## Supplemental Data 1 for "A domestic cat whole exome sequencing resource for trait discovery"

A1 90201 90250
refGene_gn=LOC101090826;refGene_tx=XM_023247174.1
A1 91101 91190
refGene_gn=LOC101090826;refGene_tx=XM_023247174.1
A1 91982 92197
refGene_gn=LOC101090826;refGene_tx=XM_023247174.1
A1 93602 93721
refGene_gn=LOC101090826;refGene_tx=XM_023247174.1
A1 94964 95117
refGene_gn=LOC101090826;refGene_tx=XM_023247174.1
A1 96654 96853
refGene_gn=LOC101090826;refGene_tx=XM_023247174.1
A1 100055 100174
refGene_gn=LOC101090826;refGene_tx=XM_023247174.1
A1 101221 101400
refGene_gn=LOC101090826;refGene_tx=XM_023247174.1
A1 157484 157597
refGene_gn=CCDC115;refGene_tx=XM_003980242.5;refGene_gn=CCDC115;refGen
e_tx=XM_011289120.3
A1 159363 159510
refGene_gn=CCDC115;refGene_tx=XM_003980242.5;refGene_gn=CCDC115;refGen
e_tx=XM_011289120.3
A1 160113 160196
refGene_gn=CCDC115;refGene_tx=XM_003980242.5;refGene_gn=CCDC115;refGen
e_tx=XM_011289120.3
A1 160284 160388
refGene_gn=CCDC115;refGene_tx=XM_003980242.5;refGene_gn=CCDC115;refGen
e_tx=XM_011289120.3
A1 160462 160636
refGene_gn=CCDC115;refGene_tx=XM_003980242.5;refGene_gn=CCDC115;refGen
e_tx=XM_011289120.3
A1 161347 161350 refGene_gn=IMP4;refGene_tx=XM_003980243.4
A1 161492 161601 refGene_gn=IMP4;refGene_tx=XM_003980243.4
A1 163599 163683 refGene_gn=IMP4;refGene_tx=XM_003980243.4
A1 164390 164500 refGene_gn=IMP4;refGene_tx=XM_003980243.4
A1 164578 164711 refGene_gn=IMP4;refGene_tx=XM_003980243.4
A1 164809 164964 refGene_gn=IMP4;refGene_tx=XM_003980243.4
A1 165046 165141 refGene_gn=IMP4;refGene_tx=XM_003980243.4
A1 165261 165335 refGene_gn=IMP4;refGene_tx=XM_003980243.4
A1 165425 165538 refGene_gn=IMP4;refGene_tx=XM_003980243.4
A1 178409 178575 refGene_gn=PTPN18;refGene_tx=XM_023250065.1
A1 179078 179155 refGene_gn=PTPN18;refGene_tx=XM_023250065.1
A1 179234 179322 refGene_gn=PTPN18;refGene_tx=XM_023250065.1
A1 179427 179474 refGene_gn=PTPN18;refGene_tx=XM_023250065.1
A1 186565 186634 refGene_gn=PTPN18;refGene_tx=XM_023250065.1
A1 187220 187283 refGene_gn=PTPN18;refGene_tx=XM_023250065.1
A1 187496 187639 refGene_gn=PTPN18;refGene_tx=XM_023250065.1
A1 188539 188580 refGene_gn=PTPN18;refGene_tx=XM_023250065.1
A1 188642 188761 refGene_gn=PTPN18;refGene_tx=XM_023250065.1
A1 188891 189004 refGene_gn=PTPN18;refGene_tx=XM_023250065.1

A1 189208 189339 refGene_gn=PTPN18;refGene_tx=XM_023250065.1
A1 189658 190210 refGene_gn=PTPN18;refGene_tx=XM_023250065.1
A1 190638 190713 refGene_gn=PTPN18;refGene_tx=XM_023250065.1
A1 190921 190989 refGene_gn=PTPN18;refGene_tx=XM_023250065.1
A1 226178 226236
refGene_gn=LOC111559468;refGene_tx=XM_023250181.1
A1 227855 227973
refGene_gn=LOC111559468;refGene_tx=XM_023250181.1
A1 228534 228622
refGene_gn=LOC111559468;refGene_tx=XM_023250181.1
A1 229404 229513
refGene_gn=LOC111559468;refGene_tx=XM_023250181.1
A1 232952 233018
refGene_gn=LOC111559468;refGene_tx=XM_023250181.1
A1 233943 234045
refGene_gn=LOC111559468;refGene_tx=XM_023250181.1
A1 237069 237262
refGene_gn=LOC111559468;refGene_tx=XM_023250181.1
A1 237633 237770
refGene_gn=LOC111559468;refGene_tx=XM_023250181.1
A1 238117 238342
refGene_gn=LOC111559468;refGene_tx=XM_023250181.1
A1 238680 238822
refGene_gn=LOC111559468;refGene_tx=XM_023250181.1
A1 241224 241348
refGene_gn=LOC111559468;refGene_tx=XM_023250181.1
A1 241715 241850
refGene_gn=LOC111559468;refGene_tx=XM_023250181.1
A1 242603 242915
refGene_gn=LOC111559468;refGene_tx=XM_023250181.1
A1 245054 245112
refGene_gn=LOC102899369;refGene_tx=XM_023247446.1
A1 246898 247025
refGene_gn=LOC102899369;refGene_tx=XM_023247446.1
A1 247722 247862
refGene_gn=LOC102899369;refGene_tx=XM_023247446.1
A1 249665 249869
refGene_gn=LOC102899369;refGene_tx=XM_023247446.1
A1 250483 250633
refGene_gn=LOC102899369;refGene_tx=XM_023247446.1
A1 250841 251021
refGene_gn=LOC102899369;refGene_tx=XM_023247446.1
A1 251731 251848
refGene_gn=LOC102899369;refGene_tx=XM_023247446.1
A1 252379 252529
refGene_gn=LOC102899369;refGene_tx=XM_023247446.1
A1 252721 252937
refGene_gn=LOC102899369;refGene_tx=XM_023247446.1
A1 253942 254047
refGene_gn=LOC102899369;refGene_tx=XM_023247446.1

A1	254606	254748
refGene_gn=LOC102899369;refGene_tx=XM_023247446.1
A1	255063	255278
refGene_gn=LOC102899369;refGene_tx=XM_023247446.1
A1	255531	255750
refGene_gn=LOC102899369;refGene_tx=XM_023247446.1
A1	257165	257272
refGene_gn=LOC102899369;refGene_tx=XM_023247446.1
A1	258438	258592
refGene_gn=LOC102899369;refGene_tx=XM_023247446.1
A1	258758	258858
refGene_gn=LOC102899369;refGene_tx=XM_023247446.1
A1	259304	259471
refGene_gn=LOC102899369;refGene_tx=XM_023247446.1
A1	259685	259859
refGene_gn=LOC102899369;refGene_tx=XM_023247446.1
A1	263362	263587
refGene_gn=LOC102899369;refGene_tx=XM_023247446.1
A1	264535	264685
refGene_gn=LOC102899369;refGene_tx=XM_023247446.1
A1	265267	265371
refGene_gn=LOC102899369;refGene_tx=XM_023247446.1
A1	266181	266393
refGene_gn=LOC102899369;refGene_tx=XM_023247446.1
A1	267134	267355
refGene_gn=LOC102899369;refGene_tx=XM_023247446.1
A1	268389	268536
refGene_gn=LOC102899369;refGene_tx=XM_023247446.1
A1	269173	269338
refGene_gn=LOC102899369;refGene_tx=XM_023247446.1
A1	269791	269943
refGene_gn=LOC102899369;refGene_tx=XM_023247446.1
A1	270269	270438
refGene_gn=LOC102899369;refGene_tx=XM_023247446.1
A1	272609	272718
refGene_gn=LOC102899369;refGene_tx=XM_023247446.1
A1	274279	274416
refGene_gn=LOC102899369;refGene_tx=XM_023247446.1
A1	274910	275036
refGene_gn=LOC102899369;refGene_tx=XM_023247446.1
A1	275139	275310
refGene_gn=LOC102899369;refGene_tx=XM_023247446.1
A1	275409	275634
refGene_gn=LOC102899369;refGene_tx=XM_023247446.1
A1	275888	276015
refGene_gn=LOC102899369;refGene_tx=XM_023247446.1
A1	276845	277045
refGene_gn=LOC102899369;refGene_tx=XM_023247446.1
A1	277707	277920
refGene_gn=LOC102899369;refGene_tx=XM_023247446.1

A1 284214 284360
refGene_gn=LOC102899369;refGene_tx=XM_023247446.1
A1 284872 285000
refGene_gn=LOC102899369;refGene_tx=XM_023247446.1
A1 285725 285870
refGene_gn=LOC102899369;refGene_tx=XM_023247446.1
A1 287008 287141
refGene_gn=LOC102899369;refGene_tx=XM_023247446.1
A1 287459 287621
refGene_gn=LOC102899369;refGene_tx=XM_023247446.1
A1 289796 289923
refGene_gn=LOC102899369;refGene_tx=XM_023247446.1
A1 290230 290442
refGene_gn=LOC102899369;refGene_tx=XM_023247446.1
A1 290983 291146
refGene_gn=LOC102899369;refGene_tx=XM_023247446.1
A1 291587 291670
refGene_gn=LOC102899369;refGene_tx=XM_023247446.1
A1 294275 294436
refGene_gn=LOC102899369;refGene_tx=XM_023247446.1
A1 294554 294720
refGene_gn=LOC102899369;refGene_tx=XM_023247446.1
A1 296840 296956
refGene_gn=LOC102899369;refGene_tx=XM_023247446.1
A1 297757 297876
refGene_gn=LOC102899369;refGene_tx=XM_023247446.1
A1 301247 301377
refGene_gn=LOC102899369;refGene_tx=XM_023247446.1
A1 308061 308199
refGene_gn=LOC102899369;refGene_tx=XM_023247446.1
A1 309560 309809
refGene_gn=LOC102899369;refGene_tx=XM_023247446.1
A1 384105 384114 refGene_gn=ATP12A;refGene_tx=XM_023247527.1
A1 384781 384937 refGene_gn=ATP12A;refGene_tx=XM_023247527.1
A1 388513 388573 refGene_gn=ATP12A;refGene_tx=XM_023247527.1
A1 390999 391203 refGene_gn=ATP12A;refGene_tx=XM_023247527.1
A1 391973 392087 refGene_gn=ATP12A;refGene_tx=XM_023247527.1
A1 393050 393185 refGene_gn=ATP12A;refGene_tx=XM_023247527.1
A1 393315 393433 refGene_gn=ATP12A;refGene_tx=XM_023247527.1
A1 393687 393955 refGene_gn=ATP12A;refGene_tx=XM_023247527.1
A1 395524 395724 refGene_gn=ATP12A;refGene_tx=XM_023247527.1
A1 395872 395982 refGene_gn=ATP12A;refGene_tx=XM_023247527.1
A1 397540 397675 refGene_gn=ATP12A;refGene_tx=XM_023247527.1
A1 401281 401474 refGene_gn=ATP12A;refGene_tx=XM_023247527.1
A1 403423 403599 refGene_gn=ATP12A;refGene_tx=XM_023247527.1
A1 404279 404416 refGene_gn=ATP12A;refGene_tx=XM_023247527.1
A1 406819 406970 refGene_gn=ATP12A;refGene_tx=XM_023247527.1
A1 408000 408169 refGene_gn=ATP12A;refGene_tx=XM_023247527.1
A1 408252 408407 refGene_gn=ATP12A;refGene_tx=XM_023247527.1
A1 410414 410538 refGene_gn=ATP12A;refGene_tx=XM_023247527.1

```
A1	410727	410873	refGene_gn=ATP12A;refGene_tx=XM_023247527.1
A1	411352	411486	refGene_gn=ATP12A;refGene_tx=XM_023247527.1
A1	411625	411727	refGene_gn=ATP12A;refGene_tx=XM_023247527.1
A1	412352	412444	refGene_gn=ATP12A;refGene_tx=XM_023247527.1
A1	412529	412558	refGene_gn=ATP12A;refGene_tx=XM_023247527.1
A1	449293	449423
refGene_gn=RNF17;refGene_tx=XM_006927145.4;refGene_gn=RNF17;refGene_tx
=XM_011281073.3
A1	451572	451667
refGene_gn=RNF17;refGene_tx=XM_006927145.4;refGene_gn=RNF17;refGene_tx
=XM_011281073.3
A1	460777	460869
refGene_gn=RNF17;refGene_tx=XM_006927145.4;refGene_gn=RNF17;refGene_tx
=XM_011281073.3
A1	470706	470818
refGene_gn=RNF17;refGene_tx=XM_006927145.4;refGene_gn=RNF17;refGene_tx
=XM_011281073.3
A1	471729	471813
refGene_gn=RNF17;refGene_tx=XM_006927145.4;refGene_gn=RNF17;refGene_tx
=XM_011281073.3
A1	473001	473102
refGene_gn=RNF17;refGene_tx=XM_006927145.4;refGene_gn=RNF17;refGene_tx
=XM_011281073.3
A1	477961	478133
refGene_gn=RNF17;refGene_tx=XM_006927145.4;refGene_gn=RNF17;refGene_tx
=XM_011281073.3
A1	479569	479646
refGene_gn=RNF17;refGene_tx=XM_006927145.4;refGene_gn=RNF17;refGene_tx
=XM_011281073.3
A1	480617	480692
refGene_gn=RNF17;refGene_tx=XM_006927145.4;refGene_gn=RNF17;refGene_tx
=XM_011281073.3
A1	485304	485612
refGene_gn=RNF17;refGene_tx=XM_006927145.4;refGene_gn=RNF17;refGene_tx
=XM_011281073.3
A1	486615	486705
refGene_gn=RNF17;refGene_tx=XM_006927145.4;refGene_gn=RNF17;refGene_tx
=XM_011281073.3
A1	487930	488089
refGene_gn=RNF17;refGene_tx=XM_006927145.4;refGene_gn=RNF17;refGene_tx
=XM_011281073.3
A1	490761	490951
refGene_gn=RNF17;refGene_tx=XM_006927145.4;refGene_gn=RNF17;refGene_tx
=XM_011281073.3
A1	492067	492236
refGene_gn=RNF17;refGene_tx=XM_006927145.4;refGene_gn=RNF17;refGene_tx
=XM_011281073.3
A1	494324	494515
refGene_gn=RNF17;refGene_tx=XM_006927145.4;refGene_gn=RNF17;refGene_tx
=XM_011281073.3
```

A1 495442 495584
refGene_gn=RNF17;refGene_tx=XM_006927145.4;refGene_gn=RNF17;refGene_tx=XM_011281073.3
A1 507833 507987
refGene_gn=RNF17;refGene_tx=XM_006927145.4;refGene_gn=RNF17;refGene_tx=XM_011281073.3
A1 508200 508316
refGene_gn=RNF17;refGene_tx=XM_006927145.4;refGene_gn=RNF17;refGene_tx=XM_011281073.3
A1 509274 509395
refGene_gn=RNF17;refGene_tx=XM_006927145.4;refGene_gn=RNF17;refGene_tx=XM_011281073.3
A1 512732 512853
refGene_gn=RNF17;refGene_tx=XM_006927145.4;refGene_gn=RNF17;refGene_tx=XM_011281073.3
A1 514890 515118
refGene_gn=RNF17;refGene_tx=XM_006927145.4;refGene_gn=RNF17;refGene_tx=XM_011281073.3
A1 515777 515928
refGene_gn=RNF17;refGene_tx=XM_006927145.4;refGene_gn=RNF17;refGene_tx=XM_011281073.3
A1 516083 516202
refGene_gn=RNF17;refGene_tx=XM_006927145.4;refGene_gn=RNF17;refGene_tx=XM_011281073.3
A1 522126 522217
refGene_gn=RNF17;refGene_tx=XM_006927145.4;refGene_gn=RNF17;refGene_tx=XM_011281073.3
A1 523378 523494
refGene_gn=RNF17;refGene_tx=XM_006927145.4;refGene_gn=RNF17;refGene_tx=XM_011281073.3
A1 525124 525411
refGene_gn=RNF17;refGene_tx=XM_006927145.4;refGene_gn=RNF17;refGene_tx=XM_011281073.3
A1 526738 526902
refGene_gn=RNF17;refGene_tx=XM_006927145.4;refGene_gn=RNF17;refGene_tx=XM_011281073.3
A1 542854 542974
refGene_gn=RNF17;refGene_tx=XM_006927145.4;refGene_gn=RNF17;refGene_tx=XM_011281073.3
A1 544144 544225
refGene_gn=RNF17;refGene_tx=XM_006927145.4;refGene_gn=RNF17;refGene_tx=XM_011281073.3
A1 546429 546555
refGene_gn=RNF17;refGene_tx=XM_006927145.4;refGene_gn=RNF17;refGene_tx=XM_011281073.3
A1 548254 548311 refGene_gn=RNF17;refGene_tx=XM_006927145.4
A1 551817 551934
refGene_gn=RNF17;refGene_tx=XM_006927145.4;refGene_gn=RNF17;refGene_tx=XM_011281073.3
A1 553174 553343

refGene_gn=RNF17;refGene_tx=XM_006927145.4;refGene_gn=RNF17;refGene_tx=XM_011281073.3
A1 555894 556030
refGene_gn=RNF17;refGene_tx=XM_006927145.4;refGene_gn=RNF17;refGene_tx=XM_011281073.3
A1 560658 560848
refGene_gn=RNF17;refGene_tx=XM_006927145.4;refGene_gn=RNF17;refGene_tx=XM_011281073.3
A1 564740 564839
refGene_gn=RNF17;refGene_tx=XM_006927145.4;refGene_gn=RNF17;refGene_tx=XM_011281073.3
A1 567168 567361
refGene_gn=CENPJ;refGene_tx=XM_003980244.4;refGene_gn=CENPJ;refGene_tx=XM_006927146.4;refGene_gn=CENPJ;refGene_tx=XM_019815063.2;refGene_gn=CENPJ;refGene_tx=XM_019815137.2;refGene_gn=CENPJ;refGene_tx=XM_019815190.2;refGene_gn=CENPJ;refGene_tx=XM_019815226.2;refGene_gn=CENPJ;refGene_tx=XM_019815312.2
A1 569678 569799
refGene_gn=CENPJ;refGene_tx=XM_003980244.4;refGene_gn=CENPJ;refGene_tx=XM_019815063.2;refGene_gn=CENPJ;refGene_tx=XM_019815137.2;refGene_gn=CENPJ;refGene_tx=XM_019815190.2;refGene_gn=CENPJ;refGene_tx=XM_019815226.2
A1 570102 570187
refGene_gn=CENPJ;refGene_tx=XM_003980244.4;refGene_gn=CENPJ;refGene_tx=XM_006927146.4;refGene_gn=CENPJ;refGene_tx=XM_019815063.2;refGene_gn=CENPJ;refGene_tx=XM_019815137.2;refGene_gn=CENPJ;refGene_tx=XM_019815190.2;refGene_gn=CENPJ;refGene_tx=XM_019815226.2
A1 570482 570623
refGene_gn=CENPJ;refGene_tx=XM_003980244.4;refGene_gn=CENPJ;refGene_tx=XM_006927146.4;refGene_gn=CENPJ;refGene_tx=XM_019815063.2;refGene_gn=CENPJ;refGene_tx=XM_019815137.2;refGene_gn=CENPJ;refGene_tx=XM_019815190.2;refGene_gn=CENPJ;refGene_tx=XM_019815226.2;refGene_gn=CENPJ;refGene_tx=XM_019815312.2
A1 573623 573743
refGene_gn=CENPJ;refGene_tx=XM_003980244.4;refGene_gn=CENPJ;refGene_tx=XM_006927146.4;refGene_gn=CENPJ;refGene_tx=XM_019815063.2;refGene_gn=CENPJ;refGene_tx=XM_019815137.2;refGene_gn=CENPJ;refGene_tx=XM_019815190.2;refGene_gn=CENPJ;refGene_tx=XM_019815226.2;refGene_gn=CENPJ;refGene_tx=XM_019815312.2
A1 574227 574292
refGene_gn=CENPJ;refGene_tx=XM_003980244.4;refGene_gn=CENPJ;refGene_tx=XM_006927146.4;refGene_gn=CENPJ;refGene_tx=XM_019815063.2;refGene_gn=CENPJ;refGene_tx=XM_019815137.2;refGene_gn=CENPJ;refGene_tx=XM_019815190.2;refGene_gn=CENPJ;refGene_tx=XM_019815226.2;refGene_gn=CENPJ;refGene_tx=XM_019815312.2
A1 583060 583145
refGene_gn=CENPJ;refGene_tx=XM_003980244.4;refGene_gn=CENPJ;refGene_tx=XM_006927146.4;refGene_gn=CENPJ;refGene_tx=XM_019815063.2;refGene_gn=CENPJ;refGene_tx=XM_019815137.2;refGene_gn=CENPJ;refGene_tx=XM_019815190.2;refGene_gn=CENPJ;refGene_tx=XM_019815226.2;refGene_gn=CENPJ;refGe

ne_tx=XM_019815312.2
A1 586885 587128
refGene_gn=CENPJ;refGene_tx=XM_003980244.4;refGene_gn=CENPJ;refGene_tx=XM_006927146.4;refGene_gn=CENPJ;refGene_tx=XM_019815063.2;refGene_gn=CENPJ;refGene_tx=XM_019815137.2;refGene_gn=CENPJ;refGene_tx=XM_019815190.2;refGene_gn=CENPJ;refGene_tx=XM_019815312.2
A1 590479 590645
refGene_gn=CENPJ;refGene_tx=XM_003980244.4;refGene_gn=CENPJ;refGene_tx=XM_006927146.4;refGene_gn=CENPJ;refGene_tx=XM_019815063.2;refGene_gn=CENPJ;refGene_tx=XM_019815137.2;refGene_gn=CENPJ;refGene_tx=XM_019815190.2;refGene_gn=CENPJ;refGene_tx=XM_019815226.2;refGene_gn=CENPJ;refGene_tx=XM_019815312.2
A1 600629 600762
refGene_gn=CENPJ;refGene_tx=XM_003980244.4;refGene_gn=CENPJ;refGene_tx=XM_006927146.4;refGene_gn=CENPJ;refGene_tx=XM_019815063.2;refGene_gn=CENPJ;refGene_tx=XM_019815137.2;refGene_gn=CENPJ;refGene_tx=XM_019815190.2;refGene_gn=CENPJ;refGene_tx=XM_019815226.2;refGene_gn=CENPJ;refGene_tx=XM_019815312.2
A1 604244 605874
refGene_gn=CENPJ;refGene_tx=XM_003980244.4;refGene_gn=CENPJ;refGene_tx=XM_006927146.4;refGene_gn=CENPJ;refGene_tx=XM_019815063.2;refGene_gn=CENPJ;refGene_tx=XM_019815137.2;refGene_gn=CENPJ;refGene_tx=XM_019815190.2;refGene_gn=CENPJ;refGene_tx=XM_019815226.2;refGene_gn=CENPJ;refGene_tx=XM_019815312.2
A1 605999 606102
refGene_gn=CENPJ;refGene_tx=XM_003980244.4;refGene_gn=CENPJ;refGene_tx=XM_006927146.4;refGene_gn=CENPJ;refGene_tx=XM_019815063.2;refGene_gn=CENPJ;refGene_tx=XM_019815137.2;refGene_gn=CENPJ;refGene_tx=XM_019815190.2;refGene_gn=CENPJ;refGene_tx=XM_019815226.2;refGene_gn=CENPJ;refGene_tx=XM_019815312.2
A1 607632 607744
refGene_gn=CENPJ;refGene_tx=XM_003980244.4;refGene_gn=CENPJ;refGene_tx=XM_006927146.4;refGene_gn=CENPJ;refGene_tx=XM_019815063.2;refGene_gn=CENPJ;refGene_tx=XM_019815137.2;refGene_gn=CENPJ;refGene_tx=XM_019815190.2;refGene_gn=CENPJ;refGene_tx=XM_019815226.2;refGene_gn=CENPJ;refGene_tx=XM_019815312.2
A1 609680 609980
refGene_gn=CENPJ;refGene_tx=XM_003980244.4;refGene_gn=CENPJ;refGene_tx=XM_006927146.4;refGene_gn=CENPJ;refGene_tx=XM_019815063.2;refGene_gn=CENPJ;refGene_tx=XM_019815137.2;refGene_gn=CENPJ;refGene_tx=XM_019815190.2;refGene_gn=CENPJ;refGene_tx=XM_019815226.2;refGene_gn=CENPJ;refGene_tx=XM_019815312.2
A1 615542 615666
refGene_gn=CENPJ;refGene_tx=XM_003980244.4;refGene_gn=CENPJ;refGene_tx=XM_006927146.4;refGene_gn=CENPJ;refGene_tx=XM_019815063.2;refGene_gn=CENPJ;refGene_tx=XM_019815137.2;refGene_gn=CENPJ;refGene_tx=XM_019815190.2;refGene_gn=CENPJ;refGene_tx=XM_019815226.2;refGene_gn=CENPJ;refGene_tx=XM_019815312.2
A1 616164 616608
refGene_gn=CENPJ;refGene_tx=XM_003980244.4;refGene_gn=CENPJ;refGene_tx

=XM_006927146.4;refGene_gn=CENPJ;refGene_tx=XM_019815063.2;refGene_gn=CENPJ;refGene_tx=XM_019815137.2;refGene_gn=CENPJ;refGene_tx=XM_019815190.2;refGene_gn=CENPJ;refGene_tx=XM_019815226.2;refGene_gn=CENPJ;refGene_tx=XM_019815312.2
A1 640579 640711
refGene_gn=PARP4;refGene_tx=XM_011282022.3;refGene_gn=PARP4;refGene_tx=XM_019814699.2;refGene_gn=PARP4;refGene_tx=XM_019814760.2;refGene_gn=PARP4;refGene_tx=XM_019814802.2;refGene_gn=PARP4;refGene_tx=XM_019814835.2
A1 643665 643867
refGene_gn=PARP4;refGene_tx=XM_011282022.3;refGene_gn=PARP4;refGene_tx=XM_019814699.2;refGene_gn=PARP4;refGene_tx=XM_019814760.2;refGene_gn=PARP4;refGene_tx=XM_019814802.2;refGene_gn=PARP4;refGene_tx=XM_019814835.2
A1 646443 646510
refGene_gn=PARP4;refGene_tx=XM_011282022.3;refGene_gn=PARP4;refGene_tx=XM_019814699.2;refGene_gn=PARP4;refGene_tx=XM_019814760.2;refGene_gn=PARP4;refGene_tx=XM_019814802.2;refGene_gn=PARP4;refGene_tx=XM_019814835.2;refGene_gn=PARP4;refGene_tx=XM_019814919.2
A1 647400 647476
refGene_gn=PARP4;refGene_tx=XM_011282022.3;refGene_gn=PARP4;refGene_tx=XM_019814699.2;refGene_gn=PARP4;refGene_tx=XM_019814760.2;refGene_gn=PARP4;refGene_tx=XM_019814802.2;refGene_gn=PARP4;refGene_tx=XM_019814835.2;refGene_gn=PARP4;refGene_tx=XM_019814919.2
A1 648212 648329
refGene_gn=PARP4;refGene_tx=XM_011282022.3;refGene_gn=PARP4;refGene_tx=XM_019814699.2;refGene_gn=PARP4;refGene_tx=XM_019814760.2;refGene_gn=PARP4;refGene_tx=XM_019814802.2;refGene_gn=PARP4;refGene_tx=XM_019814835.2;refGene_gn=PARP4;refGene_tx=XM_019814919.2
A1 650958 651108
refGene_gn=PARP4;refGene_tx=XM_011282022.3;refGene_gn=PARP4;refGene_tx=XM_019814699.2;refGene_gn=PARP4;refGene_tx=XM_019814760.2;refGene_gn=PARP4;refGene_tx=XM_019814802.2;refGene_gn=PARP4;refGene_tx=XM_019814835.2;refGene_gn=PARP4;refGene_tx=XM_019814919.2
A1 651639 651777
refGene_gn=PARP4;refGene_tx=XM_011282022.3;refGene_gn=PARP4;refGene_tx=XM_019814699.2;refGene_gn=PARP4;refGene_tx=XM_019814760.2;refGene_gn=PARP4;refGene_tx=XM_019814802.2;refGene_gn=PARP4;refGene_tx=XM_019814835.2;refGene_gn=PARP4;refGene_tx=XM_019814919.2
A1 652833 653007
refGene_gn=PARP4;refGene_tx=XM_011282022.3;refGene_gn=PARP4;refGene_tx=XM_019814699.2;refGene_gn=PARP4;refGene_tx=XM_019814760.2;refGene_gn=PARP4;refGene_tx=XM_019814802.2;refGene_gn=PARP4;refGene_tx=XM_019814835.2;refGene_gn=PARP4;refGene_tx=XM_019814919.2
A1 656468 656629
refGene_gn=PARP4;refGene_tx=XM_011282022.3;refGene_gn=PARP4;refGene_tx=XM_019814699.2;refGene_gn=PARP4;refGene_tx=XM_019814760.2;refGene_gn=PARP4;refGene_tx=XM_019814802.2;refGene_gn=PARP4;refGene_tx=XM_019814835.2;refGene_gn=PARP4;refGene_tx=XM_019814919.2
A1 660441 660579

refGene_gn=PARP4;refGene_tx=XM_011282022.3;refGene_gn=PARP4;refGene_tx=XM_019814699.2;refGene_gn=PARP4;refGene_tx=XM_019814760.2;refGene_gn=PARP4;refGene_tx=XM_019814802.2;refGene_gn=PARP4;refGene_tx=XM_019814835.2;refGene_gn=PARP4;refGene_tx=XM_019814919.2
A1 662811 662907
refGene_gn=PARP4;refGene_tx=XM_011282022.3;refGene_gn=PARP4;refGene_tx=XM_019814699.2;refGene_gn=PARP4;refGene_tx=XM_019814760.2;refGene_gn=PARP4;refGene_tx=XM_019814802.2;refGene_gn=PARP4;refGene_tx=XM_019814835.2;refGene_gn=PARP4;refGene_tx=XM_019814919.2
A1 666823 667007
refGene_gn=PARP4;refGene_tx=XM_011282022.3;refGene_gn=PARP4;refGene_tx=XM_019814699.2;refGene_gn=PARP4;refGene_tx=XM_019814760.2;refGene_gn=PARP4;refGene_tx=XM_019814802.2;refGene_gn=PARP4;refGene_tx=XM_019814835.2;refGene_gn=PARP4;refGene_tx=XM_019814919.2
A1 667198 667355
refGene_gn=PARP4;refGene_tx=XM_011282022.3;refGene_gn=PARP4;refGene_tx=XM_019814699.2;refGene_gn=PARP4;refGene_tx=XM_019814760.2;refGene_gn=PARP4;refGene_tx=XM_019814802.2;refGene_gn=PARP4;refGene_tx=XM_019814835.2;refGene_gn=PARP4;refGene_tx=XM_019814919.2
A1 668809 668934
refGene_gn=PARP4;refGene_tx=XM_011282022.3;refGene_gn=PARP4;refGene_tx=XM_019814699.2;refGene_gn=PARP4;refGene_tx=XM_019814760.2;refGene_gn=PARP4;refGene_tx=XM_019814802.2;refGene_gn=PARP4;refGene_tx=XM_019814835.2;refGene_gn=PARP4;refGene_tx=XM_019814919.2
A1 678051 678183
refGene_gn=PARP4;refGene_tx=XM_011282022.3;refGene_gn=PARP4;refGene_tx=XM_019814699.2;refGene_gn=PARP4;refGene_tx=XM_019814760.2;refGene_gn=PARP4;refGene_tx=XM_019814802.2;refGene_gn=PARP4;refGene_tx=XM_019814835.2;refGene_gn=PARP4;refGene_tx=XM_019814919.2
A1 678957 679044
refGene_gn=PARP4;refGene_tx=XM_011282022.3;refGene_gn=PARP4;refGene_tx=XM_019814699.2;refGene_gn=PARP4;refGene_tx=XM_019814760.2;refGene_gn=PARP4;refGene_tx=XM_019814802.2;refGene_gn=PARP4;refGene_tx=XM_019814835.2;refGene_gn=PARP4;refGene_tx=XM_019814919.2
A1 683166 683331
refGene_gn=PARP4;refGene_tx=XM_011282022.3;refGene_gn=PARP4;refGene_tx=XM_019814699.2;refGene_gn=PARP4;refGene_tx=XM_019814760.2;refGene_gn=PARP4;refGene_tx=XM_019814802.2;refGene_gn=PARP4;refGene_tx=XM_019814835.2;refGene_gn=PARP4;refGene_tx=XM_019814919.2
A1 683993 684040
refGene_gn=PARP4;refGene_tx=XM_011282022.3;refGene_gn=PARP4;refGene_tx=XM_019814699.2;refGene_gn=PARP4;refGene_tx=XM_019814760.2;refGene_gn=PARP4;refGene_tx=XM_019814802.2;refGene_gn=PARP4;refGene_tx=XM_019814835.2;refGene_gn=PARP4;refGene_tx=XM_019814919.2
A1 684395 684474
refGene_gn=PARP4;refGene_tx=XM_011282022.3;refGene_gn=PARP4;refGene_tx=XM_019814699.2;refGene_gn=PARP4;refGene_tx=XM_019814760.2;refGene_gn=PARP4;refGene_tx=XM_019814802.2;refGene_gn=PARP4;refGene_tx=XM_019814835.2;refGene_gn=PARP4;refGene_tx=XM_019814919.2
A1 693377 693515

refGene_gn=PARP4;refGene_tx=XM_011282022.3;refGene_gn=PARP4;refGene_tx=XM_019814699.2;refGene_gn=PARP4;refGene_tx=XM_019814760.2;refGene_gn=PARP4;refGene_tx=XM_019814802.2;refGene_gn=PARP4;refGene_tx=XM_019814835.2;refGene_gn=PARP4;refGene_tx=XM_019814919.2
A1 705400 705596
refGene_gn=PARP4;refGene_tx=XM_011282022.3;refGene_gn=PARP4;refGene_tx=XM_019814699.2;refGene_gn=PARP4;refGene_tx=XM_019814760.2;refGene_gn=PARP4;refGene_tx=XM_019814802.2;refGene_gn=PARP4;refGene_tx=XM_019814835.2;refGene_gn=PARP4;refGene_tx=XM_019814919.2
A1 706598 706666
refGene_gn=PARP4;refGene_tx=XM_011282022.3;refGene_gn=PARP4;refGene_tx=XM_019814699.2;refGene_gn=PARP4;refGene_tx=XM_019814760.2;refGene_gn=PARP4;refGene_tx=XM_019814802.2;refGene_gn=PARP4;refGene_tx=XM_019814835.2;refGene_gn=PARP4;refGene_tx=XM_019814919.2
A1 707243 707431
refGene_gn=PARP4;refGene_tx=XM_011282022.3;refGene_gn=PARP4;refGene_tx=XM_019814699.2;refGene_gn=PARP4;refGene_tx=XM_019814760.2;refGene_gn=PARP4;refGene_tx=XM_019814802.2;refGene_gn=PARP4;refGene_tx=XM_019814835.2;refGene_gn=PARP4;refGene_tx=XM_019814919.2
A1 710379 710479
refGene_gn=PARP4;refGene_tx=XM_011282022.3;refGene_gn=PARP4;refGene_tx=XM_019814699.2;refGene_gn=PARP4;refGene_tx=XM_019814760.2;refGene_gn=PARP4;refGene_tx=XM_019814802.2;refGene_gn=PARP4;refGene_tx=XM_019814835.2;refGene_gn=PARP4;refGene_tx=XM_019814919.2
A1 712148 712319
refGene_gn=PARP4;refGene_tx=XM_011282022.3;refGene_gn=PARP4;refGene_tx=XM_019814699.2;refGene_gn=PARP4;refGene_tx=XM_019814760.2;refGene_gn=PARP4;refGene_tx=XM_019814802.2;refGene_gn=PARP4;refGene_tx=XM_019814835.2;refGene_gn=PARP4;refGene_tx=XM_019814919.2
A1 712601 712682
refGene_gn=PARP4;refGene_tx=XM_011282022.3;refGene_gn=PARP4;refGene_tx=XM_019814699.2;refGene_gn=PARP4;refGene_tx=XM_019814760.2;refGene_gn=PARP4;refGene_tx=XM_019814802.2;refGene_gn=PARP4;refGene_tx=XM_019814835.2;refGene_gn=PARP4;refGene_tx=XM_019814919.2
A1 714059 714140
refGene_gn=PARP4;refGene_tx=XM_011282022.3;refGene_gn=PARP4;refGene_tx=XM_019814699.2;refGene_gn=PARP4;refGene_tx=XM_019814760.2;refGene_gn=PARP4;refGene_tx=XM_019814802.2;refGene_gn=PARP4;refGene_tx=XM_019814835.2;refGene_gn=PARP4;refGene_tx=XM_019814919.2
A1 715218 715314
refGene_gn=PARP4;refGene_tx=XM_011282022.3;refGene_gn=PARP4;refGene_tx=XM_019814699.2;refGene_gn=PARP4;refGene_tx=XM_019814760.2;refGene_gn=PARP4;refGene_tx=XM_019814802.2;refGene_gn=PARP4;refGene_tx=XM_019814835.2;refGene_gn=PARP4;refGene_tx=XM_019814919.2
A1 715902 716025
refGene_gn=PARP4;refGene_tx=XM_011282022.3;refGene_gn=PARP4;refGene_tx=XM_019814699.2;refGene_gn=PARP4;refGene_tx=XM_019814760.2;refGene_gn=PARP4;refGene_tx=XM_019814802.2;refGene_gn=PARP4;refGene_tx=XM_019814835.2;refGene_gn=PARP4;refGene_tx=XM_019814919.2
A1 723455 724460

refGene_gn=PARP4;refGene_tx=XM_011282022.3;refGene_gn=PARP4;refGene_tx=XM_019814699.2;refGene_gn=PARP4;refGene_tx=XM_019814760.2;refGene_gn=PARP4;refGene_tx=XM_019814802.2;refGene_gn=PARP4;refGene_tx=XM_019814835.2;refGene_gn=PARP4;refGene_tx=XM_019814919.2
A1 726647 726747
refGene_gn=PARP4;refGene_tx=XM_011282022.3;refGene_gn=PARP4;refGene_tx=XM_019814699.2;refGene_gn=PARP4;refGene_tx=XM_019814760.2;refGene_gn=PARP4;refGene_tx=XM_019814802.2;refGene_gn=PARP4;refGene_tx=XM_019814835.2;refGene_gn=PARP4;refGene_tx=XM_019814919.2
A1 747559 747692
refGene_gn=PARP4;refGene_tx=XM_011282022.3;refGene_gn=PARP4;refGene_tx=XM_019814699.2;refGene_gn=PARP4;refGene_tx=XM_019814760.2;refGene_gn=PARP4;refGene_tx=XM_019814802.2;refGene_gn=PARP4;refGene_tx=XM_019814835.2;refGene_gn=PARP4;refGene_tx=XM_019814919.2
A1 751369 751565
refGene_gn=PARP4;refGene_tx=XM_011282022.3;refGene_gn=PARP4;refGene_tx=XM_019814699.2;refGene_gn=PARP4;refGene_tx=XM_019814760.2;refGene_gn=PARP4;refGene_tx=XM_019814802.2;refGene_gn=PARP4;refGene_tx=XM_019814835.2;refGene_gn=PARP4;refGene_tx=XM_019814919.2
A1 757083 757305
refGene_gn=MPHOSPH8;refGene_tx=XM_011282719.3;refGene_gn=MPHOSPH8;refGene_tx=XM_019815407.2;refGene_gn=MPHOSPH8;refGene_tx=XM_019815491.2
A1 777427 777583
refGene_gn=MPHOSPH8;refGene_tx=XM_011282719.3;refGene_gn=MPHOSPH8;refGene_tx=XM_019815407.2;refGene_gn=MPHOSPH8;refGene_tx=XM_019815491.2
A1 779951 780797
refGene_gn=MPHOSPH8;refGene_tx=XM_011282719.3;refGene_gn=MPHOSPH8;refGene_tx=XM_019815407.2;refGene_gn=MPHOSPH8;refGene_tx=XM_019815491.2
A1 782106 782206
refGene_gn=MPHOSPH8;refGene_tx=XM_011282719.3;refGene_gn=MPHOSPH8;refGene_tx=XM_019815407.2;refGene_gn=MPHOSPH8;refGene_tx=XM_019815491.2
A1 783994 784252
refGene_gn=MPHOSPH8;refGene_tx=XM_011282719.3;refGene_gn=MPHOSPH8;refGene_tx=XM_019815407.2;refGene_gn=MPHOSPH8;refGene_tx=XM_019815491.2
A1 791854 791980
refGene_gn=MPHOSPH8;refGene_tx=XM_011282719.3;refGene_gn=MPHOSPH8;refGene_tx=XM_019815407.2;refGene_gn=MPHOSPH8;refGene_tx=XM_019815491.2
A1 792062 792151
refGene_gn=MPHOSPH8;refGene_tx=XM_011282719.3;refGene_gn=MPHOSPH8;refGene_tx=XM_019815407.2;refGene_gn=MPHOSPH8;refGene_tx=XM_019815491.2
A1 795619 795760
refGene_gn=MPHOSPH8;refGene_tx=XM_011282719.3;refGene_gn=MPHOSPH8;refGene_tx=XM_019815407.2;refGene_gn=MPHOSPH8;refGene_tx=XM_019815491.2
A1 796670 796757
refGene_gn=MPHOSPH8;refGene_tx=XM_011282719.3;refGene_gn=MPHOSPH8;refGene_tx=XM_019815491.2
A1 800784 800838 refGene_gn=MPHOSPH8;refGene_tx=XM_019815491.2
A1 801047 801202
refGene_gn=MPHOSPH8;refGene_tx=XM_011282719.3;refGene_gn=MPHOSPH8;refGene_tx=XM_019815407.2

```
A1	803672	803827
refGene_gn=MPHOSPH8;refGene_tx=XM_011282719.3;refGene_gn=MPHOSPH8;refGene_tx=XM_019815407.2
A1	806805	806933
refGene_gn=MPHOSPH8;refGene_tx=XM_011282719.3;refGene_gn=MPHOSPH8;refGene_tx=XM_019815407.2
A1	807789	807873
refGene_gn=MPHOSPH8;refGene_tx=XM_011282719.3;refGene_gn=MPHOSPH8;refGene_tx=XM_019815407.2
A1	808496	808538
refGene_gn=MPHOSPH8;refGene_tx=XM_011282719.3;refGene_gn=MPHOSPH8;refGene_tx=XM_019815407.2
A1	843570	843756	refGene_gn=PSPC1;refGene_tx=XM_006927147.3
A1	845791	845961	refGene_gn=PSPC1;refGene_tx=XM_006927147.3
A1	849522	849580	refGene_gn=PSPC1;refGene_tx=XM_006927147.3
A1	867776	867882	refGene_gn=PSPC1;refGene_tx=XM_006927147.3
A1	881623	881708	refGene_gn=PSPC1;refGene_tx=XM_006927147.3
A1	890767	890964	refGene_gn=PSPC1;refGene_tx=XM_006927147.3
A1	895026	895122	refGene_gn=PSPC1;refGene_tx=XM_006927147.3
A1	907342	907644	refGene_gn=PSPC1;refGene_tx=XM_006927147.3
A1	916048	916414	refGene_gn=PSPC1;refGene_tx=XM_006927147.3
A1	939267	940020
refGene_gn=ZMYM5;refGene_tx=XM_011283106.3;refGene_gn=ZMYM5;refGene_tx=XM_023248252.1;refGene_gn=ZMYM5;refGene_tx=XM_023248313.1
A1	946651	946864
refGene_gn=ZMYM5;refGene_tx=XM_011283106.3;refGene_gn=ZMYM5;refGene_tx=XM_023248252.1;refGene_gn=ZMYM5;refGene_tx=XM_023248313.1
A1	947802	947959
refGene_gn=ZMYM5;refGene_tx=XM_011283106.3;refGene_gn=ZMYM5;refGene_tx=XM_023248252.1;refGene_gn=ZMYM5;refGene_tx=XM_023248313.1;refGene_gn=ZMYM5;refGene_tx=XM_023248337.1
A1	948506	948792
refGene_gn=ZMYM5;refGene_tx=XM_011283106.3;refGene_gn=ZMYM5;refGene_tx=XM_023248252.1;refGene_gn=ZMYM5;refGene_tx=XM_023248313.1;refGene_gn=ZMYM5;refGene_tx=XM_023248401.1
A1	957601	957612	refGene_gn=ZMYM5;refGene_tx=XM_023248444.1
A1	961170	961271
refGene_gn=ZMYM5;refGene_tx=XM_011283106.3;refGene_gn=ZMYM5;refGene_tx=XM_023248252.1;refGene_gn=ZMYM5;refGene_tx=XM_023248313.1;refGene_gn=ZMYM5;refGene_tx=XM_023248337.1;refGene_gn=ZMYM5;refGene_tx=XM_023248401.1;refGene_gn=ZMYM5;refGene_tx=XM_023248444.1
A1	961508	962021
refGene_gn=ZMYM5;refGene_tx=XM_011283106.3;refGene_gn=ZMYM5;refGene_tx=XM_023248252.1;refGene_gn=ZMYM5;refGene_tx=XM_023248313.1;refGene_gn=ZMYM5;refGene_tx=XM_023248337.1;refGene_gn=ZMYM5;refGene_tx=XM_023248401.1;refGene_gn=ZMYM5;refGene_tx=XM_023248444.1
A1	1044920	1045292
refGene_gn=LOC111559482;refGene_tx=XM_023250291.1
A1	1045387	1045567
refGene_gn=LOC111559482;refGene_tx=XM_023250291.1
```

A1	1045798	1046302
refGene_gn=LOC111559482;refGene_tx=XM_023250291.1
A1	1076308	1077155
refGene_gn=ZMYM2;refGene_tx=XM_019815621.2;refGene_gn=ZMYM2;refGene_tx=XM_019815643.1;refGene_gn=ZMYM2;refGene_tx=XM_019815692.2
A1	1081550	1081836
refGene_gn=ZMYM2;refGene_tx=XM_019815621.2;refGene_gn=ZMYM2;refGene_tx=XM_019815643.1;refGene_gn=ZMYM2;refGene_tx=XM_019815692.2
A1	1083342	1083508
refGene_gn=ZMYM2;refGene_tx=XM_019815621.2;refGene_gn=ZMYM2;refGene_tx=XM_019815643.1;refGene_gn=ZMYM2;refGene_tx=XM_019815692.2
A1	1084240	1084453
refGene_gn=ZMYM2;refGene_tx=XM_019815621.2;refGene_gn=ZMYM2;refGene_tx=XM_019815643.1;refGene_gn=ZMYM2;refGene_tx=XM_019815692.2
A1	1096602	1096674
refGene_gn=ZMYM2;refGene_tx=XM_019815621.2;refGene_gn=ZMYM2;refGene_tx=XM_019815643.1;refGene_gn=ZMYM2;refGene_tx=XM_019815692.2
A1	1100566	1100717
refGene_gn=ZMYM2;refGene_tx=XM_019815621.2;refGene_gn=ZMYM2;refGene_tx=XM_019815643.1;refGene_gn=ZMYM2;refGene_tx=XM_019815692.2
A1	1101424	1101540
refGene_gn=ZMYM2;refGene_tx=XM_019815621.2;refGene_gn=ZMYM2;refGene_tx=XM_019815643.1;refGene_gn=ZMYM2;refGene_tx=XM_019815692.2
A1	1104347	1104464
refGene_gn=ZMYM2;refGene_tx=XM_019815621.2;refGene_gn=ZMYM2;refGene_tx=XM_019815643.1;refGene_gn=ZMYM2;refGene_tx=XM_019815692.2
A1	1106632	1106783
refGene_gn=ZMYM2;refGene_tx=XM_019815621.2;refGene_gn=ZMYM2;refGene_tx=XM_019815643.1;refGene_gn=ZMYM2;refGene_tx=XM_019815692.2
A1	1109312	1109485
refGene_gn=ZMYM2;refGene_tx=XM_019815621.2;refGene_gn=ZMYM2;refGene_tx=XM_019815643.1;refGene_gn=ZMYM2;refGene_tx=XM_019815692.2
A1	1116386	1116552
refGene_gn=ZMYM2;refGene_tx=XM_019815621.2;refGene_gn=ZMYM2;refGene_tx=XM_019815643.1;refGene_gn=ZMYM2;refGene_tx=XM_019815692.2
A1	1117126	1117161
refGene_gn=ZMYM2;refGene_tx=XM_019815621.2;refGene_gn=ZMYM2;refGene_tx=XM_019815643.1;refGene_gn=ZMYM2;refGene_tx=XM_019815692.2
A1	1123665	1123795
refGene_gn=ZMYM2;refGene_tx=XM_019815621.2;refGene_gn=ZMYM2;refGene_tx=XM_019815643.1;refGene_gn=ZMYM2;refGene_tx=XM_019815692.2
A1	1124482	1124598
refGene_gn=ZMYM2;refGene_tx=XM_019815621.2;refGene_gn=ZMYM2;refGene_tx=XM_019815643.1;refGene_gn=ZMYM2;refGene_tx=XM_019815692.2
A1	1125355	1125527
refGene_gn=ZMYM2;refGene_tx=XM_019815621.2;refGene_gn=ZMYM2;refGene_tx=XM_019815643.1;refGene_gn=ZMYM2;refGene_tx=XM_019815692.2
A1	1126778	1126904
refGene_gn=ZMYM2;refGene_tx=XM_019815621.2;refGene_gn=ZMYM2;refGene_tx=XM_019815643.1;refGene_gn=ZMYM2;refGene_tx=XM_019815692.2

A1	1128195	1128290
refGene_gn=ZMYM2;refGene_tx=XM_019815621.2;refGene_gn=ZMYM2;refGene_tx=XM_019815643.1;refGene_gn=ZMYM2;refGene_tx=XM_019815692.2
A1	1128833	1128860	refGene_gn=ZMYM2;refGene_tx=XM_019815692.2
A1	1129531	1129700
refGene_gn=ZMYM2;refGene_tx=XM_019815621.2;refGene_gn=ZMYM2;refGene_tx=XM_019815643.1
A1	1129951	1130103
refGene_gn=ZMYM2;refGene_tx=XM_019815621.2;refGene_gn=ZMYM2;refGene_tx=XM_019815643.1
A1	1141801	1141916
refGene_gn=ZMYM2;refGene_tx=XM_019815621.2;refGene_gn=ZMYM2;refGene_tx=XM_019815643.1
A1	1142710	1142962
refGene_gn=ZMYM2;refGene_tx=XM_019815621.2;refGene_gn=ZMYM2;refGene_tx=XM_019815643.1
A1	1143224	1143345
refGene_gn=ZMYM2;refGene_tx=XM_019815621.2;refGene_gn=ZMYM2;refGene_tx=XM_019815643.1
A1	1144769	1144962
refGene_gn=ZMYM2;refGene_tx=XM_019815621.2;refGene_gn=ZMYM2;refGene_tx=XM_019815643.1
A1	1160237	1160516
refGene_gn=LOC111558556;refGene_tx=XM_023248588.1
A1	1161554	1161617
refGene_gn=LOC111558556;refGene_tx=XM_023248588.1
A1	1196831	1198187
refGene_gn=GJA3;refGene_tx=XM_023248644.1;refGene_gn=GJA3;refGene_tx=XM_023248653.1
A1	1246105	1246786
refGene_gn=GJB2;refGene_tx=XM_003980250.5;refGene_gn=GJB2;refGene_tx=XM_019815796.2;refGene_gn=GJB2;refGene_tx=XM_023248847.1
A1	1279430	1280216
refGene_gn=GJB6;refGene_tx=XM_023248688.1;refGene_gn=GJB6;refGene_tx=XM_023248721.1;refGene_gn=GJB6;refGene_tx=XM_023248732.1;refGene_gn=GJB6;refGene_tx=XM_023248774.1;refGene_gn=GJB6;refGene_tx=XM_023248789.1;refGene_gn=GJB6;refGene_tx=XM_023248809.1
A1	1462818	1462932	refGene_gn=CRYL1;refGene_tx=XM_003980288.4
A1	1463311	1463418	refGene_gn=CRYL1;refGene_tx=XM_003980288.4
A1	1470037	1470143	refGene_gn=CRYL1;refGene_tx=XM_003980288.4
A1	1487197	1487392	refGene_gn=CRYL1;refGene_tx=XM_003980288.4
A1	1494298	1494460	refGene_gn=CRYL1;refGene_tx=XM_003980288.4
A1	1556242	1556283	refGene_gn=CRYL1;refGene_tx=XM_003980288.4
A1	1556285	1556368	refGene_gn=CRYL1;refGene_tx=XM_003980288.4
A1	1572217	1572325	refGene_gn=CRYL1;refGene_tx=XM_003980288.4
A1	1591203	1591244	refGene_gn=CRYL1;refGene_tx=XM_003980288.4
A1	1624368	1624458
refGene_gn=IFT88;refGene_tx=XM_023248958.1;refGene_gn=IFT88;refGene_tx=XM_023248971.1;refGene_gn=IFT88;refGene_tx=XM_023248988.1;refGene_gn=IFT88;refGene_tx=XM_023249092.1

A1	1628603	1628666
refGene_gn=IFT88;refGene_tx=XM_023248958.1;refGene_gn=IFT88;refGene_tx=XM_023248971.1
A1	1644672	1644729
refGene_gn=IFT88;refGene_tx=XM_023248958.1;refGene_gn=IFT88;refGene_tx=XM_023248988.1
A1	1645855	1645909
refGene_gn=IFT88;refGene_tx=XM_023248958.1;refGene_gn=IFT88;refGene_tx=XM_023248971.1;refGene_gn=IFT88;refGene_tx=XM_023248988.1;refGene_gn=IFT88;refGene_tx=XM_023249016.1;refGene_gn=IFT88;refGene_tx=XM_023249045.1;refGene_gn=IFT88;refGene_tx=XM_023249073.1
A1	1646544	1646608
refGene_gn=IFT88;refGene_tx=XM_023248958.1;refGene_gn=IFT88;refGene_tx=XM_023248971.1;refGene_gn=IFT88;refGene_tx=XM_023248988.1;refGene_gn=IFT88;refGene_tx=XM_023249016.1;refGene_gn=IFT88;refGene_tx=XM_023249045.1;refGene_gn=IFT88;refGene_tx=XM_023249073.1
A1	1647596	1647666
refGene_gn=IFT88;refGene_tx=XM_023248958.1;refGene_gn=IFT88;refGene_tx=XM_023248971.1;refGene_gn=IFT88;refGene_tx=XM_023248988.1;refGene_gn=IFT88;refGene_tx=XM_023249016.1;refGene_gn=IFT88;refGene_tx=XM_023249045.1;refGene_gn=IFT88;refGene_tx=XM_023249073.1
A1	1649865	1649956
refGene_gn=IFT88;refGene_tx=XM_023248958.1;refGene_gn=IFT88;refGene_tx=XM_023248971.1;refGene_gn=IFT88;refGene_tx=XM_023248988.1;refGene_gn=IFT88;refGene_tx=XM_023249016.1;refGene_gn=IFT88;refGene_tx=XM_023249045.1;refGene_gn=IFT88;refGene_tx=XM_023249073.1;refGene_gn=IFT88;refGene_tx=XM_023249092.1
A1	1651554	1651659
refGene_gn=IFT88;refGene_tx=XM_023248958.1;refGene_gn=IFT88;refGene_tx=XM_023248971.1;refGene_gn=IFT88;refGene_tx=XM_023248988.1;refGene_gn=IFT88;refGene_tx=XM_023249016.1;refGene_gn=IFT88;refGene_tx=XM_023249045.1;refGene_gn=IFT88;refGene_tx=XM_023249073.1;refGene_gn=IFT88;refGene_tx=XM_023249092.1
A1	1653059	1653162
refGene_gn=IFT88;refGene_tx=XM_023248958.1;refGene_gn=IFT88;refGene_tx=XM_023248971.1;refGene_gn=IFT88;refGene_tx=XM_023248988.1;refGene_gn=IFT88;refGene_tx=XM_023249016.1;refGene_gn=IFT88;refGene_tx=XM_023249045.1;refGene_gn=IFT88;refGene_tx=XM_023249073.1;refGene_gn=IFT88;refGene_tx=XM_023249092.1
A1	1653995	1654110
refGene_gn=IFT88;refGene_tx=XM_023248958.1;refGene_gn=IFT88;refGene_tx=XM_023248971.1;refGene_gn=IFT88;refGene_tx=XM_023248988.1;refGene_gn=IFT88;refGene_tx=XM_023249016.1;refGene_gn=IFT88;refGene_tx=XM_023249045.1;refGene_gn=IFT88;refGene_tx=XM_023249073.1;refGene_gn=IFT88;refGene_tx=XM_023249092.1
A1	1655910	1656142
refGene_gn=IFT88;refGene_tx=XM_023248958.1;refGene_gn=IFT88;refGene_tx=XM_023248971.1;refGene_gn=IFT88;refGene_tx=XM_023248988.1;refGene_gn=IFT88;refGene_tx=XM_023249016.1;refGene_gn=IFT88;refGene_tx=XM_023249045.1;refGene_gn=IFT88;refGene_tx=XM_023249073.1;refGene_gn=IFT88;refGe

ne_tx=XM_023249092.1
A1      1657974  1658045
refGene_gn=IFT88;refGene_tx=XM_023248958.1;refGene_gn=IFT88;refGene_tx=XM_023248971.1;refGene_gn=IFT88;refGene_tx=XM_023248988.1;refGene_gn=IFT88;refGene_tx=XM_023249016.1;refGene_gn=IFT88;refGene_tx=XM_023249045.1;refGene_gn=IFT88;refGene_tx=XM_023249073.1;refGene_gn=IFT88;refGene_tx=XM_023249092.1
A1      1667333  1667420
refGene_gn=IFT88;refGene_tx=XM_023248958.1;refGene_gn=IFT88;refGene_tx=XM_023248971.1;refGene_gn=IFT88;refGene_tx=XM_023248988.1;refGene_gn=IFT88;refGene_tx=XM_023249016.1;refGene_gn=IFT88;refGene_tx=XM_023249045.1;refGene_gn=IFT88;refGene_tx=XM_023249073.1;refGene_gn=IFT88;refGene_tx=XM_023249092.1
A1      1686228  1686328
refGene_gn=IFT88;refGene_tx=XM_023248958.1;refGene_gn=IFT88;refGene_tx=XM_023248971.1;refGene_gn=IFT88;refGene_tx=XM_023248988.1;refGene_gn=IFT88;refGene_tx=XM_023249016.1;refGene_gn=IFT88;refGene_tx=XM_023249045.1;refGene_gn=IFT88;refGene_tx=XM_023249073.1;refGene_gn=IFT88;refGene_tx=XM_023249092.1
A1      1690483  1690570
refGene_gn=IFT88;refGene_tx=XM_023248958.1;refGene_gn=IFT88;refGene_tx=XM_023248971.1;refGene_gn=IFT88;refGene_tx=XM_023248988.1;refGene_gn=IFT88;refGene_tx=XM_023249016.1;refGene_gn=IFT88;refGene_tx=XM_023249045.1;refGene_gn=IFT88;refGene_tx=XM_023249073.1;refGene_gn=IFT88;refGene_tx=XM_023249092.1
A1      1696626  1696813
refGene_gn=IFT88;refGene_tx=XM_023248958.1;refGene_gn=IFT88;refGene_tx=XM_023248971.1;refGene_gn=IFT88;refGene_tx=XM_023248988.1;refGene_gn=IFT88;refGene_tx=XM_023249016.1;refGene_gn=IFT88;refGene_tx=XM_023249045.1;refGene_gn=IFT88;refGene_tx=XM_023249073.1;refGene_gn=IFT88;refGene_tx=XM_023249092.1
A1      1698856  1698965
refGene_gn=IFT88;refGene_tx=XM_023248958.1;refGene_gn=IFT88;refGene_tx=XM_023248971.1;refGene_gn=IFT88;refGene_tx=XM_023248988.1;refGene_gn=IFT88;refGene_tx=XM_023249016.1;refGene_gn=IFT88;refGene_tx=XM_023249045.1;refGene_gn=IFT88;refGene_tx=XM_023249073.1;refGene_gn=IFT88;refGene_tx=XM_023249092.1
A1      1701274  1701425
refGene_gn=IFT88;refGene_tx=XM_023248958.1;refGene_gn=IFT88;refGene_tx=XM_023248971.1;refGene_gn=IFT88;refGene_tx=XM_023248988.1;refGene_gn=IFT88;refGene_tx=XM_023249016.1;refGene_gn=IFT88;refGene_tx=XM_023249045.1;refGene_gn=IFT88;refGene_tx=XM_023249073.1;refGene_gn=IFT88;refGene_tx=XM_023249092.1
A1      1701947  1702063
refGene_gn=IFT88;refGene_tx=XM_023248958.1;refGene_gn=IFT88;refGene_tx=XM_023248971.1;refGene_gn=IFT88;refGene_tx=XM_023248988.1;refGene_gn=IFT88;refGene_tx=XM_023249016.1;refGene_gn=IFT88;refGene_tx=XM_023249045.1;refGene_gn=IFT88;refGene_tx=XM_023249073.1;refGene_gn=IFT88;refGene_tx=XM_023249092.1
A1      1713412  1713465

refGene_gn=IFT88;refGene_tx=XM_023248958.1;refGene_gn=IFT88;refGene_tx=XM_023248971.1;refGene_gn=IFT88;refGene_tx=XM_023248988.1;refGene_gn=IFT88;refGene_tx=XM_023249016.1;refGene_gn=IFT88;refGene_tx=XM_023249045.1;refGene_gn=IFT88;refGene_tx=XM_023249073.1;refGene_gn=IFT88;refGene_tx=XM_023249092.1
A1 1715281 1715347
refGene_gn=IFT88;refGene_tx=XM_023248958.1;refGene_gn=IFT88;refGene_tx=XM_023248971.1;refGene_gn=IFT88;refGene_tx=XM_023248988.1;refGene_gn=IFT88;refGene_tx=XM_023249016.1;refGene_gn=IFT88;refGene_tx=XM_023249045.1;refGene_gn=IFT88;refGene_tx=XM_023249073.1;refGene_gn=IFT88;refGene_tx=XM_023249092.1
A1 1718792 1718899
refGene_gn=IFT88;refGene_tx=XM_023248958.1;refGene_gn=IFT88;refGene_tx=XM_023248971.1;refGene_gn=IFT88;refGene_tx=XM_023248988.1;refGene_gn=IFT88;refGene_tx=XM_023249016.1;refGene_gn=IFT88;refGene_tx=XM_023249045.1;refGene_gn=IFT88;refGene_tx=XM_023249073.1;refGene_gn=IFT88;refGene_tx=XM_023249092.1
A1 1728587 1728654
refGene_gn=IFT88;refGene_tx=XM_023248958.1;refGene_gn=IFT88;refGene_tx=XM_023248971.1;refGene_gn=IFT88;refGene_tx=XM_023248988.1;refGene_gn=IFT88;refGene_tx=XM_023249016.1;refGene_gn=IFT88;refGene_tx=XM_023249045.1;refGene_gn=IFT88;refGene_tx=XM_023249073.1;refGene_gn=IFT88;refGene_tx=XM_023249092.1
A1 1752758 1752869
refGene_gn=IFT88;refGene_tx=XM_023248958.1;refGene_gn=IFT88;refGene_tx=XM_023248971.1;refGene_gn=IFT88;refGene_tx=XM_023248988.1;refGene_gn=IFT88;refGene_tx=XM_023249016.1;refGene_gn=IFT88;refGene_tx=XM_023249045.1;refGene_gn=IFT88;refGene_tx=XM_023249073.1;refGene_gn=IFT88;refGene_tx=XM_023249092.1
A1 1753108 1753230
refGene_gn=IFT88;refGene_tx=XM_023248958.1;refGene_gn=IFT88;refGene_tx=XM_023248971.1;refGene_gn=IFT88;refGene_tx=XM_023248988.1;refGene_gn=IFT88;refGene_tx=XM_023249016.1;refGene_gn=IFT88;refGene_tx=XM_023249045.1;refGene_gn=IFT88;refGene_tx=XM_023249073.1;refGene_gn=IFT88;refGene_tx=XM_023249092.1
A1 1762593 1762608 refGene_gn=IL17D;refGene_tx=XM_023249208.1
A1 1762967 1763263 refGene_gn=IL17D;refGene_tx=XM_023249208.1
A1 1780403 1780722 refGene_gn=IL17D;refGene_tx=XM_023249208.1
A1 1786785 1786928
refGene_gn=EEF1AKMT1;refGene_tx=XM_023249167.1
A1 1788987 1789268
refGene_gn=EEF1AKMT1;refGene_tx=XM_023249167.1
A1 1793958 1794005
refGene_gn=EEF1AKMT1;refGene_tx=XM_023249167.1
A1 1794006 1794042
refGene_gn=EEF1AKMT1;refGene_tx=XM_023249167.1
A1 1797858 1797935
refGene_gn=EEF1AKMT1;refGene_tx=XM_023249167.1
A1 1797937 1798001
refGene_gn=EEF1AKMT1;refGene_tx=XM_023249167.1

A1 1820730 1820928
refGene_gn=XPO4;refGene_tx=XM_003980253.5;refGene_gn=XPO4;refGene_tx=XM_006927154.3;refGene_gn=XPO4;refGene_tx=XM_019816971.2;refGene_gn=XPO4;refGene_tx=XM_019817033.1;refGene_gn=XPO4;refGene_tx=XM_019817077.2;refGene_gn=XPO4;refGene_tx=XM_023249438.1
A1 1823975 1824068
refGene_gn=XPO4;refGene_tx=XM_003980253.5;refGene_gn=XPO4;refGene_tx=XM_006927154.3;refGene_gn=XPO4;refGene_tx=XM_019816971.2;refGene_gn=XPO4;refGene_tx=XM_019817033.1;refGene_gn=XPO4;refGene_tx=XM_019817077.2
A1 1824497 1824615
refGene_gn=XPO4;refGene_tx=XM_003980253.5;refGene_gn=XPO4;refGene_tx=XM_006927154.3;refGene_gn=XPO4;refGene_tx=XM_019816971.2;refGene_gn=XPO4;refGene_tx=XM_019817033.1;refGene_gn=XPO4;refGene_tx=XM_019817077.2
A1 1825160 1825291
refGene_gn=XPO4;refGene_tx=XM_003980253.5;refGene_gn=XPO4;refGene_tx=XM_006927154.3;refGene_gn=XPO4;refGene_tx=XM_019816971.2;refGene_gn=XPO4;refGene_tx=XM_019817033.1;refGene_gn=XPO4;refGene_tx=XM_019817077.2
A1 1826794 1826913
refGene_gn=XPO4;refGene_tx=XM_003980253.5;refGene_gn=XPO4;refGene_tx=XM_006927154.3;refGene_gn=XPO4;refGene_tx=XM_019816971.2;refGene_gn=XPO4;refGene_tx=XM_019817033.1;refGene_gn=XPO4;refGene_tx=XM_019817077.2;refGene_gn=XPO4;refGene_tx=XM_023249438.1
A1 1828320 1828501
refGene_gn=XPO4;refGene_tx=XM_003980253.5;refGene_gn=XPO4;refGene_tx=XM_006927154.3;refGene_gn=XPO4;refGene_tx=XM_019816971.2;refGene_gn=XPO4;refGene_tx=XM_019817033.1;refGene_gn=XPO4;refGene_tx=XM_019817077.2;refGene_gn=XPO4;refGene_tx=XM_023249438.1
A1 1829563 1829857
refGene_gn=XPO4;refGene_tx=XM_003980253.5;refGene_gn=XPO4;refGene_tx=XM_006927154.3;refGene_gn=XPO4;refGene_tx=XM_019816971.2;refGene_gn=XPO4;refGene_tx=XM_019817033.1;refGene_gn=XPO4;refGene_tx=XM_019817077.2;refGene_gn=XPO4;refGene_tx=XM_023249438.1
A1 1832429 1832604
refGene_gn=XPO4;refGene_tx=XM_003980253.5;refGene_gn=XPO4;refGene_tx=XM_006927154.3;refGene_gn=XPO4;refGene_tx=XM_019816971.2;refGene_gn=XPO4;refGene_tx=XM_019817033.1;refGene_gn=XPO4;refGene_tx=XM_019817077.2;refGene_gn=XPO4;refGene_tx=XM_023249438.1
A1 1834187 1834357
refGene_gn=XPO4;refGene_tx=XM_003980253.5;refGene_gn=XPO4;refGene_tx=XM_006927154.3;refGene_gn=XPO4;refGene_tx=XM_019816971.2;refGene_gn=XPO4;refGene_tx=XM_019817033.1;refGene_gn=XPO4;refGene_tx=XM_019817077.2;refGene_gn=XPO4;refGene_tx=XM_023249438.1
A1 1834836 1834996
refGene_gn=XPO4;refGene_tx=XM_003980253.5;refGene_gn=XPO4;refGene_tx=XM_006927154.3;refGene_gn=XPO4;refGene_tx=XM_019816971.2;refGene_gn=XPO4;refGene_tx=XM_019817033.1;refGene_gn=XPO4;refGene_tx=XM_019817077.2;refGene_gn=XPO4;refGene_tx=XM_023249438.1
A1 1840680 1840858
refGene_gn=XPO4;refGene_tx=XM_003980253.5;refGene_gn=XPO4;refGene_tx=XM_006927154.3;refGene_gn=XPO4;refGene_tx=XM_019816971.2;refGene_gn=XPO

4;refGene_tx=XM_019817033.1;refGene_gn=XPO4;refGene_tx=XM_019817077.2;
refGene_gn=XPO4;refGene_tx=XM_023249438.1
A1 1841661 1841807
refGene_gn=XPO4;refGene_tx=XM_003980253.5;refGene_gn=XPO4;refGene_tx=X
M_006927154.3;refGene_gn=XPO4;refGene_tx=XM_019816971.2;refGene_gn=XPO
4;refGene_tx=XM_019817033.1;refGene_gn=XPO4;refGene_tx=XM_019817077.2;
refGene_gn=XPO4;refGene_tx=XM_023249438.1
A1 1842322 1842465
refGene_gn=XPO4;refGene_tx=XM_003980253.5;refGene_gn=XPO4;refGene_tx=X
M_006927154.3;refGene_gn=XPO4;refGene_tx=XM_019816971.2;refGene_gn=XPO
4;refGene_tx=XM_019817033.1;refGene_gn=XPO4;refGene_tx=XM_019817077.2;
refGene_gn=XPO4;refGene_tx=XM_023249438.1
A1 1843081 1843258
refGene_gn=XPO4;refGene_tx=XM_003980253.5;refGene_gn=XPO4;refGene_tx=X
M_006927154.3;refGene_gn=XPO4;refGene_tx=XM_019816971.2;refGene_gn=XPO
4;refGene_tx=XM_019817033.1;refGene_gn=XPO4;refGene_tx=XM_019817077.2;
refGene_gn=XPO4;refGene_tx=XM_023249438.1
A1 1854815 1854990
refGene_gn=XPO4;refGene_tx=XM_003980253.5;refGene_gn=XPO4;refGene_tx=X
M_006927154.3;refGene_gn=XPO4;refGene_tx=XM_019816971.2;refGene_gn=XPO
4;refGene_tx=XM_019817033.1;refGene_gn=XPO4;refGene_tx=XM_019817077.2;
refGene_gn=XPO4;refGene_tx=XM_023249438.1
A1 1855240 1855398
refGene_gn=XPO4;refGene_tx=XM_003980253.5;refGene_gn=XPO4;refGene_tx=X
M_006927154.3;refGene_gn=XPO4;refGene_tx=XM_019816971.2;refGene_gn=XPO
4;refGene_tx=XM_019817033.1;refGene_gn=XPO4;refGene_tx=XM_019817077.2;
refGene_gn=XPO4;refGene_tx=XM_023249438.1
A1 1860414 1860527
refGene_gn=XPO4;refGene_tx=XM_003980253.5;refGene_gn=XPO4;refGene_tx=X
M_006927154.3;refGene_gn=XPO4;refGene_tx=XM_019817033.1;refGene_gn=XPO
4;refGene_tx=XM_019817077.2;refGene_gn=XPO4;refGene_tx=XM_023249438.1
A1 1880968 1880981 refGene_gn=XPO4;refGene_tx=XM_019816971.2
A1 1881389 1881568
refGene_gn=XPO4;refGene_tx=XM_003980253.5;refGene_gn=XPO4;refGene_tx=X
M_006927154.3;refGene_gn=XPO4;refGene_tx=XM_019816971.2;refGene_gn=XPO
4;refGene_tx=XM_019817033.1;refGene_gn=XPO4;refGene_tx=XM_019817077.2;
refGene_gn=XPO4;refGene_tx=XM_023249438.1
A1 1882307 1882424
refGene_gn=XPO4;refGene_tx=XM_003980253.5;refGene_gn=XPO4;refGene_tx=X
M_006927154.3;refGene_gn=XPO4;refGene_tx=XM_019816971.2;refGene_gn=XPO
4;refGene_tx=XM_019817033.1;refGene_gn=XPO4;refGene_tx=XM_019817077.2;
refGene_gn=XPO4;refGene_tx=XM_023249438.1
A1 1892661 1892800
refGene_gn=XPO4;refGene_tx=XM_003980253.5;refGene_gn=XPO4;refGene_tx=X
M_006927154.3;refGene_gn=XPO4;refGene_tx=XM_019816971.2;refGene_gn=XPO
4;refGene_tx=XM_019817033.1;refGene_gn=XPO4;refGene_tx=XM_019817077.2;
refGene_gn=XPO4;refGene_tx=XM_023249438.1
A1 1897526 1897668
refGene_gn=XPO4;refGene_tx=XM_003980253.5;refGene_gn=XPO4;refGene_tx=X
M_006927154.3;refGene_gn=XPO4;refGene_tx=XM_019816971.2;refGene_gn=XPO

4;refGene_tx=XM_019817033.1;refGene_gn=XPO4;refGene_tx=XM_019817077.2;
refGene_gn=XPO4;refGene_tx=XM_023249438.1
A1	1903034	1903140
refGene_gn=XPO4;refGene_tx=XM_003980253.5;refGene_gn=XPO4;refGene_tx=XM_019816971.2;refGene_gn=XPO4;refGene_tx=XM_019817077.2;refGene_gn=XPO4;refGene_tx=XM_023249438.1
A1	1932073	1932142
refGene_gn=XPO4;refGene_tx=XM_003980253.5;refGene_gn=XPO4;refGene_tx=XM_019816971.2;refGene_gn=XPO4;refGene_tx=XM_023249438.1
A1	1989623	1990118
refGene_gn=LATS2;refGene_tx=XM_023249486.1;refGene_gn=LATS2;refGene_tx=XM_023249513.1;refGene_gn=LATS2;refGene_tx=XM_023249540.1
A1	1993028	1993135
refGene_gn=LATS2;refGene_tx=XM_023249486.1;refGene_gn=LATS2;refGene_tx=XM_023249513.1;refGene_gn=LATS2;refGene_tx=XM_023249540.1
A1	1994896	1995079
refGene_gn=LATS2;refGene_tx=XM_023249486.1;refGene_gn=LATS2;refGene_tx=XM_023249513.1;refGene_gn=LATS2;refGene_tx=XM_023249540.1
A1	1996356	1996939
refGene_gn=LATS2;refGene_tx=XM_023249486.1;refGene_gn=LATS2;refGene_tx=XM_023249513.1;refGene_gn=LATS2;refGene_tx=XM_023249540.1
A1	1997984	1999432
refGene_gn=LATS2;refGene_tx=XM_023249486.1;refGene_gn=LATS2;refGene_tx=XM_023249513.1;refGene_gn=LATS2;refGene_tx=XM_023249540.1
A1	2000503	2000636
refGene_gn=LATS2;refGene_tx=XM_023249486.1;refGene_gn=LATS2;refGene_tx=XM_023249513.1;refGene_gn=LATS2;refGene_tx=XM_023249540.1
A1	2049043	2049064	refGene_gn=LATS2;refGene_tx=XM_023249513.1
A1	2051506	2051845	refGene_gn=LATS2;refGene_tx=XM_023249486.1
A1	2131940	2132037	refGene_gn=SAP18;refGene_tx=XM_023249619.1
A1	2150685	2150997
refGene_gn=SAP18;refGene_tx=XM_003980254.5;refGene_gn=SAP18;refGene_tx=XM_019817358.1;refGene_gn=SAP18;refGene_tx=XM_019817388.1;refGene_gn=SAP18;refGene_tx=XM_023249619.1
A1	2151154	2151264
refGene_gn=SAP18;refGene_tx=XM_003980254.5;refGene_gn=SAP18;refGene_tx=XM_019817358.1;refGene_gn=SAP18;refGene_tx=XM_019817388.1;refGene_gn=SAP18;refGene_tx=XM_023249619.1
A1	2156099	2156222
refGene_gn=SAP18;refGene_tx=XM_003980254.5;refGene_gn=SAP18;refGene_tx=XM_019817358.1;refGene_gn=SAP18;refGene_tx=XM_019817388.1;refGene_gn=SAP18;refGene_tx=XM_023249619.1
A1	2156481	2156638
refGene_gn=SAP18;refGene_tx=XM_003980254.5;refGene_gn=SAP18;refGene_tx=XM_019817358.1;refGene_gn=SAP18;refGene_tx=XM_019817388.1;refGene_gn=SAP18;refGene_tx=XM_023249619.1
A1	2164029	2164158
refGene_gn=SKA3;refGene_tx=XM_006927156.4;refGene_gn=SKA3;refGene_tx=XM_006927157.4
A1	2165896	2166100	refGene_gn=SKA3;refGene_tx=XM_006927156.4

A1 2167705 2167791
refGene_gn=SKA3;refGene_tx=XM_006927156.4;refGene_gn=SKA3;refGene_tx=X
M_006927157.4
A1 2172913 2172999
refGene_gn=SKA3;refGene_tx=XM_006927156.4;refGene_gn=SKA3;refGene_tx=X
M_006927157.4
A1 2175598 2176010
refGene_gn=SKA3;refGene_tx=XM_006927156.4;refGene_gn=SKA3;refGene_tx=X
M_006927157.4
A1 2179420 2179586
refGene_gn=SKA3;refGene_tx=XM_006927156.4;refGene_gn=SKA3;refGene_tx=X
M_006927157.4
A1 2179715 2179777
refGene_gn=SKA3;refGene_tx=XM_006927156.4;refGene_gn=SKA3;refGene_tx=X
M_006927157.4
A1 2183123 2183462
refGene_gn=MRPL57;refGene_tx=XM_006927155.4;refGene_gn=SKA3;refGene_tx
=XM_006927156.4;refGene_gn=SKA3;refGene_tx=XM_006927157.4
A1 2183669 2183983 refGene_gn=MRPL57;refGene_tx=XM_006927155.4
A1 2208991 2208999 refGene_gn=ZDHHC20;refGene_tx=XM_023249858.1
A1 2210758 2210796
refGene_gn=ZDHHC20;refGene_tx=XM_023249851.1;refGene_gn=ZDHHC20;refGen
e_tx=XM_023249859.1
A1 2211888 2212004
refGene_gn=ZDHHC20;refGene_tx=XM_023249851.1;refGene_gn=ZDHHC20;refGen
e_tx=XM_023249858.1;refGene_gn=ZDHHC20;refGene_tx=XM_023249859.1
A1 2212374 2212393
refGene_gn=ZDHHC20;refGene_tx=XM_023249861.1;refGene_gn=ZDHHC20;refGen
e_tx=XM_023249862.1
A1 2213438 2213528
refGene_gn=ZDHHC20;refGene_tx=XM_023249851.1;refGene_gn=ZDHHC20;refGen
e_tx=XM_023249858.1;refGene_gn=ZDHHC20;refGene_tx=XM_023249859.1;refGe
ne_gn=ZDHHC20;refGene_tx=XM_023249861.1;refGene_gn=ZDHHC20;refGene_tx=
XM_023249862.1
A1 2216608 2216735
refGene_gn=ZDHHC20;refGene_tx=XM_023249851.1;refGene_gn=ZDHHC20;refGen
e_tx=XM_023249858.1;refGene_gn=ZDHHC20;refGene_tx=XM_023249859.1;refGe
ne_gn=ZDHHC20;refGene_tx=XM_023249861.1;refGene_gn=ZDHHC20;refGene_tx=
XM_023249862.1
A1 2220702 2220835
refGene_gn=ZDHHC20;refGene_tx=XM_023249851.1;refGene_gn=ZDHHC20;refGen
e_tx=XM_023249858.1;refGene_gn=ZDHHC20;refGene_tx=XM_023249859.1;refGe
ne_gn=ZDHHC20;refGene_tx=XM_023249861.1;refGene_gn=ZDHHC20;refGene_tx=
XM_023249862.1
A1 2221797 2221833
refGene_gn=ZDHHC20;refGene_tx=XM_023249851.1;refGene_gn=ZDHHC20;refGen
e_tx=XM_023249858.1;refGene_gn=ZDHHC20;refGene_tx=XM_023249861.1
A1 2230239 2230360
refGene_gn=ZDHHC20;refGene_tx=XM_023249851.1;refGene_gn=ZDHHC20;refGen
e_tx=XM_023249858.1;refGene_gn=ZDHHC20;refGene_tx=XM_023249859.1;refGe

ne_gn=ZDHHC20;refGene_tx=XM_023249861.1;refGene_gn=ZDHHC20;refGene_tx=
XM_023249862.1
A1	2231437	2231470
refGene_gn=ZDHHC20;refGene_tx=XM_023249851.1;refGene_gn=ZDHHC20;refGen
e_tx=XM_023249858.1;refGene_gn=ZDHHC20;refGene_tx=XM_023249859.1;refGe
ne_gn=ZDHHC20;refGene_tx=XM_023249861.1;refGene_gn=ZDHHC20;refGene_tx=
XM_023249862.1
A1	2233015	2233085
refGene_gn=ZDHHC20;refGene_tx=XM_023249851.1;refGene_gn=ZDHHC20;refGen
e_tx=XM_023249858.1;refGene_gn=ZDHHC20;refGene_tx=XM_023249859.1;refGe
ne_gn=ZDHHC20;refGene_tx=XM_023249861.1;refGene_gn=ZDHHC20;refGene_tx=
XM_023249862.1
A1	2241403	2241524
refGene_gn=ZDHHC20;refGene_tx=XM_023249851.1;refGene_gn=ZDHHC20;refGen
e_tx=XM_023249858.1;refGene_gn=ZDHHC20;refGene_tx=XM_023249859.1;refGe
ne_gn=ZDHHC20;refGene_tx=XM_023249861.1;refGene_gn=ZDHHC20;refGene_tx=
XM_023249862.1
A1	2247497	2247601
refGene_gn=ZDHHC20;refGene_tx=XM_023249851.1;refGene_gn=ZDHHC20;refGen
e_tx=XM_023249858.1;refGene_gn=ZDHHC20;refGene_tx=XM_023249859.1;refGe
ne_gn=ZDHHC20;refGene_tx=XM_023249861.1;refGene_gn=ZDHHC20;refGene_tx=
XM_023249862.1
A1	2252004	2252031
refGene_gn=ZDHHC20;refGene_tx=XM_023249851.1;refGene_gn=ZDHHC20;refGen
e_tx=XM_023249858.1;refGene_gn=ZDHHC20;refGene_tx=XM_023249859.1;refGe
ne_gn=ZDHHC20;refGene_tx=XM_023249861.1;refGene_gn=ZDHHC20;refGene_tx=
XM_023249862.1
A1	2277715	2277833
refGene_gn=ZDHHC20;refGene_tx=XM_023249851.1;refGene_gn=ZDHHC20;refGen
e_tx=XM_023249858.1;refGene_gn=ZDHHC20;refGene_tx=XM_023249859.1;refGe
ne_gn=ZDHHC20;refGene_tx=XM_023249861.1;refGene_gn=ZDHHC20;refGene_tx=
XM_023249862.1
A1	2306531	2306636	refGene_gn=MICU2;refGene_tx=XM_023249868.1
A1	2308988	2309146	refGene_gn=MICU2;refGene_tx=XM_023249868.1
A1	2309609	2309718	refGene_gn=MICU2;refGene_tx=XM_023249868.1
A1	2314837	2315009	refGene_gn=MICU2;refGene_tx=XM_023249868.1
A1	2327167	2327265	refGene_gn=MICU2;refGene_tx=XM_023249868.1
A1	2338150	2338213	refGene_gn=MICU2;refGene_tx=XM_023249868.1
A1	2366344	2366427	refGene_gn=MICU2;refGene_tx=XM_023249868.1
A1	2368060	2368108	refGene_gn=MICU2;refGene_tx=XM_023249868.1
A1	2383144	2383220	refGene_gn=MICU2;refGene_tx=XM_023249868.1
A1	2383335	2383367	refGene_gn=MICU2;refGene_tx=XM_023249868.1
A1	2411378	2411523	refGene_gn=MICU2;refGene_tx=XM_023249868.1
A1	2445678	2445693	refGene_gn=MICU2;refGene_tx=XM_023249868.1
A1	2510131	2510408	refGene_gn=FGF9;refGene_tx=XM_003980257.5
A1	2519093	2519197	refGene_gn=FGF9;refGene_tx=XM_003980257.5
A1	2541988	2542234	refGene_gn=FGF9;refGene_tx=XM_003980257.5
A1	3844486	3844675
refGene_gn=SGCG;refGene_tx=XM_023249893.1;refGene_gn=SGCG;refGene_tx=X
M_023249897.1

A1 3895195 3895390
refGene_gn=SGCG;refGene_tx=XM_003980258.5;refGene_gn=SGCG;refGene_tx=XM_023249893.1;refGene_gn=SGCG;refGene_tx=XM_023249897.1
A1 3925173 3925275
refGene_gn=SGCG;refGene_tx=XM_003980258.5;refGene_gn=SGCG;refGene_tx=XM_023249893.1;refGene_gn=SGCG;refGene_tx=XM_023249897.1
A1 3930157 3930245
refGene_gn=SGCG;refGene_tx=XM_003980258.5;refGene_gn=SGCG;refGene_tx=XM_023249893.1;refGene_gn=SGCG;refGene_tx=XM_023249897.1
A1 3938514 3938634
refGene_gn=SGCG;refGene_tx=XM_003980258.5;refGene_gn=SGCG;refGene_tx=XM_023249893.1;refGene_gn=SGCG;refGene_tx=XM_023249897.1
A1 3957016 3957089
refGene_gn=SGCG;refGene_tx=XM_003980258.5;refGene_gn=SGCG;refGene_tx=XM_023249893.1;refGene_gn=SGCG;refGene_tx=XM_023249897.1
A1 3971802 3971926
refGene_gn=SGCG;refGene_tx=XM_003980258.5;refGene_gn=SGCG;refGene_tx=XM_023249893.1
A1 3974620 3974870
refGene_gn=SGCG;refGene_tx=XM_003980258.5;refGene_gn=SGCG;refGene_tx=XM_023249893.1;refGene_gn=SGCG;refGene_tx=XM_023249897.1
A1 3981198 3983985 refGene_gn=SACS;refGene_tx=XM_006927166.4
A1 3983986 3992757 refGene_gn=SACS;refGene_tx=XM_006927166.4
A1 4002698 4002790 refGene_gn=SACS;refGene_tx=XM_006927166.4
A1 4003438 4004921 refGene_gn=SACS;refGene_tx=XM_006927166.4
A1 4007544 4007691 refGene_gn=SACS;refGene_tx=XM_006927166.4
A1 4012196 4012308 refGene_gn=SACS;refGene_tx=XM_006927166.4
A1 4015818 4015904 refGene_gn=SACS;refGene_tx=XM_006927166.4
A1 4016793 4016881 refGene_gn=SACS;refGene_tx=XM_006927166.4
A1 4022470 4022621 refGene_gn=SACS;refGene_tx=XM_006927166.4
A1 4051578 4051598 refGene_gn=SACS;refGene_tx=XM_006927166.4
A1 4209454 4209493
refGene_gn=TNFRSF19;refGene_tx=XM_006927167.4;refGene_gn=TNFRSF19;refGene_tx=XM_011280485.3;refGene_gn=TNFRSF19;refGene_tx=XM_019819168.2;refGene_gn=TNFRSF19;refGene_tx=XM_023249941.1
A1 4219557 4219650
refGene_gn=TNFRSF19;refGene_tx=XM_006927167.4;refGene_gn=TNFRSF19;refGene_tx=XM_011280485.3;refGene_gn=TNFRSF19;refGene_tx=XM_019819168.2;refGene_gn=TNFRSF19;refGene_tx=XM_019819241.2;refGene_gn=TNFRSF19;refGene_tx=XM_019819279.2;refGene_gn=TNFRSF19;refGene_tx=XM_023249934.1;refGene_gn=TNFRSF19;refGene_tx=XM_023249941.1
A1 4221466 4221577
refGene_gn=TNFRSF19;refGene_tx=XM_006927167.4;refGene_gn=TNFRSF19;refGene_tx=XM_011280485.3;refGene_gn=TNFRSF19;refGene_tx=XM_019819168.2;refGene_gn=TNFRSF19;refGene_tx=XM_019819241.2;refGene_gn=TNFRSF19;refGene_tx=XM_019819279.2;refGene_gn=TNFRSF19;refGene_tx=XM_019819307.1;refGene_gn=TNFRSF19;refGene_tx=XM_023249934.1;refGene_gn=TNFRSF19;refGene_tx=XM_023249941.1
A1 4235528 4235707
refGene_gn=TNFRSF19;refGene_tx=XM_006927167.4;refGene_gn=TNFRSF19;refG

ene_tx=XM_011280485.3;refGene_gn=TNFRSF19;refGene_tx=XM_019819168.2;refGene_gn=TNFRSF19;refGene_tx=XM_019819241.2;refGene_gn=TNFRSF19;refGene_tx=XM_019819279.2;refGene_gn=TNFRSF19;refGene_tx=XM_023249934.1;refGene_gn=TNFRSF19;refGene_tx=XM_023249941.1
A1 4246257 4246343 refGene_gn=TNFRSF19;refGene_tx=XM_006927167.4;refGene_gn=TNFRSF19;refGene_tx=XM_011280485.3;refGene_gn=TNFRSF19;refGene_tx=XM_019819168.2;refGene_gn=TNFRSF19;refGene_tx=XM_019819241.2;refGene_gn=TNFRSF19;refGene_tx=XM_019819279.2;refGene_gn=TNFRSF19;refGene_tx=XM_019819307.1;refGene_gn=TNFRSF19;refGene_tx=XM_023249934.1;refGene_gn=TNFRSF19;refGene_tx=XM_023249941.1
A1 4275374 4275588 refGene_gn=TNFRSF19;refGene_tx=XM_023249943.1
A1 4280662 4280827 refGene_gn=TNFRSF19;refGene_tx=XM_006927167.4;refGene_gn=TNFRSF19;refGene_tx=XM_011280485.3;refGene_gn=TNFRSF19;refGene_tx=XM_019819168.2;refGene_gn=TNFRSF19;refGene_tx=XM_019819241.2;refGene_gn=TNFRSF19;refGene_tx=XM_019819279.2;refGene_gn=TNFRSF19;refGene_tx=XM_019819307.1;refGene_gn=TNFRSF19;refGene_tx=XM_023249934.1;refGene_gn=TNFRSF19;refGene_tx=XM_023249943.1
A1 4281273 4281399 refGene_gn=TNFRSF19;refGene_tx=XM_006927167.4;refGene_gn=TNFRSF19;refGene_tx=XM_011280485.3;refGene_gn=TNFRSF19;refGene_tx=XM_019819168.2;refGene_gn=TNFRSF19;refGene_tx=XM_019819241.2;refGene_gn=TNFRSF19;refGene_tx=XM_019819279.2;refGene_gn=TNFRSF19;refGene_tx=XM_019819307.1;refGene_gn=TNFRSF19;refGene_tx=XM_023249934.1;refGene_gn=TNFRSF19;refGene_tx=XM_023249943.1
A1 4284510 4284613 refGene_gn=TNFRSF19;refGene_tx=XM_006927167.4;refGene_gn=TNFRSF19;refGene_tx=XM_011280485.3;refGene_gn=TNFRSF19;refGene_tx=XM_019819168.2;refGene_gn=TNFRSF19;refGene_tx=XM_019819241.2;refGene_gn=TNFRSF19;refGene_tx=XM_019819279.2;refGene_gn=TNFRSF19;refGene_tx=XM_019819307.1;refGene_gn=TNFRSF19;refGene_tx=XM_023249934.1;refGene_gn=TNFRSF19;refGene_tx=XM_023249941.1;refGene_gn=TNFRSF19;refGene_tx=XM_023249943.1
A1 4285188 4285594 refGene_gn=TNFRSF19;refGene_tx=XM_006927167.4;refGene_gn=TNFRSF19;refGene_tx=XM_011280485.3;refGene_gn=TNFRSF19;refGene_tx=XM_019819168.2;refGene_gn=TNFRSF19;refGene_tx=XM_019819241.2;refGene_gn=TNFRSF19;refGene_tx=XM_019819279.2;refGene_gn=TNFRSF19;refGene_tx=XM_019819307.1;refGene_gn=TNFRSF19;refGene_tx=XM_023249934.1;refGene_gn=TNFRSF19;refGene_tx=XM_023249941.1;refGene_gn=TNFRSF19;refGene_tx=XM_023249943.1
A1 4287984 4288023 refGene_gn=TNFRSF19;refGene_tx=XM_019819168.2
A1 4289792 4289852 refGene_gn=TNFRSF19;refGene_tx=XM_006927167.4;refGene_gn=TNFRSF19;refGene_tx=XM_011280485.3;refGene_gn=TNFRSF19;refGene_tx=XM_019819241.2;refGene_gn=TNFRSF19;refGene_tx=XM_019819279.2;refGene_gn=TNFRSF19;refGene_tx=XM_019819307.1;refGene_gn=TNFRSF19;refGene_tx=XM_023249934.1;refGene_gn=TNFRSF19;refGene_tx=XM_023249941.1;refGene_gn=TNFRSF19;refGene_tx=XM_023249943.1
A1 4295419 4295517 refGene_gn=MIPEP;refGene_tx=XM_023249909.1
A1 4334333 4334431

refGene_gn=MIPEP;refGene_tx=XM_003980259.5;refGene_gn=MIPEP;refGene_tx=XM_006927168.3
A1 4361225 4361299
refGene_gn=MIPEP;refGene_tx=XM_003980259.5;refGene_gn=MIPEP;refGene_tx=XM_023249909.1
A1 4364363 4364485
refGene_gn=MIPEP;refGene_tx=XM_003980259.5;refGene_gn=MIPEP;refGene_tx=XM_023249909.1
A1 4398536 4398656
refGene_gn=MIPEP;refGene_tx=XM_003980259.5;refGene_gn=MIPEP;refGene_tx=XM_006927168.3;refGene_gn=MIPEP;refGene_tx=XM_023249909.1
A1 4401711 4401786
refGene_gn=MIPEP;refGene_tx=XM_003980259.5;refGene_gn=MIPEP;refGene_tx=XM_006927168.3;refGene_gn=MIPEP;refGene_tx=XM_023249909.1
A1 4429824 4429934
refGene_gn=MIPEP;refGene_tx=XM_003980259.5;refGene_gn=MIPEP;refGene_tx=XM_006927168.3;refGene_gn=MIPEP;refGene_tx=XM_023249909.1
A1 4432212 4432417
refGene_gn=MIPEP;refGene_tx=XM_003980259.5;refGene_gn=MIPEP;refGene_tx=XM_006927168.3;refGene_gn=MIPEP;refGene_tx=XM_023249909.1
A1 4434351 4434429
refGene_gn=MIPEP;refGene_tx=XM_003980259.5;refGene_gn=MIPEP;refGene_tx=XM_006927168.3;refGene_gn=MIPEP;refGene_tx=XM_023249909.1
A1 4435978 4436132
refGene_gn=MIPEP;refGene_tx=XM_003980259.5;refGene_gn=MIPEP;refGene_tx=XM_006927168.3;refGene_gn=MIPEP;refGene_tx=XM_023249909.1
A1 4461776 4461829
refGene_gn=MIPEP;refGene_tx=XM_003980259.5;refGene_gn=MIPEP;refGene_tx=XM_006927168.3;refGene_gn=MIPEP;refGene_tx=XM_023249909.1
A1 4465233 4465294
refGene_gn=MIPEP;refGene_tx=XM_003980259.5;refGene_gn=MIPEP;refGene_tx=XM_006927168.3;refGene_gn=MIPEP;refGene_tx=XM_023249909.1
A1 4466900 4466949
refGene_gn=MIPEP;refGene_tx=XM_003980259.5;refGene_gn=MIPEP;refGene_tx=XM_006927168.3;refGene_gn=MIPEP;refGene_tx=XM_023249909.1
A1 4472142 4472299
refGene_gn=MIPEP;refGene_tx=XM_003980259.5;refGene_gn=MIPEP;refGene_tx=XM_006927168.3;refGene_gn=MIPEP;refGene_tx=XM_023249909.1
A1 4473359 4473542
refGene_gn=MIPEP;refGene_tx=XM_003980259.5;refGene_gn=MIPEP;refGene_tx=XM_006927168.3;refGene_gn=MIPEP;refGene_tx=XM_023249909.1
A1 4477869 4477933
refGene_gn=MIPEP;refGene_tx=XM_003980259.5;refGene_gn=MIPEP;refGene_tx=XM_006927168.3;refGene_gn=MIPEP;refGene_tx=XM_023249909.1
A1 4483216 4483303
refGene_gn=MIPEP;refGene_tx=XM_003980259.5;refGene_gn=MIPEP;refGene_tx=XM_006927168.3;refGene_gn=MIPEP;refGene_tx=XM_023249909.1
A1 4486232 4486321
refGene_gn=MIPEP;refGene_tx=XM_003980259.5;refGene_gn=MIPEP;refGene_tx=XM_006927168.3;refGene_gn=MIPEP;refGene_tx=XM_023249909.1

A1 4489510 4489684
refGene_gn=MIPEP;refGene_tx=XM_003980259.5;refGene_gn=MIPEP;refGene_tx
=XM_006927168.3;refGene_gn=MIPEP;refGene_tx=XM_023249909.1
A1 4492359 4492548
refGene_gn=MIPEP;refGene_tx=XM_003980259.5;refGene_gn=MIPEP;refGene_tx
=XM_006927168.3;refGene_gn=MIPEP;refGene_tx=XM_023249909.1
A1 4498886 4499656 refGene_gn=C1QTNF9;refGene_tx=XM_011280638.2
A1 4501377 4501440 refGene_gn=C1QTNF9;refGene_tx=XM_011280638.2
A1 4504431 4504597 refGene_gn=C1QTNF9;refGene_tx=XM_011280638.2
A1 4519484 4519660
refGene_gn=SPATA13;refGene_tx=XM_006927170.4;refGene_gn=SPATA13;refGen
e_tx=XM_019819646.2;refGene_gn=SPATA13;refGene_tx=XM_019819779.2;refGe
ne_gn=SPATA13;refGene_tx=XM_019819820.2;refGene_gn=SPATA13;refGene_tx=
XM_019819853.2;refGene_gn=SPATA13;refGene_tx=XM_023249962.1;refGene_gn
=SPATA13;refGene_tx=XM_023249964.1;refGene_gn=SPATA13;refGene_tx=XM_02
3249966.1
A1 4521850 4521925
refGene_gn=SPATA13;refGene_tx=XM_006927170.4;refGene_gn=SPATA13;refGen
e_tx=XM_019819646.2;refGene_gn=SPATA13;refGene_tx=XM_019819779.2;refGe
ne_gn=SPATA13;refGene_tx=XM_019819820.2;refGene_gn=SPATA13;refGene_tx=
XM_019819853.2;refGene_gn=SPATA13;refGene_tx=XM_023249962.1;refGene_gn
=SPATA13;refGene_tx=XM_023249964.1;refGene_gn=SPATA13;refGene_tx=XM_02
3249966.1
A1 4524389 4524762
refGene_gn=SPATA13;refGene_tx=XM_006927170.4;refGene_gn=SPATA13;refGen
e_tx=XM_019819646.2;refGene_gn=SPATA13;refGene_tx=XM_019819779.2;refGe
ne_gn=SPATA13;refGene_tx=XM_019819820.2;refGene_gn=SPATA13;refGene_tx=
XM_019819853.2;refGene_gn=SPATA13;refGene_tx=XM_023249962.1;refGene_gn
=SPATA13;refGene_tx=XM_023249964.1;refGene_gn=SPATA13;refGene_tx=XM_02
3249966.1
A1 4526500 4526630
refGene_gn=SPATA13;refGene_tx=XM_006927170.4;refGene_gn=SPATA13;refGen
e_tx=XM_019819646.2;refGene_gn=SPATA13;refGene_tx=XM_019819779.2;refGe
ne_gn=SPATA13;refGene_tx=XM_019819820.2;refGene_gn=SPATA13;refGene_tx=
XM_019819853.2;refGene_gn=SPATA13;refGene_tx=XM_023249962.1;refGene_gn
=SPATA13;refGene_tx=XM_023249964.1;refGene_gn=SPATA13;refGene_tx=XM_02
3249966.1
A1 4529882 4530115
refGene_gn=SPATA13;refGene_tx=XM_006927170.4;refGene_gn=SPATA13;refGen
e_tx=XM_019819646.2;refGene_gn=SPATA13;refGene_tx=XM_019819779.2;refGe
ne_gn=SPATA13;refGene_tx=XM_019819820.2;refGene_gn=SPATA13;refGene_tx=
XM_019819853.2;refGene_gn=SPATA13;refGene_tx=XM_023249962.1;refGene_gn
=SPATA13;refGene_tx=XM_023249964.1;refGene_gn=SPATA13;refGene_tx=XM_02
3249966.1
A1 4532259 4532439
refGene_gn=SPATA13;refGene_tx=XM_006927170.4;refGene_gn=SPATA13;refGen
e_tx=XM_019819646.2;refGene_gn=SPATA13;refGene_tx=XM_019819779.2;refGe
ne_gn=SPATA13;refGene_tx=XM_019819820.2;refGene_gn=SPATA13;refGene_tx=
XM_019819853.2;refGene_gn=SPATA13;refGene_tx=XM_023249962.1;refGene_gn
=SPATA13;refGene_tx=XM_023249964.1;refGene_gn=SPATA13;refGene_tx=XM_02

3249966.1
A1 4534365 4534551
refGene_gn=SPATA13;refGene_tx=XM_006927170.4;refGene_gn=SPATA13;refGene_tx=XM_019819646.2;refGene_gn=SPATA13;refGene_tx=XM_019819779.2;refGene_gn=SPATA13;refGene_tx=XM_019819820.2;refGene_gn=SPATA13;refGene_tx=XM_019819853.2;refGene_gn=SPATA13;refGene_tx=XM_023249962.1;refGene_gn=SPATA13;refGene_tx=XM_023249964.1;refGene_gn=SPATA13;refGene_tx=XM_023249966.1
A1 4535148 4535328
refGene_gn=SPATA13;refGene_tx=XM_006927170.4;refGene_gn=SPATA13;refGene_tx=XM_019819646.2;refGene_gn=SPATA13;refGene_tx=XM_019819779.2;refGene_gn=SPATA13;refGene_tx=XM_019819820.2;refGene_gn=SPATA13;refGene_tx=XM_019819853.2;refGene_gn=SPATA13;refGene_tx=XM_023249962.1;refGene_gn=SPATA13;refGene_tx=XM_023249964.1;refGene_gn=SPATA13;refGene_tx=XM_023249966.1
A1 4536471 4536608
refGene_gn=SPATA13;refGene_tx=XM_006927170.4;refGene_gn=SPATA13;refGene_tx=XM_019819646.2;refGene_gn=SPATA13;refGene_tx=XM_019819779.2;refGene_gn=SPATA13;refGene_tx=XM_023249962.1;refGene_gn=SPATA13;refGene_tx=XM_023249964.1;refGene_gn=SPATA13;refGene_tx=XM_023249966.1
A1 4549667 4549722 refGene_gn=SPATA13;refGene_tx=XM_006927170.4
A1 4571244 4571389
refGene_gn=SPATA13;refGene_tx=XM_019819646.2;refGene_gn=SPATA13;refGene_tx=XM_019819779.2;refGene_gn=SPATA13;refGene_tx=XM_023249962.1;refGene_gn=SPATA13;refGene_tx=XM_023249964.1;refGene_gn=SPATA13;refGene_tx=XM_023249966.1
A1 4573035 4573401
refGene_gn=SPATA13;refGene_tx=XM_019819646.2;refGene_gn=SPATA13;refGene_tx=XM_023249962.1;refGene_gn=SPATA13;refGene_tx=XM_023249964.1
A1 4577296 4577359 refGene_gn=SPATA13;refGene_tx=XM_019819646.2
A1 4598397 4600140
refGene_gn=SPATA13;refGene_tx=XM_023249962.1;refGene_gn=SPATA13;refGene_tx=XM_023249964.1;refGene_gn=SPATA13;refGene_tx=XM_023249966.1
A1 4819656 4819692
refGene_gn=SPATA13;refGene_tx=XM_023249962.1;refGene_gn=SPATA13;refGene_tx=XM_023249966.1
A1 4864604 4864616
refGene_gn=SPATA13;refGene_tx=XM_023249962.1;refGene_gn=SPATA13;refGene_tx=XM_023249966.1
A1 4967931 4970028 refGene_gn=AMER2;refGene_tx=XM_023250360.1
A1 5041466 5041526 refGene_gn=MTMR6;refGene_tx=XM_019820543.1
A1 5044744 5045005
refGene_gn=MTMR6;refGene_tx=XM_003980260.5;refGene_gn=MTMR6;refGene_tx=XM_006927171.4
A1 5047208 5047335
refGene_gn=MTMR6;refGene_tx=XM_003980260.5;refGene_gn=MTMR6;refGene_tx=XM_006927171.4;refGene_gn=MTMR6;refGene_tx=XM_019820543.1
A1 5047420 5047552
refGene_gn=MTMR6;refGene_tx=XM_003980260.5;refGene_gn=MTMR6;refGene_tx=XM_006927171.4;refGene_gn=MTMR6;refGene_tx=XM_019820543.1

A1	5049730	5049931
refGene_gn=MTMR6;refGene_tx=XM_003980260.5;refGene_gn=MTMR6;refGene_tx=XM_006927171.4;refGene_gn=MTMR6;refGene_tx=XM_019820543.1
A1	5050358	5050408
refGene_gn=MTMR6;refGene_tx=XM_003980260.5;refGene_gn=MTMR6;refGene_tx=XM_006927171.4;refGene_gn=MTMR6;refGene_tx=XM_019820543.1
A1	5053749	5053875
refGene_gn=MTMR6;refGene_tx=XM_003980260.5;refGene_gn=MTMR6;refGene_tx=XM_006927171.4;refGene_gn=MTMR6;refGene_tx=XM_019820543.1
A1	5054299	5054409
refGene_gn=MTMR6;refGene_tx=XM_003980260.5;refGene_gn=MTMR6;refGene_tx=XM_006927171.4;refGene_gn=MTMR6;refGene_tx=XM_019820543.1
A1	5055373	5055506
refGene_gn=MTMR6;refGene_tx=XM_003980260.5;refGene_gn=MTMR6;refGene_tx=XM_006927171.4;refGene_gn=MTMR6;refGene_tx=XM_019820543.1
A1	5056922	5057036
refGene_gn=MTMR6;refGene_tx=XM_006927171.4;refGene_gn=MTMR6;refGene_tx=XM_019820543.1
A1	5057690	5057825
refGene_gn=MTMR6;refGene_tx=XM_003980260.5;refGene_gn=MTMR6;refGene_tx=XM_006927171.4;refGene_gn=MTMR6;refGene_tx=XM_019820543.1
A1	5060833	5060962
refGene_gn=MTMR6;refGene_tx=XM_003980260.5;refGene_gn=MTMR6;refGene_tx=XM_006927171.4;refGene_gn=MTMR6;refGene_tx=XM_019820543.1
A1	5061487	5061645
refGene_gn=MTMR6;refGene_tx=XM_003980260.5;refGene_gn=MTMR6;refGene_tx=XM_006927171.4;refGene_gn=MTMR6;refGene_tx=XM_019820543.1
A1	5063945	5064108
refGene_gn=MTMR6;refGene_tx=XM_003980260.5;refGene_gn=MTMR6;refGene_tx=XM_006927171.4;refGene_gn=MTMR6;refGene_tx=XM_019820543.1
A1	5068235	5068352
refGene_gn=MTMR6;refGene_tx=XM_003980260.5;refGene_gn=MTMR6;refGene_tx=XM_006927171.4;refGene_gn=MTMR6;refGene_tx=XM_019820543.1
A1	5079967	5079991
refGene_gn=MTMR6;refGene_tx=XM_003980260.5;refGene_gn=MTMR6;refGene_tx=XM_006927171.4;refGene_gn=MTMR6;refGene_tx=XM_019820543.1
A1	5093790	5093897
refGene_gn=NUP58;refGene_tx=XM_023250027.1;refGene_gn=NUP58;refGene_tx=XM_023250028.1
A1	5098605	5098748
refGene_gn=NUP58;refGene_tx=XM_023250027.1;refGene_gn=NUP58;refGene_tx=XM_023250028.1
A1	5102400	5102550
refGene_gn=NUP58;refGene_tx=XM_023250027.1;refGene_gn=NUP58;refGene_tx=XM_023250028.1
A1	5103235	5103373
refGene_gn=NUP58;refGene_tx=XM_023250027.1;refGene_gn=NUP58;refGene_tx=XM_023250028.1
A1	5106911	5107022
refGene_gn=NUP58;refGene_tx=XM_023250027.1;refGene_gn=NUP58;refGene_tx

=XM_023250028.1
A1 5112285 5112310
refGene_gn=NUP58;refGene_tx=XM_023250027.1;refGene_gn=NUP58;refGene_tx
=XM_023250028.1
A1 5113408 5113474
refGene_gn=NUP58;refGene_tx=XM_023250027.1;refGene_gn=NUP58;refGene_tx
=XM_023250028.1
A1 5113790 5113965
refGene_gn=NUP58;refGene_tx=XM_023250027.1;refGene_gn=NUP58;refGene_tx
=XM_023250028.1
A1 5117036 5117116
refGene_gn=NUP58;refGene_tx=XM_023250027.1;refGene_gn=NUP58;refGene_tx
=XM_023250028.1
A1 5119155 5119274
refGene_gn=NUP58;refGene_tx=XM_023250027.1;refGene_gn=NUP58;refGene_tx
=XM_023250028.1
A1 5119695 5119778
refGene_gn=NUP58;refGene_tx=XM_023250027.1;refGene_gn=NUP58;refGene_tx
=XM_023250028.1
A1 5122601 5122803
refGene_gn=NUP58;refGene_tx=XM_023250027.1;refGene_gn=NUP58;refGene_tx
=XM_023250028.1
A1 5128186 5128282
refGene_gn=NUP58;refGene_tx=XM_023250027.1;refGene_gn=NUP58;refGene_tx
=XM_023250028.1
A1 5129874 5129970
refGene_gn=NUP58;refGene_tx=XM_023250027.1;refGene_gn=NUP58;refGene_tx
=XM_023250028.1
A1 5131208 5131378 refGene_gn=NUP58;refGene_tx=XM_023250027.1
A1 5142496 5142579 refGene_gn=NUP58;refGene_tx=XM_023250028.1
A1 5155404 5155769
refGene_gn=LOC111559502;refGene_tx=XM_023250404.1
A1 5157131 5157237
refGene_gn=LOC111559502;refGene_tx=XM_023250404.1
A1 5157525 5158041
refGene_gn=LOC111559502;refGene_tx=XM_023250404.1
A1 5158523 5158547
refGene_gn=LOC111559502;refGene_tx=XM_023250404.1
A1 5169046 5169122
refGene_gn=LOC101085594;refGene_tx=XM_023250034.1;refGene_gn=LOC101085
594;refGene_tx=XM_023250035.1;refGene_gn=LOC101085594;refGene_tx=XM_02
3250041.1;refGene_gn=LOC101085594;refGene_tx=XM_023250050.1
A1 5227487 5227527
refGene_gn=LOC101085594;refGene_tx=XM_019820599.2
A1 5258484 5258629
refGene_gn=LOC101085594;refGene_tx=XM_006927173.4;refGene_gn=LOC101085
594;refGene_tx=XM_019820599.2;refGene_gn=LOC101085594;refGene_tx=XM_01
9820927.2;refGene_gn=LOC101085594;refGene_tx=XM_019820968.2;refGene_gn
=LOC101085594;refGene_tx=XM_019820987.2;refGene_gn=LOC101085594;refGen
e_tx=XM_023250034.1;refGene_gn=LOC101085594;refGene_tx=XM_023250035.1;

refGene_gn=LOC101085594;refGene_tx=XM_023250041.1;refGene_gn=LOC101085
594;refGene_tx=XM_023250050.1
A1	5309438	5309538
refGene_gn=LOC101085594;refGene_tx=XM_006927173.4;refGene_gn=LOC101085
594;refGene_tx=XM_019820599.2;refGene_gn=LOC101085594;refGene_tx=XM_01
9820927.2;refGene_gn=LOC101085594;refGene_tx=XM_019820968.2;refGene_gn
=LOC101085594;refGene_tx=XM_019820987.2;refGene_gn=LOC101085594;refGen
e_tx=XM_023250034.1;refGene_gn=LOC101085594;refGene_tx=XM_023250035.1;
refGene_gn=LOC101085594;refGene_tx=XM_023250041.1;refGene_gn=LOC101085
594;refGene_tx=XM_023250050.1
A1	5309996	5310095
refGene_gn=LOC101085594;refGene_tx=XM_006927173.4;refGene_gn=LOC101085
594;refGene_tx=XM_019820599.2;refGene_gn=LOC101085594;refGene_tx=XM_01
9820927.2;refGene_gn=LOC101085594;refGene_tx=XM_019820968.2;refGene_gn
=LOC101085594;refGene_tx=XM_019820987.2;refGene_gn=LOC101085594;refGen
e_tx=XM_023250034.1;refGene_gn=LOC101085594;refGene_tx=XM_023250035.1;
refGene_gn=LOC101085594;refGene_tx=XM_023250041.1;refGene_gn=LOC101085
594;refGene_tx=XM_023250050.1
A1	5311166	5311212
refGene_gn=LOC101085594;refGene_tx=XM_006927173.4;refGene_gn=LOC101085
594;refGene_tx=XM_019820599.2;refGene_gn=LOC101085594;refGene_tx=XM_01
9820927.2;refGene_gn=LOC101085594;refGene_tx=XM_019820968.2;refGene_gn
=LOC101085594;refGene_tx=XM_019820987.2;refGene_gn=LOC101085594;refGen
e_tx=XM_023250034.1;refGene_gn=LOC101085594;refGene_tx=XM_023250035.1;
refGene_gn=LOC101085594;refGene_tx=XM_023250041.1;refGene_gn=LOC101085
594;refGene_tx=XM_023250050.1
A1	5311832	5311873
refGene_gn=LOC101085594;refGene_tx=XM_006927173.4;refGene_gn=LOC101085
594;refGene_tx=XM_019820599.2;refGene_gn=LOC101085594;refGene_tx=XM_01
9820927.2;refGene_gn=LOC101085594;refGene_tx=XM_019820968.2;refGene_gn
=LOC101085594;refGene_tx=XM_019820987.2;refGene_gn=LOC101085594;refGen
e_tx=XM_023250034.1;refGene_gn=LOC101085594;refGene_tx=XM_023250035.1;
refGene_gn=LOC101085594;refGene_tx=XM_023250041.1;refGene_gn=LOC101085
594;refGene_tx=XM_023250050.1
A1	5316869	5316943
refGene_gn=LOC101085594;refGene_tx=XM_006927173.4;refGene_gn=LOC101085
594;refGene_tx=XM_019820599.2;refGene_gn=LOC101085594;refGene_tx=XM_01
9820927.2;refGene_gn=LOC101085594;refGene_tx=XM_019820968.2;refGene_gn
=LOC101085594;refGene_tx=XM_019820987.2;refGene_gn=LOC101085594;refGen
e_tx=XM_023250034.1;refGene_gn=LOC101085594;refGene_tx=XM_023250035.1;
refGene_gn=LOC101085594;refGene_tx=XM_023250041.1;refGene_gn=LOC101085
594;refGene_tx=XM_023250050.1
A1	5318839	5318909
refGene_gn=LOC101085594;refGene_tx=XM_006927173.4;refGene_gn=LOC101085
594;refGene_tx=XM_019820599.2;refGene_gn=LOC101085594;refGene_tx=XM_01
9820927.2;refGene_gn=LOC101085594;refGene_tx=XM_019820968.2;refGene_gn
=LOC101085594;refGene_tx=XM_019820987.2;refGene_gn=LOC101085594;refGen
e_tx=XM_023250034.1;refGene_gn=LOC101085594;refGene_tx=XM_023250035.1;
refGene_gn=LOC101085594;refGene_tx=XM_023250041.1;refGene_gn=LOC101085
594;refGene_tx=XM_023250050.1

A1 5319903 5320031
refGene_gn=LOC101085594;refGene_tx=XM_006927173.4;refGene_gn=LOC101085594;refGene_tx=XM_019820599.2;refGene_gn=LOC101085594;refGene_tx=XM_019820927.2;refGene_gn=LOC101085594;refGene_tx=XM_019820968.2;refGene_gn=LOC101085594;refGene_tx=XM_019820987.2;refGene_gn=LOC101085594;refGene_tx=XM_023250034.1;refGene_gn=LOC101085594;refGene_tx=XM_023250035.1;refGene_gn=LOC101085594;refGene_tx=XM_023250041.1;refGene_gn=LOC101085594;refGene_tx=XM_023250050.1
A1 5321253 5321365
refGene_gn=LOC101085594;refGene_tx=XM_006927173.4;refGene_gn=LOC101085594;refGene_tx=XM_019820599.2;refGene_gn=LOC101085594;refGene_tx=XM_019820927.2;refGene_gn=LOC101085594;refGene_tx=XM_019820968.2;refGene_gn=LOC101085594;refGene_tx=XM_019820987.2;refGene_gn=LOC101085594;refGene_tx=XM_023250034.1;refGene_gn=LOC101085594;refGene_tx=XM_023250035.1;refGene_gn=LOC101085594;refGene_tx=XM_023250041.1;refGene_gn=LOC101085594;refGene_tx=XM_023250050.1
A1 5327638 5327804
refGene_gn=LOC101085594;refGene_tx=XM_006927173.4;refGene_gn=LOC101085594;refGene_tx=XM_019820599.2;refGene_gn=LOC101085594;refGene_tx=XM_019820927.2;refGene_gn=LOC101085594;refGene_tx=XM_019820968.2;refGene_gn=LOC101085594;refGene_tx=XM_019820987.2;refGene_gn=LOC101085594;refGene_tx=XM_023250034.1;refGene_gn=LOC101085594;refGene_tx=XM_023250035.1;refGene_gn=LOC101085594;refGene_tx=XM_023250041.1;refGene_gn=LOC101085594;refGene_tx=XM_023250050.1
A1 5332746 5332874
refGene_gn=LOC101085594;refGene_tx=XM_006927173.4;refGene_gn=LOC101085594;refGene_tx=XM_019820599.2;refGene_gn=LOC101085594;refGene_tx=XM_019820927.2;refGene_gn=LOC101085594;refGene_tx=XM_019820968.2;refGene_gn=LOC101085594;refGene_tx=XM_019820987.2;refGene_gn=LOC101085594;refGene_tx=XM_023250034.1;refGene_gn=LOC101085594;refGene_tx=XM_023250035.1;refGene_gn=LOC101085594;refGene_tx=XM_023250041.1;refGene_gn=LOC101085594;refGene_tx=XM_023250050.1
A1 5333712 5333790
refGene_gn=LOC101085594;refGene_tx=XM_006927173.4;refGene_gn=LOC101085594;refGene_tx=XM_019820599.2;refGene_gn=LOC101085594;refGene_tx=XM_019820927.2;refGene_gn=LOC101085594;refGene_tx=XM_019820968.2;refGene_gn=LOC101085594;refGene_tx=XM_019820987.2;refGene_gn=LOC101085594;refGene_tx=XM_023250034.1;refGene_gn=LOC101085594;refGene_tx=XM_023250035.1;refGene_gn=LOC101085594;refGene_tx=XM_023250041.1;refGene_gn=LOC101085594;refGene_tx=XM_023250050.1
A1 5336967 5337056
refGene_gn=LOC101085594;refGene_tx=XM_006927173.4;refGene_gn=LOC101085594;refGene_tx=XM_019820599.2;refGene_gn=LOC101085594;refGene_tx=XM_019820927.2;refGene_gn=LOC101085594;refGene_tx=XM_019820968.2;refGene_gn=LOC101085594;refGene_tx=XM_019820987.2;refGene_gn=LOC101085594;refGene_tx=XM_023250034.1;refGene_gn=LOC101085594;refGene_tx=XM_023250035.1;refGene_gn=LOC101085594;refGene_tx=XM_023250041.1;refGene_gn=LOC101085594;refGene_tx=XM_023250050.1
A1 5341519 5341564
refGene_gn=LOC101085594;refGene_tx=XM_006927173.4;refGene_gn=LOC101085

594;refGene_tx=XM_019820599.2;refGene_gn=LOC101085594;refGene_tx=XM_019820927.2;refGene_gn=LOC101085594;refGene_tx=XM_019820968.2;refGene_gn=LOC101085594;refGene_tx=XM_019820987.2;refGene_gn=LOC101085594;refGene_tx=XM_023250034.1;refGene_gn=LOC101085594;refGene_tx=XM_023250035.1;refGene_gn=LOC101085594;refGene_tx=XM_023250041.1;refGene_gn=LOC101085594;refGene_tx=XM_023250050.1
A1 5343704 5343780
refGene_gn=LOC101085594;refGene_tx=XM_006927173.4;refGene_gn=LOC101085594;refGene_tx=XM_019820599.2;refGene_gn=LOC101085594;refGene_tx=XM_019820927.2;refGene_gn=LOC101085594;refGene_tx=XM_019820968.2;refGene_gn=LOC101085594;refGene_tx=XM_019820987.2;refGene_gn=LOC101085594;refGene_tx=XM_023250034.1;refGene_gn=LOC101085594;refGene_tx=XM_023250035.1;refGene_gn=LOC101085594;refGene_tx=XM_023250041.1;refGene_gn=LOC101085594;refGene_tx=XM_023250050.1
A1 5351216 5351322
refGene_gn=LOC101085594;refGene_tx=XM_006927173.4;refGene_gn=LOC101085594;refGene_tx=XM_019820599.2;refGene_gn=LOC101085594;refGene_tx=XM_019820927.2;refGene_gn=LOC101085594;refGene_tx=XM_019820968.2;refGene_gn=LOC101085594;refGene_tx=XM_019820987.2;refGene_gn=LOC101085594;refGene_tx=XM_023250034.1;refGene_gn=LOC101085594;refGene_tx=XM_023250035.1;refGene_gn=LOC101085594;refGene_tx=XM_023250041.1;refGene_gn=LOC101085594;refGene_tx=XM_023250050.1
A1 5352320 5352403
refGene_gn=LOC101085594;refGene_tx=XM_006927173.4;refGene_gn=LOC101085594;refGene_tx=XM_019820599.2;refGene_gn=LOC101085594;refGene_tx=XM_019820927.2;refGene_gn=LOC101085594;refGene_tx=XM_019820968.2;refGene_gn=LOC101085594;refGene_tx=XM_019820987.2;refGene_gn=LOC101085594;refGene_tx=XM_023250034.1;refGene_gn=LOC101085594;refGene_tx=XM_023250035.1;refGene_gn=LOC101085594;refGene_tx=XM_023250041.1;refGene_gn=LOC101085594;refGene_tx=XM_023250050.1
A1 5355191 5355241
refGene_gn=LOC101085594;refGene_tx=XM_006927173.4;refGene_gn=LOC101085594;refGene_tx=XM_019820599.2;refGene_gn=LOC101085594;refGene_tx=XM_019820927.2;refGene_gn=LOC101085594;refGene_tx=XM_019820968.2;refGene_gn=LOC101085594;refGene_tx=XM_019820987.2;refGene_gn=LOC101085594;refGene_tx=XM_023250034.1;refGene_gn=LOC101085594;refGene_tx=XM_023250035.1;refGene_gn=LOC101085594;refGene_tx=XM_023250041.1;refGene_gn=LOC101085594;refGene_tx=XM_023250050.1
A1 5357719 5357789
refGene_gn=LOC101085594;refGene_tx=XM_006927173.4;refGene_gn=LOC101085594;refGene_tx=XM_019820599.2;refGene_gn=LOC101085594;refGene_tx=XM_019820927.2;refGene_gn=LOC101085594;refGene_tx=XM_019820968.2;refGene_gn=LOC101085594;refGene_tx=XM_019820987.2;refGene_gn=LOC101085594;refGene_tx=XM_023250034.1;refGene_gn=LOC101085594;refGene_tx=XM_023250035.1;refGene_gn=LOC101085594;refGene_tx=XM_023250041.1;refGene_gn=LOC101085594;refGene_tx=XM_023250050.1
A1 5358696 5358781
refGene_gn=LOC101085594;refGene_tx=XM_006927173.4;refGene_gn=LOC101085594;refGene_tx=XM_019820599.2;refGene_gn=LOC101085594;refGene_tx=XM_019820927.2;refGene_gn=LOC101085594;refGene_tx=XM_019820968.2;refGene_gn

=LOC101085594;refGene_tx=XM_019820987.2;refGene_gn=LOC101085594;refGene_tx=XM_023250034.1;refGene_gn=LOC101085594;refGene_tx=XM_023250035.1;refGene_gn=LOC101085594;refGene_tx=XM_023250041.1;refGene_gn=LOC101085594;refGene_tx=XM_023250050.1
A1 5359764 5359904
refGene_gn=LOC101085594;refGene_tx=XM_006927173.4;refGene_gn=LOC101085594;refGene_tx=XM_019820599.2;refGene_gn=LOC101085594;refGene_tx=XM_019820927.2;refGene_gn=LOC101085594;refGene_tx=XM_019820968.2;refGene_gn=LOC101085594;refGene_tx=XM_019820987.2;refGene_gn=LOC101085594;refGene_tx=XM_023250034.1;refGene_gn=LOC101085594;refGene_tx=XM_023250035.1;refGene_gn=LOC101085594;refGene_tx=XM_023250041.1;refGene_gn=LOC101085594;refGene_tx=XM_023250050.1
A1 5362238 5362377
refGene_gn=LOC101085594;refGene_tx=XM_006927173.4;refGene_gn=LOC101085594;refGene_tx=XM_019820599.2;refGene_gn=LOC101085594;refGene_tx=XM_019820927.2;refGene_gn=LOC101085594;refGene_tx=XM_019820968.2;refGene_gn=LOC101085594;refGene_tx=XM_019820987.2;refGene_gn=LOC101085594;refGene_tx=XM_023250034.1;refGene_gn=LOC101085594;refGene_tx=XM_023250035.1;refGene_gn=LOC101085594;refGene_tx=XM_023250041.1;refGene_gn=LOC101085594;refGene_tx=XM_023250050.1
A1 5371645 5371710
refGene_gn=LOC101085594;refGene_tx=XM_006927173.4;refGene_gn=LOC101085594;refGene_tx=XM_019820599.2;refGene_gn=LOC101085594;refGene_tx=XM_019820927.2;refGene_gn=LOC101085594;refGene_tx=XM_019820968.2;refGene_gn=LOC101085594;refGene_tx=XM_019820987.2;refGene_gn=LOC101085594;refGene_tx=XM_023250034.1;refGene_gn=LOC101085594;refGene_tx=XM_023250035.1;refGene_gn=LOC101085594;refGene_tx=XM_023250041.1;refGene_gn=LOC101085594;refGene_tx=XM_023250050.1
A1 5466860 5467033
refGene_gn=LOC101085594;refGene_tx=XM_006927173.4;refGene_gn=LOC101085594;refGene_tx=XM_019820599.2;refGene_gn=LOC101085594;refGene_tx=XM_019820927.2;refGene_gn=LOC101085594;refGene_tx=XM_019820968.2;refGene_gn=LOC101085594;refGene_tx=XM_019820987.2;refGene_gn=LOC101085594;refGene_tx=XM_023250034.1;refGene_gn=LOC101085594;refGene_tx=XM_023250035.1;refGene_gn=LOC101085594;refGene_tx=XM_023250041.1;refGene_gn=LOC101085594;refGene_tx=XM_023250050.1
A1 5537757 5537941
refGene_gn=LOC101085594;refGene_tx=XM_006927173.4;refGene_gn=LOC101085594;refGene_tx=XM_019820599.2;refGene_gn=LOC101085594;refGene_tx=XM_019820927.2;refGene_gn=LOC101085594;refGene_tx=XM_019820968.2;refGene_gn=LOC101085594;refGene_tx=XM_019820987.2;refGene_gn=LOC101085594;refGene_tx=XM_023250034.1;refGene_gn=LOC101085594;refGene_tx=XM_023250035.1;refGene_gn=LOC101085594;refGene_tx=XM_023250041.1;refGene_gn=LOC101085594;refGene_tx=XM_023250050.1
A1 5542871 5542982
refGene_gn=LOC101085594;refGene_tx=XM_006927173.4;refGene_gn=LOC101085594;refGene_tx=XM_019820599.2;refGene_gn=LOC101085594;refGene_tx=XM_019820927.2;refGene_gn=LOC101085594;refGene_tx=XM_019820968.2;refGene_gn=LOC101085594;refGene_tx=XM_019820987.2;refGene_gn=LOC101085594;refGene_tx=XM_023250034.1;refGene_gn=LOC101085594;refGene_tx=XM_023250035.1;

refGene_gn=LOC101085594;refGene_tx=XM_023250041.1;refGene_gn=LOC101085594;refGene_tx=XM_023250050.1
A1 5595700 5595775
refGene_gn=LOC101085594;refGene_tx=XM_006927173.4;refGene_gn=LOC101085594;refGene_tx=XM_019820599.2;refGene_gn=LOC101085594;refGene_tx=XM_019820927.2;refGene_gn=LOC101085594;refGene_tx=XM_019820968.2;refGene_gn=LOC101085594;refGene_tx=XM_019820987.2;refGene_gn=LOC101085594;refGene_tx=XM_023250034.1;refGene_gn=LOC101085594;refGene_tx=XM_023250035.1;refGene_gn=LOC101085594;refGene_tx=XM_023250041.1;refGene_gn=LOC101085594;refGene_tx=XM_023250050.1
A1 5603220 5603343
refGene_gn=LOC101085594;refGene_tx=XM_006927173.4;refGene_gn=LOC101085594;refGene_tx=XM_019820599.2;refGene_gn=LOC101085594;refGene_tx=XM_019820927.2;refGene_gn=LOC101085594;refGene_tx=XM_019820968.2;refGene_gn=LOC101085594;refGene_tx=XM_019820987.2;refGene_gn=LOC101085594;refGene_tx=XM_023250034.1;refGene_gn=LOC101085594;refGene_tx=XM_023250035.1;refGene_gn=LOC101085594;refGene_tx=XM_023250041.1;refGene_gn=LOC101085594;refGene_tx=XM_023250050.1
A1 5605060 5605139
refGene_gn=LOC101085594;refGene_tx=XM_006927173.4;refGene_gn=LOC101085594;refGene_tx=XM_019820599.2;refGene_gn=LOC101085594;refGene_tx=XM_019820927.2;refGene_gn=LOC101085594;refGene_tx=XM_019820968.2;refGene_gn=LOC101085594;refGene_tx=XM_019820987.2;refGene_gn=LOC101085594;refGene_tx=XM_023250034.1;refGene_gn=LOC101085594;refGene_tx=XM_023250035.1;refGene_gn=LOC101085594;refGene_tx=XM_023250041.1;refGene_gn=LOC101085594;refGene_tx=XM_023250050.1
A1 5619908 5619970
refGene_gn=LOC101085594;refGene_tx=XM_006927173.4;refGene_gn=LOC101085594;refGene_tx=XM_019820599.2;refGene_gn=LOC101085594;refGene_tx=XM_019820927.2;refGene_gn=LOC101085594;refGene_tx=XM_019820968.2;refGene_gn=LOC101085594;refGene_tx=XM_019820987.2;refGene_gn=LOC101085594;refGene_tx=XM_023250034.1;refGene_gn=LOC101085594;refGene_tx=XM_023250035.1;refGene_gn=LOC101085594;refGene_tx=XM_023250041.1;refGene_gn=LOC101085594;refGene_tx=XM_023250050.1
A1 5620393 5620450
refGene_gn=LOC101085594;refGene_tx=XM_006927173.4;refGene_gn=LOC101085594;refGene_tx=XM_019820599.2;refGene_gn=LOC101085594;refGene_tx=XM_019820927.2;refGene_gn=LOC101085594;refGene_tx=XM_019820968.2;refGene_gn=LOC101085594;refGene_tx=XM_019820987.2;refGene_gn=LOC101085594;refGene_tx=XM_023250034.1;refGene_gn=LOC101085594;refGene_tx=XM_023250035.1;refGene_gn=LOC101085594;refGene_tx=XM_023250041.1
A1 5621838 5621946
refGene_gn=LOC101085594;refGene_tx=XM_006927173.4;refGene_gn=LOC101085594;refGene_tx=XM_019820599.2;refGene_gn=LOC101085594;refGene_tx=XM_019820927.2;refGene_gn=LOC101085594;refGene_tx=XM_019820968.2;refGene_gn=LOC101085594;refGene_tx=XM_019820987.2;refGene_gn=LOC101085594;refGene_tx=XM_023250034.1;refGene_gn=LOC101085594;refGene_tx=XM_023250035.1;refGene_gn=LOC101085594;refGene_tx=XM_023250041.1;refGene_gn=LOC101085594;refGene_tx=XM_023250050.1
A1 5714182 5714271

refGene_gn=LOC101085594;refGene_tx=XM_006927173.4;refGene_gn=LOC101085594;refGene_tx=XM_019820599.2;refGene_gn=LOC101085594;refGene_tx=XM_019820927.2;refGene_gn=LOC101085594;refGene_tx=XM_019820968.2;refGene_gn=LOC101085594;refGene_tx=XM_019820987.2;refGene_gn=LOC101085594;refGene_tx=XM_023250034.1;refGene_gn=LOC101085594;refGene_tx=XM_023250035.1;refGene_gn=LOC101085594;refGene_tx=XM_023250041.1;refGene_gn=LOC101085594;refGene_tx=XM_023250050.1
A1 5720308 5720413
refGene_gn=LOC101085594;refGene_tx=XM_006927173.4;refGene_gn=LOC101085594;refGene_tx=XM_019820599.2;refGene_gn=LOC101085594;refGene_tx=XM_019820927.2;refGene_gn=LOC101085594;refGene_tx=XM_019820968.2;refGene_gn=LOC101085594;refGene_tx=XM_019820987.2;refGene_gn=LOC101085594;refGene_tx=XM_023250034.1;refGene_gn=LOC101085594;refGene_tx=XM_023250035.1;refGene_gn=LOC101085594;refGene_tx=XM_023250041.1;refGene_gn=LOC101085594;refGene_tx=XM_023250050.1
A1 5758152 5758244
refGene_gn=LOC101085594;refGene_tx=XM_006927173.4;refGene_gn=LOC101085594;refGene_tx=XM_019820599.2;refGene_gn=LOC101085594;refGene_tx=XM_019820927.2;refGene_gn=LOC101085594;refGene_tx=XM_019820968.2;refGene_gn=LOC101085594;refGene_tx=XM_019820987.2;refGene_gn=LOC101085594;refGene_tx=XM_023250034.1;refGene_gn=LOC101085594;refGene_tx=XM_023250035.1;refGene_gn=LOC101085594;refGene_tx=XM_023250041.1;refGene_gn=LOC101085594;refGene_tx=XM_023250050.1
A1 5764217 5764315
refGene_gn=LOC101085594;refGene_tx=XM_006927173.4;refGene_gn=LOC101085594;refGene_tx=XM_019820599.2;refGene_gn=LOC101085594;refGene_tx=XM_019820927.2;refGene_gn=LOC101085594;refGene_tx=XM_019820968.2;refGene_gn=LOC101085594;refGene_tx=XM_019820987.2;refGene_gn=LOC101085594;refGene_tx=XM_023250034.1;refGene_gn=LOC101085594;refGene_tx=XM_023250035.1;refGene_gn=LOC101085594;refGene_tx=XM_023250041.1;refGene_gn=LOC101085594;refGene_tx=XM_023250050.1
A1 5783562 5784116 refGene_gn=SHISA2;refGene_tx=XM_023250064.1
A1 5787777 5788111 refGene_gn=SHISA2;refGene_tx=XM_023250064.1
A1 5874730 5874826
refGene_gn=LOC101096779;refGene_tx=XM_019821051.2;refGene_gn=LOC101096779;refGene_tx=XM_023250074.1;refGene_gn=LOC101096779;refGene_tx=XM_023250077.1;refGene_gn=LOC101096779;refGene_tx=XM_023250080.1;refGene_gn=LOC101096779;refGene_tx=XM_023250086.1;refGene_gn=LOC101096779;refGene_tx=XM_023250089.1;refGene_gn=LOC101096779;refGene_tx=XM_023250092.1;refGene_gn=LOC101096779;refGene_tx=XM_023250096.1
A1 5877358 5877415
refGene_gn=LOC101096779;refGene_tx=XM_019821051.2;refGene_gn=LOC101096779;refGene_tx=XM_023250074.1;refGene_gn=LOC101096779;refGene_tx=XM_023250077.1;refGene_gn=LOC101096779;refGene_tx=XM_023250080.1;refGene_gn=LOC101096779;refGene_tx=XM_023250086.1;refGene_gn=LOC101096779;refGene_tx=XM_023250089.1;refGene_gn=LOC101096779;refGene_tx=XM_023250092.1;refGene_gn=LOC101096779;refGene_tx=XM_023250096.1
A1 5880543 5880605
refGene_gn=LOC101096779;refGene_tx=XM_019821051.2;refGene_gn=LOC101096779;refGene_tx=XM_023250074.1;refGene_gn=LOC101096779;refGene_tx=XM_02

3250077.1;refGene_gn=LOC101096779;refGene_tx=XM_023250080.1;refGene_gn
=LOC101096779;refGene_tx=XM_023250086.1;refGene_gn=LOC101096779;refGen
e_tx=XM_023250089.1;refGene_gn=LOC101096779;refGene_tx=XM_023250092.1;
refGene_gn=LOC101096779;refGene_tx=XM_023250096.1
A1 5896384 5896463
refGene_gn=LOC101096779;refGene_tx=XM_019821051.2;refGene_gn=LOC101096
779;refGene_tx=XM_023250074.1;refGene_gn=LOC101096779;refGene_tx=XM_02
3250077.1;refGene_gn=LOC101096779;refGene_tx=XM_023250080.1;refGene_gn
=LOC101096779;refGene_tx=XM_023250086.1;refGene_gn=LOC101096779;refGen
e_tx=XM_023250089.1;refGene_gn=LOC101096779;refGene_tx=XM_023250092.1;
refGene_gn=LOC101096779;refGene_tx=XM_023250096.1
A1 5903197 5903320
refGene_gn=LOC101096779;refGene_tx=XM_019821051.2;refGene_gn=LOC101096
779;refGene_tx=XM_023250074.1;refGene_gn=LOC101096779;refGene_tx=XM_02
3250077.1;refGene_gn=LOC101096779;refGene_tx=XM_023250080.1;refGene_gn
=LOC101096779;refGene_tx=XM_023250086.1;refGene_gn=LOC101096779;refGen
e_tx=XM_023250092.1;refGene_gn=LOC101096779;refGene_tx=XM_023250096.1
A1 5923070 5923145
refGene_gn=LOC101096779;refGene_tx=XM_019821051.2;refGene_gn=LOC101096
779;refGene_tx=XM_023250074.1;refGene_gn=LOC101096779;refGene_tx=XM_02
3250077.1;refGene_gn=LOC101096779;refGene_tx=XM_023250080.1;refGene_gn
=LOC101096779;refGene_tx=XM_023250086.1;refGene_gn=LOC101096779;refGen
e_tx=XM_023250089.1;refGene_gn=LOC101096779;refGene_tx=XM_023250092.1;
refGene_gn=LOC101096779;refGene_tx=XM_023250096.1
A1 5928915 5929026
refGene_gn=LOC101096779;refGene_tx=XM_019821051.2;refGene_gn=LOC101096
779;refGene_tx=XM_023250074.1;refGene_gn=LOC101096779;refGene_tx=XM_02
3250077.1;refGene_gn=LOC101096779;refGene_tx=XM_023250080.1;refGene_gn
=LOC101096779;refGene_tx=XM_023250086.1;refGene_gn=LOC101096779;refGen
e_tx=XM_023250089.1;refGene_gn=LOC101096779;refGene_tx=XM_023250092.1;
refGene_gn=LOC101096779;refGene_tx=XM_023250096.1
A1 5935393 5935577
refGene_gn=LOC101096779;refGene_tx=XM_019821051.2;refGene_gn=LOC101096
779;refGene_tx=XM_023250074.1;refGene_gn=LOC101096779;refGene_tx=XM_02
3250077.1;refGene_gn=LOC101096779;refGene_tx=XM_023250080.1;refGene_gn
=LOC101096779;refGene_tx=XM_023250086.1;refGene_gn=LOC101096779;refGen
e_tx=XM_023250089.1;refGene_gn=LOC101096779;refGene_tx=XM_023250092.1;
refGene_gn=LOC101096779;refGene_tx=XM_023250096.1
A1 5944417 5944590
refGene_gn=LOC101096779;refGene_tx=XM_019821051.2;refGene_gn=LOC101096
779;refGene_tx=XM_023250074.1;refGene_gn=LOC101096779;refGene_tx=XM_02
3250077.1;refGene_gn=LOC101096779;refGene_tx=XM_023250080.1;refGene_gn
=LOC101096779;refGene_tx=XM_023250086.1;refGene_gn=LOC101096779;refGen
e_tx=XM_023250089.1;refGene_gn=LOC101096779;refGene_tx=XM_023250092.1;
refGene_gn=LOC101096779;refGene_tx=XM_023250096.1
A1 5953407 5953472
refGene_gn=LOC101096779;refGene_tx=XM_019821051.2;refGene_gn=LOC101096
779;refGene_tx=XM_023250074.1;refGene_gn=LOC101096779;refGene_tx=XM_02
3250077.1;refGene_gn=LOC101096779;refGene_tx=XM_023250080.1;refGene_gn
=LOC101096779;refGene_tx=XM_023250086.1;refGene_gn=LOC101096779;refGen

e_tx=XM_023250089.1;refGene_gn=LOC101096779;refGene_tx=XM_023250092.1;
refGene_gn=LOC101096779;refGene_tx=XM_023250096.1
A1 5959595 5959734
refGene_gn=LOC101096779;refGene_tx=XM_019821051.2;refGene_gn=LOC101096
779;refGene_tx=XM_023250074.1;refGene_gn=LOC101096779;refGene_tx=XM_02
3250077.1;refGene_gn=LOC101096779;refGene_tx=XM_023250080.1;refGene_gn
=LOC101096779;refGene_tx=XM_023250086.1;refGene_gn=LOC101096779;refGen
e_tx=XM_023250089.1;refGene_gn=LOC101096779;refGene_tx=XM_023250092.1;
refGene_gn=LOC101096779;refGene_tx=XM_023250096.1
A1 5964928 5965068
refGene_gn=LOC101096779;refGene_tx=XM_019821051.2;refGene_gn=LOC101096
779;refGene_tx=XM_023250074.1;refGene_gn=LOC101096779;refGene_tx=XM_02
3250077.1;refGene_gn=LOC101096779;refGene_tx=XM_023250080.1;refGene_gn
=LOC101096779;refGene_tx=XM_023250086.1;refGene_gn=LOC101096779;refGen
e_tx=XM_023250089.1;refGene_gn=LOC101096779;refGene_tx=XM_023250092.1;
refGene_gn=LOC101096779;refGene_tx=XM_023250096.1
A1 5966801 5966886
refGene_gn=LOC101096779;refGene_tx=XM_019821051.2;refGene_gn=LOC101096
779;refGene_tx=XM_023250074.1;refGene_gn=LOC101096779;refGene_tx=XM_02
3250077.1;refGene_gn=LOC101096779;refGene_tx=XM_023250080.1;refGene_gn
=LOC101096779;refGene_tx=XM_023250086.1;refGene_gn=LOC101096779;refGen
e_tx=XM_023250089.1;refGene_gn=LOC101096779;refGene_tx=XM_023250092.1;
refGene_gn=LOC101096779;refGene_tx=XM_023250096.1
A1 5970266 5970336
refGene_gn=LOC101096779;refGene_tx=XM_019821051.2;refGene_gn=LOC101096
779;refGene_tx=XM_023250074.1;refGene_gn=LOC101096779;refGene_tx=XM_02
3250077.1;refGene_gn=LOC101096779;refGene_tx=XM_023250080.1;refGene_gn
=LOC101096779;refGene_tx=XM_023250086.1;refGene_gn=LOC101096779;refGen
e_tx=XM_023250089.1;refGene_gn=LOC101096779;refGene_tx=XM_023250092.1;
refGene_gn=LOC101096779;refGene_tx=XM_023250096.1
A1 5972885 5972935
refGene_gn=LOC101096779;refGene_tx=XM_019821051.2;refGene_gn=LOC101096
779;refGene_tx=XM_023250074.1;refGene_gn=LOC101096779;refGene_tx=XM_02
3250077.1;refGene_gn=LOC101096779;refGene_tx=XM_023250080.1;refGene_gn
=LOC101096779;refGene_tx=XM_023250086.1;refGene_gn=LOC101096779;refGen
e_tx=XM_023250089.1;refGene_gn=LOC101096779;refGene_tx=XM_023250092.1;
refGene_gn=LOC101096779;refGene_tx=XM_023250096.1
A1 5974144 5974227
refGene_gn=LOC101096779;refGene_tx=XM_019821051.2;refGene_gn=LOC101096
779;refGene_tx=XM_023250074.1;refGene_gn=LOC101096779;refGene_tx=XM_02
3250077.1;refGene_gn=LOC101096779;refGene_tx=XM_023250080.1;refGene_gn
=LOC101096779;refGene_tx=XM_023250086.1;refGene_gn=LOC101096779;refGen
e_tx=XM_023250089.1;refGene_gn=LOC101096779;refGene_tx=XM_023250092.1;
refGene_gn=LOC101096779;refGene_tx=XM_023250096.1
A1 5974824 5974930
refGene_gn=LOC101096779;refGene_tx=XM_019821051.2;refGene_gn=LOC101096
779;refGene_tx=XM_023250074.1;refGene_gn=LOC101096779;refGene_tx=XM_02
3250077.1;refGene_gn=LOC101096779;refGene_tx=XM_023250080.1;refGene_gn
=LOC101096779;refGene_tx=XM_023250086.1;refGene_gn=LOC101096779;refGen
e_tx=XM_023250089.1;refGene_gn=LOC101096779;refGene_tx=XM_023250092.1;

refGene_gn=LOC101096779;refGene_tx=XM_023250096.1
A1	5976442	5976518
refGene_gn=LOC101096779;refGene_tx=XM_019821051.2;refGene_gn=LOC101096779;refGene_tx=XM_023250074.1;refGene_gn=LOC101096779;refGene_tx=XM_023250077.1;refGene_gn=LOC101096779;refGene_tx=XM_023250080.1;refGene_gn=LOC101096779;refGene_tx=XM_023250086.1;refGene_gn=LOC101096779;refGene_tx=XM_023250089.1;refGene_gn=LOC101096779;refGene_tx=XM_023250092.1;refGene_gn=LOC101096779;refGene_tx=XM_023250096.1
A1	5977304	5977346
refGene_gn=LOC101096779;refGene_tx=XM_019821051.2;refGene_gn=LOC101096779;refGene_tx=XM_023250074.1;refGene_gn=LOC101096779;refGene_tx=XM_023250077.1;refGene_gn=LOC101096779;refGene_tx=XM_023250080.1;refGene_gn=LOC101096779;refGene_tx=XM_023250086.1;refGene_gn=LOC101096779;refGene_tx=XM_023250089.1;refGene_gn=LOC101096779;refGene_tx=XM_023250092.1;refGene_gn=LOC101096779;refGene_tx=XM_023250096.1
A1	5977594	5977683
refGene_gn=LOC101096779;refGene_tx=XM_019821051.2;refGene_gn=LOC101096779;refGene_tx=XM_023250074.1;refGene_gn=LOC101096779;refGene_tx=XM_023250077.1;refGene_gn=LOC101096779;refGene_tx=XM_023250080.1;refGene_gn=LOC101096779;refGene_tx=XM_023250086.1;refGene_gn=LOC101096779;refGene_tx=XM_023250089.1;refGene_gn=LOC101096779;refGene_tx=XM_023250092.1;refGene_gn=LOC101096779;refGene_tx=XM_023250096.1
A1	5983471	5983549
refGene_gn=LOC101096779;refGene_tx=XM_019821051.2;refGene_gn=LOC101096779;refGene_tx=XM_023250074.1;refGene_gn=LOC101096779;refGene_tx=XM_023250077.1;refGene_gn=LOC101096779;refGene_tx=XM_023250080.1;refGene_gn=LOC101096779;refGene_tx=XM_023250086.1;refGene_gn=LOC101096779;refGene_tx=XM_023250089.1;refGene_gn=LOC101096779;refGene_tx=XM_023250092.1;refGene_gn=LOC101096779;refGene_tx=XM_023250096.1
A1	5984558	5984686
refGene_gn=LOC101096779;refGene_tx=XM_019821051.2;refGene_gn=LOC101096779;refGene_tx=XM_023250074.1;refGene_gn=LOC101096779;refGene_tx=XM_023250077.1;refGene_gn=LOC101096779;refGene_tx=XM_023250080.1;refGene_gn=LOC101096779;refGene_tx=XM_023250086.1;refGene_gn=LOC101096779;refGene_tx=XM_023250089.1;refGene_gn=LOC101096779;refGene_tx=XM_023250092.1;refGene_gn=LOC101096779;refGene_tx=XM_023250096.1
A1	5990126	5990292
refGene_gn=LOC101096779;refGene_tx=XM_019821051.2;refGene_gn=LOC101096779;refGene_tx=XM_023250074.1;refGene_gn=LOC101096779;refGene_tx=XM_023250077.1;refGene_gn=LOC101096779;refGene_tx=XM_023250080.1;refGene_gn=LOC101096779;refGene_tx=XM_023250086.1;refGene_gn=LOC101096779;refGene_tx=XM_023250089.1;refGene_gn=LOC101096779;refGene_tx=XM_023250092.1;refGene_gn=LOC101096779;refGene_tx=XM_023250096.1
A1	5993186	5993298
refGene_gn=LOC101096779;refGene_tx=XM_019821051.2;refGene_gn=LOC101096779;refGene_tx=XM_023250074.1;refGene_gn=LOC101096779;refGene_tx=XM_023250077.1;refGene_gn=LOC101096779;refGene_tx=XM_023250080.1;refGene_gn=LOC101096779;refGene_tx=XM_023250086.1;refGene_gn=LOC101096779;refGene_tx=XM_023250089.1;refGene_gn=LOC101096779;refGene_tx=XM_023250092.1;refGene_gn=LOC101096779;refGene_tx=XM_023250096.1

A1 5994597 5994722
refGene_gn=LOC101096779;refGene_tx=XM_019821051.2;refGene_gn=LOC101096779;refGene_tx=XM_023250074.1;refGene_gn=LOC101096779;refGene_tx=XM_023250077.1;refGene_gn=LOC101096779;refGene_tx=XM_023250080.1;refGene_gn=LOC101096779;refGene_tx=XM_023250086.1;refGene_gn=LOC101096779;refGene_tx=XM_023250089.1;refGene_gn=LOC101096779;refGene_tx=XM_023250092.1;refGene_gn=LOC101096779;refGene_tx=XM_023250096.1
A1 6003589 6003659
refGene_gn=LOC101096779;refGene_tx=XM_019821051.2;refGene_gn=LOC101096779;refGene_tx=XM_023250074.1;refGene_gn=LOC101096779;refGene_tx=XM_023250077.1;refGene_gn=LOC101096779;refGene_tx=XM_023250080.1;refGene_gn=LOC101096779;refGene_tx=XM_023250086.1;refGene_gn=LOC101096779;refGene_tx=XM_023250089.1;refGene_gn=LOC101096779;refGene_tx=XM_023250092.1;refGene_gn=LOC101096779;refGene_tx=XM_023250096.1
A1 6011241 6011315
refGene_gn=LOC101096779;refGene_tx=XM_019821051.2;refGene_gn=LOC101096779;refGene_tx=XM_023250074.1;refGene_gn=LOC101096779;refGene_tx=XM_023250077.1;refGene_gn=LOC101096779;refGene_tx=XM_023250080.1;refGene_gn=LOC101096779;refGene_tx=XM_023250086.1;refGene_gn=LOC101096779;refGene_tx=XM_023250089.1;refGene_gn=LOC101096779;refGene_tx=XM_023250092.1
A1 6013116 6013157
refGene_gn=LOC101096779;refGene_tx=XM_019821051.2;refGene_gn=LOC101096779;refGene_tx=XM_023250074.1;refGene_gn=LOC101096779;refGene_tx=XM_023250077.1;refGene_gn=LOC101096779;refGene_tx=XM_023250080.1;refGene_gn=LOC101096779;refGene_tx=XM_023250086.1;refGene_gn=LOC101096779;refGene_tx=XM_023250089.1;refGene_gn=LOC101096779;refGene_tx=XM_023250092.1
A1 6013982 6014028
refGene_gn=LOC101096779;refGene_tx=XM_019821051.2;refGene_gn=LOC101096779;refGene_tx=XM_023250074.1;refGene_gn=LOC101096779;refGene_tx=XM_023250077.1;refGene_gn=LOC101096779;refGene_tx=XM_023250080.1;refGene_gn=LOC101096779;refGene_tx=XM_023250086.1;refGene_gn=LOC101096779;refGene_tx=XM_023250089.1;refGene_gn=LOC101096779;refGene_tx=XM_023250092.1
A1 6015603 6015702
refGene_gn=LOC101096779;refGene_tx=XM_019821051.2;refGene_gn=LOC101096779;refGene_tx=XM_023250074.1;refGene_gn=LOC101096779;refGene_tx=XM_023250077.1;refGene_gn=LOC101096779;refGene_tx=XM_023250080.1;refGene_gn=LOC101096779;refGene_tx=XM_023250086.1;refGene_gn=LOC101096779;refGene_tx=XM_023250089.1;refGene_gn=LOC101096779;refGene_tx=XM_023250092.1
A1 6019639 6019739
refGene_gn=LOC101096779;refGene_tx=XM_019821051.2;refGene_gn=LOC101096779;refGene_tx=XM_023250074.1;refGene_gn=LOC101096779;refGene_tx=XM_023250077.1;refGene_gn=LOC101096779;refGene_tx=XM_023250080.1;refGene_gn=LOC101096779;refGene_tx=XM_023250086.1;refGene_gn=LOC101096779;refGene_tx=XM_023250089.1
A1 6029048 6029149
refGene_gn=LOC101096779;refGene_tx=XM_019821051.2;refGene_gn=LOC101096779;refGene_tx=XM_023250074.1;refGene_gn=LOC101096779;refGene_tx=XM_023250077.1;refGene_gn=LOC101096779;refGene_tx=XM_023250080.1;refGene_gn=LOC101096779;refGene_tx=XM_023250086.1;refGene_gn=LOC101096779;refGene_tx=XM_023250089.1;refGene_gn=LOC101096779;refGene_tx=XM_023250096.1

A1	6084641	6086404
refGene_gn=RNF6;refGene_tx=XM_003980263.5;refGene_gn=RNF6;refGene_tx=XM_011280889.3;refGene_gn=RNF6;refGene_tx=XM_011280919.3;refGene_gn=RNF6;refGene_tx=XM_019821526.2
A1	6089265	6089361
refGene_gn=RNF6;refGene_tx=XM_003980263.5;refGene_gn=RNF6;refGene_tx=XM_011280889.3;refGene_gn=RNF6;refGene_tx=XM_011280919.3;refGene_gn=RNF6;refGene_tx=XM_019821526.2
A1	6090674	6090867
refGene_gn=RNF6;refGene_tx=XM_003980263.5;refGene_gn=RNF6;refGene_tx=XM_011280889.3;refGene_gn=RNF6;refGene_tx=XM_011280919.3;refGene_gn=RNF6;refGene_tx=XM_019821526.2
A1	6114760	6114888
refGene_gn=CDK8;refGene_tx=XM_003980264.5;refGene_gn=CDK8;refGene_tx=XM_006927174.4;refGene_gn=CDK8;refGene_tx=XM_019821740.2
A1	6171940	6172016
refGene_gn=CDK8;refGene_tx=XM_003980264.5;refGene_gn=CDK8;refGene_tx=XM_006927174.4;refGene_gn=CDK8;refGene_tx=XM_019821740.2
A1	6181431	6181542
refGene_gn=CDK8;refGene_tx=XM_003980264.5;refGene_gn=CDK8;refGene_tx=XM_006927174.4;refGene_gn=CDK8;refGene_tx=XM_019821740.2
A1	6186295	6186436
refGene_gn=CDK8;refGene_tx=XM_003980264.5;refGene_gn=CDK8;refGene_tx=XM_006927174.4;refGene_gn=CDK8;refGene_tx=XM_019821740.2
A1	6206982	6207040
refGene_gn=CDK8;refGene_tx=XM_003980264.5;refGene_gn=CDK8;refGene_tx=XM_006927174.4;refGene_gn=CDK8;refGene_tx=XM_019821740.2
A1	6209409	6209541
refGene_gn=CDK8;refGene_tx=XM_003980264.5;refGene_gn=CDK8;refGene_tx=XM_006927174.4;refGene_gn=CDK8;refGene_tx=XM_019821740.2;refGene_gn=CDK8;refGene_tx=XM_019821783.1
A1	6216639	6216783
refGene_gn=CDK8;refGene_tx=XM_003980264.5;refGene_gn=CDK8;refGene_tx=XM_006927174.4;refGene_gn=CDK8;refGene_tx=XM_019821783.1
A1	6219284	6219354
refGene_gn=CDK8;refGene_tx=XM_003980264.5;refGene_gn=CDK8;refGene_tx=XM_006927174.4;refGene_gn=CDK8;refGene_tx=XM_019821740.2;refGene_gn=CDK8;refGene_tx=XM_019821783.1
A1	6220118	6220191
refGene_gn=CDK8;refGene_tx=XM_003980264.5;refGene_gn=CDK8;refGene_tx=XM_006927174.4;refGene_gn=CDK8;refGene_tx=XM_019821740.2;refGene_gn=CDK8;refGene_tx=XM_019821783.1
A1	6222989	6223087
refGene_gn=CDK8;refGene_tx=XM_003980264.5;refGene_gn=CDK8;refGene_tx=XM_006927174.4;refGene_gn=CDK8;refGene_tx=XM_019821740.2;refGene_gn=CDK8;refGene_tx=XM_019821783.1
A1	6223714	6223793
refGene_gn=CDK8;refGene_tx=XM_003980264.5;refGene_gn=CDK8;refGene_tx=XM_006927174.4;refGene_gn=CDK8;refGene_tx=XM_019821740.2;refGene_gn=CDK8;refGene_tx=XM_019821783.1

A1 6223912 6224071
refGene_gn=CDK8;refGene_tx=XM_003980264.5;refGene_gn=CDK8;refGene_tx=XM_006927174.4;refGene_gn=CDK8;refGene_tx=XM_019821740.2;refGene_gn=CDK8;refGene_tx=XM_019821783.1
A1 6228083 6228209
refGene_gn=CDK8;refGene_tx=XM_003980264.5;refGene_gn=CDK8;refGene_tx=XM_006927174.4;refGene_gn=CDK8;refGene_tx=XM_019821740.2;refGene_gn=CDK8;refGene_tx=XM_019821783.1
A1 6437412 6437545
refGene_gn=WASF3;refGene_tx=XM_006927176.4;refGene_gn=WASF3;refGene_tx=XM_019822210.2;refGene_gn=WASF3;refGene_tx=XM_019822250.2;refGene_gn=WASF3;refGene_tx=XM_019822289.2;refGene_gn=WASF3;refGene_tx=XM_019822322.2;refGene_gn=WASF3;refGene_tx=XM_019822353.2;refGene_gn=WASF3;refGene_tx=XM_019822405.2;refGene_gn=WASF3;refGene_tx=XM_019822445.2;refGene_gn=WASF3;refGene_tx=XM_023250152.1;refGene_gn=WASF3;refGene_tx=XM_023250184.1;refGene_gn=WASF3;refGene_tx=XM_023250199.1
A1 6460374 6460509
refGene_gn=WASF3;refGene_tx=XM_006927176.4;refGene_gn=WASF3;refGene_tx=XM_019822210.2;refGene_gn=WASF3;refGene_tx=XM_019822250.2;refGene_gn=WASF3;refGene_tx=XM_019822289.2;refGene_gn=WASF3;refGene_tx=XM_019822322.2;refGene_gn=WASF3;refGene_tx=XM_019822353.2;refGene_gn=WASF3;refGene_tx=XM_019822405.2;refGene_gn=WASF3;refGene_tx=XM_019822445.2;refGene_gn=WASF3;refGene_tx=XM_023250152.1;refGene_gn=WASF3;refGene_tx=XM_023250184.1;refGene_gn=WASF3;refGene_tx=XM_023250199.1
A1 6462627 6462781
refGene_gn=WASF3;refGene_tx=XM_006927176.4;refGene_gn=WASF3;refGene_tx=XM_019822210.2;refGene_gn=WASF3;refGene_tx=XM_019822250.2;refGene_gn=WASF3;refGene_tx=XM_019822289.2;refGene_gn=WASF3;refGene_tx=XM_019822322.2;refGene_gn=WASF3;refGene_tx=XM_019822353.2;refGene_gn=WASF3;refGene_tx=XM_019822405.2;refGene_gn=WASF3;refGene_tx=XM_019822445.2;refGene_gn=WASF3;refGene_tx=XM_023250152.1;refGene_gn=WASF3;refGene_tx=XM_023250184.1;refGene_gn=WASF3;refGene_tx=XM_023250199.1
A1 6466708 6466826
refGene_gn=WASF3;refGene_tx=XM_006927176.4;refGene_gn=WASF3;refGene_tx=XM_019822210.2;refGene_gn=WASF3;refGene_tx=XM_019822250.2;refGene_gn=WASF3;refGene_tx=XM_019822289.2;refGene_gn=WASF3;refGene_tx=XM_019822322.2;refGene_gn=WASF3;refGene_tx=XM_019822353.2;refGene_gn=WASF3;refGene_tx=XM_019822405.2;refGene_gn=WASF3;refGene_tx=XM_019822445.2;refGene_gn=WASF3;refGene_tx=XM_023250152.1;refGene_gn=WASF3;refGene_tx=XM_023250184.1;refGene_gn=WASF3;refGene_tx=XM_023250199.1
A1 6471785 6471961
refGene_gn=WASF3;refGene_tx=XM_006927176.4;refGene_gn=WASF3;refGene_tx=XM_019822210.2;refGene_gn=WASF3;refGene_tx=XM_019822250.2;refGene_gn=WASF3;refGene_tx=XM_019822289.2;refGene_gn=WASF3;refGene_tx=XM_019822322.2;refGene_gn=WASF3;refGene_tx=XM_019822353.2;refGene_gn=WASF3;refGene_tx=XM_019822405.2;refGene_gn=WASF3;refGene_tx=XM_019822445.2;refGene_gn=WASF3;refGene_tx=XM_023250152.1;refGene_gn=WASF3;refGene_tx=XM_023250184.1
A1 6475617 6475784 refGene_gn=WASF3;refGene_tx=XM_023250199.1
A1 6476668 6476935

refGene_gn=WASF3;refGene_tx=XM_006927176.4;refGene_gn=WASF3;refGene_tx=XM_019822210.2;refGene_gn=WASF3;refGene_tx=XM_019822250.2;refGene_gn=WASF3;refGene_tx=XM_019822289.2;refGene_gn=WASF3;refGene_tx=XM_019822322.2;refGene_gn=WASF3;refGene_tx=XM_019822353.2;refGene_gn=WASF3;refGene_tx=XM_019822405.2;refGene_gn=WASF3;refGene_tx=XM_019822445.2;refGene_gn=WASF3;refGene_tx=XM_023250152.1;refGene_gn=WASF3;refGene_tx=XM_023250184.1;refGene_gn=WASF3;refGene_tx=XM_023250199.1
A1 6478302 6478670
refGene_gn=WASF3;refGene_tx=XM_006927176.4;refGene_gn=WASF3;refGene_tx=XM_019822210.2;refGene_gn=WASF3;refGene_tx=XM_019822250.2;refGene_gn=WASF3;refGene_tx=XM_019822289.2;refGene_gn=WASF3;refGene_tx=XM_019822322.2;refGene_gn=WASF3;refGene_tx=XM_019822353.2;refGene_gn=WASF3;refGene_tx=XM_019822405.2;refGene_gn=WASF3;refGene_tx=XM_019822445.2;refGene_gn=WASF3;refGene_tx=XM_023250152.1;refGene_gn=WASF3;refGene_tx=XM_023250184.1;refGene_gn=WASF3;refGene_tx=XM_023250199.1
A1 6480968 6481126
refGene_gn=WASF3;refGene_tx=XM_006927176.4;refGene_gn=WASF3;refGene_tx=XM_019822210.2;refGene_gn=WASF3;refGene_tx=XM_019822250.2;refGene_gn=WASF3;refGene_tx=XM_019822289.2;refGene_gn=WASF3;refGene_tx=XM_019822322.2;refGene_gn=WASF3;refGene_tx=XM_019822353.2;refGene_gn=WASF3;refGene_tx=XM_019822405.2;refGene_gn=WASF3;refGene_tx=XM_019822445.2;refGene_gn=WASF3;refGene_tx=XM_023250152.1;refGene_gn=WASF3;refGene_tx=XM_023250184.1;refGene_gn=WASF3;refGene_tx=XM_023250199.1
A1 6540918 6541923 refGene_gn=GPR12;refGene_tx=XM_023250214.1
A1 6812399 6812501
refGene_gn=USP12;refGene_tx=XM_023250218.1;refGene_gn=USP12;refGene_tx=XM_023250221.1;refGene_gn=USP12;refGene_tx=XM_023250224.1;refGene_gn=USP12;refGene_tx=XM_023250229.1
A1 6813762 6813841
refGene_gn=USP12;refGene_tx=XM_023250218.1;refGene_gn=USP12;refGene_tx=XM_023250221.1;refGene_gn=USP12;refGene_tx=XM_023250224.1;refGene_gn=USP12;refGene_tx=XM_023250229.1
A1 6816330 6816528
refGene_gn=USP12;refGene_tx=XM_023250218.1;refGene_gn=USP12;refGene_tx=XM_023250221.1;refGene_gn=USP12;refGene_tx=XM_023250224.1;refGene_gn=USP12;refGene_tx=XM_023250229.1
A1 6829355 6829439
refGene_gn=USP12;refGene_tx=XM_023250218.1;refGene_gn=USP12;refGene_tx=XM_023250221.1;refGene_gn=USP12;refGene_tx=XM_023250224.1;refGene_gn=USP12;refGene_tx=XM_023250229.1
A1 6829552 6829629
refGene_gn=USP12;refGene_tx=XM_023250218.1;refGene_gn=USP12;refGene_tx=XM_023250221.1;refGene_gn=USP12;refGene_tx=XM_023250224.1;refGene_gn=USP12;refGene_tx=XM_023250229.1
A1 6835119 6835349
refGene_gn=USP12;refGene_tx=XM_023250218.1;refGene_gn=USP12;refGene_tx=XM_023250221.1;refGene_gn=USP12;refGene_tx=XM_023250224.1;refGene_gn=USP12;refGene_tx=XM_023250229.1
A1 6844322 6844536 refGene_gn=USP12;refGene_tx=XM_023250229.1
A1 6858531 6858612 refGene_gn=USP12;refGene_tx=XM_023250229.1

```
A1	6915023	6915071	refGene_gn=USP12;refGene_tx=XM_023250229.1
A1	6921986	6922643	
refGene_gn=LOC111559476;refGene_tx=XM_023250247.1
A1	6923419	6923854	
refGene_gn=LOC111559476;refGene_tx=XM_023250247.1
A1	6965762	6965829	refGene_gn=RPL21;refGene_tx=XM_003980267.5
A1	6966099	6966161	refGene_gn=RPL21;refGene_tx=XM_003980267.5
A1	6966737	6966850	refGene_gn=RPL21;refGene_tx=XM_003980267.5
A1	6967602	6967753	refGene_gn=RPL21;refGene_tx=XM_003980267.5
A1	6967937	6968027	refGene_gn=RPL21;refGene_tx=XM_003980267.5
A1	6976907	6977031	
refGene_gn=RASL11A;refGene_tx=XM_003980301.4;refGene_gn=RASL11A;refGen
e_tx=XM_023250253.1
A1	6977460	6977517	refGene_gn=RASL11A;refGene_tx=XM_003980301.4
A1	6977629	6977709	
refGene_gn=RASL11A;refGene_tx=XM_003980301.4;refGene_gn=RASL11A;refGen
e_tx=XM_023250253.1
A1	6978650	6979118	
refGene_gn=RASL11A;refGene_tx=XM_003980301.4;refGene_gn=RASL11A;refGen
e_tx=XM_023250253.1
A1	7100209	7100608	refGene_gn=GTF3A;refGene_tx=XM_011282105.2
A1	7102822	7102923	refGene_gn=GTF3A;refGene_tx=XM_011282105.2
A1	7106997	7107094	refGene_gn=GTF3A;refGene_tx=XM_011282105.2
A1	7107483	7107572	refGene_gn=GTF3A;refGene_tx=XM_011282105.2
A1	7109027	7109101	refGene_gn=GTF3A;refGene_tx=XM_011282105.2
A1	7110315	7110396	refGene_gn=GTF3A;refGene_tx=XM_011282105.2
A1	7110915	7111145	refGene_gn=GTF3A;refGene_tx=XM_011282105.2
A1	7112223	7112283	refGene_gn=GTF3A;refGene_tx=XM_011282105.2
A1	7112514	7112679	refGene_gn=GTF3A;refGene_tx=XM_011282105.2
A1	7112761	7112980	
refGene_gn=MTIF3;refGene_tx=XM_003980268.4;refGene_gn=MTIF3;refGene_tx
=XM_006927181.4;refGene_gn=MTIF3;refGene_tx=XM_006927182.4;refGene_gn=
MTIF3;refGene_tx=XM_006927183.4;refGene_gn=MTIF3;refGene_tx=XM_0232502
94.1
A1	7113924	7114079	
refGene_gn=MTIF3;refGene_tx=XM_003980268.4;refGene_gn=MTIF3;refGene_tx
=XM_006927181.4;refGene_gn=MTIF3;refGene_tx=XM_006927182.4;refGene_gn=
MTIF3;refGene_tx=XM_006927183.4
A1	7116929	7117390	
refGene_gn=MTIF3;refGene_tx=XM_003980268.4;refGene_gn=MTIF3;refGene_tx
=XM_006927181.4;refGene_gn=MTIF3;refGene_tx=XM_006927182.4;refGene_gn=
MTIF3;refGene_tx=XM_006927183.4;refGene_gn=MTIF3;refGene_tx=XM_0232502
94.1
A1	7118803	7118832	refGene_gn=MTIF3;refGene_tx=XM_023250294.1
A1	7120340	7120402	
refGene_gn=MTIF3;refGene_tx=XM_006927181.4;refGene_gn=MTIF3;refGene_tx
=XM_006927182.4
A1	7209379	7209515	
refGene_gn=LNX2;refGene_tx=XM_003980269.5;refGene_gn=LNX2;refGene_tx=X
M_023250304.1;refGene_gn=LNX2;refGene_tx=XM_023250306.1;refGene_gn=LNX
```

2;refGene_tx=XM_023250309.1
A1 7211358 7211517
refGene_gn=LNX2;refGene_tx=XM_003980269.5;refGene_gn=LNX2;refGene_tx=XM_023250304.1;refGene_gn=LNX2;refGene_tx=XM_023250306.1;refGene_gn=LNX2;refGene_tx=XM_023250309.1
A1 7214983 7215215
refGene_gn=LNX2;refGene_tx=XM_003980269.5;refGene_gn=LNX2;refGene_tx=XM_023250304.1;refGene_gn=LNX2;refGene_tx=XM_023250306.1;refGene_gn=LNX2;refGene_tx=XM_023250309.1
A1 7218249 7218427
refGene_gn=LNX2;refGene_tx=XM_003980269.5;refGene_gn=LNX2;refGene_tx=XM_023250304.1;refGene_gn=LNX2;refGene_tx=XM_023250306.1;refGene_gn=LNX2;refGene_tx=XM_023250309.1
A1 7221768 7221909
refGene_gn=LNX2;refGene_tx=XM_003980269.5;refGene_gn=LNX2;refGene_tx=XM_023250304.1;refGene_gn=LNX2;refGene_tx=XM_023250306.1
A1 7225660 7226029
refGene_gn=LNX2;refGene_tx=XM_003980269.5;refGene_gn=LNX2;refGene_tx=XM_023250304.1;refGene_gn=LNX2;refGene_tx=XM_023250306.1;refGene_gn=LNX2;refGene_tx=XM_023250309.1
A1 7231711 7231911
refGene_gn=LNX2;refGene_tx=XM_003980269.5;refGene_gn=LNX2;refGene_tx=XM_023250304.1;refGene_gn=LNX2;refGene_tx=XM_023250306.1;refGene_gn=LNX2;refGene_tx=XM_023250309.1
A1 7233101 7233349
refGene_gn=LNX2;refGene_tx=XM_003980269.5;refGene_gn=LNX2;refGene_tx=XM_023250304.1;refGene_gn=LNX2;refGene_tx=XM_023250306.1;refGene_gn=LNX2;refGene_tx=XM_023250309.1
A1 7243721 7244128
refGene_gn=LNX2;refGene_tx=XM_003980269.5;refGene_gn=LNX2;refGene_tx=XM_023250304.1;refGene_gn=LNX2;refGene_tx=XM_023250306.1;refGene_gn=LNX2;refGene_tx=XM_023250309.1
A1 7288973 7288999
refGene_gn=LOC111559488;refGene_tx=XM_023250315.1;refGene_gn=POLR1D;refGene_tx=XM_011281469.3
A1 7289961 7290337 refGene_gn=POLR1D;refGene_tx=XM_011281469.3
A1 7317069 7317144
refGene_gn=LOC111559488;refGene_tx=XM_023250315.1
A1 7336183 7336478
refGene_gn=LOC111559488;refGene_tx=XM_023250315.1
A1 7445462 7445877 refGene_gn=GSX1;refGene_tx=XM_011282206.3
A1 7446355 7446735 refGene_gn=GSX1;refGene_tx=XM_011282206.3
A1 7559519 7559922 refGene_gn=PDX1;refGene_tx=XM_011281516.2
A1 7563411 7563857 refGene_gn=PDX1;refGene_tx=XM_011281516.2
A1 7597974 7598229
refGene_gn=CDX2;refGene_tx=XM_023250575.1;refGene_gn=CDX2;refGene_tx=XM_023250591.1
A1 7599300 7599446
refGene_gn=CDX2;refGene_tx=XM_023250575.1;refGene_gn=CDX2;refGene_tx=XM_023250591.1

A1 7602940 7603481
refGene_gn=CDX2;refGene_tx=XM_023250575.1;refGene_gn=CDX2;refGene_tx=XM_023250591.1
A1 7612041 7612391 refGene_gn=URAD;refGene_tx=XM_023250321.1
A1 7619743 7619918 refGene_gn=URAD;refGene_tx=XM_023250321.1
A1 7630742 7630865 refGene_gn=FLT3;refGene_tx=XM_023250335.1
A1 7638929 7639035 refGene_gn=FLT3;refGene_tx=XM_023250335.1
A1 7639744 7639844 refGene_gn=FLT3;refGene_tx=XM_023250335.1
A1 7640626 7640738 refGene_gn=FLT3;refGene_tx=XM_023250335.1
A1 7643132 7643255 refGene_gn=FLT3;refGene_tx=XM_023250335.1
A1 7646820 7646948 refGene_gn=FLT3;refGene_tx=XM_023250335.1
A1 7647842 7647925 refGene_gn=FLT3;refGene_tx=XM_023250335.1
A1 7650020 7650174 refGene_gn=FLT3;refGene_tx=XM_023250335.1
A1 7651440 7651551 refGene_gn=FLT3;refGene_tx=XM_023250335.1
A1 7654581 7654686 refGene_gn=FLT3;refGene_tx=XM_023250335.1
A1 7654774 7654907 refGene_gn=FLT3;refGene_tx=XM_023250335.1
A1 7654994 7655101 refGene_gn=FLT3;refGene_tx=XM_023250335.1
A1 7656636 7656815 refGene_gn=FLT3;refGene_tx=XM_023250335.1
A1 7657087 7657199 refGene_gn=FLT3;refGene_tx=XM_023250335.1
A1 7658182 7658286 refGene_gn=FLT3;refGene_tx=XM_023250335.1
A1 7667400 7667569 refGene_gn=FLT3;refGene_tx=XM_023250335.1
A1 7669063 7669217 refGene_gn=FLT3;refGene_tx=XM_023250335.1
A1 7669314 7669454 refGene_gn=FLT3;refGene_tx=XM_023250335.1
A1 7669770 7669898 refGene_gn=FLT3;refGene_tx=XM_023250335.1
A1 7671519 7671649 refGene_gn=FLT3;refGene_tx=XM_023250335.1
A1 7675578 7675694 refGene_gn=FLT3;refGene_tx=XM_023250335.1
A1 7678442 7678645 refGene_gn=FLT3;refGene_tx=XM_023250335.1
A1 7685784 7685906 refGene_gn=FLT3;refGene_tx=XM_023250335.1
A1 7706716 7707519
refGene_gn=FLT3;refGene_tx=XM_023250335.1;refGene_gn=LOC109496442;refGene_tx=XM_023250336.1
A1 7754843 7755279
refGene_gn=PAN3;refGene_tx=XM_023250322.1;refGene_gn=PAN3;refGene_tx=XM_023250324.1;refGene_gn=PAN3;refGene_tx=XM_023250326.1;refGene_gn=PAN3;refGene_tx=XM_023250332.1;refGene_gn=PAN3;refGene_tx=XM_023250333.1
A1 7773726 7773848
refGene_gn=PAN3;refGene_tx=XM_023250322.1;refGene_gn=PAN3;refGene_tx=XM_023250324.1;refGene_gn=PAN3;refGene_tx=XM_023250326.1;refGene_gn=PAN3;refGene_tx=XM_023250332.1;refGene_gn=PAN3;refGene_tx=XM_023250333.1
A1 7774599 7774666
refGene_gn=PAN3;refGene_tx=XM_023250322.1;refGene_gn=PAN3;refGene_tx=XM_023250324.1;refGene_gn=PAN3;refGene_tx=XM_023250326.1;refGene_gn=PAN3;refGene_tx=XM_023250332.1;refGene_gn=PAN3;refGene_tx=XM_023250333.1
A1 7776523 7776594
refGene_gn=PAN3;refGene_tx=XM_023250322.1;refGene_gn=PAN3;refGene_tx=XM_023250324.1;refGene_gn=PAN3;refGene_tx=XM_023250326.1;refGene_gn=PAN3;refGene_tx=XM_023250332.1;refGene_gn=PAN3;refGene_tx=XM_023250333.1
A1 7792066 7792228
refGene_gn=PAN3;refGene_tx=XM_023250322.1;refGene_gn=PAN3;refGene_tx=XM_023250324.1;refGene_gn=PAN3;refGene_tx=XM_023250326.1

A1 7813464 7813612
refGene_gn=PAN3;refGene_tx=XM_023250322.1;refGene_gn=PAN3;refGene_tx=XM_023250324.1;refGene_gn=PAN3;refGene_tx=XM_023250326.1;refGene_gn=PAN3;refGene_tx=XM_023250332.1;refGene_gn=PAN3;refGene_tx=XM_023250333.1
A1 7849895 7850143
refGene_gn=PAN3;refGene_tx=XM_023250322.1;refGene_gn=PAN3;refGene_tx=XM_023250324.1;refGene_gn=PAN3;refGene_tx=XM_023250326.1;refGene_gn=PAN3;refGene_tx=XM_023250332.1;refGene_gn=PAN3;refGene_tx=XM_023250333.1
A1 7851276 7851333
refGene_gn=PAN3;refGene_tx=XM_023250322.1;refGene_gn=PAN3;refGene_tx=XM_023250324.1;refGene_gn=PAN3;refGene_tx=XM_023250332.1
A1 7852575 7852680
refGene_gn=PAN3;refGene_tx=XM_023250322.1;refGene_gn=PAN3;refGene_tx=XM_023250324.1;refGene_gn=PAN3;refGene_tx=XM_023250326.1;refGene_gn=PAN3;refGene_tx=XM_023250332.1;refGene_gn=PAN3;refGene_tx=XM_023250333.1
A1 7853764 7853822
refGene_gn=PAN3;refGene_tx=XM_023250322.1;refGene_gn=PAN3;refGene_tx=XM_023250324.1;refGene_gn=PAN3;refGene_tx=XM_023250326.1;refGene_gn=PAN3;refGene_tx=XM_023250332.1;refGene_gn=PAN3;refGene_tx=XM_023250333.1
A1 7861016 7861178
refGene_gn=PAN3;refGene_tx=XM_023250322.1;refGene_gn=PAN3;refGene_tx=XM_023250324.1;refGene_gn=PAN3;refGene_tx=XM_023250326.1;refGene_gn=PAN3;refGene_tx=XM_023250332.1;refGene_gn=PAN3;refGene_tx=XM_023250333.1
A1 7861411 7861528
refGene_gn=PAN3;refGene_tx=XM_023250322.1;refGene_gn=PAN3;refGene_tx=XM_023250324.1;refGene_gn=PAN3;refGene_tx=XM_023250326.1;refGene_gn=PAN3;refGene_tx=XM_023250332.1;refGene_gn=PAN3;refGene_tx=XM_023250333.1
A1 7861618 7861720
refGene_gn=PAN3;refGene_tx=XM_023250322.1;refGene_gn=PAN3;refGene_tx=XM_023250324.1;refGene_gn=PAN3;refGene_tx=XM_023250326.1;refGene_gn=PAN3;refGene_tx=XM_023250332.1;refGene_gn=PAN3;refGene_tx=XM_023250333.1
A1 7865763 7865929
refGene_gn=PAN3;refGene_tx=XM_023250322.1;refGene_gn=PAN3;refGene_tx=XM_023250324.1;refGene_gn=PAN3;refGene_tx=XM_023250326.1;refGene_gn=PAN3;refGene_tx=XM_023250332.1;refGene_gn=PAN3;refGene_tx=XM_023250333.1
A1 7867072 7867163
refGene_gn=PAN3;refGene_tx=XM_023250322.1;refGene_gn=PAN3;refGene_tx=XM_023250324.1;refGene_gn=PAN3;refGene_tx=XM_023250326.1;refGene_gn=PAN3;refGene_tx=XM_023250332.1;refGene_gn=PAN3;refGene_tx=XM_023250333.1
A1 7871637 7871777
refGene_gn=PAN3;refGene_tx=XM_023250322.1;refGene_gn=PAN3;refGene_tx=XM_023250324.1;refGene_gn=PAN3;refGene_tx=XM_023250326.1;refGene_gn=PAN3;refGene_tx=XM_023250332.1;refGene_gn=PAN3;refGene_tx=XM_023250333.1
A1 7874508 7874638
refGene_gn=PAN3;refGene_tx=XM_023250322.1;refGene_gn=PAN3;refGene_tx=XM_023250324.1;refGene_gn=PAN3;refGene_tx=XM_023250326.1;refGene_gn=PAN3;refGene_tx=XM_023250332.1;refGene_gn=PAN3;refGene_tx=XM_023250333.1
A1 7875069 7875134
refGene_gn=PAN3;refGene_tx=XM_023250322.1;refGene_gn=PAN3;refGene_tx=XM_023250324.1;refGene_gn=PAN3;refGene_tx=XM_023250326.1;refGene_gn=PAN

3;refGene_tx=XM_023250332.1;refGene_gn=PAN3;refGene_tx=XM_023250333.1
A1 7882213 7882352
refGene_gn=PAN3;refGene_tx=XM_023250322.1;refGene_gn=PAN3;refGene_tx=XM_023250324.1;refGene_gn=PAN3;refGene_tx=XM_023250326.1;refGene_gn=PAN3;refGene_tx=XM_023250332.1;refGene_gn=PAN3;refGene_tx=XM_023250333.1
A1 7884811 7884952
refGene_gn=PAN3;refGene_tx=XM_023250322.1;refGene_gn=PAN3;refGene_tx=XM_023250324.1;refGene_gn=PAN3;refGene_tx=XM_023250326.1;refGene_gn=PAN3;refGene_tx=XM_023250332.1;refGene_gn=PAN3;refGene_tx=XM_023250333.1
A1 7896583 7896785 refGene_gn=FLT1;refGene_tx=XM_023250349.1
A1 7900362 7900457 refGene_gn=FLT1;refGene_tx=XM_023250349.1
A1 7902634 7902719 refGene_gn=FLT1;refGene_tx=XM_023250349.1
A1 7905185 7905328 refGene_gn=FLT1;refGene_tx=XM_023250349.1
A1 7905592 7905698 refGene_gn=FLT1;refGene_tx=XM_023250349.1
A1 7911043 7911143 refGene_gn=FLT1;refGene_tx=XM_023250349.1
A1 7913000 7913112 refGene_gn=FLT1;refGene_tx=XM_023250349.1
A1 7914881 7915004 refGene_gn=FLT1;refGene_tx=XM_023250349.1
A1 7915695 7915793 refGene_gn=FLT1;refGene_tx=XM_023250349.1
A1 7916222 7916379 refGene_gn=FLT1;refGene_tx=XM_023250349.1
A1 7920041 7920130 refGene_gn=FLT1;refGene_tx=XM_023250349.1
A1 7920894 7921008 refGene_gn=FLT1;refGene_tx=XM_023250349.1
A1 7924435 7924540 refGene_gn=FLT1;refGene_tx=XM_023250349.1
A1 7929185 7929318 refGene_gn=FLT1;refGene_tx=XM_023250349.1
A1 7933655 7933762 refGene_gn=FLT1;refGene_tx=XM_023250349.1
A1 7946646 7946778 refGene_gn=FLT1;refGene_tx=XM_023250349.1
A1 7961234 7961254 refGene_gn=FLT1;refGene_tx=XM_023250353.1
A1 7965069 7965077 refGene_gn=FLT1;refGene_tx=XM_023250359.1
A1 7967077 7967224
refGene_gn=FLT1;refGene_tx=XM_023250349.1;refGene_gn=FLT1;refGene_tx=XM_023250353.1;refGene_gn=FLT1;refGene_tx=XM_023250359.1
A1 7971901 7972210
refGene_gn=FLT1;refGene_tx=XM_023250349.1;refGene_gn=FLT1;refGene_tx=XM_023250353.1;refGene_gn=FLT1;refGene_tx=XM_023250359.1
A1 7978489 7978598
refGene_gn=FLT1;refGene_tx=XM_023250349.1;refGene_gn=FLT1;refGene_tx=XM_023250353.1;refGene_gn=FLT1;refGene_tx=XM_023250359.1
A1 7987189 7987304
refGene_gn=FLT1;refGene_tx=XM_023250349.1;refGene_gn=FLT1;refGene_tx=XM_023250353.1;refGene_gn=FLT1;refGene_tx=XM_023250359.1
A1 8008384 8008544
refGene_gn=FLT1;refGene_tx=XM_023250349.1;refGene_gn=FLT1;refGene_tx=XM_023250353.1;refGene_gn=FLT1;refGene_tx=XM_023250359.1
A1 8009184 8009354
refGene_gn=FLT1;refGene_tx=XM_023250349.1;refGene_gn=FLT1;refGene_tx=XM_023250353.1;refGene_gn=FLT1;refGene_tx=XM_023250359.1
A1 8011654 8011772
refGene_gn=FLT1;refGene_tx=XM_023250349.1;refGene_gn=FLT1;refGene_tx=XM_023250353.1;refGene_gn=FLT1;refGene_tx=XM_023250359.1
A1 8013210 8013385
refGene_gn=FLT1;refGene_tx=XM_023250349.1;refGene_gn=FLT1;refGene_tx=X

M_023250353.1;refGene_gn=FLT1;refGene_tx=XM_023250359.1
A1 8015845 8015982
refGene_gn=FLT1;refGene_tx=XM_023250349.1;refGene_gn=FLT1;refGene_tx=X
M_023250353.1;refGene_gn=FLT1;refGene_tx=XM_023250359.1
A1 8016319 8016482
refGene_gn=FLT1;refGene_tx=XM_023250349.1;refGene_gn=FLT1;refGene_tx=X
M_023250353.1;refGene_gn=FLT1;refGene_tx=XM_023250359.1
A1 8020222 8020347
refGene_gn=FLT1;refGene_tx=XM_023250349.1;refGene_gn=FLT1;refGene_tx=X
M_023250353.1;refGene_gn=FLT1;refGene_tx=XM_023250359.1
A1 8043846 8044073
refGene_gn=FLT1;refGene_tx=XM_023250349.1;refGene_gn=FLT1;refGene_tx=X
M_023250353.1;refGene_gn=FLT1;refGene_tx=XM_023250359.1
A1 8044465 8044562
refGene_gn=FLT1;refGene_tx=XM_023250349.1;refGene_gn=FLT1;refGene_tx=X
M_023250353.1;refGene_gn=FLT1;refGene_tx=XM_023250359.1
A1 8071669 8071733
refGene_gn=FLT1;refGene_tx=XM_023250349.1;refGene_gn=FLT1;refGene_tx=X
M_023250353.1;refGene_gn=FLT1;refGene_tx=XM_023250359.1
A1 8216468 8216471 refGene_gn=POMP;refGene_tx=XM_023250367.1
A1 8219060 8219158 refGene_gn=POMP;refGene_tx=XM_023250367.1
A1 8221549 8221610 refGene_gn=POMP;refGene_tx=XM_023250367.1
A1 8224441 8224543 refGene_gn=POMP;refGene_tx=XM_023250367.1
A1 8228769 8228863 refGene_gn=POMP;refGene_tx=XM_023250367.1
A1 8232233 8232301 refGene_gn=POMP;refGene_tx=XM_023250367.1
A1 8242507 8242592
refGene_gn=SLC46A3;refGene_tx=XM_003980273.5;refGene_gn=SLC46A3;refGen
e_tx=XM_006927186.4
A1 8246129 8246286
refGene_gn=SLC46A3;refGene_tx=XM_003980273.5;refGene_gn=SLC46A3;refGen
e_tx=XM_006927186.4
A1 8251426 8251510
refGene_gn=SLC46A3;refGene_tx=XM_003980273.5;refGene_gn=SLC46A3;refGen
e_tx=XM_006927186.4
A1 8253093 8253964
refGene_gn=SLC46A3;refGene_tx=XM_003980273.5;refGene_gn=SLC46A3;refGen
e_tx=XM_006927186.4
A1 8260582 8260771
refGene_gn=SLC46A3;refGene_tx=XM_003980273.5;refGene_gn=SLC46A3;refGen
e_tx=XM_006927186.4
A1 8542718 8544926
refGene_gn=MTUS2;refGene_tx=XM_023250373.1;refGene_gn=MTUS2;refGene_tx
=XM_023250374.1
A1 8553603 8553844
refGene_gn=MTUS2;refGene_tx=XM_023250373.1;refGene_gn=MTUS2;refGene_tx
=XM_023250374.1
A1 8597975 8598173
refGene_gn=MTUS2;refGene_tx=XM_023250373.1;refGene_gn=MTUS2;refGene_tx
=XM_023250374.1
A1 8742166 8742328

refGene_gn=MTUS2;refGene_tx=XM_023250373.1;refGene_gn=MTUS2;refGene_tx=XM_023250374.1
A1	8771304	8771403	refGene_gn=MTUS2;refGene_tx=XM_023250373.1;refGene_gn=MTUS2;refGene_tx=XM_023250374.1
A1	8782532	8782744	refGene_gn=MTUS2;refGene_tx=XM_023250373.1;refGene_gn=MTUS2;refGene_tx=XM_023250374.1
A1	8847871	8847925	refGene_gn=MTUS2;refGene_tx=XM_003980274.5
A1	8858819	8858886	refGene_gn=MTUS2;refGene_tx=XM_003980274.5;refGene_gn=MTUS2;refGene_tx=XM_023250373.1;refGene_gn=MTUS2;refGene_tx=XM_023250374.1
A1	8893574	8893789	refGene_gn=MTUS2;refGene_tx=XM_003980274.5;refGene_gn=MTUS2;refGene_tx=XM_023250373.1;refGene_gn=MTUS2;refGene_tx=XM_023250374.1
A1	8900820	8900926	refGene_gn=MTUS2;refGene_tx=XM_003980274.5;refGene_gn=MTUS2;refGene_tx=XM_023250373.1;refGene_gn=MTUS2;refGene_tx=XM_023250374.1
A1	8903264	8903338	refGene_gn=MTUS2;refGene_tx=XM_003980274.5;refGene_gn=MTUS2;refGene_tx=XM_023250373.1;refGene_gn=MTUS2;refGene_tx=XM_023250374.1
A1	8907344	8907443	refGene_gn=MTUS2;refGene_tx=XM_003980274.5;refGene_gn=MTUS2;refGene_tx=XM_023250373.1;refGene_gn=MTUS2;refGene_tx=XM_023250374.1
A1	8908541	8908661	refGene_gn=MTUS2;refGene_tx=XM_003980274.5;refGene_gn=MTUS2;refGene_tx=XM_023250373.1;refGene_gn=MTUS2;refGene_tx=XM_023250374.1
A1	8911179	8911277	refGene_gn=MTUS2;refGene_tx=XM_003980274.5;refGene_gn=MTUS2;refGene_tx=XM_023250373.1;refGene_gn=MTUS2;refGene_tx=XM_023250374.1
A1	8915156	8915370	refGene_gn=MTUS2;refGene_tx=XM_003980274.5;refGene_gn=MTUS2;refGene_tx=XM_023250373.1;refGene_gn=MTUS2;refGene_tx=XM_023250374.1
A1	8926359	8926463	refGene_gn=SLC7A1;refGene_tx=XM_003980275.5;refGene_gn=SLC7A1;refGene_tx=XM_023250398.1
A1	8927973	8928082	refGene_gn=SLC7A1;refGene_tx=XM_003980275.5;refGene_gn=SLC7A1;refGene_tx=XM_023250398.1
A1	8928709	8928876	refGene_gn=SLC7A1;refGene_tx=XM_003980275.5;refGene_gn=SLC7A1;refGene_tx=XM_023250398.1
A1	8929158	8929376	refGene_gn=SLC7A1;refGene_tx=XM_003980275.5;refGene_gn=SLC7A1;refGene_tx=XM_023250398.1
A1	8931098	8931201	refGene_gn=SLC7A1;refGene_tx=XM_003980275.5;refGene_gn=SLC7A1;refGene_tx=XM_023250398.1
A1	8933750	8933890	refGene_gn=SLC7A1;refGene_tx=XM_003980275.5;refGene_gn=SLC7A1;refGene_

tx=XM_023250398.1
A1 8934661 8934884
refGene_gn=SLC7A1;refGene_tx=XM_003980275.5;refGene_gn=SLC7A1;refGene_tx=XM_023250398.1
A1 8935564 8935686
refGene_gn=SLC7A1;refGene_tx=XM_003980275.5;refGene_gn=SLC7A1;refGene_tx=XM_023250398.1
A1 8941465 8941640
refGene_gn=SLC7A1;refGene_tx=XM_003980275.5;refGene_gn=SLC7A1;refGene_tx=XM_023250398.1
A1 8943699 8943858
refGene_gn=SLC7A1;refGene_tx=XM_003980275.5;refGene_gn=SLC7A1;refGene_tx=XM_023250398.1
A1 8947365 8947735
refGene_gn=SLC7A1;refGene_tx=XM_003980275.5;refGene_gn=SLC7A1;refGene_tx=XM_023250398.1
A1 9121803 9121856 refGene_gn=UBL3;refGene_tx=XM_003980276.5
A1 9122173 9122251 refGene_gn=UBL3;refGene_tx=XM_003980276.5
A1 9127734 9127821 refGene_gn=UBL3;refGene_tx=XM_003980276.5
A1 9132038 9132147 refGene_gn=UBL3;refGene_tx=XM_003980276.5
A1 9186182 9186209 refGene_gn=UBL3;refGene_tx=XM_003980276.5
A1 9503520 9503719
refGene_gn=KATNAL1;refGene_tx=XM_006927188.4;refGene_gn=KATNAL1;refGene_tx=XM_023250424.1;refGene_gn=KATNAL1;refGene_tx=XM_023250425.1;refGene_gn=KATNAL1;refGene_tx=XM_023250430.1;refGene_gn=KATNAL1;refGene_tx=XM_023250438.1;refGene_gn=KATNAL1;refGene_tx=XM_023250439.1
A1 9505326 9505453
refGene_gn=KATNAL1;refGene_tx=XM_006927188.4;refGene_gn=KATNAL1;refGene_tx=XM_023250424.1;refGene_gn=KATNAL1;refGene_tx=XM_023250425.1;refGene_gn=KATNAL1;refGene_tx=XM_023250430.1;refGene_gn=KATNAL1;refGene_tx=XM_023250438.1;refGene_gn=KATNAL1;refGene_tx=XM_023250439.1
A1 9520585 9520720
refGene_gn=KATNAL1;refGene_tx=XM_006927188.4;refGene_gn=KATNAL1;refGene_tx=XM_023250424.1;refGene_gn=KATNAL1;refGene_tx=XM_023250425.1;refGene_gn=KATNAL1;refGene_tx=XM_023250430.1;refGene_gn=KATNAL1;refGene_tx=XM_023250438.1;refGene_gn=KATNAL1;refGene_tx=XM_023250439.1
A1 9523665 9523792
refGene_gn=KATNAL1;refGene_tx=XM_006927188.4;refGene_gn=KATNAL1;refGene_tx=XM_023250424.1;refGene_gn=KATNAL1;refGene_tx=XM_023250425.1;refGene_gn=KATNAL1;refGene_tx=XM_023250430.1;refGene_gn=KATNAL1;refGene_tx=XM_023250438.1;refGene_gn=KATNAL1;refGene_tx=XM_023250439.1
A1 9524517 9524676
refGene_gn=KATNAL1;refGene_tx=XM_006927188.4;refGene_gn=KATNAL1;refGene_tx=XM_023250424.1;refGene_gn=KATNAL1;refGene_tx=XM_023250425.1;refGene_gn=KATNAL1;refGene_tx=XM_023250430.1;refGene_gn=KATNAL1;refGene_tx=XM_023250438.1;refGene_gn=KATNAL1;refGene_tx=XM_023250439.1
A1 9530323 9530429
refGene_gn=KATNAL1;refGene_tx=XM_006927188.4;refGene_gn=KATNAL1;refGene_tx=XM_023250424.1;refGene_gn=KATNAL1;refGene_tx=XM_023250425.1;refGene_gn=KATNAL1;refGene_tx=XM_023250430.1;refGene_gn=KATNAL1;refGene_tx=

XM_023250438.1;refGene_gn=KATNAL1;refGene_tx=XM_023250439.1
A1 9530823 9530951
refGene_gn=KATNAL1;refGene_tx=XM_006927188.4;refGene_gn=KATNAL1;refGene_tx=XM_023250424.1;refGene_gn=KATNAL1;refGene_tx=XM_023250425.1;refGene_gn=KATNAL1;refGene_tx=XM_023250430.1;refGene_gn=KATNAL1;refGene_tx=XM_023250438.1;refGene_gn=KATNAL1;refGene_tx=XM_023250439.1
A1 9539575 9539744
refGene_gn=KATNAL1;refGene_tx=XM_006927188.4;refGene_gn=KATNAL1;refGene_tx=XM_023250424.1;refGene_gn=KATNAL1;refGene_tx=XM_023250425.1;refGene_gn=KATNAL1;refGene_tx=XM_023250430.1;refGene_gn=KATNAL1;refGene_tx=XM_023250438.1;refGene_gn=KATNAL1;refGene_tx=XM_023250439.1
A1 9562350 9562511
refGene_gn=KATNAL1;refGene_tx=XM_006927188.4;refGene_gn=KATNAL1;refGene_tx=XM_023250424.1;refGene_gn=KATNAL1;refGene_tx=XM_023250425.1;refGene_gn=KATNAL1;refGene_tx=XM_023250430.1;refGene_gn=KATNAL1;refGene_tx=XM_023250438.1;refGene_gn=KATNAL1;refGene_tx=XM_023250439.1
A1 9564503 9564665
refGene_gn=KATNAL1;refGene_tx=XM_006927188.4;refGene_gn=KATNAL1;refGene_tx=XM_023250424.1;refGene_gn=KATNAL1;refGene_tx=XM_023250425.1;refGene_gn=KATNAL1;refGene_tx=XM_023250430.1;refGene_gn=KATNAL1;refGene_tx=XM_023250438.1;refGene_gn=KATNAL1;refGene_tx=XM_023250439.1
A1 9739197 9739374 refGene_gn=HMGB1;refGene_tx=XM_003980278.5
A1 9740405 9740580 refGene_gn=HMGB1;refGene_tx=XM_003980278.5
A1 9741068 9741214 refGene_gn=HMGB1;refGene_tx=XM_003980278.5
A1 9741438 9741588 refGene_gn=HMGB1;refGene_tx=XM_003980278.5
A1 9875022 9875121
refGene_gn=USPL1;refGene_tx=XM_003980279.4;refGene_gn=USPL1;refGene_tx=XM_019824892.1;refGene_gn=USPL1;refGene_tx=XM_023250450.1
A1 9875644 9875773
refGene_gn=USPL1;refGene_tx=XM_003980279.4;refGene_gn=USPL1;refGene_tx=XM_019824892.1;refGene_gn=USPL1;refGene_tx=XM_023250450.1
A1 9882367 9883013
refGene_gn=USPL1;refGene_tx=XM_003980279.4;refGene_gn=USPL1;refGene_tx=XM_019824892.1;refGene_gn=USPL1;refGene_tx=XM_019824898.1;refGene_gn=USPL1;refGene_tx=XM_019824901.1;refGene_gn=USPL1;refGene_tx=XM_023250450.1
A1 9887018 9887132
refGene_gn=USPL1;refGene_tx=XM_003980279.4;refGene_gn=USPL1;refGene_tx=XM_019824892.1;refGene_gn=USPL1;refGene_tx=XM_019824898.1;refGene_gn=USPL1;refGene_tx=XM_019824901.1;refGene_gn=USPL1;refGene_tx=XM_023250450.1
A1 9891837 9891967
refGene_gn=USPL1;refGene_tx=XM_003980279.4;refGene_gn=USPL1;refGene_tx=XM_019824892.1;refGene_gn=USPL1;refGene_tx=XM_019824898.1;refGene_gn=USPL1;refGene_tx=XM_019824901.1;refGene_gn=USPL1;refGene_tx=XM_023250450.1
A1 9895646 9895772
refGene_gn=USPL1;refGene_tx=XM_003980279.4;refGene_gn=USPL1;refGene_tx=XM_019824892.1;refGene_gn=USPL1;refGene_tx=XM_019824898.1;refGene_gn=USPL1;refGene_tx=XM_019824901.1;refGene_gn=USPL1;refGene_tx=XM_0232504

50.1
A1	9904172	9904330
refGene_gn=USPL1;refGene_tx=XM_003980279.4;refGene_gn=USPL1;refGene_tx=XM_019824892.1;refGene_gn=USPL1;refGene_tx=XM_019824898.1;refGene_gn=USPL1;refGene_tx=XM_019824901.1;refGene_gn=USPL1;refGene_tx=XM_023250450.1
A1	9906668	9908554
refGene_gn=USPL1;refGene_tx=XM_003980279.4;refGene_gn=USPL1;refGene_tx=XM_019824892.1;refGene_gn=USPL1;refGene_tx=XM_019824898.1;refGene_gn=USPL1;refGene_tx=XM_019824901.1;refGene_gn=USPL1;refGene_tx=XM_023250450.1
A1	9961243	9961313
refGene_gn=ALOX5AP;refGene_tx=XM_006927193.4;refGene_gn=ALOX5AP;refGene_tx=XM_011282720.3;refGene_gn=ALOX5AP;refGene_tx=XM_011282769.2;refGene_gn=ALOX5AP;refGene_tx=XM_011282835.3
A1	9967564	9967664
refGene_gn=ALOX5AP;refGene_tx=XM_006927193.4;refGene_gn=ALOX5AP;refGene_tx=XM_011282720.3;refGene_gn=ALOX5AP;refGene_tx=XM_011282769.2;refGene_gn=ALOX5AP;refGene_tx=XM_011282835.3
A1	9976984	9977159
refGene_gn=ALOX5AP;refGene_tx=XM_011282720.3;refGene_gn=ALOX5AP;refGene_tx=XM_011282769.2
A1	9977831	9977902
refGene_gn=ALOX5AP;refGene_tx=XM_006927193.4;refGene_gn=ALOX5AP;refGene_tx=XM_011282720.3;refGene_gn=ALOX5AP;refGene_tx=XM_011282769.2
A1	9982135	9982217
refGene_gn=ALOX5AP;refGene_tx=XM_006927193.4;refGene_gn=ALOX5AP;refGene_tx=XM_011282720.3;refGene_gn=ALOX5AP;refGene_tx=XM_011282769.2;refGene_gn=ALOX5AP;refGene_tx=XM_011282835.3
A1	9982646	9982799
refGene_gn=ALOX5AP;refGene_tx=XM_011282720.3;refGene_gn=ALOX5AP;refGene_tx=XM_011282835.3
A1	9989877	9990040	refGene_gn=ALOX5AP;refGene_tx=XM_006927193.4
A1	9990577	9990667	refGene_gn=ALOX5AP;refGene_tx=XM_011282769.2
A1	10118229	10118516	refGene_gn=MEDAG;refGene_tx=XM_023250494.1
A1	10128261	10128371	refGene_gn=MEDAG;refGene_tx=XM_023250494.1
A1	10130958	10131071	refGene_gn=MEDAG;refGene_tx=XM_023250494.1
A1	10131499	10131785	refGene_gn=MEDAG;refGene_tx=XM_023250494.1
A1	10133865	10133990	refGene_gn=MEDAG;refGene_tx=XM_023250494.1
A1	10142213	10142365	refGene_gn=TEX26;refGene_tx=XM_023250499.1
A1	10147680	10147762	refGene_gn=TEX26;refGene_tx=XM_023250499.1
A1	10152717	10152876	refGene_gn=TEX26;refGene_tx=XM_023250499.1
A1	10166180	10166352	refGene_gn=TEX26;refGene_tx=XM_023250499.1
A1	10168046	10168208	refGene_gn=TEX26;refGene_tx=XM_023250499.1
A1	10171986	10172012	refGene_gn=TEX26;refGene_tx=XM_023250499.1
A1	10314780	10315097
refGene_gn=LOC102899076;refGene_tx=XM_023256491.1
A1	10315997	10316084
refGene_gn=LOC102899076;refGene_tx=XM_023256491.1
A1	10319103	10319250

refGene_gn=LOC102899076;refGene_tx=XM_023256491.1
A1 10321808 10321973
refGene_gn=LOC102899076;refGene_tx=XM_023256491.1
A1 10322444 10322556
refGene_gn=LOC102899076;refGene_tx=XM_023256491.1
A1 10322920 10322977
refGene_gn=LOC102899076;refGene_tx=XM_023256491.1
A1 10329390 10329597
refGene_gn=HSPH1;refGene_tx=XM_023250511.1;refGene_gn=HSPH1;refGene_tx=XM_023250512.1
A1 10330552 10330714
refGene_gn=HSPH1;refGene_tx=XM_023250511.1;refGene_gn=HSPH1;refGene_tx=XM_023250512.1
A1 10330932 10331052
refGene_gn=HSPH1;refGene_tx=XM_023250511.1;refGene_gn=HSPH1;refGene_tx=XM_023250512.1
A1 10331131 10331239
refGene_gn=HSPH1;refGene_tx=XM_023250511.1;refGene_gn=HSPH1;refGene_tx=XM_023250512.1
A1 10332689 10332815
refGene_gn=HSPH1;refGene_tx=XM_023250511.1;refGene_gn=HSPH1;refGene_tx=XM_023250512.1
A1 10333690 10333828
refGene_gn=HSPH1;refGene_tx=XM_023250511.1;refGene_gn=HSPH1;refGene_tx=XM_023250512.1
A1 10336373 10336505
refGene_gn=HSPH1;refGene_tx=XM_023250511.1;refGene_gn=HSPH1;refGene_tx=XM_023250512.1
A1 10338130 10338339
refGene_gn=HSPH1;refGene_tx=XM_023250511.1;refGene_gn=HSPH1;refGene_tx=XM_023250512.1
A1 10340147 10340281
refGene_gn=HSPH1;refGene_tx=XM_023250511.1;refGene_gn=HSPH1;refGene_tx=XM_023250512.1
A1 10340529 10340636
refGene_gn=HSPH1;refGene_tx=XM_023250511.1;refGene_gn=HSPH1;refGene_tx=XM_023250512.1
A1 10342084 10342313
refGene_gn=HSPH1;refGene_tx=XM_023250511.1;refGene_gn=HSPH1;refGene_tx=XM_023250512.1
A1 10343146 10343391
refGene_gn=HSPH1;refGene_tx=XM_023250511.1;refGene_gn=HSPH1;refGene_tx=XM_023250512.1
A1 10343825 10343959
refGene_gn=HSPH1;refGene_tx=XM_023250511.1;refGene_gn=HSPH1;refGene_tx=XM_023250512.1
A1 10345078 10345178
refGene_gn=HSPH1;refGene_tx=XM_023250511.1;refGene_gn=HSPH1;refGene_tx=XM_023250512.1
A1 10346902 10347025

refGene_gn=HSPH1;refGene_tx=XM_023250511.1;refGene_gn=HSPH1;refGene_tx=XM_023250512.1
A1 10347725 10347866 refGene_gn=HSPH1;refGene_tx=XM_023250511.1;refGene_gn=HSPH1;refGene_tx=XM_023250512.1
A1 10349717 10349775 refGene_gn=HSPH1;refGene_tx=XM_023250511.1;refGene_gn=HSPH1;refGene_tx=XM_023250512.1
A1 10351297 10351332 refGene_gn=HSPH1;refGene_tx=XM_023250511.1
A1 10352288 10352395 refGene_gn=HSPH1;refGene_tx=XM_023250512.1
A1 10383932 10384008 refGene_gn=B3GLCT;refGene_tx=XM_023250518.1
A1 10398853 10398903 refGene_gn=B3GLCT;refGene_tx=XM_023250518.1
A1 10409052 10409092 refGene_gn=B3GLCT;refGene_tx=XM_023250518.1
A1 10415012 10415122 refGene_gn=B3GLCT;refGene_tx=XM_023250518.1
A1 10430850 10430927 refGene_gn=B3GLCT;refGene_tx=XM_023250518.1;refGene_gn=B3GLCT;refGene_tx=XM_023250520.1;refGene_gn=B3GLCT;refGene_tx=XM_023250525.1
A1 10431377 10431489 refGene_gn=B3GLCT;refGene_tx=XM_023250518.1;refGene_gn=B3GLCT;refGene_tx=XM_023250520.1;refGene_gn=B3GLCT;refGene_tx=XM_023250525.1
A1 10437564 10437701 refGene_gn=B3GLCT;refGene_tx=XM_023250518.1;refGene_gn=B3GLCT;refGene_tx=XM_023250520.1;refGene_gn=B3GLCT;refGene_tx=XM_023250525.1
A1 10445312 10445376 refGene_gn=B3GLCT;refGene_tx=XM_023250518.1;refGene_gn=B3GLCT;refGene_tx=XM_023250520.1;refGene_gn=B3GLCT;refGene_tx=XM_023250525.1
A1 10450700 10450820 refGene_gn=B3GLCT;refGene_tx=XM_023250518.1;refGene_gn=B3GLCT;refGene_tx=XM_023250520.1;refGene_gn=B3GLCT;refGene_tx=XM_023250525.1
A1 10452517 10452587 refGene_gn=B3GLCT;refGene_tx=XM_023250518.1;refGene_gn=B3GLCT;refGene_tx=XM_023250520.1;refGene_gn=B3GLCT;refGene_tx=XM_023250525.1
A1 10459215 10459329 refGene_gn=B3GLCT;refGene_tx=XM_023250518.1;refGene_gn=B3GLCT;refGene_tx=XM_023250520.1;refGene_gn=B3GLCT;refGene_tx=XM_023250525.1
A1 10461231 10461331 refGene_gn=B3GLCT;refGene_tx=XM_023250518.1;refGene_gn=B3GLCT;refGene_tx=XM_023250520.1;refGene_gn=B3GLCT;refGene_tx=XM_023250525.1
A1 10481100 10481220 refGene_gn=B3GLCT;refGene_tx=XM_023250518.1;refGene_gn=B3GLCT;refGene_tx=XM_023250520.1;refGene_gn=B3GLCT;refGene_tx=XM_023250525.1
A1 10488902 10489107 refGene_gn=B3GLCT;refGene_tx=XM_023250518.1;refGene_gn=B3GLCT;refGene_tx=XM_023250520.1;refGene_gn=B3GLCT;refGene_tx=XM_023250525.1
A1 10501363 10501531 refGene_gn=B3GLCT;refGene_tx=XM_023250518.1;refGene_gn=B3GLCT;refGene_tx=XM_023250520.1;refGene_gn=B3GLCT;refGene_tx=XM_023250525.1
A1 10807666 10807718 refGene_gn=RXFP2;refGene_tx=XM_023257316.1
A1 10977449 10977629 refGene_gn=RXFP2;refGene_tx=XM_023257316.1
A1 10996012 10996159 refGene_gn=RXFP2;refGene_tx=XM_023257316.1

A1 10998602 10998680 refGene_gn=RXFP2;refGene_tx=XM_023257316.1
A1 11001501 11001607 refGene_gn=RXFP2;refGene_tx=XM_023257316.1
A1 11002443 11002515 refGene_gn=RXFP2;refGene_tx=XM_023257316.1
A1 11010036 11010108 refGene_gn=RXFP2;refGene_tx=XM_023257316.1
A1 11010687 11010759 refGene_gn=RXFP2;refGene_tx=XM_023257316.1
A1 11012325 11012397 refGene_gn=RXFP2;refGene_tx=XM_023257316.1
A1 11013438 11013510 refGene_gn=RXFP2;refGene_tx=XM_023257316.1
A1 11017349 11017421 refGene_gn=RXFP2;refGene_tx=XM_023257316.1
A1 11018865 11018937 refGene_gn=RXFP2;refGene_tx=XM_023257316.1
A1 11022254 11022326 refGene_gn=RXFP2;refGene_tx=XM_023257316.1
A1 11022437 11022509 refGene_gn=RXFP2;refGene_tx=XM_023257316.1
A1 11024421 11024493 refGene_gn=RXFP2;refGene_tx=XM_023257316.1
A1 11027044 11027274 refGene_gn=RXFP2;refGene_tx=XM_023257316.1
A1 11027902 11028313 refGene_gn=RXFP2;refGene_tx=XM_023257316.1
A1 11031317 11031536 refGene_gn=RXFP2;refGene_tx=XM_023257316.1
A1 11035990 11036250 refGene_gn=RXFP2;refGene_tx=XM_023257316.1
A1 11176854 11177059
refGene_gn=FRY;refGene_tx=XM_019825096.1;refGene_gn=FRY;refGene_tx=XM_019825104.1;refGene_gn=FRY;refGene_tx=XM_019825114.2;refGene_gn=FRY;refGene_tx=XM_023250547.1
A1 11259801 11259871
refGene_gn=FRY;refGene_tx=XM_019825116.2;refGene_gn=FRY;refGene_tx=XM_019825125.2
A1 11290312 11290373 refGene_gn=FRY;refGene_tx=XM_006927213.4
A1 11306214 11306414
refGene_gn=FRY;refGene_tx=XM_006927213.4;refGene_gn=FRY;refGene_tx=XM_011286958.3;refGene_gn=FRY;refGene_tx=XM_019825096.1;refGene_gn=FRY;refGene_tx=XM_019825104.1;refGene_gn=FRY;refGene_tx=XM_019825114.2;refGene_gn=FRY;refGene_tx=XM_019825116.2;refGene_gn=FRY;refGene_tx=XM_019825125.2;refGene_gn=FRY;refGene_tx=XM_023250547.1
A1 11329512 11329566
refGene_gn=FRY;refGene_tx=XM_006927213.4;refGene_gn=FRY;refGene_tx=XM_011286958.3;refGene_gn=FRY;refGene_tx=XM_019825096.1;refGene_gn=FRY;refGene_tx=XM_019825104.1;refGene_gn=FRY;refGene_tx=XM_019825114.2;refGene_gn=FRY;refGene_tx=XM_019825116.2;refGene_gn=FRY;refGene_tx=XM_019825125.2;refGene_gn=FRY;refGene_tx=XM_023250547.1
A1 11345856 11345996
refGene_gn=FRY;refGene_tx=XM_006927213.4;refGene_gn=FRY;refGene_tx=XM_011286958.3;refGene_gn=FRY;refGene_tx=XM_019825096.1;refGene_gn=FRY;refGene_tx=XM_019825104.1;refGene_gn=FRY;refGene_tx=XM_019825114.2;refGene_gn=FRY;refGene_tx=XM_019825116.2;refGene_gn=FRY;refGene_tx=XM_019825125.2;refGene_gn=FRY;refGene_tx=XM_023250547.1
A1 11350198 11350289
refGene_gn=FRY;refGene_tx=XM_006927213.4;refGene_gn=FRY;refGene_tx=XM_011286958.3;refGene_gn=FRY;refGene_tx=XM_019825096.1;refGene_gn=FRY;refGene_tx=XM_019825104.1;refGene_gn=FRY;refGene_tx=XM_019825114.2;refGene_gn=FRY;refGene_tx=XM_019825116.2;refGene_gn=FRY;refGene_tx=XM_019825125.2;refGene_gn=FRY;refGene_tx=XM_023250547.1
A1 11350497 11350577
refGene_gn=FRY;refGene_tx=XM_006927213.4;refGene_gn=FRY;refGene_tx=XM_

011286958.3;refGene_gn=FRY;refGene_tx=XM_019825096.1;refGene_gn=FRY;refGene_tx=XM_019825104.1;refGene_gn=FRY;refGene_tx=XM_019825114.2;refGene_gn=FRY;refGene_tx=XM_019825116.2;refGene_gn=FRY;refGene_tx=XM_019825125.2;refGene_gn=FRY;refGene_tx=XM_023250547.1
A1 11350687 11350768
refGene_gn=FRY;refGene_tx=XM_006927213.4;refGene_gn=FRY;refGene_tx=XM_011286958.3;refGene_gn=FRY;refGene_tx=XM_019825096.1;refGene_gn=FRY;refGene_tx=XM_019825104.1;refGene_gn=FRY;refGene_tx=XM_019825114.2;refGene_gn=FRY;refGene_tx=XM_019825116.2;refGene_gn=FRY;refGene_tx=XM_019825125.2;refGene_gn=FRY;refGene_tx=XM_023250547.1
A1 11356644 11356813
refGene_gn=FRY;refGene_tx=XM_006927213.4;refGene_gn=FRY;refGene_tx=XM_011286958.3;refGene_gn=FRY;refGene_tx=XM_019825096.1;refGene_gn=FRY;refGene_tx=XM_019825104.1;refGene_gn=FRY;refGene_tx=XM_019825114.2;refGene_gn=FRY;refGene_tx=XM_019825116.2;refGene_gn=FRY;refGene_tx=XM_019825125.2;refGene_gn=FRY;refGene_tx=XM_023250547.1
A1 11359864 11359957
refGene_gn=FRY;refGene_tx=XM_006927213.4;refGene_gn=FRY;refGene_tx=XM_011286958.3;refGene_gn=FRY;refGene_tx=XM_019825096.1;refGene_gn=FRY;refGene_tx=XM_019825104.1;refGene_gn=FRY;refGene_tx=XM_019825114.2;refGene_gn=FRY;refGene_tx=XM_019825116.2;refGene_gn=FRY;refGene_tx=XM_019825125.2;refGene_gn=FRY;refGene_tx=XM_023250547.1
A1 11360051 11360150
refGene_gn=FRY;refGene_tx=XM_006927213.4;refGene_gn=FRY;refGene_tx=XM_011286958.3;refGene_gn=FRY;refGene_tx=XM_019825096.1;refGene_gn=FRY;refGene_tx=XM_019825104.1;refGene_gn=FRY;refGene_tx=XM_019825114.2;refGene_gn=FRY;refGene_tx=XM_019825116.2;refGene_gn=FRY;refGene_tx=XM_019825125.2;refGene_gn=FRY;refGene_tx=XM_023250547.1
A1 11361917 11362019
refGene_gn=FRY;refGene_tx=XM_006927213.4;refGene_gn=FRY;refGene_tx=XM_011286958.3;refGene_gn=FRY;refGene_tx=XM_019825096.1;refGene_gn=FRY;refGene_tx=XM_019825104.1;refGene_gn=FRY;refGene_tx=XM_019825114.2;refGene_gn=FRY;refGene_tx=XM_019825116.2;refGene_gn=FRY;refGene_tx=XM_019825125.2;refGene_gn=FRY;refGene_tx=XM_023250547.1
A1 11379012 11379116
refGene_gn=FRY;refGene_tx=XM_006927213.4;refGene_gn=FRY;refGene_tx=XM_011286958.3;refGene_gn=FRY;refGene_tx=XM_019825096.1;refGene_gn=FRY;refGene_tx=XM_019825104.1;refGene_gn=FRY;refGene_tx=XM_019825114.2;refGene_gn=FRY;refGene_tx=XM_019825116.2;refGene_gn=FRY;refGene_tx=XM_019825125.2;refGene_gn=FRY;refGene_tx=XM_023250547.1
A1 11379593 11379702
refGene_gn=FRY;refGene_tx=XM_006927213.4;refGene_gn=FRY;refGene_tx=XM_011286958.3;refGene_gn=FRY;refGene_tx=XM_019825096.1;refGene_gn=FRY;refGene_tx=XM_019825104.1;refGene_gn=FRY;refGene_tx=XM_019825114.2;refGene_gn=FRY;refGene_tx=XM_019825116.2;refGene_gn=FRY;refGene_tx=XM_019825125.2;refGene_gn=FRY;refGene_tx=XM_023250547.1
A1 11381637 11381724
refGene_gn=FRY;refGene_tx=XM_006927213.4;refGene_gn=FRY;refGene_tx=XM_011286958.3;refGene_gn=FRY;refGene_tx=XM_019825096.1;refGene_gn=FRY;refGene_tx=XM_019825104.1;refGene_gn=FRY;refGene_tx=XM_019825114.2;refGe

ne_gn=FRY;refGene_tx=XM_019825116.2;refGene_gn=FRY;refGene_tx=XM_019825125.2;refGene_gn=FRY;refGene_tx=XM_023250547.1
A1 11390944 11391116
refGene_gn=FRY;refGene_tx=XM_006927213.4;refGene_gn=FRY;refGene_tx=XM_011286958.3;refGene_gn=FRY;refGene_tx=XM_019825096.1;refGene_gn=FRY;refGene_tx=XM_019825104.1;refGene_gn=FRY;refGene_tx=XM_019825114.2;refGene_gn=FRY;refGene_tx=XM_019825116.2;refGene_gn=FRY;refGene_tx=XM_019825125.2;refGene_gn=FRY;refGene_tx=XM_023250547.1
A1 11392479 11392612
refGene_gn=FRY;refGene_tx=XM_006927213.4;refGene_gn=FRY;refGene_tx=XM_011286958.3;refGene_gn=FRY;refGene_tx=XM_019825096.1;refGene_gn=FRY;refGene_tx=XM_019825104.1;refGene_gn=FRY;refGene_tx=XM_019825114.2;refGene_gn=FRY;refGene_tx=XM_019825116.2;refGene_gn=FRY;refGene_tx=XM_019825125.2;refGene_gn=FRY;refGene_tx=XM_023250547.1
A1 11396061 11396169
refGene_gn=FRY;refGene_tx=XM_006927213.4;refGene_gn=FRY;refGene_tx=XM_011286958.3;refGene_gn=FRY;refGene_tx=XM_019825096.1;refGene_gn=FRY;refGene_tx=XM_019825104.1;refGene_gn=FRY;refGene_tx=XM_019825114.2;refGene_gn=FRY;refGene_tx=XM_019825116.2;refGene_gn=FRY;refGene_tx=XM_019825125.2;refGene_gn=FRY;refGene_tx=XM_023250547.1
A1 11404433 11404692
refGene_gn=FRY;refGene_tx=XM_006927213.4;refGene_gn=FRY;refGene_tx=XM_011286958.3;refGene_gn=FRY;refGene_tx=XM_019825096.1;refGene_gn=FRY;refGene_tx=XM_019825104.1;refGene_gn=FRY;refGene_tx=XM_019825114.2;refGene_gn=FRY;refGene_tx=XM_019825116.2;refGene_gn=FRY;refGene_tx=XM_019825125.2;refGene_gn=FRY;refGene_tx=XM_023250547.1
A1 11405968 11406151
refGene_gn=FRY;refGene_tx=XM_006927213.4;refGene_gn=FRY;refGene_tx=XM_011286958.3;refGene_gn=FRY;refGene_tx=XM_019825096.1;refGene_gn=FRY;refGene_tx=XM_019825104.1;refGene_gn=FRY;refGene_tx=XM_019825114.2;refGene_gn=FRY;refGene_tx=XM_019825116.2;refGene_gn=FRY;refGene_tx=XM_019825125.2;refGene_gn=FRY;refGene_tx=XM_023250547.1
A1 11408697 11408784
refGene_gn=FRY;refGene_tx=XM_006927213.4;refGene_gn=FRY;refGene_tx=XM_011286958.3;refGene_gn=FRY;refGene_tx=XM_019825096.1;refGene_gn=FRY;refGene_tx=XM_019825104.1;refGene_gn=FRY;refGene_tx=XM_019825114.2;refGene_gn=FRY;refGene_tx=XM_019825116.2;refGene_gn=FRY;refGene_tx=XM_019825125.2;refGene_gn=FRY;refGene_tx=XM_023250547.1
A1 11410968 11411228
refGene_gn=FRY;refGene_tx=XM_006927213.4;refGene_gn=FRY;refGene_tx=XM_011286958.3;refGene_gn=FRY;refGene_tx=XM_019825096.1;refGene_gn=FRY;refGene_tx=XM_019825104.1;refGene_gn=FRY;refGene_tx=XM_019825114.2;refGene_gn=FRY;refGene_tx=XM_019825116.2;refGene_gn=FRY;refGene_tx=XM_019825125.2;refGene_gn=FRY;refGene_tx=XM_023250547.1
A1 11411657 11411847
refGene_gn=FRY;refGene_tx=XM_006927213.4;refGene_gn=FRY;refGene_tx=XM_011286958.3;refGene_gn=FRY;refGene_tx=XM_019825096.1;refGene_gn=FRY;refGene_tx=XM_019825104.1;refGene_gn=FRY;refGene_tx=XM_019825114.2;refGene_gn=FRY;refGene_tx=XM_019825116.2;refGene_gn=FRY;refGene_tx=XM_019825125.2;refGene_gn=FRY;refGene_tx=XM_023250547.1

A1 11412468 11412593
refGene_gn=FRY;refGene_tx=XM_006927213.4;refGene_gn=FRY;refGene_tx=XM_011286958.3;refGene_gn=FRY;refGene_tx=XM_019825096.1;refGene_gn=FRY;refGene_tx=XM_019825104.1;refGene_gn=FRY;refGene_tx=XM_019825114.2;refGene_gn=FRY;refGene_tx=XM_019825116.2;refGene_gn=FRY;refGene_tx=XM_019825125.2;refGene_gn=FRY;refGene_tx=XM_023250547.1
A1 11414562 11414620
refGene_gn=FRY;refGene_tx=XM_006927213.4;refGene_gn=FRY;refGene_tx=XM_011286958.3;refGene_gn=FRY;refGene_tx=XM_019825096.1;refGene_gn=FRY;refGene_tx=XM_019825104.1;refGene_gn=FRY;refGene_tx=XM_019825114.2;refGene_gn=FRY;refGene_tx=XM_019825116.2;refGene_gn=FRY;refGene_tx=XM_019825125.2;refGene_gn=FRY;refGene_tx=XM_023250547.1
A1 11415700 11415792
refGene_gn=FRY;refGene_tx=XM_006927213.4;refGene_gn=FRY;refGene_tx=XM_011286958.3;refGene_gn=FRY;refGene_tx=XM_019825096.1;refGene_gn=FRY;refGene_tx=XM_019825104.1;refGene_gn=FRY;refGene_tx=XM_019825114.2;refGene_gn=FRY;refGene_tx=XM_019825116.2;refGene_gn=FRY;refGene_tx=XM_019825125.2;refGene_gn=FRY;refGene_tx=XM_023250547.1
A1 11416089 11416262
refGene_gn=FRY;refGene_tx=XM_006927213.4;refGene_gn=FRY;refGene_tx=XM_011286958.3;refGene_gn=FRY;refGene_tx=XM_019825096.1;refGene_gn=FRY;refGene_tx=XM_019825104.1;refGene_gn=FRY;refGene_tx=XM_019825114.2;refGene_gn=FRY;refGene_tx=XM_019825116.2;refGene_gn=FRY;refGene_tx=XM_019825125.2;refGene_gn=FRY;refGene_tx=XM_023250547.1
A1 11417576 11417737
refGene_gn=FRY;refGene_tx=XM_006927213.4;refGene_gn=FRY;refGene_tx=XM_011286958.3;refGene_gn=FRY;refGene_tx=XM_019825096.1;refGene_gn=FRY;refGene_tx=XM_019825104.1;refGene_gn=FRY;refGene_tx=XM_019825114.2;refGene_gn=FRY;refGene_tx=XM_019825116.2;refGene_gn=FRY;refGene_tx=XM_019825125.2;refGene_gn=FRY;refGene_tx=XM_023250547.1
A1 11419061 11419172
refGene_gn=FRY;refGene_tx=XM_006927213.4;refGene_gn=FRY;refGene_tx=XM_011286958.3;refGene_gn=FRY;refGene_tx=XM_019825096.1;refGene_gn=FRY;refGene_tx=XM_019825104.1;refGene_gn=FRY;refGene_tx=XM_019825114.2;refGene_gn=FRY;refGene_tx=XM_019825116.2;refGene_gn=FRY;refGene_tx=XM_019825125.2;refGene_gn=FRY;refGene_tx=XM_023250547.1
A1 11423043 11423198
refGene_gn=FRY;refGene_tx=XM_006927213.4;refGene_gn=FRY;refGene_tx=XM_011286958.3;refGene_gn=FRY;refGene_tx=XM_019825096.1;refGene_gn=FRY;refGene_tx=XM_019825104.1;refGene_gn=FRY;refGene_tx=XM_019825114.2;refGene_gn=FRY;refGene_tx=XM_019825116.2;refGene_gn=FRY;refGene_tx=XM_019825125.2;refGene_gn=FRY;refGene_tx=XM_023250547.1
A1 11430991 11431091
refGene_gn=FRY;refGene_tx=XM_006927213.4;refGene_gn=FRY;refGene_tx=XM_011286958.3;refGene_gn=FRY;refGene_tx=XM_019825096.1;refGene_gn=FRY;refGene_tx=XM_019825104.1;refGene_gn=FRY;refGene_tx=XM_019825114.2;refGene_gn=FRY;refGene_tx=XM_019825116.2;refGene_gn=FRY;refGene_tx=XM_019825125.2;refGene_gn=FRY;refGene_tx=XM_023250547.1
A1 11431386 11431558
refGene_gn=FRY;refGene_tx=XM_006927213.4;refGene_gn=FRY;refGene_tx=XM_

011286958.3;refGene_gn=FRY;refGene_tx=XM_019825096.1;refGene_gn=FRY;refGene_tx=XM_019825104.1;refGene_gn=FRY;refGene_tx=XM_019825114.2;refGene_gn=FRY;refGene_tx=XM_019825116.2;refGene_gn=FRY;refGene_tx=XM_019825125.2;refGene_gn=FRY;refGene_tx=XM_023250547.1
A1 11436987 11437244
refGene_gn=FRY;refGene_tx=XM_006927213.4;refGene_gn=FRY;refGene_tx=XM_011286958.3;refGene_gn=FRY;refGene_tx=XM_019825096.1;refGene_gn=FRY;refGene_tx=XM_019825104.1;refGene_gn=FRY;refGene_tx=XM_019825114.2;refGene_gn=FRY;refGene_tx=XM_019825116.2;refGene_gn=FRY;refGene_tx=XM_019825125.2;refGene_gn=FRY;refGene_tx=XM_023250547.1
A1 11437689 11437836
refGene_gn=FRY;refGene_tx=XM_006927213.4;refGene_gn=FRY;refGene_tx=XM_011286958.3;refGene_gn=FRY;refGene_tx=XM_019825096.1;refGene_gn=FRY;refGene_tx=XM_019825104.1;refGene_gn=FRY;refGene_tx=XM_019825114.2;refGene_gn=FRY;refGene_tx=XM_019825116.2;refGene_gn=FRY;refGene_tx=XM_019825125.2;refGene_gn=FRY;refGene_tx=XM_023250547.1
A1 11439424 11439593
refGene_gn=FRY;refGene_tx=XM_006927213.4;refGene_gn=FRY;refGene_tx=XM_011286958.3;refGene_gn=FRY;refGene_tx=XM_019825096.1;refGene_gn=FRY;refGene_tx=XM_019825104.1;refGene_gn=FRY;refGene_tx=XM_019825114.2;refGene_gn=FRY;refGene_tx=XM_019825116.2;refGene_gn=FRY;refGene_tx=XM_019825125.2;refGene_gn=FRY;refGene_tx=XM_023250547.1
A1 11440859 11440950
refGene_gn=FRY;refGene_tx=XM_006927213.4;refGene_gn=FRY;refGene_tx=XM_011286958.3;refGene_gn=FRY;refGene_tx=XM_019825096.1;refGene_gn=FRY;refGene_tx=XM_019825104.1;refGene_gn=FRY;refGene_tx=XM_019825114.2;refGene_gn=FRY;refGene_tx=XM_019825116.2;refGene_gn=FRY;refGene_tx=XM_019825125.2;refGene_gn=FRY;refGene_tx=XM_023250547.1
A1 11446573 11446656
refGene_gn=FRY;refGene_tx=XM_006927213.4;refGene_gn=FRY;refGene_tx=XM_011286958.3;refGene_gn=FRY;refGene_tx=XM_019825096.1;refGene_gn=FRY;refGene_tx=XM_019825104.1;refGene_gn=FRY;refGene_tx=XM_019825114.2;refGene_gn=FRY;refGene_tx=XM_019825116.2;refGene_gn=FRY;refGene_tx=XM_019825125.2;refGene_gn=FRY;refGene_tx=XM_023250547.1
A1 11451793 11451944
refGene_gn=FRY;refGene_tx=XM_006927213.4;refGene_gn=FRY;refGene_tx=XM_011286958.3;refGene_gn=FRY;refGene_tx=XM_019825096.1;refGene_gn=FRY;refGene_tx=XM_019825104.1;refGene_gn=FRY;refGene_tx=XM_019825114.2;refGene_gn=FRY;refGene_tx=XM_019825116.2;refGene_gn=FRY;refGene_tx=XM_019825125.2;refGene_gn=FRY;refGene_tx=XM_023250547.1
A1 11452492 11452596
refGene_gn=FRY;refGene_tx=XM_006927213.4;refGene_gn=FRY;refGene_tx=XM_011286958.3;refGene_gn=FRY;refGene_tx=XM_019825096.1;refGene_gn=FRY;refGene_tx=XM_019825104.1;refGene_gn=FRY;refGene_tx=XM_019825114.2;refGene_gn=FRY;refGene_tx=XM_019825116.2;refGene_gn=FRY;refGene_tx=XM_019825125.2;refGene_gn=FRY;refGene_tx=XM_023250547.1
A1 11453353 11453539
refGene_gn=FRY;refGene_tx=XM_006927213.4;refGene_gn=FRY;refGene_tx=XM_011286958.3;refGene_gn=FRY;refGene_tx=XM_019825096.1;refGene_gn=FRY;refGene_tx=XM_019825104.1;refGene_gn=FRY;refGene_tx=XM_019825114.2;refGe

ne_gn=FRY;refGene_tx=XM_019825116.2;refGene_gn=FRY;refGene_tx=XM_019825125.2;refGene_gn=FRY;refGene_tx=XM_023250547.1
A1 11455731 11455930
refGene_gn=FRY;refGene_tx=XM_006927213.4;refGene_gn=FRY;refGene_tx=XM_011286958.3;refGene_gn=FRY;refGene_tx=XM_019825096.1;refGene_gn=FRY;refGene_tx=XM_019825104.1;refGene_gn=FRY;refGene_tx=XM_019825114.2;refGene_gn=FRY;refGene_tx=XM_019825116.2;refGene_gn=FRY;refGene_tx=XM_019825125.2;refGene_gn=FRY;refGene_tx=XM_023250547.1
A1 11457398 11457520
refGene_gn=FRY;refGene_tx=XM_006927213.4;refGene_gn=FRY;refGene_tx=XM_011286958.3;refGene_gn=FRY;refGene_tx=XM_019825096.1;refGene_gn=FRY;refGene_tx=XM_019825104.1;refGene_gn=FRY;refGene_tx=XM_019825114.2;refGene_gn=FRY;refGene_tx=XM_019825116.2;refGene_gn=FRY;refGene_tx=XM_019825125.2;refGene_gn=FRY;refGene_tx=XM_023250547.1
A1 11461499 11461687
refGene_gn=FRY;refGene_tx=XM_006927213.4;refGene_gn=FRY;refGene_tx=XM_011286958.3;refGene_gn=FRY;refGene_tx=XM_019825096.1;refGene_gn=FRY;refGene_tx=XM_019825104.1;refGene_gn=FRY;refGene_tx=XM_019825114.2;refGene_gn=FRY;refGene_tx=XM_019825116.2;refGene_gn=FRY;refGene_tx=XM_019825125.2;refGene_gn=FRY;refGene_tx=XM_023250547.1
A1 11463022 11463117
refGene_gn=FRY;refGene_tx=XM_006927213.4;refGene_gn=FRY;refGene_tx=XM_011286958.3;refGene_gn=FRY;refGene_tx=XM_019825096.1;refGene_gn=FRY;refGene_tx=XM_019825104.1;refGene_gn=FRY;refGene_tx=XM_019825114.2;refGene_gn=FRY;refGene_tx=XM_019825116.2;refGene_gn=FRY;refGene_tx=XM_019825125.2;refGene_gn=FRY;refGene_tx=XM_023250547.1
A1 11464357 11464965
refGene_gn=FRY;refGene_tx=XM_006927213.4;refGene_gn=FRY;refGene_tx=XM_011286958.3;refGene_gn=FRY;refGene_tx=XM_019825096.1;refGene_gn=FRY;refGene_tx=XM_019825104.1;refGene_gn=FRY;refGene_tx=XM_019825114.2;refGene_gn=FRY;refGene_tx=XM_019825116.2;refGene_gn=FRY;refGene_tx=XM_019825125.2;refGene_gn=FRY;refGene_tx=XM_023250547.1
A1 11465891 11465989
refGene_gn=FRY;refGene_tx=XM_006927213.4;refGene_gn=FRY;refGene_tx=XM_011286958.3;refGene_gn=FRY;refGene_tx=XM_019825096.1;refGene_gn=FRY;refGene_tx=XM_019825104.1;refGene_gn=FRY;refGene_tx=XM_019825114.2;refGene_gn=FRY;refGene_tx=XM_019825116.2;refGene_gn=FRY;refGene_tx=XM_019825125.2;refGene_gn=FRY;refGene_tx=XM_023250547.1
A1 11466376 11466547
refGene_gn=FRY;refGene_tx=XM_006927213.4;refGene_gn=FRY;refGene_tx=XM_011286958.3;refGene_gn=FRY;refGene_tx=XM_019825096.1;refGene_gn=FRY;refGene_tx=XM_019825104.1;refGene_gn=FRY;refGene_tx=XM_019825114.2;refGene_gn=FRY;refGene_tx=XM_019825116.2;refGene_gn=FRY;refGene_tx=XM_019825125.2;refGene_gn=FRY;refGene_tx=XM_023250547.1
A1 11470264 11470405
refGene_gn=FRY;refGene_tx=XM_006927213.4;refGene_gn=FRY;refGene_tx=XM_011286958.3;refGene_gn=FRY;refGene_tx=XM_019825096.1;refGene_gn=FRY;refGene_tx=XM_019825104.1;refGene_gn=FRY;refGene_tx=XM_019825114.2;refGene_gn=FRY;refGene_tx=XM_019825116.2;refGene_gn=FRY;refGene_tx=XM_019825125.2;refGene_gn=FRY;refGene_tx=XM_023250547.1

A1 11472193 11472373
refGene_gn=FRY;refGene_tx=XM_006927213.4;refGene_gn=FRY;refGene_tx=XM_011286958.3;refGene_gn=FRY;refGene_tx=XM_019825096.1;refGene_gn=FRY;refGene_tx=XM_019825104.1;refGene_gn=FRY;refGene_tx=XM_019825114.2;refGene_gn=FRY;refGene_tx=XM_019825116.2;refGene_gn=FRY;refGene_tx=XM_019825125.2;refGene_gn=FRY;refGene_tx=XM_023250547.1
A1 11475134 11475296
refGene_gn=FRY;refGene_tx=XM_006927213.4;refGene_gn=FRY;refGene_tx=XM_011286958.3;refGene_gn=FRY;refGene_tx=XM_019825096.1;refGene_gn=FRY;refGene_tx=XM_019825104.1;refGene_gn=FRY;refGene_tx=XM_019825114.2;refGene_gn=FRY;refGene_tx=XM_019825116.2;refGene_gn=FRY;refGene_tx=XM_019825125.2;refGene_gn=FRY;refGene_tx=XM_023250547.1
A1 11477976 11478051
refGene_gn=FRY;refGene_tx=XM_006927213.4;refGene_gn=FRY;refGene_tx=XM_011286958.3;refGene_gn=FRY;refGene_tx=XM_019825096.1;refGene_gn=FRY;refGene_tx=XM_019825104.1;refGene_gn=FRY;refGene_tx=XM_019825114.2;refGene_gn=FRY;refGene_tx=XM_019825116.2;refGene_gn=FRY;refGene_tx=XM_019825125.2;refGene_gn=FRY;refGene_tx=XM_023250547.1
A1 11480385 11480556
refGene_gn=FRY;refGene_tx=XM_006927213.4;refGene_gn=FRY;refGene_tx=XM_011286958.3;refGene_gn=FRY;refGene_tx=XM_019825096.1;refGene_gn=FRY;refGene_tx=XM_019825104.1;refGene_gn=FRY;refGene_tx=XM_019825114.2;refGene_gn=FRY;refGene_tx=XM_019825116.2;refGene_gn=FRY;refGene_tx=XM_019825125.2;refGene_gn=FRY;refGene_tx=XM_023250547.1
A1 11483891 11483900
refGene_gn=FRY;refGene_tx=XM_006927213.4;refGene_gn=FRY;refGene_tx=XM_011286958.3;refGene_gn=FRY;refGene_tx=XM_019825104.1;refGene_gn=FRY;refGene_tx=XM_019825114.2;refGene_gn=FRY;refGene_tx=XM_019825116.2;refGene_gn=FRY;refGene_tx=XM_023250547.1
A1 11486851 11487052
refGene_gn=FRY;refGene_tx=XM_006927213.4;refGene_gn=FRY;refGene_tx=XM_011286958.3;refGene_gn=FRY;refGene_tx=XM_019825096.1;refGene_gn=FRY;refGene_tx=XM_019825104.1;refGene_gn=FRY;refGene_tx=XM_019825114.2;refGene_gn=FRY;refGene_tx=XM_019825116.2;refGene_gn=FRY;refGene_tx=XM_019825125.2;refGene_gn=FRY;refGene_tx=XM_023250547.1
A1 11487604 11487766
refGene_gn=FRY;refGene_tx=XM_006927213.4;refGene_gn=FRY;refGene_tx=XM_011286958.3;refGene_gn=FRY;refGene_tx=XM_019825096.1;refGene_gn=FRY;refGene_tx=XM_019825104.1;refGene_gn=FRY;refGene_tx=XM_019825114.2;refGene_gn=FRY;refGene_tx=XM_019825116.2;refGene_gn=FRY;refGene_tx=XM_019825125.2;refGene_gn=FRY;refGene_tx=XM_023250547.1
A1 11490470 11490637
refGene_gn=FRY;refGene_tx=XM_006927213.4;refGene_gn=FRY;refGene_tx=XM_011286958.3;refGene_gn=FRY;refGene_tx=XM_019825096.1;refGene_gn=FRY;refGene_tx=XM_019825104.1;refGene_gn=FRY;refGene_tx=XM_019825114.2;refGene_gn=FRY;refGene_tx=XM_019825116.2;refGene_gn=FRY;refGene_tx=XM_019825125.2;refGene_gn=FRY;refGene_tx=XM_023250547.1
A1 11492140 11492330
refGene_gn=FRY;refGene_tx=XM_006927213.4;refGene_gn=FRY;refGene_tx=XM_011286958.3;refGene_gn=FRY;refGene_tx=XM_019825096.1;refGene_gn=FRY;re

fGene_tx=XM_019825104.1;refGene_gn=FRY;refGene_tx=XM_019825114.2;refGene_gn=FRY;refGene_tx=XM_019825116.2;refGene_gn=FRY;refGene_tx=XM_019825125.2;refGene_gn=FRY;refGene_tx=XM_023250547.1
A1 11497543 11497693
refGene_gn=FRY;refGene_tx=XM_006927213.4;refGene_gn=FRY;refGene_tx=XM_011286958.3;refGene_gn=FRY;refGene_tx=XM_019825096.1;refGene_gn=FRY;refGene_tx=XM_019825104.1;refGene_gn=FRY;refGene_tx=XM_019825114.2;refGene_gn=FRY;refGene_tx=XM_019825116.2;refGene_gn=FRY;refGene_tx=XM_019825125.2;refGene_gn=FRY;refGene_tx=XM_023250547.1
A1 11498827 11498926
refGene_gn=FRY;refGene_tx=XM_006927213.4;refGene_gn=FRY;refGene_tx=XM_011286958.3;refGene_gn=FRY;refGene_tx=XM_019825096.1;refGene_gn=FRY;refGene_tx=XM_019825104.1;refGene_gn=FRY;refGene_tx=XM_019825114.2;refGene_gn=FRY;refGene_tx=XM_019825116.2;refGene_gn=FRY;refGene_tx=XM_019825125.2;refGene_gn=FRY;refGene_tx=XM_023250547.1
A1 11500792 11500876
refGene_gn=FRY;refGene_tx=XM_006927213.4;refGene_gn=FRY;refGene_tx=XM_011286958.3;refGene_gn=FRY;refGene_tx=XM_019825096.1;refGene_gn=FRY;refGene_tx=XM_019825104.1;refGene_gn=FRY;refGene_tx=XM_019825114.2;refGene_gn=FRY;refGene_tx=XM_019825116.2;refGene_gn=FRY;refGene_tx=XM_019825125.2;refGene_gn=FRY;refGene_tx=XM_023250547.1
A1 11509352 11509370
refGene_gn=FRY;refGene_tx=XM_006927213.4;refGene_gn=FRY;refGene_tx=XM_011286958.3;refGene_gn=FRY;refGene_tx=XM_019825096.1;refGene_gn=FRY;refGene_tx=XM_019825116.2;refGene_gn=FRY;refGene_tx=XM_023250547.1
A1 11511230 11511341
refGene_gn=FRY;refGene_tx=XM_006927213.4;refGene_gn=FRY;refGene_tx=XM_011286958.3;refGene_gn=FRY;refGene_tx=XM_019825096.1;refGene_gn=FRY;refGene_tx=XM_019825104.1;refGene_gn=FRY;refGene_tx=XM_019825116.2;refGene_gn=FRY;refGene_tx=XM_019825125.2;refGene_gn=FRY;refGene_tx=XM_023250547.1
A1 11519555 11519758
refGene_gn=FRY;refGene_tx=XM_006927213.4;refGene_gn=FRY;refGene_tx=XM_011286958.3;refGene_gn=FRY;refGene_tx=XM_019825096.1;refGene_gn=FRY;refGene_tx=XM_019825104.1;refGene_gn=FRY;refGene_tx=XM_019825114.2;refGene_gn=FRY;refGene_tx=XM_019825116.2;refGene_gn=FRY;refGene_tx=XM_019825125.2;refGene_gn=FRY;refGene_tx=XM_023250547.1
A1 11520352 11520611
refGene_gn=FRY;refGene_tx=XM_006927213.4;refGene_gn=FRY;refGene_tx=XM_011286958.3;refGene_gn=FRY;refGene_tx=XM_019825096.1;refGene_gn=FRY;refGene_tx=XM_019825104.1;refGene_gn=FRY;refGene_tx=XM_019825114.2;refGene_gn=FRY;refGene_tx=XM_019825116.2;refGene_gn=FRY;refGene_tx=XM_019825125.2;refGene_gn=FRY;refGene_tx=XM_023250547.1
A1 11552973 11553117
refGene_gn=ZAR1L;refGene_tx=XM_004001338.5;refGene_gn=ZAR1L;refGene_tx=XM_011286772.3
A1 11557454 11557529
refGene_gn=ZAR1L;refGene_tx=XM_004001338.5;refGene_gn=ZAR1L;refGene_tx=XM_011286772.3
A1 11559057 11559150 refGene_gn=ZAR1L;refGene_tx=XM_004001338.5

A1 11559902 11560568
refGene_gn=ZAR1L;refGene_tx=XM_004001338.5;refGene_gn=ZAR1L;refGene_tx=XM_011286772.3
A1 11563271 11563338
refGene_gn=BRCA2;refGene_tx=NM_001009858.1;refGene_gn=BRCA2;refGene_tx=XM_011283605.3;refGene_gn=BRCA2;refGene_tx=XM_019810443.2;refGene_gn=BRCA2;refGene_tx=XM_019839322.2;refGene_gn=BRCA2;refGene_tx=XM_019839734.2
A1 11565646 11565892
refGene_gn=BRCA2;refGene_tx=NM_001009858.1;refGene_gn=BRCA2;refGene_tx=XM_011283605.3;refGene_gn=BRCA2;refGene_tx=XM_019810443.2;refGene_gn=BRCA2;refGene_tx=XM_019839322.2;refGene_gn=BRCA2;refGene_tx=XM_019839734.2
A1 11571100 11571209
refGene_gn=BRCA2;refGene_tx=NM_001009858.1;refGene_gn=BRCA2;refGene_tx=XM_011283605.3;refGene_gn=BRCA2;refGene_tx=XM_019810443.2;refGene_gn=BRCA2;refGene_tx=XM_019839322.2;refGene_gn=BRCA2;refGene_tx=XM_019839734.2
A1 11571829 11571876
refGene_gn=BRCA2;refGene_tx=NM_001009858.1;refGene_gn=BRCA2;refGene_tx=XM_011283605.3;refGene_gn=BRCA2;refGene_tx=XM_019810443.2;refGene_gn=BRCA2;refGene_tx=XM_019839322.2;refGene_gn=BRCA2;refGene_tx=XM_019839734.2
A1 11571975 11572016
refGene_gn=BRCA2;refGene_tx=NM_001009858.1;refGene_gn=BRCA2;refGene_tx=XM_011283605.3;refGene_gn=BRCA2;refGene_tx=XM_019810443.2;refGene_gn=BRCA2;refGene_tx=XM_019839322.2;refGene_gn=BRCA2;refGene_tx=XM_019839734.2
A1 11572208 11572320
refGene_gn=BRCA2;refGene_tx=NM_001009858.1;refGene_gn=BRCA2;refGene_tx=XM_011283605.3;refGene_gn=BRCA2;refGene_tx=XM_019810443.2;refGene_gn=BRCA2;refGene_tx=XM_019839322.2;refGene_gn=BRCA2;refGene_tx=XM_019839734.2
A1 11573064 11573114
refGene_gn=BRCA2;refGene_tx=NM_001009858.1;refGene_gn=BRCA2;refGene_tx=XM_011283605.3;refGene_gn=BRCA2;refGene_tx=XM_019810443.2;refGene_gn=BRCA2;refGene_tx=XM_019839322.2;refGene_gn=BRCA2;refGene_tx=XM_019839734.2
A1 11574005 11574117
refGene_gn=BRCA2;refGene_tx=NM_001009858.1;refGene_gn=BRCA2;refGene_tx=XM_011283605.3;refGene_gn=BRCA2;refGene_tx=XM_019810443.2;refGene_gn=BRCA2;refGene_tx=XM_019839322.2;refGene_gn=BRCA2;refGene_tx=XM_019839734.2
A1 11574951 11576070
refGene_gn=BRCA2;refGene_tx=NM_001009858.1;refGene_gn=BRCA2;refGene_tx=XM_011283605.3;refGene_gn=BRCA2;refGene_tx=XM_019810443.2;refGene_gn=BRCA2;refGene_tx=XM_019839322.2;refGene_gn=BRCA2;refGene_tx=XM_019839734.2
A1 11577487 11582281
refGene_gn=BRCA2;refGene_tx=NM_001009858.1;refGene_gn=BRCA2;refGene_tx

=XM_011283605.3;refGene_gn=BRCA2;refGene_tx=XM_019810443.2;refGene_gn=BRCA2;refGene_tx=XM_019839322.2;refGene_gn=BRCA2;refGene_tx=XM_019839734.2
A1	11584239	11584335
refGene_gn=BRCA2;refGene_tx=NM_001009858.1;refGene_gn=BRCA2;refGene_tx=XM_011283605.3;refGene_gn=BRCA2;refGene_tx=XM_019810443.2;refGene_gn=BRCA2;refGene_tx=XM_019839322.2;refGene_gn=BRCA2;refGene_tx=XM_019839734.2
A1	11586027	11586097
refGene_gn=BRCA2;refGene_tx=NM_001009858.1;refGene_gn=BRCA2;refGene_tx=XM_011283605.3;refGene_gn=BRCA2;refGene_tx=XM_019810443.2;refGene_gn=BRCA2;refGene_tx=XM_019839322.2;refGene_gn=BRCA2;refGene_tx=XM_019839734.2
A1	11592475	11592909
refGene_gn=BRCA2;refGene_tx=NM_001009858.1;refGene_gn=BRCA2;refGene_tx=XM_011283605.3;refGene_gn=BRCA2;refGene_tx=XM_019810443.2;refGene_gn=BRCA2;refGene_tx=XM_019839322.2;refGene_gn=BRCA2;refGene_tx=XM_019839734.2
A1	11593358	11593540
refGene_gn=BRCA2;refGene_tx=NM_001009858.1;refGene_gn=BRCA2;refGene_tx=XM_011283605.3;refGene_gn=BRCA2;refGene_tx=XM_019810443.2;refGene_gn=BRCA2;refGene_tx=XM_019839322.2;refGene_gn=BRCA2;refGene_tx=XM_019839734.2
A1	11594838	11595026
refGene_gn=BRCA2;refGene_tx=NM_001009858.1;refGene_gn=BRCA2;refGene_tx=XM_011283605.3;refGene_gn=BRCA2;refGene_tx=XM_019810443.2;refGene_gn=BRCA2;refGene_tx=XM_019839322.2;refGene_gn=BRCA2;refGene_tx=XM_019839734.2
A1	11598358	11598529
refGene_gn=BRCA2;refGene_tx=NM_001009858.1;refGene_gn=BRCA2;refGene_tx=XM_011283605.3;refGene_gn=BRCA2;refGene_tx=XM_019810443.2;refGene_gn=BRCA2;refGene_tx=XM_019839322.2;refGene_gn=BRCA2;refGene_tx=XM_019839734.2
A1	11599036	11599391
refGene_gn=BRCA2;refGene_tx=NM_001009858.1;refGene_gn=BRCA2;refGene_tx=XM_011283605.3;refGene_gn=BRCA2;refGene_tx=XM_019810443.2;refGene_gn=BRCA2;refGene_tx=XM_019839322.2;refGene_gn=BRCA2;refGene_tx=XM_019839734.2
A1	11602846	11603002
refGene_gn=BRCA2;refGene_tx=NM_001009858.1;refGene_gn=BRCA2;refGene_tx=XM_011283605.3;refGene_gn=BRCA2;refGene_tx=XM_019810443.2;refGene_gn=BRCA2;refGene_tx=XM_019839322.2;refGene_gn=BRCA2;refGene_tx=XM_019839734.2
A1	11603105	11603250
refGene_gn=BRCA2;refGene_tx=NM_001009858.1;refGene_gn=BRCA2;refGene_tx=XM_011283605.3;refGene_gn=BRCA2;refGene_tx=XM_019810443.2;refGene_gn=BRCA2;refGene_tx=XM_019839322.2;refGene_gn=BRCA2;refGene_tx=XM_019839734.2
A1	11605051	11605173
refGene_gn=BRCA2;refGene_tx=NM_001009858.1;refGene_gn=BRCA2;refGene_tx

=XM_011283605.3;refGene_gn=BRCA2;refGene_tx=XM_019810443.2;refGene_gn=BRCA2;refGene_tx=XM_019839322.2;refGene_gn=BRCA2;refGene_tx=XM_019839734.2
A1	11607537	11607736	refGene_gn=BRCA2;refGene_tx=NM_001009858.1;refGene_gn=BRCA2;refGene_tx=XM_011283605.3;refGene_gn=BRCA2;refGene_tx=XM_019810443.2;refGene_gn=BRCA2;refGene_tx=XM_019839322.2;refGene_gn=BRCA2;refGene_tx=XM_019839734.2
A1	11607955	11608119	refGene_gn=BRCA2;refGene_tx=NM_001009858.1;refGene_gn=BRCA2;refGene_tx=XM_011283605.3;refGene_gn=BRCA2;refGene_tx=XM_019810443.2;refGene_gn=BRCA2;refGene_tx=XM_019839322.2;refGene_gn=BRCA2;refGene_tx=XM_019839734.2
A1	11608206	11608345	refGene_gn=BRCA2;refGene_tx=NM_001009858.1;refGene_gn=BRCA2;refGene_tx=XM_011283605.3;refGene_gn=BRCA2;refGene_tx=XM_019839322.2;refGene_gn=BRCA2;refGene_tx=XM_019839734.2
A1	11618464	11618709	refGene_gn=BRCA2;refGene_tx=NM_001009858.1;refGene_gn=BRCA2;refGene_tx=XM_011283605.3;refGene_gn=BRCA2;refGene_tx=XM_019810443.2;refGene_gn=BRCA2;refGene_tx=XM_019839322.2;refGene_gn=BRCA2;refGene_tx=XM_019839734.2
A1	11620480	11620627	refGene_gn=BRCA2;refGene_tx=NM_001009858.1;refGene_gn=BRCA2;refGene_tx=XM_011283605.3;refGene_gn=BRCA2;refGene_tx=XM_019839322.2;refGene_gn=BRCA2;refGene_tx=XM_019839734.2
A1	11621547	11622186	refGene_gn=BRCA2;refGene_tx=NM_001009858.1;refGene_gn=BRCA2;refGene_tx=XM_011283605.3;refGene_gn=BRCA2;refGene_tx=XM_019839322.2;refGene_gn=BRCA2;refGene_tx=XM_019839734.2
A1	11625438	11625697	refGene_gn=N4BP2L1;refGene_tx=XM_023250585.1;refGene_gn=N4BP2L1;refGene_tx=XM_023250587.1;refGene_gn=N4BP2L1;refGene_tx=XM_023250588.1
A1	11626730	11626799	refGene_gn=N4BP2L1;refGene_tx=XM_023250587.1;refGene_gn=N4BP2L1;refGene_tx=XM_023250588.1
A1	11626848	11626925	refGene_gn=N4BP2L1;refGene_tx=XM_023250585.1
A1	11628825	11628914	refGene_gn=N4BP2L1;refGene_tx=XM_023250585.1;refGene_gn=N4BP2L1;refGene_tx=XM_023250587.1
A1	11629200	11629328	refGene_gn=N4BP2L1;refGene_tx=XM_023250585.1;refGene_gn=N4BP2L1;refGene_tx=XM_023250587.1;refGene_gn=N4BP2L1;refGene_tx=XM_023250588.1
A1	11655220	11655414	refGene_gn=N4BP2L1;refGene_tx=XM_023250585.1;refGene_gn=N4BP2L1;refGene_tx=XM_023250587.1;refGene_gn=N4BP2L1;refGene_tx=XM_023250588.1
A1	11663403	11663427	refGene_gn=N4BP2L2;refGene_tx=XM_006927220.4
A1	11664466	11664552	refGene_gn=N4BP2L2;refGene_tx=XM_006927220.4
A1	11667174	11668928	refGene_gn=N4BP2L2;refGene_tx=XM_006927220.4;refGene_gn=N4BP2L2;refGen

e_tx=XM_019825150.2
A1 11714997 11715144
refGene_gn=N4BP2L2;refGene_tx=XM_006927220.4;refGene_gn=N4BP2L2;refGene_tx=XM_019825150.2
A1 11718020 11718097
refGene_gn=N4BP2L2;refGene_tx=XM_006927220.4;refGene_gn=N4BP2L2;refGene_tx=XM_019825150.2
A1 11718488 11718577
refGene_gn=N4BP2L2;refGene_tx=XM_006927220.4;refGene_gn=N4BP2L2;refGene_tx=XM_019825150.2
A1 11723137 11723262
refGene_gn=N4BP2L2;refGene_tx=XM_006927220.4;refGene_gn=N4BP2L2;refGene_tx=XM_019825150.2
A1 11732836 11734095
refGene_gn=N4BP2L2;refGene_tx=XM_006927220.4;refGene_gn=N4BP2L2;refGene_tx=XM_019825150.2
A1 11837614 11837722
refGene_gn=PDS5B;refGene_tx=XM_023250592.1;refGene_gn=PDS5B;refGene_tx=XM_023250593.1;refGene_gn=PDS5B;refGene_tx=XM_023250597.1;refGene_gn=PDS5B;refGene_tx=XM_023250601.1;refGene_gn=PDS5B;refGene_tx=XM_023250605.1
A1 11841118 11841322
refGene_gn=PDS5B;refGene_tx=XM_023250592.1;refGene_gn=PDS5B;refGene_tx=XM_023250593.1;refGene_gn=PDS5B;refGene_tx=XM_023250597.1;refGene_gn=PDS5B;refGene_tx=XM_023250601.1;refGene_gn=PDS5B;refGene_tx=XM_023250605.1
A1 11843985 11844072
refGene_gn=PDS5B;refGene_tx=XM_023250592.1;refGene_gn=PDS5B;refGene_tx=XM_023250593.1;refGene_gn=PDS5B;refGene_tx=XM_023250597.1;refGene_gn=PDS5B;refGene_tx=XM_023250601.1;refGene_gn=PDS5B;refGene_tx=XM_023250605.1
A1 11844174 11844272
refGene_gn=PDS5B;refGene_tx=XM_023250592.1;refGene_gn=PDS5B;refGene_tx=XM_023250593.1;refGene_gn=PDS5B;refGene_tx=XM_023250597.1;refGene_gn=PDS5B;refGene_tx=XM_023250601.1;refGene_gn=PDS5B;refGene_tx=XM_023250605.1
A1 11845496 11845623
refGene_gn=PDS5B;refGene_tx=XM_023250592.1;refGene_gn=PDS5B;refGene_tx=XM_023250593.1;refGene_gn=PDS5B;refGene_tx=XM_023250597.1;refGene_gn=PDS5B;refGene_tx=XM_023250601.1;refGene_gn=PDS5B;refGene_tx=XM_023250605.1
A1 11852109 11852190
refGene_gn=PDS5B;refGene_tx=XM_023250592.1;refGene_gn=PDS5B;refGene_tx=XM_023250593.1;refGene_gn=PDS5B;refGene_tx=XM_023250597.1;refGene_gn=PDS5B;refGene_tx=XM_023250601.1;refGene_gn=PDS5B;refGene_tx=XM_023250605.1
A1 11859344 11859485
refGene_gn=PDS5B;refGene_tx=XM_023250592.1;refGene_gn=PDS5B;refGene_tx=XM_023250593.1;refGene_gn=PDS5B;refGene_tx=XM_023250597.1;refGene_gn=PDS5B;refGene_tx=XM_023250601.1;refGene_gn=PDS5B;refGene_tx=XM_0232506

05.1
A1 11861284 11861400
refGene_gn=PDS5B;refGene_tx=XM_023250592.1;refGene_gn=PDS5B;refGene_tx=XM_023250593.1;refGene_gn=PDS5B;refGene_tx=XM_023250597.1;refGene_gn=PDS5B;refGene_tx=XM_023250601.1;refGene_gn=PDS5B;refGene_tx=XM_023250605.1
A1 11864280 11864375
refGene_gn=PDS5B;refGene_tx=XM_023250592.1;refGene_gn=PDS5B;refGene_tx=XM_023250593.1;refGene_gn=PDS5B;refGene_tx=XM_023250597.1;refGene_gn=PDS5B;refGene_tx=XM_023250601.1;refGene_gn=PDS5B;refGene_tx=XM_023250605.1
A1 11875684 11875830
refGene_gn=PDS5B;refGene_tx=XM_023250592.1;refGene_gn=PDS5B;refGene_tx=XM_023250593.1;refGene_gn=PDS5B;refGene_tx=XM_023250597.1;refGene_gn=PDS5B;refGene_tx=XM_023250601.1;refGene_gn=PDS5B;refGene_tx=XM_023250605.1
A1 11877677 11877829
refGene_gn=PDS5B;refGene_tx=XM_023250592.1;refGene_gn=PDS5B;refGene_tx=XM_023250593.1;refGene_gn=PDS5B;refGene_tx=XM_023250597.1;refGene_gn=PDS5B;refGene_tx=XM_023250601.1;refGene_gn=PDS5B;refGene_tx=XM_023250605.1
A1 11879253 11879367
refGene_gn=PDS5B;refGene_tx=XM_023250592.1;refGene_gn=PDS5B;refGene_tx=XM_023250593.1;refGene_gn=PDS5B;refGene_tx=XM_023250597.1;refGene_gn=PDS5B;refGene_tx=XM_023250601.1;refGene_gn=PDS5B;refGene_tx=XM_023250605.1
A1 11887014 11887096
refGene_gn=PDS5B;refGene_tx=XM_023250592.1;refGene_gn=PDS5B;refGene_tx=XM_023250593.1;refGene_gn=PDS5B;refGene_tx=XM_023250597.1;refGene_gn=PDS5B;refGene_tx=XM_023250601.1;refGene_gn=PDS5B;refGene_tx=XM_023250605.1
A1 11890340 11890389
refGene_gn=PDS5B;refGene_tx=XM_023250592.1;refGene_gn=PDS5B;refGene_tx=XM_023250593.1;refGene_gn=PDS5B;refGene_tx=XM_023250597.1;refGene_gn=PDS5B;refGene_tx=XM_023250601.1;refGene_gn=PDS5B;refGene_tx=XM_023250605.1
A1 11892587 11892727
refGene_gn=PDS5B;refGene_tx=XM_023250592.1;refGene_gn=PDS5B;refGene_tx=XM_023250593.1;refGene_gn=PDS5B;refGene_tx=XM_023250597.1;refGene_gn=PDS5B;refGene_tx=XM_023250601.1;refGene_gn=PDS5B;refGene_tx=XM_023250605.1
A1 11894032 11894148
refGene_gn=PDS5B;refGene_tx=XM_023250592.1;refGene_gn=PDS5B;refGene_tx=XM_023250593.1;refGene_gn=PDS5B;refGene_tx=XM_023250597.1;refGene_gn=PDS5B;refGene_tx=XM_023250601.1;refGene_gn=PDS5B;refGene_tx=XM_023250605.1
A1 11898999 11899105
refGene_gn=PDS5B;refGene_tx=XM_023250592.1;refGene_gn=PDS5B;refGene_tx=XM_023250593.1;refGene_gn=PDS5B;refGene_tx=XM_023250597.1;refGene_gn=PDS5B;refGene_tx=XM_0232506

05.1
A1 11901898 11902059
refGene_gn=PDS5B;refGene_tx=XM_023250592.1;refGene_gn=PDS5B;refGene_tx=XM_023250593.1;refGene_gn=PDS5B;refGene_tx=XM_023250597.1;refGene_gn=PDS5B;refGene_tx=XM_023250601.1;refGene_gn=PDS5B;refGene_tx=XM_023250605.1
A1 11915055 11915179
refGene_gn=PDS5B;refGene_tx=XM_023250592.1;refGene_gn=PDS5B;refGene_tx=XM_023250593.1;refGene_gn=PDS5B;refGene_tx=XM_023250597.1;refGene_gn=PDS5B;refGene_tx=XM_023250601.1;refGene_gn=PDS5B;refGene_tx=XM_023250605.1
A1 11917848 11918007
refGene_gn=PDS5B;refGene_tx=XM_023250592.1;refGene_gn=PDS5B;refGene_tx=XM_023250593.1;refGene_gn=PDS5B;refGene_tx=XM_023250597.1;refGene_gn=PDS5B;refGene_tx=XM_023250601.1;refGene_gn=PDS5B;refGene_tx=XM_023250605.1
A1 11929685 11929754
refGene_gn=PDS5B;refGene_tx=XM_023250592.1;refGene_gn=PDS5B;refGene_tx=XM_023250593.1;refGene_gn=PDS5B;refGene_tx=XM_023250597.1;refGene_gn=PDS5B;refGene_tx=XM_023250601.1;refGene_gn=PDS5B;refGene_tx=XM_023250605.1
A1 11931319 11931456
refGene_gn=PDS5B;refGene_tx=XM_023250592.1;refGene_gn=PDS5B;refGene_tx=XM_023250593.1;refGene_gn=PDS5B;refGene_tx=XM_023250597.1;refGene_gn=PDS5B;refGene_tx=XM_023250601.1;refGene_gn=PDS5B;refGene_tx=XM_023250605.1
A1 11934102 11934226
refGene_gn=PDS5B;refGene_tx=XM_023250592.1;refGene_gn=PDS5B;refGene_tx=XM_023250593.1;refGene_gn=PDS5B;refGene_tx=XM_023250597.1;refGene_gn=PDS5B;refGene_tx=XM_023250601.1;refGene_gn=PDS5B;refGene_tx=XM_023250605.1
A1 11940413 11940618
refGene_gn=PDS5B;refGene_tx=XM_023250592.1;refGene_gn=PDS5B;refGene_tx=XM_023250593.1;refGene_gn=PDS5B;refGene_tx=XM_023250597.1;refGene_gn=PDS5B;refGene_tx=XM_023250601.1;refGene_gn=PDS5B;refGene_tx=XM_023250605.1
A1 11943355 11943470
refGene_gn=PDS5B;refGene_tx=XM_023250592.1;refGene_gn=PDS5B;refGene_tx=XM_023250593.1;refGene_gn=PDS5B;refGene_tx=XM_023250597.1;refGene_gn=PDS5B;refGene_tx=XM_023250601.1;refGene_gn=PDS5B;refGene_tx=XM_023250605.1
A1 11945241 11945374
refGene_gn=PDS5B;refGene_tx=XM_023250592.1;refGene_gn=PDS5B;refGene_tx=XM_023250593.1;refGene_gn=PDS5B;refGene_tx=XM_023250597.1;refGene_gn=PDS5B;refGene_tx=XM_023250601.1;refGene_gn=PDS5B;refGene_tx=XM_023250605.1
A1 11948227 11948347
refGene_gn=PDS5B;refGene_tx=XM_023250592.1;refGene_gn=PDS5B;refGene_tx=XM_023250593.1;refGene_gn=PDS5B;refGene_tx=XM_023250597.1;refGene_gn=PDS5B;refGene_tx=XM_023250601.1;refGene_gn=PDS5B;refGene_tx=XM_0232506

05.1
A1	11949598	11949661
refGene_gn=PDS5B;refGene_tx=XM_023250592.1;refGene_gn=PDS5B;refGene_tx
=XM_023250593.1;refGene_gn=PDS5B;refGene_tx=XM_023250597.1;refGene_gn=
PDS5B;refGene_tx=XM_023250601.1;refGene_gn=PDS5B;refGene_tx=XM_0232506
05.1
A1	11950243	11950389
refGene_gn=PDS5B;refGene_tx=XM_023250592.1;refGene_gn=PDS5B;refGene_tx
=XM_023250593.1;refGene_gn=PDS5B;refGene_tx=XM_023250597.1;refGene_gn=
PDS5B;refGene_tx=XM_023250601.1;refGene_gn=PDS5B;refGene_tx=XM_0232506
05.1
A1	11952893	11952999
refGene_gn=PDS5B;refGene_tx=XM_023250592.1;refGene_gn=PDS5B;refGene_tx
=XM_023250593.1;refGene_gn=PDS5B;refGene_tx=XM_023250597.1;refGene_gn=
PDS5B;refGene_tx=XM_023250601.1;refGene_gn=PDS5B;refGene_tx=XM_0232506
05.1
A1	11957608	11958048
refGene_gn=PDS5B;refGene_tx=XM_023250592.1;refGene_gn=PDS5B;refGene_tx
=XM_023250593.1;refGene_gn=PDS5B;refGene_tx=XM_023250597.1;refGene_gn=
PDS5B;refGene_tx=XM_023250601.1;refGene_gn=PDS5B;refGene_tx=XM_0232506
05.1
A1	11958138	11958249
refGene_gn=PDS5B;refGene_tx=XM_023250592.1;refGene_gn=PDS5B;refGene_tx
=XM_023250593.1;refGene_gn=PDS5B;refGene_tx=XM_023250597.1;refGene_gn=
PDS5B;refGene_tx=XM_023250601.1
A1	11961342	11961481
refGene_gn=PDS5B;refGene_tx=XM_023250592.1;refGene_gn=PDS5B;refGene_tx
=XM_023250593.1;refGene_gn=PDS5B;refGene_tx=XM_023250597.1;refGene_gn=
PDS5B;refGene_tx=XM_023250601.1;refGene_gn=PDS5B;refGene_tx=XM_0232506
05.1
A1	11962913	11962949
refGene_gn=PDS5B;refGene_tx=XM_023250592.1;refGene_gn=PDS5B;refGene_tx
=XM_023250593.1;refGene_gn=PDS5B;refGene_tx=XM_023250597.1;refGene_gn=
PDS5B;refGene_tx=XM_023250601.1;refGene_gn=PDS5B;refGene_tx=XM_0232506
05.1
A1	12168486	12169311	refGene_gn=KL;refGene_tx=XM_019825242.2
A1	12205483	12205994	refGene_gn=KL;refGene_tx=XM_019825242.2
A1	12206737	12207006	refGene_gn=KL;refGene_tx=XM_019825242.2
A1	12212504	12213606	refGene_gn=KL;refGene_tx=XM_019825242.2
A1	12214165	12214503	refGene_gn=KL;refGene_tx=XM_019825242.2
A1	12245350	12245480	refGene_gn=STARD13;refGene_tx=XM_006927222.3
A1	12248143	12248261
refGene_gn=STARD13;refGene_tx=XM_003980317.4;refGene_gn=STARD13;refGen
e_tx=XM_003980318.4;refGene_gn=STARD13;refGene_tx=XM_023250644.1
A1	12248546	12248736
refGene_gn=STARD13;refGene_tx=XM_019825255.1;refGene_gn=STARD13;refGen
e_tx=XM_019825258.1;refGene_gn=STARD13;refGene_tx=XM_023250634.1;refGe
ne_gn=STARD13;refGene_tx=XM_023250643.1
A1	12248980	12249157
refGene_gn=STARD13;refGene_tx=XM_003980317.4;refGene_gn=STARD13;refGen

e_tx=XM_003980318.4;refGene_gn=STARD13;refGene_tx=XM_006927222.3;refGene_gn=STARD13;refGene_tx=XM_019825255.1;refGene_gn=STARD13;refGene_tx=XM_019825258.1;refGene_gn=STARD13;refGene_tx=XM_023250634.1;refGene_gn=STARD13;refGene_tx=XM_023250643.1;refGene_gn=STARD13;refGene_tx=XM_023250644.1
A1 12252623 12252841
refGene_gn=STARD13;refGene_tx=XM_003980317.4;refGene_gn=STARD13;refGene_tx=XM_003980318.4;refGene_gn=STARD13;refGene_tx=XM_006927222.3;refGene_gn=STARD13;refGene_tx=XM_019825255.1;refGene_gn=STARD13;refGene_tx=XM_019825258.1;refGene_gn=STARD13;refGene_tx=XM_023250634.1;refGene_gn=STARD13;refGene_tx=XM_023250643.1;refGene_gn=STARD13;refGene_tx=XM_023250644.1
A1 12253836 12254058
refGene_gn=STARD13;refGene_tx=XM_003980317.4;refGene_gn=STARD13;refGene_tx=XM_003980318.4;refGene_gn=STARD13;refGene_tx=XM_006927222.3;refGene_gn=STARD13;refGene_tx=XM_019825255.1;refGene_gn=STARD13;refGene_tx=XM_019825258.1;refGene_gn=STARD13;refGene_tx=XM_023250634.1;refGene_gn=STARD13;refGene_tx=XM_023250643.1;refGene_gn=STARD13;refGene_tx=XM_023250644.1
A1 12254881 12254996
refGene_gn=STARD13;refGene_tx=XM_003980317.4;refGene_gn=STARD13;refGene_tx=XM_003980318.4;refGene_gn=STARD13;refGene_tx=XM_006927222.3;refGene_gn=STARD13;refGene_tx=XM_019825255.1;refGene_gn=STARD13;refGene_tx=XM_019825258.1;refGene_gn=STARD13;refGene_tx=XM_023250634.1;refGene_gn=STARD13;refGene_tx=XM_023250643.1;refGene_gn=STARD13;refGene_tx=XM_023250644.1
A1 12255822 12256033
refGene_gn=STARD13;refGene_tx=XM_003980317.4;refGene_gn=STARD13;refGene_tx=XM_003980318.4;refGene_gn=STARD13;refGene_tx=XM_006927222.3;refGene_gn=STARD13;refGene_tx=XM_019825255.1;refGene_gn=STARD13;refGene_tx=XM_019825258.1;refGene_gn=STARD13;refGene_tx=XM_023250634.1;refGene_gn=STARD13;refGene_tx=XM_023250643.1;refGene_gn=STARD13;refGene_tx=XM_023250644.1
A1 12261003 12261202
refGene_gn=STARD13;refGene_tx=XM_003980317.4;refGene_gn=STARD13;refGene_tx=XM_003980318.4;refGene_gn=STARD13;refGene_tx=XM_006927222.3;refGene_gn=STARD13;refGene_tx=XM_019825255.1;refGene_gn=STARD13;refGene_tx=XM_019825258.1;refGene_gn=STARD13;refGene_tx=XM_023250634.1;refGene_gn=STARD13;refGene_tx=XM_023250643.1;refGene_gn=STARD13;refGene_tx=XM_023250644.1
A1 12268777 12268994
refGene_gn=STARD13;refGene_tx=XM_003980317.4;refGene_gn=STARD13;refGene_tx=XM_003980318.4;refGene_gn=STARD13;refGene_tx=XM_006927222.3;refGene_gn=STARD13;refGene_tx=XM_019825255.1;refGene_gn=STARD13;refGene_tx=XM_019825258.1;refGene_gn=STARD13;refGene_tx=XM_023250634.1;refGene_gn=STARD13;refGene_tx=XM_023250643.1;refGene_gn=STARD13;refGene_tx=XM_023250644.1
A1 12270065 12270236
refGene_gn=STARD13;refGene_tx=XM_003980317.4;refGene_gn=STARD13;refGene_tx=XM_003980318.4;refGene_gn=STARD13;refGene_tx=XM_006927222.3;refGe

ne_gn=STARD13;refGene_tx=XM_019825255.1;refGene_gn=STARD13;refGene_tx=XM_019825258.1;refGene_gn=STARD13;refGene_tx=XM_023250634.1;refGene_gn=STARD13;refGene_tx=XM_023250643.1;refGene_gn=STARD13;refGene_tx=XM_023250644.1
A1 12271610 12273004
refGene_gn=STARD13;refGene_tx=XM_003980317.4;refGene_gn=STARD13;refGene_tx=XM_003980318.4;refGene_gn=STARD13;refGene_tx=XM_006927222.3;refGene_gn=STARD13;refGene_tx=XM_019825255.1;refGene_gn=STARD13;refGene_tx=XM_019825258.1;refGene_gn=STARD13;refGene_tx=XM_023250634.1;refGene_gn=STARD13;refGene_tx=XM_023250643.1;refGene_gn=STARD13;refGene_tx=XM_023250644.1
A1 12282323 12282387
refGene_gn=STARD13;refGene_tx=XM_003980317.4;refGene_gn=STARD13;refGene_tx=XM_003980318.4;refGene_gn=STARD13;refGene_tx=XM_006927222.3;refGene_gn=STARD13;refGene_tx=XM_019825255.1;refGene_gn=STARD13;refGene_tx=XM_019825258.1;refGene_gn=STARD13;refGene_tx=XM_023250634.1;refGene_gn=STARD13;refGene_tx=XM_023250643.1;refGene_gn=STARD13;refGene_tx=XM_023250644.1
A1 12304182 12304264
refGene_gn=STARD13;refGene_tx=XM_003980317.4;refGene_gn=STARD13;refGene_tx=XM_003980318.4;refGene_gn=STARD13;refGene_tx=XM_006927222.3;refGene_gn=STARD13;refGene_tx=XM_019825255.1;refGene_gn=STARD13;refGene_tx=XM_019825258.1;refGene_gn=STARD13;refGene_tx=XM_023250634.1;refGene_gn=STARD13;refGene_tx=XM_023250643.1;refGene_gn=STARD13;refGene_tx=XM_023250644.1
A1 12307858 12307930
refGene_gn=STARD13;refGene_tx=XM_003980317.4;refGene_gn=STARD13;refGene_tx=XM_003980318.4;refGene_gn=STARD13;refGene_tx=XM_006927222.3;refGene_gn=STARD13;refGene_tx=XM_019825255.1;refGene_gn=STARD13;refGene_tx=XM_019825258.1;refGene_gn=STARD13;refGene_tx=XM_023250634.1;refGene_gn=STARD13;refGene_tx=XM_023250643.1;refGene_gn=STARD13;refGene_tx=XM_023250644.1
A1 12325780 12325844 refGene_gn=STARD13;refGene_tx=XM_023250643.1
A1 12427559 12427728
refGene_gn=STARD13;refGene_tx=XM_003980317.4;refGene_gn=STARD13;refGene_tx=XM_003980318.4;refGene_gn=STARD13;refGene_tx=XM_006927222.3;refGene_gn=STARD13;refGene_tx=XM_019825255.1;refGene_gn=STARD13;refGene_tx=XM_019825258.1
A1 12482591 12482715
refGene_gn=STARD13;refGene_tx=XM_023250634.1;refGene_gn=STARD13;refGene_tx=XM_023250644.1
A1 12912196 12912283
refGene_gn=RFC3;refGene_tx=XM_023250694.1;refGene_gn=RFC3;refGene_tx=XM_023250701.1
A1 12915098 12915236
refGene_gn=RFC3;refGene_tx=XM_023250694.1;refGene_gn=RFC3;refGene_tx=XM_023250701.1
A1 12918371 12918439
refGene_gn=RFC3;refGene_tx=XM_023250694.1;refGene_gn=RFC3;refGene_tx=XM_023250701.1

A1 12920696 12920794
refGene_gn=RFC3;refGene_tx=XM_023250694.1;refGene_gn=RFC3;refGene_tx=XM_023250701.1
A1 12924030 12924212
refGene_gn=RFC3;refGene_tx=XM_023250694.1;refGene_gn=RFC3;refGene_tx=XM_023250701.1
A1 12925230 12925367
refGene_gn=RFC3;refGene_tx=XM_023250694.1;refGene_gn=RFC3;refGene_tx=XM_023250701.1
A1 12925452 12925551
refGene_gn=RFC3;refGene_tx=XM_023250694.1;refGene_gn=RFC3;refGene_tx=XM_023250701.1
A1 12935478 12935548
refGene_gn=RFC3;refGene_tx=XM_023250694.1;refGene_gn=RFC3;refGene_tx=XM_023250701.1
A1 12935637 12935653 refGene_gn=RFC3;refGene_tx=XM_023250701.1
A1 12937078 12937270
refGene_gn=RFC3;refGene_tx=XM_023250694.1;refGene_gn=RFC3;refGene_tx=XM_023250701.1
A1 13904732 13905032
refGene_gn=NBEA;refGene_tx=XM_011286341.3;refGene_gn=NBEA;refGene_tx=XM_019825371.2;refGene_gn=NBEA;refGene_tx=XM_019825374.2;refGene_gn=NBEA;refGene_tx=XM_019825380.2;refGene_gn=NBEA;refGene_tx=XM_023250717.1;refGene_gn=NBEA;refGene_tx=XM_023250721.1;refGene_gn=NBEA;refGene_tx=XM_023250724.1;refGene_gn=NBEA;refGene_tx=XM_023250731.1;refGene_gn=NBEA;refGene_tx=XM_023250733.1
A1 13937131 13937313
refGene_gn=LOC101083390;refGene_tx=XM_019825389.2
A1 13937315 13937436
refGene_gn=LOC101083390;refGene_tx=XM_019825389.2
A1 13982209 13982441
refGene_gn=NBEA;refGene_tx=XM_011286341.3;refGene_gn=NBEA;refGene_tx=XM_019825371.2;refGene_gn=NBEA;refGene_tx=XM_019825374.2;refGene_gn=NBEA;refGene_tx=XM_019825380.2;refGene_gn=NBEA;refGene_tx=XM_023250717.1;refGene_gn=NBEA;refGene_tx=XM_023250721.1;refGene_gn=NBEA;refGene_tx=XM_023250724.1;refGene_gn=NBEA;refGene_tx=XM_023250731.1;refGene_gn=NBEA;refGene_tx=XM_023250733.1
A1 13985845 13985946
refGene_gn=NBEA;refGene_tx=XM_011286341.3;refGene_gn=NBEA;refGene_tx=XM_019825371.2;refGene_gn=NBEA;refGene_tx=XM_019825374.2;refGene_gn=NBEA;refGene_tx=XM_019825380.2;refGene_gn=NBEA;refGene_tx=XM_023250717.1;refGene_gn=NBEA;refGene_tx=XM_023250721.1;refGene_gn=NBEA;refGene_tx=XM_023250724.1;refGene_gn=NBEA;refGene_tx=XM_023250731.1;refGene_gn=NBEA;refGene_tx=XM_023250733.1
A1 13986185 13986281
refGene_gn=NBEA;refGene_tx=XM_011286341.3;refGene_gn=NBEA;refGene_tx=XM_019825371.2;refGene_gn=NBEA;refGene_tx=XM_019825374.2;refGene_gn=NBEA;refGene_tx=XM_019825380.2;refGene_gn=NBEA;refGene_tx=XM_023250717.1;refGene_gn=NBEA;refGene_tx=XM_023250721.1;refGene_gn=NBEA;refGene_tx=XM_023250724.1;refGene_gn=NBEA;refGene_tx=XM_023250731.1;refGene_gn=NBE

A;refGene_tx=XM_023250733.1
A1 13992036 13992158
refGene_gn=NBEA;refGene_tx=XM_011286341.3;refGene_gn=NBEA;refGene_tx=X
M_019825371.2;refGene_gn=NBEA;refGene_tx=XM_019825374.2;refGene_gn=NBE
A;refGene_tx=XM_019825380.2;refGene_gn=NBEA;refGene_tx=XM_023250717.1;
refGene_gn=NBEA;refGene_tx=XM_023250721.1;refGene_gn=NBEA;refGene_tx=X
M_023250724.1;refGene_gn=NBEA;refGene_tx=XM_023250731.1;refGene_gn=NBE
A;refGene_tx=XM_023250733.1
A1 13994236 13994363
refGene_gn=NBEA;refGene_tx=XM_011286341.3;refGene_gn=NBEA;refGene_tx=X
M_019825371.2;refGene_gn=NBEA;refGene_tx=XM_019825374.2;refGene_gn=NBE
A;refGene_tx=XM_019825380.2;refGene_gn=NBEA;refGene_tx=XM_023250717.1;
refGene_gn=NBEA;refGene_tx=XM_023250721.1;refGene_gn=NBEA;refGene_tx=X
M_023250724.1;refGene_gn=NBEA;refGene_tx=XM_023250731.1;refGene_gn=NBE
A;refGene_tx=XM_023250733.1
A1 14003623 14003743
refGene_gn=NBEA;refGene_tx=XM_011286341.3;refGene_gn=NBEA;refGene_tx=X
M_019825371.2;refGene_gn=NBEA;refGene_tx=XM_019825374.2;refGene_gn=NBE
A;refGene_tx=XM_019825380.2;refGene_gn=NBEA;refGene_tx=XM_023250717.1;
refGene_gn=NBEA;refGene_tx=XM_023250721.1;refGene_gn=NBEA;refGene_tx=X
M_023250724.1;refGene_gn=NBEA;refGene_tx=XM_023250731.1;refGene_gn=NBE
A;refGene_tx=XM_023250733.1
A1 14006668 14006815
refGene_gn=NBEA;refGene_tx=XM_011286341.3;refGene_gn=NBEA;refGene_tx=X
M_019825371.2;refGene_gn=NBEA;refGene_tx=XM_019825374.2;refGene_gn=NBE
A;refGene_tx=XM_019825380.2;refGene_gn=NBEA;refGene_tx=XM_023250717.1;
refGene_gn=NBEA;refGene_tx=XM_023250721.1;refGene_gn=NBEA;refGene_tx=X
M_023250724.1;refGene_gn=NBEA;refGene_tx=XM_023250731.1;refGene_gn=NBE
A;refGene_tx=XM_023250733.1
A1 14035677 14035875
refGene_gn=NBEA;refGene_tx=XM_011286341.3;refGene_gn=NBEA;refGene_tx=X
M_019825371.2;refGene_gn=NBEA;refGene_tx=XM_019825374.2;refGene_gn=NBE
A;refGene_tx=XM_019825380.2;refGene_gn=NBEA;refGene_tx=XM_023250717.1;
refGene_gn=NBEA;refGene_tx=XM_023250721.1;refGene_gn=NBEA;refGene_tx=X
M_023250724.1;refGene_gn=NBEA;refGene_tx=XM_023250731.1;refGene_gn=NBE
A;refGene_tx=XM_023250733.1
A1 14036509 14036643
refGene_gn=NBEA;refGene_tx=XM_011286341.3;refGene_gn=NBEA;refGene_tx=X
M_019825371.2;refGene_gn=NBEA;refGene_tx=XM_019825374.2;refGene_gn=NBE
A;refGene_tx=XM_019825380.2;refGene_gn=NBEA;refGene_tx=XM_023250717.1;
refGene_gn=NBEA;refGene_tx=XM_023250721.1;refGene_gn=NBEA;refGene_tx=X
M_023250724.1;refGene_gn=NBEA;refGene_tx=XM_023250731.1;refGene_gn=NBE
A;refGene_tx=XM_023250733.1;refGene_gn=NBEA;refGene_tx=XM_023250742.1
A1 14056958 14057064 refGene_gn=NBEA;refGene_tx=XM_023250742.1
A1 14059255 14059364
refGene_gn=NBEA;refGene_tx=XM_011286341.3;refGene_gn=NBEA;refGene_tx=X
M_019825371.2;refGene_gn=NBEA;refGene_tx=XM_019825374.2;refGene_gn=NBE
A;refGene_tx=XM_019825380.2;refGene_gn=NBEA;refGene_tx=XM_023250717.1;
refGene_gn=NBEA;refGene_tx=XM_023250721.1;refGene_gn=NBEA;refGene_tx=X
M_023250724.1;refGene_gn=NBEA;refGene_tx=XM_023250731.1;refGene_gn=NBE

A;refGene_tx=XM_023250733.1;refGene_gn=NBEA;refGene_tx=XM_023250742.1
A1 14070641 14070794
refGene_gn=NBEA;refGene_tx=XM_011286341.3;refGene_gn=NBEA;refGene_tx=XM_019825371.2;refGene_gn=NBEA;refGene_tx=XM_019825374.2;refGene_gn=NBEA;refGene_tx=XM_019825380.2;refGene_gn=NBEA;refGene_tx=XM_023250717.1;refGene_gn=NBEA;refGene_tx=XM_023250721.1;refGene_gn=NBEA;refGene_tx=XM_023250724.1;refGene_gn=NBEA;refGene_tx=XM_023250731.1;refGene_gn=NBEA;refGene_tx=XM_023250733.1;refGene_gn=NBEA;refGene_tx=XM_023250742.1
A1 14071957 14072126
refGene_gn=NBEA;refGene_tx=XM_011286341.3;refGene_gn=NBEA;refGene_tx=XM_019825371.2;refGene_gn=NBEA;refGene_tx=XM_019825374.2;refGene_gn=NBEA;refGene_tx=XM_019825380.2;refGene_gn=NBEA;refGene_tx=XM_023250717.1;refGene_gn=NBEA;refGene_tx=XM_023250721.1;refGene_gn=NBEA;refGene_tx=XM_023250724.1;refGene_gn=NBEA;refGene_tx=XM_023250731.1;refGene_gn=NBEA;refGene_tx=XM_023250733.1;refGene_gn=NBEA;refGene_tx=XM_023250742.1
A1 14078643 14078723
refGene_gn=NBEA;refGene_tx=XM_011286341.3;refGene_gn=NBEA;refGene_tx=XM_019825371.2;refGene_gn=NBEA;refGene_tx=XM_019825374.2;refGene_gn=NBEA;refGene_tx=XM_019825380.2;refGene_gn=NBEA;refGene_tx=XM_023250717.1;refGene_gn=NBEA;refGene_tx=XM_023250721.1;refGene_gn=NBEA;refGene_tx=XM_023250724.1;refGene_gn=NBEA;refGene_tx=XM_023250731.1;refGene_gn=NBEA;refGene_tx=XM_023250733.1;refGene_gn=NBEA;refGene_tx=XM_023250742.1
A1 14079442 14079505
refGene_gn=NBEA;refGene_tx=XM_011286341.3;refGene_gn=NBEA;refGene_tx=XM_019825371.2;refGene_gn=NBEA;refGene_tx=XM_019825374.2;refGene_gn=NBEA;refGene_tx=XM_019825380.2;refGene_gn=NBEA;refGene_tx=XM_023250717.1;refGene_gn=NBEA;refGene_tx=XM_023250721.1;refGene_gn=NBEA;refGene_tx=XM_023250724.1;refGene_gn=NBEA;refGene_tx=XM_023250731.1;refGene_gn=NBEA;refGene_tx=XM_023250733.1;refGene_gn=NBEA;refGene_tx=XM_023250742.1
A1 14079597 14079695
refGene_gn=NBEA;refGene_tx=XM_011286341.3;refGene_gn=NBEA;refGene_tx=XM_019825371.2;refGene_gn=NBEA;refGene_tx=XM_019825374.2;refGene_gn=NBEA;refGene_tx=XM_019825380.2;refGene_gn=NBEA;refGene_tx=XM_023250717.1;refGene_gn=NBEA;refGene_tx=XM_023250721.1;refGene_gn=NBEA;refGene_tx=XM_023250724.1;refGene_gn=NBEA;refGene_tx=XM_023250731.1;refGene_gn=NBEA;refGene_tx=XM_023250733.1;refGene_gn=NBEA;refGene_tx=XM_023250742.1
A1 14083419 14083512
refGene_gn=NBEA;refGene_tx=XM_011286341.3;refGene_gn=NBEA;refGene_tx=XM_019825371.2;refGene_gn=NBEA;refGene_tx=XM_019825374.2;refGene_gn=NBEA;refGene_tx=XM_019825380.2;refGene_gn=NBEA;refGene_tx=XM_023250717.1;refGene_gn=NBEA;refGene_tx=XM_023250721.1;refGene_gn=NBEA;refGene_tx=XM_023250724.1;refGene_gn=NBEA;refGene_tx=XM_023250731.1;refGene_gn=NBEA;refGene_tx=XM_023250733.1;refGene_gn=NBEA;refGene_tx=XM_023250742.1
A1 14098330 14098439
refGene_gn=NBEA;refGene_tx=XM_011286341.3;refGene_gn=NBEA;refGene_tx=XM_019825371.2;refGene_gn=NBEA;refGene_tx=XM_019825374.2;refGene_gn=NBEA;refGene_tx=XM_019825380.2;refGene_gn=NBEA;refGene_tx=XM_023250717.1;refGene_gn=NBEA;refGene_tx=XM_023250721.1;refGene_gn=NBEA;refGene_tx=XM_023250724.1;refGene_gn=NBEA;refGene_tx=XM_023250731.1;refGene_gn=NBEA;refGene_tx=XM_023250733.1;refGene_gn=NBEA;refGene_tx=XM_023250742.1

A1 14107105 14107187
refGene_gn=NBEA;refGene_tx=XM_011286341.3;refGene_gn=NBEA;refGene_tx=XM_019825371.2;refGene_gn=NBEA;refGene_tx=XM_019825374.2;refGene_gn=NBEA;refGene_tx=XM_019825380.2;refGene_gn=NBEA;refGene_tx=XM_023250717.1;refGene_gn=NBEA;refGene_tx=XM_023250721.1;refGene_gn=NBEA;refGene_tx=XM_023250724.1;refGene_gn=NBEA;refGene_tx=XM_023250731.1;refGene_gn=NBEA;refGene_tx=XM_023250733.1;refGene_gn=NBEA;refGene_tx=XM_023250742.1
A1 14108109 14108233
refGene_gn=NBEA;refGene_tx=XM_011286341.3;refGene_gn=NBEA;refGene_tx=XM_019825371.2;refGene_gn=NBEA;refGene_tx=XM_019825374.2;refGene_gn=NBEA;refGene_tx=XM_019825380.2;refGene_gn=NBEA;refGene_tx=XM_023250717.1;refGene_gn=NBEA;refGene_tx=XM_023250721.1;refGene_gn=NBEA;refGene_tx=XM_023250724.1;refGene_gn=NBEA;refGene_tx=XM_023250731.1;refGene_gn=NBEA;refGene_tx=XM_023250733.1;refGene_gn=NBEA;refGene_tx=XM_023250742.1
A1 14109065 14109258
refGene_gn=NBEA;refGene_tx=XM_011286341.3;refGene_gn=NBEA;refGene_tx=XM_019825371.2;refGene_gn=NBEA;refGene_tx=XM_019825374.2;refGene_gn=NBEA;refGene_tx=XM_019825380.2;refGene_gn=NBEA;refGene_tx=XM_023250717.1;refGene_gn=NBEA;refGene_tx=XM_023250721.1;refGene_gn=NBEA;refGene_tx=XM_023250724.1;refGene_gn=NBEA;refGene_tx=XM_023250731.1;refGene_gn=NBEA;refGene_tx=XM_023250733.1;refGene_gn=NBEA;refGene_tx=XM_023250742.1
A1 14111043 14112060
refGene_gn=NBEA;refGene_tx=XM_011286341.3;refGene_gn=NBEA;refGene_tx=XM_019825371.2;refGene_gn=NBEA;refGene_tx=XM_019825374.2;refGene_gn=NBEA;refGene_tx=XM_019825380.2;refGene_gn=NBEA;refGene_tx=XM_023250717.1;refGene_gn=NBEA;refGene_tx=XM_023250721.1;refGene_gn=NBEA;refGene_tx=XM_023250724.1;refGene_gn=NBEA;refGene_tx=XM_023250731.1;refGene_gn=NBEA;refGene_tx=XM_023250733.1;refGene_gn=NBEA;refGene_tx=XM_023250742.1
A1 14113786 14114004
refGene_gn=NBEA;refGene_tx=XM_011286341.3;refGene_gn=NBEA;refGene_tx=XM_019825371.2;refGene_gn=NBEA;refGene_tx=XM_019825374.2;refGene_gn=NBEA;refGene_tx=XM_019825380.2;refGene_gn=NBEA;refGene_tx=XM_023250717.1;refGene_gn=NBEA;refGene_tx=XM_023250721.1;refGene_gn=NBEA;refGene_tx=XM_023250724.1;refGene_gn=NBEA;refGene_tx=XM_023250731.1;refGene_gn=NBEA;refGene_tx=XM_023250733.1;refGene_gn=NBEA;refGene_tx=XM_023250742.1
A1 14116543 14116697
refGene_gn=NBEA;refGene_tx=XM_011286341.3;refGene_gn=NBEA;refGene_tx=XM_019825371.2;refGene_gn=NBEA;refGene_tx=XM_019825374.2;refGene_gn=NBEA;refGene_tx=XM_019825380.2;refGene_gn=NBEA;refGene_tx=XM_023250717.1;refGene_gn=NBEA;refGene_tx=XM_023250721.1;refGene_gn=NBEA;refGene_tx=XM_023250724.1;refGene_gn=NBEA;refGene_tx=XM_023250731.1;refGene_gn=NBEA;refGene_tx=XM_023250733.1;refGene_gn=NBEA;refGene_tx=XM_023250742.1
A1 14122619 14122628
refGene_gn=NBEA;refGene_tx=XM_011286341.3;refGene_gn=NBEA;refGene_tx=XM_019825374.2;refGene_gn=NBEA;refGene_tx=XM_019825380.2;refGene_gn=NBEA;refGene_tx=XM_023250717.1;refGene_gn=NBEA;refGene_tx=XM_023250724.1;refGene_gn=NBEA;refGene_tx=XM_023250731.1;refGene_gn=NBEA;refGene_tx=XM_023250733.1;refGene_gn=NBEA;refGene_tx=XM_023250742.1
A1 14124952 14125133
refGene_gn=NBEA;refGene_tx=XM_011286341.3;refGene_gn=NBEA;refGene_tx=X

M_019825371.2;refGene_gn=NBEA;refGene_tx=XM_019825374.2;refGene_gn=NBE
A;refGene_tx=XM_019825380.2;refGene_gn=NBEA;refGene_tx=XM_023250717.1;
refGene_gn=NBEA;refGene_tx=XM_023250721.1;refGene_gn=NBEA;refGene_tx=X
M_023250724.1;refGene_gn=NBEA;refGene_tx=XM_023250731.1;refGene_gn=NBE
A;refGene_tx=XM_023250733.1;refGene_gn=NBEA;refGene_tx=XM_023250742.1
A1 14127343 14127474
refGene_gn=NBEA;refGene_tx=XM_011286341.3;refGene_gn=NBEA;refGene_tx=X
M_019825371.2;refGene_gn=NBEA;refGene_tx=XM_019825374.2;refGene_gn=NBE
A;refGene_tx=XM_019825380.2;refGene_gn=NBEA;refGene_tx=XM_023250717.1;
refGene_gn=NBEA;refGene_tx=XM_023250721.1;refGene_gn=NBEA;refGene_tx=X
M_023250724.1;refGene_gn=NBEA;refGene_tx=XM_023250731.1;refGene_gn=NBE
A;refGene_tx=XM_023250733.1;refGene_gn=NBEA;refGene_tx=XM_023250742.1
A1 14131551 14131659
refGene_gn=NBEA;refGene_tx=XM_011286341.3;refGene_gn=NBEA;refGene_tx=X
M_019825371.2;refGene_gn=NBEA;refGene_tx=XM_019825374.2;refGene_gn=NBE
A;refGene_tx=XM_019825380.2;refGene_gn=NBEA;refGene_tx=XM_023250717.1;
refGene_gn=NBEA;refGene_tx=XM_023250721.1;refGene_gn=NBEA;refGene_tx=X
M_023250724.1;refGene_gn=NBEA;refGene_tx=XM_023250731.1;refGene_gn=NBE
A;refGene_tx=XM_023250733.1;refGene_gn=NBEA;refGene_tx=XM_023250742.1
A1 14136741 14136910
refGene_gn=NBEA;refGene_tx=XM_011286341.3;refGene_gn=NBEA;refGene_tx=X
M_019825371.2;refGene_gn=NBEA;refGene_tx=XM_019825374.2;refGene_gn=NBE
A;refGene_tx=XM_019825380.2;refGene_gn=NBEA;refGene_tx=XM_023250717.1;
refGene_gn=NBEA;refGene_tx=XM_023250721.1;refGene_gn=NBEA;refGene_tx=X
M_023250724.1;refGene_gn=NBEA;refGene_tx=XM_023250731.1;refGene_gn=NBE
A;refGene_tx=XM_023250733.1;refGene_gn=NBEA;refGene_tx=XM_023250742.1
A1 14138352 14138448
refGene_gn=NBEA;refGene_tx=XM_011286341.3;refGene_gn=NBEA;refGene_tx=X
M_019825371.2;refGene_gn=NBEA;refGene_tx=XM_019825374.2;refGene_gn=NBE
A;refGene_tx=XM_023250717.1;refGene_gn=NBEA;refGene_tx=XM_023250721.1;
refGene_gn=NBEA;refGene_tx=XM_023250724.1;refGene_gn=NBEA;refGene_tx=X
M_023250731.1;refGene_gn=NBEA;refGene_tx=XM_023250742.1
A1 14155448 14155887
refGene_gn=NBEA;refGene_tx=XM_011286341.3;refGene_gn=NBEA;refGene_tx=X
M_019825371.2;refGene_gn=NBEA;refGene_tx=XM_019825374.2;refGene_gn=NBE
A;refGene_tx=XM_019825380.2;refGene_gn=NBEA;refGene_tx=XM_023250717.1;
refGene_gn=NBEA;refGene_tx=XM_023250721.1;refGene_gn=NBEA;refGene_tx=X
M_023250724.1;refGene_gn=NBEA;refGene_tx=XM_023250731.1;refGene_gn=NBE
A;refGene_tx=XM_023250733.1;refGene_gn=NBEA;refGene_tx=XM_023250742.1
A1 14167264 14167419
refGene_gn=NBEA;refGene_tx=XM_011286341.3;refGene_gn=NBEA;refGene_tx=X
M_019825371.2;refGene_gn=NBEA;refGene_tx=XM_019825374.2;refGene_gn=NBE
A;refGene_tx=XM_019825380.2;refGene_gn=NBEA;refGene_tx=XM_023250717.1;
refGene_gn=NBEA;refGene_tx=XM_023250721.1;refGene_gn=NBEA;refGene_tx=X
M_023250724.1;refGene_gn=NBEA;refGene_tx=XM_023250731.1;refGene_gn=NBE
A;refGene_tx=XM_023250733.1;refGene_gn=NBEA;refGene_tx=XM_023250742.1
A1 14169659 14169786
refGene_gn=NBEA;refGene_tx=XM_011286341.3;refGene_gn=NBEA;refGene_tx=X
M_019825371.2;refGene_gn=NBEA;refGene_tx=XM_019825374.2;refGene_gn=NBE
A;refGene_tx=XM_019825380.2;refGene_gn=NBEA;refGene_tx=XM_023250717.1;

refGene_gn=NBEA;refGene_tx=XM_023250721.1;refGene_gn=NBEA;refGene_tx=XM_023250724.1;refGene_gn=NBEA;refGene_tx=XM_023250731.1;refGene_gn=NBEA;refGene_tx=XM_023250733.1;refGene_gn=NBEA;refGene_tx=XM_023250742.1
A1 14190775 14190903
refGene_gn=NBEA;refGene_tx=XM_011286341.3;refGene_gn=NBEA;refGene_tx=XM_019825371.2;refGene_gn=NBEA;refGene_tx=XM_019825374.2;refGene_gn=NBEA;refGene_tx=XM_019825380.2;refGene_gn=NBEA;refGene_tx=XM_023250717.1;refGene_gn=NBEA;refGene_tx=XM_023250721.1;refGene_gn=NBEA;refGene_tx=XM_023250724.1;refGene_gn=NBEA;refGene_tx=XM_023250731.1;refGene_gn=NBEA;refGene_tx=XM_023250733.1;refGene_gn=NBEA;refGene_tx=XM_023250742.1
A1 14232774 14232836
refGene_gn=NBEA;refGene_tx=XM_011286341.3;refGene_gn=NBEA;refGene_tx=XM_019825371.2;refGene_gn=NBEA;refGene_tx=XM_019825374.2;refGene_gn=NBEA;refGene_tx=XM_019825380.2;refGene_gn=NBEA;refGene_tx=XM_023250717.1;refGene_gn=NBEA;refGene_tx=XM_023250721.1;refGene_gn=NBEA;refGene_tx=XM_023250724.1;refGene_gn=NBEA;refGene_tx=XM_023250731.1;refGene_gn=NBEA;refGene_tx=XM_023250733.1;refGene_gn=NBEA;refGene_tx=XM_023250742.1
A1 14252551 14252616
refGene_gn=NBEA;refGene_tx=XM_011286341.3;refGene_gn=NBEA;refGene_tx=XM_019825371.2;refGene_gn=NBEA;refGene_tx=XM_019825374.2;refGene_gn=NBEA;refGene_tx=XM_019825380.2;refGene_gn=NBEA;refGene_tx=XM_023250717.1;refGene_gn=NBEA;refGene_tx=XM_023250721.1;refGene_gn=NBEA;refGene_tx=XM_023250724.1;refGene_gn=NBEA;refGene_tx=XM_023250731.1;refGene_gn=NBEA;refGene_tx=XM_023250733.1;refGene_gn=NBEA;refGene_tx=XM_023250742.1
A1 14274073 14274182
refGene_gn=NBEA;refGene_tx=XM_011286341.3;refGene_gn=NBEA;refGene_tx=XM_019825371.2;refGene_gn=NBEA;refGene_tx=XM_019825374.2;refGene_gn=NBEA;refGene_tx=XM_019825380.2;refGene_gn=NBEA;refGene_tx=XM_023250717.1;refGene_gn=NBEA;refGene_tx=XM_023250721.1;refGene_gn=NBEA;refGene_tx=XM_023250724.1;refGene_gn=NBEA;refGene_tx=XM_023250731.1;refGene_gn=NBEA;refGene_tx=XM_023250733.1;refGene_gn=NBEA;refGene_tx=XM_023250742.1
A1 14277654 14277821
refGene_gn=NBEA;refGene_tx=XM_011286341.3;refGene_gn=NBEA;refGene_tx=XM_019825371.2;refGene_gn=NBEA;refGene_tx=XM_019825374.2;refGene_gn=NBEA;refGene_tx=XM_019825380.2;refGene_gn=NBEA;refGene_tx=XM_023250717.1;refGene_gn=NBEA;refGene_tx=XM_023250721.1;refGene_gn=NBEA;refGene_tx=XM_023250724.1;refGene_gn=NBEA;refGene_tx=XM_023250731.1;refGene_gn=NBEA;refGene_tx=XM_023250733.1;refGene_gn=NBEA;refGene_tx=XM_023250742.1
A1 14344095 14344220
refGene_gn=NBEA;refGene_tx=XM_011286341.3;refGene_gn=NBEA;refGene_tx=XM_019825371.2;refGene_gn=NBEA;refGene_tx=XM_019825374.2;refGene_gn=NBEA;refGene_tx=XM_019825380.2;refGene_gn=NBEA;refGene_tx=XM_023250717.1;refGene_gn=NBEA;refGene_tx=XM_023250721.1;refGene_gn=NBEA;refGene_tx=XM_023250724.1;refGene_gn=NBEA;refGene_tx=XM_023250731.1;refGene_gn=NBEA;refGene_tx=XM_023250733.1;refGene_gn=NBEA;refGene_tx=XM_023250742.1
A1 14364672 14364816
refGene_gn=NBEA;refGene_tx=XM_011286341.3;refGene_gn=NBEA;refGene_tx=XM_019825371.2;refGene_gn=NBEA;refGene_tx=XM_019825374.2;refGene_gn=NBEA;refGene_tx=XM_019825380.2;refGene_gn=NBEA;refGene_tx=XM_023250717.1;refGene_gn=NBEA;refGene_tx=XM_023250721.1;refGene_gn=NBEA;refGene_tx=X

M_023250724.1;refGene_gn=NBEA;refGene_tx=XM_023250731.1;refGene_gn=NBE
A;refGene_tx=XM_023250733.1;refGene_gn=NBEA;refGene_tx=XM_023250742.1
A1 14384483 14384620
refGene_gn=NBEA;refGene_tx=XM_011286341.3;refGene_gn=NBEA;refGene_tx=X
M_019825371.2;refGene_gn=NBEA;refGene_tx=XM_019825374.2;refGene_gn=NBE
A;refGene_tx=XM_019825380.2;refGene_gn=NBEA;refGene_tx=XM_023250717.1;
refGene_gn=NBEA;refGene_tx=XM_023250721.1;refGene_gn=NBEA;refGene_tx=X
M_023250724.1;refGene_gn=NBEA;refGene_tx=XM_023250731.1;refGene_gn=NBE
A;refGene_tx=XM_023250733.1;refGene_gn=NBEA;refGene_tx=XM_023250742.1
A1 14387209 14388289 refGene_gn=MAB21L1;refGene_tx=XM_003980322.4
A1 14464485 14464603
refGene_gn=NBEA;refGene_tx=XM_011286341.3;refGene_gn=NBEA;refGene_tx=X
M_019825371.2;refGene_gn=NBEA;refGene_tx=XM_019825374.2;refGene_gn=NBE
A;refGene_tx=XM_019825380.2;refGene_gn=NBEA;refGene_tx=XM_023250717.1;
refGene_gn=NBEA;refGene_tx=XM_023250721.1;refGene_gn=NBEA;refGene_tx=X
M_023250724.1;refGene_gn=NBEA;refGene_tx=XM_023250731.1;refGene_gn=NBE
A;refGene_tx=XM_023250733.1;refGene_gn=NBEA;refGene_tx=XM_023250742.1
A1 14464916 14465019
refGene_gn=NBEA;refGene_tx=XM_011286341.3;refGene_gn=NBEA;refGene_tx=X
M_019825371.2;refGene_gn=NBEA;refGene_tx=XM_019825374.2;refGene_gn=NBE
A;refGene_tx=XM_019825380.2;refGene_gn=NBEA;refGene_tx=XM_023250717.1;
refGene_gn=NBEA;refGene_tx=XM_023250721.1;refGene_gn=NBEA;refGene_tx=X
M_023250724.1;refGene_gn=NBEA;refGene_tx=XM_023250731.1;refGene_gn=NBE
A;refGene_tx=XM_023250733.1;refGene_gn=NBEA;refGene_tx=XM_023250742.1
A1 14472493 14472609
refGene_gn=NBEA;refGene_tx=XM_011286341.3;refGene_gn=NBEA;refGene_tx=X
M_019825371.2;refGene_gn=NBEA;refGene_tx=XM_019825374.2;refGene_gn=NBE
A;refGene_tx=XM_019825380.2;refGene_gn=NBEA;refGene_tx=XM_023250717.1;
refGene_gn=NBEA;refGene_tx=XM_023250721.1;refGene_gn=NBEA;refGene_tx=X
M_023250724.1;refGene_gn=NBEA;refGene_tx=XM_023250731.1;refGene_gn=NBE
A;refGene_tx=XM_023250733.1;refGene_gn=NBEA;refGene_tx=XM_023250742.1
A1 14484441 14484554
refGene_gn=NBEA;refGene_tx=XM_011286341.3;refGene_gn=NBEA;refGene_tx=X
M_019825371.2;refGene_gn=NBEA;refGene_tx=XM_019825374.2;refGene_gn=NBE
A;refGene_tx=XM_019825380.2;refGene_gn=NBEA;refGene_tx=XM_023250717.1;
refGene_gn=NBEA;refGene_tx=XM_023250721.1;refGene_gn=NBEA;refGene_tx=X
M_023250724.1;refGene_gn=NBEA;refGene_tx=XM_023250731.1;refGene_gn=NBE
A;refGene_tx=XM_023250733.1;refGene_gn=NBEA;refGene_tx=XM_023250742.1
A1 14505765 14505906
refGene_gn=NBEA;refGene_tx=XM_011286341.3;refGene_gn=NBEA;refGene_tx=X
M_019825371.2;refGene_gn=NBEA;refGene_tx=XM_019825374.2;refGene_gn=NBE
A;refGene_tx=XM_019825380.2;refGene_gn=NBEA;refGene_tx=XM_023250717.1;
refGene_gn=NBEA;refGene_tx=XM_023250721.1;refGene_gn=NBEA;refGene_tx=X
M_023250724.1;refGene_gn=NBEA;refGene_tx=XM_023250731.1;refGene_gn=NBE
A;refGene_tx=XM_023250733.1;refGene_gn=NBEA;refGene_tx=XM_023250742.1
A1 14516147 14516267
refGene_gn=NBEA;refGene_tx=XM_011286341.3;refGene_gn=NBEA;refGene_tx=X
M_019825371.2;refGene_gn=NBEA;refGene_tx=XM_019825374.2;refGene_gn=NBE
A;refGene_tx=XM_019825380.2;refGene_gn=NBEA;refGene_tx=XM_023250717.1;
refGene_gn=NBEA;refGene_tx=XM_023250721.1;refGene_gn=NBEA;refGene_tx=X

M_023250724.1;refGene_gn=NBEA;refGene_tx=XM_023250731.1;refGene_gn=NBEA;refGene_tx=XM_023250733.1;refGene_gn=NBEA;refGene_tx=XM_023250742.1
A1 14528698 14528851
refGene_gn=NBEA;refGene_tx=XM_011286341.3;refGene_gn=NBEA;refGene_tx=XM_019825371.2;refGene_gn=NBEA;refGene_tx=XM_019825374.2;refGene_gn=NBEA;refGene_tx=XM_019825380.2;refGene_gn=NBEA;refGene_tx=XM_023250717.1;refGene_gn=NBEA;refGene_tx=XM_023250721.1;refGene_gn=NBEA;refGene_tx=XM_023250724.1;refGene_gn=NBEA;refGene_tx=XM_023250731.1;refGene_gn=NBEA;refGene_tx=XM_023250733.1;refGene_gn=NBEA;refGene_tx=XM_023250742.1
A1 14547904 14548072
refGene_gn=NBEA;refGene_tx=XM_011286341.3;refGene_gn=NBEA;refGene_tx=XM_019825371.2;refGene_gn=NBEA;refGene_tx=XM_019825374.2;refGene_gn=NBEA;refGene_tx=XM_019825380.2;refGene_gn=NBEA;refGene_tx=XM_023250717.1;refGene_gn=NBEA;refGene_tx=XM_023250721.1;refGene_gn=NBEA;refGene_tx=XM_023250724.1;refGene_gn=NBEA;refGene_tx=XM_023250731.1;refGene_gn=NBEA;refGene_tx=XM_023250733.1;refGene_gn=NBEA;refGene_tx=XM_023250742.1
A1 14566475 14566538
refGene_gn=NBEA;refGene_tx=XM_019825374.2;refGene_gn=NBEA;refGene_tx=XM_023250717.1;refGene_gn=NBEA;refGene_tx=XM_023250721.1;refGene_gn=NBEA;refGene_tx=XM_023250724.1;refGene_gn=NBEA;refGene_tx=XM_023250733.1;refGene_gn=NBEA;refGene_tx=XM_023250742.1
A1 14566862 14566952
refGene_gn=NBEA;refGene_tx=XM_011286341.3;refGene_gn=NBEA;refGene_tx=XM_019825371.2;refGene_gn=NBEA;refGene_tx=XM_019825374.2;refGene_gn=NBEA;refGene_tx=XM_019825380.2;refGene_gn=NBEA;refGene_tx=XM_023250717.1;refGene_gn=NBEA;refGene_tx=XM_023250721.1;refGene_gn=NBEA;refGene_tx=XM_023250724.1;refGene_gn=NBEA;refGene_tx=XM_023250731.1;refGene_gn=NBEA;refGene_tx=XM_023250733.1;refGene_gn=NBEA;refGene_tx=XM_023250742.1
A1 14569603 14569796
refGene_gn=NBEA;refGene_tx=XM_011286341.3;refGene_gn=NBEA;refGene_tx=XM_019825371.2;refGene_gn=NBEA;refGene_tx=XM_019825374.2;refGene_gn=NBEA;refGene_tx=XM_019825380.2;refGene_gn=NBEA;refGene_tx=XM_023250717.1;refGene_gn=NBEA;refGene_tx=XM_023250721.1;refGene_gn=NBEA;refGene_tx=XM_023250724.1;refGene_gn=NBEA;refGene_tx=XM_023250731.1;refGene_gn=NBEA;refGene_tx=XM_023250733.1;refGene_gn=NBEA;refGene_tx=XM_023250742.1
A1 14571420 14571492
refGene_gn=NBEA;refGene_tx=XM_011286341.3;refGene_gn=NBEA;refGene_tx=XM_019825371.2;refGene_gn=NBEA;refGene_tx=XM_019825374.2;refGene_gn=NBEA;refGene_tx=XM_019825380.2;refGene_gn=NBEA;refGene_tx=XM_023250717.1;refGene_gn=NBEA;refGene_tx=XM_023250721.1;refGene_gn=NBEA;refGene_tx=XM_023250724.1;refGene_gn=NBEA;refGene_tx=XM_023250731.1;refGene_gn=NBEA;refGene_tx=XM_023250733.1;refGene_gn=NBEA;refGene_tx=XM_023250742.1
A1 14573904 14574060
refGene_gn=NBEA;refGene_tx=XM_011286341.3;refGene_gn=NBEA;refGene_tx=XM_019825371.2;refGene_gn=NBEA;refGene_tx=XM_019825374.2;refGene_gn=NBEA;refGene_tx=XM_019825380.2;refGene_gn=NBEA;refGene_tx=XM_023250717.1;refGene_gn=NBEA;refGene_tx=XM_023250721.1;refGene_gn=NBEA;refGene_tx=XM_023250724.1;refGene_gn=NBEA;refGene_tx=XM_023250731.1;refGene_gn=NBEA;refGene_tx=XM_023250733.1;refGene_gn=NBEA;refGene_tx=XM_023250742.1
A1 14574567 14574738

refGene_gn=NBEA;refGene_tx=XM_011286341.3;refGene_gn=NBEA;refGene_tx=XM_019825371.2;refGene_gn=NBEA;refGene_tx=XM_019825374.2;refGene_gn=NBEA;refGene_tx=XM_019825380.2;refGene_gn=NBEA;refGene_tx=XM_023250717.1;refGene_gn=NBEA;refGene_tx=XM_023250721.1;refGene_gn=NBEA;refGene_tx=XM_023250724.1;refGene_gn=NBEA;refGene_tx=XM_023250731.1;refGene_gn=NBEA;refGene_tx=XM_023250733.1;refGene_gn=NBEA;refGene_tx=XM_023250742.1
A1 14586027 14586129
refGene_gn=NBEA;refGene_tx=XM_011286341.3;refGene_gn=NBEA;refGene_tx=XM_019825371.2;refGene_gn=NBEA;refGene_tx=XM_019825374.2;refGene_gn=NBEA;refGene_tx=XM_019825380.2;refGene_gn=NBEA;refGene_tx=XM_023250717.1;refGene_gn=NBEA;refGene_tx=XM_023250721.1;refGene_gn=NBEA;refGene_tx=XM_023250724.1;refGene_gn=NBEA;refGene_tx=XM_023250731.1;refGene_gn=NBEA;refGene_tx=XM_023250733.1;refGene_gn=NBEA;refGene_tx=XM_023250742.1
A1 14588047 14588244
refGene_gn=NBEA;refGene_tx=XM_011286341.3;refGene_gn=NBEA;refGene_tx=XM_019825371.2;refGene_gn=NBEA;refGene_tx=XM_019825374.2;refGene_gn=NBEA;refGene_tx=XM_019825380.2;refGene_gn=NBEA;refGene_tx=XM_023250717.1;refGene_gn=NBEA;refGene_tx=XM_023250721.1;refGene_gn=NBEA;refGene_tx=XM_023250724.1;refGene_gn=NBEA;refGene_tx=XM_023250731.1;refGene_gn=NBEA;refGene_tx=XM_023250733.1;refGene_gn=NBEA;refGene_tx=XM_023250742.1
A1 14589022 14589174
refGene_gn=NBEA;refGene_tx=XM_011286341.3;refGene_gn=NBEA;refGene_tx=XM_019825371.2;refGene_gn=NBEA;refGene_tx=XM_019825374.2;refGene_gn=NBEA;refGene_tx=XM_019825380.2;refGene_gn=NBEA;refGene_tx=XM_023250717.1;refGene_gn=NBEA;refGene_tx=XM_023250721.1;refGene_gn=NBEA;refGene_tx=XM_023250724.1;refGene_gn=NBEA;refGene_tx=XM_023250731.1;refGene_gn=NBEA;refGene_tx=XM_023250733.1;refGene_gn=NBEA;refGene_tx=XM_023250742.1
A1 14591220 14591311
refGene_gn=NBEA;refGene_tx=XM_011286341.3;refGene_gn=NBEA;refGene_tx=XM_019825371.2;refGene_gn=NBEA;refGene_tx=XM_019825374.2;refGene_gn=NBEA;refGene_tx=XM_019825380.2;refGene_gn=NBEA;refGene_tx=XM_023250717.1;refGene_gn=NBEA;refGene_tx=XM_023250721.1;refGene_gn=NBEA;refGene_tx=XM_023250724.1;refGene_gn=NBEA;refGene_tx=XM_023250731.1;refGene_gn=NBEA;refGene_tx=XM_023250733.1;refGene_gn=NBEA;refGene_tx=XM_023250742.1
A1 14676979 14677144
refGene_gn=DCLK1;refGene_tx=XM_011286161.3;refGene_gn=DCLK1;refGene_tx=XM_011286192.3;refGene_gn=DCLK1;refGene_tx=XM_023250781.1;refGene_gn=DCLK1;refGene_tx=XM_023250785.1;refGene_gn=DCLK1;refGene_tx=XM_023250793.1;refGene_gn=DCLK1;refGene_tx=XM_023250802.1
A1 14690148 14690222
refGene_gn=DCLK1;refGene_tx=XM_011286192.3;refGene_gn=DCLK1;refGene_tx=XM_023250793.1;refGene_gn=DCLK1;refGene_tx=XM_023250802.1
A1 14695633 14695747
refGene_gn=DCLK1;refGene_tx=XM_011286161.3;refGene_gn=DCLK1;refGene_tx=XM_011286192.3;refGene_gn=DCLK1;refGene_tx=XM_023250781.1;refGene_gn=DCLK1;refGene_tx=XM_023250785.1;refGene_gn=DCLK1;refGene_tx=XM_023250793.1;refGene_gn=DCLK1;refGene_tx=XM_023250802.1
A1 14706856 14706937
refGene_gn=DCLK1;refGene_tx=XM_011286161.3;refGene_gn=DCLK1;refGene_tx=XM_011286192.3;refGene_gn=DCLK1;refGene_tx=XM_023250781.1;refGene_gn=

DCLK1;refGene_tx=XM_023250785.1;refGene_gn=DCLK1;refGene_tx=XM_023250793.1;refGene_gn=DCLK1;refGene_tx=XM_023250802.1
A1 14709092 14709189
refGene_gn=DCLK1;refGene_tx=XM_011286161.3;refGene_gn=DCLK1;refGene_tx=XM_011286192.3;refGene_gn=DCLK1;refGene_tx=XM_023250781.1;refGene_gn=DCLK1;refGene_tx=XM_023250785.1;refGene_gn=DCLK1;refGene_tx=XM_023250793.1;refGene_gn=DCLK1;refGene_tx=XM_023250802.1
A1 14709568 14709646
refGene_gn=DCLK1;refGene_tx=XM_011286161.3;refGene_gn=DCLK1;refGene_tx=XM_011286192.3;refGene_gn=DCLK1;refGene_tx=XM_023250781.1;refGene_gn=DCLK1;refGene_tx=XM_023250785.1;refGene_gn=DCLK1;refGene_tx=XM_023250793.1;refGene_gn=DCLK1;refGene_tx=XM_023250802.1
A1 14711652 14711786
refGene_gn=DCLK1;refGene_tx=XM_011286161.3;refGene_gn=DCLK1;refGene_tx=XM_011286192.3;refGene_gn=DCLK1;refGene_tx=XM_023250781.1;refGene_gn=DCLK1;refGene_tx=XM_023250785.1;refGene_gn=DCLK1;refGene_tx=XM_023250793.1;refGene_gn=DCLK1;refGene_tx=XM_023250802.1
A1 14723150 14723297
refGene_gn=DCLK1;refGene_tx=XM_011286161.3;refGene_gn=DCLK1;refGene_tx=XM_011286192.3;refGene_gn=DCLK1;refGene_tx=XM_023250781.1;refGene_gn=DCLK1;refGene_tx=XM_023250785.1;refGene_gn=DCLK1;refGene_tx=XM_023250793.1;refGene_gn=DCLK1;refGene_tx=XM_023250802.1
A1 14727573 14727693
refGene_gn=DCLK1;refGene_tx=XM_011286161.3;refGene_gn=DCLK1;refGene_tx=XM_011286192.3;refGene_gn=DCLK1;refGene_tx=XM_023250781.1;refGene_gn=DCLK1;refGene_tx=XM_023250785.1;refGene_gn=DCLK1;refGene_tx=XM_023250793.1;refGene_gn=DCLK1;refGene_tx=XM_023250802.1
A1 14728182 14728240
refGene_gn=DCLK1;refGene_tx=XM_011286161.3;refGene_gn=DCLK1;refGene_tx=XM_011286192.3;refGene_gn=DCLK1;refGene_tx=XM_023250781.1;refGene_gn=DCLK1;refGene_tx=XM_023250785.1;refGene_gn=DCLK1;refGene_tx=XM_023250793.1;refGene_gn=DCLK1;refGene_tx=XM_023250802.1
A1 14734858 14734967
refGene_gn=DCLK1;refGene_tx=XM_011286161.3;refGene_gn=DCLK1;refGene_tx=XM_011286192.3;refGene_gn=DCLK1;refGene_tx=XM_023250781.1;refGene_gn=DCLK1;refGene_tx=XM_023250785.1;refGene_gn=DCLK1;refGene_tx=XM_023250793.1;refGene_gn=DCLK1;refGene_tx=XM_023250802.1
A1 14737660 14737748
refGene_gn=DCLK1;refGene_tx=XM_011286161.3;refGene_gn=DCLK1;refGene_tx=XM_011286192.3;refGene_gn=DCLK1;refGene_tx=XM_023250781.1;refGene_gn=DCLK1;refGene_tx=XM_023250785.1;refGene_gn=DCLK1;refGene_tx=XM_023250793.1;refGene_gn=DCLK1;refGene_tx=XM_023250802.1
A1 14748793 14748850
refGene_gn=DCLK1;refGene_tx=XM_011286228.2;refGene_gn=DCLK1;refGene_tx=XM_023250814.1
A1 14752518 14752613
refGene_gn=DCLK1;refGene_tx=XM_011286161.3;refGene_gn=DCLK1;refGene_tx=XM_011286192.3;refGene_gn=DCLK1;refGene_tx=XM_011286228.2;refGene_gn=DCLK1;refGene_tx=XM_023250781.1;refGene_gn=DCLK1;refGene_tx=XM_023250785.1;refGene_gn=DCLK1;refGene_tx=XM_023250793.1;refGene_gn=DCLK1;refGe

ne_tx=XM_023250802.1;refGene_gn=DCLK1;refGene_tx=XM_023250814.1
A1 14753544 14753563
refGene_gn=DCLK1;refGene_tx=XM_011286161.3;refGene_gn=DCLK1;refGene_tx=XM_011286192.3;refGene_gn=DCLK1;refGene_tx=XM_011286228.2
A1 14769307 14769424
refGene_gn=DCLK1;refGene_tx=XM_023250781.1;refGene_gn=DCLK1;refGene_tx=XM_023250785.1;refGene_gn=DCLK1;refGene_tx=XM_023250793.1;refGene_gn=DCLK1;refGene_tx=XM_023250802.1;refGene_gn=DCLK1;refGene_tx=XM_023250814.1
A1 14838632 14838732
refGene_gn=DCLK1;refGene_tx=XM_023250781.1;refGene_gn=DCLK1;refGene_tx=XM_023250785.1;refGene_gn=DCLK1;refGene_tx=XM_023250793.1;refGene_gn=DCLK1;refGene_tx=XM_023250802.1;refGene_gn=DCLK1;refGene_tx=XM_023250814.1
A1 14994005 14994352
refGene_gn=DCLK1;refGene_tx=XM_023250781.1;refGene_gn=DCLK1;refGene_tx=XM_023250785.1;refGene_gn=DCLK1;refGene_tx=XM_023250793.1;refGene_gn=DCLK1;refGene_tx=XM_023250802.1;refGene_gn=DCLK1;refGene_tx=XM_023250814.1
A1 15013095 15013471
refGene_gn=DCLK1;refGene_tx=XM_023250781.1;refGene_gn=DCLK1;refGene_tx=XM_023250785.1;refGene_gn=DCLK1;refGene_tx=XM_023250793.1;refGene_gn=DCLK1;refGene_tx=XM_023250802.1;refGene_gn=DCLK1;refGene_tx=XM_023250814.1
A1 15071629 15071650 refGene_gn=SOHLH2;refGene_tx=XM_006927229.3
A1 15072230 15072290
refGene_gn=SOHLH2;refGene_tx=XM_019825441.2;refGene_gn=SOHLH2;refGene_tx=XM_019825451.2
A1 15073130 15073387
refGene_gn=SOHLH2;refGene_tx=XM_006927229.3;refGene_gn=SOHLH2;refGene_tx=XM_019825441.2;refGene_gn=SOHLH2;refGene_tx=XM_019825451.2
A1 15079684 15079806
refGene_gn=SOHLH2;refGene_tx=XM_006927229.3;refGene_gn=SOHLH2;refGene_tx=XM_019825441.2;refGene_gn=SOHLH2;refGene_tx=XM_019825451.2
A1 15080479 15080568
refGene_gn=SOHLH2;refGene_tx=XM_006927229.3;refGene_gn=SOHLH2;refGene_tx=XM_019825441.2;refGene_gn=SOHLH2;refGene_tx=XM_019825451.2
A1 15080701 15080849
refGene_gn=SOHLH2;refGene_tx=XM_006927229.3;refGene_gn=SOHLH2;refGene_tx=XM_019825441.2;refGene_gn=SOHLH2;refGene_tx=XM_019825451.2
A1 15098903 15099014
refGene_gn=SOHLH2;refGene_tx=XM_006927229.3;refGene_gn=SOHLH2;refGene_tx=XM_019825441.2;refGene_gn=SOHLH2;refGene_tx=XM_019825451.2
A1 15100644 15100744
refGene_gn=SOHLH2;refGene_tx=XM_006927229.3;refGene_gn=SOHLH2;refGene_tx=XM_019825441.2;refGene_gn=SOHLH2;refGene_tx=XM_019825451.2
A1 15102265 15102370
refGene_gn=SOHLH2;refGene_tx=XM_006927229.3;refGene_gn=SOHLH2;refGene_tx=XM_019825441.2
A1 15102448 15102510

refGene_gn=SOHLH2;refGene_tx=XM_006927229.3;refGene_gn=SOHLH2;refGene_tx=XM_019825441.2;refGene_gn=SOHLH2;refGene_tx=XM_019825451.2
A1 15113890 15114105
refGene_gn=SOHLH2;refGene_tx=XM_006927229.3;refGene_gn=SOHLH2;refGene_tx=XM_019825441.2;refGene_gn=SOHLH2;refGene_tx=XM_019825451.2
A1 15127930 15128134
refGene_gn=SOHLH2;refGene_tx=XM_006927229.3;refGene_gn=SOHLH2;refGene_tx=XM_019825441.2;refGene_gn=SOHLH2;refGene_tx=XM_019825451.2
A1 15146133 15146233
refGene_gn=CCDC169;refGene_tx=XM_006927230.3;refGene_gn=CCDC169;refGene_tx=XM_019825458.2;refGene_gn=CCDC169;refGene_tx=XM_019825465.1
A1 15156702 15156779
refGene_gn=CCDC169;refGene_tx=XM_006927230.3;refGene_gn=CCDC169;refGene_tx=XM_019825458.2;refGene_gn=CCDC169;refGene_tx=XM_019825465.1
A1 15161529 15161583
refGene_gn=CCDC169;refGene_tx=XM_006927230.3;refGene_gn=CCDC169;refGene_tx=XM_019825458.2;refGene_gn=CCDC169;refGene_tx=XM_019825465.1
A1 15161802 15161901
refGene_gn=CCDC169;refGene_tx=XM_006927230.3;refGene_gn=CCDC169;refGene_tx=XM_019825458.2;refGene_gn=CCDC169;refGene_tx=XM_019825465.1
A1 15187587 15187628
refGene_gn=CCDC169;refGene_tx=XM_006927230.3;refGene_gn=CCDC169;refGene_tx=XM_019825458.2;refGene_gn=CCDC169;refGene_tx=XM_019825465.1
A1 15187954 15188065
refGene_gn=CCDC169;refGene_tx=XM_006927230.3;refGene_gn=CCDC169;refGene_tx=XM_019825458.2
A1 15197025 15197105 refGene_gn=CCDC169;refGene_tx=XM_006927230.3
A1 15198839 15198922 refGene_gn=CCDC169;refGene_tx=XM_006927230.3
A1 15207843 15208117
refGene_gn=SPART;refGene_tx=XM_011285953.3;refGene_gn=SPART;refGene_tx=XM_019825488.2;refGene_gn=SPART;refGene_tx=XM_019825498.2;refGene_gn=SPART;refGene_tx=XM_023250855.1;refGene_gn=SPART;refGene_tx=XM_023250868.1
A1 15213770 15213864 refGene_gn=SPART;refGene_tx=XM_023250879.1
A1 15217309 15217400
refGene_gn=SPART;refGene_tx=XM_011285953.3;refGene_gn=SPART;refGene_tx=XM_019825488.2;refGene_gn=SPART;refGene_tx=XM_019825498.2;refGene_gn=SPART;refGene_tx=XM_023250855.1;refGene_gn=SPART;refGene_tx=XM_023250868.1;refGene_gn=SPART;refGene_tx=XM_023250879.1
A1 15217479 15217638
refGene_gn=SPART;refGene_tx=XM_011285953.3;refGene_gn=SPART;refGene_tx=XM_019825488.2;refGene_gn=SPART;refGene_tx=XM_019825498.2;refGene_gn=SPART;refGene_tx=XM_023250855.1;refGene_gn=SPART;refGene_tx=XM_023250868.1;refGene_gn=SPART;refGene_tx=XM_023250879.1
A1 15218675 15218870
refGene_gn=SPART;refGene_tx=XM_011285953.3;refGene_gn=SPART;refGene_tx=XM_019825488.2;refGene_gn=SPART;refGene_tx=XM_019825498.2;refGene_gn=SPART;refGene_tx=XM_023250855.1;refGene_gn=SPART;refGene_tx=XM_023250868.1;refGene_gn=SPART;refGene_tx=XM_023250879.1
A1 15223460 15223584

refGene_gn=SPART;refGene_tx=XM_011285953.3;refGene_gn=SPART;refGene_tx
=XM_019825488.2;refGene_gn=SPART;refGene_tx=XM_019825498.2;refGene_gn=
SPART;refGene_tx=XM_023250855.1;refGene_gn=SPART;refGene_tx=XM_0232508
68.1;refGene_gn=SPART;refGene_tx=XM_023250879.1
A1 15226998 15227166
refGene_gn=SPART;refGene_tx=XM_011285953.3;refGene_gn=SPART;refGene_tx
=XM_019825488.2;refGene_gn=SPART;refGene_tx=XM_019825498.2;refGene_gn=
SPART;refGene_tx=XM_023250855.1;refGene_gn=SPART;refGene_tx=XM_0232508
68.1;refGene_gn=SPART;refGene_tx=XM_023250879.1
A1 15229739 15229937
refGene_gn=SPART;refGene_tx=XM_011285953.3;refGene_gn=SPART;refGene_tx
=XM_019825488.2;refGene_gn=SPART;refGene_tx=XM_019825498.2;refGene_gn=
SPART;refGene_tx=XM_023250855.1;refGene_gn=SPART;refGene_tx=XM_0232508
68.1;refGene_gn=SPART;refGene_tx=XM_023250879.1
A1 15234084 15234894
refGene_gn=SPART;refGene_tx=XM_011285953.3;refGene_gn=SPART;refGene_tx
=XM_019825488.2;refGene_gn=SPART;refGene_tx=XM_019825498.2;refGene_gn=
SPART;refGene_tx=XM_023250855.1;refGene_gn=SPART;refGene_tx=XM_0232508
68.1;refGene_gn=SPART;refGene_tx=XM_023250879.1
A1 15299456 15299621
refGene_gn=CCNA1;refGene_tx=XM_011285894.2;refGene_gn=CCNA1;refGene_tx
=XM_011285915.3;refGene_gn=CCNA1;refGene_tx=XM_019825510.2;refGene_gn=
CCNA1;refGene_tx=XM_019825511.1;refGene_gn=CCNA1;refGene_tx=XM_0232508
86.1
A1 15302994 15303241
refGene_gn=CCNA1;refGene_tx=XM_011285894.2;refGene_gn=CCNA1;refGene_tx
=XM_011285915.3;refGene_gn=CCNA1;refGene_tx=XM_019825510.2;refGene_gn=
CCNA1;refGene_tx=XM_019825511.1;refGene_gn=CCNA1;refGene_tx=XM_0232508
86.1
A1 15303427 15303552
refGene_gn=CCNA1;refGene_tx=XM_011285894.2;refGene_gn=CCNA1;refGene_tx
=XM_011285915.3;refGene_gn=CCNA1;refGene_tx=XM_019825510.2;refGene_gn=
CCNA1;refGene_tx=XM_019825511.1;refGene_gn=CCNA1;refGene_tx=XM_0232508
86.1
A1 15303982 15304206
refGene_gn=CCNA1;refGene_tx=XM_011285894.2;refGene_gn=CCNA1;refGene_tx
=XM_011285915.3;refGene_gn=CCNA1;refGene_tx=XM_019825510.2;refGene_gn=
CCNA1;refGene_tx=XM_019825511.1;refGene_gn=CCNA1;refGene_tx=XM_0232508
86.1
A1 15305283 15305488
refGene_gn=CCNA1;refGene_tx=XM_011285894.2;refGene_gn=CCNA1;refGene_tx
=XM_011285915.3;refGene_gn=CCNA1;refGene_tx=XM_019825510.2;refGene_gn=
CCNA1;refGene_tx=XM_019825511.1;refGene_gn=CCNA1;refGene_tx=XM_0232508
86.1
A1 15306462 15306576
refGene_gn=CCNA1;refGene_tx=XM_011285894.2;refGene_gn=CCNA1;refGene_tx
=XM_011285915.3;refGene_gn=CCNA1;refGene_tx=XM_019825510.2;refGene_gn=
CCNA1;refGene_tx=XM_023250886.1
A1 15307416 15307550
refGene_gn=CCNA1;refGene_tx=XM_011285894.2;refGene_gn=CCNA1;refGene_tx

=XM_011285915.3;refGene_gn=CCNA1;refGene_tx=XM_019825510.2;refGene_gn=CCNA1;refGene_tx=XM_019825511.1;refGene_gn=CCNA1;refGene_tx=XM_023250886.1
A1 15307837 15307889
refGene_gn=CCNA1;refGene_tx=XM_011285894.2;refGene_gn=CCNA1;refGene_tx=XM_011285915.3;refGene_gn=CCNA1;refGene_tx=XM_019825510.2;refGene_gn=CCNA1;refGene_tx=XM_019825511.1;refGene_gn=CCNA1;refGene_tx=XM_023250886.1
A1 15537154 15537478
refGene_gn=SERTM1;refGene_tx=XM_023250894.1;refGene_gn=SERTM1;refGene_tx=XM_023250897.1;refGene_gn=SERTM1;refGene_tx=XM_023250899.1
A1 15572244 15572351 refGene_gn=RFXAP;refGene_tx=XM_011287640.2
A1 15575432 15575649 refGene_gn=RFXAP;refGene_tx=XM_011287640.2
A1 15601482 15601597 refGene_gn=RFXAP;refGene_tx=XM_011287640.2
A1 15643444 15643593 refGene_gn=RFXAP;refGene_tx=XM_011287640.2
A1 15643721 15644453 refGene_gn=RFXAP;refGene_tx=XM_011287640.2
A1 15648915 15649023 refGene_gn=RFXAP;refGene_tx=XM_011287640.2
A1 15651536 15651647 refGene_gn=RFXAP;refGene_tx=XM_011287640.2
A1 15666862 15667006
refGene_gn=SMAD9;refGene_tx=XM_023250900.1;refGene_gn=SMAD9;refGene_tx=XM_023250901.1;refGene_gn=SMAD9;refGene_tx=XM_023250902.1;refGene_gn=SMAD9;refGene_tx=XM_023250903.1;refGene_gn=SMAD9;refGene_tx=XM_023250904.1
A1 15678574 15678831
refGene_gn=SMAD9;refGene_tx=XM_023250900.1;refGene_gn=SMAD9;refGene_tx=XM_023250901.1;refGene_gn=SMAD9;refGene_tx=XM_023250902.1;refGene_gn=SMAD9;refGene_tx=XM_023250903.1;refGene_gn=SMAD9;refGene_tx=XM_023250904.1
A1 15685494 15685716
refGene_gn=SMAD9;refGene_tx=XM_023250900.1;refGene_gn=SMAD9;refGene_tx=XM_023250901.1;refGene_gn=SMAD9;refGene_tx=XM_023250902.1;refGene_gn=SMAD9;refGene_tx=XM_023250903.1;refGene_gn=SMAD9;refGene_tx=XM_023250904.1
A1 15687218 15687329
refGene_gn=SMAD9;refGene_tx=XM_023250900.1;refGene_gn=SMAD9;refGene_tx=XM_023250901.1;refGene_gn=SMAD9;refGene_tx=XM_023250902.1;refGene_gn=SMAD9;refGene_tx=XM_023250903.1
A1 15691170 15691428
refGene_gn=SMAD9;refGene_tx=XM_023250900.1;refGene_gn=SMAD9;refGene_tx=XM_023250901.1;refGene_gn=SMAD9;refGene_tx=XM_023250902.1;refGene_gn=SMAD9;refGene_tx=XM_023250903.1;refGene_gn=SMAD9;refGene_tx=XM_023250904.1
A1 15698284 15698696
refGene_gn=SMAD9;refGene_tx=XM_023250900.1;refGene_gn=SMAD9;refGene_tx=XM_023250901.1;refGene_gn=SMAD9;refGene_tx=XM_023250902.1;refGene_gn=SMAD9;refGene_tx=XM_023250903.1;refGene_gn=SMAD9;refGene_tx=XM_023250904.1
A1 15761559 15761675
refGene_gn=ALG5;refGene_tx=XM_003980336.5;refGene_gn=ALG5;refGene_tx=XM_011285655.3;refGene_gn=ALG5;refGene_tx=XM_019825665.2

A1 15763808 15763894
refGene_gn=ALG5;refGene_tx=XM_003980336.5;refGene_gn=ALG5;refGene_tx=XM_011285655.3;refGene_gn=ALG5;refGene_tx=XM_019825665.2
A1 15772813 15772965
refGene_gn=ALG5;refGene_tx=XM_003980336.5;refGene_gn=ALG5;refGene_tx=XM_011285655.3;refGene_gn=ALG5;refGene_tx=XM_019825665.2
A1 15777650 15777710
refGene_gn=ALG5;refGene_tx=XM_003980336.5;refGene_gn=ALG5;refGene_tx=XM_011285655.3;refGene_gn=ALG5;refGene_tx=XM_019825665.2
A1 15791954 15792068
refGene_gn=ALG5;refGene_tx=XM_003980336.5;refGene_gn=ALG5;refGene_tx=XM_011285655.3;refGene_gn=ALG5;refGene_tx=XM_019825665.2
A1 15796420 15796513
refGene_gn=ALG5;refGene_tx=XM_003980336.5;refGene_gn=ALG5;refGene_tx=XM_011285655.3;refGene_gn=ALG5;refGene_tx=XM_019825665.2
A1 15797837 15797906
refGene_gn=ALG5;refGene_tx=XM_003980336.5;refGene_gn=ALG5;refGene_tx=XM_011285655.3;refGene_gn=ALG5;refGene_tx=XM_019825665.2
A1 15799944 15799991
refGene_gn=ALG5;refGene_tx=XM_003980336.5;refGene_gn=ALG5;refGene_tx=XM_011285655.3;refGene_gn=ALG5;refGene_tx=XM_019825665.2
A1 15800398 15800570
refGene_gn=ALG5;refGene_tx=XM_003980336.5;refGene_gn=ALG5;refGene_tx=XM_011285655.3;refGene_gn=ALG5;refGene_tx=XM_019825665.2
A1 15804889 15804955 refGene_gn=ALG5;refGene_tx=XM_003980336.5
A1 15806391 15806408 refGene_gn=EXOSC8;refGene_tx=XM_011285118.3
A1 15807164 15807307 refGene_gn=EXOSC8;refGene_tx=XM_019825671.2
A1 15807505 15807542
refGene_gn=EXOSC8;refGene_tx=XM_011285118.3;refGene_gn=EXOSC8;refGene_tx=XM_019825671.2
A1 15807711 15807775
refGene_gn=EXOSC8;refGene_tx=XM_011285118.3;refGene_gn=EXOSC8;refGene_tx=XM_019825671.2
A1 15808752 15808826
refGene_gn=EXOSC8;refGene_tx=XM_011285118.3;refGene_gn=EXOSC8;refGene_tx=XM_019825671.2
A1 15809954 15810000
refGene_gn=EXOSC8;refGene_tx=XM_011285118.3;refGene_gn=EXOSC8;refGene_tx=XM_019825671.2
A1 15810716 15810822
refGene_gn=EXOSC8;refGene_tx=XM_011285118.3;refGene_gn=EXOSC8;refGene_tx=XM_019825671.2
A1 15810906 15810952
refGene_gn=EXOSC8;refGene_tx=XM_011285118.3;refGene_gn=EXOSC8;refGene_tx=XM_019825671.2
A1 15811648 15811745
refGene_gn=EXOSC8;refGene_tx=XM_011285118.3;refGene_gn=EXOSC8;refGene_tx=XM_019825671.2
A1 15812025 15812146
refGene_gn=EXOSC8;refGene_tx=XM_011285118.3;refGene_gn=EXOSC8;refGene_

tx=XM_019825671.2
A1 15812605 15812712
refGene_gn=EXOSC8;refGene_tx=XM_011285118.3;refGene_gn=EXOSC8;refGene_
tx=XM_019825671.2
A1 15813286 15813402
refGene_gn=EXOSC8;refGene_tx=XM_011285118.3;refGene_gn=EXOSC8;refGene_
tx=XM_019825671.2
A1 15813783 15813921
refGene_gn=SUPT20H;refGene_tx=XM_006927233.4;refGene_gn=SUPT20H;refGen
e_tx=XM_006927234.4;refGene_gn=SUPT20H;refGene_tx=XM_006927235.2;refGe
ne_gn=SUPT20H;refGene_tx=XM_006927236.2;refGene_gn=SUPT20H;refGene_tx=
XM_006927237.4;refGene_gn=SUPT20H;refGene_tx=XM_011285155.3;refGene_gn
=SUPT20H;refGene_tx=XM_011285320.2;refGene_gn=SUPT20H;refGene_tx=XM_01
1285350.3;refGene_gn=SUPT20H;refGene_tx=XM_011285378.3;refGene_gn=SUPT
20H;refGene_tx=XM_011285402.3;refGene_gn=SUPT20H;refGene_tx=XM_0112854
32.2;refGene_gn=SUPT20H;refGene_tx=XM_011285467.2;refGene_gn=SUPT20H;r
efGene_tx=XM_011285509.3;refGene_gn=SUPT20H;refGene_tx=XM_011285532.3;
refGene_gn=SUPT20H;refGene_tx=XM_011285553.2;refGene_gn=SUPT20H;refGen
e_tx=XM_011285584.2;refGene_gn=SUPT20H;refGene_tx=XM_019825611.2;refGe
ne_gn=SUPT20H;refGene_tx=XM_019825613.2;refGene_gn=SUPT20H;refGene_tx=
XM_019825615.2;refGene_gn=SUPT20H;refGene_tx=XM_019825616.1;refGene_gn
=SUPT20H;refGene_tx=XM_023250917.1
A1 15814681 15814785
refGene_gn=SUPT20H;refGene_tx=XM_006927233.4;refGene_gn=SUPT20H;refGen
e_tx=XM_006927234.4;refGene_gn=SUPT20H;refGene_tx=XM_006927235.2;refGe
ne_gn=SUPT20H;refGene_tx=XM_006927236.2;refGene_gn=SUPT20H;refGene_tx=
XM_006927237.4;refGene_gn=SUPT20H;refGene_tx=XM_011285155.3;refGene_gn
=SUPT20H;refGene_tx=XM_011285320.2;refGene_gn=SUPT20H;refGene_tx=XM_01
1285350.3;refGene_gn=SUPT20H;refGene_tx=XM_011285378.3;refGene_gn=SUPT
20H;refGene_tx=XM_011285402.3;refGene_gn=SUPT20H;refGene_tx=XM_0112855
09.3;refGene_gn=SUPT20H;refGene_tx=XM_011285532.3;refGene_gn=SUPT20H;r
efGene_tx=XM_019825611.2;refGene_gn=SUPT20H;refGene_tx=XM_019825613.2;
refGene_gn=SUPT20H;refGene_tx=XM_019825615.2;refGene_gn=SUPT20H;refGen
e_tx=XM_019825616.1
A1 15816250 15816356
refGene_gn=SUPT20H;refGene_tx=XM_006927233.4;refGene_gn=SUPT20H;refGen
e_tx=XM_006927234.4;refGene_gn=SUPT20H;refGene_tx=XM_006927235.2;refGe
ne_gn=SUPT20H;refGene_tx=XM_006927236.2;refGene_gn=SUPT20H;refGene_tx=
XM_006927237.4;refGene_gn=SUPT20H;refGene_tx=XM_011285155.3;refGene_gn
=SUPT20H;refGene_tx=XM_011285320.2;refGene_gn=SUPT20H;refGene_tx=XM_01
1285350.3;refGene_gn=SUPT20H;refGene_tx=XM_011285378.3;refGene_gn=SUPT
20H;refGene_tx=XM_011285402.3;refGene_gn=SUPT20H;refGene_tx=XM_0112854
32.2;refGene_gn=SUPT20H;refGene_tx=XM_011285467.2;refGene_gn=SUPT20H;r
efGene_tx=XM_011285509.3;refGene_gn=SUPT20H;refGene_tx=XM_011285532.3;
refGene_gn=SUPT20H;refGene_tx=XM_011285553.2;refGene_gn=SUPT20H;refGen
e_tx=XM_011285584.2;refGene_gn=SUPT20H;refGene_tx=XM_019825611.2;refGe
ne_gn=SUPT20H;refGene_tx=XM_019825613.2;refGene_gn=SUPT20H;refGene_tx=
XM_019825615.2;refGene_gn=SUPT20H;refGene_tx=XM_019825616.1
A1 15820696 15820816
refGene_gn=SUPT20H;refGene_tx=XM_006927233.4;refGene_gn=SUPT20H;refGen

e_tx=XM_006927234.4;refGene_gn=SUPT20H;refGene_tx=XM_006927235.2;refGene_gn=SUPT20H;refGene_tx=XM_006927236.2;refGene_gn=SUPT20H;refGene_tx=XM_006927237.4;refGene_gn=SUPT20H;refGene_tx=XM_011285155.3;refGene_gn=SUPT20H;refGene_tx=XM_011285320.2;refGene_gn=SUPT20H;refGene_tx=XM_011285350.3;refGene_gn=SUPT20H;refGene_tx=XM_011285378.3;refGene_gn=SUPT20H;refGene_tx=XM_011285402.3;refGene_gn=SUPT20H;refGene_tx=XM_011285432.2;refGene_gn=SUPT20H;refGene_tx=XM_011285467.2;refGene_gn=SUPT20H;refGene_tx=XM_011285509.3;refGene_gn=SUPT20H;refGene_tx=XM_011285532.3;refGene_gn=SUPT20H;refGene_tx=XM_011285553.2;refGene_gn=SUPT20H;refGene_tx=XM_011285584.2;refGene_gn=SUPT20H;refGene_tx=XM_019825611.2;refGene_gn=SUPT20H;refGene_tx=XM_019825613.2;refGene_gn=SUPT20H;refGene_tx=XM_019825615.2;refGene_gn=SUPT20H;refGene_tx=XM_019825616.1;refGene_gn=SUPT20H;refGene_tx=XM_023250917.1
A1 15823427 15823483
refGene_gn=SUPT20H;refGene_tx=XM_006927233.4;refGene_gn=SUPT20H;refGene_tx=XM_006927234.4;refGene_gn=SUPT20H;refGene_tx=XM_006927235.2;refGene_gn=SUPT20H;refGene_tx=XM_006927236.2;refGene_gn=SUPT20H;refGene_tx=XM_006927237.4;refGene_gn=SUPT20H;refGene_tx=XM_011285155.3;refGene_gn=SUPT20H;refGene_tx=XM_011285320.2;refGene_gn=SUPT20H;refGene_tx=XM_011285350.3;refGene_gn=SUPT20H;refGene_tx=XM_011285378.3;refGene_gn=SUPT20H;refGene_tx=XM_011285402.3;refGene_gn=SUPT20H;refGene_tx=XM_011285432.2;refGene_gn=SUPT20H;refGene_tx=XM_011285467.2;refGene_gn=SUPT20H;refGene_tx=XM_011285509.3;refGene_gn=SUPT20H;refGene_tx=XM_011285532.3;refGene_gn=SUPT20H;refGene_tx=XM_011285553.2;refGene_gn=SUPT20H;refGene_tx=XM_011285584.2;refGene_gn=SUPT20H;refGene_tx=XM_019825611.2;refGene_gn=SUPT20H;refGene_tx=XM_019825613.2;refGene_gn=SUPT20H;refGene_tx=XM_019825615.2;refGene_gn=SUPT20H;refGene_tx=XM_019825616.1;refGene_gn=SUPT20H;refGene_tx=XM_023250917.1
A1 15826078 15826233
refGene_gn=SUPT20H;refGene_tx=XM_006927233.4;refGene_gn=SUPT20H;refGene_tx=XM_006927234.4;refGene_gn=SUPT20H;refGene_tx=XM_006927235.2;refGene_gn=SUPT20H;refGene_tx=XM_006927236.2;refGene_gn=SUPT20H;refGene_tx=XM_006927237.4;refGene_gn=SUPT20H;refGene_tx=XM_011285155.3;refGene_gn=SUPT20H;refGene_tx=XM_011285320.2;refGene_gn=SUPT20H;refGene_tx=XM_011285350.3;refGene_gn=SUPT20H;refGene_tx=XM_011285378.3;refGene_gn=SUPT20H;refGene_tx=XM_011285402.3;refGene_gn=SUPT20H;refGene_tx=XM_011285432.2;refGene_gn=SUPT20H;refGene_tx=XM_011285467.2;refGene_gn=SUPT20H;refGene_tx=XM_011285509.3;refGene_gn=SUPT20H;refGene_tx=XM_011285532.3;refGene_gn=SUPT20H;refGene_tx=XM_011285553.2;refGene_gn=SUPT20H;refGene_tx=XM_011285584.2;refGene_gn=SUPT20H;refGene_tx=XM_019825611.2;refGene_gn=SUPT20H;refGene_tx=XM_019825613.2;refGene_gn=SUPT20H;refGene_tx=XM_019825615.2;refGene_gn=SUPT20H;refGene_tx=XM_019825616.1;refGene_gn=SUPT20H;refGene_tx=XM_023250917.1
A1 15826731 15827038
refGene_gn=SUPT20H;refGene_tx=XM_006927233.4;refGene_gn=SUPT20H;refGene_tx=XM_006927234.4;refGene_gn=SUPT20H;refGene_tx=XM_006927235.2;refGene_gn=SUPT20H;refGene_tx=XM_006927236.2;refGene_gn=SUPT20H;refGene_tx=XM_006927237.4;refGene_gn=SUPT20H;refGene_tx=XM_011285155.3;refGene_gn=SUPT20H;refGene_tx=XM_011285320.2;refGene_gn=SUPT20H;refGene_tx=XM_011285350.3;refGene_gn=SUPT20H;refGene_tx=XM_011285378.3;refGene_gn=SUPT

20H;refGene_tx=XM_011285402.3;refGene_gn=SUPT20H;refGene_tx=XM_0112854
32.2;refGene_gn=SUPT20H;refGene_tx=XM_011285467.2;refGene_gn=SUPT20H;r
efGene_tx=XM_011285509.3;refGene_gn=SUPT20H;refGene_tx=XM_011285532.3;
refGene_gn=SUPT20H;refGene_tx=XM_011285553.2;refGene_gn=SUPT20H;refGen
e_tx=XM_011285584.2;refGene_gn=SUPT20H;refGene_tx=XM_019825611.2;refGe
ne_gn=SUPT20H;refGene_tx=XM_019825613.2;refGene_gn=SUPT20H;refGene_tx=
XM_019825615.2;refGene_gn=SUPT20H;refGene_tx=XM_019825616.1;refGene_gn
=SUPT20H;refGene_tx=XM_023250917.1
A1 15828742 15828901
refGene_gn=SUPT20H;refGene_tx=XM_006927233.4;refGene_gn=SUPT20H;refGen
e_tx=XM_006927234.4;refGene_gn=SUPT20H;refGene_tx=XM_006927235.2;refGe
ne_gn=SUPT20H;refGene_tx=XM_006927236.2;refGene_gn=SUPT20H;refGene_tx=
XM_006927237.4;refGene_gn=SUPT20H;refGene_tx=XM_011285155.3;refGene_gn
=SUPT20H;refGene_tx=XM_011285320.2;refGene_gn=SUPT20H;refGene_tx=XM_01
1285350.3;refGene_gn=SUPT20H;refGene_tx=XM_011285378.3;refGene_gn=SUPT
20H;refGene_tx=XM_011285402.3;refGene_gn=SUPT20H;refGene_tx=XM_0112854
32.2;refGene_gn=SUPT20H;refGene_tx=XM_011285467.2;refGene_gn=SUPT20H;r
efGene_tx=XM_011285509.3;refGene_gn=SUPT20H;refGene_tx=XM_011285532.3;
refGene_gn=SUPT20H;refGene_tx=XM_011285553.2;refGene_gn=SUPT20H;refGen
e_tx=XM_011285584.2;refGene_gn=SUPT20H;refGene_tx=XM_019825611.2;refGe
ne_gn=SUPT20H;refGene_tx=XM_019825613.2;refGene_gn=SUPT20H;refGene_tx=
XM_019825615.2;refGene_gn=SUPT20H;refGene_tx=XM_019825616.1;refGene_gn
=SUPT20H;refGene_tx=XM_023250917.1
A1 15829044 15829147
refGene_gn=SUPT20H;refGene_tx=XM_006927233.4;refGene_gn=SUPT20H;refGen
e_tx=XM_006927234.4;refGene_gn=SUPT20H;refGene_tx=XM_006927235.2;refGe
ne_gn=SUPT20H;refGene_tx=XM_006927236.2;refGene_gn=SUPT20H;refGene_tx=
XM_006927237.4;refGene_gn=SUPT20H;refGene_tx=XM_011285155.3;refGene_gn
=SUPT20H;refGene_tx=XM_011285320.2;refGene_gn=SUPT20H;refGene_tx=XM_01
1285350.3;refGene_gn=SUPT20H;refGene_tx=XM_011285378.3;refGene_gn=SUPT
20H;refGene_tx=XM_011285402.3;refGene_gn=SUPT20H;refGene_tx=XM_0112854
32.2;refGene_gn=SUPT20H;refGene_tx=XM_011285467.2;refGene_gn=SUPT20H;r
efGene_tx=XM_011285509.3;refGene_gn=SUPT20H;refGene_tx=XM_011285532.3;
refGene_gn=SUPT20H;refGene_tx=XM_011285553.2;refGene_gn=SUPT20H;refGen
e_tx=XM_011285584.2;refGene_gn=SUPT20H;refGene_tx=XM_019825611.2;refGe
ne_gn=SUPT20H;refGene_tx=XM_019825613.2;refGene_gn=SUPT20H;refGene_tx=
XM_019825615.2;refGene_gn=SUPT20H;refGene_tx=XM_019825616.1;refGene_gn
=SUPT20H;refGene_tx=XM_023250917.1
A1 15829687 15829805
refGene_gn=SUPT20H;refGene_tx=XM_006927233.4;refGene_gn=SUPT20H;refGen
e_tx=XM_006927234.4;refGene_gn=SUPT20H;refGene_tx=XM_006927235.2;refGe
ne_gn=SUPT20H;refGene_tx=XM_006927236.2;refGene_gn=SUPT20H;refGene_tx=
XM_006927237.4;refGene_gn=SUPT20H;refGene_tx=XM_011285155.3;refGene_gn
=SUPT20H;refGene_tx=XM_011285320.2;refGene_gn=SUPT20H;refGene_tx=XM_01
1285350.3;refGene_gn=SUPT20H;refGene_tx=XM_011285378.3;refGene_gn=SUPT
20H;refGene_tx=XM_011285402.3;refGene_gn=SUPT20H;refGene_tx=XM_0112854
32.2;refGene_gn=SUPT20H;refGene_tx=XM_011285467.2;refGene_gn=SUPT20H;r
efGene_tx=XM_011285509.3;refGene_gn=SUPT20H;refGene_tx=XM_011285532.3;
refGene_gn=SUPT20H;refGene_tx=XM_011285553.2;refGene_gn=SUPT20H;refGen
e_tx=XM_011285584.2;refGene_gn=SUPT20H;refGene_tx=XM_019825611.2;refGe

ne_gn=SUPT20H;refGene_tx=XM_019825613.2;refGene_gn=SUPT20H;refGene_tx=XM_019825615.2;refGene_gn=SUPT20H;refGene_tx=XM_019825616.1;refGene_gn=SUPT20H;refGene_tx=XM_023250917.1
A1 15830628 15830661
refGene_gn=SUPT20H;refGene_tx=XM_006927233.4;refGene_gn=SUPT20H;refGene_tx=XM_006927234.4;refGene_gn=SUPT20H;refGene_tx=XM_006927235.2;refGene_gn=SUPT20H;refGene_tx=XM_006927236.2;refGene_gn=SUPT20H;refGene_tx=XM_006927237.4;refGene_gn=SUPT20H;refGene_tx=XM_011285155.3;refGene_gn=SUPT20H;refGene_tx=XM_011285320.2;refGene_gn=SUPT20H;refGene_tx=XM_011285350.3;refGene_gn=SUPT20H;refGene_tx=XM_011285378.3;refGene_gn=SUPT20H;refGene_tx=XM_011285402.3;refGene_gn=SUPT20H;refGene_tx=XM_011285432.2;refGene_gn=SUPT20H;refGene_tx=XM_011285467.2;refGene_gn=SUPT20H;refGene_tx=XM_011285509.3;refGene_gn=SUPT20H;refGene_tx=XM_011285532.3;refGene_gn=SUPT20H;refGene_tx=XM_011285553.2;refGene_gn=SUPT20H;refGene_tx=XM_011285584.2;refGene_gn=SUPT20H;refGene_tx=XM_019825611.2;refGene_gn=SUPT20H;refGene_tx=XM_019825613.2;refGene_gn=SUPT20H;refGene_tx=XM_019825615.2;refGene_gn=SUPT20H;refGene_tx=XM_019825616.1;refGene_gn=SUPT20H;refGene_tx=XM_023250917.1
A1 15831223 15831250
refGene_gn=SUPT20H;refGene_tx=XM_006927233.4;refGene_gn=SUPT20H;refGene_tx=XM_006927234.4;refGene_gn=SUPT20H;refGene_tx=XM_006927235.2;refGene_gn=SUPT20H;refGene_tx=XM_006927236.2;refGene_gn=SUPT20H;refGene_tx=XM_006927237.4;refGene_gn=SUPT20H;refGene_tx=XM_011285155.3;refGene_gn=SUPT20H;refGene_tx=XM_011285320.2;refGene_gn=SUPT20H;refGene_tx=XM_011285350.3;refGene_gn=SUPT20H;refGene_tx=XM_011285402.3;refGene_gn=SUPT20H;refGene_tx=XM_011285432.2;refGene_gn=SUPT20H;refGene_tx=XM_011285467.2;refGene_gn=SUPT20H;refGene_tx=XM_011285509.3;refGene_gn=SUPT20H;refGene_tx=XM_011285532.3;refGene_gn=SUPT20H;refGene_tx=XM_011285553.2;refGene_gn=SUPT20H;refGene_tx=XM_011285584.2;refGene_gn=SUPT20H;refGene_tx=XM_019825611.2;refGene_gn=SUPT20H;refGene_tx=XM_019825613.2;refGene_gn=SUPT20H;refGene_tx=XM_019825615.2;refGene_gn=SUPT20H;refGene_tx=XM_019825616.1;refGene_gn=SUPT20H;refGene_tx=XM_023250917.1
A1 15832454 15832612
refGene_gn=SUPT20H;refGene_tx=XM_006927233.4;refGene_gn=SUPT20H;refGene_tx=XM_006927234.4;refGene_gn=SUPT20H;refGene_tx=XM_006927235.2;refGene_gn=SUPT20H;refGene_tx=XM_006927236.2;refGene_gn=SUPT20H;refGene_tx=XM_006927237.4;refGene_gn=SUPT20H;refGene_tx=XM_011285155.3;refGene_gn=SUPT20H;refGene_tx=XM_011285320.2;refGene_gn=SUPT20H;refGene_tx=XM_011285350.3;refGene_gn=SUPT20H;refGene_tx=XM_011285378.3;refGene_gn=SUPT20H;refGene_tx=XM_011285402.3;refGene_gn=SUPT20H;refGene_tx=XM_011285432.2;refGene_gn=SUPT20H;refGene_tx=XM_011285467.2;refGene_gn=SUPT20H;refGene_tx=XM_011285509.3;refGene_gn=SUPT20H;refGene_tx=XM_011285532.3;refGene_gn=SUPT20H;refGene_tx=XM_011285553.2;refGene_gn=SUPT20H;refGene_tx=XM_011285584.2;refGene_gn=SUPT20H;refGene_tx=XM_019825611.2;refGene_gn=SUPT20H;refGene_tx=XM_019825613.2;refGene_gn=SUPT20H;refGene_tx=XM_019825615.2;refGene_gn=SUPT20H;refGene_tx=XM_019825616.1;refGene_gn=SUPT20H;refGene_tx=XM_023250917.1
A1 15833133 15833205
refGene_gn=SUPT20H;refGene_tx=XM_006927233.4;refGene_gn=SUPT20H;refGene_tx=XM_006927234.4;refGene_gn=SUPT20H;refGene_tx=XM_006927235.2;refGe

ne_gn=SUPT20H;refGene_tx=XM_006927236.2;refGene_gn=SUPT20H;refGene_tx=XM_006927237.4;refGene_gn=SUPT20H;refGene_tx=XM_011285155.3;refGene_gn=SUPT20H;refGene_tx=XM_011285320.2;refGene_gn=SUPT20H;refGene_tx=XM_011285350.3;refGene_gn=SUPT20H;refGene_tx=XM_011285378.3;refGene_gn=SUPT20H;refGene_tx=XM_011285402.3;refGene_gn=SUPT20H;refGene_tx=XM_011285432.2;refGene_gn=SUPT20H;refGene_tx=XM_011285467.2;refGene_gn=SUPT20H;refGene_tx=XM_011285509.3;refGene_gn=SUPT20H;refGene_tx=XM_011285532.3;refGene_gn=SUPT20H;refGene_tx=XM_011285553.2;refGene_gn=SUPT20H;refGene_tx=XM_011285584.2;refGene_gn=SUPT20H;refGene_tx=XM_019825611.2;refGene_gn=SUPT20H;refGene_tx=XM_019825613.2;refGene_gn=SUPT20H;refGene_tx=XM_019825615.2;refGene_gn=SUPT20H;refGene_tx=XM_019825616.1;refGene_gn=SUPT20H;refGene_tx=XM_023250917.1
A1 15836674 15836731
refGene_gn=SUPT20H;refGene_tx=XM_006927233.4;refGene_gn=SUPT20H;refGene_tx=XM_006927234.4;refGene_gn=SUPT20H;refGene_tx=XM_006927235.2;refGene_gn=SUPT20H;refGene_tx=XM_006927236.2;refGene_gn=SUPT20H;refGene_tx=XM_006927237.4;refGene_gn=SUPT20H;refGene_tx=XM_011285155.3;refGene_gn=SUPT20H;refGene_tx=XM_011285320.2;refGene_gn=SUPT20H;refGene_tx=XM_011285350.3;refGene_gn=SUPT20H;refGene_tx=XM_011285378.3;refGene_gn=SUPT20H;refGene_tx=XM_011285402.3;refGene_gn=SUPT20H;refGene_tx=XM_011285432.2;refGene_gn=SUPT20H;refGene_tx=XM_011285467.2;refGene_gn=SUPT20H;refGene_tx=XM_011285509.3;refGene_gn=SUPT20H;refGene_tx=XM_011285532.3;refGene_gn=SUPT20H;refGene_tx=XM_011285553.2;refGene_gn=SUPT20H;refGene_tx=XM_011285584.2;refGene_gn=SUPT20H;refGene_tx=XM_019825611.2;refGene_gn=SUPT20H;refGene_tx=XM_019825613.2;refGene_gn=SUPT20H;refGene_tx=XM_019825615.2;refGene_gn=SUPT20H;refGene_tx=XM_019825616.1;refGene_gn=SUPT20H;refGene_tx=XM_023250917.1
A1 15836822 15836979
refGene_gn=SUPT20H;refGene_tx=XM_006927233.4;refGene_gn=SUPT20H;refGene_tx=XM_006927234.4;refGene_gn=SUPT20H;refGene_tx=XM_006927235.2;refGene_gn=SUPT20H;refGene_tx=XM_006927236.2;refGene_gn=SUPT20H;refGene_tx=XM_006927237.4;refGene_gn=SUPT20H;refGene_tx=XM_011285155.3;refGene_gn=SUPT20H;refGene_tx=XM_011285320.2;refGene_gn=SUPT20H;refGene_tx=XM_011285350.3;refGene_gn=SUPT20H;refGene_tx=XM_011285378.3;refGene_gn=SUPT20H;refGene_tx=XM_011285402.3;refGene_gn=SUPT20H;refGene_tx=XM_011285432.2;refGene_gn=SUPT20H;refGene_tx=XM_011285467.2;refGene_gn=SUPT20H;refGene_tx=XM_011285509.3;refGene_gn=SUPT20H;refGene_tx=XM_011285532.3;refGene_gn=SUPT20H;refGene_tx=XM_011285553.2;refGene_gn=SUPT20H;refGene_tx=XM_011285584.2;refGene_gn=SUPT20H;refGene_tx=XM_019825611.2;refGene_gn=SUPT20H;refGene_tx=XM_019825613.2;refGene_gn=SUPT20H;refGene_tx=XM_019825615.2;refGene_gn=SUPT20H;refGene_tx=XM_019825616.1;refGene_gn=SUPT20H;refGene_tx=XM_023250917.1
A1 15838691 15838831
refGene_gn=SUPT20H;refGene_tx=XM_006927233.4;refGene_gn=SUPT20H;refGene_tx=XM_006927234.4;refGene_gn=SUPT20H;refGene_tx=XM_006927235.2;refGene_gn=SUPT20H;refGene_tx=XM_006927236.2;refGene_gn=SUPT20H;refGene_tx=XM_006927237.4;refGene_gn=SUPT20H;refGene_tx=XM_011285155.3;refGene_gn=SUPT20H;refGene_tx=XM_011285320.2;refGene_gn=SUPT20H;refGene_tx=XM_011285350.3;refGene_gn=SUPT20H;refGene_tx=XM_011285378.3;refGene_gn=SUPT20H;refGene_tx=XM_011285402.3;refGene_gn=SUPT20H;refGene_tx=XM_0112854

32.2;refGene_gn=SUPT20H;refGene_tx=XM_011285467.2;refGene_gn=SUPT20H;refGene_tx=XM_011285509.3;refGene_gn=SUPT20H;refGene_tx=XM_011285532.3;refGene_gn=SUPT20H;refGene_tx=XM_011285553.2;refGene_gn=SUPT20H;refGene_tx=XM_011285584.2;refGene_gn=SUPT20H;refGene_tx=XM_019825611.2;refGene_gn=SUPT20H;refGene_tx=XM_019825613.2;refGene_gn=SUPT20H;refGene_tx=XM_019825615.2;refGene_gn=SUPT20H;refGene_tx=XM_019825616.1;refGene_gn=SUPT20H;refGene_tx=XM_023250917.1
A1 15842155 15842209
refGene_gn=SUPT20H;refGene_tx=XM_006927233.4;refGene_gn=SUPT20H;refGene_tx=XM_006927234.4;refGene_gn=SUPT20H;refGene_tx=XM_006927235.2;refGene_gn=SUPT20H;refGene_tx=XM_006927236.2;refGene_gn=SUPT20H;refGene_tx=XM_006927237.4;refGene_gn=SUPT20H;refGene_tx=XM_011285155.3;refGene_gn=SUPT20H;refGene_tx=XM_011285320.2;refGene_gn=SUPT20H;refGene_tx=XM_011285350.3;refGene_gn=SUPT20H;refGene_tx=XM_011285378.3;refGene_gn=SUPT20H;refGene_tx=XM_011285432.2;refGene_gn=SUPT20H;refGene_tx=XM_011285467.2;refGene_gn=SUPT20H;refGene_tx=XM_011285509.3;refGene_gn=SUPT20H;refGene_tx=XM_011285532.3;refGene_gn=SUPT20H;refGene_tx=XM_011285553.2;refGene_gn=SUPT20H;refGene_tx=XM_011285584.2;refGene_gn=SUPT20H;refGene_tx=XM_019825611.2;refGene_gn=SUPT20H;refGene_tx=XM_019825613.2;refGene_gn=SUPT20H;refGene_tx=XM_019825615.2;refGene_gn=SUPT20H;refGene_tx=XM_019825616.1;refGene_gn=SUPT20H;refGene_tx=XM_023250917.1
A1 15842889 15843006
refGene_gn=SUPT20H;refGene_tx=XM_006927233.4;refGene_gn=SUPT20H;refGene_tx=XM_006927234.4;refGene_gn=SUPT20H;refGene_tx=XM_006927235.2;refGene_gn=SUPT20H;refGene_tx=XM_006927236.2;refGene_gn=SUPT20H;refGene_tx=XM_006927237.4;refGene_gn=SUPT20H;refGene_tx=XM_011285155.3;refGene_gn=SUPT20H;refGene_tx=XM_011285320.2;refGene_gn=SUPT20H;refGene_tx=XM_011285350.3;refGene_gn=SUPT20H;refGene_tx=XM_011285378.3;refGene_gn=SUPT20H;refGene_tx=XM_011285402.3;refGene_gn=SUPT20H;refGene_tx=XM_011285432.2;refGene_gn=SUPT20H;refGene_tx=XM_011285467.2;refGene_gn=SUPT20H;refGene_tx=XM_011285509.3;refGene_gn=SUPT20H;refGene_tx=XM_011285532.3;refGene_gn=SUPT20H;refGene_tx=XM_011285553.2;refGene_gn=SUPT20H;refGene_tx=XM_011285584.2;refGene_gn=SUPT20H;refGene_tx=XM_019825611.2;refGene_gn=SUPT20H;refGene_tx=XM_019825613.2;refGene_gn=SUPT20H;refGene_tx=XM_019825615.2;refGene_gn=SUPT20H;refGene_tx=XM_019825616.1;refGene_gn=SUPT20H;refGene_tx=XM_023250917.1
A1 15845938 15846042
refGene_gn=SUPT20H;refGene_tx=XM_006927233.4;refGene_gn=SUPT20H;refGene_tx=XM_006927234.4;refGene_gn=SUPT20H;refGene_tx=XM_006927235.2;refGene_gn=SUPT20H;refGene_tx=XM_006927236.2;refGene_gn=SUPT20H;refGene_tx=XM_006927237.4;refGene_gn=SUPT20H;refGene_tx=XM_011285155.3;refGene_gn=SUPT20H;refGene_tx=XM_011285320.2;refGene_gn=SUPT20H;refGene_tx=XM_011285350.3;refGene_gn=SUPT20H;refGene_tx=XM_011285378.3;refGene_gn=SUPT20H;refGene_tx=XM_011285402.3;refGene_gn=SUPT20H;refGene_tx=XM_011285432.2;refGene_gn=SUPT20H;refGene_tx=XM_011285467.2;refGene_gn=SUPT20H;refGene_tx=XM_011285509.3;refGene_gn=SUPT20H;refGene_tx=XM_011285532.3;refGene_gn=SUPT20H;refGene_tx=XM_011285553.2;refGene_gn=SUPT20H;refGene_tx=XM_011285584.2;refGene_gn=SUPT20H;refGene_tx=XM_019825611.2;refGene_gn=SUPT20H;refGene_tx=XM_019825613.2;refGene_gn=SUPT20H;refGene_tx=XM_019825615.2;refGene_gn=SUPT20H;refGene_tx=XM_019825616.1;refGene_gn

=SUPT20H;refGene_tx=XM_023250917.1
A1 15847021 15847151
refGene_gn=SUPT20H;refGene_tx=XM_006927233.4;refGene_gn=SUPT20H;refGene_tx=XM_006927234.4;refGene_gn=SUPT20H;refGene_tx=XM_006927235.2;refGene_gn=SUPT20H;refGene_tx=XM_006927236.2;refGene_gn=SUPT20H;refGene_tx=XM_006927237.4;refGene_gn=SUPT20H;refGene_tx=XM_011285155.3;refGene_gn=SUPT20H;refGene_tx=XM_011285320.2;refGene_gn=SUPT20H;refGene_tx=XM_011285350.3;refGene_gn=SUPT20H;refGene_tx=XM_011285378.3;refGene_gn=SUPT20H;refGene_tx=XM_011285402.3;refGene_gn=SUPT20H;refGene_tx=XM_011285432.2;refGene_gn=SUPT20H;refGene_tx=XM_011285467.2;refGene_gn=SUPT20H;refGene_tx=XM_011285509.3;refGene_gn=SUPT20H;refGene_tx=XM_011285532.3;refGene_gn=SUPT20H;refGene_tx=XM_011285553.2;refGene_gn=SUPT20H;refGene_tx=XM_011285584.2;refGene_gn=SUPT20H;refGene_tx=XM_019825611.2;refGene_gn=SUPT20H;refGene_tx=XM_019825613.2;refGene_gn=SUPT20H;refGene_tx=XM_019825615.2;refGene_gn=SUPT20H;refGene_tx=XM_019825616.1;refGene_gn=SUPT20H;refGene_tx=XM_023250917.1
A1 15850165 15850232
refGene_gn=SUPT20H;refGene_tx=XM_006927233.4;refGene_gn=SUPT20H;refGene_tx=XM_006927234.4;refGene_gn=SUPT20H;refGene_tx=XM_006927235.2;refGene_gn=SUPT20H;refGene_tx=XM_006927236.2;refGene_gn=SUPT20H;refGene_tx=XM_006927237.4;refGene_gn=SUPT20H;refGene_tx=XM_011285155.3;refGene_gn=SUPT20H;refGene_tx=XM_011285320.2;refGene_gn=SUPT20H;refGene_tx=XM_011285350.3;refGene_gn=SUPT20H;refGene_tx=XM_011285378.3;refGene_gn=SUPT20H;refGene_tx=XM_011285402.3;refGene_gn=SUPT20H;refGene_tx=XM_011285432.2;refGene_gn=SUPT20H;refGene_tx=XM_011285467.2;refGene_gn=SUPT20H;refGene_tx=XM_011285509.3;refGene_gn=SUPT20H;refGene_tx=XM_011285532.3;refGene_gn=SUPT20H;refGene_tx=XM_011285553.2;refGene_gn=SUPT20H;refGene_tx=XM_011285584.2;refGene_gn=SUPT20H;refGene_tx=XM_019825611.2;refGene_gn=SUPT20H;refGene_tx=XM_019825613.2;refGene_gn=SUPT20H;refGene_tx=XM_019825615.2;refGene_gn=SUPT20H;refGene_tx=XM_019825616.1;refGene_gn=SUPT20H;refGene_tx=XM_023250917.1
A1 15850491 15850550
refGene_gn=SUPT20H;refGene_tx=XM_006927233.4;refGene_gn=SUPT20H;refGene_tx=XM_006927234.4;refGene_gn=SUPT20H;refGene_tx=XM_006927235.2;refGene_gn=SUPT20H;refGene_tx=XM_006927236.2;refGene_gn=SUPT20H;refGene_tx=XM_006927237.4;refGene_gn=SUPT20H;refGene_tx=XM_011285155.3;refGene_gn=SUPT20H;refGene_tx=XM_011285320.2;refGene_gn=SUPT20H;refGene_tx=XM_011285350.3;refGene_gn=SUPT20H;refGene_tx=XM_011285378.3;refGene_gn=SUPT20H;refGene_tx=XM_011285402.3;refGene_gn=SUPT20H;refGene_tx=XM_011285432.2;refGene_gn=SUPT20H;refGene_tx=XM_011285467.2;refGene_gn=SUPT20H;refGene_tx=XM_011285509.3;refGene_gn=SUPT20H;refGene_tx=XM_011285532.3;refGene_gn=SUPT20H;refGene_tx=XM_011285553.2;refGene_gn=SUPT20H;refGene_tx=XM_011285584.2;refGene_gn=SUPT20H;refGene_tx=XM_019825611.2;refGene_gn=SUPT20H;refGene_tx=XM_019825613.2;refGene_gn=SUPT20H;refGene_tx=XM_019825615.2;refGene_gn=SUPT20H;refGene_tx=XM_019825616.1;refGene_gn=SUPT20H;refGene_tx=XM_023250917.1
A1 15851207 15851243
refGene_gn=SUPT20H;refGene_tx=XM_006927233.4;refGene_gn=SUPT20H;refGene_tx=XM_006927234.4;refGene_gn=SUPT20H;refGene_tx=XM_006927235.2;refGene_gn=SUPT20H;refGene_tx=XM_006927236.2;refGene_gn=SUPT20H;refGene_tx=

XM_006927237.4;refGene_gn=SUPT20H;refGene_tx=XM_011285155.3;refGene_gn=SUPT20H;refGene_tx=XM_011285320.2;refGene_gn=SUPT20H;refGene_tx=XM_011285350.3;refGene_gn=SUPT20H;refGene_tx=XM_011285378.3;refGene_gn=SUPT20H;refGene_tx=XM_011285402.3;refGene_gn=SUPT20H;refGene_tx=XM_011285432.2;refGene_gn=SUPT20H;refGene_tx=XM_011285467.2;refGene_gn=SUPT20H;refGene_tx=XM_011285509.3;refGene_gn=SUPT20H;refGene_tx=XM_011285532.3;refGene_gn=SUPT20H;refGene_tx=XM_011285553.2;refGene_gn=SUPT20H;refGene_tx=XM_011285584.2;refGene_gn=SUPT20H;refGene_tx=XM_023250917.1
A1 15854250 15854253
refGene_gn=SUPT20H;refGene_tx=XM_006927233.4;refGene_gn=SUPT20H;refGene_tx=XM_006927234.4;refGene_gn=SUPT20H;refGene_tx=XM_006927235.2;refGene_gn=SUPT20H;refGene_tx=XM_006927236.2;refGene_gn=SUPT20H;refGene_tx=XM_006927237.4;refGene_gn=SUPT20H;refGene_tx=XM_011285155.3;refGene_gn=SUPT20H;refGene_tx=XM_011285320.2;refGene_gn=SUPT20H;refGene_tx=XM_011285350.3;refGene_gn=SUPT20H;refGene_tx=XM_011285378.3;refGene_gn=SUPT20H;refGene_tx=XM_011285402.3;refGene_gn=SUPT20H;refGene_tx=XM_011285432.2;refGene_gn=SUPT20H;refGene_tx=XM_011285467.2;refGene_gn=SUPT20H;refGene_tx=XM_011285509.3;refGene_gn=SUPT20H;refGene_tx=XM_011285532.3;refGene_gn=SUPT20H;refGene_tx=XM_011285553.2;refGene_gn=SUPT20H;refGene_tx=XM_011285584.2;refGene_gn=SUPT20H;refGene_tx=XM_019825611.2;refGene_gn=SUPT20H;refGene_tx=XM_019825613.2;refGene_gn=SUPT20H;refGene_tx=XM_019825615.2;refGene_gn=SUPT20H;refGene_tx=XM_019825616.1;refGene_gn=SUPT20H;refGene_tx=XM_023250917.1
A1 16331461 16331499
refGene_gn=POSTN;refGene_tx=XM_011284921.3;refGene_gn=POSTN;refGene_tx=XM_011284946.3;refGene_gn=POSTN;refGene_tx=XM_011284973.3;refGene_gn=POSTN;refGene_tx=XM_011284999.3;refGene_gn=POSTN;refGene_tx=XM_011285020.3;refGene_gn=POSTN;refGene_tx=XM_011285038.3;refGene_gn=POSTN;refGene_tx=XM_011285062.3
A1 16332667 16332709
refGene_gn=POSTN;refGene_tx=XM_011284921.3;refGene_gn=POSTN;refGene_tx=XM_011284946.3;refGene_gn=POSTN;refGene_tx=XM_011284973.3;refGene_gn=POSTN;refGene_tx=XM_011284999.3;refGene_gn=POSTN;refGene_tx=XM_011285020.3;refGene_gn=POSTN;refGene_tx=XM_011285038.3;refGene_gn=POSTN;refGene_tx=XM_011285062.3
A1 16337067 16337151
refGene_gn=POSTN;refGene_tx=XM_011284921.3;refGene_gn=POSTN;refGene_tx=XM_011284946.3;refGene_gn=POSTN;refGene_tx=XM_011284999.3;refGene_gn=POSTN;refGene_tx=XM_011285038.3
A1 16337542 16337620
refGene_gn=POSTN;refGene_tx=XM_011284921.3;refGene_gn=POSTN;refGene_tx=XM_011284946.3;refGene_gn=POSTN;refGene_tx=XM_011284973.3;refGene_gn=POSTN;refGene_tx=XM_011284999.3;refGene_gn=POSTN;refGene_tx=XM_011285020.3;refGene_gn=POSTN;refGene_tx=XM_011285038.3;refGene_gn=POSTN;refGene_tx=XM_011285062.3
A1 16338599 16338689
refGene_gn=POSTN;refGene_tx=XM_011284921.3;refGene_gn=POSTN;refGene_tx=XM_011284946.3;refGene_gn=POSTN;refGene_tx=XM_011284973.3;refGene_gn=POSTN;refGene_tx=XM_011284999.3;refGene_gn=POSTN;refGene_tx=XM_011285020.3;refGene_gn=POSTN;refGene_tx=XM_011285038.3;refGene_gn=POSTN;refGe

ne_tx=XM_011285062.3
A1 16339360 16339450
refGene_gn=POSTN;refGene_tx=XM_011284921.3;refGene_gn=POSTN;refGene_tx=XM_011284946.3;refGene_gn=POSTN;refGene_tx=XM_011284973.3;refGene_gn=POSTN;refGene_tx=XM_011285020.3
A1 16342591 16342672
refGene_gn=POSTN;refGene_tx=XM_011284921.3;refGene_gn=POSTN;refGene_tx=XM_011284973.3;refGene_gn=POSTN;refGene_tx=XM_011284999.3
A1 16346818 16346864
refGene_gn=POSTN;refGene_tx=XM_011284921.3;refGene_gn=POSTN;refGene_tx=XM_011284946.3;refGene_gn=POSTN;refGene_tx=XM_011284973.3;refGene_gn=POSTN;refGene_tx=XM_011284999.3;refGene_gn=POSTN;refGene_tx=XM_011285020.3;refGene_gn=POSTN;refGene_tx=XM_011285038.3;refGene_gn=POSTN;refGene_tx=XM_011285062.3
A1 16347588 16347656
refGene_gn=POSTN;refGene_tx=XM_011284921.3;refGene_gn=POSTN;refGene_tx=XM_011284946.3;refGene_gn=POSTN;refGene_tx=XM_011284973.3;refGene_gn=POSTN;refGene_tx=XM_011284999.3;refGene_gn=POSTN;refGene_tx=XM_011285020.3;refGene_gn=POSTN;refGene_tx=XM_011285038.3;refGene_gn=POSTN;refGene_tx=XM_011285062.3
A1 16347762 16347865
refGene_gn=POSTN;refGene_tx=XM_011284921.3;refGene_gn=POSTN;refGene_tx=XM_011284946.3;refGene_gn=POSTN;refGene_tx=XM_011284973.3;refGene_gn=POSTN;refGene_tx=XM_011284999.3;refGene_gn=POSTN;refGene_tx=XM_011285020.3;refGene_gn=POSTN;refGene_tx=XM_011285038.3;refGene_gn=POSTN;refGene_tx=XM_011285062.3
A1 16347971 16348102
refGene_gn=POSTN;refGene_tx=XM_011284921.3;refGene_gn=POSTN;refGene_tx=XM_011284946.3;refGene_gn=POSTN;refGene_tx=XM_011284973.3;refGene_gn=POSTN;refGene_tx=XM_011284999.3;refGene_gn=POSTN;refGene_tx=XM_011285020.3;refGene_gn=POSTN;refGene_tx=XM_011285038.3;refGene_gn=POSTN;refGene_tx=XM_011285062.3
A1 16348641 16348772
refGene_gn=POSTN;refGene_tx=XM_011284921.3;refGene_gn=POSTN;refGene_tx=XM_011284946.3;refGene_gn=POSTN;refGene_tx=XM_011284973.3;refGene_gn=POSTN;refGene_tx=XM_011284999.3;refGene_gn=POSTN;refGene_tx=XM_011285020.3;refGene_gn=POSTN;refGene_tx=XM_011285038.3;refGene_gn=POSTN;refGene_tx=XM_011285062.3
A1 16350171 16350308
refGene_gn=POSTN;refGene_tx=XM_011284921.3;refGene_gn=POSTN;refGene_tx=XM_011284946.3;refGene_gn=POSTN;refGene_tx=XM_011284973.3;refGene_gn=POSTN;refGene_tx=XM_011284999.3;refGene_gn=POSTN;refGene_tx=XM_011285020.3;refGene_gn=POSTN;refGene_tx=XM_011285038.3;refGene_gn=POSTN;refGene_tx=XM_011285062.3
A1 16351412 16351561
refGene_gn=POSTN;refGene_tx=XM_011284921.3;refGene_gn=POSTN;refGene_tx=XM_011284946.3;refGene_gn=POSTN;refGene_tx=XM_011284973.3;refGene_gn=POSTN;refGene_tx=XM_011284999.3;refGene_gn=POSTN;refGene_tx=XM_011285020.3;refGene_gn=POSTN;refGene_tx=XM_011285038.3;refGene_gn=POSTN;refGene_tx=XM_011285062.3

A1 16353199 16353334
refGene_gn=POSTN;refGene_tx=XM_011284921.3;refGene_gn=POSTN;refGene_tx=XM_011284946.3;refGene_gn=POSTN;refGene_tx=XM_011284973.3;refGene_gn=POSTN;refGene_tx=XM_011284999.3;refGene_gn=POSTN;refGene_tx=XM_011285020.3;refGene_gn=POSTN;refGene_tx=XM_011285038.3;refGene_gn=POSTN;refGene_tx=XM_011285062.3
A1 16353914 16354127
refGene_gn=POSTN;refGene_tx=XM_011284921.3;refGene_gn=POSTN;refGene_tx=XM_011284946.3;refGene_gn=POSTN;refGene_tx=XM_011284973.3;refGene_gn=POSTN;refGene_tx=XM_011284999.3;refGene_gn=POSTN;refGene_tx=XM_011285020.3;refGene_gn=POSTN;refGene_tx=XM_011285038.3;refGene_gn=POSTN;refGene_tx=XM_011285062.3
A1 16355251 16355393
refGene_gn=POSTN;refGene_tx=XM_011284921.3;refGene_gn=POSTN;refGene_tx=XM_011284946.3;refGene_gn=POSTN;refGene_tx=XM_011284973.3;refGene_gn=POSTN;refGene_tx=XM_011284999.3;refGene_gn=POSTN;refGene_tx=XM_011285020.3;refGene_gn=POSTN;refGene_tx=XM_011285038.3;refGene_gn=POSTN;refGene_tx=XM_011285062.3
A1 16355889 16356036
refGene_gn=POSTN;refGene_tx=XM_011284921.3;refGene_gn=POSTN;refGene_tx=XM_011284946.3;refGene_gn=POSTN;refGene_tx=XM_011284973.3;refGene_gn=POSTN;refGene_tx=XM_011284999.3;refGene_gn=POSTN;refGene_tx=XM_011285020.3;refGene_gn=POSTN;refGene_tx=XM_011285038.3;refGene_gn=POSTN;refGene_tx=XM_011285062.3
A1 16356942 16357107
refGene_gn=POSTN;refGene_tx=XM_011284921.3;refGene_gn=POSTN;refGene_tx=XM_011284946.3;refGene_gn=POSTN;refGene_tx=XM_011284973.3;refGene_gn=POSTN;refGene_tx=XM_011284999.3;refGene_gn=POSTN;refGene_tx=XM_011285020.3;refGene_gn=POSTN;refGene_tx=XM_011285038.3;refGene_gn=POSTN;refGene_tx=XM_011285062.3
A1 16359484 16359642
refGene_gn=POSTN;refGene_tx=XM_011284921.3;refGene_gn=POSTN;refGene_tx=XM_011284946.3;refGene_gn=POSTN;refGene_tx=XM_011284973.3;refGene_gn=POSTN;refGene_tx=XM_011284999.3;refGene_gn=POSTN;refGene_tx=XM_011285020.3;refGene_gn=POSTN;refGene_tx=XM_011285038.3;refGene_gn=POSTN;refGene_tx=XM_011285062.3
A1 16361405 16361470
refGene_gn=POSTN;refGene_tx=XM_011284921.3;refGene_gn=POSTN;refGene_tx=XM_011284946.3;refGene_gn=POSTN;refGene_tx=XM_011284973.3;refGene_gn=POSTN;refGene_tx=XM_011284999.3;refGene_gn=POSTN;refGene_tx=XM_011285020.3;refGene_gn=POSTN;refGene_tx=XM_011285038.3;refGene_gn=POSTN;refGene_tx=XM_011285062.3
A1 16366294 16366393
refGene_gn=POSTN;refGene_tx=XM_011284921.3;refGene_gn=POSTN;refGene_tx=XM_011284946.3;refGene_gn=POSTN;refGene_tx=XM_011284973.3;refGene_gn=POSTN;refGene_tx=XM_011284999.3;refGene_gn=POSTN;refGene_tx=XM_011285020.3;refGene_gn=POSTN;refGene_tx=XM_011285038.3;refGene_gn=POSTN;refGene_tx=XM_011285062.3
A1 16367692 16367811
refGene_gn=POSTN;refGene_tx=XM_011284921.3;refGene_gn=POSTN;refGene_tx

=XM_011284946.3;refGene_gn=POSTN;refGene_tx=XM_011284973.3;refGene_gn=POSTN;refGene_tx=XM_011284999.3;refGene_gn=POSTN;refGene_tx=XM_011285020.3;refGene_gn=POSTN;refGene_tx=XM_011285038.3;refGene_gn=POSTN;refGene_tx=XM_011285062.3
A1 16409724 16410447 refGene_gn=TRPC4;refGene_tx=XM_011284874.3;refGene_gn=TRPC4;refGene_tx=XM_019825540.2;refGene_gn=TRPC4;refGene_tx=XM_019825579.2;refGene_gn=TRPC4;refGene_tx=XM_023250940.1;refGene_gn=TRPC4;refGene_tx=XM_023250949.1
A1 16412619 16412709 refGene_gn=TRPC4;refGene_tx=XM_011284874.3;refGene_gn=TRPC4;refGene_tx=XM_019825540.2;refGene_gn=TRPC4;refGene_tx=XM_019825579.2;refGene_gn=TRPC4;refGene_tx=XM_023250940.1;refGene_gn=TRPC4;refGene_tx=XM_023250949.1
A1 16412836 16412878 refGene_gn=TRPC4;refGene_tx=XM_011284874.3;refGene_gn=TRPC4;refGene_tx=XM_019825540.2;refGene_gn=TRPC4;refGene_tx=XM_019825579.2;refGene_gn=TRPC4;refGene_tx=XM_023250940.1;refGene_gn=TRPC4;refGene_tx=XM_023250949.1
A1 16426818 16427013 refGene_gn=TRPC4;refGene_tx=XM_011284874.3;refGene_gn=TRPC4;refGene_tx=XM_019825540.2;refGene_gn=TRPC4;refGene_tx=XM_019825579.2;refGene_gn=TRPC4;refGene_tx=XM_023250940.1;refGene_gn=TRPC4;refGene_tx=XM_023250949.1
A1 16430315 16430511 refGene_gn=TRPC4;refGene_tx=XM_011284874.3;refGene_gn=TRPC4;refGene_tx=XM_019825540.2;refGene_gn=TRPC4;refGene_tx=XM_019825579.2;refGene_gn=TRPC4;refGene_tx=XM_023250940.1;refGene_gn=TRPC4;refGene_tx=XM_023250949.1
A1 16438395 16438709 refGene_gn=TRPC4;refGene_tx=XM_011284874.3;refGene_gn=TRPC4;refGene_tx=XM_019825540.2;refGene_gn=TRPC4;refGene_tx=XM_019825579.2;refGene_gn=TRPC4;refGene_tx=XM_023250940.1;refGene_gn=TRPC4;refGene_tx=XM_023250949.1
A1 16448132 16448272 refGene_gn=TRPC4;refGene_tx=XM_011284874.3;refGene_gn=TRPC4;refGene_tx=XM_019825540.2;refGene_gn=TRPC4;refGene_tx=XM_019825579.2;refGene_gn=TRPC4;refGene_tx=XM_023250940.1;refGene_gn=TRPC4;refGene_tx=XM_023250949.1
A1 16467610 16467947 refGene_gn=TRPC4;refGene_tx=XM_011284874.3;refGene_gn=TRPC4;refGene_tx=XM_019825540.2;refGene_gn=TRPC4;refGene_tx=XM_019825579.2;refGene_gn=TRPC4;refGene_tx=XM_023250940.1;refGene_gn=TRPC4;refGene_tx=XM_023250949.1
A1 16479131 16479140 refGene_gn=TRPC4;refGene_tx=XM_023250949.1
A1 16519175 16519694 refGene_gn=TRPC4;refGene_tx=XM_011284874.3;refGene_gn=TRPC4;refGene_tx=XM_019825540.2;refGene_gn=TRPC4;refGene_tx=XM_019825579.2;refGene_gn=TRPC4;refGene_tx=XM_023250940.1
A1 16553455 16553833

refGene_gn=TRPC4;refGene_tx=XM_011284874.3;refGene_gn=TRPC4;refGene_tx
=XM_019825540.2;refGene_gn=TRPC4;refGene_tx=XM_019825579.2;refGene_gn=
TRPC4;refGene_tx=XM_023250940.1
A1 16980763 16980765 refGene_gn=UFM1;refGene_tx=XM_006927246.2
A1 16980855 16980912 refGene_gn=UFM1;refGene_tx=XM_006927246.2
A1 16983351 16983409 refGene_gn=UFM1;refGene_tx=XM_006927246.2
A1 16987192 16987232 refGene_gn=UFM1;refGene_tx=XM_006927246.2
A1 16988443 16988476 refGene_gn=UFM1;refGene_tx=XM_006927246.2
A1 16989954 16990022 refGene_gn=UFM1;refGene_tx=XM_006927246.2
A1 17273647 17278829
refGene_gn=FREM2;refGene_tx=XM_003980340.5;refGene_gn=FREM2;refGene_tx
=XM_023250973.1
A1 17284088 17284178
refGene_gn=FREM2;refGene_tx=XM_003980340.5;refGene_gn=FREM2;refGene_tx
=XM_023250973.1
A1 17347794 17347941
refGene_gn=FREM2;refGene_tx=XM_003980340.5;refGene_gn=FREM2;refGene_tx
=XM_023250973.1
A1 17355001 17355232
refGene_gn=FREM2;refGene_tx=XM_003980340.5;refGene_gn=FREM2;refGene_tx
=XM_023250973.1
A1 17369450 17369576
refGene_gn=FREM2;refGene_tx=XM_003980340.5;refGene_gn=FREM2;refGene_tx
=XM_023250973.1
A1 17370637 17370889
refGene_gn=FREM2;refGene_tx=XM_003980340.5;refGene_gn=FREM2;refGene_tx
=XM_023250973.1
A1 17401151 17401301
refGene_gn=FREM2;refGene_tx=XM_003980340.5;refGene_gn=FREM2;refGene_tx
=XM_023250973.1
A1 17403286 17403496
refGene_gn=FREM2;refGene_tx=XM_003980340.5;refGene_gn=FREM2;refGene_tx
=XM_023250973.1
A1 17404935 17405133
refGene_gn=FREM2;refGene_tx=XM_003980340.5;refGene_gn=FREM2;refGene_tx
=XM_023250973.1
A1 17405923 17406088
refGene_gn=FREM2;refGene_tx=XM_003980340.5;refGene_gn=FREM2;refGene_tx
=XM_023250973.1
A1 17406880 17407063
refGene_gn=FREM2;refGene_tx=XM_003980340.5;refGene_gn=FREM2;refGene_tx
=XM_023250973.1
A1 17410589 17410720
refGene_gn=FREM2;refGene_tx=XM_003980340.5;refGene_gn=FREM2;refGene_tx
=XM_023250973.1
A1 17412253 17412412
refGene_gn=FREM2;refGene_tx=XM_003980340.5;refGene_gn=FREM2;refGene_tx
=XM_023250973.1
A1 17413696 17414000
refGene_gn=FREM2;refGene_tx=XM_003980340.5;refGene_gn=FREM2;refGene_tx

=XM_023250973.1
A1 17416672 17416804
refGene_gn=FREM2;refGene_tx=XM_003980340.5;refGene_gn=FREM2;refGene_tx=XM_023250973.1
A1 17419264 17419596
refGene_gn=FREM2;refGene_tx=XM_003980340.5;refGene_gn=FREM2;refGene_tx=XM_023250973.1
A1 17428226 17428419
refGene_gn=FREM2;refGene_tx=XM_003980340.5;refGene_gn=FREM2;refGene_tx=XM_023250973.1
A1 17430436 17430541
refGene_gn=FREM2;refGene_tx=XM_003980340.5;refGene_gn=FREM2;refGene_tx=XM_023250973.1
A1 17431947 17432075
refGene_gn=FREM2;refGene_tx=XM_003980340.5;refGene_gn=FREM2;refGene_tx=XM_023250973.1
A1 17432422 17432557
refGene_gn=FREM2;refGene_tx=XM_003980340.5;refGene_gn=FREM2;refGene_tx=XM_023250973.1
A1 17433573 17433700
refGene_gn=FREM2;refGene_tx=XM_003980340.5;refGene_gn=FREM2;refGene_tx=XM_023250973.1
A1 17434540 17434728
refGene_gn=FREM2;refGene_tx=XM_003980340.5;refGene_gn=FREM2;refGene_tx=XM_023250973.1
A1 17434946 17435093
refGene_gn=FREM2;refGene_tx=XM_003980340.5;refGene_gn=FREM2;refGene_tx=XM_023250973.1
A1 17435864 17435888 refGene_gn=FREM2;refGene_tx=XM_023250973.1
A1 17436590 17437103 refGene_gn=FREM2;refGene_tx=XM_003980340.5
A1 17449882 17450107
refGene_gn=STOML3;refGene_tx=XM_019825717.2;refGene_gn=STOML3;refGene_tx=XM_019825723.2
A1 17451924 17452059
refGene_gn=STOML3;refGene_tx=XM_019825717.2;refGene_gn=STOML3;refGene_tx=XM_019825723.2
A1 17453915 17454119
refGene_gn=STOML3;refGene_tx=XM_019825717.2;refGene_gn=STOML3;refGene_tx=XM_019825723.2
A1 17455588 17455671
refGene_gn=STOML3;refGene_tx=XM_019825717.2;refGene_gn=STOML3;refGene_tx=XM_019825723.2
A1 17457812 17457885
refGene_gn=STOML3;refGene_tx=XM_019825717.2;refGene_gn=STOML3;refGene_tx=XM_019825723.2
A1 17457964 17458068
refGene_gn=STOML3;refGene_tx=XM_019825717.2;refGene_gn=STOML3;refGene_tx=XM_019825723.2
A1 17471064 17471116
refGene_gn=STOML3;refGene_tx=XM_019825717.2;refGene_gn=STOML3;refGene_

tx=XM_019825723.2
A1 17479949 17480072 refGene_gn=PROSER1;refGene_tx=XM_023251008.1
A1 17480685 17480836 refGene_gn=PROSER1;refGene_tx=XM_023251008.1
A1 17481285 17483068 refGene_gn=PROSER1;refGene_tx=XM_023251008.1
A1 17486940 17486985 refGene_gn=PROSER1;refGene_tx=XM_023251008.1
A1 17490762 17490858 refGene_gn=PROSER1;refGene_tx=XM_023251008.1
A1 17491436 17491515 refGene_gn=PROSER1;refGene_tx=XM_023251008.1
A1 17492871 17492955 refGene_gn=PROSER1;refGene_tx=XM_023251008.1
A1 17494489 17494600 refGene_gn=PROSER1;refGene_tx=XM_023251008.1
A1 17496939 17497033 refGene_gn=PROSER1;refGene_tx=XM_023251008.1
A1 17498061 17498156 refGene_gn=PROSER1;refGene_tx=XM_023251008.1
A1 17501016 17501085 refGene_gn=PROSER1;refGene_tx=XM_023251008.1
A1 17503573 17503639 refGene_gn=PROSER1;refGene_tx=XM_023251008.1
A1 17506134 17506179 refGene_gn=PROSER1;refGene_tx=XM_023251008.1
A1 17507530 17507614
refGene_gn=NHLRC3;refGene_tx=XM_003980342.5;refGene_gn=NHLRC3;refGene_
tx=XM_006927248.4;refGene_gn=NHLRC3;refGene_tx=XM_019825736.2
A1 17507826 17507855
refGene_gn=NHLRC3;refGene_tx=XM_006927248.4;refGene_gn=NHLRC3;refGene_
tx=XM_019825736.2
A1 17508292 17508445
refGene_gn=NHLRC3;refGene_tx=XM_003980342.5;refGene_gn=NHLRC3;refGene_
tx=XM_006927248.4;refGene_gn=NHLRC3;refGene_tx=XM_019825736.2
A1 17508748 17508896
refGene_gn=NHLRC3;refGene_tx=XM_003980342.5;refGene_gn=NHLRC3;refGene_
tx=XM_006927248.4;refGene_gn=NHLRC3;refGene_tx=XM_019825736.2
A1 17511060 17511261
refGene_gn=NHLRC3;refGene_tx=XM_003980342.5;refGene_gn=NHLRC3;refGene_
tx=XM_006927248.4;refGene_gn=NHLRC3;refGene_tx=XM_019825736.2
A1 17513273 17513365
refGene_gn=NHLRC3;refGene_tx=XM_003980342.5;refGene_gn=NHLRC3;refGene_
tx=XM_006927248.4;refGene_gn=NHLRC3;refGene_tx=XM_019825736.2
A1 17515971 17516084
refGene_gn=NHLRC3;refGene_tx=XM_003980342.5;refGene_gn=NHLRC3;refGene_
tx=XM_006927248.4;refGene_gn=NHLRC3;refGene_tx=XM_019825736.2
A1 17516688 17516944
refGene_gn=NHLRC3;refGene_tx=XM_003980342.5;refGene_gn=NHLRC3;refGene_
tx=XM_006927248.4;refGene_gn=NHLRC3;refGene_tx=XM_019825736.2
A1 17760418 17760537
refGene_gn=LHFPL6;refGene_tx=XM_003980348.5;refGene_gn=LHFPL6;refGene_
tx=XM_019825737.2
A1 17789384 17789483 refGene_gn=LHFPL6;refGene_tx=XM_003980348.5
A1 18009103 18009488
refGene_gn=LHFPL6;refGene_tx=XM_003980348.5;refGene_gn=LHFPL6;refGene_
tx=XM_019825737.2
A1 18049823 18049976
refGene_gn=COG6;refGene_tx=XM_003980349.5;refGene_gn=COG6;refGene_tx=X
M_019825758.2;refGene_gn=COG6;refGene_tx=XM_019825759.2
A1 18052832 18052976
refGene_gn=COG6;refGene_tx=XM_003980349.5;refGene_gn=COG6;refGene_tx=X

M_019825758.2;refGene_gn=COG6;refGene_tx=XM_019825759.2
A1 18058118 18058190
refGene_gn=COG6;refGene_tx=XM_003980349.5;refGene_gn=COG6;refGene_tx=XM_019825758.2;refGene_gn=COG6;refGene_tx=XM_019825759.2
A1 18062635 18062694
refGene_gn=COG6;refGene_tx=XM_003980349.5;refGene_gn=COG6;refGene_tx=XM_019825758.2;refGene_gn=COG6;refGene_tx=XM_019825759.2
A1 18068546 18068658
refGene_gn=COG6;refGene_tx=XM_003980349.5;refGene_gn=COG6;refGene_tx=XM_019825758.2;refGene_gn=COG6;refGene_tx=XM_019825759.2
A1 18070348 18070431
refGene_gn=COG6;refGene_tx=XM_003980349.5;refGene_gn=COG6;refGene_tx=XM_019825758.2;refGene_gn=COG6;refGene_tx=XM_019825759.2
A1 18070801 18070872
refGene_gn=COG6;refGene_tx=XM_003980349.5;refGene_gn=COG6;refGene_tx=XM_019825758.2;refGene_gn=COG6;refGene_tx=XM_019825759.2
A1 18072249 18072343
refGene_gn=COG6;refGene_tx=XM_003980349.5;refGene_gn=COG6;refGene_tx=XM_019825758.2;refGene_gn=COG6;refGene_tx=XM_019825759.2
A1 18075948 18076077
refGene_gn=COG6;refGene_tx=XM_003980349.5;refGene_gn=COG6;refGene_tx=XM_019825758.2;refGene_gn=COG6;refGene_tx=XM_019825759.2
A1 18076155 18076247
refGene_gn=COG6;refGene_tx=XM_003980349.5;refGene_gn=COG6;refGene_tx=XM_019825758.2;refGene_gn=COG6;refGene_tx=XM_019825759.2
A1 18077727 18077792
refGene_gn=COG6;refGene_tx=XM_003980349.5;refGene_gn=COG6;refGene_tx=XM_019825758.2;refGene_gn=COG6;refGene_tx=XM_019825759.2
A1 18080511 18080603
refGene_gn=COG6;refGene_tx=XM_003980349.5;refGene_gn=COG6;refGene_tx=XM_019825758.2;refGene_gn=COG6;refGene_tx=XM_019825759.2
A1 18086548 18086666
refGene_gn=COG6;refGene_tx=XM_003980349.5;refGene_gn=COG6;refGene_tx=XM_019825758.2;refGene_gn=COG6;refGene_tx=XM_019825759.2
A1 18108607 18108739
refGene_gn=COG6;refGene_tx=XM_003980349.5;refGene_gn=COG6;refGene_tx=XM_019825758.2;refGene_gn=COG6;refGene_tx=XM_019825759.2
A1 18109063 18109231
refGene_gn=COG6;refGene_tx=XM_003980349.5;refGene_gn=COG6;refGene_tx=XM_019825758.2;refGene_gn=COG6;refGene_tx=XM_019825759.2
A1 18112297 18112405
refGene_gn=COG6;refGene_tx=XM_003980349.5;refGene_gn=COG6;refGene_tx=XM_019825758.2;refGene_gn=COG6;refGene_tx=XM_019825759.2
A1 18113630 18113684
refGene_gn=COG6;refGene_tx=XM_003980349.5;refGene_gn=COG6;refGene_tx=XM_019825758.2;refGene_gn=COG6;refGene_tx=XM_019825759.2
A1 18116729 18116809
refGene_gn=COG6;refGene_tx=XM_003980349.5;refGene_gn=COG6;refGene_tx=XM_019825758.2;refGene_gn=COG6;refGene_tx=XM_019825759.2
A1 18116900 18116924

refGene_gn=COG6;refGene_tx=XM_019825758.2;refGene_gn=COG6;refGene_tx=XM_019825759.2
A1 18135168 18135316 refGene_gn=COG6;refGene_tx=XM_003980349.5
A1 18853382 18854720 refGene_gn=FOXO1;refGene_tx=XM_023251080.1
A1 18946069 18946720 refGene_gn=FOXO1;refGene_tx=XM_023251080.1
A1 18955008 18955058 refGene_gn=MRPS31;refGene_tx=XM_019825811.2
A1 18986690 18986920
refGene_gn=MRPS31;refGene_tx=XM_019825795.2;refGene_gn=MRPS31;refGene_tx=XM_019825799.2
A1 18994172 18994282
refGene_gn=MRPS31;refGene_tx=XM_006927255.3;refGene_gn=MRPS31;refGene_tx=XM_019825809.2
A1 18997734 18997878
refGene_gn=MRPS31;refGene_tx=XM_006927255.3;refGene_gn=MRPS31;refGene_tx=XM_019825795.2;refGene_gn=MRPS31;refGene_tx=XM_019825799.2;refGene_gn=MRPS31;refGene_tx=XM_019825809.2;refGene_gn=MRPS31;refGene_tx=XM_019825811.2
A1 19001311 19001391
refGene_gn=MRPS31;refGene_tx=XM_006927255.3;refGene_gn=MRPS31;refGene_tx=XM_019825795.2;refGene_gn=MRPS31;refGene_tx=XM_019825799.2;refGene_gn=MRPS31;refGene_tx=XM_019825809.2;refGene_gn=MRPS31;refGene_tx=XM_019825811.2
A1 19004321 19004462
refGene_gn=MRPS31;refGene_tx=XM_006927255.3;refGene_gn=MRPS31;refGene_tx=XM_019825795.2;refGene_gn=MRPS31;refGene_tx=XM_019825799.2;refGene_gn=MRPS31;refGene_tx=XM_019825809.2;refGene_gn=MRPS31;refGene_tx=XM_019825811.2
A1 19005456 19005615
refGene_gn=MRPS31;refGene_tx=XM_006927255.3;refGene_gn=MRPS31;refGene_tx=XM_019825795.2;refGene_gn=MRPS31;refGene_tx=XM_019825799.2;refGene_gn=MRPS31;refGene_tx=XM_019825809.2;refGene_gn=MRPS31;refGene_tx=XM_019825811.2
A1 19012984 19013269
refGene_gn=MRPS31;refGene_tx=XM_006927255.3;refGene_gn=MRPS31;refGene_tx=XM_019825795.2;refGene_gn=MRPS31;refGene_tx=XM_019825799.2;refGene_gn=MRPS31;refGene_tx=XM_019825809.2;refGene_gn=MRPS31;refGene_tx=XM_019825811.2
A1 19018083 19018208
refGene_gn=MRPS31;refGene_tx=XM_006927255.3;refGene_gn=MRPS31;refGene_tx=XM_019825795.2;refGene_gn=MRPS31;refGene_tx=XM_019825799.2;refGene_gn=MRPS31;refGene_tx=XM_019825809.2;refGene_gn=MRPS31;refGene_tx=XM_019825811.2
A1 19048568 19048623
refGene_gn=SLC25A15;refGene_tx=XM_003980351.5;refGene_gn=SLC25A15;refGene_tx=XM_019825873.2;refGene_gn=SLC25A15;refGene_tx=XM_023251112.1
A1 19069094 19069353
refGene_gn=SLC25A15;refGene_tx=XM_003980351.5;refGene_gn=SLC25A15;refGene_tx=XM_006927256.3;refGene_gn=SLC25A15;refGene_tx=XM_019825873.2;refGene_gn=SLC25A15;refGene_tx=XM_023251112.1
A1 19074241 19074379

refGene_gn=SLC25A15;refGene_tx=XM_003980351.5;refGene_gn=SLC25A15;refGene_tx=XM_006927256.3;refGene_gn=SLC25A15;refGene_tx=XM_019825873.2;refGene_gn=SLC25A15;refGene_tx=XM_023251112.1
A1 19078014 19078184
refGene_gn=SLC25A15;refGene_tx=XM_003980351.5;refGene_gn=SLC25A15;refGene_tx=XM_006927256.3;refGene_gn=SLC25A15;refGene_tx=XM_019825873.2;refGene_gn=SLC25A15;refGene_tx=XM_023251112.1
A1 19079075 19079234
refGene_gn=SLC25A15;refGene_tx=XM_003980351.5;refGene_gn=SLC25A15;refGene_tx=XM_006927256.3;refGene_gn=SLC25A15;refGene_tx=XM_019825873.2;refGene_gn=SLC25A15;refGene_tx=XM_023251112.1
A1 19080704 19080829
refGene_gn=SLC25A15;refGene_tx=XM_003980351.5;refGene_gn=SLC25A15;refGene_tx=XM_006927256.3;refGene_gn=SLC25A15;refGene_tx=XM_019825873.2;refGene_gn=SLC25A15;refGene_tx=XM_023251112.1
A1 19084858 19084973
refGene_gn=LOC101096867;refGene_tx=XM_011284412.3;refGene_gn=LOC101096867;refGene_tx=XM_019825824.2;refGene_gn=LOC101096867;refGene_tx=XM_019825849.2;refGene_gn=LOC101096867;refGene_tx=XM_023251097.1
A1 19086913 19086984
refGene_gn=LOC101096867;refGene_tx=XM_011284412.3;refGene_gn=LOC101096867;refGene_tx=XM_019825824.2;refGene_gn=LOC101096867;refGene_tx=XM_019825849.2;refGene_gn=LOC101096867;refGene_tx=XM_023251097.1
A1 19088955 19089051
refGene_gn=LOC101096867;refGene_tx=XM_011284412.3;refGene_gn=LOC101096867;refGene_tx=XM_019825824.2;refGene_gn=LOC101096867;refGene_tx=XM_019825849.2;refGene_gn=LOC101096867;refGene_tx=XM_023251097.1
A1 19091474 19091566
refGene_gn=LOC101096867;refGene_tx=XM_011284412.3;refGene_gn=LOC101096867;refGene_tx=XM_019825824.2;refGene_gn=LOC101096867;refGene_tx=XM_019825849.2;refGene_gn=LOC101096867;refGene_tx=XM_023251097.1
A1 19106833 19106939
refGene_gn=LOC101096867;refGene_tx=XM_011284412.3;refGene_gn=LOC101096867;refGene_tx=XM_019825824.2;refGene_gn=LOC101096867;refGene_tx=XM_019825849.2;refGene_gn=LOC101096867;refGene_tx=XM_023251097.1
A1 19113436 19113517
refGene_gn=LOC101096867;refGene_tx=XM_011284412.3;refGene_gn=LOC101096867;refGene_tx=XM_019825824.2;refGene_gn=LOC101096867;refGene_tx=XM_019825849.2;refGene_gn=LOC101096867;refGene_tx=XM_023251097.1
A1 19115496 19115558
refGene_gn=LOC101096867;refGene_tx=XM_011284412.3;refGene_gn=LOC101096867;refGene_tx=XM_019825824.2;refGene_gn=LOC101096867;refGene_tx=XM_019825849.2;refGene_gn=LOC101096867;refGene_tx=XM_023251097.1
A1 19147614 19147703
refGene_gn=LOC101096867;refGene_tx=XM_011284412.3;refGene_gn=LOC101096867;refGene_tx=XM_019825824.2;refGene_gn=LOC101096867;refGene_tx=XM_019825849.2;refGene_gn=LOC101096867;refGene_tx=XM_023251097.1
A1 19147805 19147887
refGene_gn=LOC101096867;refGene_tx=XM_011284412.3;refGene_gn=LOC101096867;refGene_tx=XM_019825824.2;refGene_gn=LOC101096867;refGene_tx=XM_01

9825849.2;refGene_gn=LOC101096867;refGene_tx=XM_023251097.1
A1 19148713 19148774
refGene_gn=LOC101096867;refGene_tx=XM_011284412.3;refGene_gn=LOC101096867;refGene_tx=XM_019825824.2;refGene_gn=LOC101096867;refGene_tx=XM_019825849.2;refGene_gn=LOC101096867;refGene_tx=XM_023251097.1
A1 19170352 19170417
refGene_gn=LOC101096867;refGene_tx=XM_011284412.3;refGene_gn=LOC101096867;refGene_tx=XM_019825824.2;refGene_gn=LOC101096867;refGene_tx=XM_019825849.2;refGene_gn=LOC101096867;refGene_tx=XM_023251097.1
A1 19171366 19171430
refGene_gn=LOC101096867;refGene_tx=XM_011284412.3;refGene_gn=LOC101096867;refGene_tx=XM_019825824.2;refGene_gn=LOC101096867;refGene_tx=XM_019825849.2;refGene_gn=LOC101096867;refGene_tx=XM_023251097.1
A1 19171569 19171669
refGene_gn=LOC101096867;refGene_tx=XM_011284412.3;refGene_gn=LOC101096867;refGene_tx=XM_019825824.2;refGene_gn=LOC101096867;refGene_tx=XM_019825849.2;refGene_gn=LOC101096867;refGene_tx=XM_023251097.1
A1 19173161 19173281
refGene_gn=LOC101096867;refGene_tx=XM_011284412.3;refGene_gn=LOC101096867;refGene_tx=XM_019825824.2;refGene_gn=LOC101096867;refGene_tx=XM_019825849.2;refGene_gn=LOC101096867;refGene_tx=XM_023251097.1
A1 19176032 19176194
refGene_gn=LOC101096867;refGene_tx=XM_011284412.3;refGene_gn=LOC101096867;refGene_tx=XM_019825824.2;refGene_gn=LOC101096867;refGene_tx=XM_019825849.2;refGene_gn=LOC101096867;refGene_tx=XM_023251097.1
A1 19177352 19177400
refGene_gn=LOC101096867;refGene_tx=XM_011284412.3;refGene_gn=LOC101096867;refGene_tx=XM_019825824.2;refGene_gn=LOC101096867;refGene_tx=XM_019825849.2;refGene_gn=LOC101096867;refGene_tx=XM_023251097.1
A1 19195180 19195249
refGene_gn=LOC101096867;refGene_tx=XM_011284412.3;refGene_gn=LOC101096867;refGene_tx=XM_019825824.2;refGene_gn=LOC101096867;refGene_tx=XM_019825849.2;refGene_gn=LOC101096867;refGene_tx=XM_023251097.1
A1 19196093 19196147
refGene_gn=LOC101096867;refGene_tx=XM_011284412.3;refGene_gn=LOC101096867;refGene_tx=XM_019825824.2;refGene_gn=LOC101096867;refGene_tx=XM_019825849.2;refGene_gn=LOC101096867;refGene_tx=XM_023251097.1
A1 19198810 19198864
refGene_gn=LOC101096867;refGene_tx=XM_011284412.3;refGene_gn=LOC101096867;refGene_tx=XM_019825824.2;refGene_gn=LOC101096867;refGene_tx=XM_019825849.2;refGene_gn=LOC101096867;refGene_tx=XM_023251097.1
A1 19199773 19199827
refGene_gn=LOC101096867;refGene_tx=XM_011284412.3;refGene_gn=LOC101096867;refGene_tx=XM_019825824.2;refGene_gn=LOC101096867;refGene_tx=XM_019825849.2;refGene_gn=LOC101096867;refGene_tx=XM_023251097.1
A1 19200778 19200832
refGene_gn=LOC101096867;refGene_tx=XM_011284412.3;refGene_gn=LOC101096867;refGene_tx=XM_019825824.2;refGene_gn=LOC101096867;refGene_tx=XM_019825849.2;refGene_gn=LOC101096867;refGene_tx=XM_023251097.1
A1 19208452 19208506

refGene_gn=LOC101096867;refGene_tx=XM_011284412.3;refGene_gn=LOC101096867;refGene_tx=XM_019825824.2;refGene_gn=LOC101096867;refGene_tx=XM_019825849.2;refGene_gn=LOC101096867;refGene_tx=XM_023251097.1
A1 19209486 19209540 refGene_gn=LOC101096867;refGene_tx=XM_011284412.3;refGene_gn=LOC101096867;refGene_tx=XM_019825824.2;refGene_gn=LOC101096867;refGene_tx=XM_019825849.2;refGene_gn=LOC101096867;refGene_tx=XM_023251097.1
A1 19210664 19210718 refGene_gn=LOC101096867;refGene_tx=XM_011284412.3;refGene_gn=LOC101096867;refGene_tx=XM_019825824.2;refGene_gn=LOC101096867;refGene_tx=XM_019825849.2;refGene_gn=LOC101096867;refGene_tx=XM_023251097.1
A1 19211720 19211774 refGene_gn=LOC101096867;refGene_tx=XM_011284412.3;refGene_gn=LOC101096867;refGene_tx=XM_019825824.2;refGene_gn=LOC101096867;refGene_tx=XM_019825849.2;refGene_gn=LOC101096867;refGene_tx=XM_023251097.1
A1 19219805 19219862 refGene_gn=LOC101096867;refGene_tx=XM_011284412.3;refGene_gn=LOC101096867;refGene_tx=XM_019825824.2;refGene_gn=LOC101096867;refGene_tx=XM_019825849.2;refGene_gn=LOC101096867;refGene_tx=XM_023251097.1
A1 19248426 19248480 refGene_gn=LOC101096867;refGene_tx=XM_019825824.2;refGene_gn=LOC101096867;refGene_tx=XM_023251097.1
A1 19249387 19249426 refGene_gn=LOC101096867;refGene_tx=XM_019825824.2;refGene_gn=LOC101096867;refGene_tx=XM_019825849.2;refGene_gn=LOC101096867;refGene_tx=XM_023251097.1
A1 19252861 19252920 refGene_gn=LOC101096867;refGene_tx=XM_011284412.3;refGene_gn=LOC101096867;refGene_tx=XM_019825824.2;refGene_gn=LOC101096867;refGene_tx=XM_019825849.2;refGene_gn=LOC101096867;refGene_tx=XM_023251097.1
A1 19328888 19330337 refGene_gn=THSD1;refGene_tx=XM_003980353.5;refGene_gn=THSD1;refGene_tx=XM_019825819.2;refGene_gn=THSD1;refGene_tx=XM_023251129.1;refGene_gn=THSD1;refGene_tx=XM_023251135.1
A1 19337243 19337402 refGene_gn=THSD1;refGene_tx=XM_019825819.2;refGene_gn=THSD1;refGene_tx=XM_023251129.1;refGene_gn=THSD1;refGene_tx=XM_023251135.1
A1 19349116 19350082 refGene_gn=THSD1;refGene_tx=XM_003980353.5;refGene_gn=THSD1;refGene_tx=XM_019825819.2;refGene_gn=THSD1;refGene_tx=XM_023251129.1;refGene_gn=THSD1;refGene_tx=XM_023251135.1
A1 19354623 19354681 refGene_gn=THSD1;refGene_tx=XM_003980353.5;refGene_gn=THSD1;refGene_tx=XM_019825819.2;refGene_gn=THSD1;refGene_tx=XM_023251129.1;refGene_gn=THSD1;refGene_tx=XM_023251135.1
A1 19364098 19364192 refGene_gn=VPS36;refGene_tx=XM_023251151.1
A1 19364275 19364352 refGene_gn=VPS36;refGene_tx=XM_023251151.1
A1 19365255 19365340 refGene_gn=VPS36;refGene_tx=XM_023251151.1
A1 19365952 19366017 refGene_gn=VPS36;refGene_tx=XM_023251151.1
A1 19367408 19367474 refGene_gn=VPS36;refGene_tx=XM_023251151.1

A1 19369101 19369236 refGene_gn=VPS36;refGene_tx=XM_023251151.1
A1 19369866 19369944 refGene_gn=VPS36;refGene_tx=XM_023251151.1
A1 19370065 19370098 refGene_gn=VPS36;refGene_tx=XM_023251151.1
A1 19377863 19377950 refGene_gn=VPS36;refGene_tx=XM_023251151.1
A1 19378667 19378757 refGene_gn=VPS36;refGene_tx=XM_023251151.1
A1 19379826 19379941 refGene_gn=VPS36;refGene_tx=XM_023251151.1
A1 19382696 19382767 refGene_gn=VPS36;refGene_tx=XM_023251151.1
A1 19385095 19385164 refGene_gn=VPS36;refGene_tx=XM_023251151.1
A1 19391415 19391511 refGene_gn=VPS36;refGene_tx=XM_023251151.1
A1 19409228 19409298 refGene_gn=CKAP2;refGene_tx=XM_011284205.3
A1 19410241 19410323 refGene_gn=CKAP2;refGene_tx=XM_011284205.3
A1 19416866 19416945 refGene_gn=CKAP2;refGene_tx=XM_011284205.3
A1 19417056 19417332 refGene_gn=CKAP2;refGene_tx=XM_011284205.3
A1 19417333 19417920 refGene_gn=CKAP2;refGene_tx=XM_011284205.3
A1 19418370 19418575 refGene_gn=CKAP2;refGene_tx=XM_011284205.3
A1 19419816 19419987 refGene_gn=CKAP2;refGene_tx=XM_011284205.3
A1 19422174 19422244 refGene_gn=CKAP2;refGene_tx=XM_011284205.3
A1 19423757 19424010 refGene_gn=CKAP2;refGene_tx=XM_011284205.3
A1 19424503 19424759 refGene_gn=CKAP2;refGene_tx=XM_011284205.3
A1 19434186 19434303 refGene_gn=NEK3;refGene_tx=XM_023251162.1
A1 19436416 19436510 refGene_gn=NEK3;refGene_tx=XM_023251162.1
A1 19436597 19436695 refGene_gn=NEK3;refGene_tx=XM_023251162.1
A1 19438381 19438465 refGene_gn=NEK3;refGene_tx=XM_023251162.1
A1 19439037 19439105 refGene_gn=NEK3;refGene_tx=XM_023251162.1
A1 19439189 19439276 refGene_gn=NEK3;refGene_tx=XM_023251162.1
A1 19441346 19441401 refGene_gn=NEK3;refGene_tx=XM_023251162.1
A1 19444242 19444443 refGene_gn=NEK3;refGene_tx=XM_023251162.1
A1 19445080 19445153 refGene_gn=NEK3;refGene_tx=XM_023251162.1
A1 19448393 19448443 refGene_gn=NEK3;refGene_tx=XM_023251162.1
A1 19451858 19451958 refGene_gn=NEK3;refGene_tx=XM_023251162.1
A1 19452480 19452621 refGene_gn=NEK3;refGene_tx=XM_023251162.1
A1 19452875 19453010 refGene_gn=NEK3;refGene_tx=XM_023251162.1
A1 19455163 19455290 refGene_gn=NEK3;refGene_tx=XM_023251162.1
A1 19455725 19455806 refGene_gn=NEK3;refGene_tx=XM_023251162.1
A1 19455915 19455943 refGene_gn=NEK3;refGene_tx=XM_023251162.1
A1 19459174 19459437 refGene_gn=NEK5;refGene_tx=XM_019825941.2;refGene_gn=NEK5;refGene_tx=XM_019825945.2
A1 19460947 19461086 refGene_gn=NEK5;refGene_tx=XM_019825941.2;refGene_gn=NEK5;refGene_tx=XM_019825945.2;refGene_gn=NEK5;refGene_tx=XM_019825962.1;refGene_gn=NEK5;refGene_tx=XM_023251202.1
A1 19470903 19471000 refGene_gn=NEK5;refGene_tx=XM_019825941.2;refGene_gn=NEK5;refGene_tx=XM_019825945.2;refGene_gn=NEK5;refGene_tx=XM_019825962.1;refGene_gn=NEK5;refGene_tx=XM_023251202.1
A1 19477795 19477893 refGene_gn=NEK5;refGene_tx=XM_019825941.2;refGene_gn=NEK5;refGene_tx=XM_019825945.2;refGene_gn=NEK5;refGene_tx=XM_019825962.1;refGene_gn=NEK5;refGene_tx=XM_023251202.1

A1 19479556 19479640
refGene_gn=NEK5;refGene_tx=XM_019825941.2;refGene_gn=NEK5;refGene_tx=XM_019825945.2;refGene_gn=NEK5;refGene_tx=XM_019825962.1;refGene_gn=NEK5;refGene_tx=XM_023251202.1
A1 19479730 19479801
refGene_gn=NEK5;refGene_tx=XM_019825941.2;refGene_gn=NEK5;refGene_tx=XM_019825945.2;refGene_gn=NEK5;refGene_tx=XM_019825962.1;refGene_gn=NEK5;refGene_tx=XM_023251202.1
A1 19480868 19480955
refGene_gn=NEK5;refGene_tx=XM_019825941.2;refGene_gn=NEK5;refGene_tx=XM_019825945.2;refGene_gn=NEK5;refGene_tx=XM_019825962.1;refGene_gn=NEK5;refGene_tx=XM_023251202.1
A1 19482952 19483007
refGene_gn=NEK5;refGene_tx=XM_019825941.2;refGene_gn=NEK5;refGene_tx=XM_019825945.2;refGene_gn=NEK5;refGene_tx=XM_019825962.1;refGene_gn=NEK5;refGene_tx=XM_023251202.1
A1 19484918 19485119
refGene_gn=NEK5;refGene_tx=XM_019825941.2;refGene_gn=NEK5;refGene_tx=XM_019825945.2;refGene_gn=NEK5;refGene_tx=XM_019825962.1;refGene_gn=NEK5;refGene_tx=XM_023251202.1
A1 19485198 19485280
refGene_gn=NEK5;refGene_tx=XM_019825941.2;refGene_gn=NEK5;refGene_tx=XM_019825945.2;refGene_gn=NEK5;refGene_tx=XM_019825962.1;refGene_gn=NEK5;refGene_tx=XM_023251202.1
A1 19487456 19487590
refGene_gn=NEK5;refGene_tx=XM_019825941.2;refGene_gn=NEK5;refGene_tx=XM_019825945.2;refGene_gn=NEK5;refGene_tx=XM_019825962.1;refGene_gn=NEK5;refGene_tx=XM_023251202.1
A1 19490014 19490199
refGene_gn=NEK5;refGene_tx=XM_019825941.2;refGene_gn=NEK5;refGene_tx=XM_019825945.2;refGene_gn=NEK5;refGene_tx=XM_019825962.1;refGene_gn=NEK5;refGene_tx=XM_023251202.1
A1 19491018 19491085
refGene_gn=NEK5;refGene_tx=XM_019825941.2;refGene_gn=NEK5;refGene_tx=XM_019825945.2;refGene_gn=NEK5;refGene_tx=XM_019825962.1;refGene_gn=NEK5;refGene_tx=XM_023251202.1
A1 19492615 19492732
refGene_gn=NEK5;refGene_tx=XM_019825941.2;refGene_gn=NEK5;refGene_tx=XM_019825945.2;refGene_gn=NEK5;refGene_tx=XM_019825962.1;refGene_gn=NEK5;refGene_tx=XM_023251202.1
A1 19493726 19493813
refGene_gn=NEK5;refGene_tx=XM_019825941.2;refGene_gn=NEK5;refGene_tx=XM_019825945.2;refGene_gn=NEK5;refGene_tx=XM_019825962.1;refGene_gn=NEK5;refGene_tx=XM_023251202.1
A1 19495413 19495506
refGene_gn=NEK5;refGene_tx=XM_019825941.2;refGene_gn=NEK5;refGene_tx=XM_019825962.1;refGene_gn=NEK5;refGene_tx=XM_023251202.1
A1 19497829 19497910
refGene_gn=NEK5;refGene_tx=XM_019825941.2;refGene_gn=NEK5;refGene_tx=XM_019825945.2;refGene_gn=NEK5;refGene_tx=XM_019825962.1;refGene_gn=NEK

5;refGene_tx=XM_023251202.1
A1 19498138 19498207
refGene_gn=NEK5;refGene_tx=XM_019825941.2;refGene_gn=NEK5;refGene_tx=XM_019825945.2;refGene_gn=NEK5;refGene_tx=XM_019825962.1;refGene_gn=NEK5;refGene_tx=XM_023251202.1
A1 19500189 19500298
refGene_gn=NEK5;refGene_tx=XM_019825941.2;refGene_gn=NEK5;refGene_tx=XM_019825945.2;refGene_gn=NEK5;refGene_tx=XM_019825962.1;refGene_gn=NEK5;refGene_tx=XM_023251202.1
A1 19505597 19505747
refGene_gn=NEK5;refGene_tx=XM_019825941.2;refGene_gn=NEK5;refGene_tx=XM_019825945.2;refGene_gn=NEK5;refGene_tx=XM_019825962.1;refGene_gn=NEK5;refGene_tx=XM_023251202.1
A1 19506794 19506932
refGene_gn=NEK5;refGene_tx=XM_019825941.2;refGene_gn=NEK5;refGene_tx=XM_019825945.2;refGene_gn=NEK5;refGene_tx=XM_019825962.1;refGene_gn=NEK5;refGene_tx=XM_023251202.1
A1 19513030 19513136
refGene_gn=NEK5;refGene_tx=XM_019825941.2;refGene_gn=NEK5;refGene_tx=XM_019825945.2;refGene_gn=NEK5;refGene_tx=XM_019825962.1;refGene_gn=NEK5;refGene_tx=XM_023251202.1
A1 19520754 19521022
refGene_gn=NEK5;refGene_tx=XM_019825941.2;refGene_gn=NEK5;refGene_tx=XM_019825945.2;refGene_gn=NEK5;refGene_tx=XM_019825962.1;refGene_gn=NEK5;refGene_tx=XM_023251202.1
A1 19526942 19527214 refGene_gn=ALG11;refGene_tx=XM_003980354.4
A1 19529722 19530654 refGene_gn=ALG11;refGene_tx=XM_003980354.4
A1 19534315 19534546 refGene_gn=ALG11;refGene_tx=XM_003980354.4
A1 19540477 19540521 refGene_gn=ALG11;refGene_tx=XM_003980354.4
A1 19541513 19541564 refGene_gn=ATP7B;refGene_tx=XM_023251186.1
A1 19562915 19563164
refGene_gn=ATP7B;refGene_tx=XM_023251165.1;refGene_gn=ATP7B;refGene_tx=XM_023251171.1;refGene_gn=ATP7B;refGene_tx=XM_023251176.1;refGene_gn=ATP7B;refGene_tx=XM_023251182.1
A1 19568203 19568212 refGene_gn=ATP7B;refGene_tx=XM_023251193.1
A1 19573529 19574778
refGene_gn=ATP7B;refGene_tx=XM_006927260.4;refGene_gn=ATP7B;refGene_tx=XM_019825930.2;refGene_gn=ATP7B;refGene_tx=XM_023251165.1;refGene_gn=ATP7B;refGene_tx=XM_023251171.1;refGene_gn=ATP7B;refGene_tx=XM_023251176.1;refGene_gn=ATP7B;refGene_tx=XM_023251182.1;refGene_gn=ATP7B;refGene_tx=XM_023251186.1;refGene_gn=ATP7B;refGene_tx=XM_023251193.1
A1 19581759 19582017
refGene_gn=ATP7B;refGene_tx=XM_006927260.4;refGene_gn=ATP7B;refGene_tx=XM_019825930.2;refGene_gn=ATP7B;refGene_tx=XM_023251165.1;refGene_gn=ATP7B;refGene_tx=XM_023251171.1;refGene_gn=ATP7B;refGene_tx=XM_023251176.1;refGene_gn=ATP7B;refGene_tx=XM_023251182.1;refGene_gn=ATP7B;refGene_tx=XM_023251186.1;refGene_gn=ATP7B;refGene_tx=XM_023251193.1
A1 19584071 19584235
refGene_gn=ATP7B;refGene_tx=XM_006927260.4;refGene_gn=ATP7B;refGene_tx=XM_019825930.2;refGene_gn=ATP7B;refGene_tx=XM_023251165.1;refGene_gn=

ATP7B;refGene_tx=XM_023251171.1;refGene_gn=ATP7B;refGene_tx=XM_0232511
76.1;refGene_gn=ATP7B;refGene_tx=XM_023251182.1;refGene_gn=ATP7B;refGe
ne_tx=XM_023251186.1;refGene_gn=ATP7B;refGene_tx=XM_023251193.1
A1 19586041 19586203
refGene_gn=ATP7B;refGene_tx=XM_006927260.4;refGene_gn=ATP7B;refGene_tx
=XM_019825930.2;refGene_gn=ATP7B;refGene_tx=XM_023251165.1;refGene_gn=
ATP7B;refGene_tx=XM_023251171.1;refGene_gn=ATP7B;refGene_tx=XM_0232511
76.1;refGene_gn=ATP7B;refGene_tx=XM_023251182.1;refGene_gn=ATP7B;refGe
ne_tx=XM_023251186.1;refGene_gn=ATP7B;refGene_tx=XM_023251193.1
A1 19588316 19588393
refGene_gn=ATP7B;refGene_tx=XM_006927260.4;refGene_gn=ATP7B;refGene_tx
=XM_019825930.2;refGene_gn=ATP7B;refGene_tx=XM_023251165.1;refGene_gn=
ATP7B;refGene_tx=XM_023251171.1;refGene_gn=ATP7B;refGene_tx=XM_0232511
76.1;refGene_gn=ATP7B;refGene_tx=XM_023251182.1;refGene_gn=ATP7B;refGe
ne_tx=XM_023251186.1;refGene_gn=ATP7B;refGene_tx=XM_023251193.1
A1 19590156 19590331
refGene_gn=ATP7B;refGene_tx=XM_006927260.4;refGene_gn=ATP7B;refGene_tx
=XM_019825930.2;refGene_gn=ATP7B;refGene_tx=XM_023251165.1;refGene_gn=
ATP7B;refGene_tx=XM_023251171.1;refGene_gn=ATP7B;refGene_tx=XM_0232511
76.1;refGene_gn=ATP7B;refGene_tx=XM_023251182.1;refGene_gn=ATP7B;refGe
ne_tx=XM_023251186.1;refGene_gn=ATP7B;refGene_tx=XM_023251193.1
A1 19591985 19592219
refGene_gn=ATP7B;refGene_tx=XM_006927260.4;refGene_gn=ATP7B;refGene_tx
=XM_019825930.2;refGene_gn=ATP7B;refGene_tx=XM_023251165.1;refGene_gn=
ATP7B;refGene_tx=XM_023251171.1;refGene_gn=ATP7B;refGene_tx=XM_0232511
76.1;refGene_gn=ATP7B;refGene_tx=XM_023251182.1;refGene_gn=ATP7B;refGe
ne_tx=XM_023251186.1;refGene_gn=ATP7B;refGene_tx=XM_023251193.1
A1 19592920 19593012
refGene_gn=ATP7B;refGene_tx=XM_006927260.4;refGene_gn=ATP7B;refGene_tx
=XM_019825930.2;refGene_gn=ATP7B;refGene_tx=XM_023251165.1;refGene_gn=
ATP7B;refGene_tx=XM_023251171.1;refGene_gn=ATP7B;refGene_tx=XM_0232511
76.1;refGene_gn=ATP7B;refGene_tx=XM_023251182.1;refGene_gn=ATP7B;refGe
ne_tx=XM_023251186.1;refGene_gn=ATP7B;refGene_tx=XM_023251193.1
A1 19600148 19600276
refGene_gn=ATP7B;refGene_tx=XM_006927260.4;refGene_gn=ATP7B;refGene_tx
=XM_019825930.2;refGene_gn=ATP7B;refGene_tx=XM_023251165.1;refGene_gn=
ATP7B;refGene_tx=XM_023251171.1;refGene_gn=ATP7B;refGene_tx=XM_0232511
76.1;refGene_gn=ATP7B;refGene_tx=XM_023251182.1;refGene_gn=ATP7B;refGe
ne_tx=XM_023251186.1;refGene_gn=ATP7B;refGene_tx=XM_023251193.1
A1 19600390 19600545
refGene_gn=ATP7B;refGene_tx=XM_006927260.4;refGene_gn=ATP7B;refGene_tx
=XM_019825930.2;refGene_gn=ATP7B;refGene_tx=XM_023251165.1;refGene_gn=
ATP7B;refGene_tx=XM_023251171.1;refGene_gn=ATP7B;refGene_tx=XM_0232511
76.1;refGene_gn=ATP7B;refGene_tx=XM_023251182.1;refGene_gn=ATP7B;refGe
ne_tx=XM_023251186.1;refGene_gn=ATP7B;refGene_tx=XM_023251193.1
A1 19600763 19600898
refGene_gn=ATP7B;refGene_tx=XM_006927260.4;refGene_gn=ATP7B;refGene_tx
=XM_019825930.2;refGene_gn=ATP7B;refGene_tx=XM_023251165.1;refGene_gn=
ATP7B;refGene_tx=XM_023251171.1;refGene_gn=ATP7B;refGene_tx=XM_0232511
76.1;refGene_gn=ATP7B;refGene_tx=XM_023251182.1;refGene_gn=ATP7B;refGe

ne_tx=XM_023251186.1;refGene_gn=ATP7B;refGene_tx=XM_023251193.1
A1 19604064 19604259
refGene_gn=ATP7B;refGene_tx=XM_006927260.4;refGene_gn=ATP7B;refGene_tx=XM_019825930.2;refGene_gn=ATP7B;refGene_tx=XM_023251165.1;refGene_gn=ATP7B;refGene_tx=XM_023251171.1;refGene_gn=ATP7B;refGene_tx=XM_023251176.1;refGene_gn=ATP7B;refGene_tx=XM_023251182.1;refGene_gn=ATP7B;refGene_tx=XM_023251186.1;refGene_gn=ATP7B;refGene_tx=XM_023251193.1
A1 19606185 19606368
refGene_gn=ATP7B;refGene_tx=XM_006927260.4;refGene_gn=ATP7B;refGene_tx=XM_019825930.2;refGene_gn=ATP7B;refGene_tx=XM_023251165.1;refGene_gn=ATP7B;refGene_tx=XM_023251171.1;refGene_gn=ATP7B;refGene_tx=XM_023251176.1;refGene_gn=ATP7B;refGene_tx=XM_023251186.1;refGene_gn=ATP7B;refGene_tx=XM_023251193.1
A1 19607633 19607802
refGene_gn=ATP7B;refGene_tx=XM_006927260.4;refGene_gn=ATP7B;refGene_tx=XM_019825930.2;refGene_gn=ATP7B;refGene_tx=XM_023251165.1;refGene_gn=ATP7B;refGene_tx=XM_023251171.1;refGene_gn=ATP7B;refGene_tx=XM_023251176.1;refGene_gn=ATP7B;refGene_tx=XM_023251182.1;refGene_gn=ATP7B;refGene_tx=XM_023251186.1;refGene_gn=ATP7B;refGene_tx=XM_023251193.1
A1 19609466 19609610
refGene_gn=ATP7B;refGene_tx=XM_006927260.4;refGene_gn=ATP7B;refGene_tx=XM_019825930.2;refGene_gn=ATP7B;refGene_tx=XM_023251165.1;refGene_gn=ATP7B;refGene_tx=XM_023251171.1;refGene_gn=ATP7B;refGene_tx=XM_023251176.1;refGene_gn=ATP7B;refGene_tx=XM_023251182.1;refGene_gn=ATP7B;refGene_tx=XM_023251186.1;refGene_gn=ATP7B;refGene_tx=XM_023251193.1
A1 19610881 19611024
refGene_gn=ATP7B;refGene_tx=XM_006927260.4;refGene_gn=ATP7B;refGene_tx=XM_019825930.2;refGene_gn=ATP7B;refGene_tx=XM_023251165.1;refGene_gn=ATP7B;refGene_tx=XM_023251171.1;refGene_gn=ATP7B;refGene_tx=XM_023251176.1;refGene_gn=ATP7B;refGene_tx=XM_023251182.1;refGene_gn=ATP7B;refGene_tx=XM_023251186.1;refGene_gn=ATP7B;refGene_tx=XM_023251193.1
A1 19612664 19612868
refGene_gn=ATP7B;refGene_tx=XM_006927260.4;refGene_gn=ATP7B;refGene_tx=XM_019825930.2;refGene_gn=ATP7B;refGene_tx=XM_023251165.1;refGene_gn=ATP7B;refGene_tx=XM_023251171.1;refGene_gn=ATP7B;refGene_tx=XM_023251176.1;refGene_gn=ATP7B;refGene_tx=XM_023251182.1;refGene_gn=ATP7B;refGene_tx=XM_023251186.1;refGene_gn=ATP7B;refGene_tx=XM_023251193.1
A1 19612968 19613086
refGene_gn=ATP7B;refGene_tx=XM_006927260.4;refGene_gn=ATP7B;refGene_tx=XM_019825930.2;refGene_gn=ATP7B;refGene_tx=XM_023251165.1;refGene_gn=ATP7B;refGene_tx=XM_023251171.1;refGene_gn=ATP7B;refGene_tx=XM_023251176.1;refGene_gn=ATP7B;refGene_tx=XM_023251182.1;refGene_gn=ATP7B;refGene_tx=XM_023251186.1;refGene_gn=ATP7B;refGene_tx=XM_023251193.1
A1 19613550 19613653
refGene_gn=ATP7B;refGene_tx=XM_006927260.4;refGene_gn=ATP7B;refGene_tx=XM_019825930.2;refGene_gn=ATP7B;refGene_tx=XM_023251165.1;refGene_gn=ATP7B;refGene_tx=XM_023251171.1;refGene_gn=ATP7B;refGene_tx=XM_023251176.1;refGene_gn=ATP7B;refGene_tx=XM_023251182.1;refGene_gn=ATP7B;refGene_tx=XM_023251186.1;refGene_gn=ATP7B;refGene_tx=XM_023251193.1
A1 19614274 19614548

refGene_gn=ATP7B;refGene_tx=XM_006927260.4;refGene_gn=ATP7B;refGene_tx=XM_019825930.2;refGene_gn=ATP7B;refGene_tx=XM_023251165.1;refGene_gn=ATP7B;refGene_tx=XM_023251171.1;refGene_gn=ATP7B;refGene_tx=XM_023251176.1;refGene_gn=ATP7B;refGene_tx=XM_023251182.1;refGene_gn=ATP7B;refGene_tx=XM_023251186.1;refGene_gn=ATP7B;refGene_tx=XM_023251193.1
A1 19618569 19618599 refGene_gn=ATP7B;refGene_tx=XM_023251176.1
A1 19692473 19693222 refGene_gn=CCDC70;refGene_tx=XM_023251213.1
A1 19699231 19699289 refGene_gn=CCDC70;refGene_tx=XM_023251213.1
A1 19749983 19750033 refGene_gn=DHRS12;refGene_tx=XM_023259804.1
A1 19750169 19750348 refGene_gn=DHRS12;refGene_tx=XM_023259804.1
A1 19752995 19753486 refGene_gn=DHRS12;refGene_tx=XM_023259804.1
A1 19765962 19766055 refGene_gn=DHRS12;refGene_tx=XM_023259804.1
A1 19785807 19785911 refGene_gn=DHRS12;refGene_tx=XM_023259804.1
A1 19786392 19786825 refGene_gn=DHRS12;refGene_tx=XM_023259804.1
A1 19799796 19799826
refGene_gn=WDFY2;refGene_tx=XM_003980360.5;refGene_gn=WDFY2;refGene_tx=XM_019826005.2
A1 19800883 19800992
refGene_gn=WDFY2;refGene_tx=XM_003980360.5;refGene_gn=WDFY2;refGene_tx=XM_019826005.2
A1 19803448 19803579
refGene_gn=WDFY2;refGene_tx=XM_003980360.5;refGene_gn=WDFY2;refGene_tx=XM_019826005.2
A1 19804401 19804503
refGene_gn=WDFY2;refGene_tx=XM_003980360.5;refGene_gn=WDFY2;refGene_tx=XM_019826005.2
A1 19808366 19808472
refGene_gn=WDFY2;refGene_tx=XM_003980360.5;refGene_gn=WDFY2;refGene_tx=XM_019826005.2
A1 19819825 19819952
refGene_gn=WDFY2;refGene_tx=XM_003980360.5;refGene_gn=WDFY2;refGene_tx=XM_019826005.2
A1 19834840 19834953
refGene_gn=WDFY2;refGene_tx=XM_003980360.5;refGene_gn=WDFY2;refGene_tx=XM_019826005.2
A1 19843147 19843298
refGene_gn=WDFY2;refGene_tx=XM_003980360.5;refGene_gn=WDFY2;refGene_tx=XM_019826005.2
A1 19853272 19853327
refGene_gn=WDFY2;refGene_tx=XM_003980360.5;refGene_gn=WDFY2;refGene_tx=XM_019826005.2
A1 19887339 19887413
refGene_gn=WDFY2;refGene_tx=XM_003980360.5;refGene_gn=WDFY2;refGene_tx=XM_019826005.2
A1 19898949 19899017
refGene_gn=WDFY2;refGene_tx=XM_003980360.5;refGene_gn=WDFY2;refGene_tx=XM_019826005.2
A1 19925567 19925569 refGene_gn=WDFY2;refGene_tx=XM_019826005.2
A1 19973632 19973769 refGene_gn=WDFY2;refGene_tx=XM_003980360.5
A1 20049790 20049831

refGene_gn=LOC102900676;refGene_tx=XM_019826022.2
A1 20054969 20055273
refGene_gn=LOC102900676;refGene_tx=XM_019826022.2
A1 20055442 20055534
refGene_gn=LOC102900676;refGene_tx=XM_019826022.2
A1 20056964 20057171
refGene_gn=LOC102900676;refGene_tx=XM_019826022.2
A1 20063400 20063459
refGene_gn=LOC102900676;refGene_tx=XM_019826022.2
A1 20063732 20063773
refGene_gn=LOC102900676;refGene_tx=XM_019826022.2
A1 20068534 20068645
refGene_gn=INTS6;refGene_tx=XM_023251226.1;refGene_gn=INTS6;refGene_tx=XM_023251229.1
A1 20069037 20069115
refGene_gn=INTS6;refGene_tx=XM_023251226.1;refGene_gn=INTS6;refGene_tx=XM_023251229.1
A1 20069942 20070092
refGene_gn=INTS6;refGene_tx=XM_023251226.1;refGene_gn=INTS6;refGene_tx=XM_023251229.1
A1 20089951 20090041
refGene_gn=INTS6;refGene_tx=XM_023251226.1;refGene_gn=INTS6;refGene_tx=XM_023251229.1
A1 20127706 20127890
refGene_gn=INTS6;refGene_tx=XM_023251226.1;refGene_gn=INTS6;refGene_tx=XM_023251229.1
A1 20133762 20133888
refGene_gn=INTS6;refGene_tx=XM_023251226.1;refGene_gn=INTS6;refGene_tx=XM_023251229.1
A1 20135292 20135447
refGene_gn=INTS6;refGene_tx=XM_023251226.1;refGene_gn=INTS6;refGene_tx=XM_023251229.1
A1 20140623 20140776
refGene_gn=INTS6;refGene_tx=XM_023251226.1;refGene_gn=INTS6;refGene_tx=XM_023251229.1
A1 20140918 20141051
refGene_gn=INTS6;refGene_tx=XM_023251226.1;refGene_gn=INTS6;refGene_tx=XM_023251229.1
A1 20142177 20142272
refGene_gn=INTS6;refGene_tx=XM_023251226.1;refGene_gn=INTS6;refGene_tx=XM_023251229.1
A1 20144414 20144525
refGene_gn=INTS6;refGene_tx=XM_023251226.1;refGene_gn=INTS6;refGene_tx=XM_023251229.1
A1 20145528 20145744
refGene_gn=INTS6;refGene_tx=XM_023251226.1;refGene_gn=INTS6;refGene_tx=XM_023251229.1
A1 20149394 20149521
refGene_gn=INTS6;refGene_tx=XM_023251226.1;refGene_gn=INTS6;refGene_tx=XM_023251229.1

A1	20150586	20150729
refGene_gn=INTS6;refGene_tx=XM_023251226.1;refGene_gn=INTS6;refGene_tx=XM_023251229.1
A1	20150931	20151163
refGene_gn=INTS6;refGene_tx=XM_023251226.1;refGene_gn=INTS6;refGene_tx=XM_023251229.1
A1	20155210	20155582
refGene_gn=INTS6;refGene_tx=XM_023251226.1;refGene_gn=INTS6;refGene_tx=XM_023251229.1
A1	20156605	20156699
refGene_gn=INTS6;refGene_tx=XM_023251226.1;refGene_gn=INTS6;refGene_tx=XM_023251229.1
A1	20158603	20158697
refGene_gn=INTS6;refGene_tx=XM_023251226.1;refGene_gn=INTS6;refGene_tx=XM_023251229.1
A1	20160221	20160265
refGene_gn=SERPINE3;refGene_tx=XM_019826076.2;refGene_gn=SERPINE3;refGene_tx=XM_019826081.2
A1	20162676	20162760
refGene_gn=SERPINE3;refGene_tx=XM_019826076.2;refGene_gn=SERPINE3;refGene_tx=XM_019826081.2
A1	20163258	20163345
refGene_gn=SERPINE3;refGene_tx=XM_019826076.2;refGene_gn=SERPINE3;refGene_tx=XM_019826081.2
A1	20174104	20174205
refGene_gn=SERPINE3;refGene_tx=XM_019826076.2;refGene_gn=SERPINE3;refGene_tx=XM_019826081.2
A1	20182503	20182702
refGene_gn=SERPINE3;refGene_tx=XM_019826076.2;refGene_gn=SERPINE3;refGene_tx=XM_019826081.2
A1	20184268	20184466
refGene_gn=SERPINE3;refGene_tx=XM_019826076.2;refGene_gn=SERPINE3;refGene_tx=XM_019826081.2
A1	20186248	20186482
refGene_gn=SERPINE3;refGene_tx=XM_019826076.2;refGene_gn=SERPINE3;refGene_tx=XM_019826081.2
A1	20191441	20191714
refGene_gn=SERPINE3;refGene_tx=XM_019826076.2;refGene_gn=SERPINE3;refGene_tx=XM_019826081.2
A1	20193412	20193428	refGene_gn=SERPINE3;refGene_tx=XM_019826076.2
A1	20278051	20278849
refGene_gn=FAM124A;refGene_tx=XM_011283724.3;refGene_gn=FAM124A;refGene_tx=XM_023251259.1
A1	20340707	20340785	refGene_gn=FAM124A;refGene_tx=XM_023251261.1
A1	20346640	20347374
refGene_gn=FAM124A;refGene_tx=XM_011283724.3;refGene_gn=FAM124A;refGene_tx=XM_023251259.1;refGene_gn=FAM124A;refGene_tx=XM_023251261.1
A1	20366465	20366497
refGene_gn=FAM124A;refGene_tx=XM_011283724.3;refGene_gn=FAM124A;refGene_tx=XM_023251261.1

A1 20376316 20376381
refGene_gn=FAM124A;refGene_tx=XM_011283724.3;refGene_gn=FAM124A;refGene_tx=XM_023251259.1;refGene_gn=FAM124A;refGene_tx=XM_023251261.1
A1 20450131 20450548
refGene_gn=LOC109498200;refGene_tx=XM_023251265.1
A1 20468536 20468684
refGene_gn=LOC109498200;refGene_tx=XM_023251265.1
A1 20470951 20471192
refGene_gn=LOC109498200;refGene_tx=XM_023251265.1
A1 20472354 20472529
refGene_gn=LOC109498200;refGene_tx=XM_023251265.1
A1 20529129 20529152
refGene_gn=LOC101098598;refGene_tx=XM_023259861.1
A1 20529315 20529448
refGene_gn=LOC101098598;refGene_tx=XM_023259861.1
A1 20534001 20534218
refGene_gn=LOC101098598;refGene_tx=XM_023259861.1
A1 20536194 20536580
refGene_gn=LOC101098598;refGene_tx=XM_023259861.1
A1 20590677 20590785
refGene_gn=RNASEH2B;refGene_tx=XM_019826097.2;refGene_gn=RNASEH2B;refGene_tx=XM_023251282.1;refGene_gn=RNASEH2B;refGene_tx=XM_023251287.1
A1 20592485 20592566
refGene_gn=RNASEH2B;refGene_tx=XM_019826097.2;refGene_gn=RNASEH2B;refGene_tx=XM_023251282.1
A1 20596738 20596781
refGene_gn=RNASEH2B;refGene_tx=XM_019826097.2;refGene_gn=RNASEH2B;refGene_tx=XM_023251282.1;refGene_gn=RNASEH2B;refGene_tx=XM_023251287.1
A1 20598355 20598437
refGene_gn=RNASEH2B;refGene_tx=XM_019826097.2;refGene_gn=RNASEH2B;refGene_tx=XM_023251282.1;refGene_gn=RNASEH2B;refGene_tx=XM_023251287.1
A1 20600763 20600869
refGene_gn=RNASEH2B;refGene_tx=XM_019826097.2;refGene_gn=RNASEH2B;refGene_tx=XM_023251282.1;refGene_gn=RNASEH2B;refGene_tx=XM_023251287.1
A1 20602456 20602521
refGene_gn=RNASEH2B;refGene_tx=XM_019826097.2;refGene_gn=RNASEH2B;refGene_tx=XM_023251282.1;refGene_gn=RNASEH2B;refGene_tx=XM_023251287.1
A1 20612895 20613010
refGene_gn=RNASEH2B;refGene_tx=XM_019826097.2;refGene_gn=RNASEH2B;refGene_tx=XM_023251282.1;refGene_gn=RNASEH2B;refGene_tx=XM_023251287.1
A1 20617304 20617381
refGene_gn=RNASEH2B;refGene_tx=XM_019826097.2;refGene_gn=RNASEH2B;refGene_tx=XM_023251282.1;refGene_gn=RNASEH2B;refGene_tx=XM_023251287.1
A1 20621803 20621911
refGene_gn=RNASEH2B;refGene_tx=XM_019826097.2;refGene_gn=RNASEH2B;refGene_tx=XM_023251282.1;refGene_gn=RNASEH2B;refGene_tx=XM_023251287.1
A1 20623780 20623852
refGene_gn=RNASEH2B;refGene_tx=XM_019826097.2;refGene_gn=RNASEH2B;refGene_tx=XM_023251282.1;refGene_gn=RNASEH2B;refGene_tx=XM_023251287.1
A1 20647478 20647518 refGene_gn=RNASEH2B;refGene_tx=XM_019826097.2

A1 20648099 20648151
refGene_gn=RNASEH2B;refGene_tx=XM_023251282.1;refGene_gn=RNASEH2B;refGene_tx=XM_023251287.1
A1 20701904 20702545 refGene_gn=DLEU7;refGene_tx=XM_023259966.1
A1 20702546 20702679 refGene_gn=DLEU7;refGene_tx=XM_023259966.1
A1 20719046 20719313 refGene_gn=DLEU7;refGene_tx=XM_023259966.1
A1 21549146 21549387 refGene_gn=KCNRG;refGene_tx=XM_003980362.4
A1 21553587 21554168 refGene_gn=KCNRG;refGene_tx=XM_003980362.4
A1 21556335 21557559
refGene_gn=TRIM13;refGene_tx=XM_006927263.4;refGene_gn=TRIM13;refGene_tx=XM_006927264.4
A1 21616943 21617049 refGene_gn=SPRYD7;refGene_tx=XM_003980364.5
A1 21617637 21617754 refGene_gn=SPRYD7;refGene_tx=XM_003980364.5
A1 21620286 21620453 refGene_gn=SPRYD7;refGene_tx=XM_003980364.5
A1 21627440 21627543 refGene_gn=SPRYD7;refGene_tx=XM_003980364.5
A1 21631780 21631878 refGene_gn=SPRYD7;refGene_tx=XM_003980364.5
A1 21717947 21718016 refGene_gn=KPNA3;refGene_tx=XM_023251315.1
A1 21769795 21769840 refGene_gn=KPNA3;refGene_tx=XM_023251315.1
A1 21786899 21786989 refGene_gn=KPNA3;refGene_tx=XM_023251315.1
A1 21787178 21787208 refGene_gn=KPNA3;refGene_tx=XM_023251315.1
A1 21787294 21787347 refGene_gn=KPNA3;refGene_tx=XM_023251315.1
A1 21787479 21787575 refGene_gn=KPNA3;refGene_tx=XM_023251315.1
A1 21790485 21790571 refGene_gn=KPNA3;refGene_tx=XM_023251315.1
A1 21794117 21794204 refGene_gn=KPNA3;refGene_tx=XM_023251315.1
A1 21794567 21794737 refGene_gn=KPNA3;refGene_tx=XM_023251315.1
A1 21796072 21796117 refGene_gn=KPNA3;refGene_tx=XM_023251315.1
A1 21810990 21811122 refGene_gn=KPNA3;refGene_tx=XM_023251315.1
A1 21812096 21812225 refGene_gn=KPNA3;refGene_tx=XM_023251315.1
A1 21814544 21814649 refGene_gn=KPNA3;refGene_tx=XM_023251315.1
A1 21814745 21814817 refGene_gn=KPNA3;refGene_tx=XM_023251315.1
A1 21814968 21815131 refGene_gn=KPNA3;refGene_tx=XM_023251315.1
A1 21816983 21817078 refGene_gn=KPNA3;refGene_tx=XM_023251315.1
A1 21818809 21818908 refGene_gn=KPNA3;refGene_tx=XM_023251315.1
A1 21828762 21828933 refGene_gn=EBPL;refGene_tx=XM_023251321.1
A1 21855299 21855369 refGene_gn=EBPL;refGene_tx=XM_023251321.1
A1 21860969 21861108 refGene_gn=EBPL;refGene_tx=XM_023251321.1
A1 21862611 21862852 refGene_gn=EBPL;refGene_tx=XM_023251321.1
A1 21890461 21890654 refGene_gn=ARL11;refGene_tx=XM_003980388.5
A1 21890655 21891191 refGene_gn=ARL11;refGene_tx=XM_003980388.5
A1 21932542 21932668
refGene_gn=RCBTB1;refGene_tx=XM_023251323.1;refGene_gn=RCBTB1;refGene_tx=XM_023251325.1;refGene_gn=RCBTB1;refGene_tx=XM_023251330.1;refGene_gn=RCBTB1;refGene_tx=XM_023251336.1;refGene_gn=RCBTB1;refGene_tx=XM_023251343.1;refGene_gn=RCBTB1;refGene_tx=XM_023251349.1;refGene_gn=RCBTB1;refGene_tx=XM_023251351.1;refGene_gn=RCBTB1;refGene_tx=XM_023251354.1;refGene_gn=RCBTB1;refGene_tx=XM_023251358.1
A1 21933261 21933412
refGene_gn=RCBTB1;refGene_tx=XM_023251323.1;refGene_gn=RCBTB1;refGene_tx=XM_023251325.1;refGene_gn=RCBTB1;refGene_tx=XM_023251330.1;refGene_gn=RCBTB1;refGene_tx=XM_023251336.1;refGene_gn=RCBTB1;refGene_tx=XM_02

3251343.1;refGene_gn=RCBTB1;refGene_tx=XM_023251349.1;refGene_gn=RCBTB
1;refGene_tx=XM_023251351.1;refGene_gn=RCBTB1;refGene_tx=XM_023251354.
1;refGene_gn=RCBTB1;refGene_tx=XM_023251358.1;refGene_gn=RCBTB1;refGen
e_tx=XM_023251360.1
A1 21943810 21943977
refGene_gn=RCBTB1;refGene_tx=XM_023251323.1;refGene_gn=RCBTB1;refGene_
tx=XM_023251325.1;refGene_gn=RCBTB1;refGene_tx=XM_023251330.1;refGene_
gn=RCBTB1;refGene_tx=XM_023251336.1;refGene_gn=RCBTB1;refGene_tx=XM_02
3251343.1;refGene_gn=RCBTB1;refGene_tx=XM_023251349.1;refGene_gn=RCBTB
1;refGene_tx=XM_023251351.1;refGene_gn=RCBTB1;refGene_tx=XM_023251354.
1;refGene_gn=RCBTB1;refGene_tx=XM_023251358.1
A1 21947215 21947374
refGene_gn=RCBTB1;refGene_tx=XM_023251323.1;refGene_gn=RCBTB1;refGene_
tx=XM_023251325.1;refGene_gn=RCBTB1;refGene_tx=XM_023251330.1;refGene_
gn=RCBTB1;refGene_tx=XM_023251336.1;refGene_gn=RCBTB1;refGene_tx=XM_02
3251343.1;refGene_gn=RCBTB1;refGene_tx=XM_023251349.1;refGene_gn=RCBTB
1;refGene_tx=XM_023251351.1;refGene_gn=RCBTB1;refGene_tx=XM_023251354.
1;refGene_gn=RCBTB1;refGene_tx=XM_023251358.1;refGene_gn=RCBTB1;refGen
e_tx=XM_023251360.1
A1 21950047 21950155
refGene_gn=RCBTB1;refGene_tx=XM_023251323.1;refGene_gn=RCBTB1;refGene_
tx=XM_023251325.1;refGene_gn=RCBTB1;refGene_tx=XM_023251330.1;refGene_
gn=RCBTB1;refGene_tx=XM_023251336.1;refGene_gn=RCBTB1;refGene_tx=XM_02
3251343.1;refGene_gn=RCBTB1;refGene_tx=XM_023251349.1;refGene_gn=RCBTB
1;refGene_tx=XM_023251351.1;refGene_gn=RCBTB1;refGene_tx=XM_023251354.
1;refGene_gn=RCBTB1;refGene_tx=XM_023251358.1;refGene_gn=RCBTB1;refGen
e_tx=XM_023251360.1
A1 21951226 21951369
refGene_gn=RCBTB1;refGene_tx=XM_023251323.1;refGene_gn=RCBTB1;refGene_
tx=XM_023251325.1;refGene_gn=RCBTB1;refGene_tx=XM_023251330.1;refGene_
gn=RCBTB1;refGene_tx=XM_023251336.1;refGene_gn=RCBTB1;refGene_tx=XM_02
3251343.1;refGene_gn=RCBTB1;refGene_tx=XM_023251349.1;refGene_gn=RCBTB
1;refGene_tx=XM_023251351.1;refGene_gn=RCBTB1;refGene_tx=XM_023251354.
1;refGene_gn=RCBTB1;refGene_tx=XM_023251358.1;refGene_gn=RCBTB1;refGen
e_tx=XM_023251360.1
A1 21952364 21952555
refGene_gn=RCBTB1;refGene_tx=XM_023251323.1;refGene_gn=RCBTB1;refGene_
tx=XM_023251325.1;refGene_gn=RCBTB1;refGene_tx=XM_023251330.1;refGene_
gn=RCBTB1;refGene_tx=XM_023251336.1;refGene_gn=RCBTB1;refGene_tx=XM_02
3251343.1;refGene_gn=RCBTB1;refGene_tx=XM_023251349.1;refGene_gn=RCBTB
1;refGene_tx=XM_023251351.1;refGene_gn=RCBTB1;refGene_tx=XM_023251354.
1;refGene_gn=RCBTB1;refGene_tx=XM_023251358.1;refGene_gn=RCBTB1;refGen
e_tx=XM_023251360.1
A1 21956856 21956983
refGene_gn=RCBTB1;refGene_tx=XM_023251323.1;refGene_gn=RCBTB1;refGene_
tx=XM_023251325.1;refGene_gn=RCBTB1;refGene_tx=XM_023251330.1;refGene_
gn=RCBTB1;refGene_tx=XM_023251336.1;refGene_gn=RCBTB1;refGene_tx=XM_02
3251343.1;refGene_gn=RCBTB1;refGene_tx=XM_023251349.1;refGene_gn=RCBTB
1;refGene_tx=XM_023251351.1;refGene_gn=RCBTB1;refGene_tx=XM_023251354.
1;refGene_gn=RCBTB1;refGene_tx=XM_023251358.1;refGene_gn=RCBTB1;refGen

e_tx=XM_023251360.1
A1 21959236 21959388
refGene_gn=RCBTB1;refGene_tx=XM_023251323.1;refGene_gn=RCBTB1;refGene_tx=XM_023251325.1;refGene_gn=RCBTB1;refGene_tx=XM_023251330.1;refGene_gn=RCBTB1;refGene_tx=XM_023251336.1;refGene_gn=RCBTB1;refGene_tx=XM_023251343.1;refGene_gn=RCBTB1;refGene_tx=XM_023251349.1;refGene_gn=RCBTB1;refGene_tx=XM_023251351.1;refGene_gn=RCBTB1;refGene_tx=XM_023251354.1;refGene_gn=RCBTB1;refGene_tx=XM_023251358.1;refGene_gn=RCBTB1;refGene_tx=XM_023251360.1
A1 21959927 21960067
refGene_gn=RCBTB1;refGene_tx=XM_023251323.1;refGene_gn=RCBTB1;refGene_tx=XM_023251325.1;refGene_gn=RCBTB1;refGene_tx=XM_023251330.1;refGene_gn=RCBTB1;refGene_tx=XM_023251336.1;refGene_gn=RCBTB1;refGene_tx=XM_023251343.1;refGene_gn=RCBTB1;refGene_tx=XM_023251349.1;refGene_gn=RCBTB1;refGene_tx=XM_023251351.1;refGene_gn=RCBTB1;refGene_tx=XM_023251354.1;refGene_gn=RCBTB1;refGene_tx=XM_023251358.1;refGene_gn=RCBTB1;refGene_tx=XM_023251360.1
A1 21966512 21966653
refGene_gn=RCBTB1;refGene_tx=XM_023251323.1;refGene_gn=RCBTB1;refGene_tx=XM_023251325.1;refGene_gn=RCBTB1;refGene_tx=XM_023251330.1;refGene_gn=RCBTB1;refGene_tx=XM_023251336.1;refGene_gn=RCBTB1;refGene_tx=XM_023251343.1;refGene_gn=RCBTB1;refGene_tx=XM_023251349.1;refGene_gn=RCBTB1;refGene_tx=XM_023251351.1;refGene_gn=RCBTB1;refGene_tx=XM_023251354.1;refGene_gn=RCBTB1;refGene_tx=XM_023251360.1
A1 21975822 21976004
refGene_gn=LOC101100105;refGene_tx=XM_011287848.3;refGene_gn=LOC101100105;refGene_tx=XM_019826124.2
A1 21977703 21977775
refGene_gn=LOC101100105;refGene_tx=XM_011287848.3;refGene_gn=LOC101100105;refGene_tx=XM_019826124.2
A1 21978781 21978922
refGene_gn=LOC101100105;refGene_tx=XM_011287848.3;refGene_gn=LOC101100105;refGene_tx=XM_019826124.2
A1 21979573 21979640
refGene_gn=LOC101100105;refGene_tx=XM_011287848.3;refGene_gn=LOC101100105;refGene_tx=XM_019826124.2
A1 21980315 21980371
refGene_gn=LOC101100105;refGene_tx=XM_011287848.3;refGene_gn=LOC101100105;refGene_tx=XM_019826124.2
A1 21982507 21982567
refGene_gn=LOC101100105;refGene_tx=XM_011287848.3;refGene_gn=LOC101100105;refGene_tx=XM_019826124.2
A1 21982763 21982807
refGene_gn=LOC101100105;refGene_tx=XM_011287848.3;refGene_gn=LOC101100105;refGene_tx=XM_019826124.2
A1 21986116 21986250
refGene_gn=LOC101100105;refGene_tx=XM_011287848.3;refGene_gn=LOC101100105;refGene_tx=XM_019826124.2
A1 21992146 21992254
refGene_gn=LOC101100105;refGene_tx=XM_011287848.3;refGene_gn=LOC101100

105;refGene_tx=XM_019826124.2
A1 21998279 21998401
refGene_gn=LOC101100105;refGene_tx=XM_011287848.3;refGene_gn=LOC101100105;refGene_tx=XM_019826124.2
A1 22006512 22006573
refGene_gn=LOC101100105;refGene_tx=XM_011287848.3
A1 22010777 22010895
refGene_gn=LOC101100105;refGene_tx=XM_019826101.2;refGene_gn=LOC101100105;refGene_tx=XM_019826102.2;refGene_gn=LOC101100105;refGene_tx=XM_019826104.2;refGene_gn=LOC101100105;refGene_tx=XM_019826109.2;refGene_gn=LOC101100105;refGene_tx=XM_023251371.1
A1 22011705 22011794
refGene_gn=LOC101100105;refGene_tx=XM_019826101.2;refGene_gn=LOC101100105;refGene_tx=XM_019826102.2;refGene_gn=LOC101100105;refGene_tx=XM_019826104.2;refGene_gn=LOC101100105;refGene_tx=XM_019826109.2;refGene_gn=LOC101100105;refGene_tx=XM_023251371.1
A1 22014444 22014785
refGene_gn=LOC101100105;refGene_tx=XM_019826101.2;refGene_gn=LOC101100105;refGene_tx=XM_019826102.2;refGene_gn=LOC101100105;refGene_tx=XM_019826104.2;refGene_gn=LOC101100105;refGene_tx=XM_019826109.2;refGene_gn=LOC101100105;refGene_tx=XM_023251371.1
A1 22015138 22015216
refGene_gn=LOC101100105;refGene_tx=XM_019826101.2;refGene_gn=LOC101100105;refGene_tx=XM_019826102.2;refGene_gn=LOC101100105;refGene_tx=XM_019826104.2;refGene_gn=LOC101100105;refGene_tx=XM_023251371.1
A1 22017470 22017564
refGene_gn=LOC101100105;refGene_tx=XM_019826101.2;refGene_gn=LOC101100105;refGene_tx=XM_019826102.2;refGene_gn=LOC101100105;refGene_tx=XM_019826104.2;refGene_gn=LOC101100105;refGene_tx=XM_019826109.2;refGene_gn=LOC101100105;refGene_tx=XM_019826124.2;refGene_gn=LOC101100105;refGene_tx=XM_023251371.1
A1 22018951 22019051
refGene_gn=LOC101100105;refGene_tx=XM_019826101.2;refGene_gn=LOC101100105;refGene_tx=XM_019826102.2;refGene_gn=LOC101100105;refGene_tx=XM_019826104.2;refGene_gn=LOC101100105;refGene_tx=XM_019826109.2;refGene_gn=LOC101100105;refGene_tx=XM_019826124.2;refGene_gn=LOC101100105;refGene_tx=XM_023251371.1
A1 22019795 22020012
refGene_gn=LOC101100105;refGene_tx=XM_019826101.2;refGene_gn=LOC101100105;refGene_tx=XM_019826102.2;refGene_gn=LOC101100105;refGene_tx=XM_019826104.2;refGene_gn=LOC101100105;refGene_tx=XM_019826109.2;refGene_gn=LOC101100105;refGene_tx=XM_019826124.2;refGene_gn=LOC101100105;refGene_tx=XM_023251371.1
A1 22022481 22022651
refGene_gn=LOC101100105;refGene_tx=XM_019826101.2;refGene_gn=LOC101100105;refGene_tx=XM_019826102.2;refGene_gn=LOC101100105;refGene_tx=XM_019826104.2;refGene_gn=LOC101100105;refGene_tx=XM_019826109.2;refGene_gn=LOC101100105;refGene_tx=XM_019826124.2;refGene_gn=LOC101100105;refGene_tx=XM_023251371.1
A1 22023391 22023508

refGene_gn=LOC101100105;refGene_tx=XM_019826101.2;refGene_gn=LOC101100105;refGene_tx=XM_019826102.2;refGene_gn=LOC101100105;refGene_tx=XM_019826104.2;refGene_gn=LOC101100105;refGene_tx=XM_019826109.2;refGene_gn=LOC101100105;refGene_tx=XM_019826124.2;refGene_gn=LOC101100105;refGene_tx=XM_023251371.1
A1 22037237 22037801
refGene_gn=LOC101100105;refGene_tx=XM_019826101.2;refGene_gn=LOC101100105;refGene_tx=XM_019826102.2;refGene_gn=LOC101100105;refGene_tx=XM_019826104.2;refGene_gn=LOC101100105;refGene_tx=XM_019826109.2;refGene_gn=LOC101100105;refGene_tx=XM_019826124.2;refGene_gn=LOC101100105;refGene_tx=XM_023251371.1
A1 22046009 22046106
refGene_gn=LOC101100105;refGene_tx=XM_019826101.2;refGene_gn=LOC101100105;refGene_tx=XM_019826102.2;refGene_gn=LOC101100105;refGene_tx=XM_019826104.2;refGene_gn=LOC101100105;refGene_tx=XM_019826109.2;refGene_gn=LOC101100105;refGene_tx=XM_019826124.2;refGene_gn=LOC101100105;refGene_tx=XM_023251371.1
A1 22049846 22049912
refGene_gn=LOC101100105;refGene_tx=XM_019826101.2;refGene_gn=LOC101100105;refGene_tx=XM_019826102.2;refGene_gn=LOC101100105;refGene_tx=XM_019826104.2;refGene_gn=LOC101100105;refGene_tx=XM_019826109.2;refGene_gn=LOC101100105;refGene_tx=XM_019826124.2;refGene_gn=LOC101100105;refGene_tx=XM_023251371.1
A1 22050877 22051003
refGene_gn=LOC101100105;refGene_tx=XM_019826101.2;refGene_gn=LOC101100105;refGene_tx=XM_019826102.2;refGene_gn=LOC101100105;refGene_tx=XM_019826104.2;refGene_gn=LOC101100105;refGene_tx=XM_019826109.2;refGene_gn=LOC101100105;refGene_tx=XM_019826124.2;refGene_gn=LOC101100105;refGene_tx=XM_023251371.1
A1 22057418 22057434
refGene_gn=LOC101100105;refGene_tx=XM_019826101.2;refGene_gn=LOC101100105;refGene_tx=XM_019826102.2;refGene_gn=LOC101100105;refGene_tx=XM_019826109.2;refGene_gn=LOC101100105;refGene_tx=XM_019826124.2;refGene_gn=LOC101100105;refGene_tx=XM_023251371.1
A1 22114671 22114743 refGene_gn=CAB39L;refGene_tx=XM_019826173.2
A1 22114744 22114783 refGene_gn=CAB39L;refGene_tx=XM_019826173.2
A1 22118378 22118543 refGene_gn=CAB39L;refGene_tx=XM_019826173.2
A1 22133881 22134000 refGene_gn=CAB39L;refGene_tx=XM_019826173.2
A1 22153625 22153794 refGene_gn=CAB39L;refGene_tx=XM_019826173.2
A1 22159722 22159782 refGene_gn=CAB39L;refGene_tx=XM_019826173.2
A1 22163718 22163784 refGene_gn=CAB39L;refGene_tx=XM_019826173.2
A1 22179586 22179730 refGene_gn=CAB39L;refGene_tx=XM_019826173.2
A1 22194669 22194849 refGene_gn=CAB39L;refGene_tx=XM_019826173.2
A1 22219299 22219373 refGene_gn=CDADC1;refGene_tx=XM_003980370.5
A1 22221095 22221156 refGene_gn=CDADC1;refGene_tx=XM_003980370.5
A1 22226508 22226698 refGene_gn=CDADC1;refGene_tx=XM_003980370.5
A1 22229985 22230155 refGene_gn=CDADC1;refGene_tx=XM_003980370.5
A1 22233216 22233266 refGene_gn=CDADC1;refGene_tx=XM_003980370.5
A1 22238794 22239364 refGene_gn=CDADC1;refGene_tx=XM_003980370.5
A1 22246069 22246247 refGene_gn=CDADC1;refGene_tx=XM_003980370.5

A1 22257440 22257515 refGene_gn=CDADC1;refGene_tx=XM_003980370.5
A1 22264804 22264905 refGene_gn=CDADC1;refGene_tx=XM_003980370.5
A1 22265688 22265770 refGene_gn=CDADC1;refGene_tx=XM_003980370.5
A1 22286134 22286245 refGene_gn=MLNR;refGene_tx=XM_023250919.1
A1 22286246 22286398 refGene_gn=MLNR;refGene_tx=XM_023250919.1
A1 22286400 22286523 refGene_gn=MLNR;refGene_tx=XM_023250919.1
A1 22287263 22288164 refGene_gn=MLNR;refGene_tx=XM_023250919.1
A1 22297040 22297355
refGene_gn=FNDC3A;refGene_tx=XM_023251397.1;refGene_gn=FNDC3A;refGene_tx=XM_023251404.1;refGene_gn=FNDC3A;refGene_tx=XM_023251406.1;refGene_gn=FNDC3A;refGene_tx=XM_023251409.1;refGene_gn=FNDC3A;refGene_tx=XM_023251412.1
A1 22304804 22304932
refGene_gn=FNDC3A;refGene_tx=XM_023251397.1;refGene_gn=FNDC3A;refGene_tx=XM_023251404.1;refGene_gn=FNDC3A;refGene_tx=XM_023251406.1;refGene_gn=FNDC3A;refGene_tx=XM_023251409.1
A1 22307022 22307189
refGene_gn=FNDC3A;refGene_tx=XM_023251397.1;refGene_gn=FNDC3A;refGene_tx=XM_023251404.1;refGene_gn=FNDC3A;refGene_tx=XM_023251406.1;refGene_gn=FNDC3A;refGene_tx=XM_023251409.1;refGene_gn=FNDC3A;refGene_tx=XM_023251412.1
A1 22309471 22309684
refGene_gn=FNDC3A;refGene_tx=XM_023251397.1;refGene_gn=FNDC3A;refGene_tx=XM_023251404.1;refGene_gn=FNDC3A;refGene_tx=XM_023251406.1;refGene_gn=FNDC3A;refGene_tx=XM_023251409.1;refGene_gn=FNDC3A;refGene_tx=XM_023251412.1
A1 22309784 22310068
refGene_gn=FNDC3A;refGene_tx=XM_023251397.1;refGene_gn=FNDC3A;refGene_tx=XM_023251404.1;refGene_gn=FNDC3A;refGene_tx=XM_023251406.1;refGene_gn=FNDC3A;refGene_tx=XM_023251409.1;refGene_gn=FNDC3A;refGene_tx=XM_023251412.1
A1 22310169 22310319
refGene_gn=FNDC3A;refGene_tx=XM_023251397.1;refGene_gn=FNDC3A;refGene_tx=XM_023251404.1;refGene_gn=FNDC3A;refGene_tx=XM_023251406.1;refGene_gn=FNDC3A;refGene_tx=XM_023251409.1;refGene_gn=FNDC3A;refGene_tx=XM_023251412.1
A1 22311191 22311305
refGene_gn=FNDC3A;refGene_tx=XM_023251397.1;refGene_gn=FNDC3A;refGene_tx=XM_023251404.1;refGene_gn=FNDC3A;refGene_tx=XM_023251406.1;refGene_gn=FNDC3A;refGene_tx=XM_023251409.1;refGene_gn=FNDC3A;refGene_tx=XM_023251412.1
A1 22312771 22312947
refGene_gn=FNDC3A;refGene_tx=XM_023251397.1;refGene_gn=FNDC3A;refGene_tx=XM_023251404.1;refGene_gn=FNDC3A;refGene_tx=XM_023251406.1;refGene_gn=FNDC3A;refGene_tx=XM_023251409.1;refGene_gn=FNDC3A;refGene_tx=XM_023251412.1
A1 22313034 22313140
refGene_gn=FNDC3A;refGene_tx=XM_023251397.1;refGene_gn=FNDC3A;refGene_tx=XM_023251404.1;refGene_gn=FNDC3A;refGene_tx=XM_023251406.1;refGene_gn=FNDC3A;refGene_tx=XM_023251409.1;refGene_gn=FNDC3A;refGene_tx=XM_02

3251412.1
A1 22314943 22315062
refGene_gn=FNDC3A;refGene_tx=XM_023251397.1;refGene_gn=FNDC3A;refGene_tx=XM_023251404.1;refGene_gn=FNDC3A;refGene_tx=XM_023251406.1;refGene_gn=FNDC3A;refGene_tx=XM_023251409.1;refGene_gn=FNDC3A;refGene_tx=XM_023251412.1
A1 22315878 22315947
refGene_gn=FNDC3A;refGene_tx=XM_023251397.1;refGene_gn=FNDC3A;refGene_tx=XM_023251404.1;refGene_gn=FNDC3A;refGene_tx=XM_023251406.1;refGene_gn=FNDC3A;refGene_tx=XM_023251409.1;refGene_gn=FNDC3A;refGene_tx=XM_023251412.1
A1 22316609 22316748
refGene_gn=FNDC3A;refGene_tx=XM_023251397.1;refGene_gn=FNDC3A;refGene_tx=XM_023251404.1;refGene_gn=FNDC3A;refGene_tx=XM_023251406.1;refGene_gn=FNDC3A;refGene_tx=XM_023251409.1;refGene_gn=FNDC3A;refGene_tx=XM_023251412.1
A1 22318847 22318934
refGene_gn=FNDC3A;refGene_tx=XM_023251397.1;refGene_gn=FNDC3A;refGene_tx=XM_023251404.1;refGene_gn=FNDC3A;refGene_tx=XM_023251406.1;refGene_gn=FNDC3A;refGene_tx=XM_023251409.1;refGene_gn=FNDC3A;refGene_tx=XM_023251412.1
A1 22320505 22320680
refGene_gn=FNDC3A;refGene_tx=XM_023251397.1;refGene_gn=FNDC3A;refGene_tx=XM_023251404.1;refGene_gn=FNDC3A;refGene_tx=XM_023251406.1;refGene_gn=FNDC3A;refGene_tx=XM_023251409.1;refGene_gn=FNDC3A;refGene_tx=XM_023251412.1
A1 22321048 22321173
refGene_gn=FNDC3A;refGene_tx=XM_023251397.1;refGene_gn=FNDC3A;refGene_tx=XM_023251404.1;refGene_gn=FNDC3A;refGene_tx=XM_023251406.1;refGene_gn=FNDC3A;refGene_tx=XM_023251409.1;refGene_gn=FNDC3A;refGene_tx=XM_023251412.1
A1 22324953 22325007
refGene_gn=FNDC3A;refGene_tx=XM_023251397.1;refGene_gn=FNDC3A;refGene_tx=XM_023251404.1;refGene_gn=FNDC3A;refGene_tx=XM_023251406.1;refGene_gn=FNDC3A;refGene_tx=XM_023251409.1;refGene_gn=FNDC3A;refGene_tx=XM_023251412.1
A1 22328390 22328529
refGene_gn=FNDC3A;refGene_tx=XM_023251397.1;refGene_gn=FNDC3A;refGene_tx=XM_023251404.1;refGene_gn=FNDC3A;refGene_tx=XM_023251406.1;refGene_gn=FNDC3A;refGene_tx=XM_023251409.1;refGene_gn=FNDC3A;refGene_tx=XM_023251412.1
A1 22330326 22330386
refGene_gn=FNDC3A;refGene_tx=XM_023251397.1;refGene_gn=FNDC3A;refGene_tx=XM_023251404.1;refGene_gn=FNDC3A;refGene_tx=XM_023251406.1;refGene_gn=FNDC3A;refGene_tx=XM_023251409.1;refGene_gn=FNDC3A;refGene_tx=XM_023251412.1
A1 22332273 22332431
refGene_gn=FNDC3A;refGene_tx=XM_023251397.1;refGene_gn=FNDC3A;refGene_tx=XM_023251404.1;refGene_gn=FNDC3A;refGene_tx=XM_023251406.1;refGene_gn=FNDC3A;refGene_tx=XM_023251409.1;refGene_gn=FNDC3A;refGene_tx=XM_02

3251412.1
A1 22338150 22338209
refGene_gn=FNDC3A;refGene_tx=XM_023251397.1;refGene_gn=FNDC3A;refGene_tx=XM_023251404.1;refGene_gn=FNDC3A;refGene_tx=XM_023251406.1;refGene_gn=FNDC3A;refGene_tx=XM_023251409.1;refGene_gn=FNDC3A;refGene_tx=XM_023251412.1
A1 22339336 22339606
refGene_gn=FNDC3A;refGene_tx=XM_023251397.1;refGene_gn=FNDC3A;refGene_tx=XM_023251404.1;refGene_gn=FNDC3A;refGene_tx=XM_023251406.1;refGene_gn=FNDC3A;refGene_tx=XM_023251409.1;refGene_gn=FNDC3A;refGene_tx=XM_023251412.1
A1 22343564 22343805
refGene_gn=FNDC3A;refGene_tx=XM_023251397.1;refGene_gn=FNDC3A;refGene_tx=XM_023251404.1;refGene_gn=FNDC3A;refGene_tx=XM_023251406.1;refGene_gn=FNDC3A;refGene_tx=XM_023251409.1;refGene_gn=FNDC3A;refGene_tx=XM_023251412.1
A1 22359235 22359312
refGene_gn=FNDC3A;refGene_tx=XM_023251397.1;refGene_gn=FNDC3A;refGene_tx=XM_023251404.1;refGene_gn=FNDC3A;refGene_tx=XM_023251406.1;refGene_gn=FNDC3A;refGene_tx=XM_023251409.1;refGene_gn=FNDC3A;refGene_tx=XM_023251412.1
A1 22363737 22363744 refGene_gn=FNDC3A;refGene_tx=XM_023251409.1
A1 22395559 22395635
refGene_gn=FNDC3A;refGene_tx=XM_023251397.1;refGene_gn=FNDC3A;refGene_tx=XM_023251404.1;refGene_gn=FNDC3A;refGene_tx=XM_023251406.1;refGene_gn=FNDC3A;refGene_tx=XM_023251412.1
A1 22414942 22415041
refGene_gn=FNDC3A;refGene_tx=XM_023251397.1;refGene_gn=FNDC3A;refGene_tx=XM_023251404.1;refGene_gn=FNDC3A;refGene_tx=XM_023251406.1;refGene_gn=FNDC3A;refGene_tx=XM_023251412.1
A1 22669208 22670198
refGene_gn=CYSLTR2;refGene_tx=XM_006927270.4;refGene_gn=CYSLTR2;refGene_tx=XM_006927271.4;refGene_gn=CYSLTR2;refGene_tx=XM_023251419.1
A1 22833805 22833847
refGene_gn=RCBTB2;refGene_tx=XM_019826356.2;refGene_gn=RCBTB2;refGene_tx=XM_019826359.2;refGene_gn=RCBTB2;refGene_tx=XM_019826361.2;refGene_gn=RCBTB2;refGene_tx=XM_019826365.2;refGene_gn=RCBTB2;refGene_tx=XM_019826368.2;refGene_gn=RCBTB2;refGene_tx=XM_019826370.2;refGene_gn=RCBTB2;refGene_tx=XM_019826374.2;refGene_gn=RCBTB2;refGene_tx=XM_023251448.1
A1 22838794 22838950
refGene_gn=RCBTB2;refGene_tx=XM_019826356.2;refGene_gn=RCBTB2;refGene_tx=XM_019826359.2;refGene_gn=RCBTB2;refGene_tx=XM_019826361.2;refGene_gn=RCBTB2;refGene_tx=XM_019826365.2;refGene_gn=RCBTB2;refGene_tx=XM_019826368.2;refGene_gn=RCBTB2;refGene_tx=XM_019826370.2;refGene_gn=RCBTB2;refGene_tx=XM_019826374.2;refGene_gn=RCBTB2;refGene_tx=XM_019826377.2;refGene_gn=RCBTB2;refGene_tx=XM_019826383.2;refGene_gn=RCBTB2;refGene_tx=XM_023251448.1
A1 22839142 22839293
refGene_gn=RCBTB2;refGene_tx=XM_019826356.2;refGene_gn=RCBTB2;refGene_

tx=XM_019826359.2;refGene_gn=RCBTB2;refGene_tx=XM_019826361.2;refGene_gn=RCBTB2;refGene_tx=XM_019826365.2;refGene_gn=RCBTB2;refGene_tx=XM_019826368.2;refGene_gn=RCBTB2;refGene_tx=XM_019826370.2;refGene_gn=RCBTB2;refGene_tx=XM_019826374.2;refGene_gn=RCBTB2;refGene_tx=XM_019826377.2;refGene_gn=RCBTB2;refGene_tx=XM_019826383.2;refGene_gn=RCBTB2;refGene_tx=XM_023251448.1
A1 22841465 22841632
refGene_gn=RCBTB2;refGene_tx=XM_019826356.2;refGene_gn=RCBTB2;refGene_tx=XM_019826359.2;refGene_gn=RCBTB2;refGene_tx=XM_019826361.2;refGene_gn=RCBTB2;refGene_tx=XM_019826365.2;refGene_gn=RCBTB2;refGene_tx=XM_019826368.2;refGene_gn=RCBTB2;refGene_tx=XM_019826370.2;refGene_gn=RCBTB2;refGene_tx=XM_019826374.2;refGene_gn=RCBTB2;refGene_tx=XM_019826377.2;refGene_gn=RCBTB2;refGene_tx=XM_019826383.2;refGene_gn=RCBTB2;refGene_tx=XM_023251448.1
A1 22842181 22842340
refGene_gn=RCBTB2;refGene_tx=XM_019826356.2;refGene_gn=RCBTB2;refGene_tx=XM_019826359.2;refGene_gn=RCBTB2;refGene_tx=XM_019826361.2;refGene_gn=RCBTB2;refGene_tx=XM_019826365.2;refGene_gn=RCBTB2;refGene_tx=XM_019826368.2;refGene_gn=RCBTB2;refGene_tx=XM_019826370.2;refGene_gn=RCBTB2;refGene_tx=XM_019826374.2;refGene_gn=RCBTB2;refGene_tx=XM_019826377.2;refGene_gn=RCBTB2;refGene_tx=XM_019826383.2;refGene_gn=RCBTB2;refGene_tx=XM_023251448.1
A1 22842467 22842575
refGene_gn=RCBTB2;refGene_tx=XM_019826356.2;refGene_gn=RCBTB2;refGene_tx=XM_019826359.2;refGene_gn=RCBTB2;refGene_tx=XM_019826361.2;refGene_gn=RCBTB2;refGene_tx=XM_019826365.2;refGene_gn=RCBTB2;refGene_tx=XM_019826368.2;refGene_gn=RCBTB2;refGene_tx=XM_019826370.2;refGene_gn=RCBTB2;refGene_tx=XM_019826374.2;refGene_gn=RCBTB2;refGene_tx=XM_019826377.2;refGene_gn=RCBTB2;refGene_tx=XM_019826383.2;refGene_gn=RCBTB2;refGene_tx=XM_023251448.1
A1 22843677 22843820
refGene_gn=RCBTB2;refGene_tx=XM_019826356.2;refGene_gn=RCBTB2;refGene_tx=XM_019826359.2;refGene_gn=RCBTB2;refGene_tx=XM_019826361.2;refGene_gn=RCBTB2;refGene_tx=XM_019826365.2;refGene_gn=RCBTB2;refGene_tx=XM_019826368.2;refGene_gn=RCBTB2;refGene_tx=XM_019826370.2;refGene_gn=RCBTB2;refGene_tx=XM_019826374.2;refGene_gn=RCBTB2;refGene_tx=XM_019826377.2;refGene_gn=RCBTB2;refGene_tx=XM_019826383.2;refGene_gn=RCBTB2;refGene_tx=XM_023251448.1
A1 22846657 22846723
refGene_gn=RCBTB2;refGene_tx=XM_019826356.2;refGene_gn=RCBTB2;refGene_tx=XM_019826359.2;refGene_gn=RCBTB2;refGene_tx=XM_019826361.2;refGene_gn=RCBTB2;refGene_tx=XM_019826365.2;refGene_gn=RCBTB2;refGene_tx=XM_019826368.2;refGene_gn=RCBTB2;refGene_tx=XM_019826370.2;refGene_gn=RCBTB2;refGene_tx=XM_019826377.2;refGene_gn=RCBTB2;refGene_tx=XM_019826383.2;refGene_gn=RCBTB2;refGene_tx=XM_023251448.1
A1 22849528 22849719
refGene_gn=RCBTB2;refGene_tx=XM_019826356.2;refGene_gn=RCBTB2;refGene_tx=XM_019826359.2;refGene_gn=RCBTB2;refGene_tx=XM_019826361.2;refGene_gn=RCBTB2;refGene_tx=XM_019826365.2;refGene_gn=RCBTB2;refGene_tx=XM_019826368.2;refGene_gn=RCBTB2;refGene_tx=XM_019826370.2;refGene_gn=RCBTB

2;refGene_tx=XM_019826374.2;refGene_gn=RCBTB2;refGene_tx=XM_019826377.
2;refGene_gn=RCBTB2;refGene_tx=XM_019826383.2;refGene_gn=RCBTB2;refGen
e_tx=XM_023251448.1
A1 22850222 22850349
refGene_gn=RCBTB2;refGene_tx=XM_019826356.2;refGene_gn=RCBTB2;refGene_
tx=XM_019826359.2;refGene_gn=RCBTB2;refGene_tx=XM_019826361.2;refGene_
gn=RCBTB2;refGene_tx=XM_019826365.2;refGene_gn=RCBTB2;refGene_tx=XM_01
9826368.2;refGene_gn=RCBTB2;refGene_tx=XM_019826370.2;refGene_gn=RCBTB
2;refGene_tx=XM_019826374.2;refGene_gn=RCBTB2;refGene_tx=XM_019826377.
2;refGene_gn=RCBTB2;refGene_tx=XM_019826383.2;refGene_gn=RCBTB2;refGen
e_tx=XM_023251448.1
A1 22851973 22852113
refGene_gn=RCBTB2;refGene_tx=XM_019826356.2;refGene_gn=RCBTB2;refGene_
tx=XM_019826359.2;refGene_gn=RCBTB2;refGene_tx=XM_019826361.2;refGene_
gn=RCBTB2;refGene_tx=XM_019826365.2;refGene_gn=RCBTB2;refGene_tx=XM_01
9826368.2;refGene_gn=RCBTB2;refGene_tx=XM_019826370.2;refGene_gn=RCBTB
2;refGene_tx=XM_019826374.2;refGene_gn=RCBTB2;refGene_tx=XM_019826377.
2;refGene_gn=RCBTB2;refGene_tx=XM_019826383.2;refGene_gn=RCBTB2;refGen
e_tx=XM_023251448.1
A1 22855477 22855608
refGene_gn=RCBTB2;refGene_tx=XM_019826356.2;refGene_gn=RCBTB2;refGene_
tx=XM_019826359.2;refGene_gn=RCBTB2;refGene_tx=XM_019826361.2;refGene_
gn=RCBTB2;refGene_tx=XM_019826365.2;refGene_gn=RCBTB2;refGene_tx=XM_01
9826368.2;refGene_gn=RCBTB2;refGene_tx=XM_019826370.2;refGene_gn=RCBTB
2;refGene_tx=XM_019826374.2;refGene_gn=RCBTB2;refGene_tx=XM_019826377.
2;refGene_gn=RCBTB2;refGene_tx=XM_019826383.2;refGene_gn=RCBTB2;refGen
e_tx=XM_023251448.1
A1 22857904 22858045
refGene_gn=RCBTB2;refGene_tx=XM_019826356.2;refGene_gn=RCBTB2;refGene_
tx=XM_019826359.2;refGene_gn=RCBTB2;refGene_tx=XM_019826361.2;refGene_
gn=RCBTB2;refGene_tx=XM_019826365.2;refGene_gn=RCBTB2;refGene_tx=XM_01
9826368.2;refGene_gn=RCBTB2;refGene_tx=XM_019826370.2;refGene_gn=RCBTB
2;refGene_tx=XM_019826374.2;refGene_gn=RCBTB2;refGene_tx=XM_019826377.
2;refGene_gn=RCBTB2;refGene_tx=XM_019826383.2;refGene_gn=RCBTB2;refGen
e_tx=XM_023251448.1
A1 22865202 22865276
refGene_gn=RB1;refGene_tx=XM_023251429.1;refGene_gn=RB1;refGene_tx=XM_
023251440.1
A1 22866308 22866337 refGene_gn=RB1;refGene_tx=XM_023251437.1
A1 22867893 22867943
refGene_gn=RB1;refGene_tx=XM_023251429.1;refGene_gn=RB1;refGene_tx=XM_
023251437.1;refGene_gn=RB1;refGene_tx=XM_023251440.1
A1 22868452 22868595
refGene_gn=RB1;refGene_tx=XM_023251429.1;refGene_gn=RB1;refGene_tx=XM_
023251437.1;refGene_gn=RB1;refGene_tx=XM_023251440.1
A1 22870419 22870450
refGene_gn=RB1;refGene_tx=XM_023251429.1;refGene_gn=RB1;refGene_tx=XM_
023251437.1;refGene_gn=RB1;refGene_tx=XM_023251440.1
A1 22877277 22877441
refGene_gn=RB1;refGene_tx=XM_023251429.1;refGene_gn=RB1;refGene_tx=XM_

023251437.1;refGene_gn=RB1;refGene_tx=XM_023251440.1
A1	22877541 22877655
refGene_gn=RB1;refGene_tx=XM_023251429.1;refGene_gn=RB1;refGene_tx=XM_023251437.1;refGene_gn=RB1;refGene_tx=XM_023251440.1
A1	22878380 22878485
refGene_gn=RB1;refGene_tx=XM_023251429.1;refGene_gn=RB1;refGene_tx=XM_023251437.1;refGene_gn=RB1;refGene_tx=XM_023251440.1
A1	22883316 22883462
refGene_gn=RB1;refGene_tx=XM_023251429.1;refGene_gn=RB1;refGene_tx=XM_023251437.1;refGene_gn=RB1;refGene_tx=XM_023251440.1
A1	22886673 22886816
refGene_gn=RB1;refGene_tx=XM_023251429.1;refGene_gn=RB1;refGene_tx=XM_023251437.1;refGene_gn=RB1;refGene_tx=XM_023251440.1
A1	22890132 22890251
refGene_gn=RB1;refGene_tx=XM_023251429.1;refGene_gn=RB1;refGene_tx=XM_023251437.1;refGene_gn=RB1;refGene_tx=XM_023251440.1
A1	22919057 22920092
refGene_gn=LPAR6;refGene_tx=NM_001309049.1;refGene_gn=LPAR6;refGene_tx=XM_006927279.3
A1	22949947 22950144
refGene_gn=RB1;refGene_tx=XM_023251429.1;refGene_gn=RB1;refGene_tx=XM_023251437.1;refGene_gn=RB1;refGene_tx=XM_023251440.1
A1	22951277 22951354
refGene_gn=RB1;refGene_tx=XM_023251429.1;refGene_gn=RB1;refGene_tx=XM_023251437.1;refGene_gn=RB1;refGene_tx=XM_023251440.1
A1	22951436 22951468
refGene_gn=RB1;refGene_tx=XM_023251429.1;refGene_gn=RB1;refGene_tx=XM_023251437.1;refGene_gn=RB1;refGene_tx=XM_023251440.1
A1	22951553 22951610
refGene_gn=RB1;refGene_tx=XM_023251429.1;refGene_gn=RB1;refGene_tx=XM_023251437.1;refGene_gn=RB1;refGene_tx=XM_023251440.1
A1	22952836 22952953
refGene_gn=RB1;refGene_tx=XM_023251429.1;refGene_gn=RB1;refGene_tx=XM_023251437.1;refGene_gn=RB1;refGene_tx=XM_023251440.1
A1	22970811 22970899
refGene_gn=RB1;refGene_tx=XM_023251429.1;refGene_gn=RB1;refGene_tx=XM_023251437.1;refGene_gn=RB1;refGene_tx=XM_023251440.1
A1	22973533 22973611
refGene_gn=RB1;refGene_tx=XM_023251429.1;refGene_gn=RB1;refGene_tx=XM_023251437.1;refGene_gn=RB1;refGene_tx=XM_023251440.1
A1	22974472 22974582
refGene_gn=RB1;refGene_tx=XM_023251429.1;refGene_gn=RB1;refGene_tx=XM_023251437.1;refGene_gn=RB1;refGene_tx=XM_023251440.1
A1	22977405 22977483
refGene_gn=RB1;refGene_tx=XM_023251429.1;refGene_gn=RB1;refGene_tx=XM_023251437.1;refGene_gn=RB1;refGene_tx=XM_023251440.1
A1	22979651 22979794
refGene_gn=RB1;refGene_tx=XM_023251429.1;refGene_gn=RB1;refGene_tx=XM_023251437.1;refGene_gn=RB1;refGene_tx=XM_023251440.1
A1	22981380 22981491

refGene_gn=RB1;refGene_tx=XM_023251429.1;refGene_gn=RB1;refGene_tx=XM_023251437.1;refGene_gn=RB1;refGene_tx=XM_023251440.1
A1 22984530 22984598
refGene_gn=RB1;refGene_tx=XM_023251429.1;refGene_gn=RB1;refGene_tx=XM_023251437.1;refGene_gn=RB1;refGene_tx=XM_023251440.1
A1 22985960 22985999
refGene_gn=RB1;refGene_tx=XM_023251429.1;refGene_gn=RB1;refGene_tx=XM_023251437.1;refGene_gn=RB1;refGene_tx=XM_023251440.1
A1 22992526 22992646
refGene_gn=RB1;refGene_tx=XM_023251429.1;refGene_gn=RB1;refGene_tx=XM_023251437.1;refGene_gn=RB1;refGene_tx=XM_023251440.1
A1 22996512 22996628
refGene_gn=RB1;refGene_tx=XM_023251429.1;refGene_gn=RB1;refGene_tx=XM_023251437.1
A1 23035190 23035317
refGene_gn=RB1;refGene_tx=XM_023251429.1;refGene_gn=RB1;refGene_tx=XM_023251437.1;refGene_gn=RB1;refGene_tx=XM_023251440.1
A1 23038060 23038206
refGene_gn=RB1;refGene_tx=XM_023251429.1;refGene_gn=RB1;refGene_tx=XM_023251437.1
A1 23083806 23083892 refGene_gn=ITM2B;refGene_tx=XM_011280139.3
A1 23084998 23085149 refGene_gn=ITM2B;refGene_tx=XM_011280139.3
A1 23085576 23085687 refGene_gn=ITM2B;refGene_tx=XM_011280139.3
A1 23087162 23087369 refGene_gn=ITM2B;refGene_tx=XM_011280139.3
A1 23088878 23089007 refGene_gn=ITM2B;refGene_tx=XM_011280139.3
A1 23110971 23111088 refGene_gn=ITM2B;refGene_tx=XM_011280139.3
A1 23222930 23223055
refGene_gn=MED4;refGene_tx=XM_003980375.5;refGene_gn=MED4;refGene_tx=XM_019826448.1
A1 23230572 23230639
refGene_gn=MED4;refGene_tx=XM_003980375.5;refGene_gn=MED4;refGene_tx=XM_019826448.1
A1 23235606 23235777
refGene_gn=MED4;refGene_tx=XM_003980375.5;refGene_gn=MED4;refGene_tx=XM_019826448.1
A1 23237770 23237828
refGene_gn=MED4;refGene_tx=XM_003980375.5;refGene_gn=MED4;refGene_tx=XM_019826448.1
A1 23239338 23239425
refGene_gn=MED4;refGene_tx=XM_003980375.5;refGene_gn=MED4;refGene_tx=XM_019826448.1
A1 23240638 23240770
refGene_gn=MED4;refGene_tx=XM_003980375.5;refGene_gn=MED4;refGene_tx=XM_019826448.1
A1 23242864 23243037
refGene_gn=MED4;refGene_tx=XM_003980375.5;refGene_gn=MED4;refGene_tx=XM_019826448.1
A1 23251041 23251181
refGene_gn=NUDT15;refGene_tx=XM_003980376.5;refGene_gn=NUDT15;refGene_tx=XM_019826461.2;refGene_gn=NUDT15;refGene_tx=XM_019826468.2

A1 23253981 23254077
refGene_gn=NUDT15;refGene_tx=XM_019826461.2;refGene_gn=NUDT15;refGene_
tx=XM_019826468.2
A1 23255853 23256050
refGene_gn=NUDT15;refGene_tx=XM_003980376.5;refGene_gn=NUDT15;refGene_
tx=XM_019826461.2;refGene_gn=NUDT15;refGene_tx=XM_019826468.2
A1 23259030 23259116 refGene_gn=NUDT15;refGene_tx=XM_019826468.2
A1 23259701 23259859
refGene_gn=NUDT15;refGene_tx=XM_003980376.5;refGene_gn=NUDT15;refGene_
tx=XM_019826461.2
A1 23274811 23274901 refGene_gn=SUCLA2;refGene_tx=XM_003980377.4
A1 23279717 23279898
refGene_gn=SUCLA2;refGene_tx=XM_003980377.4;refGene_gn=SUCLA2;refGene_
tx=XM_023251507.1
A1 23280162 23280235 refGene_gn=SUCLA2;refGene_tx=XM_023251510.1
A1 23291785 23291885
refGene_gn=SUCLA2;refGene_tx=XM_003980377.4;refGene_gn=SUCLA2;refGene_
tx=XM_023251507.1;refGene_gn=SUCLA2;refGene_tx=XM_023251510.1
A1 23292074 23292237
refGene_gn=SUCLA2;refGene_tx=XM_003980377.4;refGene_gn=SUCLA2;refGene_
tx=XM_023251507.1;refGene_gn=SUCLA2;refGene_tx=XM_023251510.1
A1 23302644 23302773
refGene_gn=SUCLA2;refGene_tx=XM_003980377.4;refGene_gn=SUCLA2;refGene_
tx=XM_023251507.1;refGene_gn=SUCLA2;refGene_tx=XM_023251510.1
A1 23305511 23305650
refGene_gn=SUCLA2;refGene_tx=XM_003980377.4;refGene_gn=SUCLA2;refGene_
tx=XM_023251507.1;refGene_gn=SUCLA2;refGene_tx=XM_023251510.1
A1 23309366 23309528
refGene_gn=SUCLA2;refGene_tx=XM_003980377.4;refGene_gn=SUCLA2;refGene_
tx=XM_023251507.1;refGene_gn=SUCLA2;refGene_tx=XM_023251510.1
A1 23309623 23309766
refGene_gn=SUCLA2;refGene_tx=XM_003980377.4;refGene_gn=SUCLA2;refGene_
tx=XM_023251507.1;refGene_gn=SUCLA2;refGene_tx=XM_023251510.1
A1 23314798 23314919
refGene_gn=SUCLA2;refGene_tx=XM_003980377.4;refGene_gn=SUCLA2;refGene_
tx=XM_023251507.1;refGene_gn=SUCLA2;refGene_tx=XM_023251510.1
A1 23315375 23315464
refGene_gn=SUCLA2;refGene_tx=XM_003980377.4;refGene_gn=SUCLA2;refGene_
tx=XM_023251507.1;refGene_gn=SUCLA2;refGene_tx=XM_023251510.1
A1 23317493 23317568
refGene_gn=SUCLA2;refGene_tx=XM_003980377.4;refGene_gn=SUCLA2;refGene_
tx=XM_023251507.1;refGene_gn=SUCLA2;refGene_tx=XM_023251510.1
A1 24301368 24301780
refGene_gn=HTR2A;refGene_tx=XM_003980393.5;refGene_gn=HTR2A;refGene_tx
=XM_023251524.1
A1 24304576 24304777
refGene_gn=HTR2A;refGene_tx=XM_003980393.5;refGene_gn=HTR2A;refGene_tx
=XM_023251524.1
A1 24356188 24356988
refGene_gn=HTR2A;refGene_tx=XM_003980393.5;refGene_gn=HTR2A;refGene_tx

=XM_023251524.1
A1	24401835	24401903	refGene_gn=ESD;refGene_tx=XM_006927289.3
A1	24402477	24402566	refGene_gn=ESD;refGene_tx=XM_006927289.3
A1	24406059	24406158	refGene_gn=ESD;refGene_tx=XM_006927289.3
A1	24408391	24408516	refGene_gn=ESD;refGene_tx=XM_006927289.3
A1	24409562	24409682	refGene_gn=ESD;refGene_tx=XM_006927289.3
A1	24411392	24411491	refGene_gn=ESD;refGene_tx=XM_006927289.3
A1	24413454	24413622	refGene_gn=ESD;refGene_tx=XM_006927289.3
A1	24421445	24421526	refGene_gn=ESD;refGene_tx=XM_006927289.3
A1	24437643	24437754
refGene_gn=LRCH1;refGene_tx=NM_001048155.1;refGene_gn=LRCH1;refGene_tx=XM_011288570.3;refGene_gn=LRCH1;refGene_tx=XM_023239734.1
A1	24446182	24446389
refGene_gn=LRCH1;refGene_tx=XM_011288548.3;refGene_gn=LRCH1;refGene_tx=XM_019813191.2
A1	24449228	24449238	refGene_gn=LRCH1;refGene_tx=XM_023240296.1
A1	24454590	24454592	refGene_gn=LRCH1;refGene_tx=XM_023240296.1
A1	24455532	24455610
refGene_gn=LRCH1;refGene_tx=NM_001048155.1;refGene_gn=LRCH1;refGene_tx=XM_011288548.3;refGene_gn=LRCH1;refGene_tx=XM_011288570.3;refGene_gn=LRCH1;refGene_tx=XM_019813191.2;refGene_gn=LRCH1;refGene_tx=XM_023239734.1;refGene_gn=LRCH1;refGene_tx=XM_023240296.1
A1	24460023	24460161
refGene_gn=LRCH1;refGene_tx=NM_001048155.1;refGene_gn=LRCH1;refGene_tx=XM_011288548.3;refGene_gn=LRCH1;refGene_tx=XM_011288570.3;refGene_gn=LRCH1;refGene_tx=XM_019813191.2;refGene_gn=LRCH1;refGene_tx=XM_023239734.1;refGene_gn=LRCH1;refGene_tx=XM_023240296.1
A1	24461220	24461330
refGene_gn=LRCH1;refGene_tx=NM_001048155.1;refGene_gn=LRCH1;refGene_tx=XM_011288548.3;refGene_gn=LRCH1;refGene_tx=XM_011288570.3;refGene_gn=LRCH1;refGene_tx=XM_019813191.2;refGene_gn=LRCH1;refGene_tx=XM_023239734.1;refGene_gn=LRCH1;refGene_tx=XM_023240296.1
A1	24469218	24469323
refGene_gn=LRCH1;refGene_tx=XM_019813191.2;refGene_gn=LRCH1;refGene_tx=XM_023239734.1;refGene_gn=LRCH1;refGene_tx=XM_023240296.1
A1	24472201	24472274
refGene_gn=LRCH1;refGene_tx=XM_011288548.3;refGene_gn=LRCH1;refGene_tx=XM_011288570.3;refGene_gn=LRCH1;refGene_tx=XM_019813191.2;refGene_gn=LRCH1;refGene_tx=XM_023239734.1;refGene_gn=LRCH1;refGene_tx=XM_023240296.1
A1	24473001	24473055
refGene_gn=LRCH1;refGene_tx=XM_011288548.3;refGene_gn=LRCH1;refGene_tx=XM_011288570.3;refGene_gn=LRCH1;refGene_tx=XM_019813191.2;refGene_gn=LRCH1;refGene_tx=XM_023239734.1;refGene_gn=LRCH1;refGene_tx=XM_023240296.1
A1	24476952	24476989
refGene_gn=LRCH1;refGene_tx=XM_011288548.3;refGene_gn=LRCH1;refGene_tx=XM_011288570.3;refGene_gn=LRCH1;refGene_tx=XM_019813191.2;refGene_gn=LRCH1;refGene_tx=XM_023239734.1;refGene_gn=LRCH1;refGene_tx=XM_023240296.1

A1 24477084 24477174
refGene_gn=LRCH1;refGene_tx=XM_011288548.3;refGene_gn=LRCH1;refGene_tx=XM_011288570.3;refGene_gn=LRCH1;refGene_tx=XM_019813191.2;refGene_gn=LRCH1;refGene_tx=XM_023239734.1;refGene_gn=LRCH1;refGene_tx=XM_023240296.1
A1 24481792 24481879
refGene_gn=LRCH1;refGene_tx=XM_011288548.3;refGene_gn=LRCH1;refGene_tx=XM_011288570.3;refGene_gn=LRCH1;refGene_tx=XM_019813191.2;refGene_gn=LRCH1;refGene_tx=XM_023239734.1;refGene_gn=LRCH1;refGene_tx=XM_023240296.1
A1 24484229 24484297
refGene_gn=LRCH1;refGene_tx=XM_011288548.3;refGene_gn=LRCH1;refGene_tx=XM_011288570.3;refGene_gn=LRCH1;refGene_tx=XM_019813191.2;refGene_gn=LRCH1;refGene_tx=XM_023239734.1;refGene_gn=LRCH1;refGene_tx=XM_023240296.1
A1 24485591 24485719
refGene_gn=LRCH1;refGene_tx=XM_011288548.3;refGene_gn=LRCH1;refGene_tx=XM_011288570.3;refGene_gn=LRCH1;refGene_tx=XM_019813191.2;refGene_gn=LRCH1;refGene_tx=XM_023239734.1;refGene_gn=LRCH1;refGene_tx=XM_023240296.1
A1 24487493 24487599
refGene_gn=LRCH1;refGene_tx=XM_011288548.3;refGene_gn=LRCH1;refGene_tx=XM_011288570.3;refGene_gn=LRCH1;refGene_tx=XM_019813191.2;refGene_gn=LRCH1;refGene_tx=XM_023239734.1;refGene_gn=LRCH1;refGene_tx=XM_023240296.1
A1 24491328 24491392
refGene_gn=LRCH1;refGene_tx=XM_011288548.3;refGene_gn=LRCH1;refGene_tx=XM_011288570.3;refGene_gn=LRCH1;refGene_tx=XM_019813191.2;refGene_gn=LRCH1;refGene_tx=XM_023239734.1;refGene_gn=LRCH1;refGene_tx=XM_023240296.1
A1 24492502 24492630
refGene_gn=LRCH1;refGene_tx=XM_011288548.3;refGene_gn=LRCH1;refGene_tx=XM_011288570.3;refGene_gn=LRCH1;refGene_tx=XM_019813191.2;refGene_gn=LRCH1;refGene_tx=XM_023239734.1;refGene_gn=LRCH1;refGene_tx=XM_023240296.1
A1 24493360 24493497
refGene_gn=LRCH1;refGene_tx=XM_011288548.3;refGene_gn=LRCH1;refGene_tx=XM_011288570.3;refGene_gn=LRCH1;refGene_tx=XM_019813191.2;refGene_gn=LRCH1;refGene_tx=XM_023239734.1;refGene_gn=LRCH1;refGene_tx=XM_023240296.1
A1 24498705 24498811
refGene_gn=LRCH1;refGene_tx=XM_011288548.3;refGene_gn=LRCH1;refGene_tx=XM_011288570.3;refGene_gn=LRCH1;refGene_tx=XM_019813191.2;refGene_gn=LRCH1;refGene_tx=XM_023239734.1;refGene_gn=LRCH1;refGene_tx=XM_023240296.1
A1 24515691 24515818
refGene_gn=LRCH1;refGene_tx=XM_011288548.3;refGene_gn=LRCH1;refGene_tx=XM_011288570.3;refGene_gn=LRCH1;refGene_tx=XM_019813191.2;refGene_gn=LRCH1;refGene_tx=XM_023239734.1;refGene_gn=LRCH1;refGene_tx=XM_023240296.1

A1 24538325 24538470
refGene_gn=LRCH1;refGene_tx=XM_011288548.3;refGene_gn=LRCH1;refGene_tx=XM_011288570.3;refGene_gn=LRCH1;refGene_tx=XM_019813191.2;refGene_gn=LRCH1;refGene_tx=XM_023239734.1;refGene_gn=LRCH1;refGene_tx=XM_023240296.1
A1 24637978 24638294
refGene_gn=LRCH1;refGene_tx=XM_011288548.3;refGene_gn=LRCH1;refGene_tx=XM_011288570.3;refGene_gn=LRCH1;refGene_tx=XM_019813191.2;refGene_gn=LRCH1;refGene_tx=XM_023239734.1;refGene_gn=LRCH1;refGene_tx=XM_023240296.1
A1 24738902 24739416
refGene_gn=RUBCNL;refGene_tx=XM_006927291.4;refGene_gn=RUBCNL;refGene_tx=XM_006927292.4;refGene_gn=RUBCNL;refGene_tx=XM_006927293.4;refGene_gn=RUBCNL;refGene_tx=XM_019826547.2;refGene_gn=RUBCNL;refGene_tx=XM_019826552.2;refGene_gn=RUBCNL;refGene_tx=XM_019826556.2;refGene_gn=RUBCNL;refGene_tx=XM_023251532.1;refGene_gn=RUBCNL;refGene_tx=XM_023251536.1;refGene_gn=RUBCNL;refGene_tx=XM_023251553.1;refGene_gn=RUBCNL;refGene_tx=XM_023251554.1;refGene_gn=RUBCNL;refGene_tx=XM_023251555.1
A1 24742175 24742299 refGene_gn=RUBCNL;refGene_tx=XM_019826580.2
A1 24742700 24742783
refGene_gn=RUBCNL;refGene_tx=XM_006927291.4;refGene_gn=RUBCNL;refGene_tx=XM_006927292.4;refGene_gn=RUBCNL;refGene_tx=XM_006927293.4;refGene_gn=RUBCNL;refGene_tx=XM_019826547.2;refGene_gn=RUBCNL;refGene_tx=XM_019826552.2;refGene_gn=RUBCNL;refGene_tx=XM_019826556.2;refGene_gn=RUBCNL;refGene_tx=XM_019826580.2;refGene_gn=RUBCNL;refGene_tx=XM_023251532.1;refGene_gn=RUBCNL;refGene_tx=XM_023251536.1;refGene_gn=RUBCNL;refGene_tx=XM_023251553.1;refGene_gn=RUBCNL;refGene_tx=XM_023251554.1;refGene_gn=RUBCNL;refGene_tx=XM_023251555.1
A1 24743511 24743722
refGene_gn=RUBCNL;refGene_tx=XM_006927291.4;refGene_gn=RUBCNL;refGene_tx=XM_006927292.4;refGene_gn=RUBCNL;refGene_tx=XM_006927293.4;refGene_gn=RUBCNL;refGene_tx=XM_019826547.2;refGene_gn=RUBCNL;refGene_tx=XM_019826552.2;refGene_gn=RUBCNL;refGene_tx=XM_019826556.2;refGene_gn=RUBCNL;refGene_tx=XM_019826580.2;refGene_gn=RUBCNL;refGene_tx=XM_023251532.1;refGene_gn=RUBCNL;refGene_tx=XM_023251536.1;refGene_gn=RUBCNL;refGene_tx=XM_023251553.1;refGene_gn=RUBCNL;refGene_tx=XM_023251554.1;refGene_gn=RUBCNL;refGene_tx=XM_023251555.1
A1 24748451 24748553
refGene_gn=RUBCNL;refGene_tx=XM_006927291.4;refGene_gn=RUBCNL;refGene_tx=XM_006927292.4;refGene_gn=RUBCNL;refGene_tx=XM_006927293.4;refGene_gn=RUBCNL;refGene_tx=XM_019826547.2;refGene_gn=RUBCNL;refGene_tx=XM_019826552.2;refGene_gn=RUBCNL;refGene_tx=XM_019826556.2;refGene_gn=RUBCNL;refGene_tx=XM_019826580.2;refGene_gn=RUBCNL;refGene_tx=XM_023251532.1;refGene_gn=RUBCNL;refGene_tx=XM_023251536.1;refGene_gn=RUBCNL;refGene_tx=XM_023251553.1;refGene_gn=RUBCNL;refGene_tx=XM_023251554.1;refGene_gn=RUBCNL;refGene_tx=XM_023251555.1
A1 24749034 24749095
refGene_gn=RUBCNL;refGene_tx=XM_006927291.4;refGene_gn=RUBCNL;refGene_tx=XM_006927292.4;refGene_gn=RUBCNL;refGene_tx=XM_006927293.4;refGene_gn=RUBCNL;refGene_tx=XM_019826547.2;refGene_gn=RUBCNL;refGene_tx=XM_01

9826552.2;refGene_gn=RUBCNL;refGene_tx=XM_019826556.2;refGene_gn=RUBCNL;refGene_tx=XM_019826580.2;refGene_gn=RUBCNL;refGene_tx=XM_023251532.1;refGene_gn=RUBCNL;refGene_tx=XM_023251536.1;refGene_gn=RUBCNL;refGene_tx=XM_023251553.1;refGene_gn=RUBCNL;refGene_tx=XM_023251554.1;refGene_gn=RUBCNL;refGene_tx=XM_023251555.1
A1 24749676 24749809
refGene_gn=RUBCNL;refGene_tx=XM_006927291.4;refGene_gn=RUBCNL;refGene_tx=XM_006927292.4;refGene_gn=RUBCNL;refGene_tx=XM_006927293.4;refGene_gn=RUBCNL;refGene_tx=XM_019826547.2;refGene_gn=RUBCNL;refGene_tx=XM_019826552.2;refGene_gn=RUBCNL;refGene_tx=XM_019826556.2;refGene_gn=RUBCNL;refGene_tx=XM_019826580.2;refGene_gn=RUBCNL;refGene_tx=XM_023251532.1;refGene_gn=RUBCNL;refGene_tx=XM_023251536.1;refGene_gn=RUBCNL;refGene_tx=XM_023251553.1;refGene_gn=RUBCNL;refGene_tx=XM_023251554.1;refGene_gn=RUBCNL;refGene_tx=XM_023251555.1
A1 24751498 24751644
refGene_gn=RUBCNL;refGene_tx=XM_006927291.4;refGene_gn=RUBCNL;refGene_tx=XM_006927292.4;refGene_gn=RUBCNL;refGene_tx=XM_006927293.4;refGene_gn=RUBCNL;refGene_tx=XM_019826547.2;refGene_gn=RUBCNL;refGene_tx=XM_019826552.2;refGene_gn=RUBCNL;refGene_tx=XM_019826556.2;refGene_gn=RUBCNL;refGene_tx=XM_019826580.2;refGene_gn=RUBCNL;refGene_tx=XM_023251532.1;refGene_gn=RUBCNL;refGene_tx=XM_023251536.1;refGene_gn=RUBCNL;refGene_tx=XM_023251553.1;refGene_gn=RUBCNL;refGene_tx=XM_023251554.1;refGene_gn=RUBCNL;refGene_tx=XM_023251555.1
A1 24753138 24753203
refGene_gn=RUBCNL;refGene_tx=XM_006927291.4;refGene_gn=RUBCNL;refGene_tx=XM_006927292.4;refGene_gn=RUBCNL;refGene_tx=XM_006927293.4;refGene_gn=RUBCNL;refGene_tx=XM_019826547.2;refGene_gn=RUBCNL;refGene_tx=XM_019826552.2;refGene_gn=RUBCNL;refGene_tx=XM_019826556.2;refGene_gn=RUBCNL;refGene_tx=XM_019826580.2;refGene_gn=RUBCNL;refGene_tx=XM_023251532.1;refGene_gn=RUBCNL;refGene_tx=XM_023251536.1;refGene_gn=RUBCNL;refGene_tx=XM_023251553.1;refGene_gn=RUBCNL;refGene_tx=XM_023251554.1;refGene_gn=RUBCNL;refGene_tx=XM_023251555.1
A1 24760260 24760499
refGene_gn=RUBCNL;refGene_tx=XM_006927291.4;refGene_gn=RUBCNL;refGene_tx=XM_006927292.4;refGene_gn=RUBCNL;refGene_tx=XM_006927293.4;refGene_gn=RUBCNL;refGene_tx=XM_019826547.2;refGene_gn=RUBCNL;refGene_tx=XM_019826552.2;refGene_gn=RUBCNL;refGene_tx=XM_019826556.2;refGene_gn=RUBCNL;refGene_tx=XM_019826580.2;refGene_gn=RUBCNL;refGene_tx=XM_023251532.1;refGene_gn=RUBCNL;refGene_tx=XM_023251536.1;refGene_gn=RUBCNL;refGene_tx=XM_023251553.1;refGene_gn=RUBCNL;refGene_tx=XM_023251554.1;refGene_gn=RUBCNL;refGene_tx=XM_023251555.1
A1 24761508 24761570
refGene_gn=RUBCNL;refGene_tx=XM_006927291.4;refGene_gn=RUBCNL;refGene_tx=XM_006927292.4;refGene_gn=RUBCNL;refGene_tx=XM_006927293.4;refGene_gn=RUBCNL;refGene_tx=XM_019826547.2;refGene_gn=RUBCNL;refGene_tx=XM_019826552.2;refGene_gn=RUBCNL;refGene_tx=XM_019826556.2;refGene_gn=RUBCNL;refGene_tx=XM_019826580.2;refGene_gn=RUBCNL;refGene_tx=XM_023251532.1;refGene_gn=RUBCNL;refGene_tx=XM_023251536.1;refGene_gn=RUBCNL;refGene_tx=XM_023251553.1;refGene_gn=RUBCNL;refGene_tx=XM_023251554.1;refGene_gn=RUBCNL;refGene_tx=XM_023251555.1

A1 24763950 24764104
refGene_gn=RUBCNL;refGene_tx=XM_006927291.4;refGene_gn=RUBCNL;refGene_tx=XM_006927292.4;refGene_gn=RUBCNL;refGene_tx=XM_006927293.4;refGene_gn=RUBCNL;refGene_tx=XM_019826547.2;refGene_gn=RUBCNL;refGene_tx=XM_019826552.2;refGene_gn=RUBCNL;refGene_tx=XM_019826556.2;refGene_gn=RUBCNL;refGene_tx=XM_019826580.2;refGene_gn=RUBCNL;refGene_tx=XM_023251532.1;refGene_gn=RUBCNL;refGene_tx=XM_023251536.1;refGene_gn=RUBCNL;refGene_tx=XM_023251553.1;refGene_gn=RUBCNL;refGene_tx=XM_023251554.1;refGene_gn=RUBCNL;refGene_tx=XM_023251555.1
A1 24764742 24764853
refGene_gn=RUBCNL;refGene_tx=XM_006927291.4;refGene_gn=RUBCNL;refGene_tx=XM_006927292.4;refGene_gn=RUBCNL;refGene_tx=XM_006927293.4;refGene_gn=RUBCNL;refGene_tx=XM_019826547.2;refGene_gn=RUBCNL;refGene_tx=XM_019826552.2;refGene_gn=RUBCNL;refGene_tx=XM_019826556.2;refGene_gn=RUBCNL;refGene_tx=XM_019826580.2;refGene_gn=RUBCNL;refGene_tx=XM_023251532.1;refGene_gn=RUBCNL;refGene_tx=XM_023251536.1;refGene_gn=RUBCNL;refGene_tx=XM_023251553.1;refGene_gn=RUBCNL;refGene_tx=XM_023251554.1;refGene_gn=RUBCNL;refGene_tx=XM_023251555.1
A1 24765320 24765386
refGene_gn=RUBCNL;refGene_tx=XM_006927291.4;refGene_gn=RUBCNL;refGene_tx=XM_006927292.4;refGene_gn=RUBCNL;refGene_tx=XM_006927293.4;refGene_gn=RUBCNL;refGene_tx=XM_019826547.2;refGene_gn=RUBCNL;refGene_tx=XM_019826552.2;refGene_gn=RUBCNL;refGene_tx=XM_019826556.2;refGene_gn=RUBCNL;refGene_tx=XM_019826580.2;refGene_gn=RUBCNL;refGene_tx=XM_023251532.1;refGene_gn=RUBCNL;refGene_tx=XM_023251536.1
A1 24766062 24766175
refGene_gn=RUBCNL;refGene_tx=XM_006927291.4;refGene_gn=RUBCNL;refGene_tx=XM_006927292.4;refGene_gn=RUBCNL;refGene_tx=XM_006927293.4;refGene_gn=RUBCNL;refGene_tx=XM_019826547.2;refGene_gn=RUBCNL;refGene_tx=XM_019826552.2;refGene_gn=RUBCNL;refGene_tx=XM_019826556.2;refGene_gn=RUBCNL;refGene_tx=XM_019826580.2;refGene_gn=RUBCNL;refGene_tx=XM_023251532.1;refGene_gn=RUBCNL;refGene_tx=XM_023251536.1;refGene_gn=RUBCNL;refGene_tx=XM_023251554.1
A1 24793197 24793370 refGene_gn=RUBCNL;refGene_tx=XM_023251553.1
A1 24806623 24806843
refGene_gn=LRRC63;refGene_tx=XM_019826586.2;refGene_gn=LRRC63;refGene_tx=XM_019826597.2;refGene_gn=LRRC63;refGene_tx=XM_019826605.2;refGene_gn=LRRC63;refGene_tx=XM_019826608.2;refGene_gn=LRRC63;refGene_tx=XM_019826612.2
A1 24811735 24811975
refGene_gn=LRRC63;refGene_tx=XM_019826586.2;refGene_gn=LRRC63;refGene_tx=XM_019826597.2;refGene_gn=LRRC63;refGene_tx=XM_019826605.2;refGene_gn=LRRC63;refGene_tx=XM_019826608.2;refGene_gn=LRRC63;refGene_tx=XM_019826612.2
A1 24815234 24815318
refGene_gn=LRRC63;refGene_tx=XM_019826586.2;refGene_gn=LRRC63;refGene_tx=XM_019826597.2;refGene_gn=LRRC63;refGene_tx=XM_019826608.2;refGene_gn=LRRC63;refGene_tx=XM_019826612.2
A1 24818968 24819105
refGene_gn=LRRC63;refGene_tx=XM_019826586.2;refGene_gn=LRRC63;refGene_

tx=XM_019826597.2;refGene_gn=LRRC63;refGene_tx=XM_019826605.2;refGene_gn=LRRC63;refGene_tx=XM_019826608.2;refGene_gn=LRRC63;refGene_tx=XM_019826612.2
A1 24819397 24819496
refGene_gn=LRRC63;refGene_tx=XM_019826586.2;refGene_gn=LRRC63;refGene_tx=XM_019826597.2;refGene_gn=LRRC63;refGene_tx=XM_019826605.2;refGene_gn=LRRC63;refGene_tx=XM_019826608.2
A1 24820153 24820252
refGene_gn=LRRC63;refGene_tx=XM_019826586.2;refGene_gn=LRRC63;refGene_tx=XM_019826597.2;refGene_gn=LRRC63;refGene_tx=XM_019826605.2
A1 24829715 24829876
refGene_gn=LRRC63;refGene_tx=XM_019826586.2;refGene_gn=LRRC63;refGene_tx=XM_019826597.2;refGene_gn=LRRC63;refGene_tx=XM_019826605.2;refGene_gn=LRRC63;refGene_tx=XM_019826608.2;refGene_gn=LRRC63;refGene_tx=XM_019826612.2
A1 24832317 24832386
refGene_gn=LRRC63;refGene_tx=XM_019826586.2;refGene_gn=LRRC63;refGene_tx=XM_019826597.2;refGene_gn=LRRC63;refGene_tx=XM_019826605.2;refGene_gn=LRRC63;refGene_tx=XM_019826608.2;refGene_gn=LRRC63;refGene_tx=XM_019826612.2
A1 24832930 24833620
refGene_gn=LRRC63;refGene_tx=XM_019826586.2;refGene_gn=LRRC63;refGene_tx=XM_019826597.2;refGene_gn=LRRC63;refGene_tx=XM_019826605.2;refGene_gn=LRRC63;refGene_tx=XM_019826608.2;refGene_gn=LRRC63;refGene_tx=XM_019826612.2
A1 24848742 24848827
refGene_gn=LRRC63;refGene_tx=XM_019826586.2;refGene_gn=LRRC63;refGene_tx=XM_019826597.2;refGene_gn=LRRC63;refGene_tx=XM_019826605.2;refGene_gn=LRRC63;refGene_tx=XM_019826608.2;refGene_gn=LRRC63;refGene_tx=XM_019826612.2
A1 24902127 24902191 refGene_gn=LCP1;refGene_tx=XM_003980396.5
A1 24902776 24902940 refGene_gn=LCP1;refGene_tx=XM_003980396.5
A1 24903108 24903238 refGene_gn=LCP1;refGene_tx=XM_003980396.5
A1 24904913 24905046 refGene_gn=LCP1;refGene_tx=XM_003980396.5
A1 24907488 24907570 refGene_gn=LCP1;refGene_tx=XM_003980396.5
A1 24909404 24909570 refGene_gn=LCP1;refGene_tx=XM_003980396.5
A1 24911378 24911521 refGene_gn=LCP1;refGene_tx=XM_003980396.5
A1 24913763 24913859 refGene_gn=LCP1;refGene_tx=XM_003980396.5
A1 24915430 24915626 refGene_gn=LCP1;refGene_tx=XM_003980396.5
A1 24917694 24917773 refGene_gn=LCP1;refGene_tx=XM_003980396.5
A1 24918621 24918736 refGene_gn=LCP1;refGene_tx=XM_003980396.5
A1 24919642 24919776 refGene_gn=LCP1;refGene_tx=XM_003980396.5
A1 24926580 24926704 refGene_gn=LCP1;refGene_tx=XM_003980396.5
A1 24929846 24929971 refGene_gn=LCP1;refGene_tx=XM_003980396.5
A1 24932592 24932725 refGene_gn=LCP1;refGene_tx=XM_003980396.5
A1 24950120 24950194 refGene_gn=CPB2;refGene_tx=XM_019826669.1
A1 24963942 24964018 refGene_gn=CPB2;refGene_tx=XM_019826669.1
A1 24966200 24966325 refGene_gn=CPB2;refGene_tx=XM_019826669.1
A1 24967844 24967953 refGene_gn=CPB2;refGene_tx=XM_019826669.1
A1 24971597 24971699 refGene_gn=CPB2;refGene_tx=XM_019826669.1

A1 24974865 24974970 refGene_gn=CPB2;refGene_tx=XM_019826669.1
A1 24982375 24982486 refGene_gn=CPB2;refGene_tx=XM_019826669.1
A1 24985087 24985181 refGene_gn=CPB2;refGene_tx=XM_019826669.1
A1 24991660 24991863 refGene_gn=CPB2;refGene_tx=XM_019826669.1
A1 24995351 24995439 refGene_gn=CPB2;refGene_tx=XM_019826669.1
A1 24999951 25000136 refGene_gn=CPB2;refGene_tx=XM_019826669.1
A1 25007509 25007626
refGene_gn=ZC3H13;refGene_tx=XM_006927298.4;refGene_gn=ZC3H13;refGene_tx=XM_019826627.2;refGene_gn=ZC3H13;refGene_tx=XM_019826630.2;refGene_gn=ZC3H13;refGene_tx=XM_019826634.2;refGene_gn=ZC3H13;refGene_tx=XM_019826636.2;refGene_gn=ZC3H13;refGene_tx=XM_019826640.2;refGene_gn=ZC3H13;refGene_tx=XM_019826641.2;refGene_gn=ZC3H13;refGene_tx=XM_019826649.2;refGene_gn=ZC3H13;refGene_tx=XM_019826664.2
A1 25007935 25008045
refGene_gn=ZC3H13;refGene_tx=XM_006927298.4;refGene_gn=ZC3H13;refGene_tx=XM_019826627.2;refGene_gn=ZC3H13;refGene_tx=XM_019826630.2;refGene_gn=ZC3H13;refGene_tx=XM_019826634.2;refGene_gn=ZC3H13;refGene_tx=XM_019826636.2;refGene_gn=ZC3H13;refGene_tx=XM_019826640.2;refGene_gn=ZC3H13;refGene_tx=XM_019826641.2;refGene_gn=ZC3H13;refGene_tx=XM_019826649.2;refGene_gn=ZC3H13;refGene_tx=XM_019826664.2
A1 25010604 25010716
refGene_gn=ZC3H13;refGene_tx=XM_006927298.4;refGene_gn=ZC3H13;refGene_tx=XM_019826627.2;refGene_gn=ZC3H13;refGene_tx=XM_019826630.2;refGene_gn=ZC3H13;refGene_tx=XM_019826634.2;refGene_gn=ZC3H13;refGene_tx=XM_019826636.2;refGene_gn=ZC3H13;refGene_tx=XM_019826640.2;refGene_gn=ZC3H13;refGene_tx=XM_019826641.2;refGene_gn=ZC3H13;refGene_tx=XM_019826649.2;refGene_gn=ZC3H13;refGene_tx=XM_019826664.2
A1 25022161 25022212
refGene_gn=ZC3H13;refGene_tx=XM_019826627.2;refGene_gn=ZC3H13;refGene_tx=XM_019826630.2;refGene_gn=ZC3H13;refGene_tx=XM_019826634.2;refGene_gn=ZC3H13;refGene_tx=XM_019826636.2;refGene_gn=ZC3H13;refGene_tx=XM_019826640.2;refGene_gn=ZC3H13;refGene_tx=XM_019826649.2;refGene_gn=ZC3H13;refGene_tx=XM_019826664.2
A1 25028078 25028187
refGene_gn=ZC3H13;refGene_tx=XM_006927298.4;refGene_gn=ZC3H13;refGene_tx=XM_019826627.2;refGene_gn=ZC3H13;refGene_tx=XM_019826630.2;refGene_gn=ZC3H13;refGene_tx=XM_019826634.2;refGene_gn=ZC3H13;refGene_tx=XM_019826636.2;refGene_gn=ZC3H13;refGene_tx=XM_019826640.2;refGene_gn=ZC3H13;refGene_tx=XM_019826641.2;refGene_gn=ZC3H13;refGene_tx=XM_019826649.2;refGene_gn=ZC3H13;refGene_tx=XM_019826664.2
A1 25037794 25037934
refGene_gn=ZC3H13;refGene_tx=XM_006927298.4;refGene_gn=ZC3H13;refGene_tx=XM_019826627.2;refGene_gn=ZC3H13;refGene_tx=XM_019826630.2;refGene_gn=ZC3H13;refGene_tx=XM_019826634.2;refGene_gn=ZC3H13;refGene_tx=XM_019826636.2;refGene_gn=ZC3H13;refGene_tx=XM_019826640.2;refGene_gn=ZC3H13;refGene_tx=XM_019826641.2;refGene_gn=ZC3H13;refGene_tx=XM_019826649.2;refGene_gn=ZC3H13;refGene_tx=XM_019826664.2
A1 25038535 25038696
refGene_gn=ZC3H13;refGene_tx=XM_006927298.4;refGene_gn=ZC3H13;refGene_tx=XM_019826627.2;refGene_gn=ZC3H13;refGene_tx=XM_019826630.2;refGene_

gn=ZC3H13;refGene_tx=XM_019826634.2;refGene_gn=ZC3H13;refGene_tx=XM_01
9826636.2;refGene_gn=ZC3H13;refGene_tx=XM_019826640.2;refGene_gn=ZC3H1
3;refGene_tx=XM_019826641.2;refGene_gn=ZC3H13;refGene_tx=XM_019826649.
2;refGene_gn=ZC3H13;refGene_tx=XM_019826664.2
A1 25042839 25043037
refGene_gn=ZC3H13;refGene_tx=XM_006927298.4;refGene_gn=ZC3H13;refGene_
tx=XM_019826627.2;refGene_gn=ZC3H13;refGene_tx=XM_019826630.2;refGene_
gn=ZC3H13;refGene_tx=XM_019826634.2;refGene_gn=ZC3H13;refGene_tx=XM_01
9826636.2;refGene_gn=ZC3H13;refGene_tx=XM_019826640.2;refGene_gn=ZC3H1
3;refGene_tx=XM_019826641.2;refGene_gn=ZC3H13;refGene_tx=XM_019826649.
2;refGene_gn=ZC3H13;refGene_tx=XM_019826664.2
A1 25061522 25061833
refGene_gn=ZC3H13;refGene_tx=XM_006927298.4;refGene_gn=ZC3H13;refGene_
tx=XM_019826627.2;refGene_gn=ZC3H13;refGene_tx=XM_019826630.2;refGene_
gn=ZC3H13;refGene_tx=XM_019826634.2;refGene_gn=ZC3H13;refGene_tx=XM_01
9826636.2;refGene_gn=ZC3H13;refGene_tx=XM_019826640.2;refGene_gn=ZC3H1
3;refGene_tx=XM_019826641.2;refGene_gn=ZC3H13;refGene_tx=XM_019826649.
2;refGene_gn=ZC3H13;refGene_tx=XM_019826664.2
A1 25064381 25064846
refGene_gn=ZC3H13;refGene_tx=XM_006927298.4;refGene_gn=ZC3H13;refGene_
tx=XM_019826627.2;refGene_gn=ZC3H13;refGene_tx=XM_019826630.2;refGene_
gn=ZC3H13;refGene_tx=XM_019826634.2;refGene_gn=ZC3H13;refGene_tx=XM_01
9826636.2;refGene_gn=ZC3H13;refGene_tx=XM_019826640.2;refGene_gn=ZC3H1
3;refGene_tx=XM_019826641.2;refGene_gn=ZC3H13;refGene_tx=XM_019826649.
2;refGene_gn=ZC3H13;refGene_tx=XM_019826664.2
A1 25071656 25071848
refGene_gn=ZC3H13;refGene_tx=XM_006927298.4;refGene_gn=ZC3H13;refGene_
tx=XM_019826627.2;refGene_gn=ZC3H13;refGene_tx=XM_019826630.2;refGene_
gn=ZC3H13;refGene_tx=XM_019826634.2;refGene_gn=ZC3H13;refGene_tx=XM_01
9826636.2;refGene_gn=ZC3H13;refGene_tx=XM_019826640.2;refGene_gn=ZC3H1
3;refGene_tx=XM_019826641.2;refGene_gn=ZC3H13;refGene_tx=XM_019826649.
2
A1 25075952 25076036
refGene_gn=ZC3H13;refGene_tx=XM_019826627.2;refGene_gn=ZC3H13;refGene_
tx=XM_019826630.2;refGene_gn=ZC3H13;refGene_tx=XM_019826634.2;refGene_
gn=ZC3H13;refGene_tx=XM_019826636.2;refGene_gn=ZC3H13;refGene_tx=XM_01
9826640.2;refGene_gn=ZC3H13;refGene_tx=XM_019826641.2
A1 25076379 25076941
refGene_gn=ZC3H13;refGene_tx=XM_006927298.4;refGene_gn=ZC3H13;refGene_
tx=XM_019826627.2;refGene_gn=ZC3H13;refGene_tx=XM_019826630.2;refGene_
gn=ZC3H13;refGene_tx=XM_019826634.2;refGene_gn=ZC3H13;refGene_tx=XM_01
9826636.2;refGene_gn=ZC3H13;refGene_tx=XM_019826640.2;refGene_gn=ZC3H1
3;refGene_tx=XM_019826641.2;refGene_gn=ZC3H13;refGene_tx=XM_019826649.
2;refGene_gn=ZC3H13;refGene_tx=XM_019826664.2
A1 25080287 25080391
refGene_gn=ZC3H13;refGene_tx=XM_006927298.4;refGene_gn=ZC3H13;refGene_
tx=XM_019826627.2;refGene_gn=ZC3H13;refGene_tx=XM_019826630.2;refGene_
gn=ZC3H13;refGene_tx=XM_019826634.2;refGene_gn=ZC3H13;refGene_tx=XM_01
9826636.2;refGene_gn=ZC3H13;refGene_tx=XM_019826640.2;refGene_gn=ZC3H1
3;refGene_tx=XM_019826641.2;refGene_gn=ZC3H13;refGene_tx=XM_019826649.

2;refGene_gn=ZC3H13;refGene_tx=XM_019826664.2
A1	25080731	25081925
refGene_gn=ZC3H13;refGene_tx=XM_006927298.4;refGene_gn=ZC3H13;refGene_tx=XM_019826627.2;refGene_gn=ZC3H13;refGene_tx=XM_019826630.2;refGene_gn=ZC3H13;refGene_tx=XM_019826634.2;refGene_gn=ZC3H13;refGene_tx=XM_019826636.2;refGene_gn=ZC3H13;refGene_tx=XM_019826640.2;refGene_gn=ZC3H13;refGene_tx=XM_019826641.2;refGene_gn=ZC3H13;refGene_tx=XM_019826649.2;refGene_gn=ZC3H13;refGene_tx=XM_019826664.2
A1	25082775	25083297
refGene_gn=ZC3H13;refGene_tx=XM_006927298.4;refGene_gn=ZC3H13;refGene_tx=XM_019826627.2;refGene_gn=ZC3H13;refGene_tx=XM_019826630.2;refGene_gn=ZC3H13;refGene_tx=XM_019826634.2;refGene_gn=ZC3H13;refGene_tx=XM_019826636.2;refGene_gn=ZC3H13;refGene_tx=XM_019826640.2;refGene_gn=ZC3H13;refGene_tx=XM_019826641.2;refGene_gn=ZC3H13;refGene_tx=XM_019826649.2;refGene_gn=ZC3H13;refGene_tx=XM_019826664.2
A1	25085612	25085762
refGene_gn=ZC3H13;refGene_tx=XM_006927298.4;refGene_gn=ZC3H13;refGene_tx=XM_019826627.2;refGene_gn=ZC3H13;refGene_tx=XM_019826630.2;refGene_gn=ZC3H13;refGene_tx=XM_019826634.2;refGene_gn=ZC3H13;refGene_tx=XM_019826636.2;refGene_gn=ZC3H13;refGene_tx=XM_019826640.2;refGene_gn=ZC3H13;refGene_tx=XM_019826641.2;refGene_gn=ZC3H13;refGene_tx=XM_019826649.2;refGene_gn=ZC3H13;refGene_tx=XM_019826664.2
A1	25086975	25087176
refGene_gn=ZC3H13;refGene_tx=XM_006927298.4;refGene_gn=ZC3H13;refGene_tx=XM_019826627.2;refGene_gn=ZC3H13;refGene_tx=XM_019826630.2;refGene_gn=ZC3H13;refGene_tx=XM_019826634.2;refGene_gn=ZC3H13;refGene_tx=XM_019826636.2;refGene_gn=ZC3H13;refGene_tx=XM_019826640.2;refGene_gn=ZC3H13;refGene_tx=XM_019826641.2;refGene_gn=ZC3H13;refGene_tx=XM_019826649.2;refGene_gn=ZC3H13;refGene_tx=XM_019826664.2
A1	25092542	25092706
refGene_gn=ZC3H13;refGene_tx=XM_006927298.4;refGene_gn=ZC3H13;refGene_tx=XM_019826627.2;refGene_gn=ZC3H13;refGene_tx=XM_019826630.2;refGene_gn=ZC3H13;refGene_tx=XM_019826634.2;refGene_gn=ZC3H13;refGene_tx=XM_019826636.2;refGene_gn=ZC3H13;refGene_tx=XM_019826640.2;refGene_gn=ZC3H13;refGene_tx=XM_019826641.2;refGene_gn=ZC3H13;refGene_tx=XM_019826649.2;refGene_gn=ZC3H13;refGene_tx=XM_019826664.2
A1	25097875	25098046
refGene_gn=ZC3H13;refGene_tx=XM_006927298.4;refGene_gn=ZC3H13;refGene_tx=XM_019826627.2;refGene_gn=ZC3H13;refGene_tx=XM_019826630.2;refGene_gn=ZC3H13;refGene_tx=XM_019826634.2;refGene_gn=ZC3H13;refGene_tx=XM_019826636.2;refGene_gn=ZC3H13;refGene_tx=XM_019826640.2;refGene_gn=ZC3H13;refGene_tx=XM_019826641.2;refGene_gn=ZC3H13;refGene_tx=XM_019826649.2;refGene_gn=ZC3H13;refGene_tx=XM_019826664.2
A1	25215745	25215784	refGene_gn=SIAH3;refGene_tx=XM_023251641.1
A1	25216800	25216935	refGene_gn=SIAH3;refGene_tx=XM_006927299.4
A1	25286020	25286689
refGene_gn=SIAH3;refGene_tx=XM_006927299.4;refGene_gn=SIAH3;refGene_tx=XM_023251641.1
A1	25338560	25339778
refGene_gn=SPERT;refGene_tx=XM_003980399.4;refGene_gn=SPERT;refGene_tx

=XM_006927300.3;refGene_gn=SPERT;refGene_tx=XM_019826714.2;refGene_gn=SPERT;refGene_tx=XM_019826718.2;refGene_gn=SPERT;refGene_tx=XM_023251649.1
A1 25350623 25350680 refGene_gn=SPERT;refGene_tx=XM_006927300.3
A1 25351377 25351459 refGene_gn=SPERT;refGene_tx=XM_003980399.4
A1 25351980 25352054 refGene_gn=SPERT;refGene_tx=XM_003980399.4
A1 25405212 25405534 refGene_gn=ERICH6B;refGene_tx=XM_019826676.2
A1 25411547 25411593 refGene_gn=ERICH6B;refGene_tx=XM_019826676.2
A1 25415233 25415400 refGene_gn=ERICH6B;refGene_tx=XM_019826676.2
A1 25423029 25423101 refGene_gn=ERICH6B;refGene_tx=XM_019826676.2
A1 25424144 25424183 refGene_gn=ERICH6B;refGene_tx=XM_019826676.2
A1 25427520 25427609 refGene_gn=ERICH6B;refGene_tx=XM_019826676.2
A1 25431403 25431540 refGene_gn=ERICH6B;refGene_tx=XM_019826676.2
A1 25433870 25433932 refGene_gn=ERICH6B;refGene_tx=XM_019826676.2
A1 25440516 25440683 refGene_gn=ERICH6B;refGene_tx=XM_019826676.2
A1 25455169 25455255 refGene_gn=ERICH6B;refGene_tx=XM_019826676.2
A1 25455419 25455572 refGene_gn=ERICH6B;refGene_tx=XM_019826676.2
A1 25457257 25457483 refGene_gn=ERICH6B;refGene_tx=XM_019826676.2
A1 25459745 25459961 refGene_gn=ERICH6B;refGene_tx=XM_019826676.2
A1 25466666 25466696
refGene_gn=COG3;refGene_tx=XM_003980400.5;refGene_gn=COG3;refGene_tx=XM_006927301.4
A1 25471535 25471634
refGene_gn=COG3;refGene_tx=XM_003980400.5;refGene_gn=COG3;refGene_tx=XM_006927301.4
A1 25472912 25473040
refGene_gn=COG3;refGene_tx=XM_003980400.5;refGene_gn=COG3;refGene_tx=XM_006927301.4;refGene_gn=COG3;refGene_tx=XM_019826762.2
A1 25477092 25477168
refGene_gn=COG3;refGene_tx=XM_003980400.5;refGene_gn=COG3;refGene_tx=XM_006927301.4;refGene_gn=COG3;refGene_tx=XM_019826762.2
A1 25482355 25482490
refGene_gn=COG3;refGene_tx=XM_003980400.5;refGene_gn=COG3;refGene_tx=XM_006927301.4
A1 25482599 25482688
refGene_gn=COG3;refGene_tx=XM_003980400.5;refGene_gn=COG3;refGene_tx=XM_006927301.4
A1 25485733 25485854
refGene_gn=COG3;refGene_tx=XM_003980400.5;refGene_gn=COG3;refGene_tx=XM_006927301.4;refGene_gn=COG3;refGene_tx=XM_019826762.2
A1 25489316 25489406
refGene_gn=COG3;refGene_tx=XM_003980400.5;refGene_gn=COG3;refGene_tx=XM_006927301.4;refGene_gn=COG3;refGene_tx=XM_019826762.2
A1 25491119 25491244
refGene_gn=COG3;refGene_tx=XM_003980400.5;refGene_gn=COG3;refGene_tx=XM_006927301.4;refGene_gn=COG3;refGene_tx=XM_019826762.2
A1 25496925 25497031
refGene_gn=COG3;refGene_tx=XM_003980400.5;refGene_gn=COG3;refGene_tx=XM_006927301.4;refGene_gn=COG3;refGene_tx=XM_019826762.2
A1 25511656 25511817

refGene_gn=COG3;refGene_tx=XM_003980400.5;refGene_gn=COG3;refGene_tx=XM_006927301.4;refGene_gn=COG3;refGene_tx=XM_019826762.2
A1 25513106 25513246
refGene_gn=COG3;refGene_tx=XM_003980400.5;refGene_gn=COG3;refGene_tx=XM_006927301.4;refGene_gn=COG3;refGene_tx=XM_019826762.2
A1 25515787 25515879
refGene_gn=COG3;refGene_tx=XM_003980400.5;refGene_gn=COG3;refGene_tx=XM_006927301.4;refGene_gn=COG3;refGene_tx=XM_019826762.2
A1 25516478 25516605
refGene_gn=COG3;refGene_tx=XM_003980400.5;refGene_gn=COG3;refGene_tx=XM_006927301.4;refGene_gn=COG3;refGene_tx=XM_019826762.2
A1 25517282 25517326
refGene_gn=COG3;refGene_tx=XM_003980400.5;refGene_gn=COG3;refGene_tx=XM_006927301.4;refGene_gn=COG3;refGene_tx=XM_019826762.2
A1 25520552 25520633
refGene_gn=COG3;refGene_tx=XM_003980400.5;refGene_gn=COG3;refGene_tx=XM_006927301.4;refGene_gn=COG3;refGene_tx=XM_019826762.2
A1 25521917 25522043
refGene_gn=COG3;refGene_tx=XM_003980400.5;refGene_gn=COG3;refGene_tx=XM_006927301.4;refGene_gn=COG3;refGene_tx=XM_019826762.2
A1 25523023 25523116
refGene_gn=COG3;refGene_tx=XM_003980400.5;refGene_gn=COG3;refGene_tx=XM_006927301.4;refGene_gn=COG3;refGene_tx=XM_019826762.2
A1 25524359 25524434
refGene_gn=COG3;refGene_tx=XM_003980400.5;refGene_gn=COG3;refGene_tx=XM_006927301.4;refGene_gn=COG3;refGene_tx=XM_019826762.2
A1 25525086 25525252
refGene_gn=COG3;refGene_tx=XM_003980400.5;refGene_gn=COG3;refGene_tx=XM_006927301.4;refGene_gn=COG3;refGene_tx=XM_019826762.2
A1 25526494 25526556
refGene_gn=COG3;refGene_tx=XM_003980400.5;refGene_gn=COG3;refGene_tx=XM_006927301.4;refGene_gn=COG3;refGene_tx=XM_019826762.2
A1 25528764 25528911
refGene_gn=COG3;refGene_tx=XM_003980400.5;refGene_gn=COG3;refGene_tx=XM_006927301.4;refGene_gn=COG3;refGene_tx=XM_019826762.2
A1 25537004 25537178
refGene_gn=COG3;refGene_tx=XM_003980400.5;refGene_gn=COG3;refGene_tx=XM_006927301.4;refGene_gn=COG3;refGene_tx=XM_019826762.2
A1 25546156 25546690 refGene_gn=SLC25A30;refGene_tx=XM_019826772.2
A1 25563068 25563110 refGene_gn=SLC25A30;refGene_tx=XM_006927302.3
A1 25564047 25564128 refGene_gn=SLC25A30;refGene_tx=XM_006927302.3
A1 25565663 25565802
refGene_gn=SLC25A30;refGene_tx=XM_006927302.3;refGene_gn=SLC25A30;refGene_tx=XM_019826772.2
A1 25566476 25566601
refGene_gn=SLC25A30;refGene_tx=XM_006927302.3;refGene_gn=SLC25A30;refGene_tx=XM_019826772.2
A1 25567458 25567554
refGene_gn=SLC25A30;refGene_tx=XM_006927302.3;refGene_gn=SLC25A30;refGene_tx=XM_019826772.2

A1 25569720 25569806
refGene_gn=SLC25A30;refGene_tx=XM_006927302.3;refGene_gn=SLC25A30;refGene_tx=XM_019826772.2
A1 25571254 25571349
refGene_gn=SLC25A30;refGene_tx=XM_006927302.3;refGene_gn=SLC25A30;refGene_tx=XM_019826772.2
A1 25574929 25575077
refGene_gn=SLC25A30;refGene_tx=XM_006927302.3;refGene_gn=SLC25A30;refGene_tx=XM_019826772.2
A1 25576814 25576878
refGene_gn=SLC25A30;refGene_tx=XM_006927302.3;refGene_gn=SLC25A30;refGene_tx=XM_019826772.2
A1 25637479 25637507 refGene_gn=TPT1;refGene_tx=XM_003980403.5
A1 25637776 25637850 refGene_gn=TPT1;refGene_tx=XM_003980403.5
A1 25638398 25638589 refGene_gn=TPT1;refGene_tx=XM_003980403.5
A1 25639009 25639115 refGene_gn=TPT1;refGene_tx=XM_003980403.5
A1 25639870 25639987 refGene_gn=TPT1;refGene_tx=XM_003980403.5
A1 25640977 25640980 refGene_gn=TPT1;refGene_tx=XM_003980403.5
A1 25686004 25686124
refGene_gn=GTF2F2;refGene_tx=XM_003980404.5;refGene_gn=GTF2F2;refGene_tx=XM_019826870.2
A1 25700918 25701062
refGene_gn=GTF2F2;refGene_tx=XM_003980404.5;refGene_gn=GTF2F2;refGene_tx=XM_019826870.2
A1 25714361 25714461
refGene_gn=GTF2F2;refGene_tx=XM_003980404.5;refGene_gn=GTF2F2;refGene_tx=XM_019826870.2
A1 25755970 25756052
refGene_gn=GTF2F2;refGene_tx=XM_003980404.5;refGene_gn=GTF2F2;refGene_tx=XM_019826870.2
A1 25772721 25773501
refGene_gn=KCTD4;refGene_tx=XM_003980405.5;refGene_gn=KCTD4;refGene_tx=XM_019826857.2;refGene_gn=KCTD4;refGene_tx=XM_019826861.2
A1 25817811 25817956
refGene_gn=GTF2F2;refGene_tx=XM_003980404.5;refGene_gn=GTF2F2;refGene_tx=XM_019826870.2
A1 25819307 25819326
refGene_gn=GTF2F2;refGene_tx=XM_003980404.5;refGene_gn=GTF2F2;refGene_tx=XM_019826870.2
A1 25828702 25828776
refGene_gn=GTF2F2;refGene_tx=XM_003980404.5;refGene_gn=GTF2F2;refGene_tx=XM_019826870.2
A1 25834518 25834566 refGene_gn=GTF2F2;refGene_tx=XM_019826870.2
A1 25842372 25842438 refGene_gn=GTF2F2;refGene_tx=XM_003980404.5
A1 25936784 25937003
refGene_gn=GPALPP1;refGene_tx=XM_006927303.4;refGene_gn=GPALPP1;refGene_tx=XM_019826820.2;refGene_gn=GPALPP1;refGene_tx=XM_019826822.2;refGene_gn=GPALPP1;refGene_tx=XM_019826830.2;refGene_gn=GPALPP1;refGene_tx=XM_019826833.2;refGene_gn=GPALPP1;refGene_tx=XM_019826837.2
A1 25940123 25940222

refGene_gn=GPALPP1;refGene_tx=XM_006927303.4;refGene_gn=GPALPP1;refGene_tx=XM_019826820.2;refGene_gn=GPALPP1;refGene_tx=XM_019826822.2;refGene_gn=GPALPP1;refGene_tx=XM_019826830.2;refGene_gn=GPALPP1;refGene_tx=XM_019826833.2;refGene_gn=GPALPP1;refGene_tx=XM_019826837.2
A1 25948339 25948504
refGene_gn=GPALPP1;refGene_tx=XM_006927303.4;refGene_gn=GPALPP1;refGene_tx=XM_019826820.2;refGene_gn=GPALPP1;refGene_tx=XM_019826822.2;refGene_gn=GPALPP1;refGene_tx=XM_019826830.2;refGene_gn=GPALPP1;refGene_tx=XM_019826833.2;refGene_gn=GPALPP1;refGene_tx=XM_019826837.2
A1 25948949 25949081
refGene_gn=GPALPP1;refGene_tx=XM_006927303.4;refGene_gn=GPALPP1;refGene_tx=XM_019826820.2;refGene_gn=GPALPP1;refGene_tx=XM_019826822.2;refGene_gn=GPALPP1;refGene_tx=XM_019826830.2;refGene_gn=GPALPP1;refGene_tx=XM_019826833.2;refGene_gn=GPALPP1;refGene_tx=XM_019826837.2
A1 25952590 25952699
refGene_gn=GPALPP1;refGene_tx=XM_006927303.4;refGene_gn=GPALPP1;refGene_tx=XM_019826820.2;refGene_gn=GPALPP1;refGene_tx=XM_019826822.2;refGene_gn=GPALPP1;refGene_tx=XM_019826830.2;refGene_gn=GPALPP1;refGene_tx=XM_019826833.2;refGene_gn=GPALPP1;refGene_tx=XM_019826837.2
A1 25954709 25954808
refGene_gn=GPALPP1;refGene_tx=XM_006927303.4;refGene_gn=GPALPP1;refGene_tx=XM_019826820.2;refGene_gn=GPALPP1;refGene_tx=XM_019826822.2;refGene_gn=GPALPP1;refGene_tx=XM_019826830.2;refGene_gn=GPALPP1;refGene_tx=XM_019826833.2;refGene_gn=GPALPP1;refGene_tx=XM_019826837.2
A1 25955582 25955715
refGene_gn=GPALPP1;refGene_tx=XM_006927303.4;refGene_gn=GPALPP1;refGene_tx=XM_019826820.2;refGene_gn=GPALPP1;refGene_tx=XM_019826822.2;refGene_gn=GPALPP1;refGene_tx=XM_019826830.2;refGene_gn=GPALPP1;refGene_tx=XM_019826833.2;refGene_gn=GPALPP1;refGene_tx=XM_019826837.2
A1 25959138 25959226
refGene_gn=GPALPP1;refGene_tx=XM_006927303.4;refGene_gn=GPALPP1;refGene_tx=XM_019826820.2;refGene_gn=GPALPP1;refGene_tx=XM_019826822.2;refGene_gn=GPALPP1;refGene_tx=XM_019826830.2;refGene_gn=GPALPP1;refGene_tx=XM_019826833.2;refGene_gn=GPALPP1;refGene_tx=XM_019826837.2
A1 25959448 25959851 refGene_gn=NUFIP1;refGene_tx=XM_023251699.1
A1 25961889 25961972 refGene_gn=NUFIP1;refGene_tx=XM_023251699.1
A1 25963765 25963864 refGene_gn=NUFIP1;refGene_tx=XM_023251699.1
A1 25964602 25964665 refGene_gn=NUFIP1;refGene_tx=XM_023251699.1
A1 25965445 25965522 refGene_gn=NUFIP1;refGene_tx=XM_023251699.1
A1 25978153 25978246 refGene_gn=NUFIP1;refGene_tx=XM_023251699.1
A1 25984264 25984458 refGene_gn=NUFIP1;refGene_tx=XM_023251699.1
A1 25992556 25992673 refGene_gn=NUFIP1;refGene_tx=XM_023251699.1
A1 25997445 25997663 refGene_gn=NUFIP1;refGene_tx=XM_023251699.1
A1 25998982 25999099 refGene_gn=NUFIP1;refGene_tx=XM_023251699.1
A1 26013563 26013584
refGene_gn=LOC101085341;refGene_tx=XM_019827629.2
A1 26014037 26014328
refGene_gn=LOC101085341;refGene_tx=XM_019827629.2
A1 26296978 26297188
refGene_gn=TSC22D1;refGene_tx=XM_023251721.1;refGene_gn=TSC22D1;refGen

e_tx=XM_023251724.1;refGene_gn=TSC22D1;refGene_tx=XM_023251725.1
A1 26297640 26300693
refGene_gn=TSC22D1;refGene_tx=XM_011291259.3;refGene_gn=TSC22D1;refGen
e_tx=XM_023251721.1;refGene_gn=TSC22D1;refGene_tx=XM_023251724.1;refGe
ne_gn=TSC22D1;refGene_tx=XM_023251725.1;refGene_gn=TSC22D1;refGene_tx=
XM_023251728.1
A1 26330113 26330474
refGene_gn=TSC22D1;refGene_tx=XM_023251721.1;refGene_gn=TSC22D1;refGen
e_tx=XM_023251724.1;refGene_gn=TSC22D1;refGene_tx=XM_023251725.1;refGe
ne_gn=TSC22D1;refGene_tx=XM_023251728.1
A1 26331122 26331131 refGene_gn=TSC22D1;refGene_tx=XM_023251725.1
A1 26331169 26331187 refGene_gn=TSC22D1;refGene_tx=XM_023251724.1
A1 26428252 26428377 refGene_gn=TSC22D1;refGene_tx=XM_011291310.3
A1 26428780 26428832
refGene_gn=TSC22D1;refGene_tx=XM_011291259.3;refGene_gn=TSC22D1;refGen
e_tx=XM_011291310.3;refGene_gn=TSC22D1;refGene_tx=XM_011291338.3;refGe
ne_gn=TSC22D1;refGene_tx=XM_019826959.2
A1 26430003 26430261
refGene_gn=TSC22D1;refGene_tx=XM_011291259.3;refGene_gn=TSC22D1;refGen
e_tx=XM_011291310.3;refGene_gn=TSC22D1;refGene_tx=XM_011291338.3;refGe
ne_gn=TSC22D1;refGene_tx=XM_019826959.2
A1 26466526 26466540 refGene_gn=SERP2;refGene_tx=XM_023251767.1
A1 26466860 26466898 refGene_gn=SERP2;refGene_tx=XM_023251763.1
A1 26467645 26467686 refGene_gn=SERP2;refGene_tx=XM_023251758.1
A1 26473723 26473830
refGene_gn=SERP2;refGene_tx=XM_023251748.1;refGene_gn=SERP2;refGene_tx
=XM_023251754.1;refGene_gn=SERP2;refGene_tx=XM_023251756.1
A1 26473910 26474005 refGene_gn=SERP2;refGene_tx=XM_023251757.1
A1 26483128 26483201
refGene_gn=SERP2;refGene_tx=XM_023251748.1;refGene_gn=SERP2;refGene_tx
=XM_023251754.1;refGene_gn=SERP2;refGene_tx=XM_023251756.1;refGene_gn=
SERP2;refGene_tx=XM_023251757.1;refGene_gn=SERP2;refGene_tx=XM_0232517
58.1;refGene_gn=SERP2;refGene_tx=XM_023251763.1;refGene_gn=SERP2;refGe
ne_tx=XM_023251767.1
A1 26488067 26488151
refGene_gn=SERP2;refGene_tx=XM_023251748.1;refGene_gn=SERP2;refGene_tx
=XM_023251754.1;refGene_gn=SERP2;refGene_tx=XM_023251756.1;refGene_gn=
SERP2;refGene_tx=XM_023251757.1;refGene_gn=SERP2;refGene_tx=XM_0232517
58.1;refGene_gn=SERP2;refGene_tx=XM_023251763.1;refGene_gn=SERP2;refGe
ne_tx=XM_023251767.1
A1 26680499 26680736 refGene_gn=SMIM2;refGene_tx=XM_006944347.3
A1 26680737 26680776 refGene_gn=SMIM2;refGene_tx=XM_006944347.3
A1 26939208 26939368
refGene_gn=LACC1;refGene_tx=XM_023251780.1;refGene_gn=LACC1;refGene_tx
=XM_023251785.1;refGene_gn=LACC1;refGene_tx=XM_023251786.1;refGene_gn=
LACC1;refGene_tx=XM_023251787.1;refGene_gn=LACC1;refGene_tx=XM_0232517
88.1
A1 26940776 26941002
refGene_gn=LACC1;refGene_tx=XM_023251780.1;refGene_gn=LACC1;refGene_tx
=XM_023251785.1;refGene_gn=LACC1;refGene_tx=XM_023251786.1;refGene_gn=

LACC1;refGene_tx=XM_023251787.1;refGene_gn=LACC1;refGene_tx=XM_0232517
88.1
A1 26943993 26944159
refGene_gn=LACC1;refGene_tx=XM_023251780.1;refGene_gn=LACC1;refGene_tx
=XM_023251785.1;refGene_gn=LACC1;refGene_tx=XM_023251786.1;refGene_gn=
LACC1;refGene_tx=XM_023251787.1;refGene_gn=LACC1;refGene_tx=XM_0232517
88.1
A1 26944918 26945097
refGene_gn=LACC1;refGene_tx=XM_023251780.1;refGene_gn=LACC1;refGene_tx
=XM_023251785.1;refGene_gn=LACC1;refGene_tx=XM_023251786.1;refGene_gn=
LACC1;refGene_tx=XM_023251787.1;refGene_gn=LACC1;refGene_tx=XM_0232517
88.1
A1 26945753 26946312
refGene_gn=LACC1;refGene_tx=XM_023251780.1;refGene_gn=LACC1;refGene_tx
=XM_023251785.1;refGene_gn=LACC1;refGene_tx=XM_023251786.1;refGene_gn=
LACC1;refGene_tx=XM_023251787.1;refGene_gn=LACC1;refGene_tx=XM_0232517
88.1
A1 26956920 26956966 refGene_gn=CCDC122;refGene_tx=XM_023251792.1
A1 26957476 26957592 refGene_gn=CCDC122;refGene_tx=XM_023251792.1
A1 26966776 26967175 refGene_gn=CCDC122;refGene_tx=XM_023251792.1
A1 26967940 26968057 refGene_gn=CCDC122;refGene_tx=XM_023251792.1
A1 26978873 26979023 refGene_gn=CCDC122;refGene_tx=XM_023251792.1
A1 27338243 27338331
refGene_gn=ENOX1;refGene_tx=XM_019827068.2;refGene_gn=ENOX1;refGene_tx
=XM_019827075.2;refGene_gn=ENOX1;refGene_tx=XM_023251799.1;refGene_gn=
ENOX1;refGene_tx=XM_023251807.1;refGene_gn=ENOX1;refGene_tx=XM_0232518
15.1;refGene_gn=ENOX1;refGene_tx=XM_023251820.1;refGene_gn=ENOX1;refGe
ne_tx=XM_023251825.1;refGene_gn=ENOX1;refGene_tx=XM_023251828.1;refGen
e_gn=ENOX1;refGene_tx=XM_023251831.1;refGene_gn=ENOX1;refGene_tx=XM_02
3251835.1
A1 27376516 27376660
refGene_gn=ENOX1;refGene_tx=XM_019827068.2;refGene_gn=ENOX1;refGene_tx
=XM_019827075.2;refGene_gn=ENOX1;refGene_tx=XM_023251799.1;refGene_gn=
ENOX1;refGene_tx=XM_023251807.1;refGene_gn=ENOX1;refGene_tx=XM_0232518
15.1;refGene_gn=ENOX1;refGene_tx=XM_023251820.1;refGene_gn=ENOX1;refGe
ne_tx=XM_023251825.1;refGene_gn=ENOX1;refGene_tx=XM_023251828.1;refGen
e_gn=ENOX1;refGene_tx=XM_023251831.1;refGene_gn=ENOX1;refGene_tx=XM_02
3251834.1;refGene_gn=ENOX1;refGene_tx=XM_023251835.1
A1 27377523 27377661
refGene_gn=ENOX1;refGene_tx=XM_019827068.2;refGene_gn=ENOX1;refGene_tx
=XM_019827075.2;refGene_gn=ENOX1;refGene_tx=XM_023251799.1;refGene_gn=
ENOX1;refGene_tx=XM_023251807.1;refGene_gn=ENOX1;refGene_tx=XM_0232518
15.1;refGene_gn=ENOX1;refGene_tx=XM_023251820.1;refGene_gn=ENOX1;refGe
ne_tx=XM_023251825.1;refGene_gn=ENOX1;refGene_tx=XM_023251828.1;refGen
e_gn=ENOX1;refGene_tx=XM_023251831.1;refGene_gn=ENOX1;refGene_tx=XM_02
3251834.1;refGene_gn=ENOX1;refGene_tx=XM_023251835.1
A1 27427550 27427724
refGene_gn=ENOX1;refGene_tx=XM_019827068.2;refGene_gn=ENOX1;refGene_tx
=XM_019827075.2;refGene_gn=ENOX1;refGene_tx=XM_023251799.1;refGene_gn=
ENOX1;refGene_tx=XM_023251807.1;refGene_gn=ENOX1;refGene_tx=XM_0232518

15.1;refGene_gn=ENOX1;refGene_tx=XM_023251820.1;refGene_gn=ENOX1;refGene_tx=XM_023251825.1;refGene_gn=ENOX1;refGene_tx=XM_023251828.1;refGene_gn=ENOX1;refGene_tx=XM_023251831.1;refGene_gn=ENOX1;refGene_tx=XM_023251834.1;refGene_gn=ENOX1;refGene_tx=XM_023251835.1
A1 27429318 27429525
refGene_gn=ENOX1;refGene_tx=XM_019827068.2;refGene_gn=ENOX1;refGene_tx=XM_019827075.2;refGene_gn=ENOX1;refGene_tx=XM_023251799.1;refGene_gn=ENOX1;refGene_tx=XM_023251807.1;refGene_gn=ENOX1;refGene_tx=XM_023251815.1;refGene_gn=ENOX1;refGene_tx=XM_023251820.1;refGene_gn=ENOX1;refGene_tx=XM_023251825.1;refGene_gn=ENOX1;refGene_tx=XM_023251828.1;refGene_gn=ENOX1;refGene_tx=XM_023251831.1;refGene_gn=ENOX1;refGene_tx=XM_023251834.1;refGene_gn=ENOX1;refGene_tx=XM_023251835.1
A1 27433146 27433380
refGene_gn=ENOX1;refGene_tx=XM_019827068.2;refGene_gn=ENOX1;refGene_tx=XM_019827075.2;refGene_gn=ENOX1;refGene_tx=XM_023251799.1;refGene_gn=ENOX1;refGene_tx=XM_023251807.1;refGene_gn=ENOX1;refGene_tx=XM_023251815.1;refGene_gn=ENOX1;refGene_tx=XM_023251820.1;refGene_gn=ENOX1;refGene_tx=XM_023251825.1;refGene_gn=ENOX1;refGene_tx=XM_023251828.1;refGene_gn=ENOX1;refGene_tx=XM_023251831.1;refGene_gn=ENOX1;refGene_tx=XM_023251834.1;refGene_gn=ENOX1;refGene_tx=XM_023251835.1
A1 27443669 27443882
refGene_gn=ENOX1;refGene_tx=XM_019827068.2;refGene_gn=ENOX1;refGene_tx=XM_019827075.2;refGene_gn=ENOX1;refGene_tx=XM_023251799.1;refGene_gn=ENOX1;refGene_tx=XM_023251807.1;refGene_gn=ENOX1;refGene_tx=XM_023251815.1;refGene_gn=ENOX1;refGene_tx=XM_023251820.1;refGene_gn=ENOX1;refGene_tx=XM_023251825.1;refGene_gn=ENOX1;refGene_tx=XM_023251828.1;refGene_gn=ENOX1;refGene_tx=XM_023251831.1;refGene_gn=ENOX1;refGene_tx=XM_023251834.1;refGene_gn=ENOX1;refGene_tx=XM_023251835.1
A1 27463275 27463382
refGene_gn=ENOX1;refGene_tx=XM_019827068.2;refGene_gn=ENOX1;refGene_tx=XM_019827075.2;refGene_gn=ENOX1;refGene_tx=XM_023251799.1;refGene_gn=ENOX1;refGene_tx=XM_023251807.1;refGene_gn=ENOX1;refGene_tx=XM_023251815.1;refGene_gn=ENOX1;refGene_tx=XM_023251820.1;refGene_gn=ENOX1;refGene_tx=XM_023251825.1;refGene_gn=ENOX1;refGene_tx=XM_023251828.1;refGene_gn=ENOX1;refGene_tx=XM_023251831.1;refGene_gn=ENOX1;refGene_tx=XM_023251834.1;refGene_gn=ENOX1;refGene_tx=XM_023251835.1
A1 27467585 27467703
refGene_gn=ENOX1;refGene_tx=XM_019827068.2;refGene_gn=ENOX1;refGene_tx=XM_019827075.2;refGene_gn=ENOX1;refGene_tx=XM_023251799.1;refGene_gn=ENOX1;refGene_tx=XM_023251807.1;refGene_gn=ENOX1;refGene_tx=XM_023251815.1;refGene_gn=ENOX1;refGene_tx=XM_023251820.1;refGene_gn=ENOX1;refGene_tx=XM_023251825.1;refGene_gn=ENOX1;refGene_tx=XM_023251828.1;refGene_gn=ENOX1;refGene_tx=XM_023251831.1;refGene_gn=ENOX1;refGene_tx=XM_023251834.1;refGene_gn=ENOX1;refGene_tx=XM_023251835.1
A1 27491980 27492165
refGene_gn=ENOX1;refGene_tx=XM_019827068.2;refGene_gn=ENOX1;refGene_tx=XM_019827075.2;refGene_gn=ENOX1;refGene_tx=XM_023251799.1;refGene_gn=ENOX1;refGene_tx=XM_023251807.1;refGene_gn=ENOX1;refGene_tx=XM_023251815.1;refGene_gn=ENOX1;refGene_tx=XM_023251820.1;refGene_gn=ENOX1;refGene_tx=XM_023251825.1;refGene_gn=ENOX1;refGene_tx=XM_023251828.1;refGen

e_gn=ENOX1;refGene_tx=XM_023251831.1;refGene_gn=ENOX1;refGene_tx=XM_023251834.1;refGene_gn=ENOX1;refGene_tx=XM_023251835.1
A1 27524921 27525029 refGene_gn=ENOX1;refGene_tx=XM_019827068.2;refGene_gn=ENOX1;refGene_tx=XM_019827075.2;refGene_gn=ENOX1;refGene_tx=XM_023251799.1;refGene_gn=ENOX1;refGene_tx=XM_023251807.1;refGene_gn=ENOX1;refGene_tx=XM_023251815.1;refGene_gn=ENOX1;refGene_tx=XM_023251820.1;refGene_gn=ENOX1;refGene_tx=XM_023251825.1;refGene_gn=ENOX1;refGene_tx=XM_023251828.1;refGene_gn=ENOX1;refGene_tx=XM_023251831.1;refGene_gn=ENOX1;refGene_tx=XM_023251834.1;refGene_gn=ENOX1;refGene_tx=XM_023251835.1
A1 27529311 27529368 refGene_gn=ENOX1;refGene_tx=XM_019827068.2;refGene_gn=ENOX1;refGene_tx=XM_019827075.2;refGene_gn=ENOX1;refGene_tx=XM_023251799.1;refGene_gn=ENOX1;refGene_tx=XM_023251807.1;refGene_gn=ENOX1;refGene_tx=XM_023251815.1;refGene_gn=ENOX1;refGene_tx=XM_023251820.1;refGene_gn=ENOX1;refGene_tx=XM_023251825.1;refGene_gn=ENOX1;refGene_tx=XM_023251831.1;refGene_gn=ENOX1;refGene_tx=XM_023251834.1;refGene_gn=ENOX1;refGene_tx=XM_023251835.1
A1 27565275 27565378 refGene_gn=ENOX1;refGene_tx=XM_019827068.2;refGene_gn=ENOX1;refGene_tx=XM_019827075.2;refGene_gn=ENOX1;refGene_tx=XM_023251799.1;refGene_gn=ENOX1;refGene_tx=XM_023251807.1;refGene_gn=ENOX1;refGene_tx=XM_023251815.1;refGene_gn=ENOX1;refGene_tx=XM_023251820.1;refGene_gn=ENOX1;refGene_tx=XM_023251825.1;refGene_gn=ENOX1;refGene_tx=XM_023251828.1;refGene_gn=ENOX1;refGene_tx=XM_023251831.1;refGene_gn=ENOX1;refGene_tx=XM_023251834.1;refGene_gn=ENOX1;refGene_tx=XM_023251835.1
A1 27577400 27577486 refGene_gn=ENOX1;refGene_tx=XM_019827068.2;refGene_gn=ENOX1;refGene_tx=XM_019827075.2;refGene_gn=ENOX1;refGene_tx=XM_023251799.1;refGene_gn=ENOX1;refGene_tx=XM_023251807.1;refGene_gn=ENOX1;refGene_tx=XM_023251815.1;refGene_gn=ENOX1;refGene_tx=XM_023251820.1;refGene_gn=ENOX1;refGene_tx=XM_023251825.1;refGene_gn=ENOX1;refGene_tx=XM_023251828.1;refGene_gn=ENOX1;refGene_tx=XM_023251831.1;refGene_gn=ENOX1;refGene_tx=XM_023251834.1;refGene_gn=ENOX1;refGene_tx=XM_023251835.1
A1 27581725 27581779 refGene_gn=ENOX1;refGene_tx=XM_023251831.1
A1 27588041 27588173 refGene_gn=ENOX1;refGene_tx=XM_023251825.1;refGene_gn=ENOX1;refGene_tx=XM_023251835.1
A1 27612362 27612539 refGene_gn=ENOX1;refGene_tx=XM_019827068.2;refGene_gn=ENOX1;refGene_tx=XM_019827075.2;refGene_gn=ENOX1;refGene_tx=XM_023251799.1;refGene_gn=ENOX1;refGene_tx=XM_023251807.1;refGene_gn=ENOX1;refGene_tx=XM_023251815.1;refGene_gn=ENOX1;refGene_tx=XM_023251820.1;refGene_gn=ENOX1;refGene_tx=XM_023251828.1;refGene_gn=ENOX1;refGene_tx=XM_023251834.1
A1 27717054 27717125 refGene_gn=DNAJC15;refGene_tx=XM_003980414.5
A1 27734185 27734256 refGene_gn=DNAJC15;refGene_tx=XM_003980414.5
A1 27743671 27743748 refGene_gn=DNAJC15;refGene_tx=XM_003980414.5
A1 27752928 27753002 refGene_gn=DNAJC15;refGene_tx=XM_003980414.5
A1 27755986 27756038 refGene_gn=DNAJC15;refGene_tx=XM_003980414.5
A1 27789427 27789532 refGene_gn=DNAJC15;refGene_tx=XM_003980414.5

A1 27819351 27819530
refGene_gn=EPSTI1;refGene_tx=XM_006927315.4;refGene_gn=EPSTI1;refGene_tx=XM_011290734.3;refGene_gn=EPSTI1;refGene_tx=XM_023251861.1
A1 27836414 27836473
refGene_gn=EPSTI1;refGene_tx=XM_006927315.4;refGene_gn=EPSTI1;refGene_tx=XM_011290734.3;refGene_gn=EPSTI1;refGene_tx=XM_023251861.1
A1 27838467 27838551
refGene_gn=EPSTI1;refGene_tx=XM_006927315.4;refGene_gn=EPSTI1;refGene_tx=XM_011290734.3;refGene_gn=EPSTI1;refGene_tx=XM_023251861.1
A1 27840551 27840625
refGene_gn=EPSTI1;refGene_tx=XM_006927315.4;refGene_gn=EPSTI1;refGene_tx=XM_011290734.3;refGene_gn=EPSTI1;refGene_tx=XM_023251861.1
A1 27841267 27841351
refGene_gn=EPSTI1;refGene_tx=XM_006927315.4;refGene_gn=EPSTI1;refGene_tx=XM_011290734.3;refGene_gn=EPSTI1;refGene_tx=XM_023251861.1
A1 27851457 27851531
refGene_gn=EPSTI1;refGene_tx=XM_006927315.4;refGene_gn=EPSTI1;refGene_tx=XM_011290734.3;refGene_gn=EPSTI1;refGene_tx=XM_023251861.1
A1 27871107 27871201
refGene_gn=EPSTI1;refGene_tx=XM_006927315.4;refGene_gn=EPSTI1;refGene_tx=XM_011290734.3;refGene_gn=EPSTI1;refGene_tx=XM_023251861.1
A1 27879025 27879058 refGene_gn=EPSTI1;refGene_tx=XM_011290734.3
A1 27881745 27881829
refGene_gn=EPSTI1;refGene_tx=XM_006927315.4;refGene_gn=EPSTI1;refGene_tx=XM_011290734.3
A1 27904949 27905023
refGene_gn=EPSTI1;refGene_tx=XM_006927315.4;refGene_gn=EPSTI1;refGene_tx=XM_011290734.3;refGene_gn=EPSTI1;refGene_tx=XM_023251861.1
A1 27911187 27911287
refGene_gn=EPSTI1;refGene_tx=XM_006927315.4;refGene_gn=EPSTI1;refGene_tx=XM_011290734.3;refGene_gn=EPSTI1;refGene_tx=XM_023251861.1
A1 27917599 27917608
refGene_gn=EPSTI1;refGene_tx=XM_006927315.4;refGene_gn=EPSTI1;refGene_tx=XM_011290734.3;refGene_gn=EPSTI1;refGene_tx=XM_023251861.1
A1 28014899 28015093 refGene_gn=FAM216B;refGene_tx=XM_019827152.1
A1 28016816 28016937 refGene_gn=FAM216B;refGene_tx=XM_019827152.1
A1 28019678 28019777 refGene_gn=FAM216B;refGene_tx=XM_019827152.1
A1 28177896 28178318 refGene_gn=TNFSF11;refGene_tx=XM_023251873.1
A1 28182503 28182602 refGene_gn=TNFSF11;refGene_tx=XM_023251873.1
A1 28182682 28182728 refGene_gn=TNFSF11;refGene_tx=XM_023251873.1
A1 28202186 28202354 refGene_gn=TNFSF11;refGene_tx=XM_023251873.1
A1 28209505 28209736 refGene_gn=TNFSF11;refGene_tx=XM_023251873.1
A1 28420390 28420531
refGene_gn=AKAP11;refGene_tx=XM_006927316.2;refGene_gn=AKAP11;refGene_tx=XM_006927317.2;refGene_gn=AKAP11;refGene_tx=XM_011290613.3;refGene_gn=AKAP11;refGene_tx=XM_023251876.1;refGene_gn=AKAP11;refGene_tx=XM_023251883.1;refGene_gn=AKAP11;refGene_tx=XM_023251888.1
A1 28423177 28423338
refGene_gn=AKAP11;refGene_tx=XM_006927316.2;refGene_gn=AKAP11;refGene_tx=XM_006927317.2;refGene_gn=AKAP11;refGene_tx=XM_011290613.3;refGene_

gn=AKAP11;refGene_tx=XM_023251876.1;refGene_gn=AKAP11;refGene_tx=XM_023251883.1;refGene_gn=AKAP11;refGene_tx=XM_023251888.1
A1 28426510 28426557
refGene_gn=AKAP11;refGene_tx=XM_006927316.2;refGene_gn=AKAP11;refGene_tx=XM_006927317.2;refGene_gn=AKAP11;refGene_tx=XM_011290613.3;refGene_gn=AKAP11;refGene_tx=XM_023251876.1;refGene_gn=AKAP11;refGene_tx=XM_023251883.1;refGene_gn=AKAP11;refGene_tx=XM_023251888.1
A1 28427342 28427426
refGene_gn=AKAP11;refGene_tx=XM_006927316.2;refGene_gn=AKAP11;refGene_tx=XM_011290613.3;refGene_gn=AKAP11;refGene_tx=XM_023251876.1;refGene_gn=AKAP11;refGene_tx=XM_023251883.1;refGene_gn=AKAP11;refGene_tx=XM_023251888.1
A1 28432459 28432615
refGene_gn=AKAP11;refGene_tx=XM_006927316.2;refGene_gn=AKAP11;refGene_tx=XM_006927317.2;refGene_gn=AKAP11;refGene_tx=XM_011290613.3;refGene_gn=AKAP11;refGene_tx=XM_023251876.1;refGene_gn=AKAP11;refGene_tx=XM_023251883.1;refGene_gn=AKAP11;refGene_tx=XM_023251888.1
A1 28436250 28440763
refGene_gn=AKAP11;refGene_tx=XM_006927316.2;refGene_gn=AKAP11;refGene_tx=XM_006927317.2;refGene_gn=AKAP11;refGene_tx=XM_011290613.3;refGene_gn=AKAP11;refGene_tx=XM_023251876.1;refGene_gn=AKAP11;refGene_tx=XM_023251883.1;refGene_gn=AKAP11;refGene_tx=XM_023251888.1
A1 28441349 28441614
refGene_gn=AKAP11;refGene_tx=XM_006927316.2;refGene_gn=AKAP11;refGene_tx=XM_006927317.2;refGene_gn=AKAP11;refGene_tx=XM_011290613.3;refGene_gn=AKAP11;refGene_tx=XM_023251876.1;refGene_gn=AKAP11;refGene_tx=XM_023251883.1;refGene_gn=AKAP11;refGene_tx=XM_023251888.1
A1 28442985 28443120
refGene_gn=AKAP11;refGene_tx=XM_006927316.2;refGene_gn=AKAP11;refGene_tx=XM_006927317.2;refGene_gn=AKAP11;refGene_tx=XM_011290613.3;refGene_gn=AKAP11;refGene_tx=XM_023251876.1;refGene_gn=AKAP11;refGene_tx=XM_023251883.1;refGene_gn=AKAP11;refGene_tx=XM_023251888.1
A1 28445212 28445260
refGene_gn=AKAP11;refGene_tx=XM_006927316.2;refGene_gn=AKAP11;refGene_tx=XM_006927317.2;refGene_gn=AKAP11;refGene_tx=XM_011290613.3;refGene_gn=AKAP11;refGene_tx=XM_023251876.1;refGene_gn=AKAP11;refGene_tx=XM_023251883.1;refGene_gn=AKAP11;refGene_tx=XM_023251888.1
A1 28448084 28448201
refGene_gn=AKAP11;refGene_tx=XM_006927316.2;refGene_gn=AKAP11;refGene_tx=XM_006927317.2;refGene_gn=AKAP11;refGene_tx=XM_011290613.3;refGene_gn=AKAP11;refGene_tx=XM_023251876.1;refGene_gn=AKAP11;refGene_tx=XM_023251883.1;refGene_gn=AKAP11;refGene_tx=XM_023251888.1
A1 28453670 28453730
refGene_gn=AKAP11;refGene_tx=XM_006927316.2;refGene_gn=AKAP11;refGene_tx=XM_006927317.2;refGene_gn=AKAP11;refGene_tx=XM_011290613.3;refGene_gn=AKAP11;refGene_tx=XM_023251876.1;refGene_gn=AKAP11;refGene_tx=XM_023251883.1;refGene_gn=AKAP11;refGene_tx=XM_023251888.1
A1 28501561 28501651
refGene_gn=DGKH;refGene_tx=XM_011290413.3;refGene_gn=DGKH;refGene_tx=XM_019827249.2;refGene_gn=DGKH;refGene_tx=XM_019827250.2;refGene_gn=DGK

H;refGene_tx=XM_019827254.2;refGene_gn=DGKH;refGene_tx=XM_023251900.1;
refGene_gn=DGKH;refGene_tx=XM_023251905.1;refGene_gn=DGKH;refGene_tx=X
M_023251910.1;refGene_gn=DGKH;refGene_tx=XM_023251911.1;refGene_gn=DGK
H;refGene_tx=XM_023251912.1;refGene_gn=DGKH;refGene_tx=XM_023251913.1
A1 28506964 28507095
refGene_gn=DGKH;refGene_tx=XM_011290413.3;refGene_gn=DGKH;refGene_tx=X
M_019827249.2;refGene_gn=DGKH;refGene_tx=XM_019827250.2;refGene_gn=DGK
H;refGene_tx=XM_019827254.2;refGene_gn=DGKH;refGene_tx=XM_023251900.1;
refGene_gn=DGKH;refGene_tx=XM_023251905.1;refGene_gn=DGKH;refGene_tx=X
M_023251911.1;refGene_gn=DGKH;refGene_tx=XM_023251912.1;refGene_gn=DGK
H;refGene_tx=XM_023251913.1
A1 28508533 28508642
refGene_gn=DGKH;refGene_tx=XM_011290413.3;refGene_gn=DGKH;refGene_tx=X
M_019827249.2;refGene_gn=DGKH;refGene_tx=XM_019827250.2;refGene_gn=DGK
H;refGene_tx=XM_019827254.2;refGene_gn=DGKH;refGene_tx=XM_023251900.1;
refGene_gn=DGKH;refGene_tx=XM_023251905.1;refGene_gn=DGKH;refGene_tx=X
M_023251910.1;refGene_gn=DGKH;refGene_tx=XM_023251911.1;refGene_gn=DGK
H;refGene_tx=XM_023251912.1;refGene_gn=DGKH;refGene_tx=XM_023251913.1
A1 28509001 28509121
refGene_gn=DGKH;refGene_tx=XM_011290413.3;refGene_gn=DGKH;refGene_tx=X
M_019827249.2;refGene_gn=DGKH;refGene_tx=XM_019827250.2;refGene_gn=DGK
H;refGene_tx=XM_019827254.2;refGene_gn=DGKH;refGene_tx=XM_023251900.1;
refGene_gn=DGKH;refGene_tx=XM_023251905.1;refGene_gn=DGKH;refGene_tx=X
M_023251910.1;refGene_gn=DGKH;refGene_tx=XM_023251911.1;refGene_gn=DGK
H;refGene_tx=XM_023251912.1;refGene_gn=DGKH;refGene_tx=XM_023251913.1
A1 28512025 28512118
refGene_gn=DGKH;refGene_tx=XM_011290413.3;refGene_gn=DGKH;refGene_tx=X
M_019827249.2;refGene_gn=DGKH;refGene_tx=XM_019827250.2;refGene_gn=DGK
H;refGene_tx=XM_019827254.2;refGene_gn=DGKH;refGene_tx=XM_023251900.1;
refGene_gn=DGKH;refGene_tx=XM_023251910.1;refGene_gn=DGKH;refGene_tx=X
M_023251911.1;refGene_gn=DGKH;refGene_tx=XM_023251912.1;refGene_gn=DGK
H;refGene_tx=XM_023251913.1
A1 28513207 28513313
refGene_gn=DGKH;refGene_tx=XM_011290413.3;refGene_gn=DGKH;refGene_tx=X
M_019827249.2;refGene_gn=DGKH;refGene_tx=XM_019827250.2;refGene_gn=DGK
H;refGene_tx=XM_019827254.2;refGene_gn=DGKH;refGene_tx=XM_023251900.1;
refGene_gn=DGKH;refGene_tx=XM_023251905.1;refGene_gn=DGKH;refGene_tx=X
M_023251910.1;refGene_gn=DGKH;refGene_tx=XM_023251911.1;refGene_gn=DGK
H;refGene_tx=XM_023251912.1;refGene_gn=DGKH;refGene_tx=XM_023251913.1
A1 28516911 28517075
refGene_gn=DGKH;refGene_tx=XM_011290413.3;refGene_gn=DGKH;refGene_tx=X
M_019827249.2;refGene_gn=DGKH;refGene_tx=XM_019827250.2;refGene_gn=DGK
H;refGene_tx=XM_019827254.2;refGene_gn=DGKH;refGene_tx=XM_023251900.1;
refGene_gn=DGKH;refGene_tx=XM_023251905.1;refGene_gn=DGKH;refGene_tx=X
M_023251910.1;refGene_gn=DGKH;refGene_tx=XM_023251911.1;refGene_gn=DGK
H;refGene_tx=XM_023251912.1;refGene_gn=DGKH;refGene_tx=XM_023251913.1
A1 28517957 28518092
refGene_gn=DGKH;refGene_tx=XM_011290413.3;refGene_gn=DGKH;refGene_tx=X
M_019827249.2;refGene_gn=DGKH;refGene_tx=XM_019827250.2;refGene_gn=DGK
H;refGene_tx=XM_019827254.2;refGene_gn=DGKH;refGene_tx=XM_023251900.1;

refGene_gn=DGKH;refGene_tx=XM_023251905.1;refGene_gn=DGKH;refGene_tx=XM_023251910.1;refGene_gn=DGKH;refGene_tx=XM_023251911.1;refGene_gn=DGKH;refGene_tx=XM_023251912.1;refGene_gn=DGKH;refGene_tx=XM_023251913.1
A1 28518337 28518451
refGene_gn=DGKH;refGene_tx=XM_011290413.3;refGene_gn=DGKH;refGene_tx=XM_019827249.2;refGene_gn=DGKH;refGene_tx=XM_019827250.2;refGene_gn=DGKH;refGene_tx=XM_019827254.2;refGene_gn=DGKH;refGene_tx=XM_023251900.1;refGene_gn=DGKH;refGene_tx=XM_023251905.1;refGene_gn=DGKH;refGene_tx=XM_023251910.1;refGene_gn=DGKH;refGene_tx=XM_023251911.1;refGene_gn=DGKH;refGene_tx=XM_023251912.1;refGene_gn=DGKH;refGene_tx=XM_023251913.1
A1 28520547 28520655
refGene_gn=DGKH;refGene_tx=XM_011290413.3;refGene_gn=DGKH;refGene_tx=XM_019827249.2;refGene_gn=DGKH;refGene_tx=XM_019827250.2;refGene_gn=DGKH;refGene_tx=XM_019827254.2;refGene_gn=DGKH;refGene_tx=XM_023251900.1;refGene_gn=DGKH;refGene_tx=XM_023251905.1;refGene_gn=DGKH;refGene_tx=XM_023251910.1;refGene_gn=DGKH;refGene_tx=XM_023251911.1;refGene_gn=DGKH;refGene_tx=XM_023251912.1;refGene_gn=DGKH;refGene_tx=XM_023251913.1
A1 28522872 28522969
refGene_gn=DGKH;refGene_tx=XM_011290413.3;refGene_gn=DGKH;refGene_tx=XM_019827249.2;refGene_gn=DGKH;refGene_tx=XM_019827250.2;refGene_gn=DGKH;refGene_tx=XM_019827254.2;refGene_gn=DGKH;refGene_tx=XM_023251900.1;refGene_gn=DGKH;refGene_tx=XM_023251905.1;refGene_gn=DGKH;refGene_tx=XM_023251910.1;refGene_gn=DGKH;refGene_tx=XM_023251911.1;refGene_gn=DGKH;refGene_tx=XM_023251912.1;refGene_gn=DGKH;refGene_tx=XM_023251913.1
A1 28523107 28523218
refGene_gn=DGKH;refGene_tx=XM_011290413.3;refGene_gn=DGKH;refGene_tx=XM_019827249.2;refGene_gn=DGKH;refGene_tx=XM_019827250.2;refGene_gn=DGKH;refGene_tx=XM_019827254.2;refGene_gn=DGKH;refGene_tx=XM_023251900.1;refGene_gn=DGKH;refGene_tx=XM_023251905.1;refGene_gn=DGKH;refGene_tx=XM_023251910.1;refGene_gn=DGKH;refGene_tx=XM_023251911.1;refGene_gn=DGKH;refGene_tx=XM_023251912.1;refGene_gn=DGKH;refGene_tx=XM_023251913.1
A1 28524339 28524457
refGene_gn=DGKH;refGene_tx=XM_011290413.3;refGene_gn=DGKH;refGene_tx=XM_019827249.2;refGene_gn=DGKH;refGene_tx=XM_019827250.2;refGene_gn=DGKH;refGene_tx=XM_019827254.2;refGene_gn=DGKH;refGene_tx=XM_023251900.1;refGene_gn=DGKH;refGene_tx=XM_023251905.1;refGene_gn=DGKH;refGene_tx=XM_023251910.1;refGene_gn=DGKH;refGene_tx=XM_023251911.1;refGene_gn=DGKH;refGene_tx=XM_023251912.1;refGene_gn=DGKH;refGene_tx=XM_023251913.1
A1 28530813 28530945
refGene_gn=DGKH;refGene_tx=XM_011290413.3;refGene_gn=DGKH;refGene_tx=XM_019827249.2;refGene_gn=DGKH;refGene_tx=XM_019827250.2;refGene_gn=DGKH;refGene_tx=XM_019827254.2;refGene_gn=DGKH;refGene_tx=XM_023251900.1;refGene_gn=DGKH;refGene_tx=XM_023251905.1;refGene_gn=DGKH;refGene_tx=XM_023251910.1;refGene_gn=DGKH;refGene_tx=XM_023251911.1;refGene_gn=DGKH;refGene_tx=XM_023251912.1;refGene_gn=DGKH;refGene_tx=XM_023251913.1
A1 28534658 28534781
refGene_gn=DGKH;refGene_tx=XM_011290413.3;refGene_gn=DGKH;refGene_tx=XM_019827249.2;refGene_gn=DGKH;refGene_tx=XM_019827250.2;refGene_gn=DGKH;refGene_tx=XM_019827254.2;refGene_gn=DGKH;refGene_tx=XM_023251900.1;refGene_gn=DGKH;refGene_tx=XM_023251905.1;refGene_gn=DGKH;refGene_tx=X

M_023251910.1;refGene_gn=DGKH;refGene_tx=XM_023251911.1;refGene_gn=DGK
H;refGene_tx=XM_023251912.1;refGene_gn=DGKH;refGene_tx=XM_023251913.1
A1 28535564 28535838
refGene_gn=DGKH;refGene_tx=XM_011290413.3;refGene_gn=DGKH;refGene_tx=X
M_019827249.2;refGene_gn=DGKH;refGene_tx=XM_019827250.2;refGene_gn=DGK
H;refGene_tx=XM_019827254.2;refGene_gn=DGKH;refGene_tx=XM_023251900.1;
refGene_gn=DGKH;refGene_tx=XM_023251905.1;refGene_gn=DGKH;refGene_tx=X
M_023251910.1;refGene_gn=DGKH;refGene_tx=XM_023251911.1;refGene_gn=DGK
H;refGene_tx=XM_023251912.1;refGene_gn=DGKH;refGene_tx=XM_023251913.1
A1 28537355 28537455
refGene_gn=DGKH;refGene_tx=XM_011290413.3;refGene_gn=DGKH;refGene_tx=X
M_019827249.2;refGene_gn=DGKH;refGene_tx=XM_019827250.2;refGene_gn=DGK
H;refGene_tx=XM_019827254.2;refGene_gn=DGKH;refGene_tx=XM_023251900.1;
refGene_gn=DGKH;refGene_tx=XM_023251905.1;refGene_gn=DGKH;refGene_tx=X
M_023251910.1;refGene_gn=DGKH;refGene_tx=XM_023251911.1;refGene_gn=DGK
H;refGene_tx=XM_023251912.1;refGene_gn=DGKH;refGene_tx=XM_023251913.1
A1 28545504 28545590
refGene_gn=DGKH;refGene_tx=XM_011290413.3;refGene_gn=DGKH;refGene_tx=X
M_019827249.2;refGene_gn=DGKH;refGene_tx=XM_019827250.2;refGene_gn=DGK
H;refGene_tx=XM_019827254.2;refGene_gn=DGKH;refGene_tx=XM_023251900.1;
refGene_gn=DGKH;refGene_tx=XM_023251905.1;refGene_gn=DGKH;refGene_tx=X
M_023251910.1;refGene_gn=DGKH;refGene_tx=XM_023251911.1;refGene_gn=DGK
H;refGene_tx=XM_023251912.1;refGene_gn=DGKH;refGene_tx=XM_023251913.1
A1 28552242 28552327
refGene_gn=DGKH;refGene_tx=XM_011290413.3;refGene_gn=DGKH;refGene_tx=X
M_019827249.2;refGene_gn=DGKH;refGene_tx=XM_019827250.2;refGene_gn=DGK
H;refGene_tx=XM_019827254.2;refGene_gn=DGKH;refGene_tx=XM_023251900.1;
refGene_gn=DGKH;refGene_tx=XM_023251905.1;refGene_gn=DGKH;refGene_tx=X
M_023251910.1;refGene_gn=DGKH;refGene_tx=XM_023251911.1;refGene_gn=DGK
H;refGene_tx=XM_023251912.1;refGene_gn=DGKH;refGene_tx=XM_023251913.1
A1 28553978 28554118
refGene_gn=DGKH;refGene_tx=XM_011290413.3;refGene_gn=DGKH;refGene_tx=X
M_019827249.2;refGene_gn=DGKH;refGene_tx=XM_019827250.2;refGene_gn=DGK
H;refGene_tx=XM_019827254.2;refGene_gn=DGKH;refGene_tx=XM_023251900.1;
refGene_gn=DGKH;refGene_tx=XM_023251905.1;refGene_gn=DGKH;refGene_tx=X
M_023251910.1;refGene_gn=DGKH;refGene_tx=XM_023251911.1;refGene_gn=DGK
H;refGene_tx=XM_023251912.1;refGene_gn=DGKH;refGene_tx=XM_023251913.1
A1 28554269 28554378
refGene_gn=DGKH;refGene_tx=XM_011290413.3;refGene_gn=DGKH;refGene_tx=X
M_019827249.2;refGene_gn=DGKH;refGene_tx=XM_019827250.2;refGene_gn=DGK
H;refGene_tx=XM_019827254.2;refGene_gn=DGKH;refGene_tx=XM_023251900.1;
refGene_gn=DGKH;refGene_tx=XM_023251905.1;refGene_gn=DGKH;refGene_tx=X
M_023251910.1;refGene_gn=DGKH;refGene_tx=XM_023251911.1;refGene_gn=DGK
H;refGene_tx=XM_023251912.1;refGene_gn=DGKH;refGene_tx=XM_023251913.1
A1 28556053 28556213
refGene_gn=DGKH;refGene_tx=XM_011290413.3;refGene_gn=DGKH;refGene_tx=X
M_019827249.2;refGene_gn=DGKH;refGene_tx=XM_019827250.2;refGene_gn=DGK
H;refGene_tx=XM_019827254.2;refGene_gn=DGKH;refGene_tx=XM_023251900.1;
refGene_gn=DGKH;refGene_tx=XM_023251905.1;refGene_gn=DGKH;refGene_tx=X
M_023251910.1;refGene_gn=DGKH;refGene_tx=XM_023251911.1;refGene_gn=DGK

H;refGene_tx=XM_023251912.1;refGene_gn=DGKH;refGene_tx=XM_023251913.1
A1 28557762 28557865
refGene_gn=DGKH;refGene_tx=XM_011290413.3;refGene_gn=DGKH;refGene_tx=XM_019827249.2;refGene_gn=DGKH;refGene_tx=XM_019827250.2;refGene_gn=DGKH;refGene_tx=XM_019827254.2;refGene_gn=DGKH;refGene_tx=XM_023251900.1;refGene_gn=DGKH;refGene_tx=XM_023251905.1;refGene_gn=DGKH;refGene_tx=XM_023251910.1;refGene_gn=DGKH;refGene_tx=XM_023251911.1;refGene_gn=DGKH;refGene_tx=XM_023251912.1;refGene_gn=DGKH;refGene_tx=XM_023251913.1
A1 28561707 28561833
refGene_gn=DGKH;refGene_tx=XM_011290413.3;refGene_gn=DGKH;refGene_tx=XM_019827249.2;refGene_gn=DGKH;refGene_tx=XM_019827250.2;refGene_gn=DGKH;refGene_tx=XM_019827254.2;refGene_gn=DGKH;refGene_tx=XM_023251900.1;refGene_gn=DGKH;refGene_tx=XM_023251905.1;refGene_gn=DGKH;refGene_tx=XM_023251910.1;refGene_gn=DGKH;refGene_tx=XM_023251911.1;refGene_gn=DGKH;refGene_tx=XM_023251912.1;refGene_gn=DGKH;refGene_tx=XM_023251913.1
A1 28562489 28562596
refGene_gn=DGKH;refGene_tx=XM_011290413.3;refGene_gn=DGKH;refGene_tx=XM_019827249.2;refGene_gn=DGKH;refGene_tx=XM_019827250.2;refGene_gn=DGKH;refGene_tx=XM_019827254.2;refGene_gn=DGKH;refGene_tx=XM_023251900.1;refGene_gn=DGKH;refGene_tx=XM_023251905.1;refGene_gn=DGKH;refGene_tx=XM_023251910.1;refGene_gn=DGKH;refGene_tx=XM_023251911.1;refGene_gn=DGKH;refGene_tx=XM_023251912.1;refGene_gn=DGKH;refGene_tx=XM_023251913.1
A1 28569700 28569833
refGene_gn=DGKH;refGene_tx=XM_011290413.3;refGene_gn=DGKH;refGene_tx=XM_019827249.2;refGene_gn=DGKH;refGene_tx=XM_019827250.2;refGene_gn=DGKH;refGene_tx=XM_019827254.2;refGene_gn=DGKH;refGene_tx=XM_023251900.1;refGene_gn=DGKH;refGene_tx=XM_023251905.1;refGene_gn=DGKH;refGene_tx=XM_023251910.1;refGene_gn=DGKH;refGene_tx=XM_023251911.1;refGene_gn=DGKH;refGene_tx=XM_023251912.1;refGene_gn=DGKH;refGene_tx=XM_023251913.1
A1 28570124 28570229
refGene_gn=DGKH;refGene_tx=XM_011290413.3;refGene_gn=DGKH;refGene_tx=XM_019827249.2;refGene_gn=DGKH;refGene_tx=XM_019827250.2;refGene_gn=DGKH;refGene_tx=XM_019827254.2;refGene_gn=DGKH;refGene_tx=XM_023251900.1;refGene_gn=DGKH;refGene_tx=XM_023251905.1;refGene_gn=DGKH;refGene_tx=XM_023251910.1;refGene_gn=DGKH;refGene_tx=XM_023251911.1;refGene_gn=DGKH;refGene_tx=XM_023251912.1;refGene_gn=DGKH;refGene_tx=XM_023251913.1
A1 28594882 28594963
refGene_gn=DGKH;refGene_tx=XM_023251900.1;refGene_gn=DGKH;refGene_tx=XM_023251905.1;refGene_gn=DGKH;refGene_tx=XM_023251910.1;refGene_gn=DGKH;refGene_tx=XM_023251911.1;refGene_gn=DGKH;refGene_tx=XM_023251912.1
A1 28597033 28597144
refGene_gn=DGKH;refGene_tx=XM_023251900.1;refGene_gn=DGKH;refGene_tx=XM_023251905.1;refGene_gn=DGKH;refGene_tx=XM_023251910.1;refGene_gn=DGKH;refGene_tx=XM_023251911.1;refGene_gn=DGKH;refGene_tx=XM_023251912.1
A1 28669602 28669827
refGene_gn=DGKH;refGene_tx=XM_023251900.1;refGene_gn=DGKH;refGene_tx=XM_023251905.1;refGene_gn=DGKH;refGene_tx=XM_023251910.1;refGene_gn=DGKH;refGene_tx=XM_023251911.1;refGene_gn=DGKH;refGene_tx=XM_023251912.1
A1 28673153 28673297
refGene_gn=DGKH;refGene_tx=XM_023251900.1;refGene_gn=DGKH;refGene_tx=X

M_023251905.1;refGene_gn=DGKH;refGene_tx=XM_023251910.1

| | | | |
|---|---|---|---|
| A1 | 28755465 | 28755628 | refGene_gn=VWA8;refGene_tx=XM_023260078.1 |
| A1 | 28774967 | 28775045 | refGene_gn=VWA8;refGene_tx=XM_023260078.1 |
| A1 | 28802493 | 28802624 | refGene_gn=VWA8;refGene_tx=XM_023260078.1 |
| A1 | 28806105 | 28806216 | refGene_gn=VWA8;refGene_tx=XM_023260078.1 |
| A1 | 28823852 | 28824020 | refGene_gn=VWA8;refGene_tx=XM_023260078.1 |
| A1 | 28828611 | 28828776 | refGene_gn=VWA8;refGene_tx=XM_023260078.1 |
| A1 | 28829121 | 28829171 | refGene_gn=VWA8;refGene_tx=XM_023260078.1 |
| A1 | 28830139 | 28830248 | refGene_gn=VWA8;refGene_tx=XM_023260078.1 |
| A1 | 28832453 | 28832558 | refGene_gn=VWA8;refGene_tx=XM_023260078.1 |
| A1 | 28844673 | 28844805 | refGene_gn=VWA8;refGene_tx=XM_023260078.1 |
| A1 | 28847210 | 28847345 | refGene_gn=VWA8;refGene_tx=XM_023260078.1 |
| A1 | 28847437 | 28847515 | refGene_gn=VWA8;refGene_tx=XM_023260078.1 |
| A1 | 28878860 | 28879021 | refGene_gn=VWA8;refGene_tx=XM_023260078.1 |
| A1 | 28881239 | 28881353 | refGene_gn=VWA8;refGene_tx=XM_023260078.1 |
| A1 | 28889840 | 28890009 | refGene_gn=VWA8;refGene_tx=XM_023260078.1 |
| A1 | 28892085 | 28892163 | refGene_gn=VWA8;refGene_tx=XM_023260078.1 |
| A1 | 28899887 | 28900003 | refGene_gn=VWA8;refGene_tx=XM_023260078.1 |
| A1 | 28909088 | 28909195 | refGene_gn=VWA8;refGene_tx=XM_023260078.1 |
| A1 | 28912217 | 28912324 | refGene_gn=VWA8;refGene_tx=XM_023260078.1 |
| A1 | 28923110 | 28923182 | refGene_gn=VWA8;refGene_tx=XM_023260078.1 |
| A1 | 28942766 | 28942843 | refGene_gn=VWA8;refGene_tx=XM_023260078.1 |
| A1 | 28966766 | 28966842 | refGene_gn=VWA8;refGene_tx=XM_023260078.1 |
| A1 | 28969196 | 28969332 | refGene_gn=VWA8;refGene_tx=XM_023260078.1 |
| A1 | 28972560 | 28972680 | refGene_gn=VWA8;refGene_tx=XM_023260078.1 |
| A1 | 28975594 | 28975800 | refGene_gn=VWA8;refGene_tx=XM_023260078.1 |
| A1 | 28977084 | 28977236 | refGene_gn=VWA8;refGene_tx=XM_023260078.1 |
| A1 | 28992206 | 28992315 | refGene_gn=VWA8;refGene_tx=XM_023260078.1 |
| A1 | 28994030 | 28994169 | refGene_gn=VWA8;refGene_tx=XM_023260078.1 |
| A1 | 28995822 | 28996022 | refGene_gn=VWA8;refGene_tx=XM_023260078.1 |
| A1 | 29003224 | 29003335 | refGene_gn=VWA8;refGene_tx=XM_023260078.1 |
| A1 | 29004278 | 29004343 | refGene_gn=VWA8;refGene_tx=XM_023260078.1 |
| A1 | 29004773 | 29004899 | refGene_gn=VWA8;refGene_tx=XM_023260078.1 |
| A1 | 29006053 | 29006163 | refGene_gn=VWA8;refGene_tx=XM_023260078.1 |
| A1 | 29006991 | 29007146 | refGene_gn=VWA8;refGene_tx=XM_023260078.1 |
| A1 | 29011785 | 29011999 | refGene_gn=VWA8;refGene_tx=XM_023260078.1 |
| A1 | 29017678 | 29017744 | refGene_gn=VWA8;refGene_tx=XM_023260078.1 |
| A1 | 29020252 | 29020420 | refGene_gn=VWA8;refGene_tx=XM_023260078.1 |
| A1 | 29022814 | 29022855 | refGene_gn=VWA8;refGene_tx=XM_023260078.1 |
| A1 | 29022978 | 29023227 | refGene_gn=VWA8;refGene_tx=XM_023260078.1 |
| A1 | 29070200 | 29070309 | refGene_gn=VWA8;refGene_tx=XM_023260078.1 |
| A1 | 29073371 | 29073528 | refGene_gn=VWA8;refGene_tx=XM_023260078.1 |
| A1 | 29080519 | 29080628 | refGene_gn=VWA8;refGene_tx=XM_023260078.1 |
| A1 | 29096436 | 29096562 | refGene_gn=VWA8;refGene_tx=XM_023260078.1 |
| A1 | 29098261 | 29098420 | refGene_gn=VWA8;refGene_tx=XM_023260078.1 |
| A1 | 29112242 | 29112341 | refGene_gn=VWA8;refGene_tx=XM_023260078.1 |
| A1 | 29113296 | 29113535 | refGene_gn=VWA8;refGene_tx=XM_023260078.1 |
| A1 | 29115650 | 29115759 | refGene_gn=VWA8;refGene_tx=XM_023260078.1 |
| A1 | 29170680 | 29170688 | refGene_gn=RGCC;refGene_tx=XM_023251933.1 |
| A1 | 29171972 | 29172035 | refGene_gn=RGCC;refGene_tx=XM_023251933.1 |

A1 29173780 29173888 refGene_gn=RGCC;refGene_tx=XM_023251933.1
A1 29182838 29183024 refGene_gn=RGCC;refGene_tx=XM_023251933.1
A1 29183563 29183612 refGene_gn=RGCC;refGene_tx=XM_023251933.1
A1 29229038 29229236
refGene_gn=NAA16;refGene_tx=XM_011289968.3;refGene_gn=NAA16;refGene_tx=XM_011289993.3;refGene_gn=NAA16;refGene_tx=XM_019827309.2;refGene_gn=NAA16;refGene_tx=XM_019827313.2;refGene_gn=NAA16;refGene_tx=XM_019827320.2;refGene_gn=NAA16;refGene_tx=XM_019827325.2;refGene_gn=NAA16;refGene_tx=XM_019827333.2;refGene_gn=NAA16;refGene_tx=XM_019827345.2;refGene_gn=NAA16;refGene_tx=XM_019827346.2;refGene_gn=NAA16;refGene_tx=XM_019827348.2;refGene_gn=NAA16;refGene_tx=XM_019827350.2;refGene_gn=NAA16;refGene_tx=XM_019827359.2;refGene_gn=NAA16;refGene_tx=XM_019827367.2;refGene_gn=NAA16;refGene_tx=XM_019827374.2;refGene_gn=NAA16;refGene_tx=XM_019827381.2;refGene_gn=NAA16;refGene_tx=XM_019827386.2
A1 29230360 29230458
refGene_gn=NAA16;refGene_tx=XM_011289968.3;refGene_gn=NAA16;refGene_tx=XM_011289993.3;refGene_gn=NAA16;refGene_tx=XM_019827309.2;refGene_gn=NAA16;refGene_tx=XM_019827313.2;refGene_gn=NAA16;refGene_tx=XM_019827320.2;refGene_gn=NAA16;refGene_tx=XM_019827325.2;refGene_gn=NAA16;refGene_tx=XM_019827333.2;refGene_gn=NAA16;refGene_tx=XM_019827345.2;refGene_gn=NAA16;refGene_tx=XM_019827346.2;refGene_gn=NAA16;refGene_tx=XM_019827348.2;refGene_gn=NAA16;refGene_tx=XM_019827350.2;refGene_gn=NAA16;refGene_tx=XM_019827359.2;refGene_gn=NAA16;refGene_tx=XM_019827367.2;refGene_gn=NAA16;refGene_tx=XM_019827374.2;refGene_gn=NAA16;refGene_tx=XM_019827381.2;refGene_gn=NAA16;refGene_tx=XM_019827386.2
A1 29230588 29230708
refGene_gn=NAA16;refGene_tx=XM_019827309.2;refGene_gn=NAA16;refGene_tx=XM_019827313.2;refGene_gn=NAA16;refGene_tx=XM_019827320.2;refGene_gn=NAA16;refGene_tx=XM_019827325.2;refGene_gn=NAA16;refGene_tx=XM_019827333.2;refGene_gn=NAA16;refGene_tx=XM_019827345.2;refGene_gn=NAA16;refGene_tx=XM_019827346.2;refGene_gn=NAA16;refGene_tx=XM_019827348.2;refGene_gn=NAA16;refGene_tx=XM_019827350.2;refGene_gn=NAA16;refGene_tx=XM_019827359.2;refGene_gn=NAA16;refGene_tx=XM_019827367.2;refGene_gn=NAA16;refGene_tx=XM_019827374.2;refGene_gn=NAA16;refGene_tx=XM_019827381.2;refGene_gn=NAA16;refGene_tx=XM_019827386.2
A1 29234045 29234189
refGene_gn=NAA16;refGene_tx=XM_011289968.3;refGene_gn=NAA16;refGene_tx=XM_011289993.3;refGene_gn=NAA16;refGene_tx=XM_019827309.2;refGene_gn=NAA16;refGene_tx=XM_019827313.2;refGene_gn=NAA16;refGene_tx=XM_019827320.2;refGene_gn=NAA16;refGene_tx=XM_019827325.2;refGene_gn=NAA16;refGene_tx=XM_019827333.2;refGene_gn=NAA16;refGene_tx=XM_019827345.2;refGene_gn=NAA16;refGene_tx=XM_019827346.2;refGene_gn=NAA16;refGene_tx=XM_019827348.2;refGene_gn=NAA16;refGene_tx=XM_019827350.2;refGene_gn=NAA16;refGene_tx=XM_019827359.2;refGene_gn=NAA16;refGene_tx=XM_019827367.2;refGene_gn=NAA16;refGene_tx=XM_019827374.2;refGene_gn=NAA16;refGene_tx=XM_019827381.2;refGene_gn=NAA16;refGene_tx=XM_019827386.2
A1 29234517 29234616
refGene_gn=NAA16;refGene_tx=XM_011289968.3;refGene_gn=NAA16;refGene_tx=XM_011289993.3;refGene_gn=NAA16;refGene_tx=XM_019827309.2;refGene_gn=NAA16;refGene_tx=XM_019827313.2;refGene_gn=NAA16;refGene_tx=XM_0198273

20.2;refGene_gn=NAA16;refGene_tx=XM_019827325.2;refGene_gn=NAA16;refGene_tx=XM_019827333.2;refGene_gn=NAA16;refGene_tx=XM_019827345.2;refGene_gn=NAA16;refGene_tx=XM_019827346.2;refGene_gn=NAA16;refGene_tx=XM_019827348.2;refGene_gn=NAA16;refGene_tx=XM_019827350.2;refGene_gn=NAA16;refGene_tx=XM_019827359.2;refGene_gn=NAA16;refGene_tx=XM_019827367.2;refGene_gn=NAA16;refGene_tx=XM_019827374.2;refGene_gn=NAA16;refGene_tx=XM_019827381.2;refGene_gn=NAA16;refGene_tx=XM_019827386.2
A1 29235052 29235161
refGene_gn=NAA16;refGene_tx=XM_011289968.3;refGene_gn=NAA16;refGene_tx=XM_011289993.3;refGene_gn=NAA16;refGene_tx=XM_019827309.2;refGene_gn=NAA16;refGene_tx=XM_019827313.2;refGene_gn=NAA16;refGene_tx=XM_019827320.2;refGene_gn=NAA16;refGene_tx=XM_019827325.2;refGene_gn=NAA16;refGene_tx=XM_019827333.2;refGene_gn=NAA16;refGene_tx=XM_019827345.2;refGene_gn=NAA16;refGene_tx=XM_019827346.2;refGene_gn=NAA16;refGene_tx=XM_019827348.2;refGene_gn=NAA16;refGene_tx=XM_019827350.2;refGene_gn=NAA16;refGene_tx=XM_019827359.2;refGene_gn=NAA16;refGene_tx=XM_019827367.2;refGene_gn=NAA16;refGene_tx=XM_019827374.2;refGene_gn=NAA16;refGene_tx=XM_019827381.2;refGene_gn=NAA16;refGene_tx=XM_019827386.2
A1 29236377 29236571
refGene_gn=NAA16;refGene_tx=XM_011289968.3;refGene_gn=NAA16;refGene_tx=XM_011289993.3;refGene_gn=NAA16;refGene_tx=XM_019827309.2;refGene_gn=NAA16;refGene_tx=XM_019827313.2;refGene_gn=NAA16;refGene_tx=XM_019827320.2;refGene_gn=NAA16;refGene_tx=XM_019827325.2;refGene_gn=NAA16;refGene_tx=XM_019827333.2;refGene_gn=NAA16;refGene_tx=XM_019827345.2;refGene_gn=NAA16;refGene_tx=XM_019827346.2;refGene_gn=NAA16;refGene_tx=XM_019827348.2;refGene_gn=NAA16;refGene_tx=XM_019827350.2;refGene_gn=NAA16;refGene_tx=XM_019827359.2;refGene_gn=NAA16;refGene_tx=XM_019827367.2;refGene_gn=NAA16;refGene_tx=XM_019827374.2;refGene_gn=NAA16;refGene_tx=XM_019827381.2;refGene_gn=NAA16;refGene_tx=XM_019827386.2
A1 29237651 29237865
refGene_gn=NAA16;refGene_tx=XM_011289968.3;refGene_gn=NAA16;refGene_tx=XM_011289993.3;refGene_gn=NAA16;refGene_tx=XM_019827309.2;refGene_gn=NAA16;refGene_tx=XM_019827313.2;refGene_gn=NAA16;refGene_tx=XM_019827320.2;refGene_gn=NAA16;refGene_tx=XM_019827325.2;refGene_gn=NAA16;refGene_tx=XM_019827333.2;refGene_gn=NAA16;refGene_tx=XM_019827345.2;refGene_gn=NAA16;refGene_tx=XM_019827346.2;refGene_gn=NAA16;refGene_tx=XM_019827348.2;refGene_gn=NAA16;refGene_tx=XM_019827350.2;refGene_gn=NAA16;refGene_tx=XM_019827359.2;refGene_gn=NAA16;refGene_tx=XM_019827367.2;refGene_gn=NAA16;refGene_tx=XM_019827374.2;refGene_gn=NAA16;refGene_tx=XM_019827381.2;refGene_gn=NAA16;refGene_tx=XM_019827386.2
A1 29243438 29243567
refGene_gn=NAA16;refGene_tx=XM_011289968.3;refGene_gn=NAA16;refGene_tx=XM_011289993.3;refGene_gn=NAA16;refGene_tx=XM_019827309.2;refGene_gn=NAA16;refGene_tx=XM_019827313.2;refGene_gn=NAA16;refGene_tx=XM_019827320.2;refGene_gn=NAA16;refGene_tx=XM_019827325.2;refGene_gn=NAA16;refGene_tx=XM_019827333.2;refGene_gn=NAA16;refGene_tx=XM_019827345.2;refGene_gn=NAA16;refGene_tx=XM_019827346.2;refGene_gn=NAA16;refGene_tx=XM_019827348.2;refGene_gn=NAA16;refGene_tx=XM_019827350.2;refGene_gn=NAA16;refGene_tx=XM_019827359.2;refGene_gn=NAA16;refGene_tx=XM_019827367.2;refGene_gn=NAA16;refGene_tx=XM_019827374.2;refGene_gn=NAA16;refGene_tx=

XM_019827381.2;refGene_gn=NAA16;refGene_tx=XM_019827386.2
A1 29248456 29248609
refGene_gn=NAA16;refGene_tx=XM_011289968.3;refGene_gn=NAA16;refGene_tx=XM_011289993.3;refGene_gn=NAA16;refGene_tx=XM_019827309.2;refGene_gn=NAA16;refGene_tx=XM_019827313.2;refGene_gn=NAA16;refGene_tx=XM_019827320.2;refGene_gn=NAA16;refGene_tx=XM_019827325.2;refGene_gn=NAA16;refGene_tx=XM_019827333.2;refGene_gn=NAA16;refGene_tx=XM_019827345.2;refGene_gn=NAA16;refGene_tx=XM_019827346.2;refGene_gn=NAA16;refGene_tx=XM_019827348.2;refGene_gn=NAA16;refGene_tx=XM_019827350.2;refGene_gn=NAA16;refGene_tx=XM_019827359.2;refGene_gn=NAA16;refGene_tx=XM_019827367.2;refGene_gn=NAA16;refGene_tx=XM_019827374.2;refGene_gn=NAA16;refGene_tx=XM_019827381.2;refGene_gn=NAA16;refGene_tx=XM_019827386.2
A1 29248970 29249140
refGene_gn=NAA16;refGene_tx=XM_011289968.3;refGene_gn=NAA16;refGene_tx=XM_011289993.3;refGene_gn=NAA16;refGene_tx=XM_019827309.2;refGene_gn=NAA16;refGene_tx=XM_019827313.2;refGene_gn=NAA16;refGene_tx=XM_019827320.2;refGene_gn=NAA16;refGene_tx=XM_019827325.2;refGene_gn=NAA16;refGene_tx=XM_019827333.2;refGene_gn=NAA16;refGene_tx=XM_019827345.2;refGene_gn=NAA16;refGene_tx=XM_019827346.2;refGene_gn=NAA16;refGene_tx=XM_019827348.2;refGene_gn=NAA16;refGene_tx=XM_019827350.2;refGene_gn=NAA16;refGene_tx=XM_019827359.2;refGene_gn=NAA16;refGene_tx=XM_019827367.2;refGene_gn=NAA16;refGene_tx=XM_019827374.2;refGene_gn=NAA16;refGene_tx=XM_019827381.2;refGene_gn=NAA16;refGene_tx=XM_019827386.2
A1 29252239 29252312
refGene_gn=NAA16;refGene_tx=XM_011289968.3;refGene_gn=NAA16;refGene_tx=XM_011289993.3;refGene_gn=NAA16;refGene_tx=XM_019827309.2;refGene_gn=NAA16;refGene_tx=XM_019827313.2;refGene_gn=NAA16;refGene_tx=XM_019827320.2;refGene_gn=NAA16;refGene_tx=XM_019827325.2;refGene_gn=NAA16;refGene_tx=XM_019827333.2;refGene_gn=NAA16;refGene_tx=XM_019827345.2;refGene_gn=NAA16;refGene_tx=XM_019827346.2;refGene_gn=NAA16;refGene_tx=XM_019827348.2;refGene_gn=NAA16;refGene_tx=XM_019827350.2;refGene_gn=NAA16;refGene_tx=XM_019827359.2;refGene_gn=NAA16;refGene_tx=XM_019827367.2;refGene_gn=NAA16;refGene_tx=XM_019827374.2;refGene_gn=NAA16;refGene_tx=XM_019827381.2;refGene_gn=NAA16;refGene_tx=XM_019827386.2
A1 29264731 29264838
refGene_gn=NAA16;refGene_tx=XM_011289968.3;refGene_gn=NAA16;refGene_tx=XM_011289993.3;refGene_gn=NAA16;refGene_tx=XM_019827309.2;refGene_gn=NAA16;refGene_tx=XM_019827313.2;refGene_gn=NAA16;refGene_tx=XM_019827320.2;refGene_gn=NAA16;refGene_tx=XM_019827325.2;refGene_gn=NAA16;refGene_tx=XM_019827333.2;refGene_gn=NAA16;refGene_tx=XM_019827345.2;refGene_gn=NAA16;refGene_tx=XM_019827346.2;refGene_gn=NAA16;refGene_tx=XM_019827348.2;refGene_gn=NAA16;refGene_tx=XM_019827350.2;refGene_gn=NAA16;refGene_tx=XM_019827359.2;refGene_gn=NAA16;refGene_tx=XM_019827367.2;refGene_gn=NAA16;refGene_tx=XM_019827374.2;refGene_gn=NAA16;refGene_tx=XM_019827381.2;refGene_gn=NAA16;refGene_tx=XM_019827386.2
A1 29271277 29271373
refGene_gn=NAA16;refGene_tx=XM_011289968.3;refGene_gn=NAA16;refGene_tx=XM_011289993.3;refGene_gn=NAA16;refGene_tx=XM_019827309.2;refGene_gn=NAA16;refGene_tx=XM_019827313.2;refGene_gn=NAA16;refGene_tx=XM_019827325.2;refGene_gn=NAA16;refGene_tx=XM_019827346.2;refGene_gn=NAA16;refGe

ne_tx=XM_019827348.2;refGene_gn=NAA16;refGene_tx=XM_019827350.2;refGene_gn=NAA16;refGene_tx=XM_019827359.2;refGene_gn=NAA16;refGene_tx=XM_019827367.2;refGene_gn=NAA16;refGene_tx=XM_019827374.2;refGene_gn=NAA16;refGene_tx=XM_019827381.2;refGene_gn=NAA16;refGene_tx=XM_019827386.2
A1 29272778 29272805
refGene_gn=NAA16;refGene_tx=XM_011289968.3;refGene_gn=NAA16;refGene_tx=XM_019827309.2;refGene_gn=NAA16;refGene_tx=XM_019827320.2;refGene_gn=NAA16;refGene_tx=XM_019827325.2;refGene_gn=NAA16;refGene_tx=XM_019827346.2;refGene_gn=NAA16;refGene_tx=XM_019827348.2;refGene_gn=NAA16;refGene_tx=XM_019827350.2;refGene_gn=NAA16;refGene_tx=XM_019827359.2;refGene_gn=NAA16;refGene_tx=XM_019827367.2;refGene_gn=NAA16;refGene_tx=XM_019827374.2;refGene_gn=NAA16;refGene_tx=XM_019827381.2;refGene_gn=NAA16;refGene_tx=XM_019827386.2
A1 29273871 29273991
refGene_gn=NAA16;refGene_tx=XM_011289968.3;refGene_gn=NAA16;refGene_tx=XM_011289993.3;refGene_gn=NAA16;refGene_tx=XM_019827309.2;refGene_gn=NAA16;refGene_tx=XM_019827313.2;refGene_gn=NAA16;refGene_tx=XM_019827320.2;refGene_gn=NAA16;refGene_tx=XM_019827325.2;refGene_gn=NAA16;refGene_tx=XM_019827333.2;refGene_gn=NAA16;refGene_tx=XM_019827346.2;refGene_gn=NAA16;refGene_tx=XM_019827348.2;refGene_gn=NAA16;refGene_tx=XM_019827350.2;refGene_gn=NAA16;refGene_tx=XM_019827359.2;refGene_gn=NAA16;refGene_tx=XM_019827367.2;refGene_gn=NAA16;refGene_tx=XM_019827374.2;refGene_gn=NAA16;refGene_tx=XM_019827381.2;refGene_gn=NAA16;refGene_tx=XM_019827386.2
A1 29278034 29278188
refGene_gn=NAA16;refGene_tx=XM_011289968.3;refGene_gn=NAA16;refGene_tx=XM_011289993.3;refGene_gn=NAA16;refGene_tx=XM_019827309.2;refGene_gn=NAA16;refGene_tx=XM_019827313.2;refGene_gn=NAA16;refGene_tx=XM_019827320.2;refGene_gn=NAA16;refGene_tx=XM_019827325.2;refGene_gn=NAA16;refGene_tx=XM_019827333.2;refGene_gn=NAA16;refGene_tx=XM_019827345.2;refGene_gn=NAA16;refGene_tx=XM_019827346.2;refGene_gn=NAA16;refGene_tx=XM_019827348.2;refGene_gn=NAA16;refGene_tx=XM_019827350.2;refGene_gn=NAA16;refGene_tx=XM_019827359.2;refGene_gn=NAA16;refGene_tx=XM_019827367.2;refGene_gn=NAA16;refGene_tx=XM_019827374.2;refGene_gn=NAA16;refGene_tx=XM_019827381.2;refGene_gn=NAA16;refGene_tx=XM_019827386.2
A1 29280779 29280914
refGene_gn=NAA16;refGene_tx=XM_011289968.3;refGene_gn=NAA16;refGene_tx=XM_011289993.3;refGene_gn=NAA16;refGene_tx=XM_019827309.2;refGene_gn=NAA16;refGene_tx=XM_019827313.2;refGene_gn=NAA16;refGene_tx=XM_019827320.2;refGene_gn=NAA16;refGene_tx=XM_019827325.2;refGene_gn=NAA16;refGene_tx=XM_019827333.2;refGene_gn=NAA16;refGene_tx=XM_019827345.2;refGene_gn=NAA16;refGene_tx=XM_019827346.2;refGene_gn=NAA16;refGene_tx=XM_019827348.2;refGene_gn=NAA16;refGene_tx=XM_019827350.2;refGene_gn=NAA16;refGene_tx=XM_019827359.2;refGene_gn=NAA16;refGene_tx=XM_019827367.2;refGene_gn=NAA16;refGene_tx=XM_019827374.2;refGene_gn=NAA16;refGene_tx=XM_019827381.2;refGene_gn=NAA16;refGene_tx=XM_019827386.2
A1 29282974 29283132
refGene_gn=NAA16;refGene_tx=XM_011289968.3;refGene_gn=NAA16;refGene_tx=XM_011289993.3;refGene_gn=NAA16;refGene_tx=XM_019827309.2;refGene_gn=NAA16;refGene_tx=XM_019827313.2;refGene_gn=NAA16;refGene_tx=XM_0198273

20.2;refGene_gn=NAA16;refGene_tx=XM_019827325.2;refGene_gn=NAA16;refGene_tx=XM_019827333.2;refGene_gn=NAA16;refGene_tx=XM_019827345.2;refGene_gn=NAA16;refGene_tx=XM_019827346.2;refGene_gn=NAA16;refGene_tx=XM_019827348.2;refGene_gn=NAA16;refGene_tx=XM_019827350.2;refGene_gn=NAA16;refGene_tx=XM_019827359.2;refGene_gn=NAA16;refGene_tx=XM_019827367.2
A1 29283675 29283705
refGene_gn=NAA16;refGene_tx=XM_019827325.2;refGene_gn=NAA16;refGene_tx=XM_019827346.2
A1 29285212 29285317
refGene_gn=NAA16;refGene_tx=XM_011289968.3;refGene_gn=NAA16;refGene_tx=XM_011289993.3;refGene_gn=NAA16;refGene_tx=XM_019827309.2;refGene_gn=NAA16;refGene_tx=XM_019827313.2;refGene_gn=NAA16;refGene_tx=XM_019827320.2;refGene_gn=NAA16;refGene_tx=XM_019827325.2;refGene_gn=NAA16;refGene_tx=XM_019827333.2;refGene_gn=NAA16;refGene_tx=XM_019827345.2;refGene_gn=NAA16;refGene_tx=XM_019827346.2
A1 29287132 29287217
refGene_gn=NAA16;refGene_tx=XM_011289968.3;refGene_gn=NAA16;refGene_tx=XM_011289993.3;refGene_gn=NAA16;refGene_tx=XM_019827309.2;refGene_gn=NAA16;refGene_tx=XM_019827313.2;refGene_gn=NAA16;refGene_tx=XM_019827320.2;refGene_gn=NAA16;refGene_tx=XM_019827325.2;refGene_gn=NAA16;refGene_tx=XM_019827333.2;refGene_gn=NAA16;refGene_tx=XM_019827345.2
A1 29292135 29292189
refGene_gn=NAA16;refGene_tx=XM_011289968.3;refGene_gn=NAA16;refGene_tx=XM_011289993.3;refGene_gn=NAA16;refGene_tx=XM_019827309.2;refGene_gn=NAA16;refGene_tx=XM_019827313.2;refGene_gn=NAA16;refGene_tx=XM_019827320.2;refGene_gn=NAA16;refGene_tx=XM_019827325.2;refGene_gn=NAA16;refGene_tx=XM_019827333.2;refGene_gn=NAA16;refGene_tx=XM_019827345.2
A1 29310398 29310670
refGene_gn=MTRF1;refGene_tx=XM_023251976.1;refGene_gn=MTRF1;refGene_tx=XM_023251979.1;refGene_gn=MTRF1;refGene_tx=XM_023251984.1;refGene_gn=MTRF1;refGene_tx=XM_023252005.1
A1 29310781 29310914
refGene_gn=MTRF1;refGene_tx=XM_023251976.1;refGene_gn=MTRF1;refGene_tx=XM_023251984.1;refGene_gn=MTRF1;refGene_tx=XM_023252005.1
A1 29312529 29312955
refGene_gn=MTRF1;refGene_tx=XM_023251976.1;refGene_gn=MTRF1;refGene_tx=XM_023251979.1;refGene_gn=MTRF1;refGene_tx=XM_023251984.1;refGene_gn=MTRF1;refGene_tx=XM_023251992.1;refGene_gn=MTRF1;refGene_tx=XM_023251994.1;refGene_gn=MTRF1;refGene_tx=XM_023251997.1;refGene_gn=MTRF1;refGene_tx=XM_023252002.1;refGene_gn=MTRF1;refGene_tx=XM_023252006.1;refGene_gn=MTRF1;refGene_tx=XM_023252007.1
A1 29317361 29317453
refGene_gn=MTRF1;refGene_tx=XM_023251976.1;refGene_gn=MTRF1;refGene_tx=XM_023251979.1;refGene_gn=MTRF1;refGene_tx=XM_023251984.1;refGene_gn=MTRF1;refGene_tx=XM_023251992.1;refGene_gn=MTRF1;refGene_tx=XM_023251994.1;refGene_gn=MTRF1;refGene_tx=XM_023251997.1;refGene_gn=MTRF1;refGene_tx=XM_023252002.1;refGene_gn=MTRF1;refGene_tx=XM_023252005.1;refGene_gn=MTRF1;refGene_tx=XM_023252006.1;refGene_gn=MTRF1;refGene_tx=XM_023252007.1
A1 29318833 29318915

refGene_gn=MTRF1;refGene_tx=XM_023251976.1;refGene_gn=MTRF1;refGene_tx=XM_023251979.1;refGene_gn=MTRF1;refGene_tx=XM_023251984.1;refGene_gn=MTRF1;refGene_tx=XM_023251992.1;refGene_gn=MTRF1;refGene_tx=XM_023251994.1;refGene_gn=MTRF1;refGene_tx=XM_023251997.1;refGene_gn=MTRF1;refGene_tx=XM_023252002.1;refGene_gn=MTRF1;refGene_tx=XM_023252005.1;refGene_gn=MTRF1;refGene_tx=XM_023252006.1;refGene_gn=MTRF1;refGene_tx=XM_023252007.1
A1 29319123 29319231
refGene_gn=MTRF1;refGene_tx=XM_023251976.1;refGene_gn=MTRF1;refGene_tx=XM_023251979.1;refGene_gn=MTRF1;refGene_tx=XM_023251984.1;refGene_gn=MTRF1;refGene_tx=XM_023251992.1;refGene_gn=MTRF1;refGene_tx=XM_023251994.1;refGene_gn=MTRF1;refGene_tx=XM_023251997.1;refGene_gn=MTRF1;refGene_tx=XM_023252002.1;refGene_gn=MTRF1;refGene_tx=XM_023252005.1;refGene_gn=MTRF1;refGene_tx=XM_023252006.1;refGene_gn=MTRF1;refGene_tx=XM_023252007.1
A1 29328372 29328545
refGene_gn=MTRF1;refGene_tx=XM_023251976.1;refGene_gn=MTRF1;refGene_tx=XM_023251979.1;refGene_gn=MTRF1;refGene_tx=XM_023251984.1;refGene_gn=MTRF1;refGene_tx=XM_023251992.1;refGene_gn=MTRF1;refGene_tx=XM_023251994.1;refGene_gn=MTRF1;refGene_tx=XM_023251997.1;refGene_gn=MTRF1;refGene_tx=XM_023252002.1;refGene_gn=MTRF1;refGene_tx=XM_023252005.1;refGene_gn=MTRF1;refGene_tx=XM_023252006.1;refGene_gn=MTRF1;refGene_tx=XM_023252007.1
A1 29337021 29337139
refGene_gn=MTRF1;refGene_tx=XM_023251976.1;refGene_gn=MTRF1;refGene_tx=XM_023251979.1;refGene_gn=MTRF1;refGene_tx=XM_023251984.1;refGene_gn=MTRF1;refGene_tx=XM_023251992.1;refGene_gn=MTRF1;refGene_tx=XM_023251994.1;refGene_gn=MTRF1;refGene_tx=XM_023251997.1;refGene_gn=MTRF1;refGene_tx=XM_023252002.1;refGene_gn=MTRF1;refGene_tx=XM_023252005.1;refGene_gn=MTRF1;refGene_tx=XM_023252006.1;refGene_gn=MTRF1;refGene_tx=XM_023252007.1
A1 29353067 29353204
refGene_gn=MTRF1;refGene_tx=XM_023251976.1;refGene_gn=MTRF1;refGene_tx=XM_023251979.1;refGene_gn=MTRF1;refGene_tx=XM_023251984.1;refGene_gn=MTRF1;refGene_tx=XM_023251992.1;refGene_gn=MTRF1;refGene_tx=XM_023251994.1;refGene_gn=MTRF1;refGene_tx=XM_023251997.1;refGene_gn=MTRF1;refGene_tx=XM_023252002.1;refGene_gn=MTRF1;refGene_tx=XM_023252005.1;refGene_gn=MTRF1;refGene_tx=XM_023252006.1;refGene_gn=MTRF1;refGene_tx=XM_023252007.1
A1 29356908 29357007
refGene_gn=MTRF1;refGene_tx=XM_023251976.1;refGene_gn=MTRF1;refGene_tx=XM_023251979.1;refGene_gn=MTRF1;refGene_tx=XM_023251984.1;refGene_gn=MTRF1;refGene_tx=XM_023251992.1;refGene_gn=MTRF1;refGene_tx=XM_023251994.1;refGene_gn=MTRF1;refGene_tx=XM_023251997.1;refGene_gn=MTRF1;refGene_tx=XM_023252002.1;refGene_gn=MTRF1;refGene_tx=XM_023252005.1;refGene_gn=MTRF1;refGene_tx=XM_023252006.1;refGene_gn=MTRF1;refGene_tx=XM_023252007.1
A1 29361890 29362004
refGene_gn=MTRF1;refGene_tx=XM_023251976.1;refGene_gn=MTRF1;refGene_tx=XM_023251979.1;refGene_gn=MTRF1;refGene_tx=XM_023251984.1;refGene_gn=

MTRF1;refGene_tx=XM_023251992.1;refGene_gn=MTRF1;refGene_tx=XM_023251994.1;refGene_gn=MTRF1;refGene_tx=XM_023251997.1;refGene_gn=MTRF1;refGene_tx=XM_023252002.1;refGene_gn=MTRF1;refGene_tx=XM_023252005.1;refGene_gn=MTRF1;refGene_tx=XM_023252006.1;refGene_gn=MTRF1;refGene_tx=XM_023252007.1
A1 29371828 29373868
refGene_gn=LOC101088295;refGene_tx=XM_003980419.5
A1 29446752 29448873
refGene_gn=LOC101090830;refGene_tx=XM_023252011.1
A1 29485633 29485844 refGene_gn=WBP4;refGene_tx=XM_011289664.2
A1 29533364 29533528 refGene_gn=WBP4;refGene_tx=XM_011289664.2
A1 29538279 29538473 refGene_gn=WBP4;refGene_tx=XM_011289664.2
A1 29540327 29540403 refGene_gn=WBP4;refGene_tx=XM_011289664.2
A1 29540874 29540921 refGene_gn=WBP4;refGene_tx=XM_011289664.2
A1 29544607 29544775 refGene_gn=WBP4;refGene_tx=XM_011289664.2
A1 29548166 29548290 refGene_gn=WBP4;refGene_tx=XM_011289664.2
A1 29548379 29548442 refGene_gn=WBP4;refGene_tx=XM_011289664.2
A1 29550561 29550634 refGene_gn=WBP4;refGene_tx=XM_011289664.2
A1 29551581 29551583 refGene_gn=WBP4;refGene_tx=XM_011289664.2
A1 29610095 29610167
refGene_gn=ELF1;refGene_tx=XM_003980420.5;refGene_gn=ELF1;refGene_tx=XM_011289711.3;refGene_gn=ELF1;refGene_tx=XM_011289776.3;refGene_gn=ELF1;refGene_tx=XM_019827475.2;refGene_gn=ELF1;refGene_tx=XM_019827479.2
A1 29631537 29631718
refGene_gn=ELF1;refGene_tx=XM_003980420.5;refGene_gn=ELF1;refGene_tx=XM_011289711.3;refGene_gn=ELF1;refGene_tx=XM_011289776.3;refGene_gn=ELF1;refGene_tx=XM_019827475.2;refGene_gn=ELF1;refGene_tx=XM_019827479.2
A1 29644167 29644275
refGene_gn=ELF1;refGene_tx=XM_003980420.5;refGene_gn=ELF1;refGene_tx=XM_011289711.3;refGene_gn=ELF1;refGene_tx=XM_011289776.3;refGene_gn=ELF1;refGene_tx=XM_019827475.2;refGene_gn=ELF1;refGene_tx=XM_019827479.2
A1 29645903 29646071
refGene_gn=ELF1;refGene_tx=XM_003980420.5;refGene_gn=ELF1;refGene_tx=XM_011289711.3;refGene_gn=ELF1;refGene_tx=XM_011289776.3;refGene_gn=ELF1;refGene_tx=XM_019827475.2;refGene_gn=ELF1;refGene_tx=XM_019827479.2
A1 29651160 29651244
refGene_gn=ELF1;refGene_tx=XM_003980420.5;refGene_gn=ELF1;refGene_tx=XM_011289711.3;refGene_gn=ELF1;refGene_tx=XM_011289776.3;refGene_gn=ELF1;refGene_tx=XM_019827475.2;refGene_gn=ELF1;refGene_tx=XM_019827479.2
A1 29651827 29652020
refGene_gn=ELF1;refGene_tx=XM_003980420.5;refGene_gn=ELF1;refGene_tx=XM_011289711.3;refGene_gn=ELF1;refGene_tx=XM_011289776.3;refGene_gn=ELF1;refGene_tx=XM_019827475.2;refGene_gn=ELF1;refGene_tx=XM_019827479.2
A1 29654206 29654656
refGene_gn=ELF1;refGene_tx=XM_003980420.5;refGene_gn=ELF1;refGene_tx=XM_011289711.3;refGene_gn=ELF1;refGene_tx=XM_011289776.3;refGene_gn=ELF1;refGene_tx=XM_019827475.2;refGene_gn=ELF1;refGene_tx=XM_019827479.2
A1 29661615 29662219
refGene_gn=ELF1;refGene_tx=XM_003980420.5;refGene_gn=ELF1;refGene_tx=XM_011289711.3;refGene_gn=ELF1;refGene_tx=XM_011289776.3;refGene_gn=ELF

1;refGene_tx=XM_019827475.2;refGene_gn=ELF1;refGene_tx=XM_019827479.2
A1 29671747 29671785 refGene_gn=SUGT1;refGene_tx=XM_023260101.1
A1 29671877 29671935 refGene_gn=SUGT1;refGene_tx=XM_023260101.1
A1 29679206 29679297 refGene_gn=SUGT1;refGene_tx=XM_023260101.1
A1 29679412 29679420 refGene_gn=SUGT1;refGene_tx=XM_023260101.1
A1 29679799 29679956 refGene_gn=SUGT1;refGene_tx=XM_023260101.1
A1 29682847 29682901 refGene_gn=SUGT1;refGene_tx=XM_023260101.1
A1 29683279 29683296 refGene_gn=SUGT1;refGene_tx=XM_023260101.1
A1 29686086 29686121 refGene_gn=SUGT1;refGene_tx=XM_023260101.1
A1 29687718 29687815 refGene_gn=SUGT1;refGene_tx=XM_023260101.1
A1 29688581 29688689 refGene_gn=SUGT1;refGene_tx=XM_023260101.1
A1 29698179 29698270 refGene_gn=SUGT1;refGene_tx=XM_023260101.1
A1 29702866 29703048 refGene_gn=SUGT1;refGene_tx=XM_023260101.1
A1 29713870 29713972 refGene_gn=SUGT1;refGene_tx=XM_023260101.1
A1 29739849 29740065
refGene_gn=CNMD;refGene_tx=XM_023252028.1;refGene_gn=CNMD;refGene_tx=XM_023252029.1
A1 29741333 29741393 refGene_gn=CNMD;refGene_tx=XM_023252036.1
A1 29743823 29743990
refGene_gn=CNMD;refGene_tx=XM_023252028.1;refGene_gn=CNMD;refGene_tx=XM_023252029.1;refGene_gn=CNMD;refGene_tx=XM_023252036.1
A1 29754416 29754570
refGene_gn=CNMD;refGene_tx=XM_023252028.1;refGene_gn=CNMD;refGene_tx=XM_023252029.1;refGene_gn=CNMD;refGene_tx=XM_023252036.1
A1 29758949 29759063
refGene_gn=CNMD;refGene_tx=XM_023252028.1;refGene_gn=CNMD;refGene_tx=XM_023252029.1;refGene_gn=CNMD;refGene_tx=XM_023252036.1
A1 29766042 29766183
refGene_gn=CNMD;refGene_tx=XM_023252028.1;refGene_gn=CNMD;refGene_tx=XM_023252029.1;refGene_gn=CNMD;refGene_tx=XM_023252036.1
A1 29771688 29771829
refGene_gn=CNMD;refGene_tx=XM_023252028.1;refGene_gn=CNMD;refGene_tx=XM_023252029.1;refGene_gn=CNMD;refGene_tx=XM_023252036.1
A1 29772249 29772402
refGene_gn=CNMD;refGene_tx=XM_023252028.1;refGene_gn=CNMD;refGene_tx=XM_023252029.1;refGene_gn=CNMD;refGene_tx=XM_023252036.1
A1 29867981 29868355
refGene_gn=LOC101091586;refGene_tx=XM_023252040.1
A1 29868843 29869051
refGene_gn=LOC101091586;refGene_tx=XM_023252040.1
A1 29869255 29871190
refGene_gn=LOC101091586;refGene_tx=XM_023252040.1
A1 29871191 29871896
refGene_gn=LOC101091586;refGene_tx=XM_023252040.1
A1 30076173 30076359 refGene_gn=OLFM4;refGene_tx=XM_003980432.4
A1 30081428 30081581 refGene_gn=OLFM4;refGene_tx=XM_003980432.4
A1 30087818 30088031 refGene_gn=OLFM4;refGene_tx=XM_003980432.4
A1 30089007 30089167 refGene_gn=OLFM4;refGene_tx=XM_003980432.4
A1 30096181 30096984 refGene_gn=OLFM4;refGene_tx=XM_003980432.4
A1 30232578 30233590

refGene_gn=LOC111556289;refGene_tx=XM_023252063.1
A1 30233592 30234293
refGene_gn=LOC111556289;refGene_tx=XM_023252063.1
A1 34066445 34069037
refGene_gn=PCDH17;refGene_tx=XM_003980433.5;refGene_gn=PCDH17;refGene_tx=XM_019827548.2;refGene_gn=PCDH17;refGene_tx=XM_019827553.2
A1 34100788 34100847
refGene_gn=PCDH17;refGene_tx=XM_003980433.5;refGene_gn=PCDH17;refGene_tx=XM_019827548.2;refGene_gn=PCDH17;refGene_tx=XM_019827553.2
A1 34100980 34101153
refGene_gn=PCDH17;refGene_tx=XM_003980433.5;refGene_gn=PCDH17;refGene_tx=XM_019827548.2;refGene_gn=PCDH17;refGene_tx=XM_019827553.2
A1 34143328 34143412
refGene_gn=PCDH17;refGene_tx=XM_019827548.2;refGene_gn=PCDH17;refGene_tx=XM_019827553.2
A1 34159075 34159758
refGene_gn=PCDH17;refGene_tx=XM_003980433.5;refGene_gn=PCDH17;refGene_tx=XM_019827548.2;refGene_gn=PCDH17;refGene_tx=XM_019827553.2
A1 35846590 35846853
refGene_gn=DIAPH3;refGene_tx=XM_006927330.4;refGene_gn=DIAPH3;refGene_tx=XM_019827642.2;refGene_gn=DIAPH3;refGene_tx=XM_019827647.2;refGene_gn=DIAPH3;refGene_tx=XM_023252114.1;refGene_gn=DIAPH3;refGene_tx=XM_023252126.1
A1 35972266 35972370 refGene_gn=DIAPH3;refGene_tx=XM_019827645.2
A1 35977267 35977327
refGene_gn=DIAPH3;refGene_tx=XM_006927330.4;refGene_gn=DIAPH3;refGene_tx=XM_019827642.2;refGene_gn=DIAPH3;refGene_tx=XM_019827645.2;refGene_gn=DIAPH3;refGene_tx=XM_019827647.2;refGene_gn=DIAPH3;refGene_tx=XM_023252114.1;refGene_gn=DIAPH3;refGene_tx=XM_023252126.1
A1 35977821 35977917
refGene_gn=DIAPH3;refGene_tx=XM_006927330.4;refGene_gn=DIAPH3;refGene_tx=XM_019827642.2;refGene_gn=DIAPH3;refGene_tx=XM_019827645.2;refGene_gn=DIAPH3;refGene_tx=XM_019827647.2;refGene_gn=DIAPH3;refGene_tx=XM_023252114.1;refGene_gn=DIAPH3;refGene_tx=XM_023252126.1
A1 36017698 36017834
refGene_gn=DIAPH3;refGene_tx=XM_006927330.4;refGene_gn=DIAPH3;refGene_tx=XM_019827642.2;refGene_gn=DIAPH3;refGene_tx=XM_019827645.2;refGene_gn=DIAPH3;refGene_tx=XM_019827647.2;refGene_gn=DIAPH3;refGene_tx=XM_023252114.1;refGene_gn=DIAPH3;refGene_tx=XM_023252126.1
A1 36039337 36039502
refGene_gn=DIAPH3;refGene_tx=XM_006927330.4;refGene_gn=DIAPH3;refGene_tx=XM_019827642.2;refGene_gn=DIAPH3;refGene_tx=XM_019827645.2;refGene_gn=DIAPH3;refGene_tx=XM_019827647.2;refGene_gn=DIAPH3;refGene_tx=XM_023252114.1;refGene_gn=DIAPH3;refGene_tx=XM_023252126.1
A1 36047222 36047347
refGene_gn=DIAPH3;refGene_tx=XM_006927330.4;refGene_gn=DIAPH3;refGene_tx=XM_019827642.2;refGene_gn=DIAPH3;refGene_tx=XM_019827645.2;refGene_gn=DIAPH3;refGene_tx=XM_019827647.2;refGene_gn=DIAPH3;refGene_tx=XM_023252114.1;refGene_gn=DIAPH3;refGene_tx=XM_023252126.1
A1 36065554 36065684

refGene_gn=DIAPH3;refGene_tx=XM_006927330.4;refGene_gn=DIAPH3;refGene_tx=XM_019827642.2;refGene_gn=DIAPH3;refGene_tx=XM_019827645.2;refGene_gn=DIAPH3;refGene_tx=XM_019827647.2;refGene_gn=DIAPH3;refGene_tx=XM_023252114.1;refGene_gn=DIAPH3;refGene_tx=XM_023252126.1
A1 36082614 36082854
refGene_gn=DIAPH3;refGene_tx=XM_006927330.4;refGene_gn=DIAPH3;refGene_tx=XM_019827642.2;refGene_gn=DIAPH3;refGene_tx=XM_019827645.2;refGene_gn=DIAPH3;refGene_tx=XM_019827647.2;refGene_gn=DIAPH3;refGene_tx=XM_023252114.1
A1 36117948 36118050
refGene_gn=DIAPH3;refGene_tx=XM_006927330.4;refGene_gn=DIAPH3;refGene_tx=XM_019827642.2;refGene_gn=DIAPH3;refGene_tx=XM_019827645.2;refGene_gn=DIAPH3;refGene_tx=XM_019827647.2;refGene_gn=DIAPH3;refGene_tx=XM_023252114.1;refGene_gn=DIAPH3;refGene_tx=XM_023252126.1
A1 36122407 36122502
refGene_gn=DIAPH3;refGene_tx=XM_006927330.4;refGene_gn=DIAPH3;refGene_tx=XM_019827642.2;refGene_gn=DIAPH3;refGene_tx=XM_019827645.2;refGene_gn=DIAPH3;refGene_tx=XM_019827647.2;refGene_gn=DIAPH3;refGene_tx=XM_023252114.1;refGene_gn=DIAPH3;refGene_tx=XM_023252126.1
A1 36130048 36130144
refGene_gn=DIAPH3;refGene_tx=XM_006927330.4;refGene_gn=DIAPH3;refGene_tx=XM_019827642.2;refGene_gn=DIAPH3;refGene_tx=XM_019827645.2;refGene_gn=DIAPH3;refGene_tx=XM_019827647.2;refGene_gn=DIAPH3;refGene_tx=XM_023252114.1;refGene_gn=DIAPH3;refGene_tx=XM_023252126.1
A1 36166138 36166253
refGene_gn=DIAPH3;refGene_tx=XM_006927330.4;refGene_gn=DIAPH3;refGene_tx=XM_019827642.2;refGene_gn=DIAPH3;refGene_tx=XM_019827645.2;refGene_gn=DIAPH3;refGene_tx=XM_019827647.2;refGene_gn=DIAPH3;refGene_tx=XM_023252114.1;refGene_gn=DIAPH3;refGene_tx=XM_023252126.1
A1 36167003 36167312
refGene_gn=DIAPH3;refGene_tx=XM_006927330.4;refGene_gn=DIAPH3;refGene_tx=XM_019827642.2;refGene_gn=DIAPH3;refGene_tx=XM_019827645.2;refGene_gn=DIAPH3;refGene_tx=XM_019827647.2;refGene_gn=DIAPH3;refGene_tx=XM_023252114.1;refGene_gn=DIAPH3;refGene_tx=XM_023252126.1
A1 36170792 36170897
refGene_gn=DIAPH3;refGene_tx=XM_006927330.4;refGene_gn=DIAPH3;refGene_tx=XM_019827642.2;refGene_gn=DIAPH3;refGene_tx=XM_019827645.2;refGene_gn=DIAPH3;refGene_tx=XM_019827647.2;refGene_gn=DIAPH3;refGene_tx=XM_023252114.1;refGene_gn=DIAPH3;refGene_tx=XM_023252126.1
A1 36177537 36177602
refGene_gn=DIAPH3;refGene_tx=XM_006927330.4;refGene_gn=DIAPH3;refGene_tx=XM_019827642.2;refGene_gn=DIAPH3;refGene_tx=XM_019827645.2;refGene_gn=DIAPH3;refGene_tx=XM_019827647.2;refGene_gn=DIAPH3;refGene_tx=XM_023252114.1;refGene_gn=DIAPH3;refGene_tx=XM_023252126.1
A1 36180501 36180620
refGene_gn=DIAPH3;refGene_tx=XM_006927330.4;refGene_gn=DIAPH3;refGene_tx=XM_019827642.2;refGene_gn=DIAPH3;refGene_tx=XM_019827645.2;refGene_gn=DIAPH3;refGene_tx=XM_019827647.2;refGene_gn=DIAPH3;refGene_tx=XM_023252114.1;refGene_gn=DIAPH3;refGene_tx=XM_023252126.1
A1 36185286 36185403

refGene_gn=DIAPH3;refGene_tx=XM_006927330.4;refGene_gn=DIAPH3;refGene_tx=XM_019827642.2;refGene_gn=DIAPH3;refGene_tx=XM_019827645.2;refGene_gn=DIAPH3;refGene_tx=XM_019827647.2;refGene_gn=DIAPH3;refGene_tx=XM_023252114.1;refGene_gn=DIAPH3;refGene_tx=XM_023252126.1
A1 36186206 36186325
refGene_gn=DIAPH3;refGene_tx=XM_006927330.4;refGene_gn=DIAPH3;refGene_tx=XM_019827642.2;refGene_gn=DIAPH3;refGene_tx=XM_019827645.2;refGene_gn=DIAPH3;refGene_tx=XM_019827647.2;refGene_gn=DIAPH3;refGene_tx=XM_023252114.1;refGene_gn=DIAPH3;refGene_tx=XM_023252126.1
A1 36186611 36186722
refGene_gn=DIAPH3;refGene_tx=XM_006927330.4;refGene_gn=DIAPH3;refGene_tx=XM_019827642.2;refGene_gn=DIAPH3;refGene_tx=XM_019827645.2;refGene_gn=DIAPH3;refGene_tx=XM_019827647.2;refGene_gn=DIAPH3;refGene_tx=XM_023252114.1;refGene_gn=DIAPH3;refGene_tx=XM_023252126.1
A1 36203126 36203232
refGene_gn=DIAPH3;refGene_tx=XM_006927330.4;refGene_gn=DIAPH3;refGene_tx=XM_019827642.2;refGene_gn=DIAPH3;refGene_tx=XM_019827645.2;refGene_gn=DIAPH3;refGene_tx=XM_019827647.2;refGene_gn=DIAPH3;refGene_tx=XM_023252114.1;refGene_gn=DIAPH3;refGene_tx=XM_023252126.1
A1 36205620 36205757
refGene_gn=DIAPH3;refGene_tx=XM_006927330.4;refGene_gn=DIAPH3;refGene_tx=XM_019827642.2;refGene_gn=DIAPH3;refGene_tx=XM_019827645.2;refGene_gn=DIAPH3;refGene_tx=XM_019827647.2;refGene_gn=DIAPH3;refGene_tx=XM_023252114.1;refGene_gn=DIAPH3;refGene_tx=XM_023252126.1
A1 36207902 36207914 refGene_gn=DIAPH3;refGene_tx=XM_019827647.2
A1 36210672 36210742
refGene_gn=DIAPH3;refGene_tx=XM_006927330.4;refGene_gn=DIAPH3;refGene_tx=XM_019827642.2;refGene_gn=DIAPH3;refGene_tx=XM_019827645.2;refGene_gn=DIAPH3;refGene_tx=XM_023252114.1;refGene_gn=DIAPH3;refGene_tx=XM_023252126.1
A1 36210829 36210904
refGene_gn=DIAPH3;refGene_tx=XM_006927330.4;refGene_gn=DIAPH3;refGene_tx=XM_019827642.2;refGene_gn=DIAPH3;refGene_tx=XM_019827645.2;refGene_gn=DIAPH3;refGene_tx=XM_023252114.1;refGene_gn=DIAPH3;refGene_tx=XM_023252126.1
A1 36236970 36237098
refGene_gn=DIAPH3;refGene_tx=XM_006927330.4;refGene_gn=DIAPH3;refGene_tx=XM_019827642.2;refGene_gn=DIAPH3;refGene_tx=XM_019827645.2;refGene_gn=DIAPH3;refGene_tx=XM_023252114.1;refGene_gn=DIAPH3;refGene_tx=XM_023252126.1
A1 36282484 36282589
refGene_gn=DIAPH3;refGene_tx=XM_006927330.4;refGene_gn=DIAPH3;refGene_tx=XM_019827642.2;refGene_gn=DIAPH3;refGene_tx=XM_019827645.2;refGene_gn=DIAPH3;refGene_tx=XM_023252114.1;refGene_gn=DIAPH3;refGene_tx=XM_023252126.1
A1 36302816 36302984
refGene_gn=DIAPH3;refGene_tx=XM_006927330.4;refGene_gn=DIAPH3;refGene_tx=XM_019827642.2;refGene_gn=DIAPH3;refGene_tx=XM_019827645.2;refGene_gn=DIAPH3;refGene_tx=XM_023252114.1;refGene_gn=DIAPH3;refGene_tx=XM_023252126.1

A1 36323579 36323612
refGene_gn=DIAPH3;refGene_tx=XM_006927330.4;refGene_gn=DIAPH3;refGene_tx=XM_019827645.2;refGene_gn=DIAPH3;refGene_tx=XM_023252114.1;refGene_gn=DIAPH3;refGene_tx=XM_023252126.1
A1 36349920 36350097
refGene_gn=DIAPH3;refGene_tx=XM_006927330.4;refGene_gn=DIAPH3;refGene_tx=XM_019827642.2;refGene_gn=DIAPH3;refGene_tx=XM_019827645.2;refGene_gn=DIAPH3;refGene_tx=XM_023252114.1;refGene_gn=DIAPH3;refGene_tx=XM_023252126.1
A1 36567175 36567216
refGene_gn=TDRD3;refGene_tx=XM_023252137.1;refGene_gn=TDRD3;refGene_tx=XM_023252140.1;refGene_gn=TDRD3;refGene_tx=XM_023252149.1;refGene_gn=TDRD3;refGene_tx=XM_023252153.1;refGene_gn=TDRD3;refGene_tx=XM_023252159.1;refGene_gn=TDRD3;refGene_tx=XM_023252161.1
A1 36626924 36627009
refGene_gn=TDRD3;refGene_tx=XM_023252137.1;refGene_gn=TDRD3;refGene_tx=XM_023252140.1;refGene_gn=TDRD3;refGene_tx=XM_023252149.1;refGene_gn=TDRD3;refGene_tx=XM_023252153.1;refGene_gn=TDRD3;refGene_tx=XM_023252159.1;refGene_gn=TDRD3;refGene_tx=XM_023252161.1
A1 36632292 36632358
refGene_gn=TDRD3;refGene_tx=XM_023252137.1;refGene_gn=TDRD3;refGene_tx=XM_023252140.1;refGene_gn=TDRD3;refGene_tx=XM_023252149.1;refGene_gn=TDRD3;refGene_tx=XM_023252153.1;refGene_gn=TDRD3;refGene_tx=XM_023252159.1;refGene_gn=TDRD3;refGene_tx=XM_023252161.1
A1 36646694 36646855
refGene_gn=TDRD3;refGene_tx=XM_019827665.1;refGene_gn=TDRD3;refGene_tx=XM_023252137.1;refGene_gn=TDRD3;refGene_tx=XM_023252140.1;refGene_gn=TDRD3;refGene_tx=XM_023252149.1;refGene_gn=TDRD3;refGene_tx=XM_023252153.1;refGene_gn=TDRD3;refGene_tx=XM_023252159.1;refGene_gn=TDRD3;refGene_tx=XM_023252161.1
A1 36651337 36651479
refGene_gn=TDRD3;refGene_tx=XM_019827665.1;refGene_gn=TDRD3;refGene_tx=XM_023252137.1;refGene_gn=TDRD3;refGene_tx=XM_023252140.1;refGene_gn=TDRD3;refGene_tx=XM_023252149.1;refGene_gn=TDRD3;refGene_tx=XM_023252153.1;refGene_gn=TDRD3;refGene_tx=XM_023252159.1;refGene_gn=TDRD3;refGene_tx=XM_023252161.1
A1 36657914 36657986
refGene_gn=TDRD3;refGene_tx=XM_019827665.1;refGene_gn=TDRD3;refGene_tx=XM_023252137.1;refGene_gn=TDRD3;refGene_tx=XM_023252140.1;refGene_gn=TDRD3;refGene_tx=XM_023252149.1;refGene_gn=TDRD3;refGene_tx=XM_023252153.1;refGene_gn=TDRD3;refGene_tx=XM_023252159.1;refGene_gn=TDRD3;refGene_tx=XM_023252161.1
A1 36660517 36660667
refGene_gn=TDRD3;refGene_tx=XM_019827665.1;refGene_gn=TDRD3;refGene_tx=XM_023252137.1;refGene_gn=TDRD3;refGene_tx=XM_023252140.1;refGene_gn=TDRD3;refGene_tx=XM_023252149.1;refGene_gn=TDRD3;refGene_tx=XM_023252153.1;refGene_gn=TDRD3;refGene_tx=XM_023252159.1;refGene_gn=TDRD3;refGene_tx=XM_023252161.1
A1 36665900 36666041
refGene_gn=TDRD3;refGene_tx=XM_019827665.1;refGene_gn=TDRD3;refGene_tx

=XM_023252137.1;refGene_gn=TDRD3;refGene_tx=XM_023252140.1;refGene_gn=TDRD3;refGene_tx=XM_023252149.1;refGene_gn=TDRD3;refGene_tx=XM_023252153.1;refGene_gn=TDRD3;refGene_tx=XM_023252159.1;refGene_gn=TDRD3;refGene_tx=XM_023252161.1
A1 36668820 36668977
refGene_gn=TDRD3;refGene_tx=XM_019827665.1;refGene_gn=TDRD3;refGene_tx=XM_023252137.1;refGene_gn=TDRD3;refGene_tx=XM_023252140.1;refGene_gn=TDRD3;refGene_tx=XM_023252149.1;refGene_gn=TDRD3;refGene_tx=XM_023252153.1;refGene_gn=TDRD3;refGene_tx=XM_023252159.1;refGene_gn=TDRD3;refGene_tx=XM_023252161.1
A1 36669652 36669778
refGene_gn=TDRD3;refGene_tx=XM_019827665.1;refGene_gn=TDRD3;refGene_tx=XM_023252137.1;refGene_gn=TDRD3;refGene_tx=XM_023252140.1;refGene_gn=TDRD3;refGene_tx=XM_023252149.1;refGene_gn=TDRD3;refGene_tx=XM_023252153.1;refGene_gn=TDRD3;refGene_tx=XM_023252159.1;refGene_gn=TDRD3;refGene_tx=XM_023252161.1
A1 36688697 36689548
refGene_gn=TDRD3;refGene_tx=XM_019827665.1;refGene_gn=TDRD3;refGene_tx=XM_023252137.1;refGene_gn=TDRD3;refGene_tx=XM_023252140.1;refGene_gn=TDRD3;refGene_tx=XM_023252149.1;refGene_gn=TDRD3;refGene_tx=XM_023252153.1;refGene_gn=TDRD3;refGene_tx=XM_023252159.1;refGene_gn=TDRD3;refGene_tx=XM_023252161.1
A1 36694667 36694793
refGene_gn=TDRD3;refGene_tx=XM_019827665.1;refGene_gn=TDRD3;refGene_tx=XM_023252137.1;refGene_gn=TDRD3;refGene_tx=XM_023252140.1;refGene_gn=TDRD3;refGene_tx=XM_023252149.1;refGene_gn=TDRD3;refGene_tx=XM_023252153.1;refGene_gn=TDRD3;refGene_tx=XM_023252159.1;refGene_gn=TDRD3;refGene_tx=XM_023252161.1
A1 36699840 36699900 refGene_gn=TDRD3;refGene_tx=XM_023252161.1
A1 36705652 36705738
refGene_gn=TDRD3;refGene_tx=XM_019827665.1;refGene_gn=TDRD3;refGene_tx=XM_023252137.1;refGene_gn=TDRD3;refGene_tx=XM_023252140.1;refGene_gn=TDRD3;refGene_tx=XM_023252149.1
A1 36708936 36708970 refGene_gn=TDRD3;refGene_tx=XM_023252149.1
A1 36724810 36724936
refGene_gn=TDRD3;refGene_tx=XM_019827665.1;refGene_gn=TDRD3;refGene_tx=XM_023252137.1;refGene_gn=TDRD3;refGene_tx=XM_023252140.1;refGene_gn=TDRD3;refGene_tx=XM_023252153.1;refGene_gn=TDRD3;refGene_tx=XM_023252159.1
A1 36728106 36728122 refGene_gn=TDRD3;refGene_tx=XM_023252140.1
A1 36730200 36730231
refGene_gn=TDRD3;refGene_tx=XM_019827665.1;refGene_gn=TDRD3;refGene_tx=XM_023252137.1
A1 37454863 37457590 refGene_gn=PCDH20;refGene_tx=XM_003980439.4
A1 37458628 37458760 refGene_gn=PCDH20;refGene_tx=XM_003980439.4
A1 41764301 41764675
refGene_gn=PCDH9;refGene_tx=XM_011288853.3;refGene_gn=PCDH9;refGene_tx=XM_011288878.3;refGene_gn=PCDH9;refGene_tx=XM_011288914.3;refGene_gn=PCDH9;refGene_tx=XM_019827769.2;refGene_gn=PCDH9;refGene_tx=XM_019827777.2;refGene_gn=PCDH9;refGene_tx=XM_019827782.2;refGene_gn=PCDH9;refGe

ne_tx=XM_019827804.2
A1 41900688 41900747 refGene_gn=PCDH9;refGene_tx=XM_019827787.2
A1 41994285 41994290 refGene_gn=PCDH9;refGene_tx=XM_019827791.2
A1 42076381 42076583
refGene_gn=PCDH9;refGene_tx=XM_011288853.3;refGene_gn=PCDH9;refGene_tx=XM_011288878.3;refGene_gn=PCDH9;refGene_tx=XM_011288914.3;refGene_gn=PCDH9;refGene_tx=XM_019827769.2;refGene_gn=PCDH9;refGene_tx=XM_019827777.2;refGene_gn=PCDH9;refGene_tx=XM_019827782.2;refGene_gn=PCDH9;refGene_tx=XM_019827787.2;refGene_gn=PCDH9;refGene_tx=XM_019827791.2;refGene_gn=PCDH9;refGene_tx=XM_019827804.2
A1 42338657 42338759
refGene_gn=PCDH9;refGene_tx=XM_011288853.3;refGene_gn=PCDH9;refGene_tx=XM_019827769.2;refGene_gn=PCDH9;refGene_tx=XM_019827777.2;refGene_gn=PCDH9;refGene_tx=XM_019827782.2;refGene_gn=PCDH9;refGene_tx=XM_019827787.2;refGene_gn=PCDH9;refGene_tx=XM_019827791.2;refGene_gn=PCDH9;refGene_tx=XM_019827804.2
A1 42664796 42664943 refGene_gn=PCDH9;refGene_tx=XM_019827798.2
A1 42676625 42679661
refGene_gn=PCDH9;refGene_tx=XM_011288853.3;refGene_gn=PCDH9;refGene_tx=XM_011288878.3;refGene_gn=PCDH9;refGene_tx=XM_011288914.3;refGene_gn=PCDH9;refGene_tx=XM_019827769.2;refGene_gn=PCDH9;refGene_tx=XM_019827777.2;refGene_gn=PCDH9;refGene_tx=XM_019827782.2;refGene_gn=PCDH9;refGene_tx=XM_019827787.2;refGene_gn=PCDH9;refGene_tx=XM_019827791.2;refGene_gn=PCDH9;refGene_tx=XM_019827798.2;refGene_gn=PCDH9;refGene_tx=XM_019827804.2
A1 44551319 44552039
refGene_gn=LOC101100024;refGene_tx=XM_011288689.2
A1 44552040 44552112
refGene_gn=LOC101100024;refGene_tx=XM_011288689.2
A1 44748030 44748090
refGene_gn=KLHL1;refGene_tx=XM_011288605.3;refGene_gn=KLHL1;refGene_tx=XM_019827858.2
A1 44754226 44754398 refGene_gn=KLHL1;refGene_tx=XM_011288605.3
A1 44772134 44772347
refGene_gn=KLHL1;refGene_tx=XM_011288605.3;refGene_gn=KLHL1;refGene_tx=XM_019827858.2
A1 44791942 44792105
refGene_gn=KLHL1;refGene_tx=XM_011288605.3;refGene_gn=KLHL1;refGene_tx=XM_019827858.2
A1 44840080 44840305
refGene_gn=KLHL1;refGene_tx=XM_011288605.3;refGene_gn=KLHL1;refGene_tx=XM_019827858.2
A1 44868907 44869094
refGene_gn=KLHL1;refGene_tx=XM_011288605.3;refGene_gn=KLHL1;refGene_tx=XM_019827858.2
A1 44898436 44898649
refGene_gn=KLHL1;refGene_tx=XM_011288605.3;refGene_gn=KLHL1;refGene_tx=XM_019827858.2
A1 44951700 44951897
refGene_gn=KLHL1;refGene_tx=XM_011288605.3;refGene_gn=KLHL1;refGene_tx

=XM_019827858.2
A1	44973961	44974098
refGene_gn=KLHL1;refGene_tx=XM_011288605.3;refGene_gn=KLHL1;refGene_tx
=XM_019827858.2
A1	44989967	44990150
refGene_gn=KLHL1;refGene_tx=XM_011288605.3;refGene_gn=KLHL1;refGene_tx
=XM_019827858.2
A1	45108204	45108707
refGene_gn=KLHL1;refGene_tx=XM_011288605.3;refGene_gn=KLHL1;refGene_tx
=XM_019827858.2
A1	46291720	46291758	refGene_gn=DACH1;refGene_tx=XM_019827867.2
A1	46325150	46325219	refGene_gn=DACH1;refGene_tx=XM_019827867.2
A1	46325728	46325872	refGene_gn=DACH1;refGene_tx=XM_019827867.2
A1	46328937	46329085	refGene_gn=DACH1;refGene_tx=XM_019827867.2
A1	46337715	46337867	refGene_gn=DACH1;refGene_tx=XM_019827867.2
A1	46403501	46403636	refGene_gn=DACH1;refGene_tx=XM_019827867.2
A1	46406821	46406957	refGene_gn=DACH1;refGene_tx=XM_019827867.2
A1	46422599	46422772	refGene_gn=DACH1;refGene_tx=XM_019827867.2
A1	46482192	46482354	refGene_gn=DACH1;refGene_tx=XM_019827867.2
A1	46534464	46534580	refGene_gn=DACH1;refGene_tx=XM_019827867.2
A1	46720227	46720747	refGene_gn=DACH1;refGene_tx=XM_019827867.2
A1	46720797	46721095	refGene_gn=DACH1;refGene_tx=XM_019827867.2
A1	47485786	47485810
refGene_gn=MZT1;refGene_tx=XM_019827887.2;refGene_gn=MZT1;refGene_tx=X
M_019827888.2
A1	47491627	47491773
refGene_gn=MZT1;refGene_tx=XM_019827887.2;refGene_gn=MZT1;refGene_tx=X
M_019827888.2
A1	47496136	47496287	refGene_gn=MZT1;refGene_tx=XM_019827888.2
A1	47501739	47501835	refGene_gn=MZT1;refGene_tx=XM_019827887.2
A1	47502010	47502148
refGene_gn=BORA;refGene_tx=XM_019827882.2;refGene_gn=BORA;refGene_tx=X
M_023252205.1
A1	47502174	47502247	refGene_gn=MZT1;refGene_tx=XM_019827887.2
A1	47503614	47503782
refGene_gn=BORA;refGene_tx=XM_019827882.2;refGene_gn=BORA;refGene_tx=X
M_023252205.1
A1	47505531	47505638
refGene_gn=BORA;refGene_tx=XM_019827882.2;refGene_gn=BORA;refGene_tx=X
M_023252205.1
A1	47508585	47508631
refGene_gn=BORA;refGene_tx=XM_019827882.2;refGene_gn=BORA;refGene_tx=X
M_023252205.1
A1	47511556	47511638
refGene_gn=BORA;refGene_tx=XM_019827882.2;refGene_gn=BORA;refGene_tx=X
M_023252205.1
A1	47516333	47516399
refGene_gn=BORA;refGene_tx=XM_019827882.2;refGene_gn=BORA;refGene_tx=X
M_023252205.1
A1	47517756	47517813

refGene_gn=BORA;refGene_tx=XM_019827882.2;refGene_gn=BORA;refGene_tx=XM_023252205.1
A1 47518227 47518454
refGene_gn=BORA;refGene_tx=XM_019827882.2;refGene_gn=BORA;refGene_tx=XM_023252205.1
A1 47519018 47519151
refGene_gn=BORA;refGene_tx=XM_019827882.2;refGene_gn=BORA;refGene_tx=XM_023252205.1
A1 47521779 47522387
refGene_gn=BORA;refGene_tx=XM_019827882.2;refGene_gn=BORA;refGene_tx=XM_023252205.1
A1 47529613 47529730
refGene_gn=BORA;refGene_tx=XM_019827882.2;refGene_gn=BORA;refGene_tx=XM_023252205.1
A1 47530747 47530774
refGene_gn=BORA;refGene_tx=XM_019827882.2;refGene_gn=BORA;refGene_tx=XM_023252205.1
A1 47534361 47534445
refGene_gn=DIS3;refGene_tx=XM_003980443.5;refGene_gn=DIS3;refGene_tx=XM_019827895.1;refGene_gn=DIS3;refGene_tx=XM_019827899.2
A1 47535324 47535447
refGene_gn=DIS3;refGene_tx=XM_003980443.5;refGene_gn=DIS3;refGene_tx=XM_019827895.1;refGene_gn=DIS3;refGene_tx=XM_019827899.2
A1 47536119 47536278
refGene_gn=DIS3;refGene_tx=XM_003980443.5;refGene_gn=DIS3;refGene_tx=XM_019827895.1;refGene_gn=DIS3;refGene_tx=XM_019827899.2
A1 47536413 47536582
refGene_gn=DIS3;refGene_tx=XM_003980443.5;refGene_gn=DIS3;refGene_tx=XM_019827895.1;refGene_gn=DIS3;refGene_tx=XM_019827899.2
A1 47536694 47536909
refGene_gn=DIS3;refGene_tx=XM_003980443.5;refGene_gn=DIS3;refGene_tx=XM_019827895.1;refGene_gn=DIS3;refGene_tx=XM_019827899.2
A1 47538272 47538429
refGene_gn=DIS3;refGene_tx=XM_003980443.5;refGene_gn=DIS3;refGene_tx=XM_019827895.1;refGene_gn=DIS3;refGene_tx=XM_019827899.2
A1 47540418 47540505
refGene_gn=DIS3;refGene_tx=XM_003980443.5;refGene_gn=DIS3;refGene_tx=XM_019827895.1;refGene_gn=DIS3;refGene_tx=XM_019827899.2
A1 47544024 47544152
refGene_gn=DIS3;refGene_tx=XM_003980443.5;refGene_gn=DIS3;refGene_tx=XM_019827895.1;refGene_gn=DIS3;refGene_tx=XM_019827899.2
A1 47545538 47545623
refGene_gn=DIS3;refGene_tx=XM_003980443.5;refGene_gn=DIS3;refGene_tx=XM_019827895.1;refGene_gn=DIS3;refGene_tx=XM_019827899.2
A1 47545720 47545785
refGene_gn=DIS3;refGene_tx=XM_003980443.5;refGene_gn=DIS3;refGene_tx=XM_019827895.1;refGene_gn=DIS3;refGene_tx=XM_019827899.2
A1 47546939 47547041
refGene_gn=DIS3;refGene_tx=XM_003980443.5;refGene_gn=DIS3;refGene_tx=XM_019827895.1;refGene_gn=DIS3;refGene_tx=XM_019827899.2

A1 47547298 47547415
refGene_gn=DIS3;refGene_tx=XM_003980443.5;refGene_gn=DIS3;refGene_tx=XM_019827895.1;refGene_gn=DIS3;refGene_tx=XM_019827899.2
A1 47547774 47547921
refGene_gn=DIS3;refGene_tx=XM_003980443.5;refGene_gn=DIS3;refGene_tx=XM_019827895.1;refGene_gn=DIS3;refGene_tx=XM_019827899.2
A1 47548938 47549076
refGene_gn=DIS3;refGene_tx=XM_003980443.5;refGene_gn=DIS3;refGene_tx=XM_019827895.1;refGene_gn=DIS3;refGene_tx=XM_019827899.2
A1 47549202 47549316
refGene_gn=DIS3;refGene_tx=XM_003980443.5;refGene_gn=DIS3;refGene_tx=XM_019827895.1;refGene_gn=DIS3;refGene_tx=XM_019827899.2
A1 47550644 47550809
refGene_gn=DIS3;refGene_tx=XM_003980443.5;refGene_gn=DIS3;refGene_tx=XM_019827895.1;refGene_gn=DIS3;refGene_tx=XM_019827899.2
A1 47551556 47551724
refGene_gn=DIS3;refGene_tx=XM_003980443.5;refGene_gn=DIS3;refGene_tx=XM_019827895.1;refGene_gn=DIS3;refGene_tx=XM_019827899.2
A1 47557608 47557682
refGene_gn=DIS3;refGene_tx=XM_003980443.5;refGene_gn=DIS3;refGene_tx=XM_019827895.1;refGene_gn=DIS3;refGene_tx=XM_019827899.2
A1 47558525 47558719
refGene_gn=DIS3;refGene_tx=XM_003980443.5;refGene_gn=DIS3;refGene_tx=XM_019827895.1;refGene_gn=DIS3;refGene_tx=XM_019827899.2
A1 47561179 47561337 refGene_gn=DIS3;refGene_tx=XM_003980443.5
A1 47561595 47561663 refGene_gn=DIS3;refGene_tx=XM_019827899.2
A1 47561940 47562168
refGene_gn=DIS3;refGene_tx=XM_003980443.5;refGene_gn=DIS3;refGene_tx=XM_019827899.2
A1 47563798 47564053 refGene_gn=PIBF1;refGene_tx=XM_023252216.1
A1 47568189 47568290 refGene_gn=PIBF1;refGene_tx=XM_023252216.1
A1 47571404 47571603 refGene_gn=PIBF1;refGene_tx=XM_023252216.1
A1 47574122 47574242 refGene_gn=PIBF1;refGene_tx=XM_023252216.1
A1 47600543 47600677 refGene_gn=PIBF1;refGene_tx=XM_023252216.1
A1 47606878 47606987 refGene_gn=PIBF1;refGene_tx=XM_023252216.1
A1 47607242 47607424 refGene_gn=PIBF1;refGene_tx=XM_023252216.1
A1 47608584 47608710 refGene_gn=PIBF1;refGene_tx=XM_023252216.1
A1 47623949 47624048 refGene_gn=PIBF1;refGene_tx=XM_023252216.1
A1 47677297 47677463 refGene_gn=PIBF1;refGene_tx=XM_023252216.1
A1 47683519 47683670 refGene_gn=PIBF1;refGene_tx=XM_023252216.1
A1 47690881 47690972 refGene_gn=PIBF1;refGene_tx=XM_023252216.1
A1 47706048 47706151 refGene_gn=PIBF1;refGene_tx=XM_023252216.1
A1 47733339 47733470 refGene_gn=PIBF1;refGene_tx=XM_023252216.1
A1 47741227 47741312 refGene_gn=PIBF1;refGene_tx=XM_023252216.1
A1 47762373 47762547 refGene_gn=PIBF1;refGene_tx=XM_023252216.1
A1 47771837 47771888 refGene_gn=PIBF1;refGene_tx=XM_023252216.1
A1 47809122 47809383 refGene_gn=KLF5;refGene_tx=XM_023252219.1
A1 47811728 47812602
refGene_gn=KLF5;refGene_tx=XM_011280394.3;refGene_gn=KLF5;refGene_tx=XM_023252219.1

A1	47813608	47813668
refGene_gn=KLF5;refGene_tx=XM_011280394.3;refGene_gn=KLF5;refGene_tx=XM_023252219.1
A1	47826036	47826215
refGene_gn=KLF5;refGene_tx=XM_011280394.3;refGene_gn=KLF5;refGene_tx=XM_023252219.1
A1	48378786	48378968
refGene_gn=KLF12;refGene_tx=XM_019827919.2;refGene_gn=KLF12;refGene_tx=XM_023252226.1;refGene_gn=KLF12;refGene_tx=XM_023252227.1;refGene_gn=KLF12;refGene_tx=XM_023252228.1;refGene_gn=KLF12;refGene_tx=XM_023252229.1;refGene_gn=KLF12;refGene_tx=XM_023252230.1;refGene_gn=KLF12;refGene_tx=XM_023252231.1;refGene_gn=KLF12;refGene_tx=XM_023252233.1;refGene_gn=KLF12;refGene_tx=XM_023252234.1;refGene_gn=KLF12;refGene_tx=XM_023252236.1
A1	48397013	48397171
refGene_gn=KLF12;refGene_tx=XM_019827919.2;refGene_gn=KLF12;refGene_tx=XM_023252226.1;refGene_gn=KLF12;refGene_tx=XM_023252227.1;refGene_gn=KLF12;refGene_tx=XM_023252228.1;refGene_gn=KLF12;refGene_tx=XM_023252229.1;refGene_gn=KLF12;refGene_tx=XM_023252230.1;refGene_gn=KLF12;refGene_tx=XM_023252231.1;refGene_gn=KLF12;refGene_tx=XM_023252233.1;refGene_gn=KLF12;refGene_tx=XM_023252234.1;refGene_gn=KLF12;refGene_tx=XM_023252236.1
A1	48440805	48440868
refGene_gn=KLF12;refGene_tx=XM_019827919.2;refGene_gn=KLF12;refGene_tx=XM_023252226.1;refGene_gn=KLF12;refGene_tx=XM_023252227.1;refGene_gn=KLF12;refGene_tx=XM_023252228.1;refGene_gn=KLF12;refGene_tx=XM_023252229.1;refGene_gn=KLF12;refGene_tx=XM_023252230.1;refGene_gn=KLF12;refGene_tx=XM_023252231.1;refGene_gn=KLF12;refGene_tx=XM_023252233.1;refGene_gn=KLF12;refGene_tx=XM_023252234.1;refGene_gn=KLF12;refGene_tx=XM_023252236.1
A1	48493063	48493199
refGene_gn=KLF12;refGene_tx=XM_019827919.2;refGene_gn=KLF12;refGene_tx=XM_023252226.1;refGene_gn=KLF12;refGene_tx=XM_023252227.1;refGene_gn=KLF12;refGene_tx=XM_023252228.1;refGene_gn=KLF12;refGene_tx=XM_023252229.1;refGene_gn=KLF12;refGene_tx=XM_023252230.1;refGene_gn=KLF12;refGene_tx=XM_023252231.1;refGene_gn=KLF12;refGene_tx=XM_023252233.1;refGene_gn=KLF12;refGene_tx=XM_023252234.1;refGene_gn=KLF12;refGene_tx=XM_023252236.1
A1	48522045	48522592
refGene_gn=KLF12;refGene_tx=XM_019827919.2;refGene_gn=KLF12;refGene_tx=XM_023252226.1;refGene_gn=KLF12;refGene_tx=XM_023252227.1;refGene_gn=KLF12;refGene_tx=XM_023252228.1;refGene_gn=KLF12;refGene_tx=XM_023252229.1;refGene_gn=KLF12;refGene_tx=XM_023252230.1;refGene_gn=KLF12;refGene_tx=XM_023252231.1;refGene_gn=KLF12;refGene_tx=XM_023252233.1;refGene_gn=KLF12;refGene_tx=XM_023252234.1;refGene_gn=KLF12;refGene_tx=XM_023252236.1
A1	48620062	48620152
refGene_gn=KLF12;refGene_tx=XM_019827919.2;refGene_gn=KLF12;refGene_tx=XM_023252226.1;refGene_gn=KLF12;refGene_tx=XM_023252227.1;refGene_gn=KLF12;refGene_tx=XM_023252228.1;refGene_gn=KLF12;refGene_tx=XM_0232522

29.1;refGene_gn=KLF12;refGene_tx=XM_023252230.1;refGene_gn=KLF12;refGene_tx=XM_023252231.1;refGene_gn=KLF12;refGene_tx=XM_023252233.1;refGene_gn=KLF12;refGene_tx=XM_023252234.1
A1 48672757 48672790
refGene_gn=KLF12;refGene_tx=XM_019827919.2;refGene_gn=KLF12;refGene_tx=XM_023252226.1;refGene_gn=KLF12;refGene_tx=XM_023252227.1;refGene_gn=KLF12;refGene_tx=XM_023252228.1;refGene_gn=KLF12;refGene_tx=XM_023252229.1;refGene_gn=KLF12;refGene_tx=XM_023252230.1;refGene_gn=KLF12;refGene_tx=XM_023252231.1;refGene_gn=KLF12;refGene_tx=XM_023252233.1;refGene_gn=KLF12;refGene_tx=XM_023252234.1
A1 49800882 49801116 refGene_gn=TBC1D4;refGene_tx=XM_023252267.1
A1 49803650 49803827 refGene_gn=TBC1D4;refGene_tx=XM_023252267.1
A1 49806230 49806400 refGene_gn=TBC1D4;refGene_tx=XM_023252267.1
A1 49808938 49809098 refGene_gn=TBC1D4;refGene_tx=XM_023252267.1
A1 49813438 49813683 refGene_gn=TBC1D4;refGene_tx=XM_023252267.1
A1 49816085 49816244 refGene_gn=TBC1D4;refGene_tx=XM_023252267.1
A1 49818916 49819075 refGene_gn=TBC1D4;refGene_tx=XM_023252267.1
A1 49819884 49819887 refGene_gn=TBC1D4;refGene_tx=XM_023252267.1
A1 49821772 49821985 refGene_gn=TBC1D4;refGene_tx=XM_023252267.1
A1 49823959 49824117 refGene_gn=TBC1D4;refGene_tx=XM_023252267.1
A1 49830590 49830614 refGene_gn=TBC1D4;refGene_tx=XM_023252267.1
A1 49834805 49834970 refGene_gn=TBC1D4;refGene_tx=XM_023252267.1
A1 49836509 49836736 refGene_gn=TBC1D4;refGene_tx=XM_023252267.1
A1 49838056 49838131 refGene_gn=TBC1D4;refGene_tx=XM_023252267.1
A1 49844478 49844598 refGene_gn=TBC1D4;refGene_tx=XM_023252267.1
A1 49848300 49848411 refGene_gn=TBC1D4;refGene_tx=XM_023252267.1
A1 49848698 49848790 refGene_gn=TBC1D4;refGene_tx=XM_023252267.1
A1 49854016 49854149 refGene_gn=TBC1D4;refGene_tx=XM_023252267.1
A1 49859696 49859801 refGene_gn=TBC1D4;refGene_tx=XM_023252267.1
A1 49863678 49863768 refGene_gn=TBC1D4;refGene_tx=XM_023252267.1
A1 49865205 49865784 refGene_gn=TBC1D4;refGene_tx=XM_023252267.1
A1 49987122 49987533 refGene_gn=TBC1D4;refGene_tx=XM_023252267.1
A1 49987534 49987621 refGene_gn=TBC1D4;refGene_tx=XM_023252267.1
A1 50014830 50014881 refGene_gn=COMMD6;refGene_tx=XM_003980446.3
A1 50016681 50016834 refGene_gn=COMMD6;refGene_tx=XM_003980446.3
A1 50032897 50032909 refGene_gn=COMMD6;refGene_tx=XM_003980446.3
A1 50032998 50033040 refGene_gn=COMMD6;refGene_tx=XM_003980446.3
A1 50044528 50044570
refGene_gn=UCHL3;refGene_tx=XM_003980447.5;refGene_gn=UCHL3;refGene_tx=XM_019828102.1
A1 50044687 50044699
refGene_gn=UCHL3;refGene_tx=XM_003980447.5;refGene_gn=UCHL3;refGene_tx=XM_019828102.1
A1 50052640 50052769
refGene_gn=UCHL3;refGene_tx=XM_003980447.5;refGene_gn=UCHL3;refGene_tx=XM_019828102.1;refGene_gn=UCHL3;refGene_tx=XM_019828107.1
A1 50057238 50057395
refGene_gn=UCHL3;refGene_tx=XM_003980447.5;refGene_gn=UCHL3;refGene_tx=XM_019828102.1;refGene_gn=UCHL3;refGene_tx=XM_019828107.1
A1 50057794 50057880

refGene_gn=UCHL3;refGene_tx=XM_003980447.5;refGene_gn=UCHL3;refGene_tx=XM_019828102.1;refGene_gn=UCHL3;refGene_tx=XM_019828107.1
A1 50059705 50059753
refGene_gn=UCHL3;refGene_tx=XM_003980447.5;refGene_gn=UCHL3;refGene_tx=XM_019828102.1;refGene_gn=UCHL3;refGene_tx=XM_019828107.1
A1 50077299 50077491
refGene_gn=UCHL3;refGene_tx=XM_019828102.1;refGene_gn=UCHL3;refGene_tx=XM_019828107.1
A1 50081771 50081847
refGene_gn=UCHL3;refGene_tx=XM_003980447.5;refGene_gn=UCHL3;refGene_tx=XM_019828102.1;refGene_gn=UCHL3;refGene_tx=XM_019828107.1
A1 50087855 50087914
refGene_gn=UCHL3;refGene_tx=XM_003980447.5;refGene_gn=UCHL3;refGene_tx=XM_019828102.1;refGene_gn=UCHL3;refGene_tx=XM_019828107.1
A1 50088480 50088564
refGene_gn=UCHL3;refGene_tx=XM_003980447.5;refGene_gn=UCHL3;refGene_tx=XM_019828102.1;refGene_gn=UCHL3;refGene_tx=XM_019828107.1
A1 50109562 50109622
refGene_gn=LMO7;refGene_tx=XM_006927340.4;refGene_gn=LMO7;refGene_tx=XM_019828007.2;refGene_gn=LMO7;refGene_tx=XM_019828014.2;refGene_gn=LMO7;refGene_tx=XM_019828020.2;refGene_gn=LMO7;refGene_tx=XM_019828022.2;refGene_gn=LMO7;refGene_tx=XM_019828026.2;refGene_gn=LMO7;refGene_tx=XM_019828033.2;refGene_gn=LMO7;refGene_tx=XM_019828041.2;refGene_gn=LMO7;refGene_tx=XM_019828047.2;refGene_gn=LMO7;refGene_tx=XM_019828051.2;refGene_gn=LMO7;refGene_tx=XM_019828056.2;refGene_gn=LMO7;refGene_tx=XM_019828067.2;refGene_gn=LMO7;refGene_tx=XM_023252243.1;refGene_gn=LMO7;refGene_tx=XM_023252263.1;refGene_gn=LMO7;refGene_tx=XM_023252264.1
A1 50177516 50177587
refGene_gn=LMO7;refGene_tx=XM_006927340.4;refGene_gn=LMO7;refGene_tx=XM_019828007.2;refGene_gn=LMO7;refGene_tx=XM_019828014.2;refGene_gn=LMO7;refGene_tx=XM_019828020.2;refGene_gn=LMO7;refGene_tx=XM_019828022.2;refGene_gn=LMO7;refGene_tx=XM_019828026.2;refGene_gn=LMO7;refGene_tx=XM_019828033.2;refGene_gn=LMO7;refGene_tx=XM_019828041.2;refGene_gn=LMO7;refGene_tx=XM_019828047.2;refGene_gn=LMO7;refGene_tx=XM_019828051.2;refGene_gn=LMO7;refGene_tx=XM_019828056.2;refGene_gn=LMO7;refGene_tx=XM_019828067.2;refGene_gn=LMO7;refGene_tx=XM_023252243.1;refGene_gn=LMO7;refGene_tx=XM_023252263.1;refGene_gn=LMO7;refGene_tx=XM_023252264.1
A1 50191232 50191302
refGene_gn=LMO7;refGene_tx=XM_006927340.4;refGene_gn=LMO7;refGene_tx=XM_019828007.2;refGene_gn=LMO7;refGene_tx=XM_019828014.2;refGene_gn=LMO7;refGene_tx=XM_019828020.2;refGene_gn=LMO7;refGene_tx=XM_019828022.2;refGene_gn=LMO7;refGene_tx=XM_019828026.2;refGene_gn=LMO7;refGene_tx=XM_019828033.2;refGene_gn=LMO7;refGene_tx=XM_019828041.2;refGene_gn=LMO7;refGene_tx=XM_019828047.2;refGene_gn=LMO7;refGene_tx=XM_019828051.2;refGene_gn=LMO7;refGene_tx=XM_019828056.2;refGene_gn=LMO7;refGene_tx=XM_019828067.2;refGene_gn=LMO7;refGene_tx=XM_023252243.1;refGene_gn=LMO7;refGene_tx=XM_023252263.1;refGene_gn=LMO7;refGene_tx=XM_023252264.1
A1 50210093 50210200
refGene_gn=LMO7;refGene_tx=XM_006927340.4;refGene_gn=LMO7;refGene_tx=XM_019828007.2;refGene_gn=LMO7;refGene_tx=XM_019828014.2;refGene_gn=LMO

7;refGene_tx=XM_019828020.2;refGene_gn=LMO7;refGene_tx=XM_019828022.2;
refGene_gn=LMO7;refGene_tx=XM_019828026.2;refGene_gn=LMO7;refGene_tx=X
M_019828033.2;refGene_gn=LMO7;refGene_tx=XM_019828041.2;refGene_gn=LMO
7;refGene_tx=XM_019828047.2;refGene_gn=LMO7;refGene_tx=XM_019828051.2;
refGene_gn=LMO7;refGene_tx=XM_019828056.2;refGene_gn=LMO7;refGene_tx=X
M_019828067.2;refGene_gn=LMO7;refGene_tx=XM_023252243.1;refGene_gn=LMO
7;refGene_tx=XM_023252263.1;refGene_gn=LMO7;refGene_tx=XM_023252264.1
A1 50244763 50244794
refGene_gn=LMO7;refGene_tx=XM_006927340.4;refGene_gn=LMO7;refGene_tx=X
M_019828007.2;refGene_gn=LMO7;refGene_tx=XM_019828014.2;refGene_gn=LMO
7;refGene_tx=XM_019828020.2;refGene_gn=LMO7;refGene_tx=XM_019828022.2;
refGene_gn=LMO7;refGene_tx=XM_019828026.2;refGene_gn=LMO7;refGene_tx=X
M_019828033.2;refGene_gn=LMO7;refGene_tx=XM_019828041.2;refGene_gn=LMO
7;refGene_tx=XM_019828047.2;refGene_gn=LMO7;refGene_tx=XM_019828051.2;
refGene_gn=LMO7;refGene_tx=XM_019828056.2;refGene_gn=LMO7;refGene_tx=X
M_019828067.2;refGene_gn=LMO7;refGene_tx=XM_023252243.1;refGene_gn=LMO
7;refGene_tx=XM_023252263.1;refGene_gn=LMO7;refGene_tx=XM_023252264.1
A1 50245591 50245705
refGene_gn=LMO7;refGene_tx=XM_006927340.4;refGene_gn=LMO7;refGene_tx=X
M_019828007.2;refGene_gn=LMO7;refGene_tx=XM_019828014.2;refGene_gn=LMO
7;refGene_tx=XM_019828020.2;refGene_gn=LMO7;refGene_tx=XM_019828022.2;
refGene_gn=LMO7;refGene_tx=XM_019828033.2;refGene_gn=LMO7;refGene_tx=X
M_019828041.2;refGene_gn=LMO7;refGene_tx=XM_019828047.2;refGene_gn=LMO
7;refGene_tx=XM_019828051.2;refGene_gn=LMO7;refGene_tx=XM_019828056.2;
refGene_gn=LMO7;refGene_tx=XM_019828067.2;refGene_gn=LMO7;refGene_tx=X
M_023252243.1;refGene_gn=LMO7;refGene_tx=XM_023252263.1;refGene_gn=LMO
7;refGene_tx=XM_023252264.1
A1 50249850 50250049
refGene_gn=LMO7;refGene_tx=XM_006927340.4;refGene_gn=LMO7;refGene_tx=X
M_019828007.2;refGene_gn=LMO7;refGene_tx=XM_019828014.2;refGene_gn=LMO
7;refGene_tx=XM_019828020.2;refGene_gn=LMO7;refGene_tx=XM_019828022.2;
refGene_gn=LMO7;refGene_tx=XM_019828026.2;refGene_gn=LMO7;refGene_tx=X
M_019828033.2;refGene_gn=LMO7;refGene_tx=XM_019828041.2;refGene_gn=LMO
7;refGene_tx=XM_019828047.2;refGene_gn=LMO7;refGene_tx=XM_019828051.2;
refGene_gn=LMO7;refGene_tx=XM_019828056.2;refGene_gn=LMO7;refGene_tx=X
M_019828067.2;refGene_gn=LMO7;refGene_tx=XM_023252243.1;refGene_gn=LMO
7;refGene_tx=XM_023252263.1;refGene_gn=LMO7;refGene_tx=XM_023252264.1
A1 50253436 50253689
refGene_gn=LMO7;refGene_tx=XM_006927340.4;refGene_gn=LMO7;refGene_tx=X
M_006927341.4;refGene_gn=LMO7;refGene_tx=XM_019828007.2;refGene_gn=LMO
7;refGene_tx=XM_019828014.2;refGene_gn=LMO7;refGene_tx=XM_019828020.2;
refGene_gn=LMO7;refGene_tx=XM_019828022.2;refGene_gn=LMO7;refGene_tx=X
M_019828026.2;refGene_gn=LMO7;refGene_tx=XM_019828033.2;refGene_gn=LMO
7;refGene_tx=XM_019828041.2;refGene_gn=LMO7;refGene_tx=XM_019828047.2;
refGene_gn=LMO7;refGene_tx=XM_019828051.2;refGene_gn=LMO7;refGene_tx=X
M_019828056.2;refGene_gn=LMO7;refGene_tx=XM_019828067.2;refGene_gn=LMO
7;refGene_tx=XM_023252243.1;refGene_gn=LMO7;refGene_tx=XM_023252263.1;
refGene_gn=LMO7;refGene_tx=XM_023252264.1
A1 50254026 50254311
refGene_gn=LMO7;refGene_tx=XM_006927340.4;refGene_gn=LMO7;refGene_tx=X

M_006927341.4;refGene_gn=LMO7;refGene_tx=XM_019828007.2;refGene_gn=LMO
7;refGene_tx=XM_019828014.2;refGene_gn=LMO7;refGene_tx=XM_019828020.2;
refGene_gn=LMO7;refGene_tx=XM_019828022.2;refGene_gn=LMO7;refGene_tx=X
M_019828026.2;refGene_gn=LMO7;refGene_tx=XM_019828033.2
A1 50254624 50254903
refGene_gn=LMO7;refGene_tx=XM_006927340.4;refGene_gn=LMO7;refGene_tx=X
M_006927341.4;refGene_gn=LMO7;refGene_tx=XM_019828007.2;refGene_gn=LMO
7;refGene_tx=XM_019828014.2;refGene_gn=LMO7;refGene_tx=XM_019828020.2;
refGene_gn=LMO7;refGene_tx=XM_019828022.2;refGene_gn=LMO7;refGene_tx=X
M_019828026.2;refGene_gn=LMO7;refGene_tx=XM_019828033.2;refGene_gn=LMO
7;refGene_tx=XM_019828041.2;refGene_gn=LMO7;refGene_tx=XM_019828047.2;
refGene_gn=LMO7;refGene_tx=XM_019828051.2;refGene_gn=LMO7;refGene_tx=X
M_023252243.1
A1 50255175 50255379
refGene_gn=LMO7;refGene_tx=XM_006927340.4;refGene_gn=LMO7;refGene_tx=X
M_006927341.4;refGene_gn=LMO7;refGene_tx=XM_019828007.2;refGene_gn=LMO
7;refGene_tx=XM_019828014.2;refGene_gn=LMO7;refGene_tx=XM_019828020.2;
refGene_gn=LMO7;refGene_tx=XM_019828022.2;refGene_gn=LMO7;refGene_tx=X
M_019828026.2;refGene_gn=LMO7;refGene_tx=XM_019828041.2
A1 50256698 50257418
refGene_gn=LMO7;refGene_tx=XM_006927340.4;refGene_gn=LMO7;refGene_tx=X
M_006927341.4;refGene_gn=LMO7;refGene_tx=XM_019828007.2;refGene_gn=LMO
7;refGene_tx=XM_019828014.2;refGene_gn=LMO7;refGene_tx=XM_019828020.2;
refGene_gn=LMO7;refGene_tx=XM_019828022.2;refGene_gn=LMO7;refGene_tx=X
M_019828026.2;refGene_gn=LMO7;refGene_tx=XM_019828033.2;refGene_gn=LMO
7;refGene_tx=XM_019828041.2;refGene_gn=LMO7;refGene_tx=XM_019828047.2;
refGene_gn=LMO7;refGene_tx=XM_019828051.2;refGene_gn=LMO7;refGene_tx=X
M_019828056.2;refGene_gn=LMO7;refGene_tx=XM_023252243.1
A1 50258374 50258404
refGene_gn=LMO7;refGene_tx=XM_006927340.4;refGene_gn=LMO7;refGene_tx=X
M_006927341.4;refGene_gn=LMO7;refGene_tx=XM_019828007.2;refGene_gn=LMO
7;refGene_tx=XM_019828014.2;refGene_gn=LMO7;refGene_tx=XM_019828020.2;
refGene_gn=LMO7;refGene_tx=XM_019828022.2;refGene_gn=LMO7;refGene_tx=X
M_019828026.2;refGene_gn=LMO7;refGene_tx=XM_019828033.2;refGene_gn=LMO
7;refGene_tx=XM_019828041.2;refGene_gn=LMO7;refGene_tx=XM_019828047.2;
refGene_gn=LMO7;refGene_tx=XM_019828051.2;refGene_gn=LMO7;refGene_tx=X
M_019828056.2;refGene_gn=LMO7;refGene_tx=XM_019828067.2;refGene_gn=LMO
7;refGene_tx=XM_023252243.1;refGene_gn=LMO7;refGene_tx=XM_023252263.1
A1 50265740 50265858
refGene_gn=LMO7;refGene_tx=XM_006927340.4;refGene_gn=LMO7;refGene_tx=X
M_006927341.4;refGene_gn=LMO7;refGene_tx=XM_019828007.2;refGene_gn=LMO
7;refGene_tx=XM_019828014.2;refGene_gn=LMO7;refGene_tx=XM_019828020.2;
refGene_gn=LMO7;refGene_tx=XM_019828022.2;refGene_gn=LMO7;refGene_tx=X
M_019828026.2;refGene_gn=LMO7;refGene_tx=XM_019828033.2;refGene_gn=LMO
7;refGene_tx=XM_019828041.2;refGene_gn=LMO7;refGene_tx=XM_019828047.2;
refGene_gn=LMO7;refGene_tx=XM_019828051.2;refGene_gn=LMO7;refGene_tx=X
M_019828056.2;refGene_gn=LMO7;refGene_tx=XM_019828067.2;refGene_gn=LMO
7;refGene_tx=XM_023252243.1;refGene_gn=LMO7;refGene_tx=XM_023252263.1;
refGene_gn=LMO7;refGene_tx=XM_023252264.1
A1 50267919 50268062

refGene_gn=LMO7;refGene_tx=XM_006927340.4;refGene_gn=LMO7;refGene_tx=X
M_006927341.4;refGene_gn=LMO7;refGene_tx=XM_019828007.2;refGene_gn=LMO
7;refGene_tx=XM_019828014.2;refGene_gn=LMO7;refGene_tx=XM_019828020.2;
refGene_gn=LMO7;refGene_tx=XM_019828022.2;refGene_gn=LMO7;refGene_tx=X
M_019828026.2;refGene_gn=LMO7;refGene_tx=XM_019828033.2;refGene_gn=LMO
7;refGene_tx=XM_019828041.2;refGene_gn=LMO7;refGene_tx=XM_019828047.2;
refGene_gn=LMO7;refGene_tx=XM_019828051.2;refGene_gn=LMO7;refGene_tx=X
M_019828056.2;refGene_gn=LMO7;refGene_tx=XM_019828067.2;refGene_gn=LMO
7;refGene_tx=XM_023252243.1;refGene_gn=LMO7;refGene_tx=XM_023252263.1;
refGene_gn=LMO7;refGene_tx=XM_023252264.1
A1 50269373 50269806
refGene_gn=LMO7;refGene_tx=XM_006927340.4;refGene_gn=LMO7;refGene_tx=X
M_006927341.4;refGene_gn=LMO7;refGene_tx=XM_019828007.2;refGene_gn=LMO
7;refGene_tx=XM_019828014.2;refGene_gn=LMO7;refGene_tx=XM_019828020.2;
refGene_gn=LMO7;refGene_tx=XM_019828022.2;refGene_gn=LMO7;refGene_tx=X
M_019828026.2;refGene_gn=LMO7;refGene_tx=XM_019828033.2;refGene_gn=LMO
7;refGene_tx=XM_019828041.2;refGene_gn=LMO7;refGene_tx=XM_019828047.2;
refGene_gn=LMO7;refGene_tx=XM_019828051.2;refGene_gn=LMO7;refGene_tx=X
M_019828056.2;refGene_gn=LMO7;refGene_tx=XM_019828067.2;refGene_gn=LMO
7;refGene_tx=XM_023252243.1;refGene_gn=LMO7;refGene_tx=XM_023252263.1;
refGene_gn=LMO7;refGene_tx=XM_023252264.1
A1 50271531 50271843
refGene_gn=LMO7;refGene_tx=XM_006927340.4;refGene_gn=LMO7;refGene_tx=X
M_006927341.4;refGene_gn=LMO7;refGene_tx=XM_019828007.2;refGene_gn=LMO
7;refGene_tx=XM_019828014.2;refGene_gn=LMO7;refGene_tx=XM_019828020.2;
refGene_gn=LMO7;refGene_tx=XM_019828022.2;refGene_gn=LMO7;refGene_tx=X
M_019828026.2;refGene_gn=LMO7;refGene_tx=XM_019828033.2;refGene_gn=LMO
7;refGene_tx=XM_019828041.2;refGene_gn=LMO7;refGene_tx=XM_019828047.2;
refGene_gn=LMO7;refGene_tx=XM_019828051.2;refGene_gn=LMO7;refGene_tx=X
M_019828056.2;refGene_gn=LMO7;refGene_tx=XM_019828067.2;refGene_gn=LMO
7;refGene_tx=XM_023252243.1;refGene_gn=LMO7;refGene_tx=XM_023252263.1;
refGene_gn=LMO7;refGene_tx=XM_023252264.1
A1 50280403 50280518
refGene_gn=LMO7;refGene_tx=XM_006927340.4;refGene_gn=LMO7;refGene_tx=X
M_006927341.4;refGene_gn=LMO7;refGene_tx=XM_019828007.2;refGene_gn=LMO
7;refGene_tx=XM_019828014.2;refGene_gn=LMO7;refGene_tx=XM_019828020.2;
refGene_gn=LMO7;refGene_tx=XM_019828022.2;refGene_gn=LMO7;refGene_tx=X
M_019828026.2;refGene_gn=LMO7;refGene_tx=XM_019828033.2;refGene_gn=LMO
7;refGene_tx=XM_019828041.2;refGene_gn=LMO7;refGene_tx=XM_019828047.2;
refGene_gn=LMO7;refGene_tx=XM_019828051.2;refGene_gn=LMO7;refGene_tx=X
M_019828056.2;refGene_gn=LMO7;refGene_tx=XM_019828067.2;refGene_gn=LMO
7;refGene_tx=XM_023252243.1;refGene_gn=LMO7;refGene_tx=XM_023252263.1;
refGene_gn=LMO7;refGene_tx=XM_023252264.1
A1 50281640 50281802
refGene_gn=LMO7;refGene_tx=XM_006927340.4;refGene_gn=LMO7;refGene_tx=X
M_006927341.4;refGene_gn=LMO7;refGene_tx=XM_019828007.2;refGene_gn=LMO
7;refGene_tx=XM_019828014.2;refGene_gn=LMO7;refGene_tx=XM_019828020.2;
refGene_gn=LMO7;refGene_tx=XM_019828022.2;refGene_gn=LMO7;refGene_tx=X
M_019828026.2;refGene_gn=LMO7;refGene_tx=XM_019828033.2;refGene_gn=LMO
7;refGene_tx=XM_019828041.2;refGene_gn=LMO7;refGene_tx=XM_019828047.2;

refGene_gn=LMO7;refGene_tx=XM_019828051.2;refGene_gn=LMO7;refGene_tx=XM_019828056.2;refGene_gn=LMO7;refGene_tx=XM_019828067.2;refGene_gn=LMO7;refGene_tx=XM_023252243.1;refGene_gn=LMO7;refGene_tx=XM_023252263.1;refGene_gn=LMO7;refGene_tx=XM_023252264.1
A1 50282580 50282768
refGene_gn=LMO7;refGene_tx=XM_006927340.4;refGene_gn=LMO7;refGene_tx=XM_006927341.4;refGene_gn=LMO7;refGene_tx=XM_019828007.2;refGene_gn=LMO7;refGene_tx=XM_019828014.2;refGene_gn=LMO7;refGene_tx=XM_019828020.2;refGene_gn=LMO7;refGene_tx=XM_019828022.2;refGene_gn=LMO7;refGene_tx=XM_019828026.2;refGene_gn=LMO7;refGene_tx=XM_019828033.2;refGene_gn=LMO7;refGene_tx=XM_019828041.2;refGene_gn=LMO7;refGene_tx=XM_019828047.2;refGene_gn=LMO7;refGene_tx=XM_019828051.2;refGene_gn=LMO7;refGene_tx=XM_019828056.2;refGene_gn=LMO7;refGene_tx=XM_019828067.2;refGene_gn=LMO7;refGene_tx=XM_023252243.1;refGene_gn=LMO7;refGene_tx=XM_023252263.1;refGene_gn=LMO7;refGene_tx=XM_023252264.1
A1 50283712 50283773
refGene_gn=LMO7;refGene_tx=XM_006927340.4;refGene_gn=LMO7;refGene_tx=XM_006927341.4;refGene_gn=LMO7;refGene_tx=XM_019828007.2;refGene_gn=LMO7;refGene_tx=XM_019828014.2;refGene_gn=LMO7;refGene_tx=XM_019828020.2;refGene_gn=LMO7;refGene_tx=XM_019828022.2;refGene_gn=LMO7;refGene_tx=XM_019828026.2;refGene_gn=LMO7;refGene_tx=XM_019828033.2;refGene_gn=LMO7;refGene_tx=XM_019828041.2;refGene_gn=LMO7;refGene_tx=XM_019828047.2;refGene_gn=LMO7;refGene_tx=XM_019828051.2;refGene_gn=LMO7;refGene_tx=XM_019828056.2;refGene_gn=LMO7;refGene_tx=XM_019828067.2;refGene_gn=LMO7;refGene_tx=XM_023252243.1;refGene_gn=LMO7;refGene_tx=XM_023252263.1;refGene_gn=LMO7;refGene_tx=XM_023252264.1
A1 50285688 50285745
refGene_gn=LMO7;refGene_tx=XM_006927340.4;refGene_gn=LMO7;refGene_tx=XM_006927341.4;refGene_gn=LMO7;refGene_tx=XM_019828007.2;refGene_gn=LMO7;refGene_tx=XM_019828020.2;refGene_gn=LMO7;refGene_tx=XM_019828022.2;refGene_gn=LMO7;refGene_tx=XM_019828026.2;refGene_gn=LMO7;refGene_tx=XM_019828033.2;refGene_gn=LMO7;refGene_tx=XM_019828041.2;refGene_gn=LMO7;refGene_tx=XM_019828047.2;refGene_gn=LMO7;refGene_tx=XM_019828051.2;refGene_gn=LMO7;refGene_tx=XM_019828056.2;refGene_gn=LMO7;refGene_tx=XM_019828067.2;refGene_gn=LMO7;refGene_tx=XM_023252243.1;refGene_gn=LMO7;refGene_tx=XM_023252263.1;refGene_gn=LMO7;refGene_tx=XM_023252264.1
A1 50287635 50287661
refGene_gn=LMO7;refGene_tx=XM_006927340.4;refGene_gn=LMO7;refGene_tx=XM_006927341.4;refGene_gn=LMO7;refGene_tx=XM_019828007.2;refGene_gn=LMO7;refGene_tx=XM_019828014.2;refGene_gn=LMO7;refGene_tx=XM_019828020.2;refGene_gn=LMO7;refGene_tx=XM_019828022.2;refGene_gn=LMO7;refGene_tx=XM_019828026.2;refGene_gn=LMO7;refGene_tx=XM_019828033.2;refGene_gn=LMO7;refGene_tx=XM_019828041.2;refGene_gn=LMO7;refGene_tx=XM_019828047.2;refGene_gn=LMO7;refGene_tx=XM_019828051.2;refGene_gn=LMO7;refGene_tx=XM_019828056.2;refGene_gn=LMO7;refGene_tx=XM_019828067.2;refGene_gn=LMO7;refGene_tx=XM_023252243.1;refGene_gn=LMO7;refGene_tx=XM_023252263.1;refGene_gn=LMO7;refGene_tx=XM_023252264.1
A1 50287942 50288047
refGene_gn=LMO7;refGene_tx=XM_006927340.4;refGene_gn=LMO7;refGene_tx=XM_006927341.4;refGene_gn=LMO7;refGene_tx=XM_019828007.2;refGene_gn=LMO

7;refGene_tx=XM_019828014.2;refGene_gn=LMO7;refGene_tx=XM_019828020.2;
refGene_gn=LMO7;refGene_tx=XM_019828022.2;refGene_gn=LMO7;refGene_tx=X
M_019828026.2;refGene_gn=LMO7;refGene_tx=XM_019828033.2;refGene_gn=LMO
7;refGene_tx=XM_019828041.2;refGene_gn=LMO7;refGene_tx=XM_019828047.2;
refGene_gn=LMO7;refGene_tx=XM_019828051.2;refGene_gn=LMO7;refGene_tx=X
M_019828056.2;refGene_gn=LMO7;refGene_tx=XM_019828067.2;refGene_gn=LMO
7;refGene_tx=XM_023252243.1;refGene_gn=LMO7;refGene_tx=XM_023252263.1;
refGene_gn=LMO7;refGene_tx=XM_023252264.1
A1 50288822 50288915
refGene_gn=LMO7;refGene_tx=XM_006927340.4;refGene_gn=LMO7;refGene_tx=X
M_006927341.4;refGene_gn=LMO7;refGene_tx=XM_019828007.2;refGene_gn=LMO
7;refGene_tx=XM_019828014.2;refGene_gn=LMO7;refGene_tx=XM_019828020.2;
refGene_gn=LMO7;refGene_tx=XM_019828026.2;refGene_gn=LMO7;refGene_tx=X
M_019828033.2;refGene_gn=LMO7;refGene_tx=XM_019828041.2;refGene_gn=LMO
7;refGene_tx=XM_019828047.2;refGene_gn=LMO7;refGene_tx=XM_019828051.2;
refGene_gn=LMO7;refGene_tx=XM_019828056.2;refGene_gn=LMO7;refGene_tx=X
M_019828067.2;refGene_gn=LMO7;refGene_tx=XM_023252243.1;refGene_gn=LMO
7;refGene_tx=XM_023252263.1;refGene_gn=LMO7;refGene_tx=XM_023252264.1
A1 50289320 50289658
refGene_gn=LMO7;refGene_tx=XM_006927340.4;refGene_gn=LMO7;refGene_tx=X
M_006927341.4;refGene_gn=LMO7;refGene_tx=XM_019828007.2;refGene_gn=LMO
7;refGene_tx=XM_019828014.2;refGene_gn=LMO7;refGene_tx=XM_019828020.2;
refGene_gn=LMO7;refGene_tx=XM_019828022.2;refGene_gn=LMO7;refGene_tx=X
M_019828026.2;refGene_gn=LMO7;refGene_tx=XM_019828033.2;refGene_gn=LMO
7;refGene_tx=XM_019828041.2;refGene_gn=LMO7;refGene_tx=XM_019828047.2;
refGene_gn=LMO7;refGene_tx=XM_019828051.2;refGene_gn=LMO7;refGene_tx=X
M_019828056.2;refGene_gn=LMO7;refGene_tx=XM_019828067.2;refGene_gn=LMO
7;refGene_tx=XM_023252243.1;refGene_gn=LMO7;refGene_tx=XM_023252263.1;
refGene_gn=LMO7;refGene_tx=XM_023252264.1
A1 50290506 50290572
refGene_gn=LMO7;refGene_tx=XM_006927340.4;refGene_gn=LMO7;refGene_tx=X
M_006927341.4;refGene_gn=LMO7;refGene_tx=XM_019828007.2;refGene_gn=LMO
7;refGene_tx=XM_019828014.2;refGene_gn=LMO7;refGene_tx=XM_019828020.2;
refGene_gn=LMO7;refGene_tx=XM_019828022.2;refGene_gn=LMO7;refGene_tx=X
M_019828026.2;refGene_gn=LMO7;refGene_tx=XM_019828033.2;refGene_gn=LMO
7;refGene_tx=XM_019828041.2;refGene_gn=LMO7;refGene_tx=XM_019828047.2;
refGene_gn=LMO7;refGene_tx=XM_019828051.2;refGene_gn=LMO7;refGene_tx=X
M_019828056.2;refGene_gn=LMO7;refGene_tx=XM_019828067.2;refGene_gn=LMO
7;refGene_tx=XM_023252243.1;refGene_gn=LMO7;refGene_tx=XM_023252263.1;
refGene_gn=LMO7;refGene_tx=XM_023252264.1
A1 50292644 50292697
refGene_gn=LMO7;refGene_tx=XM_006927340.4;refGene_gn=LMO7;refGene_tx=X
M_006927341.4;refGene_gn=LMO7;refGene_tx=XM_019828007.2;refGene_gn=LMO
7;refGene_tx=XM_019828014.2;refGene_gn=LMO7;refGene_tx=XM_019828020.2;
refGene_gn=LMO7;refGene_tx=XM_019828022.2;refGene_gn=LMO7;refGene_tx=X
M_019828026.2;refGene_gn=LMO7;refGene_tx=XM_019828033.2;refGene_gn=LMO
7;refGene_tx=XM_019828041.2;refGene_gn=LMO7;refGene_tx=XM_019828047.2;
refGene_gn=LMO7;refGene_tx=XM_019828051.2;refGene_gn=LMO7;refGene_tx=X
M_019828056.2;refGene_gn=LMO7;refGene_tx=XM_019828067.2;refGene_gn=LMO
7;refGene_tx=XM_023252243.1;refGene_gn=LMO7;refGene_tx=XM_023252263.1;

refGene_gn=LMO7;refGene_tx=XM_023252264.1
A1 50295630 50295844
refGene_gn=LMO7;refGene_tx=XM_006927340.4;refGene_gn=LMO7;refGene_tx=X
M_006927341.4;refGene_gn=LMO7;refGene_tx=XM_019828007.2;refGene_gn=LMO
7;refGene_tx=XM_019828014.2;refGene_gn=LMO7;refGene_tx=XM_019828020.2;
refGene_gn=LMO7;refGene_tx=XM_019828022.2;refGene_gn=LMO7;refGene_tx=X
M_019828026.2;refGene_gn=LMO7;refGene_tx=XM_019828033.2;refGene_gn=LMO
7;refGene_tx=XM_019828041.2;refGene_gn=LMO7;refGene_tx=XM_019828047.2;
refGene_gn=LMO7;refGene_tx=XM_019828051.2;refGene_gn=LMO7;refGene_tx=X
M_019828056.2;refGene_gn=LMO7;refGene_tx=XM_019828067.2;refGene_gn=LMO
7;refGene_tx=XM_023252243.1;refGene_gn=LMO7;refGene_tx=XM_023252263.1;
refGene_gn=LMO7;refGene_tx=XM_023252264.1
A1 50298797 50299079
refGene_gn=LMO7;refGene_tx=XM_006927340.4;refGene_gn=LMO7;refGene_tx=X
M_006927341.4;refGene_gn=LMO7;refGene_tx=XM_019828007.2;refGene_gn=LMO
7;refGene_tx=XM_019828014.2;refGene_gn=LMO7;refGene_tx=XM_019828020.2;
refGene_gn=LMO7;refGene_tx=XM_019828022.2;refGene_gn=LMO7;refGene_tx=X
M_019828026.2;refGene_gn=LMO7;refGene_tx=XM_019828033.2;refGene_gn=LMO
7;refGene_tx=XM_019828041.2;refGene_gn=LMO7;refGene_tx=XM_019828047.2;
refGene_gn=LMO7;refGene_tx=XM_019828051.2;refGene_gn=LMO7;refGene_tx=X
M_019828056.2;refGene_gn=LMO7;refGene_tx=XM_019828067.2;refGene_gn=LMO
7;refGene_tx=XM_023252243.1;refGene_gn=LMO7;refGene_tx=XM_023252263.1;
refGene_gn=LMO7;refGene_tx=XM_023252264.1
A1 50300712 50300821
refGene_gn=LMO7;refGene_tx=XM_006927340.4;refGene_gn=LMO7;refGene_tx=X
M_006927341.4;refGene_gn=LMO7;refGene_tx=XM_019828007.2;refGene_gn=LMO
7;refGene_tx=XM_019828014.2;refGene_gn=LMO7;refGene_tx=XM_019828020.2;
refGene_gn=LMO7;refGene_tx=XM_019828022.2;refGene_gn=LMO7;refGene_tx=X
M_019828026.2;refGene_gn=LMO7;refGene_tx=XM_019828033.2;refGene_gn=LMO
7;refGene_tx=XM_019828041.2;refGene_gn=LMO7;refGene_tx=XM_019828047.2;
refGene_gn=LMO7;refGene_tx=XM_019828056.2;refGene_gn=LMO7;refGene_tx=X
M_019828067.2;refGene_gn=LMO7;refGene_tx=XM_023252243.1;refGene_gn=LMO
7;refGene_tx=XM_023252263.1;refGene_gn=LMO7;refGene_tx=XM_023252264.1
A1 50301934 50302037
refGene_gn=LMO7;refGene_tx=XM_006927340.4;refGene_gn=LMO7;refGene_tx=X
M_006927341.4;refGene_gn=LMO7;refGene_tx=XM_019828007.2;refGene_gn=LMO
7;refGene_tx=XM_019828014.2;refGene_gn=LMO7;refGene_tx=XM_019828020.2;
refGene_gn=LMO7;refGene_tx=XM_019828022.2;refGene_gn=LMO7;refGene_tx=X
M_019828026.2;refGene_gn=LMO7;refGene_tx=XM_019828033.2;refGene_gn=LMO
7;refGene_tx=XM_019828041.2;refGene_gn=LMO7;refGene_tx=XM_019828047.2;
refGene_gn=LMO7;refGene_tx=XM_019828051.2;refGene_gn=LMO7;refGene_tx=X
M_019828056.2;refGene_gn=LMO7;refGene_tx=XM_019828067.2;refGene_gn=LMO
7;refGene_tx=XM_023252243.1;refGene_gn=LMO7;refGene_tx=XM_023252263.1;
refGene_gn=LMO7;refGene_tx=XM_023252264.1
A1 50303364 50303457
refGene_gn=LMO7;refGene_tx=XM_006927340.4;refGene_gn=LMO7;refGene_tx=X
M_006927341.4;refGene_gn=LMO7;refGene_tx=XM_019828007.2;refGene_gn=LMO
7;refGene_tx=XM_019828014.2;refGene_gn=LMO7;refGene_tx=XM_019828020.2;
refGene_gn=LMO7;refGene_tx=XM_019828022.2;refGene_gn=LMO7;refGene_tx=X
M_019828026.2;refGene_gn=LMO7;refGene_tx=XM_019828033.2;refGene_gn=LMO

7;refGene_tx=XM_019828041.2;refGene_gn=LMO7;refGene_tx=XM_019828047.2;
refGene_gn=LMO7;refGene_tx=XM_019828051.2;refGene_gn=LMO7;refGene_tx=X
M_019828056.2;refGene_gn=LMO7;refGene_tx=XM_019828067.2;refGene_gn=LMO
7;refGene_tx=XM_023252243.1;refGene_gn=LMO7;refGene_tx=XM_023252263.1;
refGene_gn=LMO7;refGene_tx=XM_023252264.1
A1 51164092 51165076 refGene_gn=KCTD12;refGene_tx=XM_011280424.3
A1 51219987 51220047 refGene_gn=ACOD1;refGene_tx=XM_023238571.1
A1 51221774 51221936 refGene_gn=ACOD1;refGene_tx=XM_023238571.1
A1 51222712 51222802 refGene_gn=ACOD1;refGene_tx=XM_023238571.1
A1 51224056 51224262 refGene_gn=ACOD1;refGene_tx=XM_023238571.1
A1 51225171 51226117 refGene_gn=ACOD1;refGene_tx=XM_023238571.1
A1 51242873 51243553 refGene_gn=CLN5;refGene_tx=XM_023252371.1
A1 51247357 51247523 refGene_gn=CLN5;refGene_tx=XM_023252371.1
A1 51248191 51248287 refGene_gn=CLN5;refGene_tx=XM_023252371.1
A1 51248288 51248418 refGene_gn=CLN5;refGene_tx=XM_023252371.1
A1 51251483 51251995 refGene_gn=CLN5;refGene_tx=XM_023252371.1
A1 51258118 51258762
refGene_gn=FBXL3;refGene_tx=XM_019828194.2;refGene_gn=FBXL3;refGene_tx
=XM_023252376.1
A1 51265212 51265384
refGene_gn=FBXL3;refGene_tx=XM_019828194.2;refGene_gn=FBXL3;refGene_tx
=XM_023252376.1
A1 51268254 51268377
refGene_gn=FBXL3;refGene_tx=XM_019828194.2;refGene_gn=FBXL3;refGene_tx
=XM_023252376.1
A1 51270003 51270351
refGene_gn=FBXL3;refGene_tx=XM_019828194.2;refGene_gn=FBXL3;refGene_tx
=XM_023252376.1
A1 51289532 51289648 refGene_gn=MYCBP2;refGene_tx=XM_011280436.3
A1 51295455 51295621 refGene_gn=MYCBP2;refGene_tx=XM_011280436.3
A1 51296262 51296370 refGene_gn=MYCBP2;refGene_tx=XM_011280436.3
A1 51297781 51297991 refGene_gn=MYCBP2;refGene_tx=XM_011280436.3
A1 51299161 51299269 refGene_gn=MYCBP2;refGene_tx=XM_011280436.3
A1 51300067 51300256 refGene_gn=MYCBP2;refGene_tx=XM_011280436.3
A1 51301484 51301588 refGene_gn=MYCBP2;refGene_tx=XM_011280436.3
A1 51302775 51302908 refGene_gn=MYCBP2;refGene_tx=XM_011280436.3
A1 51303285 51303414 refGene_gn=MYCBP2;refGene_tx=XM_011280436.3
A1 51304216 51304318 refGene_gn=MYCBP2;refGene_tx=XM_011280436.3
A1 51305878 51305998 refGene_gn=MYCBP2;refGene_tx=XM_011280436.3
A1 51307794 51307891 refGene_gn=MYCBP2;refGene_tx=XM_011280436.3
A1 51309233 51309517 refGene_gn=MYCBP2;refGene_tx=XM_011280436.3
A1 51310243 51310510 refGene_gn=MYCBP2;refGene_tx=XM_011280436.3
A1 51311512 51311593 refGene_gn=MYCBP2;refGene_tx=XM_011280436.3
A1 51317611 51317710 refGene_gn=MYCBP2;refGene_tx=XM_011280436.3
A1 51318026 51318266 refGene_gn=MYCBP2;refGene_tx=XM_011280436.3
A1 51319582 51319648 refGene_gn=MYCBP2;refGene_tx=XM_011280436.3
A1 51322063 51322288 refGene_gn=MYCBP2;refGene_tx=XM_011280436.3
A1 51322489 51322646 refGene_gn=MYCBP2;refGene_tx=XM_011280436.3
A1 51323687 51323848 refGene_gn=MYCBP2;refGene_tx=XM_011280436.3
A1 51324789 51324939 refGene_gn=MYCBP2;refGene_tx=XM_011280436.3

| | | | |
|---|---|---|---|
| A1 | 51325793 | 51325993 | refGene_gn=MYCBP2;refGene_tx=XM_011280436.3 |
| A1 | 51327052 | 51327210 | refGene_gn=MYCBP2;refGene_tx=XM_011280436.3 |
| A1 | 51332051 | 51332219 | refGene_gn=MYCBP2;refGene_tx=XM_011280436.3 |
| A1 | 51334202 | 51334447 | refGene_gn=MYCBP2;refGene_tx=XM_011280436.3 |
| A1 | 51335208 | 51335378 | refGene_gn=MYCBP2;refGene_tx=XM_011280436.3 |
| A1 | 51336261 | 51337905 | refGene_gn=MYCBP2;refGene_tx=XM_011280436.3 |
| A1 | 51354384 | 51354507 | refGene_gn=MYCBP2;refGene_tx=XM_011280436.3 |
| A1 | 51358234 | 51358367 | refGene_gn=MYCBP2;refGene_tx=XM_011280436.3 |
| A1 | 51359234 | 51359459 | refGene_gn=MYCBP2;refGene_tx=XM_011280436.3 |
| A1 | 51370641 | 51370782 | refGene_gn=MYCBP2;refGene_tx=XM_011280436.3 |
| A1 | 51371469 | 51371586 | refGene_gn=MYCBP2;refGene_tx=XM_011280436.3 |
| A1 | 51372287 | 51372385 | refGene_gn=MYCBP2;refGene_tx=XM_011280436.3 |
| A1 | 51376378 | 51376494 | refGene_gn=MYCBP2;refGene_tx=XM_011280436.3 |
| A1 | 51379837 | 51379893 | refGene_gn=MYCBP2;refGene_tx=XM_011280436.3 |
| A1 | 51384451 | 51384667 | refGene_gn=MYCBP2;refGene_tx=XM_011280436.3 |
| A1 | 51394056 | 51394201 | refGene_gn=MYCBP2;refGene_tx=XM_011280436.3 |
| A1 | 51395670 | 51395843 | refGene_gn=MYCBP2;refGene_tx=XM_011280436.3 |
| A1 | 51397945 | 51397995 | refGene_gn=MYCBP2;refGene_tx=XM_011280436.3 |
| A1 | 51400764 | 51400852 | refGene_gn=MYCBP2;refGene_tx=XM_011280436.3 |
| A1 | 51401523 | 51401642 | refGene_gn=MYCBP2;refGene_tx=XM_011280436.3 |
| A1 | 51402670 | 51402896 | refGene_gn=MYCBP2;refGene_tx=XM_011280436.3 |
| A1 | 51408842 | 51409061 | refGene_gn=MYCBP2;refGene_tx=XM_011280436.3 |
| A1 | 51409723 | 51409824 | refGene_gn=MYCBP2;refGene_tx=XM_011280436.3 |
| A1 | 51411284 | 51411427 | refGene_gn=MYCBP2;refGene_tx=XM_011280436.3 |
| A1 | 51413044 | 51413223 | refGene_gn=MYCBP2;refGene_tx=XM_011280436.3 |
| A1 | 51414802 | 51414934 | refGene_gn=MYCBP2;refGene_tx=XM_011280436.3 |
| A1 | 51415768 | 51415975 | refGene_gn=MYCBP2;refGene_tx=XM_011280436.3 |
| A1 | 51418588 | 51418780 | refGene_gn=MYCBP2;refGene_tx=XM_011280436.3 |
| A1 | 51424731 | 51424953 | refGene_gn=MYCBP2;refGene_tx=XM_011280436.3 |
| A1 | 51428428 | 51428703 | refGene_gn=MYCBP2;refGene_tx=XM_011280436.3 |
| A1 | 51429205 | 51429398 | refGene_gn=MYCBP2;refGene_tx=XM_011280436.3 |
| A1 | 51430992 | 51431089 | refGene_gn=MYCBP2;refGene_tx=XM_011280436.3 |
| A1 | 51432103 | 51432187 | refGene_gn=MYCBP2;refGene_tx=XM_011280436.3 |
| A1 | 51433541 | 51433676 | refGene_gn=MYCBP2;refGene_tx=XM_011280436.3 |
| A1 | 51437056 | 51437148 | refGene_gn=MYCBP2;refGene_tx=XM_011280436.3 |
| A1 | 51443012 | 51443140 | refGene_gn=MYCBP2;refGene_tx=XM_011280436.3 |
| A1 | 51443226 | 51443352 | refGene_gn=MYCBP2;refGene_tx=XM_011280436.3 |
| A1 | 51444538 | 51444711 | refGene_gn=MYCBP2;refGene_tx=XM_011280436.3 |
| A1 | 51459490 | 51459644 | refGene_gn=MYCBP2;refGene_tx=XM_011280436.3 |
| A1 | 51460245 | 51460450 | refGene_gn=MYCBP2;refGene_tx=XM_011280436.3 |
| A1 | 51466076 | 51466194 | refGene_gn=MYCBP2;refGene_tx=XM_011280436.3 |
| A1 | 51474258 | 51474340 | refGene_gn=MYCBP2;refGene_tx=XM_011280436.3 |
| A1 | 51475620 | 51475740 | refGene_gn=MYCBP2;refGene_tx=XM_011280436.3 |
| A1 | 51482241 | 51482349 | refGene_gn=MYCBP2;refGene_tx=XM_011280436.3 |
| A1 | 51490074 | 51490176 | refGene_gn=MYCBP2;refGene_tx=XM_011280436.3 |
| A1 | 51490510 | 51490656 | refGene_gn=MYCBP2;refGene_tx=XM_011280436.3 |
| A1 | 51493173 | 51493378 | refGene_gn=MYCBP2;refGene_tx=XM_011280436.3 |
| A1 | 51498542 | 51498701 | refGene_gn=MYCBP2;refGene_tx=XM_011280436.3 |
| A1 | 51500971 | 51501136 | refGene_gn=MYCBP2;refGene_tx=XM_011280436.3 |
| A1 | 51502617 | 51502822 | refGene_gn=MYCBP2;refGene_tx=XM_011280436.3 |

A1 51503917 51503994 refGene_gn=MYCBP2;refGene_tx=XM_011280436.3
A1 51505537 51505676 refGene_gn=MYCBP2;refGene_tx=XM_011280436.3
A1 51505809 51505883 refGene_gn=MYCBP2;refGene_tx=XM_011280436.3
A1 51509834 51509931 refGene_gn=MYCBP2;refGene_tx=XM_011280436.3
A1 51511319 51511391 refGene_gn=MYCBP2;refGene_tx=XM_011280436.3
A1 51511590 51511833 refGene_gn=MYCBP2;refGene_tx=XM_011280436.3
A1 51520786 51520983 refGene_gn=MYCBP2;refGene_tx=XM_011280436.3
A1 51525053 51525207 refGene_gn=MYCBP2;refGene_tx=XM_011280436.3
A1 51533474 51533690 refGene_gn=MYCBP2;refGene_tx=XM_011280436.3
A1 51544790 51544866 refGene_gn=MYCBP2;refGene_tx=XM_011280436.3
A1 51573559 51573861 refGene_gn=MYCBP2;refGene_tx=XM_011280436.3
A1 51750675 51750703
refGene_gn=SCEL;refGene_tx=XM_019828134.1;refGene_gn=SCEL;refGene_tx=XM_019828136.1;refGene_gn=SCEL;refGene_tx=XM_019828156.1;refGene_gn=SCEL;refGene_tx=XM_019828157.1;refGene_gn=SCEL;refGene_tx=XM_019828162.1;refGene_gn=SCEL;refGene_tx=XM_019828164.1;refGene_gn=SCEL;refGene_tx=XM_019828175.1;refGene_gn=SCEL;refGene_tx=XM_019828178.1;refGene_gn=SCEL;refGene_tx=XM_019828184.2;refGene_gn=SCEL;refGene_tx=XM_019828188.1;refGene_gn=SCEL;refGene_tx=XM_019828190.1
A1 51798075 51798193
refGene_gn=SCEL;refGene_tx=XM_019828134.1;refGene_gn=SCEL;refGene_tx=XM_019828136.1;refGene_gn=SCEL;refGene_tx=XM_019828137.1;refGene_gn=SCEL;refGene_tx=XM_019828143.1;refGene_gn=SCEL;refGene_tx=XM_019828147.1;refGene_gn=SCEL;refGene_tx=XM_019828148.1;refGene_gn=SCEL;refGene_tx=XM_019828151.2;refGene_gn=SCEL;refGene_tx=XM_019828156.1;refGene_gn=SCEL;refGene_tx=XM_019828157.1;refGene_gn=SCEL;refGene_tx=XM_019828162.1;refGene_gn=SCEL;refGene_tx=XM_019828164.1;refGene_gn=SCEL;refGene_tx=XM_019828175.1;refGene_gn=SCEL;refGene_tx=XM_019828178.1;refGene_gn=SCEL;refGene_tx=XM_019828184.2;refGene_gn=SCEL;refGene_tx=XM_019828188.1;refGene_gn=SCEL;refGene_tx=XM_019828190.1;refGene_gn=SCEL;refGene_tx=XM_019828191.1
A1 51801341 51801401
refGene_gn=SCEL;refGene_tx=XM_019828134.1;refGene_gn=SCEL;refGene_tx=XM_019828136.1;refGene_gn=SCEL;refGene_tx=XM_019828137.1;refGene_gn=SCEL;refGene_tx=XM_019828143.1;refGene_gn=SCEL;refGene_tx=XM_019828147.1;refGene_gn=SCEL;refGene_tx=XM_019828148.1;refGene_gn=SCEL;refGene_tx=XM_019828151.2;refGene_gn=SCEL;refGene_tx=XM_019828156.1;refGene_gn=SCEL;refGene_tx=XM_019828157.1;refGene_gn=SCEL;refGene_tx=XM_019828162.1;refGene_gn=SCEL;refGene_tx=XM_019828164.1;refGene_gn=SCEL;refGene_tx=XM_019828175.1;refGene_gn=SCEL;refGene_tx=XM_019828178.1;refGene_gn=SCEL;refGene_tx=XM_019828184.2;refGene_gn=SCEL;refGene_tx=XM_019828188.1;refGene_gn=SCEL;refGene_tx=XM_019828190.1;refGene_gn=SCEL;refGene_tx=XM_019828191.1
A1 51805641 51805710
refGene_gn=SCEL;refGene_tx=XM_019828134.1;refGene_gn=SCEL;refGene_tx=XM_019828136.1;refGene_gn=SCEL;refGene_tx=XM_019828137.1;refGene_gn=SCEL;refGene_tx=XM_019828143.1;refGene_gn=SCEL;refGene_tx=XM_019828147.1;refGene_gn=SCEL;refGene_tx=XM_019828148.1;refGene_gn=SCEL;refGene_tx=XM_019828151.2;refGene_gn=SCEL;refGene_tx=XM_019828156.1;refGene_gn=SCEL;refGene_tx=XM_019828157.1;refGene_gn=SCEL;refGene_tx=XM_019828162.1;

refGene_gn=SCEL;refGene_tx=XM_019828164.1;refGene_gn=SCEL;refGene_tx=XM_019828175.1;refGene_gn=SCEL;refGene_tx=XM_019828178.1;refGene_gn=SCEL;refGene_tx=XM_019828184.2;refGene_gn=SCEL;refGene_tx=XM_019828188.1;refGene_gn=SCEL;refGene_tx=XM_019828190.1;refGene_gn=SCEL;refGene_tx=XM_019828191.1
A1 51810835 51810904
refGene_gn=SCEL;refGene_tx=XM_019828134.1;refGene_gn=SCEL;refGene_tx=XM_019828136.1;refGene_gn=SCEL;refGene_tx=XM_019828137.1;refGene_gn=SCEL;refGene_tx=XM_019828143.1;refGene_gn=SCEL;refGene_tx=XM_019828147.1;refGene_gn=SCEL;refGene_tx=XM_019828148.1;refGene_gn=SCEL;refGene_tx=XM_019828151.2;refGene_gn=SCEL;refGene_tx=XM_019828156.1;refGene_gn=SCEL;refGene_tx=XM_019828157.1;refGene_gn=SCEL;refGene_tx=XM_019828162.1;refGene_gn=SCEL;refGene_tx=XM_019828164.1;refGene_gn=SCEL;refGene_tx=XM_019828178.1;refGene_gn=SCEL;refGene_tx=XM_019828184.2;refGene_gn=SCEL;refGene_tx=XM_019828188.1;refGene_gn=SCEL;refGene_tx=XM_019828190.1;refGene_gn=SCEL;refGene_tx=XM_019828191.1
A1 51811500 51811539
refGene_gn=SCEL;refGene_tx=XM_019828134.1;refGene_gn=SCEL;refGene_tx=XM_019828136.1;refGene_gn=SCEL;refGene_tx=XM_019828137.1;refGene_gn=SCEL;refGene_tx=XM_019828143.1;refGene_gn=SCEL;refGene_tx=XM_019828147.1;refGene_gn=SCEL;refGene_tx=XM_019828148.1;refGene_gn=SCEL;refGene_tx=XM_019828151.2;refGene_gn=SCEL;refGene_tx=XM_019828156.1;refGene_gn=SCEL;refGene_tx=XM_019828157.1;refGene_gn=SCEL;refGene_tx=XM_019828162.1;refGene_gn=SCEL;refGene_tx=XM_019828164.1;refGene_gn=SCEL;refGene_tx=XM_019828175.1;refGene_gn=SCEL;refGene_tx=XM_019828178.1;refGene_gn=SCEL;refGene_tx=XM_019828184.2;refGene_gn=SCEL;refGene_tx=XM_019828188.1;refGene_gn=SCEL;refGene_tx=XM_019828190.1;refGene_gn=SCEL;refGene_tx=XM_019828191.1
A1 51812830 51812911
refGene_gn=SCEL;refGene_tx=XM_019828134.1;refGene_gn=SCEL;refGene_tx=XM_019828136.1;refGene_gn=SCEL;refGene_tx=XM_019828137.1;refGene_gn=SCEL;refGene_tx=XM_019828143.1;refGene_gn=SCEL;refGene_tx=XM_019828147.1;refGene_gn=SCEL;refGene_tx=XM_019828148.1;refGene_gn=SCEL;refGene_tx=XM_019828151.2;refGene_gn=SCEL;refGene_tx=XM_019828156.1;refGene_gn=SCEL;refGene_tx=XM_019828157.1;refGene_gn=SCEL;refGene_tx=XM_019828162.1;refGene_gn=SCEL;refGene_tx=XM_019828164.1;refGene_gn=SCEL;refGene_tx=XM_019828175.1;refGene_gn=SCEL;refGene_tx=XM_019828178.1;refGene_gn=SCEL;refGene_tx=XM_019828184.2;refGene_gn=SCEL;refGene_tx=XM_019828188.1;refGene_gn=SCEL;refGene_tx=XM_019828190.1;refGene_gn=SCEL;refGene_tx=XM_019828191.1
A1 51814739 51814805
refGene_gn=SCEL;refGene_tx=XM_019828134.1;refGene_gn=SCEL;refGene_tx=XM_019828136.1;refGene_gn=SCEL;refGene_tx=XM_019828137.1;refGene_gn=SCEL;refGene_tx=XM_019828143.1;refGene_gn=SCEL;refGene_tx=XM_019828147.1;refGene_gn=SCEL;refGene_tx=XM_019828148.1;refGene_gn=SCEL;refGene_tx=XM_019828151.2;refGene_gn=SCEL;refGene_tx=XM_019828156.1;refGene_gn=SCEL;refGene_tx=XM_019828157.1;refGene_gn=SCEL;refGene_tx=XM_019828162.1;refGene_gn=SCEL;refGene_tx=XM_019828164.1;refGene_gn=SCEL;refGene_tx=XM_019828175.1;refGene_gn=SCEL;refGene_tx=XM_019828178.1;refGene_gn=SCEL;refGene_tx=XM_019828184.2;refGene_gn=SCEL;refGene_tx=XM_019828188.1;

refGene_gn=SCEL;refGene_tx=XM_019828190.1;refGene_gn=SCEL;refGene_tx=X
M_019828191.1
A1 51831880 51831961
refGene_gn=SCEL;refGene_tx=XM_019828134.1;refGene_gn=SCEL;refGene_tx=X
M_019828136.1;refGene_gn=SCEL;refGene_tx=XM_019828137.1;refGene_gn=SCE
L;refGene_tx=XM_019828143.1;refGene_gn=SCEL;refGene_tx=XM_019828147.1;
refGene_gn=SCEL;refGene_tx=XM_019828148.1;refGene_gn=SCEL;refGene_tx=X
M_019828151.2;refGene_gn=SCEL;refGene_tx=XM_019828156.1;refGene_gn=SCE
L;refGene_tx=XM_019828157.1;refGene_gn=SCEL;refGene_tx=XM_019828162.1;
refGene_gn=SCEL;refGene_tx=XM_019828164.1;refGene_gn=SCEL;refGene_tx=X
M_019828175.1;refGene_gn=SCEL;refGene_tx=XM_019828178.1;refGene_gn=SCE
L;refGene_tx=XM_019828184.2;refGene_gn=SCEL;refGene_tx=XM_019828188.1;
refGene_gn=SCEL;refGene_tx=XM_019828190.1;refGene_gn=SCEL;refGene_tx=X
M_019828191.1
A1 51834357 51834423
refGene_gn=SCEL;refGene_tx=XM_019828134.1;refGene_gn=SCEL;refGene_tx=X
M_019828136.1;refGene_gn=SCEL;refGene_tx=XM_019828137.1;refGene_gn=SCE
L;refGene_tx=XM_019828143.1;refGene_gn=SCEL;refGene_tx=XM_019828147.1;
refGene_gn=SCEL;refGene_tx=XM_019828148.1;refGene_gn=SCEL;refGene_tx=X
M_019828151.2;refGene_gn=SCEL;refGene_tx=XM_019828156.1;refGene_gn=SCE
L;refGene_tx=XM_019828157.1;refGene_gn=SCEL;refGene_tx=XM_019828162.1;
refGene_gn=SCEL;refGene_tx=XM_019828164.1;refGene_gn=SCEL;refGene_tx=X
M_019828175.1;refGene_gn=SCEL;refGene_tx=XM_019828178.1;refGene_gn=SCE
L;refGene_tx=XM_019828184.2;refGene_gn=SCEL;refGene_tx=XM_019828188.1;
refGene_gn=SCEL;refGene_tx=XM_019828190.1;refGene_gn=SCEL;refGene_tx=X
M_019828191.1
A1 51837020 51837080
refGene_gn=SCEL;refGene_tx=XM_019828134.1;refGene_gn=SCEL;refGene_tx=X
M_019828136.1;refGene_gn=SCEL;refGene_tx=XM_019828137.1;refGene_gn=SCE
L;refGene_tx=XM_019828143.1;refGene_gn=SCEL;refGene_tx=XM_019828147.1;
refGene_gn=SCEL;refGene_tx=XM_019828148.1;refGene_gn=SCEL;refGene_tx=X
M_019828151.2;refGene_gn=SCEL;refGene_tx=XM_019828156.1;refGene_gn=SCE
L;refGene_tx=XM_019828157.1;refGene_gn=SCEL;refGene_tx=XM_019828164.1;
refGene_gn=SCEL;refGene_tx=XM_019828175.1;refGene_gn=SCEL;refGene_tx=X
M_019828178.1;refGene_gn=SCEL;refGene_tx=XM_019828188.1;refGene_gn=SCE
L;refGene_tx=XM_019828190.1;refGene_gn=SCEL;refGene_tx=XM_019828191.1
A1 51841067 51841112
refGene_gn=SCEL;refGene_tx=XM_019828134.1;refGene_gn=SCEL;refGene_tx=X
M_019828136.1;refGene_gn=SCEL;refGene_tx=XM_019828137.1;refGene_gn=SCE
L;refGene_tx=XM_019828143.1;refGene_gn=SCEL;refGene_tx=XM_019828147.1;
refGene_gn=SCEL;refGene_tx=XM_019828148.1;refGene_gn=SCEL;refGene_tx=X
M_019828151.2;refGene_gn=SCEL;refGene_tx=XM_019828156.1;refGene_gn=SCE
L;refGene_tx=XM_019828157.1;refGene_gn=SCEL;refGene_tx=XM_019828162.1;
refGene_gn=SCEL;refGene_tx=XM_019828164.1;refGene_gn=SCEL;refGene_tx=X
M_019828175.1;refGene_gn=SCEL;refGene_tx=XM_019828178.1;refGene_gn=SCE
L;refGene_tx=XM_019828188.1;refGene_gn=SCEL;refGene_tx=XM_019828190.1;
refGene_gn=SCEL;refGene_tx=XM_019828191.1
A1 51842436 51842496
refGene_gn=SCEL;refGene_tx=XM_019828134.1;refGene_gn=SCEL;refGene_tx=X
M_019828136.1;refGene_gn=SCEL;refGene_tx=XM_019828137.1;refGene_gn=SCE

L;refGene_tx=XM_019828143.1;refGene_gn=SCEL;refGene_tx=XM_019828147.1;
refGene_gn=SCEL;refGene_tx=XM_019828148.1;refGene_gn=SCEL;refGene_tx=X
M_019828151.2;refGene_gn=SCEL;refGene_tx=XM_019828156.1;refGene_gn=SCE
L;refGene_tx=XM_019828157.1;refGene_gn=SCEL;refGene_tx=XM_019828162.1;
refGene_gn=SCEL;refGene_tx=XM_019828164.1;refGene_gn=SCEL;refGene_tx=X
M_019828175.1;refGene_gn=SCEL;refGene_tx=XM_019828178.1;refGene_gn=SCE
L;refGene_tx=XM_019828184.2;refGene_gn=SCEL;refGene_tx=XM_019828188.1;
refGene_gn=SCEL;refGene_tx=XM_019828190.1;refGene_gn=SCEL;refGene_tx=X
M_019828191.1
A1 51842798 51842858
refGene_gn=SCEL;refGene_tx=XM_019828134.1;refGene_gn=SCEL;refGene_tx=X
M_019828136.1;refGene_gn=SCEL;refGene_tx=XM_019828137.1;refGene_gn=SCE
L;refGene_tx=XM_019828143.1;refGene_gn=SCEL;refGene_tx=XM_019828147.1;
refGene_gn=SCEL;refGene_tx=XM_019828148.1;refGene_gn=SCEL;refGene_tx=X
M_019828151.2;refGene_gn=SCEL;refGene_tx=XM_019828156.1;refGene_gn=SCE
L;refGene_tx=XM_019828157.1;refGene_gn=SCEL;refGene_tx=XM_019828162.1;
refGene_gn=SCEL;refGene_tx=XM_019828164.1;refGene_gn=SCEL;refGene_tx=X
M_019828175.1;refGene_gn=SCEL;refGene_tx=XM_019828178.1;refGene_gn=SCE
L;refGene_tx=XM_019828184.2;refGene_gn=SCEL;refGene_tx=XM_019828188.1;
refGene_gn=SCEL;refGene_tx=XM_019828190.1;refGene_gn=SCEL;refGene_tx=X
M_019828191.1
A1 51843497 51843557
refGene_gn=SCEL;refGene_tx=XM_019828134.1;refGene_gn=SCEL;refGene_tx=X
M_019828136.1;refGene_gn=SCEL;refGene_tx=XM_019828137.1;refGene_gn=SCE
L;refGene_tx=XM_019828143.1;refGene_gn=SCEL;refGene_tx=XM_019828147.1;
refGene_gn=SCEL;refGene_tx=XM_019828148.1;refGene_gn=SCEL;refGene_tx=X
M_019828151.2;refGene_gn=SCEL;refGene_tx=XM_019828156.1;refGene_gn=SCE
L;refGene_tx=XM_019828157.1;refGene_gn=SCEL;refGene_tx=XM_019828162.1;
refGene_gn=SCEL;refGene_tx=XM_019828164.1;refGene_gn=SCEL;refGene_tx=X
M_019828175.1;refGene_gn=SCEL;refGene_tx=XM_019828178.1;refGene_gn=SCE
L;refGene_tx=XM_019828184.2;refGene_gn=SCEL;refGene_tx=XM_019828188.1;
refGene_gn=SCEL;refGene_tx=XM_019828190.1;refGene_gn=SCEL;refGene_tx=X
M_019828191.1
A1 51846121 51846181
refGene_gn=SCEL;refGene_tx=XM_019828134.1;refGene_gn=SCEL;refGene_tx=X
M_019828136.1;refGene_gn=SCEL;refGene_tx=XM_019828137.1;refGene_gn=SCE
L;refGene_tx=XM_019828143.1;refGene_gn=SCEL;refGene_tx=XM_019828147.1;
refGene_gn=SCEL;refGene_tx=XM_019828148.1;refGene_gn=SCEL;refGene_tx=X
M_019828151.2;refGene_gn=SCEL;refGene_tx=XM_019828156.1;refGene_gn=SCE
L;refGene_tx=XM_019828157.1;refGene_gn=SCEL;refGene_tx=XM_019828162.1;
refGene_gn=SCEL;refGene_tx=XM_019828164.1;refGene_gn=SCEL;refGene_tx=X
M_019828175.1;refGene_gn=SCEL;refGene_tx=XM_019828178.1;refGene_gn=SCE
L;refGene_tx=XM_019828184.2;refGene_gn=SCEL;refGene_tx=XM_019828188.1;
refGene_gn=SCEL;refGene_tx=XM_019828190.1;refGene_gn=SCEL;refGene_tx=X
M_019828191.1
A1 51847126 51847186
refGene_gn=SCEL;refGene_tx=XM_019828134.1;refGene_gn=SCEL;refGene_tx=X
M_019828136.1;refGene_gn=SCEL;refGene_tx=XM_019828137.1;refGene_gn=SCE
L;refGene_tx=XM_019828143.1;refGene_gn=SCEL;refGene_tx=XM_019828147.1;
refGene_gn=SCEL;refGene_tx=XM_019828148.1;refGene_gn=SCEL;refGene_tx=X

M_019828151.2;refGene_gn=SCEL;refGene_tx=XM_019828156.1;refGene_gn=SCE
L;refGene_tx=XM_019828157.1;refGene_gn=SCEL;refGene_tx=XM_019828162.1;
refGene_gn=SCEL;refGene_tx=XM_019828164.1;refGene_gn=SCEL;refGene_tx=X
M_019828175.1;refGene_gn=SCEL;refGene_tx=XM_019828178.1;refGene_gn=SCE
L;refGene_tx=XM_019828184.2;refGene_gn=SCEL;refGene_tx=XM_019828188.1;
refGene_gn=SCEL;refGene_tx=XM_019828190.1;refGene_gn=SCEL;refGene_tx=X
M_019828191.1
A1 51848162 51848222
refGene_gn=SCEL;refGene_tx=XM_019828134.1;refGene_gn=SCEL;refGene_tx=X
M_019828136.1;refGene_gn=SCEL;refGene_tx=XM_019828137.1;refGene_gn=SCE
L;refGene_tx=XM_019828143.1;refGene_gn=SCEL;refGene_tx=XM_019828147.1;
refGene_gn=SCEL;refGene_tx=XM_019828148.1;refGene_gn=SCEL;refGene_tx=X
M_019828151.2;refGene_gn=SCEL;refGene_tx=XM_019828156.1;refGene_gn=SCE
L;refGene_tx=XM_019828157.1;refGene_gn=SCEL;refGene_tx=XM_019828162.1;
refGene_gn=SCEL;refGene_tx=XM_019828164.1;refGene_gn=SCEL;refGene_tx=X
M_019828175.1;refGene_gn=SCEL;refGene_tx=XM_019828178.1;refGene_gn=SCE
L;refGene_tx=XM_019828184.2;refGene_gn=SCEL;refGene_tx=XM_019828188.1;
refGene_gn=SCEL;refGene_tx=XM_019828190.1;refGene_gn=SCEL;refGene_tx=X
M_019828191.1
A1 51853078 51853138
refGene_gn=SCEL;refGene_tx=XM_019828134.1;refGene_gn=SCEL;refGene_tx=X
M_019828136.1;refGene_gn=SCEL;refGene_tx=XM_019828137.1;refGene_gn=SCE
L;refGene_tx=XM_019828143.1;refGene_gn=SCEL;refGene_tx=XM_019828147.1;
refGene_gn=SCEL;refGene_tx=XM_019828148.1;refGene_gn=SCEL;refGene_tx=X
M_019828151.2;refGene_gn=SCEL;refGene_tx=XM_019828157.1;refGene_gn=SCE
L;refGene_tx=XM_019828162.1;refGene_gn=SCEL;refGene_tx=XM_019828164.1;
refGene_gn=SCEL;refGene_tx=XM_019828175.1;refGene_gn=SCEL;refGene_tx=X
M_019828178.1;refGene_gn=SCEL;refGene_tx=XM_019828184.2;refGene_gn=SCE
L;refGene_tx=XM_019828191.1
A1 51853766 51853826
refGene_gn=SCEL;refGene_tx=XM_019828134.1;refGene_gn=SCEL;refGene_tx=X
M_019828136.1;refGene_gn=SCEL;refGene_tx=XM_019828137.1;refGene_gn=SCE
L;refGene_tx=XM_019828143.1;refGene_gn=SCEL;refGene_tx=XM_019828147.1;
refGene_gn=SCEL;refGene_tx=XM_019828148.1;refGene_gn=SCEL;refGene_tx=X
M_019828151.2;refGene_gn=SCEL;refGene_tx=XM_019828156.1;refGene_gn=SCE
L;refGene_tx=XM_019828157.1;refGene_gn=SCEL;refGene_tx=XM_019828162.1;
refGene_gn=SCEL;refGene_tx=XM_019828164.1;refGene_gn=SCEL;refGene_tx=X
M_019828175.1;refGene_gn=SCEL;refGene_tx=XM_019828178.1;refGene_gn=SCE
L;refGene_tx=XM_019828184.2;refGene_gn=SCEL;refGene_tx=XM_019828188.1;
refGene_gn=SCEL;refGene_tx=XM_019828190.1;refGene_gn=SCEL;refGene_tx=X
M_019828191.1
A1 51859701 51859761
refGene_gn=SCEL;refGene_tx=XM_019828134.1;refGene_gn=SCEL;refGene_tx=X
M_019828136.1;refGene_gn=SCEL;refGene_tx=XM_019828137.1;refGene_gn=SCE
L;refGene_tx=XM_019828143.1;refGene_gn=SCEL;refGene_tx=XM_019828147.1;
refGene_gn=SCEL;refGene_tx=XM_019828148.1;refGene_gn=SCEL;refGene_tx=X
M_019828151.2;refGene_gn=SCEL;refGene_tx=XM_019828156.1;refGene_gn=SCE
L;refGene_tx=XM_019828162.1;refGene_gn=SCEL;refGene_tx=XM_019828175.1;
refGene_gn=SCEL;refGene_tx=XM_019828178.1;refGene_gn=SCEL;refGene_tx=X
M_019828184.2;refGene_gn=SCEL;refGene_tx=XM_019828191.1

A1 51861851 51861914
refGene_gn=SCEL;refGene_tx=XM_019828134.1;refGene_gn=SCEL;refGene_tx=XM_019828136.1;refGene_gn=SCEL;refGene_tx=XM_019828137.1;refGene_gn=SCEL;refGene_tx=XM_019828143.1;refGene_gn=SCEL;refGene_tx=XM_019828147.1;refGene_gn=SCEL;refGene_tx=XM_019828148.1;refGene_gn=SCEL;refGene_tx=XM_019828151.2;refGene_gn=SCEL;refGene_tx=XM_019828156.1;refGene_gn=SCEL;refGene_tx=XM_019828157.1;refGene_gn=SCEL;refGene_tx=XM_019828162.1;refGene_gn=SCEL;refGene_tx=XM_019828164.1;refGene_gn=SCEL;refGene_tx=XM_019828175.1;refGene_gn=SCEL;refGene_tx=XM_019828178.1;refGene_gn=SCEL;refGene_tx=XM_019828184.2;refGene_gn=SCEL;refGene_tx=XM_019828188.1;refGene_gn=SCEL;refGene_tx=XM_019828190.1;refGene_gn=SCEL;refGene_tx=XM_019828191.1
A1 51866033 51866093
refGene_gn=SCEL;refGene_tx=XM_019828134.1;refGene_gn=SCEL;refGene_tx=XM_019828136.1;refGene_gn=SCEL;refGene_tx=XM_019828137.1;refGene_gn=SCEL;refGene_tx=XM_019828143.1;refGene_gn=SCEL;refGene_tx=XM_019828147.1;refGene_gn=SCEL;refGene_tx=XM_019828148.1;refGene_gn=SCEL;refGene_tx=XM_019828151.2;refGene_gn=SCEL;refGene_tx=XM_019828156.1;refGene_gn=SCEL;refGene_tx=XM_019828157.1;refGene_gn=SCEL;refGene_tx=XM_019828162.1;refGene_gn=SCEL;refGene_tx=XM_019828164.1;refGene_gn=SCEL;refGene_tx=XM_019828175.1;refGene_gn=SCEL;refGene_tx=XM_019828178.1;refGene_gn=SCEL;refGene_tx=XM_019828184.2;refGene_gn=SCEL;refGene_tx=XM_019828188.1;refGene_gn=SCEL;refGene_tx=XM_019828190.1;refGene_gn=SCEL;refGene_tx=XM_019828191.1
A1 51866236 51866296
refGene_gn=SCEL;refGene_tx=XM_019828134.1;refGene_gn=SCEL;refGene_tx=XM_019828136.1;refGene_gn=SCEL;refGene_tx=XM_019828137.1;refGene_gn=SCEL;refGene_tx=XM_019828143.1;refGene_gn=SCEL;refGene_tx=XM_019828147.1;refGene_gn=SCEL;refGene_tx=XM_019828148.1;refGene_gn=SCEL;refGene_tx=XM_019828151.2;refGene_gn=SCEL;refGene_tx=XM_019828156.1;refGene_gn=SCEL;refGene_tx=XM_019828157.1;refGene_gn=SCEL;refGene_tx=XM_019828162.1;refGene_gn=SCEL;refGene_tx=XM_019828164.1;refGene_gn=SCEL;refGene_tx=XM_019828175.1;refGene_gn=SCEL;refGene_tx=XM_019828178.1;refGene_gn=SCEL;refGene_tx=XM_019828184.2;refGene_gn=SCEL;refGene_tx=XM_019828188.1;refGene_gn=SCEL;refGene_tx=XM_019828190.1;refGene_gn=SCEL;refGene_tx=XM_019828191.1
A1 51866439 51866496
refGene_gn=SCEL;refGene_tx=XM_019828134.1;refGene_gn=SCEL;refGene_tx=XM_019828136.1;refGene_gn=SCEL;refGene_tx=XM_019828137.1;refGene_gn=SCEL;refGene_tx=XM_019828143.1;refGene_gn=SCEL;refGene_tx=XM_019828147.1;refGene_gn=SCEL;refGene_tx=XM_019828148.1;refGene_gn=SCEL;refGene_tx=XM_019828151.2;refGene_gn=SCEL;refGene_tx=XM_019828156.1;refGene_gn=SCEL;refGene_tx=XM_019828157.1;refGene_gn=SCEL;refGene_tx=XM_019828162.1;refGene_gn=SCEL;refGene_tx=XM_019828164.1;refGene_gn=SCEL;refGene_tx=XM_019828175.1;refGene_gn=SCEL;refGene_tx=XM_019828178.1;refGene_gn=SCEL;refGene_tx=XM_019828184.2;refGene_gn=SCEL;refGene_tx=XM_019828188.1;refGene_gn=SCEL;refGene_tx=XM_019828190.1;refGene_gn=SCEL;refGene_tx=XM_019828191.1
A1 51874451 51874514
refGene_gn=SCEL;refGene_tx=XM_019828134.1;refGene_gn=SCEL;refGene_tx=X

M_019828136.1;refGene_gn=SCEL;refGene_tx=XM_019828137.1;refGene_gn=SCE
L;refGene_tx=XM_019828143.1;refGene_gn=SCEL;refGene_tx=XM_019828147.1;
refGene_gn=SCEL;refGene_tx=XM_019828148.1;refGene_gn=SCEL;refGene_tx=X
M_019828151.2;refGene_gn=SCEL;refGene_tx=XM_019828156.1;refGene_gn=SCE
L;refGene_tx=XM_019828157.1;refGene_gn=SCEL;refGene_tx=XM_019828162.1;
refGene_gn=SCEL;refGene_tx=XM_019828164.1;refGene_gn=SCEL;refGene_tx=X
M_019828175.1;refGene_gn=SCEL;refGene_tx=XM_019828178.1;refGene_gn=SCE
L;refGene_tx=XM_019828184.2;refGene_gn=SCEL;refGene_tx=XM_019828188.1;
refGene_gn=SCEL;refGene_tx=XM_019828190.1;refGene_gn=SCEL;refGene_tx=X
M_019828191.1
A1 51880142 51880235
refGene_gn=SCEL;refGene_tx=XM_019828134.1;refGene_gn=SCEL;refGene_tx=X
M_019828136.1;refGene_gn=SCEL;refGene_tx=XM_019828137.1;refGene_gn=SCE
L;refGene_tx=XM_019828143.1;refGene_gn=SCEL;refGene_tx=XM_019828147.1;
refGene_gn=SCEL;refGene_tx=XM_019828148.1;refGene_gn=SCEL;refGene_tx=X
M_019828151.2;refGene_gn=SCEL;refGene_tx=XM_019828156.1;refGene_gn=SCE
L;refGene_tx=XM_019828157.1;refGene_gn=SCEL;refGene_tx=XM_019828162.1;
refGene_gn=SCEL;refGene_tx=XM_019828164.1;refGene_gn=SCEL;refGene_tx=X
M_019828175.1;refGene_gn=SCEL;refGene_tx=XM_019828178.1;refGene_gn=SCE
L;refGene_tx=XM_019828184.2;refGene_gn=SCEL;refGene_tx=XM_019828188.1;
refGene_gn=SCEL;refGene_tx=XM_019828190.1;refGene_gn=SCEL;refGene_tx=X
M_019828191.1
A1 51882103 51882175
refGene_gn=SCEL;refGene_tx=XM_019828134.1;refGene_gn=SCEL;refGene_tx=X
M_019828136.1;refGene_gn=SCEL;refGene_tx=XM_019828137.1;refGene_gn=SCE
L;refGene_tx=XM_019828143.1;refGene_gn=SCEL;refGene_tx=XM_019828147.1;
refGene_gn=SCEL;refGene_tx=XM_019828148.1;refGene_gn=SCEL;refGene_tx=X
M_019828151.2;refGene_gn=SCEL;refGene_tx=XM_019828156.1;refGene_gn=SCE
L;refGene_tx=XM_019828157.1;refGene_gn=SCEL;refGene_tx=XM_019828162.1;
refGene_gn=SCEL;refGene_tx=XM_019828164.1;refGene_gn=SCEL;refGene_tx=X
M_019828175.1;refGene_gn=SCEL;refGene_tx=XM_019828178.1;refGene_gn=SCE
L;refGene_tx=XM_019828184.2;refGene_gn=SCEL;refGene_tx=XM_019828188.1;
refGene_gn=SCEL;refGene_tx=XM_019828190.1;refGene_gn=SCEL;refGene_tx=X
M_019828191.1
A1 51884145 51884220
refGene_gn=SCEL;refGene_tx=XM_019828134.1;refGene_gn=SCEL;refGene_tx=X
M_019828136.1;refGene_gn=SCEL;refGene_tx=XM_019828137.1;refGene_gn=SCE
L;refGene_tx=XM_019828143.1;refGene_gn=SCEL;refGene_tx=XM_019828147.1;
refGene_gn=SCEL;refGene_tx=XM_019828148.1;refGene_gn=SCEL;refGene_tx=X
M_019828151.2;refGene_gn=SCEL;refGene_tx=XM_019828156.1;refGene_gn=SCE
L;refGene_tx=XM_019828157.1;refGene_gn=SCEL;refGene_tx=XM_019828162.1;
refGene_gn=SCEL;refGene_tx=XM_019828164.1;refGene_gn=SCEL;refGene_tx=X
M_019828175.1;refGene_gn=SCEL;refGene_tx=XM_019828184.2;refGene_gn=SCE
L;refGene_tx=XM_019828188.1;refGene_gn=SCEL;refGene_tx=XM_019828190.1;
refGene_gn=SCEL;refGene_tx=XM_019828191.1
A1 51888385 51888494
refGene_gn=SCEL;refGene_tx=XM_019828134.1;refGene_gn=SCEL;refGene_tx=X
M_019828136.1;refGene_gn=SCEL;refGene_tx=XM_019828137.1;refGene_gn=SCE
L;refGene_tx=XM_019828143.1;refGene_gn=SCEL;refGene_tx=XM_019828147.1;
refGene_gn=SCEL;refGene_tx=XM_019828148.1;refGene_gn=SCEL;refGene_tx=X

M_019828151.2;refGene_gn=SCEL;refGene_tx=XM_019828156.1;refGene_gn=SCE
L;refGene_tx=XM_019828157.1;refGene_gn=SCEL;refGene_tx=XM_019828162.1;
refGene_gn=SCEL;refGene_tx=XM_019828164.1;refGene_gn=SCEL;refGene_tx=X
M_019828175.1;refGene_gn=SCEL;refGene_tx=XM_019828178.1;refGene_gn=SCE
L;refGene_tx=XM_019828184.2;refGene_gn=SCEL;refGene_tx=XM_019828188.1;
refGene_gn=SCEL;refGene_tx=XM_019828190.1;refGene_gn=SCEL;refGene_tx=X
M_019828191.1
A1 51890466 51890569
refGene_gn=SCEL;refGene_tx=XM_019828134.1;refGene_gn=SCEL;refGene_tx=X
M_019828136.1;refGene_gn=SCEL;refGene_tx=XM_019828137.1;refGene_gn=SCE
L;refGene_tx=XM_019828143.1;refGene_gn=SCEL;refGene_tx=XM_019828147.1;
refGene_gn=SCEL;refGene_tx=XM_019828148.1;refGene_gn=SCEL;refGene_tx=X
M_019828151.2;refGene_gn=SCEL;refGene_tx=XM_019828156.1;refGene_gn=SCE
L;refGene_tx=XM_019828157.1;refGene_gn=SCEL;refGene_tx=XM_019828162.1;
refGene_gn=SCEL;refGene_tx=XM_019828164.1;refGene_gn=SCEL;refGene_tx=X
M_019828175.1;refGene_gn=SCEL;refGene_tx=XM_019828178.1;refGene_gn=SCE
L;refGene_tx=XM_019828184.2;refGene_gn=SCEL;refGene_tx=XM_019828188.1;
refGene_gn=SCEL;refGene_tx=XM_019828190.1;refGene_gn=SCEL;refGene_tx=X
M_019828191.1
A1 51892013 51892030
refGene_gn=SCEL;refGene_tx=XM_019828134.1;refGene_gn=SCEL;refGene_tx=X
M_019828136.1;refGene_gn=SCEL;refGene_tx=XM_019828137.1;refGene_gn=SCE
L;refGene_tx=XM_019828143.1;refGene_gn=SCEL;refGene_tx=XM_019828147.1;
refGene_gn=SCEL;refGene_tx=XM_019828148.1;refGene_gn=SCEL;refGene_tx=X
M_019828151.2;refGene_gn=SCEL;refGene_tx=XM_019828156.1;refGene_gn=SCE
L;refGene_tx=XM_019828157.1;refGene_gn=SCEL;refGene_tx=XM_019828162.1;
refGene_gn=SCEL;refGene_tx=XM_019828164.1;refGene_gn=SCEL;refGene_tx=X
M_019828175.1;refGene_gn=SCEL;refGene_tx=XM_019828178.1;refGene_gn=SCE
L;refGene_tx=XM_019828184.2;refGene_gn=SCEL;refGene_tx=XM_019828188.1;
refGene_gn=SCEL;refGene_tx=XM_019828190.1;refGene_gn=SCEL;refGene_tx=X
M_019828191.1
A1 51922196 51922819
refGene_gn=SLAIN1;refGene_tx=XM_023252403.1;refGene_gn=SLAIN1;refGene_
tx=XM_023252405.1
A1 51936355 51936495
refGene_gn=SLAIN1;refGene_tx=XM_023252403.1;refGene_gn=SLAIN1;refGene_
tx=XM_023252405.1
A1 51964901 51965051 refGene_gn=SLAIN1;refGene_tx=XM_023252403.1
A1 51966190 51966340
refGene_gn=SLAIN1;refGene_tx=XM_023252403.1;refGene_gn=SLAIN1;refGene_
tx=XM_023252405.1
A1 51968406 51968748
refGene_gn=SLAIN1;refGene_tx=XM_023252403.1;refGene_gn=SLAIN1;refGene_
tx=XM_023252405.1
A1 51975398 51975554
refGene_gn=SLAIN1;refGene_tx=XM_023252403.1;refGene_gn=SLAIN1;refGene_
tx=XM_023252405.1
A1 51980763 51981046
refGene_gn=SLAIN1;refGene_tx=XM_023252403.1;refGene_gn=SLAIN1;refGene_
tx=XM_023252405.1

A1 51982749 51982825
refGene_gn=SLAIN1;refGene_tx=XM_023252403.1;refGene_gn=SLAIN1;refGene_tx=XM_023252405.1
A1 52139198 52139333
refGene_gn=EDNRB;refGene_tx=XM_003980457.4;refGene_gn=EDNRB;refGene_tx=XM_019828234.2
A1 52140772 52140881
refGene_gn=EDNRB;refGene_tx=XM_003980457.4;refGene_gn=EDNRB;refGene_tx=XM_019828234.2
A1 52141408 52141542
refGene_gn=EDNRB;refGene_tx=XM_003980457.4;refGene_gn=EDNRB;refGene_tx=XM_019828234.2
A1 52141944 52142094
refGene_gn=EDNRB;refGene_tx=XM_003980457.4;refGene_gn=EDNRB;refGene_tx=XM_019828234.2
A1 52143685 52143890
refGene_gn=EDNRB;refGene_tx=XM_003980457.4;refGene_gn=EDNRB;refGene_tx=XM_019828234.2
A1 52144024 52144137
refGene_gn=EDNRB;refGene_tx=XM_003980457.4;refGene_gn=EDNRB;refGene_tx=XM_019828234.2
A1 52159534 52160014
refGene_gn=EDNRB;refGene_tx=XM_003980457.4;refGene_gn=EDNRB;refGene_tx=XM_019828234.2
A1 52739280 52740423 refGene_gn=POU4F1;refGene_tx=XM_023252424.1
A1 52741117 52741240 refGene_gn=POU4F1;refGene_tx=XM_023252424.1
A1 52752163 52753679 refGene_gn=RNF219;refGene_tx=XM_003980460.4
A1 52772362 52772451 refGene_gn=RNF219;refGene_tx=XM_003980460.4
A1 52775786 52776035 refGene_gn=RNF219;refGene_tx=XM_003980460.4
A1 52778566 52778658 refGene_gn=RNF219;refGene_tx=XM_003980460.4
A1 52779748 52779884 refGene_gn=RNF219;refGene_tx=XM_003980460.4
A1 52792227 52792299 refGene_gn=RNF219;refGene_tx=XM_003980460.4
A1 53385333 53385357
refGene_gn=RBM26;refGene_tx=XM_023252452.1;refGene_gn=RBM26;refGene_tx=XM_023252453.1
A1 53390885 53390975
refGene_gn=RBM26;refGene_tx=XM_011280455.3;refGene_gn=RBM26;refGene_tx=XM_011280460.3;refGene_gn=RBM26;refGene_tx=XM_011280462.3;refGene_gn=RBM26;refGene_tx=XM_011280467.3;refGene_gn=RBM26;refGene_tx=XM_011280477.3;refGene_gn=RBM26;refGene_tx=XM_011280478.3;refGene_gn=RBM26;refGene_tx=XM_011280481.3;refGene_gn=RBM26;refGene_tx=XM_023252438.1;refGene_gn=RBM26;refGene_tx=XM_023252441.1;refGene_gn=RBM26;refGene_tx=XM_023252446.1;refGene_gn=RBM26;refGene_tx=XM_023252450.1;refGene_gn=RBM26;refGene_tx=XM_023252457.1
A1 53392718 53392832
refGene_gn=RBM26;refGene_tx=XM_011280455.3;refGene_gn=RBM26;refGene_tx=XM_011280460.3;refGene_gn=RBM26;refGene_tx=XM_011280462.3;refGene_gn=RBM26;refGene_tx=XM_011280467.3;refGene_gn=RBM26;refGene_tx=XM_011280477.3;refGene_gn=RBM26;refGene_tx=XM_011280478.3;refGene_gn=RBM26;refGene_tx=XM_011280481.3;refGene_gn=RBM26;refGene_tx=XM_023252438.1;refGen

e_gn=RBM26;refGene_tx=XM_023252441.1;refGene_gn=RBM26;refGene_tx=XM_02
3252446.1;refGene_gn=RBM26;refGene_tx=XM_023252450.1;refGene_gn=RBM26;
refGene_tx=XM_023252452.1;refGene_gn=RBM26;refGene_tx=XM_023252453.1;r
efGene_gn=RBM26;refGene_tx=XM_023252457.1
A1 53400981 53401068
refGene_gn=RBM26;refGene_tx=XM_011280455.3;refGene_gn=RBM26;refGene_tx
=XM_011280460.3;refGene_gn=RBM26;refGene_tx=XM_011280462.3;refGene_gn=
RBM26;refGene_tx=XM_011280467.3;refGene_gn=RBM26;refGene_tx=XM_0112804
77.3;refGene_gn=RBM26;refGene_tx=XM_011280478.3;refGene_gn=RBM26;refGe
ne_tx=XM_011280481.3;refGene_gn=RBM26;refGene_tx=XM_023252438.1;refGen
e_gn=RBM26;refGene_tx=XM_023252441.1;refGene_gn=RBM26;refGene_tx=XM_02
3252446.1;refGene_gn=RBM26;refGene_tx=XM_023252450.1;refGene_gn=RBM26;
refGene_tx=XM_023252452.1;refGene_gn=RBM26;refGene_tx=XM_023252453.1;r
efGene_gn=RBM26;refGene_tx=XM_023252457.1
A1 53403381 53403582
refGene_gn=RBM26;refGene_tx=XM_011280455.3;refGene_gn=RBM26;refGene_tx
=XM_011280460.3;refGene_gn=RBM26;refGene_tx=XM_011280462.3;refGene_gn=
RBM26;refGene_tx=XM_011280467.3;refGene_gn=RBM26;refGene_tx=XM_0112804
77.3;refGene_gn=RBM26;refGene_tx=XM_011280478.3;refGene_gn=RBM26;refGe
ne_tx=XM_011280481.3;refGene_gn=RBM26;refGene_tx=XM_023252438.1;refGen
e_gn=RBM26;refGene_tx=XM_023252441.1;refGene_gn=RBM26;refGene_tx=XM_02
3252446.1;refGene_gn=RBM26;refGene_tx=XM_023252450.1;refGene_gn=RBM26;
refGene_tx=XM_023252452.1;refGene_gn=RBM26;refGene_tx=XM_023252453.1;r
efGene_gn=RBM26;refGene_tx=XM_023252457.1
A1 53406715 53406820
refGene_gn=RBM26;refGene_tx=XM_011280455.3;refGene_gn=RBM26;refGene_tx
=XM_011280460.3;refGene_gn=RBM26;refGene_tx=XM_011280462.3;refGene_gn=
RBM26;refGene_tx=XM_011280467.3;refGene_gn=RBM26;refGene_tx=XM_0112804
77.3;refGene_gn=RBM26;refGene_tx=XM_011280478.3;refGene_gn=RBM26;refGe
ne_tx=XM_011280481.3;refGene_gn=RBM26;refGene_tx=XM_023252438.1;refGen
e_gn=RBM26;refGene_tx=XM_023252441.1;refGene_gn=RBM26;refGene_tx=XM_02
3252446.1;refGene_gn=RBM26;refGene_tx=XM_023252450.1;refGene_gn=RBM26;
refGene_tx=XM_023252452.1;refGene_gn=RBM26;refGene_tx=XM_023252453.1;r
efGene_gn=RBM26;refGene_tx=XM_023252457.1
A1 53408340 53408508
refGene_gn=RBM26;refGene_tx=XM_011280455.3;refGene_gn=RBM26;refGene_tx
=XM_011280460.3;refGene_gn=RBM26;refGene_tx=XM_011280462.3;refGene_gn=
RBM26;refGene_tx=XM_011280467.3;refGene_gn=RBM26;refGene_tx=XM_0112804
77.3;refGene_gn=RBM26;refGene_tx=XM_011280478.3;refGene_gn=RBM26;refGe
ne_tx=XM_011280481.3;refGene_gn=RBM26;refGene_tx=XM_023252438.1;refGen
e_gn=RBM26;refGene_tx=XM_023252441.1;refGene_gn=RBM26;refGene_tx=XM_02
3252446.1;refGene_gn=RBM26;refGene_tx=XM_023252450.1;refGene_gn=RBM26;
refGene_tx=XM_023252452.1;refGene_gn=RBM26;refGene_tx=XM_023252453.1;r
efGene_gn=RBM26;refGene_tx=XM_023252457.1
A1 53410157 53410232
refGene_gn=RBM26;refGene_tx=XM_011280455.3;refGene_gn=RBM26;refGene_tx
=XM_011280460.3;refGene_gn=RBM26;refGene_tx=XM_011280462.3;refGene_gn=
RBM26;refGene_tx=XM_011280467.3;refGene_gn=RBM26;refGene_tx=XM_0112804
77.3;refGene_gn=RBM26;refGene_tx=XM_011280478.3;refGene_gn=RBM26;refGe
ne_tx=XM_011280481.3;refGene_gn=RBM26;refGene_tx=XM_023252438.1;refGen

e_gn=RBM26;refGene_tx=XM_023252441.1;refGene_gn=RBM26;refGene_tx=XM_02
3252446.1;refGene_gn=RBM26;refGene_tx=XM_023252450.1;refGene_gn=RBM26;
refGene_tx=XM_023252452.1;refGene_gn=RBM26;refGene_tx=XM_023252453.1;r
efGene_gn=RBM26;refGene_tx=XM_023252457.1
A1 53410612 53410738
refGene_gn=RBM26;refGene_tx=XM_011280455.3;refGene_gn=RBM26;refGene_tx
=XM_011280460.3;refGene_gn=RBM26;refGene_tx=XM_011280462.3;refGene_gn=
RBM26;refGene_tx=XM_011280467.3;refGene_gn=RBM26;refGene_tx=XM_0112804
77.3;refGene_gn=RBM26;refGene_tx=XM_011280478.3;refGene_gn=RBM26;refGe
ne_tx=XM_011280481.3;refGene_gn=RBM26;refGene_tx=XM_023252438.1;refGen
e_gn=RBM26;refGene_tx=XM_023252441.1;refGene_gn=RBM26;refGene_tx=XM_02
3252446.1;refGene_gn=RBM26;refGene_tx=XM_023252450.1;refGene_gn=RBM26;
refGene_tx=XM_023252452.1;refGene_gn=RBM26;refGene_tx=XM_023252453.1;r
efGene_gn=RBM26;refGene_tx=XM_023252457.1
A1 53418576 53418648
refGene_gn=RBM26;refGene_tx=XM_011280455.3;refGene_gn=RBM26;refGene_tx
=XM_011280460.3;refGene_gn=RBM26;refGene_tx=XM_011280462.3;refGene_gn=
RBM26;refGene_tx=XM_011280467.3;refGene_gn=RBM26;refGene_tx=XM_0232524
38.1;refGene_gn=RBM26;refGene_tx=XM_023252441.1;refGene_gn=RBM26;refGe
ne_tx=XM_023252446.1;refGene_gn=RBM26;refGene_tx=XM_023252450.1;refGen
e_gn=RBM26;refGene_tx=XM_023252452.1;refGene_gn=RBM26;refGene_tx=XM_02
3252453.1
A1 53419849 53419981
refGene_gn=RBM26;refGene_tx=XM_011280455.3;refGene_gn=RBM26;refGene_tx
=XM_011280460.3;refGene_gn=RBM26;refGene_tx=XM_011280462.3;refGene_gn=
RBM26;refGene_tx=XM_011280467.3;refGene_gn=RBM26;refGene_tx=XM_0112804
77.3;refGene_gn=RBM26;refGene_tx=XM_011280478.3;refGene_gn=RBM26;refGe
ne_tx=XM_011280481.3;refGene_gn=RBM26;refGene_tx=XM_023252438.1;refGen
e_gn=RBM26;refGene_tx=XM_023252441.1;refGene_gn=RBM26;refGene_tx=XM_02
3252446.1;refGene_gn=RBM26;refGene_tx=XM_023252450.1;refGene_gn=RBM26;
refGene_tx=XM_023252452.1;refGene_gn=RBM26;refGene_tx=XM_023252453.1;r
efGene_gn=RBM26;refGene_tx=XM_023252457.1
A1 53420669 53420834
refGene_gn=RBM26;refGene_tx=XM_011280455.3;refGene_gn=RBM26;refGene_tx
=XM_011280460.3;refGene_gn=RBM26;refGene_tx=XM_011280462.3;refGene_gn=
RBM26;refGene_tx=XM_011280467.3;refGene_gn=RBM26;refGene_tx=XM_0112804
77.3;refGene_gn=RBM26;refGene_tx=XM_011280478.3;refGene_gn=RBM26;refGe
ne_tx=XM_011280481.3;refGene_gn=RBM26;refGene_tx=XM_023252438.1;refGen
e_gn=RBM26;refGene_tx=XM_023252441.1;refGene_gn=RBM26;refGene_tx=XM_02
3252446.1;refGene_gn=RBM26;refGene_tx=XM_023252450.1;refGene_gn=RBM26;
refGene_tx=XM_023252452.1;refGene_gn=RBM26;refGene_tx=XM_023252453.1;r
efGene_gn=RBM26;refGene_tx=XM_023252457.1
A1 53423489 53423649
refGene_gn=RBM26;refGene_tx=XM_011280455.3;refGene_gn=RBM26;refGene_tx
=XM_011280460.3;refGene_gn=RBM26;refGene_tx=XM_011280462.3;refGene_gn=
RBM26;refGene_tx=XM_011280467.3;refGene_gn=RBM26;refGene_tx=XM_0112804
77.3;refGene_gn=RBM26;refGene_tx=XM_011280478.3;refGene_gn=RBM26;refGe
ne_tx=XM_011280481.3;refGene_gn=RBM26;refGene_tx=XM_023252438.1;refGen
e_gn=RBM26;refGene_tx=XM_023252441.1;refGene_gn=RBM26;refGene_tx=XM_02
3252446.1;refGene_gn=RBM26;refGene_tx=XM_023252450.1;refGene_gn=RBM26;

refGene_tx=XM_023252452.1;refGene_gn=RBM26;refGene_tx=XM_023252453.1;refGene_gn=RBM26;refGene_tx=XM_023252457.1
A1 53424743 53424855
refGene_gn=RBM26;refGene_tx=XM_011280455.3;refGene_gn=RBM26;refGene_tx=XM_011280460.3;refGene_gn=RBM26;refGene_tx=XM_011280462.3;refGene_gn=RBM26;refGene_tx=XM_011280467.3;refGene_gn=RBM26;refGene_tx=XM_011280477.3;refGene_gn=RBM26;refGene_tx=XM_011280478.3;refGene_gn=RBM26;refGene_tx=XM_011280481.3;refGene_gn=RBM26;refGene_tx=XM_023252438.1;refGene_gn=RBM26;refGene_tx=XM_023252441.1;refGene_gn=RBM26;refGene_tx=XM_023252446.1;refGene_gn=RBM26;refGene_tx=XM_023252450.1;refGene_gn=RBM26;refGene_tx=XM_023252452.1;refGene_gn=RBM26;refGene_tx=XM_023252453.1;refGene_gn=RBM26;refGene_tx=XM_023252457.1
A1 53429733 53429889
refGene_gn=RBM26;refGene_tx=XM_011280455.3;refGene_gn=RBM26;refGene_tx=XM_011280460.3;refGene_gn=RBM26;refGene_tx=XM_011280462.3;refGene_gn=RBM26;refGene_tx=XM_011280467.3;refGene_gn=RBM26;refGene_tx=XM_011280477.3;refGene_gn=RBM26;refGene_tx=XM_011280478.3;refGene_gn=RBM26;refGene_tx=XM_011280481.3;refGene_gn=RBM26;refGene_tx=XM_023252438.1;refGene_gn=RBM26;refGene_tx=XM_023252441.1;refGene_gn=RBM26;refGene_tx=XM_023252446.1;refGene_gn=RBM26;refGene_tx=XM_023252450.1;refGene_gn=RBM26;refGene_tx=XM_023252452.1;refGene_gn=RBM26;refGene_tx=XM_023252453.1;refGene_gn=RBM26;refGene_tx=XM_023252457.1
A1 53430253 53430424
refGene_gn=RBM26;refGene_tx=XM_011280455.3;refGene_gn=RBM26;refGene_tx=XM_011280460.3;refGene_gn=RBM26;refGene_tx=XM_011280462.3;refGene_gn=RBM26;refGene_tx=XM_011280467.3;refGene_gn=RBM26;refGene_tx=XM_011280477.3;refGene_gn=RBM26;refGene_tx=XM_011280478.3;refGene_gn=RBM26;refGene_tx=XM_011280481.3;refGene_gn=RBM26;refGene_tx=XM_023252438.1;refGene_gn=RBM26;refGene_tx=XM_023252441.1;refGene_gn=RBM26;refGene_tx=XM_023252446.1;refGene_gn=RBM26;refGene_tx=XM_023252450.1;refGene_gn=RBM26;refGene_tx=XM_023252452.1;refGene_gn=RBM26;refGene_tx=XM_023252453.1;refGene_gn=RBM26;refGene_tx=XM_023252457.1
A1 53430851 53431091
refGene_gn=RBM26;refGene_tx=XM_011280455.3;refGene_gn=RBM26;refGene_tx=XM_011280460.3;refGene_gn=RBM26;refGene_tx=XM_011280462.3;refGene_gn=RBM26;refGene_tx=XM_011280467.3;refGene_gn=RBM26;refGene_tx=XM_011280477.3;refGene_gn=RBM26;refGene_tx=XM_011280478.3;refGene_gn=RBM26;refGene_tx=XM_011280481.3;refGene_gn=RBM26;refGene_tx=XM_023252438.1;refGene_gn=RBM26;refGene_tx=XM_023252441.1;refGene_gn=RBM26;refGene_tx=XM_023252446.1;refGene_gn=RBM26;refGene_tx=XM_023252450.1;refGene_gn=RBM26;refGene_tx=XM_023252452.1;refGene_gn=RBM26;refGene_tx=XM_023252453.1;refGene_gn=RBM26;refGene_tx=XM_023252457.1
A1 53431865 53432126
refGene_gn=RBM26;refGene_tx=XM_011280455.3;refGene_gn=RBM26;refGene_tx=XM_011280460.3;refGene_gn=RBM26;refGene_tx=XM_011280462.3;refGene_gn=RBM26;refGene_tx=XM_011280467.3;refGene_gn=RBM26;refGene_tx=XM_011280477.3;refGene_gn=RBM26;refGene_tx=XM_011280478.3;refGene_gn=RBM26;refGene_tx=XM_011280481.3;refGene_gn=RBM26;refGene_tx=XM_023252438.1;refGene_gn=RBM26;refGene_tx=XM_023252441.1;refGene_gn=RBM26;refGene_tx=XM_023252446.1;refGene_gn=RBM26;refGene_tx=XM_023252450.1;refGene_gn=RBM26;

refGene_tx=XM_023252452.1;refGene_gn=RBM26;refGene_tx=XM_023252453.1;refGene_gn=RBM26;refGene_tx=XM_023252457.1
A1 53433848 53434066
refGene_gn=RBM26;refGene_tx=XM_011280455.3;refGene_gn=RBM26;refGene_tx=XM_011280460.3;refGene_gn=RBM26;refGene_tx=XM_011280462.3;refGene_gn=RBM26;refGene_tx=XM_011280467.3;refGene_gn=RBM26;refGene_tx=XM_011280477.3;refGene_gn=RBM26;refGene_tx=XM_011280478.3;refGene_gn=RBM26;refGene_tx=XM_011280481.3;refGene_gn=RBM26;refGene_tx=XM_023252438.1;refGene_gn=RBM26;refGene_tx=XM_023252441.1;refGene_gn=RBM26;refGene_tx=XM_023252446.1;refGene_gn=RBM26;refGene_tx=XM_023252450.1;refGene_gn=RBM26;refGene_tx=XM_023252452.1;refGene_gn=RBM26;refGene_tx=XM_023252453.1;refGene_gn=RBM26;refGene_tx=XM_023252457.1
A1 53434736 53434825
refGene_gn=RBM26;refGene_tx=XM_011280455.3;refGene_gn=RBM26;refGene_tx=XM_011280460.3;refGene_gn=RBM26;refGene_tx=XM_011280462.3;refGene_gn=RBM26;refGene_tx=XM_011280467.3;refGene_gn=RBM26;refGene_tx=XM_011280477.3;refGene_gn=RBM26;refGene_tx=XM_011280478.3;refGene_gn=RBM26;refGene_tx=XM_011280481.3;refGene_gn=RBM26;refGene_tx=XM_023252438.1;refGene_gn=RBM26;refGene_tx=XM_023252441.1;refGene_gn=RBM26;refGene_tx=XM_023252446.1;refGene_gn=RBM26;refGene_tx=XM_023252450.1;refGene_gn=RBM26;refGene_tx=XM_023252452.1;refGene_gn=RBM26;refGene_tx=XM_023252453.1;refGene_gn=RBM26;refGene_tx=XM_023252457.1
A1 53437119 53437256
refGene_gn=RBM26;refGene_tx=XM_011280455.3;refGene_gn=RBM26;refGene_tx=XM_011280460.3;refGene_gn=RBM26;refGene_tx=XM_011280462.3;refGene_gn=RBM26;refGene_tx=XM_011280467.3;refGene_gn=RBM26;refGene_tx=XM_011280477.3;refGene_gn=RBM26;refGene_tx=XM_011280478.3;refGene_gn=RBM26;refGene_tx=XM_011280481.3;refGene_gn=RBM26;refGene_tx=XM_023252438.1;refGene_gn=RBM26;refGene_tx=XM_023252441.1;refGene_gn=RBM26;refGene_tx=XM_023252446.1;refGene_gn=RBM26;refGene_tx=XM_023252450.1;refGene_gn=RBM26;refGene_tx=XM_023252452.1;refGene_gn=RBM26;refGene_tx=XM_023252453.1;refGene_gn=RBM26;refGene_tx=XM_023252457.1
A1 53437878 53437997
refGene_gn=RBM26;refGene_tx=XM_011280455.3;refGene_gn=RBM26;refGene_tx=XM_011280460.3;refGene_gn=RBM26;refGene_tx=XM_011280462.3;refGene_gn=RBM26;refGene_tx=XM_011280467.3;refGene_gn=RBM26;refGene_tx=XM_011280477.3;refGene_gn=RBM26;refGene_tx=XM_011280478.3;refGene_gn=RBM26;refGene_tx=XM_011280481.3;refGene_gn=RBM26;refGene_tx=XM_023252438.1;refGene_gn=RBM26;refGene_tx=XM_023252441.1;refGene_gn=RBM26;refGene_tx=XM_023252446.1;refGene_gn=RBM26;refGene_tx=XM_023252450.1;refGene_gn=RBM26;refGene_tx=XM_023252452.1;refGene_gn=RBM26;refGene_tx=XM_023252453.1;refGene_gn=RBM26;refGene_tx=XM_023252457.1
A1 53459857 53459943
refGene_gn=RBM26;refGene_tx=XM_023252438.1;refGene_gn=RBM26;refGene_tx=XM_023252441.1;refGene_gn=RBM26;refGene_tx=XM_023252446.1;refGene_gn=RBM26;refGene_tx=XM_023252450.1;refGene_gn=RBM26;refGene_tx=XM_023252452.1;refGene_gn=RBM26;refGene_tx=XM_023252453.1;refGene_gn=RBM26;refGene_tx=XM_023252457.1
A1 53467787 53467858
refGene_gn=RBM26;refGene_tx=XM_011280455.3;refGene_gn=RBM26;refGene_tx

=XM_011280460.3;refGene_gn=RBM26;refGene_tx=XM_011280462.3;refGene_gn=RBM26;refGene_tx=XM_011280467.3;refGene_gn=RBM26;refGene_tx=XM_011280477.3;refGene_gn=RBM26;refGene_tx=XM_011280478.3;refGene_gn=RBM26;refGene_tx=XM_011280481.3
A1 53538998 53539271 refGene_gn=NDFIP2;refGene_tx=XM_019828338.2;refGene_gn=NDFIP2;refGene_tx=XM_023252476.1
A1 53568539 53568696 refGene_gn=NDFIP2;refGene_tx=XM_019828338.2;refGene_gn=NDFIP2;refGene_tx=XM_023252476.1
A1 53574659 53574793 refGene_gn=NDFIP2;refGene_tx=XM_019828338.2;refGene_gn=NDFIP2;refGene_tx=XM_023252476.1
A1 53580123 53580217 refGene_gn=NDFIP2;refGene_tx=XM_019828338.2;refGene_gn=NDFIP2;refGene_tx=XM_023252476.1
A1 53588373 53588498 refGene_gn=NDFIP2;refGene_tx=XM_019828338.2;refGene_gn=NDFIP2;refGene_tx=XM_023252476.1
A1 53593073 53593140 refGene_gn=NDFIP2;refGene_tx=XM_019828338.2;refGene_gn=NDFIP2;refGene_tx=XM_023252476.1
A1 53596032 53596136 refGene_gn=NDFIP2;refGene_tx=XM_019828338.2;refGene_gn=NDFIP2;refGene_tx=XM_023252476.1
A1 54188573 54189521 refGene_gn=SPRY2;refGene_tx=XM_003980464.4
A1 57338856 57340947 refGene_gn=SLITRK1;refGene_tx=XM_003980471.5
A1 59000084 59002607 refGene_gn=SLITRK6;refGene_tx=XM_006927353.4
A1 60514907 60517784 refGene_gn=SLITRK5;refGene_tx=XM_003980472.5;refGene_gn=SLITRK5;refGene_tx=XM_011280561.3;refGene_gn=SLITRK5;refGene_tx=XM_023252549.1;refGene_gn=SLITRK5;refGene_tx=XM_023252555.1
A1 63716735 63717201 refGene_gn=GPC5;refGene_tx=XM_011281029.3
A1 63759275 63759437 refGene_gn=GPC5;refGene_tx=XM_011281029.3
A1 63991012 63991707 refGene_gn=GPC5;refGene_tx=XM_011281029.3
A1 64032602 64032736 refGene_gn=GPC5;refGene_tx=XM_011281029.3
A1 64054989 64055115 refGene_gn=GPC5;refGene_tx=XM_011281029.3
A1 64194284 64194405 refGene_gn=GPC5;refGene_tx=XM_011281029.3
A1 64399433 64399590 refGene_gn=GPC5;refGene_tx=XM_011281029.3
A1 65138679 65138834 refGene_gn=GPC5;refGene_tx=XM_011281029.3
A1 65449165 65449325 refGene_gn=GPC6;refGene_tx=XM_011280573.2;refGene_gn=GPC6;refGene_tx=XM_019828517.2
A1 65737323 65737482 refGene_gn=GPC6;refGene_tx=XM_011280573.2;refGene_gn=GPC6;refGene_tx=XM_019828517.2
A1 66019176 66019568 refGene_gn=GPC6;refGene_tx=XM_011280573.2;refGene_gn=GPC6;refGene_tx=XM_019828517.2
A1 66214599 66214765

refGene_gn=GPC6;refGene_tx=XM_011280573.2;refGene_gn=GPC6;refGene_tx=XM_019828517.2
A1 66460506 66460637
refGene_gn=GPC6;refGene_tx=XM_011280573.2;refGene_gn=GPC6;refGene_tx=XM_019828517.2
A1 66478671 66478815
refGene_gn=GPC6;refGene_tx=XM_011280573.2;refGene_gn=GPC6;refGene_tx=XM_019828517.2
A1 66482866 66482896 refGene_gn=GPC6;refGene_tx=XM_019828517.2
A1 66547903 66548040
refGene_gn=GPC6;refGene_tx=XM_011280573.2;refGene_gn=GPC6;refGene_tx=XM_019828517.2
A1 66562738 66562914
refGene_gn=GPC6;refGene_tx=XM_011280573.2;refGene_gn=GPC6;refGene_tx=XM_019828517.2
A1 66566617 66566820
refGene_gn=GPC6;refGene_tx=XM_011280573.2;refGene_gn=GPC6;refGene_tx=XM_019828517.2
A1 66603912 66604085 refGene_gn=DCT;refGene_tx=XM_003980474.5
A1 66607526 66607728 refGene_gn=DCT;refGene_tx=XM_003980474.5
A1 66620450 66620586 refGene_gn=DCT;refGene_tx=XM_003980474.5
A1 66622258 66622438 refGene_gn=DCT;refGene_tx=XM_003980474.5
A1 66624666 66624833 refGene_gn=DCT;refGene_tx=XM_003980474.5
A1 66624966 66625067 refGene_gn=DCT;refGene_tx=XM_003980474.5
A1 66626910 66627210 refGene_gn=DCT;refGene_tx=XM_003980474.5
A1 66635963 66636354 refGene_gn=DCT;refGene_tx=XM_003980474.5
A1 66706600 66706686
refGene_gn=TGDS;refGene_tx=XM_003980475.4;refGene_gn=TGDS;refGene_tx=XM_006927355.3;refGene_gn=TGDS;refGene_tx=XM_019828546.2;refGene_gn=TGDS;refGene_tx=XM_019828553.2
A1 66707900 66707998
refGene_gn=TGDS;refGene_tx=XM_003980475.4;refGene_gn=TGDS;refGene_tx=XM_019828546.2;refGene_gn=TGDS;refGene_tx=XM_019828553.2
A1 66708428 66708487
refGene_gn=TGDS;refGene_tx=XM_003980475.4;refGene_gn=TGDS;refGene_tx=XM_006927355.3;refGene_gn=TGDS;refGene_tx=XM_019828546.2;refGene_gn=TGDS;refGene_tx=XM_019828553.2
A1 66709859 66710025
refGene_gn=TGDS;refGene_tx=XM_003980475.4;refGene_gn=TGDS;refGene_tx=XM_006927355.3;refGene_gn=TGDS;refGene_tx=XM_019828546.2;refGene_gn=TGDS;refGene_tx=XM_019828553.2
A1 66710659 66710703
refGene_gn=TGDS;refGene_tx=XM_003980475.4;refGene_gn=TGDS;refGene_tx=XM_006927355.3;refGene_gn=TGDS;refGene_tx=XM_019828546.2;refGene_gn=TGDS;refGene_tx=XM_019828553.2
A1 66711915 66711975
refGene_gn=TGDS;refGene_tx=XM_003980475.4;refGene_gn=TGDS;refGene_tx=XM_006927355.3;refGene_gn=TGDS;refGene_tx=XM_019828546.2;refGene_gn=TGDS;refGene_tx=XM_019828553.2
A1 66713027 66713126

refGene_gn=TGDS;refGene_tx=XM_003980475.4;refGene_gn=TGDS;refGene_tx=XM_006927355.3;refGene_gn=TGDS;refGene_tx=XM_019828546.2;refGene_gn=TGDS;refGene_tx=XM_019828553.2
A1 66714379 66714522
refGene_gn=TGDS;refGene_tx=XM_003980475.4;refGene_gn=TGDS;refGene_tx=XM_006927355.3;refGene_gn=TGDS;refGene_tx=XM_019828546.2;refGene_gn=TGDS;refGene_tx=XM_019828553.2
A1 66716903 66716994
refGene_gn=TGDS;refGene_tx=XM_003980475.4;refGene_gn=TGDS;refGene_tx=XM_006927355.3;refGene_gn=TGDS;refGene_tx=XM_019828546.2;refGene_gn=TGDS;refGene_tx=XM_019828553.2
A1 66718819 66718888
refGene_gn=TGDS;refGene_tx=XM_003980475.4;refGene_gn=TGDS;refGene_tx=XM_006927355.3;refGene_gn=TGDS;refGene_tx=XM_019828546.2;refGene_gn=TGDS;refGene_tx=XM_019828553.2
A1 66719865 66719932
refGene_gn=TGDS;refGene_tx=XM_003980475.4;refGene_gn=TGDS;refGene_tx=XM_006927355.3;refGene_gn=TGDS;refGene_tx=XM_019828546.2;refGene_gn=TGDS;refGene_tx=XM_019828553.2
A1 66722103 66722189
refGene_gn=TGDS;refGene_tx=XM_003980475.4;refGene_gn=TGDS;refGene_tx=XM_006927355.3;refGene_gn=TGDS;refGene_tx=XM_019828546.2
A1 66725788 66725933 refGene_gn=GPR180;refGene_tx=XM_023252632.1
A1 66728572 66728731 refGene_gn=GPR180;refGene_tx=XM_023252632.1
A1 66736846 66737047 refGene_gn=GPR180;refGene_tx=XM_023252632.1
A1 66743468 66743649 refGene_gn=GPR180;refGene_tx=XM_023252632.1
A1 66743790 66743840 refGene_gn=GPR180;refGene_tx=XM_023252632.1
A1 66746536 66746694 refGene_gn=GPR180;refGene_tx=XM_023252632.1
A1 66749198 66749390 refGene_gn=GPR180;refGene_tx=XM_023252632.1
A1 66751258 66751336 refGene_gn=GPR180;refGene_tx=XM_023252632.1
A1 66752591 66752750 refGene_gn=GPR180;refGene_tx=XM_023252632.1
A1 66829085 66829916 refGene_gn=SOX21;refGene_tx=XM_023252642.1
A1 66830566 66830741
refGene_gn=LOC111556548;refGene_tx=XM_023239061.1
A1 66830854 66831300
refGene_gn=LOC111556548;refGene_tx=XM_023239061.1
A1 66831540 66831750
refGene_gn=LOC111556548;refGene_tx=XM_023239061.1
A1 67085635 67085743
refGene_gn=ABCC4;refGene_tx=XM_023252652.1;refGene_gn=ABCC4;refGene_tx=XM_023252655.1;refGene_gn=ABCC4;refGene_tx=XM_023252657.1;refGene_gn=ABCC4;refGene_tx=XM_023252659.1
A1 67098164 67098299
refGene_gn=ABCC4;refGene_tx=XM_023252652.1;refGene_gn=ABCC4;refGene_tx=XM_023252655.1;refGene_gn=ABCC4;refGene_tx=XM_023252657.1;refGene_gn=ABCC4;refGene_tx=XM_023252659.1
A1 67107137 67107243
refGene_gn=ABCC4;refGene_tx=XM_023252652.1;refGene_gn=ABCC4;refGene_tx=XM_023252655.1;refGene_gn=ABCC4;refGene_tx=XM_023252657.1;refGene_gn=ABCC4;refGene_tx=XM_023252659.1

A1 67107900 67108073
refGene_gn=ABCC4;refGene_tx=XM_023252652.1;refGene_gn=ABCC4;refGene_tx=XM_023252655.1;refGene_gn=ABCC4;refGene_tx=XM_023252657.1;refGene_gn=ABCC4;refGene_tx=XM_023252659.1
A1 67115801 67115891
refGene_gn=ABCC4;refGene_tx=XM_023252652.1;refGene_gn=ABCC4;refGene_tx=XM_023252655.1;refGene_gn=ABCC4;refGene_tx=XM_023252657.1;refGene_gn=ABCC4;refGene_tx=XM_023252659.1
A1 67125555 67125711
refGene_gn=ABCC4;refGene_tx=XM_023252652.1;refGene_gn=ABCC4;refGene_tx=XM_023252655.1;refGene_gn=ABCC4;refGene_tx=XM_023252657.1;refGene_gn=ABCC4;refGene_tx=XM_023252659.1
A1 67134755 67134947
refGene_gn=ABCC4;refGene_tx=XM_023252652.1;refGene_gn=ABCC4;refGene_tx=XM_023252655.1;refGene_gn=ABCC4;refGene_tx=XM_023252657.1;refGene_gn=ABCC4;refGene_tx=XM_023252659.1
A1 67136488 67136589
refGene_gn=ABCC4;refGene_tx=XM_023252652.1;refGene_gn=ABCC4;refGene_tx=XM_023252655.1;refGene_gn=ABCC4;refGene_tx=XM_023252657.1;refGene_gn=ABCC4;refGene_tx=XM_023252659.1
A1 67137549 67137660
refGene_gn=ABCC4;refGene_tx=XM_023252652.1;refGene_gn=ABCC4;refGene_tx=XM_023252655.1;refGene_gn=ABCC4;refGene_tx=XM_023252657.1;refGene_gn=ABCC4;refGene_tx=XM_023252659.1
A1 67138941 67139061
refGene_gn=ABCC4;refGene_tx=XM_023252652.1;refGene_gn=ABCC4;refGene_tx=XM_023252655.1;refGene_gn=ABCC4;refGene_tx=XM_023252657.1;refGene_gn=ABCC4;refGene_tx=XM_023252659.1
A1 67145172 67145323
refGene_gn=ABCC4;refGene_tx=XM_023252652.1;refGene_gn=ABCC4;refGene_tx=XM_023252655.1;refGene_gn=ABCC4;refGene_tx=XM_023252657.1;refGene_gn=ABCC4;refGene_tx=XM_023252659.1
A1 67176128 67176208
refGene_gn=ABCC4;refGene_tx=XM_023252652.1;refGene_gn=ABCC4;refGene_tx=XM_023252655.1;refGene_gn=ABCC4;refGene_tx=XM_023252657.1;refGene_gn=ABCC4;refGene_tx=XM_023252659.1
A1 67219044 67219191
refGene_gn=ABCC4;refGene_tx=XM_023252652.1;refGene_gn=ABCC4;refGene_tx=XM_023252655.1;refGene_gn=ABCC4;refGene_tx=XM_023252657.1
A1 67221008 67221103
refGene_gn=ABCC4;refGene_tx=XM_023252652.1;refGene_gn=ABCC4;refGene_tx=XM_023252655.1;refGene_gn=ABCC4;refGene_tx=XM_023252657.1;refGene_gn=ABCC4;refGene_tx=XM_023252659.1
A1 67221493 67221531
refGene_gn=ABCC4;refGene_tx=XM_023252652.1;refGene_gn=ABCC4;refGene_tx=XM_023252655.1;refGene_gn=ABCC4;refGene_tx=XM_023252657.1;refGene_gn=ABCC4;refGene_tx=XM_023252659.1
A1 67222151 67222292
refGene_gn=ABCC4;refGene_tx=XM_023252652.1;refGene_gn=ABCC4;refGene_tx=XM_023252655.1;refGene_gn=ABCC4;refGene_tx=XM_023252657.1;refGene_gn=

ABCC4;refGene_tx=XM_023252659.1
A1 67224134 67224344
refGene_gn=ABCC4;refGene_tx=XM_023252652.1;refGene_gn=ABCC4;refGene_tx=XM_023252655.1;refGene_gn=ABCC4;refGene_tx=XM_023252657.1;refGene_gn=ABCC4;refGene_tx=XM_023252659.1
A1 67227536 67227633
refGene_gn=ABCC4;refGene_tx=XM_023252652.1;refGene_gn=ABCC4;refGene_tx=XM_023252655.1;refGene_gn=ABCC4;refGene_tx=XM_023252657.1;refGene_gn=ABCC4;refGene_tx=XM_023252659.1
A1 67233147 67233234
refGene_gn=ABCC4;refGene_tx=XM_023252652.1;refGene_gn=ABCC4;refGene_tx=XM_023252655.1;refGene_gn=ABCC4;refGene_tx=XM_023252657.1;refGene_gn=ABCC4;refGene_tx=XM_023252659.1
A1 67233489 67233584
refGene_gn=ABCC4;refGene_tx=XM_023252652.1;refGene_gn=ABCC4;refGene_tx=XM_023252655.1;refGene_gn=ABCC4;refGene_tx=XM_023252657.1;refGene_gn=ABCC4;refGene_tx=XM_023252659.1
A1 67243892 67244084
refGene_gn=ABCC4;refGene_tx=XM_023252652.1;refGene_gn=ABCC4;refGene_tx=XM_023252655.1;refGene_gn=ABCC4;refGene_tx=XM_023252657.1;refGene_gn=ABCC4;refGene_tx=XM_023252659.1
A1 67245429 67245519
refGene_gn=ABCC4;refGene_tx=XM_023252652.1;refGene_gn=ABCC4;refGene_tx=XM_023252655.1;refGene_gn=ABCC4;refGene_tx=XM_023252657.1;refGene_gn=ABCC4;refGene_tx=XM_023252659.1
A1 67249406 67249508
refGene_gn=ABCC4;refGene_tx=XM_023252652.1;refGene_gn=ABCC4;refGene_tx=XM_023252655.1;refGene_gn=ABCC4;refGene_tx=XM_023252657.1;refGene_gn=ABCC4;refGene_tx=XM_023252659.1
A1 67261951 67262201
refGene_gn=ABCC4;refGene_tx=XM_023252652.1;refGene_gn=ABCC4;refGene_tx=XM_023252655.1;refGene_gn=ABCC4;refGene_tx=XM_023252657.1;refGene_gn=ABCC4;refGene_tx=XM_023252659.1
A1 67263403 67263529
refGene_gn=ABCC4;refGene_tx=XM_023252652.1;refGene_gn=ABCC4;refGene_tx=XM_023252655.1;refGene_gn=ABCC4;refGene_tx=XM_023252657.1;refGene_gn=ABCC4;refGene_tx=XM_023252659.1
A1 67264879 67265043
refGene_gn=ABCC4;refGene_tx=XM_023252652.1;refGene_gn=ABCC4;refGene_tx=XM_023252655.1;refGene_gn=ABCC4;refGene_tx=XM_023252657.1;refGene_gn=ABCC4;refGene_tx=XM_023252659.1
A1 67266347 67266437
refGene_gn=ABCC4;refGene_tx=XM_023252652.1;refGene_gn=ABCC4;refGene_tx=XM_023252655.1;refGene_gn=ABCC4;refGene_tx=XM_023252657.1;refGene_gn=ABCC4;refGene_tx=XM_023252659.1
A1 67267237 67267279
refGene_gn=ABCC4;refGene_tx=XM_023252652.1;refGene_gn=ABCC4;refGene_tx=XM_023252657.1;refGene_gn=ABCC4;refGene_tx=XM_023252659.1
A1 67291014 67291239
refGene_gn=ABCC4;refGene_tx=XM_023252652.1;refGene_gn=ABCC4;refGene_tx

=XM_023252655.1;refGene_gn=ABCC4;refGene_tx=XM_023252657.1;refGene_gn=ABCC4;refGene_tx=XM_023252659.1
A1 67302074 67302195
refGene_gn=ABCC4;refGene_tx=XM_023252652.1;refGene_gn=ABCC4;refGene_tx=XM_023252655.1;refGene_gn=ABCC4;refGene_tx=XM_023252657.1;refGene_gn=ABCC4;refGene_tx=XM_023252659.1
A1 67302764 67302875
refGene_gn=ABCC4;refGene_tx=XM_023252652.1;refGene_gn=ABCC4;refGene_tx=XM_023252655.1;refGene_gn=ABCC4;refGene_tx=XM_023252657.1;refGene_gn=ABCC4;refGene_tx=XM_023252659.1
A1 67351090 67351164
refGene_gn=ABCC4;refGene_tx=XM_023252652.1;refGene_gn=ABCC4;refGene_tx=XM_023252655.1;refGene_gn=ABCC4;refGene_tx=XM_023252657.1;refGene_gn=ABCC4;refGene_tx=XM_023252659.1
A1 67431813 67432027
refGene_gn=CLDN10;refGene_tx=XM_003980480.5;refGene_gn=CLDN10;refGene_tx=XM_019828594.2
A1 67523693 67523913
refGene_gn=CLDN10;refGene_tx=XM_023252663.1;refGene_gn=CLDN10;refGene_tx=XM_023252668.1
A1 67530708 67530870
refGene_gn=CLDN10;refGene_tx=XM_003980480.5;refGene_gn=CLDN10;refGene_tx=XM_019828594.2;refGene_gn=CLDN10;refGene_tx=XM_023252663.1;refGene_gn=CLDN10;refGene_tx=XM_023252668.1
A1 67530958 67531040
refGene_gn=CLDN10;refGene_tx=XM_003980480.5;refGene_gn=CLDN10;refGene_tx=XM_019828594.2;refGene_gn=CLDN10;refGene_tx=XM_023252663.1;refGene_gn=CLDN10;refGene_tx=XM_023252668.1
A1 67545707 67545815
refGene_gn=CLDN10;refGene_tx=XM_003980480.5;refGene_gn=CLDN10;refGene_tx=XM_019828594.2;refGene_gn=CLDN10;refGene_tx=XM_023252663.1
A1 67546425 67546549
refGene_gn=CLDN10;refGene_tx=XM_003980480.5;refGene_gn=CLDN10;refGene_tx=XM_019828594.2;refGene_gn=CLDN10;refGene_tx=XM_023252663.1;refGene_gn=CLDN10;refGene_tx=XM_023252668.1
A1 67549091 67549168
refGene_gn=DZIP1;refGene_tx=XM_023252677.1;refGene_gn=DZIP1;refGene_tx=XM_023252682.1;refGene_gn=DZIP1;refGene_tx=XM_023252687.1;refGene_gn=DZIP1;refGene_tx=XM_023252689.1;refGene_gn=DZIP1;refGene_tx=XM_023252698.1
A1 67550741 67550916
refGene_gn=DZIP1;refGene_tx=XM_023252677.1;refGene_gn=DZIP1;refGene_tx=XM_023252682.1;refGene_gn=DZIP1;refGene_tx=XM_023252687.1;refGene_gn=DZIP1;refGene_tx=XM_023252689.1;refGene_gn=DZIP1;refGene_tx=XM_023252698.1
A1 67552134 67552265
refGene_gn=DZIP1;refGene_tx=XM_023252677.1;refGene_gn=DZIP1;refGene_tx=XM_023252682.1;refGene_gn=DZIP1;refGene_tx=XM_023252687.1;refGene_gn=DZIP1;refGene_tx=XM_023252689.1;refGene_gn=DZIP1;refGene_tx=XM_023252698.1

A1 67554189 67554383
refGene_gn=DZIP1;refGene_tx=XM_023252677.1;refGene_gn=DZIP1;refGene_tx=XM_023252682.1;refGene_gn=DZIP1;refGene_tx=XM_023252687.1;refGene_gn=DZIP1;refGene_tx=XM_023252689.1;refGene_gn=DZIP1;refGene_tx=XM_023252698.1
A1 67556107 67556161
refGene_gn=DZIP1;refGene_tx=XM_023252677.1;refGene_gn=DZIP1;refGene_tx=XM_023252682.1;refGene_gn=DZIP1;refGene_tx=XM_023252687.1;refGene_gn=DZIP1;refGene_tx=XM_023252689.1;refGene_gn=DZIP1;refGene_tx=XM_023252698.1
A1 67557004 67557134
refGene_gn=DZIP1;refGene_tx=XM_023252677.1;refGene_gn=DZIP1;refGene_tx=XM_023252682.1;refGene_gn=DZIP1;refGene_tx=XM_023252687.1;refGene_gn=DZIP1;refGene_tx=XM_023252689.1;refGene_gn=DZIP1;refGene_tx=XM_023252698.1
A1 67557436 67557599
refGene_gn=DZIP1;refGene_tx=XM_023252677.1;refGene_gn=DZIP1;refGene_tx=XM_023252682.1;refGene_gn=DZIP1;refGene_tx=XM_023252687.1;refGene_gn=DZIP1;refGene_tx=XM_023252689.1;refGene_gn=DZIP1;refGene_tx=XM_023252698.1
A1 67558052 67558195
refGene_gn=DZIP1;refGene_tx=XM_023252677.1;refGene_gn=DZIP1;refGene_tx=XM_023252682.1;refGene_gn=DZIP1;refGene_tx=XM_023252687.1;refGene_gn=DZIP1;refGene_tx=XM_023252689.1;refGene_gn=DZIP1;refGene_tx=XM_023252698.1
A1 67563604 67563664
refGene_gn=DZIP1;refGene_tx=XM_023252677.1;refGene_gn=DZIP1;refGene_tx=XM_023252682.1;refGene_gn=DZIP1;refGene_tx=XM_023252687.1;refGene_gn=DZIP1;refGene_tx=XM_023252698.1
A1 67568722 67568779
refGene_gn=DZIP1;refGene_tx=XM_023252677.1;refGene_gn=DZIP1;refGene_tx=XM_023252682.1;refGene_gn=DZIP1;refGene_tx=XM_023252687.1;refGene_gn=DZIP1;refGene_tx=XM_023252689.1;refGene_gn=DZIP1;refGene_tx=XM_023252698.1
A1 67571005 67571062
refGene_gn=DZIP1;refGene_tx=XM_023252677.1;refGene_gn=DZIP1;refGene_tx=XM_023252682.1;refGene_gn=DZIP1;refGene_tx=XM_023252687.1;refGene_gn=DZIP1;refGene_tx=XM_023252689.1;refGene_gn=DZIP1;refGene_tx=XM_023252698.1
A1 67572204 67572253
refGene_gn=DZIP1;refGene_tx=XM_023252677.1;refGene_gn=DZIP1;refGene_tx=XM_023252682.1;refGene_gn=DZIP1;refGene_tx=XM_023252687.1;refGene_gn=DZIP1;refGene_tx=XM_023252689.1;refGene_gn=DZIP1;refGene_tx=XM_023252698.1
A1 67572701 67572842
refGene_gn=DZIP1;refGene_tx=XM_023252677.1;refGene_gn=DZIP1;refGene_tx=XM_023252682.1;refGene_gn=DZIP1;refGene_tx=XM_023252687.1;refGene_gn=DZIP1;refGene_tx=XM_023252689.1;refGene_gn=DZIP1;refGene_tx=XM_023252698.1
A1 67578989 67579052

refGene_gn=DZIP1;refGene_tx=XM_023252677.1;refGene_gn=DZIP1;refGene_tx=XM_023252682.1;refGene_gn=DZIP1;refGene_tx=XM_023252687.1;refGene_gn=DZIP1;refGene_tx=XM_023252689.1;refGene_gn=DZIP1;refGene_tx=XM_023252698.1
A1 67581988 67582126 refGene_gn=DZIP1;refGene_tx=XM_023252677.1;refGene_gn=DZIP1;refGene_tx=XM_023252682.1;refGene_gn=DZIP1;refGene_tx=XM_023252687.1;refGene_gn=DZIP1;refGene_tx=XM_023252689.1;refGene_gn=DZIP1;refGene_tx=XM_023252698.1
A1 67587612 67587774 refGene_gn=DZIP1;refGene_tx=XM_023252677.1;refGene_gn=DZIP1;refGene_tx=XM_023252682.1;refGene_gn=DZIP1;refGene_tx=XM_023252687.1;refGene_gn=DZIP1;refGene_tx=XM_023252689.1;refGene_gn=DZIP1;refGene_tx=XM_023252698.1
A1 67590611 67590736 refGene_gn=DZIP1;refGene_tx=XM_023252677.1;refGene_gn=DZIP1;refGene_tx=XM_023252682.1;refGene_gn=DZIP1;refGene_tx=XM_023252687.1;refGene_gn=DZIP1;refGene_tx=XM_023252689.1;refGene_gn=DZIP1;refGene_tx=XM_023252698.1
A1 67593430 67593515 refGene_gn=DZIP1;refGene_tx=XM_023252677.1;refGene_gn=DZIP1;refGene_tx=XM_023252682.1;refGene_gn=DZIP1;refGene_tx=XM_023252687.1;refGene_gn=DZIP1;refGene_tx=XM_023252689.1;refGene_gn=DZIP1;refGene_tx=XM_023252698.1
A1 67600923 67601484 refGene_gn=DZIP1;refGene_tx=XM_023252677.1;refGene_gn=DZIP1;refGene_tx=XM_023252682.1;refGene_gn=DZIP1;refGene_tx=XM_023252687.1;refGene_gn=DZIP1;refGene_tx=XM_023252689.1;refGene_gn=DZIP1;refGene_tx=XM_023252698.1
A1 67601595 67601598 refGene_gn=DZIP1;refGene_tx=XM_023252677.1;refGene_gn=DZIP1;refGene_tx=XM_023252682.1;refGene_gn=DZIP1;refGene_tx=XM_023252687.1;refGene_gn=DZIP1;refGene_tx=XM_023252689.1
A1 67628306 67628388 refGene_gn=DNAJC3;refGene_tx=XM_023252703.1
A1 67647664 67647775 refGene_gn=DNAJC3;refGene_tx=XM_023252703.1
A1 67658368 67658493 refGene_gn=DNAJC3;refGene_tx=XM_023252703.1
A1 67660655 67660730 refGene_gn=DNAJC3;refGene_tx=XM_023252703.1
A1 67682000 67682153 refGene_gn=DNAJC3;refGene_tx=XM_023252703.1
A1 67683651 67683833 refGene_gn=DNAJC3;refGene_tx=XM_023252703.1
A1 67684285 67684405 refGene_gn=DNAJC3;refGene_tx=XM_023252703.1
A1 67687657 67687763 refGene_gn=DNAJC3;refGene_tx=XM_023252703.1
A1 67687847 67687968 refGene_gn=DNAJC3;refGene_tx=XM_023252703.1
A1 67712747 67712880 refGene_gn=DNAJC3;refGene_tx=XM_023252703.1
A1 67713729 67713878 refGene_gn=DNAJC3;refGene_tx=XM_023252703.1
A1 67715820 67715978 refGene_gn=DNAJC3;refGene_tx=XM_023252703.1
A1 67725491 67725514 refGene_gn=UGGT2;refGene_tx=XM_011280607.3
A1 67733205 67733332 refGene_gn=UGGT2;refGene_tx=XM_011280607.3
A1 67737852 67737969 refGene_gn=UGGT2;refGene_tx=XM_011280607.3
A1 67739215 67739330 refGene_gn=UGGT2;refGene_tx=XM_011280607.3
A1 67740460 67740621 refGene_gn=UGGT2;refGene_tx=XM_011280607.3

A1 67742265 67742448 refGene_gn=UGGT2;refGene_tx=XM_011280607.3
A1 67746685 67746770 refGene_gn=UGGT2;refGene_tx=XM_011280607.3
A1 67747831 67747924 refGene_gn=UGGT2;refGene_tx=XM_011280607.3
A1 67749791 67749877 refGene_gn=UGGT2;refGene_tx=XM_011280607.3
A1 67755174 67755259 refGene_gn=UGGT2;refGene_tx=XM_011280607.3
A1 67757156 67757242 refGene_gn=UGGT2;refGene_tx=XM_011280607.3
A1 67757594 67757753 refGene_gn=UGGT2;refGene_tx=XM_011280607.3
A1 67763038 67763222 refGene_gn=UGGT2;refGene_tx=XM_011280607.3
A1 67769379 67769459 refGene_gn=UGGT2;refGene_tx=XM_011280607.3
A1 67771832 67771935 refGene_gn=UGGT2;refGene_tx=XM_011280607.3
A1 67776196 67776292 refGene_gn=UGGT2;refGene_tx=XM_011280607.3
A1 67781990 67782115 refGene_gn=UGGT2;refGene_tx=XM_011280607.3
A1 67787027 67787159 refGene_gn=UGGT2;refGene_tx=XM_011280607.3
A1 67791947 67792154 refGene_gn=UGGT2;refGene_tx=XM_011280607.3
A1 67796889 67796984 refGene_gn=UGGT2;refGene_tx=XM_011280607.3
A1 67799067 67799166 refGene_gn=UGGT2;refGene_tx=XM_011280607.3
A1 67799257 67799381 refGene_gn=UGGT2;refGene_tx=XM_011280607.3
A1 67799775 67799940 refGene_gn=UGGT2;refGene_tx=XM_011280607.3
A1 67813471 67813603 refGene_gn=UGGT2;refGene_tx=XM_011280607.3
A1 67821753 67821889 refGene_gn=UGGT2;refGene_tx=XM_011280607.3
A1 67822447 67822533 refGene_gn=UGGT2;refGene_tx=XM_011280607.3
A1 67823360 67823480 refGene_gn=UGGT2;refGene_tx=XM_011280607.3
A1 67833627 67833778 refGene_gn=UGGT2;refGene_tx=XM_011280607.3
A1 67838515 67838607 refGene_gn=UGGT2;refGene_tx=XM_011280607.3
A1 67859220 67859281 refGene_gn=UGGT2;refGene_tx=XM_011280607.3
A1 67861502 67861602 refGene_gn=UGGT2;refGene_tx=XM_011280607.3
A1 67867593 67867694 refGene_gn=UGGT2;refGene_tx=XM_011280607.3
A1 67869979 67870052 refGene_gn=UGGT2;refGene_tx=XM_011280607.3
A1 67877154 67877251 refGene_gn=UGGT2;refGene_tx=XM_011280607.3
A1 67881719 67881894 refGene_gn=UGGT2;refGene_tx=XM_011280607.3
A1 67893328 67893441 refGene_gn=UGGT2;refGene_tx=XM_011280607.3
A1 67893902 67893952 refGene_gn=UGGT2;refGene_tx=XM_011280607.3
A1 67893954 67894032 refGene_gn=UGGT2;refGene_tx=XM_011280607.3
A1 67902231 67902314 refGene_gn=UGGT2;refGene_tx=XM_011280607.3
A1 67916904 67917056 refGene_gn=UGGT2;refGene_tx=XM_011280607.3
A1 67956857 67957561 refGene_gn=HS6ST3;refGene_tx=XM_011280614.3
A1 68612139 68612848 refGene_gn=HS6ST3;refGene_tx=XM_011280614.3
A1 68725145 68726162 refGene_gn=OXGR1;refGene_tx=XM_023252451.1
A1 68976683 68976857
refGene_gn=MBNL2;refGene_tx=XM_003980483.5;refGene_gn=MBNL2;refGene_tx=XM_006927357.4;refGene_gn=MBNL2;refGene_tx=XM_011280623.3;refGene_gn=MBNL2;refGene_tx=XM_011280630.3;refGene_gn=MBNL2;refGene_tx=XM_011280633.3;refGene_gn=MBNL2;refGene_tx=XM_011280635.3;refGene_gn=MBNL2;refGene_tx=XM_019828698.2;refGene_gn=MBNL2;refGene_tx=XM_019828713.2
A1 69029099 69029264
refGene_gn=MBNL2;refGene_tx=XM_003980483.5;refGene_gn=MBNL2;refGene_tx=XM_006927357.4;refGene_gn=MBNL2;refGene_tx=XM_011280623.3;refGene_gn=MBNL2;refGene_tx=XM_011280630.3;refGene_gn=MBNL2;refGene_tx=XM_011280633.3;refGene_gn=MBNL2;refGene_tx=XM_011280635.3;refGene_gn=MBNL2;refGene_tx=XM_019828698.2;refGene_gn=MBNL2;refGene_tx=XM_019828713.2

A1 69039598 69039799
refGene_gn=MBNL2;refGene_tx=XM_003980483.5;refGene_gn=MBNL2;refGene_tx=XM_006927357.4;refGene_gn=MBNL2;refGene_tx=XM_011280623.3;refGene_gn=MBNL2;refGene_tx=XM_011280630.3;refGene_gn=MBNL2;refGene_tx=XM_011280633.3;refGene_gn=MBNL2;refGene_tx=XM_011280635.3;refGene_gn=MBNL2;refGene_tx=XM_019828698.2;refGene_gn=MBNL2;refGene_tx=XM_019828713.2
A1 69042545 69042809
refGene_gn=MBNL2;refGene_tx=XM_003980483.5;refGene_gn=MBNL2;refGene_tx=XM_006927357.4;refGene_gn=MBNL2;refGene_tx=XM_011280623.3;refGene_gn=MBNL2;refGene_tx=XM_011280630.3;refGene_gn=MBNL2;refGene_tx=XM_011280633.3;refGene_gn=MBNL2;refGene_tx=XM_011280635.3;refGene_gn=MBNL2;refGene_tx=XM_019828698.2;refGene_gn=MBNL2;refGene_tx=XM_019828713.2
A1 69050050 69050104
refGene_gn=MBNL2;refGene_tx=XM_011280623.3;refGene_gn=MBNL2;refGene_tx=XM_011280630.3;refGene_gn=MBNL2;refGene_tx=XM_011280633.3;refGene_gn=MBNL2;refGene_tx=XM_019828698.2
A1 69050784 69050938
refGene_gn=MBNL2;refGene_tx=XM_003980483.5;refGene_gn=MBNL2;refGene_tx=XM_006927357.4;refGene_gn=MBNL2;refGene_tx=XM_011280623.3;refGene_gn=MBNL2;refGene_tx=XM_011280630.3;refGene_gn=MBNL2;refGene_tx=XM_011280633.3;refGene_gn=MBNL2;refGene_tx=XM_011280635.3;refGene_gn=MBNL2;refGene_tx=XM_019828698.2;refGene_gn=MBNL2;refGene_tx=XM_019828713.2
A1 69058084 69058120
refGene_gn=MBNL2;refGene_tx=XM_011280623.3;refGene_gn=MBNL2;refGene_tx=XM_011280635.3;refGene_gn=MBNL2;refGene_tx=XM_019828698.2;refGene_gn=MBNL2;refGene_tx=XM_019828713.2
A1 69059411 69059506
refGene_gn=MBNL2;refGene_tx=XM_006927357.4;refGene_gn=MBNL2;refGene_tx=XM_011280623.3;refGene_gn=MBNL2;refGene_tx=XM_011280633.3;refGene_gn=MBNL2;refGene_tx=XM_011280635.3
A1 69082790 69082918
refGene_gn=MBNL2;refGene_tx=XM_003980483.5;refGene_gn=MBNL2;refGene_tx=XM_006927357.4;refGene_gn=MBNL2;refGene_tx=XM_011280623.3;refGene_gn=MBNL2;refGene_tx=XM_011280630.3;refGene_gn=MBNL2;refGene_tx=XM_011280633.3;refGene_gn=MBNL2;refGene_tx=XM_011280635.3;refGene_gn=MBNL2;refGene_tx=XM_019828698.2;refGene_gn=MBNL2;refGene_tx=XM_019828713.2
A1 69128884 69129198 refGene_gn=RAP2A;refGene_tx=XM_011280649.2
A1 69160754 69160992 refGene_gn=RAP2A;refGene_tx=XM_011280649.2
A1 69515660 69515750 refGene_gn=IPO5;refGene_tx=XM_023252761.1
A1 69522117 69522198 refGene_gn=IPO5;refGene_tx=XM_023252761.1
A1 69525079 69525272 refGene_gn=IPO5;refGene_tx=XM_023252761.1
A1 69527165 69527268 refGene_gn=IPO5;refGene_tx=XM_023252761.1
A1 69528374 69528471 refGene_gn=IPO5;refGene_tx=XM_023252761.1
A1 69528688 69528793 refGene_gn=IPO5;refGene_tx=XM_023252761.1
A1 69531004 69531127 refGene_gn=IPO5;refGene_tx=XM_023252761.1
A1 69531212 69531333 refGene_gn=IPO5;refGene_tx=XM_023252761.1
A1 69533124 69533212 refGene_gn=IPO5;refGene_tx=XM_023252761.1
A1 69535002 69535109 refGene_gn=IPO5;refGene_tx=XM_023252761.1
A1 69539814 69539939 refGene_gn=IPO5;refGene_tx=XM_023252761.1
A1 69540031 69540121 refGene_gn=IPO5;refGene_tx=XM_023252761.1

A1 69540213 69540387 refGene_gn=IPO5;refGene_tx=XM_023252761.1
A1 69542735 69542954 refGene_gn=IPO5;refGene_tx=XM_023252761.1
A1 69544462 69544546 refGene_gn=IPO5;refGene_tx=XM_023252761.1
A1 69545903 69546036 refGene_gn=IPO5;refGene_tx=XM_023252761.1
A1 69546123 69546245 refGene_gn=IPO5;refGene_tx=XM_023252761.1
A1 69547827 69547924 refGene_gn=IPO5;refGene_tx=XM_023252761.1
A1 69549564 69549737 refGene_gn=IPO5;refGene_tx=XM_023252761.1
A1 69550683 69550795 refGene_gn=IPO5;refGene_tx=XM_023252761.1
A1 69550882 69550938 refGene_gn=IPO5;refGene_tx=XM_023252761.1
A1 69552092 69552215 refGene_gn=IPO5;refGene_tx=XM_023252761.1
A1 69553083 69553303 refGene_gn=IPO5;refGene_tx=XM_023252761.1
A1 69553915 69554144 refGene_gn=IPO5;refGene_tx=XM_023252761.1
A1 69556316 69556458 refGene_gn=IPO5;refGene_tx=XM_023252761.1
A1 69557061 69557148 refGene_gn=IPO5;refGene_tx=XM_023252761.1
A1 69718123 69718294
refGene_gn=FARP1;refGene_tx=XM_011280659.3;refGene_gn=FARP1;refGene_tx=XM_023252762.1;refGene_gn=FARP1;refGene_tx=XM_023252768.1
A1 69847765 69847870
refGene_gn=FARP1;refGene_tx=XM_011280659.3;refGene_gn=FARP1;refGene_tx=XM_023252762.1;refGene_gn=FARP1;refGene_tx=XM_023252768.1
A1 69867126 69867169
refGene_gn=FARP1;refGene_tx=XM_011280659.3;refGene_gn=FARP1;refGene_tx=XM_023252762.1;refGene_gn=FARP1;refGene_tx=XM_023252768.1
A1 69869667 69869746
refGene_gn=FARP1;refGene_tx=XM_011280659.3;refGene_gn=FARP1;refGene_tx=XM_023252762.1;refGene_gn=FARP1;refGene_tx=XM_023252768.1
A1 69878934 69879032
refGene_gn=FARP1;refGene_tx=XM_011280659.3;refGene_gn=FARP1;refGene_tx=XM_023252762.1;refGene_gn=FARP1;refGene_tx=XM_023252768.1
A1 69880960 69881075
refGene_gn=FARP1;refGene_tx=XM_011280659.3;refGene_gn=FARP1;refGene_tx=XM_023252762.1;refGene_gn=FARP1;refGene_tx=XM_023252768.1
A1 69881969 69882117
refGene_gn=FARP1;refGene_tx=XM_011280659.3;refGene_gn=FARP1;refGene_tx=XM_023252762.1;refGene_gn=FARP1;refGene_tx=XM_023252768.1
A1 69884661 69884757
refGene_gn=FARP1;refGene_tx=XM_011280659.3;refGene_gn=FARP1;refGene_tx=XM_023252762.1;refGene_gn=FARP1;refGene_tx=XM_023252768.1
A1 69886576 69886740
refGene_gn=FARP1;refGene_tx=XM_011280659.3;refGene_gn=FARP1;refGene_tx=XM_023252762.1;refGene_gn=FARP1;refGene_tx=XM_023252768.1
A1 69887460 69887529
refGene_gn=FARP1;refGene_tx=XM_011280659.3;refGene_gn=FARP1;refGene_tx=XM_023252762.1;refGene_gn=FARP1;refGene_tx=XM_023252768.1
A1 69890067 69890143
refGene_gn=FARP1;refGene_tx=XM_011280659.3;refGene_gn=FARP1;refGene_tx=XM_023252762.1;refGene_gn=FARP1;refGene_tx=XM_023252768.1
A1 69891210 69891451
refGene_gn=FARP1;refGene_tx=XM_011280659.3;refGene_gn=FARP1;refGene_tx=XM_023252762.1;refGene_gn=FARP1;refGene_tx=XM_023252768.1

A1 69904592 69904780
refGene_gn=FARP1;refGene_tx=XM_011280659.3;refGene_gn=FARP1;refGene_tx=XM_023252762.1;refGene_gn=FARP1;refGene_tx=XM_023252768.1
A1 69905893 69905983
refGene_gn=FARP1;refGene_tx=XM_011280659.3;refGene_gn=FARP1;refGene_tx=XM_023252762.1;refGene_gn=FARP1;refGene_tx=XM_023252768.1
A1 69907022 69907156
refGene_gn=FARP1;refGene_tx=XM_011280659.3;refGene_gn=FARP1;refGene_tx=XM_023252762.1;refGene_gn=FARP1;refGene_tx=XM_023252768.1
A1 69925714 69925796
refGene_gn=FARP1;refGene_tx=XM_011280659.3;refGene_gn=FARP1;refGene_tx=XM_023252762.1;refGene_gn=FARP1;refGene_tx=XM_023252768.1
A1 69931745 69931983
refGene_gn=FARP1;refGene_tx=XM_011280659.3;refGene_gn=FARP1;refGene_tx=XM_023252762.1;refGene_gn=FARP1;refGene_tx=XM_023252768.1
A1 69934958 69935089
refGene_gn=FARP1;refGene_tx=XM_011280659.3;refGene_gn=FARP1;refGene_tx=XM_023252762.1;refGene_gn=FARP1;refGene_tx=XM_023252768.1
A1 69939662 69939731
refGene_gn=FARP1;refGene_tx=XM_011280659.3;refGene_gn=FARP1;refGene_tx=XM_023252762.1;refGene_gn=FARP1;refGene_tx=XM_023252768.1
A1 69939927 69940017
refGene_gn=FARP1;refGene_tx=XM_011280659.3;refGene_gn=FARP1;refGene_tx=XM_023252762.1;refGene_gn=FARP1;refGene_tx=XM_023252768.1
A1 69940655 69940738
refGene_gn=FARP1;refGene_tx=XM_011280659.3;refGene_gn=FARP1;refGene_tx=XM_023252762.1;refGene_gn=FARP1;refGene_tx=XM_023252768.1
A1 69940813 69940926
refGene_gn=FARP1;refGene_tx=XM_011280659.3;refGene_gn=FARP1;refGene_tx=XM_023252762.1;refGene_gn=FARP1;refGene_tx=XM_023252768.1
A1 69941452 69941619
refGene_gn=FARP1;refGene_tx=XM_011280659.3;refGene_gn=FARP1;refGene_tx=XM_023252762.1;refGene_gn=FARP1;refGene_tx=XM_023252768.1
A1 69945327 69945435
refGene_gn=FARP1;refGene_tx=XM_011280659.3;refGene_gn=FARP1;refGene_tx=XM_023252762.1;refGene_gn=FARP1;refGene_tx=XM_023252768.1
A1 69945951 69946103
refGene_gn=FARP1;refGene_tx=XM_011280659.3;refGene_gn=FARP1;refGene_tx=XM_023252762.1;refGene_gn=FARP1;refGene_tx=XM_023252768.1
A1 69947537 69947619
refGene_gn=FARP1;refGene_tx=XM_011280659.3;refGene_gn=FARP1;refGene_tx=XM_023252762.1;refGene_gn=FARP1;refGene_tx=XM_023252768.1
A1 69953195 69953232
refGene_gn=STK24;refGene_tx=XM_003980489.5;refGene_gn=STK24;refGene_tx=XM_019828795.2;refGene_gn=STK24;refGene_tx=XM_023252773.1
A1 69957478 69957615
refGene_gn=STK24;refGene_tx=XM_003980489.5;refGene_gn=STK24;refGene_tx=XM_019828795.2;refGene_gn=STK24;refGene_tx=XM_023252773.1
A1 69960191 69960260
refGene_gn=STK24;refGene_tx=XM_003980489.5;refGene_gn=STK24;refGene_tx

=XM_019828795.2;refGene_gn=STK24;refGene_tx=XM_023252773.1
A1	69961507	69961637
refGene_gn=STK24;refGene_tx=XM_003980489.5;refGene_gn=STK24;refGene_tx
=XM_019828795.2;refGene_gn=STK24;refGene_tx=XM_023252773.1
A1	69963105	69963251
refGene_gn=STK24;refGene_tx=XM_003980489.5;refGene_gn=STK24;refGene_tx
=XM_019828795.2;refGene_gn=STK24;refGene_tx=XM_023252773.1
A1	69965536	69965722
refGene_gn=STK24;refGene_tx=XM_003980489.5;refGene_gn=STK24;refGene_tx
=XM_019828795.2;refGene_gn=STK24;refGene_tx=XM_023252773.1
A1	69975517	69975675
refGene_gn=STK24;refGene_tx=XM_003980489.5;refGene_gn=STK24;refGene_tx
=XM_023252773.1
A1	69975976	69976085
refGene_gn=STK24;refGene_tx=XM_003980489.5;refGene_gn=STK24;refGene_tx
=XM_023252773.1
A1	69986336	69986393
refGene_gn=STK24;refGene_tx=XM_003980489.5;refGene_gn=STK24;refGene_tx
=XM_019828795.2;refGene_gn=STK24;refGene_tx=XM_023252773.1
A1	70023616	70023847
refGene_gn=STK24;refGene_tx=XM_003980489.5;refGene_gn=STK24;refGene_tx
=XM_019828795.2;refGene_gn=STK24;refGene_tx=XM_023252773.1
A1	70025352	70025430
refGene_gn=STK24;refGene_tx=XM_003980489.5;refGene_gn=STK24;refGene_tx
=XM_019828795.2
A1	70074221	70074263	refGene_gn=STK24;refGene_tx=XM_023252773.1
A1	70155944	70156130	refGene_gn=SLC15A1;refGene_tx=XM_023252783.1
A1	70157047	70157155	refGene_gn=SLC15A1;refGene_tx=XM_023252783.1
A1	70158142	70158286	refGene_gn=SLC15A1;refGene_tx=XM_023252783.1
A1	70158828	70158940	refGene_gn=SLC15A1;refGene_tx=XM_023252783.1
A1	70159056	70159164	refGene_gn=SLC15A1;refGene_tx=XM_023252783.1
A1	70168721	70168771	refGene_gn=SLC15A1;refGene_tx=XM_023252783.1
A1	70169749	70169896	refGene_gn=SLC15A1;refGene_tx=XM_023252783.1
A1	70171194	70171314	refGene_gn=SLC15A1;refGene_tx=XM_023252783.1
A1	70173121	70173203	refGene_gn=SLC15A1;refGene_tx=XM_023252783.1
A1	70173704	70173793	refGene_gn=SLC15A1;refGene_tx=XM_023252783.1
A1	70173877	70173910	refGene_gn=SLC15A1;refGene_tx=XM_023252783.1
A1	70174070	70174115	refGene_gn=SLC15A1;refGene_tx=XM_023252783.1
A1	70176190	70176280	refGene_gn=SLC15A1;refGene_tx=XM_023252783.1
A1	70177325	70177412	refGene_gn=SLC15A1;refGene_tx=XM_023252783.1
A1	70180674	70180757	refGene_gn=SLC15A1;refGene_tx=XM_023252783.1
A1	70183473	70183557	refGene_gn=SLC15A1;refGene_tx=XM_023252783.1
A1	70186273	70186364	refGene_gn=SLC15A1;refGene_tx=XM_023252783.1
A1	70186540	70186640	refGene_gn=SLC15A1;refGene_tx=XM_023252783.1
A1	70193498	70193618	refGene_gn=SLC15A1;refGene_tx=XM_023252783.1
A1	70194993	70195135	refGene_gn=SLC15A1;refGene_tx=XM_023252783.1
A1	70195236	70195318	refGene_gn=SLC15A1;refGene_tx=XM_023252783.1
A1	70195925	70195942	refGene_gn=SLC15A1;refGene_tx=XM_023252783.1
A1	70213141	70213145	refGene_gn=SLC15A1;refGene_tx=XM_023252783.1
A1	70276327	70276450

refGene_gn=DOCK9;refGene_tx=XM_006927365.4;refGene_gn=DOCK9;refGene_tx
=XM_006927368.4;refGene_gn=DOCK9;refGene_tx=XM_011280686.3;refGene_gn=
DOCK9;refGene_tx=XM_019828858.2;refGene_gn=DOCK9;refGene_tx=XM_0198288
65.2;refGene_gn=DOCK9;refGene_tx=XM_019828880.2;refGene_gn=DOCK9;refGe
ne_tx=XM_019828896.2;refGene_gn=DOCK9;refGene_tx=XM_019828903.2;refGen
e_gn=DOCK9;refGene_tx=XM_019828909.2;refGene_gn=DOCK9;refGene_tx=XM_02
3252786.1;refGene_gn=DOCK9;refGene_tx=XM_023252788.1;refGene_gn=DOCK9;
refGene_tx=XM_023252805.1;refGene_gn=DOCK9;refGene_tx=XM_023252809.1;r
efGene_gn=DOCK9;refGene_tx=XM_023252810.1;refGene_gn=DOCK9;refGene_tx=
XM_023252814.1;refGene_gn=DOCK9;refGene_tx=XM_023252826.1;refGene_gn=D
OCK9;refGene_tx=XM_023252829.1;refGene_gn=DOCK9;refGene_tx=XM_02325283
1.1
A1 70277521 70277529 refGene_gn=DOCK9;refGene_tx=XM_023252814.1
A1 70278487 70278626
refGene_gn=DOCK9;refGene_tx=XM_006927365.4;refGene_gn=DOCK9;refGene_tx
=XM_006927368.4;refGene_gn=DOCK9;refGene_tx=XM_011280686.3;refGene_gn=
DOCK9;refGene_tx=XM_019828858.2;refGene_gn=DOCK9;refGene_tx=XM_0198288
65.2;refGene_gn=DOCK9;refGene_tx=XM_019828880.2;refGene_gn=DOCK9;refGe
ne_tx=XM_019828896.2;refGene_gn=DOCK9;refGene_tx=XM_019828903.2;refGen
e_gn=DOCK9;refGene_tx=XM_019828909.2;refGene_gn=DOCK9;refGene_tx=XM_02
3252786.1;refGene_gn=DOCK9;refGene_tx=XM_023252788.1;refGene_gn=DOCK9;
refGene_tx=XM_023252805.1;refGene_gn=DOCK9;refGene_tx=XM_023252809.1;r
efGene_gn=DOCK9;refGene_tx=XM_023252810.1;refGene_gn=DOCK9;refGene_tx=
XM_023252814.1;refGene_gn=DOCK9;refGene_tx=XM_023252826.1;refGene_gn=D
OCK9;refGene_tx=XM_023252829.1;refGene_gn=DOCK9;refGene_tx=XM_02325283
1.1
A1 70278761 70278862
refGene_gn=DOCK9;refGene_tx=XM_006927365.4;refGene_gn=DOCK9;refGene_tx
=XM_006927368.4;refGene_gn=DOCK9;refGene_tx=XM_011280686.3;refGene_gn=
DOCK9;refGene_tx=XM_019828858.2;refGene_gn=DOCK9;refGene_tx=XM_0198288
65.2;refGene_gn=DOCK9;refGene_tx=XM_019828880.2;refGene_gn=DOCK9;refGe
ne_tx=XM_019828896.2;refGene_gn=DOCK9;refGene_tx=XM_019828903.2;refGen
e_gn=DOCK9;refGene_tx=XM_019828909.2;refGene_gn=DOCK9;refGene_tx=XM_02
3252786.1;refGene_gn=DOCK9;refGene_tx=XM_023252788.1;refGene_gn=DOCK9;
refGene_tx=XM_023252805.1;refGene_gn=DOCK9;refGene_tx=XM_023252809.1;r
efGene_gn=DOCK9;refGene_tx=XM_023252810.1;refGene_gn=DOCK9;refGene_tx=
XM_023252814.1;refGene_gn=DOCK9;refGene_tx=XM_023252826.1;refGene_gn=D
OCK9;refGene_tx=XM_023252829.1;refGene_gn=DOCK9;refGene_tx=XM_02325283
1.1
A1 70281950 70282141
refGene_gn=DOCK9;refGene_tx=XM_006927365.4;refGene_gn=DOCK9;refGene_tx
=XM_006927368.4;refGene_gn=DOCK9;refGene_tx=XM_011280686.3;refGene_gn=
DOCK9;refGene_tx=XM_019828858.2;refGene_gn=DOCK9;refGene_tx=XM_0198288
65.2;refGene_gn=DOCK9;refGene_tx=XM_019828880.2;refGene_gn=DOCK9;refGe
ne_tx=XM_019828896.2;refGene_gn=DOCK9;refGene_tx=XM_019828903.2;refGen
e_gn=DOCK9;refGene_tx=XM_019828909.2;refGene_gn=DOCK9;refGene_tx=XM_02
3252786.1;refGene_gn=DOCK9;refGene_tx=XM_023252788.1;refGene_gn=DOCK9;
refGene_tx=XM_023252805.1;refGene_gn=DOCK9;refGene_tx=XM_023252809.1;r
efGene_gn=DOCK9;refGene_tx=XM_023252810.1;refGene_gn=DOCK9;refGene_tx=
XM_023252814.1;refGene_gn=DOCK9;refGene_tx=XM_023252826.1;refGene_gn=D

OCK9;refGene_tx=XM_023252829.1;refGene_gn=DOCK9;refGene_tx=XM_02325283
1.1
A1 70287769 70287980
refGene_gn=DOCK9;refGene_tx=XM_006927365.4;refGene_gn=DOCK9;refGene_tx
=XM_006927368.4;refGene_gn=DOCK9;refGene_tx=XM_011280686.3;refGene_gn=
DOCK9;refGene_tx=XM_019828858.2;refGene_gn=DOCK9;refGene_tx=XM_0198288
65.2;refGene_gn=DOCK9;refGene_tx=XM_019828880.2;refGene_gn=DOCK9;refGe
ne_tx=XM_019828896.2;refGene_gn=DOCK9;refGene_tx=XM_019828903.2;refGen
e_gn=DOCK9;refGene_tx=XM_019828909.2;refGene_gn=DOCK9;refGene_tx=XM_02
3252786.1;refGene_gn=DOCK9;refGene_tx=XM_023252788.1;refGene_gn=DOCK9;
refGene_tx=XM_023252805.1;refGene_gn=DOCK9;refGene_tx=XM_023252809.1;r
efGene_gn=DOCK9;refGene_tx=XM_023252810.1;refGene_gn=DOCK9;refGene_tx=
XM_023252814.1;refGene_gn=DOCK9;refGene_tx=XM_023252826.1;refGene_gn=D
OCK9;refGene_tx=XM_023252829.1;refGene_gn=DOCK9;refGene_tx=XM_02325283
1.1
A1 70290737 70290884
refGene_gn=DOCK9;refGene_tx=XM_006927365.4;refGene_gn=DOCK9;refGene_tx
=XM_006927368.4;refGene_gn=DOCK9;refGene_tx=XM_011280686.3;refGene_gn=
DOCK9;refGene_tx=XM_019828858.2;refGene_gn=DOCK9;refGene_tx=XM_0198288
65.2;refGene_gn=DOCK9;refGene_tx=XM_019828880.2;refGene_gn=DOCK9;refGe
ne_tx=XM_019828896.2;refGene_gn=DOCK9;refGene_tx=XM_019828903.2;refGen
e_gn=DOCK9;refGene_tx=XM_019828909.2;refGene_gn=DOCK9;refGene_tx=XM_02
3252786.1;refGene_gn=DOCK9;refGene_tx=XM_023252788.1;refGene_gn=DOCK9;
refGene_tx=XM_023252805.1;refGene_gn=DOCK9;refGene_tx=XM_023252809.1;r
efGene_gn=DOCK9;refGene_tx=XM_023252810.1;refGene_gn=DOCK9;refGene_tx=
XM_023252814.1;refGene_gn=DOCK9;refGene_tx=XM_023252826.1;refGene_gn=D
OCK9;refGene_tx=XM_023252829.1;refGene_gn=DOCK9;refGene_tx=XM_02325283
1.1
A1 70291724 70291760
refGene_gn=DOCK9;refGene_tx=XM_019828858.2;refGene_gn=DOCK9;refGene_tx
=XM_019828865.2;refGene_gn=DOCK9;refGene_tx=XM_019828880.2;refGene_gn=
DOCK9;refGene_tx=XM_019828896.2;refGene_gn=DOCK9;refGene_tx=XM_0198289
09.2;refGene_gn=DOCK9;refGene_tx=XM_023252786.1;refGene_gn=DOCK9;refGe
ne_tx=XM_023252788.1;refGene_gn=DOCK9;refGene_tx=XM_023252805.1;refGen
e_gn=DOCK9;refGene_tx=XM_023252809.1;refGene_gn=DOCK9;refGene_tx=XM_02
3252810.1;refGene_gn=DOCK9;refGene_tx=XM_023252814.1;refGene_gn=DOCK9;
refGene_tx=XM_023252829.1
A1 70292225 70292231
refGene_gn=DOCK9;refGene_tx=XM_019828858.2;refGene_gn=DOCK9;refGene_tx
=XM_019828865.2;refGene_gn=DOCK9;refGene_tx=XM_019828880.2;refGene_gn=
DOCK9;refGene_tx=XM_019828896.2;refGene_gn=DOCK9;refGene_tx=XM_0198289
09.2;refGene_gn=DOCK9;refGene_tx=XM_023252788.1;refGene_gn=DOCK9;refGe
ne_tx=XM_023252805.1;refGene_gn=DOCK9;refGene_tx=XM_023252809.1;refGen
e_gn=DOCK9;refGene_tx=XM_023252810.1;refGene_gn=DOCK9;refGene_tx=XM_02
3252814.1;refGene_gn=DOCK9;refGene_tx=XM_023252829.1
A1 70292446 70292560
refGene_gn=DOCK9;refGene_tx=XM_006927365.4;refGene_gn=DOCK9;refGene_tx
=XM_006927368.4;refGene_gn=DOCK9;refGene_tx=XM_011280686.3;refGene_gn=
DOCK9;refGene_tx=XM_019828858.2;refGene_gn=DOCK9;refGene_tx=XM_0198288
65.2;refGene_gn=DOCK9;refGene_tx=XM_019828880.2;refGene_gn=DOCK9;refGe

ne_tx=XM_019828896.2;refGene_gn=DOCK9;refGene_tx=XM_019828903.2;refGene_gn=DOCK9;refGene_tx=XM_019828909.2;refGene_gn=DOCK9;refGene_tx=XM_023252786.1;refGene_gn=DOCK9;refGene_tx=XM_023252788.1;refGene_gn=DOCK9;refGene_tx=XM_023252805.1;refGene_gn=DOCK9;refGene_tx=XM_023252809.1;refGene_gn=DOCK9;refGene_tx=XM_023252810.1;refGene_gn=DOCK9;refGene_tx=XM_023252814.1;refGene_gn=DOCK9;refGene_tx=XM_023252826.1;refGene_gn=DOCK9;refGene_tx=XM_023252829.1;refGene_gn=DOCK9;refGene_tx=XM_023252831.1
A1 70293288 70293411
refGene_gn=DOCK9;refGene_tx=XM_006927365.4;refGene_gn=DOCK9;refGene_tx=XM_006927368.4;refGene_gn=DOCK9;refGene_tx=XM_011280686.3;refGene_gn=DOCK9;refGene_tx=XM_019828858.2;refGene_gn=DOCK9;refGene_tx=XM_019828865.2;refGene_gn=DOCK9;refGene_tx=XM_019828880.2;refGene_gn=DOCK9;refGene_tx=XM_019828896.2;refGene_gn=DOCK9;refGene_tx=XM_019828903.2;refGene_gn=DOCK9;refGene_tx=XM_019828909.2;refGene_gn=DOCK9;refGene_tx=XM_023252786.1;refGene_gn=DOCK9;refGene_tx=XM_023252788.1;refGene_gn=DOCK9;refGene_tx=XM_023252805.1;refGene_gn=DOCK9;refGene_tx=XM_023252809.1;refGene_gn=DOCK9;refGene_tx=XM_023252810.1;refGene_gn=DOCK9;refGene_tx=XM_023252814.1;refGene_gn=DOCK9;refGene_tx=XM_023252826.1;refGene_gn=DOCK9;refGene_tx=XM_023252829.1;refGene_gn=DOCK9;refGene_tx=XM_023252831.1
A1 70303626 70303733
refGene_gn=DOCK9;refGene_tx=XM_006927365.4;refGene_gn=DOCK9;refGene_tx=XM_006927368.4;refGene_gn=DOCK9;refGene_tx=XM_011280686.3;refGene_gn=DOCK9;refGene_tx=XM_019828858.2;refGene_gn=DOCK9;refGene_tx=XM_019828865.2;refGene_gn=DOCK9;refGene_tx=XM_019828880.2;refGene_gn=DOCK9;refGene_tx=XM_019828896.2;refGene_gn=DOCK9;refGene_tx=XM_019828903.2;refGene_gn=DOCK9;refGene_tx=XM_019828909.2;refGene_gn=DOCK9;refGene_tx=XM_023252786.1;refGene_gn=DOCK9;refGene_tx=XM_023252788.1;refGene_gn=DOCK9;refGene_tx=XM_023252805.1;refGene_gn=DOCK9;refGene_tx=XM_023252809.1;refGene_gn=DOCK9;refGene_tx=XM_023252810.1;refGene_gn=DOCK9;refGene_tx=XM_023252814.1;refGene_gn=DOCK9;refGene_tx=XM_023252826.1;refGene_gn=DOCK9;refGene_tx=XM_023252829.1;refGene_gn=DOCK9;refGene_tx=XM_023252831.1
A1 70305153 70305231
refGene_gn=DOCK9;refGene_tx=XM_006927365.4;refGene_gn=DOCK9;refGene_tx=XM_011280686.3;refGene_gn=DOCK9;refGene_tx=XM_019828858.2;refGene_gn=DOCK9;refGene_tx=XM_019828865.2;refGene_gn=DOCK9;refGene_tx=XM_019828880.2;refGene_gn=DOCK9;refGene_tx=XM_019828896.2;refGene_gn=DOCK9;refGene_tx=XM_023252786.1;refGene_gn=DOCK9;refGene_tx=XM_023252788.1;refGene_gn=DOCK9;refGene_tx=XM_023252809.1;refGene_gn=DOCK9;refGene_tx=XM_023252810.1;refGene_gn=DOCK9;refGene_tx=XM_023252814.1
A1 70306161 70306219
refGene_gn=DOCK9;refGene_tx=XM_006927365.4;refGene_gn=DOCK9;refGene_tx=XM_006927368.4;refGene_gn=DOCK9;refGene_tx=XM_011280686.3;refGene_gn=DOCK9;refGene_tx=XM_019828858.2;refGene_gn=DOCK9;refGene_tx=XM_019828865.2;refGene_gn=DOCK9;refGene_tx=XM_019828880.2;refGene_gn=DOCK9;refGene_tx=XM_019828896.2;refGene_gn=DOCK9;refGene_tx=XM_019828903.2;refGene_gn=DOCK9;refGene_tx=XM_019828909.2;refGene_gn=DOCK9;refGene_tx=XM_023252786.1;refGene_gn=DOCK9;refGene_tx=XM_023252788.1;refGene_gn=DOCK9;

refGene_tx=XM_023252805.1;refGene_gn=DOCK9;refGene_tx=XM_023252809.1;refGene_gn=DOCK9;refGene_tx=XM_023252810.1;refGene_gn=DOCK9;refGene_tx=XM_023252814.1;refGene_gn=DOCK9;refGene_tx=XM_023252826.1;refGene_gn=DOCK9;refGene_tx=XM_023252829.1;refGene_gn=DOCK9;refGene_tx=XM_023252831.1

A1 70307926 70308142
refGene_gn=DOCK9;refGene_tx=XM_006927365.4;refGene_gn=DOCK9;refGene_tx=XM_006927368.4;refGene_gn=DOCK9;refGene_tx=XM_011280686.3;refGene_gn=DOCK9;refGene_tx=XM_019828858.2;refGene_gn=DOCK9;refGene_tx=XM_019828865.2;refGene_gn=DOCK9;refGene_tx=XM_019828880.2;refGene_gn=DOCK9;refGene_tx=XM_019828896.2;refGene_gn=DOCK9;refGene_tx=XM_019828903.2;refGene_gn=DOCK9;refGene_tx=XM_019828909.2;refGene_gn=DOCK9;refGene_tx=XM_023252786.1;refGene_gn=DOCK9;refGene_tx=XM_023252788.1;refGene_gn=DOCK9;refGene_tx=XM_023252805.1;refGene_gn=DOCK9;refGene_tx=XM_023252809.1;refGene_gn=DOCK9;refGene_tx=XM_023252810.1;refGene_gn=DOCK9;refGene_tx=XM_023252814.1;refGene_gn=DOCK9;refGene_tx=XM_023252826.1;refGene_gn=DOCK9;refGene_tx=XM_023252829.1;refGene_gn=DOCK9;refGene_tx=XM_023252831.1

A1 70308236 70308350
refGene_gn=DOCK9;refGene_tx=XM_006927365.4;refGene_gn=DOCK9;refGene_tx=XM_006927368.4;refGene_gn=DOCK9;refGene_tx=XM_011280686.3;refGene_gn=DOCK9;refGene_tx=XM_019828858.2;refGene_gn=DOCK9;refGene_tx=XM_019828865.2;refGene_gn=DOCK9;refGene_tx=XM_019828880.2;refGene_gn=DOCK9;refGene_tx=XM_019828896.2;refGene_gn=DOCK9;refGene_tx=XM_019828903.2;refGene_gn=DOCK9;refGene_tx=XM_019828909.2;refGene_gn=DOCK9;refGene_tx=XM_023252786.1;refGene_gn=DOCK9;refGene_tx=XM_023252788.1;refGene_gn=DOCK9;refGene_tx=XM_023252805.1;refGene_gn=DOCK9;refGene_tx=XM_023252809.1;refGene_gn=DOCK9;refGene_tx=XM_023252810.1;refGene_gn=DOCK9;refGene_tx=XM_023252814.1;refGene_gn=DOCK9;refGene_tx=XM_023252826.1;refGene_gn=DOCK9;refGene_tx=XM_023252829.1;refGene_gn=DOCK9;refGene_tx=XM_023252831.1

A1 70309935 70310118
refGene_gn=DOCK9;refGene_tx=XM_006927365.4;refGene_gn=DOCK9;refGene_tx=XM_006927368.4;refGene_gn=DOCK9;refGene_tx=XM_011280686.3;refGene_gn=DOCK9;refGene_tx=XM_019828858.2;refGene_gn=DOCK9;refGene_tx=XM_019828865.2;refGene_gn=DOCK9;refGene_tx=XM_019828880.2;refGene_gn=DOCK9;refGene_tx=XM_019828896.2;refGene_gn=DOCK9;refGene_tx=XM_019828903.2;refGene_gn=DOCK9;refGene_tx=XM_019828909.2;refGene_gn=DOCK9;refGene_tx=XM_023252786.1;refGene_gn=DOCK9;refGene_tx=XM_023252788.1;refGene_gn=DOCK9;refGene_tx=XM_023252805.1;refGene_gn=DOCK9;refGene_tx=XM_023252809.1;refGene_gn=DOCK9;refGene_tx=XM_023252810.1;refGene_gn=DOCK9;refGene_tx=XM_023252814.1;refGene_gn=DOCK9;refGene_tx=XM_023252826.1;refGene_gn=DOCK9;refGene_tx=XM_023252829.1;refGene_gn=DOCK9;refGene_tx=XM_023252831.1

A1 70310234 70310372
refGene_gn=DOCK9;refGene_tx=XM_006927365.4;refGene_gn=DOCK9;refGene_tx=XM_006927368.4;refGene_gn=DOCK9;refGene_tx=XM_011280686.3;refGene_gn=DOCK9;refGene_tx=XM_019828858.2;refGene_gn=DOCK9;refGene_tx=XM_019828865.2;refGene_gn=DOCK9;refGene_tx=XM_019828880.2;refGene_gn=DOCK9;refGene_tx=XM_019828896.2;refGene_gn=DOCK9;refGene_tx=XM_019828903.2;refGen

e_gn=DOCK9;refGene_tx=XM_019828909.2;refGene_gn=DOCK9;refGene_tx=XM_02
3252786.1;refGene_gn=DOCK9;refGene_tx=XM_023252788.1;refGene_gn=DOCK9;
refGene_tx=XM_023252805.1;refGene_gn=DOCK9;refGene_tx=XM_023252809.1;r
efGene_gn=DOCK9;refGene_tx=XM_023252810.1;refGene_gn=DOCK9;refGene_tx=
XM_023252814.1;refGene_gn=DOCK9;refGene_tx=XM_023252826.1;refGene_gn=D
OCK9;refGene_tx=XM_023252829.1;refGene_gn=DOCK9;refGene_tx=XM_02325283
1.1
A1 70316343 70316459
refGene_gn=DOCK9;refGene_tx=XM_006927365.4;refGene_gn=DOCK9;refGene_tx
=XM_006927368.4;refGene_gn=DOCK9;refGene_tx=XM_011280686.3;refGene_gn=
DOCK9;refGene_tx=XM_019828858.2;refGene_gn=DOCK9;refGene_tx=XM_0198288
65.2;refGene_gn=DOCK9;refGene_tx=XM_019828880.2;refGene_gn=DOCK9;refGe
ne_tx=XM_019828896.2;refGene_gn=DOCK9;refGene_tx=XM_019828903.2;refGen
e_gn=DOCK9;refGene_tx=XM_019828909.2;refGene_gn=DOCK9;refGene_tx=XM_02
3252786.1;refGene_gn=DOCK9;refGene_tx=XM_023252788.1;refGene_gn=DOCK9;
refGene_tx=XM_023252805.1;refGene_gn=DOCK9;refGene_tx=XM_023252809.1;r
efGene_gn=DOCK9;refGene_tx=XM_023252810.1;refGene_gn=DOCK9;refGene_tx=
XM_023252814.1;refGene_gn=DOCK9;refGene_tx=XM_023252826.1;refGene_gn=D
OCK9;refGene_tx=XM_023252829.1;refGene_gn=DOCK9;refGene_tx=XM_02325283
1.1
A1 70323871 70323915
refGene_gn=DOCK9;refGene_tx=XM_023252809.1;refGene_gn=DOCK9;refGene_tx
=XM_023252831.1
A1 70324426 70324563
refGene_gn=DOCK9;refGene_tx=XM_006927365.4;refGene_gn=DOCK9;refGene_tx
=XM_006927368.4;refGene_gn=DOCK9;refGene_tx=XM_011280686.3;refGene_gn=
DOCK9;refGene_tx=XM_019828858.2;refGene_gn=DOCK9;refGene_tx=XM_0198288
65.2;refGene_gn=DOCK9;refGene_tx=XM_019828880.2;refGene_gn=DOCK9;refGe
ne_tx=XM_019828896.2;refGene_gn=DOCK9;refGene_tx=XM_019828903.2;refGen
e_gn=DOCK9;refGene_tx=XM_019828909.2;refGene_gn=DOCK9;refGene_tx=XM_02
3252786.1;refGene_gn=DOCK9;refGene_tx=XM_023252788.1;refGene_gn=DOCK9;
refGene_tx=XM_023252805.1;refGene_gn=DOCK9;refGene_tx=XM_023252810.1;r
efGene_gn=DOCK9;refGene_tx=XM_023252814.1;refGene_gn=DOCK9;refGene_tx=
XM_023252826.1;refGene_gn=DOCK9;refGene_tx=XM_023252829.1
A1 70327396 70327444
refGene_gn=DOCK9;refGene_tx=XM_006927365.4;refGene_gn=DOCK9;refGene_tx
=XM_006927368.4;refGene_gn=DOCK9;refGene_tx=XM_011280686.3;refGene_gn=
DOCK9;refGene_tx=XM_019828858.2;refGene_gn=DOCK9;refGene_tx=XM_0198288
65.2;refGene_gn=DOCK9;refGene_tx=XM_019828880.2;refGene_gn=DOCK9;refGe
ne_tx=XM_019828896.2;refGene_gn=DOCK9;refGene_tx=XM_019828903.2;refGen
e_gn=DOCK9;refGene_tx=XM_019828909.2;refGene_gn=DOCK9;refGene_tx=XM_02
3252786.1;refGene_gn=DOCK9;refGene_tx=XM_023252788.1;refGene_gn=DOCK9;
refGene_tx=XM_023252805.1;refGene_gn=DOCK9;refGene_tx=XM_023252809.1;r
efGene_gn=DOCK9;refGene_tx=XM_023252810.1;refGene_gn=DOCK9;refGene_tx=
XM_023252814.1;refGene_gn=DOCK9;refGene_tx=XM_023252826.1;refGene_gn=D
OCK9;refGene_tx=XM_023252829.1;refGene_gn=DOCK9;refGene_tx=XM_02325283
1.1
A1 70329287 70329354
refGene_gn=DOCK9;refGene_tx=XM_006927365.4;refGene_gn=DOCK9;refGene_tx
=XM_006927368.4;refGene_gn=DOCK9;refGene_tx=XM_011280686.3;refGene_gn=

DOCK9;refGene_tx=XM_019828858.2;refGene_gn=DOCK9;refGene_tx=XM_0198288
65.2;refGene_gn=DOCK9;refGene_tx=XM_019828880.2;refGene_gn=DOCK9;refGe
ne_tx=XM_019828896.2;refGene_gn=DOCK9;refGene_tx=XM_019828903.2;refGen
e_gn=DOCK9;refGene_tx=XM_019828909.2;refGene_gn=DOCK9;refGene_tx=XM_02
3252786.1;refGene_gn=DOCK9;refGene_tx=XM_023252788.1;refGene_gn=DOCK9;
refGene_tx=XM_023252805.1;refGene_gn=DOCK9;refGene_tx=XM_023252809.1;r
efGene_gn=DOCK9;refGene_tx=XM_023252810.1;refGene_gn=DOCK9;refGene_tx=
XM_023252814.1;refGene_gn=DOCK9;refGene_tx=XM_023252826.1;refGene_gn=D
OCK9;refGene_tx=XM_023252829.1;refGene_gn=DOCK9;refGene_tx=XM_02325283
1.1
A1 70334483 70334598
refGene_gn=DOCK9;refGene_tx=XM_006927365.4;refGene_gn=DOCK9;refGene_tx
=XM_006927368.4;refGene_gn=DOCK9;refGene_tx=XM_011280686.3;refGene_gn=
DOCK9;refGene_tx=XM_019828858.2;refGene_gn=DOCK9;refGene_tx=XM_0198288
65.2;refGene_gn=DOCK9;refGene_tx=XM_019828880.2;refGene_gn=DOCK9;refGe
ne_tx=XM_019828896.2;refGene_gn=DOCK9;refGene_tx=XM_019828903.2;refGen
e_gn=DOCK9;refGene_tx=XM_019828909.2;refGene_gn=DOCK9;refGene_tx=XM_02
3252786.1;refGene_gn=DOCK9;refGene_tx=XM_023252788.1;refGene_gn=DOCK9;
refGene_tx=XM_023252805.1;refGene_gn=DOCK9;refGene_tx=XM_023252809.1;r
efGene_gn=DOCK9;refGene_tx=XM_023252810.1;refGene_gn=DOCK9;refGene_tx=
XM_023252814.1;refGene_gn=DOCK9;refGene_tx=XM_023252826.1;refGene_gn=D
OCK9;refGene_tx=XM_023252829.1;refGene_gn=DOCK9;refGene_tx=XM_02325283
1.1
A1 70336618 70336752
refGene_gn=DOCK9;refGene_tx=XM_006927365.4;refGene_gn=DOCK9;refGene_tx
=XM_006927368.4;refGene_gn=DOCK9;refGene_tx=XM_011280686.3;refGene_gn=
DOCK9;refGene_tx=XM_019828858.2;refGene_gn=DOCK9;refGene_tx=XM_0198288
65.2;refGene_gn=DOCK9;refGene_tx=XM_019828880.2;refGene_gn=DOCK9;refGe
ne_tx=XM_019828896.2;refGene_gn=DOCK9;refGene_tx=XM_019828903.2;refGen
e_gn=DOCK9;refGene_tx=XM_019828909.2;refGene_gn=DOCK9;refGene_tx=XM_02
3252786.1;refGene_gn=DOCK9;refGene_tx=XM_023252788.1;refGene_gn=DOCK9;
refGene_tx=XM_023252805.1;refGene_gn=DOCK9;refGene_tx=XM_023252809.1;r
efGene_gn=DOCK9;refGene_tx=XM_023252810.1;refGene_gn=DOCK9;refGene_tx=
XM_023252814.1;refGene_gn=DOCK9;refGene_tx=XM_023252826.1;refGene_gn=D
OCK9;refGene_tx=XM_023252829.1;refGene_gn=DOCK9;refGene_tx=XM_02325283
1.1
A1 70342444 70342565
refGene_gn=DOCK9;refGene_tx=XM_006927365.4;refGene_gn=DOCK9;refGene_tx
=XM_006927368.4;refGene_gn=DOCK9;refGene_tx=XM_011280686.3;refGene_gn=
DOCK9;refGene_tx=XM_019828858.2;refGene_gn=DOCK9;refGene_tx=XM_0198288
65.2;refGene_gn=DOCK9;refGene_tx=XM_019828880.2;refGene_gn=DOCK9;refGe
ne_tx=XM_019828896.2;refGene_gn=DOCK9;refGene_tx=XM_019828903.2;refGen
e_gn=DOCK9;refGene_tx=XM_019828909.2;refGene_gn=DOCK9;refGene_tx=XM_02
3252786.1;refGene_gn=DOCK9;refGene_tx=XM_023252788.1;refGene_gn=DOCK9;
refGene_tx=XM_023252805.1;refGene_gn=DOCK9;refGene_tx=XM_023252809.1;r
efGene_gn=DOCK9;refGene_tx=XM_023252810.1;refGene_gn=DOCK9;refGene_tx=
XM_023252814.1;refGene_gn=DOCK9;refGene_tx=XM_023252826.1;refGene_gn=D
OCK9;refGene_tx=XM_023252829.1;refGene_gn=DOCK9;refGene_tx=XM_02325283
1.1
A1 70344681 70344795

refGene_gn=DOCK9;refGene_tx=XM_006927365.4;refGene_gn=DOCK9;refGene_tx=XM_006927368.4;refGene_gn=DOCK9;refGene_tx=XM_011280686.3;refGene_gn=DOCK9;refGene_tx=XM_019828858.2;refGene_gn=DOCK9;refGene_tx=XM_019828865.2;refGene_gn=DOCK9;refGene_tx=XM_019828880.2;refGene_gn=DOCK9;refGene_tx=XM_019828896.2;refGene_gn=DOCK9;refGene_tx=XM_019828903.2;refGene_gn=DOCK9;refGene_tx=XM_019828909.2;refGene_gn=DOCK9;refGene_tx=XM_023252786.1;refGene_gn=DOCK9;refGene_tx=XM_023252788.1;refGene_gn=DOCK9;refGene_tx=XM_023252805.1;refGene_gn=DOCK9;refGene_tx=XM_023252809.1;refGene_gn=DOCK9;refGene_tx=XM_023252810.1;refGene_gn=DOCK9;refGene_tx=XM_023252814.1;refGene_gn=DOCK9;refGene_tx=XM_023252826.1;refGene_gn=DOCK9;refGene_tx=XM_023252829.1;refGene_gn=DOCK9;refGene_tx=XM_023252831.1

A1 70345038 70345217

refGene_gn=DOCK9;refGene_tx=XM_006927365.4;refGene_gn=DOCK9;refGene_tx=XM_006927368.4;refGene_gn=DOCK9;refGene_tx=XM_011280686.3;refGene_gn=DOCK9;refGene_tx=XM_019828858.2;refGene_gn=DOCK9;refGene_tx=XM_019828865.2;refGene_gn=DOCK9;refGene_tx=XM_019828880.2;refGene_gn=DOCK9;refGene_tx=XM_019828896.2;refGene_gn=DOCK9;refGene_tx=XM_019828903.2;refGene_gn=DOCK9;refGene_tx=XM_019828909.2;refGene_gn=DOCK9;refGene_tx=XM_023252786.1;refGene_gn=DOCK9;refGene_tx=XM_023252788.1;refGene_gn=DOCK9;refGene_tx=XM_023252805.1;refGene_gn=DOCK9;refGene_tx=XM_023252809.1;refGene_gn=DOCK9;refGene_tx=XM_023252810.1;refGene_gn=DOCK9;refGene_tx=XM_023252814.1;refGene_gn=DOCK9;refGene_tx=XM_023252826.1;refGene_gn=DOCK9;refGene_tx=XM_023252829.1;refGene_gn=DOCK9;refGene_tx=XM_023252831.1

A1 70349671 70349783

refGene_gn=DOCK9;refGene_tx=XM_006927365.4;refGene_gn=DOCK9;refGene_tx=XM_006927368.4;refGene_gn=DOCK9;refGene_tx=XM_011280686.3;refGene_gn=DOCK9;refGene_tx=XM_019828858.2;refGene_gn=DOCK9;refGene_tx=XM_019828865.2;refGene_gn=DOCK9;refGene_tx=XM_019828880.2;refGene_gn=DOCK9;refGene_tx=XM_019828896.2;refGene_gn=DOCK9;refGene_tx=XM_019828903.2;refGene_gn=DOCK9;refGene_tx=XM_019828909.2;refGene_gn=DOCK9;refGene_tx=XM_023252786.1;refGene_gn=DOCK9;refGene_tx=XM_023252788.1;refGene_gn=DOCK9;refGene_tx=XM_023252805.1;refGene_gn=DOCK9;refGene_tx=XM_023252809.1;refGene_gn=DOCK9;refGene_tx=XM_023252810.1;refGene_gn=DOCK9;refGene_tx=XM_023252814.1;refGene_gn=DOCK9;refGene_tx=XM_023252826.1;refGene_gn=DOCK9;refGene_tx=XM_023252829.1;refGene_gn=DOCK9;refGene_tx=XM_023252831.1

A1 70350449 70350533

refGene_gn=DOCK9;refGene_tx=XM_006927365.4;refGene_gn=DOCK9;refGene_tx=XM_006927368.4;refGene_gn=DOCK9;refGene_tx=XM_011280686.3;refGene_gn=DOCK9;refGene_tx=XM_019828858.2;refGene_gn=DOCK9;refGene_tx=XM_019828865.2;refGene_gn=DOCK9;refGene_tx=XM_019828880.2;refGene_gn=DOCK9;refGene_tx=XM_019828896.2;refGene_gn=DOCK9;refGene_tx=XM_019828903.2;refGene_gn=DOCK9;refGene_tx=XM_019828909.2;refGene_gn=DOCK9;refGene_tx=XM_023252786.1;refGene_gn=DOCK9;refGene_tx=XM_023252788.1;refGene_gn=DOCK9;refGene_tx=XM_023252805.1;refGene_gn=DOCK9;refGene_tx=XM_023252809.1;refGene_gn=DOCK9;refGene_tx=XM_023252810.1;refGene_gn=DOCK9;refGene_tx=XM_023252814.1;refGene_gn=DOCK9;refGene_tx=XM_023252826.1;refGene_gn=DOCK9;refGene_tx=XM_023252829.1;refGene_gn=DOCK9;refGene_tx=XM_02325283

1.1
A1	70350959 70351106
refGene_gn=DOCK9;refGene_tx=XM_006927365.4;refGene_gn=DOCK9;refGene_tx=XM_006927368.4;refGene_gn=DOCK9;refGene_tx=XM_011280686.3;refGene_gn=DOCK9;refGene_tx=XM_019828858.2;refGene_gn=DOCK9;refGene_tx=XM_019828865.2;refGene_gn=DOCK9;refGene_tx=XM_019828880.2;refGene_gn=DOCK9;refGene_tx=XM_019828896.2;refGene_gn=DOCK9;refGene_tx=XM_019828903.2;refGene_gn=DOCK9;refGene_tx=XM_019828909.2;refGene_gn=DOCK9;refGene_tx=XM_023252786.1;refGene_gn=DOCK9;refGene_tx=XM_023252788.1;refGene_gn=DOCK9;refGene_tx=XM_023252805.1;refGene_gn=DOCK9;refGene_tx=XM_023252809.1;refGene_gn=DOCK9;refGene_tx=XM_023252810.1;refGene_gn=DOCK9;refGene_tx=XM_023252814.1;refGene_gn=DOCK9;refGene_tx=XM_023252826.1;refGene_gn=DOCK9;refGene_tx=XM_023252829.1;refGene_gn=DOCK9;refGene_tx=XM_023252831.1
A1	70359940 70360012
refGene_gn=DOCK9;refGene_tx=XM_006927365.4;refGene_gn=DOCK9;refGene_tx=XM_006927368.4;refGene_gn=DOCK9;refGene_tx=XM_011280686.3;refGene_gn=DOCK9;refGene_tx=XM_019828858.2;refGene_gn=DOCK9;refGene_tx=XM_019828865.2;refGene_gn=DOCK9;refGene_tx=XM_019828880.2;refGene_gn=DOCK9;refGene_tx=XM_019828896.2;refGene_gn=DOCK9;refGene_tx=XM_019828903.2;refGene_gn=DOCK9;refGene_tx=XM_019828909.2;refGene_gn=DOCK9;refGene_tx=XM_023252786.1;refGene_gn=DOCK9;refGene_tx=XM_023252788.1;refGene_gn=DOCK9;refGene_tx=XM_023252805.1;refGene_gn=DOCK9;refGene_tx=XM_023252809.1;refGene_gn=DOCK9;refGene_tx=XM_023252810.1;refGene_gn=DOCK9;refGene_tx=XM_023252814.1;refGene_gn=DOCK9;refGene_tx=XM_023252826.1;refGene_gn=DOCK9;refGene_tx=XM_023252829.1;refGene_gn=DOCK9;refGene_tx=XM_023252831.1
A1	70360756 70360882
refGene_gn=DOCK9;refGene_tx=XM_006927365.4;refGene_gn=DOCK9;refGene_tx=XM_006927368.4;refGene_gn=DOCK9;refGene_tx=XM_011280686.3;refGene_gn=DOCK9;refGene_tx=XM_019828858.2;refGene_gn=DOCK9;refGene_tx=XM_019828865.2;refGene_gn=DOCK9;refGene_tx=XM_019828880.2;refGene_gn=DOCK9;refGene_tx=XM_019828896.2;refGene_gn=DOCK9;refGene_tx=XM_019828903.2;refGene_gn=DOCK9;refGene_tx=XM_019828909.2;refGene_gn=DOCK9;refGene_tx=XM_023252786.1;refGene_gn=DOCK9;refGene_tx=XM_023252788.1;refGene_gn=DOCK9;refGene_tx=XM_023252805.1;refGene_gn=DOCK9;refGene_tx=XM_023252809.1;refGene_gn=DOCK9;refGene_tx=XM_023252810.1;refGene_gn=DOCK9;refGene_tx=XM_023252814.1;refGene_gn=DOCK9;refGene_tx=XM_023252826.1;refGene_gn=DOCK9;refGene_tx=XM_023252829.1;refGene_gn=DOCK9;refGene_tx=XM_023252831.1
A1	70361693 70361763
refGene_gn=DOCK9;refGene_tx=XM_006927365.4;refGene_gn=DOCK9;refGene_tx=XM_006927368.4;refGene_gn=DOCK9;refGene_tx=XM_011280686.3;refGene_gn=DOCK9;refGene_tx=XM_019828858.2;refGene_gn=DOCK9;refGene_tx=XM_019828865.2;refGene_gn=DOCK9;refGene_tx=XM_019828880.2;refGene_gn=DOCK9;refGene_tx=XM_019828896.2;refGene_gn=DOCK9;refGene_tx=XM_019828903.2;refGene_gn=DOCK9;refGene_tx=XM_019828909.2;refGene_gn=DOCK9;refGene_tx=XM_023252786.1;refGene_gn=DOCK9;refGene_tx=XM_023252788.1;refGene_gn=DOCK9;refGene_tx=XM_023252805.1;refGene_gn=DOCK9;refGene_tx=XM_023252809.1;refGene_gn=DOCK9;refGene_tx=XM_023252810.1;refGene_gn=DOCK9;refGene_tx=

XM_023252814.1;refGene_gn=DOCK9;refGene_tx=XM_023252826.1;refGene_gn=DOCK9;refGene_tx=XM_023252829.1;refGene_gn=DOCK9;refGene_tx=XM_023252831.1
A1 70362035 70362151
refGene_gn=DOCK9;refGene_tx=XM_006927365.4;refGene_gn=DOCK9;refGene_tx=XM_006927368.4;refGene_gn=DOCK9;refGene_tx=XM_011280686.3;refGene_gn=DOCK9;refGene_tx=XM_019828858.2;refGene_gn=DOCK9;refGene_tx=XM_019828865.2;refGene_gn=DOCK9;refGene_tx=XM_019828880.2;refGene_gn=DOCK9;refGene_tx=XM_019828896.2;refGene_gn=DOCK9;refGene_tx=XM_019828903.2;refGene_gn=DOCK9;refGene_tx=XM_019828909.2;refGene_gn=DOCK9;refGene_tx=XM_023252786.1;refGene_gn=DOCK9;refGene_tx=XM_023252788.1;refGene_gn=DOCK9;refGene_tx=XM_023252805.1;refGene_gn=DOCK9;refGene_tx=XM_023252809.1;refGene_gn=DOCK9;refGene_tx=XM_023252810.1;refGene_gn=DOCK9;refGene_tx=XM_023252814.1;refGene_gn=DOCK9;refGene_tx=XM_023252826.1;refGene_gn=DOCK9;refGene_tx=XM_023252829.1;refGene_gn=DOCK9;refGene_tx=XM_023252831.1
A1 70363539 70363629
refGene_gn=DOCK9;refGene_tx=XM_006927365.4;refGene_gn=DOCK9;refGene_tx=XM_006927368.4;refGene_gn=DOCK9;refGene_tx=XM_011280686.3;refGene_gn=DOCK9;refGene_tx=XM_019828858.2;refGene_gn=DOCK9;refGene_tx=XM_019828865.2;refGene_gn=DOCK9;refGene_tx=XM_019828880.2;refGene_gn=DOCK9;refGene_tx=XM_019828896.2;refGene_gn=DOCK9;refGene_tx=XM_019828903.2;refGene_gn=DOCK9;refGene_tx=XM_019828909.2;refGene_gn=DOCK9;refGene_tx=XM_023252786.1;refGene_gn=DOCK9;refGene_tx=XM_023252788.1;refGene_gn=DOCK9;refGene_tx=XM_023252805.1;refGene_gn=DOCK9;refGene_tx=XM_023252809.1;refGene_gn=DOCK9;refGene_tx=XM_023252810.1;refGene_gn=DOCK9;refGene_tx=XM_023252814.1;refGene_gn=DOCK9;refGene_tx=XM_023252826.1;refGene_gn=DOCK9;refGene_tx=XM_023252829.1;refGene_gn=DOCK9;refGene_tx=XM_023252831.1
A1 70363988 70364075
refGene_gn=DOCK9;refGene_tx=XM_006927365.4;refGene_gn=DOCK9;refGene_tx=XM_006927368.4;refGene_gn=DOCK9;refGene_tx=XM_011280686.3;refGene_gn=DOCK9;refGene_tx=XM_019828858.2;refGene_gn=DOCK9;refGene_tx=XM_019828865.2;refGene_gn=DOCK9;refGene_tx=XM_019828880.2;refGene_gn=DOCK9;refGene_tx=XM_019828896.2;refGene_gn=DOCK9;refGene_tx=XM_019828903.2;refGene_gn=DOCK9;refGene_tx=XM_019828909.2;refGene_gn=DOCK9;refGene_tx=XM_023252786.1;refGene_gn=DOCK9;refGene_tx=XM_023252788.1;refGene_gn=DOCK9;refGene_tx=XM_023252805.1;refGene_gn=DOCK9;refGene_tx=XM_023252809.1;refGene_gn=DOCK9;refGene_tx=XM_023252810.1;refGene_gn=DOCK9;refGene_tx=XM_023252814.1;refGene_gn=DOCK9;refGene_tx=XM_023252826.1;refGene_gn=DOCK9;refGene_tx=XM_023252829.1;refGene_gn=DOCK9;refGene_tx=XM_023252831.1
A1 70365164 70365286
refGene_gn=DOCK9;refGene_tx=XM_006927365.4;refGene_gn=DOCK9;refGene_tx=XM_006927368.4;refGene_gn=DOCK9;refGene_tx=XM_011280686.3;refGene_gn=DOCK9;refGene_tx=XM_019828858.2;refGene_gn=DOCK9;refGene_tx=XM_019828865.2;refGene_gn=DOCK9;refGene_tx=XM_019828880.2;refGene_gn=DOCK9;refGene_tx=XM_019828896.2;refGene_gn=DOCK9;refGene_tx=XM_019828903.2;refGene_gn=DOCK9;refGene_tx=XM_019828909.2;refGene_gn=DOCK9;refGene_tx=XM_023252786.1;refGene_gn=DOCK9;refGene_tx=XM_023252788.1;refGene_gn=DOCK9;

refGene_tx=XM_023252805.1;refGene_gn=DOCK9;refGene_tx=XM_023252809.1;refGene_gn=DOCK9;refGene_tx=XM_023252810.1;refGene_gn=DOCK9;refGene_tx=XM_023252814.1;refGene_gn=DOCK9;refGene_tx=XM_023252826.1;refGene_gn=DOCK9;refGene_tx=XM_023252829.1;refGene_gn=DOCK9;refGene_tx=XM_023252831.1
A1 70366045 70366169
refGene_gn=DOCK9;refGene_tx=XM_006927365.4;refGene_gn=DOCK9;refGene_tx=XM_006927368.4;refGene_gn=DOCK9;refGene_tx=XM_011280686.3;refGene_gn=DOCK9;refGene_tx=XM_019828858.2;refGene_gn=DOCK9;refGene_tx=XM_019828865.2;refGene_gn=DOCK9;refGene_tx=XM_019828880.2;refGene_gn=DOCK9;refGene_tx=XM_019828896.2;refGene_gn=DOCK9;refGene_tx=XM_019828903.2;refGene_gn=DOCK9;refGene_tx=XM_019828909.2;refGene_gn=DOCK9;refGene_tx=XM_023252786.1;refGene_gn=DOCK9;refGene_tx=XM_023252788.1;refGene_gn=DOCK9;refGene_tx=XM_023252805.1;refGene_gn=DOCK9;refGene_tx=XM_023252809.1;refGene_gn=DOCK9;refGene_tx=XM_023252810.1;refGene_gn=DOCK9;refGene_tx=XM_023252814.1;refGene_gn=DOCK9;refGene_tx=XM_023252826.1;refGene_gn=DOCK9;refGene_tx=XM_023252829.1;refGene_gn=DOCK9;refGene_tx=XM_023252831.1
A1 70367447 70367540
refGene_gn=DOCK9;refGene_tx=XM_006927365.4;refGene_gn=DOCK9;refGene_tx=XM_006927368.4;refGene_gn=DOCK9;refGene_tx=XM_011280686.3;refGene_gn=DOCK9;refGene_tx=XM_019828858.2;refGene_gn=DOCK9;refGene_tx=XM_019828865.2;refGene_gn=DOCK9;refGene_tx=XM_019828880.2;refGene_gn=DOCK9;refGene_tx=XM_019828896.2;refGene_gn=DOCK9;refGene_tx=XM_019828903.2;refGene_gn=DOCK9;refGene_tx=XM_019828909.2;refGene_gn=DOCK9;refGene_tx=XM_023252786.1;refGene_gn=DOCK9;refGene_tx=XM_023252788.1;refGene_gn=DOCK9;refGene_tx=XM_023252805.1;refGene_gn=DOCK9;refGene_tx=XM_023252809.1;refGene_gn=DOCK9;refGene_tx=XM_023252810.1;refGene_gn=DOCK9;refGene_tx=XM_023252814.1;refGene_gn=DOCK9;refGene_tx=XM_023252826.1;refGene_gn=DOCK9;refGene_tx=XM_023252829.1;refGene_gn=DOCK9;refGene_tx=XM_023252831.1
A1 70368936 70369002
refGene_gn=DOCK9;refGene_tx=XM_006927365.4;refGene_gn=DOCK9;refGene_tx=XM_006927368.4;refGene_gn=DOCK9;refGene_tx=XM_011280686.3;refGene_gn=DOCK9;refGene_tx=XM_019828858.2;refGene_gn=DOCK9;refGene_tx=XM_019828865.2;refGene_gn=DOCK9;refGene_tx=XM_019828880.2;refGene_gn=DOCK9;refGene_tx=XM_019828896.2;refGene_gn=DOCK9;refGene_tx=XM_019828903.2;refGene_gn=DOCK9;refGene_tx=XM_019828909.2;refGene_gn=DOCK9;refGene_tx=XM_023252786.1;refGene_gn=DOCK9;refGene_tx=XM_023252788.1;refGene_gn=DOCK9;refGene_tx=XM_023252805.1;refGene_gn=DOCK9;refGene_tx=XM_023252809.1;refGene_gn=DOCK9;refGene_tx=XM_023252810.1;refGene_gn=DOCK9;refGene_tx=XM_023252814.1;refGene_gn=DOCK9;refGene_tx=XM_023252826.1;refGene_gn=DOCK9;refGene_tx=XM_023252829.1;refGene_gn=DOCK9;refGene_tx=XM_023252831.1
A1 70369134 70369322
refGene_gn=DOCK9;refGene_tx=XM_006927365.4;refGene_gn=DOCK9;refGene_tx=XM_006927368.4;refGene_gn=DOCK9;refGene_tx=XM_011280686.3;refGene_gn=DOCK9;refGene_tx=XM_019828858.2;refGene_gn=DOCK9;refGene_tx=XM_019828865.2;refGene_gn=DOCK9;refGene_tx=XM_019828880.2;refGene_gn=DOCK9;refGene_tx=XM_019828896.2;refGene_gn=DOCK9;refGene_tx=XM_019828903.2;refGen

e_gn=DOCK9;refGene_tx=XM_019828909.2;refGene_gn=DOCK9;refGene_tx=XM_02
3252786.1;refGene_gn=DOCK9;refGene_tx=XM_023252788.1;refGene_gn=DOCK9;
refGene_tx=XM_023252805.1;refGene_gn=DOCK9;refGene_tx=XM_023252809.1;r
efGene_gn=DOCK9;refGene_tx=XM_023252810.1;refGene_gn=DOCK9;refGene_tx=
XM_023252814.1;refGene_gn=DOCK9;refGene_tx=XM_023252826.1;refGene_gn=D
OCK9;refGene_tx=XM_023252829.1;refGene_gn=DOCK9;refGene_tx=XM_02325283
1.1
A1 70369401 70369481
refGene_gn=DOCK9;refGene_tx=XM_006927365.4;refGene_gn=DOCK9;refGene_tx
=XM_006927368.4;refGene_gn=DOCK9;refGene_tx=XM_011280686.3;refGene_gn=
DOCK9;refGene_tx=XM_019828858.2;refGene_gn=DOCK9;refGene_tx=XM_0198288
65.2;refGene_gn=DOCK9;refGene_tx=XM_019828880.2;refGene_gn=DOCK9;refGe
ne_tx=XM_019828896.2;refGene_gn=DOCK9;refGene_tx=XM_019828903.2;refGen
e_gn=DOCK9;refGene_tx=XM_019828909.2;refGene_gn=DOCK9;refGene_tx=XM_02
3252786.1;refGene_gn=DOCK9;refGene_tx=XM_023252788.1;refGene_gn=DOCK9;
refGene_tx=XM_023252805.1;refGene_gn=DOCK9;refGene_tx=XM_023252809.1;r
efGene_gn=DOCK9;refGene_tx=XM_023252810.1;refGene_gn=DOCK9;refGene_tx=
XM_023252814.1;refGene_gn=DOCK9;refGene_tx=XM_023252826.1;refGene_gn=D
OCK9;refGene_tx=XM_023252829.1;refGene_gn=DOCK9;refGene_tx=XM_02325283
1.1
A1 70380063 70380186
refGene_gn=DOCK9;refGene_tx=XM_006927365.4;refGene_gn=DOCK9;refGene_tx
=XM_006927368.4;refGene_gn=DOCK9;refGene_tx=XM_011280686.3;refGene_gn=
DOCK9;refGene_tx=XM_019828858.2;refGene_gn=DOCK9;refGene_tx=XM_0198288
65.2;refGene_gn=DOCK9;refGene_tx=XM_019828880.2;refGene_gn=DOCK9;refGe
ne_tx=XM_019828896.2;refGene_gn=DOCK9;refGene_tx=XM_019828903.2;refGen
e_gn=DOCK9;refGene_tx=XM_019828909.2;refGene_gn=DOCK9;refGene_tx=XM_02
3252786.1;refGene_gn=DOCK9;refGene_tx=XM_023252788.1;refGene_gn=DOCK9;
refGene_tx=XM_023252805.1;refGene_gn=DOCK9;refGene_tx=XM_023252809.1;r
efGene_gn=DOCK9;refGene_tx=XM_023252810.1;refGene_gn=DOCK9;refGene_tx=
XM_023252814.1;refGene_gn=DOCK9;refGene_tx=XM_023252826.1;refGene_gn=D
OCK9;refGene_tx=XM_023252829.1;refGene_gn=DOCK9;refGene_tx=XM_02325283
1.1
A1 70380794 70380877
refGene_gn=DOCK9;refGene_tx=XM_006927365.4;refGene_gn=DOCK9;refGene_tx
=XM_006927368.4;refGene_gn=DOCK9;refGene_tx=XM_011280686.3;refGene_gn=
DOCK9;refGene_tx=XM_019828858.2;refGene_gn=DOCK9;refGene_tx=XM_0198288
65.2;refGene_gn=DOCK9;refGene_tx=XM_019828880.2;refGene_gn=DOCK9;refGe
ne_tx=XM_019828896.2;refGene_gn=DOCK9;refGene_tx=XM_019828903.2;refGen
e_gn=DOCK9;refGene_tx=XM_019828909.2;refGene_gn=DOCK9;refGene_tx=XM_02
3252786.1;refGene_gn=DOCK9;refGene_tx=XM_023252788.1;refGene_gn=DOCK9;
refGene_tx=XM_023252805.1;refGene_gn=DOCK9;refGene_tx=XM_023252809.1;r
efGene_gn=DOCK9;refGene_tx=XM_023252810.1;refGene_gn=DOCK9;refGene_tx=
XM_023252814.1;refGene_gn=DOCK9;refGene_tx=XM_023252826.1;refGene_gn=D
OCK9;refGene_tx=XM_023252829.1;refGene_gn=DOCK9;refGene_tx=XM_02325283
1.1
A1 70384628 70384751
refGene_gn=DOCK9;refGene_tx=XM_006927365.4;refGene_gn=DOCK9;refGene_tx
=XM_006927368.4;refGene_gn=DOCK9;refGene_tx=XM_011280686.3;refGene_gn=
DOCK9;refGene_tx=XM_019828858.2;refGene_gn=DOCK9;refGene_tx=XM_0198288

65.2;refGene_gn=DOCK9;refGene_tx=XM_019828880.2;refGene_gn=DOCK9;refGene_tx=XM_019828896.2;refGene_gn=DOCK9;refGene_tx=XM_019828903.2;refGene_gn=DOCK9;refGene_tx=XM_019828909.2;refGene_gn=DOCK9;refGene_tx=XM_023252786.1;refGene_gn=DOCK9;refGene_tx=XM_023252788.1;refGene_gn=DOCK9;refGene_tx=XM_023252805.1;refGene_gn=DOCK9;refGene_tx=XM_023252809.1;refGene_gn=DOCK9;refGene_tx=XM_023252810.1;refGene_gn=DOCK9;refGene_tx=XM_023252814.1;refGene_gn=DOCK9;refGene_tx=XM_023252826.1;refGene_gn=DOCK9;refGene_tx=XM_023252829.1;refGene_gn=DOCK9;refGene_tx=XM_023252831.1

A1 70385138 70385342
refGene_gn=DOCK9;refGene_tx=XM_006927365.4;refGene_gn=DOCK9;refGene_tx=XM_006927368.4;refGene_gn=DOCK9;refGene_tx=XM_011280686.3;refGene_gn=DOCK9;refGene_tx=XM_019828858.2;refGene_gn=DOCK9;refGene_tx=XM_019828865.2;refGene_gn=DOCK9;refGene_tx=XM_019828880.2;refGene_gn=DOCK9;refGene_tx=XM_019828896.2;refGene_gn=DOCK9;refGene_tx=XM_019828903.2;refGene_gn=DOCK9;refGene_tx=XM_019828909.2;refGene_gn=DOCK9;refGene_tx=XM_023252786.1;refGene_gn=DOCK9;refGene_tx=XM_023252788.1;refGene_gn=DOCK9;refGene_tx=XM_023252805.1;refGene_gn=DOCK9;refGene_tx=XM_023252809.1;refGene_gn=DOCK9;refGene_tx=XM_023252810.1;refGene_gn=DOCK9;refGene_tx=XM_023252814.1;refGene_gn=DOCK9;refGene_tx=XM_023252826.1;refGene_gn=DOCK9;refGene_tx=XM_023252829.1;refGene_gn=DOCK9;refGene_tx=XM_023252831.1

A1 70385649 70385790
refGene_gn=DOCK9;refGene_tx=XM_006927365.4;refGene_gn=DOCK9;refGene_tx=XM_006927368.4;refGene_gn=DOCK9;refGene_tx=XM_011280686.3;refGene_gn=DOCK9;refGene_tx=XM_019828858.2;refGene_gn=DOCK9;refGene_tx=XM_019828865.2;refGene_gn=DOCK9;refGene_tx=XM_019828880.2;refGene_gn=DOCK9;refGene_tx=XM_019828896.2;refGene_gn=DOCK9;refGene_tx=XM_019828903.2;refGene_gn=DOCK9;refGene_tx=XM_019828909.2;refGene_gn=DOCK9;refGene_tx=XM_023252786.1;refGene_gn=DOCK9;refGene_tx=XM_023252788.1;refGene_gn=DOCK9;refGene_tx=XM_023252805.1;refGene_gn=DOCK9;refGene_tx=XM_023252809.1;refGene_gn=DOCK9;refGene_tx=XM_023252810.1;refGene_gn=DOCK9;refGene_tx=XM_023252814.1;refGene_gn=DOCK9;refGene_tx=XM_023252826.1;refGene_gn=DOCK9;refGene_tx=XM_023252829.1;refGene_gn=DOCK9;refGene_tx=XM_023252831.1

A1 70387048 70387123
refGene_gn=DOCK9;refGene_tx=XM_006927365.4;refGene_gn=DOCK9;refGene_tx=XM_006927368.4;refGene_gn=DOCK9;refGene_tx=XM_011280686.3;refGene_gn=DOCK9;refGene_tx=XM_019828858.2;refGene_gn=DOCK9;refGene_tx=XM_019828865.2;refGene_gn=DOCK9;refGene_tx=XM_019828880.2;refGene_gn=DOCK9;refGene_tx=XM_019828896.2;refGene_gn=DOCK9;refGene_tx=XM_019828903.2;refGene_gn=DOCK9;refGene_tx=XM_019828909.2;refGene_gn=DOCK9;refGene_tx=XM_023252786.1;refGene_gn=DOCK9;refGene_tx=XM_023252788.1;refGene_gn=DOCK9;refGene_tx=XM_023252805.1;refGene_gn=DOCK9;refGene_tx=XM_023252809.1;refGene_gn=DOCK9;refGene_tx=XM_023252810.1;refGene_gn=DOCK9;refGene_tx=XM_023252814.1;refGene_gn=DOCK9;refGene_tx=XM_023252826.1;refGene_gn=DOCK9;refGene_tx=XM_023252829.1;refGene_gn=DOCK9;refGene_tx=XM_023252831.1

A1 70396758 70396826
refGene_gn=DOCK9;refGene_tx=XM_006927365.4;refGene_gn=DOCK9;refGene_tx

=XM_006927368.4;refGene_gn=DOCK9;refGene_tx=XM_011280686.3;refGene_gn=DOCK9;refGene_tx=XM_019828858.2;refGene_gn=DOCK9;refGene_tx=XM_019828865.2;refGene_gn=DOCK9;refGene_tx=XM_019828880.2;refGene_gn=DOCK9;refGene_tx=XM_019828896.2;refGene_gn=DOCK9;refGene_tx=XM_019828903.2;refGene_gn=DOCK9;refGene_tx=XM_019828909.2;refGene_gn=DOCK9;refGene_tx=XM_023252786.1;refGene_gn=DOCK9;refGene_tx=XM_023252788.1;refGene_gn=DOCK9;refGene_tx=XM_023252805.1;refGene_gn=DOCK9;refGene_tx=XM_023252809.1;refGene_gn=DOCK9;refGene_tx=XM_023252810.1;refGene_gn=DOCK9;refGene_tx=XM_023252814.1;refGene_gn=DOCK9;refGene_tx=XM_023252826.1;refGene_gn=DOCK9;refGene_tx=XM_023252829.1;refGene_gn=DOCK9;refGene_tx=XM_023252831.1

A1 70397728 70397903

refGene_gn=DOCK9;refGene_tx=XM_006927365.4;refGene_gn=DOCK9;refGene_tx=XM_006927368.4;refGene_gn=DOCK9;refGene_tx=XM_011280686.3;refGene_gn=DOCK9;refGene_tx=XM_019828858.2;refGene_gn=DOCK9;refGene_tx=XM_019828865.2;refGene_gn=DOCK9;refGene_tx=XM_019828880.2;refGene_gn=DOCK9;refGene_tx=XM_019828896.2;refGene_gn=DOCK9;refGene_tx=XM_019828903.2;refGene_gn=DOCK9;refGene_tx=XM_019828909.2;refGene_gn=DOCK9;refGene_tx=XM_023252786.1;refGene_gn=DOCK9;refGene_tx=XM_023252788.1;refGene_gn=DOCK9;refGene_tx=XM_023252805.1;refGene_gn=DOCK9;refGene_tx=XM_023252809.1;refGene_gn=DOCK9;refGene_tx=XM_023252810.1;refGene_gn=DOCK9;refGene_tx=XM_023252814.1;refGene_gn=DOCK9;refGene_tx=XM_023252826.1;refGene_gn=DOCK9;refGene_tx=XM_023252829.1;refGene_gn=DOCK9;refGene_tx=XM_023252831.1

A1 70403522 70403657

refGene_gn=DOCK9;refGene_tx=XM_006927365.4;refGene_gn=DOCK9;refGene_tx=XM_006927368.4;refGene_gn=DOCK9;refGene_tx=XM_011280686.3;refGene_gn=DOCK9;refGene_tx=XM_019828858.2;refGene_gn=DOCK9;refGene_tx=XM_019828865.2;refGene_gn=DOCK9;refGene_tx=XM_019828880.2;refGene_gn=DOCK9;refGene_tx=XM_019828896.2;refGene_gn=DOCK9;refGene_tx=XM_019828903.2;refGene_gn=DOCK9;refGene_tx=XM_019828909.2;refGene_gn=DOCK9;refGene_tx=XM_023252786.1;refGene_gn=DOCK9;refGene_tx=XM_023252788.1;refGene_gn=DOCK9;refGene_tx=XM_023252805.1;refGene_gn=DOCK9;refGene_tx=XM_023252809.1;refGene_gn=DOCK9;refGene_tx=XM_023252810.1;refGene_gn=DOCK9;refGene_tx=XM_023252814.1;refGene_gn=DOCK9;refGene_tx=XM_023252826.1;refGene_gn=DOCK9;refGene_tx=XM_023252829.1;refGene_gn=DOCK9;refGene_tx=XM_023252831.1

A1 70404658 70404754

refGene_gn=DOCK9;refGene_tx=XM_006927365.4;refGene_gn=DOCK9;refGene_tx=XM_006927368.4;refGene_gn=DOCK9;refGene_tx=XM_011280686.3;refGene_gn=DOCK9;refGene_tx=XM_019828858.2;refGene_gn=DOCK9;refGene_tx=XM_019828865.2;refGene_gn=DOCK9;refGene_tx=XM_019828880.2;refGene_gn=DOCK9;refGene_tx=XM_019828896.2;refGene_gn=DOCK9;refGene_tx=XM_019828903.2;refGene_gn=DOCK9;refGene_tx=XM_019828909.2;refGene_gn=DOCK9;refGene_tx=XM_023252786.1;refGene_gn=DOCK9;refGene_tx=XM_023252788.1;refGene_gn=DOCK9;refGene_tx=XM_023252805.1;refGene_gn=DOCK9;refGene_tx=XM_023252809.1;refGene_gn=DOCK9;refGene_tx=XM_023252810.1;refGene_gn=DOCK9;refGene_tx=XM_023252814.1;refGene_gn=DOCK9;refGene_tx=XM_023252826.1;refGene_gn=DOCK9;refGene_tx=XM_023252829.1;refGene_gn=DOCK9;refGene_tx=XM_023252831.1

A1 70406589 70406659
refGene_gn=DOCK9;refGene_tx=XM_006927365.4;refGene_gn=DOCK9;refGene_tx
=XM_006927368.4;refGene_gn=DOCK9;refGene_tx=XM_011280686.3;refGene_gn=
DOCK9;refGene_tx=XM_019828858.2;refGene_gn=DOCK9;refGene_tx=XM_0198288
65.2;refGene_gn=DOCK9;refGene_tx=XM_019828880.2;refGene_gn=DOCK9;refGe
ne_tx=XM_019828896.2;refGene_gn=DOCK9;refGene_tx=XM_019828903.2;refGen
e_gn=DOCK9;refGene_tx=XM_019828909.2;refGene_gn=DOCK9;refGene_tx=XM_02
3252786.1;refGene_gn=DOCK9;refGene_tx=XM_023252788.1;refGene_gn=DOCK9;
refGene_tx=XM_023252805.1;refGene_gn=DOCK9;refGene_tx=XM_023252809.1;r
efGene_gn=DOCK9;refGene_tx=XM_023252810.1;refGene_gn=DOCK9;refGene_tx=
XM_023252814.1;refGene_gn=DOCK9;refGene_tx=XM_023252826.1;refGene_gn=D
OCK9;refGene_tx=XM_023252829.1;refGene_gn=DOCK9;refGene_tx=XM_02325283
1.1
A1 70408389 70408472
refGene_gn=DOCK9;refGene_tx=XM_006927365.4;refGene_gn=DOCK9;refGene_tx
=XM_006927368.4;refGene_gn=DOCK9;refGene_tx=XM_011280686.3;refGene_gn=
DOCK9;refGene_tx=XM_019828858.2;refGene_gn=DOCK9;refGene_tx=XM_0198288
65.2;refGene_gn=DOCK9;refGene_tx=XM_019828880.2;refGene_gn=DOCK9;refGe
ne_tx=XM_019828896.2;refGene_gn=DOCK9;refGene_tx=XM_019828903.2;refGen
e_gn=DOCK9;refGene_tx=XM_019828909.2;refGene_gn=DOCK9;refGene_tx=XM_02
3252786.1;refGene_gn=DOCK9;refGene_tx=XM_023252788.1;refGene_gn=DOCK9;
refGene_tx=XM_023252805.1;refGene_gn=DOCK9;refGene_tx=XM_023252809.1;r
efGene_gn=DOCK9;refGene_tx=XM_023252810.1;refGene_gn=DOCK9;refGene_tx=
XM_023252814.1;refGene_gn=DOCK9;refGene_tx=XM_023252826.1;refGene_gn=D
OCK9;refGene_tx=XM_023252829.1;refGene_gn=DOCK9;refGene_tx=XM_02325283
1.1
A1 70412469 70412559
refGene_gn=DOCK9;refGene_tx=XM_006927365.4;refGene_gn=DOCK9;refGene_tx
=XM_006927368.4;refGene_gn=DOCK9;refGene_tx=XM_011280686.3;refGene_gn=
DOCK9;refGene_tx=XM_019828858.2;refGene_gn=DOCK9;refGene_tx=XM_0198288
65.2;refGene_gn=DOCK9;refGene_tx=XM_019828880.2;refGene_gn=DOCK9;refGe
ne_tx=XM_019828896.2;refGene_gn=DOCK9;refGene_tx=XM_019828903.2;refGen
e_gn=DOCK9;refGene_tx=XM_019828909.2;refGene_gn=DOCK9;refGene_tx=XM_02
3252786.1;refGene_gn=DOCK9;refGene_tx=XM_023252788.1;refGene_gn=DOCK9;
refGene_tx=XM_023252805.1;refGene_gn=DOCK9;refGene_tx=XM_023252809.1;r
efGene_gn=DOCK9;refGene_tx=XM_023252810.1;refGene_gn=DOCK9;refGene_tx=
XM_023252814.1;refGene_gn=DOCK9;refGene_tx=XM_023252826.1;refGene_gn=D
OCK9;refGene_tx=XM_023252829.1;refGene_gn=DOCK9;refGene_tx=XM_02325283
1.1
A1 70434646 70434763
refGene_gn=DOCK9;refGene_tx=XM_006927365.4;refGene_gn=DOCK9;refGene_tx
=XM_006927368.4;refGene_gn=DOCK9;refGene_tx=XM_011280686.3;refGene_gn=
DOCK9;refGene_tx=XM_019828858.2;refGene_gn=DOCK9;refGene_tx=XM_0198288
65.2;refGene_gn=DOCK9;refGene_tx=XM_019828880.2;refGene_gn=DOCK9;refGe
ne_tx=XM_019828896.2;refGene_gn=DOCK9;refGene_tx=XM_019828903.2;refGen
e_gn=DOCK9;refGene_tx=XM_019828909.2;refGene_gn=DOCK9;refGene_tx=XM_02
3252786.1;refGene_gn=DOCK9;refGene_tx=XM_023252788.1;refGene_gn=DOCK9;
refGene_tx=XM_023252805.1;refGene_gn=DOCK9;refGene_tx=XM_023252809.1;r
efGene_gn=DOCK9;refGene_tx=XM_023252810.1;refGene_gn=DOCK9;refGene_tx=
XM_023252814.1;refGene_gn=DOCK9;refGene_tx=XM_023252826.1;refGene_gn=D

OCK9;refGene_tx=XM_023252829.1;refGene_gn=DOCK9;refGene_tx=XM_02325283
1.1
A1 70456352 70456478
refGene_gn=DOCK9;refGene_tx=XM_006927365.4;refGene_gn=DOCK9;refGene_tx
=XM_006927368.4;refGene_gn=DOCK9;refGene_tx=XM_019828865.2;refGene_gn=
DOCK9;refGene_tx=XM_019828896.2
A1 70499357 70499519
refGene_gn=DOCK9;refGene_tx=XM_011280686.3;refGene_gn=DOCK9;refGene_tx
=XM_019828858.2;refGene_gn=DOCK9;refGene_tx=XM_019828880.2;refGene_gn=
DOCK9;refGene_tx=XM_019828903.2;refGene_gn=DOCK9;refGene_tx=XM_0198289
09.2;refGene_gn=DOCK9;refGene_tx=XM_023252786.1;refGene_gn=DOCK9;refGe
ne_tx=XM_023252805.1;refGene_gn=DOCK9;refGene_tx=XM_023252809.1;refGen
e_gn=DOCK9;refGene_tx=XM_023252814.1
A1 70567854 70567983
refGene_gn=DOCK9;refGene_tx=XM_023252788.1;refGene_gn=DOCK9;refGene_tx
=XM_023252810.1;refGene_gn=DOCK9;refGene_tx=XM_023252826.1;refGene_gn=
DOCK9;refGene_tx=XM_023252829.1;refGene_gn=DOCK9;refGene_tx=XM_0232528
31.1
A1 70638596 70638711 refGene_gn=UBAC2;refGene_tx=XM_019828928.2
A1 70651884 70652062
refGene_gn=UBAC2;refGene_tx=XM_006927374.4;refGene_gn=UBAC2;refGene_tx
=XM_019828945.2;refGene_gn=UBAC2;refGene_tx=XM_019828949.2;refGene_gn=
UBAC2;refGene_tx=XM_019828950.2;refGene_gn=UBAC2;refGene_tx=XM_0198289
54.2
A1 70679330 70679361 refGene_gn=UBAC2;refGene_tx=XM_019828937.2
A1 70681137 70681265
refGene_gn=UBAC2;refGene_tx=XM_006927374.4;refGene_gn=UBAC2;refGene_tx
=XM_019828928.2;refGene_gn=UBAC2;refGene_tx=XM_019828937.2;refGene_gn=
UBAC2;refGene_tx=XM_019828945.2;refGene_gn=UBAC2;refGene_tx=XM_0198289
49.2;refGene_gn=UBAC2;refGene_tx=XM_019828950.2;refGene_gn=UBAC2;refGe
ne_tx=XM_019828954.2
A1 70683490 70683610
refGene_gn=UBAC2;refGene_tx=XM_006927374.4;refGene_gn=UBAC2;refGene_tx
=XM_019828928.2;refGene_gn=UBAC2;refGene_tx=XM_019828937.2;refGene_gn=
UBAC2;refGene_tx=XM_019828945.2;refGene_gn=UBAC2;refGene_tx=XM_0198289
49.2;refGene_gn=UBAC2;refGene_tx=XM_019828950.2;refGene_gn=UBAC2;refGe
ne_tx=XM_019828954.2
A1 70684327 70684437
refGene_gn=UBAC2;refGene_tx=XM_006927374.4;refGene_gn=UBAC2;refGene_tx
=XM_019828928.2;refGene_gn=UBAC2;refGene_tx=XM_019828937.2;refGene_gn=
UBAC2;refGene_tx=XM_019828945.2;refGene_gn=UBAC2;refGene_tx=XM_0198289
49.2;refGene_gn=UBAC2;refGene_tx=XM_019828950.2;refGene_gn=UBAC2;refGe
ne_tx=XM_019828954.2
A1 70690712 70691708 refGene_gn=GPR18;refGene_tx=XM_019828970.2
A1 70738613 70739702
refGene_gn=LOC101101025;refGene_tx=XM_019828961.2;refGene_gn=LOC101101
025;refGene_tx=XM_019828962.2;refGene_gn=LOC101101025;refGene_tx=XM_02
3252858.1
A1 70761467 70761591
refGene_gn=UBAC2;refGene_tx=XM_006927374.4;refGene_gn=UBAC2;refGene_tx

=XM_019828928.2;refGene_gn=UBAC2;refGene_tx=XM_019828937.2;refGene_gn=
UBAC2;refGene_tx=XM_019828945.2;refGene_gn=UBAC2;refGene_tx=XM_0198289
49.2;refGene_gn=UBAC2;refGene_tx=XM_019828950.2;refGene_gn=UBAC2;refGe
ne_tx=XM_019828954.2
A1 70765632 70765680
refGene_gn=UBAC2;refGene_tx=XM_006927374.4;refGene_gn=UBAC2;refGene_tx
=XM_019828928.2;refGene_gn=UBAC2;refGene_tx=XM_019828937.2;refGene_gn=
UBAC2;refGene_tx=XM_019828945.2;refGene_gn=UBAC2;refGene_tx=XM_0198289
49.2;refGene_gn=UBAC2;refGene_tx=XM_019828950.2;refGene_gn=UBAC2;refGe
ne_tx=XM_019828954.2
A1 70787933 70788176
refGene_gn=UBAC2;refGene_tx=XM_006927374.4;refGene_gn=UBAC2;refGene_tx
=XM_019828928.2;refGene_gn=UBAC2;refGene_tx=XM_019828937.2;refGene_gn=
UBAC2;refGene_tx=XM_019828945.2;refGene_gn=UBAC2;refGene_tx=XM_0198289
49.2;refGene_gn=UBAC2;refGene_tx=XM_019828950.2;refGene_gn=UBAC2;refGe
ne_tx=XM_019828954.2
A1 70809499 70809622
refGene_gn=UBAC2;refGene_tx=XM_006927374.4;refGene_gn=UBAC2;refGene_tx
=XM_019828928.2;refGene_gn=UBAC2;refGene_tx=XM_019828937.2;refGene_gn=
UBAC2;refGene_tx=XM_019828945.2;refGene_gn=UBAC2;refGene_tx=XM_0198289
49.2;refGene_gn=UBAC2;refGene_tx=XM_019828950.2;refGene_gn=UBAC2;refGe
ne_tx=XM_019828954.2
A1 70814727 70814781 refGene_gn=UBAC2;refGene_tx=XM_019828945.2
A1 70818944 70818965 refGene_gn=UBAC2;refGene_tx=XM_019828950.2
A1 70819009 70819195
refGene_gn=UBAC2;refGene_tx=XM_019828928.2;refGene_gn=UBAC2;refGene_tx
=XM_019828937.2;refGene_gn=UBAC2;refGene_tx=XM_019828954.2
A1 70822987 70823095 refGene_gn=UBAC2;refGene_tx=XM_006927374.4
A1 70833008 70833062 refGene_gn=UBAC2;refGene_tx=XM_019828949.2
A1 70918139 70918307 refGene_gn=TM9SF2;refGene_tx=XM_023252871.1
A1 70936754 70936822 refGene_gn=TM9SF2;refGene_tx=XM_023252871.1
A1 70940124 70940218 refGene_gn=TM9SF2;refGene_tx=XM_023252871.1
A1 70946692 70946820 refGene_gn=TM9SF2;refGene_tx=XM_023252871.1
A1 70952326 70952456 refGene_gn=TM9SF2;refGene_tx=XM_023252871.1
A1 70953473 70953598 refGene_gn=TM9SF2;refGene_tx=XM_023252871.1
A1 70954881 70954993 refGene_gn=TM9SF2;refGene_tx=XM_023252871.1
A1 70956952 70957032 refGene_gn=TM9SF2;refGene_tx=XM_023252871.1
A1 70958267 70958376 refGene_gn=TM9SF2;refGene_tx=XM_023252871.1
A1 70961024 70961157 refGene_gn=TM9SF2;refGene_tx=XM_023252871.1
A1 70964273 70964393 refGene_gn=TM9SF2;refGene_tx=XM_023252871.1
A1 70967744 70967802 refGene_gn=TM9SF2;refGene_tx=XM_023252871.1
A1 70972041 70972201 refGene_gn=TM9SF2;refGene_tx=XM_023252871.1
A1 70974345 70974497 refGene_gn=TM9SF2;refGene_tx=XM_023252871.1
A1 70975586 70975698 refGene_gn=TM9SF2;refGene_tx=XM_023252871.1
A1 70979367 70979539 refGene_gn=TM9SF2;refGene_tx=XM_023252871.1
A1 70982570 70982638 refGene_gn=TM9SF2;refGene_tx=XM_023252871.1
A1 71013864 71013926
refGene_gn=CLYBL;refGene_tx=XM_011280713.2;refGene_gn=CLYBL;refGene_tx
=XM_011280715.2;refGene_gn=CLYBL;refGene_tx=XM_011280717.2;refGene_gn=
CLYBL;refGene_tx=XM_011280722.2;refGene_gn=CLYBL;refGene_tx=XM_0112807

30.2;refGene_gn=CLYBL;refGene_tx=XM_011280732.3;refGene_gn=CLYBL;refGene_tx=XM_019829017.2;refGene_gn=CLYBL;refGene_tx=XM_019829036.1
A1 71121046 71121090 refGene_gn=CLYBL;refGene_tx=XM_019829047.2
A1 71159247 71159434
refGene_gn=CLYBL;refGene_tx=XM_011280713.2;refGene_gn=CLYBL;refGene_tx=XM_011280715.2;refGene_gn=CLYBL;refGene_tx=XM_011280717.2;refGene_gn=CLYBL;refGene_tx=XM_011280722.2;refGene_gn=CLYBL;refGene_tx=XM_011280730.2;refGene_gn=CLYBL;refGene_tx=XM_011280732.3;refGene_gn=CLYBL;refGene_tx=XM_019829017.2;refGene_gn=CLYBL;refGene_tx=XM_019829036.1;refGene_gn=CLYBL;refGene_tx=XM_019829047.2
A1 71229913 71230102
refGene_gn=CLYBL;refGene_tx=XM_011280713.2;refGene_gn=CLYBL;refGene_tx=XM_011280715.2;refGene_gn=CLYBL;refGene_tx=XM_011280717.2;refGene_gn=CLYBL;refGene_tx=XM_011280722.2;refGene_gn=CLYBL;refGene_tx=XM_011280730.2;refGene_gn=CLYBL;refGene_tx=XM_011280732.3;refGene_gn=CLYBL;refGene_tx=XM_019829017.2;refGene_gn=CLYBL;refGene_tx=XM_019829036.1;refGene_gn=CLYBL;refGene_tx=XM_019829047.2
A1 71234296 71234398
refGene_gn=CLYBL;refGene_tx=XM_011280713.2;refGene_gn=CLYBL;refGene_tx=XM_011280715.2;refGene_gn=CLYBL;refGene_tx=XM_011280717.2;refGene_gn=CLYBL;refGene_tx=XM_011280722.2;refGene_gn=CLYBL;refGene_tx=XM_011280730.2;refGene_gn=CLYBL;refGene_tx=XM_011280732.3;refGene_gn=CLYBL;refGene_tx=XM_019829017.2;refGene_gn=CLYBL;refGene_tx=XM_019829036.1;refGene_gn=CLYBL;refGene_tx=XM_019829047.2
A1 71236104 71236198
refGene_gn=CLYBL;refGene_tx=XM_011280713.2;refGene_gn=CLYBL;refGene_tx=XM_011280715.2;refGene_gn=CLYBL;refGene_tx=XM_011280717.2;refGene_gn=CLYBL;refGene_tx=XM_011280722.2;refGene_gn=CLYBL;refGene_tx=XM_011280730.2;refGene_gn=CLYBL;refGene_tx=XM_011280732.3;refGene_gn=CLYBL;refGene_tx=XM_019829017.2;refGene_gn=CLYBL;refGene_tx=XM_019829036.1;refGene_gn=CLYBL;refGene_tx=XM_019829047.2
A1 71237573 71237741
refGene_gn=CLYBL;refGene_tx=XM_011280713.2;refGene_gn=CLYBL;refGene_tx=XM_011280715.2;refGene_gn=CLYBL;refGene_tx=XM_011280717.2;refGene_gn=CLYBL;refGene_tx=XM_011280722.2;refGene_gn=CLYBL;refGene_tx=XM_011280730.2;refGene_gn=CLYBL;refGene_tx=XM_011280732.3;refGene_gn=CLYBL;refGene_tx=XM_019829017.2;refGene_gn=CLYBL;refGene_tx=XM_019829036.1;refGene_gn=CLYBL;refGene_tx=XM_019829047.2
A1 71242291 71242416
refGene_gn=CLYBL;refGene_tx=XM_011280713.2;refGene_gn=CLYBL;refGene_tx=XM_011280715.2;refGene_gn=CLYBL;refGene_tx=XM_011280717.2;refGene_gn=CLYBL;refGene_tx=XM_011280722.2;refGene_gn=CLYBL;refGene_tx=XM_011280730.2;refGene_gn=CLYBL;refGene_tx=XM_011280732.3;refGene_gn=CLYBL;refGene_tx=XM_019829017.2;refGene_gn=CLYBL;refGene_tx=XM_019829036.1;refGene_gn=CLYBL;refGene_tx=XM_019829047.2
A1 71260163 71260259
refGene_gn=CLYBL;refGene_tx=XM_011280713.2;refGene_gn=CLYBL;refGene_tx=XM_011280715.2;refGene_gn=CLYBL;refGene_tx=XM_011280717.2;refGene_gn=CLYBL;refGene_tx=XM_011280722.2;refGene_gn=CLYBL;refGene_tx=XM_011280730.2;refGene_gn=CLYBL;refGene_tx=XM_011280732.3;refGene_gn=CLYBL;refGe

ne_tx=XM_019829017.2;refGene_gn=CLYBL;refGene_tx=XM_019829036.1;refGene_gn=CLYBL;refGene_tx=XM_019829047.2
A1 71322273 71322716 refGene_gn=ZIC5;refGene_tx=XM_023252902.1
A1 71327040 71328583 refGene_gn=ZIC5;refGene_tx=XM_023252902.1
A1 71339277 71340349 refGene_gn=ZIC2;refGene_tx=XM_023252910.1
A1 71342142 71342306 refGene_gn=ZIC2;refGene_tx=XM_023252910.1
A1 71342531 71342903 refGene_gn=ZIC2;refGene_tx=XM_023252910.1
A1 71419247 71419352
refGene_gn=PCCA;refGene_tx=XM_003980503.4;refGene_gn=PCCA;refGene_tx=XM_003980504.4;refGene_gn=PCCA;refGene_tx=XM_003980505.4;refGene_gn=PCCA;refGene_tx=XM_019829079.2;refGene_gn=PCCA;refGene_tx=XM_019829094.2;refGene_gn=PCCA;refGene_tx=XM_023252917.1
A1 71433791 71433869
refGene_gn=PCCA;refGene_tx=XM_003980503.4;refGene_gn=PCCA;refGene_tx=XM_003980505.4;refGene_gn=PCCA;refGene_tx=XM_019829079.2;refGene_gn=PCCA;refGene_tx=XM_019829094.2
A1 71439816 71439864
refGene_gn=PCCA;refGene_tx=XM_003980503.4;refGene_gn=PCCA;refGene_tx=XM_003980504.4;refGene_gn=PCCA;refGene_tx=XM_003980505.4;refGene_gn=PCCA;refGene_tx=XM_019829079.2;refGene_gn=PCCA;refGene_tx=XM_019829094.2;refGene_gn=PCCA;refGene_tx=XM_023252917.1
A1 71439970 71440039
refGene_gn=PCCA;refGene_tx=XM_003980503.4;refGene_gn=PCCA;refGene_tx=XM_003980504.4;refGene_gn=PCCA;refGene_tx=XM_003980505.4;refGene_gn=PCCA;refGene_tx=XM_019829079.2;refGene_gn=PCCA;refGene_tx=XM_019829094.2;refGene_gn=PCCA;refGene_tx=XM_023252917.1;refGene_gn=PCCA;refGene_tx=XM_023252923.1
A1 71488567 71488681
refGene_gn=PCCA;refGene_tx=XM_003980503.4;refGene_gn=PCCA;refGene_tx=XM_003980504.4;refGene_gn=PCCA;refGene_tx=XM_003980505.4;refGene_gn=PCCA;refGene_tx=XM_019829079.2;refGene_gn=PCCA;refGene_tx=XM_019829094.2;refGene_gn=PCCA;refGene_tx=XM_023252917.1;refGene_gn=PCCA;refGene_tx=XM_023252923.1
A1 71490490 71490544
refGene_gn=PCCA;refGene_tx=XM_003980503.4;refGene_gn=PCCA;refGene_tx=XM_003980504.4;refGene_gn=PCCA;refGene_tx=XM_003980505.4;refGene_gn=PCCA;refGene_tx=XM_019829079.2;refGene_gn=PCCA;refGene_tx=XM_019829094.2;refGene_gn=PCCA;refGene_tx=XM_023252917.1;refGene_gn=PCCA;refGene_tx=XM_023252923.1
A1 71535947 71536079
refGene_gn=PCCA;refGene_tx=XM_003980503.4;refGene_gn=PCCA;refGene_tx=XM_003980504.4;refGene_gn=PCCA;refGene_tx=XM_003980505.4;refGene_gn=PCCA;refGene_tx=XM_019829079.2;refGene_gn=PCCA;refGene_tx=XM_019829094.2;refGene_gn=PCCA;refGene_tx=XM_023252917.1;refGene_gn=PCCA;refGene_tx=XM_023252923.1
A1 71556646 71556683
refGene_gn=PCCA;refGene_tx=XM_003980503.4;refGene_gn=PCCA;refGene_tx=XM_003980504.4;refGene_gn=PCCA;refGene_tx=XM_003980505.4;refGene_gn=PCCA;refGene_tx=XM_019829079.2;refGene_gn=PCCA;refGene_tx=XM_019829094.2;refGene_gn=PCCA;refGene_tx=XM_023252917.1;refGene_gn=PCCA;refGene_tx=X

M_023252923.1
A1 71578192 71578271
refGene_gn=PCCA;refGene_tx=XM_003980503.4;refGene_gn=PCCA;refGene_tx=XM_003980504.4;refGene_gn=PCCA;refGene_tx=XM_003980505.4;refGene_gn=PCCA;refGene_tx=XM_019829079.2;refGene_gn=PCCA;refGene_tx=XM_019829094.2;refGene_gn=PCCA;refGene_tx=XM_023252917.1;refGene_gn=PCCA;refGene_tx=XM_023252923.1
A1 71583030 71583133
refGene_gn=PCCA;refGene_tx=XM_003980503.4;refGene_gn=PCCA;refGene_tx=XM_003980504.4;refGene_gn=PCCA;refGene_tx=XM_003980505.4;refGene_gn=PCCA;refGene_tx=XM_019829079.2;refGene_gn=PCCA;refGene_tx=XM_019829094.2;refGene_gn=PCCA;refGene_tx=XM_023252917.1;refGene_gn=PCCA;refGene_tx=XM_023252923.1
A1 71590913 71591008
refGene_gn=PCCA;refGene_tx=XM_003980503.4;refGene_gn=PCCA;refGene_tx=XM_003980504.4;refGene_gn=PCCA;refGene_tx=XM_003980505.4;refGene_gn=PCCA;refGene_tx=XM_019829079.2;refGene_gn=PCCA;refGene_tx=XM_019829094.2;refGene_gn=PCCA;refGene_tx=XM_023252917.1;refGene_gn=PCCA;refGene_tx=XM_023252923.1
A1 71593906 71594057
refGene_gn=PCCA;refGene_tx=XM_003980503.4;refGene_gn=PCCA;refGene_tx=XM_003980504.4;refGene_gn=PCCA;refGene_tx=XM_003980505.4;refGene_gn=PCCA;refGene_tx=XM_019829079.2;refGene_gn=PCCA;refGene_tx=XM_019829094.2;refGene_gn=PCCA;refGene_tx=XM_023252917.1;refGene_gn=PCCA;refGene_tx=XM_023252923.1
A1 71619788 71619932
refGene_gn=PCCA;refGene_tx=XM_003980503.4;refGene_gn=PCCA;refGene_tx=XM_003980504.4;refGene_gn=PCCA;refGene_tx=XM_003980505.4;refGene_gn=PCCA;refGene_tx=XM_019829079.2;refGene_gn=PCCA;refGene_tx=XM_019829094.2;refGene_gn=PCCA;refGene_tx=XM_023252917.1;refGene_gn=PCCA;refGene_tx=XM_023252923.1
A1 71621280 71621355
refGene_gn=PCCA;refGene_tx=XM_003980503.4;refGene_gn=PCCA;refGene_tx=XM_003980504.4;refGene_gn=PCCA;refGene_tx=XM_003980505.4;refGene_gn=PCCA;refGene_tx=XM_019829079.2;refGene_gn=PCCA;refGene_tx=XM_019829094.2;refGene_gn=PCCA;refGene_tx=XM_023252917.1;refGene_gn=PCCA;refGene_tx=XM_023252923.1
A1 71625574 71625643
refGene_gn=PCCA;refGene_tx=XM_003980503.4;refGene_gn=PCCA;refGene_tx=XM_003980504.4;refGene_gn=PCCA;refGene_tx=XM_003980505.4;refGene_gn=PCCA;refGene_tx=XM_019829079.2;refGene_gn=PCCA;refGene_tx=XM_019829094.2;refGene_gn=PCCA;refGene_tx=XM_023252917.1;refGene_gn=PCCA;refGene_tx=XM_023252923.1
A1 71627542 71627618
refGene_gn=PCCA;refGene_tx=XM_003980503.4;refGene_gn=PCCA;refGene_tx=XM_003980504.4;refGene_gn=PCCA;refGene_tx=XM_003980505.4;refGene_gn=PCCA;refGene_tx=XM_019829079.2;refGene_gn=PCCA;refGene_tx=XM_019829094.2;refGene_gn=PCCA;refGene_tx=XM_023252917.1;refGene_gn=PCCA;refGene_tx=XM_023252923.1
A1 71644423 71644534

refGene_gn=PCCA;refGene_tx=XM_003980503.4;refGene_gn=PCCA;refGene_tx=XM_003980504.4;refGene_gn=PCCA;refGene_tx=XM_003980505.4;refGene_gn=PCCA;refGene_tx=XM_019829079.2;refGene_gn=PCCA;refGene_tx=XM_019829094.2;refGene_gn=PCCA;refGene_tx=XM_023252917.1;refGene_gn=PCCA;refGene_tx=XM_023252923.1
A1 71660222 71660331
refGene_gn=PCCA;refGene_tx=XM_003980503.4;refGene_gn=PCCA;refGene_tx=XM_003980504.4;refGene_gn=PCCA;refGene_tx=XM_003980505.4;refGene_gn=PCCA;refGene_tx=XM_019829079.2;refGene_gn=PCCA;refGene_tx=XM_019829094.2;refGene_gn=PCCA;refGene_tx=XM_023252917.1;refGene_gn=PCCA;refGene_tx=XM_023252923.1
A1 71696312 71696415
refGene_gn=PCCA;refGene_tx=XM_003980503.4;refGene_gn=PCCA;refGene_tx=XM_003980504.4;refGene_gn=PCCA;refGene_tx=XM_003980505.4;refGene_gn=PCCA;refGene_tx=XM_019829079.2;refGene_gn=PCCA;refGene_tx=XM_019829094.2;refGene_gn=PCCA;refGene_tx=XM_023252917.1;refGene_gn=PCCA;refGene_tx=XM_023252923.1
A1 71705202 71705229 refGene_gn=PCCA;refGene_tx=XM_019829094.2
A1 71745774 71745873
refGene_gn=PCCA;refGene_tx=XM_003980503.4;refGene_gn=PCCA;refGene_tx=XM_003980504.4;refGene_gn=PCCA;refGene_tx=XM_003980505.4;refGene_gn=PCCA;refGene_tx=XM_019829079.2;refGene_gn=PCCA;refGene_tx=XM_023252917.1;refGene_gn=PCCA;refGene_tx=XM_023252923.1
A1 71768536 71768590
refGene_gn=PCCA;refGene_tx=XM_003980503.4;refGene_gn=PCCA;refGene_tx=XM_003980504.4;refGene_gn=PCCA;refGene_tx=XM_003980505.4;refGene_gn=PCCA;refGene_tx=XM_019829079.2;refGene_gn=PCCA;refGene_tx=XM_023252917.1;refGene_gn=PCCA;refGene_tx=XM_023252923.1
A1 71826048 71826189
refGene_gn=PCCA;refGene_tx=XM_003980503.4;refGene_gn=PCCA;refGene_tx=XM_003980504.4;refGene_gn=PCCA;refGene_tx=XM_019829079.2;refGene_gn=PCCA;refGene_tx=XM_023252923.1
A1 71837306 71837384
refGene_gn=PCCA;refGene_tx=XM_003980503.4;refGene_gn=PCCA;refGene_tx=XM_003980504.4;refGene_gn=PCCA;refGene_tx=XM_003980505.4;refGene_gn=PCCA;refGene_tx=XM_019829079.2;refGene_gn=PCCA;refGene_tx=XM_023252917.1;refGene_gn=PCCA;refGene_tx=XM_023252923.1
A1 71839864 71839933 refGene_gn=PCCA;refGene_tx=XM_019829079.2
A1 71839996 71840065
refGene_gn=PCCA;refGene_tx=XM_003980503.4;refGene_gn=PCCA;refGene_tx=XM_003980504.4;refGene_gn=PCCA;refGene_tx=XM_003980505.4;refGene_gn=PCCA;refGene_tx=XM_023252917.1;refGene_gn=PCCA;refGene_tx=XM_023252923.1
A1 71842033 71842518
refGene_gn=GGACT;refGene_tx=XM_006927379.3;refGene_gn=GGACT;refGene_tx=XM_006927382.3;refGene_gn=GGACT;refGene_tx=XM_019829110.2;refGene_gn=GGACT;refGene_tx=XM_019829120.2;refGene_gn=GGACT;refGene_tx=XM_019829128.2;refGene_gn=GGACT;refGene_tx=XM_019829134.2;refGene_gn=GGACT;refGene_tx=XM_019829142.2
A1 71851624 71851748
refGene_gn=GGACT;refGene_tx=XM_006927379.3;refGene_gn=GGACT;refGene_tx

=XM_019829110.2;refGene_gn=GGACT;refGene_tx=XM_019829120.2
A1 71901816 71901965
refGene_gn=TMTC4;refGene_tx=XM_023252953.1;refGene_gn=TMTC4;refGene_tx=XM_023252959.1;refGene_gn=TMTC4;refGene_tx=XM_023252960.1
A1 71903618 71903688
refGene_gn=TMTC4;refGene_tx=XM_023252953.1;refGene_gn=TMTC4;refGene_tx=XM_023252959.1;refGene_gn=TMTC4;refGene_tx=XM_023252960.1
A1 71907586 71907699
refGene_gn=TMTC4;refGene_tx=XM_023252953.1;refGene_gn=TMTC4;refGene_tx=XM_023252959.1;refGene_gn=TMTC4;refGene_tx=XM_023252960.1
A1 71909642 71909757
refGene_gn=TMTC4;refGene_tx=XM_023252953.1;refGene_gn=TMTC4;refGene_tx=XM_023252959.1;refGene_gn=TMTC4;refGene_tx=XM_023252960.1
A1 71916560 71916702
refGene_gn=TMTC4;refGene_tx=XM_023252953.1;refGene_gn=TMTC4;refGene_tx=XM_023252959.1;refGene_gn=TMTC4;refGene_tx=XM_023252960.1
A1 71916837 71916945
refGene_gn=TMTC4;refGene_tx=XM_023252953.1;refGene_gn=TMTC4;refGene_tx=XM_023252959.1;refGene_gn=TMTC4;refGene_tx=XM_023252960.1
A1 71917086 71917166
refGene_gn=TMTC4;refGene_tx=XM_023252953.1;refGene_gn=TMTC4;refGene_tx=XM_023252959.1;refGene_gn=TMTC4;refGene_tx=XM_023252960.1
A1 71925666 71925798
refGene_gn=TMTC4;refGene_tx=XM_023252953.1;refGene_gn=TMTC4;refGene_tx=XM_023252959.1;refGene_gn=TMTC4;refGene_tx=XM_023252960.1
A1 71925886 71926058
refGene_gn=TMTC4;refGene_tx=XM_023252953.1;refGene_gn=TMTC4;refGene_tx=XM_023252959.1;refGene_gn=TMTC4;refGene_tx=XM_023252960.1
A1 71927692 71927895
refGene_gn=TMTC4;refGene_tx=XM_023252953.1;refGene_gn=TMTC4;refGene_tx=XM_023252959.1;refGene_gn=TMTC4;refGene_tx=XM_023252960.1
A1 71928778 71928943
refGene_gn=TMTC4;refGene_tx=XM_023252953.1;refGene_gn=TMTC4;refGene_tx=XM_023252959.1;refGene_gn=TMTC4;refGene_tx=XM_023252960.1
A1 71929172 71929265
refGene_gn=TMTC4;refGene_tx=XM_023252953.1;refGene_gn=TMTC4;refGene_tx=XM_023252959.1;refGene_gn=TMTC4;refGene_tx=XM_023252960.1
A1 71932085 71932186
refGene_gn=TMTC4;refGene_tx=XM_023252953.1;refGene_gn=TMTC4;refGene_tx=XM_023252959.1;refGene_gn=TMTC4;refGene_tx=XM_023252960.1
A1 71948805 71948893
refGene_gn=TMTC4;refGene_tx=XM_023252953.1;refGene_gn=TMTC4;refGene_tx=XM_023252959.1;refGene_gn=TMTC4;refGene_tx=XM_023252960.1
A1 71954945 71955162
refGene_gn=TMTC4;refGene_tx=XM_023252953.1;refGene_gn=TMTC4;refGene_tx=XM_023252959.1;refGene_gn=TMTC4;refGene_tx=XM_023252960.1
A1 71956235 71956351
refGene_gn=TMTC4;refGene_tx=XM_023252953.1;refGene_gn=TMTC4;refGene_tx=XM_023252959.1;refGene_gn=TMTC4;refGene_tx=XM_023252960.1
A1 71960837 71960999

refGene_gn=TMTC4;refGene_tx=XM_023252953.1;refGene_gn=TMTC4;refGene_tx=XM_023252959.1;refGene_gn=TMTC4;refGene_tx=XM_023252960.1
A1 72241241 72241435 refGene_gn=NALCN;refGene_tx=XM_023252962.1
A1 72241969 72242087 refGene_gn=NALCN;refGene_tx=XM_023252962.1
A1 72244399 72244549 refGene_gn=NALCN;refGene_tx=XM_023252962.1
A1 72245890 72246041 refGene_gn=NALCN;refGene_tx=XM_023252962.1
A1 72249868 72250026 refGene_gn=NALCN;refGene_tx=XM_023252962.1
A1 72251917 72252033 refGene_gn=NALCN;refGene_tx=XM_023252962.1
A1 72252706 72252839 refGene_gn=NALCN;refGene_tx=XM_023252962.1
A1 72254535 72254629 refGene_gn=NALCN;refGene_tx=XM_023252962.1
A1 72255470 72255619 refGene_gn=NALCN;refGene_tx=XM_023252962.1
A1 72256569 72256638 refGene_gn=NALCN;refGene_tx=XM_023252962.1
A1 72260121 72260241 refGene_gn=NALCN;refGene_tx=XM_023252962.1
A1 72261269 72261344 refGene_gn=NALCN;refGene_tx=XM_023252962.1
A1 72261552 72261659 refGene_gn=NALCN;refGene_tx=XM_023252962.1
A1 72262167 72262261 refGene_gn=NALCN;refGene_tx=XM_023252962.1
A1 72268142 72268241 refGene_gn=NALCN;refGene_tx=XM_023252962.1
A1 72268326 72268447 refGene_gn=NALCN;refGene_tx=XM_023252962.1
A1 72274761 72274868 refGene_gn=NALCN;refGene_tx=XM_023252962.1
A1 72278819 72278924 refGene_gn=NALCN;refGene_tx=XM_023252962.1
A1 72280600 72280768 refGene_gn=NALCN;refGene_tx=XM_023252962.1
A1 72281727 72281859 refGene_gn=NALCN;refGene_tx=XM_023252962.1
A1 72281955 72282076 refGene_gn=NALCN;refGene_tx=XM_023252962.1
A1 72282560 72282617 refGene_gn=NALCN;refGene_tx=XM_023252962.1
A1 72285418 72285541 refGene_gn=NALCN;refGene_tx=XM_023252962.1
A1 72285628 72285720 refGene_gn=NALCN;refGene_tx=XM_023252962.1
A1 72288708 72288738 refGene_gn=NALCN;refGene_tx=XM_023252962.1
A1 72288739 72288779 refGene_gn=NALCN;refGene_tx=XM_023252962.1
A1 72289215 72289317 refGene_gn=NALCN;refGene_tx=XM_023252962.1
A1 72296895 72296982 refGene_gn=NALCN;refGene_tx=XM_023252962.1
A1 72308269 72308343 refGene_gn=NALCN;refGene_tx=XM_023252962.1
A1 72326898 72327040 refGene_gn=NALCN;refGene_tx=XM_023252962.1
A1 72328978 72329115 refGene_gn=NALCN;refGene_tx=XM_023252962.1
A1 72356813 72356888 refGene_gn=NALCN;refGene_tx=XM_023252962.1
A1 72370072 72370210 refGene_gn=NALCN;refGene_tx=XM_023252962.1
A1 72397059 72397251 refGene_gn=NALCN;refGene_tx=XM_023252962.1
A1 72403742 72403910 refGene_gn=NALCN;refGene_tx=XM_023252962.1
A1 72420572 72420704 refGene_gn=NALCN;refGene_tx=XM_023252962.1
A1 72446108 72446195 refGene_gn=NALCN;refGene_tx=XM_023252962.1
A1 72452031 72452136 refGene_gn=NALCN;refGene_tx=XM_023252962.1
A1 72452258 72452401 refGene_gn=NALCN;refGene_tx=XM_023252962.1
A1 72511648 72511803 refGene_gn=NALCN;refGene_tx=XM_023252962.1
A1 72527586 72527715 refGene_gn=NALCN;refGene_tx=XM_023252962.1
A1 72527796 72527950 refGene_gn=NALCN;refGene_tx=XM_023252962.1
A1 72529098 72529168 refGene_gn=NALCN;refGene_tx=XM_023252962.1
A1 72544928 72545111 refGene_gn=NALCN;refGene_tx=XM_023252962.1
A1 72546457 72546565 refGene_gn=NALCN;refGene_tx=XM_023252962.1
A1 72585384 72585482
refGene_gn=ITGBL1;refGene_tx=XM_011280788.3;refGene_gn=ITGBL1;refGene_tx=XM_019829193.2

A1 72586637 72586855
refGene_gn=ITGBL1;refGene_tx=XM_011280788.3;refGene_gn=ITGBL1;refGene_tx=XM_019829193.2
A1 72669328 72669475
refGene_gn=ITGBL1;refGene_tx=XM_011280788.3;refGene_gn=ITGBL1;refGene_tx=XM_019829193.2
A1 72676963 72677086
refGene_gn=ITGBL1;refGene_tx=XM_011280788.3;refGene_gn=ITGBL1;refGene_tx=XM_019829193.2
A1 72680830 72680971
refGene_gn=ITGBL1;refGene_tx=XM_011280788.3;refGene_gn=ITGBL1;refGene_tx=XM_019829193.2
A1 72683872 72684013
refGene_gn=ITGBL1;refGene_tx=XM_011280788.3;refGene_gn=ITGBL1;refGene_tx=XM_019829193.2
A1 72694441 72694588
refGene_gn=ITGBL1;refGene_tx=XM_011280788.3;refGene_gn=ITGBL1;refGene_tx=XM_019829193.2
A1 72699825 72700007 refGene_gn=ITGBL1;refGene_tx=XM_019829193.2
A1 72769606 72769723 refGene_gn=ITGBL1;refGene_tx=XM_011280788.3
A1 72785276 72785423 refGene_gn=ITGBL1;refGene_tx=XM_011280788.3
A1 72792899 72793013 refGene_gn=ITGBL1;refGene_tx=XM_011280788.3
A1 72793490 72793582 refGene_gn=ITGBL1;refGene_tx=XM_011280788.3
A1 72796898 72796918
refGene_gn=FGF14;refGene_tx=XM_019829208.2;refGene_gn=FGF14;refGene_tx=XM_019829216.2;refGene_gn=FGF14;refGene_tx=XM_019829233.2;refGene_gn=FGF14;refGene_tx=XM_023252979.1
A1 72800699 72800822
refGene_gn=FGF14;refGene_tx=XM_019829208.2;refGene_gn=FGF14;refGene_tx=XM_019829216.2;refGene_gn=FGF14;refGene_tx=XM_019829233.2;refGene_gn=FGF14;refGene_tx=XM_023252979.1
A1 72803944 72804143
refGene_gn=FGF14;refGene_tx=XM_019829208.2;refGene_gn=FGF14;refGene_tx=XM_019829216.2;refGene_gn=FGF14;refGene_tx=XM_019829233.2;refGene_gn=FGF14;refGene_tx=XM_023252979.1
A1 72917703 72917807
refGene_gn=FGF14;refGene_tx=XM_019829208.2;refGene_gn=FGF14;refGene_tx=XM_019829216.2;refGene_gn=FGF14;refGene_tx=XM_019829233.2;refGene_gn=FGF14;refGene_tx=XM_023252979.1
A1 72923772 72923883
refGene_gn=FGF14;refGene_tx=XM_019829208.2;refGene_gn=FGF14;refGene_tx=XM_019829216.2;refGene_gn=FGF14;refGene_tx=XM_019829233.2;refGene_gn=FGF14;refGene_tx=XM_023252979.1
A1 72972440 72972633 refGene_gn=FGF14;refGene_tx=XM_023252979.1
A1 73417141 73417349 refGene_gn=FGF14;refGene_tx=XM_019829208.2
A1 73531419 73531584
refGene_gn=TPP2;refGene_tx=XM_023252982.1;refGene_gn=TPP2;refGene_tx=XM_023252986.1;refGene_gn=TPP2;refGene_tx=XM_023252993.1;refGene_gn=TPP2;refGene_tx=XM_023252997.1;refGene_gn=TPP2;refGene_tx=XM_023253002.1;refGene_gn=TPP2;refGene_tx=XM_023253007.1;refGene_gn=TPP2;refGene_tx=X

M_023253011.1
A1 73538702 73538831
refGene_gn=TPP2;refGene_tx=XM_023252982.1;refGene_gn=TPP2;refGene_tx=XM_023252986.1;refGene_gn=TPP2;refGene_tx=XM_023252993.1;refGene_gn=TPP2;refGene_tx=XM_023252997.1;refGene_gn=TPP2;refGene_tx=XM_023253002.1;refGene_gn=TPP2;refGene_tx=XM_023253007.1;refGene_gn=TPP2;refGene_tx=XM_023253011.1
A1 73547307 73547403
refGene_gn=TPP2;refGene_tx=XM_023252982.1;refGene_gn=TPP2;refGene_tx=XM_023252986.1;refGene_gn=TPP2;refGene_tx=XM_023252993.1;refGene_gn=TPP2;refGene_tx=XM_023252997.1;refGene_gn=TPP2;refGene_tx=XM_023253002.1;refGene_gn=TPP2;refGene_tx=XM_023253007.1;refGene_gn=TPP2;refGene_tx=XM_023253011.1
A1 73549170 73549275
refGene_gn=TPP2;refGene_tx=XM_023252982.1;refGene_gn=TPP2;refGene_tx=XM_023252986.1;refGene_gn=TPP2;refGene_tx=XM_023252993.1;refGene_gn=TPP2;refGene_tx=XM_023252997.1;refGene_gn=TPP2;refGene_tx=XM_023253002.1;refGene_gn=TPP2;refGene_tx=XM_023253007.1;refGene_gn=TPP2;refGene_tx=XM_023253011.1
A1 73551236 73551361
refGene_gn=TPP2;refGene_tx=XM_023252982.1;refGene_gn=TPP2;refGene_tx=XM_023252986.1;refGene_gn=TPP2;refGene_tx=XM_023252993.1;refGene_gn=TPP2;refGene_tx=XM_023252997.1;refGene_gn=TPP2;refGene_tx=XM_023253002.1;refGene_gn=TPP2;refGene_tx=XM_023253007.1;refGene_gn=TPP2;refGene_tx=XM_023253011.1
A1 73554263 73554427
refGene_gn=TPP2;refGene_tx=XM_023252982.1;refGene_gn=TPP2;refGene_tx=XM_023252986.1;refGene_gn=TPP2;refGene_tx=XM_023252993.1;refGene_gn=TPP2;refGene_tx=XM_023252997.1;refGene_gn=TPP2;refGene_tx=XM_023253002.1;refGene_gn=TPP2;refGene_tx=XM_023253007.1;refGene_gn=TPP2;refGene_tx=XM_023253011.1;refGene_gn=TPP2;refGene_tx=XM_023253017.1
A1 73557514 73557669
refGene_gn=TPP2;refGene_tx=XM_023252982.1;refGene_gn=TPP2;refGene_tx=XM_023252986.1;refGene_gn=TPP2;refGene_tx=XM_023252993.1;refGene_gn=TPP2;refGene_tx=XM_023252997.1;refGene_gn=TPP2;refGene_tx=XM_023253002.1;refGene_gn=TPP2;refGene_tx=XM_023253007.1;refGene_gn=TPP2;refGene_tx=XM_023253011.1;refGene_gn=TPP2;refGene_tx=XM_023253017.1
A1 73558285 73558362
refGene_gn=TPP2;refGene_tx=XM_023252982.1;refGene_gn=TPP2;refGene_tx=XM_023252986.1;refGene_gn=TPP2;refGene_tx=XM_023252993.1;refGene_gn=TPP2;refGene_tx=XM_023252997.1;refGene_gn=TPP2;refGene_tx=XM_023253002.1;refGene_gn=TPP2;refGene_tx=XM_023253007.1;refGene_gn=TPP2;refGene_tx=XM_023253011.1;refGene_gn=TPP2;refGene_tx=XM_023253017.1
A1 73560361 73560489
refGene_gn=TPP2;refGene_tx=XM_023252982.1;refGene_gn=TPP2;refGene_tx=XM_023252986.1;refGene_gn=TPP2;refGene_tx=XM_023252993.1;refGene_gn=TPP2;refGene_tx=XM_023252997.1;refGene_gn=TPP2;refGene_tx=XM_023253002.1;refGene_gn=TPP2;refGene_tx=XM_023253007.1;refGene_gn=TPP2;refGene_tx=XM_023253011.1;refGene_gn=TPP2;refGene_tx=XM_023253017.1
A1 73562546 73562646

refGene_gn=TPP2;refGene_tx=XM_023252982.1;refGene_gn=TPP2;refGene_tx=XM_023252986.1;refGene_gn=TPP2;refGene_tx=XM_023252993.1;refGene_gn=TPP2;refGene_tx=XM_023252997.1;refGene_gn=TPP2;refGene_tx=XM_023253002.1;refGene_gn=TPP2;refGene_tx=XM_023253007.1;refGene_gn=TPP2;refGene_tx=XM_023253011.1;refGene_gn=TPP2;refGene_tx=XM_023253017.1
A1 73566175 73566324
refGene_gn=TPP2;refGene_tx=XM_023252982.1;refGene_gn=TPP2;refGene_tx=XM_023252986.1;refGene_gn=TPP2;refGene_tx=XM_023252993.1;refGene_gn=TPP2;refGene_tx=XM_023252997.1;refGene_gn=TPP2;refGene_tx=XM_023253002.1;refGene_gn=TPP2;refGene_tx=XM_023253007.1;refGene_gn=TPP2;refGene_tx=XM_023253011.1;refGene_gn=TPP2;refGene_tx=XM_023253017.1
A1 73568106 73568222
refGene_gn=TPP2;refGene_tx=XM_023252982.1;refGene_gn=TPP2;refGene_tx=XM_023252986.1;refGene_gn=TPP2;refGene_tx=XM_023252993.1;refGene_gn=TPP2;refGene_tx=XM_023252997.1;refGene_gn=TPP2;refGene_tx=XM_023253002.1;refGene_gn=TPP2;refGene_tx=XM_023253007.1;refGene_gn=TPP2;refGene_tx=XM_023253011.1;refGene_gn=TPP2;refGene_tx=XM_023253017.1
A1 73568734 73568903
refGene_gn=TPP2;refGene_tx=XM_023252982.1;refGene_gn=TPP2;refGene_tx=XM_023252986.1;refGene_gn=TPP2;refGene_tx=XM_023252993.1;refGene_gn=TPP2;refGene_tx=XM_023252997.1;refGene_gn=TPP2;refGene_tx=XM_023253002.1;refGene_gn=TPP2;refGene_tx=XM_023253007.1;refGene_gn=TPP2;refGene_tx=XM_023253011.1;refGene_gn=TPP2;refGene_tx=XM_023253017.1
A1 73569586 73569744
refGene_gn=TPP2;refGene_tx=XM_023252982.1;refGene_gn=TPP2;refGene_tx=XM_023252986.1;refGene_gn=TPP2;refGene_tx=XM_023252993.1;refGene_gn=TPP2;refGene_tx=XM_023252997.1;refGene_gn=TPP2;refGene_tx=XM_023253002.1;refGene_gn=TPP2;refGene_tx=XM_023253007.1;refGene_gn=TPP2;refGene_tx=XM_023253011.1;refGene_gn=TPP2;refGene_tx=XM_023253017.1
A1 73570021 73570098
refGene_gn=TPP2;refGene_tx=XM_023252982.1;refGene_gn=TPP2;refGene_tx=XM_023252986.1;refGene_gn=TPP2;refGene_tx=XM_023252993.1;refGene_gn=TPP2;refGene_tx=XM_023252997.1;refGene_gn=TPP2;refGene_tx=XM_023253002.1;refGene_gn=TPP2;refGene_tx=XM_023253007.1;refGene_gn=TPP2;refGene_tx=XM_023253011.1;refGene_gn=TPP2;refGene_tx=XM_023253017.1
A1 73572061 73572168
refGene_gn=TPP2;refGene_tx=XM_023252982.1;refGene_gn=TPP2;refGene_tx=XM_023252986.1;refGene_gn=TPP2;refGene_tx=XM_023252993.1;refGene_gn=TPP2;refGene_tx=XM_023252997.1;refGene_gn=TPP2;refGene_tx=XM_023253002.1;refGene_gn=TPP2;refGene_tx=XM_023253007.1;refGene_gn=TPP2;refGene_tx=XM_023253011.1;refGene_gn=TPP2;refGene_tx=XM_023253017.1
A1 73574072 73574227
refGene_gn=TPP2;refGene_tx=XM_023252982.1;refGene_gn=TPP2;refGene_tx=XM_023252986.1;refGene_gn=TPP2;refGene_tx=XM_023252993.1;refGene_gn=TPP2;refGene_tx=XM_023252997.1;refGene_gn=TPP2;refGene_tx=XM_023253002.1;refGene_gn=TPP2;refGene_tx=XM_023253007.1;refGene_gn=TPP2;refGene_tx=XM_023253011.1;refGene_gn=TPP2;refGene_tx=XM_023253017.1
A1 73575342 73575459
refGene_gn=TPP2;refGene_tx=XM_023252982.1;refGene_gn=TPP2;refGene_tx=XM_023252986.1;refGene_gn=TPP2;refGene_tx=XM_023252993.1;refGene_gn=TPP

2;refGene_tx=XM_023252997.1;refGene_gn=TPP2;refGene_tx=XM_023253002.1;refGene_gn=TPP2;refGene_tx=XM_023253007.1;refGene_gn=TPP2;refGene_tx=XM_023253011.1;refGene_gn=TPP2;refGene_tx=XM_023253017.1
A1 73575700 73575801
refGene_gn=TPP2;refGene_tx=XM_023252982.1;refGene_gn=TPP2;refGene_tx=XM_023252986.1;refGene_gn=TPP2;refGene_tx=XM_023252993.1;refGene_gn=TPP2;refGene_tx=XM_023252997.1;refGene_gn=TPP2;refGene_tx=XM_023253002.1;refGene_gn=TPP2;refGene_tx=XM_023253007.1;refGene_gn=TPP2;refGene_tx=XM_023253011.1;refGene_gn=TPP2;refGene_tx=XM_023253017.1
A1 73576418 73576515
refGene_gn=TPP2;refGene_tx=XM_023252982.1;refGene_gn=TPP2;refGene_tx=XM_023252986.1;refGene_gn=TPP2;refGene_tx=XM_023252993.1;refGene_gn=TPP2;refGene_tx=XM_023252997.1;refGene_gn=TPP2;refGene_tx=XM_023253002.1;refGene_gn=TPP2;refGene_tx=XM_023253007.1;refGene_gn=TPP2;refGene_tx=XM_023253011.1;refGene_gn=TPP2;refGene_tx=XM_023253017.1
A1 73577565 73577703
refGene_gn=TPP2;refGene_tx=XM_023252982.1;refGene_gn=TPP2;refGene_tx=XM_023252986.1;refGene_gn=TPP2;refGene_tx=XM_023252993.1;refGene_gn=TPP2;refGene_tx=XM_023252997.1;refGene_gn=TPP2;refGene_tx=XM_023253002.1;refGene_gn=TPP2;refGene_tx=XM_023253007.1;refGene_gn=TPP2;refGene_tx=XM_023253011.1;refGene_gn=TPP2;refGene_tx=XM_023253017.1
A1 73579036 73579281
refGene_gn=TPP2;refGene_tx=XM_023252982.1;refGene_gn=TPP2;refGene_tx=XM_023252986.1;refGene_gn=TPP2;refGene_tx=XM_023252993.1;refGene_gn=TPP2;refGene_tx=XM_023252997.1;refGene_gn=TPP2;refGene_tx=XM_023253002.1;refGene_gn=TPP2;refGene_tx=XM_023253007.1;refGene_gn=TPP2;refGene_tx=XM_023253011.1;refGene_gn=TPP2;refGene_tx=XM_023253017.1
A1 73579537 73579616
refGene_gn=TPP2;refGene_tx=XM_023252982.1;refGene_gn=TPP2;refGene_tx=XM_023252986.1;refGene_gn=TPP2;refGene_tx=XM_023252993.1;refGene_gn=TPP2;refGene_tx=XM_023252997.1;refGene_gn=TPP2;refGene_tx=XM_023253002.1;refGene_gn=TPP2;refGene_tx=XM_023253007.1;refGene_gn=TPP2;refGene_tx=XM_023253011.1;refGene_gn=TPP2;refGene_tx=XM_023253017.1
A1 73581777 73581816
refGene_gn=TPP2;refGene_tx=XM_023252982.1;refGene_gn=TPP2;refGene_tx=XM_023252993.1;refGene_gn=TPP2;refGene_tx=XM_023252997.1;refGene_gn=TPP2;refGene_tx=XM_023253007.1;refGene_gn=TPP2;refGene_tx=XM_023253011.1;refGene_gn=TPP2;refGene_tx=XM_023253017.1
A1 73587270 73587422
refGene_gn=TPP2;refGene_tx=XM_023252982.1;refGene_gn=TPP2;refGene_tx=XM_023252986.1;refGene_gn=TPP2;refGene_tx=XM_023252993.1;refGene_gn=TPP2;refGene_tx=XM_023252997.1;refGene_gn=TPP2;refGene_tx=XM_023253002.1;refGene_gn=TPP2;refGene_tx=XM_023253007.1;refGene_gn=TPP2;refGene_tx=XM_023253011.1;refGene_gn=TPP2;refGene_tx=XM_023253017.1
A1 73588557 73588654
refGene_gn=TPP2;refGene_tx=XM_023252982.1;refGene_gn=TPP2;refGene_tx=XM_023252986.1;refGene_gn=TPP2;refGene_tx=XM_023252993.1;refGene_gn=TPP2;refGene_tx=XM_023252997.1;refGene_gn=TPP2;refGene_tx=XM_023253002.1;refGene_gn=TPP2;refGene_tx=XM_023253007.1;refGene_gn=TPP2;refGene_tx=XM_023253011.1;refGene_gn=TPP2;refGene_tx=XM_023253017.1

A1 73589943 73590085
refGene_gn=TPP2;refGene_tx=XM_023252982.1;refGene_gn=TPP2;refGene_tx=XM_023252986.1;refGene_gn=TPP2;refGene_tx=XM_023252993.1;refGene_gn=TPP2;refGene_tx=XM_023252997.1;refGene_gn=TPP2;refGene_tx=XM_023253002.1;refGene_gn=TPP2;refGene_tx=XM_023253007.1;refGene_gn=TPP2;refGene_tx=XM_023253011.1;refGene_gn=TPP2;refGene_tx=XM_023253017.1
A1 73594295 73594503
refGene_gn=TPP2;refGene_tx=XM_023252982.1;refGene_gn=TPP2;refGene_tx=XM_023252986.1;refGene_gn=TPP2;refGene_tx=XM_023252993.1;refGene_gn=TPP2;refGene_tx=XM_023252997.1;refGene_gn=TPP2;refGene_tx=XM_023253002.1;refGene_gn=TPP2;refGene_tx=XM_023253007.1;refGene_gn=TPP2;refGene_tx=XM_023253011.1;refGene_gn=TPP2;refGene_tx=XM_023253017.1
A1 73594596 73594668
refGene_gn=TPP2;refGene_tx=XM_023252982.1;refGene_gn=TPP2;refGene_tx=XM_023252986.1;refGene_gn=TPP2;refGene_tx=XM_023252993.1;refGene_gn=TPP2;refGene_tx=XM_023253017.1
A1 73596227 73596347
refGene_gn=TPP2;refGene_tx=XM_023252982.1;refGene_gn=TPP2;refGene_tx=XM_023252986.1;refGene_gn=TPP2;refGene_tx=XM_023252993.1;refGene_gn=TPP2;refGene_tx=XM_023252997.1;refGene_gn=TPP2;refGene_tx=XM_023253002.1;refGene_gn=TPP2;refGene_tx=XM_023253017.1
A1 73598422 73598512
refGene_gn=TPP2;refGene_tx=XM_023252982.1;refGene_gn=TPP2;refGene_tx=XM_023252986.1;refGene_gn=TPP2;refGene_tx=XM_023252993.1;refGene_gn=TPP2;refGene_tx=XM_023252997.1;refGene_gn=TPP2;refGene_tx=XM_023253002.1;refGene_gn=TPP2;refGene_tx=XM_023253017.1
A1 73606357 73606752 refGene_gn=METTL21C;refGene_tx=XM_006927386.4
A1 73607760 73607878 refGene_gn=METTL21C;refGene_tx=XM_006927386.4
A1 73610840 73610992 refGene_gn=METTL21C;refGene_tx=XM_006927386.4
A1 73614317 73614452 refGene_gn=METTL21C;refGene_tx=XM_006927386.4
A1 73614680 73614789 refGene_gn=METTL21C;refGene_tx=XM_006927386.4
A1 73660124 73675741 refGene_gn=CCDC168;refGene_tx=XM_023239134.1
A1 73675743 73679673 refGene_gn=CCDC168;refGene_tx=XM_023239134.1
A1 73679675 73681838 refGene_gn=CCDC168;refGene_tx=XM_023239134.1
A1 73683987 73684010 refGene_gn=CCDC168;refGene_tx=XM_023239134.1
A1 73688660 73688700 refGene_gn=CCDC168;refGene_tx=XM_023239134.1
A1 73688802 73688836 refGene_gn=CCDC168;refGene_tx=XM_023239134.1
A1 73688963 73689389 refGene_gn=CCDC168;refGene_tx=XM_023239134.1
A1 73694366 73694540
refGene_gn=TEX30;refGene_tx=XM_023253026.1;refGene_gn=TEX30;refGene_tx=XM_023253030.1;refGene_gn=TEX30;refGene_tx=XM_023253036.1
A1 73695146 73695352
refGene_gn=TEX30;refGene_tx=XM_023253026.1;refGene_gn=TEX30;refGene_tx=XM_023253030.1;refGene_gn=TEX30;refGene_tx=XM_023253036.1
A1 73695860 73695912
refGene_gn=TEX30;refGene_tx=XM_023253026.1;refGene_gn=TEX30;refGene_tx=XM_023253030.1;refGene_gn=TEX30;refGene_tx=XM_023253036.1
A1 73696675 73696906
refGene_gn=TEX30;refGene_tx=XM_023253026.1;refGene_gn=TEX30;refGene_tx=XM_023253030.1;refGene_gn=TEX30;refGene_tx=XM_023253036.1

A1 73697374 73697389
refGene_gn=TEX30;refGene_tx=XM_023253026.1;refGene_gn=TEX30;refGene_tx=XM_023253030.1;refGene_gn=TEX30;refGene_tx=XM_023253036.1
A1 73718416 73718434
refGene_gn=KDELC1;refGene_tx=XM_003980511.5;refGene_gn=KDELC1;refGene_tx=XM_019829270.2
A1 73720393 73720501
refGene_gn=KDELC1;refGene_tx=XM_003980511.5;refGene_gn=KDELC1;refGene_tx=XM_019829270.2
A1 73722318 73722408
refGene_gn=KDELC1;refGene_tx=XM_003980511.5;refGene_gn=KDELC1;refGene_tx=XM_019829270.2
A1 73723091 73723301
refGene_gn=KDELC1;refGene_tx=XM_003980511.5;refGene_gn=KDELC1;refGene_tx=XM_019829270.2
A1 73725312 73725550
refGene_gn=KDELC1;refGene_tx=XM_003980511.5;refGene_gn=KDELC1;refGene_tx=XM_019829270.2
A1 73725657 73725830
refGene_gn=KDELC1;refGene_tx=XM_003980511.5;refGene_gn=KDELC1;refGene_tx=XM_019829270.2
A1 73729428 73729506
refGene_gn=KDELC1;refGene_tx=XM_003980511.5;refGene_gn=KDELC1;refGene_tx=XM_019829270.2
A1 73730263 73730469
refGene_gn=KDELC1;refGene_tx=XM_003980511.5;refGene_gn=KDELC1;refGene_tx=XM_019829270.2
A1 73730591 73730797
refGene_gn=KDELC1;refGene_tx=XM_003980511.5;refGene_gn=KDELC1;refGene_tx=XM_019829270.2
A1 73732295 73732552 refGene_gn=KDELC1;refGene_tx=XM_019829270.2
A1 73732680 73732862 refGene_gn=KDELC1;refGene_tx=XM_003980511.5
A1 73740147 73740625
refGene_gn=LOC101085685;refGene_tx=XM_011280818.3;refGene_gn=LOC101085685;refGene_tx=XM_023253052.1;refGene_gn=LOC101085685;refGene_tx=XM_023253054.1
A1 73747714 73747841
refGene_gn=LOC101085685;refGene_tx=XM_011280818.3;refGene_gn=LOC101085685;refGene_tx=XM_023253052.1;refGene_gn=LOC101085685;refGene_tx=XM_023253054.1
A1 73757954 73758050
refGene_gn=LOC101085685;refGene_tx=XM_011280818.3;refGene_gn=LOC101085685;refGene_tx=XM_023253052.1;refGene_gn=LOC101085685;refGene_tx=XM_023253054.1
A1 73759273 73759378
refGene_gn=LOC101085685;refGene_tx=XM_011280818.3;refGene_gn=LOC101085685;refGene_tx=XM_023253052.1;refGene_gn=LOC101085685;refGene_tx=XM_023253054.1
A1 73759591 73759686
refGene_gn=LOC101085685;refGene_tx=XM_011280818.3;refGene_gn=LOC101085

685;refGene_tx=XM_023253052.1;refGene_gn=LOC101085685;refGene_tx=XM_023253054.1
A1 73770969 73771102
refGene_gn=LOC101085685;refGene_tx=XM_011280818.3;refGene_gn=LOC101085685;refGene_tx=XM_023253052.1;refGene_gn=LOC101085685;refGene_tx=XM_023253054.1
A1 73772462 73772549
refGene_gn=LOC101085685;refGene_tx=XM_011280818.3;refGene_gn=LOC101085685;refGene_tx=XM_023253052.1;refGene_gn=LOC101085685;refGene_tx=XM_023253054.1
A1 73774774 73774871
refGene_gn=LOC101085685;refGene_tx=XM_011280818.3;refGene_gn=LOC101085685;refGene_tx=XM_023253052.1;refGene_gn=LOC101085685;refGene_tx=XM_023253054.1
A1 73775738 73776032
refGene_gn=LOC101085685;refGene_tx=XM_011280818.3;refGene_gn=LOC101085685;refGene_tx=XM_023253052.1;refGene_gn=LOC101085685;refGene_tx=XM_023253054.1
A1 73780447 73780535
refGene_gn=LOC101085685;refGene_tx=XM_011280813.3
A1 73785678 73785854
refGene_gn=LOC101085685;refGene_tx=XM_011280813.3;refGene_gn=LOC101085685;refGene_tx=XM_023253054.1
A1 73788731 73788847
refGene_gn=LOC101085685;refGene_tx=XM_011280813.3;refGene_gn=LOC101085685;refGene_tx=XM_023253054.1
A1 73789318 73789402
refGene_gn=LOC101085685;refGene_tx=XM_011280813.3;refGene_gn=LOC101085685;refGene_tx=XM_023253054.1
A1 73790688 73790749
refGene_gn=LOC101085685;refGene_tx=XM_011280813.3;refGene_gn=LOC101085685;refGene_tx=XM_023253054.1
A1 73792988 73793132
refGene_gn=LOC101085685;refGene_tx=XM_011280813.3;refGene_gn=LOC101085685;refGene_tx=XM_023253054.1
A1 73794911 73795119
refGene_gn=LOC101085685;refGene_tx=XM_011280813.3;refGene_gn=LOC101085685;refGene_tx=XM_023253054.1
A1 73795502 73796603
refGene_gn=LOC101085685;refGene_tx=XM_011280813.3;refGene_gn=LOC101085685;refGene_tx=XM_023253054.1
A1 73800021 73800269
refGene_gn=LOC101085685;refGene_tx=XM_011280813.3;refGene_gn=LOC101085685;refGene_tx=XM_023253054.1
A1 73803411 73803531
refGene_gn=LOC101085685;refGene_tx=XM_011280813.3;refGene_gn=LOC101085685;refGene_tx=XM_023253054.1
A1 73803698 73803912
refGene_gn=LOC101085685;refGene_tx=XM_011280813.3;refGene_gn=LOC101085685;refGene_tx=XM_023253054.1

A1 73804667 73804812
refGene_gn=LOC101085685;refGene_tx=XM_011280813.3;refGene_gn=LOC101085685;refGene_tx=XM_023253054.1
A1 73808765 73808966
refGene_gn=LOC101085685;refGene_tx=XM_011280813.3;refGene_gn=LOC101085685;refGene_tx=XM_023253054.1
A1 73810083 73810168
refGene_gn=LOC101085685;refGene_tx=XM_011280813.3;refGene_gn=LOC101085685;refGene_tx=XM_023253054.1
A1 73811836 73812427
refGene_gn=LOC101085685;refGene_tx=XM_011280813.3;refGene_gn=LOC101085685;refGene_tx=XM_023253054.1
A1 73816894 73816964
refGene_gn=LOC101085945;refGene_tx=XM_006927390.4
A1 73827673 73827837
refGene_gn=LOC101085945;refGene_tx=XM_006927390.4;refGene_gn=LOC101085945;refGene_tx=XM_023253067.1
A1 73829215 73829333
refGene_gn=LOC101085945;refGene_tx=XM_006927390.4;refGene_gn=LOC101085945;refGene_tx=XM_023253067.1
A1 73833587 73834027
refGene_gn=LOC101085945;refGene_tx=XM_006927390.4;refGene_gn=LOC101085945;refGene_tx=XM_023253067.1
A1 73966311 73966439 refGene_gn=SLC10A2;refGene_tx=XM_003980513.4
A1 73969349 73969507 refGene_gn=SLC10A2;refGene_tx=XM_003980513.4
A1 73971501 73971677 refGene_gn=SLC10A2;refGene_tx=XM_003980513.4
A1 73972604 73972693 refGene_gn=SLC10A2;refGene_tx=XM_003980513.4
A1 73976996 73977115 refGene_gn=SLC10A2;refGene_tx=XM_003980513.4
A1 73984281 73984658 refGene_gn=SLC10A2;refGene_tx=XM_003980513.4
A1 77088366 77088755
refGene_gn=EFNB2;refGene_tx=NM_001114344.1;refGene_gn=EFNB2;refGene_tx=XM_019834389.2
A1 77090096 77090210
refGene_gn=EFNB2;refGene_tx=NM_001114344.1;refGene_gn=EFNB2;refGene_tx=XM_019834389.2
A1 77090957 77091050 refGene_gn=EFNB2;refGene_tx=NM_001114344.1
A1 77107822 77108106
refGene_gn=EFNB2;refGene_tx=NM_001114344.1;refGene_gn=EFNB2;refGene_tx=XM_019834389.2
A1 77130022 77130144
refGene_gn=EFNB2;refGene_tx=NM_001114344.1;refGene_gn=EFNB2;refGene_tx=XM_019834389.2
A1 77138312 77138477 refGene_gn=ARGLU1;refGene_tx=XM_003980522.4
A1 77152234 77152318 refGene_gn=ARGLU1;refGene_tx=XM_003980522.4
A1 77154614 77154840 refGene_gn=ARGLU1;refGene_tx=XM_003980522.4
A1 77162776 77163123 refGene_gn=ARGLU1;refGene_tx=XM_003980522.4
A1 77679941 77680231 refGene_gn=FAM155A;refGene_tx=XM_003980527.5
A1 77681933 77681956 refGene_gn=FAM155A;refGene_tx=XM_023253100.1
A1 77714196 77714368
refGene_gn=FAM155A;refGene_tx=XM_003980527.5;refGene_gn=FAM155A;refGen

e_tx=XM_023253100.1
A1 78307456 78308398
refGene_gn=FAM155A;refGene_tx=XM_003980527.5;refGene_gn=FAM155A;refGen
e_tx=XM_023253100.1
A1 78548120 78550856
refGene_gn=LIG4;refGene_tx=XM_006927396.4;refGene_gn=LIG4;refGene_tx=X
M_019829453.1;refGene_gn=LIG4;refGene_tx=XM_019829456.1
A1 78575417 78576431
refGene_gn=ABHD13;refGene_tx=XM_003980524.5;refGene_gn=ABHD13;refGene_
tx=XM_019829468.2
A1 78607153 78607705 refGene_gn=TNFSF13B;refGene_tx=XM_023253118.1
A1 78607798 78607886 refGene_gn=TNFSF13B;refGene_tx=XM_023253118.1
A1 78625065 78625122 refGene_gn=TNFSF13B;refGene_tx=XM_023253118.1
A1 78635554 78635667 refGene_gn=TNFSF13B;refGene_tx=XM_023253118.1
A1 78635752 78635903 refGene_gn=TNFSF13B;refGene_tx=XM_023253118.1
A1 78639192 78639305 refGene_gn=TNFSF13B;refGene_tx=XM_023253118.1
A1 78921485 78921711 refGene_gn=MYO16;refGene_tx=XM_023253136.1
A1 78966523 78966594 refGene_gn=MYO16;refGene_tx=XM_023253136.1
A1 78981770 78981914 refGene_gn=MYO16;refGene_tx=XM_023253136.1
A1 79010125 79010234 refGene_gn=MYO16;refGene_tx=XM_023253136.1
A1 79017327 79017452 refGene_gn=MYO16;refGene_tx=XM_023253136.1
A1 79027295 79027421 refGene_gn=MYO16;refGene_tx=XM_023253136.1
A1 79040290 79040366 refGene_gn=MYO16;refGene_tx=XM_023253136.1
A1 79044063 79044217 refGene_gn=MYO16;refGene_tx=XM_023253136.1
A1 79062201 79062352 refGene_gn=MYO16;refGene_tx=XM_023253136.1
A1 79071226 79071337 refGene_gn=MYO16;refGene_tx=XM_023253136.1
A1 79080324 79080390 refGene_gn=MYO16;refGene_tx=XM_023253136.1
A1 79089631 79089759 refGene_gn=MYO16;refGene_tx=XM_023253136.1
A1 79095612 79095718 refGene_gn=MYO16;refGene_tx=XM_023253136.1
A1 79105173 79105291 refGene_gn=MYO16;refGene_tx=XM_023253136.1
A1 79120858 79121006 refGene_gn=MYO16;refGene_tx=XM_023253136.1
A1 79163768 79163880 refGene_gn=MYO16;refGene_tx=XM_023253136.1
A1 79167054 79167172 refGene_gn=MYO16;refGene_tx=XM_023253136.1
A1 79167974 79168046 refGene_gn=MYO16;refGene_tx=XM_023253136.1
A1 79170616 79170758 refGene_gn=MYO16;refGene_tx=XM_023253136.1
A1 79195787 79195860 refGene_gn=MYO16;refGene_tx=XM_023253136.1
A1 79211990 79212143 refGene_gn=MYO16;refGene_tx=XM_023253136.1
A1 79215687 79215888 refGene_gn=MYO16;refGene_tx=XM_023253136.1
A1 79237868 79237944 refGene_gn=MYO16;refGene_tx=XM_023253136.1
A1 79243016 79243192 refGene_gn=MYO16;refGene_tx=XM_023253136.1
A1 79245622 79245703 refGene_gn=MYO16;refGene_tx=XM_023253136.1
A1 79245942 79246148 refGene_gn=MYO16;refGene_tx=XM_023253136.1
A1 79282867 79282970 refGene_gn=MYO16;refGene_tx=XM_023253136.1
A1 79301038 79301135 refGene_gn=MYO16;refGene_tx=XM_023253136.1
A1 79306385 79306632 refGene_gn=MYO16;refGene_tx=XM_023253136.1
A1 79308798 79309070 refGene_gn=MYO16;refGene_tx=XM_023253136.1
A1 79320294 79321002 refGene_gn=MYO16;refGene_tx=XM_023253136.1
A1 79321004 79321340 refGene_gn=MYO16;refGene_tx=XM_023253136.1
A1 79342215 79342374 refGene_gn=MYO16;refGene_tx=XM_023253136.1
A1 79358974 79359066 refGene_gn=MYO16;refGene_tx=XM_023253136.1

A1 79385376 79385604 refGene_gn=MYO16;refGene_tx=XM_023253136.1
A1 79812040 79812045 refGene_gn=IRS2;refGene_tx=XM_023253148.1
A1 79837762 79841765 refGene_gn=IRS2;refGene_tx=XM_023253148.1
A1 80093032 80093085
refGene_gn=LOC109492654;refGene_tx=XM_023239364.1
A1 80093619 80094239
refGene_gn=LOC109492654;refGene_tx=XM_023239364.1
A1 80095793 80095819
refGene_gn=LOC109492654;refGene_tx=XM_023239364.1
A1 80133796 80133878 refGene_gn=COL4A1;refGene_tx=XM_023253172.1
A1 80136339 80136512 refGene_gn=COL4A1;refGene_tx=XM_023253172.1
A1 80138476 80138591 refGene_gn=COL4A1;refGene_tx=XM_023253172.1
A1 80142609 80142787 refGene_gn=COL4A1;refGene_tx=XM_023253172.1
A1 80144010 80144223 refGene_gn=COL4A1;refGene_tx=XM_023253172.1
A1 80145182 80145281 refGene_gn=COL4A1;refGene_tx=XM_023253172.1
A1 80146232 80146361 refGene_gn=COL4A1;refGene_tx=XM_023253172.1
A1 80147923 80147995 refGene_gn=COL4A1;refGene_tx=XM_023253172.1
A1 80148881 80148954 refGene_gn=COL4A1;refGene_tx=XM_023253172.1
A1 80150656 80150790 refGene_gn=COL4A1;refGene_tx=XM_023253172.1
A1 80151347 80151533 refGene_gn=COL4A1;refGene_tx=XM_023253172.1
A1 80152646 80152697 refGene_gn=COL4A1;refGene_tx=XM_023253172.1
A1 80153987 80154086 refGene_gn=COL4A1;refGene_tx=XM_023253172.1
A1 80154489 80154570 refGene_gn=COL4A1;refGene_tx=XM_023253172.1
A1 80154660 80154787 refGene_gn=COL4A1;refGene_tx=XM_023253172.1
A1 80155176 80155316 refGene_gn=COL4A1;refGene_tx=XM_023253172.1
A1 80156317 80156407 refGene_gn=COL4A1;refGene_tx=XM_023253172.1
A1 80156518 80156617 refGene_gn=COL4A1;refGene_tx=XM_023253172.1
A1 80156758 80156911 refGene_gn=COL4A1;refGene_tx=XM_023253172.1
A1 80157700 80157790 refGene_gn=COL4A1;refGene_tx=XM_023253172.1
A1 80157875 80158043 refGene_gn=COL4A1;refGene_tx=XM_023253172.1
A1 80158739 80158853 refGene_gn=COL4A1;refGene_tx=XM_023253172.1
A1 80159066 80159217 refGene_gn=COL4A1;refGene_tx=XM_023253172.1
A1 80160811 80160909 refGene_gn=COL4A1;refGene_tx=XM_023253172.1
A1 80163219 80163324 refGene_gn=COL4A1;refGene_tx=XM_023253172.1
A1 80163412 80163505 refGene_gn=COL4A1;refGene_tx=XM_023253172.1
A1 80166302 80166471 refGene_gn=COL4A1;refGene_tx=XM_023253172.1
A1 80167042 80167234 refGene_gn=COL4A1;refGene_tx=XM_023253172.1
A1 80170737 80170808 refGene_gn=COL4A1;refGene_tx=XM_023253172.1
A1 80171589 80171673 refGene_gn=COL4A1;refGene_tx=XM_023253172.1
A1 80173694 80173790 refGene_gn=COL4A1;refGene_tx=XM_023253172.1
A1 80177310 80177475 refGene_gn=COL4A1;refGene_tx=XM_023253172.1
A1 80179824 80179860 refGene_gn=COL4A1;refGene_tx=XM_023253172.1
A1 80180469 80180554 refGene_gn=COL4A1;refGene_tx=XM_023253172.1
A1 80181903 80181945 refGene_gn=COL4A1;refGene_tx=XM_023253172.1
A1 80184012 80184066 refGene_gn=COL4A1;refGene_tx=XM_023253172.1
A1 80184681 80184726 refGene_gn=COL4A1;refGene_tx=XM_023253172.1
A1 80185491 80185542 refGene_gn=COL4A1;refGene_tx=XM_023253172.1
A1 80185695 80185722 refGene_gn=COL4A1;refGene_tx=XM_023253172.1
A1 80186186 80186273 refGene_gn=COL4A1;refGene_tx=XM_023253172.1
A1 80187666 80187708 refGene_gn=COL4A1;refGene_tx=XM_023253172.1

A1 80188627 80188663 refGene_gn=COL4A1;refGene_tx=XM_023253172.1
A1 80190980 80191043 refGene_gn=COL4A1;refGene_tx=XM_023253172.1
A1 80191127 80191211 refGene_gn=COL4A1;refGene_tx=XM_023253172.1
A1 80193100 80193127 refGene_gn=COL4A1;refGene_tx=XM_023253172.1
A1 80193497 80193551 refGene_gn=COL4A1;refGene_tx=XM_023253172.1
A1 80194028 80194091 refGene_gn=COL4A1;refGene_tx=XM_023253172.1
A1 80194198 80194243 refGene_gn=COL4A1;refGene_tx=XM_023253172.1
A1 80195524 80195569 refGene_gn=COL4A1;refGene_tx=XM_023253172.1
A1 80195658 80195748 refGene_gn=COL4A1;refGene_tx=XM_023253172.1
A1 80222810 80222870 refGene_gn=COL4A1;refGene_tx=XM_023253172.1
A1 80288822 80288906 refGene_gn=COL4A1;refGene_tx=XM_023253172.1
A1 80289739 80289783
refGene_gn=COL4A2;refGene_tx=XM_023253158.1;refGene_gn=COL4A2;refGene_tx=XM_023253164.1
A1 80289905 80289960
refGene_gn=COL4A2;refGene_tx=XM_023253158.1;refGene_gn=COL4A2;refGene_tx=XM_023253164.1
A1 80335661 80335742
refGene_gn=COL4A2;refGene_tx=XM_023253158.1;refGene_gn=COL4A2;refGene_tx=XM_023253164.1
A1 80390392 80390527
refGene_gn=COL4A2;refGene_tx=XM_023253158.1;refGene_gn=COL4A2;refGene_tx=XM_023253164.1
A1 80390613 80390658
refGene_gn=COL4A2;refGene_tx=XM_023253158.1;refGene_gn=COL4A2;refGene_tx=XM_023253164.1
A1 80393574 80393691
refGene_gn=COL4A2;refGene_tx=XM_023253158.1;refGene_gn=COL4A2;refGene_tx=XM_023253164.1
A1 80395384 80395456
refGene_gn=COL4A2;refGene_tx=XM_023253158.1;refGene_gn=COL4A2;refGene_tx=XM_023253164.1
A1 80396228 80396264
refGene_gn=COL4A2;refGene_tx=XM_023253158.1;refGene_gn=COL4A2;refGene_tx=XM_023253164.1
A1 80396371 80396434
refGene_gn=COL4A2;refGene_tx=XM_023253158.1;refGene_gn=COL4A2;refGene_tx=XM_023253164.1
A1 80398002 80398038
refGene_gn=COL4A2;refGene_tx=XM_023253158.1;refGene_gn=COL4A2;refGene_tx=XM_023253164.1
A1 80400485 80400527
refGene_gn=COL4A2;refGene_tx=XM_023253158.1;refGene_gn=COL4A2;refGene_tx=XM_023253164.1
A1 80402237 80402336
refGene_gn=COL4A2;refGene_tx=XM_023253158.1;refGene_gn=COL4A2;refGene_tx=XM_023253164.1
A1 80404220 80404256
refGene_gn=COL4A2;refGene_tx=XM_023253158.1;refGene_gn=COL4A2;refGene_tx=XM_023253164.1

A1 80404801 80404852
refGene_gn=COL4A2;refGene_tx=XM_023253158.1;refGene_gn=COL4A2;refGene_tx=XM_023253164.1
A1 80405765 80405810
refGene_gn=COL4A2;refGene_tx=XM_023253158.1;refGene_gn=COL4A2;refGene_tx=XM_023253164.1
A1 80408923 80408977
refGene_gn=COL4A2;refGene_tx=XM_023253158.1;refGene_gn=COL4A2;refGene_tx=XM_023253164.1
A1 80409844 80409911
refGene_gn=COL4A2;refGene_tx=XM_023253158.1;refGene_gn=COL4A2;refGene_tx=XM_023253164.1
A1 80413349 80413460
refGene_gn=COL4A2;refGene_tx=XM_023253158.1;refGene_gn=COL4A2;refGene_tx=XM_023253164.1
A1 80413962 80414112
refGene_gn=COL4A2;refGene_tx=XM_023253158.1;refGene_gn=COL4A2;refGene_tx=XM_023253164.1
A1 80419575 80419668
refGene_gn=COL4A2;refGene_tx=XM_023253158.1;refGene_gn=COL4A2;refGene_tx=XM_023253164.1
A1 80420809 80420973
refGene_gn=COL4A2;refGene_tx=XM_023253158.1;refGene_gn=COL4A2;refGene_tx=XM_023253164.1
A1 80424277 80424350
refGene_gn=COL4A2;refGene_tx=XM_023253158.1;refGene_gn=COL4A2;refGene_tx=XM_023253164.1
A1 80424441 80424548
refGene_gn=COL4A2;refGene_tx=XM_023253158.1;refGene_gn=COL4A2;refGene_tx=XM_023253164.1
A1 80426324 80426526
refGene_gn=COL4A2;refGene_tx=XM_023253158.1;refGene_gn=COL4A2;refGene_tx=XM_023253164.1
A1 80426876 80426936
refGene_gn=COL4A2;refGene_tx=XM_023253158.1;refGene_gn=COL4A2;refGene_tx=XM_023253164.1
A1 80428447 80428504
refGene_gn=COL4A2;refGene_tx=XM_023253158.1;refGene_gn=COL4A2;refGene_tx=XM_023253164.1
A1 80431721 80431829
refGene_gn=COL4A2;refGene_tx=XM_023253158.1;refGene_gn=COL4A2;refGene_tx=XM_023253164.1
A1 80435172 80435394
refGene_gn=COL4A2;refGene_tx=XM_023253158.1;refGene_gn=COL4A2;refGene_tx=XM_023253164.1
A1 80439443 80439605
refGene_gn=COL4A2;refGene_tx=XM_023253158.1;refGene_gn=COL4A2;refGene_tx=XM_023253164.1
A1 80440395 80440566
refGene_gn=COL4A2;refGene_tx=XM_023253158.1;refGene_gn=COL4A2;refGene_

tx=XM_023253164.1
A1 80441319 80441463
refGene_gn=COL4A2;refGene_tx=XM_023253158.1;refGene_gn=COL4A2;refGene_tx=XM_023253164.1
A1 80443267 80443390
refGene_gn=COL4A2;refGene_tx=XM_023253158.1;refGene_gn=COL4A2;refGene_tx=XM_023253164.1
A1 80443631 80443813
refGene_gn=COL4A2;refGene_tx=XM_023253158.1;refGene_gn=COL4A2;refGene_tx=XM_023253164.1
A1 80446128 80446192
refGene_gn=COL4A2;refGene_tx=XM_023253158.1;refGene_gn=COL4A2;refGene_tx=XM_023253164.1
A1 80446549 80446624
refGene_gn=COL4A2;refGene_tx=XM_023253158.1;refGene_gn=COL4A2;refGene_tx=XM_023253164.1
A1 80448745 80448853
refGene_gn=COL4A2;refGene_tx=XM_023253158.1;refGene_gn=COL4A2;refGene_tx=XM_023253164.1
A1 80449575 80449683
refGene_gn=COL4A2;refGene_tx=XM_023253158.1;refGene_gn=COL4A2;refGene_tx=XM_023253164.1
A1 80450392 80450464
refGene_gn=COL4A2;refGene_tx=XM_023253158.1;refGene_gn=COL4A2;refGene_tx=XM_023253164.1
A1 80450808 80450934
refGene_gn=COL4A2;refGene_tx=XM_023253158.1;refGene_gn=COL4A2;refGene_tx=XM_023253164.1
A1 80454710 80454827
refGene_gn=COL4A2;refGene_tx=XM_023253158.1;refGene_gn=COL4A2;refGene_tx=XM_023253164.1
A1 80455666 80455828
refGene_gn=COL4A2;refGene_tx=XM_023253158.1;refGene_gn=COL4A2;refGene_tx=XM_023253164.1
A1 80455970 80456069
refGene_gn=COL4A2;refGene_tx=XM_023253158.1;refGene_gn=COL4A2;refGene_tx=XM_023253164.1
A1 80456414 80456561
refGene_gn=COL4A2;refGene_tx=XM_023253158.1;refGene_gn=COL4A2;refGene_tx=XM_023253164.1
A1 80456717 80456834
refGene_gn=COL4A2;refGene_tx=XM_023253158.1;refGene_gn=COL4A2;refGene_tx=XM_023253164.1
A1 80457938 80458130
refGene_gn=COL4A2;refGene_tx=XM_023253158.1;refGene_gn=COL4A2;refGene_tx=XM_023253164.1
A1 80459273 80459560
refGene_gn=COL4A2;refGene_tx=XM_023253158.1;refGene_gn=COL4A2;refGene_tx=XM_023253164.1
A1 80461379 80461637

refGene_gn=COL4A2;refGene_tx=XM_023253158.1;refGene_gn=COL4A2;refGene_tx=XM_023253164.1
A1 80467349 80467882 refGene_gn=RAB20;refGene_tx=XM_003980528.5
A1 80492452 80492624 refGene_gn=RAB20;refGene_tx=XM_003980528.5
A1 80518793 80518839 refGene_gn=NAXD;refGene_tx=XM_023253195.1
A1 80518865 80518965 refGene_gn=NAXD;refGene_tx=XM_023253199.1
A1 80522563 80522714
refGene_gn=NAXD;refGene_tx=XM_023253195.1;refGene_gn=NAXD;refGene_tx=XM_023253199.1
A1 80525362 80525408
refGene_gn=NAXD;refGene_tx=XM_023253195.1;refGene_gn=NAXD;refGene_tx=XM_023253199.1
A1 80526101 80526190
refGene_gn=NAXD;refGene_tx=XM_023253195.1;refGene_gn=NAXD;refGene_tx=XM_023253199.1
A1 80528185 80528294
refGene_gn=NAXD;refGene_tx=XM_023253195.1;refGene_gn=NAXD;refGene_tx=XM_023253199.1
A1 80533982 80534033
refGene_gn=NAXD;refGene_tx=XM_023253195.1;refGene_gn=NAXD;refGene_tx=XM_023253199.1
A1 80534113 80534218
refGene_gn=NAXD;refGene_tx=XM_023253195.1;refGene_gn=NAXD;refGene_tx=XM_023253199.1
A1 80534774 80534895
refGene_gn=NAXD;refGene_tx=XM_023253195.1;refGene_gn=NAXD;refGene_tx=XM_023253199.1
A1 80536281 80536402
refGene_gn=NAXD;refGene_tx=XM_023253195.1;refGene_gn=NAXD;refGene_tx=XM_023253199.1
A1 80537288 80537439
refGene_gn=NAXD;refGene_tx=XM_023253195.1;refGene_gn=NAXD;refGene_tx=XM_023253199.1
A1 80539540 80539637
refGene_gn=CARS2;refGene_tx=XM_011280866.2;refGene_gn=CARS2;refGene_tx=XM_019829550.2;refGene_gn=CARS2;refGene_tx=XM_019829552.2
A1 80539952 80540159
refGene_gn=CARS2;refGene_tx=XM_011280866.2;refGene_gn=CARS2;refGene_tx=XM_019829550.2;refGene_gn=CARS2;refGene_tx=XM_019829552.2
A1 80541389 80541514
refGene_gn=CARS2;refGene_tx=XM_011280866.2;refGene_gn=CARS2;refGene_tx=XM_019829550.2;refGene_gn=CARS2;refGene_tx=XM_019829552.2
A1 80542162 80542286
refGene_gn=CARS2;refGene_tx=XM_011280866.2;refGene_gn=CARS2;refGene_tx=XM_019829550.2;refGene_gn=CARS2;refGene_tx=XM_019829552.2
A1 80544043 80544182
refGene_gn=CARS2;refGene_tx=XM_011280866.2;refGene_gn=CARS2;refGene_tx=XM_019829550.2;refGene_gn=CARS2;refGene_tx=XM_019829552.2
A1 80544735 80544802
refGene_gn=CARS2;refGene_tx=XM_011280866.2;refGene_gn=CARS2;refGene_tx

=XM_019829550.2;refGene_gn=CARS2;refGene_tx=XM_019829552.2
A1 80551217 80551285
refGene_gn=CARS2;refGene_tx=XM_011280866.2;refGene_gn=CARS2;refGene_tx
=XM_019829550.2;refGene_gn=CARS2;refGene_tx=XM_019829552.2
A1 80553609 80553743
refGene_gn=CARS2;refGene_tx=XM_011280866.2;refGene_gn=CARS2;refGene_tx
=XM_019829550.2;refGene_gn=CARS2;refGene_tx=XM_019829552.2
A1 80566310 80566440
refGene_gn=CARS2;refGene_tx=XM_011280866.2;refGene_gn=CARS2;refGene_tx
=XM_019829550.2;refGene_gn=CARS2;refGene_tx=XM_019829552.2
A1 80568608 80568692
refGene_gn=CARS2;refGene_tx=XM_011280866.2;refGene_gn=CARS2;refGene_tx
=XM_019829550.2;refGene_gn=CARS2;refGene_tx=XM_019829552.2
A1 80571389 80571495
refGene_gn=CARS2;refGene_tx=XM_011280866.2;refGene_gn=CARS2;refGene_tx
=XM_019829550.2;refGene_gn=CARS2;refGene_tx=XM_019829552.2
A1 80571612 80571684
refGene_gn=CARS2;refGene_tx=XM_011280866.2;refGene_gn=CARS2;refGene_tx
=XM_019829550.2;refGene_gn=CARS2;refGene_tx=XM_019829552.2
A1 80578800 80578918
refGene_gn=CARS2;refGene_tx=XM_011280866.2;refGene_gn=CARS2;refGene_tx
=XM_019829550.2;refGene_gn=CARS2;refGene_tx=XM_019829552.2
A1 80583815 80583866
refGene_gn=CARS2;refGene_tx=XM_011280866.2;refGene_gn=CARS2;refGene_tx
=XM_019829550.2
A1 80584123 80584335
refGene_gn=CARS2;refGene_tx=XM_011280866.2;refGene_gn=CARS2;refGene_tx
=XM_019829550.2;refGene_gn=CARS2;refGene_tx=XM_019829552.2
A1 80589021 80589163 refGene_gn=ING1;refGene_tx=XM_006927399.4
A1 80590157 80590293 refGene_gn=ING1;refGene_tx=XM_023253203.1
A1 80594976 80595680
refGene_gn=ING1;refGene_tx=XM_006927399.4;refGene_gn=ING1;refGene_tx=X
M_023253203.1
A1 80698976 80699452 refGene_gn=ANKRD10;refGene_tx=XM_023253212.1
A1 80702808 80702904 refGene_gn=ANKRD10;refGene_tx=XM_023253212.1
A1 80712209 80712445 refGene_gn=ANKRD10;refGene_tx=XM_023253212.1
A1 80723455 80723547 refGene_gn=ANKRD10;refGene_tx=XM_023253212.1
A1 80728104 80728257 refGene_gn=ANKRD10;refGene_tx=XM_023253212.1
A1 80731118 80731328 refGene_gn=ANKRD10;refGene_tx=XM_023253212.1
A1 80743452 80743538
refGene_gn=LOC109499197;refGene_tx=XM_019829602.2;refGene_gn=LOC109499
197;refGene_tx=XM_019829612.2;refGene_gn=LOC109499197;refGene_tx=XM_01
9829620.2;refGene_gn=LOC109499197;refGene_tx=XM_019829625.2
A1 80745981 80746062
refGene_gn=LOC109499197;refGene_tx=XM_019829602.2;refGene_gn=LOC109499
197;refGene_tx=XM_019829612.2;refGene_gn=LOC109499197;refGene_tx=XM_01
9829620.2;refGene_gn=LOC109499197;refGene_tx=XM_019829625.2
A1 80756688 80756784
refGene_gn=LOC109499197;refGene_tx=XM_019829602.2;refGene_gn=LOC109499
197;refGene_tx=XM_019829612.2;refGene_gn=LOC109499197;refGene_tx=XM_01

9829620.2;refGene_gn=LOC109499197;refGene_tx=XM_019829625.2
A1 80757666 80757765
refGene_gn=LOC109499197;refGene_tx=XM_019829602.2;refGene_gn=LOC109499197;refGene_tx=XM_019829612.2
A1 80759114 80759177
refGene_gn=LOC109499197;refGene_tx=XM_019829602.2;refGene_gn=LOC109499197;refGene_tx=XM_019829612.2;refGene_gn=LOC109499197;refGene_tx=XM_019829620.2;refGene_gn=LOC109499197;refGene_tx=XM_019829625.2
A1 80762667 80762977
refGene_gn=LOC109499197;refGene_tx=XM_019829602.2;refGene_gn=LOC109499197;refGene_tx=XM_019829612.2;refGene_gn=LOC109499197;refGene_tx=XM_019829620.2;refGene_gn=LOC109499197;refGene_tx=XM_019829625.2
A1 80813936 80814101
refGene_gn=ARHGEF7;refGene_tx=XM_023253244.1;refGene_gn=ARHGEF7;refGene_tx=XM_023253251.1;refGene_gn=ARHGEF7;refGene_tx=XM_023253258.1;refGene_gn=ARHGEF7;refGene_tx=XM_023253259.1
A1 80843327 80843414
refGene_gn=ARHGEF7;refGene_tx=XM_023253244.1;refGene_gn=ARHGEF7;refGene_tx=XM_023253258.1;refGene_gn=ARHGEF7;refGene_tx=XM_023253259.1
A1 80869671 80869677 refGene_gn=ARHGEF7;refGene_tx=XM_003980531.5
A1 80871888 80871973
refGene_gn=ARHGEF7;refGene_tx=XM_003980531.5;refGene_gn=ARHGEF7;refGene_tx=XM_023253244.1;refGene_gn=ARHGEF7;refGene_tx=XM_023253251.1;refGene_gn=ARHGEF7;refGene_tx=XM_023253258.1
A1 80875512 80875643
refGene_gn=ARHGEF7;refGene_tx=XM_003980531.5;refGene_gn=ARHGEF7;refGene_tx=XM_023253244.1;refGene_gn=ARHGEF7;refGene_tx=XM_023253251.1;refGene_gn=ARHGEF7;refGene_tx=XM_023253258.1
A1 80884482 80884684
refGene_gn=ARHGEF7;refGene_tx=XM_003980531.5;refGene_gn=ARHGEF7;refGene_tx=XM_019829649.1;refGene_gn=ARHGEF7;refGene_tx=XM_023253241.1;refGene_gn=ARHGEF7;refGene_tx=XM_023253244.1;refGene_gn=ARHGEF7;refGene_tx=XM_023253248.1;refGene_gn=ARHGEF7;refGene_tx=XM_023253251.1;refGene_gn=ARHGEF7;refGene_tx=XM_023253258.1;refGene_gn=ARHGEF7;refGene_tx=XM_023253259.1;refGene_gn=ARHGEF7;refGene_tx=XM_023253271.1;refGene_gn=ARHGEF7;refGene_tx=XM_023253276.1
A1 80894494 80894583
refGene_gn=ARHGEF7;refGene_tx=XM_003980531.5;refGene_gn=ARHGEF7;refGene_tx=XM_019829649.1;refGene_gn=ARHGEF7;refGene_tx=XM_023253241.1;refGene_gn=ARHGEF7;refGene_tx=XM_023253244.1;refGene_gn=ARHGEF7;refGene_tx=XM_023253248.1;refGene_gn=ARHGEF7;refGene_tx=XM_023253251.1;refGene_gn=ARHGEF7;refGene_tx=XM_023253258.1;refGene_gn=ARHGEF7;refGene_tx=XM_023253259.1;refGene_gn=ARHGEF7;refGene_tx=XM_023253271.1;refGene_gn=ARHGEF7;refGene_tx=XM_023253276.1
A1 80907203 80907298
refGene_gn=ARHGEF7;refGene_tx=XM_003980531.5;refGene_gn=ARHGEF7;refGene_tx=XM_019829649.1;refGene_gn=ARHGEF7;refGene_tx=XM_023253241.1;refGene_gn=ARHGEF7;refGene_tx=XM_023253244.1;refGene_gn=ARHGEF7;refGene_tx=XM_023253248.1;refGene_gn=ARHGEF7;refGene_tx=XM_023253251.1;refGene_gn=ARHGEF7;refGene_tx=XM_023253258.1;refGene_gn=ARHGEF7;refGene_tx=XM_02

3253259.1;refGene_gn=ARHGEF7;refGene_tx=XM_023253271.1;refGene_gn=ARHGEF7;refGene_tx=XM_023253276.1
A1 80907514 80907610
refGene_gn=ARHGEF7;refGene_tx=XM_003980531.5;refGene_gn=ARHGEF7;refGene_tx=XM_019829649.1;refGene_gn=ARHGEF7;refGene_tx=XM_023253241.1;refGene_gn=ARHGEF7;refGene_tx=XM_023253244.1;refGene_gn=ARHGEF7;refGene_tx=XM_023253248.1;refGene_gn=ARHGEF7;refGene_tx=XM_023253251.1;refGene_gn=ARHGEF7;refGene_tx=XM_023253258.1;refGene_gn=ARHGEF7;refGene_tx=XM_023253259.1;refGene_gn=ARHGEF7;refGene_tx=XM_023253271.1;refGene_gn=ARHGEF7;refGene_tx=XM_023253276.1
A1 80910098 80910221
refGene_gn=ARHGEF7;refGene_tx=XM_003980531.5;refGene_gn=ARHGEF7;refGene_tx=XM_019829649.1;refGene_gn=ARHGEF7;refGene_tx=XM_023253241.1;refGene_gn=ARHGEF7;refGene_tx=XM_023253244.1;refGene_gn=ARHGEF7;refGene_tx=XM_023253248.1;refGene_gn=ARHGEF7;refGene_tx=XM_023253251.1;refGene_gn=ARHGEF7;refGene_tx=XM_023253258.1;refGene_gn=ARHGEF7;refGene_tx=XM_023253259.1;refGene_gn=ARHGEF7;refGene_tx=XM_023253271.1;refGene_gn=ARHGEF7;refGene_tx=XM_023253276.1
A1 80919034 80919173
refGene_gn=ARHGEF7;refGene_tx=XM_003980531.5;refGene_gn=ARHGEF7;refGene_tx=XM_019829649.1;refGene_gn=ARHGEF7;refGene_tx=XM_023253241.1;refGene_gn=ARHGEF7;refGene_tx=XM_023253244.1;refGene_gn=ARHGEF7;refGene_tx=XM_023253248.1;refGene_gn=ARHGEF7;refGene_tx=XM_023253251.1;refGene_gn=ARHGEF7;refGene_tx=XM_023253258.1;refGene_gn=ARHGEF7;refGene_tx=XM_023253259.1;refGene_gn=ARHGEF7;refGene_tx=XM_023253271.1;refGene_gn=ARHGEF7;refGene_tx=XM_023253276.1
A1 80920147 80920207
refGene_gn=ARHGEF7;refGene_tx=XM_003980531.5;refGene_gn=ARHGEF7;refGene_tx=XM_019829649.1;refGene_gn=ARHGEF7;refGene_tx=XM_023253241.1;refGene_gn=ARHGEF7;refGene_tx=XM_023253244.1;refGene_gn=ARHGEF7;refGene_tx=XM_023253248.1;refGene_gn=ARHGEF7;refGene_tx=XM_023253251.1;refGene_gn=ARHGEF7;refGene_tx=XM_023253258.1;refGene_gn=ARHGEF7;refGene_tx=XM_023253259.1;refGene_gn=ARHGEF7;refGene_tx=XM_023253271.1;refGene_gn=ARHGEF7;refGene_tx=XM_023253276.1
A1 80920840 80920987
refGene_gn=ARHGEF7;refGene_tx=XM_003980531.5;refGene_gn=ARHGEF7;refGene_tx=XM_019829649.1;refGene_gn=ARHGEF7;refGene_tx=XM_023253241.1;refGene_gn=ARHGEF7;refGene_tx=XM_023253244.1;refGene_gn=ARHGEF7;refGene_tx=XM_023253248.1;refGene_gn=ARHGEF7;refGene_tx=XM_023253251.1;refGene_gn=ARHGEF7;refGene_tx=XM_023253258.1;refGene_gn=ARHGEF7;refGene_tx=XM_023253259.1;refGene_gn=ARHGEF7;refGene_tx=XM_023253271.1;refGene_gn=ARHGEF7;refGene_tx=XM_023253276.1
A1 80921830 80921917
refGene_gn=ARHGEF7;refGene_tx=XM_003980531.5;refGene_gn=ARHGEF7;refGene_tx=XM_019829649.1;refGene_gn=ARHGEF7;refGene_tx=XM_023253241.1;refGene_gn=ARHGEF7;refGene_tx=XM_023253244.1;refGene_gn=ARHGEF7;refGene_tx=XM_023253248.1;refGene_gn=ARHGEF7;refGene_tx=XM_023253251.1;refGene_gn=ARHGEF7;refGene_tx=XM_023253258.1;refGene_gn=ARHGEF7;refGene_tx=XM_023253259.1;refGene_gn=ARHGEF7;refGene_tx=XM_023253271.1;refGene_gn=ARHGEF7;refGene_tx=XM_023253276.1

A1 80923600 80923679
refGene_gn=ARHGEF7;refGene_tx=XM_003980531.5;refGene_gn=ARHGEF7;refGene_tx=XM_019829649.1;refGene_gn=ARHGEF7;refGene_tx=XM_023253241.1;refGene_gn=ARHGEF7;refGene_tx=XM_023253244.1;refGene_gn=ARHGEF7;refGene_tx=XM_023253248.1;refGene_gn=ARHGEF7;refGene_tx=XM_023253251.1;refGene_gn=ARHGEF7;refGene_tx=XM_023253258.1;refGene_gn=ARHGEF7;refGene_tx=XM_023253259.1;refGene_gn=ARHGEF7;refGene_tx=XM_023253271.1;refGene_gn=ARHGEF7;refGene_tx=XM_023253276.1
A1 80923870 80924010
refGene_gn=ARHGEF7;refGene_tx=XM_003980531.5;refGene_gn=ARHGEF7;refGene_tx=XM_019829649.1;refGene_gn=ARHGEF7;refGene_tx=XM_023253241.1;refGene_gn=ARHGEF7;refGene_tx=XM_023253244.1;refGene_gn=ARHGEF7;refGene_tx=XM_023253248.1;refGene_gn=ARHGEF7;refGene_tx=XM_023253251.1;refGene_gn=ARHGEF7;refGene_tx=XM_023253258.1;refGene_gn=ARHGEF7;refGene_tx=XM_023253259.1;refGene_gn=ARHGEF7;refGene_tx=XM_023253271.1;refGene_gn=ARHGEF7;refGene_tx=XM_023253276.1
A1 80926989 80927214
refGene_gn=ARHGEF7;refGene_tx=XM_003980531.5;refGene_gn=ARHGEF7;refGene_tx=XM_019829649.1;refGene_gn=ARHGEF7;refGene_tx=XM_023253241.1;refGene_gn=ARHGEF7;refGene_tx=XM_023253244.1;refGene_gn=ARHGEF7;refGene_tx=XM_023253248.1;refGene_gn=ARHGEF7;refGene_tx=XM_023253251.1;refGene_gn=ARHGEF7;refGene_tx=XM_023253258.1;refGene_gn=ARHGEF7;refGene_tx=XM_023253259.1;refGene_gn=ARHGEF7;refGene_tx=XM_023253271.1;refGene_gn=ARHGEF7;refGene_tx=XM_023253276.1
A1 80929270 80929364
refGene_gn=ARHGEF7;refGene_tx=XM_003980531.5;refGene_gn=ARHGEF7;refGene_tx=XM_019829649.1;refGene_gn=ARHGEF7;refGene_tx=XM_023253241.1;refGene_gn=ARHGEF7;refGene_tx=XM_023253244.1;refGene_gn=ARHGEF7;refGene_tx=XM_023253248.1;refGene_gn=ARHGEF7;refGene_tx=XM_023253251.1;refGene_gn=ARHGEF7;refGene_tx=XM_023253258.1;refGene_gn=ARHGEF7;refGene_tx=XM_023253259.1;refGene_gn=ARHGEF7;refGene_tx=XM_023253271.1;refGene_gn=ARHGEF7;refGene_tx=XM_023253276.1
A1 80932615 80932705
refGene_gn=ARHGEF7;refGene_tx=XM_003980531.5;refGene_gn=ARHGEF7;refGene_tx=XM_019829649.1;refGene_gn=ARHGEF7;refGene_tx=XM_023253241.1;refGene_gn=ARHGEF7;refGene_tx=XM_023253244.1;refGene_gn=ARHGEF7;refGene_tx=XM_023253248.1;refGene_gn=ARHGEF7;refGene_tx=XM_023253251.1;refGene_gn=ARHGEF7;refGene_tx=XM_023253258.1;refGene_gn=ARHGEF7;refGene_tx=XM_023253259.1;refGene_gn=ARHGEF7;refGene_tx=XM_023253271.1;refGene_gn=ARHGEF7;refGene_tx=XM_023253276.1
A1 80935586 80935763
refGene_gn=ARHGEF7;refGene_tx=XM_003980531.5;refGene_gn=ARHGEF7;refGene_tx=XM_019829649.1;refGene_gn=ARHGEF7;refGene_tx=XM_023253241.1;refGene_gn=ARHGEF7;refGene_tx=XM_023253244.1;refGene_gn=ARHGEF7;refGene_tx=XM_023253251.1;refGene_gn=ARHGEF7;refGene_tx=XM_023253259.1;refGene_gn=ARHGEF7;refGene_tx=XM_023253271.1;refGene_gn=ARHGEF7;refGene_tx=XM_023253276.1
A1 80943759 80943859
refGene_gn=ARHGEF7;refGene_tx=XM_003980531.5;refGene_gn=ARHGEF7;refGene_tx=XM_019829649.1;refGene_gn=ARHGEF7;refGene_tx=XM_023253241.1;refGe

ne_gn=ARHGEF7;refGene_tx=XM_023253244.1;refGene_gn=ARHGEF7;refGene_tx=XM_023253248.1;refGene_gn=ARHGEF7;refGene_tx=XM_023253251.1;refGene_gn=ARHGEF7;refGene_tx=XM_023253258.1;refGene_gn=ARHGEF7;refGene_tx=XM_023253259.1;refGene_gn=ARHGEF7;refGene_tx=XM_023253271.1;refGene_gn=ARHGEF7;refGene_tx=XM_023253276.1
A1 80944119 80944174
refGene_gn=ARHGEF7;refGene_tx=XM_003980531.5;refGene_gn=ARHGEF7;refGene_tx=XM_019829649.1;refGene_gn=ARHGEF7;refGene_tx=XM_023253241.1;refGene_gn=ARHGEF7;refGene_tx=XM_023253244.1;refGene_gn=ARHGEF7;refGene_tx=XM_023253248.1;refGene_gn=ARHGEF7;refGene_tx=XM_023253251.1;refGene_gn=ARHGEF7;refGene_tx=XM_023253258.1;refGene_gn=ARHGEF7;refGene_tx=XM_023253259.1;refGene_gn=ARHGEF7;refGene_tx=XM_023253271.1;refGene_gn=ARHGEF7;refGene_tx=XM_023253276.1
A1 80945211 80945334
refGene_gn=ARHGEF7;refGene_tx=XM_003980531.5;refGene_gn=ARHGEF7;refGene_tx=XM_019829649.1;refGene_gn=ARHGEF7;refGene_tx=XM_023253241.1;refGene_gn=ARHGEF7;refGene_tx=XM_023253244.1;refGene_gn=ARHGEF7;refGene_tx=XM_023253248.1;refGene_gn=ARHGEF7;refGene_tx=XM_023253251.1;refGene_gn=ARHGEF7;refGene_tx=XM_023253258.1;refGene_gn=ARHGEF7;refGene_tx=XM_023253259.1;refGene_gn=ARHGEF7;refGene_tx=XM_023253271.1;refGene_gn=ARHGEF7;refGene_tx=XM_023253276.1
A1 80953045 80953103 refGene_gn=TEX29;refGene_tx=XM_023253282.1
A1 80956685 80956796 refGene_gn=TEX29;refGene_tx=XM_023253282.1
A1 80967771 80967841 refGene_gn=TEX29;refGene_tx=XM_023253282.1
A1 80968527 80968697 refGene_gn=TEX29;refGene_tx=XM_023253282.1
A1 80969362 80969403 refGene_gn=TEX29;refGene_tx=XM_023253282.1
A1 81571821 81572994 refGene_gn=SOX1;refGene_tx=XM_023253292.1
A1 81773887 81774071
refGene_gn=SPACA7;refGene_tx=XM_006927404.4;refGene_gn=SPACA7;refGene_tx=XM_006927405.4;refGene_gn=SPACA7;refGene_tx=XM_019829692.2;refGene_gn=SPACA7;refGene_tx=XM_019829700.2
A1 81777549 81777609
refGene_gn=SPACA7;refGene_tx=XM_006927404.4;refGene_gn=SPACA7;refGene_tx=XM_006927405.4;refGene_gn=SPACA7;refGene_tx=XM_019829692.2;refGene_gn=SPACA7;refGene_tx=XM_019829700.2
A1 81782354 81782456
refGene_gn=SPACA7;refGene_tx=XM_006927404.4;refGene_gn=SPACA7;refGene_tx=XM_006927405.4
A1 81783341 81783449
refGene_gn=SPACA7;refGene_tx=XM_006927404.4;refGene_gn=SPACA7;refGene_tx=XM_006927405.4;refGene_gn=SPACA7;refGene_tx=XM_019829692.2;refGene_gn=SPACA7;refGene_tx=XM_019829700.2
A1 81786155 81786227
refGene_gn=SPACA7;refGene_tx=XM_006927404.4;refGene_gn=SPACA7;refGene_tx=XM_006927405.4;refGene_gn=SPACA7;refGene_tx=XM_019829692.2;refGene_gn=SPACA7;refGene_tx=XM_019829700.2
A1 81799657 81799735
refGene_gn=SPACA7;refGene_tx=XM_006927404.4;refGene_gn=SPACA7;refGene_tx=XM_006927405.4;refGene_gn=SPACA7;refGene_tx=XM_019829692.2;refGene_gn=SPACA7;refGene_tx=XM_019829700.2

A1	81801797	81801814
refGene_gn=SPACA7;refGene_tx=XM_006927404.4;refGene_gn=SPACA7;refGene_tx=XM_006927405.4;refGene_gn=SPACA7;refGene_tx=XM_019829692.2;refGene_gn=SPACA7;refGene_tx=XM_019829700.2
A1	81833563	81833775
refGene_gn=TUBGCP3;refGene_tx=XM_003980556.5;refGene_gn=TUBGCP3;refGene_tx=XM_019829712.2;refGene_gn=TUBGCP3;refGene_tx=XM_019829713.2
A1	81836553	81836731
refGene_gn=TUBGCP3;refGene_tx=XM_003980556.5;refGene_gn=TUBGCP3;refGene_tx=XM_019829712.2;refGene_gn=TUBGCP3;refGene_tx=XM_019829713.2
A1	81839234	81839375
refGene_gn=TUBGCP3;refGene_tx=XM_003980556.5;refGene_gn=TUBGCP3;refGene_tx=XM_019829712.2;refGene_gn=TUBGCP3;refGene_tx=XM_019829713.2
A1	81843127	81843259
refGene_gn=TUBGCP3;refGene_tx=XM_003980556.5;refGene_gn=TUBGCP3;refGene_tx=XM_019829712.2;refGene_gn=TUBGCP3;refGene_tx=XM_019829713.2
A1	81843380	81843469
refGene_gn=TUBGCP3;refGene_tx=XM_003980556.5;refGene_gn=TUBGCP3;refGene_tx=XM_019829712.2;refGene_gn=TUBGCP3;refGene_tx=XM_019829713.2
A1	81850402	81850538
refGene_gn=TUBGCP3;refGene_tx=XM_003980556.5;refGene_gn=TUBGCP3;refGene_tx=XM_019829712.2;refGene_gn=TUBGCP3;refGene_tx=XM_019829713.2
A1	81854159	81854228
refGene_gn=TUBGCP3;refGene_tx=XM_003980556.5;refGene_gn=TUBGCP3;refGene_tx=XM_019829712.2;refGene_gn=TUBGCP3;refGene_tx=XM_019829713.2
A1	81855559	81855695
refGene_gn=TUBGCP3;refGene_tx=XM_003980556.5;refGene_gn=TUBGCP3;refGene_tx=XM_019829712.2;refGene_gn=TUBGCP3;refGene_tx=XM_019829713.2
A1	81858044	81858234
refGene_gn=TUBGCP3;refGene_tx=XM_003980556.5;refGene_gn=TUBGCP3;refGene_tx=XM_019829712.2;refGene_gn=TUBGCP3;refGene_tx=XM_019829713.2
A1	81859974	81860083
refGene_gn=TUBGCP3;refGene_tx=XM_003980556.5;refGene_gn=TUBGCP3;refGene_tx=XM_019829712.2;refGene_gn=TUBGCP3;refGene_tx=XM_019829713.2
A1	81860599	81860710
refGene_gn=TUBGCP3;refGene_tx=XM_003980556.5;refGene_gn=TUBGCP3;refGene_tx=XM_019829712.2;refGene_gn=TUBGCP3;refGene_tx=XM_019829713.2
A1	81876098	81876265
refGene_gn=TUBGCP3;refGene_tx=XM_003980556.5;refGene_gn=TUBGCP3;refGene_tx=XM_019829712.2
A1	81877831	81877964
refGene_gn=TUBGCP3;refGene_tx=XM_003980556.5;refGene_gn=TUBGCP3;refGene_tx=XM_019829712.2
A1	81878371	81878440
refGene_gn=TUBGCP3;refGene_tx=XM_003980556.5;refGene_gn=TUBGCP3;refGene_tx=XM_019829712.2;refGene_gn=TUBGCP3;refGene_tx=XM_019829713.2
A1	81879766	81879892
refGene_gn=TUBGCP3;refGene_tx=XM_003980556.5;refGene_gn=TUBGCP3;refGene_tx=XM_019829712.2;refGene_gn=TUBGCP3;refGene_tx=XM_019829713.2
A1	81880627	81880746

refGene_gn=TUBGCP3;refGene_tx=XM_003980556.5;refGene_gn=TUBGCP3;refGene_tx=XM_019829712.2;refGene_gn=TUBGCP3;refGene_tx=XM_019829713.2
A1 81882963 81883136
refGene_gn=TUBGCP3;refGene_tx=XM_003980556.5;refGene_gn=TUBGCP3;refGene_tx=XM_019829712.2;refGene_gn=TUBGCP3;refGene_tx=XM_019829713.2
A1 81884988 81885206
refGene_gn=TUBGCP3;refGene_tx=XM_003980556.5;refGene_gn=TUBGCP3;refGene_tx=XM_019829712.2;refGene_gn=TUBGCP3;refGene_tx=XM_019829713.2
A1 81885710 81885788
refGene_gn=TUBGCP3;refGene_tx=XM_003980556.5;refGene_gn=TUBGCP3;refGene_tx=XM_019829712.2;refGene_gn=TUBGCP3;refGene_tx=XM_019829713.2
A1 81890001 81890069
refGene_gn=TUBGCP3;refGene_tx=XM_003980556.5;refGene_gn=TUBGCP3;refGene_tx=XM_019829712.2;refGene_gn=TUBGCP3;refGene_tx=XM_019829713.2
A1 81893965 81894073
refGene_gn=TUBGCP3;refGene_tx=XM_003980556.5;refGene_gn=TUBGCP3;refGene_tx=XM_019829712.2;refGene_gn=TUBGCP3;refGene_tx=XM_019829713.2
A1 81904011 81904087
refGene_gn=TUBGCP3;refGene_tx=XM_003980556.5;refGene_gn=TUBGCP3;refGene_tx=XM_019829712.2;refGene_gn=TUBGCP3;refGene_tx=XM_019829713.2
A1 81974817 81974856
refGene_gn=ATP11A;refGene_tx=XM_023253346.1;refGene_gn=ATP11A;refGene_tx=XM_023253353.1
A1 82032274 82032397
refGene_gn=ATP11A;refGene_tx=XM_023253346.1;refGene_gn=ATP11A;refGene_tx=XM_023253353.1
A1 82039492 82039582
refGene_gn=ATP11A;refGene_tx=XM_023253346.1;refGene_gn=ATP11A;refGene_tx=XM_023253353.1
A1 82040366 82040447
refGene_gn=ATP11A;refGene_tx=XM_023253346.1;refGene_gn=ATP11A;refGene_tx=XM_023253353.1
A1 82044412 82044520
refGene_gn=ATP11A;refGene_tx=XM_023253346.1;refGene_gn=ATP11A;refGene_tx=XM_023253353.1
A1 82045145 82045274
refGene_gn=ATP11A;refGene_tx=XM_023253346.1;refGene_gn=ATP11A;refGene_tx=XM_023253353.1
A1 82047292 82047396
refGene_gn=ATP11A;refGene_tx=XM_023253346.1;refGene_gn=ATP11A;refGene_tx=XM_023253353.1
A1 82047857 82047908
refGene_gn=ATP11A;refGene_tx=XM_023253346.1;refGene_gn=ATP11A;refGene_tx=XM_023253353.1
A1 82050457 82050522
refGene_gn=ATP11A;refGene_tx=XM_023253346.1;refGene_gn=ATP11A;refGene_tx=XM_023253353.1
A1 82051382 82051464
refGene_gn=ATP11A;refGene_tx=XM_023253346.1;refGene_gn=ATP11A;refGene_tx=XM_023253353.1

A1 82052772 82052923
refGene_gn=ATP11A;refGene_tx=XM_023253346.1;refGene_gn=ATP11A;refGene_tx=XM_023253353.1
A1 82054022 82054220
refGene_gn=ATP11A;refGene_tx=XM_023253346.1;refGene_gn=ATP11A;refGene_tx=XM_023253353.1
A1 82056400 82056574
refGene_gn=ATP11A;refGene_tx=XM_023253346.1;refGene_gn=ATP11A;refGene_tx=XM_023253353.1
A1 82057863 82058027
refGene_gn=ATP11A;refGene_tx=XM_023253346.1;refGene_gn=ATP11A;refGene_tx=XM_023253353.1
A1 82059783 82059855
refGene_gn=ATP11A;refGene_tx=XM_023253346.1;refGene_gn=ATP11A;refGene_tx=XM_023253353.1
A1 82061279 82061353
refGene_gn=ATP11A;refGene_tx=XM_023253346.1;refGene_gn=ATP11A;refGene_tx=XM_023253353.1
A1 82065105 82065209
refGene_gn=ATP11A;refGene_tx=XM_023253346.1;refGene_gn=ATP11A;refGene_tx=XM_023253353.1
A1 82070077 82070259
refGene_gn=ATP11A;refGene_tx=XM_023253346.1;refGene_gn=ATP11A;refGene_tx=XM_023253353.1
A1 82072189 82072441
refGene_gn=ATP11A;refGene_tx=XM_023253346.1;refGene_gn=ATP11A;refGene_tx=XM_023253353.1
A1 82073892 82074067
refGene_gn=ATP11A;refGene_tx=XM_023253346.1;refGene_gn=ATP11A;refGene_tx=XM_023253353.1
A1 82075113 82075216
refGene_gn=ATP11A;refGene_tx=XM_023253346.1;refGene_gn=ATP11A;refGene_tx=XM_023253353.1
A1 82075403 82075549
refGene_gn=ATP11A;refGene_tx=XM_023253346.1;refGene_gn=ATP11A;refGene_tx=XM_023253353.1
A1 82076037 82076097
refGene_gn=ATP11A;refGene_tx=XM_023253346.1;refGene_gn=ATP11A;refGene_tx=XM_023253353.1
A1 82076810 82076938
refGene_gn=ATP11A;refGene_tx=XM_023253346.1;refGene_gn=ATP11A;refGene_tx=XM_023253353.1
A1 82078100 82078236
refGene_gn=ATP11A;refGene_tx=XM_023253346.1;refGene_gn=ATP11A;refGene_tx=XM_023253353.1
A1 82083906 82083972
refGene_gn=ATP11A;refGene_tx=XM_023253346.1;refGene_gn=ATP11A;refGene_tx=XM_023253353.1
A1 82085205 82085309
refGene_gn=ATP11A;refGene_tx=XM_023253346.1;refGene_gn=ATP11A;refGene_

tx=XM_023253353.1
A1 82087307 82087473
refGene_gn=ATP11A;refGene_tx=XM_023253346.1;refGene_gn=ATP11A;refGene_
tx=XM_023253353.1
A1 82088895 82088973 refGene_gn=ATP11A;refGene_tx=XM_023253353.1
A1 82092174 82092384 refGene_gn=ATP11A;refGene_tx=XM_023253346.1
A1 82107730 82107899
refGene_gn=MCF2L;refGene_tx=XM_019829755.1;refGene_gn=MCF2L;refGene_tx
=XM_019829789.1
A1 82136919 82137148
refGene_gn=MCF2L;refGene_tx=XM_019829754.1;refGene_gn=MCF2L;refGene_tx
=XM_019829760.1;refGene_gn=MCF2L;refGene_tx=XM_019829766.1
A1 82137746 82137825
refGene_gn=MCF2L;refGene_tx=XM_011280921.2;refGene_gn=MCF2L;refGene_tx
=XM_019829794.1;refGene_gn=MCF2L;refGene_tx=XM_019829815.1
A1 82150274 82150419
refGene_gn=MCF2L;refGene_tx=XM_006927406.3;refGene_gn=MCF2L;refGene_tx
=XM_011280911.2
A1 82168264 82168337 refGene_gn=MCF2L;refGene_tx=XM_019829802.1
A1 82180362 82180446
refGene_gn=MCF2L;refGene_tx=XM_006927406.3;refGene_gn=MCF2L;refGene_tx
=XM_011280911.2;refGene_gn=MCF2L;refGene_tx=XM_011280921.2;refGene_gn=
MCF2L;refGene_tx=XM_019829754.1;refGene_gn=MCF2L;refGene_tx=XM_0198297
55.1;refGene_gn=MCF2L;refGene_tx=XM_019829760.1;refGene_gn=MCF2L;refGe
ne_tx=XM_019829766.1;refGene_gn=MCF2L;refGene_tx=XM_019829789.1;refGen
e_gn=MCF2L;refGene_tx=XM_019829794.1;refGene_gn=MCF2L;refGene_tx=XM_01
9829802.1;refGene_gn=MCF2L;refGene_tx=XM_019829809.1;refGene_gn=MCF2L;
refGene_tx=XM_019829815.1
A1 82183057 82183172
refGene_gn=MCF2L;refGene_tx=XM_006927406.3;refGene_gn=MCF2L;refGene_tx
=XM_011280911.2;refGene_gn=MCF2L;refGene_tx=XM_011280921.2;refGene_gn=
MCF2L;refGene_tx=XM_019829754.1;refGene_gn=MCF2L;refGene_tx=XM_0198297
55.1;refGene_gn=MCF2L;refGene_tx=XM_019829760.1;refGene_gn=MCF2L;refGe
ne_tx=XM_019829766.1;refGene_gn=MCF2L;refGene_tx=XM_019829789.1;refGen
e_gn=MCF2L;refGene_tx=XM_019829794.1;refGene_gn=MCF2L;refGene_tx=XM_01
9829802.1;refGene_gn=MCF2L;refGene_tx=XM_019829809.1;refGene_gn=MCF2L;
refGene_tx=XM_019829815.1
A1 82199028 82199119
refGene_gn=MCF2L;refGene_tx=XM_006927406.3;refGene_gn=MCF2L;refGene_tx
=XM_011280911.2;refGene_gn=MCF2L;refGene_tx=XM_011280921.2;refGene_gn=
MCF2L;refGene_tx=XM_019829754.1;refGene_gn=MCF2L;refGene_tx=XM_0198297
55.1;refGene_gn=MCF2L;refGene_tx=XM_019829760.1;refGene_gn=MCF2L;refGe
ne_tx=XM_019829766.1;refGene_gn=MCF2L;refGene_tx=XM_019829789.1;refGen
e_gn=MCF2L;refGene_tx=XM_019829794.1;refGene_gn=MCF2L;refGene_tx=XM_01
9829802.1;refGene_gn=MCF2L;refGene_tx=XM_019829809.1;refGene_gn=MCF2L;
refGene_tx=XM_019829815.1
A1 82208348 82208468
refGene_gn=MCF2L;refGene_tx=XM_006927406.3;refGene_gn=MCF2L;refGene_tx
=XM_011280911.2;refGene_gn=MCF2L;refGene_tx=XM_011280921.2;refGene_gn=
MCF2L;refGene_tx=XM_019829754.1;refGene_gn=MCF2L;refGene_tx=XM_0198297

55.1;refGene_gn=MCF2L;refGene_tx=XM_019829760.1;refGene_gn=MCF2L;refGene_tx=XM_019829766.1;refGene_gn=MCF2L;refGene_tx=XM_019829789.1;refGene_gn=MCF2L;refGene_tx=XM_019829794.1;refGene_gn=MCF2L;refGene_tx=XM_019829802.1;refGene_gn=MCF2L;refGene_tx=XM_019829809.1;refGene_gn=MCF2L;refGene_tx=XM_019829815.1
A1 82210014 82210131
refGene_gn=MCF2L;refGene_tx=XM_006927406.3;refGene_gn=MCF2L;refGene_tx=XM_011280911.2;refGene_gn=MCF2L;refGene_tx=XM_011280921.2;refGene_gn=MCF2L;refGene_tx=XM_019829754.1;refGene_gn=MCF2L;refGene_tx=XM_019829755.1;refGene_gn=MCF2L;refGene_tx=XM_019829760.1;refGene_gn=MCF2L;refGene_tx=XM_019829766.1;refGene_gn=MCF2L;refGene_tx=XM_019829789.1;refGene_gn=MCF2L;refGene_tx=XM_019829794.1;refGene_gn=MCF2L;refGene_tx=XM_019829802.1;refGene_gn=MCF2L;refGene_tx=XM_019829809.1;refGene_gn=MCF2L;refGene_tx=XM_019829815.1
A1 82210640 82210790
refGene_gn=MCF2L;refGene_tx=XM_006927406.3;refGene_gn=MCF2L;refGene_tx=XM_011280911.2;refGene_gn=MCF2L;refGene_tx=XM_011280921.2;refGene_gn=MCF2L;refGene_tx=XM_019829754.1;refGene_gn=MCF2L;refGene_tx=XM_019829755.1;refGene_gn=MCF2L;refGene_tx=XM_019829760.1;refGene_gn=MCF2L;refGene_tx=XM_019829766.1;refGene_gn=MCF2L;refGene_tx=XM_019829789.1;refGene_gn=MCF2L;refGene_tx=XM_019829794.1;refGene_gn=MCF2L;refGene_tx=XM_019829802.1;refGene_gn=MCF2L;refGene_tx=XM_019829809.1;refGene_gn=MCF2L;refGene_tx=XM_019829815.1
A1 82211497 82211622
refGene_gn=MCF2L;refGene_tx=XM_006927406.3;refGene_gn=MCF2L;refGene_tx=XM_011280911.2;refGene_gn=MCF2L;refGene_tx=XM_011280921.2;refGene_gn=MCF2L;refGene_tx=XM_019829754.1;refGene_gn=MCF2L;refGene_tx=XM_019829755.1;refGene_gn=MCF2L;refGene_tx=XM_019829760.1;refGene_gn=MCF2L;refGene_tx=XM_019829766.1;refGene_gn=MCF2L;refGene_tx=XM_019829789.1;refGene_gn=MCF2L;refGene_tx=XM_019829794.1;refGene_gn=MCF2L;refGene_tx=XM_019829802.1;refGene_gn=MCF2L;refGene_tx=XM_019829809.1;refGene_gn=MCF2L;refGene_tx=XM_019829815.1
A1 82212571 82212686
refGene_gn=MCF2L;refGene_tx=XM_006927406.3;refGene_gn=MCF2L;refGene_tx=XM_011280911.2;refGene_gn=MCF2L;refGene_tx=XM_011280921.2;refGene_gn=MCF2L;refGene_tx=XM_019829754.1;refGene_gn=MCF2L;refGene_tx=XM_019829755.1;refGene_gn=MCF2L;refGene_tx=XM_019829760.1;refGene_gn=MCF2L;refGene_tx=XM_019829766.1;refGene_gn=MCF2L;refGene_tx=XM_019829789.1;refGene_gn=MCF2L;refGene_tx=XM_019829794.1;refGene_gn=MCF2L;refGene_tx=XM_019829802.1;refGene_gn=MCF2L;refGene_tx=XM_019829809.1;refGene_gn=MCF2L;refGene_tx=XM_019829815.1
A1 82216066 82216186
refGene_gn=MCF2L;refGene_tx=XM_006927406.3;refGene_gn=MCF2L;refGene_tx=XM_011280911.2;refGene_gn=MCF2L;refGene_tx=XM_011280921.2;refGene_gn=MCF2L;refGene_tx=XM_019829754.1;refGene_gn=MCF2L;refGene_tx=XM_019829755.1;refGene_gn=MCF2L;refGene_tx=XM_019829760.1;refGene_gn=MCF2L;refGene_tx=XM_019829766.1;refGene_gn=MCF2L;refGene_tx=XM_019829789.1;refGene_gn=MCF2L;refGene_tx=XM_019829794.1;refGene_gn=MCF2L;refGene_tx=XM_019829802.1;refGene_gn=MCF2L;refGene_tx=XM_019829809.1;refGene_gn=MCF2L;refGene_tx=XM_019829815.1

A1	82216434 82216626
refGene_gn=MCF2L;refGene_tx=XM_006927406.3;refGene_gn=MCF2L;refGene_tx=XM_011280911.2;refGene_gn=MCF2L;refGene_tx=XM_011280921.2;refGene_gn=MCF2L;refGene_tx=XM_019829754.1;refGene_gn=MCF2L;refGene_tx=XM_019829755.1;refGene_gn=MCF2L;refGene_tx=XM_019829760.1;refGene_gn=MCF2L;refGene_tx=XM_019829766.1;refGene_gn=MCF2L;refGene_tx=XM_019829789.1;refGene_gn=MCF2L;refGene_tx=XM_019829794.1;refGene_gn=MCF2L;refGene_tx=XM_019829802.1;refGene_gn=MCF2L;refGene_tx=XM_019829809.1;refGene_gn=MCF2L;refGene_tx=XM_019829815.1
A1	82217045 82217237
refGene_gn=MCF2L;refGene_tx=XM_006927406.3;refGene_gn=MCF2L;refGene_tx=XM_011280911.2;refGene_gn=MCF2L;refGene_tx=XM_011280921.2;refGene_gn=MCF2L;refGene_tx=XM_019829754.1;refGene_gn=MCF2L;refGene_tx=XM_019829755.1;refGene_gn=MCF2L;refGene_tx=XM_019829760.1;refGene_gn=MCF2L;refGene_tx=XM_019829766.1;refGene_gn=MCF2L;refGene_tx=XM_019829789.1;refGene_gn=MCF2L;refGene_tx=XM_019829794.1;refGene_gn=MCF2L;refGene_tx=XM_019829802.1;refGene_gn=MCF2L;refGene_tx=XM_019829809.1;refGene_gn=MCF2L;refGene_tx=XM_019829815.1
A1	82217762 82217922
refGene_gn=MCF2L;refGene_tx=XM_006927406.3;refGene_gn=MCF2L;refGene_tx=XM_011280911.2;refGene_gn=MCF2L;refGene_tx=XM_011280921.2;refGene_gn=MCF2L;refGene_tx=XM_019829754.1;refGene_gn=MCF2L;refGene_tx=XM_019829755.1;refGene_gn=MCF2L;refGene_tx=XM_019829760.1;refGene_gn=MCF2L;refGene_tx=XM_019829766.1;refGene_gn=MCF2L;refGene_tx=XM_019829789.1;refGene_gn=MCF2L;refGene_tx=XM_019829794.1;refGene_gn=MCF2L;refGene_tx=XM_019829802.1;refGene_gn=MCF2L;refGene_tx=XM_019829809.1;refGene_gn=MCF2L;refGene_tx=XM_019829815.1
A1	82218606 82218677
refGene_gn=MCF2L;refGene_tx=XM_006927406.3;refGene_gn=MCF2L;refGene_tx=XM_011280911.2;refGene_gn=MCF2L;refGene_tx=XM_011280921.2;refGene_gn=MCF2L;refGene_tx=XM_019829754.1;refGene_gn=MCF2L;refGene_tx=XM_019829755.1;refGene_gn=MCF2L;refGene_tx=XM_019829760.1;refGene_gn=MCF2L;refGene_tx=XM_019829766.1;refGene_gn=MCF2L;refGene_tx=XM_019829789.1;refGene_gn=MCF2L;refGene_tx=XM_019829794.1;refGene_gn=MCF2L;refGene_tx=XM_019829802.1;refGene_gn=MCF2L;refGene_tx=XM_019829809.1;refGene_gn=MCF2L;refGene_tx=XM_019829815.1
A1	82218875 82218952
refGene_gn=MCF2L;refGene_tx=XM_006927406.3;refGene_gn=MCF2L;refGene_tx=XM_011280911.2;refGene_gn=MCF2L;refGene_tx=XM_011280921.2;refGene_gn=MCF2L;refGene_tx=XM_019829754.1;refGene_gn=MCF2L;refGene_tx=XM_019829755.1;refGene_gn=MCF2L;refGene_tx=XM_019829760.1;refGene_gn=MCF2L;refGene_tx=XM_019829766.1;refGene_gn=MCF2L;refGene_tx=XM_019829789.1;refGene_gn=MCF2L;refGene_tx=XM_019829794.1;refGene_gn=MCF2L;refGene_tx=XM_019829802.1;refGene_gn=MCF2L;refGene_tx=XM_019829809.1;refGene_gn=MCF2L;refGene_tx=XM_019829815.1
A1	82219405 82219472
refGene_gn=MCF2L;refGene_tx=XM_006927406.3;refGene_gn=MCF2L;refGene_tx=XM_011280911.2;refGene_gn=MCF2L;refGene_tx=XM_011280921.2;refGene_gn=MCF2L;refGene_tx=XM_019829754.1;refGene_gn=MCF2L;refGene_tx=XM_019829755.1;refGene_gn=MCF2L;refGene_tx=XM_019829760.1;refGene_gn=MCF2L;refGe

ne_tx=XM_019829766.1;refGene_gn=MCF2L;refGene_tx=XM_019829789.1;refGene_gn=MCF2L;refGene_tx=XM_019829794.1;refGene_gn=MCF2L;refGene_tx=XM_019829802.1;refGene_gn=MCF2L;refGene_tx=XM_019829809.1;refGene_gn=MCF2L;refGene_tx=XM_019829815.1
A1 82220124 82220240
refGene_gn=MCF2L;refGene_tx=XM_006927406.3;refGene_gn=MCF2L;refGene_tx=XM_011280911.2;refGene_gn=MCF2L;refGene_tx=XM_011280921.2;refGene_gn=MCF2L;refGene_tx=XM_019829754.1;refGene_gn=MCF2L;refGene_tx=XM_019829755.1;refGene_gn=MCF2L;refGene_tx=XM_019829760.1;refGene_gn=MCF2L;refGene_tx=XM_019829766.1;refGene_gn=MCF2L;refGene_tx=XM_019829789.1;refGene_gn=MCF2L;refGene_tx=XM_019829794.1;refGene_gn=MCF2L;refGene_tx=XM_019829802.1;refGene_gn=MCF2L;refGene_tx=XM_019829809.1;refGene_gn=MCF2L;refGene_tx=XM_019829815.1
A1 82221360 82221430
refGene_gn=MCF2L;refGene_tx=XM_006927406.3;refGene_gn=MCF2L;refGene_tx=XM_011280911.2;refGene_gn=MCF2L;refGene_tx=XM_011280921.2;refGene_gn=MCF2L;refGene_tx=XM_019829754.1;refGene_gn=MCF2L;refGene_tx=XM_019829755.1;refGene_gn=MCF2L;refGene_tx=XM_019829760.1;refGene_gn=MCF2L;refGene_tx=XM_019829766.1;refGene_gn=MCF2L;refGene_tx=XM_019829789.1;refGene_gn=MCF2L;refGene_tx=XM_019829794.1;refGene_gn=MCF2L;refGene_tx=XM_019829802.1;refGene_gn=MCF2L;refGene_tx=XM_019829809.1;refGene_gn=MCF2L;refGene_tx=XM_019829815.1
A1 82222751 82222844
refGene_gn=MCF2L;refGene_tx=XM_006927406.3;refGene_gn=MCF2L;refGene_tx=XM_011280911.2;refGene_gn=MCF2L;refGene_tx=XM_011280921.2;refGene_gn=MCF2L;refGene_tx=XM_019829754.1;refGene_gn=MCF2L;refGene_tx=XM_019829755.1;refGene_gn=MCF2L;refGene_tx=XM_019829760.1;refGene_gn=MCF2L;refGene_tx=XM_019829766.1;refGene_gn=MCF2L;refGene_tx=XM_019829789.1;refGene_gn=MCF2L;refGene_tx=XM_019829794.1;refGene_gn=MCF2L;refGene_tx=XM_019829802.1;refGene_gn=MCF2L;refGene_tx=XM_019829809.1;refGene_gn=MCF2L;refGene_tx=XM_019829815.1
A1 82222945 82223038
refGene_gn=MCF2L;refGene_tx=XM_006927406.3;refGene_gn=MCF2L;refGene_tx=XM_011280911.2;refGene_gn=MCF2L;refGene_tx=XM_011280921.2;refGene_gn=MCF2L;refGene_tx=XM_019829754.1;refGene_gn=MCF2L;refGene_tx=XM_019829755.1;refGene_gn=MCF2L;refGene_tx=XM_019829760.1;refGene_gn=MCF2L;refGene_tx=XM_019829766.1;refGene_gn=MCF2L;refGene_tx=XM_019829789.1;refGene_gn=MCF2L;refGene_tx=XM_019829794.1;refGene_gn=MCF2L;refGene_tx=XM_019829802.1;refGene_gn=MCF2L;refGene_tx=XM_019829809.1;refGene_gn=MCF2L;refGene_tx=XM_019829815.1
A1 82223671 82223797
refGene_gn=MCF2L;refGene_tx=XM_006927406.3;refGene_gn=MCF2L;refGene_tx=XM_011280911.2;refGene_gn=MCF2L;refGene_tx=XM_011280921.2;refGene_gn=MCF2L;refGene_tx=XM_019829754.1;refGene_gn=MCF2L;refGene_tx=XM_019829755.1;refGene_gn=MCF2L;refGene_tx=XM_019829760.1;refGene_gn=MCF2L;refGene_tx=XM_019829766.1;refGene_gn=MCF2L;refGene_tx=XM_019829789.1;refGene_gn=MCF2L;refGene_tx=XM_019829794.1;refGene_gn=MCF2L;refGene_tx=XM_019829802.1;refGene_gn=MCF2L;refGene_tx=XM_019829809.1;refGene_gn=MCF2L;refGene_tx=XM_019829815.1
A1 82224435 82224657

refGene_gn=MCF2L;refGene_tx=XM_006927406.3;refGene_gn=MCF2L;refGene_tx=XM_011280911.2;refGene_gn=MCF2L;refGene_tx=XM_011280921.2;refGene_gn=MCF2L;refGene_tx=XM_019829754.1;refGene_gn=MCF2L;refGene_tx=XM_019829755.1;refGene_gn=MCF2L;refGene_tx=XM_019829760.1;refGene_gn=MCF2L;refGene_tx=XM_019829766.1;refGene_gn=MCF2L;refGene_tx=XM_019829789.1;refGene_gn=MCF2L;refGene_tx=XM_019829794.1;refGene_gn=MCF2L;refGene_tx=XM_019829802.1;refGene_gn=MCF2L;refGene_tx=XM_019829809.1;refGene_gn=MCF2L;refGene_tx=XM_019829815.1
A1 82224823 82224916
refGene_gn=MCF2L;refGene_tx=XM_006927406.3;refGene_gn=MCF2L;refGene_tx=XM_011280911.2;refGene_gn=MCF2L;refGene_tx=XM_011280921.2;refGene_gn=MCF2L;refGene_tx=XM_019829754.1;refGene_gn=MCF2L;refGene_tx=XM_019829755.1;refGene_gn=MCF2L;refGene_tx=XM_019829760.1;refGene_gn=MCF2L;refGene_tx=XM_019829766.1;refGene_gn=MCF2L;refGene_tx=XM_019829789.1;refGene_gn=MCF2L;refGene_tx=XM_019829794.1;refGene_gn=MCF2L;refGene_tx=XM_019829802.1;refGene_gn=MCF2L;refGene_tx=XM_019829809.1;refGene_gn=MCF2L;refGene_tx=XM_019829815.1
A1 82225876 82225955
refGene_gn=MCF2L;refGene_tx=XM_006927406.3;refGene_gn=MCF2L;refGene_tx=XM_011280911.2;refGene_gn=MCF2L;refGene_tx=XM_011280921.2;refGene_gn=MCF2L;refGene_tx=XM_019829754.1;refGene_gn=MCF2L;refGene_tx=XM_019829755.1;refGene_gn=MCF2L;refGene_tx=XM_019829760.1;refGene_gn=MCF2L;refGene_tx=XM_019829766.1;refGene_gn=MCF2L;refGene_tx=XM_019829789.1;refGene_gn=MCF2L;refGene_tx=XM_019829794.1;refGene_gn=MCF2L;refGene_tx=XM_019829802.1;refGene_gn=MCF2L;refGene_tx=XM_019829809.1;refGene_gn=MCF2L;refGene_tx=XM_019829815.1
A1 82226091 82226158
refGene_gn=MCF2L;refGene_tx=XM_006927406.3;refGene_gn=MCF2L;refGene_tx=XM_011280911.2;refGene_gn=MCF2L;refGene_tx=XM_011280921.2;refGene_gn=MCF2L;refGene_tx=XM_019829754.1;refGene_gn=MCF2L;refGene_tx=XM_019829755.1;refGene_gn=MCF2L;refGene_tx=XM_019829760.1;refGene_gn=MCF2L;refGene_tx=XM_019829766.1;refGene_gn=MCF2L;refGene_tx=XM_019829789.1;refGene_gn=MCF2L;refGene_tx=XM_019829794.1;refGene_gn=MCF2L;refGene_tx=XM_019829802.1;refGene_gn=MCF2L;refGene_tx=XM_019829809.1;refGene_gn=MCF2L;refGene_tx=XM_019829815.1
A1 82226871 82226990
refGene_gn=MCF2L;refGene_tx=XM_006927406.3;refGene_gn=MCF2L;refGene_tx=XM_011280911.2;refGene_gn=MCF2L;refGene_tx=XM_011280921.2;refGene_gn=MCF2L;refGene_tx=XM_019829754.1;refGene_gn=MCF2L;refGene_tx=XM_019829755.1;refGene_gn=MCF2L;refGene_tx=XM_019829760.1;refGene_gn=MCF2L;refGene_tx=XM_019829766.1;refGene_gn=MCF2L;refGene_tx=XM_019829789.1;refGene_gn=MCF2L;refGene_tx=XM_019829794.1;refGene_gn=MCF2L;refGene_tx=XM_019829802.1;refGene_gn=MCF2L;refGene_tx=XM_019829809.1;refGene_gn=MCF2L;refGene_tx=XM_019829815.1
A1 82227137 82227230 refGene_gn=MCF2L;refGene_tx=XM_019829815.1
A1 82228469 82228544
refGene_gn=MCF2L;refGene_tx=XM_006927406.3;refGene_gn=MCF2L;refGene_tx=XM_011280911.2;refGene_gn=MCF2L;refGene_tx=XM_019829754.1;refGene_gn=MCF2L;refGene_tx=XM_019829755.1;refGene_gn=MCF2L;refGene_tx=XM_019829766.1;refGene_gn=MCF2L;refGene_tx=XM_019829794.1;refGene_gn=MCF2L;refGe

ne_tx=XM_019829802.1;refGene_gn=MCF2L;refGene_tx=XM_019829809.1;refGene_gn=MCF2L;refGene_tx=XM_019829815.1
A1 82230913 82231032
refGene_gn=MCF2L;refGene_tx=XM_006927406.3;refGene_gn=MCF2L;refGene_tx=XM_011280911.2;refGene_gn=MCF2L;refGene_tx=XM_011280921.2;refGene_gn=MCF2L;refGene_tx=XM_019829754.1;refGene_gn=MCF2L;refGene_tx=XM_019829755.1;refGene_gn=MCF2L;refGene_tx=XM_019829760.1;refGene_gn=MCF2L;refGene_tx=XM_019829766.1;refGene_gn=MCF2L;refGene_tx=XM_019829789.1;refGene_gn=MCF2L;refGene_tx=XM_019829794.1;refGene_gn=MCF2L;refGene_tx=XM_019829802.1;refGene_gn=MCF2L;refGene_tx=XM_019829809.1;refGene_gn=MCF2L;refGene_tx=XM_019829815.1
A1 82232266 82232379
refGene_gn=MCF2L;refGene_tx=XM_006927406.3;refGene_gn=MCF2L;refGene_tx=XM_011280911.2;refGene_gn=MCF2L;refGene_tx=XM_011280921.2;refGene_gn=MCF2L;refGene_tx=XM_019829754.1;refGene_gn=MCF2L;refGene_tx=XM_019829755.1;refGene_gn=MCF2L;refGene_tx=XM_019829760.1;refGene_gn=MCF2L;refGene_tx=XM_019829766.1;refGene_gn=MCF2L;refGene_tx=XM_019829789.1;refGene_gn=MCF2L;refGene_tx=XM_019829794.1;refGene_gn=MCF2L;refGene_tx=XM_019829802.1;refGene_gn=MCF2L;refGene_tx=XM_019829809.1;refGene_gn=MCF2L;refGene_tx=XM_019829815.1
A1 82232466 82232570
refGene_gn=MCF2L;refGene_tx=XM_006927406.3;refGene_gn=MCF2L;refGene_tx=XM_011280911.2;refGene_gn=MCF2L;refGene_tx=XM_011280921.2;refGene_gn=MCF2L;refGene_tx=XM_019829754.1;refGene_gn=MCF2L;refGene_tx=XM_019829755.1;refGene_gn=MCF2L;refGene_tx=XM_019829760.1;refGene_gn=MCF2L;refGene_tx=XM_019829766.1;refGene_gn=MCF2L;refGene_tx=XM_019829789.1;refGene_gn=MCF2L;refGene_tx=XM_019829794.1;refGene_gn=MCF2L;refGene_tx=XM_019829802.1;refGene_gn=MCF2L;refGene_tx=XM_019829809.1;refGene_gn=MCF2L;refGene_tx=XM_019829815.1
A1 82232680 82232766
refGene_gn=MCF2L;refGene_tx=XM_006927406.3;refGene_gn=MCF2L;refGene_tx=XM_011280911.2;refGene_gn=MCF2L;refGene_tx=XM_011280921.2;refGene_gn=MCF2L;refGene_tx=XM_019829754.1;refGene_gn=MCF2L;refGene_tx=XM_019829755.1;refGene_gn=MCF2L;refGene_tx=XM_019829760.1;refGene_gn=MCF2L;refGene_tx=XM_019829789.1;refGene_gn=MCF2L;refGene_tx=XM_019829794.1;refGene_gn=MCF2L;refGene_tx=XM_019829802.1;refGene_gn=MCF2L;refGene_tx=XM_019829809.1;refGene_gn=MCF2L;refGene_tx=XM_019829815.1
A1 82232831 82232839 refGene_gn=MCF2L;refGene_tx=XM_019829766.1
A1 82236885 82236955 refGene_gn=F7;refGene_tx=XM_003980533.3
A1 82238491 82238652 refGene_gn=F7;refGene_tx=XM_003980533.3
A1 82239928 82239953 refGene_gn=F7;refGene_tx=XM_003980533.3
A1 82240466 82240580 refGene_gn=F7;refGene_tx=XM_003980533.3
A1 82242194 82242335 refGene_gn=F7;refGene_tx=XM_003980533.3
A1 82243327 82243437 refGene_gn=F7;refGene_tx=XM_003980533.3
A1 82244905 82245029 refGene_gn=F7;refGene_tx=XM_003980533.3
A1 82245500 82246096 refGene_gn=F7;refGene_tx=XM_003980533.3
A1 82249418 82249488 refGene_gn=F10;refGene_tx=XM_003980558.2
A1 82251057 82251218 refGene_gn=F10;refGene_tx=XM_003980558.2
A1 82254350 82254375 refGene_gn=F10;refGene_tx=XM_003980558.2
A1 82255271 82255385 refGene_gn=F10;refGene_tx=XM_003980558.2

A1 82256957 82257089 refGene_gn=F10;refGene_tx=XM_003980558.2
A1 82257829 82258083 refGene_gn=F10;refGene_tx=XM_003980558.2
A1 82259898 82260016 refGene_gn=F10;refGene_tx=XM_003980558.2
A1 82260901 82261461 refGene_gn=F10;refGene_tx=XM_003980558.2
A1 82268743 82268833
refGene_gn=PCID2;refGene_tx=XM_003980534.5;refGene_gn=PCID2;refGene_tx=XM_011280926.3;refGene_gn=PCID2;refGene_tx=XM_011280927.3;refGene_gn=PCID2;refGene_tx=XM_023253400.1
A1 82269151 82269275
refGene_gn=PCID2;refGene_tx=XM_003980534.5;refGene_gn=PCID2;refGene_tx=XM_011280926.3;refGene_gn=PCID2;refGene_tx=XM_011280927.3;refGene_gn=PCID2;refGene_tx=XM_023253400.1
A1 82269769 82269895
refGene_gn=PCID2;refGene_tx=XM_003980534.5;refGene_gn=PCID2;refGene_tx=XM_011280926.3;refGene_gn=PCID2;refGene_tx=XM_011280927.3;refGene_gn=PCID2;refGene_tx=XM_023253400.1
A1 82270047 82270121
refGene_gn=PCID2;refGene_tx=XM_003980534.5;refGene_gn=PCID2;refGene_tx=XM_011280926.3;refGene_gn=PCID2;refGene_tx=XM_011280927.3;refGene_gn=PCID2;refGene_tx=XM_023253400.1
A1 82271268 82271369
refGene_gn=PCID2;refGene_tx=XM_003980534.5;refGene_gn=PCID2;refGene_tx=XM_011280926.3;refGene_gn=PCID2;refGene_tx=XM_011280927.3;refGene_gn=PCID2;refGene_tx=XM_023253400.1
A1 82272426 82272568
refGene_gn=PCID2;refGene_tx=XM_003980534.5;refGene_gn=PCID2;refGene_tx=XM_011280926.3;refGene_gn=PCID2;refGene_tx=XM_011280927.3;refGene_gn=PCID2;refGene_tx=XM_023253400.1
A1 82273218 82273294
refGene_gn=PCID2;refGene_tx=XM_003980534.5;refGene_gn=PCID2;refGene_tx=XM_011280926.3;refGene_gn=PCID2;refGene_tx=XM_011280927.3;refGene_gn=PCID2;refGene_tx=XM_023253400.1
A1 82276169 82276273
refGene_gn=PCID2;refGene_tx=XM_003980534.5;refGene_gn=PCID2;refGene_tx=XM_011280926.3;refGene_gn=PCID2;refGene_tx=XM_011280927.3;refGene_gn=PCID2;refGene_tx=XM_023253400.1
A1 82279666 82279721
refGene_gn=PCID2;refGene_tx=XM_003980534.5;refGene_gn=PCID2;refGene_tx=XM_011280926.3;refGene_gn=PCID2;refGene_tx=XM_011280927.3;refGene_gn=PCID2;refGene_tx=XM_023253400.1
A1 82280670 82280712
refGene_gn=PCID2;refGene_tx=XM_003980534.5;refGene_gn=PCID2;refGene_tx=XM_011280926.3
A1 82281584 82281650
refGene_gn=PCID2;refGene_tx=XM_003980534.5;refGene_gn=PCID2;refGene_tx=XM_011280926.3
A1 82283075 82283149
refGene_gn=PCID2;refGene_tx=XM_003980534.5;refGene_gn=PCID2;refGene_tx=XM_011280926.3;refGene_gn=PCID2;refGene_tx=XM_011280927.3
A1 82286228 82286318

refGene_gn=PCID2;refGene_tx=XM_003980534.5;refGene_gn=PCID2;refGene_tx=XM_011280926.3;refGene_gn=PCID2;refGene_tx=XM_011280927.3
A1 82292673 82292709
refGene_gn=PCID2;refGene_tx=XM_003980534.5;refGene_gn=PCID2;refGene_tx=XM_011280926.3;refGene_gn=PCID2;refGene_tx=XM_011280927.3
A1 82293222 82293376 refGene_gn=CUL4A;refGene_tx=XM_023253394.1
A1 82293704 82293820 refGene_gn=CUL4A;refGene_tx=XM_023253394.1
A1 82302214 82302318 refGene_gn=CUL4A;refGene_tx=XM_023253394.1
A1 82308158 82308228 refGene_gn=CUL4A;refGene_tx=XM_023253394.1
A1 82309372 82309446 refGene_gn=CUL4A;refGene_tx=XM_023253394.1
A1 82311960 82312123 refGene_gn=CUL4A;refGene_tx=XM_023253394.1
A1 82312614 82312704 refGene_gn=CUL4A;refGene_tx=XM_023253394.1
A1 82314030 82314113 refGene_gn=CUL4A;refGene_tx=XM_023253394.1
A1 82317095 82317163 refGene_gn=CUL4A;refGene_tx=XM_023253394.1
A1 82319237 82319356 refGene_gn=CUL4A;refGene_tx=XM_023253394.1
A1 82321730 82321923 refGene_gn=CUL4A;refGene_tx=XM_023253394.1
A1 82323174 82323279 refGene_gn=CUL4A;refGene_tx=XM_023253394.1
A1 82323588 82323699 refGene_gn=CUL4A;refGene_tx=XM_023253394.1
A1 82323782 82323868 refGene_gn=CUL4A;refGene_tx=XM_023253394.1
A1 82324288 82324396 refGene_gn=CUL4A;refGene_tx=XM_023253394.1
A1 82330637 82330751 refGene_gn=CUL4A;refGene_tx=XM_023253394.1
A1 82331776 82331882 refGene_gn=CUL4A;refGene_tx=XM_023253394.1
A1 82332041 82332214 refGene_gn=CUL4A;refGene_tx=XM_023253394.1
A1 82336279 82336432 refGene_gn=CUL4A;refGene_tx=XM_023253394.1
A1 82337969 82338065 refGene_gn=CUL4A;refGene_tx=XM_023253394.1
A1 82349964 82350016 refGene_gn=LAMP1;refGene_tx=NM_001163655.1
A1 82355010 82355132 refGene_gn=LAMP1;refGene_tx=NM_001163655.1
A1 82356955 82357175 refGene_gn=LAMP1;refGene_tx=NM_001163655.1
A1 82357292 82357451 refGene_gn=LAMP1;refGene_tx=NM_001163655.1
A1 82363801 82363989 refGene_gn=LAMP1;refGene_tx=NM_001163655.1
A1 82364528 82364654 refGene_gn=LAMP1;refGene_tx=NM_001163655.1
A1 82365408 82365475 refGene_gn=LAMP1;refGene_tx=NM_001163655.1
A1 82365544 82365715 refGene_gn=LAMP1;refGene_tx=NM_001163655.1
A1 82366135 82366275 refGene_gn=LAMP1;refGene_tx=NM_001163655.1
A1 82368525 82368615 refGene_gn=GRTP1;refGene_tx=XM_023253407.1
A1 82370588 82370774 refGene_gn=GRTP1;refGene_tx=XM_023253407.1
A1 82370850 82371023 refGene_gn=GRTP1;refGene_tx=XM_023253407.1
A1 82375830 82375927 refGene_gn=GRTP1;refGene_tx=XM_023253407.1
A1 82378366 82378491 refGene_gn=GRTP1;refGene_tx=XM_023253407.1
A1 82380692 82380851 refGene_gn=GRTP1;refGene_tx=XM_023253407.1
A1 82386425 82386577 refGene_gn=GRTP1;refGene_tx=XM_023253407.1
A1 82386835 82386873 refGene_gn=GRTP1;refGene_tx=XM_023253407.1
A1 82396408 82396472
refGene_gn=LOC105260444;refGene_tx=XM_023239413.1
A1 82398284 82398469
refGene_gn=LOC105260444;refGene_tx=XM_023239413.1
A1 82398572 82399450
refGene_gn=LOC105260444;refGene_tx=XM_023239413.1
A1 82400273 82401925
refGene_gn=LOC105260444;refGene_tx=XM_023239413.1

A1 82403856 82404356
refGene_gn=LOC105260444;refGene_tx=XM_023239413.1
A1 82406291 82406908
refGene_gn=ADPRHL1;refGene_tx=XM_003980536.4;refGene_gn=ADPRHL1;refGene_tx=XM_019829901.2
A1 82410511 82410669
refGene_gn=ADPRHL1;refGene_tx=XM_003980536.4;refGene_gn=ADPRHL1;refGene_tx=XM_019829904.1
A1 82411368 82411501
refGene_gn=ADPRHL1;refGene_tx=XM_003980536.4;refGene_gn=ADPRHL1;refGene_tx=XM_019829901.2;refGene_gn=ADPRHL1;refGene_tx=XM_019829904.1
A1 82412658 82412786
refGene_gn=ADPRHL1;refGene_tx=XM_003980536.4;refGene_gn=ADPRHL1;refGene_tx=XM_019829901.2;refGene_gn=ADPRHL1;refGene_tx=XM_019829904.1
A1 82415434 82415575
refGene_gn=ADPRHL1;refGene_tx=XM_003980536.4;refGene_gn=ADPRHL1;refGene_tx=XM_019829901.2;refGene_gn=ADPRHL1;refGene_tx=XM_019829904.1
A1 82416736 82416862
refGene_gn=ADPRHL1;refGene_tx=XM_003980536.4;refGene_gn=ADPRHL1;refGene_tx=XM_019829901.2;refGene_gn=ADPRHL1;refGene_tx=XM_019829904.1
A1 82421441 82421606
refGene_gn=ADPRHL1;refGene_tx=XM_003980536.4;refGene_gn=ADPRHL1;refGene_tx=XM_019829901.2;refGene_gn=ADPRHL1;refGene_tx=XM_019829904.1
A1 82427811 82428025
refGene_gn=ADPRHL1;refGene_tx=XM_003980536.4;refGene_gn=ADPRHL1;refGene_tx=XM_019829901.2;refGene_gn=ADPRHL1;refGene_tx=XM_019829904.1
A1 82431700 82431780
refGene_gn=DCUN1D2;refGene_tx=XM_006927413.2;refGene_gn=DCUN1D2;refGene_tx=XM_006927414.2;refGene_gn=DCUN1D2;refGene_tx=XM_006927416.3
A1 82432895 82432992
refGene_gn=DCUN1D2;refGene_tx=XM_006927413.2;refGene_gn=DCUN1D2;refGene_tx=XM_006927414.2
A1 82434559 82434642
refGene_gn=DCUN1D2;refGene_tx=XM_006927413.2;refGene_gn=DCUN1D2;refGene_tx=XM_006927414.2;refGene_gn=DCUN1D2;refGene_tx=XM_006927416.3
A1 82443930 82444061
refGene_gn=DCUN1D2;refGene_tx=XM_006927413.2;refGene_gn=DCUN1D2;refGene_tx=XM_006927414.2;refGene_gn=DCUN1D2;refGene_tx=XM_006927416.3
A1 82451506 82451675
refGene_gn=DCUN1D2;refGene_tx=XM_006927413.2;refGene_gn=DCUN1D2;refGene_tx=XM_006927414.2;refGene_gn=DCUN1D2;refGene_tx=XM_006927416.3
A1 82454884 82454939
refGene_gn=DCUN1D2;refGene_tx=XM_006927413.2;refGene_gn=DCUN1D2;refGene_tx=XM_006927414.2;refGene_gn=DCUN1D2;refGene_tx=XM_006927416.3
A1 82470716 82471160
refGene_gn=TMCO3;refGene_tx=XM_003980537.4;refGene_gn=TMCO3;refGene_tx=XM_006927417.4;refGene_gn=TMCO3;refGene_tx=XM_006927421.4;refGene_gn=TMCO3;refGene_tx=XM_019829866.2;refGene_gn=TMCO3;refGene_tx=XM_019829872.2;refGene_gn=TMCO3;refGene_tx=XM_019829877.2;refGene_gn=TMCO3;refGene_tx=XM_019829882.2

A1 82472687 82472867
refGene_gn=TMCO3;refGene_tx=XM_003980537.4;refGene_gn=TMCO3;refGene_tx=XM_006927417.4;refGene_gn=TMCO3;refGene_tx=XM_006927421.4;refGene_gn=TMCO3;refGene_tx=XM_019829866.2;refGene_gn=TMCO3;refGene_tx=XM_019829872.2;refGene_gn=TMCO3;refGene_tx=XM_019829877.2;refGene_gn=TMCO3;refGene_tx=XM_019829882.2
A1 82474358 82474550
refGene_gn=TMCO3;refGene_tx=XM_003980537.4;refGene_gn=TMCO3;refGene_tx=XM_006927417.4;refGene_gn=TMCO3;refGene_tx=XM_006927421.4;refGene_gn=TMCO3;refGene_tx=XM_019829866.2;refGene_gn=TMCO3;refGene_tx=XM_019829872.2;refGene_gn=TMCO3;refGene_tx=XM_019829877.2;refGene_gn=TMCO3;refGene_tx=XM_019829882.2
A1 82475409 82475538
refGene_gn=TMCO3;refGene_tx=XM_003980537.4;refGene_gn=TMCO3;refGene_tx=XM_006927417.4;refGene_gn=TMCO3;refGene_tx=XM_006927421.4;refGene_gn=TMCO3;refGene_tx=XM_019829866.2;refGene_gn=TMCO3;refGene_tx=XM_019829872.2;refGene_gn=TMCO3;refGene_tx=XM_019829877.2;refGene_gn=TMCO3;refGene_tx=XM_019829882.2
A1 82478121 82478214
refGene_gn=TMCO3;refGene_tx=XM_003980537.4;refGene_gn=TMCO3;refGene_tx=XM_006927417.4;refGene_gn=TMCO3;refGene_tx=XM_006927421.4;refGene_gn=TMCO3;refGene_tx=XM_019829866.2;refGene_gn=TMCO3;refGene_tx=XM_019829872.2;refGene_gn=TMCO3;refGene_tx=XM_019829877.2
A1 82482750 82482937
refGene_gn=TMCO3;refGene_tx=XM_003980537.4;refGene_gn=TMCO3;refGene_tx=XM_006927417.4;refGene_gn=TMCO3;refGene_tx=XM_006927421.4;refGene_gn=TMCO3;refGene_tx=XM_019829866.2;refGene_gn=TMCO3;refGene_tx=XM_019829872.2;refGene_gn=TMCO3;refGene_tx=XM_019829877.2;refGene_gn=TMCO3;refGene_tx=XM_019829882.2
A1 82489472 82489593
refGene_gn=TMCO3;refGene_tx=XM_003980537.4;refGene_gn=TMCO3;refGene_tx=XM_006927417.4;refGene_gn=TMCO3;refGene_tx=XM_006927421.4;refGene_gn=TMCO3;refGene_tx=XM_019829866.2;refGene_gn=TMCO3;refGene_tx=XM_019829872.2;refGene_gn=TMCO3;refGene_tx=XM_019829877.2;refGene_gn=TMCO3;refGene_tx=XM_019829882.2
A1 82500786 82500982
refGene_gn=TMCO3;refGene_tx=XM_003980537.4;refGene_gn=TMCO3;refGene_tx=XM_006927417.4;refGene_gn=TMCO3;refGene_tx=XM_006927421.4;refGene_gn=TMCO3;refGene_tx=XM_019829866.2;refGene_gn=TMCO3;refGene_tx=XM_019829872.2;refGene_gn=TMCO3;refGene_tx=XM_019829877.2;refGene_gn=TMCO3;refGene_tx=XM_019829882.2
A1 82504609 82504760
refGene_gn=TMCO3;refGene_tx=XM_003980537.4;refGene_gn=TMCO3;refGene_tx=XM_006927417.4;refGene_gn=TMCO3;refGene_tx=XM_006927421.4;refGene_gn=TMCO3;refGene_tx=XM_019829866.2;refGene_gn=TMCO3;refGene_tx=XM_019829872.2;refGene_gn=TMCO3;refGene_tx=XM_019829877.2;refGene_gn=TMCO3;refGene_tx=XM_019829882.2
A1 82508649 82508729
refGene_gn=TMCO3;refGene_tx=XM_003980537.4;refGene_gn=TMCO3;refGene_tx=XM_006927417.4;refGene_gn=TMCO3;refGene_tx=XM_006927421.4;refGene_gn=

TMCO3;refGene_tx=XM_019829866.2;refGene_gn=TMCO3;refGene_tx=XM_0198298
72.2;refGene_gn=TMCO3;refGene_tx=XM_019829877.2;refGene_gn=TMCO3;refGe
ne_tx=XM_019829882.2
A1 82509530 82509782
refGene_gn=TMCO3;refGene_tx=XM_003980537.4;refGene_gn=TMCO3;refGene_tx
=XM_006927417.4;refGene_gn=TMCO3;refGene_tx=XM_019829866.2;refGene_gn=
TMCO3;refGene_tx=XM_019829872.2;refGene_gn=TMCO3;refGene_tx=XM_0198298
77.2;refGene_gn=TMCO3;refGene_tx=XM_019829882.2
A1 82511728 82511842
refGene_gn=TMCO3;refGene_tx=XM_003980537.4;refGene_gn=TMCO3;refGene_tx
=XM_006927417.4;refGene_gn=TMCO3;refGene_tx=XM_006927421.4;refGene_gn=
TMCO3;refGene_tx=XM_019829866.2;refGene_gn=TMCO3;refGene_tx=XM_0198298
72.2;refGene_gn=TMCO3;refGene_tx=XM_019829877.2;refGene_gn=TMCO3;refGe
ne_tx=XM_019829882.2
A1 82536796 82536808
refGene_gn=TFDP1;refGene_tx=XM_023253517.1;refGene_gn=TFDP1;refGene_tx
=XM_023253524.1;refGene_gn=TFDP1;refGene_tx=XM_023253527.1;refGene_gn=
TFDP1;refGene_tx=XM_023253531.1
A1 82555166 82555233
refGene_gn=TFDP1;refGene_tx=XM_023253517.1;refGene_gn=TFDP1;refGene_tx
=XM_023253524.1;refGene_gn=TFDP1;refGene_tx=XM_023253527.1;refGene_gn=
TFDP1;refGene_tx=XM_023253531.1
A1 82563491 82563598
refGene_gn=TFDP1;refGene_tx=XM_023253517.1;refGene_gn=TFDP1;refGene_tx
=XM_023253524.1;refGene_gn=TFDP1;refGene_tx=XM_023253527.1;refGene_gn=
TFDP1;refGene_tx=XM_023253531.1
A1 82566321 82566443
refGene_gn=TFDP1;refGene_tx=XM_023253517.1;refGene_gn=TFDP1;refGene_tx
=XM_023253524.1;refGene_gn=TFDP1;refGene_tx=XM_023253527.1;refGene_gn=
TFDP1;refGene_tx=XM_023253531.1
A1 82567611 82567777
refGene_gn=TFDP1;refGene_tx=XM_023253517.1;refGene_gn=TFDP1;refGene_tx
=XM_023253524.1;refGene_gn=TFDP1;refGene_tx=XM_023253527.1;refGene_gn=
TFDP1;refGene_tx=XM_023253531.1
A1 82568124 82568268
refGene_gn=TFDP1;refGene_tx=XM_023253517.1;refGene_gn=TFDP1;refGene_tx
=XM_023253524.1;refGene_gn=TFDP1;refGene_tx=XM_023253527.1;refGene_gn=
TFDP1;refGene_tx=XM_023253531.1
A1 82568824 82568893
refGene_gn=TFDP1;refGene_tx=XM_023253517.1;refGene_gn=TFDP1;refGene_tx
=XM_023253524.1;refGene_gn=TFDP1;refGene_tx=XM_023253527.1;refGene_gn=
TFDP1;refGene_tx=XM_023253531.1
A1 82570524 82570676
refGene_gn=TFDP1;refGene_tx=XM_023253517.1;refGene_gn=TFDP1;refGene_tx
=XM_023253524.1;refGene_gn=TFDP1;refGene_tx=XM_023253527.1;refGene_gn=
TFDP1;refGene_tx=XM_023253531.1
A1 82571329 82571535
refGene_gn=TFDP1;refGene_tx=XM_023253517.1;refGene_gn=TFDP1;refGene_tx
=XM_023253524.1;refGene_gn=TFDP1;refGene_tx=XM_023253527.1;refGene_gn=
TFDP1;refGene_tx=XM_023253531.1

A1 82572209 82572285
refGene_gn=TFDP1;refGene_tx=XM_023253517.1;refGene_gn=TFDP1;refGene_tx=XM_023253524.1;refGene_gn=TFDP1;refGene_tx=XM_023253527.1;refGene_gn=TFDP1;refGene_tx=XM_023253531.1
A1 82574977 82575125
refGene_gn=TFDP1;refGene_tx=XM_023253517.1;refGene_gn=TFDP1;refGene_tx=XM_023253524.1;refGene_gn=TFDP1;refGene_tx=XM_023253527.1;refGene_gn=TFDP1;refGene_tx=XM_023253531.1
A1 82577214 82577376 refGene_gn=ATP4B;refGene_tx=XM_023253535.1
A1 82577932 82578031 refGene_gn=ATP4B;refGene_tx=XM_023253535.1
A1 82579288 82579345 refGene_gn=ATP4B;refGene_tx=XM_023253535.1
A1 82580230 82580430 refGene_gn=ATP4B;refGene_tx=XM_023253535.1
A1 82580541 82580655 refGene_gn=ATP4B;refGene_tx=XM_023253535.1
A1 82582449 82582578 refGene_gn=ATP4B;refGene_tx=XM_023253535.1
A1 82583274 82583386 refGene_gn=ATP4B;refGene_tx=XM_023253535.1
A1 82592092 82592791 refGene_gn=GRK1;refGene_tx=XM_003980540.4
A1 82594094 82594222 refGene_gn=GRK1;refGene_tx=XM_003980540.4
A1 82597035 82597193 refGene_gn=GRK1;refGene_tx=XM_003980540.4
A1 82597904 82597988 refGene_gn=GRK1;refGene_tx=XM_003980540.4
A1 82600749 82600874 refGene_gn=GRK1;refGene_tx=XM_003980540.4
A1 82602773 82602975 refGene_gn=GRK1;refGene_tx=XM_003980540.4
A1 82604006 82604305 refGene_gn=GRK1;refGene_tx=XM_003980540.4
A1 82618689 82618735 refGene_gn=TMEM255B;refGene_tx=XM_019829932.2
A1 82623176 82623313 refGene_gn=TMEM255B;refGene_tx=XM_019829932.2
A1 82625642 82625705 refGene_gn=TMEM255B;refGene_tx=XM_019829932.2
A1 82638087 82638177 refGene_gn=TMEM255B;refGene_tx=XM_019829932.2
A1 82646513 82646594 refGene_gn=TMEM255B;refGene_tx=XM_019829932.2
A1 82648849 82648935 refGene_gn=TMEM255B;refGene_tx=XM_019829932.2
A1 82650055 82650185 refGene_gn=TMEM255B;refGene_tx=XM_019829932.2
A1 82650186 82650216 refGene_gn=TMEM255B;refGene_tx=XM_019829932.2
A1 82652843 82652987 refGene_gn=TMEM255B;refGene_tx=XM_019829932.2
A1 82657335 82657470 refGene_gn=TMEM255B;refGene_tx=XM_019829932.2
A1 82664850 82665005 refGene_gn=GAS6;refGene_tx=XM_023253536.1
A1 82666036 82666265 refGene_gn=GAS6;refGene_tx=XM_023253536.1
A1 82667806 82667979 refGene_gn=GAS6;refGene_tx=XM_023253536.1
A1 82670623 82670792 refGene_gn=GAS6;refGene_tx=XM_023253536.1
A1 82671953 82672118 refGene_gn=GAS6;refGene_tx=XM_023253536.1
A1 82673807 82673997 refGene_gn=GAS6;refGene_tx=XM_023253536.1
A1 82674203 82674322 refGene_gn=GAS6;refGene_tx=XM_023253536.1
A1 82674714 82674836 refGene_gn=GAS6;refGene_tx=XM_023253536.1
A1 82675368 82675491 refGene_gn=GAS6;refGene_tx=XM_023253536.1
A1 82678329 82678452 refGene_gn=GAS6;refGene_tx=XM_023253536.1
A1 82679021 82679144 refGene_gn=GAS6;refGene_tx=XM_023253536.1
A1 82685237 82685300 refGene_gn=GAS6;refGene_tx=XM_023253536.1
A1 82686671 82686696 refGene_gn=GAS6;refGene_tx=XM_023253536.1
A1 82697316 82697483 refGene_gn=GAS6;refGene_tx=XM_023253536.1
A1 82697736 82697815 refGene_gn=GAS6;refGene_tx=XM_023253536.1
A1 82773108 82773141
refGene_gn=CA1H13orf46;refGene_tx=XM_006927424.3;refGene_gn=CA1H13orf46;refGene_tx=XM_006927425.3

A1 82773322 82773403
refGene_gn=CA1H13orf46;refGene_tx=XM_006927424.3;refGene_gn=CA1H13orf46;refGene_tx=XM_006927425.3;refGene_gn=CA1H13orf46;refGene_tx=XM_011280980.3
A1 82775598 82775665
refGene_gn=CA1H13orf46;refGene_tx=XM_006927424.3;refGene_gn=CA1H13orf46;refGene_tx=XM_006927425.3
A1 82775885 82775953
refGene_gn=CA1H13orf46;refGene_tx=XM_006927424.3;refGene_gn=CA1H13orf46;refGene_tx=XM_006927425.3;refGene_gn=CA1H13orf46;refGene_tx=XM_011280980.3
A1 82777012 82777060
refGene_gn=CA1H13orf46;refGene_tx=XM_006927424.3;refGene_gn=CA1H13orf46;refGene_tx=XM_011280980.3
A1 82777972 82778020
refGene_gn=CA1H13orf46;refGene_tx=XM_006927424.3;refGene_gn=CA1H13orf46;refGene_tx=XM_006927425.3;refGene_gn=CA1H13orf46;refGene_tx=XM_011280980.3
A1 82778141 82778229
refGene_gn=CA1H13orf46;refGene_tx=XM_006927424.3;refGene_gn=CA1H13orf46;refGene_tx=XM_006927425.3;refGene_gn=CA1H13orf46;refGene_tx=XM_011280980.3
A1 82778904 82778956
refGene_gn=CA1H13orf46;refGene_tx=XM_006927424.3;refGene_gn=CA1H13orf46;refGene_tx=XM_006927425.3;refGene_gn=CA1H13orf46;refGene_tx=XM_011280980.3
A1 82782060 82782250
refGene_gn=CA1H13orf46;refGene_tx=XM_006927424.3;refGene_gn=CA1H13orf46;refGene_tx=XM_006927425.3;refGene_gn=CA1H13orf46;refGene_tx=XM_011280980.3
A1 82786721 82786797 refGene_gn=RASA3;refGene_tx=XM_023239448.1
A1 82787969 82788153 refGene_gn=RASA3;refGene_tx=XM_023239448.1
A1 82792493 82792597 refGene_gn=RASA3;refGene_tx=XM_023239448.1
A1 82794248 82794457 refGene_gn=RASA3;refGene_tx=XM_023239448.1
A1 82800648 82800731 refGene_gn=RASA3;refGene_tx=XM_023239448.1
A1 82801168 82801275 refGene_gn=RASA3;refGene_tx=XM_023239448.1
A1 82801983 82802057 refGene_gn=RASA3;refGene_tx=XM_023239448.1
A1 82802924 82803002 refGene_gn=RASA3;refGene_tx=XM_023239448.1
A1 82804958 82805036 refGene_gn=RASA3;refGene_tx=XM_023239448.1
A1 82805840 82805947 refGene_gn=RASA3;refGene_tx=XM_023239448.1
A1 82807276 82807400 refGene_gn=RASA3;refGene_tx=XM_023239448.1
A1 82808143 82808218 refGene_gn=RASA3;refGene_tx=XM_023239448.1
A1 82809523 82809682 refGene_gn=RASA3;refGene_tx=XM_023239448.1
A1 82812231 82812478 refGene_gn=RASA3;refGene_tx=XM_023239448.1
A1 82814005 82814110 refGene_gn=RASA3;refGene_tx=XM_023239448.1
A1 82816177 82816254 refGene_gn=RASA3;refGene_tx=XM_023239448.1
A1 82819043 82819116 refGene_gn=RASA3;refGene_tx=XM_023239448.1
A1 82819630 82819711 refGene_gn=RASA3;refGene_tx=XM_023239448.1
A1 82830233 82830310 refGene_gn=RASA3;refGene_tx=XM_023239448.1
A1 82837475 82837570 refGene_gn=RASA3;refGene_tx=XM_023239448.1

A1 82846703 82846807 refGene_gn=RASA3;refGene_tx=XM_023239448.1
A1 82857964 82858082 refGene_gn=RASA3;refGene_tx=XM_023239448.1
A1 82907589 82907644 refGene_gn=RASA3;refGene_tx=XM_023239448.1
A1 82941132 82941199
refGene_gn=LOC109499896;refGene_tx=XM_023239486.1
A1 82941549 82941605
refGene_gn=LOC109499896;refGene_tx=XM_023239486.1
A1 82946334 82946463
refGene_gn=LOC109499896;refGene_tx=XM_023239486.1
A1 82960172 82960253
refGene_gn=LOC109499896;refGene_tx=XM_023239486.1
A1 82961166 82961284
refGene_gn=LOC109499896;refGene_tx=XM_023239486.1
A1 82980882 82980902
refGene_gn=LOC109499896;refGene_tx=XM_023239486.1
A1 82997602 82997650
refGene_gn=CDC16;refGene_tx=XM_003980542.4;refGene_gn=CDC16;refGene_tx
=XM_006927426.4
A1 82998639 82998694
refGene_gn=CDC16;refGene_tx=XM_003980542.4;refGene_gn=CDC16;refGene_tx
=XM_006927426.4
A1 82998796 82998894
refGene_gn=CDC16;refGene_tx=XM_003980542.4;refGene_gn=CDC16;refGene_tx
=XM_006927426.4
A1 82999839 82999878
refGene_gn=CDC16;refGene_tx=XM_003980542.4;refGene_gn=CDC16;refGene_tx
=XM_006927426.4
A1 83000219 83000360
refGene_gn=CDC16;refGene_tx=XM_003980542.4;refGene_gn=CDC16;refGene_tx
=XM_006927426.4
A1 83001853 83002013 refGene_gn=CDC16;refGene_tx=XM_003980542.4
A1 83002963 83003055 refGene_gn=CDC16;refGene_tx=XM_003980542.4
A1 83003974 83004108
refGene_gn=CDC16;refGene_tx=XM_003980542.4;refGene_gn=CDC16;refGene_tx
=XM_006927426.4
A1 83004927 83005007
refGene_gn=CDC16;refGene_tx=XM_003980542.4;refGene_gn=CDC16;refGene_tx
=XM_006927426.4
A1 83006036 83006086
refGene_gn=CDC16;refGene_tx=XM_003980542.4;refGene_gn=CDC16;refGene_tx
=XM_006927426.4
A1 83007266 83007340
refGene_gn=CDC16;refGene_tx=XM_003980542.4;refGene_gn=CDC16;refGene_tx
=XM_006927426.4
A1 83009069 83009195
refGene_gn=CDC16;refGene_tx=XM_003980542.4;refGene_gn=CDC16;refGene_tx
=XM_006927426.4
A1 83019658 83019811
refGene_gn=CDC16;refGene_tx=XM_003980542.4;refGene_gn=CDC16;refGene_tx
=XM_006927426.4

A1 83021356 83021420
refGene_gn=CDC16;refGene_tx=XM_003980542.4;refGene_gn=CDC16;refGene_tx=XM_006927426.4
A1 83023826 83023888
refGene_gn=CDC16;refGene_tx=XM_003980542.4;refGene_gn=CDC16;refGene_tx=XM_006927426.4
A1 83025001 83025137
refGene_gn=CDC16;refGene_tx=XM_003980542.4;refGene_gn=CDC16;refGene_tx=XM_006927426.4
A1 83026846 83026937
refGene_gn=CDC16;refGene_tx=XM_003980542.4;refGene_gn=CDC16;refGene_tx=XM_006927426.4
A1 83030415 83030675
refGene_gn=CDC16;refGene_tx=XM_003980542.4;refGene_gn=CDC16;refGene_tx=XM_006927426.4
A1 83042517 83042847
refGene_gn=UPF3A;refGene_tx=XM_019829987.2;refGene_gn=UPF3A;refGene_tx=XM_023253549.1;refGene_gn=UPF3A;refGene_tx=XM_023253552.1;refGene_gn=UPF3A;refGene_tx=XM_023253553.1;refGene_gn=UPF3A;refGene_tx=XM_023253554.1;refGene_gn=UPF3A;refGene_tx=XM_023253556.1;refGene_gn=UPF3A;refGene_tx=XM_023253557.1;refGene_gn=UPF3A;refGene_tx=XM_023253558.1;refGene_gn=UPF3A;refGene_tx=XM_023253560.1
A1 83043853 83043960
refGene_gn=UPF3A;refGene_tx=XM_019829987.2;refGene_gn=UPF3A;refGene_tx=XM_023253549.1;refGene_gn=UPF3A;refGene_tx=XM_023253552.1;refGene_gn=UPF3A;refGene_tx=XM_023253553.1;refGene_gn=UPF3A;refGene_tx=XM_023253554.1;refGene_gn=UPF3A;refGene_tx=XM_023253556.1;refGene_gn=UPF3A;refGene_tx=XM_023253557.1;refGene_gn=UPF3A;refGene_tx=XM_023253558.1;refGene_gn=UPF3A;refGene_tx=XM_023253560.1
A1 83045520 83045627
refGene_gn=UPF3A;refGene_tx=XM_019829987.2;refGene_gn=UPF3A;refGene_tx=XM_023253549.1;refGene_gn=UPF3A;refGene_tx=XM_023253552.1;refGene_gn=UPF3A;refGene_tx=XM_023253553.1;refGene_gn=UPF3A;refGene_tx=XM_023253554.1;refGene_gn=UPF3A;refGene_tx=XM_023253556.1;refGene_gn=UPF3A;refGene_tx=XM_023253557.1;refGene_gn=UPF3A;refGene_tx=XM_023253558.1;refGene_gn=UPF3A;refGene_tx=XM_023253560.1
A1 83048914 83049013
refGene_gn=UPF3A;refGene_tx=XM_019829987.2;refGene_gn=UPF3A;refGene_tx=XM_023253549.1;refGene_gn=UPF3A;refGene_tx=XM_023253552.1;refGene_gn=UPF3A;refGene_tx=XM_023253553.1;refGene_gn=UPF3A;refGene_tx=XM_023253554.1;refGene_gn=UPF3A;refGene_tx=XM_023253556.1;refGene_gn=UPF3A;refGene_tx=XM_023253557.1;refGene_gn=UPF3A;refGene_tx=XM_023253558.1;refGene_gn=UPF3A;refGene_tx=XM_023253560.1
A1 83049137 83049248
refGene_gn=UPF3A;refGene_tx=XM_019829987.2;refGene_gn=UPF3A;refGene_tx=XM_023253549.1;refGene_gn=UPF3A;refGene_tx=XM_023253552.1;refGene_gn=UPF3A;refGene_tx=XM_023253553.1;refGene_gn=UPF3A;refGene_tx=XM_023253554.1;refGene_gn=UPF3A;refGene_tx=XM_023253556.1;refGene_gn=UPF3A;refGene_tx=XM_023253557.1;refGene_gn=UPF3A;refGene_tx=XM_023253558.1;refGene_gn=UPF3A;refGene_tx=XM_023253560.1

A1 83053214 83053270
refGene_gn=UPF3A;refGene_tx=XM_019829987.2;refGene_gn=UPF3A;refGene_tx=XM_023253549.1;refGene_gn=UPF3A;refGene_tx=XM_023253552.1;refGene_gn=UPF3A;refGene_tx=XM_023253553.1;refGene_gn=UPF3A;refGene_tx=XM_023253554.1;refGene_gn=UPF3A;refGene_tx=XM_023253556.1;refGene_gn=UPF3A;refGene_tx=XM_023253557.1
A1 83053327 83053519
refGene_gn=UPF3A;refGene_tx=XM_019829987.2;refGene_gn=UPF3A;refGene_tx=XM_023253549.1;refGene_gn=UPF3A;refGene_tx=XM_023253552.1;refGene_gn=UPF3A;refGene_tx=XM_023253553.1;refGene_gn=UPF3A;refGene_tx=XM_023253554.1;refGene_gn=UPF3A;refGene_tx=XM_023253556.1;refGene_gn=UPF3A;refGene_tx=XM_023253557.1;refGene_gn=UPF3A;refGene_tx=XM_023253558.1;refGene_gn=UPF3A;refGene_tx=XM_023253560.1
A1 83059323 83059488
refGene_gn=UPF3A;refGene_tx=XM_019829987.2;refGene_gn=UPF3A;refGene_tx=XM_023253549.1;refGene_gn=UPF3A;refGene_tx=XM_023253552.1;refGene_gn=UPF3A;refGene_tx=XM_023253553.1;refGene_gn=UPF3A;refGene_tx=XM_023253554.1;refGene_gn=UPF3A;refGene_tx=XM_023253556.1;refGene_gn=UPF3A;refGene_tx=XM_023253557.1
A1 83061340 83061638
refGene_gn=UPF3A;refGene_tx=XM_019829987.2;refGene_gn=UPF3A;refGene_tx=XM_023253549.1;refGene_gn=UPF3A;refGene_tx=XM_023253552.1;refGene_gn=UPF3A;refGene_tx=XM_023253553.1;refGene_gn=UPF3A;refGene_tx=XM_023253554.1;refGene_gn=UPF3A;refGene_tx=XM_023253556.1
A1 83062378 83062473
refGene_gn=UPF3A;refGene_tx=XM_023253549.1;refGene_gn=UPF3A;refGene_tx=XM_023253552.1;refGene_gn=UPF3A;refGene_tx=XM_023253553.1
A1 83063694 83063847
refGene_gn=UPF3A;refGene_tx=XM_019829987.2;refGene_gn=UPF3A;refGene_tx=XM_023253549.1;refGene_gn=UPF3A;refGene_tx=XM_023253552.1;refGene_gn=UPF3A;refGene_tx=XM_023253553.1;refGene_gn=UPF3A;refGene_tx=XM_023253554.1;refGene_gn=UPF3A;refGene_tx=XM_023253556.1
A1 83065659 83065781
refGene_gn=UPF3A;refGene_tx=XM_019829987.2;refGene_gn=UPF3A;refGene_tx=XM_023253549.1;refGene_gn=UPF3A;refGene_tx=XM_023253552.1
A1 83067464 83067472 refGene_gn=UPF3A;refGene_tx=XM_023253553.1
A1 83083839 83086311
refGene_gn=CHAMP1;refGene_tx=XM_023253537.1;refGene_gn=CHAMP1;refGene_tx=XM_023253538.1
A1 83380660 83381053
refGene_gn=LOC101099772;refGene_tx=XM_019831422.1
A1 83381054 83381168
refGene_gn=LOC101099772;refGene_tx=XM_019831422.1
A1 83381170 83381433
refGene_gn=LOC101099772;refGene_tx=XM_019831422.1
A1 83384727 83384809
refGene_gn=LOC101099772;refGene_tx=XM_019831422.1
A1 83398036 83398987
refGene_gn=LOC101094075;refGene_tx=XM_003980544.1
A1 83436821 83437769

refGene_gn=LOC101100025;refGene_tx=XM_011283070.2
A1 83665030 83666002
refGene_gn=LOC101100275;refGene_tx=XM_006927434.2
A1 83691921 83692881
refGene_gn=LOC101094566;refGene_tx=XM_011281170.2
A1 83744263 83745238
refGene_gn=LOC101100535;refGene_tx=XM_003980565.1
A1 83761544 83762498
refGene_gn=LOC101094818;refGene_tx=XM_003980546.2
A1 83793161 83794127
refGene_gn=LOC101100780;refGene_tx=XM_003980566.3
A1 83809841 83810801
refGene_gn=LOC101101026;refGene_tx=XM_003980567.1
A1 83844334 83845291
refGene_gn=LOC101101283;refGene_tx=XM_003980568.1
A1 83950896 83951859
refGene_gn=LOC101095073;refGene_tx=XM_003980547.2
A1 83979281 83980241
refGene_gn=LOC101080353;refGene_tx=XM_006927435.2
A1 84123040 84124000
refGene_gn=LOC101095308;refGene_tx=XM_003980548.1
A1 84274710 84275640
refGene_gn=LOC101081111;refGene_tx=XM_003980570.1
A1 84374086 84374246
refGene_gn=LOC101084082;refGene_tx=XM_023239879.1
A1 84378044 84379011
refGene_gn=LOC101084082;refGene_tx=XM_023239879.1
A1 84401197 84401290
refGene_gn=LOC102899961;refGene_tx=XM_011283103.3
A1 84405396 84405486
refGene_gn=LOC102899961;refGene_tx=XM_011283103.3
A1 84405817 84405981
refGene_gn=LOC102899961;refGene_tx=XM_011283103.3
A1 84413488 84413644
refGene_gn=LOC102899961;refGene_tx=XM_011283103.3
A1 84413801 84413890
refGene_gn=LOC102899961;refGene_tx=XM_011283103.3
A1 84428964 84429155
refGene_gn=LOC102899961;refGene_tx=XM_011283103.3
A1 84493160 84493450 refGene_gn=LYPD8;refGene_tx=XM_019829992.2
A1 84496752 84496890 refGene_gn=LYPD8;refGene_tx=XM_019829992.2
A1 84498669 84498837 refGene_gn=LYPD8;refGene_tx=XM_019829992.2
A1 84500653 84500773 refGene_gn=LYPD8;refGene_tx=XM_019829992.2
A1 84501158 84501213 refGene_gn=LYPD8;refGene_tx=XM_019829992.2
A1 84582900 84583371 refGene_gn=SH3BP5L;refGene_tx=XM_003980571.3
A1 84583944 84584118 refGene_gn=SH3BP5L;refGene_tx=XM_003980571.3
A1 84585137 84585299 refGene_gn=SH3BP5L;refGene_tx=XM_003980571.3
A1 84586996 84587125 refGene_gn=SH3BP5L;refGene_tx=XM_003980571.3
A1 84587290 84587353 refGene_gn=SH3BP5L;refGene_tx=XM_003980571.3
A1 84594026 84594209 refGene_gn=SH3BP5L;refGene_tx=XM_003980571.3

A1 84613135 84613175
refGene_gn=ZNF672;refGene_tx=XM_006927436.4;refGene_gn=ZNF672;refGene_tx=XM_006927439.4
A1 84613967 84614058
refGene_gn=ZNF672;refGene_tx=XM_006927436.4;refGene_gn=ZNF672;refGene_tx=XM_006927439.4
A1 84614824 84616325
refGene_gn=ZNF672;refGene_tx=XM_006927436.4;refGene_gn=ZNF672;refGene_tx=XM_006927439.4;refGene_gn=ZNF672;refGene_tx=XM_019829995.2
A1 84617929 84618233
refGene_gn=ZNF692;refGene_tx=XM_006927442.3;refGene_gn=ZNF692;refGene_tx=XM_006927443.3;refGene_gn=ZNF692;refGene_tx=XM_006927444.3;refGene_gn=ZNF692;refGene_tx=XM_011281197.3;refGene_gn=ZNF692;refGene_tx=XM_019829996.2;refGene_gn=ZNF692;refGene_tx=XM_019830008.2;refGene_gn=ZNF692;refGene_tx=XM_019830009.2;refGene_gn=ZNF692;refGene_tx=XM_023253579.1;refGene_gn=ZNF692;refGene_tx=XM_023253590.1
A1 84618354 84618454
refGene_gn=ZNF692;refGene_tx=XM_006927442.3;refGene_gn=ZNF692;refGene_tx=XM_006927443.3;refGene_gn=ZNF692;refGene_tx=XM_006927444.3;refGene_gn=ZNF692;refGene_tx=XM_011281197.3;refGene_gn=ZNF692;refGene_tx=XM_019829996.2;refGene_gn=ZNF692;refGene_tx=XM_019830008.2;refGene_gn=ZNF692;refGene_tx=XM_019830009.2;refGene_gn=ZNF692;refGene_tx=XM_023253579.1;refGene_gn=ZNF692;refGene_tx=XM_023253590.1
A1 84621057 84621172
refGene_gn=ZNF692;refGene_tx=XM_006927442.3;refGene_gn=ZNF692;refGene_tx=XM_006927443.3;refGene_gn=ZNF692;refGene_tx=XM_006927444.3;refGene_gn=ZNF692;refGene_tx=XM_011281197.3;refGene_gn=ZNF692;refGene_tx=XM_019829996.2;refGene_gn=ZNF692;refGene_tx=XM_019830008.2;refGene_gn=ZNF692;refGene_tx=XM_019830009.2;refGene_gn=ZNF692;refGene_tx=XM_023253579.1;refGene_gn=ZNF692;refGene_tx=XM_023253590.1
A1 84622609 84622688
refGene_gn=ZNF692;refGene_tx=XM_006927442.3;refGene_gn=ZNF692;refGene_tx=XM_006927443.3;refGene_gn=ZNF692;refGene_tx=XM_006927444.3;refGene_gn=ZNF692;refGene_tx=XM_011281197.3;refGene_gn=ZNF692;refGene_tx=XM_019829996.2;refGene_gn=ZNF692;refGene_tx=XM_019830008.2;refGene_gn=ZNF692;refGene_tx=XM_019830009.2;refGene_gn=ZNF692;refGene_tx=XM_023253579.1;refGene_gn=ZNF692;refGene_tx=XM_023253590.1
A1 84622789 84622867
refGene_gn=ZNF692;refGene_tx=XM_006927442.3;refGene_gn=ZNF692;refGene_tx=XM_006927443.3;refGene_gn=ZNF692;refGene_tx=XM_006927444.3;refGene_gn=ZNF692;refGene_tx=XM_019829996.2;refGene_gn=ZNF692;refGene_tx=XM_019830008.2;refGene_gn=ZNF692;refGene_tx=XM_019830009.2;refGene_gn=ZNF692;refGene_tx=XM_023253579.1;refGene_gn=ZNF692;refGene_tx=XM_023253590.1
A1 84622954 84623173
refGene_gn=ZNF692;refGene_tx=XM_006927442.3;refGene_gn=ZNF692;refGene_tx=XM_006927443.3;refGene_gn=ZNF692;refGene_tx=XM_006927444.3;refGene_gn=ZNF692;refGene_tx=XM_011281197.3;refGene_gn=ZNF692;refGene_tx=XM_019829996.2;refGene_gn=ZNF692;refGene_tx=XM_019830008.2;refGene_gn=ZNF692;refGene_tx=XM_019830009.2;refGene_gn=ZNF692;refGene_tx=XM_023253579.

1;refGene_gn=ZNF692;refGene_tx=XM_023253590.1
A1 84623489 84623615
refGene_gn=ZNF692;refGene_tx=XM_006927442.3;refGene_gn=ZNF692;refGene_tx=XM_006927443.3;refGene_gn=ZNF692;refGene_tx=XM_011281197.3;refGene_gn=ZNF692;refGene_tx=XM_019829996.2;refGene_gn=ZNF692;refGene_tx=XM_019830008.2;refGene_gn=ZNF692;refGene_tx=XM_023253579.1;refGene_gn=ZNF692;refGene_tx=XM_023253590.1
A1 84623704 84623753
refGene_gn=ZNF692;refGene_tx=XM_006927442.3;refGene_gn=ZNF692;refGene_tx=XM_006927443.3;refGene_gn=ZNF692;refGene_tx=XM_006927444.3;refGene_gn=ZNF692;refGene_tx=XM_011281197.3;refGene_gn=ZNF692;refGene_tx=XM_019829996.2;refGene_gn=ZNF692;refGene_tx=XM_019830008.2;refGene_gn=ZNF692;refGene_tx=XM_019830009.2;refGene_gn=ZNF692;refGene_tx=XM_023253579.1
A1 84624275 84624539
refGene_gn=ZNF692;refGene_tx=XM_006927442.3;refGene_gn=ZNF692;refGene_tx=XM_006927443.3;refGene_gn=ZNF692;refGene_tx=XM_006927444.3;refGene_gn=ZNF692;refGene_tx=XM_011281197.3;refGene_gn=ZNF692;refGene_tx=XM_019829996.2;refGene_gn=ZNF692;refGene_tx=XM_019830008.2;refGene_gn=ZNF692;refGene_tx=XM_019830009.2;refGene_gn=ZNF692;refGene_tx=XM_023253579.1
A1 84624881 84624913
refGene_gn=ZNF692;refGene_tx=XM_006927442.3;refGene_gn=ZNF692;refGene_tx=XM_006927443.3;refGene_gn=ZNF692;refGene_tx=XM_006927444.3;refGene_gn=ZNF692;refGene_tx=XM_011281197.3;refGene_gn=ZNF692;refGene_tx=XM_019829996.2;refGene_gn=ZNF692;refGene_tx=XM_019830008.2;refGene_gn=ZNF692;refGene_tx=XM_019830009.2;refGene_gn=ZNF692;refGene_tx=XM_023253579.1
A1 84625182 84625358
refGene_gn=ZNF692;refGene_tx=XM_006927442.3;refGene_gn=ZNF692;refGene_tx=XM_006927443.3;refGene_gn=ZNF692;refGene_tx=XM_006927444.3;refGene_gn=ZNF692;refGene_tx=XM_011281197.3;refGene_gn=ZNF692;refGene_tx=XM_019829996.2;refGene_gn=ZNF692;refGene_tx=XM_019830008.2;refGene_gn=ZNF692;refGene_tx=XM_019830009.2;refGene_gn=ZNF692;refGene_tx=XM_023253579.1
A1 84740960 84740977
refGene_gn=PGBD2;refGene_tx=XM_006927446.3;refGene_gn=PGBD2;refGene_tx=XM_023253614.1
A1 84760021 84761783
refGene_gn=PGBD2;refGene_tx=XM_006927446.3;refGene_gn=PGBD2;refGene_tx=XM_019830010.1;refGene_gn=PGBD2;refGene_tx=XM_023253614.1
A1 85029832 85030816
refGene_gn=LOC101086111;refGene_tx=XM_023253606.1
A1 85085040 85085979
refGene_gn=LOC101086367;refGene_tx=XM_003980583.3
A1 85183413 85184382
refGene_gn=LOC101086869;refGene_tx=XM_003980584.1
A1 85215288 85215556
refGene_gn=LOC101087125;refGene_tx=XM_011283147.2
A1 85225562 85226511

refGene_gn=LOC101087125;refGene_tx=XM_011283147.2
A1 85243332 85243396
refGene_gn=LOC101087125;refGene_tx=XM_011283147.2
A1 85322528 85323464
refGene_gn=LOC101087376;refGene_tx=XM_011283154.1
A1 85387944 85388886
refGene_gn=LOC101087635;refGene_tx=XM_003980587.1
A1 85428977 85429937
refGene_gn=LOC101087896;refGene_tx=XM_003980588.1
A1 85454563 85455520
refGene_gn=LOC111556948;refGene_tx=XM_023240315.1
A1 85551364 85552321
refGene_gn=LOC111556985;refGene_tx=XM_023240355.1
A1 85580186 85580393
refGene_gn=LOC102900767;refGene_tx=XM_011283180.1
A1 85580395 85581148
refGene_gn=LOC102900767;refGene_tx=XM_011283180.1
A1 85631232 85632204
refGene_gn=LOC101088904;refGene_tx=XM_003980591.1
A1 85647808 85648231 refGene_gn=TRIM58;refGene_tx=XM_003980592.2
A1 85652099 85652195 refGene_gn=TRIM58;refGene_tx=XM_003980592.2
A1 85655935 85656166 refGene_gn=TRIM58;refGene_tx=XM_003980592.2
A1 85658904 85658927 refGene_gn=TRIM58;refGene_tx=XM_003980592.2
A1 85659024 85659125 refGene_gn=TRIM58;refGene_tx=XM_003980592.2
A1 85663303 85663893 refGene_gn=TRIM58;refGene_tx=XM_003980592.2
A1 85674969 85675914
refGene_gn=LOC101082541;refGene_tx=XM_003980573.1;refGene_gn=LOC101082541;refGene_tx=XM_023253727.1
A1 85679861 85679901
refGene_gn=LOC101082541;refGene_tx=XM_023253727.1
A1 85715444 85716356
refGene_gn=LOC101089671;refGene_tx=XM_003980593.2
A1 85807142 85808058
refGene_gn=LOC101089917;refGene_tx=XM_003980594.2
A1 85808096 85808134
refGene_gn=LOC101089917;refGene_tx=XM_003980594.2
A1 85821919 85822858
refGene_gn=LOC101082797;refGene_tx=XM_003980574.2
A1 85856671 85857607
refGene_gn=LOC101090164;refGene_tx=XM_003980595.1
A1 85893723 85894662
refGene_gn=LOC101083056;refGene_tx=XM_003980575.2
A1 86045102 86045674
refGene_gn=LOC109500764;refGene_tx=XM_019832822.1
A1 86047210 86047237
refGene_gn=LOC109500764;refGene_tx=XM_019832822.1
A1 86047478 86047499
refGene_gn=LOC109500764;refGene_tx=XM_019832822.1
A1 86058535 86058556
refGene_gn=LOC109500764;refGene_tx=XM_019832822.1

A1 86073992 86074013
refGene_gn=LOC109500764;refGene_tx=XM_019832822.1
A1 86104062 86104176
refGene_gn=LOC109500764;refGene_tx=XM_019832822.1
A1 86106125 86106407
refGene_gn=LOC109500764;refGene_tx=XM_019832822.1
A1 86112110 86112458
refGene_gn=LOC109500764;refGene_tx=XM_019832822.1
A1 86115566 86115663
refGene_gn=LOC109500764;refGene_tx=XM_019832822.1
A1 86172695 86173628
refGene_gn=LOC105260451;refGene_tx=XM_011281225.1
A1 86265719 86266316
refGene_gn=LOC109500775;refGene_tx=XM_023240613.1
A1 86269564 86269690
refGene_gn=LOC109500775;refGene_tx=XM_023240613.1
A1 86274893 86274914
refGene_gn=LOC109500775;refGene_tx=XM_023240613.1
A1 86343544 86343595
refGene_gn=LOC109500775;refGene_tx=XM_023240613.1
A1 86353216 86353490
refGene_gn=LOC109500775;refGene_tx=XM_023240613.1
A1 86555156 86555369
refGene_gn=LOC101080302;refGene_tx=XM_019832928.2
A1 86593223 86593373
refGene_gn=LOC101080302;refGene_tx=XM_019832928.2
A1 86615729 86616281
refGene_gn=LOC101080302;refGene_tx=XM_019832928.2
A1 86616282 86616678
refGene_gn=LOC101080302;refGene_tx=XM_019832928.2
A1 86694731 86695667
refGene_gn=LOC101101231;refGene_tx=XM_003988577.1
A1 86710763 86710778
refGene_gn=LOC101100977;refGene_tx=XM_003988576.2
A1 86711728 86711747
refGene_gn=LOC101100977;refGene_tx=XM_003988576.2
A1 86716688 86717590
refGene_gn=LOC101100977;refGene_tx=XM_003988576.2
A1 86845024 86845978
refGene_gn=LOC101099980;refGene_tx=XM_003988573.2
A1 86880124 86881066
refGene_gn=LOC101098527;refGene_tx=XM_011281230.1
A1 86902377 86903307
refGene_gn=LOC101093328;refGene_tx=XM_011283236.1
A1 86923347 86924295
refGene_gn=LOC101093577;refGene_tx=XM_003980603.1
A1 86934172 86935123
refGene_gn=LOC101093829;refGene_tx=XM_003980604.1
A1 86957999 86958944
refGene_gn=LOC101091078;refGene_tx=XM_003980597.1

A1 87003189 87004137
refGene_gn=LOC101094321;refGene_tx=XM_003980605.1
A1 87029190 87030117
refGene_gn=LOC101091338;refGene_tx=XM_003980598.1
A1 87087665 87088589
refGene_gn=LOC101091589;refGene_tx=XM_003980599.1
A1 87182110 87183049
refGene_gn=LOC101091839;refGene_tx=XM_003980600.2
A1 87212240 87212588
refGene_gn=LOC101094567;refGene_tx=XM_011283251.2
A1 87212590 87212803
refGene_gn=LOC101094567;refGene_tx=XM_011283251.2
A1 87212804 87213197
refGene_gn=LOC101094567;refGene_tx=XM_011283251.2
A1 87234416 87235349
refGene_gn=LOC101094819;refGene_tx=XM_011281246.1
A1 87299099 87299342
refGene_gn=GCSAML;refGene_tx=XM_019813897.2;refGene_gn=GCSAML;refGene_tx=XM_019813899.1
A1 87300217 87300246
refGene_gn=GCSAML;refGene_tx=XM_019813897.2;refGene_gn=GCSAML;refGene_tx=XM_019813899.1
A1 87312218 87312268
refGene_gn=GCSAML;refGene_tx=XM_019813897.2;refGene_gn=GCSAML;refGene_tx=XM_019813899.1
A1 87318214 87318274 refGene_gn=GCSAML;refGene_tx=XM_019813897.2
A1 87325949 87325978
refGene_gn=GCSAML;refGene_tx=XM_019813897.2;refGene_gn=GCSAML;refGene_tx=XM_019813899.1
A1 87345739 87346720
refGene_gn=LOC101095546;refGene_tx=XM_003980607.1
A1 87376135 87377083
refGene_gn=LOC101095792;refGene_tx=XM_003980608.2
A1 87418136 87419087
refGene_gn=LOC101096031;refGene_tx=XM_003980609.1
A1 87443097 87444042
refGene_gn=LOC101096283;refGene_tx=XM_003980610.4
A1 87473483 87474416
refGene_gn=LOC101096535;refGene_tx=XM_003980611.3
A1 87494865 87495834
refGene_gn=LOC101096784;refGene_tx=XM_003980612.2
A1 87502892 87502940
refGene_gn=LOC101097040;refGene_tx=XM_019833838.1
A1 87531645 87531713
refGene_gn=LOC101097040;refGene_tx=XM_019833838.1
A1 87535931 87536892
refGene_gn=LOC101097040;refGene_tx=XM_019833838.1
A1 87550798 87550898 refGene_gn=NLRP3;refGene_tx=XM_023253644.1
A1 87554382 87554553 refGene_gn=NLRP3;refGene_tx=XM_023253644.1
A1 87555272 87555443 refGene_gn=NLRP3;refGene_tx=XM_023253644.1

A1 87584030 87584201 refGene_gn=NLRP3;refGene_tx=XM_023253644.1
A1 87589343 87589514 refGene_gn=NLRP3;refGene_tx=XM_023253644.1
A1 87591637 87591808 refGene_gn=NLRP3;refGene_tx=XM_023253644.1
A1 87594440 87596202 refGene_gn=NLRP3;refGene_tx=XM_023253644.1
A1 87596928 87597027 refGene_gn=NLRP3;refGene_tx=XM_023253644.1
A1 87602022 87602305 refGene_gn=NLRP3;refGene_tx=XM_023253644.1
A1 87674042 87674432
refGene_gn=ZNF496;refGene_tx=XM_023253645.1;refGene_gn=ZNF496;refGene_tx=XM_023253648.1;refGene_gn=ZNF496;refGene_tx=XM_023253656.1;refGene_gn=ZNF496;refGene_tx=XM_023253661.1;refGene_gn=ZNF496;refGene_tx=XM_023253669.1;refGene_gn=ZNF496;refGene_tx=XM_023253675.1;refGene_gn=ZNF496;refGene_tx=XM_023253682.1;refGene_gn=ZNF496;refGene_tx=XM_023253683.1
A1 87674706 87674890
refGene_gn=ZNF496;refGene_tx=XM_023253645.1;refGene_gn=ZNF496;refGene_tx=XM_023253648.1;refGene_gn=ZNF496;refGene_tx=XM_023253656.1;refGene_gn=ZNF496;refGene_tx=XM_023253661.1;refGene_gn=ZNF496;refGene_tx=XM_023253669.1;refGene_gn=ZNF496;refGene_tx=XM_023253675.1;refGene_gn=ZNF496;refGene_tx=XM_023253682.1;refGene_gn=ZNF496;refGene_tx=XM_023253683.1
A1 87678622 87678699
refGene_gn=ZNF496;refGene_tx=XM_023253645.1;refGene_gn=ZNF496;refGene_tx=XM_023253648.1;refGene_gn=ZNF496;refGene_tx=XM_023253656.1;refGene_gn=ZNF496;refGene_tx=XM_023253661.1;refGene_gn=ZNF496;refGene_tx=XM_023253669.1;refGene_gn=ZNF496;refGene_tx=XM_023253675.1;refGene_gn=ZNF496;refGene_tx=XM_023253682.1;refGene_gn=ZNF496;refGene_tx=XM_023253683.1
A1 87679585 87679648
refGene_gn=ZNF496;refGene_tx=XM_023253645.1;refGene_gn=ZNF496;refGene_tx=XM_023253648.1;refGene_gn=ZNF496;refGene_tx=XM_023253656.1;refGene_gn=ZNF496;refGene_tx=XM_023253661.1
A1 87697974 87698107
refGene_gn=ZNF496;refGene_tx=XM_023253645.1;refGene_gn=ZNF496;refGene_tx=XM_023253648.1;refGene_gn=ZNF496;refGene_tx=XM_023253656.1;refGene_gn=ZNF496;refGene_tx=XM_023253661.1;refGene_gn=ZNF496;refGene_tx=XM_023253669.1;refGene_gn=ZNF496;refGene_tx=XM_023253675.1;refGene_gn=ZNF496;refGene_tx=XM_023253682.1;refGene_gn=ZNF496;refGene_tx=XM_023253683.1
A1 87698733 87698841
refGene_gn=ZNF496;refGene_tx=XM_023253645.1;refGene_gn=ZNF496;refGene_tx=XM_023253648.1;refGene_gn=ZNF496;refGene_tx=XM_023253656.1;refGene_gn=ZNF496;refGene_tx=XM_023253661.1;refGene_gn=ZNF496;refGene_tx=XM_023253669.1;refGene_gn=ZNF496;refGene_tx=XM_023253675.1;refGene_gn=ZNF496;refGene_tx=XM_023253682.1;refGene_gn=ZNF496;refGene_tx=XM_023253683.1
A1 87699828 87699942
refGene_gn=ZNF496;refGene_tx=XM_023253645.1;refGene_gn=ZNF496;refGene_tx=XM_023253648.1;refGene_gn=ZNF496;refGene_tx=XM_023253656.1;refGene_gn=ZNF496;refGene_tx=XM_023253661.1;refGene_gn=ZNF496;refGene_tx=XM_023253669.1;refGene_gn=ZNF496;refGene_tx=XM_023253675.1;refGene_gn=ZNF49

6;refGene_tx=XM_023253682.1;refGene_gn=ZNF496;refGene_tx=XM_023253683.
1
A1 87705494 87706231
refGene_gn=ZNF496;refGene_tx=XM_023253645.1;refGene_gn=ZNF496;refGene_
tx=XM_023253648.1;refGene_gn=ZNF496;refGene_tx=XM_023253656.1;refGene_
gn=ZNF496;refGene_tx=XM_023253661.1;refGene_gn=ZNF496;refGene_tx=XM_02
3253669.1;refGene_gn=ZNF496;refGene_tx=XM_023253675.1;refGene_gn=ZNF49
6;refGene_tx=XM_023253682.1;refGene_gn=ZNF496;refGene_tx=XM_023253683.
1
A1 87759881 87760829
refGene_gn=LOC101097294;refGene_tx=XM_019833844.1
A1 87858589 87858753
refGene_gn=BTNL10;refGene_tx=XM_019830106.2;refGene_gn=BTNL10;refGene_
tx=XM_019830112.2;refGene_gn=BTNL10;refGene_tx=XM_019830122.2
A1 87859839 87860121
refGene_gn=BTNL10;refGene_tx=XM_019830106.2;refGene_gn=BTNL10;refGene_
tx=XM_019830112.2;refGene_gn=BTNL10;refGene_tx=XM_019830122.2
A1 87860630 87860984
refGene_gn=BTNL10;refGene_tx=XM_019830106.2;refGene_gn=BTNL10;refGene_
tx=XM_019830112.2;refGene_gn=BTNL10;refGene_tx=XM_019830122.2
A1 87861270 87861364
refGene_gn=BTNL10;refGene_tx=XM_019830106.2;refGene_gn=BTNL10;refGene_
tx=XM_019830112.2;refGene_gn=BTNL10;refGene_tx=XM_019830122.2
A1 87898207 87898301
refGene_gn=LOC101097782;refGene_tx=XM_006927459.3;refGene_gn=LOC101097
782;refGene_tx=XM_023253695.1
A1 87898615 87898969
refGene_gn=LOC101097782;refGene_tx=XM_006927459.3;refGene_gn=LOC101097
782;refGene_tx=XM_023253695.1
A1 87899563 87899845
refGene_gn=LOC101097782;refGene_tx=XM_006927459.3;refGene_gn=LOC101097
782;refGene_tx=XM_023253695.1
A1 87900923 87901115
refGene_gn=LOC101097782;refGene_tx=XM_006927459.3;refGene_gn=LOC101097
782;refGene_tx=XM_023253695.1
A1 87901829 87901853
refGene_gn=LOC101097782;refGene_tx=XM_006927459.3;refGene_gn=LOC101097
782;refGene_tx=XM_023253695.1
A1 87906544 87907143
refGene_gn=LOC101097782;refGene_tx=XM_006927459.3;refGene_gn=LOC101097
782;refGene_tx=XM_023253695.1
A1 87924987 87924990 refGene_gn=RNF187;refGene_tx=XM_023241705.1
A1 87925131 87925353 refGene_gn=RNF187;refGene_tx=XM_023241705.1
A1 87925554 87925647 refGene_gn=RNF187;refGene_tx=XM_023241705.1
A1 87927067 87927846 refGene_gn=RNF187;refGene_tx=XM_023241705.1
A1 87928024 87928534 refGene_gn=RNF187;refGene_tx=XM_023241705.1
A1 87932741 87933078 refGene_gn=RNF187;refGene_tx=XM_023241705.1
A1 87933827 87933983 refGene_gn=RNF187;refGene_tx=XM_023241705.1
A1 87936151 87936226 refGene_gn=RNF187;refGene_tx=XM_023241705.1
A1 87963375 87963756

refGene_gn=LOC101098354;refGene_tx=XM_019830161.2
A1 87964068 87964461
refGene_gn=LOC101098601;refGene_tx=XM_006927460.3
A1 87989408 87989547
refGene_gn=LOC101101285;refGene_tx=XM_011281294.3
A1 87989549 87989806
refGene_gn=LOC101101285;refGene_tx=XM_011281294.3
A1 88004064 88004493
refGene_gn=TRIM17;refGene_tx=XM_003980620.3;refGene_gn=TRIM17;refGene_tx=XM_023253715.1
A1 88005195 88005291 refGene_gn=TRIM17;refGene_tx=XM_003980620.3
A1 88005673 88005904
refGene_gn=TRIM17;refGene_tx=XM_003980620.3;refGene_gn=TRIM17;refGene_tx=XM_023253715.1
A1 88007716 88007739
refGene_gn=TRIM17;refGene_tx=XM_003980620.3;refGene_gn=TRIM17;refGene_tx=XM_023253715.1
A1 88007829 88007930
refGene_gn=TRIM17;refGene_tx=XM_003980620.3;refGene_gn=TRIM17;refGene_tx=XM_023253715.1
A1 88008174 88008722
refGene_gn=TRIM17;refGene_tx=XM_003980620.3;refGene_gn=TRIM17;refGene_tx=XM_023253715.1
A1 88009914 88010322 refGene_gn=TRIM11;refGene_tx=XM_023253716.1
A1 88012986 88013082 refGene_gn=TRIM11;refGene_tx=XM_023253716.1
A1 88013280 88013511 refGene_gn=TRIM11;refGene_tx=XM_023253716.1
A1 88015833 88015856 refGene_gn=TRIM11;refGene_tx=XM_023253716.1
A1 88015944 88016045 refGene_gn=TRIM11;refGene_tx=XM_023253716.1
A1 88016565 88017113 refGene_gn=TRIM11;refGene_tx=XM_023253716.1
A1 88037030 88037208 refGene_gn=OBSCN;refGene_tx=XM_023253724.1
A1 88037352 88037502 refGene_gn=OBSCN;refGene_tx=XM_023253724.1
A1 88037635 88037711 refGene_gn=OBSCN;refGene_tx=XM_023253724.1
A1 88037833 88038057 refGene_gn=OBSCN;refGene_tx=XM_023253724.1
A1 88038284 88038524 refGene_gn=OBSCN;refGene_tx=XM_023253724.1
A1 88038745 88038830 refGene_gn=OBSCN;refGene_tx=XM_023253724.1
A1 88039263 88039434 refGene_gn=OBSCN;refGene_tx=XM_023253724.1
A1 88039611 88039717 refGene_gn=OBSCN;refGene_tx=XM_023253724.1
A1 88039718 88039735 refGene_gn=OBSCN;refGene_tx=XM_023253724.1
A1 88040647 88040815 refGene_gn=OBSCN;refGene_tx=XM_023253724.1
A1 88040962 88041073 refGene_gn=OBSCN;refGene_tx=XM_023253724.1
A1 88041390 88041468 refGene_gn=OBSCN;refGene_tx=XM_023253724.1
A1 88041965 88043991 refGene_gn=OBSCN;refGene_tx=XM_023253724.1
A1 88044377 88044455 refGene_gn=OBSCN;refGene_tx=XM_023253724.1
A1 88044589 88044635 refGene_gn=OBSCN;refGene_tx=XM_023253724.1
A1 88044732 88044885 refGene_gn=OBSCN;refGene_tx=XM_023253724.1
A1 88045457 88045484 refGene_gn=OBSCN;refGene_tx=XM_023253724.1
A1 88045627 88045800 refGene_gn=OBSCN;refGene_tx=XM_023253724.1
A1 88046108 88046186 refGene_gn=OBSCN;refGene_tx=XM_023253724.1
A1 88046573 88046634 refGene_gn=OBSCN;refGene_tx=XM_023253724.1
A1 88047137 88047338 refGene_gn=OBSCN;refGene_tx=XM_023253724.1

| | | | |
|---|---|---|---|
| A1 | 88048057 | 88048139 | refGene_gn=OBSCN;refGene_tx=XM_023253724.1 |
| A1 | 88048288 | 88048457 | refGene_gn=OBSCN;refGene_tx=XM_023253724.1 |
| A1 | 88048603 | 88048716 | refGene_gn=OBSCN;refGene_tx=XM_023253724.1 |
| A1 | 88049270 | 88049392 | refGene_gn=OBSCN;refGene_tx=XM_023253724.1 |
| A1 | 88049626 | 88049723 | refGene_gn=OBSCN;refGene_tx=XM_023253724.1 |
| A1 | 88050782 | 88051067 | refGene_gn=OBSCN;refGene_tx=XM_023253724.1 |
| A1 | 88058554 | 88058663 | refGene_gn=OBSCN;refGene_tx=XM_023253724.1 |
| A1 | 88059651 | 88059781 | refGene_gn=OBSCN;refGene_tx=XM_023253724.1 |
| A1 | 88059940 | 88060043 | refGene_gn=OBSCN;refGene_tx=XM_023253724.1 |
| A1 | 88060254 | 88060422 | refGene_gn=OBSCN;refGene_tx=XM_023253724.1 |
| A1 | 88065005 | 88065119 | refGene_gn=OBSCN;refGene_tx=XM_023253724.1 |
| A1 | 88065541 | 88065719 | refGene_gn=OBSCN;refGene_tx=XM_023253724.1 |
| A1 | 88065828 | 88065987 | refGene_gn=OBSCN;refGene_tx=XM_023253724.1 |
| A1 | 88066339 | 88066504 | refGene_gn=OBSCN;refGene_tx=XM_023253724.1 |
| A1 | 88066600 | 88066717 | refGene_gn=OBSCN;refGene_tx=XM_023253724.1 |
| A1 | 88066965 | 88067122 | refGene_gn=OBSCN;refGene_tx=XM_023253724.1 |
| A1 | 88067721 | 88067895 | refGene_gn=OBSCN;refGene_tx=XM_023253724.1 |
| A1 | 88068071 | 88068154 | refGene_gn=OBSCN;refGene_tx=XM_023253724.1 |
| A1 | 88068263 | 88068464 | refGene_gn=OBSCN;refGene_tx=XM_023253724.1 |
| A1 | 88068815 | 88068955 | refGene_gn=OBSCN;refGene_tx=XM_023253724.1 |
| A1 | 88069087 | 88069207 | refGene_gn=OBSCN;refGene_tx=XM_023253724.1 |
| A1 | 88069904 | 88069972 | refGene_gn=OBSCN;refGene_tx=XM_023253724.1 |
| A1 | 88070182 | 88070255 | refGene_gn=OBSCN;refGene_tx=XM_023253724.1 |
| A1 | 88070577 | 88070796 | refGene_gn=OBSCN;refGene_tx=XM_023253724.1 |
| A1 | 88070994 | 88071093 | refGene_gn=OBSCN;refGene_tx=XM_023253724.1 |
| A1 | 88071306 | 88071339 | refGene_gn=OBSCN;refGene_tx=XM_023253724.1 |
| A1 | 88071763 | 88071961 | refGene_gn=OBSCN;refGene_tx=XM_023253724.1 |
| A1 | 88072093 | 88072218 | refGene_gn=OBSCN;refGene_tx=XM_023253724.1 |
| A1 | 88072430 | 88072509 | refGene_gn=OBSCN;refGene_tx=XM_023253724.1 |
| A1 | 88080561 | 88080840 | refGene_gn=OBSCN;refGene_tx=XM_023253724.1 |
| A1 | 88081908 | 88082773 | refGene_gn=OBSCN;refGene_tx=XM_023253724.1 |
| A1 | 88083805 | 88084179 | refGene_gn=OBSCN;refGene_tx=XM_023253724.1 |
| A1 | 88085457 | 88085817 | refGene_gn=OBSCN;refGene_tx=XM_023253724.1 |
| A1 | 88085932 | 88086220 | refGene_gn=OBSCN;refGene_tx=XM_023253724.1 |
| A1 | 88086492 | 88086768 | refGene_gn=OBSCN;refGene_tx=XM_023253724.1 |
| A1 | 88087093 | 88087366 | refGene_gn=OBSCN;refGene_tx=XM_023253724.1 |
| A1 | 88089190 | 88089457 | refGene_gn=OBSCN;refGene_tx=XM_023253724.1 |
| A1 | 88090398 | 88090671 | refGene_gn=OBSCN;refGene_tx=XM_023253724.1 |
| A1 | 88093016 | 88093092 | refGene_gn=OBSCN;refGene_tx=XM_023253724.1 |
| A1 | 88093673 | 88093870 | refGene_gn=OBSCN;refGene_tx=XM_023253724.1 |
| A1 | 88094118 | 88094391 | refGene_gn=OBSCN;refGene_tx=XM_023253724.1 |
| A1 | 88094817 | 88095090 | refGene_gn=OBSCN;refGene_tx=XM_023253724.1 |
| A1 | 88095194 | 88095461 | refGene_gn=OBSCN;refGene_tx=XM_023253724.1 |
| A1 | 88096265 | 88096532 | refGene_gn=OBSCN;refGene_tx=XM_023253724.1 |
| A1 | 88096848 | 88097112 | refGene_gn=OBSCN;refGene_tx=XM_023253724.1 |
| A1 | 88097448 | 88097712 | refGene_gn=OBSCN;refGene_tx=XM_023253724.1 |
| A1 | 88107880 | 88108144 | refGene_gn=OBSCN;refGene_tx=XM_023253724.1 |
| A1 | 88109309 | 88109573 | refGene_gn=OBSCN;refGene_tx=XM_023253724.1 |
| A1 | 88110106 | 88110370 | refGene_gn=OBSCN;refGene_tx=XM_023253724.1 |
| A1 | 88113290 | 88113554 | refGene_gn=OBSCN;refGene_tx=XM_023253724.1 |

A1 88113895 88114159 refGene_gn=OBSCN;refGene_tx=XM_023253724.1
A1 88114500 88114764 refGene_gn=OBSCN;refGene_tx=XM_023253724.1
A1 88115105 88115369 refGene_gn=OBSCN;refGene_tx=XM_023253724.1
A1 88118466 88118730 refGene_gn=OBSCN;refGene_tx=XM_023253724.1
A1 88119047 88119311 refGene_gn=OBSCN;refGene_tx=XM_023253724.1
A1 88119398 88119665 refGene_gn=OBSCN;refGene_tx=XM_023253724.1
A1 88120423 88120690 refGene_gn=OBSCN;refGene_tx=XM_023253724.1
A1 88121003 88121276 refGene_gn=OBSCN;refGene_tx=XM_023253724.1
A1 88122842 88123109 refGene_gn=OBSCN;refGene_tx=XM_023253724.1
A1 88123252 88123519 refGene_gn=OBSCN;refGene_tx=XM_023253724.1
A1 88124116 88124389 refGene_gn=OBSCN;refGene_tx=XM_023253724.1
A1 88126553 88126820 refGene_gn=OBSCN;refGene_tx=XM_023253724.1
A1 88126923 88127190 refGene_gn=OBSCN;refGene_tx=XM_023253724.1
A1 88127565 88127832 refGene_gn=OBSCN;refGene_tx=XM_023253724.1
A1 88127960 88128230 refGene_gn=OBSCN;refGene_tx=XM_023253724.1
A1 88128327 88128594 refGene_gn=OBSCN;refGene_tx=XM_023253724.1
A1 88129287 88129392 refGene_gn=OBSCN;refGene_tx=XM_023253724.1
A1 88129479 88129641 refGene_gn=OBSCN;refGene_tx=XM_023253724.1
A1 88129820 88129974 refGene_gn=OBSCN;refGene_tx=XM_023253724.1
A1 88130072 88130185 refGene_gn=OBSCN;refGene_tx=XM_023253724.1
A1 88130353 88130620 refGene_gn=OBSCN;refGene_tx=XM_023253724.1
A1 88130874 88131141 refGene_gn=OBSCN;refGene_tx=XM_023253724.1
A1 88131927 88132197 refGene_gn=OBSCN;refGene_tx=XM_023253724.1
A1 88132320 88132587 refGene_gn=OBSCN;refGene_tx=XM_023253724.1
A1 88132782 88133067 refGene_gn=OBSCN;refGene_tx=XM_023253724.1
A1 88135957 88136233 refGene_gn=OBSCN;refGene_tx=XM_023253724.1
A1 88140234 88140510 refGene_gn=OBSCN;refGene_tx=XM_023253724.1
A1 88142465 88142741 refGene_gn=OBSCN;refGene_tx=XM_023253724.1
A1 88144604 88144880 refGene_gn=OBSCN;refGene_tx=XM_023253724.1
A1 88148227 88148503 refGene_gn=OBSCN;refGene_tx=XM_023253724.1
A1 88152278 88152554 refGene_gn=OBSCN;refGene_tx=XM_023253724.1
A1 88164623 88164899 refGene_gn=OBSCN;refGene_tx=XM_023253724.1
A1 88168253 88168529 refGene_gn=OBSCN;refGene_tx=XM_023253724.1
A1 88172293 88172569 refGene_gn=OBSCN;refGene_tx=XM_023253724.1
A1 88175514 88175790 refGene_gn=OBSCN;refGene_tx=XM_023253724.1
A1 88192983 88193259 refGene_gn=OBSCN;refGene_tx=XM_023253724.1
A1 88193899 88194175 refGene_gn=OBSCN;refGene_tx=XM_023253724.1
A1 88194594 88194870 refGene_gn=OBSCN;refGene_tx=XM_023253724.1
A1 88195579 88195846 refGene_gn=OBSCN;refGene_tx=XM_023253724.1
A1 88195968 88196277 refGene_gn=OBSCN;refGene_tx=XM_023253724.1
A1 88197029 88197290 refGene_gn=OBSCN;refGene_tx=XM_023253724.1
A1 88197981 88198251 refGene_gn=OBSCN;refGene_tx=XM_023253724.1
A1 88199408 88200387 refGene_gn=OBSCN;refGene_tx=XM_023253724.1
A1 88236320 88236712 refGene_gn=IBA57;refGene_tx=XM_023253725.1
A1 88236859 88237209 refGene_gn=IBA57;refGene_tx=XM_023253725.1
A1 88244194 88244535 refGene_gn=IBA57;refGene_tx=XM_023253725.1
A1 88252092 88253382
refGene_gn=GJC2;refGene_tx=XM_006927462.3;refGene_gn=GJC2;refGene_tx=X
M_019830194.2
A1 88254432 88254804 refGene_gn=GJC2;refGene_tx=XM_006927462.3

A1 88262033 88262069
refGene_gn=GUK1;refGene_tx=XM_006927463.3;refGene_gn=GUK1;refGene_tx=XM_019830200.1;refGene_gn=GUK1;refGene_tx=XM_023253733.1;refGene_gn=GUK1;refGene_tx=XM_023253735.1
A1 88262181 88262267
refGene_gn=GUK1;refGene_tx=XM_006927463.3;refGene_gn=GUK1;refGene_tx=XM_019830200.1;refGene_gn=GUK1;refGene_tx=XM_023253733.1;refGene_gn=GUK1;refGene_tx=XM_023253735.1
A1 88262382 88262467
refGene_gn=GUK1;refGene_tx=XM_006927463.3;refGene_gn=GUK1;refGene_tx=XM_019830200.1;refGene_gn=GUK1;refGene_tx=XM_023253733.1;refGene_gn=GUK1;refGene_tx=XM_023253735.1
A1 88262553 88262692
refGene_gn=GUK1;refGene_tx=XM_006927463.3;refGene_gn=GUK1;refGene_tx=XM_019830200.1;refGene_gn=GUK1;refGene_tx=XM_023253733.1;refGene_gn=GUK1;refGene_tx=XM_023253735.1
A1 88263415 88263512
refGene_gn=GUK1;refGene_tx=XM_006927463.3;refGene_gn=GUK1;refGene_tx=XM_019830200.1;refGene_gn=GUK1;refGene_tx=XM_023253733.1;refGene_gn=GUK1;refGene_tx=XM_023253735.1
A1 88264262 88264304
refGene_gn=GUK1;refGene_tx=XM_006927463.3;refGene_gn=GUK1;refGene_tx=XM_019830200.1;refGene_gn=GUK1;refGene_tx=XM_023253733.1;refGene_gn=GUK1;refGene_tx=XM_023253735.1
A1 88264541 88264655
refGene_gn=GUK1;refGene_tx=XM_006927463.3;refGene_gn=GUK1;refGene_tx=XM_019830200.1;refGene_gn=GUK1;refGene_tx=XM_023253733.1;refGene_gn=GUK1;refGene_tx=XM_023253735.1
A1 88268369 88268415 refGene_gn=GUK1;refGene_tx=XM_023253735.1
A1 88268855 88268916 refGene_gn=GUK1;refGene_tx=XM_023253733.1
A1 88270256 88270309
refGene_gn=MRPL55;refGene_tx=XM_003980624.5;refGene_gn=MRPL55;refGene_tx=XM_019830208.2
A1 88270346 88270408 refGene_gn=MRPL55;refGene_tx=XM_006927464.3
A1 88272113 88272315
refGene_gn=MRPL55;refGene_tx=XM_003980624.5;refGene_gn=MRPL55;refGene_tx=XM_006927464.3;refGene_gn=MRPL55;refGene_tx=XM_019830208.2
A1 88272665 88272818
refGene_gn=MRPL55;refGene_tx=XM_003980624.5;refGene_gn=MRPL55;refGene_tx=XM_006927464.3;refGene_gn=MRPL55;refGene_tx=XM_019830208.2
A1 88274912 88275006
refGene_gn=CA1H1orf35;refGene_tx=XM_023241736.1
A1 88275158 88275309
refGene_gn=CA1H1orf35;refGene_tx=XM_023241736.1
A1 88275427 88275473
refGene_gn=CA1H1orf35;refGene_tx=XM_023241736.1
A1 88275559 88275642
refGene_gn=CA1H1orf35;refGene_tx=XM_023241736.1
A1 88275753 88275826
refGene_gn=CA1H1orf35;refGene_tx=XM_023241736.1

A1 88275995 88276084
refGene_gn=CA1H1orf35;refGene_tx=XM_023241736.1
A1 88276502 88276631
refGene_gn=CA1H1orf35;refGene_tx=XM_023241736.1
A1 88276726 88276811
refGene_gn=CA1H1orf35;refGene_tx=XM_023241736.1
A1 88278856 88279018 refGene_gn=ARF1;refGene_tx=XM_003980625.4
A1 88279118 88279243 refGene_gn=ARF1;refGene_tx=XM_003980625.4
A1 88279319 88279430 refGene_gn=ARF1;refGene_tx=XM_003980625.4
A1 88279532 88279680 refGene_gn=ARF1;refGene_tx=XM_003980625.4
A1 88312285 88312765 refGene_gn=WNT3A;refGene_tx=XM_023253742.1
A1 88318995 88319261 refGene_gn=WNT3A;refGene_tx=XM_023253742.1
A1 88338683 88338925 refGene_gn=WNT3A;refGene_tx=XM_023253742.1
A1 88353175 88353246 refGene_gn=WNT3A;refGene_tx=XM_023253742.1
A1 88379886 88379981
refGene_gn=WNT9A;refGene_tx=XM_023253746.1;refGene_gn=WNT9A;refGene_tx
=XM_023253747.1
A1 88397178 88397435
refGene_gn=WNT9A;refGene_tx=XM_023253746.1;refGene_gn=WNT9A;refGene_tx
=XM_023253747.1
A1 88398233 88398496
refGene_gn=WNT9A;refGene_tx=XM_023253746.1;refGene_gn=WNT9A;refGene_tx
=XM_023253747.1
A1 88400435 88400918
refGene_gn=WNT9A;refGene_tx=XM_023253746.1;refGene_gn=WNT9A;refGene_tx
=XM_023253747.1
A1 88487505 88487760 refGene_gn=PRSS38;refGene_tx=XM_019813939.2
A1 88488192 88488335 refGene_gn=PRSS38;refGene_tx=XM_019813939.2
A1 88497978 88498250 refGene_gn=PRSS38;refGene_tx=XM_019813939.2
A1 88498805 88498968 refGene_gn=PRSS38;refGene_tx=XM_019813939.2
A1 88499416 88499576 refGene_gn=PRSS38;refGene_tx=XM_019813939.2
A1 88547963 88548110
refGene_gn=SNAP47;refGene_tx=XM_003980626.4;refGene_gn=SNAP47;refGene_
tx=XM_023253754.1
A1 88559990 88560115 refGene_gn=SNAP47;refGene_tx=XM_003980626.4
A1 88570779 88571270
refGene_gn=SNAP47;refGene_tx=XM_003980626.4;refGene_gn=SNAP47;refGene_
tx=XM_023253754.1
A1 88591488 88591973
refGene_gn=SNAP47;refGene_tx=XM_003980626.4;refGene_gn=SNAP47;refGene_
tx=XM_023253754.1
A1 88601687 88601952 refGene_gn=JMJD4;refGene_tx=XM_003980627.5
A1 88602052 88602218 refGene_gn=JMJD4;refGene_tx=XM_003980627.5
A1 88604214 88604340 refGene_gn=JMJD4;refGene_tx=XM_003980627.5
A1 88604548 88604816 refGene_gn=JMJD4;refGene_tx=XM_003980627.5
A1 88605205 88605352 refGene_gn=JMJD4;refGene_tx=XM_003980627.5
A1 88605807 88606116 refGene_gn=JMJD4;refGene_tx=XM_003980627.5
A1 88732632 88732711
refGene_gn=LOC109492746;refGene_tx=XM_019813951.1;refGene_gn=LOC109492
746;refGene_tx=XM_019813960.1

A1 88734752 88734876
refGene_gn=LOC109492746;refGene_tx=XM_019813951.1;refGene_gn=LOC109492
746;refGene_tx=XM_019813960.1
A1 88735234 88735318
refGene_gn=LOC109492746;refGene_tx=XM_019813951.1
A1 88736981 88737035
refGene_gn=LOC109492746;refGene_tx=XM_019813951.1;refGene_gn=LOC109492
746;refGene_tx=XM_019813960.1
A1 88749101 88749173
refGene_gn=LOC109492746;refGene_tx=XM_019813951.1;refGene_gn=LOC109492
746;refGene_tx=XM_019813960.1
A1 88750229 88750308
refGene_gn=LOC109492746;refGene_tx=XM_019813951.1;refGene_gn=LOC109492
746;refGene_tx=XM_019813960.1
A1 88750769 88750840
refGene_gn=LOC109492746;refGene_tx=XM_019813951.1;refGene_gn=LOC109492
746;refGene_tx=XM_019813960.1
A1 88754380 88754497
refGene_gn=LOC109492746;refGene_tx=XM_019813951.1;refGene_gn=LOC109492
746;refGene_tx=XM_019813960.1
A1 88756647 88756678
refGene_gn=LOC109492746;refGene_tx=XM_019813951.1;refGene_gn=LOC109492
746;refGene_tx=XM_019813960.1
A1 88849063 88849437
refGene_gn=LOC109497082;refGene_tx=XM_019824797.2;refGene_gn=LOC109497
082;refGene_tx=XM_019824799.2
A1 88849692 88849886
refGene_gn=LOC109497082;refGene_tx=XM_019824797.2;refGene_gn=LOC109497
082;refGene_tx=XM_019824799.2
A1 88850820 88850880
refGene_gn=LOC109497082;refGene_tx=XM_019824797.2;refGene_gn=LOC109497
082;refGene_tx=XM_019824799.2
A1 88851726 88851731
refGene_gn=LOC109497082;refGene_tx=XM_019824797.2;refGene_gn=LOC109497
082;refGene_tx=XM_019824799.2
A1 88857839 88858850
refGene_gn=LOC109497081;refGene_tx=XM_023241845.1
A1 90984171 90984911
refGene_gn=LOC111557322;refGene_tx=XM_023241873.1
A1 90984913 90985532
refGene_gn=LOC111557322;refGene_tx=XM_023241873.1
A1 90985672 90985836
refGene_gn=LOC111557322;refGene_tx=XM_023241873.1
A1 90985838 90986053
refGene_gn=LOC111557322;refGene_tx=XM_023241873.1
A1 90992815 90992898
refGene_gn=LOC111557322;refGene_tx=XM_023241873.1
A1 91247344 91247833 refGene_gn=MGAT1;refGene_tx=XM_023253766.1
A1 91247834 91248689 refGene_gn=MGAT1;refGene_tx=XM_023253766.1
A1 91306564 91309347

refGene_gn=ZFP62;refGene_tx=XM_003980634.4;refGene_gn=ZFP62;refGene_tx=XM_019830280.2;refGene_gn=ZFP62;refGene_tx=XM_019830289.2;refGene_gn=ZFP62;refGene_tx=XM_019830298.2;refGene_gn=ZFP62;refGene_tx=XM_019830301.1;refGene_gn=ZFP62;refGene_tx=XM_019830303.2;refGene_gn=ZFP62;refGene_tx=XM_023253799.1
A1 91310086 91310087 refGene_gn=ZFP62;refGene_tx=XM_019830301.1
A1 91310273 91310274
refGene_gn=ZFP62;refGene_tx=XM_019830280.2;refGene_gn=ZFP62;refGene_tx=XM_019830289.2;refGene_gn=ZFP62;refGene_tx=XM_019830298.2;refGene_gn=ZFP62;refGene_tx=XM_023253799.1
A1 91319549 91319826
refGene_gn=ZFP62;refGene_tx=XM_003980634.4;refGene_gn=ZFP62;refGene_tx=XM_019830303.2
A1 91373073 91373185
refGene_gn=BTNL9;refGene_tx=XM_003980676.3;refGene_gn=BTNL9;refGene_tx=XM_006927479.2;refGene_gn=BTNL9;refGene_tx=XM_019830340.1
A1 91375012 91375360
refGene_gn=BTNL9;refGene_tx=XM_003980676.3;refGene_gn=BTNL9;refGene_tx=XM_006927479.2;refGene_gn=BTNL9;refGene_tx=XM_011281396.2;refGene_gn=BTNL9;refGene_tx=XM_019830340.1
A1 91378399 91378681
refGene_gn=BTNL9;refGene_tx=XM_003980676.3;refGene_gn=BTNL9;refGene_tx=XM_006927479.2;refGene_gn=BTNL9;refGene_tx=XM_011281396.2;refGene_gn=BTNL9;refGene_tx=XM_019830340.1
A1 91383702 91383831
refGene_gn=BTNL9;refGene_tx=XM_003980676.3;refGene_gn=BTNL9;refGene_tx=XM_011281396.2;refGene_gn=BTNL9;refGene_tx=XM_019830340.1
A1 91384015 91384048
refGene_gn=BTNL9;refGene_tx=XM_003980676.3;refGene_gn=BTNL9;refGene_tx=XM_011281396.2
A1 91384348 91384369
refGene_gn=BTNL9;refGene_tx=XM_003980676.3;refGene_gn=BTNL9;refGene_tx=XM_011281396.2;refGene_gn=BTNL9;refGene_tx=XM_019830340.1
A1 91393825 91393846
refGene_gn=BTNL9;refGene_tx=XM_003980676.3;refGene_gn=BTNL9;refGene_tx=XM_006927479.2;refGene_gn=BTNL9;refGene_tx=XM_011281396.2;refGene_gn=BTNL9;refGene_tx=XM_019830340.1
A1 91394105 91394132
refGene_gn=BTNL9;refGene_tx=XM_003980676.3;refGene_gn=BTNL9;refGene_tx=XM_006927479.2;refGene_gn=BTNL9;refGene_tx=XM_011281396.2;refGene_gn=BTNL9;refGene_tx=XM_019830340.1
A1 91396520 91396547
refGene_gn=BTNL9;refGene_tx=XM_003980676.3;refGene_gn=BTNL9;refGene_tx=XM_006927479.2;refGene_gn=BTNL9;refGene_tx=XM_011281396.2;refGene_gn=BTNL9;refGene_tx=XM_019830340.1
A1 91397408 91398031
refGene_gn=BTNL9;refGene_tx=XM_003980676.3;refGene_gn=BTNL9;refGene_tx=XM_006927479.2;refGene_gn=BTNL9;refGene_tx=XM_011281396.2;refGene_gn=BTNL9;refGene_tx=XM_019830340.1
A1 91415867 91416701

refGene_gn=LOC101093739;refGene_tx=XM_011281399.2
A1	91416702	91416825
refGene_gn=LOC101093739;refGene_tx=XM_011281399.2
A1	91419438	91419447
refGene_gn=LOC101093739;refGene_tx=XM_011281399.2
A1	91430585	91430594
refGene_gn=LOC111557337;refGene_tx=XM_023242044.1
A1	91433207	91433327
refGene_gn=LOC111557337;refGene_tx=XM_023242044.1
A1	91433328	91434165
refGene_gn=LOC111557337;refGene_tx=XM_023242044.1
A1	91457504	91458455
refGene_gn=LOC101093987;refGene_tx=XM_003980678.1
A1	91503520	91503861
refGene_gn=LOC101082715;refGene_tx=XM_011281402.2
A1	91503863	91504048
refGene_gn=LOC101082715;refGene_tx=XM_011281402.2
A1	91504049	91504207
refGene_gn=LOC101082715;refGene_tx=XM_011281402.2
A1	91504208	91504463
refGene_gn=LOC101082715;refGene_tx=XM_011281402.2
A1	91544686	91545634
refGene_gn=LOC101082972;refGene_tx=XM_003980636.1
A1	91604301	91604813
refGene_gn=TRIM7;refGene_tx=XM_003980637.5;refGene_gn=TRIM7;refGene_tx=XM_006927481.3;refGene_gn=TRIM7;refGene_tx=XM_006927483.4;refGene_gn=TRIM7;refGene_tx=XM_006927484.4;refGene_gn=TRIM7;refGene_tx=XM_019830393.2;refGene_gn=TRIM7;refGene_tx=XM_019830419.2
A1	91605863	91605972
refGene_gn=TRIM7;refGene_tx=XM_006927481.3;refGene_gn=TRIM7;refGene_tx=XM_006927482.4
A1	91606210	91606246
refGene_gn=TRIM7;refGene_tx=XM_003980637.5;refGene_gn=TRIM7;refGene_tx=XM_006927481.3;refGene_gn=TRIM7;refGene_tx=XM_006927482.4;refGene_gn=TRIM7;refGene_tx=XM_006927483.4;refGene_gn=TRIM7;refGene_tx=XM_019830393.2;refGene_gn=TRIM7;refGene_tx=XM_019830419.2
A1	91606728	91606844
refGene_gn=TRIM7;refGene_tx=XM_003980637.5;refGene_gn=TRIM7;refGene_tx=XM_006927481.3;refGene_gn=TRIM7;refGene_tx=XM_019830393.2;refGene_gn=TRIM7;refGene_tx=XM_019830419.2
A1	91607136	91607159
refGene_gn=TRIM7;refGene_tx=XM_003980637.5;refGene_gn=TRIM7;refGene_tx=XM_006927481.3;refGene_gn=TRIM7;refGene_tx=XM_006927482.4;refGene_gn=TRIM7;refGene_tx=XM_006927483.4;refGene_gn=TRIM7;refGene_tx=XM_006927484.4;refGene_gn=TRIM7;refGene_tx=XM_019830393.2;refGene_gn=TRIM7;refGene_tx=XM_019830419.2
A1	91608365	91608596
refGene_gn=TRIM7;refGene_tx=XM_003980637.5;refGene_gn=TRIM7;refGene_tx=XM_006927481.3;refGene_gn=TRIM7;refGene_tx=XM_006927482.4;refGene_gn=TRIM7;refGene_tx=XM_006927483.4;refGene_gn=TRIM7;refGene_tx=XM_0069274

84.4;refGene_gn=TRIM7;refGene_tx=XM_019830393.2;refGene_gn=TRIM7;refGene_tx=XM_019830419.2
A1 91611008 91611104
refGene_gn=TRIM7;refGene_tx=XM_003980637.5;refGene_gn=TRIM7;refGene_tx=XM_006927481.3;refGene_gn=TRIM7;refGene_tx=XM_006927482.4;refGene_gn=TRIM7;refGene_tx=XM_006927483.4;refGene_gn=TRIM7;refGene_tx=XM_006927484.4
A1 91612023 91612530
refGene_gn=TRIM7;refGene_tx=XM_003980637.5;refGene_gn=TRIM7;refGene_tx=XM_006927481.3;refGene_gn=TRIM7;refGene_tx=XM_006927482.4;refGene_gn=TRIM7;refGene_tx=XM_006927483.4;refGene_gn=TRIM7;refGene_tx=XM_006927484.4
A1 91628065 91628875
refGene_gn=TRIM41;refGene_tx=XM_003980638.4;refGene_gn=TRIM41;refGene_tx=XM_006927488.3;refGene_gn=TRIM41;refGene_tx=XM_006927489.3;refGene_gn=TRIM41;refGene_tx=XM_019830363.1;refGene_gn=TRIM41;refGene_tx=XM_019830367.1;refGene_gn=TRIM41;refGene_tx=XM_019830377.1;refGene_gn=TRIM41;refGene_tx=XM_019830384.1
A1 91630931 91630979
refGene_gn=TRIM41;refGene_tx=XM_019830363.1;refGene_gn=TRIM41;refGene_tx=XM_019830367.1;refGene_gn=TRIM41;refGene_tx=XM_019830377.1;refGene_gn=TRIM41;refGene_tx=XM_019830384.1
A1 91633130 91633226
refGene_gn=TRIM41;refGene_tx=XM_003980638.4;refGene_gn=TRIM41;refGene_tx=XM_006927488.3;refGene_gn=TRIM41;refGene_tx=XM_006927489.3;refGene_gn=TRIM41;refGene_tx=XM_019830363.1;refGene_gn=TRIM41;refGene_tx=XM_019830367.1;refGene_gn=TRIM41;refGene_tx=XM_019830377.1;refGene_gn=TRIM41;refGene_tx=XM_019830384.1
A1 91634943 91635207
refGene_gn=TRIM41;refGene_tx=XM_003980638.4;refGene_gn=TRIM41;refGene_tx=XM_006927488.3;refGene_gn=TRIM41;refGene_tx=XM_006927489.3;refGene_gn=TRIM41;refGene_tx=XM_019830363.1;refGene_gn=TRIM41;refGene_tx=XM_019830367.1;refGene_gn=TRIM41;refGene_tx=XM_019830377.1;refGene_gn=TRIM41;refGene_tx=XM_019830384.1
A1 91635772 91635795
refGene_gn=TRIM41;refGene_tx=XM_003980638.4;refGene_gn=TRIM41;refGene_tx=XM_006927488.3;refGene_gn=TRIM41;refGene_tx=XM_006927489.3;refGene_gn=TRIM41;refGene_tx=XM_019830363.1;refGene_gn=TRIM41;refGene_tx=XM_019830367.1;refGene_gn=TRIM41;refGene_tx=XM_019830377.1
A1 91635995 91636123
refGene_gn=TRIM41;refGene_tx=XM_003980638.4;refGene_gn=TRIM41;refGene_tx=XM_006927488.3;refGene_gn=TRIM41;refGene_tx=XM_006927489.3;refGene_gn=TRIM41;refGene_tx=XM_019830363.1;refGene_gn=TRIM41;refGene_tx=XM_019830367.1;refGene_gn=TRIM41;refGene_tx=XM_019830377.1;refGene_gn=TRIM41;refGene_tx=XM_019830384.1
A1 91636546 91637179
refGene_gn=TRIM41;refGene_tx=XM_003980638.4;refGene_gn=TRIM41;refGene_tx=XM_006927488.3;refGene_gn=TRIM41;refGene_tx=XM_006927489.3;refGene_gn=TRIM41;refGene_tx=XM_019830363.1;refGene_gn=TRIM41;refGene_tx=XM_019830367.1;refGene_gn=TRIM41;refGene_tx=XM_019830377.1

A1 91637325 91637460 refGene_gn=TRIM41;refGene_tx=XM_006927489.3
A1 91639065 91639131 refGene_gn=RACK1;refGene_tx=XM_003980639.4
A1 91639515 91639626 refGene_gn=RACK1;refGene_tx=XM_003980639.4
A1 91640016 91640157 refGene_gn=RACK1;refGene_tx=XM_003980639.4
A1 91640570 91640681 refGene_gn=RACK1;refGene_tx=XM_003980639.4
A1 91641078 91641174 refGene_gn=RACK1;refGene_tx=XM_003980639.4
A1 91641419 91641567 refGene_gn=RACK1;refGene_tx=XM_003980639.4
A1 91642105 91642277 refGene_gn=RACK1;refGene_tx=XM_003980639.4
A1 91643298 91643407 refGene_gn=RACK1;refGene_tx=XM_003980639.4
A1 91645800 91645842 refGene_gn=TRIM52;refGene_tx=XM_006927492.4
A1 91654013 91654082 refGene_gn=TRIM52;refGene_tx=XM_006927490.4
A1 91654633 91654645 refGene_gn=TRIM52;refGene_tx=XM_019830464.2
A1 91657425 91658244
refGene_gn=TRIM52;refGene_tx=XM_006927490.4;refGene_gn=TRIM52;refGene_tx=XM_006927492.4;refGene_gn=TRIM52;refGene_tx=XM_019830464.2
A1 91668095 91668160
refGene_gn=LOC101094232;refGene_tx=XM_023253928.1
A1 91668420 91668453
refGene_gn=LOC101094232;refGene_tx=XM_023253928.1
A1 91668795 91668906
refGene_gn=LOC101094232;refGene_tx=XM_023253928.1
A1 91669019 91669295
refGene_gn=LOC101094232;refGene_tx=XM_023253928.1
A1 91671070 91671346
refGene_gn=LOC101094232;refGene_tx=XM_023253928.1
A1 91672455 91672716
refGene_gn=LOC101094232;refGene_tx=XM_023253928.1
A1 91677899 91678007
refGene_gn=LOC101094232;refGene_tx=XM_023253928.1
A1 91681373 91681422
refGene_gn=LOC101094232;refGene_tx=XM_023253928.1
A1 91725271 91726174
refGene_gn=FELCATV1R2;refGene_tx=NM_001167474.1
A1 91756345 91757278
refGene_gn=LOC111556195;refGene_tx=XM_023253937.1
A1 91852856 91852904
refGene_gn=LOC102901611;refGene_tx=XM_006927496.3;refGene_gn=LOC102901611;refGene_tx=XM_006927497.3
A1 91877676 91878893
refGene_gn=LOC102901611;refGene_tx=XM_006927496.3;refGene_gn=LOC102901611;refGene_tx=XM_006927497.3
A1 91918974 91919041
refGene_gn=FLT4;refGene_tx=XM_006927498.4;refGene_gn=FLT4;refGene_tx=XM_006927499.4
A1 91933848 91933945
refGene_gn=FLT4;refGene_tx=XM_006927498.4;refGene_gn=FLT4;refGene_tx=XM_006927499.4
A1 91934560 91934805
refGene_gn=FLT4;refGene_tx=XM_006927498.4;refGene_gn=FLT4;refGene_tx=XM_006927499.4

A1 91934988 91935101
refGene_gn=FLT4;refGene_tx=XM_006927498.4;refGene_gn=FLT4;refGene_tx=XM_006927499.4
A1 91935219 91935382
refGene_gn=FLT4;refGene_tx=XM_006927498.4;refGene_gn=FLT4;refGene_tx=XM_006927499.4
A1 91935598 91935738
refGene_gn=FLT4;refGene_tx=XM_006927498.4;refGene_gn=FLT4;refGene_tx=XM_006927499.4
A1 91936048 91936217
refGene_gn=FLT4;refGene_tx=XM_006927498.4;refGene_gn=FLT4;refGene_tx=XM_006927499.4
A1 91936495 91936613
refGene_gn=FLT4;refGene_tx=XM_006927498.4;refGene_gn=FLT4;refGene_tx=XM_006927499.4
A1 91939246 91939401
refGene_gn=FLT4;refGene_tx=XM_006927498.4;refGene_gn=FLT4;refGene_tx=XM_006927499.4
A1 91939566 91939729
refGene_gn=FLT4;refGene_tx=XM_006927498.4;refGene_gn=FLT4;refGene_tx=XM_006927499.4
A1 91941151 91941278
refGene_gn=FLT4;refGene_tx=XM_006927498.4;refGene_gn=FLT4;refGene_tx=XM_006927499.4
A1 91942345 91942454
refGene_gn=FLT4;refGene_tx=XM_006927498.4;refGene_gn=FLT4;refGene_tx=XM_006927499.4
A1 91943437 91943800
refGene_gn=FLT4;refGene_tx=XM_006927498.4;refGene_gn=FLT4;refGene_tx=XM_006927499.4
A1 91944081 91944228
refGene_gn=FLT4;refGene_tx=XM_006927498.4;refGene_gn=FLT4;refGene_tx=XM_006927499.4
A1 91944330 91944462
refGene_gn=FLT4;refGene_tx=XM_006927498.4;refGene_gn=FLT4;refGene_tx=XM_006927499.4
A1 91944622 91944729
refGene_gn=FLT4;refGene_tx=XM_006927498.4;refGene_gn=FLT4;refGene_tx=XM_006927499.4
A1 91945024 91945160
refGene_gn=FLT4;refGene_tx=XM_006927498.4;refGene_gn=FLT4;refGene_tx=XM_006927499.4
A1 91945613 91945718
refGene_gn=FLT4;refGene_tx=XM_006927498.4;refGene_gn=FLT4;refGene_tx=XM_006927499.4
A1 91946075 91946189
refGene_gn=FLT4;refGene_tx=XM_006927498.4;refGene_gn=FLT4;refGene_tx=XM_006927499.4
A1 91946365 91946454
refGene_gn=FLT4;refGene_tx=XM_006927498.4;refGene_gn=FLT4;refGene_tx=X

M_006927499.4
A1 91946587 91946735
refGene_gn=FLT4;refGene_tx=XM_006927498.4;refGene_gn=FLT4;refGene_tx=XM_006927499.4
A1 91947241 91947336
refGene_gn=FLT4;refGene_tx=XM_006927498.4;refGene_gn=FLT4;refGene_tx=XM_006927499.4
A1 91947922 91948045
refGene_gn=FLT4;refGene_tx=XM_006927498.4;refGene_gn=FLT4;refGene_tx=XM_006927499.4
A1 91949162 91949274
refGene_gn=FLT4;refGene_tx=XM_006927498.4;refGene_gn=FLT4;refGene_tx=XM_006927499.4
A1 91951105 91951205
refGene_gn=FLT4;refGene_tx=XM_006927498.4;refGene_gn=FLT4;refGene_tx=XM_006927499.4
A1 91951513 91951619
refGene_gn=FLT4;refGene_tx=XM_006927498.4;refGene_gn=FLT4;refGene_tx=XM_006927499.4
A1 91952128 91952274
refGene_gn=FLT4;refGene_tx=XM_006927498.4;refGene_gn=FLT4;refGene_tx=XM_006927499.4
A1 91953530 91953651
refGene_gn=FLT4;refGene_tx=XM_006927498.4;refGene_gn=FLT4;refGene_tx=XM_006927499.4
A1 91954762 91954848
refGene_gn=FLT4;refGene_tx=XM_006927498.4;refGene_gn=FLT4;refGene_tx=XM_006927499.4
A1 91960350 91960552
refGene_gn=FLT4;refGene_tx=XM_006927498.4;refGene_gn=FLT4;refGene_tx=XM_006927499.4
A1 91981947 91982002 refGene_gn=SCGB3A1;refGene_tx=XM_023254649.1
A1 91982654 91982944 refGene_gn=SCGB3A1;refGene_tx=XM_023254649.1
A1 91983268 91983292 refGene_gn=SCGB3A1;refGene_tx=XM_023254649.1
A1 91998271 91998484
refGene_gn=CNOT6;refGene_tx=XM_011281483.2;refGene_gn=CNOT6;refGene_tx=XM_011281491.2;refGene_gn=CNOT6;refGene_tx=XM_019830518.2
A1 92001104 92001307
refGene_gn=CNOT6;refGene_tx=XM_011281483.2;refGene_gn=CNOT6;refGene_tx=XM_011281491.2;refGene_gn=CNOT6;refGene_tx=XM_019830518.2
A1 92003109 92003340
refGene_gn=CNOT6;refGene_tx=XM_011281483.2;refGene_gn=CNOT6;refGene_tx=XM_011281491.2;refGene_gn=CNOT6;refGene_tx=XM_019830518.2
A1 92004262 92004417
refGene_gn=CNOT6;refGene_tx=XM_011281483.2;refGene_gn=CNOT6;refGene_tx=XM_011281491.2;refGene_gn=CNOT6;refGene_tx=XM_019830518.2
A1 92005009 92005164
refGene_gn=CNOT6;refGene_tx=XM_011281483.2;refGene_gn=CNOT6;refGene_tx=XM_011281491.2;refGene_gn=CNOT6;refGene_tx=XM_019830518.2
A1 92005951 92006109

refGene_gn=CNOT6;refGene_tx=XM_011281483.2;refGene_gn=CNOT6;refGene_tx=XM_011281491.2;refGene_gn=CNOT6;refGene_tx=XM_019830518.2
A1 92007421 92007490
refGene_gn=CNOT6;refGene_tx=XM_011281483.2;refGene_gn=CNOT6;refGene_tx=XM_011281491.2;refGene_gn=CNOT6;refGene_tx=XM_019830518.2
A1 92007570 92007675
refGene_gn=CNOT6;refGene_tx=XM_011281483.2;refGene_gn=CNOT6;refGene_tx=XM_011281491.2;refGene_gn=CNOT6;refGene_tx=XM_019830518.2
A1 92016087 92016173
refGene_gn=CNOT6;refGene_tx=XM_011281483.2;refGene_gn=CNOT6;refGene_tx=XM_011281491.2
A1 92018339 92018526
refGene_gn=CNOT6;refGene_tx=XM_011281483.2;refGene_gn=CNOT6;refGene_tx=XM_011281491.2
A1 92048295 92048407
refGene_gn=CNOT6;refGene_tx=XM_011281483.2;refGene_gn=CNOT6;refGene_tx=XM_011281491.2;refGene_gn=CNOT6;refGene_tx=XM_019830518.2
A1 92136582 92136891
refGene_gn=LOC105260654;refGene_tx=XM_011283345.1
A1 92183425 92183432 refGene_gn=GFPT2;refGene_tx=XM_006927503.3
A1 92198786 92198894 refGene_gn=GFPT2;refGene_tx=XM_006927503.3
A1 92200232 92200331 refGene_gn=GFPT2;refGene_tx=XM_006927503.3
A1 92200863 92200989 refGene_gn=GFPT2;refGene_tx=XM_006927503.3
A1 92204608 92204667 refGene_gn=GFPT2;refGene_tx=XM_006927503.3
A1 92205308 92205443 refGene_gn=GFPT2;refGene_tx=XM_006927503.3
A1 92208019 92208081 refGene_gn=GFPT2;refGene_tx=XM_006927503.3
A1 92210772 92210852 refGene_gn=GFPT2;refGene_tx=XM_006927503.3
A1 92211294 92211412 refGene_gn=GFPT2;refGene_tx=XM_006927503.3
A1 92218057 92218221 refGene_gn=GFPT2;refGene_tx=XM_006927503.3
A1 92220062 92220158 refGene_gn=GFPT2;refGene_tx=XM_006927503.3
A1 92220236 92220334 refGene_gn=GFPT2;refGene_tx=XM_006927503.3
A1 92220613 92220734 refGene_gn=GFPT2;refGene_tx=XM_006927503.3
A1 92222926 92223084 refGene_gn=GFPT2;refGene_tx=XM_006927503.3
A1 92223246 92223361 refGene_gn=GFPT2;refGene_tx=XM_006927503.3
A1 92226843 92226971 refGene_gn=GFPT2;refGene_tx=XM_006927503.3
A1 92228755 92228923 refGene_gn=GFPT2;refGene_tx=XM_006927503.3
A1 92230013 92230175 refGene_gn=GFPT2;refGene_tx=XM_006927503.3
A1 92232060 92232105 refGene_gn=GFPT2;refGene_tx=XM_006927503.3
A1 92259777 92259899
refGene_gn=MAPK9;refGene_tx=XM_023253983.1;refGene_gn=MAPK9;refGene_tx=XM_023253986.1;refGene_gn=MAPK9;refGene_tx=XM_023253987.1
A1 92271315 92271445
refGene_gn=MAPK9;refGene_tx=XM_023253983.1;refGene_gn=MAPK9;refGene_tx=XM_023253986.1;refGene_gn=MAPK9;refGene_tx=XM_023253987.1;refGene_gn=MAPK9;refGene_tx=XM_023253990.1
A1 92274215 92274274
refGene_gn=MAPK9;refGene_tx=XM_023253983.1;refGene_gn=MAPK9;refGene_tx=XM_023253986.1;refGene_gn=MAPK9;refGene_tx=XM_023253987.1
A1 92278400 92278539
refGene_gn=MAPK9;refGene_tx=XM_023253983.1;refGene_gn=MAPK9;refGene_tx

=XM_023253986.1;refGene_gn=MAPK9;refGene_tx=XM_023253987.1;refGene_gn=MAPK9;refGene_tx=XM_023253990.1
A1 92288000 92288166 refGene_gn=MAPK9;refGene_tx=XM_023253983.1;refGene_gn=MAPK9;refGene_tx=XM_023253986.1;refGene_gn=MAPK9;refGene_tx=XM_023253987.1;refGene_gn=MAPK9;refGene_tx=XM_023253990.1
A1 92289074 92289146 refGene_gn=MAPK9;refGene_tx=XM_023253986.1
A1 92289492 92289564 refGene_gn=MAPK9;refGene_tx=XM_023253983.1;refGene_gn=MAPK9;refGene_tx=XM_023253987.1;refGene_gn=MAPK9;refGene_tx=XM_023253990.1
A1 92292643 92292826 refGene_gn=MAPK9;refGene_tx=XM_023253983.1;refGene_gn=MAPK9;refGene_tx=XM_023253986.1;refGene_gn=MAPK9;refGene_tx=XM_023253987.1;refGene_gn=MAPK9;refGene_tx=XM_023253990.1
A1 92293977 92294102 refGene_gn=MAPK9;refGene_tx=XM_023253983.1;refGene_gn=MAPK9;refGene_tx=XM_023253986.1;refGene_gn=MAPK9;refGene_tx=XM_023253987.1;refGene_gn=MAPK9;refGene_tx=XM_023253990.1
A1 92295155 92295219 refGene_gn=MAPK9;refGene_tx=XM_023253983.1;refGene_gn=MAPK9;refGene_tx=XM_023253986.1;refGene_gn=MAPK9;refGene_tx=XM_023253987.1;refGene_gn=MAPK9;refGene_tx=XM_023253990.1
A1 92296241 92296313 refGene_gn=MAPK9;refGene_tx=XM_023253983.1;refGene_gn=MAPK9;refGene_tx=XM_023253986.1;refGene_gn=MAPK9;refGene_tx=XM_023253987.1;refGene_gn=MAPK9;refGene_tx=XM_023253990.1
A1 92298060 92298208 refGene_gn=MAPK9;refGene_tx=XM_023253983.1;refGene_gn=MAPK9;refGene_tx=XM_023253986.1;refGene_gn=MAPK9;refGene_tx=XM_023253987.1;refGene_gn=MAPK9;refGene_tx=XM_023253990.1
A1 92365975 92366152 refGene_gn=RASGEF1C;refGene_tx=XM_011281526.3;refGene_gn=RASGEF1C;refGene_tx=XM_023254000.1
A1 92366308 92366431 refGene_gn=RASGEF1C;refGene_tx=XM_011281526.3;refGene_gn=RASGEF1C;refGene_tx=XM_023254000.1
A1 92367638 92367776 refGene_gn=RASGEF1C;refGene_tx=XM_011281526.3;refGene_gn=RASGEF1C;refGene_tx=XM_023254000.1
A1 92375422 92375623 refGene_gn=RASGEF1C;refGene_tx=XM_011281526.3;refGene_gn=RASGEF1C;refGene_tx=XM_023254000.1
A1 92376694 92376769 refGene_gn=RASGEF1C;refGene_tx=XM_011281526.3;refGene_gn=RASGEF1C;refGene_tx=XM_023254000.1
A1 92387711 92387801 refGene_gn=RASGEF1C;refGene_tx=XM_011281526.3;refGene_gn=RASGEF1C;refGene_tx=XM_023254000.1
A1 92389376 92389479 refGene_gn=RASGEF1C;refGene_tx=XM_011281526.3;refGene_gn=RASGEF1C;refG

ene_tx=XM_023254000.1
A1 92389966 92390046
refGene_gn=RASGEF1C;refGene_tx=XM_011281526.3;refGene_gn=RASGEF1C;refGene_tx=XM_023254000.1
A1 92390129 92390225
refGene_gn=RASGEF1C;refGene_tx=XM_011281526.3;refGene_gn=RASGEF1C;refGene_tx=XM_023254000.1
A1 92394873 92394969
refGene_gn=RASGEF1C;refGene_tx=XM_011281526.3;refGene_gn=RASGEF1C;refGene_tx=XM_023254000.1
A1 92396804 92396928
refGene_gn=RASGEF1C;refGene_tx=XM_011281526.3;refGene_gn=RASGEF1C;refGene_tx=XM_023254000.1
A1 92403797 92403870
refGene_gn=RASGEF1C;refGene_tx=XM_011281526.3;refGene_gn=RASGEF1C;refGene_tx=XM_023254000.1
A1 92404346 92404371
refGene_gn=RASGEF1C;refGene_tx=XM_011281526.3;refGene_gn=RASGEF1C;refGene_tx=XM_023254000.1
A1 92438709 92438956
refGene_gn=RNF130;refGene_tx=XM_023254012.1;refGene_gn=RNF130;refGene_tx=XM_023254015.1;refGene_gn=RNF130;refGene_tx=XM_023254016.1
A1 92471008 92471203
refGene_gn=RNF130;refGene_tx=XM_023254012.1;refGene_gn=RNF130;refGene_tx=XM_023254015.1;refGene_gn=RNF130;refGene_tx=XM_023254016.1
A1 92497050 92497301
refGene_gn=RNF130;refGene_tx=XM_023254012.1;refGene_gn=RNF130;refGene_tx=XM_023254015.1;refGene_gn=RNF130;refGene_tx=XM_023254016.1
A1 92524585 92524657
refGene_gn=RNF130;refGene_tx=XM_023254012.1;refGene_gn=RNF130;refGene_tx=XM_023254015.1;refGene_gn=RNF130;refGene_tx=XM_023254016.1
A1 92526473 92526556
refGene_gn=RNF130;refGene_tx=XM_023254012.1;refGene_gn=RNF130;refGene_tx=XM_023254015.1;refGene_gn=RNF130;refGene_tx=XM_023254016.1
A1 92533987 92534084
refGene_gn=RNF130;refGene_tx=XM_023254012.1;refGene_gn=RNF130;refGene_tx=XM_023254015.1;refGene_gn=RNF130;refGene_tx=XM_023254016.1
A1 92536142 92536347
refGene_gn=RNF130;refGene_tx=XM_023254012.1;refGene_gn=RNF130;refGene_tx=XM_023254015.1;refGene_gn=RNF130;refGene_tx=XM_023254016.1
A1 92539470 92539564 refGene_gn=RNF130;refGene_tx=XM_023254012.1
A1 92545994 92546010
refGene_gn=RNF130;refGene_tx=XM_023254012.1;refGene_gn=RNF130;refGene_tx=XM_023254016.1
A1 92570548 92570658 refGene_gn=RNF130;refGene_tx=XM_023254015.1
A1 92581714 92581832
refGene_gn=TBC1D9B;refGene_tx=XM_023254017.1;refGene_gn=TBC1D9B;refGene_tx=XM_023254018.1
A1 92585220 92585331
refGene_gn=TBC1D9B;refGene_tx=XM_023254017.1;refGene_gn=TBC1D9B;refGen

e_tx=XM_023254018.1
A1	92591040	92591159	refGene_gn=TBC1D9B;refGene_tx=XM_023254017.1;refGene_gn=TBC1D9B;refGene_tx=XM_023254018.1
A1	92593920	92594149	refGene_gn=TBC1D9B;refGene_tx=XM_023254017.1;refGene_gn=TBC1D9B;refGene_tx=XM_023254018.1
A1	92595040	92595299	refGene_gn=TBC1D9B;refGene_tx=XM_023254017.1;refGene_gn=TBC1D9B;refGene_tx=XM_023254018.1
A1	92596375	92596583	refGene_gn=TBC1D9B;refGene_tx=XM_023254017.1;refGene_gn=TBC1D9B;refGene_tx=XM_023254018.1
A1	92599487	92599700	refGene_gn=TBC1D9B;refGene_tx=XM_023254017.1;refGene_gn=TBC1D9B;refGene_tx=XM_023254018.1
A1	92613118	92613280	refGene_gn=TBC1D9B;refGene_tx=XM_023254017.1;refGene_gn=TBC1D9B;refGene_tx=XM_023254018.1
A1	92613570	92613721	refGene_gn=TBC1D9B;refGene_tx=XM_023254017.1;refGene_gn=TBC1D9B;refGene_tx=XM_023254018.1
A1	92614212	92614427	refGene_gn=TBC1D9B;refGene_tx=XM_023254017.1;refGene_gn=TBC1D9B;refGene_tx=XM_023254018.1
A1	92615944	92616062	refGene_gn=TBC1D9B;refGene_tx=XM_023254017.1;refGene_gn=TBC1D9B;refGene_tx=XM_023254018.1
A1	92617155	92617441	refGene_gn=TBC1D9B;refGene_tx=XM_023254017.1;refGene_gn=TBC1D9B;refGene_tx=XM_023254018.1
A1	92618693	92618823	refGene_gn=TBC1D9B;refGene_tx=XM_023254017.1;refGene_gn=TBC1D9B;refGene_tx=XM_023254018.1
A1	92618952	92619051	refGene_gn=TBC1D9B;refGene_tx=XM_023254017.1;refGene_gn=TBC1D9B;refGene_tx=XM_023254018.1
A1	92619831	92619900	refGene_gn=TBC1D9B;refGene_tx=XM_023254017.1;refGene_gn=TBC1D9B;refGene_tx=XM_023254018.1
A1	92620791	92621032	refGene_gn=TBC1D9B;refGene_tx=XM_023254017.1;refGene_gn=TBC1D9B;refGene_tx=XM_023254018.1
A1	92621404	92621470	refGene_gn=TBC1D9B;refGene_tx=XM_023254017.1;refGene_gn=TBC1D9B;refGene_tx=XM_023254018.1
A1	92623015	92623087	refGene_gn=TBC1D9B;refGene_tx=XM_023254017.1;refGene_gn=TBC1D9B;refGene_tx=XM_023254018.1
A1	92623310	92623361	refGene_gn=TBC1D9B;refGene_tx=XM_023254017.1

A1 92625044 92625107
refGene_gn=TBC1D9B;refGene_tx=XM_023254017.1;refGene_gn=TBC1D9B;refGene_tx=XM_023254018.1
A1 92625684 92625791
refGene_gn=TBC1D9B;refGene_tx=XM_023254017.1;refGene_gn=TBC1D9B;refGene_tx=XM_023254018.1
A1 92626864 92627551
refGene_gn=TBC1D9B;refGene_tx=XM_023254017.1;refGene_gn=TBC1D9B;refGene_tx=XM_023254018.1
A1 92631798 92631864 refGene_gn=MRNIP;refGene_tx=XM_006927506.2
A1 92634295 92634355 refGene_gn=MRNIP;refGene_tx=XM_006927506.2
A1 92634731 92634743 refGene_gn=MRNIP;refGene_tx=XM_011281530.2
A1 92634751 92634778 refGene_gn=MRNIP;refGene_tx=XM_011281529.2
A1 92639832 92639921
refGene_gn=MRNIP;refGene_tx=XM_006927506.2;refGene_gn=MRNIP;refGene_tx=XM_011281529.2;refGene_gn=MRNIP;refGene_tx=XM_011281530.2
A1 92643419 92643495
refGene_gn=MRNIP;refGene_tx=XM_006927506.2;refGene_gn=MRNIP;refGene_tx=XM_011281529.2;refGene_gn=MRNIP;refGene_tx=XM_011281530.2
A1 92644678 92644824
refGene_gn=MRNIP;refGene_tx=XM_006927506.2;refGene_gn=MRNIP;refGene_tx=XM_011281529.2;refGene_gn=MRNIP;refGene_tx=XM_011281530.2
A1 92645510 92645598
refGene_gn=MRNIP;refGene_tx=XM_006927506.2;refGene_gn=MRNIP;refGene_tx=XM_011281529.2;refGene_gn=MRNIP;refGene_tx=XM_011281530.2
A1 92652125 92652611
refGene_gn=MRNIP;refGene_tx=XM_006927506.2;refGene_gn=MRNIP;refGene_tx=XM_011281529.2;refGene_gn=MRNIP;refGene_tx=XM_011281530.2
A1 92653731 92653889
refGene_gn=SQSTM1;refGene_tx=XM_003980646.4;refGene_gn=SQSTM1;refGene_tx=XM_023254040.1
A1 92654673 92654869
refGene_gn=SQSTM1;refGene_tx=XM_003980646.4;refGene_gn=SQSTM1;refGene_tx=XM_023254040.1
A1 92655058 92655270
refGene_gn=SQSTM1;refGene_tx=XM_003980646.4;refGene_gn=SQSTM1;refGene_tx=XM_023254040.1
A1 92660760 92660841
refGene_gn=SQSTM1;refGene_tx=XM_003980646.4;refGene_gn=SQSTM1;refGene_tx=XM_023254040.1
A1 92662220 92662362
refGene_gn=SQSTM1;refGene_tx=XM_003980646.4;refGene_gn=SQSTM1;refGene_tx=XM_023254040.1
A1 92662467 92662697
refGene_gn=SQSTM1;refGene_tx=XM_003980646.4;refGene_gn=SQSTM1;refGene_tx=XM_023254040.1
A1 92663195 92663291
refGene_gn=SQSTM1;refGene_tx=XM_003980646.4;refGene_gn=SQSTM1;refGene_tx=XM_023254040.1
A1 92665021 92665238 refGene_gn=SQSTM1;refGene_tx=XM_003980646.4

A1 92678315 92678412
refGene_gn=MGAT4B;refGene_tx=XM_023254024.1;refGene_gn=MGAT4B;refGene_tx=XM_023254032.1
A1 92683459 92683645
refGene_gn=MGAT4B;refGene_tx=XM_023254024.1;refGene_gn=MGAT4B;refGene_tx=XM_023254032.1
A1 92683730 92683871
refGene_gn=MGAT4B;refGene_tx=XM_023254024.1;refGene_gn=MGAT4B;refGene_tx=XM_023254032.1
A1 92683961 92684095
refGene_gn=MGAT4B;refGene_tx=XM_023254024.1;refGene_gn=MGAT4B;refGene_tx=XM_023254032.1
A1 92684485 92684532
refGene_gn=MGAT4B;refGene_tx=XM_023254024.1;refGene_gn=MGAT4B;refGene_tx=XM_023254032.1
A1 92684802 92684916
refGene_gn=MGAT4B;refGene_tx=XM_023254024.1;refGene_gn=MGAT4B;refGene_tx=XM_023254032.1
A1 92685141 92685217
refGene_gn=MGAT4B;refGene_tx=XM_023254024.1;refGene_gn=MGAT4B;refGene_tx=XM_023254032.1
A1 92685349 92685464
refGene_gn=MGAT4B;refGene_tx=XM_023254024.1;refGene_gn=MGAT4B;refGene_tx=XM_023254032.1
A1 92685790 92685921
refGene_gn=MGAT4B;refGene_tx=XM_023254024.1;refGene_gn=MGAT4B;refGene_tx=XM_023254032.1
A1 92686063 92686171
refGene_gn=MGAT4B;refGene_tx=XM_023254024.1;refGene_gn=MGAT4B;refGene_tx=XM_023254032.1
A1 92686290 92686484
refGene_gn=MGAT4B;refGene_tx=XM_023254024.1;refGene_gn=MGAT4B;refGene_tx=XM_023254032.1
A1 92686882 92686961
refGene_gn=MGAT4B;refGene_tx=XM_023254024.1;refGene_gn=MGAT4B;refGene_tx=XM_023254032.1
A1 92687043 92687131
refGene_gn=MGAT4B;refGene_tx=XM_023254024.1;refGene_gn=MGAT4B;refGene_tx=XM_023254032.1
A1 92687214 92687327
refGene_gn=MGAT4B;refGene_tx=XM_023254024.1;refGene_gn=MGAT4B;refGene_tx=XM_023254032.1
A1 92687422 92687446
refGene_gn=MGAT4B;refGene_tx=XM_023254024.1;refGene_gn=MGAT4B;refGene_tx=XM_023254032.1
A1 92689012 92689154
refGene_gn=LTC4S;refGene_tx=XM_003980687.5;refGene_gn=LTC4S;refGene_tx=XM_011281552.3
A1 92689387 92689469
refGene_gn=LTC4S;refGene_tx=XM_003980687.5;refGene_gn=LTC4S;refGene_tx

=XM_011281552.3
A1 92689556 92689627
refGene_gn=LTC4S;refGene_tx=XM_003980687.5;refGene_gn=LTC4S;refGene_tx
=XM_011281552.3
A1 92689733 92689833
refGene_gn=LTC4S;refGene_tx=XM_003980687.5;refGene_gn=LTC4S;refGene_tx
=XM_011281552.3
A1 92691674 92691732 refGene_gn=LTC4S;refGene_tx=XM_003980687.5
A1 92702660 92702718 refGene_gn=LTC4S;refGene_tx=XM_011281552.3
A1 92713189 92714169 refGene_gn=MAML1;refGene_tx=XM_023254056.1
A1 92716698 92716795 refGene_gn=MAML1;refGene_tx=XM_023254056.1
A1 92718952 92719192 refGene_gn=MAML1;refGene_tx=XM_023254056.1
A1 92722051 92723464 refGene_gn=MAML1;refGene_tx=XM_023254056.1
A1 92754440 92754794 refGene_gn=MAML1;refGene_tx=XM_023254056.1
A1 92758997 92759051
refGene_gn=CANX;refGene_tx=XM_003980647.4;refGene_gn=CANX;refGene_tx=X
M_023254058.1
A1 92761147 92761227
refGene_gn=CANX;refGene_tx=XM_003980647.4;refGene_gn=CANX;refGene_tx=X
M_023254058.1
A1 92761395 92761522
refGene_gn=CANX;refGene_tx=XM_003980647.4;refGene_gn=CANX;refGene_tx=X
M_023254058.1
A1 92762484 92762604
refGene_gn=CANX;refGene_tx=XM_003980647.4;refGene_gn=CANX;refGene_tx=X
M_023254058.1
A1 92763497 92763713
refGene_gn=CANX;refGene_tx=XM_003980647.4;refGene_gn=CANX;refGene_tx=X
M_023254058.1
A1 92764873 92765030
refGene_gn=CANX;refGene_tx=XM_003980647.4;refGene_gn=CANX;refGene_tx=X
M_023254058.1
A1 92765797 92765911
refGene_gn=CANX;refGene_tx=XM_003980647.4;refGene_gn=CANX;refGene_tx=X
M_023254058.1
A1 92768412 92768602
refGene_gn=CANX;refGene_tx=XM_003980647.4;refGene_gn=CANX;refGene_tx=X
M_023254058.1
A1 92769987 92770180
refGene_gn=CANX;refGene_tx=XM_003980647.4;refGene_gn=CANX;refGene_tx=X
M_023254058.1
A1 92771760 92771842
refGene_gn=CANX;refGene_tx=XM_003980647.4;refGene_gn=CANX;refGene_tx=X
M_023254058.1
A1 92772395 92772537
refGene_gn=CANX;refGene_tx=XM_003980647.4;refGene_gn=CANX;refGene_tx=X
M_023254058.1
A1 92772987 92773046
refGene_gn=CANX;refGene_tx=XM_003980647.4;refGene_gn=CANX;refGene_tx=X
M_023254058.1

A1 92773920 92773994
refGene_gn=CANX;refGene_tx=XM_003980647.4;refGene_gn=CANX;refGene_tx=XM_023254058.1
A1 92774249 92774423
refGene_gn=CANX;refGene_tx=XM_003980647.4;refGene_gn=CANX;refGene_tx=XM_023254058.1
A1 92808827 92808877
refGene_gn=CBY3;refGene_tx=XM_003980689.3;refGene_gn=CBY3;refGene_tx=XM_006927510.2;refGene_gn=CBY3;refGene_tx=XM_019830632.1;refGene_gn=CBY3;refGene_tx=XM_019830637.1;refGene_gn=CBY3;refGene_tx=XM_019830653.1;refGene_gn=CBY3;refGene_tx=XM_019830659.1;refGene_gn=CBY3;refGene_tx=XM_023254066.1;refGene_gn=CBY3;refGene_tx=XM_023254072.1;refGene_gn=CBY3;refGene_tx=XM_023254074.1;refGene_gn=CBY3;refGene_tx=XM_023254079.1
A1 92810067 92810755
refGene_gn=CBY3;refGene_tx=XM_003980689.3;refGene_gn=CBY3;refGene_tx=XM_006927510.2;refGene_gn=CBY3;refGene_tx=XM_019830632.1;refGene_gn=CBY3;refGene_tx=XM_019830637.1;refGene_gn=CBY3;refGene_tx=XM_019830653.1;refGene_gn=CBY3;refGene_tx=XM_019830659.1;refGene_gn=CBY3;refGene_tx=XM_023254066.1;refGene_gn=CBY3;refGene_tx=XM_023254072.1;refGene_gn=CBY3;refGene_tx=XM_023254074.1;refGene_gn=CBY3;refGene_tx=XM_023254079.1
A1 92851594 92851691
refGene_gn=HNRNPH1;refGene_tx=XM_003980649.4;refGene_gn=HNRNPH1;refGene_tx=XM_006927511.2;refGene_gn=HNRNPH1;refGene_tx=XM_006927512.3;refGene_gn=HNRNPH1;refGene_tx=XM_006927513.3;refGene_gn=HNRNPH1;refGene_tx=XM_019830705.2
A1 92853035 92853191
refGene_gn=HNRNPH1;refGene_tx=XM_003980649.4;refGene_gn=HNRNPH1;refGene_tx=XM_006927511.2;refGene_gn=HNRNPH1;refGene_tx=XM_006927512.3;refGene_gn=HNRNPH1;refGene_tx=XM_006927513.3;refGene_gn=HNRNPH1;refGene_tx=XM_019830705.2
A1 92853403 92853547
refGene_gn=HNRNPH1;refGene_tx=XM_003980649.4;refGene_gn=HNRNPH1;refGene_tx=XM_006927511.2;refGene_gn=HNRNPH1;refGene_tx=XM_006927512.3;refGene_gn=HNRNPH1;refGene_tx=XM_006927513.3;refGene_gn=HNRNPH1;refGene_tx=XM_019830705.2
A1 92855033 92855172
refGene_gn=HNRNPH1;refGene_tx=XM_003980649.4;refGene_gn=HNRNPH1;refGene_tx=XM_006927511.2;refGene_gn=HNRNPH1;refGene_tx=XM_006927512.3;refGene_gn=HNRNPH1;refGene_tx=XM_006927513.3;refGene_gn=HNRNPH1;refGene_tx=XM_019830705.2
A1 92856124 92856303
refGene_gn=HNRNPH1;refGene_tx=XM_003980649.4;refGene_gn=HNRNPH1;refGene_tx=XM_006927511.2;refGene_gn=HNRNPH1;refGene_tx=XM_006927512.3;refGene_gn=HNRNPH1;refGene_tx=XM_006927513.3;refGene_gn=HNRNPH1;refGene_tx=XM_019830705.2
A1 92856384 92856456
refGene_gn=HNRNPH1;refGene_tx=XM_003980649.4;refGene_gn=HNRNPH1;refGene_tx=XM_006927511.2;refGene_gn=HNRNPH1;refGene_tx=XM_006927512.3;refGene_gn=HNRNPH1;refGene_tx=XM_006927513.3;refGene_gn=HNRNPH1;refGene_tx=XM_019830705.2

A1	92856512	92856646
refGene_gn=HNRNPH1;refGene_tx=XM_003980649.4;refGene_gn=HNRNPH1;refGene_tx=XM_006927511.2;refGene_gn=HNRNPH1;refGene_tx=XM_006927512.3;refGene_gn=HNRNPH1;refGene_tx=XM_006927513.3;refGene_gn=HNRNPH1;refGene_tx=XM_019830705.2
A1	92856789	92856925
refGene_gn=HNRNPH1;refGene_tx=XM_003980649.4;refGene_gn=HNRNPH1;refGene_tx=XM_006927511.2;refGene_gn=HNRNPH1;refGene_tx=XM_006927512.3;refGene_gn=HNRNPH1;refGene_tx=XM_006927513.3;refGene_gn=HNRNPH1;refGene_tx=XM_019830705.2
A1	92857404	92857494
refGene_gn=HNRNPH1;refGene_tx=XM_003980649.4;refGene_gn=HNRNPH1;refGene_tx=XM_006927511.2;refGene_gn=HNRNPH1;refGene_tx=XM_006927512.3;refGene_gn=HNRNPH1;refGene_tx=XM_006927513.3;refGene_gn=HNRNPH1;refGene_tx=XM_019830705.2
A1	92857556	92857646
refGene_gn=HNRNPH1;refGene_tx=XM_003980649.4;refGene_gn=HNRNPH1;refGene_tx=XM_006927511.2;refGene_gn=HNRNPH1;refGene_tx=XM_006927512.3
A1	92858304	92858397
refGene_gn=HNRNPH1;refGene_tx=XM_003980649.4;refGene_gn=HNRNPH1;refGene_tx=XM_006927511.2;refGene_gn=HNRNPH1;refGene_tx=XM_006927512.3;refGene_gn=HNRNPH1;refGene_tx=XM_006927513.3;refGene_gn=HNRNPH1;refGene_tx=XM_019830705.2
A1	92858868	92858918
refGene_gn=HNRNPH1;refGene_tx=XM_003980649.4;refGene_gn=HNRNPH1;refGene_tx=XM_019830705.2
A1	92859518	92859637
refGene_gn=HNRNPH1;refGene_tx=XM_006927511.2;refGene_gn=HNRNPH1;refGene_tx=XM_006927512.3;refGene_gn=HNRNPH1;refGene_tx=XM_006927513.3
A1	92863191	92863338
refGene_gn=RUFY1;refGene_tx=XM_003980650.4;refGene_gn=RUFY1;refGene_tx=XM_006927515.3
A1	92864458	92864536
refGene_gn=RUFY1;refGene_tx=XM_003980650.4;refGene_gn=RUFY1;refGene_tx=XM_006927515.3
A1	92866036	92866085
refGene_gn=RUFY1;refGene_tx=XM_003980650.4;refGene_gn=RUFY1;refGene_tx=XM_006927515.3
A1	92868404	92868499
refGene_gn=RUFY1;refGene_tx=XM_003980650.4;refGene_gn=RUFY1;refGene_tx=XM_006927515.3
A1	92871233	92871363
refGene_gn=RUFY1;refGene_tx=XM_003980650.4;refGene_gn=RUFY1;refGene_tx=XM_006927515.3
A1	92871966	92872086
refGene_gn=RUFY1;refGene_tx=XM_003980650.4;refGene_gn=RUFY1;refGene_tx=XM_006927515.3
A1	92874166	92874264
refGene_gn=RUFY1;refGene_tx=XM_003980650.4;refGene_gn=RUFY1;refGene_tx=XM_006927515.3

A1 92887347 92887515
refGene_gn=RUFY1;refGene_tx=XM_003980650.4;refGene_gn=RUFY1;refGene_tx
=XM_006927515.3
A1 92888963 92889080
refGene_gn=RUFY1;refGene_tx=XM_003980650.4;refGene_gn=RUFY1;refGene_tx
=XM_006927515.3
A1 92889955 92890057
refGene_gn=RUFY1;refGene_tx=XM_003980650.4;refGene_gn=RUFY1;refGene_tx
=XM_006927515.3
A1 92892401 92892471
refGene_gn=RUFY1;refGene_tx=XM_003980650.4;refGene_gn=RUFY1;refGene_tx
=XM_006927515.3
A1 92895344 92895410
refGene_gn=RUFY1;refGene_tx=XM_003980650.4;refGene_gn=RUFY1;refGene_tx
=XM_006927515.3
A1 92898559 92898621
refGene_gn=RUFY1;refGene_tx=XM_003980650.4;refGene_gn=RUFY1;refGene_tx
=XM_006927515.3
A1 92907831 92907955
refGene_gn=RUFY1;refGene_tx=XM_003980650.4;refGene_gn=RUFY1;refGene_tx
=XM_006927515.3
A1 92909810 92909912 refGene_gn=RUFY1;refGene_tx=XM_003980650.4
A1 92912731 92912849
refGene_gn=RUFY1;refGene_tx=XM_003980650.4;refGene_gn=RUFY1;refGene_tx
=XM_006927515.3
A1 92914142 92914316
refGene_gn=RUFY1;refGene_tx=XM_003980650.4;refGene_gn=RUFY1;refGene_tx
=XM_006927515.3
A1 92924715 92925001
refGene_gn=RUFY1;refGene_tx=XM_003980650.4;refGene_gn=RUFY1;refGene_tx
=XM_006927515.3
A1 93100271 93100404 refGene_gn=ADAMTS2;refGene_tx=XM_023254116.1
A1 93101410 93101805 refGene_gn=ADAMTS2;refGene_tx=XM_023254116.1
A1 93178580 93178740 refGene_gn=ADAMTS2;refGene_tx=XM_023254116.1
A1 93243776 93243979 refGene_gn=ADAMTS2;refGene_tx=XM_023254116.1
A1 93267891 93267975 refGene_gn=ADAMTS2;refGene_tx=XM_023254116.1
A1 93283618 93283775 refGene_gn=ADAMTS2;refGene_tx=XM_023254116.1
A1 93294439 93294545 refGene_gn=ADAMTS2;refGene_tx=XM_023254116.1
A1 93295509 93295653 refGene_gn=ADAMTS2;refGene_tx=XM_023254116.1
A1 93296125 93296258 refGene_gn=ADAMTS2;refGene_tx=XM_023254116.1
A1 93297597 93297711 refGene_gn=ADAMTS2;refGene_tx=XM_023254116.1
A1 93314857 93315003 refGene_gn=ADAMTS2;refGene_tx=XM_023254116.1
A1 93318347 93318523 refGene_gn=ADAMTS2;refGene_tx=XM_023254116.1
A1 93319187 93319321 refGene_gn=ADAMTS2;refGene_tx=XM_023254116.1
A1 93323886 93324010 refGene_gn=ADAMTS2;refGene_tx=XM_023254116.1
A1 93324880 93324961 refGene_gn=ADAMTS2;refGene_tx=XM_023254116.1
A1 93327214 93327381 refGene_gn=ADAMTS2;refGene_tx=XM_023254116.1
A1 93329314 93329474 refGene_gn=ADAMTS2;refGene_tx=XM_023254116.1
A1 93331320 93331453 refGene_gn=ADAMTS2;refGene_tx=XM_023254116.1
A1 93332205 93332413 refGene_gn=ADAMTS2;refGene_tx=XM_023254116.1

A1 93333671 93333801 refGene_gn=ADAMTS2;refGene_tx=XM_023254116.1
A1 93334622 93334712 refGene_gn=ADAMTS2;refGene_tx=XM_023254116.1
A1 93342536 93342994 refGene_gn=ADAMTS2;refGene_tx=XM_023254116.1
A1 93366168 93367583 refGene_gn=ZNF879;refGene_tx=XM_023242170.1
A1 93373089 93373185 refGene_gn=ZNF879;refGene_tx=XM_023242170.1
A1 93373594 93373721 refGene_gn=ZNF879;refGene_tx=XM_023242170.1
A1 93379989 93380069 refGene_gn=ZNF879;refGene_tx=XM_023242170.1
A1 93381485 93381717 refGene_gn=ZNF879;refGene_tx=XM_023242170.1
A1 93391329 93391357 refGene_gn=ZNF879;refGene_tx=XM_023242170.1
A1 93391358 93392790 refGene_gn=ZNF879;refGene_tx=XM_023242170.1
A1 93398562 93398658 refGene_gn=ZNF879;refGene_tx=XM_023242170.1
A1 93399078 93399205 refGene_gn=ZNF879;refGene_tx=XM_023242170.1
A1 93399353 93399428 refGene_gn=ZNF879;refGene_tx=XM_023242170.1
A1 93403382 93403450 refGene_gn=ZNF879;refGene_tx=XM_023242170.1
A1 93403857 93403864 refGene_gn=ZNF879;refGene_tx=XM_023242170.1
A1 93420728 93421235
refGene_gn=GRM6;refGene_tx=XM_023254130.1;refGene_gn=GRM6;refGene_tx=XM_023254137.1
A1 93423830 93423941 refGene_gn=GRM6;refGene_tx=XM_019830717.1
A1 93424170 93424387
refGene_gn=GRM6;refGene_tx=XM_019830717.1;refGene_gn=GRM6;refGene_tx=XM_019830721.1;refGene_gn=GRM6;refGene_tx=XM_019830722.1;refGene_gn=GRM6;refGene_tx=XM_023254130.1;refGene_gn=GRM6;refGene_tx=XM_023254137.1
A1 93424582 93424718
refGene_gn=GRM6;refGene_tx=XM_019830717.1;refGene_gn=GRM6;refGene_tx=XM_019830721.1;refGene_gn=GRM6;refGene_tx=XM_019830722.1;refGene_gn=GRM6;refGene_tx=XM_023254130.1;refGene_gn=GRM6;refGene_tx=XM_023254137.1
A1 93425417 93425572
refGene_gn=GRM6;refGene_tx=XM_019830717.1;refGene_gn=GRM6;refGene_tx=XM_019830721.1;refGene_gn=GRM6;refGene_tx=XM_019830722.1;refGene_gn=GRM6;refGene_tx=XM_023254130.1;refGene_gn=GRM6;refGene_tx=XM_023254137.1
A1 93426700 93426841
refGene_gn=GRM6;refGene_tx=XM_019830717.1;refGene_gn=GRM6;refGene_tx=XM_019830721.1;refGene_gn=GRM6;refGene_tx=XM_019830722.1;refGene_gn=GRM6;refGene_tx=XM_023254130.1;refGene_gn=GRM6;refGene_tx=XM_023254137.1
A1 93426944 93427145
refGene_gn=GRM6;refGene_tx=XM_019830717.1;refGene_gn=GRM6;refGene_tx=XM_019830721.1;refGene_gn=GRM6;refGene_tx=XM_019830722.1;refGene_gn=GRM6;refGene_tx=XM_023254130.1;refGene_gn=GRM6;refGene_tx=XM_023254137.1
A1 93429733 93429879
refGene_gn=GRM6;refGene_tx=XM_019830717.1;refGene_gn=GRM6;refGene_tx=XM_019830721.1;refGene_gn=GRM6;refGene_tx=XM_019830722.1;refGene_gn=GRM6;refGene_tx=XM_023254130.1;refGene_gn=GRM6;refGene_tx=XM_023254137.1
A1 93429972 93430596
refGene_gn=GRM6;refGene_tx=XM_019830717.1;refGene_gn=GRM6;refGene_tx=XM_019830721.1;refGene_gn=GRM6;refGene_tx=XM_019830722.1;refGene_gn=GRM6;refGene_tx=XM_023254130.1;refGene_gn=GRM6;refGene_tx=XM_023254137.1
A1 93433108 93433420
refGene_gn=GRM6;refGene_tx=XM_019830717.1;refGene_gn=GRM6;refGene_tx=XM_019830721.1;refGene_gn=GRM6;refGene_tx=XM_019830722.1;refGene_gn=GRM

6;refGene_tx=XM_023254130.1;refGene_gn=GRM6;refGene_tx=XM_023254137.1
A1 93434135 93434333
refGene_gn=GRM6;refGene_tx=XM_019830717.1;refGene_gn=GRM6;refGene_tx=XM_019830721.1;refGene_gn=GRM6;refGene_tx=XM_019830722.1;refGene_gn=GRM6;refGene_tx=XM_023254130.1;refGene_gn=GRM6;refGene_tx=XM_023254137.1
A1 93448039 93449355
refGene_gn=ZNF454;refGene_tx=XM_006927523.3;refGene_gn=ZNF454;refGene_tx=XM_011281609.3;refGene_gn=ZNF454;refGene_tx=XM_019830786.2
A1 93465410 93465500
refGene_gn=ZNF454;refGene_tx=XM_006927523.3;refGene_gn=ZNF454;refGene_tx=XM_011281609.3
A1 93465919 93466046
refGene_gn=ZNF454;refGene_tx=XM_006927523.3;refGene_gn=ZNF454;refGene_tx=XM_011281609.3
A1 93468780 93468813
refGene_gn=ZNF454;refGene_tx=XM_006927523.3;refGene_gn=ZNF454;refGene_tx=XM_011281609.3
A1 93474706 93475011 refGene_gn=ZFP2;refGene_tx=XM_019830779.2
A1 93475013 93475182 refGene_gn=ZFP2;refGene_tx=XM_019830779.2
A1 93475183 93476101 refGene_gn=ZFP2;refGene_tx=XM_019830779.2
A1 93503447 93505030 refGene_gn=ZNF354B;refGene_tx=XM_019830725.2
A1 93519352 93519448 refGene_gn=ZNF354B;refGene_tx=XM_019830725.2
A1 93520627 93520754 refGene_gn=ZNF354B;refGene_tx=XM_019830725.2
A1 93524851 93524884 refGene_gn=ZNF354B;refGene_tx=XM_019830725.2
A1 93539324 93540257
refGene_gn=LOC101097942;refGene_tx=XM_003980693.1
A1 93570850 93570983 refGene_gn=PROP1;refGene_tx=XM_003980653.3
A1 93572433 93572544 refGene_gn=PROP1;refGene_tx=XM_003980653.3
A1 93573737 93573907 refGene_gn=PROP1;refGene_tx=XM_003980653.3
A1 93573908 93573971 refGene_gn=PROP1;refGene_tx=XM_003980653.3
A1 93574909 93575248 refGene_gn=PROP1;refGene_tx=XM_003980653.3
A1 93758577 93759519
refGene_gn=LOC111560612;refGene_tx=XM_023254753.1
A1 93784888 93784921
refGene_gn=LOC101087636;refGene_tx=XM_003980655.5
A1 93788989 93789116
refGene_gn=LOC101087636;refGene_tx=XM_003980655.5;refGene_gn=LOC101087636;refGene_tx=XM_019830822.2
A1 93790345 93790441
refGene_gn=LOC101087636;refGene_tx=XM_003980655.5;refGene_gn=LOC101087636;refGene_tx=XM_019830822.2
A1 93805454 93807007
refGene_gn=LOC101087636;refGene_tx=XM_003980655.5;refGene_gn=LOC101087636;refGene_tx=XM_019830822.2;refGene_gn=LOC101087636;refGene_tx=XM_019830823.2
A1 93836990 93837151
refGene_gn=CLK4;refGene_tx=XM_003980656.5;refGene_gn=CLK4;refGene_tx=XM_019830843.2
A1 93841902 93842125
refGene_gn=CLK4;refGene_tx=XM_003980656.5;refGene_gn=CLK4;refGene_tx=X

M_019830843.2
A1 93843238 93843329
refGene_gn=CLK4;refGene_tx=XM_003980656.5;refGene_gn=CLK4;refGene_tx=XM_019830843.2
A1 93843723 93843790
refGene_gn=CLK4;refGene_tx=XM_003980656.5;refGene_gn=CLK4;refGene_tx=XM_019830843.2;refGene_gn=CLK4;refGene_tx=XM_019830853.1
A1 93848094 93848211
refGene_gn=CLK4;refGene_tx=XM_003980656.5;refGene_gn=CLK4;refGene_tx=XM_019830843.2;refGene_gn=CLK4;refGene_tx=XM_019830853.1
A1 93848300 93848467
refGene_gn=CLK4;refGene_tx=XM_003980656.5;refGene_gn=CLK4;refGene_tx=XM_019830843.2;refGene_gn=CLK4;refGene_tx=XM_019830853.1
A1 93848665 93848760
refGene_gn=CLK4;refGene_tx=XM_003980656.5;refGene_gn=CLK4;refGene_tx=XM_019830843.2;refGene_gn=CLK4;refGene_tx=XM_019830853.1
A1 93849014 93849144
refGene_gn=CLK4;refGene_tx=XM_003980656.5;refGene_gn=CLK4;refGene_tx=XM_019830843.2;refGene_gn=CLK4;refGene_tx=XM_019830853.1
A1 93852386 93852469
refGene_gn=CLK4;refGene_tx=XM_003980656.5;refGene_gn=CLK4;refGene_tx=XM_019830843.2;refGene_gn=CLK4;refGene_tx=XM_019830853.1
A1 93856113 93856193
refGene_gn=CLK4;refGene_tx=XM_003980656.5;refGene_gn=CLK4;refGene_tx=XM_019830843.2;refGene_gn=CLK4;refGene_tx=XM_019830853.1
A1 93856848 93856939
refGene_gn=CLK4;refGene_tx=XM_003980656.5;refGene_gn=CLK4;refGene_tx=XM_019830843.2;refGene_gn=CLK4;refGene_tx=XM_019830853.1
A1 93857031 93857172
refGene_gn=CLK4;refGene_tx=XM_003980656.5;refGene_gn=CLK4;refGene_tx=XM_019830843.2;refGene_gn=CLK4;refGene_tx=XM_019830853.1
A1 93871088 93871379
refGene_gn=COL23A1;refGene_tx=XM_023254174.1;refGene_gn=COL23A1;refGene_tx=XM_023254176.1;refGene_gn=COL23A1;refGene_tx=XM_023254178.1
A1 93895926 93895993
refGene_gn=COL23A1;refGene_tx=XM_023254174.1;refGene_gn=COL23A1;refGene_tx=XM_023254176.1;refGene_gn=COL23A1;refGene_tx=XM_023254178.1
A1 94165531 94165576
refGene_gn=COL23A1;refGene_tx=XM_023254174.1;refGene_gn=COL23A1;refGene_tx=XM_023254176.1;refGene_gn=COL23A1;refGene_tx=XM_023254178.1
A1 94185228 94185236
refGene_gn=COL23A1;refGene_tx=XM_023254174.1;refGene_gn=COL23A1;refGene_tx=XM_023254176.1;refGene_gn=COL23A1;refGene_tx=XM_023254178.1
A1 94186836 94186863
refGene_gn=COL23A1;refGene_tx=XM_023254174.1;refGene_gn=COL23A1;refGene_tx=XM_023254178.1
A1 94205349 94205376
refGene_gn=COL23A1;refGene_tx=XM_023254174.1;refGene_gn=COL23A1;refGene_tx=XM_023254176.1;refGene_gn=COL23A1;refGene_tx=XM_023254178.1
A1 94206933 94206960

refGene_gn=COL23A1;refGene_tx=XM_023254174.1;refGene_gn=COL23A1;refGene_tx=XM_023254176.1;refGene_gn=COL23A1;refGene_tx=XM_023254178.1
A1 94208328 94208355
refGene_gn=COL23A1;refGene_tx=XM_023254174.1;refGene_gn=COL23A1;refGene_tx=XM_023254176.1;refGene_gn=COL23A1;refGene_tx=XM_023254178.1
A1 94212935 94213052
refGene_gn=COL23A1;refGene_tx=XM_023254174.1;refGene_gn=COL23A1;refGene_tx=XM_023254176.1;refGene_gn=COL23A1;refGene_tx=XM_023254178.1
A1 94213849 94213885
refGene_gn=COL23A1;refGene_tx=XM_023254174.1;refGene_gn=COL23A1;refGene_tx=XM_023254176.1
A1 94214269 94214296
refGene_gn=COL23A1;refGene_tx=XM_023254174.1;refGene_gn=COL23A1;refGene_tx=XM_023254176.1;refGene_gn=COL23A1;refGene_tx=XM_023254178.1
A1 94214762 94214789
refGene_gn=COL23A1;refGene_tx=XM_023254174.1;refGene_gn=COL23A1;refGene_tx=XM_023254176.1;refGene_gn=COL23A1;refGene_tx=XM_023254178.1
A1 94216632 94216686
refGene_gn=COL23A1;refGene_tx=XM_023254174.1;refGene_gn=COL23A1;refGene_tx=XM_023254176.1;refGene_gn=COL23A1;refGene_tx=XM_023254178.1
A1 94217200 94217263
refGene_gn=COL23A1;refGene_tx=XM_023254174.1;refGene_gn=COL23A1;refGene_tx=XM_023254176.1;refGene_gn=COL23A1;refGene_tx=XM_023254178.1
A1 94217697 94217742
refGene_gn=COL23A1;refGene_tx=XM_023254174.1;refGene_gn=COL23A1;refGene_tx=XM_023254176.1;refGene_gn=COL23A1;refGene_tx=XM_023254178.1
A1 94219447 94219525
refGene_gn=COL23A1;refGene_tx=XM_023254174.1;refGene_gn=COL23A1;refGene_tx=XM_023254176.1;refGene_gn=COL23A1;refGene_tx=XM_023254178.1
A1 94221824 94221878
refGene_gn=COL23A1;refGene_tx=XM_023254174.1;refGene_gn=COL23A1;refGene_tx=XM_023254176.1;refGene_gn=COL23A1;refGene_tx=XM_023254178.1
A1 94222533 94222578
refGene_gn=COL23A1;refGene_tx=XM_023254174.1;refGene_gn=COL23A1;refGene_tx=XM_023254176.1;refGene_gn=COL23A1;refGene_tx=XM_023254178.1
A1 94223437 94223527
refGene_gn=COL23A1;refGene_tx=XM_023254174.1;refGene_gn=COL23A1;refGene_tx=XM_023254176.1;refGene_gn=COL23A1;refGene_tx=XM_023254178.1
A1 94224351 94224414
refGene_gn=COL23A1;refGene_tx=XM_023254174.1;refGene_gn=COL23A1;refGene_tx=XM_023254176.1;refGene_gn=COL23A1;refGene_tx=XM_023254178.1
A1 94224757 94224814
refGene_gn=COL23A1;refGene_tx=XM_023254174.1;refGene_gn=COL23A1;refGene_tx=XM_023254176.1;refGene_gn=COL23A1;refGene_tx=XM_023254178.1
A1 94225090 94225117
refGene_gn=COL23A1;refGene_tx=XM_023254174.1;refGene_gn=COL23A1;refGene_tx=XM_023254176.1;refGene_gn=COL23A1;refGene_tx=XM_023254178.1
A1 94226073 94226136
refGene_gn=COL23A1;refGene_tx=XM_023254174.1;refGene_gn=COL23A1;refGene_tx=XM_023254176.1;refGene_gn=COL23A1;refGene_tx=XM_023254178.1

A1 94226221 94226275
refGene_gn=COL23A1;refGene_tx=XM_023254174.1;refGene_gn=COL23A1;refGene_tx=XM_023254176.1;refGene_gn=COL23A1;refGene_tx=XM_023254178.1
A1 94226479 94226506
refGene_gn=COL23A1;refGene_tx=XM_023254174.1;refGene_gn=COL23A1;refGene_tx=XM_023254176.1;refGene_gn=COL23A1;refGene_tx=XM_023254178.1
A1 94229678 94229732
refGene_gn=COL23A1;refGene_tx=XM_023254174.1;refGene_gn=COL23A1;refGene_tx=XM_023254176.1;refGene_gn=COL23A1;refGene_tx=XM_023254178.1
A1 94229871 94229958
refGene_gn=COL23A1;refGene_tx=XM_023254174.1;refGene_gn=COL23A1;refGene_tx=XM_023254176.1;refGene_gn=COL23A1;refGene_tx=XM_023254178.1
A1 94231801 94231840
refGene_gn=COL23A1;refGene_tx=XM_023254174.1;refGene_gn=COL23A1;refGene_tx=XM_023254176.1;refGene_gn=COL23A1;refGene_tx=XM_023254178.1
A1 94232260 94232263
refGene_gn=COL23A1;refGene_tx=XM_023254174.1;refGene_gn=COL23A1;refGene_tx=XM_023254176.1;refGene_gn=COL23A1;refGene_tx=XM_023254178.1
A1 94236958 94237017 refGene_gn=PHYKPL;refGene_tx=XM_023254182.1
A1 94237735 94238009
refGene_gn=PHYKPL;refGene_tx=XM_006927528.4;refGene_gn=PHYKPL;refGene_tx=XM_023254182.1
A1 94239063 94239223
refGene_gn=PHYKPL;refGene_tx=XM_006927528.4;refGene_gn=PHYKPL;refGene_tx=XM_023254182.1
A1 94243253 94243328
refGene_gn=PHYKPL;refGene_tx=XM_006927528.4;refGene_gn=PHYKPL;refGene_tx=XM_023254182.1
A1 94243894 94243982
refGene_gn=PHYKPL;refGene_tx=XM_006927528.4;refGene_gn=PHYKPL;refGene_tx=XM_023254182.1
A1 94244059 94244176
refGene_gn=PHYKPL;refGene_tx=XM_006927528.4;refGene_gn=PHYKPL;refGene_tx=XM_023254182.1
A1 94245524 94245607
refGene_gn=PHYKPL;refGene_tx=XM_006927528.4;refGene_gn=PHYKPL;refGene_tx=XM_023254182.1
A1 94245851 94246077
refGene_gn=PHYKPL;refGene_tx=XM_006927528.4;refGene_gn=PHYKPL;refGene_tx=XM_023254182.1
A1 94268597 94268752
refGene_gn=PHYKPL;refGene_tx=XM_006927528.4;refGene_gn=PHYKPL;refGene_tx=XM_023254182.1
A1 94269148 94269238
refGene_gn=PHYKPL;refGene_tx=XM_006927528.4;refGene_gn=PHYKPL;refGene_tx=XM_023254182.1
A1 94270886 94271017
refGene_gn=PHYKPL;refGene_tx=XM_006927528.4;refGene_gn=PHYKPL;refGene_tx=XM_023254182.1
A1 94271910 94271960

refGene_gn=PHYKPL;refGene_tx=XM_006927528.4;refGene_gn=PHYKPL;refGene_tx=XM_023254182.1
A1 94273355 94273426 refGene_gn=HNRNPAB;refGene_tx=XM_023254183.1;refGene_gn=HNRNPAB;refGene_tx=XM_023254184.1
A1 94273727 94273868 refGene_gn=HNRNPAB;refGene_tx=XM_023254183.1
A1 94274518 94274636 refGene_gn=HNRNPAB;refGene_tx=XM_023254183.1;refGene_gn=HNRNPAB;refGene_tx=XM_023254184.1
A1 94276511 94276643 refGene_gn=HNRNPAB;refGene_tx=XM_023254183.1;refGene_gn=HNRNPAB;refGene_tx=XM_023254184.1
A1 94276845 94277004 refGene_gn=HNRNPAB;refGene_tx=XM_023254183.1;refGene_gn=HNRNPAB;refGene_tx=XM_023254184.1
A1 94277647 94277816 refGene_gn=HNRNPAB;refGene_tx=XM_023254183.1;refGene_gn=HNRNPAB;refGene_tx=XM_023254184.1
A1 94278585 94278794 refGene_gn=HNRNPAB;refGene_tx=XM_023254183.1;refGene_gn=HNRNPAB;refGene_tx=XM_023254184.1
A1 94311597 94311757 refGene_gn=NHP2;refGene_tx=XM_003980657.5
A1 94311836 94311906 refGene_gn=NHP2;refGene_tx=XM_003980657.5
A1 94314262 94314368 refGene_gn=NHP2;refGene_tx=XM_003980657.5
A1 94314978 94315104 refGene_gn=NHP2;refGene_tx=XM_003980657.5
A1 94316581 94316645 refGene_gn=RMND5B;refGene_tx=XM_003980658.4;refGene_gn=RMND5B;refGene_tx=XM_006927529.3;refGene_gn=RMND5B;refGene_tx=XM_006927530.3;refGene_gn=RMND5B;refGene_tx=XM_011281652.2;refGene_gn=RMND5B;refGene_tx=XM_019830884.1;refGene_gn=RMND5B;refGene_tx=XM_023254191.1
A1 94316731 94316886 refGene_gn=RMND5B;refGene_tx=XM_003980658.4;refGene_gn=RMND5B;refGene_tx=XM_006927529.3;refGene_gn=RMND5B;refGene_tx=XM_006927530.3;refGene_gn=RMND5B;refGene_tx=XM_011281652.2;refGene_gn=RMND5B;refGene_tx=XM_019830884.1;refGene_gn=RMND5B;refGene_tx=XM_023254191.1
A1 94316987 94317090 refGene_gn=RMND5B;refGene_tx=XM_003980658.4;refGene_gn=RMND5B;refGene_tx=XM_006927529.3;refGene_gn=RMND5B;refGene_tx=XM_006927530.3;refGene_gn=RMND5B;refGene_tx=XM_011281652.2;refGene_gn=RMND5B;refGene_tx=XM_019830884.1;refGene_gn=RMND5B;refGene_tx=XM_023254191.1
A1 94318292 94318458 refGene_gn=RMND5B;refGene_tx=XM_003980658.4;refGene_gn=RMND5B;refGene_tx=XM_006927529.3;refGene_gn=RMND5B;refGene_tx=XM_006927530.3;refGene_gn=RMND5B;refGene_tx=XM_011281652.2;refGene_gn=RMND5B;refGene_tx=XM_019830884.1;refGene_gn=RMND5B;refGene_tx=XM_023254191.1
A1 94319362 94319529 refGene_gn=RMND5B;refGene_tx=XM_003980658.4;refGene_gn=RMND5B;refGene_tx=XM_006927529.3;refGene_gn=RMND5B;refGene_tx=XM_006927530.3;refGene_gn=RMND5B;refGene_tx=XM_011281652.2;refGene_gn=RMND5B;refGene_tx=XM_019830884.1;refGene_gn=RMND5B;refGene_tx=XM_023254191.1

A1 94319710 94319811
refGene_gn=RMND5B;refGene_tx=XM_003980658.4;refGene_gn=RMND5B;refGene_tx=XM_006927529.3;refGene_gn=RMND5B;refGene_tx=XM_006927530.3;refGene_gn=RMND5B;refGene_tx=XM_011281652.2;refGene_gn=RMND5B;refGene_tx=XM_019830884.1;refGene_gn=RMND5B;refGene_tx=XM_023254191.1
A1 94320321 94320462
refGene_gn=RMND5B;refGene_tx=XM_003980658.4;refGene_gn=RMND5B;refGene_tx=XM_006927529.3;refGene_gn=RMND5B;refGene_tx=XM_006927530.3;refGene_gn=RMND5B;refGene_tx=XM_011281652.2;refGene_gn=RMND5B;refGene_tx=XM_019830884.1;refGene_gn=RMND5B;refGene_tx=XM_023254191.1
A1 94320597 94320743
refGene_gn=RMND5B;refGene_tx=XM_003980658.4;refGene_gn=RMND5B;refGene_tx=XM_006927529.3;refGene_gn=RMND5B;refGene_tx=XM_006927530.3;refGene_gn=RMND5B;refGene_tx=XM_011281652.2;refGene_gn=RMND5B;refGene_tx=XM_019830884.1;refGene_gn=RMND5B;refGene_tx=XM_023254191.1
A1 94321882 94322018 refGene_gn=RMND5B;refGene_tx=XM_019830884.1
A1 94328662 94328801
refGene_gn=RMND5B;refGene_tx=XM_003980658.4;refGene_gn=RMND5B;refGene_tx=XM_006927529.3;refGene_gn=RMND5B;refGene_tx=XM_011281652.2;refGene_gn=RMND5B;refGene_tx=XM_023254191.1
A1 94342096 94342627
refGene_gn=N4BP3;refGene_tx=XM_006927531.4;refGene_gn=N4BP3;refGene_tx=XM_023254206.1
A1 94342758 94343013
refGene_gn=N4BP3;refGene_tx=XM_006927531.4;refGene_gn=N4BP3;refGene_tx=XM_023254206.1
A1 94343364 94343892
refGene_gn=N4BP3;refGene_tx=XM_006927531.4;refGene_gn=N4BP3;refGene_tx=XM_023254206.1
A1 94344150 94344471
refGene_gn=N4BP3;refGene_tx=XM_006927531.4;refGene_gn=N4BP3;refGene_tx=XM_023254206.1
A1 94349227 94349403
refGene_gn=LOC109499762;refGene_tx=XM_023242237.1
A1 94349517 94349648
refGene_gn=LOC109499762;refGene_tx=XM_023242237.1
A1 94349986 94350311
refGene_gn=LOC109499762;refGene_tx=XM_023242237.1
A1 94377075 94377225
refGene_gn=LOC109499762;refGene_tx=XM_023242237.1
A1 94388654 94388763
refGene_gn=LOC109499762;refGene_tx=XM_023242237.1
A1 94428317 94428473
refGene_gn=B4GALT7;refGene_tx=XM_023254211.1;refGene_gn=B4GALT7;refGene_tx=XM_023254218.1;refGene_gn=B4GALT7;refGene_tx=XM_023254221.1;refGene_gn=B4GALT7;refGene_tx=XM_023254225.1
A1 94428906 94429011
refGene_gn=B4GALT7;refGene_tx=XM_023254211.1;refGene_gn=B4GALT7;refGene_tx=XM_023254221.1
A1 94429279 94429363

refGene_gn=B4GALT7;refGene_tx=XM_023254211.1;refGene_gn=B4GALT7;refGene_tx=XM_023254218.1;refGene_gn=B4GALT7;refGene_tx=XM_023254221.1;refGene_gn=B4GALT7;refGene_tx=XM_023254225.1
A1 94430654 94430880
refGene_gn=B4GALT7;refGene_tx=XM_023254211.1;refGene_gn=B4GALT7;refGene_tx=XM_023254218.1;refGene_gn=B4GALT7;refGene_tx=XM_023254221.1;refGene_gn=B4GALT7;refGene_tx=XM_023254225.1
A1 94433589 94433952
refGene_gn=B4GALT7;refGene_tx=XM_023254211.1;refGene_gn=B4GALT7;refGene_tx=XM_023254218.1;refGene_gn=B4GALT7;refGene_tx=XM_023254221.1;refGene_gn=B4GALT7;refGene_tx=XM_023254225.1
A1 94435448 94435610
refGene_gn=B4GALT7;refGene_tx=XM_023254211.1;refGene_gn=B4GALT7;refGene_tx=XM_023254218.1
A1 94437374 94437424
refGene_gn=B4GALT7;refGene_tx=XM_023254211.1;refGene_gn=B4GALT7;refGene_tx=XM_023254218.1;refGene_gn=B4GALT7;refGene_tx=XM_023254221.1;refGene_gn=B4GALT7;refGene_tx=XM_023254225.1
A1 94440813 94440963 refGene_gn=TMED9;refGene_tx=XM_003980660.5
A1 94442029 94442176 refGene_gn=TMED9;refGene_tx=XM_003980660.5
A1 94442535 94442661 refGene_gn=TMED9;refGene_tx=XM_003980660.5
A1 94443435 94443536 refGene_gn=TMED9;refGene_tx=XM_003980660.5
A1 94443706 94443890 refGene_gn=TMED9;refGene_tx=XM_003980660.5
A1 94453703 94453833
refGene_gn=LOC101099424;refGene_tx=XM_003980699.5;refGene_gn=LOC101099424;refGene_tx=XM_019830937.2;refGene_gn=LOC101099424;refGene_tx=XM_023254236.1
A1 94454198 94454299
refGene_gn=LOC101099424;refGene_tx=XM_003980699.5;refGene_gn=LOC101099424;refGene_tx=XM_019830937.2;refGene_gn=LOC101099424;refGene_tx=XM_023254236.1
A1 94454629 94454755
refGene_gn=LOC101099424;refGene_tx=XM_003980699.5;refGene_gn=LOC101099424;refGene_tx=XM_019830937.2
A1 94455304 94455451
refGene_gn=LOC101099424;refGene_tx=XM_003980699.5;refGene_gn=LOC101099424;refGene_tx=XM_019830937.2;refGene_gn=LOC101099424;refGene_tx=XM_023254236.1
A1 94455594 94455744
refGene_gn=LOC101099424;refGene_tx=XM_003980699.5;refGene_gn=LOC101099424;refGene_tx=XM_019830937.2;refGene_gn=LOC101099424;refGene_tx=XM_023254236.1
A1 94471241 94471422 refGene_gn=YTHDC2;refGene_tx=XM_019830944.2
A1 94472604 94472695 refGene_gn=YTHDC2;refGene_tx=XM_019830944.2
A1 94480236 94480433 refGene_gn=YTHDC2;refGene_tx=XM_019830944.2
A1 94481750 94481950
refGene_gn=YTHDC2;refGene_tx=XM_019830944.2;refGene_gn=YTHDC2;refGene_tx=XM_019830946.1
A1 94485325 94485492
refGene_gn=YTHDC2;refGene_tx=XM_019830944.2;refGene_gn=YTHDC2;refGene_

tx=XM_019830946.1
A1	94486008	94486111
refGene_gn=YTHDC2;refGene_tx=XM_019830944.2;refGene_gn=YTHDC2;refGene_tx=XM_019830946.1
A1	94487001	94487158
refGene_gn=YTHDC2;refGene_tx=XM_019830944.2;refGene_gn=YTHDC2;refGene_tx=XM_019830946.1
A1	94491084	94491192
refGene_gn=YTHDC2;refGene_tx=XM_019830944.2;refGene_gn=YTHDC2;refGene_tx=XM_019830946.1
A1	94492531	94492680
refGene_gn=YTHDC2;refGene_tx=XM_019830944.2;refGene_gn=YTHDC2;refGene_tx=XM_019830946.1
A1	94494188	94494324
refGene_gn=YTHDC2;refGene_tx=XM_019830944.2;refGene_gn=YTHDC2;refGene_tx=XM_019830946.1
A1	94498429	94498556
refGene_gn=YTHDC2;refGene_tx=XM_019830944.2;refGene_gn=YTHDC2;refGene_tx=XM_019830946.1
A1	94498759	94498825
refGene_gn=YTHDC2;refGene_tx=XM_019830944.2;refGene_gn=YTHDC2;refGene_tx=XM_019830946.1
A1	94503276	94503455
refGene_gn=YTHDC2;refGene_tx=XM_019830944.2;refGene_gn=YTHDC2;refGene_tx=XM_019830946.1
A1	94503705	94503802
refGene_gn=YTHDC2;refGene_tx=XM_019830944.2;refGene_gn=YTHDC2;refGene_tx=XM_019830946.1
A1	94503881	94503969
refGene_gn=YTHDC2;refGene_tx=XM_019830944.2;refGene_gn=YTHDC2;refGene_tx=XM_019830946.1
A1	94504057	94504138
refGene_gn=YTHDC2;refGene_tx=XM_019830944.2;refGene_gn=YTHDC2;refGene_tx=XM_019830946.1
A1	94506090	94506173
refGene_gn=YTHDC2;refGene_tx=XM_019830944.2;refGene_gn=YTHDC2;refGene_tx=XM_019830946.1
A1	94509598	94509704
refGene_gn=YTHDC2;refGene_tx=XM_019830944.2;refGene_gn=YTHDC2;refGene_tx=XM_019830946.1
A1	94511162	94511282
refGene_gn=YTHDC2;refGene_tx=XM_019830944.2;refGene_gn=YTHDC2;refGene_tx=XM_019830946.1
A1	94511603	94511876
refGene_gn=YTHDC2;refGene_tx=XM_019830944.2;refGene_gn=YTHDC2;refGene_tx=XM_019830946.1
A1	94515069	94515196
refGene_gn=YTHDC2;refGene_tx=XM_019830944.2;refGene_gn=YTHDC2;refGene_tx=XM_019830946.1
A1	94516349	94516555

refGene_gn=YTHDC2;refGene_tx=XM_019830944.2;refGene_gn=YTHDC2;refGene_tx=XM_019830946.1
A1 94516899 94517095
refGene_gn=YTHDC2;refGene_tx=XM_019830944.2;refGene_gn=YTHDC2;refGene_tx=XM_019830946.1
A1 94524767 94524877
refGene_gn=YTHDC2;refGene_tx=XM_019830944.2;refGene_gn=YTHDC2;refGene_tx=XM_019830946.1
A1 94526592 94526885
refGene_gn=YTHDC2;refGene_tx=XM_019830944.2;refGene_gn=YTHDC2;refGene_tx=XM_019830946.1
A1 94530922 94531100
refGene_gn=YTHDC2;refGene_tx=XM_019830944.2;refGene_gn=YTHDC2;refGene_tx=XM_019830946.1
A1 94537004 94537208
refGene_gn=YTHDC2;refGene_tx=XM_019830944.2;refGene_gn=YTHDC2;refGene_tx=XM_019830946.1
A1 94537904 94538087
refGene_gn=YTHDC2;refGene_tx=XM_019830944.2;refGene_gn=YTHDC2;refGene_tx=XM_019830946.1
A1 94539281 94539362
refGene_gn=YTHDC2;refGene_tx=XM_019830944.2;refGene_gn=YTHDC2;refGene_tx=XM_019830946.1
A1 95252358 95252380
refGene_gn=KCNN2;refGene_tx=XM_003980701.5;refGene_gn=KCNN2;refGene_tx=XM_019830963.2
A1 95253245 95254531
refGene_gn=KCNN2;refGene_tx=XM_003980701.5;refGene_gn=KCNN2;refGene_tx=XM_019830963.2
A1 95255185 95255281
refGene_gn=KCNN2;refGene_tx=XM_003980701.5;refGene_gn=KCNN2;refGene_tx=XM_019830963.2
A1 95298330 95298749
refGene_gn=KCNN2;refGene_tx=XM_003980701.5;refGene_gn=KCNN2;refGene_tx=XM_019830963.2
A1 95372667 95372809
refGene_gn=KCNN2;refGene_tx=XM_003980701.5;refGene_gn=KCNN2;refGene_tx=XM_019830963.2
A1 95385418 95385529
refGene_gn=KCNN2;refGene_tx=XM_003980701.5;refGene_gn=KCNN2;refGene_tx=XM_019830963.2
A1 95399912 95400040
refGene_gn=KCNN2;refGene_tx=XM_003980701.5;refGene_gn=KCNN2;refGene_tx=XM_019830963.2
A1 95403564 95403573 refGene_gn=KCNN2;refGene_tx=XM_019830963.2
A1 95406343 95406413
refGene_gn=KCNN2;refGene_tx=XM_003980701.5;refGene_gn=KCNN2;refGene_tx=XM_019830963.2
A1 95409160 95409448
refGene_gn=KCNN2;refGene_tx=XM_003980701.5;refGene_gn=KCNN2;refGene_tx

=XM_019830963.2
A1 95888855 95889207
refGene_gn=TRIM36;refGene_tx=XM_003980661.5;refGene_gn=TRIM36;refGene_tx=XM_011281694.3
A1 95890955 95891253
refGene_gn=TRIM36;refGene_tx=XM_003980661.5;refGene_gn=TRIM36;refGene_tx=XM_011281694.3
A1 95894759 95895050
refGene_gn=TRIM36;refGene_tx=XM_003980661.5;refGene_gn=TRIM36;refGene_tx=XM_011281694.3
A1 95899183 95899308
refGene_gn=TRIM36;refGene_tx=XM_003980661.5;refGene_gn=TRIM36;refGene_tx=XM_011281694.3
A1 95899517 95899771
refGene_gn=TRIM36;refGene_tx=XM_003980661.5;refGene_gn=TRIM36;refGene_tx=XM_011281694.3
A1 95903324 95903420
refGene_gn=TRIM36;refGene_tx=XM_003980661.5;refGene_gn=TRIM36;refGene_tx=XM_011281694.3
A1 95905569 95905716 refGene_gn=TRIM36;refGene_tx=XM_003980661.5
A1 95909843 95910169
refGene_gn=TRIM36;refGene_tx=XM_003980661.5;refGene_gn=TRIM36;refGene_tx=XM_011281694.3
A1 95918772 95919007
refGene_gn=TRIM36;refGene_tx=XM_003980661.5;refGene_gn=TRIM36;refGene_tx=XM_011281694.3
A1 95921053 95921080
refGene_gn=TRIM36;refGene_tx=XM_003980661.5;refGene_gn=TRIM36;refGene_tx=XM_011281694.3
A1 95967609 95967791 refGene_gn=PGGT1B;refGene_tx=XM_003980663.3
A1 95973934 95974043 refGene_gn=PGGT1B;refGene_tx=XM_003980663.3
A1 95980416 95980601 refGene_gn=PGGT1B;refGene_tx=XM_003980663.3
A1 95989664 95989710 refGene_gn=PGGT1B;refGene_tx=XM_003980663.3
A1 95995993 95996126 refGene_gn=PGGT1B;refGene_tx=XM_003980663.3
A1 95997954 95998106 refGene_gn=PGGT1B;refGene_tx=XM_003980663.3
A1 96002376 96002444 refGene_gn=PGGT1B;refGene_tx=XM_003980663.3
A1 96020254 96020373 refGene_gn=PGGT1B;refGene_tx=XM_003980663.3
A1 96029044 96029184 refGene_gn=PGGT1B;refGene_tx=XM_003980663.3
A1 96033490 96033533 refGene_gn=CCDC112;refGene_tx=XM_006927533.4
A1 96033741 96033860 refGene_gn=CCDC112;refGene_tx=XM_006927533.4
A1 96035341 96035437 refGene_gn=CCDC112;refGene_tx=XM_006927533.4
A1 96036162 96036576 refGene_gn=CCDC112;refGene_tx=XM_006927533.4
A1 96041425 96041816 refGene_gn=CCDC112;refGene_tx=XM_006927533.4
A1 96042229 96042305 refGene_gn=CCDC112;refGene_tx=XM_006927533.4
A1 96044941 96045031 refGene_gn=CCDC112;refGene_tx=XM_006927533.4
A1 96048400 96048522 refGene_gn=CCDC112;refGene_tx=XM_006927533.4
A1 96053075 96053197 refGene_gn=CCDC112;refGene_tx=XM_006927533.4
A1 96063812 96063929 refGene_gn=CCDC112;refGene_tx=XM_006927533.4
A1 96291132 96292442 refGene_gn=FEM1C;refGene_tx=XM_023254280.1
A1 96313377 96313921 refGene_gn=FEM1C;refGene_tx=XM_023254280.1

A1 96344097 96344811
refGene_gn=LOC101090324;refGene_tx=XM_006927535.2;refGene_gn=LOC101090324;refGene_tx=XM_019831045.1
A1 96377405 96377642 refGene_gn=TMED7;refGene_tx=XM_006927554.4
A1 96381545 96381791 refGene_gn=TMED7;refGene_tx=XM_006927554.4
A1 96387144 96387336 refGene_gn=TMED7;refGene_tx=XM_006927554.4
A1 96563426 96563456 refGene_gn=CDO1;refGene_tx=XM_003980666.5
A1 96564859 96565029 refGene_gn=CDO1;refGene_tx=XM_003980666.5
A1 96570240 96570395 refGene_gn=CDO1;refGene_tx=XM_003980666.5
A1 96572329 96572407 refGene_gn=CDO1;refGene_tx=XM_003980666.5
A1 96575928 96576098 refGene_gn=CDO1;refGene_tx=XM_003980666.5
A1 96584829 96584889 refGene_gn=ATG12;refGene_tx=XM_006927536.4
A1 96586482 96586545
refGene_gn=ATG12;refGene_tx=XM_006927536.4;refGene_gn=ATG12;refGene_tx=XM_023254325.1
A1 96592074 96592211 refGene_gn=ATG12;refGene_tx=XM_006927536.4
A1 96596161 96596324
refGene_gn=ATG12;refGene_tx=XM_006927536.4;refGene_gn=ATG12;refGene_tx=XM_023254325.1
A1 96596810 96596879
refGene_gn=AP3S1;refGene_tx=XM_023254310.1;refGene_gn=AP3S1;refGene_tx=XM_023254311.1;refGene_gn=AP3S1;refGene_tx=XM_023254313.1
A1 96623739 96623831
refGene_gn=AP3S1;refGene_tx=XM_023254310.1;refGene_gn=AP3S1;refGene_tx=XM_023254311.1;refGene_gn=AP3S1;refGene_tx=XM_023254313.1
A1 96625707 96625819
refGene_gn=AP3S1;refGene_tx=XM_023254310.1;refGene_gn=AP3S1;refGene_tx=XM_023254311.1;refGene_gn=AP3S1;refGene_tx=XM_023254313.1
A1 96647303 96647375
refGene_gn=AP3S1;refGene_tx=XM_023254310.1;refGene_gn=AP3S1;refGene_tx=XM_023254313.1
A1 96663173 96663281
refGene_gn=AP3S1;refGene_tx=XM_023254310.1;refGene_gn=AP3S1;refGene_tx=XM_023254311.1;refGene_gn=AP3S1;refGene_tx=XM_023254313.1
A1 96666367 96666403 refGene_gn=AP3S1;refGene_tx=XM_023254313.1
A1 96672376 96672505
refGene_gn=AP3S1;refGene_tx=XM_023254310.1;refGene_gn=AP3S1;refGene_tx=XM_023254311.1
A1 96753476 96754174 refGene_gn=LVRN;refGene_tx=XM_023254329.1
A1 96773774 96773917 refGene_gn=LVRN;refGene_tx=XM_023254329.1
A1 96775780 96775920 refGene_gn=LVRN;refGene_tx=XM_023254329.1
A1 96780358 96780485 refGene_gn=LVRN;refGene_tx=XM_023254329.1
A1 96787266 96787421 refGene_gn=LVRN;refGene_tx=XM_023254329.1
A1 96789410 96789524 refGene_gn=LVRN;refGene_tx=XM_023254329.1
A1 96795259 96795403 refGene_gn=LVRN;refGene_tx=XM_023254329.1
A1 96797545 96797611 refGene_gn=LVRN;refGene_tx=XM_023254329.1
A1 96797715 96797781 refGene_gn=LVRN;refGene_tx=XM_023254329.1
A1 96798010 96798183 refGene_gn=LVRN;refGene_tx=XM_023254329.1
A1 96800246 96800323 refGene_gn=LVRN;refGene_tx=XM_023254329.1
A1 96800689 96800829 refGene_gn=LVRN;refGene_tx=XM_023254329.1

A1 96804253 96804309 refGene_gn=LVRN;refGene_tx=XM_023254329.1
A1 96807841 96807995 refGene_gn=LVRN;refGene_tx=XM_023254329.1
A1 96809887 96809982 refGene_gn=LVRN;refGene_tx=XM_023254329.1
A1 96811759 96811867 refGene_gn=LVRN;refGene_tx=XM_023254329.1
A1 96813418 96813586 refGene_gn=LVRN;refGene_tx=XM_023254329.1
A1 96814030 96814168 refGene_gn=LVRN;refGene_tx=XM_023254329.1
A1 96821854 96821930 refGene_gn=LVRN;refGene_tx=XM_023254329.1
A1 96827225 96827366 refGene_gn=LVRN;refGene_tx=XM_023254329.1
A1 96853387 96853480 refGene_gn=ARL14EPL;refGene_tx=XM_023254342.1
A1 96855388 96855528 refGene_gn=ARL14EPL;refGene_tx=XM_023254342.1
A1 96860088 96860311 refGene_gn=ARL14EPL;refGene_tx=XM_023254342.1
A1 96881312 96881353
refGene_gn=COMMD10;refGene_tx=XM_003980668.4;refGene_gn=COMMD10;refGene_tx=XM_006927537.4
A1 96881371 96881547 refGene_gn=COMMD10;refGene_tx=XM_023254378.1
A1 96882682 96882773
refGene_gn=COMMD10;refGene_tx=XM_003980668.4;refGene_gn=COMMD10;refGene_tx=XM_006927537.4;refGene_gn=COMMD10;refGene_tx=XM_019831091.2;refGene_gn=COMMD10;refGene_tx=XM_019831098.1;refGene_gn=COMMD10;refGene_tx=XM_019831100.2;refGene_gn=COMMD10;refGene_tx=XM_019831101.2;refGene_gn=COMMD10;refGene_tx=XM_019831107.2;refGene_gn=COMMD10;refGene_tx=XM_023254378.1
A1 96892452 96892563
refGene_gn=COMMD10;refGene_tx=XM_003980668.4;refGene_gn=COMMD10;refGene_tx=XM_006927537.4;refGene_gn=COMMD10;refGene_tx=XM_019831091.2;refGene_gn=COMMD10;refGene_tx=XM_019831098.1;refGene_gn=COMMD10;refGene_tx=XM_019831100.2;refGene_gn=COMMD10;refGene_tx=XM_019831101.2;refGene_gn=COMMD10;refGene_tx=XM_019831107.2
A1 96894675 96894831
refGene_gn=COMMD10;refGene_tx=XM_003980668.4;refGene_gn=COMMD10;refGene_tx=XM_006927537.4;refGene_gn=COMMD10;refGene_tx=XM_019831091.2;refGene_gn=COMMD10;refGene_tx=XM_019831098.1;refGene_gn=COMMD10;refGene_tx=XM_019831100.2;refGene_gn=COMMD10;refGene_tx=XM_019831101.2;refGene_gn=COMMD10;refGene_tx=XM_019831107.2
A1 96930789 96930900
refGene_gn=COMMD10;refGene_tx=XM_003980668.4;refGene_gn=COMMD10;refGene_tx=XM_006927537.4;refGene_gn=COMMD10;refGene_tx=XM_019831091.2;refGene_gn=COMMD10;refGene_tx=XM_019831098.1;refGene_gn=COMMD10;refGene_tx=XM_019831100.2;refGene_gn=COMMD10;refGene_tx=XM_019831101.2;refGene_gn=COMMD10;refGene_tx=XM_019831107.2;refGene_gn=COMMD10;refGene_tx=XM_023254378.1
A1 97014418 97014451 refGene_gn=COMMD10;refGene_tx=XM_019831101.2
A1 97099126 97099186
refGene_gn=COMMD10;refGene_tx=XM_003980668.4;refGene_gn=COMMD10;refGene_tx=XM_006927537.4;refGene_gn=COMMD10;refGene_tx=XM_019831091.2;refGene_gn=COMMD10;refGene_tx=XM_019831098.1;refGene_gn=COMMD10;refGene_tx=XM_019831100.2;refGene_gn=COMMD10;refGene_tx=XM_019831107.2;refGene_gn=COMMD10;refGene_tx=XM_023254378.1
A1 97099325 97099397
refGene_gn=COMMD10;refGene_tx=XM_006927537.4;refGene_gn=COMMD10;refGen

e_tx=XM_019831091.2;refGene_gn=COMMD10;refGene_tx=XM_019831100.2;refGene_gn=COMMD10;refGene_tx=XM_019831107.2;refGene_gn=COMMD10;refGene_tx=XM_023254378.1
A1 97100358 97100397
refGene_gn=COMMD10;refGene_tx=XM_003980668.4;refGene_gn=COMMD10;refGene_tx=XM_019831098.1
A1 97256065 97257264
refGene_gn=SEMA6A;refGene_tx=XM_023254418.1;refGene_gn=SEMA6A;refGene_tx=XM_023254419.1;refGene_gn=SEMA6A;refGene_tx=XM_023254423.1
A1 97276868 97277033
refGene_gn=SEMA6A;refGene_tx=XM_023254418.1;refGene_gn=SEMA6A;refGene_tx=XM_023254419.1
A1 97282706 97282757 refGene_gn=SEMA6A;refGene_tx=XM_023254418.1
A1 97282894 97282915
refGene_gn=SEMA6A;refGene_tx=XM_023254418.1;refGene_gn=SEMA6A;refGene_tx=XM_023254419.1;refGene_gn=SEMA6A;refGene_tx=XM_023254423.1
A1 97285145 97285204
refGene_gn=SEMA6A;refGene_tx=XM_023254418.1;refGene_gn=SEMA6A;refGene_tx=XM_023254419.1;refGene_gn=SEMA6A;refGene_tx=XM_023254423.1
A1 97287816 97287897
refGene_gn=SEMA6A;refGene_tx=XM_023254418.1;refGene_gn=SEMA6A;refGene_tx=XM_023254419.1;refGene_gn=SEMA6A;refGene_tx=XM_023254423.1
A1 97287982 97288123
refGene_gn=SEMA6A;refGene_tx=XM_023254418.1;refGene_gn=SEMA6A;refGene_tx=XM_023254419.1;refGene_gn=SEMA6A;refGene_tx=XM_023254423.1
A1 97288450 97288627
refGene_gn=SEMA6A;refGene_tx=XM_023254418.1;refGene_gn=SEMA6A;refGene_tx=XM_023254419.1;refGene_gn=SEMA6A;refGene_tx=XM_023254423.1
A1 97290116 97290272
refGene_gn=SEMA6A;refGene_tx=XM_023254418.1;refGene_gn=SEMA6A;refGene_tx=XM_023254419.1;refGene_gn=SEMA6A;refGene_tx=XM_023254423.1
A1 97292504 97292636
refGene_gn=SEMA6A;refGene_tx=XM_023254418.1;refGene_gn=SEMA6A;refGene_tx=XM_023254419.1;refGene_gn=SEMA6A;refGene_tx=XM_023254423.1
A1 97296812 97297030
refGene_gn=SEMA6A;refGene_tx=XM_023254418.1;refGene_gn=SEMA6A;refGene_tx=XM_023254419.1;refGene_gn=SEMA6A;refGene_tx=XM_023254423.1
A1 97298137 97298226
refGene_gn=SEMA6A;refGene_tx=XM_023254418.1;refGene_gn=SEMA6A;refGene_tx=XM_023254419.1;refGene_gn=SEMA6A;refGene_tx=XM_023254423.1
A1 97298904 97299024
refGene_gn=SEMA6A;refGene_tx=XM_023254418.1;refGene_gn=SEMA6A;refGene_tx=XM_023254419.1;refGene_gn=SEMA6A;refGene_tx=XM_023254423.1
A1 97302549 97302640
refGene_gn=SEMA6A;refGene_tx=XM_023254418.1;refGene_gn=SEMA6A;refGene_tx=XM_023254419.1;refGene_gn=SEMA6A;refGene_tx=XM_023254423.1
A1 97308425 97308527
refGene_gn=SEMA6A;refGene_tx=XM_023254418.1;refGene_gn=SEMA6A;refGene_tx=XM_023254419.1;refGene_gn=SEMA6A;refGene_tx=XM_023254423.1
A1 97309257 97309320

refGene_gn=SEMA6A;refGene_tx=XM_023254418.1;refGene_gn=SEMA6A;refGene_tx=XM_023254419.1;refGene_gn=SEMA6A;refGene_tx=XM_023254423.1
A1 97310287 97310348
refGene_gn=SEMA6A;refGene_tx=XM_023254418.1;refGene_gn=SEMA6A;refGene_tx=XM_023254419.1;refGene_gn=SEMA6A;refGene_tx=XM_023254423.1
A1 97315186 97315304
refGene_gn=SEMA6A;refGene_tx=XM_023254418.1;refGene_gn=SEMA6A;refGene_tx=XM_023254419.1;refGene_gn=SEMA6A;refGene_tx=XM_023254423.1
A1 97317846 97317946
refGene_gn=SEMA6A;refGene_tx=XM_023254418.1;refGene_gn=SEMA6A;refGene_tx=XM_023254419.1;refGene_gn=SEMA6A;refGene_tx=XM_023254423.1
A1 99555179 99555350 refGene_gn=DTWD2;refGene_tx=XM_023254474.1
A1 99556503 99556593 refGene_gn=DTWD2;refGene_tx=XM_023254475.1
A1 99560907 99561036
refGene_gn=DTWD2;refGene_tx=XM_023254474.1;refGene_gn=DTWD2;refGene_tx=XM_023254475.1
A1 99623687 99623880
refGene_gn=DTWD2;refGene_tx=XM_023254474.1;refGene_gn=DTWD2;refGene_tx=XM_023254475.1
A1 99626096 99626191
refGene_gn=DTWD2;refGene_tx=XM_023254474.1;refGene_gn=DTWD2;refGene_tx=XM_023254475.1
A1 99633127 99633218
refGene_gn=DTWD2;refGene_tx=XM_023254474.1;refGene_gn=DTWD2;refGene_tx=XM_023254475.1
A1 99655940 99656158
refGene_gn=DTWD2;refGene_tx=XM_023254474.1;refGene_gn=DTWD2;refGene_tx=XM_023254475.1
A1 99732703 99732790
refGene_gn=DMXL1;refGene_tx=XM_006927555.4;refGene_gn=DMXL1;refGene_tx=XM_011281743.3;refGene_gn=DMXL1;refGene_tx=XM_019831227.2;refGene_gn=DMXL1;refGene_tx=XM_019831232.2;refGene_gn=DMXL1;refGene_tx=XM_019831235.2;refGene_gn=DMXL1;refGene_tx=XM_019831239.2
A1 99751172 99751298
refGene_gn=DMXL1;refGene_tx=XM_006927555.4;refGene_gn=DMXL1;refGene_tx=XM_011281743.3;refGene_gn=DMXL1;refGene_tx=XM_019831227.2;refGene_gn=DMXL1;refGene_tx=XM_019831232.2;refGene_gn=DMXL1;refGene_tx=XM_019831235.2;refGene_gn=DMXL1;refGene_tx=XM_019831239.2
A1 99754209 99754281
refGene_gn=DMXL1;refGene_tx=XM_006927555.4;refGene_gn=DMXL1;refGene_tx=XM_011281743.3;refGene_gn=DMXL1;refGene_tx=XM_019831227.2;refGene_gn=DMXL1;refGene_tx=XM_019831232.2;refGene_gn=DMXL1;refGene_tx=XM_019831235.2;refGene_gn=DMXL1;refGene_tx=XM_019831239.2
A1 99757117 99757196
refGene_gn=DMXL1;refGene_tx=XM_006927555.4;refGene_gn=DMXL1;refGene_tx=XM_011281743.3;refGene_gn=DMXL1;refGene_tx=XM_019831227.2;refGene_gn=DMXL1;refGene_tx=XM_019831232.2;refGene_gn=DMXL1;refGene_tx=XM_019831235.2;refGene_gn=DMXL1;refGene_tx=XM_019831239.2
A1 99762795 99762928
refGene_gn=DMXL1;refGene_tx=XM_006927555.4;refGene_gn=DMXL1;refGene_tx

=XM_011281743.3;refGene_gn=DMXL1;refGene_tx=XM_019831227.2;refGene_gn=DMXL1;refGene_tx=XM_019831232.2;refGene_gn=DMXL1;refGene_tx=XM_019831235.2;refGene_gn=DMXL1;refGene_tx=XM_019831239.2
A1 99766772 99766839
refGene_gn=DMXL1;refGene_tx=XM_006927555.4;refGene_gn=DMXL1;refGene_tx=XM_011281743.3;refGene_gn=DMXL1;refGene_tx=XM_019831227.2;refGene_gn=DMXL1;refGene_tx=XM_019831232.2;refGene_gn=DMXL1;refGene_tx=XM_019831235.2;refGene_gn=DMXL1;refGene_tx=XM_019831239.2
A1 99768370 99768549
refGene_gn=DMXL1;refGene_tx=XM_006927555.4;refGene_gn=DMXL1;refGene_tx=XM_011281743.3;refGene_gn=DMXL1;refGene_tx=XM_019831227.2;refGene_gn=DMXL1;refGene_tx=XM_019831232.2;refGene_gn=DMXL1;refGene_tx=XM_019831235.2;refGene_gn=DMXL1;refGene_tx=XM_019831239.2
A1 99772787 99772977
refGene_gn=DMXL1;refGene_tx=XM_006927555.4;refGene_gn=DMXL1;refGene_tx=XM_011281743.3;refGene_gn=DMXL1;refGene_tx=XM_019831227.2;refGene_gn=DMXL1;refGene_tx=XM_019831232.2;refGene_gn=DMXL1;refGene_tx=XM_019831235.2;refGene_gn=DMXL1;refGene_tx=XM_019831239.2
A1 99775136 99775305
refGene_gn=DMXL1;refGene_tx=XM_006927555.4;refGene_gn=DMXL1;refGene_tx=XM_011281743.3;refGene_gn=DMXL1;refGene_tx=XM_019831227.2;refGene_gn=DMXL1;refGene_tx=XM_019831232.2;refGene_gn=DMXL1;refGene_tx=XM_019831235.2;refGene_gn=DMXL1;refGene_tx=XM_019831239.2
A1 99779828 99780041
refGene_gn=DMXL1;refGene_tx=XM_006927555.4;refGene_gn=DMXL1;refGene_tx=XM_011281743.3;refGene_gn=DMXL1;refGene_tx=XM_019831227.2;refGene_gn=DMXL1;refGene_tx=XM_019831232.2;refGene_gn=DMXL1;refGene_tx=XM_019831235.2;refGene_gn=DMXL1;refGene_tx=XM_019831239.2
A1 99782096 99782350
refGene_gn=DMXL1;refGene_tx=XM_006927555.4;refGene_gn=DMXL1;refGene_tx=XM_011281743.3;refGene_gn=DMXL1;refGene_tx=XM_019831227.2;refGene_gn=DMXL1;refGene_tx=XM_019831232.2;refGene_gn=DMXL1;refGene_tx=XM_019831235.2;refGene_gn=DMXL1;refGene_tx=XM_019831239.2
A1 99782461 99783146
refGene_gn=DMXL1;refGene_tx=XM_006927555.4;refGene_gn=DMXL1;refGene_tx=XM_011281743.3;refGene_gn=DMXL1;refGene_tx=XM_019831227.2;refGene_gn=DMXL1;refGene_tx=XM_019831232.2;refGene_gn=DMXL1;refGene_tx=XM_019831235.2;refGene_gn=DMXL1;refGene_tx=XM_019831239.2
A1 99783236 99783358
refGene_gn=DMXL1;refGene_tx=XM_006927555.4;refGene_gn=DMXL1;refGene_tx=XM_011281743.3;refGene_gn=DMXL1;refGene_tx=XM_019831227.2;refGene_gn=DMXL1;refGene_tx=XM_019831232.2;refGene_gn=DMXL1;refGene_tx=XM_019831235.2;refGene_gn=DMXL1;refGene_tx=XM_019831239.2
A1 99786808 99786898
refGene_gn=DMXL1;refGene_tx=XM_006927555.4;refGene_gn=DMXL1;refGene_tx=XM_011281743.3;refGene_gn=DMXL1;refGene_tx=XM_019831227.2;refGene_gn=DMXL1;refGene_tx=XM_019831232.2;refGene_gn=DMXL1;refGene_tx=XM_019831235.2;refGene_gn=DMXL1;refGene_tx=XM_019831239.2
A1 99786983 99787086
refGene_gn=DMXL1;refGene_tx=XM_006927555.4;refGene_gn=DMXL1;refGene_tx

=XM_011281743.3;refGene_gn=DMXL1;refGene_tx=XM_019831227.2;refGene_gn=DMXL1;refGene_tx=XM_019831232.2;refGene_gn=DMXL1;refGene_tx=XM_019831235.2;refGene_gn=DMXL1;refGene_tx=XM_019831239.2
A1 99789968 99790088
refGene_gn=DMXL1;refGene_tx=XM_006927555.4;refGene_gn=DMXL1;refGene_tx=XM_011281743.3;refGene_gn=DMXL1;refGene_tx=XM_019831227.2;refGene_gn=DMXL1;refGene_tx=XM_019831232.2;refGene_gn=DMXL1;refGene_tx=XM_019831235.2;refGene_gn=DMXL1;refGene_tx=XM_019831239.2
A1 99790380 99790602
refGene_gn=DMXL1;refGene_tx=XM_006927555.4;refGene_gn=DMXL1;refGene_tx=XM_011281743.3;refGene_gn=DMXL1;refGene_tx=XM_019831227.2;refGene_gn=DMXL1;refGene_tx=XM_019831232.2;refGene_gn=DMXL1;refGene_tx=XM_019831235.2;refGene_gn=DMXL1;refGene_tx=XM_019831239.2
A1 99792954 99794640
refGene_gn=DMXL1;refGene_tx=XM_006927555.4;refGene_gn=DMXL1;refGene_tx=XM_011281743.3;refGene_gn=DMXL1;refGene_tx=XM_019831227.2;refGene_gn=DMXL1;refGene_tx=XM_019831232.2;refGene_gn=DMXL1;refGene_tx=XM_019831235.2;refGene_gn=DMXL1;refGene_tx=XM_019831239.2
A1 99796151 99796259
refGene_gn=DMXL1;refGene_tx=XM_006927555.4;refGene_gn=DMXL1;refGene_tx=XM_011281743.3;refGene_gn=DMXL1;refGene_tx=XM_019831227.2;refGene_gn=DMXL1;refGene_tx=XM_019831232.2;refGene_gn=DMXL1;refGene_tx=XM_019831235.2;refGene_gn=DMXL1;refGene_tx=XM_019831239.2
A1 99798171 99798341
refGene_gn=DMXL1;refGene_tx=XM_006927555.4;refGene_gn=DMXL1;refGene_tx=XM_011281743.3;refGene_gn=DMXL1;refGene_tx=XM_019831227.2;refGene_gn=DMXL1;refGene_tx=XM_019831232.2;refGene_gn=DMXL1;refGene_tx=XM_019831235.2;refGene_gn=DMXL1;refGene_tx=XM_019831239.2
A1 99798899 99798997
refGene_gn=DMXL1;refGene_tx=XM_006927555.4;refGene_gn=DMXL1;refGene_tx=XM_011281743.3;refGene_gn=DMXL1;refGene_tx=XM_019831227.2;refGene_gn=DMXL1;refGene_tx=XM_019831232.2;refGene_gn=DMXL1;refGene_tx=XM_019831235.2;refGene_gn=DMXL1;refGene_tx=XM_019831239.2
A1 99799796 99799962
refGene_gn=DMXL1;refGene_tx=XM_006927555.4;refGene_gn=DMXL1;refGene_tx=XM_011281743.3;refGene_gn=DMXL1;refGene_tx=XM_019831227.2;refGene_gn=DMXL1;refGene_tx=XM_019831232.2;refGene_gn=DMXL1;refGene_tx=XM_019831235.2;refGene_gn=DMXL1;refGene_tx=XM_019831239.2
A1 99800664 99800926
refGene_gn=DMXL1;refGene_tx=XM_006927555.4;refGene_gn=DMXL1;refGene_tx=XM_011281743.3;refGene_gn=DMXL1;refGene_tx=XM_019831227.2;refGene_gn=DMXL1;refGene_tx=XM_019831232.2;refGene_gn=DMXL1;refGene_tx=XM_019831235.2;refGene_gn=DMXL1;refGene_tx=XM_019831239.2
A1 99802735 99803829
refGene_gn=DMXL1;refGene_tx=XM_006927555.4;refGene_gn=DMXL1;refGene_tx=XM_011281743.3;refGene_gn=DMXL1;refGene_tx=XM_019831227.2;refGene_gn=DMXL1;refGene_tx=XM_019831232.2;refGene_gn=DMXL1;refGene_tx=XM_019831235.2;refGene_gn=DMXL1;refGene_tx=XM_019831239.2
A1 99804288 99804480
refGene_gn=DMXL1;refGene_tx=XM_006927555.4;refGene_gn=DMXL1;refGene_tx

=XM_011281743.3;refGene_gn=DMXL1;refGene_tx=XM_019831227.2;refGene_gn=DMXL1;refGene_tx=XM_019831232.2;refGene_gn=DMXL1;refGene_tx=XM_019831235.2;refGene_gn=DMXL1;refGene_tx=XM_019831239.2
A1 99807535 99807612
refGene_gn=DMXL1;refGene_tx=XM_006927555.4;refGene_gn=DMXL1;refGene_tx=XM_011281743.3;refGene_gn=DMXL1;refGene_tx=XM_019831227.2;refGene_gn=DMXL1;refGene_tx=XM_019831232.2;refGene_gn=DMXL1;refGene_tx=XM_019831235.2;refGene_gn=DMXL1;refGene_tx=XM_019831239.2
A1 99811580 99811708
refGene_gn=DMXL1;refGene_tx=XM_006927555.4;refGene_gn=DMXL1;refGene_tx=XM_011281743.3;refGene_gn=DMXL1;refGene_tx=XM_019831227.2;refGene_gn=DMXL1;refGene_tx=XM_019831232.2;refGene_gn=DMXL1;refGene_tx=XM_019831235.2;refGene_gn=DMXL1;refGene_tx=XM_019831239.2
A1 99812233 99812482
refGene_gn=DMXL1;refGene_tx=XM_006927555.4;refGene_gn=DMXL1;refGene_tx=XM_011281743.3;refGene_gn=DMXL1;refGene_tx=XM_019831227.2;refGene_gn=DMXL1;refGene_tx=XM_019831232.2;refGene_gn=DMXL1;refGene_tx=XM_019831235.2;refGene_gn=DMXL1;refGene_tx=XM_019831239.2
A1 99812832 99812946
refGene_gn=DMXL1;refGene_tx=XM_019831227.2;refGene_gn=DMXL1;refGene_tx=XM_019831232.2;refGene_gn=DMXL1;refGene_tx=XM_019831235.2
A1 99820260 99820439
refGene_gn=DMXL1;refGene_tx=XM_006927555.4;refGene_gn=DMXL1;refGene_tx=XM_011281743.3;refGene_gn=DMXL1;refGene_tx=XM_019831227.2;refGene_gn=DMXL1;refGene_tx=XM_019831232.2;refGene_gn=DMXL1;refGene_tx=XM_019831235.2;refGene_gn=DMXL1;refGene_tx=XM_019831239.2
A1 99825483 99825620
refGene_gn=DMXL1;refGene_tx=XM_006927555.4;refGene_gn=DMXL1;refGene_tx=XM_011281743.3;refGene_gn=DMXL1;refGene_tx=XM_019831227.2;refGene_gn=DMXL1;refGene_tx=XM_019831232.2;refGene_gn=DMXL1;refGene_tx=XM_019831235.2;refGene_gn=DMXL1;refGene_tx=XM_019831239.2
A1 99827844 99827930
refGene_gn=DMXL1;refGene_tx=XM_006927555.4;refGene_gn=DMXL1;refGene_tx=XM_011281743.3;refGene_gn=DMXL1;refGene_tx=XM_019831227.2;refGene_gn=DMXL1;refGene_tx=XM_019831232.2;refGene_gn=DMXL1;refGene_tx=XM_019831235.2;refGene_gn=DMXL1;refGene_tx=XM_019831239.2
A1 99829427 99829629
refGene_gn=DMXL1;refGene_tx=XM_006927555.4;refGene_gn=DMXL1;refGene_tx=XM_011281743.3;refGene_gn=DMXL1;refGene_tx=XM_019831227.2;refGene_gn=DMXL1;refGene_tx=XM_019831232.2;refGene_gn=DMXL1;refGene_tx=XM_019831235.2;refGene_gn=DMXL1;refGene_tx=XM_019831239.2
A1 99830893 99831011
refGene_gn=DMXL1;refGene_tx=XM_006927555.4;refGene_gn=DMXL1;refGene_tx=XM_011281743.3;refGene_gn=DMXL1;refGene_tx=XM_019831227.2;refGene_gn=DMXL1;refGene_tx=XM_019831232.2;refGene_gn=DMXL1;refGene_tx=XM_019831235.2;refGene_gn=DMXL1;refGene_tx=XM_019831239.2
A1 99834235 99834298
refGene_gn=DMXL1;refGene_tx=XM_006927555.4;refGene_gn=DMXL1;refGene_tx=XM_019831227.2;refGene_gn=DMXL1;refGene_tx=XM_019831232.2;refGene_gn=DMXL1;refGene_tx=XM_019831239.2

A1	99843924	99844011
refGene_gn=DMXL1;refGene_tx=XM_006927555.4;refGene_gn=DMXL1;refGene_tx=XM_011281743.3;refGene_gn=DMXL1;refGene_tx=XM_019831227.2;refGene_gn=DMXL1;refGene_tx=XM_019831232.2;refGene_gn=DMXL1;refGene_tx=XM_019831235.2;refGene_gn=DMXL1;refGene_tx=XM_019831239.2
A1	99846372	99846494
refGene_gn=DMXL1;refGene_tx=XM_006927555.4;refGene_gn=DMXL1;refGene_tx=XM_011281743.3;refGene_gn=DMXL1;refGene_tx=XM_019831227.2;refGene_gn=DMXL1;refGene_tx=XM_019831232.2;refGene_gn=DMXL1;refGene_tx=XM_019831235.2;refGene_gn=DMXL1;refGene_tx=XM_019831239.2
A1	99846862	99847004
refGene_gn=DMXL1;refGene_tx=XM_006927555.4;refGene_gn=DMXL1;refGene_tx=XM_011281743.3;refGene_gn=DMXL1;refGene_tx=XM_019831227.2;refGene_gn=DMXL1;refGene_tx=XM_019831232.2;refGene_gn=DMXL1;refGene_tx=XM_019831235.2;refGene_gn=DMXL1;refGene_tx=XM_019831239.2
A1	99850346	99850397
refGene_gn=DMXL1;refGene_tx=XM_019831227.2;refGene_gn=DMXL1;refGene_tx=XM_019831235.2;refGene_gn=DMXL1;refGene_tx=XM_019831239.2
A1	99850660	99850721
refGene_gn=DMXL1;refGene_tx=XM_006927555.4;refGene_gn=DMXL1;refGene_tx=XM_011281743.3;refGene_gn=DMXL1;refGene_tx=XM_019831227.2;refGene_gn=DMXL1;refGene_tx=XM_019831232.2;refGene_gn=DMXL1;refGene_tx=XM_019831235.2;refGene_gn=DMXL1;refGene_tx=XM_019831239.2
A1	99852229	99852357
refGene_gn=DMXL1;refGene_tx=XM_006927555.4;refGene_gn=DMXL1;refGene_tx=XM_011281743.3;refGene_gn=DMXL1;refGene_tx=XM_019831227.2;refGene_gn=DMXL1;refGene_tx=XM_019831232.2;refGene_gn=DMXL1;refGene_tx=XM_019831235.2;refGene_gn=DMXL1;refGene_tx=XM_019831239.2
A1	99855581	99855674
refGene_gn=DMXL1;refGene_tx=XM_006927555.4;refGene_gn=DMXL1;refGene_tx=XM_011281743.3;refGene_gn=DMXL1;refGene_tx=XM_019831227.2;refGene_gn=DMXL1;refGene_tx=XM_019831232.2;refGene_gn=DMXL1;refGene_tx=XM_019831235.2;refGene_gn=DMXL1;refGene_tx=XM_019831239.2
A1	99857296	99857388
refGene_gn=DMXL1;refGene_tx=XM_006927555.4;refGene_gn=DMXL1;refGene_tx=XM_011281743.3;refGene_gn=DMXL1;refGene_tx=XM_019831227.2;refGene_gn=DMXL1;refGene_tx=XM_019831232.2;refGene_gn=DMXL1;refGene_tx=XM_019831235.2;refGene_gn=DMXL1;refGene_tx=XM_019831239.2
A1	99858470	99858523
refGene_gn=DMXL1;refGene_tx=XM_006927555.4;refGene_gn=DMXL1;refGene_tx=XM_011281743.3;refGene_gn=DMXL1;refGene_tx=XM_019831227.2;refGene_gn=DMXL1;refGene_tx=XM_019831232.2;refGene_gn=DMXL1;refGene_tx=XM_019831235.2;refGene_gn=DMXL1;refGene_tx=XM_019831239.2
A1	99862615	99862833
refGene_gn=DMXL1;refGene_tx=XM_006927555.4;refGene_gn=DMXL1;refGene_tx=XM_011281743.3;refGene_gn=DMXL1;refGene_tx=XM_019831227.2;refGene_gn=DMXL1;refGene_tx=XM_019831232.2;refGene_gn=DMXL1;refGene_tx=XM_019831235.2;refGene_gn=DMXL1;refGene_tx=XM_019831239.2
A1	99865702	99865927
refGene_gn=DMXL1;refGene_tx=XM_006927555.4;refGene_gn=DMXL1;refGene_tx

=XM_011281743.3;refGene_gn=DMXL1;refGene_tx=XM_019831227.2;refGene_gn=DMXL1;refGene_tx=XM_019831232.2;refGene_gn=DMXL1;refGene_tx=XM_019831235.2;refGene_gn=DMXL1;refGene_tx=XM_019831239.2
A1 99878165 99878166 refGene_gn=TNFAIP8;refGene_tx=XM_006927543.4
A1 99948498 99948511 refGene_gn=TNFAIP8;refGene_tx=XM_019831275.2
A1 99973137 99973177 refGene_gn=TNFAIP8;refGene_tx=XM_006927542.3
A1 99973379 99973428 refGene_gn=TNFAIP8;refGene_tx=XM_006927541.4
A1 99973864 99973937 refGene_gn=TNFAIP8;refGene_tx=XM_003980670.5
A1 99984231 99984232 refGene_gn=TNFAIP8;refGene_tx=XM_006927544.4
A1 100009777 100010343
refGene_gn=TNFAIP8;refGene_tx=XM_003980670.5;refGene_gn=TNFAIP8;refGene_tx=XM_006927541.4;refGene_gn=TNFAIP8;refGene_tx=XM_006927542.3;refGene_gn=TNFAIP8;refGene_tx=XM_006927543.4;refGene_gn=TNFAIP8;refGene_tx=XM_006927544.4;refGene_gn=TNFAIP8;refGene_tx=XM_019831275.2
A1 100046452 100046513
refGene_gn=HSD17B4;refGene_tx=XM_019831288.2
A1 100049194 100049252
refGene_gn=HSD17B4;refGene_tx=XM_006927547.4
A1 100054072 100054126
refGene_gn=HSD17B4;refGene_tx=XM_006927547.4;refGene_gn=HSD17B4;refGene_tx=XM_019831288.2
A1 100062458 100062566
refGene_gn=HSD17B4;refGene_tx=XM_006927547.4;refGene_gn=HSD17B4;refGene_tx=XM_019831288.2
A1 100064687 100064747
refGene_gn=HSD17B4;refGene_tx=XM_006927547.4;refGene_gn=HSD17B4;refGene_tx=XM_019831288.2
A1 100066516 100066538
refGene_gn=HSD17B4;refGene_tx=XM_006927547.4;refGene_gn=HSD17B4;refGene_tx=XM_019831288.2
A1 100066644 100066691
refGene_gn=HSD17B4;refGene_tx=XM_006927547.4;refGene_gn=HSD17B4;refGene_tx=XM_019831288.2
A1 100071219 100071304
refGene_gn=HSD17B4;refGene_tx=XM_006927547.4;refGene_gn=HSD17B4;refGene_tx=XM_019831288.2
A1 100072613 100072801
refGene_gn=HSD17B4;refGene_tx=XM_006927547.4;refGene_gn=HSD17B4;refGene_tx=XM_019831288.2
A1 100079546 100079638
refGene_gn=HSD17B4;refGene_tx=XM_006927547.4;refGene_gn=HSD17B4;refGene_tx=XM_019831288.2
A1 100082892 100082917
refGene_gn=HSD17B4;refGene_tx=XM_006927547.4;refGene_gn=HSD17B4;refGene_tx=XM_019831288.2
A1 100084787 100084916
refGene_gn=HSD17B4;refGene_tx=XM_006927547.4;refGene_gn=HSD17B4;refGene_tx=XM_019831288.2
A1 100087397 100087501
refGene_gn=HSD17B4;refGene_tx=XM_006927547.4;refGene_gn=HSD17B4;refGen

e_tx=XM_019831288.2
A1 100088987 100089224
refGene_gn=HSD17B4;refGene_tx=XM_006927547.4;refGene_gn=HSD17B4;refGene_tx=XM_019831288.2
A1 100092259 100092311
refGene_gn=HSD17B4;refGene_tx=XM_006927547.4;refGene_gn=HSD17B4;refGene_tx=XM_019831288.2
A1 100094494 100094566
refGene_gn=HSD17B4;refGene_tx=XM_006927547.4;refGene_gn=HSD17B4;refGene_tx=XM_019831288.2
A1 100097879 100097983
refGene_gn=HSD17B4;refGene_tx=XM_006927547.4;refGene_gn=HSD17B4;refGene_tx=XM_019831288.2
A1 100100868 100100934
refGene_gn=HSD17B4;refGene_tx=XM_006927547.4;refGene_gn=HSD17B4;refGene_tx=XM_019831288.2
A1 100110541 100110611
refGene_gn=HSD17B4;refGene_tx=XM_006927547.4;refGene_gn=HSD17B4;refGene_tx=XM_019831288.2
A1 100112002 100112109
refGene_gn=HSD17B4;refGene_tx=XM_006927547.4;refGene_gn=HSD17B4;refGene_tx=XM_019831288.2
A1 100113184 100113271
refGene_gn=HSD17B4;refGene_tx=XM_006927547.4;refGene_gn=HSD17B4;refGene_tx=XM_019831288.2
A1 100118229 100118316
refGene_gn=HSD17B4;refGene_tx=XM_006927547.4;refGene_gn=HSD17B4;refGene_tx=XM_019831288.2
A1 100119533 100119672
refGene_gn=HSD17B4;refGene_tx=XM_006927547.4;refGene_gn=HSD17B4;refGene_tx=XM_019831288.2
A1 100129152 100129280
refGene_gn=HSD17B4;refGene_tx=XM_006927547.4;refGene_gn=HSD17B4;refGene_tx=XM_019831288.2
A1 100135308 100135398
refGene_gn=HSD17B4;refGene_tx=XM_006927547.4;refGene_gn=HSD17B4;refGene_tx=XM_019831288.2
A1 100192147 100192217
refGene_gn=FAM170A;refGene_tx=XM_023254526.1;refGene_gn=FAM170A;refGene_tx=XM_023254532.1
A1 100194090 100194231
refGene_gn=FAM170A;refGene_tx=XM_023254526.1;refGene_gn=FAM170A;refGene_tx=XM_023254532.1
A1 100195032 100195754
refGene_gn=FAM170A;refGene_tx=XM_023254526.1;refGene_gn=FAM170A;refGene_tx=XM_023254532.1
A1 100936744 100936903
refGene_gn=PRR16;refGene_tx=XM_011281772.3;refGene_gn=PRR16;refGene_tx=XM_023254546.1
A1 101073632 101073644

refGene_gn=PRR16;refGene_tx=XM_023254550.1
A1 101137420 101138176
refGene_gn=PRR16;refGene_tx=XM_011281772.3;refGene_gn=PRR16;refGene_tx=XM_023254546.1;refGene_gn=PRR16;refGene_tx=XM_023254550.1
A1 101466245 101466315
refGene_gn=LOC102901339;refGene_tx=XM_019831324.2;refGene_gn=LOC102901339;refGene_tx=XM_019831334.2
A1 101468112 101468247
refGene_gn=LOC102901339;refGene_tx=XM_019831324.2
A1 101470360 101470501
refGene_gn=LOC102901339;refGene_tx=XM_019831324.2;refGene_gn=LOC102901339;refGene_tx=XM_019831334.2
A1 101475130 101475265
refGene_gn=LOC102901339;refGene_tx=XM_019831324.2;refGene_gn=LOC102901339;refGene_tx=XM_019831334.2
A1 101476264 101476414
refGene_gn=LOC102901339;refGene_tx=XM_019831324.2;refGene_gn=LOC102901339;refGene_tx=XM_019831334.2
A1 101477650 101478339
refGene_gn=LOC102901339;refGene_tx=XM_019831324.2;refGene_gn=LOC102901339;refGene_tx=XM_019831334.2
A1 102067221 102067241
refGene_gn=LOC101092483;refGene_tx=XM_011281788.1
A1 102070013 102070500
refGene_gn=LOC101092483;refGene_tx=XM_011281788.1
A1 102216304 102217033
refGene_gn=FTMT;refGene_tx=XM_023254576.1
A1 102272522 102274064
refGene_gn=LOC101092731;refGene_tx=XM_011281791.3
A1 102344923 102344959
refGene_gn=SRFBP1;refGene_tx=XM_006927549.4
A1 102355458 102355547
refGene_gn=SRFBP1;refGene_tx=XM_006927549.4
A1 102356578 102356651
refGene_gn=SRFBP1;refGene_tx=XM_006927549.4
A1 102393428 102393500
refGene_gn=SRFBP1;refGene_tx=XM_006927549.4
A1 102403929 102404011
refGene_gn=SRFBP1;refGene_tx=XM_006927549.4
A1 102404806 102405515
refGene_gn=SRFBP1;refGene_tx=XM_006927549.4
A1 102407214 102407243
refGene_gn=SRFBP1;refGene_tx=XM_006927549.4
A1 102411094 102411279
refGene_gn=SRFBP1;refGene_tx=XM_006927549.4
A1 102436804 102436811
refGene_gn=LOX;refGene_tx=XM_023254614.1
A1 102440857 102440977
refGene_gn=LOX;refGene_tx=XM_023254614.1;refGene_gn=LOX;refGene_tx=XM_023254616.1

A1 102441858 102441954
refGene_gn=LOX;refGene_tx=XM_023254614.1;refGene_gn=LOX;refGene_tx=XM_
023254616.1
A1 102445040 102445197
refGene_gn=LOX;refGene_tx=XM_023254614.1;refGene_gn=LOX;refGene_tx=XM_
023254616.1
A1 102446915 102447053
refGene_gn=LOX;refGene_tx=XM_023254614.1;refGene_gn=LOX;refGene_tx=XM_
023254616.1
A1 102448669 102448778
refGene_gn=LOX;refGene_tx=XM_023254614.1;refGene_gn=LOX;refGene_tx=XM_
023254616.1
A1 102449151 102449776
refGene_gn=LOX;refGene_tx=XM_023254614.1;refGene_gn=LOX;refGene_tx=XM_
023254616.1
A1 102530313 102531345
refGene_gn=LOC105260510;refGene_tx=XM_023254625.1
A1 102558838 102559019
refGene_gn=LOC105260510;refGene_tx=XM_023254625.1
A1 102562162 102562291
refGene_gn=LOC105260510;refGene_tx=XM_023254625.1
A1 102570824 102570922
refGene_gn=LOC105260510;refGene_tx=XM_023254625.1
A1 102730877 102730934
refGene_gn=SNCAIP;refGene_tx=XM_011281801.3;refGene_gn=SNCAIP;refGene_
tx=XM_019831430.2;refGene_gn=SNCAIP;refGene_tx=XM_019831431.2;refGene_
gn=SNCAIP;refGene_tx=XM_023254633.1;refGene_gn=SNCAIP;refGene_tx=XM_02
3254634.1;refGene_gn=SNCAIP;refGene_tx=XM_023254641.1;refGene_gn=SNCAI
P;refGene_tx=XM_023254645.1;refGene_gn=SNCAIP;refGene_tx=XM_023254650.
1
A1 102744499 102744572
refGene_gn=SNCAIP;refGene_tx=XM_011281801.3;refGene_gn=SNCAIP;refGene_
tx=XM_019831430.2;refGene_gn=SNCAIP;refGene_tx=XM_019831431.2;refGene_
gn=SNCAIP;refGene_tx=XM_023254633.1;refGene_gn=SNCAIP;refGene_tx=XM_02
3254634.1;refGene_gn=SNCAIP;refGene_tx=XM_023254641.1;refGene_gn=SNCAI
P;refGene_tx=XM_023254645.1;refGene_gn=SNCAIP;refGene_tx=XM_023254650.
1
A1 102764730 102765599
refGene_gn=SNCAIP;refGene_tx=XM_011281801.3;refGene_gn=SNCAIP;refGene_
tx=XM_019831430.2;refGene_gn=SNCAIP;refGene_tx=XM_019831431.2;refGene_
gn=SNCAIP;refGene_tx=XM_023254633.1;refGene_gn=SNCAIP;refGene_tx=XM_02
3254634.1;refGene_gn=SNCAIP;refGene_tx=XM_023254641.1;refGene_gn=SNCAI
P;refGene_tx=XM_023254645.1;refGene_gn=SNCAIP;refGene_tx=XM_023254650.
1
A1 102767467 102767647
refGene_gn=SNCAIP;refGene_tx=XM_011281801.3;refGene_gn=SNCAIP;refGene_
tx=XM_019831430.2;refGene_gn=SNCAIP;refGene_tx=XM_019831431.2;refGene_
gn=SNCAIP;refGene_tx=XM_023254633.1;refGene_gn=SNCAIP;refGene_tx=XM_02
3254634.1;refGene_gn=SNCAIP;refGene_tx=XM_023254641.1;refGene_gn=SNCAI
P;refGene_tx=XM_023254645.1;refGene_gn=SNCAIP;refGene_tx=XM_023254650.

1
A1 102774227 102774341
refGene_gn=SNCAIP;refGene_tx=XM_011281801.3;refGene_gn=SNCAIP;refGene_tx=XM_019831430.2;refGene_gn=SNCAIP;refGene_tx=XM_019831431.2;refGene_gn=SNCAIP;refGene_tx=XM_023254633.1;refGene_gn=SNCAIP;refGene_tx=XM_023254634.1;refGene_gn=SNCAIP;refGene_tx=XM_023254645.1;refGene_gn=SNCAIP;refGene_tx=XM_023254650.1
A1 102782175 102782301
refGene_gn=SNCAIP;refGene_tx=XM_011281801.3;refGene_gn=SNCAIP;refGene_tx=XM_019831430.2;refGene_gn=SNCAIP;refGene_tx=XM_019831431.2;refGene_gn=SNCAIP;refGene_tx=XM_023254633.1;refGene_gn=SNCAIP;refGene_tx=XM_023254634.1;refGene_gn=SNCAIP;refGene_tx=XM_023254641.1;refGene_gn=SNCAIP;refGene_tx=XM_023254645.1;refGene_gn=SNCAIP;refGene_tx=XM_023254650.1
A1 102787174 102787344
refGene_gn=SNCAIP;refGene_tx=XM_011281801.3;refGene_gn=SNCAIP;refGene_tx=XM_019831430.2;refGene_gn=SNCAIP;refGene_tx=XM_019831431.2;refGene_gn=SNCAIP;refGene_tx=XM_023254633.1;refGene_gn=SNCAIP;refGene_tx=XM_023254634.1;refGene_gn=SNCAIP;refGene_tx=XM_023254641.1;refGene_gn=SNCAIP;refGene_tx=XM_023254645.1;refGene_gn=SNCAIP;refGene_tx=XM_023254650.1
A1 102792752 102792845
refGene_gn=SNCAIP;refGene_tx=XM_011281801.3;refGene_gn=SNCAIP;refGene_tx=XM_019831430.2;refGene_gn=SNCAIP;refGene_tx=XM_019831431.2;refGene_gn=SNCAIP;refGene_tx=XM_023254633.1;refGene_gn=SNCAIP;refGene_tx=XM_023254634.1;refGene_gn=SNCAIP;refGene_tx=XM_023254641.1;refGene_gn=SNCAIP;refGene_tx=XM_023254645.1;refGene_gn=SNCAIP;refGene_tx=XM_023254650.1
A1 102793386 102794455
refGene_gn=SNCAIP;refGene_tx=XM_011281801.3;refGene_gn=SNCAIP;refGene_tx=XM_019831430.2;refGene_gn=SNCAIP;refGene_tx=XM_019831431.2;refGene_gn=SNCAIP;refGene_tx=XM_023254633.1;refGene_gn=SNCAIP;refGene_tx=XM_023254634.1;refGene_gn=SNCAIP;refGene_tx=XM_023254641.1;refGene_gn=SNCAIP;refGene_tx=XM_023254645.1;refGene_gn=SNCAIP;refGene_tx=XM_023254650.1
A1 102795596 102795602
refGene_gn=SNCAIP;refGene_tx=XM_023254650.1
A1 102795755 102795826
refGene_gn=SNCAIP;refGene_tx=XM_011281801.3;refGene_gn=SNCAIP;refGene_tx=XM_019831430.2;refGene_gn=SNCAIP;refGene_tx=XM_019831431.2;refGene_gn=SNCAIP;refGene_tx=XM_023254633.1;refGene_gn=SNCAIP;refGene_tx=XM_023254634.1;refGene_gn=SNCAIP;refGene_tx=XM_023254641.1
A1 102806723 102806808
refGene_gn=SNCAIP;refGene_tx=XM_011281801.3;refGene_gn=SNCAIP;refGene_tx=XM_019831430.2;refGene_gn=SNCAIP;refGene_tx=XM_019831431.2;refGene_gn=SNCAIP;refGene_tx=XM_023254633.1;refGene_gn=SNCAIP;refGene_tx=XM_023254634.1;refGene_gn=SNCAIP;refGene_tx=XM_023254641.1;refGene_gn=SNCAIP;refGene_tx=XM_023254645.1
A1 103079796 103079904
refGene_gn=SNX2;refGene_tx=XM_011281811.3;refGene_gn=SNX2;refGene_tx=X

M_019831444.2
A1	103099875	103099993
refGene_gn=SNX2;refGene_tx=XM_011281811.3;refGene_gn=SNX2;refGene_tx=X
M_019831444.2
A1	103105128	103105292
refGene_gn=SNX2;refGene_tx=XM_011281811.3;refGene_gn=SNX2;refGene_tx=X
M_019831444.2
A1	103106784	103106851
refGene_gn=SNX2;refGene_tx=XM_011281811.3;refGene_gn=SNX2;refGene_tx=X
M_019831444.2
A1	103107225	103107269
refGene_gn=SNX2;refGene_tx=XM_011281811.3;refGene_gn=SNX2;refGene_tx=X
M_019831444.2
A1	103108638	103108780
refGene_gn=SNX2;refGene_tx=XM_011281811.3;refGene_gn=SNX2;refGene_tx=X
M_019831444.2
A1	103114072	103114151
refGene_gn=SNX2;refGene_tx=XM_011281811.3;refGene_gn=SNX2;refGene_tx=X
M_019831444.2
A1	103122071	103122147
refGene_gn=SNX2;refGene_tx=XM_011281811.3;refGene_gn=SNX2;refGene_tx=X
M_019831444.2
A1	103123554	103123668
refGene_gn=SNX2;refGene_tx=XM_011281811.3;refGene_gn=SNX2;refGene_tx=X
M_019831444.2
A1	103123928	103124022
refGene_gn=SNX2;refGene_tx=XM_011281811.3;refGene_gn=SNX2;refGene_tx=X
M_019831444.2
A1	103125411	103125617
refGene_gn=SNX2;refGene_tx=XM_011281811.3;refGene_gn=SNX2;refGene_tx=X
M_019831444.2
A1	103132168	103132312
refGene_gn=SNX2;refGene_tx=XM_011281811.3;refGene_gn=SNX2;refGene_tx=X
M_019831444.2
A1	103132767	103132779
refGene_gn=SNX2;refGene_tx=XM_019831444.2
A1	103133725	103133806
refGene_gn=SNX2;refGene_tx=XM_011281811.3;refGene_gn=SNX2;refGene_tx=X
M_019831444.2
A1	103133946	103134018
refGene_gn=SNX2;refGene_tx=XM_011281811.3;refGene_gn=SNX2;refGene_tx=X
M_019831444.2
A1	103135586	103135637
refGene_gn=SNX2;refGene_tx=XM_011281811.3;refGene_gn=SNX2;refGene_tx=X
M_019831444.2
A1	103154748	103154808
refGene_gn=SNX24;refGene_tx=XM_023254692.1;refGene_gn=SNX24;refGene_tx
=XM_023254694.1
A1	103244039	103244123
refGene_gn=SNX24;refGene_tx=XM_023254692.1;refGene_gn=SNX24;refGene_tx

=XM_023254694.1
A1 103256610 103256715
refGene_gn=SNX24;refGene_tx=XM_023254692.1;refGene_gn=SNX24;refGene_tx
=XM_023254694.1
A1 103306999 103307094
refGene_gn=SNX24;refGene_tx=XM_023254692.1;refGene_gn=SNX24;refGene_tx
=XM_023254694.1
A1 103308431 103308464
refGene_gn=SNX24;refGene_tx=XM_023254692.1;refGene_gn=SNX24;refGene_tx
=XM_023254694.1
A1 103308998 103309063
refGene_gn=SNX24;refGene_tx=XM_023254692.1;refGene_gn=SNX24;refGene_tx
=XM_023254694.1
A1 103313385 103313453
refGene_gn=SNX24;refGene_tx=XM_023254692.1;refGene_gn=SNX24;refGene_tx
=XM_023254694.1
A1 103315313 103315320
refGene_gn=SNX24;refGene_tx=XM_023254692.1
A1 103327571 103327700
refGene_gn=PPIC;refGene_tx=XM_023254682.1
A1 103330437 103330622
refGene_gn=PPIC;refGene_tx=XM_023254682.1
A1 103333153 103333247
refGene_gn=PPIC;refGene_tx=XM_023254682.1
A1 103333712 103333826
refGene_gn=PPIC;refGene_tx=XM_023254682.1
A1 103339936 103340053
refGene_gn=PPIC;refGene_tx=XM_023254682.1
A1 103399336 103399931
refGene_gn=PRDM6;refGene_tx=XM_011283422.3;refGene_gn=PRDM6;refGene_tx
=XM_023254702.1
A1 103408858 103409166
refGene_gn=PRDM6;refGene_tx=XM_011283422.3;refGene_gn=PRDM6;refGene_tx
=XM_023254702.1
A1 103472383 103472511
refGene_gn=PRDM6;refGene_tx=XM_011283422.3;refGene_gn=PRDM6;refGene_tx
=XM_023254702.1
A1 103476111 103476236
refGene_gn=PRDM6;refGene_tx=XM_011283422.3;refGene_gn=PRDM6;refGene_tx
=XM_023254702.1
A1 103490089 103490432
refGene_gn=PRDM6;refGene_tx=XM_011283422.3;refGene_gn=PRDM6;refGene_tx
=XM_023254702.1
A1 103499119 103499296
refGene_gn=PRDM6;refGene_tx=XM_011283422.3;refGene_gn=PRDM6;refGene_tx
=XM_023254702.1
A1 103506805 103506920
refGene_gn=PRDM6;refGene_tx=XM_011283422.3;refGene_gn=PRDM6;refGene_tx
=XM_023254702.1
A1 103636974 103637145

refGene_gn=CEP120;refGene_tx=XM_019831472.2;refGene_gn=CEP120;refGene_tx=XM_019831483.2
A1 103637373 103637524
refGene_gn=CEP120;refGene_tx=XM_019831472.2;refGene_gn=CEP120;refGene_tx=XM_019831480.2;refGene_gn=CEP120;refGene_tx=XM_019831483.2
A1 103644084 103644319
refGene_gn=CEP120;refGene_tx=XM_011281825.2;refGene_gn=CEP120;refGene_tx=XM_019831469.2
A1 103648525 103648671
refGene_gn=CEP120;refGene_tx=XM_011281825.2;refGene_gn=CEP120;refGene_tx=XM_019831469.2;refGene_gn=CEP120;refGene_tx=XM_019831480.2;refGene_gn=CEP120;refGene_tx=XM_019831483.2
A1 103664580 103664679
refGene_gn=CEP120;refGene_tx=XM_011281825.2;refGene_gn=CEP120;refGene_tx=XM_019831469.2;refGene_gn=CEP120;refGene_tx=XM_019831472.2;refGene_gn=CEP120;refGene_tx=XM_019831480.2;refGene_gn=CEP120;refGene_tx=XM_019831483.2
A1 103671608 103671731
refGene_gn=CEP120;refGene_tx=XM_011281825.2;refGene_gn=CEP120;refGene_tx=XM_019831469.2;refGene_gn=CEP120;refGene_tx=XM_019831472.2;refGene_gn=CEP120;refGene_tx=XM_019831480.2;refGene_gn=CEP120;refGene_tx=XM_019831483.2
A1 103674025 103674187
refGene_gn=CEP120;refGene_tx=XM_011281825.2;refGene_gn=CEP120;refGene_tx=XM_019831469.2;refGene_gn=CEP120;refGene_tx=XM_019831472.2;refGene_gn=CEP120;refGene_tx=XM_019831480.2;refGene_gn=CEP120;refGene_tx=XM_019831483.2
A1 103675380 103675473
refGene_gn=CEP120;refGene_tx=XM_011281825.2;refGene_gn=CEP120;refGene_tx=XM_019831469.2;refGene_gn=CEP120;refGene_tx=XM_019831472.2;refGene_gn=CEP120;refGene_tx=XM_019831480.2;refGene_gn=CEP120;refGene_tx=XM_019831483.2
A1 103684194 103684284
refGene_gn=CEP120;refGene_tx=XM_011281825.2;refGene_gn=CEP120;refGene_tx=XM_019831469.2;refGene_gn=CEP120;refGene_tx=XM_019831472.2;refGene_gn=CEP120;refGene_tx=XM_019831480.2;refGene_gn=CEP120;refGene_tx=XM_019831483.2
A1 103685064 103685217
refGene_gn=CEP120;refGene_tx=XM_011281825.2;refGene_gn=CEP120;refGene_tx=XM_019831469.2;refGene_gn=CEP120;refGene_tx=XM_019831472.2;refGene_gn=CEP120;refGene_tx=XM_019831480.2;refGene_gn=CEP120;refGene_tx=XM_019831483.2
A1 103685315 103685412
refGene_gn=CEP120;refGene_tx=XM_011281825.2;refGene_gn=CEP120;refGene_tx=XM_019831469.2;refGene_gn=CEP120;refGene_tx=XM_019831472.2;refGene_gn=CEP120;refGene_tx=XM_019831480.2;refGene_gn=CEP120;refGene_tx=XM_019831483.2
A1 103687393 103687576
refGene_gn=CEP120;refGene_tx=XM_011281825.2;refGene_gn=CEP120;refGene_tx=XM_019831469.2;refGene_gn=CEP120;refGene_tx=XM_019831472.2;refGene_

gn=CEP120;refGene_tx=XM_019831480.2;refGene_gn=CEP120;refGene_tx=XM_01
9831483.2
A1 103688297 103688447
refGene_gn=CEP120;refGene_tx=XM_011281825.2;refGene_gn=CEP120;refGene_
tx=XM_019831469.2;refGene_gn=CEP120;refGene_tx=XM_019831472.2;refGene_
gn=CEP120;refGene_tx=XM_019831480.2;refGene_gn=CEP120;refGene_tx=XM_01
9831483.2
A1 103691717 103691895
refGene_gn=CEP120;refGene_tx=XM_011281825.2;refGene_gn=CEP120;refGene_
tx=XM_019831469.2;refGene_gn=CEP120;refGene_tx=XM_019831472.2;refGene_
gn=CEP120;refGene_tx=XM_019831480.2;refGene_gn=CEP120;refGene_tx=XM_01
9831483.2
A1 103692929 103693146
refGene_gn=CEP120;refGene_tx=XM_011281825.2;refGene_gn=CEP120;refGene_
tx=XM_019831469.2;refGene_gn=CEP120;refGene_tx=XM_019831472.2;refGene_
gn=CEP120;refGene_tx=XM_019831480.2;refGene_gn=CEP120;refGene_tx=XM_01
9831483.2
A1 103694142 103694370
refGene_gn=CEP120;refGene_tx=XM_011281825.2;refGene_gn=CEP120;refGene_
tx=XM_019831469.2;refGene_gn=CEP120;refGene_tx=XM_019831472.2;refGene_
gn=CEP120;refGene_tx=XM_019831480.2;refGene_gn=CEP120;refGene_tx=XM_01
9831483.2
A1 103697046 103697244
refGene_gn=CEP120;refGene_tx=XM_011281825.2;refGene_gn=CEP120;refGene_
tx=XM_019831469.2;refGene_gn=CEP120;refGene_tx=XM_019831472.2;refGene_
gn=CEP120;refGene_tx=XM_019831480.2;refGene_gn=CEP120;refGene_tx=XM_01
9831483.2
A1 103701100 103701249
refGene_gn=CEP120;refGene_tx=XM_011281825.2;refGene_gn=CEP120;refGene_
tx=XM_019831469.2;refGene_gn=CEP120;refGene_tx=XM_019831472.2;refGene_
gn=CEP120;refGene_tx=XM_019831480.2;refGene_gn=CEP120;refGene_tx=XM_01
9831483.2
A1 103708281 103708423
refGene_gn=CEP120;refGene_tx=XM_011281825.2;refGene_gn=CEP120;refGene_
tx=XM_019831469.2;refGene_gn=CEP120;refGene_tx=XM_019831472.2;refGene_
gn=CEP120;refGene_tx=XM_019831480.2;refGene_gn=CEP120;refGene_tx=XM_01
9831483.2
A1 103711353 103711468
refGene_gn=CEP120;refGene_tx=XM_011281825.2;refGene_gn=CEP120;refGene_
tx=XM_019831469.2;refGene_gn=CEP120;refGene_tx=XM_019831472.2;refGene_
gn=CEP120;refGene_tx=XM_019831480.2;refGene_gn=CEP120;refGene_tx=XM_01
9831483.2
A1 103713074 103713231
refGene_gn=CEP120;refGene_tx=XM_011281825.2;refGene_gn=CEP120;refGene_
tx=XM_019831469.2;refGene_gn=CEP120;refGene_tx=XM_019831472.2;refGene_
gn=CEP120;refGene_tx=XM_019831480.2;refGene_gn=CEP120;refGene_tx=XM_01
9831483.2
A1 103717988 103718037
refGene_gn=CEP120;refGene_tx=XM_011281825.2;refGene_gn=CEP120;refGene_
tx=XM_019831469.2;refGene_gn=CEP120;refGene_tx=XM_019831472.2;refGene_

gn=CEP120;refGene_tx=XM_019831480.2;refGene_gn=CEP120;refGene_tx=XM_019831483.2
A1 103838005 103838183
refGene_gn=CSNK1G3;refGene_tx=XM_011281832.3;refGene_gn=CSNK1G3;refGene_tx=XM_019831506.2;refGene_gn=CSNK1G3;refGene_tx=XM_019831514.2;refGene_gn=CSNK1G3;refGene_tx=XM_023254710.1;refGene_gn=CSNK1G3;refGene_tx=XM_023254718.1;refGene_gn=CSNK1G3;refGene_tx=XM_023254722.1;refGene_gn=CSNK1G3;refGene_tx=XM_023254726.1;refGene_gn=CSNK1G3;refGene_tx=XM_023254728.1;refGene_gn=CSNK1G3;refGene_tx=XM_023254730.1;refGene_gn=CSNK1G3;refGene_tx=XM_023254731.1
A1 103845955 103845996
refGene_gn=CSNK1G3;refGene_tx=XM_011281832.3;refGene_gn=CSNK1G3;refGene_tx=XM_019831506.2;refGene_gn=CSNK1G3;refGene_tx=XM_019831514.2;refGene_gn=CSNK1G3;refGene_tx=XM_023254710.1;refGene_gn=CSNK1G3;refGene_tx=XM_023254718.1;refGene_gn=CSNK1G3;refGene_tx=XM_023254722.1;refGene_gn=CSNK1G3;refGene_tx=XM_023254726.1;refGene_gn=CSNK1G3;refGene_tx=XM_023254728.1;refGene_gn=CSNK1G3;refGene_tx=XM_023254730.1;refGene_gn=CSNK1G3;refGene_tx=XM_023254731.1
A1 103850660 103850730
refGene_gn=CSNK1G3;refGene_tx=XM_011281832.3;refGene_gn=CSNK1G3;refGene_tx=XM_019831506.2;refGene_gn=CSNK1G3;refGene_tx=XM_019831514.2;refGene_gn=CSNK1G3;refGene_tx=XM_023254710.1;refGene_gn=CSNK1G3;refGene_tx=XM_023254718.1;refGene_gn=CSNK1G3;refGene_tx=XM_023254722.1;refGene_gn=CSNK1G3;refGene_tx=XM_023254726.1;refGene_gn=CSNK1G3;refGene_tx=XM_023254728.1;refGene_gn=CSNK1G3;refGene_tx=XM_023254730.1;refGene_gn=CSNK1G3;refGene_tx=XM_023254731.1
A1 103864249 103864398
refGene_gn=CSNK1G3;refGene_tx=XM_011281832.3;refGene_gn=CSNK1G3;refGene_tx=XM_019831506.2;refGene_gn=CSNK1G3;refGene_tx=XM_019831514.2;refGene_gn=CSNK1G3;refGene_tx=XM_023254710.1;refGene_gn=CSNK1G3;refGene_tx=XM_023254718.1;refGene_gn=CSNK1G3;refGene_tx=XM_023254722.1;refGene_gn=CSNK1G3;refGene_tx=XM_023254726.1;refGene_gn=CSNK1G3;refGene_tx=XM_023254728.1;refGene_gn=CSNK1G3;refGene_tx=XM_023254730.1;refGene_gn=CSNK1G3;refGene_tx=XM_023254731.1
A1 103865949 103866184
refGene_gn=CSNK1G3;refGene_tx=XM_011281832.3;refGene_gn=CSNK1G3;refGene_tx=XM_019831506.2;refGene_gn=CSNK1G3;refGene_tx=XM_019831514.2;refGene_gn=CSNK1G3;refGene_tx=XM_023254710.1;refGene_gn=CSNK1G3;refGene_tx=XM_023254718.1;refGene_gn=CSNK1G3;refGene_tx=XM_023254722.1;refGene_gn=CSNK1G3;refGene_tx=XM_023254726.1;refGene_gn=CSNK1G3;refGene_tx=XM_023254728.1;refGene_gn=CSNK1G3;refGene_tx=XM_023254730.1;refGene_gn=CSNK1G3;refGene_tx=XM_023254731.1
A1 103881079 103881165
refGene_gn=CSNK1G3;refGene_tx=XM_011281832.3;refGene_gn=CSNK1G3;refGene_tx=XM_019831506.2;refGene_gn=CSNK1G3;refGene_tx=XM_019831514.2;refGene_gn=CSNK1G3;refGene_tx=XM_023254710.1;refGene_gn=CSNK1G3;refGene_tx=XM_023254718.1;refGene_gn=CSNK1G3;refGene_tx=XM_023254722.1;refGene_gn=CSNK1G3;refGene_tx=XM_023254726.1;refGene_gn=CSNK1G3;refGene_tx=XM_023254728.1;refGene_gn=CSNK1G3;refGene_tx=XM_023254730.1;refGene_gn=CSNK1G3;refGene_tx=XM_023254731.1

A1 103881287 103881372
refGene_gn=CSNK1G3;refGene_tx=XM_011281832.3;refGene_gn=CSNK1G3;refGene_tx=XM_019831506.2;refGene_gn=CSNK1G3;refGene_tx=XM_019831514.2;refGene_gn=CSNK1G3;refGene_tx=XM_023254710.1;refGene_gn=CSNK1G3;refGene_tx=XM_023254718.1;refGene_gn=CSNK1G3;refGene_tx=XM_023254722.1;refGene_gn=CSNK1G3;refGene_tx=XM_023254726.1;refGene_gn=CSNK1G3;refGene_tx=XM_023254728.1;refGene_gn=CSNK1G3;refGene_tx=XM_023254730.1;refGene_gn=CSNK1G3;refGene_tx=XM_023254731.1
A1 103883697 103883846
refGene_gn=CSNK1G3;refGene_tx=XM_011281832.3;refGene_gn=CSNK1G3;refGene_tx=XM_019831506.2;refGene_gn=CSNK1G3;refGene_tx=XM_019831514.2;refGene_gn=CSNK1G3;refGene_tx=XM_023254710.1;refGene_gn=CSNK1G3;refGene_tx=XM_023254718.1;refGene_gn=CSNK1G3;refGene_tx=XM_023254722.1;refGene_gn=CSNK1G3;refGene_tx=XM_023254726.1;refGene_gn=CSNK1G3;refGene_tx=XM_023254728.1;refGene_gn=CSNK1G3;refGene_tx=XM_023254730.1;refGene_gn=CSNK1G3;refGene_tx=XM_023254731.1
A1 103885839 103885935
refGene_gn=CSNK1G3;refGene_tx=XM_011281832.3;refGene_gn=CSNK1G3;refGene_tx=XM_019831506.2;refGene_gn=CSNK1G3;refGene_tx=XM_019831514.2;refGene_gn=CSNK1G3;refGene_tx=XM_023254710.1;refGene_gn=CSNK1G3;refGene_tx=XM_023254718.1;refGene_gn=CSNK1G3;refGene_tx=XM_023254722.1;refGene_gn=CSNK1G3;refGene_tx=XM_023254726.1;refGene_gn=CSNK1G3;refGene_tx=XM_023254728.1;refGene_gn=CSNK1G3;refGene_tx=XM_023254730.1;refGene_gn=CSNK1G3;refGene_tx=XM_023254731.1
A1 103889616 103889712
refGene_gn=CSNK1G3;refGene_tx=XM_011281832.3;refGene_gn=CSNK1G3;refGene_tx=XM_019831506.2;refGene_gn=CSNK1G3;refGene_tx=XM_019831514.2;refGene_gn=CSNK1G3;refGene_tx=XM_023254710.1;refGene_gn=CSNK1G3;refGene_tx=XM_023254718.1;refGene_gn=CSNK1G3;refGene_tx=XM_023254722.1;refGene_gn=CSNK1G3;refGene_tx=XM_023254726.1
A1 103899552 103899659
refGene_gn=CSNK1G3;refGene_tx=XM_011281832.3;refGene_gn=CSNK1G3;refGene_tx=XM_019831506.2;refGene_gn=CSNK1G3;refGene_tx=XM_019831514.2;refGene_gn=CSNK1G3;refGene_tx=XM_023254710.1;refGene_gn=CSNK1G3;refGene_tx=XM_023254718.1;refGene_gn=CSNK1G3;refGene_tx=XM_023254722.1;refGene_gn=CSNK1G3;refGene_tx=XM_023254726.1;refGene_gn=CSNK1G3;refGene_tx=XM_023254728.1;refGene_gn=CSNK1G3;refGene_tx=XM_023254730.1;refGene_gn=CSNK1G3;refGene_tx=XM_023254731.1
A1 103900259 103900287
refGene_gn=CSNK1G3;refGene_tx=XM_011281832.3;refGene_gn=CSNK1G3;refGene_tx=XM_019831506.2;refGene_gn=CSNK1G3;refGene_tx=XM_019831514.2;refGene_gn=CSNK1G3;refGene_tx=XM_023254710.1;refGene_gn=CSNK1G3;refGene_tx=XM_023254718.1;refGene_gn=CSNK1G3;refGene_tx=XM_023254722.1;refGene_gn=CSNK1G3;refGene_tx=XM_023254728.1;refGene_gn=CSNK1G3;refGene_tx=XM_023254730.1;refGene_gn=CSNK1G3;refGene_tx=XM_023254731.1
A1 103910542 103910597
refGene_gn=CSNK1G3;refGene_tx=XM_011281832.3;refGene_gn=CSNK1G3;refGene_tx=XM_019831506.2;refGene_gn=CSNK1G3;refGene_tx=XM_019831514.2;refGene_gn=CSNK1G3;refGene_tx=XM_023254710.1;refGene_gn=CSNK1G3;refGene_tx=XM_023254718.1;refGene_gn=CSNK1G3;refGene_tx=XM_023254722.1;refGene_gn

=CSNK1G3;refGene_tx=XM_023254726.1;refGene_gn=CSNK1G3;refGene_tx=XM_023254728.1;refGene_gn=CSNK1G3;refGene_tx=XM_023254730.1
A1 104778849 104778935
refGene_gn=ZNF608;refGene_tx=XM_011281854.3
A1 104782449 104782603
refGene_gn=ZNF608;refGene_tx=XM_011281854.3
A1 104784489 104784662
refGene_gn=ZNF608;refGene_tx=XM_011281854.3
A1 104785248 104785417
refGene_gn=ZNF608;refGene_tx=XM_011281854.3
A1 104785419 104785668
refGene_gn=ZNF608;refGene_tx=XM_011281854.3
A1 104788557 104789850
refGene_gn=ZNF608;refGene_tx=XM_011281854.3
A1 104789851 104790004
refGene_gn=ZNF608;refGene_tx=XM_011281854.3
A1 104790005 104791011
refGene_gn=ZNF608;refGene_tx=XM_011281854.3
A1 104791511 104791599
refGene_gn=ZNF608;refGene_tx=XM_011281854.3
A1 104844737 104844993
refGene_gn=ZNF608;refGene_tx=XM_011281854.3
A1 104896628 104897534
refGene_gn=ZNF608;refGene_tx=XM_011281854.3
A1 106252713 106252838
refGene_gn=GRAMD2B;refGene_tx=XM_019831574.2
A1 106295070 106295153
refGene_gn=GRAMD2B;refGene_tx=XM_003980720.5
A1 106344299 106344419
refGene_gn=GRAMD2B;refGene_tx=XM_003980720.5;refGene_gn=GRAMD2B;refGene_tx=XM_019831574.2
A1 106347630 106347742
refGene_gn=GRAMD2B;refGene_tx=XM_003980720.5;refGene_gn=GRAMD2B;refGene_tx=XM_019831574.2
A1 106351891 106351958
refGene_gn=GRAMD2B;refGene_tx=XM_003980720.5;refGene_gn=GRAMD2B;refGene_tx=XM_019831574.2
A1 106352885 106352989
refGene_gn=GRAMD2B;refGene_tx=XM_003980720.5;refGene_gn=GRAMD2B;refGene_tx=XM_019831574.2
A1 106358254 106358350
refGene_gn=GRAMD2B;refGene_tx=XM_003980720.5;refGene_gn=GRAMD2B;refGene_tx=XM_019831574.2
A1 106360972 106361044
refGene_gn=GRAMD2B;refGene_tx=XM_003980720.5;refGene_gn=GRAMD2B;refGene_tx=XM_019831574.2
A1 106361143 106361224
refGene_gn=GRAMD2B;refGene_tx=XM_003980720.5;refGene_gn=GRAMD2B;refGene_tx=XM_019831574.2
A1 106363860 106363972

refGene_gn=GRAMD2B;refGene_tx=XM_003980720.5;refGene_gn=GRAMD2B;refGene_tx=XM_019831574.2
A1 106364947 106365070
refGene_gn=GRAMD2B;refGene_tx=XM_003980720.5;refGene_gn=GRAMD2B;refGene_tx=XM_019831574.2
A1 106366176 106366264
refGene_gn=GRAMD2B;refGene_tx=XM_003980720.5;refGene_gn=GRAMD2B;refGene_tx=XM_019831574.2
A1 106367417 106367522
refGene_gn=GRAMD2B;refGene_tx=XM_003980720.5;refGene_gn=GRAMD2B;refGene_tx=XM_019831574.2
A1 106370958 106371052
refGene_gn=GRAMD2B;refGene_tx=XM_003980720.5;refGene_gn=GRAMD2B;refGene_tx=XM_019831574.2
A1 106373311 106373353
refGene_gn=GRAMD2B;refGene_tx=XM_003980720.5;refGene_gn=GRAMD2B;refGene_tx=XM_019831574.2
A1 106419624 106419679
refGene_gn=ALDH7A1;refGene_tx=XM_003980723.5
A1 106420627 106420703
refGene_gn=ALDH7A1;refGene_tx=XM_003980723.5
A1 106423888 106423962
refGene_gn=ALDH7A1;refGene_tx=XM_003980723.5
A1 106424410 106424508
refGene_gn=ALDH7A1;refGene_tx=XM_003980723.5
A1 106425279 106425396
refGene_gn=ALDH7A1;refGene_tx=XM_003980723.5
A1 106428449 106428556
refGene_gn=ALDH7A1;refGene_tx=XM_003980723.5
A1 106429975 106430060
refGene_gn=ALDH7A1;refGene_tx=XM_003980723.5
A1 106431923 106432018
refGene_gn=ALDH7A1;refGene_tx=XM_003980723.5
A1 106433267 106433309
refGene_gn=ALDH7A1;refGene_tx=XM_003980723.5
A1 106438057 106438155
refGene_gn=ALDH7A1;refGene_tx=XM_003980723.5
A1 106440881 106440959
refGene_gn=ALDH7A1;refGene_tx=XM_003980723.5
A1 106444464 106444509
refGene_gn=ALDH7A1;refGene_tx=XM_003980723.5
A1 106445609 106445742
refGene_gn=ALDH7A1;refGene_tx=XM_003980723.5
A1 106450199 106450323
refGene_gn=ALDH7A1;refGene_tx=XM_003980723.5
A1 106450882 106450963
refGene_gn=ALDH7A1;refGene_tx=XM_003980723.5
A1 106456199 106456265
refGene_gn=ALDH7A1;refGene_tx=XM_003980723.5
A1 106457194 106457248

refGene_gn=ALDH7A1;refGene_tx=XM_003980723.5
A1 106458717 106458909
refGene_gn=ALDH7A1;refGene_tx=XM_003980723.5
A1 106464457 106464553
refGene_gn=PHAX;refGene_tx=XM_023254752.1
A1 106466482 106467096
refGene_gn=PHAX;refGene_tx=XM_023254752.1
A1 106469457 106469578
refGene_gn=PHAX;refGene_tx=XM_023254752.1
A1 106471533 106471617
refGene_gn=PHAX;refGene_tx=XM_023254752.1
A1 106477004 106477274
refGene_gn=PHAX;refGene_tx=XM_023254752.1
A1 106482660 106482773
refGene_gn=TEX43;refGene_tx=XM_006927566.3
A1 106483299 106483417
refGene_gn=TEX43;refGene_tx=XM_006927566.3
A1 106485474 106485684
refGene_gn=TEX43;refGene_tx=XM_006927566.3
A1 106573341 106573700
refGene_gn=LMNB1;refGene_tx=XM_019835509.2
A1 106600470 106600627
refGene_gn=LMNB1;refGene_tx=XM_019835509.2
A1 106601253 106601379
refGene_gn=LMNB1;refGene_tx=XM_019835509.2
A1 106604254 106604425
refGene_gn=LMNB1;refGene_tx=XM_019835509.2
A1 106606526 106606652
refGene_gn=LMNB1;refGene_tx=XM_019835509.2
A1 106613690 106613911
refGene_gn=LMNB1;refGene_tx=XM_019835509.2
A1 106615265 106615491
refGene_gn=LMNB1;refGene_tx=XM_019835509.2
A1 106617166 106617271
refGene_gn=LMNB1;refGene_tx=XM_019835509.2
A1 106621295 106621415
refGene_gn=LMNB1;refGene_tx=XM_019835509.2
A1 106626548 106626656
refGene_gn=LMNB1;refGene_tx=XM_019835509.2
A1 106629498 106629540
refGene_gn=LMNB1;refGene_tx=XM_019835509.2
A1 106662739 106662898
refGene_gn=MARCH3;refGene_tx=XM_003980726.5
A1 106669112 106669322
refGene_gn=MARCH3;refGene_tx=XM_003980726.5
A1 106695646 106695851
refGene_gn=MARCH3;refGene_tx=XM_003980726.5
A1 106699579 106699767
refGene_gn=MARCH3;refGene_tx=XM_003980726.5
A1 106818073 106818226

refGene_gn=CA1H5orf63;refGene_tx=XM_023254754.1
A1 106833330 106833357
refGene_gn=CA1H5orf63;refGene_tx=XM_023254756.1
A1 106834834 106834927
refGene_gn=CA1H5orf63;refGene_tx=XM_023254754.1;refGene_gn=CA1H5orf63;
refGene_tx=XM_023254756.1
A1 106839206 106839377
refGene_gn=CA1H5orf63;refGene_tx=XM_023254755.1
A1 106840515 106840572
refGene_gn=CA1H5orf63;refGene_tx=XM_023254754.1;refGene_gn=CA1H5orf63;
refGene_tx=XM_023254755.1;refGene_gn=CA1H5orf63;refGene_tx=XM_02325475
6.1
A1 106844572 106844686
refGene_gn=CA1H5orf63;refGene_tx=XM_023254755.1;refGene_gn=CA1H5orf63;
refGene_tx=XM_023254756.1
A1 106849925 106849979
refGene_gn=CA1H5orf63;refGene_tx=XM_023254754.1
A1 106859687 106859984
refGene_gn=CA1H5orf63;refGene_tx=XM_023254754.1
A1 107056642 107056758
refGene_gn=MEGF10;refGene_tx=XM_006927570.4
A1 107065207 107065309
refGene_gn=MEGF10;refGene_tx=XM_006927570.4
A1 107067131 107067232
refGene_gn=MEGF10;refGene_tx=XM_006927570.4
A1 107101302 107101395
refGene_gn=MEGF10;refGene_tx=XM_006927570.4
A1 107107891 107107934
refGene_gn=MEGF10;refGene_tx=XM_023254757.1
A1 107123908 107124155
refGene_gn=MEGF10;refGene_tx=XM_006927570.4;refGene_gn=MEGF10;refGene_
tx=XM_023254757.1
A1 107125544 107125665
refGene_gn=MEGF10;refGene_tx=XM_006927570.4;refGene_gn=MEGF10;refGene_
tx=XM_023254757.1
A1 107130731 107130868
refGene_gn=MEGF10;refGene_tx=XM_006927570.4;refGene_gn=MEGF10;refGene_
tx=XM_023254757.1
A1 107138453 107138666
refGene_gn=MEGF10;refGene_tx=XM_006927570.4;refGene_gn=MEGF10;refGene_
tx=XM_023254757.1
A1 107144593 107144768
refGene_gn=MEGF10;refGene_tx=XM_006927570.4;refGene_gn=MEGF10;refGene_
tx=XM_023254757.1
A1 107146100 107146221
refGene_gn=MEGF10;refGene_tx=XM_006927570.4;refGene_gn=MEGF10;refGene_
tx=XM_023254757.1
A1 107147011 107147175
refGene_gn=MEGF10;refGene_tx=XM_006927570.4;refGene_gn=MEGF10;refGene_
tx=XM_023254757.1

A1 107149819 107149922
refGene_gn=MEGF10;refGene_tx=XM_006927570.4;refGene_gn=MEGF10;refGene_tx=XM_023254757.1
A1 107163273 107163420
refGene_gn=MEGF10;refGene_tx=XM_006927570.4;refGene_gn=MEGF10;refGene_tx=XM_023254757.1
A1 107164609 107164744
refGene_gn=MEGF10;refGene_tx=XM_006927570.4;refGene_gn=MEGF10;refGene_tx=XM_023254757.1
A1 107165281 107165410
refGene_gn=MEGF10;refGene_tx=XM_006927570.4;refGene_gn=MEGF10;refGene_tx=XM_023254757.1
A1 107167253 107167382
refGene_gn=MEGF10;refGene_tx=XM_006927570.4;refGene_gn=MEGF10;refGene_tx=XM_023254757.1
A1 107172541 107172670
refGene_gn=MEGF10;refGene_tx=XM_006927570.4;refGene_gn=MEGF10;refGene_tx=XM_023254757.1
A1 107174824 107174953
refGene_gn=MEGF10;refGene_tx=XM_006927570.4;refGene_gn=MEGF10;refGene_tx=XM_023254757.1
A1 107177903 107178140
refGene_gn=MEGF10;refGene_tx=XM_006927570.4;refGene_gn=MEGF10;refGene_tx=XM_023254757.1
A1 107179404 107179532
refGene_gn=MEGF10;refGene_tx=XM_006927570.4;refGene_gn=MEGF10;refGene_tx=XM_023254757.1
A1 107181134 107181258
refGene_gn=MEGF10;refGene_tx=XM_006927570.4;refGene_gn=MEGF10;refGene_tx=XM_023254757.1
A1 107185635 107185680
refGene_gn=MEGF10;refGene_tx=XM_006927570.4;refGene_gn=MEGF10;refGene_tx=XM_023254757.1
A1 107186453 107186660
refGene_gn=MEGF10;refGene_tx=XM_006927570.4;refGene_gn=MEGF10;refGene_tx=XM_023254757.1
A1 107187922 107188107
refGene_gn=MEGF10;refGene_tx=XM_006927570.4;refGene_gn=MEGF10;refGene_tx=XM_023254757.1
A1 107246640 107246743
refGene_gn=PRRC1;refGene_tx=XM_006927571.4;refGene_gn=PRRC1;refGene_tx=XM_006927572.4;refGene_gn=PRRC1;refGene_tx=XM_023254758.1
A1 107247386 107247770
refGene_gn=PRRC1;refGene_tx=XM_006927571.4;refGene_gn=PRRC1;refGene_tx=XM_006927572.4;refGene_gn=PRRC1;refGene_tx=XM_023254758.1
A1 107249888 107250049
refGene_gn=PRRC1;refGene_tx=XM_006927571.4;refGene_gn=PRRC1;refGene_tx=XM_006927572.4;refGene_gn=PRRC1;refGene_tx=XM_023254758.1
A1 107252717 107252820
refGene_gn=PRRC1;refGene_tx=XM_006927571.4;refGene_gn=PRRC1;refGene_tx

=XM_006927572.4;refGene_gn=PRRC1;refGene_tx=XM_023254758.1
A1 107256570 107256734
refGene_gn=PRRC1;refGene_tx=XM_006927571.4;refGene_gn=PRRC1;refGene_tx
=XM_006927572.4;refGene_gn=PRRC1;refGene_tx=XM_023254758.1
A1 107262514 107262618
refGene_gn=PRRC1;refGene_tx=XM_006927571.4;refGene_gn=PRRC1;refGene_tx
=XM_006927572.4;refGene_gn=PRRC1;refGene_tx=XM_023254758.1
A1 107268612 107268715
refGene_gn=PRRC1;refGene_tx=XM_006927571.4;refGene_gn=PRRC1;refGene_tx
=XM_006927572.4;refGene_gn=PRRC1;refGene_tx=XM_023254758.1
A1 107274806 107275016
refGene_gn=PRRC1;refGene_tx=XM_006927571.4;refGene_gn=PRRC1;refGene_tx
=XM_006927572.4;refGene_gn=PRRC1;refGene_tx=XM_023254758.1
A1 107396490 107396736
refGene_gn=CTXN3;refGene_tx=XM_006927573.4;refGene_gn=CTXN3;refGene_tx
=XM_019831650.2
A1 107445814 107445918
refGene_gn=CCDC192;refGene_tx=XM_019831645.2;refGene_gn=CCDC192;refGen
e_tx=XM_019831647.2
A1 107450773 107450825
refGene_gn=CCDC192;refGene_tx=XM_019831645.2;refGene_gn=CCDC192;refGen
e_tx=XM_019831647.2
A1 107479861 107479969
refGene_gn=CCDC192;refGene_tx=XM_019831645.2;refGene_gn=CCDC192;refGen
e_tx=XM_019831647.2
A1 107504130 107504262
refGene_gn=CCDC192;refGene_tx=XM_019831645.2;refGene_gn=CCDC192;refGen
e_tx=XM_019831647.2
A1 107505375 107505432
refGene_gn=CCDC192;refGene_tx=XM_019831645.2;refGene_gn=CCDC192;refGen
e_tx=XM_019831647.2
A1 107589031 107589164
refGene_gn=CCDC192;refGene_tx=XM_019831645.2;refGene_gn=CCDC192;refGen
e_tx=XM_019831647.2
A1 107590035 107590097
refGene_gn=CCDC192;refGene_tx=XM_019831647.2
A1 107656366 107656653
refGene_gn=CCDC192;refGene_tx=XM_019831645.2
A1 107798529 107799261
refGene_gn=SLC12A2;refGene_tx=XM_011283447.2;refGene_gn=SLC12A2;refGen
e_tx=XM_023254763.1
A1 107827092 107827212
refGene_gn=SLC12A2;refGene_tx=XM_011283447.2;refGene_gn=SLC12A2;refGen
e_tx=XM_023254763.1
A1 107828537 107828613
refGene_gn=SLC12A2;refGene_tx=XM_011283447.2;refGene_gn=SLC12A2;refGen
e_tx=XM_023254763.1
A1 107828911 107829007
refGene_gn=SLC12A2;refGene_tx=XM_011283447.2;refGene_gn=SLC12A2;refGen
e_tx=XM_023254763.1

A1 107845309 107845449
refGene_gn=SLC12A2;refGene_tx=XM_011283447.2;refGene_gn=SLC12A2;refGene_tx=XM_023254763.1
A1 107847181 107847292
refGene_gn=SLC12A2;refGene_tx=XM_011283447.2;refGene_gn=SLC12A2;refGene_tx=XM_023254763.1
A1 107849314 107849423
refGene_gn=SLC12A2;refGene_tx=XM_011283447.2;refGene_gn=SLC12A2;refGene_tx=XM_023254763.1
A1 107851733 107851861
refGene_gn=SLC12A2;refGene_tx=XM_011283447.2;refGene_gn=SLC12A2;refGene_tx=XM_023254763.1
A1 107851957 107852042
refGene_gn=SLC12A2;refGene_tx=XM_011283447.2;refGene_gn=SLC12A2;refGene_tx=XM_023254763.1
A1 107855210 107855362
refGene_gn=SLC12A2;refGene_tx=XM_011283447.2;refGene_gn=SLC12A2;refGene_tx=XM_023254763.1
A1 107860567 107860675
refGene_gn=SLC12A2;refGene_tx=XM_011283447.2;refGene_gn=SLC12A2;refGene_tx=XM_023254763.1
A1 107861726 107861850
refGene_gn=SLC12A2;refGene_tx=XM_011283447.2;refGene_gn=SLC12A2;refGene_tx=XM_023254763.1
A1 107862941 107863043
refGene_gn=SLC12A2;refGene_tx=XM_011283447.2;refGene_gn=SLC12A2;refGene_tx=XM_023254763.1
A1 107864695 107864851
refGene_gn=SLC12A2;refGene_tx=XM_011283447.2;refGene_gn=SLC12A2;refGene_tx=XM_023254763.1
A1 107866134 107866234
refGene_gn=SLC12A2;refGene_tx=XM_011283447.2;refGene_gn=SLC12A2;refGene_tx=XM_023254763.1
A1 107877257 107877369
refGene_gn=SLC12A2;refGene_tx=XM_011283447.2;refGene_gn=SLC12A2;refGene_tx=XM_023254763.1
A1 107879440 107879581
refGene_gn=SLC12A2;refGene_tx=XM_011283447.2;refGene_gn=SLC12A2;refGene_tx=XM_023254763.1
A1 107884505 107884612
refGene_gn=SLC12A2;refGene_tx=XM_011283447.2;refGene_gn=SLC12A2;refGene_tx=XM_023254763.1
A1 107888548 107888628
refGene_gn=SLC12A2;refGene_tx=XM_011283447.2;refGene_gn=SLC12A2;refGene_tx=XM_023254763.1
A1 107891087 107891213
refGene_gn=SLC12A2;refGene_tx=XM_011283447.2;refGene_gn=SLC12A2;refGene_tx=XM_023254763.1
A1 107894015 107894063
refGene_gn=SLC12A2;refGene_tx=XM_011283447.2

A1 107895563 107895686
refGene_gn=SLC12A2;refGene_tx=XM_011283447.2;refGene_gn=SLC12A2;refGene_tx=XM_023254763.1
A1 107897098 107897210
refGene_gn=SLC12A2;refGene_tx=XM_011283447.2;refGene_gn=SLC12A2;refGene_tx=XM_023254763.1
A1 107899189 107899276
refGene_gn=SLC12A2;refGene_tx=XM_011283447.2;refGene_gn=SLC12A2;refGene_tx=XM_023254763.1
A1 107900535 107900671
refGene_gn=SLC12A2;refGene_tx=XM_011283447.2;refGene_gn=SLC12A2;refGene_tx=XM_023254763.1
A1 107900985 107901053
refGene_gn=SLC12A2;refGene_tx=XM_011283447.2;refGene_gn=SLC12A2;refGene_tx=XM_023254763.1
A1 107903101 107903237
refGene_gn=SLC12A2;refGene_tx=XM_011283447.2;refGene_gn=SLC12A2;refGene_tx=XM_023254763.1
A1 107964444 107964819
refGene_gn=FBN2;refGene_tx=XM_003980732.4
A1 107966653 107966825
refGene_gn=FBN2;refGene_tx=XM_003980732.4
A1 107968004 107968236
refGene_gn=FBN2;refGene_tx=XM_003980732.4
A1 107973274 107973394
refGene_gn=FBN2;refGene_tx=XM_003980732.4
A1 107974982 107975111
refGene_gn=FBN2;refGene_tx=XM_003980732.4
A1 107975716 107975833
refGene_gn=FBN2;refGene_tx=XM_003980732.4
A1 107977225 107977348
refGene_gn=FBN2;refGene_tx=XM_003980732.4
A1 107979533 107979659
refGene_gn=FBN2;refGene_tx=XM_003980732.4
A1 107980665 107980872
refGene_gn=FBN2;refGene_tx=XM_003980732.4
A1 107982196 107982322
refGene_gn=FBN2;refGene_tx=XM_003980732.4
A1 107992518 107992650
refGene_gn=FBN2;refGene_tx=XM_003980732.4
A1 107993101 107993224
refGene_gn=FBN2;refGene_tx=XM_003980732.4
A1 107994104 107994224
refGene_gn=FBN2;refGene_tx=XM_003980732.4
A1 107994816 107994942
refGene_gn=FBN2;refGene_tx=XM_003980732.4
A1 107995252 107995318
refGene_gn=FBN2;refGene_tx=XM_003980732.4
A1 107996244 107996397
refGene_gn=FBN2;refGene_tx=XM_003980732.4

A1 107997078 107997204
refGene_gn=FBN2;refGene_tx=XM_003980732.4
A1 107999884 108000004
refGene_gn=FBN2;refGene_tx=XM_003980732.4
A1 108000436 108000565
refGene_gn=FBN2;refGene_tx=XM_003980732.4
A1 108002142 108002259
refGene_gn=FBN2;refGene_tx=XM_003980732.4
A1 108004080 108004206
refGene_gn=FBN2;refGene_tx=XM_003980732.4
A1 108004667 108004793
refGene_gn=FBN2;refGene_tx=XM_003980732.4
A1 108004976 108005102
refGene_gn=FBN2;refGene_tx=XM_003980732.4
A1 108005979 108006048
refGene_gn=FBN2;refGene_tx=XM_003980732.4
A1 108008373 108008526
refGene_gn=FBN2;refGene_tx=XM_003980732.4
A1 108009151 108009277
refGene_gn=FBN2;refGene_tx=XM_003980732.4
A1 108011542 108011668
refGene_gn=FBN2;refGene_tx=XM_003980732.4
A1 108012117 108012186
refGene_gn=FBN2;refGene_tx=XM_003980732.4
A1 108012931 108013093
refGene_gn=FBN2;refGene_tx=XM_003980732.4
A1 108018455 108018578
refGene_gn=FBN2;refGene_tx=XM_003980732.4
A1 108019142 108019265
refGene_gn=FBN2;refGene_tx=XM_003980732.4
A1 108024744 108024870
refGene_gn=FBN2;refGene_tx=XM_003980732.4
A1 108026550 108026673
refGene_gn=FBN2;refGene_tx=XM_003980732.4
A1 108029352 108029475
refGene_gn=FBN2;refGene_tx=XM_003980732.4
A1 108031055 108031181
refGene_gn=FBN2;refGene_tx=XM_003980732.4
A1 108031500 108031626
refGene_gn=FBN2;refGene_tx=XM_003980732.4
A1 108031788 108031911
refGene_gn=FBN2;refGene_tx=XM_003980732.4
A1 108032384 108032510
refGene_gn=FBN2;refGene_tx=XM_003980732.4
A1 108035271 108035397
refGene_gn=FBN2;refGene_tx=XM_003980732.4
A1 108036172 108036301
refGene_gn=FBN2;refGene_tx=XM_003980732.4
A1 108041130 108041256
refGene_gn=FBN2;refGene_tx=XM_003980732.4

A1 108041784 108042012
refGene_gn=FBN2;refGene_tx=XM_003980732.4
A1 108045812 108045938
refGene_gn=FBN2;refGene_tx=XM_003980732.4
A1 108046391 108046442
refGene_gn=FBN2;refGene_tx=XM_003980732.4
A1 108047036 108047174
refGene_gn=FBN2;refGene_tx=XM_003980732.4
A1 108052095 108052215
refGene_gn=FBN2;refGene_tx=XM_003980732.4
A1 108057780 108057906
refGene_gn=FBN2;refGene_tx=XM_003980732.4
A1 108060592 108060718
refGene_gn=FBN2;refGene_tx=XM_003980732.4
A1 108062373 108062427
refGene_gn=FBN2;refGene_tx=XM_003980732.4
A1 108065434 108065587
refGene_gn=FBN2;refGene_tx=XM_003980732.4
A1 108071390 108071513
refGene_gn=FBN2;refGene_tx=XM_003980732.4
A1 108073543 108073666
refGene_gn=FBN2;refGene_tx=XM_003980732.4
A1 108074488 108074614
refGene_gn=FBN2;refGene_tx=XM_003980732.4
A1 108075517 108075637
refGene_gn=FBN2;refGene_tx=XM_003980732.4
A1 108080561 108080699
refGene_gn=FBN2;refGene_tx=XM_003980732.4
A1 108081653 108081887
refGene_gn=FBN2;refGene_tx=XM_003980732.4
A1 108083389 108083542
refGene_gn=FBN2;refGene_tx=XM_003980732.4
A1 108095704 108095830
refGene_gn=FBN2;refGene_tx=XM_003980732.4
A1 108136088 108136214
refGene_gn=FBN2;refGene_tx=XM_003980732.4
A1 108150942 108151140
refGene_gn=FBN2;refGene_tx=XM_003980732.4
A1 108204481 108204577
refGene_gn=FBN2;refGene_tx=XM_003980732.4
A1 108212952 108213048
refGene_gn=FBN2;refGene_tx=XM_003980732.4
A1 108216191 108216290
refGene_gn=FBN2;refGene_tx=XM_003980732.4
A1 108221111 108221194
refGene_gn=FBN2;refGene_tx=XM_003980732.4
A1 108222000 108222254
refGene_gn=FBN2;refGene_tx=XM_003980732.4
A1 108580173 108580654
refGene_gn=SLC27A6;refGene_tx=XM_003980733.4

A1 108599079 108599283
refGene_gn=SLC27A6;refGene_tx=XM_003980733.4
A1 108614473 108614632
refGene_gn=SLC27A6;refGene_tx=XM_003980733.4
A1 108615952 108616077
refGene_gn=SLC27A6;refGene_tx=XM_003980733.4
A1 108635277 108635472
refGene_gn=SLC27A6;refGene_tx=XM_003980733.4
A1 108643756 108643847
refGene_gn=SLC27A6;refGene_tx=XM_003980733.4
A1 108647157 108647356
refGene_gn=SLC27A6;refGene_tx=XM_003980733.4
A1 108648986 108649084
refGene_gn=SLC27A6;refGene_tx=XM_003980733.4
A1 108650249 108650380
refGene_gn=SLC27A6;refGene_tx=XM_003980733.4
A1 108654357 108654534
refGene_gn=SLC27A6;refGene_tx=XM_003980733.4
A1 108693163 108693466
refGene_gn=ISOC1;refGene_tx=XM_023254778.1
A1 108702830 108702950
refGene_gn=ISOC1;refGene_tx=XM_023254778.1
A1 108703059 108703263
refGene_gn=ISOC1;refGene_tx=XM_023254778.1
A1 108704892 108705009
refGene_gn=ISOC1;refGene_tx=XM_023254778.1
A1 108709672 108709819
refGene_gn=ISOC1;refGene_tx=XM_023254778.1
A1 108980140 108980213
refGene_gn=ADAMTS19;refGene_tx=XM_023254779.1;refGene_gn=ADAMTS19;refGene_tx=XM_023254781.1;refGene_gn=ADAMTS19;refGene_tx=XM_023254786.1;refGene_gn=ADAMTS19;refGene_tx=XM_023254790.1
A1 108980841 108981500
refGene_gn=ADAMTS19;refGene_tx=XM_023254779.1;refGene_gn=ADAMTS19;refGene_tx=XM_023254781.1;refGene_gn=ADAMTS19;refGene_tx=XM_023254786.1;refGene_gn=ADAMTS19;refGene_tx=XM_023254790.1
A1 109026978 109027144
refGene_gn=ADAMTS19;refGene_tx=XM_023254779.1;refGene_gn=ADAMTS19;refGene_tx=XM_023254781.1;refGene_gn=ADAMTS19;refGene_tx=XM_023254786.1;refGene_gn=ADAMTS19;refGene_tx=XM_023254790.1
A1 109035341 109035514
refGene_gn=ADAMTS19;refGene_tx=XM_023254779.1;refGene_gn=ADAMTS19;refGene_tx=XM_023254781.1;refGene_gn=ADAMTS19;refGene_tx=XM_023254786.1;refGene_gn=ADAMTS19;refGene_tx=XM_023254790.1
A1 109037158 109037242
refGene_gn=ADAMTS19;refGene_tx=XM_023254781.1;refGene_gn=ADAMTS19;refGene_tx=XM_023254786.1;refGene_gn=ADAMTS19;refGene_tx=XM_023254790.1
A1 109037925 109038083
refGene_gn=ADAMTS19;refGene_tx=XM_023254779.1;refGene_gn=ADAMTS19;refGene_tx=XM_023254781.1;refGene_gn=ADAMTS19;refGene_tx=XM_023254786.1;re

fGene_gn=ADAMTS19;refGene_tx=XM_023254790.1
A1 109054577 109054621
refGene_gn=ADAMTS19;refGene_tx=XM_023254779.1;refGene_gn=ADAMTS19;refGene_tx=XM_023254781.1;refGene_gn=ADAMTS19;refGene_tx=XM_023254786.1;refGene_gn=ADAMTS19;refGene_tx=XM_023254790.1
A1 109091091 109091197
refGene_gn=ADAMTS19;refGene_tx=XM_023254779.1;refGene_gn=ADAMTS19;refGene_tx=XM_023254781.1;refGene_gn=ADAMTS19;refGene_tx=XM_023254786.1;refGene_gn=ADAMTS19;refGene_tx=XM_023254790.1
A1 109114883 109115024
refGene_gn=ADAMTS19;refGene_tx=XM_023254779.1;refGene_gn=ADAMTS19;refGene_tx=XM_023254781.1;refGene_gn=ADAMTS19;refGene_tx=XM_023254786.1;refGene_gn=ADAMTS19;refGene_tx=XM_023254790.1
A1 109116478 109116629
refGene_gn=ADAMTS19;refGene_tx=XM_023254779.1;refGene_gn=ADAMTS19;refGene_tx=XM_023254781.1;refGene_gn=ADAMTS19;refGene_tx=XM_023254786.1;refGene_gn=ADAMTS19;refGene_tx=XM_023254790.1
A1 109130751 109130853
refGene_gn=ADAMTS19;refGene_tx=XM_023254779.1;refGene_gn=ADAMTS19;refGene_tx=XM_023254781.1;refGene_gn=ADAMTS19;refGene_tx=XM_023254790.1
A1 109136677 109136808
refGene_gn=ADAMTS19;refGene_tx=XM_023254779.1;refGene_gn=ADAMTS19;refGene_tx=XM_023254781.1;refGene_gn=ADAMTS19;refGene_tx=XM_023254790.1
A1 109137670 109137843
refGene_gn=ADAMTS19;refGene_tx=XM_023254779.1;refGene_gn=ADAMTS19;refGene_tx=XM_023254781.1;refGene_gn=ADAMTS19;refGene_tx=XM_023254790.1
A1 109145079 109145207
refGene_gn=ADAMTS19;refGene_tx=XM_023254779.1;refGene_gn=ADAMTS19;refGene_tx=XM_023254781.1;refGene_gn=ADAMTS19;refGene_tx=XM_023254790.1
A1 109149444 109149565
refGene_gn=ADAMTS19;refGene_tx=XM_023254779.1;refGene_gn=ADAMTS19;refGene_tx=XM_023254781.1;refGene_gn=ADAMTS19;refGene_tx=XM_023254786.1;refGene_gn=ADAMTS19;refGene_tx=XM_023254790.1
A1 109155946 109156027
refGene_gn=ADAMTS19;refGene_tx=XM_023254779.1;refGene_gn=ADAMTS19;refGene_tx=XM_023254781.1;refGene_gn=ADAMTS19;refGene_tx=XM_023254786.1;refGene_gn=ADAMTS19;refGene_tx=XM_023254790.1
A1 109169840 109169998
refGene_gn=ADAMTS19;refGene_tx=XM_023254779.1;refGene_gn=ADAMTS19;refGene_tx=XM_023254781.1;refGene_gn=ADAMTS19;refGene_tx=XM_023254786.1;refGene_gn=ADAMTS19;refGene_tx=XM_023254790.1
A1 109174242 109174396
refGene_gn=ADAMTS19;refGene_tx=XM_023254779.1;refGene_gn=ADAMTS19;refGene_tx=XM_023254781.1;refGene_gn=ADAMTS19;refGene_tx=XM_023254786.1;refGene_gn=ADAMTS19;refGene_tx=XM_023254790.1
A1 109185885 109186021
refGene_gn=ADAMTS19;refGene_tx=XM_023254779.1;refGene_gn=ADAMTS19;refGene_tx=XM_023254781.1;refGene_gn=ADAMTS19;refGene_tx=XM_023254786.1;refGene_gn=ADAMTS19;refGene_tx=XM_023254790.1
A1 109193071 109193276

refGene_gn=ADAMTS19;refGene_tx=XM_023254779.1;refGene_gn=ADAMTS19;refGene_tx=XM_023254781.1;refGene_gn=ADAMTS19;refGene_tx=XM_023254786.1
A1 109196868 109197021
refGene_gn=ADAMTS19;refGene_tx=XM_023254779.1;refGene_gn=ADAMTS19;refGene_tx=XM_023254781.1;refGene_gn=ADAMTS19;refGene_tx=XM_023254786.1;refGene_gn=ADAMTS19;refGene_tx=XM_023254790.1
A1 109216298 109216476
refGene_gn=ADAMTS19;refGene_tx=XM_023254779.1;refGene_gn=ADAMTS19;refGene_tx=XM_023254781.1;refGene_gn=ADAMTS19;refGene_tx=XM_023254786.1
A1 109220649 109220801
refGene_gn=ADAMTS19;refGene_tx=XM_023254779.1;refGene_gn=ADAMTS19;refGene_tx=XM_023254781.1;refGene_gn=ADAMTS19;refGene_tx=XM_023254786.1
A1 109229595 109229760
refGene_gn=KIAA1024L;refGene_tx=XM_003980737.2
A1 109241814 109241967
refGene_gn=KIAA1024L;refGene_tx=XM_003980737.2
A1 109241968 109242043
refGene_gn=KIAA1024L;refGene_tx=XM_003980737.2
A1 109245609 109245789
refGene_gn=KIAA1024L;refGene_tx=XM_003980737.2
A1 109349296 109350104
refGene_gn=CHSY3;refGene_tx=XM_023254802.1
A1 109352538 109352822
refGene_gn=CHSY3;refGene_tx=XM_023254802.1
A1 109668476 109668573
refGene_gn=CHSY3;refGene_tx=XM_023254802.1
A1 109668575 109668997
refGene_gn=CHSY3;refGene_tx=XM_023254802.1
A1 109668998 109669248
refGene_gn=CHSY3;refGene_tx=XM_023254802.1
A1 109669250 109669735
refGene_gn=CHSY3;refGene_tx=XM_023254802.1
A1 109669736 109670039
refGene_gn=CHSY3;refGene_tx=XM_023254802.1
A1 110397888 110398053
refGene_gn=HINT1;refGene_tx=XM_023254816.1
A1 110399237 110399342
refGene_gn=HINT1;refGene_tx=XM_023254816.1
A1 110401585 110401696
refGene_gn=HINT1;refGene_tx=XM_023254816.1
A1 110406350 110406485
refGene_gn=LYRM7;refGene_tx=XM_019831696.2
A1 110418882 110418955
refGene_gn=LYRM7;refGene_tx=XM_019831696.2
A1 110420450 110420521
refGene_gn=LYRM7;refGene_tx=XM_019831696.2
A1 110425147 110425229
refGene_gn=LYRM7;refGene_tx=XM_019831696.2
A1 110435731 110435802
refGene_gn=LYRM7;refGene_tx=XM_019831696.2

A1 110579095 110579149
refGene_gn=CDC42SE2;refGene_tx=XM_003980739.5;refGene_gn=CDC42SE2;refGene_tx=XM_006927575.4;refGene_gn=CDC42SE2;refGene_tx=XM_006927576.4;refGene_gn=CDC42SE2;refGene_tx=XM_023254838.1;refGene_gn=CDC42SE2;refGene_tx=XM_023254850.1
A1 110607478 110607580
refGene_gn=CDC42SE2;refGene_tx=XM_003980739.5;refGene_gn=CDC42SE2;refGene_tx=XM_006927575.4;refGene_gn=CDC42SE2;refGene_tx=XM_006927576.4;refGene_gn=CDC42SE2;refGene_tx=XM_023254838.1;refGene_gn=CDC42SE2;refGene_tx=XM_023254850.1
A1 110611435 110611534
refGene_gn=CDC42SE2;refGene_tx=XM_003980739.5;refGene_gn=CDC42SE2;refGene_tx=XM_006927575.4;refGene_gn=CDC42SE2;refGene_tx=XM_006927576.4;refGene_gn=CDC42SE2;refGene_tx=XM_023254838.1;refGene_gn=CDC42SE2;refGene_tx=XM_023254850.1
A1 110638505 110638531
refGene_gn=RAPGEF6;refGene_tx=XM_003980740.5;refGene_gn=RAPGEF6;refGene_tx=XM_011281964.3;refGene_gn=RAPGEF6;refGene_tx=XM_011281969.3;refGene_gn=RAPGEF6;refGene_tx=XM_019831704.2;refGene_gn=RAPGEF6;refGene_tx=XM_019831705.2;refGene_gn=RAPGEF6;refGene_tx=XM_019831706.2;refGene_gn=RAPGEF6;refGene_tx=XM_019831708.2;refGene_gn=RAPGEF6;refGene_tx=XM_019831709.2;refGene_gn=RAPGEF6;refGene_tx=XM_019831710.2;refGene_gn=RAPGEF6;refGene_tx=XM_019831711.2;refGene_gn=RAPGEF6;refGene_tx=XM_019831712.2;refGene_gn=RAPGEF6;refGene_tx=XM_019831714.2;refGene_gn=RAPGEF6;refGene_tx=XM_023254897.1
A1 110639230 110639545
refGene_gn=RAPGEF6;refGene_tx=XM_003980740.5;refGene_gn=RAPGEF6;refGene_tx=XM_011281964.3;refGene_gn=RAPGEF6;refGene_tx=XM_019831704.2;refGene_gn=RAPGEF6;refGene_tx=XM_019831705.2;refGene_gn=RAPGEF6;refGene_tx=XM_019831706.2;refGene_gn=RAPGEF6;refGene_tx=XM_019831708.2;refGene_gn=RAPGEF6;refGene_tx=XM_019831709.2;refGene_gn=RAPGEF6;refGene_tx=XM_019831710.2;refGene_gn=RAPGEF6;refGene_tx=XM_019831714.2;refGene_gn=RAPGEF6;refGene_tx=XM_023254897.1
A1 110641824 110642327
refGene_gn=RAPGEF6;refGene_tx=XM_003980740.5;refGene_gn=RAPGEF6;refGene_tx=XM_011281964.3;refGene_gn=RAPGEF6;refGene_tx=XM_011281969.3;refGene_gn=RAPGEF6;refGene_tx=XM_019831704.2;refGene_gn=RAPGEF6;refGene_tx=XM_019831705.2;refGene_gn=RAPGEF6;refGene_tx=XM_019831706.2;refGene_gn=RAPGEF6;refGene_tx=XM_019831708.2;refGene_gn=RAPGEF6;refGene_tx=XM_019831709.2;refGene_gn=RAPGEF6;refGene_tx=XM_019831710.2;refGene_gn=RAPGEF6;refGene_tx=XM_019831711.2;refGene_gn=RAPGEF6;refGene_tx=XM_019831712.2;refGene_gn=RAPGEF6;refGene_tx=XM_019831714.2;refGene_gn=RAPGEF6;refGene_tx=XM_023254897.1
A1 110643623 110643852
refGene_gn=RAPGEF6;refGene_tx=XM_003980740.5;refGene_gn=RAPGEF6;refGene_tx=XM_011281964.3;refGene_gn=RAPGEF6;refGene_tx=XM_011281969.3;refGene_gn=RAPGEF6;refGene_tx=XM_019831704.2;refGene_gn=RAPGEF6;refGene_tx=XM_019831705.2;refGene_gn=RAPGEF6;refGene_tx=XM_019831706.2;refGene_gn=RAPGEF6;refGene_tx=XM_019831708.2;refGene_gn=RAPGEF6;refGene_tx=XM_019831709.2;refGene_gn=RAPGEF6;refGene_tx=XM_019831710.2;refGene_gn=RAPG

EF6;refGene_tx=XM_019831711.2;refGene_gn=RAPGEF6;refGene_tx=XM_0198317
12.2;refGene_gn=RAPGEF6;refGene_tx=XM_019831714.2;refGene_gn=RAPGEF6;r
efGene_tx=XM_023254897.1
A1 110645927 110646409
refGene_gn=RAPGEF6;refGene_tx=XM_011281970.3
A1 110648915 110649050
refGene_gn=RAPGEF6;refGene_tx=XM_003980740.5;refGene_gn=RAPGEF6;refGen
e_tx=XM_011281964.3;refGene_gn=RAPGEF6;refGene_tx=XM_011281969.3;refGe
ne_gn=RAPGEF6;refGene_tx=XM_011281970.3;refGene_gn=RAPGEF6;refGene_tx=
XM_019831704.2;refGene_gn=RAPGEF6;refGene_tx=XM_019831705.2;refGene_gn
=RAPGEF6;refGene_tx=XM_019831706.2;refGene_gn=RAPGEF6;refGene_tx=XM_01
9831708.2;refGene_gn=RAPGEF6;refGene_tx=XM_019831709.2;refGene_gn=RAPG
EF6;refGene_tx=XM_019831710.2;refGene_gn=RAPGEF6;refGene_tx=XM_0198317
11.2;refGene_gn=RAPGEF6;refGene_tx=XM_019831712.2;refGene_gn=RAPGEF6;r
efGene_tx=XM_019831714.2;refGene_gn=RAPGEF6;refGene_tx=XM_023254897.1
A1 110651750 110651942
refGene_gn=RAPGEF6;refGene_tx=XM_003980740.5;refGene_gn=RAPGEF6;refGen
e_tx=XM_011281964.3;refGene_gn=RAPGEF6;refGene_tx=XM_011281969.3;refGe
ne_gn=RAPGEF6;refGene_tx=XM_011281970.3;refGene_gn=RAPGEF6;refGene_tx=
XM_019831704.2;refGene_gn=RAPGEF6;refGene_tx=XM_019831705.2;refGene_gn
=RAPGEF6;refGene_tx=XM_019831706.2;refGene_gn=RAPGEF6;refGene_tx=XM_01
9831708.2;refGene_gn=RAPGEF6;refGene_tx=XM_019831709.2;refGene_gn=RAPG
EF6;refGene_tx=XM_019831710.2;refGene_gn=RAPGEF6;refGene_tx=XM_0198317
11.2;refGene_gn=RAPGEF6;refGene_tx=XM_019831712.2;refGene_gn=RAPGEF6;r
efGene_tx=XM_019831714.2;refGene_gn=RAPGEF6;refGene_tx=XM_023254897.1
A1 110654150 110654371
refGene_gn=RAPGEF6;refGene_tx=XM_003980740.5;refGene_gn=RAPGEF6;refGen
e_tx=XM_011281964.3;refGene_gn=RAPGEF6;refGene_tx=XM_011281969.3;refGe
ne_gn=RAPGEF6;refGene_tx=XM_011281970.3;refGene_gn=RAPGEF6;refGene_tx=
XM_019831704.2;refGene_gn=RAPGEF6;refGene_tx=XM_019831705.2;refGene_gn
=RAPGEF6;refGene_tx=XM_019831706.2;refGene_gn=RAPGEF6;refGene_tx=XM_01
9831708.2;refGene_gn=RAPGEF6;refGene_tx=XM_019831709.2;refGene_gn=RAPG
EF6;refGene_tx=XM_019831710.2;refGene_gn=RAPGEF6;refGene_tx=XM_0198317
11.2;refGene_gn=RAPGEF6;refGene_tx=XM_019831712.2;refGene_gn=RAPGEF6;r
efGene_tx=XM_019831714.2;refGene_gn=RAPGEF6;refGene_tx=XM_023254897.1
A1 110657802 110657826
refGene_gn=RAPGEF6;refGene_tx=XM_003980740.5;refGene_gn=RAPGEF6;refGen
e_tx=XM_011281969.3;refGene_gn=RAPGEF6;refGene_tx=XM_019831704.2;refGe
ne_gn=RAPGEF6;refGene_tx=XM_019831705.2;refGene_gn=RAPGEF6;refGene_tx=
XM_019831706.2;refGene_gn=RAPGEF6;refGene_tx=XM_019831710.2;refGene_gn
=RAPGEF6;refGene_tx=XM_019831711.2;refGene_gn=RAPGEF6;refGene_tx=XM_01
9831712.2;refGene_gn=RAPGEF6;refGene_tx=XM_019831714.2;refGene_gn=RAPG
EF6;refGene_tx=XM_023254897.1
A1 110660079 110660203
refGene_gn=RAPGEF6;refGene_tx=XM_003980740.5;refGene_gn=RAPGEF6;refGen
e_tx=XM_011281964.3;refGene_gn=RAPGEF6;refGene_tx=XM_011281969.3;refGe
ne_gn=RAPGEF6;refGene_tx=XM_011281970.3;refGene_gn=RAPGEF6;refGene_tx=
XM_019831704.2;refGene_gn=RAPGEF6;refGene_tx=XM_019831705.2;refGene_gn
=RAPGEF6;refGene_tx=XM_019831706.2;refGene_gn=RAPGEF6;refGene_tx=XM_01
9831708.2;refGene_gn=RAPGEF6;refGene_tx=XM_019831709.2;refGene_gn=RAPG

EF6;refGene_tx=XM_019831710.2;refGene_gn=RAPGEF6;refGene_tx=XM_0198317
11.2;refGene_gn=RAPGEF6;refGene_tx=XM_019831712.2;refGene_gn=RAPGEF6;r
efGene_tx=XM_019831714.2;refGene_gn=RAPGEF6;refGene_tx=XM_023254897.1
A1 110664707 110664919
refGene_gn=RAPGEF6;refGene_tx=XM_003980740.5;refGene_gn=RAPGEF6;refGen
e_tx=XM_011281964.3;refGene_gn=RAPGEF6;refGene_tx=XM_011281969.3;refGe
ne_gn=RAPGEF6;refGene_tx=XM_011281970.3;refGene_gn=RAPGEF6;refGene_tx=
XM_019831704.2;refGene_gn=RAPGEF6;refGene_tx=XM_019831705.2;refGene_gn
=RAPGEF6;refGene_tx=XM_019831706.2;refGene_gn=RAPGEF6;refGene_tx=XM_01
9831708.2;refGene_gn=RAPGEF6;refGene_tx=XM_019831709.2;refGene_gn=RAPG
EF6;refGene_tx=XM_019831710.2;refGene_gn=RAPGEF6;refGene_tx=XM_0198317
11.2;refGene_gn=RAPGEF6;refGene_tx=XM_019831712.2;refGene_gn=RAPGEF6;r
efGene_tx=XM_019831714.2;refGene_gn=RAPGEF6;refGene_tx=XM_023254897.1
A1 110674840 110675224
refGene_gn=RAPGEF6;refGene_tx=XM_003980740.5;refGene_gn=RAPGEF6;refGen
e_tx=XM_011281964.3;refGene_gn=RAPGEF6;refGene_tx=XM_011281969.3;refGe
ne_gn=RAPGEF6;refGene_tx=XM_011281970.3;refGene_gn=RAPGEF6;refGene_tx=
XM_019831704.2;refGene_gn=RAPGEF6;refGene_tx=XM_019831705.2;refGene_gn
=RAPGEF6;refGene_tx=XM_019831706.2;refGene_gn=RAPGEF6;refGene_tx=XM_01
9831708.2;refGene_gn=RAPGEF6;refGene_tx=XM_019831709.2;refGene_gn=RAPG
EF6;refGene_tx=XM_019831710.2;refGene_gn=RAPGEF6;refGene_tx=XM_0198317
11.2;refGene_gn=RAPGEF6;refGene_tx=XM_019831712.2;refGene_gn=RAPGEF6;r
efGene_tx=XM_019831714.2;refGene_gn=RAPGEF6;refGene_tx=XM_023254897.1
A1 110677307 110677548
refGene_gn=RAPGEF6;refGene_tx=XM_003980740.5;refGene_gn=RAPGEF6;refGen
e_tx=XM_011281964.3;refGene_gn=RAPGEF6;refGene_tx=XM_011281969.3;refGe
ne_gn=RAPGEF6;refGene_tx=XM_011281970.3;refGene_gn=RAPGEF6;refGene_tx=
XM_019831704.2;refGene_gn=RAPGEF6;refGene_tx=XM_019831705.2;refGene_gn
=RAPGEF6;refGene_tx=XM_019831706.2;refGene_gn=RAPGEF6;refGene_tx=XM_01
9831708.2;refGene_gn=RAPGEF6;refGene_tx=XM_019831709.2;refGene_gn=RAPG
EF6;refGene_tx=XM_019831710.2;refGene_gn=RAPGEF6;refGene_tx=XM_0198317
11.2;refGene_gn=RAPGEF6;refGene_tx=XM_019831712.2;refGene_gn=RAPGEF6;r
efGene_tx=XM_019831714.2;refGene_gn=RAPGEF6;refGene_tx=XM_023254897.1
A1 110682941 110682956
refGene_gn=RAPGEF6;refGene_tx=XM_019831704.2;refGene_gn=RAPGEF6;refGen
e_tx=XM_019831705.2;refGene_gn=RAPGEF6;refGene_tx=XM_019831708.2;refGe
ne_gn=RAPGEF6;refGene_tx=XM_019831709.2;refGene_gn=RAPGEF6;refGene_tx=
XM_019831710.2;refGene_gn=RAPGEF6;refGene_tx=XM_019831711.2;refGene_gn
=RAPGEF6;refGene_tx=XM_019831712.2;refGene_gn=RAPGEF6;refGene_tx=XM_01
9831714.2;refGene_gn=RAPGEF6;refGene_tx=XM_023254897.1
A1 110685872 110686030
refGene_gn=RAPGEF6;refGene_tx=XM_003980740.5;refGene_gn=RAPGEF6;refGen
e_tx=XM_011281964.3;refGene_gn=RAPGEF6;refGene_tx=XM_011281969.3;refGe
ne_gn=RAPGEF6;refGene_tx=XM_011281970.3;refGene_gn=RAPGEF6;refGene_tx=
XM_019831704.2;refGene_gn=RAPGEF6;refGene_tx=XM_019831705.2;refGene_gn
=RAPGEF6;refGene_tx=XM_019831706.2;refGene_gn=RAPGEF6;refGene_tx=XM_01
9831708.2;refGene_gn=RAPGEF6;refGene_tx=XM_019831709.2;refGene_gn=RAPG
EF6;refGene_tx=XM_019831710.2;refGene_gn=RAPGEF6;refGene_tx=XM_0198317
11.2;refGene_gn=RAPGEF6;refGene_tx=XM_019831712.2;refGene_gn=RAPGEF6;r
efGene_tx=XM_019831714.2;refGene_gn=RAPGEF6;refGene_tx=XM_023254897.1

A1 110688214 110688226
refGene_gn=RAPGEF6;refGene_tx=XM_019831704.2;refGene_gn=RAPGEF6;refGene_tx=XM_019831706.2;refGene_gn=RAPGEF6;refGene_tx=XM_019831708.2;refGene_gn=RAPGEF6;refGene_tx=XM_019831710.2;refGene_gn=RAPGEF6;refGene_tx=XM_019831711.2;refGene_gn=RAPGEF6;refGene_tx=XM_019831714.2;refGene_gn=RAPGEF6;refGene_tx=XM_023254897.1
A1 110692564 110692805
refGene_gn=RAPGEF6;refGene_tx=XM_003980740.5;refGene_gn=RAPGEF6;refGene_tx=XM_011281964.3;refGene_gn=RAPGEF6;refGene_tx=XM_011281969.3;refGene_gn=RAPGEF6;refGene_tx=XM_011281970.3;refGene_gn=RAPGEF6;refGene_tx=XM_019831704.2;refGene_gn=RAPGEF6;refGene_tx=XM_019831705.2;refGene_gn=RAPGEF6;refGene_tx=XM_019831706.2;refGene_gn=RAPGEF6;refGene_tx=XM_019831708.2;refGene_gn=RAPGEF6;refGene_tx=XM_019831709.2;refGene_gn=RAPGEF6;refGene_tx=XM_019831710.2;refGene_gn=RAPGEF6;refGene_tx=XM_019831711.2;refGene_gn=RAPGEF6;refGene_tx=XM_019831712.2;refGene_gn=RAPGEF6;refGene_tx=XM_019831714.2;refGene_gn=RAPGEF6;refGene_tx=XM_023254897.1
A1 110702242 110702351
refGene_gn=RAPGEF6;refGene_tx=XM_003980740.5;refGene_gn=RAPGEF6;refGene_tx=XM_011281964.3;refGene_gn=RAPGEF6;refGene_tx=XM_011281969.3;refGene_gn=RAPGEF6;refGene_tx=XM_011281970.3;refGene_gn=RAPGEF6;refGene_tx=XM_019831704.2;refGene_gn=RAPGEF6;refGene_tx=XM_019831705.2;refGene_gn=RAPGEF6;refGene_tx=XM_019831706.2;refGene_gn=RAPGEF6;refGene_tx=XM_019831708.2;refGene_gn=RAPGEF6;refGene_tx=XM_019831709.2;refGene_gn=RAPGEF6;refGene_tx=XM_019831710.2;refGene_gn=RAPGEF6;refGene_tx=XM_019831711.2;refGene_gn=RAPGEF6;refGene_tx=XM_019831712.2;refGene_gn=RAPGEF6;refGene_tx=XM_019831714.2;refGene_gn=RAPGEF6;refGene_tx=XM_023254897.1
A1 110705054 110705258
refGene_gn=RAPGEF6;refGene_tx=XM_003980740.5;refGene_gn=RAPGEF6;refGene_tx=XM_011281964.3;refGene_gn=RAPGEF6;refGene_tx=XM_011281969.3;refGene_gn=RAPGEF6;refGene_tx=XM_011281970.3;refGene_gn=RAPGEF6;refGene_tx=XM_019831704.2;refGene_gn=RAPGEF6;refGene_tx=XM_019831705.2;refGene_gn=RAPGEF6;refGene_tx=XM_019831706.2;refGene_gn=RAPGEF6;refGene_tx=XM_019831708.2;refGene_gn=RAPGEF6;refGene_tx=XM_019831709.2;refGene_gn=RAPGEF6;refGene_tx=XM_019831710.2;refGene_gn=RAPGEF6;refGene_tx=XM_019831711.2;refGene_gn=RAPGEF6;refGene_tx=XM_019831712.2;refGene_gn=RAPGEF6;refGene_tx=XM_019831714.2;refGene_gn=RAPGEF6;refGene_tx=XM_023254897.1
A1 110707582 110707690
refGene_gn=RAPGEF6;refGene_tx=XM_003980740.5;refGene_gn=RAPGEF6;refGene_tx=XM_011281964.3;refGene_gn=RAPGEF6;refGene_tx=XM_011281969.3;refGene_gn=RAPGEF6;refGene_tx=XM_011281970.3;refGene_gn=RAPGEF6;refGene_tx=XM_019831704.2;refGene_gn=RAPGEF6;refGene_tx=XM_019831705.2;refGene_gn=RAPGEF6;refGene_tx=XM_019831706.2;refGene_gn=RAPGEF6;refGene_tx=XM_019831708.2;refGene_gn=RAPGEF6;refGene_tx=XM_019831709.2;refGene_gn=RAPGEF6;refGene_tx=XM_019831710.2;refGene_gn=RAPGEF6;refGene_tx=XM_019831711.2;refGene_gn=RAPGEF6;refGene_tx=XM_019831712.2;refGene_gn=RAPGEF6;refGene_tx=XM_019831714.2;refGene_gn=RAPGEF6;refGene_tx=XM_023254897.1
A1 110710919 110711084
refGene_gn=RAPGEF6;refGene_tx=XM_003980740.5;refGene_gn=RAPGEF6;refGene_tx=XM_011281964.3;refGene_gn=RAPGEF6;refGene_tx=XM_011281969.3;refGene_gn=RAPGEF6;refGene_tx=XM_011281970.3;refGene_gn=RAPGEF6;refGene_tx=

XM_019831704.2;refGene_gn=RAPGEF6;refGene_tx=XM_019831705.2;refGene_gn=RAPGEF6;refGene_tx=XM_019831706.2;refGene_gn=RAPGEF6;refGene_tx=XM_019831708.2;refGene_gn=RAPGEF6;refGene_tx=XM_019831709.2;refGene_gn=RAPGEF6;refGene_tx=XM_019831710.2;refGene_gn=RAPGEF6;refGene_tx=XM_019831711.2;refGene_gn=RAPGEF6;refGene_tx=XM_019831712.2;refGene_gn=RAPGEF6;refGene_tx=XM_019831714.2;refGene_gn=RAPGEF6;refGene_tx=XM_023254897.1
A1 110715603 110715756
refGene_gn=RAPGEF6;refGene_tx=XM_003980740.5;refGene_gn=RAPGEF6;refGene_tx=XM_011281964.3;refGene_gn=RAPGEF6;refGene_tx=XM_011281969.3;refGene_gn=RAPGEF6;refGene_tx=XM_011281970.3;refGene_gn=RAPGEF6;refGene_tx=XM_019831704.2;refGene_gn=RAPGEF6;refGene_tx=XM_019831705.2;refGene_gn=RAPGEF6;refGene_tx=XM_019831706.2;refGene_gn=RAPGEF6;refGene_tx=XM_019831708.2;refGene_gn=RAPGEF6;refGene_tx=XM_019831709.2;refGene_gn=RAPGEF6;refGene_tx=XM_019831710.2;refGene_gn=RAPGEF6;refGene_tx=XM_019831711.2;refGene_gn=RAPGEF6;refGene_tx=XM_019831712.2;refGene_gn=RAPGEF6;refGene_tx=XM_019831714.2;refGene_gn=RAPGEF6;refGene_tx=XM_023254897.1
A1 110716351 110716510
refGene_gn=RAPGEF6;refGene_tx=XM_003980740.5;refGene_gn=RAPGEF6;refGene_tx=XM_011281964.3;refGene_gn=RAPGEF6;refGene_tx=XM_011281969.3;refGene_gn=RAPGEF6;refGene_tx=XM_011281970.3;refGene_gn=RAPGEF6;refGene_tx=XM_019831704.2;refGene_gn=RAPGEF6;refGene_tx=XM_019831705.2;refGene_gn=RAPGEF6;refGene_tx=XM_019831706.2;refGene_gn=RAPGEF6;refGene_tx=XM_019831708.2;refGene_gn=RAPGEF6;refGene_tx=XM_019831709.2;refGene_gn=RAPGEF6;refGene_tx=XM_019831710.2;refGene_gn=RAPGEF6;refGene_tx=XM_019831711.2;refGene_gn=RAPGEF6;refGene_tx=XM_019831712.2;refGene_gn=RAPGEF6;refGene_tx=XM_019831714.2;refGene_gn=RAPGEF6;refGene_tx=XM_023254897.1
A1 110718604 110718741
refGene_gn=RAPGEF6;refGene_tx=XM_003980740.5;refGene_gn=RAPGEF6;refGene_tx=XM_011281964.3;refGene_gn=RAPGEF6;refGene_tx=XM_011281969.3;refGene_gn=RAPGEF6;refGene_tx=XM_011281970.3;refGene_gn=RAPGEF6;refGene_tx=XM_019831704.2;refGene_gn=RAPGEF6;refGene_tx=XM_019831705.2;refGene_gn=RAPGEF6;refGene_tx=XM_019831706.2;refGene_gn=RAPGEF6;refGene_tx=XM_019831708.2;refGene_gn=RAPGEF6;refGene_tx=XM_019831709.2;refGene_gn=RAPGEF6;refGene_tx=XM_019831710.2;refGene_gn=RAPGEF6;refGene_tx=XM_019831711.2;refGene_gn=RAPGEF6;refGene_tx=XM_019831712.2;refGene_gn=RAPGEF6;refGene_tx=XM_019831714.2;refGene_gn=RAPGEF6;refGene_tx=XM_023254897.1
A1 110720827 110721005
refGene_gn=RAPGEF6;refGene_tx=XM_003980740.5;refGene_gn=RAPGEF6;refGene_tx=XM_011281964.3;refGene_gn=RAPGEF6;refGene_tx=XM_011281969.3;refGene_gn=RAPGEF6;refGene_tx=XM_011281970.3;refGene_gn=RAPGEF6;refGene_tx=XM_019831704.2;refGene_gn=RAPGEF6;refGene_tx=XM_019831705.2;refGene_gn=RAPGEF6;refGene_tx=XM_019831706.2;refGene_gn=RAPGEF6;refGene_tx=XM_019831708.2;refGene_gn=RAPGEF6;refGene_tx=XM_019831709.2;refGene_gn=RAPGEF6;refGene_tx=XM_019831710.2;refGene_gn=RAPGEF6;refGene_tx=XM_019831711.2;refGene_gn=RAPGEF6;refGene_tx=XM_019831712.2;refGene_gn=RAPGEF6;refGene_tx=XM_019831714.2;refGene_gn=RAPGEF6;refGene_tx=XM_023254897.1
A1 110733227 110733359
refGene_gn=RAPGEF6;refGene_tx=XM_003980740.5;refGene_gn=RAPGEF6;refGene_tx=XM_011281964.3;refGene_gn=RAPGEF6;refGene_tx=XM_011281969.3;refGene_gn=RAPGEF6;refGene_tx=XM_011281970.3;refGene_gn=RAPGEF6;refGene_tx=

XM_019831704.2;refGene_gn=RAPGEF6;refGene_tx=XM_019831705.2;refGene_gn=RAPGEF6;refGene_tx=XM_019831706.2;refGene_gn=RAPGEF6;refGene_tx=XM_019831708.2;refGene_gn=RAPGEF6;refGene_tx=XM_019831709.2;refGene_gn=RAPGEF6;refGene_tx=XM_019831710.2;refGene_gn=RAPGEF6;refGene_tx=XM_019831711.2;refGene_gn=RAPGEF6;refGene_tx=XM_019831712.2;refGene_gn=RAPGEF6;refGene_tx=XM_019831714.2;refGene_gn=RAPGEF6;refGene_tx=XM_023254897.1
A1 110744609 110744663
refGene_gn=RAPGEF6;refGene_tx=XM_019831714.2;refGene_gn=RAPGEF6;refGene_tx=XM_023254897.1
A1 110757862 110758006
refGene_gn=RAPGEF6;refGene_tx=XM_003980740.5;refGene_gn=RAPGEF6;refGene_tx=XM_011281964.3;refGene_gn=RAPGEF6;refGene_tx=XM_011281969.3;refGene_gn=RAPGEF6;refGene_tx=XM_011281970.3;refGene_gn=RAPGEF6;refGene_tx=XM_019831704.2;refGene_gn=RAPGEF6;refGene_tx=XM_019831705.2;refGene_gn=RAPGEF6;refGene_tx=XM_019831706.2;refGene_gn=RAPGEF6;refGene_tx=XM_019831708.2;refGene_gn=RAPGEF6;refGene_tx=XM_019831709.2;refGene_gn=RAPGEF6;refGene_tx=XM_019831710.2;refGene_gn=RAPGEF6;refGene_tx=XM_019831711.2;refGene_gn=RAPGEF6;refGene_tx=XM_019831712.2
A1 110778201 110778271
refGene_gn=RAPGEF6;refGene_tx=XM_003980740.5;refGene_gn=RAPGEF6;refGene_tx=XM_011281964.3;refGene_gn=RAPGEF6;refGene_tx=XM_011281969.3;refGene_gn=RAPGEF6;refGene_tx=XM_011281970.3;refGene_gn=RAPGEF6;refGene_tx=XM_019831704.2;refGene_gn=RAPGEF6;refGene_tx=XM_019831705.2;refGene_gn=RAPGEF6;refGene_tx=XM_019831706.2;refGene_gn=RAPGEF6;refGene_tx=XM_019831708.2;refGene_gn=RAPGEF6;refGene_tx=XM_019831709.2;refGene_gn=RAPGEF6;refGene_tx=XM_019831710.2;refGene_gn=RAPGEF6;refGene_tx=XM_019831711.2;refGene_gn=RAPGEF6;refGene_tx=XM_019831712.2
A1 110799727 110799811
refGene_gn=RAPGEF6;refGene_tx=XM_003980740.5;refGene_gn=RAPGEF6;refGene_tx=XM_011281964.3;refGene_gn=RAPGEF6;refGene_tx=XM_011281969.3;refGene_gn=RAPGEF6;refGene_tx=XM_011281970.3;refGene_gn=RAPGEF6;refGene_tx=XM_019831704.2;refGene_gn=RAPGEF6;refGene_tx=XM_019831705.2;refGene_gn=RAPGEF6;refGene_tx=XM_019831706.2;refGene_gn=RAPGEF6;refGene_tx=XM_019831708.2;refGene_gn=RAPGEF6;refGene_tx=XM_019831709.2;refGene_gn=RAPGEF6;refGene_tx=XM_019831711.2;refGene_gn=RAPGEF6;refGene_tx=XM_019831712.2
A1 110804114 110804171
refGene_gn=RAPGEF6;refGene_tx=XM_003980740.5;refGene_gn=RAPGEF6;refGene_tx=XM_011281964.3;refGene_gn=RAPGEF6;refGene_tx=XM_011281969.3;refGene_gn=RAPGEF6;refGene_tx=XM_011281970.3;refGene_gn=RAPGEF6;refGene_tx=XM_019831704.2;refGene_gn=RAPGEF6;refGene_tx=XM_019831705.2;refGene_gn=RAPGEF6;refGene_tx=XM_019831706.2;refGene_gn=RAPGEF6;refGene_tx=XM_019831708.2;refGene_gn=RAPGEF6;refGene_tx=XM_019831709.2;refGene_gn=RAPGEF6;refGene_tx=XM_019831710.2;refGene_gn=RAPGEF6;refGene_tx=XM_019831711.2;refGene_gn=RAPGEF6;refGene_tx=XM_019831712.2
A1 110806088 110806159
refGene_gn=RAPGEF6;refGene_tx=XM_003980740.5;refGene_gn=RAPGEF6;refGene_tx=XM_011281964.3;refGene_gn=RAPGEF6;refGene_tx=XM_011281969.3;refGene_gn=RAPGEF6;refGene_tx=XM_011281970.3;refGene_gn=RAPGEF6;refGene_tx=XM_019831704.2;refGene_gn=RAPGEF6;refGene_tx=XM_019831705.2;refGene_gn

=RAPGEF6;refGene_tx=XM_019831706.2;refGene_gn=RAPGEF6;refGene_tx=XM_019831708.2;refGene_gn=RAPGEF6;refGene_tx=XM_019831709.2;refGene_gn=RAPGEF6;refGene_tx=XM_019831710.2;refGene_gn=RAPGEF6;refGene_tx=XM_019831711.2;refGene_gn=RAPGEF6;refGene_tx=XM_019831712.2
A1 110824707 110824776
refGene_gn=RAPGEF6;refGene_tx=XM_003980740.5;refGene_gn=RAPGEF6;refGene_tx=XM_011281964.3;refGene_gn=RAPGEF6;refGene_tx=XM_011281969.3;refGene_gn=RAPGEF6;refGene_tx=XM_011281970.3;refGene_gn=RAPGEF6;refGene_tx=XM_019831704.2;refGene_gn=RAPGEF6;refGene_tx=XM_019831705.2;refGene_gn=RAPGEF6;refGene_tx=XM_019831706.2;refGene_gn=RAPGEF6;refGene_tx=XM_019831708.2;refGene_gn=RAPGEF6;refGene_tx=XM_019831709.2;refGene_gn=RAPGEF6;refGene_tx=XM_019831710.2;refGene_gn=RAPGEF6;refGene_tx=XM_019831711.2;refGene_gn=RAPGEF6;refGene_tx=XM_019831712.2
A1 110833491 110833570
refGene_gn=FNIP1;refGene_tx=XM_003980741.5;refGene_gn=FNIP1;refGene_tx=XM_003980742.5
A1 110835951 110836067
refGene_gn=FNIP1;refGene_tx=XM_003980741.5;refGene_gn=FNIP1;refGene_tx=XM_003980742.5
A1 110840527 110840725
refGene_gn=FNIP1;refGene_tx=XM_003980741.5;refGene_gn=FNIP1;refGene_tx=XM_003980742.5
A1 110852885 110853054
refGene_gn=FNIP1;refGene_tx=XM_003980741.5;refGene_gn=FNIP1;refGene_tx=XM_003980742.5
A1 110853844 110855273
refGene_gn=FNIP1;refGene_tx=XM_003980741.5;refGene_gn=FNIP1;refGene_tx=XM_003980742.5
A1 110858392 110858562
refGene_gn=FNIP1;refGene_tx=XM_003980741.5;refGene_gn=FNIP1;refGene_tx=XM_003980742.5
A1 110859677 110859824
refGene_gn=FNIP1;refGene_tx=XM_003980741.5;refGene_gn=FNIP1;refGene_tx=XM_003980742.5
A1 110867668 110867754
refGene_gn=FNIP1;refGene_tx=XM_003980741.5;refGene_gn=FNIP1;refGene_tx=XM_003980742.5
A1 110871463 110871665
refGene_gn=FNIP1;refGene_tx=XM_003980741.5;refGene_gn=FNIP1;refGene_tx=XM_003980742.5
A1 110873523 110873659
refGene_gn=FNIP1;refGene_tx=XM_003980741.5;refGene_gn=FNIP1;refGene_tx=XM_003980742.5
A1 110879036 110879108
refGene_gn=FNIP1;refGene_tx=XM_003980741.5;refGene_gn=FNIP1;refGene_tx=XM_003980742.5
A1 110880264 110880348
refGene_gn=FNIP1;refGene_tx=XM_003980742.5
A1 110885649 110885741
refGene_gn=FNIP1;refGene_tx=XM_003980741.5;refGene_gn=FNIP1;refGene_tx

=XM_003980742.5
A1	110891871	110891946
refGene_gn=FNIP1;refGene_tx=XM_003980741.5;refGene_gn=FNIP1;refGene_tx
=XM_003980742.5
A1	110892198	110892299
refGene_gn=FNIP1;refGene_tx=XM_003980741.5;refGene_gn=FNIP1;refGene_tx
=XM_003980742.5
A1	110896680	110896815
refGene_gn=FNIP1;refGene_tx=XM_003980741.5;refGene_gn=FNIP1;refGene_tx
=XM_003980742.5
A1	110917111	110917238
refGene_gn=FNIP1;refGene_tx=XM_003980741.5;refGene_gn=FNIP1;refGene_tx
=XM_003980742.5
A1	110977482	110977574
refGene_gn=FNIP1;refGene_tx=XM_003980741.5;refGene_gn=FNIP1;refGene_tx
=XM_003980742.5
A1	110989002	110989034
refGene_gn=MEIKIN;refGene_tx=XM_019831716.2;refGene_gn=MEIKIN;refGene_
tx=XM_023254914.1;refGene_gn=MEIKIN;refGene_tx=XM_023254917.1;refGene_
gn=MEIKIN;refGene_tx=XM_023254922.1
A1	110998345	110998469
refGene_gn=MEIKIN;refGene_tx=XM_019831716.2;refGene_gn=MEIKIN;refGene_
tx=XM_023254914.1;refGene_gn=MEIKIN;refGene_tx=XM_023254917.1;refGene_
gn=MEIKIN;refGene_tx=XM_023254922.1
A1	111008166	111008286
refGene_gn=MEIKIN;refGene_tx=XM_019831716.2;refGene_gn=MEIKIN;refGene_
tx=XM_023254914.1;refGene_gn=MEIKIN;refGene_tx=XM_023254922.1
A1	111016206	111016287
refGene_gn=MEIKIN;refGene_tx=XM_019831716.2;refGene_gn=MEIKIN;refGene_
tx=XM_023254914.1;refGene_gn=MEIKIN;refGene_tx=XM_023254917.1;refGene_
gn=MEIKIN;refGene_tx=XM_023254922.1
A1	111018352	111018517
refGene_gn=MEIKIN;refGene_tx=XM_019831716.2;refGene_gn=MEIKIN;refGene_
tx=XM_023254914.1;refGene_gn=MEIKIN;refGene_tx=XM_023254917.1;refGene_
gn=MEIKIN;refGene_tx=XM_023254922.1
A1	111039232	111039303
refGene_gn=MEIKIN;refGene_tx=XM_019831716.2;refGene_gn=MEIKIN;refGene_
tx=XM_023254914.1;refGene_gn=MEIKIN;refGene_tx=XM_023254917.1;refGene_
gn=MEIKIN;refGene_tx=XM_023254922.1
A1	111045202	111045267
refGene_gn=MEIKIN;refGene_tx=XM_019831716.2;refGene_gn=MEIKIN;refGene_
tx=XM_023254914.1;refGene_gn=MEIKIN;refGene_tx=XM_023254917.1;refGene_
gn=MEIKIN;refGene_tx=XM_023254922.1
A1	111046977	111047017
refGene_gn=MEIKIN;refGene_tx=XM_019831716.2;refGene_gn=MEIKIN;refGene_
tx=XM_023254914.1;refGene_gn=MEIKIN;refGene_tx=XM_023254917.1;refGene_
gn=MEIKIN;refGene_tx=XM_023254922.1
A1	111056796	111056916
refGene_gn=MEIKIN;refGene_tx=XM_019831716.2;refGene_gn=MEIKIN;refGene_
tx=XM_023254914.1;refGene_gn=MEIKIN;refGene_tx=XM_023254917.1;refGene_

gn=MEIKIN;refGene_tx=XM_023254922.1
A1 111063613 111063751
refGene_gn=MEIKIN;refGene_tx=XM_019831716.2;refGene_gn=MEIKIN;refGene_tx=XM_023254914.1;refGene_gn=MEIKIN;refGene_tx=XM_023254917.1;refGene_gn=MEIKIN;refGene_tx=XM_023254922.1
A1 111067599 111067660
refGene_gn=MEIKIN;refGene_tx=XM_019831716.2;refGene_gn=MEIKIN;refGene_tx=XM_023254914.1;refGene_gn=MEIKIN;refGene_tx=XM_023254917.1;refGene_gn=MEIKIN;refGene_tx=XM_023254922.1
A1 111069292 111069380
refGene_gn=MEIKIN;refGene_tx=XM_019831716.2;refGene_gn=MEIKIN;refGene_tx=XM_023254914.1;refGene_gn=MEIKIN;refGene_tx=XM_023254917.1;refGene_gn=MEIKIN;refGene_tx=XM_023254922.1
A1 111069751 111069839
refGene_gn=MEIKIN;refGene_tx=XM_019831716.2;refGene_gn=MEIKIN;refGene_tx=XM_023254917.1
A1 111069984 111070174
refGene_gn=MEIKIN;refGene_tx=XM_019831716.2;refGene_gn=MEIKIN;refGene_tx=XM_023254917.1
A1 111070225 111070497
refGene_gn=MEIKIN;refGene_tx=XM_023254914.1
A1 111070968 111071009
refGene_gn=MEIKIN;refGene_tx=XM_023254922.1
A1 111077544 111077682
refGene_gn=ACSL6;refGene_tx=XM_011281973.3;refGene_gn=ACSL6;refGene_tx=XM_011281980.3;refGene_gn=ACSL6;refGene_tx=XM_019831731.2;refGene_gn=ACSL6;refGene_tx=XM_023254930.1;refGene_gn=ACSL6;refGene_tx=XM_023254935.1;refGene_gn=ACSL6;refGene_tx=XM_023254938.1
A1 111087452 111087524
refGene_gn=ACSL6;refGene_tx=XM_011281973.3;refGene_gn=ACSL6;refGene_tx=XM_011281980.3;refGene_gn=ACSL6;refGene_tx=XM_019831731.2;refGene_gn=ACSL6;refGene_tx=XM_023254930.1;refGene_gn=ACSL6;refGene_tx=XM_023254935.1;refGene_gn=ACSL6;refGene_tx=XM_023254938.1
A1 111088451 111088553
refGene_gn=ACSL6;refGene_tx=XM_011281973.3;refGene_gn=ACSL6;refGene_tx=XM_011281980.3;refGene_gn=ACSL6;refGene_tx=XM_019831731.2;refGene_gn=ACSL6;refGene_tx=XM_023254930.1;refGene_gn=ACSL6;refGene_tx=XM_023254935.1;refGene_gn=ACSL6;refGene_tx=XM_023254938.1
A1 111090530 111090674
refGene_gn=ACSL6;refGene_tx=XM_011281973.3;refGene_gn=ACSL6;refGene_tx=XM_011281980.3;refGene_gn=ACSL6;refGene_tx=XM_019831731.2;refGene_gn=ACSL6;refGene_tx=XM_023254930.1;refGene_gn=ACSL6;refGene_tx=XM_023254935.1;refGene_gn=ACSL6;refGene_tx=XM_023254938.1
A1 111093648 111093765
refGene_gn=ACSL6;refGene_tx=XM_011281973.3;refGene_gn=ACSL6;refGene_tx=XM_011281980.3;refGene_gn=ACSL6;refGene_tx=XM_019831731.2;refGene_gn=ACSL6;refGene_tx=XM_023254930.1;refGene_gn=ACSL6;refGene_tx=XM_023254935.1;refGene_gn=ACSL6;refGene_tx=XM_023254938.1
A1 111094493 111094582
refGene_gn=ACSL6;refGene_tx=XM_011281973.3;refGene_gn=ACSL6;refGene_tx

=XM_011281980.3;refGene_gn=ACSL6;refGene_tx=XM_019831731.2;refGene_gn=ACSL6;refGene_tx=XM_023254930.1;refGene_gn=ACSL6;refGene_tx=XM_023254935.1;refGene_gn=ACSL6;refGene_tx=XM_023254938.1
A1 111097542 111097615
refGene_gn=ACSL6;refGene_tx=XM_011281973.3;refGene_gn=ACSL6;refGene_tx=XM_011281980.3;refGene_gn=ACSL6;refGene_tx=XM_019831731.2;refGene_gn=ACSL6;refGene_tx=XM_023254930.1;refGene_gn=ACSL6;refGene_tx=XM_023254935.1;refGene_gn=ACSL6;refGene_tx=XM_023254938.1
A1 111098941 111099037
refGene_gn=ACSL6;refGene_tx=XM_011281973.3;refGene_gn=ACSL6;refGene_tx=XM_011281980.3;refGene_gn=ACSL6;refGene_tx=XM_019831731.2;refGene_gn=ACSL6;refGene_tx=XM_023254930.1;refGene_gn=ACSL6;refGene_tx=XM_023254935.1;refGene_gn=ACSL6;refGene_tx=XM_023254938.1
A1 111100087 111100222
refGene_gn=ACSL6;refGene_tx=XM_011281973.3;refGene_gn=ACSL6;refGene_tx=XM_011281980.3;refGene_gn=ACSL6;refGene_tx=XM_019831731.2;refGene_gn=ACSL6;refGene_tx=XM_023254930.1;refGene_gn=ACSL6;refGene_tx=XM_023254935.1;refGene_gn=ACSL6;refGene_tx=XM_023254938.1
A1 111100614 111100749
refGene_gn=ACSL6;refGene_tx=XM_011281973.3;refGene_gn=ACSL6;refGene_tx=XM_011281980.3;refGene_gn=ACSL6;refGene_tx=XM_019831731.2;refGene_gn=ACSL6;refGene_tx=XM_023254930.1;refGene_gn=ACSL6;refGene_tx=XM_023254935.1;refGene_gn=ACSL6;refGene_tx=XM_023254938.1
A1 111102041 111102119
refGene_gn=ACSL6;refGene_tx=XM_023254930.1
A1 111102176 111102254
refGene_gn=ACSL6;refGene_tx=XM_011281973.3;refGene_gn=ACSL6;refGene_tx=XM_011281980.3;refGene_gn=ACSL6;refGene_tx=XM_019831731.2;refGene_gn=ACSL6;refGene_tx=XM_023254935.1;refGene_gn=ACSL6;refGene_tx=XM_023254938.1
A1 111103832 111103906
refGene_gn=ACSL6;refGene_tx=XM_011281973.3;refGene_gn=ACSL6;refGene_tx=XM_011281980.3;refGene_gn=ACSL6;refGene_tx=XM_019831731.2;refGene_gn=ACSL6;refGene_tx=XM_023254930.1;refGene_gn=ACSL6;refGene_tx=XM_023254935.1;refGene_gn=ACSL6;refGene_tx=XM_023254938.1
A1 111112549 111112601
refGene_gn=ACSL6;refGene_tx=XM_011281973.3;refGene_gn=ACSL6;refGene_tx=XM_011281980.3;refGene_gn=ACSL6;refGene_tx=XM_019831731.2;refGene_gn=ACSL6;refGene_tx=XM_023254930.1;refGene_gn=ACSL6;refGene_tx=XM_023254935.1;refGene_gn=ACSL6;refGene_tx=XM_023254938.1
A1 111114107 111114140
refGene_gn=ACSL6;refGene_tx=XM_011281973.3;refGene_gn=ACSL6;refGene_tx=XM_011281980.3;refGene_gn=ACSL6;refGene_tx=XM_019831731.2;refGene_gn=ACSL6;refGene_tx=XM_023254930.1;refGene_gn=ACSL6;refGene_tx=XM_023254935.1;refGene_gn=ACSL6;refGene_tx=XM_023254938.1
A1 111115139 111115318
refGene_gn=ACSL6;refGene_tx=XM_011281973.3;refGene_gn=ACSL6;refGene_tx=XM_011281980.3;refGene_gn=ACSL6;refGene_tx=XM_019831731.2;refGene_gn=ACSL6;refGene_tx=XM_023254930.1;refGene_gn=ACSL6;refGene_tx=XM_023254935.1;refGene_gn=ACSL6;refGene_tx=XM_023254938.1

A1 111115871 111115971
refGene_gn=ACSL6;refGene_tx=XM_011281973.3;refGene_gn=ACSL6;refGene_tx=XM_011281980.3;refGene_gn=ACSL6;refGene_tx=XM_019831731.2;refGene_gn=ACSL6;refGene_tx=XM_023254930.1;refGene_gn=ACSL6;refGene_tx=XM_023254935.1;refGene_gn=ACSL6;refGene_tx=XM_023254938.1
A1 111116481 111116583
refGene_gn=ACSL6;refGene_tx=XM_011281973.3;refGene_gn=ACSL6;refGene_tx=XM_011281980.3;refGene_gn=ACSL6;refGene_tx=XM_019831731.2;refGene_gn=ACSL6;refGene_tx=XM_023254930.1;refGene_gn=ACSL6;refGene_tx=XM_023254935.1;refGene_gn=ACSL6;refGene_tx=XM_023254938.1
A1 111116847 111116912
refGene_gn=ACSL6;refGene_tx=XM_011281973.3;refGene_gn=ACSL6;refGene_tx=XM_011281980.3;refGene_gn=ACSL6;refGene_tx=XM_019831731.2;refGene_gn=ACSL6;refGene_tx=XM_023254930.1;refGene_gn=ACSL6;refGene_tx=XM_023254935.1;refGene_gn=ACSL6;refGene_tx=XM_023254938.1
A1 111117818 111117933
refGene_gn=ACSL6;refGene_tx=XM_011281973.3;refGene_gn=ACSL6;refGene_tx=XM_011281980.3;refGene_gn=ACSL6;refGene_tx=XM_019831731.2;refGene_gn=ACSL6;refGene_tx=XM_023254930.1;refGene_gn=ACSL6;refGene_tx=XM_023254935.1;refGene_gn=ACSL6;refGene_tx=XM_023254938.1
A1 111120027 111120248
refGene_gn=ACSL6;refGene_tx=XM_011281973.3;refGene_gn=ACSL6;refGene_tx=XM_011281980.3;refGene_gn=ACSL6;refGene_tx=XM_019831731.2;refGene_gn=ACSL6;refGene_tx=XM_023254930.1;refGene_gn=ACSL6;refGene_tx=XM_023254935.1;refGene_gn=ACSL6;refGene_tx=XM_023254938.1
A1 111135176 111135195
refGene_gn=ACSL6;refGene_tx=XM_011281973.3
A1 111137309 111137358
refGene_gn=ACSL6;refGene_tx=XM_011281980.3;refGene_gn=ACSL6;refGene_tx=XM_023254930.1
A1 111176459 111176618
refGene_gn=LOC111560806;refGene_tx=XM_023255164.1
A1 111176715 111176751
refGene_gn=LOC111560806;refGene_tx=XM_023255164.1
A1 111177763 111177856
refGene_gn=LOC111560806;refGene_tx=XM_023255164.1
A1 111177985 111178021
refGene_gn=LOC111560806;refGene_tx=XM_023255164.1
A1 111178130 111178238
refGene_gn=LOC111560806;refGene_tx=XM_023255164.1
A1 111190386 111190545
refGene_gn=CSF2;refGene_tx=NM_001309037.1;refGene_gn=CSF2;refGene_tx=XM_019833687.2
A1 111190641 111190683
refGene_gn=CSF2;refGene_tx=NM_001309037.1;refGene_gn=CSF2;refGene_tx=XM_019833687.2
A1 111191340 111191466
refGene_gn=CSF2;refGene_tx=NM_001309037.1;refGene_gn=CSF2;refGene_tx=XM_019833687.2
A1 111192184 111192290

refGene_gn=CSF2;refGene_tx=NM_001309037.1;refGene_gn=CSF2;refGene_tx=XM_019833687.2
A1 111333742 111333813
refGene_gn=P4HA2;refGene_tx=XM_006927582.4;refGene_gn=P4HA2;refGene_tx=XM_011282013.3;refGene_gn=P4HA2;refGene_tx=XM_019831766.2;refGene_gn=P4HA2;refGene_tx=XM_019831794.2;refGene_gn=P4HA2;refGene_tx=XM_023254950.1;refGene_gn=P4HA2;refGene_tx=XM_023254961.1;refGene_gn=P4HA2;refGene_tx=XM_023254962.1
A1 111335516 111335613
refGene_gn=P4HA2;refGene_tx=XM_006927582.4;refGene_gn=P4HA2;refGene_tx=XM_011282013.3;refGene_gn=P4HA2;refGene_tx=XM_019831766.2;refGene_gn=P4HA2;refGene_tx=XM_019831794.2;refGene_gn=P4HA2;refGene_tx=XM_023254950.1;refGene_gn=P4HA2;refGene_tx=XM_023254961.1;refGene_gn=P4HA2;refGene_tx=XM_023254962.1
A1 111336009 111336078
refGene_gn=P4HA2;refGene_tx=XM_006927582.4;refGene_gn=P4HA2;refGene_tx=XM_011282013.3;refGene_gn=P4HA2;refGene_tx=XM_019831766.2;refGene_gn=P4HA2;refGene_tx=XM_019831794.2;refGene_gn=P4HA2;refGene_tx=XM_023254950.1;refGene_gn=P4HA2;refGene_tx=XM_023254961.1;refGene_gn=P4HA2;refGene_tx=XM_023254962.1;refGene_gn=P4HA2;refGene_tx=XM_023254967.1
A1 111336981 111337073
refGene_gn=P4HA2;refGene_tx=XM_023254967.1
A1 111337842 111337908
refGene_gn=P4HA2;refGene_tx=XM_006927582.4;refGene_gn=P4HA2;refGene_tx=XM_011282013.3;refGene_gn=P4HA2;refGene_tx=XM_019831766.2;refGene_gn=P4HA2;refGene_tx=XM_019831794.2;refGene_gn=P4HA2;refGene_tx=XM_023254950.1;refGene_gn=P4HA2;refGene_tx=XM_023254961.1
A1 111337957 111338017
refGene_gn=P4HA2;refGene_tx=XM_019831794.2;refGene_gn=P4HA2;refGene_tx=XM_023254962.1;refGene_gn=P4HA2;refGene_tx=XM_023254967.1
A1 111338276 111338360
refGene_gn=P4HA2;refGene_tx=XM_023254970.1
A1 111338470 111338524
refGene_gn=P4HA2;refGene_tx=XM_006927582.4;refGene_gn=P4HA2;refGene_tx=XM_011282013.3;refGene_gn=P4HA2;refGene_tx=XM_019831766.2;refGene_gn=P4HA2;refGene_tx=XM_019831794.2;refGene_gn=P4HA2;refGene_tx=XM_023254950.1;refGene_gn=P4HA2;refGene_tx=XM_023254961.1;refGene_gn=P4HA2;refGene_tx=XM_023254962.1;refGene_gn=P4HA2;refGene_tx=XM_023254967.1;refGene_gn=P4HA2;refGene_tx=XM_023254970.1
A1 111343098 111343198
refGene_gn=P4HA2;refGene_tx=XM_006927582.4;refGene_gn=P4HA2;refGene_tx=XM_011282013.3;refGene_gn=P4HA2;refGene_tx=XM_019831766.2;refGene_gn=P4HA2;refGene_tx=XM_019831794.2;refGene_gn=P4HA2;refGene_tx=XM_023254950.1;refGene_gn=P4HA2;refGene_tx=XM_023254961.1;refGene_gn=P4HA2;refGene_tx=XM_023254962.1;refGene_gn=P4HA2;refGene_tx=XM_023254967.1;refGene_gn=P4HA2;refGene_tx=XM_023254970.1
A1 111343429 111343500
refGene_gn=P4HA2;refGene_tx=XM_006927582.4;refGene_gn=P4HA2;refGene_tx=XM_011282013.3;refGene_gn=P4HA2;refGene_tx=XM_019831766.2;refGene_gn=P4HA2;refGene_tx=XM_019831794.2;refGene_gn=P4HA2;refGene_tx=XM_0232549

50.1;refGene_gn=P4HA2;refGene_tx=XM_023254961.1;refGene_gn=P4HA2;refGene_tx=XM_023254962.1;refGene_gn=P4HA2;refGene_tx=XM_023254967.1;refGene_gn=P4HA2;refGene_tx=XM_023254970.1
A1 111346836 111347013
refGene_gn=P4HA2;refGene_tx=XM_006927582.4;refGene_gn=P4HA2;refGene_tx=XM_011282013.3;refGene_gn=P4HA2;refGene_tx=XM_019831766.2;refGene_gn=P4HA2;refGene_tx=XM_019831794.2;refGene_gn=P4HA2;refGene_tx=XM_023254950.1;refGene_gn=P4HA2;refGene_tx=XM_023254961.1;refGene_gn=P4HA2;refGene_tx=XM_023254962.1;refGene_gn=P4HA2;refGene_tx=XM_023254967.1;refGene_gn=P4HA2;refGene_tx=XM_023254970.1
A1 111348065 111348259
refGene_gn=P4HA2;refGene_tx=XM_006927582.4;refGene_gn=P4HA2;refGene_tx=XM_011282013.3;refGene_gn=P4HA2;refGene_tx=XM_019831766.2;refGene_gn=P4HA2;refGene_tx=XM_019831794.2;refGene_gn=P4HA2;refGene_tx=XM_023254950.1;refGene_gn=P4HA2;refGene_tx=XM_023254961.1;refGene_gn=P4HA2;refGene_tx=XM_023254962.1;refGene_gn=P4HA2;refGene_tx=XM_023254967.1;refGene_gn=P4HA2;refGene_tx=XM_023254970.1
A1 111349749 111349989
refGene_gn=P4HA2;refGene_tx=XM_006927582.4;refGene_gn=P4HA2;refGene_tx=XM_011282013.3;refGene_gn=P4HA2;refGene_tx=XM_019831766.2;refGene_gn=P4HA2;refGene_tx=XM_019831794.2;refGene_gn=P4HA2;refGene_tx=XM_023254950.1;refGene_gn=P4HA2;refGene_tx=XM_023254961.1;refGene_gn=P4HA2;refGene_tx=XM_023254962.1;refGene_gn=P4HA2;refGene_tx=XM_023254967.1;refGene_gn=P4HA2;refGene_tx=XM_023254970.1
A1 111353589 111353727
refGene_gn=P4HA2;refGene_tx=XM_006927582.4;refGene_gn=P4HA2;refGene_tx=XM_011282013.3;refGene_gn=P4HA2;refGene_tx=XM_019831766.2;refGene_gn=P4HA2;refGene_tx=XM_019831794.2;refGene_gn=P4HA2;refGene_tx=XM_023254950.1;refGene_gn=P4HA2;refGene_tx=XM_023254961.1;refGene_gn=P4HA2;refGene_tx=XM_023254962.1;refGene_gn=P4HA2;refGene_tx=XM_023254967.1;refGene_gn=P4HA2;refGene_tx=XM_023254970.1
A1 111357523 111357675
refGene_gn=P4HA2;refGene_tx=XM_006927582.4;refGene_gn=P4HA2;refGene_tx=XM_011282013.3;refGene_gn=P4HA2;refGene_tx=XM_019831766.2;refGene_gn=P4HA2;refGene_tx=XM_019831794.2;refGene_gn=P4HA2;refGene_tx=XM_023254950.1;refGene_gn=P4HA2;refGene_tx=XM_023254961.1;refGene_gn=P4HA2;refGene_tx=XM_023254962.1;refGene_gn=P4HA2;refGene_tx=XM_023254967.1;refGene_gn=P4HA2;refGene_tx=XM_023254970.1
A1 111358036 111358133
refGene_gn=P4HA2;refGene_tx=XM_006927582.4;refGene_gn=P4HA2;refGene_tx=XM_011282013.3;refGene_gn=P4HA2;refGene_tx=XM_019831766.2;refGene_gn=P4HA2;refGene_tx=XM_019831794.2;refGene_gn=P4HA2;refGene_tx=XM_023254950.1;refGene_gn=P4HA2;refGene_tx=XM_023254961.1;refGene_gn=P4HA2;refGene_tx=XM_023254962.1;refGene_gn=P4HA2;refGene_tx=XM_023254967.1;refGene_gn=P4HA2;refGene_tx=XM_023254970.1
A1 111358764 111358846
refGene_gn=P4HA2;refGene_tx=XM_006927582.4;refGene_gn=P4HA2;refGene_tx=XM_011282013.3;refGene_gn=P4HA2;refGene_tx=XM_019831766.2;refGene_gn=P4HA2;refGene_tx=XM_019831794.2;refGene_gn=P4HA2;refGene_tx=XM_023254950.1;refGene_gn=P4HA2;refGene_tx=XM_023254961.1;refGene_gn=P4HA2;refGe

ne_tx=XM_023254962.1;refGene_gn=P4HA2;refGene_tx=XM_023254967.1;refGen
e_gn=P4HA2;refGene_tx=XM_023254970.1
A1 111391863 111391956
refGene_gn=PDLIM4;refGene_tx=XM_003980749.4;refGene_gn=PDLIM4;refGene_
tx=XM_003980750.4
A1 111397323 111397475
refGene_gn=PDLIM4;refGene_tx=XM_003980749.4;refGene_gn=PDLIM4;refGene_
tx=XM_003980750.4
A1 111400584 111400663
refGene_gn=PDLIM4;refGene_tx=XM_003980749.4;refGene_gn=PDLIM4;refGene_
tx=XM_003980750.4
A1 111405075 111405257
refGene_gn=PDLIM4;refGene_tx=XM_003980749.4;refGene_gn=PDLIM4;refGene_
tx=XM_003980750.4
A1 111405461 111405625
refGene_gn=PDLIM4;refGene_tx=XM_003980749.4;refGene_gn=PDLIM4;refGene_
tx=XM_003980750.4
A1 111405961 111406079
refGene_gn=PDLIM4;refGene_tx=XM_003980749.4
A1 111406220 111406425
refGene_gn=PDLIM4;refGene_tx=XM_003980749.4;refGene_gn=PDLIM4;refGene_
tx=XM_003980750.4
A1 111420515 111420908
refGene_gn=LOC101095395;refGene_tx=XM_006927585.3;refGene_gn=LOC101095
395;refGene_tx=XM_019831758.2
A1 111432838 111432942
refGene_gn=LOC101095395;refGene_tx=XM_006927585.3;refGene_gn=LOC101095
395;refGene_tx=XM_019831758.2
A1 111434377 111434532
refGene_gn=LOC101095395;refGene_tx=XM_006927585.3;refGene_gn=LOC101095
395;refGene_tx=XM_019831758.2
A1 111442964 111443136
refGene_gn=LOC101095395;refGene_tx=XM_006927585.3;refGene_gn=LOC101095
395;refGene_tx=XM_019831758.2
A1 111447528 111447655
refGene_gn=LOC101095395;refGene_tx=XM_006927585.3;refGene_gn=LOC101095
395;refGene_tx=XM_019831758.2
A1 111451587 111451688
refGene_gn=LOC101095395;refGene_tx=XM_006927585.3;refGene_gn=LOC101095
395;refGene_tx=XM_019831758.2
A1 111453319 111453534
refGene_gn=LOC101095395;refGene_tx=XM_006927585.3;refGene_gn=LOC101095
395;refGene_tx=XM_019831758.2
A1 111456103 111456286
refGene_gn=LOC101095395;refGene_tx=XM_006927585.3
A1 111459314 111459450
refGene_gn=LOC101095395;refGene_tx=XM_006927585.3;refGene_gn=LOC101095
395;refGene_tx=XM_019831758.2
A1 111461580 111461656
refGene_gn=LOC101095395;refGene_tx=XM_006927585.3;refGene_gn=LOC101095

395;refGene_tx=XM_019831758.2
A1 111477057 111477450
refGene_gn=LOC101084417;refGene_tx=XM_006927692.3;refGene_gn=LOC101084417;refGene_tx=XM_023254993.1;refGene_gn=LOC101084417;refGene_tx=XM_023254997.1
A1 111484742 111484846
refGene_gn=LOC101084417;refGene_tx=XM_006927692.3;refGene_gn=LOC101084417;refGene_tx=XM_023254993.1;refGene_gn=LOC101084417;refGene_tx=XM_023254997.1
A1 111491386 111491541
refGene_gn=LOC101084417;refGene_tx=XM_006927692.3;refGene_gn=LOC101084417;refGene_tx=XM_023254993.1
A1 111492542 111492714
refGene_gn=LOC101084417;refGene_tx=XM_006927692.3;refGene_gn=LOC101084417;refGene_tx=XM_023254993.1;refGene_gn=LOC101084417;refGene_tx=XM_023254997.1
A1 111493914 111494041
refGene_gn=LOC101084417;refGene_tx=XM_006927692.3;refGene_gn=LOC101084417;refGene_tx=XM_023254993.1;refGene_gn=LOC101084417;refGene_tx=XM_023254997.1
A1 111495748 111495849
refGene_gn=LOC101084417;refGene_tx=XM_006927692.3;refGene_gn=LOC101084417;refGene_tx=XM_023254993.1;refGene_gn=LOC101084417;refGene_tx=XM_023254997.1
A1 111497514 111497729
refGene_gn=LOC101084417;refGene_tx=XM_006927692.3;refGene_gn=LOC101084417;refGene_tx=XM_023254993.1;refGene_gn=LOC101084417;refGene_tx=XM_023254997.1
A1 111499895 111500078
refGene_gn=LOC101084417;refGene_tx=XM_006927692.3;refGene_gn=LOC101084417;refGene_tx=XM_023254993.1;refGene_gn=LOC101084417;refGene_tx=XM_023254997.1
A1 111501107 111501243
refGene_gn=LOC101084417;refGene_tx=XM_006927692.3;refGene_gn=LOC101084417;refGene_tx=XM_023254993.1;refGene_gn=LOC101084417;refGene_tx=XM_023254997.1
A1 111501607 111501695
refGene_gn=LOC101084417;refGene_tx=XM_006927692.3;refGene_gn=LOC101084417;refGene_tx=XM_023254993.1;refGene_gn=LOC101084417;refGene_tx=XM_023254997.1
A1 111586721 111586834
refGene_gn=IRF1;refGene_tx=XM_003980752.5;refGene_gn=IRF1;refGene_tx=XM_019831835.2
A1 111587146 111587282
refGene_gn=IRF1;refGene_tx=XM_003980752.5;refGene_gn=IRF1;refGene_tx=XM_019831835.2
A1 111588163 111588213
refGene_gn=IRF1;refGene_tx=XM_003980752.5;refGene_gn=IRF1;refGene_tx=XM_019831835.2
A1 111588634 111588757

refGene_gn=IRF1;refGene_tx=XM_003980752.5;refGene_gn=IRF1;refGene_tx=XM_019831835.2
A1 111588942 111589075
refGene_gn=IRF1;refGene_tx=XM_003980752.5;refGene_gn=IRF1;refGene_tx=XM_019831835.2
A1 111589182 111589232
refGene_gn=IRF1;refGene_tx=XM_003980752.5;refGene_gn=IRF1;refGene_tx=XM_019831835.2
A1 111589345 111589522
refGene_gn=IRF1;refGene_tx=XM_003980752.5;refGene_gn=IRF1;refGene_tx=XM_019831835.2
A1 111590335 111590435
refGene_gn=IRF1;refGene_tx=XM_003980752.5;refGene_gn=IRF1;refGene_tx=XM_019831835.2
A1 111591748 111591835
refGene_gn=IRF1;refGene_tx=XM_003980752.5;refGene_gn=IRF1;refGene_tx=XM_019831835.2
A1 111639610 111639709
refGene_gn=IL5;refGene_tx=NM_001009845.2
A1 111639816 111639945
refGene_gn=IL5;refGene_tx=NM_001009845.2
A1 111640817 111640850
refGene_gn=IL5;refGene_tx=NM_001009845.2
A1 111641052 111641196
refGene_gn=IL5;refGene_tx=NM_001009845.2
A1 111650598 111650727
refGene_gn=RAD50;refGene_tx=XM_003980753.5
A1 111652350 111652434
refGene_gn=RAD50;refGene_tx=XM_003980753.5
A1 111662869 111663021
refGene_gn=RAD50;refGene_tx=XM_003980753.5
A1 111665966 111666152
refGene_gn=RAD50;refGene_tx=XM_003980753.5
A1 111666505 111666710
refGene_gn=RAD50;refGene_tx=XM_003980753.5
A1 111683714 111683843
refGene_gn=RAD50;refGene_tx=XM_003980753.5
A1 111684295 111684461
refGene_gn=RAD50;refGene_tx=XM_003980753.5
A1 111686816 111687010
refGene_gn=RAD50;refGene_tx=XM_003980753.5
A1 111688511 111688539
refGene_gn=RAD50;refGene_tx=XM_003980753.5
A1 111688541 111688717
refGene_gn=RAD50;refGene_tx=XM_003980753.5
A1 111689626 111689809
refGene_gn=RAD50;refGene_tx=XM_003980753.5
A1 111691352 111691510
refGene_gn=RAD50;refGene_tx=XM_003980753.5
A1 111694601 111694777

refGene_gn=RAD50;refGene_tx=XM_003980753.5
A1 111695522 111695760
refGene_gn=RAD50;refGene_tx=XM_003980753.5
A1 111702423 111702613
refGene_gn=RAD50;refGene_tx=XM_003980753.5
A1 111703024 111703151
refGene_gn=RAD50;refGene_tx=XM_003980753.5
A1 111703535 111703729
refGene_gn=RAD50;refGene_tx=XM_003980753.5
A1 111705550 111705661
refGene_gn=RAD50;refGene_tx=XM_003980753.5
A1 111707186 111707279
refGene_gn=RAD50;refGene_tx=XM_003980753.5
A1 111707360 111707474
refGene_gn=RAD50;refGene_tx=XM_003980753.5
A1 111712937 111713065
refGene_gn=RAD50;refGene_tx=XM_003980753.5
A1 111714378 111714603
refGene_gn=RAD50;refGene_tx=XM_003980753.5
A1 111734886 111734972
refGene_gn=RAD50;refGene_tx=XM_003980753.5
A1 111735478 111735621
refGene_gn=RAD50;refGene_tx=XM_003980753.5
A1 111737866 111738000
refGene_gn=RAD50;refGene_tx=XM_003980753.5
A1 111739303 111739490
refGene_gn=RAD50;refGene_tx=XM_003980753.5
A1 111762083 111762177
refGene_gn=IL13;refGene_tx=XM_019831874.2
A1 111763940 111764225
refGene_gn=IL13;refGene_tx=XM_003980880.4;refGene_gn=IL13;refGene_tx=XM_006927586.3;refGene_gn=IL13;refGene_tx=XM_006927587.3;refGene_gn=IL13;refGene_tx=XM_019831874.2
A1 111765400 111765454
refGene_gn=IL13;refGene_tx=XM_003980880.4;refGene_gn=IL13;refGene_tx=XM_006927586.3;refGene_gn=IL13;refGene_tx=XM_006927587.3;refGene_gn=IL13;refGene_tx=XM_019831874.2
A1 111765694 111765802
refGene_gn=IL13;refGene_tx=XM_003980880.4;refGene_gn=IL13;refGene_tx=XM_006927586.3;refGene_gn=IL13;refGene_tx=XM_006927587.3;refGene_gn=IL13;refGene_tx=XM_019831874.2
A1 111766165 111766273
refGene_gn=IL13;refGene_tx=XM_003980880.4;refGene_gn=IL13;refGene_tx=XM_006927586.3;refGene_gn=IL13;refGene_tx=XM_006927587.3;refGene_gn=IL13;refGene_tx=XM_019831874.2
A1 111778745 111778880
refGene_gn=IL4;refGene_tx=NM_001043339.1
A1 111779150 111779198
refGene_gn=IL4;refGene_tx=NM_001043339.1
A1 111785000 111785126

refGene_gn=IL4;refGene_tx=NM_001043339.1
A1 111787356 111787449
refGene_gn=IL4;refGene_tx=NM_001043339.1
A1 111798027 111798076
refGene_gn=KIF3A;refGene_tx=XM_003980755.5;refGene_gn=KIF3A;refGene_tx=XM_003980756.5;refGene_gn=KIF3A;refGene_tx=XM_011282050.3;refGene_gn=KIF3A;refGene_tx=XM_019831883.2;refGene_gn=KIF3A;refGene_tx=XM_019831897.2
A1 111799745 111799870
refGene_gn=KIF3A;refGene_tx=XM_003980755.5;refGene_gn=KIF3A;refGene_tx=XM_003980756.5;refGene_gn=KIF3A;refGene_tx=XM_011282050.3;refGene_gn=KIF3A;refGene_tx=XM_019831883.2;refGene_gn=KIF3A;refGene_tx=XM_019831897.2
A1 111800441 111800510
refGene_gn=KIF3A;refGene_tx=XM_003980755.5;refGene_gn=KIF3A;refGene_tx=XM_003980756.5;refGene_gn=KIF3A;refGene_tx=XM_011282050.3;refGene_gn=KIF3A;refGene_tx=XM_019831883.2;refGene_gn=KIF3A;refGene_tx=XM_019831897.2
A1 111800866 111800920
refGene_gn=KIF3A;refGene_tx=XM_003980755.5;refGene_gn=KIF3A;refGene_tx=XM_003980756.5;refGene_gn=KIF3A;refGene_tx=XM_011282050.3;refGene_gn=KIF3A;refGene_tx=XM_019831883.2;refGene_gn=KIF3A;refGene_tx=XM_019831897.2
A1 111802093 111802219
refGene_gn=KIF3A;refGene_tx=XM_003980755.5;refGene_gn=KIF3A;refGene_tx=XM_003980756.5;refGene_gn=KIF3A;refGene_tx=XM_011282050.3;refGene_gn=KIF3A;refGene_tx=XM_019831883.2;refGene_gn=KIF3A;refGene_tx=XM_019831897.2
A1 111802818 111802929
refGene_gn=KIF3A;refGene_tx=XM_003980755.5;refGene_gn=KIF3A;refGene_tx=XM_003980756.5;refGene_gn=KIF3A;refGene_tx=XM_011282050.3;refGene_gn=KIF3A;refGene_tx=XM_019831883.2;refGene_gn=KIF3A;refGene_tx=XM_019831897.2
A1 111803532 111803713
refGene_gn=KIF3A;refGene_tx=XM_003980755.5;refGene_gn=KIF3A;refGene_tx=XM_003980756.5;refGene_gn=KIF3A;refGene_tx=XM_011282050.3;refGene_gn=KIF3A;refGene_tx=XM_019831883.2;refGene_gn=KIF3A;refGene_tx=XM_019831897.2
A1 111804107 111804264
refGene_gn=KIF3A;refGene_tx=XM_003980755.5;refGene_gn=KIF3A;refGene_tx=XM_003980756.5;refGene_gn=KIF3A;refGene_tx=XM_011282050.3;refGene_gn=KIF3A;refGene_tx=XM_019831883.2;refGene_gn=KIF3A;refGene_tx=XM_019831897.2
A1 111807553 111807562
refGene_gn=KIF3A;refGene_tx=XM_003980756.5;refGene_gn=KIF3A;refGene_tx=XM_011282050.3;refGene_gn=KIF3A;refGene_tx=XM_019831897.2
A1 111809663 111809735
refGene_gn=KIF3A;refGene_tx=XM_011282050.3;refGene_gn=KIF3A;refGene_tx=XM_019831883.2;refGene_gn=KIF3A;refGene_tx=XM_019831897.2
A1 111811447 111811546

refGene_gn=KIF3A;refGene_tx=XM_003980755.5;refGene_gn=KIF3A;refGene_tx=XM_003980756.5;refGene_gn=KIF3A;refGene_tx=XM_011282050.3;refGene_gn=KIF3A;refGene_tx=XM_019831883.2;refGene_gn=KIF3A;refGene_tx=XM_019831897.2
A1 111815263 111815438
refGene_gn=KIF3A;refGene_tx=XM_003980755.5;refGene_gn=KIF3A;refGene_tx=XM_003980756.5;refGene_gn=KIF3A;refGene_tx=XM_011282050.3;refGene_gn=KIF3A;refGene_tx=XM_019831883.2;refGene_gn=KIF3A;refGene_tx=XM_019831897.2
A1 111815744 111815942
refGene_gn=KIF3A;refGene_tx=XM_003980755.5;refGene_gn=KIF3A;refGene_tx=XM_003980756.5;refGene_gn=KIF3A;refGene_tx=XM_011282050.3;refGene_gn=KIF3A;refGene_tx=XM_019831883.2;refGene_gn=KIF3A;refGene_tx=XM_019831897.2
A1 111816579 111816719
refGene_gn=KIF3A;refGene_tx=XM_003980755.5;refGene_gn=KIF3A;refGene_tx=XM_003980756.5;refGene_gn=KIF3A;refGene_tx=XM_011282050.3;refGene_gn=KIF3A;refGene_tx=XM_019831883.2;refGene_gn=KIF3A;refGene_tx=XM_019831897.2
A1 111821646 111821752
refGene_gn=KIF3A;refGene_tx=XM_003980755.5;refGene_gn=KIF3A;refGene_tx=XM_003980756.5;refGene_gn=KIF3A;refGene_tx=XM_011282050.3;refGene_gn=KIF3A;refGene_tx=XM_019831883.2;refGene_gn=KIF3A;refGene_tx=XM_019831897.2
A1 111830332 111830417
refGene_gn=KIF3A;refGene_tx=XM_003980755.5;refGene_gn=KIF3A;refGene_tx=XM_003980756.5;refGene_gn=KIF3A;refGene_tx=XM_011282050.3;refGene_gn=KIF3A;refGene_tx=XM_019831883.2;refGene_gn=KIF3A;refGene_tx=XM_019831897.2
A1 111830556 111830701
refGene_gn=KIF3A;refGene_tx=XM_003980755.5;refGene_gn=KIF3A;refGene_tx=XM_003980756.5;refGene_gn=KIF3A;refGene_tx=XM_011282050.3;refGene_gn=KIF3A;refGene_tx=XM_019831883.2;refGene_gn=KIF3A;refGene_tx=XM_019831897.2
A1 111840043 111840317
refGene_gn=KIF3A;refGene_tx=XM_003980755.5;refGene_gn=KIF3A;refGene_tx=XM_003980756.5;refGene_gn=KIF3A;refGene_tx=XM_011282050.3;refGene_gn=KIF3A;refGene_tx=XM_019831883.2;refGene_gn=KIF3A;refGene_tx=XM_019831897.2
A1 111843364 111843370
refGene_gn=KIF3A;refGene_tx=XM_003980755.5;refGene_gn=KIF3A;refGene_tx=XM_003980756.5;refGene_gn=KIF3A;refGene_tx=XM_011282050.3;refGene_gn=KIF3A;refGene_tx=XM_019831883.2
A1 111853430 111853596
refGene_gn=SEPT8;refGene_tx=XM_011282055.3;refGene_gn=SEPT8;refGene_tx=XM_011282057.3;refGene_gn=SEPT8;refGene_tx=XM_011282062.3;refGene_gn=SEPT8;refGene_tx=XM_023255052.1
A1 111860431 111860474
refGene_gn=SEPT8;refGene_tx=XM_019831914.2;refGene_gn=SEPT8;refGene_tx=XM_019831916.2

A1	111860738	111860742
refGene_gn=SEPT8;refGene_tx=XM_011282065.3
A1	111863229	111863420
refGene_gn=SEPT8;refGene_tx=XM_011282055.3;refGene_gn=SEPT8;refGene_tx=XM_011282057.3;refGene_gn=SEPT8;refGene_tx=XM_011282062.3;refGene_gn=SEPT8;refGene_tx=XM_011282065.3;refGene_gn=SEPT8;refGene_tx=XM_019831914.2;refGene_gn=SEPT8;refGene_tx=XM_019831916.2;refGene_gn=SEPT8;refGene_tx=XM_023255052.1
A1	111863514	111863647
refGene_gn=SEPT8;refGene_tx=XM_011282055.3;refGene_gn=SEPT8;refGene_tx=XM_011282057.3;refGene_gn=SEPT8;refGene_tx=XM_011282062.3;refGene_gn=SEPT8;refGene_tx=XM_011282065.3;refGene_gn=SEPT8;refGene_tx=XM_019831914.2;refGene_gn=SEPT8;refGene_tx=XM_019831916.2;refGene_gn=SEPT8;refGene_tx=XM_023255052.1
A1	111863884	111864053
refGene_gn=SEPT8;refGene_tx=XM_011282055.3;refGene_gn=SEPT8;refGene_tx=XM_011282057.3;refGene_gn=SEPT8;refGene_tx=XM_011282062.3;refGene_gn=SEPT8;refGene_tx=XM_011282065.3;refGene_gn=SEPT8;refGene_tx=XM_019831914.2;refGene_gn=SEPT8;refGene_tx=XM_019831916.2;refGene_gn=SEPT8;refGene_tx=XM_023255052.1
A1	111864258	111864355
refGene_gn=SEPT8;refGene_tx=XM_011282055.3;refGene_gn=SEPT8;refGene_tx=XM_011282057.3;refGene_gn=SEPT8;refGene_tx=XM_011282062.3;refGene_gn=SEPT8;refGene_tx=XM_011282065.3;refGene_gn=SEPT8;refGene_tx=XM_019831914.2;refGene_gn=SEPT8;refGene_tx=XM_019831916.2;refGene_gn=SEPT8;refGene_tx=XM_023255052.1
A1	111864939	111865101
refGene_gn=SEPT8;refGene_tx=XM_011282055.3;refGene_gn=SEPT8;refGene_tx=XM_011282057.3;refGene_gn=SEPT8;refGene_tx=XM_011282062.3;refGene_gn=SEPT8;refGene_tx=XM_011282065.3;refGene_gn=SEPT8;refGene_tx=XM_019831914.2;refGene_gn=SEPT8;refGene_tx=XM_019831916.2;refGene_gn=SEPT8;refGene_tx=XM_023255052.1
A1	111865618	111865805
refGene_gn=SEPT8;refGene_tx=XM_011282055.3;refGene_gn=SEPT8;refGene_tx=XM_011282057.3;refGene_gn=SEPT8;refGene_tx=XM_011282062.3;refGene_gn=SEPT8;refGene_tx=XM_011282065.3;refGene_gn=SEPT8;refGene_tx=XM_019831914.2;refGene_gn=SEPT8;refGene_tx=XM_019831916.2;refGene_gn=SEPT8;refGene_tx=XM_023255052.1
A1	111866140	111866336
refGene_gn=SEPT8;refGene_tx=XM_011282055.3;refGene_gn=SEPT8;refGene_tx=XM_011282057.3;refGene_gn=SEPT8;refGene_tx=XM_011282062.3;refGene_gn=SEPT8;refGene_tx=XM_011282065.3;refGene_gn=SEPT8;refGene_tx=XM_019831914.2;refGene_gn=SEPT8;refGene_tx=XM_019831916.2;refGene_gn=SEPT8;refGene_tx=XM_023255052.1
A1	111867280	111867401
refGene_gn=SEPT8;refGene_tx=XM_011282055.3;refGene_gn=SEPT8;refGene_tx=XM_011282057.3;refGene_gn=SEPT8;refGene_tx=XM_011282062.3;refGene_gn=SEPT8;refGene_tx=XM_011282065.3;refGene_gn=SEPT8;refGene_tx=XM_019831914.2;refGene_gn=SEPT8;refGene_tx=XM_019831916.2
A1	111877371	111877473

refGene_gn=SEPT8;refGene_tx=XM_011282055.3;refGene_gn=SEPT8;refGene_tx=XM_011282057.3;refGene_gn=SEPT8;refGene_tx=XM_011282062.3;refGene_gn=SEPT8;refGene_tx=XM_011282065.3;refGene_gn=SEPT8;refGene_tx=XM_019831914.2;refGene_gn=SEPT8;refGene_tx=XM_019831916.2
A1 111910123 111911833
refGene_gn=SOWAHA;refGene_tx=XM_023255054.1
A1 111918860 111919193
refGene_gn=SHROOM1;refGene_tx=XM_023255057.1;refGene_gn=SHROOM1;refGene_tx=XM_023255061.1;refGene_gn=SHROOM1;refGene_tx=XM_023255064.1;refGene_gn=SHROOM1;refGene_tx=XM_023255065.1;refGene_gn=SHROOM1;refGene_tx=XM_023255068.1;refGene_gn=SHROOM1;refGene_tx=XM_023255070.1;refGene_gn=SHROOM1;refGene_tx=XM_023255076.1
A1 111919312 111919585
refGene_gn=SHROOM1;refGene_tx=XM_023255057.1;refGene_gn=SHROOM1;refGene_tx=XM_023255061.1;refGene_gn=SHROOM1;refGene_tx=XM_023255064.1;refGene_gn=SHROOM1;refGene_tx=XM_023255065.1;refGene_gn=SHROOM1;refGene_tx=XM_023255068.1;refGene_gn=SHROOM1;refGene_tx=XM_023255070.1;refGene_gn=SHROOM1;refGene_tx=XM_023255076.1
A1 111919691 111919827
refGene_gn=SHROOM1;refGene_tx=XM_023255057.1;refGene_gn=SHROOM1;refGene_tx=XM_023255061.1;refGene_gn=SHROOM1;refGene_tx=XM_023255064.1;refGene_gn=SHROOM1;refGene_tx=XM_023255065.1;refGene_gn=SHROOM1;refGene_tx=XM_023255068.1;refGene_gn=SHROOM1;refGene_tx=XM_023255070.1;refGene_gn=SHROOM1;refGene_tx=XM_023255076.1
A1 111919926 111920493
refGene_gn=SHROOM1;refGene_tx=XM_023255057.1;refGene_gn=SHROOM1;refGene_tx=XM_023255061.1;refGene_gn=SHROOM1;refGene_tx=XM_023255064.1;refGene_gn=SHROOM1;refGene_tx=XM_023255065.1;refGene_gn=SHROOM1;refGene_tx=XM_023255068.1;refGene_gn=SHROOM1;refGene_tx=XM_023255070.1;refGene_gn=SHROOM1;refGene_tx=XM_023255076.1
A1 111920684 111920894
refGene_gn=SHROOM1;refGene_tx=XM_023255057.1;refGene_gn=SHROOM1;refGene_tx=XM_023255061.1;refGene_gn=SHROOM1;refGene_tx=XM_023255064.1;refGene_gn=SHROOM1;refGene_tx=XM_023255065.1;refGene_gn=SHROOM1;refGene_tx=XM_023255068.1;refGene_gn=SHROOM1;refGene_tx=XM_023255070.1;refGene_gn=SHROOM1;refGene_tx=XM_023255076.1
A1 111921123 111921179
refGene_gn=SHROOM1;refGene_tx=XM_023255057.1;refGene_gn=SHROOM1;refGene_tx=XM_023255061.1;refGene_gn=SHROOM1;refGene_tx=XM_023255064.1;refGene_gn=SHROOM1;refGene_tx=XM_023255065.1;refGene_gn=SHROOM1;refGene_tx=XM_023255068.1;refGene_gn=SHROOM1;refGene_tx=XM_023255070.1;refGene_gn=SHROOM1;refGene_tx=XM_023255076.1
A1 111921269 111922241
refGene_gn=SHROOM1;refGene_tx=XM_023255057.1;refGene_gn=SHROOM1;refGene_tx=XM_023255061.1;refGene_gn=SHROOM1;refGene_tx=XM_023255064.1;refGene_gn=SHROOM1;refGene_tx=XM_023255065.1;refGene_gn=SHROOM1;refGene_tx=XM_023255068.1;refGene_gn=SHROOM1;refGene_tx=XM_023255070.1;refGene_gn=SHROOM1;refGene_tx=XM_023255076.1
A1 111946492 111947457
refGene_gn=GDF9;refGene_tx=NM_001165900.1;refGene_gn=GDF9;refGene_tx=X

M_011281152.2;refGene_gn=GDF9;refGene_tx=XM_011281154.2;refGene_gn=GDF9;refGene_tx=XM_019825218.2;refGene_gn=GDF9;refGene_tx=XM_019826668.1;refGene_gn=GDF9;refGene_tx=XM_019827116.1
A1 111949115 111949509
refGene_gn=GDF9;refGene_tx=NM_001165900.1;refGene_gn=GDF9;refGene_tx=XM_011281152.2;refGene_gn=GDF9;refGene_tx=XM_011281154.2;refGene_gn=GDF9;refGene_tx=XM_019825218.2;refGene_gn=GDF9;refGene_tx=XM_019826668.1;refGene_gn=GDF9;refGene_tx=XM_019827116.1
A1 111951250 111951256
refGene_gn=GDF9;refGene_tx=XM_011281152.2
A1 111952073 111952227
refGene_gn=LOC101096871;refGene_tx=XM_003980758.4
A1 111952902 111952997
refGene_gn=LOC101096871;refGene_tx=XM_003980758.4
A1 111958338 111958395
refGene_gn=LEAP2;refGene_tx=XM_003980760.4;refGene_gn=LEAP2;refGene_tx=XM_019831929.2
A1 111958614 111958754
refGene_gn=LEAP2;refGene_tx=XM_003980760.4;refGene_gn=LEAP2;refGene_tx=XM_019831929.2
A1 111959045 111959082
refGene_gn=LEAP2;refGene_tx=XM_003980760.4
A1 111965615 111965743
refGene_gn=AFF4;refGene_tx=XM_011282068.3;refGene_gn=AFF4;refGene_tx=XM_011282070.3;refGene_gn=AFF4;refGene_tx=XM_023255091.1
A1 111968877 111969098
refGene_gn=AFF4;refGene_tx=XM_011282068.3;refGene_gn=AFF4;refGene_tx=XM_011282070.3;refGene_gn=AFF4;refGene_tx=XM_023255091.1;refGene_gn=LEAP2;refGene_tx=XM_019831929.2
A1 111970284 111970328
refGene_gn=AFF4;refGene_tx=XM_011282068.3;refGene_gn=AFF4;refGene_tx=XM_011282070.3;refGene_gn=AFF4;refGene_tx=XM_023255091.1
A1 111971379 111971473
refGene_gn=AFF4;refGene_tx=XM_011282068.3;refGene_gn=AFF4;refGene_tx=XM_011282070.3;refGene_gn=AFF4;refGene_tx=XM_023255091.1
A1 111972595 111972667
refGene_gn=AFF4;refGene_tx=XM_011282068.3;refGene_gn=AFF4;refGene_tx=XM_011282070.3;refGene_gn=AFF4;refGene_tx=XM_023255091.1
A1 111972927 111973064
refGene_gn=AFF4;refGene_tx=XM_011282068.3;refGene_gn=AFF4;refGene_tx=XM_011282070.3;refGene_gn=AFF4;refGene_tx=XM_023255091.1
A1 111973181 111973245
refGene_gn=AFF4;refGene_tx=XM_011282068.3;refGene_gn=AFF4;refGene_tx=XM_011282070.3;refGene_gn=AFF4;refGene_tx=XM_023255091.1
A1 111973672 111973767
refGene_gn=AFF4;refGene_tx=XM_011282068.3;refGene_gn=AFF4;refGene_tx=XM_011282070.3;refGene_gn=AFF4;refGene_tx=XM_023255091.1
A1 111978588 111978829
refGene_gn=AFF4;refGene_tx=XM_011282068.3;refGene_gn=AFF4;refGene_tx=XM_011282070.3;refGene_gn=AFF4;refGene_tx=XM_023255091.1

A1	111979435	111979524
refGene_gn=AFF4;refGene_tx=XM_011282068.3;refGene_gn=AFF4;refGene_tx=XM_011282070.3;refGene_gn=AFF4;refGene_tx=XM_023255091.1
A1	111984192	111985110
refGene_gn=AFF4;refGene_tx=XM_011282068.3;refGene_gn=AFF4;refGene_tx=XM_011282070.3;refGene_gn=AFF4;refGene_tx=XM_023255091.1
A1	111985731	111985894
refGene_gn=AFF4;refGene_tx=XM_011282068.3;refGene_gn=AFF4;refGene_tx=XM_011282070.3;refGene_gn=AFF4;refGene_tx=XM_023255091.1
A1	111986536	111986574
refGene_gn=AFF4;refGene_tx=XM_011282068.3;refGene_gn=AFF4;refGene_tx=XM_011282070.3;refGene_gn=AFF4;refGene_tx=XM_023255091.1
A1	111987016	111987071
refGene_gn=AFF4;refGene_tx=XM_011282068.3;refGene_gn=AFF4;refGene_tx=XM_011282070.3;refGene_gn=AFF4;refGene_tx=XM_023255091.1
A1	111989428	111989477
refGene_gn=AFF4;refGene_tx=XM_011282068.3;refGene_gn=AFF4;refGene_tx=XM_011282070.3;refGene_gn=AFF4;refGene_tx=XM_023255091.1
A1	111991216	111991253
refGene_gn=AFF4;refGene_tx=XM_011282068.3;refGene_gn=AFF4;refGene_tx=XM_011282070.3;refGene_gn=AFF4;refGene_tx=XM_023255091.1
A1	112015178	112015265
refGene_gn=AFF4;refGene_tx=XM_011282068.3;refGene_gn=AFF4;refGene_tx=XM_023255091.1
A1	112019195	112019240
refGene_gn=AFF4;refGene_tx=XM_011282068.3;refGene_gn=AFF4;refGene_tx=XM_011282070.3;refGene_gn=AFF4;refGene_tx=XM_023255091.1
A1	112020984	112021779
refGene_gn=AFF4;refGene_tx=XM_011282068.3;refGene_gn=AFF4;refGene_tx=XM_011282070.3;refGene_gn=AFF4;refGene_tx=XM_023255091.1
A1	112024928	112025055
refGene_gn=AFF4;refGene_tx=XM_011282068.3;refGene_gn=AFF4;refGene_tx=XM_011282070.3;refGene_gn=AFF4;refGene_tx=XM_023255091.1
A1	112034020	112034115
refGene_gn=AFF4;refGene_tx=XM_011282068.3;refGene_gn=AFF4;refGene_tx=XM_011282070.3
A1	112086677	112086945
refGene_gn=ZCCHC10;refGene_tx=XM_003980761.5
A1	112087595	112087637
refGene_gn=ZCCHC10;refGene_tx=XM_003980761.5
A1	112094077	112094242
refGene_gn=ZCCHC10;refGene_tx=XM_003980761.5
A1	112112886	112112927
refGene_gn=ZCCHC10;refGene_tx=XM_003980761.5
A1	112142315	112142422
refGene_gn=HSPA4;refGene_tx=XM_003980762.4
A1	112143907	112143909
refGene_gn=HSPA4;refGene_tx=XM_023255115.1
A1	112155392	112155450
refGene_gn=HSPA4;refGene_tx=XM_003980762.4;refGene_gn=HSPA4;refGene_tx

=XM_023255115.1
A1 112157502 112157643
refGene_gn=HSPA4;refGene_tx=XM_003980762.4;refGene_gn=HSPA4;refGene_tx
=XM_023255115.1
A1 112159509 112159632
refGene_gn=HSPA4;refGene_tx=XM_003980762.4;refGene_gn=HSPA4;refGene_tx
=XM_006927594.3;refGene_gn=HSPA4;refGene_tx=XM_023255115.1
A1 112163619 112163719
refGene_gn=HSPA4;refGene_tx=XM_003980762.4;refGene_gn=HSPA4;refGene_tx
=XM_006927594.3;refGene_gn=HSPA4;refGene_tx=XM_023255115.1
A1 112164360 112164494
refGene_gn=HSPA4;refGene_tx=XM_003980762.4;refGene_gn=HSPA4;refGene_tx
=XM_006927594.3;refGene_gn=HSPA4;refGene_tx=XM_023255115.1
A1 112166476 112166721
refGene_gn=HSPA4;refGene_tx=XM_003980762.4;refGene_gn=HSPA4;refGene_tx
=XM_006927594.3;refGene_gn=HSPA4;refGene_tx=XM_023255115.1
A1 112172133 112172210
refGene_gn=HSPA4;refGene_tx=XM_003980762.4;refGene_gn=HSPA4;refGene_tx
=XM_006927594.3;refGene_gn=HSPA4;refGene_tx=XM_023255115.1
A1 112173800 112173952
refGene_gn=HSPA4;refGene_tx=XM_003980762.4;refGene_gn=HSPA4;refGene_tx
=XM_006927594.3;refGene_gn=HSPA4;refGene_tx=XM_023255115.1
A1 112174462 112174569
refGene_gn=HSPA4;refGene_tx=XM_003980762.4;refGene_gn=HSPA4;refGene_tx
=XM_006927594.3;refGene_gn=HSPA4;refGene_tx=XM_023255115.1
A1 112175007 112175141
refGene_gn=HSPA4;refGene_tx=XM_003980762.4;refGene_gn=HSPA4;refGene_tx
=XM_006927594.3;refGene_gn=HSPA4;refGene_tx=XM_023255115.1
A1 112176139 112176321
refGene_gn=HSPA4;refGene_tx=XM_003980762.4;refGene_gn=HSPA4;refGene_tx
=XM_006927594.3;refGene_gn=HSPA4;refGene_tx=XM_023255115.1
A1 112177249 112177339
refGene_gn=HSPA4;refGene_tx=XM_003980762.4;refGene_gn=HSPA4;refGene_tx
=XM_006927594.3;refGene_gn=HSPA4;refGene_tx=XM_023255115.1
A1 112180737 112180890
refGene_gn=HSPA4;refGene_tx=XM_003980762.4;refGene_gn=HSPA4;refGene_tx
=XM_006927594.3;refGene_gn=HSPA4;refGene_tx=XM_023255115.1
A1 112182010 112182136
refGene_gn=HSPA4;refGene_tx=XM_003980762.4;refGene_gn=HSPA4;refGene_tx
=XM_006927594.3;refGene_gn=HSPA4;refGene_tx=XM_023255115.1
A1 112182851 112182959
refGene_gn=HSPA4;refGene_tx=XM_003980762.4;refGene_gn=HSPA4;refGene_tx
=XM_006927594.3;refGene_gn=HSPA4;refGene_tx=XM_023255115.1
A1 112185461 112185581
refGene_gn=HSPA4;refGene_tx=XM_003980762.4;refGene_gn=HSPA4;refGene_tx
=XM_006927594.3;refGene_gn=HSPA4;refGene_tx=XM_023255115.1
A1 112186760 112186922
refGene_gn=HSPA4;refGene_tx=XM_003980762.4;refGene_gn=HSPA4;refGene_tx
=XM_006927594.3;refGene_gn=HSPA4;refGene_tx=XM_023255115.1
A1 112187135 112187339

refGene_gn=HSPA4;refGene_tx=XM_003980762.4;refGene_gn=HSPA4;refGene_tx
=XM_006927594.3;refGene_gn=HSPA4;refGene_tx=XM_023255115.1
A1 112287411 112288114
refGene_gn=FSTL4;refGene_tx=XM_019831942.2;refGene_gn=FSTL4;refGene_tx
=XM_019831947.2;refGene_gn=FSTL4;refGene_tx=XM_023255130.1
A1 112290086 112290196
refGene_gn=FSTL4;refGene_tx=XM_019831942.2;refGene_gn=FSTL4;refGene_tx
=XM_019831947.2;refGene_gn=FSTL4;refGene_tx=XM_023255130.1
A1 112296458 112296566
refGene_gn=FSTL4;refGene_tx=XM_019831942.2;refGene_gn=FSTL4;refGene_tx
=XM_019831947.2;refGene_gn=FSTL4;refGene_tx=XM_023255130.1
A1 112303840 112303990
refGene_gn=FSTL4;refGene_tx=XM_019831942.2;refGene_gn=FSTL4;refGene_tx
=XM_019831947.2;refGene_gn=FSTL4;refGene_tx=XM_023255130.1
A1 112308951 112309070
refGene_gn=FSTL4;refGene_tx=XM_019831942.2;refGene_gn=FSTL4;refGene_tx
=XM_019831947.2;refGene_gn=FSTL4;refGene_tx=XM_023255130.1
A1 112312878 112312905
refGene_gn=FSTL4;refGene_tx=XM_019831942.2;refGene_gn=FSTL4;refGene_tx
=XM_019831947.2;refGene_gn=FSTL4;refGene_tx=XM_023255130.1
A1 112313922 112314057
refGene_gn=FSTL4;refGene_tx=XM_019831942.2;refGene_gn=FSTL4;refGene_tx
=XM_019831947.2;refGene_gn=FSTL4;refGene_tx=XM_023255130.1
A1 112314467 112314629
refGene_gn=FSTL4;refGene_tx=XM_019831942.2;refGene_gn=FSTL4;refGene_tx
=XM_019831947.2;refGene_gn=FSTL4;refGene_tx=XM_023255130.1
A1 112329579 112329700
refGene_gn=FSTL4;refGene_tx=XM_019831942.2;refGene_gn=FSTL4;refGene_tx
=XM_019831947.2;refGene_gn=FSTL4;refGene_tx=XM_023255130.1
A1 112346763 112346930
refGene_gn=FSTL4;refGene_tx=XM_019831942.2;refGene_gn=FSTL4;refGene_tx
=XM_019831947.2;refGene_gn=FSTL4;refGene_tx=XM_023255130.1
A1 112408604 112408728
refGene_gn=FSTL4;refGene_tx=XM_019831942.2;refGene_gn=FSTL4;refGene_tx
=XM_019831947.2;refGene_gn=FSTL4;refGene_tx=XM_023255130.1
A1 112412599 112412793
refGene_gn=FSTL4;refGene_tx=XM_019831942.2;refGene_gn=FSTL4;refGene_tx
=XM_019831947.2;refGene_gn=FSTL4;refGene_tx=XM_023255130.1
A1 112496793 112497042
refGene_gn=FSTL4;refGene_tx=XM_019831942.2;refGene_gn=FSTL4;refGene_tx
=XM_019831947.2;refGene_gn=FSTL4;refGene_tx=XM_023255130.1
A1 112656873 112656907
refGene_gn=FSTL4;refGene_tx=XM_019831942.2;refGene_gn=FSTL4;refGene_tx
=XM_019831947.2;refGene_gn=FSTL4;refGene_tx=XM_023255130.1
A1 112689420 112689546
refGene_gn=FSTL4;refGene_tx=XM_019831942.2;refGene_gn=FSTL4;refGene_tx
=XM_019831947.2;refGene_gn=FSTL4;refGene_tx=XM_023255130.1
A1 113042704 113042836
refGene_gn=CA1H5orf15;refGene_tx=XM_003980764.4;refGene_gn=CA1H5orf15;
refGene_tx=XM_023255148.1

A1	113043804	113044331
refGene_gn=CA1H5orf15;refGene_tx=XM_003980764.4;refGene_gn=CA1H5orf15;
refGene_tx=XM_023255148.1
A1	113047166	113047238
refGene_gn=CA1H5orf15;refGene_tx=XM_023255148.1
A1	113053860	113053999
refGene_gn=CA1H5orf15;refGene_tx=XM_003980764.4;refGene_gn=CA1H5orf15;
refGene_tx=XM_023255148.1
A1	113058960	113059052
refGene_gn=VDAC1;refGene_tx=XM_023255158.1;refGene_gn=VDAC1;refGene_tx
=XM_023255160.1
A1	113059976	113060034
refGene_gn=VDAC1;refGene_tx=XM_023255158.1;refGene_gn=VDAC1;refGene_tx
=XM_023255160.1
A1	113061907	113062058
refGene_gn=VDAC1;refGene_tx=XM_023255158.1;refGene_gn=VDAC1;refGene_tx
=XM_023255160.1
A1	113065998	113066226
refGene_gn=VDAC1;refGene_tx=XM_023255158.1;refGene_gn=VDAC1;refGene_tx
=XM_023255160.1
A1	113072768	113072821
refGene_gn=VDAC1;refGene_tx=XM_023255158.1;refGene_gn=VDAC1;refGene_tx
=XM_023255160.1
A1	113072905	113073058
refGene_gn=VDAC1;refGene_tx=XM_023255158.1;refGene_gn=VDAC1;refGene_tx
=XM_023255160.1
A1	113074466	113074516
refGene_gn=VDAC1;refGene_tx=XM_023255158.1;refGene_gn=VDAC1;refGene_tx
=XM_023255160.1
A1	113075011	113075078
refGene_gn=VDAC1;refGene_tx=XM_023255158.1;refGene_gn=VDAC1;refGene_tx
=XM_023255160.1
A1	113172706	113172958
refGene_gn=TCF7;refGene_tx=XM_023255168.1;refGene_gn=TCF7;refGene_tx=X
M_023255177.1;refGene_gn=TCF7;refGene_tx=XM_023255179.1;refGene_gn=TCF
7;refGene_tx=XM_023255196.1;refGene_gn=TCF7;refGene_tx=XM_023255200.1;
refGene_gn=TCF7;refGene_tx=XM_023255201.1;refGene_gn=TCF7;refGene_tx=X
M_023255202.1;refGene_gn=TCF7;refGene_tx=XM_023255205.1;refGene_gn=TCF
7;refGene_tx=XM_023255210.1
A1	113173123	113173190
refGene_gn=TCF7;refGene_tx=XM_023255168.1;refGene_gn=TCF7;refGene_tx=X
M_023255177.1;refGene_gn=TCF7;refGene_tx=XM_023255179.1;refGene_gn=TCF
7;refGene_tx=XM_023255196.1;refGene_gn=TCF7;refGene_tx=XM_023255200.1;
refGene_gn=TCF7;refGene_tx=XM_023255201.1;refGene_gn=TCF7;refGene_tx=X
M_023255202.1;refGene_gn=TCF7;refGene_tx=XM_023255205.1;refGene_gn=TCF
7;refGene_tx=XM_023255210.1
A1	113173693	113173818
refGene_gn=TCF7;refGene_tx=XM_023255168.1;refGene_gn=TCF7;refGene_tx=X
M_023255177.1;refGene_gn=TCF7;refGene_tx=XM_023255179.1;refGene_gn=TCF
7;refGene_tx=XM_023255183.1;refGene_gn=TCF7;refGene_tx=XM_023255193.1;

refGene_gn=TCF7;refGene_tx=XM_023255196.1;refGene_gn=TCF7;refGene_tx=XM_023255200.1;refGene_gn=TCF7;refGene_tx=XM_023255201.1;refGene_gn=TCF7;refGene_tx=XM_023255202.1;refGene_gn=TCF7;refGene_tx=XM_023255205.1;refGene_gn=TCF7;refGene_tx=XM_023255210.1
A1 113195678 113195790
refGene_gn=TCF7;refGene_tx=XM_023255168.1;refGene_gn=TCF7;refGene_tx=XM_023255177.1;refGene_gn=TCF7;refGene_tx=XM_023255179.1;refGene_gn=TCF7;refGene_tx=XM_023255183.1;refGene_gn=TCF7;refGene_tx=XM_023255193.1;refGene_gn=TCF7;refGene_tx=XM_023255196.1;refGene_gn=TCF7;refGene_tx=XM_023255200.1;refGene_gn=TCF7;refGene_tx=XM_023255201.1;refGene_gn=TCF7;refGene_tx=XM_023255202.1;refGene_gn=TCF7;refGene_tx=XM_023255205.1;refGene_gn=TCF7;refGene_tx=XM_023255210.1
A1 113196623 113196714
refGene_gn=TCF7;refGene_tx=XM_023255168.1;refGene_gn=TCF7;refGene_tx=XM_023255177.1;refGene_gn=TCF7;refGene_tx=XM_023255179.1;refGene_gn=TCF7;refGene_tx=XM_023255183.1;refGene_gn=TCF7;refGene_tx=XM_023255193.1;refGene_gn=TCF7;refGene_tx=XM_023255196.1;refGene_gn=TCF7;refGene_tx=XM_023255200.1;refGene_gn=TCF7;refGene_tx=XM_023255201.1;refGene_gn=TCF7;refGene_tx=XM_023255202.1;refGene_gn=TCF7;refGene_tx=XM_023255205.1;refGene_gn=TCF7;refGene_tx=XM_023255210.1
A1 113198142 113198235
refGene_gn=TCF7;refGene_tx=XM_023255168.1;refGene_gn=TCF7;refGene_tx=XM_023255179.1;refGene_gn=TCF7;refGene_tx=XM_023255183.1;refGene_gn=TCF7;refGene_tx=XM_023255193.1;refGene_gn=TCF7;refGene_tx=XM_023255196.1;refGene_gn=TCF7;refGene_tx=XM_023255200.1;refGene_gn=TCF7;refGene_tx=XM_023255201.1;refGene_gn=TCF7;refGene_tx=XM_023255202.1;refGene_gn=TCF7;refGene_tx=XM_023255205.1
A1 113199528 113199648
refGene_gn=TCF7;refGene_tx=XM_023255168.1;refGene_gn=TCF7;refGene_tx=XM_023255177.1;refGene_gn=TCF7;refGene_tx=XM_023255179.1;refGene_gn=TCF7;refGene_tx=XM_023255183.1;refGene_gn=TCF7;refGene_tx=XM_023255193.1;refGene_gn=TCF7;refGene_tx=XM_023255196.1;refGene_gn=TCF7;refGene_tx=XM_023255200.1;refGene_gn=TCF7;refGene_tx=XM_023255201.1;refGene_gn=TCF7;refGene_tx=XM_023255202.1;refGene_gn=TCF7;refGene_tx=XM_023255205.1;refGene_gn=TCF7;refGene_tx=XM_023255210.1
A1 113200253 113200416
refGene_gn=TCF7;refGene_tx=XM_023255168.1;refGene_gn=TCF7;refGene_tx=XM_023255177.1;refGene_gn=TCF7;refGene_tx=XM_023255179.1;refGene_gn=TCF7;refGene_tx=XM_023255183.1;refGene_gn=TCF7;refGene_tx=XM_023255193.1;refGene_gn=TCF7;refGene_tx=XM_023255196.1;refGene_gn=TCF7;refGene_tx=XM_023255200.1;refGene_gn=TCF7;refGene_tx=XM_023255201.1;refGene_gn=TCF7;refGene_tx=XM_023255202.1;refGene_gn=TCF7;refGene_tx=XM_023255205.1;refGene_gn=TCF7;refGene_tx=XM_023255210.1
A1 113200514 113200622
refGene_gn=TCF7;refGene_tx=XM_023255168.1;refGene_gn=TCF7;refGene_tx=XM_023255177.1;refGene_gn=TCF7;refGene_tx=XM_023255179.1;refGene_gn=TCF7;refGene_tx=XM_023255183.1;refGene_gn=TCF7;refGene_tx=XM_023255193.1;refGene_gn=TCF7;refGene_tx=XM_023255196.1;refGene_gn=TCF7;refGene_tx=XM_023255200.1;refGene_gn=TCF7;refGene_tx=XM_023255201.1;refGene_gn=TCF7;refGene_tx=XM_023255202.1;refGene_gn=TCF7;refGene_tx=XM_023255205.1;

refGene_gn=TCF7;refGene_tx=XM_023255210.1
A1 113201142 113201191
refGene_gn=TCF7;refGene_tx=XM_023255168.1;refGene_gn=TCF7;refGene_tx=XM_023255177.1;refGene_gn=TCF7;refGene_tx=XM_023255179.1;refGene_gn=TCF7;refGene_tx=XM_023255183.1;refGene_gn=TCF7;refGene_tx=XM_023255193.1;refGene_gn=TCF7;refGene_tx=XM_023255196.1;refGene_gn=TCF7;refGene_tx=XM_023255200.1;refGene_gn=TCF7;refGene_tx=XM_023255201.1;refGene_gn=TCF7;refGene_tx=XM_023255202.1;refGene_gn=TCF7;refGene_tx=XM_023255205.1;refGene_gn=TCF7;refGene_tx=XM_023255210.1
A1 113201986 113202059
refGene_gn=TCF7;refGene_tx=XM_023255168.1;refGene_gn=TCF7;refGene_tx=XM_023255177.1;refGene_gn=TCF7;refGene_tx=XM_023255183.1;refGene_gn=TCF7;refGene_tx=XM_023255193.1;refGene_gn=TCF7;refGene_tx=XM_023255196.1;refGene_gn=TCF7;refGene_tx=XM_023255201.1
A1 113203143 113203617
refGene_gn=TCF7;refGene_tx=XM_023255168.1;refGene_gn=TCF7;refGene_tx=XM_023255177.1;refGene_gn=TCF7;refGene_tx=XM_023255179.1;refGene_gn=TCF7;refGene_tx=XM_023255183.1;refGene_gn=TCF7;refGene_tx=XM_023255193.1;refGene_gn=TCF7;refGene_tx=XM_023255196.1;refGene_gn=TCF7;refGene_tx=XM_023255200.1;refGene_gn=TCF7;refGene_tx=XM_023255201.1;refGene_gn=TCF7;refGene_tx=XM_023255202.1;refGene_gn=TCF7;refGene_tx=XM_023255205.1;refGene_gn=TCF7;refGene_tx=XM_023255210.1
A1 113212055 113212091
refGene_gn=SKP1;refGene_tx=XM_011282104.3
A1 113212789 113212930
refGene_gn=SKP1;refGene_tx=XM_011282104.3
A1 113214385 113214529
refGene_gn=SKP1;refGene_tx=XM_011282104.3
A1 113218314 113218388
refGene_gn=SKP1;refGene_tx=XM_011282104.3
A1 113224303 113224400
refGene_gn=SKP1;refGene_tx=XM_011282104.3
A1 113241888 113241961
refGene_gn=PPP2CA;refGene_tx=XM_003980768.5
A1 113242718 113242837
refGene_gn=PPP2CA;refGene_tx=XM_003980768.5
A1 113244169 113244331
refGene_gn=PPP2CA;refGene_tx=XM_003980768.5
A1 113244846 113244936
refGene_gn=PPP2CA;refGene_tx=XM_003980768.5
A1 113245733 113245907
refGene_gn=PPP2CA;refGene_tx=XM_003980768.5
A1 113249887 113250097
refGene_gn=PPP2CA;refGene_tx=XM_003980768.5
A1 113265250 113265352
refGene_gn=PPP2CA;refGene_tx=XM_003980768.5
A1 113272934 113272966
refGene_gn=CDKL3;refGene_tx=XM_023255230.1
A1 113330247 113330267
refGene_gn=CDKL3;refGene_tx=XM_019832038.2

A1	113333756	113333919
refGene_gn=CDKL3;refGene_tx=XM_019832038.2;refGene_gn=CDKL3;refGene_tx=XM_023255230.1
A1	113336761	113337085
refGene_gn=CDKL3;refGene_tx=XM_019832038.2;refGene_gn=CDKL3;refGene_tx=XM_023255230.1
A1	113337199	113337353
refGene_gn=CDKL3;refGene_tx=XM_019832038.2;refGene_gn=CDKL3;refGene_tx=XM_023255230.1
A1	113339327	113339416
refGene_gn=CDKL3;refGene_tx=XM_019832038.2;refGene_gn=CDKL3;refGene_tx=XM_023255230.1
A1	113344743	113344883
refGene_gn=CDKL3;refGene_tx=XM_019832038.2;refGene_gn=CDKL3;refGene_tx=XM_023255230.1
A1	113346715	113346828
refGene_gn=CDKL3;refGene_tx=XM_019832038.2;refGene_gn=CDKL3;refGene_tx=XM_023255230.1
A1	113353101	113353280
refGene_gn=CDKL3;refGene_tx=XM_019832038.2;refGene_gn=CDKL3;refGene_tx=XM_023255230.1
A1	113365062	113365257
refGene_gn=CDKL3;refGene_tx=XM_019832038.2;refGene_gn=CDKL3;refGene_tx=XM_023255230.1
A1	113372652	113372835
refGene_gn=CDKL3;refGene_tx=XM_019832038.2;refGene_gn=CDKL3;refGene_tx=XM_023255230.1
A1	113373253	113373286
refGene_gn=CDKL3;refGene_tx=XM_019832038.2;refGene_gn=CDKL3;refGene_tx=XM_023255230.1
A1	113377275	113377319
refGene_gn=UBE2B;refGene_tx=XM_023255298.1
A1	113379371	113379452
refGene_gn=UBE2B;refGene_tx=XM_023255298.1
A1	113380670	113380696
refGene_gn=UBE2B;refGene_tx=XM_023255298.1
A1	113385855	113385945
refGene_gn=UBE2B;refGene_tx=XM_023255298.1
A1	113389984	113390073
refGene_gn=UBE2B;refGene_tx=XM_023255298.1
A1	113391031	113391160
refGene_gn=UBE2B;refGene_tx=XM_023255298.1
A1	113400808	113400820
refGene_gn=CDKN2AIPNL;refGene_tx=XM_003980771.5
A1	113406067	113406167
refGene_gn=CDKN2AIPNL;refGene_tx=XM_003980771.5
A1	113407435	113407674
refGene_gn=CDKN2AIPNL;refGene_tx=XM_003980771.5
A1	113498861	113498919
refGene_gn=JADE2;refGene_tx=XM_006927603.4;refGene_gn=JADE2;refGene_tx

=XM_006927604.4;refGene_gn=JADE2;refGene_tx=XM_019832025.2;refGene_gn=JADE2;refGene_tx=XM_023255268.1;refGene_gn=JADE2;refGene_tx=XM_023255270.1;refGene_gn=JADE2;refGene_tx=XM_023255273.1;refGene_gn=JADE2;refGene_tx=XM_023255281.1;refGene_gn=JADE2;refGene_tx=XM_023255283.1;refGene_gn=JADE2;refGene_tx=XM_023255285.1;refGene_gn=JADE2;refGene_tx=XM_023255291.1;refGene_gn=JADE2;refGene_tx=XM_023255296.1
A1 113500958 113501047
refGene_gn=JADE2;refGene_tx=XM_006927603.4;refGene_gn=JADE2;refGene_tx=XM_006927604.4;refGene_gn=JADE2;refGene_tx=XM_019832025.2;refGene_gn=JADE2;refGene_tx=XM_023255268.1;refGene_gn=JADE2;refGene_tx=XM_023255270.1;refGene_gn=JADE2;refGene_tx=XM_023255273.1;refGene_gn=JADE2;refGene_tx=XM_023255281.1;refGene_gn=JADE2;refGene_tx=XM_023255283.1;refGene_gn=JADE2;refGene_tx=XM_023255285.1;refGene_gn=JADE2;refGene_tx=XM_023255291.1;refGene_gn=JADE2;refGene_tx=XM_023255296.1
A1 113513879 113514037
refGene_gn=JADE2;refGene_tx=XM_006927603.4;refGene_gn=JADE2;refGene_tx=XM_006927604.4;refGene_gn=JADE2;refGene_tx=XM_019832025.2;refGene_gn=JADE2;refGene_tx=XM_023255268.1;refGene_gn=JADE2;refGene_tx=XM_023255270.1;refGene_gn=JADE2;refGene_tx=XM_023255273.1;refGene_gn=JADE2;refGene_tx=XM_023255281.1;refGene_gn=JADE2;refGene_tx=XM_023255283.1;refGene_gn=JADE2;refGene_tx=XM_023255285.1;refGene_gn=JADE2;refGene_tx=XM_023255291.1;refGene_gn=JADE2;refGene_tx=XM_023255296.1
A1 113518539 113518700
refGene_gn=JADE2;refGene_tx=XM_006927603.4;refGene_gn=JADE2;refGene_tx=XM_006927604.4;refGene_gn=JADE2;refGene_tx=XM_019832025.2;refGene_gn=JADE2;refGene_tx=XM_023255268.1;refGene_gn=JADE2;refGene_tx=XM_023255270.1;refGene_gn=JADE2;refGene_tx=XM_023255273.1;refGene_gn=JADE2;refGene_tx=XM_023255281.1;refGene_gn=JADE2;refGene_tx=XM_023255283.1;refGene_gn=JADE2;refGene_tx=XM_023255285.1;refGene_gn=JADE2;refGene_tx=XM_023255291.1;refGene_gn=JADE2;refGene_tx=XM_023255296.1
A1 113519370 113519582
refGene_gn=JADE2;refGene_tx=XM_006927603.4;refGene_gn=JADE2;refGene_tx=XM_006927604.4;refGene_gn=JADE2;refGene_tx=XM_019832025.2;refGene_gn=JADE2;refGene_tx=XM_023255268.1;refGene_gn=JADE2;refGene_tx=XM_023255270.1;refGene_gn=JADE2;refGene_tx=XM_023255273.1;refGene_gn=JADE2;refGene_tx=XM_023255281.1;refGene_gn=JADE2;refGene_tx=XM_023255283.1;refGene_gn=JADE2;refGene_tx=XM_023255285.1;refGene_gn=JADE2;refGene_tx=XM_023255291.1;refGene_gn=JADE2;refGene_tx=XM_023255296.1
A1 113520777 113520945
refGene_gn=JADE2;refGene_tx=XM_006927603.4;refGene_gn=JADE2;refGene_tx=XM_006927604.4;refGene_gn=JADE2;refGene_tx=XM_019832025.2;refGene_gn=JADE2;refGene_tx=XM_023255268.1;refGene_gn=JADE2;refGene_tx=XM_023255270.1;refGene_gn=JADE2;refGene_tx=XM_023255273.1;refGene_gn=JADE2;refGene_tx=XM_023255281.1;refGene_gn=JADE2;refGene_tx=XM_023255283.1;refGene_gn=JADE2;refGene_tx=XM_023255285.1;refGene_gn=JADE2;refGene_tx=XM_023255291.1;refGene_gn=JADE2;refGene_tx=XM_023255296.1
A1 113522410 113522527
refGene_gn=JADE2;refGene_tx=XM_006927603.4;refGene_gn=JADE2;refGene_tx=XM_006927604.4;refGene_gn=JADE2;refGene_tx=XM_019832025.2;refGene_gn=JADE2;refGene_tx=XM_023255268.1;refGene_gn=JADE2;refGene_tx=XM_0232552

70.1;refGene_gn=JADE2;refGene_tx=XM_023255273.1;refGene_gn=JADE2;refGene_tx=XM_023255281.1;refGene_gn=JADE2;refGene_tx=XM_023255283.1;refGene_gn=JADE2;refGene_tx=XM_023255285.1;refGene_gn=JADE2;refGene_tx=XM_023255291.1;refGene_gn=JADE2;refGene_tx=XM_023255296.1
A1 113523790 113524255
refGene_gn=JADE2;refGene_tx=XM_006927603.4;refGene_gn=JADE2;refGene_tx=XM_006927604.4;refGene_gn=JADE2;refGene_tx=XM_019832025.2;refGene_gn=JADE2;refGene_tx=XM_023255268.1;refGene_gn=JADE2;refGene_tx=XM_023255270.1;refGene_gn=JADE2;refGene_tx=XM_023255273.1;refGene_gn=JADE2;refGene_tx=XM_023255281.1;refGene_gn=JADE2;refGene_tx=XM_023255283.1;refGene_gn=JADE2;refGene_tx=XM_023255285.1;refGene_gn=JADE2;refGene_tx=XM_023255291.1;refGene_gn=JADE2;refGene_tx=XM_023255296.1
A1 113532921 113533039
refGene_gn=JADE2;refGene_tx=XM_006927603.4;refGene_gn=JADE2;refGene_tx=XM_006927604.4;refGene_gn=JADE2;refGene_tx=XM_019832025.2;refGene_gn=JADE2;refGene_tx=XM_023255268.1;refGene_gn=JADE2;refGene_tx=XM_023255270.1;refGene_gn=JADE2;refGene_tx=XM_023255273.1;refGene_gn=JADE2;refGene_tx=XM_023255281.1;refGene_gn=JADE2;refGene_tx=XM_023255283.1;refGene_gn=JADE2;refGene_tx=XM_023255285.1;refGene_gn=JADE2;refGene_tx=XM_023255291.1;refGene_gn=JADE2;refGene_tx=XM_023255296.1
A1 113536452 113536581
refGene_gn=JADE2;refGene_tx=XM_006927603.4;refGene_gn=JADE2;refGene_tx=XM_006927604.4;refGene_gn=JADE2;refGene_tx=XM_019832025.2;refGene_gn=JADE2;refGene_tx=XM_023255268.1;refGene_gn=JADE2;refGene_tx=XM_023255273.1;refGene_gn=JADE2;refGene_tx=XM_023255281.1;refGene_gn=JADE2;refGene_tx=XM_023255283.1;refGene_gn=JADE2;refGene_tx=XM_023255285.1;refGene_gn=JADE2;refGene_tx=XM_023255291.1;refGene_gn=JADE2;refGene_tx=XM_023255296.1
A1 113537892 113538701
refGene_gn=JADE2;refGene_tx=XM_006927603.4;refGene_gn=JADE2;refGene_tx=XM_006927604.4;refGene_gn=JADE2;refGene_tx=XM_019832025.2;refGene_gn=JADE2;refGene_tx=XM_023255268.1;refGene_gn=JADE2;refGene_tx=XM_023255270.1;refGene_gn=JADE2;refGene_tx=XM_023255273.1;refGene_gn=JADE2;refGene_tx=XM_023255281.1;refGene_gn=JADE2;refGene_tx=XM_023255283.1;refGene_gn=JADE2;refGene_tx=XM_023255285.1;refGene_gn=JADE2;refGene_tx=XM_023255291.1;refGene_gn=JADE2;refGene_tx=XM_023255296.1
A1 113561087 113561204
refGene_gn=SAR1B;refGene_tx=XM_003980773.5;refGene_gn=SAR1B;refGene_tx=XM_011282142.3
A1 113562283 113562415
refGene_gn=SAR1B;refGene_tx=XM_003980773.5;refGene_gn=SAR1B;refGene_tx=XM_011282142.3
A1 113563605 113563709
refGene_gn=SAR1B;refGene_tx=XM_003980773.5;refGene_gn=SAR1B;refGene_tx=XM_011282142.3
A1 113565201 113565267
refGene_gn=SAR1B;refGene_tx=XM_003980773.5;refGene_gn=SAR1B;refGene_tx=XM_011282142.3
A1 113569523 113569643
refGene_gn=SAR1B;refGene_tx=XM_003980773.5;refGene_gn=SAR1B;refGene_tx

=XM_011282142.3
A1 113573677 113573735
refGene_gn=SAR1B;refGene_tx=XM_003980773.5;refGene_gn=SAR1B;refGene_tx=XM_011282142.3
A1 113593136 113593233
refGene_gn=SEC24A;refGene_tx=XM_011282147.3;refGene_gn=SEC24A;refGene_tx=XM_019832106.2;refGene_gn=SEC24A;refGene_tx=XM_019832120.2
A1 113614225 113614714
refGene_gn=SEC24A;refGene_tx=XM_011282147.3;refGene_gn=SEC24A;refGene_tx=XM_019832106.2;refGene_gn=SEC24A;refGene_tx=XM_019832120.2
A1 113620507 113620678
refGene_gn=SEC24A;refGene_tx=XM_011282147.3;refGene_gn=SEC24A;refGene_tx=XM_019832106.2
A1 113622330 113622408
refGene_gn=SEC24A;refGene_tx=XM_011282147.3;refGene_gn=SEC24A;refGene_tx=XM_019832106.2;refGene_gn=SEC24A;refGene_tx=XM_019832120.2
A1 113622996 113623157
refGene_gn=SEC24A;refGene_tx=XM_011282147.3;refGene_gn=SEC24A;refGene_tx=XM_019832106.2;refGene_gn=SEC24A;refGene_tx=XM_019832120.2
A1 113623466 113623639
refGene_gn=SEC24A;refGene_tx=XM_011282147.3;refGene_gn=SEC24A;refGene_tx=XM_019832106.2;refGene_gn=SEC24A;refGene_tx=XM_019832120.2
A1 113624994 113625097
refGene_gn=SEC24A;refGene_tx=XM_011282147.3;refGene_gn=SEC24A;refGene_tx=XM_019832106.2;refGene_gn=SEC24A;refGene_tx=XM_019832120.2
A1 113626340 113626467
refGene_gn=SEC24A;refGene_tx=XM_011282147.3;refGene_gn=SEC24A;refGene_tx=XM_019832106.2;refGene_gn=SEC24A;refGene_tx=XM_019832120.2
A1 113628339 113628449
refGene_gn=SEC24A;refGene_tx=XM_011282147.3;refGene_gn=SEC24A;refGene_tx=XM_019832106.2;refGene_gn=SEC24A;refGene_tx=XM_019832120.2
A1 113632749 113632862
refGene_gn=SEC24A;refGene_tx=XM_011282147.3;refGene_gn=SEC24A;refGene_tx=XM_019832106.2;refGene_gn=SEC24A;refGene_tx=XM_019832120.2
A1 113635340 113635459
refGene_gn=SEC24A;refGene_tx=XM_011282147.3;refGene_gn=SEC24A;refGene_tx=XM_019832106.2;refGene_gn=SEC24A;refGene_tx=XM_019832120.2
A1 113640585 113640641
refGene_gn=SEC24A;refGene_tx=XM_011282147.3;refGene_gn=SEC24A;refGene_tx=XM_019832106.2;refGene_gn=SEC24A;refGene_tx=XM_019832120.2
A1 113641739 113641946
refGene_gn=SEC24A;refGene_tx=XM_011282147.3;refGene_gn=SEC24A;refGene_tx=XM_019832106.2;refGene_gn=SEC24A;refGene_tx=XM_019832120.2
A1 113643259 113643380
refGene_gn=SEC24A;refGene_tx=XM_011282147.3;refGene_gn=SEC24A;refGene_tx=XM_019832106.2;refGene_gn=SEC24A;refGene_tx=XM_019832120.2
A1 113643696 113643855
refGene_gn=SEC24A;refGene_tx=XM_011282147.3;refGene_gn=SEC24A;refGene_tx=XM_019832106.2;refGene_gn=SEC24A;refGene_tx=XM_019832120.2
A1 113646274 113646448

refGene_gn=SEC24A;refGene_tx=XM_011282147.3;refGene_gn=SEC24A;refGene_tx=XM_019832106.2;refGene_gn=SEC24A;refGene_tx=XM_019832120.2
A1 113647920 113648031
refGene_gn=SEC24A;refGene_tx=XM_011282147.3;refGene_gn=SEC24A;refGene_tx=XM_019832106.2;refGene_gn=SEC24A;refGene_tx=XM_019832120.2
A1 113652611 113652787
refGene_gn=SEC24A;refGene_tx=XM_011282147.3;refGene_gn=SEC24A;refGene_tx=XM_019832106.2;refGene_gn=SEC24A;refGene_tx=XM_019832120.2
A1 113655338 113655476
refGene_gn=SEC24A;refGene_tx=XM_011282147.3;refGene_gn=SEC24A;refGene_tx=XM_019832106.2;refGene_gn=SEC24A;refGene_tx=XM_019832120.2
A1 113658339 113658444
refGene_gn=SEC24A;refGene_tx=XM_011282147.3;refGene_gn=SEC24A;refGene_tx=XM_019832106.2;refGene_gn=SEC24A;refGene_tx=XM_019832120.2
A1 113659010 113659103
refGene_gn=SEC24A;refGene_tx=XM_011282147.3;refGene_gn=SEC24A;refGene_tx=XM_019832106.2;refGene_gn=SEC24A;refGene_tx=XM_019832120.2
A1 113662456 113662560
refGene_gn=SEC24A;refGene_tx=XM_011282147.3;refGene_gn=SEC24A;refGene_tx=XM_019832106.2;refGene_gn=SEC24A;refGene_tx=XM_019832120.2
A1 113663003 113663118
refGene_gn=SEC24A;refGene_tx=XM_011282147.3;refGene_gn=SEC24A;refGene_tx=XM_019832106.2;refGene_gn=SEC24A;refGene_tx=XM_019832120.2
A1 113672090 113672274
refGene_gn=CAMLG;refGene_tx=XM_003980774.2
A1 113673959 113674417
refGene_gn=CAMLG;refGene_tx=XM_003980774.2
A1 113676084 113676150
refGene_gn=CAMLG;refGene_tx=XM_003980774.2
A1 113681436 113681628
refGene_gn=CAMLG;refGene_tx=XM_003980774.2
A1 113686149 113686166
refGene_gn=DDX46;refGene_tx=XM_003980775.4;refGene_gn=DDX46;refGene_tx=XM_006927606.4
A1 113690283 113690469
refGene_gn=DDX46;refGene_tx=XM_003980775.4;refGene_gn=DDX46;refGene_tx=XM_006927606.4
A1 113693275 113693419
refGene_gn=DDX46;refGene_tx=XM_003980775.4;refGene_gn=DDX46;refGene_tx=XM_006927606.4
A1 113698993 113699090
refGene_gn=DDX46;refGene_tx=XM_003980775.4;refGene_gn=DDX46;refGene_tx=XM_006927606.4
A1 113700156 113700322
refGene_gn=DDX46;refGene_tx=XM_003980775.4;refGene_gn=DDX46;refGene_tx=XM_006927606.4
A1 113701951 113702103
refGene_gn=DDX46;refGene_tx=XM_003980775.4;refGene_gn=DDX46;refGene_tx=XM_006927606.4
A1 113702893 113703007

refGene_gn=DDX46;refGene_tx=XM_003980775.4;refGene_gn=DDX46;refGene_tx=XM_006927606.4
A1 113703564 113703730
refGene_gn=DDX46;refGene_tx=XM_003980775.4;refGene_gn=DDX46;refGene_tx=XM_006927606.4
A1 113705090 113705211
refGene_gn=DDX46;refGene_tx=XM_003980775.4;refGene_gn=DDX46;refGene_tx=XM_006927606.4
A1 113705793 113705969
refGene_gn=DDX46;refGene_tx=XM_003980775.4;refGene_gn=DDX46;refGene_tx=XM_006927606.4
A1 113706912 113707034
refGene_gn=DDX46;refGene_tx=XM_003980775.4;refGene_gn=DDX46;refGene_tx=XM_006927606.4
A1 113709567 113709646
refGene_gn=DDX46;refGene_tx=XM_003980775.4;refGene_gn=DDX46;refGene_tx=XM_006927606.4
A1 113711427 113711510
refGene_gn=DDX46;refGene_tx=XM_003980775.4;refGene_gn=DDX46;refGene_tx=XM_006927606.4
A1 113721071 113721236
refGene_gn=DDX46;refGene_tx=XM_003980775.4;refGene_gn=DDX46;refGene_tx=XM_006927606.4
A1 113721746 113721909
refGene_gn=DDX46;refGene_tx=XM_003980775.4;refGene_gn=DDX46;refGene_tx=XM_006927606.4
A1 113736681 113736875
refGene_gn=DDX46;refGene_tx=XM_003980775.4;refGene_gn=DDX46;refGene_tx=XM_006927606.4
A1 113739314 113739452
refGene_gn=DDX46;refGene_tx=XM_003980775.4;refGene_gn=DDX46;refGene_tx=XM_006927606.4
A1 113739774 113739924
refGene_gn=DDX46;refGene_tx=XM_003980775.4;refGene_gn=DDX46;refGene_tx=XM_006927606.4
A1 113744637 113744814
refGene_gn=DDX46;refGene_tx=XM_003980775.4;refGene_gn=DDX46;refGene_tx=XM_006927606.4
A1 113746431 113746650
refGene_gn=DDX46;refGene_tx=XM_003980775.4;refGene_gn=DDX46;refGene_tx=XM_006927606.4
A1 113746968 113747113
refGene_gn=DDX46;refGene_tx=XM_003980775.4;refGene_gn=DDX46;refGene_tx=XM_006927606.4
A1 113754840 113754914
refGene_gn=DDX46;refGene_tx=XM_003980775.4;refGene_gn=DDX46;refGene_tx=XM_006927606.4
A1 113756078 113756126
refGene_gn=DDX46;refGene_tx=XM_003980775.4;refGene_gn=DDX46;refGene_tx=XM_006927606.4

A1 113770455 113771022
refGene_gn=CA1H5orf24;refGene_tx=XM_003980776.4;refGene_gn=CA1H5orf24;
refGene_tx=XM_011282156.3
A1 113772643 113772684
refGene_gn=CA1H5orf24;refGene_tx=XM_011282156.3
A1 113788547 113788758
refGene_gn=TXNDC15;refGene_tx=XM_003980886.5
A1 113794289 113794783
refGene_gn=TXNDC15;refGene_tx=XM_003980886.5
A1 113796799 113796963
refGene_gn=TXNDC15;refGene_tx=XM_003980886.5
A1 113799832 113799963
refGene_gn=TXNDC15;refGene_tx=XM_003980886.5
A1 113802294 113802491
refGene_gn=TXNDC15;refGene_tx=XM_003980886.5
A1 113806840 113806924
refGene_gn=PCBD2;refGene_tx=XM_023255386.1;refGene_gn=PCBD2;refGene_tx
=XM_023255388.1;refGene_gn=PCBD2;refGene_tx=XM_023255395.1
A1 113809877 113810009
refGene_gn=PCBD2;refGene_tx=XM_023255386.1;refGene_gn=PCBD2;refGene_tx
=XM_023255388.1;refGene_gn=PCBD2;refGene_tx=XM_023255395.1
A1 113855469 113855565
refGene_gn=PCBD2;refGene_tx=XM_019832235.2;refGene_gn=PCBD2;refGene_tx
=XM_023255386.1;refGene_gn=PCBD2;refGene_tx=XM_023255388.1;refGene_gn=
PCBD2;refGene_tx=XM_023255395.1
A1 113856788 113856884
refGene_gn=PCBD2;refGene_tx=XM_019832235.2;refGene_gn=PCBD2;refGene_tx
=XM_023255386.1;refGene_gn=PCBD2;refGene_tx=XM_023255388.1;refGene_gn=
PCBD2;refGene_tx=XM_023255395.1
A1 113863584 113863706
refGene_gn=CATSPER3;refGene_tx=XM_003980777.5;refGene_gn=CATSPER3;refG
ene_tx=XM_023255379.1
A1 113865716 113865870
refGene_gn=CATSPER3;refGene_tx=XM_003980777.5;refGene_gn=CATSPER3;refG
ene_tx=XM_023255379.1
A1 113885778 113886018
refGene_gn=CATSPER3;refGene_tx=XM_003980777.5;refGene_gn=CATSPER3;refG
ene_tx=XM_023255379.1
A1 113891642 113891825
refGene_gn=CATSPER3;refGene_tx=XM_003980777.5;refGene_gn=CATSPER3;refG
ene_tx=XM_023255379.1
A1 113892496 113892637
refGene_gn=CATSPER3;refGene_tx=XM_003980777.5;refGene_gn=CATSPER3;refG
ene_tx=XM_023255379.1
A1 113892997 113893117
refGene_gn=CATSPER3;refGene_tx=XM_003980777.5;refGene_gn=CATSPER3;refG
ene_tx=XM_023255379.1
A1 113894322 113894544
refGene_gn=CATSPER3;refGene_tx=XM_003980777.5;refGene_gn=CATSPER3;refG
ene_tx=XM_023255379.1

A1 113895447 113895553
refGene_gn=CATSPER3;refGene_tx=XM_003980777.5;refGene_gn=CATSPER3;refGene_tx=XM_023255379.1
A1 113917872 113918415
refGene_gn=PITX1;refGene_tx=XM_023255424.1
A1 113920438 113920671
refGene_gn=PITX1;refGene_tx=XM_023255424.1
A1 113923000 113923166
refGene_gn=PITX1;refGene_tx=XM_023255424.1
A1 114225050 114225216
refGene_gn=LOC101080257;refGene_tx=XM_023255427.1;refGene_gn=LOC101080257;refGene_tx=XM_023255428.1;refGene_gn=LOC101080257;refGene_tx=XM_023255433.1;refGene_gn=LOC101080257;refGene_tx=XM_023255437.1
A1 114232676 114232851
refGene_gn=LOC101080257;refGene_tx=XM_023255427.1;refGene_gn=LOC101080257;refGene_tx=XM_023255428.1;refGene_gn=LOC101080257;refGene_tx=XM_023255433.1;refGene_gn=LOC101080257;refGene_tx=XM_023255437.1
A1 114235314 114235404
refGene_gn=LOC101080257;refGene_tx=XM_023255427.1;refGene_gn=LOC101080257;refGene_tx=XM_023255428.1;refGene_gn=LOC101080257;refGene_tx=XM_023255433.1;refGene_gn=LOC101080257;refGene_tx=XM_023255437.1
A1 114240250 114240341
refGene_gn=LOC101080257;refGene_tx=XM_023255433.1;refGene_gn=LOC101080257;refGene_tx=XM_023255437.1
A1 114242101 114242201
refGene_gn=LOC101080257;refGene_tx=XM_023255427.1;refGene_gn=LOC101080257;refGene_tx=XM_023255428.1
A1 114250100 114250211
refGene_gn=LOC101080257;refGene_tx=XM_023255427.1;refGene_gn=LOC101080257;refGene_tx=XM_023255428.1;refGene_gn=LOC101080257;refGene_tx=XM_023255433.1;refGene_gn=LOC101080257;refGene_tx=XM_023255437.1
A1 114271146 114271344
refGene_gn=LOC101080257;refGene_tx=XM_023255427.1;refGene_gn=LOC101080257;refGene_tx=XM_023255428.1;refGene_gn=LOC101080257;refGene_tx=XM_023255433.1;refGene_gn=LOC101080257;refGene_tx=XM_023255437.1
A1 114271771 114271878
refGene_gn=LOC101080257;refGene_tx=XM_023255427.1;refGene_gn=LOC101080257;refGene_tx=XM_023255428.1;refGene_gn=LOC101080257;refGene_tx=XM_023255433.1;refGene_gn=LOC101080257;refGene_tx=XM_023255437.1
A1 114293136 114293308
refGene_gn=LOC101080257;refGene_tx=XM_023255427.1;refGene_gn=LOC101080257;refGene_tx=XM_023255428.1;refGene_gn=LOC101080257;refGene_tx=XM_023255433.1;refGene_gn=LOC101080257;refGene_tx=XM_023255437.1
A1 114346305 114346909
refGene_gn=TIFAB;refGene_tx=XM_003980889.3
A1 114349569 114349610
refGene_gn=TIFAB;refGene_tx=XM_003980889.3
A1 114420990 114421722
refGene_gn=NEUROG1;refGene_tx=XM_019832334.2
A1 114456966 114456982

refGene_gn=CXCL14;refGene_tx=XM_011282200.3
A1 114459535 114459649
refGene_gn=CXCL14;refGene_tx=XM_011282200.3
A1 114463412 114463518
refGene_gn=CXCL14;refGene_tx=XM_011282200.3
A1 114463692 114463990
refGene_gn=CXCL14;refGene_tx=XM_011282200.3
A1 114623129 114623223
refGene_gn=SLC25A48;refGene_tx=XM_019832322.1;refGene_gn=SLC25A48;refGene_tx=XM_023255442.1;refGene_gn=SLC25A48;refGene_tx=XM_023255447.1;refGene_gn=SLC25A48;refGene_tx=XM_023255448.1;refGene_gn=SLC25A48;refGene_tx=XM_023255456.1;refGene_gn=SLC25A48;refGene_tx=XM_023255461.1;refGene_gn=SLC25A48;refGene_tx=XM_023255462.1;refGene_gn=SLC25A48;refGene_tx=XM_023255468.1
A1 114628921 114629010
refGene_gn=SLC25A48;refGene_tx=XM_019832322.1;refGene_gn=SLC25A48;refGene_tx=XM_023255442.1;refGene_gn=SLC25A48;refGene_tx=XM_023255447.1;refGene_gn=SLC25A48;refGene_tx=XM_023255448.1;refGene_gn=SLC25A48;refGene_tx=XM_023255456.1;refGene_gn=SLC25A48;refGene_tx=XM_023255461.1;refGene_gn=SLC25A48;refGene_tx=XM_023255462.1;refGene_gn=SLC25A48;refGene_tx=XM_023255468.1
A1 114633573 114633645
refGene_gn=SLC25A48;refGene_tx=XM_019832322.1;refGene_gn=SLC25A48;refGene_tx=XM_023255442.1;refGene_gn=SLC25A48;refGene_tx=XM_023255447.1;refGene_gn=SLC25A48;refGene_tx=XM_023255448.1;refGene_gn=SLC25A48;refGene_tx=XM_023255456.1;refGene_gn=SLC25A48;refGene_tx=XM_023255461.1;refGene_gn=SLC25A48;refGene_tx=XM_023255462.1;refGene_gn=SLC25A48;refGene_tx=XM_023255468.1
A1 114636370 114636629
refGene_gn=SLC25A48;refGene_tx=XM_019832322.1;refGene_gn=SLC25A48;refGene_tx=XM_019832331.2;refGene_gn=SLC25A48;refGene_tx=XM_023255442.1;refGene_gn=SLC25A48;refGene_tx=XM_023255447.1;refGene_gn=SLC25A48;refGene_tx=XM_023255448.1;refGene_gn=SLC25A48;refGene_tx=XM_023255456.1;refGene_gn=SLC25A48;refGene_tx=XM_023255461.1;refGene_gn=SLC25A48;refGene_tx=XM_023255462.1;refGene_gn=SLC25A48;refGene_tx=XM_023255468.1
A1 114651984 114652242
refGene_gn=SLC25A48;refGene_tx=XM_019832322.1;refGene_gn=SLC25A48;refGene_tx=XM_019832331.2;refGene_gn=SLC25A48;refGene_tx=XM_023255442.1;refGene_gn=SLC25A48;refGene_tx=XM_023255447.1;refGene_gn=SLC25A48;refGene_tx=XM_023255448.1;refGene_gn=SLC25A48;refGene_tx=XM_023255456.1;refGene_gn=SLC25A48;refGene_tx=XM_023255461.1;refGene_gn=SLC25A48;refGene_tx=XM_023255462.1;refGene_gn=SLC25A48;refGene_tx=XM_023255468.1
A1 114654704 114655138
refGene_gn=SLC25A48;refGene_tx=XM_019832322.1;refGene_gn=SLC25A48;refGene_tx=XM_019832331.2;refGene_gn=SLC25A48;refGene_tx=XM_023255442.1;refGene_gn=SLC25A48;refGene_tx=XM_023255447.1;refGene_gn=SLC25A48;refGene_tx=XM_023255448.1;refGene_gn=SLC25A48;refGene_tx=XM_023255456.1;refGene_gn=SLC25A48;refGene_tx=XM_023255461.1;refGene_gn=SLC25A48;refGene_tx=XM_023255462.1;refGene_gn=SLC25A48;refGene_tx=XM_023255468.1
A1 114662073 114662196

refGene_gn=SLC25A48;refGene_tx=XM_019832322.1;refGene_gn=SLC25A48;refGene_tx=XM_023255461.1;refGene_gn=SLC25A48;refGene_tx=XM_023255462.1;refGene_gn=SLC25A48;refGene_tx=XM_023255468.1
A1 114675341 114675452
refGene_gn=IL9;refGene_tx=XM_003980893.3
A1 114676582 114676714
refGene_gn=IL9;refGene_tx=XM_003980893.3
A1 114677924 114677957
refGene_gn=IL9;refGene_tx=XM_003980893.3
A1 114678042 114678078
refGene_gn=IL9;refGene_tx=XM_003980893.3
A1 114678383 114678497
refGene_gn=IL9;refGene_tx=XM_003980893.3
A1 114714048 114714131
refGene_gn=FBXL21;refGene_tx=XM_023255477.1
A1 114716355 114716722
refGene_gn=FBXL21;refGene_tx=XM_023255477.1
A1 114717288 114717411
refGene_gn=FBXL21;refGene_tx=XM_023255477.1
A1 114720506 114720678
refGene_gn=FBXL21;refGene_tx=XM_023255477.1
A1 114721049 114721693
refGene_gn=FBXL21;refGene_tx=XM_023255477.1
A1 114726468 114726635
refGene_gn=LECT2;refGene_tx=XM_003980781.3
A1 114733533 114733679
refGene_gn=LECT2;refGene_tx=XM_003980781.3
A1 114735842 114735939
refGene_gn=LECT2;refGene_tx=XM_003980781.3
A1 114737434 114737480
refGene_gn=LECT2;refGene_tx=XM_003980781.3
A1 114812778 114812912
refGene_gn=TGFBI;refGene_tx=XM_003980782.4
A1 114817761 114817860
refGene_gn=TGFBI;refGene_tx=XM_003980782.4
A1 114827652 114827717
refGene_gn=TGFBI;refGene_tx=XM_003980782.4
A1 114828969 114829130
refGene_gn=TGFBI;refGene_tx=XM_003980782.4
A1 114829466 114829631
refGene_gn=TGFBI;refGene_tx=XM_003980782.4
A1 114829888 114830035
refGene_gn=TGFBI;refGene_tx=XM_003980782.4
A1 114832072 114832214
refGene_gn=TGFBI;refGene_tx=XM_003980782.4
A1 114834775 114834988
refGene_gn=TGFBI;refGene_tx=XM_003980782.4
A1 114835832 114835970
refGene_gn=TGFBI;refGene_tx=XM_003980782.4
A1 114836555 114836701

refGene_gn=TGFBI;refGene_tx=XM_003980782.4
A1	114837675	114837812
refGene_gn=TGFBI;refGene_tx=XM_003980782.4
A1	114838598	114838729
refGene_gn=TGFBI;refGene_tx=XM_003980782.4
A1	114841160	114841285
refGene_gn=TGFBI;refGene_tx=XM_003980782.4
A1	114842559	114842662
refGene_gn=TGFBI;refGene_tx=XM_003980782.4
A1	114843241	114843321
refGene_gn=TGFBI;refGene_tx=XM_003980782.4
A1	114844422	114844447
refGene_gn=TGFBI;refGene_tx=XM_003980782.4
A1	114844869	114844910
refGene_gn=TGFBI;refGene_tx=XM_003980782.4
A1	114921749	114922152
refGene_gn=SMAD5;refGene_tx=XM_006927618.4
A1	114926875	114927127
refGene_gn=SMAD5;refGene_tx=XM_006927618.4
A1	114931564	114931684
refGene_gn=SMAD5;refGene_tx=XM_006927618.4
A1	114941387	114941609
refGene_gn=SMAD5;refGene_tx=XM_006927618.4
A1	114943520	114943777
refGene_gn=SMAD5;refGene_tx=XM_006927618.4
A1	114945172	114945316
refGene_gn=SMAD5;refGene_tx=XM_006927618.4
A1	114982326	114982496
refGene_gn=TRPC7;refGene_tx=XM_006927698.3
A1	114984629	114984705
refGene_gn=TRPC7;refGene_tx=XM_006927698.3
A1	114991194	114991275
refGene_gn=TRPC7;refGene_tx=XM_006927698.3
A1	114991910	114992132
refGene_gn=TRPC7;refGene_tx=XM_006927698.3
A1	114997633	114997829
refGene_gn=TRPC7;refGene_tx=XM_006927698.3
A1	115012661	115012926
refGene_gn=TRPC7;refGene_tx=XM_006927698.3
A1	115017647	115017881
refGene_gn=TRPC7;refGene_tx=XM_006927698.3
A1	115031602	115031819
refGene_gn=TRPC7;refGene_tx=XM_006927698.3
A1	115039141	115039306
refGene_gn=TRPC7;refGene_tx=XM_006927698.3
A1	115072751	115072934
refGene_gn=TRPC7;refGene_tx=XM_006927698.3
A1	115111403	115112181
refGene_gn=TRPC7;refGene_tx=XM_006927698.3
A1	115120708	115120710

refGene_gn=TRPC7;refGene_tx=XM_006927698.3
A1 115627855 115628046
refGene_gn=SPOCK1;refGene_tx=XM_023255519.1;refGene_gn=SPOCK1;refGene_tx=XM_023255527.1
A1 115628504 115628642
refGene_gn=SPOCK1;refGene_tx=XM_023255519.1;refGene_gn=SPOCK1;refGene_tx=XM_023255527.1
A1 115633896 115633959
refGene_gn=SPOCK1;refGene_tx=XM_023255519.1;refGene_gn=SPOCK1;refGene_tx=XM_023255527.1
A1 115639464 115639686
refGene_gn=SPOCK1;refGene_tx=XM_023255519.1;refGene_gn=SPOCK1;refGene_tx=XM_023255527.1
A1 115643204 115643321
refGene_gn=SPOCK1;refGene_tx=XM_023255519.1;refGene_gn=SPOCK1;refGene_tx=XM_023255527.1
A1 115713002 115713117
refGene_gn=SPOCK1;refGene_tx=XM_023255519.1;refGene_gn=SPOCK1;refGene_tx=XM_023255527.1
A1 115760786 115760913
refGene_gn=SPOCK1;refGene_tx=XM_023255519.1;refGene_gn=SPOCK1;refGene_tx=XM_023255527.1
A1 115785136 115785251
refGene_gn=SPOCK1;refGene_tx=XM_023255519.1;refGene_gn=SPOCK1;refGene_tx=XM_023255527.1
A1 115906758 115906804
refGene_gn=SPOCK1;refGene_tx=XM_023255519.1;refGene_gn=SPOCK1;refGene_tx=XM_023255527.1
A1 115913276 115913285
refGene_gn=SPOCK1;refGene_tx=XM_023255519.1
A1 116143140 116143326
refGene_gn=SPOCK1;refGene_tx=XM_023255519.1;refGene_gn=SPOCK1;refGene_tx=XM_023255527.1
A1 116259947 116259976
refGene_gn=KLHL3;refGene_tx=XM_019832555.1;refGene_gn=KLHL3;refGene_tx=XM_019832568.1;refGene_gn=KLHL3;refGene_tx=XM_019832575.1;refGene_gn=KLHL3;refGene_tx=XM_019832582.1;refGene_gn=KLHL3;refGene_tx=XM_019832593.1;refGene_gn=KLHL3;refGene_tx=XM_019832598.1;refGene_gn=KLHL3;refGene_tx=XM_019832607.1;refGene_gn=KLHL3;refGene_tx=XM_019832613.2;refGene_gn=KLHL3;refGene_tx=XM_019832617.2;refGene_gn=KLHL3;refGene_tx=XM_019832618.1;refGene_gn=KLHL3;refGene_tx=XM_023255572.1
A1 116263930 116264074
refGene_gn=KLHL3;refGene_tx=XM_019832555.1;refGene_gn=KLHL3;refGene_tx=XM_019832568.1;refGene_gn=KLHL3;refGene_tx=XM_019832575.1;refGene_gn=KLHL3;refGene_tx=XM_019832582.1;refGene_gn=KLHL3;refGene_tx=XM_019832593.1;refGene_gn=KLHL3;refGene_tx=XM_019832598.1;refGene_gn=KLHL3;refGene_tx=XM_019832607.1;refGene_gn=KLHL3;refGene_tx=XM_019832613.2;refGene_gn=KLHL3;refGene_tx=XM_019832617.2;refGene_gn=KLHL3;refGene_tx=XM_019832618.1;refGene_gn=KLHL3;refGene_tx=XM_023255572.1
A1 116266304 116266445

refGene_gn=KLHL3;refGene_tx=XM_019832555.1;refGene_gn=KLHL3;refGene_tx
=XM_019832568.1;refGene_gn=KLHL3;refGene_tx=XM_019832575.1;refGene_gn=
KLHL3;refGene_tx=XM_019832582.1;refGene_gn=KLHL3;refGene_tx=XM_0198325
93.1;refGene_gn=KLHL3;refGene_tx=XM_019832598.1;refGene_gn=KLHL3;refGe
ne_tx=XM_019832607.1;refGene_gn=KLHL3;refGene_tx=XM_019832613.2;refGen
e_gn=KLHL3;refGene_tx=XM_019832617.2;refGene_gn=KLHL3;refGene_tx=XM_01
9832618.1;refGene_gn=KLHL3;refGene_tx=XM_023255572.1
A1 116269510 116269639
refGene_gn=KLHL3;refGene_tx=XM_019832555.1;refGene_gn=KLHL3;refGene_tx
=XM_019832568.1;refGene_gn=KLHL3;refGene_tx=XM_019832575.1;refGene_gn=
KLHL3;refGene_tx=XM_019832582.1;refGene_gn=KLHL3;refGene_tx=XM_0198325
93.1;refGene_gn=KLHL3;refGene_tx=XM_019832598.1;refGene_gn=KLHL3;refGe
ne_tx=XM_019832607.1;refGene_gn=KLHL3;refGene_tx=XM_019832613.2;refGen
e_gn=KLHL3;refGene_tx=XM_019832617.2;refGene_gn=KLHL3;refGene_tx=XM_01
9832618.1;refGene_gn=KLHL3;refGene_tx=XM_023255572.1
A1 116273023 116273125
refGene_gn=KLHL3;refGene_tx=XM_019832555.1;refGene_gn=KLHL3;refGene_tx
=XM_019832575.1;refGene_gn=KLHL3;refGene_tx=XM_019832582.1;refGene_gn=
KLHL3;refGene_tx=XM_019832593.1;refGene_gn=KLHL3;refGene_tx=XM_0198325
98.1;refGene_gn=KLHL3;refGene_tx=XM_019832607.1;refGene_gn=KLHL3;refGe
ne_tx=XM_019832613.2;refGene_gn=KLHL3;refGene_tx=XM_019832617.2;refGen
e_gn=KLHL3;refGene_tx=XM_019832618.1;refGene_gn=KLHL3;refGene_tx=XM_02
3255572.1
A1 116274613 116274811
refGene_gn=KLHL3;refGene_tx=XM_019832555.1;refGene_gn=KLHL3;refGene_tx
=XM_019832568.1;refGene_gn=KLHL3;refGene_tx=XM_019832575.1;refGene_gn=
KLHL3;refGene_tx=XM_019832582.1;refGene_gn=KLHL3;refGene_tx=XM_0198325
93.1;refGene_gn=KLHL3;refGene_tx=XM_019832598.1;refGene_gn=KLHL3;refGe
ne_tx=XM_019832607.1;refGene_gn=KLHL3;refGene_tx=XM_019832613.2;refGen
e_gn=KLHL3;refGene_tx=XM_019832617.2;refGene_gn=KLHL3;refGene_tx=XM_01
9832618.1;refGene_gn=KLHL3;refGene_tx=XM_023255572.1
A1 116275196 116275314
refGene_gn=KLHL3;refGene_tx=XM_019832555.1;refGene_gn=KLHL3;refGene_tx
=XM_019832568.1;refGene_gn=KLHL3;refGene_tx=XM_019832575.1;refGene_gn=
KLHL3;refGene_tx=XM_019832582.1;refGene_gn=KLHL3;refGene_tx=XM_0198325
93.1;refGene_gn=KLHL3;refGene_tx=XM_019832598.1;refGene_gn=KLHL3;refGe
ne_tx=XM_019832607.1;refGene_gn=KLHL3;refGene_tx=XM_019832613.2;refGen
e_gn=KLHL3;refGene_tx=XM_019832617.2;refGene_gn=KLHL3;refGene_tx=XM_01
9832618.1;refGene_gn=KLHL3;refGene_tx=XM_023255572.1
A1 116289985 116290135
refGene_gn=KLHL3;refGene_tx=XM_019832555.1;refGene_gn=KLHL3;refGene_tx
=XM_019832568.1;refGene_gn=KLHL3;refGene_tx=XM_019832575.1;refGene_gn=
KLHL3;refGene_tx=XM_019832582.1;refGene_gn=KLHL3;refGene_tx=XM_0198325
93.1;refGene_gn=KLHL3;refGene_tx=XM_019832598.1;refGene_gn=KLHL3;refGe
ne_tx=XM_019832607.1;refGene_gn=KLHL3;refGene_tx=XM_019832613.2;refGen
e_gn=KLHL3;refGene_tx=XM_019832617.2;refGene_gn=KLHL3;refGene_tx=XM_01
9832618.1;refGene_gn=KLHL3;refGene_tx=XM_023255572.1
A1 116294085 116294202
refGene_gn=KLHL3;refGene_tx=XM_019832555.1;refGene_gn=KLHL3;refGene_tx
=XM_019832568.1;refGene_gn=KLHL3;refGene_tx=XM_019832575.1;refGene_gn=

KLHL3;refGene_tx=XM_019832582.1;refGene_gn=KLHL3;refGene_tx=XM_0198325
93.1;refGene_gn=KLHL3;refGene_tx=XM_019832598.1;refGene_gn=KLHL3;refGe
ne_tx=XM_019832607.1;refGene_gn=KLHL3;refGene_tx=XM_019832613.2;refGen
e_gn=KLHL3;refGene_tx=XM_019832617.2;refGene_gn=KLHL3;refGene_tx=XM_01
9832618.1;refGene_gn=KLHL3;refGene_tx=XM_023255572.1
A1	116308736	116308846
refGene_gn=KLHL3;refGene_tx=XM_019832555.1;refGene_gn=KLHL3;refGene_tx
=XM_019832568.1;refGene_gn=KLHL3;refGene_tx=XM_019832575.1;refGene_gn=
KLHL3;refGene_tx=XM_019832582.1;refGene_gn=KLHL3;refGene_tx=XM_0198325
93.1;refGene_gn=KLHL3;refGene_tx=XM_019832598.1;refGene_gn=KLHL3;refGe
ne_tx=XM_019832607.1;refGene_gn=KLHL3;refGene_tx=XM_019832613.2;refGen
e_gn=KLHL3;refGene_tx=XM_019832617.2;refGene_gn=KLHL3;refGene_tx=XM_01
9832618.1;refGene_gn=KLHL3;refGene_tx=XM_023255572.1
A1	116321841	116322004
refGene_gn=KLHL3;refGene_tx=XM_019832555.1;refGene_gn=KLHL3;refGene_tx
=XM_019832568.1;refGene_gn=KLHL3;refGene_tx=XM_019832575.1;refGene_gn=
KLHL3;refGene_tx=XM_019832582.1;refGene_gn=KLHL3;refGene_tx=XM_0198325
93.1;refGene_gn=KLHL3;refGene_tx=XM_019832598.1;refGene_gn=KLHL3;refGe
ne_tx=XM_019832607.1;refGene_gn=KLHL3;refGene_tx=XM_019832613.2;refGen
e_gn=KLHL3;refGene_tx=XM_019832617.2;refGene_gn=KLHL3;refGene_tx=XM_01
9832618.1;refGene_gn=KLHL3;refGene_tx=XM_023255572.1
A1	116331164	116331286
refGene_gn=KLHL3;refGene_tx=XM_019832555.1;refGene_gn=KLHL3;refGene_tx
=XM_019832568.1;refGene_gn=KLHL3;refGene_tx=XM_019832575.1;refGene_gn=
KLHL3;refGene_tx=XM_019832582.1;refGene_gn=KLHL3;refGene_tx=XM_0198325
93.1;refGene_gn=KLHL3;refGene_tx=XM_019832598.1;refGene_gn=KLHL3;refGe
ne_tx=XM_019832607.1;refGene_gn=KLHL3;refGene_tx=XM_019832613.2;refGen
e_gn=KLHL3;refGene_tx=XM_019832617.2;refGene_gn=KLHL3;refGene_tx=XM_01
9832618.1;refGene_gn=KLHL3;refGene_tx=XM_023255572.1
A1	116341224	116341303
refGene_gn=KLHL3;refGene_tx=XM_019832617.2
A1	116341691	116341798
refGene_gn=KLHL3;refGene_tx=XM_019832555.1;refGene_gn=KLHL3;refGene_tx
=XM_019832568.1;refGene_gn=KLHL3;refGene_tx=XM_019832575.1;refGene_gn=
KLHL3;refGene_tx=XM_019832582.1;refGene_gn=KLHL3;refGene_tx=XM_0198325
93.1;refGene_gn=KLHL3;refGene_tx=XM_019832598.1;refGene_gn=KLHL3;refGe
ne_tx=XM_019832607.1;refGene_gn=KLHL3;refGene_tx=XM_019832613.2;refGen
e_gn=KLHL3;refGene_tx=XM_023255572.1
A1	116356020	116356140
refGene_gn=KLHL3;refGene_tx=XM_019832555.1;refGene_gn=KLHL3;refGene_tx
=XM_019832568.1;refGene_gn=KLHL3;refGene_tx=XM_019832575.1;refGene_gn=
KLHL3;refGene_tx=XM_019832582.1;refGene_gn=KLHL3;refGene_tx=XM_0198325
93.1;refGene_gn=KLHL3;refGene_tx=XM_019832618.1;refGene_gn=KLHL3;refGe
ne_tx=XM_023255572.1
A1	116372498	116372608
refGene_gn=KLHL3;refGene_tx=XM_019832555.1;refGene_gn=KLHL3;refGene_tx
=XM_019832568.1;refGene_gn=KLHL3;refGene_tx=XM_023255572.1
A1	116394664	116395585
refGene_gn=HNRNPA0;refGene_tx=XM_011282239.3
A1	116402231	116402234

refGene_gn=KLHL3;refGene_tx=XM_019832555.1;refGene_gn=KLHL3;refGene_tx=XM_019832568.1
A1 116527648 116528004
refGene_gn=MYOT;refGene_tx=XM_003980790.3
A1 116532695 116532873
refGene_gn=MYOT;refGene_tx=XM_003980790.3
A1 116533762 116533867
refGene_gn=MYOT;refGene_tx=XM_003980790.3;refGene_gn=MYOT;refGene_tx=XM_006927620.2
A1 116536949 116536996
refGene_gn=MYOT;refGene_tx=XM_003980790.3;refGene_gn=MYOT;refGene_tx=XM_006927620.2
A1 116537728 116537861
refGene_gn=MYOT;refGene_tx=XM_003980790.3;refGene_gn=MYOT;refGene_tx=XM_006927620.2
A1 116538884 116539092
refGene_gn=MYOT;refGene_tx=XM_003980790.3;refGene_gn=MYOT;refGene_tx=XM_006927620.2
A1 116540985 116541151
refGene_gn=MYOT;refGene_tx=XM_003980790.3;refGene_gn=MYOT;refGene_tx=XM_006927620.2
A1 116541583 116541717
refGene_gn=MYOT;refGene_tx=XM_003980790.3;refGene_gn=MYOT;refGene_tx=XM_006927620.2
A1 116541941 116542114
refGene_gn=MYOT;refGene_tx=XM_003980790.3;refGene_gn=MYOT;refGene_tx=XM_006927620.2
A1 116544178 116544209
refGene_gn=PKD2L2;refGene_tx=XM_006927625.4;refGene_gn=PKD2L2;refGene_tx=XM_019832539.2
A1 116544950 116545052
refGene_gn=PKD2L2;refGene_tx=XM_006927625.4;refGene_gn=PKD2L2;refGene_tx=XM_019832539.2
A1 116546621 116546755
refGene_gn=PKD2L2;refGene_tx=XM_006927625.4;refGene_gn=PKD2L2;refGene_tx=XM_019832539.2
A1 116548342 116548599
refGene_gn=PKD2L2;refGene_tx=XM_006927625.4;refGene_gn=PKD2L2;refGene_tx=XM_019832539.2
A1 116551488 116551710
refGene_gn=PKD2L2;refGene_tx=XM_006927625.4;refGene_gn=PKD2L2;refGene_tx=XM_019832539.2
A1 116555497 116555726
refGene_gn=PKD2L2;refGene_tx=XM_006927625.4;refGene_gn=PKD2L2;refGene_tx=XM_019832539.2
A1 116557379 116557550
refGene_gn=PKD2L2;refGene_tx=XM_006927625.4
A1 116557837 116558019
refGene_gn=PKD2L2;refGene_tx=XM_006927625.4;refGene_gn=PKD2L2;refGene_tx=XM_019832539.2

A1 116565501 116565622
refGene_gn=PKD2L2;refGene_tx=XM_006927625.4;refGene_gn=PKD2L2;refGene_tx=XM_019832539.2
A1 116567150 116567252
refGene_gn=PKD2L2;refGene_tx=XM_006927625.4;refGene_gn=PKD2L2;refGene_tx=XM_019832539.2
A1 116569337 116569402
refGene_gn=PKD2L2;refGene_tx=XM_006927625.4;refGene_gn=PKD2L2;refGene_tx=XM_019832539.2
A1 116570016 116570068
refGene_gn=PKD2L2;refGene_tx=XM_006927625.4;refGene_gn=PKD2L2;refGene_tx=XM_019832539.2
A1 116582553 116582666
refGene_gn=PKD2L2;refGene_tx=XM_006927625.4;refGene_gn=PKD2L2;refGene_tx=XM_019832539.2
A1 116583354 116583451
refGene_gn=PKD2L2;refGene_tx=XM_006927625.4;refGene_gn=PKD2L2;refGene_tx=XM_019832539.2
A1 116586973 116587097
refGene_gn=FAM13B;refGene_tx=XM_023255534.1;refGene_gn=FAM13B;refGene_tx=XM_023255539.1;refGene_gn=FAM13B;refGene_tx=XM_023255543.1;refGene_gn=FAM13B;refGene_tx=XM_023255545.1;refGene_gn=FAM13B;refGene_tx=XM_023255548.1;refGene_gn=FAM13B;refGene_tx=XM_023255556.1;refGene_gn=FAM13B;refGene_tx=XM_023255563.1;refGene_gn=FAM13B;refGene_tx=XM_023255564.1;refGene_gn=FAM13B;refGene_tx=XM_023255566.1
A1 116588598 116588700
refGene_gn=FAM13B;refGene_tx=XM_023255534.1;refGene_gn=FAM13B;refGene_tx=XM_023255539.1;refGene_gn=FAM13B;refGene_tx=XM_023255543.1;refGene_gn=FAM13B;refGene_tx=XM_023255545.1;refGene_gn=FAM13B;refGene_tx=XM_023255548.1;refGene_gn=FAM13B;refGene_tx=XM_023255556.1;refGene_gn=FAM13B;refGene_tx=XM_023255563.1;refGene_gn=FAM13B;refGene_tx=XM_023255564.1;refGene_gn=FAM13B;refGene_tx=XM_023255566.1
A1 116590233 116590397
refGene_gn=FAM13B;refGene_tx=XM_023255534.1;refGene_gn=FAM13B;refGene_tx=XM_023255539.1;refGene_gn=FAM13B;refGene_tx=XM_023255543.1;refGene_gn=FAM13B;refGene_tx=XM_023255545.1;refGene_gn=FAM13B;refGene_tx=XM_023255548.1;refGene_gn=FAM13B;refGene_tx=XM_023255556.1;refGene_gn=FAM13B;refGene_tx=XM_023255563.1;refGene_gn=FAM13B;refGene_tx=XM_023255564.1;refGene_gn=FAM13B;refGene_tx=XM_023255566.1
A1 116590484 116590568
refGene_gn=FAM13B;refGene_tx=XM_023255534.1;refGene_gn=FAM13B;refGene_tx=XM_023255539.1;refGene_gn=FAM13B;refGene_tx=XM_023255543.1;refGene_gn=FAM13B;refGene_tx=XM_023255545.1;refGene_gn=FAM13B;refGene_tx=XM_023255548.1;refGene_gn=FAM13B;refGene_tx=XM_023255556.1;refGene_gn=FAM13B;refGene_tx=XM_023255563.1;refGene_gn=FAM13B;refGene_tx=XM_023255564.1;refGene_gn=FAM13B;refGene_tx=XM_023255566.1
A1 116596658 116596754
refGene_gn=FAM13B;refGene_tx=XM_023255534.1;refGene_gn=FAM13B;refGene_tx=XM_023255539.1;refGene_gn=FAM13B;refGene_tx=XM_023255543.1;refGene_gn=FAM13B;refGene_tx=XM_023255545.1;refGene_gn=FAM13B;refGene_tx=XM_02

3255548.1;refGene_gn=FAM13B;refGene_tx=XM_023255556.1;refGene_gn=FAM13B;refGene_tx=XM_023255563.1;refGene_gn=FAM13B;refGene_tx=XM_023255564.1;refGene_gn=FAM13B;refGene_tx=XM_023255566.1
A1 116596983 116597067
refGene_gn=FAM13B;refGene_tx=XM_023255534.1;refGene_gn=FAM13B;refGene_tx=XM_023255539.1;refGene_gn=FAM13B;refGene_tx=XM_023255543.1;refGene_gn=FAM13B;refGene_tx=XM_023255545.1;refGene_gn=FAM13B;refGene_tx=XM_023255548.1;refGene_gn=FAM13B;refGene_tx=XM_023255556.1;refGene_gn=FAM13B;refGene_tx=XM_023255563.1;refGene_gn=FAM13B;refGene_tx=XM_023255564.1;refGene_gn=FAM13B;refGene_tx=XM_023255566.1
A1 116599370 116599600
refGene_gn=FAM13B;refGene_tx=XM_023255534.1;refGene_gn=FAM13B;refGene_tx=XM_023255539.1;refGene_gn=FAM13B;refGene_tx=XM_023255543.1;refGene_gn=FAM13B;refGene_tx=XM_023255545.1;refGene_gn=FAM13B;refGene_tx=XM_023255548.1;refGene_gn=FAM13B;refGene_tx=XM_023255556.1;refGene_gn=FAM13B;refGene_tx=XM_023255563.1;refGene_gn=FAM13B;refGene_tx=XM_023255564.1;refGene_gn=FAM13B;refGene_tx=XM_023255566.1
A1 116601651 116601751
refGene_gn=FAM13B;refGene_tx=XM_023255534.1;refGene_gn=FAM13B;refGene_tx=XM_023255539.1;refGene_gn=FAM13B;refGene_tx=XM_023255543.1;refGene_gn=FAM13B;refGene_tx=XM_023255545.1;refGene_gn=FAM13B;refGene_tx=XM_023255548.1;refGene_gn=FAM13B;refGene_tx=XM_023255556.1;refGene_gn=FAM13B;refGene_tx=XM_023255563.1;refGene_gn=FAM13B;refGene_tx=XM_023255564.1;refGene_gn=FAM13B;refGene_tx=XM_023255566.1
A1 116602587 116602717
refGene_gn=FAM13B;refGene_tx=XM_023255534.1;refGene_gn=FAM13B;refGene_tx=XM_023255539.1;refGene_gn=FAM13B;refGene_tx=XM_023255543.1;refGene_gn=FAM13B;refGene_tx=XM_023255545.1;refGene_gn=FAM13B;refGene_tx=XM_023255548.1;refGene_gn=FAM13B;refGene_tx=XM_023255556.1;refGene_gn=FAM13B;refGene_tx=XM_023255563.1;refGene_gn=FAM13B;refGene_tx=XM_023255564.1;refGene_gn=FAM13B;refGene_tx=XM_023255566.1
A1 116603831 116604042
refGene_gn=FAM13B;refGene_tx=XM_023255534.1;refGene_gn=FAM13B;refGene_tx=XM_023255539.1;refGene_gn=FAM13B;refGene_tx=XM_023255543.1;refGene_gn=FAM13B;refGene_tx=XM_023255545.1;refGene_gn=FAM13B;refGene_tx=XM_023255548.1;refGene_gn=FAM13B;refGene_tx=XM_023255556.1;refGene_gn=FAM13B;refGene_tx=XM_023255563.1;refGene_gn=FAM13B;refGene_tx=XM_023255564.1;refGene_gn=FAM13B;refGene_tx=XM_023255566.1
A1 116605441 116605507
refGene_gn=FAM13B;refGene_tx=XM_023255534.1;refGene_gn=FAM13B;refGene_tx=XM_023255539.1;refGene_gn=FAM13B;refGene_tx=XM_023255545.1;refGene_gn=FAM13B;refGene_tx=XM_023255548.1;refGene_gn=FAM13B;refGene_tx=XM_023255556.1;refGene_gn=FAM13B;refGene_tx=XM_023255563.1;refGene_gn=FAM13B;refGene_tx=XM_023255564.1
A1 116608499 116608650
refGene_gn=FAM13B;refGene_tx=XM_023255534.1;refGene_gn=FAM13B;refGene_tx=XM_023255539.1;refGene_gn=FAM13B;refGene_tx=XM_023255543.1;refGene_gn=FAM13B;refGene_tx=XM_023255545.1;refGene_gn=FAM13B;refGene_tx=XM_023255548.1;refGene_gn=FAM13B;refGene_tx=XM_023255556.1;refGene_gn=FAM13B;refGene_tx=XM_023255563.1;refGene_gn=FAM13B;refGene_tx=XM_023255564.

1;refGene_gn=FAM13B;refGene_tx=XM_023255566.1
A1 116609049 116609098
refGene_gn=FAM13B;refGene_tx=XM_023255534.1;refGene_gn=FAM13B;refGene_tx=XM_023255539.1;refGene_gn=FAM13B;refGene_tx=XM_023255543.1;refGene_gn=FAM13B;refGene_tx=XM_023255545.1;refGene_gn=FAM13B;refGene_tx=XM_023255548.1;refGene_gn=FAM13B;refGene_tx=XM_023255556.1;refGene_gn=FAM13B;refGene_tx=XM_023255563.1;refGene_gn=FAM13B;refGene_tx=XM_023255564.1;refGene_gn=FAM13B;refGene_tx=XM_023255566.1
A1 116611254 116611319
refGene_gn=FAM13B;refGene_tx=XM_023255534.1;refGene_gn=FAM13B;refGene_tx=XM_023255539.1;refGene_gn=FAM13B;refGene_tx=XM_023255543.1;refGene_gn=FAM13B;refGene_tx=XM_023255545.1;refGene_gn=FAM13B;refGene_tx=XM_023255548.1;refGene_gn=FAM13B;refGene_tx=XM_023255556.1;refGene_gn=FAM13B;refGene_tx=XM_023255563.1;refGene_gn=FAM13B;refGene_tx=XM_023255564.1;refGene_gn=FAM13B;refGene_tx=XM_023255566.1
A1 116621597 116621730
refGene_gn=FAM13B;refGene_tx=XM_023255534.1;refGene_gn=FAM13B;refGene_tx=XM_023255539.1;refGene_gn=FAM13B;refGene_tx=XM_023255543.1;refGene_gn=FAM13B;refGene_tx=XM_023255545.1;refGene_gn=FAM13B;refGene_tx=XM_023255548.1;refGene_gn=FAM13B;refGene_tx=XM_023255556.1;refGene_gn=FAM13B;refGene_tx=XM_023255563.1;refGene_gn=FAM13B;refGene_tx=XM_023255564.1;refGene_gn=FAM13B;refGene_tx=XM_023255566.1
A1 116623513 116623669
refGene_gn=FAM13B;refGene_tx=XM_023255534.1;refGene_gn=FAM13B;refGene_tx=XM_023255539.1;refGene_gn=FAM13B;refGene_tx=XM_023255543.1;refGene_gn=FAM13B;refGene_tx=XM_023255545.1;refGene_gn=FAM13B;refGene_tx=XM_023255548.1;refGene_gn=FAM13B;refGene_tx=XM_023255556.1;refGene_gn=FAM13B;refGene_tx=XM_023255563.1;refGene_gn=FAM13B;refGene_tx=XM_023255564.1;refGene_gn=FAM13B;refGene_tx=XM_023255566.1
A1 116624399 116624441
refGene_gn=FAM13B;refGene_tx=XM_023255534.1;refGene_gn=FAM13B;refGene_tx=XM_023255539.1;refGene_gn=FAM13B;refGene_tx=XM_023255543.1;refGene_gn=FAM13B;refGene_tx=XM_023255545.1;refGene_gn=FAM13B;refGene_tx=XM_023255548.1;refGene_gn=FAM13B;refGene_tx=XM_023255556.1;refGene_gn=FAM13B;refGene_tx=XM_023255563.1;refGene_gn=FAM13B;refGene_tx=XM_023255564.1;refGene_gn=FAM13B;refGene_tx=XM_023255566.1
A1 116638559 116638720
refGene_gn=FAM13B;refGene_tx=XM_023255534.1;refGene_gn=FAM13B;refGene_tx=XM_023255539.1;refGene_gn=FAM13B;refGene_tx=XM_023255543.1;refGene_gn=FAM13B;refGene_tx=XM_023255545.1;refGene_gn=FAM13B;refGene_tx=XM_023255548.1;refGene_gn=FAM13B;refGene_tx=XM_023255556.1;refGene_gn=FAM13B;refGene_tx=XM_023255563.1;refGene_gn=FAM13B;refGene_tx=XM_023255564.1;refGene_gn=FAM13B;refGene_tx=XM_023255566.1
A1 116641847 116641943
refGene_gn=FAM13B;refGene_tx=XM_023255534.1;refGene_gn=FAM13B;refGene_tx=XM_023255539.1;refGene_gn=FAM13B;refGene_tx=XM_023255543.1;refGene_gn=FAM13B;refGene_tx=XM_023255545.1;refGene_gn=FAM13B;refGene_tx=XM_023255548.1;refGene_gn=FAM13B;refGene_tx=XM_023255563.1
A1 116647253 116647395
refGene_gn=FAM13B;refGene_tx=XM_023255534.1;refGene_gn=FAM13B;refGene_

tx=XM_023255539.1;refGene_gn=FAM13B;refGene_tx=XM_023255543.1;refGene_gn=FAM13B;refGene_tx=XM_023255545.1;refGene_gn=FAM13B;refGene_tx=XM_023255548.1;refGene_gn=FAM13B;refGene_tx=XM_023255556.1;refGene_gn=FAM13B;refGene_tx=XM_023255563.1;refGene_gn=FAM13B;refGene_tx=XM_023255564.1;refGene_gn=FAM13B;refGene_tx=XM_023255566.1
A1 116647700 116647878
refGene_gn=FAM13B;refGene_tx=XM_023255534.1;refGene_gn=FAM13B;refGene_tx=XM_023255539.1;refGene_gn=FAM13B;refGene_tx=XM_023255543.1;refGene_gn=FAM13B;refGene_tx=XM_023255545.1;refGene_gn=FAM13B;refGene_tx=XM_023255548.1;refGene_gn=FAM13B;refGene_tx=XM_023255556.1;refGene_gn=FAM13B;refGene_tx=XM_023255563.1;refGene_gn=FAM13B;refGene_tx=XM_023255564.1;refGene_gn=FAM13B;refGene_tx=XM_023255566.1
A1 116652954 116653167
refGene_gn=FAM13B;refGene_tx=XM_023255534.1;refGene_gn=FAM13B;refGene_tx=XM_023255539.1;refGene_gn=FAM13B;refGene_tx=XM_023255543.1;refGene_gn=FAM13B;refGene_tx=XM_023255545.1;refGene_gn=FAM13B;refGene_tx=XM_023255548.1;refGene_gn=FAM13B;refGene_tx=XM_023255556.1;refGene_gn=FAM13B;refGene_tx=XM_023255563.1;refGene_gn=FAM13B;refGene_tx=XM_023255564.1;refGene_gn=FAM13B;refGene_tx=XM_023255566.1
A1 116653736 116653893
refGene_gn=FAM13B;refGene_tx=XM_023255534.1;refGene_gn=FAM13B;refGene_tx=XM_023255539.1;refGene_gn=FAM13B;refGene_tx=XM_023255543.1;refGene_gn=FAM13B;refGene_tx=XM_023255545.1;refGene_gn=FAM13B;refGene_tx=XM_023255548.1;refGene_gn=FAM13B;refGene_tx=XM_023255556.1;refGene_gn=FAM13B;refGene_tx=XM_023255563.1;refGene_gn=FAM13B;refGene_tx=XM_023255564.1;refGene_gn=FAM13B;refGene_tx=XM_023255566.1
A1 116720769 116720837
refGene_gn=WNT8A;refGene_tx=XM_003980896.2
A1 116720939 116720973
refGene_gn=WNT8A;refGene_tx=XM_003980896.2
A1 116721186 116721325
refGene_gn=WNT8A;refGene_tx=XM_003980896.2
A1 116723190 116723316
refGene_gn=WNT8A;refGene_tx=XM_003980896.2
A1 116724014 116724157
refGene_gn=WNT8A;refGene_tx=XM_003980896.2
A1 116725278 116725800
refGene_gn=WNT8A;refGene_tx=XM_003980896.2
A1 116752556 116752637
refGene_gn=NME5;refGene_tx=XM_023255585.1;refGene_gn=NME5;refGene_tx=XM_023255586.1
A1 116757553 116757672
refGene_gn=NME5;refGene_tx=XM_023255585.1;refGene_gn=NME5;refGene_tx=XM_023255586.1
A1 116764025 116764126
refGene_gn=NME5;refGene_tx=XM_023255585.1;refGene_gn=NME5;refGene_tx=XM_023255586.1
A1 116764794 116764999
refGene_gn=NME5;refGene_tx=XM_023255585.1;refGene_gn=NME5;refGene_tx=XM_023255586.1

A1 116773754 116773883
refGene_gn=NME5;refGene_tx=XM_023255586.1
A1 116776373 116776424
refGene_gn=BRD8;refGene_tx=XM_023242937.1
A1 116776992 116777170
refGene_gn=BRD8;refGene_tx=XM_023242937.1
A1 116781142 116781211
refGene_gn=BRD8;refGene_tx=XM_023242937.1
A1 116781309 116781399
refGene_gn=BRD8;refGene_tx=XM_023242937.1
A1 116782328 116782486
refGene_gn=BRD8;refGene_tx=XM_023242937.1
A1 116783135 116783399
refGene_gn=BRD8;refGene_tx=XM_023242937.1
A1 116784670 116784949
refGene_gn=BRD8;refGene_tx=XM_023242937.1
A1 116789170 116789215
refGene_gn=BRD8;refGene_tx=XM_023242937.1
A1 116789509 116789580
refGene_gn=BRD8;refGene_tx=XM_023242937.1
A1 116789707 116789810
refGene_gn=BRD8;refGene_tx=XM_023242937.1
A1 116790753 116790931
refGene_gn=BRD8;refGene_tx=XM_023242937.1
A1 116791325 116791394
refGene_gn=BRD8;refGene_tx=XM_023242937.1
A1 116791587 116791680
refGene_gn=BRD8;refGene_tx=XM_023242937.1
A1 116791864 116792079
refGene_gn=BRD8;refGene_tx=XM_023242937.1
A1 116792847 116792894
refGene_gn=BRD8;refGene_tx=XM_023242937.1
A1 116793096 116793190
refGene_gn=BRD8;refGene_tx=XM_023242937.1
A1 116796792 116797245
refGene_gn=BRD8;refGene_tx=XM_023242937.1
A1 116797583 116797864
refGene_gn=BRD8;refGene_tx=XM_023242937.1
A1 116798273 116798483
refGene_gn=BRD8;refGene_tx=XM_023242937.1
A1 116799388 116799533
refGene_gn=BRD8;refGene_tx=XM_023242937.1
A1 116800663 116800800
refGene_gn=BRD8;refGene_tx=XM_023242937.1
A1 116801829 116801894
refGene_gn=BRD8;refGene_tx=XM_023242937.1
A1 116802300 116802381
refGene_gn=BRD8;refGene_tx=XM_023242937.1
A1 116802504 116802627
refGene_gn=BRD8;refGene_tx=XM_023242937.1

A1 116802964 116803014
refGene_gn=BRD8;refGene_tx=XM_023242937.1
A1 116804047 116804117
refGene_gn=BRD8;refGene_tx=XM_023242937.1
A1 116809205 116809302
refGene_gn=BRD8;refGene_tx=XM_023242937.1
A1 116810240 116810259
refGene_gn=BRD8;refGene_tx=XM_023242937.1
A1 116811312 116811477
refGene_gn=KIF20A;refGene_tx=XM_003980801.4;refGene_gn=KIF20A;refGene_tx=XM_006927632.3
A1 116812616 116812706
refGene_gn=KIF20A;refGene_tx=XM_003980801.4;refGene_gn=KIF20A;refGene_tx=XM_006927632.3
A1 116812809 116812929
refGene_gn=KIF20A;refGene_tx=XM_003980801.4;refGene_gn=KIF20A;refGene_tx=XM_006927632.3
A1 116813500 116813639
refGene_gn=KIF20A;refGene_tx=XM_003980801.4;refGene_gn=KIF20A;refGene_tx=XM_006927632.3
A1 116813754 116813942
refGene_gn=KIF20A;refGene_tx=XM_003980801.4;refGene_gn=KIF20A;refGene_tx=XM_006927632.3
A1 116814028 116814158
refGene_gn=KIF20A;refGene_tx=XM_003980801.4;refGene_gn=KIF20A;refGene_tx=XM_006927632.3
A1 116814340 116814535
refGene_gn=KIF20A;refGene_tx=XM_003980801.4;refGene_gn=KIF20A;refGene_tx=XM_006927632.3
A1 116814642 116814754
refGene_gn=KIF20A;refGene_tx=XM_003980801.4;refGene_gn=KIF20A;refGene_tx=XM_006927632.3
A1 116814861 116814930
refGene_gn=KIF20A;refGene_tx=XM_003980801.4;refGene_gn=KIF20A;refGene_tx=XM_006927632.3
A1 116815594 116815738
refGene_gn=KIF20A;refGene_tx=XM_003980801.4;refGene_gn=KIF20A;refGene_tx=XM_006927632.3
A1 116815879 116816045
refGene_gn=KIF20A;refGene_tx=XM_003980801.4;refGene_gn=KIF20A;refGene_tx=XM_006927632.3
A1 116816155 116816320
refGene_gn=KIF20A;refGene_tx=XM_003980801.4;refGene_gn=KIF20A;refGene_tx=XM_006927632.3
A1 116816409 116816549
refGene_gn=KIF20A;refGene_tx=XM_003980801.4;refGene_gn=KIF20A;refGene_tx=XM_006927632.3
A1 116818176 116818279
refGene_gn=KIF20A;refGene_tx=XM_003980801.4;refGene_gn=KIF20A;refGene_tx=XM_006927632.3

A1	116818577	116818776
refGene_gn=KIF20A;refGene_tx=XM_003980801.4;refGene_gn=KIF20A;refGene_tx=XM_006927632.3
A1	116819011	116819103
refGene_gn=KIF20A;refGene_tx=XM_003980801.4;refGene_gn=KIF20A;refGene_tx=XM_006927632.3
A1	116819328	116819466
refGene_gn=KIF20A;refGene_tx=XM_003980801.4;refGene_gn=KIF20A;refGene_tx=XM_006927632.3
A1	116820344	116820662
refGene_gn=KIF20A;refGene_tx=XM_003980801.4;refGene_gn=KIF20A;refGene_tx=XM_006927632.3
A1	116822220	116822391
refGene_gn=CDC23;refGene_tx=XM_023243003.1
A1	116822807	116822927
refGene_gn=CDC23;refGene_tx=XM_023243003.1
A1	116823004	116823083
refGene_gn=CDC23;refGene_tx=XM_023243003.1
A1	116823781	116823843
refGene_gn=CDC23;refGene_tx=XM_023243003.1
A1	116824149	116824225
refGene_gn=CDC23;refGene_tx=XM_023243003.1
A1	116824565	116824685
refGene_gn=CDC23;refGene_tx=XM_023243003.1
A1	116824800	116824954
refGene_gn=CDC23;refGene_tx=XM_023243003.1
A1	116833250	116833332
refGene_gn=CDC23;refGene_tx=XM_023243003.1
A1	116834780	116834913
refGene_gn=CDC23;refGene_tx=XM_023243003.1
A1	116835011	116835117
refGene_gn=CDC23;refGene_tx=XM_023243003.1
A1	116835649	116835692
refGene_gn=CDC23;refGene_tx=XM_023243003.1
A1	116837844	116837982
refGene_gn=CDC23;refGene_tx=XM_023243003.1
A1	116840131	116840204
refGene_gn=CDC23;refGene_tx=XM_023243003.1
A1	116840288	116840443
refGene_gn=CDC23;refGene_tx=XM_023243003.1
A1	116870935	116871025
refGene_gn=GFRA3;refGene_tx=XM_003980803.5
A1	116871230	116871319
refGene_gn=GFRA3;refGene_tx=XM_003980803.5
A1	116871659	116871794
refGene_gn=GFRA3;refGene_tx=XM_003980803.5
A1	116871959	116872063
refGene_gn=GFRA3;refGene_tx=XM_003980803.5
A1	116874073	116874386
refGene_gn=GFRA3;refGene_tx=XM_003980803.5

A1 116875688 116875781
refGene_gn=GFRA3;refGene_tx=XM_003980803.5
A1 116879159 116879447
refGene_gn=GFRA3;refGene_tx=XM_003980803.5
A1 116887508 116887566
refGene_gn=GFRA3;refGene_tx=XM_003980803.5
A1 116887567 116887585
refGene_gn=GFRA3;refGene_tx=XM_003980803.5
A1 116894831 116894981
refGene_gn=CDC25C;refGene_tx=XM_023255596.1;refGene_gn=CDC25C;refGene_tx=XM_023255597.1;refGene_gn=CDC25C;refGene_tx=XM_023255598.1;refGene_gn=CDC25C;refGene_tx=XM_023255601.1;refGene_gn=CDC25C;refGene_tx=XM_023255602.1
A1 116895179 116895291
refGene_gn=CDC25C;refGene_tx=XM_023255596.1;refGene_gn=CDC25C;refGene_tx=XM_023255597.1;refGene_gn=CDC25C;refGene_tx=XM_023255598.1;refGene_gn=CDC25C;refGene_tx=XM_023255601.1;refGene_gn=CDC25C;refGene_tx=XM_023255602.1
A1 116895914 116896048
refGene_gn=CDC25C;refGene_tx=XM_023255596.1;refGene_gn=CDC25C;refGene_tx=XM_023255597.1;refGene_gn=CDC25C;refGene_tx=XM_023255598.1;refGene_gn=CDC25C;refGene_tx=XM_023255601.1;refGene_gn=CDC25C;refGene_tx=XM_023255602.1
A1 116896592 116896691
refGene_gn=CDC25C;refGene_tx=XM_023255596.1;refGene_gn=CDC25C;refGene_tx=XM_023255597.1;refGene_gn=CDC25C;refGene_tx=XM_023255598.1;refGene_gn=CDC25C;refGene_tx=XM_023255601.1;refGene_gn=CDC25C;refGene_tx=XM_023255602.1
A1 116897987 116898050
refGene_gn=CDC25C;refGene_tx=XM_023255596.1;refGene_gn=CDC25C;refGene_tx=XM_023255597.1;refGene_gn=CDC25C;refGene_tx=XM_023255598.1;refGene_gn=CDC25C;refGene_tx=XM_023255601.1;refGene_gn=CDC25C;refGene_tx=XM_023255602.1
A1 116901059 116901161
refGene_gn=CDC25C;refGene_tx=XM_023255596.1;refGene_gn=CDC25C;refGene_tx=XM_023255597.1;refGene_gn=CDC25C;refGene_tx=XM_023255598.1;refGene_gn=CDC25C;refGene_tx=XM_023255601.1;refGene_gn=CDC25C;refGene_tx=XM_023255602.1
A1 116902544 116902697
refGene_gn=CDC25C;refGene_tx=XM_023255596.1;refGene_gn=CDC25C;refGene_tx=XM_023255597.1;refGene_gn=CDC25C;refGene_tx=XM_023255601.1;refGene_gn=CDC25C;refGene_tx=XM_023255602.1
A1 116914705 116914864
refGene_gn=CDC25C;refGene_tx=XM_023255596.1;refGene_gn=CDC25C;refGene_tx=XM_023255597.1;refGene_gn=CDC25C;refGene_tx=XM_023255598.1;refGene_gn=CDC25C;refGene_tx=XM_023255601.1;refGene_gn=CDC25C;refGene_tx=XM_023255602.1
A1 116918430 116918529
refGene_gn=CDC25C;refGene_tx=XM_023255596.1;refGene_gn=CDC25C;refGene_tx=XM_023255598.1;refGene_gn=CDC25C;refGene_tx=XM_023255601.1;refGene_

gn=CDC25C;refGene_tx=XM_023255602.1
A1 116919738 116919828
refGene_gn=CDC25C;refGene_tx=XM_023255596.1;refGene_gn=CDC25C;refGene_tx=XM_023255597.1;refGene_gn=CDC25C;refGene_tx=XM_023255598.1;refGene_gn=CDC25C;refGene_tx=XM_023255601.1;refGene_gn=CDC25C;refGene_tx=XM_023255602.1
A1 116919943 116919977
refGene_gn=CDC25C;refGene_tx=XM_023255596.1;refGene_gn=CDC25C;refGene_tx=XM_023255597.1;refGene_gn=CDC25C;refGene_tx=XM_023255598.1;refGene_gn=CDC25C;refGene_tx=XM_023255601.1;refGene_gn=CDC25C;refGene_tx=XM_023255602.1
A1 116921657 116921703
refGene_gn=CDC25C;refGene_tx=XM_023255596.1;refGene_gn=CDC25C;refGene_tx=XM_023255597.1;refGene_gn=CDC25C;refGene_tx=XM_023255598.1;refGene_gn=CDC25C;refGene_tx=XM_023255601.1;refGene_gn=CDC25C;refGene_tx=XM_023255602.1
A1 116923113 116923211
refGene_gn=CDC25C;refGene_tx=XM_023255596.1;refGene_gn=CDC25C;refGene_tx=XM_023255597.1;refGene_gn=CDC25C;refGene_tx=XM_023255598.1;refGene_gn=CDC25C;refGene_tx=XM_023255601.1;refGene_gn=CDC25C;refGene_tx=XM_023255602.1
A1 116924574 116924768
refGene_gn=CDC25C;refGene_tx=XM_023255596.1;refGene_gn=CDC25C;refGene_tx=XM_023255597.1;refGene_gn=CDC25C;refGene_tx=XM_023255598.1
A1 116927237 116927278
refGene_gn=LOC101087194;refGene_tx=XM_023255607.1
A1 116927389 116927494
refGene_gn=LOC101087194;refGene_tx=XM_023255607.1
A1 116928121 116928274
refGene_gn=LOC101087194;refGene_tx=XM_023255607.1
A1 116928416 116928537
refGene_gn=LOC101087194;refGene_tx=XM_023255607.1
A1 116929383 116929466
refGene_gn=LOC101087194;refGene_tx=XM_023255607.1
A1 116929467 116929513
refGene_gn=LOC101087194;refGene_tx=XM_023255607.1
A1 116933888 116933966
refGene_gn=FAM53C;refGene_tx=XM_003980805.5;refGene_gn=FAM53C;refGene_tx=XM_003980806.4
A1 116934360 116934418
refGene_gn=FAM53C;refGene_tx=XM_003980805.5;refGene_gn=FAM53C;refGene_tx=XM_003980806.4
A1 116935940 116936725
refGene_gn=FAM53C;refGene_tx=XM_003980805.5;refGene_gn=FAM53C;refGene_tx=XM_003980806.4
A1 116937634 116937892
refGene_gn=FAM53C;refGene_tx=XM_003980805.5;refGene_gn=FAM53C;refGene_tx=XM_003980806.4
A1 116943984 116944179
refGene_gn=KDM3B;refGene_tx=XM_023255610.1

A1 116960120 116960288
refGene_gn=KDM3B;refGene_tx=XM_023255610.1
A1 116965206 116965320
refGene_gn=KDM3B;refGene_tx=XM_023255610.1
A1 116966839 116966945
refGene_gn=KDM3B;refGene_tx=XM_023255610.1
A1 116968784 116968909
refGene_gn=KDM3B;refGene_tx=XM_023255610.1
A1 116970641 116970716
refGene_gn=KDM3B;refGene_tx=XM_023255610.1
A1 116973089 116973689
refGene_gn=KDM3B;refGene_tx=XM_023255610.1
A1 116977620 116978181
refGene_gn=KDM3B;refGene_tx=XM_023255610.1
A1 116978182 116978389
refGene_gn=KDM3B;refGene_tx=XM_023255610.1
A1 116978390 116978859
refGene_gn=KDM3B;refGene_tx=XM_023255610.1
A1 116979743 116979945
refGene_gn=KDM3B;refGene_tx=XM_023255610.1
A1 116984054 116984269
refGene_gn=KDM3B;refGene_tx=XM_023255610.1
A1 116984967 116985120
refGene_gn=KDM3B;refGene_tx=XM_023255610.1
A1 116995224 116995332
refGene_gn=KDM3B;refGene_tx=XM_023255610.1
A1 116998144 116998272
refGene_gn=KDM3B;refGene_tx=XM_023255610.1
A1 116999404 116999684
refGene_gn=KDM3B;refGene_tx=XM_023255610.1
A1 117000542 117000799
refGene_gn=KDM3B;refGene_tx=XM_023255610.1
A1 117004099 117004366
refGene_gn=KDM3B;refGene_tx=XM_023255610.1
A1 117005150 117005322
refGene_gn=KDM3B;refGene_tx=XM_023255610.1
A1 117005710 117005801
refGene_gn=KDM3B;refGene_tx=XM_023255610.1
A1 117005938 117006069
refGene_gn=KDM3B;refGene_tx=XM_023255610.1
A1 117007051 117007171
refGene_gn=KDM3B;refGene_tx=XM_023255610.1
A1 117008144 117008284
refGene_gn=KDM3B;refGene_tx=XM_023255610.1
A1 117008769 117008946
refGene_gn=KDM3B;refGene_tx=XM_023255610.1
A1 117009439 117009574
refGene_gn=KDM3B;refGene_tx=XM_023255610.1
A1 117012420 117012501
refGene_gn=KDM3B;refGene_tx=XM_023255610.1

A1 117015814 117015846
refGene_gn=REEP2;refGene_tx=XM_023255612.1;refGene_gn=REEP2;refGene_tx=XM_023255614.1
A1 117017699 117017772
refGene_gn=REEP2;refGene_tx=XM_023255612.1;refGene_gn=REEP2;refGene_tx=XM_023255614.1
A1 117018055 117018132
refGene_gn=REEP2;refGene_tx=XM_023255612.1;refGene_gn=REEP2;refGene_tx=XM_023255614.1;refGene_gn=REEP2;refGene_tx=XM_023255616.1
A1 117019826 117019947
refGene_gn=REEP2;refGene_tx=XM_023255612.1;refGene_gn=REEP2;refGene_tx=XM_023255614.1;refGene_gn=REEP2;refGene_tx=XM_023255616.1
A1 117020183 117020297
refGene_gn=REEP2;refGene_tx=XM_023255612.1;refGene_gn=REEP2;refGene_tx=XM_023255614.1;refGene_gn=REEP2;refGene_tx=XM_023255616.1
A1 117020645 117020793
refGene_gn=REEP2;refGene_tx=XM_023255612.1;refGene_gn=REEP2;refGene_tx=XM_023255614.1;refGene_gn=REEP2;refGene_tx=XM_023255616.1
A1 117020900 117021031
refGene_gn=REEP2;refGene_tx=XM_023255612.1;refGene_gn=REEP2;refGene_tx=XM_023255614.1;refGene_gn=REEP2;refGene_tx=XM_023255616.1
A1 117021117 117021186
refGene_gn=REEP2;refGene_tx=XM_023255612.1;refGene_gn=REEP2;refGene_tx=XM_023255614.1;refGene_gn=REEP2;refGene_tx=XM_023255616.1
A1 117036864 117037165
refGene_gn=EGR1;refGene_tx=XM_003980809.4
A1 117037870 117039201
refGene_gn=EGR1;refGene_tx=XM_003980809.4
A1 117071563 117071646
refGene_gn=ETF1;refGene_tx=XM_003980810.5;refGene_gn=ETF1;refGene_tx=XM_011282344.2
A1 117071928 117072076
refGene_gn=ETF1;refGene_tx=XM_003980810.5;refGene_gn=ETF1;refGene_tx=XM_011282344.2
A1 117072853 117072918
refGene_gn=ETF1;refGene_tx=XM_003980810.5;refGene_gn=ETF1;refGene_tx=XM_011282344.2
A1 117073290 117073446
refGene_gn=ETF1;refGene_tx=XM_003980810.5;refGene_gn=ETF1;refGene_tx=XM_011282344.2
A1 117074400 117074530
refGene_gn=ETF1;refGene_tx=XM_003980810.5;refGene_gn=ETF1;refGene_tx=XM_011282344.2
A1 117075366 117075557
refGene_gn=ETF1;refGene_tx=XM_003980810.5;refGene_gn=ETF1;refGene_tx=XM_011282344.2
A1 117076496 117076635
refGene_gn=ETF1;refGene_tx=XM_003980810.5;refGene_gn=ETF1;refGene_tx=XM_011282344.2
A1 117079234 117079374

refGene_gn=ETF1;refGene_tx=XM_003980810.5;refGene_gn=ETF1;refGene_tx=XM_011282344.2
A1 117080116 117080292
refGene_gn=ETF1;refGene_tx=XM_003980810.5;refGene_gn=ETF1;refGene_tx=XM_011282344.2
A1 117096228 117096314
refGene_gn=ETF1;refGene_tx=XM_003980810.5
A1 117108621 117108699
refGene_gn=HSPA9;refGene_tx=XM_003980812.4
A1 117109014 117109155
refGene_gn=HSPA9;refGene_tx=XM_003980812.4
A1 117109450 117109543
refGene_gn=HSPA9;refGene_tx=XM_003980812.4
A1 117109847 117109942
refGene_gn=HSPA9;refGene_tx=XM_003980812.4
A1 117110571 117110689
refGene_gn=HSPA9;refGene_tx=XM_003980812.4
A1 117111285 117111390
refGene_gn=HSPA9;refGene_tx=XM_003980812.4
A1 117112262 117112490
refGene_gn=HSPA9;refGene_tx=XM_003980812.4
A1 117113790 117114000
refGene_gn=HSPA9;refGene_tx=XM_003980812.4
A1 117118132 117118225
refGene_gn=HSPA9;refGene_tx=XM_003980812.4
A1 117118703 117118866
refGene_gn=HSPA9;refGene_tx=XM_003980812.4
A1 117119512 117119619
refGene_gn=HSPA9;refGene_tx=XM_003980812.4
A1 117119705 117119779
refGene_gn=HSPA9;refGene_tx=XM_003980812.4
A1 117120588 117120713
refGene_gn=HSPA9;refGene_tx=XM_003980812.4
A1 117121899 117122081
refGene_gn=HSPA9;refGene_tx=XM_003980812.4
A1 117122954 117123042
refGene_gn=HSPA9;refGene_tx=XM_003980812.4
A1 117123330 117123389
refGene_gn=HSPA9;refGene_tx=XM_003980812.4
A1 117124694 117124775
refGene_gn=HSPA9;refGene_tx=XM_003980812.4
A1 117290130 117290235
refGene_gn=CTNNA1;refGene_tx=XM_011282359.3;refGene_gn=CTNNA1;refGene_tx=XM_023255622.1
A1 117292210 117292406
refGene_gn=CTNNA1;refGene_tx=XM_011282359.3;refGene_gn=CTNNA1;refGene_tx=XM_023255622.1
A1 117317289 117317456
refGene_gn=CTNNA1;refGene_tx=XM_011282359.3;refGene_gn=CTNNA1;refGene_tx=XM_023255622.1

A1 117318204 117318324
refGene_gn=CTNNA1;refGene_tx=XM_011282359.3;refGene_gn=CTNNA1;refGene_tx=XM_023255622.1
A1 117326493 117326763
refGene_gn=CTNNA1;refGene_tx=XM_011282359.3;refGene_gn=CTNNA1;refGene_tx=XM_023255622.1
A1 117329209 117329413
refGene_gn=CTNNA1;refGene_tx=XM_011282359.3;refGene_gn=CTNNA1;refGene_tx=XM_023255622.1
A1 117377800 117379347
refGene_gn=LRRTM2;refGene_tx=XM_003980815.5
A1 117379692 117379696
refGene_gn=LRRTM2;refGene_tx=XM_003980815.5
A1 117391370 117391451
refGene_gn=CTNNA1;refGene_tx=XM_011282359.3;refGene_gn=CTNNA1;refGene_tx=XM_023255622.1
A1 117392763 117392916
refGene_gn=CTNNA1;refGene_tx=XM_011282359.3;refGene_gn=CTNNA1;refGene_tx=XM_023255622.1
A1 117406507 117406600
refGene_gn=CTNNA1;refGene_tx=XM_011282359.3;refGene_gn=CTNNA1;refGene_tx=XM_023255622.1
A1 117418745 117418902
refGene_gn=CTNNA1;refGene_tx=XM_011282359.3;refGene_gn=CTNNA1;refGene_tx=XM_023255622.1
A1 117426530 117426731
refGene_gn=CTNNA1;refGene_tx=XM_011282359.3;refGene_gn=CTNNA1;refGene_tx=XM_023255622.1
A1 117427307 117427459
refGene_gn=CTNNA1;refGene_tx=XM_011282359.3;refGene_gn=CTNNA1;refGene_tx=XM_023255622.1
A1 117431854 117431965
refGene_gn=CTNNA1;refGene_tx=XM_011282359.3;refGene_gn=CTNNA1;refGene_tx=XM_023255622.1
A1 117433071 117433253
refGene_gn=CTNNA1;refGene_tx=XM_011282359.3;refGene_gn=CTNNA1;refGene_tx=XM_023255622.1
A1 117433371 117433477
refGene_gn=CTNNA1;refGene_tx=XM_011282359.3;refGene_gn=CTNNA1;refGene_tx=XM_023255622.1
A1 117435478 117435613
refGene_gn=CTNNA1;refGene_tx=XM_011282359.3;refGene_gn=CTNNA1;refGene_tx=XM_023255622.1
A1 117436490 117436778
refGene_gn=CTNNA1;refGene_tx=XM_011282359.3;refGene_gn=CTNNA1;refGene_tx=XM_023255622.1
A1 117451725 117452082
refGene_gn=SIL1;refGene_tx=XM_006927639.4;refGene_gn=SIL1;refGene_tx=XM_019832833.2
A1 117455668 117455833

refGene_gn=SIL1;refGene_tx=XM_006927639.4;refGene_gn=SIL1;refGene_tx=XM_019832833.2
A1 117456265 117456362
refGene_gn=SIL1;refGene_tx=XM_006927639.4;refGene_gn=SIL1;refGene_tx=XM_019832833.2
A1 117524352 117524474
refGene_gn=SIL1;refGene_tx=XM_006927639.4;refGene_gn=SIL1;refGene_tx=XM_019832833.2
A1 117530235 117530427
refGene_gn=SIL1;refGene_tx=XM_006927639.4;refGene_gn=SIL1;refGene_tx=XM_019832833.2
A1 117542281 117542381
refGene_gn=SIL1;refGene_tx=XM_006927639.4;refGene_gn=SIL1;refGene_tx=XM_019832833.2
A1 117548568 117548689
refGene_gn=SIL1;refGene_tx=XM_006927639.4;refGene_gn=SIL1;refGene_tx=XM_019832833.2
A1 117610044 117610180
refGene_gn=SIL1;refGene_tx=XM_006927639.4;refGene_gn=SIL1;refGene_tx=XM_019832833.2
A1 117616687 117616762
refGene_gn=SIL1;refGene_tx=XM_006927639.4
A1 117667572 117667653
refGene_gn=SIL1;refGene_tx=XM_019832833.2
A1 117746870 117747782
refGene_gn=MATR3;refGene_tx=XM_011282371.3;refGene_gn=MATR3;refGene_tx=XM_011282373.3;refGene_gn=MATR3;refGene_tx=XM_011282379.3;refGene_gn=MATR3;refGene_tx=XM_023255657.1;refGene_gn=MATR3;refGene_tx=XM_023255658.1
A1 117752596 117752658
refGene_gn=MATR3;refGene_tx=XM_011282371.3;refGene_gn=MATR3;refGene_tx=XM_011282373.3;refGene_gn=MATR3;refGene_tx=XM_011282379.3;refGene_gn=MATR3;refGene_tx=XM_023255657.1;refGene_gn=MATR3;refGene_tx=XM_023255658.1
A1 117753612 117753654
refGene_gn=MATR3;refGene_tx=XM_011282371.3;refGene_gn=MATR3;refGene_tx=XM_011282373.3;refGene_gn=MATR3;refGene_tx=XM_011282379.3;refGene_gn=MATR3;refGene_tx=XM_023255657.1;refGene_gn=MATR3;refGene_tx=XM_023255658.1
A1 117753989 117754102
refGene_gn=MATR3;refGene_tx=XM_011282371.3;refGene_gn=MATR3;refGene_tx=XM_011282373.3;refGene_gn=MATR3;refGene_tx=XM_011282379.3;refGene_gn=MATR3;refGene_tx=XM_023255657.1;refGene_gn=MATR3;refGene_tx=XM_023255658.1
A1 117754324 117754377
refGene_gn=MATR3;refGene_tx=XM_011282371.3;refGene_gn=MATR3;refGene_tx=XM_011282373.3;refGene_gn=MATR3;refGene_tx=XM_011282379.3;refGene_gn=MATR3;refGene_tx=XM_023255657.1;refGene_gn=MATR3;refGene_tx=XM_023255658.1
A1 117755019 117755145

refGene_gn=MATR3;refGene_tx=XM_011282371.3;refGene_gn=MATR3;refGene_tx=XM_011282373.3;refGene_gn=MATR3;refGene_tx=XM_011282379.3;refGene_gn=MATR3;refGene_tx=XM_023255657.1;refGene_gn=MATR3;refGene_tx=XM_023255658.1
A1 117755665 117755791
refGene_gn=MATR3;refGene_tx=XM_011282371.3;refGene_gn=MATR3;refGene_tx=XM_011282373.3;refGene_gn=MATR3;refGene_tx=XM_011282379.3;refGene_gn=MATR3;refGene_tx=XM_023255657.1;refGene_gn=MATR3;refGene_tx=XM_023255658.1
A1 117755983 117756151
refGene_gn=MATR3;refGene_tx=XM_011282371.3;refGene_gn=MATR3;refGene_tx=XM_011282373.3;refGene_gn=MATR3;refGene_tx=XM_011282379.3;refGene_gn=MATR3;refGene_tx=XM_023255657.1;refGene_gn=MATR3;refGene_tx=XM_023255658.1
A1 117758171 117758303
refGene_gn=MATR3;refGene_tx=XM_011282371.3;refGene_gn=MATR3;refGene_tx=XM_011282373.3;refGene_gn=MATR3;refGene_tx=XM_011282379.3;refGene_gn=MATR3;refGene_tx=XM_023255657.1;refGene_gn=MATR3;refGene_tx=XM_023255658.1
A1 117759201 117759245
refGene_gn=MATR3;refGene_tx=XM_011282371.3;refGene_gn=MATR3;refGene_tx=XM_011282373.3;refGene_gn=MATR3;refGene_tx=XM_011282379.3;refGene_gn=MATR3;refGene_tx=XM_023255657.1;refGene_gn=MATR3;refGene_tx=XM_023255658.1
A1 117759334 117759710
refGene_gn=MATR3;refGene_tx=XM_011282371.3;refGene_gn=MATR3;refGene_tx=XM_011282373.3;refGene_gn=MATR3;refGene_tx=XM_011282379.3;refGene_gn=MATR3;refGene_tx=XM_023255657.1;refGene_gn=MATR3;refGene_tx=XM_023255658.1
A1 117761297 117761664
refGene_gn=MATR3;refGene_tx=XM_011282371.3;refGene_gn=MATR3;refGene_tx=XM_011282373.3;refGene_gn=MATR3;refGene_tx=XM_011282379.3;refGene_gn=MATR3;refGene_tx=XM_023255657.1;refGene_gn=MATR3;refGene_tx=XM_023255658.1
A1 117762141 117762263
refGene_gn=MATR3;refGene_tx=XM_011282371.3;refGene_gn=MATR3;refGene_tx=XM_011282373.3;refGene_gn=MATR3;refGene_tx=XM_011282379.3;refGene_gn=MATR3;refGene_tx=XM_023255657.1;refGene_gn=MATR3;refGene_tx=XM_023255658.1
A1 117764075 117764126
refGene_gn=MATR3;refGene_tx=XM_011282371.3;refGene_gn=MATR3;refGene_tx=XM_011282373.3;refGene_gn=MATR3;refGene_tx=XM_011282379.3;refGene_gn=MATR3;refGene_tx=XM_023255657.1;refGene_gn=MATR3;refGene_tx=XM_023255658.1
A1 117788113 117788455
refGene_gn=LOC101090836;refGene_tx=XM_003980819.5
A1 117790343 117790481
refGene_gn=PAIP2;refGene_tx=XM_003980817.5;refGene_gn=PAIP2;refGene_tx=XM_019832886.2
A1 117791866 117792046

refGene_gn=PAIP2;refGene_tx=XM_003980817.5;refGene_gn=PAIP2;refGene_tx=XM_019832886.2
A1 117793766 117793810
refGene_gn=SLC23A1;refGene_tx=XM_019832881.2
A1 117794378 117794435
refGene_gn=PAIP2;refGene_tx=XM_003980817.5;refGene_gn=PAIP2;refGene_tx=XM_019832886.2
A1 117798622 117798867
refGene_gn=SLC23A1;refGene_tx=XM_003980820.4;refGene_gn=SLC23A1;refGene_tx=XM_019832872.1;refGene_gn=SLC23A1;refGene_tx=XM_019832879.2;refGene_gn=SLC23A1;refGene_tx=XM_019832883.2
A1 117801352 117801448
refGene_gn=SLC23A1;refGene_tx=XM_003980820.4;refGene_gn=SLC23A1;refGene_tx=XM_019832872.1;refGene_gn=SLC23A1;refGene_tx=XM_019832881.2
A1 117801955 117802116
refGene_gn=SLC23A1;refGene_tx=XM_003980820.4;refGene_gn=SLC23A1;refGene_tx=XM_019832872.1;refGene_gn=SLC23A1;refGene_tx=XM_019832879.2;refGene_gn=SLC23A1;refGene_tx=XM_019832881.2;refGene_gn=SLC23A1;refGene_tx=XM_023255683.1
A1 117802198 117802328
refGene_gn=SLC23A1;refGene_tx=XM_003980820.4;refGene_gn=SLC23A1;refGene_tx=XM_019832872.1;refGene_gn=SLC23A1;refGene_tx=XM_019832879.2;refGene_gn=SLC23A1;refGene_tx=XM_019832881.2;refGene_gn=SLC23A1;refGene_tx=XM_019832883.2;refGene_gn=SLC23A1;refGene_tx=XM_023255683.1
A1 117802569 117802675
refGene_gn=SLC23A1;refGene_tx=XM_003980820.4;refGene_gn=SLC23A1;refGene_tx=XM_019832872.1;refGene_gn=SLC23A1;refGene_tx=XM_019832879.2;refGene_gn=SLC23A1;refGene_tx=XM_019832881.2;refGene_gn=SLC23A1;refGene_tx=XM_019832883.2;refGene_gn=SLC23A1;refGene_tx=XM_023255683.1
A1 117803224 117803372
refGene_gn=SLC23A1;refGene_tx=XM_003980820.4;refGene_gn=SLC23A1;refGene_tx=XM_019832872.1;refGene_gn=SLC23A1;refGene_tx=XM_019832879.2;refGene_gn=SLC23A1;refGene_tx=XM_019832881.2;refGene_gn=SLC23A1;refGene_tx=XM_019832883.2;refGene_gn=SLC23A1;refGene_tx=XM_023255683.1
A1 117803674 117803831
refGene_gn=SLC23A1;refGene_tx=XM_003980820.4;refGene_gn=SLC23A1;refGene_tx=XM_019832872.1;refGene_gn=SLC23A1;refGene_tx=XM_019832879.2;refGene_gn=SLC23A1;refGene_tx=XM_019832881.2;refGene_gn=SLC23A1;refGene_tx=XM_019832883.2;refGene_gn=SLC23A1;refGene_tx=XM_023255683.1
A1 117804127 117804248
refGene_gn=SLC23A1;refGene_tx=XM_003980820.4;refGene_gn=SLC23A1;refGene_tx=XM_019832872.1;refGene_gn=SLC23A1;refGene_tx=XM_019832879.2;refGene_gn=SLC23A1;refGene_tx=XM_019832881.2;refGene_gn=SLC23A1;refGene_tx=XM_019832883.2;refGene_gn=SLC23A1;refGene_tx=XM_023255683.1
A1 117804359 117804541
refGene_gn=SLC23A1;refGene_tx=XM_003980820.4;refGene_gn=SLC23A1;refGene_tx=XM_019832872.1;refGene_gn=SLC23A1;refGene_tx=XM_019832879.2;refGene_gn=SLC23A1;refGene_tx=XM_019832881.2;refGene_gn=SLC23A1;refGene_tx=XM_019832883.2;refGene_gn=SLC23A1;refGene_tx=XM_023255683.1
A1 117804724 117804792

refGene_gn=SLC23A1;refGene_tx=XM_003980820.4;refGene_gn=SLC23A1;refGene_tx=XM_019832872.1;refGene_gn=SLC23A1;refGene_tx=XM_019832879.2;refGene_gn=SLC23A1;refGene_tx=XM_019832881.2;refGene_gn=SLC23A1;refGene_tx=XM_019832883.2;refGene_gn=SLC23A1;refGene_tx=XM_023255683.1
A1 117804952 117805041
refGene_gn=SLC23A1;refGene_tx=XM_003980820.4;refGene_gn=SLC23A1;refGene_tx=XM_019832872.1;refGene_gn=SLC23A1;refGene_tx=XM_019832879.2;refGene_gn=SLC23A1;refGene_tx=XM_019832881.2;refGene_gn=SLC23A1;refGene_tx=XM_019832883.2;refGene_gn=SLC23A1;refGene_tx=XM_023255683.1
A1 117805783 117805941
refGene_gn=SLC23A1;refGene_tx=XM_003980820.4;refGene_gn=SLC23A1;refGene_tx=XM_019832872.1;refGene_gn=SLC23A1;refGene_tx=XM_019832879.2;refGene_gn=SLC23A1;refGene_tx=XM_019832881.2;refGene_gn=SLC23A1;refGene_tx=XM_019832883.2;refGene_gn=SLC23A1;refGene_tx=XM_023255683.1
A1 117806389 117806524
refGene_gn=SLC23A1;refGene_tx=XM_003980820.4;refGene_gn=SLC23A1;refGene_tx=XM_019832872.1;refGene_gn=SLC23A1;refGene_tx=XM_019832879.2;refGene_gn=SLC23A1;refGene_tx=XM_019832881.2;refGene_gn=SLC23A1;refGene_tx=XM_019832883.2;refGene_gn=SLC23A1;refGene_tx=XM_023255683.1
A1 117807164 117807226
refGene_gn=SLC23A1;refGene_tx=XM_003980820.4;refGene_gn=SLC23A1;refGene_tx=XM_019832872.1;refGene_gn=SLC23A1;refGene_tx=XM_019832879.2;refGene_gn=SLC23A1;refGene_tx=XM_019832881.2;refGene_gn=SLC23A1;refGene_tx=XM_019832883.2;refGene_gn=SLC23A1;refGene_tx=XM_023255683.1
A1 117808313 117808395
refGene_gn=SLC23A1;refGene_tx=XM_003980820.4;refGene_gn=SLC23A1;refGene_tx=XM_019832872.1;refGene_gn=SLC23A1;refGene_tx=XM_019832879.2;refGene_gn=SLC23A1;refGene_tx=XM_019832881.2;refGene_gn=SLC23A1;refGene_tx=XM_019832883.2;refGene_gn=SLC23A1;refGene_tx=XM_023255683.1
A1 117810238 117810395
refGene_gn=MZB1;refGene_tx=XM_003980821.4
A1 117810492 117810603
refGene_gn=MZB1;refGene_tx=XM_003980821.4
A1 117810971 117811096
refGene_gn=MZB1;refGene_tx=XM_003980821.4
A1 117812243 117812447
refGene_gn=MZB1;refGene_tx=XM_003980821.4
A1 117814604 117817670
refGene_gn=PROB1;refGene_tx=XM_006927702.4
A1 117819462 117819802
refGene_gn=SPATA24;refGene_tx=XM_003980822.4;refGene_gn=SPATA24;refGene_tx=XM_006927648.3;refGene_gn=SPATA24;refGene_tx=XM_019832934.2;refGene_gn=SPATA24;refGene_tx=XM_019832938.2;refGene_gn=SPATA24;refGene_tx=XM_023255715.1
A1 117819883 117819986
refGene_gn=SPATA24;refGene_tx=XM_003980822.4;refGene_gn=SPATA24;refGene_tx=XM_006927648.3;refGene_gn=SPATA24;refGene_tx=XM_019832934.2;refGene_gn=SPATA24;refGene_tx=XM_019832938.2;refGene_gn=SPATA24;refGene_tx=XM_023255715.1
A1 117822880 117822951

refGene_gn=SPATA24;refGene_tx=XM_003980822.4;refGene_gn=SPATA24;refGene_tx=XM_006927648.3;refGene_gn=SPATA24;refGene_tx=XM_019832934.2;refGene_gn=SPATA24;refGene_tx=XM_019832938.2;refGene_gn=SPATA24;refGene_tx=XM_023255715.1
A1 117823043 117823174
refGene_gn=SPATA24;refGene_tx=XM_003980822.4;refGene_gn=SPATA24;refGene_tx=XM_006927648.3;refGene_gn=SPATA24;refGene_tx=XM_019832934.2;refGene_gn=SPATA24;refGene_tx=XM_019832938.2;refGene_gn=SPATA24;refGene_tx=XM_023255715.1
A1 117823401 117823467
refGene_gn=SPATA24;refGene_tx=XM_003980822.4;refGene_gn=SPATA24;refGene_tx=XM_006927648.3;refGene_gn=SPATA24;refGene_tx=XM_019832934.2;refGene_gn=SPATA24;refGene_tx=XM_019832938.2;refGene_gn=SPATA24;refGene_tx=XM_023255715.1
A1 117824298 117824304
refGene_gn=SPATA24;refGene_tx=XM_019832938.2
A1 117824644 117824761
refGene_gn=SPATA24;refGene_tx=XM_003980822.4;refGene_gn=SPATA24;refGene_tx=XM_006927648.3;refGene_gn=SPATA24;refGene_tx=XM_019832934.2;refGene_gn=SPATA24;refGene_tx=XM_023255715.1
A1 117834243 117834368
refGene_gn=DNAJC18;refGene_tx=XM_006927645.4;refGene_gn=DNAJC18;refGene_tx=XM_006927646.4;refGene_gn=DNAJC18;refGene_tx=XM_019832929.2;refGene_gn=DNAJC18;refGene_tx=XM_023255711.1
A1 117837378 117837500
refGene_gn=DNAJC18;refGene_tx=XM_019832925.2
A1 117839404 117839577
refGene_gn=DNAJC18;refGene_tx=XM_006927645.4;refGene_gn=DNAJC18;refGene_tx=XM_006927646.4;refGene_gn=DNAJC18;refGene_tx=XM_019832925.2;refGene_gn=DNAJC18;refGene_tx=XM_019832929.2;refGene_gn=DNAJC18;refGene_tx=XM_023255711.1
A1 117841810 117841920
refGene_gn=DNAJC18;refGene_tx=XM_006927645.4;refGene_gn=DNAJC18;refGene_tx=XM_006927646.4;refGene_gn=DNAJC18;refGene_tx=XM_019832925.2;refGene_gn=DNAJC18;refGene_tx=XM_019832929.2;refGene_gn=DNAJC18;refGene_tx=XM_023255711.1
A1 117842915 117843031
refGene_gn=DNAJC18;refGene_tx=XM_006927645.4;refGene_gn=DNAJC18;refGene_tx=XM_006927646.4;refGene_gn=DNAJC18;refGene_tx=XM_019832925.2;refGene_gn=DNAJC18;refGene_tx=XM_019832929.2;refGene_gn=DNAJC18;refGene_tx=XM_023255711.1
A1 117844195 117844381
refGene_gn=DNAJC18;refGene_tx=XM_006927645.4;refGene_gn=DNAJC18;refGene_tx=XM_006927646.4;refGene_gn=DNAJC18;refGene_tx=XM_019832925.2;refGene_gn=DNAJC18;refGene_tx=XM_019832929.2;refGene_gn=DNAJC18;refGene_tx=XM_023255711.1
A1 117847514 117847660
refGene_gn=DNAJC18;refGene_tx=XM_006927645.4;refGene_gn=DNAJC18;refGene_tx=XM_006927646.4;refGene_gn=DNAJC18;refGene_tx=XM_019832925.2;refGene_gn=DNAJC18;refGene_tx=XM_019832929.2;refGene_gn=DNAJC18;refGene_tx=

XM_023255711.1
A1 117854317 117854504
refGene_gn=DNAJC18;refGene_tx=XM_006927645.4;refGene_gn=DNAJC18;refGene_tx=XM_006927646.4;refGene_gn=DNAJC18;refGene_tx=XM_019832925.2;refGene_gn=DNAJC18;refGene_tx=XM_019832929.2;refGene_gn=DNAJC18;refGene_tx=XM_023255711.1
A1 117855920 117855927
refGene_gn=DNAJC18;refGene_tx=XM_019832929.2
A1 117856538 117856578
refGene_gn=DNAJC18;refGene_tx=XM_006927645.4;refGene_gn=DNAJC18;refGene_tx=XM_006927646.4;refGene_gn=DNAJC18;refGene_tx=XM_019832925.2
A1 117862065 117862074
refGene_gn=ECSCR;refGene_tx=XM_006927641.2;refGene_gn=ECSCR;refGene_tx=XM_006927642.2;refGene_gn=ECSCR;refGene_tx=XM_006927643.2;refGene_gn=ECSCR;refGene_tx=XM_006927644.2
A1 117862243 117862340
refGene_gn=ECSCR;refGene_tx=XM_006927641.2;refGene_gn=ECSCR;refGene_tx=XM_006927642.2;refGene_gn=ECSCR;refGene_tx=XM_006927643.2;refGene_gn=ECSCR;refGene_tx=XM_006927644.2
A1 117863495 117863532
refGene_gn=ECSCR;refGene_tx=XM_006927641.2;refGene_gn=ECSCR;refGene_tx=XM_006927642.2;refGene_gn=ECSCR;refGene_tx=XM_006927643.2;refGene_gn=ECSCR;refGene_tx=XM_006927644.2
A1 117863751 117863850
refGene_gn=ECSCR;refGene_tx=XM_006927641.2;refGene_gn=ECSCR;refGene_tx=XM_006927642.2;refGene_gn=ECSCR;refGene_tx=XM_006927643.2;refGene_gn=ECSCR;refGene_tx=XM_006927644.2
A1 117864248 117864362
refGene_gn=ECSCR;refGene_tx=XM_006927641.2;refGene_gn=ECSCR;refGene_tx=XM_006927642.2;refGene_gn=ECSCR;refGene_tx=XM_006927643.2;refGene_gn=ECSCR;refGene_tx=XM_006927644.2
A1 117865791 117865926
refGene_gn=ECSCR;refGene_tx=XM_006927641.2;refGene_gn=ECSCR;refGene_tx=XM_006927642.2;refGene_gn=ECSCR;refGene_tx=XM_006927643.2;refGene_gn=ECSCR;refGene_tx=XM_006927644.2
A1 117866645 117866705
refGene_gn=ECSCR;refGene_tx=XM_006927641.2;refGene_gn=ECSCR;refGene_tx=XM_006927642.2
A1 117866866 117866917
refGene_gn=ECSCR;refGene_tx=XM_006927641.2;refGene_gn=ECSCR;refGene_tx=XM_006927642.2;refGene_gn=ECSCR;refGene_tx=XM_006927643.2;refGene_gn=ECSCR;refGene_tx=XM_006927644.2
A1 117867223 117867268
refGene_gn=ECSCR;refGene_tx=XM_006927641.2;refGene_gn=ECSCR;refGene_tx=XM_006927643.2
A1 117870446 117870507
refGene_gn=ECSCR;refGene_tx=XM_006927641.2;refGene_gn=ECSCR;refGene_tx=XM_006927642.2;refGene_gn=ECSCR;refGene_tx=XM_006927643.2;refGene_gn=ECSCR;refGene_tx=XM_006927644.2
A1 117879854 117880045

refGene_gn=TMEM173;refGene_tx=XM_006927649.4;refGene_gn=TMEM173;refGene_tx=XM_023255699.1
A1 117880622 117880809
refGene_gn=TMEM173;refGene_tx=XM_006927649.4;refGene_gn=TMEM173;refGene_tx=XM_023255699.1
A1 117881525 117881764
refGene_gn=TMEM173;refGene_tx=XM_006927649.4;refGene_gn=TMEM173;refGene_tx=XM_023255699.1
A1 117882930 117883039
refGene_gn=TMEM173;refGene_tx=XM_006927649.4;refGene_gn=TMEM173;refGene_tx=XM_023255699.1
A1 117883289 117883470
refGene_gn=TMEM173;refGene_tx=XM_006927649.4;refGene_gn=TMEM173;refGene_tx=XM_023255699.1
A1 117883590 117883817
refGene_gn=TMEM173;refGene_tx=XM_023255699.1
A1 117884325 117884561
refGene_gn=TMEM173;refGene_tx=XM_006927649.4
A1 117921293 117921317
refGene_gn=UBE2D2;refGene_tx=XM_023255744.1
A1 117948609 117948615
refGene_gn=UBE2D2;refGene_tx=XM_006927651.3
A1 117959199 117959263
refGene_gn=UBE2D2;refGene_tx=XM_006927651.3;refGene_gn=UBE2D2;refGene_tx=XM_023255744.1;refGene_gn=UBE2D2;refGene_tx=XM_023255748.1
A1 117966630 117966631
refGene_gn=UBE2D2;refGene_tx=XM_023255749.1
A1 117967339 117967371
refGene_gn=UBE2D2;refGene_tx=XM_006927651.3;refGene_gn=UBE2D2;refGene_tx=XM_023255744.1;refGene_gn=UBE2D2;refGene_tx=XM_023255748.1;refGene_gn=UBE2D2;refGene_tx=XM_023255749.1
A1 117967449 117967527
refGene_gn=UBE2D2;refGene_tx=XM_006927651.3;refGene_gn=UBE2D2;refGene_tx=XM_023255744.1;refGene_gn=UBE2D2;refGene_tx=XM_023255748.1;refGene_gn=UBE2D2;refGene_tx=XM_023255749.1
A1 117967615 117967721
refGene_gn=UBE2D2;refGene_tx=XM_006927651.3;refGene_gn=UBE2D2;refGene_tx=XM_023255744.1;refGene_gn=UBE2D2;refGene_tx=XM_023255748.1;refGene_gn=UBE2D2;refGene_tx=XM_023255749.1
A1 117973247 117973341
refGene_gn=UBE2D2;refGene_tx=XM_006927651.3;refGene_gn=UBE2D2;refGene_tx=XM_023255744.1;refGene_gn=UBE2D2;refGene_tx=XM_023255748.1;refGene_gn=UBE2D2;refGene_tx=XM_023255749.1
A1 117975736 117975782
refGene_gn=UBE2D2;refGene_tx=XM_006927651.3;refGene_gn=UBE2D2;refGene_tx=XM_023255744.1;refGene_gn=UBE2D2;refGene_tx=XM_023255748.1;refGene_gn=UBE2D2;refGene_tx=XM_023255749.1
A1 118028694 118029627
refGene_gn=CXXC5;refGene_tx=XM_003980825.5;refGene_gn=CXXC5;refGene_tx=XM_023255754.1;refGene_gn=CXXC5;refGene_tx=XM_023255755.1;refGene_gn=

CXXC5;refGene_tx=XM_023255761.1
A1 118031130 118031175
refGene_gn=CXXC5;refGene_tx=XM_003980825.5;refGene_gn=CXXC5;refGene_tx=XM_023255754.1;refGene_gn=CXXC5;refGene_tx=XM_023255755.1
A1 118136047 118136421
refGene_gn=PSD2;refGene_tx=XM_019833014.2
A1 118139498 118139742
refGene_gn=PSD2;refGene_tx=XM_019833014.2
A1 118139743 118139967
refGene_gn=PSD2;refGene_tx=XM_019833014.2
A1 118140393 118140588
refGene_gn=PSD2;refGene_tx=XM_019833014.2
A1 118144019 118144100
refGene_gn=PSD2;refGene_tx=XM_019833014.2
A1 118148922 118149035
refGene_gn=PSD2;refGene_tx=XM_019833014.2
A1 118149760 118149819
refGene_gn=PSD2;refGene_tx=XM_019833014.2
A1 118159738 118159828
refGene_gn=PSD2;refGene_tx=XM_019833014.2
A1 118161652 118161696
refGene_gn=PSD2;refGene_tx=XM_019833014.2
A1 118162751 118162942
refGene_gn=PSD2;refGene_tx=XM_019833014.2
A1 118163115 118163186
refGene_gn=PSD2;refGene_tx=XM_019833014.2
A1 118163612 118163770
refGene_gn=PSD2;refGene_tx=XM_019833014.2
A1 118164677 118164822
refGene_gn=PSD2;refGene_tx=XM_019833014.2
A1 118166067 118166211
refGene_gn=PSD2;refGene_tx=XM_019833014.2
A1 118169581 118169791
refGene_gn=PSD2;refGene_tx=XM_019833014.2
A1 118175269 118176023
refGene_gn=NRG2;refGene_tx=XM_023255767.1;refGene_gn=NRG2;refGene_tx=XM_023255770.1;refGene_gn=NRG2;refGene_tx=XM_023255773.1;refGene_gn=NRG2;refGene_tx=XM_023255774.1;refGene_gn=NRG2;refGene_tx=XM_023255775.1;refGene_gn=NRG2;refGene_tx=XM_023255776.1
A1 118178147 118178375
refGene_gn=NRG2;refGene_tx=XM_023255767.1;refGene_gn=NRG2;refGene_tx=XM_023255770.1;refGene_gn=NRG2;refGene_tx=XM_023255773.1;refGene_gn=NRG2;refGene_tx=XM_023255774.1;refGene_gn=NRG2;refGene_tx=XM_023255775.1;refGene_gn=NRG2;refGene_tx=XM_023255776.1
A1 118179198 118179326
refGene_gn=NRG2;refGene_tx=XM_023255767.1;refGene_gn=NRG2;refGene_tx=XM_023255770.1;refGene_gn=NRG2;refGene_tx=XM_023255773.1;refGene_gn=NRG2;refGene_tx=XM_023255774.1;refGene_gn=NRG2;refGene_tx=XM_023255775.1;refGene_gn=NRG2;refGene_tx=XM_023255776.1
A1 118179651 118179775

refGene_gn=NRG2;refGene_tx=XM_023255767.1;refGene_gn=NRG2;refGene_tx=XM_023255770.1;refGene_gn=NRG2;refGene_tx=XM_023255773.1;refGene_gn=NRG2;refGene_tx=XM_023255774.1;refGene_gn=NRG2;refGene_tx=XM_023255775.1;refGene_gn=NRG2;refGene_tx=XM_023255776.1
A1 118182153 118182256
refGene_gn=NRG2;refGene_tx=XM_023255767.1;refGene_gn=NRG2;refGene_tx=XM_023255770.1;refGene_gn=NRG2;refGene_tx=XM_023255773.1;refGene_gn=NRG2;refGene_tx=XM_023255774.1;refGene_gn=NRG2;refGene_tx=XM_023255775.1;refGene_gn=NRG2;refGene_tx=XM_023255776.1
A1 118186011 118186183
refGene_gn=NRG2;refGene_tx=XM_023255767.1;refGene_gn=NRG2;refGene_tx=XM_023255770.1;refGene_gn=NRG2;refGene_tx=XM_023255775.1;refGene_gn=NRG2;refGene_tx=XM_023255776.1
A1 118190489 118190548
refGene_gn=NRG2;refGene_tx=XM_023255767.1;refGene_gn=NRG2;refGene_tx=XM_023255770.1;refGene_gn=NRG2;refGene_tx=XM_023255774.1;refGene_gn=NRG2;refGene_tx=XM_023255775.1;refGene_gn=NRG2;refGene_tx=XM_023255776.1
A1 118190911 118190988
refGene_gn=NRG2;refGene_tx=XM_023255767.1;refGene_gn=NRG2;refGene_tx=XM_023255773.1;refGene_gn=NRG2;refGene_tx=XM_023255775.1;refGene_gn=NRG2;refGene_tx=XM_023255776.1
A1 118196993 118197114
refGene_gn=NRG2;refGene_tx=XM_023255767.1;refGene_gn=NRG2;refGene_tx=XM_023255770.1;refGene_gn=NRG2;refGene_tx=XM_023255773.1;refGene_gn=NRG2;refGene_tx=XM_023255774.1;refGene_gn=NRG2;refGene_tx=XM_023255775.1;refGene_gn=NRG2;refGene_tx=XM_023255776.1
A1 118206431 118206550
refGene_gn=NRG2;refGene_tx=XM_023255767.1;refGene_gn=NRG2;refGene_tx=XM_023255770.1;refGene_gn=NRG2;refGene_tx=XM_023255773.1;refGene_gn=NRG2;refGene_tx=XM_023255774.1;refGene_gn=NRG2;refGene_tx=XM_023255775.1;refGene_gn=NRG2;refGene_tx=XM_023255776.1
A1 118212172 118212344
refGene_gn=NRG2;refGene_tx=XM_023255767.1;refGene_gn=NRG2;refGene_tx=XM_023255770.1;refGene_gn=NRG2;refGene_tx=XM_023255773.1;refGene_gn=NRG2;refGene_tx=XM_023255774.1;refGene_gn=NRG2;refGene_tx=XM_023255775.1;refGene_gn=NRG2;refGene_tx=XM_023255776.1
A1 118231696 118231787
refGene_gn=NRG2;refGene_tx=XM_023255775.1
A1 118321412 118321431
refGene_gn=NRG2;refGene_tx=XM_023255776.1
A1 118361287 118362026
refGene_gn=NRG2;refGene_tx=XM_023255767.1;refGene_gn=NRG2;refGene_tx=XM_023255770.1;refGene_gn=NRG2;refGene_tx=XM_023255773.1;refGene_gn=NRG2;refGene_tx=XM_023255774.1
A1 118425130 118425505
refGene_gn=PURA;refGene_tx=XM_019836378.2
A1 118425506 118426100
refGene_gn=PURA;refGene_tx=XM_019836378.2
A1 118439475 118439637
refGene_gn=IGIP;refGene_tx=XM_003980827.4

A1 118467284 118467384
refGene_gn=CYSTM1;refGene_tx=XM_019833029.1
A1 118502180 118502402
refGene_gn=CYSTM1;refGene_tx=XM_019833029.1;refGene_gn=CYSTM1;refGene_tx=XM_019833033.1;refGene_gn=CYSTM1;refGene_tx=XM_019833036.2;refGene_gn=CYSTM1;refGene_tx=XM_019833038.1;refGene_gn=CYSTM1;refGene_tx=XM_023255790.1
A1 118528810 118528923
refGene_gn=CYSTM1;refGene_tx=XM_019833033.1;refGene_gn=CYSTM1;refGene_tx=XM_019833036.2;refGene_gn=CYSTM1;refGene_tx=XM_019833038.1;refGene_gn=CYSTM1;refGene_tx=XM_023255790.1
A1 118546028 118546135
refGene_gn=CYSTM1;refGene_tx=XM_019833029.1
A1 118547907 118547991
refGene_gn=PFDN1;refGene_tx=XM_003980829.4
A1 118581091 118581176
refGene_gn=PFDN1;refGene_tx=XM_003980829.4
A1 118616043 118616210
refGene_gn=PFDN1;refGene_tx=XM_003980829.4
A1 118618858 118618891
refGene_gn=PFDN1;refGene_tx=XM_003980829.4
A1 118645786 118645859
refGene_gn=HBEGF;refGene_tx=XM_003980830.4;refGene_gn=HBEGF;refGene_tx=XM_006927656.4
A1 118646658 118646814
refGene_gn=HBEGF;refGene_tx=XM_003980830.4;refGene_gn=HBEGF;refGene_tx=XM_006927656.4
A1 118652227 118652405
refGene_gn=HBEGF;refGene_tx=XM_003980830.4;refGene_gn=HBEGF;refGene_tx=XM_006927656.4
A1 118655488 118655662
refGene_gn=HBEGF;refGene_tx=XM_003980830.4;refGene_gn=HBEGF;refGene_tx=XM_006927656.4
A1 118655849 118655895
refGene_gn=HBEGF;refGene_tx=XM_003980830.4;refGene_gn=HBEGF;refGene_tx=XM_006927656.4
A1 118669663 118669881
refGene_gn=SLC4A9;refGene_tx=XM_023255813.1;refGene_gn=SLC4A9;refGene_tx=XM_023255814.1;refGene_gn=SLC4A9;refGene_tx=XM_023255816.1;refGene_gn=SLC4A9;refGene_tx=XM_023255819.1;refGene_gn=SLC4A9;refGene_tx=XM_023255827.1
A1 118670222 118670383
refGene_gn=SLC4A9;refGene_tx=XM_023255813.1;refGene_gn=SLC4A9;refGene_tx=XM_023255814.1;refGene_gn=SLC4A9;refGene_tx=XM_023255816.1;refGene_gn=SLC4A9;refGene_tx=XM_023255819.1
A1 118670653 118670767
refGene_gn=SLC4A9;refGene_tx=XM_023255813.1;refGene_gn=SLC4A9;refGene_tx=XM_023255814.1;refGene_gn=SLC4A9;refGene_tx=XM_023255816.1;refGene_gn=SLC4A9;refGene_tx=XM_023255819.1;refGene_gn=SLC4A9;refGene_tx=XM_023255827.1

A1	118671202	118671258
refGene_gn=SLC4A9;refGene_tx=XM_023255813.1;refGene_gn=SLC4A9;refGene_tx=XM_023255814.1;refGene_gn=SLC4A9;refGene_tx=XM_023255816.1;refGene_gn=SLC4A9;refGene_tx=XM_023255819.1;refGene_gn=SLC4A9;refGene_tx=XM_023255827.1
A1	118671382	118671540
refGene_gn=SLC4A9;refGene_tx=XM_023255813.1;refGene_gn=SLC4A9;refGene_tx=XM_023255814.1;refGene_gn=SLC4A9;refGene_tx=XM_023255816.1;refGene_gn=SLC4A9;refGene_tx=XM_023255819.1;refGene_gn=SLC4A9;refGene_tx=XM_023255827.1
A1	118671803	118671891
refGene_gn=SLC4A9;refGene_tx=XM_023255813.1;refGene_gn=SLC4A9;refGene_tx=XM_023255814.1;refGene_gn=SLC4A9;refGene_tx=XM_023255816.1;refGene_gn=SLC4A9;refGene_tx=XM_023255819.1;refGene_gn=SLC4A9;refGene_tx=XM_023255827.1
A1	118672230	118672385
refGene_gn=SLC4A9;refGene_tx=XM_023255813.1;refGene_gn=SLC4A9;refGene_tx=XM_023255814.1;refGene_gn=SLC4A9;refGene_tx=XM_023255816.1;refGene_gn=SLC4A9;refGene_tx=XM_023255819.1;refGene_gn=SLC4A9;refGene_tx=XM_023255827.1
A1	118672578	118672695
refGene_gn=SLC4A9;refGene_tx=XM_023255813.1;refGene_gn=SLC4A9;refGene_tx=XM_023255814.1;refGene_gn=SLC4A9;refGene_tx=XM_023255816.1;refGene_gn=SLC4A9;refGene_tx=XM_023255819.1;refGene_gn=SLC4A9;refGene_tx=XM_023255827.1
A1	118672866	118673041
refGene_gn=SLC4A9;refGene_tx=XM_023255813.1;refGene_gn=SLC4A9;refGene_tx=XM_023255814.1;refGene_gn=SLC4A9;refGene_tx=XM_023255816.1;refGene_gn=SLC4A9;refGene_tx=XM_023255819.1;refGene_gn=SLC4A9;refGene_tx=XM_023255827.1
A1	118673199	118673333
refGene_gn=SLC4A9;refGene_tx=XM_023255813.1;refGene_gn=SLC4A9;refGene_tx=XM_023255814.1;refGene_gn=SLC4A9;refGene_tx=XM_023255816.1;refGene_gn=SLC4A9;refGene_tx=XM_023255819.1;refGene_gn=SLC4A9;refGene_tx=XM_023255827.1
A1	118673506	118673769
refGene_gn=SLC4A9;refGene_tx=XM_023255813.1;refGene_gn=SLC4A9;refGene_tx=XM_023255814.1;refGene_gn=SLC4A9;refGene_tx=XM_023255816.1;refGene_gn=SLC4A9;refGene_tx=XM_023255819.1;refGene_gn=SLC4A9;refGene_tx=XM_023255827.1
A1	118674852	118674911
refGene_gn=SLC4A9;refGene_tx=XM_023255813.1;refGene_gn=SLC4A9;refGene_tx=XM_023255814.1;refGene_gn=SLC4A9;refGene_tx=XM_023255816.1;refGene_gn=SLC4A9;refGene_tx=XM_023255819.1;refGene_gn=SLC4A9;refGene_tx=XM_023255827.1
A1	118675143	118675332
refGene_gn=SLC4A9;refGene_tx=XM_023255813.1;refGene_gn=SLC4A9;refGene_tx=XM_023255814.1;refGene_gn=SLC4A9;refGene_tx=XM_023255816.1;refGene_gn=SLC4A9;refGene_tx=XM_023255819.1;refGene_gn=SLC4A9;refGene_tx=XM_023255827.1

A1 118675460 118675574
refGene_gn=SLC4A9;refGene_tx=XM_023255813.1;refGene_gn=SLC4A9;refGene_tx=XM_023255814.1;refGene_gn=SLC4A9;refGene_tx=XM_023255816.1;refGene_gn=SLC4A9;refGene_tx=XM_023255827.1
A1 118676597 118676672
refGene_gn=SLC4A9;refGene_tx=XM_023255813.1;refGene_gn=SLC4A9;refGene_tx=XM_023255814.1;refGene_gn=SLC4A9;refGene_tx=XM_023255827.1
A1 118676775 118676937
refGene_gn=SLC4A9;refGene_tx=XM_023255813.1;refGene_gn=SLC4A9;refGene_tx=XM_023255814.1;refGene_gn=SLC4A9;refGene_tx=XM_023255816.1;refGene_gn=SLC4A9;refGene_tx=XM_023255819.1;refGene_gn=SLC4A9;refGene_tx=XM_023255827.1
A1 118677084 118677263
refGene_gn=SLC4A9;refGene_tx=XM_023255813.1;refGene_gn=SLC4A9;refGene_tx=XM_023255814.1;refGene_gn=SLC4A9;refGene_tx=XM_023255816.1;refGene_gn=SLC4A9;refGene_tx=XM_023255819.1;refGene_gn=SLC4A9;refGene_tx=XM_023255827.1
A1 118678014 118678087
refGene_gn=SLC4A9;refGene_tx=XM_023255813.1;refGene_gn=SLC4A9;refGene_tx=XM_023255814.1;refGene_gn=SLC4A9;refGene_tx=XM_023255816.1;refGene_gn=SLC4A9;refGene_tx=XM_023255819.1;refGene_gn=SLC4A9;refGene_tx=XM_023255827.1
A1 118679767 118679836
refGene_gn=SLC4A9;refGene_tx=XM_023255813.1;refGene_gn=SLC4A9;refGene_tx=XM_023255814.1;refGene_gn=SLC4A9;refGene_tx=XM_023255816.1;refGene_gn=SLC4A9;refGene_tx=XM_023255819.1;refGene_gn=SLC4A9;refGene_tx=XM_023255827.1
A1 118680085 118680259
refGene_gn=SLC4A9;refGene_tx=XM_023255813.1;refGene_gn=SLC4A9;refGene_tx=XM_023255814.1;refGene_gn=SLC4A9;refGene_tx=XM_023255816.1;refGene_gn=SLC4A9;refGene_tx=XM_023255819.1;refGene_gn=SLC4A9;refGene_tx=XM_023255827.1
A1 118680863 118681016
refGene_gn=SLC4A9;refGene_tx=XM_023255813.1;refGene_gn=SLC4A9;refGene_tx=XM_023255814.1;refGene_gn=SLC4A9;refGene_tx=XM_023255816.1;refGene_gn=SLC4A9;refGene_tx=XM_023255819.1;refGene_gn=SLC4A9;refGene_tx=XM_023255827.1
A1 118681367 118681421
refGene_gn=SLC4A9;refGene_tx=XM_023255813.1;refGene_gn=SLC4A9;refGene_tx=XM_023255814.1;refGene_gn=SLC4A9;refGene_tx=XM_023255816.1;refGene_gn=SLC4A9;refGene_tx=XM_023255819.1;refGene_gn=SLC4A9;refGene_tx=XM_023255827.1
A1 118708645 118708936
refGene_gn=LOC101093912;refGene_tx=XM_003980831.3;refGene_gn=LOC101093912;refGene_tx=XM_003980833.4;refGene_gn=LOC101093912;refGene_tx=XM_006927658.2;refGene_gn=LOC101093912;refGene_tx=XM_006927659.2
A1 118737715 118737869
refGene_gn=LOC101093912;refGene_tx=XM_003980831.3;refGene_gn=LOC101093912;refGene_tx=XM_003980833.4;refGene_gn=LOC101093912;refGene_tx=XM_006927658.2;refGene_gn=LOC101093912;refGene_tx=XM_006927659.2

A1 118743526 118743683
refGene_gn=LOC101093912;refGene_tx=XM_003980831.3;refGene_gn=LOC101093912;refGene_tx=XM_003980833.4;refGene_gn=LOC101093912;refGene_tx=XM_006927658.2;refGene_gn=LOC101093912;refGene_tx=XM_006927659.2
A1 118745023 118745171
refGene_gn=LOC101093912;refGene_tx=XM_003980831.3;refGene_gn=LOC101093912;refGene_tx=XM_003980833.4;refGene_gn=LOC101093912;refGene_tx=XM_006927658.2;refGene_gn=LOC101093912;refGene_tx=XM_006927659.2
A1 118745896 118746044
refGene_gn=LOC101093912;refGene_tx=XM_003980831.3;refGene_gn=LOC101093912;refGene_tx=XM_003980833.4;refGene_gn=LOC101093912;refGene_tx=XM_006927658.2;refGene_gn=LOC101093912;refGene_tx=XM_006927659.2
A1 118747877 118748111
refGene_gn=LOC101093912;refGene_tx=XM_003980831.3;refGene_gn=LOC101093912;refGene_tx=XM_003980833.4;refGene_gn=LOC101093912;refGene_tx=XM_006927658.2;refGene_gn=LOC101093912;refGene_tx=XM_006927659.2
A1 118750479 118750574
refGene_gn=LOC101093912;refGene_tx=XM_003980831.3;refGene_gn=LOC101093912;refGene_tx=XM_003980833.4;refGene_gn=LOC101093912;refGene_tx=XM_006927658.2;refGene_gn=LOC101093912;refGene_tx=XM_006927659.2
A1 118767729 118767967
refGene_gn=LOC101093912;refGene_tx=XM_003980831.3;refGene_gn=LOC101093912;refGene_tx=XM_003980833.4;refGene_gn=LOC101093912;refGene_tx=XM_006927658.2;refGene_gn=LOC101093912;refGene_tx=XM_006927659.2
A1 118768196 118768388
refGene_gn=LOC101093912;refGene_tx=XM_003980831.3;refGene_gn=LOC101093912;refGene_tx=XM_003980833.4;refGene_gn=LOC101093912;refGene_tx=XM_006927658.2;refGene_gn=LOC101093912;refGene_tx=XM_006927659.2
A1 118773713 118773823
refGene_gn=LOC101093912;refGene_tx=XM_003980831.3;refGene_gn=LOC101093912;refGene_tx=XM_006927658.2;refGene_gn=LOC101093912;refGene_tx=XM_006927659.2
A1 118784469 118784543
refGene_gn=LOC101093912;refGene_tx=XM_003980833.4;refGene_gn=LOC101093912;refGene_tx=XM_006927658.2
A1 118793877 118793965
refGene_gn=LOC101093912;refGene_tx=XM_003980831.3;refGene_gn=LOC101093912;refGene_tx=XM_006927658.2;refGene_gn=LOC101093912;refGene_tx=XM_006927659.2
A1 118797268 118797396
refGene_gn=LOC101093912;refGene_tx=XM_003980831.3;refGene_gn=LOC101093912;refGene_tx=XM_006927658.2;refGene_gn=LOC101093912;refGene_tx=XM_006927659.2
A1 118797596 118797740
refGene_gn=LOC101093912;refGene_tx=XM_003980831.3;refGene_gn=LOC101093912;refGene_tx=XM_006927658.2;refGene_gn=LOC101093912;refGene_tx=XM_006927659.2
A1 118799034 118799137
refGene_gn=LOC101093912;refGene_tx=XM_003980831.3;refGene_gn=LOC101093912;refGene_tx=XM_006927658.2;refGene_gn=LOC101093912;refGene_tx=XM_00

6927659.2
A1 118807110 118807869
refGene_gn=LOC101093912;refGene_tx=XM_003980831.3;refGene_gn=LOC101093912;refGene_tx=XM_006927658.2;refGene_gn=LOC101093912;refGene_tx=XM_006927659.2
A1 118813515 118813661
refGene_gn=LOC101093912;refGene_tx=XM_003980831.3;refGene_gn=LOC101093912;refGene_tx=XM_006927658.2;refGene_gn=LOC101093912;refGene_tx=XM_006927659.2
A1 118813812 118813924
refGene_gn=LOC101093912;refGene_tx=XM_003980831.3;refGene_gn=LOC101093912;refGene_tx=XM_006927658.2;refGene_gn=LOC101093912;refGene_tx=XM_006927659.2
A1 118815365 118815511
refGene_gn=LOC101093912;refGene_tx=XM_003980831.3;refGene_gn=LOC101093912;refGene_tx=XM_006927658.2;refGene_gn=LOC101093912;refGene_tx=XM_006927659.2
A1 118816029 118816172
refGene_gn=LOC101093912;refGene_tx=XM_003980831.3;refGene_gn=LOC101093912;refGene_tx=XM_006927658.2;refGene_gn=LOC101093912;refGene_tx=XM_006927659.2
A1 118818715 118818929
refGene_gn=LOC101093912;refGene_tx=XM_003980831.3;refGene_gn=LOC101093912;refGene_tx=XM_006927658.2;refGene_gn=LOC101093912;refGene_tx=XM_006927659.2
A1 118820715 118820891
refGene_gn=LOC101093912;refGene_tx=XM_003980831.3;refGene_gn=LOC101093912;refGene_tx=XM_006927658.2;refGene_gn=LOC101093912;refGene_tx=XM_006927659.2
A1 118821188 118821351
refGene_gn=LOC101093912;refGene_tx=XM_003980831.3;refGene_gn=LOC101093912;refGene_tx=XM_006927658.2;refGene_gn=LOC101093912;refGene_tx=XM_006927659.2
A1 118823578 118823674
refGene_gn=LOC101093912;refGene_tx=XM_003980831.3;refGene_gn=LOC101093912;refGene_tx=XM_006927658.2;refGene_gn=LOC101093912;refGene_tx=XM_006927659.2
A1 118824146 118824263
refGene_gn=LOC101093912;refGene_tx=XM_003980831.3;refGene_gn=LOC101093912;refGene_tx=XM_006927658.2;refGene_gn=LOC101093912;refGene_tx=XM_006927659.2
A1 118828169 118828344
refGene_gn=LOC101093912;refGene_tx=XM_003980831.3;refGene_gn=LOC101093912;refGene_tx=XM_006927658.2;refGene_gn=LOC101093912;refGene_tx=XM_006927659.2
A1 118830017 118830474
refGene_gn=LOC101093912;refGene_tx=XM_003980831.3;refGene_gn=LOC101093912;refGene_tx=XM_006927658.2;refGene_gn=LOC101093912;refGene_tx=XM_006927659.2
A1 118830953 118831100

refGene_gn=LOC101093912;refGene_tx=XM_003980831.3;refGene_gn=LOC101093
912;refGene_tx=XM_006927658.2;refGene_gn=LOC101093912;refGene_tx=XM_00
6927659.2
A1 118833404 118833554
refGene_gn=LOC101093912;refGene_tx=XM_003980831.3;refGene_gn=LOC101093
912;refGene_tx=XM_006927658.2;refGene_gn=LOC101093912;refGene_tx=XM_00
6927659.2
A1 118833715 118835328
refGene_gn=LOC101093912;refGene_tx=XM_003980831.3;refGene_gn=LOC101093
912;refGene_tx=XM_006927658.2;refGene_gn=LOC101093912;refGene_tx=XM_00
6927659.2
A1 118839494 118839671
refGene_gn=LOC101093912;refGene_tx=XM_003980831.3;refGene_gn=LOC101093
912;refGene_tx=XM_006927658.2;refGene_gn=LOC101093912;refGene_tx=XM_00
6927659.2
A1 118840719 118840971
refGene_gn=LOC101093912;refGene_tx=XM_003980831.3;refGene_gn=LOC101093
912;refGene_tx=XM_006927658.2;refGene_gn=LOC101093912;refGene_tx=XM_00
6927659.2
A1 118841496 118841672
refGene_gn=LOC101093912;refGene_tx=XM_003980831.3;refGene_gn=LOC101093
912;refGene_tx=XM_006927658.2;refGene_gn=LOC101093912;refGene_tx=XM_00
6927659.2
A1 118842363 118842528
refGene_gn=LOC101093912;refGene_tx=XM_003980831.3;refGene_gn=LOC101093
912;refGene_tx=XM_006927658.2;refGene_gn=LOC101093912;refGene_tx=XM_00
6927659.2
A1 118842795 118842855
refGene_gn=LOC101093912;refGene_tx=XM_003980831.3;refGene_gn=LOC101093
912;refGene_tx=XM_006927658.2;refGene_gn=LOC101093912;refGene_tx=XM_00
6927659.2
A1 118849188 118849291
refGene_gn=EIF4EBP3;refGene_tx=XM_003980835.4
A1 118850351 118850525
refGene_gn=EIF4EBP3;refGene_tx=XM_003980835.4
A1 118850673 118850702
refGene_gn=EIF4EBP3;refGene_tx=XM_003980835.4
A1 118852104 118852304
refGene_gn=SRA1;refGene_tx=XM_003980904.3
A1 118852460 118852569
refGene_gn=SRA1;refGene_tx=XM_003980904.3
A1 118853264 118853470
refGene_gn=SRA1;refGene_tx=XM_003980904.3
A1 118857367 118857493
refGene_gn=SRA1;refGene_tx=XM_003980904.3
A1 118857618 118857756
refGene_gn=SRA1;refGene_tx=XM_003980904.3
A1 118858023 118858078
refGene_gn=SRA1;refGene_tx=XM_003980904.3
A1 118858777 118859014

refGene_gn=APBB3;refGene_tx=XM_003980837.5;refGene_gn=APBB3;refGene_tx=XM_011282508.3;refGene_gn=APBB3;refGene_tx=XM_019833145.2
A1 118860488 118860695
refGene_gn=APBB3;refGene_tx=XM_003980837.5;refGene_gn=APBB3;refGene_tx=XM_011282508.3;refGene_gn=APBB3;refGene_tx=XM_019833145.2
A1 118860813 118860929
refGene_gn=APBB3;refGene_tx=XM_003980837.5;refGene_gn=APBB3;refGene_tx=XM_011282508.3;refGene_gn=APBB3;refGene_tx=XM_019833145.2
A1 118861196 118861280
refGene_gn=APBB3;refGene_tx=XM_003980837.5;refGene_gn=APBB3;refGene_tx=XM_011282508.3;refGene_gn=APBB3;refGene_tx=XM_019833145.2
A1 118861522 118861604
refGene_gn=APBB3;refGene_tx=XM_003980837.5;refGene_gn=APBB3;refGene_tx=XM_011282508.3;refGene_gn=APBB3;refGene_tx=XM_019833145.2
A1 118861744 118861859
refGene_gn=APBB3;refGene_tx=XM_003980837.5;refGene_gn=APBB3;refGene_tx=XM_011282508.3;refGene_gn=APBB3;refGene_tx=XM_019833145.2
A1 118862033 118862039
refGene_gn=APBB3;refGene_tx=XM_011282508.3;refGene_gn=APBB3;refGene_tx=XM_019833145.2
A1 118862291 118862419
refGene_gn=APBB3;refGene_tx=XM_003980837.5;refGene_gn=APBB3;refGene_tx=XM_011282508.3;refGene_gn=APBB3;refGene_tx=XM_019833145.2
A1 118862541 118862694
refGene_gn=APBB3;refGene_tx=XM_003980837.5;refGene_gn=APBB3;refGene_tx=XM_011282508.3;refGene_gn=APBB3;refGene_tx=XM_019833145.2
A1 118862936 118862997
refGene_gn=APBB3;refGene_tx=XM_003980837.5;refGene_gn=APBB3;refGene_tx=XM_011282508.3;refGene_gn=APBB3;refGene_tx=XM_019833145.2
A1 118863559 118863636
refGene_gn=APBB3;refGene_tx=XM_003980837.5;refGene_gn=APBB3;refGene_tx=XM_011282508.3;refGene_gn=APBB3;refGene_tx=XM_019833145.2
A1 118863719 118863883
refGene_gn=APBB3;refGene_tx=XM_003980837.5;refGene_gn=APBB3;refGene_tx=XM_011282508.3;refGene_gn=APBB3;refGene_tx=XM_019833145.2
A1 118864130 118864179
refGene_gn=APBB3;refGene_tx=XM_003980837.5;refGene_gn=APBB3;refGene_tx=XM_011282508.3;refGene_gn=APBB3;refGene_tx=XM_019833145.2
A1 118864779 118864794
refGene_gn=LOC101095877;refGene_tx=XM_003980840.5
A1 118865755 118865854
refGene_gn=LOC101095877;refGene_tx=XM_003980840.5
A1 118866465 118866663
refGene_gn=LOC101095877;refGene_tx=XM_003980840.5
A1 118867090 118868065
refGene_gn=SLC35A4;refGene_tx=XM_006927660.3
A1 118869966 118870128
refGene_gn=LOC102900045;refGene_tx=XM_023255848.1
A1 118870202 118870358
refGene_gn=LOC102900045;refGene_tx=XM_023255848.1

A1 118871177 118871345
refGene_gn=LOC102900045;refGene_tx=XM_023255848.1
A1 118873766 118873850
refGene_gn=LOC102900045;refGene_tx=XM_023255848.1
A1 118873922 118874131
refGene_gn=LOC102900045;refGene_tx=XM_023255848.1
A1 118875198 118875332
refGene_gn=LOC102900045;refGene_tx=XM_023255848.1
A1 118879114 118879206
refGene_gn=LOC102900045;refGene_tx=XM_023255848.1
A1 118894490 118895615
refGene_gn=CD14;refGene_tx=XM_003980841.4
A1 118895704 118895707
refGene_gn=CD14;refGene_tx=XM_003980841.4
A1 118897289 118897457
refGene_gn=TMCO6;refGene_tx=XM_019833193.2;refGene_gn=TMCO6;refGene_tx=XM_019833198.2
A1 118899835 118899920
refGene_gn=TMCO6;refGene_tx=XM_003980905.5;refGene_gn=TMCO6;refGene_tx=XM_019833201.2
A1 118900059 118900172
refGene_gn=TMCO6;refGene_tx=XM_003980905.5;refGene_gn=TMCO6;refGene_tx=XM_011282518.3;refGene_gn=TMCO6;refGene_tx=XM_019833201.2;refGene_gn=TMCO6;refGene_tx=XM_019833205.2;refGene_gn=TMCO6;refGene_tx=XM_019833210.2
A1 118902360 118902476
refGene_gn=TMCO6;refGene_tx=XM_003980905.5;refGene_gn=TMCO6;refGene_tx=XM_011282518.3;refGene_gn=TMCO6;refGene_tx=XM_019833193.2;refGene_gn=TMCO6;refGene_tx=XM_019833198.2;refGene_gn=TMCO6;refGene_tx=XM_019833201.2;refGene_gn=TMCO6;refGene_tx=XM_019833205.2;refGene_gn=TMCO6;refGene_tx=XM_019833210.2
A1 118902581 118902765
refGene_gn=TMCO6;refGene_tx=XM_003980905.5;refGene_gn=TMCO6;refGene_tx=XM_011282518.3;refGene_gn=TMCO6;refGene_tx=XM_019833193.2;refGene_gn=TMCO6;refGene_tx=XM_019833198.2;refGene_gn=TMCO6;refGene_tx=XM_019833201.2;refGene_gn=TMCO6;refGene_tx=XM_019833205.2;refGene_gn=TMCO6;refGene_tx=XM_019833210.2;refGene_gn=TMCO6;refGene_tx=XM_019833211.2
A1 118903017 118903122
refGene_gn=TMCO6;refGene_tx=XM_003980905.5;refGene_gn=TMCO6;refGene_tx=XM_011282518.3;refGene_gn=TMCO6;refGene_tx=XM_019833193.2;refGene_gn=TMCO6;refGene_tx=XM_019833198.2;refGene_gn=TMCO6;refGene_tx=XM_019833205.2;refGene_gn=TMCO6;refGene_tx=XM_019833210.2;refGene_gn=TMCO6;refGene_tx=XM_019833211.2
A1 118903324 118903410
refGene_gn=TMCO6;refGene_tx=XM_003980905.5;refGene_gn=TMCO6;refGene_tx=XM_011282518.3;refGene_gn=TMCO6;refGene_tx=XM_011282521.3;refGene_gn=TMCO6;refGene_tx=XM_019833193.2;refGene_gn=TMCO6;refGene_tx=XM_019833198.2;refGene_gn=TMCO6;refGene_tx=XM_019833201.2;refGene_gn=TMCO6;refGene_tx=XM_019833205.2;refGene_gn=TMCO6;refGene_tx=XM_019833210.2;refGene_gn=TMCO6;refGene_tx=XM_019833211.2;refGene_gn=TMCO6;refGene_tx=XM_01

9833216.2
A1	118903587	118903760
refGene_gn=TMCO6;refGene_tx=XM_003980905.5;refGene_gn=TMCO6;refGene_tx=XM_011282518.3;refGene_gn=TMCO6;refGene_tx=XM_011282521.3;refGene_gn=TMCO6;refGene_tx=XM_019833193.2;refGene_gn=TMCO6;refGene_tx=XM_019833198.2;refGene_gn=TMCO6;refGene_tx=XM_019833201.2;refGene_gn=TMCO6;refGene_tx=XM_019833205.2;refGene_gn=TMCO6;refGene_tx=XM_019833210.2;refGene_gn=TMCO6;refGene_tx=XM_019833211.2;refGene_gn=TMCO6;refGene_tx=XM_019833213.2;refGene_gn=TMCO6;refGene_tx=XM_019833216.2
A1	118903984	118904096
refGene_gn=TMCO6;refGene_tx=XM_003980905.5;refGene_gn=TMCO6;refGene_tx=XM_011282518.3;refGene_gn=TMCO6;refGene_tx=XM_011282521.3;refGene_gn=TMCO6;refGene_tx=XM_019833193.2;refGene_gn=TMCO6;refGene_tx=XM_019833198.2;refGene_gn=TMCO6;refGene_tx=XM_019833201.2;refGene_gn=TMCO6;refGene_tx=XM_019833205.2;refGene_gn=TMCO6;refGene_tx=XM_019833210.2;refGene_gn=TMCO6;refGene_tx=XM_019833211.2;refGene_gn=TMCO6;refGene_tx=XM_019833213.2;refGene_gn=TMCO6;refGene_tx=XM_019833216.2
A1	118904213	118904400
refGene_gn=TMCO6;refGene_tx=XM_003980905.5;refGene_gn=TMCO6;refGene_tx=XM_011282518.3;refGene_gn=TMCO6;refGene_tx=XM_011282521.3;refGene_gn=TMCO6;refGene_tx=XM_019833193.2;refGene_gn=TMCO6;refGene_tx=XM_019833198.2;refGene_gn=TMCO6;refGene_tx=XM_019833201.2;refGene_gn=TMCO6;refGene_tx=XM_019833205.2;refGene_gn=TMCO6;refGene_tx=XM_019833210.2;refGene_gn=TMCO6;refGene_tx=XM_019833211.2;refGene_gn=TMCO6;refGene_tx=XM_019833213.2;refGene_gn=TMCO6;refGene_tx=XM_019833216.2
A1	118904545	118904640
refGene_gn=TMCO6;refGene_tx=XM_003980905.5;refGene_gn=TMCO6;refGene_tx=XM_011282518.3;refGene_gn=TMCO6;refGene_tx=XM_011282521.3;refGene_gn=TMCO6;refGene_tx=XM_019833193.2;refGene_gn=TMCO6;refGene_tx=XM_019833198.2;refGene_gn=TMCO6;refGene_tx=XM_019833201.2;refGene_gn=TMCO6;refGene_tx=XM_019833205.2;refGene_gn=TMCO6;refGene_tx=XM_019833210.2;refGene_gn=TMCO6;refGene_tx=XM_019833211.2;refGene_gn=TMCO6;refGene_tx=XM_019833213.2;refGene_gn=TMCO6;refGene_tx=XM_019833216.2
A1	118905023	118905191
refGene_gn=TMCO6;refGene_tx=XM_003980905.5;refGene_gn=TMCO6;refGene_tx=XM_011282518.3;refGene_gn=TMCO6;refGene_tx=XM_011282521.3;refGene_gn=TMCO6;refGene_tx=XM_019833193.2;refGene_gn=TMCO6;refGene_tx=XM_019833198.2;refGene_gn=TMCO6;refGene_tx=XM_019833201.2;refGene_gn=TMCO6;refGene_tx=XM_019833205.2;refGene_gn=TMCO6;refGene_tx=XM_019833210.2;refGene_gn=TMCO6;refGene_tx=XM_019833211.2;refGene_gn=TMCO6;refGene_tx=XM_019833213.2;refGene_gn=TMCO6;refGene_tx=XM_019833216.2
A1	118905436	118905550
refGene_gn=TMCO6;refGene_tx=XM_003980905.5;refGene_gn=TMCO6;refGene_tx=XM_011282518.3;refGene_gn=TMCO6;refGene_tx=XM_011282521.3;refGene_gn=TMCO6;refGene_tx=XM_019833193.2;refGene_gn=TMCO6;refGene_tx=XM_019833198.2;refGene_gn=TMCO6;refGene_tx=XM_019833201.2;refGene_gn=TMCO6;refGene_tx=XM_019833205.2;refGene_gn=TMCO6;refGene_tx=XM_019833210.2;refGene_gn=TMCO6;refGene_tx=XM_019833211.2;refGene_gn=TMCO6;refGene_tx=XM_019833213.2;refGene_gn=TMCO6;refGene_tx=XM_019833216.2
A1	118906261	118906353

refGene_gn=NDUFA2;refGene_tx=XM_003980842.5
A1 118907995 118908102
refGene_gn=NDUFA2;refGene_tx=XM_003980842.5
A1 118908233 118908334
refGene_gn=NDUFA2;refGene_tx=XM_003980842.5
A1 118908649 118908665
refGene_gn=IK;refGene_tx=XM_006927662.4
A1 118909212 118909279
refGene_gn=IK;refGene_tx=XM_006927662.4
A1 118911365 118911458
refGene_gn=IK;refGene_tx=XM_006927662.4
A1 118911806 118911866
refGene_gn=IK;refGene_tx=XM_006927662.4
A1 118912689 118912857
refGene_gn=IK;refGene_tx=XM_006927662.4
A1 118913027 118913142
refGene_gn=IK;refGene_tx=XM_006927662.4
A1 118913534 118913605
refGene_gn=IK;refGene_tx=XM_006927662.4
A1 118913803 118913850
refGene_gn=IK;refGene_tx=XM_006927662.4
A1 118914670 118914834
refGene_gn=IK;refGene_tx=XM_006927662.4
A1 118915358 118915467
refGene_gn=IK;refGene_tx=XM_006927662.4
A1 118915771 118915811
refGene_gn=IK;refGene_tx=XM_006927662.4
A1 118915970 118916190
refGene_gn=IK;refGene_tx=XM_006927662.4
A1 118916347 118916366
refGene_gn=IK;refGene_tx=XM_006927662.4
A1 118916839 118916918
refGene_gn=IK;refGene_tx=XM_006927662.4
A1 118917371 118917452
refGene_gn=IK;refGene_tx=XM_006927662.4
A1 118917706 118917764
refGene_gn=IK;refGene_tx=XM_006927662.4
A1 118918518 118918607
refGene_gn=IK;refGene_tx=XM_006927662.4
A1 118918769 118918878
refGene_gn=IK;refGene_tx=XM_006927662.4
A1 118919044 118919079
refGene_gn=IK;refGene_tx=XM_006927662.4
A1 118919166 118919194
refGene_gn=IK;refGene_tx=XM_006927662.4
A1 118922884 118923075
refGene_gn=WDR55;refGene_tx=XM_003980844.4
A1 118925291 118925392
refGene_gn=WDR55;refGene_tx=XM_003980844.4
A1 118925474 118925562

refGene_gn=WDR55;refGene_tx=XM_003980844.4
A1	118925653	118925833
refGene_gn=WDR55;refGene_tx=XM_003980844.4
A1	118925922	118926022
refGene_gn=WDR55;refGene_tx=XM_003980844.4
A1	118926109	118926279
refGene_gn=WDR55;refGene_tx=XM_003980844.4
A1	118926362	118926681
refGene_gn=WDR55;refGene_tx=XM_003980844.4
A1	118927387	118927845
refGene_gn=DND1;refGene_tx=XM_023255905.1
A1	118928515	118928977
refGene_gn=DND1;refGene_tx=XM_023255905.1
A1	118929365	118929483
refGene_gn=DND1;refGene_tx=XM_023255905.1
A1	118929616	118929640
refGene_gn=DND1;refGene_tx=XM_023255905.1
A1	118930361	118930438
refGene_gn=HARS;refGene_tx=XM_003980846.4;refGene_gn=HARS;refGene_tx=XM_006927663.3;refGene_gn=HARS;refGene_tx=XM_006927664.3;refGene_gn=HARS;refGene_tx=XM_006927665.3
A1	118930769	118930916
refGene_gn=HARS;refGene_tx=XM_003980846.4;refGene_gn=HARS;refGene_tx=XM_006927663.3;refGene_gn=HARS;refGene_tx=XM_006927664.3;refGene_gn=HARS;refGene_tx=XM_006927665.3
A1	118931089	118931206
refGene_gn=HARS;refGene_tx=XM_003980846.4;refGene_gn=HARS;refGene_tx=XM_006927663.3;refGene_gn=HARS;refGene_tx=XM_006927664.3;refGene_gn=HARS;refGene_tx=XM_006927665.3
A1	118932726	118932969
refGene_gn=HARS;refGene_tx=XM_003980846.4;refGene_gn=HARS;refGene_tx=XM_006927663.3;refGene_gn=HARS;refGene_tx=XM_006927664.3;refGene_gn=HARS;refGene_tx=XM_006927665.3
A1	118933062	118933190
refGene_gn=HARS;refGene_tx=XM_003980846.4;refGene_gn=HARS;refGene_tx=XM_006927663.3;refGene_gn=HARS;refGene_tx=XM_006927664.3;refGene_gn=HARS;refGene_tx=XM_006927665.3
A1	118933382	118933476
refGene_gn=HARS;refGene_tx=XM_003980846.4;refGene_gn=HARS;refGene_tx=XM_006927663.3;refGene_gn=HARS;refGene_tx=XM_006927664.3;refGene_gn=HARS;refGene_tx=XM_006927665.3
A1	118933707	118933806
refGene_gn=HARS;refGene_tx=XM_003980846.4;refGene_gn=HARS;refGene_tx=XM_006927663.3;refGene_gn=HARS;refGene_tx=XM_006927664.3;refGene_gn=HARS;refGene_tx=XM_006927665.3
A1	118933962	118934070
refGene_gn=HARS;refGene_tx=XM_003980846.4;refGene_gn=HARS;refGene_tx=XM_006927663.3;refGene_gn=HARS;refGene_tx=XM_006927664.3;refGene_gn=HARS;refGene_tx=XM_006927665.3
A1	118934967	118935093

refGene_gn=HARS;refGene_tx=XM_003980846.4;refGene_gn=HARS;refGene_tx=XM_006927663.3;refGene_gn=HARS;refGene_tx=XM_006927664.3;refGene_gn=HARS;refGene_tx=XM_006927665.3
A1 118935750 118935846
refGene_gn=HARS;refGene_tx=XM_003980846.4;refGene_gn=HARS;refGene_tx=XM_006927663.3;refGene_gn=HARS;refGene_tx=XM_006927664.3;refGene_gn=HARS;refGene_tx=XM_006927665.3
A1 118938691 118938811
refGene_gn=HARS;refGene_tx=XM_003980846.4;refGene_gn=HARS;refGene_tx=XM_006927663.3;refGene_gn=HARS;refGene_tx=XM_006927664.3;refGene_gn=HARS;refGene_tx=XM_006927665.3
A1 118943194 118943284
refGene_gn=HARS;refGene_tx=XM_003980846.4;refGene_gn=HARS;refGene_tx=XM_006927663.3;refGene_gn=HARS;refGene_tx=XM_006927664.3
A1 118943550 118943640
refGene_gn=HARS;refGene_tx=XM_003980846.4;refGene_gn=HARS;refGene_tx=XM_006927663.3;refGene_gn=HARS;refGene_tx=XM_006927664.3
A1 118943969 118944077
refGene_gn=LOC101098429;refGene_tx=XM_006927667.3
A1 118945519 118945594
refGene_gn=LOC101098429;refGene_tx=XM_006927667.3
A1 118946109 118946229
refGene_gn=LOC101098429;refGene_tx=XM_006927667.3;refGene_gn=LOC101098429;refGene_tx=XM_023255919.1
A1 118946347 118946443
refGene_gn=LOC101098429;refGene_tx=XM_006927667.3;refGene_gn=LOC101098429;refGene_tx=XM_023255919.1
A1 118947164 118947290
refGene_gn=LOC101098429;refGene_tx=XM_006927667.3;refGene_gn=LOC101098429;refGene_tx=XM_023255919.1
A1 118947397 118947505
refGene_gn=LOC101098429;refGene_tx=XM_006927667.3;refGene_gn=LOC101098429;refGene_tx=XM_023255919.1
A1 118947761 118947860
refGene_gn=LOC101098429;refGene_tx=XM_006927667.3;refGene_gn=LOC101098429;refGene_tx=XM_023255919.1
A1 118948047 118948141
refGene_gn=LOC101098429;refGene_tx=XM_006927667.3;refGene_gn=LOC101098429;refGene_tx=XM_023255919.1
A1 118948779 118948907
refGene_gn=LOC101098429;refGene_tx=XM_006927667.3;refGene_gn=LOC101098429;refGene_tx=XM_023255919.1
A1 118949000 118949240
refGene_gn=LOC101098429;refGene_tx=XM_006927667.3;refGene_gn=LOC101098429;refGene_tx=XM_023255919.1
A1 118949424 118949541
refGene_gn=LOC101098429;refGene_tx=XM_006927667.3;refGene_gn=LOC101098429;refGene_tx=XM_023255919.1
A1 118949756 118949903
refGene_gn=LOC101098429;refGene_tx=XM_006927667.3;refGene_gn=LOC101098

429;refGene_tx=XM_023255919.1
A1 118950289 118950349
refGene_gn=LOC101098429;refGene_tx=XM_006927667.3;refGene_gn=LOC101098429;refGene_tx=XM_023255919.1
A1 118951930 118951948
refGene_gn=ZMAT2;refGene_tx=XM_003980853.4
A1 118952284 118952378
refGene_gn=ZMAT2;refGene_tx=XM_003980853.4
A1 118953455 118953579
refGene_gn=ZMAT2;refGene_tx=XM_003980853.4
A1 118955006 118955080
refGene_gn=ZMAT2;refGene_tx=XM_003980853.4
A1 118955520 118955666
refGene_gn=ZMAT2;refGene_tx=XM_003980853.4
A1 118956384 118956462
refGene_gn=ZMAT2;refGene_tx=XM_003980853.4
A1 119034280 119036423
refGene_gn=LOC111560912;refGene_tx=XM_023255929.1
A1 119036425 119036697
refGene_gn=LOC111560912;refGene_tx=XM_023255929.1
A1 119041778 119044175
refGene_gn=LOC101098920;refGene_tx=XM_019833312.2
A1 119063228 119065547
refGene_gn=LOC101099699;refGene_tx=XM_023255967.1
A1 119065548 119065620
refGene_gn=LOC101099699;refGene_tx=XM_023255967.1
A1 119073097 119075494
refGene_gn=LOC101098920;refGene_tx=XM_023255945.1;refGene_gn=LOC101098920;refGene_tx=XM_023255957.1
A1 119077374 119077461
refGene_gn=LOC101098920;refGene_tx=XM_023255957.1
A1 119079834 119082111
refGene_gn=LOC101092486;refGene_tx=XM_023255965.1
A1 119082112 119082234
refGene_gn=LOC101092486;refGene_tx=XM_023255965.1
A1 119112868 119115265
refGene_gn=LOC101098920;refGene_tx=XM_023255959.1
A1 119143536 119145969
refGene_gn=LOC101098920;refGene_tx=XM_023255943.1
A1 119185364 119187923
refGene_gn=LOC101098920;refGene_tx=XM_019833319.2;refGene_gn=LOC101098920;refGene_tx=XM_023255939.1
A1 119192211 119192244
refGene_gn=LOC101098920;refGene_tx=XM_019833319.2
A1 119197577 119197636
refGene_gn=LOC101092486;refGene_tx=XM_023255965.1;refGene_gn=LOC101098920;refGene_tx=XM_019833312.2;refGene_gn=LOC101098920;refGene_tx=XM_023255939.1;refGene_gn=LOC101098920;refGene_tx=XM_023255943.1;refGene_gn=LOC101098920;refGene_tx=XM_023255945.1;refGene_gn=LOC101098920;refGene_tx=XM_023255959.1

A1	119200417	119200506	refGene_gn=LOC101098920;refGene_tx=XM_019833312.2;refGene_gn=LOC101098920;refGene_tx=XM_023255939.1;refGene_gn=LOC101098920;refGene_tx=XM_023255943.1;refGene_gn=LOC101098920;refGene_tx=XM_023255945.1;refGene_gn=LOC101098920;refGene_tx=XM_023255959.1
A1	119230641	119230952	refGene_gn=LOC101098920;refGene_tx=XM_019833312.2;refGene_gn=LOC101098920;refGene_tx=XM_023255939.1;refGene_gn=LOC101098920;refGene_tx=XM_023255943.1;refGene_gn=LOC101098920;refGene_tx=XM_023255945.1;refGene_gn=LOC101098920;refGene_tx=XM_023255959.1
A1	119264436	119266893	refGene_gn=PCDHB1;refGene_tx=XM_003980856.3
A1	119286752	119289146	refGene_gn=LOC101093742;refGene_tx=XM_011282592.3
A1	119293089	119295447	refGene_gn=PCDHB3;refGene_tx=XM_019833383.2
A1	119305502	119307783	refGene_gn=LOC101093988;refGene_tx=XM_023255991.1
A1	119316514	119318767	refGene_gn=LOC111556247;refGene_tx=XM_023255994.1
A1	119318842	119318902	refGene_gn=LOC111556247;refGene_tx=XM_023255994.1
A1	119328459	119330541	refGene_gn=LOC101094981;refGene_tx=XM_011282587.3
A1	119335605	119335971	refGene_gn=LOC101094981;refGene_tx=XM_011282587.3
A1	119344818	119347206	refGene_gn=LOC109502212;refGene_tx=XM_023256012.1
A1	119350450	119352302	refGene_gn=PCDHB8;refGene_tx=XM_023256016.1
A1	119352304	119352846	refGene_gn=PCDHB8;refGene_tx=XM_023256016.1
A1	119358628	119360261	refGene_gn=PCDHB16;refGene_tx=XM_023256043.1
A1	119360391	119361014	refGene_gn=PCDHB16;refGene_tx=XM_023256043.1
A1	119363158	119365552	refGene_gn=PCDHB10;refGene_tx=XM_023256023.1
A1	119371904	119374298	refGene_gn=PCDHB12;refGene_tx=XM_023256035.1
A1	119382451	119384848	refGene_gn=LOC109500934;refGene_tx=XM_019833357.2
A1	119392870	119395258	refGene_gn=LOC101095465;refGene_tx=XM_023256060.1
A1	119398885	119401140	refGene_gn=LOC101096200;refGene_tx=XM_023256049.1
A1	119401141	119401274	refGene_gn=LOC101096200;refGene_tx=XM_023256049.1
A1	119404268	119406668	refGene_gn=LOC111560924;refGene_tx=XM_023256044.1

A1	119437598	119438504
refGene_gn=LOC101096698;refGene_tx=XM_011282600.3
A1	119447369	119448419
refGene_gn=TAF7;refGene_tx=XM_003980857.5
A1	119449485	119449498
refGene_gn=LOC101100942;refGene_tx=XM_019833581.2
A1	119452978	119453064
refGene_gn=LOC101100942;refGene_tx=XM_019833581.2
A1	119459978	119462399
refGene_gn=LOC101100942;refGene_tx=XM_019833483.2
A1	119467240	119469748
refGene_gn=LOC101100942;refGene_tx=XM_019833431.2
A1	119472766	119475190
refGene_gn=LOC101100942;refGene_tx=XM_019833458.2
A1	119478106	119480521
refGene_gn=LOC101100942;refGene_tx=XM_019833499.2
A1	119487849	119490276
refGene_gn=LOC101100942;refGene_tx=XM_023256106.1;refGene_gn=LOC101100942;refGene_tx=XM_023256123.1
A1	119492274	119494695
refGene_gn=LOC101100942;refGene_tx=XM_023256156.1
A1	119498386	119498614
refGene_gn=LOC101100942;refGene_tx=XM_023256175.1
A1	119499748	119500807
refGene_gn=LOC101100942;refGene_tx=XM_023256106.1;refGene_gn=LOC101100942;refGene_tx=XM_023256175.1
A1	119502240	119503434
refGene_gn=LOC101098682;refGene_tx=XM_006927718.4
A1	119510334	119512755
refGene_gn=LOC101100942;refGene_tx=XM_019833476.2
A1	119516128	119518525
refGene_gn=LOC101100942;refGene_tx=XM_019833511.2
A1	119520614	119523038
refGene_gn=LOC101100942;refGene_tx=XM_019833453.2
A1	119525789	119528198
refGene_gn=LOC101100942;refGene_tx=XM_011282603.3;refGene_gn=LOC101100942;refGene_tx=XM_019833506.2
A1	119530179	119530185
refGene_gn=LOC101100942;refGene_tx=XM_011282603.3
A1	119536358	119538776
refGene_gn=LOC101100942;refGene_tx=XM_019833470.2;refGene_gn=LOC101100942;refGene_tx=XM_019833488.2
A1	119540654	119543078
refGene_gn=LOC101100942;refGene_tx=XM_019833454.2
A1	119545487	119547908
refGene_gn=LOC101100942;refGene_tx=XM_019833460.2;refGene_gn=LOC101100942;refGene_tx=XM_019833470.2
A1	119549219	119551649
refGene_gn=LOC101100942;refGene_tx=XM_019833438.2
A1	119590552	119592982

refGene_gn=LOC101100942;refGene_tx=XM_019833436.2;refGene_gn=LOC101100942;refGene_tx=XM_019833584.2
A1	119599321	119601763
refGene_gn=LOC101100942;refGene_tx=XM_019833435.2
A1	119603371	119605831
refGene_gn=LOC101100942;refGene_tx=XM_019833434.2;refGene_gn=LOC101100942;refGene_tx=XM_019833550.2;refGene_gn=LOC101100942;refGene_tx=XM_019833561.2;refGene_gn=LOC101100942;refGene_tx=XM_019833593.2
A1	119608406	119608451
refGene_gn=LOC101100942;refGene_tx=XM_019833582.2
A1	119608587	119608617
refGene_gn=LOC101100942;refGene_tx=XM_019833590.2
A1	119608823	119608882
refGene_gn=LOC101100942;refGene_tx=XM_019833431.2;refGene_gn=LOC101100942;refGene_tx=XM_019833434.2;refGene_gn=LOC101100942;refGene_tx=XM_019833435.2;refGene_gn=LOC101100942;refGene_tx=XM_019833436.2;refGene_gn=LOC101100942;refGene_tx=XM_019833438.2;refGene_gn=LOC101100942;refGene_tx=XM_019833453.2;refGene_gn=LOC101100942;refGene_tx=XM_019833454.2;refGene_gn=LOC101100942;refGene_tx=XM_019833458.2;refGene_gn=LOC101100942;refGene_tx=XM_019833460.2;refGene_gn=LOC101100942;refGene_tx=XM_019833470.2;refGene_gn=LOC101100942;refGene_tx=XM_019833476.2;refGene_gn=LOC101100942;refGene_tx=XM_019833483.2;refGene_gn=LOC101100942;refGene_tx=XM_019833488.2;refGene_gn=LOC101100942;refGene_tx=XM_019833499.2;refGene_gn=LOC101100942;refGene_tx=XM_019833506.2;refGene_gn=LOC101100942;refGene_tx=XM_019833511.2;refGene_gn=LOC101100942;refGene_tx=XM_019833550.2;refGene_gn=LOC101100942;refGene_tx=XM_019833561.2;refGene_gn=LOC101100942;refGene_tx=XM_019833581.2;refGene_gn=LOC101100942;refGene_tx=XM_019833582.2;refGene_gn=LOC101100942;refGene_tx=XM_019833584.2;refGene_gn=LOC101100942;refGene_tx=XM_019833590.2;refGene_gn=LOC101100942;refGene_tx=XM_019833593.2;refGene_gn=LOC101100942;refGene_tx=XM_023256106.1;refGene_gn=LOC101100942;refGene_tx=XM_023256123.1;refGene_gn=LOC101100942;refGene_tx=XM_023256156.1;refGene_gn=LOC101100942;refGene_tx=XM_023256175.1
A1	119616897	119616986
refGene_gn=LOC101100942;refGene_tx=XM_019833431.2;refGene_gn=LOC101100942;refGene_tx=XM_019833434.2;refGene_gn=LOC101100942;refGene_tx=XM_019833435.2;refGene_gn=LOC101100942;refGene_tx=XM_019833436.2;refGene_gn=LOC101100942;refGene_tx=XM_019833438.2;refGene_gn=LOC101100942;refGene_tx=XM_019833453.2;refGene_gn=LOC101100942;refGene_tx=XM_019833454.2;refGene_gn=LOC101100942;refGene_tx=XM_019833458.2;refGene_gn=LOC101100942;refGene_tx=XM_019833460.2;refGene_gn=LOC101100942;refGene_tx=XM_019833470.2;refGene_gn=LOC101100942;refGene_tx=XM_019833476.2;refGene_gn=LOC101100942;refGene_tx=XM_019833483.2;refGene_gn=LOC101100942;refGene_tx=XM_019833488.2;refGene_gn=LOC101100942;refGene_tx=XM_019833499.2;refGene_gn=LOC101100942;refGene_tx=XM_019833506.2;refGene_gn=LOC101100942;refGene_tx=XM_019833511.2;refGene_gn=LOC101100942;refGene_tx=XM_019833550.2;refGene_gn=LOC101100942;refGene_tx=XM_019833561.2;refGene_gn=LOC101100942;refGene_tx=XM_019833581.2;refGene_gn=LOC101100942;refGene_tx=XM_019833582.2;refGene_gn=LOC101100942;refGene_tx=XM_019833584.2;refGene_gn=LOC101100942;refGene_tx=XM_019833590.2;refGene_gn=LOC101100

942;refGene_tx=XM_019833593.2;refGene_gn=LOC101100942;refGene_tx=XM_023256106.1;refGene_gn=LOC101100942;refGene_tx=XM_023256123.1;refGene_gn=LOC101100942;refGene_tx=XM_023256156.1;refGene_gn=LOC101100942;refGene_tx=XM_023256175.1
A1 119622521 119622748
refGene_gn=LOC101100942;refGene_tx=XM_019833431.2;refGene_gn=LOC101100942;refGene_tx=XM_019833434.2;refGene_gn=LOC101100942;refGene_tx=XM_019833435.2;refGene_gn=LOC101100942;refGene_tx=XM_019833436.2;refGene_gn=LOC101100942;refGene_tx=XM_019833438.2;refGene_gn=LOC101100942;refGene_tx=XM_019833453.2;refGene_gn=LOC101100942;refGene_tx=XM_019833454.2;refGene_gn=LOC101100942;refGene_tx=XM_019833458.2;refGene_gn=LOC101100942;refGene_tx=XM_019833460.2;refGene_gn=LOC101100942;refGene_tx=XM_019833470.2;refGene_gn=LOC101100942;refGene_tx=XM_019833476.2;refGene_gn=LOC101100942;refGene_tx=XM_019833483.2;refGene_gn=LOC101100942;refGene_tx=XM_019833488.2;refGene_gn=LOC101100942;refGene_tx=XM_019833499.2;refGene_gn=LOC101100942;refGene_tx=XM_019833506.2;refGene_gn=LOC101100942;refGene_tx=XM_019833511.2;refGene_gn=LOC101100942;refGene_tx=XM_019833550.2;refGene_gn=LOC101100942;refGene_tx=XM_019833561.2;refGene_gn=LOC101100942;refGene_tx=XM_019833581.2;refGene_gn=LOC101100942;refGene_tx=XM_019833582.2;refGene_gn=LOC101100942;refGene_tx=XM_019833584.2;refGene_gn=LOC101100942;refGene_tx=XM_019833590.2;refGene_gn=LOC101100942;refGene_tx=XM_019833593.2;refGene_gn=LOC101100942;refGene_tx=XM_023256106.1;refGene_gn=LOC101100942;refGene_tx=XM_023256123.1;refGene_gn=LOC101100942;refGene_tx=XM_023256156.1;refGene_gn=LOC101100942;refGene_tx=XM_023256175.1
A1 119628341 119628499
refGene_gn=DIAPH1;refGene_tx=XM_019833607.2;refGene_gn=DIAPH1;refGene_tx=XM_023256080.1;refGene_gn=DIAPH1;refGene_tx=XM_023256084.1;refGene_gn=DIAPH1;refGene_tx=XM_023256089.1;refGene_gn=DIAPH1;refGene_tx=XM_023256092.1
A1 119630073 119630123
refGene_gn=DIAPH1;refGene_tx=XM_019833607.2
A1 119634520 119634607
refGene_gn=DIAPH1;refGene_tx=XM_019833607.2;refGene_gn=DIAPH1;refGene_tx=XM_023256080.1;refGene_gn=DIAPH1;refGene_tx=XM_023256084.1;refGene_gn=DIAPH1;refGene_tx=XM_023256089.1;refGene_gn=DIAPH1;refGene_tx=XM_023256092.1
A1 119636526 119636662
refGene_gn=DIAPH1;refGene_tx=XM_019833607.2;refGene_gn=DIAPH1;refGene_tx=XM_023256080.1;refGene_gn=DIAPH1;refGene_tx=XM_023256084.1;refGene_gn=DIAPH1;refGene_tx=XM_023256089.1;refGene_gn=DIAPH1;refGene_tx=XM_023256092.1
A1 119636801 119636966
refGene_gn=DIAPH1;refGene_tx=XM_019833607.2;refGene_gn=DIAPH1;refGene_tx=XM_023256080.1;refGene_gn=DIAPH1;refGene_tx=XM_023256084.1;refGene_gn=DIAPH1;refGene_tx=XM_023256089.1;refGene_gn=DIAPH1;refGene_tx=XM_023256092.1
A1 119637539 119637664
refGene_gn=DIAPH1;refGene_tx=XM_019833607.2;refGene_gn=DIAPH1;refGene_tx=XM_023256080.1;refGene_gn=DIAPH1;refGene_tx=XM_023256084.1;refGene_

gn=DIAPH1;refGene_tx=XM_023256089.1;refGene_gn=DIAPH1;refGene_tx=XM_023256092.1
A1 119638399 119638529
refGene_gn=DIAPH1;refGene_tx=XM_019833607.2;refGene_gn=DIAPH1;refGene_tx=XM_023256080.1;refGene_gn=DIAPH1;refGene_tx=XM_023256084.1;refGene_gn=DIAPH1;refGene_tx=XM_023256089.1;refGene_gn=DIAPH1;refGene_tx=XM_023256092.1
A1 119638662 119638902
refGene_gn=DIAPH1;refGene_tx=XM_019833607.2;refGene_gn=DIAPH1;refGene_tx=XM_023256080.1;refGene_gn=DIAPH1;refGene_tx=XM_023256084.1;refGene_gn=DIAPH1;refGene_tx=XM_023256089.1;refGene_gn=DIAPH1;refGene_tx=XM_023256092.1
A1 119639132 119639234
refGene_gn=DIAPH1;refGene_tx=XM_019833607.2;refGene_gn=DIAPH1;refGene_tx=XM_023256080.1;refGene_gn=DIAPH1;refGene_tx=XM_023256084.1;refGene_gn=DIAPH1;refGene_tx=XM_023256089.1;refGene_gn=DIAPH1;refGene_tx=XM_023256092.1
A1 119639557 119639652
refGene_gn=DIAPH1;refGene_tx=XM_019833607.2;refGene_gn=DIAPH1;refGene_tx=XM_023256080.1;refGene_gn=DIAPH1;refGene_tx=XM_023256084.1;refGene_gn=DIAPH1;refGene_tx=XM_023256089.1;refGene_gn=DIAPH1;refGene_tx=XM_023256092.1
A1 119643553 119643652
refGene_gn=DIAPH1;refGene_tx=XM_019833607.2;refGene_gn=DIAPH1;refGene_tx=XM_023256080.1;refGene_gn=DIAPH1;refGene_tx=XM_023256084.1;refGene_gn=DIAPH1;refGene_tx=XM_023256089.1;refGene_gn=DIAPH1;refGene_tx=XM_023256092.1
A1 119686766 119686775
refGene_gn=DIAPH1;refGene_tx=XM_019833607.2;refGene_gn=DIAPH1;refGene_tx=XM_023256080.1;refGene_gn=DIAPH1;refGene_tx=XM_023256089.1
A1 119687269 119687384
refGene_gn=DIAPH1;refGene_tx=XM_019833607.2;refGene_gn=DIAPH1;refGene_tx=XM_023256080.1;refGene_gn=DIAPH1;refGene_tx=XM_023256084.1;refGene_gn=DIAPH1;refGene_tx=XM_023256089.1;refGene_gn=DIAPH1;refGene_tx=XM_023256092.1
A1 119688268 119689048
refGene_gn=DIAPH1;refGene_tx=XM_019833607.2;refGene_gn=DIAPH1;refGene_tx=XM_023256080.1;refGene_gn=DIAPH1;refGene_tx=XM_023256084.1;refGene_gn=DIAPH1;refGene_tx=XM_023256089.1;refGene_gn=DIAPH1;refGene_tx=XM_023256092.1
A1 119690009 119690189
refGene_gn=DIAPH1;refGene_tx=XM_019833607.2;refGene_gn=DIAPH1;refGene_tx=XM_023256080.1;refGene_gn=DIAPH1;refGene_tx=XM_023256084.1;refGene_gn=DIAPH1;refGene_tx=XM_023256089.1;refGene_gn=DIAPH1;refGene_tx=XM_023256092.1
A1 119691299 119691364
refGene_gn=DIAPH1;refGene_tx=XM_019833607.2;refGene_gn=DIAPH1;refGene_tx=XM_023256080.1;refGene_gn=DIAPH1;refGene_tx=XM_023256084.1;refGene_gn=DIAPH1;refGene_tx=XM_023256089.1;refGene_gn=DIAPH1;refGene_tx=XM_023256092.1

A1 119691875 119691991
refGene_gn=DIAPH1;refGene_tx=XM_019833607.2;refGene_gn=DIAPH1;refGene_tx=XM_023256080.1;refGene_gn=DIAPH1;refGene_tx=XM_023256084.1;refGene_gn=DIAPH1;refGene_tx=XM_023256089.1;refGene_gn=DIAPH1;refGene_tx=XM_023256092.1
A1 119692592 119692709
refGene_gn=DIAPH1;refGene_tx=XM_019833607.2;refGene_gn=DIAPH1;refGene_tx=XM_023256080.1;refGene_gn=DIAPH1;refGene_tx=XM_023256084.1;refGene_gn=DIAPH1;refGene_tx=XM_023256089.1;refGene_gn=DIAPH1;refGene_tx=XM_023256092.1
A1 119693371 119693490
refGene_gn=DIAPH1;refGene_tx=XM_019833607.2;refGene_gn=DIAPH1;refGene_tx=XM_023256080.1;refGene_gn=DIAPH1;refGene_tx=XM_023256084.1;refGene_gn=DIAPH1;refGene_tx=XM_023256089.1;refGene_gn=DIAPH1;refGene_tx=XM_023256092.1
A1 119693663 119693774
refGene_gn=DIAPH1;refGene_tx=XM_019833607.2;refGene_gn=DIAPH1;refGene_tx=XM_023256080.1;refGene_gn=DIAPH1;refGene_tx=XM_023256084.1;refGene_gn=DIAPH1;refGene_tx=XM_023256089.1;refGene_gn=DIAPH1;refGene_tx=XM_023256092.1
A1 119694672 119694781
refGene_gn=DIAPH1;refGene_tx=XM_019833607.2;refGene_gn=DIAPH1;refGene_tx=XM_023256080.1;refGene_gn=DIAPH1;refGene_tx=XM_023256084.1;refGene_gn=DIAPH1;refGene_tx=XM_023256089.1;refGene_gn=DIAPH1;refGene_tx=XM_023256092.1
A1 119696531 119696671
refGene_gn=DIAPH1;refGene_tx=XM_019833607.2;refGene_gn=DIAPH1;refGene_tx=XM_023256080.1;refGene_gn=DIAPH1;refGene_tx=XM_023256084.1;refGene_gn=DIAPH1;refGene_tx=XM_023256089.1;refGene_gn=DIAPH1;refGene_tx=XM_023256092.1
A1 119698311 119698375
refGene_gn=DIAPH1;refGene_tx=XM_019833607.2;refGene_gn=DIAPH1;refGene_tx=XM_023256080.1;refGene_gn=DIAPH1;refGene_tx=XM_023256084.1;refGene_gn=DIAPH1;refGene_tx=XM_023256089.1;refGene_gn=DIAPH1;refGene_tx=XM_023256092.1
A1 119699196 119699283
refGene_gn=DIAPH1;refGene_tx=XM_019833607.2;refGene_gn=DIAPH1;refGene_tx=XM_023256080.1;refGene_gn=DIAPH1;refGene_tx=XM_023256084.1;refGene_gn=DIAPH1;refGene_tx=XM_023256089.1;refGene_gn=DIAPH1;refGene_tx=XM_023256092.1
A1 119699445 119699576
refGene_gn=DIAPH1;refGene_tx=XM_019833607.2;refGene_gn=DIAPH1;refGene_tx=XM_023256080.1;refGene_gn=DIAPH1;refGene_tx=XM_023256084.1;refGene_gn=DIAPH1;refGene_tx=XM_023256089.1;refGene_gn=DIAPH1;refGene_tx=XM_023256092.1
A1 119699925 119700027
refGene_gn=DIAPH1;refGene_tx=XM_019833607.2;refGene_gn=DIAPH1;refGene_tx=XM_023256080.1;refGene_gn=DIAPH1;refGene_tx=XM_023256084.1;refGene_gn=DIAPH1;refGene_tx=XM_023256089.1;refGene_gn=DIAPH1;refGene_tx=XM_023256092.1

A1 119702952 119703108
refGene_gn=DIAPH1;refGene_tx=XM_019833607.2;refGene_gn=DIAPH1;refGene_tx=XM_023256080.1;refGene_gn=DIAPH1;refGene_tx=XM_023256084.1;refGene_gn=DIAPH1;refGene_tx=XM_023256089.1;refGene_gn=DIAPH1;refGene_tx=XM_023256092.1
A1 119704138 119704165
refGene_gn=DIAPH1;refGene_tx=XM_019833607.2;refGene_gn=DIAPH1;refGene_tx=XM_023256080.1;refGene_gn=DIAPH1;refGene_tx=XM_023256084.1
A1 119730429 119730546
refGene_gn=DIAPH1;refGene_tx=XM_019833607.2;refGene_gn=DIAPH1;refGene_tx=XM_023256080.1;refGene_gn=DIAPH1;refGene_tx=XM_023256084.1;refGene_gn=DIAPH1;refGene_tx=XM_023256089.1;refGene_gn=DIAPH1;refGene_tx=XM_023256092.1
A1 119732931 119733001
refGene_gn=HDAC3;refGene_tx=XM_003980862.3;refGene_gn=HDAC3;refGene_tx=XM_011282619.2;refGene_gn=HDAC3;refGene_tx=XM_011282623.2
A1 119735815 119735973
refGene_gn=HDAC3;refGene_tx=XM_003980862.3;refGene_gn=HDAC3;refGene_tx=XM_011282619.2;refGene_gn=HDAC3;refGene_tx=XM_011282623.2
A1 119736271 119736351
refGene_gn=HDAC3;refGene_tx=XM_003980862.3;refGene_gn=HDAC3;refGene_tx=XM_011282619.2;refGene_gn=HDAC3;refGene_tx=XM_011282623.2
A1 119736631 119736690
refGene_gn=HDAC3;refGene_tx=XM_003980862.3;refGene_gn=HDAC3;refGene_tx=XM_011282619.2;refGene_gn=HDAC3;refGene_tx=XM_011282623.2
A1 119736816 119736906
refGene_gn=HDAC3;refGene_tx=XM_003980862.3;refGene_gn=HDAC3;refGene_tx=XM_011282619.2;refGene_gn=HDAC3;refGene_tx=XM_011282623.2
A1 119738242 119738307
refGene_gn=HDAC3;refGene_tx=XM_003980862.3;refGene_gn=HDAC3;refGene_tx=XM_011282619.2;refGene_gn=HDAC3;refGene_tx=XM_011282623.2
A1 119738470 119738544
refGene_gn=HDAC3;refGene_tx=XM_003980862.3;refGene_gn=HDAC3;refGene_tx=XM_011282619.2;refGene_gn=HDAC3;refGene_tx=XM_011282623.2
A1 119738919 119739000
refGene_gn=HDAC3;refGene_tx=XM_003980862.3;refGene_gn=HDAC3;refGene_tx=XM_011282619.2;refGene_gn=HDAC3;refGene_tx=XM_011282623.2
A1 119739534 119739668
refGene_gn=HDAC3;refGene_tx=XM_003980862.3;refGene_gn=HDAC3;refGene_tx=XM_011282619.2;refGene_gn=HDAC3;refGene_tx=XM_011282623.2
A1 119740053 119740109
refGene_gn=HDAC3;refGene_tx=XM_003980862.3;refGene_gn=HDAC3;refGene_tx=XM_011282619.2;refGene_gn=HDAC3;refGene_tx=XM_011282623.2
A1 119740229 119740286
refGene_gn=HDAC3;refGene_tx=XM_003980862.3;refGene_gn=HDAC3;refGene_tx=XM_011282619.2;refGene_gn=HDAC3;refGene_tx=XM_011282623.2
A1 119740415 119740497
refGene_gn=HDAC3;refGene_tx=XM_003980862.3;refGene_gn=HDAC3;refGene_tx=XM_011282619.2;refGene_gn=HDAC3;refGene_tx=XM_011282623.2
A1 119744823 119744966

refGene_gn=HDAC3;refGene_tx=XM_003980862.3;refGene_gn=HDAC3;refGene_tx=XM_011282619.2;refGene_gn=HDAC3;refGene_tx=XM_011282623.2
A1 119746295 119746378
refGene_gn=HDAC3;refGene_tx=XM_003980862.3;refGene_gn=HDAC3;refGene_tx=XM_011282619.2;refGene_gn=HDAC3;refGene_tx=XM_011282623.2
A1 119746469 119746761
refGene_gn=HDAC3;refGene_tx=XM_003980862.3;refGene_gn=HDAC3;refGene_tx=XM_011282619.2;refGene_gn=HDAC3;refGene_tx=XM_011282623.2
A1 119747996 119748180
refGene_gn=RELL2;refGene_tx=XM_019833709.1;refGene_gn=RELL2;refGene_tx=XM_019833716.1;refGene_gn=RELL2;refGene_tx=XM_019833719.2;refGene_gn=RELL2;refGene_tx=XM_019833723.1
A1 119748573 119748639
refGene_gn=RELL2;refGene_tx=XM_019833709.1;refGene_gn=RELL2;refGene_tx=XM_019833716.1;refGene_gn=RELL2;refGene_tx=XM_019833719.2;refGene_gn=RELL2;refGene_tx=XM_019833723.1
A1 119748763 119748830
refGene_gn=RELL2;refGene_tx=XM_019833709.1;refGene_gn=RELL2;refGene_tx=XM_019833716.1;refGene_gn=RELL2;refGene_tx=XM_019833719.2;refGene_gn=RELL2;refGene_tx=XM_019833723.1
A1 119748858 119748951
refGene_gn=FCHSD1;refGene_tx=XM_019833656.2;refGene_gn=FCHSD1;refGene_tx=XM_019833665.2;refGene_gn=FCHSD1;refGene_tx=XM_019833675.2
A1 119749267 119749453
refGene_gn=RELL2;refGene_tx=XM_019833709.1;refGene_gn=RELL2;refGene_tx=XM_019833716.1;refGene_gn=RELL2;refGene_tx=XM_019833719.2;refGene_gn=RELL2;refGene_tx=XM_019833723.1
A1 119749724 119750100
refGene_gn=RELL2;refGene_tx=XM_019833709.1;refGene_gn=RELL2;refGene_tx=XM_019833716.1;refGene_gn=RELL2;refGene_tx=XM_019833719.2;refGene_gn=RELL2;refGene_tx=XM_019833723.1
A1 119750227 119750260
refGene_gn=RELL2;refGene_tx=XM_019833709.1;refGene_gn=RELL2;refGene_tx=XM_019833716.1;refGene_gn=RELL2;refGene_tx=XM_019833719.2;refGene_gn=RELL2;refGene_tx=XM_019833723.1
A1 119751340 119751406
refGene_gn=FCHSD1;refGene_tx=XM_006927669.4;refGene_gn=FCHSD1;refGene_tx=XM_006927670.4
A1 119751534 119751590
refGene_gn=FCHSD1;refGene_tx=XM_006927669.4;refGene_gn=FCHSD1;refGene_tx=XM_006927670.4;refGene_gn=FCHSD1;refGene_tx=XM_019833656.2;refGene_gn=FCHSD1;refGene_tx=XM_019833665.2;refGene_gn=FCHSD1;refGene_tx=XM_019833675.2
A1 119752463 119752551
refGene_gn=FCHSD1;refGene_tx=XM_006927669.4;refGene_gn=FCHSD1;refGene_tx=XM_006927670.4;refGene_gn=FCHSD1;refGene_tx=XM_019833656.2;refGene_gn=FCHSD1;refGene_tx=XM_019833665.2;refGene_gn=FCHSD1;refGene_tx=XM_019833675.2
A1 119753334 119753555
refGene_gn=FCHSD1;refGene_tx=XM_006927669.4;refGene_gn=FCHSD1;refGene_

tx=XM_006927670.4;refGene_gn=FCHSD1;refGene_tx=XM_019833656.2;refGene_
gn=FCHSD1;refGene_tx=XM_019833665.2;refGene_gn=FCHSD1;refGene_tx=XM_01
9833675.2
A1 119753671 119753789
refGene_gn=FCHSD1;refGene_tx=XM_006927669.4;refGene_gn=FCHSD1;refGene_
tx=XM_006927670.4;refGene_gn=FCHSD1;refGene_tx=XM_019833656.2;refGene_
gn=FCHSD1;refGene_tx=XM_019833665.2;refGene_gn=FCHSD1;refGene_tx=XM_01
9833675.2
A1 119753968 119754052
refGene_gn=FCHSD1;refGene_tx=XM_006927669.4;refGene_gn=FCHSD1;refGene_
tx=XM_006927670.4;refGene_gn=FCHSD1;refGene_tx=XM_019833656.2;refGene_
gn=FCHSD1;refGene_tx=XM_019833665.2;refGene_gn=FCHSD1;refGene_tx=XM_01
9833675.2
A1 119754137 119754266
refGene_gn=FCHSD1;refGene_tx=XM_006927669.4;refGene_gn=FCHSD1;refGene_
tx=XM_006927670.4;refGene_gn=FCHSD1;refGene_tx=XM_019833656.2;refGene_
gn=FCHSD1;refGene_tx=XM_019833665.2;refGene_gn=FCHSD1;refGene_tx=XM_01
9833675.2
A1 119755129 119755291
refGene_gn=FCHSD1;refGene_tx=XM_006927669.4;refGene_gn=FCHSD1;refGene_
tx=XM_006927670.4;refGene_gn=FCHSD1;refGene_tx=XM_019833656.2;refGene_
gn=FCHSD1;refGene_tx=XM_019833665.2;refGene_gn=FCHSD1;refGene_tx=XM_01
9833675.2
A1 119755479 119755584
refGene_gn=FCHSD1;refGene_tx=XM_006927669.4;refGene_gn=FCHSD1;refGene_
tx=XM_006927670.4;refGene_gn=FCHSD1;refGene_tx=XM_019833656.2;refGene_
gn=FCHSD1;refGene_tx=XM_019833665.2;refGene_gn=FCHSD1;refGene_tx=XM_01
9833675.2
A1 119755994 119756120
refGene_gn=FCHSD1;refGene_tx=XM_006927669.4;refGene_gn=FCHSD1;refGene_
tx=XM_006927670.4;refGene_gn=FCHSD1;refGene_tx=XM_019833656.2;refGene_
gn=FCHSD1;refGene_tx=XM_019833665.2;refGene_gn=FCHSD1;refGene_tx=XM_01
9833675.2
A1 119756531 119756627
refGene_gn=FCHSD1;refGene_tx=XM_006927669.4;refGene_gn=FCHSD1;refGene_
tx=XM_006927670.4;refGene_gn=FCHSD1;refGene_tx=XM_019833656.2;refGene_
gn=FCHSD1;refGene_tx=XM_019833665.2;refGene_gn=FCHSD1;refGene_tx=XM_01
9833675.2
A1 119756792 119756915
refGene_gn=FCHSD1;refGene_tx=XM_006927669.4;refGene_gn=FCHSD1;refGene_
tx=XM_006927670.4;refGene_gn=FCHSD1;refGene_tx=XM_019833656.2;refGene_
gn=FCHSD1;refGene_tx=XM_019833665.2;refGene_gn=FCHSD1;refGene_tx=XM_01
9833675.2
A1 119757374 119757503
refGene_gn=FCHSD1;refGene_tx=XM_006927669.4;refGene_gn=FCHSD1;refGene_
tx=XM_006927670.4;refGene_gn=FCHSD1;refGene_tx=XM_019833656.2;refGene_
gn=FCHSD1;refGene_tx=XM_019833665.2;refGene_gn=FCHSD1;refGene_tx=XM_01
9833675.2
A1 119758303 119758367
refGene_gn=FCHSD1;refGene_tx=XM_006927669.4;refGene_gn=FCHSD1;refGene_

tx=XM_006927670.4;refGene_gn=FCHSD1;refGene_tx=XM_019833656.2;refGene_gn=FCHSD1;refGene_tx=XM_019833665.2;refGene_gn=FCHSD1;refGene_tx=XM_019833675.2
A1 119758462 119758599
refGene_gn=FCHSD1;refGene_tx=XM_006927669.4;refGene_gn=FCHSD1;refGene_tx=XM_006927670.4;refGene_gn=FCHSD1;refGene_tx=XM_019833656.2;refGene_gn=FCHSD1;refGene_tx=XM_019833665.2;refGene_gn=FCHSD1;refGene_tx=XM_019833675.2
A1 119758684 119758841
refGene_gn=FCHSD1;refGene_tx=XM_006927669.4;refGene_gn=FCHSD1;refGene_tx=XM_006927670.4;refGene_gn=FCHSD1;refGene_tx=XM_019833656.2;refGene_gn=FCHSD1;refGene_tx=XM_019833665.2
A1 119759164 119759232
refGene_gn=FCHSD1;refGene_tx=XM_006927669.4;refGene_gn=FCHSD1;refGene_tx=XM_006927670.4;refGene_gn=FCHSD1;refGene_tx=XM_019833656.2;refGene_gn=FCHSD1;refGene_tx=XM_019833665.2
A1 119759630 119759676
refGene_gn=FCHSD1;refGene_tx=XM_006927669.4;refGene_gn=FCHSD1;refGene_tx=XM_006927670.4;refGene_gn=FCHSD1;refGene_tx=XM_019833656.2;refGene_gn=FCHSD1;refGene_tx=XM_019833665.2;refGene_gn=FCHSD1;refGene_tx=XM_019833675.2
A1 119760286 119760384
refGene_gn=FCHSD1;refGene_tx=XM_006927669.4;refGene_gn=FCHSD1;refGene_tx=XM_006927670.4;refGene_gn=FCHSD1;refGene_tx=XM_019833656.2;refGene_gn=FCHSD1;refGene_tx=XM_019833665.2;refGene_gn=FCHSD1;refGene_tx=XM_019833675.2
A1 119760617 119760638
refGene_gn=FCHSD1;refGene_tx=XM_006927669.4;refGene_gn=FCHSD1;refGene_tx=XM_006927670.4;refGene_gn=FCHSD1;refGene_tx=XM_019833656.2;refGene_gn=FCHSD1;refGene_tx=XM_019833665.2;refGene_gn=FCHSD1;refGene_tx=XM_019833675.2
A1 119763331 119763814
refGene_gn=ARAP3;refGene_tx=XM_003980865.5;refGene_gn=ARAP3;refGene_tx=XM_019833630.2;refGene_gn=ARAP3;refGene_tx=XM_023256196.1;refGene_gn=ARAP3;refGene_tx=XM_023256204.1;refGene_gn=ARAP3;refGene_tx=XM_023256209.1;refGene_gn=ARAP3;refGene_tx=XM_023256217.1
A1 119764402 119764441
refGene_gn=ARAP3;refGene_tx=XM_003980865.5;refGene_gn=ARAP3;refGene_tx=XM_019833630.2;refGene_gn=ARAP3;refGene_tx=XM_023256196.1;refGene_gn=ARAP3;refGene_tx=XM_023256204.1;refGene_gn=ARAP3;refGene_tx=XM_023256217.1
A1 119764647 119764785
refGene_gn=ARAP3;refGene_tx=XM_003980865.5;refGene_gn=ARAP3;refGene_tx=XM_019833630.2;refGene_gn=ARAP3;refGene_tx=XM_023256196.1;refGene_gn=ARAP3;refGene_tx=XM_023256204.1;refGene_gn=ARAP3;refGene_tx=XM_023256209.1;refGene_gn=ARAP3;refGene_tx=XM_023256217.1
A1 119764883 119764947
refGene_gn=ARAP3;refGene_tx=XM_003980865.5;refGene_gn=ARAP3;refGene_tx=XM_019833630.2;refGene_gn=ARAP3;refGene_tx=XM_023256196.1;refGene_gn=ARAP3;refGene_tx=XM_023256204.1;refGene_gn=ARAP3;refGene_tx=XM_0232562

09.1;refGene_gn=ARAP3;refGene_tx=XM_023256217.1
A1	119765071	119765107
refGene_gn=ARAP3;refGene_tx=XM_003980865.5;refGene_gn=ARAP3;refGene_tx=XM_019833630.2;refGene_gn=ARAP3;refGene_tx=XM_023256196.1;refGene_gn=ARAP3;refGene_tx=XM_023256204.1;refGene_gn=ARAP3;refGene_tx=XM_023256209.1;refGene_gn=ARAP3;refGene_tx=XM_023256217.1
A1	119765203	119765286
refGene_gn=ARAP3;refGene_tx=XM_003980865.5;refGene_gn=ARAP3;refGene_tx=XM_019833630.2;refGene_gn=ARAP3;refGene_tx=XM_023256196.1;refGene_gn=ARAP3;refGene_tx=XM_023256204.1;refGene_gn=ARAP3;refGene_tx=XM_023256209.1;refGene_gn=ARAP3;refGene_tx=XM_023256217.1
A1	119765500	119765634
refGene_gn=ARAP3;refGene_tx=XM_003980865.5;refGene_gn=ARAP3;refGene_tx=XM_019833630.2;refGene_gn=ARAP3;refGene_tx=XM_023256196.1;refGene_gn=ARAP3;refGene_tx=XM_023256204.1;refGene_gn=ARAP3;refGene_tx=XM_023256209.1;refGene_gn=ARAP3;refGene_tx=XM_023256217.1
A1	119765719	119765848
refGene_gn=ARAP3;refGene_tx=XM_003980865.5;refGene_gn=ARAP3;refGene_tx=XM_019833630.2;refGene_gn=ARAP3;refGene_tx=XM_023256196.1;refGene_gn=ARAP3;refGene_tx=XM_023256204.1;refGene_gn=ARAP3;refGene_tx=XM_023256209.1;refGene_gn=ARAP3;refGene_tx=XM_023256217.1
A1	119767498	119767613
refGene_gn=ARAP3;refGene_tx=XM_003980865.5;refGene_gn=ARAP3;refGene_tx=XM_019833630.2;refGene_gn=ARAP3;refGene_tx=XM_023256196.1;refGene_gn=ARAP3;refGene_tx=XM_023256204.1;refGene_gn=ARAP3;refGene_tx=XM_023256209.1;refGene_gn=ARAP3;refGene_tx=XM_023256217.1
A1	119767711	119767786
refGene_gn=ARAP3;refGene_tx=XM_003980865.5;refGene_gn=ARAP3;refGene_tx=XM_019833630.2;refGene_gn=ARAP3;refGene_tx=XM_023256196.1;refGene_gn=ARAP3;refGene_tx=XM_023256204.1;refGene_gn=ARAP3;refGene_tx=XM_023256209.1;refGene_gn=ARAP3;refGene_tx=XM_023256217.1
A1	119768690	119768759
refGene_gn=ARAP3;refGene_tx=XM_003980865.5;refGene_gn=ARAP3;refGene_tx=XM_019833630.2;refGene_gn=ARAP3;refGene_tx=XM_023256196.1;refGene_gn=ARAP3;refGene_tx=XM_023256204.1;refGene_gn=ARAP3;refGene_tx=XM_023256209.1;refGene_gn=ARAP3;refGene_tx=XM_023256217.1
A1	119769059	119769207
refGene_gn=ARAP3;refGene_tx=XM_003980865.5;refGene_gn=ARAP3;refGene_tx=XM_019833630.2;refGene_gn=ARAP3;refGene_tx=XM_023256196.1;refGene_gn=ARAP3;refGene_tx=XM_023256204.1;refGene_gn=ARAP3;refGene_tx=XM_023256209.1;refGene_gn=ARAP3;refGene_tx=XM_023256217.1
A1	119770200	119770306
refGene_gn=ARAP3;refGene_tx=XM_003980865.5;refGene_gn=ARAP3;refGene_tx=XM_019833630.2;refGene_gn=ARAP3;refGene_tx=XM_023256196.1;refGene_gn=ARAP3;refGene_tx=XM_023256204.1;refGene_gn=ARAP3;refGene_tx=XM_023256209.1;refGene_gn=ARAP3;refGene_tx=XM_023256217.1
A1	119770592	119770805
refGene_gn=ARAP3;refGene_tx=XM_003980865.5;refGene_gn=ARAP3;refGene_tx=XM_019833630.2;refGene_gn=ARAP3;refGene_tx=XM_023256196.1;refGene_gn=ARAP3;refGene_tx=XM_023256204.1;refGene_gn=ARAP3;refGene_tx=XM_0232562

09.1
A1 119772917 119773081
refGene_gn=ARAP3;refGene_tx=XM_003980865.5;refGene_gn=ARAP3;refGene_tx=XM_019833630.2;refGene_gn=ARAP3;refGene_tx=XM_023256196.1;refGene_gn=ARAP3;refGene_tx=XM_023256204.1;refGene_gn=ARAP3;refGene_tx=XM_023256209.1;refGene_gn=ARAP3;refGene_tx=XM_023256217.1
A1 119773330 119773394
refGene_gn=ARAP3;refGene_tx=XM_003980865.5;refGene_gn=ARAP3;refGene_tx=XM_019833630.2;refGene_gn=ARAP3;refGene_tx=XM_023256196.1;refGene_gn=ARAP3;refGene_tx=XM_023256204.1;refGene_gn=ARAP3;refGene_tx=XM_023256209.1;refGene_gn=ARAP3;refGene_tx=XM_023256217.1
A1 119774097 119774317
refGene_gn=ARAP3;refGene_tx=XM_003980865.5;refGene_gn=ARAP3;refGene_tx=XM_019833630.2;refGene_gn=ARAP3;refGene_tx=XM_023256196.1;refGene_gn=ARAP3;refGene_tx=XM_023256204.1;refGene_gn=ARAP3;refGene_tx=XM_023256209.1;refGene_gn=ARAP3;refGene_tx=XM_023256217.1
A1 119776362 119776465
refGene_gn=ARAP3;refGene_tx=XM_003980865.5;refGene_gn=ARAP3;refGene_tx=XM_019833630.2;refGene_gn=ARAP3;refGene_tx=XM_023256196.1;refGene_gn=ARAP3;refGene_tx=XM_023256204.1;refGene_gn=ARAP3;refGene_tx=XM_023256209.1;refGene_gn=ARAP3;refGene_tx=XM_023256217.1
A1 119776592 119776740
refGene_gn=ARAP3;refGene_tx=XM_003980865.5;refGene_gn=ARAP3;refGene_tx=XM_019833630.2;refGene_gn=ARAP3;refGene_tx=XM_023256196.1;refGene_gn=ARAP3;refGene_tx=XM_023256204.1;refGene_gn=ARAP3;refGene_tx=XM_023256209.1;refGene_gn=ARAP3;refGene_tx=XM_023256217.1
A1 119777178 119777295
refGene_gn=ARAP3;refGene_tx=XM_003980865.5;refGene_gn=ARAP3;refGene_tx=XM_019833630.2;refGene_gn=ARAP3;refGene_tx=XM_023256196.1;refGene_gn=ARAP3;refGene_tx=XM_023256204.1;refGene_gn=ARAP3;refGene_tx=XM_023256209.1;refGene_gn=ARAP3;refGene_tx=XM_023256217.1
A1 119777859 119777995
refGene_gn=ARAP3;refGene_tx=XM_003980865.5;refGene_gn=ARAP3;refGene_tx=XM_019833630.2;refGene_gn=ARAP3;refGene_tx=XM_023256196.1;refGene_gn=ARAP3;refGene_tx=XM_023256204.1;refGene_gn=ARAP3;refGene_tx=XM_023256209.1;refGene_gn=ARAP3;refGene_tx=XM_023256217.1
A1 119778152 119778335
refGene_gn=ARAP3;refGene_tx=XM_003980865.5;refGene_gn=ARAP3;refGene_tx=XM_019833630.2;refGene_gn=ARAP3;refGene_tx=XM_023256196.1;refGene_gn=ARAP3;refGene_tx=XM_023256204.1;refGene_gn=ARAP3;refGene_tx=XM_023256209.1;refGene_gn=ARAP3;refGene_tx=XM_023256217.1
A1 119778475 119778561
refGene_gn=ARAP3;refGene_tx=XM_003980865.5;refGene_gn=ARAP3;refGene_tx=XM_019833630.2;refGene_gn=ARAP3;refGene_tx=XM_023256196.1;refGene_gn=ARAP3;refGene_tx=XM_023256204.1;refGene_gn=ARAP3;refGene_tx=XM_023256209.1;refGene_gn=ARAP3;refGene_tx=XM_023256217.1
A1 119778659 119778859
refGene_gn=ARAP3;refGene_tx=XM_003980865.5;refGene_gn=ARAP3;refGene_tx=XM_019833630.2;refGene_gn=ARAP3;refGene_tx=XM_023256196.1;refGene_gn=ARAP3;refGene_tx=XM_023256204.1;refGene_gn=ARAP3;refGene_tx=XM_0232562

09.1;refGene_gn=ARAP3;refGene_tx=XM_023256217.1
A1 119779360 119779467
refGene_gn=ARAP3;refGene_tx=XM_003980865.5;refGene_gn=ARAP3;refGene_tx=XM_019833630.2;refGene_gn=ARAP3;refGene_tx=XM_023256196.1;refGene_gn=ARAP3;refGene_tx=XM_023256204.1;refGene_gn=ARAP3;refGene_tx=XM_023256209.1;refGene_gn=ARAP3;refGene_tx=XM_023256217.1
A1 119779556 119779741
refGene_gn=ARAP3;refGene_tx=XM_003980865.5;refGene_gn=ARAP3;refGene_tx=XM_019833630.2;refGene_gn=ARAP3;refGene_tx=XM_023256196.1;refGene_gn=ARAP3;refGene_tx=XM_023256204.1;refGene_gn=ARAP3;refGene_tx=XM_023256209.1;refGene_gn=ARAP3;refGene_tx=XM_023256217.1
A1 119779835 119779956
refGene_gn=ARAP3;refGene_tx=XM_003980865.5;refGene_gn=ARAP3;refGene_tx=XM_019833630.2;refGene_gn=ARAP3;refGene_tx=XM_023256196.1;refGene_gn=ARAP3;refGene_tx=XM_023256204.1;refGene_gn=ARAP3;refGene_tx=XM_023256209.1;refGene_gn=ARAP3;refGene_tx=XM_023256217.1
A1 119780229 119780299
refGene_gn=ARAP3;refGene_tx=XM_003980865.5;refGene_gn=ARAP3;refGene_tx=XM_019833630.2;refGene_gn=ARAP3;refGene_tx=XM_023256196.1;refGene_gn=ARAP3;refGene_tx=XM_023256204.1;refGene_gn=ARAP3;refGene_tx=XM_023256209.1;refGene_gn=ARAP3;refGene_tx=XM_023256217.1
A1 119780458 119780659
refGene_gn=ARAP3;refGene_tx=XM_003980865.5;refGene_gn=ARAP3;refGene_tx=XM_019833630.2;refGene_gn=ARAP3;refGene_tx=XM_023256196.1;refGene_gn=ARAP3;refGene_tx=XM_023256204.1;refGene_gn=ARAP3;refGene_tx=XM_023256209.1;refGene_gn=ARAP3;refGene_tx=XM_023256217.1
A1 119784707 119784819
refGene_gn=ARAP3;refGene_tx=XM_003980865.5;refGene_gn=ARAP3;refGene_tx=XM_019833630.2;refGene_gn=ARAP3;refGene_tx=XM_023256196.1;refGene_gn=ARAP3;refGene_tx=XM_023256204.1;refGene_gn=ARAP3;refGene_tx=XM_023256209.1;refGene_gn=ARAP3;refGene_tx=XM_023256217.1
A1 119784926 119784988
refGene_gn=ARAP3;refGene_tx=XM_003980865.5;refGene_gn=ARAP3;refGene_tx=XM_019833630.2;refGene_gn=ARAP3;refGene_tx=XM_023256196.1;refGene_gn=ARAP3;refGene_tx=XM_023256204.1;refGene_gn=ARAP3;refGene_tx=XM_023256209.1;refGene_gn=ARAP3;refGene_tx=XM_023256217.1
A1 119785104 119785628
refGene_gn=ARAP3;refGene_tx=XM_003980865.5;refGene_gn=ARAP3;refGene_tx=XM_019833630.2;refGene_gn=ARAP3;refGene_tx=XM_023256196.1;refGene_gn=ARAP3;refGene_tx=XM_023256204.1;refGene_gn=ARAP3;refGene_tx=XM_023256209.1;refGene_gn=ARAP3;refGene_tx=XM_023256217.1
A1 119950861 119951256
refGene_gn=PCDH1;refGene_tx=XM_023256238.1
A1 119952488 119952514
refGene_gn=PCDH1;refGene_tx=XM_023256240.1
A1 119953494 119953714
refGene_gn=PCDH1;refGene_tx=XM_023256238.1;refGene_gn=PCDH1;refGene_tx=XM_023256240.1
A1 119955281 119955413
refGene_gn=PCDH1;refGene_tx=XM_023256241.1

A1 119958922 119961118
refGene_gn=PCDH1;refGene_tx=XM_023256238.1;refGene_gn=PCDH1;refGene_tx=XM_023256240.1;refGene_gn=PCDH1;refGene_tx=XM_023256241.1
A1 119964390 119965253
refGene_gn=PCDH1;refGene_tx=XM_023256238.1;refGene_gn=PCDH1;refGene_tx=XM_023256240.1;refGene_gn=PCDH1;refGene_tx=XM_023256241.1
A1 119973463 119973503
refGene_gn=PCDH1;refGene_tx=XM_023256238.1;refGene_gn=PCDH1;refGene_tx=XM_023256240.1;refGene_gn=PCDH1;refGene_tx=XM_023256241.1
A1 120000081 120000112
refGene_gn=KIAA0141;refGene_tx=XM_003980867.5;refGene_gn=KIAA0141;refGene_tx=XM_019833753.1;refGene_gn=KIAA0141;refGene_tx=XM_019833757.2
A1 120000766 120000881
refGene_gn=KIAA0141;refGene_tx=XM_003980867.5;refGene_gn=KIAA0141;refGene_tx=XM_019833753.1;refGene_gn=KIAA0141;refGene_tx=XM_019833757.2
A1 120000975 120001099
refGene_gn=KIAA0141;refGene_tx=XM_003980867.5;refGene_gn=KIAA0141;refGene_tx=XM_019833753.1;refGene_gn=KIAA0141;refGene_tx=XM_019833757.2
A1 120003056 120003204
refGene_gn=KIAA0141;refGene_tx=XM_003980867.5;refGene_gn=KIAA0141;refGene_tx=XM_019833753.1;refGene_gn=KIAA0141;refGene_tx=XM_019833757.2
A1 120004143 120004305
refGene_gn=KIAA0141;refGene_tx=XM_003980867.5;refGene_gn=KIAA0141;refGene_tx=XM_019833753.1;refGene_gn=KIAA0141;refGene_tx=XM_019833757.2
A1 120004560 120004649
refGene_gn=KIAA0141;refGene_tx=XM_003980867.5;refGene_gn=KIAA0141;refGene_tx=XM_019833753.1;refGene_gn=KIAA0141;refGene_tx=XM_019833757.2
A1 120004750 120004847
refGene_gn=KIAA0141;refGene_tx=XM_003980867.5;refGene_gn=KIAA0141;refGene_tx=XM_019833753.1;refGene_gn=KIAA0141;refGene_tx=XM_019833757.2
A1 120007525 120007668
refGene_gn=KIAA0141;refGene_tx=XM_003980867.5;refGene_gn=KIAA0141;refGene_tx=XM_019833753.1;refGene_gn=KIAA0141;refGene_tx=XM_019833757.2
A1 120008690 120008849
refGene_gn=KIAA0141;refGene_tx=XM_003980867.5;refGene_gn=KIAA0141;refGene_tx=XM_019833753.1;refGene_gn=KIAA0141;refGene_tx=XM_019833757.2
A1 120008950 120009043
refGene_gn=KIAA0141;refGene_tx=XM_003980867.5;refGene_gn=KIAA0141;refGene_tx=XM_019833753.1
A1 120011769 120011935
refGene_gn=KIAA0141;refGene_tx=XM_003980867.5;refGene_gn=KIAA0141;refGene_tx=XM_019833753.1;refGene_gn=KIAA0141;refGene_tx=XM_019833757.2
A1 120012465 120012695
refGene_gn=KIAA0141;refGene_tx=XM_003980867.5;refGene_gn=KIAA0141;refGene_tx=XM_019833753.1;refGene_gn=KIAA0141;refGene_tx=XM_019833757.2
A1 120019105 120019602
refGene_gn=PCDH12;refGene_tx=XM_003980868.4
A1 120024046 120024198
refGene_gn=PCDH12;refGene_tx=XM_003980868.4
A1 120026056 120026154

refGene_gn=PCDH12;refGene_tx=XM_003980868.4
A1 120029185 120032065
refGene_gn=PCDH12;refGene_tx=XM_003980868.4
A1 120047376 120047530
refGene_gn=RNF14;refGene_tx=XM_003980869.5;refGene_gn=RNF14;refGene_tx=XM_006927673.4;refGene_gn=RNF14;refGene_tx=XM_006927674.4;refGene_gn=RNF14;refGene_tx=XM_006927675.4;refGene_gn=RNF14;refGene_tx=XM_006927676.4;refGene_gn=RNF14;refGene_tx=XM_006927678.4;refGene_gn=RNF14;refGene_tx=XM_006927679.4;refGene_gn=RNF14;refGene_tx=XM_019833776.2
A1 120048602 120048754
refGene_gn=RNF14;refGene_tx=XM_003980869.5;refGene_gn=RNF14;refGene_tx=XM_006927673.4;refGene_gn=RNF14;refGene_tx=XM_006927674.4;refGene_gn=RNF14;refGene_tx=XM_006927675.4;refGene_gn=RNF14;refGene_tx=XM_006927676.4;refGene_gn=RNF14;refGene_tx=XM_006927678.4;refGene_gn=RNF14;refGene_tx=XM_006927679.4;refGene_gn=RNF14;refGene_tx=XM_019833776.2
A1 120051609 120052137
refGene_gn=RNF14;refGene_tx=XM_003980869.5;refGene_gn=RNF14;refGene_tx=XM_006927673.4;refGene_gn=RNF14;refGene_tx=XM_006927674.4;refGene_gn=RNF14;refGene_tx=XM_006927675.4;refGene_gn=RNF14;refGene_tx=XM_006927676.4;refGene_gn=RNF14;refGene_tx=XM_006927678.4;refGene_gn=RNF14;refGene_tx=XM_006927679.4;refGene_gn=RNF14;refGene_tx=XM_006927680.3;refGene_gn=RNF14;refGene_tx=XM_019833776.2
A1 120053006 120053235
refGene_gn=RNF14;refGene_tx=XM_003980869.5;refGene_gn=RNF14;refGene_tx=XM_006927673.4;refGene_gn=RNF14;refGene_tx=XM_006927674.4;refGene_gn=RNF14;refGene_tx=XM_006927675.4;refGene_gn=RNF14;refGene_tx=XM_006927676.4;refGene_gn=RNF14;refGene_tx=XM_006927678.4;refGene_gn=RNF14;refGene_tx=XM_006927679.4;refGene_gn=RNF14;refGene_tx=XM_006927680.3;refGene_gn=RNF14;refGene_tx=XM_019833776.2
A1 120055987 120056160
refGene_gn=RNF14;refGene_tx=XM_003980869.5;refGene_gn=RNF14;refGene_tx=XM_006927673.4;refGene_gn=RNF14;refGene_tx=XM_006927674.4;refGene_gn=RNF14;refGene_tx=XM_006927675.4;refGene_gn=RNF14;refGene_tx=XM_006927676.4;refGene_gn=RNF14;refGene_tx=XM_006927678.4;refGene_gn=RNF14;refGene_tx=XM_006927679.4;refGene_gn=RNF14;refGene_tx=XM_006927680.3;refGene_gn=RNF14;refGene_tx=XM_019833776.2
A1 120057032 120057163
refGene_gn=RNF14;refGene_tx=XM_003980869.5;refGene_gn=RNF14;refGene_tx=XM_006927673.4;refGene_gn=RNF14;refGene_tx=XM_006927674.4;refGene_gn=RNF14;refGene_tx=XM_006927675.4;refGene_gn=RNF14;refGene_tx=XM_006927676.4;refGene_gn=RNF14;refGene_tx=XM_006927678.4;refGene_gn=RNF14;refGene_tx=XM_006927679.4;refGene_gn=RNF14;refGene_tx=XM_006927680.3;refGene_gn=RNF14;refGene_tx=XM_019833776.2
A1 120059311 120059369
refGene_gn=RNF14;refGene_tx=XM_003980869.5;refGene_gn=RNF14;refGene_tx=XM_006927673.4;refGene_gn=RNF14;refGene_tx=XM_006927674.4;refGene_gn=RNF14;refGene_tx=XM_006927675.4;refGene_gn=RNF14;refGene_tx=XM_006927676.4;refGene_gn=RNF14;refGene_tx=XM_006927678.4;refGene_gn=RNF14;refGene_tx=XM_006927679.4;refGene_gn=RNF14;refGene_tx=XM_006927680.3;refGene_gn=RNF14;refGene_tx=XM_019833776.2

A1	120071552	120071653
refGene_gn=GNPDA1;refGene_tx=XM_003980870.5;refGene_gn=GNPDA1;refGene_tx=XM_006927681.4;refGene_gn=GNPDA1;refGene_tx=XM_019833786.2
A1	120073288	120073463
refGene_gn=GNPDA1;refGene_tx=XM_003980870.5;refGene_gn=GNPDA1;refGene_tx=XM_006927681.4;refGene_gn=GNPDA1;refGene_tx=XM_019833786.2
A1	120075113	120075298
refGene_gn=GNPDA1;refGene_tx=XM_003980870.5;refGene_gn=GNPDA1;refGene_tx=XM_006927681.4;refGene_gn=GNPDA1;refGene_tx=XM_019833786.2
A1	120076320	120076503
refGene_gn=GNPDA1;refGene_tx=XM_003980870.5;refGene_gn=GNPDA1;refGene_tx=XM_006927681.4;refGene_gn=GNPDA1;refGene_tx=XM_019833786.2
A1	120078148	120078250
refGene_gn=GNPDA1;refGene_tx=XM_003980870.5;refGene_gn=GNPDA1;refGene_tx=XM_006927681.4;refGene_gn=GNPDA1;refGene_tx=XM_019833786.2
A1	120082005	120082129
refGene_gn=GNPDA1;refGene_tx=XM_003980870.5;refGene_gn=GNPDA1;refGene_tx=XM_006927681.4;refGene_gn=GNPDA1;refGene_tx=XM_019833786.2
A1	120146650	120146713
refGene_gn=NDFIP1;refGene_tx=XM_023256242.1
A1	120178104	120178192
refGene_gn=NDFIP1;refGene_tx=XM_023256242.1
A1	120183212	120183343
refGene_gn=NDFIP1;refGene_tx=XM_023256242.1
A1	120187708	120187796
refGene_gn=NDFIP1;refGene_tx=XM_023256242.1
A1	120188689	120188814
refGene_gn=NDFIP1;refGene_tx=XM_023256242.1
A1	120190752	120190819
refGene_gn=NDFIP1;refGene_tx=XM_023256242.1
A1	120193716	120193820
refGene_gn=NDFIP1;refGene_tx=XM_023256242.1
A1	120337499	120338399
refGene_gn=SPRY4;refGene_tx=XM_003980872.4;refGene_gn=SPRY4;refGene_tx=XM_006927682.4
A1	120597226	120597421
refGene_gn=FGF1;refGene_tx=XM_011282701.3;refGene_gn=FGF1;refGene_tx=XM_011282703.3;refGene_gn=FGF1;refGene_tx=XM_011282706.2;refGene_gn=FGF1;refGene_tx=XM_011282709.3;refGene_gn=FGF1;refGene_tx=XM_011282711.2;refGene_gn=FGF1;refGene_tx=XM_019833811.2;refGene_gn=FGF1;refGene_tx=XM_019833813.2;refGene_gn=FGF1;refGene_tx=XM_019833819.1
A1	120602900	120603004
refGene_gn=FGF1;refGene_tx=XM_011282701.3;refGene_gn=FGF1;refGene_tx=XM_011282703.3;refGene_gn=FGF1;refGene_tx=XM_011282706.2;refGene_gn=FGF1;refGene_tx=XM_011282709.3;refGene_gn=FGF1;refGene_tx=XM_019833811.2;refGene_gn=FGF1;refGene_tx=XM_019833813.2;refGene_gn=FGF1;refGene_tx=XM_019833819.1
A1	120617551	120617720
refGene_gn=FGF1;refGene_tx=XM_011282701.3;refGene_gn=FGF1;refGene_tx=XM_011282703.3;refGene_gn=FGF1;refGene_tx=XM_011282706.2;refGene_gn=FGF

1;refGene_tx=XM_011282709.3;refGene_gn=FGF1;refGene_tx=XM_011282711.2;refGene_gn=FGF1;refGene_tx=XM_019833811.2;refGene_gn=FGF1;refGene_tx=XM_019833813.2;refGene_gn=FGF1;refGene_tx=XM_019833819.1
A1 120765950 120766104
refGene_gn=ARHGAP26;refGene_tx=XM_011282717.3;refGene_gn=ARHGAP26;refGene_tx=XM_011282723.3;refGene_gn=ARHGAP26;refGene_tx=XM_019833787.2;refGene_gn=ARHGAP26;refGene_tx=XM_019833796.2;refGene_gn=ARHGAP26;refGene_tx=XM_019833798.2;refGene_gn=ARHGAP26;refGene_tx=XM_019833799.2;refGene_gn=ARHGAP26;refGene_tx=XM_019833803.2
A1 120870088 120870184
refGene_gn=ARHGAP26;refGene_tx=XM_011282717.3;refGene_gn=ARHGAP26;refGene_tx=XM_011282723.3;refGene_gn=ARHGAP26;refGene_tx=XM_019833787.2;refGene_gn=ARHGAP26;refGene_tx=XM_019833796.2;refGene_gn=ARHGAP26;refGene_tx=XM_019833798.2;refGene_gn=ARHGAP26;refGene_tx=XM_019833799.2;refGene_gn=ARHGAP26;refGene_tx=XM_019833803.2
A1 120871744 120871806
refGene_gn=ARHGAP26;refGene_tx=XM_011282717.3;refGene_gn=ARHGAP26;refGene_tx=XM_011282723.3;refGene_gn=ARHGAP26;refGene_tx=XM_019833787.2;refGene_gn=ARHGAP26;refGene_tx=XM_019833796.2;refGene_gn=ARHGAP26;refGene_tx=XM_019833798.2;refGene_gn=ARHGAP26;refGene_tx=XM_019833799.2;refGene_gn=ARHGAP26;refGene_tx=XM_019833803.2
A1 120876241 120876313
refGene_gn=ARHGAP26;refGene_tx=XM_011282717.3;refGene_gn=ARHGAP26;refGene_tx=XM_011282723.3;refGene_gn=ARHGAP26;refGene_tx=XM_019833787.2;refGene_gn=ARHGAP26;refGene_tx=XM_019833796.2;refGene_gn=ARHGAP26;refGene_tx=XM_019833798.2;refGene_gn=ARHGAP26;refGene_tx=XM_019833799.2;refGene_gn=ARHGAP26;refGene_tx=XM_019833803.2
A1 120881822 120881924
refGene_gn=ARHGAP26;refGene_tx=XM_011282717.3;refGene_gn=ARHGAP26;refGene_tx=XM_011282723.3;refGene_gn=ARHGAP26;refGene_tx=XM_019833787.2;refGene_gn=ARHGAP26;refGene_tx=XM_019833796.2;refGene_gn=ARHGAP26;refGene_tx=XM_019833798.2;refGene_gn=ARHGAP26;refGene_tx=XM_019833799.2;refGene_gn=ARHGAP26;refGene_tx=XM_019833803.2
A1 120889631 120889742
refGene_gn=ARHGAP26;refGene_tx=XM_011282717.3;refGene_gn=ARHGAP26;refGene_tx=XM_011282723.3;refGene_gn=ARHGAP26;refGene_tx=XM_019833787.2;refGene_gn=ARHGAP26;refGene_tx=XM_019833796.2;refGene_gn=ARHGAP26;refGene_tx=XM_019833798.2;refGene_gn=ARHGAP26;refGene_tx=XM_019833799.2;refGene_gn=ARHGAP26;refGene_tx=XM_019833803.2
A1 120897718 120897823
refGene_gn=ARHGAP26;refGene_tx=XM_011282717.3;refGene_gn=ARHGAP26;refGene_tx=XM_011282723.3;refGene_gn=ARHGAP26;refGene_tx=XM_019833787.2;refGene_gn=ARHGAP26;refGene_tx=XM_019833796.2;refGene_gn=ARHGAP26;refGene_tx=XM_019833798.2;refGene_gn=ARHGAP26;refGene_tx=XM_019833799.2;refGene_gn=ARHGAP26;refGene_tx=XM_019833803.2
A1 120899524 120899654
refGene_gn=ARHGAP26;refGene_tx=XM_011282717.3;refGene_gn=ARHGAP26;refGene_tx=XM_011282723.3;refGene_gn=ARHGAP26;refGene_tx=XM_019833787.2;refGene_gn=ARHGAP26;refGene_tx=XM_019833796.2;refGene_gn=ARHGAP26;refGene_tx=XM_019833798.2;refGene_gn=ARHGAP26;refGene_tx=XM_019833799.2;refG

ene_gn=ARHGAP26;refGene_tx=XM_019833803.2
A1 120903526 120903627
refGene_gn=ARHGAP26;refGene_tx=XM_011282717.3;refGene_gn=ARHGAP26;refGene_tx=XM_011282723.3;refGene_gn=ARHGAP26;refGene_tx=XM_019833787.2;refGene_gn=ARHGAP26;refGene_tx=XM_019833796.2;refGene_gn=ARHGAP26;refGene_tx=XM_019833798.2;refGene_gn=ARHGAP26;refGene_tx=XM_019833799.2;refGene_gn=ARHGAP26;refGene_tx=XM_019833803.2
A1 120910068 120910163
refGene_gn=ARHGAP26;refGene_tx=XM_011282717.3;refGene_gn=ARHGAP26;refGene_tx=XM_011282723.3;refGene_gn=ARHGAP26;refGene_tx=XM_019833787.2;refGene_gn=ARHGAP26;refGene_tx=XM_019833796.2;refGene_gn=ARHGAP26;refGene_tx=XM_019833798.2;refGene_gn=ARHGAP26;refGene_tx=XM_019833799.2;refGene_gn=ARHGAP26;refGene_tx=XM_019833803.2
A1 120929425 120929504
refGene_gn=ARHGAP26;refGene_tx=XM_011282717.3;refGene_gn=ARHGAP26;refGene_tx=XM_011282723.3;refGene_gn=ARHGAP26;refGene_tx=XM_019833787.2;refGene_gn=ARHGAP26;refGene_tx=XM_019833796.2;refGene_gn=ARHGAP26;refGene_tx=XM_019833798.2;refGene_gn=ARHGAP26;refGene_tx=XM_019833799.2;refGene_gn=ARHGAP26;refGene_tx=XM_019833803.2
A1 120993105 120993142
refGene_gn=ARHGAP26;refGene_tx=XM_011282717.3;refGene_gn=ARHGAP26;refGene_tx=XM_011282723.3;refGene_gn=ARHGAP26;refGene_tx=XM_019833787.2;refGene_gn=ARHGAP26;refGene_tx=XM_019833796.2;refGene_gn=ARHGAP26;refGene_tx=XM_019833798.2;refGene_gn=ARHGAP26;refGene_tx=XM_019833799.2;refGene_gn=ARHGAP26;refGene_tx=XM_019833803.2
A1 121016439 121016505
refGene_gn=ARHGAP26;refGene_tx=XM_011282717.3;refGene_gn=ARHGAP26;refGene_tx=XM_011282723.3;refGene_gn=ARHGAP26;refGene_tx=XM_019833787.2;refGene_gn=ARHGAP26;refGene_tx=XM_019833796.2;refGene_gn=ARHGAP26;refGene_tx=XM_019833798.2;refGene_gn=ARHGAP26;refGene_tx=XM_019833799.2;refGene_gn=ARHGAP26;refGene_tx=XM_019833803.2
A1 121020852 121020927
refGene_gn=ARHGAP26;refGene_tx=XM_011282717.3;refGene_gn=ARHGAP26;refGene_tx=XM_011282723.3;refGene_gn=ARHGAP26;refGene_tx=XM_019833787.2;refGene_gn=ARHGAP26;refGene_tx=XM_019833796.2;refGene_gn=ARHGAP26;refGene_tx=XM_019833798.2;refGene_gn=ARHGAP26;refGene_tx=XM_019833799.2;refGene_gn=ARHGAP26;refGene_tx=XM_019833803.2
A1 121034188 121034276
refGene_gn=ARHGAP26;refGene_tx=XM_011282717.3;refGene_gn=ARHGAP26;refGene_tx=XM_011282723.3;refGene_gn=ARHGAP26;refGene_tx=XM_019833787.2;refGene_gn=ARHGAP26;refGene_tx=XM_019833796.2;refGene_gn=ARHGAP26;refGene_tx=XM_019833798.2;refGene_gn=ARHGAP26;refGene_tx=XM_019833799.2;refGene_gn=ARHGAP26;refGene_tx=XM_019833803.2
A1 121036149 121036208
refGene_gn=ARHGAP26;refGene_tx=XM_011282717.3;refGene_gn=ARHGAP26;refGene_tx=XM_011282723.3;refGene_gn=ARHGAP26;refGene_tx=XM_019833787.2;refGene_gn=ARHGAP26;refGene_tx=XM_019833796.2;refGene_gn=ARHGAP26;refGene_tx=XM_019833798.2;refGene_gn=ARHGAP26;refGene_tx=XM_019833799.2;refGene_gn=ARHGAP26;refGene_tx=XM_019833803.2
A1 121037722 121037828

refGene_gn=ARHGAP26;refGene_tx=XM_011282717.3;refGene_gn=ARHGAP26;refGene_tx=XM_011282723.3;refGene_gn=ARHGAP26;refGene_tx=XM_019833787.2;refGene_gn=ARHGAP26;refGene_tx=XM_019833796.2;refGene_gn=ARHGAP26;refGene_tx=XM_019833798.2;refGene_gn=ARHGAP26;refGene_tx=XM_019833799.2;refGene_gn=ARHGAP26;refGene_tx=XM_019833803.2
A1 121090040 121090200
refGene_gn=ARHGAP26;refGene_tx=XM_011282717.3;refGene_gn=ARHGAP26;refGene_tx=XM_011282723.3;refGene_gn=ARHGAP26;refGene_tx=XM_019833787.2;refGene_gn=ARHGAP26;refGene_tx=XM_019833796.2;refGene_gn=ARHGAP26;refGene_tx=XM_019833798.2;refGene_gn=ARHGAP26;refGene_tx=XM_019833799.2;refGene_gn=ARHGAP26;refGene_tx=XM_019833803.2
A1 121105169 121105308
refGene_gn=ARHGAP26;refGene_tx=XM_011282717.3;refGene_gn=ARHGAP26;refGene_tx=XM_011282723.3;refGene_gn=ARHGAP26;refGene_tx=XM_019833787.2;refGene_gn=ARHGAP26;refGene_tx=XM_019833796.2;refGene_gn=ARHGAP26;refGene_tx=XM_019833798.2;refGene_gn=ARHGAP26;refGene_tx=XM_019833799.2;refGene_gn=ARHGAP26;refGene_tx=XM_019833803.2
A1 121119313 121119467
refGene_gn=ARHGAP26;refGene_tx=XM_011282717.3;refGene_gn=ARHGAP26;refGene_tx=XM_011282723.3;refGene_gn=ARHGAP26;refGene_tx=XM_019833787.2;refGene_gn=ARHGAP26;refGene_tx=XM_019833796.2;refGene_gn=ARHGAP26;refGene_tx=XM_019833798.2;refGene_gn=ARHGAP26;refGene_tx=XM_019833799.2;refGene_gn=ARHGAP26;refGene_tx=XM_019833803.2
A1 121173026 121173302
refGene_gn=ARHGAP26;refGene_tx=XM_011282717.3;refGene_gn=ARHGAP26;refGene_tx=XM_011282723.3;refGene_gn=ARHGAP26;refGene_tx=XM_019833787.2;refGene_gn=ARHGAP26;refGene_tx=XM_019833798.2;refGene_gn=ARHGAP26;refGene_tx=XM_019833799.2;refGene_gn=ARHGAP26;refGene_tx=XM_019833803.2
A1 121174989 121175014
refGene_gn=ARHGAP26;refGene_tx=XM_019833799.2
A1 121179011 121179103
refGene_gn=ARHGAP26;refGene_tx=XM_011282717.3;refGene_gn=ARHGAP26;refGene_tx=XM_011282723.3;refGene_gn=ARHGAP26;refGene_tx=XM_019833787.2;refGene_gn=ARHGAP26;refGene_tx=XM_019833796.2
A1 121187531 121187620
refGene_gn=ARHGAP26;refGene_tx=XM_011282717.3;refGene_gn=ARHGAP26;refGene_tx=XM_011282723.3;refGene_gn=ARHGAP26;refGene_tx=XM_019833787.2;refGene_gn=ARHGAP26;refGene_tx=XM_019833796.2
A1 121247182 121247335
refGene_gn=NR3C1;refGene_tx=NM_001305031.1;refGene_gn=NR3C1;refGene_tx=XM_003980936.5;refGene_gn=NR3C1;refGene_tx=XM_006927722.4;refGene_gn=NR3C1;refGene_tx=XM_006927723.2;refGene_gn=NR3C1;refGene_tx=XM_011282735.2;refGene_gn=NR3C1;refGene_tx=XM_019831649.2;refGene_gn=NR3C1;refGene_tx=XM_019831899.1;refGene_gn=NR3C1;refGene_tx=XM_019832312.1;refGene_gn=NR3C1;refGene_tx=XM_023254090.1
A1 121247884 121248042
refGene_gn=NR3C1;refGene_tx=NM_001305031.1;refGene_gn=NR3C1;refGene_tx=XM_003980936.5;refGene_gn=NR3C1;refGene_tx=XM_006927722.4;refGene_gn=NR3C1;refGene_tx=XM_006927723.2;refGene_gn=NR3C1;refGene_tx=XM_011282735.2;refGene_gn=NR3C1;refGene_tx=XM_019831649.2;refGene_gn=NR3C1;refGe

ne_tx=XM_019831899.1;refGene_gn=NR3C1;refGene_tx=XM_019832312.1;refGene_gn=NR3C1;refGene_tx=XM_023254090.1
A1 121257240 121257371
refGene_gn=NR3C1;refGene_tx=NM_001305031.1;refGene_gn=NR3C1;refGene_tx=XM_003980936.5;refGene_gn=NR3C1;refGene_tx=XM_006927722.4;refGene_gn=NR3C1;refGene_tx=XM_006927723.2;refGene_gn=NR3C1;refGene_tx=XM_011282735.2;refGene_gn=NR3C1;refGene_tx=XM_019831649.2;refGene_gn=NR3C1;refGene_tx=XM_019831899.1;refGene_gn=NR3C1;refGene_tx=XM_019832312.1;refGene_gn=NR3C1;refGene_tx=XM_023254090.1
A1 121261279 121261424
refGene_gn=NR3C1;refGene_tx=NM_001305031.1;refGene_gn=NR3C1;refGene_tx=XM_003980936.5;refGene_gn=NR3C1;refGene_tx=XM_006927722.4;refGene_gn=NR3C1;refGene_tx=XM_006927723.2;refGene_gn=NR3C1;refGene_tx=XM_011282735.2;refGene_gn=NR3C1;refGene_tx=XM_019831649.2;refGene_gn=NR3C1;refGene_tx=XM_019831899.1;refGene_gn=NR3C1;refGene_tx=XM_019832312.1;refGene_gn=NR3C1;refGene_tx=XM_023254090.1
A1 121262427 121262706
refGene_gn=NR3C1;refGene_tx=NM_001305031.1;refGene_gn=NR3C1;refGene_tx=XM_003980936.5;refGene_gn=NR3C1;refGene_tx=XM_006927722.4;refGene_gn=NR3C1;refGene_tx=XM_006927723.2;refGene_gn=NR3C1;refGene_tx=XM_011282735.2;refGene_gn=NR3C1;refGene_tx=XM_019831649.2;refGene_gn=NR3C1;refGene_tx=XM_019831899.1;refGene_gn=NR3C1;refGene_tx=XM_019832312.1;refGene_gn=NR3C1;refGene_tx=XM_023254090.1
A1 121267075 121267192
refGene_gn=NR3C1;refGene_tx=NM_001305031.1;refGene_gn=NR3C1;refGene_tx=XM_003980936.5;refGene_gn=NR3C1;refGene_tx=XM_006927722.4;refGene_gn=NR3C1;refGene_tx=XM_006927723.2;refGene_gn=NR3C1;refGene_tx=XM_011282735.2;refGene_gn=NR3C1;refGene_tx=XM_019831649.2;refGene_gn=NR3C1;refGene_tx=XM_019831899.1;refGene_gn=NR3C1;refGene_tx=XM_019832312.1;refGene_gn=NR3C1;refGene_tx=XM_023254090.1
A1 121271359 121271526
refGene_gn=NR3C1;refGene_tx=NM_001305031.1;refGene_gn=NR3C1;refGene_tx=XM_003980936.5;refGene_gn=NR3C1;refGene_tx=XM_006927722.4;refGene_gn=NR3C1;refGene_tx=XM_006927723.2;refGene_gn=NR3C1;refGene_tx=XM_011282735.2;refGene_gn=NR3C1;refGene_tx=XM_019831649.2;refGene_gn=NR3C1;refGene_tx=XM_019831899.1;refGene_gn=NR3C1;refGene_tx=XM_019832312.1;refGene_gn=NR3C1;refGene_tx=XM_023254090.1
A1 121355568 121356764
refGene_gn=NR3C1;refGene_tx=NM_001305031.1;refGene_gn=NR3C1;refGene_tx=XM_003980936.5;refGene_gn=NR3C1;refGene_tx=XM_006927722.4;refGene_gn=NR3C1;refGene_tx=XM_006927723.2;refGene_gn=NR3C1;refGene_tx=XM_011282735.2;refGene_gn=NR3C1;refGene_tx=XM_019831649.2;refGene_gn=NR3C1;refGene_tx=XM_019831899.1;refGene_gn=NR3C1;refGene_tx=XM_019832312.1;refGene_gn=NR3C1;refGene_tx=XM_023254090.1
A1 122036011 122036174
refGene_gn=YIPF5;refGene_tx=XM_003980938.4;refGene_gn=YIPF5;refGene_tx=XM_019833839.2
A1 122037481 122037663
refGene_gn=YIPF5;refGene_tx=XM_003980938.4;refGene_gn=YIPF5;refGene_tx=XM_019833839.2

A1 122038699 122038845
refGene_gn=YIPF5;refGene_tx=XM_003980938.4;refGene_gn=YIPF5;refGene_tx=XM_019833839.2
A1 122039680 122039853
refGene_gn=YIPF5;refGene_tx=XM_003980938.4;refGene_gn=YIPF5;refGene_tx=XM_019833839.2
A1 122046276 122046386
refGene_gn=YIPF5;refGene_tx=XM_003980938.4;refGene_gn=YIPF5;refGene_tx=XM_019833839.2
A1 122082231 122083063
refGene_gn=KCTD16;refGene_tx=XM_003980940.5
A1 122330677 122331132
refGene_gn=KCTD16;refGene_tx=XM_003980940.5
A1 123456240 123456300
refGene_gn=PRELID2;refGene_tx=XM_023256286.1;refGene_gn=PRELID2;refGene_tx=XM_023256292.1;refGene_gn=PRELID2;refGene_tx=XM_023256294.1
A1 123481849 123481955
refGene_gn=PRELID2;refGene_tx=XM_023256286.1;refGene_gn=PRELID2;refGene_tx=XM_023256292.1;refGene_gn=PRELID2;refGene_tx=XM_023256294.1
A1 123497546 123497707
refGene_gn=PRELID2;refGene_tx=XM_023256286.1;refGene_gn=PRELID2;refGene_tx=XM_023256292.1;refGene_gn=PRELID2;refGene_tx=XM_023256294.1
A1 123500097 123500171
refGene_gn=PRELID2;refGene_tx=XM_023256286.1;refGene_gn=PRELID2;refGene_tx=XM_023256292.1;refGene_gn=PRELID2;refGene_tx=XM_023256294.1
A1 123502490 123502548
refGene_gn=PRELID2;refGene_tx=XM_023256286.1;refGene_gn=PRELID2;refGene_tx=XM_023256292.1;refGene_gn=PRELID2;refGene_tx=XM_023256294.1
A1 123516091 123516166
refGene_gn=PRELID2;refGene_tx=XM_023256286.1
A1 123542280 123542463
refGene_gn=GRXCR2;refGene_tx=XM_003980942.2
A1 123547774 123548002
refGene_gn=GRXCR2;refGene_tx=XM_003980942.2
A1 123554742 123555078
refGene_gn=GRXCR2;refGene_tx=XM_003980942.2
A1 123605481 123605859
refGene_gn=SH3RF2;refGene_tx=XM_003980943.5;refGene_gn=SH3RF2;refGene_tx=XM_019833842.2;refGene_gn=SH3RF2;refGene_tx=XM_019833843.2;refGene_gn=SH3RF2;refGene_tx=XM_019833850.2;refGene_gn=SH3RF2;refGene_tx=XM_019833855.2
A1 123656589 123656859
refGene_gn=SH3RF2;refGene_tx=XM_003980943.5;refGene_gn=SH3RF2;refGene_tx=XM_019833842.2;refGene_gn=SH3RF2;refGene_tx=XM_019833843.2;refGene_gn=SH3RF2;refGene_tx=XM_019833850.2
A1 123660039 123660135
refGene_gn=SH3RF2;refGene_tx=XM_003980943.5;refGene_gn=SH3RF2;refGene_tx=XM_019833842.2;refGene_gn=SH3RF2;refGene_tx=XM_019833843.2;refGene_gn=SH3RF2;refGene_tx=XM_019833850.2;refGene_gn=SH3RF2;refGene_tx=XM_019833855.2

A1	123668007	123668331
refGene_gn=SH3RF2;refGene_tx=XM_003980943.5;refGene_gn=SH3RF2;refGene_tx=XM_019833842.2;refGene_gn=SH3RF2;refGene_tx=XM_019833843.2;refGene_gn=SH3RF2;refGene_tx=XM_019833850.2;refGene_gn=SH3RF2;refGene_tx=XM_019833855.2
A1	123705562	123705654
refGene_gn=SH3RF2;refGene_tx=XM_003980943.5;refGene_gn=SH3RF2;refGene_tx=XM_019833842.2;refGene_gn=SH3RF2;refGene_tx=XM_019833843.2;refGene_gn=SH3RF2;refGene_tx=XM_019833850.2;refGene_gn=SH3RF2;refGene_tx=XM_019833855.2
A1	123706917	123707088
refGene_gn=SH3RF2;refGene_tx=XM_003980943.5;refGene_gn=SH3RF2;refGene_tx=XM_019833842.2;refGene_gn=SH3RF2;refGene_tx=XM_019833843.2;refGene_gn=SH3RF2;refGene_tx=XM_019833855.2
A1	123714820	123715074
refGene_gn=SH3RF2;refGene_tx=XM_003980943.5;refGene_gn=SH3RF2;refGene_tx=XM_019833842.2;refGene_gn=SH3RF2;refGene_tx=XM_019833843.2;refGene_gn=SH3RF2;refGene_tx=XM_019833850.2;refGene_gn=SH3RF2;refGene_tx=XM_019833855.2
A1	123718583	123718945
refGene_gn=SH3RF2;refGene_tx=XM_003980943.5;refGene_gn=SH3RF2;refGene_tx=XM_019833842.2;refGene_gn=SH3RF2;refGene_tx=XM_019833843.2;refGene_gn=SH3RF2;refGene_tx=XM_019833850.2;refGene_gn=SH3RF2;refGene_tx=XM_019833855.2
A1	123721490	123721766
refGene_gn=SH3RF2;refGene_tx=XM_003980943.5;refGene_gn=SH3RF2;refGene_tx=XM_019833842.2;refGene_gn=SH3RF2;refGene_tx=XM_019833843.2;refGene_gn=SH3RF2;refGene_tx=XM_019833850.2;refGene_gn=SH3RF2;refGene_tx=XM_019833855.2
A1	123741470	123741611
refGene_gn=PLAC8L1;refGene_tx=XM_003980987.4;refGene_gn=PLAC8L1;refGene_tx=XM_019833864.2
A1	123742743	123742880
refGene_gn=PLAC8L1;refGene_tx=XM_003980987.4;refGene_gn=PLAC8L1;refGene_tx=XM_019833864.2
A1	123755263	123755397
refGene_gn=PLAC8L1;refGene_tx=XM_003980987.4;refGene_gn=PLAC8L1;refGene_tx=XM_019833864.2
A1	123760366	123760491
refGene_gn=PLAC8L1;refGene_tx=XM_003980987.4;refGene_gn=PLAC8L1;refGene_tx=XM_019833864.2
A1	123771031	123771237
refGene_gn=LARS;refGene_tx=XM_023256338.1
A1	123777704	123777837
refGene_gn=LARS;refGene_tx=XM_023256338.1
A1	123781881	123781893
refGene_gn=LARS;refGene_tx=XM_023256338.1
A1	123781894	123781978
refGene_gn=LARS;refGene_tx=XM_023256338.1
A1	123784354	123784459

refGene_gn=LARS;refGene_tx=XM_023256338.1
A1 123786990 123787101
refGene_gn=LARS;refGene_tx=XM_023256338.1
A1 123790418 123790529
refGene_gn=LARS;refGene_tx=XM_023256338.1
A1 123790782 123790923
refGene_gn=LARS;refGene_tx=XM_023256338.1
A1 123791982 123792123
refGene_gn=LARS;refGene_tx=XM_023256338.1
A1 123793322 123793413
refGene_gn=LARS;refGene_tx=XM_023256338.1
A1 123795129 123795313
refGene_gn=LARS;refGene_tx=XM_023256338.1
A1 123806255 123806319
refGene_gn=LARS;refGene_tx=XM_023256338.1
A1 123808295 123808353
refGene_gn=LARS;refGene_tx=XM_023256338.1
A1 123809446 123809659
refGene_gn=LARS;refGene_tx=XM_023256338.1
A1 123810719 123810858
refGene_gn=LARS;refGene_tx=XM_023256338.1
A1 123811467 123811550
refGene_gn=LARS;refGene_tx=XM_023256338.1
A1 123811648 123811714
refGene_gn=LARS;refGene_tx=XM_023256338.1
A1 123811807 123811893
refGene_gn=LARS;refGene_tx=XM_023256338.1
A1 123817231 123817309
refGene_gn=LARS;refGene_tx=XM_023256338.1
A1 123818883 123819024
refGene_gn=LARS;refGene_tx=XM_023256338.1
A1 123819793 123819847
refGene_gn=LARS;refGene_tx=XM_023256338.1
A1 123822903 123822980
refGene_gn=LARS;refGene_tx=XM_023256338.1
A1 123823107 123823195
refGene_gn=LARS;refGene_tx=XM_023256338.1
A1 123827315 123827541
refGene_gn=LARS;refGene_tx=XM_023256338.1
A1 123827645 123827713
refGene_gn=LARS;refGene_tx=XM_023256338.1
A1 123829132 123829196
refGene_gn=LARS;refGene_tx=XM_023256338.1
A1 123829876 123829989
refGene_gn=LARS;refGene_tx=XM_023256338.1
A1 123830536 123830698
refGene_gn=LARS;refGene_tx=XM_023256338.1
A1 123834844 123834982
refGene_gn=LARS;refGene_tx=XM_023256338.1
A1 123839609 123839690

refGene_gn=LARS;refGene_tx=XM_023256338.1
A1	123840622	123840710
refGene_gn=LARS;refGene_tx=XM_023256338.1
A1	123843437	123843556
refGene_gn=LARS;refGene_tx=XM_023256338.1
A1	123854030	123854036
refGene_gn=LARS;refGene_tx=XM_023256338.1
A1	123869160	123869219
refGene_gn=RBM27;refGene_tx=XM_023256343.1;refGene_gn=RBM27;refGene_tx=XM_023256351.1;refGene_gn=RBM27;refGene_tx=XM_023256355.1;refGene_gn=RBM27;refGene_tx=XM_023256363.1
A1	123884881	123885000
refGene_gn=RBM27;refGene_tx=XM_023256343.1;refGene_gn=RBM27;refGene_tx=XM_023256351.1;refGene_gn=RBM27;refGene_tx=XM_023256355.1;refGene_gn=RBM27;refGene_tx=XM_023256363.1
A1	123888886	123889011
refGene_gn=RBM27;refGene_tx=XM_023256343.1;refGene_gn=RBM27;refGene_tx=XM_023256351.1;refGene_gn=RBM27;refGene_tx=XM_023256355.1;refGene_gn=RBM27;refGene_tx=XM_023256363.1
A1	123893743	123893835
refGene_gn=RBM27;refGene_tx=XM_023256343.1;refGene_gn=RBM27;refGene_tx=XM_023256351.1;refGene_gn=RBM27;refGene_tx=XM_023256355.1;refGene_gn=RBM27;refGene_tx=XM_023256363.1
A1	123894502	123894696
refGene_gn=RBM27;refGene_tx=XM_023256343.1;refGene_gn=RBM27;refGene_tx=XM_023256351.1;refGene_gn=RBM27;refGene_tx=XM_023256355.1;refGene_gn=RBM27;refGene_tx=XM_023256363.1
A1	123895431	123895695
refGene_gn=RBM27;refGene_tx=XM_023256343.1;refGene_gn=RBM27;refGene_tx=XM_023256351.1;refGene_gn=RBM27;refGene_tx=XM_023256355.1;refGene_gn=RBM27;refGene_tx=XM_023256363.1
A1	123897853	123898141
refGene_gn=RBM27;refGene_tx=XM_023256343.1;refGene_gn=RBM27;refGene_tx=XM_023256351.1;refGene_gn=RBM27;refGene_tx=XM_023256355.1;refGene_gn=RBM27;refGene_tx=XM_023256363.1
A1	123900731	123900866
refGene_gn=RBM27;refGene_tx=XM_023256343.1;refGene_gn=RBM27;refGene_tx=XM_023256351.1;refGene_gn=RBM27;refGene_tx=XM_023256355.1;refGene_gn=RBM27;refGene_tx=XM_023256363.1
A1	123908817	123908982
refGene_gn=RBM27;refGene_tx=XM_023256343.1;refGene_gn=RBM27;refGene_tx=XM_023256351.1
A1	123910843	123910993
refGene_gn=RBM27;refGene_tx=XM_023256343.1;refGene_gn=RBM27;refGene_tx=XM_023256351.1;refGene_gn=RBM27;refGene_tx=XM_023256355.1;refGene_gn=RBM27;refGene_tx=XM_023256363.1
A1	123914437	123914582
refGene_gn=RBM27;refGene_tx=XM_023256343.1;refGene_gn=RBM27;refGene_tx=XM_023256351.1;refGene_gn=RBM27;refGene_tx=XM_023256355.1;refGene_gn=RBM27;refGene_tx=XM_023256363.1

A1 123915178 123915332
refGene_gn=RBM27;refGene_tx=XM_023256343.1;refGene_gn=RBM27;refGene_tx=XM_023256351.1;refGene_gn=RBM27;refGene_tx=XM_023256355.1;refGene_gn=RBM27;refGene_tx=XM_023256363.1
A1 123918648 123918945
refGene_gn=RBM27;refGene_tx=XM_023256343.1;refGene_gn=RBM27;refGene_tx=XM_023256351.1;refGene_gn=RBM27;refGene_tx=XM_023256355.1;refGene_gn=RBM27;refGene_tx=XM_023256363.1
A1 123920278 123920419
refGene_gn=RBM27;refGene_tx=XM_023256343.1;refGene_gn=RBM27;refGene_tx=XM_023256351.1;refGene_gn=RBM27;refGene_tx=XM_023256355.1;refGene_gn=RBM27;refGene_tx=XM_023256363.1
A1 123925347 123925467
refGene_gn=RBM27;refGene_tx=XM_023256343.1;refGene_gn=RBM27;refGene_tx=XM_023256351.1;refGene_gn=RBM27;refGene_tx=XM_023256355.1;refGene_gn=RBM27;refGene_tx=XM_023256363.1
A1 123926397 123926472
refGene_gn=RBM27;refGene_tx=XM_023256343.1;refGene_gn=RBM27;refGene_tx=XM_023256351.1;refGene_gn=RBM27;refGene_tx=XM_023256355.1;refGene_gn=RBM27;refGene_tx=XM_023256363.1
A1 123926616 123926781
refGene_gn=RBM27;refGene_tx=XM_023256343.1;refGene_gn=RBM27;refGene_tx=XM_023256351.1;refGene_gn=RBM27;refGene_tx=XM_023256355.1;refGene_gn=RBM27;refGene_tx=XM_023256363.1
A1 123928898 123929003
refGene_gn=RBM27;refGene_tx=XM_023256343.1;refGene_gn=RBM27;refGene_tx=XM_023256351.1;refGene_gn=RBM27;refGene_tx=XM_023256355.1;refGene_gn=RBM27;refGene_tx=XM_023256363.1
A1 123929086 123929278
refGene_gn=RBM27;refGene_tx=XM_023256343.1;refGene_gn=RBM27;refGene_tx=XM_023256351.1;refGene_gn=RBM27;refGene_tx=XM_023256355.1;refGene_gn=RBM27;refGene_tx=XM_023256363.1
A1 123940054 123940165
refGene_gn=RBM27;refGene_tx=XM_023256343.1;refGene_gn=RBM27;refGene_tx=XM_023256351.1;refGene_gn=RBM27;refGene_tx=XM_023256355.1;refGene_gn=RBM27;refGene_tx=XM_023256363.1
A1 123941591 123941675
refGene_gn=RBM27;refGene_tx=XM_023256343.1;refGene_gn=RBM27;refGene_tx=XM_023256351.1;refGene_gn=RBM27;refGene_tx=XM_023256355.1;refGene_gn=RBM27;refGene_tx=XM_023256363.1
A1 123992022 123992142
refGene_gn=POU4F3;refGene_tx=XM_003980945.4
A1 123992459 123993356
refGene_gn=POU4F3;refGene_tx=XM_003980945.4
A1 124087096 124087155
refGene_gn=TCERG1;refGene_tx=XM_004001503.5
A1 124093901 124094127
refGene_gn=TCERG1;refGene_tx=XM_004001503.5
A1 124096371 124096524
refGene_gn=TCERG1;refGene_tx=XM_004001503.5

A1 124097427 124097869
refGene_gn=TCERG1;refGene_tx=XM_004001503.5
A1 124102462 124102705
refGene_gn=TCERG1;refGene_tx=XM_004001503.5
A1 124107033 124107096
refGene_gn=TCERG1;refGene_tx=XM_004001503.5
A1 124108247 124108448
refGene_gn=TCERG1;refGene_tx=XM_004001503.5
A1 124109420 124109533
refGene_gn=TCERG1;refGene_tx=XM_004001503.5
A1 124110298 124110387
refGene_gn=TCERG1;refGene_tx=XM_004001503.5
A1 124118151 124118312
refGene_gn=TCERG1;refGene_tx=XM_004001503.5
A1 124119555 124119612
refGene_gn=TCERG1;refGene_tx=XM_004001503.5
A1 124119762 124119829
refGene_gn=TCERG1;refGene_tx=XM_004001503.5
A1 124122324 124122460
refGene_gn=TCERG1;refGene_tx=XM_004001503.5
A1 124123315 124123405
refGene_gn=TCERG1;refGene_tx=XM_004001503.5
A1 124131854 124131973
refGene_gn=TCERG1;refGene_tx=XM_004001503.5
A1 124136522 124136673
refGene_gn=TCERG1;refGene_tx=XM_004001503.5
A1 124140768 124140933
refGene_gn=TCERG1;refGene_tx=XM_004001503.5
A1 124141192 124141375
refGene_gn=TCERG1;refGene_tx=XM_004001503.5
A1 124143348 124143528
refGene_gn=TCERG1;refGene_tx=XM_004001503.5
A1 124144191 124144275
refGene_gn=TCERG1;refGene_tx=XM_004001503.5
A1 124145607 124145679
refGene_gn=TCERG1;refGene_tx=XM_004001503.5
A1 124145680 124145709
refGene_gn=TCERG1;refGene_tx=XM_004001503.5
A1 124146909 124147111
refGene_gn=TCERG1;refGene_tx=XM_004001503.5
A1 124151025 124152285
refGene_gn=GPR151;refGene_tx=XM_003980946.2
A1 124250392 124250672
refGene_gn=PPP2R2B;refGene_tx=XM_011282787.2;refGene_gn=PPP2R2B;refGene_tx=XM_011282789.3;refGene_gn=PPP2R2B;refGene_tx=XM_011282798.3;refGene_gn=PPP2R2B;refGene_tx=XM_019833906.2;refGene_gn=PPP2R2B;refGene_tx=XM_019833907.1;refGene_gn=PPP2R2B;refGene_tx=XM_019833919.1;refGene_gn=PPP2R2B;refGene_tx=XM_019833921.2;refGene_gn=PPP2R2B;refGene_tx=XM_019833922.1;refGene_gn=PPP2R2B;refGene_tx=XM_019833925.1;refGene_gn=PPP2R2B;refGene_tx=XM_019833935.2;refGene_gn=PPP2R2B;refGene_tx=XM_0232564

07.1;refGene_gn=PPP2R2B;refGene_tx=XM_023256414.1
A1	124253307	124253399
refGene_gn=PPP2R2B;refGene_tx=XM_011282787.2;refGene_gn=PPP2R2B;refGene_tx=XM_011282789.3;refGene_gn=PPP2R2B;refGene_tx=XM_011282798.3;refGene_gn=PPP2R2B;refGene_tx=XM_019833906.2;refGene_gn=PPP2R2B;refGene_tx=XM_019833907.1;refGene_gn=PPP2R2B;refGene_tx=XM_019833919.1;refGene_gn=PPP2R2B;refGene_tx=XM_019833921.2;refGene_gn=PPP2R2B;refGene_tx=XM_019833922.1;refGene_gn=PPP2R2B;refGene_tx=XM_019833925.1;refGene_gn=PPP2R2B;refGene_tx=XM_019833935.2;refGene_gn=PPP2R2B;refGene_tx=XM_023256407.1;refGene_gn=PPP2R2B;refGene_tx=XM_023256414.1
A1	124262021	124262191
refGene_gn=PPP2R2B;refGene_tx=XM_011282787.2;refGene_gn=PPP2R2B;refGene_tx=XM_011282789.3;refGene_gn=PPP2R2B;refGene_tx=XM_011282798.3;refGene_gn=PPP2R2B;refGene_tx=XM_019833906.2;refGene_gn=PPP2R2B;refGene_tx=XM_019833907.1;refGene_gn=PPP2R2B;refGene_tx=XM_019833919.1;refGene_gn=PPP2R2B;refGene_tx=XM_019833921.2;refGene_gn=PPP2R2B;refGene_tx=XM_019833922.1;refGene_gn=PPP2R2B;refGene_tx=XM_019833925.1;refGene_gn=PPP2R2B;refGene_tx=XM_019833935.2;refGene_gn=PPP2R2B;refGene_tx=XM_023256407.1;refGene_gn=PPP2R2B;refGene_tx=XM_023256414.1
A1	124293374	124293539
refGene_gn=PPP2R2B;refGene_tx=XM_011282787.2;refGene_gn=PPP2R2B;refGene_tx=XM_011282789.3;refGene_gn=PPP2R2B;refGene_tx=XM_011282798.3;refGene_gn=PPP2R2B;refGene_tx=XM_019833906.2;refGene_gn=PPP2R2B;refGene_tx=XM_019833907.1;refGene_gn=PPP2R2B;refGene_tx=XM_019833919.1;refGene_gn=PPP2R2B;refGene_tx=XM_019833921.2;refGene_gn=PPP2R2B;refGene_tx=XM_019833922.1;refGene_gn=PPP2R2B;refGene_tx=XM_019833925.1;refGene_gn=PPP2R2B;refGene_tx=XM_019833935.2;refGene_gn=PPP2R2B;refGene_tx=XM_023256407.1;refGene_gn=PPP2R2B;refGene_tx=XM_023256414.1
A1	124307730	124307908
refGene_gn=PPP2R2B;refGene_tx=XM_011282787.2;refGene_gn=PPP2R2B;refGene_tx=XM_011282789.3;refGene_gn=PPP2R2B;refGene_tx=XM_011282798.3;refGene_gn=PPP2R2B;refGene_tx=XM_019833906.2;refGene_gn=PPP2R2B;refGene_tx=XM_019833907.1;refGene_gn=PPP2R2B;refGene_tx=XM_019833919.1;refGene_gn=PPP2R2B;refGene_tx=XM_019833921.2;refGene_gn=PPP2R2B;refGene_tx=XM_019833922.1;refGene_gn=PPP2R2B;refGene_tx=XM_019833925.1;refGene_gn=PPP2R2B;refGene_tx=XM_019833935.2;refGene_gn=PPP2R2B;refGene_tx=XM_023256407.1;refGene_gn=PPP2R2B;refGene_tx=XM_023256414.1
A1	124349383	124349496
refGene_gn=PPP2R2B;refGene_tx=XM_011282787.2;refGene_gn=PPP2R2B;refGene_tx=XM_011282789.3;refGene_gn=PPP2R2B;refGene_tx=XM_011282798.3;refGene_gn=PPP2R2B;refGene_tx=XM_019833906.2;refGene_gn=PPP2R2B;refGene_tx=XM_019833907.1;refGene_gn=PPP2R2B;refGene_tx=XM_019833919.1;refGene_gn=PPP2R2B;refGene_tx=XM_019833921.2;refGene_gn=PPP2R2B;refGene_tx=XM_019833922.1;refGene_gn=PPP2R2B;refGene_tx=XM_019833925.1;refGene_gn=PPP2R2B;refGene_tx=XM_019833935.2;refGene_gn=PPP2R2B;refGene_tx=XM_023256407.1;refGene_gn=PPP2R2B;refGene_tx=XM_023256414.1
A1	124355098	124355264
refGene_gn=PPP2R2B;refGene_tx=XM_011282787.2;refGene_gn=PPP2R2B;refGene_tx=XM_011282789.3;refGene_gn=PPP2R2B;refGene_tx=XM_011282798.3;refGene_gn=PPP2R2B;refGene_tx=XM_019833906.2;refGene_gn=PPP2R2B;refGene_tx=

XM_019833907.1;refGene_gn=PPP2R2B;refGene_tx=XM_019833919.1;refGene_gn=PPP2R2B;refGene_tx=XM_019833921.2;refGene_gn=PPP2R2B;refGene_tx=XM_019833922.1;refGene_gn=PPP2R2B;refGene_tx=XM_019833925.1;refGene_gn=PPP2R2B;refGene_tx=XM_019833935.2;refGene_gn=PPP2R2B;refGene_tx=XM_023256407.1;refGene_gn=PPP2R2B;refGene_tx=XM_023256414.1
A1 124358174 124358272
refGene_gn=PPP2R2B;refGene_tx=XM_011282787.2;refGene_gn=PPP2R2B;refGene_tx=XM_011282789.3;refGene_gn=PPP2R2B;refGene_tx=XM_011282798.3;refGene_gn=PPP2R2B;refGene_tx=XM_019833906.2;refGene_gn=PPP2R2B;refGene_tx=XM_019833907.1;refGene_gn=PPP2R2B;refGene_tx=XM_019833919.1;refGene_gn=PPP2R2B;refGene_tx=XM_019833921.2;refGene_gn=PPP2R2B;refGene_tx=XM_019833922.1;refGene_gn=PPP2R2B;refGene_tx=XM_019833925.1;refGene_gn=PPP2R2B;refGene_tx=XM_019833935.2;refGene_gn=PPP2R2B;refGene_tx=XM_023256407.1;refGene_gn=PPP2R2B;refGene_tx=XM_023256414.1
A1 124375695 124375717
refGene_gn=PPP2R2B;refGene_tx=XM_019833935.2
A1 124498257 124498294
refGene_gn=PPP2R2B;refGene_tx=XM_011282798.3;refGene_gn=PPP2R2B;refGene_tx=XM_019833925.1
A1 124519681 124519875
refGene_gn=PPP2R2B;refGene_tx=XM_011282787.2;refGene_gn=PPP2R2B;refGene_tx=XM_011282789.3;refGene_gn=PPP2R2B;refGene_tx=XM_019833906.2;refGene_gn=PPP2R2B;refGene_tx=XM_019833922.1;refGene_gn=PPP2R2B;refGene_tx=XM_023256407.1;refGene_gn=PPP2R2B;refGene_tx=XM_023256414.1
A1 124520267 124520449
refGene_gn=PPP2R2B;refGene_tx=XM_023256414.1
A1 124672040 124672119
refGene_gn=PPP2R2B;refGene_tx=XM_019833906.2;refGene_gn=PPP2R2B;refGene_tx=XM_019833921.2
A1 124707872 124707950
refGene_gn=PPP2R2B;refGene_tx=XM_011282787.2;refGene_gn=PPP2R2B;refGene_tx=XM_019833907.1;refGene_gn=PPP2R2B;refGene_tx=XM_019833919.1
A1 124708070 124708288
refGene_gn=PPP2R2B;refGene_tx=XM_011282787.2;refGene_gn=PPP2R2B;refGene_tx=XM_019833919.1
A1 124758186 124758220
refGene_gn=PPP2R2B;refGene_tx=XM_019833907.1
A1 124820141 124820193
refGene_gn=STK32A;refGene_tx=XM_006927743.4
A1 124849067 124849123
refGene_gn=STK32A;refGene_tx=XM_006927743.4
A1 124850376 124850528
refGene_gn=STK32A;refGene_tx=XM_006927743.4
A1 124882169 124882343
refGene_gn=STK32A;refGene_tx=XM_006927743.4
A1 124905811 124905849
refGene_gn=STK32A;refGene_tx=XM_006927743.4
A1 124914251 124914341
refGene_gn=STK32A;refGene_tx=XM_006927743.4
A1 124923664 124923762

refGene_gn=STK32A;refGene_tx=XM_006927743.4
A1 124928512 124928629
refGene_gn=STK32A;refGene_tx=XM_006927743.4
A1 124931336 124931462
refGene_gn=STK32A;refGene_tx=XM_006927743.4
A1 124932917 124933046
refGene_gn=STK32A;refGene_tx=XM_006927743.4
A1 124946159 124946224
refGene_gn=STK32A;refGene_tx=XM_006927743.4
A1 124946563 124946657
refGene_gn=STK32A;refGene_tx=XM_006927743.4
A1 124956005 124956094
refGene_gn=DPYSL3;refGene_tx=XM_023256425.1;refGene_gn=DPYSL3;refGene_tx=XM_023256432.1;refGene_gn=DPYSL3;refGene_tx=XM_023256443.1
A1 124957683 124957846
refGene_gn=DPYSL3;refGene_tx=XM_023256425.1;refGene_gn=DPYSL3;refGene_tx=XM_023256432.1;refGene_gn=DPYSL3;refGene_tx=XM_023256443.1
A1 124960519 124960699
refGene_gn=DPYSL3;refGene_tx=XM_023256425.1;refGene_gn=DPYSL3;refGene_tx=XM_023256432.1;refGene_gn=DPYSL3;refGene_tx=XM_023256443.1
A1 124961935 124962106
refGene_gn=DPYSL3;refGene_tx=XM_023256425.1;refGene_gn=DPYSL3;refGene_tx=XM_023256432.1;refGene_gn=DPYSL3;refGene_tx=XM_023256443.1
A1 124964271 124964413
refGene_gn=DPYSL3;refGene_tx=XM_023256425.1;refGene_gn=DPYSL3;refGene_tx=XM_023256432.1;refGene_gn=DPYSL3;refGene_tx=XM_023256443.1
A1 124965088 124965245
refGene_gn=DPYSL3;refGene_tx=XM_023256425.1;refGene_gn=DPYSL3;refGene_tx=XM_023256432.1;refGene_gn=DPYSL3;refGene_tx=XM_023256443.1
A1 124969116 124969237
refGene_gn=DPYSL3;refGene_tx=XM_023256425.1;refGene_gn=DPYSL3;refGene_tx=XM_023256432.1;refGene_gn=DPYSL3;refGene_tx=XM_023256443.1
A1 124972116 124972185
refGene_gn=DPYSL3;refGene_tx=XM_023256425.1;refGene_gn=DPYSL3;refGene_tx=XM_023256432.1;refGene_gn=DPYSL3;refGene_tx=XM_023256443.1
A1 124975988 124976069
refGene_gn=DPYSL3;refGene_tx=XM_023256425.1;refGene_gn=DPYSL3;refGene_tx=XM_023256432.1;refGene_gn=DPYSL3;refGene_tx=XM_023256443.1
A1 124976985 124977047
refGene_gn=DPYSL3;refGene_tx=XM_023256425.1;refGene_gn=DPYSL3;refGene_tx=XM_023256432.1;refGene_gn=DPYSL3;refGene_tx=XM_023256443.1
A1 124979088 124979253
refGene_gn=DPYSL3;refGene_tx=XM_023256425.1;refGene_gn=DPYSL3;refGene_tx=XM_023256432.1;refGene_gn=DPYSL3;refGene_tx=XM_023256443.1
A1 124981949 124982134
refGene_gn=DPYSL3;refGene_tx=XM_023256425.1;refGene_gn=DPYSL3;refGene_tx=XM_023256432.1;refGene_gn=DPYSL3;refGene_tx=XM_023256443.1
A1 124987218 124987307
refGene_gn=DPYSL3;refGene_tx=XM_023256425.1;refGene_gn=DPYSL3;refGene_tx=XM_023256432.1;refGene_gn=DPYSL3;refGene_tx=XM_023256443.1

A1 125004336 125004431
refGene_gn=DPYSL3;refGene_tx=XM_023256425.1
A1 125015925 125016340
refGene_gn=DPYSL3;refGene_tx=XM_023256425.1;refGene_gn=DPYSL3;refGene_tx=XM_023256443.1
A1 125066498 125066888
refGene_gn=DPYSL3;refGene_tx=XM_023256432.1
A1 125161650 125161671
refGene_gn=JAKMIP2;refGene_tx=XM_019833985.2
A1 125169140 125169191
refGene_gn=JAKMIP2;refGene_tx=XM_003980953.5;refGene_gn=JAKMIP2;refGene_tx=XM_003980955.5;refGene_gn=JAKMIP2;refGene_tx=XM_019833978.2
A1 125172385 125172451
refGene_gn=JAKMIP2;refGene_tx=XM_003980953.5;refGene_gn=JAKMIP2;refGene_tx=XM_003980955.5;refGene_gn=JAKMIP2;refGene_tx=XM_019833978.2;refGene_gn=JAKMIP2;refGene_tx=XM_019833985.2
A1 125178398 125178602
refGene_gn=JAKMIP2;refGene_tx=XM_003980953.5;refGene_gn=JAKMIP2;refGene_tx=XM_003980955.5;refGene_gn=JAKMIP2;refGene_tx=XM_019833978.2;refGene_gn=JAKMIP2;refGene_tx=XM_019833985.2
A1 125180569 125180647
refGene_gn=JAKMIP2;refGene_tx=XM_003980953.5;refGene_gn=JAKMIP2;refGene_tx=XM_003980955.5;refGene_gn=JAKMIP2;refGene_tx=XM_019833978.2;refGene_gn=JAKMIP2;refGene_tx=XM_019833985.2
A1 125183902 125183971
refGene_gn=JAKMIP2;refGene_tx=XM_003980953.5;refGene_gn=JAKMIP2;refGene_tx=XM_003980955.5;refGene_gn=JAKMIP2;refGene_tx=XM_019833978.2;refGene_gn=JAKMIP2;refGene_tx=XM_019833985.2
A1 125188874 125188940
refGene_gn=JAKMIP2;refGene_tx=XM_003980953.5;refGene_gn=JAKMIP2;refGene_tx=XM_003980955.5;refGene_gn=JAKMIP2;refGene_tx=XM_019833978.2;refGene_gn=JAKMIP2;refGene_tx=XM_019833985.2
A1 125189804 125189858
refGene_gn=JAKMIP2;refGene_tx=XM_003980953.5;refGene_gn=JAKMIP2;refGene_tx=XM_003980955.5;refGene_gn=JAKMIP2;refGene_tx=XM_019833978.2;refGene_gn=JAKMIP2;refGene_tx=XM_019833985.2
A1 125191646 125191745
refGene_gn=JAKMIP2;refGene_tx=XM_003980953.5;refGene_gn=JAKMIP2;refGene_tx=XM_003980955.5;refGene_gn=JAKMIP2;refGene_tx=XM_019833978.2;refGene_gn=JAKMIP2;refGene_tx=XM_019833985.2
A1 125193280 125193379
refGene_gn=JAKMIP2;refGene_tx=XM_003980953.5;refGene_gn=JAKMIP2;refGene_tx=XM_003980955.5;refGene_gn=JAKMIP2;refGene_tx=XM_019833978.2;refGene_gn=JAKMIP2;refGene_tx=XM_019833985.2
A1 125195931 125195994
refGene_gn=JAKMIP2;refGene_tx=XM_003980953.5;refGene_gn=JAKMIP2;refGene_tx=XM_019833978.2;refGene_gn=JAKMIP2;refGene_tx=XM_019833985.2
A1 125196675 125196759
refGene_gn=JAKMIP2;refGene_tx=XM_003980953.5;refGene_gn=JAKMIP2;refGene_tx=XM_003980955.5;refGene_gn=JAKMIP2;refGene_tx=XM_019833978.2;refGe

ne_gn=JAKMIP2;refGene_tx=XM_019833985.2
A1 125199878 125200007
refGene_gn=JAKMIP2;refGene_tx=XM_003980953.5;refGene_gn=JAKMIP2;refGene_tx=XM_003980955.5;refGene_gn=JAKMIP2;refGene_tx=XM_019833978.2;refGene_gn=JAKMIP2;refGene_tx=XM_019833985.2
A1 125201043 125201163
refGene_gn=JAKMIP2;refGene_tx=XM_003980953.5;refGene_gn=JAKMIP2;refGene_tx=XM_003980955.5;refGene_gn=JAKMIP2;refGene_tx=XM_019833978.2;refGene_gn=JAKMIP2;refGene_tx=XM_019833985.2
A1 125202329 125202386
refGene_gn=JAKMIP2;refGene_tx=XM_003980953.5;refGene_gn=JAKMIP2;refGene_tx=XM_003980955.5;refGene_gn=JAKMIP2;refGene_tx=XM_019833978.2;refGene_gn=JAKMIP2;refGene_tx=XM_019833985.2
A1 125204872 125205013
refGene_gn=JAKMIP2;refGene_tx=XM_003980953.5;refGene_gn=JAKMIP2;refGene_tx=XM_003980955.5;refGene_gn=JAKMIP2;refGene_tx=XM_019833978.2;refGene_gn=JAKMIP2;refGene_tx=XM_019833985.2
A1 125205895 125206042
refGene_gn=JAKMIP2;refGene_tx=XM_003980953.5;refGene_gn=JAKMIP2;refGene_tx=XM_003980955.5;refGene_gn=JAKMIP2;refGene_tx=XM_019833978.2;refGene_gn=JAKMIP2;refGene_tx=XM_019833985.2
A1 125210059 125210158
refGene_gn=JAKMIP2;refGene_tx=XM_003980953.5;refGene_gn=JAKMIP2;refGene_tx=XM_003980955.5;refGene_gn=JAKMIP2;refGene_tx=XM_019833978.2;refGene_gn=JAKMIP2;refGene_tx=XM_019833985.2
A1 125212012 125212222
refGene_gn=JAKMIP2;refGene_tx=XM_003980953.5;refGene_gn=JAKMIP2;refGene_tx=XM_003980955.5;refGene_gn=JAKMIP2;refGene_tx=XM_019833978.2;refGene_gn=JAKMIP2;refGene_tx=XM_019833985.2
A1 125216054 125216552
refGene_gn=JAKMIP2;refGene_tx=XM_003980953.5;refGene_gn=JAKMIP2;refGene_tx=XM_003980955.5;refGene_gn=JAKMIP2;refGene_tx=XM_019833978.2;refGene_gn=JAKMIP2;refGene_tx=XM_019833985.2
A1 125228959 125229088
refGene_gn=JAKMIP2;refGene_tx=XM_003980953.5;refGene_gn=JAKMIP2;refGene_tx=XM_003980955.5;refGene_gn=JAKMIP2;refGene_tx=XM_019833978.2;refGene_gn=JAKMIP2;refGene_tx=XM_019833985.2
A1 125322919 125323378
refGene_gn=LOC109501341;refGene_tx=XM_019833994.2
A1 125374281 125374327
refGene_gn=SPINK1;refGene_tx=XM_003980956.2
A1 125378150 125378257
refGene_gn=SPINK1;refGene_tx=XM_003980956.2
A1 125379745 125379777
refGene_gn=SPINK1;refGene_tx=XM_003980956.2
A1 125380829 125380884
refGene_gn=SPINK1;refGene_tx=XM_003980956.2
A1 125411278 125411333
refGene_gn=SCGB3A2;refGene_tx=XM_023256473.1
A1 125414209 125414412

refGene_gn=SCGB3A2;refGene_tx=XM_023256473.1
A1 125414771 125414795
refGene_gn=SCGB3A2;refGene_tx=XM_023256473.1
A1 125426838 125426983
refGene_gn=CA1H5orf46;refGene_tx=XM_019834030.1
A1 125431018 125431067
refGene_gn=CA1H5orf46;refGene_tx=XM_019834037.2
A1 125435850 125435986
refGene_gn=CA1H5orf46;refGene_tx=XM_019834030.1;refGene_gn=CA1H5orf46;refGene_tx=XM_019834037.2
A1 125441673 125441932
refGene_gn=CA1H5orf46;refGene_tx=XM_019834030.1;refGene_gn=CA1H5orf46;refGene_tx=XM_019834037.2
A1 125569848 125569903
refGene_gn=SPINK5;refGene_tx=XM_019834057.1
A1 125571363 125571389
refGene_gn=SPINK5;refGene_tx=XM_019834057.1
A1 125575276 125575404
refGene_gn=SPINK5;refGene_tx=XM_019834057.1;refGene_gn=SPINK5;refGene_tx=XM_019834066.1
A1 125577294 125577367
refGene_gn=SPINK5;refGene_tx=XM_019834057.1;refGene_gn=SPINK5;refGene_tx=XM_019834066.1
A1 125592882 125593010
refGene_gn=SPINK5;refGene_tx=XM_019834057.1;refGene_gn=SPINK5;refGene_tx=XM_019834066.1
A1 125595034 125595098
refGene_gn=SPINK5;refGene_tx=XM_019834057.1;refGene_gn=SPINK5;refGene_tx=XM_019834066.1
A1 125596196 125596324
refGene_gn=SPINK5;refGene_tx=XM_019834057.1;refGene_gn=SPINK5;refGene_tx=XM_019834066.1
A1 125597742 125597809
refGene_gn=SPINK5;refGene_tx=XM_019834057.1;refGene_gn=SPINK5;refGene_tx=XM_019834066.1
A1 125601837 125601965
refGene_gn=SPINK5;refGene_tx=XM_019834057.1;refGene_gn=SPINK5;refGene_tx=XM_019834066.1
A1 125603293 125603381
refGene_gn=SPINK5;refGene_tx=XM_019834057.1;refGene_gn=SPINK5;refGene_tx=XM_019834066.1
A1 125604463 125604591
refGene_gn=SPINK5;refGene_tx=XM_019834057.1;refGene_gn=SPINK5;refGene_tx=XM_019834066.1
A1 125605839 125605921
refGene_gn=SPINK5;refGene_tx=XM_019834057.1;refGene_gn=SPINK5;refGene_tx=XM_019834066.1
A1 125607328 125607456
refGene_gn=SPINK5;refGene_tx=XM_019834057.1;refGene_gn=SPINK5;refGene_tx=XM_019834066.1

A1 125608956 125609035
refGene_gn=SPINK5;refGene_tx=XM_019834057.1;refGene_gn=SPINK5;refGene_
tx=XM_019834066.1
A1 125609358 125609486
refGene_gn=SPINK5;refGene_tx=XM_019834057.1;refGene_gn=SPINK5;refGene_
tx=XM_019834066.1
A1 125612194 125612243
refGene_gn=SPINK5;refGene_tx=XM_019834057.1;refGene_gn=SPINK5;refGene_
tx=XM_019834066.1
A1 125614233 125614361
refGene_gn=SPINK5;refGene_tx=XM_019834057.1;refGene_gn=SPINK5;refGene_
tx=XM_019834066.1
A1 125616442 125616530
refGene_gn=SPINK5;refGene_tx=XM_019834057.1;refGene_gn=SPINK5;refGene_
tx=XM_019834066.1
A1 125619213 125619341
refGene_gn=SPINK5;refGene_tx=XM_019834057.1;refGene_gn=SPINK5;refGene_
tx=XM_019834066.1
A1 125619987 125620054
refGene_gn=SPINK5;refGene_tx=XM_019834057.1;refGene_gn=SPINK5;refGene_
tx=XM_019834066.1
A1 125621151 125621279
refGene_gn=SPINK5;refGene_tx=XM_019834057.1;refGene_gn=SPINK5;refGene_
tx=XM_019834066.1
A1 125622849 125622952
refGene_gn=SPINK5;refGene_tx=XM_019834057.1;refGene_gn=SPINK5;refGene_
tx=XM_019834066.1
A1 125624363 125624491
refGene_gn=SPINK5;refGene_tx=XM_019834057.1;refGene_gn=SPINK5;refGene_
tx=XM_019834066.1
A1 125627725 125627795
refGene_gn=SPINK5;refGene_tx=XM_019834057.1;refGene_gn=SPINK5;refGene_
tx=XM_019834066.1
A1 125628694 125628822
refGene_gn=SPINK5;refGene_tx=XM_019834057.1;refGene_gn=SPINK5;refGene_
tx=XM_019834066.1
A1 125629244 125629323
refGene_gn=SPINK5;refGene_tx=XM_019834057.1;refGene_gn=SPINK5;refGene_
tx=XM_019834066.1
A1 125632239 125632367
refGene_gn=SPINK5;refGene_tx=XM_019834057.1;refGene_gn=SPINK5;refGene_
tx=XM_019834066.1
A1 125633386 125633459
refGene_gn=SPINK5;refGene_tx=XM_019834057.1;refGene_gn=SPINK5;refGene_
tx=XM_019834066.1
A1 125634543 125634671
refGene_gn=SPINK5;refGene_tx=XM_019834057.1;refGene_gn=SPINK5;refGene_
tx=XM_019834066.1
A1 125635560 125635660
refGene_gn=SPINK5;refGene_tx=XM_019834057.1;refGene_gn=SPINK5;refGene_

tx=XM_019834066.1
A1 125641089 125641220
refGene_gn=SPINK5;refGene_tx=XM_019834057.1;refGene_gn=SPINK5;refGene_
tx=XM_019834066.1
A1 125643465 125643556
refGene_gn=SPINK5;refGene_tx=XM_019834057.1;refGene_gn=SPINK5;refGene_
tx=XM_019834066.1
A1 125646037 125646046
refGene_gn=SPINK5;refGene_tx=XM_019834057.1;refGene_gn=SPINK5;refGene_
tx=XM_019834066.1
A1 125667424 125667459
refGene_gn=SPINK14;refGene_tx=XM_019837481.2
A1 125668554 125668700
refGene_gn=SPINK14;refGene_tx=XM_019837481.2
A1 125669806 125669844
refGene_gn=SPINK14;refGene_tx=XM_019837481.2
A1 125670952 125671089
refGene_gn=SPINK14;refGene_tx=XM_019837481.2
A1 125672353 125672399
refGene_gn=SPINK14;refGene_tx=XM_019837481.2
A1 125695760 125695818
refGene_gn=SPINK6;refGene_tx=XM_023255739.1
A1 125698843 125698866
refGene_gn=SPINK6;refGene_tx=XM_023255739.1
A1 125706885 125707001
refGene_gn=SPINK6;refGene_tx=XM_023255739.1
A1 125708150 125708196
refGene_gn=SPINK6;refGene_tx=XM_023255739.1
A1 125750622 125750671
refGene_gn=MARCOL;refGene_tx=XM_019814099.1;refGene_gn=MARCOL;refGene_
tx=XM_019814105.1
A1 125754804 125755388
refGene_gn=MARCOL;refGene_tx=XM_019814099.1;refGene_gn=MARCOL;refGene_
tx=XM_019814105.1
A1 125757246 125757368
refGene_gn=MARCOL;refGene_tx=XM_019814099.1
A1 125759362 125759408
refGene_gn=MARCOL;refGene_tx=XM_019814099.1;refGene_gn=MARCOL;refGene_
tx=XM_019814105.1
A1 125768929 125768996
refGene_gn=LOC109492841;refGene_tx=XM_019814117.2
A1 125770019 125770057
refGene_gn=LOC109492841;refGene_tx=XM_019814117.2
A1 125771242 125771358
refGene_gn=LOC109492841;refGene_tx=XM_019814117.2
A1 125772756 125772802
refGene_gn=LOC109492841;refGene_tx=XM_019814117.2
A1 125800691 125800761
refGene_gn=GZMA;refGene_tx=XM_003980989.3
A1 125803528 125803667

refGene_gn=GZMA;refGene_tx=XM_003980989.3
A1	125805257	125805399
refGene_gn=GZMA;refGene_tx=XM_003980989.3
A1	125805598	125805868
refGene_gn=GZMA;refGene_tx=XM_003980989.3
A1	125808388	125808553
refGene_gn=GZMA;refGene_tx=XM_003980989.3
A1	125811697	125811846
refGene_gn=CDC20B;refGene_tx=XM_006927746.3;refGene_gn=CDC20B;refGene_tx=XM_019834091.2;refGene_gn=CDC20B;refGene_tx=XM_019834099.2;refGene_gn=CDC20B;refGene_tx=XM_019834107.2;refGene_gn=CDC20B;refGene_tx=XM_023256512.1;refGene_gn=CDC20B;refGene_tx=XM_023256516.1
A1	125816879	125816997
refGene_gn=CDC20B;refGene_tx=XM_006927746.3;refGene_gn=CDC20B;refGene_tx=XM_019834091.2;refGene_gn=CDC20B;refGene_tx=XM_019834099.2;refGene_gn=CDC20B;refGene_tx=XM_019834107.2;refGene_gn=CDC20B;refGene_tx=XM_023256512.1;refGene_gn=CDC20B;refGene_tx=XM_023256516.1
A1	125817473	125817599
refGene_gn=CDC20B;refGene_tx=XM_019834091.2;refGene_gn=CDC20B;refGene_tx=XM_019834099.2;refGene_gn=CDC20B;refGene_tx=XM_023256512.1;refGene_gn=CDC20B;refGene_tx=XM_023256516.1
A1	125819478	125819704
refGene_gn=CDC20B;refGene_tx=XM_006927746.3;refGene_gn=CDC20B;refGene_tx=XM_019834091.2;refGene_gn=CDC20B;refGene_tx=XM_019834099.2;refGene_gn=CDC20B;refGene_tx=XM_019834107.2;refGene_gn=CDC20B;refGene_tx=XM_023256512.1;refGene_gn=CDC20B;refGene_tx=XM_023256516.1
A1	125821961	125822056
refGene_gn=CDC20B;refGene_tx=XM_006927746.3;refGene_gn=CDC20B;refGene_tx=XM_019834091.2;refGene_gn=CDC20B;refGene_tx=XM_019834099.2;refGene_gn=CDC20B;refGene_tx=XM_019834107.2;refGene_gn=CDC20B;refGene_tx=XM_023256512.1;refGene_gn=CDC20B;refGene_tx=XM_023256516.1
A1	125823603	125823800
refGene_gn=CDC20B;refGene_tx=XM_006927746.3;refGene_gn=CDC20B;refGene_tx=XM_019834091.2;refGene_gn=CDC20B;refGene_tx=XM_019834099.2;refGene_gn=CDC20B;refGene_tx=XM_019834107.2;refGene_gn=CDC20B;refGene_tx=XM_023256512.1;refGene_gn=CDC20B;refGene_tx=XM_023256516.1
A1	125826958	125827075
refGene_gn=CDC20B;refGene_tx=XM_006927746.3;refGene_gn=CDC20B;refGene_tx=XM_019834091.2;refGene_gn=CDC20B;refGene_tx=XM_019834099.2;refGene_gn=CDC20B;refGene_tx=XM_019834107.2;refGene_gn=CDC20B;refGene_tx=XM_023256512.1;refGene_gn=CDC20B;refGene_tx=XM_023256516.1
A1	125829622	125829716
refGene_gn=CDC20B;refGene_tx=XM_006927746.3;refGene_gn=CDC20B;refGene_tx=XM_019834091.2;refGene_gn=CDC20B;refGene_tx=XM_019834099.2;refGene_gn=CDC20B;refGene_tx=XM_019834107.2;refGene_gn=CDC20B;refGene_tx=XM_023256512.1;refGene_gn=CDC20B;refGene_tx=XM_023256516.1
A1	125835124	125835255
refGene_gn=CDC20B;refGene_tx=XM_006927746.3;refGene_gn=CDC20B;refGene_tx=XM_019834091.2;refGene_gn=CDC20B;refGene_tx=XM_019834099.2;refGene_gn=CDC20B;refGene_tx=XM_019834107.2;refGene_gn=CDC20B;refGene_tx=XM_02

3256512.1;refGene_gn=CDC20B;refGene_tx=XM_023256516.1
A1 125836233 125836462
refGene_gn=CDC20B;refGene_tx=XM_006927746.3;refGene_gn=CDC20B;refGene_tx=XM_019834091.2;refGene_gn=CDC20B;refGene_tx=XM_019834099.2;refGene_gn=CDC20B;refGene_tx=XM_019834107.2;refGene_gn=CDC20B;refGene_tx=XM_023256512.1;refGene_gn=CDC20B;refGene_tx=XM_023256516.1
A1 125847641 125847845
refGene_gn=GPX8;refGene_tx=XM_003980961.5;refGene_gn=GPX8;refGene_tx=XM_006927732.4
A1 125848415 125848680
refGene_gn=GPX8;refGene_tx=XM_003980961.5;refGene_gn=GPX8;refGene_tx=XM_006927732.4
A1 125852127 125852291
refGene_gn=GPX8;refGene_tx=XM_003980961.5;refGene_gn=GPX8;refGene_tx=XM_006927732.4
A1 125859090 125859153
refGene_gn=CDC20B;refGene_tx=XM_006927746.3;refGene_gn=CDC20B;refGene_tx=XM_019834091.2;refGene_gn=CDC20B;refGene_tx=XM_019834099.2;refGene_gn=CDC20B;refGene_tx=XM_019834107.2;refGene_gn=CDC20B;refGene_tx=XM_023256512.1;refGene_gn=CDC20B;refGene_tx=XM_023256516.1
A1 125859331 125859463
refGene_gn=CDC20B;refGene_tx=XM_006927746.3;refGene_gn=CDC20B;refGene_tx=XM_019834091.2;refGene_gn=CDC20B;refGene_tx=XM_019834099.2;refGene_gn=CDC20B;refGene_tx=XM_019834107.2;refGene_gn=CDC20B;refGene_tx=XM_023256512.1;refGene_gn=CDC20B;refGene_tx=XM_023256516.1
A1 125894029 125894470
refGene_gn=MCIDAS;refGene_tx=XM_003980962.3
A1 125894714 125894825
refGene_gn=MCIDAS;refGene_tx=XM_003980962.3
A1 125895444 125895668
refGene_gn=MCIDAS;refGene_tx=XM_003980962.3
A1 125896234 125896307
refGene_gn=MCIDAS;refGene_tx=XM_003980962.3
A1 125899457 125899549
refGene_gn=MCIDAS;refGene_tx=XM_003980962.3
A1 125899727 125899824
refGene_gn=MCIDAS;refGene_tx=XM_003980962.3
A1 125899911 125900028
refGene_gn=MCIDAS;refGene_tx=XM_003980962.3
A1 125906236 125906722
refGene_gn=CCNO;refGene_tx=XM_003980991.4
A1 125907240 125907426
refGene_gn=CCNO;refGene_tx=XM_003980991.4
A1 125908041 125908425
refGene_gn=CCNO;refGene_tx=XM_003980991.4
A1 125927221 125927277
refGene_gn=DHX29;refGene_tx=XM_003980963.5;refGene_gn=DHX29;refGene_tx=XM_006927733.4
A1 125928821 125928918
refGene_gn=DHX29;refGene_tx=XM_003980963.5;refGene_gn=DHX29;refGene_tx

=XM_006927733.4
A1 125929793 125929925
refGene_gn=DHX29;refGene_tx=XM_003980963.5;refGene_gn=DHX29;refGene_tx=XM_006927733.4
A1 125930842 125931145
refGene_gn=DHX29;refGene_tx=XM_003980963.5;refGene_gn=DHX29;refGene_tx=XM_006927733.4
A1 125932764 125932858
refGene_gn=DHX29;refGene_tx=XM_003980963.5;refGene_gn=DHX29;refGene_tx=XM_006927733.4
A1 125933260 125933397
refGene_gn=DHX29;refGene_tx=XM_003980963.5;refGene_gn=DHX29;refGene_tx=XM_006927733.4
A1 125934684 125934909
refGene_gn=DHX29;refGene_tx=XM_003980963.5;refGene_gn=DHX29;refGene_tx=XM_006927733.4
A1 125935552 125935628
refGene_gn=DHX29;refGene_tx=XM_003980963.5;refGene_gn=DHX29;refGene_tx=XM_006927733.4
A1 125935715 125935844
refGene_gn=DHX29;refGene_tx=XM_003980963.5;refGene_gn=DHX29;refGene_tx=XM_006927733.4
A1 125936713 125936802
refGene_gn=DHX29;refGene_tx=XM_003980963.5;refGene_gn=DHX29;refGene_tx=XM_006927733.4
A1 125938283 125938368
refGene_gn=DHX29;refGene_tx=XM_003980963.5;refGene_gn=DHX29;refGene_tx=XM_006927733.4
A1 125938977 125939095
refGene_gn=DHX29;refGene_tx=XM_003980963.5;refGene_gn=DHX29;refGene_tx=XM_006927733.4
A1 125939231 125939376
refGene_gn=DHX29;refGene_tx=XM_003980963.5;refGene_gn=DHX29;refGene_tx=XM_006927733.4
A1 125940902 125941043
refGene_gn=DHX29;refGene_tx=XM_003980963.5;refGene_gn=DHX29;refGene_tx=XM_006927733.4
A1 125941679 125941856
refGene_gn=DHX29;refGene_tx=XM_003980963.5;refGene_gn=DHX29;refGene_tx=XM_006927733.4
A1 125949250 125949394
refGene_gn=DHX29;refGene_tx=XM_003980963.5;refGene_gn=DHX29;refGene_tx=XM_006927733.4
A1 125951075 125951681
refGene_gn=DHX29;refGene_tx=XM_003980963.5;refGene_gn=DHX29;refGene_tx=XM_006927733.4
A1 125953189 125953313
refGene_gn=DHX29;refGene_tx=XM_003980963.5;refGene_gn=DHX29;refGene_tx=XM_006927733.4
A1 125953570 125953736

refGene_gn=DHX29;refGene_tx=XM_003980963.5;refGene_gn=DHX29;refGene_tx=XM_006927733.4
A1 125955403 125955562
refGene_gn=DHX29;refGene_tx=XM_003980963.5;refGene_gn=DHX29;refGene_tx=XM_006927733.4
A1 125956316 125956443
refGene_gn=DHX29;refGene_tx=XM_003980963.5;refGene_gn=DHX29;refGene_tx=XM_006927733.4
A1 125959339 125959465
refGene_gn=DHX29;refGene_tx=XM_003980963.5;refGene_gn=DHX29;refGene_tx=XM_006927733.4
A1 125961651 125961794
refGene_gn=DHX29;refGene_tx=XM_003980963.5;refGene_gn=DHX29;refGene_tx=XM_006927733.4
A1 125962512 125962642
refGene_gn=DHX29;refGene_tx=XM_003980963.5;refGene_gn=DHX29;refGene_tx=XM_006927733.4
A1 125964603 125964717
refGene_gn=DHX29;refGene_tx=XM_003980963.5;refGene_gn=DHX29;refGene_tx=XM_006927733.4
A1 125966104 125966178
refGene_gn=DHX29;refGene_tx=XM_003980963.5;refGene_gn=DHX29;refGene_tx=XM_006927733.4
A1 125973253 125973434
refGene_gn=DHX29;refGene_tx=XM_003980963.5;refGene_gn=DHX29;refGene_tx=XM_006927733.4
A1 125973875 125974009
refGene_gn=MTREX;refGene_tx=XM_003980964.5
A1 125982931 125983069
refGene_gn=MTREX;refGene_tx=XM_003980964.5
A1 125984437 125984504
refGene_gn=MTREX;refGene_tx=XM_003980964.5
A1 125988425 125988488
refGene_gn=MTREX;refGene_tx=XM_003980964.5
A1 125989344 125989457
refGene_gn=MTREX;refGene_tx=XM_003980964.5
A1 125995120 125995295
refGene_gn=MTREX;refGene_tx=XM_003980964.5
A1 125996803 125996894
refGene_gn=MTREX;refGene_tx=XM_003980964.5
A1 125998319 125998444
refGene_gn=MTREX;refGene_tx=XM_003980964.5
A1 125999709 125999808
refGene_gn=MTREX;refGene_tx=XM_003980964.5
A1 126000471 126000574
refGene_gn=MTREX;refGene_tx=XM_003980964.5
A1 126002379 126002511
refGene_gn=MTREX;refGene_tx=XM_003980964.5
A1 126004308 126004388
refGene_gn=MTREX;refGene_tx=XM_003980964.5

A1 126005904 126006015
refGene_gn=MTREX;refGene_tx=XM_003980964.5
A1 126007953 126008055
refGene_gn=MTREX;refGene_tx=XM_003980964.5
A1 126010882 126011008
refGene_gn=MTREX;refGene_tx=XM_003980964.5
A1 126018016 126018167
refGene_gn=MTREX;refGene_tx=XM_003980964.5
A1 126029289 126029462
refGene_gn=MTREX;refGene_tx=XM_003980964.5
A1 126030268 126030337
refGene_gn=MTREX;refGene_tx=XM_003980964.5
A1 126043487 126043616
refGene_gn=MTREX;refGene_tx=XM_003980964.5
A1 126054985 126055096
refGene_gn=MTREX;refGene_tx=XM_003980964.5
A1 126056327 126056516
refGene_gn=MTREX;refGene_tx=XM_003980964.5
A1 126061260 126061424
refGene_gn=MTREX;refGene_tx=XM_003980964.5
A1 126066776 126066882
refGene_gn=MTREX;refGene_tx=XM_003980964.5
A1 126071031 126071088
refGene_gn=MTREX;refGene_tx=XM_003980964.5
A1 126072227 126072390
refGene_gn=MTREX;refGene_tx=XM_003980964.5
A1 126079527 126079632
refGene_gn=MTREX;refGene_tx=XM_003980964.5
A1 126081126 126081179
refGene_gn=MTREX;refGene_tx=XM_003980964.5
A1 126081640 126081772
refGene_gn=PLPP1;refGene_tx=XM_003980965.5;refGene_gn=PLPP1;refGene_tx=XM_003980966.5;refGene_gn=PLPP1;refGene_tx=XM_019834145.2;refGene_gn=PLPP1;refGene_tx=XM_019834155.2
A1 126082292 126082469
refGene_gn=PLPP1;refGene_tx=XM_003980965.5;refGene_gn=PLPP1;refGene_tx=XM_003980966.5;refGene_gn=PLPP1;refGene_tx=XM_019834145.2;refGene_gn=PLPP1;refGene_tx=XM_019834155.2
A1 126092872 126092930
refGene_gn=PLPP1;refGene_tx=XM_003980965.5;refGene_gn=PLPP1;refGene_tx=XM_003980966.5;refGene_gn=PLPP1;refGene_tx=XM_019834145.2;refGene_gn=PLPP1;refGene_tx=XM_019834155.2
A1 126110326 126110607
refGene_gn=PLPP1;refGene_tx=XM_003980965.5;refGene_gn=PLPP1;refGene_tx=XM_003980966.5;refGene_gn=PLPP1;refGene_tx=XM_019834145.2;refGene_gn=PLPP1;refGene_tx=XM_019834155.2
A1 126116138 126116290
refGene_gn=PLPP1;refGene_tx=XM_003980965.5
A1 126141371 126141526
refGene_gn=PLPP1;refGene_tx=XM_003980966.5;refGene_gn=PLPP1;refGene_tx

=XM_019834145.2;refGene_gn=PLPP1;refGene_tx=XM_019834155.2
A1	126171605	126171663
refGene_gn=PLPP1;refGene_tx=XM_003980965.5;refGene_gn=PLPP1;refGene_tx=XM_003980966.5
A1	126247992	126248158
refGene_gn=SLC38A9;refGene_tx=XM_003980967.4;refGene_gn=SLC38A9;refGene_tx=XM_006927734.3;refGene_gn=SLC38A9;refGene_tx=XM_011282869.3;refGene_gn=SLC38A9;refGene_tx=XM_019834283.1;refGene_gn=SLC38A9;refGene_tx=XM_023256658.1;refGene_gn=SLC38A9;refGene_tx=XM_023256670.1;refGene_gn=SLC38A9;refGene_tx=XM_023256675.1
A1	126249317	126249407
refGene_gn=SLC38A9;refGene_tx=XM_003980967.4;refGene_gn=SLC38A9;refGene_tx=XM_006927734.3;refGene_gn=SLC38A9;refGene_tx=XM_011282869.3;refGene_gn=SLC38A9;refGene_tx=XM_019834283.1;refGene_gn=SLC38A9;refGene_tx=XM_023256658.1;refGene_gn=SLC38A9;refGene_tx=XM_023256670.1;refGene_gn=SLC38A9;refGene_tx=XM_023256675.1
A1	126254291	126254440
refGene_gn=SLC38A9;refGene_tx=XM_003980967.4;refGene_gn=SLC38A9;refGene_tx=XM_006927734.3;refGene_gn=SLC38A9;refGene_tx=XM_011282869.3;refGene_gn=SLC38A9;refGene_tx=XM_019834283.1;refGene_gn=SLC38A9;refGene_tx=XM_023256658.1;refGene_gn=SLC38A9;refGene_tx=XM_023256670.1;refGene_gn=SLC38A9;refGene_tx=XM_023256675.1
A1	126255855	126255969
refGene_gn=SLC38A9;refGene_tx=XM_003980967.4;refGene_gn=SLC38A9;refGene_tx=XM_006927734.3;refGene_gn=SLC38A9;refGene_tx=XM_011282869.3;refGene_gn=SLC38A9;refGene_tx=XM_019834283.1;refGene_gn=SLC38A9;refGene_tx=XM_023256658.1;refGene_gn=SLC38A9;refGene_tx=XM_023256670.1;refGene_gn=SLC38A9;refGene_tx=XM_023256675.1
A1	126266798	126266905
refGene_gn=SLC38A9;refGene_tx=XM_003980967.4;refGene_gn=SLC38A9;refGene_tx=XM_006927734.3;refGene_gn=SLC38A9;refGene_tx=XM_011282869.3;refGene_gn=SLC38A9;refGene_tx=XM_019834283.1;refGene_gn=SLC38A9;refGene_tx=XM_023256658.1;refGene_gn=SLC38A9;refGene_tx=XM_023256670.1;refGene_gn=SLC38A9;refGene_tx=XM_023256675.1
A1	126281316	126281424
refGene_gn=SLC38A9;refGene_tx=XM_003980967.4;refGene_gn=SLC38A9;refGene_tx=XM_006927734.3;refGene_gn=SLC38A9;refGene_tx=XM_011282869.3;refGene_gn=SLC38A9;refGene_tx=XM_019834283.1;refGene_gn=SLC38A9;refGene_tx=XM_023256658.1;refGene_gn=SLC38A9;refGene_tx=XM_023256670.1;refGene_gn=SLC38A9;refGene_tx=XM_023256675.1
A1	126283431	126283626
refGene_gn=SLC38A9;refGene_tx=XM_003980967.4;refGene_gn=SLC38A9;refGene_tx=XM_006927734.3;refGene_gn=SLC38A9;refGene_tx=XM_011282869.3;refGene_gn=SLC38A9;refGene_tx=XM_019834283.1;refGene_gn=SLC38A9;refGene_tx=XM_023256658.1;refGene_gn=SLC38A9;refGene_tx=XM_023256670.1
A1	126287029	126287089
refGene_gn=SLC38A9;refGene_tx=XM_003980967.4;refGene_gn=SLC38A9;refGene_tx=XM_006927734.3;refGene_gn=SLC38A9;refGene_tx=XM_011282869.3;refGene_gn=SLC38A9;refGene_tx=XM_019834283.1;refGene_gn=SLC38A9;refGene_tx=XM_023256658.1;refGene_gn=SLC38A9;refGene_tx=XM_023256670.1;refGene_gn

=SLC38A9;refGene_tx=XM_023256675.1
A1 126289798 126289969
refGene_gn=SLC38A9;refGene_tx=XM_003980967.4;refGene_gn=SLC38A9;refGene_tx=XM_006927734.3;refGene_gn=SLC38A9;refGene_tx=XM_011282869.3;refGene_gn=SLC38A9;refGene_tx=XM_019834283.1;refGene_gn=SLC38A9;refGene_tx=XM_023256658.1;refGene_gn=SLC38A9;refGene_tx=XM_023256670.1;refGene_gn=SLC38A9;refGene_tx=XM_023256675.1
A1 126291717 126291811
refGene_gn=SLC38A9;refGene_tx=XM_003980967.4;refGene_gn=SLC38A9;refGene_tx=XM_006927734.3;refGene_gn=SLC38A9;refGene_tx=XM_011282869.3;refGene_gn=SLC38A9;refGene_tx=XM_019834283.1;refGene_gn=SLC38A9;refGene_tx=XM_023256658.1;refGene_gn=SLC38A9;refGene_tx=XM_023256670.1;refGene_gn=SLC38A9;refGene_tx=XM_023256675.1
A1 126292041 126292105
refGene_gn=SLC38A9;refGene_tx=XM_003980967.4;refGene_gn=SLC38A9;refGene_tx=XM_006927734.3;refGene_gn=SLC38A9;refGene_tx=XM_011282869.3;refGene_gn=SLC38A9;refGene_tx=XM_019834283.1;refGene_gn=SLC38A9;refGene_tx=XM_023256658.1;refGene_gn=SLC38A9;refGene_tx=XM_023256670.1;refGene_gn=SLC38A9;refGene_tx=XM_023256675.1
A1 126292242 126292364
refGene_gn=SLC38A9;refGene_tx=XM_003980967.4;refGene_gn=SLC38A9;refGene_tx=XM_006927734.3;refGene_gn=SLC38A9;refGene_tx=XM_011282869.3;refGene_gn=SLC38A9;refGene_tx=XM_019834283.1;refGene_gn=SLC38A9;refGene_tx=XM_023256658.1;refGene_gn=SLC38A9;refGene_tx=XM_023256670.1;refGene_gn=SLC38A9;refGene_tx=XM_023256675.1
A1 126296482 126296615
refGene_gn=SLC38A9;refGene_tx=XM_003980967.4;refGene_gn=SLC38A9;refGene_tx=XM_006927734.3;refGene_gn=SLC38A9;refGene_tx=XM_011282869.3;refGene_gn=SLC38A9;refGene_tx=XM_019834283.1;refGene_gn=SLC38A9;refGene_tx=XM_023256658.1;refGene_gn=SLC38A9;refGene_tx=XM_023256670.1;refGene_gn=SLC38A9;refGene_tx=XM_023256675.1
A1 126298297 126298395
refGene_gn=SLC38A9;refGene_tx=XM_023256658.1
A1 126317546 126317659
refGene_gn=SLC38A9;refGene_tx=XM_003980967.4;refGene_gn=SLC38A9;refGene_tx=XM_006927734.3;refGene_gn=SLC38A9;refGene_tx=XM_023256675.1
A1 126353967 126354039
refGene_gn=DDX4;refGene_tx=XM_006927735.4
A1 126369233 126369294
refGene_gn=DDX4;refGene_tx=XM_006927735.4
A1 126392383 126392461
refGene_gn=DDX4;refGene_tx=XM_006927735.4
A1 126396463 126396541
refGene_gn=DDX4;refGene_tx=XM_006927735.4
A1 126398186 126398237
refGene_gn=DDX4;refGene_tx=XM_006927735.4
A1 126402265 126402325
refGene_gn=DDX4;refGene_tx=XM_006927735.4
A1 126411319 126411421
refGene_gn=DDX4;refGene_tx=XM_006927735.4

A1 126412832 126412861
refGene_gn=DDX4;refGene_tx=XM_006927735.4
A1 126412862 126412914
refGene_gn=DDX4;refGene_tx=XM_006927735.4
A1 126413903 126413951
refGene_gn=DDX4;refGene_tx=XM_006927735.4
A1 126424619 126424667
refGene_gn=DDX4;refGene_tx=XM_006927735.4
A1 126424777 126424825
refGene_gn=DDX4;refGene_tx=XM_006927735.4
A1 126425044 126425187
refGene_gn=DDX4;refGene_tx=XM_006927735.4
A1 126425767 126425920
refGene_gn=DDX4;refGene_tx=XM_006927735.4
A1 126426922 126427077
refGene_gn=DDX4;refGene_tx=XM_006927735.4
A1 126432263 126432393
refGene_gn=DDX4;refGene_tx=XM_006927735.4
A1 126433664 126433831
refGene_gn=DDX4;refGene_tx=XM_006927735.4
A1 126438123 126438269
refGene_gn=DDX4;refGene_tx=XM_006927735.4
A1 126449019 126449119
refGene_gn=DDX4;refGene_tx=XM_006927735.4
A1 126450306 126450577
refGene_gn=DDX4;refGene_tx=XM_006927735.4
A1 126450726 126450837
refGene_gn=DDX4;refGene_tx=XM_006927735.4
A1 126451531 126451618
refGene_gn=DDX4;refGene_tx=XM_006927735.4
A1 126462277 126462438
refGene_gn=IL31RA;refGene_tx=XM_019834303.2
A1 126481179 126481204
refGene_gn=IL31RA;refGene_tx=XM_019834303.2
A1 126487228 126487319
refGene_gn=IL31RA;refGene_tx=XM_019834303.2
A1 126502194 126502312
refGene_gn=IL31RA;refGene_tx=XM_019834303.2
A1 126505691 126505873
refGene_gn=IL31RA;refGene_tx=XM_019834303.2
A1 126508351 126508503
refGene_gn=IL31RA;refGene_tx=XM_019834303.2
A1 126513921 126514087
refGene_gn=IL31RA;refGene_tx=XM_019834303.2
A1 126522591 126522671
refGene_gn=IL31RA;refGene_tx=XM_019834303.2
A1 126524943 126525160
refGene_gn=IL31RA;refGene_tx=XM_019834303.2
A1 126531808 126531991
refGene_gn=IL31RA;refGene_tx=XM_019834303.2

A1 126532800 126532902
refGene_gn=IL31RA;refGene_tx=XM_019834303.2
A1 126533727 126533874
refGene_gn=IL31RA;refGene_tx=XM_019834303.2
A1 126535909 126536050
refGene_gn=IL31RA;refGene_tx=XM_019834303.2
A1 126537935 126538029
refGene_gn=IL31RA;refGene_tx=XM_019834303.2
A1 126539161 126539243
refGene_gn=IL31RA;refGene_tx=XM_019834303.2
A1 126540439 126540889
refGene_gn=IL31RA;refGene_tx=XM_019834303.2
A1 126566259 126566997
refGene_gn=IL6ST;refGene_tx=XM_011282895.2;refGene_gn=IL6ST;refGene_tx=XM_011282898.2;refGene_gn=IL6ST;refGene_tx=XM_011282903.3;refGene_gn=IL6ST;refGene_tx=XM_019834335.2;refGene_gn=IL6ST;refGene_tx=XM_019834342.2;refGene_gn=IL6ST;refGene_tx=XM_019834355.2
A1 126567643 126567725
refGene_gn=IL6ST;refGene_tx=XM_011282895.2;refGene_gn=IL6ST;refGene_tx=XM_011282898.2;refGene_gn=IL6ST;refGene_tx=XM_011282903.3;refGene_gn=IL6ST;refGene_tx=XM_019834335.2;refGene_gn=IL6ST;refGene_tx=XM_019834342.2;refGene_gn=IL6ST;refGene_tx=XM_019834355.2
A1 126571420 126571517
refGene_gn=IL6ST;refGene_tx=XM_011282895.2;refGene_gn=IL6ST;refGene_tx=XM_011282898.2;refGene_gn=IL6ST;refGene_tx=XM_011282903.3;refGene_gn=IL6ST;refGene_tx=XM_019834335.2;refGene_gn=IL6ST;refGene_tx=XM_019834342.2
A1 126574374 126574515
refGene_gn=IL6ST;refGene_tx=XM_011282895.2;refGene_gn=IL6ST;refGene_tx=XM_011282898.2;refGene_gn=IL6ST;refGene_tx=XM_011282903.3;refGene_gn=IL6ST;refGene_tx=XM_019834335.2;refGene_gn=IL6ST;refGene_tx=XM_019834342.2;refGene_gn=IL6ST;refGene_tx=XM_019834355.2
A1 126574827 126574974
refGene_gn=IL6ST;refGene_tx=XM_011282895.2;refGene_gn=IL6ST;refGene_tx=XM_011282898.2;refGene_gn=IL6ST;refGene_tx=XM_011282903.3;refGene_gn=IL6ST;refGene_tx=XM_019834335.2;refGene_gn=IL6ST;refGene_tx=XM_019834342.2;refGene_gn=IL6ST;refGene_tx=XM_019834355.2
A1 126575156 126575258
refGene_gn=IL6ST;refGene_tx=XM_011282895.2;refGene_gn=IL6ST;refGene_tx=XM_011282898.2;refGene_gn=IL6ST;refGene_tx=XM_011282903.3;refGene_gn=IL6ST;refGene_tx=XM_019834335.2;refGene_gn=IL6ST;refGene_tx=XM_019834342.2;refGene_gn=IL6ST;refGene_tx=XM_019834355.2
A1 126577581 126577764
refGene_gn=IL6ST;refGene_tx=XM_011282895.2;refGene_gn=IL6ST;refGene_tx=XM_011282898.2;refGene_gn=IL6ST;refGene_tx=XM_011282903.3;refGene_gn=IL6ST;refGene_tx=XM_019834335.2;refGene_gn=IL6ST;refGene_tx=XM_019834342.2;refGene_gn=IL6ST;refGene_tx=XM_019834355.2
A1 126578298 126578509
refGene_gn=IL6ST;refGene_tx=XM_011282895.2;refGene_gn=IL6ST;refGene_tx=XM_011282898.2;refGene_gn=IL6ST;refGene_tx=XM_011282903.3;refGene_gn=

IL6ST;refGene_tx=XM_019834335.2;refGene_gn=IL6ST;refGene_tx=XM_0198343
42.2;refGene_gn=IL6ST;refGene_tx=XM_019834355.2
A1 126579161 126579244
refGene_gn=IL6ST;refGene_tx=XM_011282895.2;refGene_gn=IL6ST;refGene_tx
=XM_011282898.2;refGene_gn=IL6ST;refGene_tx=XM_011282903.3;refGene_gn=
IL6ST;refGene_tx=XM_019834335.2;refGene_gn=IL6ST;refGene_tx=XM_0198343
42.2;refGene_gn=IL6ST;refGene_tx=XM_019834355.2
A1 126581995 126582155
refGene_gn=IL6ST;refGene_tx=XM_011282895.2;refGene_gn=IL6ST;refGene_tx
=XM_011282898.2;refGene_gn=IL6ST;refGene_tx=XM_011282903.3;refGene_gn=
IL6ST;refGene_tx=XM_019834335.2;refGene_gn=IL6ST;refGene_tx=XM_0198343
42.2;refGene_gn=IL6ST;refGene_tx=XM_019834355.2
A1 126583887 126584042
refGene_gn=IL6ST;refGene_tx=XM_011282895.2;refGene_gn=IL6ST;refGene_tx
=XM_011282898.2;refGene_gn=IL6ST;refGene_tx=XM_011282903.3;refGene_gn=
IL6ST;refGene_tx=XM_019834335.2;refGene_gn=IL6ST;refGene_tx=XM_0198343
42.2;refGene_gn=IL6ST;refGene_tx=XM_019834355.2
A1 126584577 126584744
refGene_gn=IL6ST;refGene_tx=XM_011282895.2;refGene_gn=IL6ST;refGene_tx
=XM_011282898.2;refGene_gn=IL6ST;refGene_tx=XM_011282903.3;refGene_gn=
IL6ST;refGene_tx=XM_019834335.2;refGene_gn=IL6ST;refGene_tx=XM_0198343
42.2;refGene_gn=IL6ST;refGene_tx=XM_019834355.2
A1 126591471 126591592
refGene_gn=IL6ST;refGene_tx=XM_011282895.2;refGene_gn=IL6ST;refGene_tx
=XM_011282898.2;refGene_gn=IL6ST;refGene_tx=XM_011282903.3;refGene_gn=
IL6ST;refGene_tx=XM_019834335.2;refGene_gn=IL6ST;refGene_tx=XM_0198343
42.2;refGene_gn=IL6ST;refGene_tx=XM_019834355.2
A1 126592456 126592762
refGene_gn=IL6ST;refGene_tx=XM_011282895.2;refGene_gn=IL6ST;refGene_tx
=XM_011282898.2;refGene_gn=IL6ST;refGene_tx=XM_011282903.3;refGene_gn=
IL6ST;refGene_tx=XM_019834335.2;refGene_gn=IL6ST;refGene_tx=XM_0198343
42.2;refGene_gn=IL6ST;refGene_tx=XM_019834355.2
A1 126597251 126597315
refGene_gn=IL6ST;refGene_tx=XM_011282895.2;refGene_gn=IL6ST;refGene_tx
=XM_011282898.2;refGene_gn=IL6ST;refGene_tx=XM_011282903.3;refGene_gn=
IL6ST;refGene_tx=XM_019834335.2;refGene_gn=IL6ST;refGene_tx=XM_0198343
42.2;refGene_gn=IL6ST;refGene_tx=XM_019834355.2
A1 126634025 126634183
refGene_gn=LOC111557658;refGene_tx=XM_023243399.1
A1 126682785 126682871
refGene_gn=LOC111557658;refGene_tx=XM_023243399.1
A1 126682948 126683463
refGene_gn=LOC111557658;refGene_tx=XM_023243399.1
A1 126693321 126693443
refGene_gn=ANKRD55;refGene_tx=XM_019834369.2;refGene_gn=ANKRD55;refGen
e_tx=XM_019834383.2;refGene_gn=ANKRD55;refGene_tx=XM_019834384.2;refGe
ne_gn=ANKRD55;refGene_tx=XM_019834390.2
A1 126695390 126695483
refGene_gn=ANKRD55;refGene_tx=XM_019834369.2;refGene_gn=ANKRD55;refGen
e_tx=XM_019834383.2;refGene_gn=ANKRD55;refGene_tx=XM_019834384.2;refGe

ne_gn=ANKRD55;refGene_tx=XM_019834390.2
A1 126703959 126704621
refGene_gn=ANKRD55;refGene_tx=XM_019834369.2;refGene_gn=ANKRD55;refGene_tx=XM_019834383.2;refGene_gn=ANKRD55;refGene_tx=XM_019834384.2;refGene_gn=ANKRD55;refGene_tx=XM_019834390.2
A1 126707692 126707860
refGene_gn=ANKRD55;refGene_tx=XM_019834369.2;refGene_gn=ANKRD55;refGene_tx=XM_019834383.2;refGene_gn=ANKRD55;refGene_tx=XM_019834384.2
A1 126712006 126712191
refGene_gn=ANKRD55;refGene_tx=XM_019834369.2;refGene_gn=ANKRD55;refGene_tx=XM_019834383.2;refGene_gn=ANKRD55;refGene_tx=XM_019834384.2;refGene_gn=ANKRD55;refGene_tx=XM_019834390.2
A1 126724012 126724141
refGene_gn=ANKRD55;refGene_tx=XM_019834369.2;refGene_gn=ANKRD55;refGene_tx=XM_019834383.2;refGene_gn=ANKRD55;refGene_tx=XM_019834384.2;refGene_gn=ANKRD55;refGene_tx=XM_019834390.2
A1 126735644 126735705
refGene_gn=ANKRD55;refGene_tx=XM_019834369.2;refGene_gn=ANKRD55;refGene_tx=XM_019834383.2;refGene_gn=ANKRD55;refGene_tx=XM_019834384.2;refGene_gn=ANKRD55;refGene_tx=XM_019834390.2
A1 126745196 126745306
refGene_gn=ANKRD55;refGene_tx=XM_019834369.2;refGene_gn=ANKRD55;refGene_tx=XM_019834383.2;refGene_gn=ANKRD55;refGene_tx=XM_019834384.2;refGene_gn=ANKRD55;refGene_tx=XM_019834390.2
A1 126749451 126749582
refGene_gn=ANKRD55;refGene_tx=XM_019834369.2;refGene_gn=ANKRD55;refGene_tx=XM_019834383.2;refGene_gn=ANKRD55;refGene_tx=XM_019834384.2;refGene_gn=ANKRD55;refGene_tx=XM_019834390.2
A1 126759783 126759906
refGene_gn=ANKRD55;refGene_tx=XM_019834369.2;refGene_gn=ANKRD55;refGene_tx=XM_019834383.2;refGene_gn=ANKRD55;refGene_tx=XM_019834384.2;refGene_gn=ANKRD55;refGene_tx=XM_019834390.2
A1 126786269 126786327
refGene_gn=ANKRD55;refGene_tx=XM_019834369.2;refGene_gn=ANKRD55;refGene_tx=XM_019834383.2;refGene_gn=ANKRD55;refGene_tx=XM_019834384.2;refGene_gn=ANKRD55;refGene_tx=XM_019834390.2
A1 127318445 127318927
refGene_gn=MAP3K1;refGene_tx=XM_003980973.4
A1 127330552 127330584
refGene_gn=MAP3K1;refGene_tx=XM_023256730.1
A1 127334631 127334642
refGene_gn=MAP3K1;refGene_tx=XM_019834410.2
A1 127363056 127363207
refGene_gn=MAP3K1;refGene_tx=XM_003980973.4;refGene_gn=MAP3K1;refGene_tx=XM_019834410.2;refGene_gn=MAP3K1;refGene_tx=XM_019834413.2;refGene_gn=MAP3K1;refGene_tx=XM_019834423.2;refGene_gn=MAP3K1;refGene_tx=XM_023256730.1;refGene_gn=MAP3K1;refGene_tx=XM_023256741.1
A1 127366081 127366282
refGene_gn=MAP3K1;refGene_tx=XM_003980973.4;refGene_gn=MAP3K1;refGene_tx=XM_019834410.2;refGene_gn=MAP3K1;refGene_tx=XM_019834413.2;refGene_

gn=MAP3K1;refGene_tx=XM_019834423.2;refGene_gn=MAP3K1;refGene_tx=XM_023256730.1;refGene_gn=MAP3K1;refGene_tx=XM_023256741.1
A1 127370162 127370363
refGene_gn=MAP3K1;refGene_tx=XM_003980973.4;refGene_gn=MAP3K1;refGene_tx=XM_019834410.2;refGene_gn=MAP3K1;refGene_tx=XM_019834413.2;refGene_gn=MAP3K1;refGene_tx=XM_019834423.2;refGene_gn=MAP3K1;refGene_tx=XM_023256730.1;refGene_gn=MAP3K1;refGene_tx=XM_023256741.1
A1 127370729 127370846
refGene_gn=MAP3K1;refGene_tx=XM_003980973.4;refGene_gn=MAP3K1;refGene_tx=XM_019834410.2;refGene_gn=MAP3K1;refGene_tx=XM_019834413.2;refGene_gn=MAP3K1;refGene_tx=XM_019834423.2;refGene_gn=MAP3K1;refGene_tx=XM_023256730.1;refGene_gn=MAP3K1;refGene_tx=XM_023256741.1
A1 127371161 127371310
refGene_gn=MAP3K1;refGene_tx=XM_003980973.4;refGene_gn=MAP3K1;refGene_tx=XM_019834410.2;refGene_gn=MAP3K1;refGene_tx=XM_019834413.2;refGene_gn=MAP3K1;refGene_tx=XM_019834423.2;refGene_gn=MAP3K1;refGene_tx=XM_023256730.1;refGene_gn=MAP3K1;refGene_tx=XM_023256741.1
A1 127375075 127375197
refGene_gn=MAP3K1;refGene_tx=XM_003980973.4;refGene_gn=MAP3K1;refGene_tx=XM_019834410.2;refGene_gn=MAP3K1;refGene_tx=XM_019834413.2;refGene_gn=MAP3K1;refGene_tx=XM_019834423.2;refGene_gn=MAP3K1;refGene_tx=XM_023256730.1;refGene_gn=MAP3K1;refGene_tx=XM_023256741.1
A1 127375797 127375879
refGene_gn=MAP3K1;refGene_tx=XM_003980973.4;refGene_gn=MAP3K1;refGene_tx=XM_019834410.2;refGene_gn=MAP3K1;refGene_tx=XM_019834413.2;refGene_gn=MAP3K1;refGene_tx=XM_019834423.2;refGene_gn=MAP3K1;refGene_tx=XM_023256730.1;refGene_gn=MAP3K1;refGene_tx=XM_023256741.1
A1 127375993 127376177
refGene_gn=MAP3K1;refGene_tx=XM_003980973.4;refGene_gn=MAP3K1;refGene_tx=XM_019834410.2;refGene_gn=MAP3K1;refGene_tx=XM_019834413.2;refGene_gn=MAP3K1;refGene_tx=XM_019834423.2;refGene_gn=MAP3K1;refGene_tx=XM_023256730.1;refGene_gn=MAP3K1;refGene_tx=XM_023256741.1
A1 127378726 127379005
refGene_gn=MAP3K1;refGene_tx=XM_003980973.4;refGene_gn=MAP3K1;refGene_tx=XM_019834410.2;refGene_gn=MAP3K1;refGene_tx=XM_019834413.2;refGene_gn=MAP3K1;refGene_tx=XM_019834423.2;refGene_gn=MAP3K1;refGene_tx=XM_023256730.1;refGene_gn=MAP3K1;refGene_tx=XM_023256741.1
A1 127381984 127382106
refGene_gn=MAP3K1;refGene_tx=XM_003980973.4;refGene_gn=MAP3K1;refGene_tx=XM_019834410.2;refGene_gn=MAP3K1;refGene_tx=XM_019834413.2;refGene_gn=MAP3K1;refGene_tx=XM_019834423.2;refGene_gn=MAP3K1;refGene_tx=XM_023256730.1;refGene_gn=MAP3K1;refGene_tx=XM_023256741.1
A1 127383703 127383795
refGene_gn=MAP3K1;refGene_tx=XM_003980973.4;refGene_gn=MAP3K1;refGene_tx=XM_019834410.2;refGene_gn=MAP3K1;refGene_tx=XM_019834413.2;refGene_gn=MAP3K1;refGene_tx=XM_019834423.2;refGene_gn=MAP3K1;refGene_tx=XM_023256730.1;refGene_gn=MAP3K1;refGene_tx=XM_023256741.1
A1 127384051 127384241
refGene_gn=MAP3K1;refGene_tx=XM_003980973.4;refGene_gn=MAP3K1;refGene_tx=XM_019834410.2;refGene_gn=MAP3K1;refGene_tx=XM_019834413.2;refGene_

gn=MAP3K1;refGene_tx=XM_019834423.2;refGene_gn=MAP3K1;refGene_tx=XM_023256730.1;refGene_gn=MAP3K1;refGene_tx=XM_023256741.1
A1 127384544 127385829
refGene_gn=MAP3K1;refGene_tx=XM_003980973.4;refGene_gn=MAP3K1;refGene_tx=XM_019834410.2;refGene_gn=MAP3K1;refGene_tx=XM_019834413.2;refGene_gn=MAP3K1;refGene_tx=XM_019834423.2;refGene_gn=MAP3K1;refGene_tx=XM_023256730.1;refGene_gn=MAP3K1;refGene_tx=XM_023256741.1
A1 127386196 127386349
refGene_gn=MAP3K1;refGene_tx=XM_003980973.4;refGene_gn=MAP3K1;refGene_tx=XM_019834410.2;refGene_gn=MAP3K1;refGene_tx=XM_019834413.2;refGene_gn=MAP3K1;refGene_tx=XM_019834423.2;refGene_gn=MAP3K1;refGene_tx=XM_023256730.1;refGene_gn=MAP3K1;refGene_tx=XM_023256741.1
A1 127387097 127387260
refGene_gn=MAP3K1;refGene_tx=XM_003980973.4;refGene_gn=MAP3K1;refGene_tx=XM_019834410.2;refGene_gn=MAP3K1;refGene_tx=XM_019834413.2;refGene_gn=MAP3K1;refGene_tx=XM_019834423.2;refGene_gn=MAP3K1;refGene_tx=XM_023256730.1;refGene_gn=MAP3K1;refGene_tx=XM_023256741.1
A1 127389182 127389314
refGene_gn=MAP3K1;refGene_tx=XM_003980973.4;refGene_gn=MAP3K1;refGene_tx=XM_019834410.2;refGene_gn=MAP3K1;refGene_tx=XM_019834413.2;refGene_gn=MAP3K1;refGene_tx=XM_019834423.2;refGene_gn=MAP3K1;refGene_tx=XM_023256730.1;refGene_gn=MAP3K1;refGene_tx=XM_023256741.1
A1 127390145 127390288
refGene_gn=MAP3K1;refGene_tx=XM_003980973.4;refGene_gn=MAP3K1;refGene_tx=XM_019834410.2;refGene_gn=MAP3K1;refGene_tx=XM_019834413.2;refGene_gn=MAP3K1;refGene_tx=XM_019834423.2;refGene_gn=MAP3K1;refGene_tx=XM_023256730.1;refGene_gn=MAP3K1;refGene_tx=XM_023256741.1
A1 127391015 127391147
refGene_gn=MAP3K1;refGene_tx=XM_003980973.4;refGene_gn=MAP3K1;refGene_tx=XM_019834410.2;refGene_gn=MAP3K1;refGene_tx=XM_019834413.2;refGene_gn=MAP3K1;refGene_tx=XM_019834423.2;refGene_gn=MAP3K1;refGene_tx=XM_023256730.1;refGene_gn=MAP3K1;refGene_tx=XM_023256741.1
A1 127392378 127392528
refGene_gn=MAP3K1;refGene_tx=XM_003980973.4;refGene_gn=MAP3K1;refGene_tx=XM_019834410.2;refGene_gn=MAP3K1;refGene_tx=XM_019834413.2;refGene_gn=MAP3K1;refGene_tx=XM_019834423.2;refGene_gn=MAP3K1;refGene_tx=XM_023256730.1;refGene_gn=MAP3K1;refGene_tx=XM_023256741.1
A1 127420824 127420922
refGene_gn=SETD9;refGene_tx=XM_011282918.3;refGene_gn=SETD9;refGene_tx=XM_011282921.3
A1 127422367 127422735
refGene_gn=SETD9;refGene_tx=XM_011282918.3;refGene_gn=SETD9;refGene_tx=XM_011282921.3
A1 127424127 127424251
refGene_gn=SETD9;refGene_tx=XM_011282918.3;refGene_gn=SETD9;refGene_tx=XM_011282921.3
A1 127424742 127424858
refGene_gn=SETD9;refGene_tx=XM_011282918.3;refGene_gn=SETD9;refGene_tx=XM_011282921.3
A1 127425575 127425681

refGene_gn=SETD9;refGene_tx=XM_011282918.3;refGene_gn=SETD9;refGene_tx=XM_011282921.3
A1 127427731 127427819
refGene_gn=SETD9;refGene_tx=XM_011282918.3
A1 127436179 127436637
refGene_gn=MIER3;refGene_tx=XM_023256761.1;refGene_gn=MIER3;refGene_tx=XM_023256765.1;refGene_gn=MIER3;refGene_tx=XM_023256767.1;refGene_gn=MIER3;refGene_tx=XM_023256770.1
A1 127436738 127436881
refGene_gn=MIER3;refGene_tx=XM_023256761.1;refGene_gn=MIER3;refGene_tx=XM_023256765.1;refGene_gn=MIER3;refGene_tx=XM_023256767.1;refGene_gn=MIER3;refGene_tx=XM_023256770.1
A1 127436958 127437086
refGene_gn=MIER3;refGene_tx=XM_023256761.1;refGene_gn=MIER3;refGene_tx=XM_023256765.1;refGene_gn=MIER3;refGene_tx=XM_023256767.1;refGene_gn=MIER3;refGene_tx=XM_023256770.1
A1 127442188 127442283
refGene_gn=MIER3;refGene_tx=XM_023256761.1;refGene_gn=MIER3;refGene_tx=XM_023256765.1;refGene_gn=MIER3;refGene_tx=XM_023256767.1;refGene_gn=MIER3;refGene_tx=XM_023256770.1
A1 127443812 127443894
refGene_gn=MIER3;refGene_tx=XM_023256761.1;refGene_gn=MIER3;refGene_tx=XM_023256765.1;refGene_gn=MIER3;refGene_tx=XM_023256767.1;refGene_gn=MIER3;refGene_tx=XM_023256770.1
A1 127447175 127447327
refGene_gn=MIER3;refGene_tx=XM_023256761.1;refGene_gn=MIER3;refGene_tx=XM_023256765.1;refGene_gn=MIER3;refGene_tx=XM_023256767.1;refGene_gn=MIER3;refGene_tx=XM_023256770.1
A1 127449306 127449379
refGene_gn=MIER3;refGene_tx=XM_023256761.1;refGene_gn=MIER3;refGene_tx=XM_023256765.1;refGene_gn=MIER3;refGene_tx=XM_023256767.1;refGene_gn=MIER3;refGene_tx=XM_023256770.1
A1 127449574 127449660
refGene_gn=MIER3;refGene_tx=XM_023256761.1;refGene_gn=MIER3;refGene_tx=XM_023256765.1;refGene_gn=MIER3;refGene_tx=XM_023256767.1;refGene_gn=MIER3;refGene_tx=XM_023256770.1
A1 127450793 127450914
refGene_gn=MIER3;refGene_tx=XM_023256761.1;refGene_gn=MIER3;refGene_tx=XM_023256765.1;refGene_gn=MIER3;refGene_tx=XM_023256767.1;refGene_gn=MIER3;refGene_tx=XM_023256770.1
A1 127452440 127452575
refGene_gn=MIER3;refGene_tx=XM_023256761.1;refGene_gn=MIER3;refGene_tx=XM_023256765.1;refGene_gn=MIER3;refGene_tx=XM_023256767.1;refGene_gn=MIER3;refGene_tx=XM_023256770.1
A1 127462229 127462375
refGene_gn=MIER3;refGene_tx=XM_023256761.1;refGene_gn=MIER3;refGene_tx=XM_023256765.1;refGene_gn=MIER3;refGene_tx=XM_023256767.1;refGene_gn=MIER3;refGene_tx=XM_023256770.1
A1 127465398 127465423
refGene_gn=MIER3;refGene_tx=XM_023256770.1

A1 127466868 127466877
refGene_gn=MIER3;refGene_tx=XM_023256770.1
A1 127475089 127475147
refGene_gn=MIER3;refGene_tx=XM_023256761.1;refGene_gn=MIER3;refGene_tx=XM_023256765.1;refGene_gn=MIER3;refGene_tx=XM_023256767.1
A1 127656748 127656811
refGene_gn=GPBP1;refGene_tx=XM_019834432.2;refGene_gn=GPBP1;refGene_tx=XM_023256787.1;refGene_gn=GPBP1;refGene_tx=XM_023256789.1
A1 127671576 127671700
refGene_gn=GPBP1;refGene_tx=XM_019834432.2;refGene_gn=GPBP1;refGene_tx=XM_023256787.1;refGene_gn=GPBP1;refGene_tx=XM_023256789.1
A1 127671825 127672049
refGene_gn=GPBP1;refGene_tx=XM_019834432.2;refGene_gn=GPBP1;refGene_tx=XM_023256787.1;refGene_gn=GPBP1;refGene_tx=XM_023256789.1
A1 127674857 127674924
refGene_gn=GPBP1;refGene_tx=XM_019834432.2;refGene_gn=GPBP1;refGene_tx=XM_023256787.1;refGene_gn=GPBP1;refGene_tx=XM_023256789.1
A1 127675985 127676045
refGene_gn=GPBP1;refGene_tx=XM_019834432.2;refGene_gn=GPBP1;refGene_tx=XM_023256787.1
A1 127682812 127682997
refGene_gn=GPBP1;refGene_tx=XM_019834432.2;refGene_gn=GPBP1;refGene_tx=XM_023256787.1;refGene_gn=GPBP1;refGene_tx=XM_023256789.1
A1 127683571 127683712
refGene_gn=GPBP1;refGene_tx=XM_019834432.2;refGene_gn=GPBP1;refGene_tx=XM_023256787.1;refGene_gn=GPBP1;refGene_tx=XM_023256789.1
A1 127685860 127686028
refGene_gn=GPBP1;refGene_tx=XM_019834432.2;refGene_gn=GPBP1;refGene_tx=XM_023256787.1;refGene_gn=GPBP1;refGene_tx=XM_023256789.1
A1 127686475 127686663
refGene_gn=GPBP1;refGene_tx=XM_019834432.2;refGene_gn=GPBP1;refGene_tx=XM_023256787.1;refGene_gn=GPBP1;refGene_tx=XM_023256789.1
A1 127693295 127693398
refGene_gn=GPBP1;refGene_tx=XM_019834432.2;refGene_gn=GPBP1;refGene_tx=XM_023256787.1;refGene_gn=GPBP1;refGene_tx=XM_023256789.1
A1 127699498 127699657
refGene_gn=GPBP1;refGene_tx=XM_019834432.2;refGene_gn=GPBP1;refGene_tx=XM_023256787.1;refGene_gn=GPBP1;refGene_tx=XM_023256789.1
A1 127761692 127761995
refGene_gn=LOC109492844;refGene_tx=XM_019814140.2
A1 127880002 127881133
refGene_gn=ACTBL2;refGene_tx=XM_003980976.3
A1 128750612 128750804
refGene_gn=PLK2;refGene_tx=XM_003980977.4
A1 128750939 128751050
refGene_gn=PLK2;refGene_tx=XM_003980977.4
A1 128751318 128751448
refGene_gn=PLK2;refGene_tx=XM_003980977.4
A1 128751570 128751811
refGene_gn=PLK2;refGene_tx=XM_003980977.4

A1 128752053 128752183
refGene_gn=PLK2;refGene_tx=XM_003980977.4
A1 128752495 128752593
refGene_gn=PLK2;refGene_tx=XM_003980977.4
A1 128752941 128753089
refGene_gn=PLK2;refGene_tx=XM_003980977.4
A1 128753178 128753377
refGene_gn=PLK2;refGene_tx=XM_003980977.4
A1 128753486 128753582
refGene_gn=PLK2;refGene_tx=XM_003980977.4
A1 128754054 128754142
refGene_gn=PLK2;refGene_tx=XM_003980977.4
A1 128754384 128754514
refGene_gn=PLK2;refGene_tx=XM_003980977.4
A1 128754707 128754824
refGene_gn=PLK2;refGene_tx=XM_003980977.4
A1 128754968 128755076
refGene_gn=PLK2;refGene_tx=XM_003980977.4
A1 128755666 128755936
refGene_gn=PLK2;refGene_tx=XM_003980977.4
A1 128797933 128798791
refGene_gn=GAPT;refGene_tx=XM_006927739.3;refGene_gn=GAPT;refGene_tx=XM_023256793.1
A1 128863003 128863021
refGene_gn=RAB3C;refGene_tx=XM_011282948.3
A1 128863805 128863829
refGene_gn=RAB3C;refGene_tx=XM_006927740.4
A1 128884477 128884705
refGene_gn=RAB3C;refGene_tx=XM_006927740.4;refGene_gn=RAB3C;refGene_tx=XM_011282948.3
A1 128994432 128994551
refGene_gn=RAB3C;refGene_tx=XM_006927740.4;refGene_gn=RAB3C;refGene_tx=XM_011282948.3
A1 129102966 129103091
refGene_gn=RAB3C;refGene_tx=XM_006927740.4;refGene_gn=RAB3C;refGene_tx=XM_011282948.3
A1 129145356 129145544
refGene_gn=RAB3C;refGene_tx=XM_006927740.4;refGene_gn=RAB3C;refGene_tx=XM_011282948.3
A1 129270723 129271140
refGene_gn=PDE4D;refGene_tx=XM_003980985.5;refGene_gn=PDE4D;refGene_tx=XM_011282955.3;refGene_gn=PDE4D;refGene_tx=XM_019834607.2;refGene_gn=PDE4D;refGene_tx=XM_019834617.2;refGene_gn=PDE4D;refGene_tx=XM_019834628.2;refGene_gn=PDE4D;refGene_tx=XM_019834633.2;refGene_gn=PDE4D;refGene_tx=XM_023256805.1
A1 129271852 129272035
refGene_gn=PDE4D;refGene_tx=XM_003980985.5;refGene_gn=PDE4D;refGene_tx=XM_011282955.3;refGene_gn=PDE4D;refGene_tx=XM_019834607.2;refGene_gn=PDE4D;refGene_tx=XM_019834617.2;refGene_gn=PDE4D;refGene_tx=XM_019834628.2;refGene_gn=PDE4D;refGene_tx=XM_019834633.2;refGene_gn=PDE4D;refGe

ne_tx=XM_023256805.1
A1 129272741 129272864
refGene_gn=PDE4D;refGene_tx=XM_003980985.5;refGene_gn=PDE4D;refGene_tx=XM_011282955.3;refGene_gn=PDE4D;refGene_tx=XM_019834607.2;refGene_gn=PDE4D;refGene_tx=XM_019834617.2;refGene_gn=PDE4D;refGene_tx=XM_019834628.2;refGene_gn=PDE4D;refGene_tx=XM_019834633.2;refGene_gn=PDE4D;refGene_tx=XM_023256805.1
A1 129273748 129273903
refGene_gn=PDE4D;refGene_tx=XM_003980985.5;refGene_gn=PDE4D;refGene_tx=XM_011282955.3;refGene_gn=PDE4D;refGene_tx=XM_019834607.2;refGene_gn=PDE4D;refGene_tx=XM_019834617.2;refGene_gn=PDE4D;refGene_tx=XM_019834628.2;refGene_gn=PDE4D;refGene_tx=XM_019834633.2;refGene_gn=PDE4D;refGene_tx=XM_023256805.1
A1 129280345 129280445
refGene_gn=PDE4D;refGene_tx=XM_003980985.5;refGene_gn=PDE4D;refGene_tx=XM_011282955.3;refGene_gn=PDE4D;refGene_tx=XM_019834607.2;refGene_gn=PDE4D;refGene_tx=XM_019834617.2;refGene_gn=PDE4D;refGene_tx=XM_019834628.2;refGene_gn=PDE4D;refGene_tx=XM_019834633.2;refGene_gn=PDE4D;refGene_tx=XM_023256805.1
A1 129281460 129281625
refGene_gn=PDE4D;refGene_tx=XM_003980985.5;refGene_gn=PDE4D;refGene_tx=XM_011282955.3;refGene_gn=PDE4D;refGene_tx=XM_019834607.2;refGene_gn=PDE4D;refGene_tx=XM_019834617.2;refGene_gn=PDE4D;refGene_tx=XM_019834628.2;refGene_gn=PDE4D;refGene_tx=XM_019834633.2;refGene_gn=PDE4D;refGene_tx=XM_023256805.1
A1 129282468 129282567
refGene_gn=PDE4D;refGene_tx=XM_003980985.5;refGene_gn=PDE4D;refGene_tx=XM_011282955.3;refGene_gn=PDE4D;refGene_tx=XM_019834607.2;refGene_gn=PDE4D;refGene_tx=XM_019834617.2;refGene_gn=PDE4D;refGene_tx=XM_019834628.2;refGene_gn=PDE4D;refGene_tx=XM_019834633.2;refGene_gn=PDE4D;refGene_tx=XM_023256805.1
A1 129283627 129283800
refGene_gn=PDE4D;refGene_tx=XM_003980985.5;refGene_gn=PDE4D;refGene_tx=XM_011282955.3;refGene_gn=PDE4D;refGene_tx=XM_019834607.2;refGene_gn=PDE4D;refGene_tx=XM_019834617.2;refGene_gn=PDE4D;refGene_tx=XM_019834628.2;refGene_gn=PDE4D;refGene_tx=XM_019834633.2;refGene_gn=PDE4D;refGene_tx=XM_023256805.1
A1 129285180 129285274
refGene_gn=PDE4D;refGene_tx=XM_003980985.5;refGene_gn=PDE4D;refGene_tx=XM_011282955.3;refGene_gn=PDE4D;refGene_tx=XM_019834607.2;refGene_gn=PDE4D;refGene_tx=XM_019834617.2;refGene_gn=PDE4D;refGene_tx=XM_019834628.2;refGene_gn=PDE4D;refGene_tx=XM_019834633.2;refGene_gn=PDE4D;refGene_tx=XM_023256805.1
A1 129290649 129290697
refGene_gn=PDE4D;refGene_tx=XM_019834633.2
A1 129321306 129321419
refGene_gn=PDE4D;refGene_tx=XM_003980985.5;refGene_gn=PDE4D;refGene_tx=XM_011282955.3;refGene_gn=PDE4D;refGene_tx=XM_019834607.2;refGene_gn=PDE4D;refGene_tx=XM_019834617.2;refGene_gn=PDE4D;refGene_tx=XM_019834628.2;refGene_gn=PDE4D;refGene_tx=XM_023256805.1

A1 129453814 129453864
refGene_gn=PDE4D;refGene_tx=XM_011282955.3;refGene_gn=PDE4D;refGene_tx=XM_019834607.2;refGene_gn=PDE4D;refGene_tx=XM_019834617.2;refGene_gn=PDE4D;refGene_tx=XM_019834628.2;refGene_gn=PDE4D;refGene_tx=XM_023256805.1
A1 129458736 129458810
refGene_gn=PDE4D;refGene_tx=XM_011282955.3;refGene_gn=PDE4D;refGene_tx=XM_019834607.2;refGene_gn=PDE4D;refGene_tx=XM_019834617.2;refGene_gn=PDE4D;refGene_tx=XM_019834628.2;refGene_gn=PDE4D;refGene_tx=XM_023256805.1
A1 129466623 129466660
refGene_gn=PDE4D;refGene_tx=XM_011282955.3;refGene_gn=PDE4D;refGene_tx=XM_019834607.2;refGene_gn=PDE4D;refGene_tx=XM_019834617.2;refGene_gn=PDE4D;refGene_tx=XM_019834628.2;refGene_gn=PDE4D;refGene_tx=XM_023256805.1
A1 129487699 129487891
refGene_gn=PDE4D;refGene_tx=XM_011282955.3;refGene_gn=PDE4D;refGene_tx=XM_019834607.2;refGene_gn=PDE4D;refGene_tx=XM_019834617.2;refGene_gn=PDE4D;refGene_tx=XM_019834628.2;refGene_gn=PDE4D;refGene_tx=XM_023256805.1
A1 129544386 129544451
refGene_gn=PDE4D;refGene_tx=XM_019834607.2
A1 129615837 129615926
refGene_gn=PDE4D;refGene_tx=XM_019834628.2
A1 129822116 129822163
refGene_gn=PDE4D;refGene_tx=XM_019834617.2
A1 129989641 129989907
refGene_gn=PDE4D;refGene_tx=XM_011282955.3
A1 130125391 130125846
refGene_gn=PDE4D;refGene_tx=XM_023256805.1
A1 130743246 130743408
refGene_gn=DEPDC1B;refGene_tx=XM_003980996.4;refGene_gn=DEPDC1B;refGene_tx=XM_019834664.2
A1 130744687 130744873
refGene_gn=DEPDC1B;refGene_tx=XM_003980996.4;refGene_gn=DEPDC1B;refGene_tx=XM_019834664.2
A1 130752520 130752697
refGene_gn=DEPDC1B;refGene_tx=XM_003980996.4;refGene_gn=DEPDC1B;refGene_tx=XM_019834664.2
A1 130754941 130755108
refGene_gn=DEPDC1B;refGene_tx=XM_003980996.4;refGene_gn=DEPDC1B;refGene_tx=XM_019834664.2
A1 130780277 130780418
refGene_gn=DEPDC1B;refGene_tx=XM_003980996.4;refGene_gn=DEPDC1B;refGene_tx=XM_019834664.2
A1 130784218 130784266
refGene_gn=DEPDC1B;refGene_tx=XM_003980996.4;refGene_gn=DEPDC1B;refGene_tx=XM_019834664.2
A1 130786200 130786331
refGene_gn=DEPDC1B;refGene_tx=XM_003980996.4;refGene_gn=DEPDC1B;refGen

e_tx=XM_019834664.2
A1 130786968 130787096
refGene_gn=DEPDC1B;refGene_tx=XM_003980996.4;refGene_gn=DEPDC1B;refGen
e_tx=XM_019834664.2
A1 130791993 130792129
refGene_gn=DEPDC1B;refGene_tx=XM_003980996.4;refGene_gn=DEPDC1B;refGen
e_tx=XM_019834664.2
A1 130826079 130826345
refGene_gn=DEPDC1B;refGene_tx=XM_003980996.4;refGene_gn=DEPDC1B;refGen
e_tx=XM_019834664.2
A1 130833846 130833894
refGene_gn=DEPDC1B;refGene_tx=XM_003980996.4
A1 130868646 130868856
refGene_gn=ELOVL7;refGene_tx=XM_011282970.2;refGene_gn=ELOVL7;refGene_
tx=XM_023256840.1
A1 130870667 130870804
refGene_gn=ELOVL7;refGene_tx=XM_011282970.2;refGene_gn=ELOVL7;refGene_
tx=XM_023256840.1
A1 130877937 130878043
refGene_gn=ELOVL7;refGene_tx=XM_011282970.2;refGene_gn=ELOVL7;refGene_
tx=XM_023256840.1
A1 130880337 130880394
refGene_gn=ELOVL7;refGene_tx=XM_011282970.2;refGene_gn=ELOVL7;refGene_
tx=XM_023256840.1
A1 130881255 130881336
refGene_gn=ELOVL7;refGene_tx=XM_011282970.2;refGene_gn=ELOVL7;refGene_
tx=XM_023256840.1
A1 130884880 130885071
refGene_gn=ELOVL7;refGene_tx=XM_011282970.2;refGene_gn=ELOVL7;refGene_
tx=XM_023256840.1
A1 130894799 130894863
refGene_gn=ELOVL7;refGene_tx=XM_011282970.2;refGene_gn=ELOVL7;refGene_
tx=XM_023256840.1
A1 130966844 130966913
refGene_gn=ERCC8;refGene_tx=XM_003981000.5;refGene_gn=ERCC8;refGene_tx
=XM_011282979.2
A1 130975711 130975792
refGene_gn=ERCC8;refGene_tx=XM_003981000.5;refGene_gn=ERCC8;refGene_tx
=XM_011282979.2
A1 130982771 130982969
refGene_gn=ERCC8;refGene_tx=XM_003981000.5;refGene_gn=ERCC8;refGene_tx
=XM_011282979.2
A1 130992847 130992972
refGene_gn=ERCC8;refGene_tx=XM_003981000.5;refGene_gn=ERCC8;refGene_tx
=XM_011282979.2
A1 130994492 130994593
refGene_gn=ERCC8;refGene_tx=XM_003981000.5;refGene_gn=ERCC8;refGene_tx
=XM_011282979.2
A1 130996943 130997010
refGene_gn=ERCC8;refGene_tx=XM_003981000.5;refGene_gn=ERCC8;refGene_tx

=XM_011282979.2
A1 130998190 130998259
refGene_gn=ERCC8;refGene_tx=XM_003981000.5;refGene_gn=ERCC8;refGene_tx
=XM_011282979.2
A1 130999305 130999387
refGene_gn=ERCC8;refGene_tx=XM_003981000.5;refGene_gn=ERCC8;refGene_tx
=XM_011282979.2
A1 131009511 131009635
refGene_gn=ERCC8;refGene_tx=XM_003981000.5;refGene_gn=ERCC8;refGene_tx
=XM_011282979.2
A1 131013429 131013531
refGene_gn=ERCC8;refGene_tx=XM_003981000.5
A1 131018460 131018556
refGene_gn=ERCC8;refGene_tx=XM_003981000.5
A1 131029517 131029594
refGene_gn=ERCC8;refGene_tx=XM_003981000.5
A1 131029824 131029951
refGene_gn=NDUFAF2;refGene_tx=XM_019838168.2
A1 131030043 131030323
refGene_gn=LOC102900994;refGene_tx=XM_023247911.1
A1 131030434 131030611
refGene_gn=LOC102900994;refGene_tx=XM_023247911.1
A1 131116660 131117040
refGene_gn=LOC102900994;refGene_tx=XM_023247911.1
A1 131117074 131117150
refGene_gn=LOC102900994;refGene_tx=XM_023247911.1
A1 131117183 131117307
refGene_gn=LOC102900994;refGene_tx=XM_023247911.1
A1 131120639 131120718
refGene_gn=LOC102900994;refGene_tx=XM_023247911.1
A1 131129371 131129461
refGene_gn=NDUFAF2;refGene_tx=XM_019838168.2
A1 131154198 131154239
refGene_gn=NDUFAF2;refGene_tx=XM_019838168.2
A1 131203386 131203635
refGene_gn=NDUFAF2;refGene_tx=XM_019838168.2
A1 131210283 131210519
refGene_gn=SMIM15;refGene_tx=XM_006927752.3;refGene_gn=SMIM15;refGene_
tx=XM_006927755.3;refGene_gn=SMIM15;refGene_tx=XM_019834719.1;refGene_
gn=SMIM15;refGene_tx=XM_019834727.1;refGene_gn=SMIM15;refGene_tx=XM_01
9834729.1;refGene_gn=SMIM15;refGene_tx=XM_019834736.1;refGene_gn=SMIM1
5;refGene_tx=XM_019834738.1
A1 131211164 131211180
refGene_gn=SMIM15;refGene_tx=XM_019834719.1;refGene_gn=SMIM15;refGene_
tx=XM_019834727.1;refGene_gn=SMIM15;refGene_tx=XM_019834729.1;refGene_
gn=SMIM15;refGene_tx=XM_019834736.1;refGene_gn=SMIM15;refGene_tx=XM_01
9834738.1
A1 131369371 131370035
refGene_gn=ZSWIM6;refGene_tx=XM_023256902.1
A1 131444448 131444548

refGene_gn=ZSWIM6;refGene_tx=XM_019834769.2
A1 131493418 131493775
refGene_gn=ZSWIM6;refGene_tx=XM_019834769.2;refGene_gn=ZSWIM6;refGene_tx=XM_023256902.1
A1 131508736 131508885
refGene_gn=ZSWIM6;refGene_tx=XM_019834769.2;refGene_gn=ZSWIM6;refGene_tx=XM_023256902.1
A1 131512057 131512208
refGene_gn=ZSWIM6;refGene_tx=XM_019834769.2;refGene_gn=ZSWIM6;refGene_tx=XM_023256902.1
A1 131540575 131540755
refGene_gn=ZSWIM6;refGene_tx=XM_019834769.2;refGene_gn=ZSWIM6;refGene_tx=XM_023256902.1
A1 131545636 131545813
refGene_gn=ZSWIM6;refGene_tx=XM_019834769.2;refGene_gn=ZSWIM6;refGene_tx=XM_023256902.1
A1 131546088 131546235
refGene_gn=ZSWIM6;refGene_tx=XM_019834769.2;refGene_gn=ZSWIM6;refGene_tx=XM_023256902.1
A1 131550572 131550719
refGene_gn=ZSWIM6;refGene_tx=XM_019834769.2;refGene_gn=ZSWIM6;refGene_tx=XM_023256902.1
A1 131552301 131552562
refGene_gn=ZSWIM6;refGene_tx=XM_019834769.2;refGene_gn=ZSWIM6;refGene_tx=XM_023256902.1
A1 131556364 131556500
refGene_gn=ZSWIM6;refGene_tx=XM_019834769.2;refGene_gn=ZSWIM6;refGene_tx=XM_023256902.1
A1 131559663 131559821
refGene_gn=ZSWIM6;refGene_tx=XM_019834769.2;refGene_gn=ZSWIM6;refGene_tx=XM_023256902.1
A1 131560456 131560620
refGene_gn=ZSWIM6;refGene_tx=XM_019834769.2;refGene_gn=ZSWIM6;refGene_tx=XM_023256902.1
A1 131562344 131562426
refGene_gn=ZSWIM6;refGene_tx=XM_019834769.2;refGene_gn=ZSWIM6;refGene_tx=XM_023256902.1
A1 131564226 131565089
refGene_gn=ZSWIM6;refGene_tx=XM_019834769.2;refGene_gn=ZSWIM6;refGene_tx=XM_023256902.1
A1 132159794 132159858
refGene_gn=KIF2A;refGene_tx=XM_023256931.1;refGene_gn=KIF2A;refGene_tx=XM_023256941.1;refGene_gn=KIF2A;refGene_tx=XM_023256945.1;refGene_gn=KIF2A;refGene_tx=XM_023256947.1
A1 132193312 132193407
refGene_gn=KIF2A;refGene_tx=XM_023256931.1;refGene_gn=KIF2A;refGene_tx=XM_023256941.1;refGene_gn=KIF2A;refGene_tx=XM_023256945.1;refGene_gn=KIF2A;refGene_tx=XM_023256947.1
A1 132194599 132194719
refGene_gn=KIF2A;refGene_tx=XM_023256931.1;refGene_gn=KIF2A;refGene_tx

=XM_023256941.1;refGene_gn=KIF2A;refGene_tx=XM_023256945.1;refGene_gn=KIF2A;refGene_tx=XM_023256947.1
A1 132196999 132197054
refGene_gn=KIF2A;refGene_tx=XM_023256931.1;refGene_gn=KIF2A;refGene_tx=XM_023256941.1;refGene_gn=KIF2A;refGene_tx=XM_023256945.1;refGene_gn=KIF2A;refGene_tx=XM_023256947.1
A1 132198462 132198585
refGene_gn=KIF2A;refGene_tx=XM_023256931.1;refGene_gn=KIF2A;refGene_tx=XM_023256941.1;refGene_gn=KIF2A;refGene_tx=XM_023256945.1;refGene_gn=KIF2A;refGene_tx=XM_023256947.1
A1 132199155 132199256
refGene_gn=KIF2A;refGene_tx=XM_023256931.1;refGene_gn=KIF2A;refGene_tx=XM_023256941.1;refGene_gn=KIF2A;refGene_tx=XM_023256945.1;refGene_gn=KIF2A;refGene_tx=XM_023256947.1
A1 132201214 132201310
refGene_gn=KIF2A;refGene_tx=XM_023256931.1;refGene_gn=KIF2A;refGene_tx=XM_023256941.1;refGene_gn=KIF2A;refGene_tx=XM_023256945.1;refGene_gn=KIF2A;refGene_tx=XM_023256947.1
A1 132202775 132202830
refGene_gn=KIF2A;refGene_tx=XM_023256931.1;refGene_gn=KIF2A;refGene_tx=XM_023256941.1;refGene_gn=KIF2A;refGene_tx=XM_023256945.1;refGene_gn=KIF2A;refGene_tx=XM_023256947.1
A1 132203201 132203364
refGene_gn=KIF2A;refGene_tx=XM_023256931.1;refGene_gn=KIF2A;refGene_tx=XM_023256941.1;refGene_gn=KIF2A;refGene_tx=XM_023256945.1;refGene_gn=KIF2A;refGene_tx=XM_023256947.1
A1 132205705 132205796
refGene_gn=KIF2A;refGene_tx=XM_023256931.1;refGene_gn=KIF2A;refGene_tx=XM_023256941.1;refGene_gn=KIF2A;refGene_tx=XM_023256945.1;refGene_gn=KIF2A;refGene_tx=XM_023256947.1
A1 132205932 132205996
refGene_gn=KIF2A;refGene_tx=XM_023256931.1;refGene_gn=KIF2A;refGene_tx=XM_023256941.1;refGene_gn=KIF2A;refGene_tx=XM_023256945.1;refGene_gn=KIF2A;refGene_tx=XM_023256947.1
A1 132206570 132206662
refGene_gn=KIF2A;refGene_tx=XM_023256931.1;refGene_gn=KIF2A;refGene_tx=XM_023256941.1;refGene_gn=KIF2A;refGene_tx=XM_023256945.1;refGene_gn=KIF2A;refGene_tx=XM_023256947.1
A1 132207020 132207163
refGene_gn=KIF2A;refGene_tx=XM_023256931.1;refGene_gn=KIF2A;refGene_tx=XM_023256941.1;refGene_gn=KIF2A;refGene_tx=XM_023256945.1;refGene_gn=KIF2A;refGene_tx=XM_023256947.1
A1 132207530 132207735
refGene_gn=KIF2A;refGene_tx=XM_023256931.1;refGene_gn=KIF2A;refGene_tx=XM_023256941.1;refGene_gn=KIF2A;refGene_tx=XM_023256945.1;refGene_gn=KIF2A;refGene_tx=XM_023256947.1
A1 132208458 132208569
refGene_gn=KIF2A;refGene_tx=XM_023256931.1;refGene_gn=KIF2A;refGene_tx=XM_023256941.1;refGene_gn=KIF2A;refGene_tx=XM_023256945.1;refGene_gn=KIF2A;refGene_tx=XM_023256947.1

A1 132209621 132209689
refGene_gn=KIF2A;refGene_tx=XM_023256931.1;refGene_gn=KIF2A;refGene_tx=XM_023256941.1;refGene_gn=KIF2A;refGene_tx=XM_023256945.1;refGene_gn=KIF2A;refGene_tx=XM_023256947.1
A1 132212974 132213088
refGene_gn=KIF2A;refGene_tx=XM_023256931.1;refGene_gn=KIF2A;refGene_tx=XM_023256941.1
A1 132213956 132214107
refGene_gn=KIF2A;refGene_tx=XM_023256931.1;refGene_gn=KIF2A;refGene_tx=XM_023256941.1;refGene_gn=KIF2A;refGene_tx=XM_023256945.1;refGene_gn=KIF2A;refGene_tx=XM_023256947.1
A1 132217409 132217511
refGene_gn=KIF2A;refGene_tx=XM_023256931.1;refGene_gn=KIF2A;refGene_tx=XM_023256941.1;refGene_gn=KIF2A;refGene_tx=XM_023256945.1;refGene_gn=KIF2A;refGene_tx=XM_023256947.1
A1 132219451 132219587
refGene_gn=KIF2A;refGene_tx=XM_023256931.1;refGene_gn=KIF2A;refGene_tx=XM_023256941.1;refGene_gn=KIF2A;refGene_tx=XM_023256945.1;refGene_gn=KIF2A;refGene_tx=XM_023256947.1
A1 132221847 132221933
refGene_gn=KIF2A;refGene_tx=XM_023256931.1;refGene_gn=KIF2A;refGene_tx=XM_023256941.1;refGene_gn=KIF2A;refGene_tx=XM_023256945.1;refGene_gn=KIF2A;refGene_tx=XM_023256947.1
A1 132225517 132225560
refGene_gn=DIMT1;refGene_tx=XM_003981005.4;refGene_gn=DIMT1;refGene_tx=XM_006927756.3;refGene_gn=DIMT1;refGene_tx=XM_019834834.2;refGene_gn=DIMT1;refGene_tx=XM_023256962.1
A1 132227082 132227189
refGene_gn=DIMT1;refGene_tx=XM_003981005.4;refGene_gn=DIMT1;refGene_tx=XM_006927756.3;refGene_gn=DIMT1;refGene_tx=XM_019834834.2;refGene_gn=DIMT1;refGene_tx=XM_023256962.1
A1 132228366 132228430
refGene_gn=DIMT1;refGene_tx=XM_003981005.4;refGene_gn=DIMT1;refGene_tx=XM_006927756.3;refGene_gn=DIMT1;refGene_tx=XM_019834834.2;refGene_gn=DIMT1;refGene_tx=XM_023256962.1
A1 132229205 132229270
refGene_gn=DIMT1;refGene_tx=XM_003981005.4;refGene_gn=DIMT1;refGene_tx=XM_006927756.3;refGene_gn=DIMT1;refGene_tx=XM_019834834.2;refGene_gn=DIMT1;refGene_tx=XM_023256962.1
A1 132230323 132230416
refGene_gn=DIMT1;refGene_tx=XM_003981005.4;refGene_gn=DIMT1;refGene_tx=XM_006927756.3;refGene_gn=DIMT1;refGene_tx=XM_019834834.2;refGene_gn=DIMT1;refGene_tx=XM_023256962.1
A1 132230782 132230906
refGene_gn=DIMT1;refGene_tx=XM_003981005.4;refGene_gn=DIMT1;refGene_tx=XM_006927756.3;refGene_gn=DIMT1;refGene_tx=XM_019834834.2;refGene_gn=DIMT1;refGene_tx=XM_023256962.1
A1 132231982 132232032
refGene_gn=DIMT1;refGene_tx=XM_003981005.4;refGene_gn=DIMT1;refGene_tx=XM_006927756.3;refGene_gn=DIMT1;refGene_tx=XM_019834834.2;refGene_gn=

DIMT1;refGene_tx=XM_023256962.1
A1 132232118 132232212
refGene_gn=DIMT1;refGene_tx=XM_003981005.4;refGene_gn=DIMT1;refGene_tx=XM_006927756.3;refGene_gn=DIMT1;refGene_tx=XM_019834834.2;refGene_gn=DIMT1;refGene_tx=XM_023256962.1
A1 132232295 132232357
refGene_gn=DIMT1;refGene_tx=XM_003981005.4;refGene_gn=DIMT1;refGene_tx=XM_006927756.3;refGene_gn=DIMT1;refGene_tx=XM_019834834.2;refGene_gn=DIMT1;refGene_tx=XM_023256962.1
A1 132234425 132234512
refGene_gn=DIMT1;refGene_tx=XM_003981005.4;refGene_gn=DIMT1;refGene_tx=XM_006927756.3;refGene_gn=DIMT1;refGene_tx=XM_019834834.2;refGene_gn=DIMT1;refGene_tx=XM_023256962.1
A1 132236115 132236189
refGene_gn=DIMT1;refGene_tx=XM_003981005.4;refGene_gn=DIMT1;refGene_tx=XM_006927756.3
A1 132236513 132236592
refGene_gn=DIMT1;refGene_tx=XM_003981005.4;refGene_gn=DIMT1;refGene_tx=XM_019834834.2
A1 132275618 132275756
refGene_gn=IPO11;refGene_tx=XM_023256975.1;refGene_gn=IPO11;refGene_tx=XM_023256984.1;refGene_gn=IPO11;refGene_tx=XM_023256985.1;refGene_gn=IPO11;refGene_tx=XM_023256987.1
A1 132279277 132279378
refGene_gn=IPO11;refGene_tx=XM_023256975.1;refGene_gn=IPO11;refGene_tx=XM_023256984.1;refGene_gn=IPO11;refGene_tx=XM_023256985.1;refGene_gn=IPO11;refGene_tx=XM_023256987.1
A1 132287055 132287128
refGene_gn=IPO11;refGene_tx=XM_023256975.1;refGene_gn=IPO11;refGene_tx=XM_023256984.1;refGene_gn=IPO11;refGene_tx=XM_023256985.1;refGene_gn=IPO11;refGene_tx=XM_023256987.1
A1 132288749 132288953
refGene_gn=IPO11;refGene_tx=XM_023256975.1;refGene_gn=IPO11;refGene_tx=XM_023256984.1;refGene_gn=IPO11;refGene_tx=XM_023256985.1;refGene_gn=IPO11;refGene_tx=XM_023256987.1
A1 132301818 132301951
refGene_gn=IPO11;refGene_tx=XM_023256975.1;refGene_gn=IPO11;refGene_tx=XM_023256984.1;refGene_gn=IPO11;refGene_tx=XM_023256985.1;refGene_gn=IPO11;refGene_tx=XM_023256987.1
A1 132304582 132304641
refGene_gn=IPO11;refGene_tx=XM_023256975.1;refGene_gn=IPO11;refGene_tx=XM_023256984.1;refGene_gn=IPO11;refGene_tx=XM_023256985.1;refGene_gn=IPO11;refGene_tx=XM_023256987.1
A1 132308762 132308811
refGene_gn=IPO11;refGene_tx=XM_023256975.1;refGene_gn=IPO11;refGene_tx=XM_023256984.1;refGene_gn=IPO11;refGene_tx=XM_023256985.1;refGene_gn=IPO11;refGene_tx=XM_023256987.1
A1 132310725 132310796
refGene_gn=IPO11;refGene_tx=XM_023256975.1;refGene_gn=IPO11;refGene_tx=XM_023256984.1;refGene_gn=IPO11;refGene_tx=XM_023256985.1;refGene_gn=

IPO11;refGene_tx=XM_023256987.1
A1 132313216 132313409
refGene_gn=IPO11;refGene_tx=XM_023256975.1;refGene_gn=IPO11;refGene_tx=XM_023256984.1;refGene_gn=IPO11;refGene_tx=XM_023256985.1;refGene_gn=IPO11;refGene_tx=XM_023256987.1
A1 132314042 132314195
refGene_gn=IPO11;refGene_tx=XM_023256975.1;refGene_gn=IPO11;refGene_tx=XM_023256984.1;refGene_gn=IPO11;refGene_tx=XM_023256985.1;refGene_gn=IPO11;refGene_tx=XM_023256987.1
A1 132315531 132315575
refGene_gn=IPO11;refGene_tx=XM_023256975.1;refGene_gn=IPO11;refGene_tx=XM_023256984.1;refGene_gn=IPO11;refGene_tx=XM_023256985.1;refGene_gn=IPO11;refGene_tx=XM_023256987.1
A1 132317007 132317098
refGene_gn=IPO11;refGene_tx=XM_023256975.1;refGene_gn=IPO11;refGene_tx=XM_023256984.1;refGene_gn=IPO11;refGene_tx=XM_023256985.1;refGene_gn=IPO11;refGene_tx=XM_023256987.1
A1 132319922 132319970
refGene_gn=IPO11;refGene_tx=XM_023256975.1;refGene_gn=IPO11;refGene_tx=XM_023256984.1;refGene_gn=IPO11;refGene_tx=XM_023256985.1;refGene_gn=IPO11;refGene_tx=XM_023256987.1
A1 132320779 132320885
refGene_gn=IPO11;refGene_tx=XM_023256975.1;refGene_gn=IPO11;refGene_tx=XM_023256984.1;refGene_gn=IPO11;refGene_tx=XM_023256985.1;refGene_gn=IPO11;refGene_tx=XM_023256987.1
A1 132323509 132323636
refGene_gn=IPO11;refGene_tx=XM_023256975.1;refGene_gn=IPO11;refGene_tx=XM_023256984.1;refGene_gn=IPO11;refGene_tx=XM_023256985.1;refGene_gn=IPO11;refGene_tx=XM_023256987.1
A1 132327376 132327410
refGene_gn=IPO11;refGene_tx=XM_023256975.1;refGene_gn=IPO11;refGene_tx=XM_023256984.1;refGene_gn=IPO11;refGene_tx=XM_023256985.1;refGene_gn=IPO11;refGene_tx=XM_023256987.1
A1 132327721 132327762
refGene_gn=IPO11;refGene_tx=XM_023256975.1;refGene_gn=IPO11;refGene_tx=XM_023256984.1;refGene_gn=IPO11;refGene_tx=XM_023256985.1;refGene_gn=IPO11;refGene_tx=XM_023256987.1
A1 132329105 132329222
refGene_gn=IPO11;refGene_tx=XM_023256975.1;refGene_gn=IPO11;refGene_tx=XM_023256984.1;refGene_gn=IPO11;refGene_tx=XM_023256985.1;refGene_gn=IPO11;refGene_tx=XM_023256987.1
A1 132334517 132334631
refGene_gn=IPO11;refGene_tx=XM_023256975.1;refGene_gn=IPO11;refGene_tx=XM_023256984.1;refGene_gn=IPO11;refGene_tx=XM_023256985.1;refGene_gn=IPO11;refGene_tx=XM_023256987.1
A1 132339913 132340029
refGene_gn=IPO11;refGene_tx=XM_023256975.1;refGene_gn=IPO11;refGene_tx=XM_023256984.1;refGene_gn=IPO11;refGene_tx=XM_023256985.1;refGene_gn=IPO11;refGene_tx=XM_023256987.1
A1 132348653 132348730

refGene_gn=IPO11;refGene_tx=XM_023256975.1;refGene_gn=IPO11;refGene_tx=XM_023256984.1;refGene_gn=IPO11;refGene_tx=XM_023256985.1;refGene_gn=IPO11;refGene_tx=XM_023256987.1
A1 132358311 132358391
refGene_gn=IPO11;refGene_tx=XM_023256975.1;refGene_gn=IPO11;refGene_tx=XM_023256984.1;refGene_gn=IPO11;refGene_tx=XM_023256985.1;refGene_gn=IPO11;refGene_tx=XM_023256987.1
A1 132358763 132358844
refGene_gn=IPO11;refGene_tx=XM_023256975.1;refGene_gn=IPO11;refGene_tx=XM_023256984.1;refGene_gn=IPO11;refGene_tx=XM_023256985.1;refGene_gn=IPO11;refGene_tx=XM_023256987.1
A1 132370883 132370979
refGene_gn=IPO11;refGene_tx=XM_023256975.1;refGene_gn=IPO11;refGene_tx=XM_023256984.1;refGene_gn=IPO11;refGene_tx=XM_023256985.1;refGene_gn=IPO11;refGene_tx=XM_023256987.1
A1 132371947 132372061
refGene_gn=IPO11;refGene_tx=XM_023256975.1;refGene_gn=IPO11;refGene_tx=XM_023256984.1;refGene_gn=IPO11;refGene_tx=XM_023256985.1;refGene_gn=IPO11;refGene_tx=XM_023256987.1
A1 132379808 132379930
refGene_gn=IPO11;refGene_tx=XM_023256975.1;refGene_gn=IPO11;refGene_tx=XM_023256984.1;refGene_gn=IPO11;refGene_tx=XM_023256985.1;refGene_gn=IPO11;refGene_tx=XM_023256987.1
A1 132394549 132394559
refGene_gn=IPO11;refGene_tx=XM_023256987.1
A1 132395602 132395624
refGene_gn=IPO11;refGene_tx=XM_023256985.1
A1 132412276 132412372
refGene_gn=IPO11;refGene_tx=XM_023256975.1;refGene_gn=IPO11;refGene_tx=XM_023256984.1
A1 132418914 132418999
refGene_gn=IPO11;refGene_tx=XM_023256975.1;refGene_gn=IPO11;refGene_tx=XM_023256984.1
A1 132424785 132424836
refGene_gn=IPO11;refGene_tx=XM_023256975.1
A1 132449870 132450035
refGene_gn=IPO11;refGene_tx=XM_023256975.1;refGene_gn=IPO11;refGene_tx=XM_023256984.1
A1 133565224 133566493
refGene_gn=HTR1A;refGene_tx=XM_019834870.2
A1 133770431 133770566
refGene_gn=RNF180;refGene_tx=XM_023256994.1;refGene_gn=RNF180;refGene_tx=XM_023257000.1
A1 133792597 133792693
refGene_gn=RNF180;refGene_tx=XM_023256994.1;refGene_gn=RNF180;refGene_tx=XM_023257000.1;refGene_gn=RNF180;refGene_tx=XM_023257004.1
A1 133794323 133795283
refGene_gn=RNF180;refGene_tx=XM_023256994.1;refGene_gn=RNF180;refGene_tx=XM_023257000.1;refGene_gn=RNF180;refGene_tx=XM_023257004.1
A1 133883362 133883588

refGene_gn=RNF180;refGene_tx=XM_023256994.1;refGene_gn=RNF180;refGene_tx=XM_023257000.1;refGene_gn=RNF180;refGene_tx=XM_023257004.1
A1 133893285 133893411
refGene_gn=RNF180;refGene_tx=XM_023256994.1;refGene_gn=RNF180;refGene_tx=XM_023257000.1;refGene_gn=RNF180;refGene_tx=XM_023257004.1
A1 133936954 133937154
refGene_gn=RNF180;refGene_tx=XM_023256994.1;refGene_gn=RNF180;refGene_tx=XM_023257000.1;refGene_gn=RNF180;refGene_tx=XM_023257004.1
A1 134028234 134028399
refGene_gn=RGS7BP;refGene_tx=XM_003981011.5;refGene_gn=RGS7BP;refGene_tx=XM_019834877.2
A1 134029291 134029458
refGene_gn=RGS7BP;refGene_tx=XM_003981011.5;refGene_gn=RGS7BP;refGene_tx=XM_019834877.2
A1 134091681 134091812
refGene_gn=RGS7BP;refGene_tx=XM_003981011.5;refGene_gn=RGS7BP;refGene_tx=XM_019834877.2
A1 134109799 134109947
refGene_gn=RGS7BP;refGene_tx=XM_003981011.5;refGene_gn=RGS7BP;refGene_tx=XM_019834877.2
A1 134120288 134120363
refGene_gn=RGS7BP;refGene_tx=XM_003981011.5;refGene_gn=RGS7BP;refGene_tx=XM_019834877.2
A1 134132028 134132120
refGene_gn=RGS7BP;refGene_tx=XM_003981011.5
A1 134199804 134199995
refGene_gn=SHISAL2B;refGene_tx=XM_003981030.5
A1 134204274 134204432
refGene_gn=SHISAL2B;refGene_tx=XM_003981030.5
A1 134217551 134217688
refGene_gn=SHISAL2B;refGene_tx=XM_003981030.5
A1 134223473 134223663
refGene_gn=SREK1IP1;refGene_tx=XM_011284065.3
A1 134226192 134226265
refGene_gn=SREK1IP1;refGene_tx=XM_011284065.3
A1 134243715 134243859
refGene_gn=SREK1IP1;refGene_tx=XM_011284065.3
A1 134253752 134253800
refGene_gn=SREK1IP1;refGene_tx=XM_011284065.3
A1 134265097 134265302
refGene_gn=SREK1IP1;refGene_tx=XM_011284065.3
A1 134265724 134265766
refGene_gn=CWC27;refGene_tx=XM_003981012.5;refGene_gn=CWC27;refGene_tx=XM_019834889.2;refGene_gn=CWC27;refGene_tx=XM_023257015.1
A1 134273418 134273515
refGene_gn=CWC27;refGene_tx=XM_003981012.5;refGene_gn=CWC27;refGene_tx=XM_019834889.2;refGene_gn=CWC27;refGene_tx=XM_023257015.1
A1 134280522 134280635
refGene_gn=CWC27;refGene_tx=XM_003981012.5;refGene_gn=CWC27;refGene_tx=XM_019834889.2;refGene_gn=CWC27;refGene_tx=XM_023257015.1

A1 134282737 134282881
refGene_gn=CWC27;refGene_tx=XM_003981012.5;refGene_gn=CWC27;refGene_tx=XM_019834889.2;refGene_gn=CWC27;refGene_tx=XM_023257015.1
A1 134284550 134284649
refGene_gn=CWC27;refGene_tx=XM_003981012.5;refGene_gn=CWC27;refGene_tx=XM_019834889.2;refGene_gn=CWC27;refGene_tx=XM_023257015.1
A1 134285286 134285390
refGene_gn=CWC27;refGene_tx=XM_003981012.5;refGene_gn=CWC27;refGene_tx=XM_019834889.2;refGene_gn=CWC27;refGene_tx=XM_023257015.1
A1 134287100 134287170
refGene_gn=CWC27;refGene_tx=XM_003981012.5;refGene_gn=CWC27;refGene_tx=XM_019834889.2;refGene_gn=CWC27;refGene_tx=XM_023257015.1
A1 134295518 134295598
refGene_gn=CWC27;refGene_tx=XM_003981012.5;refGene_gn=CWC27;refGene_tx=XM_019834889.2;refGene_gn=CWC27;refGene_tx=XM_023257015.1
A1 134296553 134296587
refGene_gn=CWC27;refGene_tx=XM_003981012.5;refGene_gn=CWC27;refGene_tx=XM_019834889.2;refGene_gn=CWC27;refGene_tx=XM_023257015.1
A1 134297836 134297991
refGene_gn=CWC27;refGene_tx=XM_003981012.5;refGene_gn=CWC27;refGene_tx=XM_019834889.2;refGene_gn=CWC27;refGene_tx=XM_023257015.1
A1 134358924 134359028
refGene_gn=CWC27;refGene_tx=XM_003981012.5;refGene_gn=CWC27;refGene_tx=XM_019834889.2;refGene_gn=CWC27;refGene_tx=XM_023257015.1
A1 134370226 134370306
refGene_gn=CWC27;refGene_tx=XM_023257015.1
A1 134384415 134384516
refGene_gn=CWC27;refGene_tx=XM_019834889.2
A1 134424933 134425043
refGene_gn=CWC27;refGene_tx=XM_003981012.5
A1 134429101 134429205
refGene_gn=CWC27;refGene_tx=XM_003981012.5
A1 134457928 134458091
refGene_gn=CWC27;refGene_tx=XM_003981012.5
A1 134576233 134576343
refGene_gn=ADAMTS6;refGene_tx=XM_023257025.1
A1 134589638 134589795
refGene_gn=ADAMTS6;refGene_tx=XM_023257025.1
A1 134592236 134592413
refGene_gn=ADAMTS6;refGene_tx=XM_023257025.1
A1 134601548 134601753
refGene_gn=ADAMTS6;refGene_tx=XM_023257025.1
A1 134611694 134611824
refGene_gn=ADAMTS6;refGene_tx=XM_023257025.1
A1 134629305 134629444
refGene_gn=ADAMTS6;refGene_tx=XM_023257025.1
A1 134629884 134630048
refGene_gn=ADAMTS6;refGene_tx=XM_023257025.1
A1 134638343 134638424
refGene_gn=ADAMTS6;refGene_tx=XM_023257025.1

A1 134638961 134639085
refGene_gn=ADAMTS6;refGene_tx=XM_023257025.1
A1 134640667 134640801
refGene_gn=ADAMTS6;refGene_tx=XM_023257025.1
A1 134647872 134647975
refGene_gn=ADAMTS6;refGene_tx=XM_023257025.1
A1 134665671 134665735
refGene_gn=ADAMTS6;refGene_tx=XM_023257025.1
A1 134667820 134667966
refGene_gn=ADAMTS6;refGene_tx=XM_023257025.1
A1 134678975 134679083
refGene_gn=ADAMTS6;refGene_tx=XM_023257025.1
A1 134698297 134698439
refGene_gn=ADAMTS6;refGene_tx=XM_023257025.1
A1 134706808 134706955
refGene_gn=ADAMTS6;refGene_tx=XM_023257025.1
A1 134745474 134745580
refGene_gn=ADAMTS6;refGene_tx=XM_023257025.1
A1 134749706 134749750
refGene_gn=ADAMTS6;refGene_tx=XM_023257025.1
A1 134847028 134847174
refGene_gn=ADAMTS6;refGene_tx=XM_023257025.1
A1 134847673 134847757
refGene_gn=ADAMTS6;refGene_tx=XM_023257025.1
A1 134848264 134848476
refGene_gn=ADAMTS6;refGene_tx=XM_023257025.1
A1 134857967 134858136
refGene_gn=ADAMTS6;refGene_tx=XM_023257025.1
A1 134867805 134868170
refGene_gn=ADAMTS6;refGene_tx=XM_023257025.1
A1 134870366 134870463
refGene_gn=ADAMTS6;refGene_tx=XM_023257025.1
A1 134870737 134870742
refGene_gn=CENPK;refGene_tx=XM_023257048.1
A1 134882424 134882473
refGene_gn=CENPK;refGene_tx=XM_023257048.1
A1 134893525 134893684
refGene_gn=CENPK;refGene_tx=XM_019834916.2
A1 134894295 134894349
refGene_gn=CENPK;refGene_tx=XM_019834916.2;refGene_gn=CENPK;refGene_tx
=XM_023257048.1
A1 134900072 134900205
refGene_gn=CENPK;refGene_tx=XM_019834916.2;refGene_gn=CENPK;refGene_tx
=XM_023257048.1
A1 134900542 134900641
refGene_gn=CENPK;refGene_tx=XM_019834916.2;refGene_gn=CENPK;refGene_tx
=XM_023257048.1
A1 134900740 134900823
refGene_gn=CENPK;refGene_tx=XM_019834916.2;refGene_gn=CENPK;refGene_tx
=XM_023257048.1

A1 134912608 134912655
refGene_gn=CENPK;refGene_tx=XM_019834916.2;refGene_gn=CENPK;refGene_tx=XM_023257048.1
A1 134916037 134916110
refGene_gn=CENPK;refGene_tx=XM_019834916.2;refGene_gn=CENPK;refGene_tx=XM_023257048.1
A1 134919343 134919400
refGene_gn=CENPK;refGene_tx=XM_019834916.2;refGene_gn=CENPK;refGene_tx=XM_023257048.1
A1 134921464 134921581
refGene_gn=CENPK;refGene_tx=XM_019834916.2;refGene_gn=CENPK;refGene_tx=XM_023257048.1
A1 134930076 134930272
refGene_gn=PPWD1;refGene_tx=XM_003981014.5
A1 134935130 134935233
refGene_gn=PPWD1;refGene_tx=XM_003981014.5;refGene_gn=PPWD1;refGene_tx=XM_006927758.4;refGene_gn=PPWD1;refGene_tx=XM_011284050.3
A1 134935937 134936038
refGene_gn=PPWD1;refGene_tx=XM_003981014.5;refGene_gn=PPWD1;refGene_tx=XM_006927758.4;refGene_gn=PPWD1;refGene_tx=XM_011284050.3
A1 134936180 134936301
refGene_gn=PPWD1;refGene_tx=XM_003981014.5;refGene_gn=PPWD1;refGene_tx=XM_006927758.4;refGene_gn=PPWD1;refGene_tx=XM_011284050.3
A1 134938095 134938543
refGene_gn=PPWD1;refGene_tx=XM_003981014.5;refGene_gn=PPWD1;refGene_tx=XM_006927758.4;refGene_gn=PPWD1;refGene_tx=XM_011284050.3
A1 134942289 134942480
refGene_gn=PPWD1;refGene_tx=XM_003981014.5;refGene_gn=PPWD1;refGene_tx=XM_006927758.4;refGene_gn=PPWD1;refGene_tx=XM_011284050.3
A1 134943331 134943521
refGene_gn=PPWD1;refGene_tx=XM_003981014.5;refGene_gn=PPWD1;refGene_tx=XM_006927758.4;refGene_gn=PPWD1;refGene_tx=XM_011284050.3
A1 134946228 134946410
refGene_gn=PPWD1;refGene_tx=XM_003981014.5;refGene_gn=PPWD1;refGene_tx=XM_006927758.4;refGene_gn=PPWD1;refGene_tx=XM_011284050.3
A1 134948234 134948316
refGene_gn=PPWD1;refGene_tx=XM_003981014.5;refGene_gn=PPWD1;refGene_tx=XM_006927758.4;refGene_gn=PPWD1;refGene_tx=XM_011284050.3
A1 134949674 134949857
refGene_gn=PPWD1;refGene_tx=XM_003981014.5;refGene_gn=PPWD1;refGene_tx=XM_006927758.4;refGene_gn=PPWD1;refGene_tx=XM_011284050.3
A1 134950903 134951047
refGene_gn=PPWD1;refGene_tx=XM_003981014.5;refGene_gn=PPWD1;refGene_tx=XM_006927758.4;refGene_gn=PPWD1;refGene_tx=XM_011284050.3
A1 134955761 134955941
refGene_gn=TRIM23;refGene_tx=XM_006927759.4;refGene_gn=TRIM23;refGene_tx=XM_019834938.2
A1 134958117 134958242
refGene_gn=TRIM23;refGene_tx=XM_006927759.4
A1 134962969 134963080

refGene_gn=TRIM23;refGene_tx=XM_006927759.4;refGene_gn=TRIM23;refGene_tx=XM_019834938.2
A1 134963835 134963965
refGene_gn=TRIM23;refGene_tx=XM_006927759.4;refGene_gn=TRIM23;refGene_tx=XM_019834938.2
A1 134972341 134972476
refGene_gn=TRIM23;refGene_tx=XM_006927759.4;refGene_gn=TRIM23;refGene_tx=XM_019834938.2
A1 134976587 134976803
refGene_gn=TRIM23;refGene_tx=XM_006927759.4;refGene_gn=TRIM23;refGene_tx=XM_019834938.2
A1 134979526 134979709
refGene_gn=TRIM23;refGene_tx=XM_006927759.4;refGene_gn=TRIM23;refGene_tx=XM_019834938.2
A1 134980273 134980552
refGene_gn=TRIM23;refGene_tx=XM_006927759.4;refGene_gn=TRIM23;refGene_tx=XM_019834938.2
A1 134982768 134982890
refGene_gn=TRIM23;refGene_tx=XM_006927759.4;refGene_gn=TRIM23;refGene_tx=XM_019834938.2
A1 134984162 134984325
refGene_gn=TRIM23;refGene_tx=XM_006927759.4;refGene_gn=TRIM23;refGene_tx=XM_019834938.2
A1 134989696 134989843
refGene_gn=TRIM23;refGene_tx=XM_006927759.4;refGene_gn=TRIM23;refGene_tx=XM_019834938.2
A1 134990546 134990592
refGene_gn=TRAPPC13;refGene_tx=XM_003981019.5;refGene_gn=TRAPPC13;refGene_tx=XM_003981020.5;refGene_gn=TRAPPC13;refGene_tx=XM_011284034.3;refGene_gn=TRAPPC13;refGene_tx=XM_011284037.3;refGene_gn=TRAPPC13;refGene_tx=XM_019834950.2;refGene_gn=TRAPPC13;refGene_tx=XM_019834959.2;refGene_gn=TRAPPC13;refGene_tx=XM_019834971.2
A1 135001274 135001343
refGene_gn=TRAPPC13;refGene_tx=XM_003981019.5;refGene_gn=TRAPPC13;refGene_tx=XM_003981020.5;refGene_gn=TRAPPC13;refGene_tx=XM_011284034.3;refGene_gn=TRAPPC13;refGene_tx=XM_011284037.3;refGene_gn=TRAPPC13;refGene_tx=XM_019834950.2;refGene_gn=TRAPPC13;refGene_tx=XM_019834959.2;refGene_gn=TRAPPC13;refGene_tx=XM_019834971.2
A1 135001857 135001957
refGene_gn=TRAPPC13;refGene_tx=XM_003981019.5;refGene_gn=TRAPPC13;refGene_tx=XM_003981020.5;refGene_gn=TRAPPC13;refGene_tx=XM_011284034.3;refGene_gn=TRAPPC13;refGene_tx=XM_011284037.3;refGene_gn=TRAPPC13;refGene_tx=XM_019834950.2;refGene_gn=TRAPPC13;refGene_tx=XM_019834959.2;refGene_gn=TRAPPC13;refGene_tx=XM_019834971.2
A1 135003079 135003164
refGene_gn=TRAPPC13;refGene_tx=XM_003981019.5;refGene_gn=TRAPPC13;refGene_tx=XM_003981020.5;refGene_gn=TRAPPC13;refGene_tx=XM_011284034.3;refGene_gn=TRAPPC13;refGene_tx=XM_011284037.3;refGene_gn=TRAPPC13;refGene_tx=XM_019834950.2;refGene_gn=TRAPPC13;refGene_tx=XM_019834959.2;refGene_gn=TRAPPC13;refGene_tx=XM_019834971.2

A1 135009606 135009734
refGene_gn=TRAPPC13;refGene_tx=XM_003981019.5;refGene_gn=TRAPPC13;refGene_tx=XM_003981020.5;refGene_gn=TRAPPC13;refGene_tx=XM_011284034.3;refGene_gn=TRAPPC13;refGene_tx=XM_011284037.3;refGene_gn=TRAPPC13;refGene_tx=XM_019834950.2;refGene_gn=TRAPPC13;refGene_tx=XM_019834959.2;refGene_gn=TRAPPC13;refGene_tx=XM_019834971.2
A1 135012451 135012542
refGene_gn=TRAPPC13;refGene_tx=XM_003981019.5;refGene_gn=TRAPPC13;refGene_tx=XM_003981020.5;refGene_gn=TRAPPC13;refGene_tx=XM_011284034.3;refGene_gn=TRAPPC13;refGene_tx=XM_011284037.3;refGene_gn=TRAPPC13;refGene_tx=XM_019834950.2;refGene_gn=TRAPPC13;refGene_tx=XM_019834959.2;refGene_gn=TRAPPC13;refGene_tx=XM_019834971.2
A1 135013881 135013926
refGene_gn=TRAPPC13;refGene_tx=XM_003981019.5;refGene_gn=TRAPPC13;refGene_tx=XM_003981020.5;refGene_gn=TRAPPC13;refGene_tx=XM_011284034.3;refGene_gn=TRAPPC13;refGene_tx=XM_011284037.3;refGene_gn=TRAPPC13;refGene_tx=XM_019834950.2;refGene_gn=TRAPPC13;refGene_tx=XM_019834959.2;refGene_gn=TRAPPC13;refGene_tx=XM_019834971.2
A1 135017287 135017305
refGene_gn=TRAPPC13;refGene_tx=XM_003981020.5;refGene_gn=TRAPPC13;refGene_tx=XM_011284034.3;refGene_gn=TRAPPC13;refGene_tx=XM_019834950.2;refGene_gn=TRAPPC13;refGene_tx=XM_019834959.2
A1 135019262 135019396
refGene_gn=TRAPPC13;refGene_tx=XM_003981019.5;refGene_gn=TRAPPC13;refGene_tx=XM_003981020.5;refGene_gn=TRAPPC13;refGene_tx=XM_011284034.3;refGene_gn=TRAPPC13;refGene_tx=XM_011284037.3;refGene_gn=TRAPPC13;refGene_tx=XM_019834950.2;refGene_gn=TRAPPC13;refGene_tx=XM_019834959.2;refGene_gn=TRAPPC13;refGene_tx=XM_019834971.2
A1 135020668 135020867
refGene_gn=TRAPPC13;refGene_tx=XM_003981019.5;refGene_gn=TRAPPC13;refGene_tx=XM_003981020.5;refGene_gn=TRAPPC13;refGene_tx=XM_011284034.3;refGene_gn=TRAPPC13;refGene_tx=XM_011284037.3;refGene_gn=TRAPPC13;refGene_tx=XM_019834950.2;refGene_gn=TRAPPC13;refGene_tx=XM_019834959.2;refGene_gn=TRAPPC13;refGene_tx=XM_019834971.2
A1 135022296 135022397
refGene_gn=TRAPPC13;refGene_tx=XM_003981019.5;refGene_gn=TRAPPC13;refGene_tx=XM_003981020.5;refGene_gn=TRAPPC13;refGene_tx=XM_011284034.3;refGene_gn=TRAPPC13;refGene_tx=XM_011284037.3;refGene_gn=TRAPPC13;refGene_tx=XM_019834950.2;refGene_gn=TRAPPC13;refGene_tx=XM_019834959.2;refGene_gn=TRAPPC13;refGene_tx=XM_019834971.2
A1 135024562 135024713
refGene_gn=TRAPPC13;refGene_tx=XM_003981019.5;refGene_gn=TRAPPC13;refGene_tx=XM_003981020.5;refGene_gn=TRAPPC13;refGene_tx=XM_011284034.3;refGene_gn=TRAPPC13;refGene_tx=XM_011284037.3;refGene_gn=TRAPPC13;refGene_tx=XM_019834950.2;refGene_gn=TRAPPC13;refGene_tx=XM_019834959.2;refGene_gn=TRAPPC13;refGene_tx=XM_019834971.2
A1 135024832 135024940
refGene_gn=TRAPPC13;refGene_tx=XM_003981019.5;refGene_gn=TRAPPC13;refGene_tx=XM_003981020.5;refGene_gn=TRAPPC13;refGene_tx=XM_011284034.3;refGene_gn=TRAPPC13;refGene_tx=XM_011284037.3;refGene_gn=TRAPPC13;refGen

e_tx=XM_019834950.2;refGene_gn=TRAPPC13;refGene_tx=XM_019834959.2;refGene_gn=TRAPPC13;refGene_tx=XM_019834971.2
A1 135029646 135029758
refGene_gn=SGTB;refGene_tx=XM_003981021.5;refGene_gn=SGTB;refGene_tx=XM_011284029.3;refGene_gn=SGTB;refGene_tx=XM_019834981.2
A1 135031157 135031241
refGene_gn=SGTB;refGene_tx=XM_003981021.5;refGene_gn=SGTB;refGene_tx=XM_011284029.3;refGene_gn=SGTB;refGene_tx=XM_019834981.2
A1 135031465 135031503
refGene_gn=SGTB;refGene_tx=XM_003981021.5;refGene_gn=SGTB;refGene_tx=XM_011284029.3;refGene_gn=SGTB;refGene_tx=XM_019834981.2
A1 135034769 135034832
refGene_gn=SGTB;refGene_tx=XM_003981021.5;refGene_gn=SGTB;refGene_tx=XM_011284029.3;refGene_gn=SGTB;refGene_tx=XM_019834981.2
A1 135034936 135035075
refGene_gn=SGTB;refGene_tx=XM_003981021.5;refGene_gn=SGTB;refGene_tx=XM_011284029.3;refGene_gn=SGTB;refGene_tx=XM_019834981.2
A1 135039000 135039105
refGene_gn=SGTB;refGene_tx=XM_003981021.5;refGene_gn=SGTB;refGene_tx=XM_011284029.3;refGene_gn=SGTB;refGene_tx=XM_019834981.2
A1 135052140 135052240
refGene_gn=SGTB;refGene_tx=XM_003981021.5;refGene_gn=SGTB;refGene_tx=XM_011284029.3;refGene_gn=SGTB;refGene_tx=XM_019834981.2
A1 135058894 135058964
refGene_gn=SGTB;refGene_tx=XM_003981021.5;refGene_gn=SGTB;refGene_tx=XM_011284029.3;refGene_gn=SGTB;refGene_tx=XM_019834981.2
A1 135063331 135063435
refGene_gn=SGTB;refGene_tx=XM_003981021.5;refGene_gn=SGTB;refGene_tx=XM_011284029.3;refGene_gn=SGTB;refGene_tx=XM_019834981.2
A1 135073040 135073140
refGene_gn=SGTB;refGene_tx=XM_003981021.5;refGene_gn=SGTB;refGene_tx=XM_011284029.3;refGene_gn=SGTB;refGene_tx=XM_019834981.2
A1 135075099 135075140
refGene_gn=NLN;refGene_tx=XM_011284017.2
A1 135121280 135121540
refGene_gn=NLN;refGene_tx=XM_011284017.2;refGene_gn=NLN;refGene_tx=XM_011284021.3
A1 135125847 135125996
refGene_gn=NLN;refGene_tx=XM_011284017.2;refGene_gn=NLN;refGene_tx=XM_011284021.3
A1 135141044 135141152
refGene_gn=NLN;refGene_tx=XM_011284017.2;refGene_gn=NLN;refGene_tx=XM_011284021.3
A1 135144422 135144525
refGene_gn=NLN;refGene_tx=XM_011284017.2;refGene_gn=NLN;refGene_tx=XM_011284021.3
A1 135145156 135145317
refGene_gn=NLN;refGene_tx=XM_011284017.2;refGene_gn=NLN;refGene_tx=XM_011284021.3
A1 135152485 135152621

refGene_gn=NLN;refGene_tx=XM_011284017.2;refGene_gn=NLN;refGene_tx=XM_011284021.3
A1 135153564 135153931
refGene_gn=NLN;refGene_tx=XM_011284017.2;refGene_gn=NLN;refGene_tx=XM_011284021.3
A1 135157970 135158172
refGene_gn=NLN;refGene_tx=XM_011284017.2;refGene_gn=NLN;refGene_tx=XM_011284021.3
A1 135167979 135168166
refGene_gn=NLN;refGene_tx=XM_011284017.2;refGene_gn=NLN;refGene_tx=XM_011284021.3
A1 135168491 135168620
refGene_gn=NLN;refGene_tx=XM_011284017.2;refGene_gn=NLN;refGene_tx=XM_011284021.3
A1 135170103 135170240
refGene_gn=NLN;refGene_tx=XM_011284017.2;refGene_gn=NLN;refGene_tx=XM_011284021.3
A1 135180401 135180536
refGene_gn=NLN;refGene_tx=XM_011284017.2;refGene_gn=NLN;refGene_tx=XM_011284021.3
A1 135314354 135314543
refGene_gn=ERBIN;refGene_tx=XM_006927761.4;refGene_gn=ERBIN;refGene_tx=XM_011284004.3;refGene_gn=ERBIN;refGene_tx=XM_011284009.3;refGene_gn=ERBIN;refGene_tx=XM_011284013.3;refGene_gn=ERBIN;refGene_tx=XM_019834997.2;refGene_gn=ERBIN;refGene_tx=XM_019835013.2
A1 135317961 135318079
refGene_gn=ERBIN;refGene_tx=XM_006927761.4;refGene_gn=ERBIN;refGene_tx=XM_011284004.3;refGene_gn=ERBIN;refGene_tx=XM_011284009.3;refGene_gn=ERBIN;refGene_tx=XM_011284013.3;refGene_gn=ERBIN;refGene_tx=XM_019834997.2;refGene_gn=ERBIN;refGene_tx=XM_019835013.2
A1 135332748 135332827
refGene_gn=ERBIN;refGene_tx=XM_006927761.4;refGene_gn=ERBIN;refGene_tx=XM_011284004.3;refGene_gn=ERBIN;refGene_tx=XM_011284009.3;refGene_gn=ERBIN;refGene_tx=XM_011284013.3;refGene_gn=ERBIN;refGene_tx=XM_019834997.2;refGene_gn=ERBIN;refGene_tx=XM_019835013.2
A1 135334394 135334484
refGene_gn=ERBIN;refGene_tx=XM_006927761.4;refGene_gn=ERBIN;refGene_tx=XM_011284004.3;refGene_gn=ERBIN;refGene_tx=XM_011284009.3;refGene_gn=ERBIN;refGene_tx=XM_011284013.3;refGene_gn=ERBIN;refGene_tx=XM_019834997.2;refGene_gn=ERBIN;refGene_tx=XM_019835013.2
A1 135336579 135336636
refGene_gn=ERBIN;refGene_tx=XM_006927761.4;refGene_gn=ERBIN;refGene_tx=XM_011284004.3;refGene_gn=ERBIN;refGene_tx=XM_011284009.3;refGene_gn=ERBIN;refGene_tx=XM_011284013.3;refGene_gn=ERBIN;refGene_tx=XM_019834997.2;refGene_gn=ERBIN;refGene_tx=XM_019835013.2;refGene_gn=ERBIN;refGene_tx=XM_019835020.2
A1 135340733 135340797
refGene_gn=ERBIN;refGene_tx=XM_006927761.4;refGene_gn=ERBIN;refGene_tx=XM_011284004.3;refGene_gn=ERBIN;refGene_tx=XM_011284009.3;refGene_gn=ERBIN;refGene_tx=XM_011284013.3;refGene_gn=ERBIN;refGene_tx=XM_0198349

97.2;refGene_gn=ERBIN;refGene_tx=XM_019835013.2;refGene_gn=ERBIN;refGene_tx=XM_019835020.2
A1 135344062 135344137
refGene_gn=ERBIN;refGene_tx=XM_006927761.4;refGene_gn=ERBIN;refGene_tx=XM_011284004.3;refGene_gn=ERBIN;refGene_tx=XM_011284009.3;refGene_gn=ERBIN;refGene_tx=XM_011284013.3;refGene_gn=ERBIN;refGene_tx=XM_019834997.2;refGene_gn=ERBIN;refGene_tx=XM_019835013.2;refGene_gn=ERBIN;refGene_tx=XM_019835020.2
A1 135344962 135345107
refGene_gn=ERBIN;refGene_tx=XM_006927761.4;refGene_gn=ERBIN;refGene_tx=XM_011284004.3;refGene_gn=ERBIN;refGene_tx=XM_011284009.3;refGene_gn=ERBIN;refGene_tx=XM_011284013.3;refGene_gn=ERBIN;refGene_tx=XM_019834997.2;refGene_gn=ERBIN;refGene_tx=XM_019835013.2;refGene_gn=ERBIN;refGene_tx=XM_019835020.2
A1 135346344 135346417
refGene_gn=ERBIN;refGene_tx=XM_006927761.4;refGene_gn=ERBIN;refGene_tx=XM_011284004.3;refGene_gn=ERBIN;refGene_tx=XM_011284009.3;refGene_gn=ERBIN;refGene_tx=XM_011284013.3;refGene_gn=ERBIN;refGene_tx=XM_019834997.2;refGene_gn=ERBIN;refGene_tx=XM_019835013.2;refGene_gn=ERBIN;refGene_tx=XM_019835020.2
A1 135346753 135346883
refGene_gn=ERBIN;refGene_tx=XM_006927761.4;refGene_gn=ERBIN;refGene_tx=XM_011284004.3;refGene_gn=ERBIN;refGene_tx=XM_011284009.3;refGene_gn=ERBIN;refGene_tx=XM_011284013.3;refGene_gn=ERBIN;refGene_tx=XM_019834997.2;refGene_gn=ERBIN;refGene_tx=XM_019835013.2;refGene_gn=ERBIN;refGene_tx=XM_019835020.2
A1 135347174 135347290
refGene_gn=ERBIN;refGene_tx=XM_006927761.4;refGene_gn=ERBIN;refGene_tx=XM_011284004.3;refGene_gn=ERBIN;refGene_tx=XM_011284009.3;refGene_gn=ERBIN;refGene_tx=XM_011284013.3;refGene_gn=ERBIN;refGene_tx=XM_019834997.2;refGene_gn=ERBIN;refGene_tx=XM_019835013.2;refGene_gn=ERBIN;refGene_tx=XM_019835020.2
A1 135349159 135349229
refGene_gn=ERBIN;refGene_tx=XM_006927761.4;refGene_gn=ERBIN;refGene_tx=XM_011284004.3;refGene_gn=ERBIN;refGene_tx=XM_011284009.3;refGene_gn=ERBIN;refGene_tx=XM_011284013.3;refGene_gn=ERBIN;refGene_tx=XM_019834997.2;refGene_gn=ERBIN;refGene_tx=XM_019835013.2;refGene_gn=ERBIN;refGene_tx=XM_019835020.2
A1 135357602 135357702
refGene_gn=ERBIN;refGene_tx=XM_006927761.4;refGene_gn=ERBIN;refGene_tx=XM_011284004.3;refGene_gn=ERBIN;refGene_tx=XM_011284009.3;refGene_gn=ERBIN;refGene_tx=XM_011284013.3;refGene_gn=ERBIN;refGene_tx=XM_019834997.2;refGene_gn=ERBIN;refGene_tx=XM_019835013.2;refGene_gn=ERBIN;refGene_tx=XM_019835020.2
A1 135367023 135367145
refGene_gn=ERBIN;refGene_tx=XM_006927761.4;refGene_gn=ERBIN;refGene_tx=XM_011284004.3;refGene_gn=ERBIN;refGene_tx=XM_011284009.3;refGene_gn=ERBIN;refGene_tx=XM_011284013.3;refGene_gn=ERBIN;refGene_tx=XM_019834997.2;refGene_gn=ERBIN;refGene_tx=XM_019835013.2;refGene_gn=ERBIN;refGene_tx=XM_019835020.2

A1	135367764	135367938
refGene_gn=ERBIN;refGene_tx=XM_006927761.4;refGene_gn=ERBIN;refGene_tx=XM_011284004.3;refGene_gn=ERBIN;refGene_tx=XM_011284009.3;refGene_gn=ERBIN;refGene_tx=XM_011284013.3;refGene_gn=ERBIN;refGene_tx=XM_019834997.2;refGene_gn=ERBIN;refGene_tx=XM_019835013.2;refGene_gn=ERBIN;refGene_tx=XM_019835020.2
A1	135369335	135369521
refGene_gn=ERBIN;refGene_tx=XM_006927761.4;refGene_gn=ERBIN;refGene_tx=XM_011284004.3;refGene_gn=ERBIN;refGene_tx=XM_011284009.3;refGene_gn=ERBIN;refGene_tx=XM_011284013.3;refGene_gn=ERBIN;refGene_tx=XM_019834997.2;refGene_gn=ERBIN;refGene_tx=XM_019835013.2;refGene_gn=ERBIN;refGene_tx=XM_019835020.2
A1	135377192	135377307
refGene_gn=ERBIN;refGene_tx=XM_006927761.4;refGene_gn=ERBIN;refGene_tx=XM_011284004.3;refGene_gn=ERBIN;refGene_tx=XM_011284009.3;refGene_gn=ERBIN;refGene_tx=XM_011284013.3;refGene_gn=ERBIN;refGene_tx=XM_019834997.2;refGene_gn=ERBIN;refGene_tx=XM_019835013.2;refGene_gn=ERBIN;refGene_tx=XM_019835020.2
A1	135379325	135379512
refGene_gn=ERBIN;refGene_tx=XM_006927761.4;refGene_gn=ERBIN;refGene_tx=XM_011284004.3;refGene_gn=ERBIN;refGene_tx=XM_011284009.3;refGene_gn=ERBIN;refGene_tx=XM_011284013.3;refGene_gn=ERBIN;refGene_tx=XM_019834997.2;refGene_gn=ERBIN;refGene_tx=XM_019835013.2;refGene_gn=ERBIN;refGene_tx=XM_019835020.2
A1	135381505	135383051
refGene_gn=ERBIN;refGene_tx=XM_006927761.4;refGene_gn=ERBIN;refGene_tx=XM_011284004.3;refGene_gn=ERBIN;refGene_tx=XM_011284009.3;refGene_gn=ERBIN;refGene_tx=XM_011284013.3;refGene_gn=ERBIN;refGene_tx=XM_019834997.2;refGene_gn=ERBIN;refGene_tx=XM_019835013.2;refGene_gn=ERBIN;refGene_tx=XM_019835020.2
A1	135386970	135387114
refGene_gn=ERBIN;refGene_tx=XM_006927761.4;refGene_gn=ERBIN;refGene_tx=XM_011284009.3;refGene_gn=ERBIN;refGene_tx=XM_019834997.2;refGene_gn=ERBIN;refGene_tx=XM_019835020.2
A1	135389764	135389884
refGene_gn=ERBIN;refGene_tx=XM_006927761.4;refGene_gn=ERBIN;refGene_tx=XM_011284004.3;refGene_gn=ERBIN;refGene_tx=XM_011284009.3;refGene_gn=ERBIN;refGene_tx=XM_019834997.2;refGene_gn=ERBIN;refGene_tx=XM_019835020.2
A1	135392497	135392707
refGene_gn=ERBIN;refGene_tx=XM_006927761.4;refGene_gn=ERBIN;refGene_tx=XM_011284004.3;refGene_gn=ERBIN;refGene_tx=XM_019834997.2;refGene_gn=ERBIN;refGene_tx=XM_019835013.2;refGene_gn=ERBIN;refGene_tx=XM_019835020.2
A1	135393788	135393881
refGene_gn=ERBIN;refGene_tx=XM_006927761.4;refGene_gn=ERBIN;refGene_tx=XM_011284004.3;refGene_gn=ERBIN;refGene_tx=XM_011284009.3;refGene_gn=ERBIN;refGene_tx=XM_011284013.3;refGene_gn=ERBIN;refGene_tx=XM_019834997.2;refGene_gn=ERBIN;refGene_tx=XM_019835013.2;refGene_gn=ERBIN;refGene_tx=XM_019835020.2

A1 135395044 135395119
refGene_gn=ERBIN;refGene_tx=XM_006927761.4;refGene_gn=ERBIN;refGene_tx=XM_011284004.3;refGene_gn=ERBIN;refGene_tx=XM_011284009.3;refGene_gn=ERBIN;refGene_tx=XM_011284013.3;refGene_gn=ERBIN;refGene_tx=XM_019834997.2;refGene_gn=ERBIN;refGene_tx=XM_019835013.2;refGene_gn=ERBIN;refGene_tx=XM_019835020.2
A1 135395904 135396012
refGene_gn=ERBIN;refGene_tx=XM_006927761.4;refGene_gn=ERBIN;refGene_tx=XM_011284004.3;refGene_gn=ERBIN;refGene_tx=XM_011284009.3;refGene_gn=ERBIN;refGene_tx=XM_011284013.3;refGene_gn=ERBIN;refGene_tx=XM_019834997.2;refGene_gn=ERBIN;refGene_tx=XM_019835013.2;refGene_gn=ERBIN;refGene_tx=XM_019835020.2
A1 135444422 135444583
refGene_gn=SREK1;refGene_tx=XM_003981024.5;refGene_gn=SREK1;refGene_tx=XM_006927762.4;refGene_gn=SREK1;refGene_tx=XM_006927764.4
A1 135453710 135453844
refGene_gn=SREK1;refGene_tx=XM_003981024.5;refGene_gn=SREK1;refGene_tx=XM_006927762.4;refGene_gn=SREK1;refGene_tx=XM_006927764.4
A1 135459375 135459491
refGene_gn=SREK1;refGene_tx=XM_003981024.5;refGene_gn=SREK1;refGene_tx=XM_006927762.4;refGene_gn=SREK1;refGene_tx=XM_006927763.4;refGene_gn=SREK1;refGene_tx=XM_006927764.4
A1 135464876 135465041
refGene_gn=SREK1;refGene_tx=XM_003981024.5;refGene_gn=SREK1;refGene_tx=XM_006927762.4;refGene_gn=SREK1;refGene_tx=XM_006927763.4;refGene_gn=SREK1;refGene_tx=XM_006927764.4
A1 135465182 135465361
refGene_gn=SREK1;refGene_tx=XM_003981024.5;refGene_gn=SREK1;refGene_tx=XM_006927762.4;refGene_gn=SREK1;refGene_tx=XM_006927763.4;refGene_gn=SREK1;refGene_tx=XM_006927764.4
A1 135466568 135466699
refGene_gn=SREK1;refGene_tx=XM_003981024.5;refGene_gn=SREK1;refGene_tx=XM_006927762.4;refGene_gn=SREK1;refGene_tx=XM_006927763.4;refGene_gn=SREK1;refGene_tx=XM_006927764.4
A1 135467577 135467692
refGene_gn=SREK1;refGene_tx=XM_003981024.5;refGene_gn=SREK1;refGene_tx=XM_006927762.4;refGene_gn=SREK1;refGene_tx=XM_006927763.4;refGene_gn=SREK1;refGene_tx=XM_006927764.4
A1 135472682 135472802
refGene_gn=SREK1;refGene_tx=XM_003981024.5;refGene_gn=SREK1;refGene_tx=XM_006927762.4;refGene_gn=SREK1;refGene_tx=XM_006927763.4;refGene_gn=SREK1;refGene_tx=XM_006927764.4
A1 135473170 135473539
refGene_gn=SREK1;refGene_tx=XM_003981024.5;refGene_gn=SREK1;refGene_tx=XM_006927762.4;refGene_gn=SREK1;refGene_tx=XM_006927763.4
A1 135477064 135477160
refGene_gn=SREK1;refGene_tx=XM_003981024.5;refGene_gn=SREK1;refGene_tx=XM_006927762.4;refGene_gn=SREK1;refGene_tx=XM_006927763.4;refGene_gn=SREK1;refGene_tx=XM_006927764.4
A1 135488438 135488583

refGene_gn=SREK1;refGene_tx=XM_003981024.5;refGene_gn=SREK1;refGene_tx=XM_006927762.4;refGene_gn=SREK1;refGene_tx=XM_006927763.4;refGene_gn=SREK1;refGene_tx=XM_006927764.4
A1 135489550 135489700
refGene_gn=SREK1;refGene_tx=XM_003981024.5;refGene_gn=SREK1;refGene_tx=XM_006927762.4;refGene_gn=SREK1;refGene_tx=XM_006927763.4;refGene_gn=SREK1;refGene_tx=XM_006927764.4
A1 135824645 135825002
refGene_gn=MAST4;refGene_tx=XM_023257117.1
A1 135969763 135969917
refGene_gn=MAST4;refGene_tx=XM_023257117.1
A1 135998512 135998637
refGene_gn=MAST4;refGene_tx=XM_023257117.1
A1 136083011 136083086
refGene_gn=MAST4;refGene_tx=XM_023257117.1
A1 136116461 136116493
refGene_gn=MAST4;refGene_tx=XM_023257117.1
A1 136131284 136131320
refGene_gn=MAST4;refGene_tx=XM_023257117.1
A1 136275243 136275332
refGene_gn=MAST4;refGene_tx=XM_023257117.1
A1 136313994 136314064
refGene_gn=MAST4;refGene_tx=XM_023257117.1
A1 136318094 136318103
refGene_gn=MAST4;refGene_tx=XM_023257117.1
A1 136319538 136319617
refGene_gn=MAST4;refGene_tx=XM_023257117.1
A1 136325985 136326143
refGene_gn=MAST4;refGene_tx=XM_023257117.1
A1 136328141 136328217
refGene_gn=MAST4;refGene_tx=XM_023257117.1
A1 136330194 136330404
refGene_gn=MAST4;refGene_tx=XM_023257117.1
A1 136334797 136334899
refGene_gn=MAST4;refGene_tx=XM_023257117.1
A1 136337521 136337654
refGene_gn=MAST4;refGene_tx=XM_023257117.1
A1 136341366 136341434
refGene_gn=MAST4;refGene_tx=XM_023257117.1
A1 136344382 136344468
refGene_gn=MAST4;refGene_tx=XM_023257117.1
A1 136351551 136351760
refGene_gn=MAST4;refGene_tx=XM_023257117.1
A1 136353130 136353269
refGene_gn=MAST4;refGene_tx=XM_023257117.1
A1 136354831 136354964
refGene_gn=MAST4;refGene_tx=XM_023257117.1
A1 136355829 136355995
refGene_gn=MAST4;refGene_tx=XM_023257117.1
A1 136357839 136357941

refGene_gn=MAST4;refGene_tx=XM_023257117.1
A1 136362612 136362735
refGene_gn=MAST4;refGene_tx=XM_023257117.1
A1 136363165 136363278
refGene_gn=MAST4;refGene_tx=XM_023257117.1
A1 136365367 136365495
refGene_gn=MAST4;refGene_tx=XM_023257117.1
A1 136365840 136366031
refGene_gn=MAST4;refGene_tx=XM_023257117.1
A1 136370688 136370889
refGene_gn=MAST4;refGene_tx=XM_023257117.1
A1 136373193 136373423
refGene_gn=MAST4;refGene_tx=XM_023257117.1
A1 136373952 136374075
refGene_gn=MAST4;refGene_tx=XM_023257117.1
A1 136383008 136383042
refGene_gn=MAST4;refGene_tx=XM_023257117.1
A1 136383044 136383144
refGene_gn=MAST4;refGene_tx=XM_023257117.1
A1 136385078 136385260
refGene_gn=MAST4;refGene_tx=XM_023257117.1
A1 136385602 136389483
refGene_gn=MAST4;refGene_tx=XM_023257117.1
A1 136401523 136403252
refGene_gn=CD180;refGene_tx=XM_003981033.5;refGene_gn=CD180;refGene_tx
=XM_019835040.2
A1 136407193 136407360
refGene_gn=CD180;refGene_tx=XM_003981033.5;refGene_gn=CD180;refGene_tx
=XM_019835040.2
A1 136415254 136415344
refGene_gn=CD180;refGene_tx=XM_003981033.5
A1 137251072 137251409
refGene_gn=PIK3R1;refGene_tx=XM_023257120.1
A1 137297217 137297310
refGene_gn=PIK3R1;refGene_tx=XM_023257120.1
A1 137297751 137297826
refGene_gn=PIK3R1;refGene_tx=XM_023257120.1
A1 137304383 137304515
refGene_gn=PIK3R1;refGene_tx=XM_023257120.1
A1 137305330 137305532
refGene_gn=PIK3R1;refGene_tx=XM_023257120.1
A1 137305738 137305818
refGene_gn=PIK3R1;refGene_tx=XM_023257120.1
A1 137314397 137314413
refGene_gn=PIK3R1;refGene_tx=XM_011283969.3
A1 137316399 137316505
refGene_gn=PIK3R1;refGene_tx=XM_003981035.5
A1 137317891 137317994
refGene_gn=PIK3R1;refGene_tx=XM_003981035.5;refGene_gn=PIK3R1;refGene_
tx=XM_011283969.3;refGene_gn=PIK3R1;refGene_tx=XM_023257120.1

A1 137318716 137318815
refGene_gn=PIK3R1;refGene_tx=XM_003981035.5;refGene_gn=PIK3R1;refGene_tx=XM_011283969.3;refGene_gn=PIK3R1;refGene_tx=XM_023257120.1
A1 137318917 137319098
refGene_gn=PIK3R1;refGene_tx=XM_003981035.5;refGene_gn=PIK3R1;refGene_tx=XM_011283969.3;refGene_gn=PIK3R1;refGene_tx=XM_023257120.1
A1 137319304 137319430
refGene_gn=PIK3R1;refGene_tx=XM_003981035.5;refGene_gn=PIK3R1;refGene_tx=XM_011283969.3;refGene_gn=PIK3R1;refGene_tx=XM_023257120.1
A1 137320159 137320302
refGene_gn=PIK3R1;refGene_tx=XM_003981035.5;refGene_gn=PIK3R1;refGene_tx=XM_011283969.3;refGene_gn=PIK3R1;refGene_tx=XM_023257120.1
A1 137320505 137320682
refGene_gn=PIK3R1;refGene_tx=XM_003981035.5;refGene_gn=PIK3R1;refGene_tx=XM_011283969.3;refGene_gn=PIK3R1;refGene_tx=XM_023257120.1
A1 137320780 137320849
refGene_gn=PIK3R1;refGene_tx=XM_003981035.5;refGene_gn=PIK3R1;refGene_tx=XM_011283969.3;refGene_gn=PIK3R1;refGene_tx=XM_023257120.1
A1 137321488 137321659
refGene_gn=PIK3R1;refGene_tx=XM_003981035.5;refGene_gn=PIK3R1;refGene_tx=XM_011283969.3;refGene_gn=PIK3R1;refGene_tx=XM_023257120.1
A1 137322581 137322771
refGene_gn=PIK3R1;refGene_tx=XM_003981035.5;refGene_gn=PIK3R1;refGene_tx=XM_011283969.3;refGene_gn=PIK3R1;refGene_tx=XM_023257120.1
A1 138035717 138035800
refGene_gn=SLC30A5;refGene_tx=XM_011283965.2
A1 138045888 138046011
refGene_gn=SLC30A5;refGene_tx=XM_011283965.2
A1 138047185 138047252
refGene_gn=SLC30A5;refGene_tx=XM_011283965.2
A1 138049128 138049214
refGene_gn=SLC30A5;refGene_tx=XM_011283965.2
A1 138053240 138053328
refGene_gn=SLC30A5;refGene_tx=XM_011283965.2
A1 138056953 138057041
refGene_gn=SLC30A5;refGene_tx=XM_011283965.2
A1 138058221 138058298
refGene_gn=SLC30A5;refGene_tx=XM_011283965.2
A1 138058536 138058707
refGene_gn=SLC30A5;refGene_tx=XM_011283965.2
A1 138060141 138060430
refGene_gn=SLC30A5;refGene_tx=XM_011283965.2
A1 138060610 138060819
refGene_gn=SLC30A5;refGene_tx=XM_011283965.2
A1 138061359 138061517
refGene_gn=SLC30A5;refGene_tx=XM_011283965.2
A1 138068784 138068845
refGene_gn=SLC30A5;refGene_tx=XM_011283965.2
A1 138068846 138068915
refGene_gn=SLC30A5;refGene_tx=XM_011283965.2

A1 138071535 138071737
refGene_gn=SLC30A5;refGene_tx=XM_011283965.2
A1 138072399 138072626
refGene_gn=SLC30A5;refGene_tx=XM_011283965.2
A1 138075999 138076128
refGene_gn=SLC30A5;refGene_tx=XM_011283965.2
A1 138076960 138077131
refGene_gn=SLC30A5;refGene_tx=XM_011283965.2
A1 138091984 138092005
refGene_gn=CCNB1;refGene_tx=XM_003981036.5
A1 138092622 138092793
refGene_gn=CCNB1;refGene_tx=XM_003981036.5
A1 138092890 138093043
refGene_gn=CCNB1;refGene_tx=XM_003981036.5
A1 138096381 138096564
refGene_gn=CCNB1;refGene_tx=XM_003981036.5
A1 138098037 138098196
refGene_gn=CCNB1;refGene_tx=XM_003981036.5
A1 138098414 138098651
refGene_gn=CCNB1;refGene_tx=XM_003981036.5
A1 138099172 138099313
refGene_gn=CCNB1;refGene_tx=XM_003981036.5
A1 138099942 138100053
refGene_gn=CCNB1;refGene_tx=XM_003981036.5
A1 138100451 138100559
refGene_gn=CCNB1;refGene_tx=XM_003981036.5
A1 138104492 138104620
refGene_gn=CENPH;refGene_tx=XM_003981043.4
A1 138115729 138115785
refGene_gn=CENPH;refGene_tx=XM_003981043.4
A1 138120591 138120640
refGene_gn=CENPH;refGene_tx=XM_003981043.4
A1 138121245 138121320
refGene_gn=CENPH;refGene_tx=XM_003981043.4
A1 138122911 138122968
refGene_gn=CENPH;refGene_tx=XM_003981043.4
A1 138125693 138125757
refGene_gn=CENPH;refGene_tx=XM_003981043.4
A1 138126103 138126155
refGene_gn=CENPH;refGene_tx=XM_003981043.4
A1 138128147 138128311
refGene_gn=CENPH;refGene_tx=XM_003981043.4
A1 138129842 138129938
refGene_gn=CENPH;refGene_tx=XM_003981043.4
A1 138137845 138137887
refGene_gn=MRPS36;refGene_tx=XM_023257144.1
A1 138146398 138146462
refGene_gn=MRPS36;refGene_tx=XM_023257144.1
A1 138148544 138148727
refGene_gn=MRPS36;refGene_tx=XM_023257144.1

A1 138149244 138149262
refGene_gn=MRPS36;refGene_tx=XM_023257144.1
A1 138154256 138154322
refGene_gn=CDK7;refGene_tx=XM_003981038.5
A1 138154657 138154717
refGene_gn=CDK7;refGene_tx=XM_003981038.5
A1 138163492 138163526
refGene_gn=CDK7;refGene_tx=XM_003981038.5
A1 138164431 138164499
refGene_gn=CDK7;refGene_tx=XM_003981038.5
A1 138165454 138165523
refGene_gn=CDK7;refGene_tx=XM_003981038.5
A1 138169082 138169166
refGene_gn=CDK7;refGene_tx=XM_003981038.5
A1 138169167 138169194
refGene_gn=CDK7;refGene_tx=XM_003981038.5
A1 138170207 138170326
refGene_gn=CDK7;refGene_tx=XM_003981038.5
A1 138172737 138172837
refGene_gn=CDK7;refGene_tx=XM_003981038.5
A1 138183913 138184000
refGene_gn=CDK7;refGene_tx=XM_003981038.5
A1 138188105 138188255
refGene_gn=CDK7;refGene_tx=XM_003981038.5
A1 138191418 138191566
refGene_gn=CDK7;refGene_tx=XM_003981038.5
A1 138191980 138192009
refGene_gn=CDK7;refGene_tx=XM_003981038.5
A1 138195844 138196153
refGene_gn=CCDC125;refGene_tx=XM_011283948.3;refGene_gn=CCDC125;refGene_tx=XM_019835116.2;refGene_gn=CCDC125;refGene_tx=XM_019835127.1
A1 138196988 138197125
refGene_gn=CCDC125;refGene_tx=XM_011283948.3;refGene_gn=CCDC125;refGene_tx=XM_019835116.2;refGene_gn=CCDC125;refGene_tx=XM_019835127.1
A1 138201450 138201625
refGene_gn=CCDC125;refGene_tx=XM_011283948.3;refGene_gn=CCDC125;refGene_tx=XM_019835116.2;refGene_gn=CCDC125;refGene_tx=XM_019835127.1
A1 138202781 138202889
refGene_gn=CCDC125;refGene_tx=XM_011283948.3;refGene_gn=CCDC125;refGene_tx=XM_019835116.2;refGene_gn=CCDC125;refGene_tx=XM_019835127.1
A1 138203632 138203748
refGene_gn=CCDC125;refGene_tx=XM_011283948.3;refGene_gn=CCDC125;refGene_tx=XM_019835116.2;refGene_gn=CCDC125;refGene_tx=XM_019835127.1
A1 138205336 138205419
refGene_gn=CCDC125;refGene_tx=XM_011283948.3;refGene_gn=CCDC125;refGene_tx=XM_019835116.2;refGene_gn=CCDC125;refGene_tx=XM_019835127.1
A1 138212092 138212178
refGene_gn=CCDC125;refGene_tx=XM_011283948.3;refGene_gn=CCDC125;refGene_tx=XM_019835116.2;refGene_gn=CCDC125;refGene_tx=XM_019835127.1
A1 138212880 138212958

refGene_gn=CCDC125;refGene_tx=XM_011283948.3;refGene_gn=CCDC125;refGene_tx=XM_019835116.2;refGene_gn=CCDC125;refGene_tx=XM_019835127.1
A1 138213742 138213829
refGene_gn=CCDC125;refGene_tx=XM_011283948.3;refGene_gn=CCDC125;refGene_tx=XM_019835116.2;refGene_gn=CCDC125;refGene_tx=XM_019835127.1
A1 138215297 138215356
refGene_gn=CCDC125;refGene_tx=XM_011283948.3;refGene_gn=CCDC125;refGene_tx=XM_019835116.2;refGene_gn=CCDC125;refGene_tx=XM_019835127.1
A1 138216202 138216314
refGene_gn=CCDC125;refGene_tx=XM_019835127.1
A1 138217843 138218153
refGene_gn=CCDC125;refGene_tx=XM_011283948.3;refGene_gn=CCDC125;refGene_tx=XM_019835116.2
A1 138236833 138237026
refGene_gn=AK6;refGene_tx=XM_006927774.3;refGene_gn=AK6;refGene_tx=XM_023257172.1
A1 138240763 138240909
refGene_gn=AK6;refGene_tx=XM_006927774.3;refGene_gn=AK6;refGene_tx=XM_023257172.1
A1 138241008 138241067
refGene_gn=AK6;refGene_tx=XM_006927774.3;refGene_gn=AK6;refGene_tx=XM_023257172.1
A1 138249033 138249831
refGene_gn=TAF9;refGene_tx=XM_011283921.2
A1 138250663 138250756
refGene_gn=AK6;refGene_tx=XM_006927774.3;refGene_gn=AK6;refGene_tx=XM_023257172.1
A1 138252256 138252290
refGene_gn=AK6;refGene_tx=XM_023257172.1
A1 138252951 138252979
refGene_gn=AK6;refGene_tx=XM_006927774.3
A1 138253420 138253436
refGene_gn=RAD17;refGene_tx=XM_011283905.3
A1 138254693 138254797
refGene_gn=RAD17;refGene_tx=XM_006927775.4;refGene_gn=RAD17;refGene_tx=XM_011283905.3;refGene_gn=RAD17;refGene_tx=XM_011283908.3;refGene_gn=RAD17;refGene_tx=XM_019835099.2
A1 138257491 138257746
refGene_gn=RAD17;refGene_tx=XM_006927775.4;refGene_gn=RAD17;refGene_tx=XM_011283905.3;refGene_gn=RAD17;refGene_tx=XM_011283908.3;refGene_gn=RAD17;refGene_tx=XM_019835099.2
A1 138258277 138258361
refGene_gn=RAD17;refGene_tx=XM_006927775.4;refGene_gn=RAD17;refGene_tx=XM_011283905.3;refGene_gn=RAD17;refGene_tx=XM_011283908.3;refGene_gn=RAD17;refGene_tx=XM_019835099.2
A1 138262324 138262481
refGene_gn=RAD17;refGene_tx=XM_006927775.4;refGene_gn=RAD17;refGene_tx=XM_011283905.3;refGene_gn=RAD17;refGene_tx=XM_011283908.3;refGene_gn=RAD17;refGene_tx=XM_019835099.2
A1 138263412 138263549

refGene_gn=RAD17;refGene_tx=XM_006927775.4;refGene_gn=RAD17;refGene_tx=XM_011283905.3;refGene_gn=RAD17;refGene_tx=XM_011283908.3;refGene_gn=RAD17;refGene_tx=XM_019835099.2
A1 138264360 138264413
refGene_gn=RAD17;refGene_tx=XM_006927775.4;refGene_gn=RAD17;refGene_tx=XM_011283905.3;refGene_gn=RAD17;refGene_tx=XM_011283908.3;refGene_gn=RAD17;refGene_tx=XM_019835099.2
A1 138264493 138264619
refGene_gn=RAD17;refGene_tx=XM_006927775.4;refGene_gn=RAD17;refGene_tx=XM_011283905.3;refGene_gn=RAD17;refGene_tx=XM_011283908.3;refGene_gn=RAD17;refGene_tx=XM_019835099.2
A1 138264711 138264781
refGene_gn=RAD17;refGene_tx=XM_006927775.4;refGene_gn=RAD17;refGene_tx=XM_011283905.3;refGene_gn=RAD17;refGene_tx=XM_011283908.3;refGene_gn=RAD17;refGene_tx=XM_019835099.2
A1 138274670 138274779
refGene_gn=RAD17;refGene_tx=XM_006927775.4;refGene_gn=RAD17;refGene_tx=XM_011283905.3;refGene_gn=RAD17;refGene_tx=XM_011283908.3;refGene_gn=RAD17;refGene_tx=XM_019835099.2
A1 138277770 138277956
refGene_gn=RAD17;refGene_tx=XM_006927775.4;refGene_gn=RAD17;refGene_tx=XM_011283905.3;refGene_gn=RAD17;refGene_tx=XM_011283908.3;refGene_gn=RAD17;refGene_tx=XM_019835099.2
A1 138279847 138279933
refGene_gn=RAD17;refGene_tx=XM_006927775.4;refGene_gn=RAD17;refGene_tx=XM_011283905.3;refGene_gn=RAD17;refGene_tx=XM_011283908.3;refGene_gn=RAD17;refGene_tx=XM_019835099.2
A1 138280052 138280199
refGene_gn=RAD17;refGene_tx=XM_006927775.4;refGene_gn=RAD17;refGene_tx=XM_011283905.3;refGene_gn=RAD17;refGene_tx=XM_011283908.3;refGene_gn=RAD17;refGene_tx=XM_019835099.2
A1 138281696 138281846
refGene_gn=RAD17;refGene_tx=XM_006927775.4;refGene_gn=RAD17;refGene_tx=XM_011283905.3;refGene_gn=RAD17;refGene_tx=XM_011283908.3;refGene_gn=RAD17;refGene_tx=XM_019835099.2
A1 138284029 138284150
refGene_gn=RAD17;refGene_tx=XM_006927775.4;refGene_gn=RAD17;refGene_tx=XM_011283905.3;refGene_gn=RAD17;refGene_tx=XM_011283908.3;refGene_gn=RAD17;refGene_tx=XM_019835099.2
A1 138288011 138288069
refGene_gn=RAD17;refGene_tx=XM_006927775.4;refGene_gn=RAD17;refGene_tx=XM_011283905.3;refGene_gn=RAD17;refGene_tx=XM_011283908.3;refGene_gn=RAD17;refGene_tx=XM_019835099.2
A1 138290126 138290391
refGene_gn=RAD17;refGene_tx=XM_006927775.4;refGene_gn=RAD17;refGene_tx=XM_011283905.3;refGene_gn=RAD17;refGene_tx=XM_011283908.3;refGene_gn=RAD17;refGene_tx=XM_019835099.2
A1 138297018 138298167
refGene_gn=MARVELD2;refGene_tx=XM_011283900.3;refGene_gn=MARVELD2;refGene_tx=XM_019835182.2;refGene_gn=MARVELD2;refGene_tx=XM_019835189.2

A1 138301196 138301232
refGene_gn=MARVELD2;refGene_tx=XM_011283900.3;refGene_gn=MARVELD2;refGene_tx=XM_019835182.2;refGene_gn=MARVELD2;refGene_tx=XM_019835189.2
A1 138310413 138310562
refGene_gn=MARVELD2;refGene_tx=XM_011283900.3;refGene_gn=MARVELD2;refGene_tx=XM_019835182.2;refGene_gn=MARVELD2;refGene_tx=XM_019835189.2
A1 138310978 138311150
refGene_gn=MARVELD2;refGene_tx=XM_011283900.3;refGene_gn=MARVELD2;refGene_tx=XM_019835182.2;refGene_gn=MARVELD2;refGene_tx=XM_019835189.2
A1 138315354 138315405
refGene_gn=MARVELD2;refGene_tx=XM_011283900.3;refGene_gn=MARVELD2;refGene_tx=XM_019835182.2;refGene_gn=MARVELD2;refGene_tx=XM_019835189.2
A1 138316187 138316310
refGene_gn=MARVELD2;refGene_tx=XM_011283900.3;refGene_gn=MARVELD2;refGene_tx=XM_019835182.2;refGene_gn=MARVELD2;refGene_tx=XM_019835189.2
A1 138362404 138362454
refGene_gn=OCLN;refGene_tx=XM_003981047.4;refGene_gn=OCLN;refGene_tx=XM_019835192.1
A1 138367801 138368477
refGene_gn=OCLN;refGene_tx=XM_003981047.4;refGene_gn=OCLN;refGene_tx=XM_019835192.1
A1 138371133 138371295
refGene_gn=OCLN;refGene_tx=XM_003981047.4;refGene_gn=OCLN;refGene_tx=XM_019835192.1
A1 138393686 138393832
refGene_gn=OCLN;refGene_tx=XM_003981047.4;refGene_gn=OCLN;refGene_tx=XM_019835192.1
A1 138399956 138400172
refGene_gn=OCLN;refGene_tx=XM_003981047.4;refGene_gn=OCLN;refGene_tx=XM_019835192.1
A1 138403400 138403572
refGene_gn=OCLN;refGene_tx=XM_003981047.4;refGene_gn=OCLN;refGene_tx=XM_019835192.1
A1 138404974 138405016
refGene_gn=OCLN;refGene_tx=XM_003981047.4;refGene_gn=OCLN;refGene_tx=XM_019835192.1
A1 138406936 138407038
refGene_gn=OCLN;refGene_tx=XM_003981047.4;refGene_gn=OCLN;refGene_tx=XM_019835192.1
A1 138428963 138429019
refGene_gn=LOC101094730;refGene_tx=XM_011283892.3;refGene_gn=LOC101094730;refGene_tx=XM_011283896.3;refGene_gn=LOC101094730;refGene_tx=XM_019835198.2;refGene_gn=LOC101094730;refGene_tx=XM_019835202.2
A1 138429815 138429893
refGene_gn=LOC101094730;refGene_tx=XM_011283892.3;refGene_gn=LOC101094730;refGene_tx=XM_011283896.3;refGene_gn=LOC101094730;refGene_tx=XM_019835198.2;refGene_gn=LOC101094730;refGene_tx=XM_019835202.2
A1 138430220 138430257
refGene_gn=LOC101094730;refGene_tx=XM_011283892.3;refGene_gn=LOC101094730;refGene_tx=XM_011283896.3;refGene_gn=LOC101094730;refGene_tx=XM_01

9835198.2;refGene_gn=LOC101094730;refGene_tx=XM_019835202.2
A1 138430922 138431009
refGene_gn=LOC101094730;refGene_tx=XM_011283892.3;refGene_gn=LOC101094730;refGene_tx=XM_011283896.3;refGene_gn=LOC101094730;refGene_tx=XM_019835198.2;refGene_gn=LOC101094730;refGene_tx=XM_019835202.2
A1 138433190 138433241
refGene_gn=LOC101094730;refGene_tx=XM_011283892.3;refGene_gn=LOC101094730;refGene_tx=XM_011283896.3;refGene_gn=LOC101094730;refGene_tx=XM_019835198.2;refGene_gn=LOC101094730;refGene_tx=XM_019835202.2
A1 138433565 138433620
refGene_gn=LOC101094730;refGene_tx=XM_011283892.3;refGene_gn=LOC101094730;refGene_tx=XM_011283896.3;refGene_gn=LOC101094730;refGene_tx=XM_019835198.2;refGene_gn=LOC101094730;refGene_tx=XM_019835202.2
A1 138437578 138437684
refGene_gn=LOC101094730;refGene_tx=XM_011283892.3;refGene_gn=LOC101094730;refGene_tx=XM_011283896.3;refGene_gn=LOC101094730;refGene_tx=XM_019835198.2;refGene_gn=LOC101094730;refGene_tx=XM_019835202.2
A1 138441464 138441555
refGene_gn=LOC101094730;refGene_tx=XM_011283892.3;refGene_gn=LOC101094730;refGene_tx=XM_011283896.3;refGene_gn=LOC101094730;refGene_tx=XM_019835198.2;refGene_gn=LOC101094730;refGene_tx=XM_019835202.2
A1 138441709 138441797
refGene_gn=LOC101094730;refGene_tx=XM_011283892.3;refGene_gn=LOC101094730;refGene_tx=XM_011283896.3;refGene_gn=LOC101094730;refGene_tx=XM_019835198.2;refGene_gn=LOC101094730;refGene_tx=XM_019835202.2
A1 138442776 138442884
refGene_gn=LOC101094730;refGene_tx=XM_011283892.3;refGene_gn=LOC101094730;refGene_tx=XM_011283896.3;refGene_gn=LOC101094730;refGene_tx=XM_019835198.2;refGene_gn=LOC101094730;refGene_tx=XM_019835202.2
A1 138445870 138445934
refGene_gn=LOC101094730;refGene_tx=XM_011283892.3;refGene_gn=LOC101094730;refGene_tx=XM_011283896.3;refGene_gn=LOC101094730;refGene_tx=XM_019835198.2;refGene_gn=LOC101094730;refGene_tx=XM_019835202.2
A1 138447710 138447814
refGene_gn=LOC101094730;refGene_tx=XM_011283892.3;refGene_gn=LOC101094730;refGene_tx=XM_011283896.3;refGene_gn=LOC101094730;refGene_tx=XM_019835198.2;refGene_gn=LOC101094730;refGene_tx=XM_019835202.2
A1 138449131 138449234
refGene_gn=LOC101094730;refGene_tx=XM_011283892.3;refGene_gn=LOC101094730;refGene_tx=XM_011283896.3;refGene_gn=LOC101094730;refGene_tx=XM_019835198.2;refGene_gn=LOC101094730;refGene_tx=XM_019835202.2
A1 138454346 138454386
refGene_gn=LOC101094730;refGene_tx=XM_011283892.3;refGene_gn=LOC101094730;refGene_tx=XM_011283896.3;refGene_gn=LOC101094730;refGene_tx=XM_019835198.2;refGene_gn=LOC101094730;refGene_tx=XM_019835202.2
A1 138458886 138459006
refGene_gn=LOC101094730;refGene_tx=XM_011283892.3;refGene_gn=LOC101094730;refGene_tx=XM_011283896.3;refGene_gn=LOC101094730;refGene_tx=XM_019835198.2;refGene_gn=LOC101094730;refGene_tx=XM_019835202.2
A1 138474062 138474630

refGene_gn=NAIP;refGene_tx=XM_019835218.1
A1 138476115 138476215
refGene_gn=NAIP;refGene_tx=XM_019835218.1
A1 138477574 138477656
refGene_gn=NAIP;refGene_tx=XM_019835218.1
A1 138478575 138478627
refGene_gn=NAIP;refGene_tx=XM_019835218.1
A1 138482216 138482336
refGene_gn=NAIP;refGene_tx=XM_019835218.1
A1 138484709 138484809
refGene_gn=NAIP;refGene_tx=XM_019835218.1
A1 138491147 138491229
refGene_gn=NAIP;refGene_tx=XM_019835218.1
A1 138492401 138492459
refGene_gn=NAIP;refGene_tx=XM_019835218.1
A1 138492576 138494682
refGene_gn=NAIP;refGene_tx=XM_019835218.1
A1 138497743 138497911
refGene_gn=NAIP;refGene_tx=XM_019835218.1
A1 138499375 138499528
refGene_gn=NAIP;refGene_tx=XM_019835218.1
A1 138499869 138499956
refGene_gn=NAIP;refGene_tx=XM_019835218.1
A1 138502013 138502178
refGene_gn=NAIP;refGene_tx=XM_019835218.1
A1 138505493 138505858
refGene_gn=NAIP;refGene_tx=XM_019835218.1
A1 138519133 138519181
refGene_gn=SMN;refGene_tx=NM_001009328.1;refGene_gn=SMN;refGene_tx=XM_
006927779.3
A1 138527727 138527838
refGene_gn=SMN;refGene_tx=NM_001009328.1;refGene_gn=SMN;refGene_tx=XM_
006927779.3
A1 138528997 138529090
refGene_gn=SMN;refGene_tx=NM_001009328.1;refGene_gn=SMN;refGene_tx=XM_
006927779.3
A1 138531851 138532007
refGene_gn=SMN;refGene_tx=NM_001009328.1;refGene_gn=SMN;refGene_tx=XM_
006927779.3
A1 138532163 138532364
refGene_gn=SMN;refGene_tx=NM_001009328.1;refGene_gn=SMN;refGene_tx=XM_
006927779.3
A1 138533273 138533393
refGene_gn=SMN;refGene_tx=NM_001009328.1;refGene_gn=SMN;refGene_tx=XM_
006927779.3
A1 138535315 138535387
refGene_gn=SMN;refGene_tx=NM_001009328.1;refGene_gn=SMN;refGene_tx=XM_
006927779.3
A1 138551281 138551353
refGene_gn=SMN;refGene_tx=NM_001009328.1;refGene_gn=SMN;refGene_tx=XM_

006927779.3
A1	138571189	138571401
refGene_gn=BDP1;refGene_tx=XM_011283883.2;refGene_gn=BDP1;refGene_tx=XM_019835223.2;refGene_gn=BDP1;refGene_tx=XM_019835227.2;refGene_gn=BDP1;refGene_tx=XM_019835230.2;refGene_gn=BDP1;refGene_tx=XM_019835240.2;refGene_gn=BDP1;refGene_tx=XM_019835244.2;refGene_gn=BDP1;refGene_tx=XM_019835246.2
A1	138574585	138574859
refGene_gn=BDP1;refGene_tx=XM_011283883.2;refGene_gn=BDP1;refGene_tx=XM_019835223.2;refGene_gn=BDP1;refGene_tx=XM_019835227.2;refGene_gn=BDP1;refGene_tx=XM_019835230.2;refGene_gn=BDP1;refGene_tx=XM_019835240.2;refGene_gn=BDP1;refGene_tx=XM_019835244.2;refGene_gn=BDP1;refGene_tx=XM_019835246.2
A1	138576439	138576549
refGene_gn=BDP1;refGene_tx=XM_011283883.2;refGene_gn=BDP1;refGene_tx=XM_019835223.2;refGene_gn=BDP1;refGene_tx=XM_019835227.2;refGene_gn=BDP1;refGene_tx=XM_019835230.2;refGene_gn=BDP1;refGene_tx=XM_019835240.2;refGene_gn=BDP1;refGene_tx=XM_019835244.2;refGene_gn=BDP1;refGene_tx=XM_019835246.2
A1	138577658	138577718
refGene_gn=BDP1;refGene_tx=XM_011283883.2;refGene_gn=BDP1;refGene_tx=XM_019835223.2;refGene_gn=BDP1;refGene_tx=XM_019835227.2;refGene_gn=BDP1;refGene_tx=XM_019835230.2;refGene_gn=BDP1;refGene_tx=XM_019835240.2;refGene_gn=BDP1;refGene_tx=XM_019835244.2;refGene_gn=BDP1;refGene_tx=XM_019835246.2
A1	138578812	138578944
refGene_gn=BDP1;refGene_tx=XM_011283883.2;refGene_gn=BDP1;refGene_tx=XM_019835223.2;refGene_gn=BDP1;refGene_tx=XM_019835227.2;refGene_gn=BDP1;refGene_tx=XM_019835230.2;refGene_gn=BDP1;refGene_tx=XM_019835240.2;refGene_gn=BDP1;refGene_tx=XM_019835244.2;refGene_gn=BDP1;refGene_tx=XM_019835246.2
A1	138580360	138580494
refGene_gn=BDP1;refGene_tx=XM_011283883.2;refGene_gn=BDP1;refGene_tx=XM_019835223.2;refGene_gn=BDP1;refGene_tx=XM_019835227.2;refGene_gn=BDP1;refGene_tx=XM_019835230.2;refGene_gn=BDP1;refGene_tx=XM_019835240.2;refGene_gn=BDP1;refGene_tx=XM_019835244.2;refGene_gn=BDP1;refGene_tx=XM_019835246.2
A1	138581746	138581841
refGene_gn=BDP1;refGene_tx=XM_011283883.2;refGene_gn=BDP1;refGene_tx=XM_019835223.2;refGene_gn=BDP1;refGene_tx=XM_019835227.2;refGene_gn=BDP1;refGene_tx=XM_019835230.2;refGene_gn=BDP1;refGene_tx=XM_019835240.2;refGene_gn=BDP1;refGene_tx=XM_019835244.2;refGene_gn=BDP1;refGene_tx=XM_019835246.2
A1	138592855	138592910
refGene_gn=BDP1;refGene_tx=XM_011283883.2;refGene_gn=BDP1;refGene_tx=XM_019835223.2;refGene_gn=BDP1;refGene_tx=XM_019835227.2;refGene_gn=BDP1;refGene_tx=XM_019835230.2;refGene_gn=BDP1;refGene_tx=XM_019835240.2;refGene_gn=BDP1;refGene_tx=XM_019835244.2;refGene_gn=BDP1;refGene_tx=XM_019835246.2
A1	138596107	138596251

refGene_gn=BDP1;refGene_tx=XM_011283883.2;refGene_gn=BDP1;refGene_tx=XM_019835223.2;refGene_gn=BDP1;refGene_tx=XM_019835227.2;refGene_gn=BDP1;refGene_tx=XM_019835230.2;refGene_gn=BDP1;refGene_tx=XM_019835240.2;refGene_gn=BDP1;refGene_tx=XM_019835244.2;refGene_gn=BDP1;refGene_tx=XM_019835246.2
A1 138599202 138599487
refGene_gn=BDP1;refGene_tx=XM_011283883.2;refGene_gn=BDP1;refGene_tx=XM_019835223.2;refGene_gn=BDP1;refGene_tx=XM_019835227.2;refGene_gn=BDP1;refGene_tx=XM_019835230.2;refGene_gn=BDP1;refGene_tx=XM_019835240.2;refGene_gn=BDP1;refGene_tx=XM_019835244.2;refGene_gn=BDP1;refGene_tx=XM_019835246.2
A1 138599834 138599982
refGene_gn=BDP1;refGene_tx=XM_011283883.2;refGene_gn=BDP1;refGene_tx=XM_019835223.2;refGene_gn=BDP1;refGene_tx=XM_019835227.2;refGene_gn=BDP1;refGene_tx=XM_019835230.2;refGene_gn=BDP1;refGene_tx=XM_019835240.2;refGene_gn=BDP1;refGene_tx=XM_019835244.2;refGene_gn=BDP1;refGene_tx=XM_019835246.2
A1 138605514 138605673
refGene_gn=BDP1;refGene_tx=XM_011283883.2;refGene_gn=BDP1;refGene_tx=XM_019835223.2;refGene_gn=BDP1;refGene_tx=XM_019835227.2;refGene_gn=BDP1;refGene_tx=XM_019835230.2;refGene_gn=BDP1;refGene_tx=XM_019835240.2;refGene_gn=BDP1;refGene_tx=XM_019835244.2;refGene_gn=BDP1;refGene_tx=XM_019835246.2
A1 138609583 138609743
refGene_gn=BDP1;refGene_tx=XM_011283883.2;refGene_gn=BDP1;refGene_tx=XM_019835223.2;refGene_gn=BDP1;refGene_tx=XM_019835227.2;refGene_gn=BDP1;refGene_tx=XM_019835230.2;refGene_gn=BDP1;refGene_tx=XM_019835240.2;refGene_gn=BDP1;refGene_tx=XM_019835244.2;refGene_gn=BDP1;refGene_tx=XM_019835246.2
A1 138610533 138610625
refGene_gn=BDP1;refGene_tx=XM_011283883.2;refGene_gn=BDP1;refGene_tx=XM_019835223.2;refGene_gn=BDP1;refGene_tx=XM_019835227.2;refGene_gn=BDP1;refGene_tx=XM_019835230.2;refGene_gn=BDP1;refGene_tx=XM_019835240.2;refGene_gn=BDP1;refGene_tx=XM_019835244.2;refGene_gn=BDP1;refGene_tx=XM_019835246.2
A1 138611511 138611716
refGene_gn=BDP1;refGene_tx=XM_011283883.2;refGene_gn=BDP1;refGene_tx=XM_019835223.2;refGene_gn=BDP1;refGene_tx=XM_019835227.2;refGene_gn=BDP1;refGene_tx=XM_019835230.2;refGene_gn=BDP1;refGene_tx=XM_019835240.2;refGene_gn=BDP1;refGene_tx=XM_019835244.2;refGene_gn=BDP1;refGene_tx=XM_019835246.2
A1 138612478 138612591
refGene_gn=BDP1;refGene_tx=XM_011283883.2;refGene_gn=BDP1;refGene_tx=XM_019835223.2;refGene_gn=BDP1;refGene_tx=XM_019835227.2;refGene_gn=BDP1;refGene_tx=XM_019835230.2;refGene_gn=BDP1;refGene_tx=XM_019835240.2;refGene_gn=BDP1;refGene_tx=XM_019835244.2;refGene_gn=BDP1;refGene_tx=XM_019835246.2
A1 138613730 138615243
refGene_gn=BDP1;refGene_tx=XM_011283883.2;refGene_gn=BDP1;refGene_tx=XM_019835223.2;refGene_gn=BDP1;refGene_tx=XM_019835227.2;refGene_gn=BDP

1;refGene_tx=XM_019835230.2;refGene_gn=BDP1;refGene_tx=XM_019835240.2;
refGene_gn=BDP1;refGene_tx=XM_019835244.2;refGene_gn=BDP1;refGene_tx=X
M_019835246.2
A1 138616303 138616491
refGene_gn=BDP1;refGene_tx=XM_011283883.2;refGene_gn=BDP1;refGene_tx=X
M_019835223.2;refGene_gn=BDP1;refGene_tx=XM_019835227.2;refGene_gn=BDP
1;refGene_tx=XM_019835230.2;refGene_gn=BDP1;refGene_tx=XM_019835240.2;
refGene_gn=BDP1;refGene_tx=XM_019835244.2;refGene_gn=BDP1;refGene_tx=X
M_019835246.2
A1 138616936 138617159
refGene_gn=BDP1;refGene_tx=XM_011283883.2;refGene_gn=BDP1;refGene_tx=X
M_019835223.2;refGene_gn=BDP1;refGene_tx=XM_019835227.2;refGene_gn=BDP
1;refGene_tx=XM_019835230.2;refGene_gn=BDP1;refGene_tx=XM_019835240.2;
refGene_gn=BDP1;refGene_tx=XM_019835244.2;refGene_gn=BDP1;refGene_tx=X
M_019835246.2
A1 138619848 138620027
refGene_gn=BDP1;refGene_tx=XM_011283883.2;refGene_gn=BDP1;refGene_tx=X
M_019835223.2;refGene_gn=BDP1;refGene_tx=XM_019835227.2;refGene_gn=BDP
1;refGene_tx=XM_019835230.2;refGene_gn=BDP1;refGene_tx=XM_019835240.2;
refGene_gn=BDP1;refGene_tx=XM_019835244.2;refGene_gn=BDP1;refGene_tx=X
M_019835246.2
A1 138620774 138620979
refGene_gn=BDP1;refGene_tx=XM_011283883.2;refGene_gn=BDP1;refGene_tx=X
M_019835223.2;refGene_gn=BDP1;refGene_tx=XM_019835227.2;refGene_gn=BDP
1;refGene_tx=XM_019835230.2;refGene_gn=BDP1;refGene_tx=XM_019835240.2;
refGene_gn=BDP1;refGene_tx=XM_019835244.2;refGene_gn=BDP1;refGene_tx=X
M_019835246.2
A1 138622036 138622176
refGene_gn=BDP1;refGene_tx=XM_011283883.2;refGene_gn=BDP1;refGene_tx=X
M_019835223.2;refGene_gn=BDP1;refGene_tx=XM_019835227.2;refGene_gn=BDP
1;refGene_tx=XM_019835230.2;refGene_gn=BDP1;refGene_tx=XM_019835240.2;
refGene_gn=BDP1;refGene_tx=XM_019835244.2;refGene_gn=BDP1;refGene_tx=X
M_019835246.2
A1 138625062 138625261
refGene_gn=BDP1;refGene_tx=XM_011283883.2;refGene_gn=BDP1;refGene_tx=X
M_019835223.2;refGene_gn=BDP1;refGene_tx=XM_019835227.2;refGene_gn=BDP
1;refGene_tx=XM_019835230.2;refGene_gn=BDP1;refGene_tx=XM_019835240.2;
refGene_gn=BDP1;refGene_tx=XM_019835244.2;refGene_gn=BDP1;refGene_tx=X
M_019835246.2
A1 138625549 138625740
refGene_gn=BDP1;refGene_tx=XM_011283883.2;refGene_gn=BDP1;refGene_tx=X
M_019835223.2;refGene_gn=BDP1;refGene_tx=XM_019835227.2;refGene_gn=BDP
1;refGene_tx=XM_019835230.2;refGene_gn=BDP1;refGene_tx=XM_019835240.2;
refGene_gn=BDP1;refGene_tx=XM_019835244.2;refGene_gn=BDP1;refGene_tx=X
M_019835246.2
A1 138626744 138627132
refGene_gn=BDP1;refGene_tx=XM_011283883.2;refGene_gn=BDP1;refGene_tx=X
M_019835223.2;refGene_gn=BDP1;refGene_tx=XM_019835227.2;refGene_gn=BDP
1;refGene_tx=XM_019835230.2;refGene_gn=BDP1;refGene_tx=XM_019835240.2;
refGene_gn=BDP1;refGene_tx=XM_019835244.2;refGene_gn=BDP1;refGene_tx=X

M_019835246.2
A1 138628794 138628914
refGene_gn=BDP1;refGene_tx=XM_011283883.2;refGene_gn=BDP1;refGene_tx=XM_019835223.2;refGene_gn=BDP1;refGene_tx=XM_019835227.2;refGene_gn=BDP1;refGene_tx=XM_019835230.2;refGene_gn=BDP1;refGene_tx=XM_019835240.2;refGene_gn=BDP1;refGene_tx=XM_019835244.2;refGene_gn=BDP1;refGene_tx=XM_019835246.2
A1 138632692 138632729
refGene_gn=BDP1;refGene_tx=XM_011283883.2;refGene_gn=BDP1;refGene_tx=XM_019835223.2;refGene_gn=BDP1;refGene_tx=XM_019835227.2;refGene_gn=BDP1;refGene_tx=XM_019835230.2;refGene_gn=BDP1;refGene_tx=XM_019835240.2;refGene_gn=BDP1;refGene_tx=XM_019835244.2;refGene_gn=BDP1;refGene_tx=XM_019835246.2
A1 138633524 138633617
refGene_gn=BDP1;refGene_tx=XM_011283883.2;refGene_gn=BDP1;refGene_tx=XM_019835223.2;refGene_gn=BDP1;refGene_tx=XM_019835227.2;refGene_gn=BDP1;refGene_tx=XM_019835230.2;refGene_gn=BDP1;refGene_tx=XM_019835240.2;refGene_gn=BDP1;refGene_tx=XM_019835244.2;refGene_gn=BDP1;refGene_tx=XM_019835246.2
A1 138634715 138634941
refGene_gn=BDP1;refGene_tx=XM_011283883.2;refGene_gn=BDP1;refGene_tx=XM_019835223.2;refGene_gn=BDP1;refGene_tx=XM_019835227.2;refGene_gn=BDP1;refGene_tx=XM_019835230.2;refGene_gn=BDP1;refGene_tx=XM_019835240.2;refGene_gn=BDP1;refGene_tx=XM_019835244.2;refGene_gn=BDP1;refGene_tx=XM_019835246.2
A1 138635872 138636030
refGene_gn=BDP1;refGene_tx=XM_011283883.2;refGene_gn=BDP1;refGene_tx=XM_019835223.2;refGene_gn=BDP1;refGene_tx=XM_019835227.2;refGene_gn=BDP1;refGene_tx=XM_019835230.2;refGene_gn=BDP1;refGene_tx=XM_019835240.2;refGene_gn=BDP1;refGene_tx=XM_019835244.2;refGene_gn=BDP1;refGene_tx=XM_019835246.2
A1 138640705 138640859
refGene_gn=BDP1;refGene_tx=XM_011283883.2;refGene_gn=BDP1;refGene_tx=XM_019835223.2;refGene_gn=BDP1;refGene_tx=XM_019835227.2;refGene_gn=BDP1;refGene_tx=XM_019835230.2;refGene_gn=BDP1;refGene_tx=XM_019835240.2;refGene_gn=BDP1;refGene_tx=XM_019835244.2;refGene_gn=BDP1;refGene_tx=XM_019835246.2
A1 138641110 138641288
refGene_gn=BDP1;refGene_tx=XM_011283883.2;refGene_gn=BDP1;refGene_tx=XM_019835223.2;refGene_gn=BDP1;refGene_tx=XM_019835227.2;refGene_gn=BDP1;refGene_tx=XM_019835230.2;refGene_gn=BDP1;refGene_tx=XM_019835240.2;refGene_gn=BDP1;refGene_tx=XM_019835244.2;refGene_gn=BDP1;refGene_tx=XM_019835246.2
A1 138644091 138644155
refGene_gn=BDP1;refGene_tx=XM_011283883.2;refGene_gn=BDP1;refGene_tx=XM_019835223.2;refGene_gn=BDP1;refGene_tx=XM_019835227.2;refGene_gn=BDP1;refGene_tx=XM_019835230.2;refGene_gn=BDP1;refGene_tx=XM_019835240.2;refGene_gn=BDP1;refGene_tx=XM_019835244.2;refGene_gn=BDP1;refGene_tx=XM_019835246.2
A1 138644714 138644901

refGene_gn=BDP1;refGene_tx=XM_011283883.2;refGene_gn=BDP1;refGene_tx=XM_019835223.2;refGene_gn=BDP1;refGene_tx=XM_019835227.2;refGene_gn=BDP1;refGene_tx=XM_019835230.2;refGene_gn=BDP1;refGene_tx=XM_019835240.2;refGene_gn=BDP1;refGene_tx=XM_019835244.2;refGene_gn=BDP1;refGene_tx=XM_019835246.2
A1 138645891 138646096
refGene_gn=BDP1;refGene_tx=XM_011283883.2;refGene_gn=BDP1;refGene_tx=XM_019835223.2;refGene_gn=BDP1;refGene_tx=XM_019835227.2;refGene_gn=BDP1;refGene_tx=XM_019835230.2;refGene_gn=BDP1;refGene_tx=XM_019835240.2;refGene_gn=BDP1;refGene_tx=XM_019835244.2
A1 138650881 138650915
refGene_gn=BDP1;refGene_tx=XM_011283883.2;refGene_gn=BDP1;refGene_tx=XM_019835223.2;refGene_gn=BDP1;refGene_tx=XM_019835227.2;refGene_gn=BDP1;refGene_tx=XM_019835230.2;refGene_gn=BDP1;refGene_tx=XM_019835240.2;refGene_gn=BDP1;refGene_tx=XM_019835244.2;refGene_gn=BDP1;refGene_tx=XM_019835246.2
A1 138651663 138651931
refGene_gn=BDP1;refGene_tx=XM_011283883.2;refGene_gn=BDP1;refGene_tx=XM_019835223.2;refGene_gn=BDP1;refGene_tx=XM_019835227.2;refGene_gn=BDP1;refGene_tx=XM_019835230.2;refGene_gn=BDP1;refGene_tx=XM_019835240.2;refGene_gn=BDP1;refGene_tx=XM_019835244.2
A1 138652391 138652641
refGene_gn=BDP1;refGene_tx=XM_011283883.2;refGene_gn=BDP1;refGene_tx=XM_019835223.2;refGene_gn=BDP1;refGene_tx=XM_019835227.2;refGene_gn=BDP1;refGene_tx=XM_019835230.2;refGene_gn=BDP1;refGene_tx=XM_019835240.2;refGene_gn=BDP1;refGene_tx=XM_019835244.2
A1 138658236 138658365
refGene_gn=BDP1;refGene_tx=XM_011283883.2;refGene_gn=BDP1;refGene_tx=XM_019835223.2;refGene_gn=BDP1;refGene_tx=XM_019835227.2;refGene_gn=BDP1;refGene_tx=XM_019835230.2;refGene_gn=BDP1;refGene_tx=XM_019835240.2;refGene_gn=BDP1;refGene_tx=XM_019835244.2
A1 138668972 138669101
refGene_gn=MCCC2;refGene_tx=XM_003981052.4
A1 138677471 138677538
refGene_gn=MCCC2;refGene_tx=XM_003981052.4
A1 138681240 138681325
refGene_gn=MCCC2;refGene_tx=XM_003981052.4
A1 138683606 138683708
refGene_gn=MCCC2;refGene_tx=XM_003981052.4
A1 138691876 138692004
refGene_gn=MCCC2;refGene_tx=XM_003981052.4
A1 138693932 138694045
refGene_gn=MCCC2;refGene_tx=XM_003981052.4
A1 138708624 138708738
refGene_gn=MCCC2;refGene_tx=XM_003981052.4
A1 138715384 138715449
refGene_gn=MCCC2;refGene_tx=XM_003981052.4
A1 138720645 138720745
refGene_gn=MCCC2;refGene_tx=XM_003981052.4
A1 138720856 138720952

refGene_gn=MCCC2;refGene_tx=XM_003981052.4
A1 138722417 138722490
refGene_gn=MCCC2;refGene_tx=XM_003981052.4
A1 138725858 138725935
refGene_gn=MCCC2;refGene_tx=XM_003981052.4
A1 138726876 138726943
refGene_gn=MCCC2;refGene_tx=XM_003981052.4
A1 138730358 138730515
refGene_gn=MCCC2;refGene_tx=XM_003981052.4
A1 138731367 138731482
refGene_gn=MCCC2;refGene_tx=XM_003981052.4
A1 138733968 138734054
refGene_gn=MCCC2;refGene_tx=XM_003981052.4
A1 138739316 138739434
refGene_gn=MCCC2;refGene_tx=XM_003981052.4
A1 138791976 138792135
refGene_gn=CARTPT;refGene_tx=XM_003981053.2
A1 138792579 138792663
refGene_gn=CARTPT;refGene_tx=XM_003981053.2
A1 138793727 138793835
refGene_gn=CARTPT;refGene_tx=XM_003981053.2
A1 139131417 139131601
refGene_gn=MAP1B;refGene_tx=XM_006927783.4
A1 139139828 139139930
refGene_gn=MAP1B;refGene_tx=XM_006927783.4
A1 139203282 139203365
refGene_gn=MAP1B;refGene_tx=XM_006927783.4
A1 139205946 139206087
refGene_gn=MAP1B;refGene_tx=XM_006927783.4;refGene_gn=MAP1B;refGene_tx
=XM_019835294.2
A1 139216846 139223357
refGene_gn=MAP1B;refGene_tx=XM_006927783.4;refGene_gn=MAP1B;refGene_tx
=XM_019835294.2
A1 139226512 139226751
refGene_gn=MAP1B;refGene_tx=XM_006927783.4;refGene_gn=MAP1B;refGene_tx
=XM_019835294.2
A1 139227954 139228110
refGene_gn=MAP1B;refGene_tx=XM_006927783.4;refGene_gn=MAP1B;refGene_tx
=XM_019835294.2
A1 139239099 139239339
refGene_gn=MRPS27;refGene_tx=XM_003981054.4;refGene_gn=MRPS27;refGene_
tx=XM_019835313.2;refGene_gn=MRPS27;refGene_tx=XM_019835320.2
A1 139241847 139242015
refGene_gn=MRPS27;refGene_tx=XM_003981054.4;refGene_gn=MRPS27;refGene_
tx=XM_019835313.2;refGene_gn=MRPS27;refGene_tx=XM_019835320.2
A1 139243920 139244063
refGene_gn=MRPS27;refGene_tx=XM_003981054.4;refGene_gn=MRPS27;refGene_
tx=XM_019835313.2;refGene_gn=MRPS27;refGene_tx=XM_019835320.2
A1 139246660 139246763
refGene_gn=MRPS27;refGene_tx=XM_003981054.4;refGene_gn=MRPS27;refGene_

tx=XM_019835313.2;refGene_gn=MRPS27;refGene_tx=XM_019835320.2
A1 139250426 139250542
refGene_gn=MRPS27;refGene_tx=XM_003981054.4;refGene_gn=MRPS27;refGene_tx=XM_019835313.2;refGene_gn=MRPS27;refGene_tx=XM_019835320.2
A1 139253741 139253820
refGene_gn=MRPS27;refGene_tx=XM_003981054.4;refGene_gn=MRPS27;refGene_tx=XM_019835313.2;refGene_gn=MRPS27;refGene_tx=XM_019835320.2
A1 139257554 139257669
refGene_gn=MRPS27;refGene_tx=XM_003981054.4;refGene_gn=MRPS27;refGene_tx=XM_019835313.2;refGene_gn=MRPS27;refGene_tx=XM_019835320.2
A1 139306462 139306521
refGene_gn=MRPS27;refGene_tx=XM_003981054.4;refGene_gn=MRPS27;refGene_tx=XM_019835313.2;refGene_gn=MRPS27;refGene_tx=XM_019835320.2
A1 139308784 139308855
refGene_gn=MRPS27;refGene_tx=XM_003981054.4;refGene_gn=MRPS27;refGene_tx=XM_019835313.2;refGene_gn=MRPS27;refGene_tx=XM_019835320.2
A1 139320033 139320111
refGene_gn=MRPS27;refGene_tx=XM_003981054.4
A1 139324799 139324872
refGene_gn=MRPS27;refGene_tx=XM_003981054.4;refGene_gn=MRPS27;refGene_tx=XM_019835313.2
A1 139324989 139325101
refGene_gn=PTCD2;refGene_tx=XM_019835328.1;refGene_gn=PTCD2;refGene_tx=XM_019835330.1;refGene_gn=PTCD2;refGene_tx=XM_019835334.1;refGene_gn=PTCD2;refGene_tx=XM_019835337.1;refGene_gn=PTCD2;refGene_tx=XM_019835346.1;refGene_gn=PTCD2;refGene_tx=XM_023257279.1;refGene_gn=PTCD2;refGene_tx=XM_023257284.1
A1 139326082 139326175
refGene_gn=PTCD2;refGene_tx=XM_019835328.1;refGene_gn=PTCD2;refGene_tx=XM_019835334.1;refGene_gn=PTCD2;refGene_tx=XM_019835337.1;refGene_gn=PTCD2;refGene_tx=XM_019835346.1;refGene_gn=PTCD2;refGene_tx=XM_023257279.1;refGene_gn=PTCD2;refGene_tx=XM_023257284.1
A1 139329581 139329711
refGene_gn=PTCD2;refGene_tx=XM_019835328.1;refGene_gn=PTCD2;refGene_tx=XM_019835330.1;refGene_gn=PTCD2;refGene_tx=XM_019835334.1;refGene_gn=PTCD2;refGene_tx=XM_019835337.1;refGene_gn=PTCD2;refGene_tx=XM_019835346.1;refGene_gn=PTCD2;refGene_tx=XM_023257279.1;refGene_gn=PTCD2;refGene_tx=XM_023257284.1
A1 139333713 139333831
refGene_gn=PTCD2;refGene_tx=XM_006927785.3;refGene_gn=PTCD2;refGene_tx=XM_019835328.1;refGene_gn=PTCD2;refGene_tx=XM_019835330.1;refGene_gn=PTCD2;refGene_tx=XM_019835334.1;refGene_gn=PTCD2;refGene_tx=XM_019835337.1;refGene_gn=PTCD2;refGene_tx=XM_019835346.1;refGene_gn=PTCD2;refGene_tx=XM_023257279.1;refGene_gn=PTCD2;refGene_tx=XM_023257284.1
A1 139336958 139337037
refGene_gn=PTCD2;refGene_tx=XM_006927785.3;refGene_gn=PTCD2;refGene_tx=XM_019835328.1;refGene_gn=PTCD2;refGene_tx=XM_019835330.1;refGene_gn=PTCD2;refGene_tx=XM_019835334.1;refGene_gn=PTCD2;refGene_tx=XM_019835337.1;refGene_gn=PTCD2;refGene_tx=XM_019835346.1;refGene_gn=PTCD2;refGene_tx=XM_023257279.1;refGene_gn=PTCD2;refGene_tx=XM_023257284.1

A1 139337457 139337549
refGene_gn=PTCD2;refGene_tx=XM_006927785.3;refGene_gn=PTCD2;refGene_tx=XM_019835328.1;refGene_gn=PTCD2;refGene_tx=XM_019835330.1;refGene_gn=PTCD2;refGene_tx=XM_019835334.1;refGene_gn=PTCD2;refGene_tx=XM_019835337.1;refGene_gn=PTCD2;refGene_tx=XM_019835346.1;refGene_gn=PTCD2;refGene_tx=XM_023257279.1;refGene_gn=PTCD2;refGene_tx=XM_023257284.1
A1 139339764 139339878
refGene_gn=PTCD2;refGene_tx=XM_006927785.3;refGene_gn=PTCD2;refGene_tx=XM_019835328.1;refGene_gn=PTCD2;refGene_tx=XM_019835330.1;refGene_gn=PTCD2;refGene_tx=XM_019835334.1;refGene_gn=PTCD2;refGene_tx=XM_019835337.1;refGene_gn=PTCD2;refGene_tx=XM_019835346.1;refGene_gn=PTCD2;refGene_tx=XM_023257279.1;refGene_gn=PTCD2;refGene_tx=XM_023257284.1
A1 139343407 139343482
refGene_gn=PTCD2;refGene_tx=XM_006927785.3;refGene_gn=PTCD2;refGene_tx=XM_019835330.1;refGene_gn=PTCD2;refGene_tx=XM_019835334.1;refGene_gn=PTCD2;refGene_tx=XM_019835337.1;refGene_gn=PTCD2;refGene_tx=XM_019835346.1;refGene_gn=PTCD2;refGene_tx=XM_023257279.1;refGene_gn=PTCD2;refGene_tx=XM_023257284.1
A1 139351875 139351989
refGene_gn=PTCD2;refGene_tx=XM_006927785.3;refGene_gn=PTCD2;refGene_tx=XM_019835328.1;refGene_gn=PTCD2;refGene_tx=XM_019835330.1;refGene_gn=PTCD2;refGene_tx=XM_019835334.1;refGene_gn=PTCD2;refGene_tx=XM_019835337.1;refGene_gn=PTCD2;refGene_tx=XM_019835346.1;refGene_gn=PTCD2;refGene_tx=XM_023257279.1;refGene_gn=PTCD2;refGene_tx=XM_023257284.1
A1 139352071 139352188
refGene_gn=PTCD2;refGene_tx=XM_019835334.1
A1 139357399 139357624
refGene_gn=PTCD2;refGene_tx=XM_006927785.3;refGene_gn=PTCD2;refGene_tx=XM_019835328.1;refGene_gn=PTCD2;refGene_tx=XM_019835330.1
A1 139367429 139367468
refGene_gn=PTCD2;refGene_tx=XM_019835346.1
A1 139377710 139377854
refGene_gn=PTCD2;refGene_tx=XM_023257279.1
A1 139380501 139380543
refGene_gn=PTCD2;refGene_tx=XM_023257284.1
A1 139432097 139432175
refGene_gn=PTCD2;refGene_tx=XM_019835337.1
A1 139439646 139440182
refGene_gn=ZNF366;refGene_tx=XM_003981055.3;refGene_gn=ZNF366;refGene_tx=XM_019835327.2
A1 139443057 139443232
refGene_gn=ZNF366;refGene_tx=XM_003981055.3;refGene_gn=ZNF366;refGene_tx=XM_019835327.2
A1 139453424 139453616
refGene_gn=ZNF366;refGene_tx=XM_003981055.3;refGene_gn=ZNF366;refGene_tx=XM_019835327.2
A1 139457225 139458557
refGene_gn=ZNF366;refGene_tx=XM_003981055.3;refGene_gn=ZNF366;refGene_tx=XM_019835327.2
A1 139712449 139712818

refGene_gn=LOC101092488;refGene_tx=XM_003981056.4
A1 139777095 139777110
refGene_gn=TNPO1;refGene_tx=XM_023257302.1
A1 139814236 139814350
refGene_gn=TNPO1;refGene_tx=XM_023257302.1;refGene_gn=TNPO1;refGene_tx=XM_023257303.1;refGene_gn=TNPO1;refGene_tx=XM_023257308.1;refGene_gn=TNPO1;refGene_tx=XM_023257312.1
A1 139817086 139817162
refGene_gn=TNPO1;refGene_tx=XM_023257302.1;refGene_gn=TNPO1;refGene_tx=XM_023257303.1;refGene_gn=TNPO1;refGene_tx=XM_023257308.1;refGene_gn=TNPO1;refGene_tx=XM_023257312.1
A1 139821290 139821440
refGene_gn=TNPO1;refGene_tx=XM_023257302.1;refGene_gn=TNPO1;refGene_tx=XM_023257303.1;refGene_gn=TNPO1;refGene_tx=XM_023257308.1;refGene_gn=TNPO1;refGene_tx=XM_023257312.1
A1 139825217 139825324
refGene_gn=TNPO1;refGene_tx=XM_023257302.1;refGene_gn=TNPO1;refGene_tx=XM_023257303.1;refGene_gn=TNPO1;refGene_tx=XM_023257308.1;refGene_gn=TNPO1;refGene_tx=XM_023257312.1
A1 139827247 139827381
refGene_gn=TNPO1;refGene_tx=XM_023257302.1;refGene_gn=TNPO1;refGene_tx=XM_023257303.1;refGene_gn=TNPO1;refGene_tx=XM_023257308.1;refGene_gn=TNPO1;refGene_tx=XM_023257312.1
A1 139831093 139831175
refGene_gn=TNPO1;refGene_tx=XM_023257302.1;refGene_gn=TNPO1;refGene_tx=XM_023257303.1;refGene_gn=TNPO1;refGene_tx=XM_023257308.1;refGene_gn=TNPO1;refGene_tx=XM_023257312.1
A1 139834267 139834390
refGene_gn=TNPO1;refGene_tx=XM_023257302.1;refGene_gn=TNPO1;refGene_tx=XM_023257303.1;refGene_gn=TNPO1;refGene_tx=XM_023257308.1;refGene_gn=TNPO1;refGene_tx=XM_023257312.1
A1 139836523 139836642
refGene_gn=TNPO1;refGene_tx=XM_023257302.1;refGene_gn=TNPO1;refGene_tx=XM_023257303.1;refGene_gn=TNPO1;refGene_tx=XM_023257308.1;refGene_gn=TNPO1;refGene_tx=XM_023257312.1
A1 139843034 139843095
refGene_gn=TNPO1;refGene_tx=XM_023257302.1;refGene_gn=TNPO1;refGene_tx=XM_023257303.1;refGene_gn=TNPO1;refGene_tx=XM_023257308.1;refGene_gn=TNPO1;refGene_tx=XM_023257312.1
A1 139843642 139843811
refGene_gn=TNPO1;refGene_tx=XM_023257302.1;refGene_gn=TNPO1;refGene_tx=XM_023257303.1;refGene_gn=TNPO1;refGene_tx=XM_023257308.1;refGene_gn=TNPO1;refGene_tx=XM_023257312.1
A1 139847131 139847284
refGene_gn=TNPO1;refGene_tx=XM_023257302.1;refGene_gn=TNPO1;refGene_tx=XM_023257303.1;refGene_gn=TNPO1;refGene_tx=XM_023257308.1;refGene_gn=TNPO1;refGene_tx=XM_023257312.1
A1 139848001 139848227
refGene_gn=TNPO1;refGene_tx=XM_023257302.1;refGene_gn=TNPO1;refGene_tx=XM_023257303.1;refGene_gn=TNPO1;refGene_tx=XM_023257308.1;refGene_gn=

TNPO1;refGene_tx=XM_023257312.1
A1 139850319 139850491
refGene_gn=TNPO1;refGene_tx=XM_023257302.1;refGene_gn=TNPO1;refGene_tx=XM_023257303.1;refGene_gn=TNPO1;refGene_tx=XM_023257308.1;refGene_gn=TNPO1;refGene_tx=XM_023257312.1
A1 139851943 139852030
refGene_gn=TNPO1;refGene_tx=XM_023257302.1;refGene_gn=TNPO1;refGene_tx=XM_023257303.1;refGene_gn=TNPO1;refGene_tx=XM_023257308.1;refGene_gn=TNPO1;refGene_tx=XM_023257312.1
A1 139852723 139852831
refGene_gn=TNPO1;refGene_tx=XM_023257302.1;refGene_gn=TNPO1;refGene_tx=XM_023257303.1;refGene_gn=TNPO1;refGene_tx=XM_023257308.1;refGene_gn=TNPO1;refGene_tx=XM_023257312.1
A1 139852963 139853122
refGene_gn=TNPO1;refGene_tx=XM_023257302.1;refGene_gn=TNPO1;refGene_tx=XM_023257303.1;refGene_gn=TNPO1;refGene_tx=XM_023257308.1;refGene_gn=TNPO1;refGene_tx=XM_023257312.1
A1 139853202 139853290
refGene_gn=TNPO1;refGene_tx=XM_023257302.1;refGene_gn=TNPO1;refGene_tx=XM_023257303.1;refGene_gn=TNPO1;refGene_tx=XM_023257308.1;refGene_gn=TNPO1;refGene_tx=XM_023257312.1
A1 139855582 139855681
refGene_gn=TNPO1;refGene_tx=XM_023257302.1;refGene_gn=TNPO1;refGene_tx=XM_023257303.1;refGene_gn=TNPO1;refGene_tx=XM_023257308.1;refGene_gn=TNPO1;refGene_tx=XM_023257312.1
A1 139855892 139855988
refGene_gn=TNPO1;refGene_tx=XM_023257302.1;refGene_gn=TNPO1;refGene_tx=XM_023257303.1;refGene_gn=TNPO1;refGene_tx=XM_023257308.1;refGene_gn=TNPO1;refGene_tx=XM_023257312.1
A1 139858838 139858914
refGene_gn=TNPO1;refGene_tx=XM_023257302.1;refGene_gn=TNPO1;refGene_tx=XM_023257303.1;refGene_gn=TNPO1;refGene_tx=XM_023257308.1;refGene_gn=TNPO1;refGene_tx=XM_023257312.1
A1 139859794 139859894
refGene_gn=TNPO1;refGene_tx=XM_023257302.1;refGene_gn=TNPO1;refGene_tx=XM_023257303.1;refGene_gn=TNPO1;refGene_tx=XM_023257308.1;refGene_gn=TNPO1;refGene_tx=XM_023257312.1
A1 139862512 139862587
refGene_gn=TNPO1;refGene_tx=XM_023257302.1;refGene_gn=TNPO1;refGene_tx=XM_023257303.1;refGene_gn=TNPO1;refGene_tx=XM_023257308.1;refGene_gn=TNPO1;refGene_tx=XM_023257312.1
A1 139863836 139863944
refGene_gn=TNPO1;refGene_tx=XM_023257302.1;refGene_gn=TNPO1;refGene_tx=XM_023257303.1;refGene_gn=TNPO1;refGene_tx=XM_023257308.1;refGene_gn=TNPO1;refGene_tx=XM_023257312.1
A1 139903473 139903506
refGene_gn=FCHO2;refGene_tx=XM_023257315.1
A1 139920061 139920153
refGene_gn=FCHO2;refGene_tx=XM_023257315.1
A1 139937559 139937634

refGene_gn=FCHO2;refGene_tx=XM_023257315.1
A1 139940966 139941108
refGene_gn=FCHO2;refGene_tx=XM_023257315.1
A1 139941199 139941352
refGene_gn=FCHO2;refGene_tx=XM_023257315.1
A1 139951850 139951955
refGene_gn=FCHO2;refGene_tx=XM_023257315.1
A1 139955639 139955738
refGene_gn=FCHO2;refGene_tx=XM_023257315.1
A1 139956793 139956890
refGene_gn=FCHO2;refGene_tx=XM_023257315.1
A1 139969900 139969945
refGene_gn=FCHO2;refGene_tx=XM_023257315.1
A1 139971619 139971692
refGene_gn=FCHO2;refGene_tx=XM_023257315.1
A1 139975010 139975035
refGene_gn=FCHO2;refGene_tx=XM_023257315.1
A1 139986323 139986381
refGene_gn=FCHO2;refGene_tx=XM_023257315.1
A1 139986657 139986833
refGene_gn=FCHO2;refGene_tx=XM_023257315.1
A1 139988300 139988312
refGene_gn=FCHO2;refGene_tx=XM_023257315.1
A1 139988877 139988902
refGene_gn=FCHO2;refGene_tx=XM_023257315.1
A1 139990528 139990571
refGene_gn=FCHO2;refGene_tx=XM_023257315.1
A1 139992975 139993070
refGene_gn=FCHO2;refGene_tx=XM_023257315.1
A1 139997811 139997915
refGene_gn=FCHO2;refGene_tx=XM_023257315.1
A1 140002270 140002400
refGene_gn=FCHO2;refGene_tx=XM_023257315.1
A1 140006542 140006654
refGene_gn=FCHO2;refGene_tx=XM_023257315.1
A1 140010350 140010506
refGene_gn=FCHO2;refGene_tx=XM_023257315.1
A1 140010838 140010971
refGene_gn=FCHO2;refGene_tx=XM_023257315.1
A1 140018706 140018906
refGene_gn=FCHO2;refGene_tx=XM_023257315.1
A1 140019603 140019668
refGene_gn=FCHO2;refGene_tx=XM_023257315.1
A1 140023175 140023340
refGene_gn=FCHO2;refGene_tx=XM_023257315.1
A1 140023661 140023684
refGene_gn=FCHO2;refGene_tx=XM_023257315.1
A1 140057123 140057763
refGene_gn=TMEM171;refGene_tx=XM_003981071.4;refGene_gn=TMEM171;refGen
e_tx=XM_019835400.2

A1 140061529 140061671
refGene_gn=TMEM171;refGene_tx=XM_003981071.4;refGene_gn=TMEM171;refGene_tx=XM_019835400.2
A1 140063394 140063593
refGene_gn=TMEM171;refGene_tx=XM_003981071.4;refGene_gn=TMEM171;refGene_tx=XM_019835400.2
A1 140095777 140096403
refGene_gn=TMEM174;refGene_tx=XM_003981057.5
A1 140096593 140096699
refGene_gn=TMEM174;refGene_tx=XM_003981057.5
A1 140349892 140350072
refGene_gn=FOXD1;refGene_tx=XM_023257363.1
A1 140350074 140351334
refGene_gn=FOXD1;refGene_tx=XM_023257363.1
A1 140398784 140398916
refGene_gn=BTF3;refGene_tx=XM_023257367.1
A1 140399252 140399321
refGene_gn=BTF3;refGene_tx=XM_023257367.1;refGene_gn=BTF3;refGene_tx=XM_023257369.1
A1 140402393 140402507
refGene_gn=BTF3;refGene_tx=XM_023257367.1;refGene_gn=BTF3;refGene_tx=XM_023257369.1
A1 140402992 140403194
refGene_gn=BTF3;refGene_tx=XM_023257367.1;refGene_gn=BTF3;refGene_tx=XM_023257369.1
A1 140404698 140404755
refGene_gn=BTF3;refGene_tx=XM_023257367.1;refGene_gn=BTF3;refGene_tx=XM_023257369.1
A1 140405261 140405308
refGene_gn=BTF3;refGene_tx=XM_023257367.1;refGene_gn=BTF3;refGene_tx=XM_023257369.1
A1 140432728 140432784
refGene_gn=ANKRA2;refGene_tx=XM_011283822.3
A1 140434583 140434664
refGene_gn=ANKRA2;refGene_tx=XM_011283822.3;refGene_gn=ANKRA2;refGene_tx=XM_019835462.2
A1 140435185 140435252
refGene_gn=ANKRA2;refGene_tx=XM_011283822.3
A1 140435548 140435674
refGene_gn=ANKRA2;refGene_tx=XM_011283822.3;refGene_gn=ANKRA2;refGene_tx=XM_019835462.2
A1 140436137 140436235
refGene_gn=ANKRA2;refGene_tx=XM_011283822.3;refGene_gn=ANKRA2;refGene_tx=XM_019835462.2
A1 140438676 140438742
refGene_gn=ANKRA2;refGene_tx=XM_011283822.3;refGene_gn=ANKRA2;refGene_tx=XM_019835462.2
A1 140441102 140441261
refGene_gn=ANKRA2;refGene_tx=XM_011283822.3;refGene_gn=ANKRA2;refGene_tx=XM_019835462.2

A1 140441935 140442224
refGene_gn=ANKRA2;refGene_tx=XM_011283822.3;refGene_gn=ANKRA2;refGene_tx=XM_019835462.2
A1 140448330 140448420
refGene_gn=UTP15;refGene_tx=XM_003981061.4
A1 140449880 140449973
refGene_gn=UTP15;refGene_tx=XM_003981061.4
A1 140450069 140450254
refGene_gn=UTP15;refGene_tx=XM_003981061.4
A1 140451708 140451887
refGene_gn=UTP15;refGene_tx=XM_003981061.4
A1 140452468 140452594
refGene_gn=UTP15;refGene_tx=XM_003981061.4
A1 140454208 140454344
refGene_gn=UTP15;refGene_tx=XM_003981061.4
A1 140456220 140456305
refGene_gn=UTP15;refGene_tx=XM_003981061.4
A1 140457167 140457317
refGene_gn=UTP15;refGene_tx=XM_003981061.4
A1 140458265 140458367
refGene_gn=UTP15;refGene_tx=XM_003981061.4
A1 140458789 140458923
refGene_gn=UTP15;refGene_tx=XM_003981061.4
A1 140459100 140459159
refGene_gn=UTP15;refGene_tx=XM_003981061.4
A1 140459639 140459857
refGene_gn=UTP15;refGene_tx=XM_003981061.4
A1 140551149 140551182
refGene_gn=ARHGEF28;refGene_tx=XM_011283815.3
A1 140618801 140618949
refGene_gn=ARHGEF28;refGene_tx=XM_011283815.3
A1 140621838 140622132
refGene_gn=ARHGEF28;refGene_tx=XM_011283815.3
A1 140641588 140641772
refGene_gn=ARHGEF28;refGene_tx=XM_011283815.3
A1 140644517 140644698
refGene_gn=ARHGEF28;refGene_tx=XM_011283815.3
A1 140649432 140649505
refGene_gn=ARHGEF28;refGene_tx=XM_011283815.3
A1 140661418 140661474
refGene_gn=ARHGEF28;refGene_tx=XM_011283815.3
A1 140662039 140662100
refGene_gn=ARHGEF28;refGene_tx=XM_011283815.3
A1 140682221 140682306
refGene_gn=ARHGEF28;refGene_tx=XM_019835471.2
A1 140748317 140748439
refGene_gn=ARHGEF28;refGene_tx=XM_011283815.3;refGene_gn=ARHGEF28;refGene_tx=XM_019835471.2
A1 140752421 140752702
refGene_gn=ARHGEF28;refGene_tx=XM_011283815.3;refGene_gn=ARHGEF28;refG

ene_tx=XM_019835471.2
A1 140756020 140756228
refGene_gn=ARHGEF28;refGene_tx=XM_011283815.3;refGene_gn=ARHGEF28;refGene_tx=XM_019835471.2
A1 140758956 140759068
refGene_gn=ARHGEF28;refGene_tx=XM_011283815.3;refGene_gn=ARHGEF28;refGene_tx=XM_019835471.2
A1 140762784 140762827
refGene_gn=ARHGEF28;refGene_tx=XM_011283815.3;refGene_gn=ARHGEF28;refGene_tx=XM_019835471.2
A1 140768550 140768674
refGene_gn=ARHGEF28;refGene_tx=XM_011283815.3;refGene_gn=ARHGEF28;refGene_tx=XM_019835471.2
A1 140769019 140769152
refGene_gn=ARHGEF28;refGene_tx=XM_011283815.3;refGene_gn=ARHGEF28;refGene_tx=XM_019835471.2
A1 140774479 140774535
refGene_gn=ARHGEF28;refGene_tx=XM_011283815.3;refGene_gn=ARHGEF28;refGene_tx=XM_019835471.2
A1 140775575 140775624
refGene_gn=ARHGEF28;refGene_tx=XM_011283815.3;refGene_gn=ARHGEF28;refGene_tx=XM_019835471.2
A1 140777683 140777828
refGene_gn=ARHGEF28;refGene_tx=XM_011283815.3;refGene_gn=ARHGEF28;refGene_tx=XM_019835471.2
A1 140777908 140778036
refGene_gn=ARHGEF28;refGene_tx=XM_011283815.3;refGene_gn=ARHGEF28;refGene_tx=XM_019835471.2
A1 140779742 140779883
refGene_gn=ARHGEF28;refGene_tx=XM_011283815.3;refGene_gn=ARHGEF28;refGene_tx=XM_019835471.2
A1 140782654 140782902
refGene_gn=ARHGEF28;refGene_tx=XM_011283815.3;refGene_gn=ARHGEF28;refGene_tx=XM_019835471.2
A1 140788543 140788666
refGene_gn=ARHGEF28;refGene_tx=XM_011283815.3;refGene_gn=ARHGEF28;refGene_tx=XM_019835471.2
A1 140789883 140790001
refGene_gn=ARHGEF28;refGene_tx=XM_011283815.3;refGene_gn=ARHGEF28;refGene_tx=XM_019835471.2
A1 140793003 140793258
refGene_gn=ARHGEF28;refGene_tx=XM_011283815.3;refGene_gn=ARHGEF28;refGene_tx=XM_019835471.2
A1 140794814 140794891
refGene_gn=ARHGEF28;refGene_tx=XM_011283815.3;refGene_gn=ARHGEF28;refGene_tx=XM_019835471.2
A1 140799992 140800171
refGene_gn=ARHGEF28;refGene_tx=XM_011283815.3;refGene_gn=ARHGEF28;refGene_tx=XM_019835471.2
A1 140801197 140801289

refGene_gn=ARHGEF28;refGene_tx=XM_011283815.3;refGene_gn=ARHGEF28;refGene_tx=XM_019835471.2
A1 140802407 140802590
refGene_gn=ARHGEF28;refGene_tx=XM_011283815.3;refGene_gn=ARHGEF28;refGene_tx=XM_019835471.2
A1 140807256 140807388
refGene_gn=ARHGEF28;refGene_tx=XM_011283815.3;refGene_gn=ARHGEF28;refGene_tx=XM_019835471.2
A1 140810309 140810410
refGene_gn=ARHGEF28;refGene_tx=XM_011283815.3;refGene_gn=ARHGEF28;refGene_tx=XM_019835471.2
A1 140813383 140813422
refGene_gn=ARHGEF28;refGene_tx=XM_011283815.3;refGene_gn=ARHGEF28;refGene_tx=XM_019835471.2
A1 140813520 140813568
refGene_gn=ARHGEF28;refGene_tx=XM_011283815.3;refGene_gn=ARHGEF28;refGene_tx=XM_019835471.2
A1 140818390 140818879
refGene_gn=ARHGEF28;refGene_tx=XM_011283815.3;refGene_gn=ARHGEF28;refGene_tx=XM_019835471.2
A1 140820190 140820491
refGene_gn=ARHGEF28;refGene_tx=XM_011283815.3;refGene_gn=ARHGEF28;refGene_tx=XM_019835471.2
A1 140848612 140848776
refGene_gn=ARHGEF28;refGene_tx=XM_011283815.3;refGene_gn=ARHGEF28;refGene_tx=XM_019835471.2
A1 141488172 141489942
refGene_gn=ENC1;refGene_tx=XM_006927790.4
A1 141506097 141506227
refGene_gn=HEXB;refGene_tx=XM_023238898.1
A1 141506564 141506666
refGene_gn=HEXB;refGene_tx=XM_023238898.1
A1 141539513 141540096
refGene_gn=HEXB;refGene_tx=NM_001009333.2;refGene_gn=HEXB;refGene_tx=XM_023238898.1
A1 141544536 141544682
refGene_gn=HEXB;refGene_tx=NM_001009333.2;refGene_gn=HEXB;refGene_tx=XM_023238898.1
A1 141548014 141548080
refGene_gn=HEXB;refGene_tx=NM_001009333.2;refGene_gn=HEXB;refGene_tx=XM_023238898.1
A1 141555109 141555156
refGene_gn=HEXB;refGene_tx=NM_001009333.2;refGene_gn=HEXB;refGene_tx=XM_023238898.1
A1 141555406 141555517
refGene_gn=HEXB;refGene_tx=NM_001009333.2;refGene_gn=HEXB;refGene_tx=XM_023238898.1
A1 141563187 141563289
refGene_gn=HEXB;refGene_tx=NM_001009333.2;refGene_gn=HEXB;refGene_tx=XM_023238898.1

A1 141565278 141565408
refGene_gn=HEXB;refGene_tx=NM_001009333.2;refGene_gn=HEXB;refGene_tx=XM_023238898.1
A1 141566813 141566994
refGene_gn=HEXB;refGene_tx=NM_001009333.2;refGene_gn=HEXB;refGene_tx=XM_023238898.1
A1 141567651 141567738
refGene_gn=HEXB;refGene_tx=NM_001009333.2;refGene_gn=HEXB;refGene_tx=XM_023238898.1
A1 141568872 141568945
refGene_gn=HEXB;refGene_tx=NM_001009333.2;refGene_gn=HEXB;refGene_tx=XM_023238898.1
A1 141569904 141570079
refGene_gn=HEXB;refGene_tx=NM_001009333.2;refGene_gn=HEXB;refGene_tx=XM_023238898.1
A1 141571037 141571128
refGene_gn=HEXB;refGene_tx=NM_001009333.2;refGene_gn=HEXB;refGene_tx=XM_023238898.1
A1 141571260 141571365
refGene_gn=HEXB;refGene_tx=NM_001009333.2;refGene_gn=HEXB;refGene_tx=XM_023238898.1
A1 141571760 141571821
refGene_gn=HEXB;refGene_tx=NM_001009333.2;refGene_gn=HEXB;refGene_tx=XM_023238898.1
A1 141572076 141572214
refGene_gn=GFM2;refGene_tx=XM_019835513.2;refGene_gn=GFM2;refGene_tx=XM_023257405.1;refGene_gn=GFM2;refGene_tx=XM_023257412.1
A1 141572588 141572771
refGene_gn=GFM2;refGene_tx=XM_019835513.2;refGene_gn=GFM2;refGene_tx=XM_023257405.1;refGene_gn=GFM2;refGene_tx=XM_023257412.1
A1 141576391 141576507
refGene_gn=GFM2;refGene_tx=XM_019835513.2;refGene_gn=GFM2;refGene_tx=XM_023257405.1;refGene_gn=GFM2;refGene_tx=XM_023257412.1
A1 141577829 141578009
refGene_gn=GFM2;refGene_tx=XM_019835513.2;refGene_gn=GFM2;refGene_tx=XM_023257405.1;refGene_gn=GFM2;refGene_tx=XM_023257412.1
A1 141579680 141579819
refGene_gn=GFM2;refGene_tx=XM_019835513.2;refGene_gn=GFM2;refGene_tx=XM_023257405.1;refGene_gn=GFM2;refGene_tx=XM_023257412.1
A1 141582249 141582326
refGene_gn=GFM2;refGene_tx=XM_019835513.2;refGene_gn=GFM2;refGene_tx=XM_023257405.1;refGene_gn=GFM2;refGene_tx=XM_023257412.1
A1 141586378 141586568
refGene_gn=GFM2;refGene_tx=XM_019835513.2;refGene_gn=GFM2;refGene_tx=XM_023257405.1;refGene_gn=GFM2;refGene_tx=XM_023257412.1
A1 141587982 141588082
refGene_gn=GFM2;refGene_tx=XM_019835513.2;refGene_gn=GFM2;refGene_tx=XM_023257405.1;refGene_gn=GFM2;refGene_tx=XM_023257412.1
A1 141588386 141588527
refGene_gn=GFM2;refGene_tx=XM_019835513.2;refGene_gn=GFM2;refGene_tx=X

M_023257412.1
A1 141589876 141590025
refGene_gn=GFM2;refGene_tx=XM_019835513.2;refGene_gn=GFM2;refGene_tx=XM_023257405.1;refGene_gn=GFM2;refGene_tx=XM_023257412.1
A1 141591861 141591942
refGene_gn=GFM2;refGene_tx=XM_019835513.2;refGene_gn=GFM2;refGene_tx=XM_023257405.1;refGene_gn=GFM2;refGene_tx=XM_023257412.1
A1 141596165 141596345
refGene_gn=GFM2;refGene_tx=XM_019835513.2;refGene_gn=GFM2;refGene_tx=XM_023257405.1;refGene_gn=GFM2;refGene_tx=XM_023257412.1
A1 141596594 141596655
refGene_gn=GFM2;refGene_tx=XM_019835513.2;refGene_gn=GFM2;refGene_tx=XM_023257405.1;refGene_gn=GFM2;refGene_tx=XM_023257412.1
A1 141598001 141598090
refGene_gn=GFM2;refGene_tx=XM_019835513.2;refGene_gn=GFM2;refGene_tx=XM_023257405.1;refGene_gn=GFM2;refGene_tx=XM_023257412.1
A1 141603568 141603657
refGene_gn=GFM2;refGene_tx=XM_019835513.2;refGene_gn=GFM2;refGene_tx=XM_023257405.1;refGene_gn=GFM2;refGene_tx=XM_023257412.1
A1 141604580 141604706
refGene_gn=GFM2;refGene_tx=XM_019835513.2;refGene_gn=GFM2;refGene_tx=XM_023257405.1;refGene_gn=GFM2;refGene_tx=XM_023257412.1
A1 141617683 141617781
refGene_gn=GFM2;refGene_tx=XM_019835513.2;refGene_gn=GFM2;refGene_tx=XM_023257405.1
A1 141618206 141618264
refGene_gn=GFM2;refGene_tx=XM_019835513.2;refGene_gn=GFM2;refGene_tx=XM_023257405.1
A1 141619880 141619965
refGene_gn=GFM2;refGene_tx=XM_019835513.2;refGene_gn=GFM2;refGene_tx=XM_023257405.1
A1 141621300 141621363
refGene_gn=GFM2;refGene_tx=XM_019835513.2;refGene_gn=GFM2;refGene_tx=XM_023257405.1
A1 141632428 141632431
refGene_gn=NSA2;refGene_tx=XM_003981073.5
A1 141633881 141634069
refGene_gn=NSA2;refGene_tx=XM_003981073.5
A1 141634161 141634312
refGene_gn=NSA2;refGene_tx=XM_003981073.5
A1 141635826 141636006
refGene_gn=NSA2;refGene_tx=XM_003981073.5
A1 141638171 141638364
refGene_gn=NSA2;refGene_tx=XM_003981073.5
A1 141639241 141639309
refGene_gn=NSA2;refGene_tx=XM_003981073.5
A1 141643862 141644411
refGene_gn=FAM169A;refGene_tx=XM_003981074.5;refGene_gn=FAM169A;refGene_tx=XM_019835558.2;refGene_gn=FAM169A;refGene_tx=XM_019835567.2;refGene_gn=FAM169A;refGene_tx=XM_019835576.2;refGene_gn=FAM169A;refGene_tx=

XM_023257417.1;refGene_gn=FAM169A;refGene_tx=XM_023257424.1;refGene_gn=FAM169A;refGene_tx=XM_023257441.1;refGene_gn=FAM169A;refGene_tx=XM_023257449.1;refGene_gn=FAM169A;refGene_tx=XM_023257456.1
A1 141645532 141645736
refGene_gn=FAM169A;refGene_tx=XM_003981074.5;refGene_gn=FAM169A;refGene_tx=XM_019835558.2;refGene_gn=FAM169A;refGene_tx=XM_019835567.2;refGene_gn=FAM169A;refGene_tx=XM_019835576.2;refGene_gn=FAM169A;refGene_tx=XM_023257417.1;refGene_gn=FAM169A;refGene_tx=XM_023257424.1;refGene_gn=FAM169A;refGene_tx=XM_023257441.1;refGene_gn=FAM169A;refGene_tx=XM_023257449.1;refGene_gn=FAM169A;refGene_tx=XM_023257456.1
A1 141649293 141649450
refGene_gn=FAM169A;refGene_tx=XM_003981074.5;refGene_gn=FAM169A;refGene_tx=XM_019835558.2;refGene_gn=FAM169A;refGene_tx=XM_019835567.2;refGene_gn=FAM169A;refGene_tx=XM_019835576.2;refGene_gn=FAM169A;refGene_tx=XM_023257417.1;refGene_gn=FAM169A;refGene_tx=XM_023257424.1;refGene_gn=FAM169A;refGene_tx=XM_023257441.1;refGene_gn=FAM169A;refGene_tx=XM_023257449.1;refGene_gn=FAM169A;refGene_tx=XM_023257456.1
A1 141653585 141653736
refGene_gn=FAM169A;refGene_tx=XM_003981074.5;refGene_gn=FAM169A;refGene_tx=XM_019835558.2;refGene_gn=FAM169A;refGene_tx=XM_019835567.2;refGene_gn=FAM169A;refGene_tx=XM_019835576.2;refGene_gn=FAM169A;refGene_tx=XM_023257417.1;refGene_gn=FAM169A;refGene_tx=XM_023257424.1;refGene_gn=FAM169A;refGene_tx=XM_023257441.1;refGene_gn=FAM169A;refGene_tx=XM_023257449.1;refGene_gn=FAM169A;refGene_tx=XM_023257456.1
A1 141654522 141654562
refGene_gn=FAM169A;refGene_tx=XM_003981074.5;refGene_gn=FAM169A;refGene_tx=XM_019835558.2;refGene_gn=FAM169A;refGene_tx=XM_019835567.2;refGene_gn=FAM169A;refGene_tx=XM_019835576.2;refGene_gn=FAM169A;refGene_tx=XM_023257417.1;refGene_gn=FAM169A;refGene_tx=XM_023257424.1;refGene_gn=FAM169A;refGene_tx=XM_023257441.1;refGene_gn=FAM169A;refGene_tx=XM_023257449.1;refGene_gn=FAM169A;refGene_tx=XM_023257456.1
A1 141657969 141658082
refGene_gn=FAM169A;refGene_tx=XM_003981074.5;refGene_gn=FAM169A;refGene_tx=XM_019835558.2;refGene_gn=FAM169A;refGene_tx=XM_019835567.2;refGene_gn=FAM169A;refGene_tx=XM_019835576.2;refGene_gn=FAM169A;refGene_tx=XM_023257417.1;refGene_gn=FAM169A;refGene_tx=XM_023257424.1;refGene_gn=FAM169A;refGene_tx=XM_023257441.1;refGene_gn=FAM169A;refGene_tx=XM_023257449.1;refGene_gn=FAM169A;refGene_tx=XM_023257456.1
A1 141658630 141658759
refGene_gn=FAM169A;refGene_tx=XM_003981074.5;refGene_gn=FAM169A;refGene_tx=XM_019835558.2;refGene_gn=FAM169A;refGene_tx=XM_019835567.2;refGene_gn=FAM169A;refGene_tx=XM_019835576.2;refGene_gn=FAM169A;refGene_tx=XM_023257417.1;refGene_gn=FAM169A;refGene_tx=XM_023257424.1;refGene_gn=FAM169A;refGene_tx=XM_023257441.1;refGene_gn=FAM169A;refGene_tx=XM_023257449.1;refGene_gn=FAM169A;refGene_tx=XM_023257456.1
A1 141670553 141670733
refGene_gn=FAM169A;refGene_tx=XM_003981074.5;refGene_gn=FAM169A;refGene_tx=XM_019835558.2;refGene_gn=FAM169A;refGene_tx=XM_019835567.2;refGene_gn=FAM169A;refGene_tx=XM_019835576.2;refGene_gn=FAM169A;refGene_tx=XM_023257417.1;refGene_gn=FAM169A;refGene_tx=XM_023257424.1;refGene_gn

=FAM169A;refGene_tx=XM_023257441.1;refGene_gn=FAM169A;refGene_tx=XM_023257449.1;refGene_gn=FAM169A;refGene_tx=XM_023257456.1
A1	141690023	141690195
refGene_gn=FAM169A;refGene_tx=XM_003981074.5;refGene_gn=FAM169A;refGene_tx=XM_019835558.2;refGene_gn=FAM169A;refGene_tx=XM_019835567.2;refGene_gn=FAM169A;refGene_tx=XM_019835576.2;refGene_gn=FAM169A;refGene_tx=XM_023257417.1;refGene_gn=FAM169A;refGene_tx=XM_023257424.1;refGene_gn=FAM169A;refGene_tx=XM_023257441.1;refGene_gn=FAM169A;refGene_tx=XM_023257449.1;refGene_gn=FAM169A;refGene_tx=XM_023257456.1
A1	141692288	141692374
refGene_gn=FAM169A;refGene_tx=XM_003981074.5;refGene_gn=FAM169A;refGene_tx=XM_019835558.2;refGene_gn=FAM169A;refGene_tx=XM_019835567.2;refGene_gn=FAM169A;refGene_tx=XM_019835576.2;refGene_gn=FAM169A;refGene_tx=XM_023257417.1;refGene_gn=FAM169A;refGene_tx=XM_023257424.1;refGene_gn=FAM169A;refGene_tx=XM_023257441.1;refGene_gn=FAM169A;refGene_tx=XM_023257449.1
A1	141692523	141692623
refGene_gn=FAM169A;refGene_tx=XM_003981074.5;refGene_gn=FAM169A;refGene_tx=XM_019835558.2;refGene_gn=FAM169A;refGene_tx=XM_019835567.2;refGene_gn=FAM169A;refGene_tx=XM_019835576.2;refGene_gn=FAM169A;refGene_tx=XM_023257417.1;refGene_gn=FAM169A;refGene_tx=XM_023257424.1;refGene_gn=FAM169A;refGene_tx=XM_023257441.1;refGene_gn=FAM169A;refGene_tx=XM_023257449.1
A1	141693681	141693813
refGene_gn=FAM169A;refGene_tx=XM_003981074.5;refGene_gn=FAM169A;refGene_tx=XM_019835558.2;refGene_gn=FAM169A;refGene_tx=XM_019835567.2;refGene_gn=FAM169A;refGene_tx=XM_019835576.2;refGene_gn=FAM169A;refGene_tx=XM_023257417.1;refGene_gn=FAM169A;refGene_tx=XM_023257424.1;refGene_gn=FAM169A;refGene_tx=XM_023257441.1;refGene_gn=FAM169A;refGene_tx=XM_023257449.1
A1	141862354	141863719
refGene_gn=GCNT4;refGene_tx=XM_003981090.5;refGene_gn=GCNT4;refGene_tx=XM_019835587.2;refGene_gn=GCNT4;refGene_tx=XM_019835593.2;refGene_gn=GCNT4;refGene_tx=XM_023257501.1
A1	141902174	141902320
refGene_gn=ANKRD31;refGene_tx=XM_023257507.1
A1	141909045	141909148
refGene_gn=ANKRD31;refGene_tx=XM_023257507.1
A1	141915389	141915530
refGene_gn=ANKRD31;refGene_tx=XM_023257507.1
A1	141932321	141933324
refGene_gn=ANKRD31;refGene_tx=XM_023257507.1
A1	141936218	141936315
refGene_gn=ANKRD31;refGene_tx=XM_023257507.1
A1	141941126	141941214
refGene_gn=ANKRD31;refGene_tx=XM_023257507.1
A1	141946422	141946538
refGene_gn=ANKRD31;refGene_tx=XM_023257507.1
A1	141948220	141948383
refGene_gn=ANKRD31;refGene_tx=XM_023257507.1

A1 141956749 141956892
refGene_gn=ANKRD31;refGene_tx=XM_023257507.1
A1 141958203 141958341
refGene_gn=ANKRD31;refGene_tx=XM_023257507.1
A1 141962469 141962634
refGene_gn=ANKRD31;refGene_tx=XM_023257507.1
A1 141963263 141964743
refGene_gn=ANKRD31;refGene_tx=XM_023257507.1
A1 141966148 141966201
refGene_gn=ANKRD31;refGene_tx=XM_023257507.1
A1 141970520 141970665
refGene_gn=ANKRD31;refGene_tx=XM_023257507.1
A1 141979852 141979995
refGene_gn=ANKRD31;refGene_tx=XM_023257507.1
A1 141987280 141987300
refGene_gn=ANKRD31;refGene_tx=XM_023257507.1
A1 141987301 141987428
refGene_gn=ANKRD31;refGene_tx=XM_023257507.1
A1 141998328 141998438
refGene_gn=ANKRD31;refGene_tx=XM_023257507.1
A1 142001357 142001632
refGene_gn=ANKRD31;refGene_tx=XM_023257507.1
A1 142013224 142013268
refGene_gn=ANKRD31;refGene_tx=XM_023257507.1
A1 142015411 142015488
refGene_gn=ANKRD31;refGene_tx=XM_023257507.1
A1 142020081 142020122
refGene_gn=ANKRD31;refGene_tx=XM_023257507.1
A1 142021265 142021372
refGene_gn=ANKRD31;refGene_tx=XM_023257507.1
A1 142025789 142025863
refGene_gn=ANKRD31;refGene_tx=XM_023257507.1
A1 142040042 142040146
refGene_gn=ANKRD31;refGene_tx=XM_023257507.1
A1 142091436 142091601
refGene_gn=HMGCR;refGene_tx=XM_003981075.5;refGene_gn=HMGCR;refGene_tx
=XM_019835641.2
A1 142092073 142092185
refGene_gn=HMGCR;refGene_tx=XM_003981075.5;refGene_gn=HMGCR;refGene_tx
=XM_019835641.2
A1 142092461 142092549
refGene_gn=HMGCR;refGene_tx=XM_003981075.5;refGene_gn=HMGCR;refGene_tx
=XM_019835641.2
A1 142093679 142093764
refGene_gn=HMGCR;refGene_tx=XM_003981075.5;refGene_gn=HMGCR;refGene_tx
=XM_019835641.2
A1 142094642 142094748
refGene_gn=HMGCR;refGene_tx=XM_003981075.5;refGene_gn=HMGCR;refGene_tx
=XM_019835641.2
A1 142097206 142097313

refGene_gn=HMGCR;refGene_tx=XM_003981075.5;refGene_gn=HMGCR;refGene_tx=XM_019835641.2
A1 142097419 142097536
refGene_gn=HMGCR;refGene_tx=XM_003981075.5;refGene_gn=HMGCR;refGene_tx=XM_019835641.2
A1 142097960 142098121
refGene_gn=HMGCR;refGene_tx=XM_003981075.5;refGene_gn=HMGCR;refGene_tx=XM_019835641.2
A1 142098241 142098489
refGene_gn=HMGCR;refGene_tx=XM_003981075.5;refGene_gn=HMGCR;refGene_tx=XM_019835641.2
A1 142098604 142098783
refGene_gn=HMGCR;refGene_tx=XM_003981075.5;refGene_gn=HMGCR;refGene_tx=XM_019835641.2
A1 142099792 142099987
refGene_gn=HMGCR;refGene_tx=XM_003981075.5;refGene_gn=HMGCR;refGene_tx=XM_019835641.2
A1 142100575 142100734
refGene_gn=HMGCR;refGene_tx=XM_003981075.5;refGene_gn=HMGCR;refGene_tx=XM_019835641.2
A1 142100858 142101016
refGene_gn=HMGCR;refGene_tx=XM_003981075.5;refGene_gn=HMGCR;refGene_tx=XM_019835641.2
A1 142102379 142102485
refGene_gn=HMGCR;refGene_tx=XM_003981075.5;refGene_gn=HMGCR;refGene_tx=XM_019835641.2
A1 142104503 142104674
refGene_gn=HMGCR;refGene_tx=XM_003981075.5;refGene_gn=HMGCR;refGene_tx=XM_019835641.2
A1 142104984 142105125
refGene_gn=HMGCR;refGene_tx=XM_003981075.5;refGene_gn=HMGCR;refGene_tx=XM_019835641.2
A1 142105209 142105368
refGene_gn=HMGCR;refGene_tx=XM_003981075.5;refGene_gn=HMGCR;refGene_tx=XM_019835641.2
A1 142105805 142105960
refGene_gn=HMGCR;refGene_tx=XM_003981075.5;refGene_gn=HMGCR;refGene_tx=XM_019835641.2
A1 142106105 142106160
refGene_gn=HMGCR;refGene_tx=XM_003981075.5;refGene_gn=HMGCR;refGene_tx=XM_019835641.2
A1 142122466 142122594
refGene_gn=COL4A3BP;refGene_tx=XM_019835672.2;refGene_gn=COL4A3BP;refGene_tx=XM_019835677.2
A1 142123750 142123880
refGene_gn=COL4A3BP;refGene_tx=XM_019835672.2;refGene_gn=COL4A3BP;refGene_tx=XM_019835677.2
A1 142124434 142124563
refGene_gn=COL4A3BP;refGene_tx=XM_019835672.2;refGene_gn=COL4A3BP;refGene_tx=XM_019835677.2

A1 142127677 142127748
refGene_gn=COL4A3BP;refGene_tx=XM_019835672.2;refGene_gn=COL4A3BP;refGene_tx=XM_019835677.2
A1 142128674 142128807
refGene_gn=COL4A3BP;refGene_tx=XM_019835672.2;refGene_gn=COL4A3BP;refGene_tx=XM_019835677.2
A1 142131532 142131628
refGene_gn=COL4A3BP;refGene_tx=XM_019835672.2;refGene_gn=COL4A3BP;refGene_tx=XM_019835677.2
A1 142139298 142139376
refGene_gn=COL4A3BP;refGene_tx=XM_019835677.2
A1 142139904 142139997
refGene_gn=COL4A3BP;refGene_tx=XM_019835672.2;refGene_gn=COL4A3BP;refGene_tx=XM_019835677.2
A1 142142405 142142492
refGene_gn=COL4A3BP;refGene_tx=XM_019835672.2;refGene_gn=COL4A3BP;refGene_tx=XM_019835677.2
A1 142147004 142147097
refGene_gn=COL4A3BP;refGene_tx=XM_019835672.2;refGene_gn=COL4A3BP;refGene_tx=XM_019835677.2
A1 142151968 142152126
refGene_gn=COL4A3BP;refGene_tx=XM_019835672.2;refGene_gn=COL4A3BP;refGene_tx=XM_019835677.2
A1 142157124 142157208
refGene_gn=COL4A3BP;refGene_tx=XM_019835672.2;refGene_gn=COL4A3BP;refGene_tx=XM_019835677.2
A1 142162596 142162735
refGene_gn=COL4A3BP;refGene_tx=XM_019835672.2;refGene_gn=COL4A3BP;refGene_tx=XM_019835677.2
A1 142163511 142163619
refGene_gn=COL4A3BP;refGene_tx=XM_019835672.2;refGene_gn=COL4A3BP;refGene_tx=XM_019835677.2
A1 142183254 142183371
refGene_gn=COL4A3BP;refGene_tx=XM_019835672.2;refGene_gn=COL4A3BP;refGene_tx=XM_019835677.2
A1 142214904 142215039
refGene_gn=COL4A3BP;refGene_tx=XM_019835672.2;refGene_gn=COL4A3BP;refGene_tx=XM_019835677.2
A1 142220665 142221071
refGene_gn=COL4A3BP;refGene_tx=XM_019835672.2;refGene_gn=COL4A3BP;refGene_tx=XM_019835677.2
A1 142221298 142221381
refGene_gn=COL4A3BP;refGene_tx=XM_019835672.2;refGene_gn=COL4A3BP;refGene_tx=XM_019835677.2
A1 142247840 142247975
refGene_gn=POLK;refGene_tx=XM_006927792.4;refGene_gn=POLK;refGene_tx=XM_006927793.4;refGene_gn=POLK;refGene_tx=XM_019835644.2;refGene_gn=POLK;refGene_tx=XM_019835651.2;refGene_gn=POLK;refGene_tx=XM_019835655.2;refGene_gn=POLK;refGene_tx=XM_019835662.2;refGene_gn=POLK;refGene_tx=XM_019835667.2;refGene_gn=POLK;refGene_tx=XM_019835670.2

A1 142254510 142254630
refGene_gn=POLK;refGene_tx=XM_006927792.4;refGene_gn=POLK;refGene_tx=XM_006927793.4;refGene_gn=POLK;refGene_tx=XM_019835644.2;refGene_gn=POLK;refGene_tx=XM_019835651.2;refGene_gn=POLK;refGene_tx=XM_019835655.2;refGene_gn=POLK;refGene_tx=XM_019835667.2;refGene_gn=POLK;refGene_tx=XM_019835670.2
A1 142261746 142261899
refGene_gn=POLK;refGene_tx=XM_006927792.4;refGene_gn=POLK;refGene_tx=XM_006927793.4;refGene_gn=POLK;refGene_tx=XM_019835644.2;refGene_gn=POLK;refGene_tx=XM_019835651.2;refGene_gn=POLK;refGene_tx=XM_019835655.2;refGene_gn=POLK;refGene_tx=XM_019835662.2;refGene_gn=POLK;refGene_tx=XM_019835670.2
A1 142266628 142266760
refGene_gn=POLK;refGene_tx=XM_006927792.4;refGene_gn=POLK;refGene_tx=XM_006927793.4;refGene_gn=POLK;refGene_tx=XM_019835644.2;refGene_gn=POLK;refGene_tx=XM_019835651.2;refGene_gn=POLK;refGene_tx=XM_019835655.2;refGene_gn=POLK;refGene_tx=XM_019835662.2;refGene_gn=POLK;refGene_tx=XM_019835667.2;refGene_gn=POLK;refGene_tx=XM_019835670.2
A1 142270077 142270231
refGene_gn=POLK;refGene_tx=XM_006927792.4;refGene_gn=POLK;refGene_tx=XM_006927793.4;refGene_gn=POLK;refGene_tx=XM_019835644.2;refGene_gn=POLK;refGene_tx=XM_019835651.2;refGene_gn=POLK;refGene_tx=XM_019835655.2;refGene_gn=POLK;refGene_tx=XM_019835662.2;refGene_gn=POLK;refGene_tx=XM_019835667.2;refGene_gn=POLK;refGene_tx=XM_019835670.2
A1 142274934 142275174
refGene_gn=POLK;refGene_tx=XM_006927792.4;refGene_gn=POLK;refGene_tx=XM_006927793.4;refGene_gn=POLK;refGene_tx=XM_019835644.2;refGene_gn=POLK;refGene_tx=XM_019835651.2;refGene_gn=POLK;refGene_tx=XM_019835655.2;refGene_gn=POLK;refGene_tx=XM_019835662.2;refGene_gn=POLK;refGene_tx=XM_019835667.2;refGene_gn=POLK;refGene_tx=XM_019835670.2
A1 142276797 142276922
refGene_gn=POLK;refGene_tx=XM_006927792.4;refGene_gn=POLK;refGene_tx=XM_006927793.4;refGene_gn=POLK;refGene_tx=XM_019835644.2;refGene_gn=POLK;refGene_tx=XM_019835651.2;refGene_gn=POLK;refGene_tx=XM_019835655.2;refGene_gn=POLK;refGene_tx=XM_019835662.2;refGene_gn=POLK;refGene_tx=XM_019835667.2;refGene_gn=POLK;refGene_tx=XM_019835670.2
A1 142278099 142278266
refGene_gn=POLK;refGene_tx=XM_006927792.4;refGene_gn=POLK;refGene_tx=XM_006927793.4;refGene_gn=POLK;refGene_tx=XM_019835644.2;refGene_gn=POLK;refGene_tx=XM_019835651.2;refGene_gn=POLK;refGene_tx=XM_019835655.2;refGene_gn=POLK;refGene_tx=XM_019835662.2;refGene_gn=POLK;refGene_tx=XM_019835667.2;refGene_gn=POLK;refGene_tx=XM_019835670.2
A1 142280557 142280590
refGene_gn=POLK;refGene_tx=XM_006927792.4;refGene_gn=POLK;refGene_tx=XM_006927793.4;refGene_gn=POLK;refGene_tx=XM_019835644.2;refGene_gn=POLK;refGene_tx=XM_019835651.2;refGene_gn=POLK;refGene_tx=XM_019835655.2;refGene_gn=POLK;refGene_tx=XM_019835662.2;refGene_gn=POLK;refGene_tx=XM_019835667.2;refGene_gn=POLK;refGene_tx=XM_019835670.2
A1 142287352 142287449
refGene_gn=POLK;refGene_tx=XM_006927792.4;refGene_gn=POLK;refGene_tx=X

M_006927793.4;refGene_gn=POLK;refGene_tx=XM_019835644.2;refGene_gn=POLK;refGene_tx=XM_019835651.2;refGene_gn=POLK;refGene_tx=XM_019835655.2;refGene_gn=POLK;refGene_tx=XM_019835662.2;refGene_gn=POLK;refGene_tx=XM_019835667.2;refGene_gn=POLK;refGene_tx=XM_019835670.2
A1 142289276 142289448 refGene_gn=POLK;refGene_tx=XM_006927792.4;refGene_gn=POLK;refGene_tx=XM_006927793.4;refGene_gn=POLK;refGene_tx=XM_019835644.2;refGene_gn=POLK;refGene_tx=XM_019835651.2;refGene_gn=POLK;refGene_tx=XM_019835655.2;refGene_gn=POLK;refGene_tx=XM_019835662.2;refGene_gn=POLK;refGene_tx=XM_019835667.2;refGene_gn=POLK;refGene_tx=XM_019835670.2
A1 142291292 142292237 refGene_gn=POLK;refGene_tx=XM_006927792.4;refGene_gn=POLK;refGene_tx=XM_006927793.4;refGene_gn=POLK;refGene_tx=XM_019835644.2;refGene_gn=POLK;refGene_tx=XM_019835651.2;refGene_gn=POLK;refGene_tx=XM_019835655.2;refGene_gn=POLK;refGene_tx=XM_019835662.2;refGene_gn=POLK;refGene_tx=XM_019835667.2
A1 142293318 142293361 refGene_gn=POLK;refGene_tx=XM_006927792.4;refGene_gn=POLK;refGene_tx=XM_006927793.4;refGene_gn=POLK;refGene_tx=XM_019835644.2;refGene_gn=POLK;refGene_tx=XM_019835651.2;refGene_gn=POLK;refGene_tx=XM_019835655.2;refGene_gn=POLK;refGene_tx=XM_019835662.2;refGene_gn=POLK;refGene_tx=XM_019835667.2;refGene_gn=POLK;refGene_tx=XM_019835670.2
A1 142293541 142293617 refGene_gn=POLK;refGene_tx=XM_006927792.4;refGene_gn=POLK;refGene_tx=XM_006927793.4;refGene_gn=POLK;refGene_tx=XM_019835644.2;refGene_gn=POLK;refGene_tx=XM_019835651.2;refGene_gn=POLK;refGene_tx=XM_019835655.2;refGene_gn=POLK;refGene_tx=XM_019835662.2;refGene_gn=POLK;refGene_tx=XM_019835667.2;refGene_gn=POLK;refGene_tx=XM_019835670.2
A1 142299666 142299841 refGene_gn=ANKDD1B;refGene_tx=XM_023257559.1
A1 142308745 142308849 refGene_gn=ANKDD1B;refGene_tx=XM_023257559.1
A1 142311885 142311984 refGene_gn=ANKDD1B;refGene_tx=XM_023257559.1
A1 142322195 142322294 refGene_gn=ANKDD1B;refGene_tx=XM_023257559.1
A1 142322399 142322504 refGene_gn=ANKDD1B;refGene_tx=XM_023257559.1
A1 142331144 142331243 refGene_gn=ANKDD1B;refGene_tx=XM_023257559.1
A1 142332001 142332100 refGene_gn=ANKDD1B;refGene_tx=XM_023257559.1
A1 142337569 142337668 refGene_gn=ANKDD1B;refGene_tx=XM_023257559.1
A1 142343551 142343650 refGene_gn=ANKDD1B;refGene_tx=XM_023257559.1
A1 142346457 142346556 refGene_gn=ANKDD1B;refGene_tx=XM_023257559.1
A1 142349063 142349162 refGene_gn=ANKDD1B;refGene_tx=XM_023257559.1

A1 142353866 142353962
refGene_gn=ANKDD1B;refGene_tx=XM_023257559.1
A1 142357895 142358100
refGene_gn=ANKDD1B;refGene_tx=XM_023257559.1
A1 142360537 142360669
refGene_gn=ANKDD1B;refGene_tx=XM_023257559.1
A1 142361737 142361799
refGene_gn=ANKDD1B;refGene_tx=XM_023257559.1
A1 142364956 142365100
refGene_gn=POC5;refGene_tx=XM_003981080.5;refGene_gn=POC5;refGene_tx=XM_006927796.4;refGene_gn=POC5;refGene_tx=XM_006927797.4;refGene_gn=POC5;refGene_tx=XM_019835708.2;refGene_gn=POC5;refGene_tx=XM_019835721.2
A1 142368102 142368279
refGene_gn=POC5;refGene_tx=XM_003981080.5;refGene_gn=POC5;refGene_tx=XM_006927796.4;refGene_gn=POC5;refGene_tx=XM_006927797.4;refGene_gn=POC5;refGene_tx=XM_019835708.2;refGene_gn=POC5;refGene_tx=XM_019835721.2
A1 142374878 142375153
refGene_gn=POC5;refGene_tx=XM_003981080.5;refGene_gn=POC5;refGene_tx=XM_006927796.4;refGene_gn=POC5;refGene_tx=XM_006927797.4;refGene_gn=POC5;refGene_tx=XM_019835708.2;refGene_gn=POC5;refGene_tx=XM_019835721.2
A1 142380393 142380547
refGene_gn=POC5;refGene_tx=XM_003981080.5;refGene_gn=POC5;refGene_tx=XM_006927796.4;refGene_gn=POC5;refGene_tx=XM_006927797.4;refGene_gn=POC5;refGene_tx=XM_019835708.2;refGene_gn=POC5;refGene_tx=XM_019835721.2
A1 142382855 142383035
refGene_gn=POC5;refGene_tx=XM_003981080.5;refGene_gn=POC5;refGene_tx=XM_006927796.4;refGene_gn=POC5;refGene_tx=XM_006927797.4;refGene_gn=POC5;refGene_tx=XM_019835708.2;refGene_gn=POC5;refGene_tx=XM_019835721.2
A1 142386378 142386483
refGene_gn=POC5;refGene_tx=XM_003981080.5;refGene_gn=POC5;refGene_tx=XM_006927796.4;refGene_gn=POC5;refGene_tx=XM_006927797.4;refGene_gn=POC5;refGene_tx=XM_019835708.2;refGene_gn=POC5;refGene_tx=XM_019835721.2
A1 142389759 142389936
refGene_gn=POC5;refGene_tx=XM_003981080.5;refGene_gn=POC5;refGene_tx=XM_006927796.4;refGene_gn=POC5;refGene_tx=XM_006927797.4;refGene_gn=POC5;refGene_tx=XM_019835708.2;refGene_gn=POC5;refGene_tx=XM_019835721.2
A1 142391854 142392060
refGene_gn=POC5;refGene_tx=XM_003981080.5;refGene_gn=POC5;refGene_tx=XM_006927796.4;refGene_gn=POC5;refGene_tx=XM_006927797.4;refGene_gn=POC5;refGene_tx=XM_019835708.2;refGene_gn=POC5;refGene_tx=XM_019835721.2
A1 142394286 142394373
refGene_gn=POC5;refGene_tx=XM_003981080.5;refGene_gn=POC5;refGene_tx=XM_006927796.4;refGene_gn=POC5;refGene_tx=XM_006927797.4;refGene_gn=POC5;refGene_tx=XM_019835708.2;refGene_gn=POC5;refGene_tx=XM_019835721.2
A1 142396314 142396456
refGene_gn=POC5;refGene_tx=XM_003981080.5;refGene_gn=POC5;refGene_tx=XM_006927796.4;refGene_gn=POC5;refGene_tx=XM_019835708.2
A1 142401828 142401912
refGene_gn=POC5;refGene_tx=XM_003981080.5;refGene_gn=POC5;refGene_tx=XM_006927796.4;refGene_gn=POC5;refGene_tx=XM_019835708.2

A1 142405437 142405468
refGene_gn=LOC111561429;refGene_tx=XM_023257620.1
A1 142413134 142413273
refGene_gn=LOC111561429;refGene_tx=XM_023257610.1
A1 142422271 142422434
refGene_gn=LOC111561429;refGene_tx=XM_023257593.1;refGene_gn=LOC111561429;refGene_tx=XM_023257600.1;refGene_gn=LOC111561429;refGene_tx=XM_023257604.1;refGene_gn=LOC111561429;refGene_tx=XM_023257610.1;refGene_gn=LOC111561429;refGene_tx=XM_023257620.1
A1 142427349 142427401
refGene_gn=LOC111561429;refGene_tx=XM_023257610.1;refGene_gn=LOC111561429;refGene_tx=XM_023257620.1
A1 142659626 142659931
refGene_gn=LOC111561429;refGene_tx=XM_023257593.1;refGene_gn=LOC111561429;refGene_tx=XM_023257600.1;refGene_gn=LOC111561429;refGene_tx=XM_023257604.1
A1 142665075 142665411
refGene_gn=LOC111561429;refGene_tx=XM_023257593.1;refGene_gn=LOC111561429;refGene_tx=XM_023257600.1;refGene_gn=LOC111561429;refGene_tx=XM_023257604.1
A1 142732511 142733091
refGene_gn=SV2C;refGene_tx=XM_003981081.5;refGene_gn=SV2C;refGene_tx=XM_019835700.2
A1 142779694 142779875
refGene_gn=SV2C;refGene_tx=XM_003981081.5;refGene_gn=SV2C;refGene_tx=XM_019835700.2
A1 142795432 142795584
refGene_gn=SV2C;refGene_tx=XM_003981081.5;refGene_gn=SV2C;refGene_tx=XM_019835700.2
A1 142856563 142856697
refGene_gn=SV2C;refGene_tx=XM_003981081.5;refGene_gn=SV2C;refGene_tx=XM_019835700.2
A1 142857166 142857256
refGene_gn=SV2C;refGene_tx=XM_003981081.5;refGene_gn=SV2C;refGene_tx=XM_019835700.2
A1 142859776 142859887
refGene_gn=SV2C;refGene_tx=XM_003981081.5;refGene_gn=SV2C;refGene_tx=XM_019835700.2
A1 142860337 142860426
refGene_gn=SV2C;refGene_tx=XM_003981081.5;refGene_gn=SV2C;refGene_tx=XM_019835700.2
A1 142864691 142864850
refGene_gn=SV2C;refGene_tx=XM_003981081.5;refGene_gn=SV2C;refGene_tx=XM_019835700.2
A1 142867697 142867831
refGene_gn=SV2C;refGene_tx=XM_003981081.5;refGene_gn=SV2C;refGene_tx=XM_019835700.2
A1 142869280 142869484
refGene_gn=SV2C;refGene_tx=XM_003981081.5;refGene_gn=SV2C;refGene_tx=XM_019835700.2

A1	142869949	142870109
refGene_gn=SV2C;refGene_tx=XM_003981081.5;refGene_gn=SV2C;refGene_tx=XM_019835700.2
A1	142890770	142890954
refGene_gn=SV2C;refGene_tx=XM_003981081.5;refGene_gn=SV2C;refGene_tx=XM_019835700.2
A1	142935750	142935796
refGene_gn=IQGAP2;refGene_tx=XM_011284171.3
A1	142986677	142986777
refGene_gn=IQGAP2;refGene_tx=XM_011284171.3;refGene_gn=IQGAP2;refGene_tx=XM_019835759.2;refGene_gn=IQGAP2;refGene_tx=XM_019835765.2
A1	143082753	143082910
refGene_gn=IQGAP2;refGene_tx=XM_011284171.3;refGene_gn=IQGAP2;refGene_tx=XM_019835759.2;refGene_gn=IQGAP2;refGene_tx=XM_019835765.2
A1	143092905	143092983
refGene_gn=IQGAP2;refGene_tx=XM_011284171.3;refGene_gn=IQGAP2;refGene_tx=XM_019835759.2;refGene_gn=IQGAP2;refGene_tx=XM_019835765.2
A1	143096332	143096409
refGene_gn=IQGAP2;refGene_tx=XM_011284171.3;refGene_gn=IQGAP2;refGene_tx=XM_019835759.2;refGene_gn=IQGAP2;refGene_tx=XM_019835765.2
A1	143112310	143112378
refGene_gn=IQGAP2;refGene_tx=XM_011284171.3;refGene_gn=IQGAP2;refGene_tx=XM_019835759.2;refGene_gn=IQGAP2;refGene_tx=XM_019835765.2
A1	143112797	143112911
refGene_gn=IQGAP2;refGene_tx=XM_011284171.3;refGene_gn=IQGAP2;refGene_tx=XM_019835765.2
A1	143113611	143113790
refGene_gn=IQGAP2;refGene_tx=XM_011284171.3;refGene_gn=IQGAP2;refGene_tx=XM_019835759.2;refGene_gn=IQGAP2;refGene_tx=XM_019835765.2
A1	143115887	143115975
refGene_gn=IQGAP2;refGene_tx=XM_011284171.3;refGene_gn=IQGAP2;refGene_tx=XM_019835759.2;refGene_gn=IQGAP2;refGene_tx=XM_019835765.2
A1	143120029	143120193
refGene_gn=IQGAP2;refGene_tx=XM_011284171.3;refGene_gn=IQGAP2;refGene_tx=XM_019835759.2;refGene_gn=IQGAP2;refGene_tx=XM_019835765.2
A1	143123338	143123499
refGene_gn=IQGAP2;refGene_tx=XM_011284171.3;refGene_gn=IQGAP2;refGene_tx=XM_019835759.2;refGene_gn=IQGAP2;refGene_tx=XM_019835765.2
A1	143129015	143129140
refGene_gn=IQGAP2;refGene_tx=XM_011284171.3;refGene_gn=IQGAP2;refGene_tx=XM_019835759.2;refGene_gn=IQGAP2;refGene_tx=XM_019835765.2
A1	143132487	143132503
refGene_gn=IQGAP2;refGene_tx=XM_006927800.4
A1	143141960	143142124
refGene_gn=IQGAP2;refGene_tx=XM_006927800.4;refGene_gn=IQGAP2;refGene_tx=XM_011284171.3;refGene_gn=IQGAP2;refGene_tx=XM_019835759.2;refGene_gn=IQGAP2;refGene_tx=XM_019835765.2
A1	143147855	143148934
refGene_gn=F2RL2;refGene_tx=XM_003981082.5;refGene_gn=F2RL2;refGene_tx=XM_006927798.4;refGene_gn=F2RL2;refGene_tx=XM_019835776.2

A1 143152993 143153057
refGene_gn=F2RL2;refGene_tx=XM_003981082.5;refGene_gn=F2RL2;refGene_tx
=XM_006927798.4
A1 143157151 143157242
refGene_gn=IQGAP2;refGene_tx=XM_006927800.4;refGene_gn=IQGAP2;refGene_
tx=XM_011284171.3;refGene_gn=IQGAP2;refGene_tx=XM_019835759.2;refGene_
gn=IQGAP2;refGene_tx=XM_019835765.2
A1 143162976 143163144
refGene_gn=IQGAP2;refGene_tx=XM_006927800.4;refGene_gn=IQGAP2;refGene_
tx=XM_011284171.3;refGene_gn=IQGAP2;refGene_tx=XM_019835759.2;refGene_
gn=IQGAP2;refGene_tx=XM_019835765.2
A1 143165698 143165841
refGene_gn=IQGAP2;refGene_tx=XM_006927800.4;refGene_gn=IQGAP2;refGene_
tx=XM_011284171.3;refGene_gn=IQGAP2;refGene_tx=XM_019835759.2;refGene_
gn=IQGAP2;refGene_tx=XM_019835765.2
A1 143167699 143167870
refGene_gn=IQGAP2;refGene_tx=XM_011284171.3;refGene_gn=IQGAP2;refGene_
tx=XM_019835759.2
A1 143179147 143179231
refGene_gn=IQGAP2;refGene_tx=XM_006927800.4;refGene_gn=IQGAP2;refGene_
tx=XM_011284171.3;refGene_gn=IQGAP2;refGene_tx=XM_019835759.2;refGene_
gn=IQGAP2;refGene_tx=XM_019835765.2
A1 143180806 143180878
refGene_gn=IQGAP2;refGene_tx=XM_006927800.4;refGene_gn=IQGAP2;refGene_
tx=XM_011284171.3;refGene_gn=IQGAP2;refGene_tx=XM_019835759.2;refGene_
gn=IQGAP2;refGene_tx=XM_019835765.2
A1 143181911 143181981
refGene_gn=IQGAP2;refGene_tx=XM_006927800.4;refGene_gn=IQGAP2;refGene_
tx=XM_011284171.3;refGene_gn=IQGAP2;refGene_tx=XM_019835759.2;refGene_
gn=IQGAP2;refGene_tx=XM_019835765.2
A1 143184189 143184398
refGene_gn=IQGAP2;refGene_tx=XM_006927800.4;refGene_gn=IQGAP2;refGene_
tx=XM_011284171.3;refGene_gn=IQGAP2;refGene_tx=XM_019835759.2;refGene_
gn=IQGAP2;refGene_tx=XM_019835765.2
A1 143192609 143192759
refGene_gn=IQGAP2;refGene_tx=XM_006927800.4;refGene_gn=IQGAP2;refGene_
tx=XM_011284171.3;refGene_gn=IQGAP2;refGene_tx=XM_019835759.2;refGene_
gn=IQGAP2;refGene_tx=XM_019835765.2
A1 143195523 143195687
refGene_gn=IQGAP2;refGene_tx=XM_006927800.4;refGene_gn=IQGAP2;refGene_
tx=XM_011284171.3;refGene_gn=IQGAP2;refGene_tx=XM_019835759.2;refGene_
gn=IQGAP2;refGene_tx=XM_019835765.2
A1 143197803 143198028
refGene_gn=IQGAP2;refGene_tx=XM_006927800.4;refGene_gn=IQGAP2;refGene_
tx=XM_011284171.3;refGene_gn=IQGAP2;refGene_tx=XM_019835759.2;refGene_
gn=IQGAP2;refGene_tx=XM_019835765.2
A1 143199718 143199859
refGene_gn=IQGAP2;refGene_tx=XM_006927800.4;refGene_gn=IQGAP2;refGene_
tx=XM_011284171.3;refGene_gn=IQGAP2;refGene_tx=XM_019835759.2;refGene_
gn=IQGAP2;refGene_tx=XM_019835765.2

A1 143200199 143200284
refGene_gn=IQGAP2;refGene_tx=XM_006927800.4;refGene_gn=IQGAP2;refGene_tx=XM_011284171.3;refGene_gn=IQGAP2;refGene_tx=XM_019835759.2;refGene_gn=IQGAP2;refGene_tx=XM_019835765.2
A1 143200729 143200962
refGene_gn=IQGAP2;refGene_tx=XM_006927800.4;refGene_gn=IQGAP2;refGene_tx=XM_011284171.3;refGene_gn=IQGAP2;refGene_tx=XM_019835759.2;refGene_gn=IQGAP2;refGene_tx=XM_019835765.2
A1 143203224 143203357
refGene_gn=IQGAP2;refGene_tx=XM_006927800.4;refGene_gn=IQGAP2;refGene_tx=XM_011284171.3;refGene_gn=IQGAP2;refGene_tx=XM_019835759.2;refGene_gn=IQGAP2;refGene_tx=XM_019835765.2
A1 143207016 143207119
refGene_gn=IQGAP2;refGene_tx=XM_006927800.4;refGene_gn=IQGAP2;refGene_tx=XM_011284171.3;refGene_gn=IQGAP2;refGene_tx=XM_019835759.2;refGene_gn=IQGAP2;refGene_tx=XM_019835765.2
A1 143207628 143207770
refGene_gn=IQGAP2;refGene_tx=XM_006927800.4;refGene_gn=IQGAP2;refGene_tx=XM_011284171.3;refGene_gn=IQGAP2;refGene_tx=XM_019835759.2;refGene_gn=IQGAP2;refGene_tx=XM_019835765.2
A1 143218607 143218695
refGene_gn=IQGAP2;refGene_tx=XM_006927800.4;refGene_gn=IQGAP2;refGene_tx=XM_011284171.3;refGene_gn=IQGAP2;refGene_tx=XM_019835759.2;refGene_gn=IQGAP2;refGene_tx=XM_019835765.2
A1 143220845 143221058
refGene_gn=IQGAP2;refGene_tx=XM_006927800.4;refGene_gn=IQGAP2;refGene_tx=XM_011284171.3;refGene_gn=IQGAP2;refGene_tx=XM_019835759.2;refGene_gn=IQGAP2;refGene_tx=XM_019835765.2
A1 143223901 143224062
refGene_gn=IQGAP2;refGene_tx=XM_006927800.4;refGene_gn=IQGAP2;refGene_tx=XM_011284171.3;refGene_gn=IQGAP2;refGene_tx=XM_019835759.2;refGene_gn=IQGAP2;refGene_tx=XM_019835765.2
A1 143226033 143226171
refGene_gn=IQGAP2;refGene_tx=XM_006927800.4;refGene_gn=IQGAP2;refGene_tx=XM_011284171.3;refGene_gn=IQGAP2;refGene_tx=XM_019835759.2;refGene_gn=IQGAP2;refGene_tx=XM_019835765.2
A1 143227232 143227341
refGene_gn=IQGAP2;refGene_tx=XM_006927800.4;refGene_gn=IQGAP2;refGene_tx=XM_011284171.3;refGene_gn=IQGAP2;refGene_tx=XM_019835759.2;refGene_gn=IQGAP2;refGene_tx=XM_019835765.2
A1 143233371 143233485
refGene_gn=IQGAP2;refGene_tx=XM_006927800.4;refGene_gn=IQGAP2;refGene_tx=XM_011284171.3;refGene_gn=IQGAP2;refGene_tx=XM_019835759.2;refGene_gn=IQGAP2;refGene_tx=XM_019835765.2
A1 143246861 143246949
refGene_gn=F2R;refGene_tx=XM_003981083.5
A1 143260667 143261863
refGene_gn=F2R;refGene_tx=XM_003981083.5
A1 143333507 143333625
refGene_gn=F2RL1;refGene_tx=XM_006927801.4

A1 143347567 143348679
refGene_gn=F2RL1;refGene_tx=XM_006927801.4
A1 143379173 143379314
refGene_gn=S100Z;refGene_tx=XM_023257685.1;refGene_gn=S100Z;refGene_tx=XM_023257687.1
A1 143382976 143383135
refGene_gn=S100Z;refGene_tx=XM_023257685.1;refGene_gn=S100Z;refGene_tx=XM_023257687.1
A1 143416359 143416437
refGene_gn=CRHBP;refGene_tx=XM_023257693.1;refGene_gn=CRHBP;refGene_tx=XM_023257696.1
A1 143416795 143416889
refGene_gn=CRHBP;refGene_tx=XM_023257693.1;refGene_gn=CRHBP;refGene_tx=XM_023257696.1
A1 143417170 143417328
refGene_gn=CRHBP;refGene_tx=XM_023257693.1;refGene_gn=CRHBP;refGene_tx=XM_023257696.1
A1 143418696 143418907
refGene_gn=CRHBP;refGene_tx=XM_023257693.1;refGene_gn=CRHBP;refGene_tx=XM_023257696.1
A1 143421120 143421269
refGene_gn=CRHBP;refGene_tx=XM_023257693.1;refGene_gn=CRHBP;refGene_tx=XM_023257696.1
A1 143425322 143425440
refGene_gn=CRHBP;refGene_tx=XM_023257693.1;refGene_gn=CRHBP;refGene_tx=XM_023257696.1
A1 143427180 143427188
refGene_gn=CRHBP;refGene_tx=XM_023257696.1
A1 143428073 143428231
refGene_gn=CRHBP;refGene_tx=XM_023257693.1
A1 143475105 143475315
refGene_gn=AGGF1;refGene_tx=XM_011284145.3
A1 143477980 143478083
refGene_gn=AGGF1;refGene_tx=XM_011284145.3
A1 143479164 143479364
refGene_gn=AGGF1;refGene_tx=XM_011284145.3
A1 143480319 143480484
refGene_gn=AGGF1;refGene_tx=XM_011284145.3
A1 143483496 143483685
refGene_gn=AGGF1;refGene_tx=XM_011284145.3
A1 143485422 143485753
refGene_gn=AGGF1;refGene_tx=XM_011284145.3
A1 143486749 143486861
refGene_gn=AGGF1;refGene_tx=XM_011284145.3
A1 143488065 143488117
refGene_gn=AGGF1;refGene_tx=XM_011284145.3
A1 143491039 143491141
refGene_gn=AGGF1;refGene_tx=XM_011284145.3
A1 143492004 143492170
refGene_gn=AGGF1;refGene_tx=XM_011284145.3

A1 143493948 143494031
refGene_gn=AGGF1;refGene_tx=XM_011284145.3
A1 143497603 143497731
refGene_gn=AGGF1;refGene_tx=XM_011284145.3
A1 143499590 143499690
refGene_gn=AGGF1;refGene_tx=XM_011284145.3
A1 143501510 143501711
refGene_gn=AGGF1;refGene_tx=XM_011284145.3
A1 143518223 143518934
refGene_gn=ZBED3;refGene_tx=XM_023257717.1;refGene_gn=ZBED3;refGene_tx=XM_023257719.1;refGene_gn=ZBED3;refGene_tx=XM_023257724.1;refGene_gn=ZBED3;refGene_tx=XM_023257729.1;refGene_gn=ZBED3;refGene_tx=XM_023257732.1;refGene_gn=ZBED3;refGene_tx=XM_023257734.1;refGene_gn=ZBED3;refGene_tx=XM_023257736.1;refGene_gn=ZBED3;refGene_tx=XM_023257738.1;refGene_gn=ZBED3;refGene_tx=XM_023257741.1;refGene_gn=ZBED3;refGene_tx=XM_023257743.1;refGene_gn=ZBED3;refGene_tx=XM_023257747.1;refGene_gn=ZBED3;refGene_tx=XM_023257749.1
A1 143637029 143637383
refGene_gn=PDE8B;refGene_tx=XM_019835832.2
A1 143734262 143734322
refGene_gn=PDE8B;refGene_tx=XM_019835832.2
A1 143754233 143754424
refGene_gn=PDE8B;refGene_tx=XM_019835832.2
A1 143757935 143757995
refGene_gn=PDE8B;refGene_tx=XM_019835832.2
A1 143760291 143760349
refGene_gn=PDE8B;refGene_tx=XM_019835832.2
A1 143766631 143766720
refGene_gn=PDE8B;refGene_tx=XM_019835832.2
A1 143773132 143773211
refGene_gn=PDE8B;refGene_tx=XM_019835832.2
A1 143776739 143776880
refGene_gn=PDE8B;refGene_tx=XM_019835832.2
A1 143778418 143778507
refGene_gn=PDE8B;refGene_tx=XM_019835832.2
A1 143780600 143780661
refGene_gn=PDE8B;refGene_tx=XM_019835832.2
A1 143843506 143843549
refGene_gn=PDE8B;refGene_tx=XM_019835832.2
A1 143847412 143847490
refGene_gn=PDE8B;refGene_tx=XM_019835832.2
A1 143856143 143856220
refGene_gn=PDE8B;refGene_tx=XM_019835832.2
A1 143857650 143857815
refGene_gn=PDE8B;refGene_tx=XM_019835832.2
A1 143859865 143859911
refGene_gn=PDE8B;refGene_tx=XM_019835832.2
A1 143860331 143860384
refGene_gn=PDE8B;refGene_tx=XM_019835832.2
A1 143860385 143860468

refGene_gn=PDE8B;refGene_tx=XM_019835832.2
A1 143861363 143861562
refGene_gn=PDE8B;refGene_tx=XM_019835832.2
A1 143865638 143865856
refGene_gn=PDE8B;refGene_tx=XM_019835832.2
A1 143867454 143867575
refGene_gn=PDE8B;refGene_tx=XM_019835832.2
A1 143870632 143870800
refGene_gn=PDE8B;refGene_tx=XM_019835832.2
A1 143872852 143872982
refGene_gn=PDE8B;refGene_tx=XM_019835832.2
A1 143873550 143873660
refGene_gn=PDE8B;refGene_tx=XM_019835832.2
A1 143877503 143877656
refGene_gn=WDR41;refGene_tx=XM_019835839.2;refGene_gn=WDR41;refGene_tx=XM_019835843.2;refGene_gn=WDR41;refGene_tx=XM_019835844.2;refGene_gn=WDR41;refGene_tx=XM_019835848.2;refGene_gn=WDR41;refGene_tx=XM_019835852.2
A1 143879899 143880033
refGene_gn=WDR41;refGene_tx=XM_019835839.2;refGene_gn=WDR41;refGene_tx=XM_019835843.2;refGene_gn=WDR41;refGene_tx=XM_019835844.2;refGene_gn=WDR41;refGene_tx=XM_019835848.2;refGene_gn=WDR41;refGene_tx=XM_019835852.2
A1 143880864 143880953
refGene_gn=WDR41;refGene_tx=XM_019835839.2;refGene_gn=WDR41;refGene_tx=XM_019835843.2;refGene_gn=WDR41;refGene_tx=XM_019835844.2;refGene_gn=WDR41;refGene_tx=XM_019835848.2;refGene_gn=WDR41;refGene_tx=XM_019835852.2
A1 143882122 143882244
refGene_gn=WDR41;refGene_tx=XM_019835839.2;refGene_gn=WDR41;refGene_tx=XM_019835843.2;refGene_gn=WDR41;refGene_tx=XM_019835844.2;refGene_gn=WDR41;refGene_tx=XM_019835848.2;refGene_gn=WDR41;refGene_tx=XM_019835852.2
A1 143885661 143885846
refGene_gn=WDR41;refGene_tx=XM_019835839.2;refGene_gn=WDR41;refGene_tx=XM_019835843.2;refGene_gn=WDR41;refGene_tx=XM_019835844.2;refGene_gn=WDR41;refGene_tx=XM_019835848.2;refGene_gn=WDR41;refGene_tx=XM_019835852.2
A1 143891919 143892030
refGene_gn=WDR41;refGene_tx=XM_019835839.2;refGene_gn=WDR41;refGene_tx=XM_019835843.2;refGene_gn=WDR41;refGene_tx=XM_019835844.2;refGene_gn=WDR41;refGene_tx=XM_019835848.2;refGene_gn=WDR41;refGene_tx=XM_019835852.2
A1 143899264 143899327
refGene_gn=WDR41;refGene_tx=XM_019835839.2;refGene_gn=WDR41;refGene_tx=XM_019835843.2;refGene_gn=WDR41;refGene_tx=XM_019835844.2;refGene_gn=WDR41;refGene_tx=XM_019835848.2;refGene_gn=WDR41;refGene_tx=XM_019835852.2
A1 143901829 143901941
refGene_gn=WDR41;refGene_tx=XM_019835839.2;refGene_gn=WDR41;refGene_tx

=XM_019835843.2;refGene_gn=WDR41;refGene_tx=XM_019835844.2;refGene_gn=WDR41;refGene_tx=XM_019835848.2;refGene_gn=WDR41;refGene_tx=XM_019835852.2
A1 143907446 143907509
refGene_gn=WDR41;refGene_tx=XM_019835839.2;refGene_gn=WDR41;refGene_tx=XM_019835843.2;refGene_gn=WDR41;refGene_tx=XM_019835844.2;refGene_gn=WDR41;refGene_tx=XM_019835848.2;refGene_gn=WDR41;refGene_tx=XM_019835852.2
A1 143910023 143910155
refGene_gn=WDR41;refGene_tx=XM_019835839.2;refGene_gn=WDR41;refGene_tx=XM_019835843.2;refGene_gn=WDR41;refGene_tx=XM_019835844.2;refGene_gn=WDR41;refGene_tx=XM_019835848.2;refGene_gn=WDR41;refGene_tx=XM_019835852.2
A1 143911599 143911648
refGene_gn=WDR41;refGene_tx=XM_019835839.2;refGene_gn=WDR41;refGene_tx=XM_019835843.2;refGene_gn=WDR41;refGene_tx=XM_019835844.2;refGene_gn=WDR41;refGene_tx=XM_019835848.2;refGene_gn=WDR41;refGene_tx=XM_019835852.2
A1 143923588 143923704
refGene_gn=WDR41;refGene_tx=XM_019835839.2;refGene_gn=WDR41;refGene_tx=XM_019835843.2;refGene_gn=WDR41;refGene_tx=XM_019835844.2;refGene_gn=WDR41;refGene_tx=XM_019835848.2;refGene_gn=WDR41;refGene_tx=XM_019835852.2
A1 143925963 143926014
refGene_gn=WDR41;refGene_tx=XM_019835848.2
A1 143950370 143950397
refGene_gn=WDR41;refGene_tx=XM_019835852.2
A1 144052013 144052127
refGene_gn=WDR41;refGene_tx=XM_019835839.2;refGene_gn=WDR41;refGene_tx=XM_019835843.2;refGene_gn=WDR41;refGene_tx=XM_019835844.2
A1 144062163 144062200
refGene_gn=WDR41;refGene_tx=XM_019835839.2;refGene_gn=WDR41;refGene_tx=XM_019835843.2;refGene_gn=WDR41;refGene_tx=XM_019835844.2
A1 144070342 144070873
refGene_gn=OTP;refGene_tx=XM_003981087.3
A1 144076591 144077001
refGene_gn=OTP;refGene_tx=XM_003981087.3
A1 144078291 144078328
refGene_gn=OTP;refGene_tx=XM_003981087.3
A1 144119341 144119422
refGene_gn=TBCA;refGene_tx=XM_023257765.1;refGene_gn=TBCA;refGene_tx=XM_023257768.1
A1 144121515 144121602
refGene_gn=TBCA;refGene_tx=XM_023257765.1;refGene_gn=TBCA;refGene_tx=XM_023257768.1
A1 144135401 144135507
refGene_gn=TBCA;refGene_tx=XM_023257765.1;refGene_gn=TBCA;refGene_tx=XM_023257768.1
A1 144151106 144151153
refGene_gn=TBCA;refGene_tx=XM_023257765.1

A1 144196878 144196931
refGene_gn=TBCA;refGene_tx=XM_023257768.1
A1 144418020 144418174
refGene_gn=AP3B1;refGene_tx=XM_011284118.3;refGene_gn=AP3B1;refGene_tx=XM_023257772.1
A1 144429586 144429725
refGene_gn=AP3B1;refGene_tx=XM_011284118.3;refGene_gn=AP3B1;refGene_tx=XM_023257772.1;refGene_gn=AP3B1;refGene_tx=XM_023257774.1
A1 144433637 144433678
refGene_gn=AP3B1;refGene_tx=XM_023257774.1
A1 144433814 144433834
refGene_gn=AP3B1;refGene_tx=XM_023257780.1
A1 144434938 144435036
refGene_gn=AP3B1;refGene_tx=XM_011284118.3;refGene_gn=AP3B1;refGene_tx=XM_023257772.1;refGene_gn=AP3B1;refGene_tx=XM_023257774.1;refGene_gn=AP3B1;refGene_tx=XM_023257780.1
A1 144448088 144448173
refGene_gn=AP3B1;refGene_tx=XM_011284118.3;refGene_gn=AP3B1;refGene_tx=XM_023257772.1;refGene_gn=AP3B1;refGene_tx=XM_023257774.1;refGene_gn=AP3B1;refGene_tx=XM_023257780.1
A1 144453340 144453572
refGene_gn=AP3B1;refGene_tx=XM_011284118.3;refGene_gn=AP3B1;refGene_tx=XM_023257772.1;refGene_gn=AP3B1;refGene_tx=XM_023257774.1;refGene_gn=AP3B1;refGene_tx=XM_023257780.1
A1 144483351 144483423
refGene_gn=AP3B1;refGene_tx=XM_023257772.1;refGene_gn=AP3B1;refGene_tx=XM_023257774.1;refGene_gn=AP3B1;refGene_tx=XM_023257780.1
A1 144485934 144486041
refGene_gn=AP3B1;refGene_tx=XM_011284118.3;refGene_gn=AP3B1;refGene_tx=XM_023257772.1;refGene_gn=AP3B1;refGene_tx=XM_023257774.1;refGene_gn=AP3B1;refGene_tx=XM_023257780.1
A1 144497826 144497899
refGene_gn=AP3B1;refGene_tx=XM_011284118.3;refGene_gn=AP3B1;refGene_tx=XM_023257772.1;refGene_gn=AP3B1;refGene_tx=XM_023257774.1;refGene_gn=AP3B1;refGene_tx=XM_023257780.1
A1 144508644 144508792
refGene_gn=AP3B1;refGene_tx=XM_011284118.3;refGene_gn=AP3B1;refGene_tx=XM_023257772.1;refGene_gn=AP3B1;refGene_tx=XM_023257774.1;refGene_gn=AP3B1;refGene_tx=XM_023257780.1
A1 144512295 144512458
refGene_gn=AP3B1;refGene_tx=XM_011284118.3;refGene_gn=AP3B1;refGene_tx=XM_023257772.1;refGene_gn=AP3B1;refGene_tx=XM_023257774.1;refGene_gn=AP3B1;refGene_tx=XM_023257780.1
A1 144516228 144516337
refGene_gn=AP3B1;refGene_tx=XM_011284118.3;refGene_gn=AP3B1;refGene_tx=XM_023257772.1;refGene_gn=AP3B1;refGene_tx=XM_023257774.1;refGene_gn=AP3B1;refGene_tx=XM_023257780.1
A1 144521641 144521772
refGene_gn=AP3B1;refGene_tx=XM_011284118.3;refGene_gn=AP3B1;refGene_tx=XM_023257772.1;refGene_gn=AP3B1;refGene_tx=XM_023257774.1;refGene_gn=

AP3B1;refGene_tx=XM_023257780.1
A1 144522727 144522914
refGene_gn=AP3B1;refGene_tx=XM_011284118.3;refGene_gn=AP3B1;refGene_tx
=XM_023257772.1;refGene_gn=AP3B1;refGene_tx=XM_023257774.1;refGene_gn=
AP3B1;refGene_tx=XM_023257780.1
A1 144533066 144533243
refGene_gn=AP3B1;refGene_tx=XM_011284118.3;refGene_gn=AP3B1;refGene_tx
=XM_023257772.1;refGene_gn=AP3B1;refGene_tx=XM_023257774.1;refGene_gn=
AP3B1;refGene_tx=XM_023257780.1
A1 144544456 144544566
refGene_gn=AP3B1;refGene_tx=XM_011284118.3;refGene_gn=AP3B1;refGene_tx
=XM_023257772.1;refGene_gn=AP3B1;refGene_tx=XM_023257774.1;refGene_gn=
AP3B1;refGene_tx=XM_023257780.1
A1 144551616 144551749
refGene_gn=AP3B1;refGene_tx=XM_011284118.3;refGene_gn=AP3B1;refGene_tx
=XM_023257772.1;refGene_gn=AP3B1;refGene_tx=XM_023257774.1;refGene_gn=
AP3B1;refGene_tx=XM_023257780.1
A1 144554854 144554917
refGene_gn=AP3B1;refGene_tx=XM_011284118.3;refGene_gn=AP3B1;refGene_tx
=XM_023257772.1;refGene_gn=AP3B1;refGene_tx=XM_023257774.1;refGene_gn=
AP3B1;refGene_tx=XM_023257780.1
A1 144557840 144557912
refGene_gn=AP3B1;refGene_tx=XM_011284118.3;refGene_gn=AP3B1;refGene_tx
=XM_023257772.1;refGene_gn=AP3B1;refGene_tx=XM_023257774.1;refGene_gn=
AP3B1;refGene_tx=XM_023257780.1
A1 144558000 144558055
refGene_gn=AP3B1;refGene_tx=XM_011284118.3;refGene_gn=AP3B1;refGene_tx
=XM_023257772.1;refGene_gn=AP3B1;refGene_tx=XM_023257774.1;refGene_gn=
AP3B1;refGene_tx=XM_023257780.1
A1 144559138 144559236
refGene_gn=AP3B1;refGene_tx=XM_011284118.3;refGene_gn=AP3B1;refGene_tx
=XM_023257772.1;refGene_gn=AP3B1;refGene_tx=XM_023257774.1;refGene_gn=
AP3B1;refGene_tx=XM_023257780.1
A1 144563183 144563339
refGene_gn=AP3B1;refGene_tx=XM_011284118.3;refGene_gn=AP3B1;refGene_tx
=XM_023257772.1;refGene_gn=AP3B1;refGene_tx=XM_023257774.1;refGene_gn=
AP3B1;refGene_tx=XM_023257780.1
A1 144598671 144598854
refGene_gn=AP3B1;refGene_tx=XM_011284118.3;refGene_gn=AP3B1;refGene_tx
=XM_023257772.1;refGene_gn=AP3B1;refGene_tx=XM_023257774.1;refGene_gn=
AP3B1;refGene_tx=XM_023257780.1
A1 144608352 144608419
refGene_gn=AP3B1;refGene_tx=XM_011284118.3;refGene_gn=AP3B1;refGene_tx
=XM_023257772.1;refGene_gn=AP3B1;refGene_tx=XM_023257774.1;refGene_gn=
AP3B1;refGene_tx=XM_023257780.1
A1 144611209 144611370
refGene_gn=AP3B1;refGene_tx=XM_011284118.3;refGene_gn=AP3B1;refGene_tx
=XM_023257772.1;refGene_gn=AP3B1;refGene_tx=XM_023257774.1;refGene_gn=
AP3B1;refGene_tx=XM_023257780.1
A1 144612597 144612693

refGene_gn=AP3B1;refGene_tx=XM_011284118.3;refGene_gn=AP3B1;refGene_tx=XM_023257772.1;refGene_gn=AP3B1;refGene_tx=XM_023257774.1;refGene_gn=AP3B1;refGene_tx=XM_023257780.1
A1 144633772 144633847
refGene_gn=AP3B1;refGene_tx=XM_011284118.3;refGene_gn=AP3B1;refGene_tx=XM_023257772.1;refGene_gn=AP3B1;refGene_tx=XM_023257774.1;refGene_gn=AP3B1;refGene_tx=XM_023257780.1
A1 144644226 144644302
refGene_gn=AP3B1;refGene_tx=XM_011284118.3;refGene_gn=AP3B1;refGene_tx=XM_023257772.1;refGene_gn=AP3B1;refGene_tx=XM_023257774.1;refGene_gn=AP3B1;refGene_tx=XM_023257780.1
A1 144680459 144680587
refGene_gn=AP3B1;refGene_tx=XM_011284118.3;refGene_gn=AP3B1;refGene_tx=XM_023257772.1;refGene_gn=AP3B1;refGene_tx=XM_023257774.1;refGene_gn=AP3B1;refGene_tx=XM_023257780.1
A1 144735337 144735394
refGene_gn=SCAMP1;refGene_tx=XM_003981088.5;refGene_gn=SCAMP1;refGene_tx=XM_019835860.2
A1 144735648 144735899
refGene_gn=LOC109501924;refGene_tx=XM_019835867.2
A1 144736062 144736093
refGene_gn=LOC109501924;refGene_tx=XM_019835867.2
A1 144758576 144758654
refGene_gn=SCAMP1;refGene_tx=XM_003981088.5
A1 144788649 144788748
refGene_gn=SCAMP1;refGene_tx=XM_003981088.5;refGene_gn=SCAMP1;refGene_tx=XM_019835860.2
A1 144789772 144789881
refGene_gn=SCAMP1;refGene_tx=XM_003981088.5;refGene_gn=SCAMP1;refGene_tx=XM_019835860.2
A1 144792736 144792865
refGene_gn=SCAMP1;refGene_tx=XM_003981088.5;refGene_gn=SCAMP1;refGene_tx=XM_019835860.2
A1 144798090 144798250
refGene_gn=SCAMP1;refGene_tx=XM_003981088.5;refGene_gn=SCAMP1;refGene_tx=XM_019835860.2
A1 144843473 144843575
refGene_gn=SCAMP1;refGene_tx=XM_003981088.5;refGene_gn=SCAMP1;refGene_tx=XM_019835860.2
A1 144846937 144847055
refGene_gn=SCAMP1;refGene_tx=XM_003981088.5;refGene_gn=SCAMP1;refGene_tx=XM_019835860.2
A1 144862063 144862228
refGene_gn=SCAMP1;refGene_tx=XM_003981088.5;refGene_gn=SCAMP1;refGene_tx=XM_019835860.2
A1 144877972 144878229
refGene_gn=LHFPL2;refGene_tx=XM_023257792.1;refGene_gn=LHFPL2;refGene_tx=XM_023257793.1;refGene_gn=LHFPL2;refGene_tx=XM_023257794.1;refGene_gn=LHFPL2;refGene_tx=XM_023257795.1
A1 144900002 144900432

refGene_gn=LHFPL2;refGene_tx=XM_023257792.1;refGene_gn=LHFPL2;refGene_tx=XM_023257793.1;refGene_gn=LHFPL2;refGene_tx=XM_023257794.1;refGene_gn=LHFPL2;refGene_tx=XM_023257795.1
A1 145138687 145138953
refGene_gn=ARSB;refGene_tx=NM_001142259.1
A1 145139995 145140118
refGene_gn=ARSB;refGene_tx=NM_001142259.1
A1 145201308 145201379
refGene_gn=ARSB;refGene_tx=NM_001142259.1
A1 145222998 145223242
refGene_gn=ARSB;refGene_tx=NM_001142259.1
A1 145277804 145278012
refGene_gn=ARSB;refGene_tx=NM_001142259.1
A1 145285355 145285546
refGene_gn=ARSB;refGene_tx=NM_001142259.1
A1 145289821 145290008
refGene_gn=ARSB;refGene_tx=NM_001142259.1
A1 145306017 145306335
refGene_gn=ARSB;refGene_tx=NM_001142259.1
A1 145323961 145324177
refGene_gn=DMGDH;refGene_tx=XM_023257800.1;refGene_gn=DMGDH;refGene_tx=XM_023257803.1
A1 145329870 145330005
refGene_gn=DMGDH;refGene_tx=XM_023257800.1;refGene_gn=DMGDH;refGene_tx=XM_023257803.1
A1 145350063 145350123
refGene_gn=DMGDH;refGene_tx=XM_023257800.1;refGene_gn=DMGDH;refGene_tx=XM_023257803.1
A1 145353100 145353258
refGene_gn=DMGDH;refGene_tx=XM_023257800.1;refGene_gn=DMGDH;refGene_tx=XM_023257803.1
A1 145354918 145355136
refGene_gn=DMGDH;refGene_tx=XM_023257800.1;refGene_gn=DMGDH;refGene_tx=XM_023257803.1
A1 145356387 145356518
refGene_gn=DMGDH;refGene_tx=XM_023257800.1;refGene_gn=DMGDH;refGene_tx=XM_023257803.1
A1 145357456 145357622
refGene_gn=DMGDH;refGene_tx=XM_023257800.1;refGene_gn=DMGDH;refGene_tx=XM_023257803.1
A1 145359326 145359480
refGene_gn=DMGDH;refGene_tx=XM_023257800.1;refGene_gn=DMGDH;refGene_tx=XM_023257803.1
A1 145360535 145360729
refGene_gn=DMGDH;refGene_tx=XM_023257800.1;refGene_gn=DMGDH;refGene_tx=XM_023257803.1
A1 145367602 145367801
refGene_gn=DMGDH;refGene_tx=XM_023257800.1;refGene_gn=DMGDH;refGene_tx=XM_023257803.1
A1 145370125 145370374

refGene_gn=DMGDH;refGene_tx=XM_023257800.1;refGene_gn=DMGDH;refGene_tx=XM_023257803.1
A1 145377821 145378026
refGene_gn=DMGDH;refGene_tx=XM_023257800.1;refGene_gn=DMGDH;refGene_tx=XM_023257803.1
A1 145380686 145380851
refGene_gn=DMGDH;refGene_tx=XM_023257800.1;refGene_gn=DMGDH;refGene_tx=XM_023257803.1
A1 145382322 145382421
refGene_gn=DMGDH;refGene_tx=XM_023257800.1;refGene_gn=DMGDH;refGene_tx=XM_023257803.1
A1 145391460 145391635
refGene_gn=DMGDH;refGene_tx=XM_023257800.1;refGene_gn=DMGDH;refGene_tx=XM_023257803.1
A1 145392650 145392745
refGene_gn=DMGDH;refGene_tx=XM_023257800.1;refGene_gn=DMGDH;refGene_tx=XM_023257803.1
A1 145392910 145392943
refGene_gn=BHMT2;refGene_tx=XM_003981099.4
A1 145397920 145398053
refGene_gn=BHMT2;refGene_tx=XM_003981099.4
A1 145399843 145399935
refGene_gn=BHMT2;refGene_tx=XM_003981099.4;refGene_gn=BHMT2;refGene_tx=XM_019835891.2;refGene_gn=BHMT2;refGene_tx=XM_019835898.2;refGene_gn=BHMT2;refGene_tx=XM_019835908.2;refGene_gn=BHMT2;refGene_tx=XM_019835909.2;refGene_gn=BHMT2;refGene_tx=XM_019835910.2
A1 145401109 145401301
refGene_gn=BHMT2;refGene_tx=XM_003981099.4;refGene_gn=BHMT2;refGene_tx=XM_019835891.2;refGene_gn=BHMT2;refGene_tx=XM_019835898.2;refGene_gn=BHMT2;refGene_tx=XM_019835908.2;refGene_gn=BHMT2;refGene_tx=XM_019835909.2;refGene_gn=BHMT2;refGene_tx=XM_019835910.2
A1 145403342 145403490
refGene_gn=BHMT2;refGene_tx=XM_003981099.4;refGene_gn=BHMT2;refGene_tx=XM_019835891.2;refGene_gn=BHMT2;refGene_tx=XM_019835898.2;refGene_gn=BHMT2;refGene_tx=XM_019835908.2;refGene_gn=BHMT2;refGene_tx=XM_019835909.2;refGene_gn=BHMT2;refGene_tx=XM_019835910.2
A1 145403717 145403900
refGene_gn=BHMT2;refGene_tx=XM_003981099.4;refGene_gn=BHMT2;refGene_tx=XM_019835891.2;refGene_gn=BHMT2;refGene_tx=XM_019835898.2;refGene_gn=BHMT2;refGene_tx=XM_019835908.2;refGene_gn=BHMT2;refGene_tx=XM_019835909.2;refGene_gn=BHMT2;refGene_tx=XM_019835910.2
A1 145404146 145404375
refGene_gn=BHMT2;refGene_tx=XM_003981099.4;refGene_gn=BHMT2;refGene_tx=XM_019835891.2;refGene_gn=BHMT2;refGene_tx=XM_019835898.2;refGene_gn=BHMT2;refGene_tx=XM_019835908.2;refGene_gn=BHMT2;refGene_tx=XM_019835909.2;refGene_gn=BHMT2;refGene_tx=XM_019835910.2
A1 145408788 145408870
refGene_gn=BHMT2;refGene_tx=XM_003981099.4;refGene_gn=BHMT2;refGene_tx=XM_019835891.2;refGene_gn=BHMT2;refGene_tx=XM_019835898.2;refGene_gn=BHMT2;refGene_tx=XM_019835908.2;refGene_gn=BHMT2;refGene_tx=XM_0198359

09.2;refGene_gn=BHMT2;refGene_tx=XM_019835910.2
A1 145438346 145438379
refGene_gn=BHMT;refGene_tx=XM_003981100.4
A1 145438970 145438973
refGene_gn=BHMT;refGene_tx=XM_023257805.1
A1 145439781 145439914
refGene_gn=BHMT;refGene_tx=XM_003981100.4;refGene_gn=BHMT;refGene_tx=XM_023257805.1
A1 145446387 145446506
refGene_gn=BHMT;refGene_tx=XM_003981100.4;refGene_gn=BHMT;refGene_tx=XM_023257805.1
A1 145447453 145447645
refGene_gn=BHMT;refGene_tx=XM_003981100.4;refGene_gn=BHMT;refGene_tx=XM_023257805.1
A1 145448004 145448152
refGene_gn=BHMT;refGene_tx=XM_003981100.4;refGene_gn=BHMT;refGene_tx=XM_023257805.1
A1 145451719 145451902
refGene_gn=BHMT;refGene_tx=XM_003981100.4;refGene_gn=BHMT;refGene_tx=XM_023257805.1
A1 145453289 145453518
refGene_gn=BHMT;refGene_tx=XM_003981100.4;refGene_gn=BHMT;refGene_tx=XM_023257805.1
A1 145455238 145455425
refGene_gn=BHMT;refGene_tx=XM_003981100.4;refGene_gn=BHMT;refGene_tx=XM_023257805.1
A1 145544683 145545712
refGene_gn=JMY;refGene_tx=XM_011284222.3
A1 145584070 145584244
refGene_gn=JMY;refGene_tx=XM_011284222.3
A1 145602711 145602862
refGene_gn=JMY;refGene_tx=XM_011284222.3
A1 145611035 145611205
refGene_gn=JMY;refGene_tx=XM_011284222.3
A1 145626195 145626361
refGene_gn=JMY;refGene_tx=XM_011284222.3
A1 145627078 145627266
refGene_gn=JMY;refGene_tx=XM_011284222.3
A1 145630223 145630310
refGene_gn=JMY;refGene_tx=XM_011284222.3
A1 145635999 145636095
refGene_gn=JMY;refGene_tx=XM_011284222.3
A1 145637452 145638050
refGene_gn=JMY;refGene_tx=XM_011284222.3
A1 145639928 145640236
refGene_gn=JMY;refGene_tx=XM_011284222.3
A1 145683258 145683447
refGene_gn=HOMER1;refGene_tx=XM_019835915.2;refGene_gn=HOMER1;refGene_tx=XM_019835916.2
A1 145698156 145698237

refGene_gn=HOMER1;refGene_tx=XM_019835915.2;refGene_gn=HOMER1;refGene_tx=XM_019835916.2
A1 145698859 145698970
refGene_gn=HOMER1;refGene_tx=XM_019835915.2;refGene_gn=HOMER1;refGene_tx=XM_019835916.2
A1 145703301 145703458
refGene_gn=HOMER1;refGene_tx=XM_019835915.2;refGene_gn=HOMER1;refGene_tx=XM_019835916.2
A1 145730428 145730441
refGene_gn=HOMER1;refGene_tx=XM_019835918.2
A1 145750446 145750586
refGene_gn=HOMER1;refGene_tx=XM_019835915.2;refGene_gn=HOMER1;refGene_tx=XM_019835916.2;refGene_gn=HOMER1;refGene_tx=XM_019835918.2
A1 145760805 145760898
refGene_gn=HOMER1;refGene_tx=XM_019835915.2;refGene_gn=HOMER1;refGene_tx=XM_019835916.2;refGene_gn=HOMER1;refGene_tx=XM_019835918.2
A1 145764392 145764524
refGene_gn=HOMER1;refGene_tx=XM_019835915.2;refGene_gn=HOMER1;refGene_tx=XM_019835916.2;refGene_gn=HOMER1;refGene_tx=XM_019835918.2
A1 145768517 145768674
refGene_gn=HOMER1;refGene_tx=XM_019835915.2;refGene_gn=HOMER1;refGene_tx=XM_019835918.2
A1 145815757 145815762
refGene_gn=HOMER1;refGene_tx=XM_019835915.2;refGene_gn=HOMER1;refGene_tx=XM_019835918.2
A1 145815906 145816082
refGene_gn=LOC111557751;refGene_tx=XM_023243709.1
A1 145816985 145817568
refGene_gn=LOC111557751;refGene_tx=XM_023243709.1
A1 145817666 145817837
refGene_gn=LOC111557751;refGene_tx=XM_023243709.1
A1 145888513 145888650
refGene_gn=PAPD4;refGene_tx=XM_006927811.4;refGene_gn=PAPD4;refGene_tx=XM_006927812.4;refGene_gn=PAPD4;refGene_tx=XM_006927813.4;refGene_gn=PAPD4;refGene_tx=XM_006927814.4
A1 145889080 145889170
refGene_gn=PAPD4;refGene_tx=XM_006927811.4;refGene_gn=PAPD4;refGene_tx=XM_006927812.4;refGene_gn=PAPD4;refGene_tx=XM_006927813.4;refGene_gn=PAPD4;refGene_tx=XM_006927814.4
A1 145891638 145891876
refGene_gn=PAPD4;refGene_tx=XM_006927811.4;refGene_gn=PAPD4;refGene_tx=XM_006927812.4;refGene_gn=PAPD4;refGene_tx=XM_006927813.4;refGene_gn=PAPD4;refGene_tx=XM_006927814.4
A1 145908374 145908489
refGene_gn=PAPD4;refGene_tx=XM_006927811.4;refGene_gn=PAPD4;refGene_tx=XM_006927812.4;refGene_gn=PAPD4;refGene_tx=XM_006927813.4;refGene_gn=PAPD4;refGene_tx=XM_006927814.4
A1 145908579 145908671
refGene_gn=PAPD4;refGene_tx=XM_006927811.4;refGene_gn=PAPD4;refGene_tx=XM_006927812.4;refGene_gn=PAPD4;refGene_tx=XM_006927813.4;refGene_gn=

PAPD4;refGene_tx=XM_006927814.4
A1 145910472 145910551
refGene_gn=PAPD4;refGene_tx=XM_006927811.4;refGene_gn=PAPD4;refGene_tx=XM_006927812.4;refGene_gn=PAPD4;refGene_tx=XM_006927813.4;refGene_gn=PAPD4;refGene_tx=XM_006927814.4
A1 145912267 145912337
refGene_gn=PAPD4;refGene_tx=XM_006927811.4;refGene_gn=PAPD4;refGene_tx=XM_006927812.4;refGene_gn=PAPD4;refGene_tx=XM_006927813.4;refGene_gn=PAPD4;refGene_tx=XM_006927814.4
A1 145919789 145919866
refGene_gn=PAPD4;refGene_tx=XM_006927811.4;refGene_gn=PAPD4;refGene_tx=XM_006927812.4;refGene_gn=PAPD4;refGene_tx=XM_006927813.4;refGene_gn=PAPD4;refGene_tx=XM_006927814.4
A1 145920239 145920368
refGene_gn=PAPD4;refGene_tx=XM_006927811.4;refGene_gn=PAPD4;refGene_tx=XM_006927812.4;refGene_gn=PAPD4;refGene_tx=XM_006927813.4;refGene_gn=PAPD4;refGene_tx=XM_006927814.4
A1 145925370 145925414
refGene_gn=PAPD4;refGene_tx=XM_006927811.4;refGene_gn=PAPD4;refGene_tx=XM_006927812.4;refGene_gn=PAPD4;refGene_tx=XM_006927813.4;refGene_gn=PAPD4;refGene_tx=XM_006927814.4
A1 145934414 145934551
refGene_gn=PAPD4;refGene_tx=XM_006927811.4;refGene_gn=PAPD4;refGene_tx=XM_006927812.4;refGene_gn=PAPD4;refGene_tx=XM_006927813.4;refGene_gn=PAPD4;refGene_tx=XM_006927814.4
A1 145945131 145945223
refGene_gn=PAPD4;refGene_tx=XM_006927811.4;refGene_gn=PAPD4;refGene_tx=XM_006927812.4;refGene_gn=PAPD4;refGene_tx=XM_006927813.4;refGene_gn=PAPD4;refGene_tx=XM_006927814.4
A1 145950332 145950412
refGene_gn=PAPD4;refGene_tx=XM_006927811.4;refGene_gn=PAPD4;refGene_tx=XM_006927812.4;refGene_gn=PAPD4;refGene_tx=XM_006927813.4;refGene_gn=PAPD4;refGene_tx=XM_006927814.4
A1 145952003 145952078
refGene_gn=PAPD4;refGene_tx=XM_006927811.4;refGene_gn=PAPD4;refGene_tx=XM_006927812.4;refGene_gn=PAPD4;refGene_tx=XM_006927813.4;refGene_gn=PAPD4;refGene_tx=XM_006927814.4
A1 145956325 145956486
refGene_gn=CMYA5;refGene_tx=XM_011284243.3
A1 145988541 145999363
refGene_gn=CMYA5;refGene_tx=XM_011284243.3
A1 146002477 146002573
refGene_gn=CMYA5;refGene_tx=XM_011284243.3
A1 146003960 146004194
refGene_gn=CMYA5;refGene_tx=XM_011284243.3
A1 146005794 146005817
refGene_gn=CMYA5;refGene_tx=XM_011284243.3
A1 146010706 146010825
refGene_gn=CMYA5;refGene_tx=XM_011284243.3
A1 146017558 146017708

refGene_gn=CMYA5;refGene_tx=XM_011284243.3
A1 146020756 146020903
refGene_gn=CMYA5;refGene_tx=XM_011284243.3
A1 146021994 146022142
refGene_gn=CMYA5;refGene_tx=XM_011284243.3
A1 146036375 146036509
refGene_gn=CMYA5;refGene_tx=XM_011284243.3
A1 146037779 146037879
refGene_gn=CMYA5;refGene_tx=XM_011284243.3
A1 146039973 146040147
refGene_gn=CMYA5;refGene_tx=XM_011284243.3
A1 146044676 146044923
refGene_gn=CMYA5;refGene_tx=XM_011284243.3
A1 146197247 146197358
refGene_gn=MTX3;refGene_tx=XM_003981106.4;refGene_gn=MTX3;refGene_tx=XM_006927815.3;refGene_gn=MTX3;refGene_tx=XM_019835955.2
A1 146201132 146201221
refGene_gn=MTX3;refGene_tx=XM_003981106.4;refGene_gn=MTX3;refGene_tx=XM_019835955.2
A1 146202472 146202630
refGene_gn=MTX3;refGene_tx=XM_003981106.4;refGene_gn=MTX3;refGene_tx=XM_006927815.3;refGene_gn=MTX3;refGene_tx=XM_019835955.2
A1 146203134 146203211
refGene_gn=MTX3;refGene_tx=XM_003981106.4;refGene_gn=MTX3;refGene_tx=XM_006927815.3;refGene_gn=MTX3;refGene_tx=XM_019835955.2
A1 146203353 146203536
refGene_gn=MTX3;refGene_tx=XM_003981106.4;refGene_gn=MTX3;refGene_tx=XM_006927815.3;refGene_gn=MTX3;refGene_tx=XM_019835955.2
A1 146204048 146204141
refGene_gn=MTX3;refGene_tx=XM_003981106.4;refGene_gn=MTX3;refGene_tx=XM_006927815.3
A1 146205102 146205179
refGene_gn=MTX3;refGene_tx=XM_003981106.4;refGene_gn=MTX3;refGene_tx=XM_006927815.3;refGene_gn=MTX3;refGene_tx=XM_019835955.2
A1 146205556 146205626
refGene_gn=MTX3;refGene_tx=XM_003981106.4;refGene_gn=MTX3;refGene_tx=XM_006927815.3;refGene_gn=MTX3;refGene_tx=XM_019835955.2
A1 146206134 146206215
refGene_gn=MTX3;refGene_tx=XM_003981106.4;refGene_gn=MTX3;refGene_tx=XM_006927815.3;refGene_gn=MTX3;refGene_tx=XM_019835955.2
A1 146252676 146252764
refGene_gn=THBS4;refGene_tx=XM_003981108.5
A1 146258919 146259123
refGene_gn=THBS4;refGene_tx=XM_003981108.5
A1 146270210 146270458
refGene_gn=THBS4;refGene_tx=XM_003981108.5
A1 146273875 146273984
refGene_gn=THBS4;refGene_tx=XM_003981108.5
A1 146274365 146274448
refGene_gn=THBS4;refGene_tx=XM_003981108.5

A1 146274776 146274828
refGene_gn=THBS4;refGene_tx=XM_003981108.5
A1 146275042 146275254
refGene_gn=THBS4;refGene_tx=XM_003981108.5
A1 146276753 146276891
refGene_gn=THBS4;refGene_tx=XM_003981108.5
A1 146279081 146279150
refGene_gn=THBS4;refGene_tx=XM_003981108.5
A1 146281089 146281242
refGene_gn=THBS4;refGene_tx=XM_003981108.5
A1 146284100 146284205
refGene_gn=THBS4;refGene_tx=XM_003981108.5
A1 146284767 146284875
refGene_gn=THBS4;refGene_tx=XM_003981108.5
A1 146285126 146285286
refGene_gn=THBS4;refGene_tx=XM_003981108.5
A1 146286612 146286731
refGene_gn=THBS4;refGene_tx=XM_003981108.5
A1 146287613 146287666
refGene_gn=THBS4;refGene_tx=XM_003981108.5
A1 146290175 146290369
refGene_gn=THBS4;refGene_tx=XM_003981108.5
A1 146291279 146291458
refGene_gn=THBS4;refGene_tx=XM_003981108.5
A1 146292095 146292144
refGene_gn=THBS4;refGene_tx=XM_003981108.5
A1 146292241 146292438
refGene_gn=THBS4;refGene_tx=XM_003981108.5
A1 146293037 146293210
refGene_gn=THBS4;refGene_tx=XM_003981108.5
A1 146295224 146295364
refGene_gn=THBS4;refGene_tx=XM_003981108.5
A1 146295896 146295958
refGene_gn=THBS4;refGene_tx=XM_003981108.5
A1 146362382 146362530
refGene_gn=SERINC5;refGene_tx=XM_011284262.3
A1 146365793 146365938
refGene_gn=SERINC5;refGene_tx=XM_011284262.3
A1 146366553 146366593
refGene_gn=SERINC5;refGene_tx=XM_011284262.3
A1 146369554 146369621
refGene_gn=SERINC5;refGene_tx=XM_011284262.3
A1 146376774 146376901
refGene_gn=SERINC5;refGene_tx=XM_011284262.3
A1 146380681 146380777
refGene_gn=SERINC5;refGene_tx=XM_011284262.3
A1 146382987 146383199
refGene_gn=SERINC5;refGene_tx=XM_011284262.3
A1 146389567 146389661
refGene_gn=SERINC5;refGene_tx=XM_011284262.3

A1 146391345 146391428
refGene_gn=SERINC5;refGene_tx=XM_011284262.3
A1 146392344 146392523
refGene_gn=SERINC5;refGene_tx=XM_011284262.3
A1 146420837 146421005
refGene_gn=SERINC5;refGene_tx=XM_011284262.3
A1 146462901 146462928
refGene_gn=SERINC5;refGene_tx=XM_011284262.3
A1 146511801 146512917
refGene_gn=SPZ1;refGene_tx=XM_003981119.4
A1 146567875 146567945
refGene_gn=ZFYVE16;refGene_tx=XM_003981109.5;refGene_gn=ZFYVE16;refGene_tx=XM_006927816.4;refGene_gn=ZFYVE16;refGene_tx=XM_019836033.2;refGene_gn=ZFYVE16;refGene_tx=XM_019836035.2;refGene_gn=ZFYVE16;refGene_tx=XM_023257833.1
A1 146569244 146571490
refGene_gn=ZFYVE16;refGene_tx=XM_003981109.5;refGene_gn=ZFYVE16;refGene_tx=XM_006927816.4;refGene_gn=ZFYVE16;refGene_tx=XM_019836033.2;refGene_gn=ZFYVE16;refGene_tx=XM_019836035.2;refGene_gn=ZFYVE16;refGene_tx=XM_023257833.1
A1 146573514 146573622
refGene_gn=ZFYVE16;refGene_tx=XM_003981109.5;refGene_gn=ZFYVE16;refGene_tx=XM_006927816.4;refGene_gn=ZFYVE16;refGene_tx=XM_019836033.2;refGene_gn=ZFYVE16;refGene_tx=XM_019836035.2;refGene_gn=ZFYVE16;refGene_tx=XM_023257833.1
A1 146576446 146576608
refGene_gn=ZFYVE16;refGene_tx=XM_003981109.5;refGene_gn=ZFYVE16;refGene_tx=XM_006927816.4;refGene_gn=ZFYVE16;refGene_tx=XM_019836033.2;refGene_gn=ZFYVE16;refGene_tx=XM_019836035.2
A1 146579471 146579614
refGene_gn=ZFYVE16;refGene_tx=XM_003981109.5;refGene_gn=ZFYVE16;refGene_tx=XM_006927816.4;refGene_gn=ZFYVE16;refGene_tx=XM_019836033.2;refGene_gn=ZFYVE16;refGene_tx=XM_019836035.2
A1 146581920 146582296
refGene_gn=ZFYVE16;refGene_tx=XM_003981109.5;refGene_gn=ZFYVE16;refGene_tx=XM_006927816.4;refGene_gn=ZFYVE16;refGene_tx=XM_019836033.2;refGene_gn=ZFYVE16;refGene_tx=XM_019836035.2
A1 146583519 146583642
refGene_gn=ZFYVE16;refGene_tx=XM_003981109.5;refGene_gn=ZFYVE16;refGene_tx=XM_006927816.4;refGene_gn=ZFYVE16;refGene_tx=XM_019836033.2;refGene_gn=ZFYVE16;refGene_tx=XM_019836035.2
A1 146584344 146584500
refGene_gn=ZFYVE16;refGene_tx=XM_003981109.5;refGene_gn=ZFYVE16;refGene_tx=XM_006927816.4;refGene_gn=ZFYVE16;refGene_tx=XM_019836033.2;refGene_gn=ZFYVE16;refGene_tx=XM_019836035.2
A1 146585081 146585306
refGene_gn=ZFYVE16;refGene_tx=XM_003981109.5;refGene_gn=ZFYVE16;refGene_tx=XM_006927816.4;refGene_gn=ZFYVE16;refGene_tx=XM_019836033.2;refGene_gn=ZFYVE16;refGene_tx=XM_019836035.2
A1 146588339 146588422

refGene_gn=ZFYVE16;refGene_tx=XM_003981109.5;refGene_gn=ZFYVE16;refGene_tx=XM_006927816.4;refGene_gn=ZFYVE16;refGene_tx=XM_019836033.2;refGene_gn=ZFYVE16;refGene_tx=XM_019836035.2
A1 146589185 146589290
refGene_gn=ZFYVE16;refGene_tx=XM_003981109.5;refGene_gn=ZFYVE16;refGene_tx=XM_006927816.4;refGene_gn=ZFYVE16;refGene_tx=XM_019836033.2;refGene_gn=ZFYVE16;refGene_tx=XM_019836035.2
A1 146590094 146590242
refGene_gn=ZFYVE16;refGene_tx=XM_003981109.5;refGene_gn=ZFYVE16;refGene_tx=XM_006927816.4;refGene_gn=ZFYVE16;refGene_tx=XM_019836033.2;refGene_gn=ZFYVE16;refGene_tx=XM_019836035.2
A1 146594649 146594730
refGene_gn=ZFYVE16;refGene_tx=XM_003981109.5;refGene_gn=ZFYVE16;refGene_tx=XM_006927816.4;refGene_gn=ZFYVE16;refGene_tx=XM_019836033.2;refGene_gn=ZFYVE16;refGene_tx=XM_019836035.2
A1 146601964 146602127
refGene_gn=ZFYVE16;refGene_tx=XM_003981109.5;refGene_gn=ZFYVE16;refGene_tx=XM_006927816.4;refGene_gn=ZFYVE16;refGene_tx=XM_019836033.2;refGene_gn=ZFYVE16;refGene_tx=XM_019836035.2
A1 146602967 146603073
refGene_gn=ZFYVE16;refGene_tx=XM_003981109.5;refGene_gn=ZFYVE16;refGene_tx=XM_006927816.4;refGene_gn=ZFYVE16;refGene_tx=XM_019836033.2;refGene_gn=ZFYVE16;refGene_tx=XM_019836035.2
A1 146603967 146604135
refGene_gn=ZFYVE16;refGene_tx=XM_003981109.5;refGene_gn=ZFYVE16;refGene_tx=XM_006927816.4;refGene_gn=ZFYVE16;refGene_tx=XM_019836033.2;refGene_gn=ZFYVE16;refGene_tx=XM_019836035.2
A1 146607866 146608028
refGene_gn=ZFYVE16;refGene_tx=XM_003981109.5;refGene_gn=ZFYVE16;refGene_tx=XM_006927816.4;refGene_gn=ZFYVE16;refGene_tx=XM_019836033.2;refGene_gn=ZFYVE16;refGene_tx=XM_019836035.2
A1 146622508 146622533
refGene_gn=FAM151B;refGene_tx=XM_023257854.1;refGene_gn=FAM151B;refGene_tx=XM_023257858.1;refGene_gn=FAM151B;refGene_tx=XM_023257859.1;refGene_gn=FAM151B;refGene_tx=XM_023257863.1;refGene_gn=FAM151B;refGene_tx=XM_023257864.1
A1 146628310 146628436
refGene_gn=FAM151B;refGene_tx=XM_023257854.1;refGene_gn=FAM151B;refGene_tx=XM_023257858.1;refGene_gn=FAM151B;refGene_tx=XM_023257859.1;refGene_gn=FAM151B;refGene_tx=XM_023257863.1;refGene_gn=FAM151B;refGene_tx=XM_023257864.1
A1 146630914 146631080
refGene_gn=FAM151B;refGene_tx=XM_023257854.1;refGene_gn=FAM151B;refGene_tx=XM_023257858.1;refGene_gn=FAM151B;refGene_tx=XM_023257859.1;refGene_gn=FAM151B;refGene_tx=XM_023257863.1;refGene_gn=FAM151B;refGene_tx=XM_023257864.1
A1 146639241 146639459
refGene_gn=FAM151B;refGene_tx=XM_023257854.1;refGene_gn=FAM151B;refGene_tx=XM_023257858.1;refGene_gn=FAM151B;refGene_tx=XM_023257859.1;refGene_gn=FAM151B;refGene_tx=XM_023257863.1;refGene_gn=FAM151B;refGene_tx=

XM_023257864.1
A1 146642685 146642821
refGene_gn=FAM151B;refGene_tx=XM_023257854.1;refGene_gn=FAM151B;refGene_tx=XM_023257858.1;refGene_gn=FAM151B;refGene_tx=XM_023257859.1;refGene_gn=FAM151B;refGene_tx=XM_023257863.1;refGene_gn=FAM151B;refGene_tx=XM_023257864.1
A1 146652508 146652527
refGene_gn=FAM151B;refGene_tx=XM_023257864.1
A1 146656305 146656364
refGene_gn=FAM151B;refGene_tx=XM_023257863.1
A1 146657487 146657647
refGene_gn=FAM151B;refGene_tx=XM_023257854.1;refGene_gn=FAM151B;refGene_tx=XM_023257863.1
A1 146658228 146658332
refGene_gn=FAM151B;refGene_tx=XM_023257858.1;refGene_gn=FAM151B;refGene_tx=XM_023257859.1
A1 146662582 146662596
refGene_gn=FAM151B;refGene_tx=XM_023257859.1
A1 146672303 146673848
refGene_gn=ANKRD34B;refGene_tx=XM_003981110.4;refGene_gn=ANKRD34B;refGene_tx=XM_019836089.1;refGene_gn=ANKRD34B;refGene_tx=XM_023257845.1
A1 146723402 146723481
refGene_gn=DHFR;refGene_tx=XM_003981111.4;refGene_gn=DHFR;refGene_tx=XM_011284300.2;refGene_gn=DHFR;refGene_tx=XM_019836103.1;refGene_gn=DHFR;refGene_tx=XM_019836107.1;refGene_gn=DHFR;refGene_tx=XM_023257865.1
A1 146729042 146729197
refGene_gn=DHFR;refGene_tx=XM_003981111.4;refGene_gn=DHFR;refGene_tx=XM_019836103.1;refGene_gn=DHFR;refGene_tx=XM_019836107.1;refGene_gn=DHFR;refGene_tx=XM_023257865.1
A1 146732923 146733050
refGene_gn=DHFR;refGene_tx=XM_003981111.4;refGene_gn=DHFR;refGene_tx=XM_011284300.2;refGene_gn=DHFR;refGene_tx=XM_019836103.1;refGene_gn=DHFR;refGene_tx=XM_019836107.1;refGene_gn=DHFR;refGene_tx=XM_023257865.1
A1 146739112 146739218
refGene_gn=DHFR;refGene_tx=XM_003981111.4;refGene_gn=DHFR;refGene_tx=XM_011284300.2;refGene_gn=DHFR;refGene_tx=XM_019836103.1;refGene_gn=DHFR;refGene_tx=XM_019836107.1;refGene_gn=DHFR;refGene_tx=XM_023257865.1
A1 146741431 146741481
refGene_gn=DHFR;refGene_tx=XM_003981111.4;refGene_gn=DHFR;refGene_tx=XM_011284300.2;refGene_gn=DHFR;refGene_tx=XM_019836103.1;refGene_gn=DHFR;refGene_tx=XM_019836107.1;refGene_gn=DHFR;refGene_tx=XM_023257865.1
A1 146741829 146741915
refGene_gn=DHFR;refGene_tx=XM_003981111.4;refGene_gn=DHFR;refGene_tx=XM_011284300.2;refGene_gn=DHFR;refGene_tx=XM_019836103.1;refGene_gn=DHFR;refGene_tx=XM_019836107.1;refGene_gn=DHFR;refGene_tx=XM_023257865.1
A1 146742153 146742360
refGene_gn=MSH3;refGene_tx=XM_003981121.5;refGene_gn=MSH3;refGene_tx=XM_006927817.4;refGene_gn=MSH3;refGene_tx=XM_019836078.2
A1 146744054 146744175
refGene_gn=MSH3;refGene_tx=XM_003981121.5;refGene_gn=MSH3;refGene_tx=X

M_006927817.4;refGene_gn=MSH3;refGene_tx=XM_019836078.2
A1 146755439 146755660
refGene_gn=MSH3;refGene_tx=XM_003981121.5;refGene_gn=MSH3;refGene_tx=XM_006927817.4;refGene_gn=MSH3;refGene_tx=XM_019836078.2
A1 146759132 146759348
refGene_gn=MSH3;refGene_tx=XM_003981121.5;refGene_gn=MSH3;refGene_tx=XM_006927817.4;refGene_gn=MSH3;refGene_tx=XM_019836078.2
A1 146761003 146761120
refGene_gn=MSH3;refGene_tx=XM_003981121.5;refGene_gn=MSH3;refGene_tx=XM_006927817.4;refGene_gn=MSH3;refGene_tx=XM_019836078.2
A1 146761884 146762002
refGene_gn=MSH3;refGene_tx=XM_003981121.5;refGene_gn=MSH3;refGene_tx=XM_006927817.4;refGene_gn=MSH3;refGene_tx=XM_019836078.2
A1 146763617 146763763
refGene_gn=MSH3;refGene_tx=XM_003981121.5;refGene_gn=MSH3;refGene_tx=XM_006927817.4;refGene_gn=MSH3;refGene_tx=XM_019836078.2
A1 146767927 146768094
refGene_gn=MSH3;refGene_tx=XM_003981121.5;refGene_gn=MSH3;refGene_tx=XM_006927817.4;refGene_gn=MSH3;refGene_tx=XM_019836078.2
A1 146799198 146799311
refGene_gn=MSH3;refGene_tx=XM_003981121.5;refGene_gn=MSH3;refGene_tx=XM_006927817.4;refGene_gn=MSH3;refGene_tx=XM_019836078.2
A1 146801933 146802048
refGene_gn=MSH3;refGene_tx=XM_003981121.5;refGene_gn=MSH3;refGene_tx=XM_006927817.4;refGene_gn=MSH3;refGene_tx=XM_019836078.2
A1 146814271 146814356
refGene_gn=MSH3;refGene_tx=XM_003981121.5;refGene_gn=MSH3;refGene_tx=XM_006927817.4;refGene_gn=MSH3;refGene_tx=XM_019836078.2
A1 146817412 146817522
refGene_gn=MSH3;refGene_tx=XM_003981121.5;refGene_gn=MSH3;refGene_tx=XM_006927817.4;refGene_gn=MSH3;refGene_tx=XM_019836078.2
A1 146835064 146835197
refGene_gn=MSH3;refGene_tx=XM_003981121.5;refGene_gn=MSH3;refGene_tx=XM_006927817.4;refGene_gn=MSH3;refGene_tx=XM_019836078.2
A1 146848826 146849014
refGene_gn=MSH3;refGene_tx=XM_003981121.5;refGene_gn=MSH3;refGene_tx=XM_006927817.4;refGene_gn=MSH3;refGene_tx=XM_019836078.2
A1 146849744 146849913
refGene_gn=MSH3;refGene_tx=XM_003981121.5;refGene_gn=MSH3;refGene_tx=XM_006927817.4;refGene_gn=MSH3;refGene_tx=XM_019836078.2
A1 146855483 146855548
refGene_gn=MSH3;refGene_tx=XM_003981121.5;refGene_gn=MSH3;refGene_tx=XM_006927817.4;refGene_gn=MSH3;refGene_tx=XM_019836078.2
A1 146858133 146858250
refGene_gn=MSH3;refGene_tx=XM_003981121.5;refGene_gn=MSH3;refGene_tx=XM_006927817.4;refGene_gn=MSH3;refGene_tx=XM_019836078.2
A1 146863591 146863699
refGene_gn=MSH3;refGene_tx=XM_003981121.5;refGene_gn=MSH3;refGene_tx=XM_006927817.4;refGene_gn=MSH3;refGene_tx=XM_019836078.2
A1 146871645 146871757

refGene_gn=MSH3;refGene_tx=XM_003981121.5;refGene_gn=MSH3;refGene_tx=XM_006927817.4;refGene_gn=MSH3;refGene_tx=XM_019836078.2
A1 146889070 146889228
refGene_gn=MSH3;refGene_tx=XM_003981121.5;refGene_gn=MSH3;refGene_tx=XM_006927817.4;refGene_gn=MSH3;refGene_tx=XM_019836078.2
A1 146914794 146914981
refGene_gn=MSH3;refGene_tx=XM_003981121.5;refGene_gn=MSH3;refGene_tx=XM_006927817.4;refGene_gn=MSH3;refGene_tx=XM_019836078.2
A1 146919407 146919537
refGene_gn=MSH3;refGene_tx=XM_003981121.5;refGene_gn=MSH3;refGene_tx=XM_006927817.4;refGene_gn=MSH3;refGene_tx=XM_019836078.2
A1 146926751 146926920
refGene_gn=MSH3;refGene_tx=XM_003981121.5;refGene_gn=MSH3;refGene_tx=XM_006927817.4
A1 146929367 146929476
refGene_gn=MSH3;refGene_tx=XM_003981121.5;refGene_gn=MSH3;refGene_tx=XM_006927817.4;refGene_gn=MSH3;refGene_tx=XM_019836078.2
A1 146976838 146977126
refGene_gn=RASGRF2;refGene_tx=XM_023257887.1;refGene_gn=RASGRF2;refGene_tx=XM_023257891.1;refGene_gn=RASGRF2;refGene_tx=XM_023257899.1;refGene_gn=RASGRF2;refGene_tx=XM_023257908.1;refGene_gn=RASGRF2;refGene_tx=XM_023257915.1
A1 147051469 147051576
refGene_gn=RASGRF2;refGene_tx=XM_023257887.1;refGene_gn=RASGRF2;refGene_tx=XM_023257891.1;refGene_gn=RASGRF2;refGene_tx=XM_023257899.1;refGene_gn=RASGRF2;refGene_tx=XM_023257908.1;refGene_gn=RASGRF2;refGene_tx=XM_023257915.1
A1 147070065 147070213
refGene_gn=RASGRF2;refGene_tx=XM_023257887.1;refGene_gn=RASGRF2;refGene_tx=XM_023257891.1;refGene_gn=RASGRF2;refGene_tx=XM_023257899.1;refGene_gn=RASGRF2;refGene_tx=XM_023257908.1;refGene_gn=RASGRF2;refGene_tx=XM_023257915.1
A1 147072541 147072631
refGene_gn=RASGRF2;refGene_tx=XM_023257887.1;refGene_gn=RASGRF2;refGene_tx=XM_023257891.1;refGene_gn=RASGRF2;refGene_tx=XM_023257899.1;refGene_gn=RASGRF2;refGene_tx=XM_023257908.1;refGene_gn=RASGRF2;refGene_tx=XM_023257915.1
A1 147077148 147077402
refGene_gn=RASGRF2;refGene_tx=XM_023257887.1;refGene_gn=RASGRF2;refGene_tx=XM_023257891.1;refGene_gn=RASGRF2;refGene_tx=XM_023257899.1;refGene_gn=RASGRF2;refGene_tx=XM_023257908.1;refGene_gn=RASGRF2;refGene_tx=XM_023257915.1
A1 147080490 147080570
refGene_gn=RASGRF2;refGene_tx=XM_023257887.1;refGene_gn=RASGRF2;refGene_tx=XM_023257891.1;refGene_gn=RASGRF2;refGene_tx=XM_023257899.1;refGene_gn=RASGRF2;refGene_tx=XM_023257908.1;refGene_gn=RASGRF2;refGene_tx=XM_023257915.1
A1 147080972 147081166
refGene_gn=RASGRF2;refGene_tx=XM_023257887.1;refGene_gn=RASGRF2;refGene_tx=XM_023257891.1;refGene_gn=RASGRF2;refGene_tx=XM_023257899.1;refGe

ne_gn=RASGRF2;refGene_tx=XM_023257908.1;refGene_gn=RASGRF2;refGene_tx=XM_023257915.1
A1 147086216 147086326
refGene_gn=RASGRF2;refGene_tx=XM_023257887.1;refGene_gn=RASGRF2;refGene_tx=XM_023257891.1;refGene_gn=RASGRF2;refGene_tx=XM_023257899.1;refGene_gn=RASGRF2;refGene_tx=XM_023257908.1;refGene_gn=RASGRF2;refGene_tx=XM_023257915.1
A1 147087071 147087190
refGene_gn=RASGRF2;refGene_tx=XM_023257887.1;refGene_gn=RASGRF2;refGene_tx=XM_023257891.1;refGene_gn=RASGRF2;refGene_tx=XM_023257899.1;refGene_gn=RASGRF2;refGene_tx=XM_023257908.1;refGene_gn=RASGRF2;refGene_tx=XM_023257915.1
A1 147093381 147093542
refGene_gn=RASGRF2;refGene_tx=XM_023257887.1;refGene_gn=RASGRF2;refGene_tx=XM_023257891.1;refGene_gn=RASGRF2;refGene_tx=XM_023257899.1;refGene_gn=RASGRF2;refGene_tx=XM_023257908.1;refGene_gn=RASGRF2;refGene_tx=XM_023257915.1
A1 147094453 147094520
refGene_gn=RASGRF2;refGene_tx=XM_023257887.1;refGene_gn=RASGRF2;refGene_tx=XM_023257891.1;refGene_gn=RASGRF2;refGene_tx=XM_023257899.1;refGene_gn=RASGRF2;refGene_tx=XM_023257908.1;refGene_gn=RASGRF2;refGene_tx=XM_023257915.1
A1 147095059 147095196
refGene_gn=RASGRF2;refGene_tx=XM_023257887.1;refGene_gn=RASGRF2;refGene_tx=XM_023257891.1;refGene_gn=RASGRF2;refGene_tx=XM_023257899.1;refGene_gn=RASGRF2;refGene_tx=XM_023257908.1;refGene_gn=RASGRF2;refGene_tx=XM_023257915.1
A1 147104013 147104096
refGene_gn=RASGRF2;refGene_tx=XM_023257887.1;refGene_gn=RASGRF2;refGene_tx=XM_023257891.1;refGene_gn=RASGRF2;refGene_tx=XM_023257899.1;refGene_gn=RASGRF2;refGene_tx=XM_023257908.1;refGene_gn=RASGRF2;refGene_tx=XM_023257915.1
A1 147109330 147109579
refGene_gn=RASGRF2;refGene_tx=XM_023257887.1;refGene_gn=RASGRF2;refGene_tx=XM_023257891.1;refGene_gn=RASGRF2;refGene_tx=XM_023257899.1;refGene_gn=RASGRF2;refGene_tx=XM_023257908.1;refGene_gn=RASGRF2;refGene_tx=XM_023257915.1
A1 147110204 147110593
refGene_gn=RASGRF2;refGene_tx=XM_023257887.1;refGene_gn=RASGRF2;refGene_tx=XM_023257891.1;refGene_gn=RASGRF2;refGene_tx=XM_023257899.1;refGene_gn=RASGRF2;refGene_tx=XM_023257908.1;refGene_gn=RASGRF2;refGene_tx=XM_023257915.1
A1 147116799 147116925
refGene_gn=RASGRF2;refGene_tx=XM_023257887.1;refGene_gn=RASGRF2;refGene_tx=XM_023257891.1;refGene_gn=RASGRF2;refGene_tx=XM_023257899.1;refGene_gn=RASGRF2;refGene_tx=XM_023257908.1;refGene_gn=RASGRF2;refGene_tx=XM_023257915.1
A1 147119598 147119688
refGene_gn=RASGRF2;refGene_tx=XM_023257887.1;refGene_gn=RASGRF2;refGene_tx=XM_023257891.1;refGene_gn=RASGRF2;refGene_tx=XM_023257899.1;refGe

ne_gn=RASGRF2;refGene_tx=XM_023257908.1;refGene_gn=RASGRF2;refGene_tx=XM_023257915.1
A1 147149625 147149670
refGene_gn=RASGRF2;refGene_tx=XM_023257887.1;refGene_gn=RASGRF2;refGene_tx=XM_023257891.1;refGene_gn=RASGRF2;refGene_tx=XM_023257908.1
A1 147171554 147171661
refGene_gn=RASGRF2;refGene_tx=XM_023257887.1;refGene_gn=RASGRF2;refGene_tx=XM_023257891.1;refGene_gn=RASGRF2;refGene_tx=XM_023257899.1;refGene_gn=RASGRF2;refGene_tx=XM_023257908.1;refGene_gn=RASGRF2;refGene_tx=XM_023257915.1
A1 147189684 147189797
refGene_gn=RASGRF2;refGene_tx=XM_023257887.1;refGene_gn=RASGRF2;refGene_tx=XM_023257891.1;refGene_gn=RASGRF2;refGene_tx=XM_023257899.1;refGene_gn=RASGRF2;refGene_tx=XM_023257908.1;refGene_gn=RASGRF2;refGene_tx=XM_023257915.1
A1 147197743 147197804
refGene_gn=RASGRF2;refGene_tx=XM_023257887.1;refGene_gn=RASGRF2;refGene_tx=XM_023257891.1;refGene_gn=RASGRF2;refGene_tx=XM_023257899.1;refGene_gn=RASGRF2;refGene_tx=XM_023257908.1;refGene_gn=RASGRF2;refGene_tx=XM_023257915.1
A1 147198111 147198215
refGene_gn=RASGRF2;refGene_tx=XM_023257887.1;refGene_gn=RASGRF2;refGene_tx=XM_023257891.1;refGene_gn=RASGRF2;refGene_tx=XM_023257899.1;refGene_gn=RASGRF2;refGene_tx=XM_023257908.1;refGene_gn=RASGRF2;refGene_tx=XM_023257915.1
A1 147200628 147200713
refGene_gn=RASGRF2;refGene_tx=XM_023257887.1;refGene_gn=RASGRF2;refGene_tx=XM_023257891.1;refGene_gn=RASGRF2;refGene_tx=XM_023257899.1;refGene_gn=RASGRF2;refGene_tx=XM_023257908.1;refGene_gn=RASGRF2;refGene_tx=XM_023257915.1
A1 147203464 147203662
refGene_gn=RASGRF2;refGene_tx=XM_023257887.1;refGene_gn=RASGRF2;refGene_tx=XM_023257891.1;refGene_gn=RASGRF2;refGene_tx=XM_023257899.1;refGene_gn=RASGRF2;refGene_tx=XM_023257908.1;refGene_gn=RASGRF2;refGene_tx=XM_023257915.1
A1 147208797 147208877
refGene_gn=RASGRF2;refGene_tx=XM_023257887.1;refGene_gn=RASGRF2;refGene_tx=XM_023257891.1;refGene_gn=RASGRF2;refGene_tx=XM_023257899.1;refGene_gn=RASGRF2;refGene_tx=XM_023257908.1;refGene_gn=RASGRF2;refGene_tx=XM_023257915.1
A1 147210198 147210316
refGene_gn=RASGRF2;refGene_tx=XM_023257887.1;refGene_gn=RASGRF2;refGene_tx=XM_023257891.1;refGene_gn=RASGRF2;refGene_tx=XM_023257899.1;refGene_gn=RASGRF2;refGene_tx=XM_023257908.1;refGene_gn=RASGRF2;refGene_tx=XM_023257915.1
A1 147210869 147210938
refGene_gn=RASGRF2;refGene_tx=XM_023257887.1;refGene_gn=RASGRF2;refGene_tx=XM_023257891.1;refGene_gn=RASGRF2;refGene_tx=XM_023257899.1;refGene_gn=RASGRF2;refGene_tx=XM_023257908.1;refGene_gn=RASGRF2;refGene_tx=XM_023257915.1

A1 147213749 147213842
refGene_gn=RASGRF2;refGene_tx=XM_023257887.1;refGene_gn=RASGRF2;refGene_tx=XM_023257891.1;refGene_gn=RASGRF2;refGene_tx=XM_023257899.1;refGene_gn=RASGRF2;refGene_tx=XM_023257908.1;refGene_gn=RASGRF2;refGene_tx=XM_023257915.1
A1 147239013 147239165
refGene_gn=CKMT2;refGene_tx=XM_003981112.3
A1 147240584 147240783
refGene_gn=CKMT2;refGene_tx=XM_003981112.3
A1 147242137 147242233
refGene_gn=CKMT2;refGene_tx=XM_003981112.3
A1 147242600 147242822
refGene_gn=CKMT2;refGene_tx=XM_003981112.3
A1 147244942 147245028
refGene_gn=CKMT2;refGene_tx=XM_003981112.3
A1 147245898 147246022
refGene_gn=CKMT2;refGene_tx=XM_003981112.3
A1 147246703 147246838
refGene_gn=CKMT2;refGene_tx=XM_003981112.3
A1 147247945 147248071
refGene_gn=CKMT2;refGene_tx=XM_003981112.3
A1 147258051 147258171
refGene_gn=CKMT2;refGene_tx=XM_003981112.3
A1 147293285 147293669
refGene_gn=ZCCHC9;refGene_tx=XM_003981113.4
A1 147294347 147294498
refGene_gn=ZCCHC9;refGene_tx=XM_003981113.4
A1 147294731 147294824
refGene_gn=ZCCHC9;refGene_tx=XM_003981113.4
A1 147296551 147296559
refGene_gn=ACOT12;refGene_tx=XM_023257929.1
A1 147296742 147296811
refGene_gn=ZCCHC9;refGene_tx=XM_003981113.4
A1 147298077 147298196
refGene_gn=ZCCHC9;refGene_tx=XM_003981113.4
A1 147301157 147301307
refGene_gn=ACOT12;refGene_tx=XM_023257923.1;refGene_gn=ACOT12;refGene_tx=XM_023257927.1;refGene_gn=ACOT12;refGene_tx=XM_023257929.1;refGene_gn=ACOT12;refGene_tx=XM_023257934.1;refGene_gn=ACOT12;refGene_tx=XM_023257940.1
A1 147301567 147301694
refGene_gn=ACOT12;refGene_tx=XM_023257923.1;refGene_gn=ACOT12;refGene_tx=XM_023257927.1;refGene_gn=ACOT12;refGene_tx=XM_023257929.1;refGene_gn=ACOT12;refGene_tx=XM_023257934.1;refGene_gn=ACOT12;refGene_tx=XM_023257940.1
A1 147303270 147303399
refGene_gn=ACOT12;refGene_tx=XM_023257923.1;refGene_gn=ACOT12;refGene_tx=XM_023257927.1;refGene_gn=ACOT12;refGene_tx=XM_023257929.1;refGene_gn=ACOT12;refGene_tx=XM_023257940.1
A1 147305600 147305734

refGene_gn=ACOT12;refGene_tx=XM_023257923.1;refGene_gn=ACOT12;refGene_tx=XM_023257927.1;refGene_gn=ACOT12;refGene_tx=XM_023257929.1;refGene_gn=ACOT12;refGene_tx=XM_023257934.1;refGene_gn=ACOT12;refGene_tx=XM_023257940.1
A1 147310677 147310761
refGene_gn=ACOT12;refGene_tx=XM_023257923.1;refGene_gn=ACOT12;refGene_tx=XM_023257927.1;refGene_gn=ACOT12;refGene_tx=XM_023257929.1;refGene_gn=ACOT12;refGene_tx=XM_023257934.1;refGene_gn=ACOT12;refGene_tx=XM_023257940.1
A1 147311439 147311503
refGene_gn=ACOT12;refGene_tx=XM_023257923.1;refGene_gn=ACOT12;refGene_tx=XM_023257927.1;refGene_gn=ACOT12;refGene_tx=XM_023257929.1;refGene_gn=ACOT12;refGene_tx=XM_023257934.1;refGene_gn=ACOT12;refGene_tx=XM_023257940.1
A1 147311791 147311847
refGene_gn=ACOT12;refGene_tx=XM_023257923.1;refGene_gn=ACOT12;refGene_tx=XM_023257927.1;refGene_gn=ACOT12;refGene_tx=XM_023257929.1;refGene_gn=ACOT12;refGene_tx=XM_023257934.1;refGene_gn=ACOT12;refGene_tx=XM_023257940.1
A1 147312501 147312652
refGene_gn=ACOT12;refGene_tx=XM_023257923.1;refGene_gn=ACOT12;refGene_tx=XM_023257927.1;refGene_gn=ACOT12;refGene_tx=XM_023257929.1;refGene_gn=ACOT12;refGene_tx=XM_023257934.1;refGene_gn=ACOT12;refGene_tx=XM_023257940.1
A1 147313592 147313712
refGene_gn=ACOT12;refGene_tx=XM_023257923.1;refGene_gn=ACOT12;refGene_tx=XM_023257927.1;refGene_gn=ACOT12;refGene_tx=XM_023257929.1;refGene_gn=ACOT12;refGene_tx=XM_023257934.1;refGene_gn=ACOT12;refGene_tx=XM_023257940.1
A1 147315721 147315878
refGene_gn=ACOT12;refGene_tx=XM_023257923.1;refGene_gn=ACOT12;refGene_tx=XM_023257927.1;refGene_gn=ACOT12;refGene_tx=XM_023257929.1;refGene_gn=ACOT12;refGene_tx=XM_023257934.1;refGene_gn=ACOT12;refGene_tx=XM_023257940.1
A1 147324291 147324427
refGene_gn=ACOT12;refGene_tx=XM_023257923.1;refGene_gn=ACOT12;refGene_tx=XM_023257927.1;refGene_gn=ACOT12;refGene_tx=XM_023257929.1;refGene_gn=ACOT12;refGene_tx=XM_023257934.1;refGene_gn=ACOT12;refGene_tx=XM_023257940.1
A1 147330185 147330287
refGene_gn=ACOT12;refGene_tx=XM_023257923.1;refGene_gn=ACOT12;refGene_tx=XM_023257927.1;refGene_gn=ACOT12;refGene_tx=XM_023257929.1;refGene_gn=ACOT12;refGene_tx=XM_023257934.1;refGene_gn=ACOT12;refGene_tx=XM_023257940.1
A1 147333818 147333879
refGene_gn=ACOT12;refGene_tx=XM_023257923.1;refGene_gn=ACOT12;refGene_tx=XM_023257927.1;refGene_gn=ACOT12;refGene_tx=XM_023257929.1;refGene_gn=ACOT12;refGene_tx=XM_023257934.1;refGene_gn=ACOT12;refGene_tx=XM_023257940.1
A1 147343751 147343821

refGene_gn=ACOT12;refGene_tx=XM_023257923.1;refGene_gn=ACOT12;refGene_tx=XM_023257927.1;refGene_gn=ACOT12;refGene_tx=XM_023257929.1;refGene_gn=ACOT12;refGene_tx=XM_023257934.1
A1 147350707 147350834
refGene_gn=ACOT12;refGene_tx=XM_023257923.1;refGene_gn=ACOT12;refGene_tx=XM_023257927.1;refGene_gn=ACOT12;refGene_tx=XM_023257929.1;refGene_gn=ACOT12;refGene_tx=XM_023257934.1
A1 147379500 147379530
refGene_gn=SSBP2;refGene_tx=XM_011284322.2;refGene_gn=SSBP2;refGene_tx=XM_011284323.3;refGene_gn=SSBP2;refGene_tx=XM_011284326.3;refGene_gn=SSBP2;refGene_tx=XM_011284330.2;refGene_gn=SSBP2;refGene_tx=XM_019836180.1
A1 147386297 147386396
refGene_gn=SSBP2;refGene_tx=XM_011284322.2;refGene_gn=SSBP2;refGene_tx=XM_011284323.3;refGene_gn=SSBP2;refGene_tx=XM_011284326.3;refGene_gn=SSBP2;refGene_tx=XM_011284330.2;refGene_gn=SSBP2;refGene_tx=XM_019836180.1
A1 147400395 147400424
refGene_gn=SSBP2;refGene_tx=XM_011284322.2;refGene_gn=SSBP2;refGene_tx=XM_011284323.3;refGene_gn=SSBP2;refGene_tx=XM_011284326.3;refGene_gn=SSBP2;refGene_tx=XM_011284330.2;refGene_gn=SSBP2;refGene_tx=XM_019836180.1
A1 147402940 147403019
refGene_gn=SSBP2;refGene_tx=XM_011284322.2;refGene_gn=SSBP2;refGene_tx=XM_011284323.3;refGene_gn=SSBP2;refGene_tx=XM_011284326.3;refGene_gn=SSBP2;refGene_tx=XM_011284330.2;refGene_gn=SSBP2;refGene_tx=XM_019836180.1
A1 147406000 147406071
refGene_gn=SSBP2;refGene_tx=XM_011284322.2;refGene_gn=SSBP2;refGene_tx=XM_011284323.3;refGene_gn=SSBP2;refGene_tx=XM_011284326.3;refGene_gn=SSBP2;refGene_tx=XM_011284330.2;refGene_gn=SSBP2;refGene_tx=XM_019836180.1
A1 147409341 147409396
refGene_gn=SSBP2;refGene_tx=XM_011284322.2;refGene_gn=SSBP2;refGene_tx=XM_011284323.3;refGene_gn=SSBP2;refGene_tx=XM_011284326.3;refGene_gn=SSBP2;refGene_tx=XM_011284330.2;refGene_gn=SSBP2;refGene_tx=XM_019836180.1
A1 147411211 147411247
refGene_gn=SSBP2;refGene_tx=XM_011284322.2;refGene_gn=SSBP2;refGene_tx=XM_011284323.3;refGene_gn=SSBP2;refGene_tx=XM_011284326.3;refGene_gn=SSBP2;refGene_tx=XM_011284330.2;refGene_gn=SSBP2;refGene_tx=XM_019836180.1
A1 147422850 147422899
refGene_gn=SSBP2;refGene_tx=XM_011284322.2;refGene_gn=SSBP2;refGene_tx=XM_011284323.3;refGene_gn=SSBP2;refGene_tx=XM_011284326.3;refGene_gn=SSBP2;refGene_tx=XM_011284330.2;refGene_gn=SSBP2;refGene_tx=XM_019836180.1
A1 147429061 147429153
refGene_gn=SSBP2;refGene_tx=XM_011284322.2;refGene_gn=SSBP2;refGene_tx=XM_011284323.3;refGene_gn=SSBP2;refGene_tx=XM_011284326.3;refGene_gn=

SSBP2;refGene_tx=XM_011284330.2;refGene_gn=SSBP2;refGene_tx=XM_0198361
80.1
A1 147435859 147435930
refGene_gn=SSBP2;refGene_tx=XM_011284322.2;refGene_gn=SSBP2;refGene_tx
=XM_011284323.3;refGene_gn=SSBP2;refGene_tx=XM_011284326.3;refGene_gn=
SSBP2;refGene_tx=XM_011284330.2;refGene_gn=SSBP2;refGene_tx=XM_0198361
80.1
A1 147436662 147436729
refGene_gn=SSBP2;refGene_tx=XM_011284322.2;refGene_gn=SSBP2;refGene_tx
=XM_011284323.3;refGene_gn=SSBP2;refGene_tx=XM_011284326.3;refGene_gn=
SSBP2;refGene_tx=XM_011284330.2;refGene_gn=SSBP2;refGene_tx=XM_0198361
80.1
A1 147448461 147448521
refGene_gn=SSBP2;refGene_tx=XM_011284322.2;refGene_gn=SSBP2;refGene_tx
=XM_011284323.3;refGene_gn=SSBP2;refGene_tx=XM_011284326.3;refGene_gn=
SSBP2;refGene_tx=XM_011284330.2;refGene_gn=SSBP2;refGene_tx=XM_0198361
80.1
A1 147469877 147469967
refGene_gn=SSBP2;refGene_tx=XM_011284322.2;refGene_gn=SSBP2;refGene_tx
=XM_011284323.3;refGene_gn=SSBP2;refGene_tx=XM_019836180.1
A1 147575091 147575176
refGene_gn=SSBP2;refGene_tx=XM_011284322.2;refGene_gn=SSBP2;refGene_tx
=XM_011284323.3;refGene_gn=SSBP2;refGene_tx=XM_011284326.3;refGene_gn=
SSBP2;refGene_tx=XM_011284330.2;refGene_gn=SSBP2;refGene_tx=XM_0198361
80.1
A1 147589442 147589504
refGene_gn=SSBP2;refGene_tx=XM_011284322.2;refGene_gn=SSBP2;refGene_tx
=XM_011284323.3;refGene_gn=SSBP2;refGene_tx=XM_011284326.3;refGene_gn=
SSBP2;refGene_tx=XM_011284330.2;refGene_gn=SSBP2;refGene_tx=XM_0198361
80.1
A1 147602228 147602301
refGene_gn=SSBP2;refGene_tx=XM_011284322.2;refGene_gn=SSBP2;refGene_tx
=XM_011284323.3;refGene_gn=SSBP2;refGene_tx=XM_011284326.3;refGene_gn=
SSBP2;refGene_tx=XM_011284330.2;refGene_gn=SSBP2;refGene_tx=XM_0198361
80.1
A1 147695967 147696029
refGene_gn=SSBP2;refGene_tx=XM_011284322.2;refGene_gn=SSBP2;refGene_tx
=XM_011284323.3;refGene_gn=SSBP2;refGene_tx=XM_011284326.3;refGene_gn=
SSBP2;refGene_tx=XM_011284330.2
A1 147697091 147697132
refGene_gn=SSBP2;refGene_tx=XM_019836180.1
A1 147876701 147876791
refGene_gn=ATG10;refGene_tx=XM_006927827.4;refGene_gn=ATG10;refGene_tx
=XM_006927828.4;refGene_gn=ATG10;refGene_tx=XM_019836218.2;refGene_gn=
ATG10;refGene_tx=XM_019836227.2;refGene_gn=ATG10;refGene_tx=XM_0198362
34.2;refGene_gn=ATG10;refGene_tx=XM_019836266.2;refGene_gn=ATG10;refGe
ne_tx=XM_019836280.2
A1 147890583 147890703
refGene_gn=ATG10;refGene_tx=XM_006927827.4;refGene_gn=ATG10;refGene_tx
=XM_006927828.4;refGene_gn=ATG10;refGene_tx=XM_006927829.4;refGene_gn=

ATG10;refGene_tx=XM_006927830.4;refGene_gn=ATG10;refGene_tx=XM_0198362
18.2;refGene_gn=ATG10;refGene_tx=XM_019836227.2;refGene_gn=ATG10;refGe
ne_tx=XM_019836234.2;refGene_gn=ATG10;refGene_tx=XM_019836263.2;refGen
e_gn=ATG10;refGene_tx=XM_019836266.2;refGene_gn=ATG10;refGene_tx=XM_01
9836280.2;refGene_gn=ATG10;refGene_tx=XM_023258050.1
A1 147925304 147925388
refGene_gn=ATG10;refGene_tx=XM_006927827.4;refGene_gn=ATG10;refGene_tx
=XM_006927829.4;refGene_gn=ATG10;refGene_tx=XM_006927830.4;refGene_gn=
ATG10;refGene_tx=XM_019836218.2;refGene_gn=ATG10;refGene_tx=XM_0198362
27.2;refGene_gn=ATG10;refGene_tx=XM_019836234.2;refGene_gn=ATG10;refGe
ne_tx=XM_019836263.2;refGene_gn=ATG10;refGene_tx=XM_019836266.2;refGen
e_gn=ATG10;refGene_tx=XM_019836280.2;refGene_gn=ATG10;refGene_tx=XM_02
3258050.1
A1 147939697 147939805
refGene_gn=ATG10;refGene_tx=XM_006927827.4;refGene_gn=ATG10;refGene_tx
=XM_006927828.4;refGene_gn=ATG10;refGene_tx=XM_006927829.4;refGene_gn=
ATG10;refGene_tx=XM_006927830.4;refGene_gn=ATG10;refGene_tx=XM_0198362
18.2;refGene_gn=ATG10;refGene_tx=XM_019836227.2;refGene_gn=ATG10;refGe
ne_tx=XM_019836234.2;refGene_gn=ATG10;refGene_tx=XM_019836263.2;refGen
e_gn=ATG10;refGene_tx=XM_019836266.2;refGene_gn=ATG10;refGene_tx=XM_01
9836280.2;refGene_gn=ATG10;refGene_tx=XM_023258050.1
A1 148038660 148038799
refGene_gn=ATG10;refGene_tx=XM_006927827.4;refGene_gn=ATG10;refGene_tx
=XM_006927828.4;refGene_gn=ATG10;refGene_tx=XM_006927829.4;refGene_gn=
ATG10;refGene_tx=XM_006927830.4;refGene_gn=ATG10;refGene_tx=XM_0198362
18.2;refGene_gn=ATG10;refGene_tx=XM_019836227.2;refGene_gn=ATG10;refGe
ne_tx=XM_019836234.2;refGene_gn=ATG10;refGene_tx=XM_019836263.2;refGen
e_gn=ATG10;refGene_tx=XM_019836266.2;refGene_gn=ATG10;refGene_tx=XM_01
9836280.2;refGene_gn=ATG10;refGene_tx=XM_023258050.1
A1 148053567 148053665
refGene_gn=ATG10;refGene_tx=XM_006927827.4;refGene_gn=ATG10;refGene_tx
=XM_006927828.4;refGene_gn=ATG10;refGene_tx=XM_006927829.4;refGene_gn=
ATG10;refGene_tx=XM_006927830.4;refGene_gn=ATG10;refGene_tx=XM_0198362
18.2;refGene_gn=ATG10;refGene_tx=XM_019836227.2;refGene_gn=ATG10;refGe
ne_tx=XM_019836234.2;refGene_gn=ATG10;refGene_tx=XM_019836263.2;refGen
e_gn=ATG10;refGene_tx=XM_019836266.2;refGene_gn=ATG10;refGene_tx=XM_01
9836280.2;refGene_gn=ATG10;refGene_tx=XM_023258050.1
A1 148092122 148093031
refGene_gn=LOC109502094;refGene_tx=XM_023258008.1
A1 148126037 148126135
refGene_gn=ATG10;refGene_tx=XM_006927827.4;refGene_gn=ATG10;refGene_tx
=XM_006927828.4;refGene_gn=ATG10;refGene_tx=XM_006927829.4;refGene_gn=
ATG10;refGene_tx=XM_006927830.4;refGene_gn=ATG10;refGene_tx=XM_0198362
18.2;refGene_gn=ATG10;refGene_tx=XM_019836227.2;refGene_gn=ATG10;refGe
ne_tx=XM_019836234.2;refGene_gn=ATG10;refGene_tx=XM_019836263.2;refGen
e_gn=ATG10;refGene_tx=XM_023258050.1
A1 148126790 148126890
refGene_gn=ATG10;refGene_tx=XM_006927827.4;refGene_gn=ATG10;refGene_tx
=XM_006927828.4;refGene_gn=ATG10;refGene_tx=XM_006927829.4;refGene_gn=
ATG10;refGene_tx=XM_006927830.4;refGene_gn=ATG10;refGene_tx=XM_0198362

18.2;refGene_gn=ATG10;refGene_tx=XM_019836227.2;refGene_gn=ATG10;refGene_tx=XM_019836234.2;refGene_gn=ATG10;refGene_tx=XM_019836263.2;refGene_gn=ATG10;refGene_tx=XM_023258050.1
A1 148138718 148138760
refGene_gn=ATG10;refGene_tx=XM_019836266.2
A1 148141157 148141304
refGene_gn=RPS23;refGene_tx=XM_003981125.3
A1 148141429 148141550
refGene_gn=RPS23;refGene_tx=XM_003981125.3
A1 148142166 148142326
refGene_gn=RPS23;refGene_tx=XM_003981125.3
A1 148142803 148143023
refGene_gn=RPS23;refGene_tx=XM_003981125.3
A1 148146566 148146605
refGene_gn=ATG10;refGene_tx=XM_019836266.2;refGene_gn=ATG10;refGene_tx=XM_019836280.2
A1 148160054 148160149
refGene_gn=ATP6AP1L;refGene_tx=XM_011284337.3
A1 148167462 148167506
refGene_gn=ATP6AP1L;refGene_tx=XM_011284337.3
A1 148167762 148167824
refGene_gn=ATP6AP1L;refGene_tx=XM_011284337.3
A1 148169368 148169604
refGene_gn=ATP6AP1L;refGene_tx=XM_011284337.3
A1 148175578 148175629
refGene_gn=ATP6AP1L;refGene_tx=XM_011284337.3
A1 148177978 148178219
refGene_gn=ATP6AP1L;refGene_tx=XM_011284337.3
A1 148182835 148183138
refGene_gn=ATP6AP1L;refGene_tx=XM_011284337.3
A1 148806011 148806082
refGene_gn=TMEM167A;refGene_tx=XM_003981126.4
A1 148810806 148810841
refGene_gn=TMEM167A;refGene_tx=XM_003981126.4
A1 148815840 148815950
refGene_gn=TMEM167A;refGene_tx=XM_003981126.4
A1 148826262 148826265
refGene_gn=TMEM167A;refGene_tx=XM_003981126.4
A1 148840226 148840365
refGene_gn=XRCC4;refGene_tx=XM_011284381.3;refGene_gn=XRCC4;refGene_tx=XM_011284386.2;refGene_gn=XRCC4;refGene_tx=XM_019836335.1
A1 148847370 148847546
refGene_gn=XRCC4;refGene_tx=XM_011284381.3;refGene_gn=XRCC4;refGene_tx=XM_011284386.2;refGene_gn=XRCC4;refGene_tx=XM_019836335.1
A1 148915632 148915799
refGene_gn=XRCC4;refGene_tx=XM_011284381.3;refGene_gn=XRCC4;refGene_tx=XM_011284386.2;refGene_gn=XRCC4;refGene_tx=XM_019836335.1
A1 148922278 148922431
refGene_gn=XRCC4;refGene_tx=XM_011284381.3;refGene_gn=XRCC4;refGene_tx=XM_011284386.2;refGene_gn=XRCC4;refGene_tx=XM_019836335.1

A1 148923519 148923626
refGene_gn=XRCC4;refGene_tx=XM_011284381.3;refGene_gn=XRCC4;refGene_tx=XM_011284386.2;refGene_gn=XRCC4;refGene_tx=XM_019836335.1
A1 148964266 148964411
refGene_gn=XRCC4;refGene_tx=XM_011284381.3;refGene_gn=XRCC4;refGene_tx=XM_011284386.2;refGene_gn=XRCC4;refGene_tx=XM_019836335.1
A1 149054919 149055031
refGene_gn=XRCC4;refGene_tx=XM_011284381.3;refGene_gn=XRCC4;refGene_tx=XM_011284386.2;refGene_gn=XRCC4;refGene_tx=XM_019836335.1
A1 149162404 149162474
refGene_gn=VCAN;refGene_tx=XM_003981130.5;refGene_gn=VCAN;refGene_tx=XM_003981131.5;refGene_gn=VCAN;refGene_tx=XM_003981132.4;refGene_gn=VCAN;refGene_tx=XM_003981133.5
A1 149167534 149167912
refGene_gn=VCAN;refGene_tx=XM_003981130.5;refGene_gn=VCAN;refGene_tx=XM_003981131.5;refGene_gn=VCAN;refGene_tx=XM_003981132.4;refGene_gn=VCAN;refGene_tx=XM_003981133.5
A1 149171091 149171266
refGene_gn=VCAN;refGene_tx=XM_003981130.5;refGene_gn=VCAN;refGene_tx=XM_003981131.5;refGene_gn=VCAN;refGene_tx=XM_003981132.4;refGene_gn=VCAN;refGene_tx=XM_003981133.5
A1 149171351 149171479
refGene_gn=VCAN;refGene_tx=XM_003981130.5;refGene_gn=VCAN;refGene_tx=XM_003981131.5;refGene_gn=VCAN;refGene_tx=XM_003981132.4;refGene_gn=VCAN;refGene_tx=XM_003981133.5
A1 149189886 149190180
refGene_gn=VCAN;refGene_tx=XM_003981130.5;refGene_gn=VCAN;refGene_tx=XM_003981131.5;refGene_gn=VCAN;refGene_tx=XM_003981132.4;refGene_gn=VCAN;refGene_tx=XM_003981133.5
A1 149196802 149199802
refGene_gn=VCAN;refGene_tx=XM_003981130.5;refGene_gn=VCAN;refGene_tx=XM_003981133.5
A1 149218112 149223329
refGene_gn=VCAN;refGene_tx=XM_003981130.5;refGene_gn=VCAN;refGene_tx=XM_003981131.5
A1 149226626 149226740
refGene_gn=VCAN;refGene_tx=XM_003981130.5;refGene_gn=VCAN;refGene_tx=XM_003981131.5;refGene_gn=VCAN;refGene_tx=XM_003981132.4;refGene_gn=VCAN;refGene_tx=XM_003981133.5
A1 149229405 149229519
refGene_gn=VCAN;refGene_tx=XM_003981130.5;refGene_gn=VCAN;refGene_tx=XM_003981131.5;refGene_gn=VCAN;refGene_tx=XM_003981132.4;refGene_gn=VCAN;refGene_tx=XM_003981133.5
A1 149234505 149234664
refGene_gn=VCAN;refGene_tx=XM_003981130.5;refGene_gn=VCAN;refGene_tx=XM_003981131.5;refGene_gn=VCAN;refGene_tx=XM_003981132.4;refGene_gn=VCAN;refGene_tx=XM_003981133.5
A1 149235925 149236008
refGene_gn=VCAN;refGene_tx=XM_003981130.5;refGene_gn=VCAN;refGene_tx=XM_003981131.5;refGene_gn=VCAN;refGene_tx=XM_003981132.4;refGene_gn=VCA

N;refGene_tx=XM_003981133.5
A1 149254051 149254196
refGene_gn=VCAN;refGene_tx=XM_003981130.5;refGene_gn=VCAN;refGene_tx=XM_003981131.5;refGene_gn=VCAN;refGene_tx=XM_003981132.4;refGene_gn=VCAN;refGene_tx=XM_003981133.5
A1 149259596 149259779
refGene_gn=VCAN;refGene_tx=XM_003981130.5;refGene_gn=VCAN;refGene_tx=XM_003981131.5;refGene_gn=VCAN;refGene_tx=XM_003981132.4;refGene_gn=VCAN;refGene_tx=XM_003981133.5
A1 149259924 149260052
refGene_gn=VCAN;refGene_tx=XM_003981130.5;refGene_gn=VCAN;refGene_tx=XM_003981131.5;refGene_gn=VCAN;refGene_tx=XM_003981132.4;refGene_gn=VCAN;refGene_tx=XM_003981133.5
A1 149320570 149320860
refGene_gn=HAPLN1;refGene_tx=XM_006927834.3;refGene_gn=HAPLN1;refGene_tx=XM_019836372.2;refGene_gn=HAPLN1;refGene_tx=XM_019836379.1;refGene_gn=HAPLN1;refGene_tx=XM_019836382.2;refGene_gn=HAPLN1;refGene_tx=XM_019836384.2
A1 149323075 149323378
refGene_gn=HAPLN1;refGene_tx=XM_006927834.3;refGene_gn=HAPLN1;refGene_tx=XM_019836372.2;refGene_gn=HAPLN1;refGene_tx=XM_019836379.1;refGene_gn=HAPLN1;refGene_tx=XM_019836382.2;refGene_gn=HAPLN1;refGene_tx=XM_019836384.2
A1 149331328 149331700
refGene_gn=HAPLN1;refGene_tx=XM_006927834.3;refGene_gn=HAPLN1;refGene_tx=XM_019836372.2;refGene_gn=HAPLN1;refGene_tx=XM_019836379.1;refGene_gn=HAPLN1;refGene_tx=XM_019836382.2;refGene_gn=HAPLN1;refGene_tx=XM_019836384.2
A1 149352021 149352124
refGene_gn=HAPLN1;refGene_tx=XM_006927834.3;refGene_gn=HAPLN1;refGene_tx=XM_019836372.2;refGene_gn=HAPLN1;refGene_tx=XM_019836379.1;refGene_gn=HAPLN1;refGene_tx=XM_019836382.2;refGene_gn=HAPLN1;refGene_tx=XM_019836384.2
A1 149582262 149582412
refGene_gn=EDIL3;refGene_tx=XM_011284414.2;refGene_gn=EDIL3;refGene_tx=XM_011284418.2
A1 149603100 149603256
refGene_gn=EDIL3;refGene_tx=XM_011284414.2;refGene_gn=EDIL3;refGene_tx=XM_011284418.2
A1 149678707 149678892
refGene_gn=EDIL3;refGene_tx=XM_011284414.2;refGene_gn=EDIL3;refGene_tx=XM_011284418.2
A1 149683374 149683519
refGene_gn=EDIL3;refGene_tx=XM_011284414.2;refGene_gn=EDIL3;refGene_tx=XM_011284418.2
A1 149685131 149685287
refGene_gn=EDIL3;refGene_tx=XM_011284414.2;refGene_gn=EDIL3;refGene_tx=XM_011284418.2
A1 149722510 149722692
refGene_gn=EDIL3;refGene_tx=XM_011284414.2;refGene_gn=EDIL3;refGene_tx

=XM_011284418.2
A1 149758858 149758972
refGene_gn=EDIL3;refGene_tx=XM_011284414.2;refGene_gn=EDIL3;refGene_tx
=XM_011284418.2
A1 149800601 149800730
refGene_gn=EDIL3;refGene_tx=XM_011284414.2;refGene_gn=EDIL3;refGene_tx
=XM_011284418.2
A1 149842298 149842328
refGene_gn=EDIL3;refGene_tx=XM_011284414.2
A1 149868083 149868212
refGene_gn=EDIL3;refGene_tx=XM_011284414.2;refGene_gn=EDIL3;refGene_tx
=XM_011284418.2
A1 149992030 149992100
refGene_gn=EDIL3;refGene_tx=XM_011284414.2;refGene_gn=EDIL3;refGene_tx
=XM_011284418.2
A1 151811886 151811961
refGene_gn=LOC102899183;refGene_tx=XM_011284438.3
A1 151812946 151813063
refGene_gn=LOC102899183;refGene_tx=XM_011284438.3
A1 152325629 152326144
refGene_gn=RASA1;refGene_tx=XM_003981135.4
A1 152377608 152377761
refGene_gn=RASA1;refGene_tx=XM_003981135.4
A1 152379006 152379142
refGene_gn=RASA1;refGene_tx=XM_003981135.4
A1 152379784 152379855
refGene_gn=RASA1;refGene_tx=XM_003981135.4
A1 152383849 152383967
refGene_gn=RASA1;refGene_tx=XM_003981135.4
A1 152387063 152387095
refGene_gn=RASA1;refGene_tx=XM_003981135.4
A1 152396095 152396148
refGene_gn=RASA1;refGene_tx=XM_003981135.4
A1 152398506 152398657
refGene_gn=RASA1;refGene_tx=XM_003981135.4
A1 152403423 152403502
refGene_gn=RASA1;refGene_tx=XM_003981135.4
A1 152409476 152409597
refGene_gn=RASA1;refGene_tx=XM_003981135.4
A1 152410808 152410965
refGene_gn=RASA1;refGene_tx=XM_003981135.4
A1 152418828 152418916
refGene_gn=RASA1;refGene_tx=XM_003981135.4
A1 152420864 152420942
refGene_gn=RASA1;refGene_tx=XM_003981135.4
A1 152422790 152422948
refGene_gn=RASA1;refGene_tx=XM_003981135.4
A1 152423459 152423536
refGene_gn=RASA1;refGene_tx=XM_003981135.4
A1 152424868 152425041

refGene_gn=RASA1;refGene_tx=XM_003981135.4
A1	152425365	152425525
refGene_gn=RASA1;refGene_tx=XM_003981135.4
A1	152426606	152426749
refGene_gn=RASA1;refGene_tx=XM_003981135.4
A1	152428163	152428279
refGene_gn=RASA1;refGene_tx=XM_003981135.4
A1	152428995	152429082
refGene_gn=RASA1;refGene_tx=XM_003981135.4
A1	152431327	152431395
refGene_gn=RASA1;refGene_tx=XM_003981135.4
A1	152433169	152433258
refGene_gn=RASA1;refGene_tx=XM_003981135.4
A1	152434613	152434691
refGene_gn=RASA1;refGene_tx=XM_003981135.4
A1	152437283	152437418
refGene_gn=RASA1;refGene_tx=XM_003981135.4
A1	152438466	152438550
refGene_gn=RASA1;refGene_tx=XM_003981135.4
A1	152441588	152441627
refGene_gn=CCNH;refGene_tx=XM_003981136.5;refGene_gn=CCNH;refGene_tx=XM_006927835.4;refGene_gn=CCNH;refGene_tx=XM_019836430.1
A1	152442170	152442246
refGene_gn=CCNH;refGene_tx=XM_003981136.5;refGene_gn=CCNH;refGene_tx=XM_006927835.4;refGene_gn=CCNH;refGene_tx=XM_019836430.1
A1	152445062	152445174
refGene_gn=CCNH;refGene_tx=XM_003981136.5;refGene_gn=CCNH;refGene_tx=XM_006927835.4;refGene_gn=CCNH;refGene_tx=XM_019836430.1
A1	152447616	152447687
refGene_gn=CCNH;refGene_tx=XM_003981136.5;refGene_gn=CCNH;refGene_tx=XM_006927835.4;refGene_gn=CCNH;refGene_tx=XM_019836430.1
A1	152450618	152450782
refGene_gn=CCNH;refGene_tx=XM_003981136.5;refGene_gn=CCNH;refGene_tx=XM_006927835.4;refGene_gn=CCNH;refGene_tx=XM_019836430.1
A1	152454461	152454672
refGene_gn=CCNH;refGene_tx=XM_003981136.5;refGene_gn=CCNH;refGene_tx=XM_006927835.4;refGene_gn=CCNH;refGene_tx=XM_019836430.1
A1	152455688	152455762
refGene_gn=CCNH;refGene_tx=XM_003981136.5;refGene_gn=CCNH;refGene_tx=XM_006927835.4;refGene_gn=CCNH;refGene_tx=XM_019836430.1
A1	152458438	152458561
refGene_gn=CCNH;refGene_tx=XM_003981136.5;refGene_gn=CCNH;refGene_tx=XM_006927835.4;refGene_gn=CCNH;refGene_tx=XM_019836430.1
A1	152459130	152459145
refGene_gn=CCNH;refGene_tx=XM_019836430.1
A1	152460637	152460754
refGene_gn=CCNH;refGene_tx=XM_003981136.5;refGene_gn=CCNH;refGene_tx=XM_006927835.4
A1	153136911	153137189
refGene_gn=TMEM161B;refGene_tx=XM_003981137.5;refGene_gn=TMEM161B;refG

ene_tx=XM_023258178.1;refGene_gn=TMEM161B;refGene_tx=XM_023258183.1
A1 153138384 153138481
refGene_gn=TMEM161B;refGene_tx=XM_003981137.5;refGene_gn=TMEM161B;refGene_tx=XM_023258178.1;refGene_gn=TMEM161B;refGene_tx=XM_023258183.1
A1 153139548 153139723
refGene_gn=TMEM161B;refGene_tx=XM_003981137.5;refGene_gn=TMEM161B;refGene_tx=XM_023258178.1;refGene_gn=TMEM161B;refGene_tx=XM_023258183.1
A1 153142007 153142121
refGene_gn=TMEM161B;refGene_tx=XM_003981137.5;refGene_gn=TMEM161B;refGene_tx=XM_023258178.1;refGene_gn=TMEM161B;refGene_tx=XM_023258183.1
A1 153144815 153144956
refGene_gn=TMEM161B;refGene_tx=XM_003981137.5;refGene_gn=TMEM161B;refGene_tx=XM_023258178.1;refGene_gn=TMEM161B;refGene_tx=XM_023258183.1
A1 153145445 153145506
refGene_gn=TMEM161B;refGene_tx=XM_003981137.5;refGene_gn=TMEM161B;refGene_tx=XM_023258178.1;refGene_gn=TMEM161B;refGene_tx=XM_023258183.1
A1 153146027 153146179
refGene_gn=TMEM161B;refGene_tx=XM_003981137.5;refGene_gn=TMEM161B;refGene_tx=XM_023258178.1;refGene_gn=TMEM161B;refGene_tx=XM_023258183.1
A1 153165508 153165665
refGene_gn=TMEM161B;refGene_tx=XM_003981137.5;refGene_gn=TMEM161B;refGene_tx=XM_023258183.1
A1 153169561 153169659
refGene_gn=TMEM161B;refGene_tx=XM_003981137.5
A1 153171741 153171825
refGene_gn=TMEM161B;refGene_tx=XM_003981137.5
A1 153183264 153183368
refGene_gn=TMEM161B;refGene_tx=XM_003981137.5
A1 153205890 153205893
refGene_gn=TMEM161B;refGene_tx=XM_003981137.5
A1 153608523 153608845
refGene_gn=MEF2C;refGene_tx=XM_011284522.3;refGene_gn=MEF2C;refGene_tx=XM_011284523.3;refGene_gn=MEF2C;refGene_tx=XM_011284527.2;refGene_gn=MEF2C;refGene_tx=XM_011284529.3;refGene_gn=MEF2C;refGene_tx=XM_011284535.2;refGene_gn=MEF2C;refGene_tx=XM_011284541.2;refGene_gn=MEF2C;refGene_tx=XM_019836497.2;refGene_gn=MEF2C;refGene_tx=XM_019836507.2;refGene_gn=MEF2C;refGene_tx=XM_019836518.1;refGene_gn=MEF2C;refGene_tx=XM_019836530.1;refGene_gn=MEF2C;refGene_tx=XM_019836537.1;refGene_gn=MEF2C;refGene_tx=XM_019836547.1;refGene_gn=MEF2C;refGene_tx=XM_019836553.2;refGene_gn=MEF2C;refGene_tx=XM_019836558.2;refGene_gn=MEF2C;refGene_tx=XM_019836564.2;refGene_gn=MEF2C;refGene_tx=XM_019836572.1;refGene_gn=MEF2C;refGene_tx=XM_019836581.2;refGene_gn=MEF2C;refGene_tx=XM_019836585.2
A1 153614396 153614532
refGene_gn=MEF2C;refGene_tx=XM_011284522.3;refGene_gn=MEF2C;refGene_tx=XM_011284523.3;refGene_gn=MEF2C;refGene_tx=XM_011284527.2;refGene_gn=MEF2C;refGene_tx=XM_011284529.3;refGene_gn=MEF2C;refGene_tx=XM_011284535.2;refGene_gn=MEF2C;refGene_tx=XM_011284541.2;refGene_gn=MEF2C;refGene_tx=XM_019836497.2;refGene_gn=MEF2C;refGene_tx=XM_019836507.2;refGene_gn=MEF2C;refGene_tx=XM_019836518.1;refGene_gn=MEF2C;refGene_tx=XM_01

9836530.1;refGene_gn=MEF2C;refGene_tx=XM_019836537.1;refGene_gn=MEF2C;refGene_tx=XM_019836547.1;refGene_gn=MEF2C;refGene_tx=XM_019836553.2;refGene_gn=MEF2C;refGene_tx=XM_019836558.2;refGene_gn=MEF2C;refGene_tx=XM_019836564.2;refGene_gn=MEF2C;refGene_tx=XM_019836572.1;refGene_gn=MEF2C;refGene_tx=XM_019836581.2;refGene_gn=MEF2C;refGene_tx=XM_019836585.2

A1 153615126 153615256
refGene_gn=MEF2C;refGene_tx=XM_011284522.3;refGene_gn=MEF2C;refGene_tx=XM_011284523.3;refGene_gn=MEF2C;refGene_tx=XM_011284527.2;refGene_gn=MEF2C;refGene_tx=XM_011284529.3;refGene_gn=MEF2C;refGene_tx=XM_011284535.2;refGene_gn=MEF2C;refGene_tx=XM_011284541.2;refGene_gn=MEF2C;refGene_tx=XM_019836497.2;refGene_gn=MEF2C;refGene_tx=XM_019836507.2;refGene_gn=MEF2C;refGene_tx=XM_019836518.1;refGene_gn=MEF2C;refGene_tx=XM_019836530.1;refGene_gn=MEF2C;refGene_tx=XM_019836537.1;refGene_gn=MEF2C;refGene_tx=XM_019836547.1;refGene_gn=MEF2C;refGene_tx=XM_019836553.2;refGene_gn=MEF2C;refGene_tx=XM_019836558.2;refGene_gn=MEF2C;refGene_tx=XM_019836564.2;refGene_gn=MEF2C;refGene_tx=XM_019836572.1;refGene_gn=MEF2C;refGene_tx=XM_019836581.2;refGene_gn=MEF2C;refGene_tx=XM_019836585.2

A1 153616145 153616169
refGene_gn=MEF2C;refGene_tx=XM_011284522.3;refGene_gn=MEF2C;refGene_tx=XM_011284523.3;refGene_gn=MEF2C;refGene_tx=XM_011284527.2;refGene_gn=MEF2C;refGene_tx=XM_011284529.3;refGene_gn=MEF2C;refGene_tx=XM_011284535.2;refGene_gn=MEF2C;refGene_tx=XM_011284541.2;refGene_gn=MEF2C;refGene_tx=XM_019836497.2;refGene_gn=MEF2C;refGene_tx=XM_019836507.2;refGene_gn=MEF2C;refGene_tx=XM_019836518.1;refGene_gn=MEF2C;refGene_tx=XM_019836530.1;refGene_gn=MEF2C;refGene_tx=XM_019836537.1;refGene_gn=MEF2C;refGene_tx=XM_019836547.1;refGene_gn=MEF2C;refGene_tx=XM_019836553.2;refGene_gn=MEF2C;refGene_tx=XM_019836572.1

A1 153617722 153617895
refGene_gn=MEF2C;refGene_tx=XM_011284522.3;refGene_gn=MEF2C;refGene_tx=XM_011284523.3;refGene_gn=MEF2C;refGene_tx=XM_011284527.2;refGene_gn=MEF2C;refGene_tx=XM_011284529.3;refGene_gn=MEF2C;refGene_tx=XM_011284535.2;refGene_gn=MEF2C;refGene_tx=XM_011284541.2;refGene_gn=MEF2C;refGene_tx=XM_019836497.2;refGene_gn=MEF2C;refGene_tx=XM_019836507.2;refGene_gn=MEF2C;refGene_tx=XM_019836518.1;refGene_gn=MEF2C;refGene_tx=XM_019836530.1;refGene_gn=MEF2C;refGene_tx=XM_019836537.1;refGene_gn=MEF2C;refGene_tx=XM_019836547.1;refGene_gn=MEF2C;refGene_tx=XM_019836553.2;refGene_gn=MEF2C;refGene_tx=XM_019836558.2;refGene_gn=MEF2C;refGene_tx=XM_019836564.2;refGene_gn=MEF2C;refGene_tx=XM_019836572.1;refGene_gn=MEF2C;refGene_tx=XM_019836581.2;refGene_gn=MEF2C;refGene_tx=XM_019836585.2

A1 153634677 153634725
refGene_gn=MEF2C;refGene_tx=XM_011284522.3;refGene_gn=MEF2C;refGene_tx=XM_011284523.3;refGene_gn=MEF2C;refGene_tx=XM_011284527.2;refGene_gn=MEF2C;refGene_tx=XM_011284529.3;refGene_gn=MEF2C;refGene_tx=XM_011284535.2;refGene_gn=MEF2C;refGene_tx=XM_011284541.2;refGene_gn=MEF2C;refGene_tx=XM_019836497.2;refGene_gn=MEF2C;refGene_tx=XM_019836507.2;refGene_gn=MEF2C;refGene_tx=XM_019836518.1;refGene_gn=MEF2C;refGene_tx=XM_019836530.1;refGene_gn=MEF2C;refGene_tx=XM_019836537.1;refGene_gn=MEF2C;

refGene_tx=XM_019836547.1;refGene_gn=MEF2C;refGene_tx=XM_019836553.2;refGene_gn=MEF2C;refGene_tx=XM_019836558.2;refGene_gn=MEF2C;refGene_tx=XM_019836564.2;refGene_gn=MEF2C;refGene_tx=XM_019836572.1;refGene_gn=MEF2C;refGene_tx=XM_019836581.2;refGene_gn=MEF2C;refGene_tx=XM_019836585.2
A1 153636792 153636979
refGene_gn=MEF2C;refGene_tx=XM_011284522.3;refGene_gn=MEF2C;refGene_tx=XM_011284523.3;refGene_gn=MEF2C;refGene_tx=XM_011284527.2;refGene_gn=MEF2C;refGene_tx=XM_011284529.3;refGene_gn=MEF2C;refGene_tx=XM_011284535.2;refGene_gn=MEF2C;refGene_tx=XM_011284541.2;refGene_gn=MEF2C;refGene_tx=XM_019836497.2;refGene_gn=MEF2C;refGene_tx=XM_019836507.2;refGene_gn=MEF2C;refGene_tx=XM_019836518.1;refGene_gn=MEF2C;refGene_tx=XM_019836530.1;refGene_gn=MEF2C;refGene_tx=XM_019836537.1;refGene_gn=MEF2C;refGene_tx=XM_019836547.1;refGene_gn=MEF2C;refGene_tx=XM_019836553.2;refGene_gn=MEF2C;refGene_tx=XM_019836558.2;refGene_gn=MEF2C;refGene_tx=XM_019836564.2;refGene_gn=MEF2C;refGene_tx=XM_019836572.1;refGene_gn=MEF2C;refGene_tx=XM_019836581.2;refGene_gn=MEF2C;refGene_tx=XM_019836585.2
A1 153645989 153646127
refGene_gn=MEF2C;refGene_tx=XM_019836547.1;refGene_gn=MEF2C;refGene_tx=XM_019836564.2;refGene_gn=MEF2C;refGene_tx=XM_019836585.2
A1 153646186 153646330
refGene_gn=MEF2C;refGene_tx=XM_011284522.3;refGene_gn=MEF2C;refGene_tx=XM_011284523.3;refGene_gn=MEF2C;refGene_tx=XM_011284527.2;refGene_gn=MEF2C;refGene_tx=XM_011284529.3;refGene_gn=MEF2C;refGene_tx=XM_011284535.2;refGene_gn=MEF2C;refGene_tx=XM_011284541.2;refGene_gn=MEF2C;refGene_tx=XM_019836497.2;refGene_gn=MEF2C;refGene_tx=XM_019836507.2;refGene_gn=MEF2C;refGene_tx=XM_019836518.1;refGene_gn=MEF2C;refGene_tx=XM_019836530.1;refGene_gn=MEF2C;refGene_tx=XM_019836537.1;refGene_gn=MEF2C;refGene_tx=XM_019836553.2;refGene_gn=MEF2C;refGene_tx=XM_019836558.2;refGene_gn=MEF2C;refGene_tx=XM_019836572.1;refGene_gn=MEF2C;refGene_tx=XM_019836581.2;refGene_gn=MEF2C;refGene_tx=XM_019836585.2
A1 153680555 153680759
refGene_gn=MEF2C;refGene_tx=XM_011284522.3;refGene_gn=MEF2C;refGene_tx=XM_011284523.3;refGene_gn=MEF2C;refGene_tx=XM_011284527.2;refGene_gn=MEF2C;refGene_tx=XM_011284529.3;refGene_gn=MEF2C;refGene_tx=XM_011284535.2;refGene_gn=MEF2C;refGene_tx=XM_011284541.2;refGene_gn=MEF2C;refGene_tx=XM_019836497.2;refGene_gn=MEF2C;refGene_tx=XM_019836507.2;refGene_gn=MEF2C;refGene_tx=XM_019836518.1;refGene_gn=MEF2C;refGene_tx=XM_019836530.1;refGene_gn=MEF2C;refGene_tx=XM_019836537.1;refGene_gn=MEF2C;refGene_tx=XM_019836547.1;refGene_gn=MEF2C;refGene_tx=XM_019836553.2;refGene_gn=MEF2C;refGene_tx=XM_019836558.2;refGene_gn=MEF2C;refGene_tx=XM_019836564.2;refGene_gn=MEF2C;refGene_tx=XM_019836572.1;refGene_gn=MEF2C;refGene_tx=XM_019836581.2;refGene_gn=MEF2C;refGene_tx=XM_019836585.2
A1 153699946 153700000
refGene_gn=MEF2C;refGene_tx=XM_011284522.3;refGene_gn=MEF2C;refGene_tx=XM_011284523.3;refGene_gn=MEF2C;refGene_tx=XM_011284527.2;refGene_gn=MEF2C;refGene_tx=XM_011284529.3;refGene_gn=MEF2C;refGene_tx=XM_011284535.2;refGene_gn=MEF2C;refGene_tx=XM_011284541.2;refGene_gn=MEF2C;refGe

ne_tx=XM_019836497.2;refGene_gn=MEF2C;refGene_tx=XM_019836507.2;refGene_gn=MEF2C;refGene_tx=XM_019836518.1;refGene_gn=MEF2C;refGene_tx=XM_019836530.1;refGene_gn=MEF2C;refGene_tx=XM_019836537.1;refGene_gn=MEF2C;refGene_tx=XM_019836547.1;refGene_gn=MEF2C;refGene_tx=XM_019836553.2;refGene_gn=MEF2C;refGene_tx=XM_019836558.2;refGene_gn=MEF2C;refGene_tx=XM_019836564.2;refGene_gn=MEF2C;refGene_tx=XM_019836572.1;refGene_gn=MEF2C;refGene_tx=XM_019836581.2;refGene_gn=MEF2C;refGene_tx=XM_019836585.2
A1 155040044 155040088
refGene_gn=CETN3;refGene_tx=XM_003981145.5
A1 155046828 155047020
refGene_gn=CETN3;refGene_tx=XM_003981145.5
A1 155052469 155052584
refGene_gn=CETN3;refGene_tx=XM_003981145.5
A1 155054348 155054484
refGene_gn=CETN3;refGene_tx=XM_003981145.5
A1 155056259 155056276
refGene_gn=CETN3;refGene_tx=XM_003981145.5
A1 155100479 155100865
refGene_gn=MBLAC2;refGene_tx=XM_003981146.5
A1 155114138 155114592
refGene_gn=MBLAC2;refGene_tx=XM_003981146.5
A1 155123356 155123473
refGene_gn=POLR3G;refGene_tx=XM_019836669.2;refGene_gn=POLR3G;refGene_tx=XM_019836704.2
A1 155125977 155126107
refGene_gn=POLR3G;refGene_tx=XM_019836669.2;refGene_gn=POLR3G;refGene_tx=XM_019836704.2
A1 155133716 155133773
refGene_gn=POLR3G;refGene_tx=XM_019836669.2;refGene_gn=POLR3G;refGene_tx=XM_019836704.2
A1 155136181 155136232
refGene_gn=POLR3G;refGene_tx=XM_019836669.2;refGene_gn=POLR3G;refGene_tx=XM_019836704.2
A1 155147456 155147539
refGene_gn=POLR3G;refGene_tx=XM_019836669.2;refGene_gn=POLR3G;refGene_tx=XM_019836704.2
A1 155151827 155151974
refGene_gn=POLR3G;refGene_tx=XM_019836669.2;refGene_gn=POLR3G;refGene_tx=XM_019836704.2
A1 155157539 155157626
refGene_gn=POLR3G;refGene_tx=XM_019836669.2;refGene_gn=POLR3G;refGene_tx=XM_019836704.2
A1 155163669 155164335
refGene_gn=LYSMD3;refGene_tx=XM_003981147.5
A1 155170479 155170734
refGene_gn=LYSMD3;refGene_tx=XM_003981147.5
A1 155202960 155202982
refGene_gn=ADGRV1;refGene_tx=XM_019836773.1;refGene_gn=ADGRV1;refGene_tx=XM_019836778.1;refGene_gn=ADGRV1;refGene_tx=XM_023258351.1;refGene_

gn=ADGRV1;refGene_tx=XM_023258353.1;refGene_gn=ADGRV1;refGene_tx=XM_023258360.1
A1 155243238 155243423
refGene_gn=ADGRV1;refGene_tx=XM_019836773.1;refGene_gn=ADGRV1;refGene_tx=XM_019836778.1;refGene_gn=ADGRV1;refGene_tx=XM_023258340.1;refGene_gn=ADGRV1;refGene_tx=XM_023258343.1;refGene_gn=ADGRV1;refGene_tx=XM_023258347.1;refGene_gn=ADGRV1;refGene_tx=XM_023258351.1;refGene_gn=ADGRV1;refGene_tx=XM_023258353.1;refGene_gn=ADGRV1;refGene_tx=XM_023258360.1
A1 155245458 155245608
refGene_gn=ADGRV1;refGene_tx=XM_019836773.1;refGene_gn=ADGRV1;refGene_tx=XM_019836778.1;refGene_gn=ADGRV1;refGene_tx=XM_023258340.1;refGene_gn=ADGRV1;refGene_tx=XM_023258343.1;refGene_gn=ADGRV1;refGene_tx=XM_023258347.1;refGene_gn=ADGRV1;refGene_tx=XM_023258351.1;refGene_gn=ADGRV1;refGene_tx=XM_023258353.1;refGene_gn=ADGRV1;refGene_tx=XM_023258360.1
A1 155246715 155246811
refGene_gn=ADGRV1;refGene_tx=XM_019836773.1;refGene_gn=ADGRV1;refGene_tx=XM_019836778.1;refGene_gn=ADGRV1;refGene_tx=XM_023258340.1;refGene_gn=ADGRV1;refGene_tx=XM_023258343.1;refGene_gn=ADGRV1;refGene_tx=XM_023258347.1;refGene_gn=ADGRV1;refGene_tx=XM_023258351.1;refGene_gn=ADGRV1;refGene_tx=XM_023258353.1;refGene_gn=ADGRV1;refGene_tx=XM_023258360.1
A1 155249551 155249656
refGene_gn=ADGRV1;refGene_tx=XM_019836773.1;refGene_gn=ADGRV1;refGene_tx=XM_019836778.1;refGene_gn=ADGRV1;refGene_tx=XM_023258340.1;refGene_gn=ADGRV1;refGene_tx=XM_023258343.1;refGene_gn=ADGRV1;refGene_tx=XM_023258347.1;refGene_gn=ADGRV1;refGene_tx=XM_023258351.1;refGene_gn=ADGRV1;refGene_tx=XM_023258353.1;refGene_gn=ADGRV1;refGene_tx=XM_023258360.1
A1 155250448 155250562
refGene_gn=ADGRV1;refGene_tx=XM_019836773.1;refGene_gn=ADGRV1;refGene_tx=XM_019836778.1;refGene_gn=ADGRV1;refGene_tx=XM_023258340.1;refGene_gn=ADGRV1;refGene_tx=XM_023258343.1;refGene_gn=ADGRV1;refGene_tx=XM_023258347.1;refGene_gn=ADGRV1;refGene_tx=XM_023258351.1;refGene_gn=ADGRV1;refGene_tx=XM_023258353.1;refGene_gn=ADGRV1;refGene_tx=XM_023258360.1
A1 155252872 155253438
refGene_gn=ADGRV1;refGene_tx=XM_019836773.1;refGene_gn=ADGRV1;refGene_tx=XM_019836778.1;refGene_gn=ADGRV1;refGene_tx=XM_023258340.1;refGene_gn=ADGRV1;refGene_tx=XM_023258343.1;refGene_gn=ADGRV1;refGene_tx=XM_023258347.1;refGene_gn=ADGRV1;refGene_tx=XM_023258351.1;refGene_gn=ADGRV1;refGene_tx=XM_023258353.1;refGene_gn=ADGRV1;refGene_tx=XM_023258360.1
A1 155253987 155254258
refGene_gn=ADGRV1;refGene_tx=XM_019836773.1;refGene_gn=ADGRV1;refGene_tx=XM_019836778.1;refGene_gn=ADGRV1;refGene_tx=XM_023258340.1;refGene_gn=ADGRV1;refGene_tx=XM_023258343.1;refGene_gn=ADGRV1;refGene_tx=XM_023258347.1;refGene_gn=ADGRV1;refGene_tx=XM_023258351.1;refGene_gn=ADGRV1;refGene_tx=XM_023258353.1;refGene_gn=ADGRV1;refGene_tx=XM_023258360.

1
A1 155254897 155255227
refGene_gn=ADGRV1;refGene_tx=XM_019836773.1;refGene_gn=ADGRV1;refGene_tx=XM_019836778.1;refGene_gn=ADGRV1;refGene_tx=XM_023258340.1;refGene_gn=ADGRV1;refGene_tx=XM_023258343.1;refGene_gn=ADGRV1;refGene_tx=XM_023258347.1;refGene_gn=ADGRV1;refGene_tx=XM_023258351.1;refGene_gn=ADGRV1;refGene_tx=XM_023258353.1;refGene_gn=ADGRV1;refGene_tx=XM_023258360.1
A1 155262001 155262178
refGene_gn=ADGRV1;refGene_tx=XM_019836773.1;refGene_gn=ADGRV1;refGene_tx=XM_019836778.1;refGene_gn=ADGRV1;refGene_tx=XM_023258340.1;refGene_gn=ADGRV1;refGene_tx=XM_023258343.1;refGene_gn=ADGRV1;refGene_tx=XM_023258347.1;refGene_gn=ADGRV1;refGene_tx=XM_023258351.1;refGene_gn=ADGRV1;refGene_tx=XM_023258353.1;refGene_gn=ADGRV1;refGene_tx=XM_023258360.1
A1 155264301 155264525
refGene_gn=ADGRV1;refGene_tx=XM_019836773.1;refGene_gn=ADGRV1;refGene_tx=XM_019836778.1;refGene_gn=ADGRV1;refGene_tx=XM_023258340.1;refGene_gn=ADGRV1;refGene_tx=XM_023258343.1;refGene_gn=ADGRV1;refGene_tx=XM_023258347.1;refGene_gn=ADGRV1;refGene_tx=XM_023258351.1;refGene_gn=ADGRV1;refGene_tx=XM_023258353.1;refGene_gn=ADGRV1;refGene_tx=XM_023258360.1
A1 155268655 155268782
refGene_gn=ADGRV1;refGene_tx=XM_019836773.1;refGene_gn=ADGRV1;refGene_tx=XM_019836778.1;refGene_gn=ADGRV1;refGene_tx=XM_023258340.1;refGene_gn=ADGRV1;refGene_tx=XM_023258343.1;refGene_gn=ADGRV1;refGene_tx=XM_023258347.1;refGene_gn=ADGRV1;refGene_tx=XM_023258351.1;refGene_gn=ADGRV1;refGene_tx=XM_023258353.1;refGene_gn=ADGRV1;refGene_tx=XM_023258360.1
A1 155268879 155269065
refGene_gn=ADGRV1;refGene_tx=XM_019836773.1;refGene_gn=ADGRV1;refGene_tx=XM_019836778.1;refGene_gn=ADGRV1;refGene_tx=XM_023258340.1;refGene_gn=ADGRV1;refGene_tx=XM_023258343.1;refGene_gn=ADGRV1;refGene_tx=XM_023258347.1;refGene_gn=ADGRV1;refGene_tx=XM_023258351.1;refGene_gn=ADGRV1;refGene_tx=XM_023258353.1;refGene_gn=ADGRV1;refGene_tx=XM_023258360.1
A1 155269815 155269996
refGene_gn=ADGRV1;refGene_tx=XM_019836773.1;refGene_gn=ADGRV1;refGene_tx=XM_019836778.1;refGene_gn=ADGRV1;refGene_tx=XM_023258340.1;refGene_gn=ADGRV1;refGene_tx=XM_023258343.1;refGene_gn=ADGRV1;refGene_tx=XM_023258347.1;refGene_gn=ADGRV1;refGene_tx=XM_023258351.1;refGene_gn=ADGRV1;refGene_tx=XM_023258353.1;refGene_gn=ADGRV1;refGene_tx=XM_023258360.1
A1 155270728 155270892
refGene_gn=ADGRV1;refGene_tx=XM_019836773.1;refGene_gn=ADGRV1;refGene_tx=XM_019836778.1;refGene_gn=ADGRV1;refGene_tx=XM_023258340.1;refGene_gn=ADGRV1;refGene_tx=XM_023258343.1;refGene_gn=ADGRV1;refGene_tx=XM_023258347.1;refGene_gn=ADGRV1;refGene_tx=XM_023258351.1;refGene_gn=ADGRV1;refGene_tx=XM_023258353.1;refGene_gn=ADGRV1;refGene_tx=XM_023258360.1

A1 155271971 155272095
refGene_gn=ADGRV1;refGene_tx=XM_019836773.1;refGene_gn=ADGRV1;refGene_tx=XM_019836778.1;refGene_gn=ADGRV1;refGene_tx=XM_023258340.1;refGene_gn=ADGRV1;refGene_tx=XM_023258343.1;refGene_gn=ADGRV1;refGene_tx=XM_023258347.1;refGene_gn=ADGRV1;refGene_tx=XM_023258351.1;refGene_gn=ADGRV1;refGene_tx=XM_023258353.1;refGene_gn=ADGRV1;refGene_tx=XM_023258360.1
A1 155273314 155273581
refGene_gn=ADGRV1;refGene_tx=XM_019836773.1;refGene_gn=ADGRV1;refGene_tx=XM_019836778.1;refGene_gn=ADGRV1;refGene_tx=XM_023258340.1;refGene_gn=ADGRV1;refGene_tx=XM_023258343.1;refGene_gn=ADGRV1;refGene_tx=XM_023258347.1;refGene_gn=ADGRV1;refGene_tx=XM_023258351.1;refGene_gn=ADGRV1;refGene_tx=XM_023258353.1;refGene_gn=ADGRV1;refGene_tx=XM_023258360.1
A1 155278885 155279012
refGene_gn=ADGRV1;refGene_tx=XM_019836773.1;refGene_gn=ADGRV1;refGene_tx=XM_019836778.1;refGene_gn=ADGRV1;refGene_tx=XM_023258340.1;refGene_gn=ADGRV1;refGene_tx=XM_023258343.1;refGene_gn=ADGRV1;refGene_tx=XM_023258347.1;refGene_gn=ADGRV1;refGene_tx=XM_023258351.1;refGene_gn=ADGRV1;refGene_tx=XM_023258353.1;refGene_gn=ADGRV1;refGene_tx=XM_023258360.1
A1 155279847 155280065
refGene_gn=ADGRV1;refGene_tx=XM_019836773.1;refGene_gn=ADGRV1;refGene_tx=XM_019836778.1;refGene_gn=ADGRV1;refGene_tx=XM_023258340.1;refGene_gn=ADGRV1;refGene_tx=XM_023258343.1;refGene_gn=ADGRV1;refGene_tx=XM_023258347.1;refGene_gn=ADGRV1;refGene_tx=XM_023258351.1;refGene_gn=ADGRV1;refGene_tx=XM_023258353.1;refGene_gn=ADGRV1;refGene_tx=XM_023258360.1
A1 155280686 155281430
refGene_gn=ADGRV1;refGene_tx=XM_019836773.1;refGene_gn=ADGRV1;refGene_tx=XM_019836778.1;refGene_gn=ADGRV1;refGene_tx=XM_023258340.1;refGene_gn=ADGRV1;refGene_tx=XM_023258343.1;refGene_gn=ADGRV1;refGene_tx=XM_023258347.1;refGene_gn=ADGRV1;refGene_tx=XM_023258351.1;refGene_gn=ADGRV1;refGene_tx=XM_023258353.1;refGene_gn=ADGRV1;refGene_tx=XM_023258360.1
A1 155283719 155284093
refGene_gn=ADGRV1;refGene_tx=XM_019836773.1;refGene_gn=ADGRV1;refGene_tx=XM_019836778.1;refGene_gn=ADGRV1;refGene_tx=XM_023258340.1;refGene_gn=ADGRV1;refGene_tx=XM_023258343.1;refGene_gn=ADGRV1;refGene_tx=XM_023258347.1;refGene_gn=ADGRV1;refGene_tx=XM_023258351.1;refGene_gn=ADGRV1;refGene_tx=XM_023258353.1;refGene_gn=ADGRV1;refGene_tx=XM_023258360.1
A1 155293634 155293811
refGene_gn=ADGRV1;refGene_tx=XM_019836773.1;refGene_gn=ADGRV1;refGene_tx=XM_019836778.1;refGene_gn=ADGRV1;refGene_tx=XM_023258340.1;refGene_gn=ADGRV1;refGene_tx=XM_023258343.1;refGene_gn=ADGRV1;refGene_tx=XM_023258347.1;refGene_gn=ADGRV1;refGene_tx=XM_023258351.1;refGene_gn=ADGRV1;refGene_tx=XM_023258353.1;refGene_gn=ADGRV1;refGene_tx=XM_023258360.1
A1 155299177 155299358

refGene_gn=ADGRV1;refGene_tx=XM_019836773.1;refGene_gn=ADGRV1;refGene_tx=XM_019836778.1;refGene_gn=ADGRV1;refGene_tx=XM_023258340.1;refGene_gn=ADGRV1;refGene_tx=XM_023258343.1;refGene_gn=ADGRV1;refGene_tx=XM_023258347.1;refGene_gn=ADGRV1;refGene_tx=XM_023258351.1;refGene_gn=ADGRV1;refGene_tx=XM_023258353.1;refGene_gn=ADGRV1;refGene_tx=XM_023258360.1
A1 155300270 155300473
refGene_gn=ADGRV1;refGene_tx=XM_019836773.1;refGene_gn=ADGRV1;refGene_tx=XM_019836778.1;refGene_gn=ADGRV1;refGene_tx=XM_023258340.1;refGene_gn=ADGRV1;refGene_tx=XM_023258343.1;refGene_gn=ADGRV1;refGene_tx=XM_023258347.1;refGene_gn=ADGRV1;refGene_tx=XM_023258351.1;refGene_gn=ADGRV1;refGene_tx=XM_023258353.1;refGene_gn=ADGRV1;refGene_tx=XM_023258360.1
A1 155304871 155305001
refGene_gn=ADGRV1;refGene_tx=XM_019836773.1;refGene_gn=ADGRV1;refGene_tx=XM_019836778.1;refGene_gn=ADGRV1;refGene_tx=XM_023258340.1;refGene_gn=ADGRV1;refGene_tx=XM_023258343.1;refGene_gn=ADGRV1;refGene_tx=XM_023258347.1;refGene_gn=ADGRV1;refGene_tx=XM_023258351.1;refGene_gn=ADGRV1;refGene_tx=XM_023258353.1;refGene_gn=ADGRV1;refGene_tx=XM_023258360.1
A1 155307828 155307909
refGene_gn=ADGRV1;refGene_tx=XM_019836773.1;refGene_gn=ADGRV1;refGene_tx=XM_019836778.1;refGene_gn=ADGRV1;refGene_tx=XM_023258340.1;refGene_gn=ADGRV1;refGene_tx=XM_023258343.1;refGene_gn=ADGRV1;refGene_tx=XM_023258347.1;refGene_gn=ADGRV1;refGene_tx=XM_023258351.1;refGene_gn=ADGRV1;refGene_tx=XM_023258353.1;refGene_gn=ADGRV1;refGene_tx=XM_023258360.1
A1 155309279 155309419
refGene_gn=ADGRV1;refGene_tx=XM_019836773.1;refGene_gn=ADGRV1;refGene_tx=XM_019836778.1;refGene_gn=ADGRV1;refGene_tx=XM_023258340.1;refGene_gn=ADGRV1;refGene_tx=XM_023258343.1;refGene_gn=ADGRV1;refGene_tx=XM_023258347.1;refGene_gn=ADGRV1;refGene_tx=XM_023258351.1;refGene_gn=ADGRV1;refGene_tx=XM_023258353.1;refGene_gn=ADGRV1;refGene_tx=XM_023258360.1
A1 155311707 155312317
refGene_gn=ADGRV1;refGene_tx=XM_019836773.1;refGene_gn=ADGRV1;refGene_tx=XM_019836778.1;refGene_gn=ADGRV1;refGene_tx=XM_023258340.1;refGene_gn=ADGRV1;refGene_tx=XM_023258343.1;refGene_gn=ADGRV1;refGene_tx=XM_023258347.1;refGene_gn=ADGRV1;refGene_tx=XM_023258351.1;refGene_gn=ADGRV1;refGene_tx=XM_023258353.1;refGene_gn=ADGRV1;refGene_tx=XM_023258360.1
A1 155315437 155315653
refGene_gn=ADGRV1;refGene_tx=XM_019836773.1;refGene_gn=ADGRV1;refGene_tx=XM_019836778.1;refGene_gn=ADGRV1;refGene_tx=XM_023258340.1;refGene_gn=ADGRV1;refGene_tx=XM_023258343.1;refGene_gn=ADGRV1;refGene_tx=XM_023258347.1;refGene_gn=ADGRV1;refGene_tx=XM_023258353.1;refGene_gn=ADGRV1;refGene_tx=XM_023258360.1
A1 155317177 155317393
refGene_gn=ADGRV1;refGene_tx=XM_019836773.1;refGene_gn=ADGRV1;refGene_tx=XM_019836778.1;refGene_gn=ADGRV1;refGene_tx=XM_023258340.1;refGene_

gn=ADGRV1;refGene_tx=XM_023258343.1;refGene_gn=ADGRV1;refGene_tx=XM_02
3258347.1;refGene_gn=ADGRV1;refGene_tx=XM_023258351.1;refGene_gn=ADGRV
1;refGene_tx=XM_023258353.1;refGene_gn=ADGRV1;refGene_tx=XM_023258360.
1
A1 155318032 155318277
refGene_gn=ADGRV1;refGene_tx=XM_019836773.1;refGene_gn=ADGRV1;refGene_
tx=XM_019836778.1;refGene_gn=ADGRV1;refGene_tx=XM_023258340.1;refGene_
gn=ADGRV1;refGene_tx=XM_023258343.1;refGene_gn=ADGRV1;refGene_tx=XM_02
3258347.1;refGene_gn=ADGRV1;refGene_tx=XM_023258351.1;refGene_gn=ADGRV
1;refGene_tx=XM_023258353.1;refGene_gn=ADGRV1;refGene_tx=XM_023258360.
1
A1 155319513 155319695
refGene_gn=ADGRV1;refGene_tx=XM_019836773.1;refGene_gn=ADGRV1;refGene_
tx=XM_019836778.1;refGene_gn=ADGRV1;refGene_tx=XM_023258340.1;refGene_
gn=ADGRV1;refGene_tx=XM_023258343.1;refGene_gn=ADGRV1;refGene_tx=XM_02
3258347.1;refGene_gn=ADGRV1;refGene_tx=XM_023258351.1;refGene_gn=ADGRV
1;refGene_tx=XM_023258353.1;refGene_gn=ADGRV1;refGene_tx=XM_023258360.
1
A1 155320482 155321294
refGene_gn=ADGRV1;refGene_tx=XM_019836773.1;refGene_gn=ADGRV1;refGene_
tx=XM_019836778.1;refGene_gn=ADGRV1;refGene_tx=XM_023258340.1;refGene_
gn=ADGRV1;refGene_tx=XM_023258343.1;refGene_gn=ADGRV1;refGene_tx=XM_02
3258347.1;refGene_gn=ADGRV1;refGene_tx=XM_023258351.1;refGene_gn=ADGRV
1;refGene_tx=XM_023258353.1;refGene_gn=ADGRV1;refGene_tx=XM_023258360.
1
A1 155323995 155324205
refGene_gn=ADGRV1;refGene_tx=XM_019836773.1;refGene_gn=ADGRV1;refGene_
tx=XM_019836778.1;refGene_gn=ADGRV1;refGene_tx=XM_023258340.1;refGene_
gn=ADGRV1;refGene_tx=XM_023258343.1;refGene_gn=ADGRV1;refGene_tx=XM_02
3258347.1;refGene_gn=ADGRV1;refGene_tx=XM_023258351.1;refGene_gn=ADGRV
1;refGene_tx=XM_023258353.1;refGene_gn=ADGRV1;refGene_tx=XM_023258360.
1
A1 155331718 155331849
refGene_gn=ADGRV1;refGene_tx=XM_019836773.1;refGene_gn=ADGRV1;refGene_
tx=XM_019836778.1;refGene_gn=ADGRV1;refGene_tx=XM_023258340.1;refGene_
gn=ADGRV1;refGene_tx=XM_023258343.1;refGene_gn=ADGRV1;refGene_tx=XM_02
3258347.1;refGene_gn=ADGRV1;refGene_tx=XM_023258351.1;refGene_gn=ADGRV
1;refGene_tx=XM_023258353.1;refGene_gn=ADGRV1;refGene_tx=XM_023258360.
1
A1 155333348 155333448
refGene_gn=ADGRV1;refGene_tx=XM_019836773.1;refGene_gn=ADGRV1;refGene_
tx=XM_019836778.1;refGene_gn=ADGRV1;refGene_tx=XM_023258340.1;refGene_
gn=ADGRV1;refGene_tx=XM_023258343.1;refGene_gn=ADGRV1;refGene_tx=XM_02
3258347.1;refGene_gn=ADGRV1;refGene_tx=XM_023258351.1;refGene_gn=ADGRV
1;refGene_tx=XM_023258353.1;refGene_gn=ADGRV1;refGene_tx=XM_023258360.
1
A1 155334100 155334280
refGene_gn=ADGRV1;refGene_tx=XM_019836773.1;refGene_gn=ADGRV1;refGene_
tx=XM_019836778.1;refGene_gn=ADGRV1;refGene_tx=XM_023258340.1;refGene_
gn=ADGRV1;refGene_tx=XM_023258343.1;refGene_gn=ADGRV1;refGene_tx=XM_02

3258347.1;refGene_gn=ADGRV1;refGene_tx=XM_023258351.1;refGene_gn=ADGRV1;refGene_tx=XM_023258353.1;refGene_gn=ADGRV1;refGene_tx=XM_023258360.1
A1 155338246 155338410
refGene_gn=ADGRV1;refGene_tx=XM_019836773.1;refGene_gn=ADGRV1;refGene_tx=XM_019836778.1;refGene_gn=ADGRV1;refGene_tx=XM_023258340.1;refGene_gn=ADGRV1;refGene_tx=XM_023258343.1;refGene_gn=ADGRV1;refGene_tx=XM_023258347.1;refGene_gn=ADGRV1;refGene_tx=XM_023258351.1;refGene_gn=ADGRV1;refGene_tx=XM_023258353.1;refGene_gn=ADGRV1;refGene_tx=XM_023258360.1
A1 155342247 155342341
refGene_gn=ADGRV1;refGene_tx=XM_019836773.1;refGene_gn=ADGRV1;refGene_tx=XM_019836778.1;refGene_gn=ADGRV1;refGene_tx=XM_023258340.1;refGene_gn=ADGRV1;refGene_tx=XM_023258343.1;refGene_gn=ADGRV1;refGene_tx=XM_023258347.1;refGene_gn=ADGRV1;refGene_tx=XM_023258351.1;refGene_gn=ADGRV1;refGene_tx=XM_023258353.1;refGene_gn=ADGRV1;refGene_tx=XM_023258360.1
A1 155342773 155342806
refGene_gn=ADGRV1;refGene_tx=XM_019836773.1;refGene_gn=ADGRV1;refGene_tx=XM_023258340.1;refGene_gn=ADGRV1;refGene_tx=XM_023258343.1;refGene_gn=ADGRV1;refGene_tx=XM_023258347.1;refGene_gn=ADGRV1;refGene_tx=XM_023258351.1;refGene_gn=ADGRV1;refGene_tx=XM_023258353.1;refGene_gn=ADGRV1;refGene_tx=XM_023258360.1
A1 155344527 155344606
refGene_gn=ADGRV1;refGene_tx=XM_019836773.1;refGene_gn=ADGRV1;refGene_tx=XM_019836778.1;refGene_gn=ADGRV1;refGene_tx=XM_023258340.1;refGene_gn=ADGRV1;refGene_tx=XM_023258343.1;refGene_gn=ADGRV1;refGene_tx=XM_023258347.1;refGene_gn=ADGRV1;refGene_tx=XM_023258351.1;refGene_gn=ADGRV1;refGene_tx=XM_023258353.1;refGene_gn=ADGRV1;refGene_tx=XM_023258360.1
A1 155344708 155344847
refGene_gn=ADGRV1;refGene_tx=XM_019836773.1;refGene_gn=ADGRV1;refGene_tx=XM_019836778.1;refGene_gn=ADGRV1;refGene_tx=XM_023258340.1;refGene_gn=ADGRV1;refGene_tx=XM_023258343.1;refGene_gn=ADGRV1;refGene_tx=XM_023258347.1;refGene_gn=ADGRV1;refGene_tx=XM_023258351.1;refGene_gn=ADGRV1;refGene_tx=XM_023258353.1;refGene_gn=ADGRV1;refGene_tx=XM_023258360.1
A1 155345633 155345775
refGene_gn=ADGRV1;refGene_tx=XM_019836773.1;refGene_gn=ADGRV1;refGene_tx=XM_019836778.1;refGene_gn=ADGRV1;refGene_tx=XM_023258340.1;refGene_gn=ADGRV1;refGene_tx=XM_023258343.1;refGene_gn=ADGRV1;refGene_tx=XM_023258347.1;refGene_gn=ADGRV1;refGene_tx=XM_023258351.1;refGene_gn=ADGRV1;refGene_tx=XM_023258353.1;refGene_gn=ADGRV1;refGene_tx=XM_023258360.1
A1 155349413 155349676
refGene_gn=ADGRV1;refGene_tx=XM_019836773.1;refGene_gn=ADGRV1;refGene_tx=XM_019836778.1;refGene_gn=ADGRV1;refGene_tx=XM_023258340.1;refGene_gn=ADGRV1;refGene_tx=XM_023258343.1;refGene_gn=ADGRV1;refGene_tx=XM_023258347.1;refGene_gn=ADGRV1;refGene_tx=XM_023258351.1;refGene_gn=ADGRV1;refGene_tx=XM_023258353.1;refGene_gn=ADGRV1;refGene_tx=XM_023258360.

1
A1 155352424 155352600
refGene_gn=ADGRV1;refGene_tx=XM_019836773.1;refGene_gn=ADGRV1;refGene_tx=XM_019836778.1;refGene_gn=ADGRV1;refGene_tx=XM_023258340.1;refGene_gn=ADGRV1;refGene_tx=XM_023258343.1;refGene_gn=ADGRV1;refGene_tx=XM_023258347.1;refGene_gn=ADGRV1;refGene_tx=XM_023258351.1;refGene_gn=ADGRV1;refGene_tx=XM_023258353.1;refGene_gn=ADGRV1;refGene_tx=XM_023258360.1
A1 155353664 155353789
refGene_gn=ADGRV1;refGene_tx=XM_019836773.1;refGene_gn=ADGRV1;refGene_tx=XM_019836778.1;refGene_gn=ADGRV1;refGene_tx=XM_023258340.1;refGene_gn=ADGRV1;refGene_tx=XM_023258343.1;refGene_gn=ADGRV1;refGene_tx=XM_023258347.1;refGene_gn=ADGRV1;refGene_tx=XM_023258351.1;refGene_gn=ADGRV1;refGene_tx=XM_023258353.1;refGene_gn=ADGRV1;refGene_tx=XM_023258360.1
A1 155356674 155356832
refGene_gn=ADGRV1;refGene_tx=XM_019836773.1;refGene_gn=ADGRV1;refGene_tx=XM_019836778.1;refGene_gn=ADGRV1;refGene_tx=XM_023258340.1;refGene_gn=ADGRV1;refGene_tx=XM_023258343.1;refGene_gn=ADGRV1;refGene_tx=XM_023258347.1;refGene_gn=ADGRV1;refGene_tx=XM_023258351.1;refGene_gn=ADGRV1;refGene_tx=XM_023258353.1;refGene_gn=ADGRV1;refGene_tx=XM_023258360.1
A1 155356922 155357069
refGene_gn=ADGRV1;refGene_tx=XM_019836773.1;refGene_gn=ADGRV1;refGene_tx=XM_019836778.1;refGene_gn=ADGRV1;refGene_tx=XM_023258340.1;refGene_gn=ADGRV1;refGene_tx=XM_023258343.1;refGene_gn=ADGRV1;refGene_tx=XM_023258347.1;refGene_gn=ADGRV1;refGene_tx=XM_023258351.1;refGene_gn=ADGRV1;refGene_tx=XM_023258353.1;refGene_gn=ADGRV1;refGene_tx=XM_023258360.1
A1 155357390 155357498
refGene_gn=ADGRV1;refGene_tx=XM_019836773.1;refGene_gn=ADGRV1;refGene_tx=XM_019836778.1;refGene_gn=ADGRV1;refGene_tx=XM_023258340.1;refGene_gn=ADGRV1;refGene_tx=XM_023258343.1;refGene_gn=ADGRV1;refGene_tx=XM_023258347.1;refGene_gn=ADGRV1;refGene_tx=XM_023258351.1;refGene_gn=ADGRV1;refGene_tx=XM_023258353.1;refGene_gn=ADGRV1;refGene_tx=XM_023258360.1
A1 155360426 155360691
refGene_gn=ADGRV1;refGene_tx=XM_019836773.1;refGene_gn=ADGRV1;refGene_tx=XM_019836778.1;refGene_gn=ADGRV1;refGene_tx=XM_023258340.1;refGene_gn=ADGRV1;refGene_tx=XM_023258343.1;refGene_gn=ADGRV1;refGene_tx=XM_023258347.1;refGene_gn=ADGRV1;refGene_tx=XM_023258351.1;refGene_gn=ADGRV1;refGene_tx=XM_023258353.1;refGene_gn=ADGRV1;refGene_tx=XM_023258360.1
A1 155361650 155361773
refGene_gn=ADGRV1;refGene_tx=XM_019836773.1;refGene_gn=ADGRV1;refGene_tx=XM_019836778.1;refGene_gn=ADGRV1;refGene_tx=XM_023258340.1;refGene_gn=ADGRV1;refGene_tx=XM_023258343.1;refGene_gn=ADGRV1;refGene_tx=XM_023258347.1;refGene_gn=ADGRV1;refGene_tx=XM_023258351.1;refGene_gn=ADGRV1;refGene_tx=XM_023258353.1;refGene_gn=ADGRV1;refGene_tx=XM_023258360.1

A1 155367911 155368134
refGene_gn=ADGRV1;refGene_tx=XM_019836773.1;refGene_gn=ADGRV1;refGene_tx=XM_019836778.1;refGene_gn=ADGRV1;refGene_tx=XM_023258340.1;refGene_gn=ADGRV1;refGene_tx=XM_023258343.1;refGene_gn=ADGRV1;refGene_tx=XM_023258347.1;refGene_gn=ADGRV1;refGene_tx=XM_023258351.1;refGene_gn=ADGRV1;refGene_tx=XM_023258353.1;refGene_gn=ADGRV1;refGene_tx=XM_023258360.1
A1 155368463 155368668
refGene_gn=ADGRV1;refGene_tx=XM_019836773.1;refGene_gn=ADGRV1;refGene_tx=XM_019836778.1;refGene_gn=ADGRV1;refGene_tx=XM_023258340.1;refGene_gn=ADGRV1;refGene_tx=XM_023258343.1;refGene_gn=ADGRV1;refGene_tx=XM_023258347.1;refGene_gn=ADGRV1;refGene_tx=XM_023258351.1;refGene_gn=ADGRV1;refGene_tx=XM_023258353.1;refGene_gn=ADGRV1;refGene_tx=XM_023258360.1
A1 155375284 155375431
refGene_gn=ADGRV1;refGene_tx=XM_019836773.1;refGene_gn=ADGRV1;refGene_tx=XM_019836778.1;refGene_gn=ADGRV1;refGene_tx=XM_023258340.1;refGene_gn=ADGRV1;refGene_tx=XM_023258343.1;refGene_gn=ADGRV1;refGene_tx=XM_023258347.1;refGene_gn=ADGRV1;refGene_tx=XM_023258351.1;refGene_gn=ADGRV1;refGene_tx=XM_023258353.1;refGene_gn=ADGRV1;refGene_tx=XM_023258360.1
A1 155377243 155377499
refGene_gn=ADGRV1;refGene_tx=XM_019836773.1;refGene_gn=ADGRV1;refGene_tx=XM_019836778.1;refGene_gn=ADGRV1;refGene_tx=XM_023258340.1;refGene_gn=ADGRV1;refGene_tx=XM_023258343.1;refGene_gn=ADGRV1;refGene_tx=XM_023258347.1;refGene_gn=ADGRV1;refGene_tx=XM_023258351.1;refGene_gn=ADGRV1;refGene_tx=XM_023258353.1;refGene_gn=ADGRV1;refGene_tx=XM_023258360.1
A1 155379203 155379406
refGene_gn=ADGRV1;refGene_tx=XM_019836773.1;refGene_gn=ADGRV1;refGene_tx=XM_019836778.1;refGene_gn=ADGRV1;refGene_tx=XM_023258340.1;refGene_gn=ADGRV1;refGene_tx=XM_023258343.1;refGene_gn=ADGRV1;refGene_tx=XM_023258347.1;refGene_gn=ADGRV1;refGene_tx=XM_023258351.1;refGene_gn=ADGRV1;refGene_tx=XM_023258353.1;refGene_gn=ADGRV1;refGene_tx=XM_023258360.1
A1 155380634 155380811
refGene_gn=ADGRV1;refGene_tx=XM_019836773.1;refGene_gn=ADGRV1;refGene_tx=XM_019836778.1;refGene_gn=ADGRV1;refGene_tx=XM_023258340.1;refGene_gn=ADGRV1;refGene_tx=XM_023258343.1;refGene_gn=ADGRV1;refGene_tx=XM_023258347.1;refGene_gn=ADGRV1;refGene_tx=XM_023258351.1;refGene_gn=ADGRV1;refGene_tx=XM_023258353.1;refGene_gn=ADGRV1;refGene_tx=XM_023258360.1
A1 155381675 155381858
refGene_gn=ADGRV1;refGene_tx=XM_019836773.1;refGene_gn=ADGRV1;refGene_tx=XM_019836778.1;refGene_gn=ADGRV1;refGene_tx=XM_023258340.1;refGene_gn=ADGRV1;refGene_tx=XM_023258343.1;refGene_gn=ADGRV1;refGene_tx=XM_023258347.1;refGene_gn=ADGRV1;refGene_tx=XM_023258351.1;refGene_gn=ADGRV1;refGene_tx=XM_023258353.1;refGene_gn=ADGRV1;refGene_tx=XM_023258360.1
A1 155383731 155383911

refGene_gn=ADGRV1;refGene_tx=XM_019836773.1;refGene_gn=ADGRV1;refGene_tx=XM_019836778.1;refGene_gn=ADGRV1;refGene_tx=XM_023258340.1;refGene_gn=ADGRV1;refGene_tx=XM_023258343.1;refGene_gn=ADGRV1;refGene_tx=XM_023258347.1;refGene_gn=ADGRV1;refGene_tx=XM_023258351.1;refGene_gn=ADGRV1;refGene_tx=XM_023258353.1;refGene_gn=ADGRV1;refGene_tx=XM_023258360.1
A1 155387543 155387708
refGene_gn=ADGRV1;refGene_tx=XM_019836773.1;refGene_gn=ADGRV1;refGene_tx=XM_019836778.1;refGene_gn=ADGRV1;refGene_tx=XM_023258340.1;refGene_gn=ADGRV1;refGene_tx=XM_023258343.1;refGene_gn=ADGRV1;refGene_tx=XM_023258347.1;refGene_gn=ADGRV1;refGene_tx=XM_023258351.1;refGene_gn=ADGRV1;refGene_tx=XM_023258353.1;refGene_gn=ADGRV1;refGene_tx=XM_023258360.1
A1 155388259 155388377
refGene_gn=ADGRV1;refGene_tx=XM_019836773.1;refGene_gn=ADGRV1;refGene_tx=XM_019836778.1;refGene_gn=ADGRV1;refGene_tx=XM_023258340.1;refGene_gn=ADGRV1;refGene_tx=XM_023258343.1;refGene_gn=ADGRV1;refGene_tx=XM_023258347.1;refGene_gn=ADGRV1;refGene_tx=XM_023258351.1;refGene_gn=ADGRV1;refGene_tx=XM_023258353.1;refGene_gn=ADGRV1;refGene_tx=XM_023258360.1
A1 155390431 155390555
refGene_gn=ADGRV1;refGene_tx=XM_019836773.1;refGene_gn=ADGRV1;refGene_tx=XM_019836778.1;refGene_gn=ADGRV1;refGene_tx=XM_023258340.1;refGene_gn=ADGRV1;refGene_tx=XM_023258343.1;refGene_gn=ADGRV1;refGene_tx=XM_023258347.1;refGene_gn=ADGRV1;refGene_tx=XM_023258351.1;refGene_gn=ADGRV1;refGene_tx=XM_023258353.1;refGene_gn=ADGRV1;refGene_tx=XM_023258360.1
A1 155391433 155391572
refGene_gn=ADGRV1;refGene_tx=XM_019836773.1;refGene_gn=ADGRV1;refGene_tx=XM_019836778.1;refGene_gn=ADGRV1;refGene_tx=XM_023258340.1;refGene_gn=ADGRV1;refGene_tx=XM_023258343.1;refGene_gn=ADGRV1;refGene_tx=XM_023258347.1;refGene_gn=ADGRV1;refGene_tx=XM_023258351.1;refGene_gn=ADGRV1;refGene_tx=XM_023258353.1;refGene_gn=ADGRV1;refGene_tx=XM_023258360.1
A1 155391746 155391932
refGene_gn=ADGRV1;refGene_tx=XM_019836773.1;refGene_gn=ADGRV1;refGene_tx=XM_019836778.1;refGene_gn=ADGRV1;refGene_tx=XM_023258340.1;refGene_gn=ADGRV1;refGene_tx=XM_023258343.1;refGene_gn=ADGRV1;refGene_tx=XM_023258347.1;refGene_gn=ADGRV1;refGene_tx=XM_023258351.1;refGene_gn=ADGRV1;refGene_tx=XM_023258353.1;refGene_gn=ADGRV1;refGene_tx=XM_023258360.1
A1 155392281 155392514
refGene_gn=ADGRV1;refGene_tx=XM_019836773.1;refGene_gn=ADGRV1;refGene_tx=XM_019836778.1;refGene_gn=ADGRV1;refGene_tx=XM_023258340.1;refGene_gn=ADGRV1;refGene_tx=XM_023258343.1;refGene_gn=ADGRV1;refGene_tx=XM_023258347.1;refGene_gn=ADGRV1;refGene_tx=XM_023258351.1;refGene_gn=ADGRV1;refGene_tx=XM_023258353.1;refGene_gn=ADGRV1;refGene_tx=XM_023258360.1
A1 155395686 155395835
refGene_gn=ADGRV1;refGene_tx=XM_019836773.1;refGene_gn=ADGRV1;refGene_

tx=XM_019836778.1;refGene_gn=ADGRV1;refGene_tx=XM_023258340.1;refGene_gn=ADGRV1;refGene_tx=XM_023258343.1;refGene_gn=ADGRV1;refGene_tx=XM_023258347.1;refGene_gn=ADGRV1;refGene_tx=XM_023258351.1;refGene_gn=ADGRV1;refGene_tx=XM_023258353.1;refGene_gn=ADGRV1;refGene_tx=XM_023258360.1
A1 155397587 155397789
refGene_gn=ADGRV1;refGene_tx=XM_019836773.1;refGene_gn=ADGRV1;refGene_tx=XM_019836778.1;refGene_gn=ADGRV1;refGene_tx=XM_023258340.1;refGene_gn=ADGRV1;refGene_tx=XM_023258343.1;refGene_gn=ADGRV1;refGene_tx=XM_023258347.1;refGene_gn=ADGRV1;refGene_tx=XM_023258351.1;refGene_gn=ADGRV1;refGene_tx=XM_023258353.1;refGene_gn=ADGRV1;refGene_tx=XM_023258360.1
A1 155398308 155398528
refGene_gn=ADGRV1;refGene_tx=XM_019836773.1;refGene_gn=ADGRV1;refGene_tx=XM_019836778.1;refGene_gn=ADGRV1;refGene_tx=XM_023258340.1;refGene_gn=ADGRV1;refGene_tx=XM_023258343.1;refGene_gn=ADGRV1;refGene_tx=XM_023258347.1;refGene_gn=ADGRV1;refGene_tx=XM_023258351.1;refGene_gn=ADGRV1;refGene_tx=XM_023258353.1;refGene_gn=ADGRV1;refGene_tx=XM_023258360.1
A1 155400990 155401230
refGene_gn=ADGRV1;refGene_tx=XM_019836773.1;refGene_gn=ADGRV1;refGene_tx=XM_019836778.1;refGene_gn=ADGRV1;refGene_tx=XM_023258340.1;refGene_gn=ADGRV1;refGene_tx=XM_023258343.1;refGene_gn=ADGRV1;refGene_tx=XM_023258347.1;refGene_gn=ADGRV1;refGene_tx=XM_023258351.1;refGene_gn=ADGRV1;refGene_tx=XM_023258353.1;refGene_gn=ADGRV1;refGene_tx=XM_023258360.1
A1 155402516 155402666
refGene_gn=ADGRV1;refGene_tx=XM_019836773.1;refGene_gn=ADGRV1;refGene_tx=XM_019836778.1;refGene_gn=ADGRV1;refGene_tx=XM_023258340.1;refGene_gn=ADGRV1;refGene_tx=XM_023258343.1;refGene_gn=ADGRV1;refGene_tx=XM_023258347.1;refGene_gn=ADGRV1;refGene_tx=XM_023258351.1;refGene_gn=ADGRV1;refGene_tx=XM_023258353.1;refGene_gn=ADGRV1;refGene_tx=XM_023258360.1
A1 155403742 155404216
refGene_gn=ADGRV1;refGene_tx=XM_019836773.1;refGene_gn=ADGRV1;refGene_tx=XM_019836778.1;refGene_gn=ADGRV1;refGene_tx=XM_023258340.1;refGene_gn=ADGRV1;refGene_tx=XM_023258343.1;refGene_gn=ADGRV1;refGene_tx=XM_023258347.1;refGene_gn=ADGRV1;refGene_tx=XM_023258351.1;refGene_gn=ADGRV1;refGene_tx=XM_023258353.1;refGene_gn=ADGRV1;refGene_tx=XM_023258360.1
A1 155406417 155406561
refGene_gn=ADGRV1;refGene_tx=XM_019836773.1;refGene_gn=ADGRV1;refGene_tx=XM_019836778.1;refGene_gn=ADGRV1;refGene_tx=XM_023258340.1;refGene_gn=ADGRV1;refGene_tx=XM_023258343.1;refGene_gn=ADGRV1;refGene_tx=XM_023258347.1;refGene_gn=ADGRV1;refGene_tx=XM_023258351.1;refGene_gn=ADGRV1;refGene_tx=XM_023258353.1;refGene_gn=ADGRV1;refGene_tx=XM_023258360.1
A1 155409458 155409633
refGene_gn=ADGRV1;refGene_tx=XM_019836773.1;refGene_gn=ADGRV1;refGene_tx=XM_019836778.1;refGene_gn=ADGRV1;refGene_tx=XM_023258340.1;refGene_

gn=ADGRV1;refGene_tx=XM_023258343.1;refGene_gn=ADGRV1;refGene_tx=XM_023258347.1;refGene_gn=ADGRV1;refGene_tx=XM_023258351.1;refGene_gn=ADGRV1;refGene_tx=XM_023258353.1;refGene_gn=ADGRV1;refGene_tx=XM_023258360.1
A1 155411163 155411299
refGene_gn=ADGRV1;refGene_tx=XM_019836773.1;refGene_gn=ADGRV1;refGene_tx=XM_019836778.1;refGene_gn=ADGRV1;refGene_tx=XM_023258340.1;refGene_gn=ADGRV1;refGene_tx=XM_023258343.1;refGene_gn=ADGRV1;refGene_tx=XM_023258347.1;refGene_gn=ADGRV1;refGene_tx=XM_023258351.1;refGene_gn=ADGRV1;refGene_tx=XM_023258353.1;refGene_gn=ADGRV1;refGene_tx=XM_023258360.1
A1 155413829 155414938
refGene_gn=ADGRV1;refGene_tx=XM_019836773.1;refGene_gn=ADGRV1;refGene_tx=XM_019836778.1;refGene_gn=ADGRV1;refGene_tx=XM_023258340.1;refGene_gn=ADGRV1;refGene_tx=XM_023258343.1;refGene_gn=ADGRV1;refGene_tx=XM_023258347.1;refGene_gn=ADGRV1;refGene_tx=XM_023258351.1;refGene_gn=ADGRV1;refGene_tx=XM_023258353.1;refGene_gn=ADGRV1;refGene_tx=XM_023258360.1
A1 155418423 155418541
refGene_gn=ADGRV1;refGene_tx=XM_019836773.1;refGene_gn=ADGRV1;refGene_tx=XM_019836778.1;refGene_gn=ADGRV1;refGene_tx=XM_023258340.1;refGene_gn=ADGRV1;refGene_tx=XM_023258343.1;refGene_gn=ADGRV1;refGene_tx=XM_023258347.1;refGene_gn=ADGRV1;refGene_tx=XM_023258351.1;refGene_gn=ADGRV1;refGene_tx=XM_023258353.1;refGene_gn=ADGRV1;refGene_tx=XM_023258360.1
A1 155419413 155419585
refGene_gn=ADGRV1;refGene_tx=XM_019836773.1;refGene_gn=ADGRV1;refGene_tx=XM_019836778.1;refGene_gn=ADGRV1;refGene_tx=XM_023258340.1;refGene_gn=ADGRV1;refGene_tx=XM_023258343.1;refGene_gn=ADGRV1;refGene_tx=XM_023258347.1;refGene_gn=ADGRV1;refGene_tx=XM_023258351.1;refGene_gn=ADGRV1;refGene_tx=XM_023258353.1;refGene_gn=ADGRV1;refGene_tx=XM_023258360.1
A1 155428855 155429098
refGene_gn=ADGRV1;refGene_tx=XM_019836773.1;refGene_gn=ADGRV1;refGene_tx=XM_019836778.1;refGene_gn=ADGRV1;refGene_tx=XM_023258340.1;refGene_gn=ADGRV1;refGene_tx=XM_023258343.1;refGene_gn=ADGRV1;refGene_tx=XM_023258347.1;refGene_gn=ADGRV1;refGene_tx=XM_023258351.1;refGene_gn=ADGRV1;refGene_tx=XM_023258353.1;refGene_gn=ADGRV1;refGene_tx=XM_023258360.1
A1 155432875 155433283
refGene_gn=ADGRV1;refGene_tx=XM_019836773.1;refGene_gn=ADGRV1;refGene_tx=XM_019836778.1;refGene_gn=ADGRV1;refGene_tx=XM_023258340.1;refGene_gn=ADGRV1;refGene_tx=XM_023258343.1;refGene_gn=ADGRV1;refGene_tx=XM_023258347.1;refGene_gn=ADGRV1;refGene_tx=XM_023258351.1;refGene_gn=ADGRV1;refGene_tx=XM_023258353.1;refGene_gn=ADGRV1;refGene_tx=XM_023258360.1
A1 155437553 155437738
refGene_gn=ADGRV1;refGene_tx=XM_019836773.1;refGene_gn=ADGRV1;refGene_tx=XM_019836778.1;refGene_gn=ADGRV1;refGene_tx=XM_023258340.1;refGene_gn=ADGRV1;refGene_tx=XM_023258343.1;refGene_gn=ADGRV1;refGene_tx=XM_02

3258347.1;refGene_gn=ADGRV1;refGene_tx=XM_023258351.1;refGene_gn=ADGRV1;refGene_tx=XM_023258353.1
A1	155442765	155443015
refGene_gn=ADGRV1;refGene_tx=XM_019836773.1;refGene_gn=ADGRV1;refGene_tx=XM_019836778.1;refGene_gn=ADGRV1;refGene_tx=XM_023258340.1;refGene_gn=ADGRV1;refGene_tx=XM_023258343.1;refGene_gn=ADGRV1;refGene_tx=XM_023258347.1;refGene_gn=ADGRV1;refGene_tx=XM_023258351.1;refGene_gn=ADGRV1;refGene_tx=XM_023258353.1;refGene_gn=ADGRV1;refGene_tx=XM_023258360.1
A1	155443534	155443674
refGene_gn=ADGRV1;refGene_tx=XM_019836773.1;refGene_gn=ADGRV1;refGene_tx=XM_019836778.1;refGene_gn=ADGRV1;refGene_tx=XM_023258340.1;refGene_gn=ADGRV1;refGene_tx=XM_023258343.1;refGene_gn=ADGRV1;refGene_tx=XM_023258347.1;refGene_gn=ADGRV1;refGene_tx=XM_023258351.1;refGene_gn=ADGRV1;refGene_tx=XM_023258353.1
A1	155445423	155445584
refGene_gn=ADGRV1;refGene_tx=XM_019836773.1;refGene_gn=ADGRV1;refGene_tx=XM_019836778.1;refGene_gn=ADGRV1;refGene_tx=XM_023258340.1;refGene_gn=ADGRV1;refGene_tx=XM_023258343.1;refGene_gn=ADGRV1;refGene_tx=XM_023258347.1;refGene_gn=ADGRV1;refGene_tx=XM_023258351.1;refGene_gn=ADGRV1;refGene_tx=XM_023258353.1
A1	155455482	155455583
refGene_gn=ADGRV1;refGene_tx=XM_019836773.1;refGene_gn=ADGRV1;refGene_tx=XM_019836778.1;refGene_gn=ADGRV1;refGene_tx=XM_023258340.1;refGene_gn=ADGRV1;refGene_tx=XM_023258343.1;refGene_gn=ADGRV1;refGene_tx=XM_023258347.1;refGene_gn=ADGRV1;refGene_tx=XM_023258351.1;refGene_gn=ADGRV1;refGene_tx=XM_023258353.1
A1	155561061	155561178
refGene_gn=ADGRV1;refGene_tx=XM_019836773.1;refGene_gn=ADGRV1;refGene_tx=XM_019836778.1;refGene_gn=ADGRV1;refGene_tx=XM_023258340.1;refGene_gn=ADGRV1;refGene_tx=XM_023258343.1;refGene_gn=ADGRV1;refGene_tx=XM_023258347.1;refGene_gn=ADGRV1;refGene_tx=XM_023258351.1;refGene_gn=ADGRV1;refGene_tx=XM_023258353.1
A1	155573281	155573460
refGene_gn=ADGRV1;refGene_tx=XM_019836773.1;refGene_gn=ADGRV1;refGene_tx=XM_019836778.1;refGene_gn=ADGRV1;refGene_tx=XM_023258340.1;refGene_gn=ADGRV1;refGene_tx=XM_023258343.1;refGene_gn=ADGRV1;refGene_tx=XM_023258347.1;refGene_gn=ADGRV1;refGene_tx=XM_023258351.1
A1	155670771	155670929
refGene_gn=ADGRV1;refGene_tx=XM_019836773.1;refGene_gn=ADGRV1;refGene_tx=XM_019836778.1;refGene_gn=ADGRV1;refGene_tx=XM_023258340.1;refGene_gn=ADGRV1;refGene_tx=XM_023258343.1;refGene_gn=ADGRV1;refGene_tx=XM_023258347.1;refGene_gn=ADGRV1;refGene_tx=XM_023258351.1;refGene_gn=ADGRV1;refGene_tx=XM_023258353.1
A1	155699350	155699472
refGene_gn=ADGRV1;refGene_tx=XM_019836773.1;refGene_gn=ADGRV1;refGene_tx=XM_019836778.1;refGene_gn=ADGRV1;refGene_tx=XM_023258340.1;refGene_gn=ADGRV1;refGene_tx=XM_023258343.1;refGene_gn=ADGRV1;refGene_tx=XM_023258347.1;refGene_gn=ADGRV1;refGene_tx=XM_023258351.1
A1	155756189	155756381

refGene_gn=ADGRV1;refGene_tx=XM_019836773.1;refGene_gn=ADGRV1;refGene_tx=XM_019836778.1;refGene_gn=ADGRV1;refGene_tx=XM_023258340.1;refGene_gn=ADGRV1;refGene_tx=XM_023258343.1;refGene_gn=ADGRV1;refGene_tx=XM_023258347.1;refGene_gn=ADGRV1;refGene_tx=XM_023258351.1
A1 155759219 155759397
refGene_gn=ADGRV1;refGene_tx=XM_019836773.1;refGene_gn=ADGRV1;refGene_tx=XM_019836778.1;refGene_gn=ADGRV1;refGene_tx=XM_023258340.1;refGene_gn=ADGRV1;refGene_tx=XM_023258343.1;refGene_gn=ADGRV1;refGene_tx=XM_023258347.1;refGene_gn=ADGRV1;refGene_tx=XM_023258351.1
A1 155768168 155768287
refGene_gn=ADGRV1;refGene_tx=XM_019836773.1;refGene_gn=ADGRV1;refGene_tx=XM_019836778.1;refGene_gn=ADGRV1;refGene_tx=XM_023258340.1;refGene_gn=ADGRV1;refGene_tx=XM_023258343.1;refGene_gn=ADGRV1;refGene_tx=XM_023258347.1;refGene_gn=ADGRV1;refGene_tx=XM_023258351.1
A1 155936237 155936294
refGene_gn=ARRDC3;refGene_tx=XM_011284590.3;refGene_gn=ARRDC3;refGene_tx=XM_019836866.2;refGene_gn=ARRDC3;refGene_tx=XM_019836868.2
A1 155938879 155939072
refGene_gn=ARRDC3;refGene_tx=XM_011284590.3;refGene_gn=ARRDC3;refGene_tx=XM_011284591.2;refGene_gn=ARRDC3;refGene_tx=XM_019836866.2;refGene_gn=ARRDC3;refGene_tx=XM_019836868.2
A1 155939306 155939469
refGene_gn=ARRDC3;refGene_tx=XM_011284590.3;refGene_gn=ARRDC3;refGene_tx=XM_011284591.2;refGene_gn=ARRDC3;refGene_tx=XM_019836866.2;refGene_gn=ARRDC3;refGene_tx=XM_019836868.2
A1 155939805 155940062
refGene_gn=ARRDC3;refGene_tx=XM_011284590.3;refGene_gn=ARRDC3;refGene_tx=XM_011284591.2;refGene_gn=ARRDC3;refGene_tx=XM_019836866.2;refGene_gn=ARRDC3;refGene_tx=XM_019836868.2
A1 155940403 155940506
refGene_gn=ARRDC3;refGene_tx=XM_011284590.3;refGene_gn=ARRDC3;refGene_tx=XM_011284591.2;refGene_gn=ARRDC3;refGene_tx=XM_019836866.2
A1 155941493 155941641
refGene_gn=ARRDC3;refGene_tx=XM_011284590.3;refGene_gn=ARRDC3;refGene_tx=XM_011284591.2;refGene_gn=ARRDC3;refGene_tx=XM_019836866.2
A1 155943554 155943636
refGene_gn=ARRDC3;refGene_tx=XM_011284590.3;refGene_gn=ARRDC3;refGene_tx=XM_011284591.2;refGene_gn=ARRDC3;refGene_tx=XM_019836866.2
A1 155944526 155944548
refGene_gn=ARRDC3;refGene_tx=XM_019836866.2
A1 155947604 155947884
refGene_gn=ARRDC3;refGene_tx=XM_011284590.3;refGene_gn=ARRDC3;refGene_tx=XM_011284591.2
A1 158001439 158001902
refGene_gn=NR2F1;refGene_tx=XM_019836924.2;refGene_gn=NR2F1;refGene_tx=XM_023258405.1
A1 158003638 158003651
refGene_gn=NR2F1;refGene_tx=XM_019836922.2
A1 158004331 158004859
refGene_gn=NR2F1;refGene_tx=XM_019836922.2;refGene_gn=NR2F1;refGene_tx

=XM_019836924.2;refGene_gn=NR2F1;refGene_tx=XM_023258405.1
A1 158009947 158010228
refGene_gn=NR2F1;refGene_tx=XM_019836922.2;refGene_gn=NR2F1;refGene_tx=XM_019836924.2;refGene_gn=NR2F1;refGene_tx=XM_023258405.1
A1 158037416 158037559
refGene_gn=FAM172A;refGene_tx=XM_023258433.1;refGene_gn=FAM172A;refGene_tx=XM_023258436.1;refGene_gn=FAM172A;refGene_tx=XM_023258451.1;refGene_gn=FAM172A;refGene_tx=XM_023258460.1;refGene_gn=FAM172A;refGene_tx=XM_023258464.1
A1 158124998 158125051
refGene_gn=FAM172A;refGene_tx=XM_019837023.1
A1 158162649 158163663
refGene_gn=POU5F2;refGene_tx=XM_019837078.2
A1 158178978 158178986
refGene_gn=FAM172A;refGene_tx=XM_019837058.2
A1 158179090 158179134
refGene_gn=FAM172A;refGene_tx=XM_019837033.2
A1 158179167 158179217
refGene_gn=FAM172A;refGene_tx=XM_019837029.2
A1 158183619 158183704
refGene_gn=FAM172A;refGene_tx=XM_019837023.1;refGene_gn=FAM172A;refGene_tx=XM_019837029.2;refGene_gn=FAM172A;refGene_tx=XM_019837033.2;refGene_gn=FAM172A;refGene_tx=XM_019837058.2;refGene_gn=FAM172A;refGene_tx=XM_023258433.1;refGene_gn=FAM172A;refGene_tx=XM_023258436.1;refGene_gn=FAM172A;refGene_tx=XM_023258451.1;refGene_gn=FAM172A;refGene_tx=XM_023258460.1;refGene_gn=FAM172A;refGene_tx=XM_023258464.1
A1 158191598 158191715
refGene_gn=FAM172A;refGene_tx=XM_019837023.1;refGene_gn=FAM172A;refGene_tx=XM_019837029.2;refGene_gn=FAM172A;refGene_tx=XM_019837033.2;refGene_gn=FAM172A;refGene_tx=XM_019837058.2;refGene_gn=FAM172A;refGene_tx=XM_023258433.1;refGene_gn=FAM172A;refGene_tx=XM_023258436.1;refGene_gn=FAM172A;refGene_tx=XM_023258451.1;refGene_gn=FAM172A;refGene_tx=XM_023258460.1;refGene_gn=FAM172A;refGene_tx=XM_023258464.1
A1 158231059 158231179
refGene_gn=FAM172A;refGene_tx=XM_019837023.1;refGene_gn=FAM172A;refGene_tx=XM_019837029.2;refGene_gn=FAM172A;refGene_tx=XM_019837033.2;refGene_gn=FAM172A;refGene_tx=XM_019837058.2;refGene_gn=FAM172A;refGene_tx=XM_023258433.1;refGene_gn=FAM172A;refGene_tx=XM_023258436.1;refGene_gn=FAM172A;refGene_tx=XM_023258451.1;refGene_gn=FAM172A;refGene_tx=XM_023258460.1;refGene_gn=FAM172A;refGene_tx=XM_023258464.1
A1 158285702 158285921
refGene_gn=FAM172A;refGene_tx=XM_019837023.1;refGene_gn=FAM172A;refGene_tx=XM_019837029.2;refGene_gn=FAM172A;refGene_tx=XM_019837033.2;refGene_gn=FAM172A;refGene_tx=XM_019837058.2;refGene_gn=FAM172A;refGene_tx=XM_023258433.1;refGene_gn=FAM172A;refGene_tx=XM_023258436.1;refGene_gn=FAM172A;refGene_tx=XM_023258451.1;refGene_gn=FAM172A;refGene_tx=XM_023258460.1;refGene_gn=FAM172A;refGene_tx=XM_023258464.1
A1 158352849 158353041
refGene_gn=FAM172A;refGene_tx=XM_019837023.1;refGene_gn=FAM172A;refGene_tx=XM_019837029.2;refGene_gn=FAM172A;refGene_tx=XM_019837033.2;refGe

ne_gn=FAM172A;refGene_tx=XM_019837058.2;refGene_gn=FAM172A;refGene_tx=XM_023258433.1;refGene_gn=FAM172A;refGene_tx=XM_023258436.1;refGene_gn=FAM172A;refGene_tx=XM_023258460.1;refGene_gn=FAM172A;refGene_tx=XM_023258464.1
A1 158358012 158358078
refGene_gn=FAM172A;refGene_tx=XM_019837023.1;refGene_gn=FAM172A;refGene_tx=XM_019837029.2;refGene_gn=FAM172A;refGene_tx=XM_019837033.2;refGene_gn=FAM172A;refGene_tx=XM_019837058.2;refGene_gn=FAM172A;refGene_tx=XM_023258433.1;refGene_gn=FAM172A;refGene_tx=XM_023258436.1;refGene_gn=FAM172A;refGene_tx=XM_023258451.1;refGene_gn=FAM172A;refGene_tx=XM_023258460.1;refGene_gn=FAM172A;refGene_tx=XM_023258464.1
A1 158408802 158408903
refGene_gn=FAM172A;refGene_tx=XM_019837023.1;refGene_gn=FAM172A;refGene_tx=XM_019837029.2;refGene_gn=FAM172A;refGene_tx=XM_019837033.2;refGene_gn=FAM172A;refGene_tx=XM_019837058.2;refGene_gn=FAM172A;refGene_tx=XM_023258433.1;refGene_gn=FAM172A;refGene_tx=XM_023258436.1;refGene_gn=FAM172A;refGene_tx=XM_023258451.1
A1 158413913 158414015
refGene_gn=FAM172A;refGene_tx=XM_019837023.1;refGene_gn=FAM172A;refGene_tx=XM_019837029.2;refGene_gn=FAM172A;refGene_tx=XM_019837033.2;refGene_gn=FAM172A;refGene_tx=XM_019837058.2;refGene_gn=FAM172A;refGene_tx=XM_023258433.1;refGene_gn=FAM172A;refGene_tx=XM_023258436.1;refGene_gn=FAM172A;refGene_tx=XM_023258451.1;refGene_gn=FAM172A;refGene_tx=XM_023258460.1;refGene_gn=FAM172A;refGene_tx=XM_023258464.1
A1 158426126 158426232
refGene_gn=FAM172A;refGene_tx=XM_023258436.1
A1 158482102 158482220
refGene_gn=KIAA0825;refGene_tx=XM_019836952.2;refGene_gn=KIAA0825;refGene_tx=XM_019836968.2;refGene_gn=KIAA0825;refGene_tx=XM_023258417.1
A1 158679608 158679703
refGene_gn=KIAA0825;refGene_tx=XM_019836952.2;refGene_gn=KIAA0825;refGene_tx=XM_019836968.2;refGene_gn=KIAA0825;refGene_tx=XM_023258417.1
A1 158681478 158681637
refGene_gn=KIAA0825;refGene_tx=XM_019836952.2;refGene_gn=KIAA0825;refGene_tx=XM_019836968.2;refGene_gn=KIAA0825;refGene_tx=XM_023258417.1
A1 158685731 158685891
refGene_gn=KIAA0825;refGene_tx=XM_019836952.2;refGene_gn=KIAA0825;refGene_tx=XM_019836968.2;refGene_gn=KIAA0825;refGene_tx=XM_023258417.1
A1 158690179 158690588
refGene_gn=KIAA0825;refGene_tx=XM_019836952.2;refGene_gn=KIAA0825;refGene_tx=XM_019836968.2;refGene_gn=KIAA0825;refGene_tx=XM_023258417.1
A1 158699138 158699363
refGene_gn=KIAA0825;refGene_tx=XM_019836952.2;refGene_gn=KIAA0825;refGene_tx=XM_019836968.2;refGene_gn=KIAA0825;refGene_tx=XM_023258417.1
A1 158704804 158704984
refGene_gn=KIAA0825;refGene_tx=XM_019836952.2;refGene_gn=KIAA0825;refGene_tx=XM_019836968.2;refGene_gn=KIAA0825;refGene_tx=XM_023258417.1
A1 158720321 158720461
refGene_gn=KIAA0825;refGene_tx=XM_019836952.2;refGene_gn=KIAA0825;refGene_tx=XM_019836968.2;refGene_gn=KIAA0825;refGene_tx=XM_023258417.1

A1	158734140	158734251
refGene_gn=KIAA0825;refGene_tx=XM_019836952.2;refGene_gn=KIAA0825;refGene_tx=XM_019836968.2;refGene_gn=KIAA0825;refGene_tx=XM_023258417.1
A1	158745869	158746052
refGene_gn=KIAA0825;refGene_tx=XM_019836952.2;refGene_gn=KIAA0825;refGene_tx=XM_019836968.2;refGene_gn=KIAA0825;refGene_tx=XM_023258417.1
A1	158748366	158748557
refGene_gn=KIAA0825;refGene_tx=XM_019836952.2;refGene_gn=KIAA0825;refGene_tx=XM_019836968.2;refGene_gn=KIAA0825;refGene_tx=XM_023258417.1
A1	158753021	158753172
refGene_gn=KIAA0825;refGene_tx=XM_019836952.2;refGene_gn=KIAA0825;refGene_tx=XM_019836968.2;refGene_gn=KIAA0825;refGene_tx=XM_023258417.1
A1	158754580	158754846
refGene_gn=KIAA0825;refGene_tx=XM_019836952.2;refGene_gn=KIAA0825;refGene_tx=XM_019836968.2;refGene_gn=KIAA0825;refGene_tx=XM_023258417.1
A1	158756272	158756500
refGene_gn=KIAA0825;refGene_tx=XM_019836952.2;refGene_gn=KIAA0825;refGene_tx=XM_019836968.2;refGene_gn=KIAA0825;refGene_tx=XM_023258417.1
A1	158758087	158758203
refGene_gn=KIAA0825;refGene_tx=XM_019836952.2;refGene_gn=KIAA0825;refGene_tx=XM_019836968.2;refGene_gn=KIAA0825;refGene_tx=XM_023258417.1
A1	158768184	158768346
refGene_gn=KIAA0825;refGene_tx=XM_019836952.2;refGene_gn=KIAA0825;refGene_tx=XM_019836968.2;refGene_gn=KIAA0825;refGene_tx=XM_023258417.1
A1	158804970	158805640
refGene_gn=KIAA0825;refGene_tx=XM_019836952.2;refGene_gn=KIAA0825;refGene_tx=XM_019836968.2;refGene_gn=KIAA0825;refGene_tx=XM_023258417.1
A1	158808629	158808798
refGene_gn=KIAA0825;refGene_tx=XM_019836952.2;refGene_gn=KIAA0825;refGene_tx=XM_019836968.2;refGene_gn=KIAA0825;refGene_tx=XM_023258417.1
A1	158823186	158823317
refGene_gn=KIAA0825;refGene_tx=XM_019836952.2;refGene_gn=KIAA0825;refGene_tx=XM_019836968.2;refGene_gn=KIAA0825;refGene_tx=XM_023258417.1
A1	158882273	158882387
refGene_gn=SLF1;refGene_tx=XM_003981155.4;refGene_gn=SLF1;refGene_tx=XM_019836978.2
A1	158882547	158882623
refGene_gn=SLF1;refGene_tx=XM_003981155.4;refGene_gn=SLF1;refGene_tx=XM_019836978.2
A1	158883759	158884000
refGene_gn=SLF1;refGene_tx=XM_003981155.4;refGene_gn=SLF1;refGene_tx=XM_019836978.2
A1	158895934	158896097
refGene_gn=SLF1;refGene_tx=XM_003981155.4;refGene_gn=SLF1;refGene_tx=XM_019836978.2
A1	158901899	158902043
refGene_gn=SLF1;refGene_tx=XM_003981155.4
A1	158904137	158904284
refGene_gn=SLF1;refGene_tx=XM_003981155.4;refGene_gn=SLF1;refGene_tx=XM_019836978.2

A1 158905587 158905734
refGene_gn=SLF1;refGene_tx=XM_003981155.4;refGene_gn=SLF1;refGene_tx=XM_019836978.2
A1 158906592 158906715
refGene_gn=SLF1;refGene_tx=XM_003981155.4;refGene_gn=SLF1;refGene_tx=XM_019836978.2
A1 158907815 158907869
refGene_gn=SLF1;refGene_tx=XM_003981155.4;refGene_gn=SLF1;refGene_tx=XM_019836978.2
A1 158908805 158908964
refGene_gn=SLF1;refGene_tx=XM_003981155.4;refGene_gn=SLF1;refGene_tx=XM_019836978.2
A1 158910764 158910928
refGene_gn=SLF1;refGene_tx=XM_003981155.4;refGene_gn=SLF1;refGene_tx=XM_019836978.2
A1 158918706 158918838
refGene_gn=SLF1;refGene_tx=XM_003981155.4;refGene_gn=SLF1;refGene_tx=XM_019836978.2
A1 158919670 158919836
refGene_gn=SLF1;refGene_tx=XM_003981155.4;refGene_gn=SLF1;refGene_tx=XM_019836978.2
A1 158926672 158926820
refGene_gn=SLF1;refGene_tx=XM_003981155.4;refGene_gn=SLF1;refGene_tx=XM_019836978.2
A1 158943970 158944116
refGene_gn=SLF1;refGene_tx=XM_003981155.4;refGene_gn=SLF1;refGene_tx=XM_019836978.2
A1 158945142 158945306
refGene_gn=SLF1;refGene_tx=XM_003981155.4;refGene_gn=SLF1;refGene_tx=XM_019836978.2
A1 158946111 158946245
refGene_gn=SLF1;refGene_tx=XM_003981155.4;refGene_gn=SLF1;refGene_tx=XM_019836978.2
A1 158948158 158948251
refGene_gn=SLF1;refGene_tx=XM_003981155.4;refGene_gn=SLF1;refGene_tx=XM_019836978.2
A1 158948640 158948823
refGene_gn=SLF1;refGene_tx=XM_003981155.4;refGene_gn=SLF1;refGene_tx=XM_019836978.2
A1 158951447 158951929
refGene_gn=SLF1;refGene_tx=XM_003981155.4;refGene_gn=SLF1;refGene_tx=XM_019836978.2
A1 158964229 158964301
refGene_gn=MCTP1;refGene_tx=XM_023258485.1;refGene_gn=MCTP1;refGene_tx=XM_023258486.1;refGene_gn=MCTP1;refGene_tx=XM_023258491.1;refGene_gn=MCTP1;refGene_tx=XM_023258493.1;refGene_gn=MCTP1;refGene_tx=XM_023258495.1;refGene_gn=MCTP1;refGene_tx=XM_023258501.1;refGene_gn=MCTP1;refGene_tx=XM_023258502.1;refGene_gn=MCTP1;refGene_tx=XM_023258511.1
A1 158965279 158965377
refGene_gn=MCTP1;refGene_tx=XM_023258485.1;refGene_gn=MCTP1;refGene_tx

=XM_023258486.1;refGene_gn=MCTP1;refGene_tx=XM_023258491.1;refGene_gn=MCTP1;refGene_tx=XM_023258493.1;refGene_gn=MCTP1;refGene_tx=XM_023258495.1;refGene_gn=MCTP1;refGene_tx=XM_023258501.1;refGene_gn=MCTP1;refGene_tx=XM_023258502.1;refGene_gn=MCTP1;refGene_tx=XM_023258511.1
A1 158973889 158973999
refGene_gn=MCTP1;refGene_tx=XM_023258485.1;refGene_gn=MCTP1;refGene_tx=XM_023258486.1;refGene_gn=MCTP1;refGene_tx=XM_023258491.1;refGene_gn=MCTP1;refGene_tx=XM_023258493.1;refGene_gn=MCTP1;refGene_tx=XM_023258495.1;refGene_gn=MCTP1;refGene_tx=XM_023258501.1;refGene_gn=MCTP1;refGene_tx=XM_023258511.1
A1 158977960 158978070
refGene_gn=MCTP1;refGene_tx=XM_023258485.1;refGene_gn=MCTP1;refGene_tx=XM_023258486.1;refGene_gn=MCTP1;refGene_tx=XM_023258491.1;refGene_gn=MCTP1;refGene_tx=XM_023258493.1;refGene_gn=MCTP1;refGene_tx=XM_023258495.1;refGene_gn=MCTP1;refGene_tx=XM_023258501.1;refGene_gn=MCTP1;refGene_tx=XM_023258511.1
A1 159026396 159026417
refGene_gn=MCTP1;refGene_tx=XM_023258504.1
A1 159034857 159034911
refGene_gn=MCTP1;refGene_tx=XM_023258485.1;refGene_gn=MCTP1;refGene_tx=XM_023258486.1;refGene_gn=MCTP1;refGene_tx=XM_023258491.1;refGene_gn=MCTP1;refGene_tx=XM_023258493.1;refGene_gn=MCTP1;refGene_tx=XM_023258495.1;refGene_gn=MCTP1;refGene_tx=XM_023258501.1;refGene_gn=MCTP1;refGene_tx=XM_023258502.1;refGene_gn=MCTP1;refGene_tx=XM_023258504.1;refGene_gn=MCTP1;refGene_tx=XM_023258511.1
A1 159056795 159056915
refGene_gn=MCTP1;refGene_tx=XM_023258485.1;refGene_gn=MCTP1;refGene_tx=XM_023258486.1;refGene_gn=MCTP1;refGene_tx=XM_023258491.1;refGene_gn=MCTP1;refGene_tx=XM_023258493.1;refGene_gn=MCTP1;refGene_tx=XM_023258495.1;refGene_gn=MCTP1;refGene_tx=XM_023258501.1;refGene_gn=MCTP1;refGene_tx=XM_023258502.1;refGene_gn=MCTP1;refGene_tx=XM_023258504.1;refGene_gn=MCTP1;refGene_tx=XM_023258511.1
A1 159108229 159108292
refGene_gn=MCTP1;refGene_tx=XM_023258508.1
A1 159114786 159114906
refGene_gn=MCTP1;refGene_tx=XM_023258485.1;refGene_gn=MCTP1;refGene_tx=XM_023258486.1;refGene_gn=MCTP1;refGene_tx=XM_023258491.1;refGene_gn=MCTP1;refGene_tx=XM_023258493.1;refGene_gn=MCTP1;refGene_tx=XM_023258495.1;refGene_gn=MCTP1;refGene_tx=XM_023258501.1;refGene_gn=MCTP1;refGene_tx=XM_023258502.1;refGene_gn=MCTP1;refGene_tx=XM_023258504.1;refGene_gn=MCTP1;refGene_tx=XM_023258508.1;refGene_gn=MCTP1;refGene_tx=XM_023258511.1
A1 159116960 159117035
refGene_gn=MCTP1;refGene_tx=XM_023258485.1;refGene_gn=MCTP1;refGene_tx=XM_023258486.1;refGene_gn=MCTP1;refGene_tx=XM_023258491.1;refGene_gn=MCTP1;refGene_tx=XM_023258493.1;refGene_gn=MCTP1;refGene_tx=XM_023258495.1;refGene_gn=MCTP1;refGene_tx=XM_023258501.1;refGene_gn=MCTP1;refGene_tx=XM_023258502.1;refGene_gn=MCTP1;refGene_tx=XM_023258504.1;refGene_gn=MCTP1;refGene_tx=XM_023258508.1;refGene_gn=MCTP1;refGene_tx=XM_023258511.1

A1 159117415 159117517
refGene_gn=MCTP1;refGene_tx=XM_023258485.1;refGene_gn=MCTP1;refGene_tx=XM_023258486.1;refGene_gn=MCTP1;refGene_tx=XM_023258491.1;refGene_gn=MCTP1;refGene_tx=XM_023258493.1;refGene_gn=MCTP1;refGene_tx=XM_023258495.1;refGene_gn=MCTP1;refGene_tx=XM_023258501.1;refGene_gn=MCTP1;refGene_tx=XM_023258502.1;refGene_gn=MCTP1;refGene_tx=XM_023258504.1;refGene_gn=MCTP1;refGene_tx=XM_023258508.1;refGene_gn=MCTP1;refGene_tx=XM_023258511.1
A1 159117846 159117949
refGene_gn=MCTP1;refGene_tx=XM_023258485.1;refGene_gn=MCTP1;refGene_tx=XM_023258486.1;refGene_gn=MCTP1;refGene_tx=XM_023258491.1;refGene_gn=MCTP1;refGene_tx=XM_023258493.1;refGene_gn=MCTP1;refGene_tx=XM_023258495.1;refGene_gn=MCTP1;refGene_tx=XM_023258501.1;refGene_gn=MCTP1;refGene_tx=XM_023258502.1;refGene_gn=MCTP1;refGene_tx=XM_023258504.1;refGene_gn=MCTP1;refGene_tx=XM_023258508.1;refGene_gn=MCTP1;refGene_tx=XM_023258511.1
A1 159119722 159119825
refGene_gn=MCTP1;refGene_tx=XM_023258485.1;refGene_gn=MCTP1;refGene_tx=XM_023258486.1;refGene_gn=MCTP1;refGene_tx=XM_023258491.1;refGene_gn=MCTP1;refGene_tx=XM_023258493.1;refGene_gn=MCTP1;refGene_tx=XM_023258495.1;refGene_gn=MCTP1;refGene_tx=XM_023258501.1;refGene_gn=MCTP1;refGene_tx=XM_023258502.1;refGene_gn=MCTP1;refGene_tx=XM_023258504.1;refGene_gn=MCTP1;refGene_tx=XM_023258508.1;refGene_gn=MCTP1;refGene_tx=XM_023258511.1
A1 159137055 159137149
refGene_gn=MCTP1;refGene_tx=XM_023258485.1;refGene_gn=MCTP1;refGene_tx=XM_023258486.1;refGene_gn=MCTP1;refGene_tx=XM_023258491.1;refGene_gn=MCTP1;refGene_tx=XM_023258493.1;refGene_gn=MCTP1;refGene_tx=XM_023258495.1;refGene_gn=MCTP1;refGene_tx=XM_023258501.1;refGene_gn=MCTP1;refGene_tx=XM_023258502.1;refGene_gn=MCTP1;refGene_tx=XM_023258504.1;refGene_gn=MCTP1;refGene_tx=XM_023258508.1;refGene_gn=MCTP1;refGene_tx=XM_023258511.1
A1 159141883 159142070
refGene_gn=MCTP1;refGene_tx=XM_023258485.1;refGene_gn=MCTP1;refGene_tx=XM_023258486.1;refGene_gn=MCTP1;refGene_tx=XM_023258491.1;refGene_gn=MCTP1;refGene_tx=XM_023258493.1;refGene_gn=MCTP1;refGene_tx=XM_023258495.1;refGene_gn=MCTP1;refGene_tx=XM_023258501.1;refGene_gn=MCTP1;refGene_tx=XM_023258502.1;refGene_gn=MCTP1;refGene_tx=XM_023258504.1;refGene_gn=MCTP1;refGene_tx=XM_023258508.1;refGene_gn=MCTP1;refGene_tx=XM_023258511.1
A1 159157644 159157775
refGene_gn=MCTP1;refGene_tx=XM_023258485.1;refGene_gn=MCTP1;refGene_tx=XM_023258486.1;refGene_gn=MCTP1;refGene_tx=XM_023258491.1;refGene_gn=MCTP1;refGene_tx=XM_023258493.1;refGene_gn=MCTP1;refGene_tx=XM_023258495.1;refGene_gn=MCTP1;refGene_tx=XM_023258501.1;refGene_gn=MCTP1;refGene_tx=XM_023258502.1;refGene_gn=MCTP1;refGene_tx=XM_023258504.1;refGene_gn=MCTP1;refGene_tx=XM_023258508.1;refGene_gn=MCTP1;refGene_tx=XM_023258511.1
A1 159160609 159160780
refGene_gn=MCTP1;refGene_tx=XM_023258485.1;refGene_gn=MCTP1;refGene_tx

=XM_023258486.1;refGene_gn=MCTP1;refGene_tx=XM_023258491.1;refGene_gn=
MCTP1;refGene_tx=XM_023258493.1;refGene_gn=MCTP1;refGene_tx=XM_0232584
95.1;refGene_gn=MCTP1;refGene_tx=XM_023258501.1;refGene_gn=MCTP1;refGe
ne_tx=XM_023258502.1;refGene_gn=MCTP1;refGene_tx=XM_023258504.1;refGen
e_gn=MCTP1;refGene_tx=XM_023258508.1;refGene_gn=MCTP1;refGene_tx=XM_02
3258511.1
A1 159165985 159166063
refGene_gn=MCTP1;refGene_tx=XM_023258485.1;refGene_gn=MCTP1;refGene_tx
=XM_023258486.1;refGene_gn=MCTP1;refGene_tx=XM_023258491.1;refGene_gn=
MCTP1;refGene_tx=XM_023258502.1;refGene_gn=MCTP1;refGene_tx=XM_0232585
04.1;refGene_gn=MCTP1;refGene_tx=XM_023258508.1;refGene_gn=MCTP1;refGe
ne_tx=XM_023258511.1
A1 159171417 159171477
refGene_gn=MCTP1;refGene_tx=XM_023258485.1;refGene_gn=MCTP1;refGene_tx
=XM_023258486.1;refGene_gn=MCTP1;refGene_tx=XM_023258491.1;refGene_gn=
MCTP1;refGene_tx=XM_023258502.1;refGene_gn=MCTP1;refGene_tx=XM_0232585
04.1;refGene_gn=MCTP1;refGene_tx=XM_023258508.1;refGene_gn=MCTP1;refGe
ne_tx=XM_023258511.1
A1 159181421 159181460
refGene_gn=MCTP1;refGene_tx=XM_023258485.1;refGene_gn=MCTP1;refGene_tx
=XM_023258486.1;refGene_gn=MCTP1;refGene_tx=XM_023258493.1;refGene_gn=
MCTP1;refGene_tx=XM_023258495.1;refGene_gn=MCTP1;refGene_tx=XM_0232585
02.1;refGene_gn=MCTP1;refGene_tx=XM_023258504.1;refGene_gn=MCTP1;refGe
ne_tx=XM_023258508.1;refGene_gn=MCTP1;refGene_tx=XM_023258511.1
A1 159185456 159185504
refGene_gn=MCTP1;refGene_tx=XM_023258485.1;refGene_gn=MCTP1;refGene_tx
=XM_023258493.1;refGene_gn=MCTP1;refGene_tx=XM_023258502.1;refGene_gn=
MCTP1;refGene_tx=XM_023258504.1;refGene_gn=MCTP1;refGene_tx=XM_0232585
08.1;refGene_gn=MCTP1;refGene_tx=XM_023258511.1
A1 159190680 159190792
refGene_gn=MCTP1;refGene_tx=XM_023258485.1;refGene_gn=MCTP1;refGene_tx
=XM_023258486.1;refGene_gn=MCTP1;refGene_tx=XM_023258491.1;refGene_gn=
MCTP1;refGene_tx=XM_023258493.1;refGene_gn=MCTP1;refGene_tx=XM_0232584
95.1;refGene_gn=MCTP1;refGene_tx=XM_023258501.1;refGene_gn=MCTP1;refGe
ne_tx=XM_023258502.1;refGene_gn=MCTP1;refGene_tx=XM_023258504.1;refGen
e_gn=MCTP1;refGene_tx=XM_023258508.1;refGene_gn=MCTP1;refGene_tx=XM_02
3258511.1
A1 159194736 159194816
refGene_gn=MCTP1;refGene_tx=XM_023258485.1;refGene_gn=MCTP1;refGene_tx
=XM_023258486.1;refGene_gn=MCTP1;refGene_tx=XM_023258491.1;refGene_gn=
MCTP1;refGene_tx=XM_023258493.1;refGene_gn=MCTP1;refGene_tx=XM_0232584
95.1;refGene_gn=MCTP1;refGene_tx=XM_023258501.1;refGene_gn=MCTP1;refGe
ne_tx=XM_023258502.1;refGene_gn=MCTP1;refGene_tx=XM_023258504.1;refGen
e_gn=MCTP1;refGene_tx=XM_023258508.1;refGene_gn=MCTP1;refGene_tx=XM_02
3258511.1
A1 159204011 159204154
refGene_gn=MCTP1;refGene_tx=XM_023258485.1;refGene_gn=MCTP1;refGene_tx
=XM_023258486.1;refGene_gn=MCTP1;refGene_tx=XM_023258491.1;refGene_gn=
MCTP1;refGene_tx=XM_023258493.1;refGene_gn=MCTP1;refGene_tx=XM_0232584
95.1;refGene_gn=MCTP1;refGene_tx=XM_023258501.1;refGene_gn=MCTP1;refGe

ne_tx=XM_023258502.1;refGene_gn=MCTP1;refGene_tx=XM_023258504.1;refGene_gn=MCTP1;refGene_tx=XM_023258508.1;refGene_gn=MCTP1;refGene_tx=XM_023258511.1
A1 159245643 159245758
refGene_gn=MCTP1;refGene_tx=XM_023258485.1;refGene_gn=MCTP1;refGene_tx=XM_023258486.1;refGene_gn=MCTP1;refGene_tx=XM_023258491.1;refGene_gn=MCTP1;refGene_tx=XM_023258493.1;refGene_gn=MCTP1;refGene_tx=XM_023258495.1;refGene_gn=MCTP1;refGene_tx=XM_023258501.1;refGene_gn=MCTP1;refGene_tx=XM_023258502.1;refGene_gn=MCTP1;refGene_tx=XM_023258504.1;refGene_gn=MCTP1;refGene_tx=XM_023258508.1;refGene_gn=MCTP1;refGene_tx=XM_023258511.1
A1 159297116 159297173
refGene_gn=MCTP1;refGene_tx=XM_023258511.1
A1 159484639 159485362
refGene_gn=MCTP1;refGene_tx=XM_023258485.1;refGene_gn=MCTP1;refGene_tx=XM_023258486.1;refGene_gn=MCTP1;refGene_tx=XM_023258491.1;refGene_gn=MCTP1;refGene_tx=XM_023258493.1;refGene_gn=MCTP1;refGene_tx=XM_023258495.1;refGene_gn=MCTP1;refGene_tx=XM_023258501.1;refGene_gn=MCTP1;refGene_tx=XM_023258502.1;refGene_gn=MCTP1;refGene_tx=XM_023258504.1;refGene_gn=MCTP1;refGene_tx=XM_023258508.1
A1 159570779 159570789
refGene_gn=FAM81B;refGene_tx=XM_003981156.5;refGene_gn=FAM81B;refGene_tx=XM_019837109.2;refGene_gn=FAM81B;refGene_tx=XM_019837119.2
A1 159571421 159571552
refGene_gn=FAM81B;refGene_tx=XM_003981156.5;refGene_gn=FAM81B;refGene_tx=XM_019837109.2;refGene_gn=FAM81B;refGene_tx=XM_019837119.2
A1 159574889 159574954
refGene_gn=FAM81B;refGene_tx=XM_003981156.5;refGene_gn=FAM81B;refGene_tx=XM_011284659.3;refGene_gn=FAM81B;refGene_tx=XM_019837109.2;refGene_gn=FAM81B;refGene_tx=XM_019837119.2
A1 159588019 159588263
refGene_gn=FAM81B;refGene_tx=XM_003981156.5;refGene_gn=FAM81B;refGene_tx=XM_011284659.3;refGene_gn=FAM81B;refGene_tx=XM_019837109.2;refGene_gn=FAM81B;refGene_tx=XM_019837119.2
A1 159594178 159594297
refGene_gn=FAM81B;refGene_tx=XM_003981156.5;refGene_gn=FAM81B;refGene_tx=XM_011284659.3;refGene_gn=FAM81B;refGene_tx=XM_019837109.2;refGene_gn=FAM81B;refGene_tx=XM_019837119.2
A1 159603961 159604091
refGene_gn=FAM81B;refGene_tx=XM_003981156.5;refGene_gn=FAM81B;refGene_tx=XM_011284659.3;refGene_gn=FAM81B;refGene_tx=XM_019837109.2;refGene_gn=FAM81B;refGene_tx=XM_019837119.2
A1 159613218 159613325
refGene_gn=FAM81B;refGene_tx=XM_003981156.5;refGene_gn=FAM81B;refGene_tx=XM_011284659.3;refGene_gn=FAM81B;refGene_tx=XM_019837109.2;refGene_gn=FAM81B;refGene_tx=XM_019837119.2
A1 159621228 159621364
refGene_gn=FAM81B;refGene_tx=XM_003981156.5;refGene_gn=FAM81B;refGene_tx=XM_011284659.3;refGene_gn=FAM81B;refGene_tx=XM_019837109.2;refGene_gn=FAM81B;refGene_tx=XM_019837119.2

A1 159623911 159624107
refGene_gn=FAM81B;refGene_tx=XM_003981156.5;refGene_gn=FAM81B;refGene_tx=XM_011284659.3;refGene_gn=FAM81B;refGene_tx=XM_019837109.2;refGene_gn=FAM81B;refGene_tx=XM_019837119.2
A1 159625813 159625947
refGene_gn=FAM81B;refGene_tx=XM_003981156.5;refGene_gn=FAM81B;refGene_tx=XM_011284659.3;refGene_gn=FAM81B;refGene_tx=XM_019837109.2;refGene_gn=FAM81B;refGene_tx=XM_019837119.2
A1 159637049 159637124
refGene_gn=TTC37;refGene_tx=XM_011284663.3;refGene_gn=TTC37;refGene_tx=XM_019837129.2;refGene_gn=TTC37;refGene_tx=XM_019837133.2;refGene_gn=TTC37;refGene_tx=XM_019837140.2;refGene_gn=TTC37;refGene_tx=XM_023258551.1
A1 159639319 159639446
refGene_gn=TTC37;refGene_tx=XM_011284663.3;refGene_gn=TTC37;refGene_tx=XM_019837129.2;refGene_gn=TTC37;refGene_tx=XM_019837133.2;refGene_gn=TTC37;refGene_tx=XM_019837140.2;refGene_gn=TTC37;refGene_tx=XM_023258551.1
A1 159641135 159641266
refGene_gn=TTC37;refGene_tx=XM_011284663.3;refGene_gn=TTC37;refGene_tx=XM_019837129.2;refGene_gn=TTC37;refGene_tx=XM_019837133.2;refGene_gn=TTC37;refGene_tx=XM_019837140.2;refGene_gn=TTC37;refGene_tx=XM_023258551.1
A1 159648572 159648725
refGene_gn=TTC37;refGene_tx=XM_011284663.3;refGene_gn=TTC37;refGene_tx=XM_019837129.2;refGene_gn=TTC37;refGene_tx=XM_019837133.2;refGene_gn=TTC37;refGene_tx=XM_019837140.2;refGene_gn=TTC37;refGene_tx=XM_023258551.1
A1 159653818 159653977
refGene_gn=TTC37;refGene_tx=XM_011284663.3;refGene_gn=TTC37;refGene_tx=XM_019837129.2;refGene_gn=TTC37;refGene_tx=XM_019837133.2;refGene_gn=TTC37;refGene_tx=XM_019837140.2;refGene_gn=TTC37;refGene_tx=XM_023258551.1
A1 159655722 159655844
refGene_gn=TTC37;refGene_tx=XM_011284663.3;refGene_gn=TTC37;refGene_tx=XM_019837129.2;refGene_gn=TTC37;refGene_tx=XM_019837133.2;refGene_gn=TTC37;refGene_tx=XM_019837140.2;refGene_gn=TTC37;refGene_tx=XM_023258551.1
A1 159659087 159659216
refGene_gn=TTC37;refGene_tx=XM_011284663.3;refGene_gn=TTC37;refGene_tx=XM_019837129.2;refGene_gn=TTC37;refGene_tx=XM_019837133.2;refGene_gn=TTC37;refGene_tx=XM_019837140.2;refGene_gn=TTC37;refGene_tx=XM_023258551.1
A1 159662892 159663004
refGene_gn=TTC37;refGene_tx=XM_011284663.3;refGene_gn=TTC37;refGene_tx=XM_019837129.2;refGene_gn=TTC37;refGene_tx=XM_019837133.2;refGene_gn=TTC37;refGene_tx=XM_019837140.2;refGene_gn=TTC37;refGene_tx=XM_023258551.1
A1 159663190 159663244
refGene_gn=TTC37;refGene_tx=XM_011284663.3;refGene_gn=TTC37;refGene_tx

=XM_019837129.2;refGene_gn=TTC37;refGene_tx=XM_019837133.2;refGene_gn=TTC37;refGene_tx=XM_019837140.2;refGene_gn=TTC37;refGene_tx=XM_023258551.1
A1 159666033 159666106
refGene_gn=TTC37;refGene_tx=XM_011284663.3;refGene_gn=TTC37;refGene_tx=XM_019837129.2;refGene_gn=TTC37;refGene_tx=XM_019837133.2;refGene_gn=TTC37;refGene_tx=XM_019837140.2;refGene_gn=TTC37;refGene_tx=XM_023258551.1
A1 159666919 159667141
refGene_gn=TTC37;refGene_tx=XM_011284663.3;refGene_gn=TTC37;refGene_tx=XM_019837129.2;refGene_gn=TTC37;refGene_tx=XM_019837133.2;refGene_gn=TTC37;refGene_tx=XM_019837140.2;refGene_gn=TTC37;refGene_tx=XM_023258551.1
A1 159668396 159668520
refGene_gn=TTC37;refGene_tx=XM_011284663.3;refGene_gn=TTC37;refGene_tx=XM_019837129.2;refGene_gn=TTC37;refGene_tx=XM_019837133.2;refGene_gn=TTC37;refGene_tx=XM_019837140.2;refGene_gn=TTC37;refGene_tx=XM_023258551.1
A1 159669318 159669443
refGene_gn=TTC37;refGene_tx=XM_011284663.3;refGene_gn=TTC37;refGene_tx=XM_019837129.2;refGene_gn=TTC37;refGene_tx=XM_019837133.2;refGene_gn=TTC37;refGene_tx=XM_019837140.2;refGene_gn=TTC37;refGene_tx=XM_023258551.1
A1 159672010 159672088
refGene_gn=TTC37;refGene_tx=XM_011284663.3;refGene_gn=TTC37;refGene_tx=XM_019837129.2;refGene_gn=TTC37;refGene_tx=XM_019837133.2;refGene_gn=TTC37;refGene_tx=XM_019837140.2;refGene_gn=TTC37;refGene_tx=XM_023258551.1
A1 159674512 159674606
refGene_gn=TTC37;refGene_tx=XM_011284663.3;refGene_gn=TTC37;refGene_tx=XM_019837129.2;refGene_gn=TTC37;refGene_tx=XM_019837133.2;refGene_gn=TTC37;refGene_tx=XM_019837140.2;refGene_gn=TTC37;refGene_tx=XM_023258551.1
A1 159677199 159677341
refGene_gn=TTC37;refGene_tx=XM_011284663.3;refGene_gn=TTC37;refGene_tx=XM_019837129.2;refGene_gn=TTC37;refGene_tx=XM_019837133.2;refGene_gn=TTC37;refGene_tx=XM_019837140.2;refGene_gn=TTC37;refGene_tx=XM_023258551.1
A1 159677951 159678026
refGene_gn=TTC37;refGene_tx=XM_011284663.3;refGene_gn=TTC37;refGene_tx=XM_019837129.2;refGene_gn=TTC37;refGene_tx=XM_019837133.2;refGene_gn=TTC37;refGene_tx=XM_019837140.2;refGene_gn=TTC37;refGene_tx=XM_023258551.1
A1 159679919 159679988
refGene_gn=TTC37;refGene_tx=XM_011284663.3;refGene_gn=TTC37;refGene_tx=XM_019837129.2;refGene_gn=TTC37;refGene_tx=XM_019837133.2;refGene_gn=TTC37;refGene_tx=XM_019837140.2;refGene_gn=TTC37;refGene_tx=XM_023258551.1
A1 159681453 159681510
refGene_gn=TTC37;refGene_tx=XM_011284663.3;refGene_gn=TTC37;refGene_tx

=XM_019837129.2;refGene_gn=TTC37;refGene_tx=XM_019837133.2;refGene_gn=TTC37;refGene_tx=XM_019837140.2;refGene_gn=TTC37;refGene_tx=XM_023258551.1
A1 159681620 159681682
refGene_gn=TTC37;refGene_tx=XM_011284663.3;refGene_gn=TTC37;refGene_tx=XM_019837129.2;refGene_gn=TTC37;refGene_tx=XM_019837133.2;refGene_gn=TTC37;refGene_tx=XM_019837140.2;refGene_gn=TTC37;refGene_tx=XM_023258551.1
A1 159681770 159681849
refGene_gn=TTC37;refGene_tx=XM_011284663.3;refGene_gn=TTC37;refGene_tx=XM_019837129.2;refGene_gn=TTC37;refGene_tx=XM_019837133.2;refGene_gn=TTC37;refGene_tx=XM_019837140.2;refGene_gn=TTC37;refGene_tx=XM_023258551.1
A1 159682008 159682153
refGene_gn=TTC37;refGene_tx=XM_011284663.3;refGene_gn=TTC37;refGene_tx=XM_019837129.2;refGene_gn=TTC37;refGene_tx=XM_019837133.2;refGene_gn=TTC37;refGene_tx=XM_019837140.2;refGene_gn=TTC37;refGene_tx=XM_023258551.1
A1 159682241 159682418
refGene_gn=TTC37;refGene_tx=XM_011284663.3;refGene_gn=TTC37;refGene_tx=XM_019837129.2;refGene_gn=TTC37;refGene_tx=XM_019837133.2;refGene_gn=TTC37;refGene_tx=XM_019837140.2;refGene_gn=TTC37;refGene_tx=XM_023258551.1
A1 159684859 159684962
refGene_gn=TTC37;refGene_tx=XM_011284663.3;refGene_gn=TTC37;refGene_tx=XM_019837129.2;refGene_gn=TTC37;refGene_tx=XM_019837133.2;refGene_gn=TTC37;refGene_tx=XM_019837140.2;refGene_gn=TTC37;refGene_tx=XM_023258551.1
A1 159686320 159686574
refGene_gn=TTC37;refGene_tx=XM_011284663.3;refGene_gn=TTC37;refGene_tx=XM_019837129.2;refGene_gn=TTC37;refGene_tx=XM_019837133.2;refGene_gn=TTC37;refGene_tx=XM_019837140.2;refGene_gn=TTC37;refGene_tx=XM_023258551.1
A1 159688073 159688198
refGene_gn=TTC37;refGene_tx=XM_011284663.3;refGene_gn=TTC37;refGene_tx=XM_019837129.2;refGene_gn=TTC37;refGene_tx=XM_019837133.2;refGene_gn=TTC37;refGene_tx=XM_019837140.2;refGene_gn=TTC37;refGene_tx=XM_023258551.1
A1 159688546 159688726
refGene_gn=TTC37;refGene_tx=XM_011284663.3;refGene_gn=TTC37;refGene_tx=XM_019837129.2;refGene_gn=TTC37;refGene_tx=XM_019837133.2;refGene_gn=TTC37;refGene_tx=XM_019837140.2;refGene_gn=TTC37;refGene_tx=XM_023258551.1
A1 159689624 159689751
refGene_gn=TTC37;refGene_tx=XM_011284663.3;refGene_gn=TTC37;refGene_tx=XM_019837129.2;refGene_gn=TTC37;refGene_tx=XM_019837133.2;refGene_gn=TTC37;refGene_tx=XM_019837140.2;refGene_gn=TTC37;refGene_tx=XM_023258551.1
A1 159690206 159690310
refGene_gn=TTC37;refGene_tx=XM_011284663.3;refGene_gn=TTC37;refGene_tx

=XM_019837129.2;refGene_gn=TTC37;refGene_tx=XM_019837133.2;refGene_gn=TTC37;refGene_tx=XM_019837140.2;refGene_gn=TTC37;refGene_tx=XM_023258551.1
A1 159690398 159690485
refGene_gn=TTC37;refGene_tx=XM_011284663.3;refGene_gn=TTC37;refGene_tx=XM_019837129.2;refGene_gn=TTC37;refGene_tx=XM_019837133.2;refGene_gn=TTC37;refGene_tx=XM_019837140.2;refGene_gn=TTC37;refGene_tx=XM_023258551.1
A1 159692623 159692763
refGene_gn=TTC37;refGene_tx=XM_011284663.3;refGene_gn=TTC37;refGene_tx=XM_019837129.2;refGene_gn=TTC37;refGene_tx=XM_019837133.2;refGene_gn=TTC37;refGene_tx=XM_019837140.2;refGene_gn=TTC37;refGene_tx=XM_023258551.1
A1 159693583 159693685
refGene_gn=TTC37;refGene_tx=XM_011284663.3;refGene_gn=TTC37;refGene_tx=XM_019837129.2;refGene_gn=TTC37;refGene_tx=XM_019837133.2;refGene_gn=TTC37;refGene_tx=XM_019837140.2;refGene_gn=TTC37;refGene_tx=XM_023258551.1
A1 159695529 159695670
refGene_gn=TTC37;refGene_tx=XM_011284663.3;refGene_gn=TTC37;refGene_tx=XM_019837129.2;refGene_gn=TTC37;refGene_tx=XM_019837133.2;refGene_gn=TTC37;refGene_tx=XM_019837140.2;refGene_gn=TTC37;refGene_tx=XM_023258551.1
A1 159703154 159703263
refGene_gn=TTC37;refGene_tx=XM_011284663.3;refGene_gn=TTC37;refGene_tx=XM_019837129.2;refGene_gn=TTC37;refGene_tx=XM_019837133.2;refGene_gn=TTC37;refGene_tx=XM_019837140.2;refGene_gn=TTC37;refGene_tx=XM_023258551.1
A1 159703371 159703470
refGene_gn=TTC37;refGene_tx=XM_011284663.3;refGene_gn=TTC37;refGene_tx=XM_019837129.2;refGene_gn=TTC37;refGene_tx=XM_019837133.2;refGene_gn=TTC37;refGene_tx=XM_019837140.2
A1 159706218 159706359
refGene_gn=TTC37;refGene_tx=XM_011284663.3;refGene_gn=TTC37;refGene_tx=XM_019837129.2;refGene_gn=TTC37;refGene_tx=XM_019837133.2;refGene_gn=TTC37;refGene_tx=XM_019837140.2
A1 159707105 159707181
refGene_gn=TTC37;refGene_tx=XM_011284663.3;refGene_gn=TTC37;refGene_tx=XM_019837129.2;refGene_gn=TTC37;refGene_tx=XM_019837133.2;refGene_gn=TTC37;refGene_tx=XM_019837140.2
A1 159707630 159707722
refGene_gn=TTC37;refGene_tx=XM_011284663.3;refGene_gn=TTC37;refGene_tx=XM_019837129.2;refGene_gn=TTC37;refGene_tx=XM_019837133.2;refGene_gn=TTC37;refGene_tx=XM_019837140.2
A1 159708786 159708930
refGene_gn=TTC37;refGene_tx=XM_011284663.3;refGene_gn=TTC37;refGene_tx=XM_019837129.2;refGene_gn=TTC37;refGene_tx=XM_019837133.2;refGene_gn=TTC37;refGene_tx=XM_019837140.2
A1 159710854 159710944
refGene_gn=TTC37;refGene_tx=XM_011284663.3;refGene_gn=TTC37;refGene_tx

=XM_019837129.2;refGene_gn=TTC37;refGene_tx=XM_019837133.2;refGene_gn=TTC37;refGene_tx=XM_019837140.2
A1 159721444 159721582
refGene_gn=ARSK;refGene_tx=XM_023258567.1
A1 159727359 159727489
refGene_gn=ARSK;refGene_tx=XM_023258567.1;refGene_gn=ARSK;refGene_tx=XM_023258569.1
A1 159733270 159733430
refGene_gn=ARSK;refGene_tx=XM_023258567.1;refGene_gn=ARSK;refGene_tx=XM_023258569.1;refGene_gn=ARSK;refGene_tx=XM_023258570.1
A1 159749539 159749822
refGene_gn=ARSK;refGene_tx=XM_023258567.1;refGene_gn=ARSK;refGene_tx=XM_023258569.1;refGene_gn=ARSK;refGene_tx=XM_023258570.1
A1 159752515 159752687
refGene_gn=ARSK;refGene_tx=XM_023258567.1;refGene_gn=ARSK;refGene_tx=XM_023258569.1;refGene_gn=ARSK;refGene_tx=XM_023258570.1
A1 159756163 159756388
refGene_gn=ARSK;refGene_tx=XM_023258567.1;refGene_gn=ARSK;refGene_tx=XM_023258569.1;refGene_gn=ARSK;refGene_tx=XM_023258570.1
A1 159764230 159764455
refGene_gn=ARSK;refGene_tx=XM_023258567.1;refGene_gn=ARSK;refGene_tx=XM_023258569.1;refGene_gn=ARSK;refGene_tx=XM_023258570.1
A1 159766513 159766815
refGene_gn=ARSK;refGene_tx=XM_023258567.1;refGene_gn=ARSK;refGene_tx=XM_023258569.1;refGene_gn=ARSK;refGene_tx=XM_023258570.1
A1 159776917 159778222
refGene_gn=GPR150;refGene_tx=XM_023258575.1
A1 159809449 159809624
refGene_gn=RFESD;refGene_tx=XM_006927855.4;refGene_gn=RFESD;refGene_tx=XM_019837197.2;refGene_gn=RFESD;refGene_tx=XM_023258582.1
A1 159809768 159809803
refGene_gn=RFESD;refGene_tx=XM_006927855.4;refGene_gn=RFESD;refGene_tx=XM_019837197.2;refGene_gn=RFESD;refGene_tx=XM_023258582.1
A1 159811430 159811694
refGene_gn=RFESD;refGene_tx=XM_006927855.4;refGene_gn=RFESD;refGene_tx=XM_019837197.2;refGene_gn=RFESD;refGene_tx=XM_023258582.1
A1 159814624 159814912
refGene_gn=SPATA9;refGene_tx=XM_003981157.4;refGene_gn=SPATA9;refGene_tx=XM_019837176.2
A1 159819711 159819807
refGene_gn=SPATA9;refGene_tx=XM_003981157.4;refGene_gn=SPATA9;refGene_tx=XM_019837176.2
A1 159829053 159829281
refGene_gn=SPATA9;refGene_tx=XM_003981157.4;refGene_gn=SPATA9;refGene_tx=XM_019837176.2
A1 159837562 159837651
refGene_gn=SPATA9;refGene_tx=XM_003981157.4;refGene_gn=SPATA9;refGene_tx=XM_019837176.2
A1 159837841 159837902
refGene_gn=SPATA9;refGene_tx=XM_003981157.4;refGene_gn=SPATA9;refGene_

tx=XM_019837176.2
A1 159875204 159875956
refGene_gn=RHOBTB3;refGene_tx=XM_011284680.3;refGene_gn=RHOBTB3;refGene_tx=XM_023258589.1;refGene_gn=RHOBTB3;refGene_tx=XM_023258594.1
A1 159876136 159876362
refGene_gn=RHOBTB3;refGene_tx=XM_011284680.3;refGene_gn=RHOBTB3;refGene_tx=XM_023258589.1;refGene_gn=RHOBTB3;refGene_tx=XM_023258594.1
A1 159881089 159881276
refGene_gn=RHOBTB3;refGene_tx=XM_011284680.3;refGene_gn=RHOBTB3;refGene_tx=XM_023258589.1;refGene_gn=RHOBTB3;refGene_tx=XM_023258594.1
A1 159890159 159890314
refGene_gn=RHOBTB3;refGene_tx=XM_011284680.3;refGene_gn=RHOBTB3;refGene_tx=XM_023258589.1;refGene_gn=RHOBTB3;refGene_tx=XM_023258594.1
A1 159893591 159893703
refGene_gn=RHOBTB3;refGene_tx=XM_011284680.3;refGene_gn=RHOBTB3;refGene_tx=XM_023258589.1;refGene_gn=RHOBTB3;refGene_tx=XM_023258594.1
A1 159896847 159897213
refGene_gn=RHOBTB3;refGene_tx=XM_011284680.3;refGene_gn=RHOBTB3;refGene_tx=XM_023258589.1;refGene_gn=RHOBTB3;refGene_tx=XM_023258594.1
A1 159904355 159904468
refGene_gn=RHOBTB3;refGene_tx=XM_011284680.3;refGene_gn=RHOBTB3;refGene_tx=XM_023258589.1;refGene_gn=RHOBTB3;refGene_tx=XM_023258594.1
A1 159907471 159907592
refGene_gn=RHOBTB3;refGene_tx=XM_011284680.3;refGene_gn=RHOBTB3;refGene_tx=XM_023258589.1;refGene_gn=RHOBTB3;refGene_tx=XM_023258594.1
A1 159912259 159912433
refGene_gn=RHOBTB3;refGene_tx=XM_011284680.3;refGene_gn=RHOBTB3;refGene_tx=XM_023258589.1;refGene_gn=RHOBTB3;refGene_tx=XM_023258594.1
A1 159916270 159916437
refGene_gn=RHOBTB3;refGene_tx=XM_011284680.3;refGene_gn=RHOBTB3;refGene_tx=XM_023258589.1
A1 159921455 159921552
refGene_gn=RHOBTB3;refGene_tx=XM_011284680.3;refGene_gn=RHOBTB3;refGene_tx=XM_023258589.1
A1 159927005 159927121
refGene_gn=RHOBTB3;refGene_tx=XM_011284680.3;refGene_gn=RHOBTB3;refGene_tx=XM_023258589.1
A1 159957267 159957381
refGene_gn=GLRX;refGene_tx=XM_006927856.4
A1 159962023 159962230
refGene_gn=GLRX;refGene_tx=XM_006927856.4
A1 160026502 160026619
refGene_gn=ELL2;refGene_tx=XM_004001508.5
A1 160026709 160026754
refGene_gn=ELL2;refGene_tx=XM_004001508.5
A1 160028820 160028992
refGene_gn=ELL2;refGene_tx=XM_004001508.5
A1 160032413 160032480
refGene_gn=ELL2;refGene_tx=XM_004001508.5
A1 160035291 160035865

refGene_gn=ELL2;refGene_tx=XM_004001508.5
A1 160037464 160037552
refGene_gn=ELL2;refGene_tx=XM_004001508.5
A1 160037708 160037833
refGene_gn=ELL2;refGene_tx=XM_004001508.5
A1 160041190 160041450
refGene_gn=ELL2;refGene_tx=XM_004001508.5
A1 160048016 160048180
refGene_gn=ELL2;refGene_tx=XM_004001508.5
A1 160052957 160053079
refGene_gn=ELL2;refGene_tx=XM_004001508.5
A1 160078765 160078813
refGene_gn=ELL2;refGene_tx=XM_004001508.5
A1 160096390 160096537
refGene_gn=ELL2;refGene_tx=XM_004001508.5
A1 160494155 160494533
refGene_gn=PCSK1;refGene_tx=XM_003981159.5
A1 160496792 160496954
refGene_gn=PCSK1;refGene_tx=XM_003981159.5
A1 160498492 160498626
refGene_gn=PCSK1;refGene_tx=XM_003981159.5
A1 160499952 160500110
refGene_gn=PCSK1;refGene_tx=XM_003981159.5
A1 160502819 160503053
refGene_gn=PCSK1;refGene_tx=XM_003981159.5
A1 160512702 160512803
refGene_gn=PCSK1;refGene_tx=XM_003981159.5
A1 160515299 160515512
refGene_gn=PCSK1;refGene_tx=XM_003981159.5
A1 160516659 160516832
refGene_gn=PCSK1;refGene_tx=XM_003981159.5
A1 160519764 160519853
refGene_gn=PCSK1;refGene_tx=XM_003981159.5
A1 160525295 160525372
refGene_gn=PCSK1;refGene_tx=XM_003981159.5
A1 160526665 160526812
refGene_gn=PCSK1;refGene_tx=XM_003981159.5
A1 160528764 160528875
refGene_gn=PCSK1;refGene_tx=XM_003981159.5
A1 160532219 160532324
refGene_gn=PCSK1;refGene_tx=XM_003981159.5
A1 160535189 160535369
refGene_gn=PCSK1;refGene_tx=XM_003981159.5
A1 160739032 160739107
refGene_gn=CAST;refGene_tx=XM_011284752.3;refGene_gn=CAST;refGene_tx=XM_023258666.1;refGene_gn=CAST;refGene_tx=XM_023258673.1;refGene_gn=CAST;refGene_tx=XM_023258676.1;refGene_gn=CAST;refGene_tx=XM_023258679.1;refGene_gn=CAST;refGene_tx=XM_023258685.1;refGene_gn=CAST;refGene_tx=XM_023258688.1
A1 160739750 160739780

refGene_gn=CAST;refGene_tx=XM_011284727.2
A1 160752056 160752119
refGene_gn=CAST;refGene_tx=XM_011284727.2;refGene_gn=CAST;refGene_tx=XM_011284752.3;refGene_gn=CAST;refGene_tx=XM_023258666.1;refGene_gn=CAST;refGene_tx=XM_023258673.1;refGene_gn=CAST;refGene_tx=XM_023258676.1;refGene_gn=CAST;refGene_tx=XM_023258679.1;refGene_gn=CAST;refGene_tx=XM_023258685.1;refGene_gn=CAST;refGene_tx=XM_023258688.1
A1 160771874 160771946
refGene_gn=CAST;refGene_tx=XM_011284727.2;refGene_gn=CAST;refGene_tx=XM_011284752.3;refGene_gn=CAST;refGene_tx=XM_023258666.1;refGene_gn=CAST;refGene_tx=XM_023258673.1;refGene_gn=CAST;refGene_tx=XM_023258676.1;refGene_gn=CAST;refGene_tx=XM_023258679.1;refGene_gn=CAST;refGene_tx=XM_023258685.1;refGene_gn=CAST;refGene_tx=XM_023258688.1
A1 160797053 160797113
refGene_gn=CAST;refGene_tx=XM_011284727.2;refGene_gn=CAST;refGene_tx=XM_011284750.2;refGene_gn=CAST;refGene_tx=XM_011284751.2;refGene_gn=CAST;refGene_tx=XM_011284752.3;refGene_gn=CAST;refGene_tx=XM_023258666.1;refGene_gn=CAST;refGene_tx=XM_023258673.1;refGene_gn=CAST;refGene_tx=XM_023258676.1;refGene_gn=CAST;refGene_tx=XM_023258679.1;refGene_gn=CAST;refGene_tx=XM_023258685.1;refGene_gn=CAST;refGene_tx=XM_023258688.1
A1 160801164 160801230
refGene_gn=CAST;refGene_tx=XM_011284727.2;refGene_gn=CAST;refGene_tx=XM_011284750.2;refGene_gn=CAST;refGene_tx=XM_011284752.3;refGene_gn=CAST;refGene_tx=XM_023258666.1;refGene_gn=CAST;refGene_tx=XM_023258673.1;refGene_gn=CAST;refGene_tx=XM_023258676.1
A1 160801840 160801882
refGene_gn=CAST;refGene_tx=XM_011284727.2;refGene_gn=CAST;refGene_tx=XM_011284750.2;refGene_gn=CAST;refGene_tx=XM_011284751.2;refGene_gn=CAST;refGene_tx=XM_011284752.3;refGene_gn=CAST;refGene_tx=XM_023258666.1;refGene_gn=CAST;refGene_tx=XM_023258673.1;refGene_gn=CAST;refGene_tx=XM_023258676.1;refGene_gn=CAST;refGene_tx=XM_023258679.1;refGene_gn=CAST;refGene_tx=XM_023258685.1;refGene_gn=CAST;refGene_tx=XM_023258688.1
A1 160803499 160803556
refGene_gn=CAST;refGene_tx=XM_011284727.2;refGene_gn=CAST;refGene_tx=XM_011284750.2;refGene_gn=CAST;refGene_tx=XM_011284751.2;refGene_gn=CAST;refGene_tx=XM_011284752.3;refGene_gn=CAST;refGene_tx=XM_023258666.1;refGene_gn=CAST;refGene_tx=XM_023258673.1;refGene_gn=CAST;refGene_tx=XM_023258676.1;refGene_gn=CAST;refGene_tx=XM_023258679.1;refGene_gn=CAST;refGene_tx=XM_023258685.1;refGene_gn=CAST;refGene_tx=XM_023258688.1
A1 160804251 160804365
refGene_gn=CAST;refGene_tx=XM_011284727.2;refGene_gn=CAST;refGene_tx=XM_011284750.2;refGene_gn=CAST;refGene_tx=XM_011284751.2;refGene_gn=CAST;refGene_tx=XM_011284752.3;refGene_gn=CAST;refGene_tx=XM_023258666.1;refGene_gn=CAST;refGene_tx=XM_023258673.1;refGene_gn=CAST;refGene_tx=XM_023258676.1;refGene_gn=CAST;refGene_tx=XM_023258679.1;refGene_gn=CAST;refGene_tx=XM_023258685.1;refGene_gn=CAST;refGene_tx=XM_023258688.1
A1 160805391 160805472
refGene_gn=CAST;refGene_tx=XM_011284727.2;refGene_gn=CAST;refGene_tx=XM_011284750.2;refGene_gn=CAST;refGene_tx=XM_011284751.2;refGene_gn=CAST;refGene_tx=XM_011284752.3;refGene_gn=CAST;refGene_tx=XM_023258666.1;

refGene_gn=CAST;refGene_tx=XM_023258673.1;refGene_gn=CAST;refGene_tx=XM_023258676.1;refGene_gn=CAST;refGene_tx=XM_023258679.1;refGene_gn=CAST;refGene_tx=XM_023258685.1;refGene_gn=CAST;refGene_tx=XM_023258688.1
A1 160809547 160809616
refGene_gn=CAST;refGene_tx=XM_011284727.2;refGene_gn=CAST;refGene_tx=XM_011284750.2;refGene_gn=CAST;refGene_tx=XM_011284751.2;refGene_gn=CAST;refGene_tx=XM_011284752.3;refGene_gn=CAST;refGene_tx=XM_023258666.1;refGene_gn=CAST;refGene_tx=XM_023258673.1;refGene_gn=CAST;refGene_tx=XM_023258676.1;refGene_gn=CAST;refGene_tx=XM_023258679.1;refGene_gn=CAST;refGene_tx=XM_023258685.1;refGene_gn=CAST;refGene_tx=XM_023258688.1
A1 160811007 160811106
refGene_gn=CAST;refGene_tx=XM_011284727.2;refGene_gn=CAST;refGene_tx=XM_011284750.2;refGene_gn=CAST;refGene_tx=XM_011284751.2;refGene_gn=CAST;refGene_tx=XM_011284752.3;refGene_gn=CAST;refGene_tx=XM_023258666.1;refGene_gn=CAST;refGene_tx=XM_023258673.1;refGene_gn=CAST;refGene_tx=XM_023258676.1;refGene_gn=CAST;refGene_tx=XM_023258679.1;refGene_gn=CAST;refGene_tx=XM_023258685.1;refGene_gn=CAST;refGene_tx=XM_023258688.1
A1 160812731 160812812
refGene_gn=CAST;refGene_tx=XM_011284727.2;refGene_gn=CAST;refGene_tx=XM_011284750.2;refGene_gn=CAST;refGene_tx=XM_011284751.2;refGene_gn=CAST;refGene_tx=XM_011284752.3;refGene_gn=CAST;refGene_tx=XM_023258666.1;refGene_gn=CAST;refGene_tx=XM_023258673.1;refGene_gn=CAST;refGene_tx=XM_023258676.1;refGene_gn=CAST;refGene_tx=XM_023258679.1;refGene_gn=CAST;refGene_tx=XM_023258685.1;refGene_gn=CAST;refGene_tx=XM_023258688.1
A1 160813641 160813680
refGene_gn=CAST;refGene_tx=XM_011284727.2;refGene_gn=CAST;refGene_tx=XM_011284750.2;refGene_gn=CAST;refGene_tx=XM_011284751.2;refGene_gn=CAST;refGene_tx=XM_011284752.3;refGene_gn=CAST;refGene_tx=XM_023258666.1;refGene_gn=CAST;refGene_tx=XM_023258673.1;refGene_gn=CAST;refGene_tx=XM_023258676.1;refGene_gn=CAST;refGene_tx=XM_023258679.1;refGene_gn=CAST;refGene_tx=XM_023258685.1;refGene_gn=CAST;refGene_tx=XM_023258688.1
A1 160814292 160814385
refGene_gn=CAST;refGene_tx=XM_011284727.2;refGene_gn=CAST;refGene_tx=XM_011284750.2;refGene_gn=CAST;refGene_tx=XM_011284751.2;refGene_gn=CAST;refGene_tx=XM_011284752.3;refGene_gn=CAST;refGene_tx=XM_023258666.1;refGene_gn=CAST;refGene_tx=XM_023258673.1;refGene_gn=CAST;refGene_tx=XM_023258676.1;refGene_gn=CAST;refGene_tx=XM_023258679.1;refGene_gn=CAST;refGene_tx=XM_023258685.1;refGene_gn=CAST;refGene_tx=XM_023258688.1
A1 160814520 160814610
refGene_gn=CAST;refGene_tx=XM_011284727.2;refGene_gn=CAST;refGene_tx=XM_011284750.2;refGene_gn=CAST;refGene_tx=XM_011284751.2;refGene_gn=CAST;refGene_tx=XM_011284752.3;refGene_gn=CAST;refGene_tx=XM_023258666.1;refGene_gn=CAST;refGene_tx=XM_023258673.1;refGene_gn=CAST;refGene_tx=XM_023258676.1;refGene_gn=CAST;refGene_tx=XM_023258679.1;refGene_gn=CAST;refGene_tx=XM_023258685.1;refGene_gn=CAST;refGene_tx=XM_023258688.1
A1 160815924 160816029
refGene_gn=CAST;refGene_tx=XM_011284727.2;refGene_gn=CAST;refGene_tx=XM_011284750.2;refGene_gn=CAST;refGene_tx=XM_011284751.2;refGene_gn=CAST;refGene_tx=XM_011284752.3;refGene_gn=CAST;refGene_tx=XM_023258666.1;refGene_gn=CAST;refGene_tx=XM_023258673.1;refGene_gn=CAST;refGene_tx=X

M_023258676.1;refGene_gn=CAST;refGene_tx=XM_023258679.1;refGene_gn=CAS
T;refGene_tx=XM_023258685.1;refGene_gn=CAST;refGene_tx=XM_023258688.1
A1 160820134 160820218
refGene_gn=CAST;refGene_tx=XM_011284727.2;refGene_gn=CAST;refGene_tx=X
M_011284750.2;refGene_gn=CAST;refGene_tx=XM_011284751.2;refGene_gn=CAS
T;refGene_tx=XM_011284752.3;refGene_gn=CAST;refGene_tx=XM_023258666.1;
refGene_gn=CAST;refGene_tx=XM_023258673.1;refGene_gn=CAST;refGene_tx=X
M_023258676.1;refGene_gn=CAST;refGene_tx=XM_023258679.1;refGene_gn=CAS
T;refGene_tx=XM_023258685.1;refGene_gn=CAST;refGene_tx=XM_023258688.1
A1 160821086 160821134
refGene_gn=CAST;refGene_tx=XM_011284727.2;refGene_gn=CAST;refGene_tx=X
M_011284750.2;refGene_gn=CAST;refGene_tx=XM_011284751.2;refGene_gn=CAS
T;refGene_tx=XM_011284752.3;refGene_gn=CAST;refGene_tx=XM_023258666.1;
refGene_gn=CAST;refGene_tx=XM_023258673.1;refGene_gn=CAST;refGene_tx=X
M_023258676.1;refGene_gn=CAST;refGene_tx=XM_023258679.1;refGene_gn=CAS
T;refGene_tx=XM_023258685.1;refGene_gn=CAST;refGene_tx=XM_023258688.1
A1 160822162 160822258
refGene_gn=CAST;refGene_tx=XM_011284727.2;refGene_gn=CAST;refGene_tx=X
M_011284750.2;refGene_gn=CAST;refGene_tx=XM_011284751.2;refGene_gn=CAS
T;refGene_tx=XM_011284752.3;refGene_gn=CAST;refGene_tx=XM_023258666.1;
refGene_gn=CAST;refGene_tx=XM_023258673.1;refGene_gn=CAST;refGene_tx=X
M_023258676.1;refGene_gn=CAST;refGene_tx=XM_023258679.1;refGene_gn=CAS
T;refGene_tx=XM_023258685.1;refGene_gn=CAST;refGene_tx=XM_023258688.1
A1 160823899 160823995
refGene_gn=CAST;refGene_tx=XM_011284727.2;refGene_gn=CAST;refGene_tx=X
M_011284750.2;refGene_gn=CAST;refGene_tx=XM_011284751.2;refGene_gn=CAS
T;refGene_tx=XM_011284752.3;refGene_gn=CAST;refGene_tx=XM_023258666.1;
refGene_gn=CAST;refGene_tx=XM_023258673.1;refGene_gn=CAST;refGene_tx=X
M_023258676.1;refGene_gn=CAST;refGene_tx=XM_023258679.1;refGene_gn=CAS
T;refGene_tx=XM_023258685.1;refGene_gn=CAST;refGene_tx=XM_023258688.1
A1 160827738 160827840
refGene_gn=CAST;refGene_tx=XM_011284727.2;refGene_gn=CAST;refGene_tx=X
M_011284750.2;refGene_gn=CAST;refGene_tx=XM_011284751.2;refGene_gn=CAS
T;refGene_tx=XM_011284752.3;refGene_gn=CAST;refGene_tx=XM_023258666.1;
refGene_gn=CAST;refGene_tx=XM_023258673.1;refGene_gn=CAST;refGene_tx=X
M_023258676.1;refGene_gn=CAST;refGene_tx=XM_023258679.1;refGene_gn=CAS
T;refGene_tx=XM_023258685.1;refGene_gn=CAST;refGene_tx=XM_023258688.1
A1 160828346 160828430
refGene_gn=CAST;refGene_tx=XM_011284727.2;refGene_gn=CAST;refGene_tx=X
M_011284750.2;refGene_gn=CAST;refGene_tx=XM_011284751.2;refGene_gn=CAS
T;refGene_tx=XM_011284752.3;refGene_gn=CAST;refGene_tx=XM_023258666.1;
refGene_gn=CAST;refGene_tx=XM_023258673.1;refGene_gn=CAST;refGene_tx=X
M_023258676.1;refGene_gn=CAST;refGene_tx=XM_023258679.1;refGene_gn=CAS
T;refGene_tx=XM_023258685.1;refGene_gn=CAST;refGene_tx=XM_023258688.1
A1 160830057 160830108
refGene_gn=CAST;refGene_tx=XM_011284727.2;refGene_gn=CAST;refGene_tx=X
M_011284750.2;refGene_gn=CAST;refGene_tx=XM_011284751.2;refGene_gn=CAS
T;refGene_tx=XM_011284752.3;refGene_gn=CAST;refGene_tx=XM_023258666.1;
refGene_gn=CAST;refGene_tx=XM_023258673.1;refGene_gn=CAST;refGene_tx=X
M_023258676.1;refGene_gn=CAST;refGene_tx=XM_023258679.1;refGene_gn=CAS

T;refGene_tx=XM_023258685.1;refGene_gn=CAST;refGene_tx=XM_023258688.1
A1 160830281 160830353
refGene_gn=CAST;refGene_tx=XM_011284727.2;refGene_gn=CAST;refGene_tx=X
M_011284750.2;refGene_gn=CAST;refGene_tx=XM_011284751.2;refGene_gn=CAS
T;refGene_tx=XM_011284752.3;refGene_gn=CAST;refGene_tx=XM_023258666.1;
refGene_gn=CAST;refGene_tx=XM_023258673.1;refGene_gn=CAST;refGene_tx=X
M_023258676.1;refGene_gn=CAST;refGene_tx=XM_023258679.1;refGene_gn=CAS
T;refGene_tx=XM_023258685.1;refGene_gn=CAST;refGene_tx=XM_023258688.1
A1 160836049 160836145
refGene_gn=CAST;refGene_tx=XM_011284727.2;refGene_gn=CAST;refGene_tx=X
M_011284750.2;refGene_gn=CAST;refGene_tx=XM_011284751.2;refGene_gn=CAS
T;refGene_tx=XM_011284752.3;refGene_gn=CAST;refGene_tx=XM_023258666.1;
refGene_gn=CAST;refGene_tx=XM_023258673.1;refGene_gn=CAST;refGene_tx=X
M_023258676.1;refGene_gn=CAST;refGene_tx=XM_023258679.1;refGene_gn=CAS
T;refGene_tx=XM_023258685.1;refGene_gn=CAST;refGene_tx=XM_023258688.1
A1 160837788 160837896
refGene_gn=CAST;refGene_tx=XM_011284727.2;refGene_gn=CAST;refGene_tx=X
M_011284750.2;refGene_gn=CAST;refGene_tx=XM_011284751.2;refGene_gn=CAS
T;refGene_tx=XM_011284752.3;refGene_gn=CAST;refGene_tx=XM_023258666.1;
refGene_gn=CAST;refGene_tx=XM_023258673.1;refGene_gn=CAST;refGene_tx=X
M_023258676.1;refGene_gn=CAST;refGene_tx=XM_023258679.1;refGene_gn=CAS
T;refGene_tx=XM_023258685.1;refGene_gn=CAST;refGene_tx=XM_023258688.1
A1 160839179 160839272
refGene_gn=CAST;refGene_tx=XM_011284727.2;refGene_gn=CAST;refGene_tx=X
M_011284750.2;refGene_gn=CAST;refGene_tx=XM_011284751.2;refGene_gn=CAS
T;refGene_tx=XM_011284752.3;refGene_gn=CAST;refGene_tx=XM_023258666.1;
refGene_gn=CAST;refGene_tx=XM_023258673.1;refGene_gn=CAST;refGene_tx=X
M_023258676.1;refGene_gn=CAST;refGene_tx=XM_023258679.1;refGene_gn=CAS
T;refGene_tx=XM_023258685.1;refGene_gn=CAST;refGene_tx=XM_023258688.1
A1 160840741 160840786
refGene_gn=CAST;refGene_tx=XM_011284727.2;refGene_gn=CAST;refGene_tx=X
M_011284750.2;refGene_gn=CAST;refGene_tx=XM_011284751.2;refGene_gn=CAS
T;refGene_tx=XM_011284752.3;refGene_gn=CAST;refGene_tx=XM_023258666.1;
refGene_gn=CAST;refGene_tx=XM_023258673.1;refGene_gn=CAST;refGene_tx=X
M_023258676.1;refGene_gn=CAST;refGene_tx=XM_023258679.1;refGene_gn=CAS
T;refGene_tx=XM_023258685.1;refGene_gn=CAST;refGene_tx=XM_023258688.1
A1 160841206 160841302
refGene_gn=CAST;refGene_tx=XM_011284727.2;refGene_gn=CAST;refGene_tx=X
M_011284750.2;refGene_gn=CAST;refGene_tx=XM_011284751.2;refGene_gn=CAS
T;refGene_tx=XM_011284752.3;refGene_gn=CAST;refGene_tx=XM_023258666.1;
refGene_gn=CAST;refGene_tx=XM_023258673.1;refGene_gn=CAST;refGene_tx=X
M_023258676.1;refGene_gn=CAST;refGene_tx=XM_023258679.1;refGene_gn=CAS
T;refGene_tx=XM_023258685.1;refGene_gn=CAST;refGene_tx=XM_023258688.1
A1 160842494 160842566
refGene_gn=CAST;refGene_tx=XM_011284727.2;refGene_gn=CAST;refGene_tx=X
M_011284750.2;refGene_gn=CAST;refGene_tx=XM_011284751.2;refGene_gn=CAS
T;refGene_tx=XM_011284752.3;refGene_gn=CAST;refGene_tx=XM_023258666.1;
refGene_gn=CAST;refGene_tx=XM_023258673.1;refGene_gn=CAST;refGene_tx=X
M_023258676.1;refGene_gn=CAST;refGene_tx=XM_023258679.1;refGene_gn=CAS
T;refGene_tx=XM_023258685.1;refGene_gn=CAST;refGene_tx=XM_023258688.1

A1 160846623 160846674
refGene_gn=CAST;refGene_tx=XM_011284727.2;refGene_gn=CAST;refGene_tx=XM_011284750.2;refGene_gn=CAST;refGene_tx=XM_011284751.2;refGene_gn=CAST;refGene_tx=XM_011284752.3;refGene_gn=CAST;refGene_tx=XM_023258666.1;refGene_gn=CAST;refGene_tx=XM_023258673.1;refGene_gn=CAST;refGene_tx=XM_023258676.1;refGene_gn=CAST;refGene_tx=XM_023258679.1;refGene_gn=CAST;refGene_tx=XM_023258685.1;refGene_gn=CAST;refGene_tx=XM_023258688.1
A1 160850925 160851081
refGene_gn=ERAP1;refGene_tx=XM_011284724.3;refGene_gn=ERAP1;refGene_tx=XM_023258646.1;refGene_gn=ERAP1;refGene_tx=XM_023258647.1;refGene_gn=ERAP1;refGene_tx=XM_023258652.1;refGene_gn=ERAP1;refGene_tx=XM_023258660.1
A1 160855349 160855431
refGene_gn=ERAP1;refGene_tx=XM_011284724.3;refGene_gn=ERAP1;refGene_tx=XM_023258646.1;refGene_gn=ERAP1;refGene_tx=XM_023258647.1;refGene_gn=ERAP1;refGene_tx=XM_023258652.1;refGene_gn=ERAP1;refGene_tx=XM_023258660.1
A1 160856251 160856392
refGene_gn=ERAP1;refGene_tx=XM_011284724.3;refGene_gn=ERAP1;refGene_tx=XM_023258646.1;refGene_gn=ERAP1;refGene_tx=XM_023258647.1;refGene_gn=ERAP1;refGene_tx=XM_023258652.1;refGene_gn=ERAP1;refGene_tx=XM_023258660.1
A1 160856871 160857033
refGene_gn=ERAP1;refGene_tx=XM_011284724.3;refGene_gn=ERAP1;refGene_tx=XM_023258646.1;refGene_gn=ERAP1;refGene_tx=XM_023258647.1;refGene_gn=ERAP1;refGene_tx=XM_023258652.1;refGene_gn=ERAP1;refGene_tx=XM_023258660.1
A1 160857984 160858169
refGene_gn=ERAP1;refGene_tx=XM_011284724.3;refGene_gn=ERAP1;refGene_tx=XM_023258646.1;refGene_gn=ERAP1;refGene_tx=XM_023258647.1;refGene_gn=ERAP1;refGene_tx=XM_023258652.1;refGene_gn=ERAP1;refGene_tx=XM_023258660.1
A1 160858818 160858975
refGene_gn=ERAP1;refGene_tx=XM_011284724.3;refGene_gn=ERAP1;refGene_tx=XM_023258646.1;refGene_gn=ERAP1;refGene_tx=XM_023258647.1;refGene_gn=ERAP1;refGene_tx=XM_023258652.1;refGene_gn=ERAP1;refGene_tx=XM_023258660.1
A1 160860749 160860933
refGene_gn=ERAP1;refGene_tx=XM_011284724.3;refGene_gn=ERAP1;refGene_tx=XM_023258646.1;refGene_gn=ERAP1;refGene_tx=XM_023258647.1;refGene_gn=ERAP1;refGene_tx=XM_023258652.1;refGene_gn=ERAP1;refGene_tx=XM_023258660.1
A1 160861528 160861608
refGene_gn=ERAP1;refGene_tx=XM_011284724.3;refGene_gn=ERAP1;refGene_tx=XM_023258646.1;refGene_gn=ERAP1;refGene_tx=XM_023258647.1;refGene_gn=ERAP1;refGene_tx=XM_023258652.1;refGene_gn=ERAP1;refGene_tx=XM_023258660.1
A1 160868327 160868482
refGene_gn=ERAP1;refGene_tx=XM_011284724.3;refGene_gn=ERAP1;refGene_tx=XM_023258646.1;refGene_gn=ERAP1;refGene_tx=XM_023258647.1;refGene_gn=

ERAP1;refGene_tx=XM_023258652.1;refGene_gn=ERAP1;refGene_tx=XM_0232586
60.1
A1 160870271 160870343
refGene_gn=ERAP1;refGene_tx=XM_011284724.3;refGene_gn=ERAP1;refGene_tx
=XM_023258646.1;refGene_gn=ERAP1;refGene_tx=XM_023258647.1;refGene_gn=
ERAP1;refGene_tx=XM_023258652.1;refGene_gn=ERAP1;refGene_tx=XM_0232586
60.1
A1 160870495 160870627
refGene_gn=ERAP1;refGene_tx=XM_011284724.3;refGene_gn=ERAP1;refGene_tx
=XM_023258646.1;refGene_gn=ERAP1;refGene_tx=XM_023258647.1;refGene_gn=
ERAP1;refGene_tx=XM_023258652.1;refGene_gn=ERAP1;refGene_tx=XM_0232586
60.1
A1 160875253 160875385
refGene_gn=ERAP1;refGene_tx=XM_011284724.3;refGene_gn=ERAP1;refGene_tx
=XM_023258646.1;refGene_gn=ERAP1;refGene_tx=XM_023258647.1;refGene_gn=
ERAP1;refGene_tx=XM_023258652.1;refGene_gn=ERAP1;refGene_tx=XM_0232586
60.1
A1 160876699 160876813
refGene_gn=ERAP1;refGene_tx=XM_011284724.3;refGene_gn=ERAP1;refGene_tx
=XM_023258646.1;refGene_gn=ERAP1;refGene_tx=XM_023258647.1;refGene_gn=
ERAP1;refGene_tx=XM_023258652.1;refGene_gn=ERAP1;refGene_tx=XM_0232586
60.1
A1 160877156 160877311
refGene_gn=ERAP1;refGene_tx=XM_011284724.3;refGene_gn=ERAP1;refGene_tx
=XM_023258646.1;refGene_gn=ERAP1;refGene_tx=XM_023258647.1;refGene_gn=
ERAP1;refGene_tx=XM_023258652.1;refGene_gn=ERAP1;refGene_tx=XM_0232586
60.1
A1 160878040 160878161
refGene_gn=ERAP1;refGene_tx=XM_011284724.3;refGene_gn=ERAP1;refGene_tx
=XM_023258646.1;refGene_gn=ERAP1;refGene_tx=XM_023258647.1;refGene_gn=
ERAP1;refGene_tx=XM_023258652.1;refGene_gn=ERAP1;refGene_tx=XM_0232586
60.1
A1 160885222 160885357
refGene_gn=ERAP1;refGene_tx=XM_011284724.3;refGene_gn=ERAP1;refGene_tx
=XM_023258646.1;refGene_gn=ERAP1;refGene_tx=XM_023258647.1;refGene_gn=
ERAP1;refGene_tx=XM_023258652.1;refGene_gn=ERAP1;refGene_tx=XM_0232586
60.1
A1 160888255 160888394
refGene_gn=ERAP1;refGene_tx=XM_011284724.3;refGene_gn=ERAP1;refGene_tx
=XM_023258646.1;refGene_gn=ERAP1;refGene_tx=XM_023258647.1;refGene_gn=
ERAP1;refGene_tx=XM_023258652.1;refGene_gn=ERAP1;refGene_tx=XM_0232586
60.1
A1 160890150 160890694
refGene_gn=ERAP1;refGene_tx=XM_011284724.3;refGene_gn=ERAP1;refGene_tx
=XM_023258646.1;refGene_gn=ERAP1;refGene_tx=XM_023258647.1;refGene_gn=
ERAP1;refGene_tx=XM_023258652.1;refGene_gn=ERAP1;refGene_tx=XM_0232586
60.1
A1 160948345 160948423
refGene_gn=ERAP1;refGene_tx=XM_011284724.3
A1 160978828 160978838

refGene_gn=ERAP1;refGene_tx=XM_011284724.3
A1 161022723 161022742
refGene_gn=LNPEP;refGene_tx=XM_003981162.5
A1 161057311 161058152
refGene_gn=LNPEP;refGene_tx=XM_003981162.5;refGene_gn=LNPEP;refGene_tx=XM_019837388.2
A1 161063242 161063381
refGene_gn=LNPEP;refGene_tx=XM_003981162.5;refGene_gn=LNPEP;refGene_tx=XM_019837388.2
A1 161065123 161065255
refGene_gn=LNPEP;refGene_tx=XM_003981162.5;refGene_gn=LNPEP;refGene_tx=XM_019837388.2
A1 161076952 161077073
refGene_gn=LNPEP;refGene_tx=XM_003981162.5;refGene_gn=LNPEP;refGene_tx=XM_019837388.2
A1 161077721 161077876
refGene_gn=LNPEP;refGene_tx=XM_003981162.5;refGene_gn=LNPEP;refGene_tx=XM_019837388.2
A1 161079883 161079997
refGene_gn=LNPEP;refGene_tx=XM_003981162.5;refGene_gn=LNPEP;refGene_tx=XM_019837388.2
A1 161081409 161081544
refGene_gn=LNPEP;refGene_tx=XM_003981162.5;refGene_gn=LNPEP;refGene_tx=XM_019837388.2
A1 161086446 161086578
refGene_gn=LNPEP;refGene_tx=XM_003981162.5;refGene_gn=LNPEP;refGene_tx=XM_019837388.2
A1 161088470 161088631
refGene_gn=LNPEP;refGene_tx=XM_003981162.5;refGene_gn=LNPEP;refGene_tx=XM_019837388.2
A1 161088873 161088962
refGene_gn=LNPEP;refGene_tx=XM_003981162.5;refGene_gn=LNPEP;refGene_tx=XM_019837388.2
A1 161094678 161094862
refGene_gn=LNPEP;refGene_tx=XM_003981162.5;refGene_gn=LNPEP;refGene_tx=XM_019837388.2
A1 161095989 161096146
refGene_gn=LNPEP;refGene_tx=XM_003981162.5;refGene_gn=LNPEP;refGene_tx=XM_019837388.2
A1 161102373 161102558
refGene_gn=LNPEP;refGene_tx=XM_003981162.5;refGene_gn=LNPEP;refGene_tx=XM_019837388.2
A1 161103985 161104147
refGene_gn=LNPEP;refGene_tx=XM_003981162.5;refGene_gn=LNPEP;refGene_tx=XM_019837388.2
A1 161106474 161106615
refGene_gn=LNPEP;refGene_tx=XM_003981162.5;refGene_gn=LNPEP;refGene_tx=XM_019837388.2
A1 161107265 161107347
refGene_gn=LNPEP;refGene_tx=XM_003981162.5;refGene_gn=LNPEP;refGene_tx

=XM_019837388.2
A1 161108215 161108347
refGene_gn=LNPEP;refGene_tx=XM_003981162.5;refGene_gn=LNPEP;refGene_tx
=XM_019837388.2
A1 161133949 161134108
refGene_gn=LOC109502451;refGene_tx=XM_023258705.1
A1 161135088 161135275
refGene_gn=LOC109502451;refGene_tx=XM_023258705.1
A1 161138004 161138195
refGene_gn=LOC109502451;refGene_tx=XM_023258705.1
A1 161140572 161140733
refGene_gn=LOC109502451;refGene_tx=XM_023258705.1
A1 161145160 161145285
refGene_gn=LOC109502451;refGene_tx=XM_023258705.1
A1 161151294 161151481
refGene_gn=LOC109502451;refGene_tx=XM_023258705.1
A1 161154661 161154871
refGene_gn=LOC109502451;refGene_tx=XM_023258705.1
A1 161155364 161155521
refGene_gn=LOC109502451;refGene_tx=XM_023258705.1
A1 161163734 161164022
refGene_gn=LIX1;refGene_tx=NM_001048162.1;refGene_gn=LIX1;refGene_tx=X
M_019816992.1;refGene_gn=LIX1;refGene_tx=XM_023241803.1;refGene_gn=LIX
1;refGene_tx=XM_023242373.1
A1 161166170 161166248
refGene_gn=LIX1;refGene_tx=NM_001048162.1;refGene_gn=LIX1;refGene_tx=X
M_019816992.1;refGene_gn=LIX1;refGene_tx=XM_023241803.1;refGene_gn=LIX
1;refGene_tx=XM_023242373.1
A1 161174535 161174631
refGene_gn=LIX1;refGene_tx=NM_001048162.1;refGene_gn=LIX1;refGene_tx=X
M_019816992.1;refGene_gn=LIX1;refGene_tx=XM_023241803.1;refGene_gn=LIX
1;refGene_tx=XM_023242373.1
A1 161176928 161177069
refGene_gn=LIX1;refGene_tx=NM_001048162.1;refGene_gn=LIX1;refGene_tx=X
M_019816992.1;refGene_gn=LIX1;refGene_tx=XM_023241803.1;refGene_gn=LIX
1;refGene_tx=XM_023242373.1
A1 161194846 161195010
refGene_gn=LIX1;refGene_tx=NM_001048162.1;refGene_gn=LIX1;refGene_tx=X
M_019816992.1;refGene_gn=LIX1;refGene_tx=XM_023241803.1;refGene_gn=LIX
1;refGene_tx=XM_023242373.1
A1 161213477 161213623
refGene_gn=LIX1;refGene_tx=NM_001048162.1;refGene_gn=LIX1;refGene_tx=X
M_019816992.1;refGene_gn=LIX1;refGene_tx=XM_023241803.1;refGene_gn=LIX
1;refGene_tx=XM_023242373.1
A1 161213827 161213859
refGene_gn=LIX1;refGene_tx=XM_019816992.1;refGene_gn=LIX1;refGene_tx=X
M_023241803.1;refGene_gn=LIX1;refGene_tx=XM_023242373.1
A1 161224053 161224218
refGene_gn=RIOK2;refGene_tx=XM_003981163.5;refGene_gn=RIOK2;refGene_tx
=XM_019837403.2

A1 161228501 161228598
refGene_gn=RIOK2;refGene_tx=XM_003981163.5;refGene_gn=RIOK2;refGene_tx
=XM_019837403.2
A1 161230091 161230616
refGene_gn=RIOK2;refGene_tx=XM_003981163.5;refGene_gn=RIOK2;refGene_tx
=XM_019837403.2
A1 161232042 161232135
refGene_gn=RIOK2;refGene_tx=XM_003981163.5;refGene_gn=RIOK2;refGene_tx
=XM_019837403.2
A1 161234349 161234541
refGene_gn=RIOK2;refGene_tx=XM_003981163.5;refGene_gn=RIOK2;refGene_tx
=XM_019837403.2
A1 161236727 161236816
refGene_gn=RIOK2;refGene_tx=XM_003981163.5;refGene_gn=RIOK2;refGene_tx
=XM_019837403.2
A1 161239960 161240136
refGene_gn=RIOK2;refGene_tx=XM_003981163.5;refGene_gn=RIOK2;refGene_tx
=XM_019837403.2
A1 161242196 161242337
refGene_gn=RIOK2;refGene_tx=XM_003981163.5;refGene_gn=RIOK2;refGene_tx
=XM_019837403.2
A1 161243136 161243275
refGene_gn=RIOK2;refGene_tx=XM_003981163.5;refGene_gn=RIOK2;refGene_tx
=XM_019837403.2
A1 161245598 161245664
refGene_gn=RIOK2;refGene_tx=XM_003981163.5;refGene_gn=RIOK2;refGene_tx
=XM_019837403.2
A1 162370153 162370274
refGene_gn=RGMB;refGene_tx=XM_003981183.3
A1 162370385 162370628
refGene_gn=RGMB;refGene_tx=XM_003981183.3
A1 162371594 162371670
refGene_gn=RGMB;refGene_tx=XM_003981183.3
A1 162374881 162375147
refGene_gn=RGMB;refGene_tx=XM_003981183.3
A1 162380431 162380940
refGene_gn=RGMB;refGene_tx=XM_003981183.3
A1 162396279 162396948
refGene_gn=RGMB;refGene_tx=XM_003981183.3
A1 162453348 162453694
refGene_gn=CHD1;refGene_tx=XM_023258732.1;refGene_gn=CHD1;refGene_tx=X
M_023258736.1
A1 162455457 162455668
refGene_gn=CHD1;refGene_tx=XM_023258732.1;refGene_gn=CHD1;refGene_tx=X
M_023258736.1
A1 162456172 162456224
refGene_gn=CHD1;refGene_tx=XM_023258732.1;refGene_gn=CHD1;refGene_tx=X
M_023258736.1
A1 162457213 162457310
refGene_gn=CHD1;refGene_tx=XM_023258732.1;refGene_gn=CHD1;refGene_tx=X

M_023258736.1
A1	162459006	162459185
refGene_gn=CHD1;refGene_tx=XM_023258732.1;refGene_gn=CHD1;refGene_tx=X
M_023258736.1
A1	162464312	162464453
refGene_gn=CHD1;refGene_tx=XM_023258732.1;refGene_gn=CHD1;refGene_tx=X
M_023258736.1
A1	162465522	162465651
refGene_gn=CHD1;refGene_tx=XM_023258732.1;refGene_gn=CHD1;refGene_tx=X
M_023258736.1
A1	162466491	162466614
refGene_gn=CHD1;refGene_tx=XM_023258732.1;refGene_gn=CHD1;refGene_tx=X
M_023258736.1
A1	162467600	162467751
refGene_gn=CHD1;refGene_tx=XM_023258732.1;refGene_gn=CHD1;refGene_tx=X
M_023258736.1
A1	162467973	162468112
refGene_gn=CHD1;refGene_tx=XM_023258732.1;refGene_gn=CHD1;refGene_tx=X
M_023258736.1
A1	162468819	162468950
refGene_gn=CHD1;refGene_tx=XM_023258732.1;refGene_gn=CHD1;refGene_tx=X
M_023258736.1
A1	162470265	162470307
refGene_gn=CHD1;refGene_tx=XM_023258732.1;refGene_gn=CHD1;refGene_tx=X
M_023258736.1
A1	162471603	162471764
refGene_gn=CHD1;refGene_tx=XM_023258732.1;refGene_gn=CHD1;refGene_tx=X
M_023258736.1
A1	162474256	162474433
refGene_gn=CHD1;refGene_tx=XM_023258732.1;refGene_gn=CHD1;refGene_tx=X
M_023258736.1
A1	162476356	162476452
refGene_gn=CHD1;refGene_tx=XM_023258732.1;refGene_gn=CHD1;refGene_tx=X
M_023258736.1
A1	162476563	162476660
refGene_gn=CHD1;refGene_tx=XM_023258732.1;refGene_gn=CHD1;refGene_tx=X
M_023258736.1
A1	162477045	162477194
refGene_gn=CHD1;refGene_tx=XM_023258732.1;refGene_gn=CHD1;refGene_tx=X
M_023258736.1
A1	162483209	162483359
refGene_gn=CHD1;refGene_tx=XM_023258732.1;refGene_gn=CHD1;refGene_tx=X
M_023258736.1
A1	162483813	162483885
refGene_gn=CHD1;refGene_tx=XM_023258732.1;refGene_gn=CHD1;refGene_tx=X
M_023258736.1
A1	162487228	162487381
refGene_gn=CHD1;refGene_tx=XM_023258732.1;refGene_gn=CHD1;refGene_tx=X
M_023258736.1
A1	162488453	162488616

refGene_gn=CHD1;refGene_tx=XM_023258732.1;refGene_gn=CHD1;refGene_tx=XM_023258736.1
A1 162490371 162490560
refGene_gn=CHD1;refGene_tx=XM_023258732.1;refGene_gn=CHD1;refGene_tx=XM_023258736.1
A1 162491278 162491469
refGene_gn=CHD1;refGene_tx=XM_023258732.1;refGene_gn=CHD1;refGene_tx=XM_023258736.1
A1 162492525 162492615
refGene_gn=CHD1;refGene_tx=XM_023258732.1;refGene_gn=CHD1;refGene_tx=XM_023258736.1
A1 162493306 162493523
refGene_gn=CHD1;refGene_tx=XM_023258732.1;refGene_gn=CHD1;refGene_tx=XM_023258736.1
A1 162494281 162494409
refGene_gn=CHD1;refGene_tx=XM_023258732.1;refGene_gn=CHD1;refGene_tx=XM_023258736.1
A1 162495354 162495533
refGene_gn=CHD1;refGene_tx=XM_023258732.1;refGene_gn=CHD1;refGene_tx=XM_023258736.1
A1 162495758 162495859
refGene_gn=CHD1;refGene_tx=XM_023258732.1;refGene_gn=CHD1;refGene_tx=XM_023258736.1
A1 162496312 162496538
refGene_gn=CHD1;refGene_tx=XM_023258732.1;refGene_gn=CHD1;refGene_tx=XM_023258736.1
A1 162498236 162498508
refGene_gn=CHD1;refGene_tx=XM_023258732.1;refGene_gn=CHD1;refGene_tx=XM_023258736.1
A1 162498637 162498787
refGene_gn=CHD1;refGene_tx=XM_023258732.1;refGene_gn=CHD1;refGene_tx=XM_023258736.1
A1 162501461 162501526
refGene_gn=CHD1;refGene_tx=XM_023258732.1;refGene_gn=CHD1;refGene_tx=XM_023258736.1
A1 162502573 162502687
refGene_gn=CHD1;refGene_tx=XM_023258732.1;refGene_gn=CHD1;refGene_tx=XM_023258736.1
A1 162503646 162503848
refGene_gn=CHD1;refGene_tx=XM_023258732.1;refGene_gn=CHD1;refGene_tx=XM_023258736.1
A1 162519594 162519652
refGene_gn=CHD1;refGene_tx=XM_023258732.1
A1 162525921 162525974
refGene_gn=CHD1;refGene_tx=XM_023258732.1;refGene_gn=CHD1;refGene_tx=XM_023258736.1
A1 163944104 163944544
refGene_gn=FAM174A;refGene_tx=XM_003981164.5;refGene_gn=FAM174A;refGene_tx=XM_011284785.3
A1 163967371 163967510

refGene_gn=FAM174A;refGene_tx=XM_003981164.5;refGene_gn=FAM174A;refGene_tx=XM_011284785.3
A1 163981524 163981528
refGene_gn=FAM174A;refGene_tx=XM_011284785.3
A1 164201338 164201621
refGene_gn=ST8SIA4;refGene_tx=XM_003981165.5
A1 164252063 164252357
refGene_gn=ST8SIA4;refGene_tx=XM_003981165.5
A1 164280283 164280541
refGene_gn=ST8SIA4;refGene_tx=XM_003981165.5
A1 164290034 164290166
refGene_gn=ST8SIA4;refGene_tx=XM_003981165.5
A1 164297024 164297143
refGene_gn=ST8SIA4;refGene_tx=XM_003981165.5
A1 164297144 164297216
refGene_gn=ST8SIA4;refGene_tx=XM_003981165.5
A1 164297217 164297304
refGene_gn=ST8SIA4;refGene_tx=XM_003981165.5
A1 164297305 164297425
refGene_gn=ST8SIA4;refGene_tx=XM_003981165.5
A1 165301553 165301711
refGene_gn=SLCO4C1;refGene_tx=XM_011284799.3;refGene_gn=SLCO4C1;refGene_tx=XM_019837484.2;refGene_gn=SLCO4C1;refGene_tx=XM_019837491.2;refGene_gn=SLCO4C1;refGene_tx=XM_019837499.2
A1 165310836 165310974
refGene_gn=SLCO4C1;refGene_tx=XM_011284799.3;refGene_gn=SLCO4C1;refGene_tx=XM_019837484.2;refGene_gn=SLCO4C1;refGene_tx=XM_019837491.2;refGene_gn=SLCO4C1;refGene_tx=XM_019837499.2
A1 165312463 165312528
refGene_gn=SLCO4C1;refGene_tx=XM_011284799.3;refGene_gn=SLCO4C1;refGene_tx=XM_019837484.2;refGene_gn=SLCO4C1;refGene_tx=XM_019837491.2;refGene_gn=SLCO4C1;refGene_tx=XM_019837499.2
A1 165313871 165314062
refGene_gn=SLCO4C1;refGene_tx=XM_011284799.3;refGene_gn=SLCO4C1;refGene_tx=XM_019837484.2;refGene_gn=SLCO4C1;refGene_tx=XM_019837491.2;refGene_gn=SLCO4C1;refGene_tx=XM_019837499.2
A1 165317634 165317788
refGene_gn=SLCO4C1;refGene_tx=XM_011284799.3;refGene_gn=SLCO4C1;refGene_tx=XM_019837484.2;refGene_gn=SLCO4C1;refGene_tx=XM_019837491.2;refGene_gn=SLCO4C1;refGene_tx=XM_019837499.2
A1 165323579 165323775
refGene_gn=SLCO4C1;refGene_tx=XM_011284799.3;refGene_gn=SLCO4C1;refGene_tx=XM_019837484.2;refGene_gn=SLCO4C1;refGene_tx=XM_019837491.2;refGene_gn=SLCO4C1;refGene_tx=XM_019837499.2
A1 165324131 165324276
refGene_gn=SLCO4C1;refGene_tx=XM_011284799.3;refGene_gn=SLCO4C1;refGene_tx=XM_019837484.2;refGene_gn=SLCO4C1;refGene_tx=XM_019837491.2;refGene_gn=SLCO4C1;refGene_tx=XM_019837499.2
A1 165326343 165326462
refGene_gn=SLCO4C1;refGene_tx=XM_011284799.3;refGene_gn=SLCO4C1;refGen

e_tx=XM_019837484.2;refGene_gn=SLCO4C1;refGene_tx=XM_019837491.2;refGene_gn=SLCO4C1;refGene_tx=XM_019837499.2
A1 165329118 165329240
refGene_gn=SLCO4C1;refGene_tx=XM_011284799.3;refGene_gn=SLCO4C1;refGene_tx=XM_019837484.2;refGene_gn=SLCO4C1;refGene_tx=XM_019837491.2;refGene_gn=SLCO4C1;refGene_tx=XM_019837499.2
A1 165331248 165331345
refGene_gn=SLCO4C1;refGene_tx=XM_011284799.3;refGene_gn=SLCO4C1;refGene_tx=XM_019837484.2;refGene_gn=SLCO4C1;refGene_tx=XM_019837491.2;refGene_gn=SLCO4C1;refGene_tx=XM_019837499.2
A1 165335881 165336064
refGene_gn=SLCO4C1;refGene_tx=XM_011284799.3;refGene_gn=SLCO4C1;refGene_tx=XM_019837484.2;refGene_gn=SLCO4C1;refGene_tx=XM_019837491.2;refGene_gn=SLCO4C1;refGene_tx=XM_019837499.2
A1 165371629 165371893
refGene_gn=SLCO4C1;refGene_tx=XM_011284799.3;refGene_gn=SLCO4C1;refGene_tx=XM_019837484.2;refGene_gn=SLCO4C1;refGene_tx=XM_019837491.2;refGene_gn=SLCO4C1;refGene_tx=XM_019837499.2
A1 165375073 165375419
refGene_gn=SLCO4C1;refGene_tx=XM_011284799.3
A1 165419450 165419587
refGene_gn=SLCO6A1;refGene_tx=XM_011284804.3
A1 165422288 165422426
refGene_gn=SLCO6A1;refGene_tx=XM_011284804.3;refGene_gn=SLCO6A1;refGene_tx=XM_019837512.2
A1 165424346 165424411
refGene_gn=SLCO6A1;refGene_tx=XM_011284804.3
A1 165432309 165432445
refGene_gn=SLCO6A1;refGene_tx=XM_019837505.2
A1 165441964 165442152
refGene_gn=SLCO6A1;refGene_tx=XM_011284804.3;refGene_gn=SLCO6A1;refGene_tx=XM_019837505.2;refGene_gn=SLCO6A1;refGene_tx=XM_019837512.2
A1 165449442 165449596
refGene_gn=SLCO6A1;refGene_tx=XM_011284804.3;refGene_gn=SLCO6A1;refGene_tx=XM_019837505.2;refGene_gn=SLCO6A1;refGene_tx=XM_019837512.2
A1 165464391 165464587
refGene_gn=SLCO6A1;refGene_tx=XM_011284804.3;refGene_gn=SLCO6A1;refGene_tx=XM_019837505.2;refGene_gn=SLCO6A1;refGene_tx=XM_019837512.2
A1 165483423 165483568
refGene_gn=SLCO6A1;refGene_tx=XM_011284804.3;refGene_gn=SLCO6A1;refGene_tx=XM_019837505.2;refGene_gn=SLCO6A1;refGene_tx=XM_019837512.2
A1 165493694 165493804
refGene_gn=SLCO6A1;refGene_tx=XM_011284804.3;refGene_gn=SLCO6A1;refGene_tx=XM_019837505.2;refGene_gn=SLCO6A1;refGene_tx=XM_019837512.2
A1 165494621 165494743
refGene_gn=SLCO6A1;refGene_tx=XM_011284804.3;refGene_gn=SLCO6A1;refGene_tx=XM_019837505.2;refGene_gn=SLCO6A1;refGene_tx=XM_019837512.2
A1 165498297 165498394
refGene_gn=SLCO6A1;refGene_tx=XM_011284804.3;refGene_gn=SLCO6A1;refGene_tx=XM_019837505.2;refGene_gn=SLCO6A1;refGene_tx=XM_019837512.2

A1 165499961 165500147
refGene_gn=SLCO6A1;refGene_tx=XM_011284804.3;refGene_gn=SLCO6A1;refGene_tx=XM_019837505.2;refGene_gn=SLCO6A1;refGene_tx=XM_019837512.2
A1 165501536 165501797
refGene_gn=SLCO6A1;refGene_tx=XM_011284804.3;refGene_gn=SLCO6A1;refGene_tx=XM_019837505.2;refGene_gn=SLCO6A1;refGene_tx=XM_019837512.2
A1 165511529 165511854
refGene_gn=SLCO6A1;refGene_tx=XM_011284804.3;refGene_gn=SLCO6A1;refGene_tx=XM_019837505.2;refGene_gn=SLCO6A1;refGene_tx=XM_019837512.2
A1 165869333 165869422
refGene_gn=PAM;refGene_tx=XM_023258790.1;refGene_gn=PAM;refGene_tx=XM_023258791.1;refGene_gn=PAM;refGene_tx=XM_023258793.1;refGene_gn=PAM;refGene_tx=XM_023258797.1;refGene_gn=PAM;refGene_tx=XM_023258807.1;refGene_gn=PAM;refGene_tx=XM_023258812.1;refGene_gn=PAM;refGene_tx=XM_023258813.1;refGene_gn=PAM;refGene_tx=XM_023258814.1;refGene_gn=PAM;refGene_tx=XM_023258815.1;refGene_gn=PAM;refGene_tx=XM_023258817.1;refGene_gn=PAM;refGene_tx=XM_023258823.1;refGene_gn=PAM;refGene_tx=XM_023258824.1;refGene_gn=PAM;refGene_tx=XM_023258826.1;refGene_gn=PAM;refGene_tx=XM_023258827.1;refGene_gn=PAM;refGene_tx=XM_023258830.1;refGene_gn=PAM;refGene_tx=XM_023258836.1
A1 165870380 165870501
refGene_gn=PAM;refGene_tx=XM_023258790.1;refGene_gn=PAM;refGene_tx=XM_023258791.1;refGene_gn=PAM;refGene_tx=XM_023258793.1;refGene_gn=PAM;refGene_tx=XM_023258797.1;refGene_gn=PAM;refGene_tx=XM_023258807.1;refGene_gn=PAM;refGene_tx=XM_023258812.1;refGene_gn=PAM;refGene_tx=XM_023258813.1;refGene_gn=PAM;refGene_tx=XM_023258814.1;refGene_gn=PAM;refGene_tx=XM_023258815.1;refGene_gn=PAM;refGene_tx=XM_023258817.1;refGene_gn=PAM;refGene_tx=XM_023258823.1;refGene_gn=PAM;refGene_tx=XM_023258824.1;refGene_gn=PAM;refGene_tx=XM_023258826.1;refGene_gn=PAM;refGene_tx=XM_023258827.1;refGene_gn=PAM;refGene_tx=XM_023258830.1;refGene_gn=PAM;refGene_tx=XM_023258836.1
A1 165901532 165901590
refGene_gn=PAM;refGene_tx=XM_023258790.1;refGene_gn=PAM;refGene_tx=XM_023258791.1;refGene_gn=PAM;refGene_tx=XM_023258793.1;refGene_gn=PAM;refGene_tx=XM_023258797.1;refGene_gn=PAM;refGene_tx=XM_023258807.1;refGene_gn=PAM;refGene_tx=XM_023258812.1;refGene_gn=PAM;refGene_tx=XM_023258813.1;refGene_gn=PAM;refGene_tx=XM_023258814.1;refGene_gn=PAM;refGene_tx=XM_023258815.1;refGene_gn=PAM;refGene_tx=XM_023258817.1;refGene_gn=PAM;refGene_tx=XM_023258823.1;refGene_gn=PAM;refGene_tx=XM_023258824.1;refGene_gn=PAM;refGene_tx=XM_023258826.1;refGene_gn=PAM;refGene_tx=XM_023258827.1;refGene_gn=PAM;refGene_tx=XM_023258830.1;refGene_gn=PAM;refGene_tx=XM_023258836.1
A1 165912480 165912568
refGene_gn=PAM;refGene_tx=XM_023258790.1;refGene_gn=PAM;refGene_tx=XM_023258791.1;refGene_gn=PAM;refGene_tx=XM_023258793.1;refGene_gn=PAM;refGene_tx=XM_023258797.1;refGene_gn=PAM;refGene_tx=XM_023258807.1;refGene_gn=PAM;refGene_tx=XM_023258812.1;refGene_gn=PAM;refGene_tx=XM_023258813.1;refGene_gn=PAM;refGene_tx=XM_023258814.1;refGene_gn=PAM;refGene_tx=XM_023258815.1;refGene_gn=PAM;refGene_tx=XM_023258817.1;refGene_gn=PAM;refGene_tx=XM_023258823.1;refGene_gn=PAM;refGene_tx=XM_023258824.

1;refGene_gn=PAM;refGene_tx=XM_023258826.1;refGene_gn=PAM;refGene_tx=XM_023258827.1;refGene_gn=PAM;refGene_tx=XM_023258830.1;refGene_gn=PAM;refGene_tx=XM_023258836.1
A1 165923224 165923310
refGene_gn=PAM;refGene_tx=XM_023258790.1;refGene_gn=PAM;refGene_tx=XM_023258791.1;refGene_gn=PAM;refGene_tx=XM_023258793.1;refGene_gn=PAM;refGene_tx=XM_023258797.1;refGene_gn=PAM;refGene_tx=XM_023258807.1;refGene_gn=PAM;refGene_tx=XM_023258812.1;refGene_gn=PAM;refGene_tx=XM_023258813.1;refGene_gn=PAM;refGene_tx=XM_023258814.1;refGene_gn=PAM;refGene_tx=XM_023258815.1;refGene_gn=PAM;refGene_tx=XM_023258817.1;refGene_gn=PAM;refGene_tx=XM_023258823.1;refGene_gn=PAM;refGene_tx=XM_023258824.1;refGene_gn=PAM;refGene_tx=XM_023258826.1;refGene_gn=PAM;refGene_tx=XM_023258827.1;refGene_gn=PAM;refGene_tx=XM_023258830.1;refGene_gn=PAM;refGene_tx=XM_023258836.1
A1 165924703 165924787
refGene_gn=PAM;refGene_tx=XM_023258790.1;refGene_gn=PAM;refGene_tx=XM_023258791.1;refGene_gn=PAM;refGene_tx=XM_023258793.1;refGene_gn=PAM;refGene_tx=XM_023258797.1;refGene_gn=PAM;refGene_tx=XM_023258807.1;refGene_gn=PAM;refGene_tx=XM_023258812.1;refGene_gn=PAM;refGene_tx=XM_023258813.1;refGene_gn=PAM;refGene_tx=XM_023258814.1;refGene_gn=PAM;refGene_tx=XM_023258815.1;refGene_gn=PAM;refGene_tx=XM_023258817.1;refGene_gn=PAM;refGene_tx=XM_023258823.1;refGene_gn=PAM;refGene_tx=XM_023258824.1;refGene_gn=PAM;refGene_tx=XM_023258826.1;refGene_gn=PAM;refGene_tx=XM_023258827.1;refGene_gn=PAM;refGene_tx=XM_023258830.1;refGene_gn=PAM;refGene_tx=XM_023258836.1
A1 165949369 165949418
refGene_gn=PAM;refGene_tx=XM_023258790.1;refGene_gn=PAM;refGene_tx=XM_023258791.1;refGene_gn=PAM;refGene_tx=XM_023258793.1;refGene_gn=PAM;refGene_tx=XM_023258797.1;refGene_gn=PAM;refGene_tx=XM_023258807.1;refGene_gn=PAM;refGene_tx=XM_023258812.1;refGene_gn=PAM;refGene_tx=XM_023258813.1;refGene_gn=PAM;refGene_tx=XM_023258814.1;refGene_gn=PAM;refGene_tx=XM_023258815.1;refGene_gn=PAM;refGene_tx=XM_023258817.1;refGene_gn=PAM;refGene_tx=XM_023258823.1;refGene_gn=PAM;refGene_tx=XM_023258824.1;refGene_gn=PAM;refGene_tx=XM_023258826.1;refGene_gn=PAM;refGene_tx=XM_023258827.1;refGene_gn=PAM;refGene_tx=XM_023258830.1;refGene_gn=PAM;refGene_tx=XM_023258836.1
A1 165950822 165950890
refGene_gn=PAM;refGene_tx=XM_023258790.1;refGene_gn=PAM;refGene_tx=XM_023258791.1;refGene_gn=PAM;refGene_tx=XM_023258793.1;refGene_gn=PAM;refGene_tx=XM_023258797.1;refGene_gn=PAM;refGene_tx=XM_023258807.1;refGene_gn=PAM;refGene_tx=XM_023258812.1;refGene_gn=PAM;refGene_tx=XM_023258813.1;refGene_gn=PAM;refGene_tx=XM_023258814.1;refGene_gn=PAM;refGene_tx=XM_023258815.1;refGene_gn=PAM;refGene_tx=XM_023258817.1;refGene_gn=PAM;refGene_tx=XM_023258823.1;refGene_gn=PAM;refGene_tx=XM_023258824.1;refGene_gn=PAM;refGene_tx=XM_023258826.1;refGene_gn=PAM;refGene_tx=XM_023258827.1;refGene_gn=PAM;refGene_tx=XM_023258830.1;refGene_gn=PAM;refGene_tx=XM_023258836.1
A1 165952011 165952092
refGene_gn=PAM;refGene_tx=XM_023258790.1;refGene_gn=PAM;refGene_tx=XM_023258791.1;refGene_gn=PAM;refGene_tx=XM_023258793.1;refGene_gn=PAM;re

fGene_tx=XM_023258797.1;refGene_gn=PAM;refGene_tx=XM_023258807.1;refGene_gn=PAM;refGene_tx=XM_023258812.1;refGene_gn=PAM;refGene_tx=XM_023258813.1;refGene_gn=PAM;refGene_tx=XM_023258814.1;refGene_gn=PAM;refGene_tx=XM_023258815.1;refGene_gn=PAM;refGene_tx=XM_023258817.1;refGene_gn=PAM;refGene_tx=XM_023258823.1;refGene_gn=PAM;refGene_tx=XM_023258824.1;refGene_gn=PAM;refGene_tx=XM_023258826.1;refGene_gn=PAM;refGene_tx=XM_023258827.1;refGene_gn=PAM;refGene_tx=XM_023258830.1;refGene_gn=PAM;refGene_tx=XM_023258836.1
A1 165952375 165952452
refGene_gn=PAM;refGene_tx=XM_023258790.1;refGene_gn=PAM;refGene_tx=XM_023258791.1;refGene_gn=PAM;refGene_tx=XM_023258793.1;refGene_gn=PAM;refGene_tx=XM_023258797.1;refGene_gn=PAM;refGene_tx=XM_023258807.1;refGene_gn=PAM;refGene_tx=XM_023258812.1;refGene_gn=PAM;refGene_tx=XM_023258813.1;refGene_gn=PAM;refGene_tx=XM_023258814.1;refGene_gn=PAM;refGene_tx=XM_023258815.1;refGene_gn=PAM;refGene_tx=XM_023258817.1;refGene_gn=PAM;refGene_tx=XM_023258823.1;refGene_gn=PAM;refGene_tx=XM_023258824.1;refGene_gn=PAM;refGene_tx=XM_023258826.1;refGene_gn=PAM;refGene_tx=XM_023258827.1;refGene_gn=PAM;refGene_tx=XM_023258830.1;refGene_gn=PAM;refGene_tx=XM_023258836.1
A1 165952769 165952873
refGene_gn=PAM;refGene_tx=XM_023258790.1;refGene_gn=PAM;refGene_tx=XM_023258791.1;refGene_gn=PAM;refGene_tx=XM_023258793.1;refGene_gn=PAM;refGene_tx=XM_023258797.1;refGene_gn=PAM;refGene_tx=XM_023258807.1;refGene_gn=PAM;refGene_tx=XM_023258812.1;refGene_gn=PAM;refGene_tx=XM_023258813.1;refGene_gn=PAM;refGene_tx=XM_023258814.1;refGene_gn=PAM;refGene_tx=XM_023258815.1;refGene_gn=PAM;refGene_tx=XM_023258817.1;refGene_gn=PAM;refGene_tx=XM_023258823.1;refGene_gn=PAM;refGene_tx=XM_023258824.1;refGene_gn=PAM;refGene_tx=XM_023258826.1;refGene_gn=PAM;refGene_tx=XM_023258827.1;refGene_gn=PAM;refGene_tx=XM_023258830.1;refGene_gn=PAM;refGene_tx=XM_023258836.1
A1 165960490 165960675
refGene_gn=PAM;refGene_tx=XM_023258790.1;refGene_gn=PAM;refGene_tx=XM_023258791.1;refGene_gn=PAM;refGene_tx=XM_023258793.1;refGene_gn=PAM;refGene_tx=XM_023258797.1;refGene_gn=PAM;refGene_tx=XM_023258807.1;refGene_gn=PAM;refGene_tx=XM_023258812.1;refGene_gn=PAM;refGene_tx=XM_023258813.1;refGene_gn=PAM;refGene_tx=XM_023258814.1;refGene_gn=PAM;refGene_tx=XM_023258815.1;refGene_gn=PAM;refGene_tx=XM_023258817.1;refGene_gn=PAM;refGene_tx=XM_023258823.1;refGene_gn=PAM;refGene_tx=XM_023258824.1;refGene_gn=PAM;refGene_tx=XM_023258826.1;refGene_gn=PAM;refGene_tx=XM_023258827.1;refGene_gn=PAM;refGene_tx=XM_023258830.1;refGene_gn=PAM;refGene_tx=XM_023258836.1
A1 165961724 165961793
refGene_gn=PAM;refGene_tx=XM_023258790.1;refGene_gn=PAM;refGene_tx=XM_023258791.1;refGene_gn=PAM;refGene_tx=XM_023258793.1;refGene_gn=PAM;refGene_tx=XM_023258797.1;refGene_gn=PAM;refGene_tx=XM_023258807.1;refGene_gn=PAM;refGene_tx=XM_023258812.1;refGene_gn=PAM;refGene_tx=XM_023258813.1;refGene_gn=PAM;refGene_tx=XM_023258814.1;refGene_gn=PAM;refGene_tx=XM_023258815.1;refGene_gn=PAM;refGene_tx=XM_023258817.1;refGene_gn=PAM;refGene_tx=XM_023258823.1;refGene_gn=PAM;refGene_tx=XM_023258824.1;refGene_gn=PAM;refGene_tx=XM_023258826.1;refGene_gn=PAM;refGene_tx=X

M_023258827.1;refGene_gn=PAM;refGene_tx=XM_023258830.1;refGene_gn=PAM;
refGene_tx=XM_023258836.1
A1 165973107 165973431
refGene_gn=PAM;refGene_tx=XM_023258790.1;refGene_gn=PAM;refGene_tx=XM_
023258791.1;refGene_gn=PAM;refGene_tx=XM_023258793.1;refGene_gn=PAM;re
fGene_tx=XM_023258797.1;refGene_gn=PAM;refGene_tx=XM_023258807.1;refGe
ne_gn=PAM;refGene_tx=XM_023258812.1;refGene_gn=PAM;refGene_tx=XM_02325
8813.1;refGene_gn=PAM;refGene_tx=XM_023258814.1;refGene_gn=PAM;refGene
_tx=XM_023258815.1;refGene_gn=PAM;refGene_tx=XM_023258817.1;refGene_gn
=PAM;refGene_tx=XM_023258826.1;refGene_gn=PAM;refGene_tx=XM_023258827.
1
A1 165982849 165982979
refGene_gn=PAM;refGene_tx=XM_023258790.1;refGene_gn=PAM;refGene_tx=XM_
023258791.1;refGene_gn=PAM;refGene_tx=XM_023258793.1;refGene_gn=PAM;re
fGene_tx=XM_023258797.1;refGene_gn=PAM;refGene_tx=XM_023258807.1;refGe
ne_gn=PAM;refGene_tx=XM_023258812.1;refGene_gn=PAM;refGene_tx=XM_02325
8813.1;refGene_gn=PAM;refGene_tx=XM_023258814.1;refGene_gn=PAM;refGene
_tx=XM_023258815.1;refGene_gn=PAM;refGene_tx=XM_023258817.1;refGene_gn
=PAM;refGene_tx=XM_023258823.1;refGene_gn=PAM;refGene_tx=XM_023258824.
1;refGene_gn=PAM;refGene_tx=XM_023258826.1;refGene_gn=PAM;refGene_tx=X
M_023258827.1;refGene_gn=PAM;refGene_tx=XM_023258830.1;refGene_gn=PAM;
refGene_tx=XM_023258836.1
A1 165992782 165992899
refGene_gn=PAM;refGene_tx=XM_023258790.1;refGene_gn=PAM;refGene_tx=XM_
023258791.1;refGene_gn=PAM;refGene_tx=XM_023258793.1;refGene_gn=PAM;re
fGene_tx=XM_023258797.1;refGene_gn=PAM;refGene_tx=XM_023258807.1;refGe
ne_gn=PAM;refGene_tx=XM_023258812.1;refGene_gn=PAM;refGene_tx=XM_02325
8813.1;refGene_gn=PAM;refGene_tx=XM_023258814.1;refGene_gn=PAM;refGene
_tx=XM_023258815.1;refGene_gn=PAM;refGene_tx=XM_023258817.1;refGene_gn
=PAM;refGene_tx=XM_023258823.1;refGene_gn=PAM;refGene_tx=XM_023258824.
1;refGene_gn=PAM;refGene_tx=XM_023258826.1;refGene_gn=PAM;refGene_tx=X
M_023258827.1;refGene_gn=PAM;refGene_tx=XM_023258830.1;refGene_gn=PAM;
refGene_tx=XM_023258836.1
A1 165994922 165994995
refGene_gn=PAM;refGene_tx=XM_023258790.1;refGene_gn=PAM;refGene_tx=XM_
023258791.1;refGene_gn=PAM;refGene_tx=XM_023258793.1;refGene_gn=PAM;re
fGene_tx=XM_023258797.1;refGene_gn=PAM;refGene_tx=XM_023258807.1;refGe
ne_gn=PAM;refGene_tx=XM_023258812.1;refGene_gn=PAM;refGene_tx=XM_02325
8813.1;refGene_gn=PAM;refGene_tx=XM_023258814.1;refGene_gn=PAM;refGene
_tx=XM_023258815.1;refGene_gn=PAM;refGene_tx=XM_023258817.1;refGene_gn
=PAM;refGene_tx=XM_023258823.1;refGene_gn=PAM;refGene_tx=XM_023258824.
1;refGene_gn=PAM;refGene_tx=XM_023258826.1;refGene_gn=PAM;refGene_tx=X
M_023258827.1;refGene_gn=PAM;refGene_tx=XM_023258830.1;refGene_gn=PAM;
refGene_tx=XM_023258836.1
A1 165995945 165996156
refGene_gn=PAM;refGene_tx=XM_023258790.1;refGene_gn=PAM;refGene_tx=XM_
023258791.1;refGene_gn=PAM;refGene_tx=XM_023258793.1;refGene_gn=PAM;re
fGene_tx=XM_023258797.1;refGene_gn=PAM;refGene_tx=XM_023258807.1;refGe
ne_gn=PAM;refGene_tx=XM_023258812.1;refGene_gn=PAM;refGene_tx=XM_02325
8813.1;refGene_gn=PAM;refGene_tx=XM_023258814.1;refGene_gn=PAM;refGene

_tx=XM_023258815.1;refGene_gn=PAM;refGene_tx=XM_023258817.1;refGene_gn=PAM;refGene_tx=XM_023258823.1;refGene_gn=PAM;refGene_tx=XM_023258824.1;refGene_gn=PAM;refGene_tx=XM_023258826.1;refGene_gn=PAM;refGene_tx=XM_023258827.1;refGene_gn=PAM;refGene_tx=XM_023258830.1;refGene_gn=PAM;refGene_tx=XM_023258836.1
A1 165996579 165996780
refGene_gn=PAM;refGene_tx=XM_023258790.1;refGene_gn=PAM;refGene_tx=XM_023258791.1;refGene_gn=PAM;refGene_tx=XM_023258793.1;refGene_gn=PAM;refGene_tx=XM_023258797.1;refGene_gn=PAM;refGene_tx=XM_023258807.1;refGene_gn=PAM;refGene_tx=XM_023258812.1;refGene_gn=PAM;refGene_tx=XM_023258813.1;refGene_gn=PAM;refGene_tx=XM_023258814.1;refGene_gn=PAM;refGene_tx=XM_023258815.1;refGene_gn=PAM;refGene_tx=XM_023258817.1;refGene_gn=PAM;refGene_tx=XM_023258823.1;refGene_gn=PAM;refGene_tx=XM_023258824.1;refGene_gn=PAM;refGene_tx=XM_023258826.1;refGene_gn=PAM;refGene_tx=XM_023258827.1;refGene_gn=PAM;refGene_tx=XM_023258830.1;refGene_gn=PAM;refGene_tx=XM_023258836.1
A1 165998979 165999095
refGene_gn=PAM;refGene_tx=XM_023258790.1;refGene_gn=PAM;refGene_tx=XM_023258791.1;refGene_gn=PAM;refGene_tx=XM_023258793.1;refGene_gn=PAM;refGene_tx=XM_023258797.1;refGene_gn=PAM;refGene_tx=XM_023258807.1;refGene_gn=PAM;refGene_tx=XM_023258812.1;refGene_gn=PAM;refGene_tx=XM_023258813.1;refGene_gn=PAM;refGene_tx=XM_023258814.1;refGene_gn=PAM;refGene_tx=XM_023258815.1;refGene_gn=PAM;refGene_tx=XM_023258817.1;refGene_gn=PAM;refGene_tx=XM_023258823.1;refGene_gn=PAM;refGene_tx=XM_023258824.1;refGene_gn=PAM;refGene_tx=XM_023258826.1;refGene_gn=PAM;refGene_tx=XM_023258827.1;refGene_gn=PAM;refGene_tx=XM_023258830.1;refGene_gn=PAM;refGene_tx=XM_023258836.1
A1 166004104 166004204
refGene_gn=PAM;refGene_tx=XM_023258790.1;refGene_gn=PAM;refGene_tx=XM_023258791.1;refGene_gn=PAM;refGene_tx=XM_023258793.1;refGene_gn=PAM;refGene_tx=XM_023258797.1;refGene_gn=PAM;refGene_tx=XM_023258807.1;refGene_gn=PAM;refGene_tx=XM_023258812.1;refGene_gn=PAM;refGene_tx=XM_023258813.1;refGene_gn=PAM;refGene_tx=XM_023258814.1;refGene_gn=PAM;refGene_tx=XM_023258815.1;refGene_gn=PAM;refGene_tx=XM_023258817.1;refGene_gn=PAM;refGene_tx=XM_023258823.1;refGene_gn=PAM;refGene_tx=XM_023258824.1;refGene_gn=PAM;refGene_tx=XM_023258826.1;refGene_gn=PAM;refGene_tx=XM_023258827.1;refGene_gn=PAM;refGene_tx=XM_023258830.1;refGene_gn=PAM;refGene_tx=XM_023258836.1
A1 166006441 166006495
refGene_gn=PAM;refGene_tx=XM_023258790.1;refGene_gn=PAM;refGene_tx=XM_023258791.1;refGene_gn=PAM;refGene_tx=XM_023258793.1;refGene_gn=PAM;refGene_tx=XM_023258797.1;refGene_gn=PAM;refGene_tx=XM_023258807.1;refGene_gn=PAM;refGene_tx=XM_023258812.1;refGene_gn=PAM;refGene_tx=XM_023258813.1;refGene_gn=PAM;refGene_tx=XM_023258814.1;refGene_gn=PAM;refGene_tx=XM_023258815.1;refGene_gn=PAM;refGene_tx=XM_023258817.1;refGene_gn=PAM;refGene_tx=XM_023258823.1;refGene_gn=PAM;refGene_tx=XM_023258824.1;refGene_gn=PAM;refGene_tx=XM_023258826.1;refGene_gn=PAM;refGene_tx=XM_023258827.1;refGene_gn=PAM;refGene_tx=XM_023258830.1;refGene_gn=PAM;refGene_tx=XM_023258836.1
A1 166011736 166011940

refGene_gn=PAM;refGene_tx=XM_023258790.1;refGene_gn=PAM;refGene_tx=XM_023258791.1;refGene_gn=PAM;refGene_tx=XM_023258793.1;refGene_gn=PAM;refGene_tx=XM_023258797.1;refGene_gn=PAM;refGene_tx=XM_023258807.1;refGene_gn=PAM;refGene_tx=XM_023258812.1;refGene_gn=PAM;refGene_tx=XM_023258813.1;refGene_gn=PAM;refGene_tx=XM_023258817.1;refGene_gn=PAM;refGene_tx=XM_023258823.1;refGene_gn=PAM;refGene_tx=XM_023258824.1;refGene_gn=PAM;refGene_tx=XM_023258830.1
A1 166014851 166014908
refGene_gn=PAM;refGene_tx=XM_023258790.1;refGene_gn=PAM;refGene_tx=XM_023258791.1;refGene_gn=PAM;refGene_tx=XM_023258793.1;refGene_gn=PAM;refGene_tx=XM_023258797.1;refGene_gn=PAM;refGene_tx=XM_023258812.1;refGene_gn=PAM;refGene_tx=XM_023258813.1;refGene_gn=PAM;refGene_tx=XM_023258814.1;refGene_gn=PAM;refGene_tx=XM_023258815.1;refGene_gn=PAM;refGene_tx=XM_023258823.1;refGene_gn=PAM;refGene_tx=XM_023258824.1;refGene_gn=PAM;refGene_tx=XM_023258826.1;refGene_gn=PAM;refGene_tx=XM_023258827.1;refGene_gn=PAM;refGene_tx=XM_023258836.1
A1 166015565 166015747
refGene_gn=PAM;refGene_tx=XM_023258790.1;refGene_gn=PAM;refGene_tx=XM_023258791.1;refGene_gn=PAM;refGene_tx=XM_023258793.1;refGene_gn=PAM;refGene_tx=XM_023258797.1;refGene_gn=PAM;refGene_tx=XM_023258807.1;refGene_gn=PAM;refGene_tx=XM_023258814.1;refGene_gn=PAM;refGene_tx=XM_023258815.1;refGene_gn=PAM;refGene_tx=XM_023258823.1;refGene_gn=PAM;refGene_tx=XM_023258824.1;refGene_gn=PAM;refGene_tx=XM_023258830.1;refGene_gn=PAM;refGene_tx=XM_023258836.1
A1 166026241 166026258
refGene_gn=PAM;refGene_tx=XM_023258812.1;refGene_gn=PAM;refGene_tx=XM_023258813.1;refGene_gn=PAM;refGene_tx=XM_023258817.1;refGene_gn=PAM;refGene_tx=XM_023258826.1;refGene_gn=PAM;refGene_tx=XM_023258827.1
A1 166032137 166032214
refGene_gn=GIN1;refGene_tx=XM_011284816.3;refGene_gn=GIN1;refGene_tx=XM_019837573.2;refGene_gn=GIN1;refGene_tx=XM_019837576.2;refGene_gn=GIN1;refGene_tx=XM_019837580.2
A1 166035493 166035768
refGene_gn=GIN1;refGene_tx=XM_003981167.5;refGene_gn=GIN1;refGene_tx=XM_011284816.3;refGene_gn=GIN1;refGene_tx=XM_019837573.2;refGene_gn=GIN1;refGene_tx=XM_019837576.2;refGene_gn=GIN1;refGene_tx=XM_019837580.2
A1 166042352 166042638
refGene_gn=GIN1;refGene_tx=XM_003981167.5;refGene_gn=GIN1;refGene_tx=XM_011284816.3;refGene_gn=GIN1;refGene_tx=XM_019837573.2;refGene_gn=GIN1;refGene_tx=XM_019837576.2;refGene_gn=GIN1;refGene_tx=XM_019837580.2
A1 166043365 166043548
refGene_gn=GIN1;refGene_tx=XM_003981167.5;refGene_gn=GIN1;refGene_tx=XM_011284816.3;refGene_gn=GIN1;refGene_tx=XM_019837573.2;refGene_gn=GIN1;refGene_tx=XM_019837576.2;refGene_gn=GIN1;refGene_tx=XM_019837580.2
A1 166043650 166043842
refGene_gn=GIN1;refGene_tx=XM_003981167.5;refGene_gn=GIN1;refGene_tx=XM_011284816.3;refGene_gn=GIN1;refGene_tx=XM_019837573.2;refGene_gn=GIN1;refGene_tx=XM_019837576.2;refGene_gn=GIN1;refGene_tx=XM_019837580.2
A1 166046834 166047140
refGene_gn=GIN1;refGene_tx=XM_003981167.5;refGene_gn=GIN1;refGene_tx=X

M_011284816.3;refGene_gn=GIN1;refGene_tx=XM_019837573.2;refGene_gn=GIN1;refGene_tx=XM_019837576.2;refGene_gn=GIN1;refGene_tx=XM_019837580.2
A1 166048922 166049116
refGene_gn=GIN1;refGene_tx=XM_003981167.5;refGene_gn=GIN1;refGene_tx=XM_011284816.3;refGene_gn=GIN1;refGene_tx=XM_019837573.2;refGene_gn=GIN1;refGene_tx=XM_019837576.2;refGene_gn=GIN1;refGene_tx=XM_019837580.2
A1 166050693 166050835
refGene_gn=GIN1;refGene_tx=XM_003981167.5;refGene_gn=GIN1;refGene_tx=XM_011284816.3;refGene_gn=GIN1;refGene_tx=XM_019837573.2;refGene_gn=GIN1;refGene_tx=XM_019837576.2;refGene_gn=GIN1;refGene_tx=XM_019837580.2
A1 166063483 166063597
refGene_gn=PPIP5K2;refGene_tx=XM_011284879.3
A1 166068078 166068274
refGene_gn=PPIP5K2;refGene_tx=XM_011284879.3
A1 166070786 166070877
refGene_gn=PPIP5K2;refGene_tx=XM_011284879.3
A1 166072077 166072163
refGene_gn=PPIP5K2;refGene_tx=XM_011284879.3
A1 166073243 166073398
refGene_gn=PPIP5K2;refGene_tx=XM_011284879.3
A1 166075673 166075775
refGene_gn=PPIP5K2;refGene_tx=XM_011284879.3
A1 166076907 166077069
refGene_gn=PPIP5K2;refGene_tx=XM_011284879.3
A1 166078621 166078743
refGene_gn=PPIP5K2;refGene_tx=XM_011284879.3
A1 166080552 166080654
refGene_gn=PPIP5K2;refGene_tx=XM_011284879.3
A1 166081787 166081874
refGene_gn=PPIP5K2;refGene_tx=XM_011284879.3
A1 166082611 166082687
refGene_gn=PPIP5K2;refGene_tx=XM_011284879.3
A1 166082768 166082878
refGene_gn=PPIP5K2;refGene_tx=XM_011284879.3
A1 166083592 166083678
refGene_gn=PPIP5K2;refGene_tx=XM_011284879.3
A1 166086542 166086577
refGene_gn=PPIP5K2;refGene_tx=XM_011284879.3
A1 166086578 166086669
refGene_gn=PPIP5K2;refGene_tx=XM_011284879.3
A1 166086811 166086933
refGene_gn=PPIP5K2;refGene_tx=XM_011284879.3
A1 166087206 166087389
refGene_gn=PPIP5K2;refGene_tx=XM_011284879.3
A1 166091994 166092136
refGene_gn=PPIP5K2;refGene_tx=XM_011284879.3
A1 166092904 166093128
refGene_gn=PPIP5K2;refGene_tx=XM_011284879.3
A1 166098467 166098595
refGene_gn=PPIP5K2;refGene_tx=XM_011284879.3

A1 166099172 166099287
refGene_gn=PPIP5K2;refGene_tx=XM_011284879.3
A1 166104894 166105002
refGene_gn=PPIP5K2;refGene_tx=XM_011284879.3
A1 166105093 166105210
refGene_gn=PPIP5K2;refGene_tx=XM_011284879.3
A1 166108616 166108784
refGene_gn=PPIP5K2;refGene_tx=XM_011284879.3
A1 166111139 166111313
refGene_gn=PPIP5K2;refGene_tx=XM_011284879.3
A1 166112597 166112670
refGene_gn=PPIP5K2;refGene_tx=XM_011284879.3
A1 166114378 166114498
refGene_gn=PPIP5K2;refGene_tx=XM_011284879.3
A1 166115377 166115440
refGene_gn=PPIP5K2;refGene_tx=XM_011284879.3
A1 166117198 166117258
refGene_gn=PPIP5K2;refGene_tx=XM_011284879.3
A1 166118875 166119016
refGene_gn=PPIP5K2;refGene_tx=XM_011284879.3
A1 166122562 166122688
refGene_gn=PPIP5K2;refGene_tx=XM_011284879.3
A1 166129010 166129123
refGene_gn=PPIP5K2;refGene_tx=XM_011284879.3
A1 166189474 166190098
refGene_gn=CA1H5orf30;refGene_tx=XM_006927862.4;refGene_gn=CA1H5orf30;
refGene_tx=XM_019837694.2;refGene_gn=CA1H5orf30;refGene_tx=XM_01983769
8.2;refGene_gn=CA1H5orf30;refGene_tx=XM_019837708.2;refGene_gn=CA1H5or
f30;refGene_tx=XM_023258847.1;refGene_gn=CA1H5orf30;refGene_tx=XM_0232
58851.1;refGene_gn=CA1H5orf30;refGene_tx=XM_023258857.1
A1 166341033 166341133
refGene_gn=LOC109502572;refGene_tx=XM_019837711.2;refGene_gn=LOC109502
572;refGene_tx=XM_019837714.2
A1 166343075 166343211
refGene_gn=LOC109502572;refGene_tx=XM_019837711.2;refGene_gn=LOC109502
572;refGene_tx=XM_019837714.2
A1 166351906 166352093
refGene_gn=LOC109502572;refGene_tx=XM_019837711.2;refGene_gn=LOC109502
572;refGene_tx=XM_019837714.2
A1 166353788 166353831
refGene_gn=LOC109502572;refGene_tx=XM_019837711.2;refGene_gn=LOC109502
572;refGene_tx=XM_019837714.2
A1 166355686 166355844
refGene_gn=LOC109502572;refGene_tx=XM_019837711.2;refGene_gn=LOC109502
572;refGene_tx=XM_019837714.2
A1 166401460 166401639
refGene_gn=LOC109502572;refGene_tx=XM_019837711.2;refGene_gn=LOC109502
572;refGene_tx=XM_019837714.2
A1 166410426 166410537
refGene_gn=LOC109502572;refGene_tx=XM_019837711.2;refGene_gn=LOC109502

572;refGene_tx=XM_019837714.2
A1 166423443 166423545
refGene_gn=LOC109502572;refGene_tx=XM_019837711.2;refGene_gn=LOC109502
572;refGene_tx=XM_019837714.2
A1 166429538 166429706
refGene_gn=LOC109502572;refGene_tx=XM_019837711.2;refGene_gn=LOC109502
572;refGene_tx=XM_019837714.2
A1 166439886 166440078
refGene_gn=LOC109502572;refGene_tx=XM_019837711.2;refGene_gn=LOC109502
572;refGene_tx=XM_019837714.2
A1 166447143 166449483
refGene_gn=LOC109502572;refGene_tx=XM_019837711.2;refGene_gn=LOC109502
572;refGene_tx=XM_019837714.2
A1 166451980 166452072
refGene_gn=LOC109502572;refGene_tx=XM_019837711.2;refGene_gn=LOC109502
572;refGene_tx=XM_019837714.2
A1 166464687 166464704
refGene_gn=LOC109502572;refGene_tx=XM_019837711.2
A1 166471972 166472083
refGene_gn=NUDT12;refGene_tx=XM_011284887.2;refGene_gn=NUDT12;refGene_
tx=XM_011284889.2
A1 166473060 166473260
refGene_gn=NUDT12;refGene_tx=XM_011284887.2;refGene_gn=NUDT12;refGene_
tx=XM_011284889.2
A1 166475928 166476042
refGene_gn=NUDT12;refGene_tx=XM_011284887.2;refGene_gn=NUDT12;refGene_
tx=XM_011284889.2
A1 166477093 166477261
refGene_gn=NUDT12;refGene_tx=XM_011284887.2;refGene_gn=NUDT12;refGene_
tx=XM_011284889.2
A1 166480574 166481161
refGene_gn=NUDT12;refGene_tx=XM_011284887.2;refGene_gn=NUDT12;refGene_
tx=XM_011284889.2
A1 166481722 166481928
refGene_gn=NUDT12;refGene_tx=XM_011284887.2;refGene_gn=NUDT12;refGene_
tx=XM_011284889.2
A1 168383564 168383780
refGene_gn=LOC102901462;refGene_tx=XM_006927863.4
A1 168383873 168384221
refGene_gn=LOC101085346;refGene_tx=XM_023258871.1
A1 169923909 169924031
refGene_gn=EFNA5;refGene_tx=XM_003981172.5;refGene_gn=EFNA5;refGene_tx
=XM_006927864.4
A1 169929214 169929295
refGene_gn=EFNA5;refGene_tx=XM_003981172.5
A1 169929712 169929778
refGene_gn=EFNA5;refGene_tx=XM_003981172.5;refGene_gn=EFNA5;refGene_tx
=XM_006927864.4
A1 169967286 169967579
refGene_gn=EFNA5;refGene_tx=XM_003981172.5;refGene_gn=EFNA5;refGene_tx

=XM_006927864.4
A1 170193400 170193525
refGene_gn=EFNA5;refGene_tx=XM_003981172.5;refGene_gn=EFNA5;refGene_tx=XM_006927864.4
A1 170375546 170375687
refGene_gn=FBXL17;refGene_tx=XM_003981190.4;refGene_gn=FBXL17;refGene_tx=XM_023258890.1
A1 170394652 170394795
refGene_gn=FBXL17;refGene_tx=XM_003981190.4;refGene_gn=FBXL17;refGene_tx=XM_023258890.1
A1 170536847 170536924
refGene_gn=FBXL17;refGene_tx=XM_003981190.4;refGene_gn=FBXL17;refGene_tx=XM_023258890.1
A1 170689116 170689247
refGene_gn=FBXL17;refGene_tx=XM_003981190.4;refGene_gn=FBXL17;refGene_tx=XM_023258890.1
A1 170714605 170714713
refGene_gn=FBXL17;refGene_tx=XM_003981190.4
A1 170823971 170824103
refGene_gn=FBXL17;refGene_tx=XM_003981190.4;refGene_gn=FBXL17;refGene_tx=XM_023258890.1
A1 170844402 170844660
refGene_gn=FBXL17;refGene_tx=XM_003981190.4;refGene_gn=FBXL17;refGene_tx=XM_023258890.1
A1 170848921 170849044
refGene_gn=FBXL17;refGene_tx=XM_003981190.4;refGene_gn=FBXL17;refGene_tx=XM_023258890.1
A1 170868209 170869196
refGene_gn=FBXL17;refGene_tx=XM_003981190.4;refGene_gn=FBXL17;refGene_tx=XM_023258890.1
A1 171222824 171223031
refGene_gn=FER;refGene_tx=XM_023258902.1
A1 171242848 171242965
refGene_gn=FER;refGene_tx=XM_023258902.1
A1 171242966 171243023
refGene_gn=FER;refGene_tx=XM_023258902.1
A1 171248421 171248521
refGene_gn=FER;refGene_tx=XM_023258902.1
A1 171268001 171268185
refGene_gn=FER;refGene_tx=XM_023258902.1
A1 171271332 171271470
refGene_gn=FER;refGene_tx=XM_023258902.1
A1 171272268 171272388
refGene_gn=FER;refGene_tx=XM_023258902.1
A1 171288176 171288299
refGene_gn=FER;refGene_tx=XM_023258902.1
A1 171302077 171302267
refGene_gn=FER;refGene_tx=XM_023258902.1
A1 171367408 171367501
refGene_gn=FER;refGene_tx=XM_023258902.1

A1 171378346 171378550
refGene_gn=FER;refGene_tx=XM_023258902.1
A1 171387861 171387984
refGene_gn=FER;refGene_tx=XM_023258902.1
A1 171452512 171452569
refGene_gn=FER;refGene_tx=XM_023258902.1
A1 171456683 171456799
refGene_gn=FER;refGene_tx=XM_023258902.1
A1 171459345 171459440
refGene_gn=FER;refGene_tx=XM_023258902.1
A1 171501675 171501799
refGene_gn=FER;refGene_tx=XM_023258902.1
A1 171589012 171589167
refGene_gn=FER;refGene_tx=XM_023258902.1
A1 171607712 171607835
refGene_gn=FER;refGene_tx=XM_023258902.1
A1 171609227 171609278
refGene_gn=FER;refGene_tx=XM_023258902.1
A1 171609280 171609372
refGene_gn=FER;refGene_tx=XM_023258902.1
A1 171726495 171726621
refGene_gn=PJA2;refGene_tx=XM_003981173.5
A1 171728443 171728565
refGene_gn=PJA2;refGene_tx=XM_003981173.5
A1 171728947 171729062
refGene_gn=PJA2;refGene_tx=XM_003981173.5
A1 171742746 171742858
refGene_gn=PJA2;refGene_tx=XM_003981173.5
A1 171748084 171748270
refGene_gn=PJA2;refGene_tx=XM_003981173.5
A1 171759606 171759792
refGene_gn=PJA2;refGene_tx=XM_003981173.5
A1 171769058 171770112
refGene_gn=PJA2;refGene_tx=XM_003981173.5
A1 171771714 171771915
refGene_gn=PJA2;refGene_tx=XM_003981173.5
A1 171773879 171773910
refGene_gn=PJA2;refGene_tx=XM_003981173.5
A1 172026300 172026435
refGene_gn=MAN2A1;refGene_tx=XM_003981174.5
A1 172050982 172051240
refGene_gn=MAN2A1;refGene_tx=XM_003981174.5
A1 172053580 172053725
refGene_gn=MAN2A1;refGene_tx=XM_003981174.5
A1 172066759 172066931
refGene_gn=MAN2A1;refGene_tx=XM_003981174.5
A1 172086162 172086290
refGene_gn=MAN2A1;refGene_tx=XM_003981174.5
A1 172090393 172090567
refGene_gn=MAN2A1;refGene_tx=XM_003981174.5

A1 172093521 172093708
refGene_gn=MAN2A1;refGene_tx=XM_003981174.5
A1 172098324 172098502
refGene_gn=MAN2A1;refGene_tx=XM_003981174.5
A1 172104490 172104693
refGene_gn=MAN2A1;refGene_tx=XM_003981174.5
A1 172107477 172107660
refGene_gn=MAN2A1;refGene_tx=XM_003981174.5
A1 172112311 172112426
refGene_gn=MAN2A1;refGene_tx=XM_003981174.5
A1 172113716 172113784
refGene_gn=MAN2A1;refGene_tx=XM_003981174.5
A1 172141882 172142048
refGene_gn=MAN2A1;refGene_tx=XM_003981174.5
A1 172143807 172144026
refGene_gn=MAN2A1;refGene_tx=XM_003981174.5
A1 172144777 172144900
refGene_gn=MAN2A1;refGene_tx=XM_003981174.5
A1 172148107 172148222
refGene_gn=MAN2A1;refGene_tx=XM_003981174.5
A1 172163156 172163290
refGene_gn=MAN2A1;refGene_tx=XM_003981174.5
A1 172166334 172166476
refGene_gn=MAN2A1;refGene_tx=XM_003981174.5
A1 172168968 172169102
refGene_gn=MAN2A1;refGene_tx=XM_003981174.5
A1 172175495 172175690
refGene_gn=MAN2A1;refGene_tx=XM_003981174.5
A1 172188067 172188178
refGene_gn=MAN2A1;refGene_tx=XM_003981174.5
A1 172189773 172189926
refGene_gn=MAN2A1;refGene_tx=XM_003981174.5
A1 172624690 172624882
refGene_gn=TMEM232;refGene_tx=XM_023258913.1;refGene_gn=TMEM232;refGene_tx=XM_023258918.1;refGene_gn=TMEM232;refGene_tx=XM_023258921.1;refGene_gn=TMEM232;refGene_tx=XM_023258928.1
A1 172628177 172628271
refGene_gn=TMEM232;refGene_tx=XM_023258913.1;refGene_gn=TMEM232;refGene_tx=XM_023258918.1;refGene_gn=TMEM232;refGene_tx=XM_023258921.1;refGene_gn=TMEM232;refGene_tx=XM_023258928.1
A1 172731944 172732189
refGene_gn=TMEM232;refGene_tx=XM_023258913.1;refGene_gn=TMEM232;refGene_tx=XM_023258918.1;refGene_gn=TMEM232;refGene_tx=XM_023258921.1;refGene_gn=TMEM232;refGene_tx=XM_023258928.1
A1 172747297 172747476
refGene_gn=TMEM232;refGene_tx=XM_023258913.1;refGene_gn=TMEM232;refGene_tx=XM_023258918.1;refGene_gn=TMEM232;refGene_tx=XM_023258921.1;refGene_gn=TMEM232;refGene_tx=XM_023258928.1
A1 172770132 172770385
refGene_gn=TMEM232;refGene_tx=XM_023258913.1;refGene_gn=TMEM232;refGen

e_tx=XM_023258918.1;refGene_gn=TMEM232;refGene_tx=XM_023258921.1;refGene_gn=TMEM232;refGene_tx=XM_023258928.1
A1 172771893 172772017
refGene_gn=TMEM232;refGene_tx=XM_023258913.1;refGene_gn=TMEM232;refGene_tx=XM_023258918.1;refGene_gn=TMEM232;refGene_tx=XM_023258921.1;refGene_gn=TMEM232;refGene_tx=XM_023258928.1
A1 172783676 172783831
refGene_gn=TMEM232;refGene_tx=XM_023258913.1;refGene_gn=TMEM232;refGene_tx=XM_023258918.1;refGene_gn=TMEM232;refGene_tx=XM_023258921.1;refGene_gn=TMEM232;refGene_tx=XM_023258928.1
A1 172791359 172791526
refGene_gn=TMEM232;refGene_tx=XM_023258913.1;refGene_gn=TMEM232;refGene_tx=XM_023258918.1;refGene_gn=TMEM232;refGene_tx=XM_023258921.1;refGene_gn=TMEM232;refGene_tx=XM_023258928.1
A1 172795757 172795857
refGene_gn=TMEM232;refGene_tx=XM_023258913.1;refGene_gn=TMEM232;refGene_tx=XM_023258918.1;refGene_gn=TMEM232;refGene_tx=XM_023258921.1;refGene_gn=TMEM232;refGene_tx=XM_023258928.1
A1 172799983 172800141
refGene_gn=TMEM232;refGene_tx=XM_023258913.1;refGene_gn=TMEM232;refGene_tx=XM_023258918.1;refGene_gn=TMEM232;refGene_tx=XM_023258921.1;refGene_gn=TMEM232;refGene_tx=XM_023258928.1
A1 172802454 172802560
refGene_gn=TMEM232;refGene_tx=XM_023258913.1;refGene_gn=TMEM232;refGene_tx=XM_023258918.1;refGene_gn=TMEM232;refGene_tx=XM_023258921.1
A1 172803898 172804010
refGene_gn=TMEM232;refGene_tx=XM_023258913.1;refGene_gn=TMEM232;refGene_tx=XM_023258918.1;refGene_gn=TMEM232;refGene_tx=XM_023258921.1
A1 172815284 172815403
refGene_gn=TMEM232;refGene_tx=XM_023258913.1;refGene_gn=TMEM232;refGene_tx=XM_023258918.1;refGene_gn=TMEM232;refGene_tx=XM_023258921.1
A1 172839650 172839669
refGene_gn=SLC25A46;refGene_tx=XM_011284939.3
A1 172840544 172840830
refGene_gn=SLC25A46;refGene_tx=XM_011284932.3
A1 172843418 172843461
refGene_gn=SLC25A46;refGene_tx=XM_011284932.3;refGene_gn=SLC25A46;refGene_tx=XM_011284939.3
A1 172844754 172844812
refGene_gn=SLC25A46;refGene_tx=XM_011284932.3;refGene_gn=SLC25A46;refGene_tx=XM_011284939.3
A1 172847662 172847740
refGene_gn=SLC25A46;refGene_tx=XM_011284932.3;refGene_gn=SLC25A46;refGene_tx=XM_011284939.3
A1 172848194 172848295
refGene_gn=SLC25A46;refGene_tx=XM_011284932.3;refGene_gn=SLC25A46;refGene_tx=XM_011284939.3
A1 172854086 172854143
refGene_gn=SLC25A46;refGene_tx=XM_011284932.3;refGene_gn=SLC25A46;refGene_tx=XM_011284939.3

A1 172855387 172855445
refGene_gn=SLC25A46;refGene_tx=XM_011284932.3;refGene_gn=SLC25A46;refGene_tx=XM_011284939.3
A1 172865574 172866153
refGene_gn=SLC25A46;refGene_tx=XM_011284932.3;refGene_gn=SLC25A46;refGene_tx=XM_011284939.3
A1 173082228 173082249
refGene_gn=TSLP;refGene_tx=XM_003981241.4
A1 173082634 173082868
refGene_gn=TSLP;refGene_tx=XM_003981241.4
A1 173083748 173083793
refGene_gn=TSLP;refGene_tx=XM_003981241.4
A1 173084651 173084786
refGene_gn=TSLP;refGene_tx=XM_003981241.4
A1 173086716 173086833
refGene_gn=TSLP;refGene_tx=XM_003981241.4
A1 173098362 173098385
refGene_gn=WDR36;refGene_tx=XM_023244671.1
A1 173098750 173098989
refGene_gn=WDR36;refGene_tx=XM_023244671.1
A1 173099063 173099207
refGene_gn=WDR36;refGene_tx=XM_023244671.1
A1 173099209 173099328
refGene_gn=WDR36;refGene_tx=XM_023244671.1
A1 173101804 173101832
refGene_gn=WDR36;refGene_tx=XM_023244671.1
A1 173103928 173104029
refGene_gn=WDR36;refGene_tx=XM_023244671.1
A1 173108815 173108933
refGene_gn=WDR36;refGene_tx=XM_023244671.1
A1 173109915 173110048
refGene_gn=WDR36;refGene_tx=XM_023244671.1
A1 173111530 173111585
refGene_gn=WDR36;refGene_tx=XM_023244671.1
A1 173113019 173113152
refGene_gn=WDR36;refGene_tx=XM_023244671.1
A1 173113435 173113611
refGene_gn=WDR36;refGene_tx=XM_023244671.1
A1 173113920 173114041
refGene_gn=WDR36;refGene_tx=XM_023244671.1
A1 173114518 173114584
refGene_gn=WDR36;refGene_tx=XM_023244671.1
A1 173115869 173115956
refGene_gn=WDR36;refGene_tx=XM_023244671.1
A1 173117248 173117394
refGene_gn=WDR36;refGene_tx=XM_023244671.1
A1 173121910 173122025
refGene_gn=WDR36;refGene_tx=XM_023244671.1
A1 173122508 173122674
refGene_gn=WDR36;refGene_tx=XM_023244671.1

A1 173122913 173123022
refGene_gn=WDR36;refGene_tx=XM_023244671.1
A1 173125212 173125292
refGene_gn=WDR36;refGene_tx=XM_023244671.1
A1 173130892 173131000
refGene_gn=WDR36;refGene_tx=XM_023244671.1
A1 173133070 173133168
refGene_gn=WDR36;refGene_tx=XM_023244671.1
A1 173133564 173133710
refGene_gn=WDR36;refGene_tx=XM_023244671.1
A1 173136251 173136371
refGene_gn=WDR36;refGene_tx=XM_023244671.1
A1 173136547 173136629
refGene_gn=WDR36;refGene_tx=XM_023244671.1
A1 173138274 173138462
refGene_gn=WDR36;refGene_tx=XM_023244671.1
A1 173139608 173139758
refGene_gn=WDR36;refGene_tx=XM_023244671.1
A1 173241621 173241782
refGene_gn=CAMK4;refGene_tx=XM_003981193.5
A1 173344759 173344838
refGene_gn=CAMK4;refGene_tx=XM_003981193.5
A1 173371390 173371453
refGene_gn=CAMK4;refGene_tx=XM_003981193.5
A1 173373607 173373690
refGene_gn=CAMK4;refGene_tx=XM_003981193.5
A1 173395848 173395921
refGene_gn=CAMK4;refGene_tx=XM_003981193.5
A1 173429606 173429697
refGene_gn=CAMK4;refGene_tx=XM_003981193.5
A1 173431786 173431861
refGene_gn=CAMK4;refGene_tx=XM_003981193.5;refGene_gn=CAMK4;refGene_tx
=XM_019837934.2
A1 173445064 173445140
refGene_gn=CAMK4;refGene_tx=XM_003981193.5;refGene_gn=CAMK4;refGene_tx
=XM_019837934.2
A1 173451677 173451804
refGene_gn=CAMK4;refGene_tx=XM_003981193.5;refGene_gn=CAMK4;refGene_tx
=XM_019837934.2
A1 173456424 173456577
refGene_gn=CAMK4;refGene_tx=XM_003981193.5;refGene_gn=CAMK4;refGene_tx
=XM_019837934.2
A1 173458192 173458633
refGene_gn=CAMK4;refGene_tx=XM_003981193.5;refGene_gn=CAMK4;refGene_tx
=XM_019837934.2
A1 173476039 173476269
refGene_gn=STARD4;refGene_tx=XM_003981243.5;refGene_gn=STARD4;refGene_
tx=XM_006927876.4;refGene_gn=STARD4;refGene_tx=XM_019837950.2
A1 173477171 173477286
refGene_gn=STARD4;refGene_tx=XM_003981243.5;refGene_gn=STARD4;refGene_

tx=XM_006927876.4;refGene_gn=STARD4;refGene_tx=XM_019837950.2
A1 173478121 173478248
refGene_gn=STARD4;refGene_tx=XM_003981243.5;refGene_gn=STARD4;refGene_
tx=XM_006927876.4;refGene_gn=STARD4;refGene_tx=XM_019837950.2
A1 173482587 173482637
refGene_gn=STARD4;refGene_tx=XM_003981243.5;refGene_gn=STARD4;refGene_
tx=XM_006927876.4;refGene_gn=STARD4;refGene_tx=XM_019837950.2
A1 173483836 173483941
refGene_gn=STARD4;refGene_tx=XM_003981243.5;refGene_gn=STARD4;refGene_
tx=XM_006927876.4;refGene_gn=STARD4;refGene_tx=XM_019837950.2
A1 173657213 173657339
refGene_gn=NREP;refGene_tx=XM_011284960.3
A1 173661601 173661679
refGene_gn=NREP;refGene_tx=XM_011284960.3
A1 173682139 173682142
refGene_gn=NREP;refGene_tx=XM_011284960.3
A1 174016423 174016558
refGene_gn=EPB41L4A;refGene_tx=XM_019838056.2;refGene_gn=EPB41L4A;refG
ene_tx=XM_019838060.2;refGene_gn=EPB41L4A;refGene_tx=XM_019838062.2;re
fGene_gn=EPB41L4A;refGene_tx=XM_019838064.2
A1 174019661 174019743
refGene_gn=EPB41L4A;refGene_tx=XM_019838056.2;refGene_gn=EPB41L4A;refG
ene_tx=XM_019838060.2;refGene_gn=EPB41L4A;refGene_tx=XM_019838062.2
A1 174020773 174020884
refGene_gn=EPB41L4A;refGene_tx=XM_019838056.2;refGene_gn=EPB41L4A;refG
ene_tx=XM_019838060.2;refGene_gn=EPB41L4A;refGene_tx=XM_019838062.2;re
fGene_gn=EPB41L4A;refGene_tx=XM_019838064.2
A1 174022135 174022204
refGene_gn=EPB41L4A;refGene_tx=XM_019838056.2;refGene_gn=EPB41L4A;refG
ene_tx=XM_019838060.2;refGene_gn=EPB41L4A;refGene_tx=XM_019838062.2;re
fGene_gn=EPB41L4A;refGene_tx=XM_019838064.2
A1 174022743 174022791
refGene_gn=EPB41L4A;refGene_tx=XM_019838056.2;refGene_gn=EPB41L4A;refG
ene_tx=XM_019838060.2;refGene_gn=EPB41L4A;refGene_tx=XM_019838062.2;re
fGene_gn=EPB41L4A;refGene_tx=XM_019838064.2
A1 174037174 174037235
refGene_gn=EPB41L4A;refGene_tx=XM_019838070.2
A1 174038406 174038526
refGene_gn=EPB41L4A;refGene_tx=XM_019838056.2;refGene_gn=EPB41L4A;refG
ene_tx=XM_019838060.2;refGene_gn=EPB41L4A;refGene_tx=XM_019838062.2;re
fGene_gn=EPB41L4A;refGene_tx=XM_019838064.2;refGene_gn=EPB41L4A;refGen
e_tx=XM_019838070.2
A1 174050828 174050906
refGene_gn=EPB41L4A;refGene_tx=XM_019838056.2;refGene_gn=EPB41L4A;refG
ene_tx=XM_019838060.2;refGene_gn=EPB41L4A;refGene_tx=XM_019838064.2;re
fGene_gn=EPB41L4A;refGene_tx=XM_019838070.2
A1 174051907 174051955
refGene_gn=EPB41L4A;refGene_tx=XM_019838056.2;refGene_gn=EPB41L4A;refG
ene_tx=XM_019838064.2;refGene_gn=EPB41L4A;refGene_tx=XM_019838070.2
A1 174059821 174059932

refGene_gn=EPB41L4A;refGene_tx=XM_019838056.2;refGene_gn=EPB41L4A;refGene_tx=XM_019838060.2;refGene_gn=EPB41L4A;refGene_tx=XM_019838062.2;refGene_gn=EPB41L4A;refGene_tx=XM_019838064.2;refGene_gn=EPB41L4A;refGene_tx=XM_019838070.2
A1 174060687 174060771
refGene_gn=EPB41L4A;refGene_tx=XM_019838056.2;refGene_gn=EPB41L4A;refGene_tx=XM_019838060.2;refGene_gn=EPB41L4A;refGene_tx=XM_019838062.2;refGene_gn=EPB41L4A;refGene_tx=XM_019838064.2;refGene_gn=EPB41L4A;refGene_tx=XM_019838070.2
A1 174063914 174064005
refGene_gn=EPB41L4A;refGene_tx=XM_019838056.2;refGene_gn=EPB41L4A;refGene_tx=XM_019838060.2;refGene_gn=EPB41L4A;refGene_tx=XM_019838062.2;refGene_gn=EPB41L4A;refGene_tx=XM_019838064.2;refGene_gn=EPB41L4A;refGene_tx=XM_019838070.2
A1 174084880 174085002
refGene_gn=EPB41L4A;refGene_tx=XM_019838056.2;refGene_gn=EPB41L4A;refGene_tx=XM_019838060.2;refGene_gn=EPB41L4A;refGene_tx=XM_019838062.2;refGene_gn=EPB41L4A;refGene_tx=XM_019838064.2;refGene_gn=EPB41L4A;refGene_tx=XM_019838070.2
A1 174090400 174090478
refGene_gn=EPB41L4A;refGene_tx=XM_019838056.2;refGene_gn=EPB41L4A;refGene_tx=XM_019838060.2;refGene_gn=EPB41L4A;refGene_tx=XM_019838062.2;refGene_gn=EPB41L4A;refGene_tx=XM_019838064.2;refGene_gn=EPB41L4A;refGene_tx=XM_019838070.2
A1 174091878 174091970
refGene_gn=EPB41L4A;refGene_tx=XM_019838056.2;refGene_gn=EPB41L4A;refGene_tx=XM_019838060.2;refGene_gn=EPB41L4A;refGene_tx=XM_019838062.2;refGene_gn=EPB41L4A;refGene_tx=XM_019838064.2;refGene_gn=EPB41L4A;refGene_tx=XM_019838070.2
A1 174107942 174108006
refGene_gn=EPB41L4A;refGene_tx=XM_019838056.2;refGene_gn=EPB41L4A;refGene_tx=XM_019838060.2;refGene_gn=EPB41L4A;refGene_tx=XM_019838062.2;refGene_gn=EPB41L4A;refGene_tx=XM_019838064.2;refGene_gn=EPB41L4A;refGene_tx=XM_019838070.2
A1 174108608 174108697
refGene_gn=EPB41L4A;refGene_tx=XM_019838056.2;refGene_gn=EPB41L4A;refGene_tx=XM_019838060.2;refGene_gn=EPB41L4A;refGene_tx=XM_019838062.2;refGene_gn=EPB41L4A;refGene_tx=XM_019838064.2;refGene_gn=EPB41L4A;refGene_tx=XM_019838070.2
A1 174110924 174111012
refGene_gn=EPB41L4A;refGene_tx=XM_019838056.2;refGene_gn=EPB41L4A;refGene_tx=XM_019838060.2;refGene_gn=EPB41L4A;refGene_tx=XM_019838062.2;refGene_gn=EPB41L4A;refGene_tx=XM_019838064.2;refGene_gn=EPB41L4A;refGene_tx=XM_019838070.2
A1 174113472 174113593
refGene_gn=EPB41L4A;refGene_tx=XM_019838056.2;refGene_gn=EPB41L4A;refGene_tx=XM_019838060.2;refGene_gn=EPB41L4A;refGene_tx=XM_019838062.2;refGene_gn=EPB41L4A;refGene_tx=XM_019838064.2;refGene_gn=EPB41L4A;refGene_tx=XM_019838070.2
A1 174114744 174114842

refGene_gn=EPB41L4A;refGene_tx=XM_019838056.2;refGene_gn=EPB41L4A;refGene_tx=XM_019838060.2;refGene_gn=EPB41L4A;refGene_tx=XM_019838062.2;refGene_gn=EPB41L4A;refGene_tx=XM_019838064.2;refGene_gn=EPB41L4A;refGene_tx=XM_019838070.2
A1 174122888 174122967
refGene_gn=EPB41L4A;refGene_tx=XM_019838056.2;refGene_gn=EPB41L4A;refGene_tx=XM_019838060.2;refGene_gn=EPB41L4A;refGene_tx=XM_019838062.2;refGene_gn=EPB41L4A;refGene_tx=XM_019838064.2;refGene_gn=EPB41L4A;refGene_tx=XM_019838070.2
A1 174130335 174130387
refGene_gn=EPB41L4A;refGene_tx=XM_019838056.2;refGene_gn=EPB41L4A;refGene_tx=XM_019838060.2;refGene_gn=EPB41L4A;refGene_tx=XM_019838062.2;refGene_gn=EPB41L4A;refGene_tx=XM_019838064.2;refGene_gn=EPB41L4A;refGene_tx=XM_019838070.2
A1 174156168 174156273
refGene_gn=EPB41L4A;refGene_tx=XM_019838056.2;refGene_gn=EPB41L4A;refGene_tx=XM_019838060.2;refGene_gn=EPB41L4A;refGene_tx=XM_019838062.2;refGene_gn=EPB41L4A;refGene_tx=XM_019838064.2;refGene_gn=EPB41L4A;refGene_tx=XM_019838070.2
A1 174268268 174268367
refGene_gn=EPB41L4A;refGene_tx=XM_019838056.2;refGene_gn=EPB41L4A;refGene_tx=XM_019838060.2;refGene_gn=EPB41L4A;refGene_tx=XM_019838062.2;refGene_gn=EPB41L4A;refGene_tx=XM_019838064.2;refGene_gn=EPB41L4A;refGene_tx=XM_019838070.2
A1 174269160 174269949
refGene_gn=LOC111557869;refGene_tx=XM_023244716.1
A1 174275792 174275818
refGene_gn=LOC111557869;refGene_tx=XM_023244716.1
A1 174321139 174321278
refGene_gn=LOC111557869;refGene_tx=XM_023244716.1
A1 174325069 174325133
refGene_gn=LOC111557869;refGene_tx=XM_023244716.1
A1 174329111 174329189
refGene_gn=LOC111557869;refGene_tx=XM_023244716.1
A1 174338250 174338343
refGene_gn=LOC111557869;refGene_tx=XM_023244716.1
A1 174340146 174340202
refGene_gn=LOC111557869;refGene_tx=XM_023244716.1
A1 174340652 174340742
refGene_gn=LOC111557869;refGene_tx=XM_023244716.1
A1 174343488 174343544
refGene_gn=LOC111557869;refGene_tx=XM_023244716.1
A1 174348443 174348525
refGene_gn=LOC111557869;refGene_tx=XM_023244716.1
A1 174443123 174443291
refGene_gn=APC;refGene_tx=XM_011285189.3;refGene_gn=APC;refGene_tx=XM_019838005.2;refGene_gn=APC;refGene_tx=XM_019838027.2;refGene_gn=APC;refGene_tx=XM_019838035.2;refGene_gn=APC;refGene_tx=XM_019838047.2;refGene_gn=APC;refGene_tx=XM_023258973.1
A1 174497661 174497808

refGene_gn=APC;refGene_tx=XM_019838007.2
A1	174516983	174517118
refGene_gn=APC;refGene_tx=XM_006927878.4;refGene_gn=APC;refGene_tx=XM_019838018.2;refGene_gn=APC;refGene_tx=XM_019838020.2;refGene_gn=APC;refGene_tx=XM_019838040.2
A1	174526694	174526779
refGene_gn=APC;refGene_tx=XM_006927878.4;refGene_gn=APC;refGene_tx=XM_011285189.3;refGene_gn=APC;refGene_tx=XM_019838005.2;refGene_gn=APC;refGene_tx=XM_019838007.2;refGene_gn=APC;refGene_tx=XM_019838018.2;refGene_gn=APC;refGene_tx=XM_019838020.2;refGene_gn=APC;refGene_tx=XM_019838027.2;refGene_gn=APC;refGene_tx=XM_019838035.2;refGene_gn=APC;refGene_tx=XM_019838040.2;refGene_gn=APC;refGene_tx=XM_019838047.2;refGene_gn=APC;refGene_tx=XM_023258973.1
A1	174527778	174527980
refGene_gn=APC;refGene_tx=XM_006927878.4;refGene_gn=APC;refGene_tx=XM_011285189.3;refGene_gn=APC;refGene_tx=XM_019838005.2;refGene_gn=APC;refGene_tx=XM_019838007.2;refGene_gn=APC;refGene_tx=XM_019838018.2;refGene_gn=APC;refGene_tx=XM_019838020.2;refGene_gn=APC;refGene_tx=XM_019838027.2;refGene_gn=APC;refGene_tx=XM_019838035.2;refGene_gn=APC;refGene_tx=XM_019838040.2;refGene_gn=APC;refGene_tx=XM_019838047.2;refGene_gn=APC;refGene_tx=XM_023258973.1
A1	174534584	174534693
refGene_gn=APC;refGene_tx=XM_006927878.4;refGene_gn=APC;refGene_tx=XM_011285189.3;refGene_gn=APC;refGene_tx=XM_019838005.2;refGene_gn=APC;refGene_tx=XM_019838007.2;refGene_gn=APC;refGene_tx=XM_019838018.2;refGene_gn=APC;refGene_tx=XM_019838020.2;refGene_gn=APC;refGene_tx=XM_019838027.2;refGene_gn=APC;refGene_tx=XM_019838035.2;refGene_gn=APC;refGene_tx=XM_019838040.2;refGene_gn=APC;refGene_tx=XM_019838047.2;refGene_gn=APC;refGene_tx=XM_023258973.1
A1	174538139	174538253
refGene_gn=APC;refGene_tx=XM_006927878.4;refGene_gn=APC;refGene_tx=XM_011285189.3;refGene_gn=APC;refGene_tx=XM_019838005.2;refGene_gn=APC;refGene_tx=XM_019838007.2;refGene_gn=APC;refGene_tx=XM_019838018.2;refGene_gn=APC;refGene_tx=XM_019838020.2;refGene_gn=APC;refGene_tx=XM_019838027.2;refGene_gn=APC;refGene_tx=XM_019838035.2;refGene_gn=APC;refGene_tx=XM_019838040.2;refGene_gn=APC;refGene_tx=XM_019838047.2;refGene_gn=APC;refGene_tx=XM_023258973.1
A1	174552413	174552497
refGene_gn=APC;refGene_tx=XM_006927878.4;refGene_gn=APC;refGene_tx=XM_011285189.3;refGene_gn=APC;refGene_tx=XM_019838005.2;refGene_gn=APC;refGene_tx=XM_019838007.2;refGene_gn=APC;refGene_tx=XM_019838018.2;refGene_gn=APC;refGene_tx=XM_019838020.2;refGene_gn=APC;refGene_tx=XM_019838027.2;refGene_gn=APC;refGene_tx=XM_019838035.2;refGene_gn=APC;refGene_tx=XM_019838040.2;refGene_gn=APC;refGene_tx=XM_019838047.2;refGene_gn=APC;refGene_tx=XM_023258973.1
A1	174560837	174560942
refGene_gn=APC;refGene_tx=XM_006927878.4;refGene_gn=APC;refGene_tx=XM_011285189.3;refGene_gn=APC;refGene_tx=XM_019838005.2;refGene_gn=APC;refGene_tx=XM_019838007.2;refGene_gn=APC;refGene_tx=XM_019838018.2;refGene_gn=APC;refGene_tx=XM_019838020.2;refGene_gn=APC;refGene_tx=XM_01983

8035.2;refGene_gn=APC;refGene_tx=XM_019838040.2;refGene_gn=APC;refGene_tx=XM_023258973.1
A1 174568611 174568710
refGene_gn=APC;refGene_tx=XM_006927878.4;refGene_gn=APC;refGene_tx=XM_011285189.3;refGene_gn=APC;refGene_tx=XM_019838005.2;refGene_gn=APC;refGene_tx=XM_019838007.2;refGene_gn=APC;refGene_tx=XM_019838018.2;refGene_gn=APC;refGene_tx=XM_019838020.2;refGene_gn=APC;refGene_tx=XM_019838027.2;refGene_gn=APC;refGene_tx=XM_019838035.2;refGene_gn=APC;refGene_tx=XM_019838040.2;refGene_gn=APC;refGene_tx=XM_019838047.2;refGene_gn=APC;refGene_tx=XM_023258973.1
A1 174571952 174572331
refGene_gn=APC;refGene_tx=XM_006927878.4;refGene_gn=APC;refGene_tx=XM_011285189.3;refGene_gn=APC;refGene_tx=XM_019838005.2;refGene_gn=APC;refGene_tx=XM_019838007.2;refGene_gn=APC;refGene_tx=XM_019838018.2;refGene_gn=APC;refGene_tx=XM_019838020.2;refGene_gn=APC;refGene_tx=XM_019838027.2;refGene_gn=APC;refGene_tx=XM_019838035.2;refGene_gn=APC;refGene_tx=XM_019838040.2;refGene_gn=APC;refGene_tx=XM_019838047.2;refGene_gn=APC;refGene_tx=XM_023258973.1
A1 174574566 174574662
refGene_gn=APC;refGene_tx=XM_006927878.4;refGene_gn=APC;refGene_tx=XM_011285189.3;refGene_gn=APC;refGene_tx=XM_019838005.2;refGene_gn=APC;refGene_tx=XM_019838007.2;refGene_gn=APC;refGene_tx=XM_019838018.2;refGene_gn=APC;refGene_tx=XM_019838020.2;refGene_gn=APC;refGene_tx=XM_019838027.2;refGene_gn=APC;refGene_tx=XM_019838035.2;refGene_gn=APC;refGene_tx=XM_019838040.2;refGene_gn=APC;refGene_tx=XM_019838047.2;refGene_gn=APC;refGene_tx=XM_023258973.1
A1 174575962 174576016
refGene_gn=APC;refGene_tx=XM_019838005.2;refGene_gn=APC;refGene_tx=XM_019838007.2;refGene_gn=APC;refGene_tx=XM_019838018.2;refGene_gn=APC;refGene_tx=XM_019838020.2;refGene_gn=APC;refGene_tx=XM_019838027.2;refGene_gn=APC;refGene_tx=XM_019838035.2;refGene_gn=APC;refGene_tx=XM_019838047.2
A1 174579682 174579822
refGene_gn=APC;refGene_tx=XM_006927878.4;refGene_gn=APC;refGene_tx=XM_011285189.3;refGene_gn=APC;refGene_tx=XM_019838005.2;refGene_gn=APC;refGene_tx=XM_019838007.2;refGene_gn=APC;refGene_tx=XM_019838018.2;refGene_gn=APC;refGene_tx=XM_019838020.2;refGene_gn=APC;refGene_tx=XM_019838027.2;refGene_gn=APC;refGene_tx=XM_019838035.2;refGene_gn=APC;refGene_tx=XM_019838040.2;refGene_gn=APC;refGene_tx=XM_019838047.2;refGene_gn=APC;refGene_tx=XM_023258973.1
A1 174580372 174580450
refGene_gn=APC;refGene_tx=XM_006927878.4;refGene_gn=APC;refGene_tx=XM_011285189.3;refGene_gn=APC;refGene_tx=XM_019838005.2;refGene_gn=APC;refGene_tx=XM_019838007.2;refGene_gn=APC;refGene_tx=XM_019838018.2;refGene_gn=APC;refGene_tx=XM_019838020.2;refGene_gn=APC;refGene_tx=XM_019838027.2;refGene_gn=APC;refGene_tx=XM_019838035.2;refGene_gn=APC;refGene_tx=XM_019838040.2;refGene_gn=APC;refGene_tx=XM_019838047.2;refGene_gn=APC;refGene_tx=XM_023258973.1
A1 174581630 174581747
refGene_gn=APC;refGene_tx=XM_006927878.4;refGene_gn=APC;refGene_tx=XM_

011285189.3;refGene_gn=APC;refGene_tx=XM_019838005.2;refGene_gn=APC;refGene_tx=XM_019838007.2;refGene_gn=APC;refGene_tx=XM_019838018.2;refGene_gn=APC;refGene_tx=XM_019838020.2;refGene_gn=APC;refGene_tx=XM_019838027.2;refGene_gn=APC;refGene_tx=XM_019838035.2;refGene_gn=APC;refGene_tx=XM_019838040.2;refGene_gn=APC;refGene_tx=XM_019838047.2;refGene_gn=APC;refGene_tx=XM_023258973.1
A1 174584292 174584507
refGene_gn=APC;refGene_tx=XM_006927878.4;refGene_gn=APC;refGene_tx=XM_011285189.3;refGene_gn=APC;refGene_tx=XM_019838005.2;refGene_gn=APC;refGene_tx=XM_019838007.2;refGene_gn=APC;refGene_tx=XM_019838018.2;refGene_gn=APC;refGene_tx=XM_019838020.2;refGene_gn=APC;refGene_tx=XM_019838027.2;refGene_gn=APC;refGene_tx=XM_019838035.2;refGene_gn=APC;refGene_tx=XM_019838040.2;refGene_gn=APC;refGene_tx=XM_019838047.2;refGene_gn=APC;refGene_tx=XM_023258973.1
A1 174588524 174595104
refGene_gn=APC;refGene_tx=XM_006927878.4;refGene_gn=APC;refGene_tx=XM_011285189.3;refGene_gn=APC;refGene_tx=XM_019838005.2;refGene_gn=APC;refGene_tx=XM_019838007.2;refGene_gn=APC;refGene_tx=XM_019838018.2;refGene_gn=APC;refGene_tx=XM_019838020.2;refGene_gn=APC;refGene_tx=XM_019838027.2;refGene_gn=APC;refGene_tx=XM_019838035.2;refGene_gn=APC;refGene_tx=XM_019838040.2;refGene_gn=APC;refGene_tx=XM_019838047.2;refGene_gn=APC;refGene_tx=XM_023258973.1
A1 174606861 174606902
refGene_gn=SRP19;refGene_tx=XM_003981198.4
A1 174607541 174607617
refGene_gn=SRP19;refGene_tx=XM_003981198.4
A1 174610282 174610354
refGene_gn=SRP19;refGene_tx=XM_003981198.4
A1 174610483 174610595
refGene_gn=SRP19;refGene_tx=XM_003981198.4
A1 174612178 174612312
refGene_gn=SRP19;refGene_tx=XM_003981198.4
A1 174621753 174621803
refGene_gn=REEP5;refGene_tx=XM_003981201.4
A1 174626575 174626744
refGene_gn=REEP5;refGene_tx=XM_003981201.4
A1 174639059 174639198
refGene_gn=REEP5;refGene_tx=XM_003981201.4
A1 174658471 174658565
refGene_gn=REEP5;refGene_tx=XM_003981201.4
A1 174659293 174659411
refGene_gn=REEP5;refGene_tx=XM_003981201.4
A1 174707710 174707763
refGene_gn=DCP2;refGene_tx=XM_003981202.3
A1 174724953 174725105
refGene_gn=DCP2;refGene_tx=XM_003981202.3
A1 174730416 174730544
refGene_gn=DCP2;refGene_tx=XM_003981202.3
A1 174730947 174731046
refGene_gn=DCP2;refGene_tx=XM_003981202.3

A1 174740903 174741056
refGene_gn=DCP2;refGene_tx=XM_003981202.3
A1 174741178 174741291
refGene_gn=DCP2;refGene_tx=XM_003981202.3
A1 174741390 174741498
refGene_gn=DCP2;refGene_tx=XM_003981202.3
A1 174743826 174743962
refGene_gn=DCP2;refGene_tx=XM_003981202.3
A1 174747019 174747130
refGene_gn=DCP2;refGene_tx=XM_003981202.3
A1 174748521 174748573
refGene_gn=DCP2;refGene_tx=XM_003981202.3
A1 174750899 174751063
refGene_gn=DCP2;refGene_tx=XM_003981202.3
A1 174764650 174764831
refGene_gn=MCC;refGene_tx=XM_003981204.5;refGene_gn=MCC;refGene_tx=XM_019838097.2;refGene_gn=MCC;refGene_tx=XM_019838108.2
A1 174766605 174766728
refGene_gn=MCC;refGene_tx=XM_003981204.5;refGene_gn=MCC;refGene_tx=XM_019838097.2;refGene_gn=MCC;refGene_tx=XM_019838108.2
A1 174780601 174780702
refGene_gn=MCC;refGene_tx=XM_003981204.5;refGene_gn=MCC;refGene_tx=XM_019838097.2;refGene_gn=MCC;refGene_tx=XM_019838108.2
A1 174785190 174785397
refGene_gn=MCC;refGene_tx=XM_003981204.5;refGene_gn=MCC;refGene_tx=XM_019838097.2;refGene_gn=MCC;refGene_tx=XM_019838108.2
A1 174790221 174790456
refGene_gn=MCC;refGene_tx=XM_003981204.5;refGene_gn=MCC;refGene_tx=XM_019838097.2;refGene_gn=MCC;refGene_tx=XM_019838108.2
A1 174800136 174800320
refGene_gn=MCC;refGene_tx=XM_003981204.5;refGene_gn=MCC;refGene_tx=XM_019838097.2;refGene_gn=MCC;refGene_tx=XM_019838108.2
A1 174803640 174803744
refGene_gn=MCC;refGene_tx=XM_003981204.5;refGene_gn=MCC;refGene_tx=XM_019838097.2;refGene_gn=MCC;refGene_tx=XM_019838108.2
A1 174806542 174806683
refGene_gn=MCC;refGene_tx=XM_003981204.5;refGene_gn=MCC;refGene_tx=XM_019838097.2;refGene_gn=MCC;refGene_tx=XM_019838108.2
A1 174811302 174811451
refGene_gn=MCC;refGene_tx=XM_003981204.5;refGene_gn=MCC;refGene_tx=XM_019838097.2;refGene_gn=MCC;refGene_tx=XM_019838108.2
A1 174812555 174812645
refGene_gn=MCC;refGene_tx=XM_003981204.5;refGene_gn=MCC;refGene_tx=XM_019838097.2;refGene_gn=MCC;refGene_tx=XM_019838108.2
A1 174813589 174813736
refGene_gn=MCC;refGene_tx=XM_003981204.5;refGene_gn=MCC;refGene_tx=XM_019838097.2;refGene_gn=MCC;refGene_tx=XM_019838108.2
A1 174833196 174833403
refGene_gn=MCC;refGene_tx=XM_003981204.5;refGene_gn=MCC;refGene_tx=XM_019838097.2;refGene_gn=MCC;refGene_tx=XM_019838108.2

A1 174836497 174836661
refGene_gn=MCC;refGene_tx=XM_003981204.5;refGene_gn=MCC;refGene_tx=XM_019838097.2;refGene_gn=MCC;refGene_tx=XM_019838108.2
A1 174853255 174853398
refGene_gn=MCC;refGene_tx=XM_003981204.5;refGene_gn=MCC;refGene_tx=XM_019838097.2;refGene_gn=MCC;refGene_tx=XM_019838108.2
A1 174862635 174862778
refGene_gn=MCC;refGene_tx=XM_003981204.5;refGene_gn=MCC;refGene_tx=XM_019838097.2;refGene_gn=MCC;refGene_tx=XM_019838108.2
A1 174876153 174876267
refGene_gn=MCC;refGene_tx=XM_003981204.5;refGene_gn=MCC;refGene_tx=XM_019838097.2;refGene_gn=MCC;refGene_tx=XM_019838108.2
A1 174918992 174919019
refGene_gn=MCC;refGene_tx=XM_019838108.2
A1 175037200 175037257
refGene_gn=MCC;refGene_tx=XM_003981204.5
A1 175077647 175077862
refGene_gn=MCC;refGene_tx=XM_019838097.2
A1 175107625 175107873
refGene_gn=MCC;refGene_tx=XM_019838097.2
A1 175196463 175196615
refGene_gn=MCC;refGene_tx=XM_019838097.2
A1 175220115 175220271
refGene_gn=LOC102900222;refGene_tx=XM_023244761.1
A1 175220272 175220383
refGene_gn=LOC102900222;refGene_tx=XM_023244761.1
A1 175245280 175245490
refGene_gn=FAM193B;refGene_tx=XM_011285137.2;refGene_gn=FAM193B;refGene_tx=XM_011285138.2;refGene_gn=FAM193B;refGene_tx=XM_011285140.2;refGene_gn=FAM193B;refGene_tx=XM_011285141.2;refGene_gn=FAM193B;refGene_tx=XM_019838139.2
A1 175259630 175259873
refGene_gn=FAM193B;refGene_tx=XM_011285137.2;refGene_gn=FAM193B;refGene_tx=XM_011285138.2;refGene_gn=FAM193B;refGene_tx=XM_011285140.2;refGene_gn=FAM193B;refGene_tx=XM_011285141.2;refGene_gn=FAM193B;refGene_tx=XM_011285151.3;refGene_gn=FAM193B;refGene_tx=XM_019838139.2;refGene_gn=FAM193B;refGene_tx=XM_019838155.2;refGene_gn=FAM193B;refGene_tx=XM_019838157.2;refGene_gn=FAM193B;refGene_tx=XM_019838160.2;refGene_gn=FAM193B;refGene_tx=XM_019838164.2;refGene_gn=FAM193B;refGene_tx=XM_019838174.2;refGene_gn=FAM193B;refGene_tx=XM_019838178.2;refGene_gn=FAM193B;refGene_tx=XM_019838182.2
A1 175260673 175260902
refGene_gn=FAM193B;refGene_tx=XM_011285137.2;refGene_gn=FAM193B;refGene_tx=XM_011285138.2;refGene_gn=FAM193B;refGene_tx=XM_011285140.2;refGene_gn=FAM193B;refGene_tx=XM_011285141.2;refGene_gn=FAM193B;refGene_tx=XM_011285151.3;refGene_gn=FAM193B;refGene_tx=XM_019838139.2;refGene_gn=FAM193B;refGene_tx=XM_019838155.2;refGene_gn=FAM193B;refGene_tx=XM_019838157.2;refGene_gn=FAM193B;refGene_tx=XM_019838160.2;refGene_gn=FAM193B;refGene_tx=XM_019838164.2;refGene_gn=FAM193B;refGene_tx=XM_019838174.2;refGene_gn=FAM193B;refGene_tx=XM_019838178.2;refGene_gn=FAM193B;r

efGene_tx=XM_019838182.2
A1 175262124 175262512
refGene_gn=FAM193B;refGene_tx=XM_011285137.2;refGene_gn=FAM193B;refGene_tx=XM_011285138.2;refGene_gn=FAM193B;refGene_tx=XM_011285140.2;refGene_gn=FAM193B;refGene_tx=XM_011285141.2;refGene_gn=FAM193B;refGene_tx=XM_011285151.3;refGene_gn=FAM193B;refGene_tx=XM_019838139.2;refGene_gn=FAM193B;refGene_tx=XM_019838155.2;refGene_gn=FAM193B;refGene_tx=XM_019838157.2;refGene_gn=FAM193B;refGene_tx=XM_019838160.2;refGene_gn=FAM193B;refGene_tx=XM_019838164.2;refGene_gn=FAM193B;refGene_tx=XM_019838174.2;refGene_gn=FAM193B;refGene_tx=XM_019838178.2;refGene_gn=FAM193B;refGene_tx=XM_019838182.2
A1 175264562 175264764
refGene_gn=FAM193B;refGene_tx=XM_011285137.2;refGene_gn=FAM193B;refGene_tx=XM_011285138.2;refGene_gn=FAM193B;refGene_tx=XM_011285140.2;refGene_gn=FAM193B;refGene_tx=XM_011285141.2;refGene_gn=FAM193B;refGene_tx=XM_011285151.3;refGene_gn=FAM193B;refGene_tx=XM_019838139.2;refGene_gn=FAM193B;refGene_tx=XM_019838155.2;refGene_gn=FAM193B;refGene_tx=XM_019838157.2;refGene_gn=FAM193B;refGene_tx=XM_019838160.2;refGene_gn=FAM193B;refGene_tx=XM_019838164.2;refGene_gn=FAM193B;refGene_tx=XM_019838174.2;refGene_gn=FAM193B;refGene_tx=XM_019838178.2;refGene_gn=FAM193B;refGene_tx=XM_019838182.2
A1 175265683 175265923
refGene_gn=FAM193B;refGene_tx=XM_011285137.2;refGene_gn=FAM193B;refGene_tx=XM_011285138.2;refGene_gn=FAM193B;refGene_tx=XM_011285151.3;refGene_gn=FAM193B;refGene_tx=XM_019838139.2;refGene_gn=FAM193B;refGene_tx=XM_019838155.2;refGene_gn=FAM193B;refGene_tx=XM_019838157.2;refGene_gn=FAM193B;refGene_tx=XM_019838160.2;refGene_gn=FAM193B;refGene_tx=XM_019838164.2;refGene_gn=FAM193B;refGene_tx=XM_019838174.2;refGene_gn=FAM193B;refGene_tx=XM_019838178.2;refGene_gn=FAM193B;refGene_tx=XM_019838182.2
A1 175272076 175273097
refGene_gn=FAM193B;refGene_tx=XM_011285137.2;refGene_gn=FAM193B;refGene_tx=XM_011285138.2;refGene_gn=FAM193B;refGene_tx=XM_011285140.2;refGene_gn=FAM193B;refGene_tx=XM_011285141.2;refGene_gn=FAM193B;refGene_tx=XM_011285151.3;refGene_gn=FAM193B;refGene_tx=XM_019838139.2;refGene_gn=FAM193B;refGene_tx=XM_019838155.2;refGene_gn=FAM193B;refGene_tx=XM_019838157.2;refGene_gn=FAM193B;refGene_tx=XM_019838160.2;refGene_gn=FAM193B;refGene_tx=XM_019838164.2;refGene_gn=FAM193B;refGene_tx=XM_019838174.2;refGene_gn=FAM193B;refGene_tx=XM_019838178.2;refGene_gn=FAM193B;refGene_tx=XM_019838182.2
A1 175273254 175273330
refGene_gn=FAM193B;refGene_tx=XM_011285137.2;refGene_gn=FAM193B;refGene_tx=XM_011285138.2;refGene_gn=FAM193B;refGene_tx=XM_011285140.2;refGene_gn=FAM193B;refGene_tx=XM_011285141.2;refGene_gn=FAM193B;refGene_tx=XM_011285151.3;refGene_gn=FAM193B;refGene_tx=XM_019838139.2;refGene_gn=FAM193B;refGene_tx=XM_019838155.2;refGene_gn=FAM193B;refGene_tx=XM_019838157.2;refGene_gn=FAM193B;refGene_tx=XM_019838160.2;refGene_gn=FAM193B;refGene_tx=XM_019838164.2;refGene_gn=FAM193B;refGene_tx=XM_019838174.2;refGene_gn=FAM193B;refGene_tx=XM_019838178.2;refGene_gn=FAM193B;refGene_tx=XM_019838182.2

A1 175274169 175274266
refGene_gn=FAM193B;refGene_tx=XM_011285137.2;refGene_gn=FAM193B;refGene_tx=XM_011285138.2;refGene_gn=FAM193B;refGene_tx=XM_011285140.2;refGene_gn=FAM193B;refGene_tx=XM_011285141.2;refGene_gn=FAM193B;refGene_tx=XM_011285151.3;refGene_gn=FAM193B;refGene_tx=XM_019838139.2;refGene_gn=FAM193B;refGene_tx=XM_019838155.2;refGene_gn=FAM193B;refGene_tx=XM_019838157.2;refGene_gn=FAM193B;refGene_tx=XM_019838160.2;refGene_gn=FAM193B;refGene_tx=XM_019838164.2;refGene_gn=FAM193B;refGene_tx=XM_019838174.2;refGene_gn=FAM193B;refGene_tx=XM_019838178.2;refGene_gn=FAM193B;refGene_tx=XM_019838182.2
A1 175279382 175279409
refGene_gn=DDX41;refGene_tx=XM_019838189.2
A1 175279490 175279601
refGene_gn=DDX41;refGene_tx=XM_019838189.2
A1 175279855 175280015
refGene_gn=DDX41;refGene_tx=XM_019838189.2
A1 175280087 175280162
refGene_gn=DDX41;refGene_tx=XM_019838189.2
A1 175280288 175280349
refGene_gn=DDX41;refGene_tx=XM_019838189.2
A1 175280476 175280613
refGene_gn=DDX41;refGene_tx=XM_019838189.2
A1 175281000 175281073
refGene_gn=DDX41;refGene_tx=XM_019838189.2
A1 175281194 175281348
refGene_gn=DDX41;refGene_tx=XM_019838189.2
A1 175281424 175281561
refGene_gn=DDX41;refGene_tx=XM_019838189.2
A1 175282414 175282577
refGene_gn=DDX41;refGene_tx=XM_019838189.2
A1 175282748 175282880
refGene_gn=DDX41;refGene_tx=XM_019838189.2
A1 175283145 175283217
refGene_gn=DDX41;refGene_tx=XM_019838189.2
A1 175283352 175283449
refGene_gn=DDX41;refGene_tx=XM_019838189.2
A1 175283588 175283738
refGene_gn=DDX41;refGene_tx=XM_019838189.2
A1 175283821 175283893
refGene_gn=DDX41;refGene_tx=XM_019838189.2
A1 175284009 175284120
refGene_gn=DDX41;refGene_tx=XM_019838189.2
A1 175284301 175284438
refGene_gn=DDX41;refGene_tx=XM_019838189.2
A1 175286696 175286762
refGene_gn=DOK3;refGene_tx=XM_006927879.4;refGene_gn=DOK3;refGene_tx=XM_019838202.2
A1 175287719 175288025
refGene_gn=DOK3;refGene_tx=XM_006927879.4;refGene_gn=DOK3;refGene_tx=XM_019838202.2

A1 175289807 175289907
refGene_gn=DOK3;refGene_tx=XM_006927879.4;refGene_gn=DOK3;refGene_tx=X
M_019838202.2
A1 175290017 175290190
refGene_gn=DOK3;refGene_tx=XM_006927879.4;refGene_gn=DOK3;refGene_tx=X
M_019838202.2
A1 175290310 175291009
refGene_gn=DOK3;refGene_tx=XM_006927879.4;refGene_gn=DOK3;refGene_tx=X
M_019838202.2
A1 175298999 175299095
refGene_gn=PDLIM7;refGene_tx=XM_023259069.1;refGene_gn=PDLIM7;refGene_
tx=XM_023259070.1;refGene_gn=PDLIM7;refGene_tx=XM_023259073.1;refGene_
gn=PDLIM7;refGene_tx=XM_023259076.1;refGene_gn=PDLIM7;refGene_tx=XM_02
3259081.1
A1 175302862 175303014
refGene_gn=PDLIM7;refGene_tx=XM_023259069.1;refGene_gn=PDLIM7;refGene_
tx=XM_023259070.1;refGene_gn=PDLIM7;refGene_tx=XM_023259073.1;refGene_
gn=PDLIM7;refGene_tx=XM_023259076.1;refGene_gn=PDLIM7;refGene_tx=XM_02
3259080.1;refGene_gn=PDLIM7;refGene_tx=XM_023259081.1
A1 175303094 175303125
refGene_gn=PDLIM7;refGene_tx=XM_023259069.1;refGene_gn=PDLIM7;refGene_
tx=XM_023259070.1;refGene_gn=PDLIM7;refGene_tx=XM_023259073.1;refGene_
gn=PDLIM7;refGene_tx=XM_023259076.1;refGene_gn=PDLIM7;refGene_tx=XM_02
3259080.1;refGene_gn=PDLIM7;refGene_tx=XM_023259081.1
A1 175303611 175303730
refGene_gn=PDLIM7;refGene_tx=XM_023259069.1;refGene_gn=PDLIM7;refGene_
tx=XM_023259070.1;refGene_gn=PDLIM7;refGene_tx=XM_023259081.1
A1 175304132 175304149
refGene_gn=PDLIM7;refGene_tx=XM_023259073.1;refGene_gn=PDLIM7;refGene_
tx=XM_023259076.1
A1 175304396 175304533
refGene_gn=PDLIM7;refGene_tx=XM_023259069.1;refGene_gn=PDLIM7;refGene_
tx=XM_023259070.1;refGene_gn=PDLIM7;refGene_tx=XM_023259073.1;refGene_
gn=PDLIM7;refGene_tx=XM_023259076.1;refGene_gn=PDLIM7;refGene_tx=XM_02
3259080.1;refGene_gn=PDLIM7;refGene_tx=XM_023259081.1
A1 175304694 175304731
refGene_gn=PDLIM7;refGene_tx=XM_023259069.1;refGene_gn=PDLIM7;refGene_
tx=XM_023259070.1;refGene_gn=PDLIM7;refGene_tx=XM_023259073.1;refGene_
gn=PDLIM7;refGene_tx=XM_023259076.1;refGene_gn=PDLIM7;refGene_tx=XM_02
3259080.1;refGene_gn=PDLIM7;refGene_tx=XM_023259081.1
A1 175305017 175305114
refGene_gn=PDLIM7;refGene_tx=XM_023259081.1
A1 175305737 175305808
refGene_gn=PDLIM7;refGene_tx=XM_023259069.1;refGene_gn=PDLIM7;refGene_
tx=XM_023259070.1;refGene_gn=PDLIM7;refGene_tx=XM_023259073.1;refGene_
gn=PDLIM7;refGene_tx=XM_023259076.1;refGene_gn=PDLIM7;refGene_tx=XM_02
3259080.1
A1 175305964 175306202
refGene_gn=PDLIM7;refGene_tx=XM_023259069.1;refGene_gn=PDLIM7;refGene_
tx=XM_023259070.1;refGene_gn=PDLIM7;refGene_tx=XM_023259073.1;refGene_

gn=PDLIM7;refGene_tx=XM_023259076.1;refGene_gn=PDLIM7;refGene_tx=XM_02
3259080.1
A1 175307201 175307382
refGene_gn=PDLIM7;refGene_tx=XM_023259069.1;refGene_gn=PDLIM7;refGene_
tx=XM_023259070.1;refGene_gn=PDLIM7;refGene_tx=XM_023259073.1;refGene_
gn=PDLIM7;refGene_tx=XM_023259076.1;refGene_gn=PDLIM7;refGene_tx=XM_02
3259080.1
A1 175315288 175315409
refGene_gn=PDLIM7;refGene_tx=XM_023259069.1;refGene_gn=PDLIM7;refGene_
tx=XM_023259070.1;refGene_gn=PDLIM7;refGene_tx=XM_023259073.1;refGene_
gn=PDLIM7;refGene_tx=XM_023259076.1;refGene_gn=PDLIM7;refGene_tx=XM_02
3259080.1
A1 175315500 175315616
refGene_gn=PDLIM7;refGene_tx=XM_023259069.1;refGene_gn=PDLIM7;refGene_
tx=XM_023259070.1;refGene_gn=PDLIM7;refGene_tx=XM_023259073.1;refGene_
gn=PDLIM7;refGene_tx=XM_023259076.1;refGene_gn=PDLIM7;refGene_tx=XM_02
3259080.1
A1 175315771 175315858
refGene_gn=PDLIM7;refGene_tx=XM_023259069.1;refGene_gn=PDLIM7;refGene_
tx=XM_023259070.1;refGene_gn=PDLIM7;refGene_tx=XM_023259073.1;refGene_
gn=PDLIM7;refGene_tx=XM_023259076.1;refGene_gn=PDLIM7;refGene_tx=XM_02
3259080.1
A1 175323980 175324066
refGene_gn=DBN1;refGene_tx=XM_023259103.1;refGene_gn=DBN1;refGene_tx=X
M_023259105.1
A1 175328781 175328837
refGene_gn=DBN1;refGene_tx=XM_023259103.1;refGene_gn=DBN1;refGene_tx=X
M_023259105.1
A1 175329448 175329561
refGene_gn=DBN1;refGene_tx=XM_023259103.1;refGene_gn=DBN1;refGene_tx=X
M_023259105.1
A1 175329719 175329794
refGene_gn=DBN1;refGene_tx=XM_023259103.1;refGene_gn=DBN1;refGene_tx=X
M_023259105.1
A1 175329894 175330041
refGene_gn=DBN1;refGene_tx=XM_023259103.1;refGene_gn=DBN1;refGene_tx=X
M_023259105.1
A1 175330179 175330257
refGene_gn=DBN1;refGene_tx=XM_023259103.1;refGene_gn=DBN1;refGene_tx=X
M_023259105.1
A1 175330531 175330683
refGene_gn=DBN1;refGene_tx=XM_023259103.1;refGene_gn=DBN1;refGene_tx=X
M_023259105.1
A1 175330767 175330831
refGene_gn=DBN1;refGene_tx=XM_023259103.1;refGene_gn=DBN1;refGene_tx=X
M_023259105.1
A1 175334842 175334899
refGene_gn=DBN1;refGene_tx=XM_023259103.1;refGene_gn=DBN1;refGene_tx=X
M_023259105.1
A1 175334985 175335109

refGene_gn=DBN1;refGene_tx=XM_023259103.1;refGene_gn=DBN1;refGene_tx=XM_023259105.1
A1 175335910 175336048
refGene_gn=DBN1;refGene_tx=XM_023259103.1
A1 175336384 175336549
refGene_gn=DBN1;refGene_tx=XM_023259103.1;refGene_gn=DBN1;refGene_tx=XM_023259105.1
A1 175336978 175337658
refGene_gn=DBN1;refGene_tx=XM_023259103.1;refGene_gn=DBN1;refGene_tx=XM_023259105.1
A1 175337923 175338026
refGene_gn=DBN1;refGene_tx=XM_023259103.1;refGene_gn=DBN1;refGene_tx=XM_023259105.1
A1 175338253 175338324
refGene_gn=DBN1;refGene_tx=XM_023259103.1;refGene_gn=DBN1;refGene_tx=XM_023259105.1
A1 175339587 175339985
refGene_gn=PRR7;refGene_tx=XM_023259115.1
A1 175340174 175340586
refGene_gn=PRR7;refGene_tx=XM_023259115.1
A1 175353727 175353799
refGene_gn=GRK6;refGene_tx=XM_023259108.1
A1 175354271 175354277
refGene_gn=GRK6;refGene_tx=XM_023259109.1
A1 175354469 175354604
refGene_gn=GRK6;refGene_tx=XM_023259108.1;refGene_gn=GRK6;refGene_tx=XM_023259109.1
A1 175354685 175354823
refGene_gn=GRK6;refGene_tx=XM_023259108.1;refGene_gn=GRK6;refGene_tx=XM_023259109.1
A1 175358912 175359050
refGene_gn=GRK6;refGene_tx=XM_023259108.1;refGene_gn=GRK6;refGene_tx=XM_023259109.1
A1 175359141 175359350
refGene_gn=GRK6;refGene_tx=XM_023259108.1;refGene_gn=GRK6;refGene_tx=XM_023259109.1
A1 175360506 175360596
refGene_gn=GRK6;refGene_tx=XM_023259108.1;refGene_gn=GRK6;refGene_tx=XM_023259109.1
A1 175360692 175360730
refGene_gn=GRK6;refGene_tx=XM_023259108.1;refGene_gn=GRK6;refGene_tx=XM_023259109.1
A1 175361443 175361634
refGene_gn=GRK6;refGene_tx=XM_023259108.1;refGene_gn=GRK6;refGene_tx=XM_023259109.1
A1 175361905 175362046
refGene_gn=GRK6;refGene_tx=XM_023259108.1;refGene_gn=GRK6;refGene_tx=XM_023259109.1
A1 175362185 175362249
refGene_gn=GRK6;refGene_tx=XM_023259108.1;refGene_gn=GRK6;refGene_tx=X

M_023259109.1
A1 175362354 175362447
refGene_gn=GRK6;refGene_tx=XM_023259108.1;refGene_gn=GRK6;refGene_tx=X
M_023259109.1
A1 175362784 175362885
refGene_gn=GRK6;refGene_tx=XM_023259108.1;refGene_gn=GRK6;refGene_tx=X
M_023259109.1
A1 175363229 175363307
refGene_gn=GRK6;refGene_tx=XM_023259108.1;refGene_gn=GRK6;refGene_tx=X
M_023259109.1
A1 175363426 175363539
refGene_gn=GRK6;refGene_tx=XM_023259108.1;refGene_gn=GRK6;refGene_tx=X
M_023259109.1
A1 175364535 175364631
refGene_gn=GRK6;refGene_tx=XM_023259108.1;refGene_gn=GRK6;refGene_tx=X
M_023259109.1
A1 175368319 175368371
refGene_gn=GRK6;refGene_tx=XM_023259108.1;refGene_gn=GRK6;refGene_tx=X
M_023259109.1
A1 175374295 175374352
refGene_gn=F12;refGene_tx=NM_001168212.2
A1 175374721 175374779
refGene_gn=F12;refGene_tx=NM_001168212.2
A1 175378614 175378714
refGene_gn=F12;refGene_tx=NM_001168212.2
A1 175378871 175378942
refGene_gn=F12;refGene_tx=NM_001168212.2
A1 175379235 175379346
refGene_gn=F12;refGene_tx=NM_001168212.2
A1 175379490 175379622
refGene_gn=F12;refGene_tx=NM_001168212.2
A1 175379773 175379878
refGene_gn=F12;refGene_tx=NM_001168212.2
A1 175380026 175380192
refGene_gn=F12;refGene_tx=NM_001168212.2
A1 175380285 175380488
refGene_gn=F12;refGene_tx=NM_001168212.2
A1 175380584 175380807
refGene_gn=F12;refGene_tx=NM_001168212.2
A1 175381019 175381156
refGene_gn=F12;refGene_tx=NM_001168212.2
A1 175381256 175381406
refGene_gn=F12;refGene_tx=NM_001168212.2
A1 175381947 175382096
refGene_gn=F12;refGene_tx=NM_001168212.2
A1 175382186 175382354
refGene_gn=F12;refGene_tx=NM_001168212.2
A1 175383789 175384203
refGene_gn=PFN3;refGene_tx=XM_003981209.4
A1 175386365 175386869

refGene_gn=SLC34A1;refGene_tx=XM_003981210.5;refGene_gn=SLC34A1;refGene_tx=XM_019838251.2;refGene_gn=SLC34A1;refGene_tx=XM_019838255.2;refGene_gn=SLC34A1;refGene_tx=XM_019838264.2
A1 175387551 175387676
refGene_gn=SLC34A1;refGene_tx=XM_003981210.5;refGene_gn=SLC34A1;refGene_tx=XM_019838251.2;refGene_gn=SLC34A1;refGene_tx=XM_019838255.2;refGene_gn=SLC34A1;refGene_tx=XM_019838264.2
A1 175387776 175387893
refGene_gn=SLC34A1;refGene_tx=XM_003981210.5;refGene_gn=SLC34A1;refGene_tx=XM_019838251.2;refGene_gn=SLC34A1;refGene_tx=XM_019838255.2
A1 175388913 175389081
refGene_gn=SLC34A1;refGene_tx=XM_003981210.5;refGene_gn=SLC34A1;refGene_tx=XM_019838251.2;refGene_gn=SLC34A1;refGene_tx=XM_019838255.2;refGene_gn=SLC34A1;refGene_tx=XM_019838264.2
A1 175389354 175389424
refGene_gn=SLC34A1;refGene_tx=XM_003981210.5;refGene_gn=SLC34A1;refGene_tx=XM_019838251.2;refGene_gn=SLC34A1;refGene_tx=XM_019838255.2;refGene_gn=SLC34A1;refGene_tx=XM_019838264.2
A1 175394328 175394424
refGene_gn=SLC34A1;refGene_tx=XM_003981210.5;refGene_gn=SLC34A1;refGene_tx=XM_019838251.2;refGene_gn=SLC34A1;refGene_tx=XM_019838255.2;refGene_gn=SLC34A1;refGene_tx=XM_019838264.2
A1 175394508 175394704
refGene_gn=SLC34A1;refGene_tx=XM_003981210.5;refGene_gn=SLC34A1;refGene_tx=XM_019838251.2;refGene_gn=SLC34A1;refGene_tx=XM_019838255.2;refGene_gn=SLC34A1;refGene_tx=XM_019838264.2
A1 175394827 175394939
refGene_gn=SLC34A1;refGene_tx=XM_003981210.5;refGene_gn=SLC34A1;refGene_tx=XM_019838251.2;refGene_gn=SLC34A1;refGene_tx=XM_019838255.2;refGene_gn=SLC34A1;refGene_tx=XM_019838264.2
A1 175395863 175396007
refGene_gn=SLC34A1;refGene_tx=XM_003981210.5;refGene_gn=SLC34A1;refGene_tx=XM_019838251.2;refGene_gn=SLC34A1;refGene_tx=XM_019838255.2;refGene_gn=SLC34A1;refGene_tx=XM_019838264.2
A1 175396078 175396207
refGene_gn=SLC34A1;refGene_tx=XM_003981210.5;refGene_gn=SLC34A1;refGene_tx=XM_019838251.2;refGene_gn=SLC34A1;refGene_tx=XM_019838255.2;refGene_gn=SLC34A1;refGene_tx=XM_019838264.2
A1 175396296 175396446
refGene_gn=SLC34A1;refGene_tx=XM_003981210.5;refGene_gn=SLC34A1;refGene_tx=XM_019838251.2;refGene_gn=SLC34A1;refGene_tx=XM_019838255.2;refGene_gn=SLC34A1;refGene_tx=XM_019838264.2
A1 175396599 175396708
refGene_gn=SLC34A1;refGene_tx=XM_003981210.5;refGene_gn=SLC34A1;refGene_tx=XM_019838251.2;refGene_gn=SLC34A1;refGene_tx=XM_019838255.2;refGene_gn=SLC34A1;refGene_tx=XM_019838264.2
A1 175409499 175409702
refGene_gn=RGS14;refGene_tx=XM_006927882.3
A1 175409961 175410076
refGene_gn=RGS14;refGene_tx=XM_006927882.3

A1 175410155 175410202
refGene_gn=RGS14;refGene_tx=XM_006927882.3
A1 175410303 175410385
refGene_gn=RGS14;refGene_tx=XM_006927882.3
A1 175410467 175410594
refGene_gn=RGS14;refGene_tx=XM_006927882.3
A1 175410817 175410894
refGene_gn=RGS14;refGene_tx=XM_006927882.3
A1 175412605 175412809
refGene_gn=RGS14;refGene_tx=XM_006927882.3
A1 175413306 175413419
refGene_gn=RGS14;refGene_tx=XM_006927882.3
A1 175413719 175413831
refGene_gn=RGS14;refGene_tx=XM_006927882.3
A1 175413948 175414092
refGene_gn=RGS14;refGene_tx=XM_006927882.3
A1 175414515 175414659
refGene_gn=RGS14;refGene_tx=XM_006927882.3
A1 175414744 175414837
refGene_gn=RGS14;refGene_tx=XM_006927882.3
A1 175415191 175415370
refGene_gn=RGS14;refGene_tx=XM_006927882.3
A1 175415538 175415560
refGene_gn=RGS14;refGene_tx=XM_006927882.3
A1 175422430 175422475
refGene_gn=RGS14;refGene_tx=XM_006927882.3
A1 175427619 175427815
refGene_gn=LMAN2;refGene_tx=XM_003981212.5;refGene_gn=LMAN2;refGene_tx=XM_006927883.4;refGene_gn=LMAN2;refGene_tx=XM_023259161.1
A1 175427983 175428102
refGene_gn=LMAN2;refGene_tx=XM_003981212.5;refGene_gn=LMAN2;refGene_tx=XM_006927883.4;refGene_gn=LMAN2;refGene_tx=XM_023259161.1
A1 175439452 175439570
refGene_gn=LMAN2;refGene_tx=XM_003981212.5;refGene_gn=LMAN2;refGene_tx=XM_006927883.4;refGene_gn=LMAN2;refGene_tx=XM_023259161.1
A1 175440165 175440245
refGene_gn=LMAN2;refGene_tx=XM_003981212.5;refGene_gn=LMAN2;refGene_tx=XM_006927883.4;refGene_gn=LMAN2;refGene_tx=XM_023259161.1
A1 175440426 175440588
refGene_gn=LMAN2;refGene_tx=XM_003981212.5;refGene_gn=LMAN2;refGene_tx=XM_006927883.4;refGene_gn=LMAN2;refGene_tx=XM_023259161.1
A1 175440696 175440811
refGene_gn=LMAN2;refGene_tx=XM_003981212.5;refGene_gn=LMAN2;refGene_tx=XM_006927883.4;refGene_gn=LMAN2;refGene_tx=XM_023259161.1
A1 175443766 175443886
refGene_gn=LMAN2;refGene_tx=XM_003981212.5;refGene_gn=LMAN2;refGene_tx=XM_006927883.4;refGene_gn=LMAN2;refGene_tx=XM_023259161.1
A1 175445627 175445810
refGene_gn=LMAN2;refGene_tx=XM_003981212.5;refGene_gn=LMAN2;refGene_tx=XM_006927883.4

A1 175450415 175450456
refGene_gn=LMAN2;refGene_tx=XM_023259161.1
A1 175470580 175470650
refGene_gn=MXD3;refGene_tx=XM_003981248.5
A1 175470903 175471009
refGene_gn=MXD3;refGene_tx=XM_003981248.5
A1 175471659 175471689
refGene_gn=MXD3;refGene_tx=XM_003981248.5
A1 175471816 175471931
refGene_gn=MXD3;refGene_tx=XM_003981248.5
A1 175474010 175474194
refGene_gn=MXD3;refGene_tx=XM_003981248.5
A1 175474268 175474384
refGene_gn=MXD3;refGene_tx=XM_003981248.5
A1 175475432 175475581
refGene_gn=PRELID1;refGene_tx=XM_003981213.5
A1 175475812 175475891
refGene_gn=PRELID1;refGene_tx=XM_003981213.5
A1 175476004 175476118
refGene_gn=PRELID1;refGene_tx=XM_003981213.5
A1 175477356 175477582
refGene_gn=PRELID1;refGene_tx=XM_003981213.5
A1 175478146 175478238
refGene_gn=PRELID1;refGene_tx=XM_003981213.5
A1 175478881 175478998
refGene_gn=RAB24;refGene_tx=XM_006927886.4;refGene_gn=RAB24;refGene_tx=XM_006927887.3
A1 175479081 175479150
refGene_gn=RAB24;refGene_tx=XM_006927886.4;refGene_gn=RAB24;refGene_tx=XM_006927887.3
A1 175479349 175479428
refGene_gn=RAB24;refGene_tx=XM_006927886.4;refGene_gn=RAB24;refGene_tx=XM_006927887.3
A1 175479536 175479604
refGene_gn=RAB24;refGene_tx=XM_006927886.4;refGene_gn=RAB24;refGene_tx=XM_006927887.3
A1 175479680 175479780
refGene_gn=RAB24;refGene_tx=XM_006927886.4;refGene_gn=RAB24;refGene_tx=XM_006927887.3
A1 175479994 175480045
refGene_gn=RAB24;refGene_tx=XM_006927886.4;refGene_gn=RAB24;refGene_tx=XM_006927887.3
A1 175480184 175480247
refGene_gn=RAB24;refGene_tx=XM_006927886.4;refGene_gn=RAB24;refGene_tx=XM_006927887.3
A1 175480364 175480429
refGene_gn=RAB24;refGene_tx=XM_006927886.4
A1 175483360 175483413
refGene_gn=RAB24;refGene_tx=XM_006927887.3
A1 175486702 175488333

refGene_gn=NSD1;refGene_tx=XM_006927884.4;refGene_gn=NSD1;refGene_tx=XM_019838275.2;refGene_gn=NSD1;refGene_tx=XM_023259129.1;refGene_gn=NSD1;refGene_tx=XM_023259132.1;refGene_gn=NSD1;refGene_tx=XM_023259133.1;refGene_gn=NSD1;refGene_tx=XM_023259134.1;refGene_gn=NSD1;refGene_tx=XM_023259150.1;refGene_gn=NSD1;refGene_tx=XM_023259154.1
A1 175489930 175490135
refGene_gn=NSD1;refGene_tx=XM_006927884.4;refGene_gn=NSD1;refGene_tx=XM_019838275.2;refGene_gn=NSD1;refGene_tx=XM_023259129.1;refGene_gn=NSD1;refGene_tx=XM_023259132.1;refGene_gn=NSD1;refGene_tx=XM_023259133.1;refGene_gn=NSD1;refGene_tx=XM_023259134.1;refGene_gn=NSD1;refGene_tx=XM_023259150.1;refGene_gn=NSD1;refGene_tx=XM_023259154.1
A1 175491891 175491998
refGene_gn=NSD1;refGene_tx=XM_006927884.4;refGene_gn=NSD1;refGene_tx=XM_019838275.2;refGene_gn=NSD1;refGene_tx=XM_023259129.1;refGene_gn=NSD1;refGene_tx=XM_023259132.1;refGene_gn=NSD1;refGene_tx=XM_023259133.1;refGene_gn=NSD1;refGene_tx=XM_023259134.1;refGene_gn=NSD1;refGene_tx=XM_023259150.1;refGene_gn=NSD1;refGene_tx=XM_023259154.1
A1 175494826 175494968
refGene_gn=NSD1;refGene_tx=XM_006927884.4;refGene_gn=NSD1;refGene_tx=XM_019838275.2;refGene_gn=NSD1;refGene_tx=XM_023259129.1;refGene_gn=NSD1;refGene_tx=XM_023259132.1;refGene_gn=NSD1;refGene_tx=XM_023259133.1;refGene_gn=NSD1;refGene_tx=XM_023259134.1;refGene_gn=NSD1;refGene_tx=XM_023259150.1;refGene_gn=NSD1;refGene_tx=XM_023259154.1
A1 175496227 175496344
refGene_gn=NSD1;refGene_tx=XM_006927884.4;refGene_gn=NSD1;refGene_tx=XM_019838275.2;refGene_gn=NSD1;refGene_tx=XM_023259129.1;refGene_gn=NSD1;refGene_tx=XM_023259132.1;refGene_gn=NSD1;refGene_tx=XM_023259133.1;refGene_gn=NSD1;refGene_tx=XM_023259134.1;refGene_gn=NSD1;refGene_tx=XM_023259150.1;refGene_gn=NSD1;refGene_tx=XM_023259154.1
A1 175497687 175497957
refGene_gn=NSD1;refGene_tx=XM_006927884.4;refGene_gn=NSD1;refGene_tx=XM_019838275.2;refGene_gn=NSD1;refGene_tx=XM_023259129.1;refGene_gn=NSD1;refGene_tx=XM_023259132.1;refGene_gn=NSD1;refGene_tx=XM_023259133.1;refGene_gn=NSD1;refGene_tx=XM_023259134.1;refGene_gn=NSD1;refGene_tx=XM_023259150.1;refGene_gn=NSD1;refGene_tx=XM_023259154.1
A1 175502413 175502526
refGene_gn=NSD1;refGene_tx=XM_006927884.4;refGene_gn=NSD1;refGene_tx=XM_019838275.2;refGene_gn=NSD1;refGene_tx=XM_023259129.1;refGene_gn=NSD1;refGene_tx=XM_023259132.1;refGene_gn=NSD1;refGene_tx=XM_023259133.1;refGene_gn=NSD1;refGene_tx=XM_023259134.1;refGene_gn=NSD1;refGene_tx=XM_023259150.1;refGene_gn=NSD1;refGene_tx=XM_023259154.1
A1 175506687 175506893
refGene_gn=NSD1;refGene_tx=XM_006927884.4;refGene_gn=NSD1;refGene_tx=XM_019838275.2;refGene_gn=NSD1;refGene_tx=XM_023259129.1;refGene_gn=NSD1;refGene_tx=XM_023259132.1;refGene_gn=NSD1;refGene_tx=XM_023259133.1;refGene_gn=NSD1;refGene_tx=XM_023259134.1;refGene_gn=NSD1;refGene_tx=XM_023259150.1;refGene_gn=NSD1;refGene_tx=XM_023259154.1
A1 175508417 175508574
refGene_gn=NSD1;refGene_tx=XM_006927884.4;refGene_gn=NSD1;refGene_tx=XM_019838275.2;refGene_gn=NSD1;refGene_tx=XM_023259129.1;refGene_gn=NSD

1;refGene_tx=XM_023259132.1;refGene_gn=NSD1;refGene_tx=XM_023259133.1;
refGene_gn=NSD1;refGene_tx=XM_023259134.1;refGene_gn=NSD1;refGene_tx=X
M_023259150.1;refGene_gn=NSD1;refGene_tx=XM_023259154.1
A1 175513975 175514155
refGene_gn=NSD1;refGene_tx=XM_006927884.4;refGene_gn=NSD1;refGene_tx=X
M_019838275.2;refGene_gn=NSD1;refGene_tx=XM_023259129.1;refGene_gn=NSD
1;refGene_tx=XM_023259132.1;refGene_gn=NSD1;refGene_tx=XM_023259133.1;
refGene_gn=NSD1;refGene_tx=XM_023259134.1;refGene_gn=NSD1;refGene_tx=X
M_023259150.1;refGene_gn=NSD1;refGene_tx=XM_023259154.1
A1 175515467 175515668
refGene_gn=NSD1;refGene_tx=XM_006927884.4;refGene_gn=NSD1;refGene_tx=X
M_019838275.2;refGene_gn=NSD1;refGene_tx=XM_023259129.1;refGene_gn=NSD
1;refGene_tx=XM_023259132.1;refGene_gn=NSD1;refGene_tx=XM_023259133.1;
refGene_gn=NSD1;refGene_tx=XM_023259134.1;refGene_gn=NSD1;refGene_tx=X
M_023259150.1;refGene_gn=NSD1;refGene_tx=XM_023259154.1
A1 175523226 175523350
refGene_gn=NSD1;refGene_tx=XM_006927884.4;refGene_gn=NSD1;refGene_tx=X
M_019838275.2;refGene_gn=NSD1;refGene_tx=XM_023259129.1;refGene_gn=NSD
1;refGene_tx=XM_023259132.1;refGene_gn=NSD1;refGene_tx=XM_023259133.1;
refGene_gn=NSD1;refGene_tx=XM_023259134.1;refGene_gn=NSD1;refGene_tx=X
M_023259150.1;refGene_gn=NSD1;refGene_tx=XM_023259154.1
A1 175525779 175525923
refGene_gn=NSD1;refGene_tx=XM_006927884.4;refGene_gn=NSD1;refGene_tx=X
M_019838275.2;refGene_gn=NSD1;refGene_tx=XM_023259129.1;refGene_gn=NSD
1;refGene_tx=XM_023259132.1;refGene_gn=NSD1;refGene_tx=XM_023259133.1;
refGene_gn=NSD1;refGene_tx=XM_023259134.1;refGene_gn=NSD1;refGene_tx=X
M_023259150.1;refGene_gn=NSD1;refGene_tx=XM_023259154.1
A1 175526869 175526988
refGene_gn=NSD1;refGene_tx=XM_006927884.4;refGene_gn=NSD1;refGene_tx=X
M_019838275.2;refGene_gn=NSD1;refGene_tx=XM_023259129.1;refGene_gn=NSD
1;refGene_tx=XM_023259132.1;refGene_gn=NSD1;refGene_tx=XM_023259133.1;
refGene_gn=NSD1;refGene_tx=XM_023259134.1;refGene_gn=NSD1;refGene_tx=X
M_023259150.1;refGene_gn=NSD1;refGene_tx=XM_023259154.1
A1 175529264 175529340
refGene_gn=NSD1;refGene_tx=XM_006927884.4;refGene_gn=NSD1;refGene_tx=X
M_019838275.2;refGene_gn=NSD1;refGene_tx=XM_023259129.1;refGene_gn=NSD
1;refGene_tx=XM_023259132.1;refGene_gn=NSD1;refGene_tx=XM_023259133.1;
refGene_gn=NSD1;refGene_tx=XM_023259134.1;refGene_gn=NSD1;refGene_tx=X
M_023259150.1;refGene_gn=NSD1;refGene_tx=XM_023259154.1
A1 175531783 175531893
refGene_gn=NSD1;refGene_tx=XM_006927884.4;refGene_gn=NSD1;refGene_tx=X
M_019838275.2;refGene_gn=NSD1;refGene_tx=XM_023259129.1;refGene_gn=NSD
1;refGene_tx=XM_023259132.1;refGene_gn=NSD1;refGene_tx=XM_023259133.1;
refGene_gn=NSD1;refGene_tx=XM_023259134.1;refGene_gn=NSD1;refGene_tx=X
M_023259150.1;refGene_gn=NSD1;refGene_tx=XM_023259154.1
A1 175533094 175533365
refGene_gn=NSD1;refGene_tx=XM_006927884.4;refGene_gn=NSD1;refGene_tx=X
M_019838275.2;refGene_gn=NSD1;refGene_tx=XM_023259129.1;refGene_gn=NSD
1;refGene_tx=XM_023259132.1;refGene_gn=NSD1;refGene_tx=XM_023259133.1;
refGene_gn=NSD1;refGene_tx=XM_023259134.1;refGene_gn=NSD1;refGene_tx=X

M_023259150.1;refGene_gn=NSD1;refGene_tx=XM_023259154.1
A1 175535453 175535578
refGene_gn=NSD1;refGene_tx=XM_006927884.4;refGene_gn=NSD1;refGene_tx=XM_019838275.2;refGene_gn=NSD1;refGene_tx=XM_023259129.1;refGene_gn=NSD1;refGene_tx=XM_023259132.1;refGene_gn=NSD1;refGene_tx=XM_023259133.1;refGene_gn=NSD1;refGene_tx=XM_023259134.1;refGene_gn=NSD1;refGene_tx=XM_023259150.1;refGene_gn=NSD1;refGene_tx=XM_023259154.1
A1 175550489 175553061
refGene_gn=NSD1;refGene_tx=XM_006927884.4;refGene_gn=NSD1;refGene_tx=XM_019838275.2;refGene_gn=NSD1;refGene_tx=XM_023259129.1;refGene_gn=NSD1;refGene_tx=XM_023259132.1;refGene_gn=NSD1;refGene_tx=XM_023259133.1;refGene_gn=NSD1;refGene_tx=XM_023259134.1;refGene_gn=NSD1;refGene_tx=XM_023259150.1;refGene_gn=NSD1;refGene_tx=XM_023259154.1
A1 175555361 175555534
refGene_gn=NSD1;refGene_tx=XM_006927884.4;refGene_gn=NSD1;refGene_tx=XM_019838275.2;refGene_gn=NSD1;refGene_tx=XM_023259129.1;refGene_gn=NSD1;refGene_tx=XM_023259132.1;refGene_gn=NSD1;refGene_tx=XM_023259133.1;refGene_gn=NSD1;refGene_tx=XM_023259134.1;refGene_gn=NSD1;refGene_tx=XM_023259150.1;refGene_gn=NSD1;refGene_tx=XM_023259154.1
A1 175564083 175564219
refGene_gn=NSD1;refGene_tx=XM_006927884.4;refGene_gn=NSD1;refGene_tx=XM_019838275.2;refGene_gn=NSD1;refGene_tx=XM_023259129.1;refGene_gn=NSD1;refGene_tx=XM_023259132.1;refGene_gn=NSD1;refGene_tx=XM_023259133.1;refGene_gn=NSD1;refGene_tx=XM_023259134.1;refGene_gn=NSD1;refGene_tx=XM_023259150.1;refGene_gn=NSD1;refGene_tx=XM_023259154.1
A1 175625245 175626172
refGene_gn=NSD1;refGene_tx=XM_006927884.4;refGene_gn=NSD1;refGene_tx=XM_019838275.2;refGene_gn=NSD1;refGene_tx=XM_023259129.1;refGene_gn=NSD1;refGene_tx=XM_023259132.1;refGene_gn=NSD1;refGene_tx=XM_023259133.1;refGene_gn=NSD1;refGene_tx=XM_023259134.1;refGene_gn=NSD1;refGene_tx=XM_023259150.1;refGene_gn=NSD1;refGene_tx=XM_023259154.1
A1 175654348 175654495
refGene_gn=FGFR4;refGene_tx=XM_003981217.5;refGene_gn=FGFR4;refGene_tx=XM_011285066.3;refGene_gn=FGFR4;refGene_tx=XM_023259201.1;refGene_gn=FGFR4;refGene_tx=XM_023259202.1
A1 175654593 175654699
refGene_gn=FGFR4;refGene_tx=XM_003981217.5;refGene_gn=FGFR4;refGene_tx=XM_011285066.3;refGene_gn=FGFR4;refGene_tx=XM_023259201.1;refGene_gn=FGFR4;refGene_tx=XM_023259202.1
A1 175654947 175655085
refGene_gn=FGFR4;refGene_tx=XM_003981217.5;refGene_gn=FGFR4;refGene_tx=XM_011285066.3;refGene_gn=FGFR4;refGene_tx=XM_023259201.1;refGene_gn=FGFR4;refGene_tx=XM_023259202.1
A1 175655308 175655379
refGene_gn=FGFR4;refGene_tx=XM_003981217.5;refGene_gn=FGFR4;refGene_tx=XM_011285066.3;refGene_gn=FGFR4;refGene_tx=XM_023259201.1;refGene_gn=FGFR4;refGene_tx=XM_023259202.1
A1 175655468 175655591
refGene_gn=FGFR4;refGene_tx=XM_003981217.5;refGene_gn=FGFR4;refGene_tx=XM_011285066.3;refGene_gn=FGFR4;refGene_tx=XM_023259201.1;refGene_gn=

FGFR4;refGene_tx=XM_023259202.1
A1 175655990 175656181
refGene_gn=FGFR4;refGene_tx=XM_003981217.5;refGene_gn=FGFR4;refGene_tx=XM_011285066.3;refGene_gn=FGFR4;refGene_tx=XM_023259201.1;refGene_gn=FGFR4;refGene_tx=XM_023259202.1
A1 175656267 175656378
refGene_gn=FGFR4;refGene_tx=XM_003981217.5;refGene_gn=FGFR4;refGene_tx=XM_011285066.3;refGene_gn=FGFR4;refGene_tx=XM_023259201.1;refGene_gn=FGFR4;refGene_tx=XM_023259202.1
A1 175657544 175657666
refGene_gn=FGFR4;refGene_tx=XM_003981217.5;refGene_gn=FGFR4;refGene_tx=XM_011285066.3;refGene_gn=FGFR4;refGene_tx=XM_023259201.1;refGene_gn=FGFR4;refGene_tx=XM_023259202.1
A1 175657752 175657898
refGene_gn=FGFR4;refGene_tx=XM_003981217.5;refGene_gn=FGFR4;refGene_tx=XM_011285066.3;refGene_gn=FGFR4;refGene_tx=XM_023259201.1;refGene_gn=FGFR4;refGene_tx=XM_023259202.1
A1 175658006 175658200
refGene_gn=FGFR4;refGene_tx=XM_003981217.5;refGene_gn=FGFR4;refGene_tx=XM_011285066.3;refGene_gn=FGFR4;refGene_tx=XM_023259201.1;refGene_gn=FGFR4;refGene_tx=XM_023259202.1
A1 175658523 175658662
refGene_gn=FGFR4;refGene_tx=XM_003981217.5;refGene_gn=FGFR4;refGene_tx=XM_011285066.3;refGene_gn=FGFR4;refGene_tx=XM_023259201.1;refGene_gn=FGFR4;refGene_tx=XM_023259202.1
A1 175658794 175658985
refGene_gn=FGFR4;refGene_tx=XM_003981217.5;refGene_gn=FGFR4;refGene_tx=XM_011285066.3;refGene_gn=FGFR4;refGene_tx=XM_023259201.1;refGene_gn=FGFR4;refGene_tx=XM_023259202.1
A1 175659495 175659619
refGene_gn=FGFR4;refGene_tx=XM_003981217.5;refGene_gn=FGFR4;refGene_tx=XM_011285066.3;refGene_gn=FGFR4;refGene_tx=XM_023259201.1;refGene_gn=FGFR4;refGene_tx=XM_023259202.1
A1 175660186 175660353
refGene_gn=FGFR4;refGene_tx=XM_003981217.5;refGene_gn=FGFR4;refGene_tx=XM_011285066.3;refGene_gn=FGFR4;refGene_tx=XM_023259201.1;refGene_gn=FGFR4;refGene_tx=XM_023259202.1
A1 175660476 175660557
refGene_gn=FGFR4;refGene_tx=XM_003981217.5;refGene_gn=FGFR4;refGene_tx=XM_011285066.3;refGene_gn=FGFR4;refGene_tx=XM_023259201.1;refGene_gn=FGFR4;refGene_tx=XM_023259202.1
A1 175660648 175660912
refGene_gn=FGFR4;refGene_tx=XM_003981217.5;refGene_gn=FGFR4;refGene_tx=XM_011285066.3;refGene_gn=FGFR4;refGene_tx=XM_023259201.1
A1 175661592 175661770
refGene_gn=FGFR4;refGene_tx=XM_003981217.5;refGene_gn=FGFR4;refGene_tx=XM_011285066.3;refGene_gn=FGFR4;refGene_tx=XM_023259201.1;refGene_gn=FGFR4;refGene_tx=XM_023259202.1
A1 175680568 175680656
refGene_gn=ZNF346;refGene_tx=XM_003981219.3;refGene_gn=ZNF346;refGene_

tx=XM_006927889.3;refGene_gn=ZNF346;refGene_tx=XM_019838336.1;refGene_gn=ZNF346;refGene_tx=XM_019838339.2
A1 175683149 175683243
refGene_gn=ZNF346;refGene_tx=XM_003981219.3;refGene_gn=ZNF346;refGene_tx=XM_006927889.3;refGene_gn=ZNF346;refGene_tx=XM_019838336.1;refGene_gn=ZNF346;refGene_tx=XM_019838339.2
A1 175684269 175684337
refGene_gn=ZNF346;refGene_tx=XM_019838327.2
A1 175689938 175690172
refGene_gn=ZNF346;refGene_tx=XM_003981219.3;refGene_gn=ZNF346;refGene_tx=XM_006927889.3;refGene_gn=ZNF346;refGene_tx=XM_019838327.2;refGene_gn=ZNF346;refGene_tx=XM_019838339.2
A1 175694000 175694145
refGene_gn=ZNF346;refGene_tx=XM_003981219.3;refGene_gn=ZNF346;refGene_tx=XM_006927889.3;refGene_gn=ZNF346;refGene_tx=XM_019838327.2;refGene_gn=ZNF346;refGene_tx=XM_019838336.1
A1 175695900 175695993
refGene_gn=ZNF346;refGene_tx=XM_003981219.3;refGene_gn=ZNF346;refGene_tx=XM_006927889.3;refGene_gn=ZNF346;refGene_tx=XM_019838327.2;refGene_gn=ZNF346;refGene_tx=XM_019838336.1
A1 175696565 175696669
refGene_gn=ZNF346;refGene_tx=XM_003981219.3;refGene_gn=ZNF346;refGene_tx=XM_006927889.3;refGene_gn=ZNF346;refGene_tx=XM_019838327.2;refGene_gn=ZNF346;refGene_tx=XM_019838336.1
A1 175709400 175709575
refGene_gn=ZNF346;refGene_tx=XM_003981219.3;refGene_gn=ZNF346;refGene_tx=XM_006927889.3;refGene_gn=ZNF346;refGene_tx=XM_019838327.2;refGene_gn=ZNF346;refGene_tx=XM_019838336.1;refGene_gn=ZNF346;refGene_tx=XM_019838339.2
A1 175760243 175760390
refGene_gn=UIMC1;refGene_tx=XM_023259245.1;refGene_gn=UIMC1;refGene_tx=XM_023259248.1;refGene_gn=UIMC1;refGene_tx=XM_023259260.1
A1 175787189 175787274
refGene_gn=UIMC1;refGene_tx=XM_023259245.1;refGene_gn=UIMC1;refGene_tx=XM_023259248.1;refGene_gn=UIMC1;refGene_tx=XM_023259260.1
A1 175805644 175805769
refGene_gn=UIMC1;refGene_tx=XM_019838358.1;refGene_gn=UIMC1;refGene_tx=XM_023259245.1;refGene_gn=UIMC1;refGene_tx=XM_023259248.1;refGene_gn=UIMC1;refGene_tx=XM_023259260.1
A1 175806521 175806627
refGene_gn=UIMC1;refGene_tx=XM_019838358.1;refGene_gn=UIMC1;refGene_tx=XM_023259245.1;refGene_gn=UIMC1;refGene_tx=XM_023259248.1;refGene_gn=UIMC1;refGene_tx=XM_023259260.1
A1 175806936 175807670
refGene_gn=UIMC1;refGene_tx=XM_019838358.1;refGene_gn=UIMC1;refGene_tx=XM_023259245.1;refGene_gn=UIMC1;refGene_tx=XM_023259248.1;refGene_gn=UIMC1;refGene_tx=XM_023259260.1
A1 175815255 175815320
refGene_gn=UIMC1;refGene_tx=XM_019838358.1;refGene_gn=UIMC1;refGene_tx=XM_023259245.1;refGene_gn=UIMC1;refGene_tx=XM_023259248.1;refGene_gn=

UIMC1;refGene_tx=XM_023259260.1
A1 175817331 175817408
refGene_gn=UIMC1;refGene_tx=XM_019838358.1;refGene_gn=UIMC1;refGene_tx=XM_023259245.1;refGene_gn=UIMC1;refGene_tx=XM_023259248.1;refGene_gn=UIMC1;refGene_tx=XM_023259260.1
A1 175821945 175822049
refGene_gn=UIMC1;refGene_tx=XM_019838358.1;refGene_gn=UIMC1;refGene_tx=XM_023259245.1;refGene_gn=UIMC1;refGene_tx=XM_023259248.1;refGene_gn=UIMC1;refGene_tx=XM_023259260.1
A1 175827164 175827315
refGene_gn=UIMC1;refGene_tx=XM_019838358.1;refGene_gn=UIMC1;refGene_tx=XM_023259245.1;refGene_gn=UIMC1;refGene_tx=XM_023259248.1;refGene_gn=UIMC1;refGene_tx=XM_023259260.1
A1 175862451 175862536
refGene_gn=UIMC1;refGene_tx=XM_019838358.1;refGene_gn=UIMC1;refGene_tx=XM_023259245.1;refGene_gn=UIMC1;refGene_tx=XM_023259248.1;refGene_gn=UIMC1;refGene_tx=XM_023259260.1
A1 175864623 175864795
refGene_gn=UIMC1;refGene_tx=XM_019838358.1;refGene_gn=UIMC1;refGene_tx=XM_023259245.1;refGene_gn=UIMC1;refGene_tx=XM_023259248.1;refGene_gn=UIMC1;refGene_tx=XM_023259260.1
A1 175866118 175866182
refGene_gn=UIMC1;refGene_tx=XM_019838358.1;refGene_gn=UIMC1;refGene_tx=XM_023259245.1;refGene_gn=UIMC1;refGene_tx=XM_023259248.1;refGene_gn=UIMC1;refGene_tx=XM_023259260.1
A1 175867228 175867271
refGene_gn=UIMC1;refGene_tx=XM_019838358.1;refGene_gn=UIMC1;refGene_tx=XM_023259245.1;refGene_gn=UIMC1;refGene_tx=XM_023259248.1;refGene_gn=UIMC1;refGene_tx=XM_023259260.1
A1 175867824 175868035
refGene_gn=UIMC1;refGene_tx=XM_019838358.1;refGene_gn=UIMC1;refGene_tx=XM_023259245.1;refGene_gn=UIMC1;refGene_tx=XM_023259248.1;refGene_gn=UIMC1;refGene_tx=XM_023259260.1
A1 175874346 175874407
refGene_gn=HK3;refGene_tx=XM_019838343.2
A1 175878243 175878365
refGene_gn=HK3;refGene_tx=XM_019838343.2
A1 175881628 175881791
refGene_gn=HK3;refGene_tx=XM_019838343.2
A1 175881994 175882149
refGene_gn=HK3;refGene_tx=XM_019838343.2
A1 175882235 175882355
refGene_gn=HK3;refGene_tx=XM_019838343.2
A1 175882444 175882540
refGene_gn=HK3;refGene_tx=XM_019838343.2
A1 175883461 175883561
refGene_gn=HK3;refGene_tx=XM_019838343.2
A1 175883642 175883826
refGene_gn=HK3;refGene_tx=XM_019838343.2
A1 175884272 175884428

refGene_gn=HK3;refGene_tx=XM_019838343.2
A1 175884573 175884807
refGene_gn=HK3;refGene_tx=XM_019838343.2
A1 175885185 175885481
refGene_gn=HK3;refGene_tx=XM_019838343.2
A1 175885592 175885732
refGene_gn=HK3;refGene_tx=XM_019838343.2
A1 175885811 175885931
refGene_gn=HK3;refGene_tx=XM_019838343.2
A1 175888328 175888424
refGene_gn=HK3;refGene_tx=XM_019838343.2
A1 175888608 175888708
refGene_gn=HK3;refGene_tx=XM_019838343.2
A1 175890468 175890652
refGene_gn=HK3;refGene_tx=XM_019838343.2
A1 175890751 175890907
refGene_gn=HK3;refGene_tx=XM_019838343.2
A1 175891072 175891306
refGene_gn=HK3;refGene_tx=XM_019838343.2
A1 175891393 175891538
refGene_gn=HK3;refGene_tx=XM_019838343.2
A1 175892711 175892877
refGene_gn=UNC5A;refGene_tx=XM_011285307.2;refGene_gn=UNC5A;refGene_tx
=XM_023259242.1
A1 175893123 175893302
refGene_gn=UNC5A;refGene_tx=XM_011285307.2;refGene_gn=UNC5A;refGene_tx
=XM_023259242.1
A1 175893977 175894142
refGene_gn=UNC5A;refGene_tx=XM_011285307.2;refGene_gn=UNC5A;refGene_tx
=XM_023259242.1
A1 175894214 175894364
refGene_gn=UNC5A;refGene_tx=XM_011285307.2;refGene_gn=UNC5A;refGene_tx
=XM_023259242.1
A1 175894470 175894704
refGene_gn=UNC5A;refGene_tx=XM_011285307.2;refGene_gn=UNC5A;refGene_tx
=XM_023259242.1
A1 175894859 175895028
refGene_gn=UNC5A;refGene_tx=XM_011285307.2;refGene_gn=UNC5A;refGene_tx
=XM_023259242.1
A1 175895278 175895366
refGene_gn=UNC5A;refGene_tx=XM_011285307.2;refGene_gn=UNC5A;refGene_tx
=XM_023259242.1
A1 175898168 175898471
refGene_gn=UNC5A;refGene_tx=XM_011285307.2;refGene_gn=UNC5A;refGene_tx
=XM_023259242.1
A1 175898572 175898761
refGene_gn=UNC5A;refGene_tx=XM_011285307.2;refGene_gn=UNC5A;refGene_tx
=XM_023259242.1
A1 175899713 175899878
refGene_gn=UNC5A;refGene_tx=XM_011285307.2;refGene_gn=UNC5A;refGene_tx

=XM_023259242.1
A1 175900448 175900616
refGene_gn=UNC5A;refGene_tx=XM_011285307.2
A1 175901264 175901445
refGene_gn=UNC5A;refGene_tx=XM_011285307.2;refGene_gn=UNC5A;refGene_tx
=XM_023259242.1
A1 175901591 175901695
refGene_gn=UNC5A;refGene_tx=XM_011285307.2;refGene_gn=UNC5A;refGene_tx
=XM_023259242.1
A1 175901959 175902103
refGene_gn=UNC5A;refGene_tx=XM_011285307.2;refGene_gn=UNC5A;refGene_tx
=XM_023259242.1
A1 175907266 175907488
refGene_gn=UNC5A;refGene_tx=XM_011285307.2;refGene_gn=UNC5A;refGene_tx
=XM_023259242.1
A1 175953337 175953407
refGene_gn=UNC5A;refGene_tx=XM_011285307.2;refGene_gn=UNC5A;refGene_tx
=XM_023259242.1
A1 176084915 176084952
refGene_gn=TSPAN17;refGene_tx=XM_003981222.5;refGene_gn=TSPAN17;refGen
e_tx=XM_011285205.3;refGene_gn=TSPAN17;refGene_tx=XM_019838374.2
A1 176085498 176085560
refGene_gn=TSPAN17;refGene_tx=XM_003981222.5;refGene_gn=TSPAN17;refGen
e_tx=XM_011285205.3;refGene_gn=TSPAN17;refGene_tx=XM_019838374.2
A1 176085640 176085757
refGene_gn=TSPAN17;refGene_tx=XM_003981222.5;refGene_gn=TSPAN17;refGen
e_tx=XM_019838374.2
A1 176086397 176086445
refGene_gn=TSPAN17;refGene_tx=XM_003981222.5;refGene_gn=TSPAN17;refGen
e_tx=XM_011285205.3;refGene_gn=TSPAN17;refGene_tx=XM_019838374.2
A1 176087077 176087203
refGene_gn=TSPAN17;refGene_tx=XM_003981222.5;refGene_gn=TSPAN17;refGen
e_tx=XM_011285205.3;refGene_gn=TSPAN17;refGene_tx=XM_019838374.2
A1 176088983 176089154
refGene_gn=TSPAN17;refGene_tx=XM_003981222.5;refGene_gn=TSPAN17;refGen
e_tx=XM_011285205.3;refGene_gn=TSPAN17;refGene_tx=XM_019838374.2
A1 176090243 176090390
refGene_gn=TSPAN17;refGene_tx=XM_003981222.5;refGene_gn=TSPAN17;refGen
e_tx=XM_011285205.3;refGene_gn=TSPAN17;refGene_tx=XM_019838374.2
A1 176090479 176090530
refGene_gn=TSPAN17;refGene_tx=XM_003981222.5;refGene_gn=TSPAN17;refGen
e_tx=XM_011285205.3
A1 176093978 176094065
refGene_gn=TSPAN17;refGene_tx=XM_003981222.5;refGene_gn=TSPAN17;refGen
e_tx=XM_011285205.3;refGene_gn=TSPAN17;refGene_tx=XM_019838374.2
A1 176095954 176096069
refGene_gn=EIF4E1B;refGene_tx=XM_023244820.1
A1 176096402 176096542
refGene_gn=EIF4E1B;refGene_tx=XM_023244820.1
A1 176096683 176096797

refGene_gn=EIF4E1B;refGene_tx=XM_023244820.1
A1 176097578 176097642
refGene_gn=EIF4E1B;refGene_tx=XM_023244820.1
A1 176098417 176098513
refGene_gn=EIF4E1B;refGene_tx=XM_023244820.1
A1 176098829 176099106
refGene_gn=EIF4E1B;refGene_tx=XM_023244820.1
A1 176099378 176099382
refGene_gn=EIF4E1B;refGene_tx=XM_023244820.1
A1 176112750 176112871
refGene_gn=SNCB;refGene_tx=XM_003981223.5;refGene_gn=SNCB;refGene_tx=XM_011285213.3
A1 176115814 176115856
refGene_gn=SNCB;refGene_tx=XM_003981223.5;refGene_gn=SNCB;refGene_tx=XM_011285213.3
A1 176116079 176116198
refGene_gn=SNCB;refGene_tx=XM_003981223.5;refGene_gn=SNCB;refGene_tx=XM_011285213.3
A1 176120544 176120634
refGene_gn=SNCB;refGene_tx=XM_003981223.5;refGene_gn=SNCB;refGene_tx=XM_011285213.3
A1 176120995 176121028
refGene_gn=SNCB;refGene_tx=XM_003981223.5;refGene_gn=SNCB;refGene_tx=XM_011285213.3
A1 176139139 176142028
refGene_gn=GPRIN1;refGene_tx=XM_023259289.1;refGene_gn=GPRIN1;refGene_tx=XM_023259290.1;refGene_gn=GPRIN1;refGene_tx=XM_023259295.1
A1 176143182 176143317
refGene_gn=CDHR2;refGene_tx=XM_019838400.2;refGene_gn=CDHR2;refGene_tx=XM_019838405.2
A1 176144378 176144436
refGene_gn=CDHR2;refGene_tx=XM_019838400.2;refGene_gn=CDHR2;refGene_tx=XM_019838405.2
A1 176144561 176144642
refGene_gn=CDHR2;refGene_tx=XM_019838400.2;refGene_gn=CDHR2;refGene_tx=XM_019838405.2
A1 176144723 176144837
refGene_gn=CDHR2;refGene_tx=XM_019838400.2;refGene_gn=CDHR2;refGene_tx=XM_019838405.2
A1 176145369 176145494
refGene_gn=CDHR2;refGene_tx=XM_019838400.2;refGene_gn=CDHR2;refGene_tx=XM_019838405.2
A1 176145576 176145637
refGene_gn=CDHR2;refGene_tx=XM_019838400.2;refGene_gn=CDHR2;refGene_tx=XM_019838405.2
A1 176145731 176145809
refGene_gn=CDHR2;refGene_tx=XM_019838400.2;refGene_gn=CDHR2;refGene_tx=XM_019838405.2
A1 176145912 176145981
refGene_gn=CDHR2;refGene_tx=XM_019838400.2;refGene_gn=CDHR2;refGene_tx

=XM_019838405.2
A1	176146431	176146520
refGene_gn=CDHR2;refGene_tx=XM_019838400.2;refGene_gn=CDHR2;refGene_tx
=XM_019838405.2
A1	176146604	176146713
refGene_gn=CDHR2;refGene_tx=XM_019838400.2;refGene_gn=CDHR2;refGene_tx
=XM_019838405.2
A1	176146859	176147011
refGene_gn=CDHR2;refGene_tx=XM_019838400.2;refGene_gn=CDHR2;refGene_tx
=XM_019838405.2
A1	176148378	176148428
refGene_gn=CDHR2;refGene_tx=XM_019838400.2;refGene_gn=CDHR2;refGene_tx
=XM_019838405.2
A1	176148851	176148923
refGene_gn=CDHR2;refGene_tx=XM_019838400.2;refGene_gn=CDHR2;refGene_tx
=XM_019838405.2
A1	176149695	176150301
refGene_gn=CDHR2;refGene_tx=XM_019838400.2;refGene_gn=CDHR2;refGene_tx
=XM_019838405.2
A1	176150449	176150519
refGene_gn=CDHR2;refGene_tx=XM_019838400.2;refGene_gn=CDHR2;refGene_tx
=XM_019838405.2
A1	176152896	176153136
refGene_gn=CDHR2;refGene_tx=XM_019838400.2;refGene_gn=CDHR2;refGene_tx
=XM_019838405.2
A1	176155348	176155592
refGene_gn=CDHR2;refGene_tx=XM_019838400.2;refGene_gn=CDHR2;refGene_tx
=XM_019838405.2
A1	176155860	176155922
refGene_gn=CDHR2;refGene_tx=XM_019838400.2;refGene_gn=CDHR2;refGene_tx
=XM_019838405.2
A1	176156200	176156362
refGene_gn=CDHR2;refGene_tx=XM_019838400.2;refGene_gn=CDHR2;refGene_tx
=XM_019838405.2
A1	176156443	176156599
refGene_gn=CDHR2;refGene_tx=XM_019838400.2;refGene_gn=CDHR2;refGene_tx
=XM_019838405.2
A1	176157630	176157864
refGene_gn=CDHR2;refGene_tx=XM_019838400.2;refGene_gn=CDHR2;refGene_tx
=XM_019838405.2
A1	176157985	176158101
refGene_gn=CDHR2;refGene_tx=XM_019838400.2;refGene_gn=CDHR2;refGene_tx
=XM_019838405.2
A1	176158189	176158265
refGene_gn=CDHR2;refGene_tx=XM_019838400.2;refGene_gn=CDHR2;refGene_tx
=XM_019838405.2
A1	176158353	176158500
refGene_gn=CDHR2;refGene_tx=XM_019838400.2;refGene_gn=CDHR2;refGene_tx
=XM_019838405.2
A1	176158571	176158697

refGene_gn=CDHR2;refGene_tx=XM_019838400.2;refGene_gn=CDHR2;refGene_tx
=XM_019838405.2
A1 176159259 176159349
refGene_gn=CDHR2;refGene_tx=XM_019838400.2;refGene_gn=CDHR2;refGene_tx
=XM_019838405.2
A1 176161153 176161243
refGene_gn=CDHR2;refGene_tx=XM_019838400.2;refGene_gn=CDHR2;refGene_tx
=XM_019838405.2
A1 176162506 176162560
refGene_gn=CDHR2;refGene_tx=XM_019838400.2
A1 176162677 176162817
refGene_gn=CDHR2;refGene_tx=XM_019838400.2;refGene_gn=CDHR2;refGene_tx
=XM_019838405.2
A1 176168506 176168578
refGene_gn=CDHR2;refGene_tx=XM_019838400.2;refGene_gn=CDHR2;refGene_tx
=XM_019838405.2
A1 176168849 176168916
refGene_gn=CDHR2;refGene_tx=XM_019838400.2;refGene_gn=CDHR2;refGene_tx
=XM_019838405.2
A1 176179243 176179315
refGene_gn=CDHR2;refGene_tx=XM_019838400.2;refGene_gn=CDHR2;refGene_tx
=XM_019838405.2
A1 176192452 176192460
refGene_gn=RNF44;refGene_tx=XM_019838437.2;refGene_gn=RNF44;refGene_tx
=XM_019838440.2;refGene_gn=RNF44;refGene_tx=XM_019838444.2
A1 176197753 176197860
refGene_gn=RNF44;refGene_tx=XM_006927895.4;refGene_gn=RNF44;refGene_tx
=XM_011285233.2;refGene_gn=RNF44;refGene_tx=XM_011285236.2;refGene_gn=
RNF44;refGene_tx=XM_011285240.2;refGene_gn=RNF44;refGene_tx=XM_0112852
43.2;refGene_gn=RNF44;refGene_tx=XM_011285245.3
A1 176198024 176198254
refGene_gn=RNF44;refGene_tx=XM_006927895.4;refGene_gn=RNF44;refGene_tx
=XM_011285233.2;refGene_gn=RNF44;refGene_tx=XM_011285236.2;refGene_gn=
RNF44;refGene_tx=XM_011285240.2;refGene_gn=RNF44;refGene_tx=XM_0112852
43.2;refGene_gn=RNF44;refGene_tx=XM_011285245.3;refGene_gn=RNF44;refGe
ne_tx=XM_011285251.2;refGene_gn=RNF44;refGene_tx=XM_011285255.2;refGen
e_gn=RNF44;refGene_tx=XM_011285261.2;refGene_gn=RNF44;refGene_tx=XM_01
9838437.2;refGene_gn=RNF44;refGene_tx=XM_019838440.2;refGene_gn=RNF44;
refGene_tx=XM_019838444.2;refGene_gn=RNF44;refGene_tx=XM_019838461.2;r
efGene_gn=RNF44;refGene_tx=XM_019838467.2;refGene_gn=RNF44;refGene_tx=
XM_019838474.1
A1 176198676 176198844
refGene_gn=RNF44;refGene_tx=XM_006927895.4;refGene_gn=RNF44;refGene_tx
=XM_011285233.2;refGene_gn=RNF44;refGene_tx=XM_011285236.2;refGene_gn=
RNF44;refGene_tx=XM_011285240.2;refGene_gn=RNF44;refGene_tx=XM_0112852
43.2;refGene_gn=RNF44;refGene_tx=XM_011285245.3;refGene_gn=RNF44;refGe
ne_tx=XM_011285251.2;refGene_gn=RNF44;refGene_tx=XM_011285255.2;refGen
e_gn=RNF44;refGene_tx=XM_011285261.2;refGene_gn=RNF44;refGene_tx=XM_01
9838437.2;refGene_gn=RNF44;refGene_tx=XM_019838440.2;refGene_gn=RNF44;
refGene_tx=XM_019838444.2;refGene_gn=RNF44;refGene_tx=XM_019838461.2;r

efGene_gn=RNF44;refGene_tx=XM_019838467.2;refGene_gn=RNF44;refGene_tx=XM_019838474.1
A1 176199261 176199435
refGene_gn=RNF44;refGene_tx=XM_006927895.4;refGene_gn=RNF44;refGene_tx=XM_011285233.2;refGene_gn=RNF44;refGene_tx=XM_011285236.2;refGene_gn=RNF44;refGene_tx=XM_011285240.2;refGene_gn=RNF44;refGene_tx=XM_011285243.2;refGene_gn=RNF44;refGene_tx=XM_011285245.3;refGene_gn=RNF44;refGene_tx=XM_011285251.2;refGene_gn=RNF44;refGene_tx=XM_011285255.2;refGene_gn=RNF44;refGene_tx=XM_011285261.2;refGene_gn=RNF44;refGene_tx=XM_019838437.2;refGene_gn=RNF44;refGene_tx=XM_019838440.2;refGene_gn=RNF44;refGene_tx=XM_019838444.2;refGene_gn=RNF44;refGene_tx=XM_019838461.2;refGene_gn=RNF44;refGene_tx=XM_019838467.2;refGene_gn=RNF44;refGene_tx=XM_019838474.1
A1 176199533 176199695
refGene_gn=RNF44;refGene_tx=XM_006927895.4;refGene_gn=RNF44;refGene_tx=XM_011285233.2;refGene_gn=RNF44;refGene_tx=XM_011285236.2;refGene_gn=RNF44;refGene_tx=XM_011285240.2;refGene_gn=RNF44;refGene_tx=XM_011285243.2;refGene_gn=RNF44;refGene_tx=XM_011285245.3;refGene_gn=RNF44;refGene_tx=XM_011285251.2;refGene_gn=RNF44;refGene_tx=XM_011285255.2;refGene_gn=RNF44;refGene_tx=XM_011285261.2;refGene_gn=RNF44;refGene_tx=XM_019838437.2;refGene_gn=RNF44;refGene_tx=XM_019838440.2;refGene_gn=RNF44;refGene_tx=XM_019838444.2;refGene_gn=RNF44;refGene_tx=XM_019838461.2;refGene_gn=RNF44;refGene_tx=XM_019838467.2;refGene_gn=RNF44;refGene_tx=XM_019838474.1
A1 176199779 176199904
refGene_gn=RNF44;refGene_tx=XM_006927895.4;refGene_gn=RNF44;refGene_tx=XM_011285233.2;refGene_gn=RNF44;refGene_tx=XM_011285236.2;refGene_gn=RNF44;refGene_tx=XM_011285240.2;refGene_gn=RNF44;refGene_tx=XM_011285243.2;refGene_gn=RNF44;refGene_tx=XM_011285245.3;refGene_gn=RNF44;refGene_tx=XM_011285251.2;refGene_gn=RNF44;refGene_tx=XM_011285255.2;refGene_gn=RNF44;refGene_tx=XM_011285261.2;refGene_gn=RNF44;refGene_tx=XM_019838437.2;refGene_gn=RNF44;refGene_tx=XM_019838440.2;refGene_gn=RNF44;refGene_tx=XM_019838444.2;refGene_gn=RNF44;refGene_tx=XM_019838461.2;refGene_gn=RNF44;refGene_tx=XM_019838467.2;refGene_gn=RNF44;refGene_tx=XM_019838474.1
A1 176200219 176200307
refGene_gn=RNF44;refGene_tx=XM_006927895.4;refGene_gn=RNF44;refGene_tx=XM_011285233.2;refGene_gn=RNF44;refGene_tx=XM_011285236.2;refGene_gn=RNF44;refGene_tx=XM_011285240.2;refGene_gn=RNF44;refGene_tx=XM_011285243.2;refGene_gn=RNF44;refGene_tx=XM_011285245.3;refGene_gn=RNF44;refGene_tx=XM_011285251.2;refGene_gn=RNF44;refGene_tx=XM_011285255.2;refGene_gn=RNF44;refGene_tx=XM_011285261.2;refGene_gn=RNF44;refGene_tx=XM_019838437.2;refGene_gn=RNF44;refGene_tx=XM_019838440.2;refGene_gn=RNF44;refGene_tx=XM_019838444.2;refGene_gn=RNF44;refGene_tx=XM_019838461.2;refGene_gn=RNF44;refGene_tx=XM_019838467.2;refGene_gn=RNF44;refGene_tx=XM_019838474.1
A1 176200395 176200517
refGene_gn=RNF44;refGene_tx=XM_006927895.4;refGene_gn=RNF44;refGene_tx=XM_011285233.2;refGene_gn=RNF44;refGene_tx=XM_011285236.2;refGene_gn=RNF44;refGene_tx=XM_011285240.2;refGene_gn=RNF44;refGene_tx=XM_0112852

43.2;refGene_gn=RNF44;refGene_tx=XM_011285245.3;refGene_gn=RNF44;refGene_tx=XM_011285251.2;refGene_gn=RNF44;refGene_tx=XM_011285255.2;refGene_gn=RNF44;refGene_tx=XM_011285261.2;refGene_gn=RNF44;refGene_tx=XM_019838437.2;refGene_gn=RNF44;refGene_tx=XM_019838440.2;refGene_gn=RNF44;refGene_tx=XM_019838444.2;refGene_gn=RNF44;refGene_tx=XM_019838461.2;refGene_gn=RNF44;refGene_tx=XM_019838467.2;refGene_gn=RNF44;refGene_tx=XM_019838474.1
A1 176200617 176200717
refGene_gn=RNF44;refGene_tx=XM_006927895.4;refGene_gn=RNF44;refGene_tx=XM_011285233.2;refGene_gn=RNF44;refGene_tx=XM_011285236.2;refGene_gn=RNF44;refGene_tx=XM_011285240.2;refGene_gn=RNF44;refGene_tx=XM_011285243.2;refGene_gn=RNF44;refGene_tx=XM_011285245.3;refGene_gn=RNF44;refGene_tx=XM_011285251.2;refGene_gn=RNF44;refGene_tx=XM_011285255.2;refGene_gn=RNF44;refGene_tx=XM_011285261.2;refGene_gn=RNF44;refGene_tx=XM_019838437.2;refGene_gn=RNF44;refGene_tx=XM_019838440.2;refGene_gn=RNF44;refGene_tx=XM_019838444.2;refGene_gn=RNF44;refGene_tx=XM_019838461.2;refGene_gn=RNF44;refGene_tx=XM_019838467.2;refGene_gn=RNF44;refGene_tx=XM_019838474.1
A1 176200916 176200979
refGene_gn=RNF44;refGene_tx=XM_011285236.2;refGene_gn=RNF44;refGene_tx=XM_011285243.2
A1 176202428 176202521
refGene_gn=RNF44;refGene_tx=XM_006927895.4;refGene_gn=RNF44;refGene_tx=XM_011285233.2;refGene_gn=RNF44;refGene_tx=XM_011285240.2;refGene_gn=RNF44;refGene_tx=XM_011285245.3;refGene_gn=RNF44;refGene_tx=XM_011285251.2;refGene_gn=RNF44;refGene_tx=XM_011285255.2;refGene_gn=RNF44;refGene_tx=XM_011285261.2;refGene_gn=RNF44;refGene_tx=XM_019838437.2;refGene_gn=RNF44;refGene_tx=XM_019838440.2;refGene_gn=RNF44;refGene_tx=XM_019838444.2;refGene_gn=RNF44;refGene_tx=XM_019838461.2;refGene_gn=RNF44;refGene_tx=XM_019838467.2;refGene_gn=RNF44;refGene_tx=XM_019838474.1
A1 176222109 176222292
refGene_gn=FAF2;refGene_tx=XM_003981228.4
A1 176227047 176227191
refGene_gn=FAF2;refGene_tx=XM_003981228.4
A1 176228135 176228307
refGene_gn=FAF2;refGene_tx=XM_003981228.4
A1 176229299 176229477
refGene_gn=FAF2;refGene_tx=XM_003981228.4
A1 176231124 176231216
refGene_gn=FAF2;refGene_tx=XM_003981228.4
A1 176231332 176231418
refGene_gn=FAF2;refGene_tx=XM_003981228.4
A1 176233017 176233156
refGene_gn=FAF2;refGene_tx=XM_003981228.4
A1 176235689 176235766
refGene_gn=FAF2;refGene_tx=XM_003981228.4
A1 176237986 176238121
refGene_gn=FAF2;refGene_tx=XM_003981228.4
A1 176250529 176250598
refGene_gn=FAF2;refGene_tx=XM_003981228.4

A1 176297461 176297524
refGene_gn=FAF2;refGene_tx=XM_003981228.4
A1 176315951 176316138
refGene_gn=CLTB;refGene_tx=XM_006927908.4;refGene_gn=CLTB;refGene_tx=XM_019838511.2
A1 176321424 176321471
refGene_gn=CLTB;refGene_tx=XM_006927908.4;refGene_gn=CLTB;refGene_tx=XM_019838511.2
A1 176331041 176331159
refGene_gn=CLTB;refGene_tx=XM_006927908.4;refGene_gn=CLTB;refGene_tx=XM_019838511.2
A1 176331399 176331511
refGene_gn=CLTB;refGene_tx=XM_006927908.4;refGene_gn=CLTB;refGene_tx=XM_019838511.2
A1 176333503 176333557
refGene_gn=CLTB;refGene_tx=XM_019838511.2
A1 176336787 176336959
refGene_gn=CLTB;refGene_tx=XM_006927908.4;refGene_gn=CLTB;refGene_tx=XM_019838511.2
A1 176339413 176339580
refGene_gn=HIGD2A;refGene_tx=XM_003981229.5
A1 176339938 176340092
refGene_gn=HIGD2A;refGene_tx=XM_003981229.5
A1 176340376 176340483
refGene_gn=NOP16;refGene_tx=XM_003981230.5
A1 176340570 176340679
refGene_gn=NOP16;refGene_tx=XM_003981230.5
A1 176343675 176343745
refGene_gn=NOP16;refGene_tx=XM_003981230.5
A1 176345531 176345638
refGene_gn=NOP16;refGene_tx=XM_003981230.5
A1 176346474 176346618
refGene_gn=NOP16;refGene_tx=XM_003981230.5
A1 176359009 176359183
refGene_gn=ARL10;refGene_tx=XM_003981231.5
A1 176360882 176361058
refGene_gn=ARL10;refGene_tx=XM_003981231.5
A1 176363843 176364045
refGene_gn=ARL10;refGene_tx=XM_003981231.5
A1 176364727 176364910
refGene_gn=ARL10;refGene_tx=XM_003981231.5
A1 176390592 176390620
refGene_gn=KIAA1191;refGene_tx=XM_003981232.4;refGene_gn=KIAA1191;refGene_tx=XM_019838541.1
A1 176393248 176393427
refGene_gn=KIAA1191;refGene_tx=XM_003981232.4;refGene_gn=KIAA1191;refGene_tx=XM_003981233.4;refGene_gn=KIAA1191;refGene_tx=XM_019838541.1
A1 176396717 176396844
refGene_gn=KIAA1191;refGene_tx=XM_003981232.4;refGene_gn=KIAA1191;refGene_tx=XM_003981233.4;refGene_gn=KIAA1191;refGene_tx=XM_019838541.1

A1 176398977 176399102
refGene_gn=KIAA1191;refGene_tx=XM_003981232.4;refGene_gn=KIAA1191;refGene_tx=XM_003981233.4;refGene_gn=KIAA1191;refGene_tx=XM_019838541.1
A1 176401103 176401210
refGene_gn=KIAA1191;refGene_tx=XM_003981232.4;refGene_gn=KIAA1191;refGene_tx=XM_003981233.4;refGene_gn=KIAA1191;refGene_tx=XM_019838541.1
A1 176401401 176401544
refGene_gn=KIAA1191;refGene_tx=XM_003981232.4;refGene_gn=KIAA1191;refGene_tx=XM_003981233.4;refGene_gn=KIAA1191;refGene_tx=XM_019838541.1
A1 176401678 176401887
refGene_gn=KIAA1191;refGene_tx=XM_003981232.4;refGene_gn=KIAA1191;refGene_tx=XM_003981233.4;refGene_gn=KIAA1191;refGene_tx=XM_019838541.1
A1 176403809 176404072
refGene_gn=SIMC1;refGene_tx=XM_023259336.1
A1 176411074 176411159
refGene_gn=SIMC1;refGene_tx=XM_023259336.1
A1 176411264 176411421
refGene_gn=SIMC1;refGene_tx=XM_023259336.1
A1 176422496 176422625
refGene_gn=SIMC1;refGene_tx=XM_023259336.1
A1 176424791 176424944
refGene_gn=SIMC1;refGene_tx=XM_023259336.1
A1 176432273 176432428
refGene_gn=SIMC1;refGene_tx=XM_023259336.1
A1 176444303 176444373
refGene_gn=SIMC1;refGene_tx=XM_023259336.1
A1 176445117 176445350
refGene_gn=SIMC1;refGene_tx=XM_023259336.1
A1 176449185 176450599
refGene_gn=SIMC1;refGene_tx=XM_023259336.1
A1 176450600 176450668
refGene_gn=SIMC1;refGene_tx=XM_023259336.1
A1 176486474 176486603
refGene_gn=SIMC1;refGene_tx=XM_023259336.1
A1 176558633 176558900
refGene_gn=THOC3;refGene_tx=XM_003981234.5
A1 176559662 176559819
refGene_gn=THOC3;refGene_tx=XM_003981234.5
A1 176561242 176561447
refGene_gn=THOC3;refGene_tx=XM_003981234.5
A1 176567595 176567757
refGene_gn=THOC3;refGene_tx=XM_003981234.5
A1 176567876 176567977
refGene_gn=THOC3;refGene_tx=XM_003981234.5
A1 176574805 176574969
refGene_gn=THOC3;refGene_tx=XM_003981234.5
A1 176649439 176649637
refGene_gn=CPLX2;refGene_tx=XM_006927899.4;refGene_gn=CPLX2;refGene_tx=XM_019838643.2;refGene_gn=CPLX2;refGene_tx=XM_023259421.1
A1 176650398 176650574

refGene_gn=CPLX2;refGene_tx=XM_006927899.4;refGene_gn=CPLX2;refGene_tx=XM_019838643.2;refGene_gn=CPLX2;refGene_tx=XM_023259421.1
A1 176650710 176650741
refGene_gn=CPLX2;refGene_tx=XM_006927899.4;refGene_gn=CPLX2;refGene_tx=XM_019838643.2;refGene_gn=CPLX2;refGene_tx=XM_023259421.1
A1 176769320 176769324
refGene_gn=HRH2;refGene_tx=XM_023259434.1
A1 176797578 176797771
refGene_gn=HRH2;refGene_tx=XM_023259432.1;refGene_gn=HRH2;refGene_tx=XM_023259433.1
A1 176819238 176819429
refGene_gn=HRH2;refGene_tx=XM_023259432.1
A1 176825153 176826233
refGene_gn=HRH2;refGene_tx=XM_023259432.1;refGene_gn=HRH2;refGene_tx=XM_023259433.1;refGene_gn=HRH2;refGene_tx=XM_023259434.1;refGene_gn=HRH2;refGene_tx=XM_023259441.1
A1 176930746 176930843
refGene_gn=SFXN1;refGene_tx=XM_011285368.3;refGene_gn=SFXN1;refGene_tx=XM_019838650.2;refGene_gn=SFXN1;refGene_tx=XM_019838651.2;refGene_gn=SFXN1;refGene_tx=XM_023259452.1
A1 176935198 176935246
refGene_gn=SFXN1;refGene_tx=XM_011285368.3;refGene_gn=SFXN1;refGene_tx=XM_019838650.2;refGene_gn=SFXN1;refGene_tx=XM_019838651.2;refGene_gn=SFXN1;refGene_tx=XM_023259452.1
A1 176935877 176935927
refGene_gn=SFXN1;refGene_tx=XM_011285368.3;refGene_gn=SFXN1;refGene_tx=XM_019838650.2;refGene_gn=SFXN1;refGene_tx=XM_019838651.2;refGene_gn=SFXN1;refGene_tx=XM_023259452.1
A1 176936866 176936902
refGene_gn=SFXN1;refGene_tx=XM_019838657.2
A1 176943119 176943169
refGene_gn=SFXN1;refGene_tx=XM_011285368.3;refGene_gn=SFXN1;refGene_tx=XM_019838650.2;refGene_gn=SFXN1;refGene_tx=XM_019838651.2;refGene_gn=SFXN1;refGene_tx=XM_019838657.2;refGene_gn=SFXN1;refGene_tx=XM_023259452.1
A1 176945052 176945180
refGene_gn=SFXN1;refGene_tx=XM_011285368.3;refGene_gn=SFXN1;refGene_tx=XM_019838650.2;refGene_gn=SFXN1;refGene_tx=XM_019838651.2;refGene_gn=SFXN1;refGene_tx=XM_019838657.2;refGene_gn=SFXN1;refGene_tx=XM_023259452.1
A1 176946143 176946229
refGene_gn=SFXN1;refGene_tx=XM_011285368.3;refGene_gn=SFXN1;refGene_tx=XM_019838650.2;refGene_gn=SFXN1;refGene_tx=XM_019838651.2;refGene_gn=SFXN1;refGene_tx=XM_019838657.2;refGene_gn=SFXN1;refGene_tx=XM_023259452.1
A1 176946749 176946825
refGene_gn=SFXN1;refGene_tx=XM_011285368.3;refGene_gn=SFXN1;refGene_tx=XM_019838650.2;refGene_gn=SFXN1;refGene_tx=XM_019838651.2;refGene_gn=SFXN1;refGene_tx=XM_019838657.2;refGene_gn=SFXN1;refGene_tx=XM_023259452.1

A1 176948124 176948223
refGene_gn=SFXN1;refGene_tx=XM_011285368.3;refGene_gn=SFXN1;refGene_tx=XM_019838650.2;refGene_gn=SFXN1;refGene_tx=XM_019838651.2;refGene_gn=SFXN1;refGene_tx=XM_019838657.2;refGene_gn=SFXN1;refGene_tx=XM_023259452.1
A1 176948964 176949135
refGene_gn=SFXN1;refGene_tx=XM_011285368.3;refGene_gn=SFXN1;refGene_tx=XM_019838650.2;refGene_gn=SFXN1;refGene_tx=XM_019838651.2;refGene_gn=SFXN1;refGene_tx=XM_019838657.2;refGene_gn=SFXN1;refGene_tx=XM_023259452.1
A1 176955552 176955716
refGene_gn=SFXN1;refGene_tx=XM_011285368.3;refGene_gn=SFXN1;refGene_tx=XM_019838650.2;refGene_gn=SFXN1;refGene_tx=XM_019838651.2;refGene_gn=SFXN1;refGene_tx=XM_019838657.2;refGene_gn=SFXN1;refGene_tx=XM_023259452.1
A1 176996366 176997707
refGene_gn=DRD1;refGene_tx=XM_003981239.5;refGene_gn=DRD1;refGene_tx=XM_019838667.2;refGene_gn=DRD1;refGene_tx=XM_023259460.1
A1 177460382 177460807
refGene_gn=MSX2;refGene_tx=XM_006927909.4
A1 177464668 177465047
refGene_gn=MSX2;refGene_tx=XM_006927909.4
A1 177976059 177976251
refGene_gn=NSG2;refGene_tx=XM_003981254.5
A1 177977170 177977281
refGene_gn=NSG2;refGene_tx=XM_003981254.5
A1 178013149 178013233
refGene_gn=NSG2;refGene_tx=XM_003981254.5
A1 178029827 178029956
refGene_gn=NSG2;refGene_tx=XM_003981254.5
A1 178092773 178093001
refGene_gn=CPEB4;refGene_tx=XM_003981256.5;refGene_gn=CPEB4;refGene_tx=XM_006927910.4;refGene_gn=CPEB4;refGene_tx=XM_011285394.3;refGene_gn=CPEB4;refGene_tx=XM_019838717.2
A1 178095497 178095679
refGene_gn=CPEB4;refGene_tx=XM_003981256.5;refGene_gn=CPEB4;refGene_tx=XM_006927910.4;refGene_gn=CPEB4;refGene_tx=XM_011285394.3;refGene_gn=CPEB4;refGene_tx=XM_019838717.2
A1 178097050 178097165
refGene_gn=CPEB4;refGene_tx=XM_003981256.5;refGene_gn=CPEB4;refGene_tx=XM_006927910.4;refGene_gn=CPEB4;refGene_tx=XM_011285394.3;refGene_gn=CPEB4;refGene_tx=XM_019838717.2
A1 178099073 178099192
refGene_gn=CPEB4;refGene_tx=XM_003981256.5;refGene_gn=CPEB4;refGene_tx=XM_006927910.4;refGene_gn=CPEB4;refGene_tx=XM_011285394.3;refGene_gn=CPEB4;refGene_tx=XM_019838717.2
A1 178099573 178099663
refGene_gn=CPEB4;refGene_tx=XM_003981256.5;refGene_gn=CPEB4;refGene_tx=XM_006927910.4;refGene_gn=CPEB4;refGene_tx=XM_011285394.3;refGene_gn=CPEB4;refGene_tx=XM_019838717.2

A1 178102795 178102969
refGene_gn=CPEB4;refGene_tx=XM_003981256.5;refGene_gn=CPEB4;refGene_tx=XM_006927910.4;refGene_gn=CPEB4;refGene_tx=XM_011285394.3;refGene_gn=CPEB4;refGene_tx=XM_019838717.2
A1 178104690 178104714
refGene_gn=CPEB4;refGene_tx=XM_011285394.3;refGene_gn=CPEB4;refGene_tx=XM_019838717.2
A1 178114538 178114589
refGene_gn=CPEB4;refGene_tx=XM_006927910.4;refGene_gn=CPEB4;refGene_tx=XM_019838717.2
A1 178133711 178133793
refGene_gn=CPEB4;refGene_tx=XM_003981256.5;refGene_gn=CPEB4;refGene_tx=XM_006927910.4;refGene_gn=CPEB4;refGene_tx=XM_011285394.3;refGene_gn=CPEB4;refGene_tx=XM_019838717.2
A1 178154955 178156080
refGene_gn=CPEB4;refGene_tx=XM_003981256.5;refGene_gn=CPEB4;refGene_tx=XM_006927910.4;refGene_gn=CPEB4;refGene_tx=XM_011285394.3;refGene_gn=CPEB4;refGene_tx=XM_019838717.2
A1 178382416 178382992
refGene_gn=BOD1;refGene_tx=XM_006927914.3;refGene_gn=BOD1;refGene_tx=XM_023259485.1
A1 178387400 178387525
refGene_gn=BOD1;refGene_tx=XM_006927914.3
A1 178388674 178388870
refGene_gn=BOD1;refGene_tx=XM_006927914.3
A1 178647788 178647939
refGene_gn=STC2;refGene_tx=XM_003981258.4
A1 178650026 178650169
refGene_gn=STC2;refGene_tx=XM_003981258.4
A1 178652686 178652898
refGene_gn=STC2;refGene_tx=XM_003981258.4
A1 178657981 178658384
refGene_gn=STC2;refGene_tx=XM_003981258.4
A1 178674213 178674267
refGene_gn=LOC102899379;refGene_tx=XM_019839788.2
A1 178674322 178674611
refGene_gn=LOC102899379;refGene_tx=XM_019839788.2
A1 178674911 178675262
refGene_gn=LOC102899379;refGene_tx=XM_019839788.2
A1 178678366 178678452
refGene_gn=LOC102899379;refGene_tx=XM_019839788.2
A1 178678545 178679031
refGene_gn=LOC102899379;refGene_tx=XM_019839788.2
A1 178735702 178736033
refGene_gn=NKX2-5;refGene_tx=XM_003981259.3
A1 178737617 178738258
refGene_gn=NKX2-5;refGene_tx=XM_003981259.3
A1 178788125 178788322
refGene_gn=BNIP1;refGene_tx=XM_003981260.4;refGene_gn=BNIP1;refGene_tx=XM_019838821.2

A1	178790601	178790693
refGene_gn=BNIP1;refGene_tx=XM_019838815.2;refGene_gn=BNIP1;refGene_tx=XM_019838821.2
A1	178791647	178791766
refGene_gn=BNIP1;refGene_tx=XM_003981260.4;refGene_gn=BNIP1;refGene_tx=XM_019838815.2;refGene_gn=BNIP1;refGene_tx=XM_019838821.2
A1	178792757	178792859
refGene_gn=BNIP1;refGene_tx=XM_003981260.4;refGene_gn=BNIP1;refGene_tx=XM_019838815.2;refGene_gn=BNIP1;refGene_tx=XM_019838821.2
A1	178795186	178795278
refGene_gn=BNIP1;refGene_tx=XM_003981260.4;refGene_gn=BNIP1;refGene_tx=XM_019838815.2;refGene_gn=BNIP1;refGene_tx=XM_019838821.2
A1	178798752	178798845
refGene_gn=BNIP1;refGene_tx=XM_003981260.4;refGene_gn=BNIP1;refGene_tx=XM_019838815.2;refGene_gn=BNIP1;refGene_tx=XM_019838821.2
A1	178800740	178800824
refGene_gn=BNIP1;refGene_tx=XM_003981260.4;refGene_gn=BNIP1;refGene_tx=XM_019838815.2;refGene_gn=BNIP1;refGene_tx=XM_019838821.2
A1	178810882	178810998
refGene_gn=CREBRF;refGene_tx=XM_003981262.5;refGene_gn=CREBRF;refGene_tx=XM_019838830.2;refGene_gn=CREBRF;refGene_tx=XM_023259550.1
A1	178821014	178821137
refGene_gn=CREBRF;refGene_tx=XM_003981262.5;refGene_gn=CREBRF;refGene_tx=XM_019838830.2;refGene_gn=CREBRF;refGene_tx=XM_023259550.1
A1	178827067	178827141
refGene_gn=CREBRF;refGene_tx=XM_003981262.5;refGene_gn=CREBRF;refGene_tx=XM_019838830.2;refGene_gn=CREBRF;refGene_tx=XM_023259550.1
A1	178828333	178828523
refGene_gn=CREBRF;refGene_tx=XM_003981262.5;refGene_gn=CREBRF;refGene_tx=XM_019838830.2;refGene_gn=CREBRF;refGene_tx=XM_023259550.1
A1	178830390	178830585
refGene_gn=CREBRF;refGene_tx=XM_003981262.5;refGene_gn=CREBRF;refGene_tx=XM_019838830.2;refGene_gn=CREBRF;refGene_tx=XM_023259550.1
A1	178837060	178838147
refGene_gn=CREBRF;refGene_tx=XM_003981262.5;refGene_gn=CREBRF;refGene_tx=XM_019838830.2;refGene_gn=CREBRF;refGene_tx=XM_023259550.1
A1	178839823	178839949
refGene_gn=CREBRF;refGene_tx=XM_003981262.5;refGene_gn=CREBRF;refGene_tx=XM_019838830.2;refGene_gn=CREBRF;refGene_tx=XM_023259550.1
A1	178842155	178842164
refGene_gn=CREBRF;refGene_tx=XM_003981262.5;refGene_gn=CREBRF;refGene_tx=XM_019838830.2;refGene_gn=CREBRF;refGene_tx=XM_023259550.1
A1	178884590	178884684
refGene_gn=ATP6V0E1;refGene_tx=XM_011285428.3
A1	178903028	178903076
refGene_gn=ATP6V0E1;refGene_tx=XM_011285428.3
A1	178909518	178909622
refGene_gn=ATP6V0E1;refGene_tx=XM_011285428.3
A1	178918869	178918998
refGene_gn=RPL26L1;refGene_tx=XM_019838863.2;refGene_gn=RPL26L1;refGen

e_tx=XM_019838865.2;refGene_gn=RPL26L1;refGene_tx=XM_019838869.2
A1 178919743 178919884
refGene_gn=RPL26L1;refGene_tx=XM_019838863.2;refGene_gn=RPL26L1;refGen
e_tx=XM_019838865.2;refGene_gn=RPL26L1;refGene_tx=XM_019838869.2
A1 178926997 178927165
refGene_gn=RPL26L1;refGene_tx=XM_019838863.2;refGene_gn=RPL26L1;refGen
e_tx=XM_019838865.2;refGene_gn=RPL26L1;refGene_tx=XM_019838869.2
A1 178935159 178935446
refGene_gn=ERGIC1;refGene_tx=XM_023259579.1;refGene_gn=ERGIC1;refGene_
tx=XM_023259581.1
A1 178945499 178945560
refGene_gn=ERGIC1;refGene_tx=XM_023259581.1
A1 178946380 178946503
refGene_gn=ERGIC1;refGene_tx=XM_023259579.1;refGene_gn=ERGIC1;refGene_
tx=XM_023259581.1
A1 178949312 178949413
refGene_gn=ERGIC1;refGene_tx=XM_023259579.1;refGene_gn=ERGIC1;refGene_
tx=XM_023259581.1
A1 178953620 178953681
refGene_gn=ERGIC1;refGene_tx=XM_023259579.1;refGene_gn=ERGIC1;refGene_
tx=XM_023259581.1
A1 178958471 178958576
refGene_gn=ERGIC1;refGene_tx=XM_023259579.1;refGene_gn=ERGIC1;refGene_
tx=XM_023259581.1
A1 178967021 178967146
refGene_gn=ERGIC1;refGene_tx=XM_023259579.1;refGene_gn=ERGIC1;refGene_
tx=XM_023259581.1
A1 178970528 178970623
refGene_gn=ERGIC1;refGene_tx=XM_023259579.1
A1 178981250 178981323
refGene_gn=ERGIC1;refGene_tx=XM_023259579.1
A1 178988309 178988371
refGene_gn=ERGIC1;refGene_tx=XM_023259579.1
A1 179036501 179036521
refGene_gn=ERGIC1;refGene_tx=XM_023259579.1
A1 179089671 179090038
refGene_gn=DUSP1;refGene_tx=XM_023259591.1
A1 179090321 179090467
refGene_gn=DUSP1;refGene_tx=XM_023259591.1
A1 179090819 179091039
refGene_gn=DUSP1;refGene_tx=XM_023259591.1
A1 179091459 179091830
refGene_gn=DUSP1;refGene_tx=XM_023259591.1
A1 179164386 179164631
refGene_gn=NEURL1B;refGene_tx=XM_023259599.1
A1 179165016 179165142
refGene_gn=NEURL1B;refGene_tx=XM_023259599.1
A1 179167276 179167996
refGene_gn=NEURL1B;refGene_tx=XM_023259599.1
A1 179180107 179180653

refGene_gn=NEURL1B;refGene_tx=XM_023259599.1
A1 179198492 179198523
refGene_gn=NEURL1B;refGene_tx=XM_023259599.1
A1 179362610 179362685
refGene_gn=SH3PXD2B;refGene_tx=XM_023259607.1
A1 179396284 179396365
refGene_gn=SH3PXD2B;refGene_tx=XM_023259607.1
A1 179407885 179407961
refGene_gn=SH3PXD2B;refGene_tx=XM_023259607.1
A1 179414305 179414382
refGene_gn=SH3PXD2B;refGene_tx=XM_023259607.1
A1 179426622 179426714
refGene_gn=SH3PXD2B;refGene_tx=XM_023259607.1
A1 179430687 179430713
refGene_gn=SH3PXD2B;refGene_tx=XM_023259607.1
A1 179438689 179438824
refGene_gn=SH3PXD2B;refGene_tx=XM_023259607.1
A1 179442924 179443029
refGene_gn=SH3PXD2B;refGene_tx=XM_023259607.1
A1 179447254 179447372
refGene_gn=SH3PXD2B;refGene_tx=XM_023259607.1
A1 179450400 179450627
refGene_gn=SH3PXD2B;refGene_tx=XM_023259607.1
A1 179452511 179452561
refGene_gn=SH3PXD2B;refGene_tx=XM_023259607.1
A1 179453474 179453600
refGene_gn=SH3PXD2B;refGene_tx=XM_023259607.1
A1 179460653 179462198
refGene_gn=SH3PXD2B;refGene_tx=XM_023259607.1
A1 179500588 179500658
refGene_gn=UBTD2;refGene_tx=XM_023259612.1
A1 179516185 179516222
refGene_gn=UBTD2;refGene_tx=XM_019838991.2
A1 179540732 179540969
refGene_gn=UBTD2;refGene_tx=XM_019838991.2;refGene_gn=UBTD2;refGene_tx
=XM_023259612.1
A1 179558593 179558991
refGene_gn=UBTD2;refGene_tx=XM_019838991.2;refGene_gn=UBTD2;refGene_tx
=XM_023259612.1
A1 179568334 179568532
refGene_gn=EFCAB9;refGene_tx=XM_023259610.1
A1 179569261 179569438
refGene_gn=EFCAB9;refGene_tx=XM_023259610.1
A1 179570383 179570532
refGene_gn=EFCAB9;refGene_tx=XM_023259610.1
A1 179573965 179574203
refGene_gn=EFCAB9;refGene_tx=XM_023259610.1
A1 179585031 179585187
refGene_gn=STK10;refGene_tx=XM_011285468.3
A1 179610560 179610725

refGene_gn=STK10;refGene_tx=XM_011285468.3
A1 179638600 179638649
refGene_gn=STK10;refGene_tx=XM_011285468.3
A1 179644355 179644505
refGene_gn=STK10;refGene_tx=XM_011285468.3
A1 179660539 179660612
refGene_gn=STK10;refGene_tx=XM_011285468.3
A1 179661870 179662065
refGene_gn=STK10;refGene_tx=XM_011285468.3
A1 179662645 179662727
refGene_gn=STK10;refGene_tx=XM_011285468.3
A1 179670102 179670237
refGene_gn=STK10;refGene_tx=XM_011285468.3
A1 179672335 179672881
refGene_gn=STK10;refGene_tx=XM_011285468.3
A1 179675223 179675354
refGene_gn=STK10;refGene_tx=XM_011285468.3
A1 179679617 179679741
refGene_gn=STK10;refGene_tx=XM_011285468.3
A1 179680159 179680339
refGene_gn=STK10;refGene_tx=XM_011285468.3
A1 179684406 179684499
refGene_gn=STK10;refGene_tx=XM_011285468.3
A1 179687314 179687444
refGene_gn=STK10;refGene_tx=XM_011285468.3
A1 179689217 179689342
refGene_gn=STK10;refGene_tx=XM_011285468.3
A1 179690339 179690528
refGene_gn=STK10;refGene_tx=XM_011285468.3
A1 179691120 179691246
refGene_gn=STK10;refGene_tx=XM_011285468.3
A1 179694160 179694274
refGene_gn=STK10;refGene_tx=XM_011285468.3
A1 179701293 179701434
refGene_gn=STK10;refGene_tx=XM_011285468.3
A1 179751087 179751132
refGene_gn=FBXW11;refGene_tx=XM_003981273.5;refGene_gn=FBXW11;refGene_tx=XM_003981275.5;refGene_gn=FBXW11;refGene_tx=XM_006927920.4
A1 179773129 179773174
refGene_gn=FBXW11;refGene_tx=XM_019838940.2;refGene_gn=FBXW11;refGene_tx=XM_019838963.2;refGene_gn=FBXW11;refGene_tx=XM_023259633.1;refGene_gn=FBXW11;refGene_tx=XM_023259634.1
A1 179788601 179788703
refGene_gn=FBXW11;refGene_tx=XM_006927920.4;refGene_gn=FBXW11;refGene_tx=XM_019838940.2;refGene_gn=FBXW11;refGene_tx=XM_023259633.1;refGene_gn=FBXW11;refGene_tx=XM_023259634.1
A1 179829141 179829144
refGene_gn=FBXW11;refGene_tx=XM_019838966.1
A1 179830690 179830753
refGene_gn=FBXW11;refGene_tx=XM_003981273.5;refGene_gn=FBXW11;refGene_

tx=XM_006927920.4;refGene_gn=FBXW11;refGene_tx=XM_006927921.3;refGene_gn=FBXW11;refGene_tx=XM_019838940.2;refGene_gn=FBXW11;refGene_tx=XM_019838963.2;refGene_gn=FBXW11;refGene_tx=XM_019838966.1;refGene_gn=FBXW11;refGene_tx=XM_023259633.1
A1 179833802 179834028
refGene_gn=FBXW11;refGene_tx=XM_003981273.5;refGene_gn=FBXW11;refGene_tx=XM_003981275.5;refGene_gn=FBXW11;refGene_tx=XM_006927920.4;refGene_gn=FBXW11;refGene_tx=XM_006927921.3;refGene_gn=FBXW11;refGene_tx=XM_019838940.2;refGene_gn=FBXW11;refGene_tx=XM_019838963.2;refGene_gn=FBXW11;refGene_tx=XM_019838966.1;refGene_gn=FBXW11;refGene_tx=XM_023259633.1;refGene_gn=FBXW11;refGene_tx=XM_023259634.1
A1 179843870 179844057
refGene_gn=FBXW11;refGene_tx=XM_003981273.5;refGene_gn=FBXW11;refGene_tx=XM_003981275.5;refGene_gn=FBXW11;refGene_tx=XM_006927920.4;refGene_gn=FBXW11;refGene_tx=XM_006927921.3;refGene_gn=FBXW11;refGene_tx=XM_019838940.2;refGene_gn=FBXW11;refGene_tx=XM_019838963.2;refGene_gn=FBXW11;refGene_tx=XM_019838966.1;refGene_gn=FBXW11;refGene_tx=XM_023259633.1;refGene_gn=FBXW11;refGene_tx=XM_023259634.1
A1 179844925 179845016
refGene_gn=FBXW11;refGene_tx=XM_003981273.5;refGene_gn=FBXW11;refGene_tx=XM_003981275.5;refGene_gn=FBXW11;refGene_tx=XM_006927920.4;refGene_gn=FBXW11;refGene_tx=XM_006927921.3;refGene_gn=FBXW11;refGene_tx=XM_019838940.2;refGene_gn=FBXW11;refGene_tx=XM_019838963.2;refGene_gn=FBXW11;refGene_tx=XM_019838966.1;refGene_gn=FBXW11;refGene_tx=XM_023259633.1;refGene_gn=FBXW11;refGene_tx=XM_023259634.1
A1 179852511 179852649
refGene_gn=FBXW11;refGene_tx=XM_003981273.5;refGene_gn=FBXW11;refGene_tx=XM_003981275.5;refGene_gn=FBXW11;refGene_tx=XM_006927920.4;refGene_gn=FBXW11;refGene_tx=XM_006927921.3;refGene_gn=FBXW11;refGene_tx=XM_019838940.2;refGene_gn=FBXW11;refGene_tx=XM_019838963.2;refGene_gn=FBXW11;refGene_tx=XM_019838966.1;refGene_gn=FBXW11;refGene_tx=XM_023259633.1;refGene_gn=FBXW11;refGene_tx=XM_023259634.1
A1 179856962 179857081
refGene_gn=FBXW11;refGene_tx=XM_003981273.5;refGene_gn=FBXW11;refGene_tx=XM_003981275.5;refGene_gn=FBXW11;refGene_tx=XM_006927920.4;refGene_gn=FBXW11;refGene_tx=XM_006927921.3;refGene_gn=FBXW11;refGene_tx=XM_019838940.2;refGene_gn=FBXW11;refGene_tx=XM_019838963.2;refGene_gn=FBXW11;refGene_tx=XM_019838966.1;refGene_gn=FBXW11;refGene_tx=XM_023259633.1;refGene_gn=FBXW11;refGene_tx=XM_023259634.1
A1 179858312 179858562
refGene_gn=FBXW11;refGene_tx=XM_003981273.5;refGene_gn=FBXW11;refGene_tx=XM_003981275.5;refGene_gn=FBXW11;refGene_tx=XM_006927920.4;refGene_gn=FBXW11;refGene_tx=XM_006927921.3;refGene_gn=FBXW11;refGene_tx=XM_019838940.2;refGene_gn=FBXW11;refGene_tx=XM_019838963.2;refGene_gn=FBXW11;refGene_tx=XM_019838966.1;refGene_gn=FBXW11;refGene_tx=XM_023259633.1;refGene_gn=FBXW11;refGene_tx=XM_023259634.1
A1 179861289 179861408
refGene_gn=FBXW11;refGene_tx=XM_003981273.5;refGene_gn=FBXW11;refGene_tx=XM_003981275.5;refGene_gn=FBXW11;refGene_tx=XM_006927920.4;refGene_gn=FBXW11;refGene_tx=XM_006927921.3;refGene_gn=FBXW11;refGene_tx=XM_01

9838940.2;refGene_gn=FBXW11;refGene_tx=XM_019838963.2;refGene_gn=FBXW1
1;refGene_tx=XM_019838966.1;refGene_gn=FBXW11;refGene_tx=XM_023259633.
1;refGene_gn=FBXW11;refGene_tx=XM_023259634.1
A1 179863236 179863347
refGene_gn=FBXW11;refGene_tx=XM_003981273.5;refGene_gn=FBXW11;refGene_
tx=XM_003981275.5;refGene_gn=FBXW11;refGene_tx=XM_006927920.4;refGene_
gn=FBXW11;refGene_tx=XM_006927921.3;refGene_gn=FBXW11;refGene_tx=XM_01
9838940.2;refGene_gn=FBXW11;refGene_tx=XM_019838963.2;refGene_gn=FBXW1
1;refGene_tx=XM_019838966.1;refGene_gn=FBXW11;refGene_tx=XM_023259633.
1;refGene_gn=FBXW11;refGene_tx=XM_023259634.1
A1 179864516 179864595
refGene_gn=FBXW11;refGene_tx=XM_003981273.5;refGene_gn=FBXW11;refGene_
tx=XM_003981275.5;refGene_gn=FBXW11;refGene_tx=XM_006927920.4;refGene_
gn=FBXW11;refGene_tx=XM_006927921.3;refGene_gn=FBXW11;refGene_tx=XM_01
9838940.2;refGene_gn=FBXW11;refGene_tx=XM_019838963.2;refGene_gn=FBXW1
1;refGene_tx=XM_019838966.1;refGene_gn=FBXW11;refGene_tx=XM_023259633.
1;refGene_gn=FBXW11;refGene_tx=XM_023259634.1
A1 179865897 179866059
refGene_gn=FBXW11;refGene_tx=XM_003981273.5;refGene_gn=FBXW11;refGene_
tx=XM_003981275.5;refGene_gn=FBXW11;refGene_tx=XM_006927920.4;refGene_
gn=FBXW11;refGene_tx=XM_006927921.3;refGene_gn=FBXW11;refGene_tx=XM_01
9838940.2;refGene_gn=FBXW11;refGene_tx=XM_019838963.2;refGene_gn=FBXW1
1;refGene_tx=XM_019838966.1;refGene_gn=FBXW11;refGene_tx=XM_023259633.
1;refGene_gn=FBXW11;refGene_tx=XM_023259634.1
A1 179942611 179942824
refGene_gn=SMIM23;refGene_tx=XM_023259648.1
A1 179943090 179943158
refGene_gn=SMIM23;refGene_tx=XM_023259648.1
A1 179943382 179943434
refGene_gn=SMIM23;refGene_tx=XM_023259648.1
A1 179948985 179949075
refGene_gn=SMIM23;refGene_tx=XM_023259648.1
A1 180245642 180245909
refGene_gn=FGF18;refGene_tx=XM_011285502.3
A1 180252177 180252284
refGene_gn=FGF18;refGene_tx=XM_011285502.3
A1 180263198 180263379
refGene_gn=FGF18;refGene_tx=XM_011285502.3
A1 180276756 180276793
refGene_gn=FGF18;refGene_tx=XM_011285502.3
A1 180276966 180276998
refGene_gn=FGF18;refGene_tx=XM_011285502.3
A1 180284627 180284666
refGene_gn=NPM1;refGene_tx=XM_003981277.3
A1 180287641 180287716
refGene_gn=NPM1;refGene_tx=XM_003981277.3
A1 180288947 180288956
refGene_gn=NPM1;refGene_tx=XM_023259650.1
A1 180289939 180290041
refGene_gn=NPM1;refGene_tx=XM_003981277.3;refGene_gn=NPM1;refGene_tx=X

M_023259650.1
A1 180290741 180290825
refGene_gn=NPM1;refGene_tx=XM_003981277.3;refGene_gn=NPM1;refGene_tx=XM_023259650.1
A1 180291389 180291447
refGene_gn=NPM1;refGene_tx=XM_003981277.3;refGene_gn=NPM1;refGene_tx=XM_023259650.1
A1 180296252 180296320
refGene_gn=NPM1;refGene_tx=XM_003981277.3;refGene_gn=NPM1;refGene_tx=XM_023259650.1
A1 180296433 180296540
refGene_gn=NPM1;refGene_tx=XM_003981277.3;refGene_gn=NPM1;refGene_tx=XM_023259650.1
A1 180296880 180296974
refGene_gn=NPM1;refGene_tx=XM_003981277.3;refGene_gn=NPM1;refGene_tx=XM_023259650.1
A1 180297306 180297426
refGene_gn=NPM1;refGene_tx=XM_003981277.3;refGene_gn=NPM1;refGene_tx=XM_023259650.1
A1 180298007 180298087
refGene_gn=NPM1;refGene_tx=XM_003981277.3;refGene_gn=NPM1;refGene_tx=XM_023259650.1
A1 180300155 180300213
refGene_gn=NPM1;refGene_tx=XM_003981277.3;refGene_gn=NPM1;refGene_tx=XM_023259650.1
A1 180351646 180351857
refGene_gn=TLX3;refGene_tx=XM_011285513.3
A1 180352806 180353050
refGene_gn=TLX3;refGene_tx=XM_011285513.3
A1 180353404 180353966
refGene_gn=TLX3;refGene_tx=XM_011285513.3
A1 180365297 180365394
refGene_gn=RANBP17;refGene_tx=XM_023259662.1;refGene_gn=RANBP17;refGene_tx=XM_023259663.1;refGene_gn=RANBP17;refGene_tx=XM_023259664.1
A1 180367229 180367357
refGene_gn=RANBP17;refGene_tx=XM_023259662.1;refGene_gn=RANBP17;refGene_tx=XM_023259663.1;refGene_gn=RANBP17;refGene_tx=XM_023259664.1
A1 180369241 180369340
refGene_gn=RANBP17;refGene_tx=XM_023259662.1;refGene_gn=RANBP17;refGene_tx=XM_023259663.1;refGene_gn=RANBP17;refGene_tx=XM_023259664.1
A1 180394683 180394850
refGene_gn=RANBP17;refGene_tx=XM_023259662.1;refGene_gn=RANBP17;refGene_tx=XM_023259663.1;refGene_gn=RANBP17;refGene_tx=XM_023259664.1;refGene_gn=RANBP17;refGene_tx=XM_023259665.1
A1 180406783 180406922
refGene_gn=RANBP17;refGene_tx=XM_023259662.1;refGene_gn=RANBP17;refGene_tx=XM_023259663.1;refGene_gn=RANBP17;refGene_tx=XM_023259664.1;refGene_gn=RANBP17;refGene_tx=XM_023259665.1
A1 180409702 180409917
refGene_gn=RANBP17;refGene_tx=XM_023259662.1;refGene_gn=RANBP17;refGen

e_tx=XM_023259663.1;refGene_gn=RANBP17;refGene_tx=XM_023259664.1
A1 180422505 180422588
refGene_gn=RANBP17;refGene_tx=XM_023259662.1;refGene_gn=RANBP17;refGene_tx=XM_023259663.1;refGene_gn=RANBP17;refGene_tx=XM_023259664.1;refGene_gn=RANBP17;refGene_tx=XM_023259665.1
A1 180425231 180425339
refGene_gn=RANBP17;refGene_tx=XM_023259662.1;refGene_gn=RANBP17;refGene_tx=XM_023259663.1;refGene_gn=RANBP17;refGene_tx=XM_023259664.1;refGene_gn=RANBP17;refGene_tx=XM_023259665.1
A1 180433045 180433134
refGene_gn=RANBP17;refGene_tx=XM_023259662.1;refGene_gn=RANBP17;refGene_tx=XM_023259663.1;refGene_gn=RANBP17;refGene_tx=XM_023259664.1;refGene_gn=RANBP17;refGene_tx=XM_023259665.1
A1 180438940 180439044
refGene_gn=RANBP17;refGene_tx=XM_023259662.1;refGene_gn=RANBP17;refGene_tx=XM_023259663.1;refGene_gn=RANBP17;refGene_tx=XM_023259664.1;refGene_gn=RANBP17;refGene_tx=XM_023259665.1
A1 180451494 180451603
refGene_gn=RANBP17;refGene_tx=XM_023259662.1;refGene_gn=RANBP17;refGene_tx=XM_023259663.1;refGene_gn=RANBP17;refGene_tx=XM_023259664.1;refGene_gn=RANBP17;refGene_tx=XM_023259665.1
A1 180451695 180451759
refGene_gn=RANBP17;refGene_tx=XM_023259662.1;refGene_gn=RANBP17;refGene_tx=XM_023259663.1;refGene_gn=RANBP17;refGene_tx=XM_023259664.1;refGene_gn=RANBP17;refGene_tx=XM_023259665.1
A1 180462026 180462107
refGene_gn=RANBP17;refGene_tx=XM_023259662.1;refGene_gn=RANBP17;refGene_tx=XM_023259663.1;refGene_gn=RANBP17;refGene_tx=XM_023259664.1;refGene_gn=RANBP17;refGene_tx=XM_023259665.1
A1 180463160 180463234
refGene_gn=RANBP17;refGene_tx=XM_023259662.1;refGene_gn=RANBP17;refGene_tx=XM_023259663.1;refGene_gn=RANBP17;refGene_tx=XM_023259664.1;refGene_gn=RANBP17;refGene_tx=XM_023259665.1
A1 180470811 180470871
refGene_gn=RANBP17;refGene_tx=XM_023259662.1;refGene_gn=RANBP17;refGene_tx=XM_023259664.1;refGene_gn=RANBP17;refGene_tx=XM_023259665.1
A1 180624008 180624144
refGene_gn=RANBP17;refGene_tx=XM_023259662.1;refGene_gn=RANBP17;refGene_tx=XM_023259663.1;refGene_gn=RANBP17;refGene_tx=XM_023259664.1;refGene_gn=RANBP17;refGene_tx=XM_023259665.1
A1 180634265 180634371
refGene_gn=RANBP17;refGene_tx=XM_023259662.1;refGene_gn=RANBP17;refGene_tx=XM_023259663.1;refGene_gn=RANBP17;refGene_tx=XM_023259664.1;refGene_gn=RANBP17;refGene_tx=XM_023259665.1
A1 180656950 180657144
refGene_gn=RANBP17;refGene_tx=XM_023259662.1;refGene_gn=RANBP17;refGene_tx=XM_023259663.1;refGene_gn=RANBP17;refGene_tx=XM_023259664.1;refGene_gn=RANBP17;refGene_tx=XM_023259665.1
A1 180659150 180659323
refGene_gn=RANBP17;refGene_tx=XM_023259662.1;refGene_gn=RANBP17;refGen

e_tx=XM_023259663.1;refGene_gn=RANBP17;refGene_tx=XM_023259664.1;refGene_gn=RANBP17;refGene_tx=XM_023259665.1
A1 180659927 180660074
refGene_gn=RANBP17;refGene_tx=XM_023259662.1;refGene_gn=RANBP17;refGene_tx=XM_023259663.1;refGene_gn=RANBP17;refGene_tx=XM_023259664.1;refGene_gn=RANBP17;refGene_tx=XM_023259665.1
A1 180661969 180662089
refGene_gn=RANBP17;refGene_tx=XM_023259662.1;refGene_gn=RANBP17;refGene_tx=XM_023259663.1;refGene_gn=RANBP17;refGene_tx=XM_023259665.1
A1 180664535 180664609
refGene_gn=RANBP17;refGene_tx=XM_023259662.1;refGene_gn=RANBP17;refGene_tx=XM_023259663.1;refGene_gn=RANBP17;refGene_tx=XM_023259664.1;refGene_gn=RANBP17;refGene_tx=XM_023259665.1
A1 180667374 180667540
refGene_gn=RANBP17;refGene_tx=XM_023259662.1;refGene_gn=RANBP17;refGene_tx=XM_023259663.1;refGene_gn=RANBP17;refGene_tx=XM_023259664.1;refGene_gn=RANBP17;refGene_tx=XM_023259665.1
A1 180669195 180669300
refGene_gn=RANBP17;refGene_tx=XM_023259662.1;refGene_gn=RANBP17;refGene_tx=XM_023259663.1;refGene_gn=RANBP17;refGene_tx=XM_023259664.1;refGene_gn=RANBP17;refGene_tx=XM_023259665.1
A1 180673815 180673881
refGene_gn=RANBP17;refGene_tx=XM_023259662.1;refGene_gn=RANBP17;refGene_tx=XM_023259663.1;refGene_gn=RANBP17;refGene_tx=XM_023259664.1;refGene_gn=RANBP17;refGene_tx=XM_023259665.1
A1 180676914 180677081
refGene_gn=RANBP17;refGene_tx=XM_023259662.1;refGene_gn=RANBP17;refGene_tx=XM_023259663.1;refGene_gn=RANBP17;refGene_tx=XM_023259664.1;refGene_gn=RANBP17;refGene_tx=XM_023259665.1
A1 180685212 180685303
refGene_gn=RANBP17;refGene_tx=XM_023259662.1;refGene_gn=RANBP17;refGene_tx=XM_023259663.1;refGene_gn=RANBP17;refGene_tx=XM_023259664.1;refGene_gn=RANBP17;refGene_tx=XM_023259665.1
A1 180688825 180688972
refGene_gn=RANBP17;refGene_tx=XM_023259662.1;refGene_gn=RANBP17;refGene_tx=XM_023259663.1;refGene_gn=RANBP17;refGene_tx=XM_023259664.1;refGene_gn=RANBP17;refGene_tx=XM_023259665.1
A1 180698294 180698312
refGene_gn=RANBP17;refGene_tx=XM_023259662.1;refGene_gn=RANBP17;refGene_tx=XM_023259663.1;refGene_gn=RANBP17;refGene_tx=XM_023259664.1;refGene_gn=RANBP17;refGene_tx=XM_023259665.1
A1 180725395 180725404
refGene_gn=GABRP;refGene_tx=XM_019839066.2
A1 180740844 180741147
refGene_gn=GABRP;refGene_tx=XM_006927927.4;refGene_gn=GABRP;refGene_tx=XM_019839020.2;refGene_gn=GABRP;refGene_tx=XM_019839025.2;refGene_gn=GABRP;refGene_tx=XM_019839035.2;refGene_gn=GABRP;refGene_tx=XM_019839043.2;refGene_gn=GABRP;refGene_tx=XM_019839050.2;refGene_gn=GABRP;refGene_tx=XM_019839052.2;refGene_gn=GABRP;refGene_tx=XM_019839056.2;refGene_gn=GABRP;refGene_tx=XM_019839061.2

A1 180742851 180743039
refGene_gn=GABRP;refGene_tx=XM_006927927.4;refGene_gn=GABRP;refGene_tx=XM_019839020.2;refGene_gn=GABRP;refGene_tx=XM_019839025.2;refGene_gn=GABRP;refGene_tx=XM_019839035.2;refGene_gn=GABRP;refGene_tx=XM_019839043.2;refGene_gn=GABRP;refGene_tx=XM_019839050.2;refGene_gn=GABRP;refGene_tx=XM_019839052.2;refGene_gn=GABRP;refGene_tx=XM_019839056.2;refGene_gn=GABRP;refGene_tx=XM_019839061.2;refGene_gn=GABRP;refGene_tx=XM_019839066.2
A1 180743562 180743715
refGene_gn=GABRP;refGene_tx=XM_006927927.4;refGene_gn=GABRP;refGene_tx=XM_019839020.2;refGene_gn=GABRP;refGene_tx=XM_019839025.2;refGene_gn=GABRP;refGene_tx=XM_019839035.2;refGene_gn=GABRP;refGene_tx=XM_019839043.2;refGene_gn=GABRP;refGene_tx=XM_019839050.2;refGene_gn=GABRP;refGene_tx=XM_019839052.2;refGene_gn=GABRP;refGene_tx=XM_019839056.2;refGene_gn=GABRP;refGene_tx=XM_019839061.2;refGene_gn=GABRP;refGene_tx=XM_019839066.2
A1 180744424 180744562
refGene_gn=GABRP;refGene_tx=XM_006927927.4;refGene_gn=GABRP;refGene_tx=XM_019839020.2;refGene_gn=GABRP;refGene_tx=XM_019839025.2;refGene_gn=GABRP;refGene_tx=XM_019839035.2;refGene_gn=GABRP;refGene_tx=XM_019839043.2;refGene_gn=GABRP;refGene_tx=XM_019839050.2;refGene_gn=GABRP;refGene_tx=XM_019839052.2;refGene_gn=GABRP;refGene_tx=XM_019839056.2;refGene_gn=GABRP;refGene_tx=XM_019839061.2;refGene_gn=GABRP;refGene_tx=XM_019839066.2
A1 180748218 180748301
refGene_gn=GABRP;refGene_tx=XM_006927927.4;refGene_gn=GABRP;refGene_tx=XM_019839020.2;refGene_gn=GABRP;refGene_tx=XM_019839025.2;refGene_gn=GABRP;refGene_tx=XM_019839035.2;refGene_gn=GABRP;refGene_tx=XM_019839043.2;refGene_gn=GABRP;refGene_tx=XM_019839050.2;refGene_gn=GABRP;refGene_tx=XM_019839052.2;refGene_gn=GABRP;refGene_tx=XM_019839056.2;refGene_gn=GABRP;refGene_tx=XM_019839061.2;refGene_gn=GABRP;refGene_tx=XM_019839066.2
A1 180750318 180750581
refGene_gn=GABRP;refGene_tx=XM_006927927.4;refGene_gn=GABRP;refGene_tx=XM_019839020.2;refGene_gn=GABRP;refGene_tx=XM_019839025.2;refGene_gn=GABRP;refGene_tx=XM_019839035.2;refGene_gn=GABRP;refGene_tx=XM_019839043.2;refGene_gn=GABRP;refGene_tx=XM_019839050.2;refGene_gn=GABRP;refGene_tx=XM_019839052.2;refGene_gn=GABRP;refGene_tx=XM_019839056.2;refGene_gn=GABRP;refGene_tx=XM_019839061.2;refGene_gn=GABRP;refGene_tx=XM_019839066.2
A1 180751331 180751399
refGene_gn=GABRP;refGene_tx=XM_006927927.4;refGene_gn=GABRP;refGene_tx=XM_019839020.2;refGene_gn=GABRP;refGene_tx=XM_019839025.2;refGene_gn=GABRP;refGene_tx=XM_019839035.2;refGene_gn=GABRP;refGene_tx=XM_019839043.2;refGene_gn=GABRP;refGene_tx=XM_019839050.2;refGene_gn=GABRP;refGene_tx=XM_019839052.2;refGene_gn=GABRP;refGene_tx=XM_019839056.2;refGene_gn=GABRP;refGene_tx=XM_019839061.2;refGene_gn=GABRP;refGene_tx=XM_019839066.2
A1 180755388 180755507
refGene_gn=GABRP;refGene_tx=XM_006927927.4;refGene_gn=GABRP;refGene_tx

=XM_019839020.2;refGene_gn=GABRP;refGene_tx=XM_019839025.2;refGene_gn=GABRP;refGene_tx=XM_019839066.2
A1	180756194	180756247
refGene_gn=GABRP;refGene_tx=XM_006927927.4;refGene_gn=GABRP;refGene_tx=XM_019839020.2;refGene_gn=GABRP;refGene_tx=XM_019839025.2;refGene_gn=GABRP;refGene_tx=XM_019839066.2
A1	180803477	180805517
refGene_gn=LOC109503008;refGene_tx=XM_019839074.2;refGene_gn=LOC109503008;refGene_tx=XM_019839076.2
A1	180821380	180821428
refGene_gn=KCNIP1;refGene_tx=XM_011285529.3;refGene_gn=KCNIP1;refGene_tx=XM_011285533.2;refGene_gn=KCNIP1;refGene_tx=XM_019839082.2;refGene_gn=KCNIP1;refGene_tx=XM_019839086.2
A1	180825106	180825169
refGene_gn=KCNIP1;refGene_tx=XM_011285529.3;refGene_gn=KCNIP1;refGene_tx=XM_011285533.2;refGene_gn=KCNIP1;refGene_tx=XM_019839082.2;refGene_gn=KCNIP1;refGene_tx=XM_019839086.2
A1	180826068	180826173
refGene_gn=KCNIP1;refGene_tx=XM_011285529.3;refGene_gn=KCNIP1;refGene_tx=XM_011285533.2;refGene_gn=KCNIP1;refGene_tx=XM_019839082.2;refGene_gn=KCNIP1;refGene_tx=XM_019839086.2
A1	180831337	180831445
refGene_gn=KCNIP1;refGene_tx=XM_011285529.3;refGene_gn=KCNIP1;refGene_tx=XM_011285533.2;refGene_gn=KCNIP1;refGene_tx=XM_019839082.2;refGene_gn=KCNIP1;refGene_tx=XM_019839086.2
A1	180832144	180832215
refGene_gn=KCNIP1;refGene_tx=XM_011285529.3;refGene_gn=KCNIP1;refGene_tx=XM_011285533.2;refGene_gn=KCNIP1;refGene_tx=XM_019839082.2;refGene_gn=KCNIP1;refGene_tx=XM_019839086.2
A1	180833701	180833771
refGene_gn=KCNIP1;refGene_tx=XM_011285529.3;refGene_gn=KCNIP1;refGene_tx=XM_011285533.2;refGene_gn=KCNIP1;refGene_tx=XM_019839082.2;refGene_gn=KCNIP1;refGene_tx=XM_019839086.2
A1	180834578	180834703
refGene_gn=KCNIP1;refGene_tx=XM_011285529.3;refGene_gn=KCNIP1;refGene_tx=XM_011285533.2;refGene_gn=KCNIP1;refGene_tx=XM_019839082.2;refGene_gn=KCNIP1;refGene_tx=XM_019839086.2
A1	180839628	180839661
refGene_gn=KCNIP1;refGene_tx=XM_019839082.2
A1	181032660	181032721
refGene_gn=KCNIP1;refGene_tx=XM_011285533.2;refGene_gn=KCNIP1;refGene_tx=XM_019839082.2
A1	181133139	181133273
refGene_gn=KCNMB1;refGene_tx=XM_003981286.4;refGene_gn=KCNMB1;refGene_tx=XM_019839091.2
A1	181134652	181134853
refGene_gn=KCNMB1;refGene_tx=XM_003981286.4;refGene_gn=KCNMB1;refGene_tx=XM_019839091.2;refGene_gn=KCNMB1;refGene_tx=XM_023259685.1
A1	181139756	181140026
refGene_gn=KCNMB1;refGene_tx=XM_003981286.4;refGene_gn=KCNMB1;refGene_

tx=XM_019839091.2;refGene_gn=KCNMB1;refGene_tx=XM_023259685.1
A1 181163419 181163507
refGene_gn=KCNIP1;refGene_tx=XM_011285529.3
A1 181211357 181211435
refGene_gn=LCP2;refGene_tx=XM_023259696.1
A1 181216000 181216063
refGene_gn=LCP2;refGene_tx=XM_023259696.1
A1 181221873 181221920
refGene_gn=LCP2;refGene_tx=XM_023259696.1
A1 181231275 181231341
refGene_gn=LCP2;refGene_tx=XM_023259688.1;refGene_gn=LCP2;refGene_tx=XM_023259696.1
A1 181231797 181231829
refGene_gn=LCP2;refGene_tx=XM_023259688.1;refGene_gn=LCP2;refGene_tx=XM_023259696.1
A1 181232440 181232478
refGene_gn=LCP2;refGene_tx=XM_023259688.1;refGene_gn=LCP2;refGene_tx=XM_023259696.1
A1 181235339 181235541
refGene_gn=LCP2;refGene_tx=XM_023259688.1;refGene_gn=LCP2;refGene_tx=XM_023259696.1
A1 181237828 181237926
refGene_gn=LCP2;refGene_tx=XM_023259688.1;refGene_gn=LCP2;refGene_tx=XM_023259696.1
A1 181239416 181239487
refGene_gn=LCP2;refGene_tx=XM_023259688.1;refGene_gn=LCP2;refGene_tx=XM_023259696.1
A1 181239572 181239673
refGene_gn=LCP2;refGene_tx=XM_023259688.1;refGene_gn=LCP2;refGene_tx=XM_023259696.1
A1 181243078 181243104
refGene_gn=LCP2;refGene_tx=XM_023259688.1;refGene_gn=LCP2;refGene_tx=XM_023259696.1
A1 181243218 181243238
refGene_gn=LCP2;refGene_tx=XM_023259688.1;refGene_gn=LCP2;refGene_tx=XM_023259696.1
A1 181243334 181243442
refGene_gn=LCP2;refGene_tx=XM_023259688.1;refGene_gn=LCP2;refGene_tx=XM_023259696.1
A1 181245290 181245321
refGene_gn=LCP2;refGene_tx=XM_023259688.1;refGene_gn=LCP2;refGene_tx=XM_023259696.1
A1 181247528 181247541
refGene_gn=LCP2;refGene_tx=XM_023259688.1;refGene_gn=LCP2;refGene_tx=XM_023259696.1
A1 181248220 181248350
refGene_gn=LCP2;refGene_tx=XM_023259688.1;refGene_gn=LCP2;refGene_tx=XM_023259696.1
A1 181249132 181249182
refGene_gn=LCP2;refGene_tx=XM_023259688.1;refGene_gn=LCP2;refGene_tx=X

M_023259696.1
A1 181252654 181252749
refGene_gn=LCP2;refGene_tx=XM_023259688.1;refGene_gn=LCP2;refGene_tx=X
M_023259696.1
A1 181253354 181253432
refGene_gn=LCP2;refGene_tx=XM_023259688.1;refGene_gn=LCP2;refGene_tx=X
M_023259696.1
A1 181254939 181255095
refGene_gn=LCP2;refGene_tx=XM_023259688.1;refGene_gn=LCP2;refGene_tx=X
M_023259696.1
A1 181256618 181256741
refGene_gn=LCP2;refGene_tx=XM_023259688.1;refGene_gn=LCP2;refGene_tx=X
M_023259696.1
A1 181258880 181259032
refGene_gn=CA1H5orf58;refGene_tx=XM_023259707.1
A1 181264537 181264631
refGene_gn=CA1H5orf58;refGene_tx=XM_023259707.1
A1 181265383 181265413
refGene_gn=CA1H5orf58;refGene_tx=XM_023259707.1
A1 181368178 181368738
refGene_gn=FOXI1;refGene_tx=XM_003981288.3
A1 181370218 181370795
refGene_gn=FOXI1;refGene_tx=XM_003981288.3
A1 181388957 181389020
refGene_gn=DOCK2;refGene_tx=XM_003981290.3
A1 181389915 181390058
refGene_gn=DOCK2;refGene_tx=XM_003981290.3
A1 181391544 181391665
refGene_gn=DOCK2;refGene_tx=XM_003981290.3
A1 181392656 181392828
refGene_gn=DOCK2;refGene_tx=XM_003981290.3
A1 181393677 181393805
refGene_gn=DOCK2;refGene_tx=XM_003981290.3
A1 181395343 181395481
refGene_gn=DOCK2;refGene_tx=XM_003981290.3
A1 181401964 181402048
refGene_gn=DOCK2;refGene_tx=XM_003981290.3
A1 181403488 181403665
refGene_gn=DOCK2;refGene_tx=XM_003981290.3
A1 181409104 181409191
refGene_gn=DOCK2;refGene_tx=XM_003981290.3
A1 181410685 181410770
refGene_gn=DOCK2;refGene_tx=XM_003981290.3
A1 181412214 181412296
refGene_gn=DOCK2;refGene_tx=XM_003981290.3
A1 181416933 181417075
refGene_gn=DOCK2;refGene_tx=XM_003981290.3
A1 181419355 181419460
refGene_gn=DOCK2;refGene_tx=XM_003981290.3
A1 181421520 181421610

refGene_gn=DOCK2;refGene_tx=XM_003981290.3
A1 181425001 181425121
refGene_gn=DOCK2;refGene_tx=XM_003981290.3
A1 181425838 181425929
refGene_gn=DOCK2;refGene_tx=XM_003981290.3
A1 181429304 181429345
refGene_gn=DOCK2;refGene_tx=XM_003981290.3
A1 181431249 181431406
refGene_gn=DOCK2;refGene_tx=XM_003981290.3
A1 181438202 181438288
refGene_gn=DOCK2;refGene_tx=XM_003981290.3
A1 181447680 181447829
refGene_gn=DOCK2;refGene_tx=XM_003981290.3
A1 181458797 181458856
refGene_gn=DOCK2;refGene_tx=XM_003981290.3
A1 181458946 181459047
refGene_gn=DOCK2;refGene_tx=XM_003981290.3
A1 181469345 181469424
refGene_gn=DOCK2;refGene_tx=XM_003981290.3
A1 181479880 181479975
refGene_gn=DOCK2;refGene_tx=XM_003981290.3
A1 181482974 181483073
refGene_gn=DOCK2;refGene_tx=XM_003981290.3
A1 181585295 181586641
refGene_gn=FAM196B;refGene_tx=XM_003981291.4;refGene_gn=FAM196B;refGene_tx=XM_019839124.1;refGene_gn=FAM196B;refGene_tx=XM_019839132.1;refGene_gn=FAM196B;refGene_tx=XM_019839135.2;refGene_gn=FAM196B;refGene_tx=XM_023259736.1
A1 181587764 181587839
refGene_gn=FAM196B;refGene_tx=XM_003981291.4;refGene_gn=FAM196B;refGene_tx=XM_019839124.1;refGene_gn=FAM196B;refGene_tx=XM_019839132.1;refGene_gn=FAM196B;refGene_tx=XM_019839135.2;refGene_gn=FAM196B;refGene_tx=XM_023259736.1
A1 181603039 181603226
refGene_gn=FAM196B;refGene_tx=XM_003981291.4;refGene_gn=FAM196B;refGene_tx=XM_019839124.1;refGene_gn=FAM196B;refGene_tx=XM_019839135.2;refGene_gn=FAM196B;refGene_tx=XM_023259736.1
A1 181605223 181605365
refGene_gn=FAM196B;refGene_tx=XM_019839132.1
A1 181623241 181623337
refGene_gn=DOCK2;refGene_tx=XM_003981290.3
A1 181649756 181649905
refGene_gn=DOCK2;refGene_tx=XM_003981290.3
A1 181683273 181683380
refGene_gn=DOCK2;refGene_tx=XM_003981290.3
A1 181685173 181685244
refGene_gn=DOCK2;refGene_tx=XM_003981290.3
A1 181696508 181696617
refGene_gn=DOCK2;refGene_tx=XM_003981290.3
A1 181725434 181725569

refGene_gn=DOCK2;refGene_tx=XM_003981290.3
A1 181726408 181726509
refGene_gn=DOCK2;refGene_tx=XM_003981290.3
A1 181727507 181727597
refGene_gn=DOCK2;refGene_tx=XM_003981290.3
A1 181729195 181729293
refGene_gn=DOCK2;refGene_tx=XM_003981290.3
A1 181729415 181729599
refGene_gn=DOCK2;refGene_tx=XM_003981290.3
A1 181731360 181731464
refGene_gn=DOCK2;refGene_tx=XM_003981290.3
A1 181731575 181731648
refGene_gn=DOCK2;refGene_tx=XM_003981290.3
A1 181733441 181733540
refGene_gn=DOCK2;refGene_tx=XM_003981290.3
A1 181738654 181738779
refGene_gn=DOCK2;refGene_tx=XM_003981290.3
A1 181739982 181740108
refGene_gn=DOCK2;refGene_tx=XM_003981290.3
A1 181740701 181740778
refGene_gn=DOCK2;refGene_tx=XM_003981290.3
A1 181741742 181741818
refGene_gn=DOCK2;refGene_tx=XM_003981290.3
A1 181744383 181744519
refGene_gn=DOCK2;refGene_tx=XM_003981290.3
A1 181750149 181750231
refGene_gn=DOCK2;refGene_tx=XM_003981290.3
A1 181754710 181754865
refGene_gn=DOCK2;refGene_tx=XM_003981290.3
A1 181757146 181757282
refGene_gn=DOCK2;refGene_tx=XM_003981290.3
A1 181769799 181769948
refGene_gn=DOCK2;refGene_tx=XM_003981290.3
A1 181772583 181772680
refGene_gn=DOCK2;refGene_tx=XM_003981290.3
A1 181773174 181773230
refGene_gn=DOCK2;refGene_tx=XM_003981290.3
A1 181774311 181774352
refGene_gn=DOCK2;refGene_tx=XM_003981290.3
A1 181784633 181784717
refGene_gn=DOCK2;refGene_tx=XM_003981290.3
A1 181805620 181805663
refGene_gn=DOCK2;refGene_tx=XM_003981290.3
A1 181839885 181840033
refGene_gn=SPDL1;refGene_tx=XM_023259759.1
A1 181842403 181842945
refGene_gn=SPDL1;refGene_tx=XM_023259750.1;refGene_gn=SPDL1;refGene_tx
=XM_023259754.1;refGene_gn=SPDL1;refGene_tx=XM_023259759.1
A1 181845085 181845273
refGene_gn=SPDL1;refGene_tx=XM_023259750.1;refGene_gn=SPDL1;refGene_tx

=XM_023259754.1;refGene_gn=SPDL1;refGene_tx=XM_023259759.1
A1	181845900	181846004
refGene_gn=SPDL1;refGene_tx=XM_023259750.1;refGene_gn=SPDL1;refGene_tx=XM_023259754.1;refGene_gn=SPDL1;refGene_tx=XM_023259759.1
A1	181846434	181846575
refGene_gn=SPDL1;refGene_tx=XM_023259750.1;refGene_gn=SPDL1;refGene_tx=XM_023259754.1;refGene_gn=SPDL1;refGene_tx=XM_023259759.1
A1	181848755	181848866
refGene_gn=SPDL1;refGene_tx=XM_023259750.1;refGene_gn=SPDL1;refGene_tx=XM_023259754.1;refGene_gn=SPDL1;refGene_tx=XM_023259759.1
A1	181848944	181849043
refGene_gn=SPDL1;refGene_tx=XM_023259750.1;refGene_gn=SPDL1;refGene_tx=XM_023259754.1;refGene_gn=SPDL1;refGene_tx=XM_023259759.1
A1	181849142	181849292
refGene_gn=SPDL1;refGene_tx=XM_023259750.1;refGene_gn=SPDL1;refGene_tx=XM_023259754.1;refGene_gn=SPDL1;refGene_tx=XM_023259759.1
A1	181849853	181850048
refGene_gn=SPDL1;refGene_tx=XM_023259750.1;refGene_gn=SPDL1;refGene_tx=XM_023259754.1;refGene_gn=SPDL1;refGene_tx=XM_023259759.1
A1	181852334	181852511
refGene_gn=SPDL1;refGene_tx=XM_023259750.1;refGene_gn=SPDL1;refGene_tx=XM_023259754.1;refGene_gn=SPDL1;refGene_tx=XM_023259759.1
A1	181854618	181854777
refGene_gn=SPDL1;refGene_tx=XM_023259750.1;refGene_gn=SPDL1;refGene_tx=XM_023259754.1;refGene_gn=SPDL1;refGene_tx=XM_023259759.1
A1	182130948	182131145
refGene_gn=SLIT3;refGene_tx=XM_023259788.1
A1	182176415	182176487
refGene_gn=SLIT3;refGene_tx=XM_023259788.1
A1	182183152	182183224
refGene_gn=SLIT3;refGene_tx=XM_023259788.1
A1	182232944	182233016
refGene_gn=SLIT3;refGene_tx=XM_023259788.1
A1	182536516	182536588
refGene_gn=SLIT3;refGene_tx=XM_023259788.1
A1	182573145	182573217
refGene_gn=SLIT3;refGene_tx=XM_023259788.1
A1	182593678	182593750
refGene_gn=SLIT3;refGene_tx=XM_023259788.1
A1	182599543	182599707
refGene_gn=SLIT3;refGene_tx=XM_023259788.1
A1	182608786	182608928
refGene_gn=SLIT3;refGene_tx=XM_023259788.1
A1	182619331	182619403
refGene_gn=SLIT3;refGene_tx=XM_023259788.1
A1	182624222	182624294
refGene_gn=SLIT3;refGene_tx=XM_023259788.1
A1	182627928	182628000
refGene_gn=SLIT3;refGene_tx=XM_023259788.1
A1	182639698	182639842

refGene_gn=SLIT3;refGene_tx=XM_023259788.1
A1	182640984	182641148
refGene_gn=SLIT3;refGene_tx=XM_023259788.1
A1	182651299	182651450
refGene_gn=SLIT3;refGene_tx=XM_023259788.1
A1	182652418	182652493
refGene_gn=SLIT3;refGene_tx=XM_023259788.1
A1	182659837	182659981
refGene_gn=SLIT3;refGene_tx=XM_023259788.1
A1	182660756	182660900
refGene_gn=SLIT3;refGene_tx=XM_023259788.1
A1	182663691	182663855
refGene_gn=SLIT3;refGene_tx=XM_023259788.1
A1	182664959	182665092
refGene_gn=SLIT3;refGene_tx=XM_023259788.1
A1	182676499	182676568
refGene_gn=SLIT3;refGene_tx=XM_023259788.1
A1	182678252	182678324
refGene_gn=SLIT3;refGene_tx=XM_023259788.1
A1	182678946	182679018
refGene_gn=SLIT3;refGene_tx=XM_023259788.1
A1	182687087	182687159
refGene_gn=SLIT3;refGene_tx=XM_023259788.1
A1	182688757	182688921
refGene_gn=SLIT3;refGene_tx=XM_023259788.1
A1	182691147	182691272
refGene_gn=SLIT3;refGene_tx=XM_023259788.1
A1	182697166	182697264
refGene_gn=SLIT3;refGene_tx=XM_023259788.1
A1	182701517	182701657
refGene_gn=SLIT3;refGene_tx=XM_023259788.1
A1	182704615	182704709
refGene_gn=SLIT3;refGene_tx=XM_023259788.1
A1	182710022	182710160
refGene_gn=SLIT3;refGene_tx=XM_023259788.1
A1	182711708	182711949
refGene_gn=SLIT3;refGene_tx=XM_023259788.1
A1	182713847	182713978
refGene_gn=SLIT3;refGene_tx=XM_023259788.1
A1	182721572	182721727
refGene_gn=SLIT3;refGene_tx=XM_023259788.1
A1	182722875	182723161
refGene_gn=SLIT3;refGene_tx=XM_023259788.1
A1	182723966	182724175
refGene_gn=SLIT3;refGene_tx=XM_023259788.1
A1	182725920	182726156
refGene_gn=SLIT3;refGene_tx=XM_023259788.1
A1	182822815	182822843
refGene_gn=PANK3;refGene_tx=XM_003981327.5
A1	182829711	182829790

refGene_gn=PANK3;refGene_tx=XM_023259796.1
A1 182829884 182830237
refGene_gn=PANK3;refGene_tx=XM_003981327.5;refGene_gn=PANK3;refGene_tx=XM_023259796.1;refGene_gn=PANK3;refGene_tx=XM_023259803.1
A1 182830958 182830967
refGene_gn=PANK3;refGene_tx=XM_019839170.2
A1 182832474 182832728
refGene_gn=PANK3;refGene_tx=XM_003981327.5;refGene_gn=PANK3;refGene_tx=XM_019839170.2;refGene_gn=PANK3;refGene_tx=XM_023259796.1;refGene_gn=PANK3;refGene_tx=XM_023259803.1
A1 182834294 182834471
refGene_gn=PANK3;refGene_tx=XM_003981327.5;refGene_gn=PANK3;refGene_tx=XM_019839170.2;refGene_gn=PANK3;refGene_tx=XM_023259796.1;refGene_gn=PANK3;refGene_tx=XM_023259803.1
A1 182835101 182835225
refGene_gn=PANK3;refGene_tx=XM_003981327.5;refGene_gn=PANK3;refGene_tx=XM_019839170.2;refGene_gn=PANK3;refGene_tx=XM_023259796.1;refGene_gn=PANK3;refGene_tx=XM_023259803.1
A1 182837178 182837304
refGene_gn=PANK3;refGene_tx=XM_003981327.5;refGene_gn=PANK3;refGene_tx=XM_019839170.2;refGene_gn=PANK3;refGene_tx=XM_023259796.1;refGene_gn=PANK3;refGene_tx=XM_023259803.1
A1 182838501 182838552
refGene_gn=PANK3;refGene_tx=XM_003981327.5;refGene_gn=PANK3;refGene_tx=XM_019839170.2;refGene_gn=PANK3;refGene_tx=XM_023259796.1;refGene_gn=PANK3;refGene_tx=XM_023259803.1
A1 182866405 182866918
refGene_gn=LOC101085349;refGene_tx=XM_019839171.2
A1 182866919 182867402
refGene_gn=LOC101085349;refGene_tx=XM_019839171.2
A1 182895507 182895617
refGene_gn=RARS;refGene_tx=XM_003981329.5
A1 182896477 182896725
refGene_gn=RARS;refGene_tx=XM_003981329.5
A1 182896863 182897036
refGene_gn=RARS;refGene_tx=XM_003981329.5
A1 182902315 182902421
refGene_gn=RARS;refGene_tx=XM_003981329.5
A1 182907414 182907524
refGene_gn=RARS;refGene_tx=XM_003981329.5
A1 182908044 182908223
refGene_gn=RARS;refGene_tx=XM_003981329.5
A1 182909679 182909784
refGene_gn=RARS;refGene_tx=XM_003981329.5
A1 182910586 182910716
refGene_gn=RARS;refGene_tx=XM_003981329.5
A1 182912543 182912664
refGene_gn=RARS;refGene_tx=XM_003981329.5
A1 182913950 182914072
refGene_gn=RARS;refGene_tx=XM_003981329.5

A1 182914626 182914727
refGene_gn=RARS;refGene_tx=XM_003981329.5
A1 182915279 182915388
refGene_gn=RARS;refGene_tx=XM_003981329.5
A1 182916330 182916519
refGene_gn=RARS;refGene_tx=XM_003981329.5
A1 182917959 182918094
refGene_gn=RARS;refGene_tx=XM_003981329.5
A1 182919976 182920021
refGene_gn=RARS;refGene_tx=XM_003981329.5
A1 182940659 182940726
refGene_gn=WWC1;refGene_tx=XM_023259830.1;refGene_gn=WWC1;refGene_tx=XM_023259831.1
A1 182941900 182942022
refGene_gn=WWC1;refGene_tx=XM_023259830.1;refGene_gn=WWC1;refGene_tx=XM_023259831.1
A1 182944622 182944856
refGene_gn=WWC1;refGene_tx=XM_023259830.1;refGene_gn=WWC1;refGene_tx=XM_023259831.1
A1 182948389 182948482
refGene_gn=WWC1;refGene_tx=XM_023259830.1;refGene_gn=WWC1;refGene_tx=XM_023259831.1
A1 182953924 182954089
refGene_gn=WWC1;refGene_tx=XM_023259830.1;refGene_gn=WWC1;refGene_tx=XM_023259831.1
A1 182955325 182955440
refGene_gn=WWC1;refGene_tx=XM_023259830.1;refGene_gn=WWC1;refGene_tx=XM_023259831.1
A1 182961647 182961739
refGene_gn=WWC1;refGene_tx=XM_023259830.1;refGene_gn=WWC1;refGene_tx=XM_023259831.1
A1 182964315 182964468
refGene_gn=WWC1;refGene_tx=XM_023259830.1;refGene_gn=WWC1;refGene_tx=XM_023259831.1
A1 182971948 182972141
refGene_gn=WWC1;refGene_tx=XM_023259830.1;refGene_gn=WWC1;refGene_tx=XM_023259831.1
A1 182973329 182973416
refGene_gn=WWC1;refGene_tx=XM_023259830.1;refGene_gn=WWC1;refGene_tx=XM_023259831.1
A1 182974334 182974415
refGene_gn=WWC1;refGene_tx=XM_023259830.1;refGene_gn=WWC1;refGene_tx=XM_023259831.1
A1 182974853 182974962
refGene_gn=WWC1;refGene_tx=XM_023259830.1;refGene_gn=WWC1;refGene_tx=XM_023259831.1
A1 182977974 182978510
refGene_gn=WWC1;refGene_tx=XM_023259830.1;refGene_gn=WWC1;refGene_tx=XM_023259831.1
A1 182979332 182979422

refGene_gn=WWC1;refGene_tx=XM_023259830.1;refGene_gn=WWC1;refGene_tx=XM_023259831.1
A1 182985846 182986089
refGene_gn=WWC1;refGene_tx=XM_023259830.1;refGene_gn=WWC1;refGene_tx=XM_023259831.1
A1 182989835 182989909
refGene_gn=WWC1;refGene_tx=XM_023259830.1;refGene_gn=WWC1;refGene_tx=XM_023259831.1
A1 182991110 182991257
refGene_gn=WWC1;refGene_tx=XM_023259830.1;refGene_gn=WWC1;refGene_tx=XM_023259831.1
A1 182993114 182993244
refGene_gn=WWC1;refGene_tx=XM_023259830.1;refGene_gn=WWC1;refGene_tx=XM_023259831.1
A1 182998063 182998143
refGene_gn=WWC1;refGene_tx=XM_023259830.1;refGene_gn=WWC1;refGene_tx=XM_023259831.1
A1 182999480 182999557
refGene_gn=WWC1;refGene_tx=XM_023259830.1;refGene_gn=WWC1;refGene_tx=XM_023259831.1
A1 183010550 183010754
refGene_gn=WWC1;refGene_tx=XM_023259830.1;refGene_gn=WWC1;refGene_tx=XM_023259831.1
A1 183022130 183022240
refGene_gn=WWC1;refGene_tx=XM_023259830.1
A1 183093471 183093590
refGene_gn=WWC1;refGene_tx=XM_023259830.1
A1 183116414 183117176
refGene_gn=TENM2;refGene_tx=XM_019839174.2;refGene_gn=TENM2;refGene_tx=XM_019839189.2;refGene_gn=TENM2;refGene_tx=XM_019839191.2;refGene_gn=TENM2;refGene_tx=XM_019839194.2;refGene_gn=TENM2;refGene_tx=XM_019839195.2;refGene_gn=TENM2;refGene_tx=XM_019839196.2;refGene_gn=TENM2;refGene_tx=XM_019839198.2;refGene_gn=TENM2;refGene_tx=XM_019839204.2;refGene_gn=TENM2;refGene_tx=XM_023259843.1;refGene_gn=TENM2;refGene_tx=XM_023259850.1;refGene_gn=TENM2;refGene_tx=XM_023259852.1;refGene_gn=TENM2;refGene_tx=XM_023259862.1;refGene_gn=TENM2;refGene_tx=XM_023259868.1;refGene_gn=TENM2;refGene_tx=XM_023259877.1;refGene_gn=TENM2;refGene_tx=XM_023259887.1;refGene_gn=TENM2;refGene_tx=XM_023259913.1;refGene_gn=TENM2;refGene_tx=XM_023259942.1
A1 183118819 183118950
refGene_gn=TENM2;refGene_tx=XM_019839174.2;refGene_gn=TENM2;refGene_tx=XM_019839189.2;refGene_gn=TENM2;refGene_tx=XM_019839191.2;refGene_gn=TENM2;refGene_tx=XM_019839194.2;refGene_gn=TENM2;refGene_tx=XM_019839195.2;refGene_gn=TENM2;refGene_tx=XM_019839196.2;refGene_gn=TENM2;refGene_tx=XM_019839198.2;refGene_gn=TENM2;refGene_tx=XM_019839204.2;refGene_gn=TENM2;refGene_tx=XM_023259843.1;refGene_gn=TENM2;refGene_tx=XM_023259850.1;refGene_gn=TENM2;refGene_tx=XM_023259852.1;refGene_gn=TENM2;refGene_tx=XM_023259862.1;refGene_gn=TENM2;refGene_tx=XM_023259868.1;refGene_gn=TENM2;refGene_tx=XM_023259877.1;refGene_gn=TENM2;refGene_tx=XM_023259887.1;refGene_gn=TENM2;refGene_tx=XM_023259913.1

A1 183129471 183131086
refGene_gn=TENM2;refGene_tx=XM_019839174.2;refGene_gn=TENM2;refGene_tx=XM_019839189.2;refGene_gn=TENM2;refGene_tx=XM_019839191.2;refGene_gn=TENM2;refGene_tx=XM_019839194.2;refGene_gn=TENM2;refGene_tx=XM_019839195.2;refGene_gn=TENM2;refGene_tx=XM_019839196.2;refGene_gn=TENM2;refGene_tx=XM_019839198.2;refGene_gn=TENM2;refGene_tx=XM_019839204.2;refGene_gn=TENM2;refGene_tx=XM_023259843.1;refGene_gn=TENM2;refGene_tx=XM_023259850.1;refGene_gn=TENM2;refGene_tx=XM_023259852.1;refGene_gn=TENM2;refGene_tx=XM_023259862.1;refGene_gn=TENM2;refGene_tx=XM_023259868.1;refGene_gn=TENM2;refGene_tx=XM_023259877.1;refGene_gn=TENM2;refGene_tx=XM_023259887.1;refGene_gn=TENM2;refGene_tx=XM_023259913.1;refGene_gn=TENM2;refGene_tx=XM_023259942.1
A1 183132676 183132973
refGene_gn=TENM2;refGene_tx=XM_019839174.2;refGene_gn=TENM2;refGene_tx=XM_019839189.2;refGene_gn=TENM2;refGene_tx=XM_019839191.2;refGene_gn=TENM2;refGene_tx=XM_019839194.2;refGene_gn=TENM2;refGene_tx=XM_019839195.2;refGene_gn=TENM2;refGene_tx=XM_019839196.2;refGene_gn=TENM2;refGene_tx=XM_019839198.2;refGene_gn=TENM2;refGene_tx=XM_019839204.2;refGene_gn=TENM2;refGene_tx=XM_023259843.1;refGene_gn=TENM2;refGene_tx=XM_023259850.1;refGene_gn=TENM2;refGene_tx=XM_023259852.1;refGene_gn=TENM2;refGene_tx=XM_023259862.1;refGene_gn=TENM2;refGene_tx=XM_023259868.1;refGene_gn=TENM2;refGene_tx=XM_023259877.1;refGene_gn=TENM2;refGene_tx=XM_023259887.1;refGene_gn=TENM2;refGene_tx=XM_023259913.1;refGene_gn=TENM2;refGene_tx=XM_023259942.1
A1 183162610 183162846
refGene_gn=TENM2;refGene_tx=XM_019839174.2;refGene_gn=TENM2;refGene_tx=XM_019839189.2;refGene_gn=TENM2;refGene_tx=XM_019839191.2;refGene_gn=TENM2;refGene_tx=XM_019839194.2;refGene_gn=TENM2;refGene_tx=XM_019839195.2;refGene_gn=TENM2;refGene_tx=XM_019839196.2;refGene_gn=TENM2;refGene_tx=XM_019839198.2;refGene_gn=TENM2;refGene_tx=XM_019839204.2;refGene_gn=TENM2;refGene_tx=XM_023259843.1;refGene_gn=TENM2;refGene_tx=XM_023259850.1;refGene_gn=TENM2;refGene_tx=XM_023259852.1;refGene_gn=TENM2;refGene_tx=XM_023259862.1;refGene_gn=TENM2;refGene_tx=XM_023259868.1;refGene_gn=TENM2;refGene_tx=XM_023259877.1;refGene_gn=TENM2;refGene_tx=XM_023259887.1;refGene_gn=TENM2;refGene_tx=XM_023259913.1;refGene_gn=TENM2;refGene_tx=XM_023259942.1
A1 183164334 183164510
refGene_gn=TENM2;refGene_tx=XM_019839174.2;refGene_gn=TENM2;refGene_tx=XM_019839189.2;refGene_gn=TENM2;refGene_tx=XM_019839191.2;refGene_gn=TENM2;refGene_tx=XM_019839194.2;refGene_gn=TENM2;refGene_tx=XM_019839195.2;refGene_gn=TENM2;refGene_tx=XM_019839196.2;refGene_gn=TENM2;refGene_tx=XM_019839198.2;refGene_gn=TENM2;refGene_tx=XM_019839204.2;refGene_gn=TENM2;refGene_tx=XM_023259843.1;refGene_gn=TENM2;refGene_tx=XM_023259850.1;refGene_gn=TENM2;refGene_tx=XM_023259852.1;refGene_gn=TENM2;refGene_tx=XM_023259862.1;refGene_gn=TENM2;refGene_tx=XM_023259868.1;refGene_gn=TENM2;refGene_tx=XM_023259877.1;refGene_gn=TENM2;refGene_tx=XM_023259887.1;refGene_gn=TENM2;refGene_tx=XM_023259913.1;refGene_gn=TENM2;refGene_tx=XM_023259942.1
A1 183171004 183171879
refGene_gn=TENM2;refGene_tx=XM_019839174.2;refGene_gn=TENM2;refGene_tx

=XM_019839189.2;refGene_gn=TENM2;refGene_tx=XM_019839191.2;refGene_gn=TENM2;refGene_tx=XM_019839194.2;refGene_gn=TENM2;refGene_tx=XM_019839195.2;refGene_gn=TENM2;refGene_tx=XM_019839196.2;refGene_gn=TENM2;refGene_tx=XM_019839198.2;refGene_gn=TENM2;refGene_tx=XM_019839204.2;refGene_gn=TENM2;refGene_tx=XM_023259843.1;refGene_gn=TENM2;refGene_tx=XM_023259850.1;refGene_gn=TENM2;refGene_tx=XM_023259852.1;refGene_gn=TENM2;refGene_tx=XM_023259862.1;refGene_gn=TENM2;refGene_tx=XM_023259868.1;refGene_gn=TENM2;refGene_tx=XM_023259877.1;refGene_gn=TENM2;refGene_tx=XM_023259887.1;refGene_gn=TENM2;refGene_tx=XM_023259913.1;refGene_gn=TENM2;refGene_tx=XM_023259942.1
A1 183173091 183173246
refGene_gn=TENM2;refGene_tx=XM_019839174.2;refGene_gn=TENM2;refGene_tx=XM_019839189.2;refGene_gn=TENM2;refGene_tx=XM_019839191.2;refGene_gn=TENM2;refGene_tx=XM_019839194.2;refGene_gn=TENM2;refGene_tx=XM_019839195.2;refGene_gn=TENM2;refGene_tx=XM_019839196.2;refGene_gn=TENM2;refGene_tx=XM_019839198.2;refGene_gn=TENM2;refGene_tx=XM_019839204.2;refGene_gn=TENM2;refGene_tx=XM_023259843.1;refGene_gn=TENM2;refGene_tx=XM_023259850.1;refGene_gn=TENM2;refGene_tx=XM_023259852.1;refGene_gn=TENM2;refGene_tx=XM_023259862.1;refGene_gn=TENM2;refGene_tx=XM_023259868.1;refGene_gn=TENM2;refGene_tx=XM_023259877.1;refGene_gn=TENM2;refGene_tx=XM_023259887.1;refGene_gn=TENM2;refGene_tx=XM_023259913.1;refGene_gn=TENM2;refGene_tx=XM_023259942.1
A1 183174473 183174706
refGene_gn=TENM2;refGene_tx=XM_019839174.2;refGene_gn=TENM2;refGene_tx=XM_019839189.2;refGene_gn=TENM2;refGene_tx=XM_019839191.2;refGene_gn=TENM2;refGene_tx=XM_019839194.2;refGene_gn=TENM2;refGene_tx=XM_019839195.2;refGene_gn=TENM2;refGene_tx=XM_019839196.2;refGene_gn=TENM2;refGene_tx=XM_019839198.2;refGene_gn=TENM2;refGene_tx=XM_019839204.2;refGene_gn=TENM2;refGene_tx=XM_023259843.1;refGene_gn=TENM2;refGene_tx=XM_023259850.1;refGene_gn=TENM2;refGene_tx=XM_023259852.1;refGene_gn=TENM2;refGene_tx=XM_023259862.1;refGene_gn=TENM2;refGene_tx=XM_023259868.1;refGene_gn=TENM2;refGene_tx=XM_023259877.1;refGene_gn=TENM2;refGene_tx=XM_023259887.1;refGene_gn=TENM2;refGene_tx=XM_023259913.1;refGene_gn=TENM2;refGene_tx=XM_023259942.1
A1 183177722 183177743
refGene_gn=TENM2;refGene_tx=XM_019839174.2;refGene_gn=TENM2;refGene_tx=XM_019839189.2;refGene_gn=TENM2;refGene_tx=XM_019839191.2;refGene_gn=TENM2;refGene_tx=XM_019839194.2;refGene_gn=TENM2;refGene_tx=XM_019839195.2;refGene_gn=TENM2;refGene_tx=XM_019839196.2;refGene_gn=TENM2;refGene_tx=XM_019839198.2;refGene_gn=TENM2;refGene_tx=XM_019839204.2;refGene_gn=TENM2;refGene_tx=XM_023259850.1;refGene_gn=TENM2;refGene_tx=XM_023259852.1;refGene_gn=TENM2;refGene_tx=XM_023259862.1;refGene_gn=TENM2;refGene_tx=XM_023259868.1;refGene_gn=TENM2;refGene_tx=XM_023259877.1;refGene_gn=TENM2;refGene_tx=XM_023259887.1;refGene_gn=TENM2;refGene_tx=XM_023259913.1;refGene_gn=TENM2;refGene_tx=XM_023259942.1
A1 183187811 183188061
refGene_gn=TENM2;refGene_tx=XM_019839174.2;refGene_gn=TENM2;refGene_tx=XM_019839189.2;refGene_gn=TENM2;refGene_tx=XM_019839191.2;refGene_gn=TENM2;refGene_tx=XM_019839194.2;refGene_gn=TENM2;refGene_tx=XM_019839195.2;refGene_gn=TENM2;refGene_tx=XM_019839196.2;refGene_gn=TENM2;refGe

ne_tx=XM_019839198.2;refGene_gn=TENM2;refGene_tx=XM_019839204.2;refGen
e_gn=TENM2;refGene_tx=XM_023259843.1;refGene_gn=TENM2;refGene_tx=XM_02
3259850.1;refGene_gn=TENM2;refGene_tx=XM_023259852.1;refGene_gn=TENM2;
refGene_tx=XM_023259862.1;refGene_gn=TENM2;refGene_tx=XM_023259868.1;r
efGene_gn=TENM2;refGene_tx=XM_023259877.1;refGene_gn=TENM2;refGene_tx=
XM_023259887.1;refGene_gn=TENM2;refGene_tx=XM_023259913.1;refGene_gn=T
ENM2;refGene_tx=XM_023259942.1
A1 183188482 183188626
refGene_gn=TENM2;refGene_tx=XM_019839174.2;refGene_gn=TENM2;refGene_tx
=XM_019839189.2;refGene_gn=TENM2;refGene_tx=XM_019839191.2;refGene_gn=
TENM2;refGene_tx=XM_019839194.2;refGene_gn=TENM2;refGene_tx=XM_0198391
95.2;refGene_gn=TENM2;refGene_tx=XM_019839196.2;refGene_gn=TENM2;refGe
ne_tx=XM_019839198.2;refGene_gn=TENM2;refGene_tx=XM_019839204.2;refGen
e_gn=TENM2;refGene_tx=XM_023259843.1;refGene_gn=TENM2;refGene_tx=XM_02
3259850.1;refGene_gn=TENM2;refGene_tx=XM_023259852.1;refGene_gn=TENM2;
refGene_tx=XM_023259862.1;refGene_gn=TENM2;refGene_tx=XM_023259868.1;r
efGene_gn=TENM2;refGene_tx=XM_023259877.1;refGene_gn=TENM2;refGene_tx=
XM_023259887.1;refGene_gn=TENM2;refGene_tx=XM_023259913.1;refGene_gn=T
ENM2;refGene_tx=XM_023259942.1
A1 183194228 183194496
refGene_gn=TENM2;refGene_tx=XM_019839174.2;refGene_gn=TENM2;refGene_tx
=XM_019839189.2;refGene_gn=TENM2;refGene_tx=XM_019839191.2;refGene_gn=
TENM2;refGene_tx=XM_019839194.2;refGene_gn=TENM2;refGene_tx=XM_0198391
95.2;refGene_gn=TENM2;refGene_tx=XM_019839196.2;refGene_gn=TENM2;refGe
ne_tx=XM_019839198.2;refGene_gn=TENM2;refGene_tx=XM_019839204.2;refGen
e_gn=TENM2;refGene_tx=XM_023259843.1;refGene_gn=TENM2;refGene_tx=XM_02
3259850.1;refGene_gn=TENM2;refGene_tx=XM_023259852.1;refGene_gn=TENM2;
refGene_tx=XM_023259862.1;refGene_gn=TENM2;refGene_tx=XM_023259868.1;r
efGene_gn=TENM2;refGene_tx=XM_023259877.1;refGene_gn=TENM2;refGene_tx=
XM_023259887.1;refGene_gn=TENM2;refGene_tx=XM_023259913.1;refGene_gn=T
ENM2;refGene_tx=XM_023259942.1
A1 183195281 183195543
refGene_gn=TENM2;refGene_tx=XM_019839174.2;refGene_gn=TENM2;refGene_tx
=XM_019839189.2;refGene_gn=TENM2;refGene_tx=XM_019839191.2;refGene_gn=
TENM2;refGene_tx=XM_019839194.2;refGene_gn=TENM2;refGene_tx=XM_0198391
95.2;refGene_gn=TENM2;refGene_tx=XM_019839196.2;refGene_gn=TENM2;refGe
ne_tx=XM_019839198.2;refGene_gn=TENM2;refGene_tx=XM_019839204.2;refGen
e_gn=TENM2;refGene_tx=XM_023259843.1;refGene_gn=TENM2;refGene_tx=XM_02
3259850.1;refGene_gn=TENM2;refGene_tx=XM_023259852.1;refGene_gn=TENM2;
refGene_tx=XM_023259862.1;refGene_gn=TENM2;refGene_tx=XM_023259868.1;r
efGene_gn=TENM2;refGene_tx=XM_023259877.1;refGene_gn=TENM2;refGene_tx=
XM_023259887.1;refGene_gn=TENM2;refGene_tx=XM_023259913.1;refGene_gn=T
ENM2;refGene_tx=XM_023259942.1
A1 183198955 183199075
refGene_gn=TENM2;refGene_tx=XM_019839174.2;refGene_gn=TENM2;refGene_tx
=XM_019839189.2;refGene_gn=TENM2;refGene_tx=XM_019839191.2;refGene_gn=
TENM2;refGene_tx=XM_019839194.2;refGene_gn=TENM2;refGene_tx=XM_0198391
95.2;refGene_gn=TENM2;refGene_tx=XM_019839196.2;refGene_gn=TENM2;refGe
ne_tx=XM_019839198.2;refGene_gn=TENM2;refGene_tx=XM_019839204.2;refGen
e_gn=TENM2;refGene_tx=XM_023259843.1;refGene_gn=TENM2;refGene_tx=XM_02

3259850.1;refGene_gn=TENM2;refGene_tx=XM_023259852.1;refGene_gn=TENM2;refGene_tx=XM_023259862.1;refGene_gn=TENM2;refGene_tx=XM_023259868.1;refGene_gn=TENM2;refGene_tx=XM_023259877.1;refGene_gn=TENM2;refGene_tx=XM_023259887.1;refGene_gn=TENM2;refGene_tx=XM_023259913.1;refGene_gn=TENM2;refGene_tx=XM_023259942.1
A1 183203352 183203563
refGene_gn=TENM2;refGene_tx=XM_019839174.2;refGene_gn=TENM2;refGene_tx=XM_019839189.2;refGene_gn=TENM2;refGene_tx=XM_019839191.2;refGene_gn=TENM2;refGene_tx=XM_019839194.2;refGene_gn=TENM2;refGene_tx=XM_019839195.2;refGene_gn=TENM2;refGene_tx=XM_019839196.2;refGene_gn=TENM2;refGene_tx=XM_019839198.2;refGene_gn=TENM2;refGene_tx=XM_019839204.2;refGene_gn=TENM2;refGene_tx=XM_023259843.1;refGene_gn=TENM2;refGene_tx=XM_023259850.1;refGene_gn=TENM2;refGene_tx=XM_023259852.1;refGene_gn=TENM2;refGene_tx=XM_023259862.1;refGene_gn=TENM2;refGene_tx=XM_023259868.1;refGene_gn=TENM2;refGene_tx=XM_023259877.1;refGene_gn=TENM2;refGene_tx=XM_023259887.1;refGene_gn=TENM2;refGene_tx=XM_023259913.1;refGene_gn=TENM2;refGene_tx=XM_023259942.1
A1 183229913 183230060
refGene_gn=TENM2;refGene_tx=XM_019839174.2;refGene_gn=TENM2;refGene_tx=XM_019839189.2;refGene_gn=TENM2;refGene_tx=XM_019839191.2;refGene_gn=TENM2;refGene_tx=XM_019839194.2;refGene_gn=TENM2;refGene_tx=XM_019839195.2;refGene_gn=TENM2;refGene_tx=XM_019839196.2;refGene_gn=TENM2;refGene_tx=XM_019839198.2;refGene_gn=TENM2;refGene_tx=XM_019839204.2;refGene_gn=TENM2;refGene_tx=XM_023259843.1;refGene_gn=TENM2;refGene_tx=XM_023259850.1;refGene_gn=TENM2;refGene_tx=XM_023259852.1;refGene_gn=TENM2;refGene_tx=XM_023259862.1;refGene_gn=TENM2;refGene_tx=XM_023259868.1;refGene_gn=TENM2;refGene_tx=XM_023259877.1;refGene_gn=TENM2;refGene_tx=XM_023259887.1;refGene_gn=TENM2;refGene_tx=XM_023259913.1;refGene_gn=TENM2;refGene_tx=XM_023259942.1
A1 183260291 183260504
refGene_gn=TENM2;refGene_tx=XM_019839174.2;refGene_gn=TENM2;refGene_tx=XM_019839189.2;refGene_gn=TENM2;refGene_tx=XM_019839191.2;refGene_gn=TENM2;refGene_tx=XM_019839194.2;refGene_gn=TENM2;refGene_tx=XM_019839195.2;refGene_gn=TENM2;refGene_tx=XM_019839196.2;refGene_gn=TENM2;refGene_tx=XM_019839198.2;refGene_gn=TENM2;refGene_tx=XM_019839204.2;refGene_gn=TENM2;refGene_tx=XM_023259843.1;refGene_gn=TENM2;refGene_tx=XM_023259850.1;refGene_gn=TENM2;refGene_tx=XM_023259852.1;refGene_gn=TENM2;refGene_tx=XM_023259862.1;refGene_gn=TENM2;refGene_tx=XM_023259868.1;refGene_gn=TENM2;refGene_tx=XM_023259887.1;refGene_gn=TENM2;refGene_tx=XM_023259913.1;refGene_gn=TENM2;refGene_tx=XM_023259942.1
A1 183262224 183262425
refGene_gn=TENM2;refGene_tx=XM_019839174.2;refGene_gn=TENM2;refGene_tx=XM_019839189.2;refGene_gn=TENM2;refGene_tx=XM_019839191.2;refGene_gn=TENM2;refGene_tx=XM_019839194.2;refGene_gn=TENM2;refGene_tx=XM_019839195.2;refGene_gn=TENM2;refGene_tx=XM_019839196.2;refGene_gn=TENM2;refGene_tx=XM_019839198.2;refGene_gn=TENM2;refGene_tx=XM_019839204.2;refGene_gn=TENM2;refGene_tx=XM_023259843.1;refGene_gn=TENM2;refGene_tx=XM_023259850.1;refGene_gn=TENM2;refGene_tx=XM_023259852.1;refGene_gn=TENM2;refGene_tx=XM_023259862.1;refGene_gn=TENM2;refGene_tx=XM_023259868.1;refGene_gn=TENM2;refGene_tx=XM_023259877.1;refGene_gn=TENM2;refGene_tx=

XM_023259887.1;refGene_gn=TENM2;refGene_tx=XM_023259913.1;refGene_gn=TENM2;refGene_tx=XM_023259942.1
A1 183268336 183268531
refGene_gn=TENM2;refGene_tx=XM_019839174.2;refGene_gn=TENM2;refGene_tx=XM_019839189.2;refGene_gn=TENM2;refGene_tx=XM_019839191.2;refGene_gn=TENM2;refGene_tx=XM_019839194.2;refGene_gn=TENM2;refGene_tx=XM_019839195.2;refGene_gn=TENM2;refGene_tx=XM_019839196.2;refGene_gn=TENM2;refGene_tx=XM_019839198.2;refGene_gn=TENM2;refGene_tx=XM_019839204.2;refGene_gn=TENM2;refGene_tx=XM_023259843.1;refGene_gn=TENM2;refGene_tx=XM_023259850.1;refGene_gn=TENM2;refGene_tx=XM_023259852.1;refGene_gn=TENM2;refGene_tx=XM_023259862.1;refGene_gn=TENM2;refGene_tx=XM_023259868.1;refGene_gn=TENM2;refGene_tx=XM_023259877.1;refGene_gn=TENM2;refGene_tx=XM_023259887.1;refGene_gn=TENM2;refGene_tx=XM_023259913.1;refGene_gn=TENM2;refGene_tx=XM_023259942.1
A1 183292107 183292209
refGene_gn=TENM2;refGene_tx=XM_019839174.2;refGene_gn=TENM2;refGene_tx=XM_019839189.2;refGene_gn=TENM2;refGene_tx=XM_019839191.2;refGene_gn=TENM2;refGene_tx=XM_019839194.2;refGene_gn=TENM2;refGene_tx=XM_019839195.2;refGene_gn=TENM2;refGene_tx=XM_019839196.2;refGene_gn=TENM2;refGene_tx=XM_019839198.2;refGene_gn=TENM2;refGene_tx=XM_019839204.2;refGene_gn=TENM2;refGene_tx=XM_023259843.1;refGene_gn=TENM2;refGene_tx=XM_023259850.1;refGene_gn=TENM2;refGene_tx=XM_023259852.1;refGene_gn=TENM2;refGene_tx=XM_023259862.1;refGene_gn=TENM2;refGene_tx=XM_023259868.1;refGene_gn=TENM2;refGene_tx=XM_023259877.1;refGene_gn=TENM2;refGene_tx=XM_023259887.1;refGene_gn=TENM2;refGene_tx=XM_023259913.1;refGene_gn=TENM2;refGene_tx=XM_023259942.1
A1 183299056 183299252
refGene_gn=TENM2;refGene_tx=XM_019839174.2;refGene_gn=TENM2;refGene_tx=XM_019839189.2;refGene_gn=TENM2;refGene_tx=XM_019839191.2;refGene_gn=TENM2;refGene_tx=XM_019839194.2;refGene_gn=TENM2;refGene_tx=XM_019839195.2;refGene_gn=TENM2;refGene_tx=XM_019839196.2;refGene_gn=TENM2;refGene_tx=XM_019839198.2;refGene_gn=TENM2;refGene_tx=XM_019839204.2;refGene_gn=TENM2;refGene_tx=XM_023259843.1;refGene_gn=TENM2;refGene_tx=XM_023259850.1;refGene_gn=TENM2;refGene_tx=XM_023259852.1;refGene_gn=TENM2;refGene_tx=XM_023259862.1;refGene_gn=TENM2;refGene_tx=XM_023259868.1;refGene_gn=TENM2;refGene_tx=XM_023259877.1;refGene_gn=TENM2;refGene_tx=XM_023259887.1;refGene_gn=TENM2;refGene_tx=XM_023259913.1;refGene_gn=TENM2;refGene_tx=XM_023259942.1
A1 183321570 183321776
refGene_gn=TENM2;refGene_tx=XM_019839174.2;refGene_gn=TENM2;refGene_tx=XM_019839189.2;refGene_gn=TENM2;refGene_tx=XM_019839191.2;refGene_gn=TENM2;refGene_tx=XM_019839194.2;refGene_gn=TENM2;refGene_tx=XM_019839195.2;refGene_gn=TENM2;refGene_tx=XM_019839196.2;refGene_gn=TENM2;refGene_tx=XM_019839198.2;refGene_gn=TENM2;refGene_tx=XM_019839204.2;refGene_gn=TENM2;refGene_tx=XM_023259843.1;refGene_gn=TENM2;refGene_tx=XM_023259850.1;refGene_gn=TENM2;refGene_tx=XM_023259852.1;refGene_gn=TENM2;refGene_tx=XM_023259862.1;refGene_gn=TENM2;refGene_tx=XM_023259868.1;refGene_gn=TENM2;refGene_tx=XM_023259877.1;refGene_gn=TENM2;refGene_tx=XM_023259887.1;refGene_gn=TENM2;refGene_tx=XM_023259913.1;refGene_gn=TENM2;refGene_tx=XM_023259942.1

A1 183335068 183335191
refGene_gn=TENM2;refGene_tx=XM_019839174.2;refGene_gn=TENM2;refGene_tx=XM_019839189.2;refGene_gn=TENM2;refGene_tx=XM_019839191.2;refGene_gn=TENM2;refGene_tx=XM_019839194.2;refGene_gn=TENM2;refGene_tx=XM_019839195.2;refGene_gn=TENM2;refGene_tx=XM_019839196.2;refGene_gn=TENM2;refGene_tx=XM_019839198.2;refGene_gn=TENM2;refGene_tx=XM_019839204.2;refGene_gn=TENM2;refGene_tx=XM_023259843.1;refGene_gn=TENM2;refGene_tx=XM_023259850.1;refGene_gn=TENM2;refGene_tx=XM_023259852.1;refGene_gn=TENM2;refGene_tx=XM_023259862.1;refGene_gn=TENM2;refGene_tx=XM_023259868.1;refGene_gn=TENM2;refGene_tx=XM_023259877.1;refGene_gn=TENM2;refGene_tx=XM_023259887.1;refGene_gn=TENM2;refGene_tx=XM_023259913.1;refGene_gn=TENM2;refGene_tx=XM_023259942.1
A1 183386493 183386732
refGene_gn=TENM2;refGene_tx=XM_019839174.2;refGene_gn=TENM2;refGene_tx=XM_019839189.2;refGene_gn=TENM2;refGene_tx=XM_019839191.2;refGene_gn=TENM2;refGene_tx=XM_019839194.2;refGene_gn=TENM2;refGene_tx=XM_019839195.2;refGene_gn=TENM2;refGene_tx=XM_019839196.2;refGene_gn=TENM2;refGene_tx=XM_019839198.2;refGene_gn=TENM2;refGene_tx=XM_019839204.2;refGene_gn=TENM2;refGene_tx=XM_023259843.1;refGene_gn=TENM2;refGene_tx=XM_023259850.1;refGene_gn=TENM2;refGene_tx=XM_023259852.1;refGene_gn=TENM2;refGene_tx=XM_023259862.1;refGene_gn=TENM2;refGene_tx=XM_023259868.1;refGene_gn=TENM2;refGene_tx=XM_023259877.1;refGene_gn=TENM2;refGene_tx=XM_023259887.1;refGene_gn=TENM2;refGene_tx=XM_023259913.1;refGene_gn=TENM2;refGene_tx=XM_023259942.1
A1 183418171 183418406
refGene_gn=TENM2;refGene_tx=XM_019839174.2;refGene_gn=TENM2;refGene_tx=XM_019839189.2;refGene_gn=TENM2;refGene_tx=XM_019839191.2;refGene_gn=TENM2;refGene_tx=XM_019839194.2;refGene_gn=TENM2;refGene_tx=XM_019839195.2;refGene_gn=TENM2;refGene_tx=XM_019839196.2;refGene_gn=TENM2;refGene_tx=XM_019839198.2;refGene_gn=TENM2;refGene_tx=XM_019839204.2;refGene_gn=TENM2;refGene_tx=XM_023259843.1;refGene_gn=TENM2;refGene_tx=XM_023259850.1;refGene_gn=TENM2;refGene_tx=XM_023259852.1;refGene_gn=TENM2;refGene_tx=XM_023259862.1;refGene_gn=TENM2;refGene_tx=XM_023259868.1;refGene_gn=TENM2;refGene_tx=XM_023259877.1;refGene_gn=TENM2;refGene_tx=XM_023259887.1;refGene_gn=TENM2;refGene_tx=XM_023259913.1;refGene_gn=TENM2;refGene_tx=XM_023259942.1
A1 183493883 183494093
refGene_gn=TENM2;refGene_tx=XM_019839174.2;refGene_gn=TENM2;refGene_tx=XM_019839189.2;refGene_gn=TENM2;refGene_tx=XM_019839191.2;refGene_gn=TENM2;refGene_tx=XM_019839194.2;refGene_gn=TENM2;refGene_tx=XM_019839195.2;refGene_gn=TENM2;refGene_tx=XM_019839196.2;refGene_gn=TENM2;refGene_tx=XM_023259843.1;refGene_gn=TENM2;refGene_tx=XM_023259850.1;refGene_gn=TENM2;refGene_tx=XM_023259852.1;refGene_gn=TENM2;refGene_tx=XM_023259862.1;refGene_gn=TENM2;refGene_tx=XM_023259877.1;refGene_gn=TENM2;refGene_tx=XM_023259887.1;refGene_gn=TENM2;refGene_tx=XM_023259942.1
A1 183542061 183542077
refGene_gn=TENM2;refGene_tx=XM_019839191.2;refGene_gn=TENM2;refGene_tx=XM_019839204.2
A1 183781218 183781264
refGene_gn=TENM2;refGene_tx=XM_019839189.2;refGene_gn=TENM2;refGene_tx

=XM_019839198.2
A1 183981161 183981437
refGene_gn=TENM2;refGene_tx=XM_019839174.2;refGene_gn=TENM2;refGene_tx=XM_023259843.1;refGene_gn=TENM2;refGene_tx=XM_023259850.1;refGene_gn=TENM2;refGene_tx=XM_023259852.1;refGene_gn=TENM2;refGene_tx=XM_023259862.1;refGene_gn=TENM2;refGene_tx=XM_023259868.1;refGene_gn=TENM2;refGene_tx=XM_023259877.1;refGene_gn=TENM2;refGene_tx=XM_023259942.1
A1 183984869 183984944
refGene_gn=TENM2;refGene_tx=XM_019839174.2;refGene_gn=TENM2;refGene_tx=XM_023259843.1;refGene_gn=TENM2;refGene_tx=XM_023259850.1;refGene_gn=TENM2;refGene_tx=XM_023259868.1;refGene_gn=TENM2;refGene_tx=XM_023259877.1;refGene_gn=TENM2;refGene_tx=XM_023259942.1
A1 184059272 184059498
refGene_gn=TENM2;refGene_tx=XM_019839174.2;refGene_gn=TENM2;refGene_tx=XM_023259843.1;refGene_gn=TENM2;refGene_tx=XM_023259850.1;refGene_gn=TENM2;refGene_tx=XM_023259852.1;refGene_gn=TENM2;refGene_tx=XM_023259862.1;refGene_gn=TENM2;refGene_tx=XM_023259868.1;refGene_gn=TENM2;refGene_tx=XM_023259877.1;refGene_gn=TENM2;refGene_tx=XM_023259887.1;refGene_gn=TENM2;refGene_tx=XM_023259913.1;refGene_gn=TENM2;refGene_tx=XM_023259942.1
A1 187494473 187494644
refGene_gn=MAT2B;refGene_tx=XM_003981335.5;refGene_gn=MAT2B;refGene_tx=XM_003981336.5
A1 187495263 187495377
refGene_gn=MAT2B;refGene_tx=XM_003981335.5;refGene_gn=MAT2B;refGene_tx=XM_003981336.5
A1 187496719 187496913
refGene_gn=MAT2B;refGene_tx=XM_003981335.5;refGene_gn=MAT2B;refGene_tx=XM_003981336.5
A1 187499807 187499960
refGene_gn=MAT2B;refGene_tx=XM_003981335.5;refGene_gn=MAT2B;refGene_tx=XM_003981336.5
A1 187500147 187500262
refGene_gn=MAT2B;refGene_tx=XM_003981335.5;refGene_gn=MAT2B;refGene_tx=XM_003981336.5
A1 187502610 187502805
refGene_gn=MAT2B;refGene_tx=XM_003981335.5;refGene_gn=MAT2B;refGene_tx=XM_003981336.5
A1 187509073 187509136
refGene_gn=MAT2B;refGene_tx=XM_003981335.5
A1 187511488 187511518
refGene_gn=MAT2B;refGene_tx=XM_003981336.5
A1 187528109 187528129
refGene_gn=HMMR;refGene_tx=XM_023260007.1;refGene_gn=HMMR;refGene_tx=XM_023260013.1
A1 187528652 187528815
refGene_gn=HMMR;refGene_tx=XM_023260007.1;refGene_gn=HMMR;refGene_tx=XM_023260013.1
A1 187534433 187534610
refGene_gn=HMMR;refGene_tx=XM_023260007.1;refGene_gn=HMMR;refGene_tx=X

M_023260013.1
A1 187535293 187535393
refGene_gn=HMMR;refGene_tx=XM_023260007.1;refGene_gn=HMMR;refGene_tx=XM_023260013.1
A1 187535483 187535636
refGene_gn=HMMR;refGene_tx=XM_023260007.1;refGene_gn=HMMR;refGene_tx=XM_023260013.1
A1 187535922 187536069
refGene_gn=HMMR;refGene_tx=XM_023260007.1;refGene_gn=HMMR;refGene_tx=XM_023260013.1
A1 187542933 187543050
refGene_gn=HMMR;refGene_tx=XM_023260007.1;refGene_gn=HMMR;refGene_tx=XM_023260013.1
A1 187547876 187548091
refGene_gn=HMMR;refGene_tx=XM_023260007.1;refGene_gn=HMMR;refGene_tx=XM_023260013.1
A1 187551430 187551579
refGene_gn=HMMR;refGene_tx=XM_023260007.1;refGene_gn=HMMR;refGene_tx=XM_023260013.1
A1 187552207 187552386
refGene_gn=HMMR;refGene_tx=XM_023260007.1;refGene_gn=HMMR;refGene_tx=XM_023260013.1
A1 187552751 187552826
refGene_gn=HMMR;refGene_tx=XM_023260007.1;refGene_gn=HMMR;refGene_tx=XM_023260013.1
A1 187554389 187554490
refGene_gn=HMMR;refGene_tx=XM_023260007.1;refGene_gn=HMMR;refGene_tx=XM_023260013.1
A1 187554577 187554664
refGene_gn=HMMR;refGene_tx=XM_023260007.1;refGene_gn=HMMR;refGene_tx=XM_023260013.1
A1 187556119 187556308
refGene_gn=HMMR;refGene_tx=XM_023260007.1;refGene_gn=HMMR;refGene_tx=XM_023260013.1
A1 187559648 187559696
refGene_gn=HMMR;refGene_tx=XM_023260007.1;refGene_gn=HMMR;refGene_tx=XM_023260013.1
A1 187563140 187563220
refGene_gn=HMMR;refGene_tx=XM_023260007.1;refGene_gn=HMMR;refGene_tx=XM_023260013.1
A1 187563688 187563787
refGene_gn=HMMR;refGene_tx=XM_023260007.1;refGene_gn=HMMR;refGene_tx=XM_023260013.1
A1 187567827 187567873
refGene_gn=HMMR;refGene_tx=XM_023260007.1;refGene_gn=HMMR;refGene_tx=XM_023260013.1
A1 187568538 187568727
refGene_gn=NUDCD2;refGene_tx=XM_003981338.5
A1 187570923 187570972
refGene_gn=NUDCD2;refGene_tx=XM_003981338.5

A1 187571177 187571329
refGene_gn=NUDCD2;refGene_tx=XM_003981338.5
A1 187573674 187573758
refGene_gn=NUDCD2;refGene_tx=XM_003981338.5
A1 187584119 187584311
refGene_gn=CCNG1;refGene_tx=XM_003981339.5
A1 187584539 187584638
refGene_gn=CCNG1;refGene_tx=XM_003981339.5
A1 187584717 187584796
refGene_gn=CCNG1;refGene_tx=XM_003981339.5
A1 187585365 187585619
refGene_gn=CCNG1;refGene_tx=XM_003981339.5
A1 187587330 187587594
refGene_gn=CCNG1;refGene_tx=XM_003981339.5
A1 188550454 188550730
refGene_gn=GABRG2;refGene_tx=XM_003981341.5;refGene_gn=GABRG2;refGene_tx=XM_003981342.5
A1 188552142 188552166
refGene_gn=GABRG2;refGene_tx=XM_003981341.5
A1 188554143 188554349
refGene_gn=GABRG2;refGene_tx=XM_003981341.5;refGene_gn=GABRG2;refGene_tx=XM_003981342.5
A1 188567358 188567511
refGene_gn=GABRG2;refGene_tx=XM_003981341.5;refGene_gn=GABRG2;refGene_tx=XM_003981342.5
A1 188617582 188617720
refGene_gn=GABRG2;refGene_tx=XM_003981341.5;refGene_gn=GABRG2;refGene_tx=XM_003981342.5
A1 188621113 188621196
refGene_gn=GABRG2;refGene_tx=XM_003981341.5;refGene_gn=GABRG2;refGene_tx=XM_003981342.5
A1 188624402 188624623
refGene_gn=GABRG2;refGene_tx=XM_003981341.5;refGene_gn=GABRG2;refGene_tx=XM_003981342.5
A1 188627170 188627238
refGene_gn=GABRG2;refGene_tx=XM_003981341.5;refGene_gn=GABRG2;refGene_tx=XM_003981342.5
A1 188628792 188628944
refGene_gn=GABRG2;refGene_tx=XM_003981341.5;refGene_gn=GABRG2;refGene_tx=XM_003981342.5
A1 188662124 188662231
refGene_gn=GABRG2;refGene_tx=XM_003981341.5;refGene_gn=GABRG2;refGene_tx=XM_003981342.5
A1 188772096 188772408
refGene_gn=GABRA1;refGene_tx=XM_006927959.4;refGene_gn=GABRA1;refGene_tx=XM_019839343.2;refGene_gn=GABRA1;refGene_tx=XM_019839347.2;refGene_gn=GABRA1;refGene_tx=XM_019839350.2
A1 188777606 188777809
refGene_gn=GABRA1;refGene_tx=XM_006927959.4;refGene_gn=GABRA1;refGene_tx=XM_019839343.2;refGene_gn=GABRA1;refGene_tx=XM_019839347.2;refGene_

gn=GABRA1;refGene_tx=XM_019839350.2
A1 188783550 188783703
refGene_gn=GABRA1;refGene_tx=XM_006927959.4;refGene_gn=GABRA1;refGene_tx=XM_019839343.2;refGene_gn=GABRA1;refGene_tx=XM_019839347.2;refGene_gn=GABRA1;refGene_tx=XM_019839350.2
A1 188792969 188793113
refGene_gn=GABRA1;refGene_tx=XM_006927959.4;refGene_gn=GABRA1;refGene_tx=XM_019839343.2;refGene_gn=GABRA1;refGene_tx=XM_019839347.2;refGene_gn=GABRA1;refGene_tx=XM_019839350.2
A1 188800226 188800309
refGene_gn=GABRA1;refGene_tx=XM_006927959.4;refGene_gn=GABRA1;refGene_tx=XM_019839343.2;refGene_gn=GABRA1;refGene_tx=XM_019839347.2;refGene_gn=GABRA1;refGene_tx=XM_019839350.2
A1 188802557 188802778
refGene_gn=GABRA1;refGene_tx=XM_006927959.4;refGene_gn=GABRA1;refGene_tx=XM_019839343.2;refGene_gn=GABRA1;refGene_tx=XM_019839347.2;refGene_gn=GABRA1;refGene_tx=XM_019839350.2
A1 188809732 188809800
refGene_gn=GABRA1;refGene_tx=XM_006927959.4;refGene_gn=GABRA1;refGene_tx=XM_019839343.2
A1 188824272 188824385
refGene_gn=GABRA1;refGene_tx=XM_006927959.4;refGene_gn=GABRA1;refGene_tx=XM_019839343.2;refGene_gn=GABRA1;refGene_tx=XM_019839347.2;refGene_gn=GABRA1;refGene_tx=XM_019839350.2
A1 188827828 188827902
refGene_gn=GABRA1;refGene_tx=XM_006927959.4;refGene_gn=GABRA1;refGene_tx=XM_019839343.2
A1 188965998 188966274
refGene_gn=GABRA6;refGene_tx=XM_006927962.3;refGene_gn=GABRA6;refGene_tx=XM_019839361.2
A1 188975330 188975590
refGene_gn=GABRA6;refGene_tx=XM_006927962.3;refGene_gn=GABRA6;refGene_tx=XM_019839361.2
A1 188977576 188977729
refGene_gn=GABRA6;refGene_tx=XM_006927962.3;refGene_gn=GABRA6;refGene_tx=XM_019839361.2
A1 188978117 188978261
refGene_gn=GABRA6;refGene_tx=XM_006927962.3;refGene_gn=GABRA6;refGene_tx=XM_019839361.2
A1 188978562 188978645
refGene_gn=GABRA6;refGene_tx=XM_006927962.3;refGene_gn=GABRA6;refGene_tx=XM_019839361.2
A1 188978729 188978950
refGene_gn=GABRA6;refGene_tx=XM_006927962.3;refGene_gn=GABRA6;refGene_tx=XM_019839361.2
A1 188980898 188980966
refGene_gn=GABRA6;refGene_tx=XM_006927962.3
A1 188981558 188981677
refGene_gn=GABRA6;refGene_tx=XM_006927962.3
A1 188981871 188981909

refGene_gn=GABRA6;refGene_tx=XM_006927962.3
A1 189108826 189108903
refGene_gn=GABRB2;refGene_tx=XM_011285648.3;refGene_gn=GABRB2;refGene_tx=XM_011285651.3;refGene_gn=GABRB2;refGene_tx=XM_019839372.2
A1 189109067 189109159
refGene_gn=GABRB2;refGene_tx=XM_011285648.3;refGene_gn=GABRB2;refGene_tx=XM_011285651.3;refGene_gn=GABRB2;refGene_tx=XM_019839372.2
A1 189110153 189110221
refGene_gn=GABRB2;refGene_tx=XM_011285648.3;refGene_gn=GABRB2;refGene_tx=XM_011285651.3;refGene_gn=GABRB2;refGene_tx=XM_019839372.2
A1 189185441 189185662
refGene_gn=GABRB2;refGene_tx=XM_011285648.3;refGene_gn=GABRB2;refGene_tx=XM_011285651.3;refGene_gn=GABRB2;refGene_tx=XM_019839372.2
A1 189224824 189224907
refGene_gn=GABRB2;refGene_tx=XM_011285648.3;refGene_gn=GABRB2;refGene_tx=XM_011285651.3;refGene_gn=GABRB2;refGene_tx=XM_019839372.2
A1 189298539 189298677
refGene_gn=GABRB2;refGene_tx=XM_011285648.3;refGene_gn=GABRB2;refGene_tx=XM_011285651.3;refGene_gn=GABRB2;refGene_tx=XM_019839372.2
A1 189300413 189300566
refGene_gn=GABRB2;refGene_tx=XM_011285648.3;refGene_gn=GABRB2;refGene_tx=XM_011285651.3;refGene_gn=GABRB2;refGene_tx=XM_019839372.2
A1 189303923 189304168
refGene_gn=GABRB2;refGene_tx=XM_011285648.3;refGene_gn=GABRB2;refGene_tx=XM_011285651.3;refGene_gn=GABRB2;refGene_tx=XM_019839372.2
A1 189309316 189309430
refGene_gn=GABRB2;refGene_tx=XM_011285651.3;refGene_gn=GABRB2;refGene_tx=XM_019839372.2
A1 189341098 189341446
refGene_gn=GABRB2;refGene_tx=XM_011285648.3;refGene_gn=GABRB2;refGene_tx=XM_011285651.3;refGene_gn=GABRB2;refGene_tx=XM_019839372.2
A1 189737268 189737305
refGene_gn=ATP10B;refGene_tx=XM_023260052.1
A1 189899424 189899719
refGene_gn=ATP10B;refGene_tx=XM_023260052.1
A1 189900727 189900922
refGene_gn=ATP10B;refGene_tx=XM_023260052.1
A1 189918298 189918503
refGene_gn=ATP10B;refGene_tx=XM_023260052.1
A1 189936482 189936568
refGene_gn=ATP10B;refGene_tx=XM_023260052.1
A1 189940965 189941072
refGene_gn=ATP10B;refGene_tx=XM_023260052.1
A1 189944491 189944623
refGene_gn=ATP10B;refGene_tx=XM_023260052.1
A1 189947956 189948084
refGene_gn=ATP10B;refGene_tx=XM_023260052.1
A1 189950034 189950287
refGene_gn=ATP10B;refGene_tx=XM_023260052.1
A1 189951386 189951625

refGene_gn=ATP10B;refGene_tx=XM_023260052.1
A1	189960926	189961118
refGene_gn=ATP10B;refGene_tx=XM_023260052.1
A1	189962632	189963233
refGene_gn=ATP10B;refGene_tx=XM_023260052.1
A1	189965599	189965709
refGene_gn=ATP10B;refGene_tx=XM_023260052.1
A1	189967538	189967665
refGene_gn=ATP10B;refGene_tx=XM_023260052.1
A1	189970583	189970768
refGene_gn=ATP10B;refGene_tx=XM_023260052.1
A1	189974783	189975105
refGene_gn=ATP10B;refGene_tx=XM_023260052.1
A1	189977878	189977955
refGene_gn=ATP10B;refGene_tx=XM_023260052.1
A1	189979322	189979448
refGene_gn=ATP10B;refGene_tx=XM_023260052.1
A1	189986667	189986868
refGene_gn=ATP10B;refGene_tx=XM_023260052.1
A1	189987938	189988019
refGene_gn=ATP10B;refGene_tx=XM_023260052.1
A1	189989401	189989506
refGene_gn=ATP10B;refGene_tx=XM_023260052.1
A1	189998582	189998770
refGene_gn=ATP10B;refGene_tx=XM_023260052.1
A1	190004058	190004296
refGene_gn=ATP10B;refGene_tx=XM_023260052.1
A1	190191759	190191836
refGene_gn=PTTG1;refGene_tx=XM_003981324.5;refGene_gn=PTTG1;refGene_tx
=XM_006927954.3;refGene_gn=PTTG1;refGene_tx=XM_006927956.3;refGene_gn=
PTTG1;refGene_tx=XM_006927957.3;refGene_gn=PTTG1;refGene_tx=XM_0198394
24.2;refGene_gn=PTTG1;refGene_tx=XM_019839430.2
A1	190192599	190192758
refGene_gn=PTTG1;refGene_tx=XM_003981324.5;refGene_gn=PTTG1;refGene_tx
=XM_006927954.3;refGene_gn=PTTG1;refGene_tx=XM_006927956.3;refGene_gn=
PTTG1;refGene_tx=XM_006927957.3;refGene_gn=PTTG1;refGene_tx=XM_0198394
24.2;refGene_gn=PTTG1;refGene_tx=XM_019839430.2
A1	190196330	190196424
refGene_gn=PTTG1;refGene_tx=XM_003981324.5;refGene_gn=PTTG1;refGene_tx
=XM_006927954.3;refGene_gn=PTTG1;refGene_tx=XM_006927956.3;refGene_gn=
PTTG1;refGene_tx=XM_006927957.3;refGene_gn=PTTG1;refGene_tx=XM_0198394
24.2;refGene_gn=PTTG1;refGene_tx=XM_019839430.2
A1	190198074	190198272
refGene_gn=PTTG1;refGene_tx=XM_003981324.5;refGene_gn=PTTG1;refGene_tx
=XM_006927954.3;refGene_gn=PTTG1;refGene_tx=XM_006927956.3;refGene_gn=
PTTG1;refGene_tx=XM_006927957.3;refGene_gn=PTTG1;refGene_tx=XM_0198394
24.2;refGene_gn=PTTG1;refGene_tx=XM_019839430.2
A1	190198636	190198727
refGene_gn=PTTG1;refGene_tx=XM_003981324.5;refGene_gn=PTTG1;refGene_tx
=XM_006927954.3;refGene_gn=PTTG1;refGene_tx=XM_006927956.3;refGene_gn=

PTTG1;refGene_tx=XM_006927957.3;refGene_gn=PTTG1;refGene_tx=XM_0198394
24.2;refGene_gn=PTTG1;refGene_tx=XM_019839430.2
A1 190204853 190205023
refGene_gn=SLU7;refGene_tx=XM_023260055.1
A1 190206380 190206534
refGene_gn=SLU7;refGene_tx=XM_023260055.1
A1 190206875 190206956
refGene_gn=SLU7;refGene_tx=XM_023260055.1
A1 190207283 190207448
refGene_gn=SLU7;refGene_tx=XM_023260055.1
A1 190208339 190208408
refGene_gn=SLU7;refGene_tx=XM_023260055.1
A1 190212863 190212911
refGene_gn=SLU7;refGene_tx=XM_023260055.1
A1 190213095 190213227
refGene_gn=SLU7;refGene_tx=XM_023260055.1
A1 190213483 190213581
refGene_gn=SLU7;refGene_tx=XM_023260055.1
A1 190214000 190214068
refGene_gn=SLU7;refGene_tx=XM_023260055.1
A1 190214201 190214341
refGene_gn=SLU7;refGene_tx=XM_023260055.1
A1 190215228 190215390
refGene_gn=SLU7;refGene_tx=XM_023260055.1
A1 190216686 190216791
refGene_gn=SLU7;refGene_tx=XM_023260055.1
A1 190216948 190217020
refGene_gn=SLU7;refGene_tx=XM_023260055.1
A1 190217339 190217456
refGene_gn=SLU7;refGene_tx=XM_023260055.1
A1 190218069 190218249
refGene_gn=SLU7;refGene_tx=XM_023260055.1
A1 190225107 190226892
refGene_gn=ZBED8;refGene_tx=XM_003981321.5
A1 190243452 190243656
refGene_gn=C1QTNF2;refGene_tx=XM_003981320.4
A1 190254085 190254130
refGene_gn=C1QTNF2;refGene_tx=XM_006927949.3;refGene_gn=C1QTNF2;refGen
e_tx=XM_006927952.4;refGene_gn=C1QTNF2;refGene_tx=XM_019839471.2;refGe
ne_gn=C1QTNF2;refGene_tx=XM_019839478.2;refGene_gn=C1QTNF2;refGene_tx=
XM_019839483.2;refGene_gn=C1QTNF2;refGene_tx=XM_023260059.1
A1 190260340 190260593
refGene_gn=C1QTNF2;refGene_tx=XM_003981320.4;refGene_gn=C1QTNF2;refGen
e_tx=XM_006927949.3;refGene_gn=C1QTNF2;refGene_tx=XM_006927952.4;refGe
ne_gn=C1QTNF2;refGene_tx=XM_006927953.3;refGene_gn=C1QTNF2;refGene_tx=
XM_019839471.2;refGene_gn=C1QTNF2;refGene_tx=XM_019839478.2;refGene_gn
=C1QTNF2;refGene_tx=XM_019839483.2;refGene_gn=C1QTNF2;refGene_tx=XM_02
3260059.1
A1 190264517 190265131
refGene_gn=C1QTNF2;refGene_tx=XM_003981320.4;refGene_gn=C1QTNF2;refGen

e_tx=XM_006927949.3;refGene_gn=C1QTNF2;refGene_tx=XM_006927952.4;refGene_gn=C1QTNF2;refGene_tx=XM_006927953.3;refGene_gn=C1QTNF2;refGene_tx=XM_019839471.2;refGene_gn=C1QTNF2;refGene_tx=XM_019839478.2;refGene_gn=C1QTNF2;refGene_tx=XM_019839483.2;refGene_gn=C1QTNF2;refGene_tx=XM_023260059.1
A1 190306179 190306245
refGene_gn=CCNJL;refGene_tx=XM_011285695.3;refGene_gn=CCNJL;refGene_tx=XM_019839458.2
A1 190336987 190337201
refGene_gn=CCNJL;refGene_tx=XM_011285695.3;refGene_gn=CCNJL;refGene_tx=XM_019839458.2
A1 190359125 190359428
refGene_gn=CCNJL;refGene_tx=XM_011285695.3;refGene_gn=CCNJL;refGene_tx=XM_019839458.2
A1 190361250 190361410
refGene_gn=CCNJL;refGene_tx=XM_011285695.3;refGene_gn=CCNJL;refGene_tx=XM_019839458.2
A1 190362897 190363321
refGene_gn=CCNJL;refGene_tx=XM_011285695.3;refGene_gn=CCNJL;refGene_tx=XM_019839458.2
A1 190373041 190373095
refGene_gn=FABP6;refGene_tx=XM_003981318.4;refGene_gn=FABP6;refGene_tx=XM_019839490.2
A1 190375490 190375580
refGene_gn=FABP6;refGene_tx=XM_003981318.4;refGene_gn=FABP6;refGene_tx=XM_019839490.2
A1 190376859 190377035
refGene_gn=FABP6;refGene_tx=XM_003981318.4;refGene_gn=FABP6;refGene_tx=XM_019839490.2
A1 190378238 190378317
refGene_gn=FABP6;refGene_tx=XM_003981318.4;refGene_gn=FABP6;refGene_tx=XM_019839490.2
A1 190411505 190411508
refGene_gn=FABP6;refGene_tx=XM_019839490.2
A1 190452660 190453244
refGene_gn=PWWP2A;refGene_tx=XM_023260064.1;refGene_gn=PWWP2A;refGene_tx=XM_023260065.1;refGene_gn=PWWP2A;refGene_tx=XM_023260066.1;refGene_gn=PWWP2A;refGene_tx=XM_023260067.1;refGene_gn=PWWP2A;refGene_tx=XM_023260068.1;refGene_gn=PWWP2A;refGene_tx=XM_023260069.1;refGene_gn=PWWP2A;refGene_tx=XM_023260070.1;refGene_gn=PWWP2A;refGene_tx=XM_023260071.1;refGene_gn=PWWP2A;refGene_tx=XM_023260072.1;refGene_gn=PWWP2A;refGene_tx=XM_023260073.1;refGene_gn=PWWP2A;refGene_tx=XM_023260074.1;refGene_gn=PWWP2A;refGene_tx=XM_023260075.1;refGene_gn=PWWP2A;refGene_tx=XM_023260077.1
A1 190466447 190466497
refGene_gn=PWWP2A;refGene_tx=XM_011285712.3
A1 190471104 190472788
refGene_gn=PWWP2A;refGene_tx=XM_011285712.3;refGene_gn=PWWP2A;refGene_tx=XM_023260064.1;refGene_gn=PWWP2A;refGene_tx=XM_023260065.1;refGene_gn=PWWP2A;refGene_tx=XM_023260066.1;refGene_gn=PWWP2A;refGene_tx=XM_02

3260067.1;refGene_gn=PWWP2A;refGene_tx=XM_023260068.1;refGene_gn=PWWP2
A;refGene_tx=XM_023260069.1;refGene_gn=PWWP2A;refGene_tx=XM_023260070.
1;refGene_gn=PWWP2A;refGene_tx=XM_023260071.1;refGene_gn=PWWP2A;refGen
e_tx=XM_023260072.1;refGene_gn=PWWP2A;refGene_tx=XM_023260073.1;refGen
e_gn=PWWP2A;refGene_tx=XM_023260074.1;refGene_gn=PWWP2A;refGene_tx=XM_
023260075.1;refGene_gn=PWWP2A;refGene_tx=XM_023260076.1;refGene_gn=PWW
P2A;refGene_tx=XM_023260077.1
A1 190476887 190476975
refGene_gn=PWWP2A;refGene_tx=XM_023260068.1;refGene_gn=PWWP2A;refGene_
tx=XM_023260070.1;refGene_gn=PWWP2A;refGene_tx=XM_023260071.1;refGene_
gn=PWWP2A;refGene_tx=XM_023260073.1
A1 190483459 190483645
refGene_gn=PWWP2A;refGene_tx=XM_023260068.1;refGene_gn=PWWP2A;refGene_
tx=XM_023260069.1;refGene_gn=PWWP2A;refGene_tx=XM_023260070.1;refGene_
gn=PWWP2A;refGene_tx=XM_023260071.1;refGene_gn=PWWP2A;refGene_tx=XM_02
3260072.1;refGene_gn=PWWP2A;refGene_tx=XM_023260074.1
A1 190484747 190484836
refGene_gn=PWWP2A;refGene_tx=XM_023260069.1;refGene_gn=PWWP2A;refGene_
tx=XM_023260071.1
A1 190485987 190486089
refGene_gn=PWWP2A;refGene_tx=XM_023260068.1;refGene_gn=PWWP2A;refGene_
tx=XM_023260069.1;refGene_gn=PWWP2A;refGene_tx=XM_023260070.1;refGene_
gn=PWWP2A;refGene_tx=XM_023260073.1;refGene_gn=PWWP2A;refGene_tx=XM_02
3260074.1;refGene_gn=PWWP2A;refGene_tx=XM_023260077.1
A1 190496900 190497034
refGene_gn=TTC1;refGene_tx=XM_003981316.5;refGene_gn=TTC1;refGene_tx=X
M_006927944.4;refGene_gn=TTC1;refGene_tx=XM_019839575.1
A1 190504650 190504705
refGene_gn=TTC1;refGene_tx=XM_003981316.5;refGene_gn=TTC1;refGene_tx=X
M_006927944.4;refGene_gn=TTC1;refGene_tx=XM_019839575.1
A1 190507214 190507363
refGene_gn=TTC1;refGene_tx=XM_003981316.5;refGene_gn=TTC1;refGene_tx=X
M_006927944.4;refGene_gn=TTC1;refGene_tx=XM_019839575.1
A1 190510411 190510448
refGene_gn=TTC1;refGene_tx=XM_003981316.5;refGene_gn=TTC1;refGene_tx=X
M_006927944.4;refGene_gn=TTC1;refGene_tx=XM_019839575.1
A1 190515318 190515431
refGene_gn=TTC1;refGene_tx=XM_003981316.5;refGene_gn=TTC1;refGene_tx=X
M_006927944.4;refGene_gn=TTC1;refGene_tx=XM_019839575.1
A1 190517086 190517147
refGene_gn=TTC1;refGene_tx=XM_003981316.5;refGene_gn=TTC1;refGene_tx=X
M_006927944.4;refGene_gn=TTC1;refGene_tx=XM_019839575.1
A1 190541720 190542053
refGene_gn=TTC1;refGene_tx=XM_003981316.5;refGene_gn=TTC1;refGene_tx=X
M_006927944.4;refGene_gn=TTC1;refGene_tx=XM_019839575.1
A1 190575210 190575818
refGene_gn=ADRA1B;refGene_tx=XM_023260079.1
A1 190625446 190626395
refGene_gn=ADRA1B;refGene_tx=XM_023260079.1
A1 190925169 190925424

refGene_gn=LOC109492996;refGene_tx=XM_019814310.2
A1 191099915 191100003
refGene_gn=IL12B;refGene_tx=NM_001077413.1
A1 191103416 191103692
refGene_gn=IL12B;refGene_tx=NM_001077413.1
A1 191104195 191104313
refGene_gn=IL12B;refGene_tx=NM_001077413.1
A1 191105836 191106054
refGene_gn=IL12B;refGene_tx=NM_001077413.1
A1 191107717 191107875
refGene_gn=IL12B;refGene_tx=NM_001077413.1
A1 191109243 191109375
refGene_gn=IL12B;refGene_tx=NM_001077413.1
A1 191134945 191135176
refGene_gn=UBLCP1;refGene_tx=XM_003981314.5;refGene_gn=UBLCP1;refGene_tx=XM_023260088.1
A1 191135315 191135499
refGene_gn=UBLCP1;refGene_tx=XM_023260088.1
A1 191137473 191137601
refGene_gn=UBLCP1;refGene_tx=XM_003981314.5;refGene_gn=UBLCP1;refGene_tx=XM_023260088.1
A1 191139423 191139447
refGene_gn=UBLCP1;refGene_tx=XM_019839638.2
A1 191141820 191141937
refGene_gn=UBLCP1;refGene_tx=XM_003981314.5;refGene_gn=UBLCP1;refGene_tx=XM_019839638.2;refGene_gn=UBLCP1;refGene_tx=XM_023260088.1
A1 191145472 191145571
refGene_gn=UBLCP1;refGene_tx=XM_003981314.5;refGene_gn=UBLCP1;refGene_tx=XM_019839638.2;refGene_gn=UBLCP1;refGene_tx=XM_023260088.1
A1 191146478 191146516
refGene_gn=UBLCP1;refGene_tx=XM_003981314.5;refGene_gn=UBLCP1;refGene_tx=XM_019839638.2;refGene_gn=UBLCP1;refGene_tx=XM_023260088.1
A1 191149064 191149163
refGene_gn=UBLCP1;refGene_tx=XM_003981314.5;refGene_gn=UBLCP1;refGene_tx=XM_019839638.2;refGene_gn=UBLCP1;refGene_tx=XM_023260088.1
A1 191150413 191150529
refGene_gn=UBLCP1;refGene_tx=XM_003981314.5;refGene_gn=UBLCP1;refGene_tx=XM_019839638.2;refGene_gn=UBLCP1;refGene_tx=XM_023260088.1
A1 191150617 191150703
refGene_gn=UBLCP1;refGene_tx=XM_003981314.5;refGene_gn=UBLCP1;refGene_tx=XM_019839638.2;refGene_gn=UBLCP1;refGene_tx=XM_023260088.1
A1 191151105 191151197
refGene_gn=UBLCP1;refGene_tx=XM_003981314.5;refGene_gn=UBLCP1;refGene_tx=XM_019839638.2;refGene_gn=UBLCP1;refGene_tx=XM_023260088.1
A1 191151973 191152127
refGene_gn=UBLCP1;refGene_tx=XM_003981314.5;refGene_gn=UBLCP1;refGene_tx=XM_019839638.2;refGene_gn=UBLCP1;refGene_tx=XM_023260088.1
A1 191197125 191197309
refGene_gn=RNF145;refGene_tx=XM_023260095.1
A1 191208319 191208428

refGene_gn=RNF145;refGene_tx=XM_023260095.1
A1 191222328 191222420
refGene_gn=RNF145;refGene_tx=XM_023260095.1
A1 191229607 191229843
refGene_gn=RNF145;refGene_tx=XM_023260095.1
A1 191232570 191232746
refGene_gn=RNF145;refGene_tx=XM_023260095.1
A1 191236343 191236484
refGene_gn=RNF145;refGene_tx=XM_023260095.1
A1 191239125 191239308
refGene_gn=RNF145;refGene_tx=XM_023260095.1
A1 191248358 191248506
refGene_gn=RNF145;refGene_tx=XM_023260095.1
A1 191249386 191249743
refGene_gn=RNF145;refGene_tx=XM_023260095.1
A1 191251975 191252362
refGene_gn=RNF145;refGene_tx=XM_023260095.1
A1 191302082 191302216
refGene_gn=EBF1;refGene_tx=XM_023260105.1
A1 191304351 191304508
refGene_gn=EBF1;refGene_tx=XM_023260105.1
A1 191305046 191305110
refGene_gn=EBF1;refGene_tx=XM_023260105.1
A1 191305735 191305791
refGene_gn=EBF1;refGene_tx=XM_023260105.1
A1 191316417 191316444
refGene_gn=EBF1;refGene_tx=XM_023260105.1
A1 191316445 191316492
refGene_gn=EBF1;refGene_tx=XM_023260105.1
A1 191326282 191326351
refGene_gn=EBF1;refGene_tx=XM_023260105.1
A1 191548755 191548837
refGene_gn=EBF1;refGene_tx=XM_023260105.1
A1 191565438 191565556
refGene_gn=EBF1;refGene_tx=XM_023260105.1
A1 191592048 191592179
refGene_gn=EBF1;refGene_tx=XM_023260105.1
A1 191612930 191613057
refGene_gn=EBF1;refGene_tx=XM_023260105.1
A1 191659025 191659114
refGene_gn=EBF1;refGene_tx=XM_023260105.1
A1 191675667 191675733
refGene_gn=EBF1;refGene_tx=XM_023260105.1
A1 191676714 191676892
refGene_gn=EBF1;refGene_tx=XM_023260105.1
A1 191677533 191677713
refGene_gn=EBF1;refGene_tx=XM_023260105.1
A1 191681378 191681573
refGene_gn=EBF1;refGene_tx=XM_023260105.1
A1 191690034 191690066

refGene_gn=EBF1;refGene_tx=XM_023260105.1
A1 192420656 192420697
refGene_gn=CLINT1;refGene_tx=XM_003981309.5;refGene_gn=CLINT1;refGene_tx=XM_003981310.5
A1 192451470 192451575
refGene_gn=CLINT1;refGene_tx=XM_003981309.5;refGene_gn=CLINT1;refGene_tx=XM_003981310.5
A1 192452112 192452209
refGene_gn=CLINT1;refGene_tx=XM_003981309.5;refGene_gn=CLINT1;refGene_tx=XM_003981310.5
A1 192453644 192453753
refGene_gn=CLINT1;refGene_tx=XM_003981309.5;refGene_gn=CLINT1;refGene_tx=XM_003981310.5
A1 192454706 192454871
refGene_gn=CLINT1;refGene_tx=XM_003981309.5;refGene_gn=CLINT1;refGene_tx=XM_003981310.5
A1 192457669 192457847
refGene_gn=CLINT1;refGene_tx=XM_003981309.5;refGene_gn=CLINT1;refGene_tx=XM_003981310.5
A1 192461245 192461492
refGene_gn=CLINT1;refGene_tx=XM_003981309.5;refGene_gn=CLINT1;refGene_tx=XM_003981310.5
A1 192463673 192463743
refGene_gn=CLINT1;refGene_tx=XM_003981309.5;refGene_gn=CLINT1;refGene_tx=XM_003981310.5
A1 192470874 192470949
refGene_gn=CLINT1;refGene_tx=XM_003981309.5;refGene_gn=CLINT1;refGene_tx=XM_003981310.5
A1 192474714 192475007
refGene_gn=CLINT1;refGene_tx=XM_003981309.5;refGene_gn=CLINT1;refGene_tx=XM_003981310.5
A1 192476584 192476789
refGene_gn=CLINT1;refGene_tx=XM_003981309.5;refGene_gn=CLINT1;refGene_tx=XM_003981310.5
A1 192478022 192478369
refGene_gn=CLINT1;refGene_tx=XM_003981309.5;refGene_gn=CLINT1;refGene_tx=XM_003981310.5
A1 192511420 192511831
refGene_gn=LSM11;refGene_tx=XM_023260127.1
A1 192512295 192512379
refGene_gn=LSM11;refGene_tx=XM_023260127.1
A1 192514755 192514895
refGene_gn=LSM11;refGene_tx=XM_023260127.1
A1 192520398 192520843
refGene_gn=LSM11;refGene_tx=XM_023260127.1
A1 192524188 192524350
refGene_gn=THG1L;refGene_tx=XM_003981308.5;refGene_gn=THG1L;refGene_tx=XM_019839760.2;refGene_gn=THG1L;refGene_tx=XM_019839771.2;refGene_gn=THG1L;refGene_tx=XM_019839775.2;refGene_gn=THG1L;refGene_tx=XM_019839776.2

A1	192525634	192525742
refGene_gn=THG1L;refGene_tx=XM_003981308.5;refGene_gn=THG1L;refGene_tx=XM_019839771.2;refGene_gn=THG1L;refGene_tx=XM_019839775.2;refGene_gn=THG1L;refGene_tx=XM_019839776.2
A1	192526872	192526961
refGene_gn=THG1L;refGene_tx=XM_003981308.5;refGene_gn=THG1L;refGene_tx=XM_019839760.2;refGene_gn=THG1L;refGene_tx=XM_019839771.2;refGene_gn=THG1L;refGene_tx=XM_019839775.2;refGene_gn=THG1L;refGene_tx=XM_019839776.2
A1	192527976	192528146
refGene_gn=THG1L;refGene_tx=XM_003981308.5;refGene_gn=THG1L;refGene_tx=XM_019839760.2;refGene_gn=THG1L;refGene_tx=XM_019839771.2;refGene_gn=THG1L;refGene_tx=XM_019839775.2;refGene_gn=THG1L;refGene_tx=XM_019839776.2
A1	192529119	192529296
refGene_gn=THG1L;refGene_tx=XM_003981308.5;refGene_gn=THG1L;refGene_tx=XM_019839760.2;refGene_gn=THG1L;refGene_tx=XM_019839771.2;refGene_gn=THG1L;refGene_tx=XM_019839775.2
A1	192530018	192530299
refGene_gn=THG1L;refGene_tx=XM_003981308.5;refGene_gn=THG1L;refGene_tx=XM_019839760.2
A1	192556185	192557197
refGene_gn=SOX30;refGene_tx=XM_011285779.3;refGene_gn=SOX30;refGene_tx=XM_023260132.1
A1	192558796	192559036
refGene_gn=SOX30;refGene_tx=XM_011285779.3;refGene_gn=SOX30;refGene_tx=XM_023260132.1
A1	192560226	192560406
refGene_gn=SOX30;refGene_tx=XM_011285779.3;refGene_gn=SOX30;refGene_tx=XM_023260132.1
A1	192566941	192567434
refGene_gn=SOX30;refGene_tx=XM_011285779.3
A1	192592097	192592479
refGene_gn=SOX30;refGene_tx=XM_011285779.3;refGene_gn=SOX30;refGene_tx=XM_023260132.1
A1	192632210	192632277
refGene_gn=ADAM19;refGene_tx=XM_011286766.2
A1	192635276	192635362
refGene_gn=ADAM19;refGene_tx=XM_011286766.2
A1	192639654	192639725
refGene_gn=ADAM19;refGene_tx=XM_011286766.2
A1	192664250	192664329
refGene_gn=ADAM19;refGene_tx=XM_011286766.2
A1	192670219	192670296
refGene_gn=ADAM19;refGene_tx=XM_011286766.2
A1	192679255	192679448
refGene_gn=ADAM19;refGene_tx=XM_011286766.2
A1	192680209	192680275
refGene_gn=ADAM19;refGene_tx=XM_011286766.2
A1	192684794	192684866

refGene_gn=ADAM19;refGene_tx=XM_011286766.2
A1 192688401 192688568
refGene_gn=ADAM19;refGene_tx=XM_011286766.2
A1 192690951 192691036
refGene_gn=ADAM19;refGene_tx=XM_011286766.2
A1 192692162 192692302
refGene_gn=ADAM19;refGene_tx=XM_011286766.2
A1 192694056 192694234
refGene_gn=ADAM19;refGene_tx=XM_011286766.2
A1 192696696 192696786
refGene_gn=ADAM19;refGene_tx=XM_011286766.2
A1 192698937 192699133
refGene_gn=ADAM19;refGene_tx=XM_011286766.2
A1 192700325 192700434
refGene_gn=ADAM19;refGene_tx=XM_011286766.2
A1 192701745 192701950
refGene_gn=ADAM19;refGene_tx=XM_011286766.2
A1 192702689 192702767
refGene_gn=ADAM19;refGene_tx=XM_011286766.2
A1 192702902 192703011
refGene_gn=ADAM19;refGene_tx=XM_011286766.2
A1 192704077 192704222
refGene_gn=ADAM19;refGene_tx=XM_011286766.2
A1 192705052 192705137
refGene_gn=ADAM19;refGene_tx=XM_011286766.2
A1 192705458 192705683
refGene_gn=ADAM19;refGene_tx=XM_011286766.2
A1 192708181 192708334
refGene_gn=ADAM19;refGene_tx=XM_011286766.2
A1 192709102 192709156
refGene_gn=ADAM19;refGene_tx=XM_011286766.2
A1 192716510 192717139
refGene_gn=NIPAL4;refGene_tx=XM_023260156.1
A1 192718164 192718325
refGene_gn=NIPAL4;refGene_tx=XM_023260156.1
A1 192720913 192721004
refGene_gn=NIPAL4;refGene_tx=XM_023260156.1
A1 192722170 192722227
refGene_gn=NIPAL4;refGene_tx=XM_023260156.1
A1 192725415 192725655
refGene_gn=NIPAL4;refGene_tx=XM_023260156.1
A1 192729022 192729059
refGene_gn=NIPAL4;refGene_tx=XM_023260156.1
A1 192754829 192754997
refGene_gn=CYFIP2;refGene_tx=XM_019839827.2;refGene_gn=CYFIP2;refGene_tx=XM_023260157.1;refGene_gn=CYFIP2;refGene_tx=XM_023260160.1;refGene_gn=CYFIP2;refGene_tx=XM_023260164.1;refGene_gn=CYFIP2;refGene_tx=XM_023260171.1
A1 192758139 192758287
refGene_gn=CYFIP2;refGene_tx=XM_019839827.2;refGene_gn=CYFIP2;refGene_

tx=XM_023260157.1;refGene_gn=CYFIP2;refGene_tx=XM_023260160.1;refGene_gn=CYFIP2;refGene_tx=XM_023260164.1;refGene_gn=CYFIP2;refGene_tx=XM_023260171.1
A1 192759830 192760069
refGene_gn=CYFIP2;refGene_tx=XM_019839827.2;refGene_gn=CYFIP2;refGene_tx=XM_023260157.1;refGene_gn=CYFIP2;refGene_tx=XM_023260160.1;refGene_gn=CYFIP2;refGene_tx=XM_023260164.1;refGene_gn=CYFIP2;refGene_tx=XM_023260171.1
A1 192765723 192765818
refGene_gn=CYFIP2;refGene_tx=XM_019839827.2;refGene_gn=CYFIP2;refGene_tx=XM_023260157.1;refGene_gn=CYFIP2;refGene_tx=XM_023260160.1;refGene_gn=CYFIP2;refGene_tx=XM_023260164.1;refGene_gn=CYFIP2;refGene_tx=XM_023260171.1
A1 192766282 192766355
refGene_gn=CYFIP2;refGene_tx=XM_019839827.2;refGene_gn=CYFIP2;refGene_tx=XM_023260157.1;refGene_gn=CYFIP2;refGene_tx=XM_023260160.1;refGene_gn=CYFIP2;refGene_tx=XM_023260164.1;refGene_gn=CYFIP2;refGene_tx=XM_023260171.1
A1 192774784 192774808
refGene_gn=CYFIP2;refGene_tx=XM_023260174.1
A1 192788310 192788441
refGene_gn=CYFIP2;refGene_tx=XM_019839827.2;refGene_gn=CYFIP2;refGene_tx=XM_023260157.1;refGene_gn=CYFIP2;refGene_tx=XM_023260160.1;refGene_gn=CYFIP2;refGene_tx=XM_023260164.1;refGene_gn=CYFIP2;refGene_tx=XM_023260171.1;refGene_gn=CYFIP2;refGene_tx=XM_023260174.1
A1 192789854 192789945
refGene_gn=CYFIP2;refGene_tx=XM_019839827.2;refGene_gn=CYFIP2;refGene_tx=XM_023260157.1;refGene_gn=CYFIP2;refGene_tx=XM_023260160.1;refGene_gn=CYFIP2;refGene_tx=XM_023260164.1;refGene_gn=CYFIP2;refGene_tx=XM_023260171.1;refGene_gn=CYFIP2;refGene_tx=XM_023260174.1
A1 192791410 192791554
refGene_gn=CYFIP2;refGene_tx=XM_019839827.2;refGene_gn=CYFIP2;refGene_tx=XM_023260157.1;refGene_gn=CYFIP2;refGene_tx=XM_023260160.1;refGene_gn=CYFIP2;refGene_tx=XM_023260164.1;refGene_gn=CYFIP2;refGene_tx=XM_023260171.1;refGene_gn=CYFIP2;refGene_tx=XM_023260174.1
A1 192804123 192804837
refGene_gn=FNDC9;refGene_tx=XM_006927960.4
A1 192806854 192806942
refGene_gn=CYFIP2;refGene_tx=XM_019839827.2;refGene_gn=CYFIP2;refGene_tx=XM_023260157.1;refGene_gn=CYFIP2;refGene_tx=XM_023260160.1;refGene_gn=CYFIP2;refGene_tx=XM_023260164.1;refGene_gn=CYFIP2;refGene_tx=XM_023260171.1;refGene_gn=CYFIP2;refGene_tx=XM_023260174.1
A1 192808766 192808966
refGene_gn=CYFIP2;refGene_tx=XM_019839827.2;refGene_gn=CYFIP2;refGene_tx=XM_023260157.1;refGene_gn=CYFIP2;refGene_tx=XM_023260160.1;refGene_gn=CYFIP2;refGene_tx=XM_023260164.1;refGene_gn=CYFIP2;refGene_tx=XM_023260171.1;refGene_gn=CYFIP2;refGene_tx=XM_023260174.1
A1 192815227 192815347
refGene_gn=CYFIP2;refGene_tx=XM_019839827.2;refGene_gn=CYFIP2;refGene_tx=XM_023260157.1;refGene_gn=CYFIP2;refGene_tx=XM_023260160.1;refGene_

gn=CYFIP2;refGene_tx=XM_023260164.1;refGene_gn=CYFIP2;refGene_tx=XM_02
3260171.1;refGene_gn=CYFIP2;refGene_tx=XM_023260174.1
A1 192817483 192817592
refGene_gn=CYFIP2;refGene_tx=XM_019839827.2;refGene_gn=CYFIP2;refGene_
tx=XM_023260157.1;refGene_gn=CYFIP2;refGene_tx=XM_023260160.1;refGene_
gn=CYFIP2;refGene_tx=XM_023260164.1;refGene_gn=CYFIP2;refGene_tx=XM_02
3260171.1;refGene_gn=CYFIP2;refGene_tx=XM_023260174.1
A1 192820726 192820803
refGene_gn=CYFIP2;refGene_tx=XM_019839827.2;refGene_gn=CYFIP2;refGene_
tx=XM_023260157.1;refGene_gn=CYFIP2;refGene_tx=XM_023260160.1;refGene_
gn=CYFIP2;refGene_tx=XM_023260164.1;refGene_gn=CYFIP2;refGene_tx=XM_02
3260171.1;refGene_gn=CYFIP2;refGene_tx=XM_023260174.1
A1 192824127 192824224
refGene_gn=CYFIP2;refGene_tx=XM_019839827.2;refGene_gn=CYFIP2;refGene_
tx=XM_023260157.1;refGene_gn=CYFIP2;refGene_tx=XM_023260160.1;refGene_
gn=CYFIP2;refGene_tx=XM_023260164.1;refGene_gn=CYFIP2;refGene_tx=XM_02
3260171.1;refGene_gn=CYFIP2;refGene_tx=XM_023260174.1
A1 192824761 192824918
refGene_gn=CYFIP2;refGene_tx=XM_019839827.2;refGene_gn=CYFIP2;refGene_
tx=XM_023260157.1;refGene_gn=CYFIP2;refGene_tx=XM_023260160.1;refGene_
gn=CYFIP2;refGene_tx=XM_023260164.1;refGene_gn=CYFIP2;refGene_tx=XM_02
3260171.1;refGene_gn=CYFIP2;refGene_tx=XM_023260174.1
A1 192825869 192826023
refGene_gn=CYFIP2;refGene_tx=XM_019839827.2;refGene_gn=CYFIP2;refGene_
tx=XM_023260157.1;refGene_gn=CYFIP2;refGene_tx=XM_023260160.1;refGene_
gn=CYFIP2;refGene_tx=XM_023260164.1;refGene_gn=CYFIP2;refGene_tx=XM_02
3260171.1;refGene_gn=CYFIP2;refGene_tx=XM_023260174.1
A1 192826951 192827026
refGene_gn=CYFIP2;refGene_tx=XM_019839827.2;refGene_gn=CYFIP2;refGene_
tx=XM_023260157.1;refGene_gn=CYFIP2;refGene_tx=XM_023260164.1;refGene_
gn=CYFIP2;refGene_tx=XM_023260171.1;refGene_gn=CYFIP2;refGene_tx=XM_02
3260174.1
A1 192829183 192829331
refGene_gn=CYFIP2;refGene_tx=XM_019839827.2;refGene_gn=CYFIP2;refGene_
tx=XM_023260157.1;refGene_gn=CYFIP2;refGene_tx=XM_023260160.1;refGene_
gn=CYFIP2;refGene_tx=XM_023260164.1;refGene_gn=CYFIP2;refGene_tx=XM_02
3260171.1;refGene_gn=CYFIP2;refGene_tx=XM_023260174.1
A1 192830002 192830169
refGene_gn=CYFIP2;refGene_tx=XM_019839827.2;refGene_gn=CYFIP2;refGene_
tx=XM_023260157.1;refGene_gn=CYFIP2;refGene_tx=XM_023260160.1;refGene_
gn=CYFIP2;refGene_tx=XM_023260164.1;refGene_gn=CYFIP2;refGene_tx=XM_02
3260171.1;refGene_gn=CYFIP2;refGene_tx=XM_023260174.1
A1 192833789 192833915
refGene_gn=CYFIP2;refGene_tx=XM_019839827.2;refGene_gn=CYFIP2;refGene_
tx=XM_023260157.1;refGene_gn=CYFIP2;refGene_tx=XM_023260160.1;refGene_
gn=CYFIP2;refGene_tx=XM_023260164.1;refGene_gn=CYFIP2;refGene_tx=XM_02
3260171.1;refGene_gn=CYFIP2;refGene_tx=XM_023260174.1
A1 192834458 192834578
refGene_gn=CYFIP2;refGene_tx=XM_019839827.2;refGene_gn=CYFIP2;refGene_
tx=XM_023260157.1;refGene_gn=CYFIP2;refGene_tx=XM_023260160.1;refGene_

gn=CYFIP2;refGene_tx=XM_023260164.1;refGene_gn=CYFIP2;refGene_tx=XM_02
3260171.1;refGene_gn=CYFIP2;refGene_tx=XM_023260174.1
A1 192836891 192837009
refGene_gn=CYFIP2;refGene_tx=XM_019839827.2;refGene_gn=CYFIP2;refGene_
tx=XM_023260157.1;refGene_gn=CYFIP2;refGene_tx=XM_023260160.1;refGene_
gn=CYFIP2;refGene_tx=XM_023260164.1;refGene_gn=CYFIP2;refGene_tx=XM_02
3260171.1;refGene_gn=CYFIP2;refGene_tx=XM_023260174.1
A1 192838820 192838912
refGene_gn=CYFIP2;refGene_tx=XM_019839827.2;refGene_gn=CYFIP2;refGene_
tx=XM_023260157.1;refGene_gn=CYFIP2;refGene_tx=XM_023260160.1;refGene_
gn=CYFIP2;refGene_tx=XM_023260164.1;refGene_gn=CYFIP2;refGene_tx=XM_02
3260171.1;refGene_gn=CYFIP2;refGene_tx=XM_023260174.1
A1 192841044 192841149
refGene_gn=CYFIP2;refGene_tx=XM_019839827.2;refGene_gn=CYFIP2;refGene_
tx=XM_023260157.1;refGene_gn=CYFIP2;refGene_tx=XM_023260160.1;refGene_
gn=CYFIP2;refGene_tx=XM_023260164.1;refGene_gn=CYFIP2;refGene_tx=XM_02
3260171.1;refGene_gn=CYFIP2;refGene_tx=XM_023260174.1
A1 192843582 192843711
refGene_gn=CYFIP2;refGene_tx=XM_019839827.2;refGene_gn=CYFIP2;refGene_
tx=XM_023260157.1;refGene_gn=CYFIP2;refGene_tx=XM_023260160.1;refGene_
gn=CYFIP2;refGene_tx=XM_023260164.1;refGene_gn=CYFIP2;refGene_tx=XM_02
3260171.1;refGene_gn=CYFIP2;refGene_tx=XM_023260174.1
A1 192845031 192845128
refGene_gn=CYFIP2;refGene_tx=XM_019839827.2;refGene_gn=CYFIP2;refGene_
tx=XM_023260157.1;refGene_gn=CYFIP2;refGene_tx=XM_023260160.1;refGene_
gn=CYFIP2;refGene_tx=XM_023260164.1;refGene_gn=CYFIP2;refGene_tx=XM_02
3260171.1;refGene_gn=CYFIP2;refGene_tx=XM_023260174.1
A1 192846935 192847117
refGene_gn=CYFIP2;refGene_tx=XM_019839827.2;refGene_gn=CYFIP2;refGene_
tx=XM_023260157.1;refGene_gn=CYFIP2;refGene_tx=XM_023260160.1;refGene_
gn=CYFIP2;refGene_tx=XM_023260164.1;refGene_gn=CYFIP2;refGene_tx=XM_02
3260171.1;refGene_gn=CYFIP2;refGene_tx=XM_023260174.1
A1 192850495 192850597
refGene_gn=CYFIP2;refGene_tx=XM_019839827.2;refGene_gn=CYFIP2;refGene_
tx=XM_023260157.1;refGene_gn=CYFIP2;refGene_tx=XM_023260160.1;refGene_
gn=CYFIP2;refGene_tx=XM_023260164.1;refGene_gn=CYFIP2;refGene_tx=XM_02
3260171.1;refGene_gn=CYFIP2;refGene_tx=XM_023260174.1
A1 192852288 192852366
refGene_gn=CYFIP2;refGene_tx=XM_019839827.2;refGene_gn=CYFIP2;refGene_
tx=XM_023260157.1;refGene_gn=CYFIP2;refGene_tx=XM_023260160.1;refGene_
gn=CYFIP2;refGene_tx=XM_023260164.1;refGene_gn=CYFIP2;refGene_tx=XM_02
3260171.1;refGene_gn=CYFIP2;refGene_tx=XM_023260174.1
A1 192857627 192857717
refGene_gn=CYFIP2;refGene_tx=XM_019839827.2;refGene_gn=CYFIP2;refGene_
tx=XM_023260157.1;refGene_gn=CYFIP2;refGene_tx=XM_023260160.1;refGene_
gn=CYFIP2;refGene_tx=XM_023260164.1;refGene_gn=CYFIP2;refGene_tx=XM_02
3260171.1;refGene_gn=CYFIP2;refGene_tx=XM_023260174.1
A1 192859221 192859361
refGene_gn=CYFIP2;refGene_tx=XM_019839827.2;refGene_gn=CYFIP2;refGene_
tx=XM_023260157.1;refGene_gn=CYFIP2;refGene_tx=XM_023260160.1;refGene_

gn=CYFIP2;refGene_tx=XM_023260164.1;refGene_gn=CYFIP2;refGene_tx=XM_023260171.1;refGene_gn=CYFIP2;refGene_tx=XM_023260174.1
A1 192865979 192866046
refGene_gn=CYFIP2;refGene_tx=XM_023260157.1;refGene_gn=CYFIP2;refGene_tx=XM_023260160.1;refGene_gn=CYFIP2;refGene_tx=XM_023260174.1
A1 192891290 192891362
refGene_gn=ITK;refGene_tx=XM_023260178.1
A1 192894027 192894185
refGene_gn=ITK;refGene_tx=XM_023260178.1
A1 192897641 192897760
refGene_gn=ITK;refGene_tx=XM_023260178.1
A1 192897850 192897915
refGene_gn=ITK;refGene_tx=XM_023260178.1
A1 192898831 192899048
refGene_gn=ITK;refGene_tx=XM_023260178.1
A1 192899428 192899600
refGene_gn=ITK;refGene_tx=XM_023260178.1
A1 192900175 192900250
refGene_gn=ITK;refGene_tx=XM_023260178.1
A1 192900665 192900799
refGene_gn=ITK;refGene_tx=XM_023260178.1
A1 192901635 192901718
refGene_gn=ITK;refGene_tx=XM_023260178.1
A1 192909209 192909264
refGene_gn=ITK;refGene_tx=XM_023260178.1
A1 192913537 192913603
refGene_gn=ITK;refGene_tx=XM_023260178.1
A1 192917873 192918025
refGene_gn=ITK;refGene_tx=XM_023260178.1
A1 192920331 192920372
refGene_gn=ITK;refGene_tx=XM_023260178.1
A1 192923629 192923758
refGene_gn=ITK;refGene_tx=XM_023260178.1
A1 192925555 192925637
refGene_gn=ITK;refGene_tx=XM_023260178.1
A1 192927581 192927686
refGene_gn=ITK;refGene_tx=XM_023260178.1
A1 192947177 192947315
refGene_gn=ITK;refGene_tx=XM_023260178.1
A1 192959942 192960563
refGene_gn=FAM71B;refGene_tx=XM_003981348.5
A1 192961354 192962539
refGene_gn=FAM71B;refGene_tx=XM_003981348.5
A1 192983066 192983768
refGene_gn=MED7;refGene_tx=XM_003981300.4;refGene_gn=MED7;refGene_tx=XM_019839853.2
A1 193007115 193007170
refGene_gn=HAVCR2;refGene_tx=XM_003981347.3
A1 193008423 193008762
refGene_gn=HAVCR2;refGene_tx=XM_003981347.3

```
A1	193009665	193009749
refGene_gn=HAVCR2;refGene_tx=XM_003981347.3
A1	193013460	193013501
refGene_gn=HAVCR2;refGene_tx=XM_003981347.3
A1	193015217	193015344
refGene_gn=HAVCR2;refGene_tx=XM_003981347.3
A1	193018907	193018944
refGene_gn=HAVCR2;refGene_tx=XM_003981347.3
A1	193020810	193020970
refGene_gn=HAVCR2;refGene_tx=XM_003981347.3
A1	193037229	193037275
refGene_gn=HAVCR1;refGene_tx=NM_001305024.1
A1	193040167	193040506
refGene_gn=HAVCR1;refGene_tx=NM_001305024.1
A1	193043463	193043547
refGene_gn=HAVCR1;refGene_tx=NM_001305024.1
A1	193046593	193046698
refGene_gn=HAVCR1;refGene_tx=NM_001305024.1
A1	193049503	193049559
refGene_gn=HAVCR1;refGene_tx=NM_001305024.1
A1	193054714	193054749
refGene_gn=HAVCR1;refGene_tx=NM_001305024.1
A1	193056967	193056972
refGene_gn=HAVCR1;refGene_tx=NM_001305024.1
A1	193056974	193057081
refGene_gn=HAVCR1;refGene_tx=NM_001305024.1
A1	193094256	193094314
refGene_gn=TIMD4;refGene_tx=XM_006927929.4;refGene_gn=TIMD4;refGene_tx
=XM_006927930.4;refGene_gn=TIMD4;refGene_tx=XM_019839877.2
A1	193098516	193098855
refGene_gn=TIMD4;refGene_tx=XM_006927929.4;refGene_gn=TIMD4;refGene_tx
=XM_006927930.4;refGene_gn=TIMD4;refGene_tx=XM_019839877.2;refGene_gn=
TIMD4;refGene_tx=XM_019839889.2;refGene_gn=TIMD4;refGene_tx=XM_0232601
91.1
A1	193101415	193101643
refGene_gn=TIMD4;refGene_tx=XM_006927929.4;refGene_gn=TIMD4;refGene_tx
=XM_006927930.4;refGene_gn=TIMD4;refGene_tx=XM_019839877.2;refGene_gn=
TIMD4;refGene_tx=XM_019839889.2;refGene_gn=TIMD4;refGene_tx=XM_0232601
91.1
A1	193103027	193103111
refGene_gn=TIMD4;refGene_tx=XM_006927929.4;refGene_gn=TIMD4;refGene_tx
=XM_006927930.4;refGene_gn=TIMD4;refGene_tx=XM_019839877.2;refGene_gn=
TIMD4;refGene_tx=XM_019839889.2;refGene_gn=TIMD4;refGene_tx=XM_0232601
91.1
A1	193104450	193104534
refGene_gn=TIMD4;refGene_tx=XM_006927929.4;refGene_gn=TIMD4;refGene_tx
=XM_006927930.4;refGene_gn=TIMD4;refGene_tx=XM_019839877.2;refGene_gn=
TIMD4;refGene_tx=XM_019839889.2;refGene_gn=TIMD4;refGene_tx=XM_0232601
91.1
A1	193114464	193114538
```

refGene_gn=TIMD4;refGene_tx=XM_006927930.4
A1 193121399 193121482
refGene_gn=TIMD4;refGene_tx=XM_006927929.4;refGene_gn=TIMD4;refGene_tx=XM_019839877.2;refGene_gn=TIMD4;refGene_tx=XM_019839889.2;refGene_gn=TIMD4;refGene_tx=XM_023260191.1
A1 193122952 193123046
refGene_gn=TIMD4;refGene_tx=XM_006927929.4;refGene_gn=TIMD4;refGene_tx=XM_019839877.2;refGene_gn=TIMD4;refGene_tx=XM_019839889.2;refGene_gn=TIMD4;refGene_tx=XM_023260191.1
A1 193124323 193124363
refGene_gn=TIMD4;refGene_tx=XM_006927929.4;refGene_gn=TIMD4;refGene_tx=XM_019839877.2;refGene_gn=TIMD4;refGene_tx=XM_019839889.2;refGene_gn=TIMD4;refGene_tx=XM_023260191.1
A1 193125271 193125356
refGene_gn=TIMD4;refGene_tx=XM_006927929.4;refGene_gn=TIMD4;refGene_tx=XM_019839877.2;refGene_gn=TIMD4;refGene_tx=XM_019839889.2;refGene_gn=TIMD4;refGene_tx=XM_023260191.1
A1 193285757 193285931
refGene_gn=SGCD;refGene_tx=XM_011285858.3;refGene_gn=SGCD;refGene_tx=XM_019839908.2;refGene_gn=SGCD;refGene_tx=XM_019839916.2;refGene_gn=SGCD;refGene_tx=XM_019839924.2
A1 193287277 193287401
refGene_gn=SGCD;refGene_tx=XM_011285858.3;refGene_gn=SGCD;refGene_tx=XM_019839908.2;refGene_gn=SGCD;refGene_tx=XM_019839916.2;refGene_gn=SGCD;refGene_tx=XM_019839924.2
A1 193385379 193385452
refGene_gn=SGCD;refGene_tx=XM_011285858.3;refGene_gn=SGCD;refGene_tx=XM_019839908.2;refGene_gn=SGCD;refGene_tx=XM_019839916.2;refGene_gn=SGCD;refGene_tx=XM_019839924.2
A1 193421485 193421605
refGene_gn=SGCD;refGene_tx=XM_011285858.3;refGene_gn=SGCD;refGene_tx=XM_019839908.2;refGene_gn=SGCD;refGene_tx=XM_019839916.2
A1 193427324 193427412
refGene_gn=SGCD;refGene_tx=XM_011285858.3;refGene_gn=SGCD;refGene_tx=XM_019839908.2;refGene_gn=SGCD;refGene_tx=XM_019839916.2;refGene_gn=SGCD;refGene_tx=XM_019839924.2
A1 193494748 193494850
refGene_gn=SGCD;refGene_tx=XM_011285858.3;refGene_gn=SGCD;refGene_tx=XM_019839908.2;refGene_gn=SGCD;refGene_tx=XM_019839916.2;refGene_gn=SGCD;refGene_tx=XM_019839924.2
A1 193645233 193645422
refGene_gn=SGCD;refGene_tx=XM_011285858.3;refGene_gn=SGCD;refGene_tx=XM_019839908.2;refGene_gn=SGCD;refGene_tx=XM_019839916.2;refGene_gn=SGCD;refGene_tx=XM_019839924.2
A1 194883212 194883424
refGene_gn=MRPL22;refGene_tx=XM_003981358.5;refGene_gn=MRPL22;refGene_tx=XM_006927964.4;refGene_gn=MRPL22;refGene_tx=XM_006927965.2
A1 194889406 194889476
refGene_gn=MRPL22;refGene_tx=XM_003981358.5;refGene_gn=MRPL22;refGene_tx=XM_006927964.4;refGene_gn=MRPL22;refGene_tx=XM_006927965.2

A1 194896151 194896229
refGene_gn=MRPL22;refGene_tx=XM_003981358.5;refGene_gn=MRPL22;refGene_tx=XM_006927964.4;refGene_gn=MRPL22;refGene_tx=XM_006927965.2
A1 194897467 194897533
refGene_gn=MRPL22;refGene_tx=XM_003981358.5;refGene_gn=MRPL22;refGene_tx=XM_006927964.4;refGene_gn=MRPL22;refGene_tx=XM_006927965.2
A1 194902938 194903074
refGene_gn=MRPL22;refGene_tx=XM_003981358.5;refGene_gn=MRPL22;refGene_tx=XM_006927964.4
A1 194917895 194917938
refGene_gn=MRPL22;refGene_tx=XM_003981358.5;refGene_gn=MRPL22;refGene_tx=XM_006927964.4
A1 194918017 194918045
refGene_gn=MRPL22;refGene_tx=XM_003981358.5;refGene_gn=MRPL22;refGene_tx=XM_006927964.4
A1 194919502 194919668
refGene_gn=GEMIN5;refGene_tx=XM_003981475.5
A1 194920477 194920638
refGene_gn=GEMIN5;refGene_tx=XM_003981475.5
A1 194921319 194921501
refGene_gn=GEMIN5;refGene_tx=XM_003981475.5
A1 194924315 194924467
refGene_gn=GEMIN5;refGene_tx=XM_003981475.5
A1 194924661 194924781
refGene_gn=GEMIN5;refGene_tx=XM_003981475.5
A1 194925881 194926014
refGene_gn=GEMIN5;refGene_tx=XM_003981475.5
A1 194926916 194927082
refGene_gn=GEMIN5;refGene_tx=XM_003981475.5
A1 194928037 194928250
refGene_gn=GEMIN5;refGene_tx=XM_003981475.5
A1 194929160 194929246
refGene_gn=GEMIN5;refGene_tx=XM_003981475.5
A1 194931828 194931911
refGene_gn=GEMIN5;refGene_tx=XM_003981475.5
A1 194933358 194933495
refGene_gn=GEMIN5;refGene_tx=XM_003981475.5
A1 194935060 194935134
refGene_gn=GEMIN5;refGene_tx=XM_003981475.5
A1 194935814 194935996
refGene_gn=GEMIN5;refGene_tx=XM_003981475.5
A1 194939100 194939240
refGene_gn=GEMIN5;refGene_tx=XM_003981475.5
A1 194940123 194940295
refGene_gn=GEMIN5;refGene_tx=XM_003981475.5
A1 194942825 194943053
refGene_gn=GEMIN5;refGene_tx=XM_003981475.5
A1 194944080 194944194
refGene_gn=GEMIN5;refGene_tx=XM_003981475.5
A1 194944654 194944777

refGene_gn=GEMIN5;refGene_tx=XM_003981475.5
A1 194946072 194946168
refGene_gn=GEMIN5;refGene_tx=XM_003981475.5
A1 194946579 194946717
refGene_gn=GEMIN5;refGene_tx=XM_003981475.5
A1 194947718 194947866
refGene_gn=GEMIN5;refGene_tx=XM_003981475.5
A1 194950990 194951110
refGene_gn=GEMIN5;refGene_tx=XM_003981475.5
A1 194951537 194951748
refGene_gn=GEMIN5;refGene_tx=XM_003981475.5
A1 194953257 194953506
refGene_gn=GEMIN5;refGene_tx=XM_003981475.5
A1 194955038 194955201
refGene_gn=GEMIN5;refGene_tx=XM_003981475.5
A1 194955974 194956479
refGene_gn=GEMIN5;refGene_tx=XM_003981475.5
A1 194957990 194958087
refGene_gn=GEMIN5;refGene_tx=XM_003981475.5
A1 194958666 194958834
refGene_gn=GEMIN5;refGene_tx=XM_003981475.5
A1 194963113 194963263
refGene_gn=CNOT8;refGene_tx=XM_011285873.3;refGene_gn=CNOT8;refGene_tx=XM_011285880.3;refGene_gn=CNOT8;refGene_tx=XM_011285882.3;refGene_gn=CNOT8;refGene_tx=XM_019839959.2
A1 194964938 194965049
refGene_gn=CNOT8;refGene_tx=XM_011285873.3;refGene_gn=CNOT8;refGene_tx=XM_011285880.3;refGene_gn=CNOT8;refGene_tx=XM_011285882.3;refGene_gn=CNOT8;refGene_tx=XM_019839959.2
A1 194966011 194966156
refGene_gn=CNOT8;refGene_tx=XM_011285873.3;refGene_gn=CNOT8;refGene_tx=XM_011285880.3;refGene_gn=CNOT8;refGene_tx=XM_011285882.3;refGene_gn=CNOT8;refGene_tx=XM_019839959.2
A1 194966627 194966641
refGene_gn=CNOT8;refGene_tx=XM_011285882.3
A1 194966752 194966914
refGene_gn=CNOT8;refGene_tx=XM_011285873.3;refGene_gn=CNOT8;refGene_tx=XM_011285880.3;refGene_gn=CNOT8;refGene_tx=XM_019839959.2
A1 194969588 194969670
refGene_gn=CNOT8;refGene_tx=XM_019839959.2
A1 194969776 194969970
refGene_gn=CNOT8;refGene_tx=XM_011285873.3;refGene_gn=CNOT8;refGene_tx=XM_011285880.3;refGene_gn=CNOT8;refGene_tx=XM_019839959.2
A1 194971352 194971469
refGene_gn=CNOT8;refGene_tx=XM_011285873.3;refGene_gn=CNOT8;refGene_tx=XM_011285880.3
A1 194996833 194996878
refGene_gn=FAXDC2;refGene_tx=XM_011285891.2
A1 194999741 194999833
refGene_gn=FAXDC2;refGene_tx=XM_011285891.2

A1	194999948	195000052
refGene_gn=FAXDC2;refGene_tx=XM_011285891.2
A1	195002904	195003026
refGene_gn=FAXDC2;refGene_tx=XM_011285891.2
A1	195007532	195007738
refGene_gn=FAXDC2;refGene_tx=XM_011285891.2
A1	195008221	195008327
refGene_gn=FAXDC2;refGene_tx=XM_011285891.2
A1	195008717	195008884
refGene_gn=FAXDC2;refGene_tx=XM_011285891.2
A1	195009820	195009977
refGene_gn=FAXDC2;refGene_tx=XM_011285891.2
A1	195017043	195017253
refGene_gn=LARP1;refGene_tx=XM_023260226.1
A1	195018771	195018899
refGene_gn=LARP1;refGene_tx=XM_023260226.1
A1	195019094	195019204
refGene_gn=LARP1;refGene_tx=XM_023260226.1
A1	195020737	195020882
refGene_gn=LARP1;refGene_tx=XM_023260226.1
A1	195024287	195024439
refGene_gn=LARP1;refGene_tx=XM_023260226.1
A1	195025327	195025434
refGene_gn=LARP1;refGene_tx=XM_023260226.1
A1	195025887	195026093
refGene_gn=LARP1;refGene_tx=XM_023260226.1
A1	195026216	195026340
refGene_gn=LARP1;refGene_tx=XM_023260226.1
A1	195027154	195027547
refGene_gn=LARP1;refGene_tx=XM_023260226.1
A1	195029420	195029590
refGene_gn=LARP1;refGene_tx=XM_023260226.1
A1	195029694	195029869
refGene_gn=LARP1;refGene_tx=XM_023260226.1
A1	195032603	195032748
refGene_gn=LARP1;refGene_tx=XM_023260226.1
A1	195033650	195033813
refGene_gn=LARP1;refGene_tx=XM_023260226.1
A1	195033912	195034119
refGene_gn=LARP1;refGene_tx=XM_023260226.1
A1	195034195	195034324
refGene_gn=LARP1;refGene_tx=XM_023260226.1
A1	195035249	195035424
refGene_gn=LARP1;refGene_tx=XM_023260226.1
A1	195036373	195036439
refGene_gn=LARP1;refGene_tx=XM_023260226.1
A1	195036689	195036751
refGene_gn=LARP1;refGene_tx=XM_023260226.1
A1	195069108	195069550
refGene_gn=LARP1;refGene_tx=XM_023260226.1

A1 195246334 195246886
refGene_gn=HAND1;refGene_tx=XM_003981360.3;refGene_gn=HAND1;refGene_tx
=XM_019839968.1
A1 195248407 195248512
refGene_gn=HAND1;refGene_tx=XM_003981360.3;refGene_gn=HAND1;refGene_tx
=XM_019839968.1
A1 195268624 195268753
refGene_gn=SAP30L;refGene_tx=XM_003981361.4
A1 195270534 195270633
refGene_gn=SAP30L;refGene_tx=XM_003981361.4
A1 195272880 195273003
refGene_gn=SAP30L;refGene_tx=XM_003981361.4
A1 195278204 195278402
refGene_gn=SAP30L;refGene_tx=XM_003981361.4
A1 195295609 195295768
refGene_gn=GALNT10;refGene_tx=XM_023260250.1;refGene_gn=GALNT10;refGen
e_tx=XM_023260253.1
A1 195296296 195296446
refGene_gn=GALNT10;refGene_tx=XM_023260250.1;refGene_gn=GALNT10;refGen
e_tx=XM_023260253.1
A1 195299583 195299700
refGene_gn=GALNT10;refGene_tx=XM_023260250.1;refGene_gn=GALNT10;refGen
e_tx=XM_023260253.1
A1 195303500 195303722
refGene_gn=GALNT10;refGene_tx=XM_023260250.1;refGene_gn=GALNT10;refGen
e_tx=XM_023260253.1
A1 195307926 195308034
refGene_gn=GALNT10;refGene_tx=XM_023260250.1;refGene_gn=GALNT10;refGen
e_tx=XM_023260253.1
A1 195322907 195323025
refGene_gn=GALNT10;refGene_tx=XM_023260250.1;refGene_gn=GALNT10;refGen
e_tx=XM_023260253.1
A1 195328592 195328776
refGene_gn=GALNT10;refGene_tx=XM_023260250.1;refGene_gn=GALNT10;refGen
e_tx=XM_023260253.1
A1 195332619 195332805
refGene_gn=GALNT10;refGene_tx=XM_023260250.1;refGene_gn=GALNT10;refGen
e_tx=XM_023260253.1
A1 195393205 195393372
refGene_gn=GALNT10;refGene_tx=XM_023260250.1;refGene_gn=GALNT10;refGen
e_tx=XM_023260253.1
A1 195420230 195420369
refGene_gn=GALNT10;refGene_tx=XM_023260250.1;refGene_gn=GALNT10;refGen
e_tx=XM_023260253.1
A1 195422830 195422933
refGene_gn=GALNT10;refGene_tx=XM_023260250.1;refGene_gn=GALNT10;refGen
e_tx=XM_023260253.1
A1 195440753 195440798
refGene_gn=GALNT10;refGene_tx=XM_023260253.1
A1 195514715 195514874

refGene_gn=GALNT10;refGene_tx=XM_023260250.1
A1	195647423	195648217
refGene_gn=MFAP3;refGene_tx=XM_003981364.4;refGene_gn=MFAP3;refGene_tx=XM_006927966.3;refGene_gn=MFAP3;refGene_tx=XM_006927967.4
A1	195650468	195650763
refGene_gn=MFAP3;refGene_tx=XM_003981364.4;refGene_gn=MFAP3;refGene_tx=XM_006927966.3;refGene_gn=MFAP3;refGene_tx=XM_006927967.4
A1	195666700	195666931
refGene_gn=FAM114A2;refGene_tx=XM_011285908.2;refGene_gn=FAM114A2;refGene_tx=XM_011285913.3;refGene_gn=FAM114A2;refGene_tx=XM_019840021.2
A1	195667948	195668048
refGene_gn=FAM114A2;refGene_tx=XM_011285908.2;refGene_gn=FAM114A2;refGene_tx=XM_011285913.3;refGene_gn=FAM114A2;refGene_tx=XM_019840021.2
A1	195668486	195668579
refGene_gn=FAM114A2;refGene_tx=XM_011285908.2;refGene_gn=FAM114A2;refGene_tx=XM_011285913.3;refGene_gn=FAM114A2;refGene_tx=XM_019840021.2
A1	195669825	195669917
refGene_gn=FAM114A2;refGene_tx=XM_011285908.2;refGene_gn=FAM114A2;refGene_tx=XM_011285913.3;refGene_gn=FAM114A2;refGene_tx=XM_019840021.2
A1	195670973	195671108
refGene_gn=FAM114A2;refGene_tx=XM_011285908.2;refGene_gn=FAM114A2;refGene_tx=XM_011285913.3;refGene_gn=FAM114A2;refGene_tx=XM_019840021.2
A1	195671623	195671782
refGene_gn=FAM114A2;refGene_tx=XM_011285908.2;refGene_gn=FAM114A2;refGene_tx=XM_011285913.3;refGene_gn=FAM114A2;refGene_tx=XM_019840021.2
A1	195672393	195672517
refGene_gn=FAM114A2;refGene_tx=XM_011285908.2;refGene_gn=FAM114A2;refGene_tx=XM_011285913.3;refGene_gn=FAM114A2;refGene_tx=XM_019840021.2
A1	195677916	195677996
refGene_gn=FAM114A2;refGene_tx=XM_011285908.2;refGene_gn=FAM114A2;refGene_tx=XM_011285913.3;refGene_gn=FAM114A2;refGene_tx=XM_019840021.2
A1	195685875	195685998
refGene_gn=FAM114A2;refGene_tx=XM_011285908.2;refGene_gn=FAM114A2;refGene_tx=XM_011285913.3;refGene_gn=FAM114A2;refGene_tx=XM_019840021.2
A1	195686501	195686641
refGene_gn=FAM114A2;refGene_tx=XM_011285908.2;refGene_gn=FAM114A2;refGene_tx=XM_011285913.3;refGene_gn=FAM114A2;refGene_tx=XM_019840021.2
A1	195688142	195688215
refGene_gn=FAM114A2;refGene_tx=XM_011285908.2;refGene_gn=FAM114A2;refGene_tx=XM_011285913.3;refGene_gn=FAM114A2;refGene_tx=XM_019840021.2
A1	195691717	195691771
refGene_gn=FAM114A2;refGene_tx=XM_011285908.2;refGene_gn=FAM114A2;refGene_tx=XM_011285913.3;refGene_gn=FAM114A2;refGene_tx=XM_019840021.2
A1	195693653	195693755
refGene_gn=FAM114A2;refGene_tx=XM_011285908.2;refGene_gn=FAM114A2;refGene_tx=XM_011285913.3;refGene_gn=FAM114A2;refGene_tx=XM_019840021.2
A1	195862685	195862886
refGene_gn=GRIA1;refGene_tx=XM_003981365.4;refGene_gn=GRIA1;refGene_tx=XM_003981366.4;refGene_gn=GRIA1;refGene_tx=XM_019840051.2
A1	195870770	195870905

refGene_gn=GRIA1;refGene_tx=XM_003981365.4;refGene_gn=GRIA1;refGene_tx=XM_003981366.4;refGene_gn=GRIA1;refGene_tx=XM_019840051.2
A1	195877571	195877686
refGene_gn=GRIA1;refGene_tx=XM_003981365.4
A1	195878427	195878542
refGene_gn=GRIA1;refGene_tx=XM_003981366.4;refGene_gn=GRIA1;refGene_tx=XM_019840051.2
A1	195900060	195900308
refGene_gn=GRIA1;refGene_tx=XM_003981365.4;refGene_gn=GRIA1;refGene_tx=XM_003981366.4;refGene_gn=GRIA1;refGene_tx=XM_019840051.2
A1	195905542	195905741
refGene_gn=GRIA1;refGene_tx=XM_003981365.4;refGene_gn=GRIA1;refGene_tx=XM_003981366.4;refGene_gn=GRIA1;refGene_tx=XM_019840051.2
A1	195949477	195949848
refGene_gn=GRIA1;refGene_tx=XM_003981365.4;refGene_gn=GRIA1;refGene_tx=XM_003981366.4;refGene_gn=GRIA1;refGene_tx=XM_019840051.2
A1	195956158	195956365
refGene_gn=GRIA1;refGene_tx=XM_003981365.4;refGene_gn=GRIA1;refGene_tx=XM_003981366.4;refGene_gn=GRIA1;refGene_tx=XM_019840051.2
A1	195957076	195957187
refGene_gn=GRIA1;refGene_tx=XM_003981365.4;refGene_gn=GRIA1;refGene_tx=XM_003981366.4;refGene_gn=GRIA1;refGene_tx=XM_019840051.2
A1	195969170	195969275
refGene_gn=GRIA1;refGene_tx=XM_003981365.4;refGene_gn=GRIA1;refGene_tx=XM_003981366.4;refGene_gn=GRIA1;refGene_tx=XM_019840051.2
A1	195977469	195977637
refGene_gn=GRIA1;refGene_tx=XM_003981365.4;refGene_gn=GRIA1;refGene_tx=XM_003981366.4;refGene_gn=GRIA1;refGene_tx=XM_019840051.2
A1	195980152	195980314
refGene_gn=GRIA1;refGene_tx=XM_003981365.4;refGene_gn=GRIA1;refGene_tx=XM_003981366.4;refGene_gn=GRIA1;refGene_tx=XM_019840051.2
A1	196000145	196000199
refGene_gn=GRIA1;refGene_tx=XM_003981365.4;refGene_gn=GRIA1;refGene_tx=XM_003981366.4;refGene_gn=GRIA1;refGene_tx=XM_019840051.2
A1	196005300	196005485
refGene_gn=GRIA1;refGene_tx=XM_003981365.4;refGene_gn=GRIA1;refGene_tx=XM_003981366.4;refGene_gn=GRIA1;refGene_tx=XM_019840051.2
A1	196008414	196008654
refGene_gn=GRIA1;refGene_tx=XM_003981365.4;refGene_gn=GRIA1;refGene_tx=XM_003981366.4;refGene_gn=GRIA1;refGene_tx=XM_019840051.2
A1	196164112	196164250
refGene_gn=GRIA1;refGene_tx=XM_003981365.4;refGene_gn=GRIA1;refGene_tx=XM_003981366.4
A1	196167283	196167365
refGene_gn=GRIA1;refGene_tx=XM_003981365.4;refGene_gn=GRIA1;refGene_tx=XM_003981366.4
A1	197118086	197118803
refGene_gn=NMUR2;refGene_tx=XM_003981478.5;refGene_gn=NMUR2;refGene_tx=XM_019840059.2
A1	197125319	197125404

refGene_gn=NMUR2;refGene_tx=XM_003981478.5;refGene_gn=NMUR2;refGene_tx=XM_019840059.2
A1 197127834 197127960
refGene_gn=NMUR2;refGene_tx=XM_003981478.5
A1 197130967 197131239
refGene_gn=NMUR2;refGene_tx=XM_003981478.5;refGene_gn=NMUR2;refGene_tx=XM_019840059.2
A1 197558630 197558686
refGene_gn=GLRA1;refGene_tx=XM_003981370.4;refGene_gn=GLRA1;refGene_tx=XM_003981371.4
A1 197593036 197593164
refGene_gn=GLRA1;refGene_tx=XM_003981370.4;refGene_gn=GLRA1;refGene_tx=XM_003981371.4
A1 197599195 197599263
refGene_gn=GLRA1;refGene_tx=XM_003981370.4;refGene_gn=GLRA1;refGene_tx=XM_003981371.4
A1 197619631 197619855
refGene_gn=GLRA1;refGene_tx=XM_003981370.4;refGene_gn=GLRA1;refGene_tx=XM_003981371.4
A1 197622589 197622672
refGene_gn=GLRA1;refGene_tx=XM_003981370.4;refGene_gn=GLRA1;refGene_tx=XM_003981371.4
A1 197624551 197624689
refGene_gn=GLRA1;refGene_tx=XM_003981370.4;refGene_gn=GLRA1;refGene_tx=XM_003981371.4
A1 197627015 197627230
refGene_gn=GLRA1;refGene_tx=XM_003981370.4;refGene_gn=GLRA1;refGene_tx=XM_003981371.4
A1 197636028 197636175
refGene_gn=GLRA1;refGene_tx=XM_003981370.4;refGene_gn=GLRA1;refGene_tx=XM_003981371.4
A1 197642579 197642894
refGene_gn=GLRA1;refGene_tx=XM_003981370.4;refGene_gn=GLRA1;refGene_tx=XM_003981371.4
A1 197663470 197663677
refGene_gn=G3BP1;refGene_tx=XM_003981373.5;refGene_gn=G3BP1;refGene_tx=XM_006927968.4;refGene_gn=G3BP1;refGene_tx=XM_006927969.4
A1 197666806 197666916
refGene_gn=G3BP1;refGene_tx=XM_003981373.5;refGene_gn=G3BP1;refGene_tx=XM_006927968.4;refGene_gn=G3BP1;refGene_tx=XM_006927969.4
A1 197667594 197667723
refGene_gn=G3BP1;refGene_tx=XM_003981373.5;refGene_gn=G3BP1;refGene_tx=XM_006927968.4;refGene_gn=G3BP1;refGene_tx=XM_006927969.4
A1 197667927 197668039
refGene_gn=G3BP1;refGene_tx=XM_003981373.5;refGene_gn=G3BP1;refGene_tx=XM_006927968.4;refGene_gn=G3BP1;refGene_tx=XM_006927969.4
A1 197668663 197668765
refGene_gn=G3BP1;refGene_tx=XM_003981373.5;refGene_gn=G3BP1;refGene_tx=XM_006927968.4;refGene_gn=G3BP1;refGene_tx=XM_006927969.4
A1 197670028 197670230

refGene_gn=G3BP1;refGene_tx=XM_003981373.5;refGene_gn=G3BP1;refGene_tx=XM_006927968.4;refGene_gn=G3BP1;refGene_tx=XM_006927969.4
A1	197672124	197672218
refGene_gn=G3BP1;refGene_tx=XM_003981373.5;refGene_gn=G3BP1;refGene_tx=XM_006927968.4;refGene_gn=G3BP1;refGene_tx=XM_006927969.4
A1	197673629	197673720
refGene_gn=G3BP1;refGene_tx=XM_003981373.5;refGene_gn=G3BP1;refGene_tx=XM_006927968.4;refGene_gn=G3BP1;refGene_tx=XM_006927969.4
A1	197675725	197675899
refGene_gn=G3BP1;refGene_tx=XM_003981373.5;refGene_gn=G3BP1;refGene_tx=XM_006927968.4;refGene_gn=G3BP1;refGene_tx=XM_006927969.4
A1	197676357	197676439
refGene_gn=G3BP1;refGene_tx=XM_003981373.5;refGene_gn=G3BP1;refGene_tx=XM_006927968.4;refGene_gn=G3BP1;refGene_tx=XM_006927969.4
A1	197681488	197681583
refGene_gn=G3BP1;refGene_tx=XM_003981373.5;refGene_gn=G3BP1;refGene_tx=XM_006927968.4;refGene_gn=G3BP1;refGene_tx=XM_006927969.4
A1	197707523	197707529
refGene_gn=ATOX1;refGene_tx=XM_011285958.3
A1	197714503	197714579
refGene_gn=ATOX1;refGene_tx=XM_011285958.3
A1	197718239	197718364
refGene_gn=ATOX1;refGene_tx=XM_011285958.3
A1	197771692	197771749
refGene_gn=SPARC;refGene_tx=XM_003981374.5
A1	197773046	197773109
refGene_gn=SPARC;refGene_tx=XM_003981374.5
A1	197773660	197773748
refGene_gn=SPARC;refGene_tx=XM_003981374.5
A1	197775190	197775312
refGene_gn=SPARC;refGene_tx=XM_003981374.5
A1	197777128	197777249
refGene_gn=SPARC;refGene_tx=XM_003981374.5
A1	197780053	197780187
refGene_gn=SPARC;refGene_tx=XM_003981374.5
A1	197780853	197781002
refGene_gn=SPARC;refGene_tx=XM_003981374.5
A1	197783072	197783221
refGene_gn=SPARC;refGene_tx=XM_003981374.5
A1	197783688	197783717
refGene_gn=SPARC;refGene_tx=XM_003981374.5
A1	197827014	197827121
refGene_gn=SLC36A1;refGene_tx=XM_019840113.2
A1	197833623	197833652
refGene_gn=SLC36A1;refGene_tx=XM_019840121.2
A1	197871649	197874911
refGene_gn=FAT2;refGene_tx=XM_006927970.4
A1	197877380	197877695
refGene_gn=FAT2;refGene_tx=XM_006927970.4
A1	197882901	197882960

refGene_gn=FAT2;refGene_tx=XM_006927970.4
A1 197884333 197884645
refGene_gn=FAT2;refGene_tx=XM_006927970.4
A1 197885646 197885857
refGene_gn=FAT2;refGene_tx=XM_006927970.4
A1 197887673 197887813
refGene_gn=FAT2;refGene_tx=XM_006927970.4
A1 197888645 197888927
refGene_gn=FAT2;refGene_tx=XM_006927970.4
A1 197890247 197890458
refGene_gn=FAT2;refGene_tx=XM_006927970.4
A1 197894767 197898826
refGene_gn=FAT2;refGene_tx=XM_006927970.4
A1 197900047 197900244
refGene_gn=FAT2;refGene_tx=XM_006927970.4
A1 197902600 197902754
refGene_gn=FAT2;refGene_tx=XM_006927970.4
A1 197904790 197905024
refGene_gn=FAT2;refGene_tx=XM_006927970.4
A1 197907517 197907901
refGene_gn=FAT2;refGene_tx=XM_006927970.4
A1 197909726 197909941
refGene_gn=FAT2;refGene_tx=XM_006927970.4
A1 197910935 197911073
refGene_gn=FAT2;refGene_tx=XM_006927970.4
A1 197911668 197911812
refGene_gn=FAT2;refGene_tx=XM_006927970.4
A1 197914192 197914390
refGene_gn=FAT2;refGene_tx=XM_006927970.4
A1 197917859 197918670
refGene_gn=FAT2;refGene_tx=XM_006927970.4
A1 197923337 197923483
refGene_gn=FAT2;refGene_tx=XM_006927970.4
A1 197927449 197927891
refGene_gn=FAT2;refGene_tx=XM_006927970.4
A1 197929777 197929931
refGene_gn=FAT2;refGene_tx=XM_006927970.4
A1 197931579 197932037
refGene_gn=FAT2;refGene_tx=XM_006927970.4
A1 197933119 197933652
refGene_gn=FAT2;refGene_tx=XM_006927970.4
A1 197947360 197947632
refGene_gn=SLC36A1;refGene_tx=XM_006927971.4
A1 197956832 197957002
refGene_gn=SLC36A1;refGene_tx=XM_006927971.4;refGene_gn=SLC36A1;refGen
e_tx=XM_019840113.2;refGene_gn=SLC36A1;refGene_tx=XM_019840121.2
A1 197960331 197960498
refGene_gn=SLC36A1;refGene_tx=XM_006927971.4;refGene_gn=SLC36A1;refGen
e_tx=XM_019840113.2;refGene_gn=SLC36A1;refGene_tx=XM_019840121.2
A1 197962700 197962799

refGene_gn=SLC36A1;refGene_tx=XM_006927971.4;refGene_gn=SLC36A1;refGene_tx=XM_019840113.2;refGene_gn=SLC36A1;refGene_tx=XM_019840121.2
A1 197967976 197968195
refGene_gn=SLC36A1;refGene_tx=XM_006927971.4;refGene_gn=SLC36A1;refGene_tx=XM_019840113.2;refGene_gn=SLC36A1;refGene_tx=XM_019840121.2
A1 197968612 197968697
refGene_gn=SLC36A1;refGene_tx=XM_006927971.4;refGene_gn=SLC36A1;refGene_tx=XM_019840113.2;refGene_gn=SLC36A1;refGene_tx=XM_019840121.2
A1 197970708 197970804
refGene_gn=SLC36A1;refGene_tx=XM_006927971.4;refGene_gn=SLC36A1;refGene_tx=XM_019840113.2;refGene_gn=SLC36A1;refGene_tx=XM_019840121.2
A1 197971254 197971343
refGene_gn=SLC36A1;refGene_tx=XM_006927971.4;refGene_gn=SLC36A1;refGene_tx=XM_019840113.2;refGene_gn=SLC36A1;refGene_tx=XM_019840121.2
A1 197972258 197972349
refGene_gn=SLC36A1;refGene_tx=XM_006927971.4;refGene_gn=SLC36A1;refGene_tx=XM_019840113.2;refGene_gn=SLC36A1;refGene_tx=XM_019840121.2
A1 197979665 197979808
refGene_gn=SLC36A1;refGene_tx=XM_006927971.4;refGene_gn=SLC36A1;refGene_tx=XM_019840113.2;refGene_gn=SLC36A1;refGene_tx=XM_019840121.2
A1 198048125 198048289
refGene_gn=SLC36A2;refGene_tx=XM_003981376.5;refGene_gn=SLC36A2;refGene_tx=XM_006927972.4
A1 198051250 198051341
refGene_gn=SLC36A2;refGene_tx=XM_003981376.5;refGene_gn=SLC36A2;refGene_tx=XM_006927972.4
A1 198051901 198051990
refGene_gn=SLC36A2;refGene_tx=XM_003981376.5;refGene_gn=SLC36A2;refGene_tx=XM_006927972.4
A1 198052535 198052631
refGene_gn=SLC36A2;refGene_tx=XM_003981376.5;refGene_gn=SLC36A2;refGene_tx=XM_006927972.4
A1 198057664 198057749
refGene_gn=SLC36A2;refGene_tx=XM_003981376.5;refGene_gn=SLC36A2;refGene_tx=XM_006927972.4
A1 198060779 198060998
refGene_gn=SLC36A2;refGene_tx=XM_003981376.5;refGene_gn=SLC36A2;refGene_tx=XM_006927972.4
A1 198062264 198062363
refGene_gn=SLC36A2;refGene_tx=XM_003981376.5;refGene_gn=SLC36A2;refGene_tx=XM_006927972.4
A1 198067173 198067340
refGene_gn=SLC36A2;refGene_tx=XM_003981376.5;refGene_gn=SLC36A2;refGene_tx=XM_006927972.4
A1 198069323 198069493
refGene_gn=SLC36A2;refGene_tx=XM_003981376.5;refGene_gn=SLC36A2;refGene_tx=XM_006927972.4
A1 198072695 198072967
refGene_gn=SLC36A2;refGene_tx=XM_003981376.5;refGene_gn=SLC36A2;refGene_tx=XM_006927972.4

A1	198079216	198079338
refGene_gn=SLC36A3;refGene_tx=XM_003981377.5;refGene_gn=SLC36A3;refGene_tx=XM_019810447.2;refGene_gn=SLC36A3;refGene_tx=XM_023260328.1
A1	198082675	198082766
refGene_gn=SLC36A3;refGene_tx=XM_003981377.5;refGene_gn=SLC36A3;refGene_tx=XM_019810447.2;refGene_gn=SLC36A3;refGene_tx=XM_023260328.1
A1	198085223	198085312
refGene_gn=SLC36A3;refGene_tx=XM_003981377.5;refGene_gn=SLC36A3;refGene_tx=XM_019810447.2;refGene_gn=SLC36A3;refGene_tx=XM_023260328.1
A1	198088163	198088259
refGene_gn=SLC36A3;refGene_tx=XM_003981377.5;refGene_gn=SLC36A3;refGene_tx=XM_019810447.2;refGene_gn=SLC36A3;refGene_tx=XM_023260328.1
A1	198093061	198093146
refGene_gn=SLC36A3;refGene_tx=XM_003981377.5;refGene_gn=SLC36A3;refGene_tx=XM_019810447.2;refGene_gn=SLC36A3;refGene_tx=XM_023260328.1
A1	198094065	198094284
refGene_gn=SLC36A3;refGene_tx=XM_003981377.5;refGene_gn=SLC36A3;refGene_tx=XM_019810447.2;refGene_gn=SLC36A3;refGene_tx=XM_023260328.1
A1	198096414	198096513
refGene_gn=SLC36A3;refGene_tx=XM_003981377.5;refGene_gn=SLC36A3;refGene_tx=XM_019810447.2;refGene_gn=SLC36A3;refGene_tx=XM_023260328.1
A1	198096892	198097059
refGene_gn=SLC36A3;refGene_tx=XM_003981377.5;refGene_gn=SLC36A3;refGene_tx=XM_019810447.2;refGene_gn=SLC36A3;refGene_tx=XM_023260328.1
A1	198100215	198100385
refGene_gn=SLC36A3;refGene_tx=XM_003981377.5;refGene_gn=SLC36A3;refGene_tx=XM_019810447.2;refGene_gn=SLC36A3;refGene_tx=XM_023260328.1
A1	198103551	198103820
refGene_gn=SLC36A3;refGene_tx=XM_003981377.5;refGene_gn=SLC36A3;refGene_tx=XM_019810447.2;refGene_gn=SLC36A3;refGene_tx=XM_023260328.1
A1	198103947	198104039
refGene_gn=SLC36A3;refGene_tx=XM_019810447.2;refGene_gn=SLC36A3;refGene_tx=XM_023260328.1
A1	198114600	198114756
refGene_gn=GM2A;refGene_tx=XM_003981379.5
A1	198115140	198115323
refGene_gn=GM2A;refGene_tx=XM_003981379.5
A1	198121704	198121857
refGene_gn=GM2A;refGene_tx=XM_003981379.5
A1	198127283	198127373
refGene_gn=GM2A;refGene_tx=XM_003981379.5
A1	198146868	198146943
refGene_gn=CCDC69;refGene_tx=XM_006928069.2
A1	198148091	198148502
refGene_gn=CCDC69;refGene_tx=XM_006928069.2
A1	198162998	198163068
refGene_gn=CCDC69;refGene_tx=XM_006928069.2
A1	198165590	198165697
refGene_gn=CCDC69;refGene_tx=XM_006928069.2
A1	198166222	198166310

refGene_gn=CCDC69;refGene_tx=XM_006928069.2
A1 198178027 198178101
refGene_gn=CCDC69;refGene_tx=XM_006928069.2
A1 198179087 198179189
refGene_gn=CCDC69;refGene_tx=XM_006928069.2
A1 198179659 198179779
refGene_gn=CCDC69;refGene_tx=XM_006928069.2
A1 198180699 198180797
refGene_gn=CCDC69;refGene_tx=XM_006928069.2
A1 198181523 198181701
refGene_gn=CCDC69;refGene_tx=XM_006928069.2
A1 198182198 198182241
refGene_gn=LOC101084936;refGene_tx=XM_019811055.1
A1 198183992 198184044
refGene_gn=LOC101084936;refGene_tx=XM_019811055.1
A1 198184133 198184260
refGene_gn=LOC101084936;refGene_tx=XM_019811055.1
A1 198184938 198184954
refGene_gn=LOC101084936;refGene_tx=XM_019811055.1
A1 198185779 198185930
refGene_gn=LOC101084936;refGene_tx=XM_019811055.1
A1 198186384 198186678
refGene_gn=LOC101084936;refGene_tx=XM_019811055.1
A1 198186926 198187078
refGene_gn=LOC101084936;refGene_tx=XM_019811055.1
A1 198187849 198188192
refGene_gn=LOC101084936;refGene_tx=XM_019811055.1
A1 198193783 198193841
refGene_gn=LOC101084936;refGene_tx=XM_019811055.1
A1 198216698 198216716
refGene_gn=ANXA6;refGene_tx=XM_003981380.5;refGene_gn=ANXA6;refGene_tx=XM_003981381.5;refGene_gn=ANXA6;refGene_tx=XM_019810500.1
A1 198221827 198221918
refGene_gn=ANXA6;refGene_tx=XM_003981380.5;refGene_gn=ANXA6;refGene_tx=XM_003981381.5;refGene_gn=ANXA6;refGene_tx=XM_019810500.1
A1 198222618 198222713
refGene_gn=ANXA6;refGene_tx=XM_003981380.5;refGene_gn=ANXA6;refGene_tx=XM_003981381.5;refGene_gn=ANXA6;refGene_tx=XM_019810500.1
A1 198223260 198223374
refGene_gn=ANXA6;refGene_tx=XM_003981380.5;refGene_gn=ANXA6;refGene_tx=XM_003981381.5;refGene_gn=ANXA6;refGene_tx=XM_019810500.1
A1 198224429 198224520
refGene_gn=ANXA6;refGene_tx=XM_003981380.5;refGene_gn=ANXA6;refGene_tx=XM_003981381.5;refGene_gn=ANXA6;refGene_tx=XM_019810500.1
A1 198225455 198225535
refGene_gn=ANXA6;refGene_tx=XM_003981380.5;refGene_gn=ANXA6;refGene_tx=XM_003981381.5;refGene_gn=ANXA6;refGene_tx=XM_019810500.1
A1 198227621 198227678
refGene_gn=ANXA6;refGene_tx=XM_003981380.5;refGene_gn=ANXA6;refGene_tx=XM_003981381.5;refGene_gn=ANXA6;refGene_tx=XM_019810500.1

A1	198228365	198228459
refGene_gn=ANXA6;refGene_tx=XM_003981380.5;refGene_gn=ANXA6;refGene_tx=XM_003981381.5;refGene_gn=ANXA6;refGene_tx=XM_019810500.1
A1	198228889	198228985
refGene_gn=ANXA6;refGene_tx=XM_003981380.5;refGene_gn=ANXA6;refGene_tx=XM_003981381.5;refGene_gn=ANXA6;refGene_tx=XM_019810500.1
A1	198230117	198230176
refGene_gn=ANXA6;refGene_tx=XM_003981380.5;refGene_gn=ANXA6;refGene_tx=XM_003981381.5;refGene_gn=ANXA6;refGene_tx=XM_019810500.1
A1	198231258	198231381
refGene_gn=ANXA6;refGene_tx=XM_003981380.5;refGene_gn=ANXA6;refGene_tx=XM_003981381.5;refGene_gn=ANXA6;refGene_tx=XM_019810500.1
A1	198232483	198232542
refGene_gn=ANXA6;refGene_tx=XM_003981380.5;refGene_gn=ANXA6;refGene_tx=XM_003981381.5;refGene_gn=ANXA6;refGene_tx=XM_019810500.1
A1	198233884	198233963
refGene_gn=ANXA6;refGene_tx=XM_003981380.5;refGene_gn=ANXA6;refGene_tx=XM_003981381.5;refGene_gn=ANXA6;refGene_tx=XM_019810500.1
A1	198235578	198235660
refGene_gn=ANXA6;refGene_tx=XM_003981380.5;refGene_gn=ANXA6;refGene_tx=XM_003981381.5;refGene_gn=ANXA6;refGene_tx=XM_019810500.1
A1	198236879	198236974
refGene_gn=ANXA6;refGene_tx=XM_003981380.5;refGene_gn=ANXA6;refGene_tx=XM_003981381.5;refGene_gn=ANXA6;refGene_tx=XM_019810500.1
A1	198237609	198237723
refGene_gn=ANXA6;refGene_tx=XM_003981380.5;refGene_gn=ANXA6;refGene_tx=XM_003981381.5;refGene_gn=ANXA6;refGene_tx=XM_019810500.1
A1	198240354	198240445
refGene_gn=ANXA6;refGene_tx=XM_003981380.5;refGene_gn=ANXA6;refGene_tx=XM_003981381.5;refGene_gn=ANXA6;refGene_tx=XM_019810500.1
A1	198241162	198241242
refGene_gn=ANXA6;refGene_tx=XM_003981380.5;refGene_gn=ANXA6;refGene_tx=XM_003981381.5;refGene_gn=ANXA6;refGene_tx=XM_019810500.1
A1	198242022	198242076
refGene_gn=ANXA6;refGene_tx=XM_003981380.5;refGene_gn=ANXA6;refGene_tx=XM_003981381.5;refGene_gn=ANXA6;refGene_tx=XM_019810500.1
A1	198243286	198243517
refGene_gn=ANXA6;refGene_tx=XM_019810500.1
A1	198248372	198248390
refGene_gn=ANXA6;refGene_tx=XM_003981380.5
A1	198249151	198249245
refGene_gn=ANXA6;refGene_tx=XM_003981380.5;refGene_gn=ANXA6;refGene_tx=XM_003981381.5
A1	198250372	198250468
refGene_gn=ANXA6;refGene_tx=XM_003981380.5;refGene_gn=ANXA6;refGene_tx=XM_003981381.5
A1	198251855	198251914
refGene_gn=ANXA6;refGene_tx=XM_003981380.5;refGene_gn=ANXA6;refGene_tx=XM_003981381.5
A1	198253264	198253387

refGene_gn=ANXA6;refGene_tx=XM_003981380.5;refGene_gn=ANXA6;refGene_tx=XM_003981381.5
A1 198255593 198255653
refGene_gn=ANXA6;refGene_tx=XM_003981380.5;refGene_gn=ANXA6;refGene_tx=XM_003981381.5
A1 198289297 198289433
refGene_gn=TNIP1;refGene_tx=XM_003981382.4;refGene_gn=TNIP1;refGene_tx=XM_006927975.4
A1 198290586 198290721
refGene_gn=TNIP1;refGene_tx=XM_003981382.4;refGene_gn=TNIP1;refGene_tx=XM_006927975.4;refGene_gn=TNIP1;refGene_tx=XM_019810511.2
A1 198291995 198292081
refGene_gn=TNIP1;refGene_tx=XM_003981382.4;refGene_gn=TNIP1;refGene_tx=XM_006927975.4;refGene_gn=TNIP1;refGene_tx=XM_019810511.2
A1 198293508 198293592
refGene_gn=TNIP1;refGene_tx=XM_003981382.4;refGene_gn=TNIP1;refGene_tx=XM_006927975.4;refGene_gn=TNIP1;refGene_tx=XM_019810511.2
A1 198296027 198296219
refGene_gn=TNIP1;refGene_tx=XM_003981382.4;refGene_gn=TNIP1;refGene_tx=XM_006927975.4;refGene_gn=TNIP1;refGene_tx=XM_019810511.2
A1 198299999 198300094
refGene_gn=TNIP1;refGene_tx=XM_003981382.4;refGene_gn=TNIP1;refGene_tx=XM_006927975.4;refGene_gn=TNIP1;refGene_tx=XM_019810511.2
A1 198301724 198301845
refGene_gn=TNIP1;refGene_tx=XM_003981382.4;refGene_gn=TNIP1;refGene_tx=XM_006927975.4;refGene_gn=TNIP1;refGene_tx=XM_019810511.2
A1 198305977 198306067
refGene_gn=TNIP1;refGene_tx=XM_003981382.4;refGene_gn=TNIP1;refGene_tx=XM_006927975.4;refGene_gn=TNIP1;refGene_tx=XM_019810511.2
A1 198308322 198308388
refGene_gn=TNIP1;refGene_tx=XM_003981382.4;refGene_gn=TNIP1;refGene_tx=XM_006927975.4;refGene_gn=TNIP1;refGene_tx=XM_019810511.2
A1 198308552 198308684
refGene_gn=TNIP1;refGene_tx=XM_003981382.4;refGene_gn=TNIP1;refGene_tx=XM_006927975.4;refGene_gn=TNIP1;refGene_tx=XM_019810511.2
A1 198311384 198311513
refGene_gn=TNIP1;refGene_tx=XM_003981382.4;refGene_gn=TNIP1;refGene_tx=XM_006927975.4;refGene_gn=TNIP1;refGene_tx=XM_019810511.2
A1 198314049 198314181
refGene_gn=TNIP1;refGene_tx=XM_003981382.4;refGene_gn=TNIP1;refGene_tx=XM_006927975.4;refGene_gn=TNIP1;refGene_tx=XM_019810511.2
A1 198315276 198315402
refGene_gn=TNIP1;refGene_tx=XM_003981382.4;refGene_gn=TNIP1;refGene_tx=XM_006927975.4;refGene_gn=TNIP1;refGene_tx=XM_019810511.2
A1 198315903 198315969
refGene_gn=TNIP1;refGene_tx=XM_003981382.4;refGene_gn=TNIP1;refGene_tx=XM_006927975.4;refGene_gn=TNIP1;refGene_tx=XM_019810511.2
A1 198316308 198316500
refGene_gn=TNIP1;refGene_tx=XM_003981382.4;refGene_gn=TNIP1;refGene_tx=XM_006927975.4;refGene_gn=TNIP1;refGene_tx=XM_019810511.2

A1 198317560 198317651
refGene_gn=TNIP1;refGene_tx=XM_003981382.4;refGene_gn=TNIP1;refGene_tx=XM_006927975.4;refGene_gn=TNIP1;refGene_tx=XM_019810511.2
A1 198319361 198319396
refGene_gn=TNIP1;refGene_tx=XM_003981382.4;refGene_gn=TNIP1;refGene_tx=XM_019810511.2
A1 198321539 198321761
refGene_gn=GPX3;refGene_tx=XM_003981387.5
A1 198322175 198322275
refGene_gn=GPX3;refGene_tx=XM_003981387.5
A1 198322729 198322847
refGene_gn=GPX3;refGene_tx=XM_003981387.5
A1 198324323 198324477
refGene_gn=GPX3;refGene_tx=XM_003981387.5
A1 198329104 198329191
refGene_gn=GPX3;refGene_tx=XM_003981387.5
A1 198352272 198353201
refGene_gn=LOC101085183;refGene_tx=XM_019810527.2
A1 198356631 198356727
refGene_gn=LOC101085183;refGene_tx=XM_019810527.2
A1 198357018 198357145
refGene_gn=LOC101085183;refGene_tx=XM_019810527.2
A1 198357915 198357960
refGene_gn=LOC101085183;refGene_tx=XM_019810527.2
A1 198359988 198360078
refGene_gn=LOC101085183;refGene_tx=XM_019810527.2
A1 198373840 198373937
refGene_gn=ZNF300;refGene_tx=XM_023260378.1;refGene_gn=ZNF300;refGene_tx=XM_023260382.1;refGene_gn=ZNF300;refGene_tx=XM_023260385.1;refGene_gn=ZNF300;refGene_tx=XM_023260392.1;refGene_gn=ZNF300;refGene_tx=XM_023260399.1
A1 198375184 198375311
refGene_gn=ZNF300;refGene_tx=XM_023260378.1;refGene_gn=ZNF300;refGene_tx=XM_023260382.1;refGene_gn=ZNF300;refGene_tx=XM_023260385.1;refGene_gn=ZNF300;refGene_tx=XM_023260392.1;refGene_gn=ZNF300;refGene_tx=XM_023260393.1;refGene_gn=ZNF300;refGene_tx=XM_023260399.1
A1 198375569 198375665
refGene_gn=ZNF300;refGene_tx=XM_023260378.1;refGene_gn=ZNF300;refGene_tx=XM_023260382.1;refGene_gn=ZNF300;refGene_tx=XM_023260385.1;refGene_gn=ZNF300;refGene_tx=XM_023260392.1;refGene_gn=ZNF300;refGene_tx=XM_023260393.1;refGene_gn=ZNF300;refGene_tx=XM_023260399.1
A1 198391001 198392545
refGene_gn=ZNF300;refGene_tx=XM_023260378.1;refGene_gn=ZNF300;refGene_tx=XM_023260382.1;refGene_gn=ZNF300;refGene_tx=XM_023260385.1;refGene_gn=ZNF300;refGene_tx=XM_023260392.1;refGene_gn=ZNF300;refGene_tx=XM_023260393.1;refGene_gn=ZNF300;refGene_tx=XM_023260399.1
A1 198424120 198425140
refGene_gn=ZNF300;refGene_tx=XM_023260399.1
A1 198470150 198470333
refGene_gn=SMIM3;refGene_tx=XM_006927980.4;refGene_gn=SMIM3;refGene_tx

=XM_006927981.4;refGene_gn=SMIM3;refGene_tx=XM_006927983.4;refGene_gn=SMIM3;refGene_tx=XM_011286028.2;refGene_gn=SMIM3;refGene_tx=XM_019810578.2
A1 198521835 198521970
refGene_gn=DCTN4;refGene_tx=XM_003981390.4;refGene_gn=DCTN4;refGene_tx=XM_019810590.2
A1 198524394 198524465
refGene_gn=DCTN4;refGene_tx=XM_003981390.4;refGene_gn=DCTN4;refGene_tx=XM_019810590.2
A1 198526917 198527096
refGene_gn=DCTN4;refGene_tx=XM_003981390.4;refGene_gn=DCTN4;refGene_tx=XM_019810590.2
A1 198532364 198532408
refGene_gn=DCTN4;refGene_tx=XM_003981390.4;refGene_gn=DCTN4;refGene_tx=XM_019810590.2
A1 198534734 198534842
refGene_gn=DCTN4;refGene_tx=XM_003981390.4;refGene_gn=DCTN4;refGene_tx=XM_019810590.2
A1 198536069 198536090
refGene_gn=DCTN4;refGene_tx=XM_019810590.2
A1 198536516 198536590
refGene_gn=DCTN4;refGene_tx=XM_003981390.4;refGene_gn=DCTN4;refGene_tx=XM_019810590.2
A1 198536818 198536931
refGene_gn=DCTN4;refGene_tx=XM_003981390.4;refGene_gn=DCTN4;refGene_tx=XM_019810590.2
A1 198537646 198537756
refGene_gn=DCTN4;refGene_tx=XM_003981390.4;refGene_gn=DCTN4;refGene_tx=XM_019810590.2
A1 198543068 198543142
refGene_gn=DCTN4;refGene_tx=XM_003981390.4;refGene_gn=DCTN4;refGene_tx=XM_019810590.2
A1 198545285 198545340
refGene_gn=DCTN4;refGene_tx=XM_003981390.4;refGene_gn=DCTN4;refGene_tx=XM_019810590.2
A1 198546321 198546429
refGene_gn=DCTN4;refGene_tx=XM_003981390.4;refGene_gn=DCTN4;refGene_tx=XM_019810590.2
A1 198548662 198548760
refGene_gn=DCTN4;refGene_tx=XM_003981390.4;refGene_gn=DCTN4;refGene_tx=XM_019810590.2
A1 198551022 198551236
refGene_gn=DCTN4;refGene_tx=XM_003981390.4;refGene_gn=DCTN4;refGene_tx=XM_019810590.2
A1 198560526 198560580
refGene_gn=RBM22;refGene_tx=XM_003981392.5
A1 198561024 198561078
refGene_gn=RBM22;refGene_tx=XM_003981392.5
A1 198562034 198562064
refGene_gn=RBM22;refGene_tx=XM_003981392.5

A1	198562899	198563032
refGene_gn=RBM22;refGene_tx=XM_003981392.5
A1	198564518	198564619
refGene_gn=RBM22;refGene_tx=XM_003981392.5
A1	198564720	198564893
refGene_gn=RBM22;refGene_tx=XM_003981392.5
A1	198566189	198566390
refGene_gn=RBM22;refGene_tx=XM_003981392.5
A1	198567849	198568014
refGene_gn=RBM22;refGene_tx=XM_003981392.5
A1	198568770	198568859
refGene_gn=RBM22;refGene_tx=XM_003981392.5
A1	198569073	198569205
refGene_gn=RBM22;refGene_tx=XM_003981392.5
A1	198570055	198570186
refGene_gn=RBM22;refGene_tx=XM_003981392.5
A1	198582758	198582927
refGene_gn=MYOZ3;refGene_tx=XM_006927985.4;refGene_gn=MYOZ3;refGene_tx=XM_019810611.2
A1	198585935	198586098
refGene_gn=MYOZ3;refGene_tx=XM_006927985.4;refGene_gn=MYOZ3;refGene_tx=XM_019810611.2
A1	198586516	198586652
refGene_gn=MYOZ3;refGene_tx=XM_006927985.4;refGene_gn=MYOZ3;refGene_tx=XM_019810611.2
A1	198586753	198586834
refGene_gn=MYOZ3;refGene_tx=XM_006927985.4;refGene_gn=MYOZ3;refGene_tx=XM_019810611.2
A1	198588037	198588192
refGene_gn=MYOZ3;refGene_tx=XM_006927985.4;refGene_gn=MYOZ3;refGene_tx=XM_019810611.2
A1	198594640	198594701
refGene_gn=MYOZ3;refGene_tx=XM_006927985.4;refGene_gn=MYOZ3;refGene_tx=XM_019810611.2
A1	198600701	198601049
refGene_gn=SYNPO;refGene_tx=XM_023260430.1
A1	198601050	198601395
refGene_gn=SYNPO;refGene_tx=XM_023260430.1
A1	198606660	198609017
refGene_gn=SYNPO;refGene_tx=XM_023260430.1
A1	198635607	198636013
refGene_gn=SYNPO;refGene_tx=XM_023260430.1
A1	198689326	198689446
refGene_gn=NDST1;refGene_tx=XM_023260438.1
A1	198690699	198690802
refGene_gn=NDST1;refGene_tx=XM_023260438.1
A1	198692496	198692606
refGene_gn=NDST1;refGene_tx=XM_023260438.1
A1	198693264	198693435
refGene_gn=NDST1;refGene_tx=XM_023260438.1

A1 198695492 198695631
refGene_gn=NDST1;refGene_tx=XM_023260438.1
A1 198695632 198695665
refGene_gn=NDST1;refGene_tx=XM_023260438.1
A1 198698014 198698138
refGene_gn=NDST1;refGene_tx=XM_023260438.1
A1 198699366 198699463
refGene_gn=NDST1;refGene_tx=XM_023260438.1
A1 198700580 198700763
refGene_gn=NDST1;refGene_tx=XM_023260438.1
A1 198701591 198701720
refGene_gn=NDST1;refGene_tx=XM_023260438.1
A1 198703570 198703756
refGene_gn=NDST1;refGene_tx=XM_023260438.1
A1 198704431 198704586
refGene_gn=NDST1;refGene_tx=XM_023260438.1
A1 198705753 198705841
refGene_gn=NDST1;refGene_tx=XM_023260438.1
A1 198708767 198709262
refGene_gn=NDST1;refGene_tx=XM_023260438.1
A1 198715567 198716080
refGene_gn=NDST1;refGene_tx=XM_023260438.1
A1 198716185 198716466
refGene_gn=LOC111558042;refGene_tx=XM_023245487.1
A1 198744966 198745038
refGene_gn=LOC111558042;refGene_tx=XM_023245487.1
A1 198746672 198747198
refGene_gn=LOC111558042;refGene_tx=XM_023245487.1
A1 198749718 198749805
refGene_gn=LOC111558042;refGene_tx=XM_023245487.1
A1 198777788 198777937
refGene_gn=RPS14;refGene_tx=XM_006927988.2
A1 198778541 198778703
refGene_gn=RPS14;refGene_tx=XM_006927988.2
A1 198780575 198780652
refGene_gn=RPS14;refGene_tx=XM_006927988.2
A1 198781678 198781746
refGene_gn=RPS14;refGene_tx=XM_006927988.2
A1 198782939 198782940
refGene_gn=CD74;refGene_tx=XM_003981485.5;refGene_gn=CD74;refGene_tx=XM_006927989.4
A1 198804483 198804721
refGene_gn=CD74;refGene_tx=XM_003981485.5;refGene_gn=CD74;refGene_tx=XM_006927989.4
A1 198808592 198808765
refGene_gn=CD74;refGene_tx=XM_003981485.5;refGene_gn=CD74;refGene_tx=XM_006927989.4
A1 198808934 198809014
refGene_gn=CD74;refGene_tx=XM_003981485.5;refGene_gn=CD74;refGene_tx=XM_006927989.4

A1 198809553 198809616
refGene_gn=CD74;refGene_tx=XM_003981485.5;refGene_gn=CD74;refGene_tx=XM_006927989.4
A1 198810337 198810433
refGene_gn=CD74;refGene_tx=XM_003981485.5;refGene_gn=CD74;refGene_tx=XM_006927989.4
A1 198810754 198810842
refGene_gn=CD74;refGene_tx=XM_003981485.5;refGene_gn=CD74;refGene_tx=XM_006927989.4
A1 198811880 198812072
refGene_gn=CD74;refGene_tx=XM_003981485.5
A1 198812529 198812592
refGene_gn=CD74;refGene_tx=XM_003981485.5;refGene_gn=CD74;refGene_tx=XM_006927989.4
A1 198812850 198812861
refGene_gn=CD74;refGene_tx=XM_003981485.5;refGene_gn=CD74;refGene_tx=XM_006927989.4
A1 198815850 198815877
refGene_gn=TCOF1;refGene_tx=XM_006927991.4
A1 198816554 198816652
refGene_gn=TCOF1;refGene_tx=XM_006927991.4
A1 198817282 198817831
refGene_gn=TCOF1;refGene_tx=XM_006927991.4
A1 198819729 198819910
refGene_gn=TCOF1;refGene_tx=XM_006927991.4
A1 198820420 198820503
refGene_gn=TCOF1;refGene_tx=XM_006927991.4
A1 198821023 198821270
refGene_gn=TCOF1;refGene_tx=XM_006927991.4
A1 198821599 198821713
refGene_gn=TCOF1;refGene_tx=XM_006927991.4
A1 198823159 198823278
refGene_gn=TCOF1;refGene_tx=XM_006927991.4
A1 198824548 198824735
refGene_gn=TCOF1;refGene_tx=XM_006927991.4
A1 198828897 198829008
refGene_gn=TCOF1;refGene_tx=XM_006927991.4
A1 198831696 198831837
refGene_gn=TCOF1;refGene_tx=XM_006927991.4
A1 198831997 198832201
refGene_gn=TCOF1;refGene_tx=XM_006927991.4
A1 198832278 198832464
refGene_gn=TCOF1;refGene_tx=XM_006927991.4
A1 198832659 198832797
refGene_gn=TCOF1;refGene_tx=XM_006927991.4
A1 198838738 198838936
refGene_gn=TCOF1;refGene_tx=XM_006927991.4
A1 198839026 198839278
refGene_gn=TCOF1;refGene_tx=XM_006927991.4
A1 198839437 198839620

refGene_gn=TCOF1;refGene_tx=XM_006927991.4
A1 198839786 198839951
refGene_gn=TCOF1;refGene_tx=XM_006927991.4
A1 198840133 198840367
refGene_gn=TCOF1;refGene_tx=XM_006927991.4
A1 198840584 198840797
refGene_gn=TCOF1;refGene_tx=XM_006927991.4
A1 198842033 198842264
refGene_gn=TCOF1;refGene_tx=XM_006927991.4
A1 198843777 198843851
refGene_gn=TCOF1;refGene_tx=XM_006927991.4
A1 198844456 198844469
refGene_gn=TCOF1;refGene_tx=XM_006927991.4
A1 198844470 198844644
refGene_gn=TCOF1;refGene_tx=XM_006927991.4
A1 198845391 198845465
refGene_gn=TCOF1;refGene_tx=XM_006927991.4
A1 198847491 198847631
refGene_gn=TCOF1;refGene_tx=XM_006927991.4
A1 198850878 198850934
refGene_gn=TCOF1;refGene_tx=XM_006927991.4
A1 198853305 198853413
refGene_gn=TCOF1;refGene_tx=XM_006927991.4
A1 198904718 198905038
refGene_gn=ARSI;refGene_tx=XM_003981486.5
A1 198908383 198909791
refGene_gn=ARSI;refGene_tx=XM_003981486.5
A1 198917376 198917438
refGene_gn=CAMK2A;refGene_tx=XM_011286079.3;refGene_gn=CAMK2A;refGene_tx=XM_011286088.3;refGene_gn=CAMK2A;refGene_tx=XM_023260459.1
A1 198930852 198930947
refGene_gn=CAMK2A;refGene_tx=XM_011286079.3;refGene_gn=CAMK2A;refGene_tx=XM_011286088.3;refGene_gn=CAMK2A;refGene_tx=XM_023260459.1
A1 198938176 198938236
refGene_gn=CAMK2A;refGene_tx=XM_011286079.3;refGene_gn=CAMK2A;refGene_tx=XM_011286088.3;refGene_gn=CAMK2A;refGene_tx=XM_023260459.1
A1 198944779 198944834
refGene_gn=CAMK2A;refGene_tx=XM_011286079.3;refGene_gn=CAMK2A;refGene_tx=XM_011286088.3;refGene_gn=CAMK2A;refGene_tx=XM_023260459.1
A1 198945706 198945772
refGene_gn=CAMK2A;refGene_tx=XM_011286079.3;refGene_gn=CAMK2A;refGene_tx=XM_011286088.3;refGene_gn=CAMK2A;refGene_tx=XM_023260459.1
A1 198945892 198945965
refGene_gn=CAMK2A;refGene_tx=XM_011286079.3;refGene_gn=CAMK2A;refGene_tx=XM_011286088.3;refGene_gn=CAMK2A;refGene_tx=XM_023260459.1
A1 198948994 198949097
refGene_gn=CAMK2A;refGene_tx=XM_011286079.3;refGene_gn=CAMK2A;refGene_tx=XM_011286088.3;refGene_gn=CAMK2A;refGene_tx=XM_023260459.1
A1 198950406 198950490
refGene_gn=CAMK2A;refGene_tx=XM_011286079.3;refGene_gn=CAMK2A;refGene_

tx=XM_011286088.3;refGene_gn=CAMK2A;refGene_tx=XM_023260459.1
A1 198950620 198950715
refGene_gn=CAMK2A;refGene_tx=XM_011286079.3;refGene_gn=CAMK2A;refGene_
tx=XM_011286088.3;refGene_gn=CAMK2A;refGene_tx=XM_023260459.1
A1 198951948 198952071
refGene_gn=CAMK2A;refGene_tx=XM_011286079.3;refGene_gn=CAMK2A;refGene_
tx=XM_011286088.3;refGene_gn=CAMK2A;refGene_tx=XM_023260459.1
A1 198952435 198952519
refGene_gn=CAMK2A;refGene_tx=XM_011286079.3;refGene_gn=CAMK2A;refGene_
tx=XM_011286088.3;refGene_gn=CAMK2A;refGene_tx=XM_023260459.1
A1 198954019 198954062
refGene_gn=CAMK2A;refGene_tx=XM_011286079.3;refGene_gn=CAMK2A;refGene_
tx=XM_011286088.3;refGene_gn=CAMK2A;refGene_tx=XM_023260459.1
A1 198956492 198956533
refGene_gn=CAMK2A;refGene_tx=XM_011286079.3;refGene_gn=CAMK2A;refGene_
tx=XM_011286088.3;refGene_gn=CAMK2A;refGene_tx=XM_023260459.1
A1 198961141 198961174
refGene_gn=CAMK2A;refGene_tx=XM_011286079.3
A1 198962157 198962206
refGene_gn=CAMK2A;refGene_tx=XM_011286079.3;refGene_gn=CAMK2A;refGene_
tx=XM_011286088.3;refGene_gn=CAMK2A;refGene_tx=XM_023260459.1
A1 198969516 198969592
refGene_gn=CAMK2A;refGene_tx=XM_011286079.3;refGene_gn=CAMK2A;refGene_
tx=XM_011286088.3;refGene_gn=CAMK2A;refGene_tx=XM_023260459.1
A1 198972207 198972302
refGene_gn=CAMK2A;refGene_tx=XM_011286079.3;refGene_gn=CAMK2A;refGene_
tx=XM_011286088.3;refGene_gn=CAMK2A;refGene_tx=XM_023260459.1
A1 198976386 198976619
refGene_gn=CAMK2A;refGene_tx=XM_011286079.3;refGene_gn=CAMK2A;refGene_
tx=XM_011286088.3;refGene_gn=CAMK2A;refGene_tx=XM_023260459.1
A1 198976925 198976929
refGene_gn=CAMK2A;refGene_tx=XM_011286079.3;refGene_gn=CAMK2A;refGene_
tx=XM_011286088.3
A1 198989039 198989249
refGene_gn=SLC6A7;refGene_tx=XM_011286099.3;refGene_gn=SLC6A7;refGene_
tx=XM_019810690.2
A1 198992537 198992705
refGene_gn=SLC6A7;refGene_tx=XM_011286099.3;refGene_gn=SLC6A7;refGene_
tx=XM_019810690.2
A1 198993251 198993352
refGene_gn=SLC6A7;refGene_tx=XM_011286099.3;refGene_gn=SLC6A7;refGene_
tx=XM_019810690.2
A1 198993527 198993627
refGene_gn=SLC6A7;refGene_tx=XM_011286099.3;refGene_gn=SLC6A7;refGene_
tx=XM_019810690.2
A1 198994109 198994241
refGene_gn=SLC6A7;refGene_tx=XM_011286099.3;refGene_gn=SLC6A7;refGene_
tx=XM_019810690.2
A1 198994490 198994603
refGene_gn=SLC6A7;refGene_tx=XM_011286099.3

A1 198995518 198995643
refGene_gn=SLC6A7;refGene_tx=XM_011286099.3;refGene_gn=SLC6A7;refGene_tx=XM_019810690.2
A1 198995773 198995877
refGene_gn=SLC6A7;refGene_tx=XM_011286099.3;refGene_gn=SLC6A7;refGene_tx=XM_019810690.2
A1 198997375 198997510
refGene_gn=SLC6A7;refGene_tx=XM_011286099.3;refGene_gn=SLC6A7;refGene_tx=XM_019810690.2
A1 198998949 198999088
refGene_gn=SLC6A7;refGene_tx=XM_011286099.3;refGene_gn=SLC6A7;refGene_tx=XM_019810690.2
A1 198999509 198999744
refGene_gn=SLC6A7;refGene_tx=XM_011286099.3;refGene_gn=SLC6A7;refGene_tx=XM_019810690.2
A1 198999925 199000057
refGene_gn=SLC6A7;refGene_tx=XM_011286099.3;refGene_gn=SLC6A7;refGene_tx=XM_019810690.2
A1 199001720 199001904
refGene_gn=SLC6A7;refGene_tx=XM_011286099.3;refGene_gn=SLC6A7;refGene_tx=XM_019810690.2
A1 199006936 199006969
refGene_gn=SLC6A7;refGene_tx=XM_011286099.3;refGene_gn=SLC6A7;refGene_tx=XM_019810690.2
A1 199012851 199013052
refGene_gn=CDX1;refGene_tx=XM_023260468.1
A1 199013575 199013721
refGene_gn=CDX1;refGene_tx=XM_023260468.1
A1 199028518 199028963
refGene_gn=CDX1;refGene_tx=XM_023260468.1
A1 199054874 199054896
refGene_gn=PDGFRB;refGene_tx=XM_011286103.3;refGene_gn=PDGFRB;refGene_tx=XM_019810730.2
A1 199056079 199056403
refGene_gn=PDGFRB;refGene_tx=XM_011286103.3;refGene_gn=PDGFRB;refGene_tx=XM_019810730.2
A1 199056925 199057226
refGene_gn=PDGFRB;refGene_tx=XM_011286103.3;refGene_gn=PDGFRB;refGene_tx=XM_019810730.2;refGene_gn=PDGFRB;refGene_tx=XM_023260465.1
A1 199057903 199058031
refGene_gn=PDGFRB;refGene_tx=XM_011286103.3;refGene_gn=PDGFRB;refGene_tx=XM_019810730.2;refGene_gn=PDGFRB;refGene_tx=XM_023260465.1
A1 199058145 199058320
refGene_gn=PDGFRB;refGene_tx=XM_011286103.3;refGene_gn=PDGFRB;refGene_tx=XM_019810730.2;refGene_gn=PDGFRB;refGene_tx=XM_023260465.1
A1 199058932 199059125
refGene_gn=PDGFRB;refGene_tx=XM_011286103.3;refGene_gn=PDGFRB;refGene_tx=XM_019810730.2;refGene_gn=PDGFRB;refGene_tx=XM_023260465.1
A1 199059817 199059933
refGene_gn=PDGFRB;refGene_tx=XM_011286103.3;refGene_gn=PDGFRB;refGene_

tx=XM_019810730.2;refGene_gn=PDGFRB;refGene_tx=XM_023260465.1
A1 199061067 199061191
refGene_gn=PDGFRB;refGene_tx=XM_011286103.3;refGene_gn=PDGFRB;refGene_tx=XM_019810730.2;refGene_gn=PDGFRB;refGene_tx=XM_023260465.1
A1 199061666 199061878
refGene_gn=PDGFRB;refGene_tx=XM_011286103.3;refGene_gn=PDGFRB;refGene_tx=XM_019810730.2;refGene_gn=PDGFRB;refGene_tx=XM_023260465.1
A1 199064278 199064373
refGene_gn=PDGFRB;refGene_tx=XM_011286103.3;refGene_gn=PDGFRB;refGene_tx=XM_019810730.2;refGene_gn=PDGFRB;refGene_tx=XM_023260465.1
A1 199065327 199065460
refGene_gn=PDGFRB;refGene_tx=XM_011286103.3;refGene_gn=PDGFRB;refGene_tx=XM_019810730.2;refGene_gn=PDGFRB;refGene_tx=XM_023260465.1
A1 199066009 199066114
refGene_gn=PDGFRB;refGene_tx=XM_011286103.3;refGene_gn=PDGFRB;refGene_tx=XM_019810730.2;refGene_gn=PDGFRB;refGene_tx=XM_023260465.1
A1 199066464 199066575
refGene_gn=PDGFRB;refGene_tx=XM_011286103.3;refGene_gn=PDGFRB;refGene_tx=XM_019810730.2;refGene_gn=PDGFRB;refGene_tx=XM_023260465.1
A1 199067855 199068015
refGene_gn=PDGFRB;refGene_tx=XM_011286103.3;refGene_gn=PDGFRB;refGene_tx=XM_019810730.2;refGene_gn=PDGFRB;refGene_tx=XM_023260465.1
A1 199069074 199069235
refGene_gn=PDGFRB;refGene_tx=XM_011286103.3;refGene_gn=PDGFRB;refGene_tx=XM_019810730.2;refGene_gn=PDGFRB;refGene_tx=XM_023260465.1
A1 199069758 199069877
refGene_gn=PDGFRB;refGene_tx=XM_011286103.3;refGene_gn=PDGFRB;refGene_tx=XM_019810730.2;refGene_gn=PDGFRB;refGene_tx=XM_023260465.1
A1 199070132 199070255
refGene_gn=PDGFRB;refGene_tx=XM_011286103.3;refGene_gn=PDGFRB;refGene_tx=XM_019810730.2;refGene_gn=PDGFRB;refGene_tx=XM_023260465.1
A1 199070988 199071100
refGene_gn=PDGFRB;refGene_tx=XM_011286103.3;refGene_gn=PDGFRB;refGene_tx=XM_019810730.2;refGene_gn=PDGFRB;refGene_tx=XM_023260465.1
A1 199071542 199071642
refGene_gn=PDGFRB;refGene_tx=XM_011286103.3;refGene_gn=PDGFRB;refGene_tx=XM_019810730.2;refGene_gn=PDGFRB;refGene_tx=XM_023260465.1
A1 199072161 199072267
refGene_gn=PDGFRB;refGene_tx=XM_011286103.3;refGene_gn=PDGFRB;refGene_tx=XM_019810730.2;refGene_gn=PDGFRB;refGene_tx=XM_023260465.1
A1 199073105 199073338
refGene_gn=PDGFRB;refGene_tx=XM_011286103.3;refGene_gn=PDGFRB;refGene_tx=XM_019810730.2;refGene_gn=PDGFRB;refGene_tx=XM_023260465.1
A1 199074862 199075034
refGene_gn=PDGFRB;refGene_tx=XM_011286103.3;refGene_gn=PDGFRB;refGene_tx=XM_019810730.2;refGene_gn=PDGFRB;refGene_tx=XM_023260465.1
A1 199097079 199097128
refGene_gn=CSF1R;refGene_tx=NM_001009231.1
A1 199100989 199101247
refGene_gn=CSF1R;refGene_tx=NM_001009231.1

A1 199101685 199101970
refGene_gn=CSF1R;refGene_tx=NM_001009231.1
A1 199103618 199103755
refGene_gn=CSF1R;refGene_tx=NM_001009231.1
A1 199104405 199104565
refGene_gn=CSF1R;refGene_tx=NM_001009231.1
A1 199108345 199108532
refGene_gn=CSF1R;refGene_tx=NM_001009231.1
A1 199110852 199110968
refGene_gn=CSF1R;refGene_tx=NM_001009231.1
A1 199111121 199111242
refGene_gn=CSF1R;refGene_tx=NM_001009231.1
A1 199111335 199111523
refGene_gn=CSF1R;refGene_tx=NM_001009231.1
A1 199113257 199113373
refGene_gn=CSF1R;refGene_tx=NM_001009231.1
A1 199118328 199118455
refGene_gn=CSF1R;refGene_tx=NM_001009231.1
A1 199118576 199118681
refGene_gn=CSF1R;refGene_tx=NM_001009231.1
A1 199119170 199119281
refGene_gn=CSF1R;refGene_tx=NM_001009231.1
A1 199119874 199120037
refGene_gn=CSF1R;refGene_tx=NM_001009231.1
A1 199122115 199122207
refGene_gn=CSF1R;refGene_tx=NM_001009231.1
A1 199122305 199122403
refGene_gn=CSF1R;refGene_tx=NM_001009231.1
A1 199123334 199123457
refGene_gn=CSF1R;refGene_tx=NM_001009231.1
A1 199123537 199123649
refGene_gn=CSF1R;refGene_tx=NM_001009231.1
A1 199124191 199124291
refGene_gn=CSF1R;refGene_tx=NM_001009231.1
A1 199125156 199125265
refGene_gn=CSF1R;refGene_tx=NM_001009231.1
A1 199125363 199125567
refGene_gn=CSF1R;refGene_tx=NM_001009231.1
A1 199127466 199127934
refGene_gn=HMGXB3;refGene_tx=XM_006927997.4;refGene_gn=HMGXB3;refGene_tx=XM_019810700.2;refGene_gn=HMGXB3;refGene_tx=XM_019810706.2;refGene_gn=HMGXB3;refGene_tx=XM_023260488.1;refGene_gn=HMGXB3;refGene_tx=XM_023260490.1;refGene_gn=HMGXB3;refGene_tx=XM_023260494.1;refGene_gn=HMGXB3;refGene_tx=XM_023260501.1
A1 199128944 199129154
refGene_gn=HMGXB3;refGene_tx=XM_006927997.4;refGene_gn=HMGXB3;refGene_tx=XM_019810700.2;refGene_gn=HMGXB3;refGene_tx=XM_019810706.2;refGene_gn=HMGXB3;refGene_tx=XM_023260488.1;refGene_gn=HMGXB3;refGene_tx=XM_023260490.1;refGene_gn=HMGXB3;refGene_tx=XM_023260501.1
A1 199130762 199130879

refGene_gn=HMGXB3;refGene_tx=XM_006927997.4;refGene_gn=HMGXB3;refGene_tx=XM_019810700.2;refGene_gn=HMGXB3;refGene_tx=XM_019810706.2;refGene_gn=HMGXB3;refGene_tx=XM_023260488.1;refGene_gn=HMGXB3;refGene_tx=XM_023260490.1;refGene_gn=HMGXB3;refGene_tx=XM_023260494.1;refGene_gn=HMGXB3;refGene_tx=XM_023260501.1
A1 199131698 199131832
refGene_gn=HMGXB3;refGene_tx=XM_006927997.4;refGene_gn=HMGXB3;refGene_tx=XM_019810700.2;refGene_gn=HMGXB3;refGene_tx=XM_019810706.2;refGene_gn=HMGXB3;refGene_tx=XM_023260488.1;refGene_gn=HMGXB3;refGene_tx=XM_023260490.1;refGene_gn=HMGXB3;refGene_tx=XM_023260494.1;refGene_gn=HMGXB3;refGene_tx=XM_023260501.1
A1 199132671 199132891
refGene_gn=HMGXB3;refGene_tx=XM_006927997.4;refGene_gn=HMGXB3;refGene_tx=XM_019810700.2;refGene_gn=HMGXB3;refGene_tx=XM_019810706.2;refGene_gn=HMGXB3;refGene_tx=XM_023260488.1;refGene_gn=HMGXB3;refGene_tx=XM_023260490.1;refGene_gn=HMGXB3;refGene_tx=XM_023260494.1;refGene_gn=HMGXB3;refGene_tx=XM_023260501.1
A1 199136149 199136334
refGene_gn=HMGXB3;refGene_tx=XM_006927997.4;refGene_gn=HMGXB3;refGene_tx=XM_019810700.2;refGene_gn=HMGXB3;refGene_tx=XM_019810706.2;refGene_gn=HMGXB3;refGene_tx=XM_023260488.1;refGene_gn=HMGXB3;refGene_tx=XM_023260490.1;refGene_gn=HMGXB3;refGene_tx=XM_023260494.1;refGene_gn=HMGXB3;refGene_tx=XM_023260501.1
A1 199137240 199137372
refGene_gn=HMGXB3;refGene_tx=XM_006927997.4;refGene_gn=HMGXB3;refGene_tx=XM_019810700.2;refGene_gn=HMGXB3;refGene_tx=XM_019810706.2;refGene_gn=HMGXB3;refGene_tx=XM_023260488.1;refGene_gn=HMGXB3;refGene_tx=XM_023260490.1;refGene_gn=HMGXB3;refGene_tx=XM_023260494.1;refGene_gn=HMGXB3;refGene_tx=XM_023260501.1
A1 199141740 199141868
refGene_gn=HMGXB3;refGene_tx=XM_006927997.4;refGene_gn=HMGXB3;refGene_tx=XM_019810700.2;refGene_gn=HMGXB3;refGene_tx=XM_019810706.2;refGene_gn=HMGXB3;refGene_tx=XM_023260488.1;refGene_gn=HMGXB3;refGene_tx=XM_023260490.1;refGene_gn=HMGXB3;refGene_tx=XM_023260494.1;refGene_gn=HMGXB3;refGene_tx=XM_023260501.1
A1 199142384 199142686
refGene_gn=HMGXB3;refGene_tx=XM_006927997.4;refGene_gn=HMGXB3;refGene_tx=XM_019810700.2;refGene_gn=HMGXB3;refGene_tx=XM_019810706.2;refGene_gn=HMGXB3;refGene_tx=XM_023260488.1;refGene_gn=HMGXB3;refGene_tx=XM_023260490.1;refGene_gn=HMGXB3;refGene_tx=XM_023260494.1;refGene_gn=HMGXB3;refGene_tx=XM_023260501.1
A1 199146495 199146645
refGene_gn=HMGXB3;refGene_tx=XM_006927997.4;refGene_gn=HMGXB3;refGene_tx=XM_019810700.2;refGene_gn=HMGXB3;refGene_tx=XM_019810706.2;refGene_gn=HMGXB3;refGene_tx=XM_023260488.1;refGene_gn=HMGXB3;refGene_tx=XM_023260490.1;refGene_gn=HMGXB3;refGene_tx=XM_023260494.1;refGene_gn=HMGXB3;refGene_tx=XM_023260501.1
A1 199149240 199149339
refGene_gn=HMGXB3;refGene_tx=XM_006927997.4;refGene_gn=HMGXB3;refGene_tx=XM_019810700.2;refGene_gn=HMGXB3;refGene_tx=XM_019810706.2;refGene_

gn=HMGXB3;refGene_tx=XM_023260488.1;refGene_gn=HMGXB3;refGene_tx=XM_02
3260490.1;refGene_gn=HMGXB3;refGene_tx=XM_023260494.1;refGene_gn=HMGXB
3;refGene_tx=XM_023260501.1
A1 199151540 199151638
refGene_gn=HMGXB3;refGene_tx=XM_006927997.4;refGene_gn=HMGXB3;refGene_
tx=XM_019810700.2;refGene_gn=HMGXB3;refGene_tx=XM_019810706.2;refGene_
gn=HMGXB3;refGene_tx=XM_023260488.1;refGene_gn=HMGXB3;refGene_tx=XM_02
3260490.1;refGene_gn=HMGXB3;refGene_tx=XM_023260494.1;refGene_gn=HMGXB
3;refGene_tx=XM_023260501.1
A1 199151778 199151954
refGene_gn=HMGXB3;refGene_tx=XM_006927997.4;refGene_gn=HMGXB3;refGene_
tx=XM_019810700.2;refGene_gn=HMGXB3;refGene_tx=XM_019810706.2;refGene_
gn=HMGXB3;refGene_tx=XM_023260488.1;refGene_gn=HMGXB3;refGene_tx=XM_02
3260490.1;refGene_gn=HMGXB3;refGene_tx=XM_023260494.1;refGene_gn=HMGXB
3;refGene_tx=XM_023260501.1
A1 199153566 199153985
refGene_gn=HMGXB3;refGene_tx=XM_006927997.4;refGene_gn=HMGXB3;refGene_
tx=XM_019810700.2;refGene_gn=HMGXB3;refGene_tx=XM_019810706.2;refGene_
gn=HMGXB3;refGene_tx=XM_023260488.1;refGene_gn=HMGXB3;refGene_tx=XM_02
3260490.1;refGene_gn=HMGXB3;refGene_tx=XM_023260494.1;refGene_gn=HMGXB
3;refGene_tx=XM_023260501.1
A1 199158368 199158500
refGene_gn=HMGXB3;refGene_tx=XM_006927997.4;refGene_gn=HMGXB3;refGene_
tx=XM_019810700.2;refGene_gn=HMGXB3;refGene_tx=XM_019810706.2;refGene_
gn=HMGXB3;refGene_tx=XM_023260488.1;refGene_gn=HMGXB3;refGene_tx=XM_02
3260494.1;refGene_gn=HMGXB3;refGene_tx=XM_023260501.1
A1 199164424 199164523
refGene_gn=HMGXB3;refGene_tx=XM_006927997.4;refGene_gn=HMGXB3;refGene_
tx=XM_019810700.2;refGene_gn=HMGXB3;refGene_tx=XM_019810706.2;refGene_
gn=HMGXB3;refGene_tx=XM_023260488.1;refGene_gn=HMGXB3;refGene_tx=XM_02
3260490.1;refGene_gn=HMGXB3;refGene_tx=XM_023260494.1
A1 199165568 199166066
refGene_gn=HMGXB3;refGene_tx=XM_006927997.4;refGene_gn=HMGXB3;refGene_
tx=XM_019810700.2;refGene_gn=HMGXB3;refGene_tx=XM_019810706.2;refGene_
gn=HMGXB3;refGene_tx=XM_023260488.1;refGene_gn=HMGXB3;refGene_tx=XM_02
3260490.1;refGene_gn=HMGXB3;refGene_tx=XM_023260494.1
A1 199169612 199169787
refGene_gn=HMGXB3;refGene_tx=XM_006927997.4;refGene_gn=HMGXB3;refGene_
tx=XM_019810700.2;refGene_gn=HMGXB3;refGene_tx=XM_019810706.2;refGene_
gn=HMGXB3;refGene_tx=XM_023260488.1;refGene_gn=HMGXB3;refGene_tx=XM_02
3260490.1;refGene_gn=HMGXB3;refGene_tx=XM_023260494.1;refGene_gn=HMGXB
3;refGene_tx=XM_023260501.1
A1 199170662 199170799
refGene_gn=HMGXB3;refGene_tx=XM_006927997.4;refGene_gn=HMGXB3;refGene_
tx=XM_019810700.2;refGene_gn=HMGXB3;refGene_tx=XM_019810706.2;refGene_
gn=HMGXB3;refGene_tx=XM_023260488.1;refGene_gn=HMGXB3;refGene_tx=XM_02
3260490.1;refGene_gn=HMGXB3;refGene_tx=XM_023260494.1;refGene_gn=HMGXB
3;refGene_tx=XM_023260501.1
A1 199177378 199178953
refGene_gn=TIGD6;refGene_tx=XM_003981401.5

A1 199190908 199192429
refGene_gn=SLC26A2;refGene_tx=XM_011286115.3;refGene_gn=SLC26A2;refGene_tx=XM_019810737.2
A1 199193913 199194618
refGene_gn=SLC26A2;refGene_tx=XM_011286115.3;refGene_gn=SLC26A2;refGene_tx=XM_019810737.2
A1 199217747 199218221
refGene_gn=PDE6A;refGene_tx=XM_023260520.1;refGene_gn=PDE6A;refGene_tx=XM_023260521.1;refGene_gn=PDE6A;refGene_tx=XM_023260522.1
A1 199226054 199226207
refGene_gn=PDE6A;refGene_tx=XM_023260520.1;refGene_gn=PDE6A;refGene_tx=XM_023260521.1;refGene_gn=PDE6A;refGene_tx=XM_023260522.1
A1 199226811 199226901
refGene_gn=PDE6A;refGene_tx=XM_023260520.1;refGene_gn=PDE6A;refGene_tx=XM_023260521.1;refGene_gn=PDE6A;refGene_tx=XM_023260522.1
A1 199227508 199227649
refGene_gn=PDE6A;refGene_tx=XM_023260520.1;refGene_gn=PDE6A;refGene_tx=XM_023260521.1;refGene_gn=PDE6A;refGene_tx=XM_023260522.1
A1 199232710 199232785
refGene_gn=PDE6A;refGene_tx=XM_023260520.1;refGene_gn=PDE6A;refGene_tx=XM_023260521.1;refGene_gn=PDE6A;refGene_tx=XM_023260522.1
A1 199239031 199239096
refGene_gn=PDE6A;refGene_tx=XM_023260520.1;refGene_gn=PDE6A;refGene_tx=XM_023260521.1;refGene_gn=PDE6A;refGene_tx=XM_023260522.1
A1 199247275 199247342
refGene_gn=PDE6A;refGene_tx=XM_023260520.1;refGene_gn=PDE6A;refGene_tx=XM_023260521.1;refGene_gn=PDE6A;refGene_tx=XM_023260522.1
A1 199249996 199250044
refGene_gn=PDE6A;refGene_tx=XM_023260520.1;refGene_gn=PDE6A;refGene_tx=XM_023260521.1;refGene_gn=PDE6A;refGene_tx=XM_023260522.1
A1 199253213 199253363
refGene_gn=PDE6A;refGene_tx=XM_023260520.1;refGene_gn=PDE6A;refGene_tx=XM_023260521.1;refGene_gn=PDE6A;refGene_tx=XM_023260522.1
A1 199253862 199254006
refGene_gn=PDE6A;refGene_tx=XM_023260520.1;refGene_gn=PDE6A;refGene_tx=XM_023260521.1;refGene_gn=PDE6A;refGene_tx=XM_023260522.1
A1 199255638 199255704
refGene_gn=PDE6A;refGene_tx=XM_023260520.1;refGene_gn=PDE6A;refGene_tx=XM_023260521.1;refGene_gn=PDE6A;refGene_tx=XM_023260522.1
A1 199255917 199256064
refGene_gn=PDE6A;refGene_tx=XM_023260520.1;refGene_gn=PDE6A;refGene_tx=XM_023260521.1;refGene_gn=PDE6A;refGene_tx=XM_023260522.1
A1 199257294 199257402
refGene_gn=PDE6A;refGene_tx=XM_023260520.1;refGene_gn=PDE6A;refGene_tx=XM_023260521.1;refGene_gn=PDE6A;refGene_tx=XM_023260522.1
A1 199259069 199259179
refGene_gn=PDE6A;refGene_tx=XM_023260520.1;refGene_gn=PDE6A;refGene_tx=XM_023260521.1;refGene_gn=PDE6A;refGene_tx=XM_023260522.1
A1 199260658 199260746
refGene_gn=PDE6A;refGene_tx=XM_023260520.1;refGene_gn=PDE6A;refGene_tx

=XM_023260521.1;refGene_gn=PDE6A;refGene_tx=XM_023260522.1
A1 199260929 199261030
refGene_gn=PDE6A;refGene_tx=XM_023260520.1;refGene_gn=PDE6A;refGene_tx
=XM_023260521.1;refGene_gn=PDE6A;refGene_tx=XM_023260522.1
A1 199261452 199261560
refGene_gn=PDE6A;refGene_tx=XM_023260520.1;refGene_gn=PDE6A;refGene_tx
=XM_023260521.1;refGene_gn=PDE6A;refGene_tx=XM_023260522.1
A1 199261919 199262137
refGene_gn=PDE6A;refGene_tx=XM_023260521.1;refGene_gn=PDE6A;refGene_tx
=XM_023260522.1
A1 199275335 199275399
refGene_gn=PDE6A;refGene_tx=XM_023260520.1
A1 199275685 199275760
refGene_gn=PDE6A;refGene_tx=XM_023260520.1
A1 199277562 199277646
refGene_gn=PDE6A;refGene_tx=XM_023260520.1
A1 199279222 199279370
refGene_gn=PDE6A;refGene_tx=XM_023260520.1
A1 199281052 199281132
refGene_gn=PDE6A;refGene_tx=XM_023260520.1
A1 199292030 199292131
refGene_gn=PPARGC1B;refGene_tx=XM_011286128.3
A1 199293106 199293261
refGene_gn=PPARGC1B;refGene_tx=XM_011286128.3
A1 199296269 199296391
refGene_gn=PPARGC1B;refGene_tx=XM_011286128.3
A1 199298726 199298802
refGene_gn=PPARGC1B;refGene_tx=XM_011286128.3
A1 199301922 199302721
refGene_gn=PPARGC1B;refGene_tx=XM_011286128.3
A1 199303879 199303944
refGene_gn=PPARGC1B;refGene_tx=XM_011286128.3
A1 199304539 199304576
refGene_gn=PPARGC1B;refGene_tx=XM_011286128.3
A1 199305764 199306020
refGene_gn=PPARGC1B;refGene_tx=XM_011286128.3
A1 199306021 199306492
refGene_gn=PPARGC1B;refGene_tx=XM_011286128.3
A1 199306493 199306922
refGene_gn=PPARGC1B;refGene_tx=XM_011286128.3
A1 199308132 199308249
refGene_gn=PPARGC1B;refGene_tx=XM_011286128.3
A1 199311563 199311785
refGene_gn=PPARGC1B;refGene_tx=XM_011286128.3
A1 199317396 199317570
refGene_gn=PPARGC1B;refGene_tx=XM_011286128.3
A1 199396946 199397024
refGene_gn=PPARGC1B;refGene_tx=XM_011286128.3
A1 199473579 199473789
refGene_gn=ARHGEF37;refGene_tx=XM_006927999.2

A1 199476270 199476428
refGene_gn=ARHGEF37;refGene_tx=XM_006927999.2
A1 199477935 199478131
refGene_gn=ARHGEF37;refGene_tx=XM_006927999.2
A1 199480431 199480560
refGene_gn=ARHGEF37;refGene_tx=XM_006927999.2
A1 199482979 199483309
refGene_gn=ARHGEF37;refGene_tx=XM_006927999.2
A1 199484117 199484228
refGene_gn=ARHGEF37;refGene_tx=XM_006927999.2
A1 199486450 199486555
refGene_gn=ARHGEF37;refGene_tx=XM_006927999.2
A1 199487257 199487388
refGene_gn=ARHGEF37;refGene_tx=XM_006927999.2
A1 199488701 199488901
refGene_gn=ARHGEF37;refGene_tx=XM_006927999.2
A1 199498347 199498495
refGene_gn=ARHGEF37;refGene_tx=XM_006927999.2
A1 199507785 199507909
refGene_gn=ARHGEF37;refGene_tx=XM_006927999.2
A1 199509404 199509590
refGene_gn=ARHGEF37;refGene_tx=XM_006927999.2
A1 199562086 199562209
refGene_gn=CSNK1A1;refGene_tx=XM_003981404.5;refGene_gn=CSNK1A1;refGen
e_tx=XM_003981405.5;refGene_gn=CSNK1A1;refGene_tx=XM_006928000.4;refGe
ne_gn=CSNK1A1;refGene_tx=XM_006928002.4
A1 199562825 199562932
refGene_gn=CSNK1A1;refGene_tx=XM_003981404.5;refGene_gn=CSNK1A1;refGen
e_tx=XM_003981405.5;refGene_gn=CSNK1A1;refGene_tx=XM_006928000.4;refGe
ne_gn=CSNK1A1;refGene_tx=XM_006928002.4
A1 199581489 199581616
refGene_gn=CSNK1A1;refGene_tx=XM_003981404.5;refGene_gn=CSNK1A1;refGen
e_tx=XM_003981405.5;refGene_gn=CSNK1A1;refGene_tx=XM_006928000.4;refGe
ne_gn=CSNK1A1;refGene_tx=XM_006928002.4
A1 199585808 199585907
refGene_gn=CSNK1A1;refGene_tx=XM_003981404.5;refGene_gn=CSNK1A1;refGen
e_tx=XM_003981405.5;refGene_gn=CSNK1A1;refGene_tx=XM_006928000.4;refGe
ne_gn=CSNK1A1;refGene_tx=XM_006928002.4
A1 199588304 199588388
refGene_gn=CSNK1A1;refGene_tx=XM_003981405.5;refGene_gn=CSNK1A1;refGen
e_tx=XM_006928000.4;refGene_gn=CSNK1A1;refGene_tx=XM_006928002.4
A1 199592645 199592785
refGene_gn=CSNK1A1;refGene_tx=XM_003981404.5;refGene_gn=CSNK1A1;refGen
e_tx=XM_003981405.5;refGene_gn=CSNK1A1;refGene_tx=XM_006928000.4;refGe
ne_gn=CSNK1A1;refGene_tx=XM_006928002.4
A1 199594142 199594221
refGene_gn=CSNK1A1;refGene_tx=XM_003981404.5;refGene_gn=CSNK1A1;refGen
e_tx=XM_003981405.5;refGene_gn=CSNK1A1;refGene_tx=XM_006928000.4;refGe
ne_gn=CSNK1A1;refGene_tx=XM_006928002.4
A1 199595220 199595295

refGene_gn=CSNK1A1;refGene_tx=XM_003981404.5;refGene_gn=CSNK1A1;refGene_tx=XM_003981405.5;refGene_gn=CSNK1A1;refGene_tx=XM_006928000.4;refGene_gn=CSNK1A1;refGene_tx=XM_006928002.4
A1 199598367 199598474
refGene_gn=CSNK1A1;refGene_tx=XM_003981404.5;refGene_gn=CSNK1A1;refGene_tx=XM_003981405.5;refGene_gn=CSNK1A1;refGene_tx=XM_006928000.4;refGene_gn=CSNK1A1;refGene_tx=XM_006928002.4
A1 199599466 199599620
refGene_gn=CSNK1A1;refGene_tx=XM_003981404.5;refGene_gn=CSNK1A1;refGene_tx=XM_003981405.5;refGene_gn=CSNK1A1;refGene_tx=XM_006928000.4;refGene_gn=CSNK1A1;refGene_tx=XM_006928002.4
A1 199606753 199606761
refGene_gn=CSNK1A1;refGene_tx=XM_003981404.5;refGene_gn=CSNK1A1;refGene_tx=XM_003981405.5;refGene_gn=CSNK1A1;refGene_tx=XM_006928002.4
A1 199687740 199687806
refGene_gn=IL17B;refGene_tx=XM_019810830.2
A1 199706820 199706841
refGene_gn=IL17B;refGene_tx=XM_003981407.4
A1 199709015 199709305
refGene_gn=IL17B;refGene_tx=XM_003981407.4;refGene_gn=IL17B;refGene_tx=XM_019810830.2
A1 199711091 199711323
refGene_gn=IL17B;refGene_tx=XM_003981407.4;refGene_gn=IL17B;refGene_tx=XM_019810830.2
A1 199716015 199716677
refGene_gn=PCYOX1L;refGene_tx=XM_006928072.3
A1 199717161 199717302
refGene_gn=PCYOX1L;refGene_tx=XM_006928072.3
A1 199719362 199719574
refGene_gn=PCYOX1L;refGene_tx=XM_006928072.3
A1 199721156 199721331
refGene_gn=PCYOX1L;refGene_tx=XM_006928072.3
A1 199723011 199723218
refGene_gn=PCYOX1L;refGene_tx=XM_006928072.3
A1 199726887 199726975
refGene_gn=PCYOX1L;refGene_tx=XM_006928072.3
A1 199733579 199733944
refGene_gn=GRPEL2;refGene_tx=XM_003981408.5
A1 199737725 199737807
refGene_gn=GRPEL2;refGene_tx=XM_003981408.5
A1 199738804 199738958
refGene_gn=GRPEL2;refGene_tx=XM_003981408.5
A1 199741570 199741647
refGene_gn=GRPEL2;refGene_tx=XM_003981408.5
A1 199746101 199746125
refGene_gn=AFAP1L1;refGene_tx=XM_023260562.1
A1 199748163 199748292
refGene_gn=AFAP1L1;refGene_tx=XM_023260562.1
A1 199750730 199750909
refGene_gn=AFAP1L1;refGene_tx=XM_023260562.1

A1 199753366 199753531
refGene_gn=AFAP1L1;refGene_tx=XM_023260562.1
A1 199757127 199757239
refGene_gn=AFAP1L1;refGene_tx=XM_023260562.1
A1 199759059 199759132
refGene_gn=AFAP1L1;refGene_tx=XM_023260562.1
A1 199759754 199759900
refGene_gn=AFAP1L1;refGene_tx=XM_023260562.1
A1 199762790 199763002
refGene_gn=AFAP1L1;refGene_tx=XM_023260562.1
A1 199764458 199764611
refGene_gn=AFAP1L1;refGene_tx=XM_023260562.1
A1 199764853 199764947
refGene_gn=AFAP1L1;refGene_tx=XM_023260562.1
A1 199767320 199767413
refGene_gn=AFAP1L1;refGene_tx=XM_023260562.1
A1 199769363 199769543
refGene_gn=AFAP1L1;refGene_tx=XM_023260562.1
A1 199771489 199771701
refGene_gn=AFAP1L1;refGene_tx=XM_023260562.1
A1 199772371 199772497
refGene_gn=AFAP1L1;refGene_tx=XM_023260562.1
A1 199776452 199776561
refGene_gn=AFAP1L1;refGene_tx=XM_023260562.1
A1 199778134 199778232
refGene_gn=AFAP1L1;refGene_tx=XM_023260562.1
A1 199779018 199779102
refGene_gn=AFAP1L1;refGene_tx=XM_023260562.1
A1 199779864 199779993
refGene_gn=AFAP1L1;refGene_tx=XM_023260562.1
A1 199807537 199807553
refGene_gn=AFAP1L1;refGene_tx=XM_023260562.1
A1 199821195 199821309
refGene_gn=ABLIM3;refGene_tx=XM_006928009.3;refGene_gn=ABLIM3;refGene_tx=XM_006928010.3;refGene_gn=ABLIM3;refGene_tx=XM_011286180.2;refGene_gn=ABLIM3;refGene_tx=XM_019810867.1;refGene_gn=ABLIM3;refGene_tx=XM_019810880.1;refGene_gn=ABLIM3;refGene_tx=XM_019810883.1;refGene_gn=ABLIM3;refGene_tx=XM_019810886.1
A1 199825703 199825784
refGene_gn=ABLIM3;refGene_tx=XM_006928009.3;refGene_gn=ABLIM3;refGene_tx=XM_006928010.3;refGene_gn=ABLIM3;refGene_tx=XM_011286180.2;refGene_gn=ABLIM3;refGene_tx=XM_019810867.1;refGene_gn=ABLIM3;refGene_tx=XM_019810880.1;refGene_gn=ABLIM3;refGene_tx=XM_019810883.1;refGene_gn=ABLIM3;refGene_tx=XM_019810886.1
A1 199826326 199826334
refGene_gn=ABLIM3;refGene_tx=XM_006928009.3;refGene_gn=ABLIM3;refGene_tx=XM_006928010.3;refGene_gn=ABLIM3;refGene_tx=XM_011286180.2;refGene_gn=ABLIM3;refGene_tx=XM_019810867.1;refGene_gn=ABLIM3;refGene_tx=XM_019810880.1;refGene_gn=ABLIM3;refGene_tx=XM_019810883.1;refGene_gn=ABLIM3;refGene_tx=XM_019810886.1

A1	199827056	199827117
refGene_gn=ABLIM3;refGene_tx=XM_006928009.3;refGene_gn=ABLIM3;refGene_tx=XM_006928010.3;refGene_gn=ABLIM3;refGene_tx=XM_011286180.2;refGene_gn=ABLIM3;refGene_tx=XM_019810867.1;refGene_gn=ABLIM3;refGene_tx=XM_019810880.1;refGene_gn=ABLIM3;refGene_tx=XM_019810883.1;refGene_gn=ABLIM3;refGene_tx=XM_019810886.1
A1	199827730	199827789
refGene_gn=ABLIM3;refGene_tx=XM_006928009.3;refGene_gn=ABLIM3;refGene_tx=XM_006928010.3;refGene_gn=ABLIM3;refGene_tx=XM_011286180.2;refGene_gn=ABLIM3;refGene_tx=XM_019810867.1;refGene_gn=ABLIM3;refGene_tx=XM_019810880.1;refGene_gn=ABLIM3;refGene_tx=XM_019810883.1;refGene_gn=ABLIM3;refGene_tx=XM_019810886.1
A1	199828370	199828400
refGene_gn=ABLIM3;refGene_tx=XM_006928009.3;refGene_gn=ABLIM3;refGene_tx=XM_006928010.3;refGene_gn=ABLIM3;refGene_tx=XM_011286180.2;refGene_gn=ABLIM3;refGene_tx=XM_019810867.1;refGene_gn=ABLIM3;refGene_tx=XM_019810880.1;refGene_gn=ABLIM3;refGene_tx=XM_019810883.1;refGene_gn=ABLIM3;refGene_tx=XM_019810886.1
A1	199829667	199829815
refGene_gn=ABLIM3;refGene_tx=XM_006928009.3;refGene_gn=ABLIM3;refGene_tx=XM_006928010.3;refGene_gn=ABLIM3;refGene_tx=XM_011286180.2;refGene_gn=ABLIM3;refGene_tx=XM_019810867.1;refGene_gn=ABLIM3;refGene_tx=XM_019810880.1;refGene_gn=ABLIM3;refGene_tx=XM_019810883.1;refGene_gn=ABLIM3;refGene_tx=XM_019810886.1
A1	199830942	199831007
refGene_gn=ABLIM3;refGene_tx=XM_006928009.3;refGene_gn=ABLIM3;refGene_tx=XM_006928010.3;refGene_gn=ABLIM3;refGene_tx=XM_011286180.2;refGene_gn=ABLIM3;refGene_tx=XM_019810867.1;refGene_gn=ABLIM3;refGene_tx=XM_019810880.1;refGene_gn=ABLIM3;refGene_tx=XM_019810883.1;refGene_gn=ABLIM3;refGene_tx=XM_019810886.1
A1	199832493	199832628
refGene_gn=ABLIM3;refGene_tx=XM_006928009.3;refGene_gn=ABLIM3;refGene_tx=XM_006928010.3;refGene_gn=ABLIM3;refGene_tx=XM_011286180.2;refGene_gn=ABLIM3;refGene_tx=XM_019810867.1;refGene_gn=ABLIM3;refGene_tx=XM_019810880.1;refGene_gn=ABLIM3;refGene_tx=XM_019810883.1;refGene_gn=ABLIM3;refGene_tx=XM_019810886.1
A1	199834639	199834687
refGene_gn=ABLIM3;refGene_tx=XM_006928009.3;refGene_gn=ABLIM3;refGene_tx=XM_006928010.3;refGene_gn=ABLIM3;refGene_tx=XM_019810867.1;refGene_gn=ABLIM3;refGene_tx=XM_019810880.1;refGene_gn=ABLIM3;refGene_tx=XM_019810886.1
A1	199836022	199836121
refGene_gn=ABLIM3;refGene_tx=XM_006928010.3;refGene_gn=ABLIM3;refGene_tx=XM_019810867.1;refGene_gn=ABLIM3;refGene_tx=XM_019810886.1
A1	199836812	199836942
refGene_gn=ABLIM3;refGene_tx=XM_006928009.3;refGene_gn=ABLIM3;refGene_tx=XM_006928010.3;refGene_gn=ABLIM3;refGene_tx=XM_011286180.2;refGene_gn=ABLIM3;refGene_tx=XM_019810867.1;refGene_gn=ABLIM3;refGene_tx=XM_019810880.1;refGene_gn=ABLIM3;refGene_tx=XM_019810883.1;refGene_gn=ABLIM3;refGene_tx=XM_019810886.1

A1 199837417 199837447
refGene_gn=ABLIM3;refGene_tx=XM_019810867.1;refGene_gn=ABLIM3;refGene_tx=XM_019810880.1;refGene_gn=ABLIM3;refGene_tx=XM_019810883.1
A1 199838976 199839132
refGene_gn=ABLIM3;refGene_tx=XM_006928010.3;refGene_gn=ABLIM3;refGene_tx=XM_019810867.1;refGene_gn=ABLIM3;refGene_tx=XM_019810880.1;refGene_gn=ABLIM3;refGene_tx=XM_019810883.1
A1 199843224 199843296
refGene_gn=ABLIM3;refGene_tx=XM_006928009.3;refGene_gn=ABLIM3;refGene_tx=XM_006928010.3;refGene_gn=ABLIM3;refGene_tx=XM_011286180.2;refGene_gn=ABLIM3;refGene_tx=XM_019810867.1;refGene_gn=ABLIM3;refGene_tx=XM_019810880.1;refGene_gn=ABLIM3;refGene_tx=XM_019810883.1;refGene_gn=ABLIM3;refGene_tx=XM_019810886.1
A1 199845565 199845624
refGene_gn=ABLIM3;refGene_tx=XM_006928009.3;refGene_gn=ABLIM3;refGene_tx=XM_006928010.3;refGene_gn=ABLIM3;refGene_tx=XM_011286180.2;refGene_gn=ABLIM3;refGene_tx=XM_019810867.1;refGene_gn=ABLIM3;refGene_tx=XM_019810880.1;refGene_gn=ABLIM3;refGene_tx=XM_019810883.1;refGene_gn=ABLIM3;refGene_tx=XM_019810886.1
A1 199857577 199857665
refGene_gn=ABLIM3;refGene_tx=XM_006928009.3;refGene_gn=ABLIM3;refGene_tx=XM_006928010.3;refGene_gn=ABLIM3;refGene_tx=XM_011286180.2;refGene_gn=ABLIM3;refGene_tx=XM_019810867.1;refGene_gn=ABLIM3;refGene_tx=XM_019810880.1;refGene_gn=ABLIM3;refGene_tx=XM_019810883.1;refGene_gn=ABLIM3;refGene_tx=XM_019810886.1
A1 199865646 199865740
refGene_gn=ABLIM3;refGene_tx=XM_006928009.3;refGene_gn=ABLIM3;refGene_tx=XM_006928010.3;refGene_gn=ABLIM3;refGene_tx=XM_011286180.2;refGene_gn=ABLIM3;refGene_tx=XM_019810867.1;refGene_gn=ABLIM3;refGene_tx=XM_019810880.1;refGene_gn=ABLIM3;refGene_tx=XM_019810883.1;refGene_gn=ABLIM3;refGene_tx=XM_019810886.1
A1 199869531 199869658
refGene_gn=ABLIM3;refGene_tx=XM_006928009.3;refGene_gn=ABLIM3;refGene_tx=XM_006928010.3;refGene_gn=ABLIM3;refGene_tx=XM_011286180.2;refGene_gn=ABLIM3;refGene_tx=XM_019810867.1;refGene_gn=ABLIM3;refGene_tx=XM_019810880.1;refGene_gn=ABLIM3;refGene_tx=XM_019810883.1;refGene_gn=ABLIM3;refGene_tx=XM_019810886.1
A1 199877593 199877706
refGene_gn=ABLIM3;refGene_tx=XM_006928009.3;refGene_gn=ABLIM3;refGene_tx=XM_006928010.3;refGene_gn=ABLIM3;refGene_tx=XM_011286180.2;refGene_gn=ABLIM3;refGene_tx=XM_019810867.1;refGene_gn=ABLIM3;refGene_tx=XM_019810880.1;refGene_gn=ABLIM3;refGene_tx=XM_019810883.1;refGene_gn=ABLIM3;refGene_tx=XM_019810886.1
A1 199879773 199879957
refGene_gn=ABLIM3;refGene_tx=XM_006928009.3;refGene_gn=ABLIM3;refGene_tx=XM_006928010.3;refGene_gn=ABLIM3;refGene_tx=XM_011286180.2;refGene_gn=ABLIM3;refGene_tx=XM_019810867.1;refGene_gn=ABLIM3;refGene_tx=XM_019810880.1;refGene_gn=ABLIM3;refGene_tx=XM_019810883.1;refGene_gn=ABLIM3;refGene_tx=XM_019810886.1
A1 199890528 199890666

refGene_gn=ABLIM3;refGene_tx=XM_006928009.3;refGene_gn=ABLIM3;refGene_tx=XM_006928010.3;refGene_gn=ABLIM3;refGene_tx=XM_011286180.2;refGene_gn=ABLIM3;refGene_tx=XM_019810867.1;refGene_gn=ABLIM3;refGene_tx=XM_019810880.1;refGene_gn=ABLIM3;refGene_tx=XM_019810883.1;refGene_gn=ABLIM3;refGene_tx=XM_019810886.1
A1 199928986 199928999
refGene_gn=ABLIM3;refGene_tx=XM_006928009.3;refGene_gn=ABLIM3;refGene_tx=XM_006928010.3;refGene_gn=ABLIM3;refGene_tx=XM_011286180.2;refGene_gn=ABLIM3;refGene_tx=XM_019810867.1;refGene_gn=ABLIM3;refGene_tx=XM_019810880.1;refGene_gn=ABLIM3;refGene_tx=XM_019810883.1;refGene_gn=ABLIM3;refGene_tx=XM_019810886.1
A1 199979556 199979608
refGene_gn=SH3TC2;refGene_tx=XM_003981410.5;refGene_gn=SH3TC2;refGene_tx=XM_006928011.4
A1 199987033 199987229
refGene_gn=SH3TC2;refGene_tx=XM_023260595.1;refGene_gn=SH3TC2;refGene_tx=XM_023260598.1
A1 199988385 199988484
refGene_gn=SH3TC2;refGene_tx=XM_003981410.5;refGene_gn=SH3TC2;refGene_tx=XM_006928011.4;refGene_gn=SH3TC2;refGene_tx=XM_019810918.2;refGene_gn=SH3TC2;refGene_tx=XM_023260595.1;refGene_gn=SH3TC2;refGene_tx=XM_023260598.1
A1 199990275 199990360
refGene_gn=SH3TC2;refGene_tx=XM_019810911.2
A1 199991478 199991606
refGene_gn=SH3TC2;refGene_tx=XM_003981410.5;refGene_gn=SH3TC2;refGene_tx=XM_006928011.4;refGene_gn=SH3TC2;refGene_tx=XM_019810911.2;refGene_gn=SH3TC2;refGene_tx=XM_019810918.2;refGene_gn=SH3TC2;refGene_tx=XM_023260595.1;refGene_gn=SH3TC2;refGene_tx=XM_023260598.1
A1 199994672 199994878
refGene_gn=SH3TC2;refGene_tx=XM_003981410.5;refGene_gn=SH3TC2;refGene_tx=XM_006928011.4;refGene_gn=SH3TC2;refGene_tx=XM_006928013.4;refGene_gn=SH3TC2;refGene_tx=XM_019810911.2;refGene_gn=SH3TC2;refGene_tx=XM_019810918.2;refGene_gn=SH3TC2;refGene_tx=XM_023260595.1;refGene_gn=SH3TC2;refGene_tx=XM_023260598.1
A1 199996340 199996484
refGene_gn=SH3TC2;refGene_tx=XM_003981410.5;refGene_gn=SH3TC2;refGene_tx=XM_006928011.4;refGene_gn=SH3TC2;refGene_tx=XM_006928013.4;refGene_gn=SH3TC2;refGene_tx=XM_019810911.2;refGene_gn=SH3TC2;refGene_tx=XM_019810918.2;refGene_gn=SH3TC2;refGene_tx=XM_023260595.1;refGene_gn=SH3TC2;refGene_tx=XM_023260598.1
A1 199997538 199997740
refGene_gn=SH3TC2;refGene_tx=XM_003981410.5;refGene_gn=SH3TC2;refGene_tx=XM_006928011.4;refGene_gn=SH3TC2;refGene_tx=XM_006928013.4;refGene_gn=SH3TC2;refGene_tx=XM_019810911.2;refGene_gn=SH3TC2;refGene_tx=XM_019810918.2;refGene_gn=SH3TC2;refGene_tx=XM_023260595.1;refGene_gn=SH3TC2;refGene_tx=XM_023260598.1
A1 199998518 199998592
refGene_gn=SH3TC2;refGene_tx=XM_003981410.5;refGene_gn=SH3TC2;refGene_tx=XM_006928011.4;refGene_gn=SH3TC2;refGene_tx=XM_006928013.4;refGene_

gn=SH3TC2;refGene_tx=XM_019810911.2;refGene_gn=SH3TC2;refGene_tx=XM_01
9810918.2;refGene_gn=SH3TC2;refGene_tx=XM_023260595.1;refGene_gn=SH3TC
2;refGene_tx=XM_023260598.1
A1 200000862 200001058
refGene_gn=SH3TC2;refGene_tx=XM_003981410.5;refGene_gn=SH3TC2;refGene_
tx=XM_006928011.4;refGene_gn=SH3TC2;refGene_tx=XM_006928013.4;refGene_
gn=SH3TC2;refGene_tx=XM_019810911.2;refGene_gn=SH3TC2;refGene_tx=XM_01
9810918.2;refGene_gn=SH3TC2;refGene_tx=XM_023260595.1;refGene_gn=SH3TC
2;refGene_tx=XM_023260598.1
A1 200009212 200009346
refGene_gn=SH3TC2;refGene_tx=XM_003981410.5;refGene_gn=SH3TC2;refGene_
tx=XM_006928011.4;refGene_gn=SH3TC2;refGene_tx=XM_006928013.4;refGene_
gn=SH3TC2;refGene_tx=XM_019810911.2;refGene_gn=SH3TC2;refGene_tx=XM_01
9810918.2;refGene_gn=SH3TC2;refGene_tx=XM_023260595.1;refGene_gn=SH3TC
2;refGene_tx=XM_023260598.1
A1 200011928 200011970
refGene_gn=SH3TC2;refGene_tx=XM_003981410.5;refGene_gn=SH3TC2;refGene_
tx=XM_006928011.4;refGene_gn=SH3TC2;refGene_tx=XM_006928013.4;refGene_
gn=SH3TC2;refGene_tx=XM_019810911.2;refGene_gn=SH3TC2;refGene_tx=XM_01
9810918.2;refGene_gn=SH3TC2;refGene_tx=XM_023260595.1;refGene_gn=SH3TC
2;refGene_tx=XM_023260598.1
A1 200012094 200013795
refGene_gn=SH3TC2;refGene_tx=XM_003981410.5;refGene_gn=SH3TC2;refGene_
tx=XM_006928011.4;refGene_gn=SH3TC2;refGene_tx=XM_006928013.4;refGene_
gn=SH3TC2;refGene_tx=XM_019810911.2;refGene_gn=SH3TC2;refGene_tx=XM_01
9810918.2;refGene_gn=SH3TC2;refGene_tx=XM_023260595.1;refGene_gn=SH3TC
2;refGene_tx=XM_023260598.1
A1 200013901 200014082
refGene_gn=SH3TC2;refGene_tx=XM_003981410.5;refGene_gn=SH3TC2;refGene_
tx=XM_006928011.4;refGene_gn=SH3TC2;refGene_tx=XM_006928013.4;refGene_
gn=SH3TC2;refGene_tx=XM_019810911.2;refGene_gn=SH3TC2;refGene_tx=XM_01
9810918.2;refGene_gn=SH3TC2;refGene_tx=XM_023260595.1;refGene_gn=SH3TC
2;refGene_tx=XM_023260598.1
A1 200021580 200021731
refGene_gn=SH3TC2;refGene_tx=XM_003981410.5;refGene_gn=SH3TC2;refGene_
tx=XM_006928011.4;refGene_gn=SH3TC2;refGene_tx=XM_006928013.4;refGene_
gn=SH3TC2;refGene_tx=XM_019810911.2;refGene_gn=SH3TC2;refGene_tx=XM_01
9810918.2;refGene_gn=SH3TC2;refGene_tx=XM_023260595.1;refGene_gn=SH3TC
2;refGene_tx=XM_023260598.1
A1 200023784 200023907
refGene_gn=SH3TC2;refGene_tx=XM_003981410.5;refGene_gn=SH3TC2;refGene_
tx=XM_006928011.4;refGene_gn=SH3TC2;refGene_tx=XM_006928013.4;refGene_
gn=SH3TC2;refGene_tx=XM_019810911.2;refGene_gn=SH3TC2;refGene_tx=XM_01
9810918.2;refGene_gn=SH3TC2;refGene_tx=XM_023260595.1;refGene_gn=SH3TC
2;refGene_tx=XM_023260598.1
A1 200025056 200025207
refGene_gn=SH3TC2;refGene_tx=XM_003981410.5;refGene_gn=SH3TC2;refGene_
tx=XM_006928011.4;refGene_gn=SH3TC2;refGene_tx=XM_006928013.4;refGene_
gn=SH3TC2;refGene_tx=XM_019810911.2;refGene_gn=SH3TC2;refGene_tx=XM_01
9810918.2;refGene_gn=SH3TC2;refGene_tx=XM_023260595.1;refGene_gn=SH3TC

2;refGene_tx=XM_023260598.1
A1 200027058 200027270
refGene_gn=SH3TC2;refGene_tx=XM_003981410.5;refGene_gn=SH3TC2;refGene_tx=XM_006928011.4;refGene_gn=SH3TC2;refGene_tx=XM_006928013.4;refGene_gn=SH3TC2;refGene_tx=XM_019810911.2;refGene_gn=SH3TC2;refGene_tx=XM_019810918.2;refGene_gn=SH3TC2;refGene_tx=XM_023260595.1;refGene_gn=SH3TC2;refGene_tx=XM_023260598.1
A1 200029144 200029336
refGene_gn=SH3TC2;refGene_tx=XM_003981410.5;refGene_gn=SH3TC2;refGene_tx=XM_006928011.4;refGene_gn=SH3TC2;refGene_tx=XM_006928013.4;refGene_gn=SH3TC2;refGene_tx=XM_019810911.2;refGene_gn=SH3TC2;refGene_tx=XM_019810918.2;refGene_gn=SH3TC2;refGene_tx=XM_023260595.1;refGene_gn=SH3TC2;refGene_tx=XM_023260598.1
A1 200182785 200184042
refGene_gn=ADRB2;refGene_tx=NM_001009247.1
A1 200341775 200341801
refGene_gn=HTR4;refGene_tx=XM_023260627.1;refGene_gn=HTR4;refGene_tx=XM_023260628.1;refGene_gn=HTR4;refGene_tx=XM_023260630.1
A1 200418872 200418998
refGene_gn=HTR4;refGene_tx=XM_023260627.1;refGene_gn=HTR4;refGene_tx=XM_023260628.1;refGene_gn=HTR4;refGene_tx=XM_023260630.1
A1 200420257 200420458
refGene_gn=HTR4;refGene_tx=XM_023260627.1;refGene_gn=HTR4;refGene_tx=XM_023260628.1;refGene_gn=HTR4;refGene_tx=XM_023260630.1
A1 200440148 200440302
refGene_gn=HTR4;refGene_tx=XM_023260627.1;refGene_gn=HTR4;refGene_tx=XM_023260628.1;refGene_gn=HTR4;refGene_tx=XM_023260630.1
A1 200441436 200441478
refGene_gn=HTR4;refGene_tx=XM_023260627.1
A1 200444194 200444763
refGene_gn=HTR4;refGene_tx=XM_023260627.1;refGene_gn=HTR4;refGene_tx=XM_023260628.1;refGene_gn=HTR4;refGene_tx=XM_023260630.1
A1 200468866 200468957
refGene_gn=HTR4;refGene_tx=XM_023260627.1;refGene_gn=HTR4;refGene_tx=XM_023260628.1
A1 200496594 200496682
refGene_gn=HTR4;refGene_tx=XM_023260630.1
A1 200505412 200505591
refGene_gn=FBXO38;refGene_tx=XM_003981411.4;refGene_gn=FBXO38;refGene_tx=XM_019810975.2;refGene_gn=FBXO38;refGene_tx=XM_019810982.2;refGene_gn=FBXO38;refGene_tx=XM_019810989.2;refGene_gn=FBXO38;refGene_tx=XM_019810994.2;refGene_gn=FBXO38;refGene_tx=XM_019810998.2;refGene_gn=FBXO38;refGene_tx=XM_019811006.2
A1 200506574 200506688
refGene_gn=FBXO38;refGene_tx=XM_003981411.4;refGene_gn=FBXO38;refGene_tx=XM_019810975.2;refGene_gn=FBXO38;refGene_tx=XM_019810982.2;refGene_gn=FBXO38;refGene_tx=XM_019810989.2;refGene_gn=FBXO38;refGene_tx=XM_019810994.2;refGene_gn=FBXO38;refGene_tx=XM_019810998.2;refGene_gn=FBXO38;refGene_tx=XM_019811006.2
A1 200507216 200507320

refGene_gn=FBXO38;refGene_tx=XM_003981411.4;refGene_gn=FBXO38;refGene_
tx=XM_019810975.2;refGene_gn=FBXO38;refGene_tx=XM_019810982.2;refGene_
gn=FBXO38;refGene_tx=XM_019810989.2;refGene_gn=FBXO38;refGene_tx=XM_01
9810994.2;refGene_gn=FBXO38;refGene_tx=XM_019810998.2;refGene_gn=FBXO3
8;refGene_tx=XM_019811006.2
A1 200507954 200508100
refGene_gn=FBXO38;refGene_tx=XM_003981411.4;refGene_gn=FBXO38;refGene_
tx=XM_019810975.2;refGene_gn=FBXO38;refGene_tx=XM_019810982.2;refGene_
gn=FBXO38;refGene_tx=XM_019810989.2;refGene_gn=FBXO38;refGene_tx=XM_01
9810994.2;refGene_gn=FBXO38;refGene_tx=XM_019810998.2;refGene_gn=FBXO3
8;refGene_tx=XM_019811006.2
A1 200509457 200509624
refGene_gn=FBXO38;refGene_tx=XM_003981411.4;refGene_gn=FBXO38;refGene_
tx=XM_019810975.2;refGene_gn=FBXO38;refGene_tx=XM_019810982.2;refGene_
gn=FBXO38;refGene_tx=XM_019810989.2;refGene_gn=FBXO38;refGene_tx=XM_01
9810994.2;refGene_gn=FBXO38;refGene_tx=XM_019810998.2;refGene_gn=FBXO3
8;refGene_tx=XM_019811006.2
A1 200512384 200512487
refGene_gn=FBXO38;refGene_tx=XM_003981411.4;refGene_gn=FBXO38;refGene_
tx=XM_019810975.2;refGene_gn=FBXO38;refGene_tx=XM_019810982.2;refGene_
gn=FBXO38;refGene_tx=XM_019810989.2;refGene_gn=FBXO38;refGene_tx=XM_01
9810994.2;refGene_gn=FBXO38;refGene_tx=XM_019810998.2;refGene_gn=FBXO3
8;refGene_tx=XM_019811006.2
A1 200512694 200512795
refGene_gn=FBXO38;refGene_tx=XM_003981411.4;refGene_gn=FBXO38;refGene_
tx=XM_019810975.2;refGene_gn=FBXO38;refGene_tx=XM_019810982.2;refGene_
gn=FBXO38;refGene_tx=XM_019810989.2;refGene_gn=FBXO38;refGene_tx=XM_01
9810994.2;refGene_gn=FBXO38;refGene_tx=XM_019810998.2;refGene_gn=FBXO3
8;refGene_tx=XM_019811006.2
A1 200517825 200518560
refGene_gn=FBXO38;refGene_tx=XM_003981411.4;refGene_gn=FBXO38;refGene_
tx=XM_019810975.2;refGene_gn=FBXO38;refGene_tx=XM_019810982.2;refGene_
gn=FBXO38;refGene_tx=XM_019810989.2;refGene_gn=FBXO38;refGene_tx=XM_01
9810994.2;refGene_gn=FBXO38;refGene_tx=XM_019810998.2
A1 200520065 200520245
refGene_gn=FBXO38;refGene_tx=XM_003981411.4;refGene_gn=FBXO38;refGene_
tx=XM_019810975.2;refGene_gn=FBXO38;refGene_tx=XM_019810982.2;refGene_
gn=FBXO38;refGene_tx=XM_019810989.2;refGene_gn=FBXO38;refGene_tx=XM_01
9810994.2;refGene_gn=FBXO38;refGene_tx=XM_019810998.2;refGene_gn=FBXO3
8;refGene_tx=XM_019811006.2
A1 200521648 200521768
refGene_gn=FBXO38;refGene_tx=XM_003981411.4;refGene_gn=FBXO38;refGene_
tx=XM_019810975.2;refGene_gn=FBXO38;refGene_tx=XM_019810982.2;refGene_
gn=FBXO38;refGene_tx=XM_019810989.2;refGene_gn=FBXO38;refGene_tx=XM_01
9810994.2;refGene_gn=FBXO38;refGene_tx=XM_019810998.2;refGene_gn=FBXO3
8;refGene_tx=XM_019811006.2
A1 200528762 200528973
refGene_gn=FBXO38;refGene_tx=XM_003981411.4;refGene_gn=FBXO38;refGene_
tx=XM_019810975.2;refGene_gn=FBXO38;refGene_tx=XM_019810982.2;refGene_
gn=FBXO38;refGene_tx=XM_019810989.2;refGene_gn=FBXO38;refGene_tx=XM_01

9810994.2;refGene_gn=FBXO38;refGene_tx=XM_019810998.2;refGene_gn=FBXO3
8;refGene_tx=XM_019811006.2
A1	200530282	200530425
refGene_gn=FBXO38;refGene_tx=XM_003981411.4;refGene_gn=FBXO38;refGene_
tx=XM_019810975.2;refGene_gn=FBXO38;refGene_tx=XM_019810982.2;refGene_
gn=FBXO38;refGene_tx=XM_019810989.2;refGene_gn=FBXO38;refGene_tx=XM_01
9810994.2;refGene_gn=FBXO38;refGene_tx=XM_019810998.2;refGene_gn=FBXO3
8;refGene_tx=XM_019811006.2
A1	200532066	200532237
refGene_gn=FBXO38;refGene_tx=XM_003981411.4;refGene_gn=FBXO38;refGene_
tx=XM_019810975.2;refGene_gn=FBXO38;refGene_tx=XM_019810982.2;refGene_
gn=FBXO38;refGene_tx=XM_019810989.2;refGene_gn=FBXO38;refGene_tx=XM_01
9810994.2;refGene_gn=FBXO38;refGene_tx=XM_019810998.2;refGene_gn=FBXO3
8;refGene_tx=XM_019811006.2
A1	200535807	200535938
refGene_gn=FBXO38;refGene_tx=XM_003981411.4;refGene_gn=FBXO38;refGene_
tx=XM_019810975.2;refGene_gn=FBXO38;refGene_tx=XM_019810982.2;refGene_
gn=FBXO38;refGene_tx=XM_019810989.2;refGene_gn=FBXO38;refGene_tx=XM_01
9810994.2;refGene_gn=FBXO38;refGene_tx=XM_019810998.2;refGene_gn=FBXO3
8;refGene_tx=XM_019811006.2
A1	200537397	200537491
refGene_gn=FBXO38;refGene_tx=XM_003981411.4;refGene_gn=FBXO38;refGene_
tx=XM_019810975.2;refGene_gn=FBXO38;refGene_tx=XM_019810982.2;refGene_
gn=FBXO38;refGene_tx=XM_019810989.2;refGene_gn=FBXO38;refGene_tx=XM_01
9810994.2;refGene_gn=FBXO38;refGene_tx=XM_019810998.2;refGene_gn=FBXO3
8;refGene_tx=XM_019811006.2
A1	200540407	200540545
refGene_gn=FBXO38;refGene_tx=XM_003981411.4;refGene_gn=FBXO38;refGene_
tx=XM_019810975.2;refGene_gn=FBXO38;refGene_tx=XM_019810982.2;refGene_
gn=FBXO38;refGene_tx=XM_019810989.2;refGene_gn=FBXO38;refGene_tx=XM_01
9810994.2;refGene_gn=FBXO38;refGene_tx=XM_019810998.2;refGene_gn=FBXO3
8;refGene_tx=XM_019811006.2
A1	200541744	200541882
refGene_gn=FBXO38;refGene_tx=XM_003981411.4;refGene_gn=FBXO38;refGene_
tx=XM_019810975.2;refGene_gn=FBXO38;refGene_tx=XM_019810982.2;refGene_
gn=FBXO38;refGene_tx=XM_019810989.2;refGene_gn=FBXO38;refGene_tx=XM_01
9810994.2;refGene_gn=FBXO38;refGene_tx=XM_019810998.2;refGene_gn=FBXO3
8;refGene_tx=XM_019811006.2
A1	200544800	200544966
refGene_gn=FBXO38;refGene_tx=XM_003981411.4;refGene_gn=FBXO38;refGene_
tx=XM_019810975.2;refGene_gn=FBXO38;refGene_tx=XM_019810982.2;refGene_
gn=FBXO38;refGene_tx=XM_019810989.2;refGene_gn=FBXO38;refGene_tx=XM_01
9810994.2;refGene_gn=FBXO38;refGene_tx=XM_019811006.2
A1	200545164	200545328
refGene_gn=FBXO38;refGene_tx=XM_003981411.4;refGene_gn=FBXO38;refGene_
tx=XM_019810975.2;refGene_gn=FBXO38;refGene_tx=XM_019810982.2;refGene_
gn=FBXO38;refGene_tx=XM_019810989.2;refGene_gn=FBXO38;refGene_tx=XM_01
9810994.2;refGene_gn=FBXO38;refGene_tx=XM_019811006.2
A1	200548108	200548242
refGene_gn=FBXO38;refGene_tx=XM_003981411.4;refGene_gn=FBXO38;refGene_

tx=XM_019810975.2;refGene_gn=FBXO38;refGene_tx=XM_019810982.2;refGene_gn=FBXO38;refGene_tx=XM_019810989.2;refGene_gn=FBXO38;refGene_tx=XM_019810994.2;refGene_gn=FBXO38;refGene_tx=XM_019811006.2
A1 200552207 200552209
refGene_gn=FBXO38;refGene_tx=XM_019810989.2
A1 200552609 200552737
refGene_gn=FBXO38;refGene_tx=XM_003981411.4;refGene_gn=FBXO38;refGene_tx=XM_019810975.2;refGene_gn=FBXO38;refGene_tx=XM_019810982.2;refGene_gn=FBXO38;refGene_tx=XM_019810994.2;refGene_gn=FBXO38;refGene_tx=XM_019810998.2;refGene_gn=FBXO38;refGene_tx=XM_019811006.2
A1 200580728 200580804
refGene_gn=SPINK9;refGene_tx=XM_006928017.3
A1 200586568 200586699
refGene_gn=SPINK9;refGene_tx=XM_006928017.3
A1 200588655 200588681
refGene_gn=SPINK9;refGene_tx=XM_006928017.3
A1 200589838 200589884
refGene_gn=SPINK9;refGene_tx=XM_006928017.3
A1 200605406 200605452
refGene_gn=LOC101087814;refGene_tx=XM_003981414.5
A1 200606927 200607055
refGene_gn=LOC101087814;refGene_tx=XM_003981414.5
A1 200609045 200609074
refGene_gn=LOC101087814;refGene_tx=XM_003981414.5
A1 200609836 200609891
refGene_gn=LOC101087814;refGene_tx=XM_003981414.5
A1 200620816 200620862
refGene_gn=SPINK7;refGene_tx=XM_003981415.3
A1 200622219 200622344
refGene_gn=SPINK7;refGene_tx=XM_003981415.3
A1 200623040 200623066
refGene_gn=SPINK7;refGene_tx=XM_003981415.3
A1 200624360 200624421
refGene_gn=SPINK7;refGene_tx=XM_003981415.3
A1 200631720 200631769
refGene_gn=SPINK13;refGene_tx=XM_023260666.1
A1 200642090 200642218
refGene_gn=SPINK13;refGene_tx=XM_023260666.1
A1 200642880 200642918
refGene_gn=SPINK13;refGene_tx=XM_023260666.1
A1 200645435 200645505
refGene_gn=SPINK13;refGene_tx=XM_023260666.1
A1 200736493 200736667
refGene_gn=GZMK;refGene_tx=XM_003981494.4
A1 200738141 200738411
refGene_gn=GZMK;refGene_tx=XM_003981494.4
A1 200739189 200739340
refGene_gn=GZMK;refGene_tx=XM_003981494.4
A1 200745708 200745856
refGene_gn=GZMK;refGene_tx=XM_003981494.4

A1	200746217	200746278
refGene_gn=GZMK;refGene_tx=XM_003981494.4
A1	200772061	200772362
refGene_gn=ESM1;refGene_tx=XM_003981416.5
A1	200775852	200776002
refGene_gn=ESM1;refGene_tx=XM_003981416.5
A1	200779632	200779730
refGene_gn=ESM1;refGene_tx=XM_003981416.5
A1	200846836	200846915
refGene_gn=LOC101089242;refGene_tx=XM_023260693.1;refGene_gn=LOC101089
242;refGene_tx=XM_023260694.1;refGene_gn=LOC101089242;refGene_tx=XM_02
3260695.1;refGene_gn=LOC101089242;refGene_tx=XM_023260699.1
A1	200865121	200865285
refGene_gn=LOC101089242;refGene_tx=XM_023260693.1;refGene_gn=LOC101089
242;refGene_tx=XM_023260694.1;refGene_gn=LOC101089242;refGene_tx=XM_02
3260695.1;refGene_gn=LOC101089242;refGene_tx=XM_023260699.1
A1	200865626	200869193
refGene_gn=LOC101089242;refGene_tx=XM_023260693.1;refGene_gn=LOC101089
242;refGene_tx=XM_023260694.1;refGene_gn=LOC101089242;refGene_tx=XM_02
3260695.1;refGene_gn=LOC101089242;refGene_tx=XM_023260699.1
A1	200870632	200871103
refGene_gn=LOC101089242;refGene_tx=XM_023260693.1;refGene_gn=LOC101089
242;refGene_tx=XM_023260694.1;refGene_gn=LOC101089242;refGene_tx=XM_02
3260695.1;refGene_gn=LOC101089242;refGene_tx=XM_023260699.1
A1	200875417	200875594
refGene_gn=LOC101089242;refGene_tx=XM_023260693.1;refGene_gn=LOC101089
242;refGene_tx=XM_023260694.1;refGene_gn=LOC101089242;refGene_tx=XM_02
3260695.1;refGene_gn=LOC101089242;refGene_tx=XM_023260699.1
A1	200883422	200883621
refGene_gn=LOC101089242;refGene_tx=XM_023260693.1;refGene_gn=LOC101089
242;refGene_tx=XM_023260694.1;refGene_gn=LOC101089242;refGene_tx=XM_02
3260695.1;refGene_gn=LOC101089242;refGene_tx=XM_023260699.1
A1	200887267	200887399
refGene_gn=LOC101089242;refGene_tx=XM_023260693.1;refGene_gn=LOC101089
242;refGene_tx=XM_023260694.1;refGene_gn=LOC101089242;refGene_tx=XM_02
3260699.1
A1	200889780	200889846
refGene_gn=LOC101089242;refGene_tx=XM_023260693.1;refGene_gn=LOC101089
242;refGene_tx=XM_023260699.1
A1	200897110	200897283
refGene_gn=LOC101089242;refGene_tx=XM_023260693.1;refGene_gn=LOC101089
242;refGene_tx=XM_023260694.1;refGene_gn=LOC101089242;refGene_tx=XM_02
3260695.1;refGene_gn=LOC101089242;refGene_tx=XM_023260699.1
A1	200911141	200911325
refGene_gn=LOC101089242;refGene_tx=XM_023260693.1;refGene_gn=LOC101089
242;refGene_tx=XM_023260694.1;refGene_gn=LOC101089242;refGene_tx=XM_02
3260695.1;refGene_gn=LOC101089242;refGene_tx=XM_023260699.1
A1	200912214	200914262
refGene_gn=LOC101089242;refGene_tx=XM_023260693.1;refGene_gn=LOC101089
242;refGene_tx=XM_023260694.1;refGene_gn=LOC101089242;refGene_tx=XM_02

3260695.1;refGene_gn=LOC101089242;refGene_tx=XM_023260699.1
A1 201106482 201106736
refGene_gn=SNX18;refGene_tx=XM_023260708.1
A1 201130548 201130667
refGene_gn=SNX18;refGene_tx=XM_023260712.1
A1 201132063 201133684
refGene_gn=SNX18;refGene_tx=XM_023260708.1;refGene_gn=SNX18;refGene_tx=XM_023260712.1
A1 201173700 201174174
refGene_gn=HSPB3;refGene_tx=XM_006928019.4
A1 201285526 201285574
refGene_gn=ARL15;refGene_tx=XM_003981419.4;refGene_gn=ARL15;refGene_tx=XM_023260732.1
A1 201408048 201408096
refGene_gn=ARL15;refGene_tx=XM_019811096.2
A1 201428956 201429101
refGene_gn=ARL15;refGene_tx=XM_003981419.4;refGene_gn=ARL15;refGene_tx=XM_019811096.2;refGene_gn=ARL15;refGene_tx=XM_023260732.1
A1 201444391 201444451
refGene_gn=ARL15;refGene_tx=XM_003981419.4;refGene_gn=ARL15;refGene_tx=XM_019811096.2;refGene_gn=ARL15;refGene_tx=XM_023260732.1
A1 201484828 201485037
refGene_gn=ARL15;refGene_tx=XM_003981419.4;refGene_gn=ARL15;refGene_tx=XM_019811096.2;refGene_gn=ARL15;refGene_tx=XM_023260732.1
A1 201682849 201683002
refGene_gn=ARL15;refGene_tx=XM_003981419.4;refGene_gn=ARL15;refGene_tx=XM_019811096.2;refGene_gn=ARL15;refGene_tx=XM_023260732.1
A1 201832797 201832901
refGene_gn=NDUFS4;refGene_tx=XM_003981420.3;refGene_gn=NDUFS4;refGene_tx=XM_019811113.1;refGene_gn=NDUFS4;refGene_tx=XM_019811115.1
A1 201854272 201854346
refGene_gn=NDUFS4;refGene_tx=XM_003981420.3;refGene_gn=NDUFS4;refGene_tx=XM_019811113.1;refGene_gn=NDUFS4;refGene_tx=XM_019811115.1
A1 201868965 201869138
refGene_gn=NDUFS4;refGene_tx=XM_003981420.3;refGene_gn=NDUFS4;refGene_tx=XM_019811113.1;refGene_gn=NDUFS4;refGene_tx=XM_019811115.1
A1 201905902 201905981
refGene_gn=NDUFS4;refGene_tx=XM_003981420.3;refGene_gn=NDUFS4;refGene_tx=XM_019811115.1
A1 201934432 201934452
refGene_gn=NDUFS4;refGene_tx=XM_019811115.1
A1 201945269 201945443
refGene_gn=NDUFS4;refGene_tx=XM_003981420.3;refGene_gn=NDUFS4;refGene_tx=XM_019811113.1
A1 202006318 202006401
refGene_gn=FST;refGene_tx=XM_023260781.1;refGene_gn=FST;refGene_tx=XM_023260782.1
A1 202006662 202006664
refGene_gn=FST;refGene_tx=XM_023260784.1
A1 202007360 202007591

refGene_gn=FST;refGene_tx=XM_023260781.1;refGene_gn=FST;refGene_tx=XM_023260782.1;refGene_gn=FST;refGene_tx=XM_023260784.1
A1 202008257 202008482
refGene_gn=FST;refGene_tx=XM_023260781.1;refGene_gn=FST;refGene_tx=XM_023260782.1;refGene_gn=FST;refGene_tx=XM_023260784.1
A1 202008818 202009037
refGene_gn=FST;refGene_tx=XM_023260781.1;refGene_gn=FST;refGene_tx=XM_023260782.1;refGene_gn=FST;refGene_tx=XM_023260784.1
A1 202009474 202009666
refGene_gn=FST;refGene_tx=XM_023260781.1;refGene_gn=FST;refGene_tx=XM_023260782.1;refGene_gn=FST;refGene_tx=XM_023260784.1
A1 202011531 202011616
refGene_gn=FST;refGene_tx=XM_023260781.1;refGene_gn=FST;refGene_tx=XM_023260782.1;refGene_gn=FST;refGene_tx=XM_023260784.1
A1 202387800 202387907
refGene_gn=MOCS2;refGene_tx=XM_006928021.2;refGene_gn=MOCS2;refGene_tx=XM_006928022.2
A1 202401574 202401702
refGene_gn=MOCS2;refGene_tx=XM_006928021.2;refGene_gn=MOCS2;refGene_tx=XM_006928022.2
A1 202402227 202402378
refGene_gn=MOCS2;refGene_tx=XM_006928021.2;refGene_gn=MOCS2;refGene_tx=XM_006928022.2
A1 202403123 202403247
refGene_gn=MOCS2;refGene_tx=XM_006928021.2;refGene_gn=MOCS2;refGene_tx=XM_006928022.2
A1 202405807 202405876
refGene_gn=MOCS2;refGene_tx=XM_006928021.2;refGene_gn=MOCS2;refGene_tx=XM_006928022.2
A1 202412174 202412255
refGene_gn=ITGA2;refGene_tx=XM_023260785.1
A1 202412753 202412870
refGene_gn=ITGA2;refGene_tx=XM_023260785.1
A1 202415652 202415742
refGene_gn=ITGA2;refGene_tx=XM_023260785.1
A1 202418909 202419023
refGene_gn=ITGA2;refGene_tx=XM_023260785.1
A1 202420946 202421051
refGene_gn=ITGA2;refGene_tx=XM_023260785.1
A1 202422010 202422121
refGene_gn=ITGA2;refGene_tx=XM_023260785.1
A1 202423736 202423839
refGene_gn=ITGA2;refGene_tx=XM_023260785.1
A1 202427332 202427416
refGene_gn=ITGA2;refGene_tx=XM_023260785.1
A1 202427502 202427579
refGene_gn=ITGA2;refGene_tx=XM_023260785.1
A1 202428296 202428389
refGene_gn=ITGA2;refGene_tx=XM_023260785.1
A1 202429513 202429655

refGene_gn=ITGA2;refGene_tx=XM_023260785.1
A1 202430082 202430165
refGene_gn=ITGA2;refGene_tx=XM_023260785.1
A1 202430704 202430815
refGene_gn=ITGA2;refGene_tx=XM_023260785.1
A1 202433111 202433263
refGene_gn=ITGA2;refGene_tx=XM_023260785.1
A1 202437293 202437433
refGene_gn=ITGA2;refGene_tx=XM_023260785.1
A1 202440116 202440253
refGene_gn=ITGA2;refGene_tx=XM_023260785.1
A1 202440916 202441120
refGene_gn=ITGA2;refGene_tx=XM_023260785.1
A1 202450199 202450343
refGene_gn=ITGA2;refGene_tx=XM_023260785.1
A1 202451949 202452095
refGene_gn=ITGA2;refGene_tx=XM_023260785.1
A1 202453900 202454039
refGene_gn=ITGA2;refGene_tx=XM_023260785.1
A1 202455806 202455883
refGene_gn=ITGA2;refGene_tx=XM_023260785.1
A1 202458278 202458444
refGene_gn=ITGA2;refGene_tx=XM_023260785.1
A1 202458730 202458881
refGene_gn=ITGA2;refGene_tx=XM_023260785.1
A1 202463713 202463862
refGene_gn=ITGA2;refGene_tx=XM_023260785.1
A1 202466454 202466582
refGene_gn=ITGA2;refGene_tx=XM_023260785.1
A1 202466756 202466871
refGene_gn=ITGA2;refGene_tx=XM_023260785.1
A1 202470162 202470254
refGene_gn=ITGA2;refGene_tx=XM_023260785.1
A1 202475796 202475906
refGene_gn=ITGA2;refGene_tx=XM_023260785.1
A1 202490540 202490658
refGene_gn=ITGA2;refGene_tx=XM_023260785.1
A1 202517352 202517407
refGene_gn=ITGA2;refGene_tx=XM_023260785.1
A1 202544236 202544281
refGene_gn=ITGA1;refGene_tx=XM_003981422.4
A1 202549826 202549943
refGene_gn=ITGA1;refGene_tx=XM_003981422.4
A1 202553084 202553177
refGene_gn=ITGA1;refGene_tx=XM_003981422.4
A1 202554319 202554424
refGene_gn=ITGA1;refGene_tx=XM_003981422.4
A1 202554586 202554691
refGene_gn=ITGA1;refGene_tx=XM_003981422.4
A1 202556733 202556847

refGene_gn=ITGA1;refGene_tx=XM_003981422.4
A1 202560256 202560359
refGene_gn=ITGA1;refGene_tx=XM_003981422.4
A1 202561718 202561808
refGene_gn=ITGA1;refGene_tx=XM_003981422.4
A1 202564692 202564769
refGene_gn=ITGA1;refGene_tx=XM_003981422.4
A1 202569110 202569191
refGene_gn=ITGA1;refGene_tx=XM_003981422.4
A1 202570947 202571157
refGene_gn=ITGA1;refGene_tx=XM_003981422.4
A1 202572999 202573110
refGene_gn=ITGA1;refGene_tx=XM_003981422.4
A1 202575026 202575163
refGene_gn=ITGA1;refGene_tx=XM_003981422.4
A1 202576587 202576754
refGene_gn=ITGA1;refGene_tx=XM_003981422.4
A1 202581290 202581421
refGene_gn=ITGA1;refGene_tx=XM_003981422.4
A1 202584258 202584516
refGene_gn=ITGA1;refGene_tx=XM_003981422.4
A1 202585740 202585884
refGene_gn=ITGA1;refGene_tx=XM_003981422.4
A1 202588741 202588884
refGene_gn=ITGA1;refGene_tx=XM_003981422.4
A1 202593729 202593874
refGene_gn=ITGA1;refGene_tx=XM_003981422.4
A1 202594528 202594602
refGene_gn=ITGA1;refGene_tx=XM_003981422.4
A1 202599163 202599329
refGene_gn=ITGA1;refGene_tx=XM_003981422.4
A1 202607665 202607816
refGene_gn=ITGA1;refGene_tx=XM_003981422.4
A1 202611129 202611278
refGene_gn=ITGA1;refGene_tx=XM_003981422.4
A1 202627226 202627354
refGene_gn=ITGA1;refGene_tx=XM_003981422.4
A1 202627971 202628083
refGene_gn=ITGA1;refGene_tx=XM_003981422.4
A1 202628188 202628277
refGene_gn=ITGA1;refGene_tx=XM_003981422.4
A1 202630890 202631003
refGene_gn=ITGA1;refGene_tx=XM_003981422.4
A1 202644108 202644229
refGene_gn=ITGA1;refGene_tx=XM_003981422.4
A1 202686838 202687270
refGene_gn=PELO;refGene_tx=XM_003981423.5
A1 202687553 202688279
refGene_gn=PELO;refGene_tx=XM_003981423.5
A1 202701523 202701584

refGene_gn=ITGA1;refGene_tx=XM_003981422.4
A1 203889703 203889820
refGene_gn=ISL1;refGene_tx=XM_003981424.4
A1 203891892 203892060
refGene_gn=ISL1;refGene_tx=XM_003981424.4
A1 203893261 203893548
refGene_gn=ISL1;refGene_tx=XM_003981424.4
A1 203895366 203895626
refGene_gn=ISL1;refGene_tx=XM_003981424.4
A1 203898401 203898591
refGene_gn=ISL1;refGene_tx=XM_003981424.4
A1 203899450 203899478
refGene_gn=ISL1;refGene_tx=XM_003981424.4
A1 204369631 204369734
refGene_gn=PARP8;refGene_tx=XM_011286265.3;refGene_gn=PARP8;refGene_tx=XM_019811146.2
A1 204381000 204381085
refGene_gn=PARP8;refGene_tx=XM_011286265.3;refGene_gn=PARP8;refGene_tx=XM_019811146.2
A1 204381940 204382010
refGene_gn=PARP8;refGene_tx=XM_011286265.3;refGene_gn=PARP8;refGene_tx=XM_019811146.2
A1 204383167 204383241
refGene_gn=PARP8;refGene_tx=XM_011286265.3;refGene_gn=PARP8;refGene_tx=XM_019811146.2
A1 204383787 204383857
refGene_gn=PARP8;refGene_tx=XM_011286265.3;refGene_gn=PARP8;refGene_tx=XM_019811146.2
A1 204385077 204385150
refGene_gn=PARP8;refGene_tx=XM_011286265.3;refGene_gn=PARP8;refGene_tx=XM_019811146.2
A1 204385385 204385498
refGene_gn=PARP8;refGene_tx=XM_011286265.3;refGene_gn=PARP8;refGene_tx=XM_019811146.2
A1 204386590 204386639
refGene_gn=PARP8;refGene_tx=XM_011286265.3;refGene_gn=PARP8;refGene_tx=XM_019811146.2
A1 204388381 204388449
refGene_gn=PARP8;refGene_tx=XM_011286265.3;refGene_gn=PARP8;refGene_tx=XM_019811146.2
A1 204390959 204391025
refGene_gn=PARP8;refGene_tx=XM_011286265.3;refGene_gn=PARP8;refGene_tx=XM_019811146.2
A1 204392296 204392422
refGene_gn=PARP8;refGene_tx=XM_011286265.3;refGene_gn=PARP8;refGene_tx=XM_019811146.2
A1 204399349 204399442
refGene_gn=PARP8;refGene_tx=XM_011286265.3;refGene_gn=PARP8;refGene_tx=XM_019811146.2
A1 204409261 204409357

refGene_gn=PARP8;refGene_tx=XM_011286265.3;refGene_gn=PARP8;refGene_tx=XM_019811146.2
A1 204409459 204409510
refGene_gn=PARP8;refGene_tx=XM_011286265.3;refGene_gn=PARP8;refGene_tx=XM_019811146.2
A1 204411425 204411990
refGene_gn=PARP8;refGene_tx=XM_011286265.3;refGene_gn=PARP8;refGene_tx=XM_019811146.2
A1 204412518 204412644
refGene_gn=PARP8;refGene_tx=XM_011286265.3;refGene_gn=PARP8;refGene_tx=XM_019811146.2
A1 204418798 204418865
refGene_gn=PARP8;refGene_tx=XM_011286265.3;refGene_gn=PARP8;refGene_tx=XM_019811146.2
A1 204424434 204424525
refGene_gn=PARP8;refGene_tx=XM_011286265.3;refGene_gn=PARP8;refGene_tx=XM_019811146.2
A1 204425000 204425061
refGene_gn=PARP8;refGene_tx=XM_011286265.3;refGene_gn=PARP8;refGene_tx=XM_019811146.2
A1 204434355 204434450
refGene_gn=PARP8;refGene_tx=XM_011286265.3;refGene_gn=PARP8;refGene_tx=XM_019811146.2
A1 204436727 204436805
refGene_gn=PARP8;refGene_tx=XM_011286265.3;refGene_gn=PARP8;refGene_tx=XM_019811146.2
A1 204438924 204438995
refGene_gn=PARP8;refGene_tx=XM_011286265.3;refGene_gn=PARP8;refGene_tx=XM_019811146.2
A1 204439589 204439679
refGene_gn=PARP8;refGene_tx=XM_011286265.3;refGene_gn=PARP8;refGene_tx=XM_019811146.2
A1 204441737 204441741
refGene_gn=PARP8;refGene_tx=XM_019811146.2
A1 204446875 204446913
refGene_gn=PARP8;refGene_tx=XM_011286265.3
A1 204542368 204542423
refGene_gn=PARP8;refGene_tx=XM_011286265.3
A1 204543394 204543496
refGene_gn=PARP8;refGene_tx=XM_011286265.3
A1 204543583 204543689
refGene_gn=PARP8;refGene_tx=XM_011286265.3
A1 204685334 204685440
refGene_gn=EMB;refGene_tx=XM_023260829.1;refGene_gn=EMB;refGene_tx=XM_023260839.1
A1 204697449 204697539
refGene_gn=EMB;refGene_tx=XM_023260829.1;refGene_gn=EMB;refGene_tx=XM_023260839.1
A1 204697837 204697927
refGene_gn=EMB;refGene_tx=XM_023260829.1;refGene_gn=EMB;refGene_tx=XM_

023260839.1;refGene_gn=EMB;refGene_tx=XM_023260842.1
A1 204698663 204698753
refGene_gn=EMB;refGene_tx=XM_023260829.1;refGene_gn=EMB;refGene_tx=XM_023260839.1;refGene_gn=EMB;refGene_tx=XM_023260842.1
A1 204699152 204699242
refGene_gn=EMB;refGene_tx=XM_023260829.1;refGene_gn=EMB;refGene_tx=XM_023260839.1;refGene_gn=EMB;refGene_tx=XM_023260842.1
A1 204699722 204699812
refGene_gn=EMB;refGene_tx=XM_023260829.1;refGene_gn=EMB;refGene_tx=XM_023260842.1
A1 204716640 204716824
refGene_gn=EMB;refGene_tx=XM_023260829.1;refGene_gn=EMB;refGene_tx=XM_023260839.1;refGene_gn=EMB;refGene_tx=XM_023260842.1
A1 204717056 204717145
refGene_gn=EMB;refGene_tx=XM_023260829.1;refGene_gn=EMB;refGene_tx=XM_023260839.1;refGene_gn=EMB;refGene_tx=XM_023260842.1
A1 204725480 204725608
refGene_gn=EMB;refGene_tx=XM_023260829.1;refGene_gn=EMB;refGene_tx=XM_023260839.1;refGene_gn=EMB;refGene_tx=XM_023260842.1
A1 204727780 204728060
refGene_gn=EMB;refGene_tx=XM_023260829.1;refGene_gn=EMB;refGene_tx=XM_023260839.1;refGene_gn=EMB;refGene_tx=XM_023260842.1
A1 204729098 204729132
refGene_gn=EMB;refGene_tx=XM_023260829.1;refGene_gn=EMB;refGene_tx=XM_023260839.1;refGene_gn=EMB;refGene_tx=XM_023260842.1
A1 204731876 204731931
refGene_gn=EMB;refGene_tx=XM_023260829.1;refGene_gn=EMB;refGene_tx=XM_023260839.1;refGene_gn=EMB;refGene_tx=XM_023260842.1
A1 204732520 204732538
refGene_gn=EMB;refGene_tx=XM_023260829.1;refGene_gn=EMB;refGene_tx=XM_023260839.1;refGene_gn=EMB;refGene_tx=XM_023260842.1
A1 205065933 205066340
refGene_gn=HCN1;refGene_tx=XM_011286847.3
A1 205114070 205114494
refGene_gn=HCN1;refGene_tx=XM_011286847.3
A1 205282952 205283114
refGene_gn=HCN1;refGene_tx=XM_011286847.3
A1 205345760 205345979
refGene_gn=HCN1;refGene_tx=XM_011286847.3
A1 205384427 205384574
refGene_gn=HCN1;refGene_tx=XM_011286847.3
A1 205404100 205404341
refGene_gn=HCN1;refGene_tx=XM_011286847.3
A1 205448473 205448638
refGene_gn=HCN1;refGene_tx=XM_011286847.3
A1 205451823 205452707
refGene_gn=HCN1;refGene_tx=XM_011286847.3
A1 205537035 205538982
refGene_gn=LOC105260991;refGene_tx=XM_019811152.2
A1 205538984 205540579

refGene_gn=LOC105260991;refGene_tx=XM_019811152.2
A1	205540580	205540962
refGene_gn=LOC105260991;refGene_tx=XM_019811152.2
A1	205540964	205542071
refGene_gn=LOC105260991;refGene_tx=XM_019811152.2
A1	205855567	205855857
refGene_gn=MRPS30;refGene_tx=XM_023260853.1
A1	205858025	205858202
refGene_gn=MRPS30;refGene_tx=XM_023260853.1
A1	205859654	205859760
refGene_gn=MRPS30;refGene_tx=XM_023260853.1
A1	205860492	205860638
refGene_gn=MRPS30;refGene_tx=XM_023260853.1
A1	205862230	205862828
refGene_gn=MRPS30;refGene_tx=XM_023260853.1
A1	206200213	206200580
refGene_gn=FGF10;refGene_tx=XM_023260856.1
A1	206276424	206276528
refGene_gn=FGF10;refGene_tx=XM_023260856.1
A1	206280564	206280762
refGene_gn=FGF10;refGene_tx=XM_023260856.1
A1	206716916	206717066
refGene_gn=NNT;refGene_tx=XM_003981426.5;refGene_gn=NNT;refGene_tx=XM_006928023.3;refGene_gn=NNT;refGene_tx=XM_011286285.3;refGene_gn=NNT;refGene_tx=XM_023260859.1
A1	206718754	206718870
refGene_gn=NNT;refGene_tx=XM_003981426.5;refGene_gn=NNT;refGene_tx=XM_006928023.3;refGene_gn=NNT;refGene_tx=XM_011286285.3;refGene_gn=NNT;refGene_tx=XM_023260859.1
A1	206720113	206720232
refGene_gn=NNT;refGene_tx=XM_003981426.5;refGene_gn=NNT;refGene_tx=XM_006928023.3;refGene_gn=NNT;refGene_tx=XM_011286285.3;refGene_gn=NNT;refGene_tx=XM_023260859.1
A1	206732400	206732419
refGene_gn=NNT;refGene_tx=XM_023260861.1
A1	206738245	206738327
refGene_gn=NNT;refGene_tx=XM_003981426.5;refGene_gn=NNT;refGene_tx=XM_006928023.3;refGene_gn=NNT;refGene_tx=XM_011286285.3;refGene_gn=NNT;refGene_tx=XM_023260859.1;refGene_gn=NNT;refGene_tx=XM_023260861.1
A1	206739241	206739401
refGene_gn=NNT;refGene_tx=XM_003981426.5;refGene_gn=NNT;refGene_tx=XM_006928023.3;refGene_gn=NNT;refGene_tx=XM_011286285.3;refGene_gn=NNT;refGene_tx=XM_023260859.1;refGene_gn=NNT;refGene_tx=XM_023260861.1
A1	206747183	206747363
refGene_gn=NNT;refGene_tx=XM_003981426.5;refGene_gn=NNT;refGene_tx=XM_006928023.3;refGene_gn=NNT;refGene_tx=XM_011286285.3;refGene_gn=NNT;refGene_tx=XM_023260859.1;refGene_gn=NNT;refGene_tx=XM_023260861.1
A1	206749370	206749531
refGene_gn=NNT;refGene_tx=XM_003981426.5;refGene_gn=NNT;refGene_tx=XM_006928023.3;refGene_gn=NNT;refGene_tx=XM_011286285.3;refGene_gn=NNT;re

fGene_tx=XM_023260859.1;refGene_gn=NNT;refGene_tx=XM_023260861.1
A1 206750683 206750917
refGene_gn=NNT;refGene_tx=XM_003981426.5;refGene_gn=NNT;refGene_tx=XM_006928023.3;refGene_gn=NNT;refGene_tx=XM_011286285.3;refGene_gn=NNT;refGene_tx=XM_023260859.1;refGene_gn=NNT;refGene_tx=XM_023260861.1
A1 206753223 206753419
refGene_gn=NNT;refGene_tx=XM_003981426.5;refGene_gn=NNT;refGene_tx=XM_006928023.3;refGene_gn=NNT;refGene_tx=XM_011286285.3;refGene_gn=NNT;refGene_tx=XM_023260859.1;refGene_gn=NNT;refGene_tx=XM_023260861.1
A1 206754662 206754808
refGene_gn=NNT;refGene_tx=XM_003981426.5;refGene_gn=NNT;refGene_tx=XM_006928023.3;refGene_gn=NNT;refGene_tx=XM_011286285.3;refGene_gn=NNT;refGene_tx=XM_023260859.1;refGene_gn=NNT;refGene_tx=XM_023260861.1
A1 206755811 206755922
refGene_gn=NNT;refGene_tx=XM_003981426.5;refGene_gn=NNT;refGene_tx=XM_006928023.3;refGene_gn=NNT;refGene_tx=XM_011286285.3;refGene_gn=NNT;refGene_tx=XM_023260859.1;refGene_gn=NNT;refGene_tx=XM_023260861.1
A1 206757441 206757603
refGene_gn=NNT;refGene_tx=XM_003981426.5;refGene_gn=NNT;refGene_tx=XM_006928023.3;refGene_gn=NNT;refGene_tx=XM_011286285.3;refGene_gn=NNT;refGene_tx=XM_023260859.1;refGene_gn=NNT;refGene_tx=XM_023260861.1
A1 206760644 206760798
refGene_gn=NNT;refGene_tx=XM_003981426.5;refGene_gn=NNT;refGene_tx=XM_006928023.3;refGene_gn=NNT;refGene_tx=XM_011286285.3;refGene_gn=NNT;refGene_tx=XM_023260859.1;refGene_gn=NNT;refGene_tx=XM_023260861.1
A1 206761419 206761611
refGene_gn=NNT;refGene_tx=XM_003981426.5;refGene_gn=NNT;refGene_tx=XM_006928023.3;refGene_gn=NNT;refGene_tx=XM_011286285.3;refGene_gn=NNT;refGene_tx=XM_023260859.1;refGene_gn=NNT;refGene_tx=XM_023260861.1
A1 206761911 206762045
refGene_gn=NNT;refGene_tx=XM_003981426.5;refGene_gn=NNT;refGene_tx=XM_006928023.3;refGene_gn=NNT;refGene_tx=XM_011286285.3;refGene_gn=NNT;refGene_tx=XM_023260859.1;refGene_gn=NNT;refGene_tx=XM_023260861.1
A1 206763229 206763646
refGene_gn=LOC101090993;refGene_tx=XM_019811190.2
A1 206780584 206780772
refGene_gn=NNT;refGene_tx=XM_003981426.5;refGene_gn=NNT;refGene_tx=XM_006928023.3;refGene_gn=NNT;refGene_tx=XM_011286285.3;refGene_gn=NNT;refGene_tx=XM_023260859.1;refGene_gn=NNT;refGene_tx=XM_023260861.1
A1 206785720 206785809
refGene_gn=NNT;refGene_tx=XM_003981426.5;refGene_gn=NNT;refGene_tx=XM_006928023.3;refGene_gn=NNT;refGene_tx=XM_011286285.3;refGene_gn=NNT;refGene_tx=XM_023260859.1;refGene_gn=NNT;refGene_tx=XM_023260861.1
A1 206792246 206792334
refGene_gn=NNT;refGene_tx=XM_003981426.5;refGene_gn=NNT;refGene_tx=XM_006928023.3;refGene_gn=NNT;refGene_tx=XM_011286285.3;refGene_gn=NNT;refGene_tx=XM_023260859.1;refGene_gn=NNT;refGene_tx=XM_023260861.1
A1 206796979 206797197
refGene_gn=NNT;refGene_tx=XM_003981426.5;refGene_gn=NNT;refGene_tx=XM_006928023.3;refGene_gn=NNT;refGene_tx=XM_011286285.3;refGene_gn=NNT;re

fGene_tx=XM_023260859.1;refGene_gn=NNT;refGene_tx=XM_023260861.1
A1 206799980 206800210
refGene_gn=NNT;refGene_tx=XM_003981426.5;refGene_gn=NNT;refGene_tx=XM_006928023.3;refGene_gn=NNT;refGene_tx=XM_011286285.3;refGene_gn=NNT;refGene_tx=XM_023260859.1;refGene_gn=NNT;refGene_tx=XM_023260861.1
A1 206805196 206805353
refGene_gn=NNT;refGene_tx=XM_003981426.5;refGene_gn=NNT;refGene_tx=XM_006928023.3;refGene_gn=NNT;refGene_tx=XM_011286285.3;refGene_gn=NNT;refGene_tx=XM_023260859.1;refGene_gn=NNT;refGene_tx=XM_023260861.1
A1 206834385 206834653
refGene_gn=PAIP1;refGene_tx=XM_023260866.1;refGene_gn=PAIP1;refGene_tx=XM_023260868.1
A1 206835244 206835414
refGene_gn=PAIP1;refGene_tx=XM_011286290.3;refGene_gn=PAIP1;refGene_tx=XM_023260866.1;refGene_gn=PAIP1;refGene_tx=XM_023260868.1;refGene_gn=PAIP1;refGene_tx=XM_023260871.1
A1 206844309 206844495
refGene_gn=PAIP1;refGene_tx=XM_011286290.3;refGene_gn=PAIP1;refGene_tx=XM_023260866.1;refGene_gn=PAIP1;refGene_tx=XM_023260868.1;refGene_gn=PAIP1;refGene_tx=XM_023260871.1
A1 206847904 206848017
refGene_gn=PAIP1;refGene_tx=XM_011286290.3;refGene_gn=PAIP1;refGene_tx=XM_023260866.1;refGene_gn=PAIP1;refGene_tx=XM_023260868.1;refGene_gn=PAIP1;refGene_tx=XM_023260871.1
A1 206851511 206851623
refGene_gn=PAIP1;refGene_tx=XM_011286290.3;refGene_gn=PAIP1;refGene_tx=XM_023260866.1;refGene_gn=PAIP1;refGene_tx=XM_023260868.1;refGene_gn=PAIP1;refGene_tx=XM_023260871.1
A1 206853565 206853691
refGene_gn=PAIP1;refGene_tx=XM_011286290.3;refGene_gn=PAIP1;refGene_tx=XM_023260866.1;refGene_gn=PAIP1;refGene_tx=XM_023260868.1;refGene_gn=PAIP1;refGene_tx=XM_023260871.1
A1 206854726 206854833
refGene_gn=PAIP1;refGene_tx=XM_011286290.3;refGene_gn=PAIP1;refGene_tx=XM_023260866.1;refGene_gn=PAIP1;refGene_tx=XM_023260868.1;refGene_gn=PAIP1;refGene_tx=XM_023260871.1
A1 206855347 206855465
refGene_gn=PAIP1;refGene_tx=XM_011286290.3;refGene_gn=PAIP1;refGene_tx=XM_023260866.1;refGene_gn=PAIP1;refGene_tx=XM_023260868.1;refGene_gn=PAIP1;refGene_tx=XM_023260871.1
A1 206856230 206856285
refGene_gn=PAIP1;refGene_tx=XM_011286290.3;refGene_gn=PAIP1;refGene_tx=XM_023260866.1;refGene_gn=PAIP1;refGene_tx=XM_023260868.1;refGene_gn=PAIP1;refGene_tx=XM_023260871.1
A1 206863432 206863526
refGene_gn=PAIP1;refGene_tx=XM_011286290.3;refGene_gn=PAIP1;refGene_tx=XM_023260866.1;refGene_gn=PAIP1;refGene_tx=XM_023260868.1;refGene_gn=PAIP1;refGene_tx=XM_023260871.1
A1 206865698 206865792
refGene_gn=PAIP1;refGene_tx=XM_011286290.3;refGene_gn=PAIP1;refGene_tx

=XM_023260866.1;refGene_gn=PAIP1;refGene_tx=XM_023260868.1;refGene_gn=PAIP1;refGene_tx=XM_023260871.1
A1 206917310 206917622
refGene_gn=TMEM267;refGene_tx=XM_003981428.4;refGene_gn=TMEM267;refGene_tx=XM_006928024.4;refGene_gn=TMEM267;refGene_tx=XM_006928025.3
A1 206922360 206922696
refGene_gn=TMEM267;refGene_tx=XM_003981428.4;refGene_gn=TMEM267;refGene_tx=XM_006928024.4;refGene_gn=TMEM267;refGene_tx=XM_006928025.3
A1 206943444 206943481
refGene_gn=CCL28;refGene_tx=XM_019811565.2
A1 206950685 206950831
refGene_gn=CCL28;refGene_tx=XM_019811565.2
A1 206950865 206951202
refGene_gn=CCL28;refGene_tx=XM_019811565.2
A1 206961642 206961769
refGene_gn=CCL28;refGene_tx=XM_019811565.2
A1 206968801 206968994
refGene_gn=CCL28;refGene_tx=XM_019811565.2
A1 207009207 207009655
refGene_gn=HMGCS1;refGene_tx=XM_003981430.5;refGene_gn=HMGCS1;refGene_tx=XM_006928026.4
A1 207009980 207010106
refGene_gn=HMGCS1;refGene_tx=XM_003981430.5;refGene_gn=HMGCS1;refGene_tx=XM_006928026.4
A1 207010638 207010803
refGene_gn=HMGCS1;refGene_tx=XM_003981430.5;refGene_gn=HMGCS1;refGene_tx=XM_006928026.4
A1 207012191 207012357
refGene_gn=HMGCS1;refGene_tx=XM_003981430.5;refGene_gn=HMGCS1;refGene_tx=XM_006928026.4
A1 207013153 207013324
refGene_gn=HMGCS1;refGene_tx=XM_003981430.5;refGene_gn=HMGCS1;refGene_tx=XM_006928026.4
A1 207013798 207013905
refGene_gn=HMGCS1;refGene_tx=XM_003981430.5;refGene_gn=HMGCS1;refGene_tx=XM_006928026.4
A1 207015245 207015371
refGene_gn=HMGCS1;refGene_tx=XM_003981430.5;refGene_gn=HMGCS1;refGene_tx=XM_006928026.4
A1 207015597 207015761
refGene_gn=HMGCS1;refGene_tx=XM_003981430.5;refGene_gn=HMGCS1;refGene_tx=XM_006928026.4
A1 207016893 207016983
refGene_gn=HMGCS1;refGene_tx=XM_003981430.5;refGene_gn=HMGCS1;refGene_tx=XM_006928026.4
A1 207026828 207027578
refGene_gn=NIM1K;refGene_tx=XM_003981431.5;refGene_gn=NIM1K;refGene_tx=XM_019811263.2;refGene_gn=NIM1K;refGene_tx=XM_023238605.1
A1 207029538 207029807
refGene_gn=NIM1K;refGene_tx=XM_003981431.5;refGene_gn=NIM1K;refGene_tx

=XM_019811263.2;refGene_gn=NIM1K;refGene_tx=XM_023238605.1
A1 207044194 207044486
refGene_gn=NIM1K;refGene_tx=XM_003981431.5;refGene_gn=NIM1K;refGene_tx
=XM_019811263.2;refGene_gn=NIM1K;refGene_tx=XM_023238605.1
A1 207088835 207088924
refGene_gn=LOC109493037;refGene_tx=XM_023245683.1
A1 207092501 207093167
refGene_gn=LOC109493037;refGene_tx=XM_023245683.1
A1 207093688 207093803
refGene_gn=LOC109493037;refGene_tx=XM_023245683.1
A1 207104472 207105159
refGene_gn=ZNF131;refGene_tx=XM_011286332.3;refGene_gn=ZNF131;refGene_
tx=XM_019811284.2;refGene_gn=ZNF131;refGene_tx=XM_023238635.1;refGene_
gn=ZNF131;refGene_tx=XM_023238642.1;refGene_gn=ZNF131;refGene_tx=XM_02
3238647.1;refGene_gn=ZNF131;refGene_tx=XM_023238649.1;refGene_gn=ZNF13
1;refGene_tx=XM_023238650.1;refGene_gn=ZNF131;refGene_tx=XM_023238656.
1;refGene_gn=ZNF131;refGene_tx=XM_023238661.1;refGene_gn=ZNF131;refGen
e_tx=XM_023238670.1
A1 207105848 207105979
refGene_gn=ZNF131;refGene_tx=XM_011286332.3;refGene_gn=ZNF131;refGene_
tx=XM_019811284.2;refGene_gn=ZNF131;refGene_tx=XM_023238635.1;refGene_
gn=ZNF131;refGene_tx=XM_023238642.1;refGene_gn=ZNF131;refGene_tx=XM_02
3238647.1;refGene_gn=ZNF131;refGene_tx=XM_023238649.1;refGene_gn=ZNF13
1;refGene_tx=XM_023238650.1;refGene_gn=ZNF131;refGene_tx=XM_023238656.
1;refGene_gn=ZNF131;refGene_tx=XM_023238661.1;refGene_gn=ZNF131;refGen
e_tx=XM_023238670.1
A1 207112879 207113565
refGene_gn=ZNF131;refGene_tx=XM_011286332.3;refGene_gn=ZNF131;refGene_
tx=XM_019811284.2;refGene_gn=ZNF131;refGene_tx=XM_023238635.1;refGene_
gn=ZNF131;refGene_tx=XM_023238642.1;refGene_gn=ZNF131;refGene_tx=XM_02
3238647.1;refGene_gn=ZNF131;refGene_tx=XM_023238649.1;refGene_gn=ZNF13
1;refGene_tx=XM_023238650.1;refGene_gn=ZNF131;refGene_tx=XM_023238656.
1;refGene_gn=ZNF131;refGene_tx=XM_023238661.1;refGene_gn=ZNF131;refGen
e_tx=XM_023238670.1
A1 207124678 207124823
refGene_gn=ZNF131;refGene_tx=XM_011286332.3;refGene_gn=ZNF131;refGene_
tx=XM_023238635.1;refGene_gn=ZNF131;refGene_tx=XM_023238642.1;refGene_
gn=ZNF131;refGene_tx=XM_023238647.1;refGene_gn=ZNF131;refGene_tx=XM_02
3238649.1;refGene_gn=ZNF131;refGene_tx=XM_023238650.1;refGene_gn=ZNF13
1;refGene_tx=XM_023238656.1;refGene_gn=ZNF131;refGene_tx=XM_023238661.
1;refGene_gn=ZNF131;refGene_tx=XM_023238670.1
A1 207131473 207131575
refGene_gn=ZNF131;refGene_tx=XM_011286332.3;refGene_gn=ZNF131;refGene_
tx=XM_023238635.1;refGene_gn=ZNF131;refGene_tx=XM_023238642.1;refGene_
gn=ZNF131;refGene_tx=XM_023238647.1;refGene_gn=ZNF131;refGene_tx=XM_02
3238649.1;refGene_gn=ZNF131;refGene_tx=XM_023238650.1;refGene_gn=ZNF13
1;refGene_tx=XM_023238656.1;refGene_gn=ZNF131;refGene_tx=XM_023238661.
1;refGene_gn=ZNF131;refGene_tx=XM_023238670.1
A1 207132597 207132721
refGene_gn=ZNF131;refGene_tx=XM_011286332.3;refGene_gn=ZNF131;refGene_

tx=XM_023238635.1;refGene_gn=ZNF131;refGene_tx=XM_023238642.1;refGene_gn=ZNF131;refGene_tx=XM_023238647.1;refGene_gn=ZNF131;refGene_tx=XM_023238649.1;refGene_gn=ZNF131;refGene_tx=XM_023238650.1;refGene_gn=ZNF131;refGene_tx=XM_023238656.1;refGene_gn=ZNF131;refGene_tx=XM_023238661.1;refGene_gn=ZNF131;refGene_tx=XM_023238670.1
A1	207316545	207316748
refGene_gn=SELENOP;refGene_tx=XM_011286344.3;refGene_gn=SELENOP;refGene_tx=XM_019811303.2
A1	207316994	207317207
refGene_gn=SELENOP;refGene_tx=XM_011286344.3;refGene_gn=SELENOP;refGene_tx=XM_019811303.2
A1	207318480	207318598
refGene_gn=SELENOP;refGene_tx=XM_011286344.3;refGene_gn=SELENOP;refGene_tx=XM_019811303.2
A1	207322967	207323609
refGene_gn=SELENOP;refGene_tx=XM_011286344.3;refGene_gn=SELENOP;refGene_tx=XM_019811303.2
A1	207324570	207324720
refGene_gn=CCDC152;refGene_tx=XM_006928031.4
A1	207325458	207325542
refGene_gn=CCDC152;refGene_tx=XM_006928031.4
A1	207327445	207327573
refGene_gn=CCDC152;refGene_tx=XM_006928031.4
A1	207329856	207329959
refGene_gn=CCDC152;refGene_tx=XM_006928031.4
A1	207331801	207331866
refGene_gn=CCDC152;refGene_tx=XM_006928031.4
A1	207344208	207344277
refGene_gn=CCDC152;refGene_tx=XM_006928031.4
A1	207351135	207351241
refGene_gn=CCDC152;refGene_tx=XM_006928031.4
A1	207353155	207353249
refGene_gn=CCDC152;refGene_tx=XM_006928031.4
A1	207355186	207355512
refGene_gn=CCDC152;refGene_tx=XM_006928031.4
A1	207383458	207384430
refGene_gn=GHR;refGene_tx=XM_011286348.3;refGene_gn=GHR;refGene_tx=XM_023238712.1;refGene_gn=GHR;refGene_tx=XM_023238718.1
A1	207384761	207384831
refGene_gn=GHR;refGene_tx=XM_011286348.3;refGene_gn=GHR;refGene_tx=XM_023238712.1;refGene_gn=GHR;refGene_tx=XM_023238718.1
A1	207389102	207389193
refGene_gn=GHR;refGene_tx=XM_011286348.3;refGene_gn=GHR;refGene_tx=XM_023238712.1;refGene_gn=GHR;refGene_tx=XM_023238718.1
A1	207392368	207392534
refGene_gn=GHR;refGene_tx=XM_011286348.3;refGene_gn=GHR;refGene_tx=XM_023238712.1;refGene_gn=GHR;refGene_tx=XM_023238718.1
A1	207400680	207400859
refGene_gn=GHR;refGene_tx=XM_011286348.3;refGene_gn=GHR;refGene_tx=XM_023238712.1;refGene_gn=GHR;refGene_tx=XM_023238718.1

A1	207405872	207406045
refGene_gn=GHR;refGene_tx=XM_011286348.3;refGene_gn=GHR;refGene_tx=XM_023238712.1;refGene_gn=GHR;refGene_tx=XM_023238718.1
A1	207412772	207412902
refGene_gn=GHR;refGene_tx=XM_011286348.3;refGene_gn=GHR;refGene_tx=XM_023238712.1;refGene_gn=GHR;refGene_tx=XM_023238718.1
A1	207457831	207457897
refGene_gn=GHR;refGene_tx=XM_011286348.3;refGene_gn=GHR;refGene_tx=XM_023238712.1;refGene_gn=GHR;refGene_tx=XM_023238718.1
A1	207526371	207526462
refGene_gn=GHR;refGene_tx=XM_011286348.3;refGene_gn=GHR;refGene_tx=XM_023238712.1;refGene_gn=GHR;refGene_tx=XM_023238718.1
A1	207977329	207977466
refGene_gn=FBXO4;refGene_tx=XM_023238755.1
A1	208019426	208019516
refGene_gn=FBXO4;refGene_tx=XM_003981435.5;refGene_gn=FBXO4;refGene_tx=XM_019811354.2;refGene_gn=FBXO4;refGene_tx=XM_023238737.1;refGene_gn=FBXO4;refGene_tx=XM_023238744.1;refGene_gn=FBXO4;refGene_tx=XM_023238759.1;refGene_gn=FBXO4;refGene_tx=XM_023238762.1;refGene_gn=FBXO4;refGene_tx=XM_023238768.1
A1	208020710	208020906
refGene_gn=FBXO4;refGene_tx=XM_003981435.5;refGene_gn=FBXO4;refGene_tx=XM_019811354.2;refGene_gn=FBXO4;refGene_tx=XM_023238737.1;refGene_gn=FBXO4;refGene_tx=XM_023238744.1;refGene_gn=FBXO4;refGene_tx=XM_023238759.1;refGene_gn=FBXO4;refGene_tx=XM_023238762.1
A1	208021463	208021645
refGene_gn=FBXO4;refGene_tx=XM_023238741.1
A1	208024675	208024851
refGene_gn=FBXO4;refGene_tx=XM_003981435.5;refGene_gn=FBXO4;refGene_tx=XM_019811354.2;refGene_gn=FBXO4;refGene_tx=XM_023238737.1;refGene_gn=FBXO4;refGene_tx=XM_023238741.1;refGene_gn=FBXO4;refGene_tx=XM_023238744.1;refGene_gn=FBXO4;refGene_tx=XM_023238755.1;refGene_gn=FBXO4;refGene_tx=XM_023238759.1;refGene_gn=FBXO4;refGene_tx=XM_023238762.1;refGene_gn=FBXO4;refGene_tx=XM_023238768.1
A1	208024961	208025037
refGene_gn=FBXO4;refGene_tx=XM_003981435.5;refGene_gn=FBXO4;refGene_tx=XM_019811354.2;refGene_gn=FBXO4;refGene_tx=XM_023238737.1;refGene_gn=FBXO4;refGene_tx=XM_023238741.1;refGene_gn=FBXO4;refGene_tx=XM_023238744.1;refGene_gn=FBXO4;refGene_tx=XM_023238755.1;refGene_gn=FBXO4;refGene_tx=XM_023238759.1;refGene_gn=FBXO4;refGene_tx=XM_023238762.1;refGene_gn=FBXO4;refGene_tx=XM_023238768.1
A1	208028568	208028789
refGene_gn=FBXO4;refGene_tx=XM_003981435.5;refGene_gn=FBXO4;refGene_tx=XM_019811354.2;refGene_gn=FBXO4;refGene_tx=XM_023238737.1;refGene_gn=FBXO4;refGene_tx=XM_023238741.1;refGene_gn=FBXO4;refGene_tx=XM_023238744.1;refGene_gn=FBXO4;refGene_tx=XM_023238755.1;refGene_gn=FBXO4;refGene_tx=XM_023238759.1;refGene_gn=FBXO4;refGene_tx=XM_023238762.1;refGene_gn=FBXO4;refGene_tx=XM_023238768.1
A1	208031373	208031609
refGene_gn=FBXO4;refGene_tx=XM_003981435.5;refGene_gn=FBXO4;refGene_tx

=XM_019811354.2;refGene_gn=FBXO4;refGene_tx=XM_023238737.1;refGene_gn=FBXO4;refGene_tx=XM_023238741.1;refGene_gn=FBXO4;refGene_tx=XM_023238744.1;refGene_gn=FBXO4;refGene_tx=XM_023238755.1;refGene_gn=FBXO4;refGene_tx=XM_023238759.1;refGene_gn=FBXO4;refGene_tx=XM_023238762.1;refGene_gn=FBXO4;refGene_tx=XM_023238768.1
A1 208032841 208033030
refGene_gn=FBXO4;refGene_tx=XM_003981435.5;refGene_gn=FBXO4;refGene_tx=XM_019811354.2;refGene_gn=FBXO4;refGene_tx=XM_023238737.1;refGene_gn=FBXO4;refGene_tx=XM_023238741.1;refGene_gn=FBXO4;refGene_tx=XM_023238744.1;refGene_gn=FBXO4;refGene_tx=XM_023238755.1;refGene_gn=FBXO4;refGene_tx=XM_023238759.1;refGene_gn=FBXO4;refGene_tx=XM_023238762.1;refGene_gn=FBXO4;refGene_tx=XM_023238768.1
A1 208042395 208042682
refGene_gn=CA1H5orf51;refGene_tx=XM_003981436.5
A1 208046945 208047060
refGene_gn=CA1H5orf51;refGene_tx=XM_003981436.5
A1 208047955 208048087
refGene_gn=CA1H5orf51;refGene_tx=XM_003981436.5
A1 208050813 208050957
refGene_gn=CA1H5orf51;refGene_tx=XM_003981436.5
A1 208058067 208058174
refGene_gn=CA1H5orf51;refGene_tx=XM_003981436.5
A1 208061810 208061910
refGene_gn=CA1H5orf51;refGene_tx=XM_003981436.5
A1 208086834 208086912
refGene_gn=OXCT1;refGene_tx=XM_003981437.5
A1 208095752 208095861
refGene_gn=OXCT1;refGene_tx=XM_003981437.5
A1 208096939 208097030
refGene_gn=OXCT1;refGene_tx=XM_003981437.5
A1 208107065 208107201
refGene_gn=OXCT1;refGene_tx=XM_003981437.5
A1 208110779 208110929
refGene_gn=OXCT1;refGene_tx=XM_003981437.5
A1 208115411 208115518
refGene_gn=OXCT1;refGene_tx=XM_003981437.5
A1 208118749 208118810
refGene_gn=OXCT1;refGene_tx=XM_003981437.5
A1 208151283 208151391
refGene_gn=OXCT1;refGene_tx=XM_003981437.5
A1 208152652 208152767
refGene_gn=OXCT1;refGene_tx=XM_003981437.5
A1 208156790 208156885
refGene_gn=OXCT1;refGene_tx=XM_003981437.5
A1 208158617 208158666
refGene_gn=OXCT1;refGene_tx=XM_003981437.5
A1 208164950 208164977
refGene_gn=OXCT1;refGene_tx=XM_003981437.5
A1 208164979 208165022
refGene_gn=OXCT1;refGene_tx=XM_003981437.5

A1 208165635 208165711
refGene_gn=OXCT1;refGene_tx=XM_003981437.5
A1 208198980 208199070
refGene_gn=OXCT1;refGene_tx=XM_003981437.5
A1 208208783 208208864
refGene_gn=OXCT1;refGene_tx=XM_003981437.5
A1 208220222 208220324
refGene_gn=OXCT1;refGene_tx=XM_003981437.5
A1 208225289 208225331
refGene_gn=OXCT1;refGene_tx=XM_003981437.5
A1 208392595 208392698
refGene_gn=PLCXD3;refGene_tx=XM_019811398.2;refGene_gn=PLCXD3;refGene_tx=XM_019811404.2
A1 208490386 208490453
refGene_gn=PLCXD3;refGene_tx=XM_019811410.2
A1 208506627 208507336
refGene_gn=PLCXD3;refGene_tx=XM_019811398.2;refGene_gn=PLCXD3;refGene_tx=XM_019811404.2;refGene_gn=PLCXD3;refGene_tx=XM_019811410.2
A1 208512979 208513121
refGene_gn=PLCXD3;refGene_tx=XM_019811404.2
A1 208556558 208556712
refGene_gn=PLCXD3;refGene_tx=XM_019811398.2;refGene_gn=PLCXD3;refGene_tx=XM_019811410.2
A1 208645106 208645249
refGene_gn=C6;refGene_tx=XM_019811437.2
A1 208646682 208646839
refGene_gn=C6;refGene_tx=XM_019811437.2
A1 208648270 208648415
refGene_gn=C6;refGene_tx=XM_019811437.2
A1 208652950 208653089
refGene_gn=C6;refGene_tx=XM_019811437.2
A1 208659716 208659855
refGene_gn=C6;refGene_tx=XM_019811437.2
A1 208665730 208665931
refGene_gn=C6;refGene_tx=XM_019811437.2
A1 208671424 208671665
refGene_gn=C6;refGene_tx=XM_019811437.2
A1 208676193 208676252
refGene_gn=C6;refGene_tx=XM_019811437.2
A1 208676253 208676317
refGene_gn=C6;refGene_tx=XM_019811437.2
A1 208683596 208683763
refGene_gn=C6;refGene_tx=XM_019811437.2
A1 208685175 208685401
refGene_gn=C6;refGene_tx=XM_019811437.2
A1 208686300 208686472
refGene_gn=C6;refGene_tx=XM_019811437.2
A1 208686667 208686779
refGene_gn=C6;refGene_tx=XM_019811437.2
A1 208689365 208689498

refGene_gn=C6;refGene_tx=XM_019811437.2
A1 208690422 208690611
refGene_gn=C6;refGene_tx=XM_019811437.2
A1 208696020 208696111
refGene_gn=C6;refGene_tx=XM_019811437.2
A1 208699020 208699262
refGene_gn=C6;refGene_tx=XM_019811437.2
A1 208704821 208705003
refGene_gn=C6;refGene_tx=XM_019811437.2
A1 208750650 208750678
refGene_gn=MROH2B;refGene_tx=XM_019811443.2;refGene_gn=MROH2B;refGene_tx=XM_019811451.2;refGene_gn=MROH2B;refGene_tx=XM_023238843.1
A1 208752310 208752372
refGene_gn=MROH2B;refGene_tx=XM_019811443.2;refGene_gn=MROH2B;refGene_tx=XM_019811451.2;refGene_gn=MROH2B;refGene_tx=XM_023238843.1
A1 208753789 208753900
refGene_gn=MROH2B;refGene_tx=XM_019811443.2;refGene_gn=MROH2B;refGene_tx=XM_019811451.2;refGene_gn=MROH2B;refGene_tx=XM_023238843.1
A1 208755425 208755585
refGene_gn=MROH2B;refGene_tx=XM_019811443.2;refGene_gn=MROH2B;refGene_tx=XM_019811451.2;refGene_gn=MROH2B;refGene_tx=XM_023238843.1
A1 208756300 208756399
refGene_gn=MROH2B;refGene_tx=XM_019811443.2;refGene_gn=MROH2B;refGene_tx=XM_019811451.2;refGene_gn=MROH2B;refGene_tx=XM_023238843.1
A1 208759119 208759274
refGene_gn=MROH2B;refGene_tx=XM_019811443.2;refGene_gn=MROH2B;refGene_tx=XM_019811451.2;refGene_gn=MROH2B;refGene_tx=XM_023238843.1;refGene_gn=MROH2B;refGene_tx=XM_023238844.1
A1 208762045 208762186
refGene_gn=MROH2B;refGene_tx=XM_019811443.2;refGene_gn=MROH2B;refGene_tx=XM_019811451.2;refGene_gn=MROH2B;refGene_tx=XM_023238843.1;refGene_gn=MROH2B;refGene_tx=XM_023238844.1
A1 208762533 208762626
refGene_gn=MROH2B;refGene_tx=XM_019811443.2;refGene_gn=MROH2B;refGene_tx=XM_019811451.2;refGene_gn=MROH2B;refGene_tx=XM_023238843.1;refGene_gn=MROH2B;refGene_tx=XM_023238844.1
A1 208762717 208762787
refGene_gn=MROH2B;refGene_tx=XM_019811443.2;refGene_gn=MROH2B;refGene_tx=XM_019811451.2;refGene_gn=MROH2B;refGene_tx=XM_023238843.1;refGene_gn=MROH2B;refGene_tx=XM_023238844.1
A1 208764678 208764792
refGene_gn=MROH2B;refGene_tx=XM_019811443.2;refGene_gn=MROH2B;refGene_tx=XM_019811451.2;refGene_gn=MROH2B;refGene_tx=XM_023238843.1;refGene_gn=MROH2B;refGene_tx=XM_023238844.1
A1 208765674 208765748
refGene_gn=MROH2B;refGene_tx=XM_019811443.2;refGene_gn=MROH2B;refGene_tx=XM_019811451.2;refGene_gn=MROH2B;refGene_tx=XM_023238843.1;refGene_gn=MROH2B;refGene_tx=XM_023238844.1
A1 208768192 208768315
refGene_gn=MROH2B;refGene_tx=XM_019811443.2;refGene_gn=MROH2B;refGene_

tx=XM_019811451.2;refGene_gn=MROH2B;refGene_tx=XM_023238843.1;refGene_gn=MROH2B;refGene_tx=XM_023238844.1
A1 208769049 208769163
refGene_gn=MROH2B;refGene_tx=XM_019811443.2;refGene_gn=MROH2B;refGene_tx=XM_019811451.2;refGene_gn=MROH2B;refGene_tx=XM_023238843.1;refGene_gn=MROH2B;refGene_tx=XM_023238844.1
A1 208771222 208771379
refGene_gn=MROH2B;refGene_tx=XM_019811443.2;refGene_gn=MROH2B;refGene_tx=XM_019811451.2;refGene_gn=MROH2B;refGene_tx=XM_023238843.1;refGene_gn=MROH2B;refGene_tx=XM_023238844.1
A1 208771525 208771566
refGene_gn=MROH2B;refGene_tx=XM_019811443.2;refGene_gn=MROH2B;refGene_tx=XM_019811451.2;refGene_gn=MROH2B;refGene_tx=XM_023238843.1;refGene_gn=MROH2B;refGene_tx=XM_023238844.1
A1 208772456 208772598
refGene_gn=MROH2B;refGene_tx=XM_019811443.2;refGene_gn=MROH2B;refGene_tx=XM_019811451.2;refGene_gn=MROH2B;refGene_tx=XM_023238843.1;refGene_gn=MROH2B;refGene_tx=XM_023238844.1
A1 208773155 208773199
refGene_gn=MROH2B;refGene_tx=XM_019811443.2;refGene_gn=MROH2B;refGene_tx=XM_019811451.2;refGene_gn=MROH2B;refGene_tx=XM_023238843.1;refGene_gn=MROH2B;refGene_tx=XM_023238844.1
A1 208775466 208775574
refGene_gn=MROH2B;refGene_tx=XM_019811443.2;refGene_gn=MROH2B;refGene_tx=XM_019811451.2;refGene_gn=MROH2B;refGene_tx=XM_023238843.1;refGene_gn=MROH2B;refGene_tx=XM_023238844.1
A1 208779047 208779164
refGene_gn=MROH2B;refGene_tx=XM_019811443.2;refGene_gn=MROH2B;refGene_tx=XM_019811451.2;refGene_gn=MROH2B;refGene_tx=XM_023238843.1;refGene_gn=MROH2B;refGene_tx=XM_023238844.1
A1 208790800 208790908
refGene_gn=MROH2B;refGene_tx=XM_019811443.2;refGene_gn=MROH2B;refGene_tx=XM_019811451.2;refGene_gn=MROH2B;refGene_tx=XM_023238843.1;refGene_gn=MROH2B;refGene_tx=XM_023238844.1
A1 208791438 208791591
refGene_gn=MROH2B;refGene_tx=XM_019811443.2;refGene_gn=MROH2B;refGene_tx=XM_019811451.2;refGene_gn=MROH2B;refGene_tx=XM_023238843.1;refGene_gn=MROH2B;refGene_tx=XM_023238844.1
A1 208794335 208794362
refGene_gn=MROH2B;refGene_tx=XM_019811443.2;refGene_gn=MROH2B;refGene_tx=XM_019811451.2;refGene_gn=MROH2B;refGene_tx=XM_023238843.1;refGene_gn=MROH2B;refGene_tx=XM_023238844.1
A1 208795152 208795272
refGene_gn=MROH2B;refGene_tx=XM_019811443.2;refGene_gn=MROH2B;refGene_tx=XM_019811451.2;refGene_gn=MROH2B;refGene_tx=XM_023238843.1;refGene_gn=MROH2B;refGene_tx=XM_023238844.1
A1 208795488 208795568
refGene_gn=MROH2B;refGene_tx=XM_019811443.2;refGene_gn=MROH2B;refGene_tx=XM_019811451.2;refGene_gn=MROH2B;refGene_tx=XM_023238843.1;refGene_gn=MROH2B;refGene_tx=XM_023238844.1

A1	208801582	208801718
refGene_gn=MROH2B;refGene_tx=XM_019811443.2;refGene_gn=MROH2B;refGene_tx=XM_019811451.2;refGene_gn=MROH2B;refGene_tx=XM_023238843.1;refGene_gn=MROH2B;refGene_tx=XM_023238844.1
A1	208801812	208801908
refGene_gn=MROH2B;refGene_tx=XM_019811443.2;refGene_gn=MROH2B;refGene_tx=XM_019811451.2;refGene_gn=MROH2B;refGene_tx=XM_023238843.1;refGene_gn=MROH2B;refGene_tx=XM_023238844.1
A1	208802169	208802259
refGene_gn=MROH2B;refGene_tx=XM_019811443.2;refGene_gn=MROH2B;refGene_tx=XM_019811451.2;refGene_gn=MROH2B;refGene_tx=XM_023238843.1;refGene_gn=MROH2B;refGene_tx=XM_023238844.1
A1	208802627	208802748
refGene_gn=MROH2B;refGene_tx=XM_019811443.2;refGene_gn=MROH2B;refGene_tx=XM_019811451.2;refGene_gn=MROH2B;refGene_tx=XM_023238843.1;refGene_gn=MROH2B;refGene_tx=XM_023238844.1
A1	208805125	208805223
refGene_gn=MROH2B;refGene_tx=XM_019811443.2;refGene_gn=MROH2B;refGene_tx=XM_019811451.2;refGene_gn=MROH2B;refGene_tx=XM_023238843.1;refGene_gn=MROH2B;refGene_tx=XM_023238844.1
A1	208807823	208807976
refGene_gn=MROH2B;refGene_tx=XM_019811443.2;refGene_gn=MROH2B;refGene_tx=XM_019811451.2;refGene_gn=MROH2B;refGene_tx=XM_023238843.1;refGene_gn=MROH2B;refGene_tx=XM_023238844.1
A1	208810138	208810296
refGene_gn=MROH2B;refGene_tx=XM_019811443.2;refGene_gn=MROH2B;refGene_tx=XM_019811451.2;refGene_gn=MROH2B;refGene_tx=XM_023238843.1;refGene_gn=MROH2B;refGene_tx=XM_023238844.1
A1	208810806	208810933
refGene_gn=MROH2B;refGene_tx=XM_019811443.2;refGene_gn=MROH2B;refGene_tx=XM_019811451.2;refGene_gn=MROH2B;refGene_tx=XM_023238843.1;refGene_gn=MROH2B;refGene_tx=XM_023238844.1
A1	208811527	208811715
refGene_gn=MROH2B;refGene_tx=XM_019811443.2;refGene_gn=MROH2B;refGene_tx=XM_019811451.2;refGene_gn=MROH2B;refGene_tx=XM_023238843.1;refGene_gn=MROH2B;refGene_tx=XM_023238844.1
A1	208813017	208813158
refGene_gn=MROH2B;refGene_tx=XM_019811443.2;refGene_gn=MROH2B;refGene_tx=XM_019811451.2;refGene_gn=MROH2B;refGene_tx=XM_023238843.1;refGene_gn=MROH2B;refGene_tx=XM_023238844.1
A1	208813477	208813592
refGene_gn=MROH2B;refGene_tx=XM_019811443.2;refGene_gn=MROH2B;refGene_tx=XM_019811451.2;refGene_gn=MROH2B;refGene_tx=XM_023238843.1;refGene_gn=MROH2B;refGene_tx=XM_023238844.1
A1	208814225	208814372
refGene_gn=MROH2B;refGene_tx=XM_019811443.2;refGene_gn=MROH2B;refGene_tx=XM_019811451.2;refGene_gn=MROH2B;refGene_tx=XM_023238843.1;refGene_gn=MROH2B;refGene_tx=XM_023238844.1
A1	208814614	208814797
refGene_gn=MROH2B;refGene_tx=XM_019811443.2;refGene_gn=MROH2B;refGene_

tx=XM_019811451.2;refGene_gn=MROH2B;refGene_tx=XM_023238843.1;refGene_gn=MROH2B;refGene_tx=XM_023238844.1
A1 208816330 208816399
refGene_gn=MROH2B;refGene_tx=XM_023238843.1
A1 208818712 208818868
refGene_gn=MROH2B;refGene_tx=XM_019811443.2;refGene_gn=MROH2B;refGene_tx=XM_019811451.2;refGene_gn=MROH2B;refGene_tx=XM_023238844.1
A1 208819211 208819346
refGene_gn=MROH2B;refGene_tx=XM_019811443.2;refGene_gn=MROH2B;refGene_tx=XM_023238844.1
A1 208819989 208820092
refGene_gn=MROH2B;refGene_tx=XM_019811443.2;refGene_gn=MROH2B;refGene_tx=XM_019811451.2;refGene_gn=MROH2B;refGene_tx=XM_023238844.1
A1 208821099 208821165
refGene_gn=MROH2B;refGene_tx=XM_019811443.2;refGene_gn=MROH2B;refGene_tx=XM_019811451.2;refGene_gn=MROH2B;refGene_tx=XM_023238844.1
A1 208821618 208821725
refGene_gn=MROH2B;refGene_tx=XM_019811443.2;refGene_gn=MROH2B;refGene_tx=XM_019811451.2;refGene_gn=MROH2B;refGene_tx=XM_023238844.1
A1 208835217 208835399
refGene_gn=C7;refGene_tx=XM_003981508.5;refGene_gn=C7;refGene_tx=XM_019811479.2;refGene_gn=C7;refGene_tx=XM_019811484.2
A1 208837576 208837761
refGene_gn=C7;refGene_tx=XM_003981508.5;refGene_gn=C7;refGene_tx=XM_019811479.2;refGene_gn=C7;refGene_tx=XM_019811484.2
A1 208841098 208841189
refGene_gn=C7;refGene_tx=XM_003981508.5;refGene_gn=C7;refGene_tx=XM_019811479.2;refGene_gn=C7;refGene_tx=XM_019811484.2
A1 208845812 208846004
refGene_gn=C7;refGene_tx=XM_003981508.5;refGene_gn=C7;refGene_tx=XM_019811479.2;refGene_gn=C7;refGene_tx=XM_019811484.2
A1 208849224 208849357
refGene_gn=C7;refGene_tx=XM_003981508.5;refGene_gn=C7;refGene_tx=XM_019811479.2;refGene_gn=C7;refGene_tx=XM_019811484.2
A1 208851762 208851850
refGene_gn=C7;refGene_tx=XM_003981508.5;refGene_gn=C7;refGene_tx=XM_019811479.2;refGene_gn=C7;refGene_tx=XM_019811484.2
A1 208853587 208853759
refGene_gn=C7;refGene_tx=XM_003981508.5;refGene_gn=C7;refGene_tx=XM_019811479.2;refGene_gn=C7;refGene_tx=XM_019811484.2
A1 208855653 208855882
refGene_gn=C7;refGene_tx=XM_003981508.5;refGene_gn=C7;refGene_tx=XM_019811479.2;refGene_gn=C7;refGene_tx=XM_019811484.2
A1 208858126 208858293
refGene_gn=C7;refGene_tx=XM_003981508.5;refGene_gn=C7;refGene_tx=XM_019811479.2;refGene_gn=C7;refGene_tx=XM_019811484.2
A1 208861861 208861972
refGene_gn=C7;refGene_tx=XM_003981508.5;refGene_gn=C7;refGene_tx=XM_019811479.2;refGene_gn=C7;refGene_tx=XM_019811484.2
A1 208864107 208864351

refGene_gn=C7;refGene_tx=XM_003981508.5;refGene_gn=C7;refGene_tx=XM_019811479.2;refGene_gn=C7;refGene_tx=XM_019811484.2
A1 208866792 208866963
refGene_gn=C7;refGene_tx=XM_003981508.5;refGene_gn=C7;refGene_tx=XM_019811479.2;refGene_gn=C7;refGene_tx=XM_019811484.2
A1 208875459 208875598
refGene_gn=C7;refGene_tx=XM_003981508.5;refGene_gn=C7;refGene_tx=XM_019811479.2;refGene_gn=C7;refGene_tx=XM_019811484.2
A1 208876654 208876802
refGene_gn=C7;refGene_tx=XM_003981508.5;refGene_gn=C7;refGene_tx=XM_019811479.2;refGene_gn=C7;refGene_tx=XM_019811484.2
A1 208879177 208879319
refGene_gn=C7;refGene_tx=XM_003981508.5;refGene_gn=C7;refGene_tx=XM_019811479.2;refGene_gn=C7;refGene_tx=XM_019811484.2
A1 208884777 208884853
refGene_gn=C7;refGene_tx=XM_003981508.5
A1 208886807 208886863
refGene_gn=C7;refGene_tx=XM_003981508.5;refGene_gn=C7;refGene_tx=XM_019811479.2
A1 208890490 208890635
refGene_gn=C7;refGene_tx=XM_003981508.5;refGene_gn=C7;refGene_tx=XM_019811479.2;refGene_gn=C7;refGene_tx=XM_019811484.2
A1 208923532 208925778
refGene_gn=CARD6;refGene_tx=XM_003981440.5
A1 208931890 208932448
refGene_gn=CARD6;refGene_tx=XM_003981440.5
A1 208933541 208933824
refGene_gn=CARD6;refGene_tx=XM_003981440.5
A1 208937902 208937905
refGene_gn=RPL37;refGene_tx=XM_003981441.4
A1 208938521 208938657
refGene_gn=RPL37;refGene_tx=XM_003981441.4
A1 208938899 208938984
refGene_gn=RPL37;refGene_tx=XM_003981441.4
A1 208940304 208940374
refGene_gn=RPL37;refGene_tx=XM_003981441.4
A1 208961064 208961191
refGene_gn=PRKAA1;refGene_tx=XM_023238864.1;refGene_gn=PRKAA1;refGene_tx=XM_023238865.1;refGene_gn=PRKAA1;refGene_tx=XM_023238876.1
A1 208963065 208963078
refGene_gn=PRKAA1;refGene_tx=XM_011286402.3
A1 208978733 208978875
refGene_gn=PRKAA1;refGene_tx=XM_011286402.3;refGene_gn=PRKAA1;refGene_tx=XM_023238864.1;refGene_gn=PRKAA1;refGene_tx=XM_023238865.1;refGene_gn=PRKAA1;refGene_tx=XM_023238876.1
A1 208980485 208980579
refGene_gn=PRKAA1;refGene_tx=XM_011286402.3;refGene_gn=PRKAA1;refGene_tx=XM_023238864.1;refGene_gn=PRKAA1;refGene_tx=XM_023238865.1;refGene_gn=PRKAA1;refGene_tx=XM_023238866.1;refGene_gn=PRKAA1;refGene_tx=XM_023238869.1;refGene_gn=PRKAA1;refGene_tx=XM_023238872.1;refGene_gn=PRKAA

1;refGene_tx=XM_023238876.1
A1	208981012	208981057
refGene_gn=PRKAA1;refGene_tx=XM_011286402.3;refGene_gn=PRKAA1;refGene_tx=XM_023238864.1;refGene_gn=PRKAA1;refGene_tx=XM_023238866.1;refGene_gn=PRKAA1;refGene_tx=XM_023238869.1;refGene_gn=PRKAA1;refGene_tx=XM_023238872.1
A1	208984692	208984837
refGene_gn=PRKAA1;refGene_tx=XM_011286402.3;refGene_gn=PRKAA1;refGene_tx=XM_023238864.1;refGene_gn=PRKAA1;refGene_tx=XM_023238865.1;refGene_gn=PRKAA1;refGene_tx=XM_023238866.1;refGene_gn=PRKAA1;refGene_tx=XM_023238869.1;refGene_gn=PRKAA1;refGene_tx=XM_023238872.1;refGene_gn=PRKAA1;refGene_tx=XM_023238874.1;refGene_gn=PRKAA1;refGene_tx=XM_023238876.1
A1	208986852	208986940
refGene_gn=PRKAA1;refGene_tx=XM_011286402.3;refGene_gn=PRKAA1;refGene_tx=XM_023238864.1;refGene_gn=PRKAA1;refGene_tx=XM_023238865.1;refGene_gn=PRKAA1;refGene_tx=XM_023238866.1;refGene_gn=PRKAA1;refGene_tx=XM_023238869.1;refGene_gn=PRKAA1;refGene_tx=XM_023238872.1;refGene_gn=PRKAA1;refGene_tx=XM_023238874.1;refGene_gn=PRKAA1;refGene_tx=XM_023238876.1
A1	208987373	208987446
refGene_gn=PRKAA1;refGene_tx=XM_023238876.1
A1	208988528	208988753
refGene_gn=PRKAA1;refGene_tx=XM_011286402.3;refGene_gn=PRKAA1;refGene_tx=XM_023238864.1;refGene_gn=PRKAA1;refGene_tx=XM_023238865.1;refGene_gn=PRKAA1;refGene_tx=XM_023238866.1;refGene_gn=PRKAA1;refGene_tx=XM_023238869.1;refGene_gn=PRKAA1;refGene_tx=XM_023238872.1;refGene_gn=PRKAA1;refGene_tx=XM_023238874.1
A1	208990075	208990562
refGene_gn=PRKAA1;refGene_tx=XM_011286402.3;refGene_gn=PRKAA1;refGene_tx=XM_023238864.1;refGene_gn=PRKAA1;refGene_tx=XM_023238865.1;refGene_gn=PRKAA1;refGene_tx=XM_023238866.1;refGene_gn=PRKAA1;refGene_tx=XM_023238869.1;refGene_gn=PRKAA1;refGene_tx=XM_023238872.1;refGene_gn=PRKAA1;refGene_tx=XM_023238874.1
A1	208990675	208990802
refGene_gn=PRKAA1;refGene_tx=XM_011286402.3;refGene_gn=PRKAA1;refGene_tx=XM_023238864.1;refGene_gn=PRKAA1;refGene_tx=XM_023238865.1;refGene_gn=PRKAA1;refGene_tx=XM_023238866.1;refGene_gn=PRKAA1;refGene_tx=XM_023238869.1;refGene_gn=PRKAA1;refGene_tx=XM_023238872.1;refGene_gn=PRKAA1;refGene_tx=XM_023238874.1
A1	208994965	208995207
refGene_gn=PRKAA1;refGene_tx=XM_011286402.3;refGene_gn=PRKAA1;refGene_tx=XM_023238864.1;refGene_gn=PRKAA1;refGene_tx=XM_023238865.1;refGene_gn=PRKAA1;refGene_tx=XM_023238866.1;refGene_gn=PRKAA1;refGene_tx=XM_023238869.1;refGene_gn=PRKAA1;refGene_tx=XM_023238872.1;refGene_gn=PRKAA1;refGene_tx=XM_023238874.1
A1	209005719	209005940
refGene_gn=TTC33;refGene_tx=XM_003981444.5;refGene_gn=TTC33;refGene_tx=XM_006928037.4;refGene_gn=TTC33;refGene_tx=XM_019811535.2;refGene_gn=TTC33;refGene_tx=XM_019811540.2;refGene_gn=TTC33;refGene_tx=XM_0198115

41.2;refGene_gn=TTC33;refGene_tx=XM_019811547.1;refGene_gn=TTC33;refGene_tx=XM_023238897.1
A1	209027161	209027261
refGene_gn=TTC33;refGene_tx=XM_019811547.1
A1	209033337	209033419
refGene_gn=TTC33;refGene_tx=XM_003981444.5;refGene_gn=TTC33;refGene_tx=XM_006928037.4;refGene_gn=TTC33;refGene_tx=XM_019811535.2;refGene_gn=TTC33;refGene_tx=XM_019811540.2;refGene_gn=TTC33;refGene_tx=XM_019811541.2;refGene_gn=TTC33;refGene_tx=XM_019811547.1;refGene_gn=TTC33;refGene_tx=XM_023238897.1
A1	209034870	209035002
refGene_gn=TTC33;refGene_tx=XM_003981444.5;refGene_gn=TTC33;refGene_tx=XM_006928037.4;refGene_gn=TTC33;refGene_tx=XM_019811535.2;refGene_gn=TTC33;refGene_tx=XM_019811540.2;refGene_gn=TTC33;refGene_tx=XM_019811541.2;refGene_gn=TTC33;refGene_tx=XM_019811547.1;refGene_gn=TTC33;refGene_tx=XM_023238897.1
A1	209039301	209039655
refGene_gn=TTC33;refGene_tx=XM_003981444.5;refGene_gn=TTC33;refGene_tx=XM_006928037.4;refGene_gn=TTC33;refGene_tx=XM_019811535.2;refGene_gn=TTC33;refGene_tx=XM_019811540.2;refGene_gn=TTC33;refGene_tx=XM_019811541.2;refGene_gn=TTC33;refGene_tx=XM_019811547.1;refGene_gn=TTC33;refGene_tx=XM_023238897.1
A1	209056489	209057080
refGene_gn=PTGER4;refGene_tx=XM_011286412.3
A1	209067419	209068298
refGene_gn=PTGER4;refGene_tx=XM_011286412.3
A1	210112487	210112578
refGene_gn=DAB2;refGene_tx=XM_011286417.2;refGene_gn=DAB2;refGene_tx=XM_019811579.1;refGene_gn=DAB2;refGene_tx=XM_019811583.1;refGene_gn=DAB2;refGene_tx=XM_023238899.1;refGene_gn=DAB2;refGene_tx=XM_023238900.1;refGene_gn=DAB2;refGene_tx=XM_023238901.1
A1	210113587	210113727
refGene_gn=DAB2;refGene_tx=XM_011286417.2;refGene_gn=DAB2;refGene_tx=XM_019811579.1;refGene_gn=DAB2;refGene_tx=XM_019811583.1;refGene_gn=DAB2;refGene_tx=XM_023238899.1;refGene_gn=DAB2;refGene_tx=XM_023238900.1;refGene_gn=DAB2;refGene_tx=XM_023238901.1
A1	210114660	210114759
refGene_gn=DAB2;refGene_tx=XM_011286417.2;refGene_gn=DAB2;refGene_tx=XM_019811579.1;refGene_gn=DAB2;refGene_tx=XM_019811583.1;refGene_gn=DAB2;refGene_tx=XM_023238899.1;refGene_gn=DAB2;refGene_tx=XM_023238900.1;refGene_gn=DAB2;refGene_tx=XM_023238901.1
A1	210116442	210116574
refGene_gn=DAB2;refGene_tx=XM_011286417.2;refGene_gn=DAB2;refGene_tx=XM_019811579.1;refGene_gn=DAB2;refGene_tx=XM_019811583.1;refGene_gn=DAB2;refGene_tx=XM_023238899.1;refGene_gn=DAB2;refGene_tx=XM_023238900.1;refGene_gn=DAB2;refGene_tx=XM_023238901.1
A1	210117292	210117373
refGene_gn=DAB2;refGene_tx=XM_011286417.2;refGene_gn=DAB2;refGene_tx=XM_019811579.1;refGene_gn=DAB2;refGene_tx=XM_019811583.1;refGene_gn=DAB2;refGene_tx=XM_023238899.1;refGene_gn=DAB2;refGene_tx=XM_023238900.1;

refGene_gn=DAB2;refGene_tx=XM_023238901.1
A1 210117894 210117921
refGene_gn=DAB2;refGene_tx=XM_011286417.2;refGene_gn=DAB2;refGene_tx=XM_019811579.1;refGene_gn=DAB2;refGene_tx=XM_019811583.1;refGene_gn=DAB2;refGene_tx=XM_023238899.1;refGene_gn=DAB2;refGene_tx=XM_023238900.1;refGene_gn=DAB2;refGene_tx=XM_023238901.1
A1 210118159 210118213
refGene_gn=DAB2;refGene_tx=XM_011286417.2;refGene_gn=DAB2;refGene_tx=XM_019811579.1;refGene_gn=DAB2;refGene_tx=XM_019811583.1;refGene_gn=DAB2;refGene_tx=XM_023238899.1;refGene_gn=DAB2;refGene_tx=XM_023238900.1;refGene_gn=DAB2;refGene_tx=XM_023238901.1
A1 210118621 210118684
refGene_gn=DAB2;refGene_tx=XM_011286417.2;refGene_gn=DAB2;refGene_tx=XM_019811579.1;refGene_gn=DAB2;refGene_tx=XM_023238899.1;refGene_gn=DAB2;refGene_tx=XM_023238900.1
A1 210123368 210124022
refGene_gn=DAB2;refGene_tx=XM_011286417.2;refGene_gn=DAB2;refGene_tx=XM_019811579.1;refGene_gn=DAB2;refGene_tx=XM_019811583.1;refGene_gn=DAB2;refGene_tx=XM_023238899.1
A1 210125019 210125182
refGene_gn=DAB2;refGene_tx=XM_011286417.2;refGene_gn=DAB2;refGene_tx=XM_019811579.1;refGene_gn=DAB2;refGene_tx=XM_019811583.1;refGene_gn=DAB2;refGene_tx=XM_023238899.1;refGene_gn=DAB2;refGene_tx=XM_023238900.1;refGene_gn=DAB2;refGene_tx=XM_023238901.1
A1 210129140 210129773
refGene_gn=DAB2;refGene_tx=XM_011286417.2;refGene_gn=DAB2;refGene_tx=XM_019811579.1;refGene_gn=DAB2;refGene_tx=XM_019811583.1;refGene_gn=DAB2;refGene_tx=XM_023238899.1;refGene_gn=DAB2;refGene_tx=XM_023238900.1;refGene_gn=DAB2;refGene_tx=XM_023238901.1
A1 210130323 210130433
refGene_gn=DAB2;refGene_tx=XM_011286417.2;refGene_gn=DAB2;refGene_tx=XM_019811579.1;refGene_gn=DAB2;refGene_tx=XM_019811583.1;refGene_gn=DAB2;refGene_tx=XM_023238899.1;refGene_gn=DAB2;refGene_tx=XM_023238900.1;refGene_gn=DAB2;refGene_tx=XM_023238901.1
A1 210131043 210131106
refGene_gn=DAB2;refGene_tx=XM_011286417.2;refGene_gn=DAB2;refGene_tx=XM_019811579.1;refGene_gn=DAB2;refGene_tx=XM_019811583.1;refGene_gn=DAB2;refGene_tx=XM_023238899.1;refGene_gn=DAB2;refGene_tx=XM_023238900.1;refGene_gn=DAB2;refGene_tx=XM_023238901.1
A1 210147430 210147507
refGene_gn=C9;refGene_tx=XM_019811604.1
A1 210161345 210161451
refGene_gn=C9;refGene_tx=XM_019811604.1
A1 210161851 210161996
refGene_gn=C9;refGene_tx=XM_019811604.1
A1 210162783 210162931
refGene_gn=C9;refGene_tx=XM_019811604.1
A1 210170316 210170455
refGene_gn=C9;refGene_tx=XM_019811604.1
A1 210178974 210179244

refGene_gn=C9;refGene_tx=XM_019811604.1
A1	210183812	210184053
refGene_gn=C9;refGene_tx=XM_019811604.1
A1	210187166	210187292
refGene_gn=C9;refGene_tx=XM_019811604.1
A1	210189002	210189181
refGene_gn=C9;refGene_tx=XM_019811604.1
A1	210199053	210199285
refGene_gn=C9;refGene_tx=XM_019811604.1
A1	210200253	210200279
refGene_gn=C9;refGene_tx=XM_019811604.1
A1	210219341	210219344
refGene_gn=FYB1;refGene_tx=XM_011286424.2
A1	210277722	210278884
refGene_gn=FYB1;refGene_tx=XM_011286424.2
A1	210327700	210327857
refGene_gn=FYB1;refGene_tx=XM_011286424.2
A1	210340637	210340684
refGene_gn=FYB1;refGene_tx=XM_011286424.2
A1	210342939	210342959
refGene_gn=FYB1;refGene_tx=XM_011286424.2
A1	210343401	210343436
refGene_gn=FYB1;refGene_tx=XM_011286424.2
A1	210344394	210344515
refGene_gn=FYB1;refGene_tx=XM_011286424.2
A1	210348220	210348380
refGene_gn=FYB1;refGene_tx=XM_011286424.2
A1	210348859	210349004
refGene_gn=FYB1;refGene_tx=XM_011286424.2
A1	210354660	210354683
refGene_gn=FYB1;refGene_tx=XM_011286424.2
A1	210355699	210355766
refGene_gn=FYB1;refGene_tx=XM_011286424.2
A1	210356549	210356687
refGene_gn=FYB1;refGene_tx=XM_011286424.2
A1	210358164	210358190
refGene_gn=FYB1;refGene_tx=XM_011286424.2
A1	210360529	210360596
refGene_gn=FYB1;refGene_tx=XM_011286424.2
A1	210363401	210363447
refGene_gn=FYB1;refGene_tx=XM_011286424.2
A1	210363448	210363493
refGene_gn=FYB1;refGene_tx=XM_011286424.2
A1	210364852	210365015
refGene_gn=FYB1;refGene_tx=XM_011286424.2
A1	210374154	210374188
refGene_gn=FYB1;refGene_tx=XM_011286424.2
A1	210376892	210376924
refGene_gn=FYB1;refGene_tx=XM_011286424.2
A1	210377702	210377725

refGene_gn=FYB1;refGene_tx=XM_011286424.2
A1 210401182 210401231
refGene_gn=RICTOR;refGene_tx=XM_023238910.1;refGene_gn=RICTOR;refGene_tx=XM_023238911.1;refGene_gn=RICTOR;refGene_tx=XM_023238916.1;refGene_gn=RICTOR;refGene_tx=XM_023238949.1
A1 210401402 210401450
refGene_gn=RICTOR;refGene_tx=XM_023238910.1;refGene_gn=RICTOR;refGene_tx=XM_023238911.1;refGene_gn=RICTOR;refGene_tx=XM_023238916.1;refGene_gn=RICTOR;refGene_tx=XM_023238949.1
A1 210441585 210441683
refGene_gn=RICTOR;refGene_tx=XM_023238910.1;refGene_gn=RICTOR;refGene_tx=XM_023238911.1;refGene_gn=RICTOR;refGene_tx=XM_023238916.1;refGene_gn=RICTOR;refGene_tx=XM_023238925.1;refGene_gn=RICTOR;refGene_tx=XM_023238926.1;refGene_gn=RICTOR;refGene_tx=XM_023238932.1;refGene_gn=RICTOR;refGene_tx=XM_023238938.1;refGene_gn=RICTOR;refGene_tx=XM_023238949.1
A1 210454889 210454954
refGene_gn=RICTOR;refGene_tx=XM_023238910.1;refGene_gn=RICTOR;refGene_tx=XM_023238911.1;refGene_gn=RICTOR;refGene_tx=XM_023238916.1;refGene_gn=RICTOR;refGene_tx=XM_023238925.1;refGene_gn=RICTOR;refGene_tx=XM_023238926.1;refGene_gn=RICTOR;refGene_tx=XM_023238932.1;refGene_gn=RICTOR;refGene_tx=XM_023238938.1;refGene_gn=RICTOR;refGene_tx=XM_023238949.1
A1 210456232 210456364
refGene_gn=RICTOR;refGene_tx=XM_023238910.1;refGene_gn=RICTOR;refGene_tx=XM_023238911.1;refGene_gn=RICTOR;refGene_tx=XM_023238916.1;refGene_gn=RICTOR;refGene_tx=XM_023238925.1;refGene_gn=RICTOR;refGene_tx=XM_023238926.1;refGene_gn=RICTOR;refGene_tx=XM_023238932.1;refGene_gn=RICTOR;refGene_tx=XM_023238938.1;refGene_gn=RICTOR;refGene_tx=XM_023238949.1
A1 210461934 210461998
refGene_gn=RICTOR;refGene_tx=XM_023238910.1;refGene_gn=RICTOR;refGene_tx=XM_023238911.1;refGene_gn=RICTOR;refGene_tx=XM_023238916.1;refGene_gn=RICTOR;refGene_tx=XM_023238925.1;refGene_gn=RICTOR;refGene_tx=XM_023238926.1;refGene_gn=RICTOR;refGene_tx=XM_023238932.1;refGene_gn=RICTOR;refGene_tx=XM_023238938.1;refGene_gn=RICTOR;refGene_tx=XM_023238949.1
A1 210466179 210466306
refGene_gn=RICTOR;refGene_tx=XM_023238910.1;refGene_gn=RICTOR;refGene_tx=XM_023238911.1;refGene_gn=RICTOR;refGene_tx=XM_023238916.1;refGene_gn=RICTOR;refGene_tx=XM_023238925.1;refGene_gn=RICTOR;refGene_tx=XM_023238926.1;refGene_gn=RICTOR;refGene_tx=XM_023238932.1;refGene_gn=RICTOR;refGene_tx=XM_023238938.1;refGene_gn=RICTOR;refGene_tx=XM_023238949.1
A1 210471403 210471573
refGene_gn=RICTOR;refGene_tx=XM_023238910.1;refGene_gn=RICTOR;refGene_tx=XM_023238911.1;refGene_gn=RICTOR;refGene_tx=XM_023238916.1;refGene_gn=RICTOR;refGene_tx=XM_023238925.1;refGene_gn=RICTOR;refGene_tx=XM_023238926.1;refGene_gn=RICTOR;refGene_tx=XM_023238932.1;refGene_gn=RICTOR;refGene_tx=XM_023238938.1;refGene_gn=RICTOR;refGene_tx=XM_023238949.

1
A1 210476103 210476171
refGene_gn=RICTOR;refGene_tx=XM_023238910.1;refGene_gn=RICTOR;refGene_tx=XM_023238911.1;refGene_gn=RICTOR;refGene_tx=XM_023238916.1;refGene_gn=RICTOR;refGene_tx=XM_023238925.1;refGene_gn=RICTOR;refGene_tx=XM_023238926.1;refGene_gn=RICTOR;refGene_tx=XM_023238932.1;refGene_gn=RICTOR;refGene_tx=XM_023238938.1;refGene_gn=RICTOR;refGene_tx=XM_023238949.1
A1 210479502 210479570
refGene_gn=RICTOR;refGene_tx=XM_023238910.1;refGene_gn=RICTOR;refGene_tx=XM_023238911.1;refGene_gn=RICTOR;refGene_tx=XM_023238916.1;refGene_gn=RICTOR;refGene_tx=XM_023238925.1;refGene_gn=RICTOR;refGene_tx=XM_023238926.1;refGene_gn=RICTOR;refGene_tx=XM_023238932.1;refGene_gn=RICTOR;refGene_tx=XM_023238938.1;refGene_gn=RICTOR;refGene_tx=XM_023238949.1
A1 210480584 210480667
refGene_gn=RICTOR;refGene_tx=XM_023238910.1;refGene_gn=RICTOR;refGene_tx=XM_023238911.1;refGene_gn=RICTOR;refGene_tx=XM_023238916.1;refGene_gn=RICTOR;refGene_tx=XM_023238925.1;refGene_gn=RICTOR;refGene_tx=XM_023238926.1;refGene_gn=RICTOR;refGene_tx=XM_023238932.1;refGene_gn=RICTOR;refGene_tx=XM_023238938.1;refGene_gn=RICTOR;refGene_tx=XM_023238949.1
A1 210482075 210482163
refGene_gn=RICTOR;refGene_tx=XM_023238910.1;refGene_gn=RICTOR;refGene_tx=XM_023238911.1;refGene_gn=RICTOR;refGene_tx=XM_023238916.1;refGene_gn=RICTOR;refGene_tx=XM_023238925.1;refGene_gn=RICTOR;refGene_tx=XM_023238926.1;refGene_gn=RICTOR;refGene_tx=XM_023238932.1;refGene_gn=RICTOR;refGene_tx=XM_023238938.1;refGene_gn=RICTOR;refGene_tx=XM_023238949.1
A1 210482688 210482779
refGene_gn=RICTOR;refGene_tx=XM_023238910.1;refGene_gn=RICTOR;refGene_tx=XM_023238911.1;refGene_gn=RICTOR;refGene_tx=XM_023238916.1;refGene_gn=RICTOR;refGene_tx=XM_023238925.1;refGene_gn=RICTOR;refGene_tx=XM_023238926.1;refGene_gn=RICTOR;refGene_tx=XM_023238932.1;refGene_gn=RICTOR;refGene_tx=XM_023238938.1;refGene_gn=RICTOR;refGene_tx=XM_023238949.1
A1 210482886 210482953
refGene_gn=RICTOR;refGene_tx=XM_023238910.1;refGene_gn=RICTOR;refGene_tx=XM_023238911.1;refGene_gn=RICTOR;refGene_tx=XM_023238916.1;refGene_gn=RICTOR;refGene_tx=XM_023238925.1;refGene_gn=RICTOR;refGene_tx=XM_023238926.1;refGene_gn=RICTOR;refGene_tx=XM_023238932.1;refGene_gn=RICTOR;refGene_tx=XM_023238938.1;refGene_gn=RICTOR;refGene_tx=XM_023238949.1
A1 210483098 210483179
refGene_gn=RICTOR;refGene_tx=XM_023238910.1;refGene_gn=RICTOR;refGene_tx=XM_023238911.1;refGene_gn=RICTOR;refGene_tx=XM_023238916.1;refGene_gn=RICTOR;refGene_tx=XM_023238925.1;refGene_gn=RICTOR;refGene_tx=XM_023238926.1;refGene_gn=RICTOR;refGene_tx=XM_023238932.1;refGene_gn=RICTOR;refGene_tx=XM_023238938.1;refGene_gn=RICTOR;refGene_tx=XM_023238949.1

A1 210485631 210485732
refGene_gn=RICTOR;refGene_tx=XM_023238910.1;refGene_gn=RICTOR;refGene_tx=XM_023238911.1;refGene_gn=RICTOR;refGene_tx=XM_023238916.1;refGene_gn=RICTOR;refGene_tx=XM_023238925.1;refGene_gn=RICTOR;refGene_tx=XM_023238926.1;refGene_gn=RICTOR;refGene_tx=XM_023238932.1;refGene_gn=RICTOR;refGene_tx=XM_023238938.1;refGene_gn=RICTOR;refGene_tx=XM_023238949.1
A1 210488435 210488601
refGene_gn=RICTOR;refGene_tx=XM_023238910.1;refGene_gn=RICTOR;refGene_tx=XM_023238911.1;refGene_gn=RICTOR;refGene_tx=XM_023238916.1;refGene_gn=RICTOR;refGene_tx=XM_023238925.1;refGene_gn=RICTOR;refGene_tx=XM_023238926.1;refGene_gn=RICTOR;refGene_tx=XM_023238932.1;refGene_gn=RICTOR;refGene_tx=XM_023238938.1;refGene_gn=RICTOR;refGene_tx=XM_023238949.1
A1 210488907 210489009
refGene_gn=RICTOR;refGene_tx=XM_023238910.1;refGene_gn=RICTOR;refGene_tx=XM_023238911.1;refGene_gn=RICTOR;refGene_tx=XM_023238916.1;refGene_gn=RICTOR;refGene_tx=XM_023238925.1;refGene_gn=RICTOR;refGene_tx=XM_023238926.1;refGene_gn=RICTOR;refGene_tx=XM_023238932.1;refGene_gn=RICTOR;refGene_tx=XM_023238938.1;refGene_gn=RICTOR;refGene_tx=XM_023238949.1
A1 210489131 210489178
refGene_gn=RICTOR;refGene_tx=XM_023238910.1;refGene_gn=RICTOR;refGene_tx=XM_023238911.1;refGene_gn=RICTOR;refGene_tx=XM_023238916.1;refGene_gn=RICTOR;refGene_tx=XM_023238925.1;refGene_gn=RICTOR;refGene_tx=XM_023238926.1;refGene_gn=RICTOR;refGene_tx=XM_023238932.1;refGene_gn=RICTOR;refGene_tx=XM_023238938.1;refGene_gn=RICTOR;refGene_tx=XM_023238949.1
A1 210490629 210490765
refGene_gn=RICTOR;refGene_tx=XM_023238910.1;refGene_gn=RICTOR;refGene_tx=XM_023238911.1;refGene_gn=RICTOR;refGene_tx=XM_023238916.1;refGene_gn=RICTOR;refGene_tx=XM_023238925.1;refGene_gn=RICTOR;refGene_tx=XM_023238926.1;refGene_gn=RICTOR;refGene_tx=XM_023238932.1;refGene_gn=RICTOR;refGene_tx=XM_023238938.1;refGene_gn=RICTOR;refGene_tx=XM_023238949.1
A1 210491155 210491355
refGene_gn=RICTOR;refGene_tx=XM_023238910.1;refGene_gn=RICTOR;refGene_tx=XM_023238911.1;refGene_gn=RICTOR;refGene_tx=XM_023238916.1;refGene_gn=RICTOR;refGene_tx=XM_023238925.1;refGene_gn=RICTOR;refGene_tx=XM_023238926.1;refGene_gn=RICTOR;refGene_tx=XM_023238932.1;refGene_gn=RICTOR;refGene_tx=XM_023238938.1;refGene_gn=RICTOR;refGene_tx=XM_023238949.1
A1 210491588 210491715
refGene_gn=RICTOR;refGene_tx=XM_023238910.1;refGene_gn=RICTOR;refGene_tx=XM_023238911.1;refGene_gn=RICTOR;refGene_tx=XM_023238916.1;refGene_gn=RICTOR;refGene_tx=XM_023238925.1;refGene_gn=RICTOR;refGene_tx=XM_023238926.1;refGene_gn=RICTOR;refGene_tx=XM_023238932.1;refGene_gn=RICTOR;refGene_tx=XM_023238938.1;refGene_gn=RICTOR;refGene_tx=XM_023238949.1
A1 210492076 210492241

refGene_gn=RICTOR;refGene_tx=XM_023238910.1;refGene_gn=RICTOR;refGene_tx=XM_023238911.1;refGene_gn=RICTOR;refGene_tx=XM_023238916.1;refGene_gn=RICTOR;refGene_tx=XM_023238925.1;refGene_gn=RICTOR;refGene_tx=XM_023238926.1;refGene_gn=RICTOR;refGene_tx=XM_023238932.1;refGene_gn=RICTOR;refGene_tx=XM_023238938.1;refGene_gn=RICTOR;refGene_tx=XM_023238949.1
A1 210492435 210492512
refGene_gn=RICTOR;refGene_tx=XM_023238910.1;refGene_gn=RICTOR;refGene_tx=XM_023238911.1;refGene_gn=RICTOR;refGene_tx=XM_023238916.1;refGene_gn=RICTOR;refGene_tx=XM_023238925.1;refGene_gn=RICTOR;refGene_tx=XM_023238926.1;refGene_gn=RICTOR;refGene_tx=XM_023238932.1;refGene_gn=RICTOR;refGene_tx=XM_023238938.1;refGene_gn=RICTOR;refGene_tx=XM_023238949.1
A1 210492961 210493040
refGene_gn=RICTOR;refGene_tx=XM_023238910.1;refGene_gn=RICTOR;refGene_tx=XM_023238911.1;refGene_gn=RICTOR;refGene_tx=XM_023238916.1;refGene_gn=RICTOR;refGene_tx=XM_023238925.1;refGene_gn=RICTOR;refGene_tx=XM_023238926.1;refGene_gn=RICTOR;refGene_tx=XM_023238932.1;refGene_gn=RICTOR;refGene_tx=XM_023238938.1;refGene_gn=RICTOR;refGene_tx=XM_023238949.1
A1 210498646 210498756
refGene_gn=RICTOR;refGene_tx=XM_023238910.1;refGene_gn=RICTOR;refGene_tx=XM_023238911.1;refGene_gn=RICTOR;refGene_tx=XM_023238916.1;refGene_gn=RICTOR;refGene_tx=XM_023238925.1;refGene_gn=RICTOR;refGene_tx=XM_023238926.1;refGene_gn=RICTOR;refGene_tx=XM_023238932.1;refGene_gn=RICTOR;refGene_tx=XM_023238938.1;refGene_gn=RICTOR;refGene_tx=XM_023238949.1
A1 210499243 210499331
refGene_gn=RICTOR;refGene_tx=XM_023238910.1;refGene_gn=RICTOR;refGene_tx=XM_023238911.1;refGene_gn=RICTOR;refGene_tx=XM_023238916.1;refGene_gn=RICTOR;refGene_tx=XM_023238925.1;refGene_gn=RICTOR;refGene_tx=XM_023238926.1;refGene_gn=RICTOR;refGene_tx=XM_023238932.1;refGene_gn=RICTOR;refGene_tx=XM_023238938.1;refGene_gn=RICTOR;refGene_tx=XM_023238949.1
A1 210500480 210500573
refGene_gn=RICTOR;refGene_tx=XM_023238910.1;refGene_gn=RICTOR;refGene_tx=XM_023238911.1;refGene_gn=RICTOR;refGene_tx=XM_023238916.1;refGene_gn=RICTOR;refGene_tx=XM_023238925.1;refGene_gn=RICTOR;refGene_tx=XM_023238926.1;refGene_gn=RICTOR;refGene_tx=XM_023238932.1;refGene_gn=RICTOR;refGene_tx=XM_023238938.1;refGene_gn=RICTOR;refGene_tx=XM_023238949.1
A1 210500927 210501034
refGene_gn=RICTOR;refGene_tx=XM_023238910.1;refGene_gn=RICTOR;refGene_tx=XM_023238911.1;refGene_gn=RICTOR;refGene_tx=XM_023238916.1;refGene_gn=RICTOR;refGene_tx=XM_023238925.1;refGene_gn=RICTOR;refGene_tx=XM_023238926.1;refGene_gn=RICTOR;refGene_tx=XM_023238932.1;refGene_gn=RICTOR;refGene_tx=XM_023238938.1;refGene_gn=RICTOR;refGene_tx=XM_023238949.1
A1 210502158 210502388
refGene_gn=RICTOR;refGene_tx=XM_023238910.1;refGene_gn=RICTOR;refGene_

tx=XM_023238911.1;refGene_gn=RICTOR;refGene_tx=XM_023238916.1;refGene_gn=RICTOR;refGene_tx=XM_023238925.1;refGene_gn=RICTOR;refGene_tx=XM_023238926.1;refGene_gn=RICTOR;refGene_tx=XM_023238932.1;refGene_gn=RICTOR;refGene_tx=XM_023238938.1;refGene_gn=RICTOR;refGene_tx=XM_023238949.1
A1 210504584 210505593
refGene_gn=RICTOR;refGene_tx=XM_023238910.1;refGene_gn=RICTOR;refGene_tx=XM_023238911.1;refGene_gn=RICTOR;refGene_tx=XM_023238916.1;refGene_gn=RICTOR;refGene_tx=XM_023238925.1;refGene_gn=RICTOR;refGene_tx=XM_023238926.1;refGene_gn=RICTOR;refGene_tx=XM_023238932.1;refGene_gn=RICTOR;refGene_tx=XM_023238938.1;refGene_gn=RICTOR;refGene_tx=XM_023238949.1
A1 210505875 210505947
refGene_gn=RICTOR;refGene_tx=XM_023238910.1;refGene_gn=RICTOR;refGene_tx=XM_023238916.1;refGene_gn=RICTOR;refGene_tx=XM_023238925.1;refGene_gn=RICTOR;refGene_tx=XM_023238926.1;refGene_gn=RICTOR;refGene_tx=XM_023238932.1;refGene_gn=RICTOR;refGene_tx=XM_023238938.1
A1 210509106 210509284
refGene_gn=RICTOR;refGene_tx=XM_023238910.1;refGene_gn=RICTOR;refGene_tx=XM_023238911.1;refGene_gn=RICTOR;refGene_tx=XM_023238916.1;refGene_gn=RICTOR;refGene_tx=XM_023238925.1;refGene_gn=RICTOR;refGene_tx=XM_023238926.1;refGene_gn=RICTOR;refGene_tx=XM_023238932.1;refGene_gn=RICTOR;refGene_tx=XM_023238938.1;refGene_gn=RICTOR;refGene_tx=XM_023238949.1
A1 210510029 210510114
refGene_gn=RICTOR;refGene_tx=XM_023238910.1;refGene_gn=RICTOR;refGene_tx=XM_023238911.1;refGene_gn=RICTOR;refGene_tx=XM_023238916.1;refGene_gn=RICTOR;refGene_tx=XM_023238925.1;refGene_gn=RICTOR;refGene_tx=XM_023238926.1;refGene_gn=RICTOR;refGene_tx=XM_023238932.1;refGene_gn=RICTOR;refGene_tx=XM_023238938.1;refGene_gn=RICTOR;refGene_tx=XM_023238949.1
A1 210510867 210511101
refGene_gn=RICTOR;refGene_tx=XM_023238910.1;refGene_gn=RICTOR;refGene_tx=XM_023238911.1;refGene_gn=RICTOR;refGene_tx=XM_023238916.1;refGene_gn=RICTOR;refGene_tx=XM_023238925.1;refGene_gn=RICTOR;refGene_tx=XM_023238926.1;refGene_gn=RICTOR;refGene_tx=XM_023238932.1;refGene_gn=RICTOR;refGene_tx=XM_023238938.1;refGene_gn=RICTOR;refGene_tx=XM_023238949.1
A1 210517222 210517378
refGene_gn=RICTOR;refGene_tx=XM_023238910.1;refGene_gn=RICTOR;refGene_tx=XM_023238911.1;refGene_gn=RICTOR;refGene_tx=XM_023238916.1;refGene_gn=RICTOR;refGene_tx=XM_023238925.1;refGene_gn=RICTOR;refGene_tx=XM_023238926.1;refGene_gn=RICTOR;refGene_tx=XM_023238932.1;refGene_gn=RICTOR;refGene_tx=XM_023238938.1;refGene_gn=RICTOR;refGene_tx=XM_023238949.1
A1 210517711 210517835
refGene_gn=RICTOR;refGene_tx=XM_023238910.1;refGene_gn=RICTOR;refGene_tx=XM_023238911.1;refGene_gn=RICTOR;refGene_tx=XM_023238916.1;refGene_gn=RICTOR;refGene_tx=XM_023238925.1;refGene_gn=RICTOR;refGene_tx=XM_023238926.1;refGene_gn=RICTOR;refGene_tx=XM_023238932.1;refGene_gn=RICTO

R;refGene_tx=XM_023238938.1;refGene_gn=RICTOR;refGene_tx=XM_023238949.
1
A1 210519368 210519507
refGene_gn=RICTOR;refGene_tx=XM_023238910.1;refGene_gn=RICTOR;refGene_
tx=XM_023238911.1;refGene_gn=RICTOR;refGene_tx=XM_023238916.1;refGene_
gn=RICTOR;refGene_tx=XM_023238925.1;refGene_gn=RICTOR;refGene_tx=XM_02
3238926.1;refGene_gn=RICTOR;refGene_tx=XM_023238932.1;refGene_gn=RICTO
R;refGene_tx=XM_023238938.1;refGene_gn=RICTOR;refGene_tx=XM_023238949.
1
A1 210519622 210519697
refGene_gn=RICTOR;refGene_tx=XM_023238910.1;refGene_gn=RICTOR;refGene_
tx=XM_023238911.1;refGene_gn=RICTOR;refGene_tx=XM_023238916.1;refGene_
gn=RICTOR;refGene_tx=XM_023238925.1;refGene_gn=RICTOR;refGene_tx=XM_02
3238926.1;refGene_gn=RICTOR;refGene_tx=XM_023238932.1;refGene_gn=RICTO
R;refGene_tx=XM_023238938.1;refGene_gn=RICTOR;refGene_tx=XM_023238949.
1
A1 210528189 210528720
refGene_gn=OSMR;refGene_tx=XM_023238963.1;refGene_gn=OSMR;refGene_tx=X
M_023238966.1;refGene_gn=OSMR;refGene_tx=XM_023238974.1
A1 210529057 210529130
refGene_gn=OSMR;refGene_tx=XM_023238963.1;refGene_gn=OSMR;refGene_tx=X
M_023238966.1;refGene_gn=OSMR;refGene_tx=XM_023238974.1
A1 210529638 210529720
refGene_gn=OSMR;refGene_tx=XM_023238963.1;refGene_gn=OSMR;refGene_tx=X
M_023238966.1;refGene_gn=OSMR;refGene_tx=XM_023238974.1
A1 210532729 210532897
refGene_gn=OSMR;refGene_tx=XM_023238963.1;refGene_gn=OSMR;refGene_tx=X
M_023238966.1;refGene_gn=OSMR;refGene_tx=XM_023238974.1
A1 210533477 210533651
refGene_gn=OSMR;refGene_tx=XM_023238963.1;refGene_gn=OSMR;refGene_tx=X
M_023238966.1;refGene_gn=OSMR;refGene_tx=XM_023238974.1
A1 210534974 210535079
refGene_gn=OSMR;refGene_tx=XM_023238963.1;refGene_gn=OSMR;refGene_tx=X
M_023238966.1;refGene_gn=OSMR;refGene_tx=XM_023238974.1
A1 210536371 210536551
refGene_gn=OSMR;refGene_tx=XM_023238963.1;refGene_gn=OSMR;refGene_tx=X
M_023238966.1;refGene_gn=OSMR;refGene_tx=XM_023238974.1
A1 210539961 210540184
refGene_gn=OSMR;refGene_tx=XM_023238963.1;refGene_gn=OSMR;refGene_tx=X
M_023238966.1;refGene_gn=OSMR;refGene_tx=XM_023238974.1
A1 210541629 210541706
refGene_gn=OSMR;refGene_tx=XM_023238963.1;refGene_gn=OSMR;refGene_tx=X
M_023238966.1;refGene_gn=OSMR;refGene_tx=XM_023238974.1
A1 210545293 210545447
refGene_gn=OSMR;refGene_tx=XM_023238963.1;refGene_gn=OSMR;refGene_tx=X
M_023238966.1;refGene_gn=OSMR;refGene_tx=XM_023238974.1
A1 210545811 210545954
refGene_gn=OSMR;refGene_tx=XM_023238963.1;refGene_gn=OSMR;refGene_tx=X
M_023238966.1;refGene_gn=OSMR;refGene_tx=XM_023238974.1
A1 210549485 210549637

refGene_gn=OSMR;refGene_tx=XM_023238963.1;refGene_gn=OSMR;refGene_tx=XM_023238966.1;refGene_gn=OSMR;refGene_tx=XM_023238974.1
A1 210550195 210550325
refGene_gn=OSMR;refGene_tx=XM_023238963.1;refGene_gn=OSMR;refGene_tx=XM_023238966.1;refGene_gn=OSMR;refGene_tx=XM_023238974.1
A1 210551655 210551931
refGene_gn=OSMR;refGene_tx=XM_023238963.1;refGene_gn=OSMR;refGene_tx=XM_023238966.1;refGene_gn=OSMR;refGene_tx=XM_023238974.1
A1 210554059 210554231
refGene_gn=OSMR;refGene_tx=XM_023238963.1;refGene_gn=OSMR;refGene_tx=XM_023238966.1;refGene_gn=OSMR;refGene_tx=XM_023238974.1
A1 210560019 210560192
refGene_gn=OSMR;refGene_tx=XM_023238963.1;refGene_gn=OSMR;refGene_tx=XM_023238966.1;refGene_gn=OSMR;refGene_tx=XM_023238974.1
A1 210565295 210565368
refGene_gn=OSMR;refGene_tx=XM_023238963.1;refGene_gn=OSMR;refGene_tx=XM_023238966.1;refGene_gn=OSMR;refGene_tx=XM_023238974.1
A1 210845170 210845312
refGene_gn=LIFR;refGene_tx=XM_023238979.1;refGene_gn=LIFR;refGene_tx=XM_023238982.1
A1 210846978 210847093
refGene_gn=LIFR;refGene_tx=XM_023238979.1;refGene_gn=LIFR;refGene_tx=XM_023238982.1
A1 210849540 210849680
refGene_gn=LIFR;refGene_tx=XM_023238979.1;refGene_gn=LIFR;refGene_tx=XM_023238982.1
A1 210855682 210855846
refGene_gn=LIFR;refGene_tx=XM_023238979.1;refGene_gn=LIFR;refGene_tx=XM_023238982.1
A1 210861334 210861509
refGene_gn=LIFR;refGene_tx=XM_023238979.1;refGene_gn=LIFR;refGene_tx=XM_023238982.1
A1 210863302 210863557
refGene_gn=LIFR;refGene_tx=XM_023238979.1;refGene_gn=LIFR;refGene_tx=XM_023238982.1
A1 210868554 210868684
refGene_gn=LIFR;refGene_tx=XM_023238979.1;refGene_gn=LIFR;refGene_tx=XM_023238982.1
A1 210869112 210869282
refGene_gn=LIFR;refGene_tx=XM_023238979.1;refGene_gn=LIFR;refGene_tx=XM_023238982.1
A1 210871160 210871306
refGene_gn=LIFR;refGene_tx=XM_023238979.1;refGene_gn=LIFR;refGene_tx=XM_023238982.1
A1 210872942 210873105
refGene_gn=LIFR;refGene_tx=XM_023238979.1;refGene_gn=LIFR;refGene_tx=XM_023238982.1
A1 210875261 210875332
refGene_gn=LIFR;refGene_tx=XM_023238979.1;refGene_gn=LIFR;refGene_tx=XM_023238982.1

A1 210878626 210878840
refGene_gn=LIFR;refGene_tx=XM_023238979.1;refGene_gn=LIFR;refGene_tx=XM_023238982.1
A1 210881512 210881692
refGene_gn=LIFR;refGene_tx=XM_023238979.1;refGene_gn=LIFR;refGene_tx=XM_023238982.1
A1 210884291 210884393
refGene_gn=LIFR;refGene_tx=XM_023238979.1;refGene_gn=LIFR;refGene_tx=XM_023238982.1
A1 210884941 210885109
refGene_gn=LIFR;refGene_tx=XM_023238979.1;refGene_gn=LIFR;refGene_tx=XM_023238982.1
A1 210888762 210888924
refGene_gn=LIFR;refGene_tx=XM_023238979.1;refGene_gn=LIFR;refGene_tx=XM_023238982.1
A1 210889838 210889932
refGene_gn=LIFR;refGene_tx=XM_023238979.1;refGene_gn=LIFR;refGene_tx=XM_023238982.1
A1 210891791 210891870
refGene_gn=LIFR;refGene_tx=XM_023238979.1;refGene_gn=LIFR;refGene_tx=XM_023238982.1
A1 210892252 210892876
refGene_gn=LIFR;refGene_tx=XM_023238979.1;refGene_gn=LIFR;refGene_tx=XM_023238982.1
A1 210910379 210910534
refGene_gn=EGFLAM;refGene_tx=XM_011286450.3;refGene_gn=EGFLAM;refGene_tx=XM_011286454.3;refGene_gn=EGFLAM;refGene_tx=XM_019811656.2
A1 210911329 210911433
refGene_gn=EGFLAM;refGene_tx=XM_011286450.3;refGene_gn=EGFLAM;refGene_tx=XM_011286454.3;refGene_gn=EGFLAM;refGene_tx=XM_019811656.2
A1 210915846 210915930
refGene_gn=EGFLAM;refGene_tx=XM_011286450.3;refGene_gn=EGFLAM;refGene_tx=XM_011286454.3;refGene_gn=EGFLAM;refGene_tx=XM_019811656.2
A1 210922378 210922522
refGene_gn=EGFLAM;refGene_tx=XM_011286450.3;refGene_gn=EGFLAM;refGene_tx=XM_011286454.3;refGene_gn=EGFLAM;refGene_tx=XM_019811656.2
A1 210926129 210926208
refGene_gn=EGFLAM;refGene_tx=XM_011286450.3;refGene_gn=EGFLAM;refGene_tx=XM_019811656.2
A1 210929161 210929185
refGene_gn=EGFLAM;refGene_tx=XM_019811656.2
A1 210934979 210935160
refGene_gn=EGFLAM;refGene_tx=XM_011286450.3;refGene_gn=EGFLAM;refGene_tx=XM_019811656.2
A1 210937663 210937780
refGene_gn=EGFLAM;refGene_tx=XM_011286450.3;refGene_gn=EGFLAM;refGene_tx=XM_019811656.2
A1 210942295 210942407
refGene_gn=EGFLAM;refGene_tx=XM_011286450.3;refGene_gn=EGFLAM;refGene_tx=XM_019811656.2

A1 210945445 210945689
refGene_gn=EGFLAM;refGene_tx=XM_011286450.3;refGene_gn=EGFLAM;refGene_tx=XM_019811656.2
A1 210947028 210947154
refGene_gn=EGFLAM;refGene_tx=XM_011286450.3;refGene_gn=EGFLAM;refGene_tx=XM_019811656.2
A1 210953726 210953916
refGene_gn=EGFLAM;refGene_tx=XM_011286450.3;refGene_gn=EGFLAM;refGene_tx=XM_019811656.2
A1 210957222 210957367
refGene_gn=EGFLAM;refGene_tx=XM_011286450.3;refGene_gn=EGFLAM;refGene_tx=XM_019811656.2
A1 210959286 210959387
refGene_gn=EGFLAM;refGene_tx=XM_011286450.3;refGene_gn=EGFLAM;refGene_tx=XM_019811656.2
A1 210960524 210960625
refGene_gn=EGFLAM;refGene_tx=XM_011286450.3;refGene_gn=EGFLAM;refGene_tx=XM_019811656.2
A1 210961270 210961589
refGene_gn=EGFLAM;refGene_tx=XM_011286450.3;refGene_gn=EGFLAM;refGene_tx=XM_019811656.2
A1 210962157 210962273
refGene_gn=EGFLAM;refGene_tx=XM_011286450.3;refGene_gn=EGFLAM;refGene_tx=XM_019811656.2
A1 210993832 210993999
refGene_gn=EGFLAM;refGene_tx=XM_011286450.3;refGene_gn=EGFLAM;refGene_tx=XM_019811656.2
A1 211013331 211013467
refGene_gn=EGFLAM;refGene_tx=XM_011286450.3;refGene_gn=EGFLAM;refGene_tx=XM_019811656.2
A1 211014714 211014832
refGene_gn=EGFLAM;refGene_tx=XM_011286450.3;refGene_gn=EGFLAM;refGene_tx=XM_019811656.2
A1 211022807 211022834
refGene_gn=EGFLAM;refGene_tx=XM_011286450.3;refGene_gn=EGFLAM;refGene_tx=XM_019811656.2
A1 211025934 211026018
refGene_gn=EGFLAM;refGene_tx=XM_011286450.3;refGene_gn=EGFLAM;refGene_tx=XM_019811656.2
A1 211027120 211027230
refGene_gn=EGFLAM;refGene_tx=XM_011286450.3;refGene_gn=EGFLAM;refGene_tx=XM_019811656.2
A1 211088218 211088315
refGene_gn=EGFLAM;refGene_tx=XM_011286450.3;refGene_gn=EGFLAM;refGene_tx=XM_019811656.2
A1 211503311 211503462
refGene_gn=GDNF;refGene_tx=XM_006928045.4
A1 211522596 211523081
refGene_gn=GDNF;refGene_tx=XM_006928045.4
A1 211561196 211561284

refGene_gn=WDR70;refGene_tx=XM_019811747.1
A1 211582450 211582613
refGene_gn=WDR70;refGene_tx=XM_019811747.1
A1 211584859 211584976
refGene_gn=WDR70;refGene_tx=XM_019811747.1
A1 211586976 211587056
refGene_gn=WDR70;refGene_tx=XM_019811747.1
A1 211588704 211588805
refGene_gn=WDR70;refGene_tx=XM_019811747.1
A1 211611978 211612117
refGene_gn=WDR70;refGene_tx=XM_019811747.1
A1 211613583 211613668
refGene_gn=WDR70;refGene_tx=XM_019811747.1
A1 211616338 211616438
refGene_gn=WDR70;refGene_tx=XM_019811747.1
A1 211675806 211675981
refGene_gn=WDR70;refGene_tx=XM_019811747.1
A1 211750054 211750131
refGene_gn=WDR70;refGene_tx=XM_019811747.1
A1 211778062 211778216
refGene_gn=WDR70;refGene_tx=XM_019811747.1
A1 211801282 211801416
refGene_gn=WDR70;refGene_tx=XM_019811747.1
A1 211805336 211805396
refGene_gn=WDR70;refGene_tx=XM_019811747.1
A1 211839896 211840095
refGene_gn=WDR70;refGene_tx=XM_019811747.1
A1 211850088 211850209
refGene_gn=WDR70;refGene_tx=XM_019811747.1
A1 211859236 211859320
refGene_gn=WDR70;refGene_tx=XM_019811747.1
A1 211861354 211861420
refGene_gn=WDR70;refGene_tx=XM_019811747.1
A1 211861517 211861542
refGene_gn=WDR70;refGene_tx=XM_019811747.1
A1 211870352 211870509
refGene_gn=NUP155;refGene_tx=XM_011286474.3
A1 211873369 211873507
refGene_gn=NUP155;refGene_tx=XM_011286474.3
A1 211879446 211879543
refGene_gn=NUP155;refGene_tx=XM_011286474.3
A1 211882346 211882417
refGene_gn=NUP155;refGene_tx=XM_011286474.3
A1 211888066 211888159
refGene_gn=NUP155;refGene_tx=XM_011286474.3
A1 211888924 211889091
refGene_gn=NUP155;refGene_tx=XM_011286474.3
A1 211891417 211891523
refGene_gn=NUP155;refGene_tx=XM_011286474.3
A1 211892298 211892333

refGene_gn=NUP155;refGene_tx=XM_011286474.3
A1 211892334 211892373
refGene_gn=NUP155;refGene_tx=XM_011286474.3
A1 211892906 211892998
refGene_gn=NUP155;refGene_tx=XM_011286474.3
A1 211894708 211894806
refGene_gn=NUP155;refGene_tx=XM_011286474.3
A1 211895302 211895455
refGene_gn=NUP155;refGene_tx=XM_011286474.3
A1 211898808 211898909
refGene_gn=NUP155;refGene_tx=XM_011286474.3
A1 211901382 211901553
refGene_gn=NUP155;refGene_tx=XM_011286474.3
A1 211903282 211903393
refGene_gn=NUP155;refGene_tx=XM_011286474.3
A1 211905014 211905109
refGene_gn=NUP155;refGene_tx=XM_011286474.3
A1 211905841 211905930
refGene_gn=NUP155;refGene_tx=XM_011286474.3
A1 211906833 211906896
refGene_gn=NUP155;refGene_tx=XM_011286474.3
A1 211907409 211907557
refGene_gn=NUP155;refGene_tx=XM_011286474.3
A1 211908914 211908981
refGene_gn=NUP155;refGene_tx=XM_011286474.3
A1 211909992 211910108
refGene_gn=NUP155;refGene_tx=XM_011286474.3
A1 211919607 211919705
refGene_gn=NUP155;refGene_tx=XM_011286474.3
A1 211922773 211922889
refGene_gn=NUP155;refGene_tx=XM_011286474.3
A1 211922890 211922905
refGene_gn=NUP155;refGene_tx=XM_011286474.3
A1 211927861 211928053
refGene_gn=NUP155;refGene_tx=XM_011286474.3
A1 211928708 211928847
refGene_gn=NUP155;refGene_tx=XM_011286474.3
A1 211929785 211929921
refGene_gn=NUP155;refGene_tx=XM_011286474.3
A1 211931209 211931363
refGene_gn=NUP155;refGene_tx=XM_011286474.3
A1 211931565 211931670
refGene_gn=NUP155;refGene_tx=XM_011286474.3
A1 211932581 211932736
refGene_gn=NUP155;refGene_tx=XM_011286474.3
A1 211935824 211935954
refGene_gn=NUP155;refGene_tx=XM_011286474.3
A1 211937344 211937458
refGene_gn=NUP155;refGene_tx=XM_011286474.3
A1 211938181 211938302

refGene_gn=NUP155;refGene_tx=XM_011286474.3
A1 211938949 211939060
refGene_gn=NUP155;refGene_tx=XM_011286474.3
A1 211941460 211941597
refGene_gn=NUP155;refGene_tx=XM_011286474.3
A1 211943302 211943409
refGene_gn=NUP155;refGene_tx=XM_011286474.3
A1 211944025 211944164
refGene_gn=NUP155;refGene_tx=XM_011286474.3
A1 211977416 211977497
refGene_gn=CA1H5orf42;refGene_tx=XM_023245967.1
A1 211981754 211981890
refGene_gn=CA1H5orf42;refGene_tx=XM_023245967.1
A1 211982003 211982123
refGene_gn=CA1H5orf42;refGene_tx=XM_023245967.1
A1 211982834 211983064
refGene_gn=CA1H5orf42;refGene_tx=XM_023245967.1
A1 211988487 211988594
refGene_gn=CA1H5orf42;refGene_tx=XM_023245967.1
A1 211991217 211991374
refGene_gn=CA1H5orf42;refGene_tx=XM_023245967.1
A1 211991806 211991910
refGene_gn=CA1H5orf42;refGene_tx=XM_023245967.1
A1 211993223 211993406
refGene_gn=CA1H5orf42;refGene_tx=XM_023245967.1
A1 211996201 211996451
refGene_gn=CA1H5orf42;refGene_tx=XM_023245967.1
A1 211996624 211996780
refGene_gn=CA1H5orf42;refGene_tx=XM_023245967.1
A1 211996943 211997713
refGene_gn=CA1H5orf42;refGene_tx=XM_023245967.1
A1 211998259 211998468
refGene_gn=CA1H5orf42;refGene_tx=XM_023245967.1
A1 211998669 211998750
refGene_gn=CA1H5orf42;refGene_tx=XM_023245967.1
A1 212001680 212001845
refGene_gn=CA1H5orf42;refGene_tx=XM_023245967.1
A1 212005766 212005940
refGene_gn=CA1H5orf42;refGene_tx=XM_023245967.1
A1 212009586 212009815
refGene_gn=CA1H5orf42;refGene_tx=XM_023245967.1
A1 212010985 212011125
refGene_gn=CA1H5orf42;refGene_tx=XM_023245967.1
A1 212013000 212013218
refGene_gn=CA1H5orf42;refGene_tx=XM_023245967.1
A1 212015407 212015572
refGene_gn=CA1H5orf42;refGene_tx=XM_023245967.1
A1 212020048 212020187
refGene_gn=CA1H5orf42;refGene_tx=XM_023245967.1
A1 212020913 212021023

refGene_gn=CA1H5orf42;refGene_tx=XM_023245967.1
A1 212021182 212021341
refGene_gn=CA1H5orf42;refGene_tx=XM_023245967.1
A1 212021458 212021482
refGene_gn=CA1H5orf42;refGene_tx=XM_023245967.1
A1 212021483 212021568
refGene_gn=CA1H5orf42;refGene_tx=XM_023245967.1
A1 212023247 212023528
refGene_gn=CA1H5orf42;refGene_tx=XM_023245967.1
A1 212023791 212023802
refGene_gn=CA1H5orf42;refGene_tx=XM_023245967.1
A1 212027218 212028131
refGene_gn=CA1H5orf42;refGene_tx=XM_023245967.1
A1 212039421 212039570
refGene_gn=CA1H5orf42;refGene_tx=XM_023245967.1
A1 212040171 212040338
refGene_gn=CA1H5orf42;refGene_tx=XM_023245967.1
A1 212040853 212040936
refGene_gn=CA1H5orf42;refGene_tx=XM_023245967.1
A1 212041665 212041745
refGene_gn=CA1H5orf42;refGene_tx=XM_023245967.1
A1 212042834 212042912
refGene_gn=CA1H5orf42;refGene_tx=XM_023245967.1
A1 212047556 212047749
refGene_gn=CA1H5orf42;refGene_tx=XM_023245967.1
A1 212049027 212049321
refGene_gn=CA1H5orf42;refGene_tx=XM_023245967.1
A1 212049509 212050280
refGene_gn=CA1H5orf42;refGene_tx=XM_023245967.1
A1 212051710 212051877
refGene_gn=CA1H5orf42;refGene_tx=XM_023245967.1
A1 212053194 212053327
refGene_gn=CA1H5orf42;refGene_tx=XM_023245967.1
A1 212054200 212054252
refGene_gn=CA1H5orf42;refGene_tx=XM_023245967.1
A1 212058484 212058589
refGene_gn=CA1H5orf42;refGene_tx=XM_023245967.1
A1 212062396 212062506
refGene_gn=CA1H5orf42;refGene_tx=XM_023245967.1
A1 212062863 212063059
refGene_gn=CA1H5orf42;refGene_tx=XM_023245967.1
A1 212063274 212063424
refGene_gn=CA1H5orf42;refGene_tx=XM_023245967.1
A1 212068191 212068445
refGene_gn=CA1H5orf42;refGene_tx=XM_023245967.1
A1 212071887 212071972
refGene_gn=CA1H5orf42;refGene_tx=XM_023245967.1
A1 212075397 212075562
refGene_gn=CA1H5orf42;refGene_tx=XM_023245967.1
A1 212079304 212079335

refGene_gn=CA1H5orf42;refGene_tx=XM_023245967.1
A1 212079873 212080005
refGene_gn=CA1H5orf42;refGene_tx=XM_023245967.1
A1 212089302 212089468
refGene_gn=CA1H5orf42;refGene_tx=XM_023245967.1
A1 212091713 212091772
refGene_gn=CA1H5orf42;refGene_tx=XM_023245967.1
A1 212092225 212092393
refGene_gn=CA1H5orf42;refGene_tx=XM_023245967.1
A1 212093169 212093294
refGene_gn=CA1H5orf42;refGene_tx=XM_023245967.1
A1 212097795 212097852
refGene_gn=CA1H5orf42;refGene_tx=XM_023245967.1
A1 212097854 212097884
refGene_gn=CA1H5orf42;refGene_tx=XM_023245967.1
A1 212107892 212108047
refGene_gn=CA1H5orf42;refGene_tx=XM_023245967.1
A1 212108510 212108684
refGene_gn=CA1H5orf42;refGene_tx=XM_023245967.1
A1 212130622 212130988
refGene_gn=NIPBL;refGene_tx=XM_023239001.1;refGene_gn=NIPBL;refGene_tx=XM_023239002.1
A1 212131507 212131737
refGene_gn=NIPBL;refGene_tx=XM_023239001.1;refGene_gn=NIPBL;refGene_tx=XM_023239002.1
A1 212133780 212133955
refGene_gn=NIPBL;refGene_tx=XM_023239001.1;refGene_gn=NIPBL;refGene_tx=XM_023239002.1
A1 212135839 212136114
refGene_gn=NIPBL;refGene_tx=XM_023239001.1;refGene_gn=NIPBL;refGene_tx=XM_023239002.1
A1 212137901 212138048
refGene_gn=NIPBL;refGene_tx=XM_023239001.1;refGene_gn=NIPBL;refGene_tx=XM_023239002.1
A1 212141122 212141323
refGene_gn=NIPBL;refGene_tx=XM_023239001.1;refGene_gn=NIPBL;refGene_tx=XM_023239002.1
A1 212143053 212143161
refGene_gn=NIPBL;refGene_tx=XM_023239001.1;refGene_gn=NIPBL;refGene_tx=XM_023239002.1
A1 212144921 212145112
refGene_gn=NIPBL;refGene_tx=XM_023239001.1;refGene_gn=NIPBL;refGene_tx=XM_023239002.1
A1 212145529 212145703
refGene_gn=NIPBL;refGene_tx=XM_023239001.1;refGene_gn=NIPBL;refGene_tx=XM_023239002.1
A1 212147152 212147243
refGene_gn=NIPBL;refGene_tx=XM_023239001.1;refGene_gn=NIPBL;refGene_tx=XM_023239002.1
A1 212147760 212147915

refGene_gn=NIPBL;refGene_tx=XM_023239001.1;refGene_gn=NIPBL;refGene_tx=XM_023239002.1
A1 212148552 212148646
refGene_gn=NIPBL;refGene_tx=XM_023239001.1;refGene_gn=NIPBL;refGene_tx=XM_023239002.1
A1 212148766 212148907
refGene_gn=NIPBL;refGene_tx=XM_023239001.1;refGene_gn=NIPBL;refGene_tx=XM_023239002.1
A1 212152920 212153057
refGene_gn=NIPBL;refGene_tx=XM_023239001.1;refGene_gn=NIPBL;refGene_tx=XM_023239002.1
A1 212154582 212154691
refGene_gn=NIPBL;refGene_tx=XM_023239001.1;refGene_gn=NIPBL;refGene_tx=XM_023239002.1
A1 212169651 212169705
refGene_gn=NIPBL;refGene_tx=XM_023239001.1;refGene_gn=NIPBL;refGene_tx=XM_023239002.1
A1 212171020 212171119
refGene_gn=NIPBL;refGene_tx=XM_023239001.1;refGene_gn=NIPBL;refGene_tx=XM_023239002.1
A1 212172389 212172524
refGene_gn=NIPBL;refGene_tx=XM_023239001.1;refGene_gn=NIPBL;refGene_tx=XM_023239002.1
A1 212175303 212175450
refGene_gn=NIPBL;refGene_tx=XM_023239001.1;refGene_gn=NIPBL;refGene_tx=XM_023239002.1
A1 212175554 212175653
refGene_gn=NIPBL;refGene_tx=XM_023239001.1;refGene_gn=NIPBL;refGene_tx=XM_023239002.1
A1 212176270 212176373
refGene_gn=NIPBL;refGene_tx=XM_023239001.1;refGene_gn=NIPBL;refGene_tx=XM_023239002.1
A1 212176477 212176692
refGene_gn=NIPBL;refGene_tx=XM_023239001.1;refGene_gn=NIPBL;refGene_tx=XM_023239002.1
A1 212178154 212178244
refGene_gn=NIPBL;refGene_tx=XM_023239001.1;refGene_gn=NIPBL;refGene_tx=XM_023239002.1
A1 212180553 212180697
refGene_gn=NIPBL;refGene_tx=XM_023239001.1;refGene_gn=NIPBL;refGene_tx=XM_023239002.1
A1 212181443 212181576
refGene_gn=NIPBL;refGene_tx=XM_023239001.1;refGene_gn=NIPBL;refGene_tx=XM_023239002.1
A1 212182172 212182255
refGene_gn=NIPBL;refGene_tx=XM_023239001.1;refGene_gn=NIPBL;refGene_tx=XM_023239002.1
A1 212184061 212184200
refGene_gn=NIPBL;refGene_tx=XM_023239001.1;refGene_gn=NIPBL;refGene_tx=XM_023239002.1

A1	212185513	212185614
refGene_gn=NIPBL;refGene_tx=XM_023239001.1;refGene_gn=NIPBL;refGene_tx
=XM_023239002.1
A1	212186066	212186147
refGene_gn=NIPBL;refGene_tx=XM_023239001.1;refGene_gn=NIPBL;refGene_tx
=XM_023239002.1
A1	212186708	212186860
refGene_gn=NIPBL;refGene_tx=XM_023239001.1;refGene_gn=NIPBL;refGene_tx
=XM_023239002.1
A1	212188269	212188501
refGene_gn=NIPBL;refGene_tx=XM_023239001.1;refGene_gn=NIPBL;refGene_tx
=XM_023239002.1
A1	212190815	212190902
refGene_gn=NIPBL;refGene_tx=XM_023239001.1;refGene_gn=NIPBL;refGene_tx
=XM_023239002.1
A1	212193657	212193761
refGene_gn=NIPBL;refGene_tx=XM_023239001.1;refGene_gn=NIPBL;refGene_tx
=XM_023239002.1
A1	212195320	212195410
refGene_gn=NIPBL;refGene_tx=XM_023239001.1;refGene_gn=NIPBL;refGene_tx
=XM_023239002.1
A1	212195498	212195570
refGene_gn=NIPBL;refGene_tx=XM_023239001.1;refGene_gn=NIPBL;refGene_tx
=XM_023239002.1
A1	212195825	212196023
refGene_gn=NIPBL;refGene_tx=XM_023239001.1;refGene_gn=NIPBL;refGene_tx
=XM_023239002.1
A1	212200412	212200595
refGene_gn=NIPBL;refGene_tx=XM_023239001.1;refGene_gn=NIPBL;refGene_tx
=XM_023239002.1
A1	212216946	212218572
refGene_gn=NIPBL;refGene_tx=XM_023239001.1;refGene_gn=NIPBL;refGene_tx
=XM_023239002.1
A1	212224617	212225244
refGene_gn=NIPBL;refGene_tx=XM_023239001.1;refGene_gn=NIPBL;refGene_tx
=XM_023239002.1
A1	212228728	212228828
refGene_gn=NIPBL;refGene_tx=XM_023239001.1;refGene_gn=NIPBL;refGene_tx
=XM_023239002.1
A1	212229751	212229912
refGene_gn=NIPBL;refGene_tx=XM_023239001.1;refGene_gn=NIPBL;refGene_tx
=XM_023239002.1
A1	212234808	212234960
refGene_gn=NIPBL;refGene_tx=XM_023239001.1;refGene_gn=NIPBL;refGene_tx
=XM_023239002.1
A1	212235786	212235886
refGene_gn=NIPBL;refGene_tx=XM_023239001.1;refGene_gn=NIPBL;refGene_tx
=XM_023239002.1
A1	212238727	212238855
refGene_gn=NIPBL;refGene_tx=XM_023239001.1;refGene_gn=NIPBL;refGene_tx

=XM_023239002.1
A1 212242307 212242473
refGene_gn=NIPBL;refGene_tx=XM_023239001.1;refGene_gn=NIPBL;refGene_tx
=XM_023239002.1
A1 212244773 212244837
refGene_gn=NIPBL;refGene_tx=XM_023239001.1;refGene_gn=NIPBL;refGene_tx
=XM_023239002.1
A1 212494239 212494444
refGene_gn=SLC1A3;refGene_tx=XM_003981454.5
A1 212496696 212496831
refGene_gn=SLC1A3;refGene_tx=XM_003981454.5
A1 212500228 212500423
refGene_gn=SLC1A3;refGene_tx=XM_003981454.5
A1 212500986 212501220
refGene_gn=SLC1A3;refGene_tx=XM_003981454.5
A1 212503947 212504240
refGene_gn=SLC1A3;refGene_tx=XM_003981454.5
A1 212507434 212507477
refGene_gn=SLC1A3;refGene_tx=XM_003981454.5
A1 212510328 212510533
refGene_gn=SLC1A3;refGene_tx=XM_003981454.5
A1 212552135 212552273
refGene_gn=SLC1A3;refGene_tx=XM_003981454.5
A1 212570312 212570493
refGene_gn=SLC1A3;refGene_tx=XM_003981454.5
A1 212614648 212614707
refGene_gn=RANBP3L;refGene_tx=XM_019811844.1;refGene_gn=RANBP3L;refGen
e_tx=XM_019811853.2
A1 212843730 212843821
refGene_gn=RANBP3L;refGene_tx=XM_011286498.3
A1 212866105 212866164
refGene_gn=RANBP3L;refGene_tx=XM_011286498.3;refGene_gn=RANBP3L;refGen
e_tx=XM_019811844.1;refGene_gn=RANBP3L;refGene_tx=XM_019811853.2
A1 212867043 212867083
refGene_gn=RANBP3L;refGene_tx=XM_011286498.3;refGene_gn=RANBP3L;refGen
e_tx=XM_019811844.1;refGene_gn=RANBP3L;refGene_tx=XM_019811853.2
A1 212867580 212867658
refGene_gn=RANBP3L;refGene_tx=XM_011286498.3;refGene_gn=RANBP3L;refGen
e_tx=XM_019811844.1;refGene_gn=RANBP3L;refGene_tx=XM_019811853.2
A1 212868701 212868776
refGene_gn=RANBP3L;refGene_tx=XM_011286498.3;refGene_gn=RANBP3L;refGen
e_tx=XM_019811844.1;refGene_gn=RANBP3L;refGene_tx=XM_019811853.2;refGe
ne_gn=RANBP3L;refGene_tx=XM_019811859.2
A1 212870504 212870576
refGene_gn=RANBP3L;refGene_tx=XM_011286498.3;refGene_gn=RANBP3L;refGen
e_tx=XM_019811844.1;refGene_gn=RANBP3L;refGene_tx=XM_019811853.2;refGe
ne_gn=RANBP3L;refGene_tx=XM_019811859.2
A1 212870936 212871076
refGene_gn=RANBP3L;refGene_tx=XM_011286498.3;refGene_gn=RANBP3L;refGen
e_tx=XM_019811844.1;refGene_gn=RANBP3L;refGene_tx=XM_019811853.2;refGe

ne_gn=RANBP3L;refGene_tx=XM_019811859.2
A1	212874431	212874535
refGene_gn=RANBP3L;refGene_tx=XM_011286498.3;refGene_gn=RANBP3L;refGene_tx=XM_019811844.1;refGene_gn=RANBP3L;refGene_tx=XM_019811853.2;refGene_gn=RANBP3L;refGene_tx=XM_019811859.2
A1	212876162	212876247
refGene_gn=RANBP3L;refGene_tx=XM_011286498.3;refGene_gn=RANBP3L;refGene_tx=XM_019811844.1;refGene_gn=RANBP3L;refGene_tx=XM_019811853.2;refGene_gn=RANBP3L;refGene_tx=XM_019811859.2
A1	212879693	212879796
refGene_gn=RANBP3L;refGene_tx=XM_011286498.3;refGene_gn=RANBP3L;refGene_tx=XM_019811844.1;refGene_gn=RANBP3L;refGene_tx=XM_019811853.2;refGene_gn=RANBP3L;refGene_tx=XM_019811859.2
A1	212880157	212880282
refGene_gn=RANBP3L;refGene_tx=XM_011286498.3;refGene_gn=RANBP3L;refGene_tx=XM_019811844.1;refGene_gn=RANBP3L;refGene_tx=XM_019811853.2;refGene_gn=RANBP3L;refGene_tx=XM_019811859.2
A1	212881667	212881788
refGene_gn=RANBP3L;refGene_tx=XM_011286498.3;refGene_gn=RANBP3L;refGene_tx=XM_019811844.1;refGene_gn=RANBP3L;refGene_tx=XM_019811853.2;refGene_gn=RANBP3L;refGene_tx=XM_019811859.2
A1	212883218	212883361
refGene_gn=RANBP3L;refGene_tx=XM_011286498.3;refGene_gn=RANBP3L;refGene_tx=XM_019811844.1;refGene_gn=RANBP3L;refGene_tx=XM_019811853.2;refGene_gn=RANBP3L;refGene_tx=XM_019811859.2
A1	212886629	212886819
refGene_gn=RANBP3L;refGene_tx=XM_011286498.3;refGene_gn=RANBP3L;refGene_tx=XM_019811844.1;refGene_gn=RANBP3L;refGene_tx=XM_019811853.2;refGene_gn=RANBP3L;refGene_tx=XM_019811859.2
A1	212889103	212889147
refGene_gn=RANBP3L;refGene_tx=XM_011286498.3;refGene_gn=RANBP3L;refGene_tx=XM_019811844.1;refGene_gn=RANBP3L;refGene_tx=XM_019811853.2;refGene_gn=RANBP3L;refGene_tx=XM_019811859.2
A1	212896771	212897071
refGene_gn=NADK2;refGene_tx=XM_011286501.3;refGene_gn=NADK2;refGene_tx=XM_023239012.1
A1	212908219	212908308
refGene_gn=NADK2;refGene_tx=XM_011286501.3;refGene_gn=NADK2;refGene_tx=XM_023239012.1
A1	212909180	212909269
refGene_gn=NADK2;refGene_tx=XM_011286501.3;refGene_gn=NADK2;refGene_tx=XM_023239012.1
A1	212910110	212910192
refGene_gn=NADK2;refGene_tx=XM_011286501.3;refGene_gn=NADK2;refGene_tx=XM_019811877.2;refGene_gn=NADK2;refGene_tx=XM_023239012.1
A1	212917258	212917342
refGene_gn=NADK2;refGene_tx=XM_011286501.3;refGene_gn=NADK2;refGene_tx=XM_019811877.2;refGene_gn=NADK2;refGene_tx=XM_023239012.1
A1	212918752	212918889
refGene_gn=NADK2;refGene_tx=XM_011286501.3;refGene_gn=NADK2;refGene_tx

=XM_019811877.2;refGene_gn=NADK2;refGene_tx=XM_023239012.1
A1 212926159 212926238
refGene_gn=NADK2;refGene_tx=XM_011286501.3;refGene_gn=NADK2;refGene_tx
=XM_019811877.2;refGene_gn=NADK2;refGene_tx=XM_023239012.1
A1 212929063 212929129
refGene_gn=NADK2;refGene_tx=XM_011286501.3;refGene_gn=NADK2;refGene_tx
=XM_019811877.2
A1 212930466 212930562
refGene_gn=NADK2;refGene_tx=XM_011286501.3;refGene_gn=NADK2;refGene_tx
=XM_019811877.2;refGene_gn=NADK2;refGene_tx=XM_023239012.1
A1 212940782 212940838
refGene_gn=NADK2;refGene_tx=XM_011286501.3;refGene_gn=NADK2;refGene_tx
=XM_019811877.2;refGene_gn=NADK2;refGene_tx=XM_023239012.1
A1 212940962 212941016
refGene_gn=NADK2;refGene_tx=XM_011286501.3;refGene_gn=NADK2;refGene_tx
=XM_019811877.2;refGene_gn=NADK2;refGene_tx=XM_023239012.1
A1 212943552 212943676
refGene_gn=NADK2;refGene_tx=XM_011286501.3;refGene_gn=NADK2;refGene_tx
=XM_019811877.2;refGene_gn=NADK2;refGene_tx=XM_023239012.1
A1 212946189 212946325
refGene_gn=NADK2;refGene_tx=XM_011286501.3;refGene_gn=NADK2;refGene_tx
=XM_019811877.2;refGene_gn=NADK2;refGene_tx=XM_023239012.1
A1 212964652 212964746
refGene_gn=SKP2;refGene_tx=XM_023239018.1
A1 212966275 212966489
refGene_gn=SKP2;refGene_tx=XM_023239016.1
A1 212971013 212971121
refGene_gn=SKP2;refGene_tx=XM_023239016.1;refGene_gn=SKP2;refGene_tx=X
M_023239018.1
A1 212971286 212971338
refGene_gn=SKP2;refGene_tx=XM_023239016.1;refGene_gn=SKP2;refGene_tx=X
M_023239018.1
A1 212977815 212977946
refGene_gn=SKP2;refGene_tx=XM_023239016.1;refGene_gn=SKP2;refGene_tx=X
M_023239018.1
A1 212980338 212980437
refGene_gn=SKP2;refGene_tx=XM_023239016.1;refGene_gn=SKP2;refGene_tx=X
M_023239018.1
A1 212982031 212982166
refGene_gn=SKP2;refGene_tx=XM_023239016.1;refGene_gn=SKP2;refGene_tx=X
M_023239018.1
A1 212983464 212983608
refGene_gn=SKP2;refGene_tx=XM_023239016.1;refGene_gn=SKP2;refGene_tx=X
M_023239018.1
A1 212986078 212986190
refGene_gn=SKP2;refGene_tx=XM_023239016.1;refGene_gn=SKP2;refGene_tx=X
M_023239018.1
A1 212997396 212997668
refGene_gn=SKP2;refGene_tx=XM_023239016.1;refGene_gn=SKP2;refGene_tx=X
M_023239018.1

A1 212998167 212998211
refGene_gn=SKP2;refGene_tx=XM_023239016.1;refGene_gn=SKP2;refGene_tx=XM_023239018.1
A1 213011530 213011704
refGene_gn=LMBRD2;refGene_tx=XM_003981457.5;refGene_gn=LMBRD2;refGene_tx=XM_006928052.4;refGene_gn=LMBRD2;refGene_tx=XM_019811891.2;refGene_gn=LMBRD2;refGene_tx=XM_019811896.2;refGene_gn=LMBRD2;refGene_tx=XM_019811915.2
A1 213011956 213012054
refGene_gn=LMBRD2;refGene_tx=XM_003981457.5;refGene_gn=LMBRD2;refGene_tx=XM_006928052.4;refGene_gn=LMBRD2;refGene_tx=XM_019811891.2;refGene_gn=LMBRD2;refGene_tx=XM_019811896.2;refGene_gn=LMBRD2;refGene_tx=XM_019811915.2
A1 213013384 213013480
refGene_gn=LMBRD2;refGene_tx=XM_003981457.5;refGene_gn=LMBRD2;refGene_tx=XM_006928052.4;refGene_gn=LMBRD2;refGene_tx=XM_019811891.2;refGene_gn=LMBRD2;refGene_tx=XM_019811896.2;refGene_gn=LMBRD2;refGene_tx=XM_019811915.2
A1 213016294 213016462
refGene_gn=LMBRD2;refGene_tx=XM_003981457.5;refGene_gn=LMBRD2;refGene_tx=XM_006928052.4;refGene_gn=LMBRD2;refGene_tx=XM_019811891.2;refGene_gn=LMBRD2;refGene_tx=XM_019811896.2;refGene_gn=LMBRD2;refGene_tx=XM_019811915.2
A1 213017705 213017916
refGene_gn=LMBRD2;refGene_tx=XM_003981457.5;refGene_gn=LMBRD2;refGene_tx=XM_006928052.4;refGene_gn=LMBRD2;refGene_tx=XM_019811891.2;refGene_gn=LMBRD2;refGene_tx=XM_019811896.2;refGene_gn=LMBRD2;refGene_tx=XM_019811915.2
A1 213020731 213020806
refGene_gn=LMBRD2;refGene_tx=XM_003981457.5;refGene_gn=LMBRD2;refGene_tx=XM_006928052.4;refGene_gn=LMBRD2;refGene_tx=XM_019811891.2;refGene_gn=LMBRD2;refGene_tx=XM_019811896.2;refGene_gn=LMBRD2;refGene_tx=XM_019811915.2
A1 213021954 213022068
refGene_gn=LMBRD2;refGene_tx=XM_003981457.5;refGene_gn=LMBRD2;refGene_tx=XM_006928052.4;refGene_gn=LMBRD2;refGene_tx=XM_019811891.2;refGene_gn=LMBRD2;refGene_tx=XM_019811896.2;refGene_gn=LMBRD2;refGene_tx=XM_019811915.2
A1 213022436 213022620
refGene_gn=LMBRD2;refGene_tx=XM_003981457.5;refGene_gn=LMBRD2;refGene_tx=XM_006928052.4;refGene_gn=LMBRD2;refGene_tx=XM_019811891.2;refGene_gn=LMBRD2;refGene_tx=XM_019811896.2;refGene_gn=LMBRD2;refGene_tx=XM_019811915.2
A1 213025486 213025668
refGene_gn=LMBRD2;refGene_tx=XM_003981457.5;refGene_gn=LMBRD2;refGene_tx=XM_006928052.4;refGene_gn=LMBRD2;refGene_tx=XM_019811891.2;refGene_gn=LMBRD2;refGene_tx=XM_019811896.2;refGene_gn=LMBRD2;refGene_tx=XM_019811915.2
A1 213026262 213026396
refGene_gn=LMBRD2;refGene_tx=XM_003981457.5;refGene_gn=LMBRD2;refGene_

tx=XM_006928052.4;refGene_gn=LMBRD2;refGene_tx=XM_019811891.2;refGene_gn=LMBRD2;refGene_tx=XM_019811896.2;refGene_gn=LMBRD2;refGene_tx=XM_019811915.2
A1 213027761 213027867
refGene_gn=LMBRD2;refGene_tx=XM_003981457.5;refGene_gn=LMBRD2;refGene_tx=XM_006928052.4;refGene_gn=LMBRD2;refGene_tx=XM_019811891.2;refGene_gn=LMBRD2;refGene_tx=XM_019811896.2;refGene_gn=LMBRD2;refGene_tx=XM_019811915.2
A1 213028526 213028624
refGene_gn=LMBRD2;refGene_tx=XM_003981457.5;refGene_gn=LMBRD2;refGene_tx=XM_006928052.4;refGene_gn=LMBRD2;refGene_tx=XM_019811891.2;refGene_gn=LMBRD2;refGene_tx=XM_019811896.2;refGene_gn=LMBRD2;refGene_tx=XM_019811915.2
A1 213033744 213033848
refGene_gn=LMBRD2;refGene_tx=XM_003981457.5;refGene_gn=LMBRD2;refGene_tx=XM_006928052.4;refGene_gn=LMBRD2;refGene_tx=XM_019811891.2;refGene_gn=LMBRD2;refGene_tx=XM_019811896.2;refGene_gn=LMBRD2;refGene_tx=XM_019811915.2
A1 213034920 213034928
refGene_gn=LMBRD2;refGene_tx=XM_019811915.2
A1 213035023 213035070
refGene_gn=LMBRD2;refGene_tx=XM_003981457.5;refGene_gn=LMBRD2;refGene_tx=XM_006928052.4;refGene_gn=LMBRD2;refGene_tx=XM_019811891.2;refGene_gn=LMBRD2;refGene_tx=XM_019811896.2
A1 213036601 213036707
refGene_gn=LMBRD2;refGene_tx=XM_003981457.5;refGene_gn=LMBRD2;refGene_tx=XM_006928052.4;refGene_gn=LMBRD2;refGene_tx=XM_019811891.2;refGene_gn=LMBRD2;refGene_tx=XM_019811896.2
A1 213039336 213039466
refGene_gn=LMBRD2;refGene_tx=XM_003981457.5;refGene_gn=LMBRD2;refGene_tx=XM_006928052.4;refGene_gn=LMBRD2;refGene_tx=XM_019811891.2;refGene_gn=LMBRD2;refGene_tx=XM_019811896.2
A1 213040345 213040430
refGene_gn=LMBRD2;refGene_tx=XM_003981457.5;refGene_gn=LMBRD2;refGene_tx=XM_006928052.4;refGene_gn=LMBRD2;refGene_tx=XM_019811891.2;refGene_gn=LMBRD2;refGene_tx=XM_019811896.2
A1 213066931 213067097
refGene_gn=LOC101099779;refGene_tx=XM_019811923.2
A1 213067191 213067285
refGene_gn=LOC101099779;refGene_tx=XM_019811925.2
A1 213069729 213069831
refGene_gn=LOC101099779;refGene_tx=XM_019811923.2;refGene_gn=LOC101099779;refGene_tx=XM_019811925.2
A1 213083966 213084081
refGene_gn=LOC101099779;refGene_tx=XM_019811923.2;refGene_gn=LOC101099779;refGene_tx=XM_019811925.2
A1 213086413 213086945
refGene_gn=LOC101099779;refGene_tx=XM_019811923.2;refGene_gn=LOC101099779;refGene_tx=XM_019811925.2
A1 213089431 213089663

refGene_gn=LOC101099779;refGene_tx=XM_019811923.2;refGene_gn=LOC101099779;refGene_tx=XM_019811925.2
A1 213091201 213091421
refGene_gn=LOC101099779;refGene_tx=XM_019811923.2;refGene_gn=LOC101099779;refGene_tx=XM_019811925.2
A1 213093769 213094046
refGene_gn=LOC101099779;refGene_tx=XM_019811923.2;refGene_gn=LOC101099779;refGene_tx=XM_019811925.2
A1 213097186 213097280
refGene_gn=LOC101081204;refGene_tx=XM_011286527.3
A1 213099514 213099616
refGene_gn=LOC101081204;refGene_tx=XM_011286527.3
A1 213110908 213111017
refGene_gn=LOC101081204;refGene_tx=XM_011286527.3
A1 213112819 213113351
refGene_gn=LOC101081204;refGene_tx=XM_011286527.3
A1 213119996 213120228
refGene_gn=LOC101081204;refGene_tx=XM_011286527.3
A1 213121691 213121911
refGene_gn=LOC101081204;refGene_tx=XM_011286527.3
A1 213123020 213123297
refGene_gn=LOC101081204;refGene_tx=XM_011286527.3
A1 213170532 213170626
refGene_gn=LOC101097701;refGene_tx=XM_019811937.2
A1 213180189 213180291
refGene_gn=LOC101097701;refGene_tx=XM_019811937.2
A1 213199896 213200005
refGene_gn=LOC101097701;refGene_tx=XM_019811937.2
A1 213205529 213206061
refGene_gn=LOC101097701;refGene_tx=XM_019811937.2
A1 213211501 213211733
refGene_gn=LOC101097701;refGene_tx=XM_019811937.2
A1 213213052 213213272
refGene_gn=LOC101097701;refGene_tx=XM_019811937.2
A1 213214387 213214664
refGene_gn=LOC101097701;refGene_tx=XM_019811937.2
A1 213237533 213237790
refGene_gn=CAPSL;refGene_tx=XM_023239042.1;refGene_gn=CAPSL;refGene_tx=XM_023239044.1
A1 213240654 213240832
refGene_gn=CAPSL;refGene_tx=XM_023239042.1;refGene_gn=CAPSL;refGene_tx=XM_023239044.1
A1 213241117 213241327
refGene_gn=CAPSL;refGene_tx=XM_023239042.1;refGene_gn=CAPSL;refGene_tx=XM_023239044.1
A1 213246127 213246229
refGene_gn=CAPSL;refGene_tx=XM_023239044.1
A1 213260136 213260250
refGene_gn=CAPSL;refGene_tx=XM_023239042.1
A1 213270789 213271293

refGene_gn=IL7R;refGene_tx=XM_006928055.3
A1	213271692	213271768
refGene_gn=IL7R;refGene_tx=XM_006928055.3
A1	213272885	213272979
refGene_gn=IL7R;refGene_tx=XM_006928055.3
A1	213273880	213274043
refGene_gn=IL7R;refGene_tx=XM_006928055.3
A1	213275852	213276010
refGene_gn=IL7R;refGene_tx=XM_006928055.3
A1	213280228	213280386
refGene_gn=IL7R;refGene_tx=XM_006928055.3
A1	213291417	213291556
refGene_gn=IL7R;refGene_tx=XM_006928055.3
A1	213296054	213296136
refGene_gn=IL7R;refGene_tx=XM_006928055.3
A1	213325991	213326081
refGene_gn=SPEF2;refGene_tx=XM_011286553.3
A1	213331852	213331975
refGene_gn=SPEF2;refGene_tx=XM_011286553.3
A1	213332172	213332412
refGene_gn=SPEF2;refGene_tx=XM_011286553.3
A1	213338496	213338676
refGene_gn=SPEF2;refGene_tx=XM_011286553.3
A1	213345251	213345344
refGene_gn=SPEF2;refGene_tx=XM_011286553.3
A1	213358790	213358973
refGene_gn=SPEF2;refGene_tx=XM_011286553.3
A1	213359915	213360022
refGene_gn=SPEF2;refGene_tx=XM_011286553.3
A1	213367496	213367848
refGene_gn=LOC111556553;refGene_tx=XM_023239064.1
A1	213367930	213368054
refGene_gn=LOC111556553;refGene_tx=XM_023239064.1
A1	213368144	213368353
refGene_gn=LOC111556553;refGene_tx=XM_023239064.1
A1	213368473	213368700
refGene_gn=LOC111556553;refGene_tx=XM_023239064.1
A1	213379254	213379484
refGene_gn=SPEF2;refGene_tx=XM_011286553.3
A1	213382226	213382365
refGene_gn=SPEF2;refGene_tx=XM_011286553.3
A1	213383462	213383591
refGene_gn=SPEF2;refGene_tx=XM_011286553.3
A1	213385444	213385592
refGene_gn=SPEF2;refGene_tx=XM_011286553.3
A1	213392548	213392729
refGene_gn=SPEF2;refGene_tx=XM_011286553.3
A1	213396432	213396584
refGene_gn=SPEF2;refGene_tx=XM_011286553.3
A1	213400718	213400856

refGene_gn=SPEF2;refGene_tx=XM_011286553.3
A1 213408337 213408476
refGene_gn=SPEF2;refGene_tx=XM_011286553.3
A1 213408559 213408687
refGene_gn=SPEF2;refGene_tx=XM_011286553.3
A1 213413658 213413807
refGene_gn=SPEF2;refGene_tx=XM_011286553.3
A1 213418836 213418911
refGene_gn=SPEF2;refGene_tx=XM_011286553.3
A1 213422403 213422568
refGene_gn=SPEF2;refGene_tx=XM_011286553.3
A1 213425421 213425579
refGene_gn=SPEF2;refGene_tx=XM_011286553.3
A1 213428339 213428448
refGene_gn=SPEF2;refGene_tx=XM_011286553.3
A1 213435895 213436152
refGene_gn=SPEF2;refGene_tx=XM_011286553.3
A1 213437455 213437559
refGene_gn=SPEF2;refGene_tx=XM_011286553.3
A1 213439771 213439833
refGene_gn=SPEF2;refGene_tx=XM_011286553.3
A1 213441204 213441280
refGene_gn=SPEF2;refGene_tx=XM_011286553.3
A1 213442941 213443096
refGene_gn=SPEF2;refGene_tx=XM_011286553.3
A1 213444611 213444831
refGene_gn=SPEF2;refGene_tx=XM_011286553.3
A1 213463084 213463253
refGene_gn=SPEF2;refGene_tx=XM_011286553.3
A1 213466296 213466484
refGene_gn=SPEF2;refGene_tx=XM_011286553.3
A1 213472499 213472688
refGene_gn=SPEF2;refGene_tx=XM_011286553.3
A1 213480445 213480632
refGene_gn=SPEF2;refGene_tx=XM_011286553.3
A1 213485858 213485923
refGene_gn=SPEF2;refGene_tx=XM_011286553.3
A1 213487389 213487530
refGene_gn=SPEF2;refGene_tx=XM_011286553.3
A1 213489949 213490120
refGene_gn=SPEF2;refGene_tx=XM_011286553.3
A1 213493200 213493453
refGene_gn=SPEF2;refGene_tx=XM_011286553.3
A1 213499778 213499881
refGene_gn=SPEF2;refGene_tx=XM_011286553.3
A1 213511997 213512055
refGene_gn=SPEF2;refGene_tx=XM_011286553.3
A1 213917951 213918018
refGene_gn=PRLR;refGene_tx=XM_003981463.4
A1 213920768 213920901

refGene_gn=PRLR;refGene_tx=XM_003981463.4
A1 213922381 213922551
refGene_gn=PRLR;refGene_tx=XM_003981463.4
A1 213926778 213926948
refGene_gn=PRLR;refGene_tx=XM_003981463.4
A1 213928657 213928799
refGene_gn=PRLR;refGene_tx=XM_003981463.4
A1 213930348 213930448
refGene_gn=PRLR;refGene_tx=XM_003981463.4
A1 213930958 213931028
refGene_gn=PRLR;refGene_tx=XM_003981463.4
A1 213932682 213933705
refGene_gn=PRLR;refGene_tx=XM_003981463.4
A1 213934419 213934441
refGene_gn=AGXT2;refGene_tx=XM_003981518.2
A1 213935395 213935530
refGene_gn=AGXT2;refGene_tx=XM_003981518.2
A1 213948982 213949184
refGene_gn=AGXT2;refGene_tx=XM_003981518.2
A1 213950463 213950729
refGene_gn=AGXT2;refGene_tx=XM_003981518.2
A1 213953365 213953451
refGene_gn=AGXT2;refGene_tx=XM_003981518.2
A1 213955181 213955366
refGene_gn=AGXT2;refGene_tx=XM_003981518.2
A1 213958140 213958264
refGene_gn=AGXT2;refGene_tx=XM_003981518.2
A1 213959839 213959934
refGene_gn=AGXT2;refGene_tx=XM_003981518.2
A1 213961980 213962074
refGene_gn=AGXT2;refGene_tx=XM_003981518.2
A1 213962721 213962815
refGene_gn=AGXT2;refGene_tx=XM_003981518.2
A1 213972871 213972972
refGene_gn=AGXT2;refGene_tx=XM_003981518.2
A1 213973555 213973648
refGene_gn=AGXT2;refGene_tx=XM_003981518.2
A1 213974252 213974335
refGene_gn=AGXT2;refGene_tx=XM_003981518.2
A1 213974336 213974374
refGene_gn=AGXT2;refGene_tx=XM_003981518.2
A1 213976417 213976509
refGene_gn=AGXT2;refGene_tx=XM_003981518.2
A1 213978523 213978673
refGene_gn=AGXT2;refGene_tx=XM_003981518.2
A1 213989774 213989873
refGene_gn=AGXT2;refGene_tx=XM_003981518.2
A1 213997894 213998011
refGene_gn=AGXT2;refGene_tx=XM_003981518.2
A1 214021857 214022022

refGene_gn=DNAJC21;refGene_tx=XM_023239071.1
A1 214022754 214022830
refGene_gn=DNAJC21;refGene_tx=XM_023239071.1
A1 214028079 214028252
refGene_gn=DNAJC21;refGene_tx=XM_023239071.1
A1 214031938 214031981
refGene_gn=DNAJC21;refGene_tx=XM_023239071.1
A1 214032409 214032568
refGene_gn=DNAJC21;refGene_tx=XM_023239071.1
A1 214035660 214035748
refGene_gn=DNAJC21;refGene_tx=XM_023239071.1
A1 214038163 214038315
refGene_gn=DNAJC21;refGene_tx=XM_023239071.1
A1 214039188 214039493
refGene_gn=DNAJC21;refGene_tx=XM_023239071.1
A1 214039876 214039999
refGene_gn=DNAJC21;refGene_tx=XM_023239071.1
A1 214040757 214040881
refGene_gn=DNAJC21;refGene_tx=XM_023239071.1
A1 214042331 214042425
refGene_gn=DNAJC21;refGene_tx=XM_023239071.1
A1 214050362 214050459
refGene_gn=DNAJC21;refGene_tx=XM_023239071.1
A1 214053740 214054010
refGene_gn=BRIX1;refGene_tx=XM_003981464.4
A1 214054257 214054386
refGene_gn=BRIX1;refGene_tx=XM_003981464.4
A1 214056514 214056616
refGene_gn=BRIX1;refGene_tx=XM_003981464.4
A1 214056755 214056806
refGene_gn=BRIX1;refGene_tx=XM_003981464.4
A1 214056965 214057039
refGene_gn=BRIX1;refGene_tx=XM_003981464.4
A1 214057141 214057191
refGene_gn=BRIX1;refGene_tx=XM_003981464.4
A1 214057465 214057536
refGene_gn=BRIX1;refGene_tx=XM_003981464.4
A1 214059438 214059482
refGene_gn=BRIX1;refGene_tx=XM_003981464.4
A1 214060520 214060632
refGene_gn=BRIX1;refGene_tx=XM_003981464.4
A1 214063434 214063593
refGene_gn=BRIX1;refGene_tx=XM_003981464.4
A1 214064393 214064588
refGene_gn=RAD1;refGene_tx=XM_003981465.4;refGene_gn=RAD1;refGene_tx=XM_003981466.4;refGene_gn=RAD1;refGene_tx=XM_006928058.3;refGene_gn=RAD1;refGene_tx=XM_006928059.3;refGene_gn=RAD1;refGene_tx=XM_019812044.1
A1 214065531 214065640
refGene_gn=RAD1;refGene_tx=XM_003981465.4;refGene_gn=RAD1;refGene_tx=XM_003981466.4;refGene_gn=RAD1;refGene_tx=XM_006928058.3;refGene_gn=RAD

1;refGene_tx=XM_006928059.3;refGene_gn=RAD1;refGene_tx=XM_019812044.1
A1 214067242 214067501
refGene_gn=RAD1;refGene_tx=XM_003981465.4;refGene_gn=RAD1;refGene_tx=XM_003981466.4;refGene_gn=RAD1;refGene_tx=XM_006928058.3;refGene_gn=RAD1;refGene_tx=XM_006928059.3;refGene_gn=RAD1;refGene_tx=XM_019812044.1
A1 214069569 214069668
refGene_gn=RAD1;refGene_tx=XM_003981465.4;refGene_gn=RAD1;refGene_tx=XM_003981466.4;refGene_gn=RAD1;refGene_tx=XM_006928058.3;refGene_gn=RAD1;refGene_tx=XM_006928059.3;refGene_gn=RAD1;refGene_tx=XM_019812044.1
A1 214069984 214070168
refGene_gn=RAD1;refGene_tx=XM_003981465.4;refGene_gn=RAD1;refGene_tx=XM_003981466.4;refGene_gn=RAD1;refGene_tx=XM_006928058.3;refGene_gn=RAD1;refGene_tx=XM_006928059.3;refGene_gn=RAD1;refGene_tx=XM_019812044.1
A1 214077964 214078046
refGene_gn=TTC23L;refGene_tx=XM_019812058.2;refGene_gn=TTC23L;refGene_tx=XM_019812076.2;refGene_gn=TTC23L;refGene_tx=XM_023239093.1;refGene_gn=TTC23L;refGene_tx=XM_023239095.1;refGene_gn=TTC23L;refGene_tx=XM_023239098.1;refGene_gn=TTC23L;refGene_tx=XM_023239109.1
A1 214085915 214085998
refGene_gn=TTC23L;refGene_tx=XM_011286575.3;refGene_gn=TTC23L;refGene_tx=XM_019812058.2;refGene_gn=TTC23L;refGene_tx=XM_019812076.2;refGene_gn=TTC23L;refGene_tx=XM_023239093.1;refGene_gn=TTC23L;refGene_tx=XM_023239095.1;refGene_gn=TTC23L;refGene_tx=XM_023239098.1;refGene_gn=TTC23L;refGene_tx=XM_023239109.1
A1 214094627 214094787
refGene_gn=TTC23L;refGene_tx=XM_011286575.3;refGene_gn=TTC23L;refGene_tx=XM_019812058.2;refGene_gn=TTC23L;refGene_tx=XM_019812076.2;refGene_gn=TTC23L;refGene_tx=XM_023239093.1;refGene_gn=TTC23L;refGene_tx=XM_023239095.1;refGene_gn=TTC23L;refGene_tx=XM_023239098.1;refGene_gn=TTC23L;refGene_tx=XM_023239109.1
A1 214095132 214095260
refGene_gn=TTC23L;refGene_tx=XM_011286575.3;refGene_gn=TTC23L;refGene_tx=XM_019812058.2;refGene_gn=TTC23L;refGene_tx=XM_019812076.2;refGene_gn=TTC23L;refGene_tx=XM_023239093.1;refGene_gn=TTC23L;refGene_tx=XM_023239095.1;refGene_gn=TTC23L;refGene_tx=XM_023239098.1;refGene_gn=TTC23L;refGene_tx=XM_023239109.1
A1 214105810 214105919
refGene_gn=TTC23L;refGene_tx=XM_011286575.3;refGene_gn=TTC23L;refGene_tx=XM_019812058.2;refGene_gn=TTC23L;refGene_tx=XM_019812076.2;refGene_gn=TTC23L;refGene_tx=XM_023239093.1;refGene_gn=TTC23L;refGene_tx=XM_023239095.1;refGene_gn=TTC23L;refGene_tx=XM_023239098.1;refGene_gn=TTC23L;refGene_tx=XM_023239109.1
A1 214107856 214108034
refGene_gn=TTC23L;refGene_tx=XM_011286575.3;refGene_gn=TTC23L;refGene_tx=XM_019812058.2;refGene_gn=TTC23L;refGene_tx=XM_019812076.2;refGene_gn=TTC23L;refGene_tx=XM_023239093.1;refGene_gn=TTC23L;refGene_tx=XM_023239095.1;refGene_gn=TTC23L;refGene_tx=XM_023239098.1;refGene_gn=TTC23L;refGene_tx=XM_023239109.1
A1 214111165 214111291
refGene_gn=TTC23L;refGene_tx=XM_011286575.3;refGene_gn=TTC23L;refGene_

tx=XM_019812058.2;refGene_gn=TTC23L;refGene_tx=XM_019812076.2;refGene_
gn=TTC23L;refGene_tx=XM_023239093.1;refGene_gn=TTC23L;refGene_tx=XM_02
3239095.1;refGene_gn=TTC23L;refGene_tx=XM_023239098.1;refGene_gn=TTC23
L;refGene_tx=XM_023239109.1
A1 214112761 214112918
refGene_gn=TTC23L;refGene_tx=XM_011286575.3;refGene_gn=TTC23L;refGene_
tx=XM_019812058.2;refGene_gn=TTC23L;refGene_tx=XM_019812076.2;refGene_
gn=TTC23L;refGene_tx=XM_023239093.1;refGene_gn=TTC23L;refGene_tx=XM_02
3239095.1;refGene_gn=TTC23L;refGene_tx=XM_023239098.1;refGene_gn=TTC23
L;refGene_tx=XM_023239109.1
A1 214123616 214123740
refGene_gn=TTC23L;refGene_tx=XM_011286575.3;refGene_gn=TTC23L;refGene_
tx=XM_019812058.2;refGene_gn=TTC23L;refGene_tx=XM_023239093.1;refGene_
gn=TTC23L;refGene_tx=XM_023239095.1;refGene_gn=TTC23L;refGene_tx=XM_02
3239098.1;refGene_gn=TTC23L;refGene_tx=XM_023239109.1
A1 214127530 214127714
refGene_gn=TTC23L;refGene_tx=XM_011286575.3;refGene_gn=TTC23L;refGene_
tx=XM_019812058.2;refGene_gn=TTC23L;refGene_tx=XM_023239093.1;refGene_
gn=TTC23L;refGene_tx=XM_023239095.1;refGene_gn=TTC23L;refGene_tx=XM_02
3239098.1;refGene_gn=TTC23L;refGene_tx=XM_023239109.1
A1 214132865 214132940
refGene_gn=TTC23L;refGene_tx=XM_011286575.3;refGene_gn=TTC23L;refGene_
tx=XM_019812058.2;refGene_gn=TTC23L;refGene_tx=XM_019812076.2;refGene_
gn=TTC23L;refGene_tx=XM_023239093.1;refGene_gn=TTC23L;refGene_tx=XM_02
3239095.1;refGene_gn=TTC23L;refGene_tx=XM_023239098.1;refGene_gn=TTC23
L;refGene_tx=XM_023239109.1
A1 214133792 214133865
refGene_gn=TTC23L;refGene_tx=XM_019812076.2
A1 214141612 214141690
refGene_gn=RAI14;refGene_tx=XM_019812089.2;refGene_gn=RAI14;refGene_tx
=XM_019812093.2;refGene_gn=RAI14;refGene_tx=XM_019812095.2;refGene_gn=
RAI14;refGene_tx=XM_023239127.1
A1 214142285 214142351
refGene_gn=RAI14;refGene_tx=XM_019812089.2;refGene_gn=RAI14;refGene_tx
=XM_019812093.2;refGene_gn=RAI14;refGene_tx=XM_019812095.2;refGene_gn=
RAI14;refGene_tx=XM_023239127.1
A1 214145909 214146059
refGene_gn=RAI14;refGene_tx=XM_019812089.2;refGene_gn=RAI14;refGene_tx
=XM_019812093.2;refGene_gn=RAI14;refGene_tx=XM_019812095.2;refGene_gn=
RAI14;refGene_tx=XM_023239127.1
A1 214147223 214148759
refGene_gn=RAI14;refGene_tx=XM_019812089.2;refGene_gn=RAI14;refGene_tx
=XM_019812093.2;refGene_gn=RAI14;refGene_tx=XM_019812095.2;refGene_gn=
RAI14;refGene_tx=XM_023239127.1
A1 214149399 214149518
refGene_gn=RAI14;refGene_tx=XM_019812089.2;refGene_gn=RAI14;refGene_tx
=XM_019812093.2;refGene_gn=RAI14;refGene_tx=XM_019812095.2;refGene_gn=
RAI14;refGene_tx=XM_023239127.1
A1 214153052 214153107
refGene_gn=RAI14;refGene_tx=XM_019812089.2;refGene_gn=RAI14;refGene_tx

=XM_019812093.2;refGene_gn=RAI14;refGene_tx=XM_019812095.2;refGene_gn=RAI14;refGene_tx=XM_023239127.1
A1 214155662 214155749
refGene_gn=RAI14;refGene_tx=XM_019812089.2;refGene_gn=RAI14;refGene_tx=XM_019812093.2;refGene_gn=RAI14;refGene_tx=XM_019812095.2;refGene_gn=RAI14;refGene_tx=XM_023239127.1
A1 214156717 214156804
refGene_gn=RAI14;refGene_tx=XM_019812089.2;refGene_gn=RAI14;refGene_tx=XM_019812093.2;refGene_gn=RAI14;refGene_tx=XM_023239127.1
A1 214157873 214157902
refGene_gn=RAI14;refGene_tx=XM_019812089.2;refGene_gn=RAI14;refGene_tx=XM_019812093.2;refGene_gn=RAI14;refGene_tx=XM_019812095.2;refGene_gn=RAI14;refGene_tx=XM_023239127.1
A1 214158158 214158337
refGene_gn=RAI14;refGene_tx=XM_019812089.2;refGene_gn=RAI14;refGene_tx=XM_019812093.2;refGene_gn=RAI14;refGene_tx=XM_019812095.2;refGene_gn=RAI14;refGene_tx=XM_023239127.1
A1 214159016 214159123
refGene_gn=RAI14;refGene_tx=XM_019812089.2;refGene_gn=RAI14;refGene_tx=XM_019812093.2;refGene_gn=RAI14;refGene_tx=XM_019812095.2;refGene_gn=RAI14;refGene_tx=XM_023239127.1
A1 214161596 214161667
refGene_gn=RAI14;refGene_tx=XM_019812089.2;refGene_gn=RAI14;refGene_tx=XM_019812093.2;refGene_gn=RAI14;refGene_tx=XM_019812095.2;refGene_gn=RAI14;refGene_tx=XM_023239127.1
A1 214162399 214162457
refGene_gn=RAI14;refGene_tx=XM_019812089.2;refGene_gn=RAI14;refGene_tx=XM_019812093.2;refGene_gn=RAI14;refGene_tx=XM_019812095.2;refGene_gn=RAI14;refGene_tx=XM_023239127.1
A1 214166250 214166315
refGene_gn=RAI14;refGene_tx=XM_019812089.2;refGene_gn=RAI14;refGene_tx=XM_019812093.2;refGene_gn=RAI14;refGene_tx=XM_019812095.2;refGene_gn=RAI14;refGene_tx=XM_023239127.1
A1 214176127 214176216
refGene_gn=RAI14;refGene_tx=XM_019812089.2;refGene_gn=RAI14;refGene_tx=XM_019812093.2;refGene_gn=RAI14;refGene_tx=XM_019812095.2;refGene_gn=RAI14;refGene_tx=XM_023239127.1
A1 214197603 214197734
refGene_gn=RAI14;refGene_tx=XM_019812089.2;refGene_gn=RAI14;refGene_tx=XM_019812093.2;refGene_gn=RAI14;refGene_tx=XM_019812095.2;refGene_gn=RAI14;refGene_tx=XM_023239127.1
A1 214197850 214197904
refGene_gn=RAI14;refGene_tx=XM_019812089.2;refGene_gn=RAI14;refGene_tx=XM_019812095.2
A1 214259672 214259708
refGene_gn=RAI14;refGene_tx=XM_019812093.2;refGene_gn=RAI14;refGene_tx=XM_023239127.1
A1 214471797 214472100
refGene_gn=C1QTNF3;refGene_tx=XM_003981468.4;refGene_gn=C1QTNF3;refGene_tx=XM_003981469.4;refGene_gn=C1QTNF3;refGene_tx=XM_006928062.3

A1 214477990 214478102
refGene_gn=C1QTNF3;refGene_tx=XM_003981468.4;refGene_gn=C1QTNF3;refGene_tx=XM_003981469.4;refGene_gn=C1QTNF3;refGene_tx=XM_006928062.3
A1 214480362 214480517
refGene_gn=C1QTNF3;refGene_tx=XM_003981468.4;refGene_gn=C1QTNF3;refGene_tx=XM_003981469.4;refGene_gn=C1QTNF3;refGene_tx=XM_006928062.3
A1 214485257 214485387
refGene_gn=C1QTNF3;refGene_tx=XM_003981468.4;refGene_gn=C1QTNF3;refGene_tx=XM_003981469.4;refGene_gn=C1QTNF3;refGene_tx=XM_006928062.3
A1 214490675 214490775
refGene_gn=C1QTNF3;refGene_tx=XM_003981468.4;refGene_gn=C1QTNF3;refGene_tx=XM_003981469.4;refGene_gn=C1QTNF3;refGene_tx=XM_006928062.3
A1 214494261 214494421
refGene_gn=C1QTNF3;refGene_tx=XM_003981468.4;refGene_gn=C1QTNF3;refGene_tx=XM_003981469.4;refGene_gn=C1QTNF3;refGene_tx=XM_006928062.3
A1 214501151 214501398
refGene_gn=AMACR;refGene_tx=XM_011286594.2
A1 214504319 214504463
refGene_gn=AMACR;refGene_tx=XM_011286594.2
A1 214505476 214505637
refGene_gn=AMACR;refGene_tx=XM_011286594.2
A1 214511248 214511435
refGene_gn=AMACR;refGene_tx=XM_011286594.2
A1 214521633 214522043
refGene_gn=AMACR;refGene_tx=XM_011286594.2
A1 214528405 214528790
refGene_gn=SLC45A2;refGene_tx=NM_001127661.1;refGene_gn=SLC45A2;refGene_tx=XM_019814042.1
A1 214531344 214531521
refGene_gn=SLC45A2;refGene_tx=NM_001127661.1;refGene_gn=SLC45A2;refGene_tx=XM_019814042.1
A1 214547225 214547551
refGene_gn=SLC45A2;refGene_tx=NM_001127661.1;refGene_gn=SLC45A2;refGene_tx=XM_019814042.1
A1 214552094 214552238
refGene_gn=SLC45A2;refGene_tx=NM_001127661.1;refGene_gn=SLC45A2;refGene_tx=XM_019814042.1
A1 214552923 214553047
refGene_gn=SLC45A2;refGene_tx=NM_001127661.1;refGene_gn=SLC45A2;refGene_tx=XM_019814042.1
A1 214557652 214557867
refGene_gn=SLC45A2;refGene_tx=NM_001127661.1;refGene_gn=SLC45A2;refGene_tx=XM_019814042.1
A1 214560189 214560411
refGene_gn=SLC45A2;refGene_tx=NM_001127661.1;refGene_gn=SLC45A2;refGene_tx=XM_019814042.1
A1 214565116 214566493
refGene_gn=RXFP3;refGene_tx=XM_003981471.4
A1 214582746 214582873
refGene_gn=ADAMTS12;refGene_tx=XM_023239155.1;refGene_gn=ADAMTS12;refG

ene_tx=XM_023239156.1;refGene_gn=ADAMTS12;refGene_tx=XM_023239159.1
A1	214590590	214590952
refGene_gn=ADAMTS12;refGene_tx=XM_023239155.1;refGene_gn=ADAMTS12;refGene_tx=XM_023239156.1;refGene_gn=ADAMTS12;refGene_tx=XM_023239159.1
A1	214705217	214705368
refGene_gn=ADAMTS12;refGene_tx=XM_023239155.1;refGene_gn=ADAMTS12;refGene_tx=XM_023239156.1;refGene_gn=ADAMTS12;refGene_tx=XM_023239159.1
A1	214771345	214771542
refGene_gn=ADAMTS12;refGene_tx=XM_023239155.1;refGene_gn=ADAMTS12;refGene_tx=XM_023239156.1;refGene_gn=ADAMTS12;refGene_tx=XM_023239159.1
A1	214772469	214772553
refGene_gn=ADAMTS12;refGene_tx=XM_023239155.1;refGene_gn=ADAMTS12;refGene_tx=XM_023239156.1;refGene_gn=ADAMTS12;refGene_tx=XM_023239159.1
A1	214787356	214787481
refGene_gn=ADAMTS12;refGene_tx=XM_023239155.1;refGene_gn=ADAMTS12;refGene_tx=XM_023239156.1;refGene_gn=ADAMTS12;refGene_tx=XM_023239159.1
A1	214791740	214791890
refGene_gn=ADAMTS12;refGene_tx=XM_023239155.1;refGene_gn=ADAMTS12;refGene_tx=XM_023239156.1;refGene_gn=ADAMTS12;refGene_tx=XM_023239159.1
A1	214800328	214800472
refGene_gn=ADAMTS12;refGene_tx=XM_023239155.1;refGene_gn=ADAMTS12;refGene_tx=XM_023239156.1;refGene_gn=ADAMTS12;refGene_tx=XM_023239159.1
A1	214800832	214800977
refGene_gn=ADAMTS12;refGene_tx=XM_023239155.1;refGene_gn=ADAMTS12;refGene_tx=XM_023239156.1;refGene_gn=ADAMTS12;refGene_tx=XM_023239159.1
A1	214806104	214806197
refGene_gn=ADAMTS12;refGene_tx=XM_023239155.1;refGene_gn=ADAMTS12;refGene_tx=XM_023239156.1;refGene_gn=ADAMTS12;refGene_tx=XM_023239159.1
A1	214807858	214808004
refGene_gn=ADAMTS12;refGene_tx=XM_023239155.1;refGene_gn=ADAMTS12;refGene_tx=XM_023239156.1;refGene_gn=ADAMTS12;refGene_tx=XM_023239159.1
A1	214812630	214812800
refGene_gn=ADAMTS12;refGene_tx=XM_023239155.1;refGene_gn=ADAMTS12;refGene_tx=XM_023239156.1;refGene_gn=ADAMTS12;refGene_tx=XM_023239159.1
A1	214824544	214824685
refGene_gn=ADAMTS12;refGene_tx=XM_023239155.1;refGene_gn=ADAMTS12;refGene_tx=XM_023239156.1;refGene_gn=ADAMTS12;refGene_tx=XM_023239159.1
A1	214832744	214832865
refGene_gn=ADAMTS12;refGene_tx=XM_023239155.1;refGene_gn=ADAMTS12;refGene_tx=XM_023239156.1;refGene_gn=ADAMTS12;refGene_tx=XM_023239159.1
A1	214843468	214843713
refGene_gn=ADAMTS12;refGene_tx=XM_023239155.1;refGene_gn=ADAMTS12;refGene_tx=XM_023239156.1
A1	214845361	214845500
refGene_gn=ADAMTS12;refGene_tx=XM_023239155.1;refGene_gn=ADAMTS12;refGene_tx=XM_023239156.1
A1	214852485	214852612
refGene_gn=ADAMTS12;refGene_tx=XM_023239155.1;refGene_gn=ADAMTS12;refGene_tx=XM_023239156.1
A1	214866808	214867019

refGene_gn=ADAMTS12;refGene_tx=XM_023239155.1;refGene_gn=ADAMTS12;refGene_tx=XM_023239156.1
A1 214877875 214879000
refGene_gn=ADAMTS12;refGene_tx=XM_023239155.1;refGene_gn=ADAMTS12;refGene_tx=XM_023239156.1
A1 214885051 214885204
refGene_gn=ADAMTS12;refGene_tx=XM_023239155.1;refGene_gn=ADAMTS12;refGene_tx=XM_023239156.1
A1 214896120 214896297
refGene_gn=ADAMTS12;refGene_tx=XM_023239155.1
A1 214900651 214900795
refGene_gn=ADAMTS12;refGene_tx=XM_023239155.1;refGene_gn=ADAMTS12;refGene_tx=XM_023239156.1
A1 214907612 214907766
refGene_gn=ADAMTS12;refGene_tx=XM_023239155.1;refGene_gn=ADAMTS12;refGene_tx=XM_023239156.1
A1 214915489 214915668
refGene_gn=ADAMTS12;refGene_tx=XM_023239155.1;refGene_gn=ADAMTS12;refGene_tx=XM_023239156.1
A1 214950838 214950987
refGene_gn=TARS;refGene_tx=XM_003981472.4
A1 214952012 214952127
refGene_gn=TARS;refGene_tx=XM_003981472.4
A1 214955225 214955298
refGene_gn=TARS;refGene_tx=XM_003981472.4
A1 214956580 214956685
refGene_gn=TARS;refGene_tx=XM_003981472.4
A1 214956789 214956890
refGene_gn=TARS;refGene_tx=XM_003981472.4
A1 214957095 214957173
refGene_gn=TARS;refGene_tx=XM_003981472.4
A1 214957855 214957993
refGene_gn=TARS;refGene_tx=XM_003981472.4
A1 214958088 214958251
refGene_gn=TARS;refGene_tx=XM_003981472.4
A1 214959144 214959311
refGene_gn=TARS;refGene_tx=XM_003981472.4
A1 214960297 214960396
refGene_gn=TARS;refGene_tx=XM_003981472.4
A1 214961511 214961658
refGene_gn=TARS;refGene_tx=XM_003981472.4
A1 214962389 214962468
refGene_gn=TARS;refGene_tx=XM_003981472.4
A1 214962552 214962617
refGene_gn=TARS;refGene_tx=XM_003981472.4
A1 214962883 214963001
refGene_gn=TARS;refGene_tx=XM_003981472.4
A1 214963483 214963605
refGene_gn=TARS;refGene_tx=XM_003981472.4
A1 214965370 214965494

refGene_gn=TARS;refGene_tx=XM_003981472.4
A1 214968328 214968519
refGene_gn=TARS;refGene_tx=XM_003981472.4
A1 214970331 214970412
refGene_gn=TARS;refGene_tx=XM_003981472.4
A1 214975655 214975712
refGene_gn=TARS;refGene_tx=XM_003981472.4
A1 215361961 215362067
refGene_gn=LOC101099435;refGene_tx=XM_019812202.1
A1 215362260 215362369
refGene_gn=LOC101099435;refGene_tx=XM_019812202.1
A1 215362437 215362498
refGene_gn=LOC101099435;refGene_tx=XM_019812202.1
A1 215603717 215603829
refGene_gn=NPR3;refGene_tx=XM_003981473.5;refGene_gn=NPR3;refGene_tx=XM_006928064.4
A1 215604864 215604952
refGene_gn=NPR3;refGene_tx=XM_003981473.5;refGene_gn=NPR3;refGene_tx=XM_006928064.4
A1 215606845 215606981
refGene_gn=NPR3;refGene_tx=XM_003981473.5;refGene_gn=NPR3;refGene_tx=XM_006928064.4
A1 215608352 215608447
refGene_gn=NPR3;refGene_tx=XM_003981473.5;refGene_gn=NPR3;refGene_tx=XM_006928064.4
A1 215614126 215614262
refGene_gn=NPR3;refGene_tx=XM_003981473.5;refGene_gn=NPR3;refGene_tx=XM_006928064.4
A1 215650717 215650884
refGene_gn=NPR3;refGene_tx=XM_003981473.5;refGene_gn=NPR3;refGene_tx=XM_006928064.4
A1 215661622 215661745
refGene_gn=NPR3;refGene_tx=XM_003981473.5;refGene_gn=NPR3;refGene_tx=XM_006928064.4
A1 215675224 215675981
refGene_gn=NPR3;refGene_tx=XM_003981473.5;refGene_gn=NPR3;refGene_tx=XM_006928064.4
A1 215747804 215747884
refGene_gn=SUB1;refGene_tx=XM_003981522.5;refGene_gn=SUB1;refGene_tx=XM_023239178.1
A1 215749830 215749939
refGene_gn=SUB1;refGene_tx=XM_003981522.5;refGene_gn=SUB1;refGene_tx=XM_023239178.1
A1 215755385 215755508
refGene_gn=SUB1;refGene_tx=XM_003981522.5;refGene_gn=SUB1;refGene_tx=XM_023239178.1
A1 215757851 215757923
refGene_gn=SUB1;refGene_tx=XM_003981522.5;refGene_gn=SUB1;refGene_tx=XM_023239178.1
A1 215854592 215854629

refGene_gn=ZFR;refGene_tx=XM_023239182.1;refGene_gn=ZFR;refGene_tx=XM_023239184.1;refGene_gn=ZFR;refGene_tx=XM_023239190.1;refGene_gn=ZFR;refGene_tx=XM_023239191.1
A1 215854927 215855027
refGene_gn=ZFR;refGene_tx=XM_023239182.1;refGene_gn=ZFR;refGene_tx=XM_023239184.1;refGene_gn=ZFR;refGene_tx=XM_023239189.1;refGene_gn=ZFR;refGene_tx=XM_023239190.1;refGene_gn=ZFR;refGene_tx=XM_023239191.1
A1 215874069 215874093
refGene_gn=ZFR;refGene_tx=XM_023239182.1;refGene_gn=ZFR;refGene_tx=XM_023239189.1;refGene_gn=ZFR;refGene_tx=XM_023239190.1;refGene_gn=ZFR;refGene_tx=XM_023239191.1
A1 215875565 215875848
refGene_gn=ZFR;refGene_tx=XM_023239182.1;refGene_gn=ZFR;refGene_tx=XM_023239184.1;refGene_gn=ZFR;refGene_tx=XM_023239189.1;refGene_gn=ZFR;refGene_tx=XM_023239190.1;refGene_gn=ZFR;refGene_tx=XM_023239191.1
A1 215877466 215877611
refGene_gn=ZFR;refGene_tx=XM_023239182.1;refGene_gn=ZFR;refGene_tx=XM_023239184.1;refGene_gn=ZFR;refGene_tx=XM_023239189.1;refGene_gn=ZFR;refGene_tx=XM_023239190.1;refGene_gn=ZFR;refGene_tx=XM_023239191.1
A1 215879497 215879716
refGene_gn=ZFR;refGene_tx=XM_023239182.1;refGene_gn=ZFR;refGene_tx=XM_023239184.1;refGene_gn=ZFR;refGene_tx=XM_023239189.1;refGene_gn=ZFR;refGene_tx=XM_023239190.1;refGene_gn=ZFR;refGene_tx=XM_023239191.1
A1 215886904 215887152
refGene_gn=ZFR;refGene_tx=XM_023239182.1;refGene_gn=ZFR;refGene_tx=XM_023239184.1;refGene_gn=ZFR;refGene_tx=XM_023239189.1;refGene_gn=ZFR;refGene_tx=XM_023239190.1
A1 215889872 215890064
refGene_gn=ZFR;refGene_tx=XM_023239182.1;refGene_gn=ZFR;refGene_tx=XM_023239184.1;refGene_gn=ZFR;refGene_tx=XM_023239189.1;refGene_gn=ZFR;refGene_tx=XM_023239190.1
A1 215890581 215890873
refGene_gn=ZFR;refGene_tx=XM_023239182.1;refGene_gn=ZFR;refGene_tx=XM_023239184.1;refGene_gn=ZFR;refGene_tx=XM_023239189.1;refGene_gn=ZFR;refGene_tx=XM_023239190.1
A1 215892733 215892927
refGene_gn=ZFR;refGene_tx=XM_023239182.1;refGene_gn=ZFR;refGene_tx=XM_023239184.1;refGene_gn=ZFR;refGene_tx=XM_023239189.1;refGene_gn=ZFR;refGene_tx=XM_023239190.1;refGene_gn=ZFR;refGene_tx=XM_023239191.1
A1 215894318 215894438
refGene_gn=ZFR;refGene_tx=XM_023239182.1;refGene_gn=ZFR;refGene_tx=XM_023239184.1;refGene_gn=ZFR;refGene_tx=XM_023239189.1;refGene_gn=ZFR;refGene_tx=XM_023239190.1;refGene_gn=ZFR;refGene_tx=XM_023239191.1
A1 215895502 215895648
refGene_gn=ZFR;refGene_tx=XM_023239182.1;refGene_gn=ZFR;refGene_tx=XM_023239184.1;refGene_gn=ZFR;refGene_tx=XM_023239189.1;refGene_gn=ZFR;refGene_tx=XM_023239190.1;refGene_gn=ZFR;refGene_tx=XM_023239191.1
A1 215900734 215900897
refGene_gn=ZFR;refGene_tx=XM_023239182.1;refGene_gn=ZFR;refGene_tx=XM_023239184.1;refGene_gn=ZFR;refGene_tx=XM_023239189.1;refGene_gn=ZFR;re

fGene_tx=XM_023239190.1;refGene_gn=ZFR;refGene_tx=XM_023239191.1
A1 215902179 215902385
refGene_gn=ZFR;refGene_tx=XM_023239182.1;refGene_gn=ZFR;refGene_tx=XM_023239184.1;refGene_gn=ZFR;refGene_tx=XM_023239189.1;refGene_gn=ZFR;refGene_tx=XM_023239190.1;refGene_gn=ZFR;refGene_tx=XM_023239191.1
A1 215903115 215903266
refGene_gn=ZFR;refGene_tx=XM_023239182.1;refGene_gn=ZFR;refGene_tx=XM_023239184.1;refGene_gn=ZFR;refGene_tx=XM_023239189.1;refGene_gn=ZFR;refGene_tx=XM_023239190.1;refGene_gn=ZFR;refGene_tx=XM_023239191.1
A1 215905278 215905420
refGene_gn=ZFR;refGene_tx=XM_023239182.1;refGene_gn=ZFR;refGene_tx=XM_023239184.1;refGene_gn=ZFR;refGene_tx=XM_023239189.1;refGene_gn=ZFR;refGene_tx=XM_023239190.1;refGene_gn=ZFR;refGene_tx=XM_023239191.1
A1 215910719 215910817
refGene_gn=ZFR;refGene_tx=XM_023239182.1;refGene_gn=ZFR;refGene_tx=XM_023239184.1;refGene_gn=ZFR;refGene_tx=XM_023239189.1;refGene_gn=ZFR;refGene_tx=XM_023239191.1
A1 215911676 215911772
refGene_gn=ZFR;refGene_tx=XM_023239182.1;refGene_gn=ZFR;refGene_tx=XM_023239184.1;refGene_gn=ZFR;refGene_tx=XM_023239189.1;refGene_gn=ZFR;refGene_tx=XM_023239190.1;refGene_gn=ZFR;refGene_tx=XM_023239191.1
A1 215932129 215932241
refGene_gn=ZFR;refGene_tx=XM_023239182.1;refGene_gn=ZFR;refGene_tx=XM_023239184.1;refGene_gn=ZFR;refGene_tx=XM_023239189.1;refGene_gn=ZFR;refGene_tx=XM_023239191.1
A1 215932349 215932447
refGene_gn=ZFR;refGene_tx=XM_023239182.1;refGene_gn=ZFR;refGene_tx=XM_023239184.1;refGene_gn=ZFR;refGene_tx=XM_023239189.1;refGene_gn=ZFR;refGene_tx=XM_023239191.1
A1 215938210 215938390
refGene_gn=ZFR;refGene_tx=XM_023239182.1;refGene_gn=ZFR;refGene_tx=XM_023239184.1;refGene_gn=ZFR;refGene_tx=XM_023239189.1;refGene_gn=ZFR;refGene_tx=XM_023239191.1
A1 215963707 215963788
refGene_gn=MTMR12;refGene_tx=XM_023239209.1;refGene_gn=MTMR12;refGene_tx=XM_023239221.1
A1 215985095 215985119
refGene_gn=MTMR12;refGene_tx=XM_006928088.4
A1 215988563 215988624
refGene_gn=MTMR12;refGene_tx=XM_006928088.4;refGene_gn=MTMR12;refGene_tx=XM_023239209.1;refGene_gn=MTMR12;refGene_tx=XM_023239221.1
A1 215991213 215991356
refGene_gn=MTMR12;refGene_tx=XM_006928088.4;refGene_gn=MTMR12;refGene_tx=XM_023239209.1;refGene_gn=MTMR12;refGene_tx=XM_023239221.1
A1 215992679 215992752
refGene_gn=MTMR12;refGene_tx=XM_006928088.4;refGene_gn=MTMR12;refGene_tx=XM_023239209.1;refGene_gn=MTMR12;refGene_tx=XM_023239221.1
A1 215993733 215993864
refGene_gn=MTMR12;refGene_tx=XM_006928088.4;refGene_gn=MTMR12;refGene_tx=XM_023239209.1;refGene_gn=MTMR12;refGene_tx=XM_023239221.1

A1 215995122 215995216
refGene_gn=MTMR12;refGene_tx=XM_006928088.4;refGene_gn=MTMR12;refGene_tx=XM_023239209.1;refGene_gn=MTMR12;refGene_tx=XM_023239221.1
A1 215998643 215998773
refGene_gn=MTMR12;refGene_tx=XM_006928088.4;refGene_gn=MTMR12;refGene_tx=XM_023239209.1;refGene_gn=MTMR12;refGene_tx=XM_023239221.1
A1 216005104 216005180
refGene_gn=MTMR12;refGene_tx=XM_006928088.4;refGene_gn=MTMR12;refGene_tx=XM_023239209.1;refGene_gn=MTMR12;refGene_tx=XM_023239221.1
A1 216013120 216013230
refGene_gn=MTMR12;refGene_tx=XM_006928088.4;refGene_gn=MTMR12;refGene_tx=XM_023239209.1;refGene_gn=MTMR12;refGene_tx=XM_023239221.1
A1 216013956 216014081
refGene_gn=MTMR12;refGene_tx=XM_006928088.4;refGene_gn=MTMR12;refGene_tx=XM_023239209.1;refGene_gn=MTMR12;refGene_tx=XM_023239221.1
A1 216015696 216015775
refGene_gn=MTMR12;refGene_tx=XM_006928088.4;refGene_gn=MTMR12;refGene_tx=XM_023239209.1;refGene_gn=MTMR12;refGene_tx=XM_023239221.1
A1 216017199 216017270
refGene_gn=MTMR12;refGene_tx=XM_006928088.4;refGene_gn=MTMR12;refGene_tx=XM_023239209.1;refGene_gn=MTMR12;refGene_tx=XM_023239221.1
A1 216019705 216019878
refGene_gn=MTMR12;refGene_tx=XM_006928088.4;refGene_gn=MTMR12;refGene_tx=XM_023239209.1;refGene_gn=MTMR12;refGene_tx=XM_023239221.1
A1 216022212 216022377
refGene_gn=MTMR12;refGene_tx=XM_006928088.4;refGene_gn=MTMR12;refGene_tx=XM_023239209.1;refGene_gn=MTMR12;refGene_tx=XM_023239221.1
A1 216023290 216023452
refGene_gn=MTMR12;refGene_tx=XM_006928088.4;refGene_gn=MTMR12;refGene_tx=XM_023239209.1;refGene_gn=MTMR12;refGene_tx=XM_023239221.1
A1 216024341 216024440
refGene_gn=MTMR12;refGene_tx=XM_006928088.4;refGene_gn=MTMR12;refGene_tx=XM_023239209.1
A1 216026309 216026879
refGene_gn=MTMR12;refGene_tx=XM_006928088.4;refGene_gn=MTMR12;refGene_tx=XM_023239209.1;refGene_gn=MTMR12;refGene_tx=XM_023239221.1
A1 216072936 216073161
refGene_gn=GOLPH3;refGene_tx=XM_023239237.1;refGene_gn=GOLPH3;refGene_tx=XM_023239239.1
A1 216106030 216106162
refGene_gn=GOLPH3;refGene_tx=XM_023239237.1
A1 216112604 216112719
refGene_gn=GOLPH3;refGene_tx=XM_023239237.1;refGene_gn=GOLPH3;refGene_tx=XM_023239239.1
A1 216121077 216121502
refGene_gn=GOLPH3;refGene_tx=XM_023239237.1;refGene_gn=GOLPH3;refGene_tx=XM_023239239.1
A1 216135683 216135850
refGene_gn=PDZD2;refGene_tx=XM_003981524.5;refGene_gn=PDZD2;refGene_tx=XM_006928089.4;refGene_gn=PDZD2;refGene_tx=XM_019812262.2;refGene_gn=

PDZD2;refGene_tx=XM_019812272.2;refGene_gn=PDZD2;refGene_tx=XM_0198122
81.2
A1 216137079 216137214
refGene_gn=PDZD2;refGene_tx=XM_003981524.5;refGene_gn=PDZD2;refGene_tx
=XM_006928089.4;refGene_gn=PDZD2;refGene_tx=XM_019812262.2;refGene_gn=
PDZD2;refGene_tx=XM_019812272.2;refGene_gn=PDZD2;refGene_tx=XM_0198122
81.2
A1 216139722 216139984
refGene_gn=PDZD2;refGene_tx=XM_003981524.5;refGene_gn=PDZD2;refGene_tx
=XM_006928089.4;refGene_gn=PDZD2;refGene_tx=XM_019812262.2;refGene_gn=
PDZD2;refGene_tx=XM_019812272.2;refGene_gn=PDZD2;refGene_tx=XM_0198122
81.2
A1 216140791 216140893
refGene_gn=PDZD2;refGene_tx=XM_003981524.5;refGene_gn=PDZD2;refGene_tx
=XM_006928089.4;refGene_gn=PDZD2;refGene_tx=XM_019812262.2;refGene_gn=
PDZD2;refGene_tx=XM_019812272.2;refGene_gn=PDZD2;refGene_tx=XM_0198122
81.2
A1 216145247 216145365
refGene_gn=PDZD2;refGene_tx=XM_003981524.5;refGene_gn=PDZD2;refGene_tx
=XM_006928089.4;refGene_gn=PDZD2;refGene_tx=XM_019812262.2;refGene_gn=
PDZD2;refGene_tx=XM_019812272.2;refGene_gn=PDZD2;refGene_tx=XM_0198122
81.2
A1 216146605 216150662
refGene_gn=PDZD2;refGene_tx=XM_003981524.5;refGene_gn=PDZD2;refGene_tx
=XM_006928089.4;refGene_gn=PDZD2;refGene_tx=XM_019812262.2;refGene_gn=
PDZD2;refGene_tx=XM_019812272.2;refGene_gn=PDZD2;refGene_tx=XM_0198122
81.2
A1 216153285 216153421
refGene_gn=PDZD2;refGene_tx=XM_003981524.5;refGene_gn=PDZD2;refGene_tx
=XM_006928089.4;refGene_gn=PDZD2;refGene_tx=XM_019812262.2;refGene_gn=
PDZD2;refGene_tx=XM_019812272.2;refGene_gn=PDZD2;refGene_tx=XM_0198122
81.2
A1 216157034 216157831
refGene_gn=PDZD2;refGene_tx=XM_003981524.5;refGene_gn=PDZD2;refGene_tx
=XM_006928089.4;refGene_gn=PDZD2;refGene_tx=XM_019812262.2;refGene_gn=
PDZD2;refGene_tx=XM_019812272.2;refGene_gn=PDZD2;refGene_tx=XM_0198122
81.2
A1 216159790 216159947
refGene_gn=PDZD2;refGene_tx=XM_003981524.5;refGene_gn=PDZD2;refGene_tx
=XM_006928089.4;refGene_gn=PDZD2;refGene_tx=XM_019812262.2;refGene_gn=
PDZD2;refGene_tx=XM_019812272.2;refGene_gn=PDZD2;refGene_tx=XM_0198122
81.2
A1 216160886 216160921
refGene_gn=PDZD2;refGene_tx=XM_003981524.5;refGene_gn=PDZD2;refGene_tx
=XM_006928089.4;refGene_gn=PDZD2;refGene_tx=XM_019812262.2;refGene_gn=
PDZD2;refGene_tx=XM_019812272.2;refGene_gn=PDZD2;refGene_tx=XM_0198122
81.2
A1 216162456 216162538
refGene_gn=PDZD2;refGene_tx=XM_003981524.5;refGene_gn=PDZD2;refGene_tx
=XM_006928089.4;refGene_gn=PDZD2;refGene_tx=XM_019812262.2;refGene_gn=

PDZD2;refGene_tx=XM_019812272.2;refGene_gn=PDZD2;refGene_tx=XM_0198122
81.2
A1 216167117 216167250
refGene_gn=PDZD2;refGene_tx=XM_003981524.5;refGene_gn=PDZD2;refGene_tx
=XM_006928089.4;refGene_gn=PDZD2;refGene_tx=XM_019812262.2;refGene_gn=
PDZD2;refGene_tx=XM_019812272.2;refGene_gn=PDZD2;refGene_tx=XM_0198122
81.2
A1 216168873 216168991
refGene_gn=PDZD2;refGene_tx=XM_003981524.5;refGene_gn=PDZD2;refGene_tx
=XM_006928089.4;refGene_gn=PDZD2;refGene_tx=XM_019812262.2;refGene_gn=
PDZD2;refGene_tx=XM_019812272.2;refGene_gn=PDZD2;refGene_tx=XM_0198122
81.2
A1 216169898 216170124
refGene_gn=PDZD2;refGene_tx=XM_003981524.5;refGene_gn=PDZD2;refGene_tx
=XM_006928089.4;refGene_gn=PDZD2;refGene_tx=XM_019812262.2;refGene_gn=
PDZD2;refGene_tx=XM_019812272.2;refGene_gn=PDZD2;refGene_tx=XM_0198122
81.2
A1 216170285 216170359
refGene_gn=PDZD2;refGene_tx=XM_003981524.5;refGene_gn=PDZD2;refGene_tx
=XM_006928089.4;refGene_gn=PDZD2;refGene_tx=XM_019812262.2;refGene_gn=
PDZD2;refGene_tx=XM_019812272.2;refGene_gn=PDZD2;refGene_tx=XM_0198122
81.2
A1 216172864 216172979
refGene_gn=PDZD2;refGene_tx=XM_003981524.5;refGene_gn=PDZD2;refGene_tx
=XM_006928089.4;refGene_gn=PDZD2;refGene_tx=XM_019812262.2;refGene_gn=
PDZD2;refGene_tx=XM_019812272.2;refGene_gn=PDZD2;refGene_tx=XM_0198122
81.2
A1 216174008 216174128
refGene_gn=PDZD2;refGene_tx=XM_003981524.5;refGene_gn=PDZD2;refGene_tx
=XM_006928089.4;refGene_gn=PDZD2;refGene_tx=XM_019812262.2;refGene_gn=
PDZD2;refGene_tx=XM_019812272.2;refGene_gn=PDZD2;refGene_tx=XM_0198122
81.2
A1 216177170 216177316
refGene_gn=PDZD2;refGene_tx=XM_003981524.5;refGene_gn=PDZD2;refGene_tx
=XM_006928089.4;refGene_gn=PDZD2;refGene_tx=XM_019812262.2;refGene_gn=
PDZD2;refGene_tx=XM_019812272.2;refGene_gn=PDZD2;refGene_tx=XM_0198122
81.2
A1 216183514 216183626
refGene_gn=PDZD2;refGene_tx=XM_003981524.5;refGene_gn=PDZD2;refGene_tx
=XM_006928089.4;refGene_gn=PDZD2;refGene_tx=XM_019812262.2;refGene_gn=
PDZD2;refGene_tx=XM_019812272.2;refGene_gn=PDZD2;refGene_tx=XM_0198122
81.2
A1 216215375 216215531
refGene_gn=PDZD2;refGene_tx=XM_003981524.5;refGene_gn=PDZD2;refGene_tx
=XM_006928089.4;refGene_gn=PDZD2;refGene_tx=XM_019812262.2;refGene_gn=
PDZD2;refGene_tx=XM_019812272.2;refGene_gn=PDZD2;refGene_tx=XM_0198122
81.2
A1 216220835 216220968
refGene_gn=PDZD2;refGene_tx=XM_003981524.5;refGene_gn=PDZD2;refGene_tx
=XM_006928089.4;refGene_gn=PDZD2;refGene_tx=XM_019812281.2

A1 216223846 216223989
refGene_gn=PDZD2;refGene_tx=XM_003981524.5;refGene_gn=PDZD2;refGene_tx=XM_006928089.4;refGene_gn=PDZD2;refGene_tx=XM_019812262.2;refGene_gn=PDZD2;refGene_tx=XM_019812281.2
A1 216233588 216234090
refGene_gn=PDZD2;refGene_tx=XM_003981524.5;refGene_gn=PDZD2;refGene_tx=XM_006928089.4;refGene_gn=PDZD2;refGene_tx=XM_019812262.2;refGene_gn=PDZD2;refGene_tx=XM_019812272.2;refGene_gn=PDZD2;refGene_tx=XM_019812281.2
A1 216365292 216365768
refGene_gn=PDZD2;refGene_tx=XM_003981524.5;refGene_gn=PDZD2;refGene_tx=XM_019812262.2;refGene_gn=PDZD2;refGene_tx=XM_019812272.2
A1 216573967 216574100
refGene_gn=CA1H5orf22;refGene_tx=XM_003981531.4;refGene_gn=CA1H5orf22;refGene_tx=XM_006928091.4;refGene_gn=CA1H5orf22;refGene_tx=XM_019812312.2;refGene_gn=CA1H5orf22;refGene_tx=XM_019812314.1
A1 216577612 216577752
refGene_gn=CA1H5orf22;refGene_tx=XM_003981531.4;refGene_gn=CA1H5orf22;refGene_tx=XM_006928091.4;refGene_gn=CA1H5orf22;refGene_tx=XM_019812312.2;refGene_gn=CA1H5orf22;refGene_tx=XM_019812314.1
A1 216579530 216579597
refGene_gn=CA1H5orf22;refGene_tx=XM_003981531.4;refGene_gn=CA1H5orf22;refGene_tx=XM_006928091.4;refGene_gn=CA1H5orf22;refGene_tx=XM_019812312.2;refGene_gn=CA1H5orf22;refGene_tx=XM_019812314.1
A1 216583303 216583425
refGene_gn=CA1H5orf22;refGene_tx=XM_003981531.4;refGene_gn=CA1H5orf22;refGene_tx=XM_006928091.4;refGene_gn=CA1H5orf22;refGene_tx=XM_019812312.2;refGene_gn=CA1H5orf22;refGene_tx=XM_019812314.1
A1 216583656 216583719
refGene_gn=CA1H5orf22;refGene_tx=XM_003981531.4;refGene_gn=CA1H5orf22;refGene_tx=XM_006928091.4;refGene_gn=CA1H5orf22;refGene_tx=XM_019812312.2;refGene_gn=CA1H5orf22;refGene_tx=XM_019812314.1
A1 216585812 216586242
refGene_gn=CA1H5orf22;refGene_tx=XM_003981531.4;refGene_gn=CA1H5orf22;refGene_tx=XM_006928091.4;refGene_gn=CA1H5orf22;refGene_tx=XM_019812312.2;refGene_gn=CA1H5orf22;refGene_tx=XM_019812314.1
A1 216588184 216588334
refGene_gn=CA1H5orf22;refGene_tx=XM_003981531.4;refGene_gn=CA1H5orf22;refGene_tx=XM_006928091.4;refGene_gn=CA1H5orf22;refGene_tx=XM_019812312.2;refGene_gn=CA1H5orf22;refGene_tx=XM_019812314.1
A1 216590838 216590984
refGene_gn=CA1H5orf22;refGene_tx=XM_003981531.4;refGene_gn=CA1H5orf22;refGene_tx=XM_019812312.2
A1 216592446 216592530
refGene_gn=CA1H5orf22;refGene_tx=XM_003981531.4
A1 216595029 216595049
refGene_gn=DROSHA;refGene_tx=XM_003981525.5;refGene_gn=DROSHA;refGene_tx=XM_011286642.2;refGene_gn=DROSHA;refGene_tx=XM_011286647.3;refGene_gn=DROSHA;refGene_tx=XM_011286648.3;refGene_gn=DROSHA;refGene_tx=XM_011286651.2;refGene_gn=DROSHA;refGene_tx=XM_011286654.2;refGene_gn=DROSH

A;refGene_tx=XM_019812340.1
A1 216596046 216596880
refGene_gn=DROSHA;refGene_tx=XM_003981525.5;refGene_gn=DROSHA;refGene_tx=XM_011286642.2;refGene_gn=DROSHA;refGene_tx=XM_011286647.3;refGene_gn=DROSHA;refGene_tx=XM_011286648.3;refGene_gn=DROSHA;refGene_tx=XM_011286651.2;refGene_gn=DROSHA;refGene_tx=XM_011286654.2;refGene_gn=DROSHA;refGene_tx=XM_019812340.1
A1 216601246 216601339
refGene_gn=DROSHA;refGene_tx=XM_003981525.5;refGene_gn=DROSHA;refGene_tx=XM_011286642.2;refGene_gn=DROSHA;refGene_tx=XM_011286647.3;refGene_gn=DROSHA;refGene_tx=XM_011286648.3;refGene_gn=DROSHA;refGene_tx=XM_011286651.2;refGene_gn=DROSHA;refGene_tx=XM_011286654.2
A1 216607304 216607415
refGene_gn=DROSHA;refGene_tx=XM_003981525.5;refGene_gn=DROSHA;refGene_tx=XM_011286642.2;refGene_gn=DROSHA;refGene_tx=XM_011286647.3;refGene_gn=DROSHA;refGene_tx=XM_011286648.3;refGene_gn=DROSHA;refGene_tx=XM_011286651.2;refGene_gn=DROSHA;refGene_tx=XM_019812340.1
A1 216607648 216607880
refGene_gn=DROSHA;refGene_tx=XM_003981525.5;refGene_gn=DROSHA;refGene_tx=XM_011286642.2;refGene_gn=DROSHA;refGene_tx=XM_011286647.3;refGene_gn=DROSHA;refGene_tx=XM_011286648.3;refGene_gn=DROSHA;refGene_tx=XM_011286651.2;refGene_gn=DROSHA;refGene_tx=XM_011286654.2;refGene_gn=DROSHA;refGene_tx=XM_019812340.1
A1 216611162 216611304
refGene_gn=DROSHA;refGene_tx=XM_003981525.5;refGene_gn=DROSHA;refGene_tx=XM_011286642.2;refGene_gn=DROSHA;refGene_tx=XM_011286647.3;refGene_gn=DROSHA;refGene_tx=XM_011286648.3;refGene_gn=DROSHA;refGene_tx=XM_011286651.2;refGene_gn=DROSHA;refGene_tx=XM_011286654.2;refGene_gn=DROSHA;refGene_tx=XM_019812340.1
A1 216613385 216613540
refGene_gn=DROSHA;refGene_tx=XM_003981525.5;refGene_gn=DROSHA;refGene_tx=XM_011286642.2;refGene_gn=DROSHA;refGene_tx=XM_011286647.3;refGene_gn=DROSHA;refGene_tx=XM_011286648.3;refGene_gn=DROSHA;refGene_tx=XM_011286651.2;refGene_gn=DROSHA;refGene_tx=XM_011286654.2;refGene_gn=DROSHA;refGene_tx=XM_019812340.1
A1 216616468 216616549
refGene_gn=DROSHA;refGene_tx=XM_003981525.5;refGene_gn=DROSHA;refGene_tx=XM_011286642.2;refGene_gn=DROSHA;refGene_tx=XM_011286647.3;refGene_gn=DROSHA;refGene_tx=XM_011286648.3;refGene_gn=DROSHA;refGene_tx=XM_011286651.2;refGene_gn=DROSHA;refGene_tx=XM_011286654.2;refGene_gn=DROSHA;refGene_tx=XM_019812340.1
A1 216622644 216622731
refGene_gn=DROSHA;refGene_tx=XM_003981525.5;refGene_gn=DROSHA;refGene_tx=XM_011286642.2;refGene_gn=DROSHA;refGene_tx=XM_011286647.3;refGene_gn=DROSHA;refGene_tx=XM_011286648.3;refGene_gn=DROSHA;refGene_tx=XM_011286651.2;refGene_gn=DROSHA;refGene_tx=XM_011286654.2;refGene_gn=DROSHA;refGene_tx=XM_019812340.1
A1 216625189 216625276
refGene_gn=DROSHA;refGene_tx=XM_003981525.5;refGene_gn=DROSHA;refGene_tx=XM_011286642.2;refGene_gn=DROSHA;refGene_tx=XM_011286647.3;refGene_

gn=DROSHA;refGene_tx=XM_011286648.3;refGene_gn=DROSHA;refGene_tx=XM_01
1286651.2;refGene_gn=DROSHA;refGene_tx=XM_011286654.2;refGene_gn=DROSH
A;refGene_tx=XM_019812340.1
A1 216632301 216632373
refGene_gn=DROSHA;refGene_tx=XM_003981525.5;refGene_gn=DROSHA;refGene_
tx=XM_011286642.2;refGene_gn=DROSHA;refGene_tx=XM_011286647.3;refGene_
gn=DROSHA;refGene_tx=XM_011286648.3;refGene_gn=DROSHA;refGene_tx=XM_01
1286651.2;refGene_gn=DROSHA;refGene_tx=XM_011286654.2;refGene_gn=DROSH
A;refGene_tx=XM_019812340.1
A1 216633928 216634010
refGene_gn=DROSHA;refGene_tx=XM_003981525.5;refGene_gn=DROSHA;refGene_
tx=XM_011286642.2;refGene_gn=DROSHA;refGene_tx=XM_011286647.3;refGene_
gn=DROSHA;refGene_tx=XM_011286648.3;refGene_gn=DROSHA;refGene_tx=XM_01
1286651.2;refGene_gn=DROSHA;refGene_tx=XM_011286654.2;refGene_gn=DROSH
A;refGene_tx=XM_019812340.1
A1 216635176 216635251
refGene_gn=DROSHA;refGene_tx=XM_003981525.5;refGene_gn=DROSHA;refGene_
tx=XM_011286642.2;refGene_gn=DROSHA;refGene_tx=XM_011286647.3;refGene_
gn=DROSHA;refGene_tx=XM_011286648.3;refGene_gn=DROSHA;refGene_tx=XM_01
1286651.2;refGene_gn=DROSHA;refGene_tx=XM_011286654.2;refGene_gn=DROSH
A;refGene_tx=XM_019812340.1
A1 216646823 216646993
refGene_gn=DROSHA;refGene_tx=XM_003981525.5;refGene_gn=DROSHA;refGene_
tx=XM_011286642.2;refGene_gn=DROSHA;refGene_tx=XM_011286647.3;refGene_
gn=DROSHA;refGene_tx=XM_011286648.3;refGene_gn=DROSHA;refGene_tx=XM_01
1286651.2;refGene_gn=DROSHA;refGene_tx=XM_011286654.2;refGene_gn=DROSH
A;refGene_tx=XM_019812340.1
A1 216653932 216654057
refGene_gn=DROSHA;refGene_tx=XM_003981525.5;refGene_gn=DROSHA;refGene_
tx=XM_011286642.2;refGene_gn=DROSHA;refGene_tx=XM_011286647.3;refGene_
gn=DROSHA;refGene_tx=XM_011286648.3;refGene_gn=DROSHA;refGene_tx=XM_01
1286651.2;refGene_gn=DROSHA;refGene_tx=XM_011286654.2;refGene_gn=DROSH
A;refGene_tx=XM_019812340.1
A1 216655692 216655792
refGene_gn=DROSHA;refGene_tx=XM_003981525.5;refGene_gn=DROSHA;refGene_
tx=XM_011286642.2;refGene_gn=DROSHA;refGene_tx=XM_011286647.3;refGene_
gn=DROSHA;refGene_tx=XM_011286648.3;refGene_gn=DROSHA;refGene_tx=XM_01
1286651.2;refGene_gn=DROSHA;refGene_tx=XM_011286654.2;refGene_gn=DROSH
A;refGene_tx=XM_019812340.1
A1 216657214 216657322
refGene_gn=DROSHA;refGene_tx=XM_003981525.5;refGene_gn=DROSHA;refGene_
tx=XM_011286642.2;refGene_gn=DROSHA;refGene_tx=XM_011286647.3;refGene_
gn=DROSHA;refGene_tx=XM_011286648.3;refGene_gn=DROSHA;refGene_tx=XM_01
1286651.2;refGene_gn=DROSHA;refGene_tx=XM_011286654.2;refGene_gn=DROSH
A;refGene_tx=XM_019812340.1
A1 216669251 216669359
refGene_gn=DROSHA;refGene_tx=XM_003981525.5;refGene_gn=DROSHA;refGene_
tx=XM_011286642.2;refGene_gn=DROSHA;refGene_tx=XM_011286647.3;refGene_
gn=DROSHA;refGene_tx=XM_011286648.3;refGene_gn=DROSHA;refGene_tx=XM_01
1286651.2;refGene_gn=DROSHA;refGene_tx=XM_011286654.2;refGene_gn=DROSH

A;refGene_tx=XM_019812340.1
A1 216669864 216670003
refGene_gn=DROSHA;refGene_tx=XM_003981525.5;refGene_gn=DROSHA;refGene_tx=XM_011286642.2;refGene_gn=DROSHA;refGene_tx=XM_011286647.3;refGene_gn=DROSHA;refGene_tx=XM_011286648.3;refGene_gn=DROSHA;refGene_tx=XM_011286651.2;refGene_gn=DROSHA;refGene_tx=XM_011286654.2;refGene_gn=DROSHA;refGene_tx=XM_019812340.1
A1 216670339 216670400
refGene_gn=DROSHA;refGene_tx=XM_003981525.5;refGene_gn=DROSHA;refGene_tx=XM_011286642.2;refGene_gn=DROSHA;refGene_tx=XM_011286647.3;refGene_gn=DROSHA;refGene_tx=XM_011286648.3;refGene_gn=DROSHA;refGene_tx=XM_011286651.2;refGene_gn=DROSHA;refGene_tx=XM_011286654.2;refGene_gn=DROSHA;refGene_tx=XM_019812340.1
A1 216681622 216681682
refGene_gn=DROSHA;refGene_tx=XM_003981525.5;refGene_gn=DROSHA;refGene_tx=XM_011286642.2;refGene_gn=DROSHA;refGene_tx=XM_011286647.3;refGene_gn=DROSHA;refGene_tx=XM_011286648.3;refGene_gn=DROSHA;refGene_tx=XM_011286651.2;refGene_gn=DROSHA;refGene_tx=XM_011286654.2;refGene_gn=DROSHA;refGene_tx=XM_019812340.1
A1 216682973 216683073
refGene_gn=DROSHA;refGene_tx=XM_003981525.5;refGene_gn=DROSHA;refGene_tx=XM_011286642.2;refGene_gn=DROSHA;refGene_tx=XM_011286647.3;refGene_gn=DROSHA;refGene_tx=XM_011286648.3;refGene_gn=DROSHA;refGene_tx=XM_011286651.2;refGene_gn=DROSHA;refGene_tx=XM_011286654.2;refGene_gn=DROSHA;refGene_tx=XM_019812340.1
A1 216686700 216686803
refGene_gn=DROSHA;refGene_tx=XM_003981525.5;refGene_gn=DROSHA;refGene_tx=XM_011286642.2;refGene_gn=DROSHA;refGene_tx=XM_011286647.3;refGene_gn=DROSHA;refGene_tx=XM_011286648.3;refGene_gn=DROSHA;refGene_tx=XM_011286651.2;refGene_gn=DROSHA;refGene_tx=XM_011286654.2;refGene_gn=DROSHA;refGene_tx=XM_019812340.1
A1 216689884 216689955
refGene_gn=DROSHA;refGene_tx=XM_003981525.5;refGene_gn=DROSHA;refGene_tx=XM_011286642.2;refGene_gn=DROSHA;refGene_tx=XM_011286647.3;refGene_gn=DROSHA;refGene_tx=XM_011286648.3;refGene_gn=DROSHA;refGene_tx=XM_011286651.2;refGene_gn=DROSHA;refGene_tx=XM_011286654.2;refGene_gn=DROSHA;refGene_tx=XM_019812340.1
A1 216696388 216696433
refGene_gn=DROSHA;refGene_tx=XM_003981525.5;refGene_gn=DROSHA;refGene_tx=XM_011286642.2;refGene_gn=DROSHA;refGene_tx=XM_011286647.3;refGene_gn=DROSHA;refGene_tx=XM_011286648.3;refGene_gn=DROSHA;refGene_tx=XM_011286651.2;refGene_gn=DROSHA;refGene_tx=XM_011286654.2;refGene_gn=DROSHA;refGene_tx=XM_019812340.1
A1 216697939 216698097
refGene_gn=DROSHA;refGene_tx=XM_003981525.5;refGene_gn=DROSHA;refGene_tx=XM_011286642.2;refGene_gn=DROSHA;refGene_tx=XM_011286647.3;refGene_gn=DROSHA;refGene_tx=XM_011286648.3;refGene_gn=DROSHA;refGene_tx=XM_011286651.2;refGene_gn=DROSHA;refGene_tx=XM_011286654.2;refGene_gn=DROSHA;refGene_tx=XM_019812340.1
A1 216699114 216699220

refGene_gn=DROSHA;refGene_tx=XM_003981525.5;refGene_gn=DROSHA;refGene_tx=XM_011286642.2;refGene_gn=DROSHA;refGene_tx=XM_011286647.3;refGene_gn=DROSHA;refGene_tx=XM_011286648.3;refGene_gn=DROSHA;refGene_tx=XM_011286651.2;refGene_gn=DROSHA;refGene_tx=XM_011286654.2;refGene_gn=DROSHA;refGene_tx=XM_019812340.1
A1 216710258 216710400
refGene_gn=DROSHA;refGene_tx=XM_003981525.5;refGene_gn=DROSHA;refGene_tx=XM_011286642.2;refGene_gn=DROSHA;refGene_tx=XM_011286647.3;refGene_gn=DROSHA;refGene_tx=XM_011286648.3;refGene_gn=DROSHA;refGene_tx=XM_011286651.2;refGene_gn=DROSHA;refGene_tx=XM_011286654.2;refGene_gn=DROSHA;refGene_tx=XM_019812340.1
A1 216711857 216711940
refGene_gn=DROSHA;refGene_tx=XM_003981525.5;refGene_gn=DROSHA;refGene_tx=XM_011286642.2;refGene_gn=DROSHA;refGene_tx=XM_011286647.3;refGene_gn=DROSHA;refGene_tx=XM_011286648.3;refGene_gn=DROSHA;refGene_tx=XM_011286651.2;refGene_gn=DROSHA;refGene_tx=XM_011286654.2;refGene_gn=DROSHA;refGene_tx=XM_019812340.1
A1 216712031 216712135
refGene_gn=DROSHA;refGene_tx=XM_003981525.5;refGene_gn=DROSHA;refGene_tx=XM_011286642.2;refGene_gn=DROSHA;refGene_tx=XM_011286647.3;refGene_gn=DROSHA;refGene_tx=XM_011286648.3;refGene_gn=DROSHA;refGene_tx=XM_011286651.2;refGene_gn=DROSHA;refGene_tx=XM_011286654.2;refGene_gn=DROSHA;refGene_tx=XM_019812340.1
A1 216714159 216714252
refGene_gn=DROSHA;refGene_tx=XM_003981525.5;refGene_gn=DROSHA;refGene_tx=XM_011286642.2;refGene_gn=DROSHA;refGene_tx=XM_011286647.3;refGene_gn=DROSHA;refGene_tx=XM_011286648.3;refGene_gn=DROSHA;refGene_tx=XM_011286651.2;refGene_gn=DROSHA;refGene_tx=XM_011286654.2;refGene_gn=DROSHA;refGene_tx=XM_019812340.1
A1 216715090 216715137
refGene_gn=DROSHA;refGene_tx=XM_003981525.5;refGene_gn=DROSHA;refGene_tx=XM_011286642.2;refGene_gn=DROSHA;refGene_tx=XM_011286647.3;refGene_gn=DROSHA;refGene_tx=XM_011286648.3;refGene_gn=DROSHA;refGene_tx=XM_011286651.2;refGene_gn=DROSHA;refGene_tx=XM_011286654.2;refGene_gn=DROSHA;refGene_tx=XM_019812340.1
A1 216718817 216718948
refGene_gn=DROSHA;refGene_tx=XM_003981525.5;refGene_gn=DROSHA;refGene_tx=XM_011286642.2;refGene_gn=DROSHA;refGene_tx=XM_011286647.3;refGene_gn=DROSHA;refGene_tx=XM_011286648.3;refGene_gn=DROSHA;refGene_tx=XM_011286651.2;refGene_gn=DROSHA;refGene_tx=XM_011286654.2;refGene_gn=DROSHA;refGene_tx=XM_019812340.1
A1 216778988 216779479
refGene_gn=CDH6;refGene_tx=XM_019812349.2;refGene_gn=CDH6;refGene_tx=XM_019812356.2;refGene_gn=CDH6;refGene_tx=XM_019812360.2;refGene_gn=CDH6;refGene_tx=XM_023239304.1
A1 216784494 216784853
refGene_gn=CDH6;refGene_tx=XM_019812349.2;refGene_gn=CDH6;refGene_tx=XM_019812356.2;refGene_gn=CDH6;refGene_tx=XM_019812360.2;refGene_gn=CDH6;refGene_tx=XM_019812377.1;refGene_gn=CDH6;refGene_tx=XM_023239304.1
A1 216784984 216785102

refGene_gn=CDH6;refGene_tx=XM_019812349.2;refGene_gn=CDH6;refGene_tx=XM_019812356.2;refGene_gn=CDH6;refGene_tx=XM_019812360.2;refGene_gn=CDH6;refGene_tx=XM_019812377.1;refGene_gn=CDH6;refGene_tx=XM_023239304.1
A1 216786151 216786273
refGene_gn=CDH6;refGene_tx=XM_019812349.2;refGene_gn=CDH6;refGene_tx=XM_019812356.2;refGene_gn=CDH6;refGene_tx=XM_019812360.2;refGene_gn=CDH6;refGene_tx=XM_019812377.1;refGene_gn=CDH6;refGene_tx=XM_023239304.1
A1 216788786 216788923
refGene_gn=CDH6;refGene_tx=XM_019812349.2;refGene_gn=CDH6;refGene_tx=XM_019812356.2;refGene_gn=CDH6;refGene_tx=XM_019812360.2;refGene_gn=CDH6;refGene_tx=XM_019812377.1;refGene_gn=CDH6;refGene_tx=XM_023239304.1
A1 216794360 216794614
refGene_gn=CDH6;refGene_tx=XM_019812349.2;refGene_gn=CDH6;refGene_tx=XM_019812356.2;refGene_gn=CDH6;refGene_tx=XM_019812360.2;refGene_gn=CDH6;refGene_tx=XM_019812377.1;refGene_gn=CDH6;refGene_tx=XM_023239304.1
A1 216796692 216796880
refGene_gn=CDH6;refGene_tx=XM_019812349.2;refGene_gn=CDH6;refGene_tx=XM_019812356.2;refGene_gn=CDH6;refGene_tx=XM_019812360.2;refGene_gn=CDH6;refGene_tx=XM_019812377.1;refGene_gn=CDH6;refGene_tx=XM_023239304.1
A1 216799233 216799401
refGene_gn=CDH6;refGene_tx=XM_019812349.2;refGene_gn=CDH6;refGene_tx=XM_019812356.2;refGene_gn=CDH6;refGene_tx=XM_019812360.2;refGene_gn=CDH6;refGene_tx=XM_019812377.1;refGene_gn=CDH6;refGene_tx=XM_023239304.1
A1 216801386 216801506
refGene_gn=CDH6;refGene_tx=XM_019812349.2;refGene_gn=CDH6;refGene_tx=XM_019812356.2;refGene_gn=CDH6;refGene_tx=XM_019812360.2;refGene_gn=CDH6;refGene_tx=XM_019812377.1;refGene_gn=CDH6;refGene_tx=XM_023239304.1
A1 216804735 216805030
refGene_gn=CDH6;refGene_tx=XM_019812349.2;refGene_gn=CDH6;refGene_tx=XM_019812356.2;refGene_gn=CDH6;refGene_tx=XM_019812360.2;refGene_gn=CDH6;refGene_tx=XM_019812377.1;refGene_gn=CDH6;refGene_tx=XM_023239304.1
A1 216829607 216829835
refGene_gn=CDH6;refGene_tx=XM_019812349.2;refGene_gn=CDH6;refGene_tx=XM_019812356.2;refGene_gn=CDH6;refGene_tx=XM_019812360.2;refGene_gn=CDH6;refGene_tx=XM_019812377.1;refGene_gn=CDH6;refGene_tx=XM_023239304.1
A1 220302219 220302447
refGene_gn=CDH9;refGene_tx=XM_003981527.4;refGene_gn=CDH9;refGene_tx=XM_019812404.2
A1 220356731 220357026
refGene_gn=CDH9;refGene_tx=XM_003981527.4;refGene_gn=CDH9;refGene_tx=XM_019812404.2
A1 220363579 220363699
refGene_gn=CDH9;refGene_tx=XM_003981527.4;refGene_gn=CDH9;refGene_tx=XM_019812404.2
A1 220364288 220364456
refGene_gn=CDH9;refGene_tx=XM_003981527.4;refGene_gn=CDH9;refGene_tx=XM_019812404.2
A1 220366506 220366694
refGene_gn=CDH9;refGene_tx=XM_003981527.4;refGene_gn=CDH9;refGene_tx=XM_019812404.2

A1	220367636	220367890
refGene_gn=CDH9;refGene_tx=XM_003981527.4;refGene_gn=CDH9;refGene_tx=XM_019812404.2
A1	220379046	220379219
refGene_gn=CDH9;refGene_tx=XM_003981527.4;refGene_gn=CDH9;refGene_tx=XM_019812404.2
A1	220379632	220379754
refGene_gn=CDH9;refGene_tx=XM_003981527.4;refGene_gn=CDH9;refGene_tx=XM_019812404.2
A1	220385529	220385647
refGene_gn=CDH9;refGene_tx=XM_003981527.4;refGene_gn=CDH9;refGene_tx=XM_019812404.2
A1	220385746	220385998
refGene_gn=CDH9;refGene_tx=XM_003981527.4;refGene_gn=CDH9;refGene_tx=XM_019812404.2
A1	220391000	220391488
refGene_gn=CDH9;refGene_tx=XM_003981527.4;refGene_gn=CDH9;refGene_tx=XM_019812404.2
A1	220676504	220676569
refGene_gn=LOC101084261;refGene_tx=XM_011286959.3
A1	220676571	220676723
refGene_gn=LOC101084261;refGene_tx=XM_011286959.3
A1	220676724	220677186
refGene_gn=LOC101084261;refGene_tx=XM_011286959.3
A1	220749032	220749222
refGene_gn=LOC101084261;refGene_tx=XM_011286959.3
A1	220774908	220775095
refGene_gn=LOC101084261;refGene_tx=XM_011286959.3
A1	220794245	220794251
refGene_gn=LOC101084261;refGene_tx=XM_011286959.3
A1	222227703	222227934
refGene_gn=LOC101086288;refGene_tx=XM_011286675.3;refGene_gn=LOC101086288;refGene_tx=XM_019812419.2
A1	222288084	222288379
refGene_gn=LOC101086288;refGene_tx=XM_011286675.3;refGene_gn=LOC101086288;refGene_tx=XM_019812419.2
A1	222289882	222290002
refGene_gn=LOC101086288;refGene_tx=XM_011286675.3;refGene_gn=LOC101086288;refGene_tx=XM_019812419.2
A1	222290421	222290589
refGene_gn=LOC101086288;refGene_tx=XM_011286675.3;refGene_gn=LOC101086288;refGene_tx=XM_019812419.2
A1	222315448	222315636
refGene_gn=LOC101086288;refGene_tx=XM_011286675.3;refGene_gn=LOC101086288;refGene_tx=XM_019812419.2
A1	222318009	222318263
refGene_gn=LOC101086288;refGene_tx=XM_011286675.3;refGene_gn=LOC101086288;refGene_tx=XM_019812419.2
A1	222322967	222323104
refGene_gn=LOC101086288;refGene_tx=XM_011286675.3;refGene_gn=LOC101086

288;refGene_tx=XM_019812419.2
A1 222335126 222335248
refGene_gn=LOC101086288;refGene_tx=XM_011286675.3;refGene_gn=LOC101086
288;refGene_tx=XM_019812419.2
A1 222341494 222341603
refGene_gn=LOC101086288;refGene_tx=XM_011286675.3;refGene_gn=LOC101086
288;refGene_tx=XM_019812419.2
A1 222346975 222347227
refGene_gn=LOC101086288;refGene_tx=XM_011286675.3;refGene_gn=LOC101086
288;refGene_tx=XM_019812419.2
A1 222350171 222350662
refGene_gn=LOC101086288;refGene_tx=XM_011286675.3;refGene_gn=LOC101086
288;refGene_tx=XM_019812419.2
A1 224284294 224284525
refGene_gn=CDH12;refGene_tx=XM_006928094.3;refGene_gn=CDH12;refGene_tx
=XM_019812450.2;refGene_gn=CDH12;refGene_tx=XM_019812457.2
A1 224357696 224357991
refGene_gn=CDH12;refGene_tx=XM_006928094.3;refGene_gn=CDH12;refGene_tx
=XM_019812450.2;refGene_gn=CDH12;refGene_tx=XM_019812457.2
A1 224459160 224459280
refGene_gn=CDH12;refGene_tx=XM_006928094.3;refGene_gn=CDH12;refGene_tx
=XM_019812450.2;refGene_gn=CDH12;refGene_tx=XM_019812457.2
A1 224471377 224471545
refGene_gn=CDH12;refGene_tx=XM_006928094.3;refGene_gn=CDH12;refGene_tx
=XM_019812450.2;refGene_gn=CDH12;refGene_tx=XM_019812457.2
A1 224493308 224493496
refGene_gn=CDH12;refGene_tx=XM_006928094.3;refGene_gn=CDH12;refGene_tx
=XM_019812450.2;refGene_gn=CDH12;refGene_tx=XM_019812457.2
A1 224506382 224506636
refGene_gn=CDH12;refGene_tx=XM_006928094.3;refGene_gn=CDH12;refGene_tx
=XM_019812450.2;refGene_gn=CDH12;refGene_tx=XM_019812457.2
A1 224523778 224523915
refGene_gn=CDH12;refGene_tx=XM_006928094.3;refGene_gn=CDH12;refGene_tx
=XM_019812450.2;refGene_gn=CDH12;refGene_tx=XM_019812457.2
A1 224540552 224540674
refGene_gn=CDH12;refGene_tx=XM_006928094.3;refGene_gn=CDH12;refGene_tx
=XM_019812450.2;refGene_gn=CDH12;refGene_tx=XM_019812457.2
A1 224545016 224545134
refGene_gn=CDH12;refGene_tx=XM_006928094.3;refGene_gn=CDH12;refGene_tx
=XM_019812450.2;refGene_gn=CDH12;refGene_tx=XM_019812457.2
A1 224549863 224550115
refGene_gn=CDH12;refGene_tx=XM_006928094.3;refGene_gn=CDH12;refGene_tx
=XM_019812450.2;refGene_gn=CDH12;refGene_tx=XM_019812457.2
A1 224552875 224553375
refGene_gn=CDH12;refGene_tx=XM_006928094.3;refGene_gn=CDH12;refGene_tx
=XM_019812450.2;refGene_gn=CDH12;refGene_tx=XM_019812457.2
A1 226056723 226056951
refGene_gn=CDH18;refGene_tx=XM_019812487.2;refGene_gn=CDH18;refGene_tx
=XM_023239357.1;refGene_gn=CDH18;refGene_tx=XM_023239358.1
A1 226142297 226142592

refGene_gn=CDH18;refGene_tx=XM_019812487.2;refGene_gn=CDH18;refGene_tx=XM_023239357.1;refGene_gn=CDH18;refGene_tx=XM_023239358.1
A1 226166601 226166721
refGene_gn=CDH18;refGene_tx=XM_019812487.2;refGene_gn=CDH18;refGene_tx=XM_023239357.1;refGene_gn=CDH18;refGene_tx=XM_023239358.1
A1 226278085 226278253
refGene_gn=CDH18;refGene_tx=XM_019812487.2;refGene_gn=CDH18;refGene_tx=XM_023239357.1;refGene_gn=CDH18;refGene_tx=XM_023239358.1
A1 226305293 226305481
refGene_gn=CDH18;refGene_tx=XM_019812487.2;refGene_gn=CDH18;refGene_tx=XM_023239357.1;refGene_gn=CDH18;refGene_tx=XM_023239358.1
A1 226316930 226317184
refGene_gn=CDH18;refGene_tx=XM_019812487.2;refGene_gn=CDH18;refGene_tx=XM_023239357.1;refGene_gn=CDH18;refGene_tx=XM_023239358.1
A1 226333781 226333918
refGene_gn=CDH18;refGene_tx=XM_019812487.2;refGene_gn=CDH18;refGene_tx=XM_023239357.1;refGene_gn=CDH18;refGene_tx=XM_023239358.1
A1 226356534 226356656
refGene_gn=CDH18;refGene_tx=XM_019812487.2;refGene_gn=CDH18;refGene_tx=XM_023239357.1;refGene_gn=CDH18;refGene_tx=XM_023239358.1
A1 226367039 226367157
refGene_gn=CDH18;refGene_tx=XM_019812487.2;refGene_gn=CDH18;refGene_tx=XM_023239357.1;refGene_gn=CDH18;refGene_tx=XM_023239358.1
A1 226386913 226387165
refGene_gn=CDH18;refGene_tx=XM_019812487.2;refGene_gn=CDH18;refGene_tx=XM_023239357.1;refGene_gn=CDH18;refGene_tx=XM_023239358.1
A1 226388104 226388369
refGene_gn=CDH18;refGene_tx=XM_023239360.1
A1 226396116 226396607
refGene_gn=CDH18;refGene_tx=XM_019812487.2;refGene_gn=CDH18;refGene_tx=XM_023239360.1
A1 227994872 227995538
refGene_gn=BASP1;refGene_tx=XM_019812583.2;refGene_gn=BASP1;refGene_tx=XM_019812588.2;refGene_gn=BASP1;refGene_tx=XM_023239400.1
A1 228254198 228254219
refGene_gn=MYO10;refGene_tx=XM_023239408.1;refGene_gn=MYO10;refGene_tx=XM_023239414.1;refGene_gn=MYO10;refGene_tx=XM_023239417.1
A1 228303324 228303423
refGene_gn=MYO10;refGene_tx=XM_023239408.1;refGene_gn=MYO10;refGene_tx=XM_023239417.1
A1 228352314 228352473
refGene_gn=MYO10;refGene_tx=XM_023239408.1;refGene_gn=MYO10;refGene_tx=XM_023239414.1;refGene_gn=MYO10;refGene_tx=XM_023239418.1
A1 228370115 228370303
refGene_gn=MYO10;refGene_tx=XM_023239408.1;refGene_gn=MYO10;refGene_tx=XM_023239414.1;refGene_gn=MYO10;refGene_tx=XM_023239417.1;refGene_gn=MYO10;refGene_tx=XM_023239418.1
A1 228382674 228382809
refGene_gn=MYO10;refGene_tx=XM_023239408.1;refGene_gn=MYO10;refGene_tx=XM_023239414.1;refGene_gn=MYO10;refGene_tx=XM_023239417.1;refGene_gn=

MYO10;refGene_tx=XM_023239418.1
A1 228385149 228385274
refGene_gn=MYO10;refGene_tx=XM_023239408.1;refGene_gn=MYO10;refGene_tx=XM_023239414.1;refGene_gn=MYO10;refGene_tx=XM_023239417.1;refGene_gn=MYO10;refGene_tx=XM_023239418.1
A1 228385991 228386005
refGene_gn=MYO10;refGene_tx=XM_023239408.1;refGene_gn=MYO10;refGene_tx=XM_023239414.1;refGene_gn=MYO10;refGene_tx=XM_023239417.1;refGene_gn=MYO10;refGene_tx=XM_023239418.1
A1 228386121 228386206
refGene_gn=MYO10;refGene_tx=XM_023239408.1;refGene_gn=MYO10;refGene_tx=XM_023239414.1;refGene_gn=MYO10;refGene_tx=XM_023239417.1;refGene_gn=MYO10;refGene_tx=XM_023239418.1
A1 228386800 228386904
refGene_gn=MYO10;refGene_tx=XM_023239408.1;refGene_gn=MYO10;refGene_tx=XM_023239414.1;refGene_gn=MYO10;refGene_tx=XM_023239417.1;refGene_gn=MYO10;refGene_tx=XM_023239418.1
A1 228393932 228394062
refGene_gn=MYO10;refGene_tx=XM_023239408.1;refGene_gn=MYO10;refGene_tx=XM_023239414.1;refGene_gn=MYO10;refGene_tx=XM_023239417.1;refGene_gn=MYO10;refGene_tx=XM_023239418.1
A1 228395449 228395568
refGene_gn=MYO10;refGene_tx=XM_023239408.1;refGene_gn=MYO10;refGene_tx=XM_023239414.1;refGene_gn=MYO10;refGene_tx=XM_023239417.1;refGene_gn=MYO10;refGene_tx=XM_023239418.1
A1 228396671 228396818
refGene_gn=MYO10;refGene_tx=XM_023239408.1;refGene_gn=MYO10;refGene_tx=XM_023239414.1;refGene_gn=MYO10;refGene_tx=XM_023239417.1;refGene_gn=MYO10;refGene_tx=XM_023239418.1
A1 228396998 228397099
refGene_gn=MYO10;refGene_tx=XM_023239408.1;refGene_gn=MYO10;refGene_tx=XM_023239414.1;refGene_gn=MYO10;refGene_tx=XM_023239417.1;refGene_gn=MYO10;refGene_tx=XM_023239418.1
A1 228397207 228397274
refGene_gn=MYO10;refGene_tx=XM_023239408.1;refGene_gn=MYO10;refGene_tx=XM_023239414.1;refGene_gn=MYO10;refGene_tx=XM_023239417.1;refGene_gn=MYO10;refGene_tx=XM_023239418.1
A1 228397998 228398091
refGene_gn=MYO10;refGene_tx=XM_023239408.1;refGene_gn=MYO10;refGene_tx=XM_023239414.1;refGene_gn=MYO10;refGene_tx=XM_023239417.1;refGene_gn=MYO10;refGene_tx=XM_023239418.1
A1 228398516 228398585
refGene_gn=MYO10;refGene_tx=XM_023239408.1;refGene_gn=MYO10;refGene_tx=XM_023239414.1;refGene_gn=MYO10;refGene_tx=XM_023239417.1;refGene_gn=MYO10;refGene_tx=XM_023239418.1
A1 228398761 228398844
refGene_gn=MYO10;refGene_tx=XM_023239408.1;refGene_gn=MYO10;refGene_tx=XM_023239414.1;refGene_gn=MYO10;refGene_tx=XM_023239417.1;refGene_gn=MYO10;refGene_tx=XM_023239418.1
A1 228401581 228401690

refGene_gn=MYO10;refGene_tx=XM_023239408.1;refGene_gn=MYO10;refGene_tx=XM_023239414.1;refGene_gn=MYO10;refGene_tx=XM_023239417.1;refGene_gn=MYO10;refGene_tx=XM_023239418.1
A1 228404519 228404600
refGene_gn=MYO10;refGene_tx=XM_023239408.1;refGene_gn=MYO10;refGene_tx=XM_023239414.1;refGene_gn=MYO10;refGene_tx=XM_023239417.1;refGene_gn=MYO10;refGene_tx=XM_023239418.1
A1 228440690 228440815
refGene_gn=MYO10;refGene_tx=XM_023239408.1;refGene_gn=MYO10;refGene_tx=XM_023239414.1;refGene_gn=MYO10;refGene_tx=XM_023239417.1;refGene_gn=MYO10;refGene_tx=XM_023239418.1
A1 228440908 228441023
refGene_gn=MYO10;refGene_tx=XM_023239408.1;refGene_gn=MYO10;refGene_tx=XM_023239414.1;refGene_gn=MYO10;refGene_tx=XM_023239417.1;refGene_gn=MYO10;refGene_tx=XM_023239418.1
A1 228447865 228447972
refGene_gn=MYO10;refGene_tx=XM_023239408.1;refGene_gn=MYO10;refGene_tx=XM_023239414.1;refGene_gn=MYO10;refGene_tx=XM_023239417.1;refGene_gn=MYO10;refGene_tx=XM_023239418.1
A1 228448819 228449029
refGene_gn=MYO10;refGene_tx=XM_023239408.1;refGene_gn=MYO10;refGene_tx=XM_023239414.1;refGene_gn=MYO10;refGene_tx=XM_023239417.1;refGene_gn=MYO10;refGene_tx=XM_023239418.1
A1 228449382 228449428
refGene_gn=MYO10;refGene_tx=XM_023239408.1;refGene_gn=MYO10;refGene_tx=XM_023239414.1;refGene_gn=MYO10;refGene_tx=XM_023239417.1;refGene_gn=MYO10;refGene_tx=XM_023239418.1
A1 228450321 228451188
refGene_gn=MYO10;refGene_tx=XM_023239408.1;refGene_gn=MYO10;refGene_tx=XM_023239414.1;refGene_gn=MYO10;refGene_tx=XM_023239417.1;refGene_gn=MYO10;refGene_tx=XM_023239418.1
A1 228452035 228452159
refGene_gn=MYO10;refGene_tx=XM_023239408.1;refGene_gn=MYO10;refGene_tx=XM_023239414.1;refGene_gn=MYO10;refGene_tx=XM_023239417.1;refGene_gn=MYO10;refGene_tx=XM_023239418.1
A1 228457653 228457897
refGene_gn=MYO10;refGene_tx=XM_023239408.1;refGene_gn=MYO10;refGene_tx=XM_023239414.1;refGene_gn=MYO10;refGene_tx=XM_023239417.1;refGene_gn=MYO10;refGene_tx=XM_023239418.1
A1 228461251 228461347
refGene_gn=MYO10;refGene_tx=XM_023239408.1;refGene_gn=MYO10;refGene_tx=XM_023239414.1;refGene_gn=MYO10;refGene_tx=XM_023239417.1;refGene_gn=MYO10;refGene_tx=XM_023239418.1
A1 228464369 228464463
refGene_gn=MYO10;refGene_tx=XM_023239408.1;refGene_gn=MYO10;refGene_tx=XM_023239414.1;refGene_gn=MYO10;refGene_tx=XM_023239417.1;refGene_gn=MYO10;refGene_tx=XM_023239418.1
A1 228465778 228465834
refGene_gn=MYO10;refGene_tx=XM_023239408.1;refGene_gn=MYO10;refGene_tx=XM_023239414.1;refGene_gn=MYO10;refGene_tx=XM_023239417.1;refGene_gn=

MYO10;refGene_tx=XM_023239418.1
A1 228467519 228467662
refGene_gn=MYO10;refGene_tx=XM_023239408.1;refGene_gn=MYO10;refGene_tx
=XM_023239414.1;refGene_gn=MYO10;refGene_tx=XM_023239417.1;refGene_gn=
MYO10;refGene_tx=XM_023239418.1
A1 228467896 228468091
refGene_gn=MYO10;refGene_tx=XM_023239408.1;refGene_gn=MYO10;refGene_tx
=XM_023239414.1;refGene_gn=MYO10;refGene_tx=XM_023239417.1;refGene_gn=
MYO10;refGene_tx=XM_023239418.1
A1 228468993 228469151
refGene_gn=MYO10;refGene_tx=XM_023239408.1;refGene_gn=MYO10;refGene_tx
=XM_023239414.1;refGene_gn=MYO10;refGene_tx=XM_023239417.1;refGene_gn=
MYO10;refGene_tx=XM_023239418.1
A1 228471564 228471688
refGene_gn=MYO10;refGene_tx=XM_023239408.1;refGene_gn=MYO10;refGene_tx
=XM_023239414.1;refGene_gn=MYO10;refGene_tx=XM_023239417.1;refGene_gn=
MYO10;refGene_tx=XM_023239418.1
A1 228472162 228472460
refGene_gn=MYO10;refGene_tx=XM_023239408.1;refGene_gn=MYO10;refGene_tx
=XM_023239414.1;refGene_gn=MYO10;refGene_tx=XM_023239417.1;refGene_gn=
MYO10;refGene_tx=XM_023239418.1
A1 228473325 228473533
refGene_gn=MYO10;refGene_tx=XM_023239408.1;refGene_gn=MYO10;refGene_tx
=XM_023239414.1;refGene_gn=MYO10;refGene_tx=XM_023239417.1;refGene_gn=
MYO10;refGene_tx=XM_023239418.1
A1 228474387 228474524
refGene_gn=MYO10;refGene_tx=XM_023239408.1;refGene_gn=MYO10;refGene_tx
=XM_023239414.1;refGene_gn=MYO10;refGene_tx=XM_023239417.1;refGene_gn=
MYO10;refGene_tx=XM_023239418.1
A1 228475746 228475867
refGene_gn=MYO10;refGene_tx=XM_023239408.1;refGene_gn=MYO10;refGene_tx
=XM_023239414.1;refGene_gn=MYO10;refGene_tx=XM_023239417.1;refGene_gn=
MYO10;refGene_tx=XM_023239418.1
A1 228476242 228476695
refGene_gn=MYO10;refGene_tx=XM_023239408.1;refGene_gn=MYO10;refGene_tx
=XM_023239414.1;refGene_gn=MYO10;refGene_tx=XM_023239417.1;refGene_gn=
MYO10;refGene_tx=XM_023239418.1
A1 228479224 228479416
refGene_gn=MYO10;refGene_tx=XM_023239408.1;refGene_gn=MYO10;refGene_tx
=XM_023239414.1;refGene_gn=MYO10;refGene_tx=XM_023239417.1;refGene_gn=
MYO10;refGene_tx=XM_023239418.1
A1 228481087 228481189
refGene_gn=MYO10;refGene_tx=XM_023239408.1;refGene_gn=MYO10;refGene_tx
=XM_023239414.1;refGene_gn=MYO10;refGene_tx=XM_023239417.1;refGene_gn=
MYO10;refGene_tx=XM_023239418.1
A1 228508825 228509118
refGene_gn=RETREG1;refGene_tx=XM_023239424.1
A1 228542157 228542264
refGene_gn=RETREG1;refGene_tx=XM_023239424.1
A1 228549620 228549651

refGene_gn=RETREG1;refGene_tx=XM_023239424.1
A1 228601596 228601631
refGene_gn=RETREG1;refGene_tx=XM_006928098.4
A1 228628358 228628485
refGene_gn=RETREG1;refGene_tx=XM_006928098.4;refGene_gn=RETREG1;refGene_tx=XM_023239424.1
A1 228630891 228630976
refGene_gn=RETREG1;refGene_tx=XM_006928098.4;refGene_gn=RETREG1;refGene_tx=XM_023239424.1
A1 228632663 228632801
refGene_gn=RETREG1;refGene_tx=XM_006928098.4;refGene_gn=RETREG1;refGene_tx=XM_023239424.1
A1 228633601 228633666
refGene_gn=RETREG1;refGene_tx=XM_006928098.4;refGene_gn=RETREG1;refGene_tx=XM_023239424.1
A1 228633913 228634040
refGene_gn=RETREG1;refGene_tx=XM_006928098.4;refGene_gn=RETREG1;refGene_tx=XM_023239424.1
A1 228636776 228637276
refGene_gn=RETREG1;refGene_tx=XM_006928098.4;refGene_gn=RETREG1;refGene_tx=XM_023239424.1
A1 228645573 228646198
refGene_gn=ZNF622;refGene_tx=XM_006928099.4
A1 228646926 228647187
refGene_gn=ZNF622;refGene_tx=XM_006928099.4
A1 228647416 228647579
refGene_gn=ZNF622;refGene_tx=XM_006928099.4
A1 228650346 228650459
refGene_gn=ZNF622;refGene_tx=XM_006928099.4
A1 228656844 228656988
refGene_gn=ZNF622;refGene_tx=XM_006928099.4
A1 228658434 228658568
refGene_gn=ZNF622;refGene_tx=XM_006928099.4
A1 228881929 228882055
refGene_gn=LOC109492397;refGene_tx=XM_023246845.1
A1 228882298 228882412
refGene_gn=LOC109492397;refGene_tx=XM_023246845.1
A1 228890106 228890755
refGene_gn=LOC109492397;refGene_tx=XM_023246845.1
A1 228890987 228891617
refGene_gn=LOC109492397;refGene_tx=XM_023246845.1;refGene_gn=MARCH11;refGene_tx=XM_023239442.1
A1 228892847 228893003
refGene_gn=MARCH11;refGene_tx=XM_023239442.1
A1 228970821 228971014
refGene_gn=MARCH11;refGene_tx=XM_023239442.1
A1 228994509 228994832
refGene_gn=MARCH11;refGene_tx=XM_023239442.1
A1 229098938 229099675
refGene_gn=FBXL7;refGene_tx=XM_006928101.4;refGene_gn=FBXL7;refGene_tx

=XM_019812685.1
A1 229108311 229108923
refGene_gn=FBXL7;refGene_tx=XM_006928101.4;refGene_gn=FBXL7;refGene_tx
=XM_019812685.1
A1 229388818 229388908
refGene_gn=FBXL7;refGene_tx=XM_006928101.4;refGene_gn=FBXL7;refGene_tx
=XM_019812685.1
A1 229434695 229434753
refGene_gn=FBXL7;refGene_tx=XM_019812685.1
A1 229493643 229493680
refGene_gn=FBXL7;refGene_tx=XM_006928101.4
A1 229953750 229953846
refGene_gn=ANKH;refGene_tx=XM_011287018.2;refGene_gn=ANKH;refGene_tx=X
M_019812703.2
A1 230036480 230036697
refGene_gn=ANKH;refGene_tx=XM_011287018.2;refGene_gn=ANKH;refGene_tx=X
M_019812703.2
A1 230044009 230044128
refGene_gn=ANKH;refGene_tx=XM_011287018.2;refGene_gn=ANKH;refGene_tx=X
M_019812703.2
A1 230047275 230047359
refGene_gn=ANKH;refGene_tx=XM_011287018.2;refGene_gn=ANKH;refGene_tx=X
M_019812703.2
A1 230055064 230055235
refGene_gn=ANKH;refGene_tx=XM_011287018.2
A1 230057057 230057192
refGene_gn=ANKH;refGene_tx=XM_011287018.2;refGene_gn=ANKH;refGene_tx=X
M_019812703.2
A1 230059390 230059483
refGene_gn=ANKH;refGene_tx=XM_011287018.2;refGene_gn=ANKH;refGene_tx=X
M_019812703.2
A1 230063778 230063874
refGene_gn=ANKH;refGene_tx=XM_011287018.2;refGene_gn=ANKH;refGene_tx=X
M_019812703.2
A1 230086836 230086966
refGene_gn=ANKH;refGene_tx=XM_011287018.2;refGene_gn=ANKH;refGene_tx=X
M_019812703.2
A1 230089406 230089530
refGene_gn=ANKH;refGene_tx=XM_011287018.2;refGene_gn=ANKH;refGene_tx=X
M_019812703.2
A1 230090102 230090202
refGene_gn=ANKH;refGene_tx=XM_011287018.2;refGene_gn=ANKH;refGene_tx=X
M_019812703.2
A1 230091759 230091873
refGene_gn=ANKH;refGene_tx=XM_011287018.2;refGene_gn=ANKH;refGene_tx=X
M_019812703.2
A1 230109431 230109626
refGene_gn=OTULIN;refGene_tx=XM_023239457.1;refGene_gn=OTULIN;refGene_
tx=XM_023239458.1;refGene_gn=OTULIN;refGene_tx=XM_023239460.1
A1 230112105 230112375

refGene_gn=OTULIN;refGene_tx=XM_023239457.1;refGene_gn=OTULIN;refGene_tx=XM_023239458.1;refGene_gn=OTULIN;refGene_tx=XM_023239460.1
A1 230115694 230115820
refGene_gn=OTULIN;refGene_tx=XM_023239457.1;refGene_gn=OTULIN;refGene_tx=XM_023239458.1;refGene_gn=OTULIN;refGene_tx=XM_023239460.1
A1 230120424 230120568
refGene_gn=OTULIN;refGene_tx=XM_023239457.1;refGene_gn=OTULIN;refGene_tx=XM_023239458.1;refGene_gn=OTULIN;refGene_tx=XM_023239460.1
A1 230123929 230124024
refGene_gn=OTULIN;refGene_tx=XM_023239457.1;refGene_gn=OTULIN;refGene_tx=XM_023239458.1;refGene_gn=OTULIN;refGene_tx=XM_023239460.1
A1 230129238 230129315
refGene_gn=OTULIN;refGene_tx=XM_023239457.1;refGene_gn=OTULIN;refGene_tx=XM_023239458.1;refGene_gn=OTULIN;refGene_tx=XM_023239460.1
A1 230131453 230131488
refGene_gn=OTULIN;refGene_tx=XM_023239458.1
A1 230135889 230136065
refGene_gn=OTULIN;refGene_tx=XM_023239457.1
A1 230179454 230179628
refGene_gn=FAM105A;refGene_tx=XM_023246883.1
A1 230180547 230180817
refGene_gn=FAM105A;refGene_tx=XM_023246883.1
A1 230182154 230182353
refGene_gn=FAM105A;refGene_tx=XM_023246883.1
A1 230186111 230186307
refGene_gn=FAM105A;refGene_tx=XM_023246883.1
A1 230186408 230186440
refGene_gn=FAM105A;refGene_tx=XM_023246883.1
A1 230186528 230186688
refGene_gn=FAM105A;refGene_tx=XM_023246883.1
A1 230207870 230208342
refGene_gn=FAM105A;refGene_tx=XM_023246883.1
A1 230252104 230252653
refGene_gn=TRIO;refGene_tx=XM_023239465.1
A1 230253199 230253338
refGene_gn=TRIO;refGene_tx=XM_023239465.1
A1 230256067 230256268
refGene_gn=TRIO;refGene_tx=XM_023239465.1
A1 230257965 230258044
refGene_gn=TRIO;refGene_tx=XM_023239465.1
A1 230259566 230259688
refGene_gn=TRIO;refGene_tx=XM_023239465.1
A1 230259954 230260117
refGene_gn=TRIO;refGene_tx=XM_023239465.1
A1 230260323 230260351
refGene_gn=TRIO;refGene_tx=XM_023239465.1
A1 230261099 230261238
refGene_gn=TRIO;refGene_tx=XM_023239465.1
A1 230262919 230263167
refGene_gn=TRIO;refGene_tx=XM_023239465.1

A1 230267157 230268002
refGene_gn=TRIO;refGene_tx=XM_023239465.1
A1 230270218 230270396
refGene_gn=TRIO;refGene_tx=XM_023239465.1
A1 230272450 230272642
refGene_gn=TRIO;refGene_tx=XM_023239465.1
A1 230273578 230273656
refGene_gn=TRIO;refGene_tx=XM_023239465.1
A1 230273955 230274006
refGene_gn=TRIO;refGene_tx=XM_023239465.1
A1 230274736 230274829
refGene_gn=TRIO;refGene_tx=XM_023239465.1
A1 230275399 230275489
refGene_gn=TRIO;refGene_tx=XM_023239465.1
A1 230277472 230277542
refGene_gn=TRIO;refGene_tx=XM_023239465.1
A1 230280264 230280368
refGene_gn=TRIO;refGene_tx=XM_023239465.1
A1 230281928 230281995
refGene_gn=TRIO;refGene_tx=XM_023239465.1
A1 230283185 230283334
refGene_gn=TRIO;refGene_tx=XM_023239465.1
A1 230288852 230288948
refGene_gn=TRIO;refGene_tx=XM_023239465.1
A1 230291449 230291620
refGene_gn=TRIO;refGene_tx=XM_023239465.1
A1 230293020 230293313
refGene_gn=TRIO;refGene_tx=XM_023239465.1
A1 230329149 230329393
refGene_gn=TRIO;refGene_tx=XM_023239465.1
A1 230341525 230341625
refGene_gn=TRIO;refGene_tx=XM_023239465.1
A1 230342202 230342345
refGene_gn=TRIO;refGene_tx=XM_023239465.1
A1 230346342 230346444
refGene_gn=TRIO;refGene_tx=XM_023239465.1
A1 230348449 230348640
refGene_gn=TRIO;refGene_tx=XM_023239465.1
A1 230350384 230350496
refGene_gn=TRIO;refGene_tx=XM_023239465.1
A1 230353170 230353263
refGene_gn=TRIO;refGene_tx=XM_023239465.1
A1 230357862 230357952
refGene_gn=TRIO;refGene_tx=XM_023239465.1
A1 230358586 230358656
refGene_gn=TRIO;refGene_tx=XM_023239465.1
A1 230359611 230359721
refGene_gn=TRIO;refGene_tx=XM_023239465.1
A1 230360335 230360402
refGene_gn=TRIO;refGene_tx=XM_023239465.1

A1 230361171 230361287
refGene_gn=TRIO;refGene_tx=XM_023239465.1
A1 230361385 230361580
refGene_gn=TRIO;refGene_tx=XM_023239465.1
A1 230368503 230368626
refGene_gn=TRIO;refGene_tx=XM_023239465.1
A1 230372391 230372507
refGene_gn=TRIO;refGene_tx=XM_023239465.1
A1 230375738 230375853
refGene_gn=TRIO;refGene_tx=XM_023239465.1
A1 230379510 230379660
refGene_gn=TRIO;refGene_tx=XM_023239465.1
A1 230380140 230380332
refGene_gn=TRIO;refGene_tx=XM_023239465.1
A1 230382048 230382168
refGene_gn=TRIO;refGene_tx=XM_023239465.1
A1 230384410 230384577
refGene_gn=TRIO;refGene_tx=XM_023239465.1
A1 230385376 230385572
refGene_gn=TRIO;refGene_tx=XM_023239465.1
A1 230389847 230390022
refGene_gn=TRIO;refGene_tx=XM_023239465.1
A1 230390923 230391093
refGene_gn=TRIO;refGene_tx=XM_023239465.1
A1 230408257 230408449
refGene_gn=TRIO;refGene_tx=XM_023239465.1
A1 230422757 230422880
refGene_gn=TRIO;refGene_tx=XM_023239465.1
A1 230437920 230438151
refGene_gn=TRIO;refGene_tx=XM_023239465.1
A1 230449423 230449555
refGene_gn=TRIO;refGene_tx=XM_023239465.1
A1 230457859 230458051
refGene_gn=TRIO;refGene_tx=XM_023239465.1
A1 230462172 230462258
refGene_gn=TRIO;refGene_tx=XM_023239465.1
A1 230462260 230462294
refGene_gn=TRIO;refGene_tx=XM_023239465.1
A1 230463779 230464292
refGene_gn=TRIO;refGene_tx=XM_023239465.1
A1 230466770 230466963
refGene_gn=TRIO;refGene_tx=XM_023239465.1
A1 230473305 230473420
refGene_gn=TRIO;refGene_tx=XM_023239465.1
A1 230483367 230483442
refGene_gn=TRIO;refGene_tx=XM_023239465.1
A1 230604398 230604471
refGene_gn=TRIO;refGene_tx=XM_023239465.1
A1 230604472 230604556
refGene_gn=TRIO;refGene_tx=XM_023239465.1

A1 230723795 230723807
refGene_gn=DNAH5;refGene_tx=XM_023239471.1
A1 230786795 230786930
refGene_gn=DNAH5;refGene_tx=XM_023239471.1
A1 230790307 230790392
refGene_gn=DNAH5;refGene_tx=XM_023239471.1
A1 230793864 230793926
refGene_gn=DNAH5;refGene_tx=XM_023239471.1
A1 230793927 230794026
refGene_gn=DNAH5;refGene_tx=XM_023239471.1
A1 230795229 230795451
refGene_gn=DNAH5;refGene_tx=XM_023239471.1
A1 230798304 230798442
refGene_gn=DNAH5;refGene_tx=XM_023239471.1
A1 230799235 230799412
refGene_gn=DNAH5;refGene_tx=XM_023239471.1
A1 230800455 230800569
refGene_gn=DNAH5;refGene_tx=XM_023239471.1
A1 230801511 230801619
refGene_gn=DNAH5;refGene_tx=XM_023239471.1
A1 230804245 230804368
refGene_gn=DNAH5;refGene_tx=XM_023239471.1
A1 230804925 230805141
refGene_gn=DNAH5;refGene_tx=XM_023239471.1
A1 230808860 230808968
refGene_gn=DNAH5;refGene_tx=XM_023239471.1
A1 230814458 230814544
refGene_gn=DNAH5;refGene_tx=XM_023239471.1
A1 230815589 230815911
refGene_gn=DNAH5;refGene_tx=XM_023239471.1
A1 230816852 230817059
refGene_gn=DNAH5;refGene_tx=XM_023239471.1
A1 230821922 230822094
refGene_gn=DNAH5;refGene_tx=XM_023239471.1
A1 230824499 230824645
refGene_gn=DNAH5;refGene_tx=XM_023239471.1
A1 230830316 230830482
refGene_gn=DNAH5;refGene_tx=XM_023239471.1
A1 230831094 230831331
refGene_gn=DNAH5;refGene_tx=XM_023239471.1
A1 230832615 230832806
refGene_gn=DNAH5;refGene_tx=XM_023239471.1
A1 230832893 230832981
refGene_gn=DNAH5;refGene_tx=XM_023239471.1
A1 230839435 230839569
refGene_gn=DNAH5;refGene_tx=XM_023239471.1
A1 230844277 230844479
refGene_gn=DNAH5;refGene_tx=XM_023239471.1
A1 230845030 230845266
refGene_gn=DNAH5;refGene_tx=XM_023239471.1

A1 230848383 230848602
refGene_gn=DNAH5;refGene_tx=XM_023239471.1
A1 230849612 230849675
refGene_gn=DNAH5;refGene_tx=XM_023239471.1
A1 230849952 230850191
refGene_gn=DNAH5;refGene_tx=XM_023239471.1
A1 230850932 230851173
refGene_gn=DNAH5;refGene_tx=XM_023239471.1
A1 230853239 230853439
refGene_gn=DNAH5;refGene_tx=XM_023239471.1
A1 230856544 230856698
refGene_gn=DNAH5;refGene_tx=XM_023239471.1
A1 230858187 230858351
refGene_gn=DNAH5;refGene_tx=XM_023239471.1
A1 230864695 230864852
refGene_gn=DNAH5;refGene_tx=XM_023239471.1
A1 230868433 230868646
refGene_gn=DNAH5;refGene_tx=XM_023239471.1
A1 230869401 230869626
refGene_gn=DNAH5;refGene_tx=XM_023239471.1
A1 230871229 230871402
refGene_gn=DNAH5;refGene_tx=XM_023239471.1
A1 230873592 230873771
refGene_gn=DNAH5;refGene_tx=XM_023239471.1
A1 230874385 230874573
refGene_gn=DNAH5;refGene_tx=XM_023239471.1
A1 230874905 230875100
refGene_gn=DNAH5;refGene_tx=XM_023239471.1
A1 230878513 230878648
refGene_gn=DNAH5;refGene_tx=XM_023239471.1
A1 230879489 230879597
refGene_gn=DNAH5;refGene_tx=XM_023239471.1
A1 230881814 230881968
refGene_gn=DNAH5;refGene_tx=XM_023239471.1
A1 230888015 230888162
refGene_gn=DNAH5;refGene_tx=XM_023239471.1
A1 230890573 230890815
refGene_gn=DNAH5;refGene_tx=XM_023239471.1
A1 230894637 230894814
refGene_gn=DNAH5;refGene_tx=XM_023239471.1
A1 230895223 230895425
refGene_gn=DNAH5;refGene_tx=XM_023239471.1
A1 230896969 230897112
refGene_gn=DNAH5;refGene_tx=XM_023239471.1
A1 230897829 230897964
refGene_gn=DNAH5;refGene_tx=XM_023239471.1
A1 230907891 230908014
refGene_gn=DNAH5;refGene_tx=XM_023239471.1
A1 230908191 230908405
refGene_gn=DNAH5;refGene_tx=XM_023239471.1

A1	230910084	230910308
refGene_gn=DNAH5;refGene_tx=XM_023239471.1
A1	230912623	230912822
refGene_gn=DNAH5;refGene_tx=XM_023239471.1
A1	230915212	230915385
refGene_gn=DNAH5;refGene_tx=XM_023239471.1
A1	230920600	230920731
refGene_gn=DNAH5;refGene_tx=XM_023239471.1
A1	230924833	230924987
refGene_gn=DNAH5;refGene_tx=XM_023239471.1
A1	230926357	230926625
refGene_gn=DNAH5;refGene_tx=XM_023239471.1
A1	230936646	230936878
refGene_gn=DNAH5;refGene_tx=XM_023239471.1
A1	230937995	230938110
refGene_gn=DNAH5;refGene_tx=XM_023239471.1
A1	230938352	230938529
refGene_gn=DNAH5;refGene_tx=XM_023239471.1
A1	230942235	230942439
refGene_gn=DNAH5;refGene_tx=XM_023239471.1
A1	230947242	230947422
refGene_gn=DNAH5;refGene_tx=XM_023239471.1
A1	230950926	230951064
refGene_gn=DNAH5;refGene_tx=XM_023239471.1
A1	230955417	230955553
refGene_gn=DNAH5;refGene_tx=XM_023239471.1
A1	230956367	230956684
refGene_gn=DNAH5;refGene_tx=XM_023239471.1
A1	230957247	230957403
refGene_gn=DNAH5;refGene_tx=XM_023239471.1
A1	230958573	230958756
refGene_gn=DNAH5;refGene_tx=XM_023239471.1
A1	230971086	230971330
refGene_gn=DNAH5;refGene_tx=XM_023239471.1
A1	230972331	230972446
refGene_gn=DNAH5;refGene_tx=XM_023239471.1
A1	230972941	230973132
refGene_gn=DNAH5;refGene_tx=XM_023239471.1
A1	230977410	230977532
refGene_gn=DNAH5;refGene_tx=XM_023239471.1
A1	230979319	230979469
refGene_gn=DNAH5;refGene_tx=XM_023239471.1
A1	230983422	230983668
refGene_gn=DNAH5;refGene_tx=XM_023239471.1
A1	230987866	230988086
refGene_gn=DNAH5;refGene_tx=XM_023239471.1
A1	230989411	230989617
refGene_gn=DNAH5;refGene_tx=XM_023239471.1
A1	230990234	230990438
refGene_gn=DNAH5;refGene_tx=XM_023239471.1

A1 230992568 230992784
refGene_gn=DNAH5;refGene_tx=XM_023239471.1
A1 230994874 230995087
refGene_gn=DNAH5;refGene_tx=XM_023239471.1
A1 230997700 230997853
refGene_gn=DNAH5;refGene_tx=XM_023239471.1
A1 230998270 230998502
refGene_gn=DNAH5;refGene_tx=XM_023239471.1
A1 231005126 231005278
refGene_gn=DNAH5;refGene_tx=XM_023239471.1
A1 232489640 232489692
refGene_gn=CTNND2;refGene_tx=XM_023239481.1;refGene_gn=CTNND2;refGene_tx=XM_023239488.1
A1 232648130 232648267
refGene_gn=CTNND2;refGene_tx=XM_023239481.1;refGene_gn=CTNND2;refGene_tx=XM_023239488.1
A1 232842620 232842733
refGene_gn=CTNND2;refGene_tx=XM_023239481.1;refGene_gn=CTNND2;refGene_tx=XM_023239488.1
A1 232998878 232998913
refGene_gn=CTNND2;refGene_tx=XM_023239481.1;refGene_gn=CTNND2;refGene_tx=XM_023239488.1
A1 232999303 232999420
refGene_gn=CTNND2;refGene_tx=XM_023239481.1;refGene_gn=CTNND2;refGene_tx=XM_023239488.1
A1 233014812 233014985
refGene_gn=CTNND2;refGene_tx=XM_023239481.1;refGene_gn=CTNND2;refGene_tx=XM_023239488.1
A1 233029439 233030013
refGene_gn=CTNND2;refGene_tx=XM_023239481.1;refGene_gn=CTNND2;refGene_tx=XM_023239488.1
A1 233052589 233052784
refGene_gn=CTNND2;refGene_tx=XM_023239481.1;refGene_gn=CTNND2;refGene_tx=XM_023239488.1
A1 233063956 233064212
refGene_gn=CTNND2;refGene_tx=XM_023239481.1;refGene_gn=CTNND2;refGene_tx=XM_023239488.1
A1 233176839 233176972
refGene_gn=CTNND2;refGene_tx=XM_023239481.1;refGene_gn=CTNND2;refGene_tx=XM_023239488.1
A1 233211091 233211305
refGene_gn=CTNND2;refGene_tx=XM_023239481.1;refGene_gn=CTNND2;refGene_tx=XM_023239488.1
A1 233256878 233257062
refGene_gn=CTNND2;refGene_tx=XM_023239481.1;refGene_gn=CTNND2;refGene_tx=XM_023239488.1
A1 233291206 233291324
refGene_gn=CTNND2;refGene_tx=XM_023239481.1;refGene_gn=CTNND2;refGene_tx=XM_023239488.1
A1 233296912 233297098

refGene_gn=CTNND2;refGene_tx=XM_023239481.1;refGene_gn=CTNND2;refGene_tx=XM_023239488.1
A1 233312034 233312208
refGene_gn=CTNND2;refGene_tx=XM_023239481.1;refGene_gn=CTNND2;refGene_tx=XM_023239488.1
A1 233325441 233325516
refGene_gn=CTNND2;refGene_tx=XM_023239481.1
A1 233326569 233326720
refGene_gn=CTNND2;refGene_tx=XM_023239481.1;refGene_gn=CTNND2;refGene_tx=XM_023239488.1
A1 233379095 233379306
refGene_gn=CTNND2;refGene_tx=XM_023239481.1;refGene_gn=CTNND2;refGene_tx=XM_023239488.1
A1 233384418 233384503
refGene_gn=CTNND2;refGene_tx=XM_023239481.1;refGene_gn=CTNND2;refGene_tx=XM_023239488.1
A1 233409851 233409978
refGene_gn=CTNND2;refGene_tx=XM_023239481.1;refGene_gn=CTNND2;refGene_tx=XM_023239488.1
A1 233414162 233414294
refGene_gn=CTNND2;refGene_tx=XM_023239481.1;refGene_gn=CTNND2;refGene_tx=XM_023239488.1
A1 233420204 233420278
refGene_gn=CTNND2;refGene_tx=XM_023239481.1;refGene_gn=CTNND2;refGene_tx=XM_023239488.1
A1 233427626 233427887
refGene_gn=CTNND2;refGene_tx=XM_023239481.1;refGene_gn=CTNND2;refGene_tx=XM_023239488.1
A1 233606486 233606541
refGene_gn=DAP;refGene_tx=XM_003981544.5;refGene_gn=DAP;refGene_tx=XM_019812830.2
A1 233618413 233618510
refGene_gn=DAP;refGene_tx=XM_003981544.5;refGene_gn=DAP;refGene_tx=XM_019812830.2
A1 233638163 233638296
refGene_gn=DAP;refGene_tx=XM_019812830.2
A1 233666991 233667034
refGene_gn=DAP;refGene_tx=XM_003981544.5
A1 233668612 233668726
refGene_gn=DAP;refGene_tx=XM_003981544.5
A1 233694254 233695087
refGene_gn=ANKRD33B;refGene_tx=XM_011287289.3
A1 233703692 233703833
refGene_gn=ANKRD33B;refGene_tx=XM_011287289.3
A1 233721708 233721838
refGene_gn=ANKRD33B;refGene_tx=XM_011287289.3
A1 233774565 233774919
refGene_gn=ANKRD33B;refGene_tx=XM_011287289.3
A1 233807620 233807896
refGene_gn=LOC102899551;refGene_tx=XM_023259200.1

A1 233808589 233809027
refGene_gn=LOC102899551;refGene_tx=XM_023259200.1
A1 233844798 233844862
refGene_gn=ROPN1L;refGene_tx=XM_003981552.5
A1 233848921 233849097
refGene_gn=ROPN1L;refGene_tx=XM_003981552.5
A1 233856395 233856557
refGene_gn=ROPN1L;refGene_tx=XM_003981552.5
A1 233857993 233858117
refGene_gn=ROPN1L;refGene_tx=XM_003981552.5
A1 233862295 233862426
refGene_gn=ROPN1L;refGene_tx=XM_003981552.5
A1 233872808 233872899
refGene_gn=MARCH6;refGene_tx=XM_023239515.1;refGene_gn=MARCH6;refGene_
tx=XM_023239519.1
A1 233876698 233876834
refGene_gn=MARCH6;refGene_tx=XM_023239515.1;refGene_gn=MARCH6;refGene_
tx=XM_023239519.1
A1 233878220 233878353
refGene_gn=MARCH6;refGene_tx=XM_023239515.1;refGene_gn=MARCH6;refGene_
tx=XM_023239519.1
A1 233880771 233880861
refGene_gn=MARCH6;refGene_tx=XM_023239515.1;refGene_gn=MARCH6;refGene_
tx=XM_023239519.1
A1 233885992 233886127
refGene_gn=MARCH6;refGene_tx=XM_023239515.1;refGene_gn=MARCH6;refGene_
tx=XM_023239519.1
A1 233887794 233887991
refGene_gn=MARCH6;refGene_tx=XM_023239515.1;refGene_gn=MARCH6;refGene_
tx=XM_023239519.1
A1 233890895 233890965
refGene_gn=MARCH6;refGene_tx=XM_023239515.1;refGene_gn=MARCH6;refGene_
tx=XM_023239519.1
A1 233893976 233894181
refGene_gn=MARCH6;refGene_tx=XM_023239515.1;refGene_gn=MARCH6;refGene_
tx=XM_023239519.1
A1 233894781 233894919
refGene_gn=MARCH6;refGene_tx=XM_023239515.1;refGene_gn=MARCH6;refGene_
tx=XM_023239519.1
A1 233897363 233897464
refGene_gn=MARCH6;refGene_tx=XM_023239515.1;refGene_gn=MARCH6;refGene_
tx=XM_023239519.1
A1 233898257 233898377
refGene_gn=MARCH6;refGene_tx=XM_023239515.1;refGene_gn=MARCH6;refGene_
tx=XM_023239519.1
A1 233900342 233900477
refGene_gn=MARCH6;refGene_tx=XM_023239515.1;refGene_gn=MARCH6;refGene_
tx=XM_023239519.1
A1 233901769 233901844
refGene_gn=MARCH6;refGene_tx=XM_023239515.1;refGene_gn=MARCH6;refGene_

tx=XM_023239519.1
A1 233901923 233901992
refGene_gn=MARCH6;refGene_tx=XM_023239515.1;refGene_gn=MARCH6;refGene_tx=XM_023239519.1
A1 233902220 233902301
refGene_gn=MARCH6;refGene_tx=XM_023239515.1;refGene_gn=MARCH6;refGene_tx=XM_023239519.1
A1 233903584 233903643
refGene_gn=MARCH6;refGene_tx=XM_023239515.1;refGene_gn=MARCH6;refGene_tx=XM_023239519.1
A1 233906782 233906834
refGene_gn=MARCH6;refGene_tx=XM_023239515.1;refGene_gn=MARCH6;refGene_tx=XM_023239519.1
A1 233907997 233908030
refGene_gn=MARCH6;refGene_tx=XM_023239515.1;refGene_gn=MARCH6;refGene_tx=XM_023239519.1
A1 233908768 233908830
refGene_gn=MARCH6;refGene_tx=XM_023239515.1;refGene_gn=MARCH6;refGene_tx=XM_023239519.1
A1 233910444 233910634
refGene_gn=MARCH6;refGene_tx=XM_023239515.1;refGene_gn=MARCH6;refGene_tx=XM_023239519.1
A1 233913236 233913405
refGene_gn=MARCH6;refGene_tx=XM_023239515.1;refGene_gn=MARCH6;refGene_tx=XM_023239519.1
A1 233917359 233917432
refGene_gn=MARCH6;refGene_tx=XM_023239515.1;refGene_gn=MARCH6;refGene_tx=XM_023239519.1
A1 233922470 233922614
refGene_gn=MARCH6;refGene_tx=XM_023239515.1;refGene_gn=MARCH6;refGene_tx=XM_023239519.1
A1 233925013 233925087
refGene_gn=MARCH6;refGene_tx=XM_023239515.1;refGene_gn=MARCH6;refGene_tx=XM_023239519.1
A1 233926118 233926215
refGene_gn=MARCH6;refGene_tx=XM_023239515.1;refGene_gn=MARCH6;refGene_tx=XM_023239519.1
A1 233944081 233944100
refGene_gn=MARCH6;refGene_tx=XM_023239515.1;refGene_gn=MARCH6;refGene_tx=XM_023239519.1
A1 233997781 233997996
refGene_gn=CMBL;refGene_tx=XM_011287042.2;refGene_gn=CMBL;refGene_tx=XM_019812899.2;refGene_gn=CMBL;refGene_tx=XM_019812904.2;refGene_gn=CMBL;refGene_tx=XM_019812906.2;refGene_gn=CMBL;refGene_tx=XM_019812908.2
A1 233999153 233999261
refGene_gn=CMBL;refGene_tx=XM_011287042.2;refGene_gn=CMBL;refGene_tx=XM_019812899.2;refGene_gn=CMBL;refGene_tx=XM_019812904.2;refGene_gn=CMBL;refGene_tx=XM_019812906.2;refGene_gn=CMBL;refGene_tx=XM_019812908.2
A1 234003843 234003986
refGene_gn=CMBL;refGene_tx=XM_011287042.2;refGene_gn=CMBL;refGene_tx=X

M_019812899.2;refGene_gn=CMBL;refGene_tx=XM_019812904.2;refGene_gn=CMB
L;refGene_tx=XM_019812906.2;refGene_gn=CMBL;refGene_tx=XM_019812908.2
A1 234006038 234006130
refGene_gn=CMBL;refGene_tx=XM_011287042.2;refGene_gn=CMBL;refGene_tx=X
M_019812899.2;refGene_gn=CMBL;refGene_tx=XM_019812904.2;refGene_gn=CMB
L;refGene_tx=XM_019812906.2;refGene_gn=CMBL;refGene_tx=XM_019812908.2
A1 234007192 234007372
refGene_gn=CMBL;refGene_tx=XM_011287042.2;refGene_gn=CMBL;refGene_tx=X
M_019812899.2;refGene_gn=CMBL;refGene_tx=XM_019812904.2;refGene_gn=CMB
L;refGene_tx=XM_019812906.2;refGene_gn=CMBL;refGene_tx=XM_019812908.2
A1 234011951 234012079
refGene_gn=CCT5;refGene_tx=XM_003981555.5
A1 234013531 234013712
refGene_gn=CCT5;refGene_tx=XM_003981555.5
A1 234014189 234014327
refGene_gn=CCT5;refGene_tx=XM_003981555.5
A1 234014923 234015109
refGene_gn=CCT5;refGene_tx=XM_003981555.5
A1 234015701 234015821
refGene_gn=CCT5;refGene_tx=XM_003981555.5
A1 234017542 234017692
refGene_gn=CCT5;refGene_tx=XM_003981555.5
A1 234017772 234017965
refGene_gn=CCT5;refGene_tx=XM_003981555.5
A1 234019547 234019746
refGene_gn=CCT5;refGene_tx=XM_003981555.5
A1 234020974 234021139
refGene_gn=CCT5;refGene_tx=XM_003981555.5
A1 234021659 234021720
refGene_gn=CCT5;refGene_tx=XM_003981555.5
A1 234024694 234024799
refGene_gn=CCT5;refGene_tx=XM_003981555.5
A1 234025251 234025267
refGene_gn=FAM173B;refGene_tx=XM_006928110.4;refGene_gn=FAM173B;refGen
e_tx=XM_019812917.2
A1 234029456 234029743
refGene_gn=FAM173B;refGene_tx=XM_006928110.4;refGene_gn=FAM173B;refGen
e_tx=XM_019812917.2
A1 234032963 234033101
refGene_gn=FAM173B;refGene_tx=XM_006928110.4;refGene_gn=FAM173B;refGen
e_tx=XM_019812917.2
A1 234034477 234034528
refGene_gn=FAM173B;refGene_tx=XM_006928110.4;refGene_gn=FAM173B;refGen
e_tx=XM_019812917.2
A1 234041347 234041512
refGene_gn=FAM173B;refGene_tx=XM_006928110.4
A1 234162249 234162510
refGene_gn=FAM173B;refGene_tx=XM_019812917.2
A1 234525304 234526201
refGene_gn=TAS2R1;refGene_tx=XM_003981557.2

A1 234772194 234772315
refGene_gn=SEMA5A;refGene_tx=XM_023239573.1;refGene_gn=SEMA5A;refGene_tx=XM_023239575.1;refGene_gn=SEMA5A;refGene_tx=XM_023239577.1;refGene_gn=SEMA5A;refGene_tx=XM_023239585.1;refGene_gn=SEMA5A;refGene_tx=XM_023239592.1;refGene_gn=SEMA5A;refGene_tx=XM_023239595.1;refGene_gn=SEMA5A;refGene_tx=XM_023239598.1;refGene_gn=SEMA5A;refGene_tx=XM_023239601.1;refGene_gn=SEMA5A;refGene_tx=XM_023239602.1
A1 234818611 234818711
refGene_gn=SEMA5A;refGene_tx=XM_023239573.1;refGene_gn=SEMA5A;refGene_tx=XM_023239575.1;refGene_gn=SEMA5A;refGene_tx=XM_023239577.1;refGene_gn=SEMA5A;refGene_tx=XM_023239585.1;refGene_gn=SEMA5A;refGene_tx=XM_023239592.1;refGene_gn=SEMA5A;refGene_tx=XM_023239595.1;refGene_gn=SEMA5A;refGene_tx=XM_023239598.1;refGene_gn=SEMA5A;refGene_tx=XM_023239601.1;refGene_gn=SEMA5A;refGene_tx=XM_023239602.1
A1 234834738 234834784
refGene_gn=SEMA5A;refGene_tx=XM_023239573.1;refGene_gn=SEMA5A;refGene_tx=XM_023239575.1;refGene_gn=SEMA5A;refGene_tx=XM_023239577.1;refGene_gn=SEMA5A;refGene_tx=XM_023239585.1;refGene_gn=SEMA5A;refGene_tx=XM_023239592.1;refGene_gn=SEMA5A;refGene_tx=XM_023239595.1;refGene_gn=SEMA5A;refGene_tx=XM_023239598.1;refGene_gn=SEMA5A;refGene_tx=XM_023239601.1;refGene_gn=SEMA5A;refGene_tx=XM_023239602.1
A1 234902929 234902992
refGene_gn=SEMA5A;refGene_tx=XM_023239573.1;refGene_gn=SEMA5A;refGene_tx=XM_023239575.1;refGene_gn=SEMA5A;refGene_tx=XM_023239577.1;refGene_gn=SEMA5A;refGene_tx=XM_023239585.1;refGene_gn=SEMA5A;refGene_tx=XM_023239592.1;refGene_gn=SEMA5A;refGene_tx=XM_023239595.1;refGene_gn=SEMA5A;refGene_tx=XM_023239598.1;refGene_gn=SEMA5A;refGene_tx=XM_023239601.1;refGene_gn=SEMA5A;refGene_tx=XM_023239602.1
A1 234913377 234913476
refGene_gn=SEMA5A;refGene_tx=XM_023239573.1;refGene_gn=SEMA5A;refGene_tx=XM_023239575.1;refGene_gn=SEMA5A;refGene_tx=XM_023239577.1;refGene_gn=SEMA5A;refGene_tx=XM_023239585.1;refGene_gn=SEMA5A;refGene_tx=XM_023239592.1;refGene_gn=SEMA5A;refGene_tx=XM_023239595.1;refGene_gn=SEMA5A;refGene_tx=XM_023239598.1;refGene_gn=SEMA5A;refGene_tx=XM_023239601.1;refGene_gn=SEMA5A;refGene_tx=XM_023239602.1
A1 234914908 234915122
refGene_gn=SEMA5A;refGene_tx=XM_023239573.1;refGene_gn=SEMA5A;refGene_tx=XM_023239575.1;refGene_gn=SEMA5A;refGene_tx=XM_023239577.1;refGene_gn=SEMA5A;refGene_tx=XM_023239585.1;refGene_gn=SEMA5A;refGene_tx=XM_023239592.1;refGene_gn=SEMA5A;refGene_tx=XM_023239595.1;refGene_gn=SEMA5A;refGene_tx=XM_023239598.1;refGene_gn=SEMA5A;refGene_tx=XM_023239601.1;refGene_gn=SEMA5A;refGene_tx=XM_023239602.1
A1 234932756 234933042
refGene_gn=SEMA5A;refGene_tx=XM_023239573.1;refGene_gn=SEMA5A;refGene_tx=XM_023239575.1;refGene_gn=SEMA5A;refGene_tx=XM_023239577.1;refGene_gn=SEMA5A;refGene_tx=XM_023239585.1;refGene_gn=SEMA5A;refGene_tx=XM_023239592.1;refGene_gn=SEMA5A;refGene_tx=XM_023239595.1;refGene_gn=SEMA5A;refGene_tx=XM_023239598.1;refGene_gn=SEMA5A;refGene_tx=XM_023239601.1;refGene_gn=SEMA5A;refGene_tx=XM_023239602.1
A1 234936155 234936291

refGene_gn=SEMA5A;refGene_tx=XM_023239573.1;refGene_gn=SEMA5A;refGene_
tx=XM_023239575.1;refGene_gn=SEMA5A;refGene_tx=XM_023239577.1;refGene_
gn=SEMA5A;refGene_tx=XM_023239585.1;refGene_gn=SEMA5A;refGene_tx=XM_02
3239592.1;refGene_gn=SEMA5A;refGene_tx=XM_023239595.1;refGene_gn=SEMA5
A;refGene_tx=XM_023239598.1;refGene_gn=SEMA5A;refGene_tx=XM_023239601.
1;refGene_gn=SEMA5A;refGene_tx=XM_023239602.1
A1 234943342 234943547
refGene_gn=SEMA5A;refGene_tx=XM_023239573.1;refGene_gn=SEMA5A;refGene_
tx=XM_023239575.1;refGene_gn=SEMA5A;refGene_tx=XM_023239577.1;refGene_
gn=SEMA5A;refGene_tx=XM_023239585.1;refGene_gn=SEMA5A;refGene_tx=XM_02
3239592.1;refGene_gn=SEMA5A;refGene_tx=XM_023239595.1;refGene_gn=SEMA5
A;refGene_tx=XM_023239598.1;refGene_gn=SEMA5A;refGene_tx=XM_023239601.
1;refGene_gn=SEMA5A;refGene_tx=XM_023239602.1
A1 234974784 234974992
refGene_gn=SEMA5A;refGene_tx=XM_023239573.1;refGene_gn=SEMA5A;refGene_
tx=XM_023239575.1;refGene_gn=SEMA5A;refGene_tx=XM_023239577.1;refGene_
gn=SEMA5A;refGene_tx=XM_023239585.1;refGene_gn=SEMA5A;refGene_tx=XM_02
3239592.1;refGene_gn=SEMA5A;refGene_tx=XM_023239595.1;refGene_gn=SEMA5
A;refGene_tx=XM_023239598.1;refGene_gn=SEMA5A;refGene_tx=XM_023239601.
1;refGene_gn=SEMA5A;refGene_tx=XM_023239602.1
A1 234992594 234992712
refGene_gn=SEMA5A;refGene_tx=XM_023239573.1;refGene_gn=SEMA5A;refGene_
tx=XM_023239575.1;refGene_gn=SEMA5A;refGene_tx=XM_023239577.1;refGene_
gn=SEMA5A;refGene_tx=XM_023239585.1;refGene_gn=SEMA5A;refGene_tx=XM_02
3239592.1;refGene_gn=SEMA5A;refGene_tx=XM_023239595.1;refGene_gn=SEMA5
A;refGene_tx=XM_023239598.1;refGene_gn=SEMA5A;refGene_tx=XM_023239601.
1;refGene_gn=SEMA5A;refGene_tx=XM_023239602.1
A1 235003220 235003402
refGene_gn=SEMA5A;refGene_tx=XM_023239573.1;refGene_gn=SEMA5A;refGene_
tx=XM_023239575.1;refGene_gn=SEMA5A;refGene_tx=XM_023239577.1;refGene_
gn=SEMA5A;refGene_tx=XM_023239585.1;refGene_gn=SEMA5A;refGene_tx=XM_02
3239592.1;refGene_gn=SEMA5A;refGene_tx=XM_023239595.1;refGene_gn=SEMA5
A;refGene_tx=XM_023239598.1;refGene_gn=SEMA5A;refGene_tx=XM_023239601.
1;refGene_gn=SEMA5A;refGene_tx=XM_023239602.1
A1 235010113 235010257
refGene_gn=SEMA5A;refGene_tx=XM_023239573.1;refGene_gn=SEMA5A;refGene_
tx=XM_023239575.1;refGene_gn=SEMA5A;refGene_tx=XM_023239577.1;refGene_
gn=SEMA5A;refGene_tx=XM_023239585.1;refGene_gn=SEMA5A;refGene_tx=XM_02
3239592.1;refGene_gn=SEMA5A;refGene_tx=XM_023239595.1;refGene_gn=SEMA5
A;refGene_tx=XM_023239598.1;refGene_gn=SEMA5A;refGene_tx=XM_023239601.
1;refGene_gn=SEMA5A;refGene_tx=XM_023239602.1
A1 235018807 235018825
refGene_gn=SEMA5A;refGene_tx=XM_023239573.1;refGene_gn=SEMA5A;refGene_
tx=XM_023239575.1;refGene_gn=SEMA5A;refGene_tx=XM_023239577.1;refGene_
gn=SEMA5A;refGene_tx=XM_023239585.1;refGene_gn=SEMA5A;refGene_tx=XM_02
3239592.1;refGene_gn=SEMA5A;refGene_tx=XM_023239595.1;refGene_gn=SEMA5
A;refGene_tx=XM_023239598.1;refGene_gn=SEMA5A;refGene_tx=XM_023239601.
1
A1 235019058 235019206
refGene_gn=SEMA5A;refGene_tx=XM_023239573.1;refGene_gn=SEMA5A;refGene_

tx=XM_023239575.1;refGene_gn=SEMA5A;refGene_tx=XM_023239577.1;refGene_gn=SEMA5A;refGene_tx=XM_023239585.1;refGene_gn=SEMA5A;refGene_tx=XM_023239592.1;refGene_gn=SEMA5A;refGene_tx=XM_023239595.1;refGene_gn=SEMA5A;refGene_tx=XM_023239598.1;refGene_gn=SEMA5A;refGene_tx=XM_023239601.1;refGene_gn=SEMA5A;refGene_tx=XM_023239602.1
A1 235059026 235059252
refGene_gn=SEMA5A;refGene_tx=XM_023239573.1;refGene_gn=SEMA5A;refGene_tx=XM_023239575.1;refGene_gn=SEMA5A;refGene_tx=XM_023239577.1;refGene_gn=SEMA5A;refGene_tx=XM_023239585.1;refGene_gn=SEMA5A;refGene_tx=XM_023239592.1;refGene_gn=SEMA5A;refGene_tx=XM_023239595.1;refGene_gn=SEMA5A;refGene_tx=XM_023239598.1;refGene_gn=SEMA5A;refGene_tx=XM_023239601.1;refGene_gn=SEMA5A;refGene_tx=XM_023239602.1
A1 235062510 235062729
refGene_gn=SEMA5A;refGene_tx=XM_023239573.1;refGene_gn=SEMA5A;refGene_tx=XM_023239575.1;refGene_gn=SEMA5A;refGene_tx=XM_023239577.1;refGene_gn=SEMA5A;refGene_tx=XM_023239585.1;refGene_gn=SEMA5A;refGene_tx=XM_023239592.1;refGene_gn=SEMA5A;refGene_tx=XM_023239595.1;refGene_gn=SEMA5A;refGene_tx=XM_023239598.1;refGene_gn=SEMA5A;refGene_tx=XM_023239601.1;refGene_gn=SEMA5A;refGene_tx=XM_023239602.1
A1 235070137 235070308
refGene_gn=SEMA5A;refGene_tx=XM_023239573.1;refGene_gn=SEMA5A;refGene_tx=XM_023239575.1;refGene_gn=SEMA5A;refGene_tx=XM_023239577.1;refGene_gn=SEMA5A;refGene_tx=XM_023239585.1;refGene_gn=SEMA5A;refGene_tx=XM_023239592.1;refGene_gn=SEMA5A;refGene_tx=XM_023239595.1;refGene_gn=SEMA5A;refGene_tx=XM_023239598.1;refGene_gn=SEMA5A;refGene_tx=XM_023239601.1;refGene_gn=SEMA5A;refGene_tx=XM_023239602.1
A1 235071829 235071985
refGene_gn=SEMA5A;refGene_tx=XM_023239573.1;refGene_gn=SEMA5A;refGene_tx=XM_023239575.1;refGene_gn=SEMA5A;refGene_tx=XM_023239577.1;refGene_gn=SEMA5A;refGene_tx=XM_023239585.1;refGene_gn=SEMA5A;refGene_tx=XM_023239592.1;refGene_gn=SEMA5A;refGene_tx=XM_023239595.1;refGene_gn=SEMA5A;refGene_tx=XM_023239598.1;refGene_gn=SEMA5A;refGene_tx=XM_023239601.1;refGene_gn=SEMA5A;refGene_tx=XM_023239602.1
A1 235073404 235073452
refGene_gn=SEMA5A;refGene_tx=XM_023239573.1;refGene_gn=SEMA5A;refGene_tx=XM_023239575.1;refGene_gn=SEMA5A;refGene_tx=XM_023239577.1;refGene_gn=SEMA5A;refGene_tx=XM_023239585.1;refGene_gn=SEMA5A;refGene_tx=XM_023239592.1;refGene_gn=SEMA5A;refGene_tx=XM_023239595.1;refGene_gn=SEMA5A;refGene_tx=XM_023239598.1;refGene_gn=SEMA5A;refGene_tx=XM_023239601.1;refGene_gn=SEMA5A;refGene_tx=XM_023239602.1
A1 235077835 235078047
refGene_gn=SEMA5A;refGene_tx=XM_023239573.1;refGene_gn=SEMA5A;refGene_tx=XM_023239575.1;refGene_gn=SEMA5A;refGene_tx=XM_023239577.1;refGene_gn=SEMA5A;refGene_tx=XM_023239585.1;refGene_gn=SEMA5A;refGene_tx=XM_023239592.1;refGene_gn=SEMA5A;refGene_tx=XM_023239595.1;refGene_gn=SEMA5A;refGene_tx=XM_023239598.1;refGene_gn=SEMA5A;refGene_tx=XM_023239601.1;refGene_gn=SEMA5A;refGene_tx=XM_023239602.1
A1 235080714 235080834
refGene_gn=SEMA5A;refGene_tx=XM_023239573.1;refGene_gn=SEMA5A;refGene_tx=XM_023239575.1;refGene_gn=SEMA5A;refGene_tx=XM_023239577.1;refGene_

gn=SEMA5A;refGene_tx=XM_023239585.1;refGene_gn=SEMA5A;refGene_tx=XM_02
3239592.1;refGene_gn=SEMA5A;refGene_tx=XM_023239595.1;refGene_gn=SEMA5
A;refGene_tx=XM_023239598.1;refGene_gn=SEMA5A;refGene_tx=XM_023239601.
1;refGene_gn=SEMA5A;refGene_tx=XM_023239602.1
A1 236035165 236035310
refGene_gn=MTRR;refGene_tx=XM_006928111.4;refGene_gn=MTRR;refGene_tx=X
M_019812938.2;refGene_gn=MTRR;refGene_tx=XM_019812941.2;refGene_gn=MTR
R;refGene_tx=XM_019812943.2
A1 236038199 236038382
refGene_gn=MTRR;refGene_tx=XM_006928111.4;refGene_gn=MTRR;refGene_tx=X
M_019812938.2;refGene_gn=MTRR;refGene_tx=XM_019812941.2
A1 236038467 236038560
refGene_gn=MTRR;refGene_tx=XM_006928111.4;refGene_gn=MTRR;refGene_tx=X
M_019812938.2;refGene_gn=MTRR;refGene_tx=XM_019812941.2;refGene_gn=MTR
R;refGene_tx=XM_019812943.2
A1 236039841 236039960
refGene_gn=MTRR;refGene_tx=XM_006928111.4;refGene_gn=MTRR;refGene_tx=X
M_019812938.2;refGene_gn=MTRR;refGene_tx=XM_019812941.2;refGene_gn=MTR
R;refGene_tx=XM_019812943.2
A1 236043142 236043347
refGene_gn=MTRR;refGene_tx=XM_006928111.4;refGene_gn=MTRR;refGene_tx=X
M_019812938.2;refGene_gn=MTRR;refGene_tx=XM_019812941.2;refGene_gn=MTR
R;refGene_tx=XM_019812943.2
A1 236043871 236043914
refGene_gn=MTRR;refGene_tx=XM_006928111.4;refGene_gn=MTRR;refGene_tx=X
M_019812938.2;refGene_gn=MTRR;refGene_tx=XM_019812941.2;refGene_gn=MTR
R;refGene_tx=XM_019812943.2
A1 236048762 236048940
refGene_gn=MTRR;refGene_tx=XM_006928111.4;refGene_gn=MTRR;refGene_tx=X
M_019812938.2;refGene_gn=MTRR;refGene_tx=XM_019812941.2;refGene_gn=MTR
R;refGene_tx=XM_019812943.2
A1 236051133 236051222
refGene_gn=MTRR;refGene_tx=XM_006928111.4;refGene_gn=MTRR;refGene_tx=X
M_019812938.2;refGene_gn=MTRR;refGene_tx=XM_019812941.2;refGene_gn=MTR
R;refGene_tx=XM_019812943.2
A1 236052098 236052252
refGene_gn=MTRR;refGene_tx=XM_006928111.4;refGene_gn=MTRR;refGene_tx=X
M_019812938.2;refGene_gn=MTRR;refGene_tx=XM_019812941.2;refGene_gn=MTR
R;refGene_tx=XM_019812943.2
A1 236053845 236053968
refGene_gn=MTRR;refGene_tx=XM_006928111.4;refGene_gn=MTRR;refGene_tx=X
M_019812938.2;refGene_gn=MTRR;refGene_tx=XM_019812941.2;refGene_gn=MTR
R;refGene_tx=XM_019812943.2
A1 236058377 236058756
refGene_gn=MTRR;refGene_tx=XM_006928111.4;refGene_gn=MTRR;refGene_tx=X
M_019812938.2;refGene_gn=MTRR;refGene_tx=XM_019812941.2;refGene_gn=MTR
R;refGene_tx=XM_019812943.2
A1 236061501 236061619
refGene_gn=MTRR;refGene_tx=XM_006928111.4;refGene_gn=MTRR;refGene_tx=X
M_019812938.2;refGene_gn=MTRR;refGene_tx=XM_019812941.2;refGene_gn=MTR

R;refGene_tx=XM_019812943.2
A1 236063602 236063756
refGene_gn=MTRR;refGene_tx=XM_006928111.4;refGene_gn=MTRR;refGene_tx=XM_019812938.2;refGene_gn=MTRR;refGene_tx=XM_019812941.2;refGene_gn=MTRR;refGene_tx=XM_019812943.2
A1 236065714 236065843
refGene_gn=MTRR;refGene_tx=XM_006928111.4;refGene_gn=MTRR;refGene_tx=XM_019812938.2;refGene_gn=MTRR;refGene_tx=XM_019812941.2;refGene_gn=MTRR;refGene_tx=XM_019812943.2
A1 236068840 236069884
refGene_gn=FASTKD3;refGene_tx=XM_006928112.4
A1 236069885 236070107
refGene_gn=FASTKD3;refGene_tx=XM_006928112.4
A1 236070108 236070280
refGene_gn=FASTKD3;refGene_tx=XM_006928112.4
A1 236071371 236071457
refGene_gn=FASTKD3;refGene_tx=XM_006928112.4
A1 236074004 236074179
refGene_gn=FASTKD3;refGene_tx=XM_006928112.4
A1 236075342 236075412
refGene_gn=FASTKD3;refGene_tx=XM_006928112.4
A1 236076394 236076509
refGene_gn=FASTKD3;refGene_tx=XM_006928112.4
A1 236077555 236077654
refGene_gn=FASTKD3;refGene_tx=XM_006928112.4
A1 236090246 236090393
refGene_gn=CA1H5orf49;refGene_tx=XM_023239643.1;refGene_gn=CA1H5orf49;refGene_tx=XM_023239645.1
A1 236091377 236091752
refGene_gn=CA1H5orf49;refGene_tx=XM_023239643.1
A1 236103099 236103216
refGene_gn=CA1H5orf49;refGene_tx=XM_023239645.1
A1 236104852 236105032
refGene_gn=CA1H5orf49;refGene_tx=XM_023239645.1
A1 236109173 236109326
refGene_gn=ADCY2;refGene_tx=XM_023239649.1;refGene_gn=ADCY2;refGene_tx=XM_023239653.1;refGene_gn=ADCY2;refGene_tx=XM_023239655.1;refGene_gn=ADCY2;refGene_tx=XM_023239661.1
A1 236113759 236113884
refGene_gn=ADCY2;refGene_tx=XM_023239649.1;refGene_gn=ADCY2;refGene_tx=XM_023239653.1;refGene_gn=ADCY2;refGene_tx=XM_023239655.1;refGene_gn=ADCY2;refGene_tx=XM_023239661.1
A1 236118261 236118376
refGene_gn=ADCY2;refGene_tx=XM_023239649.1;refGene_gn=ADCY2;refGene_tx=XM_023239653.1;refGene_gn=ADCY2;refGene_tx=XM_023239655.1;refGene_gn=ADCY2;refGene_tx=XM_023239661.1
A1 236131869 236131977
refGene_gn=ADCY2;refGene_tx=XM_023239649.1;refGene_gn=ADCY2;refGene_tx=XM_023239653.1;refGene_gn=ADCY2;refGene_tx=XM_023239655.1;refGene_gn=ADCY2;refGene_tx=XM_023239661.1

A1 236133750 236133897
refGene_gn=ADCY2;refGene_tx=XM_023239649.1;refGene_gn=ADCY2;refGene_tx
=XM_023239653.1;refGene_gn=ADCY2;refGene_tx=XM_023239655.1;refGene_gn=
ADCY2;refGene_tx=XM_023239662.1
A1 236142753 236142912
refGene_gn=ADCY2;refGene_tx=XM_023239649.1;refGene_gn=ADCY2;refGene_tx
=XM_023239653.1;refGene_gn=ADCY2;refGene_tx=XM_023239655.1;refGene_gn=
ADCY2;refGene_tx=XM_023239662.1
A1 236147781 236147866
refGene_gn=ADCY2;refGene_tx=XM_023239649.1;refGene_gn=ADCY2;refGene_tx
=XM_023239655.1;refGene_gn=ADCY2;refGene_tx=XM_023239661.1;refGene_gn=
ADCY2;refGene_tx=XM_023239662.1
A1 236159773 236159943
refGene_gn=ADCY2;refGene_tx=XM_023239649.1;refGene_gn=ADCY2;refGene_tx
=XM_023239653.1;refGene_gn=ADCY2;refGene_tx=XM_023239655.1;refGene_gn=
ADCY2;refGene_tx=XM_023239661.1;refGene_gn=ADCY2;refGene_tx=XM_0232396
62.1
A1 236166413 236166533
refGene_gn=ADCY2;refGene_tx=XM_023239649.1;refGene_gn=ADCY2;refGene_tx
=XM_023239653.1;refGene_gn=ADCY2;refGene_tx=XM_023239655.1;refGene_gn=
ADCY2;refGene_tx=XM_023239661.1;refGene_gn=ADCY2;refGene_tx=XM_0232396
62.1
A1 236175280 236175418
refGene_gn=ADCY2;refGene_tx=XM_023239649.1;refGene_gn=ADCY2;refGene_tx
=XM_023239653.1;refGene_gn=ADCY2;refGene_tx=XM_023239655.1;refGene_gn=
ADCY2;refGene_tx=XM_023239661.1;refGene_gn=ADCY2;refGene_tx=XM_0232396
62.1
A1 236188708 236188793
refGene_gn=ADCY2;refGene_tx=XM_023239649.1;refGene_gn=ADCY2;refGene_tx
=XM_023239653.1;refGene_gn=ADCY2;refGene_tx=XM_023239655.1;refGene_gn=
ADCY2;refGene_tx=XM_023239661.1;refGene_gn=ADCY2;refGene_tx=XM_0232396
62.1
A1 236207083 236207181
refGene_gn=ADCY2;refGene_tx=XM_023239649.1;refGene_gn=ADCY2;refGene_tx
=XM_023239653.1;refGene_gn=ADCY2;refGene_tx=XM_023239655.1;refGene_gn=
ADCY2;refGene_tx=XM_023239661.1;refGene_gn=ADCY2;refGene_tx=XM_0232396
62.1
A1 236210257 236210327
refGene_gn=ADCY2;refGene_tx=XM_023239649.1;refGene_gn=ADCY2;refGene_tx
=XM_023239653.1;refGene_gn=ADCY2;refGene_tx=XM_023239655.1;refGene_gn=
ADCY2;refGene_tx=XM_023239661.1;refGene_gn=ADCY2;refGene_tx=XM_0232396
62.1
A1 236216943 236217024
refGene_gn=ADCY2;refGene_tx=XM_023239649.1;refGene_gn=ADCY2;refGene_tx
=XM_023239653.1;refGene_gn=ADCY2;refGene_tx=XM_023239655.1;refGene_gn=
ADCY2;refGene_tx=XM_023239661.1;refGene_gn=ADCY2;refGene_tx=XM_0232396
62.1
A1 236220533 236220577
refGene_gn=ADCY2;refGene_tx=XM_023239649.1;refGene_gn=ADCY2;refGene_tx
=XM_023239653.1;refGene_gn=ADCY2;refGene_tx=XM_023239655.1;refGene_gn=

ADCY2;refGene_tx=XM_023239661.1;refGene_gn=ADCY2;refGene_tx=XM_0232396
62.1
A1 236223053 236223227
refGene_gn=ADCY2;refGene_tx=XM_023239649.1;refGene_gn=ADCY2;refGene_tx
=XM_023239653.1;refGene_gn=ADCY2;refGene_tx=XM_023239655.1;refGene_gn=
ADCY2;refGene_tx=XM_023239661.1;refGene_gn=ADCY2;refGene_tx=XM_0232396
62.1
A1 236224598 236224731
refGene_gn=ADCY2;refGene_tx=XM_023239649.1;refGene_gn=ADCY2;refGene_tx
=XM_023239653.1;refGene_gn=ADCY2;refGene_tx=XM_023239655.1;refGene_gn=
ADCY2;refGene_tx=XM_023239661.1;refGene_gn=ADCY2;refGene_tx=XM_0232396
62.1
A1 236225574 236225733
refGene_gn=ADCY2;refGene_tx=XM_023239649.1;refGene_gn=ADCY2;refGene_tx
=XM_023239653.1;refGene_gn=ADCY2;refGene_tx=XM_023239655.1;refGene_gn=
ADCY2;refGene_tx=XM_023239661.1;refGene_gn=ADCY2;refGene_tx=XM_0232396
62.1
A1 236231493 236231621
refGene_gn=ADCY2;refGene_tx=XM_023239649.1;refGene_gn=ADCY2;refGene_tx
=XM_023239653.1;refGene_gn=ADCY2;refGene_tx=XM_023239655.1;refGene_gn=
ADCY2;refGene_tx=XM_023239661.1;refGene_gn=ADCY2;refGene_tx=XM_0232396
62.1
A1 236237702 236237814
refGene_gn=ADCY2;refGene_tx=XM_023239649.1;refGene_gn=ADCY2;refGene_tx
=XM_023239653.1;refGene_gn=ADCY2;refGene_tx=XM_023239655.1;refGene_gn=
ADCY2;refGene_tx=XM_023239661.1;refGene_gn=ADCY2;refGene_tx=XM_0232396
62.1
A1 236241068 236241217
refGene_gn=ADCY2;refGene_tx=XM_023239649.1;refGene_gn=ADCY2;refGene_tx
=XM_023239653.1;refGene_gn=ADCY2;refGene_tx=XM_023239655.1;refGene_gn=
ADCY2;refGene_tx=XM_023239661.1;refGene_gn=ADCY2;refGene_tx=XM_0232396
62.1
A1 236311634 236311784
refGene_gn=ADCY2;refGene_tx=XM_023239649.1;refGene_gn=ADCY2;refGene_tx
=XM_023239653.1;refGene_gn=ADCY2;refGene_tx=XM_023239661.1;refGene_gn=
ADCY2;refGene_tx=XM_023239662.1
A1 236407937 236408099
refGene_gn=ADCY2;refGene_tx=XM_023239649.1;refGene_gn=ADCY2;refGene_tx
=XM_023239653.1;refGene_gn=ADCY2;refGene_tx=XM_023239655.1;refGene_gn=
ADCY2;refGene_tx=XM_023239661.1;refGene_gn=ADCY2;refGene_tx=XM_0232396
62.1
A1 236500908 236501106
refGene_gn=ADCY2;refGene_tx=XM_023239649.1;refGene_gn=ADCY2;refGene_tx
=XM_023239653.1;refGene_gn=ADCY2;refGene_tx=XM_023239655.1;refGene_gn=
ADCY2;refGene_tx=XM_023239661.1;refGene_gn=ADCY2;refGene_tx=XM_0232396
62.1
A1 236523805 236524036
refGene_gn=ADCY2;refGene_tx=XM_023239649.1;refGene_gn=ADCY2;refGene_tx
=XM_023239653.1;refGene_gn=ADCY2;refGene_tx=XM_023239655.1;refGene_gn=
ADCY2;refGene_tx=XM_023239661.1;refGene_gn=ADCY2;refGene_tx=XM_0232396

62.1
A1 237056415 237056610
refGene_gn=PAPD7;refGene_tx=XM_023239675.1;refGene_gn=PAPD7;refGene_tx=XM_023239676.1
A1 237058447 237058612
refGene_gn=PAPD7;refGene_tx=XM_023239675.1;refGene_gn=PAPD7;refGene_tx=XM_023239676.1
A1 237060889 237061048
refGene_gn=PAPD7;refGene_tx=XM_023239675.1;refGene_gn=PAPD7;refGene_tx=XM_023239676.1
A1 237061460 237061633
refGene_gn=PAPD7;refGene_tx=XM_023239675.1;refGene_gn=PAPD7;refGene_tx=XM_023239676.1
A1 237062235 237062336
refGene_gn=PAPD7;refGene_tx=XM_023239675.1;refGene_gn=PAPD7;refGene_tx=XM_023239676.1
A1 237063360 237063487
refGene_gn=PAPD7;refGene_tx=XM_023239675.1;refGene_gn=PAPD7;refGene_tx=XM_023239676.1
A1 237065644 237065858
refGene_gn=PAPD7;refGene_tx=XM_023239675.1;refGene_gn=PAPD7;refGene_tx=XM_023239676.1
A1 237068253 237068382
refGene_gn=PAPD7;refGene_tx=XM_023239675.1;refGene_gn=PAPD7;refGene_tx=XM_023239676.1
A1 237069543 237069651
refGene_gn=PAPD7;refGene_tx=XM_023239675.1;refGene_gn=PAPD7;refGene_tx=XM_023239676.1
A1 237071852 237071973
refGene_gn=PAPD7;refGene_tx=XM_023239675.1;refGene_gn=PAPD7;refGene_tx=XM_023239676.1
A1 237072898 237072945
refGene_gn=PAPD7;refGene_tx=XM_023239675.1;refGene_gn=PAPD7;refGene_tx=XM_023239676.1
A1 237073496 237073586
refGene_gn=PAPD7;refGene_tx=XM_023239675.1;refGene_gn=PAPD7;refGene_tx=XM_023239676.1
A1 237160748 237160815
refGene_gn=SRD5A1;refGene_tx=XM_023239691.1
A1 237166770 237166921
refGene_gn=SRD5A1;refGene_tx=XM_023239691.1
A1 237168374 237168476
refGene_gn=SRD5A1;refGene_tx=XM_023239691.1
A1 237172335 237172502
refGene_gn=SRD5A1;refGene_tx=XM_023239691.1
A1 237183562 237183843
refGene_gn=SRD5A1;refGene_tx=XM_023239691.1
A1 237184356 237184446
refGene_gn=NSUN2;refGene_tx=XM_023239687.1;refGene_gn=NSUN2;refGene_tx=XM_023239689.1

A1 237184563 237184721
refGene_gn=NSUN2;refGene_tx=XM_023239687.1;refGene_gn=NSUN2;refGene_tx=XM_023239689.1
A1 237185315 237185420
refGene_gn=NSUN2;refGene_tx=XM_023239687.1;refGene_gn=NSUN2;refGene_tx=XM_023239689.1
A1 237190739 237190845
refGene_gn=NSUN2;refGene_tx=XM_023239687.1;refGene_gn=NSUN2;refGene_tx=XM_023239689.1
A1 237193139 237193211
refGene_gn=NSUN2;refGene_tx=XM_023239687.1;refGene_gn=NSUN2;refGene_tx=XM_023239689.1
A1 237193951 237194036
refGene_gn=NSUN2;refGene_tx=XM_023239687.1;refGene_gn=NSUN2;refGene_tx=XM_023239689.1
A1 237195148 237195341
refGene_gn=NSUN2;refGene_tx=XM_023239687.1;refGene_gn=NSUN2;refGene_tx=XM_023239689.1
A1 237197440 237197515
refGene_gn=NSUN2;refGene_tx=XM_023239687.1;refGene_gn=NSUN2;refGene_tx=XM_023239689.1
A1 237198585 237198716
refGene_gn=NSUN2;refGene_tx=XM_023239687.1;refGene_gn=NSUN2;refGene_tx=XM_023239689.1
A1 237200797 237200871
refGene_gn=NSUN2;refGene_tx=XM_023239687.1;refGene_gn=NSUN2;refGene_tx=XM_023239689.1
A1 237201463 237201594
refGene_gn=NSUN2;refGene_tx=XM_023239687.1;refGene_gn=NSUN2;refGene_tx=XM_023239689.1
A1 237203632 237203729
refGene_gn=NSUN2;refGene_tx=XM_023239687.1;refGene_gn=NSUN2;refGene_tx=XM_023239689.1
A1 237208201 237208353
refGene_gn=NSUN2;refGene_tx=XM_023239687.1;refGene_gn=NSUN2;refGene_tx=XM_023239689.1
A1 237208606 237208699
refGene_gn=NSUN2;refGene_tx=XM_023239687.1;refGene_gn=NSUN2;refGene_tx=XM_023239689.1
A1 237209453 237209589
refGene_gn=NSUN2;refGene_tx=XM_023239687.1;refGene_gn=NSUN2;refGene_tx=XM_023239689.1
A1 237210269 237210350
refGene_gn=NSUN2;refGene_tx=XM_023239687.1;refGene_gn=NSUN2;refGene_tx=XM_023239689.1
A1 237210615 237210754
refGene_gn=NSUN2;refGene_tx=XM_023239687.1;refGene_gn=NSUN2;refGene_tx=XM_023239689.1
A1 237211568 237211816
refGene_gn=NSUN2;refGene_tx=XM_023239687.1

A1 237211854 237211894
refGene_gn=NSUN2;refGene_tx=XM_023239689.1
A1 237215049 237215341
refGene_gn=NSUN2;refGene_tx=XM_023239689.1
A1 237275037 237275169
refGene_gn=UBE2QL1;refGene_tx=XM_011287320.3
A1 237312081 237312879
refGene_gn=UBE2QL1;refGene_tx=XM_011287320.3
A1 237360489 237360611
refGene_gn=MED10;refGene_tx=XM_023239698.1;refGene_gn=MED10;refGene_tx=XM_023239703.1;refGene_gn=MED10;refGene_tx=XM_023239705.1;refGene_gn=MED10;refGene_tx=XM_023239707.1
A1 237361936 237362020
refGene_gn=MED10;refGene_tx=XM_023239698.1;refGene_gn=MED10;refGene_tx=XM_023239703.1;refGene_gn=MED10;refGene_tx=XM_023239705.1;refGene_gn=MED10;refGene_tx=XM_023239707.1
A1 237364686 237364789
refGene_gn=MED10;refGene_tx=XM_023239698.1;refGene_gn=MED10;refGene_tx=XM_023239703.1;refGene_gn=MED10;refGene_tx=XM_023239705.1;refGene_gn=MED10;refGene_tx=XM_023239707.1
A1 237366935 237367034
refGene_gn=MED10;refGene_tx=XM_023239705.1
A1 237377582 237377726
refGene_gn=MED10;refGene_tx=XM_023239703.1
A1 237384559 237384580
refGene_gn=MED10;refGene_tx=XM_023239707.1
A1 237521513 237521621
refGene_gn=MED10;refGene_tx=XM_023239698.1
A1 237531609 237531834
refGene_gn=MED10;refGene_tx=XM_023239698.1
A1 238069148 238069551
refGene_gn=ICE1;refGene_tx=XM_023239718.1;refGene_gn=ICE1;refGene_tx=XM_023239720.1
A1 238070574 238070673
refGene_gn=ICE1;refGene_tx=XM_023239718.1;refGene_gn=ICE1;refGene_tx=XM_023239720.1
A1 238075642 238075701
refGene_gn=ICE1;refGene_tx=XM_023239723.1
A1 238076361 238076468
refGene_gn=ICE1;refGene_tx=XM_023239720.1;refGene_gn=ICE1;refGene_tx=XM_023239723.1
A1 238078723 238078914
refGene_gn=ICE1;refGene_tx=XM_023239718.1;refGene_gn=ICE1;refGene_tx=XM_023239720.1;refGene_gn=ICE1;refGene_tx=XM_023239723.1;refGene_gn=ICE1;refGene_tx=XM_023239724.1
A1 238081838 238081999
refGene_gn=ICE1;refGene_tx=XM_023239718.1;refGene_gn=ICE1;refGene_tx=XM_023239720.1;refGene_gn=ICE1;refGene_tx=XM_023239723.1
A1 238084591 238084760
refGene_gn=ICE1;refGene_tx=XM_023239718.1;refGene_gn=ICE1;refGene_tx=X

M_023239720.1;refGene_gn=ICE1;refGene_tx=XM_023239723.1;refGene_gn=ICE1;refGene_tx=XM_023239724.1
A1 238085899 238090663 refGene_gn=ICE1;refGene_tx=XM_023239718.1;refGene_gn=ICE1;refGene_tx=XM_023239720.1;refGene_gn=ICE1;refGene_tx=XM_023239723.1;refGene_gn=ICE1;refGene_tx=XM_023239724.1
A1 238092947 238093360 refGene_gn=ICE1;refGene_tx=XM_023239718.1;refGene_gn=ICE1;refGene_tx=XM_023239720.1;refGene_gn=ICE1;refGene_tx=XM_023239723.1;refGene_gn=ICE1;refGene_tx=XM_023239724.1
A1 238096072 238096159 refGene_gn=ICE1;refGene_tx=XM_023239718.1;refGene_gn=ICE1;refGene_tx=XM_023239720.1;refGene_gn=ICE1;refGene_tx=XM_023239723.1;refGene_gn=ICE1;refGene_tx=XM_023239724.1
A1 238105505 238105562 refGene_gn=ICE1;refGene_tx=XM_023239718.1;refGene_gn=ICE1;refGene_tx=XM_023239720.1;refGene_gn=ICE1;refGene_tx=XM_023239723.1;refGene_gn=ICE1;refGene_tx=XM_023239724.1
A1 238105647 238105687 refGene_gn=ICE1;refGene_tx=XM_023239718.1;refGene_gn=ICE1;refGene_tx=XM_023239720.1;refGene_gn=ICE1;refGene_tx=XM_023239723.1;refGene_gn=ICE1;refGene_tx=XM_023239724.1
A1 238105909 238105992 refGene_gn=ICE1;refGene_tx=XM_023239718.1;refGene_gn=ICE1;refGene_tx=XM_023239720.1;refGene_gn=ICE1;refGene_tx=XM_023239723.1;refGene_gn=ICE1;refGene_tx=XM_023239724.1
A1 238108118 238108156 refGene_gn=ICE1;refGene_tx=XM_023239718.1;refGene_gn=ICE1;refGene_tx=XM_023239720.1;refGene_gn=ICE1;refGene_tx=XM_023239723.1;refGene_gn=ICE1;refGene_tx=XM_023239724.1
A1 238109111 238109188 refGene_gn=ICE1;refGene_tx=XM_023239718.1;refGene_gn=ICE1;refGene_tx=XM_023239720.1;refGene_gn=ICE1;refGene_tx=XM_023239723.1;refGene_gn=ICE1;refGene_tx=XM_023239724.1
A1 238110695 238110807 refGene_gn=ICE1;refGene_tx=XM_023239718.1;refGene_gn=ICE1;refGene_tx=XM_023239720.1;refGene_gn=ICE1;refGene_tx=XM_023239723.1;refGene_gn=ICE1;refGene_tx=XM_023239724.1
A1 238111552 238111571 refGene_gn=ICE1;refGene_tx=XM_023239718.1;refGene_gn=ICE1;refGene_tx=XM_023239720.1;refGene_gn=ICE1;refGene_tx=XM_023239723.1;refGene_gn=ICE1;refGene_tx=XM_023239724.1
A1 238114003 238114038 refGene_gn=ICE1;refGene_tx=XM_023239718.1;refGene_gn=ICE1;refGene_tx=XM_023239720.1;refGene_gn=ICE1;refGene_tx=XM_023239723.1;refGene_gn=ICE1;refGene_tx=XM_023239724.1
A1 238114685 238114744 refGene_gn=ICE1;refGene_tx=XM_023239718.1;refGene_gn=ICE1;refGene_tx=XM_023239720.1;refGene_gn=ICE1;refGene_tx=XM_023239723.1;refGene_gn=ICE1;refGene_tx=XM_023239724.1

A1 238126990 238127074
refGene_gn=ICE1;refGene_tx=XM_023239718.1;refGene_gn=ICE1;refGene_tx=XM_023239720.1;refGene_gn=ICE1;refGene_tx=XM_023239723.1;refGene_gn=ICE1;refGene_tx=XM_023239724.1
A1 238128479 238128838
refGene_gn=LOC102900202;refGene_tx=XM_019813024.2;refGene_gn=LOC102900202;refGene_tx=XM_019813027.2;refGene_gn=LOC102900202;refGene_tx=XM_019813031.2;refGene_gn=LOC102900202;refGene_tx=XM_019813034.2;refGene_gn=LOC102900202;refGene_tx=XM_019813035.2;refGene_gn=LOC102900202;refGene_tx=XM_019813036.2;refGene_gn=LOC102900202;refGene_tx=XM_019813038.2
A1 238130013 238130119
refGene_gn=LOC102900202;refGene_tx=XM_019813024.2;refGene_gn=LOC102900202;refGene_tx=XM_019813027.2;refGene_gn=LOC102900202;refGene_tx=XM_019813031.2;refGene_gn=LOC102900202;refGene_tx=XM_019813034.2;refGene_gn=LOC102900202;refGene_tx=XM_019813035.2;refGene_gn=LOC102900202;refGene_tx=XM_019813036.2;refGene_gn=LOC102900202;refGene_tx=XM_019813038.2
A1 238131115 238131251
refGene_gn=LOC102900202;refGene_tx=XM_019813024.2;refGene_gn=LOC102900202;refGene_tx=XM_019813027.2;refGene_gn=LOC102900202;refGene_tx=XM_019813031.2;refGene_gn=LOC102900202;refGene_tx=XM_019813034.2;refGene_gn=LOC102900202;refGene_tx=XM_019813035.2;refGene_gn=LOC102900202;refGene_tx=XM_019813036.2;refGene_gn=LOC102900202;refGene_tx=XM_019813038.2
A1 238132676 238133648
refGene_gn=LOC102900202;refGene_tx=XM_019813024.2;refGene_gn=LOC102900202;refGene_tx=XM_019813027.2;refGene_gn=LOC102900202;refGene_tx=XM_019813031.2;refGene_gn=LOC102900202;refGene_tx=XM_019813034.2;refGene_gn=LOC102900202;refGene_tx=XM_019813035.2;refGene_gn=LOC102900202;refGene_tx=XM_019813036.2;refGene_gn=LOC102900202;refGene_tx=XM_019813038.2
A1 238209223 238209339
refGene_gn=ADAMTS16;refGene_tx=XM_023239735.1
A1 238209595 238209743
refGene_gn=ADAMTS16;refGene_tx=XM_023239735.1
A1 238217169 238217394
refGene_gn=ADAMTS16;refGene_tx=XM_023239735.1
A1 238219173 238219368
refGene_gn=ADAMTS16;refGene_tx=XM_023239735.1
A1 238219518 238219720
refGene_gn=ADAMTS16;refGene_tx=XM_023239735.1
A1 238252789 238252916
refGene_gn=ADAMTS16;refGene_tx=XM_023239735.1
A1 238273897 238274680
refGene_gn=LOC101094823;refGene_tx=XM_011287129.2
A1 238276965 238277104
refGene_gn=ADAMTS16;refGene_tx=XM_023239735.1
A1 238278536 238278781
refGene_gn=ADAMTS16;refGene_tx=XM_023239735.1
A1 238279629 238279753
refGene_gn=ADAMTS16;refGene_tx=XM_023239735.1
A1 238281942 238282073
refGene_gn=ADAMTS16;refGene_tx=XM_023239735.1

A1 238284065 238284238
refGene_gn=ADAMTS16;refGene_tx=XM_023239735.1
A1 238286096 238286245
refGene_gn=ADAMTS16;refGene_tx=XM_023239735.1
A1 238291378 238291474
refGene_gn=ADAMTS16;refGene_tx=XM_023239735.1
A1 238300957 238301111
refGene_gn=ADAMTS16;refGene_tx=XM_023239735.1
A1 238310156 238310294
refGene_gn=ADAMTS16;refGene_tx=XM_023239735.1
A1 238317906 238318012
refGene_gn=ADAMTS16;refGene_tx=XM_023239735.1
A1 238319089 238319249
refGene_gn=ADAMTS16;refGene_tx=XM_023239735.1
A1 238321717 238321801
refGene_gn=ADAMTS16;refGene_tx=XM_023239735.1
A1 238323361 238323561
refGene_gn=ADAMTS16;refGene_tx=XM_023239735.1
A1 238326633 238326892
refGene_gn=ADAMTS16;refGene_tx=XM_023239735.1
A1 238365417 238365740
refGene_gn=ADAMTS16;refGene_tx=XM_023239735.1
A1 238372109 238372215
refGene_gn=ADAMTS16;refGene_tx=XM_023239735.1
A1 238372301 238372361
refGene_gn=ADAMTS16;refGene_tx=XM_023239735.1
A1 239588258 239588316
refGene_gn=IRX1;refGene_tx=XM_023239744.1
A1 239588593 239588666
refGene_gn=IRX1;refGene_tx=XM_023239744.1
A1 239589043 239590076
refGene_gn=IRX1;refGene_tx=XM_023239744.1
A1 239593052 239593328
refGene_gn=IRX1;refGene_tx=XM_023239744.1
A1 239594765 239594852
refGene_gn=LOC111556665;refGene_tx=XM_023239749.1
A1 239595352 239595464
refGene_gn=LOC111556665;refGene_tx=XM_023239749.1
A1 239597554 239598132
refGene_gn=LOC111556665;refGene_tx=XM_023239749.1
A1 240272944 240273253
refGene_gn=CA1H5orf38;refGene_tx=XM_023246998.1
A1 240274795 240274876
refGene_gn=CA1H5orf38;refGene_tx=XM_023246998.1
A1 240275353 240275444
refGene_gn=CA1H5orf38;refGene_tx=XM_023246998.1
A1 240275657 240276047
refGene_gn=CA1H5orf38;refGene_tx=XM_023246998.1
A1 240276616 240276985
refGene_gn=CA1H5orf38;refGene_tx=XM_023246998.1;refGene_gn=IRX2;refGen

e_tx=XM_023239766.1
A1 240278289 240278695
refGene_gn=IRX2;refGene_tx=XM_023239766.1
A1 240278954 240278983
refGene_gn=IRX2;refGene_tx=XM_023239766.1
A1 240278984 240279197
refGene_gn=IRX2;refGene_tx=XM_023239766.1
A1 240279198 240279660
refGene_gn=IRX2;refGene_tx=XM_023239766.1
A1 240279661 240279686
refGene_gn=IRX2;refGene_tx=XM_023239766.1
A1 240280478 240280531
refGene_gn=IRX2;refGene_tx=XM_023239766.1
A1 240955181 240955226
refGene_gn=IRX4;refGene_tx=XM_023239777.1;refGene_gn=IRX4;refGene_tx=XM_023239781.1;refGene_gn=IRX4;refGene_tx=XM_023239783.1
A1 240955709 240955961
refGene_gn=IRX4;refGene_tx=XM_023239777.1;refGene_gn=IRX4;refGene_tx=XM_023239781.1;refGene_gn=IRX4;refGene_tx=XM_023239783.1
A1 240956602 240956745
refGene_gn=IRX4;refGene_tx=XM_023239777.1;refGene_gn=IRX4;refGene_tx=XM_023239781.1;refGene_gn=IRX4;refGene_tx=XM_023239783.1
A1 240957625 240957954
refGene_gn=IRX4;refGene_tx=XM_023239777.1;refGene_gn=IRX4;refGene_tx=XM_023239781.1;refGene_gn=IRX4;refGene_tx=XM_023239783.1
A1 240958409 240959233
refGene_gn=IRX4;refGene_tx=XM_023239777.1;refGene_gn=IRX4;refGene_tx=XM_023239781.1;refGene_gn=IRX4;refGene_tx=XM_023239783.1
A1 241011381 241011450
refGene_gn=NDUFS6;refGene_tx=XM_023239793.1
A1 241012295 241012418
refGene_gn=NDUFS6;refGene_tx=XM_023239793.1
A1 241017754 241017808
refGene_gn=NDUFS6;refGene_tx=XM_023239793.1
A1 241018547 241018679
refGene_gn=NDUFS6;refGene_tx=XM_023239793.1
A1 241019898 241020143
refGene_gn=MRPL36;refGene_tx=XM_023239789.1
A1 241020442 241020758
refGene_gn=MRPL36;refGene_tx=XM_023239789.1
A1 241167155 241167290
refGene_gn=LPCAT1;refGene_tx=XM_023239800.1
A1 241169494 241169536
refGene_gn=LPCAT1;refGene_tx=XM_019813061.2
A1 241184857 241185000
refGene_gn=LPCAT1;refGene_tx=XM_019813061.2;refGene_gn=LPCAT1;refGene_tx=XM_019813062.2;refGene_gn=LPCAT1;refGene_tx=XM_023239800.1
A1 241190270 241190485
refGene_gn=LPCAT1;refGene_tx=XM_019813061.2;refGene_gn=LPCAT1;refGene_tx=XM_019813062.2;refGene_gn=LPCAT1;refGene_tx=XM_023239800.1

A1 241193680 241193793
refGene_gn=LPCAT1;refGene_tx=XM_019813061.2;refGene_gn=LPCAT1;refGene_tx=XM_019813062.2;refGene_gn=LPCAT1;refGene_tx=XM_023239800.1
A1 241195346 241195407
refGene_gn=LPCAT1;refGene_tx=XM_019813061.2;refGene_gn=LPCAT1;refGene_tx=XM_019813062.2;refGene_gn=LPCAT1;refGene_tx=XM_023239800.1
A1 241195955 241196014
refGene_gn=LPCAT1;refGene_tx=XM_019813061.2;refGene_gn=LPCAT1;refGene_tx=XM_019813062.2;refGene_gn=LPCAT1;refGene_tx=XM_023239800.1
A1 241197760 241197795
refGene_gn=LPCAT1;refGene_tx=XM_019813061.2;refGene_gn=LPCAT1;refGene_tx=XM_019813062.2;refGene_gn=LPCAT1;refGene_tx=XM_023239800.1
A1 241198945 241199000
refGene_gn=LPCAT1;refGene_tx=XM_019813061.2;refGene_gn=LPCAT1;refGene_tx=XM_019813062.2;refGene_gn=LPCAT1;refGene_tx=XM_023239800.1
A1 241200949 241201032
refGene_gn=LPCAT1;refGene_tx=XM_019813061.2;refGene_gn=LPCAT1;refGene_tx=XM_019813062.2;refGene_gn=LPCAT1;refGene_tx=XM_023239800.1
A1 241204819 241204945
refGene_gn=LPCAT1;refGene_tx=XM_019813061.2;refGene_gn=LPCAT1;refGene_tx=XM_019813062.2;refGene_gn=LPCAT1;refGene_tx=XM_023239800.1
A1 241205552 241205706
refGene_gn=LPCAT1;refGene_tx=XM_019813061.2;refGene_gn=LPCAT1;refGene_tx=XM_019813062.2;refGene_gn=LPCAT1;refGene_tx=XM_023239800.1
A1 241207990 241208089
refGene_gn=LPCAT1;refGene_tx=XM_019813061.2;refGene_gn=LPCAT1;refGene_tx=XM_019813062.2;refGene_gn=LPCAT1;refGene_tx=XM_023239800.1
A1 241209328 241209467
refGene_gn=LPCAT1;refGene_tx=XM_019813061.2;refGene_gn=LPCAT1;refGene_tx=XM_019813062.2;refGene_gn=LPCAT1;refGene_tx=XM_023239800.1
A1 241211418 241211603
refGene_gn=LPCAT1;refGene_tx=XM_019813061.2;refGene_gn=LPCAT1;refGene_tx=XM_019813062.2;refGene_gn=LPCAT1;refGene_tx=XM_023239800.1
A1 241225143 241225150
refGene_gn=SLC6A3;refGene_tx=XM_023247045.1
A1 241226682 241227546
refGene_gn=SLC6A3;refGene_tx=XM_023247045.1
A1 241229898 241230225
refGene_gn=SLC6A3;refGene_tx=XM_023247045.1
A1 241231378 241231510
refGene_gn=SLC6A3;refGene_tx=XM_023247045.1
A1 241241085 241241311
refGene_gn=SLC6A3;refGene_tx=XM_023247045.1
A1 241248792 241248931
refGene_gn=SLC6A3;refGene_tx=XM_023247045.1
A1 241249655 241249790
refGene_gn=SLC6A3;refGene_tx=XM_023247045.1
A1 241254882 241254986
refGene_gn=SLC6A3;refGene_tx=XM_023247045.1
A1 241256161 241256286

refGene_gn=SLC6A3;refGene_tx=XM_023247045.1
A1 241258930 241259043
refGene_gn=SLC6A3;refGene_tx=XM_023247045.1
A1 241260207 241260336
refGene_gn=SLC6A3;refGene_tx=XM_023247045.1
A1 241261087 241261187
refGene_gn=SLC6A3;refGene_tx=XM_023247045.1
A1 241264362 241264463
refGene_gn=SLC6A3;refGene_tx=XM_023247045.1
A1 241265680 241265848
refGene_gn=SLC6A3;refGene_tx=XM_023247045.1
A1 241266994 241267066
refGene_gn=SLC6A3;refGene_tx=XM_023247045.1
A1 241270737 241270761
refGene_gn=SLC6A3;refGene_tx=XM_023247045.1
A1 241293707 241293869
refGene_gn=CLPTM1L;refGene_tx=XM_011287175.3;refGene_gn=CLPTM1L;refGen
e_tx=XM_023239814.1;refGene_gn=CLPTM1L;refGene_tx=XM_023239815.1;refGe
ne_gn=CLPTM1L;refGene_tx=XM_023239816.1
A1 241294270 241294371
refGene_gn=CLPTM1L;refGene_tx=XM_011287175.3;refGene_gn=CLPTM1L;refGen
e_tx=XM_023239814.1;refGene_gn=CLPTM1L;refGene_tx=XM_023239815.1;refGe
ne_gn=CLPTM1L;refGene_tx=XM_023239816.1
A1 241295299 241295489
refGene_gn=CLPTM1L;refGene_tx=XM_011287175.3;refGene_gn=CLPTM1L;refGen
e_tx=XM_023239814.1;refGene_gn=CLPTM1L;refGene_tx=XM_023239815.1;refGe
ne_gn=CLPTM1L;refGene_tx=XM_023239816.1
A1 241296943 241297092
refGene_gn=CLPTM1L;refGene_tx=XM_011287175.3;refGene_gn=CLPTM1L;refGen
e_tx=XM_023239814.1;refGene_gn=CLPTM1L;refGene_tx=XM_023239815.1;refGe
ne_gn=CLPTM1L;refGene_tx=XM_023239816.1
A1 241298097 241298176
refGene_gn=CLPTM1L;refGene_tx=XM_011287175.3;refGene_gn=CLPTM1L;refGen
e_tx=XM_023239814.1;refGene_gn=CLPTM1L;refGene_tx=XM_023239815.1;refGe
ne_gn=CLPTM1L;refGene_tx=XM_023239816.1
A1 241300503 241300621
refGene_gn=CLPTM1L;refGene_tx=XM_011287175.3;refGene_gn=CLPTM1L;refGen
e_tx=XM_023239814.1;refGene_gn=CLPTM1L;refGene_tx=XM_023239815.1;refGe
ne_gn=CLPTM1L;refGene_tx=XM_023239816.1
A1 241300846 241300941
refGene_gn=CLPTM1L;refGene_tx=XM_011287175.3;refGene_gn=CLPTM1L;refGen
e_tx=XM_023239814.1;refGene_gn=CLPTM1L;refGene_tx=XM_023239815.1;refGe
ne_gn=CLPTM1L;refGene_tx=XM_023239816.1
A1 241301680 241301765
refGene_gn=CLPTM1L;refGene_tx=XM_011287175.3;refGene_gn=CLPTM1L;refGen
e_tx=XM_023239814.1;refGene_gn=CLPTM1L;refGene_tx=XM_023239815.1;refGe
ne_gn=CLPTM1L;refGene_tx=XM_023239816.1
A1 241302785 241302889
refGene_gn=CLPTM1L;refGene_tx=XM_011287175.3;refGene_gn=CLPTM1L;refGen
e_tx=XM_023239814.1;refGene_gn=CLPTM1L;refGene_tx=XM_023239815.1;refGe

ne_gn=CLPTM1L;refGene_tx=XM_023239816.1
A1 241303784 241303850
refGene_gn=CLPTM1L;refGene_tx=XM_011287175.3;refGene_gn=CLPTM1L;refGen
e_tx=XM_023239814.1;refGene_gn=CLPTM1L;refGene_tx=XM_023239815.1;refGe
ne_gn=CLPTM1L;refGene_tx=XM_023239816.1
A1 241305014 241305065
refGene_gn=CLPTM1L;refGene_tx=XM_011287175.3;refGene_gn=CLPTM1L;refGen
e_tx=XM_023239814.1;refGene_gn=CLPTM1L;refGene_tx=XM_023239815.1;refGe
ne_gn=CLPTM1L;refGene_tx=XM_023239816.1
A1 241305767 241305850
refGene_gn=CLPTM1L;refGene_tx=XM_011287175.3;refGene_gn=CLPTM1L;refGen
e_tx=XM_023239814.1;refGene_gn=CLPTM1L;refGene_tx=XM_023239815.1;refGe
ne_gn=CLPTM1L;refGene_tx=XM_023239816.1
A1 241306311 241306346
refGene_gn=CLPTM1L;refGene_tx=XM_011287175.3;refGene_gn=CLPTM1L;refGen
e_tx=XM_023239814.1;refGene_gn=CLPTM1L;refGene_tx=XM_023239815.1;refGe
ne_gn=CLPTM1L;refGene_tx=XM_023239816.1
A1 241306823 241306879
refGene_gn=CLPTM1L;refGene_tx=XM_011287175.3;refGene_gn=CLPTM1L;refGen
e_tx=XM_023239814.1;refGene_gn=CLPTM1L;refGene_tx=XM_023239815.1;refGe
ne_gn=CLPTM1L;refGene_tx=XM_023239816.1
A1 241306952 241306997
refGene_gn=CLPTM1L;refGene_tx=XM_011287175.3;refGene_gn=CLPTM1L;refGen
e_tx=XM_023239814.1;refGene_gn=CLPTM1L;refGene_tx=XM_023239815.1;refGe
ne_gn=CLPTM1L;refGene_tx=XM_023239816.1
A1 241307571 241307687
refGene_gn=CLPTM1L;refGene_tx=XM_011287175.3;refGene_gn=CLPTM1L;refGen
e_tx=XM_023239814.1;refGene_gn=CLPTM1L;refGene_tx=XM_023239815.1;refGe
ne_gn=CLPTM1L;refGene_tx=XM_023239816.1
A1 241307854 241308020
refGene_gn=CLPTM1L;refGene_tx=XM_023239814.1;refGene_gn=CLPTM1L;refGen
e_tx=XM_023239815.1
A1 241308079 241308164
refGene_gn=CLPTM1L;refGene_tx=XM_011287175.3;refGene_gn=CLPTM1L;refGen
e_tx=XM_023239816.1
A1 241322079 241323700
refGene_gn=TERT;refGene_tx=XM_023239817.1;refGene_gn=TERT;refGene_tx=X
M_023239818.1;refGene_gn=TERT;refGene_tx=XM_023239819.1;refGene_gn=TER
T;refGene_tx=XM_023239824.1;refGene_gn=TERT;refGene_tx=XM_023239829.1;
refGene_gn=TERT;refGene_tx=XM_023239833.1;refGene_gn=TERT;refGene_tx=X
M_023239836.1;refGene_gn=TERT;refGene_tx=XM_023239837.1;refGene_gn=TER
T;refGene_tx=XM_023239842.1;refGene_gn=TERT;refGene_tx=XM_023239846.1;
refGene_gn=TERT;refGene_tx=XM_023239847.1;refGene_gn=TERT;refGene_tx=X
M_023239848.1;refGene_gn=TERT;refGene_tx=XM_023239853.1;refGene_gn=TER
T;refGene_tx=XM_023239855.1;refGene_gn=TERT;refGene_tx=XM_023239857.1;
refGene_gn=TERT;refGene_tx=XM_023239861.1
A1 241327307 241327503
refGene_gn=TERT;refGene_tx=XM_023239817.1;refGene_gn=TERT;refGene_tx=X
M_023239818.1;refGene_gn=TERT;refGene_tx=XM_023239819.1;refGene_gn=TER
T;refGene_tx=XM_023239824.1;refGene_gn=TERT;refGene_tx=XM_023239829.1;

refGene_gn=TERT;refGene_tx=XM_023239833.1;refGene_gn=TERT;refGene_tx=X
M_023239836.1;refGene_gn=TERT;refGene_tx=XM_023239837.1;refGene_gn=TER
T;refGene_tx=XM_023239842.1;refGene_gn=TERT;refGene_tx=XM_023239846.1;
refGene_gn=TERT;refGene_tx=XM_023239847.1;refGene_gn=TERT;refGene_tx=X
M_023239848.1;refGene_gn=TERT;refGene_tx=XM_023239853.1;refGene_gn=TER
T;refGene_tx=XM_023239855.1;refGene_gn=TERT;refGene_tx=XM_023239857.1;
refGene_gn=TERT;refGene_tx=XM_023239861.1
A1 241328829 241329010
refGene_gn=TERT;refGene_tx=XM_023239817.1;refGene_gn=TERT;refGene_tx=X
M_023239818.1;refGene_gn=TERT;refGene_tx=XM_023239819.1;refGene_gn=TER
T;refGene_tx=XM_023239824.1;refGene_gn=TERT;refGene_tx=XM_023239829.1;
refGene_gn=TERT;refGene_tx=XM_023239833.1;refGene_gn=TERT;refGene_tx=X
M_023239836.1;refGene_gn=TERT;refGene_tx=XM_023239837.1;refGene_gn=TER
T;refGene_tx=XM_023239842.1;refGene_gn=TERT;refGene_tx=XM_023239846.1;
refGene_gn=TERT;refGene_tx=XM_023239847.1;refGene_gn=TERT;refGene_tx=X
M_023239848.1;refGene_gn=TERT;refGene_tx=XM_023239853.1;refGene_gn=TER
T;refGene_tx=XM_023239855.1;refGene_gn=TERT;refGene_tx=XM_023239857.1;
refGene_gn=TERT;refGene_tx=XM_023239861.1
A1 241329512 241329692
refGene_gn=TERT;refGene_tx=XM_023239817.1;refGene_gn=TERT;refGene_tx=X
M_023239818.1;refGene_gn=TERT;refGene_tx=XM_023239819.1;refGene_gn=TER
T;refGene_tx=XM_023239824.1;refGene_gn=TERT;refGene_tx=XM_023239829.1;
refGene_gn=TERT;refGene_tx=XM_023239833.1;refGene_gn=TERT;refGene_tx=X
M_023239836.1;refGene_gn=TERT;refGene_tx=XM_023239837.1;refGene_gn=TER
T;refGene_tx=XM_023239842.1;refGene_gn=TERT;refGene_tx=XM_023239846.1;
refGene_gn=TERT;refGene_tx=XM_023239847.1;refGene_gn=TERT;refGene_tx=X
M_023239848.1;refGene_gn=TERT;refGene_tx=XM_023239853.1;refGene_gn=TER
T;refGene_tx=XM_023239855.1;refGene_gn=TERT;refGene_tx=XM_023239857.1;
refGene_gn=TERT;refGene_tx=XM_023239861.1
A1 241330076 241330235
refGene_gn=TERT;refGene_tx=XM_023239817.1;refGene_gn=TERT;refGene_tx=X
M_023239818.1;refGene_gn=TERT;refGene_tx=XM_023239819.1;refGene_gn=TER
T;refGene_tx=XM_023239824.1;refGene_gn=TERT;refGene_tx=XM_023239829.1;
refGene_gn=TERT;refGene_tx=XM_023239833.1;refGene_gn=TERT;refGene_tx=X
M_023239836.1;refGene_gn=TERT;refGene_tx=XM_023239837.1;refGene_gn=TER
T;refGene_tx=XM_023239842.1;refGene_gn=TERT;refGene_tx=XM_023239846.1;
refGene_gn=TERT;refGene_tx=XM_023239847.1;refGene_gn=TERT;refGene_tx=X
M_023239848.1;refGene_gn=TERT;refGene_tx=XM_023239853.1;refGene_gn=TER
T;refGene_tx=XM_023239855.1;refGene_gn=TERT;refGene_tx=XM_023239857.1;
refGene_gn=TERT;refGene_tx=XM_023239861.1
A1 241330828 241330927
refGene_gn=TERT;refGene_tx=XM_023239817.1;refGene_gn=TERT;refGene_tx=X
M_023239818.1;refGene_gn=TERT;refGene_tx=XM_023239819.1;refGene_gn=TER
T;refGene_tx=XM_023239824.1;refGene_gn=TERT;refGene_tx=XM_023239829.1;
refGene_gn=TERT;refGene_tx=XM_023239833.1;refGene_gn=TERT;refGene_tx=X
M_023239836.1;refGene_gn=TERT;refGene_tx=XM_023239837.1;refGene_gn=TER
T;refGene_tx=XM_023239842.1;refGene_gn=TERT;refGene_tx=XM_023239846.1;
refGene_gn=TERT;refGene_tx=XM_023239847.1;refGene_gn=TERT;refGene_tx=X
M_023239848.1;refGene_gn=TERT;refGene_tx=XM_023239853.1;refGene_gn=TER
T;refGene_tx=XM_023239855.1;refGene_gn=TERT;refGene_tx=XM_023239857.1;

refGene_gn=TERT;refGene_tx=XM_023239861.1
A1 241331548 241331634
refGene_gn=TERT;refGene_tx=XM_023239817.1;refGene_gn=TERT;refGene_tx=XM_023239818.1;refGene_gn=TERT;refGene_tx=XM_023239819.1;refGene_gn=TERT;refGene_tx=XM_023239824.1;refGene_gn=TERT;refGene_tx=XM_023239829.1;refGene_gn=TERT;refGene_tx=XM_023239833.1;refGene_gn=TERT;refGene_tx=XM_023239836.1;refGene_gn=TERT;refGene_tx=XM_023239837.1;refGene_gn=TERT;refGene_tx=XM_023239842.1;refGene_gn=TERT;refGene_tx=XM_023239846.1;refGene_gn=TERT;refGene_tx=XM_023239847.1;refGene_gn=TERT;refGene_tx=XM_023239848.1;refGene_gn=TERT;refGene_tx=XM_023239853.1;refGene_gn=TERT;refGene_tx=XM_023239855.1;refGene_gn=TERT;refGene_tx=XM_023239857.1;refGene_gn=TERT;refGene_tx=XM_023239861.1
A1 241332356 241332470
refGene_gn=TERT;refGene_tx=XM_023239817.1;refGene_gn=TERT;refGene_tx=XM_023239818.1;refGene_gn=TERT;refGene_tx=XM_023239819.1;refGene_gn=TERT;refGene_tx=XM_023239824.1;refGene_gn=TERT;refGene_tx=XM_023239829.1;refGene_gn=TERT;refGene_tx=XM_023239833.1;refGene_gn=TERT;refGene_tx=XM_023239836.1;refGene_gn=TERT;refGene_tx=XM_023239842.1;refGene_gn=TERT;refGene_tx=XM_023239846.1;refGene_gn=TERT;refGene_tx=XM_023239847.1;refGene_gn=TERT;refGene_tx=XM_023239848.1;refGene_gn=TERT;refGene_tx=XM_023239853.1;refGene_gn=TERT;refGene_tx=XM_023239855.1
A1 241332829 241332901
refGene_gn=TERT;refGene_tx=XM_023239817.1;refGene_gn=TERT;refGene_tx=XM_023239818.1;refGene_gn=TERT;refGene_tx=XM_023239819.1;refGene_gn=TERT;refGene_tx=XM_023239833.1;refGene_gn=TERT;refGene_tx=XM_023239836.1;refGene_gn=TERT;refGene_tx=XM_023239842.1;refGene_gn=TERT;refGene_tx=XM_023239846.1;refGene_gn=TERT;refGene_tx=XM_023239853.1;refGene_gn=TERT;refGene_tx=XM_023239855.1
A1 241334509 241334698
refGene_gn=TERT;refGene_tx=XM_023239817.1;refGene_gn=TERT;refGene_tx=XM_023239818.1;refGene_gn=TERT;refGene_tx=XM_023239819.1;refGene_gn=TERT;refGene_tx=XM_023239824.1;refGene_gn=TERT;refGene_tx=XM_023239829.1;refGene_gn=TERT;refGene_tx=XM_023239833.1;refGene_gn=TERT;refGene_tx=XM_023239836.1;refGene_gn=TERT;refGene_tx=XM_023239837.1;refGene_gn=TERT;refGene_tx=XM_023239842.1;refGene_gn=TERT;refGene_tx=XM_023239846.1;refGene_gn=TERT;refGene_tx=XM_023239847.1;refGene_gn=TERT;refGene_tx=XM_023239848.1;refGene_gn=TERT;refGene_tx=XM_023239853.1;refGene_gn=TERT;refGene_tx=XM_023239855.1;refGene_gn=TERT;refGene_tx=XM_023239857.1;refGene_gn=TERT;refGene_tx=XM_023239861.1
A1 241337899 241338026
refGene_gn=TERT;refGene_tx=XM_023239817.1;refGene_gn=TERT;refGene_tx=XM_023239818.1;refGene_gn=TERT;refGene_tx=XM_023239819.1;refGene_gn=TERT;refGene_tx=XM_023239824.1;refGene_gn=TERT;refGene_tx=XM_023239829.1;refGene_gn=TERT;refGene_tx=XM_023239833.1;refGene_gn=TERT;refGene_tx=XM_023239836.1;refGene_gn=TERT;refGene_tx=XM_023239837.1;refGene_gn=TERT;refGene_tx=XM_023239842.1;refGene_gn=TERT;refGene_tx=XM_023239846.1;refGene_gn=TERT;refGene_tx=XM_023239847.1;refGene_gn=TERT;refGene_tx=XM_023239848.1;refGene_gn=TERT;refGene_tx=XM_023239853.1;refGene_gn=TERT;refGene_tx=XM_023239855.1;refGene_gn=TERT;refGene_tx=XM_023239857.1;refGene_gn=TERT;refGene_tx=XM_023239861.1

A1 241338411 241338473
refGene_gn=TERT;refGene_tx=XM_023239819.1;refGene_gn=TERT;refGene_tx=XM_023239846.1;refGene_gn=TERT;refGene_tx=XM_023239848.1;refGene_gn=TERT;refGene_tx=XM_023239853.1;refGene_gn=TERT;refGene_tx=XM_023239855.1;refGene_gn=TERT;refGene_tx=XM_023239861.1
A1 241341947 241342072
refGene_gn=TERT;refGene_tx=XM_023239817.1;refGene_gn=TERT;refGene_tx=XM_023239818.1;refGene_gn=TERT;refGene_tx=XM_023239819.1;refGene_gn=TERT;refGene_tx=XM_023239824.1;refGene_gn=TERT;refGene_tx=XM_023239829.1;refGene_gn=TERT;refGene_tx=XM_023239833.1;refGene_gn=TERT;refGene_tx=XM_023239836.1;refGene_gn=TERT;refGene_tx=XM_023239837.1;refGene_gn=TERT;refGene_tx=XM_023239842.1;refGene_gn=TERT;refGene_tx=XM_023239846.1;refGene_gn=TERT;refGene_tx=XM_023239847.1;refGene_gn=TERT;refGene_tx=XM_023239848.1;refGene_gn=TERT;refGene_tx=XM_023239853.1;refGene_gn=TERT;refGene_tx=XM_023239855.1;refGene_gn=TERT;refGene_tx=XM_023239857.1;refGene_gn=TERT;refGene_tx=XM_023239861.1
A1 241342906 241343361
refGene_gn=TERT;refGene_tx=XM_023239817.1;refGene_gn=TERT;refGene_tx=XM_023239818.1;refGene_gn=TERT;refGene_tx=XM_023239819.1;refGene_gn=TERT;refGene_tx=XM_023239824.1;refGene_gn=TERT;refGene_tx=XM_023239829.1;refGene_gn=TERT;refGene_tx=XM_023239833.1;refGene_gn=TERT;refGene_tx=XM_023239836.1;refGene_gn=TERT;refGene_tx=XM_023239837.1;refGene_gn=TERT;refGene_tx=XM_023239842.1;refGene_gn=TERT;refGene_tx=XM_023239846.1;refGene_gn=TERT;refGene_tx=XM_023239847.1;refGene_gn=TERT;refGene_tx=XM_023239848.1;refGene_gn=TERT;refGene_tx=XM_023239853.1;refGene_gn=TERT;refGene_tx=XM_023239855.1;refGene_gn=TERT;refGene_tx=XM_023239857.1;refGene_gn=TERT;refGene_tx=XM_023239861.1
A1 241343506 241343681
refGene_gn=TERT;refGene_tx=XM_023239842.1;refGene_gn=TERT;refGene_tx=XM_023239846.1;refGene_gn=TERT;refGene_tx=XM_023239847.1;refGene_gn=TERT;refGene_tx=XM_023239848.1;refGene_gn=TERT;refGene_tx=XM_023239855.1;refGene_gn=TERT;refGene_tx=XM_023239857.1;refGene_gn=TERT;refGene_tx=XM_023239861.1
A1 241345984 241346378
refGene_gn=TERT;refGene_tx=XM_023239817.1;refGene_gn=TERT;refGene_tx=XM_023239818.1;refGene_gn=TERT;refGene_tx=XM_023239824.1;refGene_gn=TERT;refGene_tx=XM_023239829.1;refGene_gn=TERT;refGene_tx=XM_023239833.1;refGene_gn=TERT;refGene_tx=XM_023239836.1;refGene_gn=TERT;refGene_tx=XM_023239837.1;refGene_gn=TERT;refGene_tx=XM_023239853.1
A1 241348636 241348807
refGene_gn=LOC101081196;refGene_tx=XM_019813133.1;refGene_gn=LOC101081196;refGene_tx=XM_019813136.1
A1 241349641 241349804
refGene_gn=LOC101081196;refGene_tx=XM_019813133.1;refGene_gn=LOC101081196;refGene_tx=XM_019813136.1
A1 241350889 241351049
refGene_gn=LOC101081196;refGene_tx=XM_019813133.1;refGene_gn=LOC101081196;refGene_tx=XM_019813136.1
A1 241351443 241351648
refGene_gn=LOC101081196;refGene_tx=XM_019813133.1;refGene_gn=LOC101081

196;refGene_tx=XM_019813136.1
A1 241352120 241352277
refGene_gn=LOC101081196;refGene_tx=XM_019813133.1;refGene_gn=LOC101081196;refGene_tx=XM_019813136.1
A1 241353813 241353942
refGene_gn=LOC101081196;refGene_tx=XM_019813133.1;refGene_gn=LOC101081196;refGene_tx=XM_019813136.1
A1 241355355 241355468
refGene_gn=LOC101081196;refGene_tx=XM_019813133.1;refGene_gn=LOC101081196;refGene_tx=XM_019813136.1
A1 241356221 241356332
refGene_gn=LOC101081196;refGene_tx=XM_019813133.1;refGene_gn=LOC101081196;refGene_tx=XM_019813136.1
A1 241357760 241357942
refGene_gn=LOC101081196;refGene_tx=XM_019813133.1;refGene_gn=LOC101081196;refGene_tx=XM_019813136.1
A1 241358539 241358677
refGene_gn=LOC101081196;refGene_tx=XM_019813133.1;refGene_gn=LOC101081196;refGene_tx=XM_019813136.1
A1 241359413 241359554
refGene_gn=LOC101081196;refGene_tx=XM_019813133.1;refGene_gn=LOC101081196;refGene_tx=XM_019813136.1
A1 241362905 241362999
refGene_gn=LOC101081196;refGene_tx=XM_019813136.1
A1 241363486 241363646
refGene_gn=LOC101081196;refGene_tx=XM_019813133.1
A1 241419631 241419821
refGene_gn=SLC12A7;refGene_tx=XM_023239871.1;refGene_gn=SLC12A7;refGene_tx=XM_023239873.1
A1 241448781 241448911
refGene_gn=SLC12A7;refGene_tx=XM_023239878.1;refGene_gn=SLC12A7;refGene_tx=XM_023239880.1
A1 241463150 241463245
refGene_gn=SLC12A7;refGene_tx=XM_006928123.4;refGene_gn=SLC12A7;refGene_tx=XM_023239871.1;refGene_gn=SLC12A7;refGene_tx=XM_023239873.1;refGene_gn=SLC12A7;refGene_tx=XM_023239878.1;refGene_gn=SLC12A7;refGene_tx=XM_023239880.1
A1 241463603 241463726
refGene_gn=SLC12A7;refGene_tx=XM_006928123.4;refGene_gn=SLC12A7;refGene_tx=XM_023239871.1;refGene_gn=SLC12A7;refGene_tx=XM_023239873.1;refGene_gn=SLC12A7;refGene_tx=XM_023239878.1;refGene_gn=SLC12A7;refGene_tx=XM_023239880.1
A1 241466113 241466260
refGene_gn=SLC12A7;refGene_tx=XM_006928123.4;refGene_gn=SLC12A7;refGene_tx=XM_023239871.1;refGene_gn=SLC12A7;refGene_tx=XM_023239873.1;refGene_gn=SLC12A7;refGene_tx=XM_023239878.1;refGene_gn=SLC12A7;refGene_tx=XM_023239880.1
A1 241467281 241467336
refGene_gn=SLC12A7;refGene_tx=XM_006928123.4;refGene_gn=SLC12A7;refGene_tx=XM_023239871.1;refGene_gn=SLC12A7;refGene_tx=XM_023239873.1;refGe

ne_gn=SLC12A7;refGene_tx=XM_023239878.1;refGene_gn=SLC12A7;refGene_tx=XM_023239880.1;refGene_gn=SLC12A7;refGene_tx=XM_023239886.1
A1 241468200 241468331
refGene_gn=SLC12A7;refGene_tx=XM_006928123.4;refGene_gn=SLC12A7;refGene_tx=XM_023239871.1;refGene_gn=SLC12A7;refGene_tx=XM_023239873.1;refGene_gn=SLC12A7;refGene_tx=XM_023239878.1;refGene_gn=SLC12A7;refGene_tx=XM_023239880.1;refGene_gn=SLC12A7;refGene_tx=XM_023239886.1
A1 241469089 241469331
refGene_gn=SLC12A7;refGene_tx=XM_006928123.4;refGene_gn=SLC12A7;refGene_tx=XM_023239871.1;refGene_gn=SLC12A7;refGene_tx=XM_023239873.1;refGene_gn=SLC12A7;refGene_tx=XM_023239878.1;refGene_gn=SLC12A7;refGene_tx=XM_023239880.1;refGene_gn=SLC12A7;refGene_tx=XM_023239886.1
A1 241470044 241470256
refGene_gn=SLC12A7;refGene_tx=XM_006928123.4;refGene_gn=SLC12A7;refGene_tx=XM_023239871.1;refGene_gn=SLC12A7;refGene_tx=XM_023239873.1;refGene_gn=SLC12A7;refGene_tx=XM_023239878.1;refGene_gn=SLC12A7;refGene_tx=XM_023239880.1;refGene_gn=SLC12A7;refGene_tx=XM_023239886.1
A1 241470639 241470807
refGene_gn=SLC12A7;refGene_tx=XM_006928123.4;refGene_gn=SLC12A7;refGene_tx=XM_023239871.1;refGene_gn=SLC12A7;refGene_tx=XM_023239873.1;refGene_gn=SLC12A7;refGene_tx=XM_023239878.1;refGene_gn=SLC12A7;refGene_tx=XM_023239880.1;refGene_gn=SLC12A7;refGene_tx=XM_023239886.1
A1 241471374 241471473
refGene_gn=SLC12A7;refGene_tx=XM_006928123.4;refGene_gn=SLC12A7;refGene_tx=XM_023239871.1;refGene_gn=SLC12A7;refGene_tx=XM_023239873.1;refGene_gn=SLC12A7;refGene_tx=XM_023239878.1;refGene_gn=SLC12A7;refGene_tx=XM_023239880.1;refGene_gn=SLC12A7;refGene_tx=XM_023239886.1
A1 241471927 241471985
refGene_gn=SLC12A7;refGene_tx=XM_006928123.4;refGene_gn=SLC12A7;refGene_tx=XM_023239871.1;refGene_gn=SLC12A7;refGene_tx=XM_023239873.1;refGene_gn=SLC12A7;refGene_tx=XM_023239878.1;refGene_gn=SLC12A7;refGene_tx=XM_023239880.1;refGene_gn=SLC12A7;refGene_tx=XM_023239886.1
A1 241472349 241472524
refGene_gn=SLC12A7;refGene_tx=XM_006928123.4;refGene_gn=SLC12A7;refGene_tx=XM_023239871.1;refGene_gn=SLC12A7;refGene_tx=XM_023239873.1;refGene_gn=SLC12A7;refGene_tx=XM_023239878.1;refGene_gn=SLC12A7;refGene_tx=XM_023239880.1;refGene_gn=SLC12A7;refGene_tx=XM_023239886.1
A1 241473348 241473467
refGene_gn=SLC12A7;refGene_tx=XM_006928123.4;refGene_gn=SLC12A7;refGene_tx=XM_023239871.1;refGene_gn=SLC12A7;refGene_tx=XM_023239873.1;refGene_gn=SLC12A7;refGene_tx=XM_023239878.1;refGene_gn=SLC12A7;refGene_tx=XM_023239880.1;refGene_gn=SLC12A7;refGene_tx=XM_023239886.1
A1 241473729 241473828
refGene_gn=SLC12A7;refGene_tx=XM_006928123.4;refGene_gn=SLC12A7;refGene_tx=XM_023239871.1;refGene_gn=SLC12A7;refGene_tx=XM_023239873.1;refGene_gn=SLC12A7;refGene_tx=XM_023239878.1;refGene_gn=SLC12A7;refGene_tx=XM_023239880.1;refGene_gn=SLC12A7;refGene_tx=XM_023239886.1
A1 241474174 241474294
refGene_gn=SLC12A7;refGene_tx=XM_006928123.4;refGene_gn=SLC12A7;refGene_tx=XM_023239871.1;refGene_gn=SLC12A7;refGene_tx=XM_023239873.1;refGe

ne_gn=SLC12A7;refGene_tx=XM_023239878.1;refGene_gn=SLC12A7;refGene_tx=XM_023239880.1;refGene_gn=SLC12A7;refGene_tx=XM_023239886.1
A1 241474817 241474922
refGene_gn=SLC12A7;refGene_tx=XM_006928123.4;refGene_gn=SLC12A7;refGene_tx=XM_023239871.1;refGene_gn=SLC12A7;refGene_tx=XM_023239873.1;refGene_gn=SLC12A7;refGene_tx=XM_023239878.1;refGene_gn=SLC12A7;refGene_tx=XM_023239880.1;refGene_gn=SLC12A7;refGene_tx=XM_023239886.1
A1 241475341 241475510
refGene_gn=SLC12A7;refGene_tx=XM_006928123.4;refGene_gn=SLC12A7;refGene_tx=XM_023239871.1;refGene_gn=SLC12A7;refGene_tx=XM_023239873.1;refGene_gn=SLC12A7;refGene_tx=XM_023239878.1;refGene_gn=SLC12A7;refGene_tx=XM_023239880.1;refGene_gn=SLC12A7;refGene_tx=XM_023239886.1
A1 241477258 241477454
refGene_gn=SLC12A7;refGene_tx=XM_006928123.4;refGene_gn=SLC12A7;refGene_tx=XM_023239871.1;refGene_gn=SLC12A7;refGene_tx=XM_023239873.1;refGene_gn=SLC12A7;refGene_tx=XM_023239878.1;refGene_gn=SLC12A7;refGene_tx=XM_023239880.1;refGene_gn=SLC12A7;refGene_tx=XM_023239886.1
A1 241477715 241477885
refGene_gn=SLC12A7;refGene_tx=XM_006928123.4;refGene_gn=SLC12A7;refGene_tx=XM_023239871.1;refGene_gn=SLC12A7;refGene_tx=XM_023239873.1;refGene_gn=SLC12A7;refGene_tx=XM_023239878.1;refGene_gn=SLC12A7;refGene_tx=XM_023239880.1;refGene_gn=SLC12A7;refGene_tx=XM_023239886.1
A1 241478524 241478656
refGene_gn=SLC12A7;refGene_tx=XM_006928123.4;refGene_gn=SLC12A7;refGene_tx=XM_023239871.1;refGene_gn=SLC12A7;refGene_tx=XM_023239873.1;refGene_gn=SLC12A7;refGene_tx=XM_023239878.1;refGene_gn=SLC12A7;refGene_tx=XM_023239880.1;refGene_gn=SLC12A7;refGene_tx=XM_023239886.1
A1 241479449 241479557
refGene_gn=SLC12A7;refGene_tx=XM_006928123.4;refGene_gn=SLC12A7;refGene_tx=XM_023239871.1;refGene_gn=SLC12A7;refGene_tx=XM_023239873.1;refGene_gn=SLC12A7;refGene_tx=XM_023239878.1;refGene_gn=SLC12A7;refGene_tx=XM_023239880.1;refGene_gn=SLC12A7;refGene_tx=XM_023239886.1
A1 241480006 241480185
refGene_gn=SLC12A7;refGene_tx=XM_006928123.4;refGene_gn=SLC12A7;refGene_tx=XM_023239871.1;refGene_gn=SLC12A7;refGene_tx=XM_023239873.1;refGene_gn=SLC12A7;refGene_tx=XM_023239878.1;refGene_gn=SLC12A7;refGene_tx=XM_023239880.1;refGene_gn=SLC12A7;refGene_tx=XM_023239886.1
A1 241480671 241480686
refGene_gn=SLC12A7;refGene_tx=XM_006928123.4;refGene_gn=SLC12A7;refGene_tx=XM_023239871.1;refGene_gn=SLC12A7;refGene_tx=XM_023239878.1;refGene_gn=SLC12A7;refGene_tx=XM_023239886.1
A1 241483509 241483643
refGene_gn=SLC12A7;refGene_tx=XM_006928123.4;refGene_gn=SLC12A7;refGene_tx=XM_023239871.1;refGene_gn=SLC12A7;refGene_tx=XM_023239873.1;refGene_gn=SLC12A7;refGene_tx=XM_023239878.1;refGene_gn=SLC12A7;refGene_tx=XM_023239880.1;refGene_gn=SLC12A7;refGene_tx=XM_023239886.1
A1 241484595 241484687
refGene_gn=SLC12A7;refGene_tx=XM_006928123.4;refGene_gn=SLC12A7;refGene_tx=XM_023239871.1;refGene_gn=SLC12A7;refGene_tx=XM_023239873.1;refGene_gn=SLC12A7;refGene_tx=XM_023239878.1;refGene_gn=SLC12A7;refGene_tx=

XM_023239880.1;refGene_gn=SLC12A7;refGene_tx=XM_023239886.1
A1	241491906	241492442
refGene_gn=NKD2;refGene_tx=XM_023239890.1
A1	241492712	241492840
refGene_gn=NKD2;refGene_tx=XM_023239890.1
A1	241493459	241493544
refGene_gn=NKD2;refGene_tx=XM_023239890.1
A1	241493809	241493957
refGene_gn=NKD2;refGene_tx=XM_023239890.1
A1	241494458	241494554
refGene_gn=NKD2;refGene_tx=XM_023239890.1
A1	241494782	241494910
refGene_gn=NKD2;refGene_tx=XM_023239890.1
A1	241496063	241496124
refGene_gn=NKD2;refGene_tx=XM_023239890.1
A1	241513517	241513597
refGene_gn=NKD2;refGene_tx=XM_023239890.1
A1	241513861	241513967
refGene_gn=NKD2;refGene_tx=XM_023239890.1
A1	241519330	241520178
refGene_gn=LOC111561902;refGene_tx=XM_023259682.1
A1	241520300	241520361
refGene_gn=LOC111561902;refGene_tx=XM_023259682.1
A1	241556279	241556390
refGene_gn=TRIP13;refGene_tx=XM_023239947.1
A1	241564102	241564198
refGene_gn=TRIP13;refGene_tx=XM_023239942.1
A1	241564576	241564646
refGene_gn=TRIP13;refGene_tx=XM_023239942.1
A1	241565689	241565802
refGene_gn=TRIP13;refGene_tx=XM_023239942.1
A1	241568073	241568227
refGene_gn=TRIP13;refGene_tx=XM_023239942.1;refGene_gn=TRIP13;refGene_
tx=XM_023239947.1
A1	241570565	241570672
refGene_gn=TRIP13;refGene_tx=XM_023239942.1;refGene_gn=TRIP13;refGene_
tx=XM_023239947.1
A1	241570914	241571001
refGene_gn=TRIP13;refGene_tx=XM_023239942.1;refGene_gn=TRIP13;refGene_
tx=XM_023239947.1
A1	241571814	241571878
refGene_gn=TRIP13;refGene_tx=XM_023239942.1;refGene_gn=TRIP13;refGene_
tx=XM_023239947.1
A1	241574410	241574483
refGene_gn=TRIP13;refGene_tx=XM_023239942.1;refGene_gn=TRIP13;refGene_
tx=XM_023239947.1
A1	241576020	241576111
refGene_gn=TRIP13;refGene_tx=XM_023239942.1;refGene_gn=TRIP13;refGene_
tx=XM_023239947.1
A1	241576819	241576875

refGene_gn=TRIP13;refGene_tx=XM_023239942.1;refGene_gn=TRIP13;refGene_tx=XM_023239947.1
A1 241577858 241577988
refGene_gn=TRIP13;refGene_tx=XM_023239942.1;refGene_gn=TRIP13;refGene_tx=XM_023239947.1
A1 241579656 241579822
refGene_gn=TRIP13;refGene_tx=XM_023239942.1;refGene_gn=TRIP13;refGene_tx=XM_023239947.1
A1 241580519 241580611
refGene_gn=TRIP13;refGene_tx=XM_023239942.1;refGene_gn=TRIP13;refGene_tx=XM_023239947.1
A1 241580959 241581011
refGene_gn=BRD9;refGene_tx=XM_023239899.1;refGene_gn=BRD9;refGene_tx=XM_023239901.1;refGene_gn=BRD9;refGene_tx=XM_023239903.1;refGene_gn=BRD9;refGene_tx=XM_023239912.1;refGene_gn=BRD9;refGene_tx=XM_023239920.1;refGene_gn=BRD9;refGene_tx=XM_023239926.1;refGene_gn=BRD9;refGene_tx=XM_023239928.1;refGene_gn=BRD9;refGene_tx=XM_023239933.1
A1 241581664 241581879
refGene_gn=BRD9;refGene_tx=XM_023239899.1;refGene_gn=BRD9;refGene_tx=XM_023239901.1;refGene_gn=BRD9;refGene_tx=XM_023239903.1;refGene_gn=BRD9;refGene_tx=XM_023239912.1;refGene_gn=BRD9;refGene_tx=XM_023239920.1;refGene_gn=BRD9;refGene_tx=XM_023239926.1;refGene_gn=BRD9;refGene_tx=XM_023239928.1;refGene_gn=BRD9;refGene_tx=XM_023239933.1
A1 241582132 241582265
refGene_gn=BRD9;refGene_tx=XM_023239899.1;refGene_gn=BRD9;refGene_tx=XM_023239901.1;refGene_gn=BRD9;refGene_tx=XM_023239903.1;refGene_gn=BRD9;refGene_tx=XM_023239912.1;refGene_gn=BRD9;refGene_tx=XM_023239920.1;refGene_gn=BRD9;refGene_tx=XM_023239926.1;refGene_gn=BRD9;refGene_tx=XM_023239928.1;refGene_gn=BRD9;refGene_tx=XM_023239933.1
A1 241583744 241583805
refGene_gn=BRD9;refGene_tx=XM_023239899.1;refGene_gn=BRD9;refGene_tx=XM_023239901.1;refGene_gn=BRD9;refGene_tx=XM_023239903.1;refGene_gn=BRD9;refGene_tx=XM_023239912.1;refGene_gn=BRD9;refGene_tx=XM_023239920.1;refGene_gn=BRD9;refGene_tx=XM_023239926.1;refGene_gn=BRD9;refGene_tx=XM_023239928.1;refGene_gn=BRD9;refGene_tx=XM_023239933.1
A1 241584233 241584378
refGene_gn=BRD9;refGene_tx=XM_023239899.1;refGene_gn=BRD9;refGene_tx=XM_023239901.1;refGene_gn=BRD9;refGene_tx=XM_023239903.1;refGene_gn=BRD9;refGene_tx=XM_023239912.1;refGene_gn=BRD9;refGene_tx=XM_023239920.1;refGene_gn=BRD9;refGene_tx=XM_023239926.1;refGene_gn=BRD9;refGene_tx=XM_023239928.1;refGene_gn=BRD9;refGene_tx=XM_023239933.1
A1 241585964 241586075
refGene_gn=BRD9;refGene_tx=XM_023239899.1;refGene_gn=BRD9;refGene_tx=XM_023239901.1;refGene_gn=BRD9;refGene_tx=XM_023239903.1;refGene_gn=BRD9;refGene_tx=XM_023239912.1;refGene_gn=BRD9;refGene_tx=XM_023239920.1;refGene_gn=BRD9;refGene_tx=XM_023239926.1;refGene_gn=BRD9;refGene_tx=XM_023239928.1;refGene_gn=BRD9;refGene_tx=XM_023239933.1
A1 241586733 241586909
refGene_gn=BRD9;refGene_tx=XM_023239899.1;refGene_gn=BRD9;refGene_tx=XM_023239901.1;refGene_gn=BRD9;refGene_tx=XM_023239903.1;refGene_gn=BRD

9;refGene_tx=XM_023239912.1;refGene_gn=BRD9;refGene_tx=XM_023239920.1;
refGene_gn=BRD9;refGene_tx=XM_023239926.1;refGene_gn=BRD9;refGene_tx=X
M_023239928.1;refGene_gn=BRD9;refGene_tx=XM_023239933.1
A1 241587957 241588090
refGene_gn=BRD9;refGene_tx=XM_023239899.1;refGene_gn=BRD9;refGene_tx=X
M_023239901.1;refGene_gn=BRD9;refGene_tx=XM_023239903.1;refGene_gn=BRD
9;refGene_tx=XM_023239912.1;refGene_gn=BRD9;refGene_tx=XM_023239920.1;
refGene_gn=BRD9;refGene_tx=XM_023239926.1;refGene_gn=BRD9;refGene_tx=X
M_023239928.1
A1 241590063 241590139
refGene_gn=BRD9;refGene_tx=XM_023239899.1;refGene_gn=BRD9;refGene_tx=X
M_023239901.1;refGene_gn=BRD9;refGene_tx=XM_023239903.1;refGene_gn=BRD
9;refGene_tx=XM_023239912.1;refGene_gn=BRD9;refGene_tx=XM_023239920.1;
refGene_gn=BRD9;refGene_tx=XM_023239926.1;refGene_gn=BRD9;refGene_tx=X
M_023239928.1
A1 241591040 241591136
refGene_gn=BRD9;refGene_tx=XM_023239899.1;refGene_gn=BRD9;refGene_tx=X
M_023239901.1;refGene_gn=BRD9;refGene_tx=XM_023239903.1;refGene_gn=BRD
9;refGene_tx=XM_023239912.1;refGene_gn=BRD9;refGene_tx=XM_023239920.1;
refGene_gn=BRD9;refGene_tx=XM_023239926.1;refGene_gn=BRD9;refGene_tx=X
M_023239928.1
A1 241592041 241592174
refGene_gn=BRD9;refGene_tx=XM_023239899.1;refGene_gn=BRD9;refGene_tx=X
M_023239901.1;refGene_gn=BRD9;refGene_tx=XM_023239903.1;refGene_gn=BRD
9;refGene_tx=XM_023239912.1;refGene_gn=BRD9;refGene_tx=XM_023239920.1;
refGene_gn=BRD9;refGene_tx=XM_023239926.1;refGene_gn=BRD9;refGene_tx=X
M_023239928.1
A1 241594713 241594825
refGene_gn=BRD9;refGene_tx=XM_023239899.1;refGene_gn=BRD9;refGene_tx=X
M_023239901.1;refGene_gn=BRD9;refGene_tx=XM_023239903.1;refGene_gn=BRD
9;refGene_tx=XM_023239912.1;refGene_gn=BRD9;refGene_tx=XM_023239920.1;
refGene_gn=BRD9;refGene_tx=XM_023239926.1;refGene_gn=BRD9;refGene_tx=X
M_023239928.1;refGene_gn=BRD9;refGene_tx=XM_023239933.1
A1 241595632 241595716
refGene_gn=BRD9;refGene_tx=XM_023239899.1;refGene_gn=BRD9;refGene_tx=X
M_023239901.1;refGene_gn=BRD9;refGene_tx=XM_023239903.1;refGene_gn=BRD
9;refGene_tx=XM_023239912.1;refGene_gn=BRD9;refGene_tx=XM_023239933.1
A1 241597831 241597870
refGene_gn=BRD9;refGene_tx=XM_023239899.1;refGene_gn=BRD9;refGene_tx=X
M_023239901.1;refGene_gn=BRD9;refGene_tx=XM_023239903.1;refGene_gn=BRD
9;refGene_tx=XM_023239912.1;refGene_gn=BRD9;refGene_tx=XM_023239920.1;
refGene_gn=BRD9;refGene_tx=XM_023239926.1;refGene_gn=BRD9;refGene_tx=X
M_023239928.1;refGene_gn=BRD9;refGene_tx=XM_023239933.1
A1 241599994 241600094
refGene_gn=BRD9;refGene_tx=XM_023239899.1;refGene_gn=BRD9;refGene_tx=X
M_023239901.1;refGene_gn=BRD9;refGene_tx=XM_023239903.1;refGene_gn=BRD
9;refGene_tx=XM_023239912.1;refGene_gn=BRD9;refGene_tx=XM_023239920.1;
refGene_gn=BRD9;refGene_tx=XM_023239926.1;refGene_gn=BRD9;refGene_tx=X
M_023239928.1;refGene_gn=BRD9;refGene_tx=XM_023239933.1
A1 241602402 241602576

refGene_gn=BRD9;refGene_tx=XM_023239899.1;refGene_gn=BRD9;refGene_tx=XM_023239901.1;refGene_gn=BRD9;refGene_tx=XM_023239903.1;refGene_gn=BRD9;refGene_tx=XM_023239912.1;refGene_gn=BRD9;refGene_tx=XM_023239920.1;refGene_gn=BRD9;refGene_tx=XM_023239926.1;refGene_gn=BRD9;refGene_tx=XM_023239928.1;refGene_gn=BRD9;refGene_tx=XM_023239933.1
A1 241603279 241603380
refGene_gn=BRD9;refGene_tx=XM_023239899.1;refGene_gn=BRD9;refGene_tx=XM_023239901.1;refGene_gn=BRD9;refGene_tx=XM_023239903.1;refGene_gn=BRD9;refGene_tx=XM_023239912.1;refGene_gn=BRD9;refGene_tx=XM_023239920.1;refGene_gn=BRD9;refGene_tx=XM_023239926.1;refGene_gn=BRD9;refGene_tx=XM_023239928.1;refGene_gn=BRD9;refGene_tx=XM_023239933.1
A1 241607848 241608070
refGene_gn=LOC105259694;refGene_tx=XM_023247103.1
A1 241608690 241608864
refGene_gn=LOC105259694;refGene_tx=XM_023247103.1
A1 241612205 241612381
refGene_gn=LOC105259694;refGene_tx=XM_023247103.1
A1 241612824 241612926
refGene_gn=LOC105259694;refGene_tx=XM_023247103.1
A1 241615530 241615655
refGene_gn=LOC105259694;refGene_tx=XM_023247103.1
A1 241617799 241617952
refGene_gn=LOC105259694;refGene_tx=XM_023247103.1
A1 241620391 241620624
refGene_gn=LOC105259694;refGene_tx=XM_023247103.1
A1 241631200 241631517
refGene_gn=TPPP;refGene_tx=XM_023239951.1
A1 241643620 241643774
refGene_gn=TPPP;refGene_tx=XM_023239951.1
A1 241643943 241644138
refGene_gn=TPPP;refGene_tx=XM_023239951.1
A1 241653874 241654016
refGene_gn=CEP72;refGene_tx=XM_003981592.4;refGene_gn=CEP72;refGene_tx=XM_019813202.2;refGene_gn=CEP72;refGene_tx=XM_019813205.2;refGene_gn=CEP72;refGene_tx=XM_023239967.1
A1 241655501 241655613
refGene_gn=CEP72;refGene_tx=XM_003981592.4;refGene_gn=CEP72;refGene_tx=XM_019813202.2;refGene_gn=CEP72;refGene_tx=XM_019813205.2;refGene_gn=CEP72;refGene_tx=XM_023239967.1
A1 241658394 241658521
refGene_gn=CEP72;refGene_tx=XM_003981592.4;refGene_gn=CEP72;refGene_tx=XM_019813202.2;refGene_gn=CEP72;refGene_tx=XM_019813205.2;refGene_gn=CEP72;refGene_tx=XM_023239967.1
A1 241665395 241665592
refGene_gn=CEP72;refGene_tx=XM_003981592.4;refGene_gn=CEP72;refGene_tx=XM_019813202.2;refGene_gn=CEP72;refGene_tx=XM_019813205.2;refGene_gn=CEP72;refGene_tx=XM_023239967.1
A1 241666663 241666805
refGene_gn=CEP72;refGene_tx=XM_003981592.4;refGene_gn=CEP72;refGene_tx=XM_019813202.2;refGene_gn=CEP72;refGene_tx=XM_019813205.2;refGene_gn=

CEP72;refGene_tx=XM_023239967.1
A1	241668934	241669236
refGene_gn=CEP72;refGene_tx=XM_003981592.4;refGene_gn=CEP72;refGene_tx=XM_019813202.2;refGene_gn=CEP72;refGene_tx=XM_019813205.2;refGene_gn=CEP72;refGene_tx=XM_023239967.1
A1	241669902	241670085
refGene_gn=CEP72;refGene_tx=XM_003981592.4;refGene_gn=CEP72;refGene_tx=XM_019813202.2;refGene_gn=CEP72;refGene_tx=XM_019813205.2;refGene_gn=CEP72;refGene_tx=XM_023239967.1
A1	241671407	241671586
refGene_gn=CEP72;refGene_tx=XM_003981592.4;refGene_gn=CEP72;refGene_tx=XM_019813202.2;refGene_gn=CEP72;refGene_tx=XM_019813205.2;refGene_gn=CEP72;refGene_tx=XM_023239967.1
A1	241679272	241679381
refGene_gn=CEP72;refGene_tx=XM_003981592.4;refGene_gn=CEP72;refGene_tx=XM_019813202.2;refGene_gn=CEP72;refGene_tx=XM_019813205.2;refGene_gn=CEP72;refGene_tx=XM_023239967.1
A1	241681082	241681275
refGene_gn=CEP72;refGene_tx=XM_003981592.4;refGene_gn=CEP72;refGene_tx=XM_019813202.2;refGene_gn=CEP72;refGene_tx=XM_019813205.2;refGene_gn=CEP72;refGene_tx=XM_023239967.1
A1	241682183	241682311
refGene_gn=CEP72;refGene_tx=XM_003981592.4;refGene_gn=CEP72;refGene_tx=XM_019813202.2;refGene_gn=CEP72;refGene_tx=XM_019813205.2;refGene_gn=CEP72;refGene_tx=XM_023239967.1
A1	241683498	241683583
refGene_gn=CEP72;refGene_tx=XM_019813202.2;refGene_gn=CEP72;refGene_tx=XM_019813205.2
A1	241687922	241688004
refGene_gn=CEP72;refGene_tx=XM_003981592.4
A1	241757268	241757476
refGene_gn=SLC9A3;refGene_tx=XM_023239980.1;refGene_gn=SLC9A3;refGene_tx=XM_023239985.1
A1	241782897	241783200
refGene_gn=SLC9A3;refGene_tx=XM_023239980.1;refGene_gn=SLC9A3;refGene_tx=XM_023239985.1
A1	241786374	241786535
refGene_gn=SLC9A3;refGene_tx=XM_023239980.1;refGene_gn=SLC9A3;refGene_tx=XM_023239985.1
A1	241788390	241788469
refGene_gn=SLC9A3;refGene_tx=XM_023239980.1;refGene_gn=SLC9A3;refGene_tx=XM_023239985.1
A1	241789077	241789255
refGene_gn=SLC9A3;refGene_tx=XM_023239980.1;refGene_gn=SLC9A3;refGene_tx=XM_023239985.1
A1	241789864	241790085
refGene_gn=SLC9A3;refGene_tx=XM_023239980.1;refGene_gn=SLC9A3;refGene_tx=XM_023239985.1
A1	241791006	241791209
refGene_gn=SLC9A3;refGene_tx=XM_023239980.1;refGene_gn=SLC9A3;refGene_

tx=XM_023239985.1
A1 241791668 241791758
refGene_gn=SLC9A3;refGene_tx=XM_023239980.1;refGene_gn=SLC9A3;refGene_
tx=XM_023239985.1
A1 241791856 241791927
refGene_gn=SLC9A3;refGene_tx=XM_023239980.1;refGene_gn=SLC9A3;refGene_
tx=XM_023239985.1
A1 241792903 241793033
refGene_gn=SLC9A3;refGene_tx=XM_023239980.1;refGene_gn=SLC9A3;refGene_
tx=XM_023239985.1
A1 241795150 241795266
refGene_gn=SLC9A3;refGene_tx=XM_023239980.1;refGene_gn=SLC9A3;refGene_
tx=XM_023239985.1
A1 241795909 241796039
refGene_gn=SLC9A3;refGene_tx=XM_023239980.1;refGene_gn=SLC9A3;refGene_
tx=XM_023239985.1
A1 241796249 241796426
refGene_gn=SLC9A3;refGene_tx=XM_023239980.1;refGene_gn=SLC9A3;refGene_
tx=XM_023239985.1
A1 241796663 241796736
refGene_gn=SLC9A3;refGene_tx=XM_023239980.1;refGene_gn=SLC9A3;refGene_
tx=XM_023239985.1
A1 241797313 241797427
refGene_gn=SLC9A3;refGene_tx=XM_023239980.1;refGene_gn=SLC9A3;refGene_
tx=XM_023239985.1
A1 241797747 241797995
refGene_gn=SLC9A3;refGene_tx=XM_023239980.1;refGene_gn=SLC9A3;refGene_
tx=XM_023239985.1
A1 241799606 241799610
refGene_gn=SLC9A3;refGene_tx=XM_023239980.1
A1 241806758 241806930
refGene_gn=EXOC3;refGene_tx=XM_003981565.5;refGene_gn=EXOC3;refGene_tx
=XM_006928127.3;refGene_gn=EXOC3;refGene_tx=XM_019813260.2
A1 241807697 241807825
refGene_gn=EXOC3;refGene_tx=XM_003981565.5;refGene_gn=EXOC3;refGene_tx
=XM_006928127.3;refGene_gn=EXOC3;refGene_tx=XM_019813260.2
A1 241808242 241808404
refGene_gn=EXOC3;refGene_tx=XM_003981565.5;refGene_gn=EXOC3;refGene_tx
=XM_006928127.3;refGene_gn=EXOC3;refGene_tx=XM_019813260.2
A1 241809738 241809861
refGene_gn=EXOC3;refGene_tx=XM_003981565.5;refGene_gn=EXOC3;refGene_tx
=XM_006928127.3;refGene_gn=EXOC3;refGene_tx=XM_019813260.2
A1 241811135 241811286
refGene_gn=EXOC3;refGene_tx=XM_003981565.5;refGene_gn=EXOC3;refGene_tx
=XM_006928127.3;refGene_gn=EXOC3;refGene_tx=XM_019813260.2
A1 241811378 241811489
refGene_gn=EXOC3;refGene_tx=XM_003981565.5;refGene_gn=EXOC3;refGene_tx
=XM_006928127.3;refGene_gn=EXOC3;refGene_tx=XM_019813260.2
A1 241813186 241813287
refGene_gn=EXOC3;refGene_tx=XM_003981565.5;refGene_gn=EXOC3;refGene_tx

=XM_006928127.3;refGene_gn=EXOC3;refGene_tx=XM_019813260.2
A1 241814324 241814450
refGene_gn=EXOC3;refGene_tx=XM_003981565.5;refGene_gn=EXOC3;refGene_tx=XM_006928127.3;refGene_gn=EXOC3;refGene_tx=XM_019813260.2
A1 241815285 241815403
refGene_gn=EXOC3;refGene_tx=XM_003981565.5;refGene_gn=EXOC3;refGene_tx=XM_006928127.3;refGene_gn=EXOC3;refGene_tx=XM_019813260.2
A1 241817157 241817839
refGene_gn=EXOC3;refGene_tx=XM_003981565.5;refGene_gn=EXOC3;refGene_tx=XM_006928127.3;refGene_gn=EXOC3;refGene_tx=XM_019813260.2
A1 241825871 241826091
refGene_gn=EXOC3;refGene_tx=XM_003981565.5;refGene_gn=EXOC3;refGene_tx=XM_006928127.3;refGene_gn=EXOC3;refGene_tx=XM_019813260.2
A1 241826968 241827112
refGene_gn=EXOC3;refGene_tx=XM_003981565.5;refGene_gn=EXOC3;refGene_tx=XM_006928127.3;refGene_gn=EXOC3;refGene_tx=XM_019813260.2
A1 241835727 241836781
refGene_gn=AHRR;refGene_tx=XM_023240021.1
A1 241837649 241837788
refGene_gn=AHRR;refGene_tx=XM_023240021.1
A1 241837965 241838027
refGene_gn=AHRR;refGene_tx=XM_023240021.1
A1 241841569 241841769
refGene_gn=AHRR;refGene_tx=XM_023240021.1
A1 241845702 241845818
refGene_gn=AHRR;refGene_tx=XM_023240021.1
A1 241847539 241847669
refGene_gn=AHRR;refGene_tx=XM_023240021.1
A1 241855036 241855050
refGene_gn=AHRR;refGene_tx=XM_023240021.1
A1 241855052 241855125
refGene_gn=AHRR;refGene_tx=XM_023240021.1
A1 241885125 241885229
refGene_gn=AHRR;refGene_tx=XM_023240021.1
A1 241903673 241903855
refGene_gn=AHRR;refGene_tx=XM_023240021.1
A1 241909903 241909968
refGene_gn=AHRR;refGene_tx=XM_023240021.1
A1 241931669 241931768
refGene_gn=PDCD6;refGene_tx=XM_003981566.4;refGene_gn=PDCD6;refGene_tx=XM_006928128.3;refGene_gn=PDCD6;refGene_tx=XM_006928129.4;refGene_gn=PDCD6;refGene_tx=XM_019813303.1;refGene_gn=PDCD6;refGene_tx=XM_019813318.2;refGene_gn=PDCD6;refGene_tx=XM_019813325.2
A1 241935406 241935516
refGene_gn=PDCD6;refGene_tx=XM_003981566.4;refGene_gn=PDCD6;refGene_tx=XM_006928128.3;refGene_gn=PDCD6;refGene_tx=XM_006928129.4;refGene_gn=PDCD6;refGene_tx=XM_019813303.1;refGene_gn=PDCD6;refGene_tx=XM_019813318.2;refGene_gn=PDCD6;refGene_tx=XM_019813325.2
A1 241941252 241941411
refGene_gn=PDCD6;refGene_tx=XM_003981566.4;refGene_gn=PDCD6;refGene_tx

=XM_006928128.3;refGene_gn=PDCD6;refGene_tx=XM_006928129.4;refGene_gn=PDCD6;refGene_tx=XM_019813303.1;refGene_gn=PDCD6;refGene_tx=XM_019813318.2
A1 241942430 241942475
refGene_gn=PDCD6;refGene_tx=XM_003981566.4;refGene_gn=PDCD6;refGene_tx=XM_006928128.3
A1 241951544 241951606
refGene_gn=PDCD6;refGene_tx=XM_003981566.4;refGene_gn=PDCD6;refGene_tx=XM_006928128.3;refGene_gn=PDCD6;refGene_tx=XM_019813303.1;refGene_gn=PDCD6;refGene_tx=XM_019813325.2
A1 241952555 241952650
refGene_gn=PDCD6;refGene_tx=XM_003981566.4;refGene_gn=PDCD6;refGene_tx=XM_006928128.3;refGene_gn=PDCD6;refGene_tx=XM_019813303.1;refGene_gn=PDCD6;refGene_tx=XM_019813325.2
A1 241958159 241958246
refGene_gn=SDHA;refGene_tx=XM_023240036.1;refGene_gn=SDHA;refGene_tx=XM_023240041.1
A1 241960134 241960248
refGene_gn=SDHA;refGene_tx=XM_023240036.1;refGene_gn=SDHA;refGene_tx=XM_023240041.1
A1 241962478 241962609
refGene_gn=SDHA;refGene_tx=XM_023240036.1;refGene_gn=SDHA;refGene_tx=XM_023240041.1
A1 241962810 241962922
refGene_gn=SDHA;refGene_tx=XM_023240036.1;refGene_gn=SDHA;refGene_tx=XM_023240041.1
A1 241967602 241967721
refGene_gn=SDHA;refGene_tx=XM_023240036.1;refGene_gn=SDHA;refGene_tx=XM_023240041.1
A1 241970506 241970678
refGene_gn=SDHA;refGene_tx=XM_023240036.1;refGene_gn=SDHA;refGene_tx=XM_023240041.1
A1 241971487 241971683
refGene_gn=SDHA;refGene_tx=XM_023240036.1;refGene_gn=SDHA;refGene_tx=XM_023240041.1
A1 241972473 241972642
refGene_gn=SDHA;refGene_tx=XM_023240036.1;refGene_gn=SDHA;refGene_tx=XM_023240041.1
A1 241974881 241975006
refGene_gn=SDHA;refGene_tx=XM_023240036.1;refGene_gn=SDHA;refGene_tx=XM_023240041.1
A1 241975660 241975809
refGene_gn=SDHA;refGene_tx=XM_023240036.1;refGene_gn=SDHA;refGene_tx=XM_023240041.1
A1 241976686 241976851
refGene_gn=SDHA;refGene_tx=XM_023240036.1;refGene_gn=SDHA;refGene_tx=XM_023240041.1
A1 241977131 241977275
refGene_gn=SDHA;refGene_tx=XM_023240036.1;refGene_gn=SDHA;refGene_tx=XM_023240041.1

A1	241979470	241979632
refGene_gn=SDHA;refGene_tx=XM_023240036.1;refGene_gn=SDHA;refGene_tx=XM_023240041.1
A1	241980865	241980955
refGene_gn=SDHA;refGene_tx=XM_023240036.1;refGene_gn=SDHA;refGene_tx=XM_023240041.1
A1	241987855	241987918
refGene_gn=SDHA;refGene_tx=XM_023240036.1;refGene_gn=SDHA;refGene_tx=XM_023240041.1
A1	241989880	241990001
refGene_gn=CCDC127;refGene_tx=XM_003981569.5;refGene_gn=CCDC127;refGene_tx=XM_011287225.3
A1	241998108	241998770
refGene_gn=CCDC127;refGene_tx=XM_003981569.5;refGene_gn=CCDC127;refGene_tx=XM_011287225.3
A1	242005328	242005971
refGene_gn=LRRC14B;refGene_tx=XM_003981570.5;refGene_gn=LRRC14B;refGene_tx=XM_019813383.2
A1	242008601	242009491
refGene_gn=LRRC14B;refGene_tx=XM_003981570.5;refGene_gn=LRRC14B;refGene_tx=XM_019813383.2
A1	242010933	242010939
refGene_gn=PLEKHG4B;refGene_tx=XM_006928146.4;refGene_gn=PLEKHG4B;refGene_tx=XM_023240052.1;refGene_gn=PLEKHG4B;refGene_tx=XM_023240054.1
A1	242012990	242013127
refGene_gn=PLEKHG4B;refGene_tx=XM_006928146.4;refGene_gn=PLEKHG4B;refGene_tx=XM_023240052.1;refGene_gn=PLEKHG4B;refGene_tx=XM_023240054.1
A1	242016275	242016536
refGene_gn=PLEKHG4B;refGene_tx=XM_006928146.4;refGene_gn=PLEKHG4B;refGene_tx=XM_023240052.1;refGene_gn=PLEKHG4B;refGene_tx=XM_023240054.1
A1	242016748	242016856
refGene_gn=PLEKHG4B;refGene_tx=XM_006928146.4;refGene_gn=PLEKHG4B;refGene_tx=XM_023240052.1;refGene_gn=PLEKHG4B;refGene_tx=XM_023240054.1
A1	242017418	242017599
refGene_gn=PLEKHG4B;refGene_tx=XM_006928146.4;refGene_gn=PLEKHG4B;refGene_tx=XM_023240052.1;refGene_gn=PLEKHG4B;refGene_tx=XM_023240054.1
A1	242018180	242018351
refGene_gn=PLEKHG4B;refGene_tx=XM_006928146.4;refGene_gn=PLEKHG4B;refGene_tx=XM_023240052.1;refGene_gn=PLEKHG4B;refGene_tx=XM_023240054.1
A1	242019539	242019770
refGene_gn=PLEKHG4B;refGene_tx=XM_006928146.4;refGene_gn=PLEKHG4B;refGene_tx=XM_023240052.1;refGene_gn=PLEKHG4B;refGene_tx=XM_023240054.1
A1	242019846	242019936
refGene_gn=PLEKHG4B;refGene_tx=XM_006928146.4;refGene_gn=PLEKHG4B;refGene_tx=XM_023240052.1;refGene_gn=PLEKHG4B;refGene_tx=XM_023240054.1
A1	242021669	242021922
refGene_gn=PLEKHG4B;refGene_tx=XM_006928146.4;refGene_gn=PLEKHG4B;refGene_tx=XM_023240052.1;refGene_gn=PLEKHG4B;refGene_tx=XM_023240054.1
A1	242024404	242025150
refGene_gn=PLEKHG4B;refGene_tx=XM_006928146.4;refGene_gn=PLEKHG4B;refG

ene_tx=XM_023240052.1;refGene_gn=PLEKHG4B;refGene_tx=XM_023240054.1
A1 242028834 242028973
refGene_gn=PLEKHG4B;refGene_tx=XM_006928146.4;refGene_gn=PLEKHG4B;refGene_tx=XM_023240052.1;refGene_gn=PLEKHG4B;refGene_tx=XM_023240054.1
A1 242029678 242029821
refGene_gn=PLEKHG4B;refGene_tx=XM_006928146.4;refGene_gn=PLEKHG4B;refGene_tx=XM_023240052.1;refGene_gn=PLEKHG4B;refGene_tx=XM_023240054.1
A1 242030450 242030549
refGene_gn=PLEKHG4B;refGene_tx=XM_006928146.4;refGene_gn=PLEKHG4B;refGene_tx=XM_023240052.1;refGene_gn=PLEKHG4B;refGene_tx=XM_023240054.1
A1 242030813 242030930
refGene_gn=PLEKHG4B;refGene_tx=XM_006928146.4;refGene_gn=PLEKHG4B;refGene_tx=XM_023240052.1;refGene_gn=PLEKHG4B;refGene_tx=XM_023240054.1
A1 242032904 242032991
refGene_gn=PLEKHG4B;refGene_tx=XM_006928146.4;refGene_gn=PLEKHG4B;refGene_tx=XM_023240052.1
A1 242038049 242038143
refGene_gn=PLEKHG4B;refGene_tx=XM_006928146.4;refGene_gn=PLEKHG4B;refGene_tx=XM_023240052.1;refGene_gn=PLEKHG4B;refGene_tx=XM_023240054.1
A1 242042901 242043025
refGene_gn=PLEKHG4B;refGene_tx=XM_006928146.4;refGene_gn=PLEKHG4B;refGene_tx=XM_023240052.1;refGene_gn=PLEKHG4B;refGene_tx=XM_023240054.1
A1 242043556 242043748
refGene_gn=PLEKHG4B;refGene_tx=XM_006928146.4;refGene_gn=PLEKHG4B;refGene_tx=XM_023240052.1;refGene_gn=PLEKHG4B;refGene_tx=XM_023240054.1
A1 242048285 242049462
refGene_gn=PLEKHG4B;refGene_tx=XM_006928146.4;refGene_gn=PLEKHG4B;refGene_tx=XM_023240052.1;refGene_gn=PLEKHG4B;refGene_tx=XM_023240054.1
A1 242069496 242069694
refGene_gn=PLEKHG4B;refGene_tx=XM_006928146.4;refGene_gn=PLEKHG4B;refGene_tx=XM_023240052.1;refGene_gn=PLEKHG4B;refGene_tx=XM_023240054.1
A1 242091682 242091721
refGene_gn=PLEKHG4B;refGene_tx=XM_006928146.4;refGene_gn=PLEKHG4B;refGene_tx=XM_023240052.1;refGene_gn=PLEKHG4B;refGene_tx=XM_023240054.1
A2 9044 9479
refGene_gn=LOC111559571;refGene_tx=XM_023250837.1
A2 23699 23826
refGene_gn=LOC111559571;refGene_tx=XM_023250837.1
A2 24635 24819
refGene_gn=LOC111559571;refGene_tx=XM_023250837.1
A2 25255 26018
refGene_gn=LOC111559571;refGene_tx=XM_023250837.1
A2 68690 68840
refGene_gn=PLPP2;refGene_tx=XM_019815034.2;refGene_gn=PLPP2;refGene_tx=XM_019815042.2;refGene_gn=PLPP2;refGene_tx=XM_023242378.1;refGene_gn=PLPP2;refGene_tx=XM_023242384.1;refGene_gn=PLPP2;refGene_tx=XM_023242385.1
A2 68984 69161
refGene_gn=PLPP2;refGene_tx=XM_019815034.2;refGene_gn=PLPP2;refGene_tx=XM_019815042.2;refGene_gn=PLPP2;refGene_tx=XM_023242378.1;refGene_gn=

PLPP2;refGene_tx=XM_023242384.1;refGene_gn=PLPP2;refGene_tx=XM_0232423
85.1
A2 69386 69444
refGene_gn=PLPP2;refGene_tx=XM_019815034.2;refGene_gn=PLPP2;refGene_tx
=XM_019815042.2;refGene_gn=PLPP2;refGene_tx=XM_023242378.1;refGene_gn=
PLPP2;refGene_tx=XM_023242384.1;refGene_gn=PLPP2;refGene_tx=XM_0232423
85.1
A2 73279 73593
refGene_gn=PLPP2;refGene_tx=XM_019815034.2;refGene_gn=PLPP2;refGene_tx
=XM_019815042.2;refGene_gn=PLPP2;refGene_tx=XM_023242378.1;refGene_gn=
PLPP2;refGene_tx=XM_023242384.1;refGene_gn=PLPP2;refGene_tx=XM_0232423
85.1
A2 73807 73959
refGene_gn=PLPP2;refGene_tx=XM_019815034.2;refGene_gn=PLPP2;refGene_tx
=XM_019815042.2;refGene_gn=PLPP2;refGene_tx=XM_023242378.1;refGene_gn=
PLPP2;refGene_tx=XM_023242384.1;refGene_gn=PLPP2;refGene_tx=XM_0232423
85.1
A2 76761 76813
refGene_gn=PLPP2;refGene_tx=XM_023242378.1;refGene_gn=PLPP2;refGene_tx
=XM_023242384.1;refGene_gn=PLPP2;refGene_tx=XM_023242385.1
A2 84440 84462
refGene_gn=MIER2;refGene_tx=XM_019815052.1;refGene_gn=MIER2;refGene_tx
=XM_023242388.1;refGene_gn=MIER2;refGene_tx=XM_023242393.1;refGene_gn=
MIER2;refGene_tx=XM_023242397.1
A2 84719 84723
refGene_gn=MIER2;refGene_tx=XM_023242390.1;refGene_gn=MIER2;refGene_tx
=XM_023242391.1
A2 84912 85342
refGene_gn=MIER2;refGene_tx=XM_019815052.1;refGene_gn=MIER2;refGene_tx
=XM_023242388.1;refGene_gn=MIER2;refGene_tx=XM_023242390.1;refGene_gn=
MIER2;refGene_tx=XM_023242391.1;refGene_gn=MIER2;refGene_tx=XM_0232423
93.1;refGene_gn=MIER2;refGene_tx=XM_023242397.1
A2 86221 86343
refGene_gn=MIER2;refGene_tx=XM_019815052.1;refGene_gn=MIER2;refGene_tx
=XM_023242388.1;refGene_gn=MIER2;refGene_tx=XM_023242390.1;refGene_gn=
MIER2;refGene_tx=XM_023242391.1;refGene_gn=MIER2;refGene_tx=XM_0232423
93.1;refGene_gn=MIER2;refGene_tx=XM_023242397.1
A2 86431 86556
refGene_gn=MIER2;refGene_tx=XM_019815052.1;refGene_gn=MIER2;refGene_tx
=XM_023242388.1;refGene_gn=MIER2;refGene_tx=XM_023242390.1;refGene_gn=
MIER2;refGene_tx=XM_023242391.1;refGene_gn=MIER2;refGene_tx=XM_0232423
93.1;refGene_gn=MIER2;refGene_tx=XM_023242397.1
A2 88740 88835
refGene_gn=MIER2;refGene_tx=XM_019815052.1;refGene_gn=MIER2;refGene_tx
=XM_023242388.1;refGene_gn=MIER2;refGene_tx=XM_023242390.1;refGene_gn=
MIER2;refGene_tx=XM_023242391.1;refGene_gn=MIER2;refGene_tx=XM_0232423
93.1;refGene_gn=MIER2;refGene_tx=XM_023242397.1
A2 88924 89006
refGene_gn=MIER2;refGene_tx=XM_019815052.1;refGene_gn=MIER2;refGene_tx
=XM_023242388.1;refGene_gn=MIER2;refGene_tx=XM_023242390.1;refGene_gn=

MIER2;refGene_tx=XM_023242391.1;refGene_gn=MIER2;refGene_tx=XM_0232423
93.1;refGene_gn=MIER2;refGene_tx=XM_023242397.1
A2 90117 90269
refGene_gn=MIER2;refGene_tx=XM_019815052.1;refGene_gn=MIER2;refGene_tx
=XM_023242388.1;refGene_gn=MIER2;refGene_tx=XM_023242390.1;refGene_gn=
MIER2;refGene_tx=XM_023242391.1;refGene_gn=MIER2;refGene_tx=XM_0232423
93.1;refGene_gn=MIER2;refGene_tx=XM_023242397.1
A2 93298 93368
refGene_gn=MIER2;refGene_tx=XM_019815052.1;refGene_gn=MIER2;refGene_tx
=XM_023242388.1;refGene_gn=MIER2;refGene_tx=XM_023242390.1;refGene_gn=
MIER2;refGene_tx=XM_023242391.1;refGene_gn=MIER2;refGene_tx=XM_0232423
93.1;refGene_gn=MIER2;refGene_tx=XM_023242397.1
A2 93888 93980
refGene_gn=MIER2;refGene_tx=XM_019815052.1;refGene_gn=MIER2;refGene_tx
=XM_023242388.1;refGene_gn=MIER2;refGene_tx=XM_023242390.1;refGene_gn=
MIER2;refGene_tx=XM_023242391.1;refGene_gn=MIER2;refGene_tx=XM_0232423
93.1;refGene_gn=MIER2;refGene_tx=XM_023242397.1
A2 94471 94595
refGene_gn=MIER2;refGene_tx=XM_019815052.1;refGene_gn=MIER2;refGene_tx
=XM_023242388.1;refGene_gn=MIER2;refGene_tx=XM_023242390.1;refGene_gn=
MIER2;refGene_tx=XM_023242391.1;refGene_gn=MIER2;refGene_tx=XM_0232423
93.1;refGene_gn=MIER2;refGene_tx=XM_023242397.1
A2 95095 95221
refGene_gn=MIER2;refGene_tx=XM_019815052.1;refGene_gn=MIER2;refGene_tx
=XM_023242388.1;refGene_gn=MIER2;refGene_tx=XM_023242390.1;refGene_gn=
MIER2;refGene_tx=XM_023242391.1;refGene_gn=MIER2;refGene_tx=XM_0232423
93.1;refGene_gn=MIER2;refGene_tx=XM_023242397.1
A2 98766 98909
refGene_gn=MIER2;refGene_tx=XM_023242388.1;refGene_gn=MIER2;refGene_tx
=XM_023242390.1;refGene_gn=MIER2;refGene_tx=XM_023242391.1;refGene_gn=
MIER2;refGene_tx=XM_023242393.1;refGene_gn=MIER2;refGene_tx=XM_0232423
97.1
A2 100095 100186
refGene_gn=MIER2;refGene_tx=XM_019815052.1;refGene_gn=MIER2;refGene_tx
=XM_023242388.1;refGene_gn=MIER2;refGene_tx=XM_023242390.1;refGene_gn=
MIER2;refGene_tx=XM_023242391.1;refGene_gn=MIER2;refGene_tx=XM_0232423
93.1
A2 105692 105709 refGene_gn=MIER2;refGene_tx=XM_019815052.1
A2 106954 106963
refGene_gn=MIER2;refGene_tx=XM_023242388.1;refGene_gn=MIER2;refGene_tx
=XM_023242390.1;refGene_gn=MIER2;refGene_tx=XM_023242391.1;refGene_gn=
MIER2;refGene_tx=XM_023242393.1
A2 114651 114912
refGene_gn=THEG;refGene_tx=XM_006928149.3;refGene_gn=THEG;refGene_tx=X
M_006928151.3;refGene_gn=THEG;refGene_tx=XM_006928152.3
A2 118346 118506
refGene_gn=THEG;refGene_tx=XM_006928149.3;refGene_gn=THEG;refGene_tx=X
M_006928151.3;refGene_gn=THEG;refGene_tx=XM_006928152.3
A2 121121 121517 refGene_gn=THEG;refGene_tx=XM_006928150.3
A2 121622 121740

refGene_gn=THEG;refGene_tx=XM_006928149.3;refGene_gn=THEG;refGene_tx=XM_006928150.3;refGene_gn=THEG;refGene_tx=XM_006928151.3
A2 123219 123289
refGene_gn=THEG;refGene_tx=XM_006928149.3;refGene_gn=THEG;refGene_tx=XM_006928150.3;refGene_gn=THEG;refGene_tx=XM_006928151.3;refGene_gn=THEG;refGene_tx=XM_006928152.3
A2 124214 124276
refGene_gn=THEG;refGene_tx=XM_006928149.3;refGene_gn=THEG;refGene_tx=XM_006928150.3;refGene_gn=THEG;refGene_tx=XM_006928151.3;refGene_gn=THEG;refGene_tx=XM_006928152.3
A2 124701 124773
refGene_gn=THEG;refGene_tx=XM_006928149.3;refGene_gn=THEG;refGene_tx=XM_006928150.3;refGene_gn=THEG;refGene_tx=XM_006928152.3
A2 125064 125184
refGene_gn=THEG;refGene_tx=XM_006928149.3;refGene_gn=THEG;refGene_tx=XM_006928150.3;refGene_gn=THEG;refGene_tx=XM_006928151.3;refGene_gn=THEG;refGene_tx=XM_006928152.3
A2 126152 126451
refGene_gn=THEG;refGene_tx=XM_006928149.3;refGene_gn=THEG;refGene_tx=XM_006928150.3;refGene_gn=THEG;refGene_tx=XM_006928151.3;refGene_gn=THEG;refGene_tx=XM_006928152.3
A2 143836 145171
refGene_gn=C2CD4C;refGene_tx=XM_023242412.1;refGene_gn=C2CD4C;refGene_tx=XM_023242415.1
A2 145699 145838 refGene_gn=C2CD4C;refGene_tx=XM_023242412.1
A2 152811 152940 refGene_gn=SHC2;refGene_tx=XM_023242418.1
A2 154838 155149 refGene_gn=SHC2;refGene_tx=XM_023242418.1
A2 156083 156307 refGene_gn=SHC2;refGene_tx=XM_023242419.1
A2 156962 157106
refGene_gn=SHC2;refGene_tx=XM_023242418.1;refGene_gn=SHC2;refGene_tx=XM_023242419.1
A2 159687 159745
refGene_gn=SHC2;refGene_tx=XM_023242418.1;refGene_gn=SHC2;refGene_tx=XM_023242419.1
A2 160374 160528
refGene_gn=SHC2;refGene_tx=XM_023242418.1;refGene_gn=SHC2;refGene_tx=XM_023242419.1
A2 161770 161897
refGene_gn=SHC2;refGene_tx=XM_023242418.1;refGene_gn=SHC2;refGene_tx=XM_023242419.1
A2 161991 162043
refGene_gn=SHC2;refGene_tx=XM_023242418.1;refGene_gn=SHC2;refGene_tx=XM_023242419.1
A2 162182 162236
refGene_gn=SHC2;refGene_tx=XM_023242418.1;refGene_gn=SHC2;refGene_tx=XM_023242419.1
A2 163625 163745
refGene_gn=SHC2;refGene_tx=XM_023242418.1;refGene_gn=SHC2;refGene_tx=XM_023242419.1
A2 163924 163985

refGene_gn=SHC2;refGene_tx=XM_023242418.1;refGene_gn=SHC2;refGene_tx=XM_023242419.1
A2 165927 165998
refGene_gn=SHC2;refGene_tx=XM_023242418.1;refGene_gn=SHC2;refGene_tx=XM_023242419.1
A2 179018 179483
refGene_gn=SHC2;refGene_tx=XM_023242418.1;refGene_gn=SHC2;refGene_tx=XM_023242419.1
A2 182138 182623 refGene_gn=ODF3L2;refGene_tx=XM_023242420.1
A2 184201 184315 refGene_gn=ODF3L2;refGene_tx=XM_023242420.1
A2 186109 186217 refGene_gn=ODF3L2;refGene_tx=XM_023242420.1
A2 188645 188748 refGene_gn=ODF3L2;refGene_tx=XM_023242420.1
A2 203142 203203
refGene_gn=MADCAM1;refGene_tx=XM_003981624.4;refGene_gn=MADCAM1;refGene_tx=XM_019815108.2
A2 203395 203668
refGene_gn=MADCAM1;refGene_tx=XM_003981624.4;refGene_gn=MADCAM1;refGene_tx=XM_019815108.2
A2 204186 204510
refGene_gn=MADCAM1;refGene_tx=XM_003981624.4;refGene_gn=MADCAM1;refGene_tx=XM_019815108.2
A2 206188 206563
refGene_gn=MADCAM1;refGene_tx=XM_003981624.4;refGene_gn=MADCAM1;refGene_tx=XM_019815108.2
A2 207223 207333
refGene_gn=MADCAM1;refGene_tx=XM_003981624.4;refGene_gn=MADCAM1;refGene_tx=XM_019815108.2
A2 208743 209081 refGene_gn=TPGS1;refGene_tx=XM_023242431.1
A2 214941 215479 refGene_gn=TPGS1;refGene_tx=XM_023242431.1
A2 221806 221983
refGene_gn=CDC34;refGene_tx=XM_023242434.1;refGene_gn=CDC34;refGene_tx=XM_023242436.1
A2 224783 224870
refGene_gn=CDC34;refGene_tx=XM_023242434.1;refGene_gn=CDC34;refGene_tx=XM_023242436.1
A2 224978 225076
refGene_gn=CDC34;refGene_tx=XM_023242434.1;refGene_gn=CDC34;refGene_tx=XM_023242436.1
A2 225854 225989
refGene_gn=CDC34;refGene_tx=XM_023242434.1;refGene_gn=CDC34;refGene_tx=XM_023242436.1
A2 229060 229322
refGene_gn=CDC34;refGene_tx=XM_023242434.1;refGene_gn=CDC34;refGene_tx=XM_023242436.1
A2 238421 238557
refGene_gn=BSG;refGene_tx=XM_023242437.1;refGene_gn=BSG;refGene_tx=XM_023242446.1
A2 244357 244514
refGene_gn=BSG;refGene_tx=XM_023242437.1;refGene_gn=BSG;refGene_tx=XM_023242446.1

A2	245231	245314	refGene_gn=BSG;refGene_tx=XM_023242437.1;refGene_gn=BSG;refGene_tx=XM_023242446.1
A2	245435	245572	refGene_gn=BSG;refGene_tx=XM_023242437.1;refGene_gn=BSG;refGene_tx=XM_023242446.1
A2	245654	245931	refGene_gn=BSG;refGene_tx=XM_023242437.1;refGene_gn=BSG;refGene_tx=XM_023242446.1
A2	246457	246482	refGene_gn=BSG;refGene_tx=XM_023242437.1;refGene_gn=BSG;refGene_tx=XM_023242446.1
A2	246642	246703	refGene_gn=BSG;refGene_tx=XM_023242437.1;refGene_gn=BSG;refGene_tx=XM_023242446.1
A2	252515	253135	refGene_gn=HCN2;refGene_tx=XM_023242496.1
A2	263552	263976	refGene_gn=HCN2;refGene_tx=XM_023242496.1
A2	264517	264679	refGene_gn=HCN2;refGene_tx=XM_023242496.1
A2	265125	265344	refGene_gn=HCN2;refGene_tx=XM_023242496.1
A2	266759	266906	refGene_gn=HCN2;refGene_tx=XM_023242496.1
A2	270285	270526	refGene_gn=HCN2;refGene_tx=XM_023242496.1
A2	270766	270931	refGene_gn=HCN2;refGene_tx=XM_023242496.1
A2	272760	273470	refGene_gn=HCN2;refGene_tx=XM_023242496.1
A2	274336	274386	refGene_gn=POLRMT;refGene_tx=XM_023242462.1;refGene_gn=POLRMT;refGene_tx=XM_023242468.1;refGene_gn=POLRMT;refGene_tx=XM_023242485.1;refGene_gn=POLRMT;refGene_tx=XM_023242487.1
A2	274485	274732	refGene_gn=POLRMT;refGene_tx=XM_023242462.1;refGene_gn=POLRMT;refGene_tx=XM_023242468.1;refGene_gn=POLRMT;refGene_tx=XM_023242471.1;refGene_gn=POLRMT;refGene_tx=XM_023242478.1;refGene_gn=POLRMT;refGene_tx=XM_023242485.1;refGene_gn=POLRMT;refGene_tx=XM_023242487.1
A2	274941	275014	refGene_gn=POLRMT;refGene_tx=XM_023242462.1;refGene_gn=POLRMT;refGene_tx=XM_023242468.1;refGene_gn=POLRMT;refGene_tx=XM_023242471.1;refGene_gn=POLRMT;refGene_tx=XM_023242478.1;refGene_gn=POLRMT;refGene_tx=XM_023242485.1;refGene_gn=POLRMT;refGene_tx=XM_023242487.1
A2	275904	276003	refGene_gn=POLRMT;refGene_tx=XM_023242462.1;refGene_gn=POLRMT;refGene_tx=XM_023242468.1;refGene_gn=POLRMT;refGene_tx=XM_023242471.1;refGene_gn=POLRMT;refGene_tx=XM_023242478.1;refGene_gn=POLRMT;refGene_tx=XM_023242485.1;refGene_gn=POLRMT;refGene_tx=XM_023242487.1
A2	276086	276139	refGene_gn=POLRMT;refGene_tx=XM_023242462.1;refGene_gn=POLRMT;refGene_tx=XM_023242468.1;refGene_gn=POLRMT;refGene_tx=XM_023242471.1;refGene_gn=POLRMT;refGene_tx=XM_023242478.1;refGene_gn=POLRMT;refGene_tx=XM_023242485.1;refGene_gn=POLRMT;refGene_tx=XM_023242487.1
A2	276223	276337	refGene_gn=POLRMT;refGene_tx=XM_023242462.1;refGene_gn=POLRMT;refGene_tx=XM_023242468.1;refGene_gn=POLRMT;refGene_tx=XM_023242471.1;refGene_

gn=POLRMT;refGene_tx=XM_023242478.1;refGene_gn=POLRMT;refGene_tx=XM_023242485.1;refGene_gn=POLRMT;refGene_tx=XM_023242487.1
A2 276421 276508
refGene_gn=POLRMT;refGene_tx=XM_023242462.1;refGene_gn=POLRMT;refGene_tx=XM_023242468.1;refGene_gn=POLRMT;refGene_tx=XM_023242471.1;refGene_gn=POLRMT;refGene_tx=XM_023242478.1;refGene_gn=POLRMT;refGene_tx=XM_023242485.1;refGene_gn=POLRMT;refGene_tx=XM_023242487.1
A2 276801 276981
refGene_gn=POLRMT;refGene_tx=XM_023242462.1;refGene_gn=POLRMT;refGene_tx=XM_023242468.1;refGene_gn=POLRMT;refGene_tx=XM_023242471.1;refGene_gn=POLRMT;refGene_tx=XM_023242478.1;refGene_gn=POLRMT;refGene_tx=XM_023242485.1;refGene_gn=POLRMT;refGene_tx=XM_023242487.1
A2 277227 277350
refGene_gn=POLRMT;refGene_tx=XM_023242462.1;refGene_gn=POLRMT;refGene_tx=XM_023242468.1;refGene_gn=POLRMT;refGene_tx=XM_023242471.1;refGene_gn=POLRMT;refGene_tx=XM_023242478.1;refGene_gn=POLRMT;refGene_tx=XM_023242485.1;refGene_gn=POLRMT;refGene_tx=XM_023242487.1
A2 277613 277736
refGene_gn=POLRMT;refGene_tx=XM_023242462.1;refGene_gn=POLRMT;refGene_tx=XM_023242468.1;refGene_gn=POLRMT;refGene_tx=XM_023242471.1;refGene_gn=POLRMT;refGene_tx=XM_023242478.1;refGene_gn=POLRMT;refGene_tx=XM_023242485.1;refGene_gn=POLRMT;refGene_tx=XM_023242487.1
A2 278049 278838
refGene_gn=POLRMT;refGene_tx=XM_023242462.1;refGene_gn=POLRMT;refGene_tx=XM_023242468.1;refGene_gn=POLRMT;refGene_tx=XM_023242471.1;refGene_gn=POLRMT;refGene_tx=XM_023242478.1;refGene_gn=POLRMT;refGene_tx=XM_023242485.1;refGene_gn=POLRMT;refGene_tx=XM_023242487.1
A2 279020 279245
refGene_gn=POLRMT;refGene_tx=XM_023242462.1;refGene_gn=POLRMT;refGene_tx=XM_023242468.1;refGene_gn=POLRMT;refGene_tx=XM_023242471.1;refGene_gn=POLRMT;refGene_tx=XM_023242478.1;refGene_gn=POLRMT;refGene_tx=XM_023242485.1;refGene_gn=POLRMT;refGene_tx=XM_023242487.1
A2 279554 279722
refGene_gn=POLRMT;refGene_tx=XM_023242462.1;refGene_gn=POLRMT;refGene_tx=XM_023242468.1;refGene_gn=POLRMT;refGene_tx=XM_023242471.1;refGene_gn=POLRMT;refGene_tx=XM_023242478.1;refGene_gn=POLRMT;refGene_tx=XM_023242485.1;refGene_gn=POLRMT;refGene_tx=XM_023242487.1
A2 279803 279968
refGene_gn=POLRMT;refGene_tx=XM_023242462.1;refGene_gn=POLRMT;refGene_tx=XM_023242468.1;refGene_gn=POLRMT;refGene_tx=XM_023242471.1;refGene_gn=POLRMT;refGene_tx=XM_023242478.1;refGene_gn=POLRMT;refGene_tx=XM_023242485.1;refGene_gn=POLRMT;refGene_tx=XM_023242487.1
A2 280169 280353
refGene_gn=POLRMT;refGene_tx=XM_023242462.1;refGene_gn=POLRMT;refGene_tx=XM_023242468.1;refGene_gn=POLRMT;refGene_tx=XM_023242471.1;refGene_gn=POLRMT;refGene_tx=XM_023242478.1;refGene_gn=POLRMT;refGene_tx=XM_023242485.1;refGene_gn=POLRMT;refGene_tx=XM_023242487.1
A2 281162 281355
refGene_gn=POLRMT;refGene_tx=XM_023242462.1;refGene_gn=POLRMT;refGene_tx=XM_023242468.1;refGene_gn=POLRMT;refGene_tx=XM_023242471.1;refGene_

gn=POLRMT;refGene_tx=XM_023242478.1;refGene_gn=POLRMT;refGene_tx=XM_023242485.1;refGene_gn=POLRMT;refGene_tx=XM_023242487.1
A2 281478 281609
refGene_gn=POLRMT;refGene_tx=XM_023242462.1;refGene_gn=POLRMT;refGene_tx=XM_023242468.1;refGene_gn=POLRMT;refGene_tx=XM_023242471.1;refGene_gn=POLRMT;refGene_tx=XM_023242478.1;refGene_gn=POLRMT;refGene_tx=XM_023242485.1;refGene_gn=POLRMT;refGene_tx=XM_023242487.1
A2 284975 285583
refGene_gn=POLRMT;refGene_tx=XM_023242462.1;refGene_gn=POLRMT;refGene_tx=XM_023242468.1;refGene_gn=POLRMT;refGene_tx=XM_023242471.1;refGene_gn=POLRMT;refGene_tx=XM_023242478.1
A2 288067 288166
refGene_gn=POLRMT;refGene_tx=XM_023242462.1;refGene_gn=POLRMT;refGene_tx=XM_023242468.1;refGene_gn=POLRMT;refGene_tx=XM_023242471.1;refGene_gn=POLRMT;refGene_tx=XM_023242478.1
A2 288694 288782
refGene_gn=POLRMT;refGene_tx=XM_023242462.1;refGene_gn=POLRMT;refGene_tx=XM_023242468.1;refGene_gn=POLRMT;refGene_tx=XM_023242471.1;refGene_gn=POLRMT;refGene_tx=XM_023242478.1
A2 295171 295385 refGene_gn=FGF22;refGene_tx=XM_023242667.1
A2 297791 297895 refGene_gn=FGF22;refGene_tx=XM_023242667.1
A2 298068 298263 refGene_gn=FGF22;refGene_tx=XM_023242667.1
A2 300013 300163
refGene_gn=RNF126;refGene_tx=XM_023242595.1;refGene_gn=RNF126;refGene_tx=XM_023242596.1;refGene_gn=RNF126;refGene_tx=XM_023242598.1;refGene_gn=RNF126;refGene_tx=XM_023242609.1
A2 300266 300382
refGene_gn=RNF126;refGene_tx=XM_023242595.1;refGene_gn=RNF126;refGene_tx=XM_023242596.1;refGene_gn=RNF126;refGene_tx=XM_023242598.1;refGene_gn=RNF126;refGene_tx=XM_023242609.1
A2 300463 300557
refGene_gn=RNF126;refGene_tx=XM_023242595.1;refGene_gn=RNF126;refGene_tx=XM_023242596.1;refGene_gn=RNF126;refGene_tx=XM_023242598.1;refGene_gn=RNF126;refGene_tx=XM_023242609.1
A2 301095 301165
refGene_gn=RNF126;refGene_tx=XM_023242595.1;refGene_gn=RNF126;refGene_tx=XM_023242596.1;refGene_gn=RNF126;refGene_tx=XM_023242598.1;refGene_gn=RNF126;refGene_tx=XM_023242609.1
A2 301239 301308
refGene_gn=RNF126;refGene_tx=XM_023242595.1;refGene_gn=RNF126;refGene_tx=XM_023242596.1;refGene_gn=RNF126;refGene_tx=XM_023242598.1;refGene_gn=RNF126;refGene_tx=XM_023242609.1
A2 302361 302606
refGene_gn=RNF126;refGene_tx=XM_023242595.1;refGene_gn=RNF126;refGene_tx=XM_023242596.1;refGene_gn=RNF126;refGene_tx=XM_023242598.1;refGene_gn=RNF126;refGene_tx=XM_023242609.1
A2 302957 303039
refGene_gn=RNF126;refGene_tx=XM_023242595.1;refGene_gn=RNF126;refGene_tx=XM_023242596.1;refGene_gn=RNF126;refGene_tx=XM_023242598.1;refGene_gn=RNF126;refGene_tx=XM_023242609.1

A2	303173	303232
refGene_gn=RNF126;refGene_tx=XM_023242595.1;refGene_gn=RNF126;refGene_tx=XM_023242596.1;refGene_gn=RNF126;refGene_tx=XM_023242598.1;refGene_gn=RNF126;refGene_tx=XM_023242609.1
A2	310724	310799
refGene_gn=RNF126;refGene_tx=XM_023242595.1;refGene_gn=RNF126;refGene_tx=XM_023242596.1;refGene_gn=RNF126;refGene_tx=XM_023242598.1;refGene_gn=RNF126;refGene_tx=XM_023242609.1
A2	319427	319530
refGene_gn=FSTL3;refGene_tx=XM_023242624.1;refGene_gn=FSTL3;refGene_tx=XM_023242630.1;refGene_gn=FSTL3;refGene_tx=XM_023242634.1;refGene_gn=FSTL3;refGene_tx=XM_023242637.1;refGene_gn=FSTL3;refGene_tx=XM_023242641.1
A2	320882	321068
refGene_gn=FSTL3;refGene_tx=XM_023242624.1;refGene_gn=FSTL3;refGene_tx=XM_023242630.1;refGene_gn=FSTL3;refGene_tx=XM_023242634.1;refGene_gn=FSTL3;refGene_tx=XM_023242637.1;refGene_gn=FSTL3;refGene_tx=XM_023242641.1
A2	323230	323446
refGene_gn=FSTL3;refGene_tx=XM_023242624.1;refGene_gn=FSTL3;refGene_tx=XM_023242630.1;refGene_gn=FSTL3;refGene_tx=XM_023242634.1;refGene_gn=FSTL3;refGene_tx=XM_023242637.1;refGene_gn=FSTL3;refGene_tx=XM_023242641.1
A2	323807	324181
refGene_gn=FSTL3;refGene_tx=XM_023242624.1;refGene_gn=FSTL3;refGene_tx=XM_023242630.1;refGene_gn=FSTL3;refGene_tx=XM_023242634.1;refGene_gn=FSTL3;refGene_tx=XM_023242637.1;refGene_gn=FSTL3;refGene_tx=XM_023242641.1
A2	324499	324551
refGene_gn=FSTL3;refGene_tx=XM_023242624.1;refGene_gn=FSTL3;refGene_tx=XM_023242630.1;refGene_gn=FSTL3;refGene_tx=XM_023242637.1
A2	326315	326402	refGene_gn=PRSS57;refGene_tx=XM_003981632.3
A2	326403	326553	refGene_gn=PRSS57;refGene_tx=XM_003981632.3
A2	327615	327879	refGene_gn=PRSS57;refGene_tx=XM_003981632.3
A2	329557	329702	refGene_gn=PRSS57;refGene_tx=XM_003981632.3
A2	330124	330278	refGene_gn=PRSS57;refGene_tx=XM_003981632.3
A2	330508	330587	refGene_gn=PRSS57;refGene_tx=XM_003981632.3
A2	337638	337643	refGene_gn=PALM;refGene_tx=XM_023242580.1
A2	345585	345746
refGene_gn=PALM;refGene_tx=XM_023242568.1;refGene_gn=PALM;refGene_tx=XM_023242570.1;refGene_gn=PALM;refGene_tx=XM_023242573.1;refGene_gn=PALM;refGene_tx=XM_023242577.1;refGene_gn=PALM;refGene_tx=XM_023242583.1
A2	348097	348149
refGene_gn=PALM;refGene_tx=XM_023242568.1;refGene_gn=PALM;refGene_tx=XM_023242570.1;refGene_gn=PALM;refGene_tx=XM_023242573.1;refGene_gn=PALM;refGene_tx=XM_023242577.1;refGene_gn=PALM;refGene_tx=XM_023242580.1;refGene_gn=PALM;refGene_tx=XM_023242583.1
A2	348556	348637
refGene_gn=PALM;refGene_tx=XM_023242568.1;refGene_gn=PALM;refGene_tx=XM_023242570.1;refGene_gn=PALM;refGene_tx=XM_023242573.1;refGene_gn=PAL

M;refGene_tx=XM_023242577.1;refGene_gn=PALM;refGene_tx=XM_023242580.1;
refGene_gn=PALM;refGene_tx=XM_023242583.1;refGene_gn=PALM;refGene_tx=X
M_023242586.1
A2 350126 350254
refGene_gn=PALM;refGene_tx=XM_023242568.1;refGene_gn=PALM;refGene_tx=X
M_023242570.1;refGene_gn=PALM;refGene_tx=XM_023242573.1;refGene_gn=PAL
M;refGene_tx=XM_023242577.1;refGene_gn=PALM;refGene_tx=XM_023242580.1;
refGene_gn=PALM;refGene_tx=XM_023242583.1;refGene_gn=PALM;refGene_tx=X
M_023242586.1
A2 351532 351656
refGene_gn=PALM;refGene_tx=XM_023242568.1;refGene_gn=PALM;refGene_tx=X
M_023242570.1;refGene_gn=PALM;refGene_tx=XM_023242573.1;refGene_gn=PAL
M;refGene_tx=XM_023242577.1;refGene_gn=PALM;refGene_tx=XM_023242580.1;
refGene_gn=PALM;refGene_tx=XM_023242583.1;refGene_gn=PALM;refGene_tx=X
M_023242586.1
A2 353056 353078
refGene_gn=PALM;refGene_tx=XM_023242568.1;refGene_gn=PALM;refGene_tx=X
M_023242570.1;refGene_gn=PALM;refGene_tx=XM_023242573.1;refGene_gn=PAL
M;refGene_tx=XM_023242577.1;refGene_gn=PALM;refGene_tx=XM_023242580.1;
refGene_gn=PALM;refGene_tx=XM_023242583.1;refGene_gn=PALM;refGene_tx=X
M_023242586.1
A2 353601 353664
refGene_gn=PALM;refGene_tx=XM_023242568.1;refGene_gn=PALM;refGene_tx=X
M_023242570.1;refGene_gn=PALM;refGene_tx=XM_023242577.1;refGene_gn=PAL
M;refGene_tx=XM_023242580.1;refGene_gn=PALM;refGene_tx=XM_023242586.1
A2 354227 354287
refGene_gn=PALM;refGene_tx=XM_023242568.1;refGene_gn=PALM;refGene_tx=X
M_023242570.1;refGene_gn=PALM;refGene_tx=XM_023242573.1;refGene_gn=PAL
M;refGene_tx=XM_023242577.1;refGene_gn=PALM;refGene_tx=XM_023242580.1;
refGene_gn=PALM;refGene_tx=XM_023242583.1;refGene_gn=PALM;refGene_tx=X
M_023242586.1
A2 356730 356862
refGene_gn=PALM;refGene_tx=XM_023242568.1;refGene_gn=PALM;refGene_tx=X
M_023242570.1;refGene_gn=PALM;refGene_tx=XM_023242573.1;refGene_gn=PAL
M;refGene_tx=XM_023242580.1;refGene_gn=PALM;refGene_tx=XM_023242586.1
A2 360493 361023
refGene_gn=PALM;refGene_tx=XM_023242568.1;refGene_gn=PALM;refGene_tx=X
M_023242570.1;refGene_gn=PALM;refGene_tx=XM_023242573.1;refGene_gn=PAL
M;refGene_tx=XM_023242577.1;refGene_gn=PALM;refGene_tx=XM_023242580.1;
refGene_gn=PALM;refGene_tx=XM_023242583.1;refGene_gn=PALM;refGene_tx=X
M_023242586.1
A2 367248 368962
refGene_gn=MISP;refGene_tx=XM_003981634.4;refGene_gn=MISP;refGene_tx=X
M_019815183.2
A2 369263 369367
refGene_gn=MISP;refGene_tx=XM_003981634.4;refGene_gn=MISP;refGene_tx=X
M_019815183.2
A2 370274 370313
refGene_gn=MISP;refGene_tx=XM_003981634.4;refGene_gn=MISP;refGene_tx=X
M_019815183.2

A2 370777 370873
refGene_gn=MISP;refGene_tx=XM_003981634.4;refGene_gn=MISP;refGene_tx=XM_019815183.2
A2 400814 400822
refGene_gn=PTBP1;refGene_tx=XM_023242548.1;refGene_gn=PTBP1;refGene_tx=XM_023242551.1
A2 403467 403498
refGene_gn=PTBP1;refGene_tx=XM_023242548.1;refGene_gn=PTBP1;refGene_tx=XM_023242551.1
A2 406617 406693
refGene_gn=PTBP1;refGene_tx=XM_023242548.1;refGene_gn=PTBP1;refGene_tx=XM_023242551.1
A2 406846 407019
refGene_gn=PTBP1;refGene_tx=XM_023242548.1;refGene_gn=PTBP1;refGene_tx=XM_023242551.1
A2 407099 407246
refGene_gn=PTBP1;refGene_tx=XM_023242548.1;refGene_gn=PTBP1;refGene_tx=XM_023242551.1
A2 407334 407505
refGene_gn=PTBP1;refGene_tx=XM_023242548.1;refGene_gn=PTBP1;refGene_tx=XM_023242551.1
A2 407598 407709
refGene_gn=PTBP1;refGene_tx=XM_023242548.1;refGene_gn=PTBP1;refGene_tx=XM_023242551.1
A2 407783 407958
refGene_gn=PTBP1;refGene_tx=XM_023242548.1;refGene_gn=PTBP1;refGene_tx=XM_023242551.1
A2 408213 408291 refGene_gn=PTBP1;refGene_tx=XM_023242548.1
A2 409011 409160
refGene_gn=PTBP1;refGene_tx=XM_023242548.1;refGene_gn=PTBP1;refGene_tx=XM_023242551.1
A2 410273 410307
refGene_gn=PTBP1;refGene_tx=XM_023242548.1;refGene_gn=PTBP1;refGene_tx=XM_023242551.1
A2 410740 410833
refGene_gn=PTBP1;refGene_tx=XM_023242548.1;refGene_gn=PTBP1;refGene_tx=XM_023242551.1
A2 410923 411140
refGene_gn=PTBP1;refGene_tx=XM_023242548.1;refGene_gn=PTBP1;refGene_tx=XM_023242551.1
A2 411238 411316
refGene_gn=PTBP1;refGene_tx=XM_023242548.1;refGene_gn=PTBP1;refGene_tx=XM_023242551.1
A2 411403 411536
refGene_gn=PTBP1;refGene_tx=XM_023242548.1;refGene_gn=PTBP1;refGene_tx=XM_023242551.1
A2 413440 414817
refGene_gn=PLPPR3;refGene_tx=XM_023242509.1;refGene_gn=PLPPR3;refGene_tx=XM_023242515.1;refGene_gn=PLPPR3;refGene_tx=XM_023242518.1
A2 415301 415620

refGene_gn=PLPPR3;refGene_tx=XM_023242509.1;refGene_gn=PLPPR3;refGene_tx=XM_023242515.1;refGene_gn=PLPPR3;refGene_tx=XM_023242518.1
A2 415718 415914
refGene_gn=PLPPR3;refGene_tx=XM_023242509.1;refGene_gn=PLPPR3;refGene_tx=XM_023242515.1;refGene_gn=PLPPR3;refGene_tx=XM_023242518.1
A2 416019 416161
refGene_gn=PLPPR3;refGene_tx=XM_023242509.1;refGene_gn=PLPPR3;refGene_tx=XM_023242515.1;refGene_gn=PLPPR3;refGene_tx=XM_023242518.1
A2 416247 416433
refGene_gn=PLPPR3;refGene_tx=XM_023242509.1;refGene_gn=PLPPR3;refGene_tx=XM_023242515.1;refGene_gn=PLPPR3;refGene_tx=XM_023242518.1
A2 418921 418996
refGene_gn=PLPPR3;refGene_tx=XM_023242509.1;refGene_gn=PLPPR3;refGene_tx=XM_023242515.1;refGene_gn=PLPPR3;refGene_tx=XM_023242518.1
A2 435723 435790
refGene_gn=LOC100301482;refGene_tx=XM_003981637.3
A2 436151 436308
refGene_gn=LOC100301482;refGene_tx=XM_003981637.3
A2 436989 437131
refGene_gn=LOC100301482;refGene_tx=XM_003981637.3
A2 437941 438172
refGene_gn=LOC100301482;refGene_tx=XM_003981637.3
A2 438296 438497
refGene_gn=LOC100301482;refGene_tx=XM_003981637.3
A2 440403 440461 refGene_gn=CFD;refGene_tx=XM_023242657.1
A2 441041 441198 refGene_gn=CFD;refGene_tx=XM_023242657.1
A2 441281 441426 refGene_gn=CFD;refGene_tx=XM_023242657.1
A2 441519 441777 refGene_gn=CFD;refGene_tx=XM_023242657.1
A2 442059 442221 refGene_gn=CFD;refGene_tx=XM_023242657.1
A2 446151 446278
refGene_gn=MED16;refGene_tx=XM_023242498.1;refGene_gn=MED16;refGene_tx=XM_023242503.1;refGene_gn=MED16;refGene_tx=XM_023242505.1
A2 446819 446903
refGene_gn=MED16;refGene_tx=XM_023242498.1;refGene_gn=MED16;refGene_tx=XM_023242503.1;refGene_gn=MED16;refGene_tx=XM_023242505.1
A2 447689 447903
refGene_gn=MED16;refGene_tx=XM_023242498.1;refGene_gn=MED16;refGene_tx=XM_023242503.1;refGene_gn=MED16;refGene_tx=XM_023242505.1
A2 448297 448490
refGene_gn=MED16;refGene_tx=XM_023242498.1;refGene_gn=MED16;refGene_tx=XM_023242503.1;refGene_gn=MED16;refGene_tx=XM_023242505.1
A2 448789 448923
refGene_gn=MED16;refGene_tx=XM_023242498.1;refGene_gn=MED16;refGene_tx=XM_023242503.1;refGene_gn=MED16;refGene_tx=XM_023242505.1
A2 449132 449343
refGene_gn=MED16;refGene_tx=XM_023242498.1;refGene_gn=MED16;refGene_tx=XM_023242503.1;refGene_gn=MED16;refGene_tx=XM_023242505.1
A2 449506 449713
refGene_gn=MED16;refGene_tx=XM_023242498.1;refGene_gn=MED16;refGene_tx=XM_023242503.1;refGene_gn=MED16;refGene_tx=XM_023242505.1

A2	450404	450616	refGene_gn=MED16;refGene_tx=XM_023242498.1;refGene_gn=MED16;refGene_tx=XM_023242503.1;refGene_gn=MED16;refGene_tx=XM_023242505.1
A2	451272	451428	refGene_gn=MED16;refGene_tx=XM_023242498.1;refGene_gn=MED16;refGene_tx=XM_023242503.1;refGene_gn=MED16;refGene_tx=XM_023242505.1
A2	453395	453501	refGene_gn=MED16;refGene_tx=XM_023242498.1;refGene_gn=MED16;refGene_tx=XM_023242503.1;refGene_gn=MED16;refGene_tx=XM_023242505.1
A2	453934	454366	refGene_gn=MED16;refGene_tx=XM_023242498.1;refGene_gn=MED16;refGene_tx=XM_023242503.1;refGene_gn=MED16;refGene_tx=XM_023242505.1
A2	454810	454980	refGene_gn=MED16;refGene_tx=XM_023242498.1;refGene_gn=MED16;refGene_tx=XM_023242503.1;refGene_gn=MED16;refGene_tx=XM_023242505.1
A2	455272	455380	refGene_gn=MED16;refGene_tx=XM_023242498.1;refGene_gn=MED16;refGene_tx=XM_023242503.1;refGene_gn=MED16;refGene_tx=XM_023242505.1
A2	456060	456193	refGene_gn=MED16;refGene_tx=XM_023242498.1;refGene_gn=MED16;refGene_tx=XM_023242503.1;refGene_gn=MED16;refGene_tx=XM_023242505.1
A2	460296	460400	refGene_gn=R3HDM4;refGene_tx=XM_023242652.1
A2	461169	461225	refGene_gn=R3HDM4;refGene_tx=XM_023242652.1
A2	461307	461393	refGene_gn=R3HDM4;refGene_tx=XM_023242652.1
A2	461849	461935	refGene_gn=R3HDM4;refGene_tx=XM_023242652.1
A2	462862	462986	refGene_gn=R3HDM4;refGene_tx=XM_023242652.1
A2	463424	463549	refGene_gn=R3HDM4;refGene_tx=XM_023242652.1
A2	463951	464106	refGene_gn=R3HDM4;refGene_tx=XM_023242652.1
A2	468919	468996	refGene_gn=R3HDM4;refGene_tx=XM_023242652.1
A2	471718	471962	refGene_gn=KISS1R;refGene_tx=XM_023242671.1
A2	473119	473244	refGene_gn=KISS1R;refGene_tx=XM_023242671.1
A2	473627	473763	refGene_gn=KISS1R;refGene_tx=XM_023242671.1
A2	474066	474299	refGene_gn=KISS1R;refGene_tx=XM_023242671.1
A2	474597	475014	refGene_gn=KISS1R;refGene_tx=XM_023242671.1
A2	485640	486038	refGene_gn=ARID3A;refGene_tx=XM_023242525.1;refGene_gn=ARID3A;refGene_tx=XM_023242526.1;refGene_gn=ARID3A;refGene_tx=XM_023242531.1;refGene_gn=ARID3A;refGene_tx=XM_023242535.1;refGene_gn=ARID3A;refGene_tx=XM_023242540.1;refGene_gn=ARID3A;refGene_tx=XM_023242542.1;refGene_gn=ARID3A;refGene_tx=XM_023242543.1
A2	486871	487223	refGene_gn=ARID3A;refGene_tx=XM_023242525.1;refGene_gn=ARID3A;refGene_tx=XM_023242526.1;refGene_gn=ARID3A;refGene_tx=XM_023242531.1;refGene_gn=ARID3A;refGene_tx=XM_023242535.1;refGene_gn=ARID3A;refGene_tx=XM_023242540.1;refGene_gn=ARID3A;refGene_tx=XM_023242542.1;refGene_gn=ARID3A;refGene_tx=XM_023242543.1
A2	503705	503778	refGene_gn=ARID3A;refGene_tx=XM_023242525.1;refGene_gn=ARID3A;refGene_tx=XM_023242526.1;refGene_gn=ARID3A;refGene_tx=XM_023242531.1;refGene_gn=ARID3A;refGene_tx=XM_023242535.1;refGene_gn=ARID3A;refGene_tx=XM_02

3242540.1;refGene_gn=ARID3A;refGene_tx=XM_023242542.1;refGene_gn=ARID3A;refGene_tx=XM_023242543.1
A2 506939 507123
refGene_gn=ARID3A;refGene_tx=XM_023242525.1;refGene_gn=ARID3A;refGene_tx=XM_023242526.1;refGene_gn=ARID3A;refGene_tx=XM_023242531.1;refGene_gn=ARID3A;refGene_tx=XM_023242535.1;refGene_gn=ARID3A;refGene_tx=XM_023242540.1;refGene_gn=ARID3A;refGene_tx=XM_023242542.1;refGene_gn=ARID3A;refGene_tx=XM_023242543.1
A2 507359 507607
refGene_gn=ARID3A;refGene_tx=XM_023242525.1;refGene_gn=ARID3A;refGene_tx=XM_023242526.1;refGene_gn=ARID3A;refGene_tx=XM_023242531.1;refGene_gn=ARID3A;refGene_tx=XM_023242535.1;refGene_gn=ARID3A;refGene_tx=XM_023242540.1;refGene_gn=ARID3A;refGene_tx=XM_023242542.1;refGene_gn=ARID3A;refGene_tx=XM_023242543.1
A2 509383 509680
refGene_gn=ARID3A;refGene_tx=XM_023242525.1;refGene_gn=ARID3A;refGene_tx=XM_023242526.1;refGene_gn=ARID3A;refGene_tx=XM_023242531.1;refGene_gn=ARID3A;refGene_tx=XM_023242535.1;refGene_gn=ARID3A;refGene_tx=XM_023242540.1;refGene_gn=ARID3A;refGene_tx=XM_023242542.1;refGene_gn=ARID3A;refGene_tx=XM_023242543.1
A2 510394 510493
refGene_gn=ARID3A;refGene_tx=XM_023242525.1;refGene_gn=ARID3A;refGene_tx=XM_023242526.1;refGene_gn=ARID3A;refGene_tx=XM_023242531.1;refGene_gn=ARID3A;refGene_tx=XM_023242535.1;refGene_gn=ARID3A;refGene_tx=XM_023242540.1;refGene_gn=ARID3A;refGene_tx=XM_023242542.1;refGene_gn=ARID3A;refGene_tx=XM_023242543.1
A2 510941 511117
refGene_gn=ARID3A;refGene_tx=XM_023242525.1;refGene_gn=ARID3A;refGene_tx=XM_023242526.1;refGene_gn=ARID3A;refGene_tx=XM_023242531.1;refGene_gn=ARID3A;refGene_tx=XM_023242535.1;refGene_gn=ARID3A;refGene_tx=XM_023242540.1;refGene_gn=ARID3A;refGene_tx=XM_023242542.1;refGene_gn=ARID3A;refGene_tx=XM_023242543.1
A2 519181 519391
refGene_gn=WDR18;refGene_tx=XM_011288240.3;refGene_gn=WDR18;refGene_tx=XM_011288241.3
A2 520441 520552
refGene_gn=WDR18;refGene_tx=XM_011288240.3;refGene_gn=WDR18;refGene_tx=XM_011288241.3;refGene_gn=WDR18;refGene_tx=XM_023242562.1
A2 521990 522124
refGene_gn=WDR18;refGene_tx=XM_011288240.3;refGene_gn=WDR18;refGene_tx=XM_011288241.3;refGene_gn=WDR18;refGene_tx=XM_023242562.1
A2 522279 522421
refGene_gn=WDR18;refGene_tx=XM_011288240.3;refGene_gn=WDR18;refGene_tx=XM_011288241.3;refGene_gn=WDR18;refGene_tx=XM_023242562.1
A2 522877 523021
refGene_gn=WDR18;refGene_tx=XM_011288240.3;refGene_gn=WDR18;refGene_tx=XM_011288241.3;refGene_gn=WDR18;refGene_tx=XM_023242562.1
A2 523161 523226
refGene_gn=WDR18;refGene_tx=XM_011288240.3;refGene_gn=WDR18;refGene_tx=XM_011288241.3;refGene_gn=WDR18;refGene_tx=XM_023242562.1

A2	523320	523445	refGene_gn=WDR18;refGene_tx=XM_011288240.3;refGene_gn=WDR18;refGene_tx=XM_011288241.3;refGene_gn=WDR18;refGene_tx=XM_023242562.1
A2	523698	523865	refGene_gn=WDR18;refGene_tx=XM_011288240.3;refGene_gn=WDR18;refGene_tx=XM_011288241.3;refGene_gn=WDR18;refGene_tx=XM_023242562.1
A2	524712	524781	refGene_gn=WDR18;refGene_tx=XM_011288240.3;refGene_gn=WDR18;refGene_tx=XM_011288241.3;refGene_gn=WDR18;refGene_tx=XM_023242562.1
A2	524872	525004	refGene_gn=WDR18;refGene_tx=XM_011288240.3;refGene_gn=WDR18;refGene_tx=XM_011288241.3;refGene_gn=WDR18;refGene_tx=XM_023242562.1
A2	527083	527147	refGene_gn=WDR18;refGene_tx=XM_011288240.3;refGene_gn=WDR18;refGene_tx=XM_023242562.1
A2	529717	530149	refGene_gn=GRIN3B;refGene_tx=XM_023242691.1
A2	531694	532287	refGene_gn=GRIN3B;refGene_tx=XM_023242691.1
A2	532641	533674	refGene_gn=GRIN3B;refGene_tx=XM_023242691.1
A2	535424	535570	refGene_gn=GRIN3B;refGene_tx=XM_023242691.1
A2	535992	536108	refGene_gn=GRIN3B;refGene_tx=XM_023242691.1
A2	536214	536366	refGene_gn=GRIN3B;refGene_tx=XM_023242691.1
A2	536589	536754	refGene_gn=GRIN3B;refGene_tx=XM_023242691.1
A2	536845	536904	refGene_gn=GRIN3B;refGene_tx=XM_023242691.1
A2	537089	537130	refGene_gn=GRIN3B;refGene_tx=XM_023242691.1
A2	537295	537591	refGene_gn=GRIN3B;refGene_tx=XM_023242691.1
A2	538312	538858	refGene_gn=TMEM259;refGene_tx=XM_023242706.1;refGene_gn=TMEM259;refGene_tx=XM_023242709.1;refGene_gn=TMEM259;refGene_tx=XM_023242711.1;refGene_gn=TMEM259;refGene_tx=XM_023242713.1;refGene_gn=TMEM259;refGene_tx=XM_023242716.1;refGene_gn=TMEM259;refGene_tx=XM_023242722.1;refGene_gn=TMEM259;refGene_tx=XM_023242726.1;refGene_gn=TMEM259;refGene_tx=XM_023242729.1;refGene_gn=TMEM259;refGene_tx=XM_023242732.1
A2	538959	539059	refGene_gn=TMEM259;refGene_tx=XM_023242706.1;refGene_gn=TMEM259;refGene_tx=XM_023242709.1;refGene_gn=TMEM259;refGene_tx=XM_023242711.1;refGene_gn=TMEM259;refGene_tx=XM_023242713.1;refGene_gn=TMEM259;refGene_tx=XM_023242716.1;refGene_gn=TMEM259;refGene_tx=XM_023242722.1;refGene_gn=TMEM259;refGene_tx=XM_023242726.1;refGene_gn=TMEM259;refGene_tx=XM_023242729.1;refGene_gn=TMEM259;refGene_tx=XM_023242732.1
A2	539128	539285	refGene_gn=TMEM259;refGene_tx=XM_023242706.1;refGene_gn=TMEM259;refGene_tx=XM_023242709.1;refGene_gn=TMEM259;refGene_tx=XM_023242711.1;refGene_gn=TMEM259;refGene_tx=XM_023242713.1;refGene_gn=TMEM259;refGene_tx=XM_023242716.1;refGene_gn=TMEM259;refGene_tx=XM_023242722.1;refGene_gn=TMEM259;refGene_tx=XM_023242726.1;refGene_gn=TMEM259;refGene_tx=XM_023242729.1;refGene_gn=TMEM259;refGene_tx=XM_023242732.1
A2	539370	539454	refGene_gn=TMEM259;refGene_tx=XM_023242706.1;refGene_gn=TMEM259;refGene_tx=XM_023242709.1;refGene_gn=TMEM259;refGene_tx=XM_023242711.1;refGene_gn=TMEM259;refGene_tx=XM_023242713.1;refGene_gn=TMEM259;refGene_tx=

XM_023242716.1;refGene_gn=TMEM259;refGene_tx=XM_023242722.1;refGene_gn=TMEM259;refGene_tx=XM_023242726.1;refGene_gn=TMEM259;refGene_tx=XM_023242729.1;refGene_gn=TMEM259;refGene_tx=XM_023242732.1
A2 539587 539645
refGene_gn=TMEM259;refGene_tx=XM_023242706.1;refGene_gn=TMEM259;refGene_tx=XM_023242709.1;refGene_gn=TMEM259;refGene_tx=XM_023242711.1;refGene_gn=TMEM259;refGene_tx=XM_023242713.1;refGene_gn=TMEM259;refGene_tx=XM_023242716.1;refGene_gn=TMEM259;refGene_tx=XM_023242722.1;refGene_gn=TMEM259;refGene_tx=XM_023242726.1
A2 539746 539847
refGene_gn=TMEM259;refGene_tx=XM_023242706.1;refGene_gn=TMEM259;refGene_tx=XM_023242709.1;refGene_gn=TMEM259;refGene_tx=XM_023242711.1;refGene_gn=TMEM259;refGene_tx=XM_023242713.1;refGene_gn=TMEM259;refGene_tx=XM_023242716.1;refGene_gn=TMEM259;refGene_tx=XM_023242722.1;refGene_gn=TMEM259;refGene_tx=XM_023242726.1;refGene_gn=TMEM259;refGene_tx=XM_023242729.1;refGene_gn=TMEM259;refGene_tx=XM_023242732.1
A2 539941 540064
refGene_gn=TMEM259;refGene_tx=XM_023242706.1;refGene_gn=TMEM259;refGene_tx=XM_023242709.1;refGene_gn=TMEM259;refGene_tx=XM_023242711.1;refGene_gn=TMEM259;refGene_tx=XM_023242713.1;refGene_gn=TMEM259;refGene_tx=XM_023242716.1;refGene_gn=TMEM259;refGene_tx=XM_023242722.1;refGene_gn=TMEM259;refGene_tx=XM_023242726.1;refGene_gn=TMEM259;refGene_tx=XM_023242729.1;refGene_gn=TMEM259;refGene_tx=XM_023242732.1
A2 540260 540371
refGene_gn=TMEM259;refGene_tx=XM_023242706.1;refGene_gn=TMEM259;refGene_tx=XM_023242709.1;refGene_gn=TMEM259;refGene_tx=XM_023242711.1;refGene_gn=TMEM259;refGene_tx=XM_023242713.1;refGene_gn=TMEM259;refGene_tx=XM_023242716.1;refGene_gn=TMEM259;refGene_tx=XM_023242722.1;refGene_gn=TMEM259;refGene_tx=XM_023242726.1;refGene_gn=TMEM259;refGene_tx=XM_023242729.1;refGene_gn=TMEM259;refGene_tx=XM_023242732.1
A2 540821 540921
refGene_gn=TMEM259;refGene_tx=XM_023242706.1;refGene_gn=TMEM259;refGene_tx=XM_023242709.1;refGene_gn=TMEM259;refGene_tx=XM_023242711.1;refGene_gn=TMEM259;refGene_tx=XM_023242713.1;refGene_gn=TMEM259;refGene_tx=XM_023242716.1;refGene_gn=TMEM259;refGene_tx=XM_023242722.1;refGene_gn=TMEM259;refGene_tx=XM_023242726.1;refGene_gn=TMEM259;refGene_tx=XM_023242729.1;refGene_gn=TMEM259;refGene_tx=XM_023242732.1
A2 541611 541887
refGene_gn=TMEM259;refGene_tx=XM_023242706.1;refGene_gn=TMEM259;refGene_tx=XM_023242709.1;refGene_gn=TMEM259;refGene_tx=XM_023242711.1;refGene_gn=TMEM259;refGene_tx=XM_023242713.1;refGene_gn=TMEM259;refGene_tx=XM_023242716.1;refGene_gn=TMEM259;refGene_tx=XM_023242722.1;refGene_gn=TMEM259;refGene_tx=XM_023242726.1;refGene_gn=TMEM259;refGene_tx=XM_023242729.1;refGene_gn=TMEM259;refGene_tx=XM_023242732.1
A2 545371 545593
refGene_gn=TMEM259;refGene_tx=XM_023242706.1;refGene_gn=TMEM259;refGene_tx=XM_023242709.1;refGene_gn=TMEM259;refGene_tx=XM_023242711.1;refGene_gn=TMEM259;refGene_tx=XM_023242713.1;refGene_gn=TMEM259;refGene_tx=XM_023242716.1;refGene_gn=TMEM259;refGene_tx=XM_023242722.1;refGene_gn=TMEM259;refGene_tx=XM_023242726.1;refGene_gn=TMEM259;refGene_tx=XM_02

3242729.1;refGene_gn=TMEM259;refGene_tx=XM_023242732.1
A2	550313	552686
refGene_gn=LOC111557541;refGene_tx=XM_023242701.1
A2	553125	553362
refGene_gn=LOC111557541;refGene_tx=XM_023242701.1
A2	560354	560417
refGene_gn=CNN2;refGene_tx=XM_003981601.5;refGene_gn=CNN2;refGene_tx=XM_019815238.2
A2	564522	564644
refGene_gn=CNN2;refGene_tx=XM_003981601.5;refGene_gn=CNN2;refGene_tx=XM_019815238.2
A2	565102	565169
refGene_gn=CNN2;refGene_tx=XM_003981601.5;refGene_gn=CNN2;refGene_tx=XM_019815238.2
A2	565273	565411
refGene_gn=CNN2;refGene_tx=XM_003981601.5;refGene_gn=CNN2;refGene_tx=XM_019815238.2
A2	566086	566203
refGene_gn=CNN2;refGene_tx=XM_003981601.5;refGene_gn=CNN2;refGene_tx=XM_019815238.2
A2	566357	566504
refGene_gn=CNN2;refGene_tx=XM_003981601.5;refGene_gn=CNN2;refGene_tx=XM_019815238.2
A2	567006	567276
refGene_gn=CNN2;refGene_tx=XM_003981601.5;refGene_gn=CNN2;refGene_tx=XM_019815238.2
A2	586152	586242	refGene_gn=ARHGAP45;refGene_tx=XM_023242738.1
A2	586349	586448	refGene_gn=ARHGAP45;refGene_tx=XM_023242736.1
A2	587150	587478
refGene_gn=ARHGAP45;refGene_tx=XM_023242736.1;refGene_gn=ARHGAP45;refGene_tx=XM_023242738.1
A2	589991	590135
refGene_gn=ARHGAP45;refGene_tx=XM_023242736.1;refGene_gn=ARHGAP45;refGene_tx=XM_023242738.1
A2	590356	590441
refGene_gn=ARHGAP45;refGene_tx=XM_023242736.1;refGene_gn=ARHGAP45;refGene_tx=XM_023242738.1
A2	590529	590602
refGene_gn=ARHGAP45;refGene_tx=XM_023242736.1;refGene_gn=ARHGAP45;refGene_tx=XM_023242738.1
A2	590703	590767
refGene_gn=ARHGAP45;refGene_tx=XM_023242736.1;refGene_gn=ARHGAP45;refGene_tx=XM_023242738.1
A2	590852	590990
refGene_gn=ARHGAP45;refGene_tx=XM_023242736.1;refGene_gn=ARHGAP45;refGene_tx=XM_023242738.1
A2	591068	591133
refGene_gn=ARHGAP45;refGene_tx=XM_023242736.1;refGene_gn=ARHGAP45;refGene_tx=XM_023242738.1
A2	591291	591402

refGene_gn=ARHGAP45;refGene_tx=XM_023242736.1;refGene_gn=ARHGAP45;refGene_tx=XM_023242738.1
A2	591488	591569	refGene_gn=ARHGAP45;refGene_tx=XM_023242736.1;refGene_gn=ARHGAP45;refGene_tx=XM_023242738.1
A2	593022	593205	refGene_gn=ARHGAP45;refGene_tx=XM_023242736.1;refGene_gn=ARHGAP45;refGene_tx=XM_023242738.1
A2	593387	593525	refGene_gn=ARHGAP45;refGene_tx=XM_023242736.1;refGene_gn=ARHGAP45;refGene_tx=XM_023242738.1
A2	593614	593805	refGene_gn=ARHGAP45;refGene_tx=XM_023242736.1;refGene_gn=ARHGAP45;refGene_tx=XM_023242738.1
A2	593909	594034	refGene_gn=ARHGAP45;refGene_tx=XM_023242736.1;refGene_gn=ARHGAP45;refGene_tx=XM_023242738.1
A2	594106	594190	refGene_gn=ARHGAP45;refGene_tx=XM_023242736.1;refGene_gn=ARHGAP45;refGene_tx=XM_023242738.1
A2	594323	594410	refGene_gn=ARHGAP45;refGene_tx=XM_023242736.1;refGene_gn=ARHGAP45;refGene_tx=XM_023242738.1
A2	594502	594675	refGene_gn=ARHGAP45;refGene_tx=XM_023242736.1;refGene_gn=ARHGAP45;refGene_tx=XM_023242738.1
A2	594997	595186	refGene_gn=ARHGAP45;refGene_tx=XM_023242736.1;refGene_gn=ARHGAP45;refGene_tx=XM_023242738.1
A2	595281	595419	refGene_gn=ARHGAP45;refGene_tx=XM_023242736.1;refGene_gn=ARHGAP45;refGene_tx=XM_023242738.1
A2	595515	595742	refGene_gn=ARHGAP45;refGene_tx=XM_023242736.1;refGene_gn=ARHGAP45;refGene_tx=XM_023242738.1
A2	595827	596020	refGene_gn=ARHGAP45;refGene_tx=XM_023242736.1;refGene_gn=ARHGAP45;refGene_tx=XM_023242738.1
A2	596708	596817	refGene_gn=ARHGAP45;refGene_tx=XM_023242736.1;refGene_gn=ARHGAP45;refGene_tx=XM_023242738.1
A2	597136	597459	refGene_gn=ARHGAP45;refGene_tx=XM_023242736.1;refGene_gn=ARHGAP45;refGene_tx=XM_023242738.1
A2	602594	602660	refGene_gn=POLR2E;refGene_tx=XM_023242745.1
A2	602826	602905	refGene_gn=POLR2E;refGene_tx=XM_023242745.1
A2	602999	603058	refGene_gn=POLR2E;refGene_tx=XM_023242745.1
A2	603617	603698	refGene_gn=POLR2E;refGene_tx=XM_023242745.1
A2	604230	604346	refGene_gn=POLR2E;refGene_tx=XM_023242745.1
A2	606125	606300	refGene_gn=POLR2E;refGene_tx=XM_023242745.1

A2 607320 607377 refGene_gn=POLR2E;refGene_tx=XM_023242745.1
A2 612746 612830 refGene_gn=GPX4;refGene_tx=XM_023242756.1
A2 613212 613425 refGene_gn=GPX4;refGene_tx=XM_011288229.3
A2 613692 613787
refGene_gn=GPX4;refGene_tx=XM_011288229.3;refGene_gn=GPX4;refGene_tx=X
M_023242756.1
A2 613871 614016
refGene_gn=GPX4;refGene_tx=XM_011288229.3;refGene_gn=GPX4;refGene_tx=X
M_023242756.1
A2 614178 614330
refGene_gn=GPX4;refGene_tx=XM_011288229.3;refGene_gn=GPX4;refGene_tx=X
M_023242756.1
A2 614428 614453
refGene_gn=GPX4;refGene_tx=XM_011288229.3;refGene_gn=GPX4;refGene_tx=X
M_023242756.1
A2 614633 614693
refGene_gn=GPX4;refGene_tx=XM_011288229.3;refGene_gn=GPX4;refGene_tx=X
M_023242756.1
A2 614770 614803
refGene_gn=GPX4;refGene_tx=XM_011288229.3;refGene_gn=GPX4;refGene_tx=X
M_023242756.1
A2 617256 617723 refGene_gn=SBNO2;refGene_tx=XM_023242762.1
A2 617801 617992 refGene_gn=SBNO2;refGene_tx=XM_023242762.1
A2 618218 618295 refGene_gn=SBNO2;refGene_tx=XM_023242762.1
A2 618522 618654 refGene_gn=SBNO2;refGene_tx=XM_023242762.1
A2 618787 618880 refGene_gn=SBNO2;refGene_tx=XM_023242762.1
A2 618963 619058 refGene_gn=SBNO2;refGene_tx=XM_023242762.1
A2 619923 620067 refGene_gn=SBNO2;refGene_tx=XM_023242762.1
A2 620314 620389 refGene_gn=SBNO2;refGene_tx=XM_023242762.1
A2 620605 620714 refGene_gn=SBNO2;refGene_tx=XM_023242762.1
A2 620794 620866 refGene_gn=SBNO2;refGene_tx=XM_023242762.1
A2 620975 621088 refGene_gn=SBNO2;refGene_tx=XM_023242762.1
A2 621170 621299 refGene_gn=SBNO2;refGene_tx=XM_023242762.1
A2 621781 621920 refGene_gn=SBNO2;refGene_tx=XM_023242762.1
A2 622484 622654 refGene_gn=SBNO2;refGene_tx=XM_023242762.1
A2 622901 623093 refGene_gn=SBNO2;refGene_tx=XM_023242762.1
A2 624150 624233 refGene_gn=SBNO2;refGene_tx=XM_023242762.1
A2 624894 624992 refGene_gn=SBNO2;refGene_tx=XM_023242762.1
A2 625491 625668 refGene_gn=SBNO2;refGene_tx=XM_023242762.1
A2 626508 626662 refGene_gn=SBNO2;refGene_tx=XM_023242762.1
A2 627067 627173 refGene_gn=SBNO2;refGene_tx=XM_023242762.1
A2 627412 627530 refGene_gn=SBNO2;refGene_tx=XM_023242762.1
A2 629762 629906 refGene_gn=SBNO2;refGene_tx=XM_023242762.1
A2 629985 630076 refGene_gn=SBNO2;refGene_tx=XM_023242762.1
A2 630155 630289 refGene_gn=SBNO2;refGene_tx=XM_023242762.1
A2 630369 630524 refGene_gn=SBNO2;refGene_tx=XM_023242762.1
A2 630857 630954 refGene_gn=SBNO2;refGene_tx=XM_023242762.1
A2 631208 631292 refGene_gn=SBNO2;refGene_tx=XM_023242762.1
A2 633557 633719 refGene_gn=SBNO2;refGene_tx=XM_023242762.1
A2 647490 647602 refGene_gn=SBNO2;refGene_tx=XM_023242762.1

A2	648204	648278	refGene_gn=SBNO2;refGene_tx=XM_023242762.1
A2	652794	652914	refGene_gn=SBNO2;refGene_tx=XM_023242762.1
A2	679475	679765
refGene_gn=STK11;refGene_tx=XM_003981604.4;refGene_gn=STK11;refGene_tx=XM_011288228.3;refGene_gn=STK11;refGene_tx=XM_019815258.2
A2	688357	688441
refGene_gn=STK11;refGene_tx=XM_003981604.4;refGene_gn=STK11;refGene_tx=XM_011288228.3;refGene_gn=STK11;refGene_tx=XM_019815258.2
A2	689532	689622
refGene_gn=STK11;refGene_tx=XM_003981604.4;refGene_gn=STK11;refGene_tx=XM_011288228.3;refGene_gn=STK11;refGene_tx=XM_019815258.2
A2	690323	690456
refGene_gn=STK11;refGene_tx=XM_003981604.4;refGene_gn=STK11;refGene_tx=XM_011288228.3;refGene_gn=STK11;refGene_tx=XM_019815258.2
A2	690544	690681
refGene_gn=STK11;refGene_tx=XM_003981604.4;refGene_gn=STK11;refGene_tx=XM_011288228.3;refGene_gn=STK11;refGene_tx=XM_019815258.2
A2	691743	691871
refGene_gn=STK11;refGene_tx=XM_003981604.4;refGene_gn=STK11;refGene_tx=XM_011288228.3;refGene_gn=STK11;refGene_tx=XM_019815258.2
A2	693972	694030
refGene_gn=STK11;refGene_tx=XM_003981604.4;refGene_gn=STK11;refGene_tx=XM_011288228.3;refGene_gn=STK11;refGene_tx=XM_019815258.2
A2	694301	694498
refGene_gn=STK11;refGene_tx=XM_003981604.4;refGene_gn=STK11;refGene_tx=XM_011288228.3;refGene_gn=STK11;refGene_tx=XM_019815258.2
A2	694715	694757	refGene_gn=STK11;refGene_tx=XM_019815258.2
A2	696959	697162
refGene_gn=STK11;refGene_tx=XM_003981604.4;refGene_gn=STK11;refGene_tx=XM_011288228.3;refGene_gn=STK11;refGene_tx=XM_019815258.2
A2	699830	700541	refGene_gn=CBARP;refGene_tx=XM_023242819.1
A2	700894	700899
refGene_gn=LOC101085272;refGene_tx=XM_023242792.1;refGene_gn=LOC101085272;refGene_tx=XM_023242795.1
A2	701329	701989
refGene_gn=LOC101085272;refGene_tx=XM_023242763.1;refGene_gn=LOC101085272;refGene_tx=XM_023242773.1;refGene_gn=LOC101085272;refGene_tx=XM_023242779.1;refGene_gn=LOC101085272;refGene_tx=XM_023242789.1;refGene_gn=LOC101085272;refGene_tx=XM_023242792.1;refGene_gn=LOC101085272;refGene_tx=XM_023242795.1
A2	703178	703389
refGene_gn=LOC101085272;refGene_tx=XM_023242763.1;refGene_gn=LOC101085272;refGene_tx=XM_023242773.1;refGene_gn=LOC101085272;refGene_tx=XM_023242779.1;refGene_gn=LOC101085272;refGene_tx=XM_023242789.1;refGene_gn=LOC101085272;refGene_tx=XM_023242792.1;refGene_gn=LOC101085272;refGene_tx=XM_023242795.1
A2	703744	703933
refGene_gn=LOC101085272;refGene_tx=XM_023242763.1;refGene_gn=LOC101085272;refGene_tx=XM_023242773.1;refGene_gn=LOC101085272;refGene_tx=XM_023242779.1;refGene_gn=LOC101085272;refGene_tx=XM_023242789.1;refGene_gn

=LOC101085272;refGene_tx=XM_023242792.1;refGene_gn=LOC101085272;refGene_tx=XM_023242795.1
A2	704112	704284	refGene_gn=LOC101085272;refGene_tx=XM_023242763.1;refGene_gn=LOC101085272;refGene_tx=XM_023242773.1;refGene_gn=LOC101085272;refGene_tx=XM_023242779.1;refGene_gn=LOC101085272;refGene_tx=XM_023242789.1;refGene_gn=LOC101085272;refGene_tx=XM_023242792.1;refGene_gn=LOC101085272;refGene_tx=XM_023242795.1
A2	704488	704633	refGene_gn=LOC101085272;refGene_tx=XM_023242763.1;refGene_gn=LOC101085272;refGene_tx=XM_023242773.1;refGene_gn=LOC101085272;refGene_tx=XM_023242779.1;refGene_gn=LOC101085272;refGene_tx=XM_023242789.1;refGene_gn=LOC101085272;refGene_tx=XM_023242792.1;refGene_gn=LOC101085272;refGene_tx=XM_023242795.1
A2	704977	705042	refGene_gn=LOC101085272;refGene_tx=XM_023242763.1;refGene_gn=LOC101085272;refGene_tx=XM_023242773.1;refGene_gn=LOC101085272;refGene_tx=XM_023242779.1;refGene_gn=LOC101085272;refGene_tx=XM_023242789.1;refGene_gn=LOC101085272;refGene_tx=XM_023242792.1;refGene_gn=LOC101085272;refGene_tx=XM_023242795.1
A2	705269	705409	refGene_gn=LOC101085272;refGene_tx=XM_023242763.1;refGene_gn=LOC101085272;refGene_tx=XM_023242773.1;refGene_gn=LOC101085272;refGene_tx=XM_023242779.1;refGene_gn=LOC101085272;refGene_tx=XM_023242789.1;refGene_gn=LOC101085272;refGene_tx=XM_023242792.1;refGene_gn=LOC101085272;refGene_tx=XM_023242795.1
A2	705502	705619	refGene_gn=LOC101085272;refGene_tx=XM_023242763.1;refGene_gn=LOC101085272;refGene_tx=XM_023242773.1;refGene_gn=LOC101085272;refGene_tx=XM_023242779.1;refGene_gn=LOC101085272;refGene_tx=XM_023242789.1;refGene_gn=LOC101085272;refGene_tx=XM_023242792.1;refGene_gn=LOC101085272;refGene_tx=XM_023242795.1
A2	707406	707970	refGene_gn=LOC101085272;refGene_tx=XM_023242763.1;refGene_gn=LOC101085272;refGene_tx=XM_023242773.1;refGene_gn=LOC101085272;refGene_tx=XM_023242779.1;refGene_gn=LOC101085272;refGene_tx=XM_023242789.1;refGene_gn=LOC101085272;refGene_tx=XM_023242792.1;refGene_gn=LOC101085272;refGene_tx=XM_023242795.1
A2	710328	710469	refGene_gn=ATP5F1D;refGene_tx=XM_003981606.5
A2	711230	711384	refGene_gn=ATP5F1D;refGene_tx=XM_003981606.5
A2	711885	711974	refGene_gn=ATP5F1D;refGene_tx=XM_003981606.5
A2	712077	712200	refGene_gn=ATP5F1D;refGene_tx=XM_003981606.5
A2	717406	717639	refGene_gn=MIDN;refGene_tx=XM_023242804.1;refGene_gn=MIDN;refGene_tx=XM_023242807.1;refGene_gn=MIDN;refGene_tx=XM_023242810.1
A2	718677	718765	refGene_gn=MIDN;refGene_tx=XM_023242804.1;refGene_gn=MIDN;refGene_tx=XM_023242807.1;refGene_gn=MIDN;refGene_tx=XM_023242810.1;refGene_gn=MIDN;refGene_tx=XM_023242811.1
A2	718945	719008

refGene_gn=MIDN;refGene_tx=XM_023242804.1;refGene_gn=MIDN;refGene_tx=XM_023242807.1;refGene_gn=MIDN;refGene_tx=XM_023242810.1;refGene_gn=MIDN;refGene_tx=XM_023242811.1
A2 720929 721058
refGene_gn=MIDN;refGene_tx=XM_023242804.1;refGene_gn=MIDN;refGene_tx=XM_023242807.1;refGene_gn=MIDN;refGene_tx=XM_023242811.1
A2 721143 721455
refGene_gn=MIDN;refGene_tx=XM_023242804.1;refGene_gn=MIDN;refGene_tx=XM_023242807.1;refGene_gn=MIDN;refGene_tx=XM_023242810.1;refGene_gn=MIDN;refGene_tx=XM_023242811.1
A2 721610 721779
refGene_gn=MIDN;refGene_tx=XM_023242804.1;refGene_gn=MIDN;refGene_tx=XM_023242807.1;refGene_gn=MIDN;refGene_tx=XM_023242810.1;refGene_gn=MIDN;refGene_tx=XM_023242811.1
A2 721936 722197
refGene_gn=MIDN;refGene_tx=XM_023242804.1;refGene_gn=MIDN;refGene_tx=XM_023242807.1;refGene_gn=MIDN;refGene_tx=XM_023242810.1;refGene_gn=MIDN;refGene_tx=XM_023242811.1
A2 722899 723177
refGene_gn=MIDN;refGene_tx=XM_023242804.1;refGene_gn=MIDN;refGene_tx=XM_023242807.1;refGene_gn=MIDN;refGene_tx=XM_023242810.1;refGene_gn=MIDN;refGene_tx=XM_023242811.1
A2 731956 731980
refGene_gn=CIRBP;refGene_tx=XM_006928167.4;refGene_gn=CIRBP;refGene_tx=XM_019815319.2
A2 735077 735186
refGene_gn=CIRBP;refGene_tx=XM_006928167.4;refGene_gn=CIRBP;refGene_tx=XM_019815319.2;refGene_gn=CIRBP;refGene_tx=XM_023242850.1
A2 735266 735373
refGene_gn=CIRBP;refGene_tx=XM_006928167.4;refGene_gn=CIRBP;refGene_tx=XM_019815319.2;refGene_gn=CIRBP;refGene_tx=XM_023242850.1
A2 735477 735631
refGene_gn=CIRBP;refGene_tx=XM_006928167.4;refGene_gn=CIRBP;refGene_tx=XM_019815319.2;refGene_gn=CIRBP;refGene_tx=XM_023242850.1
A2 735723 735805
refGene_gn=CIRBP;refGene_tx=XM_006928167.4;refGene_gn=CIRBP;refGene_tx=XM_019815319.2;refGene_gn=CIRBP;refGene_tx=XM_023242850.1
A2 735891 736018
refGene_gn=CIRBP;refGene_tx=XM_006928167.4;refGene_gn=CIRBP;refGene_tx=XM_019815319.2;refGene_gn=CIRBP;refGene_tx=XM_023242850.1
A2 736361 736378 refGene_gn=CIRBP;refGene_tx=XM_019815319.2
A2 739598 739903
refGene_gn=CA2H19orf24;refGene_tx=XM_023242828.1;refGene_gn=CA2H19orf24;refGene_tx=XM_023242829.1
A2 740852 740969
refGene_gn=CA2H19orf24;refGene_tx=XM_023242828.1;refGene_gn=CA2H19orf24;refGene_tx=XM_023242829.1
A2 741133 741248
refGene_gn=CA2H19orf24;refGene_tx=XM_023242828.1
A2 741616 741724

refGene_gn=CA2H19orf24;refGene_tx=XM_023242828.1;refGene_gn=CA2H19orf2
4;refGene_tx=XM_023242829.1
A2 747317 747460 refGene_gn=EFNA2;refGene_tx=XM_023242863.1
A2 753704 753923
refGene_gn=LOC111557553;refGene_tx=XM_023242855.1
A2 754003 754097
refGene_gn=LOC111557553;refGene_tx=XM_023242855.1
A2 755078 755280
refGene_gn=LOC111557553;refGene_tx=XM_023242855.1
A2 756149 756387
refGene_gn=LOC111557553;refGene_tx=XM_023242855.1
A2 756653 756967 refGene_gn=EFNA2;refGene_tx=XM_023242863.1
A2 759160 759226 refGene_gn=EFNA2;refGene_tx=XM_023242863.1
A2 759792 759914 refGene_gn=EFNA2;refGene_tx=XM_023242863.1
A2 812885 812942
refGene_gn=LOC101086040;refGene_tx=XM_023242871.1
A2 813568 813654
refGene_gn=LOC101086040;refGene_tx=XM_023242871.1
A2 814775 814846
refGene_gn=LOC101086040;refGene_tx=XM_023242871.1
A2 816359 817186
refGene_gn=LOC101086040;refGene_tx=XM_023242871.1
A2 817187 817203
refGene_gn=LOC101086040;refGene_tx=XM_023242871.1
A2 818357 818459
refGene_gn=LOC101086040;refGene_tx=XM_023242871.1
A2 820348 820419
refGene_gn=LOC101086040;refGene_tx=XM_023242871.1
A2 821772 821849
refGene_gn=LOC101086040;refGene_tx=XM_023242871.1
A2 822529 822590
refGene_gn=LOC101086040;refGene_tx=XM_023242871.1
A2 824311 824387
refGene_gn=LOC101086040;refGene_tx=XM_023242871.1
A2 824758 824809
refGene_gn=LOC101086040;refGene_tx=XM_023242871.1
A2 825347 825784
refGene_gn=LOC101086040;refGene_tx=XM_023242871.1
A2 827296 827385
refGene_gn=LOC101086040;refGene_tx=XM_023242871.1
A2 829153 829208
refGene_gn=LOC101086040;refGene_tx=XM_023242871.1
A2 832254 832270 refGene_gn=NDUFS7;refGene_tx=XM_023242902.1
A2 833294 833328 refGene_gn=NDUFS7;refGene_tx=XM_023242902.1
A2 834203 834272 refGene_gn=NDUFS7;refGene_tx=XM_023242902.1
A2 834615 834733 refGene_gn=NDUFS7;refGene_tx=XM_023242902.1
A2 835809 835989 refGene_gn=NDUFS7;refGene_tx=XM_023242902.1
A2 836059 836106 refGene_gn=NDUFS7;refGene_tx=XM_023242902.1
A2 836802 836891 refGene_gn=NDUFS7;refGene_tx=XM_023242902.1
A2 838462 838560 refGene_gn=NDUFS7;refGene_tx=XM_023242902.1

A2 839385 839526 refGene_gn=GAMT;refGene_tx=XM_011288196.3
A2 840056 840167 refGene_gn=GAMT;refGene_tx=XM_011288196.3
A2 840323 840391 refGene_gn=GAMT;refGene_tx=XM_011288196.3
A2 840699 840763 refGene_gn=GAMT;refGene_tx=XM_011288196.3
A2 840972 841118 refGene_gn=GAMT;refGene_tx=XM_011288196.3
A2 842119 842300 refGene_gn=GAMT;refGene_tx=XM_011288196.3
A2 848316 848345
refGene_gn=DAZAP1;refGene_tx=XM_023242994.1;refGene_gn=DAZAP1;refGene_tx=XM_023242999.1
A2 856040 856081
refGene_gn=DAZAP1;refGene_tx=XM_023242994.1;refGene_gn=DAZAP1;refGene_tx=XM_023242999.1
A2 856707 856874
refGene_gn=DAZAP1;refGene_tx=XM_023242987.1;refGene_gn=DAZAP1;refGene_tx=XM_023242991.1;refGene_gn=DAZAP1;refGene_tx=XM_023242993.1;refGene_gn=DAZAP1;refGene_tx=XM_023242994.1;refGene_gn=DAZAP1;refGene_tx=XM_023242999.1;refGene_gn=DAZAP1;refGene_tx=XM_023243002.1;refGene_gn=DAZAP1;refGene_tx=XM_023243006.1;refGene_gn=DAZAP1;refGene_tx=XM_023243016.1
A2 857164 857230
refGene_gn=DAZAP1;refGene_tx=XM_023242987.1;refGene_gn=DAZAP1;refGene_tx=XM_023242991.1;refGene_gn=DAZAP1;refGene_tx=XM_023242993.1;refGene_gn=DAZAP1;refGene_tx=XM_023242994.1;refGene_gn=DAZAP1;refGene_tx=XM_023242999.1;refGene_gn=DAZAP1;refGene_tx=XM_023243002.1;refGene_gn=DAZAP1;refGene_tx=XM_023243006.1;refGene_gn=DAZAP1;refGene_tx=XM_023243016.1
A2 859100 859211
refGene_gn=DAZAP1;refGene_tx=XM_023242987.1;refGene_gn=DAZAP1;refGene_tx=XM_023242991.1;refGene_gn=DAZAP1;refGene_tx=XM_023242993.1;refGene_gn=DAZAP1;refGene_tx=XM_023242994.1;refGene_gn=DAZAP1;refGene_tx=XM_023242999.1;refGene_gn=DAZAP1;refGene_tx=XM_023243002.1;refGene_gn=DAZAP1;refGene_tx=XM_023243006.1;refGene_gn=DAZAP1;refGene_tx=XM_023243016.1
A2 860393 860442
refGene_gn=DAZAP1;refGene_tx=XM_023242987.1;refGene_gn=DAZAP1;refGene_tx=XM_023242991.1;refGene_gn=DAZAP1;refGene_tx=XM_023242993.1;refGene_gn=DAZAP1;refGene_tx=XM_023242994.1;refGene_gn=DAZAP1;refGene_tx=XM_023242999.1;refGene_gn=DAZAP1;refGene_tx=XM_023243002.1;refGene_gn=DAZAP1;refGene_tx=XM_023243006.1;refGene_gn=DAZAP1;refGene_tx=XM_023243016.1
A2 863670 863753
refGene_gn=DAZAP1;refGene_tx=XM_023242987.1;refGene_gn=DAZAP1;refGene_tx=XM_023242991.1;refGene_gn=DAZAP1;refGene_tx=XM_023242993.1;refGene_gn=DAZAP1;refGene_tx=XM_023242994.1;refGene_gn=DAZAP1;refGene_tx=XM_023242999.1;refGene_gn=DAZAP1;refGene_tx=XM_023243002.1;refGene_gn=DAZAP1;refGene_tx=XM_023243006.1;refGene_gn=DAZAP1;refGene_tx=XM_023243016.1
A2 866545 866699
refGene_gn=DAZAP1;refGene_tx=XM_023242987.1;refGene_gn=DAZAP1;refGene_tx=XM_023242991.1;refGene_gn=DAZAP1;refGene_tx=XM_023242993.1;refGene_

gn=DAZAP1;refGene_tx=XM_023242994.1;refGene_gn=DAZAP1;refGene_tx=XM_02
3242999.1;refGene_gn=DAZAP1;refGene_tx=XM_023243002.1;refGene_gn=DAZAP
1;refGene_tx=XM_023243006.1;refGene_gn=DAZAP1;refGene_tx=XM_023243016.
1
A2 868060 868090
refGene_gn=DAZAP1;refGene_tx=XM_023242987.1;refGene_gn=DAZAP1;refGene_
tx=XM_023242991.1;refGene_gn=DAZAP1;refGene_tx=XM_023242993.1;refGene_
gn=DAZAP1;refGene_tx=XM_023242994.1;refGene_gn=DAZAP1;refGene_tx=XM_02
3242999.1;refGene_gn=DAZAP1;refGene_tx=XM_023243002.1;refGene_gn=DAZAP
1;refGene_tx=XM_023243006.1;refGene_gn=DAZAP1;refGene_tx=XM_023243016.
1
A2 868217 868358
refGene_gn=DAZAP1;refGene_tx=XM_023242987.1;refGene_gn=DAZAP1;refGene_
tx=XM_023242991.1;refGene_gn=DAZAP1;refGene_tx=XM_023242993.1;refGene_
gn=DAZAP1;refGene_tx=XM_023242994.1;refGene_gn=DAZAP1;refGene_tx=XM_02
3242999.1;refGene_gn=DAZAP1;refGene_tx=XM_023243002.1;refGene_gn=DAZAP
1;refGene_tx=XM_023243006.1;refGene_gn=DAZAP1;refGene_tx=XM_023243016.
1
A2 869697 869874
refGene_gn=DAZAP1;refGene_tx=XM_023242987.1;refGene_gn=DAZAP1;refGene_
tx=XM_023242991.1;refGene_gn=DAZAP1;refGene_tx=XM_023242993.1;refGene_
gn=DAZAP1;refGene_tx=XM_023242994.1;refGene_gn=DAZAP1;refGene_tx=XM_02
3242999.1;refGene_gn=DAZAP1;refGene_tx=XM_023243002.1;refGene_gn=DAZAP
1;refGene_tx=XM_023243006.1;refGene_gn=DAZAP1;refGene_tx=XM_023243016.
1
A2 870879 870959 refGene_gn=DAZAP1;refGene_tx=XM_023242993.1
A2 871905 872078
refGene_gn=DAZAP1;refGene_tx=XM_023242987.1;refGene_gn=DAZAP1;refGene_
tx=XM_023242991.1;refGene_gn=DAZAP1;refGene_tx=XM_023242994.1;refGene_
gn=DAZAP1;refGene_tx=XM_023242999.1;refGene_gn=DAZAP1;refGene_tx=XM_02
3243002.1;refGene_gn=DAZAP1;refGene_tx=XM_023243006.1;refGene_gn=DAZAP
1;refGene_tx=XM_023243016.1
A2 876132 876135 refGene_gn=RPS15;refGene_tx=XM_006928176.4
A2 876496 876582 refGene_gn=RPS15;refGene_tx=XM_006928176.4
A2 877092 877327 refGene_gn=RPS15;refGene_tx=XM_006928176.4
A2 877417 877531 refGene_gn=RPS15;refGene_tx=XM_006928176.4
A2 881526 881580
refGene_gn=APC2;refGene_tx=XM_023242913.1;refGene_gn=APC2;refGene_tx=X
M_023242922.1
A2 887470 887629
refGene_gn=APC2;refGene_tx=XM_023242913.1;refGene_gn=APC2;refGene_tx=X
M_023242922.1;refGene_gn=APC2;refGene_tx=XM_023242923.1
A2 887768 887859
refGene_gn=APC2;refGene_tx=XM_023242913.1;refGene_gn=APC2;refGene_tx=X
M_023242922.1;refGene_gn=APC2;refGene_tx=XM_023242923.1
A2 887964 888145
refGene_gn=APC2;refGene_tx=XM_023242913.1;refGene_gn=APC2;refGene_tx=X
M_023242922.1;refGene_gn=APC2;refGene_tx=XM_023242923.1
A2 889108 889217
refGene_gn=APC2;refGene_tx=XM_023242913.1;refGene_gn=APC2;refGene_tx=X

M_023242922.1;refGene_gn=APC2;refGene_tx=XM_023242923.1
A2	889297	889414
refGene_gn=APC2;refGene_tx=XM_023242913.1;refGene_gn=APC2;refGene_tx=XM_023242922.1;refGene_gn=APC2;refGene_tx=XM_023242923.1
A2	889934	890012
refGene_gn=APC2;refGene_tx=XM_023242913.1;refGene_gn=APC2;refGene_tx=XM_023242922.1;refGene_gn=APC2;refGene_tx=XM_023242923.1
A2	890212	890311
refGene_gn=APC2;refGene_tx=XM_023242913.1;refGene_gn=APC2;refGene_tx=XM_023242922.1;refGene_gn=APC2;refGene_tx=XM_023242923.1
A2	890848	891236
refGene_gn=APC2;refGene_tx=XM_023242913.1;refGene_gn=APC2;refGene_tx=XM_023242922.1;refGene_gn=APC2;refGene_tx=XM_023242923.1
A2	891683	891779
refGene_gn=APC2;refGene_tx=XM_023242913.1;refGene_gn=APC2;refGene_tx=XM_023242922.1;refGene_gn=APC2;refGene_tx=XM_023242923.1
A2	894468	894608
refGene_gn=APC2;refGene_tx=XM_023242913.1;refGene_gn=APC2;refGene_tx=XM_023242922.1;refGene_gn=APC2;refGene_tx=XM_023242923.1
A2	894942	895020
refGene_gn=APC2;refGene_tx=XM_023242913.1;refGene_gn=APC2;refGene_tx=XM_023242922.1;refGene_gn=APC2;refGene_tx=XM_023242923.1
A2	895243	895360
refGene_gn=APC2;refGene_tx=XM_023242913.1;refGene_gn=APC2;refGene_tx=XM_023242922.1;refGene_gn=APC2;refGene_tx=XM_023242923.1
A2	895673	895888
refGene_gn=APC2;refGene_tx=XM_023242913.1;refGene_gn=APC2;refGene_tx=XM_023242922.1;refGene_gn=APC2;refGene_tx=XM_023242923.1
A2	897706	902657
refGene_gn=APC2;refGene_tx=XM_023242913.1;refGene_gn=APC2;refGene_tx=XM_023242922.1;refGene_gn=APC2;refGene_tx=XM_023242923.1
A2	906334	906561
refGene_gn=CA2H19orf25;refGene_tx=XM_019815471.2;refGene_gn=CA2H19orf25;refGene_tx=XM_023243083.1;refGene_gn=CA2H19orf25;refGene_tx=XM_023243086.1
A2	908371	908501
refGene_gn=CA2H19orf25;refGene_tx=XM_019815471.2;refGene_gn=CA2H19orf25;refGene_tx=XM_023243083.1;refGene_gn=CA2H19orf25;refGene_tx=XM_023243086.1
A2	911132	911604
refGene_gn=PCSK4;refGene_tx=XM_023242932.1;refGene_gn=PCSK4;refGene_tx=XM_023242936.1;refGene_gn=PCSK4;refGene_tx=XM_023242938.1;refGene_gn=PCSK4;refGene_tx=XM_023242943.1;refGene_gn=PCSK4;refGene_tx=XM_023242946.1;refGene_gn=PCSK4;refGene_tx=XM_023242952.1;refGene_gn=PCSK4;refGene_tx=XM_023242954.1
A2	911741	911864
refGene_gn=PCSK4;refGene_tx=XM_023242932.1;refGene_gn=PCSK4;refGene_tx=XM_023242936.1;refGene_gn=PCSK4;refGene_tx=XM_023242938.1;refGene_gn=PCSK4;refGene_tx=XM_023242943.1;refGene_gn=PCSK4;refGene_tx=XM_023242946.1;refGene_gn=PCSK4;refGene_tx=XM_023242952.1;refGene_gn=PCSK4;refGe

ne_tx=XM_023242954.1
A2 912143 912268
refGene_gn=PCSK4;refGene_tx=XM_023242932.1;refGene_gn=PCSK4;refGene_tx=XM_023242936.1;refGene_gn=PCSK4;refGene_tx=XM_023242938.1;refGene_gn=PCSK4;refGene_tx=XM_023242943.1;refGene_gn=PCSK4;refGene_tx=XM_023242946.1;refGene_gn=PCSK4;refGene_tx=XM_023242952.1;refGene_gn=PCSK4;refGene_tx=XM_023242954.1
A2 912439 912709
refGene_gn=PCSK4;refGene_tx=XM_023242932.1;refGene_gn=PCSK4;refGene_tx=XM_023242936.1;refGene_gn=PCSK4;refGene_tx=XM_023242938.1;refGene_gn=PCSK4;refGene_tx=XM_023242943.1;refGene_gn=PCSK4;refGene_tx=XM_023242946.1;refGene_gn=PCSK4;refGene_tx=XM_023242952.1;refGene_gn=PCSK4;refGene_tx=XM_023242954.1
A2 912917 913035
refGene_gn=PCSK4;refGene_tx=XM_023242932.1;refGene_gn=PCSK4;refGene_tx=XM_023242936.1;refGene_gn=PCSK4;refGene_tx=XM_023242938.1;refGene_gn=PCSK4;refGene_tx=XM_023242943.1;refGene_gn=PCSK4;refGene_tx=XM_023242946.1;refGene_gn=PCSK4;refGene_tx=XM_023242952.1;refGene_gn=PCSK4;refGene_tx=XM_023242954.1
A2 913106 913210
refGene_gn=PCSK4;refGene_tx=XM_023242932.1;refGene_gn=PCSK4;refGene_tx=XM_023242936.1;refGene_gn=PCSK4;refGene_tx=XM_023242938.1;refGene_gn=PCSK4;refGene_tx=XM_023242943.1;refGene_gn=PCSK4;refGene_tx=XM_023242946.1;refGene_gn=PCSK4;refGene_tx=XM_023242952.1;refGene_gn=PCSK4;refGene_tx=XM_023242954.1
A2 913302 913403
refGene_gn=PCSK4;refGene_tx=XM_023242932.1;refGene_gn=PCSK4;refGene_tx=XM_023242936.1;refGene_gn=PCSK4;refGene_tx=XM_023242938.1;refGene_gn=PCSK4;refGene_tx=XM_023242943.1;refGene_gn=PCSK4;refGene_tx=XM_023242946.1;refGene_gn=PCSK4;refGene_tx=XM_023242952.1;refGene_gn=PCSK4;refGene_tx=XM_023242954.1
A2 913906 914119
refGene_gn=PCSK4;refGene_tx=XM_023242932.1;refGene_gn=PCSK4;refGene_tx=XM_023242936.1;refGene_gn=PCSK4;refGene_tx=XM_023242938.1;refGene_gn=PCSK4;refGene_tx=XM_023242943.1;refGene_gn=PCSK4;refGene_tx=XM_023242946.1;refGene_gn=PCSK4;refGene_tx=XM_023242952.1;refGene_gn=PCSK4;refGene_tx=XM_023242954.1
A2 914203 914376
refGene_gn=PCSK4;refGene_tx=XM_023242932.1;refGene_gn=PCSK4;refGene_tx=XM_023242936.1;refGene_gn=PCSK4;refGene_tx=XM_023242938.1;refGene_gn=PCSK4;refGene_tx=XM_023242943.1;refGene_gn=PCSK4;refGene_tx=XM_023242946.1;refGene_gn=PCSK4;refGene_tx=XM_023242952.1;refGene_gn=PCSK4;refGene_tx=XM_023242954.1
A2 914597 914686
refGene_gn=PCSK4;refGene_tx=XM_023242932.1;refGene_gn=PCSK4;refGene_tx=XM_023242936.1;refGene_gn=PCSK4;refGene_tx=XM_023242938.1;refGene_gn=PCSK4;refGene_tx=XM_023242943.1;refGene_gn=PCSK4;refGene_tx=XM_023242946.1;refGene_gn=PCSK4;refGene_tx=XM_023242952.1;refGene_gn=PCSK4;refGene_tx=XM_023242954.1
A2 914785 914862

refGene_gn=PCSK4;refGene_tx=XM_023242932.1;refGene_gn=PCSK4;refGene_tx
=XM_023242936.1;refGene_gn=PCSK4;refGene_tx=XM_023242938.1;refGene_gn=
PCSK4;refGene_tx=XM_023242943.1;refGene_gn=PCSK4;refGene_tx=XM_0232429
46.1;refGene_gn=PCSK4;refGene_tx=XM_023242952.1;refGene_gn=PCSK4;refGe
ne_tx=XM_023242954.1
A2 914945 915074
refGene_gn=PCSK4;refGene_tx=XM_023242932.1;refGene_gn=PCSK4;refGene_tx
=XM_023242936.1;refGene_gn=PCSK4;refGene_tx=XM_023242938.1;refGene_gn=
PCSK4;refGene_tx=XM_023242943.1;refGene_gn=PCSK4;refGene_tx=XM_0232429
46.1;refGene_gn=PCSK4;refGene_tx=XM_023242952.1;refGene_gn=PCSK4;refGe
ne_tx=XM_023242954.1
A2 915148 915241
refGene_gn=PCSK4;refGene_tx=XM_023242932.1;refGene_gn=PCSK4;refGene_tx
=XM_023242936.1;refGene_gn=PCSK4;refGene_tx=XM_023242938.1;refGene_gn=
PCSK4;refGene_tx=XM_023242943.1;refGene_gn=PCSK4;refGene_tx=XM_0232429
46.1;refGene_gn=PCSK4;refGene_tx=XM_023242952.1;refGene_gn=PCSK4;refGe
ne_tx=XM_023242954.1
A2 916718 916823
refGene_gn=PCSK4;refGene_tx=XM_023242932.1;refGene_gn=PCSK4;refGene_tx
=XM_023242936.1;refGene_gn=PCSK4;refGene_tx=XM_023242938.1;refGene_gn=
PCSK4;refGene_tx=XM_023242943.1;refGene_gn=PCSK4;refGene_tx=XM_0232429
46.1;refGene_gn=PCSK4;refGene_tx=XM_023242952.1;refGene_gn=PCSK4;refGe
ne_tx=XM_023242954.1
A2 917070 917493
refGene_gn=PCSK4;refGene_tx=XM_023242932.1;refGene_gn=PCSK4;refGene_tx
=XM_023242936.1;refGene_gn=PCSK4;refGene_tx=XM_023242938.1;refGene_gn=
PCSK4;refGene_tx=XM_023242943.1;refGene_gn=PCSK4;refGene_tx=XM_0232429
46.1;refGene_gn=PCSK4;refGene_tx=XM_023242952.1;refGene_gn=PCSK4;refGe
ne_tx=XM_023242954.1
A2 918241 918356
refGene_gn=REEP6;refGene_tx=XM_023243058.1;refGene_gn=REEP6;refGene_tx
=XM_023243067.1
A2 921375 921469
refGene_gn=REEP6;refGene_tx=XM_023243058.1;refGene_gn=REEP6;refGene_tx
=XM_023243067.1
A2 921555 921694
refGene_gn=REEP6;refGene_tx=XM_023243058.1;refGene_gn=REEP6;refGene_tx
=XM_023243067.1
A2 921878 922047
refGene_gn=REEP6;refGene_tx=XM_023243058.1;refGene_gn=REEP6;refGene_tx
=XM_023243067.1
A2 922222 922285 refGene_gn=REEP6;refGene_tx=XM_023243058.1
A2 922445 922483
refGene_gn=REEP6;refGene_tx=XM_023243058.1;refGene_gn=REEP6;refGene_tx
=XM_023243067.1
A2 928982 929284 refGene_gn=ADAMTSL5;refGene_tx=XM_006928180.4
A2 929620 929692 refGene_gn=ADAMTSL5;refGene_tx=XM_006928180.4
A2 929765 929964 refGene_gn=ADAMTSL5;refGene_tx=XM_006928180.4
A2 930105 930269 refGene_gn=ADAMTSL5;refGene_tx=XM_006928180.4
A2 930405 930457 refGene_gn=ADAMTSL5;refGene_tx=XM_006928180.4

A2 930458 930493 refGene_gn=ADAMTSL5;refGene_tx=XM_006928180.4
A2 930845 930957 refGene_gn=ADAMTSL5;refGene_tx=XM_006928180.4
A2 931221 931349 refGene_gn=ADAMTSL5;refGene_tx=XM_006928180.4
A2 932672 932781 refGene_gn=ADAMTSL5;refGene_tx=XM_006928180.4
A2 932933 932994 refGene_gn=ADAMTSL5;refGene_tx=XM_006928180.4
A2 933238 933330 refGene_gn=ADAMTSL5;refGene_tx=XM_006928180.4
A2 933439 933635 refGene_gn=ADAMTSL5;refGene_tx=XM_006928180.4
A2 933909 933965 refGene_gn=ADAMTSL5;refGene_tx=XM_006928180.4
A2 943366 943480
refGene_gn=PLK5;refGene_tx=XM_023242966.1;refGene_gn=PLK5;refGene_tx=X
M_023242973.1
A2 944100 944214
refGene_gn=PLK5;refGene_tx=XM_023242966.1;refGene_gn=PLK5;refGene_tx=X
M_023242973.1
A2 944327 944444
refGene_gn=PLK5;refGene_tx=XM_023242966.1;refGene_gn=PLK5;refGene_tx=X
M_023242973.1
A2 946035 946165
refGene_gn=PLK5;refGene_tx=XM_023242966.1;refGene_gn=PLK5;refGene_tx=X
M_023242973.1
A2 946277 946365
refGene_gn=PLK5;refGene_tx=XM_023242966.1;refGene_gn=PLK5;refGene_tx=X
M_023242973.1
A2 946450 946546
refGene_gn=PLK5;refGene_tx=XM_023242966.1;refGene_gn=PLK5;refGene_tx=X
M_023242973.1
A2 947049 947248
refGene_gn=PLK5;refGene_tx=XM_023242966.1;refGene_gn=PLK5;refGene_tx=X
M_023242973.1
A2 947325 947452
refGene_gn=PLK5;refGene_tx=XM_023242966.1;refGene_gn=PLK5;refGene_tx=X
M_023242973.1
A2 947609 947686
refGene_gn=PLK5;refGene_tx=XM_023242966.1;refGene_gn=PLK5;refGene_tx=X
M_023242973.1
A2 947837 947922
refGene_gn=PLK5;refGene_tx=XM_023242966.1;refGene_gn=PLK5;refGene_tx=X
M_023242973.1
A2 948197 948275
refGene_gn=PLK5;refGene_tx=XM_023242966.1;refGene_gn=PLK5;refGene_tx=X
M_023242973.1
A2 948983 949140
refGene_gn=PLK5;refGene_tx=XM_023242966.1;refGene_gn=PLK5;refGene_tx=X
M_023242973.1
A2 949865 949995
refGene_gn=PLK5;refGene_tx=XM_023242966.1;refGene_gn=PLK5;refGene_tx=X
M_023242973.1
A2 950078 950426
refGene_gn=PLK5;refGene_tx=XM_023242966.1;refGene_gn=PLK5;refGene_tx=X
M_023242973.1

A2 951115 951301 refGene_gn=PLK5;refGene_tx=XM_023242973.1
A2 962644 962682 refGene_gn=MEX3D;refGene_tx=XM_023250887.1
A2 962786 964157 refGene_gn=MEX3D;refGene_tx=XM_023250887.1
A2 970298 970747 refGene_gn=MEX3D;refGene_tx=XM_023250887.1
A2 970892 971088 refGene_gn=MEX3D;refGene_tx=XM_023250887.1
A2 972303 972340 refGene_gn=MEX3D;refGene_tx=XM_023250887.1
A2 981674 981873
refGene_gn=MBD3;refGene_tx=XM_023243028.1;refGene_gn=MBD3;refGene_tx=XM_023243038.1;refGene_gn=MBD3;refGene_tx=XM_023243045.1
A2 983426 983604
refGene_gn=MBD3;refGene_tx=XM_023243028.1;refGene_gn=MBD3;refGene_tx=XM_023243038.1;refGene_gn=MBD3;refGene_tx=XM_023243045.1
A2 983914 984005
refGene_gn=MBD3;refGene_tx=XM_023243028.1;refGene_gn=MBD3;refGene_tx=XM_023243038.1;refGene_gn=MBD3;refGene_tx=XM_023243045.1
A2 984810 984948
refGene_gn=MBD3;refGene_tx=XM_023243028.1;refGene_gn=MBD3;refGene_tx=XM_023243038.1;refGene_gn=MBD3;refGene_tx=XM_023243045.1
A2 985575 985735
refGene_gn=MBD3;refGene_tx=XM_023243028.1;refGene_gn=MBD3;refGene_tx=XM_023243038.1;refGene_gn=MBD3;refGene_tx=XM_023243045.1
A2 990598 990708
refGene_gn=MBD3;refGene_tx=XM_023243028.1;refGene_gn=MBD3;refGene_tx=XM_023243038.1
A2 996444 996565
refGene_gn=LOC101086703;refGene_tx=XM_003981611.5;refGene_gn=LOC101086703;refGene_tx=XM_006928183.4;refGene_gn=LOC101086703;refGene_tx=XM_006928184.4
A2 997965 998015
refGene_gn=LOC101086703;refGene_tx=XM_003981611.5;refGene_gn=LOC101086703;refGene_tx=XM_006928183.4;refGene_gn=LOC101086703;refGene_tx=XM_006928184.4
A2 1004421 1004564
refGene_gn=TCF3;refGene_tx=XM_023243101.1;refGene_gn=TCF3;refGene_tx=XM_023243110.1;refGene_gn=TCF3;refGene_tx=XM_023243112.1;refGene_gn=TCF3;refGene_tx=XM_023243143.1;refGene_gn=TCF3;refGene_tx=XM_023243151.1
A2 1004717 1004944
refGene_gn=TCF3;refGene_tx=XM_023243110.1;refGene_gn=TCF3;refGene_tx=XM_023243112.1;refGene_gn=TCF3;refGene_tx=XM_023243113.1;refGene_gn=TCF3;refGene_tx=XM_023243119.1;refGene_gn=TCF3;refGene_tx=XM_023243127.1;refGene_gn=TCF3;refGene_tx=XM_023243134.1;refGene_gn=TCF3;refGene_tx=XM_023243151.1
A2 1007036 1007272
refGene_gn=TCF3;refGene_tx=XM_023243101.1;refGene_gn=TCF3;refGene_tx=XM_023243113.1;refGene_gn=TCF3;refGene_tx=XM_023243119.1;refGene_gn=TCF3;refGene_tx=XM_023243127.1;refGene_gn=TCF3;refGene_tx=XM_023243134.1;refGene_gn=TCF3;refGene_tx=XM_023243143.1;refGene_gn=TCF3;refGene_tx=XM_023243160.1;refGene_gn=TCF3;refGene_tx=XM_023243171.1;refGene_gn=TCF3;refGene_tx=XM_023243174.1;refGene_gn=TCF3;refGene_tx=XM_023243175.1;refGene_gn=TCF3;refGene_tx=XM_023243179.1;refGene_gn=TCF3;refGene_tx=X

M_023243184.1;refGene_gn=TCF3;refGene_tx=XM_023243187.1
A2	1007439	1007575
refGene_gn=TCF3;refGene_tx=XM_023243101.1;refGene_gn=TCF3;refGene_tx=XM_023243110.1;refGene_gn=TCF3;refGene_tx=XM_023243112.1;refGene_gn=TCF3;refGene_tx=XM_023243113.1;refGene_gn=TCF3;refGene_tx=XM_023243119.1;refGene_gn=TCF3;refGene_tx=XM_023243127.1;refGene_gn=TCF3;refGene_tx=XM_023243134.1;refGene_gn=TCF3;refGene_tx=XM_023243143.1;refGene_gn=TCF3;refGene_tx=XM_023243151.1;refGene_gn=TCF3;refGene_tx=XM_023243160.1;refGene_gn=TCF3;refGene_tx=XM_023243171.1;refGene_gn=TCF3;refGene_tx=XM_023243174.1;refGene_gn=TCF3;refGene_tx=XM_023243175.1;refGene_gn=TCF3;refGene_tx=XM_023243179.1;refGene_gn=TCF3;refGene_tx=XM_023243184.1;refGene_gn=TCF3;refGene_tx=XM_023243187.1
A2	1010753	1010877
refGene_gn=TCF3;refGene_tx=XM_023243101.1;refGene_gn=TCF3;refGene_tx=XM_023243110.1;refGene_gn=TCF3;refGene_tx=XM_023243112.1;refGene_gn=TCF3;refGene_tx=XM_023243113.1;refGene_gn=TCF3;refGene_tx=XM_023243119.1;refGene_gn=TCF3;refGene_tx=XM_023243127.1;refGene_gn=TCF3;refGene_tx=XM_023243134.1;refGene_gn=TCF3;refGene_tx=XM_023243143.1;refGene_gn=TCF3;refGene_tx=XM_023243151.1;refGene_gn=TCF3;refGene_tx=XM_023243160.1;refGene_gn=TCF3;refGene_tx=XM_023243171.1;refGene_gn=TCF3;refGene_tx=XM_023243174.1;refGene_gn=TCF3;refGene_tx=XM_023243175.1;refGene_gn=TCF3;refGene_tx=XM_023243179.1;refGene_gn=TCF3;refGene_tx=XM_023243184.1;refGene_gn=TCF3;refGene_tx=XM_023243187.1
A2	1010967	1011126
refGene_gn=TCF3;refGene_tx=XM_023243101.1;refGene_gn=TCF3;refGene_tx=XM_023243110.1;refGene_gn=TCF3;refGene_tx=XM_023243112.1;refGene_gn=TCF3;refGene_tx=XM_023243113.1;refGene_gn=TCF3;refGene_tx=XM_023243119.1;refGene_gn=TCF3;refGene_tx=XM_023243127.1;refGene_gn=TCF3;refGene_tx=XM_023243134.1;refGene_gn=TCF3;refGene_tx=XM_023243143.1;refGene_gn=TCF3;refGene_tx=XM_023243151.1;refGene_gn=TCF3;refGene_tx=XM_023243160.1;refGene_gn=TCF3;refGene_tx=XM_023243171.1;refGene_gn=TCF3;refGene_tx=XM_023243174.1;refGene_gn=TCF3;refGene_tx=XM_023243175.1;refGene_gn=TCF3;refGene_tx=XM_023243179.1;refGene_gn=TCF3;refGene_tx=XM_023243184.1;refGene_gn=TCF3;refGene_tx=XM_023243187.1
A2	1011476	1011550
refGene_gn=TCF3;refGene_tx=XM_023243101.1;refGene_gn=TCF3;refGene_tx=XM_023243110.1;refGene_gn=TCF3;refGene_tx=XM_023243112.1;refGene_gn=TCF3;refGene_tx=XM_023243113.1;refGene_gn=TCF3;refGene_tx=XM_023243119.1;refGene_gn=TCF3;refGene_tx=XM_023243127.1;refGene_gn=TCF3;refGene_tx=XM_023243134.1;refGene_gn=TCF3;refGene_tx=XM_023243143.1;refGene_gn=TCF3;refGene_tx=XM_023243151.1;refGene_gn=TCF3;refGene_tx=XM_023243160.1;refGene_gn=TCF3;refGene_tx=XM_023243171.1;refGene_gn=TCF3;refGene_tx=XM_023243174.1;refGene_gn=TCF3;refGene_tx=XM_023243175.1;refGene_gn=TCF3;refGene_tx=XM_023243179.1;refGene_gn=TCF3;refGene_tx=XM_023243184.1;refGene_gn=TCF3;refGene_tx=XM_023243187.1
A2	1012325	1012404
refGene_gn=TCF3;refGene_tx=XM_023243101.1;refGene_gn=TCF3;refGene_tx=XM_023243110.1;refGene_gn=TCF3;refGene_tx=XM_023243112.1;refGene_gn=TCF3;refGene_tx=XM_023243113.1;refGene_gn=TCF3;refGene_tx=XM_023243119.1;refGene_gn=TCF3;refGene_tx=XM_023243127.1;refGene_gn=TCF3;refGene_tx=X

M_023243134.1;refGene_gn=TCF3;refGene_tx=XM_023243143.1;refGene_gn=TCF3;refGene_tx=XM_023243151.1;refGene_gn=TCF3;refGene_tx=XM_023243160.1;refGene_gn=TCF3;refGene_tx=XM_023243171.1;refGene_gn=TCF3;refGene_tx=XM_023243174.1;refGene_gn=TCF3;refGene_tx=XM_023243175.1;refGene_gn=TCF3;refGene_tx=XM_023243179.1;refGene_gn=TCF3;refGene_tx=XM_023243184.1;refGene_gn=TCF3;refGene_tx=XM_023243187.1
A2 1012510 1012569
refGene_gn=TCF3;refGene_tx=XM_023243101.1;refGene_gn=TCF3;refGene_tx=XM_023243110.1;refGene_gn=TCF3;refGene_tx=XM_023243112.1;refGene_gn=TCF3;refGene_tx=XM_023243113.1;refGene_gn=TCF3;refGene_tx=XM_023243119.1;refGene_gn=TCF3;refGene_tx=XM_023243127.1;refGene_gn=TCF3;refGene_tx=XM_023243134.1;refGene_gn=TCF3;refGene_tx=XM_023243143.1;refGene_gn=TCF3;refGene_tx=XM_023243151.1;refGene_gn=TCF3;refGene_tx=XM_023243160.1;refGene_gn=TCF3;refGene_tx=XM_023243171.1;refGene_gn=TCF3;refGene_tx=XM_023243174.1;refGene_gn=TCF3;refGene_tx=XM_023243175.1;refGene_gn=TCF3;refGene_tx=XM_023243179.1;refGene_gn=TCF3;refGene_tx=XM_023243184.1;refGene_gn=TCF3;refGene_tx=XM_023243187.1
A2 1012865 1012998
refGene_gn=TCF3;refGene_tx=XM_023243101.1;refGene_gn=TCF3;refGene_tx=XM_023243110.1;refGene_gn=TCF3;refGene_tx=XM_023243112.1;refGene_gn=TCF3;refGene_tx=XM_023243113.1;refGene_gn=TCF3;refGene_tx=XM_023243119.1;refGene_gn=TCF3;refGene_tx=XM_023243127.1;refGene_gn=TCF3;refGene_tx=XM_023243134.1;refGene_gn=TCF3;refGene_tx=XM_023243143.1;refGene_gn=TCF3;refGene_tx=XM_023243151.1;refGene_gn=TCF3;refGene_tx=XM_023243160.1;refGene_gn=TCF3;refGene_tx=XM_023243171.1;refGene_gn=TCF3;refGene_tx=XM_023243174.1;refGene_gn=TCF3;refGene_tx=XM_023243175.1;refGene_gn=TCF3;refGene_tx=XM_023243179.1;refGene_gn=TCF3;refGene_tx=XM_023243184.1;refGene_gn=TCF3;refGene_tx=XM_023243187.1
A2 1013082 1013189
refGene_gn=TCF3;refGene_tx=XM_023243101.1;refGene_gn=TCF3;refGene_tx=XM_023243110.1;refGene_gn=TCF3;refGene_tx=XM_023243112.1;refGene_gn=TCF3;refGene_tx=XM_023243113.1;refGene_gn=TCF3;refGene_tx=XM_023243119.1;refGene_gn=TCF3;refGene_tx=XM_023243127.1;refGene_gn=TCF3;refGene_tx=XM_023243134.1;refGene_gn=TCF3;refGene_tx=XM_023243143.1;refGene_gn=TCF3;refGene_tx=XM_023243151.1;refGene_gn=TCF3;refGene_tx=XM_023243160.1;refGene_gn=TCF3;refGene_tx=XM_023243171.1;refGene_gn=TCF3;refGene_tx=XM_023243174.1;refGene_gn=TCF3;refGene_tx=XM_023243175.1;refGene_gn=TCF3;refGene_tx=XM_023243179.1;refGene_gn=TCF3;refGene_tx=XM_023243184.1;refGene_gn=TCF3;refGene_tx=XM_023243187.1
A2 1013269 1013372
refGene_gn=TCF3;refGene_tx=XM_023243101.1;refGene_gn=TCF3;refGene_tx=XM_023243110.1;refGene_gn=TCF3;refGene_tx=XM_023243112.1;refGene_gn=TCF3;refGene_tx=XM_023243113.1;refGene_gn=TCF3;refGene_tx=XM_023243119.1;refGene_gn=TCF3;refGene_tx=XM_023243127.1;refGene_gn=TCF3;refGene_tx=XM_023243134.1;refGene_gn=TCF3;refGene_tx=XM_023243143.1;refGene_gn=TCF3;refGene_tx=XM_023243151.1;refGene_gn=TCF3;refGene_tx=XM_023243160.1;refGene_gn=TCF3;refGene_tx=XM_023243171.1;refGene_gn=TCF3;refGene_tx=XM_023243174.1;refGene_gn=TCF3;refGene_tx=XM_023243175.1;refGene_gn=TCF3;refGene_tx=XM_023243179.1;refGene_gn=TCF3;refGene_tx=XM_023243184.1;refGene_gn=TCF3;refGene_tx=XM_023243187.1

A2 1014463 1014513
refGene_gn=TCF3;refGene_tx=XM_023243101.1;refGene_gn=TCF3;refGene_tx=XM_023243110.1;refGene_gn=TCF3;refGene_tx=XM_023243112.1;refGene_gn=TCF3;refGene_tx=XM_023243113.1;refGene_gn=TCF3;refGene_tx=XM_023243119.1;refGene_gn=TCF3;refGene_tx=XM_023243127.1;refGene_gn=TCF3;refGene_tx=XM_023243134.1;refGene_gn=TCF3;refGene_tx=XM_023243143.1;refGene_gn=TCF3;refGene_tx=XM_023243151.1;refGene_gn=TCF3;refGene_tx=XM_023243160.1;refGene_gn=TCF3;refGene_tx=XM_023243171.1;refGene_gn=TCF3;refGene_tx=XM_023243174.1;refGene_gn=TCF3;refGene_tx=XM_023243175.1;refGene_gn=TCF3;refGene_tx=XM_023243179.1;refGene_gn=TCF3;refGene_tx=XM_023243184.1;refGene_gn=TCF3;refGene_tx=XM_023243187.1
A2 1015104 1015234
refGene_gn=TCF3;refGene_tx=XM_023243101.1;refGene_gn=TCF3;refGene_tx=XM_023243110.1;refGene_gn=TCF3;refGene_tx=XM_023243112.1;refGene_gn=TCF3;refGene_tx=XM_023243113.1;refGene_gn=TCF3;refGene_tx=XM_023243119.1;refGene_gn=TCF3;refGene_tx=XM_023243127.1;refGene_gn=TCF3;refGene_tx=XM_023243134.1;refGene_gn=TCF3;refGene_tx=XM_023243143.1;refGene_gn=TCF3;refGene_tx=XM_023243151.1;refGene_gn=TCF3;refGene_tx=XM_023243160.1;refGene_gn=TCF3;refGene_tx=XM_023243171.1;refGene_gn=TCF3;refGene_tx=XM_023243174.1;refGene_gn=TCF3;refGene_tx=XM_023243175.1;refGene_gn=TCF3;refGene_tx=XM_023243179.1;refGene_gn=TCF3;refGene_tx=XM_023243184.1;refGene_gn=TCF3;refGene_tx=XM_023243187.1
A2 1016395 1016463
refGene_gn=TCF3;refGene_tx=XM_023243101.1;refGene_gn=TCF3;refGene_tx=XM_023243110.1;refGene_gn=TCF3;refGene_tx=XM_023243112.1;refGene_gn=TCF3;refGene_tx=XM_023243113.1;refGene_gn=TCF3;refGene_tx=XM_023243119.1;refGene_gn=TCF3;refGene_tx=XM_023243127.1;refGene_gn=TCF3;refGene_tx=XM_023243134.1;refGene_gn=TCF3;refGene_tx=XM_023243143.1;refGene_gn=TCF3;refGene_tx=XM_023243151.1;refGene_gn=TCF3;refGene_tx=XM_023243160.1;refGene_gn=TCF3;refGene_tx=XM_023243171.1;refGene_gn=TCF3;refGene_tx=XM_023243174.1;refGene_gn=TCF3;refGene_tx=XM_023243175.1;refGene_gn=TCF3;refGene_tx=XM_023243179.1;refGene_gn=TCF3;refGene_tx=XM_023243184.1;refGene_gn=TCF3;refGene_tx=XM_023243187.1
A2 1018198 1018277
refGene_gn=TCF3;refGene_tx=XM_023243101.1;refGene_gn=TCF3;refGene_tx=XM_023243110.1;refGene_gn=TCF3;refGene_tx=XM_023243112.1;refGene_gn=TCF3;refGene_tx=XM_023243113.1;refGene_gn=TCF3;refGene_tx=XM_023243119.1;refGene_gn=TCF3;refGene_tx=XM_023243127.1;refGene_gn=TCF3;refGene_tx=XM_023243134.1;refGene_gn=TCF3;refGene_tx=XM_023243143.1;refGene_gn=TCF3;refGene_tx=XM_023243151.1;refGene_gn=TCF3;refGene_tx=XM_023243160.1;refGene_gn=TCF3;refGene_tx=XM_023243171.1;refGene_gn=TCF3;refGene_tx=XM_023243174.1;refGene_gn=TCF3;refGene_tx=XM_023243175.1;refGene_gn=TCF3;refGene_tx=XM_023243179.1;refGene_gn=TCF3;refGene_tx=XM_023243184.1;refGene_gn=TCF3;refGene_tx=XM_023243187.1
A2 1018474 1018548
refGene_gn=TCF3;refGene_tx=XM_023243101.1;refGene_gn=TCF3;refGene_tx=XM_023243110.1;refGene_gn=TCF3;refGene_tx=XM_023243112.1;refGene_gn=TCF3;refGene_tx=XM_023243113.1;refGene_gn=TCF3;refGene_tx=XM_023243119.1;refGene_gn=TCF3;refGene_tx=XM_023243127.1;refGene_gn=TCF3;refGene_tx=XM_023243134.1;refGene_gn=TCF3;refGene_tx=XM_023243143.1;refGene_gn=TCF

3;refGene_tx=XM_023243151.1;refGene_gn=TCF3;refGene_tx=XM_023243160.1;
refGene_gn=TCF3;refGene_tx=XM_023243171.1;refGene_gn=TCF3;refGene_tx=X
M_023243174.1;refGene_gn=TCF3;refGene_tx=XM_023243175.1;refGene_gn=TCF
3;refGene_tx=XM_023243179.1;refGene_gn=TCF3;refGene_tx=XM_023243184.1;
refGene_gn=TCF3;refGene_tx=XM_023243187.1
A2 1029500 1029573
refGene_gn=TCF3;refGene_tx=XM_023243101.1;refGene_gn=TCF3;refGene_tx=X
M_023243110.1;refGene_gn=TCF3;refGene_tx=XM_023243112.1;refGene_gn=TCF
3;refGene_tx=XM_023243113.1;refGene_gn=TCF3;refGene_tx=XM_023243119.1;
refGene_gn=TCF3;refGene_tx=XM_023243127.1;refGene_gn=TCF3;refGene_tx=X
M_023243134.1;refGene_gn=TCF3;refGene_tx=XM_023243143.1;refGene_gn=TCF
3;refGene_tx=XM_023243151.1;refGene_gn=TCF3;refGene_tx=XM_023243160.1;
refGene_gn=TCF3;refGene_tx=XM_023243171.1;refGene_gn=TCF3;refGene_tx=X
M_023243174.1;refGene_gn=TCF3;refGene_tx=XM_023243175.1;refGene_gn=TCF
3;refGene_tx=XM_023243179.1;refGene_gn=TCF3;refGene_tx=XM_023243184.1;
refGene_gn=TCF3;refGene_tx=XM_023243187.1
A2 1033208 1033280
refGene_gn=TCF3;refGene_tx=XM_023243101.1;refGene_gn=TCF3;refGene_tx=X
M_023243110.1;refGene_gn=TCF3;refGene_tx=XM_023243112.1;refGene_gn=TCF
3;refGene_tx=XM_023243113.1;refGene_gn=TCF3;refGene_tx=XM_023243119.1;
refGene_gn=TCF3;refGene_tx=XM_023243127.1;refGene_gn=TCF3;refGene_tx=X
M_023243134.1;refGene_gn=TCF3;refGene_tx=XM_023243143.1;refGene_gn=TCF
3;refGene_tx=XM_023243151.1;refGene_gn=TCF3;refGene_tx=XM_023243160.1;
refGene_gn=TCF3;refGene_tx=XM_023243171.1;refGene_gn=TCF3;refGene_tx=X
M_023243174.1;refGene_gn=TCF3;refGene_tx=XM_023243175.1;refGene_gn=TCF
3;refGene_tx=XM_023243179.1;refGene_gn=TCF3;refGene_tx=XM_023243184.1;
refGene_gn=TCF3;refGene_tx=XM_023243187.1
A2 1113249 1114450 refGene_gn=ONECUT3;refGene_tx=XM_023241230.1
A2 1133769 1134062 refGene_gn=ONECUT3;refGene_tx=XM_023241230.1
A2 1138432 1139017
refGene_gn=ATP8B3;refGene_tx=XM_023243212.1;refGene_gn=ATP8B3;refGene_
tx=XM_023243213.1;refGene_gn=ATP8B3;refGene_tx=XM_023243214.1;refGene_
gn=ATP8B3;refGene_tx=XM_023243219.1;refGene_gn=ATP8B3;refGene_tx=XM_02
3243221.1;refGene_gn=ATP8B3;refGene_tx=XM_023243229.1;refGene_gn=ATP8B
3;refGene_tx=XM_023243238.1;refGene_gn=ATP8B3;refGene_tx=XM_023243240.
1;refGene_gn=ATP8B3;refGene_tx=XM_023243241.1;refGene_gn=ATP8B3;refGen
e_tx=XM_023243246.1;refGene_gn=ATP8B3;refGene_tx=XM_023243251.1
A2 1140171 1140299
refGene_gn=ATP8B3;refGene_tx=XM_023243212.1;refGene_gn=ATP8B3;refGene_
tx=XM_023243213.1;refGene_gn=ATP8B3;refGene_tx=XM_023243214.1;refGene_
gn=ATP8B3;refGene_tx=XM_023243219.1;refGene_gn=ATP8B3;refGene_tx=XM_02
3243221.1;refGene_gn=ATP8B3;refGene_tx=XM_023243229.1;refGene_gn=ATP8B
3;refGene_tx=XM_023243238.1;refGene_gn=ATP8B3;refGene_tx=XM_023243240.
1;refGene_gn=ATP8B3;refGene_tx=XM_023243241.1;refGene_gn=ATP8B3;refGen
e_tx=XM_023243246.1;refGene_gn=ATP8B3;refGene_tx=XM_023243251.1
A2 1140588 1140727
refGene_gn=ATP8B3;refGene_tx=XM_023243212.1;refGene_gn=ATP8B3;refGene_
tx=XM_023243213.1;refGene_gn=ATP8B3;refGene_tx=XM_023243214.1;refGene_
gn=ATP8B3;refGene_tx=XM_023243219.1;refGene_gn=ATP8B3;refGene_tx=XM_02
3243221.1;refGene_gn=ATP8B3;refGene_tx=XM_023243229.1;refGene_gn=ATP8B

3;refGene_tx=XM_023243238.1;refGene_gn=ATP8B3;refGene_tx=XM_023243240.
1;refGene_gn=ATP8B3;refGene_tx=XM_023243241.1;refGene_gn=ATP8B3;refGen
e_tx=XM_023243246.1;refGene_gn=ATP8B3;refGene_tx=XM_023243251.1
A2 1140920 1141160
refGene_gn=ATP8B3;refGene_tx=XM_023243212.1;refGene_gn=ATP8B3;refGene_
tx=XM_023243213.1;refGene_gn=ATP8B3;refGene_tx=XM_023243214.1;refGene_
gn=ATP8B3;refGene_tx=XM_023243219.1;refGene_gn=ATP8B3;refGene_tx=XM_02
3243221.1;refGene_gn=ATP8B3;refGene_tx=XM_023243229.1;refGene_gn=ATP8B
3;refGene_tx=XM_023243238.1;refGene_gn=ATP8B3;refGene_tx=XM_023243240.
1;refGene_gn=ATP8B3;refGene_tx=XM_023243241.1
A2 1141317 1141401
refGene_gn=ATP8B3;refGene_tx=XM_023243212.1;refGene_gn=ATP8B3;refGene_
tx=XM_023243213.1;refGene_gn=ATP8B3;refGene_tx=XM_023243214.1;refGene_
gn=ATP8B3;refGene_tx=XM_023243219.1;refGene_gn=ATP8B3;refGene_tx=XM_02
3243221.1;refGene_gn=ATP8B3;refGene_tx=XM_023243229.1;refGene_gn=ATP8B
3;refGene_tx=XM_023243238.1;refGene_gn=ATP8B3;refGene_tx=XM_023243240.
1;refGene_gn=ATP8B3;refGene_tx=XM_023243241.1;refGene_gn=ATP8B3;refGen
e_tx=XM_023243246.1;refGene_gn=ATP8B3;refGene_tx=XM_023243251.1
A2 1143272 1143496
refGene_gn=ATP8B3;refGene_tx=XM_023243212.1;refGene_gn=ATP8B3;refGene_
tx=XM_023243213.1;refGene_gn=ATP8B3;refGene_tx=XM_023243214.1;refGene_
gn=ATP8B3;refGene_tx=XM_023243219.1;refGene_gn=ATP8B3;refGene_tx=XM_02
3243221.1;refGene_gn=ATP8B3;refGene_tx=XM_023243229.1;refGene_gn=ATP8B
3;refGene_tx=XM_023243238.1;refGene_gn=ATP8B3;refGene_tx=XM_023243240.
1;refGene_gn=ATP8B3;refGene_tx=XM_023243241.1;refGene_gn=ATP8B3;refGen
e_tx=XM_023243246.1;refGene_gn=ATP8B3;refGene_tx=XM_023243251.1
A2 1143583 1143977
refGene_gn=ATP8B3;refGene_tx=XM_023243212.1;refGene_gn=ATP8B3;refGene_
tx=XM_023243213.1;refGene_gn=ATP8B3;refGene_tx=XM_023243214.1;refGene_
gn=ATP8B3;refGene_tx=XM_023243219.1;refGene_gn=ATP8B3;refGene_tx=XM_02
3243221.1;refGene_gn=ATP8B3;refGene_tx=XM_023243229.1;refGene_gn=ATP8B
3;refGene_tx=XM_023243238.1;refGene_gn=ATP8B3;refGene_tx=XM_023243240.
1;refGene_gn=ATP8B3;refGene_tx=XM_023243241.1;refGene_gn=ATP8B3;refGen
e_tx=XM_023243246.1;refGene_gn=ATP8B3;refGene_tx=XM_023243251.1
A2 1144200 1144318
refGene_gn=ATP8B3;refGene_tx=XM_023243212.1;refGene_gn=ATP8B3;refGene_
tx=XM_023243213.1;refGene_gn=ATP8B3;refGene_tx=XM_023243214.1;refGene_
gn=ATP8B3;refGene_tx=XM_023243219.1;refGene_gn=ATP8B3;refGene_tx=XM_02
3243221.1;refGene_gn=ATP8B3;refGene_tx=XM_023243229.1;refGene_gn=ATP8B
3;refGene_tx=XM_023243238.1;refGene_gn=ATP8B3;refGene_tx=XM_023243240.
1;refGene_gn=ATP8B3;refGene_tx=XM_023243241.1;refGene_gn=ATP8B3;refGen
e_tx=XM_023243246.1;refGene_gn=ATP8B3;refGene_tx=XM_023243251.1
A2 1144527 1144603
refGene_gn=ATP8B3;refGene_tx=XM_023243212.1;refGene_gn=ATP8B3;refGene_
tx=XM_023243213.1;refGene_gn=ATP8B3;refGene_tx=XM_023243214.1;refGene_
gn=ATP8B3;refGene_tx=XM_023243219.1;refGene_gn=ATP8B3;refGene_tx=XM_02
3243221.1;refGene_gn=ATP8B3;refGene_tx=XM_023243229.1;refGene_gn=ATP8B
3;refGene_tx=XM_023243238.1;refGene_gn=ATP8B3;refGene_tx=XM_023243240.
1;refGene_gn=ATP8B3;refGene_tx=XM_023243241.1;refGene_gn=ATP8B3;refGen
e_tx=XM_023243246.1;refGene_gn=ATP8B3;refGene_tx=XM_023243251.1

A2	1145681	1145793
refGene_gn=ATP8B3;refGene_tx=XM_023243212.1;refGene_gn=ATP8B3;refGene_tx=XM_023243213.1;refGene_gn=ATP8B3;refGene_tx=XM_023243214.1;refGene_gn=ATP8B3;refGene_tx=XM_023243219.1;refGene_gn=ATP8B3;refGene_tx=XM_023243221.1;refGene_gn=ATP8B3;refGene_tx=XM_023243229.1;refGene_gn=ATP8B3;refGene_tx=XM_023243238.1;refGene_gn=ATP8B3;refGene_tx=XM_023243240.1;refGene_gn=ATP8B3;refGene_tx=XM_023243241.1;refGene_gn=ATP8B3;refGene_tx=XM_023243246.1;refGene_gn=ATP8B3;refGene_tx=XM_023243251.1
A2	1145877	1146042
refGene_gn=ATP8B3;refGene_tx=XM_023243212.1;refGene_gn=ATP8B3;refGene_tx=XM_023243213.1;refGene_gn=ATP8B3;refGene_tx=XM_023243214.1;refGene_gn=ATP8B3;refGene_tx=XM_023243219.1;refGene_gn=ATP8B3;refGene_tx=XM_023243221.1;refGene_gn=ATP8B3;refGene_tx=XM_023243229.1;refGene_gn=ATP8B3;refGene_tx=XM_023243238.1;refGene_gn=ATP8B3;refGene_tx=XM_023243240.1;refGene_gn=ATP8B3;refGene_tx=XM_023243241.1;refGene_gn=ATP8B3;refGene_tx=XM_023243246.1;refGene_gn=ATP8B3;refGene_tx=XM_023243251.1
A2	1146533	1146646
refGene_gn=ATP8B3;refGene_tx=XM_023243212.1;refGene_gn=ATP8B3;refGene_tx=XM_023243213.1;refGene_gn=ATP8B3;refGene_tx=XM_023243214.1;refGene_gn=ATP8B3;refGene_tx=XM_023243219.1;refGene_gn=ATP8B3;refGene_tx=XM_023243221.1;refGene_gn=ATP8B3;refGene_tx=XM_023243229.1;refGene_gn=ATP8B3;refGene_tx=XM_023243238.1;refGene_gn=ATP8B3;refGene_tx=XM_023243240.1;refGene_gn=ATP8B3;refGene_tx=XM_023243241.1;refGene_gn=ATP8B3;refGene_tx=XM_023243246.1;refGene_gn=ATP8B3;refGene_tx=XM_023243251.1
A2	1146724	1146916
refGene_gn=ATP8B3;refGene_tx=XM_023243212.1;refGene_gn=ATP8B3;refGene_tx=XM_023243213.1;refGene_gn=ATP8B3;refGene_tx=XM_023243214.1;refGene_gn=ATP8B3;refGene_tx=XM_023243219.1;refGene_gn=ATP8B3;refGene_tx=XM_023243221.1;refGene_gn=ATP8B3;refGene_tx=XM_023243229.1;refGene_gn=ATP8B3;refGene_tx=XM_023243238.1;refGene_gn=ATP8B3;refGene_tx=XM_023243240.1;refGene_gn=ATP8B3;refGene_tx=XM_023243241.1;refGene_gn=ATP8B3;refGene_tx=XM_023243246.1;refGene_gn=ATP8B3;refGene_tx=XM_023243251.1
A2	1147297	1147454
refGene_gn=ATP8B3;refGene_tx=XM_023243212.1;refGene_gn=ATP8B3;refGene_tx=XM_023243213.1;refGene_gn=ATP8B3;refGene_tx=XM_023243214.1;refGene_gn=ATP8B3;refGene_tx=XM_023243219.1;refGene_gn=ATP8B3;refGene_tx=XM_023243221.1;refGene_gn=ATP8B3;refGene_tx=XM_023243229.1;refGene_gn=ATP8B3;refGene_tx=XM_023243238.1;refGene_gn=ATP8B3;refGene_tx=XM_023243240.1;refGene_gn=ATP8B3;refGene_tx=XM_023243241.1;refGene_gn=ATP8B3;refGene_tx=XM_023243246.1;refGene_gn=ATP8B3;refGene_tx=XM_023243251.1
A2	1147543	1147575
refGene_gn=ATP8B3;refGene_tx=XM_023243212.1;refGene_gn=ATP8B3;refGene_tx=XM_023243213.1;refGene_gn=ATP8B3;refGene_tx=XM_023243214.1;refGene_gn=ATP8B3;refGene_tx=XM_023243219.1;refGene_gn=ATP8B3;refGene_tx=XM_023243221.1;refGene_gn=ATP8B3;refGene_tx=XM_023243229.1;refGene_gn=ATP8B3;refGene_tx=XM_023243238.1;refGene_gn=ATP8B3;refGene_tx=XM_023243240.1;refGene_gn=ATP8B3;refGene_tx=XM_023243241.1;refGene_gn=ATP8B3;refGene_tx=XM_023243246.1;refGene_gn=ATP8B3;refGene_tx=XM_023243251.1
A2	1149732	1149941
refGene_gn=ATP8B3;refGene_tx=XM_023243212.1;refGene_gn=ATP8B3;refGene_

tx=XM_023243213.1;refGene_gn=ATP8B3;refGene_tx=XM_023243214.1;refGene_gn=ATP8B3;refGene_tx=XM_023243219.1;refGene_gn=ATP8B3;refGene_tx=XM_023243221.1;refGene_gn=ATP8B3;refGene_tx=XM_023243229.1;refGene_gn=ATP8B3;refGene_tx=XM_023243238.1;refGene_gn=ATP8B3;refGene_tx=XM_023243240.1;refGene_gn=ATP8B3;refGene_tx=XM_023243241.1;refGene_gn=ATP8B3;refGene_tx=XM_023243246.1;refGene_gn=ATP8B3;refGene_tx=XM_023243251.1
A2 1150055 1150246
refGene_gn=ATP8B3;refGene_tx=XM_023243212.1;refGene_gn=ATP8B3;refGene_tx=XM_023243213.1;refGene_gn=ATP8B3;refGene_tx=XM_023243214.1;refGene_gn=ATP8B3;refGene_tx=XM_023243219.1;refGene_gn=ATP8B3;refGene_tx=XM_023243221.1;refGene_gn=ATP8B3;refGene_tx=XM_023243229.1;refGene_gn=ATP8B3;refGene_tx=XM_023243238.1;refGene_gn=ATP8B3;refGene_tx=XM_023243240.1;refGene_gn=ATP8B3;refGene_tx=XM_023243241.1;refGene_gn=ATP8B3;refGene_tx=XM_023243246.1;refGene_gn=ATP8B3;refGene_tx=XM_023243251.1
A2 1150704 1150793
refGene_gn=ATP8B3;refGene_tx=XM_023243212.1;refGene_gn=ATP8B3;refGene_tx=XM_023243213.1;refGene_gn=ATP8B3;refGene_tx=XM_023243214.1;refGene_gn=ATP8B3;refGene_tx=XM_023243219.1;refGene_gn=ATP8B3;refGene_tx=XM_023243221.1;refGene_gn=ATP8B3;refGene_tx=XM_023243229.1;refGene_gn=ATP8B3;refGene_tx=XM_023243238.1;refGene_gn=ATP8B3;refGene_tx=XM_023243240.1;refGene_gn=ATP8B3;refGene_tx=XM_023243241.1;refGene_gn=ATP8B3;refGene_tx=XM_023243246.1;refGene_gn=ATP8B3;refGene_tx=XM_023243251.1
A2 1150878 1151037
refGene_gn=ATP8B3;refGene_tx=XM_023243212.1;refGene_gn=ATP8B3;refGene_tx=XM_023243213.1;refGene_gn=ATP8B3;refGene_tx=XM_023243214.1;refGene_gn=ATP8B3;refGene_tx=XM_023243219.1;refGene_gn=ATP8B3;refGene_tx=XM_023243221.1;refGene_gn=ATP8B3;refGene_tx=XM_023243229.1;refGene_gn=ATP8B3;refGene_tx=XM_023243238.1;refGene_gn=ATP8B3;refGene_tx=XM_023243240.1;refGene_gn=ATP8B3;refGene_tx=XM_023243241.1;refGene_gn=ATP8B3;refGene_tx=XM_023243246.1;refGene_gn=ATP8B3;refGene_tx=XM_023243251.1
A2 1152299 1152382
refGene_gn=ATP8B3;refGene_tx=XM_023243212.1;refGene_gn=ATP8B3;refGene_tx=XM_023243213.1;refGene_gn=ATP8B3;refGene_tx=XM_023243214.1;refGene_gn=ATP8B3;refGene_tx=XM_023243219.1;refGene_gn=ATP8B3;refGene_tx=XM_023243221.1;refGene_gn=ATP8B3;refGene_tx=XM_023243229.1;refGene_gn=ATP8B3;refGene_tx=XM_023243238.1;refGene_gn=ATP8B3;refGene_tx=XM_023243240.1;refGene_gn=ATP8B3;refGene_tx=XM_023243241.1;refGene_gn=ATP8B3;refGene_tx=XM_023243246.1;refGene_gn=ATP8B3;refGene_tx=XM_023243251.1
A2 1152776 1152847
refGene_gn=ATP8B3;refGene_tx=XM_023243212.1;refGene_gn=ATP8B3;refGene_tx=XM_023243213.1;refGene_gn=ATP8B3;refGene_tx=XM_023243214.1;refGene_gn=ATP8B3;refGene_tx=XM_023243219.1;refGene_gn=ATP8B3;refGene_tx=XM_023243221.1;refGene_gn=ATP8B3;refGene_tx=XM_023243229.1;refGene_gn=ATP8B3;refGene_tx=XM_023243238.1;refGene_gn=ATP8B3;refGene_tx=XM_023243240.1;refGene_gn=ATP8B3;refGene_tx=XM_023243241.1;refGene_gn=ATP8B3;refGene_tx=XM_023243246.1;refGene_gn=ATP8B3;refGene_tx=XM_023243251.1
A2 1152950 1153023
refGene_gn=ATP8B3;refGene_tx=XM_023243212.1;refGene_gn=ATP8B3;refGene_tx=XM_023243213.1;refGene_gn=ATP8B3;refGene_tx=XM_023243214.1;refGene_gn=ATP8B3;refGene_tx=XM_023243219.1;refGene_gn=ATP8B3;refGene_tx=XM_02

3243221.1;refGene_gn=ATP8B3;refGene_tx=XM_023243229.1;refGene_gn=ATP8B3;refGene_tx=XM_023243238.1;refGene_gn=ATP8B3;refGene_tx=XM_023243240.1;refGene_gn=ATP8B3;refGene_tx=XM_023243241.1;refGene_gn=ATP8B3;refGene_tx=XM_023243246.1;refGene_gn=ATP8B3;refGene_tx=XM_023243251.1
A2 1153445 1153507
refGene_gn=ATP8B3;refGene_tx=XM_023243212.1;refGene_gn=ATP8B3;refGene_tx=XM_023243213.1;refGene_gn=ATP8B3;refGene_tx=XM_023243214.1;refGene_gn=ATP8B3;refGene_tx=XM_023243219.1;refGene_gn=ATP8B3;refGene_tx=XM_023243221.1;refGene_gn=ATP8B3;refGene_tx=XM_023243229.1;refGene_gn=ATP8B3;refGene_tx=XM_023243238.1;refGene_gn=ATP8B3;refGene_tx=XM_023243240.1;refGene_gn=ATP8B3;refGene_tx=XM_023243241.1;refGene_gn=ATP8B3;refGene_tx=XM_023243246.1;refGene_gn=ATP8B3;refGene_tx=XM_023243251.1
A2 1153730 1153829
refGene_gn=ATP8B3;refGene_tx=XM_023243212.1;refGene_gn=ATP8B3;refGene_tx=XM_023243213.1;refGene_gn=ATP8B3;refGene_tx=XM_023243214.1;refGene_gn=ATP8B3;refGene_tx=XM_023243219.1;refGene_gn=ATP8B3;refGene_tx=XM_023243221.1;refGene_gn=ATP8B3;refGene_tx=XM_023243229.1;refGene_gn=ATP8B3;refGene_tx=XM_023243238.1;refGene_gn=ATP8B3;refGene_tx=XM_023243240.1;refGene_gn=ATP8B3;refGene_tx=XM_023243241.1;refGene_gn=ATP8B3;refGene_tx=XM_023243246.1;refGene_gn=ATP8B3;refGene_tx=XM_023243251.1
A2 1154676 1154790
refGene_gn=ATP8B3;refGene_tx=XM_023243212.1;refGene_gn=ATP8B3;refGene_tx=XM_023243213.1;refGene_gn=ATP8B3;refGene_tx=XM_023243214.1;refGene_gn=ATP8B3;refGene_tx=XM_023243219.1;refGene_gn=ATP8B3;refGene_tx=XM_023243221.1;refGene_gn=ATP8B3;refGene_tx=XM_023243240.1;refGene_gn=ATP8B3;refGene_tx=XM_023243241.1;refGene_gn=ATP8B3;refGene_tx=XM_023243246.1;refGene_gn=ATP8B3;refGene_tx=XM_023243251.1
A2 1155209 1155301
refGene_gn=ATP8B3;refGene_tx=XM_023243212.1;refGene_gn=ATP8B3;refGene_tx=XM_023243213.1;refGene_gn=ATP8B3;refGene_tx=XM_023243214.1;refGene_gn=ATP8B3;refGene_tx=XM_023243219.1;refGene_gn=ATP8B3;refGene_tx=XM_023243221.1;refGene_gn=ATP8B3;refGene_tx=XM_023243229.1;refGene_gn=ATP8B3;refGene_tx=XM_023243238.1;refGene_gn=ATP8B3;refGene_tx=XM_023243240.1;refGene_gn=ATP8B3;refGene_tx=XM_023243246.1;refGene_gn=ATP8B3;refGene_tx=XM_023243251.1
A2 1155701 1155760
refGene_gn=ATP8B3;refGene_tx=XM_023243212.1;refGene_gn=ATP8B3;refGene_tx=XM_023243213.1;refGene_gn=ATP8B3;refGene_tx=XM_023243214.1;refGene_gn=ATP8B3;refGene_tx=XM_023243219.1;refGene_gn=ATP8B3;refGene_tx=XM_023243221.1;refGene_gn=ATP8B3;refGene_tx=XM_023243229.1;refGene_gn=ATP8B3;refGene_tx=XM_023243238.1;refGene_gn=ATP8B3;refGene_tx=XM_023243240.1;refGene_gn=ATP8B3;refGene_tx=XM_023243246.1;refGene_gn=ATP8B3;refGene_tx=XM_023243251.1
A2 1156933 1156949 refGene_gn=ATP8B3;refGene_tx=XM_023243229.1
A2 1160458 1160547 refGene_gn=REXO1;refGene_tx=XM_023243259.1
A2 1160637 1160758 refGene_gn=REXO1;refGene_tx=XM_023243259.1
A2 1160888 1161027 refGene_gn=REXO1;refGene_tx=XM_023243259.1
A2 1161121 1161237 refGene_gn=REXO1;refGene_tx=XM_023243259.1
A2 1161667 1161778 refGene_gn=REXO1;refGene_tx=XM_023243259.1
A2 1162038 1162112 refGene_gn=REXO1;refGene_tx=XM_023243259.1

| | | | |
|---|---|---|---|
| A2 | 1162220 | 1162334 | refGene_gn=REXO1;refGene_tx=XM_023243259.1 |
| A2 | 1162449 | 1162587 | refGene_gn=REXO1;refGene_tx=XM_023243259.1 |
| A2 | 1162671 | 1162785 | refGene_gn=REXO1;refGene_tx=XM_023243259.1 |
| A2 | 1163516 | 1163640 | refGene_gn=REXO1;refGene_tx=XM_023243259.1 |
| A2 | 1163799 | 1163931 | refGene_gn=REXO1;refGene_tx=XM_023243259.1 |
| A2 | 1164332 | 1164496 | refGene_gn=REXO1;refGene_tx=XM_023243259.1 |
| A2 | 1166549 | 1166763 | refGene_gn=REXO1;refGene_tx=XM_023243259.1 |
| A2 | 1167540 | 1167645 | refGene_gn=REXO1;refGene_tx=XM_023243259.1 |
| A2 | 1168679 | 1170406 | refGene_gn=REXO1;refGene_tx=XM_023243259.1 |
| A2 | 1181751 | 1181908 | refGene_gn=REXO1;refGene_tx=XM_023243259.1 |
| A2 | 1189541 | 1189843 | refGene_gn=KLF16;refGene_tx=XM_023243264.1 |
| A2 | 1198137 | 1198603 | refGene_gn=KLF16;refGene_tx=XM_023243264.1 |
| A2 | 1208836 | 1209062 | refGene_gn=ABHD17A;refGene_tx=XM_003981612.5 |
| A2 | 1209160 | 1209340 | refGene_gn=ABHD17A;refGene_tx=XM_003981612.5 |
| A2 | 1211836 | 1212031 | refGene_gn=ABHD17A;refGene_tx=XM_003981612.5 |
| A2 | 1213236 | 1213568 | refGene_gn=ABHD17A;refGene_tx=XM_003981612.5 |
| A2 | 1237791 | 1238862 | refGene_gn=ADAT3;refGene_tx=XM_023243276.1;refGene_gn=ADAT3;refGene_tx=XM_023243277.1;refGene_gn=ADAT3;refGene_tx=XM_023243280.1;refGene_gn=ADAT3;refGene_tx=XM_023243283.1 |
| A2 | 1240576 | 1240583 | refGene_gn=SCAMP4;refGene_tx=XM_003981660.5 |
| A2 | 1242531 | 1242660 | refGene_gn=SCAMP4;refGene_tx=XM_003981660.5 |
| A2 | 1242923 | 1243080 | refGene_gn=SCAMP4;refGene_tx=XM_003981660.5 |
| A2 | 1243831 | 1243933 | refGene_gn=SCAMP4;refGene_tx=XM_003981660.5 |
| A2 | 1250000 | 1250118 | refGene_gn=SCAMP4;refGene_tx=XM_003981660.5 |
| A2 | 1252121 | 1252301 | refGene_gn=SCAMP4;refGene_tx=XM_003981660.5 |
| A2 | 1276653 | 1276837 | refGene_gn=CSNK1G2;refGene_tx=XM_023243302.1 |
| A2 | 1282473 | 1282514 | refGene_gn=CSNK1G2;refGene_tx=XM_023243302.1 |
| A2 | 1282645 | 1282715 | refGene_gn=CSNK1G2;refGene_tx=XM_023243302.1 |
| A2 | 1282806 | 1282955 | refGene_gn=CSNK1G2;refGene_tx=XM_023243302.1 |
| A2 | 1283078 | 1283313 | refGene_gn=CSNK1G2;refGene_tx=XM_023243302.1 |
| A2 | 1283377 | 1283463 | refGene_gn=CSNK1G2;refGene_tx=XM_023243302.1 |
| A2 | 1283534 | 1283619 | refGene_gn=CSNK1G2;refGene_tx=XM_023243302.1 |
| A2 | 1283704 | 1283853 | refGene_gn=CSNK1G2;refGene_tx=XM_023243302.1 |
| A2 | 1284011 | 1284095 | refGene_gn=CSNK1G2;refGene_tx=XM_023243302.1 |
| A2 | 1284194 | 1284301 | refGene_gn=CSNK1G2;refGene_tx=XM_023243302.1 |
| A2 | 1284427 | 1284482 | refGene_gn=CSNK1G2;refGene_tx=XM_023243302.1 |
| A2 | 1292760 | 1292922 | refGene_gn=BTBD2;refGene_tx=XM_023243308.1 |
| A2 | 1293820 | 1293967 | refGene_gn=BTBD2;refGene_tx=XM_023243308.1 |
| A2 | 1294126 | 1294214 | refGene_gn=BTBD2;refGene_tx=XM_023243308.1 |
| A2 | 1294338 | 1294531 | refGene_gn=BTBD2;refGene_tx=XM_023243308.1 |
| A2 | 1296391 | 1296589 | refGene_gn=BTBD2;refGene_tx=XM_023243308.1 |
| A2 | 1296674 | 1296780 | refGene_gn=BTBD2;refGene_tx=XM_023243308.1 |
| A2 | 1297923 | 1298080 | refGene_gn=BTBD2;refGene_tx=XM_023243308.1 |
| A2 | 1301263 | 1301383 | refGene_gn=BTBD2;refGene_tx=XM_023243308.1 |
| A2 | 1310675 | 1311106 | refGene_gn=BTBD2;refGene_tx=XM_023243308.1 |
| A2 | 1333612 | 1333847 | refGene_gn=MKNK2;refGene_tx=XM_003981613.5 |
| A2 | 1334172 | 1334216 | refGene_gn=MKNK2;refGene_tx=XM_003981613.5 |
| A2 | 1334958 | 1335123 | refGene_gn=MKNK2;refGene_tx=XM_003981613.5 |
| A2 | 1335773 | 1335968 | refGene_gn=MKNK2;refGene_tx=XM_003981613.5 |

A2	1336411	1336507	refGene_gn=MKNK2;refGene_tx=XM_003981613.5
A2	1336610	1336666	refGene_gn=MKNK2;refGene_tx=XM_003981613.5
A2	1336782	1336887	refGene_gn=MKNK2;refGene_tx=XM_003981613.5
A2	1337168	1337242	refGene_gn=MKNK2;refGene_tx=XM_003981613.5
A2	1337567	1337647	refGene_gn=MKNK2;refGene_tx=XM_003981613.5
A2	1339640	1339738	refGene_gn=MKNK2;refGene_tx=XM_003981613.5
A2	1339816	1339918	refGene_gn=MKNK2;refGene_tx=XM_003981613.5
A2	1340024	1340112	refGene_gn=MKNK2;refGene_tx=XM_003981613.5
A2	1343346	1343397	refGene_gn=MKNK2;refGene_tx=XM_003981613.5
A2	1361393	1361423	refGene_gn=MOB3A;refGene_tx=XM_011288089.3
A2	1361645	1361834
refGene_gn=MOB3A;refGene_tx=XM_011288078.3;refGene_gn=MOB3A;refGene_tx
=XM_011288082.3;refGene_gn=MOB3A;refGene_tx=XM_011288084.3
A2	1362692	1362895
refGene_gn=MOB3A;refGene_tx=XM_011288078.3;refGene_gn=MOB3A;refGene_tx
=XM_011288082.3;refGene_gn=MOB3A;refGene_tx=XM_011288084.3;refGene_gn=
MOB3A;refGene_tx=XM_011288089.3
A2	1365616	1366037
refGene_gn=MOB3A;refGene_tx=XM_011288078.3;refGene_gn=MOB3A;refGene_tx
=XM_011288082.3;refGene_gn=MOB3A;refGene_tx=XM_011288084.3;refGene_gn=
MOB3A;refGene_tx=XM_011288089.3
A2	1375814	1376028
refGene_gn=IZUMO4;refGene_tx=XM_011288567.2;refGene_gn=IZUMO4;refGene_
tx=XM_023243340.1
A2	1376138	1376219
refGene_gn=IZUMO4;refGene_tx=XM_011288567.2;refGene_gn=IZUMO4;refGene_
tx=XM_023243340.1
A2	1376312	1376384
refGene_gn=IZUMO4;refGene_tx=XM_011288567.2;refGene_gn=IZUMO4;refGene_
tx=XM_023243340.1
A2	1377016	1377043
refGene_gn=IZUMO4;refGene_tx=XM_011288567.2;refGene_gn=IZUMO4;refGene_
tx=XM_023243340.1
A2	1377139	1377215
refGene_gn=IZUMO4;refGene_tx=XM_011288567.2;refGene_gn=IZUMO4;refGene_
tx=XM_023243340.1
A2	1377398	1377446
refGene_gn=IZUMO4;refGene_tx=XM_011288567.2;refGene_gn=IZUMO4;refGene_
tx=XM_023243340.1
A2	1377528	1377543	refGene_gn=IZUMO4;refGene_tx=XM_011288567.2
A2	1377902	1377920
refGene_gn=IZUMO4;refGene_tx=XM_011288567.2;refGene_gn=IZUMO4;refGene_
tx=XM_023243340.1
A2	1378171	1378186
refGene_gn=IZUMO4;refGene_tx=XM_011288567.2;refGene_gn=IZUMO4;refGene_
tx=XM_023243340.1
A2	1378508	1378599
refGene_gn=IZUMO4;refGene_tx=XM_011288567.2;refGene_gn=IZUMO4;refGene_
tx=XM_023243340.1
A2	1380671	1380767

refGene_gn=AP3D1;refGene_tx=XM_023243317.1;refGene_gn=AP3D1;refGene_tx
=XM_023243329.1
A2	1383734	1383814
refGene_gn=AP3D1;refGene_tx=XM_023243317.1;refGene_gn=AP3D1;refGene_tx
=XM_023243329.1
A2	1383926	1384048
refGene_gn=AP3D1;refGene_tx=XM_023243317.1;refGene_gn=AP3D1;refGene_tx
=XM_023243329.1
A2	1384605	1384691
refGene_gn=AP3D1;refGene_tx=XM_023243317.1;refGene_gn=AP3D1;refGene_tx
=XM_023243329.1
A2	1384800	1384889
refGene_gn=AP3D1;refGene_tx=XM_023243317.1;refGene_gn=AP3D1;refGene_tx
=XM_023243329.1
A2	1385097	1385287
refGene_gn=AP3D1;refGene_tx=XM_023243317.1;refGene_gn=AP3D1;refGene_tx
=XM_023243322.1;refGene_gn=AP3D1;refGene_tx=XM_023243325.1;refGene_gn=
AP3D1;refGene_tx=XM_023243329.1
A2	1385663	1385711
refGene_gn=AP3D1;refGene_tx=XM_023243317.1;refGene_gn=AP3D1;refGene_tx
=XM_023243322.1;refGene_gn=AP3D1;refGene_tx=XM_023243325.1;refGene_gn=
AP3D1;refGene_tx=XM_023243329.1
A2	1386093	1386243
refGene_gn=AP3D1;refGene_tx=XM_023243317.1;refGene_gn=AP3D1;refGene_tx
=XM_023243322.1;refGene_gn=AP3D1;refGene_tx=XM_023243325.1;refGene_gn=
AP3D1;refGene_tx=XM_023243329.1
A2	1387076	1387193
refGene_gn=AP3D1;refGene_tx=XM_023243317.1;refGene_gn=AP3D1;refGene_tx
=XM_023243322.1;refGene_gn=AP3D1;refGene_tx=XM_023243325.1;refGene_gn=
AP3D1;refGene_tx=XM_023243329.1
A2	1387516	1387593
refGene_gn=AP3D1;refGene_tx=XM_023243317.1;refGene_gn=AP3D1;refGene_tx
=XM_023243322.1;refGene_gn=AP3D1;refGene_tx=XM_023243325.1;refGene_gn=
AP3D1;refGene_tx=XM_023243329.1
A2	1387891	1388072
refGene_gn=AP3D1;refGene_tx=XM_023243317.1;refGene_gn=AP3D1;refGene_tx
=XM_023243322.1;refGene_gn=AP3D1;refGene_tx=XM_023243325.1;refGene_gn=
AP3D1;refGene_tx=XM_023243329.1
A2	1388167	1388241
refGene_gn=AP3D1;refGene_tx=XM_023243317.1;refGene_gn=AP3D1;refGene_tx
=XM_023243322.1;refGene_gn=AP3D1;refGene_tx=XM_023243325.1;refGene_gn=
AP3D1;refGene_tx=XM_023243329.1
A2	1388478	1388681
refGene_gn=AP3D1;refGene_tx=XM_023243317.1;refGene_gn=AP3D1;refGene_tx
=XM_023243322.1;refGene_gn=AP3D1;refGene_tx=XM_023243325.1;refGene_gn=
AP3D1;refGene_tx=XM_023243329.1
A2	1389148	1389224
refGene_gn=AP3D1;refGene_tx=XM_023243317.1;refGene_gn=AP3D1;refGene_tx
=XM_023243322.1;refGene_gn=AP3D1;refGene_tx=XM_023243325.1;refGene_gn=
AP3D1;refGene_tx=XM_023243329.1

A2 1389925 1389997
refGene_gn=AP3D1;refGene_tx=XM_023243317.1;refGene_gn=AP3D1;refGene_tx=XM_023243322.1;refGene_gn=AP3D1;refGene_tx=XM_023243325.1;refGene_gn=AP3D1;refGene_tx=XM_023243329.1
A2 1391485 1391627
refGene_gn=AP3D1;refGene_tx=XM_023243317.1;refGene_gn=AP3D1;refGene_tx=XM_023243322.1;refGene_gn=AP3D1;refGene_tx=XM_023243325.1;refGene_gn=AP3D1;refGene_tx=XM_023243329.1
A2 1391981 1392127
refGene_gn=AP3D1;refGene_tx=XM_023243317.1;refGene_gn=AP3D1;refGene_tx=XM_023243322.1;refGene_gn=AP3D1;refGene_tx=XM_023243325.1;refGene_gn=AP3D1;refGene_tx=XM_023243329.1
A2 1393215 1393447
refGene_gn=AP3D1;refGene_tx=XM_023243317.1;refGene_gn=AP3D1;refGene_tx=XM_023243322.1;refGene_gn=AP3D1;refGene_tx=XM_023243325.1;refGene_gn=AP3D1;refGene_tx=XM_023243329.1
A2 1393879 1394110
refGene_gn=AP3D1;refGene_tx=XM_023243317.1;refGene_gn=AP3D1;refGene_tx=XM_023243322.1;refGene_gn=AP3D1;refGene_tx=XM_023243325.1;refGene_gn=AP3D1;refGene_tx=XM_023243329.1
A2 1394180 1394329
refGene_gn=AP3D1;refGene_tx=XM_023243317.1;refGene_gn=AP3D1;refGene_tx=XM_023243322.1;refGene_gn=AP3D1;refGene_tx=XM_023243325.1;refGene_gn=AP3D1;refGene_tx=XM_023243329.1
A2 1394414 1394560
refGene_gn=AP3D1;refGene_tx=XM_023243317.1;refGene_gn=AP3D1;refGene_tx=XM_023243322.1;refGene_gn=AP3D1;refGene_tx=XM_023243325.1;refGene_gn=AP3D1;refGene_tx=XM_023243329.1
A2 1395530 1395579
refGene_gn=AP3D1;refGene_tx=XM_023243317.1;refGene_gn=AP3D1;refGene_tx=XM_023243322.1;refGene_gn=AP3D1;refGene_tx=XM_023243325.1;refGene_gn=AP3D1;refGene_tx=XM_023243329.1
A2 1395898 1395948
refGene_gn=AP3D1;refGene_tx=XM_023243317.1;refGene_gn=AP3D1;refGene_tx=XM_023243322.1;refGene_gn=AP3D1;refGene_tx=XM_023243325.1;refGene_gn=AP3D1;refGene_tx=XM_023243329.1
A2 1397138 1397188
refGene_gn=AP3D1;refGene_tx=XM_023243317.1;refGene_gn=AP3D1;refGene_tx=XM_023243322.1;refGene_gn=AP3D1;refGene_tx=XM_023243325.1;refGene_gn=AP3D1;refGene_tx=XM_023243329.1
A2 1398208 1398282
refGene_gn=AP3D1;refGene_tx=XM_023243317.1;refGene_gn=AP3D1;refGene_tx=XM_023243322.1;refGene_gn=AP3D1;refGene_tx=XM_023243325.1;refGene_gn=AP3D1;refGene_tx=XM_023243329.1
A2 1398428 1398568
refGene_gn=AP3D1;refGene_tx=XM_023243317.1;refGene_gn=AP3D1;refGene_tx=XM_023243322.1;refGene_gn=AP3D1;refGene_tx=XM_023243325.1;refGene_gn=AP3D1;refGene_tx=XM_023243329.1
A2 1399427 1399557
refGene_gn=AP3D1;refGene_tx=XM_023243317.1;refGene_gn=AP3D1;refGene_tx

=XM_023243322.1;refGene_gn=AP3D1;refGene_tx=XM_023243325.1;refGene_gn=AP3D1;refGene_tx=XM_023243329.1
A2	1401014	1401122
refGene_gn=AP3D1;refGene_tx=XM_023243317.1;refGene_gn=AP3D1;refGene_tx=XM_023243322.1;refGene_gn=AP3D1;refGene_tx=XM_023243325.1;refGene_gn=AP3D1;refGene_tx=XM_023243329.1
A2	1403065	1403146
refGene_gn=AP3D1;refGene_tx=XM_023243317.1;refGene_gn=AP3D1;refGene_tx=XM_023243322.1;refGene_gn=AP3D1;refGene_tx=XM_023243325.1;refGene_gn=AP3D1;refGene_tx=XM_023243329.1
A2	1404454	1404535
refGene_gn=AP3D1;refGene_tx=XM_023243317.1;refGene_gn=AP3D1;refGene_tx=XM_023243322.1;refGene_gn=AP3D1;refGene_tx=XM_023243325.1;refGene_gn=AP3D1;refGene_tx=XM_023243329.1
A2	1405131	1405227
refGene_gn=AP3D1;refGene_tx=XM_023243317.1;refGene_gn=AP3D1;refGene_tx=XM_023243322.1;refGene_gn=AP3D1;refGene_tx=XM_023243325.1;refGene_gn=AP3D1;refGene_tx=XM_023243329.1
A2	1416486	1416582
refGene_gn=AP3D1;refGene_tx=XM_023243317.1;refGene_gn=AP3D1;refGene_tx=XM_023243322.1;refGene_gn=AP3D1;refGene_tx=XM_023243325.1;refGene_gn=AP3D1;refGene_tx=XM_023243329.1
A2	1443882	1443963
refGene_gn=DOT1L;refGene_tx=XM_023243349.1;refGene_gn=DOT1L;refGene_tx=XM_023243350.1;refGene_gn=DOT1L;refGene_tx=XM_023243353.1
A2	1460856	1460900
refGene_gn=DOT1L;refGene_tx=XM_023243349.1;refGene_gn=DOT1L;refGene_tx=XM_023243350.1;refGene_gn=DOT1L;refGene_tx=XM_023243353.1
A2	1469549	1469624
refGene_gn=DOT1L;refGene_tx=XM_023243349.1;refGene_gn=DOT1L;refGene_tx=XM_023243350.1;refGene_gn=DOT1L;refGene_tx=XM_023243353.1
A2	1471662	1471726
refGene_gn=DOT1L;refGene_tx=XM_023243349.1;refGene_gn=DOT1L;refGene_tx=XM_023243350.1;refGene_gn=DOT1L;refGene_tx=XM_023243353.1
A2	1473063	1473292
refGene_gn=DOT1L;refGene_tx=XM_023243349.1;refGene_gn=DOT1L;refGene_tx=XM_023243350.1;refGene_gn=DOT1L;refGene_tx=XM_023243353.1
A2	1474042	1474137
refGene_gn=DOT1L;refGene_tx=XM_023243349.1;refGene_gn=DOT1L;refGene_tx=XM_023243350.1;refGene_gn=DOT1L;refGene_tx=XM_023243353.1
A2	1474494	1474557
refGene_gn=DOT1L;refGene_tx=XM_023243349.1;refGene_gn=DOT1L;refGene_tx=XM_023243350.1;refGene_gn=DOT1L;refGene_tx=XM_023243353.1
A2	1481405	1481461
refGene_gn=DOT1L;refGene_tx=XM_023243349.1;refGene_gn=DOT1L;refGene_tx=XM_023243350.1;refGene_gn=DOT1L;refGene_tx=XM_023243353.1
A2	1483500	1483580
refGene_gn=DOT1L;refGene_tx=XM_023243349.1;refGene_gn=DOT1L;refGene_tx=XM_023243350.1;refGene_gn=DOT1L;refGene_tx=XM_023243353.1
A2	1487187	1487256

```
refGene_gn=DOT1L;refGene_tx=XM_023243349.1;refGene_gn=DOT1L;refGene_tx
=XM_023243350.1;refGene_gn=DOT1L;refGene_tx=XM_023243353.1
A2      1487890 1487997
refGene_gn=DOT1L;refGene_tx=XM_023243349.1;refGene_gn=DOT1L;refGene_tx
=XM_023243350.1;refGene_gn=DOT1L;refGene_tx=XM_023243353.1
A2      1488273 1488315
refGene_gn=DOT1L;refGene_tx=XM_023243349.1;refGene_gn=DOT1L;refGene_tx
=XM_023243350.1;refGene_gn=DOT1L;refGene_tx=XM_023243353.1
A2      1489188 1489299
refGene_gn=DOT1L;refGene_tx=XM_023243349.1;refGene_gn=DOT1L;refGene_tx
=XM_023243350.1;refGene_gn=DOT1L;refGene_tx=XM_023243353.1
A2      1489443 1489678
refGene_gn=DOT1L;refGene_tx=XM_023243349.1;refGene_gn=DOT1L;refGene_tx
=XM_023243350.1;refGene_gn=DOT1L;refGene_tx=XM_023243353.1
A2      1490000 1490114
refGene_gn=DOT1L;refGene_tx=XM_023243349.1;refGene_gn=DOT1L;refGene_tx
=XM_023243350.1;refGene_gn=DOT1L;refGene_tx=XM_023243353.1
A2      1490740 1490832
refGene_gn=DOT1L;refGene_tx=XM_023243349.1;refGene_gn=DOT1L;refGene_tx
=XM_023243350.1;refGene_gn=DOT1L;refGene_tx=XM_023243353.1
A2      1491147 1491249
refGene_gn=DOT1L;refGene_tx=XM_023243349.1;refGene_gn=DOT1L;refGene_tx
=XM_023243350.1;refGene_gn=DOT1L;refGene_tx=XM_023243353.1
A2      1491513 1491651
refGene_gn=DOT1L;refGene_tx=XM_023243349.1;refGene_gn=DOT1L;refGene_tx
=XM_023243350.1;refGene_gn=DOT1L;refGene_tx=XM_023243353.1
A2      1492318 1492444
refGene_gn=DOT1L;refGene_tx=XM_023243349.1;refGene_gn=DOT1L;refGene_tx
=XM_023243350.1;refGene_gn=DOT1L;refGene_tx=XM_023243353.1
A2      1493432 1493917
refGene_gn=DOT1L;refGene_tx=XM_023243349.1;refGene_gn=DOT1L;refGene_tx
=XM_023243350.1;refGene_gn=DOT1L;refGene_tx=XM_023243353.1
A2      1494107 1494243
refGene_gn=DOT1L;refGene_tx=XM_023243349.1;refGene_gn=DOT1L;refGene_tx
=XM_023243350.1;refGene_gn=DOT1L;refGene_tx=XM_023243353.1
A2      1494638 1494785
refGene_gn=DOT1L;refGene_tx=XM_023243349.1;refGene_gn=DOT1L;refGene_tx
=XM_023243350.1;refGene_gn=DOT1L;refGene_tx=XM_023243353.1
A2      1495893 1496008
refGene_gn=DOT1L;refGene_tx=XM_023243349.1;refGene_gn=DOT1L;refGene_tx
=XM_023243350.1;refGene_gn=DOT1L;refGene_tx=XM_023243353.1
A2      1496954 1497517
refGene_gn=DOT1L;refGene_tx=XM_023243349.1;refGene_gn=DOT1L;refGene_tx
=XM_023243350.1;refGene_gn=DOT1L;refGene_tx=XM_023243353.1
A2      1497752 1497955
refGene_gn=DOT1L;refGene_tx=XM_023243349.1;refGene_gn=DOT1L;refGene_tx
=XM_023243350.1;refGene_gn=DOT1L;refGene_tx=XM_023243353.1
A2      1500274 1500339
refGene_gn=DOT1L;refGene_tx=XM_023243349.1;refGene_gn=DOT1L;refGene_tx
=XM_023243350.1;refGene_gn=DOT1L;refGene_tx=XM_023243353.1
```

A2 1500818 1501742
refGene_gn=DOT1L;refGene_tx=XM_023243349.1;refGene_gn=DOT1L;refGene_tx
=XM_023243350.1;refGene_gn=DOT1L;refGene_tx=XM_023243353.1
A2 1502509 1503027
refGene_gn=DOT1L;refGene_tx=XM_023243349.1;refGene_gn=DOT1L;refGene_tx
=XM_023243350.1
A2 1504535 1504543 refGene_gn=DOT1L;refGene_tx=XM_023243353.1
A2 1508610 1508676 refGene_gn=PLEKHJ1;refGene_tx=XM_023243362.1
A2 1508757 1508821 refGene_gn=PLEKHJ1;refGene_tx=XM_023243362.1
A2 1508907 1508998 refGene_gn=PLEKHJ1;refGene_tx=XM_023243362.1
A2 1509861 1509928 refGene_gn=PLEKHJ1;refGene_tx=XM_023243362.1
A2 1510024 1510092 refGene_gn=PLEKHJ1;refGene_tx=XM_023243362.1
A2 1510196 1510404 refGene_gn=PLEKHJ1;refGene_tx=XM_023243362.1
A2 1515573 1515699 refGene_gn=SF3A2;refGene_tx=XM_023243361.1
A2 1516904 1516976 refGene_gn=SF3A2;refGene_tx=XM_023243361.1
A2 1518004 1518051 refGene_gn=SF3A2;refGene_tx=XM_023243361.1
A2 1518479 1518589 refGene_gn=SF3A2;refGene_tx=XM_023243361.1
A2 1519734 1519784 refGene_gn=SF3A2;refGene_tx=XM_023243361.1
A2 1519869 1520010 refGene_gn=SF3A2;refGene_tx=XM_023243361.1
A2 1520334 1520403 refGene_gn=SF3A2;refGene_tx=XM_023243361.1
A2 1520507 1520865 refGene_gn=SF3A2;refGene_tx=XM_023243361.1
A2 1520867 1521307 refGene_gn=SF3A2;refGene_tx=XM_023243361.1
A2 1522172 1522611 refGene_gn=AMH;refGene_tx=XM_011288073.3
A2 1523188 1523331 refGene_gn=AMH;refGene_tx=XM_011288073.3
A2 1523548 1523657 refGene_gn=AMH;refGene_tx=XM_011288073.3
A2 1523747 1523907 refGene_gn=AMH;refGene_tx=XM_011288073.3
A2 1523983 1524899 refGene_gn=AMH;refGene_tx=XM_011288073.3
A2 1525274 1525744
refGene_gn=JSRP1;refGene_tx=XM_023243371.1;refGene_gn=JSRP1;refGene_tx
=XM_023243375.1
A2 1525846 1525948
refGene_gn=JSRP1;refGene_tx=XM_023243371.1;refGene_gn=JSRP1;refGene_tx
=XM_023243375.1
A2 1526491 1526668
refGene_gn=JSRP1;refGene_tx=XM_023243371.1;refGene_gn=JSRP1;refGene_tx
=XM_023243375.1
A2 1527018 1527148
refGene_gn=JSRP1;refGene_tx=XM_023243371.1;refGene_gn=JSRP1;refGene_tx
=XM_023243375.1
A2 1527308 1527346
refGene_gn=JSRP1;refGene_tx=XM_023243371.1;refGene_gn=JSRP1;refGene_tx
=XM_023243375.1
A2 1527735 1527853
refGene_gn=JSRP1;refGene_tx=XM_023243371.1;refGene_gn=JSRP1;refGene_tx
=XM_023243375.1
A2 1528897 1529094 refGene_gn=JSRP1;refGene_tx=XM_023243371.1
A2 1537446 1537592 refGene_gn=OAZ1;refGene_tx=XM_023243386.1
A2 1538507 1538574 refGene_gn=OAZ1;refGene_tx=XM_023243386.1
A2 1538575 1538662 refGene_gn=OAZ1;refGene_tx=XM_023243386.1
A2 1538881 1539054 refGene_gn=OAZ1;refGene_tx=XM_023243386.1

A2	1539151	1539237	refGene_gn=OAZ1;refGene_tx=XM_023243386.1
A2	1539388	1539522	refGene_gn=OAZ1;refGene_tx=XM_023243386.1
A2	1542381	1543164
refGene_gn=PEAK3;refGene_tx=XM_023243392.1;refGene_gn=PEAK3;refGene_tx=XM_023243393.1;refGene_gn=PEAK3;refGene_tx=XM_023243395.1
A2	1544467	1545009
refGene_gn=PEAK3;refGene_tx=XM_023243392.1;refGene_gn=PEAK3;refGene_tx=XM_023243393.1;refGene_gn=PEAK3;refGene_tx=XM_023243395.1
A2	1545502	1545584
refGene_gn=PEAK3;refGene_tx=XM_023243392.1;refGene_gn=PEAK3;refGene_tx=XM_023243393.1;refGene_gn=PEAK3;refGene_tx=XM_023243395.1
A2	1552624	1554403	refGene_gn=LINGO3;refGene_tx=XM_023243396.1
A2	1578022	1578165	refGene_gn=LSM7;refGene_tx=XM_003981615.5
A2	1579383	1579455	refGene_gn=LSM7;refGene_tx=XM_003981615.5
A2	1583069	1583160	refGene_gn=LSM7;refGene_tx=XM_003981615.5
A2	1583243	1583375	refGene_gn=LSM7;refGene_tx=XM_003981615.5
A2	1583401	1583482
refGene_gn=SPPL2B;refGene_tx=XM_006928195.3;refGene_gn=SPPL2B;refGene_tx=XM_006928198.3;refGene_gn=SPPL2B;refGene_tx=XM_023243397.1;refGene_gn=SPPL2B;refGene_tx=XM_023243398.1
A2	1591028	1591148
refGene_gn=SPPL2B;refGene_tx=XM_006928195.3;refGene_gn=SPPL2B;refGene_tx=XM_006928198.3;refGene_gn=SPPL2B;refGene_tx=XM_023243397.1;refGene_gn=SPPL2B;refGene_tx=XM_023243398.1
A2	1591710	1591923
refGene_gn=SPPL2B;refGene_tx=XM_006928195.3;refGene_gn=SPPL2B;refGene_tx=XM_006928198.3;refGene_gn=SPPL2B;refGene_tx=XM_023243397.1;refGene_gn=SPPL2B;refGene_tx=XM_023243398.1
A2	1592267	1592357
refGene_gn=SPPL2B;refGene_tx=XM_006928195.3;refGene_gn=SPPL2B;refGene_tx=XM_006928198.3;refGene_gn=SPPL2B;refGene_tx=XM_023243397.1;refGene_gn=SPPL2B;refGene_tx=XM_023243398.1
A2	1593081	1593221
refGene_gn=SPPL2B;refGene_tx=XM_006928195.3;refGene_gn=SPPL2B;refGene_tx=XM_006928198.3;refGene_gn=SPPL2B;refGene_tx=XM_023243397.1;refGene_gn=SPPL2B;refGene_tx=XM_023243398.1
A2	1593586	1593729
refGene_gn=SPPL2B;refGene_tx=XM_006928195.3;refGene_gn=SPPL2B;refGene_tx=XM_006928198.3;refGene_gn=SPPL2B;refGene_tx=XM_023243397.1;refGene_gn=SPPL2B;refGene_tx=XM_023243398.1
A2	1594026	1594123
refGene_gn=SPPL2B;refGene_tx=XM_006928195.3;refGene_gn=SPPL2B;refGene_tx=XM_006928198.3;refGene_gn=SPPL2B;refGene_tx=XM_023243397.1;refGene_gn=SPPL2B;refGene_tx=XM_023243398.1
A2	1594871	1594988
refGene_gn=SPPL2B;refGene_tx=XM_006928195.3;refGene_gn=SPPL2B;refGene_tx=XM_006928198.3;refGene_gn=SPPL2B;refGene_tx=XM_023243397.1;refGene_gn=SPPL2B;refGene_tx=XM_023243398.1
A2	1596102	1596184
refGene_gn=SPPL2B;refGene_tx=XM_006928195.3;refGene_gn=SPPL2B;refGene_

tx=XM_006928198.3;refGene_gn=SPPL2B;refGene_tx=XM_023243397.1;refGene_gn=SPPL2B;refGene_tx=XM_023243398.1
A2	1596876	1596951	refGene_gn=SPPL2B;refGene_tx=XM_006928195.3;refGene_gn=SPPL2B;refGene_tx=XM_006928198.3;refGene_gn=SPPL2B;refGene_tx=XM_023243397.1;refGene_gn=SPPL2B;refGene_tx=XM_023243398.1
A2	1597027	1597090	refGene_gn=SPPL2B;refGene_tx=XM_006928195.3;refGene_gn=SPPL2B;refGene_tx=XM_006928198.3;refGene_gn=SPPL2B;refGene_tx=XM_023243397.1;refGene_gn=SPPL2B;refGene_tx=XM_023243398.1
A2	1597181	1597281	refGene_gn=SPPL2B;refGene_tx=XM_006928195.3;refGene_gn=SPPL2B;refGene_tx=XM_006928198.3;refGene_gn=SPPL2B;refGene_tx=XM_023243397.1;refGene_gn=SPPL2B;refGene_tx=XM_023243398.1
A2	1597448	1597526	refGene_gn=SPPL2B;refGene_tx=XM_006928195.3;refGene_gn=SPPL2B;refGene_tx=XM_006928198.3;refGene_gn=SPPL2B;refGene_tx=XM_023243398.1
A2	1597599	1597982	refGene_gn=SPPL2B;refGene_tx=XM_023243397.1
A2	1598886	1599068	refGene_gn=SPPL2B;refGene_tx=XM_006928195.3;refGene_gn=SPPL2B;refGene_tx=XM_006928198.3;refGene_gn=SPPL2B;refGene_tx=XM_023243398.1
A2	1600264	1600510	refGene_gn=SPPL2B;refGene_tx=XM_006928195.3
A2	1624721	1624863	refGene_gn=TMPRSS9;refGene_tx=XM_023243401.1
A2	1628663	1628791	refGene_gn=TMPRSS9;refGene_tx=XM_023243401.1
A2	1631839	1631907	refGene_gn=TMPRSS9;refGene_tx=XM_023243401.1
A2	1632016	1632192	refGene_gn=TMPRSS9;refGene_tx=XM_023243401.1
A2	1632979	1633021	refGene_gn=TMPRSS9;refGene_tx=XM_023243401.1
A2	1634119	1634233	refGene_gn=TMPRSS9;refGene_tx=XM_023243401.1
A2	1636454	1636626	refGene_gn=TMPRSS9;refGene_tx=XM_023243401.1
A2	1639034	1639309	refGene_gn=TMPRSS9;refGene_tx=XM_023243401.1
A2	1641272	1641409	refGene_gn=TMPRSS9;refGene_tx=XM_023243401.1
A2	1642128	1642432	refGene_gn=TMPRSS9;refGene_tx=XM_023243401.1
A2	1642569	1642741	refGene_gn=TMPRSS9;refGene_tx=XM_023243401.1
A2	1643264	1643536	refGene_gn=TMPRSS9;refGene_tx=XM_023243401.1
A2	1644103	1644240	refGene_gn=TMPRSS9;refGene_tx=XM_023243401.1
A2	1647961	1648343	refGene_gn=TMPRSS9;refGene_tx=XM_023243401.1
A2	1651375	1651612	refGene_gn=TIMM13;refGene_tx=XM_011288043.3
A2	1652264	1652436	refGene_gn=TMPRSS9;refGene_tx=XM_023243401.1
A2	1653094	1653360	refGene_gn=TMPRSS9;refGene_tx=XM_023243401.1
A2	1653434	1653571	refGene_gn=TMPRSS9;refGene_tx=XM_023243401.1
A2	1654041	1654203	refGene_gn=TMPRSS9;refGene_tx=XM_023243401.1
A2	1655061	1655160	refGene_gn=TIMM13;refGene_tx=XM_023243409.1
A2	1655311	1655380	refGene_gn=TIMM13;refGene_tx=XM_011288043.3;refGene_gn=TIMM13;refGene_tx=XM_023243409.1
A2	1655466	1655586	refGene_gn=TIMM13;refGene_tx=XM_011288043.3;refGene_gn=TIMM13;refGene_tx=XM_023243409.1
A2	1658511	1658553	refGene_gn=LMNB2;refGene_tx=XM_023243403.1;refGene_gn=LMNB2;refGene_tx

=XM_023243404.1
A2	1659206	1659320
refGene_gn=LMNB2;refGene_tx=XM_023243403.1;refGene_gn=LMNB2;refGene_tx=XM_023243404.1
A2	1659581	1659701
refGene_gn=LMNB2;refGene_tx=XM_023243403.1;refGene_gn=LMNB2;refGene_tx=XM_023243404.1
A2	1660232	1660340
refGene_gn=LMNB2;refGene_tx=XM_023243403.1;refGene_gn=LMNB2;refGene_tx=XM_023243404.1
A2	1660428	1660723
refGene_gn=LMNB2;refGene_tx=XM_023243403.1;refGene_gn=LMNB2;refGene_tx=XM_023243404.1
A2	1660900	1661121
refGene_gn=LMNB2;refGene_tx=XM_023243403.1;refGene_gn=LMNB2;refGene_tx=XM_023243404.1
A2	1661448	1661574
refGene_gn=LMNB2;refGene_tx=XM_023243403.1;refGene_gn=LMNB2;refGene_tx=XM_023243404.1
A2	1661690	1661861
refGene_gn=LMNB2;refGene_tx=XM_023243403.1;refGene_gn=LMNB2;refGene_tx=XM_023243404.1
A2	1662629	1662992	refGene_gn=LMNB2;refGene_tx=XM_023243404.1
A2	1663297	1663423	refGene_gn=LMNB2;refGene_tx=XM_023243403.1
A2	1663642	1663799	refGene_gn=LMNB2;refGene_tx=XM_023243403.1
A2	1668160	1668297	refGene_gn=LMNB2;refGene_tx=XM_023243403.1
A2	1675154	1675358	refGene_gn=LMNB2;refGene_tx=XM_023243403.1
A2	1690193	1690237	refGene_gn=GADD45B;refGene_tx=XM_003981616.5
A2	1690328	1690430	refGene_gn=GADD45B;refGene_tx=XM_003981616.5
A2	1690838	1691061	refGene_gn=GADD45B;refGene_tx=XM_003981616.5
A2	1691319	1691433	refGene_gn=GADD45B;refGene_tx=XM_003981616.5
A2	1717625	1717751	refGene_gn=GNG7;refGene_tx=XM_023243428.1
A2	1721255	1721336	refGene_gn=GNG7;refGene_tx=XM_023243428.1
A2	1839689	1840286
refGene_gn=DIRAS1;refGene_tx=XM_006928199.4;refGene_gn=DIRAS1;refGene_tx=XM_023243432.1
A2	1850790	1851525
refGene_gn=SLC39A3;refGene_tx=XM_003981619.5;refGene_gn=SLC39A3;refGene_tx=XM_006928200.4;refGene_gn=SLC39A3;refGene_tx=XM_023243433.1
A2	1854571	1854903
refGene_gn=SLC39A3;refGene_tx=XM_003981619.5;refGene_gn=SLC39A3;refGene_tx=XM_006928200.4;refGene_gn=SLC39A3;refGene_tx=XM_023243433.1
A2	1856391	1856527	refGene_gn=SLC39A3;refGene_tx=XM_023243433.1
A2	1864221	1864336
refGene_gn=SGTA;refGene_tx=XM_011288012.3;refGene_gn=SGTA;refGene_tx=XM_011288017.3;refGene_gn=SGTA;refGene_tx=XM_019815620.2;refGene_gn=SGTA;refGene_tx=XM_023243458.1
A2	1864543	1864633
refGene_gn=SGTA;refGene_tx=XM_011288012.3;refGene_gn=SGTA;refGene_tx=XM_011288017.3;refGene_gn=SGTA;refGene_tx=XM_019815620.2;refGene_gn=SGT

A;refGene_tx=XM_023243458.1
A2 1865479 1865517 refGene_gn=SGTA;refGene_tx=XM_023243458.1
A2 1867149 1867187
refGene_gn=SGTA;refGene_tx=XM_011288012.3;refGene_gn=SGTA;refGene_tx=XM_011288017.3;refGene_gn=SGTA;refGene_tx=XM_019815620.2
A2 1868297 1868360
refGene_gn=SGTA;refGene_tx=XM_011288012.3;refGene_gn=SGTA;refGene_tx=XM_011288017.3;refGene_gn=SGTA;refGene_tx=XM_019815620.2;refGene_gn=SGTA;refGene_tx=XM_023243458.1
A2 1869036 1869175
refGene_gn=SGTA;refGene_tx=XM_011288012.3;refGene_gn=SGTA;refGene_tx=XM_011288017.3;refGene_gn=SGTA;refGene_tx=XM_019815620.2;refGene_gn=SGTA;refGene_tx=XM_023243458.1
A2 1870317 1870422
refGene_gn=SGTA;refGene_tx=XM_011288012.3;refGene_gn=SGTA;refGene_tx=XM_011288017.3;refGene_gn=SGTA;refGene_tx=XM_019815620.2;refGene_gn=SGTA;refGene_tx=XM_023243458.1
A2 1871893 1871993
refGene_gn=SGTA;refGene_tx=XM_011288012.3;refGene_gn=SGTA;refGene_tx=XM_011288017.3;refGene_gn=SGTA;refGene_tx=XM_019815620.2;refGene_gn=SGTA;refGene_tx=XM_023243458.1
A2 1872969 1873054
refGene_gn=SGTA;refGene_tx=XM_011288012.3;refGene_gn=SGTA;refGene_tx=XM_011288017.3;refGene_gn=SGTA;refGene_tx=XM_019815620.2;refGene_gn=SGTA;refGene_tx=XM_023243458.1
A2 1873277 1873384
refGene_gn=SGTA;refGene_tx=XM_011288012.3;refGene_gn=SGTA;refGene_tx=XM_011288017.3;refGene_gn=SGTA;refGene_tx=XM_019815620.2;refGene_gn=SGTA;refGene_tx=XM_023243458.1
A2 1873672 1873772
refGene_gn=SGTA;refGene_tx=XM_011288012.3;refGene_gn=SGTA;refGene_tx=XM_011288017.3;refGene_gn=SGTA;refGene_tx=XM_019815620.2;refGene_gn=SGTA;refGene_tx=XM_023243458.1
A2 1882794 1882810
refGene_gn=THOP1;refGene_tx=XM_023243465.1;refGene_gn=THOP1;refGene_tx=XM_023243468.1
A2 1885528 1885741
refGene_gn=THOP1;refGene_tx=XM_023243465.1;refGene_gn=THOP1;refGene_tx=XM_023243468.1
A2 1888505 1888654
refGene_gn=THOP1;refGene_tx=XM_023243465.1;refGene_gn=THOP1;refGene_tx=XM_023243468.1
A2 1889957 1890065
refGene_gn=THOP1;refGene_tx=XM_023243465.1;refGene_gn=THOP1;refGene_tx=XM_023243468.1
A2 1892570 1892673
refGene_gn=THOP1;refGene_tx=XM_023243465.1;refGene_gn=THOP1;refGene_tx=XM_023243468.1
A2 1896162 1896323
refGene_gn=THOP1;refGene_tx=XM_023243465.1;refGene_gn=THOP1;refGene_tx

=XM_023243468.1
A2 1897773 1897909
refGene_gn=THOP1;refGene_tx=XM_023243465.1;refGene_gn=THOP1;refGene_tx
=XM_023243468.1
A2 1898358 1898725
refGene_gn=THOP1;refGene_tx=XM_023243465.1;refGene_gn=THOP1;refGene_tx
=XM_023243468.1
A2 1899073 1899275
refGene_gn=THOP1;refGene_tx=XM_023243465.1;refGene_gn=THOP1;refGene_tx
=XM_023243468.1
A2 1901780 1901967
refGene_gn=THOP1;refGene_tx=XM_023243465.1;refGene_gn=THOP1;refGene_tx
=XM_023243468.1
A2 1902134 1902263
refGene_gn=THOP1;refGene_tx=XM_023243465.1;refGene_gn=THOP1;refGene_tx
=XM_023243468.1
A2 1903325 1903462
refGene_gn=THOP1;refGene_tx=XM_023243465.1;refGene_gn=THOP1;refGene_tx
=XM_023243468.1
A2 1903953 1904112
refGene_gn=THOP1;refGene_tx=XM_023243465.1;refGene_gn=THOP1;refGene_tx
=XM_023243468.1
A2 1925399 1925617 refGene_gn=ZNF554;refGene_tx=XM_023243484.1
A2 1934225 1934298 refGene_gn=ZNF554;refGene_tx=XM_023243484.1
A2 1936599 1936726
refGene_gn=ZNF554;refGene_tx=XM_023243484.1;refGene_gn=ZNF554;refGene_
tx=XM_023243491.1
A2 1939553 1939745
refGene_gn=ZNF554;refGene_tx=XM_023243484.1;refGene_gn=ZNF554;refGene_
tx=XM_023243491.1
A2 1940532 1941692
refGene_gn=ZNF554;refGene_tx=XM_023243484.1;refGene_gn=ZNF554;refGene_
tx=XM_023243491.1
A2 1949165 1949168
refGene_gn=LOC102900656;refGene_tx=XM_023243475.1
A2 1960641 1960768
refGene_gn=LOC102900656;refGene_tx=XM_023243475.1
A2 1961202 1961386
refGene_gn=LOC102900656;refGene_tx=XM_023243475.1
A2 1962228 1963789
refGene_gn=LOC102900656;refGene_tx=XM_023243475.1
A2 1982211 1982646
refGene_gn=ZNF556;refGene_tx=XM_011287994.3;refGene_gn=ZNF556;refGene_
tx=XM_011288005.3
A2 1996631 1996758
refGene_gn=ZNF556;refGene_tx=XM_011287994.3;refGene_gn=ZNF556;refGene_
tx=XM_011288005.3
A2 1997558 1997742
refGene_gn=ZNF556;refGene_tx=XM_011287994.3;refGene_gn=ZNF556;refGene_
tx=XM_011288005.3

A2 1998166 1999088
refGene_gn=ZNF556;refGene_tx=XM_011287994.3;refGene_gn=ZNF556;refGene_tx=XM_011288005.3
A2 2035620 2035623
refGene_gn=LOC101084518;refGene_tx=XM_011287974.3;refGene_gn=LOC101084518;refGene_tx=XM_019815671.2;refGene_gn=LOC101084518;refGene_tx=XM_023243516.1;refGene_gn=LOC101084518;refGene_tx=XM_023243521.1
A2 2055346 2055509
refGene_gn=LOC101084518;refGene_tx=XM_011287974.3;refGene_gn=LOC101084518;refGene_tx=XM_019815671.2;refGene_gn=LOC101084518;refGene_tx=XM_019815680.2;refGene_gn=LOC101084518;refGene_tx=XM_023243516.1;refGene_gn=LOC101084518;refGene_tx=XM_023243521.1
A2 2058690 2058748
refGene_gn=LOC101084518;refGene_tx=XM_011287974.3;refGene_gn=LOC101084518;refGene_tx=XM_019815671.2;refGene_gn=LOC101084518;refGene_tx=XM_019815680.2;refGene_gn=LOC101084518;refGene_tx=XM_023243516.1;refGene_gn=LOC101084518;refGene_tx=XM_023243521.1
A2 2059167 2060866
refGene_gn=LOC101084518;refGene_tx=XM_011287974.3;refGene_gn=LOC101084518;refGene_tx=XM_019815671.2;refGene_gn=LOC101084518;refGene_tx=XM_019815680.2;refGene_gn=LOC101084518;refGene_tx=XM_023243516.1;refGene_gn=LOC101084518;refGene_tx=XM_023243519.1;refGene_gn=LOC101084518;refGene_tx=XM_023243520.1;refGene_gn=LOC101084518;refGene_tx=XM_023243521.1
A2 2079265 2079268
refGene_gn=LOC101084776;refGene_tx=XM_019815693.2
A2 2087728 2087762
refGene_gn=LOC101084776;refGene_tx=XM_019815703.2;refGene_gn=LOC101084776;refGene_tx=XM_019815706.2
A2 2095999 2096002
refGene_gn=LOC101084776;refGene_tx=XM_019815688.2
A2 2110915 2111042
refGene_gn=LOC101084776;refGene_tx=XM_019815688.2;refGene_gn=LOC101084776;refGene_tx=XM_019815693.2;refGene_gn=LOC101084776;refGene_tx=XM_023243544.1
A2 2118595 2118755
refGene_gn=LOC101084776;refGene_tx=XM_019815688.2;refGene_gn=LOC101084776;refGene_tx=XM_019815693.2;refGene_gn=LOC101084776;refGene_tx=XM_019815703.2;refGene_gn=LOC101084776;refGene_tx=XM_019815706.2;refGene_gn=LOC101084776;refGene_tx=XM_023243544.1
A2 2119400 2120670
refGene_gn=LOC101084776;refGene_tx=XM_019815688.2;refGene_gn=LOC101084776;refGene_tx=XM_019815693.2;refGene_gn=LOC101084776;refGene_tx=XM_019815703.2;refGene_gn=LOC101084776;refGene_tx=XM_019815706.2;refGene_gn=LOC101084776;refGene_tx=XM_023243544.1
A2 2150216 2150227
refGene_gn=LOC109497257;refGene_tx=XM_023250888.1
A2 2211764 2213363
refGene_gn=LOC109497811;refGene_tx=XM_019826123.2;refGene_gn=LOC109497811;refGene_tx=XM_023243557.1
A2 2213759 2214028

refGene_gn=LOC109497811;refGene_tx=XM_019826123.2
A2 2216738 2216865
refGene_gn=LOC109497257;refGene_tx=XM_023250888.1
A2 2225217 2225541
refGene_gn=LOC109497257;refGene_tx=XM_023250888.1
A2 2226533 2226575
refGene_gn=LOC109497257;refGene_tx=XM_023250888.1
A2 2250986 2252853
refGene_gn=LOC105261065;refGene_tx=XM_011287956.3;refGene_gn=LOC105261065;refGene_tx=XM_019826111.2
A2 2253093 2253113
refGene_gn=LOC105261065;refGene_tx=XM_019826111.2
A2 2253287 2253345
refGene_gn=LOC105261065;refGene_tx=XM_011287956.3
A2 2256327 2256454
refGene_gn=LOC105261065;refGene_tx=XM_011287956.3
A2 2262205 2262208
refGene_gn=LOC105261065;refGene_tx=XM_011287956.3
A2 2280687 2281699
refGene_gn=LOC101085783;refGene_tx=XM_006928244.4
A2 2282096 2282277
refGene_gn=LOC101085783;refGene_tx=XM_006928244.4
A2 2289617 2289744
refGene_gn=LOC101085783;refGene_tx=XM_006928244.4
A2 2296863 2296866
refGene_gn=LOC101085783;refGene_tx=XM_006928244.4
A2 2314472 2316168
refGene_gn=LOC101086041;refGene_tx=XM_006928243.4
A2 2316575 2316756
refGene_gn=LOC101086041;refGene_tx=XM_006928243.4
A2 2318008 2318135
refGene_gn=LOC101086041;refGene_tx=XM_006928243.4
A2 2336807 2336810
refGene_gn=LOC101086041;refGene_tx=XM_006928243.4
A2 2371084 2371153
refGene_gn=TLE6;refGene_tx=XM_019826112.2;refGene_gn=TLE6;refGene_tx=XM_019826113.2
A2 2371495 2371728
refGene_gn=TLE6;refGene_tx=XM_019826112.2;refGene_gn=TLE6;refGene_tx=XM_019826113.2;refGene_gn=TLE6;refGene_tx=XM_023243578.1;refGene_gn=TLE6;refGene_tx=XM_023243579.1;refGene_gn=TLE6;refGene_tx=XM_023243584.1
A2 2372083 2372129
refGene_gn=TLE6;refGene_tx=XM_019826112.2;refGene_gn=TLE6;refGene_tx=XM_019826113.2;refGene_gn=TLE6;refGene_tx=XM_023243578.1;refGene_gn=TLE6;refGene_tx=XM_023243579.1;refGene_gn=TLE6;refGene_tx=XM_023243584.1
A2 2373132 2373174
refGene_gn=TLE6;refGene_tx=XM_019826112.2;refGene_gn=TLE6;refGene_tx=XM_019826113.2;refGene_gn=TLE6;refGene_tx=XM_023243578.1;refGene_gn=TLE6;refGene_tx=XM_023243579.1;refGene_gn=TLE6;refGene_tx=XM_023243584.1
A2 2376379 2376442

refGene_gn=TLE6;refGene_tx=XM_019826112.2;refGene_gn=TLE6;refGene_tx=XM_019826113.2;refGene_gn=TLE6;refGene_tx=XM_019826118.2;refGene_gn=TLE6;refGene_tx=XM_023243578.1;refGene_gn=TLE6;refGene_tx=XM_023243579.1;refGene_gn=TLE6;refGene_tx=XM_023243584.1
A2 2376535 2376854
refGene_gn=TLE6;refGene_tx=XM_019826112.2;refGene_gn=TLE6;refGene_tx=XM_019826113.2;refGene_gn=TLE6;refGene_tx=XM_019826118.2;refGene_gn=TLE6;refGene_tx=XM_023243578.1;refGene_gn=TLE6;refGene_tx=XM_023243579.1;refGene_gn=TLE6;refGene_tx=XM_023243584.1
A2 2376963 2376986
refGene_gn=TLE6;refGene_tx=XM_019826112.2;refGene_gn=TLE6;refGene_tx=XM_019826113.2;refGene_gn=TLE6;refGene_tx=XM_019826118.2;refGene_gn=TLE6;refGene_tx=XM_023243578.1;refGene_gn=TLE6;refGene_tx=XM_023243579.1;refGene_gn=TLE6;refGene_tx=XM_023243584.1
A2 2377270 2377331
refGene_gn=TLE6;refGene_tx=XM_019826112.2;refGene_gn=TLE6;refGene_tx=XM_019826113.2;refGene_gn=TLE6;refGene_tx=XM_019826118.2;refGene_gn=TLE6;refGene_tx=XM_023243578.1;refGene_gn=TLE6;refGene_tx=XM_023243579.1;refGene_gn=TLE6;refGene_tx=XM_023243584.1
A2 2377410 2377514
refGene_gn=TLE6;refGene_tx=XM_019826112.2;refGene_gn=TLE6;refGene_tx=XM_019826113.2;refGene_gn=TLE6;refGene_tx=XM_019826118.2;refGene_gn=TLE6;refGene_tx=XM_023243578.1;refGene_gn=TLE6;refGene_tx=XM_023243579.1;refGene_gn=TLE6;refGene_tx=XM_023243584.1
A2 2377599 2377637
refGene_gn=TLE6;refGene_tx=XM_019826112.2;refGene_gn=TLE6;refGene_tx=XM_019826113.2;refGene_gn=TLE6;refGene_tx=XM_019826118.2;refGene_gn=TLE6;refGene_tx=XM_023243578.1;refGene_gn=TLE6;refGene_tx=XM_023243579.1;refGene_gn=TLE6;refGene_tx=XM_023243584.1
A2 2378207 2378804
refGene_gn=TLE6;refGene_tx=XM_019826112.2;refGene_gn=TLE6;refGene_tx=XM_019826113.2;refGene_gn=TLE6;refGene_tx=XM_019826118.2;refGene_gn=TLE6;refGene_tx=XM_023243578.1;refGene_gn=TLE6;refGene_tx=XM_023243579.1;refGene_gn=TLE6;refGene_tx=XM_023243584.1
A2 2379096 2379238
refGene_gn=TLE6;refGene_tx=XM_019826112.2;refGene_gn=TLE6;refGene_tx=XM_019826113.2;refGene_gn=TLE6;refGene_tx=XM_019826118.2;refGene_gn=TLE6;refGene_tx=XM_023243578.1;refGene_gn=TLE6;refGene_tx=XM_023243579.1;refGene_gn=TLE6;refGene_tx=XM_023243584.1
A2 2379656 2379807
refGene_gn=TLE6;refGene_tx=XM_019826112.2;refGene_gn=TLE6;refGene_tx=XM_019826113.2;refGene_gn=TLE6;refGene_tx=XM_019826118.2;refGene_gn=TLE6;refGene_tx=XM_023243578.1;refGene_gn=TLE6;refGene_tx=XM_023243579.1;refGene_gn=TLE6;refGene_tx=XM_023243584.1
A2 2380047 2380124
refGene_gn=TLE6;refGene_tx=XM_019826112.2;refGene_gn=TLE6;refGene_tx=XM_019826113.2;refGene_gn=TLE6;refGene_tx=XM_019826118.2;refGene_gn=TLE6;refGene_tx=XM_023243578.1;refGene_gn=TLE6;refGene_tx=XM_023243579.1
A2 2380280 2380385
refGene_gn=TLE6;refGene_tx=XM_019826112.2;refGene_gn=TLE6;refGene_tx=X

M_019826113.2;refGene_gn=TLE6;refGene_tx=XM_019826118.2;refGene_gn=TLE6;refGene_tx=XM_023243578.1;refGene_gn=TLE6;refGene_tx=XM_023243579.1
A2 2382584 2382692
refGene_gn=TLE2;refGene_tx=XM_023243595.1;refGene_gn=TLE2;refGene_tx=XM_023243601.1
A2 2384002 2384079
refGene_gn=TLE2;refGene_tx=XM_023243595.1;refGene_gn=TLE2;refGene_tx=XM_023243601.1
A2 2384952 2385103
refGene_gn=TLE2;refGene_tx=XM_023243595.1;refGene_gn=TLE2;refGene_tx=XM_023243601.1
A2 2386649 2386797
refGene_gn=TLE2;refGene_tx=XM_023243595.1;refGene_gn=TLE2;refGene_tx=XM_023243601.1
A2 2386886 2387134
refGene_gn=TLE2;refGene_tx=XM_023243595.1;refGene_gn=TLE2;refGene_tx=XM_023243601.1
A2 2387335 2387594
refGene_gn=TLE2;refGene_tx=XM_023243595.1;refGene_gn=TLE2;refGene_tx=XM_023243601.1
A2 2388311 2388388
refGene_gn=TLE2;refGene_tx=XM_023243595.1;refGene_gn=TLE2;refGene_tx=XM_023243601.1
A2 2389435 2389596
refGene_gn=TLE2;refGene_tx=XM_023243595.1;refGene_gn=TLE2;refGene_tx=XM_023243601.1
A2 2390369 2390508
refGene_gn=TLE2;refGene_tx=XM_023243595.1;refGene_gn=TLE2;refGene_tx=XM_023243601.1
A2 2391968 2392118
refGene_gn=TLE2;refGene_tx=XM_023243595.1;refGene_gn=TLE2;refGene_tx=XM_023243601.1
A2 2392661 2392706
refGene_gn=TLE2;refGene_tx=XM_023243595.1;refGene_gn=TLE2;refGene_tx=XM_023243601.1
A2 2393924 2394032
refGene_gn=TLE2;refGene_tx=XM_023243595.1;refGene_gn=TLE2;refGene_tx=XM_023243601.1
A2 2399449 2399469
refGene_gn=TLE2;refGene_tx=XM_023243595.1;refGene_gn=TLE2;refGene_tx=XM_023243601.1
A2 2399828 2400012
refGene_gn=TLE2;refGene_tx=XM_023243595.1;refGene_gn=TLE2;refGene_tx=XM_023243601.1
A2 2400227 2400302
refGene_gn=TLE2;refGene_tx=XM_023243595.1;refGene_gn=TLE2;refGene_tx=XM_023243601.1
A2 2403485 2403548
refGene_gn=TLE2;refGene_tx=XM_023243595.1;refGene_gn=TLE2;refGene_tx=XM_023243601.1

A2 2405576 2405621
refGene_gn=TLE2;refGene_tx=XM_023243595.1;refGene_gn=TLE2;refGene_tx=X
M_023243601.1
A2 2405986 2406050
refGene_gn=TLE2;refGene_tx=XM_023243595.1;refGene_gn=TLE2;refGene_tx=X
M_023243601.1
A2 2406503 2406601
refGene_gn=TLE2;refGene_tx=XM_023243595.1;refGene_gn=TLE2;refGene_tx=X
M_023243601.1
A2 2406675 2406699
refGene_gn=TLE2;refGene_tx=XM_023243595.1;refGene_gn=TLE2;refGene_tx=X
M_023243601.1
A2 2420672 2420894
refGene_gn=AES;refGene_tx=XM_023243608.1;refGene_gn=AES;refGene_tx=XM_
023243612.1
A2 2420976 2421051
refGene_gn=AES;refGene_tx=XM_023243608.1;refGene_gn=AES;refGene_tx=XM_
023243612.1
A2 2422439 2422502
refGene_gn=AES;refGene_tx=XM_023243608.1;refGene_gn=AES;refGene_tx=XM_
023243612.1
A2 2423124 2423169
refGene_gn=AES;refGene_tx=XM_023243608.1;refGene_gn=AES;refGene_tx=XM_
023243612.1
A2 2424503 2424567
refGene_gn=AES;refGene_tx=XM_023243608.1;refGene_gn=AES;refGene_tx=XM_
023243612.1
A2 2426929 2427027
refGene_gn=AES;refGene_tx=XM_023243608.1;refGene_gn=AES;refGene_tx=XM_
023243612.1
A2 2428138 2428165
refGene_gn=AES;refGene_tx=XM_023243608.1;refGene_gn=AES;refGene_tx=XM_
023243612.1
A2 2459749 2459885 refGene_gn=GNA11;refGene_tx=XM_023243621.1
A2 2471212 2471397 refGene_gn=GNA11;refGene_tx=XM_023243621.1
A2 2474276 2474431 refGene_gn=GNA11;refGene_tx=XM_023243621.1
A2 2474807 2474936 refGene_gn=GNA11;refGene_tx=XM_023243621.1
A2 2477569 2477699 refGene_gn=GNA11;refGene_tx=XM_023243621.1
A2 2477775 2477929 refGene_gn=GNA11;refGene_tx=XM_023243621.1
A2 2478905 2479096 refGene_gn=GNA11;refGene_tx=XM_023243621.1
A2 2490274 2490419 refGene_gn=GNA15;refGene_tx=XM_023243623.1
A2 2496893 2497078 refGene_gn=GNA15;refGene_tx=XM_023243623.1
A2 2501585 2501740 refGene_gn=GNA15;refGene_tx=XM_023243623.1
A2 2503000 2503129 refGene_gn=GNA15;refGene_tx=XM_023243623.1
A2 2508558 2508688 refGene_gn=GNA15;refGene_tx=XM_023243623.1
A2 2509786 2509940 refGene_gn=GNA15;refGene_tx=XM_023243623.1
A2 2512880 2513107 refGene_gn=GNA15;refGene_tx=XM_023243623.1
A2 2524190 2525348 refGene_gn=S1PR4;refGene_tx=XM_006928247.4
A2 2530310 2530494
refGene_gn=NCLN;refGene_tx=XM_023243634.1;refGene_gn=NCLN;refGene_tx=X

M_023243639.1
A2	2534855	2535046
refGene_gn=NCLN;refGene_tx=XM_023243634.1;refGene_gn=NCLN;refGene_tx=X
M_023243639.1
A2	2535926	2536071
refGene_gn=NCLN;refGene_tx=XM_023243634.1;refGene_gn=NCLN;refGene_tx=X
M_023243639.1
A2	2536774	2536869
refGene_gn=NCLN;refGene_tx=XM_023243634.1;refGene_gn=NCLN;refGene_tx=X
M_023243639.1
A2	2539537	2539618
refGene_gn=NCLN;refGene_tx=XM_023243634.1;refGene_gn=NCLN;refGene_tx=X
M_023243639.1
A2	2541507	2541611
refGene_gn=NCLN;refGene_tx=XM_023243634.1;refGene_gn=NCLN;refGene_tx=X
M_023243639.1
A2	2541958	2542047
refGene_gn=NCLN;refGene_tx=XM_023243634.1;refGene_gn=NCLN;refGene_tx=X
M_023243639.1
A2	2542130	2542270
refGene_gn=NCLN;refGene_tx=XM_023243634.1;refGene_gn=NCLN;refGene_tx=X
M_023243639.1
A2	2542527	2542706
refGene_gn=NCLN;refGene_tx=XM_023243634.1;refGene_gn=NCLN;refGene_tx=X
M_023243639.1
A2	2544027	2544115
refGene_gn=NCLN;refGene_tx=XM_023243634.1;refGene_gn=NCLN;refGene_tx=X
M_023243639.1
A2	2544204	2544243
refGene_gn=NCLN;refGene_tx=XM_023243634.1;refGene_gn=NCLN;refGene_tx=X
M_023243639.1
A2	2544424	2544588
refGene_gn=NCLN;refGene_tx=XM_023243634.1;refGene_gn=NCLN;refGene_tx=X
M_023243639.1
A2	2545557	2545611
refGene_gn=NCLN;refGene_tx=XM_023243634.1;refGene_gn=NCLN;refGene_tx=X
M_023243639.1
A2	2545709	2545788
refGene_gn=NCLN;refGene_tx=XM_023243634.1;refGene_gn=NCLN;refGene_tx=X
M_023243639.1
A2	2545922	2545982
refGene_gn=NCLN;refGene_tx=XM_023243634.1;refGene_gn=NCLN;refGene_tx=X
M_023243639.1
A2	2560869	2561128
refGene_gn=CELF5;refGene_tx=XM_023243647.1;refGene_gn=CELF5;refGene_tx
=XM_023243649.1;refGene_gn=CELF5;refGene_tx=XM_023243654.1;refGene_gn=
CELF5;refGene_tx=XM_023243656.1;refGene_gn=CELF5;refGene_tx=XM_0232436
59.1;refGene_gn=CELF5;refGene_tx=XM_023243661.1
A2	2581270	2581353
refGene_gn=CELF5;refGene_tx=XM_023243647.1;refGene_gn=CELF5;refGene_tx

=XM_023243649.1;refGene_gn=CELF5;refGene_tx=XM_023243654.1;refGene_gn=
CELF5;refGene_tx=XM_023243656.1;refGene_gn=CELF5;refGene_tx=XM_0232436
59.1;refGene_gn=CELF5;refGene_tx=XM_023243661.1
A2 2591462 2591514
refGene_gn=CELF5;refGene_tx=XM_023243647.1;refGene_gn=CELF5;refGene_tx
=XM_023243649.1;refGene_gn=CELF5;refGene_tx=XM_023243654.1;refGene_gn=
CELF5;refGene_tx=XM_023243656.1;refGene_gn=CELF5;refGene_tx=XM_0232436
59.1;refGene_gn=CELF5;refGene_tx=XM_023243661.1
A2 2594323 2594452
refGene_gn=CELF5;refGene_tx=XM_023243647.1;refGene_gn=CELF5;refGene_tx
=XM_023243649.1;refGene_gn=CELF5;refGene_tx=XM_023243654.1;refGene_gn=
CELF5;refGene_tx=XM_023243656.1;refGene_gn=CELF5;refGene_tx=XM_0232436
59.1;refGene_gn=CELF5;refGene_tx=XM_023243661.1
A2 2596432 2596512
refGene_gn=CELF5;refGene_tx=XM_023243647.1;refGene_gn=CELF5;refGene_tx
=XM_023243649.1;refGene_gn=CELF5;refGene_tx=XM_023243654.1;refGene_gn=
CELF5;refGene_tx=XM_023243656.1;refGene_gn=CELF5;refGene_tx=XM_0232436
59.1;refGene_gn=CELF5;refGene_tx=XM_023243661.1
A2 2598415 2598562
refGene_gn=CELF5;refGene_tx=XM_023243647.1;refGene_gn=CELF5;refGene_tx
=XM_023243649.1;refGene_gn=CELF5;refGene_tx=XM_023243654.1;refGene_gn=
CELF5;refGene_tx=XM_023243656.1;refGene_gn=CELF5;refGene_tx=XM_0232436
59.1;refGene_gn=CELF5;refGene_tx=XM_023243661.1
A2 2599169 2599311
refGene_gn=CELF5;refGene_tx=XM_023243647.1;refGene_gn=CELF5;refGene_tx
=XM_023243649.1;refGene_gn=CELF5;refGene_tx=XM_023243654.1;refGene_gn=
CELF5;refGene_tx=XM_023243656.1;refGene_gn=CELF5;refGene_tx=XM_0232436
59.1;refGene_gn=CELF5;refGene_tx=XM_023243661.1
A2 2599409 2599556
refGene_gn=CELF5;refGene_tx=XM_023243647.1;refGene_gn=CELF5;refGene_tx
=XM_023243649.1;refGene_gn=CELF5;refGene_tx=XM_023243654.1;refGene_gn=
CELF5;refGene_tx=XM_023243656.1;refGene_gn=CELF5;refGene_tx=XM_0232436
59.1;refGene_gn=CELF5;refGene_tx=XM_023243661.1
A2 2601090 2601153
refGene_gn=CELF5;refGene_tx=XM_023243647.1;refGene_gn=CELF5;refGene_tx
=XM_023243649.1;refGene_gn=CELF5;refGene_tx=XM_023243654.1;refGene_gn=
CELF5;refGene_tx=XM_023243656.1;refGene_gn=CELF5;refGene_tx=XM_0232436
59.1;refGene_gn=CELF5;refGene_tx=XM_023243661.1
A2 2601733 2601817
refGene_gn=CELF5;refGene_tx=XM_023243647.1;refGene_gn=CELF5;refGene_tx
=XM_023243649.1;refGene_gn=CELF5;refGene_tx=XM_023243654.1;refGene_gn=
CELF5;refGene_tx=XM_023243659.1
A2 2604701 2604845
refGene_gn=CELF5;refGene_tx=XM_023243647.1;refGene_gn=CELF5;refGene_tx
=XM_023243649.1;refGene_gn=CELF5;refGene_tx=XM_023243654.1;refGene_gn=
CELF5;refGene_tx=XM_023243656.1;refGene_gn=CELF5;refGene_tx=XM_0232436
59.1;refGene_gn=CELF5;refGene_tx=XM_023243661.1
A2 2607010 2607138
refGene_gn=CELF5;refGene_tx=XM_023243647.1;refGene_gn=CELF5;refGene_tx
=XM_023243649.1;refGene_gn=CELF5;refGene_tx=XM_023243654.1;refGene_gn=

CELF5;refGene_tx=XM_023243656.1;refGene_gn=CELF5;refGene_tx=XM_0232436
59.1;refGene_gn=CELF5;refGene_tx=XM_023243661.1
A2 2660512 2660542 refGene_gn=NFIC;refGene_tx=XM_023250898.1
A2 2668758 2668912 refGene_gn=NFIC;refGene_tx=XM_023250898.1
A2 2670878 2670985 refGene_gn=NFIC;refGene_tx=XM_023250898.1
A2 2687449 2687972 refGene_gn=NFIC;refGene_tx=XM_023250898.1
A2 2694727 2694799 refGene_gn=NFIC;refGene_tx=XM_023250898.1
A2 2697744 2697819 refGene_gn=NFIC;refGene_tx=XM_023250898.1
A2 2698465 2698589 refGene_gn=NFIC;refGene_tx=XM_023250898.1
A2 2699227 2699352 refGene_gn=NFIC;refGene_tx=XM_023250898.1
A2 2705736 2705862 refGene_gn=NFIC;refGene_tx=XM_023250898.1
A2 2707546 2707731 refGene_gn=NFIC;refGene_tx=XM_023250898.1
A2 2708402 2708550 refGene_gn=NFIC;refGene_tx=XM_023250898.1
A2 2710525 2710611 refGene_gn=NFIC;refGene_tx=XM_023250898.1
A2 2714798 2714816 refGene_gn=NFIC;refGene_tx=XM_023250898.1
A2 2724182 2724309
refGene_gn=LOC102900455;refGene_tx=XM_006928248.4
A2 2724913 2725025
refGene_gn=LOC102900455;refGene_tx=XM_006928248.4
A2 2725110 2725573
refGene_gn=LOC102900455;refGene_tx=XM_006928248.4
A2 2726867 2727012
refGene_gn=LOC102901059;refGene_tx=XM_006928261.3
A2 2729457 2729517
refGene_gn=LOC102901059;refGene_tx=XM_006928261.3
A2 2729783 2729895
refGene_gn=LOC102901059;refGene_tx=XM_006928261.3
A2 2730295 2730362
refGene_gn=LOC102901059;refGene_tx=XM_006928261.3
A2 2731805 2732137
refGene_gn=LOC102901059;refGene_tx=XM_019826138.2;refGene_gn=LOC102901
059;refGene_tx=XM_023243673.1
A2 2732288 2732589
refGene_gn=LOC102901059;refGene_tx=XM_019826138.2;refGene_gn=LOC102901
059;refGene_tx=XM_023243673.1
A2 2740058 2740332
refGene_gn=DOHH;refGene_tx=XM_023243662.1;refGene_gn=DOHH;refGene_tx=X
M_023243663.1;refGene_gn=DOHH;refGene_tx=XM_023243664.1
A2 2742556 2742633
refGene_gn=DOHH;refGene_tx=XM_023243662.1;refGene_gn=DOHH;refGene_tx=X
M_023243663.1;refGene_gn=DOHH;refGene_tx=XM_023243664.1
A2 2743801 2744042
refGene_gn=DOHH;refGene_tx=XM_023243662.1;refGene_gn=DOHH;refGene_tx=X
M_023243663.1;refGene_gn=DOHH;refGene_tx=XM_023243664.1
A2 2745001 2745321
refGene_gn=DOHH;refGene_tx=XM_023243662.1;refGene_gn=DOHH;refGene_tx=X
M_023243663.1;refGene_gn=DOHH;refGene_tx=XM_023243664.1
A2 2768401 2768470
refGene_gn=FZR1;refGene_tx=XM_023243685.1;refGene_gn=FZR1;refGene_tx=X
M_023243687.1;refGene_gn=FZR1;refGene_tx=XM_023243692.1

A2 2769766 2769892
refGene_gn=FZR1;refGene_tx=XM_023243685.1;refGene_gn=FZR1;refGene_tx=XM_023243687.1;refGene_gn=FZR1;refGene_tx=XM_023243692.1
A2 2769988 2770052
refGene_gn=FZR1;refGene_tx=XM_023243685.1;refGene_gn=FZR1;refGene_tx=XM_023243687.1;refGene_gn=FZR1;refGene_tx=XM_023243692.1
A2 2770144 2770272
refGene_gn=FZR1;refGene_tx=XM_023243685.1;refGene_gn=FZR1;refGene_tx=XM_023243687.1;refGene_gn=FZR1;refGene_tx=XM_023243692.1
A2 2770763 2770846
refGene_gn=FZR1;refGene_tx=XM_023243685.1;refGene_gn=FZR1;refGene_tx=XM_023243687.1;refGene_gn=FZR1;refGene_tx=XM_023243692.1
A2 2771197 2771381
refGene_gn=FZR1;refGene_tx=XM_023243685.1;refGene_gn=FZR1;refGene_tx=XM_023243687.1;refGene_gn=FZR1;refGene_tx=XM_023243692.1
A2 2771961 2772027
refGene_gn=FZR1;refGene_tx=XM_023243685.1;refGene_gn=FZR1;refGene_tx=XM_023243687.1;refGene_gn=FZR1;refGene_tx=XM_023243692.1
A2 2772518 2772621
refGene_gn=FZR1;refGene_tx=XM_023243685.1;refGene_gn=FZR1;refGene_tx=XM_023243687.1;refGene_gn=FZR1;refGene_tx=XM_023243692.1
A2 2772706 2772891
refGene_gn=FZR1;refGene_tx=XM_023243685.1;refGene_gn=FZR1;refGene_tx=XM_023243687.1;refGene_gn=FZR1;refGene_tx=XM_023243692.1
A2 2773009 2773243
refGene_gn=FZR1;refGene_tx=XM_023243685.1;refGene_gn=FZR1;refGene_tx=XM_023243687.1;refGene_gn=FZR1;refGene_tx=XM_023243692.1
A2 2773776 2773881
refGene_gn=FZR1;refGene_tx=XM_023243685.1;refGene_gn=FZR1;refGene_tx=XM_023243687.1;refGene_gn=FZR1;refGene_tx=XM_023243692.1
A2 2774497 2774599
refGene_gn=FZR1;refGene_tx=XM_023243685.1;refGene_gn=FZR1;refGene_tx=XM_023243687.1;refGene_gn=FZR1;refGene_tx=XM_023243692.1
A2 2775052 2775094
refGene_gn=FZR1;refGene_tx=XM_023243685.1;refGene_gn=FZR1;refGene_tx=XM_023243687.1;refGene_gn=FZR1;refGene_tx=XM_023243692.1
A2 2778633 2778725
refGene_gn=CA2H19orf71;refGene_tx=XM_006928249.4
A2 2780199 2780438
refGene_gn=CA2H19orf71;refGene_tx=XM_006928249.4
A2 2780538 2780664
refGene_gn=CA2H19orf71;refGene_tx=XM_006928249.4
A2 2780777 2780953
refGene_gn=CA2H19orf71;refGene_tx=XM_006928249.4
A2 2781457 2781572
refGene_gn=MFSD12;refGene_tx=XM_011287915.3;refGene_gn=MFSD12;refGene_tx=XM_023243702.1;refGene_gn=MFSD12;refGene_tx=XM_023243703.1
A2 2781671 2781802
refGene_gn=MFSD12;refGene_tx=XM_011287915.3;refGene_gn=MFSD12;refGene_tx=XM_023243702.1

A2 2782652 2782747
refGene_gn=MFSD12;refGene_tx=XM_011287915.3;refGene_gn=MFSD12;refGene_tx=XM_023243702.1;refGene_gn=MFSD12;refGene_tx=XM_023243703.1
A2 2782824 2782995
refGene_gn=MFSD12;refGene_tx=XM_011287915.3;refGene_gn=MFSD12;refGene_tx=XM_023243702.1;refGene_gn=MFSD12;refGene_tx=XM_023243703.1
A2 2783468 2783561
refGene_gn=MFSD12;refGene_tx=XM_011287915.3;refGene_gn=MFSD12;refGene_tx=XM_023243702.1;refGene_gn=MFSD12;refGene_tx=XM_023243703.1
A2 2783643 2783736
refGene_gn=MFSD12;refGene_tx=XM_011287915.3;refGene_gn=MFSD12;refGene_tx=XM_023243702.1;refGene_gn=MFSD12;refGene_tx=XM_023243703.1
A2 2783954 2784137
refGene_gn=MFSD12;refGene_tx=XM_011287915.3;refGene_gn=MFSD12;refGene_tx=XM_023243702.1;refGene_gn=MFSD12;refGene_tx=XM_023243703.1
A2 2784223 2784362
refGene_gn=MFSD12;refGene_tx=XM_011287915.3;refGene_gn=MFSD12;refGene_tx=XM_023243702.1;refGene_gn=MFSD12;refGene_tx=XM_023243703.1
A2 2785922 2786133
refGene_gn=MFSD12;refGene_tx=XM_011287915.3;refGene_gn=MFSD12;refGene_tx=XM_023243702.1;refGene_gn=MFSD12;refGene_tx=XM_023243703.1
A2 2787570 2787754 refGene_gn=MFSD12;refGene_tx=XM_011287915.3
A2 2790115 2790413
refGene_gn=MFSD12;refGene_tx=XM_023243702.1;refGene_gn=MFSD12;refGene_tx=XM_023243703.1
A2 2801961 2801999 refGene_gn=HMG20B;refGene_tx=XM_023243715.1
A2 2802351 2802460 refGene_gn=HMG20B;refGene_tx=XM_023243715.1
A2 2802995 2803199 refGene_gn=HMG20B;refGene_tx=XM_023243715.1
A2 2803816 2803937 refGene_gn=HMG20B;refGene_tx=XM_023243715.1
A2 2804254 2804301 refGene_gn=HMG20B;refGene_tx=XM_023243715.1
A2 2804560 2804633 refGene_gn=HMG20B;refGene_tx=XM_023243715.1
A2 2804910 2805126 refGene_gn=HMG20B;refGene_tx=XM_023243715.1
A2 2805816 2805949 refGene_gn=HMG20B;refGene_tx=XM_023243715.1
A2 2806523 2806536 refGene_gn=HMG20B;refGene_tx=XM_023243715.1
A2 2809995 2810220 refGene_gn=GIPC3;refGene_tx=XM_023243724.1
A2 2810963 2811149 refGene_gn=GIPC3;refGene_tx=XM_023243724.1
A2 2811296 2811477 refGene_gn=GIPC3;refGene_tx=XM_023243724.1
A2 2812995 2813108 refGene_gn=GIPC3;refGene_tx=XM_023243724.1
A2 2813417 2813499 refGene_gn=GIPC3;refGene_tx=XM_023243724.1
A2 2813610 2813762 refGene_gn=GIPC3;refGene_tx=XM_023243724.1
A2 2817590 2817746 refGene_gn=TBXA2R;refGene_tx=XM_023243718.1
A2 2817747 2817837 refGene_gn=TBXA2R;refGene_tx=XM_023243718.1
A2 2821992 2822778 refGene_gn=TBXA2R;refGene_tx=XM_023243718.1
A2 2831580 2832071 refGene_gn=CACTIN;refGene_tx=XM_023243732.1
A2 2832279 2832566 refGene_gn=CACTIN;refGene_tx=XM_023243732.1
A2 2832693 2832816 refGene_gn=CACTIN;refGene_tx=XM_023243732.1
A2 2833395 2833588 refGene_gn=CACTIN;refGene_tx=XM_023243732.1
A2 2836545 2836660 refGene_gn=CACTIN;refGene_tx=XM_023243732.1
A2 2836752 2836915 refGene_gn=CACTIN;refGene_tx=XM_023243732.1
A2 2837749 2837895 refGene_gn=CACTIN;refGene_tx=XM_023243732.1

A2 2838346 2838442 refGene_gn=CACTIN;refGene_tx=XM_023243732.1
A2 2841203 2841687 refGene_gn=CACTIN;refGene_tx=XM_023243732.1
A2 2844166 2844333 refGene_gn=CACTIN;refGene_tx=XM_023243732.1
A2 2849411 2849414
refGene_gn=PIP5K1C;refGene_tx=XM_023243747.1;refGene_gn=PIP5K1C;refGene_tx=XM_023243753.1;refGene_gn=PIP5K1C;refGene_tx=XM_023243758.1;refGene_gn=PIP5K1C;refGene_tx=XM_023243761.1;refGene_gn=PIP5K1C;refGene_tx=XM_023243765.1;refGene_gn=PIP5K1C;refGene_tx=XM_023243767.1;refGene_gn=PIP5K1C;refGene_tx=XM_023243772.1
A2 2849569 2849653
refGene_gn=PIP5K1C;refGene_tx=XM_023243747.1;refGene_gn=PIP5K1C;refGene_tx=XM_023243753.1;refGene_gn=PIP5K1C;refGene_tx=XM_023243758.1;refGene_gn=PIP5K1C;refGene_tx=XM_023243761.1;refGene_gn=PIP5K1C;refGene_tx=XM_023243765.1;refGene_gn=PIP5K1C;refGene_tx=XM_023243767.1
A2 2852301 2852379
refGene_gn=PIP5K1C;refGene_tx=XM_023243747.1;refGene_gn=PIP5K1C;refGene_tx=XM_023243753.1;refGene_gn=PIP5K1C;refGene_tx=XM_023243758.1;refGene_gn=PIP5K1C;refGene_tx=XM_023243767.1
A2 2853503 2853624
refGene_gn=PIP5K1C;refGene_tx=XM_023243747.1;refGene_gn=PIP5K1C;refGene_tx=XM_023243753.1;refGene_gn=PIP5K1C;refGene_tx=XM_023243758.1;refGene_gn=PIP5K1C;refGene_tx=XM_023243761.1;refGene_gn=PIP5K1C;refGene_tx=XM_023243765.1;refGene_gn=PIP5K1C;refGene_tx=XM_023243767.1;refGene_gn=PIP5K1C;refGene_tx=XM_023243772.1
A2 2855925 2856030
refGene_gn=PIP5K1C;refGene_tx=XM_023243747.1;refGene_gn=PIP5K1C;refGene_tx=XM_023243753.1;refGene_gn=PIP5K1C;refGene_tx=XM_023243758.1;refGene_gn=PIP5K1C;refGene_tx=XM_023243761.1;refGene_gn=PIP5K1C;refGene_tx=XM_023243765.1;refGene_gn=PIP5K1C;refGene_tx=XM_023243767.1;refGene_gn=PIP5K1C;refGene_tx=XM_023243772.1
A2 2857178 2857211
refGene_gn=PIP5K1C;refGene_tx=XM_023243747.1;refGene_gn=PIP5K1C;refGene_tx=XM_023243753.1;refGene_gn=PIP5K1C;refGene_tx=XM_023243758.1;refGene_gn=PIP5K1C;refGene_tx=XM_023243761.1;refGene_gn=PIP5K1C;refGene_tx=XM_023243765.1;refGene_gn=PIP5K1C;refGene_tx=XM_023243767.1;refGene_gn=PIP5K1C;refGene_tx=XM_023243772.1
A2 2857324 2857487
refGene_gn=PIP5K1C;refGene_tx=XM_023243747.1;refGene_gn=PIP5K1C;refGene_tx=XM_023243753.1;refGene_gn=PIP5K1C;refGene_tx=XM_023243758.1;refGene_gn=PIP5K1C;refGene_tx=XM_023243761.1;refGene_gn=PIP5K1C;refGene_tx=XM_023243765.1;refGene_gn=PIP5K1C;refGene_tx=XM_023243767.1;refGene_gn=PIP5K1C;refGene_tx=XM_023243772.1
A2 2858187 2858349
refGene_gn=PIP5K1C;refGene_tx=XM_023243747.1;refGene_gn=PIP5K1C;refGene_tx=XM_023243753.1;refGene_gn=PIP5K1C;refGene_tx=XM_023243758.1;refGene_gn=PIP5K1C;refGene_tx=XM_023243761.1;refGene_gn=PIP5K1C;refGene_tx=XM_023243765.1;refGene_gn=PIP5K1C;refGene_tx=XM_023243767.1;refGene_gn=PIP5K1C;refGene_tx=XM_023243772.1
A2 2859836 2859921
refGene_gn=PIP5K1C;refGene_tx=XM_023243747.1;refGene_gn=PIP5K1C;refGen

e_tx=XM_023243753.1;refGene_gn=PIP5K1C;refGene_tx=XM_023243758.1;refGene_gn=PIP5K1C;refGene_tx=XM_023243761.1;refGene_gn=PIP5K1C;refGene_tx=XM_023243765.1;refGene_gn=PIP5K1C;refGene_tx=XM_023243767.1;refGene_gn=PIP5K1C;refGene_tx=XM_023243772.1
A2 2861527 2861576
refGene_gn=PIP5K1C;refGene_tx=XM_023243747.1;refGene_gn=PIP5K1C;refGene_tx=XM_023243753.1;refGene_gn=PIP5K1C;refGene_tx=XM_023243758.1;refGene_gn=PIP5K1C;refGene_tx=XM_023243761.1;refGene_gn=PIP5K1C;refGene_tx=XM_023243765.1;refGene_gn=PIP5K1C;refGene_tx=XM_023243767.1;refGene_gn=PIP5K1C;refGene_tx=XM_023243772.1
A2 2863110 2863194
refGene_gn=PIP5K1C;refGene_tx=XM_023243747.1;refGene_gn=PIP5K1C;refGene_tx=XM_023243753.1;refGene_gn=PIP5K1C;refGene_tx=XM_023243758.1;refGene_gn=PIP5K1C;refGene_tx=XM_023243761.1;refGene_gn=PIP5K1C;refGene_tx=XM_023243765.1;refGene_gn=PIP5K1C;refGene_tx=XM_023243767.1;refGene_gn=PIP5K1C;refGene_tx=XM_023243772.1
A2 2864619 2864825
refGene_gn=PIP5K1C;refGene_tx=XM_023243747.1;refGene_gn=PIP5K1C;refGene_tx=XM_023243753.1;refGene_gn=PIP5K1C;refGene_tx=XM_023243758.1;refGene_gn=PIP5K1C;refGene_tx=XM_023243761.1;refGene_gn=PIP5K1C;refGene_tx=XM_023243765.1;refGene_gn=PIP5K1C;refGene_tx=XM_023243767.1;refGene_gn=PIP5K1C;refGene_tx=XM_023243772.1
A2 2865530 2865830
refGene_gn=PIP5K1C;refGene_tx=XM_023243747.1;refGene_gn=PIP5K1C;refGene_tx=XM_023243753.1;refGene_gn=PIP5K1C;refGene_tx=XM_023243758.1;refGene_gn=PIP5K1C;refGene_tx=XM_023243761.1;refGene_gn=PIP5K1C;refGene_tx=XM_023243765.1;refGene_gn=PIP5K1C;refGene_tx=XM_023243767.1;refGene_gn=PIP5K1C;refGene_tx=XM_023243772.1
A2 2867784 2867937
refGene_gn=PIP5K1C;refGene_tx=XM_023243747.1;refGene_gn=PIP5K1C;refGene_tx=XM_023243753.1;refGene_gn=PIP5K1C;refGene_tx=XM_023243758.1;refGene_gn=PIP5K1C;refGene_tx=XM_023243761.1;refGene_gn=PIP5K1C;refGene_tx=XM_023243765.1;refGene_gn=PIP5K1C;refGene_tx=XM_023243767.1;refGene_gn=PIP5K1C;refGene_tx=XM_023243772.1
A2 2872031 2872149
refGene_gn=PIP5K1C;refGene_tx=XM_023243747.1;refGene_gn=PIP5K1C;refGene_tx=XM_023243753.1;refGene_gn=PIP5K1C;refGene_tx=XM_023243758.1;refGene_gn=PIP5K1C;refGene_tx=XM_023243761.1;refGene_gn=PIP5K1C;refGene_tx=XM_023243765.1;refGene_gn=PIP5K1C;refGene_tx=XM_023243767.1;refGene_gn=PIP5K1C;refGene_tx=XM_023243772.1
A2 2872943 2873074
refGene_gn=PIP5K1C;refGene_tx=XM_023243747.1;refGene_gn=PIP5K1C;refGene_tx=XM_023243753.1;refGene_gn=PIP5K1C;refGene_tx=XM_023243758.1;refGene_gn=PIP5K1C;refGene_tx=XM_023243761.1;refGene_gn=PIP5K1C;refGene_tx=XM_023243765.1;refGene_gn=PIP5K1C;refGene_tx=XM_023243767.1;refGene_gn=PIP5K1C;refGene_tx=XM_023243772.1
A2 2873981 2874074
refGene_gn=PIP5K1C;refGene_tx=XM_023243747.1;refGene_gn=PIP5K1C;refGene_tx=XM_023243753.1;refGene_gn=PIP5K1C;refGene_tx=XM_023243758.1;refGene_gn=PIP5K1C;refGene_tx=XM_023243761.1;refGene_gn=PIP5K1C;refGene_tx=

XM_023243765.1;refGene_gn=PIP5K1C;refGene_tx=XM_023243767.1;refGene_gn=PIP5K1C;refGene_tx=XM_023243772.1
A2 2875509 2875541
refGene_gn=PIP5K1C;refGene_tx=XM_023243747.1;refGene_gn=PIP5K1C;refGene_tx=XM_023243753.1;refGene_gn=PIP5K1C;refGene_tx=XM_023243758.1;refGene_gn=PIP5K1C;refGene_tx=XM_023243761.1;refGene_gn=PIP5K1C;refGene_tx=XM_023243765.1;refGene_gn=PIP5K1C;refGene_tx=XM_023243767.1;refGene_gn=PIP5K1C;refGene_tx=XM_023243772.1
A2 2880149 2880553
refGene_gn=LOC109497815;refGene_tx=XM_023243776.1
A2 2880994 2881141
refGene_gn=LOC109497815;refGene_tx=XM_023243776.1
A2 2882612 2882625
refGene_gn=LOC109497815;refGene_tx=XM_023243776.1
A2 2896005 2896099
refGene_gn=PIP5K1C;refGene_tx=XM_023243747.1;refGene_gn=PIP5K1C;refGene_tx=XM_023243753.1;refGene_gn=PIP5K1C;refGene_tx=XM_023243758.1;refGene_gn=PIP5K1C;refGene_tx=XM_023243761.1;refGene_gn=PIP5K1C;refGene_tx=XM_023243765.1;refGene_gn=PIP5K1C;refGene_tx=XM_023243772.1
A2 2906912 2906930
refGene_gn=TJP3;refGene_tx=XM_023243796.1;refGene_gn=TJP3;refGene_tx=XM_023243803.1
A2 2911028 2911085
refGene_gn=TJP3;refGene_tx=XM_023243796.1;refGene_gn=TJP3;refGene_tx=XM_023243803.1;refGene_gn=TJP3;refGene_tx=XM_023243807.1;refGene_gn=TJP3;refGene_tx=XM_023243810.1
A2 2911192 2911302
refGene_gn=TJP3;refGene_tx=XM_023243796.1;refGene_gn=TJP3;refGene_tx=XM_023243803.1;refGene_gn=TJP3;refGene_tx=XM_023243807.1;refGene_gn=TJP3;refGene_tx=XM_023243810.1
A2 2912498 2912601
refGene_gn=TJP3;refGene_tx=XM_023243796.1;refGene_gn=TJP3;refGene_tx=XM_023243803.1;refGene_gn=TJP3;refGene_tx=XM_023243807.1;refGene_gn=TJP3;refGene_tx=XM_023243810.1
A2 2912842 2913164
refGene_gn=TJP3;refGene_tx=XM_023243796.1;refGene_gn=TJP3;refGene_tx=XM_023243803.1;refGene_gn=TJP3;refGene_tx=XM_023243807.1;refGene_gn=TJP3;refGene_tx=XM_023243810.1
A2 2914195 2914299
refGene_gn=TJP3;refGene_tx=XM_023243796.1;refGene_gn=TJP3;refGene_tx=XM_023243803.1;refGene_gn=TJP3;refGene_tx=XM_023243807.1;refGene_gn=TJP3;refGene_tx=XM_023243810.1
A2 2915523 2915683
refGene_gn=TJP3;refGene_tx=XM_023243796.1;refGene_gn=TJP3;refGene_tx=XM_023243803.1;refGene_gn=TJP3;refGene_tx=XM_023243807.1;refGene_gn=TJP3;refGene_tx=XM_023243810.1
A2 2915941 2916050
refGene_gn=TJP3;refGene_tx=XM_023243796.1;refGene_gn=TJP3;refGene_tx=XM_023243803.1;refGene_gn=TJP3;refGene_tx=XM_023243807.1;refGene_gn=TJP3;refGene_tx=XM_023243810.1

A2 2919160 2919219
refGene_gn=TJP3;refGene_tx=XM_023243796.1;refGene_gn=TJP3;refGene_tx=XM_023243803.1;refGene_gn=TJP3;refGene_tx=XM_023243807.1;refGene_gn=TJP3;refGene_tx=XM_023243810.1
A2 2919459 2919526
refGene_gn=TJP3;refGene_tx=XM_023243796.1;refGene_gn=TJP3;refGene_tx=XM_023243803.1;refGene_gn=TJP3;refGene_tx=XM_023243807.1;refGene_gn=TJP3;refGene_tx=XM_023243810.1
A2 2919723 2919880
refGene_gn=TJP3;refGene_tx=XM_023243796.1;refGene_gn=TJP3;refGene_tx=XM_023243803.1;refGene_gn=TJP3;refGene_tx=XM_023243807.1;refGene_gn=TJP3;refGene_tx=XM_023243810.1
A2 2920386 2920495
refGene_gn=TJP3;refGene_tx=XM_023243796.1;refGene_gn=TJP3;refGene_tx=XM_023243803.1;refGene_gn=TJP3;refGene_tx=XM_023243807.1;refGene_gn=TJP3;refGene_tx=XM_023243810.1
A2 2920571 2920809
refGene_gn=TJP3;refGene_tx=XM_023243796.1;refGene_gn=TJP3;refGene_tx=XM_023243803.1;refGene_gn=TJP3;refGene_tx=XM_023243807.1;refGene_gn=TJP3;refGene_tx=XM_023243810.1
A2 2920918 2921130
refGene_gn=TJP3;refGene_tx=XM_023243796.1;refGene_gn=TJP3;refGene_tx=XM_023243803.1;refGene_gn=TJP3;refGene_tx=XM_023243807.1;refGene_gn=TJP3;refGene_tx=XM_023243810.1
A2 2922985 2923081
refGene_gn=TJP3;refGene_tx=XM_023243796.1;refGene_gn=TJP3;refGene_tx=XM_023243803.1;refGene_gn=TJP3;refGene_tx=XM_023243807.1;refGene_gn=TJP3;refGene_tx=XM_023243810.1
A2 2925681 2925752
refGene_gn=TJP3;refGene_tx=XM_023243796.1;refGene_gn=TJP3;refGene_tx=XM_023243803.1;refGene_gn=TJP3;refGene_tx=XM_023243807.1;refGene_gn=TJP3;refGene_tx=XM_023243810.1
A2 2926128 2926339
refGene_gn=TJP3;refGene_tx=XM_023243796.1;refGene_gn=TJP3;refGene_tx=XM_023243803.1;refGene_gn=TJP3;refGene_tx=XM_023243807.1;refGene_gn=TJP3;refGene_tx=XM_023243810.1
A2 2926428 2926529
refGene_gn=TJP3;refGene_tx=XM_023243796.1;refGene_gn=TJP3;refGene_tx=XM_023243803.1;refGene_gn=TJP3;refGene_tx=XM_023243807.1;refGene_gn=TJP3;refGene_tx=XM_023243810.1
A2 2926681 2926945
refGene_gn=TJP3;refGene_tx=XM_023243796.1;refGene_gn=TJP3;refGene_tx=XM_023243803.1;refGene_gn=TJP3;refGene_tx=XM_023243807.1;refGene_gn=TJP3;refGene_tx=XM_023243810.1
A2 2927787 2927834
refGene_gn=TJP3;refGene_tx=XM_023243796.1;refGene_gn=TJP3;refGene_tx=XM_023243803.1;refGene_gn=TJP3;refGene_tx=XM_023243807.1;refGene_gn=TJP3;refGene_tx=XM_023243810.1
A2 2928246 2928349
refGene_gn=TJP3;refGene_tx=XM_023243796.1;refGene_gn=TJP3;refGene_tx=X

M_023243803.1;refGene_gn=TJP3;refGene_tx=XM_023243807.1;refGene_gn=TJP3;refGene_tx=XM_023243810.1
A2	2928694	2928766	refGene_gn=APBA3;refGene_tx=XM_023243812.1
A2	2928877	2929018	refGene_gn=APBA3;refGene_tx=XM_023243812.1
A2	2929115	2929235	refGene_gn=APBA3;refGene_tx=XM_023243812.1
A2	2929859	2930072	refGene_gn=APBA3;refGene_tx=XM_023243812.1
A2	2930156	2930336	refGene_gn=APBA3;refGene_tx=XM_023243812.1
A2	2930772	2930934	refGene_gn=APBA3;refGene_tx=XM_023243812.1
A2	2931030	2931117	refGene_gn=APBA3;refGene_tx=XM_023243812.1
A2	2931283	2931429	refGene_gn=APBA3;refGene_tx=XM_023243812.1
A2	2934380	2934420	refGene_gn=APBA3;refGene_tx=XM_023243812.1
A2	2934508	2935084	refGene_gn=APBA3;refGene_tx=XM_023243812.1
A2	2936644	2936762	refGene_gn=MRPL54;refGene_tx=XM_006928269.4
A2	2937601	2937767	refGene_gn=MRPL54;refGene_tx=XM_006928269.4
A2	2938059	2938192	refGene_gn=MRPL54;refGene_tx=XM_006928269.4
A2	2940898	2941237	refGene_gn=RAX2;refGene_tx=XM_023243834.1
A2	2941348	2941621	refGene_gn=RAX2;refGene_tx=XM_023243840.1
A2	2941928	2942144
refGene_gn=RAX2;refGene_tx=XM_023243834.1;refGene_gn=RAX2;refGene_tx=XM_023243840.1
A2	2945629	2945869
refGene_gn=MATK;refGene_tx=XM_023243813.1;refGene_gn=MATK;refGene_tx=XM_023243816.1;refGene_gn=MATK;refGene_tx=XM_023243818.1;refGene_gn=MATK;refGene_tx=XM_023243826.1
A2	2945962	2946049
refGene_gn=MATK;refGene_tx=XM_023243813.1;refGene_gn=MATK;refGene_tx=XM_023243816.1;refGene_gn=MATK;refGene_tx=XM_023243818.1;refGene_gn=MATK;refGene_tx=XM_023243826.1
A2	2946343	2946539
refGene_gn=MATK;refGene_tx=XM_023243813.1;refGene_gn=MATK;refGene_tx=XM_023243816.1;refGene_gn=MATK;refGene_tx=XM_023243818.1;refGene_gn=MATK;refGene_tx=XM_023243826.1
A2	2946695	2946769
refGene_gn=MATK;refGene_tx=XM_023243813.1;refGene_gn=MATK;refGene_tx=XM_023243816.1;refGene_gn=MATK;refGene_tx=XM_023243818.1;refGene_gn=MATK;refGene_tx=XM_023243826.1
A2	2946857	2946942
refGene_gn=MATK;refGene_tx=XM_023243813.1;refGene_gn=MATK;refGene_tx=XM_023243816.1;refGene_gn=MATK;refGene_tx=XM_023243818.1;refGene_gn=MATK;refGene_tx=XM_023243826.1
A2	2947033	2947133
refGene_gn=MATK;refGene_tx=XM_023243813.1;refGene_gn=MATK;refGene_tx=XM_023243816.1;refGene_gn=MATK;refGene_tx=XM_023243818.1;refGene_gn=MATK;refGene_tx=XM_023243826.1
A2	2947846	2947912
refGene_gn=MATK;refGene_tx=XM_023243813.1;refGene_gn=MATK;refGene_tx=XM_023243816.1;refGene_gn=MATK;refGene_tx=XM_023243818.1;refGene_gn=MATK;refGene_tx=XM_023243826.1
A2	2949212	2949306
refGene_gn=MATK;refGene_tx=XM_023243813.1;refGene_gn=MATK;refGene_tx=X

M_023243816.1;refGene_gn=MATK;refGene_tx=XM_023243818.1;refGene_gn=MATK;refGene_tx=XM_023243826.1
A2 2949751 2949971
refGene_gn=MATK;refGene_tx=XM_023243813.1;refGene_gn=MATK;refGene_tx=XM_023243816.1;refGene_gn=MATK;refGene_tx=XM_023243818.1;refGene_gn=MATK;refGene_tx=XM_023243826.1
A2 2950063 2950179
refGene_gn=MATK;refGene_tx=XM_023243813.1;refGene_gn=MATK;refGene_tx=XM_023243816.1;refGene_gn=MATK;refGene_tx=XM_023243818.1;refGene_gn=MATK;refGene_tx=XM_023243826.1
A2 2950283 2950397
refGene_gn=MATK;refGene_tx=XM_023243813.1;refGene_gn=MATK;refGene_tx=XM_023243816.1;refGene_gn=MATK;refGene_tx=XM_023243818.1;refGene_gn=MATK;refGene_tx=XM_023243826.1
A2 2950702 2950762
refGene_gn=MATK;refGene_tx=XM_023243813.1;refGene_gn=MATK;refGene_tx=XM_023243816.1;refGene_gn=MATK;refGene_tx=XM_023243818.1;refGene_gn=MATK;refGene_tx=XM_023243826.1
A2 2951506 2951512
refGene_gn=MATK;refGene_tx=XM_023243813.1;refGene_gn=MATK;refGene_tx=XM_023243816.1
A2 2964351 2964534
refGene_gn=ZFR2;refGene_tx=XM_023243780.1;refGene_gn=ZFR2;refGene_tx=XM_023243787.1;refGene_gn=ZFR2;refGene_tx=XM_023243792.1
A2 2965271 2965369
refGene_gn=ZFR2;refGene_tx=XM_023243780.1;refGene_gn=ZFR2;refGene_tx=XM_023243787.1;refGene_gn=ZFR2;refGene_tx=XM_023243792.1
A2 2965923 2966035
refGene_gn=ZFR2;refGene_tx=XM_023243780.1;refGene_gn=ZFR2;refGene_tx=XM_023243787.1;refGene_gn=ZFR2;refGene_tx=XM_023243792.1
A2 2967287 2967383
refGene_gn=ZFR2;refGene_tx=XM_023243780.1;refGene_gn=ZFR2;refGene_tx=XM_023243787.1;refGene_gn=ZFR2;refGene_tx=XM_023243792.1;refGene_gn=ZFR2;refGene_tx=XM_023243795.1
A2 2967710 2967808
refGene_gn=ZFR2;refGene_tx=XM_023243780.1;refGene_gn=ZFR2;refGene_tx=XM_023243787.1;refGene_gn=ZFR2;refGene_tx=XM_023243792.1
A2 2969543 2969682
refGene_gn=ZFR2;refGene_tx=XM_023243780.1;refGene_gn=ZFR2;refGene_tx=XM_023243787.1;refGene_gn=ZFR2;refGene_tx=XM_023243792.1;refGene_gn=ZFR2;refGene_tx=XM_023243795.1
A2 2970885 2971057
refGene_gn=ZFR2;refGene_tx=XM_023243780.1;refGene_gn=ZFR2;refGene_tx=XM_023243787.1;refGene_gn=ZFR2;refGene_tx=XM_023243792.1;refGene_gn=ZFR2;refGene_tx=XM_023243795.1
A2 2974029 2974220
refGene_gn=ZFR2;refGene_tx=XM_023243780.1;refGene_gn=ZFR2;refGene_tx=XM_023243787.1;refGene_gn=ZFR2;refGene_tx=XM_023243792.1;refGene_gn=ZFR2;refGene_tx=XM_023243795.1
A2 2974885 2975018

refGene_gn=ZFR2;refGene_tx=XM_023243780.1;refGene_gn=ZFR2;refGene_tx=XM_023243787.1;refGene_gn=ZFR2;refGene_tx=XM_023243792.1;refGene_gn=ZFR2;refGene_tx=XM_023243795.1
A2 2976101 2976241
refGene_gn=ZFR2;refGene_tx=XM_023243780.1;refGene_gn=ZFR2;refGene_tx=XM_023243787.1;refGene_gn=ZFR2;refGene_tx=XM_023243792.1;refGene_gn=ZFR2;refGene_tx=XM_023243795.1
A2 2977690 2977810
refGene_gn=ZFR2;refGene_tx=XM_023243780.1;refGene_gn=ZFR2;refGene_tx=XM_023243787.1;refGene_gn=ZFR2;refGene_tx=XM_023243792.1;refGene_gn=ZFR2;refGene_tx=XM_023243795.1
A2 2978766 2978918
refGene_gn=ZFR2;refGene_tx=XM_023243780.1;refGene_gn=ZFR2;refGene_tx=XM_023243787.1;refGene_gn=ZFR2;refGene_tx=XM_023243792.1;refGene_gn=ZFR2;refGene_tx=XM_023243795.1
A2 2980022 2980251
refGene_gn=ZFR2;refGene_tx=XM_023243780.1;refGene_gn=ZFR2;refGene_tx=XM_023243787.1;refGene_gn=ZFR2;refGene_tx=XM_023243792.1;refGene_gn=ZFR2;refGene_tx=XM_023243795.1
A2 2982404 2982587
refGene_gn=ZFR2;refGene_tx=XM_023243780.1;refGene_gn=ZFR2;refGene_tx=XM_023243787.1;refGene_gn=ZFR2;refGene_tx=XM_023243792.1;refGene_gn=ZFR2;refGene_tx=XM_023243795.1
A2 2983571 2983822
refGene_gn=ZFR2;refGene_tx=XM_023243780.1;refGene_gn=ZFR2;refGene_tx=XM_023243787.1;refGene_gn=ZFR2;refGene_tx=XM_023243792.1;refGene_gn=ZFR2;refGene_tx=XM_023243795.1
A2 2983914 2984139
refGene_gn=ZFR2;refGene_tx=XM_023243780.1;refGene_gn=ZFR2;refGene_tx=XM_023243787.1;refGene_gn=ZFR2;refGene_tx=XM_023243792.1;refGene_gn=ZFR2;refGene_tx=XM_023243795.1
A2 2984596 2984711
refGene_gn=ZFR2;refGene_tx=XM_023243780.1;refGene_gn=ZFR2;refGene_tx=XM_023243787.1;refGene_gn=ZFR2;refGene_tx=XM_023243792.1;refGene_gn=ZFR2;refGene_tx=XM_023243795.1
A2 2985650 2985846
refGene_gn=ZFR2;refGene_tx=XM_023243780.1;refGene_gn=ZFR2;refGene_tx=XM_023243787.1;refGene_gn=ZFR2;refGene_tx=XM_023243792.1;refGene_gn=ZFR2;refGene_tx=XM_023243795.1
A2 3004347 3004400
refGene_gn=ZFR2;refGene_tx=XM_023243780.1;refGene_gn=ZFR2;refGene_tx=XM_023243787.1;refGene_gn=ZFR2;refGene_tx=XM_023243792.1;refGene_gn=ZFR2;refGene_tx=XM_023243795.1
A2 3013038 3013115
refGene_gn=ATCAY;refGene_tx=XM_003981738.5;refGene_gn=ATCAY;refGene_tx=XM_019826131.2
A2 3023212 3023271
refGene_gn=ATCAY;refGene_tx=XM_003981738.5;refGene_gn=ATCAY;refGene_tx=XM_019826131.2
A2 3025972 3026194

refGene_gn=ATCAY;refGene_tx=XM_003981738.5;refGene_gn=ATCAY;refGene_tx=XM_019826131.2
A2 3027750 3027936
refGene_gn=ATCAY;refGene_tx=XM_003981738.5;refGene_gn=ATCAY;refGene_tx=XM_019826131.2
A2 3028221 3028324
refGene_gn=ATCAY;refGene_tx=XM_003981738.5;refGene_gn=ATCAY;refGene_tx=XM_019826131.2
A2 3029647 3029779
refGene_gn=ATCAY;refGene_tx=XM_003981738.5;refGene_gn=ATCAY;refGene_tx=XM_019826131.2
A2 3030474 3030561
refGene_gn=ATCAY;refGene_tx=XM_003981738.5;refGene_gn=ATCAY;refGene_tx=XM_019826131.2
A2 3032078 3032157 refGene_gn=ATCAY;refGene_tx=XM_019826131.2
A2 3032776 3032875 refGene_gn=ATCAY;refGene_tx=XM_003981738.5
A2 3033564 3033600 refGene_gn=ATCAY;refGene_tx=XM_003981738.5
A2 3035568 3035640 refGene_gn=ATCAY;refGene_tx=XM_003981738.5
A2 3036940 3036970 refGene_gn=ATCAY;refGene_tx=XM_003981738.5
A2 3038823 3038833 refGene_gn=ATCAY;refGene_tx=XM_003981738.5
A2 3050228 3050765 refGene_gn=DAPK3;refGene_tx=XM_023243856.1
A2 3051163 3051209 refGene_gn=DAPK3;refGene_tx=XM_023243856.1
A2 3051956 3052109 refGene_gn=DAPK3;refGene_tx=XM_023243856.1
A2 3053140 3053167 refGene_gn=DAPK3;refGene_tx=XM_023243856.1
A2 3053741 3053790 refGene_gn=DAPK3;refGene_tx=XM_023243856.1
A2 3053994 3054124 refGene_gn=DAPK3;refGene_tx=XM_023243856.1
A2 3054306 3054667 refGene_gn=DAPK3;refGene_tx=XM_023243856.1
A2 3060615 3060677 refGene_gn=DAPK3;refGene_tx=XM_023243856.1
A2 3066207 3066401 refGene_gn=EEF2;refGene_tx=XM_003981696.5
A2 3066905 3067038 refGene_gn=EEF2;refGene_tx=XM_003981696.5
A2 3067119 3067302 refGene_gn=EEF2;refGene_tx=XM_003981696.5
A2 3067596 3067950 refGene_gn=EEF2;refGene_tx=XM_003981696.5
A2 3068402 3068510 refGene_gn=EEF2;refGene_tx=XM_003981696.5
A2 3068950 3069209 refGene_gn=EEF2;refGene_tx=XM_003981696.5
A2 3069805 3070001 refGene_gn=EEF2;refGene_tx=XM_003981696.5
A2 3070141 3070280 refGene_gn=EEF2;refGene_tx=XM_003981696.5
A2 3070501 3070615 refGene_gn=EEF2;refGene_tx=XM_003981696.5
A2 3071131 3071237 refGene_gn=EEF2;refGene_tx=XM_003981696.5
A2 3071420 3071599 refGene_gn=EEF2;refGene_tx=XM_003981696.5
A2 3072110 3072322 refGene_gn=EEF2;refGene_tx=XM_003981696.5
A2 3072404 3072586 refGene_gn=EEF2;refGene_tx=XM_003981696.5
A2 3073462 3073677 refGene_gn=EEF2;refGene_tx=XM_003981696.5
A2 3074692 3074695 refGene_gn=EEF2;refGene_tx=XM_003981696.5
A2 3090104 3090131
refGene_gn=PIAS4;refGene_tx=XM_023243883.1;refGene_gn=PIAS4;refGene_tx=XM_023243894.1
A2 3094863 3095290
refGene_gn=PIAS4;refGene_tx=XM_023243883.1;refGene_gn=PIAS4;refGene_tx=XM_023243894.1;refGene_gn=PIAS4;refGene_tx=XM_023243899.1
A2 3105953 3106038

refGene_gn=PIAS4;refGene_tx=XM_023243883.1;refGene_gn=PIAS4;refGene_tx=XM_023243894.1;refGene_gn=PIAS4;refGene_tx=XM_023243899.1
A2 3106760 3106802
refGene_gn=PIAS4;refGene_tx=XM_023243883.1;refGene_gn=PIAS4;refGene_tx=XM_023243894.1;refGene_gn=PIAS4;refGene_tx=XM_023243899.1
A2 3107167 3107258
refGene_gn=PIAS4;refGene_tx=XM_023243883.1;refGene_gn=PIAS4;refGene_tx=XM_023243894.1;refGene_gn=PIAS4;refGene_tx=XM_023243899.1
A2 3107360 3107489
refGene_gn=PIAS4;refGene_tx=XM_023243883.1;refGene_gn=PIAS4;refGene_tx=XM_023243894.1;refGene_gn=PIAS4;refGene_tx=XM_023243899.1
A2 3107600 3107706
refGene_gn=PIAS4;refGene_tx=XM_023243883.1;refGene_gn=PIAS4;refGene_tx=XM_023243894.1;refGene_gn=PIAS4;refGene_tx=XM_023243899.1
A2 3109701 3109775
refGene_gn=PIAS4;refGene_tx=XM_023243883.1;refGene_gn=PIAS4;refGene_tx=XM_023243894.1;refGene_gn=PIAS4;refGene_tx=XM_023243899.1
A2 3109873 3110034
refGene_gn=PIAS4;refGene_tx=XM_023243883.1;refGene_gn=PIAS4;refGene_tx=XM_023243894.1;refGene_gn=PIAS4;refGene_tx=XM_023243899.1
A2 3112432 3112563
refGene_gn=PIAS4;refGene_tx=XM_023243883.1;refGene_gn=PIAS4;refGene_tx=XM_023243894.1;refGene_gn=PIAS4;refGene_tx=XM_023243899.1
A2 3112689 3112961
refGene_gn=PIAS4;refGene_tx=XM_023243883.1;refGene_gn=PIAS4;refGene_tx=XM_023243894.1;refGene_gn=PIAS4;refGene_tx=XM_023243899.1
A2 3120990 3121489
refGene_gn=ZBTB7A;refGene_tx=XM_023243861.1;refGene_gn=ZBTB7A;refGene_tx=XM_023243870.1;refGene_gn=ZBTB7A;refGene_tx=XM_023243878.1;refGene_gn=ZBTB7A;refGene_tx=XM_023243880.1
A2 3125022 3126278
refGene_gn=ZBTB7A;refGene_tx=XM_023243861.1;refGene_gn=ZBTB7A;refGene_tx=XM_023243870.1;refGene_gn=ZBTB7A;refGene_tx=XM_023243878.1;refGene_gn=ZBTB7A;refGene_tx=XM_023243880.1
A2 3155995 3156106
refGene_gn=MAP2K2;refGene_tx=XM_023243905.1;refGene_gn=MAP2K2;refGene_tx=XM_023243907.1
A2 3157159 3157289 refGene_gn=MAP2K2;refGene_tx=XM_023243905.1
A2 3157703 3157811
refGene_gn=MAP2K2;refGene_tx=XM_023243905.1;refGene_gn=MAP2K2;refGene_tx=XM_023243907.1;refGene_gn=MAP2K2;refGene_tx=XM_023243914.1
A2 3158538 3158539 refGene_gn=MAP2K2;refGene_tx=XM_023243917.1
A2 3158826 3158888
refGene_gn=MAP2K2;refGene_tx=XM_023243905.1;refGene_gn=MAP2K2;refGene_tx=XM_023243907.1;refGene_gn=MAP2K2;refGene_tx=XM_023243914.1;refGene_gn=MAP2K2;refGene_tx=XM_023243917.1
A2 3159885 3159950
refGene_gn=MAP2K2;refGene_tx=XM_023243905.1;refGene_gn=MAP2K2;refGene_tx=XM_023243907.1;refGene_gn=MAP2K2;refGene_tx=XM_023243914.1;refGene_gn=MAP2K2;refGene_tx=XM_023243917.1

A2	3161778	3161992
refGene_gn=MAP2K2;refGene_tx=XM_023243905.1;refGene_gn=MAP2K2;refGene_tx=XM_023243907.1;refGene_gn=MAP2K2;refGene_tx=XM_023243914.1;refGene_gn=MAP2K2;refGene_tx=XM_023243917.1
A2	3162549	3162674
refGene_gn=MAP2K2;refGene_tx=XM_023243905.1;refGene_gn=MAP2K2;refGene_tx=XM_023243907.1;refGene_gn=MAP2K2;refGene_tx=XM_023243914.1;refGene_gn=MAP2K2;refGene_tx=XM_023243917.1
A2	3162754	3162806
refGene_gn=MAP2K2;refGene_tx=XM_023243905.1;refGene_gn=MAP2K2;refGene_tx=XM_023243907.1;refGene_gn=MAP2K2;refGene_tx=XM_023243914.1;refGene_gn=MAP2K2;refGene_tx=XM_023243917.1
A2	3163263	3163341
refGene_gn=MAP2K2;refGene_tx=XM_023243905.1;refGene_gn=MAP2K2;refGene_tx=XM_023243907.1;refGene_gn=MAP2K2;refGene_tx=XM_023243914.1;refGene_gn=MAP2K2;refGene_tx=XM_023243917.1
A2	3167154	3167301
refGene_gn=MAP2K2;refGene_tx=XM_023243905.1;refGene_gn=MAP2K2;refGene_tx=XM_023243907.1;refGene_gn=MAP2K2;refGene_tx=XM_023243914.1;refGene_gn=MAP2K2;refGene_tx=XM_023243917.1
A2	3172218	3172429
refGene_gn=MAP2K2;refGene_tx=XM_023243905.1;refGene_gn=MAP2K2;refGene_tx=XM_023243907.1;refGene_gn=MAP2K2;refGene_tx=XM_023243914.1;refGene_gn=MAP2K2;refGene_tx=XM_023243917.1
A2	3176112	3176204
refGene_gn=MAP2K2;refGene_tx=XM_023243905.1;refGene_gn=MAP2K2;refGene_tx=XM_023243907.1;refGene_gn=MAP2K2;refGene_tx=XM_023243914.1;refGene_gn=MAP2K2;refGene_tx=XM_023243917.1
A2	3196695	3196722
refGene_gn=CREB3L3;refGene_tx=XM_003981697.5;refGene_gn=CREB3L3;refGene_tx=XM_006928270.4;refGene_gn=CREB3L3;refGene_tx=XM_019815746.2
A2	3197235	3197364
refGene_gn=CREB3L3;refGene_tx=XM_003981697.5;refGene_gn=CREB3L3;refGene_tx=XM_006928270.4;refGene_gn=CREB3L3;refGene_tx=XM_019815746.2
A2	3198019	3198290
refGene_gn=CREB3L3;refGene_tx=XM_003981697.5;refGene_gn=CREB3L3;refGene_tx=XM_006928270.4;refGene_gn=CREB3L3;refGene_tx=XM_019815746.2
A2	3199164	3199277
refGene_gn=CREB3L3;refGene_tx=XM_003981697.5;refGene_gn=CREB3L3;refGene_tx=XM_006928270.4;refGene_gn=CREB3L3;refGene_tx=XM_019815746.2
A2	3199864	3200002
refGene_gn=CREB3L3;refGene_tx=XM_003981697.5;refGene_gn=CREB3L3;refGene_tx=XM_006928270.4;refGene_gn=CREB3L3;refGene_tx=XM_019815746.2
A2	3200782	3200889
refGene_gn=CREB3L3;refGene_tx=XM_003981697.5;refGene_gn=CREB3L3;refGene_tx=XM_006928270.4;refGene_gn=CREB3L3;refGene_tx=XM_019815746.2
A2	3202790	3202859
refGene_gn=CREB3L3;refGene_tx=XM_003981697.5;refGene_gn=CREB3L3;refGene_tx=XM_006928270.4;refGene_gn=CREB3L3;refGene_tx=XM_019815746.2
A2	3204043	3204128

refGene_gn=CREB3L3;refGene_tx=XM_003981697.5;refGene_gn=CREB3L3;refGene_tx=XM_006928270.4;refGene_gn=CREB3L3;refGene_tx=XM_019815746.2
A2 3204287 3204384
refGene_gn=CREB3L3;refGene_tx=XM_003981697.5;refGene_gn=CREB3L3;refGene_tx=XM_006928270.4;refGene_gn=CREB3L3;refGene_tx=XM_019815746.2
A2 3204589 3204903
refGene_gn=CREB3L3;refGene_tx=XM_003981697.5;refGene_gn=CREB3L3;refGene_tx=XM_006928270.4;refGene_gn=CREB3L3;refGene_tx=XM_019815746.2
A2 3206620 3206962 refGene_gn=SIRT6;refGene_tx=XM_023243942.1
A2 3207078 3207202 refGene_gn=SIRT6;refGene_tx=XM_023243942.1
A2 3207708 3207789 refGene_gn=SIRT6;refGene_tx=XM_023243942.1
A2 3207880 3207976 refGene_gn=SIRT6;refGene_tx=XM_023243942.1
A2 3209793 3209853 refGene_gn=SIRT6;refGene_tx=XM_023243942.1
A2 3212057 3212240 refGene_gn=SIRT6;refGene_tx=XM_023243942.1
A2 3212597 3212725 refGene_gn=SIRT6;refGene_tx=XM_023243942.1
A2 3214513 3214579 refGene_gn=SIRT6;refGene_tx=XM_023243942.1
A2 3223632 3223671 refGene_gn=ANKRD24;refGene_tx=XM_023243955.1
A2 3224497 3224584 refGene_gn=ANKRD24;refGene_tx=XM_023243955.1
A2 3224689 3224820 refGene_gn=ANKRD24;refGene_tx=XM_023243955.1
A2 3224896 3224985 refGene_gn=ANKRD24;refGene_tx=XM_023243955.1
A2 3225915 3225980 refGene_gn=ANKRD24;refGene_tx=XM_023243955.1
A2 3226509 3226567 refGene_gn=ANKRD24;refGene_tx=XM_023243955.1
A2 3228085 3228156 refGene_gn=ANKRD24;refGene_tx=XM_023243955.1
A2 3228344 3228451 refGene_gn=ANKRD24;refGene_tx=XM_023243955.1
A2 3228630 3228818 refGene_gn=ANKRD24;refGene_tx=XM_023243955.1
A2 3229303 3229332 refGene_gn=ANKRD24;refGene_tx=XM_023243955.1
A2 3229945 3230026 refGene_gn=ANKRD24;refGene_tx=XM_023243955.1
A2 3230165 3230273 refGene_gn=ANKRD24;refGene_tx=XM_023243955.1
A2 3231467 3231512 refGene_gn=ANKRD24;refGene_tx=XM_023243955.1
A2 3231611 3231710 refGene_gn=ANKRD24;refGene_tx=XM_023243955.1
A2 3232627 3232700 refGene_gn=ANKRD24;refGene_tx=XM_023243955.1
A2 3232889 3233008 refGene_gn=ANKRD24;refGene_tx=XM_023243955.1
A2 3233194 3233713 refGene_gn=ANKRD24;refGene_tx=XM_023243955.1
A2 3233747 3234557 refGene_gn=ANKRD24;refGene_tx=XM_023243955.1
A2 3234558 3234759 refGene_gn=ANKRD24;refGene_tx=XM_023243955.1
A2 3237286 3237454 refGene_gn=ANKRD24;refGene_tx=XM_023243955.1
A2 3238788 3238914 refGene_gn=ANKRD24;refGene_tx=XM_023243955.1
A2 3239095 3239161 refGene_gn=ANKRD24;refGene_tx=XM_023243955.1
A2 3239298 3239376 refGene_gn=ANKRD24;refGene_tx=XM_023243955.1
A2 3240733 3240800
refGene_gn=EBI3;refGene_tx=XM_003981744.5;refGene_gn=EBI3;refGene_tx=XM_006928272.4
A2 3241619 3241743
refGene_gn=EBI3;refGene_tx=XM_003981744.5;refGene_gn=EBI3;refGene_tx=XM_006928272.4
A2 3242807 3242986
refGene_gn=EBI3;refGene_tx=XM_003981744.5;refGene_gn=EBI3;refGene_tx=XM_006928272.4
A2 3243768 3243926
refGene_gn=EBI3;refGene_tx=XM_003981744.5;refGene_gn=EBI3;refGene_tx=X

M_006928272.4
A2 3246141 3246294
refGene_gn=EBI3;refGene_tx=XM_003981744.5;refGene_gn=EBI3;refGene_tx=XM_006928272.4
A2 3248290 3248314 refGene_gn=CCDC94;refGene_tx=XM_023243963.1
A2 3250107 3250208 refGene_gn=CCDC94;refGene_tx=XM_023243963.1
A2 3251960 3252105 refGene_gn=CCDC94;refGene_tx=XM_023243963.1
A2 3254161 3254296 refGene_gn=CCDC94;refGene_tx=XM_023243963.1
A2 3256332 3256514 refGene_gn=CCDC94;refGene_tx=XM_023243963.1
A2 3257489 3257613 refGene_gn=CCDC94;refGene_tx=XM_023243963.1
A2 3259971 3260116 refGene_gn=CCDC94;refGene_tx=XM_023243963.1
A2 3260269 3260382 refGene_gn=CCDC94;refGene_tx=XM_023243963.1
A2 3267807 3268104 refGene_gn=SHD;refGene_tx=XM_019815762.2
A2 3269085 3269191 refGene_gn=SHD;refGene_tx=XM_019815762.2
A2 3269282 3269471 refGene_gn=SHD;refGene_tx=XM_019815762.2
A2 3270471 3270595 refGene_gn=SHD;refGene_tx=XM_019815762.2
A2 3271476 3271596 refGene_gn=SHD;refGene_tx=XM_019815762.2
A2 3272453 3272640 refGene_gn=SHD;refGene_tx=XM_019815762.2
A2 3281041 3281056 refGene_gn=FSD1;refGene_tx=XM_023243966.1
A2 3281960 3282056 refGene_gn=FSD1;refGene_tx=XM_023243966.1
A2 3282272 3282404 refGene_gn=FSD1;refGene_tx=XM_023243966.1
A2 3283111 3283213 refGene_gn=FSD1;refGene_tx=XM_023243966.1
A2 3284235 3284258 refGene_gn=FSD1;refGene_tx=XM_023243966.1
A2 3284435 3284557 refGene_gn=FSD1;refGene_tx=XM_023243966.1
A2 3285603 3285813 refGene_gn=FSD1;refGene_tx=XM_023243966.1
A2 3288960 3289059 refGene_gn=FSD1;refGene_tx=XM_023243966.1
A2 3289842 3290002 refGene_gn=FSD1;refGene_tx=XM_023243966.1
A2 3290553 3290633 refGene_gn=FSD1;refGene_tx=XM_023243966.1
A2 3291874 3292126 refGene_gn=FSD1;refGene_tx=XM_023243966.1
A2 3292227 3292316 refGene_gn=FSD1;refGene_tx=XM_023243966.1
A2 3292416 3292527 refGene_gn=FSD1;refGene_tx=XM_023243966.1
A2 3292973 3293062 refGene_gn=STAP2;refGene_tx=XM_023243977.1
A2 3293294 3293369 refGene_gn=STAP2;refGene_tx=XM_023243977.1
A2 3293446 3293548 refGene_gn=STAP2;refGene_tx=XM_023243977.1
A2 3293634 3293798 refGene_gn=STAP2;refGene_tx=XM_023243977.1
A2 3294723 3294781 refGene_gn=STAP2;refGene_tx=XM_023243977.1
A2 3294904 3295007 refGene_gn=STAP2;refGene_tx=XM_023243977.1
A2 3295099 3295169 refGene_gn=STAP2;refGene_tx=XM_023243977.1
A2 3296086 3296221 refGene_gn=STAP2;refGene_tx=XM_023243977.1
A2 3296418 3296519 refGene_gn=STAP2;refGene_tx=XM_023243977.1
A2 3297001 3297058 refGene_gn=STAP2;refGene_tx=XM_023243977.1
A2 3298002 3298125 refGene_gn=STAP2;refGene_tx=XM_023243977.1
A2 3298275 3298347 refGene_gn=STAP2;refGene_tx=XM_023243977.1
A2 3300606 3300708 refGene_gn=STAP2;refGene_tx=XM_023243977.1
A2 3306898 3306905
refGene_gn=MPND;refGene_tx=XM_023243995.1;refGene_gn=MPND;refGene_tx=XM_023244002.1;refGene_gn=MPND;refGene_tx=XM_023244003.1;refGene_gn=MPND;refGene_tx=XM_023244006.1;refGene_gn=MPND;refGene_tx=XM_023244009.1;refGene_gn=MPND;refGene_tx=XM_023244010.1;refGene_gn=MPND;refGene_tx=XM_023244013.1

A2 3307007 3307300
refGene_gn=MPND;refGene_tx=XM_023243995.1;refGene_gn=MPND;refGene_tx=XM_023244002.1;refGene_gn=MPND;refGene_tx=XM_023244003.1;refGene_gn=MPND;refGene_tx=XM_023244006.1;refGene_gn=MPND;refGene_tx=XM_023244009.1;refGene_gn=MPND;refGene_tx=XM_023244010.1;refGene_gn=MPND;refGene_tx=XM_023244013.1
A2 3308035 3308272
refGene_gn=MPND;refGene_tx=XM_023243995.1;refGene_gn=MPND;refGene_tx=XM_023244002.1;refGene_gn=MPND;refGene_tx=XM_023244003.1;refGene_gn=MPND;refGene_tx=XM_023244006.1;refGene_gn=MPND;refGene_tx=XM_023244009.1;refGene_gn=MPND;refGene_tx=XM_023244010.1;refGene_gn=MPND;refGene_tx=XM_023244013.1
A2 3309955 3310088
refGene_gn=MPND;refGene_tx=XM_023243995.1;refGene_gn=MPND;refGene_tx=XM_023244002.1;refGene_gn=MPND;refGene_tx=XM_023244003.1;refGene_gn=MPND;refGene_tx=XM_023244006.1;refGene_gn=MPND;refGene_tx=XM_023244009.1;refGene_gn=MPND;refGene_tx=XM_023244010.1;refGene_gn=MPND;refGene_tx=XM_023244013.1
A2 3310809 3310894
refGene_gn=MPND;refGene_tx=XM_023243995.1;refGene_gn=MPND;refGene_tx=XM_023244002.1;refGene_gn=MPND;refGene_tx=XM_023244003.1;refGene_gn=MPND;refGene_tx=XM_023244006.1;refGene_gn=MPND;refGene_tx=XM_023244009.1;refGene_gn=MPND;refGene_tx=XM_023244010.1;refGene_gn=MPND;refGene_tx=XM_023244013.1
A2 3311065 3311162
refGene_gn=MPND;refGene_tx=XM_023243995.1;refGene_gn=MPND;refGene_tx=XM_023244002.1;refGene_gn=MPND;refGene_tx=XM_023244003.1;refGene_gn=MPND;refGene_tx=XM_023244006.1;refGene_gn=MPND;refGene_tx=XM_023244009.1;refGene_gn=MPND;refGene_tx=XM_023244010.1;refGene_gn=MPND;refGene_tx=XM_023244013.1
A2 3311444 3311517
refGene_gn=MPND;refGene_tx=XM_023243995.1;refGene_gn=MPND;refGene_tx=XM_023244002.1;refGene_gn=MPND;refGene_tx=XM_023244003.1;refGene_gn=MPND;refGene_tx=XM_023244006.1;refGene_gn=MPND;refGene_tx=XM_023244009.1;refGene_gn=MPND;refGene_tx=XM_023244010.1;refGene_gn=MPND;refGene_tx=XM_023244013.1
A2 3311597 3311674
refGene_gn=MPND;refGene_tx=XM_023243995.1;refGene_gn=MPND;refGene_tx=XM_023244002.1;refGene_gn=MPND;refGene_tx=XM_023244003.1;refGene_gn=MPND;refGene_tx=XM_023244006.1;refGene_gn=MPND;refGene_tx=XM_023244009.1;refGene_gn=MPND;refGene_tx=XM_023244010.1;refGene_gn=MPND;refGene_tx=XM_023244013.1
A2 3311841 3312222 refGene_gn=MPND;refGene_tx=XM_023244010.1
A2 3312454 3312641
refGene_gn=MPND;refGene_tx=XM_023243995.1;refGene_gn=MPND;refGene_tx=XM_023244002.1;refGene_gn=MPND;refGene_tx=XM_023244003.1;refGene_gn=MPND;refGene_tx=XM_023244006.1;refGene_gn=MPND;refGene_tx=XM_023244009.1;refGene_gn=MPND;refGene_tx=XM_023244013.1
A2 3312766 3312837
refGene_gn=MPND;refGene_tx=XM_023243995.1;refGene_gn=MPND;refGene_tx=X

M_023244002.1;refGene_gn=MPND;refGene_tx=XM_023244003.1;refGene_gn=MPND;refGene_tx=XM_023244006.1;refGene_gn=MPND;refGene_tx=XM_023244009.1;refGene_gn=MPND;refGene_tx=XM_023244013.1
A2 3313254 3313425
refGene_gn=MPND;refGene_tx=XM_023243995.1;refGene_gn=MPND;refGene_tx=XM_023244002.1;refGene_gn=MPND;refGene_tx=XM_023244003.1;refGene_gn=MPND;refGene_tx=XM_023244006.1;refGene_gn=MPND;refGene_tx=XM_023244009.1;refGene_gn=MPND;refGene_tx=XM_023244013.1
A2 3314105 3314198
refGene_gn=MPND;refGene_tx=XM_023243995.1;refGene_gn=MPND;refGene_tx=XM_023244002.1;refGene_gn=MPND;refGene_tx=XM_023244003.1;refGene_gn=MPND;refGene_tx=XM_023244006.1;refGene_gn=MPND;refGene_tx=XM_023244009.1
A2 3315004 3315091
refGene_gn=MPND;refGene_tx=XM_023243995.1;refGene_gn=MPND;refGene_tx=XM_023244002.1;refGene_gn=MPND;refGene_tx=XM_023244003.1;refGene_gn=MPND;refGene_tx=XM_023244006.1;refGene_gn=MPND;refGene_tx=XM_023244009.1
A2 3316683 3316880 refGene_gn=SH3GL1;refGene_tx=XM_023244015.1
A2 3317300 3317357 refGene_gn=SH3GL1;refGene_tx=XM_023244015.1
A2 3317463 3317588 refGene_gn=SH3GL1;refGene_tx=XM_023244015.1
A2 3318053 3318157 refGene_gn=SH3GL1;refGene_tx=XM_023244015.1
A2 3318464 3318623 refGene_gn=SH3GL1;refGene_tx=XM_023244015.1
A2 3318717 3318851 refGene_gn=SH3GL1;refGene_tx=XM_023244015.1
A2 3319193 3319337 refGene_gn=SH3GL1;refGene_tx=XM_023244015.1
A2 3320236 3320309 refGene_gn=SH3GL1;refGene_tx=XM_023244015.1
A2 3320614 3320683 refGene_gn=SH3GL1;refGene_tx=XM_023244015.1
A2 3346247 3346292 refGene_gn=SH3GL1;refGene_tx=XM_023244015.1
A2 3348981 3349033
refGene_gn=CHAF1A;refGene_tx=XM_023244018.1;refGene_gn=CHAF1A;refGene_tx=XM_023244019.1;refGene_gn=CHAF1A;refGene_tx=XM_023244020.1
A2 3352286 3352337
refGene_gn=CHAF1A;refGene_tx=XM_011287806.3;refGene_gn=CHAF1A;refGene_tx=XM_023244018.1;refGene_gn=CHAF1A;refGene_tx=XM_023244019.1;refGene_gn=CHAF1A;refGene_tx=XM_023244020.1
A2 3353132 3354001
refGene_gn=CHAF1A;refGene_tx=XM_011287806.3;refGene_gn=CHAF1A;refGene_tx=XM_023244018.1;refGene_gn=CHAF1A;refGene_tx=XM_023244019.1;refGene_gn=CHAF1A;refGene_tx=XM_023244020.1
A2 3358882 3358939
refGene_gn=CHAF1A;refGene_tx=XM_011287806.3;refGene_gn=CHAF1A;refGene_tx=XM_023244018.1;refGene_gn=CHAF1A;refGene_tx=XM_023244019.1
A2 3361194 3361424
refGene_gn=CHAF1A;refGene_tx=XM_011287806.3;refGene_gn=CHAF1A;refGene_tx=XM_023244018.1;refGene_gn=CHAF1A;refGene_tx=XM_023244019.1;refGene_gn=CHAF1A;refGene_tx=XM_023244020.1
A2 3361554 3361615
refGene_gn=CHAF1A;refGene_tx=XM_011287806.3;refGene_gn=CHAF1A;refGene_tx=XM_023244018.1;refGene_gn=CHAF1A;refGene_tx=XM_023244019.1;refGene_gn=CHAF1A;refGene_tx=XM_023244020.1
A2 3361930 3361999
refGene_gn=CHAF1A;refGene_tx=XM_011287806.3;refGene_gn=CHAF1A;refGene_

tx=XM_023244018.1;refGene_gn=CHAF1A;refGene_tx=XM_023244019.1;refGene_gn=CHAF1A;refGene_tx=XM_023244020.1
A2 3364794 3365021
refGene_gn=CHAF1A;refGene_tx=XM_011287806.3;refGene_gn=CHAF1A;refGene_tx=XM_023244018.1;refGene_gn=CHAF1A;refGene_tx=XM_023244019.1;refGene_gn=CHAF1A;refGene_tx=XM_023244020.1
A2 3365574 3365743
refGene_gn=CHAF1A;refGene_tx=XM_011287806.3;refGene_gn=CHAF1A;refGene_tx=XM_023244018.1;refGene_gn=CHAF1A;refGene_tx=XM_023244019.1;refGene_gn=CHAF1A;refGene_tx=XM_023244020.1
A2 3365887 3365968
refGene_gn=CHAF1A;refGene_tx=XM_011287806.3;refGene_gn=CHAF1A;refGene_tx=XM_023244018.1;refGene_gn=CHAF1A;refGene_tx=XM_023244019.1;refGene_gn=CHAF1A;refGene_tx=XM_023244020.1
A2 3366429 3366522
refGene_gn=CHAF1A;refGene_tx=XM_011287806.3;refGene_gn=CHAF1A;refGene_tx=XM_023244018.1;refGene_gn=CHAF1A;refGene_tx=XM_023244019.1;refGene_gn=CHAF1A;refGene_tx=XM_023244020.1
A2 3367447 3367700
refGene_gn=CHAF1A;refGene_tx=XM_011287806.3;refGene_gn=CHAF1A;refGene_tx=XM_023244018.1;refGene_gn=CHAF1A;refGene_tx=XM_023244019.1;refGene_gn=CHAF1A;refGene_tx=XM_023244020.1
A2 3368358 3368849
refGene_gn=CHAF1A;refGene_tx=XM_011287806.3;refGene_gn=CHAF1A;refGene_tx=XM_023244018.1;refGene_gn=CHAF1A;refGene_tx=XM_023244019.1;refGene_gn=CHAF1A;refGene_tx=XM_023244020.1
A2 3373740 3373837
refGene_gn=CHAF1A;refGene_tx=XM_011287806.3;refGene_gn=CHAF1A;refGene_tx=XM_023244018.1;refGene_gn=CHAF1A;refGene_tx=XM_023244019.1;refGene_gn=CHAF1A;refGene_tx=XM_023244020.1
A2 3374528 3374629
refGene_gn=CHAF1A;refGene_tx=XM_011287806.3;refGene_gn=CHAF1A;refGene_tx=XM_023244018.1;refGene_gn=CHAF1A;refGene_tx=XM_023244019.1;refGene_gn=CHAF1A;refGene_tx=XM_023244020.1
A2 3377347 3377476
refGene_gn=UBXN6;refGene_tx=XM_023244031.1;refGene_gn=UBXN6;refGene_tx=XM_023244036.1
A2 3377803 3377952
refGene_gn=UBXN6;refGene_tx=XM_023244031.1;refGene_gn=UBXN6;refGene_tx=XM_023244036.1
A2 3378163 3378514
refGene_gn=UBXN6;refGene_tx=XM_023244031.1;refGene_gn=UBXN6;refGene_tx=XM_023244036.1
A2 3378639 3378724
refGene_gn=UBXN6;refGene_tx=XM_023244031.1;refGene_gn=UBXN6;refGene_tx=XM_023244036.1
A2 3379293 3379369
refGene_gn=UBXN6;refGene_tx=XM_023244031.1;refGene_gn=UBXN6;refGene_tx=XM_023244036.1
A2 3379895 3379993

refGene_gn=UBXN6;refGene_tx=XM_023244031.1;refGene_gn=UBXN6;refGene_tx=XM_023244036.1
A2 3382718 3382847
refGene_gn=UBXN6;refGene_tx=XM_023244031.1;refGene_gn=UBXN6;refGene_tx=XM_023244036.1
A2 3384131 3384196
refGene_gn=UBXN6;refGene_tx=XM_023244031.1;refGene_gn=UBXN6;refGene_tx=XM_023244036.1
A2 3384521 3384685
refGene_gn=UBXN6;refGene_tx=XM_023244031.1;refGene_gn=UBXN6;refGene_tx=XM_023244036.1
A2 3385603 3385608 refGene_gn=UBXN6;refGene_tx=XM_023244036.1
A2 3387175 3387258 refGene_gn=UBXN6;refGene_tx=XM_023244031.1
A2 3395255 3395327
refGene_gn=HDGFL2;refGene_tx=XM_023244045.1;refGene_gn=HDGFL2;refGene_tx=XM_023244050.1
A2 3397724 3397801
refGene_gn=HDGFL2;refGene_tx=XM_023244045.1;refGene_gn=HDGFL2;refGene_tx=XM_023244050.1
A2 3397894 3398033
refGene_gn=HDGFL2;refGene_tx=XM_023244045.1;refGene_gn=HDGFL2;refGene_tx=XM_023244050.1
A2 3404310 3404502
refGene_gn=HDGFL2;refGene_tx=XM_023244045.1;refGene_gn=HDGFL2;refGene_tx=XM_023244050.1
A2 3410126 3410243
refGene_gn=HDGFL2;refGene_tx=XM_023244045.1;refGene_gn=HDGFL2;refGene_tx=XM_023244050.1
A2 3410355 3410427
refGene_gn=HDGFL2;refGene_tx=XM_023244045.1;refGene_gn=HDGFL2;refGene_tx=XM_023244050.1
A2 3411362 3411528
refGene_gn=HDGFL2;refGene_tx=XM_023244045.1;refGene_gn=HDGFL2;refGene_tx=XM_023244050.1
A2 3411623 3411699
refGene_gn=HDGFL2;refGene_tx=XM_023244045.1;refGene_gn=HDGFL2;refGene_tx=XM_023244050.1
A2 3411776 3412086
refGene_gn=HDGFL2;refGene_tx=XM_023244045.1;refGene_gn=HDGFL2;refGene_tx=XM_023244050.1
A2 3413207 3413305
refGene_gn=HDGFL2;refGene_tx=XM_023244045.1;refGene_gn=HDGFL2;refGene_tx=XM_023244050.1
A2 3413924 3413998
refGene_gn=HDGFL2;refGene_tx=XM_023244045.1;refGene_gn=HDGFL2;refGene_tx=XM_023244050.1
A2 3414142 3414213
refGene_gn=HDGFL2;refGene_tx=XM_023244045.1;refGene_gn=HDGFL2;refGene_tx=XM_023244050.1
A2 3414520 3414622

refGene_gn=HDGFL2;refGene_tx=XM_023244045.1;refGene_gn=HDGFL2;refGene_tx=XM_023244050.1
A2 3414773 3414987
refGene_gn=HDGFL2;refGene_tx=XM_023244045.1;refGene_gn=HDGFL2;refGene_tx=XM_023244050.1
A2 3415589 3415722
refGene_gn=HDGFL2;refGene_tx=XM_023244045.1;refGene_gn=HDGFL2;refGene_tx=XM_023244050.1
A2 3416070 3416173
refGene_gn=HDGFL2;refGene_tx=XM_023244045.1;refGene_gn=HDGFL2;refGene_tx=XM_023244050.1
A2 3418264 3418597 refGene_gn=PLIN4;refGene_tx=XM_023244052.1
A2 3418669 3418756 refGene_gn=PLIN4;refGene_tx=XM_023244052.1
A2 3420089 3420277 refGene_gn=PLIN4;refGene_tx=XM_023244052.1
A2 3420375 3421305 refGene_gn=PLIN4;refGene_tx=XM_023244052.1
A2 3424287 3425254 refGene_gn=PLIN4;refGene_tx=XM_023244052.1
A2 3427154 3427213 refGene_gn=PLIN4;refGene_tx=XM_023244052.1
A2 3428048 3428184 refGene_gn=PLIN4;refGene_tx=XM_023244052.1
A2 3428687 3428738 refGene_gn=PLIN4;refGene_tx=XM_023244052.1
A2 3432343 3432874
refGene_gn=PLIN5;refGene_tx=XM_011287787.2;refGene_gn=PLIN5;refGene_tx=XM_011287794.3;refGene_gn=PLIN5;refGene_tx=XM_019815807.2
A2 3433126 3433243
refGene_gn=PLIN5;refGene_tx=XM_011287787.2;refGene_gn=PLIN5;refGene_tx=XM_011287794.3;refGene_gn=PLIN5;refGene_tx=XM_019815807.2
A2 3434653 3434853
refGene_gn=PLIN5;refGene_tx=XM_011287787.2;refGene_gn=PLIN5;refGene_tx=XM_011287794.3;refGene_gn=PLIN5;refGene_tx=XM_019815807.2
A2 3436185 3436366
refGene_gn=PLIN5;refGene_tx=XM_011287787.2;refGene_gn=PLIN5;refGene_tx=XM_011287794.3;refGene_gn=PLIN5;refGene_tx=XM_019815807.2
A2 3436866 3436949
refGene_gn=PLIN5;refGene_tx=XM_011287787.2;refGene_gn=PLIN5;refGene_tx=XM_011287794.3;refGene_gn=PLIN5;refGene_tx=XM_019815807.2
A2 3437518 3437714
refGene_gn=PLIN5;refGene_tx=XM_011287787.2;refGene_gn=PLIN5;refGene_tx=XM_011287794.3;refGene_gn=PLIN5;refGene_tx=XM_019815807.2
A2 3438246 3438321
refGene_gn=PLIN5;refGene_tx=XM_011287787.2;refGene_gn=PLIN5;refGene_tx=XM_011287794.3;refGene_gn=PLIN5;refGene_tx=XM_019815807.2
A2 3440595 3440790 refGene_gn=LRG1;refGene_tx=XM_003981754.5
A2 3440791 3441605 refGene_gn=LRG1;refGene_tx=XM_003981754.5
A2 3442536 3442589 refGene_gn=LRG1;refGene_tx=XM_003981754.5
A2 3447247 3447481 refGene_gn=SEMA6B;refGene_tx=XM_023244067.1
A2 3447483 3448208 refGene_gn=SEMA6B;refGene_tx=XM_023244067.1
A2 3448555 3448614 refGene_gn=SEMA6B;refGene_tx=XM_023244067.1
A2 3448722 3448800 refGene_gn=SEMA6B;refGene_tx=XM_023244067.1
A2 3449407 3449554 refGene_gn=SEMA6B;refGene_tx=XM_023244067.1
A2 3449631 3449814 refGene_gn=SEMA6B;refGene_tx=XM_023244067.1
A2 3449986 3450136 refGene_gn=SEMA6B;refGene_tx=XM_023244067.1

A2 3450361 3450493 refGene_gn=SEMA6B;refGene_tx=XM_023244067.1
A2 3450971 3451189 refGene_gn=SEMA6B;refGene_tx=XM_023244067.1
A2 3451813 3451902 refGene_gn=SEMA6B;refGene_tx=XM_023244067.1
A2 3452382 3452502 refGene_gn=SEMA6B;refGene_tx=XM_023244067.1
A2 3452849 3452940 refGene_gn=SEMA6B;refGene_tx=XM_023244067.1
A2 3453350 3453452 refGene_gn=SEMA6B;refGene_tx=XM_023244067.1
A2 3454034 3454097 refGene_gn=SEMA6B;refGene_tx=XM_023244067.1
A2 3454252 3454313 refGene_gn=SEMA6B;refGene_tx=XM_023244067.1
A2 3455875 3455999 refGene_gn=SEMA6B;refGene_tx=XM_023244067.1
A2 3456143 3456264 refGene_gn=SEMA6B;refGene_tx=XM_023244067.1
A2 3506674 3506765
refGene_gn=TNFAIP8L1;refGene_tx=XM_019815907.2
A2 3517787 3518353
refGene_gn=TNFAIP8L1;refGene_tx=XM_006928283.4;refGene_gn=TNFAIP8L1;refGene_tx=XM_011287779.3;refGene_gn=TNFAIP8L1;refGene_tx=XM_019815907.2
A2 3522030 3522110 refGene_gn=MYDGF;refGene_tx=XM_019815898.2
A2 3523550 3523623 refGene_gn=MYDGF;refGene_tx=XM_019815898.2
A2 3524887 3524969 refGene_gn=MYDGF;refGene_tx=XM_019815898.2
A2 3525407 3525675 refGene_gn=MYDGF;refGene_tx=XM_019815896.2
A2 3526397 3526459
refGene_gn=MYDGF;refGene_tx=XM_019815896.2;refGene_gn=MYDGF;refGene_tx=XM_019815898.2
A2 3532161 3532212
refGene_gn=MYDGF;refGene_tx=XM_019815896.2;refGene_gn=MYDGF;refGene_tx=XM_019815898.2
A2 3533523 3533838
refGene_gn=MYDGF;refGene_tx=XM_019815896.2;refGene_gn=MYDGF;refGene_tx=XM_019815898.2
A2 3533964 3533991
refGene_gn=LOC102902301;refGene_tx=XM_006928282.4
A2 3534405 3534474
refGene_gn=LOC102902301;refGene_tx=XM_019815850.2
A2 3534875 3535037
refGene_gn=LOC102902301;refGene_tx=XM_019815849.2;refGene_gn=LOC102902301;refGene_tx=XM_019815856.2;refGene_gn=LOC102902301;refGene_tx=XM_019815870.2;refGene_gn=LOC102902301;refGene_tx=XM_019815873.2
A2 3535845 3536072
refGene_gn=LOC102902301;refGene_tx=XM_006928282.4;refGene_gn=LOC102902301;refGene_tx=XM_019815849.2;refGene_gn=LOC102902301;refGene_tx=XM_019815850.2;refGene_gn=LOC102902301;refGene_tx=XM_019815856.2;refGene_gn=LOC102902301;refGene_tx=XM_019815870.2;refGene_gn=LOC102902301;refGene_tx=XM_019815873.2
A2 3536668 3537029
refGene_gn=LOC102902301;refGene_tx=XM_006928282.4;refGene_gn=LOC102902301;refGene_tx=XM_019815849.2;refGene_gn=LOC102902301;refGene_tx=XM_019815850.2;refGene_gn=LOC102902301;refGene_tx=XM_019815856.2
A2 3539661 3539754
refGene_gn=DPP9;refGene_tx=XM_011287768.3;refGene_gn=DPP9;refGene_tx=XM_023250867.1;refGene_gn=DPP9;refGene_tx=XM_023250869.1;refGene_gn=DPP9;refGene_tx=XM_023250870.1;refGene_gn=DPP9;refGene_tx=XM_023250871.1

A2 3542533 3542645
refGene_gn=DPP9;refGene_tx=XM_011287768.3;refGene_gn=DPP9;refGene_tx=XM_023250867.1;refGene_gn=DPP9;refGene_tx=XM_023250869.1;refGene_gn=DPP9;refGene_tx=XM_023250870.1;refGene_gn=DPP9;refGene_tx=XM_023250871.1
A2 3543957 3544100
refGene_gn=DPP9;refGene_tx=XM_011287768.3;refGene_gn=DPP9;refGene_tx=XM_023250867.1;refGene_gn=DPP9;refGene_tx=XM_023250869.1;refGene_gn=DPP9;refGene_tx=XM_023250870.1;refGene_gn=DPP9;refGene_tx=XM_023250871.1
A2 3544582 3544687
refGene_gn=DPP9;refGene_tx=XM_011287771.3;refGene_gn=DPP9;refGene_tx=XM_011287773.3
A2 3544807 3544960
refGene_gn=DPP9;refGene_tx=XM_011287768.3;refGene_gn=DPP9;refGene_tx=XM_011287771.3;refGene_gn=DPP9;refGene_tx=XM_011287773.3;refGene_gn=DPP9;refGene_tx=XM_023250867.1;refGene_gn=DPP9;refGene_tx=XM_023250869.1;refGene_gn=DPP9;refGene_tx=XM_023250870.1;refGene_gn=DPP9;refGene_tx=XM_023250871.1
A2 3545836 3545983
refGene_gn=DPP9;refGene_tx=XM_011287768.3;refGene_gn=DPP9;refGene_tx=XM_011287771.3;refGene_gn=DPP9;refGene_tx=XM_011287773.3;refGene_gn=DPP9;refGene_tx=XM_023250867.1;refGene_gn=DPP9;refGene_tx=XM_023250869.1;refGene_gn=DPP9;refGene_tx=XM_023250870.1;refGene_gn=DPP9;refGene_tx=XM_023250871.1
A2 3546958 3547104
refGene_gn=DPP9;refGene_tx=XM_011287768.3;refGene_gn=DPP9;refGene_tx=XM_011287771.3;refGene_gn=DPP9;refGene_tx=XM_011287773.3;refGene_gn=DPP9;refGene_tx=XM_023250867.1;refGene_gn=DPP9;refGene_tx=XM_023250869.1;refGene_gn=DPP9;refGene_tx=XM_023250870.1;refGene_gn=DPP9;refGene_tx=XM_023250871.1
A2 3548694 3548830
refGene_gn=DPP9;refGene_tx=XM_011287768.3;refGene_gn=DPP9;refGene_tx=XM_011287771.3;refGene_gn=DPP9;refGene_tx=XM_011287773.3;refGene_gn=DPP9;refGene_tx=XM_023250867.1;refGene_gn=DPP9;refGene_tx=XM_023250869.1;refGene_gn=DPP9;refGene_tx=XM_023250870.1;refGene_gn=DPP9;refGene_tx=XM_023250871.1
A2 3549360 3549513
refGene_gn=DPP9;refGene_tx=XM_011287768.3;refGene_gn=DPP9;refGene_tx=XM_011287771.3;refGene_gn=DPP9;refGene_tx=XM_011287773.3;refGene_gn=DPP9;refGene_tx=XM_023250867.1;refGene_gn=DPP9;refGene_tx=XM_023250869.1;refGene_gn=DPP9;refGene_tx=XM_023250870.1;refGene_gn=DPP9;refGene_tx=XM_023250871.1
A2 3552469 3552549
refGene_gn=DPP9;refGene_tx=XM_011287768.3;refGene_gn=DPP9;refGene_tx=XM_011287771.3;refGene_gn=DPP9;refGene_tx=XM_011287773.3;refGene_gn=DPP9;refGene_tx=XM_023250867.1;refGene_gn=DPP9;refGene_tx=XM_023250869.1;refGene_gn=DPP9;refGene_tx=XM_023250870.1;refGene_gn=DPP9;refGene_tx=XM_023250871.1
A2 3553540 3553589
refGene_gn=LOC102902301;refGene_tx=XM_019815873.2
A2 3554541 3554635

refGene_gn=LOC102902301;refGene_tx=XM_019815870.2
A2	3555601	3555764
refGene_gn=DPP9;refGene_tx=XM_011287768.3;refGene_gn=DPP9;refGene_tx=XM_011287771.3;refGene_gn=DPP9;refGene_tx=XM_011287773.3;refGene_gn=DPP9;refGene_tx=XM_023250867.1;refGene_gn=DPP9;refGene_tx=XM_023250869.1;refGene_gn=DPP9;refGene_tx=XM_023250870.1;refGene_gn=DPP9;refGene_tx=XM_023250871.1
A2	3555965	3556143
refGene_gn=DPP9;refGene_tx=XM_011287768.3;refGene_gn=DPP9;refGene_tx=XM_011287771.3;refGene_gn=DPP9;refGene_tx=XM_011287773.3;refGene_gn=DPP9;refGene_tx=XM_023250867.1;refGene_gn=DPP9;refGene_tx=XM_023250869.1;refGene_gn=DPP9;refGene_tx=XM_023250870.1;refGene_gn=DPP9;refGene_tx=XM_023250871.1
A2	3557873	3557974
refGene_gn=DPP9;refGene_tx=XM_011287768.3;refGene_gn=DPP9;refGene_tx=XM_011287771.3;refGene_gn=DPP9;refGene_tx=XM_011287773.3;refGene_gn=DPP9;refGene_tx=XM_023250867.1;refGene_gn=DPP9;refGene_tx=XM_023250869.1;refGene_gn=DPP9;refGene_tx=XM_023250870.1;refGene_gn=DPP9;refGene_tx=XM_023250871.1
A2	3559369	3559431
refGene_gn=DPP9;refGene_tx=XM_011287768.3;refGene_gn=DPP9;refGene_tx=XM_011287771.3;refGene_gn=DPP9;refGene_tx=XM_011287773.3;refGene_gn=DPP9;refGene_tx=XM_023250867.1;refGene_gn=DPP9;refGene_tx=XM_023250869.1;refGene_gn=DPP9;refGene_tx=XM_023250870.1;refGene_gn=DPP9;refGene_tx=XM_023250871.1
A2	3560512	3560641
refGene_gn=DPP9;refGene_tx=XM_011287768.3;refGene_gn=DPP9;refGene_tx=XM_011287771.3;refGene_gn=DPP9;refGene_tx=XM_011287773.3;refGene_gn=DPP9;refGene_tx=XM_023250867.1;refGene_gn=DPP9;refGene_tx=XM_023250869.1;refGene_gn=DPP9;refGene_tx=XM_023250870.1;refGene_gn=DPP9;refGene_tx=XM_023250871.1
A2	3563896	3564010
refGene_gn=DPP9;refGene_tx=XM_011287768.3;refGene_gn=DPP9;refGene_tx=XM_011287771.3;refGene_gn=DPP9;refGene_tx=XM_011287773.3;refGene_gn=DPP9;refGene_tx=XM_023250867.1;refGene_gn=DPP9;refGene_tx=XM_023250869.1;refGene_gn=DPP9;refGene_tx=XM_023250870.1;refGene_gn=DPP9;refGene_tx=XM_023250871.1
A2	3564720	3564889
refGene_gn=DPP9;refGene_tx=XM_011287768.3;refGene_gn=DPP9;refGene_tx=XM_011287771.3;refGene_gn=DPP9;refGene_tx=XM_011287773.3;refGene_gn=DPP9;refGene_tx=XM_023250867.1;refGene_gn=DPP9;refGene_tx=XM_023250869.1;refGene_gn=DPP9;refGene_tx=XM_023250870.1;refGene_gn=DPP9;refGene_tx=XM_023250871.1
A2	3564971	3565145
refGene_gn=DPP9;refGene_tx=XM_011287768.3;refGene_gn=DPP9;refGene_tx=XM_011287771.3;refGene_gn=DPP9;refGene_tx=XM_011287773.3;refGene_gn=DPP9;refGene_tx=XM_023250867.1;refGene_gn=DPP9;refGene_tx=XM_023250869.1;refGene_gn=DPP9;refGene_tx=XM_023250870.1;refGene_gn=DPP9;refGene_tx=XM_023250871.1
A2	3566195	3566308

refGene_gn=DPP9;refGene_tx=XM_011287768.3;refGene_gn=DPP9;refGene_tx=XM_011287771.3;refGene_gn=DPP9;refGene_tx=XM_011287773.3;refGene_gn=DPP9;refGene_tx=XM_023250867.1;refGene_gn=DPP9;refGene_tx=XM_023250869.1;refGene_gn=DPP9;refGene_tx=XM_023250870.1;refGene_gn=DPP9;refGene_tx=XM_023250871.1
A2 3571336 3571590
refGene_gn=DPP9;refGene_tx=XM_011287768.3;refGene_gn=DPP9;refGene_tx=XM_011287771.3;refGene_gn=DPP9;refGene_tx=XM_011287773.3;refGene_gn=DPP9;refGene_tx=XM_023250867.1;refGene_gn=DPP9;refGene_tx=XM_023250869.1;refGene_gn=DPP9;refGene_tx=XM_023250870.1;refGene_gn=DPP9;refGene_tx=XM_023250871.1
A2 3573782 3573850
refGene_gn=DPP9;refGene_tx=XM_011287768.3;refGene_gn=DPP9;refGene_tx=XM_011287771.3;refGene_gn=DPP9;refGene_tx=XM_011287773.3
A2 3575459 3575515
refGene_gn=DPP9;refGene_tx=XM_023250867.1;refGene_gn=DPP9;refGene_tx=XM_023250869.1
A2 3635221 3637189 refGene_gn=FEM1A;refGene_tx=XM_019815938.2
A2 3649610 3651773 refGene_gn=TICAM1;refGene_tx=XM_023250878.1
A2 3669020 3669365
refGene_gn=PLIN3;refGene_tx=XM_023250880.1;refGene_gn=PLIN3;refGene_tx=XM_023250881.1;refGene_gn=PLIN3;refGene_tx=XM_023250882.1
A2 3671188 3671314
refGene_gn=PLIN3;refGene_tx=XM_023250880.1;refGene_gn=PLIN3;refGene_tx=XM_023250881.1;refGene_gn=PLIN3;refGene_tx=XM_023250882.1
A2 3672673 3672873
refGene_gn=PLIN3;refGene_tx=XM_023250880.1;refGene_gn=PLIN3;refGene_tx=XM_023250881.1;refGene_gn=PLIN3;refGene_tx=XM_023250882.1
A2 3677125 3677423
refGene_gn=PLIN3;refGene_tx=XM_023250880.1;refGene_gn=PLIN3;refGene_tx=XM_023250881.1;refGene_gn=PLIN3;refGene_tx=XM_023250882.1
A2 3680654 3680737
refGene_gn=PLIN3;refGene_tx=XM_023250880.1;refGene_gn=PLIN3;refGene_tx=XM_023250881.1;refGene_gn=PLIN3;refGene_tx=XM_023250882.1
A2 3680876 3681075
refGene_gn=PLIN3;refGene_tx=XM_023250880.1;refGene_gn=PLIN3;refGene_tx=XM_023250881.1;refGene_gn=PLIN3;refGene_tx=XM_023250882.1
A2 3684625 3684709
refGene_gn=PLIN3;refGene_tx=XM_023250880.1;refGene_gn=PLIN3;refGene_tx=XM_023250881.1;refGene_gn=PLIN3;refGene_tx=XM_023250882.1
A2 3700247 3700775
refGene_gn=ARRDC5;refGene_tx=XM_003981758.5;refGene_gn=ARRDC5;refGene_tx=XM_019816023.2;refGene_gn=ARRDC5;refGene_tx=XM_019816027.2
A2 3710948 3711154
refGene_gn=ARRDC5;refGene_tx=XM_003981758.5;refGene_gn=ARRDC5;refGene_tx=XM_019816023.2;refGene_gn=ARRDC5;refGene_tx=XM_019816027.2
A2 3712813 3713165 refGene_gn=ARRDC5;refGene_tx=XM_003981758.5
A2 3716701 3716858
refGene_gn=ARRDC5;refGene_tx=XM_019816023.2;refGene_gn=ARRDC5;refGene_tx=XM_019816027.2

A2 3717072 3717225
refGene_gn=LOC101091764;refGene_tx=XM_023250857.1;refGene_gn=LOC101091764;refGene_tx=XM_023250858.1;refGene_gn=LOC101091764;refGene_tx=XM_023250859.1;refGene_gn=LOC101091764;refGene_tx=XM_023250860.1;refGene_gn=LOC101091764;refGene_tx=XM_023250861.1;refGene_gn=LOC101091764;refGene_tx=XM_023250862.1;refGene_gn=LOC101091764;refGene_tx=XM_023250863.1;refGene_gn=LOC101091764;refGene_tx=XM_023250866.1
A2 3725459 3725723
refGene_gn=LOC101091764;refGene_tx=XM_023250857.1;refGene_gn=LOC101091764;refGene_tx=XM_023250858.1;refGene_gn=LOC101091764;refGene_tx=XM_023250859.1;refGene_gn=LOC101091764;refGene_tx=XM_023250860.1;refGene_gn=LOC101091764;refGene_tx=XM_023250861.1;refGene_gn=LOC101091764;refGene_tx=XM_023250862.1;refGene_gn=LOC101091764;refGene_tx=XM_023250863.1;refGene_gn=LOC101091764;refGene_tx=XM_023250866.1
A2 3726474 3726635
refGene_gn=LOC101091764;refGene_tx=XM_023250857.1;refGene_gn=LOC101091764;refGene_tx=XM_023250858.1;refGene_gn=LOC101091764;refGene_tx=XM_023250859.1;refGene_gn=LOC101091764;refGene_tx=XM_023250860.1;refGene_gn=LOC101091764;refGene_tx=XM_023250861.1;refGene_gn=LOC101091764;refGene_tx=XM_023250862.1;refGene_gn=LOC101091764;refGene_tx=XM_023250863.1;refGene_gn=LOC101091764;refGene_tx=XM_023250866.1
A2 3730227 3730443
refGene_gn=LOC101091764;refGene_tx=XM_023250857.1;refGene_gn=LOC101091764;refGene_tx=XM_023250858.1;refGene_gn=LOC101091764;refGene_tx=XM_023250859.1;refGene_gn=LOC101091764;refGene_tx=XM_023250860.1;refGene_gn=LOC101091764;refGene_tx=XM_023250861.1;refGene_gn=LOC101091764;refGene_tx=XM_023250862.1;refGene_gn=LOC101091764;refGene_tx=XM_023250863.1;refGene_gn=LOC101091764;refGene_tx=XM_023250866.1
A2 3736368 3736469
refGene_gn=LOC101091764;refGene_tx=XM_023250857.1;refGene_gn=LOC101091764;refGene_tx=XM_023250858.1;refGene_gn=LOC101091764;refGene_tx=XM_023250859.1;refGene_gn=LOC101091764;refGene_tx=XM_023250860.1;refGene_gn=LOC101091764;refGene_tx=XM_023250861.1;refGene_gn=LOC101091764;refGene_tx=XM_023250862.1;refGene_gn=LOC101091764;refGene_tx=XM_023250863.1;refGene_gn=LOC101091764;refGene_tx=XM_023250866.1
A2 3736588 3736775
refGene_gn=LOC101091764;refGene_tx=XM_023250857.1;refGene_gn=LOC101091764;refGene_tx=XM_023250858.1;refGene_gn=LOC101091764;refGene_tx=XM_023250859.1;refGene_gn=LOC101091764;refGene_tx=XM_023250860.1;refGene_gn=LOC101091764;refGene_tx=XM_023250861.1;refGene_gn=LOC101091764;refGene_tx=XM_023250862.1;refGene_gn=LOC101091764;refGene_tx=XM_023250863.1;refGene_gn=LOC101091764;refGene_tx=XM_023250866.1
A2 3737382 3737506
refGene_gn=LOC101091764;refGene_tx=XM_023250857.1;refGene_gn=LOC101091764;refGene_tx=XM_023250858.1;refGene_gn=LOC101091764;refGene_tx=XM_023250859.1;refGene_gn=LOC101091764;refGene_tx=XM_023250860.1;refGene_gn=LOC101091764;refGene_tx=XM_023250861.1;refGene_gn=LOC101091764;refGene_tx=XM_023250862.1;refGene_gn=LOC101091764;refGene_tx=XM_023250863.1;refGene_gn=LOC101091764;refGene_tx=XM_023250866.1
A2 3737736 3737844

refGene_gn=LOC101091764;refGene_tx=XM_023250857.1;refGene_gn=LOC101091764;refGene_tx=XM_023250858.1;refGene_gn=LOC101091764;refGene_tx=XM_023250859.1;refGene_gn=LOC101091764;refGene_tx=XM_023250860.1;refGene_gn=LOC101091764;refGene_tx=XM_023250861.1;refGene_gn=LOC101091764;refGene_tx=XM_023250862.1;refGene_gn=LOC101091764;refGene_tx=XM_023250863.1;refGene_gn=LOC101091764;refGene_tx=XM_023250866.1
A2 3738126 3738231
refGene_gn=LOC101091764;refGene_tx=XM_023250857.1;refGene_gn=LOC101091764;refGene_tx=XM_023250858.1;refGene_gn=LOC101091764;refGene_tx=XM_023250859.1;refGene_gn=LOC101091764;refGene_tx=XM_023250860.1;refGene_gn=LOC101091764;refGene_tx=XM_023250861.1;refGene_gn=LOC101091764;refGene_tx=XM_023250862.1;refGene_gn=LOC101091764;refGene_tx=XM_023250863.1;refGene_gn=LOC101091764;refGene_tx=XM_023250866.1
A2 3738984 3739091
refGene_gn=LOC101091764;refGene_tx=XM_023250857.1;refGene_gn=LOC101091764;refGene_tx=XM_023250858.1;refGene_gn=LOC101091764;refGene_tx=XM_023250859.1;refGene_gn=LOC101091764;refGene_tx=XM_023250860.1;refGene_gn=LOC101091764;refGene_tx=XM_023250861.1;refGene_gn=LOC101091764;refGene_tx=XM_023250862.1;refGene_gn=LOC101091764;refGene_tx=XM_023250863.1;refGene_gn=LOC101091764;refGene_tx=XM_023250866.1
A2 3742282 3742445
refGene_gn=LOC101091764;refGene_tx=XM_023250857.1;refGene_gn=LOC101091764;refGene_tx=XM_023250858.1;refGene_gn=LOC101091764;refGene_tx=XM_023250859.1;refGene_gn=LOC101091764;refGene_tx=XM_023250860.1;refGene_gn=LOC101091764;refGene_tx=XM_023250861.1;refGene_gn=LOC101091764;refGene_tx=XM_023250862.1;refGene_gn=LOC101091764;refGene_tx=XM_023250863.1;refGene_gn=LOC101091764;refGene_tx=XM_023250866.1
A2 3742526 3742664
refGene_gn=LOC101091764;refGene_tx=XM_023250857.1;refGene_gn=LOC101091764;refGene_tx=XM_023250858.1;refGene_gn=LOC101091764;refGene_tx=XM_023250859.1;refGene_gn=LOC101091764;refGene_tx=XM_023250860.1;refGene_gn=LOC101091764;refGene_tx=XM_023250861.1;refGene_gn=LOC101091764;refGene_tx=XM_023250862.1;refGene_gn=LOC101091764;refGene_tx=XM_023250863.1;refGene_gn=LOC101091764;refGene_tx=XM_023250866.1
A2 3744360 3744526
refGene_gn=LOC101091764;refGene_tx=XM_023250857.1;refGene_gn=LOC101091764;refGene_tx=XM_023250858.1;refGene_gn=LOC101091764;refGene_tx=XM_023250859.1;refGene_gn=LOC101091764;refGene_tx=XM_023250860.1;refGene_gn=LOC101091764;refGene_tx=XM_023250861.1;refGene_gn=LOC101091764;refGene_tx=XM_023250862.1;refGene_gn=LOC101091764;refGene_tx=XM_023250863.1;refGene_gn=LOC101091764;refGene_tx=XM_023250866.1
A2 3744601 3744756
refGene_gn=LOC101091764;refGene_tx=XM_023250857.1;refGene_gn=LOC101091764;refGene_tx=XM_023250858.1;refGene_gn=LOC101091764;refGene_tx=XM_023250859.1;refGene_gn=LOC101091764;refGene_tx=XM_023250860.1;refGene_gn=LOC101091764;refGene_tx=XM_023250861.1;refGene_gn=LOC101091764;refGene_tx=XM_023250862.1;refGene_gn=LOC101091764;refGene_tx=XM_023250863.1;refGene_gn=LOC101091764;refGene_tx=XM_023250866.1
A2 3746438 3746543
refGene_gn=LOC101091764;refGene_tx=XM_023250857.1;refGene_gn=LOC101091

764;refGene_tx=XM_023250858.1;refGene_gn=LOC101091764;refGene_tx=XM_02
3250859.1;refGene_gn=LOC101091764;refGene_tx=XM_023250860.1;refGene_gn
=LOC101091764;refGene_tx=XM_023250861.1;refGene_gn=LOC101091764;refGen
e_tx=XM_023250862.1;refGene_gn=LOC101091764;refGene_tx=XM_023250863.1;
refGene_gn=LOC101091764;refGene_tx=XM_023250866.1
A2 3749676 3749823
refGene_gn=LOC101091764;refGene_tx=XM_023250866.1
A2 3790712 3790810
refGene_gn=LOC101091764;refGene_tx=XM_023250857.1;refGene_gn=LOC101091
764;refGene_tx=XM_023250858.1;refGene_gn=LOC101091764;refGene_tx=XM_02
3250859.1;refGene_gn=LOC101091764;refGene_tx=XM_023250860.1;refGene_gn
=LOC101091764;refGene_tx=XM_023250861.1;refGene_gn=LOC101091764;refGen
e_tx=XM_023250862.1;refGene_gn=LOC101091764;refGene_tx=XM_023250863.1
A2 3803583 3803749
refGene_gn=LOC101091764;refGene_tx=XM_023250857.1;refGene_gn=LOC101091
764;refGene_tx=XM_023250858.1;refGene_gn=LOC101091764;refGene_tx=XM_02
3250859.1;refGene_gn=LOC101091764;refGene_tx=XM_023250860.1;refGene_gn
=LOC101091764;refGene_tx=XM_023250861.1;refGene_gn=LOC101091764;refGen
e_tx=XM_023250862.1;refGene_gn=LOC101091764;refGene_tx=XM_023250863.1;
refGene_gn=LOC101091764;refGene_tx=XM_023250864.1;refGene_gn=LOC101091
764;refGene_tx=XM_023250865.1
A2 3808912 3809088
refGene_gn=LOC101091764;refGene_tx=XM_023250857.1;refGene_gn=LOC101091
764;refGene_tx=XM_023250858.1;refGene_gn=LOC101091764;refGene_tx=XM_02
3250859.1;refGene_gn=LOC101091764;refGene_tx=XM_023250860.1;refGene_gn
=LOC101091764;refGene_tx=XM_023250861.1;refGene_gn=LOC101091764;refGen
e_tx=XM_023250862.1;refGene_gn=LOC101091764;refGene_tx=XM_023250863.1;
refGene_gn=LOC101091764;refGene_tx=XM_023250864.1;refGene_gn=LOC101091
764;refGene_tx=XM_023250865.1
A2 3809592 3809707
refGene_gn=LOC101091764;refGene_tx=XM_023250857.1;refGene_gn=LOC101091
764;refGene_tx=XM_023250858.1;refGene_gn=LOC101091764;refGene_tx=XM_02
3250859.1;refGene_gn=LOC101091764;refGene_tx=XM_023250860.1;refGene_gn
=LOC101091764;refGene_tx=XM_023250861.1;refGene_gn=LOC101091764;refGen
e_tx=XM_023250862.1;refGene_gn=LOC101091764;refGene_tx=XM_023250863.1;
refGene_gn=LOC101091764;refGene_tx=XM_023250864.1;refGene_gn=LOC101091
764;refGene_tx=XM_023250865.1
A2 3812540 3812734
refGene_gn=LOC101091764;refGene_tx=XM_023250857.1;refGene_gn=LOC101091
764;refGene_tx=XM_023250858.1;refGene_gn=LOC101091764;refGene_tx=XM_02
3250859.1;refGene_gn=LOC101091764;refGene_tx=XM_023250860.1;refGene_gn
=LOC101091764;refGene_tx=XM_023250861.1;refGene_gn=LOC101091764;refGen
e_tx=XM_023250862.1;refGene_gn=LOC101091764;refGene_tx=XM_023250863.1;
refGene_gn=LOC101091764;refGene_tx=XM_023250864.1;refGene_gn=LOC101091
764;refGene_tx=XM_023250865.1
A2 3836125 3836175
refGene_gn=LOC101091764;refGene_tx=XM_023250857.1;refGene_gn=LOC101091
764;refGene_tx=XM_023250858.1;refGene_gn=LOC101091764;refGene_tx=XM_02
3250859.1;refGene_gn=LOC101091764;refGene_tx=XM_023250860.1;refGene_gn
=LOC101091764;refGene_tx=XM_023250861.1;refGene_gn=LOC101091764;refGen

e_tx=XM_023250862.1;refGene_gn=LOC101091764;refGene_tx=XM_023250863.1;refGene_gn=LOC101091764;refGene_tx=XM_023250864.1;refGene_gn=LOC101091764;refGene_tx=XM_023250865.1
A2 3840264 3840368 refGene_gn=LOC101091764;refGene_tx=XM_023250857.1;refGene_gn=LOC101091764;refGene_tx=XM_023250858.1;refGene_gn=LOC101091764;refGene_tx=XM_023250859.1;refGene_gn=LOC101091764;refGene_tx=XM_023250860.1;refGene_gn=LOC101091764;refGene_tx=XM_023250861.1;refGene_gn=LOC101091764;refGene_tx=XM_023250862.1;refGene_gn=LOC101091764;refGene_tx=XM_023250863.1;refGene_gn=LOC101091764;refGene_tx=XM_023250864.1;refGene_gn=LOC101091764;refGene_tx=XM_023250865.1
A2 3844814 3844952 refGene_gn=LOC101091764;refGene_tx=XM_023250857.1;refGene_gn=LOC101091764;refGene_tx=XM_023250858.1;refGene_gn=LOC101091764;refGene_tx=XM_023250859.1;refGene_gn=LOC101091764;refGene_tx=XM_023250860.1;refGene_gn=LOC101091764;refGene_tx=XM_023250861.1;refGene_gn=LOC101091764;refGene_tx=XM_023250862.1;refGene_gn=LOC101091764;refGene_tx=XM_023250863.1;refGene_gn=LOC101091764;refGene_tx=XM_023250864.1;refGene_gn=LOC101091764;refGene_tx=XM_023250865.1
A2 3860834 3861031 refGene_gn=LOC101091764;refGene_tx=XM_023250857.1;refGene_gn=LOC101091764;refGene_tx=XM_023250858.1;refGene_gn=LOC101091764;refGene_tx=XM_023250859.1;refGene_gn=LOC101091764;refGene_tx=XM_023250860.1;refGene_gn=LOC101091764;refGene_tx=XM_023250861.1;refGene_gn=LOC101091764;refGene_tx=XM_023250862.1;refGene_gn=LOC101091764;refGene_tx=XM_023250863.1;refGene_gn=LOC101091764;refGene_tx=XM_023250864.1;refGene_gn=LOC101091764;refGene_tx=XM_023250865.1
A2 3870129 3870231 refGene_gn=LOC101091764;refGene_tx=XM_023250857.1;refGene_gn=LOC101091764;refGene_tx=XM_023250858.1;refGene_gn=LOC101091764;refGene_tx=XM_023250859.1;refGene_gn=LOC101091764;refGene_tx=XM_023250860.1;refGene_gn=LOC101091764;refGene_tx=XM_023250861.1;refGene_gn=LOC101091764;refGene_tx=XM_023250863.1;refGene_gn=LOC101091764;refGene_tx=XM_023250864.1;refGene_gn=LOC101091764;refGene_tx=XM_023250865.1
A2 3870616 3870816 refGene_gn=LOC101091764;refGene_tx=XM_023250857.1;refGene_gn=LOC101091764;refGene_tx=XM_023250858.1;refGene_gn=LOC101091764;refGene_tx=XM_023250859.1;refGene_gn=LOC101091764;refGene_tx=XM_023250860.1;refGene_gn=LOC101091764;refGene_tx=XM_023250861.1;refGene_gn=LOC101091764;refGene_tx=XM_023250862.1;refGene_gn=LOC101091764;refGene_tx=XM_023250863.1;refGene_gn=LOC101091764;refGene_tx=XM_023250864.1;refGene_gn=LOC101091764;refGene_tx=XM_023250865.1
A2 3878332 3878784 refGene_gn=LOC101091764;refGene_tx=XM_023250857.1;refGene_gn=LOC101091764;refGene_tx=XM_023250858.1;refGene_gn=LOC101091764;refGene_tx=XM_023250859.1;refGene_gn=LOC101091764;refGene_tx=XM_023250860.1;refGene_gn=LOC101091764;refGene_tx=XM_023250861.1;refGene_gn=LOC101091764;refGene_tx=XM_023250862.1;refGene_gn=LOC101091764;refGene_tx=XM_023250863.1;refGene_gn=LOC101091764;refGene_tx=XM_023250864.1;refGene_gn=LOC101091764;refGene_tx=XM_023250865.1

A2 3879005 3879126
refGene_gn=LOC101091764;refGene_tx=XM_023250857.1;refGene_gn=LOC101091764;refGene_tx=XM_023250858.1;refGene_gn=LOC101091764;refGene_tx=XM_023250859.1;refGene_gn=LOC101091764;refGene_tx=XM_023250860.1;refGene_gn=LOC101091764;refGene_tx=XM_023250861.1;refGene_gn=LOC101091764;refGene_tx=XM_023250862.1;refGene_gn=LOC101091764;refGene_tx=XM_023250863.1;refGene_gn=LOC101091764;refGene_tx=XM_023250864.1;refGene_gn=LOC101091764;refGene_tx=XM_023250865.1
A2 3881136 3881315
refGene_gn=LOC101091764;refGene_tx=XM_023250857.1;refGene_gn=LOC101091764;refGene_tx=XM_023250858.1;refGene_gn=LOC101091764;refGene_tx=XM_023250859.1;refGene_gn=LOC101091764;refGene_tx=XM_023250860.1;refGene_gn=LOC101091764;refGene_tx=XM_023250861.1;refGene_gn=LOC101091764;refGene_tx=XM_023250862.1;refGene_gn=LOC101091764;refGene_tx=XM_023250863.1;refGene_gn=LOC101091764;refGene_tx=XM_023250864.1;refGene_gn=LOC101091764;refGene_tx=XM_023250865.1
A2 3883294 3883517
refGene_gn=LOC101091764;refGene_tx=XM_023250857.1;refGene_gn=LOC101091764;refGene_tx=XM_023250858.1;refGene_gn=LOC101091764;refGene_tx=XM_023250859.1;refGene_gn=LOC101091764;refGene_tx=XM_023250860.1;refGene_gn=LOC101091764;refGene_tx=XM_023250861.1;refGene_gn=LOC101091764;refGene_tx=XM_023250862.1;refGene_gn=LOC101091764;refGene_tx=XM_023250863.1;refGene_gn=LOC101091764;refGene_tx=XM_023250864.1;refGene_gn=LOC101091764;refGene_tx=XM_023250865.1
A2 3884776 3884853
refGene_gn=LOC101091764;refGene_tx=XM_023250857.1;refGene_gn=LOC101091764;refGene_tx=XM_023250858.1;refGene_gn=LOC101091764;refGene_tx=XM_023250859.1;refGene_gn=LOC101091764;refGene_tx=XM_023250860.1;refGene_gn=LOC101091764;refGene_tx=XM_023250861.1;refGene_gn=LOC101091764;refGene_tx=XM_023250862.1;refGene_gn=LOC101091764;refGene_tx=XM_023250863.1;refGene_gn=LOC101091764;refGene_tx=XM_023250864.1;refGene_gn=LOC101091764;refGene_tx=XM_023250865.1
A2 3885059 3885115
refGene_gn=LOC101091764;refGene_tx=XM_023250857.1;refGene_gn=LOC101091764;refGene_tx=XM_023250858.1;refGene_gn=LOC101091764;refGene_tx=XM_023250859.1;refGene_gn=LOC101091764;refGene_tx=XM_023250860.1;refGene_gn=LOC101091764;refGene_tx=XM_023250861.1;refGene_gn=LOC101091764;refGene_tx=XM_023250862.1;refGene_gn=LOC101091764;refGene_tx=XM_023250863.1;refGene_gn=LOC101091764;refGene_tx=XM_023250864.1;refGene_gn=LOC101091764;refGene_tx=XM_023250865.1
A2 3885298 3885407
refGene_gn=LOC101091764;refGene_tx=XM_023250857.1;refGene_gn=LOC101091764;refGene_tx=XM_023250858.1;refGene_gn=LOC101091764;refGene_tx=XM_023250859.1;refGene_gn=LOC101091764;refGene_tx=XM_023250860.1;refGene_gn=LOC101091764;refGene_tx=XM_023250861.1;refGene_gn=LOC101091764;refGene_tx=XM_023250862.1;refGene_gn=LOC101091764;refGene_tx=XM_023250863.1;refGene_gn=LOC101091764;refGene_tx=XM_023250864.1;refGene_gn=LOC101091764;refGene_tx=XM_023250865.1
A2 3889051 3889237
refGene_gn=LOC101091764;refGene_tx=XM_023250857.1;refGene_gn=LOC101091

764;refGene_tx=XM_023250858.1;refGene_gn=LOC101091764;refGene_tx=XM_023250859.1;refGene_gn=LOC101091764;refGene_tx=XM_023250860.1;refGene_gn=LOC101091764;refGene_tx=XM_023250861.1;refGene_gn=LOC101091764;refGene_tx=XM_023250862.1;refGene_gn=LOC101091764;refGene_tx=XM_023250863.1;refGene_gn=LOC101091764;refGene_tx=XM_023250864.1;refGene_gn=LOC101091764;refGene_tx=XM_023250865.1
A2 3889325 3889490
refGene_gn=LOC101091764;refGene_tx=XM_023250857.1;refGene_gn=LOC101091764;refGene_tx=XM_023250858.1;refGene_gn=LOC101091764;refGene_tx=XM_023250859.1;refGene_gn=LOC101091764;refGene_tx=XM_023250860.1;refGene_gn=LOC101091764;refGene_tx=XM_023250861.1;refGene_gn=LOC101091764;refGene_tx=XM_023250862.1;refGene_gn=LOC101091764;refGene_tx=XM_023250863.1;refGene_gn=LOC101091764;refGene_tx=XM_023250864.1;refGene_gn=LOC101091764;refGene_tx=XM_023250865.1
A2 3890336 3890456
refGene_gn=LOC101091764;refGene_tx=XM_023250857.1;refGene_gn=LOC101091764;refGene_tx=XM_023250858.1;refGene_gn=LOC101091764;refGene_tx=XM_023250859.1;refGene_gn=LOC101091764;refGene_tx=XM_023250860.1;refGene_gn=LOC101091764;refGene_tx=XM_023250861.1;refGene_gn=LOC101091764;refGene_tx=XM_023250862.1;refGene_gn=LOC101091764;refGene_tx=XM_023250863.1;refGene_gn=LOC101091764;refGene_tx=XM_023250864.1;refGene_gn=LOC101091764;refGene_tx=XM_023250865.1
A2 3890951 3891044
refGene_gn=LOC101091764;refGene_tx=XM_023250857.1;refGene_gn=LOC101091764;refGene_tx=XM_023250858.1;refGene_gn=LOC101091764;refGene_tx=XM_023250859.1;refGene_gn=LOC101091764;refGene_tx=XM_023250860.1;refGene_gn=LOC101091764;refGene_tx=XM_023250861.1;refGene_gn=LOC101091764;refGene_tx=XM_023250862.1;refGene_gn=LOC101091764;refGene_tx=XM_023250863.1;refGene_gn=LOC101091764;refGene_tx=XM_023250864.1;refGene_gn=LOC101091764;refGene_tx=XM_023250865.1
A2 3892028 3892205
refGene_gn=LOC101091764;refGene_tx=XM_023250857.1;refGene_gn=LOC101091764;refGene_tx=XM_023250858.1;refGene_gn=LOC101091764;refGene_tx=XM_023250859.1;refGene_gn=LOC101091764;refGene_tx=XM_023250860.1;refGene_gn=LOC101091764;refGene_tx=XM_023250861.1;refGene_gn=LOC101091764;refGene_tx=XM_023250862.1;refGene_gn=LOC101091764;refGene_tx=XM_023250863.1;refGene_gn=LOC101091764;refGene_tx=XM_023250864.1;refGene_gn=LOC101091764;refGene_tx=XM_023250865.1
A2 3928811 3928880
refGene_gn=PTPRS;refGene_tx=XM_023250838.1;refGene_gn=PTPRS;refGene_tx=XM_023250839.1;refGene_gn=PTPRS;refGene_tx=XM_023250840.1;refGene_gn=PTPRS;refGene_tx=XM_023250841.1;refGene_gn=PTPRS;refGene_tx=XM_023250842.1;refGene_gn=PTPRS;refGene_tx=XM_023250843.1;refGene_gn=PTPRS;refGene_tx=XM_023250844.1;refGene_gn=PTPRS;refGene_tx=XM_023250845.1;refGene_gn=PTPRS;refGene_tx=XM_023250846.1;refGene_gn=PTPRS;refGene_tx=XM_023250847.1;refGene_gn=PTPRS;refGene_tx=XM_023250848.1;refGene_gn=PTPRS;refGene_tx=XM_023250849.1;refGene_gn=PTPRS;refGene_tx=XM_023250850.1;refGene_gn=PTPRS;refGene_tx=XM_023250851.1;refGene_gn=PTPRS;refGene_tx=XM_023250852.1;refGene_gn=PTPRS;refGene_tx=XM_023250853.1;refGene_gn=PTPRS;refGene_tx=XM_023250854.1;refGene_gn=PTPRS;refGene_tx=XM_02325085

6.1
A2 3929433 3929569
refGene_gn=PTPRS;refGene_tx=XM_023250838.1;refGene_gn=PTPRS;refGene_tx=XM_023250839.1;refGene_gn=PTPRS;refGene_tx=XM_023250840.1;refGene_gn=PTPRS;refGene_tx=XM_023250841.1;refGene_gn=PTPRS;refGene_tx=XM_023250842.1;refGene_gn=PTPRS;refGene_tx=XM_023250843.1;refGene_gn=PTPRS;refGene_tx=XM_023250844.1;refGene_gn=PTPRS;refGene_tx=XM_023250845.1;refGene_gn=PTPRS;refGene_tx=XM_023250846.1;refGene_gn=PTPRS;refGene_tx=XM_023250847.1;refGene_gn=PTPRS;refGene_tx=XM_023250848.1;refGene_gn=PTPRS;refGene_tx=XM_023250849.1;refGene_gn=PTPRS;refGene_tx=XM_023250850.1;refGene_gn=PTPRS;refGene_tx=XM_023250851.1;refGene_gn=PTPRS;refGene_tx=XM_023250852.1;refGene_gn=PTPRS;refGene_tx=XM_023250853.1;refGene_gn=PTPRS;refGene_tx=XM_023250854.1;refGene_gn=PTPRS;refGene_tx=XM_023250856.1
A2 3929764 3929919
refGene_gn=PTPRS;refGene_tx=XM_023250838.1;refGene_gn=PTPRS;refGene_tx=XM_023250839.1;refGene_gn=PTPRS;refGene_tx=XM_023250840.1;refGene_gn=PTPRS;refGene_tx=XM_023250841.1;refGene_gn=PTPRS;refGene_tx=XM_023250842.1;refGene_gn=PTPRS;refGene_tx=XM_023250843.1;refGene_gn=PTPRS;refGene_tx=XM_023250844.1;refGene_gn=PTPRS;refGene_tx=XM_023250845.1;refGene_gn=PTPRS;refGene_tx=XM_023250846.1;refGene_gn=PTPRS;refGene_tx=XM_023250847.1;refGene_gn=PTPRS;refGene_tx=XM_023250848.1;refGene_gn=PTPRS;refGene_tx=XM_023250849.1;refGene_gn=PTPRS;refGene_tx=XM_023250850.1;refGene_gn=PTPRS;refGene_tx=XM_023250851.1;refGene_gn=PTPRS;refGene_tx=XM_023250852.1;refGene_gn=PTPRS;refGene_tx=XM_023250853.1;refGene_gn=PTPRS;refGene_tx=XM_023250854.1;refGene_gn=PTPRS;refGene_tx=XM_023250856.1
A2 3930823 3930949
refGene_gn=PTPRS;refGene_tx=XM_023250838.1;refGene_gn=PTPRS;refGene_tx=XM_023250839.1;refGene_gn=PTPRS;refGene_tx=XM_023250840.1;refGene_gn=PTPRS;refGene_tx=XM_023250841.1;refGene_gn=PTPRS;refGene_tx=XM_023250842.1;refGene_gn=PTPRS;refGene_tx=XM_023250843.1;refGene_gn=PTPRS;refGene_tx=XM_023250844.1;refGene_gn=PTPRS;refGene_tx=XM_023250845.1;refGene_gn=PTPRS;refGene_tx=XM_023250846.1;refGene_gn=PTPRS;refGene_tx=XM_023250847.1;refGene_gn=PTPRS;refGene_tx=XM_023250848.1;refGene_gn=PTPRS;refGene_tx=XM_023250849.1;refGene_gn=PTPRS;refGene_tx=XM_023250850.1;refGene_gn=PTPRS;refGene_tx=XM_023250851.1;refGene_gn=PTPRS;refGene_tx=XM_023250852.1;refGene_gn=PTPRS;refGene_tx=XM_023250853.1;refGene_gn=PTPRS;refGene_tx=XM_023250854.1;refGene_gn=PTPRS;refGene_tx=XM_023250856.1
A2 3931111 3931238
refGene_gn=PTPRS;refGene_tx=XM_023250838.1;refGene_gn=PTPRS;refGene_tx=XM_023250839.1;refGene_gn=PTPRS;refGene_tx=XM_023250840.1;refGene_gn=PTPRS;refGene_tx=XM_023250841.1;refGene_gn=PTPRS;refGene_tx=XM_023250842.1;refGene_gn=PTPRS;refGene_tx=XM_023250843.1;refGene_gn=PTPRS;refGene_tx=XM_023250844.1;refGene_gn=PTPRS;refGene_tx=XM_023250845.1;refGene_gn=PTPRS;refGene_tx=XM_023250846.1;refGene_gn=PTPRS;refGene_tx=XM_023250847.1;refGene_gn=PTPRS;refGene_tx=XM_023250848.1;refGene_gn=PTPRS;refGene_tx=XM_023250849.1;refGene_gn=PTPRS;refGene_tx=XM_023250850.1;refGene_gn=PTPRS;refGene_tx=XM_023250851.1;refGene_gn=PTPRS;refGene_tx=

XM_023250852.1;refGene_gn=PTPRS;refGene_tx=XM_023250853.1;refGene_gn=PTPRS;refGene_tx=XM_023250854.1;refGene_gn=PTPRS;refGene_tx=XM_023250856.1
A2 3931875 3932054
refGene_gn=PTPRS;refGene_tx=XM_023250838.1;refGene_gn=PTPRS;refGene_tx=XM_023250839.1;refGene_gn=PTPRS;refGene_tx=XM_023250840.1;refGene_gn=PTPRS;refGene_tx=XM_023250841.1;refGene_gn=PTPRS;refGene_tx=XM_023250842.1;refGene_gn=PTPRS;refGene_tx=XM_023250843.1;refGene_gn=PTPRS;refGene_tx=XM_023250844.1;refGene_gn=PTPRS;refGene_tx=XM_023250845.1;refGene_gn=PTPRS;refGene_tx=XM_023250846.1;refGene_gn=PTPRS;refGene_tx=XM_023250847.1;refGene_gn=PTPRS;refGene_tx=XM_023250848.1;refGene_gn=PTPRS;refGene_tx=XM_023250849.1;refGene_gn=PTPRS;refGene_tx=XM_023250850.1;refGene_gn=PTPRS;refGene_tx=XM_023250851.1;refGene_gn=PTPRS;refGene_tx=XM_023250852.1;refGene_gn=PTPRS;refGene_tx=XM_023250853.1;refGene_gn=PTPRS;refGene_tx=XM_023250854.1;refGene_gn=PTPRS;refGene_tx=XM_023250856.1
A2 3932207 3932493
refGene_gn=PTPRS;refGene_tx=XM_023250838.1;refGene_gn=PTPRS;refGene_tx=XM_023250839.1;refGene_gn=PTPRS;refGene_tx=XM_023250840.1;refGene_gn=PTPRS;refGene_tx=XM_023250841.1;refGene_gn=PTPRS;refGene_tx=XM_023250842.1;refGene_gn=PTPRS;refGene_tx=XM_023250843.1;refGene_gn=PTPRS;refGene_tx=XM_023250844.1;refGene_gn=PTPRS;refGene_tx=XM_023250845.1;refGene_gn=PTPRS;refGene_tx=XM_023250846.1;refGene_gn=PTPRS;refGene_tx=XM_023250847.1;refGene_gn=PTPRS;refGene_tx=XM_023250848.1;refGene_gn=PTPRS;refGene_tx=XM_023250849.1;refGene_gn=PTPRS;refGene_tx=XM_023250850.1;refGene_gn=PTPRS;refGene_tx=XM_023250851.1;refGene_gn=PTPRS;refGene_tx=XM_023250852.1;refGene_gn=PTPRS;refGene_tx=XM_023250853.1;refGene_gn=PTPRS;refGene_tx=XM_023250854.1;refGene_gn=PTPRS;refGene_tx=XM_023250856.1
A2 3932569 3932724
refGene_gn=PTPRS;refGene_tx=XM_023250838.1;refGene_gn=PTPRS;refGene_tx=XM_023250839.1;refGene_gn=PTPRS;refGene_tx=XM_023250840.1;refGene_gn=PTPRS;refGene_tx=XM_023250841.1;refGene_gn=PTPRS;refGene_tx=XM_023250842.1;refGene_gn=PTPRS;refGene_tx=XM_023250843.1;refGene_gn=PTPRS;refGene_tx=XM_023250844.1;refGene_gn=PTPRS;refGene_tx=XM_023250845.1;refGene_gn=PTPRS;refGene_tx=XM_023250846.1;refGene_gn=PTPRS;refGene_tx=XM_023250847.1;refGene_gn=PTPRS;refGene_tx=XM_023250848.1;refGene_gn=PTPRS;refGene_tx=XM_023250849.1;refGene_gn=PTPRS;refGene_tx=XM_023250850.1;refGene_gn=PTPRS;refGene_tx=XM_023250851.1;refGene_gn=PTPRS;refGene_tx=XM_023250852.1;refGene_gn=PTPRS;refGene_tx=XM_023250853.1;refGene_gn=PTPRS;refGene_tx=XM_023250854.1;refGene_gn=PTPRS;refGene_tx=XM_023250856.1
A2 3934058 3934178
refGene_gn=PTPRS;refGene_tx=XM_023250838.1;refGene_gn=PTPRS;refGene_tx=XM_023250839.1;refGene_gn=PTPRS;refGene_tx=XM_023250840.1;refGene_gn=PTPRS;refGene_tx=XM_023250841.1;refGene_gn=PTPRS;refGene_tx=XM_023250842.1;refGene_gn=PTPRS;refGene_tx=XM_023250843.1;refGene_gn=PTPRS;refGene_tx=XM_023250844.1;refGene_gn=PTPRS;refGene_tx=XM_023250845.1;refGene_gn=PTPRS;refGene_tx=XM_023250846.1;refGene_gn=PTPRS;refGene_tx=XM_023250847.1;refGene_gn=PTPRS;refGene_tx=XM_023250848.1;refGene_gn=PTPRS;

refGene_tx=XM_023250849.1;refGene_gn=PTPRS;refGene_tx=XM_023250850.1;r
efGene_gn=PTPRS;refGene_tx=XM_023250851.1;refGene_gn=PTPRS;refGene_tx=
XM_023250852.1;refGene_gn=PTPRS;refGene_tx=XM_023250853.1;refGene_gn=P
TPRS;refGene_tx=XM_023250854.1;refGene_gn=PTPRS;refGene_tx=XM_02325085
6.1
A2 3934256 3934432
refGene_gn=PTPRS;refGene_tx=XM_023250838.1;refGene_gn=PTPRS;refGene_tx
=XM_023250839.1;refGene_gn=PTPRS;refGene_tx=XM_023250840.1;refGene_gn=
PTPRS;refGene_tx=XM_023250841.1;refGene_gn=PTPRS;refGene_tx=XM_0232508
42.1;refGene_gn=PTPRS;refGene_tx=XM_023250843.1;refGene_gn=PTPRS;refGe
ne_tx=XM_023250844.1;refGene_gn=PTPRS;refGene_tx=XM_023250845.1;refGen
e_gn=PTPRS;refGene_tx=XM_023250846.1;refGene_gn=PTPRS;refGene_tx=XM_02
3250847.1;refGene_gn=PTPRS;refGene_tx=XM_023250848.1;refGene_gn=PTPRS;
refGene_tx=XM_023250849.1;refGene_gn=PTPRS;refGene_tx=XM_023250850.1;r
efGene_gn=PTPRS;refGene_tx=XM_023250851.1;refGene_gn=PTPRS;refGene_tx=
XM_023250852.1;refGene_gn=PTPRS;refGene_tx=XM_023250853.1;refGene_gn=P
TPRS;refGene_tx=XM_023250854.1;refGene_gn=PTPRS;refGene_tx=XM_02325085
6.1
A2 3934916 3935040
refGene_gn=PTPRS;refGene_tx=XM_023250838.1;refGene_gn=PTPRS;refGene_tx
=XM_023250839.1;refGene_gn=PTPRS;refGene_tx=XM_023250840.1;refGene_gn=
PTPRS;refGene_tx=XM_023250841.1;refGene_gn=PTPRS;refGene_tx=XM_0232508
42.1;refGene_gn=PTPRS;refGene_tx=XM_023250843.1;refGene_gn=PTPRS;refGe
ne_tx=XM_023250844.1;refGene_gn=PTPRS;refGene_tx=XM_023250845.1;refGen
e_gn=PTPRS;refGene_tx=XM_023250846.1;refGene_gn=PTPRS;refGene_tx=XM_02
3250847.1;refGene_gn=PTPRS;refGene_tx=XM_023250848.1;refGene_gn=PTPRS;
refGene_tx=XM_023250849.1;refGene_gn=PTPRS;refGene_tx=XM_023250850.1;r
efGene_gn=PTPRS;refGene_tx=XM_023250851.1;refGene_gn=PTPRS;refGene_tx=
XM_023250852.1;refGene_gn=PTPRS;refGene_tx=XM_023250853.1;refGene_gn=P
TPRS;refGene_tx=XM_023250854.1;refGene_gn=PTPRS;refGene_tx=XM_02325085
6.1
A2 3935126 3935224
refGene_gn=PTPRS;refGene_tx=XM_023250838.1;refGene_gn=PTPRS;refGene_tx
=XM_023250839.1;refGene_gn=PTPRS;refGene_tx=XM_023250840.1;refGene_gn=
PTPRS;refGene_tx=XM_023250841.1;refGene_gn=PTPRS;refGene_tx=XM_0232508
42.1;refGene_gn=PTPRS;refGene_tx=XM_023250843.1;refGene_gn=PTPRS;refGe
ne_tx=XM_023250844.1;refGene_gn=PTPRS;refGene_tx=XM_023250845.1;refGen
e_gn=PTPRS;refGene_tx=XM_023250846.1;refGene_gn=PTPRS;refGene_tx=XM_02
3250847.1;refGene_gn=PTPRS;refGene_tx=XM_023250848.1;refGene_gn=PTPRS;
refGene_tx=XM_023250849.1;refGene_gn=PTPRS;refGene_tx=XM_023250850.1;r
efGene_gn=PTPRS;refGene_tx=XM_023250851.1;refGene_gn=PTPRS;refGene_tx=
XM_023250852.1;refGene_gn=PTPRS;refGene_tx=XM_023250853.1;refGene_gn=P
TPRS;refGene_tx=XM_023250854.1;refGene_gn=PTPRS;refGene_tx=XM_02325085
6.1
A2 3936312 3936360
refGene_gn=PTPRS;refGene_tx=XM_023250838.1;refGene_gn=PTPRS;refGene_tx
=XM_023250839.1;refGene_gn=PTPRS;refGene_tx=XM_023250840.1;refGene_gn=
PTPRS;refGene_tx=XM_023250841.1;refGene_gn=PTPRS;refGene_tx=XM_0232508
42.1;refGene_gn=PTPRS;refGene_tx=XM_023250843.1;refGene_gn=PTPRS;refGe
ne_tx=XM_023250844.1;refGene_gn=PTPRS;refGene_tx=XM_023250845.1;refGen

e_gn=PTPRS;refGene_tx=XM_023250847.1;refGene_gn=PTPRS;refGene_tx=XM_02
3250849.1;refGene_gn=PTPRS;refGene_tx=XM_023250850.1;refGene_gn=PTPRS;
refGene_tx=XM_023250851.1;refGene_gn=PTPRS;refGene_tx=XM_023250856.1
A2 3937753 3937866
refGene_gn=PTPRS;refGene_tx=XM_023250838.1;refGene_gn=PTPRS;refGene_tx
=XM_023250839.1;refGene_gn=PTPRS;refGene_tx=XM_023250840.1;refGene_gn=
PTPRS;refGene_tx=XM_023250841.1;refGene_gn=PTPRS;refGene_tx=XM_0232508
42.1;refGene_gn=PTPRS;refGene_tx=XM_023250843.1;refGene_gn=PTPRS;refGe
ne_tx=XM_023250844.1;refGene_gn=PTPRS;refGene_tx=XM_023250845.1;refGen
e_gn=PTPRS;refGene_tx=XM_023250846.1;refGene_gn=PTPRS;refGene_tx=XM_02
3250847.1;refGene_gn=PTPRS;refGene_tx=XM_023250848.1;refGene_gn=PTPRS;
refGene_tx=XM_023250849.1;refGene_gn=PTPRS;refGene_tx=XM_023250850.1;r
efGene_gn=PTPRS;refGene_tx=XM_023250851.1;refGene_gn=PTPRS;refGene_tx=
XM_023250852.1;refGene_gn=PTPRS;refGene_tx=XM_023250853.1;refGene_gn=P
TPRS;refGene_tx=XM_023250854.1;refGene_gn=PTPRS;refGene_tx=XM_02325085
6.1
A2 3938137 3938149
refGene_gn=PTPRS;refGene_tx=XM_023250838.1;refGene_gn=PTPRS;refGene_tx
=XM_023250839.1;refGene_gn=PTPRS;refGene_tx=XM_023250840.1;refGene_gn=
PTPRS;refGene_tx=XM_023250841.1;refGene_gn=PTPRS;refGene_tx=XM_0232508
42.1;refGene_gn=PTPRS;refGene_tx=XM_023250843.1;refGene_gn=PTPRS;refGe
ne_tx=XM_023250844.1;refGene_gn=PTPRS;refGene_tx=XM_023250845.1;refGen
e_gn=PTPRS;refGene_tx=XM_023250846.1;refGene_gn=PTPRS;refGene_tx=XM_02
3250847.1;refGene_gn=PTPRS;refGene_tx=XM_023250848.1;refGene_gn=PTPRS;
refGene_tx=XM_023250849.1;refGene_gn=PTPRS;refGene_tx=XM_023250850.1;r
efGene_gn=PTPRS;refGene_tx=XM_023250851.1;refGene_gn=PTPRS;refGene_tx=
XM_023250852.1;refGene_gn=PTPRS;refGene_tx=XM_023250853.1;refGene_gn=P
TPRS;refGene_tx=XM_023250854.1;refGene_gn=PTPRS;refGene_tx=XM_02325085
6.1
A2 3938654 3938812
refGene_gn=PTPRS;refGene_tx=XM_023250838.1;refGene_gn=PTPRS;refGene_tx
=XM_023250839.1;refGene_gn=PTPRS;refGene_tx=XM_023250840.1;refGene_gn=
PTPRS;refGene_tx=XM_023250841.1;refGene_gn=PTPRS;refGene_tx=XM_0232508
42.1;refGene_gn=PTPRS;refGene_tx=XM_023250843.1;refGene_gn=PTPRS;refGe
ne_tx=XM_023250844.1;refGene_gn=PTPRS;refGene_tx=XM_023250845.1;refGen
e_gn=PTPRS;refGene_tx=XM_023250846.1;refGene_gn=PTPRS;refGene_tx=XM_02
3250847.1;refGene_gn=PTPRS;refGene_tx=XM_023250848.1;refGene_gn=PTPRS;
refGene_tx=XM_023250849.1;refGene_gn=PTPRS;refGene_tx=XM_023250850.1;r
efGene_gn=PTPRS;refGene_tx=XM_023250851.1;refGene_gn=PTPRS;refGene_tx=
XM_023250852.1;refGene_gn=PTPRS;refGene_tx=XM_023250853.1;refGene_gn=P
TPRS;refGene_tx=XM_023250854.1;refGene_gn=PTPRS;refGene_tx=XM_02325085
6.1
A2 3939164 3939380
refGene_gn=PTPRS;refGene_tx=XM_023250838.1;refGene_gn=PTPRS;refGene_tx
=XM_023250839.1;refGene_gn=PTPRS;refGene_tx=XM_023250840.1;refGene_gn=
PTPRS;refGene_tx=XM_023250841.1;refGene_gn=PTPRS;refGene_tx=XM_0232508
42.1;refGene_gn=PTPRS;refGene_tx=XM_023250843.1;refGene_gn=PTPRS;refGe
ne_tx=XM_023250844.1;refGene_gn=PTPRS;refGene_tx=XM_023250845.1;refGen
e_gn=PTPRS;refGene_tx=XM_023250846.1;refGene_gn=PTPRS;refGene_tx=XM_02
3250847.1;refGene_gn=PTPRS;refGene_tx=XM_023250848.1;refGene_gn=PTPRS;

refGene_tx=XM_023250849.1;refGene_gn=PTPRS;refGene_tx=XM_023250850.1;refGene_gn=PTPRS;refGene_tx=XM_023250851.1;refGene_gn=PTPRS;refGene_tx=XM_023250852.1;refGene_gn=PTPRS;refGene_tx=XM_023250853.1;refGene_gn=PTPRS;refGene_tx=XM_023250854.1;refGene_gn=PTPRS;refGene_tx=XM_023250856.1
A2 3939471 3939565
refGene_gn=PTPRS;refGene_tx=XM_023250838.1;refGene_gn=PTPRS;refGene_tx=XM_023250839.1;refGene_gn=PTPRS;refGene_tx=XM_023250840.1;refGene_gn=PTPRS;refGene_tx=XM_023250841.1;refGene_gn=PTPRS;refGene_tx=XM_023250842.1;refGene_gn=PTPRS;refGene_tx=XM_023250843.1;refGene_gn=PTPRS;refGene_tx=XM_023250844.1;refGene_gn=PTPRS;refGene_tx=XM_023250845.1;refGene_gn=PTPRS;refGene_tx=XM_023250846.1;refGene_gn=PTPRS;refGene_tx=XM_023250847.1;refGene_gn=PTPRS;refGene_tx=XM_023250848.1;refGene_gn=PTPRS;refGene_tx=XM_023250849.1;refGene_gn=PTPRS;refGene_tx=XM_023250850.1;refGene_gn=PTPRS;refGene_tx=XM_023250851.1;refGene_gn=PTPRS;refGene_tx=XM_023250852.1;refGene_gn=PTPRS;refGene_tx=XM_023250853.1;refGene_gn=PTPRS;refGene_tx=XM_023250854.1;refGene_gn=PTPRS;refGene_tx=XM_023250856.1
A2 3939988 3940242
refGene_gn=PTPRS;refGene_tx=XM_023250838.1;refGene_gn=PTPRS;refGene_tx=XM_023250839.1;refGene_gn=PTPRS;refGene_tx=XM_023250840.1;refGene_gn=PTPRS;refGene_tx=XM_023250841.1;refGene_gn=PTPRS;refGene_tx=XM_023250842.1;refGene_gn=PTPRS;refGene_tx=XM_023250843.1;refGene_gn=PTPRS;refGene_tx=XM_023250844.1;refGene_gn=PTPRS;refGene_tx=XM_023250845.1;refGene_gn=PTPRS;refGene_tx=XM_023250846.1;refGene_gn=PTPRS;refGene_tx=XM_023250847.1;refGene_gn=PTPRS;refGene_tx=XM_023250848.1;refGene_gn=PTPRS;refGene_tx=XM_023250849.1;refGene_gn=PTPRS;refGene_tx=XM_023250850.1;refGene_gn=PTPRS;refGene_tx=XM_023250851.1;refGene_gn=PTPRS;refGene_tx=XM_023250852.1;refGene_gn=PTPRS;refGene_tx=XM_023250853.1;refGene_gn=PTPRS;refGene_tx=XM_023250854.1;refGene_gn=PTPRS;refGene_tx=XM_023250856.1
A2 3940888 3940986
refGene_gn=PTPRS;refGene_tx=XM_023250838.1;refGene_gn=PTPRS;refGene_tx=XM_023250839.1;refGene_gn=PTPRS;refGene_tx=XM_023250840.1;refGene_gn=PTPRS;refGene_tx=XM_023250841.1;refGene_gn=PTPRS;refGene_tx=XM_023250842.1;refGene_gn=PTPRS;refGene_tx=XM_023250843.1;refGene_gn=PTPRS;refGene_tx=XM_023250844.1;refGene_gn=PTPRS;refGene_tx=XM_023250845.1;refGene_gn=PTPRS;refGene_tx=XM_023250846.1;refGene_gn=PTPRS;refGene_tx=XM_023250847.1;refGene_gn=PTPRS;refGene_tx=XM_023250848.1;refGene_gn=PTPRS;refGene_tx=XM_023250849.1;refGene_gn=PTPRS;refGene_tx=XM_023250850.1;refGene_gn=PTPRS;refGene_tx=XM_023250851.1;refGene_gn=PTPRS;refGene_tx=XM_023250852.1;refGene_gn=PTPRS;refGene_tx=XM_023250853.1;refGene_gn=PTPRS;refGene_tx=XM_023250854.1;refGene_gn=PTPRS;refGene_tx=XM_023250856.1
A2 3941427 3942033
refGene_gn=PTPRS;refGene_tx=XM_023250838.1;refGene_gn=PTPRS;refGene_tx=XM_023250839.1;refGene_gn=PTPRS;refGene_tx=XM_023250840.1;refGene_gn=PTPRS;refGene_tx=XM_023250841.1;refGene_gn=PTPRS;refGene_tx=XM_023250842.1;refGene_gn=PTPRS;refGene_tx=XM_023250843.1;refGene_gn=PTPRS;refGene_tx=XM_023250844.1;refGene_gn=PTPRS;refGene_tx=XM_023250845.1;refGen

e_gn=PTPRS;refGene_tx=XM_023250846.1;refGene_gn=PTPRS;refGene_tx=XM_02
3250847.1;refGene_gn=PTPRS;refGene_tx=XM_023250848.1;refGene_gn=PTPRS;
refGene_tx=XM_023250849.1
A2 3943775 3943893
refGene_gn=PTPRS;refGene_tx=XM_023250838.1;refGene_gn=PTPRS;refGene_tx
=XM_023250839.1;refGene_gn=PTPRS;refGene_tx=XM_023250840.1;refGene_gn=
PTPRS;refGene_tx=XM_023250841.1;refGene_gn=PTPRS;refGene_tx=XM_0232508
42.1;refGene_gn=PTPRS;refGene_tx=XM_023250843.1;refGene_gn=PTPRS;refGe
ne_tx=XM_023250844.1;refGene_gn=PTPRS;refGene_tx=XM_023250845.1;refGen
e_gn=PTPRS;refGene_tx=XM_023250846.1;refGene_gn=PTPRS;refGene_tx=XM_02
3250847.1;refGene_gn=PTPRS;refGene_tx=XM_023250848.1;refGene_gn=PTPRS;
refGene_tx=XM_023250849.1
A2 3946183 3946210
refGene_gn=PTPRS;refGene_tx=XM_023250838.1;refGene_gn=PTPRS;refGene_tx
=XM_023250839.1;refGene_gn=PTPRS;refGene_tx=XM_023250840.1;refGene_gn=
PTPRS;refGene_tx=XM_023250841.1;refGene_gn=PTPRS;refGene_tx=XM_0232508
42.1;refGene_gn=PTPRS;refGene_tx=XM_023250843.1;refGene_gn=PTPRS;refGe
ne_tx=XM_023250845.1;refGene_gn=PTPRS;refGene_tx=XM_023250846.1;refGen
e_gn=PTPRS;refGene_tx=XM_023250849.1
A2 3946372 3946566
refGene_gn=PTPRS;refGene_tx=XM_023250838.1;refGene_gn=PTPRS;refGene_tx
=XM_023250839.1;refGene_gn=PTPRS;refGene_tx=XM_023250840.1;refGene_gn=
PTPRS;refGene_tx=XM_023250841.1;refGene_gn=PTPRS;refGene_tx=XM_0232508
42.1;refGene_gn=PTPRS;refGene_tx=XM_023250843.1;refGene_gn=PTPRS;refGe
ne_tx=XM_023250844.1;refGene_gn=PTPRS;refGene_tx=XM_023250845.1;refGen
e_gn=PTPRS;refGene_tx=XM_023250846.1;refGene_gn=PTPRS;refGene_tx=XM_02
3250847.1;refGene_gn=PTPRS;refGene_tx=XM_023250848.1;refGene_gn=PTPRS;
refGene_tx=XM_023250849.1
A2 3947720 3948026
refGene_gn=PTPRS;refGene_tx=XM_023250838.1;refGene_gn=PTPRS;refGene_tx
=XM_023250839.1;refGene_gn=PTPRS;refGene_tx=XM_023250840.1;refGene_gn=
PTPRS;refGene_tx=XM_023250841.1;refGene_gn=PTPRS;refGene_tx=XM_0232508
42.1;refGene_gn=PTPRS;refGene_tx=XM_023250843.1;refGene_gn=PTPRS;refGe
ne_tx=XM_023250844.1;refGene_gn=PTPRS;refGene_tx=XM_023250845.1;refGen
e_gn=PTPRS;refGene_tx=XM_023250846.1;refGene_gn=PTPRS;refGene_tx=XM_02
3250847.1;refGene_gn=PTPRS;refGene_tx=XM_023250848.1;refGene_gn=PTPRS;
refGene_tx=XM_023250849.1
A2 3956754 3956899
refGene_gn=PTPRS;refGene_tx=XM_023250838.1;refGene_gn=PTPRS;refGene_tx
=XM_023250839.1;refGene_gn=PTPRS;refGene_tx=XM_023250840.1;refGene_gn=
PTPRS;refGene_tx=XM_023250841.1;refGene_gn=PTPRS;refGene_tx=XM_0232508
42.1;refGene_gn=PTPRS;refGene_tx=XM_023250843.1;refGene_gn=PTPRS;refGe
ne_tx=XM_023250844.1;refGene_gn=PTPRS;refGene_tx=XM_023250845.1;refGen
e_gn=PTPRS;refGene_tx=XM_023250846.1;refGene_gn=PTPRS;refGene_tx=XM_02
3250847.1;refGene_gn=PTPRS;refGene_tx=XM_023250848.1;refGene_gn=PTPRS;
refGene_tx=XM_023250849.1;refGene_gn=PTPRS;refGene_tx=XM_023250850.1;r
efGene_gn=PTPRS;refGene_tx=XM_023250851.1;refGene_gn=PTPRS;refGene_tx=
XM_023250852.1;refGene_gn=PTPRS;refGene_tx=XM_023250853.1;refGene_gn=P
TPRS;refGene_tx=XM_023250854.1;refGene_gn=PTPRS;refGene_tx=XM_02325085
6.1

A2	3957159	3957293
refGene_gn=PTPRS;refGene_tx=XM_023250838.1;refGene_gn=PTPRS;refGene_tx=XM_023250839.1;refGene_gn=PTPRS;refGene_tx=XM_023250840.1;refGene_gn=PTPRS;refGene_tx=XM_023250841.1;refGene_gn=PTPRS;refGene_tx=XM_023250842.1;refGene_gn=PTPRS;refGene_tx=XM_023250843.1;refGene_gn=PTPRS;refGene_tx=XM_023250844.1;refGene_gn=PTPRS;refGene_tx=XM_023250845.1;refGene_gn=PTPRS;refGene_tx=XM_023250846.1;refGene_gn=PTPRS;refGene_tx=XM_023250847.1;refGene_gn=PTPRS;refGene_tx=XM_023250848.1;refGene_gn=PTPRS;refGene_tx=XM_023250849.1;refGene_gn=PTPRS;refGene_tx=XM_023250850.1;refGene_gn=PTPRS;refGene_tx=XM_023250851.1;refGene_gn=PTPRS;refGene_tx=XM_023250852.1;refGene_gn=PTPRS;refGene_tx=XM_023250853.1;refGene_gn=PTPRS;refGene_tx=XM_023250854.1;refGene_gn=PTPRS;refGene_tx=XM_023250856.1
A2	3958260	3958842
refGene_gn=PTPRS;refGene_tx=XM_023250838.1;refGene_gn=PTPRS;refGene_tx=XM_023250839.1;refGene_gn=PTPRS;refGene_tx=XM_023250840.1;refGene_gn=PTPRS;refGene_tx=XM_023250841.1;refGene_gn=PTPRS;refGene_tx=XM_023250842.1;refGene_gn=PTPRS;refGene_tx=XM_023250843.1;refGene_gn=PTPRS;refGene_tx=XM_023250844.1;refGene_gn=PTPRS;refGene_tx=XM_023250845.1;refGene_gn=PTPRS;refGene_tx=XM_023250846.1;refGene_gn=PTPRS;refGene_tx=XM_023250847.1;refGene_gn=PTPRS;refGene_tx=XM_023250848.1;refGene_gn=PTPRS;refGene_tx=XM_023250849.1;refGene_gn=PTPRS;refGene_tx=XM_023250850.1;refGene_gn=PTPRS;refGene_tx=XM_023250851.1;refGene_gn=PTPRS;refGene_tx=XM_023250852.1;refGene_gn=PTPRS;refGene_tx=XM_023250853.1;refGene_gn=PTPRS;refGene_tx=XM_023250854.1;refGene_gn=PTPRS;refGene_tx=XM_023250856.1
A2	3960308	3960578
refGene_gn=PTPRS;refGene_tx=XM_023250838.1;refGene_gn=PTPRS;refGene_tx=XM_023250839.1;refGene_gn=PTPRS;refGene_tx=XM_023250840.1;refGene_gn=PTPRS;refGene_tx=XM_023250841.1;refGene_gn=PTPRS;refGene_tx=XM_023250842.1;refGene_gn=PTPRS;refGene_tx=XM_023250843.1;refGene_gn=PTPRS;refGene_tx=XM_023250844.1;refGene_gn=PTPRS;refGene_tx=XM_023250845.1;refGene_gn=PTPRS;refGene_tx=XM_023250846.1;refGene_gn=PTPRS;refGene_tx=XM_023250847.1;refGene_gn=PTPRS;refGene_tx=XM_023250848.1;refGene_gn=PTPRS;refGene_tx=XM_023250849.1;refGene_gn=PTPRS;refGene_tx=XM_023250850.1;refGene_gn=PTPRS;refGene_tx=XM_023250851.1;refGene_gn=PTPRS;refGene_tx=XM_023250852.1;refGene_gn=PTPRS;refGene_tx=XM_023250853.1;refGene_gn=PTPRS;refGene_tx=XM_023250854.1;refGene_gn=PTPRS;refGene_tx=XM_023250856.1
A2	3969403	3969415
refGene_gn=PTPRS;refGene_tx=XM_023250838.1;refGene_gn=PTPRS;refGene_tx=XM_023250839.1;refGene_gn=PTPRS;refGene_tx=XM_023250840.1;refGene_gn=PTPRS;refGene_tx=XM_023250841.1;refGene_gn=PTPRS;refGene_tx=XM_023250843.1;refGene_gn=PTPRS;refGene_tx=XM_023250844.1;refGene_gn=PTPRS;refGene_tx=XM_023250846.1;refGene_gn=PTPRS;refGene_tx=XM_023250849.1;refGene_gn=PTPRS;refGene_tx=XM_023250850.1;refGene_gn=PTPRS;refGene_tx=XM_023250852.1;refGene_gn=PTPRS;refGene_tx=XM_023250853.1
A2	3971307	3971418
refGene_gn=PTPRS;refGene_tx=XM_023250838.1;refGene_gn=PTPRS;refGene_tx=XM_023250839.1;refGene_gn=PTPRS;refGene_tx=XM_023250840.1;refGene_gn=

PTPRS;refGene_tx=XM_023250841.1;refGene_gn=PTPRS;refGene_tx=XM_0232508
42.1;refGene_gn=PTPRS;refGene_tx=XM_023250843.1;refGene_gn=PTPRS;refGe
ne_tx=XM_023250844.1;refGene_gn=PTPRS;refGene_tx=XM_023250845.1;refGen
e_gn=PTPRS;refGene_tx=XM_023250846.1;refGene_gn=PTPRS;refGene_tx=XM_02
3250847.1;refGene_gn=PTPRS;refGene_tx=XM_023250848.1;refGene_gn=PTPRS;
refGene_tx=XM_023250849.1;refGene_gn=PTPRS;refGene_tx=XM_023250850.1;r
efGene_gn=PTPRS;refGene_tx=XM_023250851.1;refGene_gn=PTPRS;refGene_tx=
XM_023250852.1;refGene_gn=PTPRS;refGene_tx=XM_023250853.1;refGene_gn=P
TPRS;refGene_tx=XM_023250854.1;refGene_gn=PTPRS;refGene_tx=XM_02325085
6.1
A2 3973839 3973857
refGene_gn=PTPRS;refGene_tx=XM_023250838.1;refGene_gn=PTPRS;refGene_tx
=XM_023250839.1;refGene_gn=PTPRS;refGene_tx=XM_023250840.1;refGene_gn=
PTPRS;refGene_tx=XM_023250841.1;refGene_gn=PTPRS;refGene_tx=XM_0232508
42.1;refGene_gn=PTPRS;refGene_tx=XM_023250844.1;refGene_gn=PTPRS;refGe
ne_tx=XM_023250846.1;refGene_gn=PTPRS;refGene_tx=XM_023250849.1;refGen
e_gn=PTPRS;refGene_tx=XM_023250850.1;refGene_gn=PTPRS;refGene_tx=XM_02
3250851.1;refGene_gn=PTPRS;refGene_tx=XM_023250852.1
A2 3975532 3975541
refGene_gn=PTPRS;refGene_tx=XM_023250838.1;refGene_gn=PTPRS;refGene_tx
=XM_023250839.1;refGene_gn=PTPRS;refGene_tx=XM_023250840.1;refGene_gn=
PTPRS;refGene_tx=XM_023250842.1;refGene_gn=PTPRS;refGene_tx=XM_0232508
44.1;refGene_gn=PTPRS;refGene_tx=XM_023250846.1;refGene_gn=PTPRS;refGe
ne_tx=XM_023250849.1;refGene_gn=PTPRS;refGene_tx=XM_023250850.1;refGen
e_gn=PTPRS;refGene_tx=XM_023250851.1;refGene_gn=PTPRS;refGene_tx=XM_02
3250852.1
A2 3976866 3977055
refGene_gn=PTPRS;refGene_tx=XM_023250838.1;refGene_gn=PTPRS;refGene_tx
=XM_023250839.1;refGene_gn=PTPRS;refGene_tx=XM_023250840.1;refGene_gn=
PTPRS;refGene_tx=XM_023250841.1;refGene_gn=PTPRS;refGene_tx=XM_0232508
42.1;refGene_gn=PTPRS;refGene_tx=XM_023250843.1;refGene_gn=PTPRS;refGe
ne_tx=XM_023250844.1;refGene_gn=PTPRS;refGene_tx=XM_023250845.1;refGen
e_gn=PTPRS;refGene_tx=XM_023250846.1;refGene_gn=PTPRS;refGene_tx=XM_02
3250847.1;refGene_gn=PTPRS;refGene_tx=XM_023250848.1;refGene_gn=PTPRS;
refGene_tx=XM_023250849.1;refGene_gn=PTPRS;refGene_tx=XM_023250850.1;r
efGene_gn=PTPRS;refGene_tx=XM_023250851.1;refGene_gn=PTPRS;refGene_tx=
XM_023250852.1;refGene_gn=PTPRS;refGene_tx=XM_023250853.1;refGene_gn=P
TPRS;refGene_tx=XM_023250854.1;refGene_gn=PTPRS;refGene_tx=XM_02325085
6.1
A2 3982252 3982394
refGene_gn=PTPRS;refGene_tx=XM_023250838.1;refGene_gn=PTPRS;refGene_tx
=XM_023250839.1;refGene_gn=PTPRS;refGene_tx=XM_023250840.1;refGene_gn=
PTPRS;refGene_tx=XM_023250841.1;refGene_gn=PTPRS;refGene_tx=XM_0232508
42.1;refGene_gn=PTPRS;refGene_tx=XM_023250843.1;refGene_gn=PTPRS;refGe
ne_tx=XM_023250844.1;refGene_gn=PTPRS;refGene_tx=XM_023250845.1;refGen
e_gn=PTPRS;refGene_tx=XM_023250846.1;refGene_gn=PTPRS;refGene_tx=XM_02
3250847.1;refGene_gn=PTPRS;refGene_tx=XM_023250848.1;refGene_gn=PTPRS;
refGene_tx=XM_023250849.1;refGene_gn=PTPRS;refGene_tx=XM_023250850.1;r
efGene_gn=PTPRS;refGene_tx=XM_023250851.1;refGene_gn=PTPRS;refGene_tx=
XM_023250852.1;refGene_gn=PTPRS;refGene_tx=XM_023250853.1;refGene_gn=P

TPRS;refGene_tx=XM_023250854.1;refGene_gn=PTPRS;refGene_tx=XM_02325085
6.1
A2 3982859 3983005
refGene_gn=PTPRS;refGene_tx=XM_023250838.1;refGene_gn=PTPRS;refGene_tx
=XM_023250839.1;refGene_gn=PTPRS;refGene_tx=XM_023250840.1;refGene_gn=
PTPRS;refGene_tx=XM_023250841.1;refGene_gn=PTPRS;refGene_tx=XM_0232508
42.1;refGene_gn=PTPRS;refGene_tx=XM_023250843.1;refGene_gn=PTPRS;refGe
ne_tx=XM_023250844.1;refGene_gn=PTPRS;refGene_tx=XM_023250845.1;refGen
e_gn=PTPRS;refGene_tx=XM_023250846.1;refGene_gn=PTPRS;refGene_tx=XM_02
3250847.1;refGene_gn=PTPRS;refGene_tx=XM_023250848.1;refGene_gn=PTPRS;
refGene_tx=XM_023250849.1;refGene_gn=PTPRS;refGene_tx=XM_023250850.1;r
efGene_gn=PTPRS;refGene_tx=XM_023250851.1;refGene_gn=PTPRS;refGene_tx=
XM_023250852.1;refGene_gn=PTPRS;refGene_tx=XM_023250853.1;refGene_gn=P
TPRS;refGene_tx=XM_023250854.1;refGene_gn=PTPRS;refGene_tx=XM_02325085
6.1
A2 3989383 3989474
refGene_gn=PTPRS;refGene_tx=XM_023250838.1;refGene_gn=PTPRS;refGene_tx
=XM_023250839.1;refGene_gn=PTPRS;refGene_tx=XM_023250840.1;refGene_gn=
PTPRS;refGene_tx=XM_023250841.1;refGene_gn=PTPRS;refGene_tx=XM_0232508
42.1;refGene_gn=PTPRS;refGene_tx=XM_023250843.1;refGene_gn=PTPRS;refGe
ne_tx=XM_023250844.1;refGene_gn=PTPRS;refGene_tx=XM_023250845.1;refGen
e_gn=PTPRS;refGene_tx=XM_023250846.1;refGene_gn=PTPRS;refGene_tx=XM_02
3250847.1;refGene_gn=PTPRS;refGene_tx=XM_023250848.1;refGene_gn=PTPRS;
refGene_tx=XM_023250849.1;refGene_gn=PTPRS;refGene_tx=XM_023250850.1;r
efGene_gn=PTPRS;refGene_tx=XM_023250851.1;refGene_gn=PTPRS;refGene_tx=
XM_023250852.1;refGene_gn=PTPRS;refGene_tx=XM_023250853.1;refGene_gn=P
TPRS;refGene_tx=XM_023250854.1;refGene_gn=PTPRS;refGene_tx=XM_02325085
6.1
A2 4099394 4100756 refGene_gn=ZNRF4;refGene_tx=XM_011288395.3
A2 4178875 4179029 refGene_gn=SAFB2;refGene_tx=XM_003981706.5
A2 4179322 4179389 refGene_gn=SAFB2;refGene_tx=XM_003981706.5
A2 4179525 4179617 refGene_gn=SAFB2;refGene_tx=XM_003981706.5
A2 4181068 4181199 refGene_gn=SAFB2;refGene_tx=XM_003981706.5
A2 4182182 4182225 refGene_gn=SAFB2;refGene_tx=XM_003981706.5
A2 4183416 4183557 refGene_gn=SAFB2;refGene_tx=XM_003981706.5
A2 4184435 4184726 refGene_gn=SAFB2;refGene_tx=XM_003981706.5
A2 4186219 4186356 refGene_gn=SAFB2;refGene_tx=XM_003981706.5
A2 4189027 4189119 refGene_gn=SAFB2;refGene_tx=XM_003981706.5
A2 4190327 4190458 refGene_gn=SAFB2;refGene_tx=XM_003981706.5
A2 4194850 4194963 refGene_gn=SAFB2;refGene_tx=XM_003981706.5
A2 4195052 4195202 refGene_gn=SAFB2;refGene_tx=XM_003981706.5
A2 4199834 4199935 refGene_gn=SAFB2;refGene_tx=XM_003981706.5
A2 4200476 4200526 refGene_gn=SAFB2;refGene_tx=XM_003981706.5
A2 4200952 4201463 refGene_gn=SAFB2;refGene_tx=XM_003981706.5
A2 4202380 4202408 refGene_gn=SAFB2;refGene_tx=XM_003981706.5
A2 4203337 4203400 refGene_gn=SAFB2;refGene_tx=XM_003981706.5
A2 4205135 4205339 refGene_gn=SAFB2;refGene_tx=XM_003981706.5
A2 4205422 4205487 refGene_gn=SAFB2;refGene_tx=XM_003981706.5
A2 4210252 4210340 refGene_gn=SAFB2;refGene_tx=XM_003981706.5
A2 4211350 4211536 refGene_gn=SAFB2;refGene_tx=XM_003981706.5

A2 4212017 4212206
refGene_gn=SAFB;refGene_tx=XM_011287719.3;refGene_gn=SAFB;refGene_tx=XM_011287720.3
A2 4215336 4215421
refGene_gn=SAFB;refGene_tx=XM_011287719.3;refGene_gn=SAFB;refGene_tx=XM_011287720.3
A2 4227123 4227188
refGene_gn=SAFB;refGene_tx=XM_011287719.3;refGene_gn=SAFB;refGene_tx=XM_011287720.3
A2 4227270 4227477
refGene_gn=SAFB;refGene_tx=XM_011287719.3;refGene_gn=SAFB;refGene_tx=XM_011287720.3
A2 4230116 4230179
refGene_gn=SAFB;refGene_tx=XM_011287719.3;refGene_gn=SAFB;refGene_tx=XM_011287720.3
A2 4232846 4232874
refGene_gn=SAFB;refGene_tx=XM_011287719.3;refGene_gn=SAFB;refGene_tx=XM_011287720.3
A2 4233815 4234326
refGene_gn=SAFB;refGene_tx=XM_011287719.3;refGene_gn=SAFB;refGene_tx=XM_011287720.3
A2 4234745 4234795
refGene_gn=SAFB;refGene_tx=XM_011287719.3;refGene_gn=SAFB;refGene_tx=XM_011287720.3
A2 4235633 4235728
refGene_gn=SAFB;refGene_tx=XM_011287719.3;refGene_gn=SAFB;refGene_tx=XM_011287720.3
A2 4237446 4237596
refGene_gn=SAFB;refGene_tx=XM_011287719.3;refGene_gn=SAFB;refGene_tx=XM_011287720.3
A2 4237673 4237756
refGene_gn=SAFB;refGene_tx=XM_011287719.3;refGene_gn=SAFB;refGene_tx=XM_011287720.3
A2 4238103 4238243
refGene_gn=SAFB;refGene_tx=XM_011287719.3;refGene_gn=SAFB;refGene_tx=XM_011287720.3
A2 4238407 4238496
refGene_gn=SAFB;refGene_tx=XM_011287719.3;refGene_gn=SAFB;refGene_tx=XM_011287720.3
A2 4240929 4241036
refGene_gn=SAFB;refGene_tx=XM_011287719.3;refGene_gn=SAFB;refGene_tx=XM_011287720.3
A2 4243226 4243517
refGene_gn=SAFB;refGene_tx=XM_011287719.3;refGene_gn=SAFB;refGene_tx=XM_011287720.3
A2 4244669 4244807
refGene_gn=SAFB;refGene_tx=XM_011287719.3;refGene_gn=SAFB;refGene_tx=XM_011287720.3
A2 4245022 4245065
refGene_gn=SAFB;refGene_tx=XM_011287719.3;refGene_gn=SAFB;refGene_tx=X

M_011287720.3
A2 4247384 4247500
refGene_gn=SAFB;refGene_tx=XM_011287719.3;refGene_gn=SAFB;refGene_tx=XM_011287720.3
A2 4247705 4247812
refGene_gn=SAFB;refGene_tx=XM_011287719.3;refGene_gn=SAFB;refGene_tx=XM_011287720.3
A2 4248236 4248303
refGene_gn=SAFB;refGene_tx=XM_011287719.3;refGene_gn=SAFB;refGene_tx=XM_011287720.3
A2 4248510 4248640
refGene_gn=SAFB;refGene_tx=XM_011287719.3;refGene_gn=SAFB;refGene_tx=XM_011287720.3
A2 4253541 4253639
refGene_gn=CA2H19orf70;refGene_tx=XM_003981708.4;refGene_gn=CA2H19orf70;refGene_tx=XM_006928288.3
A2 4254026 4254078
refGene_gn=CA2H19orf70;refGene_tx=XM_003981708.4;refGene_gn=CA2H19orf70;refGene_tx=XM_006928288.3
A2 4254303 4254481
refGene_gn=CA2H19orf70;refGene_tx=XM_003981708.4;refGene_gn=CA2H19orf70;refGene_tx=XM_006928288.3
A2 4255212 4255241
refGene_gn=CA2H19orf70;refGene_tx=XM_003981708.4;refGene_gn=CA2H19orf70;refGene_tx=XM_006928288.3
A2 4255695 4255717
refGene_gn=HSD11B1L;refGene_tx=XM_006928286.4;refGene_gn=HSD11B1L;refGene_tx=XM_006928287.4
A2 4257645 4257741
refGene_gn=HSD11B1L;refGene_tx=XM_003981709.5;refGene_gn=HSD11B1L;refGene_tx=XM_006928286.4;refGene_gn=HSD11B1L;refGene_tx=XM_006928287.4
A2 4257825 4257956
refGene_gn=HSD11B1L;refGene_tx=XM_003981709.5;refGene_gn=HSD11B1L;refGene_tx=XM_006928286.4;refGene_gn=HSD11B1L;refGene_tx=XM_006928287.4
A2 4258380 4258492
refGene_gn=HSD11B1L;refGene_tx=XM_003981709.5;refGene_gn=HSD11B1L;refGene_tx=XM_006928286.4;refGene_gn=HSD11B1L;refGene_tx=XM_006928287.4
A2 4258802 4258913
refGene_gn=HSD11B1L;refGene_tx=XM_003981709.5;refGene_gn=HSD11B1L;refGene_tx=XM_006928286.4;refGene_gn=HSD11B1L;refGene_tx=XM_006928287.4;refGene_gn=HSD11B1L;refGene_tx=XM_019816119.2
A2 4259089 4259183
refGene_gn=HSD11B1L;refGene_tx=XM_003981709.5;refGene_gn=HSD11B1L;refGene_tx=XM_006928286.4;refGene_gn=HSD11B1L;refGene_tx=XM_006928287.4;refGene_gn=HSD11B1L;refGene_tx=XM_019816119.2
A2 4259312 4259478
refGene_gn=HSD11B1L;refGene_tx=XM_003981709.5;refGene_gn=HSD11B1L;refGene_tx=XM_006928286.4;refGene_gn=HSD11B1L;refGene_tx=XM_006928287.4;refGene_gn=HSD11B1L;refGene_tx=XM_019816119.2
A2 4259557 4259753

```
refGene_gn=HSD11B1L;refGene_tx=XM_003981709.5;refGene_gn=HSD11B1L;refG
ene_tx=XM_006928286.4;refGene_gn=HSD11B1L;refGene_tx=XM_006928287.4;re
fGene_gn=HSD11B1L;refGene_tx=XM_019816119.2
A2	4261743	4261836	refGene_gn=RPL36;refGene_tx=XM_011287701.3
A2	4262371	4262506	refGene_gn=RPL36;refGene_tx=XM_011287701.3
A2	4263446	4263536	refGene_gn=RPL36;refGene_tx=XM_011287701.3
A2	4263839	4264016	refGene_gn=LONP1;refGene_tx=XM_003981712.5
A2	4264723	4264888	refGene_gn=LONP1;refGene_tx=XM_003981712.5
A2	4264982	4265200	refGene_gn=LONP1;refGene_tx=XM_003981712.5
A2	4265779	4265945	refGene_gn=LONP1;refGene_tx=XM_003981712.5
A2	4266340	4266481	refGene_gn=LONP1;refGene_tx=XM_003981712.5
A2	4268970	4269087	refGene_gn=LONP1;refGene_tx=XM_003981712.5
A2	4269177	4269300	refGene_gn=LONP1;refGene_tx=XM_003981712.5
A2	4269642	4269730	refGene_gn=LONP1;refGene_tx=XM_003981712.5
A2	4271772	4271951	refGene_gn=LONP1;refGene_tx=XM_003981712.5
A2	4272887	4273026	refGene_gn=LONP1;refGene_tx=XM_003981712.5
A2	4273762	4273983	refGene_gn=LONP1;refGene_tx=XM_003981712.5
A2	4275087	4275171	refGene_gn=LONP1;refGene_tx=XM_003981712.5
A2	4275563	4275693	refGene_gn=LONP1;refGene_tx=XM_003981712.5
A2	4276245	4276307	refGene_gn=LONP1;refGene_tx=XM_003981712.5
A2	4277844	4278076	refGene_gn=LONP1;refGene_tx=XM_003981712.5
A2	4279797	4279917	refGene_gn=LONP1;refGene_tx=XM_003981712.5
A2	4280534	4280623	refGene_gn=LONP1;refGene_tx=XM_003981712.5
A2	4283235	4283667	refGene_gn=LONP1;refGene_tx=XM_003981712.5
A2	4284358	4284432
refGene_gn=CATSPERD;refGene_tx=XM_011287677.3;refGene_gn=CATSPERD;refG
ene_tx=XM_019816085.2;refGene_gn=CATSPERD;refGene_tx=XM_023250872.1
A2	4286592	4286647
refGene_gn=CATSPERD;refGene_tx=XM_011287677.3;refGene_gn=CATSPERD;refG
ene_tx=XM_019816085.2;refGene_gn=CATSPERD;refGene_tx=XM_023250872.1
A2	4287475	4287552
refGene_gn=CATSPERD;refGene_tx=XM_011287677.3;refGene_gn=CATSPERD;refG
ene_tx=XM_019816085.2;refGene_gn=CATSPERD;refGene_tx=XM_023250872.1
A2	4291289	4291362
refGene_gn=CATSPERD;refGene_tx=XM_011287677.3;refGene_gn=CATSPERD;refG
ene_tx=XM_019816085.2;refGene_gn=CATSPERD;refGene_tx=XM_023250872.1
A2	4292091	4292206
refGene_gn=CATSPERD;refGene_tx=XM_011287677.3;refGene_gn=CATSPERD;refG
ene_tx=XM_019816085.2;refGene_gn=CATSPERD;refGene_tx=XM_023250872.1
A2	4296209	4296277
refGene_gn=CATSPERD;refGene_tx=XM_011287677.3;refGene_gn=CATSPERD;refG
ene_tx=XM_019816085.2;refGene_gn=CATSPERD;refGene_tx=XM_023250872.1
A2	4298085	4298199
refGene_gn=CATSPERD;refGene_tx=XM_011287677.3;refGene_gn=CATSPERD;refG
ene_tx=XM_019816085.2;refGene_gn=CATSPERD;refGene_tx=XM_023250872.1
A2	4300098	4300317	refGene_gn=CATSPERD;refGene_tx=XM_023250873.1
A2	4301534	4301618
refGene_gn=CATSPERD;refGene_tx=XM_011287677.3;refGene_gn=CATSPERD;refG
ene_tx=XM_019816085.2;refGene_gn=CATSPERD;refGene_tx=XM_023250872.1;re
fGene_gn=CATSPERD;refGene_tx=XM_023250873.1
```

A2	4303654	4303805
refGene_gn=CATSPERD;refGene_tx=XM_011287677.3;refGene_gn=CATSPERD;refGene_tx=XM_019816085.2;refGene_gn=CATSPERD;refGene_tx=XM_023250872.1;refGene_gn=CATSPERD;refGene_tx=XM_023250873.1
A2	4304697	4304793
refGene_gn=CATSPERD;refGene_tx=XM_011287677.3;refGene_gn=CATSPERD;refGene_tx=XM_019816085.2;refGene_gn=CATSPERD;refGene_tx=XM_023250872.1;refGene_gn=CATSPERD;refGene_tx=XM_023250873.1
A2	4306934	4307017
refGene_gn=CATSPERD;refGene_tx=XM_011287677.3;refGene_gn=CATSPERD;refGene_tx=XM_019816085.2;refGene_gn=CATSPERD;refGene_tx=XM_023250872.1;refGene_gn=CATSPERD;refGene_tx=XM_023250873.1
A2	4308237	4308414
refGene_gn=CATSPERD;refGene_tx=XM_011287677.3;refGene_gn=CATSPERD;refGene_tx=XM_019816085.2;refGene_gn=CATSPERD;refGene_tx=XM_023250872.1;refGene_gn=CATSPERD;refGene_tx=XM_023250873.1
A2	4309840	4309954
refGene_gn=CATSPERD;refGene_tx=XM_011287677.3;refGene_gn=CATSPERD;refGene_tx=XM_019816085.2;refGene_gn=CATSPERD;refGene_tx=XM_023250872.1;refGene_gn=CATSPERD;refGene_tx=XM_023250873.1
A2	4310714	4310804
refGene_gn=CATSPERD;refGene_tx=XM_011287677.3;refGene_gn=CATSPERD;refGene_tx=XM_019816085.2;refGene_gn=CATSPERD;refGene_tx=XM_023250872.1;refGene_gn=CATSPERD;refGene_tx=XM_023250873.1
A2	4311564	4311623
refGene_gn=CATSPERD;refGene_tx=XM_011287677.3;refGene_gn=CATSPERD;refGene_tx=XM_019816085.2;refGene_gn=CATSPERD;refGene_tx=XM_023250872.1;refGene_gn=CATSPERD;refGene_tx=XM_023250873.1
A2	4314862	4314941
refGene_gn=CATSPERD;refGene_tx=XM_011287677.3;refGene_gn=CATSPERD;refGene_tx=XM_019816085.2;refGene_gn=CATSPERD;refGene_tx=XM_023250872.1;refGene_gn=CATSPERD;refGene_tx=XM_023250873.1
A2	4316358	4316411
refGene_gn=CATSPERD;refGene_tx=XM_011287677.3;refGene_gn=CATSPERD;refGene_tx=XM_019816085.2;refGene_gn=CATSPERD;refGene_tx=XM_023250872.1;refGene_gn=CATSPERD;refGene_tx=XM_023250873.1
A2	4318835	4318910
refGene_gn=CATSPERD;refGene_tx=XM_011287677.3;refGene_gn=CATSPERD;refGene_tx=XM_023250872.1;refGene_gn=CATSPERD;refGene_tx=XM_023250873.1
A2	4319724	4319853
refGene_gn=CATSPERD;refGene_tx=XM_011287677.3;refGene_gn=CATSPERD;refGene_tx=XM_023250872.1;refGene_gn=CATSPERD;refGene_tx=XM_023250873.1
A2	4321453	4321637
refGene_gn=CATSPERD;refGene_tx=XM_011287677.3;refGene_gn=CATSPERD;refGene_tx=XM_019816085.2;refGene_gn=CATSPERD;refGene_tx=XM_023250872.1;refGene_gn=CATSPERD;refGene_tx=XM_023250873.1
A2	4325721	4325879
refGene_gn=CATSPERD;refGene_tx=XM_011287677.3;refGene_gn=CATSPERD;refGene_tx=XM_019816085.2;refGene_gn=CATSPERD;refGene_tx=XM_023250872.1;refGene_gn=CATSPERD;refGene_tx=XM_023250873.1

A2	4326761	4326876	refGene_gn=CATSPERD;refGene_tx=XM_023250872.1
A2	4327000	4327238
refGene_gn=CATSPERD;refGene_tx=XM_011287677.3;refGene_gn=CATSPERD;refGene_tx=XM_019816085.2;refGene_gn=CATSPERD;refGene_tx=XM_023250873.1
A2	4328551	4329997
refGene_gn=PRR22;refGene_tx=XM_019816099.2;refGene_gn=PRR22;refGene_tx=XM_023250883.1
A2	4330350	4330413
refGene_gn=PRR22;refGene_tx=XM_019816099.2;refGene_gn=PRR22;refGene_tx=XM_023250883.1
A2	4330516	4330646
refGene_gn=PRR22;refGene_tx=XM_019816099.2;refGene_gn=PRR22;refGene_tx=XM_023250883.1
A2	4331256	4331329	refGene_gn=DUS3L;refGene_tx=XM_011287689.3
A2	4331423	4331552	refGene_gn=DUS3L;refGene_tx=XM_011287689.3
A2	4331627	4331816	refGene_gn=DUS3L;refGene_tx=XM_011287689.3
A2	4332132	4332208	refGene_gn=DUS3L;refGene_tx=XM_011287689.3
A2	4332409	4332506	refGene_gn=DUS3L;refGene_tx=XM_011287689.3
A2	4332756	4332867	refGene_gn=DUS3L;refGene_tx=XM_011287689.3
A2	4332949	4333015	refGene_gn=DUS3L;refGene_tx=XM_011287689.3
A2	4333215	4333332	refGene_gn=DUS3L;refGene_tx=XM_011287689.3
A2	4333593	4333746	refGene_gn=DUS3L;refGene_tx=XM_011287689.3
A2	4333870	4333912	refGene_gn=DUS3L;refGene_tx=XM_011287689.3
A2	4334262	4334775	refGene_gn=DUS3L;refGene_tx=XM_011287689.3
A2	4335029	4335303	refGene_gn=DUS3L;refGene_tx=XM_011287689.3
A2	4335988	4336086	refGene_gn=DUS3L;refGene_tx=XM_011287689.3
A2	4356108	4356277	refGene_gn=NRTN;refGene_tx=XM_023250885.1
A2	4358898	4359323	refGene_gn=NRTN;refGene_tx=XM_023250885.1
A2	4362212	4363364
refGene_gn=LOC101093666;refGene_tx=XM_023250884.1
A2	4384726	4384839	refGene_gn=NDUFA11;refGene_tx=XM_003981714.5
A2	4386159	4386282	refGene_gn=NDUFA11;refGene_tx=XM_003981714.5
A2	4386374	4386467	refGene_gn=NDUFA11;refGene_tx=XM_003981714.5
A2	4390318	4390415	refGene_gn=NDUFA11;refGene_tx=XM_003981714.5
A2	4391563	4391754	refGene_gn=VMAC;refGene_tx=XM_003981715.5
A2	4392979	4393298	refGene_gn=VMAC;refGene_tx=XM_003981715.5
A2	4396581	4396664	refGene_gn=CAPS;refGene_tx=XM_003981716.4
A2	4396737	4396915	refGene_gn=CAPS;refGene_tx=XM_003981716.4
A2	4397032	4397239	refGene_gn=CAPS;refGene_tx=XM_003981716.4
A2	4397312	4397414	refGene_gn=CAPS;refGene_tx=XM_003981716.4
A2	4399320	4399364
refGene_gn=RANBP3;refGene_tx=XM_003981718.5;refGene_gn=RANBP3;refGene_tx=XM_006928289.4;refGene_gn=RANBP3;refGene_tx=XM_006928290.4;refGene_gn=RANBP3;refGene_tx=XM_011287630.3;refGene_gn=RANBP3;refGene_tx=XM_011287639.3;refGene_gn=RANBP3;refGene_tx=XM_011287643.3;refGene_gn=RANBP3;refGene_tx=XM_011287647.3;refGene_gn=RANBP3;refGene_tx=XM_019816198.2
A2	4399470	4399651
refGene_gn=RANBP3;refGene_tx=XM_003981718.5;refGene_gn=RANBP3;refGene_tx=XM_006928289.4;refGene_gn=RANBP3;refGene_tx=XM_006928290.4;refGene_

gn=RANBP3;refGene_tx=XM_011287630.3;refGene_gn=RANBP3;refGene_tx=XM_01
1287639.3;refGene_gn=RANBP3;refGene_tx=XM_011287643.3;refGene_gn=RANBP
3;refGene_tx=XM_011287647.3;refGene_gn=RANBP3;refGene_tx=XM_019816198.
2
A2 4400113 4400256
refGene_gn=RANBP3;refGene_tx=XM_003981718.5;refGene_gn=RANBP3;refGene_
tx=XM_006928289.4;refGene_gn=RANBP3;refGene_tx=XM_006928290.4;refGene_
gn=RANBP3;refGene_tx=XM_011287630.3;refGene_gn=RANBP3;refGene_tx=XM_01
1287639.3;refGene_gn=RANBP3;refGene_tx=XM_011287643.3;refGene_gn=RANBP
3;refGene_tx=XM_011287647.3;refGene_gn=RANBP3;refGene_tx=XM_019816198.
2
A2 4401386 4401507
refGene_gn=RANBP3;refGene_tx=XM_003981718.5;refGene_gn=RANBP3;refGene_
tx=XM_006928289.4;refGene_gn=RANBP3;refGene_tx=XM_006928290.4;refGene_
gn=RANBP3;refGene_tx=XM_011287630.3;refGene_gn=RANBP3;refGene_tx=XM_01
1287639.3;refGene_gn=RANBP3;refGene_tx=XM_011287643.3;refGene_gn=RANBP
3;refGene_tx=XM_011287647.3;refGene_gn=RANBP3;refGene_tx=XM_019816198.
2
A2 4403517 4403642
refGene_gn=RANBP3;refGene_tx=XM_003981718.5;refGene_gn=RANBP3;refGene_
tx=XM_006928289.4;refGene_gn=RANBP3;refGene_tx=XM_006928290.4;refGene_
gn=RANBP3;refGene_tx=XM_011287630.3;refGene_gn=RANBP3;refGene_tx=XM_01
1287639.3;refGene_gn=RANBP3;refGene_tx=XM_011287643.3;refGene_gn=RANBP
3;refGene_tx=XM_011287647.3;refGene_gn=RANBP3;refGene_tx=XM_019816198.
2
A2 4404179 4404282
refGene_gn=RANBP3;refGene_tx=XM_003981718.5;refGene_gn=RANBP3;refGene_
tx=XM_006928289.4;refGene_gn=RANBP3;refGene_tx=XM_006928290.4;refGene_
gn=RANBP3;refGene_tx=XM_011287630.3;refGene_gn=RANBP3;refGene_tx=XM_01
1287639.3;refGene_gn=RANBP3;refGene_tx=XM_011287643.3;refGene_gn=RANBP
3;refGene_tx=XM_011287647.3;refGene_gn=RANBP3;refGene_tx=XM_019816198.
2
A2 4405454 4405533
refGene_gn=RANBP3;refGene_tx=XM_003981718.5;refGene_gn=RANBP3;refGene_
tx=XM_006928289.4;refGene_gn=RANBP3;refGene_tx=XM_006928290.4;refGene_
gn=RANBP3;refGene_tx=XM_011287630.3;refGene_gn=RANBP3;refGene_tx=XM_01
1287639.3;refGene_gn=RANBP3;refGene_tx=XM_011287643.3;refGene_gn=RANBP
3;refGene_tx=XM_011287647.3;refGene_gn=RANBP3;refGene_tx=XM_019816198.
2
A2 4407042 4407146
refGene_gn=RANBP3;refGene_tx=XM_003981718.5;refGene_gn=RANBP3;refGene_
tx=XM_006928289.4;refGene_gn=RANBP3;refGene_tx=XM_006928290.4;refGene_
gn=RANBP3;refGene_tx=XM_011287630.3;refGene_gn=RANBP3;refGene_tx=XM_01
1287639.3;refGene_gn=RANBP3;refGene_tx=XM_011287643.3;refGene_gn=RANBP
3;refGene_tx=XM_011287647.3;refGene_gn=RANBP3;refGene_tx=XM_019816198.
2
A2 4409044 4409158
refGene_gn=RANBP3;refGene_tx=XM_003981718.5;refGene_gn=RANBP3;refGene_
tx=XM_006928289.4;refGene_gn=RANBP3;refGene_tx=XM_006928290.4;refGene_
gn=RANBP3;refGene_tx=XM_011287630.3;refGene_gn=RANBP3;refGene_tx=XM_01

1287639.3;refGene_gn=RANBP3;refGene_tx=XM_011287643.3;refGene_gn=RANBP3;refGene_tx=XM_011287647.3;refGene_gn=RANBP3;refGene_tx=XM_019816198.2
A2 4410152 4410271
refGene_gn=RANBP3;refGene_tx=XM_003981718.5;refGene_gn=RANBP3;refGene_tx=XM_006928289.4;refGene_gn=RANBP3;refGene_tx=XM_006928290.4;refGene_gn=RANBP3;refGene_tx=XM_011287630.3;refGene_gn=RANBP3;refGene_tx=XM_011287639.3;refGene_gn=RANBP3;refGene_tx=XM_011287643.3;refGene_gn=RANBP3;refGene_tx=XM_011287647.3;refGene_gn=RANBP3;refGene_tx=XM_019816198.2
A2 4411654 4411747
refGene_gn=RANBP3;refGene_tx=XM_003981718.5;refGene_gn=RANBP3;refGene_tx=XM_006928289.4;refGene_gn=RANBP3;refGene_tx=XM_006928290.4;refGene_gn=RANBP3;refGene_tx=XM_011287630.3;refGene_gn=RANBP3;refGene_tx=XM_011287639.3;refGene_gn=RANBP3;refGene_tx=XM_011287643.3;refGene_gn=RANBP3;refGene_tx=XM_011287647.3;refGene_gn=RANBP3;refGene_tx=XM_019816198.2
A2 4412499 4412565
refGene_gn=RANBP3;refGene_tx=XM_003981718.5;refGene_gn=RANBP3;refGene_tx=XM_006928289.4;refGene_gn=RANBP3;refGene_tx=XM_006928290.4;refGene_gn=RANBP3;refGene_tx=XM_011287630.3;refGene_gn=RANBP3;refGene_tx=XM_011287639.3;refGene_gn=RANBP3;refGene_tx=XM_011287643.3;refGene_gn=RANBP3;refGene_tx=XM_011287647.3;refGene_gn=RANBP3;refGene_tx=XM_019816198.2
A2 4417810 4417882
refGene_gn=RANBP3;refGene_tx=XM_006928289.4;refGene_gn=RANBP3;refGene_tx=XM_011287630.3;refGene_gn=RANBP3;refGene_tx=XM_011287643.3;refGene_gn=RANBP3;refGene_tx=XM_011287647.3;refGene_gn=RANBP3;refGene_tx=XM_019816198.2
A2 4419720 4419807
refGene_gn=RANBP3;refGene_tx=XM_003981718.5;refGene_gn=RANBP3;refGene_tx=XM_006928289.4;refGene_gn=RANBP3;refGene_tx=XM_006928290.4;refGene_gn=RANBP3;refGene_tx=XM_011287630.3;refGene_gn=RANBP3;refGene_tx=XM_011287639.3;refGene_gn=RANBP3;refGene_tx=XM_011287643.3;refGene_gn=RANBP3;refGene_tx=XM_011287647.3;refGene_gn=RANBP3;refGene_tx=XM_019816198.2
A2 4419904 4419941
refGene_gn=RANBP3;refGene_tx=XM_003981718.5;refGene_gn=RANBP3;refGene_tx=XM_006928289.4;refGene_gn=RANBP3;refGene_tx=XM_006928290.4;refGene_gn=RANBP3;refGene_tx=XM_011287630.3;refGene_gn=RANBP3;refGene_tx=XM_011287639.3;refGene_gn=RANBP3;refGene_tx=XM_011287643.3;refGene_gn=RANBP3;refGene_tx=XM_011287647.3;refGene_gn=RANBP3;refGene_tx=XM_019816198.2
A2 4427124 4427133
refGene_gn=RANBP3;refGene_tx=XM_011287647.3;refGene_gn=RANBP3;refGene_tx=XM_019816198.2
A2 4427795 4427984
refGene_gn=RANBP3;refGene_tx=XM_003981718.5;refGene_gn=RANBP3;refGene_tx=XM_006928289.4;refGene_gn=RANBP3;refGene_tx=XM_011287630.3;refGene_gn=RANBP3;refGene_tx=XM_011287639.3

A2 4433659 4433715
refGene_gn=RANBP3;refGene_tx=XM_003981718.5;refGene_gn=RANBP3;refGene_tx=XM_006928289.4;refGene_gn=RANBP3;refGene_tx=XM_006928290.4;refGene_gn=RANBP3;refGene_tx=XM_011287630.3;refGene_gn=RANBP3;refGene_tx=XM_011287639.3;refGene_gn=RANBP3;refGene_tx=XM_011287643.3
A2 4453190 4453212
refGene_gn=RANBP3;refGene_tx=XM_003981718.5;refGene_gn=RANBP3;refGene_tx=XM_006928289.4;refGene_gn=RANBP3;refGene_tx=XM_006928290.4;refGene_gn=RANBP3;refGene_tx=XM_011287630.3;refGene_gn=RANBP3;refGene_tx=XM_011287639.3;refGene_gn=RANBP3;refGene_tx=XM_011287643.3
A2 4466501 4466614
refGene_gn=RFX2;refGene_tx=XM_011287614.3;refGene_gn=RFX2;refGene_tx=XM_023250874.1;refGene_gn=RFX2;refGene_tx=XM_023250875.1;refGene_gn=RFX2;refGene_tx=XM_023250876.1;refGene_gn=RFX2;refGene_tx=XM_023250877.1
A2 4467139 4467182
refGene_gn=RFX2;refGene_tx=XM_011287614.3;refGene_gn=RFX2;refGene_tx=XM_023250874.1;refGene_gn=RFX2;refGene_tx=XM_023250875.1;refGene_gn=RFX2;refGene_tx=XM_023250876.1;refGene_gn=RFX2;refGene_tx=XM_023250877.1
A2 4468660 4468844
refGene_gn=RFX2;refGene_tx=XM_011287614.3;refGene_gn=RFX2;refGene_tx=XM_023250874.1;refGene_gn=RFX2;refGene_tx=XM_023250875.1;refGene_gn=RFX2;refGene_tx=XM_023250876.1;refGene_gn=RFX2;refGene_tx=XM_023250877.1
A2 4469248 4469457
refGene_gn=RFX2;refGene_tx=XM_011287614.3;refGene_gn=RFX2;refGene_tx=XM_023250874.1;refGene_gn=RFX2;refGene_tx=XM_023250875.1;refGene_gn=RFX2;refGene_tx=XM_023250876.1;refGene_gn=RFX2;refGene_tx=XM_023250877.1
A2 4469992 4470142
refGene_gn=RFX2;refGene_tx=XM_011287614.3;refGene_gn=RFX2;refGene_tx=XM_023250874.1;refGene_gn=RFX2;refGene_tx=XM_023250875.1;refGene_gn=RFX2;refGene_tx=XM_023250876.1;refGene_gn=RFX2;refGene_tx=XM_023250877.1
A2 4470933 4471031
refGene_gn=RFX2;refGene_tx=XM_011287614.3;refGene_gn=RFX2;refGene_tx=XM_023250874.1;refGene_gn=RFX2;refGene_tx=XM_023250875.1;refGene_gn=RFX2;refGene_tx=XM_023250876.1;refGene_gn=RFX2;refGene_tx=XM_023250877.1
A2 4472306 4472461
refGene_gn=RFX2;refGene_tx=XM_011287614.3;refGene_gn=RFX2;refGene_tx=XM_023250874.1;refGene_gn=RFX2;refGene_tx=XM_023250875.1;refGene_gn=RFX2;refGene_tx=XM_023250876.1;refGene_gn=RFX2;refGene_tx=XM_023250877.1
A2 4472881 4472994
refGene_gn=RFX2;refGene_tx=XM_011287614.3;refGene_gn=RFX2;refGene_tx=XM_023250874.1;refGene_gn=RFX2;refGene_tx=XM_023250875.1;refGene_gn=RFX2;refGene_tx=XM_023250876.1;refGene_gn=RFX2;refGene_tx=XM_023250877.1
A2 4473612 4473731
refGene_gn=RFX2;refGene_tx=XM_011287614.3;refGene_gn=RFX2;refGene_tx=XM_023250874.1;refGene_gn=RFX2;refGene_tx=XM_023250875.1;refGene_gn=RFX2;refGene_tx=XM_023250876.1;refGene_gn=RFX2;refGene_tx=XM_023250877.1
A2 4474629 4474745
refGene_gn=RFX2;refGene_tx=XM_011287614.3;refGene_gn=RFX2;refGene_tx=XM_023250874.1;refGene_gn=RFX2;refGene_tx=XM_023250875.1;refGene_gn=RFX2;refGene_tx=XM_023250876.1;refGene_gn=RFX2;refGene_tx=XM_023250877.1

A2	4477448	4477568
refGene_gn=RFX2;refGene_tx=XM_011287614.3;refGene_gn=RFX2;refGene_tx=XM_023250874.1;refGene_gn=RFX2;refGene_tx=XM_023250875.1;refGene_gn=RFX2;refGene_tx=XM_023250876.1;refGene_gn=RFX2;refGene_tx=XM_023250877.1
A2	4478797	4478979
refGene_gn=RFX2;refGene_tx=XM_011287614.3;refGene_gn=RFX2;refGene_tx=XM_023250874.1;refGene_gn=RFX2;refGene_tx=XM_023250875.1;refGene_gn=RFX2;refGene_tx=XM_023250876.1;refGene_gn=RFX2;refGene_tx=XM_023250877.1
A2	4484495	4484570
refGene_gn=RFX2;refGene_tx=XM_011287614.3;refGene_gn=RFX2;refGene_tx=XM_023250874.1;refGene_gn=RFX2;refGene_tx=XM_023250875.1
A2	4498378	4498640
refGene_gn=RFX2;refGene_tx=XM_011287614.3;refGene_gn=RFX2;refGene_tx=XM_023250874.1;refGene_gn=RFX2;refGene_tx=XM_023250875.1;refGene_gn=RFX2;refGene_tx=XM_023250876.1;refGene_gn=RFX2;refGene_tx=XM_023250877.1
A2	4500055	4500135
refGene_gn=RFX2;refGene_tx=XM_011287614.3;refGene_gn=RFX2;refGene_tx=XM_023250874.1;refGene_gn=RFX2;refGene_tx=XM_023250875.1;refGene_gn=RFX2;refGene_tx=XM_023250876.1;refGene_gn=RFX2;refGene_tx=XM_023250877.1
A2	4502029	4502119
refGene_gn=RFX2;refGene_tx=XM_011287614.3;refGene_gn=RFX2;refGene_tx=XM_023250874.1;refGene_gn=RFX2;refGene_tx=XM_023250875.1;refGene_gn=RFX2;refGene_tx=XM_023250876.1;refGene_gn=RFX2;refGene_tx=XM_023250877.1
A2	4503822	4503927
refGene_gn=RFX2;refGene_tx=XM_011287614.3;refGene_gn=RFX2;refGene_tx=XM_023250874.1;refGene_gn=RFX2;refGene_tx=XM_023250875.1;refGene_gn=RFX2;refGene_tx=XM_023250876.1;refGene_gn=RFX2;refGene_tx=XM_023250877.1
A2	4570000	4570014
refGene_gn=LOC101085442;refGene_tx=XM_006928292.3;refGene_gn=LOC101085442;refGene_tx=XM_006928293.3;refGene_gn=LOC101085442;refGene_tx=XM_019816229.2;refGene_gn=LOC101085442;refGene_tx=XM_019816243.2
A2	4574766	4574867
refGene_gn=LOC101085442;refGene_tx=XM_006928292.3;refGene_gn=LOC101085442;refGene_tx=XM_006928293.3;refGene_gn=LOC101085442;refGene_tx=XM_019816229.2;refGene_gn=LOC101085442;refGene_tx=XM_019816243.2
A2	4581111	4581341
refGene_gn=LOC101085442;refGene_tx=XM_006928292.3;refGene_gn=LOC101085442;refGene_tx=XM_006928293.3;refGene_gn=LOC101085442;refGene_tx=XM_019816229.2;refGene_gn=LOC101085442;refGene_tx=XM_019816243.2
A2	4584843	4584932
refGene_gn=LOC101085442;refGene_tx=XM_006928292.3;refGene_gn=LOC101085442;refGene_tx=XM_006928293.3;refGene_gn=LOC101085442;refGene_tx=XM_019816229.2;refGene_gn=LOC101085442;refGene_tx=XM_019816243.2
A2	4585764	4585885
refGene_gn=LOC101085442;refGene_tx=XM_006928292.3;refGene_gn=LOC101085442;refGene_tx=XM_006928293.3;refGene_gn=LOC101085442;refGene_tx=XM_019816229.2;refGene_gn=LOC101085442;refGene_tx=XM_019816243.2
A2	4588081	4588165
refGene_gn=LOC101085442;refGene_tx=XM_006928292.3;refGene_gn=LOC101085442;refGene_tx=XM_006928293.3;refGene_gn=LOC101085442;refGene_tx=XM_01

9816229.2;refGene_gn=LOC101085442;refGene_tx=XM_019816243.2
A2	4590417	4590567
refGene_gn=LOC101085442;refGene_tx=XM_006928292.3;refGene_gn=LOC101085442;refGene_tx=XM_006928293.3;refGene_gn=LOC101085442;refGene_tx=XM_019816229.2;refGene_gn=LOC101085442;refGene_tx=XM_019816243.2
A2	4593978	4594155
refGene_gn=LOC101085442;refGene_tx=XM_006928292.3;refGene_gn=LOC101085442;refGene_tx=XM_006928293.3;refGene_gn=LOC101085442;refGene_tx=XM_019816229.2;refGene_gn=LOC101085442;refGene_tx=XM_019816243.2
A2	4596725	4596907
refGene_gn=LOC101085442;refGene_tx=XM_006928292.3;refGene_gn=LOC101085442;refGene_tx=XM_006928293.3;refGene_gn=LOC101085442;refGene_tx=XM_019816229.2;refGene_gn=LOC101085442;refGene_tx=XM_019816243.2
A2	4597060	4597300
refGene_gn=LOC101085442;refGene_tx=XM_006928292.3;refGene_gn=LOC101085442;refGene_tx=XM_006928293.3;refGene_gn=LOC101085442;refGene_tx=XM_019816229.2;refGene_gn=LOC101085442;refGene_tx=XM_019816243.2
A2	4599406	4599627
refGene_gn=LOC101085442;refGene_tx=XM_006928292.3;refGene_gn=LOC101085442;refGene_tx=XM_006928293.3;refGene_gn=LOC101085442;refGene_tx=XM_019816229.2;refGene_gn=LOC101085442;refGene_tx=XM_019816243.2
A2	4601060	4601267
refGene_gn=LOC101085442;refGene_tx=XM_006928292.3;refGene_gn=LOC101085442;refGene_tx=XM_019816229.2
A2	4601876	4602016
refGene_gn=LOC101085442;refGene_tx=XM_006928292.3;refGene_gn=LOC101085442;refGene_tx=XM_006928293.3;refGene_gn=LOC101085442;refGene_tx=XM_019816229.2;refGene_gn=LOC101085442;refGene_tx=XM_019816243.2
A2	4602165	4602412
refGene_gn=LOC101085442;refGene_tx=XM_006928292.3;refGene_gn=LOC101085442;refGene_tx=XM_006928293.3;refGene_gn=LOC101085442;refGene_tx=XM_019816229.2;refGene_gn=LOC101085442;refGene_tx=XM_019816243.2
A2	4603678	4603752
refGene_gn=LOC101085442;refGene_tx=XM_006928292.3;refGene_gn=LOC101085442;refGene_tx=XM_006928293.3;refGene_gn=LOC101085442;refGene_tx=XM_019816229.2;refGene_gn=LOC101085442;refGene_tx=XM_019816243.2
A2	4615099	4615160
refGene_gn=LOC102900010;refGene_tx=XM_023244096.1;refGene_gn=LOC102900010;refGene_tx=XM_023244097.1;refGene_gn=LOC102900010;refGene_tx=XM_023244098.1
A2	4615639	4615863
refGene_gn=LOC102900010;refGene_tx=XM_023244096.1;refGene_gn=LOC102900010;refGene_tx=XM_023244097.1;refGene_gn=LOC102900010;refGene_tx=XM_023244098.1
A2	4618698	4618787
refGene_gn=LOC102900010;refGene_tx=XM_023244096.1;refGene_gn=LOC102900010;refGene_tx=XM_023244097.1;refGene_gn=LOC102900010;refGene_tx=XM_023244098.1
A2	4619329	4619450
refGene_gn=LOC102900010;refGene_tx=XM_023244096.1;refGene_gn=LOC102900

010;refGene_tx=XM_023244097.1;refGene_gn=LOC102900010;refGene_tx=XM_023244098.1
A2	4624694	4624775	refGene_gn=LOC102900010;refGene_tx=XM_023244096.1;refGene_gn=LOC102900010;refGene_tx=XM_023244097.1;refGene_gn=LOC102900010;refGene_tx=XM_023244098.1
A2	4624885	4625035	refGene_gn=LOC102900010;refGene_tx=XM_023244096.1;refGene_gn=LOC102900010;refGene_tx=XM_023244097.1;refGene_gn=LOC102900010;refGene_tx=XM_023244098.1
A2	4625154	4625331	refGene_gn=LOC102900010;refGene_tx=XM_023244096.1;refGene_gn=LOC102900010;refGene_tx=XM_023244097.1;refGene_gn=LOC102900010;refGene_tx=XM_023244098.1
A2	4627105	4627287	refGene_gn=LOC102900010;refGene_tx=XM_023244096.1;refGene_gn=LOC102900010;refGene_tx=XM_023244097.1;refGene_gn=LOC102900010;refGene_tx=XM_023244098.1
A2	4627456	4627690	refGene_gn=LOC102900010;refGene_tx=XM_023244096.1;refGene_gn=LOC102900010;refGene_tx=XM_023244097.1;refGene_gn=LOC102900010;refGene_tx=XM_023244098.1
A2	4631012	4631230	refGene_gn=LOC102900010;refGene_tx=XM_023244096.1;refGene_gn=LOC102900010;refGene_tx=XM_023244097.1;refGene_gn=LOC102900010;refGene_tx=XM_023244098.1
A2	4631485	4631625	refGene_gn=LOC102900010;refGene_tx=XM_023244096.1;refGene_gn=LOC102900010;refGene_tx=XM_023244097.1;refGene_gn=LOC102900010;refGene_tx=XM_023244098.1
A2	4632706	4632953	refGene_gn=LOC102900010;refGene_tx=XM_023244096.1;refGene_gn=LOC102900010;refGene_tx=XM_023244097.1;refGene_gn=LOC102900010;refGene_tx=XM_023244098.1
A2	4633281	4633370	refGene_gn=LOC102900010;refGene_tx=XM_023244096.1;refGene_gn=LOC102900010;refGene_tx=XM_023244097.1;refGene_gn=LOC102900010;refGene_tx=XM_023244098.1
A2	4639317	4639446	refGene_gn=MLLT1;refGene_tx=XM_023244104.1;refGene_gn=MLLT1;refGene_tx=XM_023244105.1
A2	4639647	4639719	refGene_gn=MLLT1;refGene_tx=XM_023244104.1;refGene_gn=MLLT1;refGene_tx=XM_023244105.1
A2	4639923	4639995	refGene_gn=MLLT1;refGene_tx=XM_023244104.1;refGene_gn=MLLT1;refGene_tx=XM_023244105.1
A2	4640365	4640465	refGene_gn=MLLT1;refGene_tx=XM_023244104.1;refGene_gn=MLLT1;refGene_tx=XM_023244105.1

A2 4642556 4642665
refGene_gn=MLLT1;refGene_tx=XM_023244104.1;refGene_gn=MLLT1;refGene_tx
=XM_023244105.1
A2 4645094 4645182
refGene_gn=MLLT1;refGene_tx=XM_023244104.1;refGene_gn=MLLT1;refGene_tx
=XM_023244105.1
A2 4650061 4650616
refGene_gn=MLLT1;refGene_tx=XM_023244104.1;refGene_gn=MLLT1;refGene_tx
=XM_023244105.1
A2 4654524 4654650 refGene_gn=MLLT1;refGene_tx=XM_023244104.1
A2 4657965 4658109
refGene_gn=MLLT1;refGene_tx=XM_023244104.1;refGene_gn=MLLT1;refGene_tx
=XM_023244105.1
A2 4688032 4688115
refGene_gn=MLLT1;refGene_tx=XM_023244104.1;refGene_gn=MLLT1;refGene_tx
=XM_023244105.1
A2 4694823 4695004
refGene_gn=MLLT1;refGene_tx=XM_023244104.1;refGene_gn=MLLT1;refGene_tx
=XM_023244105.1
A2 4704609 4704621
refGene_gn=MLLT1;refGene_tx=XM_023244104.1;refGene_gn=MLLT1;refGene_tx
=XM_023244105.1
A2 4713253 4713419 refGene_gn=ACER1;refGene_tx=XM_003981719.5
A2 4713751 4713889 refGene_gn=ACER1;refGene_tx=XM_003981719.5
A2 4716773 4716911 refGene_gn=ACER1;refGene_tx=XM_003981719.5
A2 4717624 4717745 refGene_gn=ACER1;refGene_tx=XM_003981719.5
A2 4717747 4717765 refGene_gn=ACER1;refGene_tx=XM_003981719.5
A2 4717847 4717899 refGene_gn=ACER1;refGene_tx=XM_003981719.5
A2 4717900 4717963 refGene_gn=ACER1;refGene_tx=XM_003981719.5
A2 4732410 4732503 refGene_gn=ACER1;refGene_tx=XM_003981719.5
A2 4758527 4758610 refGene_gn=CLPP;refGene_tx=XM_003981720.5
A2 4758611 4758714 refGene_gn=CLPP;refGene_tx=XM_003981720.5
A2 4758795 4758867 refGene_gn=CLPP;refGene_tx=XM_003981720.5
A2 4759408 4759505 refGene_gn=CLPP;refGene_tx=XM_003981720.5
A2 4760317 4760505 refGene_gn=CLPP;refGene_tx=XM_003981720.5
A2 4761273 4761379 refGene_gn=CLPP;refGene_tx=XM_003981720.5
A2 4763593 4763766 refGene_gn=CLPP;refGene_tx=XM_003981720.5
A2 4766456 4766660 refGene_gn=ALKBH7;refGene_tx=XM_003981721.5
A2 4767301 4767475 refGene_gn=ALKBH7;refGene_tx=XM_003981721.5
A2 4767563 4767688 refGene_gn=ALKBH7;refGene_tx=XM_003981721.5
A2 4767900 4768063 refGene_gn=ALKBH7;refGene_tx=XM_003981721.5
A2 4768405 4768794 refGene_gn=PSPN;refGene_tx=XM_023240908.1
A2 4768859 4768932 refGene_gn=PSPN;refGene_tx=XM_023240908.1
A2 4771538 4771743
refGene_gn=GTF2F1;refGene_tx=XM_019816291.2;refGene_gn=GTF2F1;refGene_
tx=XM_019816300.2
A2 4771848 4771966
refGene_gn=GTF2F1;refGene_tx=XM_019816291.2;refGene_gn=GTF2F1;refGene_
tx=XM_019816300.2
A2 4772176 4772315

refGene_gn=GTF2F1;refGene_tx=XM_019816291.2;refGene_gn=GTF2F1;refGene_tx=XM_019816300.2
A2 4772391 4772465
refGene_gn=GTF2F1;refGene_tx=XM_019816291.2;refGene_gn=GTF2F1;refGene_tx=XM_019816300.2
A2 4772572 4772754
refGene_gn=GTF2F1;refGene_tx=XM_019816291.2;refGene_gn=GTF2F1;refGene_tx=XM_019816300.2
A2 4772851 4773005
refGene_gn=GTF2F1;refGene_tx=XM_019816291.2;refGene_gn=GTF2F1;refGene_tx=XM_019816300.2
A2 4773563 4773748
refGene_gn=GTF2F1;refGene_tx=XM_019816291.2;refGene_gn=GTF2F1;refGene_tx=XM_019816300.2
A2 4775506 4775677
refGene_gn=GTF2F1;refGene_tx=XM_019816291.2;refGene_gn=GTF2F1;refGene_tx=XM_019816300.2
A2 4776482 4776676
refGene_gn=GTF2F1;refGene_tx=XM_019816291.2;refGene_gn=GTF2F1;refGene_tx=XM_019816300.2
A2 4781307 4781380
refGene_gn=GTF2F1;refGene_tx=XM_019816291.2;refGene_gn=GTF2F1;refGene_tx=XM_019816300.2
A2 4781952 4781999
refGene_gn=GTF2F1;refGene_tx=XM_019816291.2;refGene_gn=GTF2F1;refGene_tx=XM_019816300.2
A2 4782086 4782098
refGene_gn=GTF2F1;refGene_tx=XM_019816291.2;refGene_gn=GTF2F1;refGene_tx=XM_019816300.2
A2 4782815 4782875
refGene_gn=LOC102900269;refGene_tx=XM_023244133.1;refGene_gn=LOC102900269;refGene_tx=XM_023244139.1
A2 4784917 4785031
refGene_gn=LOC102900269;refGene_tx=XM_023244133.1;refGene_gn=LOC102900269;refGene_tx=XM_023244139.1;refGene_gn=LOC102900269;refGene_tx=XM_023244149.1;refGene_gn=LOC102900269;refGene_tx=XM_023244154.1
A2 4795538 4795734
refGene_gn=LOC102900269;refGene_tx=XM_023244133.1;refGene_gn=LOC102900269;refGene_tx=XM_023244149.1;refGene_gn=LOC102900269;refGene_tx=XM_023244154.1
A2 4796716 4796863
refGene_gn=LOC102900269;refGene_tx=XM_023244133.1;refGene_gn=LOC102900269;refGene_tx=XM_023244139.1;refGene_gn=LOC102900269;refGene_tx=XM_023244149.1;refGene_gn=LOC102900269;refGene_tx=XM_023244154.1
A2 4797282 4797318
refGene_gn=LOC102900269;refGene_tx=XM_023244139.1
A2 4798719 4798728 refGene_gn=KHSRP;refGene_tx=XM_023244163.1
A2 4799738 4800016
refGene_gn=KHSRP;refGene_tx=XM_023244162.1;refGene_gn=KHSRP;refGene_tx=XM_023244163.1

A2 4800107 4800185
refGene_gn=KHSRP;refGene_tx=XM_023244162.1;refGene_gn=KHSRP;refGene_tx
=XM_023244163.1
A2 4800289 4800487
refGene_gn=KHSRP;refGene_tx=XM_023244162.1;refGene_gn=KHSRP;refGene_tx
=XM_023244163.1
A2 4800594 4800683
refGene_gn=KHSRP;refGene_tx=XM_023244162.1;refGene_gn=KHSRP;refGene_tx
=XM_023244163.1
A2 4801062 4801172
refGene_gn=KHSRP;refGene_tx=XM_023244162.1;refGene_gn=KHSRP;refGene_tx
=XM_023244163.1
A2 4801269 4801430
refGene_gn=KHSRP;refGene_tx=XM_023244162.1;refGene_gn=KHSRP;refGene_tx
=XM_023244163.1
A2 4801517 4801662
refGene_gn=KHSRP;refGene_tx=XM_023244162.1;refGene_gn=KHSRP;refGene_tx
=XM_023244163.1
A2 4801789 4801890
refGene_gn=KHSRP;refGene_tx=XM_023244162.1;refGene_gn=KHSRP;refGene_tx
=XM_023244163.1
A2 4802576 4802679
refGene_gn=KHSRP;refGene_tx=XM_023244162.1;refGene_gn=KHSRP;refGene_tx
=XM_023244163.1
A2 4802792 4802891
refGene_gn=KHSRP;refGene_tx=XM_023244162.1;refGene_gn=KHSRP;refGene_tx
=XM_023244163.1
A2 4803218 4803317
refGene_gn=KHSRP;refGene_tx=XM_023244162.1;refGene_gn=KHSRP;refGene_tx
=XM_023244163.1
A2 4803503 4803678
refGene_gn=KHSRP;refGene_tx=XM_023244162.1;refGene_gn=KHSRP;refGene_tx
=XM_023244163.1
A2 4803973 4804031
refGene_gn=KHSRP;refGene_tx=XM_023244162.1;refGene_gn=KHSRP;refGene_tx
=XM_023244163.1
A2 4804909 4804981
refGene_gn=KHSRP;refGene_tx=XM_023244162.1;refGene_gn=KHSRP;refGene_tx
=XM_023244163.1
A2 4805227 4805277
refGene_gn=KHSRP;refGene_tx=XM_023244162.1;refGene_gn=KHSRP;refGene_tx
=XM_023244163.1
A2 4806203 4806243
refGene_gn=KHSRP;refGene_tx=XM_023244162.1;refGene_gn=KHSRP;refGene_tx
=XM_023244163.1
A2 4806601 4806640
refGene_gn=KHSRP;refGene_tx=XM_023244162.1;refGene_gn=KHSRP;refGene_tx
=XM_023244163.1
A2 4807255 4807352
refGene_gn=KHSRP;refGene_tx=XM_023244162.1;refGene_gn=KHSRP;refGene_tx

=XM_023244163.1
A2	4808950	4809208
refGene_gn=KHSRP;refGene_tx=XM_023244162.1;refGene_gn=KHSRP;refGene_tx
=XM_023244163.1
A2	4810965	4811138	refGene_gn=SLC25A41;refGene_tx=XM_003981723.4
A2	4811952	4812100	refGene_gn=SLC25A41;refGene_tx=XM_003981723.4
A2	4812191	4812359	refGene_gn=SLC25A41;refGene_tx=XM_003981723.4
A2	4815742	4815850	refGene_gn=SLC25A41;refGene_tx=XM_003981723.4
A2	4816028	4816181	refGene_gn=SLC25A41;refGene_tx=XM_003981723.4
A2	4816661	4816817	refGene_gn=SLC25A41;refGene_tx=XM_003981723.4
A2	4817058	4817259	refGene_gn=SLC25A41;refGene_tx=XM_003981723.4
A2	4822432	4822808
refGene_gn=SLC25A23;refGene_tx=XM_023244217.1;refGene_gn=SLC25A23;refG
ene_tx=XM_023244223.1
A2	4822816	4822838	refGene_gn=SLC25A23;refGene_tx=XM_023244225.1
A2	4823102	4823283
refGene_gn=SLC25A23;refGene_tx=XM_023244217.1;refGene_gn=SLC25A23;refG
ene_tx=XM_023244223.1;refGene_gn=SLC25A23;refGene_tx=XM_023244225.1
A2	4824287	4824438
refGene_gn=SLC25A23;refGene_tx=XM_023244217.1;refGene_gn=SLC25A23;refG
ene_tx=XM_023244223.1;refGene_gn=SLC25A23;refGene_tx=XM_023244225.1
A2	4827909	4828077
refGene_gn=SLC25A23;refGene_tx=XM_023244217.1;refGene_gn=SLC25A23;refG
ene_tx=XM_023244223.1;refGene_gn=SLC25A23;refGene_tx=XM_023244225.1
A2	4829074	4829182
refGene_gn=SLC25A23;refGene_tx=XM_023244217.1;refGene_gn=SLC25A23;refG
ene_tx=XM_023244223.1;refGene_gn=SLC25A23;refGene_tx=XM_023244225.1
A2	4829350	4829503
refGene_gn=SLC25A23;refGene_tx=XM_023244217.1;refGene_gn=SLC25A23;refG
ene_tx=XM_023244223.1;refGene_gn=SLC25A23;refGene_tx=XM_023244225.1
A2	4829593	4829752
refGene_gn=SLC25A23;refGene_tx=XM_023244217.1;refGene_gn=SLC25A23;refG
ene_tx=XM_023244223.1;refGene_gn=SLC25A23;refGene_tx=XM_023244225.1
A2	4830799	4830911
refGene_gn=SLC25A23;refGene_tx=XM_023244217.1;refGene_gn=SLC25A23;refG
ene_tx=XM_023244223.1;refGene_gn=SLC25A23;refGene_tx=XM_023244225.1
A2	4831205	4831293
refGene_gn=SLC25A23;refGene_tx=XM_023244217.1;refGene_gn=SLC25A23;refG
ene_tx=XM_023244223.1;refGene_gn=SLC25A23;refGene_tx=XM_023244225.1
A2	4831871	4831998
refGene_gn=SLC25A23;refGene_tx=XM_023244217.1;refGene_gn=SLC25A23;refG
ene_tx=XM_023244223.1;refGene_gn=SLC25A23;refGene_tx=XM_023244225.1
A2	4833126	4833282
refGene_gn=SLC25A23;refGene_tx=XM_023244217.1;refGene_gn=SLC25A23;refG
ene_tx=XM_023244223.1;refGene_gn=SLC25A23;refGene_tx=XM_023244225.1
A2	4836070	4836152
refGene_gn=CRB3;refGene_tx=XM_023244235.1;refGene_gn=CRB3;refGene_tx=X
M_023244245.1;refGene_gn=CRB3;refGene_tx=XM_023244249.1
A2	4836862	4836936
refGene_gn=CRB3;refGene_tx=XM_023244235.1;refGene_gn=CRB3;refGene_tx=X

M_023244245.1;refGene_gn=CRB3;refGene_tx=XM_023244249.1
A2 4837229 4837373
refGene_gn=CRB3;refGene_tx=XM_023244235.1;refGene_gn=CRB3;refGene_tx=XM_023244245.1;refGene_gn=CRB3;refGene_tx=XM_023244249.1
A2 4837723 4837795
refGene_gn=CRB3;refGene_tx=XM_023244235.1;refGene_gn=CRB3;refGene_tx=XM_023244245.1;refGene_gn=CRB3;refGene_tx=XM_023244249.1
A2 4838335 4838959
refGene_gn=DENND1C;refGene_tx=XM_003981724.5;refGene_gn=DENND1C;refGene_tx=XM_019816370.2;refGene_gn=DENND1C;refGene_tx=XM_023244196.1
A2 4839068 4839276
refGene_gn=DENND1C;refGene_tx=XM_003981724.5;refGene_gn=DENND1C;refGene_tx=XM_019816370.2;refGene_gn=DENND1C;refGene_tx=XM_023244196.1
A2 4839396 4839468
refGene_gn=DENND1C;refGene_tx=XM_003981724.5;refGene_gn=DENND1C;refGene_tx=XM_019816370.2;refGene_gn=DENND1C;refGene_tx=XM_023244196.1;refGene_gn=DENND1C;refGene_tx=XM_023244209.1
A2 4839652 4839760
refGene_gn=DENND1C;refGene_tx=XM_003981724.5;refGene_gn=DENND1C;refGene_tx=XM_019816370.2;refGene_gn=DENND1C;refGene_tx=XM_023244196.1;refGene_gn=DENND1C;refGene_tx=XM_023244209.1
A2 4839832 4839877
refGene_gn=DENND1C;refGene_tx=XM_003981724.5;refGene_gn=DENND1C;refGene_tx=XM_019816370.2;refGene_gn=DENND1C;refGene_tx=XM_023244196.1;refGene_gn=DENND1C;refGene_tx=XM_023244209.1
A2 4840496 4840568
refGene_gn=DENND1C;refGene_tx=XM_003981724.5;refGene_gn=DENND1C;refGene_tx=XM_019816370.2;refGene_gn=DENND1C;refGene_tx=XM_023244196.1;refGene_gn=DENND1C;refGene_tx=XM_023244209.1
A2 4840784 4840825
refGene_gn=DENND1C;refGene_tx=XM_003981724.5;refGene_gn=DENND1C;refGene_tx=XM_019816370.2;refGene_gn=DENND1C;refGene_tx=XM_023244196.1;refGene_gn=DENND1C;refGene_tx=XM_023244209.1
A2 4840916 4841007
refGene_gn=DENND1C;refGene_tx=XM_003981724.5;refGene_gn=DENND1C;refGene_tx=XM_019816370.2;refGene_gn=DENND1C;refGene_tx=XM_023244196.1;refGene_gn=DENND1C;refGene_tx=XM_023244209.1
A2 4841808 4841913
refGene_gn=DENND1C;refGene_tx=XM_003981724.5;refGene_gn=DENND1C;refGene_tx=XM_019816370.2;refGene_gn=DENND1C;refGene_tx=XM_023244196.1;refGene_gn=DENND1C;refGene_tx=XM_023244209.1
A2 4842754 4842880
refGene_gn=DENND1C;refGene_tx=XM_003981724.5;refGene_gn=DENND1C;refGene_tx=XM_019816370.2;refGene_gn=DENND1C;refGene_tx=XM_023244196.1;refGene_gn=DENND1C;refGene_tx=XM_023244209.1
A2 4842971 4843073
refGene_gn=DENND1C;refGene_tx=XM_003981724.5;refGene_gn=DENND1C;refGene_tx=XM_019816370.2;refGene_gn=DENND1C;refGene_tx=XM_023244196.1;refGene_gn=DENND1C;refGene_tx=XM_023244209.1
A2 4843200 4843246

refGene_gn=DENND1C;refGene_tx=XM_003981724.5;refGene_gn=DENND1C;refGene_tx=XM_019816370.2;refGene_gn=DENND1C;refGene_tx=XM_023244196.1;refGene_gn=DENND1C;refGene_tx=XM_023244209.1
A2 4843340 4843441
refGene_gn=DENND1C;refGene_tx=XM_003981724.5;refGene_gn=DENND1C;refGene_tx=XM_019816370.2;refGene_gn=DENND1C;refGene_tx=XM_023244196.1;refGene_gn=DENND1C;refGene_tx=XM_023244209.1
A2 4844195 4844306
refGene_gn=DENND1C;refGene_tx=XM_003981724.5;refGene_gn=DENND1C;refGene_tx=XM_019816370.2;refGene_gn=DENND1C;refGene_tx=XM_023244196.1;refGene_gn=DENND1C;refGene_tx=XM_023244209.1
A2 4844417 4844471
refGene_gn=DENND1C;refGene_tx=XM_003981724.5;refGene_gn=DENND1C;refGene_tx=XM_019816370.2;refGene_gn=DENND1C;refGene_tx=XM_023244196.1;refGene_gn=DENND1C;refGene_tx=XM_023244209.1
A2 4844558 4844624
refGene_gn=DENND1C;refGene_tx=XM_003981724.5;refGene_gn=DENND1C;refGene_tx=XM_019816370.2;refGene_gn=DENND1C;refGene_tx=XM_023244196.1;refGene_gn=DENND1C;refGene_tx=XM_023244209.1
A2 4844718 4844793
refGene_gn=DENND1C;refGene_tx=XM_003981724.5;refGene_gn=DENND1C;refGene_tx=XM_019816370.2;refGene_gn=DENND1C;refGene_tx=XM_023244196.1;refGene_gn=DENND1C;refGene_tx=XM_023244209.1
A2 4844971 4845041
refGene_gn=DENND1C;refGene_tx=XM_003981724.5;refGene_gn=DENND1C;refGene_tx=XM_019816370.2;refGene_gn=DENND1C;refGene_tx=XM_023244196.1;refGene_gn=DENND1C;refGene_tx=XM_023244209.1
A2 4845131 4845251
refGene_gn=DENND1C;refGene_tx=XM_003981724.5;refGene_gn=DENND1C;refGene_tx=XM_019816370.2;refGene_gn=DENND1C;refGene_tx=XM_023244196.1;refGene_gn=DENND1C;refGene_tx=XM_023244209.1
A2 4845476 4845526
refGene_gn=DENND1C;refGene_tx=XM_003981724.5;refGene_gn=DENND1C;refGene_tx=XM_019816370.2;refGene_gn=DENND1C;refGene_tx=XM_023244196.1;refGene_gn=DENND1C;refGene_tx=XM_023244209.1
A2 4845650 4845694
refGene_gn=DENND1C;refGene_tx=XM_003981724.5;refGene_gn=DENND1C;refGene_tx=XM_023244209.1
A2 4845775 4845840
refGene_gn=DENND1C;refGene_tx=XM_003981724.5;refGene_gn=DENND1C;refGene_tx=XM_023244209.1
A2 4846944 4846961 refGene_gn=DENND1C;refGene_tx=XM_003981724.5
A2 4847284 4847349 refGene_gn=DENND1C;refGene_tx=XM_023244209.1
A2 4852552 4853610 refGene_gn=TUBB4A;refGene_tx=XM_023244250.1
A2 4856090 4856201 refGene_gn=TUBB4A;refGene_tx=XM_023244250.1
A2 4856313 4856422 refGene_gn=TUBB4A;refGene_tx=XM_023244250.1
A2 4856976 4857033 refGene_gn=TUBB4A;refGene_tx=XM_023244250.1
A2 4872543 4872789 refGene_gn=TNFSF9;refGene_tx=XM_023244260.1
A2 4873261 4873292 refGene_gn=TNFSF9;refGene_tx=XM_023244260.1
A2 4874020 4874520 refGene_gn=TNFSF9;refGene_tx=XM_023244260.1

A2 4914123 4914515
refGene_gn=CD70;refGene_tx=XM_003981772.5;refGene_gn=CD70;refGene_tx=XM_019826091.2
A2 4916345 4916379
refGene_gn=CD70;refGene_tx=XM_003981772.5;refGene_gn=CD70;refGene_tx=XM_019826091.2
A2 4916938 4917019 refGene_gn=CD70;refGene_tx=XM_019826091.2
A2 4917236 4917404 refGene_gn=CD70;refGene_tx=XM_003981772.5
A2 4944513 4944941 refGene_gn=TNFSF14;refGene_tx=XM_003981725.3
A2 4945528 4945570 refGene_gn=TNFSF14;refGene_tx=XM_003981725.3
A2 4945834 4945871 refGene_gn=TNFSF14;refGene_tx=XM_003981725.3
A2 4947121 4947343 refGene_gn=TNFSF14;refGene_tx=XM_003981725.3
A2 4960403 4960517
refGene_gn=LOC101088468;refGene_tx=XM_023244292.1
A2 4961992 4962044
refGene_gn=LOC101088468;refGene_tx=XM_023244292.1
A2 4962160 4962251
refGene_gn=LOC101088468;refGene_tx=XM_023244292.1
A2 4962367 4962427
refGene_gn=LOC101088468;refGene_tx=XM_023244292.1
A2 4962570 4962729
refGene_gn=LOC101088468;refGene_tx=XM_023244292.1
A2 4963796 4963960
refGene_gn=LOC101088468;refGene_tx=XM_023244292.1
A2 4965627 4965784
refGene_gn=LOC101088468;refGene_tx=XM_023244292.1
A2 4967677 4967776
refGene_gn=LOC101088468;refGene_tx=XM_023244292.1
A2 4968687 4968847
refGene_gn=LOC101088468;refGene_tx=XM_023244292.1
A2 4969133 4969209
refGene_gn=LOC101088468;refGene_tx=XM_023244292.1
A2 4969702 4969906
refGene_gn=LOC101088468;refGene_tx=XM_023244292.1
A2 4970663 4970750
refGene_gn=LOC101088468;refGene_tx=XM_023244292.1
A2 4970858 4970925
refGene_gn=LOC101088468;refGene_tx=XM_023244292.1
A2 4972976 4973189
refGene_gn=LOC101088468;refGene_tx=XM_023244292.1
A2 4973293 4973436
refGene_gn=LOC101088468;refGene_tx=XM_023244292.1
A2 4993921 4993938
refGene_gn=LOC111557813;refGene_tx=XM_023244300.1;refGene_gn=LOC111557813;refGene_tx=XM_023244303.1
A2 4994019 4994102
refGene_gn=LOC111557813;refGene_tx=XM_023244300.1;refGene_gn=LOC111557813;refGene_tx=XM_023244303.1
A2 4994182 4994277
refGene_gn=LOC111557813;refGene_tx=XM_023244300.1;refGene_gn=LOC111557

813;refGene_tx=XM_023244303.1
A2 4996042 4996113
refGene_gn=LOC111557813;refGene_tx=XM_023244300.1;refGene_gn=LOC111557
813;refGene_tx=XM_023244303.1
A2 4996209 4996372
refGene_gn=LOC111557813;refGene_tx=XM_023244300.1;refGene_gn=LOC111557
813;refGene_tx=XM_023244303.1
A2 4996761 4996954
refGene_gn=LOC111557813;refGene_tx=XM_023244300.1;refGene_gn=LOC111557
813;refGene_tx=XM_023244303.1
A2 4997346 4997417
refGene_gn=LOC111557813;refGene_tx=XM_023244303.1
A2 5054282 5054362
refGene_gn=LOC111557813;refGene_tx=XM_023244300.1
A2 5065949 5066035
refGene_gn=LOC101088468;refGene_tx=XM_023244292.1
A2 5066302 5066411
refGene_gn=LOC101088468;refGene_tx=XM_023244292.1
A2 5074468 5074666
refGene_gn=LOC101088468;refGene_tx=XM_023244292.1
A2 5074828 5074900
refGene_gn=LOC101088468;refGene_tx=XM_023244292.1
A2 5075166 5075296
refGene_gn=LOC101088468;refGene_tx=XM_023244292.1
A2 5075583 5075742
refGene_gn=LOC101088468;refGene_tx=XM_023244292.1
A2 5076072 5076279
refGene_gn=LOC101088468;refGene_tx=XM_023244292.1
A2 5076398 5076614
refGene_gn=LOC101088468;refGene_tx=XM_023244292.1
A2 5077008 5077155
refGene_gn=LOC101088468;refGene_tx=XM_023244292.1
A2 5077260 5077376
refGene_gn=LOC101088468;refGene_tx=XM_023244292.1
A2 5079673 5079800
refGene_gn=LOC101088468;refGene_tx=XM_023244292.1
A2 5079891 5079994
refGene_gn=LOC101088468;refGene_tx=XM_023244292.1
A2 5080212 5080303
refGene_gn=LOC101088468;refGene_tx=XM_023244292.1
A2 5080385 5080468
refGene_gn=LOC101088468;refGene_tx=XM_023244292.1
A2 5080548 5080643
refGene_gn=LOC101088468;refGene_tx=XM_023244292.1
A2 5082411 5082482
refGene_gn=LOC101088468;refGene_tx=XM_023244292.1
A2 5082580 5082743
refGene_gn=LOC101088468;refGene_tx=XM_023244292.1
A2 5083080 5083273
refGene_gn=LOC101088468;refGene_tx=XM_023244292.1

A2 5083664 5083741
refGene_gn=LOC101088468;refGene_tx=XM_023244292.1
A2 5089561 5089634
refGene_gn=GPR108;refGene_tx=XM_006928300.4;refGene_gn=GPR108;refGene_tx=XM_011287531.3;refGene_gn=GPR108;refGene_tx=XM_019816438.2
A2 5089815 5089940
refGene_gn=GPR108;refGene_tx=XM_006928300.4;refGene_gn=GPR108;refGene_tx=XM_011287531.3;refGene_gn=GPR108;refGene_tx=XM_019816438.2
A2 5090042 5090126
refGene_gn=GPR108;refGene_tx=XM_006928300.4;refGene_gn=GPR108;refGene_tx=XM_011287531.3;refGene_gn=GPR108;refGene_tx=XM_019816438.2
A2 5090292 5090342
refGene_gn=GPR108;refGene_tx=XM_006928300.4;refGene_gn=GPR108;refGene_tx=XM_011287531.3;refGene_gn=GPR108;refGene_tx=XM_019816438.2
A2 5090628 5090672
refGene_gn=GPR108;refGene_tx=XM_006928300.4;refGene_gn=GPR108;refGene_tx=XM_011287531.3;refGene_gn=GPR108;refGene_tx=XM_019816438.2
A2 5090785 5090916
refGene_gn=GPR108;refGene_tx=XM_006928300.4;refGene_gn=GPR108;refGene_tx=XM_011287531.3;refGene_gn=GPR108;refGene_tx=XM_019816438.2
A2 5091007 5091125
refGene_gn=GPR108;refGene_tx=XM_006928300.4;refGene_gn=GPR108;refGene_tx=XM_011287531.3;refGene_gn=GPR108;refGene_tx=XM_019816438.2
A2 5091205 5091279
refGene_gn=GPR108;refGene_tx=XM_006928300.4;refGene_gn=GPR108;refGene_tx=XM_011287531.3;refGene_gn=GPR108;refGene_tx=XM_019816438.2
A2 5091374 5091449 refGene_gn=GPR108;refGene_tx=XM_019816455.2
A2 5091802 5091878
refGene_gn=GPR108;refGene_tx=XM_006928300.4;refGene_gn=GPR108;refGene_tx=XM_011287531.3;refGene_gn=GPR108;refGene_tx=XM_019816438.2;refGene_gn=GPR108;refGene_tx=XM_019816455.2
A2 5091992 5092126
refGene_gn=GPR108;refGene_tx=XM_006928300.4;refGene_gn=GPR108;refGene_tx=XM_011287531.3;refGene_gn=GPR108;refGene_tx=XM_019816438.2;refGene_gn=GPR108;refGene_tx=XM_019816455.2
A2 5092322 5092427
refGene_gn=GPR108;refGene_tx=XM_006928300.4;refGene_gn=GPR108;refGene_tx=XM_011287531.3;refGene_gn=GPR108;refGene_tx=XM_019816438.2;refGene_gn=GPR108;refGene_tx=XM_019816455.2
A2 5092523 5092592
refGene_gn=GPR108;refGene_tx=XM_006928300.4;refGene_gn=GPR108;refGene_tx=XM_011287531.3;refGene_gn=GPR108;refGene_tx=XM_019816438.2;refGene_gn=GPR108;refGene_tx=XM_019816455.2
A2 5092675 5092737
refGene_gn=GPR108;refGene_tx=XM_006928300.4;refGene_gn=GPR108;refGene_tx=XM_011287531.3;refGene_gn=GPR108;refGene_tx=XM_019816438.2;refGene_gn=GPR108;refGene_tx=XM_019816455.2
A2 5092861 5092986
refGene_gn=GPR108;refGene_tx=XM_006928300.4;refGene_gn=GPR108;refGene_tx=XM_011287531.3;refGene_gn=GPR108;refGene_tx=XM_019816438.2;refGene_

gn=GPR108;refGene_tx=XM_019816455.2
A2 5093705 5093788
refGene_gn=GPR108;refGene_tx=XM_006928300.4;refGene_gn=GPR108;refGene_tx=XM_011287531.3;refGene_gn=GPR108;refGene_tx=XM_019816438.2;refGene_gn=GPR108;refGene_tx=XM_019816455.2
A2 5093988 5094039
refGene_gn=GPR108;refGene_tx=XM_006928300.4;refGene_gn=GPR108;refGene_tx=XM_011287531.3;refGene_gn=GPR108;refGene_tx=XM_019816438.2;refGene_gn=GPR108;refGene_tx=XM_019816455.2
A2 5094542 5094878
refGene_gn=GPR108;refGene_tx=XM_006928300.4;refGene_gn=GPR108;refGene_tx=XM_011287531.3;refGene_gn=GPR108;refGene_tx=XM_019816438.2;refGene_gn=GPR108;refGene_tx=XM_019816455.2
A2 5095414 5095766
refGene_gn=GPR108;refGene_tx=XM_006928300.4;refGene_gn=GPR108;refGene_tx=XM_019816438.2;refGene_gn=GPR108;refGene_tx=XM_019816455.2
A2 5097706 5097730
refGene_gn=TRIP10;refGene_tx=XM_023244360.1;refGene_gn=TRIP10;refGene_tx=XM_023244366.1;refGene_gn=TRIP10;refGene_tx=XM_023244374.1;refGene_gn=TRIP10;refGene_tx=XM_023244381.1
A2 5098919 5099035
refGene_gn=TRIP10;refGene_tx=XM_023244360.1;refGene_gn=TRIP10;refGene_tx=XM_023244366.1;refGene_gn=TRIP10;refGene_tx=XM_023244374.1;refGene_gn=TRIP10;refGene_tx=XM_023244381.1
A2 5099120 5099177
refGene_gn=TRIP10;refGene_tx=XM_023244360.1;refGene_gn=TRIP10;refGene_tx=XM_023244366.1;refGene_gn=TRIP10;refGene_tx=XM_023244374.1;refGene_gn=TRIP10;refGene_tx=XM_023244381.1
A2 5100021 5100169
refGene_gn=TRIP10;refGene_tx=XM_023244360.1;refGene_gn=TRIP10;refGene_tx=XM_023244366.1;refGene_gn=TRIP10;refGene_tx=XM_023244374.1;refGene_gn=TRIP10;refGene_tx=XM_023244381.1
A2 5100250 5100313
refGene_gn=TRIP10;refGene_tx=XM_023244360.1;refGene_gn=TRIP10;refGene_tx=XM_023244366.1;refGene_gn=TRIP10;refGene_tx=XM_023244374.1;refGene_gn=TRIP10;refGene_tx=XM_023244381.1
A2 5100530 5100635
refGene_gn=TRIP10;refGene_tx=XM_023244360.1;refGene_gn=TRIP10;refGene_tx=XM_023244366.1;refGene_gn=TRIP10;refGene_tx=XM_023244374.1;refGene_gn=TRIP10;refGene_tx=XM_023244381.1
A2 5100757 5100886
refGene_gn=TRIP10;refGene_tx=XM_023244360.1;refGene_gn=TRIP10;refGene_tx=XM_023244366.1;refGene_gn=TRIP10;refGene_tx=XM_023244374.1;refGene_gn=TRIP10;refGene_tx=XM_023244381.1
A2 5101473 5101620
refGene_gn=TRIP10;refGene_tx=XM_023244360.1;refGene_gn=TRIP10;refGene_tx=XM_023244366.1;refGene_gn=TRIP10;refGene_tx=XM_023244374.1;refGene_gn=TRIP10;refGene_tx=XM_023244381.1
A2 5101735 5101930
refGene_gn=TRIP10;refGene_tx=XM_023244360.1;refGene_gn=TRIP10;refGene_

tx=XM_023244366.1;refGene_gn=TRIP10;refGene_tx=XM_023244374.1;refGene_gn=TRIP10;refGene_tx=XM_023244381.1
A2 5103000 5103168
refGene_gn=TRIP10;refGene_tx=XM_023244360.1;refGene_gn=TRIP10;refGene_tx=XM_023244366.1
A2 5103436 5103546
refGene_gn=TRIP10;refGene_tx=XM_023244360.1;refGene_gn=TRIP10;refGene_tx=XM_023244366.1;refGene_gn=TRIP10;refGene_tx=XM_023244374.1;refGene_gn=TRIP10;refGene_tx=XM_023244381.1
A2 5105308 5105441
refGene_gn=TRIP10;refGene_tx=XM_023244360.1;refGene_gn=TRIP10;refGene_tx=XM_023244366.1;refGene_gn=TRIP10;refGene_tx=XM_023244374.1;refGene_gn=TRIP10;refGene_tx=XM_023244381.1
A2 5105668 5105814
refGene_gn=TRIP10;refGene_tx=XM_023244360.1;refGene_gn=TRIP10;refGene_tx=XM_023244366.1;refGene_gn=TRIP10;refGene_tx=XM_023244374.1;refGene_gn=TRIP10;refGene_tx=XM_023244381.1
A2 5105892 5106014
refGene_gn=TRIP10;refGene_tx=XM_023244360.1;refGene_gn=TRIP10;refGene_tx=XM_023244366.1;refGene_gn=TRIP10;refGene_tx=XM_023244374.1;refGene_gn=TRIP10;refGene_tx=XM_023244381.1
A2 5106091 5106282
refGene_gn=TRIP10;refGene_tx=XM_023244360.1;refGene_gn=TRIP10;refGene_tx=XM_023244366.1;refGene_gn=TRIP10;refGene_tx=XM_023244374.1;refGene_gn=TRIP10;refGene_tx=XM_023244381.1
A2 5107247 5107408
refGene_gn=SH2D3A;refGene_tx=XM_003981777.5;refGene_gn=SH2D3A;refGene_tx=XM_011287519.3
A2 5108067 5108456
refGene_gn=SH2D3A;refGene_tx=XM_003981777.5;refGene_gn=SH2D3A;refGene_tx=XM_011287511.3;refGene_gn=SH2D3A;refGene_tx=XM_011287519.3;refGene_gn=SH2D3A;refGene_tx=XM_011287525.3;refGene_gn=SH2D3A;refGene_tx=XM_019816471.2;refGene_gn=SH2D3A;refGene_tx=XM_023244346.1
A2 5108738 5108850
refGene_gn=SH2D3A;refGene_tx=XM_003981777.5;refGene_gn=SH2D3A;refGene_tx=XM_011287511.3;refGene_gn=SH2D3A;refGene_tx=XM_011287519.3;refGene_gn=SH2D3A;refGene_tx=XM_011287525.3;refGene_gn=SH2D3A;refGene_tx=XM_019816471.2;refGene_gn=SH2D3A;refGene_tx=XM_023244346.1
A2 5108928 5109103
refGene_gn=SH2D3A;refGene_tx=XM_003981777.5;refGene_gn=SH2D3A;refGene_tx=XM_011287511.3;refGene_gn=SH2D3A;refGene_tx=XM_011287519.3;refGene_gn=SH2D3A;refGene_tx=XM_011287525.3;refGene_gn=SH2D3A;refGene_tx=XM_019816471.2;refGene_gn=SH2D3A;refGene_tx=XM_023244346.1
A2 5109312 5110036
refGene_gn=SH2D3A;refGene_tx=XM_003981777.5;refGene_gn=SH2D3A;refGene_tx=XM_011287511.3;refGene_gn=SH2D3A;refGene_tx=XM_011287519.3;refGene_gn=SH2D3A;refGene_tx=XM_011287525.3;refGene_gn=SH2D3A;refGene_tx=XM_019816471.2;refGene_gn=SH2D3A;refGene_tx=XM_023244346.1
A2 5111613 5111690
refGene_gn=SH2D3A;refGene_tx=XM_003981777.5;refGene_gn=SH2D3A;refGene_

tx=XM_011287511.3;refGene_gn=SH2D3A;refGene_tx=XM_011287519.3;refGene_gn=SH2D3A;refGene_tx=XM_011287525.3;refGene_gn=SH2D3A;refGene_tx=XM_019816471.2;refGene_gn=SH2D3A;refGene_tx=XM_023244346.1
A2 5112052 5112402
refGene_gn=SH2D3A;refGene_tx=XM_003981777.5;refGene_gn=SH2D3A;refGene_tx=XM_011287511.3;refGene_gn=SH2D3A;refGene_tx=XM_011287519.3;refGene_gn=SH2D3A;refGene_tx=XM_011287525.3;refGene_gn=SH2D3A;refGene_tx=XM_019816471.2;refGene_gn=SH2D3A;refGene_tx=XM_023244346.1
A2 5113905 5113974
refGene_gn=SH2D3A;refGene_tx=XM_003981777.5;refGene_gn=SH2D3A;refGene_tx=XM_011287511.3;refGene_gn=SH2D3A;refGene_tx=XM_011287519.3;refGene_gn=SH2D3A;refGene_tx=XM_011287525.3;refGene_gn=SH2D3A;refGene_tx=XM_019816471.2;refGene_gn=SH2D3A;refGene_tx=XM_023244346.1
A2 5120607 5120811
refGene_gn=VAV1;refGene_tx=XM_003981728.4;refGene_gn=VAV1;refGene_tx=XM_006928302.3;refGene_gn=VAV1;refGene_tx=XM_006928303.3;refGene_gn=VAV1;refGene_tx=XM_019816508.2;refGene_gn=VAV1;refGene_tx=XM_023244394.1
A2 5141599 5141716
refGene_gn=VAV1;refGene_tx=XM_003981728.4;refGene_gn=VAV1;refGene_tx=XM_006928302.3;refGene_gn=VAV1;refGene_tx=XM_006928303.3;refGene_gn=VAV1;refGene_tx=XM_019816508.2;refGene_gn=VAV1;refGene_tx=XM_023244394.1
A2 5142238 5142297
refGene_gn=VAV1;refGene_tx=XM_003981728.4;refGene_gn=VAV1;refGene_tx=XM_006928302.3;refGene_gn=VAV1;refGene_tx=XM_006928303.3;refGene_gn=VAV1;refGene_tx=XM_019816508.2;refGene_gn=VAV1;refGene_tx=XM_023244394.1
A2 5142407 5142476
refGene_gn=VAV1;refGene_tx=XM_003981728.4;refGene_gn=VAV1;refGene_tx=XM_006928302.3;refGene_gn=VAV1;refGene_tx=XM_006928303.3;refGene_gn=VAV1;refGene_tx=XM_019816508.2;refGene_gn=VAV1;refGene_tx=XM_023244394.1
A2 5143364 5143476
refGene_gn=VAV1;refGene_tx=XM_003981728.4;refGene_gn=VAV1;refGene_tx=XM_006928302.3;refGene_gn=VAV1;refGene_tx=XM_006928303.3;refGene_gn=VAV1;refGene_tx=XM_019816508.2;refGene_gn=VAV1;refGene_tx=XM_023244394.1
A2 5143583 5143679
refGene_gn=VAV1;refGene_tx=XM_003981728.4;refGene_gn=VAV1;refGene_tx=XM_006928302.3;refGene_gn=VAV1;refGene_tx=XM_006928303.3;refGene_gn=VAV1;refGene_tx=XM_019816508.2;refGene_gn=VAV1;refGene_tx=XM_023244394.1
A2 5144658 5144727
refGene_gn=VAV1;refGene_tx=XM_003981728.4;refGene_gn=VAV1;refGene_tx=XM_006928302.3;refGene_gn=VAV1;refGene_tx=XM_006928303.3;refGene_gn=VAV1;refGene_tx=XM_019816508.2;refGene_gn=VAV1;refGene_tx=XM_023244394.1
A2 5144875 5144979
refGene_gn=VAV1;refGene_tx=XM_003981728.4;refGene_gn=VAV1;refGene_tx=XM_006928302.3;refGene_gn=VAV1;refGene_tx=XM_006928303.3;refGene_gn=VAV1;refGene_tx=XM_019816508.2;refGene_gn=VAV1;refGene_tx=XM_023244394.1
A2 5145553 5145653
refGene_gn=VAV1;refGene_tx=XM_003981728.4;refGene_gn=VAV1;refGene_tx=XM_006928302.3;refGene_gn=VAV1;refGene_tx=XM_006928303.3;refGene_gn=VAV1;refGene_tx=XM_019816508.2;refGene_gn=VAV1;refGene_tx=XM_023244394.1
A2 5146305 5146401

refGene_gn=VAV1;refGene_tx=XM_003981728.4;refGene_gn=VAV1;refGene_tx=XM_006928302.3;refGene_gn=VAV1;refGene_tx=XM_006928303.3;refGene_gn=VAV1;refGene_tx=XM_019816508.2;refGene_gn=VAV1;refGene_tx=XM_023244394.1
A2 5146601 5146670
refGene_gn=VAV1;refGene_tx=XM_003981728.4;refGene_gn=VAV1;refGene_tx=XM_006928302.3;refGene_gn=VAV1;refGene_tx=XM_006928303.3;refGene_gn=VAV1;refGene_tx=XM_019816508.2;refGene_gn=VAV1;refGene_tx=XM_023244394.1
A2 5146943 5147030
refGene_gn=VAV1;refGene_tx=XM_003981728.4;refGene_gn=VAV1;refGene_tx=XM_006928302.3;refGene_gn=VAV1;refGene_tx=XM_006928303.3;refGene_gn=VAV1;refGene_tx=XM_019816508.2;refGene_gn=VAV1;refGene_tx=XM_023244394.1
A2 5147140 5147226
refGene_gn=VAV1;refGene_tx=XM_003981728.4;refGene_gn=VAV1;refGene_tx=XM_006928302.3;refGene_gn=VAV1;refGene_tx=XM_006928303.3;refGene_gn=VAV1;refGene_tx=XM_019816508.2;refGene_gn=VAV1;refGene_tx=XM_023244394.1
A2 5147556 5147689
refGene_gn=VAV1;refGene_tx=XM_003981728.4;refGene_gn=VAV1;refGene_tx=XM_006928302.3;refGene_gn=VAV1;refGene_tx=XM_006928303.3;refGene_gn=VAV1;refGene_tx=XM_019816508.2;refGene_gn=VAV1;refGene_tx=XM_023244394.1
A2 5148008 5148118
refGene_gn=VAV1;refGene_tx=XM_003981728.4;refGene_gn=VAV1;refGene_tx=XM_006928302.3;refGene_gn=VAV1;refGene_tx=XM_006928303.3;refGene_gn=VAV1;refGene_tx=XM_019816508.2;refGene_gn=VAV1;refGene_tx=XM_023244394.1
A2 5149345 5149447
refGene_gn=VAV1;refGene_tx=XM_003981728.4;refGene_gn=VAV1;refGene_tx=XM_006928302.3;refGene_gn=VAV1;refGene_tx=XM_006928303.3;refGene_gn=VAV1;refGene_tx=XM_019816508.2;refGene_gn=VAV1;refGene_tx=XM_023244394.1
A2 5149687 5149785
refGene_gn=VAV1;refGene_tx=XM_003981728.4;refGene_gn=VAV1;refGene_tx=XM_006928302.3;refGene_gn=VAV1;refGene_tx=XM_006928303.3;refGene_gn=VAV1;refGene_tx=XM_019816508.2;refGene_gn=VAV1;refGene_tx=XM_023244394.1
A2 5149871 5149894
refGene_gn=VAV1;refGene_tx=XM_003981728.4;refGene_gn=VAV1;refGene_tx=XM_006928302.3;refGene_gn=VAV1;refGene_tx=XM_006928303.3;refGene_gn=VAV1;refGene_tx=XM_019816508.2;refGene_gn=VAV1;refGene_tx=XM_023244394.1
A2 5150066 5150112
refGene_gn=VAV1;refGene_tx=XM_003981728.4;refGene_gn=VAV1;refGene_tx=XM_006928302.3;refGene_gn=VAV1;refGene_tx=XM_006928303.3;refGene_gn=VAV1;refGene_tx=XM_019816508.2;refGene_gn=VAV1;refGene_tx=XM_023244394.1
A2 5152686 5152823
refGene_gn=VAV1;refGene_tx=XM_003981728.4;refGene_gn=VAV1;refGene_tx=XM_006928302.3;refGene_gn=VAV1;refGene_tx=XM_006928303.3;refGene_gn=VAV1;refGene_tx=XM_019816508.2;refGene_gn=VAV1;refGene_tx=XM_023244394.1
A2 5153157 5153223
refGene_gn=VAV1;refGene_tx=XM_003981728.4;refGene_gn=VAV1;refGene_tx=XM_006928302.3;refGene_gn=VAV1;refGene_tx=XM_006928303.3;refGene_gn=VAV1;refGene_tx=XM_019816508.2;refGene_gn=VAV1;refGene_tx=XM_023244394.1
A2 5157056 5157088
refGene_gn=VAV1;refGene_tx=XM_003981728.4;refGene_gn=VAV1;refGene_tx=XM_006928302.3;refGene_gn=VAV1;refGene_tx=XM_006928303.3;refGene_gn=VAV

1;refGene_tx=XM_019816508.2;refGene_gn=VAV1;refGene_tx=XM_023244394.1
A2 5161830 5161947
refGene_gn=VAV1;refGene_tx=XM_003981728.4;refGene_gn=VAV1;refGene_tx=XM_006928302.3;refGene_gn=VAV1;refGene_tx=XM_006928303.3;refGene_gn=VAV1;refGene_tx=XM_019816508.2;refGene_gn=VAV1;refGene_tx=XM_023244394.1
A2 5163503 5163591
refGene_gn=VAV1;refGene_tx=XM_003981728.4;refGene_gn=VAV1;refGene_tx=XM_006928302.3;refGene_gn=VAV1;refGene_tx=XM_006928303.3;refGene_gn=VAV1;refGene_tx=XM_019816508.2;refGene_gn=VAV1;refGene_tx=XM_023244394.1
A2 5165384 5165505
refGene_gn=VAV1;refGene_tx=XM_003981728.4;refGene_gn=VAV1;refGene_tx=XM_006928302.3;refGene_gn=VAV1;refGene_tx=XM_006928303.3;refGene_gn=VAV1;refGene_tx=XM_019816508.2;refGene_gn=VAV1;refGene_tx=XM_023244394.1
A2 5166688 5166840
refGene_gn=VAV1;refGene_tx=XM_003981728.4;refGene_gn=VAV1;refGene_tx=XM_006928302.3;refGene_gn=VAV1;refGene_tx=XM_006928303.3;refGene_gn=VAV1;refGene_tx=XM_019816508.2;refGene_gn=VAV1;refGene_tx=XM_023244394.1
A2 5168770 5168824
refGene_gn=VAV1;refGene_tx=XM_003981728.4;refGene_gn=VAV1;refGene_tx=XM_006928302.3;refGene_gn=VAV1;refGene_tx=XM_006928303.3;refGene_gn=VAV1;refGene_tx=XM_019816508.2;refGene_gn=VAV1;refGene_tx=XM_023244394.1
A2 5187089 5187120 refGene_gn=ADGRE1;refGene_tx=XM_023240916.1
A2 5189307 5189361 refGene_gn=ADGRE1;refGene_tx=XM_023240916.1
A2 5201056 5201203 refGene_gn=ADGRE1;refGene_tx=XM_023240916.1
A2 5203415 5203571 refGene_gn=ADGRE1;refGene_tx=XM_023240916.1
A2 5203681 5203801 refGene_gn=ADGRE1;refGene_tx=XM_023240916.1
A2 5205692 5205839 refGene_gn=ADGRE1;refGene_tx=XM_023240916.1
A2 5208646 5208796 refGene_gn=ADGRE1;refGene_tx=XM_023240916.1
A2 5223918 5224074 refGene_gn=ADGRE1;refGene_tx=XM_023240916.1
A2 5224184 5224304 refGene_gn=ADGRE1;refGene_tx=XM_023240916.1
A2 5226193 5226340 refGene_gn=ADGRE1;refGene_tx=XM_023240916.1
A2 5230866 5231007 refGene_gn=ADGRE1;refGene_tx=XM_023240916.1
A2 5231092 5231230 refGene_gn=ADGRE1;refGene_tx=XM_023240916.1
A2 5233082 5233171 refGene_gn=ADGRE1;refGene_tx=XM_023240916.1
A2 5234767 5234851 refGene_gn=ADGRE1;refGene_tx=XM_023240916.1
A2 5239466 5239605 refGene_gn=ADGRE1;refGene_tx=XM_023240916.1
A2 5240808 5241020 refGene_gn=ADGRE1;refGene_tx=XM_023240916.1
A2 5241756 5241927 refGene_gn=ADGRE1;refGene_tx=XM_023240916.1
A2 5243411 5243606 refGene_gn=ADGRE1;refGene_tx=XM_023240916.1
A2 5244848 5245084 refGene_gn=ADGRE1;refGene_tx=XM_023240916.1
A2 5248076 5248143 refGene_gn=ADGRE1;refGene_tx=XM_023240916.1
A2 5249710 5249802 refGene_gn=ADGRE1;refGene_tx=XM_023240916.1
A2 5252602 5252771 refGene_gn=ADGRE1;refGene_tx=XM_023240916.1
A2 5252880 5252985 refGene_gn=ADGRE1;refGene_tx=XM_023240916.1
A2 5255038 5255044 refGene_gn=ADGRE1;refGene_tx=XM_023240916.1
A2 5271945 5271976
refGene_gn=LOC101091924;refGene_tx=XM_019816557.2;refGene_gn=LOC101091924;refGene_tx=XM_023244396.1
A2 5275636 5275687
refGene_gn=LOC101091924;refGene_tx=XM_019816557.2;refGene_gn=LOC101091

924;refGene_tx=XM_023244396.1
A2	5278133	5278253
refGene_gn=LOC101091924;refGene_tx=XM_019816557.2;refGene_gn=LOC101091
924;refGene_tx=XM_023244396.1
A2	5283187	5283337
refGene_gn=LOC101091924;refGene_tx=XM_019816557.2;refGene_gn=LOC101091
924;refGene_tx=XM_023244396.1
A2	5286247	5286288
refGene_gn=LOC101091924;refGene_tx=XM_019816557.2;refGene_gn=LOC101091
924;refGene_tx=XM_023244396.1
A2	5287143	5287279
refGene_gn=LOC101091924;refGene_tx=XM_019816557.2;refGene_gn=LOC101091
924;refGene_tx=XM_023244396.1
A2	5287881	5287989
refGene_gn=LOC101091924;refGene_tx=XM_019816557.2;refGene_gn=LOC101091
924;refGene_tx=XM_023244396.1
A2	5290826	5291011
refGene_gn=LOC101091924;refGene_tx=XM_019816557.2;refGene_gn=LOC101091
924;refGene_tx=XM_023244396.1
A2	5291599	5291767
refGene_gn=LOC101091924;refGene_tx=XM_019816557.2;refGene_gn=LOC101091
924;refGene_tx=XM_023244396.1
A2	5294352	5294547
refGene_gn=LOC101091924;refGene_tx=XM_019816557.2;refGene_gn=LOC101091
924;refGene_tx=XM_023244396.1
A2	5303564	5303800
refGene_gn=LOC101091924;refGene_tx=XM_019816557.2;refGene_gn=LOC101091
924;refGene_tx=XM_023244396.1
A2	5305764	5305831
refGene_gn=LOC101091924;refGene_tx=XM_019816557.2;refGene_gn=LOC101091
924;refGene_tx=XM_023244396.1
A2	5306645	5306737
refGene_gn=LOC101091924;refGene_tx=XM_019816557.2;refGene_gn=LOC101091
924;refGene_tx=XM_023244396.1
A2	5307887	5308068
refGene_gn=LOC101091924;refGene_tx=XM_019816557.2;refGene_gn=LOC101091
924;refGene_tx=XM_023244396.1
A2	5310240	5310336
refGene_gn=LOC101091924;refGene_tx=XM_019816557.2;refGene_gn=LOC101091
924;refGene_tx=XM_023244396.1
A2	5312488	5312596
refGene_gn=LOC101091924;refGene_tx=XM_019816557.2
A2	5344704	5344944
refGene_gn=LOC101091924;refGene_tx=XM_023244396.1
A2	5406327	5407269
refGene_gn=LOC101092166;refGene_tx=XM_003981789.2
A2	5423185	5424436	refGene_gn=ACTL9;refGene_tx=XM_023244415.1
A2	5492812	5492900	refGene_gn=ADAMTS10;refGene_tx=XM_023244418.1
A2	5493060	5493407	refGene_gn=ADAMTS10;refGene_tx=XM_023244418.1
A2	5493769	5493926	refGene_gn=ADAMTS10;refGene_tx=XM_023244418.1

A2 5494972 5495190 refGene_gn=ADAMTS10;refGene_tx=XM_023244418.1
A2 5496819 5496903 refGene_gn=ADAMTS10;refGene_tx=XM_023244418.1
A2 5497008 5497154 refGene_gn=ADAMTS10;refGene_tx=XM_023244418.1
A2 5497570 5497614 refGene_gn=ADAMTS10;refGene_tx=XM_023244418.1
A2 5497768 5497874 refGene_gn=ADAMTS10;refGene_tx=XM_023244418.1
A2 5497961 5498108 refGene_gn=ADAMTS10;refGene_tx=XM_023244418.1
A2 5498313 5498455 refGene_gn=ADAMTS10;refGene_tx=XM_023244418.1
A2 5501091 5501199 refGene_gn=ADAMTS10;refGene_tx=XM_023244418.1
A2 5501508 5501654 refGene_gn=ADAMTS10;refGene_tx=XM_023244418.1
A2 5501750 5501814 refGene_gn=ADAMTS10;refGene_tx=XM_023244418.1
A2 5502902 5503005 refGene_gn=ADAMTS10;refGene_tx=XM_023244418.1
A2 5503315 5503449 refGene_gn=ADAMTS10;refGene_tx=XM_023244418.1
A2 5503555 5503679 refGene_gn=ADAMTS10;refGene_tx=XM_023244418.1
A2 5504621 5504702 refGene_gn=ADAMTS10;refGene_tx=XM_023244418.1
A2 5504788 5504952 refGene_gn=ADAMTS10;refGene_tx=XM_023244418.1
A2 5505046 5505173 refGene_gn=ADAMTS10;refGene_tx=XM_023244418.1
A2 5505266 5505320 refGene_gn=ADAMTS10;refGene_tx=XM_023244418.1
A2 5505322 5505395 refGene_gn=ADAMTS10;refGene_tx=XM_023244418.1
A2 5505761 5505966 refGene_gn=ADAMTS10;refGene_tx=XM_023244418.1
A2 5506211 5506388 refGene_gn=ADAMTS10;refGene_tx=XM_023244418.1
A2 5506468 5506628 refGene_gn=ADAMTS10;refGene_tx=XM_023244418.1
A2 5507854 5507964 refGene_gn=ADAMTS10;refGene_tx=XM_023244418.1
A2 5510070 5510073 refGene_gn=MYO1F;refGene_tx=XM_023244421.1
A2 5511117 5512050
refGene_gn=LOC102902512;refGene_tx=XM_023244426.1;refGene_gn=LOC102902512;refGene_tx=XM_023244431.1;refGene_gn=LOC102902512;refGene_tx=XM_023244438.1;refGene_gn=LOC102902512;refGene_tx=XM_023244451.1;refGene_gn=LOC102902512;refGene_tx=XM_023244455.1;refGene_gn=LOC102902512;refGene_tx=XM_023244456.1;refGene_gn=LOC102902512;refGene_tx=XM_023244459.1;refGene_gn=LOC102902512;refGene_tx=XM_023244464.1;refGene_gn=LOC102902512;refGene_tx=XM_023244465.1;refGene_gn=LOC102902512;refGene_tx=XM_023244467.1
A2 5518877 5519015 refGene_gn=MYO1F;refGene_tx=XM_023244421.1
A2 5519481 5519571 refGene_gn=MYO1F;refGene_tx=XM_023244421.1
A2 5519654 5519749 refGene_gn=MYO1F;refGene_tx=XM_023244421.1
A2 5520175 5520263 refGene_gn=MYO1F;refGene_tx=XM_023244421.1
A2 5520376 5520466 refGene_gn=MYO1F;refGene_tx=XM_023244421.1
A2 5523541 5523673 refGene_gn=MYO1F;refGene_tx=XM_023244421.1
A2 5523835 5523967 refGene_gn=MYO1F;refGene_tx=XM_023244421.1
A2 5524174 5524307 refGene_gn=MYO1F;refGene_tx=XM_023244421.1
A2 5524482 5524679 refGene_gn=MYO1F;refGene_tx=XM_023244421.1
A2 5525373 5525454 refGene_gn=MYO1F;refGene_tx=XM_023244421.1
A2 5525573 5525660 refGene_gn=MYO1F;refGene_tx=XM_023244421.1
A2 5526688 5526775 refGene_gn=MYO1F;refGene_tx=XM_023244421.1
A2 5527462 5527630 refGene_gn=MYO1F;refGene_tx=XM_023244421.1
A2 5528318 5528404 refGene_gn=MYO1F;refGene_tx=XM_023244421.1
A2 5531302 5531384 refGene_gn=MYO1F;refGene_tx=XM_023244421.1
A2 5532726 5532833 refGene_gn=MYO1F;refGene_tx=XM_023244421.1
A2 5533353 5533452 refGene_gn=MYO1F;refGene_tx=XM_023244421.1
A2 5533543 5533688 refGene_gn=MYO1F;refGene_tx=XM_023244421.1

```
A2      5538721 5538836 refGene_gn=MYO1F;refGene_tx=XM_023244421.1
A2      5538929 5539099 refGene_gn=MYO1F;refGene_tx=XM_023244421.1
A2      5539810 5539956 refGene_gn=MYO1F;refGene_tx=XM_023244421.1
A2      5540337 5540484 refGene_gn=MYO1F;refGene_tx=XM_023244421.1
A2      5540595 5540744 refGene_gn=MYO1F;refGene_tx=XM_023244421.1
A2      5541214 5541298 refGene_gn=MYO1F;refGene_tx=XM_023244421.1
A2      5541846 5542042 refGene_gn=MYO1F;refGene_tx=XM_023244421.1
A2      5542128 5542298 refGene_gn=MYO1F;refGene_tx=XM_023244421.1
A2      5542956 5543033 refGene_gn=MYO1F;refGene_tx=XM_023244421.1
A2      5546693 5546696
refGene_gn=ZNF414;refGene_tx=XM_003981792.5;refGene_gn=ZNF414;refGene_
tx=XM_011287456.3
A2      5547419 5547732
refGene_gn=ZNF414;refGene_tx=XM_003981792.5;refGene_gn=ZNF414;refGene_
tx=XM_011287456.3;refGene_gn=ZNF414;refGene_tx=XM_019816653.2
A2      5548063 5548171
refGene_gn=ZNF414;refGene_tx=XM_003981792.5;refGene_gn=ZNF414;refGene_
tx=XM_011287456.3;refGene_gn=ZNF414;refGene_tx=XM_019816653.2
A2      5548284 5548390
refGene_gn=ZNF414;refGene_tx=XM_003981792.5;refGene_gn=ZNF414;refGene_
tx=XM_011287456.3;refGene_gn=ZNF414;refGene_tx=XM_019816653.2
A2      5548938 5549282
refGene_gn=ZNF414;refGene_tx=XM_003981792.5;refGene_gn=ZNF414;refGene_
tx=XM_011287456.3;refGene_gn=ZNF414;refGene_tx=XM_019816653.2
A2      5549362 5549423
refGene_gn=ZNF414;refGene_tx=XM_003981792.5;refGene_gn=ZNF414;refGene_
tx=XM_011287456.3;refGene_gn=ZNF414;refGene_tx=XM_019816653.2
A2      5549865 5550039
refGene_gn=ZNF414;refGene_tx=XM_003981792.5;refGene_gn=ZNF414;refGene_
tx=XM_011287456.3;refGene_gn=ZNF414;refGene_tx=XM_019816653.2
A2      5550132 5550262
refGene_gn=ZNF414;refGene_tx=XM_003981792.5;refGene_gn=ZNF414;refGene_
tx=XM_011287456.3;refGene_gn=ZNF414;refGene_tx=XM_019816653.2
A2      5554255 5554258
refGene_gn=PRAM1;refGene_tx=XM_006928332.4;refGene_gn=PRAM1;refGene_tx
=XM_023244490.1
A2      5556335 5558016
refGene_gn=PRAM1;refGene_tx=XM_006928332.4;refGene_gn=PRAM1;refGene_tx
=XM_023244490.1
A2      5558138 5558205
refGene_gn=PRAM1;refGene_tx=XM_006928332.4;refGene_gn=PRAM1;refGene_tx
=XM_023244490.1
A2      5558279 5558356
refGene_gn=PRAM1;refGene_tx=XM_006928332.4;refGene_gn=PRAM1;refGene_tx
=XM_023244490.1
A2      5560765 5560820
refGene_gn=PRAM1;refGene_tx=XM_006928332.4;refGene_gn=PRAM1;refGene_tx
=XM_023244490.1
A2      5560936 5561045
refGene_gn=PRAM1;refGene_tx=XM_006928332.4;refGene_gn=PRAM1;refGene_tx
```

```
=XM_023244490.1
A2	5561165	5561328
refGene_gn=PRAM1;refGene_tx=XM_006928332.4;refGene_gn=PRAM1;refGene_tx
=XM_023244490.1
A2	5561405	5561439
refGene_gn=PRAM1;refGene_tx=XM_006928332.4;refGene_gn=PRAM1;refGene_tx
=XM_023244490.1
A2	5561533	5561567
refGene_gn=PRAM1;refGene_tx=XM_006928332.4;refGene_gn=PRAM1;refGene_tx
=XM_023244490.1
A2	5563187	5563351	refGene_gn=HNRNPM;refGene_tx=XM_003981779.5
A2	5565026	5565078	refGene_gn=HNRNPM;refGene_tx=XM_003981779.5
A2	5565684	5566221	refGene_gn=HNRNPM;refGene_tx=XM_003981779.5
A2	5566222	5566488	refGene_gn=HNRNPM;refGene_tx=XM_003981779.5
A2	5568483	5568537	refGene_gn=HNRNPM;refGene_tx=XM_003981779.5
A2	5575731	5575809	refGene_gn=HNRNPM;refGene_tx=XM_003981779.5
A2	5576266	5576311	refGene_gn=HNRNPM;refGene_tx=XM_003981779.5
A2	5577563	5577665	refGene_gn=HNRNPM;refGene_tx=XM_003981779.5
A2	5580644	5580705	refGene_gn=HNRNPM;refGene_tx=XM_003981779.5
A2	5581531	5581581	refGene_gn=HNRNPM;refGene_tx=XM_003981779.5
A2	5582401	5582555	refGene_gn=HNRNPM;refGene_tx=XM_003981779.5
A2	5583797	5583833	refGene_gn=HNRNPM;refGene_tx=XM_003981779.5
A2	5583950	5583989	refGene_gn=HNRNPM;refGene_tx=XM_003981779.5
A2	5585090	5585184	refGene_gn=HNRNPM;refGene_tx=XM_003981779.5
A2	5585269	5585277	refGene_gn=HNRNPM;refGene_tx=XM_003981779.5
A2	5586189	5586242	refGene_gn=HNRNPM;refGene_tx=XM_003981779.5
A2	5593345	5593515	refGene_gn=HNRNPM;refGene_tx=XM_003981779.5
A2	5602799	5602912	refGene_gn=HNRNPM;refGene_tx=XM_003981779.5
A2	5607184	5607343
refGene_gn=MARCH2;refGene_tx=XM_023244497.1;refGene_gn=MARCH2;refGene_
tx=XM_023244498.1
A2	5609237	5609447
refGene_gn=MARCH2;refGene_tx=XM_023244497.1;refGene_gn=MARCH2;refGene_
tx=XM_023244498.1
A2	5610972	5611168
refGene_gn=MARCH2;refGene_tx=XM_023244497.1;refGene_gn=MARCH2;refGene_
tx=XM_023244498.1
A2	5611857	5611928	refGene_gn=MARCH2;refGene_tx=XM_023244498.1
A2	5616004	5616180	refGene_gn=MARCH2;refGene_tx=XM_023244497.1
A2	5628087	5628233	refGene_gn=RAB11B;refGene_tx=XM_003981782.5
A2	5628893	5628974	refGene_gn=RAB11B;refGene_tx=XM_003981782.5
A2	5629301	5629495	refGene_gn=RAB11B;refGene_tx=XM_003981782.5
A2	5629995	5630191	refGene_gn=RAB11B;refGene_tx=XM_003981782.5
A2	5637429	5637469	refGene_gn=RAB11B;refGene_tx=XM_003981782.5
A2	5645110	5645292	refGene_gn=ANGPTL4;refGene_tx=XM_003981783.5
A2	5646665	5646947	refGene_gn=ANGPTL4;refGene_tx=XM_003981783.5
A2	5647043	5647139	refGene_gn=ANGPTL4;refGene_tx=XM_003981783.5
A2	5647509	5647623	refGene_gn=ANGPTL4;refGene_tx=XM_003981783.5
A2	5649851	5649969	refGene_gn=ANGPTL4;refGene_tx=XM_003981783.5
A2	5650105	5650216	refGene_gn=ANGPTL4;refGene_tx=XM_003981783.5
```

```
A2	5650412	5650754	refGene_gn=ANGPTL4;refGene_tx=XM_003981783.5
A2	5681745	5681779	refGene_gn=KANK3;refGene_tx=XM_023244508.1
A2	5682601	5683897	refGene_gn=KANK3;refGene_tx=XM_023244508.1
A2	5683975	5684075	refGene_gn=KANK3;refGene_tx=XM_023244508.1
A2	5684281	5684443	refGene_gn=KANK3;refGene_tx=XM_023244508.1
A2	5684560	5684645	refGene_gn=KANK3;refGene_tx=XM_023244508.1
A2	5685138	5685355	refGene_gn=KANK3;refGene_tx=XM_023244508.1
A2	5688679	5688822	refGene_gn=KANK3;refGene_tx=XM_023244508.1
A2	5688940	5689141	refGene_gn=KANK3;refGene_tx=XM_023244508.1
A2	5689224	5689323	refGene_gn=KANK3;refGene_tx=XM_023244508.1
A2	5690101	5690179	refGene_gn=KANK3;refGene_tx=XM_023244508.1
A2	5691800	5691851	refGene_gn=NDUFA7;refGene_tx=XM_003981785.3
A2	5692271	5692321	refGene_gn=NDUFA7;refGene_tx=XM_003981785.3
A2	5695118	5695268	refGene_gn=NDUFA7;refGene_tx=XM_003981785.3
A2	5698411	5698502	refGene_gn=NDUFA7;refGene_tx=XM_003981785.3
A2	5701117	5701244
refGene_gn=CD320;refGene_tx=XM_019824900.2;refGene_gn=CD320;refGene_tx
=XM_023244513.1
A2	5703738	5703852
refGene_gn=CD320;refGene_tx=XM_019824900.2;refGene_gn=CD320;refGene_tx
=XM_023244513.1
A2	5704119	5704457
refGene_gn=CD320;refGene_tx=XM_019824900.2;refGene_gn=CD320;refGene_tx
=XM_023244513.1
A2	5704740	5704899
refGene_gn=CD320;refGene_tx=XM_019824900.2;refGene_gn=CD320;refGene_tx
=XM_023244513.1
A2	5705025	5705168
refGene_gn=CD320;refGene_tx=XM_019824900.2;refGene_gn=CD320;refGene_tx
=XM_023244513.1
A2	5730298	5730457
refGene_gn=CERS4;refGene_tx=XM_023244528.1;refGene_gn=CERS4;refGene_tx
=XM_023244530.1;refGene_gn=CERS4;refGene_tx=XM_023244535.1
A2	5730539	5730696
refGene_gn=CERS4;refGene_tx=XM_023244528.1;refGene_gn=CERS4;refGene_tx
=XM_023244530.1;refGene_gn=CERS4;refGene_tx=XM_023244535.1
A2	5731963	5732070
refGene_gn=CERS4;refGene_tx=XM_023244528.1;refGene_gn=CERS4;refGene_tx
=XM_023244530.1;refGene_gn=CERS4;refGene_tx=XM_023244535.1
A2	5737799	5737928
refGene_gn=CERS4;refGene_tx=XM_023244528.1;refGene_gn=CERS4;refGene_tx
=XM_023244530.1;refGene_gn=CERS4;refGene_tx=XM_023244535.1
A2	5738142	5738235
refGene_gn=CERS4;refGene_tx=XM_023244528.1;refGene_gn=CERS4;refGene_tx
=XM_023244530.1;refGene_gn=CERS4;refGene_tx=XM_023244535.1
A2	5738553	5738604
refGene_gn=CERS4;refGene_tx=XM_023244528.1;refGene_gn=CERS4;refGene_tx
=XM_023244530.1;refGene_gn=CERS4;refGene_tx=XM_023244535.1
A2	5738928	5738986
refGene_gn=CERS4;refGene_tx=XM_023244528.1;refGene_gn=CERS4;refGene_tx
```

=XM_023244530.1;refGene_gn=CERS4;refGene_tx=XM_023244535.1
A2	5739086	5739205
refGene_gn=CERS4;refGene_tx=XM_023244528.1;refGene_gn=CERS4;refGene_tx=XM_023244530.1;refGene_gn=CERS4;refGene_tx=XM_023244535.1
A2	5739868	5739986
refGene_gn=CERS4;refGene_tx=XM_023244528.1;refGene_gn=CERS4;refGene_tx=XM_023244530.1
A2	5742422	5742595
refGene_gn=CERS4;refGene_tx=XM_023244528.1;refGene_gn=CERS4;refGene_tx=XM_023244530.1;refGene_gn=CERS4;refGene_tx=XM_023244535.1
A2	5815372	5815569	refGene_gn=FBN3;refGene_tx=XM_023244548.1
A2	5815669	5815752	refGene_gn=FBN3;refGene_tx=XM_023244548.1
A2	5816282	5816381	refGene_gn=FBN3;refGene_tx=XM_023244548.1
A2	5816571	5816661	refGene_gn=FBN3;refGene_tx=XM_023244548.1
A2	5816914	5817010	refGene_gn=FBN3;refGene_tx=XM_023244548.1
A2	5818562	5818760	refGene_gn=FBN3;refGene_tx=XM_023244548.1
A2	5818864	5818990	refGene_gn=FBN3;refGene_tx=XM_023244548.1
A2	5819123	5819249	refGene_gn=FBN3;refGene_tx=XM_023244548.1
A2	5821617	5821770	refGene_gn=FBN3;refGene_tx=XM_023244548.1
A2	5821894	5822077	refGene_gn=FBN3;refGene_tx=XM_023244548.1
A2	5822470	5822614	refGene_gn=FBN3;refGene_tx=XM_023244548.1
A2	5822683	5822803	refGene_gn=FBN3;refGene_tx=XM_023244548.1
A2	5822925	5823051	refGene_gn=FBN3;refGene_tx=XM_023244548.1
A2	5825470	5825593	refGene_gn=FBN3;refGene_tx=XM_023244548.1
A2	5825694	5825970	refGene_gn=FBN3;refGene_tx=XM_023244548.1
A2	5826259	5826313	refGene_gn=FBN3;refGene_tx=XM_023244548.1
A2	5827798	5827924	refGene_gn=FBN3;refGene_tx=XM_023244548.1
A2	5828025	5828151	refGene_gn=FBN3;refGene_tx=XM_023244548.1
A2	5829486	5829606	refGene_gn=FBN3;refGene_tx=XM_023244548.1
A2	5829745	5829883	refGene_gn=FBN3;refGene_tx=XM_023244548.1
A2	5830027	5830078	refGene_gn=FBN3;refGene_tx=XM_023244548.1
A2	5830338	5830464	refGene_gn=FBN3;refGene_tx=XM_023244548.1
A2	5832264	5832489	refGene_gn=FBN3;refGene_tx=XM_023244548.1
A2	5832669	5832795	refGene_gn=FBN3;refGene_tx=XM_023244548.1
A2	5833080	5833209	refGene_gn=FBN3;refGene_tx=XM_023244548.1
A2	5834473	5834599	refGene_gn=FBN3;refGene_tx=XM_023244548.1
A2	5834778	5834904	refGene_gn=FBN3;refGene_tx=XM_023244548.1
A2	5835060	5835183	refGene_gn=FBN3;refGene_tx=XM_023244548.1
A2	5835330	5835456	refGene_gn=FBN3;refGene_tx=XM_023244548.1
A2	5836431	5836557	refGene_gn=FBN3;refGene_tx=XM_023244548.1
A2	5837883	5838005	refGene_gn=FBN3;refGene_tx=XM_023244548.1
A2	5838127	5838128	refGene_gn=FBN3;refGene_tx=XM_023244548.1
A2	5838191	5838314	refGene_gn=FBN3;refGene_tx=XM_023244548.1
A2	5838716	5838842	refGene_gn=FBN3;refGene_tx=XM_023244548.1
A2	5838923	5839046	refGene_gn=FBN3;refGene_tx=XM_023244548.1
A2	5839825	5839948	refGene_gn=FBN3;refGene_tx=XM_023244548.1
A2	5840445	5840607	refGene_gn=FBN3;refGene_tx=XM_023244548.1
A2	5841576	5841645	refGene_gn=FBN3;refGene_tx=XM_023244548.1
A2	5843074	5843200	refGene_gn=FBN3;refGene_tx=XM_023244548.1
A2	5848621	5848747	refGene_gn=FBN3;refGene_tx=XM_023244548.1

| | | | |
|---|---|---|---|
| A2 | 5849157 | 5849307 | refGene_gn=FBN3;refGene_tx=XM_023244548.1 |
| A2 | 5851721 | 5851793 | refGene_gn=FBN3;refGene_tx=XM_023244548.1 |
| A2 | 5854383 | 5854509 | refGene_gn=FBN3;refGene_tx=XM_023244548.1 |
| A2 | 5854783 | 5854909 | refGene_gn=FBN3;refGene_tx=XM_023244548.1 |
| A2 | 5855284 | 5855410 | refGene_gn=FBN3;refGene_tx=XM_023244548.1 |
| A2 | 5855789 | 5855906 | refGene_gn=FBN3;refGene_tx=XM_023244548.1 |
| A2 | 5857195 | 5857324 | refGene_gn=FBN3;refGene_tx=XM_023244548.1 |
| A2 | 5858306 | 5858426 | refGene_gn=FBN3;refGene_tx=XM_023244548.1 |
| A2 | 5859176 | 5859302 | refGene_gn=FBN3;refGene_tx=XM_023244548.1 |
| A2 | 5860131 | 5860284 | refGene_gn=FBN3;refGene_tx=XM_023244548.1 |
| A2 | 5860423 | 5860489 | refGene_gn=FBN3;refGene_tx=XM_023244548.1 |
| A2 | 5860691 | 5860714 | refGene_gn=FBN3;refGene_tx=XM_023244548.1 |
| A2 | 5860715 | 5860818 | refGene_gn=FBN3;refGene_tx=XM_023244548.1 |
| A2 | 5861055 | 5861175 | refGene_gn=FBN3;refGene_tx=XM_023244548.1 |
| A2 | 5861294 | 5861417 | refGene_gn=FBN3;refGene_tx=XM_023244548.1 |
| A2 | 5862766 | 5862901 | refGene_gn=FBN3;refGene_tx=XM_023244548.1 |
| A2 | 5863046 | 5863172 | refGene_gn=FBN3;refGene_tx=XM_023244548.1 |
| A2 | 5863700 | 5863907 | refGene_gn=FBN3;refGene_tx=XM_023244548.1 |
| A2 | 5864733 | 5864859 | refGene_gn=FBN3;refGene_tx=XM_023244548.1 |
| A2 | 5865434 | 5865557 | refGene_gn=FBN3;refGene_tx=XM_023244548.1 |
| A2 | 5865714 | 5865831 | refGene_gn=FBN3;refGene_tx=XM_023244548.1 |
| A2 | 5869082 | 5869211 | refGene_gn=FBN3;refGene_tx=XM_023244548.1 |
| A2 | 5869300 | 5869420 | refGene_gn=FBN3;refGene_tx=XM_023244548.1 |
| A2 | 5870795 | 5871030 | refGene_gn=FBN3;refGene_tx=XM_023244548.1 |
| A2 | 5871573 | 5871721 | refGene_gn=FBN3;refGene_tx=XM_023244548.1 |
| A2 | 5872731 | 5873070 | refGene_gn=FBN3;refGene_tx=XM_023244548.1 |
| A2 | 5875561 | 5875569 | refGene_gn=CCL25;refGene_tx=XM_019816732.2 |
| A2 | 5875888 | 5876076 | refGene_gn=CCL25;refGene_tx=XM_019816732.2 |
| A2 | 5876179 | 5876297 | refGene_gn=CCL25;refGene_tx=XM_019816732.2 |
| A2 | 5878221 | 5878328 | refGene_gn=CCL25;refGene_tx=XM_019816732.2 |
| A2 | 5882989 | 5883033 | refGene_gn=CCL25;refGene_tx=XM_019816732.2 |
| A2 | 5883690 | 5883726 | refGene_gn=CCL25;refGene_tx=XM_019816732.2 |
| A2 | 5921574 | 5921746 | refGene_gn=ELAVL1;refGene_tx=XM_023244561.1 |
| A2 | 5932232 | 5932336 | refGene_gn=ELAVL1;refGene_tx=XM_023244561.1 |
| A2 | 5938153 | 5938307 | refGene_gn=ELAVL1;refGene_tx=XM_023244561.1 |
| A2 | 5942943 | 5943169 | refGene_gn=ELAVL1;refGene_tx=XM_023244561.1 |
| A2 | 5945928 | 5946253 | refGene_gn=ELAVL1;refGene_tx=XM_023244561.1 |
| A2 | 5970151 | 5970196 | refGene_gn=TIMM44;refGene_tx=XM_003981797.3 |
| A2 | 5972377 | 5972479 | refGene_gn=TIMM44;refGene_tx=XM_003981797.3 |
| A2 | 5974890 | 5975061 | refGene_gn=TIMM44;refGene_tx=XM_003981797.3 |
| A2 | 5976026 | 5976107 | refGene_gn=TIMM44;refGene_tx=XM_003981797.3 |
| A2 | 5976912 | 5977062 | refGene_gn=TIMM44;refGene_tx=XM_003981797.3 |
| A2 | 5977837 | 5977977 | refGene_gn=TIMM44;refGene_tx=XM_003981797.3 |
| A2 | 5978289 | 5978375 | refGene_gn=TIMM44;refGene_tx=XM_003981797.3 |
| A2 | 5979015 | 5979108 | refGene_gn=TIMM44;refGene_tx=XM_003981797.3 |
| A2 | 5979196 | 5979321 | refGene_gn=TIMM44;refGene_tx=XM_003981797.3 |
| A2 | 5980637 | 5980688 | refGene_gn=TIMM44;refGene_tx=XM_003981797.3 |
| A2 | 5982699 | 5982789 | refGene_gn=TIMM44;refGene_tx=XM_003981797.3 |
| A2 | 5983356 | 5983467 | refGene_gn=TIMM44;refGene_tx=XM_003981797.3 |
| A2 | 5983797 | 5983917 | refGene_gn=TIMM44;refGene_tx=XM_003981797.3 |

A2	5985489	5985738	refGene_gn=CTXN1;refGene_tx=XM_023244574.1
A2	5988125	5988410	refGene_gn=SNAPC2;refGene_tx=XM_003981798.5
A2	5988411	5988458	refGene_gn=SNAPC2;refGene_tx=XM_003981798.5
A2	5988546	5988847	refGene_gn=SNAPC2;refGene_tx=XM_003981798.5
A2	5989126	5989195	refGene_gn=SNAPC2;refGene_tx=XM_003981798.5
A2	5989380	5989500	refGene_gn=SNAPC2;refGene_tx=XM_003981798.5
A2	5990091	5990274	refGene_gn=SNAPC2;refGene_tx=XM_003981798.5
A2	5992005	5992424	refGene_gn=TGFBR3L;refGene_tx=XM_023244615.1
A2	5992527	5992654	refGene_gn=TGFBR3L;refGene_tx=XM_023244615.1
A2	5993242	5993563	refGene_gn=TGFBR3L;refGene_tx=XM_023244615.1
A2	5993645	5993793	refGene_gn=TGFBR3L;refGene_tx=XM_023244615.1
A2	5993794	5994387	refGene_gn=TGFBR3L;refGene_tx=XM_023244615.1
A2	5997179	5997314
refGene_gn=MAP2K7;refGene_tx=XM_011288743.3;refGene_gn=MAP2K7;refGene_tx=XM_011288745.3;refGene_gn=MAP2K7;refGene_tx=XM_019816856.2
A2	5997432	5997478
refGene_gn=MAP2K7;refGene_tx=XM_011288743.3;refGene_gn=MAP2K7;refGene_tx=XM_011288745.3;refGene_gn=MAP2K7;refGene_tx=XM_019816856.2
A2	5998065	5998208
refGene_gn=MAP2K7;refGene_tx=XM_011288743.3;refGene_gn=MAP2K7;refGene_tx=XM_011288745.3;refGene_gn=MAP2K7;refGene_tx=XM_019816856.2
A2	5998315	5998396
refGene_gn=MAP2K7;refGene_tx=XM_011288743.3;refGene_gn=MAP2K7;refGene_tx=XM_011288745.3;refGene_gn=MAP2K7;refGene_tx=XM_019816856.2
A2	5998499	5998679
refGene_gn=MAP2K7;refGene_tx=XM_011288743.3;refGene_gn=MAP2K7;refGene_tx=XM_011288745.3;refGene_gn=MAP2K7;refGene_tx=XM_019816856.2
A2	5998768	5998876
refGene_gn=MAP2K7;refGene_tx=XM_011288743.3;refGene_gn=MAP2K7;refGene_tx=XM_011288745.3
A2	5999028	5999148
refGene_gn=MAP2K7;refGene_tx=XM_011288743.3;refGene_gn=MAP2K7;refGene_tx=XM_011288745.3;refGene_gn=MAP2K7;refGene_tx=XM_019816856.2
A2	5999224	5999338
refGene_gn=MAP2K7;refGene_tx=XM_011288743.3;refGene_gn=MAP2K7;refGene_tx=XM_011288745.3;refGene_gn=MAP2K7;refGene_tx=XM_019816856.2
A2	5999453	5999520
refGene_gn=MAP2K7;refGene_tx=XM_011288743.3;refGene_gn=MAP2K7;refGene_tx=XM_011288745.3;refGene_gn=MAP2K7;refGene_tx=XM_019816856.2
A2	5999705	5999847
refGene_gn=MAP2K7;refGene_tx=XM_011288743.3;refGene_gn=MAP2K7;refGene_tx=XM_011288745.3;refGene_gn=MAP2K7;refGene_tx=XM_019816856.2
A2	6002731	6002779
refGene_gn=MAP2K7;refGene_tx=XM_011288743.3;refGene_gn=MAP2K7;refGene_tx=XM_019816856.2
A2	6004479	6004603
refGene_gn=MAP2K7;refGene_tx=XM_011288743.3;refGene_gn=MAP2K7;refGene_tx=XM_011288745.3;refGene_gn=MAP2K7;refGene_tx=XM_019816856.2
A2	6006977	6009221
refGene_gn=LRRC8E;refGene_tx=XM_003981801.4;refGene_gn=LRRC8E;refGene_

tx=XM_019816801.2;refGene_gn=LRRC8E;refGene_tx=XM_019816810.2;refGene_gn=LRRC8E;refGene_tx=XM_019816814.2;refGene_gn=LRRC8E;refGene_tx=XM_019816818.2;refGene_gn=LRRC8E;refGene_tx=XM_019816821.2;refGene_gn=LRRC8E;refGene_tx=XM_019816825.2;refGene_gn=LRRC8E;refGene_tx=XM_019816831.2;refGene_gn=LRRC8E;refGene_tx=XM_019816836.2;refGene_gn=LRRC8E;refGene_tx=XM_019816844.2;refGene_gn=LRRC8E;refGene_tx=XM_023244583.1;refGene_gn=LRRC8E;refGene_tx=XM_023244585.1;refGene_gn=LRRC8E;refGene_tx=XM_023244591.1
A2 6009966 6010109
refGene_gn=LRRC8E;refGene_tx=XM_003981801.4;refGene_gn=LRRC8E;refGene_tx=XM_019816801.2;refGene_gn=LRRC8E;refGene_tx=XM_019816810.2;refGene_gn=LRRC8E;refGene_tx=XM_019816814.2;refGene_gn=LRRC8E;refGene_tx=XM_023244583.1;refGene_gn=LRRC8E;refGene_tx=XM_023244585.1;refGene_gn=LRRC8E;refGene_tx=XM_023244591.1
A2 6011688 6011902 refGene_gn=LRRC8E;refGene_tx=XM_023244583.1
A2 6017632 6017982
refGene_gn=PRR36;refGene_tx=XM_023244641.1;refGene_gn=PRR36;refGene_tx=XM_023244644.1
A2 6018532 6018807
refGene_gn=PRR36;refGene_tx=XM_023244641.1;refGene_gn=PRR36;refGene_tx=XM_023244644.1
A2 6018927 6019027
refGene_gn=PRR36;refGene_tx=XM_023244641.1;refGene_gn=PRR36;refGene_tx=XM_023244644.1
A2 6019230 6019343
refGene_gn=PRR36;refGene_tx=XM_023244641.1;refGene_gn=PRR36;refGene_tx=XM_023244644.1
A2 6019447 6021601
refGene_gn=PRR36;refGene_tx=XM_023244641.1;refGene_gn=PRR36;refGene_tx=XM_023244644.1
A2 6021784 6022311
refGene_gn=PRR36;refGene_tx=XM_023244641.1;refGene_gn=PRR36;refGene_tx=XM_023244644.1
A2 6026749 6027034
refGene_gn=EVI5L;refGene_tx=XM_023244653.1;refGene_gn=EVI5L;refGene_tx=XM_023244658.1;refGene_gn=EVI5L;refGene_tx=XM_023244660.1;refGene_gn=EVI5L;refGene_tx=XM_023244662.1;refGene_gn=EVI5L;refGene_tx=XM_023244666.1;refGene_gn=EVI5L;refGene_tx=XM_023244672.1
A2 6027208 6027304
refGene_gn=EVI5L;refGene_tx=XM_023244653.1;refGene_gn=EVI5L;refGene_tx=XM_023244658.1;refGene_gn=EVI5L;refGene_tx=XM_023244660.1;refGene_gn=EVI5L;refGene_tx=XM_023244662.1;refGene_gn=EVI5L;refGene_tx=XM_023244666.1;refGene_gn=EVI5L;refGene_tx=XM_023244672.1
A2 6027384 6027480
refGene_gn=EVI5L;refGene_tx=XM_023244653.1;refGene_gn=EVI5L;refGene_tx=XM_023244658.1;refGene_gn=EVI5L;refGene_tx=XM_023244660.1;refGene_gn=EVI5L;refGene_tx=XM_023244662.1;refGene_gn=EVI5L;refGene_tx=XM_023244666.1;refGene_gn=EVI5L;refGene_tx=XM_023244672.1
A2 6027566 6027713
refGene_gn=EVI5L;refGene_tx=XM_023244653.1;refGene_gn=EVI5L;refGene_tx

=XM_023244658.1;refGene_gn=EVI5L;refGene_tx=XM_023244660.1;refGene_gn=EVI5L;refGene_tx=XM_023244662.1;refGene_gn=EVI5L;refGene_tx=XM_023244666.1;refGene_gn=EVI5L;refGene_tx=XM_023244672.1
A2 6027815 6027971
refGene_gn=EVI5L;refGene_tx=XM_023244653.1;refGene_gn=EVI5L;refGene_tx=XM_023244658.1;refGene_gn=EVI5L;refGene_tx=XM_023244660.1;refGene_gn=EVI5L;refGene_tx=XM_023244662.1;refGene_gn=EVI5L;refGene_tx=XM_023244666.1;refGene_gn=EVI5L;refGene_tx=XM_023244672.1
A2 6028059 6028200
refGene_gn=EVI5L;refGene_tx=XM_023244653.1;refGene_gn=EVI5L;refGene_tx=XM_023244658.1;refGene_gn=EVI5L;refGene_tx=XM_023244660.1;refGene_gn=EVI5L;refGene_tx=XM_023244662.1;refGene_gn=EVI5L;refGene_tx=XM_023244666.1;refGene_gn=EVI5L;refGene_tx=XM_023244672.1
A2 6029514 6029661
refGene_gn=EVI5L;refGene_tx=XM_023244653.1;refGene_gn=EVI5L;refGene_tx=XM_023244658.1;refGene_gn=EVI5L;refGene_tx=XM_023244660.1;refGene_gn=EVI5L;refGene_tx=XM_023244662.1;refGene_gn=EVI5L;refGene_tx=XM_023244666.1;refGene_gn=EVI5L;refGene_tx=XM_023244672.1
A2 6031224 6031365
refGene_gn=EVI5L;refGene_tx=XM_023244653.1;refGene_gn=EVI5L;refGene_tx=XM_023244658.1;refGene_gn=EVI5L;refGene_tx=XM_023244660.1;refGene_gn=EVI5L;refGene_tx=XM_023244662.1;refGene_gn=EVI5L;refGene_tx=XM_023244666.1;refGene_gn=EVI5L;refGene_tx=XM_023244672.1
A2 6032496 6032529
refGene_gn=EVI5L;refGene_tx=XM_023244653.1;refGene_gn=EVI5L;refGene_tx=XM_023244662.1;refGene_gn=EVI5L;refGene_tx=XM_023244666.1;refGene_gn=EVI5L;refGene_tx=XM_023244672.1
A2 6033489 6033543
refGene_gn=EVI5L;refGene_tx=XM_023244653.1;refGene_gn=EVI5L;refGene_tx=XM_023244658.1;refGene_gn=EVI5L;refGene_tx=XM_023244660.1;refGene_gn=EVI5L;refGene_tx=XM_023244662.1;refGene_gn=EVI5L;refGene_tx=XM_023244666.1;refGene_gn=EVI5L;refGene_tx=XM_023244672.1
A2 6036559 6036620
refGene_gn=EVI5L;refGene_tx=XM_023244653.1;refGene_gn=EVI5L;refGene_tx=XM_023244658.1;refGene_gn=EVI5L;refGene_tx=XM_023244660.1;refGene_gn=EVI5L;refGene_tx=XM_023244662.1;refGene_gn=EVI5L;refGene_tx=XM_023244666.1;refGene_gn=EVI5L;refGene_tx=XM_023244672.1
A2 6036701 6036799
refGene_gn=EVI5L;refGene_tx=XM_023244653.1;refGene_gn=EVI5L;refGene_tx=XM_023244658.1;refGene_gn=EVI5L;refGene_tx=XM_023244660.1;refGene_gn=EVI5L;refGene_tx=XM_023244662.1;refGene_gn=EVI5L;refGene_tx=XM_023244666.1;refGene_gn=EVI5L;refGene_tx=XM_023244672.1
A2 6037937 6038027
refGene_gn=EVI5L;refGene_tx=XM_023244653.1;refGene_gn=EVI5L;refGene_tx=XM_023244658.1;refGene_gn=EVI5L;refGene_tx=XM_023244660.1;refGene_gn=EVI5L;refGene_tx=XM_023244662.1;refGene_gn=EVI5L;refGene_tx=XM_023244666.1;refGene_gn=EVI5L;refGene_tx=XM_023244672.1
A2 6038163 6038307
refGene_gn=EVI5L;refGene_tx=XM_023244653.1;refGene_gn=EVI5L;refGene_tx=XM_023244658.1;refGene_gn=EVI5L;refGene_tx=XM_023244660.1;refGene_gn=

EVI5L;refGene_tx=XM_023244662.1;refGene_gn=EVI5L;refGene_tx=XM_0232446
66.1;refGene_gn=EVI5L;refGene_tx=XM_023244672.1
A2	6039664	6039790
refGene_gn=EVI5L;refGene_tx=XM_023244653.1;refGene_gn=EVI5L;refGene_tx
=XM_023244658.1;refGene_gn=EVI5L;refGene_tx=XM_023244660.1;refGene_gn=
EVI5L;refGene_tx=XM_023244662.1;refGene_gn=EVI5L;refGene_tx=XM_0232446
66.1;refGene_gn=EVI5L;refGene_tx=XM_023244672.1
A2	6040392	6040467
refGene_gn=EVI5L;refGene_tx=XM_023244653.1;refGene_gn=EVI5L;refGene_tx
=XM_023244658.1;refGene_gn=EVI5L;refGene_tx=XM_023244660.1;refGene_gn=
EVI5L;refGene_tx=XM_023244662.1;refGene_gn=EVI5L;refGene_tx=XM_0232446
66.1;refGene_gn=EVI5L;refGene_tx=XM_023244672.1
A2	6040570	6040795
refGene_gn=EVI5L;refGene_tx=XM_023244653.1;refGene_gn=EVI5L;refGene_tx
=XM_023244658.1;refGene_gn=EVI5L;refGene_tx=XM_023244660.1;refGene_gn=
EVI5L;refGene_tx=XM_023244662.1;refGene_gn=EVI5L;refGene_tx=XM_0232446
66.1;refGene_gn=EVI5L;refGene_tx=XM_023244672.1
A2	6041229	6041419
refGene_gn=EVI5L;refGene_tx=XM_023244653.1;refGene_gn=EVI5L;refGene_tx
=XM_023244658.1;refGene_gn=EVI5L;refGene_tx=XM_023244660.1;refGene_gn=
EVI5L;refGene_tx=XM_023244662.1;refGene_gn=EVI5L;refGene_tx=XM_0232446
66.1;refGene_gn=EVI5L;refGene_tx=XM_023244672.1
A2	6042590	6042727
refGene_gn=EVI5L;refGene_tx=XM_023244653.1;refGene_gn=EVI5L;refGene_tx
=XM_023244658.1;refGene_gn=EVI5L;refGene_tx=XM_023244660.1;refGene_gn=
EVI5L;refGene_tx=XM_023244662.1;refGene_gn=EVI5L;refGene_tx=XM_0232446
66.1;refGene_gn=EVI5L;refGene_tx=XM_023244672.1
A2	6076132	6076172	refGene_gn=CD209;refGene_tx=XM_003981802.4
A2	6076333	6076408	refGene_gn=CD209;refGene_tx=XM_003981802.4
A2	6076541	6076628	refGene_gn=CD209;refGene_tx=XM_003981802.4
A2	6077339	6077417	refGene_gn=CD209;refGene_tx=XM_003981802.4
A2	6077721	6077799	refGene_gn=CD209;refGene_tx=XM_003981802.4
A2	6078015	6078167	refGene_gn=CD209;refGene_tx=XM_003981802.4
A2	6078613	6078726	refGene_gn=CD209;refGene_tx=XM_003981802.4
A2	6079110	6079252	refGene_gn=CD209;refGene_tx=XM_003981802.4
A2	6106285	6106367
refGene_gn=CLEC4G;refGene_tx=XM_003981803.4;refGene_gn=CLEC4G;refGene_
tx=XM_019816863.1
A2	6106697	6106799
refGene_gn=CLEC4G;refGene_tx=XM_003981803.4;refGene_gn=CLEC4G;refGene_
tx=XM_019816863.1
A2	6107149	6107203
refGene_gn=CLEC4G;refGene_tx=XM_003981803.4;refGene_gn=CLEC4G;refGene_
tx=XM_019816863.1
A2	6107418	6107481
refGene_gn=CLEC4G;refGene_tx=XM_003981803.4;refGene_gn=CLEC4G;refGene_
tx=XM_019816863.1
A2	6107613	6107718
refGene_gn=CLEC4G;refGene_tx=XM_003981803.4;refGene_gn=CLEC4G;refGene_
tx=XM_019816863.1

A2	6108039	6108129
refGene_gn=CLEC4G;refGene_tx=XM_003981803.4;refGene_gn=CLEC4G;refGene_tx=XM_019816863.1
A2	6108332	6108481
refGene_gn=CLEC4G;refGene_tx=XM_003981803.4;refGene_gn=CLEC4G;refGene_tx=XM_019816863.1
A2	6108600	6108716
refGene_gn=CLEC4G;refGene_tx=XM_003981803.4;refGene_gn=CLEC4G;refGene_tx=XM_019816863.1
A2	6108855	6108889	refGene_gn=CLEC4G;refGene_tx=XM_019816863.1
A2	6108984	6109123	refGene_gn=CLEC4G;refGene_tx=XM_003981803.4
A2	6125431	6125453
refGene_gn=FCER2;refGene_tx=XM_023244681.1;refGene_gn=FCER2;refGene_tx=XM_023244692.1
A2	6125754	6125812
refGene_gn=FCER2;refGene_tx=XM_011288763.3;refGene_gn=FCER2;refGene_tx=XM_023244676.1
A2	6126470	6126590
refGene_gn=FCER2;refGene_tx=XM_011288763.3;refGene_gn=FCER2;refGene_tx=XM_023244676.1;refGene_gn=FCER2;refGene_tx=XM_023244681.1;refGene_gn=FCER2;refGene_tx=XM_023244692.1
A2	6126912	6126966
refGene_gn=FCER2;refGene_tx=XM_011288763.3;refGene_gn=FCER2;refGene_tx=XM_023244676.1;refGene_gn=FCER2;refGene_tx=XM_023244681.1;refGene_gn=FCER2;refGene_tx=XM_023244692.1
A2	6127501	6127564
refGene_gn=FCER2;refGene_tx=XM_011288763.3;refGene_gn=FCER2;refGene_tx=XM_023244676.1;refGene_gn=FCER2;refGene_tx=XM_023244681.1;refGene_gn=FCER2;refGene_tx=XM_023244692.1
A2	6127791	6127854
refGene_gn=FCER2;refGene_tx=XM_011288763.3;refGene_gn=FCER2;refGene_tx=XM_023244676.1;refGene_gn=FCER2;refGene_tx=XM_023244681.1;refGene_gn=FCER2;refGene_tx=XM_023244692.1
A2	6127957	6128020
refGene_gn=FCER2;refGene_tx=XM_011288763.3;refGene_gn=FCER2;refGene_tx=XM_023244676.1;refGene_gn=FCER2;refGene_tx=XM_023244681.1;refGene_gn=FCER2;refGene_tx=XM_023244692.1
A2	6128111	6128201
refGene_gn=FCER2;refGene_tx=XM_011288763.3;refGene_gn=FCER2;refGene_tx=XM_023244676.1;refGene_gn=FCER2;refGene_tx=XM_023244681.1;refGene_gn=FCER2;refGene_tx=XM_023244692.1
A2	6130012	6130497
refGene_gn=FCER2;refGene_tx=XM_011288763.3;refGene_gn=FCER2;refGene_tx=XM_023244676.1;refGene_gn=FCER2;refGene_tx=XM_023244681.1;refGene_gn=FCER2;refGene_tx=XM_023244692.1
A2	6130623	6130730
refGene_gn=FCER2;refGene_tx=XM_011288763.3;refGene_gn=FCER2;refGene_tx=XM_023244676.1;refGene_gn=FCER2;refGene_tx=XM_023244681.1;refGene_gn=FCER2;refGene_tx=XM_023244692.1
A2	6130955	6131094

refGene_gn=FCER2;refGene_tx=XM_011288763.3;refGene_gn=FCER2;refGene_tx=XM_023244676.1;refGene_gn=FCER2;refGene_tx=XM_023244681.1;refGene_gn=FCER2;refGene_tx=XM_023244692.1
A2	6135481	6136048
refGene_gn=TRAPPC5;refGene_tx=XM_006928366.3;refGene_gn=TRAPPC5;refGene_tx=XM_023244707.1
A2	6139236	6139253
refGene_gn=MCEMP1;refGene_tx=XM_006928347.3;refGene_gn=MCEMP1;refGene_tx=XM_023244712.1;refGene_gn=MCEMP1;refGene_tx=XM_023244717.1
A2	6139369	6139435
refGene_gn=MCEMP1;refGene_tx=XM_006928347.3;refGene_gn=MCEMP1;refGene_tx=XM_023244712.1;refGene_gn=MCEMP1;refGene_tx=XM_023244717.1
A2	6139714	6139861
refGene_gn=MCEMP1;refGene_tx=XM_006928347.3;refGene_gn=MCEMP1;refGene_tx=XM_023244712.1;refGene_gn=MCEMP1;refGene_tx=XM_023244717.1
A2	6139939	6139990
refGene_gn=MCEMP1;refGene_tx=XM_006928347.3;refGene_gn=MCEMP1;refGene_tx=XM_023244712.1;refGene_gn=MCEMP1;refGene_tx=XM_023244717.1
A2	6140217	6140346
refGene_gn=MCEMP1;refGene_tx=XM_006928347.3;refGene_gn=MCEMP1;refGene_tx=XM_023244712.1;refGene_gn=MCEMP1;refGene_tx=XM_023244717.1
A2	6140876	6141014
refGene_gn=MCEMP1;refGene_tx=XM_006928347.3;refGene_gn=MCEMP1;refGene_tx=XM_023244712.1;refGene_gn=MCEMP1;refGene_tx=XM_023244717.1
A2	6141316	6141425
refGene_gn=MCEMP1;refGene_tx=XM_006928347.3;refGene_gn=MCEMP1;refGene_tx=XM_023244712.1;refGene_gn=MCEMP1;refGene_tx=XM_023244717.1
A2	6146840	6146977
refGene_gn=RETN;refGene_tx=XM_023244723.1;refGene_gn=RETN;refGene_tx=XM_023244724.1
A2	6147489	6147567
refGene_gn=RETN;refGene_tx=XM_023244723.1;refGene_gn=RETN;refGene_tx=XM_023244724.1
A2	6147937	6148055
refGene_gn=RETN;refGene_tx=XM_023244723.1;refGene_gn=RETN;refGene_tx=XM_023244724.1
A2	6161246	6161332
refGene_gn=STXBP2;refGene_tx=XM_003981805.4;refGene_gn=STXBP2;refGene_tx=XM_023244735.1
A2	6161656	6161814
refGene_gn=STXBP2;refGene_tx=XM_003981805.4;refGene_gn=STXBP2;refGene_tx=XM_023244735.1
A2	6161928	6162014
refGene_gn=STXBP2;refGene_tx=XM_003981805.4;refGene_gn=STXBP2;refGene_tx=XM_023244735.1
A2	6162658	6162754
refGene_gn=STXBP2;refGene_tx=XM_003981805.4;refGene_gn=STXBP2;refGene_tx=XM_023244735.1
A2	6163661	6163771
refGene_gn=STXBP2;refGene_tx=XM_003981805.4;refGene_gn=STXBP2;refGene_

tx=XM_023244735.1
A2	6164295	6164434
refGene_gn=STXBP2;refGene_tx=XM_003981805.4;refGene_gn=STXBP2;refGene_tx=XM_023244735.1
A2	6164777	6164858
refGene_gn=STXBP2;refGene_tx=XM_003981805.4;refGene_gn=STXBP2;refGene_tx=XM_023244735.1
A2	6164967	6165033
refGene_gn=STXBP2;refGene_tx=XM_003981805.4;refGene_gn=STXBP2;refGene_tx=XM_023244735.1
A2	6165171	6165229
refGene_gn=STXBP2;refGene_tx=XM_003981805.4;refGene_gn=STXBP2;refGene_tx=XM_023244735.1
A2	6165541	6165649
refGene_gn=STXBP2;refGene_tx=XM_003981805.4;refGene_gn=STXBP2;refGene_tx=XM_023244735.1
A2	6165736	6165867
refGene_gn=STXBP2;refGene_tx=XM_003981805.4;refGene_gn=STXBP2;refGene_tx=XM_023244735.1
A2	6165956	6166041
refGene_gn=STXBP2;refGene_tx=XM_003981805.4;refGene_gn=STXBP2;refGene_tx=XM_023244735.1
A2	6166230	6166379
refGene_gn=STXBP2;refGene_tx=XM_003981805.4;refGene_gn=STXBP2;refGene_tx=XM_023244735.1
A2	6166783	6166887
refGene_gn=STXBP2;refGene_tx=XM_003981805.4;refGene_gn=STXBP2;refGene_tx=XM_023244735.1
A2	6166974	6167053
refGene_gn=STXBP2;refGene_tx=XM_003981805.4;refGene_gn=STXBP2;refGene_tx=XM_023244735.1
A2	6167256	6167333
refGene_gn=STXBP2;refGene_tx=XM_003981805.4;refGene_gn=STXBP2;refGene_tx=XM_023244735.1
A2	6167838	6167945
refGene_gn=STXBP2;refGene_tx=XM_003981805.4;refGene_gn=STXBP2;refGene_tx=XM_023244735.1
A2	6168187	6168237
refGene_gn=STXBP2;refGene_tx=XM_003981805.4;refGene_gn=STXBP2;refGene_tx=XM_023244735.1
A2	6169470	6169507	refGene_gn=STXBP2;refGene_tx=XM_003981805.4
A2	6171696	6171755	refGene_gn=PCP2;refGene_tx=XM_019816911.2
A2	6172676	6172780
refGene_gn=PCP2;refGene_tx=XM_006928351.3;refGene_gn=PCP2;refGene_tx=XM_019816911.2
A2	6173211	6173234	refGene_gn=PCP2;refGene_tx=XM_019816911.2
A2	6173307	6173422
refGene_gn=PCP2;refGene_tx=XM_006928351.3;refGene_gn=PCP2;refGene_tx=XM_019816911.2
A2	6173711	6173836

refGene_gn=PCP2;refGene_tx=XM_006928351.3;refGene_gn=PCP2;refGene_tx=XM_019816911.2
A2 6175020 6175140
refGene_gn=PCP2;refGene_tx=XM_006928351.3;refGene_gn=PCP2;refGene_tx=XM_019816911.2
A2 6175251 6175335 refGene_gn=PET100;refGene_tx=XM_006928353.3
A2 6175661 6175694 refGene_gn=PET100;refGene_tx=XM_006928353.3
A2 6175863 6175950 refGene_gn=PET100;refGene_tx=XM_006928353.3
A2 6176697 6176724 refGene_gn=PET100;refGene_tx=XM_006928353.3
A2 6177034 6177085 refGene_gn=XAB2;refGene_tx=XM_003981809.4
A2 6181747 6181896 refGene_gn=XAB2;refGene_tx=XM_003981809.4
A2 6182188 6182312 refGene_gn=XAB2;refGene_tx=XM_003981809.4
A2 6182540 6182738 refGene_gn=XAB2;refGene_tx=XM_003981809.4
A2 6184161 6184296 refGene_gn=XAB2;refGene_tx=XM_003981809.4
A2 6184398 6184563 refGene_gn=XAB2;refGene_tx=XM_003981809.4
A2 6185422 6185567 refGene_gn=XAB2;refGene_tx=XM_003981809.4
A2 6185849 6186001 refGene_gn=XAB2;refGene_tx=XM_003981809.4
A2 6186082 6186202 refGene_gn=XAB2;refGene_tx=XM_003981809.4
A2 6186353 6186485 refGene_gn=XAB2;refGene_tx=XM_003981809.4
A2 6186580 6186712 refGene_gn=XAB2;refGene_tx=XM_003981809.4
A2 6186808 6186922 refGene_gn=XAB2;refGene_tx=XM_003981809.4
A2 6187848 6188011 refGene_gn=XAB2;refGene_tx=XM_003981809.4
A2 6188099 6188290 refGene_gn=XAB2;refGene_tx=XM_003981809.4
A2 6188434 6188557 refGene_gn=XAB2;refGene_tx=XM_003981809.4
A2 6188641 6188813 refGene_gn=XAB2;refGene_tx=XM_003981809.4
A2 6189066 6189196 refGene_gn=XAB2;refGene_tx=XM_003981809.4
A2 6189393 6189503 refGene_gn=XAB2;refGene_tx=XM_003981809.4
A2 6189595 6189657 refGene_gn=XAB2;refGene_tx=XM_003981809.4
A2 6191069 6191375 refGene_gn=CAMSAP3;refGene_tx=XM_023244762.1
A2 6191464 6191583 refGene_gn=CAMSAP3;refGene_tx=XM_023244762.1
A2 6191666 6191779 refGene_gn=CAMSAP3;refGene_tx=XM_023244762.1
A2 6192034 6192134 refGene_gn=CAMSAP3;refGene_tx=XM_023244762.1
A2 6192614 6192855 refGene_gn=CAMSAP3;refGene_tx=XM_023244762.1
A2 6192856 6192914 refGene_gn=CAMSAP3;refGene_tx=XM_023244762.1
A2 6192994 6193134 refGene_gn=CAMSAP3;refGene_tx=XM_023244762.1
A2 6194384 6195822 refGene_gn=CAMSAP3;refGene_tx=XM_023244762.1
A2 6195918 6195988 refGene_gn=CAMSAP3;refGene_tx=XM_023244762.1
A2 6196240 6196314 refGene_gn=CAMSAP3;refGene_tx=XM_023244762.1
A2 6196457 6196513 refGene_gn=CAMSAP3;refGene_tx=XM_023244762.1
A2 6196569 6196680 refGene_gn=CAMSAP3;refGene_tx=XM_023244762.1
A2 6196764 6196779 refGene_gn=CAMSAP3;refGene_tx=XM_023244762.1
A2 6196780 6196905 refGene_gn=CAMSAP3;refGene_tx=XM_023244762.1
A2 6197512 6197651 refGene_gn=CAMSAP3;refGene_tx=XM_023244762.1
A2 6197902 6197950 refGene_gn=CAMSAP3;refGene_tx=XM_023244762.1
A2 6198835 6198868 refGene_gn=CAMSAP3;refGene_tx=XM_023244762.1
A2 6199035 6199131 refGene_gn=CAMSAP3;refGene_tx=XM_023244762.1
A2 6199210 6199333 refGene_gn=CAMSAP3;refGene_tx=XM_023244762.1
A2 6199655 6199909 refGene_gn=CAMSAP3;refGene_tx=XM_023244762.1
A2 6205331 6205479 refGene_gn=CAMSAP3;refGene_tx=XM_023244762.1
A2 6229186 6229261

refGene_gn=PNPLA6;refGene_tx=XM_011288813.3;refGene_gn=PNPLA6;refGene_tx=XM_023244804.1;refGene_gn=PNPLA6;refGene_tx=XM_023244809.1;refGene_gn=PNPLA6;refGene_tx=XM_023244814.1;refGene_gn=PNPLA6;refGene_tx=XM_023244817.1;refGene_gn=PNPLA6;refGene_tx=XM_023244821.1
A2 6229391 6229501
refGene_gn=PNPLA6;refGene_tx=XM_011288813.3;refGene_gn=PNPLA6;refGene_tx=XM_023244804.1;refGene_gn=PNPLA6;refGene_tx=XM_023244809.1;refGene_gn=PNPLA6;refGene_tx=XM_023244814.1;refGene_gn=PNPLA6;refGene_tx=XM_023244817.1;refGene_gn=PNPLA6;refGene_tx=XM_023244821.1
A2 6229593 6229690
refGene_gn=PNPLA6;refGene_tx=XM_011288813.3;refGene_gn=PNPLA6;refGene_tx=XM_023244804.1;refGene_gn=PNPLA6;refGene_tx=XM_023244809.1;refGene_gn=PNPLA6;refGene_tx=XM_023244814.1;refGene_gn=PNPLA6;refGene_tx=XM_023244817.1;refGene_gn=PNPLA6;refGene_tx=XM_023244821.1
A2 6229900 6230017
refGene_gn=PNPLA6;refGene_tx=XM_011288813.3;refGene_gn=PNPLA6;refGene_tx=XM_023244804.1;refGene_gn=PNPLA6;refGene_tx=XM_023244809.1;refGene_gn=PNPLA6;refGene_tx=XM_023244814.1;refGene_gn=PNPLA6;refGene_tx=XM_023244817.1;refGene_gn=PNPLA6;refGene_tx=XM_023244821.1
A2 6230911 6231213
refGene_gn=PNPLA6;refGene_tx=XM_011288813.3;refGene_gn=PNPLA6;refGene_tx=XM_023244804.1;refGene_gn=PNPLA6;refGene_tx=XM_023244809.1;refGene_gn=PNPLA6;refGene_tx=XM_023244814.1;refGene_gn=PNPLA6;refGene_tx=XM_023244817.1;refGene_gn=PNPLA6;refGene_tx=XM_023244821.1
A2 6231476 6231922 refGene_gn=PNPLA6;refGene_tx=XM_023244822.1
A2 6232773 6232890
refGene_gn=PNPLA6;refGene_tx=XM_011288813.3;refGene_gn=PNPLA6;refGene_tx=XM_023244804.1;refGene_gn=PNPLA6;refGene_tx=XM_023244814.1;refGene_gn=PNPLA6;refGene_tx=XM_023244817.1;refGene_gn=PNPLA6;refGene_tx=XM_023244821.1;refGene_gn=PNPLA6;refGene_tx=XM_023244822.1
A2 6232976 6233093 refGene_gn=PNPLA6;refGene_tx=XM_023244809.1
A2 6233329 6233399
refGene_gn=PNPLA6;refGene_tx=XM_011288813.3;refGene_gn=PNPLA6;refGene_tx=XM_023244804.1;refGene_gn=PNPLA6;refGene_tx=XM_023244809.1;refGene_gn=PNPLA6;refGene_tx=XM_023244814.1;refGene_gn=PNPLA6;refGene_tx=XM_023244817.1;refGene_gn=PNPLA6;refGene_tx=XM_023244821.1;refGene_gn=PNPLA6;refGene_tx=XM_023244822.1
A2 6233486 6233603
refGene_gn=PNPLA6;refGene_tx=XM_011288813.3;refGene_gn=PNPLA6;refGene_tx=XM_023244804.1;refGene_gn=PNPLA6;refGene_tx=XM_023244809.1;refGene_gn=PNPLA6;refGene_tx=XM_023244814.1;refGene_gn=PNPLA6;refGene_tx=XM_023244817.1;refGene_gn=PNPLA6;refGene_tx=XM_023244821.1;refGene_gn=PNPLA6;refGene_tx=XM_023244822.1
A2 6233916 6234073
refGene_gn=PNPLA6;refGene_tx=XM_011288813.3;refGene_gn=PNPLA6;refGene_tx=XM_023244804.1;refGene_gn=PNPLA6;refGene_tx=XM_023244809.1;refGene_gn=PNPLA6;refGene_tx=XM_023244814.1;refGene_gn=PNPLA6;refGene_tx=XM_023244817.1;refGene_gn=PNPLA6;refGene_tx=XM_023244821.1;refGene_gn=PNPLA6;refGene_tx=XM_023244822.1
A2 6234268 6234387

refGene_gn=PNPLA6;refGene_tx=XM_011288813.3;refGene_gn=PNPLA6;refGene_tx=XM_023244804.1;refGene_gn=PNPLA6;refGene_tx=XM_023244809.1;refGene_gn=PNPLA6;refGene_tx=XM_023244814.1;refGene_gn=PNPLA6;refGene_tx=XM_023244817.1;refGene_gn=PNPLA6;refGene_tx=XM_023244821.1;refGene_gn=PNPLA6;refGene_tx=XM_023244822.1
A2 6234579 6234762
refGene_gn=PNPLA6;refGene_tx=XM_011288813.3;refGene_gn=PNPLA6;refGene_tx=XM_023244804.1;refGene_gn=PNPLA6;refGene_tx=XM_023244809.1;refGene_gn=PNPLA6;refGene_tx=XM_023244814.1;refGene_gn=PNPLA6;refGene_tx=XM_023244817.1;refGene_gn=PNPLA6;refGene_tx=XM_023244821.1;refGene_gn=PNPLA6;refGene_tx=XM_023244822.1
A2 6234932 6235101
refGene_gn=PNPLA6;refGene_tx=XM_011288813.3;refGene_gn=PNPLA6;refGene_tx=XM_023244804.1;refGene_gn=PNPLA6;refGene_tx=XM_023244809.1;refGene_gn=PNPLA6;refGene_tx=XM_023244814.1;refGene_gn=PNPLA6;refGene_tx=XM_023244817.1;refGene_gn=PNPLA6;refGene_tx=XM_023244821.1;refGene_gn=PNPLA6;refGene_tx=XM_023244822.1
A2 6235370 6235434
refGene_gn=PNPLA6;refGene_tx=XM_011288813.3;refGene_gn=PNPLA6;refGene_tx=XM_023244804.1;refGene_gn=PNPLA6;refGene_tx=XM_023244809.1;refGene_gn=PNPLA6;refGene_tx=XM_023244814.1;refGene_gn=PNPLA6;refGene_tx=XM_023244817.1;refGene_gn=PNPLA6;refGene_tx=XM_023244821.1;refGene_gn=PNPLA6;refGene_tx=XM_023244822.1
A2 6235646 6235790
refGene_gn=PNPLA6;refGene_tx=XM_011288813.3;refGene_gn=PNPLA6;refGene_tx=XM_023244804.1;refGene_gn=PNPLA6;refGene_tx=XM_023244809.1;refGene_gn=PNPLA6;refGene_tx=XM_023244814.1;refGene_gn=PNPLA6;refGene_tx=XM_023244817.1;refGene_gn=PNPLA6;refGene_tx=XM_023244821.1;refGene_gn=PNPLA6;refGene_tx=XM_023244822.1
A2 6237749 6237825
refGene_gn=PNPLA6;refGene_tx=XM_011288813.3;refGene_gn=PNPLA6;refGene_tx=XM_023244804.1;refGene_gn=PNPLA6;refGene_tx=XM_023244809.1;refGene_gn=PNPLA6;refGene_tx=XM_023244814.1;refGene_gn=PNPLA6;refGene_tx=XM_023244817.1;refGene_gn=PNPLA6;refGene_tx=XM_023244821.1;refGene_gn=PNPLA6;refGene_tx=XM_023244822.1
A2 6238054 6238168
refGene_gn=PNPLA6;refGene_tx=XM_011288813.3;refGene_gn=PNPLA6;refGene_tx=XM_023244804.1;refGene_gn=PNPLA6;refGene_tx=XM_023244809.1;refGene_gn=PNPLA6;refGene_tx=XM_023244814.1;refGene_gn=PNPLA6;refGene_tx=XM_023244817.1;refGene_gn=PNPLA6;refGene_tx=XM_023244821.1;refGene_gn=PNPLA6;refGene_tx=XM_023244822.1
A2 6238532 6238656
refGene_gn=PNPLA6;refGene_tx=XM_011288813.3;refGene_gn=PNPLA6;refGene_tx=XM_023244804.1;refGene_gn=PNPLA6;refGene_tx=XM_023244809.1;refGene_gn=PNPLA6;refGene_tx=XM_023244814.1;refGene_gn=PNPLA6;refGene_tx=XM_023244817.1;refGene_gn=PNPLA6;refGene_tx=XM_023244821.1;refGene_gn=PNPLA6;refGene_tx=XM_023244822.1
A2 6238735 6238867
refGene_gn=PNPLA6;refGene_tx=XM_011288813.3;refGene_gn=PNPLA6;refGene_tx=XM_023244804.1;refGene_gn=PNPLA6;refGene_tx=XM_023244809.1;refGene_

gn=PNPLA6;refGene_tx=XM_023244814.1;refGene_gn=PNPLA6;refGene_tx=XM_023244817.1;refGene_gn=PNPLA6;refGene_tx=XM_023244821.1;refGene_gn=PNPLA6;refGene_tx=XM_023244822.1
A2 6239106 6239312
refGene_gn=PNPLA6;refGene_tx=XM_011288813.3;refGene_gn=PNPLA6;refGene_tx=XM_023244804.1;refGene_gn=PNPLA6;refGene_tx=XM_023244809.1;refGene_gn=PNPLA6;refGene_tx=XM_023244814.1;refGene_gn=PNPLA6;refGene_tx=XM_023244817.1;refGene_gn=PNPLA6;refGene_tx=XM_023244821.1;refGene_gn=PNPLA6;refGene_tx=XM_023244822.1
A2 6243131 6243209
refGene_gn=PNPLA6;refGene_tx=XM_011288813.3;refGene_gn=PNPLA6;refGene_tx=XM_023244804.1;refGene_gn=PNPLA6;refGene_tx=XM_023244809.1;refGene_gn=PNPLA6;refGene_tx=XM_023244814.1;refGene_gn=PNPLA6;refGene_tx=XM_023244817.1;refGene_gn=PNPLA6;refGene_tx=XM_023244821.1;refGene_gn=PNPLA6;refGene_tx=XM_023244822.1
A2 6243301 6243469
refGene_gn=PNPLA6;refGene_tx=XM_011288813.3;refGene_gn=PNPLA6;refGene_tx=XM_023244804.1;refGene_gn=PNPLA6;refGene_tx=XM_023244809.1;refGene_gn=PNPLA6;refGene_tx=XM_023244814.1;refGene_gn=PNPLA6;refGene_tx=XM_023244817.1;refGene_gn=PNPLA6;refGene_tx=XM_023244821.1;refGene_gn=PNPLA6;refGene_tx=XM_023244822.1
A2 6243594 6243704
refGene_gn=PNPLA6;refGene_tx=XM_011288813.3;refGene_gn=PNPLA6;refGene_tx=XM_023244804.1;refGene_gn=PNPLA6;refGene_tx=XM_023244809.1;refGene_gn=PNPLA6;refGene_tx=XM_023244814.1;refGene_gn=PNPLA6;refGene_tx=XM_023244817.1;refGene_gn=PNPLA6;refGene_tx=XM_023244821.1;refGene_gn=PNPLA6;refGene_tx=XM_023244822.1
A2 6243978 6244062
refGene_gn=PNPLA6;refGene_tx=XM_011288813.3;refGene_gn=PNPLA6;refGene_tx=XM_023244804.1;refGene_gn=PNPLA6;refGene_tx=XM_023244809.1;refGene_gn=PNPLA6;refGene_tx=XM_023244814.1;refGene_gn=PNPLA6;refGene_tx=XM_023244817.1;refGene_gn=PNPLA6;refGene_tx=XM_023244821.1;refGene_gn=PNPLA6;refGene_tx=XM_023244822.1
A2 6244255 6244418
refGene_gn=PNPLA6;refGene_tx=XM_011288813.3;refGene_gn=PNPLA6;refGene_tx=XM_023244804.1;refGene_gn=PNPLA6;refGene_tx=XM_023244809.1;refGene_gn=PNPLA6;refGene_tx=XM_023244814.1;refGene_gn=PNPLA6;refGene_tx=XM_023244817.1;refGene_gn=PNPLA6;refGene_tx=XM_023244821.1;refGene_gn=PNPLA6;refGene_tx=XM_023244822.1
A2 6244509 6244590
refGene_gn=PNPLA6;refGene_tx=XM_011288813.3;refGene_gn=PNPLA6;refGene_tx=XM_023244804.1;refGene_gn=PNPLA6;refGene_tx=XM_023244809.1;refGene_gn=PNPLA6;refGene_tx=XM_023244814.1;refGene_gn=PNPLA6;refGene_tx=XM_023244817.1;refGene_gn=PNPLA6;refGene_tx=XM_023244821.1;refGene_gn=PNPLA6;refGene_tx=XM_023244822.1
A2 6244968 6245097
refGene_gn=PNPLA6;refGene_tx=XM_011288813.3;refGene_gn=PNPLA6;refGene_tx=XM_023244804.1;refGene_gn=PNPLA6;refGene_tx=XM_023244809.1;refGene_gn=PNPLA6;refGene_tx=XM_023244814.1;refGene_gn=PNPLA6;refGene_tx=XM_023244817.1;refGene_gn=PNPLA6;refGene_tx=XM_023244821.1;refGene_gn=PNPLA

6;refGene_tx=XM_023244822.1
A2 6245310 6245391
refGene_gn=PNPLA6;refGene_tx=XM_011288813.3;refGene_gn=PNPLA6;refGene_tx=XM_023244804.1;refGene_gn=PNPLA6;refGene_tx=XM_023244809.1;refGene_gn=PNPLA6;refGene_tx=XM_023244814.1;refGene_gn=PNPLA6;refGene_tx=XM_023244817.1;refGene_gn=PNPLA6;refGene_tx=XM_023244821.1;refGene_gn=PNPLA6;refGene_tx=XM_023244822.1
A2 6245743 6245903
refGene_gn=PNPLA6;refGene_tx=XM_011288813.3;refGene_gn=PNPLA6;refGene_tx=XM_023244804.1;refGene_gn=PNPLA6;refGene_tx=XM_023244809.1;refGene_gn=PNPLA6;refGene_tx=XM_023244814.1;refGene_gn=PNPLA6;refGene_tx=XM_023244817.1;refGene_gn=PNPLA6;refGene_tx=XM_023244821.1;refGene_gn=PNPLA6;refGene_tx=XM_023244822.1
A2 6245994 6246135
refGene_gn=PNPLA6;refGene_tx=XM_011288813.3;refGene_gn=PNPLA6;refGene_tx=XM_023244804.1;refGene_gn=PNPLA6;refGene_tx=XM_023244809.1;refGene_gn=PNPLA6;refGene_tx=XM_023244814.1;refGene_gn=PNPLA6;refGene_tx=XM_023244817.1;refGene_gn=PNPLA6;refGene_tx=XM_023244821.1;refGene_gn=PNPLA6;refGene_tx=XM_023244822.1
A2 6248120 6248218
refGene_gn=PNPLA6;refGene_tx=XM_011288813.3;refGene_gn=PNPLA6;refGene_tx=XM_023244804.1;refGene_gn=PNPLA6;refGene_tx=XM_023244809.1;refGene_gn=PNPLA6;refGene_tx=XM_023244814.1;refGene_gn=PNPLA6;refGene_tx=XM_023244817.1;refGene_gn=PNPLA6;refGene_tx=XM_023244821.1;refGene_gn=PNPLA6;refGene_tx=XM_023244822.1
A2 6248389 6248472
refGene_gn=PNPLA6;refGene_tx=XM_011288813.3;refGene_gn=PNPLA6;refGene_tx=XM_023244804.1;refGene_gn=PNPLA6;refGene_tx=XM_023244809.1;refGene_gn=PNPLA6;refGene_tx=XM_023244814.1;refGene_gn=PNPLA6;refGene_tx=XM_023244817.1;refGene_gn=PNPLA6;refGene_tx=XM_023244821.1;refGene_gn=PNPLA6;refGene_tx=XM_023244822.1
A2 6248643 6248878
refGene_gn=PNPLA6;refGene_tx=XM_011288813.3;refGene_gn=PNPLA6;refGene_tx=XM_023244804.1;refGene_gn=PNPLA6;refGene_tx=XM_023244809.1;refGene_gn=PNPLA6;refGene_tx=XM_023244814.1;refGene_gn=PNPLA6;refGene_tx=XM_023244817.1;refGene_gn=PNPLA6;refGene_tx=XM_023244821.1;refGene_gn=PNPLA6;refGene_tx=XM_023244822.1
A2 6249069 6249138
refGene_gn=PNPLA6;refGene_tx=XM_011288813.3;refGene_gn=PNPLA6;refGene_tx=XM_023244817.1;refGene_gn=PNPLA6;refGene_tx=XM_023244821.1
A2 6249220 6249223
refGene_gn=PNPLA6;refGene_tx=XM_011288813.3;refGene_gn=PNPLA6;refGene_tx=XM_023244817.1;refGene_gn=PNPLA6;refGene_tx=XM_023244821.1
A2 6250777 6250814 refGene_gn=MCOLN1;refGene_tx=XM_003981811.5
A2 6250910 6251041 refGene_gn=MCOLN1;refGene_tx=XM_003981811.5
A2 6252263 6252731
refGene_gn=MCOLN1;refGene_tx=XM_003981811.5;refGene_gn=MCOLN1;refGene_tx=XM_006928355.4;refGene_gn=MCOLN1;refGene_tx=XM_006928356.4
A2 6252989 6253112
refGene_gn=MCOLN1;refGene_tx=XM_003981811.5;refGene_gn=MCOLN1;refGene_

tx=XM_006928355.4;refGene_gn=MCOLN1;refGene_tx=XM_006928356.4
A2	6253529	6253631
refGene_gn=MCOLN1;refGene_tx=XM_003981811.5;refGene_gn=MCOLN1;refGene_tx=XM_006928355.4;refGene_gn=MCOLN1;refGene_tx=XM_006928356.4
A2	6253774	6253924
refGene_gn=MCOLN1;refGene_tx=XM_003981811.5;refGene_gn=MCOLN1;refGene_tx=XM_006928355.4;refGene_gn=MCOLN1;refGene_tx=XM_006928356.4
A2	6254050	6254157
refGene_gn=MCOLN1;refGene_tx=XM_003981811.5;refGene_gn=MCOLN1;refGene_tx=XM_006928355.4;refGene_gn=MCOLN1;refGene_tx=XM_006928356.4
A2	6254385	6254485
refGene_gn=MCOLN1;refGene_tx=XM_003981811.5;refGene_gn=MCOLN1;refGene_tx=XM_006928355.4;refGene_gn=MCOLN1;refGene_tx=XM_006928356.4
A2	6254698	6254795
refGene_gn=MCOLN1;refGene_tx=XM_003981811.5;refGene_gn=MCOLN1;refGene_tx=XM_006928355.4;refGene_gn=MCOLN1;refGene_tx=XM_006928356.4
A2	6255041	6255150
refGene_gn=MCOLN1;refGene_tx=XM_003981811.5;refGene_gn=MCOLN1;refGene_tx=XM_006928355.4;refGene_gn=MCOLN1;refGene_tx=XM_006928356.4
A2	6255564	6255730
refGene_gn=MCOLN1;refGene_tx=XM_003981811.5;refGene_gn=MCOLN1;refGene_tx=XM_006928355.4;refGene_gn=MCOLN1;refGene_tx=XM_006928356.4
A2	6255825	6255993
refGene_gn=MCOLN1;refGene_tx=XM_003981811.5;refGene_gn=MCOLN1;refGene_tx=XM_006928355.4;refGene_gn=MCOLN1;refGene_tx=XM_006928356.4
A2	6256692	6256889
refGene_gn=MCOLN1;refGene_tx=XM_003981811.5;refGene_gn=MCOLN1;refGene_tx=XM_006928355.4;refGene_gn=MCOLN1;refGene_tx=XM_006928356.4
A2	6258867	6258892	refGene_gn=MCOLN1;refGene_tx=XM_006928356.4
A2	6259039	6259070
refGene_gn=MCOLN1;refGene_tx=XM_003981811.5;refGene_gn=MCOLN1;refGene_tx=XM_006928355.4
A2	6260917	6260973	refGene_gn=ZNF358;refGene_tx=XM_006928357.4
A2	6260975	6262622	refGene_gn=ZNF358;refGene_tx=XM_006928357.4
A2	6262865	6263157	refGene_gn=ZNF358;refGene_tx=XM_006928357.4
A2	6271143	6271371
refGene_gn=TEX45;refGene_tx=XM_003981812.4;refGene_gn=TEX45;refGene_tx=XM_023244824.1;refGene_gn=TEX45;refGene_tx=XM_023244825.1
A2	6271875	6271961
refGene_gn=TEX45;refGene_tx=XM_003981812.4;refGene_gn=TEX45;refGene_tx=XM_023244824.1
A2	6272474	6272679
refGene_gn=TEX45;refGene_tx=XM_003981812.4;refGene_gn=TEX45;refGene_tx=XM_023244824.1;refGene_gn=TEX45;refGene_tx=XM_023244825.1
A2	6273043	6273153
refGene_gn=TEX45;refGene_tx=XM_003981812.4;refGene_gn=TEX45;refGene_tx=XM_023244824.1;refGene_gn=TEX45;refGene_tx=XM_023244825.1
A2	6273234	6273347
refGene_gn=TEX45;refGene_tx=XM_003981812.4;refGene_gn=TEX45;refGene_tx=XM_023244824.1;refGene_gn=TEX45;refGene_tx=XM_023244825.1

A2 6273597 6273698
refGene_gn=TEX45;refGene_tx=XM_003981812.4;refGene_gn=TEX45;refGene_tx
=XM_023244824.1;refGene_gn=TEX45;refGene_tx=XM_023244825.1
A2 6273786 6273893
refGene_gn=TEX45;refGene_tx=XM_003981812.4;refGene_gn=TEX45;refGene_tx
=XM_023244824.1;refGene_gn=TEX45;refGene_tx=XM_023244825.1
A2 6274914 6275500
refGene_gn=TEX45;refGene_tx=XM_003981812.4;refGene_gn=TEX45;refGene_tx
=XM_023244824.1;refGene_gn=TEX45;refGene_tx=XM_023244825.1
A2 6277166 6277181
refGene_gn=TEX45;refGene_tx=XM_003981812.4;refGene_gn=TEX45;refGene_tx
=XM_023244825.1
A2 6282688 6282748
refGene_gn=PEX11G;refGene_tx=XM_003981813.5;refGene_gn=PEX11G;refGene_
tx=XM_011288822.2;refGene_gn=PEX11G;refGene_tx=XM_011288825.3
A2 6284044 6284233
refGene_gn=PEX11G;refGene_tx=XM_003981813.5;refGene_gn=PEX11G;refGene_
tx=XM_011288822.2;refGene_gn=PEX11G;refGene_tx=XM_019817027.2
A2 6286957 6287683
refGene_gn=PEX11G;refGene_tx=XM_003981813.5;refGene_gn=PEX11G;refGene_
tx=XM_011288822.2;refGene_gn=PEX11G;refGene_tx=XM_011288825.3;refGene_
gn=PEX11G;refGene_tx=XM_019817027.2
A2 6288017 6288080 refGene_gn=PEX11G;refGene_tx=XM_003981813.5
A2 6288679 6288914 refGene_gn=PEX11G;refGene_tx=XM_003981813.5
A2 6293216 6293359
refGene_gn=ARHGEF18;refGene_tx=XM_011288831.3;refGene_gn=ARHGEF18;refG
ene_tx=XM_019816930.2
A2 6293439 6293565
refGene_gn=ARHGEF18;refGene_tx=XM_011288831.3;refGene_gn=ARHGEF18;refG
ene_tx=XM_019816930.2
A2 6294417 6294751
refGene_gn=ARHGEF18;refGene_tx=XM_011288831.3;refGene_gn=ARHGEF18;refG
ene_tx=XM_019816930.2
A2 6295066 6295537
refGene_gn=ARHGEF18;refGene_tx=XM_011288831.3;refGene_gn=ARHGEF18;refG
ene_tx=XM_019816930.2
A2 6295648 6295696
refGene_gn=ARHGEF18;refGene_tx=XM_011288831.3;refGene_gn=ARHGEF18;refG
ene_tx=XM_019816930.2
A2 6295789 6295846
refGene_gn=ARHGEF18;refGene_tx=XM_011288831.3;refGene_gn=ARHGEF18;refG
ene_tx=XM_019816930.2
A2 6296357 6296479
refGene_gn=ARHGEF18;refGene_tx=XM_011288831.3;refGene_gn=ARHGEF18;refG
ene_tx=XM_019816930.2
A2 6296817 6296955
refGene_gn=ARHGEF18;refGene_tx=XM_011288831.3;refGene_gn=ARHGEF18;refG
ene_tx=XM_019816930.2
A2 6298152 6298335
refGene_gn=ARHGEF18;refGene_tx=XM_011288831.3;refGene_gn=ARHGEF18;refG

ene_tx=XM_019816930.2
A2 6300372 6300464
refGene_gn=ARHGEF18;refGene_tx=XM_011288831.3;refGene_gn=ARHGEF18;refGene_tx=XM_019816930.2
A2 6301253 6301432
refGene_gn=ARHGEF18;refGene_tx=XM_011288831.3;refGene_gn=ARHGEF18;refGene_tx=XM_019816930.2
A2 6302538 6302615
refGene_gn=ARHGEF18;refGene_tx=XM_011288831.3;refGene_gn=ARHGEF18;refGene_tx=XM_019816930.2
A2 6303617 6303866
refGene_gn=ARHGEF18;refGene_tx=XM_011288831.3;refGene_gn=ARHGEF18;refGene_tx=XM_019816930.2
A2 6304717 6304835
refGene_gn=ARHGEF18;refGene_tx=XM_011288831.3;refGene_gn=ARHGEF18;refGene_tx=XM_019816930.2
A2 6306958 6307084
refGene_gn=ARHGEF18;refGene_tx=XM_011288831.3;refGene_gn=ARHGEF18;refGene_tx=XM_019816930.2
A2 6307782 6308033
refGene_gn=ARHGEF18;refGene_tx=XM_011288831.3;refGene_gn=ARHGEF18;refGene_tx=XM_019816930.2
A2 6310009 6310150
refGene_gn=ARHGEF18;refGene_tx=XM_011288831.3;refGene_gn=ARHGEF18;refGene_tx=XM_019816930.2
A2 6310280 6310393
refGene_gn=ARHGEF18;refGene_tx=XM_011288831.3;refGene_gn=ARHGEF18;refGene_tx=XM_019816930.2
A2 6311431 6311570
refGene_gn=ARHGEF18;refGene_tx=XM_011288831.3;refGene_gn=ARHGEF18;refGene_tx=XM_019816930.2
A2 6350885 6351027 refGene_gn=ARHGEF18;refGene_tx=XM_019816930.2
A2 6351171 6351274 refGene_gn=ARHGEF18;refGene_tx=XM_019816930.2
A2 6352776 6352854 refGene_gn=ARHGEF18;refGene_tx=XM_019816930.2
A2 6353348 6353393 refGene_gn=ARHGEF18;refGene_tx=XM_019816930.2
A2 6353701 6353759 refGene_gn=ARHGEF18;refGene_tx=XM_019816930.2
A2 6354688 6354803 refGene_gn=ARHGEF18;refGene_tx=XM_019816930.2
A2 6355400 6355551 refGene_gn=ARHGEF18;refGene_tx=XM_019816930.2
A2 6356311 6356571 refGene_gn=ARHGEF18;refGene_tx=XM_019816930.2
A2 6364522 6364647 refGene_gn=ARHGEF18;refGene_tx=XM_019816930.2
A2 6374402 6374478 refGene_gn=ARHGEF18;refGene_tx=XM_019816930.2
A2 6422206 6422306
refGene_gn=INSR;refGene_tx=XM_023244792.1;refGene_gn=INSR;refGene_tx=XM_023244793.1;refGene_gn=INSR;refGene_tx=XM_023244794.1;refGene_gn=INSR;refGene_tx=XM_023244797.1
A2 6441113 6441665
refGene_gn=INSR;refGene_tx=XM_023244792.1;refGene_gn=INSR;refGene_tx=XM_023244793.1;refGene_gn=INSR;refGene_tx=XM_023244794.1;refGene_gn=INSR;refGene_tx=XM_023244797.1
A2 6510958 6511280

refGene_gn=INSR;refGene_tx=XM_023244792.1;refGene_gn=INSR;refGene_tx=XM_023244793.1;refGene_gn=INSR;refGene_tx=XM_023244794.1;refGene_gn=INSR;refGene_tx=XM_023244797.1
A2 6517523 6517672
refGene_gn=INSR;refGene_tx=XM_023244792.1;refGene_gn=INSR;refGene_tx=XM_023244793.1;refGene_gn=INSR;refGene_tx=XM_023244794.1;refGene_gn=INSR;refGene_tx=XM_023244797.1
A2 6518717 6518862
refGene_gn=INSR;refGene_tx=XM_023244792.1;refGene_gn=INSR;refGene_tx=XM_023244793.1;refGene_gn=INSR;refGene_tx=XM_023244794.1;refGene_gn=INSR;refGene_tx=XM_023244797.1
A2 6522191 6522406
refGene_gn=INSR;refGene_tx=XM_023244792.1;refGene_gn=INSR;refGene_tx=XM_023244793.1;refGene_gn=INSR;refGene_tx=XM_023244794.1;refGene_gn=INSR;refGene_tx=XM_023244797.1
A2 6523367 6523494
refGene_gn=INSR;refGene_tx=XM_023244792.1;refGene_gn=INSR;refGene_tx=XM_023244793.1;refGene_gn=INSR;refGene_tx=XM_023244794.1;refGene_gn=INSR;refGene_tx=XM_023244797.1
A2 6525329 6525580
refGene_gn=INSR;refGene_tx=XM_023244792.1;refGene_gn=INSR;refGene_tx=XM_023244793.1;refGene_gn=INSR;refGene_tx=XM_023244794.1;refGene_gn=INSR;refGene_tx=XM_023244797.1
A2 6527091 6527259
refGene_gn=INSR;refGene_tx=XM_023244792.1;refGene_gn=INSR;refGene_tx=XM_023244793.1;refGene_gn=INSR;refGene_tx=XM_023244794.1;refGene_gn=INSR;refGene_tx=XM_023244797.1
A2 6532054 6532256
refGene_gn=INSR;refGene_tx=XM_023244792.1;refGene_gn=INSR;refGene_tx=XM_023244793.1;refGene_gn=INSR;refGene_tx=XM_023244794.1;refGene_gn=INSR;refGene_tx=XM_023244797.1
A2 6533939 6533975
refGene_gn=INSR;refGene_tx=XM_023244792.1;refGene_gn=INSR;refGene_tx=XM_023244793.1
A2 6538822 6539094
refGene_gn=INSR;refGene_tx=XM_023244792.1;refGene_gn=INSR;refGene_tx=XM_023244793.1;refGene_gn=INSR;refGene_tx=XM_023244794.1;refGene_gn=INSR;refGene_tx=XM_023244797.1
A2 6540738 6540878
refGene_gn=INSR;refGene_tx=XM_023244792.1;refGene_gn=INSR;refGene_tx=XM_023244793.1;refGene_gn=INSR;refGene_tx=XM_023244794.1;refGene_gn=INSR;refGene_tx=XM_023244797.1
A2 6547231 6547391
refGene_gn=INSR;refGene_tx=XM_023244792.1;refGene_gn=INSR;refGene_tx=XM_023244793.1;refGene_gn=INSR;refGene_tx=XM_023244794.1;refGene_gn=INSR;refGene_tx=XM_023244797.1
A2 6551544 6551647
refGene_gn=INSR;refGene_tx=XM_023244792.1;refGene_gn=INSR;refGene_tx=XM_023244793.1;refGene_gn=INSR;refGene_tx=XM_023244794.1;refGene_gn=INSR;refGene_tx=XM_023244797.1

A2	6553367	6553435
refGene_gn=INSR;refGene_tx=XM_023244792.1;refGene_gn=INSR;refGene_tx=XM_023244793.1;refGene_gn=INSR;refGene_tx=XM_023244794.1;refGene_gn=INSR;refGene_tx=XM_023244797.1
A2	6555398	6555643
refGene_gn=INSR;refGene_tx=XM_023244792.1;refGene_gn=INSR;refGene_tx=XM_023244793.1;refGene_gn=INSR;refGene_tx=XM_023244794.1;refGene_gn=INSR;refGene_tx=XM_023244797.1
A2	6556581	6556692
refGene_gn=INSR;refGene_tx=XM_023244792.1;refGene_gn=INSR;refGene_tx=XM_023244793.1;refGene_gn=INSR;refGene_tx=XM_023244794.1;refGene_gn=INSR;refGene_tx=XM_023244797.1
A2	6556783	6556943
refGene_gn=INSR;refGene_tx=XM_023244792.1;refGene_gn=INSR;refGene_tx=XM_023244793.1;refGene_gn=INSR;refGene_tx=XM_023244794.1;refGene_gn=INSR;refGene_tx=XM_023244797.1
A2	6558479	6558609
refGene_gn=INSR;refGene_tx=XM_023244792.1;refGene_gn=INSR;refGene_tx=XM_023244793.1;refGene_gn=INSR;refGene_tx=XM_023244794.1;refGene_gn=INSR;refGene_tx=XM_023244797.1
A2	6558948	6559083
refGene_gn=INSR;refGene_tx=XM_023244792.1;refGene_gn=INSR;refGene_tx=XM_023244793.1;refGene_gn=INSR;refGene_tx=XM_023244794.1;refGene_gn=INSR;refGene_tx=XM_023244797.1
A2	6561428	6561783
refGene_gn=INSR;refGene_tx=XM_023244792.1;refGene_gn=INSR;refGene_tx=XM_023244793.1;refGene_gn=INSR;refGene_tx=XM_023244794.1;refGene_gn=INSR;refGene_tx=XM_023244797.1
A2	6583386	6584163
refGene_gn=ZNF557;refGene_tx=XM_006928358.4;refGene_gn=ZNF557;refGene_tx=XM_023244827.1;refGene_gn=ZNF557;refGene_tx=XM_023244831.1;refGene_gn=ZNF557;refGene_tx=XM_023244832.1;refGene_gn=ZNF557;refGene_tx=XM_023244835.1;refGene_gn=ZNF557;refGene_tx=XM_023244838.1
A2	6585554	6585637
refGene_gn=ZNF557;refGene_tx=XM_006928358.4;refGene_gn=ZNF557;refGene_tx=XM_023244827.1;refGene_gn=ZNF557;refGene_tx=XM_023244831.1;refGene_gn=ZNF557;refGene_tx=XM_023244832.1;refGene_gn=ZNF557;refGene_tx=XM_023244835.1;refGene_gn=ZNF557;refGene_tx=XM_023244838.1
A2	6586149	6586245
refGene_gn=ZNF557;refGene_tx=XM_006928358.4;refGene_gn=ZNF557;refGene_tx=XM_023244827.1;refGene_gn=ZNF557;refGene_tx=XM_023244832.1;refGene_gn=ZNF557;refGene_tx=XM_023244835.1;refGene_gn=ZNF557;refGene_tx=XM_023244838.1
A2	6589393	6589520
refGene_gn=ZNF557;refGene_tx=XM_006928358.4;refGene_gn=ZNF557;refGene_tx=XM_023244827.1;refGene_gn=ZNF557;refGene_tx=XM_023244831.1;refGene_gn=ZNF557;refGene_tx=XM_023244832.1;refGene_gn=ZNF557;refGene_tx=XM_023244835.1;refGene_gn=ZNF557;refGene_tx=XM_023244838.1
A2	6590400	6590489
refGene_gn=ZNF557;refGene_tx=XM_006928358.4;refGene_gn=ZNF557;refGene_

tx=XM_023244827.1;refGene_gn=ZNF557;refGene_tx=XM_023244831.1
A2	6591060	6591091
refGene_gn=ZNF557;refGene_tx=XM_023244827.1;refGene_gn=ZNF557;refGene_tx=XM_023244831.1
A2	6658348	6659131
refGene_gn=ZNF557;refGene_tx=XM_019816993.2;refGene_gn=ZNF557;refGene_tx=XM_023244826.1
A2	6659612	6659695
refGene_gn=ZNF557;refGene_tx=XM_019816993.2;refGene_gn=ZNF557;refGene_tx=XM_023244826.1
A2	6660330	6660426
refGene_gn=ZNF557;refGene_tx=XM_019816993.2;refGene_gn=ZNF557;refGene_tx=XM_023244826.1
A2	6675758	6675885
refGene_gn=ZNF557;refGene_tx=XM_019816993.2;refGene_gn=ZNF557;refGene_tx=XM_023244826.1
A2	6676526	6676615	refGene_gn=ZNF557;refGene_tx=XM_023244826.1
A2	6677182	6677213
refGene_gn=ZNF557;refGene_tx=XM_006928358.4;refGene_gn=ZNF557;refGene_tx=XM_023244826.1
A2	6692975	6693566
refGene_gn=LOC101081786;refGene_tx=XM_003981829.5;refGene_gn=LOC101081786;refGene_tx=XM_019817065.2
A2	6695912	6696008
refGene_gn=LOC101082044;refGene_tx=XM_023244823.1
A2	6698649	6698729
refGene_gn=LOC101082044;refGene_tx=XM_023244823.1
A2	6698931	6698973
refGene_gn=LOC101082044;refGene_tx=XM_023244823.1
A2	6700583	6700750
refGene_gn=LOC101082044;refGene_tx=XM_023244823.1
A2	6705267	6705335
refGene_gn=LOC101082044;refGene_tx=XM_023244823.1
A2	6705656	6705820
refGene_gn=LOC101082044;refGene_tx=XM_023244823.1
A2	6706905	6707054
refGene_gn=LOC101082044;refGene_tx=XM_023244823.1
A2	6707835	6707903
refGene_gn=LOC101082044;refGene_tx=XM_023244823.1
A2	6708242	6708373
refGene_gn=LOC101082044;refGene_tx=XM_023244823.1
A2	6709763	6709936
refGene_gn=LOC101082044;refGene_tx=XM_023244823.1
A2	6710094	6710162
refGene_gn=LOC101082044;refGene_tx=XM_023244823.1
A2	6710245	6710367
refGene_gn=LOC101082044;refGene_tx=XM_023244823.1
A2	6712016	6712082
refGene_gn=LOC101082044;refGene_tx=XM_023244823.1
A2	6713221	6713257

refGene_gn=LOC101082044;refGene_tx=XM_023244823.1
A2 6714622 6714795
refGene_gn=LOC101082044;refGene_tx=XM_023244823.1
A2 6716691 6716782
refGene_gn=LOC101082044;refGene_tx=XM_023244823.1
A2 6716784 6716976
refGene_gn=LOC101082044;refGene_tx=XM_023244823.1
A2 6720536 6720572
refGene_gn=LOC101082044;refGene_tx=XM_023244823.1
A2 6720858 6721031
refGene_gn=LOC101082044;refGene_tx=XM_023244823.1
A2 6721487 6721555
refGene_gn=LOC101082044;refGene_tx=XM_023244823.1
A2 6721649 6721774
refGene_gn=LOC101082044;refGene_tx=XM_023244823.1
A2 6723336 6723372
refGene_gn=LOC101082044;refGene_tx=XM_023244823.1
A2 6723885 6723921
refGene_gn=LOC101082044;refGene_tx=XM_023244823.1
A2 6724037 6724210
refGene_gn=LOC101082044;refGene_tx=XM_023244823.1
A2 6724797 6724865
refGene_gn=LOC101082044;refGene_tx=XM_023244823.1
A2 6725297 6725422
refGene_gn=LOC101082044;refGene_tx=XM_023244823.1
A2 6725896 6725932
refGene_gn=LOC101082044;refGene_tx=XM_023244823.1
A2 6726061 6726234
refGene_gn=LOC101082044;refGene_tx=XM_023244823.1
A2 6728321 6728389
refGene_gn=LOC101082044;refGene_tx=XM_023244823.1
A2 6728601 6728726
refGene_gn=LOC101082044;refGene_tx=XM_023244823.1
A2 6730233 6730296
refGene_gn=LOC101082044;refGene_tx=XM_023244823.1
A2 6730526 6730562
refGene_gn=LOC101082044;refGene_tx=XM_023244823.1
A2 6730896 6731069
refGene_gn=LOC101082044;refGene_tx=XM_023244823.1
A2 6732177 6732260
refGene_gn=LOC101082044;refGene_tx=XM_023244823.1
A2 6749917 6749961
refGene_gn=LOC111557120;refGene_tx=XM_023240926.1
A2 6750345 6750381
refGene_gn=LOC111557120;refGene_tx=XM_023240926.1
A2 6753473 6753646
refGene_gn=LOC111557120;refGene_tx=XM_023240926.1
A2 6755564 6755632
refGene_gn=LOC111557120;refGene_tx=XM_023240926.1
A2 6755726 6755851

refGene_gn=LOC111557120;refGene_tx=XM_023240926.1
A2 6760412 6760540
refGene_gn=LOC111557120;refGene_tx=XM_023240926.1
A2 6764028 6764146
refGene_gn=LOC111557120;refGene_tx=XM_023240926.1
A2 6766042 6766588
refGene_gn=LOC111557120;refGene_tx=XM_023240926.1
A2 6767722 6780137
refGene_gn=LOC111557120;refGene_tx=XM_023240926.1
A2 6786468 6787058
refGene_gn=LOC111557870;refGene_tx=XM_023244770.1
A2 6788101 6788212
refGene_gn=LOC111557870;refGene_tx=XM_023244770.1
A2 6791464 6791569
refGene_gn=LOC111557870;refGene_tx=XM_023244770.1
A2 6793541 6797985
refGene_gn=LOC111557870;refGene_tx=XM_023244770.1
A2 6854638 6855625
refGene_gn=LOC101082294;refGene_tx=XM_011288866.3
A2 6884523 6885474
refGene_gn=LOC101098693;refGene_tx=XM_003981814.3
A2 6905619 6906558
refGene_gn=LOC101082548;refGene_tx=XM_011288858.1
A2 6912250 6913171
refGene_gn=LOC101082803;refGene_tx=XM_011288856.2
A2 6924391 6925309
refGene_gn=LOC101083065;refGene_tx=XM_003981831.5
A2 6934586 6935525
refGene_gn=LOC101083324;refGene_tx=XM_003981832.4
A2 6943305 6944244
refGene_gn=LOC101083572;refGene_tx=XM_003981833.4
A2 6954478 6955432
refGene_gn=LOC101083828;refGene_tx=XM_011289502.2
A2 6968335 6969217
refGene_gn=LOC109491371;refGene_tx=XM_019817043.2
A2 6972850 6973793
refGene_gn=LOC101084344;refGene_tx=XM_023240954.1
A2 6975453 6975603
refGene_gn=LOC101084344;refGene_tx=XM_023240954.1
A2 6977275 6977565
refGene_gn=LOC101084344;refGene_tx=XM_023240954.1
A2 6990822 6991866
refGene_gn=LOC101084602;refGene_tx=XM_019824907.1
A2 7013474 7014413
refGene_gn=LOC101084856;refGene_tx=XM_006928381.4
A2 7035253 7035284
refGene_gn=ZNF317;refGene_tx=XM_006928360.2;refGene_gn=ZNF317;refGene_
tx=XM_006928361.2
A2 7036010 7036147
refGene_gn=ZNF317;refGene_tx=XM_006928360.2;refGene_gn=ZNF317;refGene_

tx=XM_006928361.2
A2 7036573 7036700
refGene_gn=ZNF317;refGene_tx=XM_006928360.2;refGene_gn=ZNF317;refGene_
tx=XM_006928361.2
A2 7037148 7037244
refGene_gn=ZNF317;refGene_tx=XM_006928360.2;refGene_gn=ZNF317;refGene_
tx=XM_006928361.2
A2 7038716 7038799
refGene_gn=ZNF317;refGene_tx=XM_006928360.2;refGene_gn=ZNF317;refGene_
tx=XM_006928361.2
A2 7039231 7040554
refGene_gn=ZNF317;refGene_tx=XM_006928360.2;refGene_gn=ZNF317;refGene_
tx=XM_006928361.2
A2 7055531 7056560
refGene_gn=LOC101085352;refGene_tx=XM_003981839.5
A2 7072050 7072989
refGene_gn=LOC101098931;refGene_tx=XM_003981815.3
A2 7086750 7087689
refGene_gn=LOC101099185;refGene_tx=XM_003981816.2
A2 7104691 7105624
refGene_gn=LOC105261207;refGene_tx=XM_011289531.2
A2 7112832 7113825
refGene_gn=LOC101082221;refGene_tx=XM_004001087.3
A2 7121898 7122841
refGene_gn=LOC101082475;refGene_tx=XM_023240976.1
A2 7125850 7125936
refGene_gn=LOC101082475;refGene_tx=XM_023240976.1
A2 7136547 7137537
refGene_gn=LOC101085604;refGene_tx=XM_023240979.1
A2 7145113 7146106
refGene_gn=LOC101099440;refGene_tx=XM_003981817.1
A2 7184244 7184368
refGene_gn=LOC105261156;refGene_tx=XM_011288921.3
A2 7184369 7184803
refGene_gn=LOC105261156;refGene_tx=XM_011288921.3
A2 7184804 7185813
refGene_gn=LOC105261156;refGene_tx=XM_011288921.3
A2 7186156 7186340
refGene_gn=LOC105261156;refGene_tx=XM_011288921.3
A2 7186735 7186834
refGene_gn=LOC105261156;refGene_tx=XM_011288921.3
A2 7190617 7190744
refGene_gn=LOC105261156;refGene_tx=XM_011288921.3
A2 7191626 7191671
refGene_gn=LOC105261156;refGene_tx=XM_011288921.3
A2 7211498 7211661
refGene_gn=LOC102899257;refGene_tx=XM_019817136.2
A2 7212277 7212366
refGene_gn=LOC102899257;refGene_tx=XM_019817136.2
A2 7213032 7213159

```
refGene_gn=LOC102899257;refGene_tx=XM_019817136.2
A2      7214410 7214487
refGene_gn=LOC102899257;refGene_tx=XM_019817136.2
A2      7241020 7241053
refGene_gn=LOC101086373;refGene_tx=XM_006928362.3
A2      7241679 7241806
refGene_gn=LOC101086373;refGene_tx=XM_006928362.3
A2      7242254 7242347
refGene_gn=LOC101086373;refGene_tx=XM_006928362.3
A2      7243050 7243133
refGene_gn=LOC101086373;refGene_tx=XM_006928362.3
A2      7243658 7244758
refGene_gn=LOC101086373;refGene_tx=XM_006928362.3
A2      7244759 7244769
refGene_gn=LOC101086373;refGene_tx=XM_006928362.3
A2      7257750 7258935
refGene_gn=LOC101086629;refGene_tx=XM_023244963.1;refGene_gn=LOC101086
629;refGene_tx=XM_023244967.1;refGene_gn=LOC101086629;refGene_tx=XM_02
3244974.1;refGene_gn=LOC101086629;refGene_tx=XM_023244980.1
A2      7259386 7259469
refGene_gn=LOC101086629;refGene_tx=XM_023244963.1;refGene_gn=LOC101086
629;refGene_tx=XM_023244967.1;refGene_gn=LOC101086629;refGene_tx=XM_02
3244974.1;refGene_gn=LOC101086629;refGene_tx=XM_023244980.1
A2      7260385 7260466
refGene_gn=LOC101086629;refGene_tx=XM_023244963.1;refGene_gn=LOC101086
629;refGene_tx=XM_023244967.1;refGene_gn=LOC101086629;refGene_tx=XM_02
3244974.1;refGene_gn=LOC101086629;refGene_tx=XM_023244980.1
A2      7261281 7261408
refGene_gn=LOC101086629;refGene_tx=XM_023244963.1;refGene_gn=LOC101086
629;refGene_tx=XM_023244967.1;refGene_gn=LOC101086629;refGene_tx=XM_02
3244974.1;refGene_gn=LOC101086629;refGene_tx=XM_023244980.1
A2      7262020 7262110
refGene_gn=LOC101086629;refGene_tx=XM_023244963.1;refGene_gn=LOC101086
629;refGene_tx=XM_023244967.1;refGene_gn=LOC101086629;refGene_tx=XM_02
3244974.1;refGene_gn=LOC101086629;refGene_tx=XM_023244980.1
A2      7262598 7262601
refGene_gn=LOC101086629;refGene_tx=XM_023244963.1;refGene_gn=LOC101086
629;refGene_tx=XM_023244967.1;refGene_gn=LOC101086629;refGene_tx=XM_02
3244974.1
A2      7279942 7281208
refGene_gn=LOC101086878;refGene_tx=XM_023244958.1
A2      7282373 7282456
refGene_gn=LOC101086878;refGene_tx=XM_023244958.1
A2      7282979 7283060
refGene_gn=LOC101086878;refGene_tx=XM_023244958.1
A2      7284272 7284399
refGene_gn=LOC101086878;refGene_tx=XM_023244958.1
A2      7285028 7285114
refGene_gn=LOC101086878;refGene_tx=XM_023244958.1
A2      7285610 7285708
```

refGene_gn=LOC101086878;refGene_tx=XM_023244958.1
A2 7289526 7289593
refGene_gn=LOC101086878;refGene_tx=XM_023244958.1
A2 7293378 7293388
refGene_gn=LOC101086878;refGene_tx=XM_023244958.1
A2 7305183 7306122
refGene_gn=LOC101099693;refGene_tx=XM_003981818.3
A2 7334656 7334726
refGene_gn=LOC111557991;refGene_tx=XM_023244990.1;refGene_gn=LOC111557991;refGene_tx=XM_023244994.1;refGene_gn=LOC111557991;refGene_tx=XM_023245000.1;refGene_gn=LOC111557991;refGene_tx=XM_023245003.1;refGene_gn=LOC111557991;refGene_tx=XM_023245005.1;refGene_gn=LOC111557991;refGene_tx=XM_023245007.1
A2 7370596 7370690
refGene_gn=LOC111557991;refGene_tx=XM_023244990.1;refGene_gn=LOC111557991;refGene_tx=XM_023244994.1;refGene_gn=LOC111557991;refGene_tx=XM_023245000.1;refGene_gn=LOC111557991;refGene_tx=XM_023245003.1;refGene_gn=LOC111557991;refGene_tx=XM_023245005.1
A2 7384701 7384785
refGene_gn=LOC111557991;refGene_tx=XM_023244990.1;refGene_gn=LOC111557991;refGene_tx=XM_023244994.1;refGene_gn=LOC111557991;refGene_tx=XM_023245000.1;refGene_gn=LOC111557991;refGene_tx=XM_023245003.1;refGene_gn=LOC111557991;refGene_tx=XM_023245007.1
A2 7387057 7387184
refGene_gn=LOC111557991;refGene_tx=XM_023244990.1;refGene_gn=LOC111557991;refGene_tx=XM_023244994.1;refGene_gn=LOC111557991;refGene_tx=XM_023245000.1;refGene_gn=LOC111557991;refGene_tx=XM_023245003.1;refGene_gn=LOC111557991;refGene_tx=XM_023245005.1;refGene_gn=LOC111557991;refGene_tx=XM_023245007.1
A2 7387823 7387856
refGene_gn=LOC111557991;refGene_tx=XM_023244990.1;refGene_gn=LOC111557991;refGene_tx=XM_023244994.1;refGene_gn=LOC111557991;refGene_tx=XM_023245000.1;refGene_gn=LOC111557991;refGene_tx=XM_023245003.1;refGene_gn=LOC111557991;refGene_tx=XM_023245005.1;refGene_gn=LOC111557991;refGene_tx=XM_023245007.1
A2 7447982 7448804
refGene_gn=FBXL12;refGene_tx=XM_019817191.1;refGene_gn=FBXL12;refGene_tx=XM_023245052.1
A2 7451433 7451506 refGene_gn=FBXL12;refGene_tx=XM_023245052.1
A2 7451623 7451709 refGene_gn=FBXL12;refGene_tx=XM_023245052.1
A2 7458337 7458393
refGene_gn=UBL5;refGene_tx=XM_003981843.4;refGene_gn=UBL5;refGene_tx=XM_006928384.4;refGene_gn=UBL5;refGene_tx=XM_006928385.3
A2 7458567 7458651
refGene_gn=UBL5;refGene_tx=XM_003981843.4;refGene_gn=UBL5;refGene_tx=XM_006928384.4;refGene_gn=UBL5;refGene_tx=XM_006928385.3
A2 7458810 7458848
refGene_gn=UBL5;refGene_tx=XM_003981843.4;refGene_gn=UBL5;refGene_tx=XM_006928384.4;refGene_gn=UBL5;refGene_tx=XM_006928385.3
A2 7459477 7459521

refGene_gn=UBL5;refGene_tx=XM_003981843.4;refGene_gn=UBL5;refGene_tx=XM_006928384.4;refGene_gn=UBL5;refGene_tx=XM_006928385.3
A2 7466387 7466445 refGene_gn=PIN1;refGene_tx=XM_023245073.1
A2 7468493 7468706 refGene_gn=PIN1;refGene_tx=XM_023245073.1
A2 7477151 7477262 refGene_gn=PIN1;refGene_tx=XM_023245073.1
A2 7478150 7478260 refGene_gn=PIN1;refGene_tx=XM_023245073.1
A2 7482884 7483562 refGene_gn=OLFM2;refGene_tx=XM_006928398.4;refGene_gn=OLFM2;refGene_tx=XM_023245077.1
A2 7484540 7484647 refGene_gn=OLFM2;refGene_tx=XM_006928398.4;refGene_gn=OLFM2;refGene_tx=XM_023245077.1
A2 7484979 7485199 refGene_gn=OLFM2;refGene_tx=XM_006928398.4;refGene_gn=OLFM2;refGene_tx=XM_023245077.1
A2 7485672 7485819 refGene_gn=OLFM2;refGene_tx=XM_006928398.4;refGene_gn=OLFM2;refGene_tx=XM_023245077.1
A2 7487135 7487285 refGene_gn=OLFM2;refGene_tx=XM_006928398.4;refGene_gn=OLFM2;refGene_tx=XM_023245077.1
A2 7525441 7525576 refGene_gn=OLFM2;refGene_tx=XM_023245077.1
A2 7542504 7542585 refGene_gn=OLFM2;refGene_tx=XM_006928398.4
A2 7556909 7557056 refGene_gn=COL5A3;refGene_tx=XM_011288971.3;refGene_gn=COL5A3;refGene_tx=XM_023245083.1;refGene_gn=COL5A3;refGene_tx=XM_023245087.1;refGene_gn=COL5A3;refGene_tx=XM_023245105.1
A2 7557137 7557377 refGene_gn=COL5A3;refGene_tx=XM_011288971.3;refGene_gn=COL5A3;refGene_tx=XM_023245083.1;refGene_gn=COL5A3;refGene_tx=XM_023245087.1;refGene_gn=COL5A3;refGene_tx=XM_023245105.1
A2 7559098 7559167 refGene_gn=COL5A3;refGene_tx=XM_011288971.3;refGene_gn=COL5A3;refGene_tx=XM_023245083.1;refGene_gn=COL5A3;refGene_tx=XM_023245087.1;refGene_gn=COL5A3;refGene_tx=XM_023245105.1
A2 7560706 7560819 refGene_gn=COL5A3;refGene_tx=XM_011288971.3;refGene_gn=COL5A3;refGene_tx=XM_023245083.1;refGene_gn=COL5A3;refGene_tx=XM_023245087.1;refGene_gn=COL5A3;refGene_tx=XM_023245105.1
A2 7560907 7561115 refGene_gn=COL5A3;refGene_tx=XM_011288971.3;refGene_gn=COL5A3;refGene_tx=XM_023245083.1;refGene_gn=COL5A3;refGene_tx=XM_023245087.1;refGene_gn=COL5A3;refGene_tx=XM_023245105.1
A2 7561710 7561764 refGene_gn=COL5A3;refGene_tx=XM_011288971.3;refGene_gn=COL5A3;refGene_tx=XM_023245083.1;refGene_gn=COL5A3;refGene_tx=XM_023245087.1;refGene_gn=COL5A3;refGene_tx=XM_023245105.1
A2 7562208 7562244 refGene_gn=COL5A3;refGene_tx=XM_011288971.3;refGene_gn=COL5A3;refGene_tx=XM_023245083.1;refGene_gn=COL5A3;refGene_tx=XM_023245087.1;refGene_

gn=COL5A3;refGene_tx=XM_023245105.1
A2	7562324	7562378
refGene_gn=COL5A3;refGene_tx=XM_011288971.3;refGene_gn=COL5A3;refGene_tx=XM_023245083.1;refGene_gn=COL5A3;refGene_tx=XM_023245087.1;refGene_gn=COL5A3;refGene_tx=XM_023245105.1
A2	7562689	7562797
refGene_gn=COL5A3;refGene_tx=XM_011288971.3;refGene_gn=COL5A3;refGene_tx=XM_023245083.1;refGene_gn=COL5A3;refGene_tx=XM_023245087.1;refGene_gn=COL5A3;refGene_tx=XM_023245105.1
A2	7562994	7563048
refGene_gn=COL5A3;refGene_tx=XM_011288971.3;refGene_gn=COL5A3;refGene_tx=XM_023245083.1;refGene_gn=COL5A3;refGene_tx=XM_023245087.1;refGene_gn=COL5A3;refGene_tx=XM_023245105.1
A2	7563500	7563554
refGene_gn=COL5A3;refGene_tx=XM_011288971.3;refGene_gn=COL5A3;refGene_tx=XM_023245083.1;refGene_gn=COL5A3;refGene_tx=XM_023245087.1;refGene_gn=COL5A3;refGene_tx=XM_023245105.1
A2	7563700	7563808
refGene_gn=COL5A3;refGene_tx=XM_011288971.3;refGene_gn=COL5A3;refGene_tx=XM_023245083.1;refGene_gn=COL5A3;refGene_tx=XM_023245087.1;refGene_gn=COL5A3;refGene_tx=XM_023245105.1
A2	7563980	7564034
refGene_gn=COL5A3;refGene_tx=XM_011288971.3;refGene_gn=COL5A3;refGene_tx=XM_023245083.1;refGene_gn=COL5A3;refGene_tx=XM_023245087.1;refGene_gn=COL5A3;refGene_tx=XM_023245105.1
A2	7564170	7564224
refGene_gn=COL5A3;refGene_tx=XM_011288971.3;refGene_gn=COL5A3;refGene_tx=XM_023245083.1;refGene_gn=COL5A3;refGene_tx=XM_023245087.1;refGene_gn=COL5A3;refGene_tx=XM_023245105.1
A2	7564521	7564575
refGene_gn=COL5A3;refGene_tx=XM_011288971.3;refGene_gn=COL5A3;refGene_tx=XM_023245083.1;refGene_gn=COL5A3;refGene_tx=XM_023245087.1;refGene_gn=COL5A3;refGene_tx=XM_023245105.1
A2	7564737	7564791
refGene_gn=COL5A3;refGene_tx=XM_011288971.3;refGene_gn=COL5A3;refGene_tx=XM_023245083.1;refGene_gn=COL5A3;refGene_tx=XM_023245087.1;refGene_gn=COL5A3;refGene_tx=XM_023245105.1
A2	7565392	7565500
refGene_gn=COL5A3;refGene_tx=XM_011288971.3;refGene_gn=COL5A3;refGene_tx=XM_023245083.1;refGene_gn=COL5A3;refGene_tx=XM_023245087.1;refGene_gn=COL5A3;refGene_tx=XM_023245105.1
A2	7566051	7566105
refGene_gn=COL5A3;refGene_tx=XM_011288971.3;refGene_gn=COL5A3;refGene_tx=XM_023245083.1;refGene_gn=COL5A3;refGene_tx=XM_023245087.1;refGene_gn=COL5A3;refGene_tx=XM_023245105.1
A2	7566241	7566295
refGene_gn=COL5A3;refGene_tx=XM_011288971.3;refGene_gn=COL5A3;refGene_tx=XM_023245083.1;refGene_gn=COL5A3;refGene_tx=XM_023245087.1;refGene_gn=COL5A3;refGene_tx=XM_023245105.1
A2	7566410	7566464

refGene_gn=COL5A3;refGene_tx=XM_011288971.3;refGene_gn=COL5A3;refGene_tx=XM_023245083.1;refGene_gn=COL5A3;refGene_tx=XM_023245087.1;refGene_gn=COL5A3;refGene_tx=XM_023245105.1
A2 7566667 7566721
refGene_gn=COL5A3;refGene_tx=XM_011288971.3;refGene_gn=COL5A3;refGene_tx=XM_023245083.1;refGene_gn=COL5A3;refGene_tx=XM_023245087.1;refGene_gn=COL5A3;refGene_tx=XM_023245105.1
A2 7566795 7566903
refGene_gn=COL5A3;refGene_tx=XM_011288971.3;refGene_gn=COL5A3;refGene_tx=XM_023245083.1;refGene_gn=COL5A3;refGene_tx=XM_023245087.1;refGene_gn=COL5A3;refGene_tx=XM_023245105.1
A2 7567544 7567598
refGene_gn=COL5A3;refGene_tx=XM_011288971.3;refGene_gn=COL5A3;refGene_tx=XM_023245083.1;refGene_gn=COL5A3;refGene_tx=XM_023245087.1;refGene_gn=COL5A3;refGene_tx=XM_023245105.1
A2 7568045 7568099
refGene_gn=COL5A3;refGene_tx=XM_011288971.3;refGene_gn=COL5A3;refGene_tx=XM_023245083.1;refGene_gn=COL5A3;refGene_tx=XM_023245087.1;refGene_gn=COL5A3;refGene_tx=XM_023245105.1
A2 7568195 7568303
refGene_gn=COL5A3;refGene_tx=XM_011288971.3;refGene_gn=COL5A3;refGene_tx=XM_023245083.1;refGene_gn=COL5A3;refGene_tx=XM_023245087.1;refGene_gn=COL5A3;refGene_tx=XM_023245105.1
A2 7568419 7568527
refGene_gn=COL5A3;refGene_tx=XM_011288971.3;refGene_gn=COL5A3;refGene_tx=XM_023245083.1;refGene_gn=COL5A3;refGene_tx=XM_023245087.1;refGene_gn=COL5A3;refGene_tx=XM_023245105.1
A2 7569337 7569391
refGene_gn=COL5A3;refGene_tx=XM_011288971.3;refGene_gn=COL5A3;refGene_tx=XM_023245083.1;refGene_gn=COL5A3;refGene_tx=XM_023245087.1;refGene_gn=COL5A3;refGene_tx=XM_023245105.1
A2 7569650 7569740
refGene_gn=COL5A3;refGene_tx=XM_011288971.3;refGene_gn=COL5A3;refGene_tx=XM_023245083.1;refGene_gn=COL5A3;refGene_tx=XM_023245087.1;refGene_gn=COL5A3;refGene_tx=XM_023245105.1
A2 7569904 7570012
refGene_gn=COL5A3;refGene_tx=XM_011288971.3;refGene_gn=COL5A3;refGene_tx=XM_023245083.1;refGene_gn=COL5A3;refGene_tx=XM_023245087.1;refGene_gn=COL5A3;refGene_tx=XM_023245105.1
A2 7570146 7570200
refGene_gn=COL5A3;refGene_tx=XM_011288971.3;refGene_gn=COL5A3;refGene_tx=XM_023245083.1;refGene_gn=COL5A3;refGene_tx=XM_023245087.1;refGene_gn=COL5A3;refGene_tx=XM_023245105.1
A2 7570289 7570343
refGene_gn=COL5A3;refGene_tx=XM_011288971.3;refGene_gn=COL5A3;refGene_tx=XM_023245083.1;refGene_gn=COL5A3;refGene_tx=XM_023245087.1;refGene_gn=COL5A3;refGene_tx=XM_023245105.1
A2 7570442 7570496
refGene_gn=COL5A3;refGene_tx=XM_011288971.3;refGene_gn=COL5A3;refGene_tx=XM_023245083.1;refGene_gn=COL5A3;refGene_tx=XM_023245087.1;refGene_

gn=COL5A3;refGene_tx=XM_023245105.1
A2	7570849	7570894
refGene_gn=COL5A3;refGene_tx=XM_011288971.3;refGene_gn=COL5A3;refGene_tx=XM_023245083.1;refGene_gn=COL5A3;refGene_tx=XM_023245087.1;refGene_gn=COL5A3;refGene_tx=XM_023245105.1
A2	7571005	7571059
refGene_gn=COL5A3;refGene_tx=XM_011288971.3;refGene_gn=COL5A3;refGene_tx=XM_023245083.1;refGene_gn=COL5A3;refGene_tx=XM_023245087.1;refGene_gn=COL5A3;refGene_tx=XM_023245105.1
A2	7571313	7571358
refGene_gn=COL5A3;refGene_tx=XM_011288971.3;refGene_gn=COL5A3;refGene_tx=XM_023245083.1;refGene_gn=COL5A3;refGene_tx=XM_023245087.1;refGene_gn=COL5A3;refGene_tx=XM_023245105.1
A2	7571818	7571872
refGene_gn=COL5A3;refGene_tx=XM_011288971.3;refGene_gn=COL5A3;refGene_tx=XM_023245083.1;refGene_gn=COL5A3;refGene_tx=XM_023245087.1;refGene_gn=COL5A3;refGene_tx=XM_023245105.1
A2	7574824	7574878
refGene_gn=COL5A3;refGene_tx=XM_011288971.3;refGene_gn=COL5A3;refGene_tx=XM_023245083.1;refGene_gn=COL5A3;refGene_tx=XM_023245087.1;refGene_gn=COL5A3;refGene_tx=XM_023245105.1
A2	7575319	7575427
refGene_gn=COL5A3;refGene_tx=XM_023245083.1;refGene_gn=COL5A3;refGene_tx=XM_023245087.1;refGene_gn=COL5A3;refGene_tx=XM_023245105.1
A2	7575569	7575623
refGene_gn=COL5A3;refGene_tx=XM_023245083.1;refGene_gn=COL5A3;refGene_tx=XM_023245087.1;refGene_gn=COL5A3;refGene_tx=XM_023245105.1
A2	7575739	7575784
refGene_gn=COL5A3;refGene_tx=XM_023245083.1;refGene_gn=COL5A3;refGene_tx=XM_023245087.1;refGene_gn=COL5A3;refGene_tx=XM_023245105.1
A2	7577272	7577326
refGene_gn=COL5A3;refGene_tx=XM_023245083.1;refGene_gn=COL5A3;refGene_tx=XM_023245087.1;refGene_gn=COL5A3;refGene_tx=XM_023245105.1
A2	7577470	7577515
refGene_gn=COL5A3;refGene_tx=XM_023245083.1;refGene_gn=COL5A3;refGene_tx=XM_023245087.1;refGene_gn=COL5A3;refGene_tx=XM_023245105.1
A2	7577609	7577663
refGene_gn=COL5A3;refGene_tx=XM_023245083.1;refGene_gn=COL5A3;refGene_tx=XM_023245087.1;refGene_gn=COL5A3;refGene_tx=XM_023245105.1
A2	7578584	7578629
refGene_gn=COL5A3;refGene_tx=XM_011288971.3;refGene_gn=COL5A3;refGene_tx=XM_023245083.1;refGene_gn=COL5A3;refGene_tx=XM_023245087.1;refGene_gn=COL5A3;refGene_tx=XM_023245105.1
A2	7578774	7578828
refGene_gn=COL5A3;refGene_tx=XM_011288971.3;refGene_gn=COL5A3;refGene_tx=XM_023245083.1;refGene_gn=COL5A3;refGene_tx=XM_023245087.1;refGene_gn=COL5A3;refGene_tx=XM_023245105.1
A2	7578988	7579033
refGene_gn=COL5A3;refGene_tx=XM_011288971.3;refGene_gn=COL5A3;refGene_tx=XM_023245083.1;refGene_gn=COL5A3;refGene_tx=XM_023245087.1;refGene_

gn=COL5A3;refGene_tx=XM_023245105.1
A2	7579215	7579269
refGene_gn=COL5A3;refGene_tx=XM_011288971.3;refGene_gn=COL5A3;refGene_tx=XM_023245083.1;refGene_gn=COL5A3;refGene_tx=XM_023245087.1;refGene_gn=COL5A3;refGene_tx=XM_023245105.1
A2	7579376	7579421
refGene_gn=COL5A3;refGene_tx=XM_011288971.3;refGene_gn=COL5A3;refGene_tx=XM_023245083.1;refGene_gn=COL5A3;refGene_tx=XM_023245087.1;refGene_gn=COL5A3;refGene_tx=XM_023245105.1
A2	7579764	7579818
refGene_gn=COL5A3;refGene_tx=XM_011288971.3;refGene_gn=COL5A3;refGene_tx=XM_023245083.1;refGene_gn=COL5A3;refGene_tx=XM_023245087.1;refGene_gn=COL5A3;refGene_tx=XM_023245105.1
A2	7580004	7580058
refGene_gn=COL5A3;refGene_tx=XM_011288971.3;refGene_gn=COL5A3;refGene_tx=XM_023245083.1;refGene_gn=COL5A3;refGene_tx=XM_023245087.1;refGene_gn=COL5A3;refGene_tx=XM_023245105.1
A2	7580139	7580193
refGene_gn=COL5A3;refGene_tx=XM_011288971.3;refGene_gn=COL5A3;refGene_tx=XM_023245083.1;refGene_gn=COL5A3;refGene_tx=XM_023245087.1;refGene_gn=COL5A3;refGene_tx=XM_023245105.1
A2	7581023	7581077
refGene_gn=COL5A3;refGene_tx=XM_011288971.3;refGene_gn=COL5A3;refGene_tx=XM_023245083.1;refGene_gn=COL5A3;refGene_tx=XM_023245087.1;refGene_gn=COL5A3;refGene_tx=XM_023245105.1
A2	7581271	7581325
refGene_gn=COL5A3;refGene_tx=XM_011288971.3;refGene_gn=COL5A3;refGene_tx=XM_023245083.1;refGene_gn=COL5A3;refGene_tx=XM_023245087.1;refGene_gn=COL5A3;refGene_tx=XM_023245105.1
A2	7581416	7581473
refGene_gn=COL5A3;refGene_tx=XM_011288971.3;refGene_gn=COL5A3;refGene_tx=XM_023245083.1;refGene_gn=COL5A3;refGene_tx=XM_023245087.1;refGene_gn=COL5A3;refGene_tx=XM_023245105.1
A2	7581655	7581739
refGene_gn=COL5A3;refGene_tx=XM_011288971.3;refGene_gn=COL5A3;refGene_tx=XM_023245083.1;refGene_gn=COL5A3;refGene_tx=XM_023245087.1;refGene_gn=COL5A3;refGene_tx=XM_023245105.1
A2	7581835	7581910
refGene_gn=COL5A3;refGene_tx=XM_011288971.3;refGene_gn=COL5A3;refGene_tx=XM_023245083.1;refGene_gn=COL5A3;refGene_tx=XM_023245087.1;refGene_gn=COL5A3;refGene_tx=XM_023245105.1
A2	7582273	7582336
refGene_gn=COL5A3;refGene_tx=XM_011288971.3;refGene_gn=COL5A3;refGene_tx=XM_023245083.1;refGene_gn=COL5A3;refGene_tx=XM_023245087.1;refGene_gn=COL5A3;refGene_tx=XM_023245105.1
A2	7582650	7582692
refGene_gn=COL5A3;refGene_tx=XM_011288971.3;refGene_gn=COL5A3;refGene_tx=XM_023245083.1;refGene_gn=COL5A3;refGene_tx=XM_023245087.1;refGene_gn=COL5A3;refGene_tx=XM_023245105.1
A2	7582765	7582813

refGene_gn=COL5A3;refGene_tx=XM_011288971.3;refGene_gn=COL5A3;refGene_tx=XM_023245083.1;refGene_gn=COL5A3;refGene_tx=XM_023245087.1;refGene_gn=COL5A3;refGene_tx=XM_023245105.1
A2 7583865 7584012
refGene_gn=COL5A3;refGene_tx=XM_011288971.3;refGene_gn=COL5A3;refGene_tx=XM_023245083.1;refGene_gn=COL5A3;refGene_tx=XM_023245087.1;refGene_gn=COL5A3;refGene_tx=XM_023245105.1
A2 7584131 7584245
refGene_gn=COL5A3;refGene_tx=XM_011288971.3;refGene_gn=COL5A3;refGene_tx=XM_023245083.1;refGene_gn=COL5A3;refGene_tx=XM_023245087.1;refGene_gn=COL5A3;refGene_tx=XM_023245105.1
A2 7586050 7586209
refGene_gn=COL5A3;refGene_tx=XM_011288971.3;refGene_gn=COL5A3;refGene_tx=XM_023245083.1;refGene_gn=COL5A3;refGene_tx=XM_023245087.1;refGene_gn=COL5A3;refGene_tx=XM_023245105.1
A2 7586528 7586633
refGene_gn=COL5A3;refGene_tx=XM_011288971.3;refGene_gn=COL5A3;refGene_tx=XM_023245083.1;refGene_gn=COL5A3;refGene_tx=XM_023245087.1;refGene_gn=COL5A3;refGene_tx=XM_023245105.1
A2 7587225 7587382
refGene_gn=COL5A3;refGene_tx=XM_011288971.3;refGene_gn=COL5A3;refGene_tx=XM_023245083.1;refGene_gn=COL5A3;refGene_tx=XM_023245087.1;refGene_gn=COL5A3;refGene_tx=XM_023245105.1
A2 7587457 7587647
refGene_gn=COL5A3;refGene_tx=XM_011288971.3;refGene_gn=COL5A3;refGene_tx=XM_023245083.1;refGene_gn=COL5A3;refGene_tx=XM_023245087.1
A2 7587741 7587903
refGene_gn=COL5A3;refGene_tx=XM_011288971.3;refGene_gn=COL5A3;refGene_tx=XM_023245083.1;refGene_gn=COL5A3;refGene_tx=XM_023245087.1
A2 7591752 7591840
refGene_gn=COL5A3;refGene_tx=XM_011288971.3;refGene_gn=COL5A3;refGene_tx=XM_023245083.1;refGene_gn=COL5A3;refGene_tx=XM_023245087.1
A2 7593889 7593941 refGene_gn=RDH8;refGene_tx=XM_003981846.5
A2 7593943 7594051 refGene_gn=RDH8;refGene_tx=XM_003981846.5
A2 7596525 7596687 refGene_gn=RDH8;refGene_tx=XM_003981846.5
A2 7597323 7597503 refGene_gn=RDH8;refGene_tx=XM_003981846.5
A2 7598472 7598566 refGene_gn=RDH8;refGene_tx=XM_003981846.5
A2 7598853 7599037 refGene_gn=RDH8;refGene_tx=XM_003981846.5
A2 7599160 7599376 refGene_gn=RDH8;refGene_tx=XM_003981846.5
A2 7635986 7636049
refGene_gn=CA2H19orf66;refGene_tx=XM_011289617.2
A2 7636389 7636476
refGene_gn=CA2H19orf66;refGene_tx=XM_011289617.2
A2 7636566 7636732
refGene_gn=CA2H19orf66;refGene_tx=XM_011289617.2
A2 7637160 7637250
refGene_gn=CA2H19orf66;refGene_tx=XM_011289617.2
A2 7637567 7637654
refGene_gn=CA2H19orf66;refGene_tx=XM_011289617.2
A2 7637742 7637862

refGene_gn=CA2H19orf66;refGene_tx=XM_011289617.2
A2 7637943 7638076
refGene_gn=CA2H19orf66;refGene_tx=XM_011289617.2
A2 7638511 7638633
refGene_gn=CA2H19orf66;refGene_tx=XM_011289617.2
A2 7657269 7657392
refGene_gn=CA2H19orf66;refGene_tx=XM_011289617.2
A2 7657586 7657636
refGene_gn=CA2H19orf66;refGene_tx=XM_011289617.2
A2 7658902 7658941
refGene_gn=CA2H19orf66;refGene_tx=XM_011289617.2
A2 7659131 7659281
refGene_gn=CA2H19orf66;refGene_tx=XM_011289617.2
A2 7659610 7659714
refGene_gn=CA2H19orf66;refGene_tx=XM_011289617.2
A2 7659841 7659996
refGene_gn=CA2H19orf66;refGene_tx=XM_011289617.2
A2 7660566 7660796
refGene_gn=CA2H19orf66;refGene_tx=XM_011289617.2
A2 7661082 7661273
refGene_gn=ANGPTL6;refGene_tx=XM_011288979.3;refGene_gn=ANGPTL6;refGene_tx=XM_011288981.3;refGene_gn=ANGPTL6;refGene_tx=XM_019817244.2;refGene_gn=ANGPTL6;refGene_tx=XM_019817247.2;refGene_gn=ANGPTL6;refGene_tx=XM_019817253.2;refGene_gn=ANGPTL6;refGene_tx=XM_019817263.1;refGene_gn=ANGPTL6;refGene_tx=XM_019817269.2;refGene_gn=ANGPTL6;refGene_tx=XM_019817274.2
A2 7661861 7662132
refGene_gn=ANGPTL6;refGene_tx=XM_011288979.3;refGene_gn=ANGPTL6;refGene_tx=XM_011288981.3;refGene_gn=ANGPTL6;refGene_tx=XM_019817244.2;refGene_gn=ANGPTL6;refGene_tx=XM_019817247.2;refGene_gn=ANGPTL6;refGene_tx=XM_019817253.2;refGene_gn=ANGPTL6;refGene_tx=XM_019817263.1;refGene_gn=ANGPTL6;refGene_tx=XM_019817269.2;refGene_gn=ANGPTL6;refGene_tx=XM_019817274.2
A2 7662211 7662399
refGene_gn=ANGPTL6;refGene_tx=XM_011288979.3;refGene_gn=ANGPTL6;refGene_tx=XM_011288981.3;refGene_gn=ANGPTL6;refGene_tx=XM_019817244.2;refGene_gn=ANGPTL6;refGene_tx=XM_019817247.2;refGene_gn=ANGPTL6;refGene_tx=XM_019817253.2;refGene_gn=ANGPTL6;refGene_tx=XM_019817263.1;refGene_gn=ANGPTL6;refGene_tx=XM_019817269.2;refGene_gn=ANGPTL6;refGene_tx=XM_019817274.2
A2 7662955 7663133
refGene_gn=ANGPTL6;refGene_tx=XM_011288979.3;refGene_gn=ANGPTL6;refGene_tx=XM_011288981.3;refGene_gn=ANGPTL6;refGene_tx=XM_019817244.2;refGene_gn=ANGPTL6;refGene_tx=XM_019817247.2;refGene_gn=ANGPTL6;refGene_tx=XM_019817253.2;refGene_gn=ANGPTL6;refGene_tx=XM_019817263.1;refGene_gn=ANGPTL6;refGene_tx=XM_019817269.2;refGene_gn=ANGPTL6;refGene_tx=XM_019817274.2
A2 7663829 7664594
refGene_gn=ANGPTL6;refGene_tx=XM_011288979.3;refGene_gn=ANGPTL6;refGene_tx=XM_011288981.3;refGene_gn=ANGPTL6;refGene_tx=XM_019817244.2;refGe

ne_gn=ANGPTL6;refGene_tx=XM_019817247.2;refGene_gn=ANGPTL6;refGene_tx=XM_019817253.2;refGene_gn=ANGPTL6;refGene_tx=XM_019817263.1;refGene_gn=ANGPTL6;refGene_tx=XM_019817269.2;refGene_gn=ANGPTL6;refGene_tx=XM_019817274.2
A2 7667043 7667273 refGene_gn=ANGPTL6;refGene_tx=XM_019817244.2
A2 7669440 7669458 refGene_gn=LOC101093001;refGene_tx=XM_023245152.1
A2 7669575 7669746 refGene_gn=LOC101093001;refGene_tx=XM_023245152.1
A2 7670247 7670323 refGene_gn=LOC101093001;refGene_tx=XM_023245152.1
A2 7670325 7670348 refGene_gn=LOC101093001;refGene_tx=XM_023245152.1
A2 7670551 7670602 refGene_gn=LOC101093001;refGene_tx=XM_023245152.1
A2 7670706 7670877 refGene_gn=LOC101093001;refGene_tx=XM_023245152.1
A2 7671533 7671610 refGene_gn=LOC101093001;refGene_tx=XM_023245152.1
A2 7671812 7671920 refGene_gn=LOC101093001;refGene_tx=XM_023245152.1
A2 7672021 7672145 refGene_gn=LOC101093001;refGene_tx=XM_023245152.1
A2 7672256 7672335 refGene_gn=LOC101093001;refGene_tx=XM_023245152.1
A2 7672445 7672575 refGene_gn=LOC101093001;refGene_tx=XM_023245152.1
A2 7672651 7672812 refGene_gn=LOC101093001;refGene_tx=XM_023245152.1
A2 7672894 7672981 refGene_gn=LOC101093001;refGene_tx=XM_023245152.1
A2 7675074 7676198 refGene_gn=LOC101093001;refGene_tx=XM_023245152.1
A2 7676498 7676514 refGene_gn=EIF3G;refGene_tx=XM_011288997.3
A2 7676800 7676907 refGene_gn=EIF3G;refGene_tx=XM_011288997.3
A2 7676986 7677123 refGene_gn=EIF3G;refGene_tx=XM_011288997.3
A2 7677226 7677334 refGene_gn=EIF3G;refGene_tx=XM_011288997.3
A2 7678046 7678236 refGene_gn=EIF3G;refGene_tx=XM_011288997.3
A2 7678329 7678434 refGene_gn=EIF3G;refGene_tx=XM_011288997.3
A2 7679257 7679317 refGene_gn=EIF3G;refGene_tx=XM_011288997.3
A2 7679465 7679554 refGene_gn=EIF3G;refGene_tx=XM_011288997.3
A2 7679659 7679743 refGene_gn=EIF3G;refGene_tx=XM_011288997.3
A2 7680233 7680280 refGene_gn=EIF3G;refGene_tx=XM_011288997.3
A2 7680415 7680435 refGene_gn=EIF3G;refGene_tx=XM_011288997.3
A2 7686718 7686753 refGene_gn=DNMT1;refGene_tx=XM_006928387.4;refGene_gn=DNMT1;refGene_tx=XM_019817312.1;refGene_gn=DNMT1;refGene_tx=XM_023245156.1;refGene_gn=DNMT1;refGene_tx=XM_023245159.1
A2 7687233 7687324 refGene_gn=DNMT1;refGene_tx=XM_006928387.4;refGene_gn=DNMT1;refGene_tx

=XM_019817312.1;refGene_gn=DNMT1;refGene_tx=XM_023245156.1;refGene_gn=DNMT1;refGene_tx=XM_023245159.1
A2 7688126 7688243
refGene_gn=DNMT1;refGene_tx=XM_006928387.4;refGene_gn=DNMT1;refGene_tx=XM_019817312.1;refGene_gn=DNMT1;refGene_tx=XM_023245156.1;refGene_gn=DNMT1;refGene_tx=XM_023245159.1
A2 7688484 7688651
refGene_gn=DNMT1;refGene_tx=XM_006928387.4;refGene_gn=DNMT1;refGene_tx=XM_019817312.1;refGene_gn=DNMT1;refGene_tx=XM_023245156.1;refGene_gn=DNMT1;refGene_tx=XM_023245159.1
A2 7688984 7689180
refGene_gn=DNMT1;refGene_tx=XM_006928387.4;refGene_gn=DNMT1;refGene_tx=XM_019817312.1;refGene_gn=DNMT1;refGene_tx=XM_023245156.1;refGene_gn=DNMT1;refGene_tx=XM_023245159.1
A2 7689865 7690043
refGene_gn=DNMT1;refGene_tx=XM_006928387.4;refGene_gn=DNMT1;refGene_tx=XM_019817312.1;refGene_gn=DNMT1;refGene_tx=XM_023245156.1;refGene_gn=DNMT1;refGene_tx=XM_023245159.1
A2 7690454 7690621
refGene_gn=DNMT1;refGene_tx=XM_006928387.4;refGene_gn=DNMT1;refGene_tx=XM_019817312.1;refGene_gn=DNMT1;refGene_tx=XM_023245156.1;refGene_gn=DNMT1;refGene_tx=XM_023245159.1
A2 7691798 7691940
refGene_gn=DNMT1;refGene_tx=XM_006928387.4;refGene_gn=DNMT1;refGene_tx=XM_019817312.1;refGene_gn=DNMT1;refGene_tx=XM_023245156.1;refGene_gn=DNMT1;refGene_tx=XM_023245159.1
A2 7692251 7692534
refGene_gn=DNMT1;refGene_tx=XM_006928387.4;refGene_gn=DNMT1;refGene_tx=XM_019817312.1;refGene_gn=DNMT1;refGene_tx=XM_023245156.1;refGene_gn=DNMT1;refGene_tx=XM_023245159.1
A2 7692681 7692810
refGene_gn=DNMT1;refGene_tx=XM_006928387.4;refGene_gn=DNMT1;refGene_tx=XM_019817312.1;refGene_gn=DNMT1;refGene_tx=XM_023245156.1;refGene_gn=DNMT1;refGene_tx=XM_023245159.1
A2 7692985 7693070
refGene_gn=DNMT1;refGene_tx=XM_006928387.4;refGene_gn=DNMT1;refGene_tx=XM_019817312.1;refGene_gn=DNMT1;refGene_tx=XM_023245156.1;refGene_gn=DNMT1;refGene_tx=XM_023245159.1
A2 7694010 7694203
refGene_gn=DNMT1;refGene_tx=XM_006928387.4;refGene_gn=DNMT1;refGene_tx=XM_019817312.1;refGene_gn=DNMT1;refGene_tx=XM_023245156.1;refGene_gn=DNMT1;refGene_tx=XM_023245159.1
A2 7695583 7695805
refGene_gn=DNMT1;refGene_tx=XM_006928387.4;refGene_gn=DNMT1;refGene_tx=XM_019817312.1;refGene_gn=DNMT1;refGene_tx=XM_023245156.1;refGene_gn=DNMT1;refGene_tx=XM_023245159.1
A2 7697274 7697448
refGene_gn=DNMT1;refGene_tx=XM_006928387.4;refGene_gn=DNMT1;refGene_tx=XM_019817312.1;refGene_gn=DNMT1;refGene_tx=XM_023245156.1;refGene_gn=DNMT1;refGene_tx=XM_023245159.1

A2	7698944	7699078
refGene_gn=DNMT1;refGene_tx=XM_006928387.4;refGene_gn=DNMT1;refGene_tx=XM_019817312.1;refGene_gn=DNMT1;refGene_tx=XM_023245156.1;refGene_gn=DNMT1;refGene_tx=XM_023245159.1
A2	7699158	7699363
refGene_gn=DNMT1;refGene_tx=XM_006928387.4;refGene_gn=DNMT1;refGene_tx=XM_019817312.1;refGene_gn=DNMT1;refGene_tx=XM_023245156.1;refGene_gn=DNMT1;refGene_tx=XM_023245159.1
A2	7699545	7699661
refGene_gn=DNMT1;refGene_tx=XM_006928387.4;refGene_gn=DNMT1;refGene_tx=XM_019817312.1;refGene_gn=DNMT1;refGene_tx=XM_023245156.1;refGene_gn=DNMT1;refGene_tx=XM_023245159.1
A2	7700692	7700840
refGene_gn=DNMT1;refGene_tx=XM_006928387.4;refGene_gn=DNMT1;refGene_tx=XM_019817312.1;refGene_gn=DNMT1;refGene_tx=XM_023245156.1;refGene_gn=DNMT1;refGene_tx=XM_023245159.1
A2	7701034	7701132
refGene_gn=DNMT1;refGene_tx=XM_006928387.4;refGene_gn=DNMT1;refGene_tx=XM_019817312.1;refGene_gn=DNMT1;refGene_tx=XM_023245156.1;refGene_gn=DNMT1;refGene_tx=XM_023245159.1
A2	7703156	7703340
refGene_gn=DNMT1;refGene_tx=XM_006928387.4;refGene_gn=DNMT1;refGene_tx=XM_019817312.1;refGene_gn=DNMT1;refGene_tx=XM_023245156.1;refGene_gn=DNMT1;refGene_tx=XM_023245159.1
A2	7703445	7703633
refGene_gn=DNMT1;refGene_tx=XM_006928387.4;refGene_gn=DNMT1;refGene_tx=XM_019817312.1;refGene_gn=DNMT1;refGene_tx=XM_023245156.1;refGene_gn=DNMT1;refGene_tx=XM_023245159.1
A2	7703792	7703944
refGene_gn=DNMT1;refGene_tx=XM_006928387.4;refGene_gn=DNMT1;refGene_tx=XM_019817312.1;refGene_gn=DNMT1;refGene_tx=XM_023245156.1;refGene_gn=DNMT1;refGene_tx=XM_023245159.1
A2	7704681	7704774
refGene_gn=DNMT1;refGene_tx=XM_006928387.4;refGene_gn=DNMT1;refGene_tx=XM_019817312.1;refGene_gn=DNMT1;refGene_tx=XM_023245156.1;refGene_gn=DNMT1;refGene_tx=XM_023245159.1
A2	7705055	7705174
refGene_gn=DNMT1;refGene_tx=XM_006928387.4;refGene_gn=DNMT1;refGene_tx=XM_019817312.1;refGene_gn=DNMT1;refGene_tx=XM_023245156.1;refGene_gn=DNMT1;refGene_tx=XM_023245159.1
A2	7707863	7707973
refGene_gn=DNMT1;refGene_tx=XM_006928387.4;refGene_gn=DNMT1;refGene_tx=XM_019817312.1;refGene_gn=DNMT1;refGene_tx=XM_023245156.1;refGene_gn=DNMT1;refGene_tx=XM_023245159.1
A2	7708068	7708149
refGene_gn=DNMT1;refGene_tx=XM_006928387.4;refGene_gn=DNMT1;refGene_tx=XM_019817312.1;refGene_gn=DNMT1;refGene_tx=XM_023245156.1;refGene_gn=DNMT1;refGene_tx=XM_023245159.1
A2	7708243	7708289
refGene_gn=DNMT1;refGene_tx=XM_006928387.4;refGene_gn=DNMT1;refGene_tx

=XM_019817312.1;refGene_gn=DNMT1;refGene_tx=XM_023245156.1;refGene_gn=DNMT1;refGene_tx=XM_023245159.1
A2 7708619 7708654
refGene_gn=DNMT1;refGene_tx=XM_006928387.4;refGene_gn=DNMT1;refGene_tx=XM_019817312.1;refGene_gn=DNMT1;refGene_tx=XM_023245156.1;refGene_gn=DNMT1;refGene_tx=XM_023245159.1
A2 7710422 7710504
refGene_gn=DNMT1;refGene_tx=XM_006928387.4;refGene_gn=DNMT1;refGene_tx=XM_019817312.1;refGene_gn=DNMT1;refGene_tx=XM_023245156.1;refGene_gn=DNMT1;refGene_tx=XM_023245159.1
A2 7710982 7711017
refGene_gn=DNMT1;refGene_tx=XM_006928387.4;refGene_gn=DNMT1;refGene_tx=XM_019817312.1;refGene_gn=DNMT1;refGene_tx=XM_023245156.1;refGene_gn=DNMT1;refGene_tx=XM_023245159.1
A2 7712913 7712998
refGene_gn=DNMT1;refGene_tx=XM_006928387.4;refGene_gn=DNMT1;refGene_tx=XM_019817312.1;refGene_gn=DNMT1;refGene_tx=XM_023245156.1;refGene_gn=DNMT1;refGene_tx=XM_023245159.1
A2 7714013 7714048
refGene_gn=DNMT1;refGene_tx=XM_006928387.4;refGene_gn=DNMT1;refGene_tx=XM_019817312.1;refGene_gn=DNMT1;refGene_tx=XM_023245156.1;refGene_gn=DNMT1;refGene_tx=XM_023245159.1
A2 7715882 7715967
refGene_gn=DNMT1;refGene_tx=XM_006928387.4;refGene_gn=DNMT1;refGene_tx=XM_019817312.1;refGene_gn=DNMT1;refGene_tx=XM_023245156.1;refGene_gn=DNMT1;refGene_tx=XM_023245159.1
A2 7716344 7716379
refGene_gn=DNMT1;refGene_tx=XM_006928387.4;refGene_gn=DNMT1;refGene_tx=XM_019817312.1;refGene_gn=DNMT1;refGene_tx=XM_023245156.1;refGene_gn=DNMT1;refGene_tx=XM_023245159.1
A2 7719830 7719906
refGene_gn=DNMT1;refGene_tx=XM_006928387.4;refGene_gn=DNMT1;refGene_tx=XM_019817312.1;refGene_gn=DNMT1;refGene_tx=XM_023245156.1;refGene_gn=DNMT1;refGene_tx=XM_023245159.1
A2 7721414 7721490
refGene_gn=DNMT1;refGene_tx=XM_006928387.4;refGene_gn=DNMT1;refGene_tx=XM_019817312.1;refGene_gn=DNMT1;refGene_tx=XM_023245156.1;refGene_gn=DNMT1;refGene_tx=XM_023245159.1
A2 7722132 7722352
refGene_gn=DNMT1;refGene_tx=XM_006928387.4;refGene_gn=DNMT1;refGene_tx=XM_019817312.1;refGene_gn=DNMT1;refGene_tx=XM_023245156.1;refGene_gn=DNMT1;refGene_tx=XM_023245159.1
A2 7722545 7722653
refGene_gn=DNMT1;refGene_tx=XM_006928387.4;refGene_gn=DNMT1;refGene_tx=XM_023245156.1;refGene_gn=DNMT1;refGene_tx=XM_023245159.1
A2 7723602 7723639
refGene_gn=DNMT1;refGene_tx=XM_006928387.4;refGene_gn=DNMT1;refGene_tx=XM_023245156.1;refGene_gn=DNMT1;refGene_tx=XM_023245159.1
A2 7731928 7731972 refGene_gn=DNMT1;refGene_tx=XM_006928387.4
A2 7732340 7732396 refGene_gn=DNMT1;refGene_tx=XM_023245159.1

A2 7732905 7732985 refGene_gn=DNMT1;refGene_tx=XM_023245156.1
A2 7751228 7752287 refGene_gn=S1PR2;refGene_tx=XM_023245167.1
A2 7769793 7769868 refGene_gn=MRPL4;refGene_tx=XM_003981849.5
A2 7769953 7770020 refGene_gn=MRPL4;refGene_tx=XM_003981849.5
A2 7770106 7770257 refGene_gn=MRPL4;refGene_tx=XM_003981849.5
A2 7772137 7772189 refGene_gn=MRPL4;refGene_tx=XM_003981849.5
A2 7773171 7773289 refGene_gn=MRPL4;refGene_tx=XM_003981849.5
A2 7774171 7774278 refGene_gn=MRPL4;refGene_tx=XM_003981849.5
A2 7774375 7774485 refGene_gn=MRPL4;refGene_tx=XM_003981849.5
A2 7774563 7774640 refGene_gn=MRPL4;refGene_tx=XM_003981849.5
A2 7775764 7775910 refGene_gn=MRPL4;refGene_tx=XM_003981849.5
A2 7786462 7786529
refGene_gn=ICAM1;refGene_tx=XM_003981850.5;refGene_gn=ICAM1;refGene_tx
=XM_006928389.4
A2 7788165 7788423
refGene_gn=ICAM1;refGene_tx=XM_003981850.5;refGene_gn=ICAM1;refGene_tx
=XM_006928389.4
A2 7792743 7793049
refGene_gn=ICAM1;refGene_tx=XM_003981850.5;refGene_gn=ICAM1;refGene_tx
=XM_006928389.4
A2 7793435 7793723
refGene_gn=ICAM1;refGene_tx=XM_003981850.5;refGene_gn=ICAM1;refGene_tx
=XM_006928389.4
A2 7793802 7794060
refGene_gn=ICAM1;refGene_tx=XM_003981850.5;refGene_gn=ICAM1;refGene_tx
=XM_006928389.4
A2 7794242 7794491
refGene_gn=ICAM1;refGene_tx=XM_003981850.5;refGene_gn=ICAM1;refGene_tx
=XM_006928389.4
A2 7794577 7794750
refGene_gn=ICAM1;refGene_tx=XM_003981850.5;refGene_gn=ICAM1;refGene_tx
=XM_006928389.4
A2 7796268 7796662 refGene_gn=ICAM4;refGene_tx=XM_003981865.2
A2 7796781 7797087 refGene_gn=ICAM4;refGene_tx=XM_003981865.2
A2 7797232 7797375 refGene_gn=ICAM4;refGene_tx=XM_003981865.2
A2 7798995 7799077 refGene_gn=ICAM5;refGene_tx=XM_003981851.5
A2 7800005 7800275 refGene_gn=ICAM5;refGene_tx=XM_003981851.5
A2 7800465 7800786 refGene_gn=ICAM5;refGene_tx=XM_003981851.5
A2 7801007 7801295 refGene_gn=ICAM5;refGene_tx=XM_003981851.5
A2 7801677 7801932 refGene_gn=ICAM5;refGene_tx=XM_003981851.5
A2 7802065 7802314 refGene_gn=ICAM5;refGene_tx=XM_003981851.5
A2 7802777 7803020 refGene_gn=ICAM5;refGene_tx=XM_003981851.5
A2 7803106 7803385 refGene_gn=ICAM5;refGene_tx=XM_003981851.5
A2 7803467 7803707 refGene_gn=ICAM5;refGene_tx=XM_003981851.5
A2 7804121 7804388 refGene_gn=ICAM5;refGene_tx=XM_003981851.5
A2 7805107 7805385 refGene_gn=ICAM5;refGene_tx=XM_003981851.5
A2 7808535 7808665 refGene_gn=ZGLP1;refGene_tx=XM_019817389.2
A2 7808811 7808937 refGene_gn=ZGLP1;refGene_tx=XM_019817389.2
A2 7809082 7809176 refGene_gn=ZGLP1;refGene_tx=XM_019817389.2
A2 7809524 7809610 refGene_gn=ZGLP1;refGene_tx=XM_019817389.2

A2 7810463 7810838 refGene_gn=ZGLP1;refGene_tx=XM_019817389.2
A2 7811075 7811212 refGene_gn=ZGLP1;refGene_tx=XM_019817389.2
A2 7811984 7812141
refGene_gn=FDX1L;refGene_tx=XM_003981852.4;refGene_gn=FDX1L;refGene_tx
=XM_006928390.4;refGene_gn=FDX1L;refGene_tx=XM_019817410.2
A2 7812393 7812481
refGene_gn=FDX1L;refGene_tx=XM_003981852.4;refGene_gn=FDX1L;refGene_tx
=XM_006928390.4
A2 7815092 7815238 refGene_gn=FDX1L;refGene_tx=XM_006928390.4
A2 7815501 7815608
refGene_gn=FDX1L;refGene_tx=XM_003981852.4;refGene_gn=FDX1L;refGene_tx
=XM_006928390.4;refGene_gn=FDX1L;refGene_tx=XM_019817410.2
A2 7815811 7815866
refGene_gn=FDX1L;refGene_tx=XM_003981852.4;refGene_gn=FDX1L;refGene_tx
=XM_006928390.4;refGene_gn=FDX1L;refGene_tx=XM_019817410.2
A2 7815952 7816128
refGene_gn=FDX1L;refGene_tx=XM_003981852.4;refGene_gn=FDX1L;refGene_tx
=XM_006928390.4;refGene_gn=FDX1L;refGene_tx=XM_019817410.2
A2 7817554 7817701
refGene_gn=RAVER1;refGene_tx=XM_011289029.3;refGene_gn=RAVER1;refGene_
tx=XM_011289030.3
A2 7817798 7817882
refGene_gn=RAVER1;refGene_tx=XM_011289029.3;refGene_gn=RAVER1;refGene_
tx=XM_011289030.3
A2 7818291 7818432
refGene_gn=RAVER1;refGene_tx=XM_011289029.3;refGene_gn=RAVER1;refGene_
tx=XM_011289030.3
A2 7818932 7819007
refGene_gn=RAVER1;refGene_tx=XM_011289029.3;refGene_gn=RAVER1;refGene_
tx=XM_011289030.3
A2 7819779 7820085
refGene_gn=RAVER1;refGene_tx=XM_011289029.3;refGene_gn=RAVER1;refGene_
tx=XM_011289030.3
A2 7820188 7820400
refGene_gn=RAVER1;refGene_tx=XM_011289029.3;refGene_gn=RAVER1;refGene_
tx=XM_011289030.3
A2 7820693 7820781
refGene_gn=RAVER1;refGene_tx=XM_011289029.3;refGene_gn=RAVER1;refGene_
tx=XM_011289030.3
A2 7821682 7821777
refGene_gn=RAVER1;refGene_tx=XM_011289029.3;refGene_gn=RAVER1;refGene_
tx=XM_011289030.3
A2 7822120 7822250
refGene_gn=RAVER1;refGene_tx=XM_011289029.3;refGene_gn=RAVER1;refGene_
tx=XM_011289030.3
A2 7822428 7822620
refGene_gn=RAVER1;refGene_tx=XM_011289029.3;refGene_gn=RAVER1;refGene_
tx=XM_011289030.3
A2 7826593 7827063
refGene_gn=RAVER1;refGene_tx=XM_011289029.3;refGene_gn=RAVER1;refGene_

tx=XM_011289030.3
A2	7829610	7829677
refGene_gn=RAVER1;refGene_tx=XM_011289029.3;refGene_gn=RAVER1;refGene_tx=XM_011289030.3
A2	7831248	7831518
refGene_gn=RAVER1;refGene_tx=XM_011289029.3;refGene_gn=RAVER1;refGene_tx=XM_011289030.3
A2	7832289	7832483	refGene_gn=ICAM3;refGene_tx=XM_023245246.1
A2	7832568	7832817	refGene_gn=ICAM3;refGene_tx=XM_023245246.1
A2	7832931	7833186	refGene_gn=ICAM3;refGene_tx=XM_023245246.1
A2	7833528	7833816	refGene_gn=ICAM3;refGene_tx=XM_023245246.1
A2	7834155	7834461	refGene_gn=ICAM3;refGene_tx=XM_023245246.1
A2	7838734	7839001	refGene_gn=ICAM3;refGene_tx=XM_023245246.1
A2	7839571	7839653	refGene_gn=ICAM3;refGene_tx=XM_023245246.1
A2	7847536	7847671	refGene_gn=TYK2;refGene_tx=XM_023245251.1
A2	7847753	7847864	refGene_gn=TYK2;refGene_tx=XM_023245251.1
A2	7848291	7848409	refGene_gn=TYK2;refGene_tx=XM_023245251.1
A2	7848623	7848796	refGene_gn=TYK2;refGene_tx=XM_023245251.1
A2	7849128	7849247	refGene_gn=TYK2;refGene_tx=XM_023245251.1
A2	7849575	7849768	refGene_gn=TYK2;refGene_tx=XM_023245251.1
A2	7850035	7850133	refGene_gn=TYK2;refGene_tx=XM_023245251.1
A2	7850557	7850708	refGene_gn=TYK2;refGene_tx=XM_023245251.1
A2	7851447	7851602	refGene_gn=TYK2;refGene_tx=XM_023245251.1
A2	7851685	7851821	refGene_gn=TYK2;refGene_tx=XM_023245251.1
A2	7852705	7852833	refGene_gn=TYK2;refGene_tx=XM_023245251.1
A2	7853759	7853847	refGene_gn=TYK2;refGene_tx=XM_023245251.1
A2	7853933	7854116	refGene_gn=TYK2;refGene_tx=XM_023245251.1
A2	7854200	7854304	refGene_gn=TYK2;refGene_tx=XM_023245251.1
A2	7854383	7854573	refGene_gn=TYK2;refGene_tx=XM_023245251.1
A2	7854664	7854773	refGene_gn=TYK2;refGene_tx=XM_023245251.1
A2	7856633	7856791	refGene_gn=TYK2;refGene_tx=XM_023245251.1
A2	7856889	7857087	refGene_gn=TYK2;refGene_tx=XM_023245251.1
A2	7857493	7857860	refGene_gn=TYK2;refGene_tx=XM_023245251.1
A2	7859440	7859604	refGene_gn=TYK2;refGene_tx=XM_023245251.1
A2	7859954	7860102	refGene_gn=TYK2;refGene_tx=XM_023245251.1
A2	7860234	7860358	refGene_gn=TYK2;refGene_tx=XM_023245251.1
A2	7864300	7864535	refGene_gn=TYK2;refGene_tx=XM_023245251.1
A2	7874157	7874313	refGene_gn=CDC37;refGene_tx=XM_023245261.1
A2	7875391	7875464	refGene_gn=CDC37;refGene_tx=XM_023245261.1
A2	7875552	7875734	refGene_gn=CDC37;refGene_tx=XM_023245261.1
A2	7876279	7876402	refGene_gn=CDC37;refGene_tx=XM_023245261.1
A2	7876482	7876598	refGene_gn=CDC37;refGene_tx=XM_023245261.1
A2	7876709	7876818	refGene_gn=CDC37;refGene_tx=XM_023245261.1
A2	7877163	7877445	refGene_gn=CDC37;refGene_tx=XM_023245261.1
A2	7882983	7883085	refGene_gn=CDC37;refGene_tx=XM_023245261.1
A2	7890207	7890255
refGene_gn=PDE4A;refGene_tx=XM_019817453.2;refGene_gn=PDE4A;refGene_tx=XM_023245276.1
A2	7890888	7891088
refGene_gn=PDE4A;refGene_tx=XM_019817453.2;refGene_gn=PDE4A;refGene_tx

=XM_023245276.1
A2	7894027	7894347	refGene_gn=PDE4A;refGene_tx=XM_011289052.3
A2	7906096	7906332	refGene_gn=PDE4A;refGene_tx=XM_023245278.1
A2	7907676	7907813	refGene_gn=PDE4A;refGene_tx=XM_011289058.3
A2	7914250	7914442
refGene_gn=PDE4A;refGene_tx=XM_011289052.3;refGene_gn=PDE4A;refGene_tx
=XM_011289058.3;refGene_gn=PDE4A;refGene_tx=XM_019817453.2;refGene_gn=
PDE4A;refGene_tx=XM_023245276.1;refGene_gn=PDE4A;refGene_tx=XM_0232452
78.1
A2	7915326	7915363
refGene_gn=PDE4A;refGene_tx=XM_011289052.3;refGene_gn=PDE4A;refGene_tx
=XM_011289058.3;refGene_gn=PDE4A;refGene_tx=XM_019817453.2;refGene_gn=
PDE4A;refGene_tx=XM_023245276.1;refGene_gn=PDE4A;refGene_tx=XM_0232452
78.1
A2	7915486	7915557
refGene_gn=PDE4A;refGene_tx=XM_011289052.3;refGene_gn=PDE4A;refGene_tx
=XM_011289058.3;refGene_gn=PDE4A;refGene_tx=XM_019817453.2;refGene_gn=
PDE4A;refGene_tx=XM_023245276.1;refGene_gn=PDE4A;refGene_tx=XM_0232452
78.1
A2	7916310	7916360
refGene_gn=PDE4A;refGene_tx=XM_011289052.3;refGene_gn=PDE4A;refGene_tx
=XM_011289058.3;refGene_gn=PDE4A;refGene_tx=XM_019817453.2;refGene_gn=
PDE4A;refGene_tx=XM_023245276.1;refGene_gn=PDE4A;refGene_tx=XM_0232452
78.1
A2	7916530	7916643
refGene_gn=PDE4A;refGene_tx=XM_011289052.3;refGene_gn=PDE4A;refGene_tx
=XM_011289058.3;refGene_gn=PDE4A;refGene_tx=XM_019817453.2;refGene_gn=
PDE4A;refGene_tx=XM_019817462.2;refGene_gn=PDE4A;refGene_tx=XM_0232452
76.1;refGene_gn=PDE4A;refGene_tx=XM_023245278.1
A2	7918616	7918682	refGene_gn=PDE4A;refGene_tx=XM_003981857.5
A2	7920184	7920278
refGene_gn=PDE4A;refGene_tx=XM_003981857.5;refGene_gn=PDE4A;refGene_tx
=XM_011289052.3;refGene_gn=PDE4A;refGene_tx=XM_011289058.3;refGene_gn=
PDE4A;refGene_tx=XM_019817453.2;refGene_gn=PDE4A;refGene_tx=XM_0198174
62.2;refGene_gn=PDE4A;refGene_tx=XM_023245276.1;refGene_gn=PDE4A;refGe
ne_tx=XM_023245278.1
A2	7921760	7921978
refGene_gn=PDE4A;refGene_tx=XM_003981857.5;refGene_gn=PDE4A;refGene_tx
=XM_011289052.3;refGene_gn=PDE4A;refGene_tx=XM_011289058.3;refGene_gn=
PDE4A;refGene_tx=XM_019817453.2;refGene_gn=PDE4A;refGene_tx=XM_0198174
62.2;refGene_gn=PDE4A;refGene_tx=XM_023245276.1;refGene_gn=PDE4A;refGe
ne_tx=XM_023245278.1
A2	7923361	7923460
refGene_gn=PDE4A;refGene_tx=XM_003981857.5;refGene_gn=PDE4A;refGene_tx
=XM_011289052.3;refGene_gn=PDE4A;refGene_tx=XM_011289058.3;refGene_gn=
PDE4A;refGene_tx=XM_019817453.2;refGene_gn=PDE4A;refGene_tx=XM_0198174
62.2;refGene_gn=PDE4A;refGene_tx=XM_023245276.1;refGene_gn=PDE4A;refGe
ne_tx=XM_023245278.1
A2	7923552	7923717
refGene_gn=PDE4A;refGene_tx=XM_003981857.5;refGene_gn=PDE4A;refGene_tx

=XM_011289052.3;refGene_gn=PDE4A;refGene_tx=XM_011289058.3;refGene_gn=PDE4A;refGene_tx=XM_019817453.2;refGene_gn=PDE4A;refGene_tx=XM_019817462.2;refGene_gn=PDE4A;refGene_tx=XM_023245276.1;refGene_gn=PDE4A;refGene_tx=XM_023245278.1
A2 7923937 7924037
refGene_gn=PDE4A;refGene_tx=XM_003981857.5;refGene_gn=PDE4A;refGene_tx=XM_011289052.3;refGene_gn=PDE4A;refGene_tx=XM_011289058.3;refGene_gn=PDE4A;refGene_tx=XM_019817453.2;refGene_gn=PDE4A;refGene_tx=XM_019817462.2;refGene_gn=PDE4A;refGene_tx=XM_023245276.1;refGene_gn=PDE4A;refGene_tx=XM_023245278.1
A2 7924160 7924315
refGene_gn=PDE4A;refGene_tx=XM_003981857.5;refGene_gn=PDE4A;refGene_tx=XM_011289052.3;refGene_gn=PDE4A;refGene_tx=XM_011289058.3;refGene_gn=PDE4A;refGene_tx=XM_019817453.2;refGene_gn=PDE4A;refGene_tx=XM_019817462.2;refGene_gn=PDE4A;refGene_tx=XM_023245276.1;refGene_gn=PDE4A;refGene_tx=XM_023245278.1
A2 7924518 7924641
refGene_gn=PDE4A;refGene_tx=XM_003981857.5;refGene_gn=PDE4A;refGene_tx=XM_011289052.3;refGene_gn=PDE4A;refGene_tx=XM_011289058.3;refGene_gn=PDE4A;refGene_tx=XM_019817453.2;refGene_gn=PDE4A;refGene_tx=XM_019817462.2;refGene_gn=PDE4A;refGene_tx=XM_023245276.1;refGene_gn=PDE4A;refGene_tx=XM_023245278.1
A2 7925082 7925265
refGene_gn=PDE4A;refGene_tx=XM_003981857.5;refGene_gn=PDE4A;refGene_tx=XM_011289052.3;refGene_gn=PDE4A;refGene_tx=XM_011289058.3;refGene_gn=PDE4A;refGene_tx=XM_019817453.2;refGene_gn=PDE4A;refGene_tx=XM_019817462.2;refGene_gn=PDE4A;refGene_tx=XM_023245276.1;refGene_gn=PDE4A;refGene_tx=XM_023245278.1
A2 7928227 7928917
refGene_gn=PDE4A;refGene_tx=XM_003981857.5;refGene_gn=PDE4A;refGene_tx=XM_011289052.3;refGene_gn=PDE4A;refGene_tx=XM_011289058.3;refGene_gn=PDE4A;refGene_tx=XM_019817453.2;refGene_gn=PDE4A;refGene_tx=XM_019817462.2;refGene_gn=PDE4A;refGene_tx=XM_023245276.1;refGene_gn=PDE4A;refGene_tx=XM_023245278.1
A2 7945644 7945811
refGene_gn=KEAP1;refGene_tx=XM_023245299.1;refGene_gn=KEAP1;refGene_tx=XM_023245303.1
A2 7945971 7946148
refGene_gn=KEAP1;refGene_tx=XM_023245299.1;refGene_gn=KEAP1;refGene_tx=XM_023245303.1
A2 7946424 7946630
refGene_gn=KEAP1;refGene_tx=XM_023245299.1;refGene_gn=KEAP1;refGene_tx=XM_023245303.1
A2 7947709 7948395
refGene_gn=KEAP1;refGene_tx=XM_023245299.1;refGene_gn=KEAP1;refGene_tx=XM_023245303.1
A2 7950618 7951257
refGene_gn=KEAP1;refGene_tx=XM_023245299.1;refGene_gn=KEAP1;refGene_tx=XM_023245303.1
A2 7959609 7960806 refGene_gn=S1PR5;refGene_tx=XM_023245305.1

A2 7981995 7982227
refGene_gn=ATG4D;refGene_tx=XM_003981869.5;refGene_gn=ATG4D;refGene_tx=XM_023245326.1
A2 7982755 7982839
refGene_gn=ATG4D;refGene_tx=XM_003981869.5;refGene_gn=ATG4D;refGene_tx=XM_023245326.1
A2 7982920 7983094
refGene_gn=ATG4D;refGene_tx=XM_003981869.5;refGene_gn=ATG4D;refGene_tx=XM_023245326.1
A2 7983390 7983667
refGene_gn=ATG4D;refGene_tx=XM_003981869.5;refGene_gn=ATG4D;refGene_tx=XM_023245326.1
A2 7983741 7983806
refGene_gn=ATG4D;refGene_tx=XM_003981869.5;refGene_gn=ATG4D;refGene_tx=XM_023245326.1
A2 7985718 7985849
refGene_gn=ATG4D;refGene_tx=XM_003981869.5;refGene_gn=ATG4D;refGene_tx=XM_023245326.1
A2 7986725 7986804 refGene_gn=ATG4D;refGene_tx=XM_003981869.5
A2 7986880 7986957
refGene_gn=ATG4D;refGene_tx=XM_003981869.5;refGene_gn=ATG4D;refGene_tx=XM_023245326.1
A2 7987040 7987160
refGene_gn=ATG4D;refGene_tx=XM_003981869.5;refGene_gn=ATG4D;refGene_tx=XM_023245326.1
A2 7987767 7987950 refGene_gn=ATG4D;refGene_tx=XM_003981869.5
A2 7988523 7988863
refGene_gn=KRI1;refGene_tx=XM_011289073.3;refGene_gn=KRI1;refGene_tx=XM_019817497.2
A2 7989461 7989560
refGene_gn=KRI1;refGene_tx=XM_011289073.3;refGene_gn=KRI1;refGene_tx=XM_019817497.2
A2 7989683 7989748
refGene_gn=KRI1;refGene_tx=XM_011289073.3;refGene_gn=KRI1;refGene_tx=XM_019817497.2
A2 7990658 7990789
refGene_gn=KRI1;refGene_tx=XM_011289073.3;refGene_gn=KRI1;refGene_tx=XM_019817497.2
A2 7990874 7991001
refGene_gn=KRI1;refGene_tx=XM_011289073.3;refGene_gn=KRI1;refGene_tx=XM_019817497.2
A2 7991080 7991172
refGene_gn=KRI1;refGene_tx=XM_011289073.3;refGene_gn=KRI1;refGene_tx=XM_019817497.2
A2 7991272 7991348
refGene_gn=KRI1;refGene_tx=XM_011289073.3;refGene_gn=KRI1;refGene_tx=XM_019817497.2
A2 7992405 7992576
refGene_gn=KRI1;refGene_tx=XM_011289073.3;refGene_gn=KRI1;refGene_tx=XM_019817497.2

A2	7992665	7992761
refGene_gn=KRI1;refGene_tx=XM_011289073.3;refGene_gn=KRI1;refGene_tx=XM_019817497.2
A2	7992868	7992995
refGene_gn=KRI1;refGene_tx=XM_011289073.3;refGene_gn=KRI1;refGene_tx=XM_019817497.2
A2	7993542	7993673
refGene_gn=KRI1;refGene_tx=XM_011289073.3;refGene_gn=KRI1;refGene_tx=XM_019817497.2
A2	7993937	7994015
refGene_gn=KRI1;refGene_tx=XM_011289073.3;refGene_gn=KRI1;refGene_tx=XM_019817497.2
A2	7994103	7994200
refGene_gn=KRI1;refGene_tx=XM_011289073.3;refGene_gn=KRI1;refGene_tx=XM_019817497.2
A2	7994495	7994545
refGene_gn=KRI1;refGene_tx=XM_011289073.3;refGene_gn=KRI1;refGene_tx=XM_019817497.2
A2	7994621	7994676
refGene_gn=KRI1;refGene_tx=XM_011289073.3;refGene_gn=KRI1;refGene_tx=XM_019817497.2
A2	7994936	7995045
refGene_gn=KRI1;refGene_tx=XM_011289073.3;refGene_gn=KRI1;refGene_tx=XM_019817497.2
A2	7995413	7995519
refGene_gn=KRI1;refGene_tx=XM_011289073.3;refGene_gn=KRI1;refGene_tx=XM_019817497.2
A2	7996295	7996369
refGene_gn=KRI1;refGene_tx=XM_011289073.3;refGene_gn=KRI1;refGene_tx=XM_019817497.2
A2	7996481	7996575
refGene_gn=KRI1;refGene_tx=XM_011289073.3;refGene_gn=KRI1;refGene_tx=XM_019817497.2
A2	7997649	7998009	refGene_gn=CDKN2D;refGene_tx=XM_003981859.5
A2	7999041	7999182	refGene_gn=CDKN2D;refGene_tx=XM_003981859.5
A2	8001355	8001378
refGene_gn=AP1M2;refGene_tx=XM_011289085.3;refGene_gn=AP1M2;refGene_tx=XM_011289091.3
A2	8001803	8001879
refGene_gn=AP1M2;refGene_tx=XM_011289085.3;refGene_gn=AP1M2;refGene_tx=XM_011289091.3
A2	8002278	8002404
refGene_gn=AP1M2;refGene_tx=XM_011289085.3;refGene_gn=AP1M2;refGene_tx=XM_011289091.3
A2	8003197	8003356
refGene_gn=AP1M2;refGene_tx=XM_011289085.3;refGene_gn=AP1M2;refGene_tx=XM_011289091.3
A2	8005168	8005240
refGene_gn=AP1M2;refGene_tx=XM_011289085.3;refGene_gn=AP1M2;refGene_tx=XM_011289091.3

A2 8005321 8005470
refGene_gn=AP1M2;refGene_tx=XM_011289085.3;refGene_gn=AP1M2;refGene_tx=XM_011289091.3
A2 8006146 8006273
refGene_gn=AP1M2;refGene_tx=XM_011289085.3;refGene_gn=AP1M2;refGene_tx=XM_011289091.3
A2 8006375 8006523
refGene_gn=AP1M2;refGene_tx=XM_011289085.3;refGene_gn=AP1M2;refGene_tx=XM_011289091.3
A2 8006611 8006742
refGene_gn=AP1M2;refGene_tx=XM_011289085.3;refGene_gn=AP1M2;refGene_tx=XM_011289091.3
A2 8008210 8008278
refGene_gn=AP1M2;refGene_tx=XM_011289085.3;refGene_gn=AP1M2;refGene_tx=XM_011289091.3
A2 8008507 8008664
refGene_gn=AP1M2;refGene_tx=XM_011289085.3;refGene_gn=AP1M2;refGene_tx=XM_011289091.3
A2 8009841 8009883
refGene_gn=AP1M2;refGene_tx=XM_011289085.3;refGene_gn=AP1M2;refGene_tx=XM_011289091.3
A2 8018467 8018498 refGene_gn=SLC44A2;refGene_tx=XM_023245347.1
A2 8034194 8034231
refGene_gn=SLC44A2;refGene_tx=XM_003981872.5;refGene_gn=SLC44A2;refGene_tx=XM_003981873.5
A2 8034808 8034857
refGene_gn=SLC44A2;refGene_tx=XM_003981872.5;refGene_gn=SLC44A2;refGene_tx=XM_003981873.5;refGene_gn=SLC44A2;refGene_tx=XM_023245347.1
A2 8036932 8037006
refGene_gn=SLC44A2;refGene_tx=XM_003981872.5;refGene_gn=SLC44A2;refGene_tx=XM_003981873.5;refGene_gn=SLC44A2;refGene_tx=XM_023245347.1
A2 8037132 8037217
refGene_gn=SLC44A2;refGene_tx=XM_003981872.5;refGene_gn=SLC44A2;refGene_tx=XM_003981873.5;refGene_gn=SLC44A2;refGene_tx=XM_023245347.1
A2 8037752 8037837
refGene_gn=SLC44A2;refGene_tx=XM_003981872.5;refGene_gn=SLC44A2;refGene_tx=XM_003981873.5;refGene_gn=SLC44A2;refGene_tx=XM_023245347.1
A2 8038026 8038137
refGene_gn=SLC44A2;refGene_tx=XM_003981872.5;refGene_gn=SLC44A2;refGene_tx=XM_003981873.5;refGene_gn=SLC44A2;refGene_tx=XM_023245347.1
A2 8038213 8038274
refGene_gn=SLC44A2;refGene_tx=XM_003981872.5;refGene_gn=SLC44A2;refGene_tx=XM_003981873.5;refGene_gn=SLC44A2;refGene_tx=XM_023245347.1
A2 8038360 8038484
refGene_gn=SLC44A2;refGene_tx=XM_003981872.5;refGene_gn=SLC44A2;refGene_tx=XM_003981873.5;refGene_gn=SLC44A2;refGene_tx=XM_023245347.1
A2 8038604 8038688
refGene_gn=SLC44A2;refGene_tx=XM_003981872.5;refGene_gn=SLC44A2;refGene_tx=XM_003981873.5;refGene_gn=SLC44A2;refGene_tx=XM_023245347.1
A2 8038778 8038891

refGene_gn=SLC44A2;refGene_tx=XM_003981872.5;refGene_gn=SLC44A2;refGene_tx=XM_003981873.5;refGene_gn=SLC44A2;refGene_tx=XM_023245347.1
A2 8040504 8040636
refGene_gn=SLC44A2;refGene_tx=XM_003981872.5;refGene_gn=SLC44A2;refGene_tx=XM_003981873.5;refGene_gn=SLC44A2;refGene_tx=XM_023245347.1
A2 8040731 8040831
refGene_gn=SLC44A2;refGene_tx=XM_003981872.5;refGene_gn=SLC44A2;refGene_tx=XM_003981873.5;refGene_gn=SLC44A2;refGene_tx=XM_023245347.1
A2 8040914 8041007
refGene_gn=SLC44A2;refGene_tx=XM_003981872.5;refGene_gn=SLC44A2;refGene_tx=XM_003981873.5;refGene_gn=SLC44A2;refGene_tx=XM_023245347.1
A2 8041191 8041276
refGene_gn=SLC44A2;refGene_tx=XM_003981872.5;refGene_gn=SLC44A2;refGene_tx=XM_003981873.5;refGene_gn=SLC44A2;refGene_tx=XM_023245347.1
A2 8041880 8042143
refGene_gn=SLC44A2;refGene_tx=XM_003981872.5;refGene_gn=SLC44A2;refGene_tx=XM_003981873.5;refGene_gn=SLC44A2;refGene_tx=XM_023245347.1
A2 8042228 8042323
refGene_gn=SLC44A2;refGene_tx=XM_003981872.5;refGene_gn=SLC44A2;refGene_tx=XM_003981873.5;refGene_gn=SLC44A2;refGene_tx=XM_023245347.1
A2 8042945 8043049
refGene_gn=SLC44A2;refGene_tx=XM_003981872.5;refGene_gn=SLC44A2;refGene_tx=XM_003981873.5;refGene_gn=SLC44A2;refGene_tx=XM_023245347.1
A2 8043176 8043250
refGene_gn=SLC44A2;refGene_tx=XM_003981872.5;refGene_gn=SLC44A2;refGene_tx=XM_003981873.5;refGene_gn=SLC44A2;refGene_tx=XM_023245347.1
A2 8043339 8043410
refGene_gn=SLC44A2;refGene_tx=XM_003981872.5;refGene_gn=SLC44A2;refGene_tx=XM_003981873.5;refGene_gn=SLC44A2;refGene_tx=XM_023245347.1
A2 8043587 8043676
refGene_gn=SLC44A2;refGene_tx=XM_003981872.5;refGene_gn=SLC44A2;refGene_tx=XM_003981873.5;refGene_gn=SLC44A2;refGene_tx=XM_023245347.1
A2 8044751 8044836
refGene_gn=SLC44A2;refGene_tx=XM_003981872.5;refGene_gn=SLC44A2;refGene_tx=XM_003981873.5;refGene_gn=SLC44A2;refGene_tx=XM_023245347.1
A2 8045274 8045396
refGene_gn=SLC44A2;refGene_tx=XM_003981873.5;refGene_gn=SLC44A2;refGene_tx=XM_023245347.1
A2 8045679 8045786 refGene_gn=SLC44A2;refGene_tx=XM_003981872.5
A2 8069929 8069932
refGene_gn=ILF3;refGene_tx=XM_023245352.1;refGene_gn=ILF3;refGene_tx=XM_023245354.1;refGene_gn=ILF3;refGene_tx=XM_023245358.1;refGene_gn=ILF3;refGene_tx=XM_023245363.1
A2 8070313 8070543
refGene_gn=ILF3;refGene_tx=XM_023245352.1;refGene_gn=ILF3;refGene_tx=XM_023245354.1;refGene_gn=ILF3;refGene_tx=XM_023245358.1;refGene_gn=ILF3;refGene_tx=XM_023245363.1
A2 8070701 8070879
refGene_gn=ILF3;refGene_tx=XM_023245352.1;refGene_gn=ILF3;refGene_tx=XM_023245354.1;refGene_gn=ILF3;refGene_tx=XM_023245358.1;refGene_gn=ILF

3;refGene_tx=XM_023245363.1
A2 8072996 8073144
refGene_gn=ILF3;refGene_tx=XM_023245352.1;refGene_gn=ILF3;refGene_tx=XM_023245354.1;refGene_gn=ILF3;refGene_tx=XM_023245358.1;refGene_gn=ILF3;refGene_tx=XM_023245363.1
A2 8074589 8074681
refGene_gn=ILF3;refGene_tx=XM_023245352.1;refGene_gn=ILF3;refGene_tx=XM_023245354.1;refGene_gn=ILF3;refGene_tx=XM_023245358.1;refGene_gn=ILF3;refGene_tx=XM_023245363.1
A2 8075069 8075165
refGene_gn=ILF3;refGene_tx=XM_023245352.1;refGene_gn=ILF3;refGene_tx=XM_023245354.1;refGene_gn=ILF3;refGene_tx=XM_023245358.1;refGene_gn=ILF3;refGene_tx=XM_023245363.1
A2 8075836 8075948
refGene_gn=ILF3;refGene_tx=XM_023245352.1;refGene_gn=ILF3;refGene_tx=XM_023245354.1;refGene_gn=ILF3;refGene_tx=XM_023245358.1;refGene_gn=ILF3;refGene_tx=XM_023245363.1
A2 8076361 8076459
refGene_gn=ILF3;refGene_tx=XM_023245352.1;refGene_gn=ILF3;refGene_tx=XM_023245354.1;refGene_gn=ILF3;refGene_tx=XM_023245358.1;refGene_gn=ILF3;refGene_tx=XM_023245363.1
A2 8077027 8077139
refGene_gn=ILF3;refGene_tx=XM_023245352.1;refGene_gn=ILF3;refGene_tx=XM_023245354.1;refGene_gn=ILF3;refGene_tx=XM_023245358.1;refGene_gn=ILF3;refGene_tx=XM_023245363.1
A2 8077247 8077358
refGene_gn=ILF3;refGene_tx=XM_023245352.1;refGene_gn=ILF3;refGene_tx=XM_023245354.1;refGene_gn=ILF3;refGene_tx=XM_023245358.1;refGene_gn=ILF3;refGene_tx=XM_023245363.1
A2 8077667 8077867
refGene_gn=ILF3;refGene_tx=XM_023245352.1;refGene_gn=ILF3;refGene_tx=XM_023245354.1;refGene_gn=ILF3;refGene_tx=XM_023245358.1;refGene_gn=ILF3;refGene_tx=XM_023245363.1
A2 8078285 8078453
refGene_gn=ILF3;refGene_tx=XM_023245352.1;refGene_gn=ILF3;refGene_tx=XM_023245354.1;refGene_gn=ILF3;refGene_tx=XM_023245358.1;refGene_gn=ILF3;refGene_tx=XM_023245363.1
A2 8078911 8079049
refGene_gn=ILF3;refGene_tx=XM_023245352.1;refGene_gn=ILF3;refGene_tx=XM_023245354.1;refGene_gn=ILF3;refGene_tx=XM_023245358.1;refGene_gn=ILF3;refGene_tx=XM_023245363.1
A2 8079135 8079312
refGene_gn=ILF3;refGene_tx=XM_023245352.1;refGene_gn=ILF3;refGene_tx=XM_023245354.1;refGene_gn=ILF3;refGene_tx=XM_023245358.1;refGene_gn=ILF3;refGene_tx=XM_023245363.1
A2 8079409 8079563
refGene_gn=ILF3;refGene_tx=XM_023245352.1;refGene_gn=ILF3;refGene_tx=XM_023245354.1;refGene_gn=ILF3;refGene_tx=XM_023245358.1;refGene_gn=ILF3;refGene_tx=XM_023245363.1
A2 8079716 8079770

refGene_gn=ILF3;refGene_tx=XM_023245352.1;refGene_gn=ILF3;refGene_tx=XM_023245354.1;refGene_gn=ILF3;refGene_tx=XM_023245358.1;refGene_gn=ILF3;refGene_tx=XM_023245363.1
A2 8080220 8080270
refGene_gn=ILF3;refGene_tx=XM_023245358.1;refGene_gn=ILF3;refGene_tx=XM_023245363.1
A2 8083183 8083549
refGene_gn=ILF3;refGene_tx=XM_023245352.1;refGene_gn=ILF3;refGene_tx=XM_023245354.1
A2 8084656 8084785
refGene_gn=ILF3;refGene_tx=XM_023245352.1;refGene_gn=ILF3;refGene_tx=XM_023245354.1
A2 8085456 8085593
refGene_gn=ILF3;refGene_tx=XM_023245352.1;refGene_gn=ILF3;refGene_tx=XM_023245354.1
A2 8101200 8101443 refGene_gn=QTRT1;refGene_tx=XM_023241003.1
A2 8101676 8101745 refGene_gn=QTRT1;refGene_tx=XM_023241003.1
A2 8101843 8102022 refGene_gn=QTRT1;refGene_tx=XM_023241003.1
A2 8104432 8104471 refGene_gn=QTRT1;refGene_tx=XM_023241003.1
A2 8104590 8104706 refGene_gn=QTRT1;refGene_tx=XM_023241003.1
A2 8107177 8107316 refGene_gn=QTRT1;refGene_tx=XM_023241003.1
A2 8107566 8107642 refGene_gn=QTRT1;refGene_tx=XM_023241003.1
A2 8107771 8107881 refGene_gn=QTRT1;refGene_tx=XM_023241003.1
A2 8107969 8108057 refGene_gn=QTRT1;refGene_tx=XM_023241003.1
A2 8108162 8108315 refGene_gn=QTRT1;refGene_tx=XM_023241003.1
A2 8110826 8110987
refGene_gn=DNM2;refGene_tx=XM_023245368.1;refGene_gn=DNM2;refGene_tx=XM_023245369.1;refGene_gn=DNM2;refGene_tx=XM_023245370.1;refGene_gn=DNM2;refGene_tx=XM_023245377.1;refGene_gn=DNM2;refGene_tx=XM_023245382.1;refGene_gn=DNM2;refGene_tx=XM_023245386.1;refGene_gn=DNM2;refGene_tx=XM_023245389.1;refGene_gn=DNM2;refGene_tx=XM_023245392.1;refGene_gn=DNM2;refGene_tx=XM_023245398.1;refGene_gn=DNM2;refGene_tx=XM_023245403.1;refGene_gn=DNM2;refGene_tx=XM_023245404.1;refGene_gn=DNM2;refGene_tx=XM_023245405.1
A2 8133693 8133794 refGene_gn=DNM2;refGene_tx=XM_023245409.1
A2 8143803 8143877
refGene_gn=DNM2;refGene_tx=XM_023245368.1;refGene_gn=DNM2;refGene_tx=XM_023245369.1;refGene_gn=DNM2;refGene_tx=XM_023245370.1;refGene_gn=DNM2;refGene_tx=XM_023245377.1;refGene_gn=DNM2;refGene_tx=XM_023245382.1;refGene_gn=DNM2;refGene_tx=XM_023245386.1;refGene_gn=DNM2;refGene_tx=XM_023245389.1;refGene_gn=DNM2;refGene_tx=XM_023245392.1;refGene_gn=DNM2;refGene_tx=XM_023245398.1;refGene_gn=DNM2;refGene_tx=XM_023245403.1;refGene_gn=DNM2;refGene_tx=XM_023245404.1;refGene_gn=DNM2;refGene_tx=XM_023245405.1;refGene_gn=DNM2;refGene_tx=XM_023245409.1
A2 8155765 8155915
refGene_gn=DNM2;refGene_tx=XM_023245368.1;refGene_gn=DNM2;refGene_tx=XM_023245369.1;refGene_gn=DNM2;refGene_tx=XM_023245370.1;refGene_gn=DNM2;refGene_tx=XM_023245377.1;refGene_gn=DNM2;refGene_tx=XM_023245382.1;refGene_gn=DNM2;refGene_tx=XM_023245386.1;refGene_gn=DNM2;refGene_tx=XM_023245389.1;refGene_gn=DNM2;refGene_tx=XM_023245392.1;refGene_gn=DNM

2;refGene_tx=XM_023245398.1;refGene_gn=DNM2;refGene_tx=XM_023245403.1;
refGene_gn=DNM2;refGene_tx=XM_023245404.1;refGene_gn=DNM2;refGene_tx=X
M_023245405.1;refGene_gn=DNM2;refGene_tx=XM_023245409.1
A2 8157228 8157432
refGene_gn=DNM2;refGene_tx=XM_023245368.1;refGene_gn=DNM2;refGene_tx=X
M_023245369.1;refGene_gn=DNM2;refGene_tx=XM_023245370.1;refGene_gn=DNM
2;refGene_tx=XM_023245377.1;refGene_gn=DNM2;refGene_tx=XM_023245382.1;
refGene_gn=DNM2;refGene_tx=XM_023245386.1;refGene_gn=DNM2;refGene_tx=X
M_023245389.1;refGene_gn=DNM2;refGene_tx=XM_023245392.1;refGene_gn=DNM
2;refGene_tx=XM_023245398.1;refGene_gn=DNM2;refGene_tx=XM_023245403.1;
refGene_gn=DNM2;refGene_tx=XM_023245404.1;refGene_gn=DNM2;refGene_tx=X
M_023245405.1;refGene_gn=DNM2;refGene_tx=XM_023245409.1;refGene_gn=DNM
2;refGene_tx=XM_023245411.1
A2 8159987 8160086
refGene_gn=DNM2;refGene_tx=XM_023245368.1;refGene_gn=DNM2;refGene_tx=X
M_023245369.1;refGene_gn=DNM2;refGene_tx=XM_023245370.1;refGene_gn=DNM
2;refGene_tx=XM_023245377.1;refGene_gn=DNM2;refGene_tx=XM_023245382.1;
refGene_gn=DNM2;refGene_tx=XM_023245386.1;refGene_gn=DNM2;refGene_tx=X
M_023245389.1;refGene_gn=DNM2;refGene_tx=XM_023245392.1;refGene_gn=DNM
2;refGene_tx=XM_023245398.1;refGene_gn=DNM2;refGene_tx=XM_023245403.1;
refGene_gn=DNM2;refGene_tx=XM_023245404.1;refGene_gn=DNM2;refGene_tx=X
M_023245405.1;refGene_gn=DNM2;refGene_tx=XM_023245409.1;refGene_gn=DNM
2;refGene_tx=XM_023245411.1
A2 8165997 8166158
refGene_gn=DNM2;refGene_tx=XM_023245368.1;refGene_gn=DNM2;refGene_tx=X
M_023245369.1;refGene_gn=DNM2;refGene_tx=XM_023245370.1;refGene_gn=DNM
2;refGene_tx=XM_023245377.1;refGene_gn=DNM2;refGene_tx=XM_023245382.1;
refGene_gn=DNM2;refGene_tx=XM_023245386.1;refGene_gn=DNM2;refGene_tx=X
M_023245389.1;refGene_gn=DNM2;refGene_tx=XM_023245392.1;refGene_gn=DNM
2;refGene_tx=XM_023245398.1;refGene_gn=DNM2;refGene_tx=XM_023245403.1;
refGene_gn=DNM2;refGene_tx=XM_023245404.1;refGene_gn=DNM2;refGene_tx=X
M_023245405.1;refGene_gn=DNM2;refGene_tx=XM_023245409.1;refGene_gn=DNM
2;refGene_tx=XM_023245411.1
A2 8169091 8169234
refGene_gn=DNM2;refGene_tx=XM_023245368.1;refGene_gn=DNM2;refGene_tx=X
M_023245369.1;refGene_gn=DNM2;refGene_tx=XM_023245370.1;refGene_gn=DNM
2;refGene_tx=XM_023245377.1;refGene_gn=DNM2;refGene_tx=XM_023245382.1;
refGene_gn=DNM2;refGene_tx=XM_023245386.1;refGene_gn=DNM2;refGene_tx=X
M_023245389.1;refGene_gn=DNM2;refGene_tx=XM_023245392.1;refGene_gn=DNM
2;refGene_tx=XM_023245398.1;refGene_gn=DNM2;refGene_tx=XM_023245403.1;
refGene_gn=DNM2;refGene_tx=XM_023245404.1;refGene_gn=DNM2;refGene_tx=X
M_023245405.1;refGene_gn=DNM2;refGene_tx=XM_023245409.1;refGene_gn=DNM
2;refGene_tx=XM_023245411.1
A2 8172155 8172291
refGene_gn=DNM2;refGene_tx=XM_023245368.1;refGene_gn=DNM2;refGene_tx=X
M_023245369.1;refGene_gn=DNM2;refGene_tx=XM_023245370.1;refGene_gn=DNM
2;refGene_tx=XM_023245377.1;refGene_gn=DNM2;refGene_tx=XM_023245382.1;
refGene_gn=DNM2;refGene_tx=XM_023245386.1;refGene_gn=DNM2;refGene_tx=X
M_023245389.1;refGene_gn=DNM2;refGene_tx=XM_023245392.1;refGene_gn=DNM
2;refGene_tx=XM_023245398.1;refGene_gn=DNM2;refGene_tx=XM_023245403.1;

refGene_gn=DNM2;refGene_tx=XM_023245404.1;refGene_gn=DNM2;refGene_tx=XM_023245405.1;refGene_gn=DNM2;refGene_tx=XM_023245409.1;refGene_gn=DNM2;refGene_tx=XM_023245411.1
A2 8173541 8173609
refGene_gn=DNM2;refGene_tx=XM_023245368.1;refGene_gn=DNM2;refGene_tx=XM_023245369.1;refGene_gn=DNM2;refGene_tx=XM_023245370.1;refGene_gn=DNM2;refGene_tx=XM_023245377.1;refGene_gn=DNM2;refGene_tx=XM_023245382.1;refGene_gn=DNM2;refGene_tx=XM_023245386.1;refGene_gn=DNM2;refGene_tx=XM_023245389.1;refGene_gn=DNM2;refGene_tx=XM_023245392.1;refGene_gn=DNM2;refGene_tx=XM_023245398.1;refGene_gn=DNM2;refGene_tx=XM_023245403.1;refGene_gn=DNM2;refGene_tx=XM_023245404.1;refGene_gn=DNM2;refGene_tx=XM_023245405.1;refGene_gn=DNM2;refGene_tx=XM_023245409.1;refGene_gn=DNM2;refGene_tx=XM_023245411.1
A2 8174235 8174374
refGene_gn=DNM2;refGene_tx=XM_023245368.1;refGene_gn=DNM2;refGene_tx=XM_023245370.1;refGene_gn=DNM2;refGene_tx=XM_023245377.1;refGene_gn=DNM2;refGene_tx=XM_023245382.1;refGene_gn=DNM2;refGene_tx=XM_023245389.1;refGene_gn=DNM2;refGene_tx=XM_023245392.1;refGene_gn=DNM2;refGene_tx=XM_023245403.1;refGene_gn=DNM2;refGene_tx=XM_023245405.1;refGene_gn=DNM2;refGene_tx=XM_023245409.1;refGene_gn=DNM2;refGene_tx=XM_023245411.1
A2 8175591 8175730
refGene_gn=DNM2;refGene_tx=XM_023245369.1;refGene_gn=DNM2;refGene_tx=XM_023245386.1;refGene_gn=DNM2;refGene_tx=XM_023245398.1;refGene_gn=DNM2;refGene_tx=XM_023245404.1
A2 8176712 8176799
refGene_gn=DNM2;refGene_tx=XM_023245368.1;refGene_gn=DNM2;refGene_tx=XM_023245369.1;refGene_gn=DNM2;refGene_tx=XM_023245370.1;refGene_gn=DNM2;refGene_tx=XM_023245377.1;refGene_gn=DNM2;refGene_tx=XM_023245382.1;refGene_gn=DNM2;refGene_tx=XM_023245386.1;refGene_gn=DNM2;refGene_tx=XM_023245389.1;refGene_gn=DNM2;refGene_tx=XM_023245392.1;refGene_gn=DNM2;refGene_tx=XM_023245398.1;refGene_gn=DNM2;refGene_tx=XM_023245403.1;refGene_gn=DNM2;refGene_tx=XM_023245404.1;refGene_gn=DNM2;refGene_tx=XM_023245405.1;refGene_gn=DNM2;refGene_tx=XM_023245409.1;refGene_gn=DNM2;refGene_tx=XM_023245411.1
A2 8178434 8178505
refGene_gn=DNM2;refGene_tx=XM_023245368.1;refGene_gn=DNM2;refGene_tx=XM_023245369.1;refGene_gn=DNM2;refGene_tx=XM_023245370.1;refGene_gn=DNM2;refGene_tx=XM_023245377.1;refGene_gn=DNM2;refGene_tx=XM_023245382.1;refGene_gn=DNM2;refGene_tx=XM_023245386.1;refGene_gn=DNM2;refGene_tx=XM_023245389.1;refGene_gn=DNM2;refGene_tx=XM_023245392.1;refGene_gn=DNM2;refGene_tx=XM_023245398.1;refGene_gn=DNM2;refGene_tx=XM_023245403.1;refGene_gn=DNM2;refGene_tx=XM_023245404.1;refGene_gn=DNM2;refGene_tx=XM_023245405.1;refGene_gn=DNM2;refGene_tx=XM_023245409.1;refGene_gn=DNM2;refGene_tx=XM_023245411.1
A2 8179882 8179912
refGene_gn=DNM2;refGene_tx=XM_023245368.1;refGene_gn=DNM2;refGene_tx=XM_023245369.1;refGene_gn=DNM2;refGene_tx=XM_023245370.1;refGene_gn=DNM2;refGene_tx=XM_023245377.1;refGene_gn=DNM2;refGene_tx=XM_023245389.1;refGene_gn=DNM2;refGene_tx=XM_023245409.1;refGene_gn=DNM2;refGene_tx=XM_023245411.1

A2	8181965	8182017
refGene_gn=DNM2;refGene_tx=XM_023245368.1;refGene_gn=DNM2;refGene_tx=XM_023245369.1;refGene_gn=DNM2;refGene_tx=XM_023245370.1;refGene_gn=DNM2;refGene_tx=XM_023245377.1;refGene_gn=DNM2;refGene_tx=XM_023245382.1;refGene_gn=DNM2;refGene_tx=XM_023245386.1;refGene_gn=DNM2;refGene_tx=XM_023245389.1;refGene_gn=DNM2;refGene_tx=XM_023245392.1;refGene_gn=DNM2;refGene_tx=XM_023245398.1;refGene_gn=DNM2;refGene_tx=XM_023245403.1;refGene_gn=DNM2;refGene_tx=XM_023245404.1;refGene_gn=DNM2;refGene_tx=XM_023245405.1;refGene_gn=DNM2;refGene_tx=XM_023245409.1;refGene_gn=DNM2;refGene_tx=XM_023245411.1
A2	8182958	8182970
refGene_gn=DNM2;refGene_tx=XM_023245368.1;refGene_gn=DNM2;refGene_tx=XM_023245369.1;refGene_gn=DNM2;refGene_tx=XM_023245370.1;refGene_gn=DNM2;refGene_tx=XM_023245382.1;refGene_gn=DNM2;refGene_tx=XM_023245386.1;refGene_gn=DNM2;refGene_tx=XM_023245389.1;refGene_gn=DNM2;refGene_tx=XM_023245392.1;refGene_gn=DNM2;refGene_tx=XM_023245398.1;refGene_gn=DNM2;refGene_tx=XM_023245405.1;refGene_gn=DNM2;refGene_tx=XM_023245409.1;refGene_gn=DNM2;refGene_tx=XM_023245411.1
A2	8186504	8186618
refGene_gn=DNM2;refGene_tx=XM_023245368.1;refGene_gn=DNM2;refGene_tx=XM_023245369.1;refGene_gn=DNM2;refGene_tx=XM_023245370.1;refGene_gn=DNM2;refGene_tx=XM_023245377.1;refGene_gn=DNM2;refGene_tx=XM_023245382.1;refGene_gn=DNM2;refGene_tx=XM_023245386.1;refGene_gn=DNM2;refGene_tx=XM_023245389.1;refGene_gn=DNM2;refGene_tx=XM_023245392.1;refGene_gn=DNM2;refGene_tx=XM_023245398.1;refGene_gn=DNM2;refGene_tx=XM_023245403.1;refGene_gn=DNM2;refGene_tx=XM_023245404.1;refGene_gn=DNM2;refGene_tx=XM_023245405.1;refGene_gn=DNM2;refGene_tx=XM_023245409.1;refGene_gn=DNM2;refGene_tx=XM_023245411.1
A2	8191129	8191239
refGene_gn=DNM2;refGene_tx=XM_023245368.1;refGene_gn=DNM2;refGene_tx=XM_023245369.1;refGene_gn=DNM2;refGene_tx=XM_023245370.1;refGene_gn=DNM2;refGene_tx=XM_023245377.1;refGene_gn=DNM2;refGene_tx=XM_023245382.1;refGene_gn=DNM2;refGene_tx=XM_023245386.1;refGene_gn=DNM2;refGene_tx=XM_023245389.1;refGene_gn=DNM2;refGene_tx=XM_023245392.1;refGene_gn=DNM2;refGene_tx=XM_023245398.1;refGene_gn=DNM2;refGene_tx=XM_023245403.1;refGene_gn=DNM2;refGene_tx=XM_023245404.1;refGene_gn=DNM2;refGene_tx=XM_023245405.1;refGene_gn=DNM2;refGene_tx=XM_023245409.1;refGene_gn=DNM2;refGene_tx=XM_023245411.1
A2	8193935	8194047
refGene_gn=DNM2;refGene_tx=XM_023245368.1;refGene_gn=DNM2;refGene_tx=XM_023245369.1;refGene_gn=DNM2;refGene_tx=XM_023245370.1;refGene_gn=DNM2;refGene_tx=XM_023245377.1;refGene_gn=DNM2;refGene_tx=XM_023245382.1;refGene_gn=DNM2;refGene_tx=XM_023245386.1;refGene_gn=DNM2;refGene_tx=XM_023245389.1;refGene_gn=DNM2;refGene_tx=XM_023245392.1;refGene_gn=DNM2;refGene_tx=XM_023245398.1;refGene_gn=DNM2;refGene_tx=XM_023245403.1;refGene_gn=DNM2;refGene_tx=XM_023245404.1;refGene_gn=DNM2;refGene_tx=XM_023245405.1;refGene_gn=DNM2;refGene_tx=XM_023245409.1;refGene_gn=DNM2;refGene_tx=XM_023245411.1
A2	8194569	8194734
refGene_gn=DNM2;refGene_tx=XM_023245368.1;refGene_gn=DNM2;refGene_tx=X

M_023245369.1;refGene_gn=DNM2;refGene_tx=XM_023245370.1;refGene_gn=DNM2;refGene_tx=XM_023245377.1;refGene_gn=DNM2;refGene_tx=XM_023245382.1;refGene_gn=DNM2;refGene_tx=XM_023245386.1;refGene_gn=DNM2;refGene_tx=XM_023245389.1;refGene_gn=DNM2;refGene_tx=XM_023245392.1;refGene_gn=DNM2;refGene_tx=XM_023245398.1;refGene_gn=DNM2;refGene_tx=XM_023245403.1;refGene_gn=DNM2;refGene_tx=XM_023245404.1;refGene_gn=DNM2;refGene_tx=XM_023245405.1;refGene_gn=DNM2;refGene_tx=XM_023245409.1;refGene_gn=DNM2;refGene_tx=XM_023245411.1
A2 8196952 8197185
refGene_gn=DNM2;refGene_tx=XM_023245368.1;refGene_gn=DNM2;refGene_tx=XM_023245369.1;refGene_gn=DNM2;refGene_tx=XM_023245370.1;refGene_gn=DNM2;refGene_tx=XM_023245377.1;refGene_gn=DNM2;refGene_tx=XM_023245382.1;refGene_gn=DNM2;refGene_tx=XM_023245386.1;refGene_gn=DNM2;refGene_tx=XM_023245389.1;refGene_gn=DNM2;refGene_tx=XM_023245392.1;refGene_gn=DNM2;refGene_tx=XM_023245398.1;refGene_gn=DNM2;refGene_tx=XM_023245403.1;refGene_gn=DNM2;refGene_tx=XM_023245404.1;refGene_gn=DNM2;refGene_tx=XM_023245405.1;refGene_gn=DNM2;refGene_tx=XM_023245409.1;refGene_gn=DNM2;refGene_tx=XM_023245411.1
A2 8198190 8198433
refGene_gn=DNM2;refGene_tx=XM_023245368.1;refGene_gn=DNM2;refGene_tx=XM_023245369.1;refGene_gn=DNM2;refGene_tx=XM_023245370.1;refGene_gn=DNM2;refGene_tx=XM_023245377.1;refGene_gn=DNM2;refGene_tx=XM_023245382.1;refGene_gn=DNM2;refGene_tx=XM_023245386.1;refGene_gn=DNM2;refGene_tx=XM_023245389.1;refGene_gn=DNM2;refGene_tx=XM_023245392.1;refGene_gn=DNM2;refGene_tx=XM_023245398.1;refGene_gn=DNM2;refGene_tx=XM_023245403.1;refGene_gn=DNM2;refGene_tx=XM_023245404.1;refGene_gn=DNM2;refGene_tx=XM_023245405.1;refGene_gn=DNM2;refGene_tx=XM_023245409.1;refGene_gn=DNM2;refGene_tx=XM_023245411.1
A2 8199066 8199136
refGene_gn=DNM2;refGene_tx=XM_023245368.1;refGene_gn=DNM2;refGene_tx=XM_023245369.1;refGene_gn=DNM2;refGene_tx=XM_023245370.1;refGene_gn=DNM2;refGene_tx=XM_023245377.1;refGene_gn=DNM2;refGene_tx=XM_023245382.1;refGene_gn=DNM2;refGene_tx=XM_023245386.1;refGene_gn=DNM2;refGene_tx=XM_023245392.1;refGene_gn=DNM2;refGene_tx=XM_023245398.1;refGene_gn=DNM2;refGene_tx=XM_023245403.1;refGene_gn=DNM2;refGene_tx=XM_023245404.1;refGene_gn=DNM2;refGene_tx=XM_023245409.1;refGene_gn=DNM2;refGene_tx=XM_023245411.1
A2 8200430 8200470
refGene_gn=DNM2;refGene_tx=XM_023245389.1;refGene_gn=DNM2;refGene_tx=XM_023245405.1
A2 8201395 8201614
refGene_gn=TMED1;refGene_tx=XM_003981876.4;refGene_gn=TMED1;refGene_tx=XM_023245422.1
A2 8202205 8202389
refGene_gn=TMED1;refGene_tx=XM_003981876.4;refGene_gn=TMED1;refGene_tx=XM_023245422.1
A2 8202531 8202629
refGene_gn=TMED1;refGene_tx=XM_003981876.4;refGene_gn=TMED1;refGene_tx=XM_023245422.1
A2 8202743 8202803 refGene_gn=TMED1;refGene_tx=XM_023245422.1

A2 8203303 8203486 refGene_gn=TMED1;refGene_tx=XM_003981876.4
A2 8211691 8211722
refGene_gn=CA2H19orf38;refGene_tx=XM_003981877.4
A2 8212842 8213145
refGene_gn=CA2H19orf38;refGene_tx=XM_003981877.4
A2 8215404 8215497
refGene_gn=CA2H19orf38;refGene_tx=XM_003981877.4
A2 8216650 8216678
refGene_gn=CA2H19orf38;refGene_tx=XM_003981877.4
A2 8217643 8217687
refGene_gn=CA2H19orf38;refGene_tx=XM_003981877.4
A2 8218847 8218885
refGene_gn=CA2H19orf38;refGene_tx=XM_003981877.4
A2 8223839 8223986
refGene_gn=CA2H19orf38;refGene_tx=XM_003981877.4
A2 8225912 8226132 refGene_gn=CARM1;refGene_tx=XM_023245429.1
A2 8226300 8226385
refGene_gn=CARM1;refGene_tx=XM_019824912.2;refGene_gn=CARM1;refGene_tx
=XM_023245441.1
A2 8250587 8250713
refGene_gn=CARM1;refGene_tx=XM_019824912.2;refGene_gn=CARM1;refGene_tx
=XM_023245429.1;refGene_gn=CARM1;refGene_tx=XM_023245441.1
A2 8252268 8252375
refGene_gn=CARM1;refGene_tx=XM_019824912.2;refGene_gn=CARM1;refGene_tx
=XM_023245429.1;refGene_gn=CARM1;refGene_tx=XM_023245441.1
A2 8253302 8253407
refGene_gn=CARM1;refGene_tx=XM_019824912.2;refGene_gn=CARM1;refGene_tx
=XM_023245429.1;refGene_gn=CARM1;refGene_tx=XM_023245441.1
A2 8254263 8254374
refGene_gn=CARM1;refGene_tx=XM_019824912.2;refGene_gn=CARM1;refGene_tx
=XM_023245429.1;refGene_gn=CARM1;refGene_tx=XM_023245441.1
A2 8255564 8255742
refGene_gn=CARM1;refGene_tx=XM_019824912.2;refGene_gn=CARM1;refGene_tx
=XM_023245429.1;refGene_gn=CARM1;refGene_tx=XM_023245441.1
A2 8257888 8257979
refGene_gn=CARM1;refGene_tx=XM_019824912.2;refGene_gn=CARM1;refGene_tx
=XM_023245429.1;refGene_gn=CARM1;refGene_tx=XM_023245441.1
A2 8258158 8258240
refGene_gn=CARM1;refGene_tx=XM_019824912.2;refGene_gn=CARM1;refGene_tx
=XM_023245429.1;refGene_gn=CARM1;refGene_tx=XM_023245441.1
A2 8261772 8261858
refGene_gn=CARM1;refGene_tx=XM_019824912.2;refGene_gn=CARM1;refGene_tx
=XM_023245429.1;refGene_gn=CARM1;refGene_tx=XM_023245441.1
A2 8262045 8262135
refGene_gn=CARM1;refGene_tx=XM_019824912.2;refGene_gn=CARM1;refGene_tx
=XM_023245429.1;refGene_gn=CARM1;refGene_tx=XM_023245441.1
A2 8262581 8262719
refGene_gn=CARM1;refGene_tx=XM_019824912.2;refGene_gn=CARM1;refGene_tx
=XM_023245429.1;refGene_gn=CARM1;refGene_tx=XM_023245441.1
A2 8262805 8262895

refGene_gn=CARM1;refGene_tx=XM_019824912.2;refGene_gn=CARM1;refGene_tx
=XM_023245429.1;refGene_gn=CARM1;refGene_tx=XM_023245441.1
A2 8262979 8263092
refGene_gn=CARM1;refGene_tx=XM_019824912.2;refGene_gn=CARM1;refGene_tx
=XM_023245429.1;refGene_gn=CARM1;refGene_tx=XM_023245441.1
A2 8263187 8263265
refGene_gn=CARM1;refGene_tx=XM_019824912.2;refGene_gn=CARM1;refGene_tx
=XM_023245429.1;refGene_gn=CARM1;refGene_tx=XM_023245441.1
A2 8263541 8263610
refGene_gn=CARM1;refGene_tx=XM_019824912.2;refGene_gn=CARM1;refGene_tx
=XM_023245429.1
A2 8263773 8263916
refGene_gn=CARM1;refGene_tx=XM_019824912.2;refGene_gn=CARM1;refGene_tx
=XM_023245429.1;refGene_gn=CARM1;refGene_tx=XM_023245441.1
A2 8265559 8265670
refGene_gn=YIPF2;refGene_tx=XM_003981896.5;refGene_gn=YIPF2;refGene_tx
=XM_011289124.3
A2 8265758 8265941
refGene_gn=YIPF2;refGene_tx=XM_003981896.5;refGene_gn=YIPF2;refGene_tx
=XM_011289124.3
A2 8266018 8266185
refGene_gn=YIPF2;refGene_tx=XM_003981896.5;refGene_gn=YIPF2;refGene_tx
=XM_011289124.3
A2 8266298 8266415
refGene_gn=YIPF2;refGene_tx=XM_003981896.5;refGene_gn=YIPF2;refGene_tx
=XM_011289124.3
A2 8267704 8267792
refGene_gn=YIPF2;refGene_tx=XM_003981896.5;refGene_gn=YIPF2;refGene_tx
=XM_011289124.3
A2 8268741 8268819
refGene_gn=YIPF2;refGene_tx=XM_003981896.5;refGene_gn=YIPF2;refGene_tx
=XM_011289124.3
A2 8268917 8269075
refGene_gn=YIPF2;refGene_tx=XM_003981896.5;refGene_gn=YIPF2;refGene_tx
=XM_011289124.3
A2 8269475 8269506
refGene_gn=YIPF2;refGene_tx=XM_003981896.5;refGene_gn=YIPF2;refGene_tx
=XM_011289124.3
A2 8269957 8270051 refGene_gn=TIMM29;refGene_tx=XM_023245448.1
A2 8270132 8270821 refGene_gn=TIMM29;refGene_tx=XM_023245448.1
A2 8322513 8323452
refGene_gn=LOC102901712;refGene_tx=XM_023245489.1
A2 8340857 8341079
refGene_gn=SMARCA4;refGene_tx=XM_006928407.4;refGene_gn=SMARCA4;refGen
e_tx=XM_006928408.4;refGene_gn=SMARCA4;refGene_tx=XM_019817549.2;refGe
ne_gn=SMARCA4;refGene_tx=XM_019817553.2;refGene_gn=SMARCA4;refGene_tx=
XM_019817559.2;refGene_gn=SMARCA4;refGene_tx=XM_019817561.2;refGene_gn
=SMARCA4;refGene_tx=XM_019817567.2
A2 8343179 8343312
refGene_gn=SMARCA4;refGene_tx=XM_006928407.4;refGene_gn=SMARCA4;refGen

e_tx=XM_006928408.4;refGene_gn=SMARCA4;refGene_tx=XM_019817549.2;refGene_gn=SMARCA4;refGene_tx=XM_019817553.2;refGene_gn=SMARCA4;refGene_tx=XM_019817559.2;refGene_gn=SMARCA4;refGene_tx=XM_019817561.2;refGene_gn=SMARCA4;refGene_tx=XM_019817567.2
A2 8344122 8344527
refGene_gn=SMARCA4;refGene_tx=XM_006928407.4;refGene_gn=SMARCA4;refGene_tx=XM_006928408.4;refGene_gn=SMARCA4;refGene_tx=XM_019817549.2;refGene_gn=SMARCA4;refGene_tx=XM_019817553.2;refGene_gn=SMARCA4;refGene_tx=XM_019817559.2;refGene_gn=SMARCA4;refGene_tx=XM_019817561.2;refGene_gn=SMARCA4;refGene_tx=XM_019817567.2
A2 8344861 8344960
refGene_gn=SMARCA4;refGene_tx=XM_006928407.4;refGene_gn=SMARCA4;refGene_tx=XM_006928408.4;refGene_gn=SMARCA4;refGene_tx=XM_019817549.2;refGene_gn=SMARCA4;refGene_tx=XM_019817553.2;refGene_gn=SMARCA4;refGene_tx=XM_019817559.2;refGene_gn=SMARCA4;refGene_tx=XM_019817561.2;refGene_gn=SMARCA4;refGene_tx=XM_019817567.2
A2 8346090 8346349
refGene_gn=SMARCA4;refGene_tx=XM_006928407.4;refGene_gn=SMARCA4;refGene_tx=XM_006928408.4;refGene_gn=SMARCA4;refGene_tx=XM_019817549.2;refGene_gn=SMARCA4;refGene_tx=XM_019817553.2;refGene_gn=SMARCA4;refGene_tx=XM_019817559.2;refGene_gn=SMARCA4;refGene_tx=XM_019817561.2;refGene_gn=SMARCA4;refGene_tx=XM_019817567.2
A2 8348647 8348774
refGene_gn=SMARCA4;refGene_tx=XM_006928407.4;refGene_gn=SMARCA4;refGene_tx=XM_006928408.4;refGene_gn=SMARCA4;refGene_tx=XM_019817549.2;refGene_gn=SMARCA4;refGene_tx=XM_019817553.2;refGene_gn=SMARCA4;refGene_tx=XM_019817559.2;refGene_gn=SMARCA4;refGene_tx=XM_019817561.2;refGene_gn=SMARCA4;refGene_tx=XM_019817567.2
A2 8349295 8349469
refGene_gn=SMARCA4;refGene_tx=XM_006928407.4;refGene_gn=SMARCA4;refGene_tx=XM_006928408.4;refGene_gn=SMARCA4;refGene_tx=XM_019817549.2;refGene_gn=SMARCA4;refGene_tx=XM_019817553.2;refGene_gn=SMARCA4;refGene_tx=XM_019817559.2;refGene_gn=SMARCA4;refGene_tx=XM_019817561.2;refGene_gn=SMARCA4;refGene_tx=XM_019817567.2
A2 8350965 8351139
refGene_gn=SMARCA4;refGene_tx=XM_006928407.4;refGene_gn=SMARCA4;refGene_tx=XM_006928408.4;refGene_gn=SMARCA4;refGene_tx=XM_019817549.2;refGene_gn=SMARCA4;refGene_tx=XM_019817553.2;refGene_gn=SMARCA4;refGene_tx=XM_019817559.2;refGene_gn=SMARCA4;refGene_tx=XM_019817561.2;refGene_gn=SMARCA4;refGene_tx=XM_019817567.2
A2 8352341 8352509
refGene_gn=SMARCA4;refGene_tx=XM_006928407.4;refGene_gn=SMARCA4;refGene_tx=XM_006928408.4;refGene_gn=SMARCA4;refGene_tx=XM_019817549.2;refGene_gn=SMARCA4;refGene_tx=XM_019817553.2;refGene_gn=SMARCA4;refGene_tx=XM_019817559.2;refGene_gn=SMARCA4;refGene_tx=XM_019817561.2;refGene_gn=SMARCA4;refGene_tx=XM_019817567.2
A2 8352626 8352677
refGene_gn=SMARCA4;refGene_tx=XM_006928407.4;refGene_gn=SMARCA4;refGene_tx=XM_006928408.4;refGene_gn=SMARCA4;refGene_tx=XM_019817549.2;refGene_gn=SMARCA4;refGene_tx=XM_019817553.2;refGene_gn=SMARCA4;refGene_tx=

XM_019817559.2;refGene_gn=SMARCA4;refGene_tx=XM_019817561.2;refGene_gn=SMARCA4;refGene_tx=XM_019817567.2
A2 8357409 8357540 refGene_gn=SMARCA4;refGene_tx=XM_006928407.4;refGene_gn=SMARCA4;refGene_tx=XM_006928408.4;refGene_gn=SMARCA4;refGene_tx=XM_019817549.2;refGene_gn=SMARCA4;refGene_tx=XM_019817553.2;refGene_gn=SMARCA4;refGene_tx=XM_019817559.2;refGene_gn=SMARCA4;refGene_tx=XM_019817561.2;refGene_gn=SMARCA4;refGene_tx=XM_019817567.2
A2 8357704 8357762 refGene_gn=SMARCA4;refGene_tx=XM_006928407.4;refGene_gn=SMARCA4;refGene_tx=XM_006928408.4;refGene_gn=SMARCA4;refGene_tx=XM_019817549.2;refGene_gn=SMARCA4;refGene_tx=XM_019817553.2;refGene_gn=SMARCA4;refGene_tx=XM_019817559.2;refGene_gn=SMARCA4;refGene_tx=XM_019817561.2;refGene_gn=SMARCA4;refGene_tx=XM_019817567.2
A2 8365583 8365705 refGene_gn=SMARCA4;refGene_tx=XM_006928407.4;refGene_gn=SMARCA4;refGene_tx=XM_006928408.4;refGene_gn=SMARCA4;refGene_tx=XM_019817549.2;refGene_gn=SMARCA4;refGene_tx=XM_019817553.2;refGene_gn=SMARCA4;refGene_tx=XM_019817559.2;refGene_gn=SMARCA4;refGene_tx=XM_019817561.2;refGene_gn=SMARCA4;refGene_tx=XM_019817567.2
A2 8367277 8367428 refGene_gn=SMARCA4;refGene_tx=XM_006928407.4;refGene_gn=SMARCA4;refGene_tx=XM_006928408.4;refGene_gn=SMARCA4;refGene_tx=XM_019817549.2;refGene_gn=SMARCA4;refGene_tx=XM_019817553.2;refGene_gn=SMARCA4;refGene_tx=XM_019817559.2;refGene_gn=SMARCA4;refGene_tx=XM_019817561.2;refGene_gn=SMARCA4;refGene_tx=XM_019817567.2
A2 8368196 8368360 refGene_gn=SMARCA4;refGene_tx=XM_006928407.4;refGene_gn=SMARCA4;refGene_tx=XM_006928408.4;refGene_gn=SMARCA4;refGene_tx=XM_019817549.2;refGene_gn=SMARCA4;refGene_tx=XM_019817553.2;refGene_gn=SMARCA4;refGene_tx=XM_019817559.2;refGene_gn=SMARCA4;refGene_tx=XM_019817561.2;refGene_gn=SMARCA4;refGene_tx=XM_019817567.2
A2 8373887 8373954 refGene_gn=SMARCA4;refGene_tx=XM_006928407.4;refGene_gn=SMARCA4;refGene_tx=XM_006928408.4;refGene_gn=SMARCA4;refGene_tx=XM_019817549.2;refGene_gn=SMARCA4;refGene_tx=XM_019817553.2;refGene_gn=SMARCA4;refGene_tx=XM_019817559.2;refGene_gn=SMARCA4;refGene_tx=XM_019817561.2;refGene_gn=SMARCA4;refGene_tx=XM_019817567.2
A2 8374945 8375056 refGene_gn=SMARCA4;refGene_tx=XM_006928407.4;refGene_gn=SMARCA4;refGene_tx=XM_006928408.4;refGene_gn=SMARCA4;refGene_tx=XM_019817549.2;refGene_gn=SMARCA4;refGene_tx=XM_019817553.2;refGene_gn=SMARCA4;refGene_tx=XM_019817559.2;refGene_gn=SMARCA4;refGene_tx=XM_019817561.2;refGene_gn=SMARCA4;refGene_tx=XM_019817567.2
A2 8376581 8376824 refGene_gn=SMARCA4;refGene_tx=XM_006928407.4;refGene_gn=SMARCA4;refGene_tx=XM_006928408.4;refGene_gn=SMARCA4;refGene_tx=XM_019817549.2;refGene_gn=SMARCA4;refGene_tx=XM_019817553.2;refGene_gn=SMARCA4;refGene_tx=XM_019817559.2;refGene_gn=SMARCA4;refGene_tx=XM_019817561.2;refGene_gn=SMARCA4;refGene_tx=XM_019817567.2

A2 8378799 8378913
refGene_gn=SMARCA4;refGene_tx=XM_006928407.4;refGene_gn=SMARCA4;refGene_tx=XM_006928408.4;refGene_gn=SMARCA4;refGene_tx=XM_019817549.2;refGene_gn=SMARCA4;refGene_tx=XM_019817553.2;refGene_gn=SMARCA4;refGene_tx=XM_019817559.2;refGene_gn=SMARCA4;refGene_tx=XM_019817561.2;refGene_gn=SMARCA4;refGene_tx=XM_019817567.2
A2 8379598 8379706
refGene_gn=SMARCA4;refGene_tx=XM_006928407.4;refGene_gn=SMARCA4;refGene_tx=XM_006928408.4;refGene_gn=SMARCA4;refGene_tx=XM_019817549.2;refGene_gn=SMARCA4;refGene_tx=XM_019817553.2;refGene_gn=SMARCA4;refGene_tx=XM_019817559.2;refGene_gn=SMARCA4;refGene_tx=XM_019817561.2;refGene_gn=SMARCA4;refGene_tx=XM_019817567.2
A2 8380744 8380831
refGene_gn=SMARCA4;refGene_tx=XM_006928407.4;refGene_gn=SMARCA4;refGene_tx=XM_006928408.4;refGene_gn=SMARCA4;refGene_tx=XM_019817549.2;refGene_gn=SMARCA4;refGene_tx=XM_019817553.2;refGene_gn=SMARCA4;refGene_tx=XM_019817559.2;refGene_gn=SMARCA4;refGene_tx=XM_019817561.2;refGene_gn=SMARCA4;refGene_tx=XM_019817567.2
A2 8381821 8381868
refGene_gn=SMARCA4;refGene_tx=XM_006928407.4;refGene_gn=SMARCA4;refGene_tx=XM_006928408.4;refGene_gn=SMARCA4;refGene_tx=XM_019817549.2;refGene_gn=SMARCA4;refGene_tx=XM_019817553.2;refGene_gn=SMARCA4;refGene_tx=XM_019817559.2;refGene_gn=SMARCA4;refGene_tx=XM_019817561.2;refGene_gn=SMARCA4;refGene_tx=XM_019817567.2
A2 8382969 8383136
refGene_gn=SMARCA4;refGene_tx=XM_006928407.4;refGene_gn=SMARCA4;refGene_tx=XM_006928408.4;refGene_gn=SMARCA4;refGene_tx=XM_019817549.2;refGene_gn=SMARCA4;refGene_tx=XM_019817553.2;refGene_gn=SMARCA4;refGene_tx=XM_019817559.2;refGene_gn=SMARCA4;refGene_tx=XM_019817561.2;refGene_gn=SMARCA4;refGene_tx=XM_019817567.2
A2 8385505 8385669
refGene_gn=SMARCA4;refGene_tx=XM_006928407.4;refGene_gn=SMARCA4;refGene_tx=XM_006928408.4;refGene_gn=SMARCA4;refGene_tx=XM_019817549.2;refGene_gn=SMARCA4;refGene_tx=XM_019817553.2;refGene_gn=SMARCA4;refGene_tx=XM_019817559.2;refGene_gn=SMARCA4;refGene_tx=XM_019817561.2;refGene_gn=SMARCA4;refGene_tx=XM_019817567.2
A2 8388023 8388251
refGene_gn=SMARCA4;refGene_tx=XM_006928407.4;refGene_gn=SMARCA4;refGene_tx=XM_006928408.4;refGene_gn=SMARCA4;refGene_tx=XM_019817549.2;refGene_gn=SMARCA4;refGene_tx=XM_019817553.2;refGene_gn=SMARCA4;refGene_tx=XM_019817559.2;refGene_gn=SMARCA4;refGene_tx=XM_019817561.2;refGene_gn=SMARCA4;refGene_tx=XM_019817567.2
A2 8388509 8388608
refGene_gn=SMARCA4;refGene_tx=XM_006928407.4;refGene_gn=SMARCA4;refGene_tx=XM_006928408.4;refGene_gn=SMARCA4;refGene_tx=XM_019817549.2;refGene_gn=SMARCA4;refGene_tx=XM_019817553.2
A2 8388842 8388920
refGene_gn=SMARCA4;refGene_tx=XM_006928407.4;refGene_gn=SMARCA4;refGene_tx=XM_006928408.4;refGene_gn=SMARCA4;refGene_tx=XM_019817549.2;refGene_gn=SMARCA4;refGene_tx=XM_019817553.2;refGene_gn=SMARCA4;refGene_tx=

XM_019817559.2;refGene_gn=SMARCA4;refGene_tx=XM_019817561.2;refGene_gn=SMARCA4;refGene_tx=XM_019817567.2
A2 8389589 8389808
refGene_gn=SMARCA4;refGene_tx=XM_006928407.4;refGene_gn=SMARCA4;refGene_tx=XM_006928408.4;refGene_gn=SMARCA4;refGene_tx=XM_019817549.2;refGene_gn=SMARCA4;refGene_tx=XM_019817553.2;refGene_gn=SMARCA4;refGene_tx=XM_019817559.2;refGene_gn=SMARCA4;refGene_tx=XM_019817561.2;refGene_gn=SMARCA4;refGene_tx=XM_019817567.2
A2 8394401 8394664
refGene_gn=SMARCA4;refGene_tx=XM_006928407.4;refGene_gn=SMARCA4;refGene_tx=XM_006928408.4;refGene_gn=SMARCA4;refGene_tx=XM_019817549.2;refGene_gn=SMARCA4;refGene_tx=XM_019817553.2;refGene_gn=SMARCA4;refGene_tx=XM_019817559.2;refGene_gn=SMARCA4;refGene_tx=XM_019817561.2;refGene_gn=SMARCA4;refGene_tx=XM_019817567.2
A2 8409079 8409188
refGene_gn=SMARCA4;refGene_tx=XM_006928407.4;refGene_gn=SMARCA4;refGene_tx=XM_006928408.4;refGene_gn=SMARCA4;refGene_tx=XM_019817549.2;refGene_gn=SMARCA4;refGene_tx=XM_019817553.2;refGene_gn=SMARCA4;refGene_tx=XM_019817559.2;refGene_gn=SMARCA4;refGene_tx=XM_019817561.2;refGene_gn=SMARCA4;refGene_tx=XM_019817567.2
A2 8409663 8409765
refGene_gn=SMARCA4;refGene_tx=XM_006928407.4;refGene_gn=SMARCA4;refGene_tx=XM_006928408.4;refGene_gn=SMARCA4;refGene_tx=XM_019817549.2;refGene_gn=SMARCA4;refGene_tx=XM_019817553.2;refGene_gn=SMARCA4;refGene_tx=XM_019817559.2;refGene_gn=SMARCA4;refGene_tx=XM_019817561.2;refGene_gn=SMARCA4;refGene_tx=XM_019817567.2
A2 8410580 8410713
refGene_gn=SMARCA4;refGene_tx=XM_006928407.4;refGene_gn=SMARCA4;refGene_tx=XM_006928408.4;refGene_gn=SMARCA4;refGene_tx=XM_019817549.2;refGene_gn=SMARCA4;refGene_tx=XM_019817553.2;refGene_gn=SMARCA4;refGene_tx=XM_019817559.2;refGene_gn=SMARCA4;refGene_tx=XM_019817561.2;refGene_gn=SMARCA4;refGene_tx=XM_019817567.2
A2 8410851 8410994
refGene_gn=SMARCA4;refGene_tx=XM_006928407.4;refGene_gn=SMARCA4;refGene_tx=XM_006928408.4;refGene_gn=SMARCA4;refGene_tx=XM_019817549.2;refGene_gn=SMARCA4;refGene_tx=XM_019817553.2;refGene_gn=SMARCA4;refGene_tx=XM_019817559.2;refGene_gn=SMARCA4;refGene_tx=XM_019817561.2;refGene_gn=SMARCA4;refGene_tx=XM_019817567.2
A2 8412380 8412413
refGene_gn=SMARCA4;refGene_tx=XM_006928407.4;refGene_gn=SMARCA4;refGene_tx=XM_006928408.4;refGene_gn=SMARCA4;refGene_tx=XM_019817549.2;refGene_gn=SMARCA4;refGene_tx=XM_019817553.2;refGene_gn=SMARCA4;refGene_tx=XM_019817559.2;refGene_gn=SMARCA4;refGene_tx=XM_019817561.2;refGene_gn=SMARCA4;refGene_tx=XM_019817567.2
A2 8428853 8428920 refGene_gn=LDLR;refGene_tx=XM_003981898.5
A2 8429520 8429689 refGene_gn=LDLR;refGene_tx=XM_023245495.1
A2 8436046 8436169
refGene_gn=LDLR;refGene_tx=XM_003981898.5;refGene_gn=LDLR;refGene_tx=XM_023245494.1;refGene_gn=LDLR;refGene_tx=XM_023245495.1
A2 8436618 8436741

refGene_gn=LDLR;refGene_tx=XM_003981898.5;refGene_gn=LDLR;refGene_tx=XM_023245494.1;refGene_gn=LDLR;refGene_tx=XM_023245495.1
A2 8438282 8438669
refGene_gn=LDLR;refGene_tx=XM_003981898.5;refGene_gn=LDLR;refGene_tx=XM_023245494.1;refGene_gn=LDLR;refGene_tx=XM_023245495.1
A2 8442063 8442186
refGene_gn=LDLR;refGene_tx=XM_003981898.5;refGene_gn=LDLR;refGene_tx=XM_023245494.1;refGene_gn=LDLR;refGene_tx=XM_023245495.1
A2 8443950 8444073
refGene_gn=LDLR;refGene_tx=XM_003981898.5;refGene_gn=LDLR;refGene_tx=XM_023245494.1;refGene_gn=LDLR;refGene_tx=XM_023245495.1
A2 8445326 8445446
refGene_gn=LDLR;refGene_tx=XM_003981898.5;refGene_gn=LDLR;refGene_tx=XM_023245494.1;refGene_gn=LDLR;refGene_tx=XM_023245495.1
A2 8445894 8446020
refGene_gn=LDLR;refGene_tx=XM_003981898.5;refGene_gn=LDLR;refGene_tx=XM_023245494.1;refGene_gn=LDLR;refGene_tx=XM_023245495.1
A2 8448033 8448205
refGene_gn=LDLR;refGene_tx=XM_003981898.5;refGene_gn=LDLR;refGene_tx=XM_023245494.1;refGene_gn=LDLR;refGene_tx=XM_023245495.1
A2 8448297 8448525
refGene_gn=LDLR;refGene_tx=XM_003981898.5;refGene_gn=LDLR;refGene_tx=XM_023245494.1;refGene_gn=LDLR;refGene_tx=XM_023245495.1
A2 8451087 8451206
refGene_gn=LDLR;refGene_tx=XM_003981898.5;refGene_gn=LDLR;refGene_tx=XM_023245494.1;refGene_gn=LDLR;refGene_tx=XM_023245495.1
A2 8451415 8451555
refGene_gn=LDLR;refGene_tx=XM_003981898.5;refGene_gn=LDLR;refGene_tx=XM_023245494.1;refGene_gn=LDLR;refGene_tx=XM_023245495.1
A2 8453572 8453714
refGene_gn=LDLR;refGene_tx=XM_003981898.5;refGene_gn=LDLR;refGene_tx=XM_023245494.1;refGene_gn=LDLR;refGene_tx=XM_023245495.1
A2 8453825 8453978
refGene_gn=LDLR;refGene_tx=XM_003981898.5;refGene_gn=LDLR;refGene_tx=XM_023245494.1;refGene_gn=LDLR;refGene_tx=XM_023245495.1
A2 8454228 8454411
refGene_gn=LDLR;refGene_tx=XM_003981898.5;refGene_gn=LDLR;refGene_tx=XM_023245494.1;refGene_gn=LDLR;refGene_tx=XM_023245495.1
A2 8456799 8456877
refGene_gn=LDLR;refGene_tx=XM_003981898.5;refGene_gn=LDLR;refGene_tx=XM_023245494.1;refGene_gn=LDLR;refGene_tx=XM_023245495.1
A2 8457916 8458074
refGene_gn=LDLR;refGene_tx=XM_003981898.5;refGene_gn=LDLR;refGene_tx=XM_023245494.1;refGene_gn=LDLR;refGene_tx=XM_023245495.1
A2 8458704 8458740
refGene_gn=LDLR;refGene_tx=XM_003981898.5;refGene_gn=LDLR;refGene_tx=XM_023245494.1;refGene_gn=LDLR;refGene_tx=XM_023245495.1
A2 8468800 8468907
refGene_gn=SPC24;refGene_tx=XM_003981879.5;refGene_gn=SPC24;refGene_tx=XM_011289140.3

A2 8470872 8470949
refGene_gn=SPC24;refGene_tx=XM_003981879.5;refGene_gn=SPC24;refGene_tx=XM_011289140.3
A2 8471084 8471189
refGene_gn=SPC24;refGene_tx=XM_003981879.5;refGene_gn=SPC24;refGene_tx=XM_011289140.3
A2 8471333 8471478
refGene_gn=SPC24;refGene_tx=XM_003981879.5;refGene_gn=SPC24;refGene_tx=XM_011289140.3
A2 8474357 8474466 refGene_gn=SPC24;refGene_tx=XM_011289140.3
A2 8474625 8474785 refGene_gn=SPC24;refGene_tx=XM_003981879.5
A2 8483012 8483036 refGene_gn=KANK2;refGene_tx=XM_006928418.4
A2 8483189 8483243 refGene_gn=KANK2;refGene_tx=XM_003981880.5
A2 8484813 8484894
refGene_gn=KANK2;refGene_tx=XM_006928415.4;refGene_gn=KANK2;refGene_tx=XM_006928416.4;refGene_gn=KANK2;refGene_tx=XM_006928417.4;refGene_gn=KANK2;refGene_tx=XM_011289143.3;refGene_gn=KANK2;refGene_tx=XM_023245505.1
A2 8485230 8485320
refGene_gn=KANK2;refGene_tx=XM_003981880.5;refGene_gn=KANK2;refGene_tx=XM_006928415.4;refGene_gn=KANK2;refGene_tx=XM_006928416.4;refGene_gn=KANK2;refGene_tx=XM_006928417.4;refGene_gn=KANK2;refGene_tx=XM_006928418.4;refGene_gn=KANK2;refGene_tx=XM_011289143.3;refGene_gn=KANK2;refGene_tx=XM_023245505.1
A2 8485398 8485599
refGene_gn=KANK2;refGene_tx=XM_003981880.5;refGene_gn=KANK2;refGene_tx=XM_006928415.4;refGene_gn=KANK2;refGene_tx=XM_006928416.4;refGene_gn=KANK2;refGene_tx=XM_006928417.4;refGene_gn=KANK2;refGene_tx=XM_006928418.4;refGene_gn=KANK2;refGene_tx=XM_011289143.3;refGene_gn=KANK2;refGene_tx=XM_023245505.1
A2 8487597 8487740
refGene_gn=KANK2;refGene_tx=XM_003981880.5;refGene_gn=KANK2;refGene_tx=XM_006928415.4;refGene_gn=KANK2;refGene_tx=XM_006928416.4;refGene_gn=KANK2;refGene_tx=XM_006928417.4;refGene_gn=KANK2;refGene_tx=XM_006928418.4;refGene_gn=KANK2;refGene_tx=XM_011289143.3;refGene_gn=KANK2;refGene_tx=XM_023245505.1
A2 8488127 8488347
refGene_gn=KANK2;refGene_tx=XM_003981880.5;refGene_gn=KANK2;refGene_tx=XM_006928415.4;refGene_gn=KANK2;refGene_tx=XM_006928416.4;refGene_gn=KANK2;refGene_tx=XM_006928417.4;refGene_gn=KANK2;refGene_tx=XM_006928418.4;refGene_gn=KANK2;refGene_tx=XM_011289143.3;refGene_gn=KANK2;refGene_tx=XM_023245505.1
A2 8488910 8488998
refGene_gn=KANK2;refGene_tx=XM_003981880.5;refGene_gn=KANK2;refGene_tx=XM_006928415.4;refGene_gn=KANK2;refGene_tx=XM_006928416.4;refGene_gn=KANK2;refGene_tx=XM_006928417.4;refGene_gn=KANK2;refGene_tx=XM_006928418.4;refGene_gn=KANK2;refGene_tx=XM_011289143.3;refGene_gn=KANK2;refGene_tx=XM_023245505.1
A2 8489504 8489744
refGene_gn=KANK2;refGene_tx=XM_003981880.5;refGene_gn=KANK2;refGene_tx

=XM_006928415.4;refGene_gn=KANK2;refGene_tx=XM_006928416.4;refGene_gn=KANK2;refGene_tx=XM_006928417.4;refGene_gn=KANK2;refGene_tx=XM_006928418.4;refGene_gn=KANK2;refGene_tx=XM_011289143.3;refGene_gn=KANK2;refGene_tx=XM_023245505.1
A2 8490991 8491094 refGene_gn=KANK2;refGene_tx=XM_003981880.5;refGene_gn=KANK2;refGene_tx=XM_006928415.4;refGene_gn=KANK2;refGene_tx=XM_006928416.4;refGene_gn=KANK2;refGene_tx=XM_006928417.4;refGene_gn=KANK2;refGene_tx=XM_006928418.4;refGene_gn=KANK2;refGene_tx=XM_011289143.3;refGene_gn=KANK2;refGene_tx=XM_023245505.1
A2 8491192 8491360 refGene_gn=KANK2;refGene_tx=XM_003981880.5;refGene_gn=KANK2;refGene_tx=XM_006928415.4;refGene_gn=KANK2;refGene_tx=XM_006928416.4;refGene_gn=KANK2;refGene_tx=XM_006928417.4;refGene_gn=KANK2;refGene_tx=XM_006928418.4;refGene_gn=KANK2;refGene_tx=XM_011289143.3;refGene_gn=KANK2;refGene_tx=XM_023245505.1
A2 8501567 8502779 refGene_gn=KANK2;refGene_tx=XM_003981880.5;refGene_gn=KANK2;refGene_tx=XM_006928415.4;refGene_gn=KANK2;refGene_tx=XM_006928416.4;refGene_gn=KANK2;refGene_tx=XM_006928417.4;refGene_gn=KANK2;refGene_tx=XM_006928418.4;refGene_gn=KANK2;refGene_tx=XM_011289143.3;refGene_gn=KANK2;refGene_tx=XM_023245505.1
A2 8503208 8503245 refGene_gn=KANK2;refGene_tx=XM_003981880.5;refGene_gn=KANK2;refGene_tx=XM_006928415.4;refGene_gn=KANK2;refGene_tx=XM_006928416.4;refGene_gn=KANK2;refGene_tx=XM_006928417.4;refGene_gn=KANK2;refGene_tx=XM_006928418.4;refGene_gn=KANK2;refGene_tx=XM_011289143.3;refGene_gn=KANK2;refGene_tx=XM_023245505.1
A2 8506932 8506975 refGene_gn=DOCK6;refGene_tx=XM_023245498.1
A2 8507355 8507517 refGene_gn=DOCK6;refGene_tx=XM_023245498.1
A2 8507738 8507845 refGene_gn=DOCK6;refGene_tx=XM_023245498.1
A2 8507928 8508072 refGene_gn=DOCK6;refGene_tx=XM_023245498.1
A2 8508462 8508699 refGene_gn=DOCK6;refGene_tx=XM_023245498.1
A2 8508953 8509043 refGene_gn=DOCK6;refGene_tx=XM_023245498.1
A2 8509139 8509265 refGene_gn=DOCK6;refGene_tx=XM_023245498.1
A2 8510582 8510597 refGene_gn=DOCK6;refGene_tx=XM_023245498.1
A2 8510689 8510821 refGene_gn=DOCK6;refGene_tx=XM_023245498.1
A2 8513723 8513867 refGene_gn=DOCK6;refGene_tx=XM_023245498.1
A2 8513953 8514146 refGene_gn=DOCK6;refGene_tx=XM_023245498.1
A2 8515520 8515621 refGene_gn=DOCK6;refGene_tx=XM_023245498.1
A2 8515747 8515906 refGene_gn=DOCK6;refGene_tx=XM_023245498.1
A2 8516045 8516198 refGene_gn=DOCK6;refGene_tx=XM_023245498.1
A2 8516512 8516647 refGene_gn=DOCK6;refGene_tx=XM_023245498.1
A2 8516779 8516876 refGene_gn=DOCK6;refGene_tx=XM_023245498.1
A2 8516975 8517060 refGene_gn=DOCK6;refGene_tx=XM_023245498.1
A2 8517393 8517520 refGene_gn=DOCK6;refGene_tx=XM_023245498.1
A2 8517878 8518061 refGene_gn=DOCK6;refGene_tx=XM_023245498.1
A2 8518212 8518373 refGene_gn=DOCK6;refGene_tx=XM_023245498.1
A2 8520212 8520382 refGene_gn=DOCK6;refGene_tx=XM_023245498.1
A2 8520471 8520611 refGene_gn=DOCK6;refGene_tx=XM_023245498.1

A2	8520964	8521135	refGene_gn=DOCK6;refGene_tx=XM_023245498.1
A2	8521237	8521351	refGene_gn=DOCK6;refGene_tx=XM_023245498.1
A2	8524576	8524717	refGene_gn=DOCK6;refGene_tx=XM_023245498.1
A2	8525387	8525483	refGene_gn=DOCK6;refGene_tx=XM_023245498.1
A2	8527852	8527957	refGene_gn=DOCK6;refGene_tx=XM_023245498.1
A2	8528510	8528674	refGene_gn=DOCK6;refGene_tx=XM_023245498.1
A2	8529347	8529509	refGene_gn=DOCK6;refGene_tx=XM_023245498.1
A2	8529770	8530002	refGene_gn=DOCK6;refGene_tx=XM_023245498.1
A2	8530367	8530454	refGene_gn=DOCK6;refGene_tx=XM_023245498.1
A2	8531136	8531238	refGene_gn=DOCK6;refGene_tx=XM_023245498.1
A2	8531325	8531464	refGene_gn=DOCK6;refGene_tx=XM_023245498.1
A2	8531654	8531725	refGene_gn=DOCK6;refGene_tx=XM_023245498.1
A2	8531809	8531927	refGene_gn=DOCK6;refGene_tx=XM_023245498.1
A2	8533014	8533311	refGene_gn=ANGPTL8;refGene_tx=XM_006928419.4
A2	8533506	8533668	refGene_gn=ANGPTL8;refGene_tx=XM_006928419.4
A2	8534208	8534318	refGene_gn=ANGPTL8;refGene_tx=XM_006928419.4
A2	8534429	8534457	refGene_gn=ANGPTL8;refGene_tx=XM_006928419.4
A2	8534817	8534980	refGene_gn=DOCK6;refGene_tx=XM_023245498.1
A2	8535387	8535481	refGene_gn=DOCK6;refGene_tx=XM_023245498.1
A2	8535594	8535722	refGene_gn=DOCK6;refGene_tx=XM_023245498.1
A2	8535885	8536039	refGene_gn=DOCK6;refGene_tx=XM_023245498.1
A2	8536155	8536236	refGene_gn=DOCK6;refGene_tx=XM_023245498.1
A2	8537120	8537270	refGene_gn=DOCK6;refGene_tx=XM_023245498.1
A2	8537370	8537437	refGene_gn=DOCK6;refGene_tx=XM_023245498.1
A2	8538966	8539052	refGene_gn=DOCK6;refGene_tx=XM_023245498.1
A2	8541026	8541239	refGene_gn=DOCK6;refGene_tx=XM_023245498.1
A2	8542378	8542480	refGene_gn=DOCK6;refGene_tx=XM_023245498.1
A2	8542481	8542509	refGene_gn=DOCK6;refGene_tx=XM_023245498.1
A2	8542709	8542778	refGene_gn=DOCK6;refGene_tx=XM_023245498.1
A2	8543038	8543214	refGene_gn=DOCK6;refGene_tx=XM_023245498.1
A2	8543773	8543861	refGene_gn=DOCK6;refGene_tx=XM_023245498.1
A2	8550934	8550978	refGene_gn=DOCK6;refGene_tx=XM_023245498.1
A2	8565778	8565847	refGene_gn=TSPAN16;refGene_tx=XM_023241013.1
A2	8567364	8567603	refGene_gn=TSPAN16;refGene_tx=XM_023241013.1
A2	8569980	8570086	refGene_gn=TSPAN16;refGene_tx=XM_023241013.1
A2	8571988	8572038	refGene_gn=TSPAN16;refGene_tx=XM_023241013.1
A2	8572039	8572094	refGene_gn=TSPAN16;refGene_tx=XM_023241013.1
A2	8573953	8574055	refGene_gn=TSPAN16;refGene_tx=XM_023241013.1
A2	8580173	8580454
refGene_gn=RAB3D;refGene_tx=XM_019817741.2;refGene_gn=RAB3D;refGene_tx
=XM_019817745.2
A2	8582610	8582798
refGene_gn=RAB3D;refGene_tx=XM_003981881.5;refGene_gn=RAB3D;refGene_tx
=XM_019817741.2;refGene_gn=RAB3D;refGene_tx=XM_019817745.2
A2	8588161	8588286
refGene_gn=RAB3D;refGene_tx=XM_003981881.5;refGene_gn=RAB3D;refGene_tx
=XM_019817741.2;refGene_gn=RAB3D;refGene_tx=XM_019817745.2
A2	8588374	8588493
refGene_gn=RAB3D;refGene_tx=XM_003981881.5;refGene_gn=RAB3D;refGene_tx
=XM_019817741.2;refGene_gn=RAB3D;refGene_tx=XM_019817745.2

A2 8589139 8589367
refGene_gn=RAB3D;refGene_tx=XM_003981881.5;refGene_gn=RAB3D;refGene_tx=XM_019817741.2;refGene_gn=RAB3D;refGene_tx=XM_019817745.2
A2 8592187 8592493
refGene_gn=TMEM205;refGene_tx=XM_003981882.5;refGene_gn=TMEM205;refGene_tx=XM_006928425.4;refGene_gn=TMEM205;refGene_tx=XM_006928426.4;refGene_gn=TMEM205;refGene_tx=XM_006928427.3;refGene_gn=TMEM205;refGene_tx=XM_019817816.2;refGene_gn=TMEM205;refGene_tx=XM_023245574.1
A2 8593528 8593692
refGene_gn=TMEM205;refGene_tx=XM_003981882.5;refGene_gn=TMEM205;refGene_tx=XM_006928425.4;refGene_gn=TMEM205;refGene_tx=XM_006928426.4;refGene_gn=TMEM205;refGene_tx=XM_006928427.3;refGene_gn=TMEM205;refGene_tx=XM_019817816.2;refGene_gn=TMEM205;refGene_tx=XM_023245574.1
A2 8593797 8593906
refGene_gn=TMEM205;refGene_tx=XM_003981882.5;refGene_gn=TMEM205;refGene_tx=XM_006928425.4;refGene_gn=TMEM205;refGene_tx=XM_006928426.4;refGene_gn=TMEM205;refGene_tx=XM_006928427.3;refGene_gn=TMEM205;refGene_tx=XM_019817816.2;refGene_gn=TMEM205;refGene_tx=XM_023245574.1
A2 8594067 8594079 refGene_gn=TMEM205;refGene_tx=XM_003981882.5
A2 8594248 8594269 refGene_gn=TMEM205;refGene_tx=XM_006928425.4
A2 8594790 8594884
refGene_gn=CCDC159;refGene_tx=XM_019817766.1;refGene_gn=CCDC159;refGene_tx=XM_019817768.1;refGene_gn=CCDC159;refGene_tx=XM_019817771.1;refGene_gn=CCDC159;refGene_tx=XM_019817777.1;refGene_gn=CCDC159;refGene_tx=XM_019817782.1;refGene_gn=CCDC159;refGene_tx=XM_019817793.1;refGene_gn=CCDC159;refGene_tx=XM_019817796.1
A2 8596311 8596427
refGene_gn=CCDC159;refGene_tx=XM_019817766.1;refGene_gn=CCDC159;refGene_tx=XM_019817768.1;refGene_gn=CCDC159;refGene_tx=XM_019817771.1;refGene_gn=CCDC159;refGene_tx=XM_019817777.1;refGene_gn=CCDC159;refGene_tx=XM_019817782.1;refGene_gn=CCDC159;refGene_tx=XM_019817793.1
A2 8596538 8596762
refGene_gn=CCDC159;refGene_tx=XM_019817766.1;refGene_gn=CCDC159;refGene_tx=XM_019817768.1;refGene_gn=CCDC159;refGene_tx=XM_019817771.1;refGene_gn=CCDC159;refGene_tx=XM_019817777.1;refGene_gn=CCDC159;refGene_tx=XM_019817782.1;refGene_gn=CCDC159;refGene_tx=XM_019817793.1;refGene_gn=CCDC159;refGene_tx=XM_019817796.1
A2 8597735 8597769
refGene_gn=CCDC159;refGene_tx=XM_019817766.1;refGene_gn=CCDC159;refGene_tx=XM_019817768.1;refGene_gn=CCDC159;refGene_tx=XM_019817771.1;refGene_gn=CCDC159;refGene_tx=XM_019817777.1;refGene_gn=CCDC159;refGene_tx=XM_019817782.1;refGene_gn=CCDC159;refGene_tx=XM_019817793.1;refGene_gn=CCDC159;refGene_tx=XM_019817796.1
A2 8598059 8598148
refGene_gn=CCDC159;refGene_tx=XM_019817766.1;refGene_gn=CCDC159;refGene_tx=XM_019817768.1;refGene_gn=CCDC159;refGene_tx=XM_019817771.1;refGene_gn=CCDC159;refGene_tx=XM_019817777.1;refGene_gn=CCDC159;refGene_tx=XM_019817782.1;refGene_gn=CCDC159;refGene_tx=XM_019817793.1;refGene_gn=CCDC159;refGene_tx=XM_019817796.1
A2 8598239 8598321

refGene_gn=CCDC159;refGene_tx=XM_019817766.1;refGene_gn=CCDC159;refGene_tx=XM_019817768.1;refGene_gn=CCDC159;refGene_tx=XM_019817771.1;refGene_gn=CCDC159;refGene_tx=XM_019817777.1;refGene_gn=CCDC159;refGene_tx=XM_019817782.1;refGene_gn=CCDC159;refGene_tx=XM_019817793.1;refGene_gn=CCDC159;refGene_tx=XM_019817796.1
A2 8598501 8598709
refGene_gn=CCDC159;refGene_tx=XM_019817766.1;refGene_gn=CCDC159;refGene_tx=XM_019817768.1;refGene_gn=CCDC159;refGene_tx=XM_019817771.1;refGene_gn=CCDC159;refGene_tx=XM_019817777.1;refGene_gn=CCDC159;refGene_tx=XM_019817782.1;refGene_gn=CCDC159;refGene_tx=XM_019817793.1;refGene_gn=CCDC159;refGene_tx=XM_019817796.1
A2 8599332 8599418
refGene_gn=CCDC159;refGene_tx=XM_019817766.1;refGene_gn=CCDC159;refGene_tx=XM_019817768.1;refGene_gn=CCDC159;refGene_tx=XM_019817771.1;refGene_gn=CCDC159;refGene_tx=XM_019817777.1;refGene_gn=CCDC159;refGene_tx=XM_019817782.1;refGene_gn=CCDC159;refGene_tx=XM_019817793.1;refGene_gn=CCDC159;refGene_tx=XM_019817796.1
A2 8599496 8599573
refGene_gn=CCDC159;refGene_tx=XM_019817766.1;refGene_gn=CCDC159;refGene_tx=XM_019817768.1;refGene_gn=CCDC159;refGene_tx=XM_019817771.1;refGene_gn=CCDC159;refGene_tx=XM_019817777.1;refGene_gn=CCDC159;refGene_tx=XM_019817782.1;refGene_gn=CCDC159;refGene_tx=XM_019817793.1;refGene_gn=CCDC159;refGene_tx=XM_019817796.1
A2 8599916 8600038
refGene_gn=CCDC159;refGene_tx=XM_019817766.1;refGene_gn=CCDC159;refGene_tx=XM_019817768.1;refGene_gn=CCDC159;refGene_tx=XM_019817771.1;refGene_gn=CCDC159;refGene_tx=XM_019817777.1;refGene_gn=CCDC159;refGene_tx=XM_019817782.1;refGene_gn=CCDC159;refGene_tx=XM_019817793.1;refGene_gn=CCDC159;refGene_tx=XM_019817796.1
A2 8600266 8600349
refGene_gn=CCDC159;refGene_tx=XM_019817766.1;refGene_gn=CCDC159;refGene_tx=XM_019817768.1;refGene_gn=CCDC159;refGene_tx=XM_019817771.1;refGene_gn=CCDC159;refGene_tx=XM_019817777.1;refGene_gn=CCDC159;refGene_tx=XM_019817782.1;refGene_gn=CCDC159;refGene_tx=XM_019817793.1;refGene_gn=CCDC159;refGene_tx=XM_019817796.1
A2 8600700 8600814
refGene_gn=CCDC159;refGene_tx=XM_019817766.1;refGene_gn=CCDC159;refGene_tx=XM_019817768.1;refGene_gn=CCDC159;refGene_tx=XM_019817771.1;refGene_gn=CCDC159;refGene_tx=XM_019817777.1;refGene_gn=CCDC159;refGene_tx=XM_019817782.1;refGene_gn=CCDC159;refGene_tx=XM_019817793.1;refGene_gn=CCDC159;refGene_tx=XM_019817796.1
A2 8600967 8600972
refGene_gn=CCDC159;refGene_tx=XM_019817766.1;refGene_gn=CCDC159;refGene_tx=XM_019817768.1;refGene_gn=CCDC159;refGene_tx=XM_019817771.1;refGene_gn=CCDC159;refGene_tx=XM_019817777.1;refGene_gn=CCDC159;refGene_tx=XM_019817782.1;refGene_gn=CCDC159;refGene_tx=XM_019817793.1;refGene_gn=CCDC159;refGene_tx=XM_019817796.1
A2 8601530 8601752
refGene_gn=PLPPR2;refGene_tx=XM_006928431.4;refGene_gn=PLPPR2;refGene_tx=XM_006928435.4

A2	8602945	8603066
refGene_gn=PLPPR2;refGene_tx=XM_006928431.4;refGene_gn=PLPPR2;refGene_tx=XM_006928433.3;refGene_gn=PLPPR2;refGene_tx=XM_006928435.4
A2	8603873	8603953
refGene_gn=PLPPR2;refGene_tx=XM_006928431.4;refGene_gn=PLPPR2;refGene_tx=XM_006928433.3;refGene_gn=PLPPR2;refGene_tx=XM_006928435.4;refGene_gn=PLPPR2;refGene_tx=XM_006928436.3
A2	8604622	8604811
refGene_gn=PLPPR2;refGene_tx=XM_006928431.4;refGene_gn=PLPPR2;refGene_tx=XM_006928433.3;refGene_gn=PLPPR2;refGene_tx=XM_006928435.4;refGene_gn=PLPPR2;refGene_tx=XM_006928436.3
A2	8604930	8605066
refGene_gn=PLPPR2;refGene_tx=XM_006928431.4;refGene_gn=PLPPR2;refGene_tx=XM_006928433.3;refGene_gn=PLPPR2;refGene_tx=XM_006928435.4;refGene_gn=PLPPR2;refGene_tx=XM_006928436.3
A2	8605900	8606172
refGene_gn=PLPPR2;refGene_tx=XM_006928431.4;refGene_gn=PLPPR2;refGene_tx=XM_006928433.3;refGene_gn=PLPPR2;refGene_tx=XM_006928435.4;refGene_gn=PLPPR2;refGene_tx=XM_006928436.3
A2	8607096	8607273
refGene_gn=PLPPR2;refGene_tx=XM_006928431.4;refGene_gn=PLPPR2;refGene_tx=XM_006928433.3;refGene_gn=PLPPR2;refGene_tx=XM_006928435.4;refGene_gn=PLPPR2;refGene_tx=XM_006928436.3
A2	8607845	8607968
refGene_gn=PLPPR2;refGene_tx=XM_006928431.4;refGene_gn=PLPPR2;refGene_tx=XM_006928433.3;refGene_gn=PLPPR2;refGene_tx=XM_006928435.4;refGene_gn=PLPPR2;refGene_tx=XM_006928436.3
A2	8608268	8608320
refGene_gn=PLPPR2;refGene_tx=XM_006928431.4;refGene_gn=PLPPR2;refGene_tx=XM_006928433.3;refGene_gn=PLPPR2;refGene_tx=XM_006928435.4;refGene_gn=PLPPR2;refGene_tx=XM_006928436.3
A2	8608453	8608797
refGene_gn=PLPPR2;refGene_tx=XM_006928431.4;refGene_gn=PLPPR2;refGene_tx=XM_006928433.3;refGene_gn=PLPPR2;refGene_tx=XM_006928435.4;refGene_gn=PLPPR2;refGene_tx=XM_006928436.3
A2	8614655	8614904
refGene_gn=SWSAP1;refGene_tx=XM_003981885.4;refGene_gn=SWSAP1;refGene_tx=XM_019817826.2
A2	8615230	8615290	refGene_gn=SWSAP1;refGene_tx=XM_019817826.2
A2	8615631	8616135
refGene_gn=SWSAP1;refGene_tx=XM_003981885.4;refGene_gn=SWSAP1;refGene_tx=XM_019817826.2
A2	8617766	8618375	refGene_gn=EPOR;refGene_tx=XM_023245578.1
A2	8618468	8618556	refGene_gn=EPOR;refGene_tx=XM_023245578.1
A2	8620503	8620591	refGene_gn=EPOR;refGene_tx=XM_023245578.1
A2	8620674	8620828	refGene_gn=EPOR;refGene_tx=XM_023245578.1
A2	8621334	8621492	refGene_gn=EPOR;refGene_tx=XM_023245578.1
A2	8621569	8621748	refGene_gn=EPOR;refGene_tx=XM_023245578.1
A2	8622147	8622263	refGene_gn=EPOR;refGene_tx=XM_023245578.1
A2	8622264	8622284	refGene_gn=EPOR;refGene_tx=XM_023245578.1

A2	8623155	8623270	refGene_gn=EPOR;refGene_tx=XM_023245578.1
A2	8628067	8628186	refGene_gn=RGL3;refGene_tx=XM_003981902.5
A2	8629169	8629284	refGene_gn=RGL3;refGene_tx=XM_003981902.5
A2	8629370	8629523
refGene_gn=RGL3;refGene_tx=XM_003981902.5;refGene_gn=RGL3;refGene_tx=XM_019817837.2
A2	8630160	8630257
refGene_gn=RGL3;refGene_tx=XM_003981902.5;refGene_gn=RGL3;refGene_tx=XM_019817837.2
A2	8630345	8630414
refGene_gn=RGL3;refGene_tx=XM_003981902.5;refGene_gn=RGL3;refGene_tx=XM_019817837.2
A2	8630515	8630611
refGene_gn=RGL3;refGene_tx=XM_003981902.5;refGene_gn=RGL3;refGene_tx=XM_019817837.2
A2	8631311	8631433
refGene_gn=RGL3;refGene_tx=XM_003981902.5;refGene_gn=RGL3;refGene_tx=XM_019817837.2
A2	8631513	8631546
refGene_gn=RGL3;refGene_tx=XM_003981902.5;refGene_gn=RGL3;refGene_tx=XM_019817837.2
A2	8631748	8631835
refGene_gn=RGL3;refGene_tx=XM_003981902.5;refGene_gn=RGL3;refGene_tx=XM_019817837.2
A2	8631943	8632000
refGene_gn=RGL3;refGene_tx=XM_003981902.5;refGene_gn=RGL3;refGene_tx=XM_019817837.2
A2	8633139	8633224
refGene_gn=RGL3;refGene_tx=XM_003981902.5;refGene_gn=RGL3;refGene_tx=XM_019817837.2
A2	8633357	8633461
refGene_gn=RGL3;refGene_tx=XM_003981902.5;refGene_gn=RGL3;refGene_tx=XM_019817837.2
A2	8634172	8634388
refGene_gn=RGL3;refGene_tx=XM_003981902.5;refGene_gn=RGL3;refGene_tx=XM_019817837.2
A2	8634475	8634618
refGene_gn=RGL3;refGene_tx=XM_003981902.5;refGene_gn=RGL3;refGene_tx=XM_019817837.2
A2	8638411	8638623
refGene_gn=RGL3;refGene_tx=XM_003981902.5;refGene_gn=RGL3;refGene_tx=XM_019817837.2
A2	8638740	8638794
refGene_gn=RGL3;refGene_tx=XM_003981902.5;refGene_gn=RGL3;refGene_tx=XM_019817837.2
A2	8638922	8639146
refGene_gn=RGL3;refGene_tx=XM_003981902.5;refGene_gn=RGL3;refGene_tx=XM_019817837.2
A2	8639790	8639904
refGene_gn=RGL3;refGene_tx=XM_003981902.5;refGene_gn=RGL3;refGene_tx=X

M_019817837.2
A2	8640371	8640404
refGene_gn=RGL3;refGene_tx=XM_003981902.5;refGene_gn=RGL3;refGene_tx=X
M_019817837.2
A2	8641208	8641327	refGene_gn=CCDC151;refGene_tx=XM_006928437.4
A2	8641494	8641579	refGene_gn=CCDC151;refGene_tx=XM_006928437.4
A2	8641989	8642145	refGene_gn=CCDC151;refGene_tx=XM_006928437.4
A2	8642754	8642911	refGene_gn=CCDC151;refGene_tx=XM_006928437.4
A2	8642992	8643153	refGene_gn=CCDC151;refGene_tx=XM_006928437.4
A2	8644111	8644264	refGene_gn=CCDC151;refGene_tx=XM_006928437.4
A2	8644752	8644875	refGene_gn=CCDC151;refGene_tx=XM_006928437.4
A2	8645030	8645156	refGene_gn=CCDC151;refGene_tx=XM_006928437.4
A2	8645257	8645359	refGene_gn=CCDC151;refGene_tx=XM_006928437.4
A2	8645457	8645628	refGene_gn=CCDC151;refGene_tx=XM_006928437.4
A2	8646820	8646898	refGene_gn=CCDC151;refGene_tx=XM_006928437.4
A2	8646998	8647072	refGene_gn=CCDC151;refGene_tx=XM_006928437.4
A2	8647074	8647119	refGene_gn=CCDC151;refGene_tx=XM_006928437.4
A2	8648353	8648594	refGene_gn=CCDC151;refGene_tx=XM_006928437.4
A2	8649867	8649943
refGene_gn=PRKCSH;refGene_tx=XM_023245611.1;refGene_gn=PRKCSH;refGene_
tx=XM_023245613.1
A2	8650127	8650244
refGene_gn=PRKCSH;refGene_tx=XM_023245611.1;refGene_gn=PRKCSH;refGene_
tx=XM_023245613.1
A2	8650867	8650963
refGene_gn=PRKCSH;refGene_tx=XM_023245611.1;refGene_gn=PRKCSH;refGene_
tx=XM_023245613.1
A2	8651075	8651133
refGene_gn=PRKCSH;refGene_tx=XM_023245611.1;refGene_gn=PRKCSH;refGene_
tx=XM_023245613.1
A2	8652016	8652134
refGene_gn=PRKCSH;refGene_tx=XM_023245611.1;refGene_gn=PRKCSH;refGene_
tx=XM_023245613.1
A2	8653604	8653734
refGene_gn=PRKCSH;refGene_tx=XM_023245611.1;refGene_gn=PRKCSH;refGene_
tx=XM_023245613.1
A2	8656299	8656384
refGene_gn=PRKCSH;refGene_tx=XM_023245611.1;refGene_gn=PRKCSH;refGene_
tx=XM_023245613.1
A2	8657001	8657080
refGene_gn=PRKCSH;refGene_tx=XM_023245611.1;refGene_gn=PRKCSH;refGene_
tx=XM_023245613.1
A2	8657756	8657843
refGene_gn=PRKCSH;refGene_tx=XM_023245611.1;refGene_gn=PRKCSH;refGene_
tx=XM_023245613.1
A2	8658143	8658332
refGene_gn=PRKCSH;refGene_tx=XM_023245611.1;refGene_gn=PRKCSH;refGene_
tx=XM_023245613.1
A2	8658404	8658504
refGene_gn=PRKCSH;refGene_tx=XM_023245611.1;refGene_gn=PRKCSH;refGene_

tx=XM_023245613.1
A2 8658962 8659032
refGene_gn=PRKCSH;refGene_tx=XM_023245611.1;refGene_gn=PRKCSH;refGene_tx=XM_023245613.1
A2 8659273 8659363
refGene_gn=PRKCSH;refGene_tx=XM_023245611.1;refGene_gn=PRKCSH;refGene_tx=XM_023245613.1
A2 8659545 8659620
refGene_gn=PRKCSH;refGene_tx=XM_023245611.1;refGene_gn=PRKCSH;refGene_tx=XM_023245613.1
A2 8659712 8659812
refGene_gn=PRKCSH;refGene_tx=XM_023245611.1;refGene_gn=PRKCSH;refGene_tx=XM_023245613.1
A2 8659895 8660042
refGene_gn=PRKCSH;refGene_tx=XM_023245611.1;refGene_gn=PRKCSH;refGene_tx=XM_023245613.1
A2 8664118 8664470
refGene_gn=ELAVL3;refGene_tx=XM_023245678.1;refGene_gn=ELAVL3;refGene_tx=XM_023245687.1;refGene_gn=ELAVL3;refGene_tx=XM_023245692.1;refGene_gn=ELAVL3;refGene_tx=XM_023245693.1
A2 8666075 8666114
refGene_gn=ELAVL3;refGene_tx=XM_023245678.1;refGene_gn=ELAVL3;refGene_tx=XM_023245687.1;refGene_gn=ELAVL3;refGene_tx=XM_023245692.1;refGene_gn=ELAVL3;refGene_tx=XM_023245693.1
A2 8666788 8667014
refGene_gn=ELAVL3;refGene_tx=XM_023245678.1;refGene_gn=ELAVL3;refGene_tx=XM_023245687.1;refGene_gn=ELAVL3;refGene_tx=XM_023245692.1;refGene_gn=ELAVL3;refGene_tx=XM_023245693.1
A2 8667157 8667311
refGene_gn=ELAVL3;refGene_tx=XM_023245678.1;refGene_gn=ELAVL3;refGene_tx=XM_023245687.1;refGene_gn=ELAVL3;refGene_tx=XM_023245692.1;refGene_gn=ELAVL3;refGene_tx=XM_023245693.1
A2 8670831 8670935
refGene_gn=ELAVL3;refGene_tx=XM_023245678.1;refGene_gn=ELAVL3;refGene_tx=XM_023245687.1;refGene_gn=ELAVL3;refGene_tx=XM_023245692.1;refGene_gn=ELAVL3;refGene_tx=XM_023245693.1
A2 8671258 8671478
refGene_gn=ELAVL3;refGene_tx=XM_023245678.1;refGene_gn=ELAVL3;refGene_tx=XM_023245687.1;refGene_gn=ELAVL3;refGene_tx=XM_023245692.1;refGene_gn=ELAVL3;refGene_tx=XM_023245693.1
A2 8677495 8677504
refGene_gn=ELAVL3;refGene_tx=XM_023245678.1;refGene_gn=ELAVL3;refGene_tx=XM_023245687.1;refGene_gn=ELAVL3;refGene_tx=XM_023245692.1
A2 8680621 8680799
refGene_gn=ZNF653;refGene_tx=XM_023245623.1;refGene_gn=ZNF653;refGene_tx=XM_023245624.1
A2 8681036 8681136
refGene_gn=ZNF653;refGene_tx=XM_023245623.1;refGene_gn=ZNF653;refGene_tx=XM_023245624.1
A2 8681661 8681776

refGene_gn=ZNF653;refGene_tx=XM_023245623.1;refGene_gn=ZNF653;refGene_tx=XM_023245624.1
A2 8682634 8682709 refGene_gn=ZNF653;refGene_tx=XM_023245626.1
A2 8682859 8682971
refGene_gn=ZNF653;refGene_tx=XM_023245623.1;refGene_gn=ZNF653;refGene_tx=XM_023245624.1;refGene_gn=ZNF653;refGene_tx=XM_023245626.1
A2 8683072 8683244
refGene_gn=ZNF653;refGene_tx=XM_023245623.1;refGene_gn=ZNF653;refGene_tx=XM_023245624.1;refGene_gn=ZNF653;refGene_tx=XM_023245626.1
A2 8683384 8683999
refGene_gn=ZNF653;refGene_tx=XM_023245623.1;refGene_gn=ZNF653;refGene_tx=XM_023245624.1;refGene_gn=ZNF653;refGene_tx=XM_023245626.1
A2 8688410 8688626
refGene_gn=ZNF653;refGene_tx=XM_023245623.1;refGene_gn=ZNF653;refGene_tx=XM_023245624.1;refGene_gn=ZNF653;refGene_tx=XM_023245626.1
A2 8690408 8690452
refGene_gn=ZNF653;refGene_tx=XM_023245623.1;refGene_gn=ZNF653;refGene_tx=XM_023245624.1;refGene_gn=ZNF653;refGene_tx=XM_023245626.1
A2 8691913 8691993 refGene_gn=ZNF653;refGene_tx=XM_023245624.1
A2 8696312 8696611
refGene_gn=ZNF653;refGene_tx=XM_023245623.1;refGene_gn=ZNF653;refGene_tx=XM_023245626.1
A2 8697193 8697444
refGene_gn=ECSIT;refGene_tx=XM_003981889.4;refGene_gn=ECSIT;refGene_tx=XM_006928438.4;refGene_gn=ECSIT;refGene_tx=XM_006928439.3;refGene_gn=ECSIT;refGene_tx=XM_006928440.4
A2 8698004 8698110
refGene_gn=ECSIT;refGene_tx=XM_003981889.4;refGene_gn=ECSIT;refGene_tx=XM_006928438.4;refGene_gn=ECSIT;refGene_tx=XM_006928439.3;refGene_gn=ECSIT;refGene_tx=XM_006928440.4
A2 8698309 8698458
refGene_gn=ECSIT;refGene_tx=XM_003981889.4;refGene_gn=ECSIT;refGene_tx=XM_006928438.4;refGene_gn=ECSIT;refGene_tx=XM_006928439.3;refGene_gn=ECSIT;refGene_tx=XM_006928440.4
A2 8698574 8698632
refGene_gn=ECSIT;refGene_tx=XM_003981889.4;refGene_gn=ECSIT;refGene_tx=XM_006928438.4;refGene_gn=ECSIT;refGene_tx=XM_006928439.3;refGene_gn=ECSIT;refGene_tx=XM_006928440.4
A2 8702204 8702428
refGene_gn=ECSIT;refGene_tx=XM_003981889.4;refGene_gn=ECSIT;refGene_tx=XM_006928438.4;refGene_gn=ECSIT;refGene_tx=XM_006928439.3;refGene_gn=ECSIT;refGene_tx=XM_006928440.4
A2 8702821 8703239
refGene_gn=ECSIT;refGene_tx=XM_003981889.4;refGene_gn=ECSIT;refGene_tx=XM_006928438.4;refGene_gn=ECSIT;refGene_tx=XM_006928439.3;refGene_gn=ECSIT;refGene_tx=XM_006928440.4
A2 8706906 8707002
refGene_gn=ECSIT;refGene_tx=XM_003981889.4;refGene_gn=ECSIT;refGene_tx=XM_006928438.4;refGene_gn=ECSIT;refGene_tx=XM_006928439.3;refGene_gn=ECSIT;refGene_tx=XM_006928440.4

A2 8723918 8723981
refGene_gn=CNN1;refGene_tx=XM_023245709.1;refGene_gn=CNN1;refGene_tx=X
M_023245711.1;refGene_gn=CNN1;refGene_tx=XM_023245717.1
A2 8726043 8726165
refGene_gn=CNN1;refGene_tx=XM_023245709.1;refGene_gn=CNN1;refGene_tx=X
M_023245711.1;refGene_gn=CNN1;refGene_tx=XM_023245717.1
A2 8728517 8728584
refGene_gn=CNN1;refGene_tx=XM_023245709.1;refGene_gn=CNN1;refGene_tx=X
M_023245711.1;refGene_gn=CNN1;refGene_tx=XM_023245717.1
A2 8728696 8728834
refGene_gn=CNN1;refGene_tx=XM_023245709.1;refGene_gn=CNN1;refGene_tx=X
M_023245711.1;refGene_gn=CNN1;refGene_tx=XM_023245717.1
A2 8729014 8729125
refGene_gn=CNN1;refGene_tx=XM_023245709.1;refGene_gn=CNN1;refGene_tx=X
M_023245711.1;refGene_gn=CNN1;refGene_tx=XM_023245717.1
A2 8729593 8729740
refGene_gn=CNN1;refGene_tx=XM_023245709.1;refGene_gn=CNN1;refGene_tx=X
M_023245711.1;refGene_gn=CNN1;refGene_tx=XM_023245717.1
A2 8729837 8730083
refGene_gn=CNN1;refGene_tx=XM_023245709.1;refGene_gn=CNN1;refGene_tx=X
M_023245711.1;refGene_gn=CNN1;refGene_tx=XM_023245717.1
A2 8730284 8730285 refGene_gn=CNN1;refGene_tx=XM_023245717.1
A2 8734301 8734371
refGene_gn=ELOF1;refGene_tx=XM_006928443.3;refGene_gn=ELOF1;refGene_tx
=XM_006928444.4;refGene_gn=ELOF1;refGene_tx=XM_011289233.3;refGene_gn=
ELOF1;refGene_tx=XM_011289243.3;refGene_gn=ELOF1;refGene_tx=XM_0198179
10.1
A2 8734678 8734749
refGene_gn=ELOF1;refGene_tx=XM_006928443.3;refGene_gn=ELOF1;refGene_tx
=XM_006928444.4;refGene_gn=ELOF1;refGene_tx=XM_011289233.3;refGene_gn=
ELOF1;refGene_tx=XM_011289243.3;refGene_gn=ELOF1;refGene_tx=XM_0198179
10.1
A2 8734906 8735022
refGene_gn=ELOF1;refGene_tx=XM_006928443.3;refGene_gn=ELOF1;refGene_tx
=XM_006928444.4;refGene_gn=ELOF1;refGene_tx=XM_011289233.3;refGene_gn=
ELOF1;refGene_tx=XM_011289243.3;refGene_gn=ELOF1;refGene_tx=XM_0198179
10.1
A2 8744786 8745059
refGene_gn=ACP5;refGene_tx=XM_003981893.5;refGene_gn=ACP5;refGene_tx=X
M_023245702.1;refGene_gn=ACP5;refGene_tx=XM_023245705.1
A2 8745455 8745801
refGene_gn=ACP5;refGene_tx=XM_003981893.5;refGene_gn=ACP5;refGene_tx=X
M_023245702.1;refGene_gn=ACP5;refGene_tx=XM_023245705.1
A2 8745944 8746072
refGene_gn=ACP5;refGene_tx=XM_003981893.5;refGene_gn=ACP5;refGene_tx=X
M_023245702.1;refGene_gn=ACP5;refGene_tx=XM_023245705.1
A2 8746280 8746541
refGene_gn=ACP5;refGene_tx=XM_003981893.5;refGene_gn=ACP5;refGene_tx=X
M_023245702.1;refGene_gn=ACP5;refGene_tx=XM_023245705.1
A2 8809971 8811625

refGene_gn=LOC102899880;refGene_tx=XM_023245634.1
A2	8812426	8812487
refGene_gn=LOC102899880;refGene_tx=XM_023245634.1
A2	8812906	8813033
refGene_gn=LOC102899880;refGene_tx=XM_023245634.1
A2	8821145	8821148
refGene_gn=LOC102899880;refGene_tx=XM_023245634.1
A2	8864183	8865921
refGene_gn=LOC101082380;refGene_tx=XM_019817923.2
A2	8866857	8866915
refGene_gn=LOC101082380;refGene_tx=XM_019817923.2
A2	8867137	8867264
refGene_gn=LOC101082380;refGene_tx=XM_019817923.2
A2	8883174	8883177
refGene_gn=LOC101082380;refGene_tx=XM_019817923.2
A2	8892007	8892078
refGene_gn=LOC101082633;refGene_tx=XM_023245661.1
A2	8914237	8914257
refGene_gn=LOC101082633;refGene_tx=XM_019817964.2;refGene_gn=LOC101082633;refGene_tx=XM_019817967.2;refGene_gn=LOC101082633;refGene_tx=XM_019817973.2;refGene_gn=LOC101082633;refGene_tx=XM_019817977.2
A2	8914570	8916134
refGene_gn=LOC101082633;refGene_tx=XM_011289265.3;refGene_gn=LOC101082633;refGene_tx=XM_019817952.2;refGene_gn=LOC101082633;refGene_tx=XM_019817959.2;refGene_gn=LOC101082633;refGene_tx=XM_019817964.2;refGene_gn=LOC101082633;refGene_tx=XM_019817967.2;refGene_gn=LOC101082633;refGene_tx=XM_019817973.2;refGene_gn=LOC101082633;refGene_tx=XM_019817977.2;refGene_gn=LOC101082633;refGene_tx=XM_023245647.1;refGene_gn=LOC101082633;refGene_tx=XM_023245661.1
A2	8916650	8916723
refGene_gn=LOC101082633;refGene_tx=XM_011289265.3;refGene_gn=LOC101082633;refGene_tx=XM_019817952.2;refGene_gn=LOC101082633;refGene_tx=XM_019817959.2;refGene_gn=LOC101082633;refGene_tx=XM_019817964.2;refGene_gn=LOC101082633;refGene_tx=XM_019817967.2;refGene_gn=LOC101082633;refGene_tx=XM_019817973.2;refGene_gn=LOC101082633;refGene_tx=XM_019817977.2;refGene_gn=LOC101082633;refGene_tx=XM_023245647.1;refGene_gn=LOC101082633;refGene_tx=XM_023245661.1
A2	8916917	8917044
refGene_gn=LOC101082633;refGene_tx=XM_011289265.3;refGene_gn=LOC101082633;refGene_tx=XM_019817952.2;refGene_gn=LOC101082633;refGene_tx=XM_019817959.2;refGene_gn=LOC101082633;refGene_tx=XM_019817964.2;refGene_gn=LOC101082633;refGene_tx=XM_019817967.2;refGene_gn=LOC101082633;refGene_tx=XM_019817973.2;refGene_gn=LOC101082633;refGene_tx=XM_019817977.2;refGene_gn=LOC101082633;refGene_tx=XM_023245647.1;refGene_gn=LOC101082633;refGene_tx=XM_023245661.1
A2	8930536	8930539
refGene_gn=LOC101082633;refGene_tx=XM_011289265.3;refGene_gn=LOC101082633;refGene_tx=XM_019817952.2;refGene_gn=LOC101082633;refGene_tx=XM_019817964.2;refGene_gn=LOC101082633;refGene_tx=XM_019817967.2;refGene_gn=LOC101082633;refGene_tx=XM_019817973.2;refGene_gn=LOC101082633;refGen

e_tx=XM_019817977.2;refGene_gn=LOC101082633;refGene_tx=XM_023245661.1
A2	8941466	8941469
refGene_gn=LOC101083149;refGene_tx=XM_023245737.1
A2	8962317	8962320
refGene_gn=LOC101083149;refGene_tx=XM_023245736.1
A2	8982413	8982540
refGene_gn=LOC101083149;refGene_tx=XM_023245736.1;refGene_gn=LOC101083149;refGene_tx=XM_023245737.1;refGene_gn=LOC101083149;refGene_tx=XM_023245738.1
A2	8982725	8982786
refGene_gn=LOC101083149;refGene_tx=XM_023245736.1;refGene_gn=LOC101083149;refGene_tx=XM_023245737.1;refGene_gn=LOC101083149;refGene_tx=XM_023245738.1
A2	8983387	8985041
refGene_gn=LOC101083149;refGene_tx=XM_023245736.1;refGene_gn=LOC101083149;refGene_tx=XM_023245737.1;refGene_gn=LOC101083149;refGene_tx=XM_023245738.1
A2	9005320	9005430
refGene_gn=MAN2B1;refGene_tx=NM_001009222.1;refGene_gn=MAN2B1;refGene_tx=XM_006928455.3
A2	9005903	9006006
refGene_gn=MAN2B1;refGene_tx=NM_001009222.1;refGene_gn=MAN2B1;refGene_tx=XM_006928455.3
A2	9006114	9006267
refGene_gn=MAN2B1;refGene_tx=NM_001009222.1;refGene_gn=MAN2B1;refGene_tx=XM_006928455.3
A2	9006767	9006995
refGene_gn=MAN2B1;refGene_tx=NM_001009222.1;refGene_gn=MAN2B1;refGene_tx=XM_006928455.3
A2	9007398	9007479
refGene_gn=MAN2B1;refGene_tx=NM_001009222.1;refGene_gn=MAN2B1;refGene_tx=XM_006928455.3
A2	9007662	9007750
refGene_gn=MAN2B1;refGene_tx=NM_001009222.1;refGene_gn=MAN2B1;refGene_tx=XM_006928455.3
A2	9007928	9008030
refGene_gn=MAN2B1;refGene_tx=NM_001009222.1;refGene_gn=MAN2B1;refGene_tx=XM_006928455.3
A2	9008120	9008239
refGene_gn=MAN2B1;refGene_tx=NM_001009222.1;refGene_gn=MAN2B1;refGene_tx=XM_006928455.3
A2	9009730	9009848
refGene_gn=MAN2B1;refGene_tx=NM_001009222.1;refGene_gn=MAN2B1;refGene_tx=XM_006928455.3
A2	9009941	9010039
refGene_gn=MAN2B1;refGene_tx=NM_001009222.1;refGene_gn=MAN2B1;refGene_tx=XM_006928455.3
A2	9011045	9011219
refGene_gn=MAN2B1;refGene_tx=NM_001009222.1;refGene_gn=MAN2B1;refGene_tx=XM_006928455.3

A2 9012486 9012603
refGene_gn=MAN2B1;refGene_tx=NM_001009222.1;refGene_gn=MAN2B1;refGene_tx=XM_006928455.3
A2 9012922 9013030
refGene_gn=MAN2B1;refGene_tx=NM_001009222.1;refGene_gn=MAN2B1;refGene_tx=XM_006928455.3
A2 9013307 9013417
refGene_gn=MAN2B1;refGene_tx=NM_001009222.1;refGene_gn=MAN2B1;refGene_tx=XM_006928455.3
A2 9013603 9013682
refGene_gn=MAN2B1;refGene_tx=NM_001009222.1;refGene_gn=MAN2B1;refGene_tx=XM_006928455.3
A2 9013762 9013883
refGene_gn=MAN2B1;refGene_tx=NM_001009222.1;refGene_gn=MAN2B1;refGene_tx=XM_006928455.3
A2 9013984 9014070
refGene_gn=MAN2B1;refGene_tx=NM_001009222.1;refGene_gn=MAN2B1;refGene_tx=XM_006928455.3
A2 9016923 9017040
refGene_gn=MAN2B1;refGene_tx=NM_001009222.1;refGene_gn=MAN2B1;refGene_tx=XM_006928455.3
A2 9017946 9018092
refGene_gn=MAN2B1;refGene_tx=NM_001009222.1;refGene_gn=MAN2B1;refGene_tx=XM_006928455.3
A2 9018351 9018484
refGene_gn=MAN2B1;refGene_tx=NM_001009222.1;refGene_gn=MAN2B1;refGene_tx=XM_006928455.3
A2 9019152 9019346
refGene_gn=MAN2B1;refGene_tx=NM_001009222.1;refGene_gn=MAN2B1;refGene_tx=XM_006928455.3
A2 9019707 9019881
refGene_gn=MAN2B1;refGene_tx=NM_001009222.1;refGene_gn=MAN2B1;refGene_tx=XM_006928455.3
A2 9020057 9020160
refGene_gn=MAN2B1;refGene_tx=NM_001009222.1;refGene_gn=MAN2B1;refGene_tx=XM_006928455.3
A2 9020874 9021036
refGene_gn=MAN2B1;refGene_tx=NM_001009222.1;refGene_gn=MAN2B1;refGene_tx=XM_006928455.3
A2 9022287 9022354 refGene_gn=WDR83OS;refGene_tx=XM_003981906.4
A2 9022449 9022547 refGene_gn=WDR83OS;refGene_tx=XM_003981906.4
A2 9023059 9023165 refGene_gn=WDR83OS;refGene_tx=XM_003981906.4
A2 9023280 9023330 refGene_gn=WDR83OS;refGene_tx=XM_003981906.4
A2 9023715 9023818 refGene_gn=WDR83;refGene_tx=XM_003981909.4
A2 9023895 9024016 refGene_gn=WDR83;refGene_tx=XM_003981909.4
A2 9024098 9024204 refGene_gn=WDR83;refGene_tx=XM_003981909.4
A2 9024477 9024526 refGene_gn=WDR83;refGene_tx=XM_003981909.4
A2 9024599 9024726 refGene_gn=WDR83;refGene_tx=XM_003981909.4
A2 9025644 9025712 refGene_gn=WDR83;refGene_tx=XM_003981909.4
A2 9025814 9025923 refGene_gn=WDR83;refGene_tx=XM_003981909.4

A2 9026010 9026125 refGene_gn=WDR83;refGene_tx=XM_003981909.4
A2 9027095 9027245 refGene_gn=WDR83;refGene_tx=XM_003981909.4
A2 9027389 9027498
refGene_gn=DHPS;refGene_tx=XM_003981907.5;refGene_gn=DHPS;refGene_tx=XM_019818021.1;refGene_gn=DHPS;refGene_tx=XM_023245761.1
A2 9027569 9027695
refGene_gn=DHPS;refGene_tx=XM_003981907.5;refGene_gn=DHPS;refGene_tx=XM_019818021.1;refGene_gn=DHPS;refGene_tx=XM_023245761.1
A2 9028425 9028529
refGene_gn=DHPS;refGene_tx=XM_003981907.5;refGene_gn=DHPS;refGene_tx=XM_019818021.1;refGene_gn=DHPS;refGene_tx=XM_023245761.1
A2 9028608 9028714
refGene_gn=DHPS;refGene_tx=XM_003981907.5;refGene_gn=DHPS;refGene_tx=XM_019818021.1;refGene_gn=DHPS;refGene_tx=XM_023245761.1
A2 9029245 9029332
refGene_gn=DHPS;refGene_tx=XM_003981907.5;refGene_gn=DHPS;refGene_tx=XM_019818021.1;refGene_gn=DHPS;refGene_tx=XM_023245761.1
A2 9029412 9029509
refGene_gn=DHPS;refGene_tx=XM_003981907.5;refGene_gn=DHPS;refGene_tx=XM_019818021.1;refGene_gn=DHPS;refGene_tx=XM_023245761.1
A2 9029590 9029712
refGene_gn=DHPS;refGene_tx=XM_003981907.5;refGene_gn=DHPS;refGene_tx=XM_019818021.1;refGene_gn=DHPS;refGene_tx=XM_023245761.1
A2 9030149 9030314
refGene_gn=DHPS;refGene_tx=XM_003981907.5;refGene_gn=DHPS;refGene_tx=XM_019818021.1;refGene_gn=DHPS;refGene_tx=XM_023245761.1
A2 9030451 9030472 refGene_gn=DHPS;refGene_tx=XM_019818021.1
A2 9030891 9031098
refGene_gn=DHPS;refGene_tx=XM_003981907.5;refGene_gn=DHPS;refGene_tx=XM_023245761.1
A2 9034327 9034420
refGene_gn=GNG14;refGene_tx=XM_003981910.4;refGene_gn=GNG14;refGene_tx=XM_006928461.4;refGene_gn=GNG14;refGene_tx=XM_006928462.4;refGene_gn=GNG14;refGene_tx=XM_006928464.4;refGene_gn=GNG14;refGene_tx=XM_019818051.2
A2 9034502 9034628
refGene_gn=GNG14;refGene_tx=XM_003981910.4;refGene_gn=GNG14;refGene_tx=XM_006928461.4;refGene_gn=GNG14;refGene_tx=XM_006928462.4;refGene_gn=GNG14;refGene_tx=XM_006928464.4;refGene_gn=GNG14;refGene_tx=XM_019818051.2
A2 9035515 9035590
refGene_gn=FBXW9;refGene_tx=XM_003981911.5;refGene_gn=FBXW9;refGene_tx=XM_019817999.2
A2 9035663 9035729
refGene_gn=FBXW9;refGene_tx=XM_003981911.5;refGene_gn=FBXW9;refGene_tx=XM_019817999.2
A2 9035840 9035930
refGene_gn=FBXW9;refGene_tx=XM_003981911.5;refGene_gn=FBXW9;refGene_tx=XM_019817999.2
A2 9036060 9036174

refGene_gn=FBXW9;refGene_tx=XM_003981911.5;refGene_gn=FBXW9;refGene_tx=XM_019817999.2
A2 9036257 9036406
refGene_gn=FBXW9;refGene_tx=XM_003981911.5;refGene_gn=FBXW9;refGene_tx=XM_019817999.2
A2 9037247 9037339
refGene_gn=FBXW9;refGene_tx=XM_003981911.5;refGene_gn=FBXW9;refGene_tx=XM_019817999.2
A2 9037425 9037565
refGene_gn=FBXW9;refGene_tx=XM_003981911.5;refGene_gn=FBXW9;refGene_tx=XM_019817999.2
A2 9038988 9039117
refGene_gn=FBXW9;refGene_tx=XM_003981911.5;refGene_gn=FBXW9;refGene_tx=XM_019817999.2
A2 9039193 9039333
refGene_gn=FBXW9;refGene_tx=XM_003981911.5;refGene_gn=FBXW9;refGene_tx=XM_019817999.2
A2 9040485 9040894
refGene_gn=FBXW9;refGene_tx=XM_003981911.5;refGene_gn=FBXW9;refGene_tx=XM_019817999.2
A2 9044632 9044740 refGene_gn=TNPO2;refGene_tx=XM_023245800.1
A2 9044857 9044932 refGene_gn=TNPO2;refGene_tx=XM_023245800.1
A2 9045010 9045110 refGene_gn=TNPO2;refGene_tx=XM_023245800.1
A2 9045378 9045454 refGene_gn=TNPO2;refGene_tx=XM_023245800.1
A2 9045734 9045830 refGene_gn=TNPO2;refGene_tx=XM_023245800.1
A2 9046403 9046502 refGene_gn=TNPO2;refGene_tx=XM_023245800.1
A2 9046687 9046775 refGene_gn=TNPO2;refGene_tx=XM_023245800.1
A2 9047331 9047490 refGene_gn=TNPO2;refGene_tx=XM_023245800.1
A2 9047586 9047694 refGene_gn=TNPO2;refGene_tx=XM_023245800.1
A2 9047773 9047860 refGene_gn=TNPO2;refGene_tx=XM_023245800.1
A2 9048315 9048487 refGene_gn=TNPO2;refGene_tx=XM_023245800.1
A2 9048691 9048917 refGene_gn=TNPO2;refGene_tx=XM_023245800.1
A2 9050240 9050393 refGene_gn=TNPO2;refGene_tx=XM_023245800.1
A2 9050599 9050765 refGene_gn=TNPO2;refGene_tx=XM_023245800.1
A2 9050867 9050928 refGene_gn=TNPO2;refGene_tx=XM_023245800.1
A2 9052000 9052088 refGene_gn=TNPO2;refGene_tx=XM_023245800.1
A2 9052090 9052118 refGene_gn=TNPO2;refGene_tx=XM_023245800.1
A2 9052216 9052339 refGene_gn=TNPO2;refGene_tx=XM_023245800.1
A2 9052413 9052495 refGene_gn=TNPO2;refGene_tx=XM_023245800.1
A2 9052588 9052722 refGene_gn=TNPO2;refGene_tx=XM_023245800.1
A2 9052808 9052915 refGene_gn=TNPO2;refGene_tx=XM_023245800.1
A2 9055399 9055549 refGene_gn=TNPO2;refGene_tx=XM_023245800.1
A2 9055631 9055707 refGene_gn=TNPO2;refGene_tx=XM_023245800.1
A2 9056324 9056423 refGene_gn=TNPO2;refGene_tx=XM_023245800.1
A2 9063508 9063616 refGene_gn=TRIR;refGene_tx=XM_023245807.1
A2 9063966 9064044
refGene_gn=TRIR;refGene_tx=XM_023245807.1;refGene_gn=TRIR;refGene_tx=XM_023245815.1
A2 9066227 9066578
refGene_gn=TRIR;refGene_tx=XM_023245807.1;refGene_gn=TRIR;refGene_tx=X

M_023245815.1
A2	9068079	9068240	refGene_gn=ASNA1;refGene_tx=XM_003981913.4
A2	9069081	9069229	refGene_gn=ASNA1;refGene_tx=XM_003981913.4
A2	9072797	9072946	refGene_gn=ASNA1;refGene_tx=XM_003981913.4
A2	9073033	9073184	refGene_gn=ASNA1;refGene_tx=XM_003981913.4
A2	9073687	9073795	refGene_gn=ASNA1;refGene_tx=XM_003981913.4
A2	9073884	9074082	refGene_gn=ASNA1;refGene_tx=XM_003981913.4
A2	9074490	9074622	refGene_gn=ASNA1;refGene_tx=XM_003981913.4
A2	9078724	9078876
refGene_gn=BEST2;refGene_tx=XM_019818105.2;refGene_gn=BEST2;refGene_tx=XM_019818107.2;refGene_gn=BEST2;refGene_tx=XM_023245829.1
A2	9078992	9079087
refGene_gn=BEST2;refGene_tx=XM_019818105.2;refGene_gn=BEST2;refGene_tx=XM_019818107.2;refGene_gn=BEST2;refGene_tx=XM_019818111.2;refGene_gn=BEST2;refGene_tx=XM_023245829.1
A2	9079226	9079269	refGene_gn=BEST2;refGene_tx=XM_019818111.2
A2	9079817	9080051
refGene_gn=BEST2;refGene_tx=XM_019818105.2;refGene_gn=BEST2;refGene_tx=XM_019818107.2;refGene_gn=BEST2;refGene_tx=XM_019818111.2;refGene_gn=BEST2;refGene_tx=XM_023245829.1
A2	9080157	9080312
refGene_gn=BEST2;refGene_tx=XM_019818105.2;refGene_gn=BEST2;refGene_tx=XM_019818107.2;refGene_gn=BEST2;refGene_tx=XM_019818111.2;refGene_gn=BEST2;refGene_tx=XM_023245829.1
A2	9080883	9080961
refGene_gn=BEST2;refGene_tx=XM_019818105.2;refGene_gn=BEST2;refGene_tx=XM_019818107.2;refGene_gn=BEST2;refGene_tx=XM_019818111.2
A2	9081136	9081289
refGene_gn=BEST2;refGene_tx=XM_019818105.2;refGene_gn=BEST2;refGene_tx=XM_019818107.2;refGene_gn=BEST2;refGene_tx=XM_019818111.2;refGene_gn=BEST2;refGene_tx=XM_023245829.1
A2	9081382	9081463
refGene_gn=BEST2;refGene_tx=XM_019818105.2;refGene_gn=BEST2;refGene_tx=XM_019818107.2;refGene_gn=BEST2;refGene_tx=XM_019818111.2;refGene_gn=BEST2;refGene_tx=XM_023245829.1
A2	9081659	9081814
refGene_gn=BEST2;refGene_tx=XM_019818105.2;refGene_gn=BEST2;refGene_tx=XM_019818107.2;refGene_gn=BEST2;refGene_tx=XM_019818111.2;refGene_gn=BEST2;refGene_tx=XM_023245829.1
A2	9082294	9082715
refGene_gn=BEST2;refGene_tx=XM_019818105.2;refGene_gn=BEST2;refGene_tx=XM_019818107.2;refGene_gn=BEST2;refGene_tx=XM_019818111.2;refGene_gn=BEST2;refGene_tx=XM_023245829.1
A2	9084829	9084979
refGene_gn=HOOK2;refGene_tx=XM_023245835.1;refGene_gn=HOOK2;refGene_tx=XM_023245839.1;refGene_gn=HOOK2;refGene_tx=XM_023245843.1;refGene_gn=HOOK2;refGene_tx=XM_023245847.1;refGene_gn=HOOK2;refGene_tx=XM_023245849.1;refGene_gn=HOOK2;refGene_tx=XM_023245850.1;refGene_gn=HOOK2;refGene_tx=XM_023245852.1;refGene_gn=HOOK2;refGene_tx=XM_023245854.1;refGene_gn=HOOK2;refGene_tx=XM_023245857.1;refGene_gn=HOOK2;refGene_tx=XM_02

3245866.1;refGene_gn=HOOK2;refGene_tx=XM_023245872.1
A2	9085062	9085134
refGene_gn=HOOK2;refGene_tx=XM_023245835.1;refGene_gn=HOOK2;refGene_tx=XM_023245839.1;refGene_gn=HOOK2;refGene_tx=XM_023245843.1;refGene_gn=HOOK2;refGene_tx=XM_023245847.1;refGene_gn=HOOK2;refGene_tx=XM_023245849.1;refGene_gn=HOOK2;refGene_tx=XM_023245850.1;refGene_gn=HOOK2;refGene_tx=XM_023245852.1;refGene_gn=HOOK2;refGene_tx=XM_023245854.1;refGene_gn=HOOK2;refGene_tx=XM_023245857.1;refGene_gn=HOOK2;refGene_tx=XM_023245866.1;refGene_gn=HOOK2;refGene_tx=XM_023245872.1
A2	9085239	9085350
refGene_gn=HOOK2;refGene_tx=XM_023245835.1;refGene_gn=HOOK2;refGene_tx=XM_023245839.1;refGene_gn=HOOK2;refGene_tx=XM_023245843.1;refGene_gn=HOOK2;refGene_tx=XM_023245847.1;refGene_gn=HOOK2;refGene_tx=XM_023245849.1;refGene_gn=HOOK2;refGene_tx=XM_023245850.1;refGene_gn=HOOK2;refGene_tx=XM_023245852.1;refGene_gn=HOOK2;refGene_tx=XM_023245854.1;refGene_gn=HOOK2;refGene_tx=XM_023245857.1;refGene_gn=HOOK2;refGene_tx=XM_023245861.1;refGene_gn=HOOK2;refGene_tx=XM_023245866.1;refGene_gn=HOOK2;refGene_tx=XM_023245872.1;refGene_gn=HOOK2;refGene_tx=XM_023245877.1;refGene_gn=HOOK2;refGene_tx=XM_023245882.1
A2	9085564	9085668
refGene_gn=HOOK2;refGene_tx=XM_023245835.1;refGene_gn=HOOK2;refGene_tx=XM_023245839.1;refGene_gn=HOOK2;refGene_tx=XM_023245843.1;refGene_gn=HOOK2;refGene_tx=XM_023245847.1;refGene_gn=HOOK2;refGene_tx=XM_023245849.1;refGene_gn=HOOK2;refGene_tx=XM_023245850.1;refGene_gn=HOOK2;refGene_tx=XM_023245852.1;refGene_gn=HOOK2;refGene_tx=XM_023245854.1;refGene_gn=HOOK2;refGene_tx=XM_023245857.1;refGene_gn=HOOK2;refGene_tx=XM_023245861.1;refGene_gn=HOOK2;refGene_tx=XM_023245866.1;refGene_gn=HOOK2;refGene_tx=XM_023245872.1;refGene_gn=HOOK2;refGene_tx=XM_023245877.1;refGene_gn=HOOK2;refGene_tx=XM_023245882.1
A2	9085753	9085836
refGene_gn=HOOK2;refGene_tx=XM_023245835.1;refGene_gn=HOOK2;refGene_tx=XM_023245839.1;refGene_gn=HOOK2;refGene_tx=XM_023245843.1;refGene_gn=HOOK2;refGene_tx=XM_023245847.1;refGene_gn=HOOK2;refGene_tx=XM_023245849.1;refGene_gn=HOOK2;refGene_tx=XM_023245850.1;refGene_gn=HOOK2;refGene_tx=XM_023245852.1;refGene_gn=HOOK2;refGene_tx=XM_023245854.1;refGene_gn=HOOK2;refGene_tx=XM_023245857.1;refGene_gn=HOOK2;refGene_tx=XM_023245861.1;refGene_gn=HOOK2;refGene_tx=XM_023245866.1;refGene_gn=HOOK2;refGene_tx=XM_023245872.1;refGene_gn=HOOK2;refGene_tx=XM_023245877.1;refGene_gn=HOOK2;refGene_tx=XM_023245882.1
A2	9086860	9086871
refGene_gn=HOOK2;refGene_tx=XM_023245847.1;refGene_gn=HOOK2;refGene_tx=XM_023245849.1;refGene_gn=HOOK2;refGene_tx=XM_023245877.1;refGene_gn=HOOK2;refGene_tx=XM_023245882.1
A2	9088331	9088652
refGene_gn=HOOK2;refGene_tx=XM_023245835.1;refGene_gn=HOOK2;refGene_tx=XM_023245839.1;refGene_gn=HOOK2;refGene_tx=XM_023245843.1;refGene_gn=HOOK2;refGene_tx=XM_023245847.1;refGene_gn=HOOK2;refGene_tx=XM_023245849.1;refGene_gn=HOOK2;refGene_tx=XM_023245850.1;refGene_gn=HOOK2;refGene_tx=XM_023245852.1;refGene_gn=HOOK2;refGene_tx=XM_023245854.1;refGene_gn=HOOK2;refGene_tx=XM_023245857.1;refGene_gn=HOOK2;refGene_tx=XM_02

3245861.1;refGene_gn=HOOK2;refGene_tx=XM_023245866.1;refGene_gn=HOOK2;
refGene_tx=XM_023245872.1;refGene_gn=HOOK2;refGene_tx=XM_023245877.1;r
efGene_gn=HOOK2;refGene_tx=XM_023245882.1
A2 9088747 9088909
refGene_gn=HOOK2;refGene_tx=XM_023245835.1;refGene_gn=HOOK2;refGene_tx
=XM_023245839.1;refGene_gn=HOOK2;refGene_tx=XM_023245843.1;refGene_gn=
HOOK2;refGene_tx=XM_023245847.1;refGene_gn=HOOK2;refGene_tx=XM_0232458
49.1;refGene_gn=HOOK2;refGene_tx=XM_023245850.1;refGene_gn=HOOK2;refGe
ne_tx=XM_023245852.1;refGene_gn=HOOK2;refGene_tx=XM_023245854.1;refGen
e_gn=HOOK2;refGene_tx=XM_023245857.1;refGene_gn=HOOK2;refGene_tx=XM_02
3245861.1;refGene_gn=HOOK2;refGene_tx=XM_023245866.1;refGene_gn=HOOK2;
refGene_tx=XM_023245872.1;refGene_gn=HOOK2;refGene_tx=XM_023245877.1;r
efGene_gn=HOOK2;refGene_tx=XM_023245882.1
A2 9089596 9089666
refGene_gn=HOOK2;refGene_tx=XM_023245835.1;refGene_gn=HOOK2;refGene_tx
=XM_023245839.1;refGene_gn=HOOK2;refGene_tx=XM_023245843.1;refGene_gn=
HOOK2;refGene_tx=XM_023245847.1;refGene_gn=HOOK2;refGene_tx=XM_0232458
49.1;refGene_gn=HOOK2;refGene_tx=XM_023245850.1;refGene_gn=HOOK2;refGe
ne_tx=XM_023245852.1;refGene_gn=HOOK2;refGene_tx=XM_023245854.1;refGen
e_gn=HOOK2;refGene_tx=XM_023245857.1;refGene_gn=HOOK2;refGene_tx=XM_02
3245861.1;refGene_gn=HOOK2;refGene_tx=XM_023245866.1;refGene_gn=HOOK2;
refGene_tx=XM_023245872.1;refGene_gn=HOOK2;refGene_tx=XM_023245877.1;r
efGene_gn=HOOK2;refGene_tx=XM_023245882.1
A2 9090007 9090095
refGene_gn=HOOK2;refGene_tx=XM_023245835.1;refGene_gn=HOOK2;refGene_tx
=XM_023245839.1;refGene_gn=HOOK2;refGene_tx=XM_023245843.1;refGene_gn=
HOOK2;refGene_tx=XM_023245847.1;refGene_gn=HOOK2;refGene_tx=XM_0232458
49.1;refGene_gn=HOOK2;refGene_tx=XM_023245850.1;refGene_gn=HOOK2;refGe
ne_tx=XM_023245852.1;refGene_gn=HOOK2;refGene_tx=XM_023245854.1;refGen
e_gn=HOOK2;refGene_tx=XM_023245857.1;refGene_gn=HOOK2;refGene_tx=XM_02
3245861.1;refGene_gn=HOOK2;refGene_tx=XM_023245866.1;refGene_gn=HOOK2;
refGene_tx=XM_023245872.1;refGene_gn=HOOK2;refGene_tx=XM_023245877.1;r
efGene_gn=HOOK2;refGene_tx=XM_023245882.1
A2 9090202 9090313
refGene_gn=HOOK2;refGene_tx=XM_023245835.1;refGene_gn=HOOK2;refGene_tx
=XM_023245839.1;refGene_gn=HOOK2;refGene_tx=XM_023245843.1;refGene_gn=
HOOK2;refGene_tx=XM_023245847.1;refGene_gn=HOOK2;refGene_tx=XM_0232458
49.1;refGene_gn=HOOK2;refGene_tx=XM_023245850.1;refGene_gn=HOOK2;refGe
ne_tx=XM_023245852.1;refGene_gn=HOOK2;refGene_tx=XM_023245854.1;refGen
e_gn=HOOK2;refGene_tx=XM_023245857.1;refGene_gn=HOOK2;refGene_tx=XM_02
3245861.1;refGene_gn=HOOK2;refGene_tx=XM_023245866.1;refGene_gn=HOOK2;
refGene_tx=XM_023245872.1;refGene_gn=HOOK2;refGene_tx=XM_023245877.1;r
efGene_gn=HOOK2;refGene_tx=XM_023245882.1
A2 9090600 9090802
refGene_gn=HOOK2;refGene_tx=XM_023245835.1;refGene_gn=HOOK2;refGene_tx
=XM_023245839.1;refGene_gn=HOOK2;refGene_tx=XM_023245843.1;refGene_gn=
HOOK2;refGene_tx=XM_023245847.1;refGene_gn=HOOK2;refGene_tx=XM_0232458
49.1;refGene_gn=HOOK2;refGene_tx=XM_023245850.1;refGene_gn=HOOK2;refGe
ne_tx=XM_023245852.1;refGene_gn=HOOK2;refGene_tx=XM_023245854.1;refGen
e_gn=HOOK2;refGene_tx=XM_023245857.1;refGene_gn=HOOK2;refGene_tx=XM_02

3245861.1;refGene_gn=HOOK2;refGene_tx=XM_023245866.1;refGene_gn=HOOK2;
refGene_tx=XM_023245872.1;refGene_gn=HOOK2;refGene_tx=XM_023245877.1;r
efGene_gn=HOOK2;refGene_tx=XM_023245882.1
A2	9092029	9092170
refGene_gn=HOOK2;refGene_tx=XM_023245835.1;refGene_gn=HOOK2;refGene_tx
=XM_023245839.1;refGene_gn=HOOK2;refGene_tx=XM_023245843.1;refGene_gn=
HOOK2;refGene_tx=XM_023245847.1;refGene_gn=HOOK2;refGene_tx=XM_0232458
49.1;refGene_gn=HOOK2;refGene_tx=XM_023245850.1;refGene_gn=HOOK2;refGe
ne_tx=XM_023245852.1;refGene_gn=HOOK2;refGene_tx=XM_023245854.1;refGen
e_gn=HOOK2;refGene_tx=XM_023245857.1;refGene_gn=HOOK2;refGene_tx=XM_02
3245861.1;refGene_gn=HOOK2;refGene_tx=XM_023245866.1;refGene_gn=HOOK2;
refGene_tx=XM_023245872.1;refGene_gn=HOOK2;refGene_tx=XM_023245877.1;r
efGene_gn=HOOK2;refGene_tx=XM_023245882.1
A2	9092258	9092419
refGene_gn=HOOK2;refGene_tx=XM_023245835.1;refGene_gn=HOOK2;refGene_tx
=XM_023245839.1;refGene_gn=HOOK2;refGene_tx=XM_023245843.1;refGene_gn=
HOOK2;refGene_tx=XM_023245847.1;refGene_gn=HOOK2;refGene_tx=XM_0232458
49.1;refGene_gn=HOOK2;refGene_tx=XM_023245850.1;refGene_gn=HOOK2;refGe
ne_tx=XM_023245852.1;refGene_gn=HOOK2;refGene_tx=XM_023245854.1;refGen
e_gn=HOOK2;refGene_tx=XM_023245857.1;refGene_gn=HOOK2;refGene_tx=XM_02
3245861.1;refGene_gn=HOOK2;refGene_tx=XM_023245866.1;refGene_gn=HOOK2;
refGene_tx=XM_023245872.1;refGene_gn=HOOK2;refGene_tx=XM_023245877.1;r
efGene_gn=HOOK2;refGene_tx=XM_023245882.1
A2	9092506	9092587
refGene_gn=HOOK2;refGene_tx=XM_023245835.1;refGene_gn=HOOK2;refGene_tx
=XM_023245839.1;refGene_gn=HOOK2;refGene_tx=XM_023245843.1;refGene_gn=
HOOK2;refGene_tx=XM_023245847.1;refGene_gn=HOOK2;refGene_tx=XM_0232458
49.1;refGene_gn=HOOK2;refGene_tx=XM_023245850.1;refGene_gn=HOOK2;refGe
ne_tx=XM_023245852.1;refGene_gn=HOOK2;refGene_tx=XM_023245854.1;refGen
e_gn=HOOK2;refGene_tx=XM_023245857.1;refGene_gn=HOOK2;refGene_tx=XM_02
3245861.1;refGene_gn=HOOK2;refGene_tx=XM_023245866.1;refGene_gn=HOOK2;
refGene_tx=XM_023245872.1;refGene_gn=HOOK2;refGene_tx=XM_023245877.1;r
efGene_gn=HOOK2;refGene_tx=XM_023245882.1
A2	9092910	9092973
refGene_gn=HOOK2;refGene_tx=XM_023245835.1;refGene_gn=HOOK2;refGene_tx
=XM_023245839.1;refGene_gn=HOOK2;refGene_tx=XM_023245843.1;refGene_gn=
HOOK2;refGene_tx=XM_023245847.1;refGene_gn=HOOK2;refGene_tx=XM_0232458
49.1;refGene_gn=HOOK2;refGene_tx=XM_023245850.1;refGene_gn=HOOK2;refGe
ne_tx=XM_023245852.1;refGene_gn=HOOK2;refGene_tx=XM_023245854.1;refGen
e_gn=HOOK2;refGene_tx=XM_023245857.1;refGene_gn=HOOK2;refGene_tx=XM_02
3245861.1;refGene_gn=HOOK2;refGene_tx=XM_023245866.1;refGene_gn=HOOK2;
refGene_tx=XM_023245872.1;refGene_gn=HOOK2;refGene_tx=XM_023245877.1;r
efGene_gn=HOOK2;refGene_tx=XM_023245882.1
A2	9093326	9093394
refGene_gn=HOOK2;refGene_tx=XM_023245835.1;refGene_gn=HOOK2;refGene_tx
=XM_023245839.1;refGene_gn=HOOK2;refGene_tx=XM_023245843.1;refGene_gn=
HOOK2;refGene_tx=XM_023245847.1;refGene_gn=HOOK2;refGene_tx=XM_0232458
49.1;refGene_gn=HOOK2;refGene_tx=XM_023245850.1;refGene_gn=HOOK2;refGe
ne_tx=XM_023245852.1;refGene_gn=HOOK2;refGene_tx=XM_023245854.1;refGen
e_gn=HOOK2;refGene_tx=XM_023245857.1;refGene_gn=HOOK2;refGene_tx=XM_02

3245861.1;refGene_gn=HOOK2;refGene_tx=XM_023245866.1;refGene_gn=HOOK2;
refGene_tx=XM_023245872.1;refGene_gn=HOOK2;refGene_tx=XM_023245877.1;r
efGene_gn=HOOK2;refGene_tx=XM_023245882.1
A2 9093500 9093633
refGene_gn=HOOK2;refGene_tx=XM_023245835.1;refGene_gn=HOOK2;refGene_tx
=XM_023245839.1;refGene_gn=HOOK2;refGene_tx=XM_023245843.1;refGene_gn=
HOOK2;refGene_tx=XM_023245847.1;refGene_gn=HOOK2;refGene_tx=XM_0232458
49.1;refGene_gn=HOOK2;refGene_tx=XM_023245850.1;refGene_gn=HOOK2;refGe
ne_tx=XM_023245852.1;refGene_gn=HOOK2;refGene_tx=XM_023245854.1;refGen
e_gn=HOOK2;refGene_tx=XM_023245857.1;refGene_gn=HOOK2;refGene_tx=XM_02
3245861.1;refGene_gn=HOOK2;refGene_tx=XM_023245877.1;refGene_gn=HOOK2;
refGene_tx=XM_023245882.1
A2 9093740 9093791
refGene_gn=HOOK2;refGene_tx=XM_023245835.1;refGene_gn=HOOK2;refGene_tx
=XM_023245839.1;refGene_gn=HOOK2;refGene_tx=XM_023245843.1;refGene_gn=
HOOK2;refGene_tx=XM_023245847.1;refGene_gn=HOOK2;refGene_tx=XM_0232458
49.1;refGene_gn=HOOK2;refGene_tx=XM_023245850.1;refGene_gn=HOOK2;refGe
ne_tx=XM_023245852.1;refGene_gn=HOOK2;refGene_tx=XM_023245854.1;refGen
e_gn=HOOK2;refGene_tx=XM_023245857.1;refGene_gn=HOOK2;refGene_tx=XM_02
3245861.1;refGene_gn=HOOK2;refGene_tx=XM_023245877.1;refGene_gn=HOOK2;
refGene_tx=XM_023245882.1
A2 9094726 9094799
refGene_gn=HOOK2;refGene_tx=XM_023245835.1;refGene_gn=HOOK2;refGene_tx
=XM_023245839.1;refGene_gn=HOOK2;refGene_tx=XM_023245843.1;refGene_gn=
HOOK2;refGene_tx=XM_023245847.1;refGene_gn=HOOK2;refGene_tx=XM_0232458
49.1;refGene_gn=HOOK2;refGene_tx=XM_023245850.1;refGene_gn=HOOK2;refGe
ne_tx=XM_023245852.1;refGene_gn=HOOK2;refGene_tx=XM_023245854.1;refGen
e_gn=HOOK2;refGene_tx=XM_023245857.1;refGene_gn=HOOK2;refGene_tx=XM_02
3245861.1;refGene_gn=HOOK2;refGene_tx=XM_023245877.1;refGene_gn=HOOK2;
refGene_tx=XM_023245882.1
A2 9094868 9094954
refGene_gn=HOOK2;refGene_tx=XM_023245835.1;refGene_gn=HOOK2;refGene_tx
=XM_023245839.1;refGene_gn=HOOK2;refGene_tx=XM_023245843.1;refGene_gn=
HOOK2;refGene_tx=XM_023245847.1;refGene_gn=HOOK2;refGene_tx=XM_0232458
49.1;refGene_gn=HOOK2;refGene_tx=XM_023245850.1;refGene_gn=HOOK2;refGe
ne_tx=XM_023245852.1;refGene_gn=HOOK2;refGene_tx=XM_023245854.1;refGen
e_gn=HOOK2;refGene_tx=XM_023245857.1;refGene_gn=HOOK2;refGene_tx=XM_02
3245861.1;refGene_gn=HOOK2;refGene_tx=XM_023245877.1;refGene_gn=HOOK2;
refGene_tx=XM_023245882.1
A2 9095464 9095509
refGene_gn=HOOK2;refGene_tx=XM_023245835.1;refGene_gn=HOOK2;refGene_tx
=XM_023245839.1;refGene_gn=HOOK2;refGene_tx=XM_023245843.1;refGene_gn=
HOOK2;refGene_tx=XM_023245847.1;refGene_gn=HOOK2;refGene_tx=XM_0232458
49.1;refGene_gn=HOOK2;refGene_tx=XM_023245850.1;refGene_gn=HOOK2;refGe
ne_tx=XM_023245852.1;refGene_gn=HOOK2;refGene_tx=XM_023245854.1;refGen
e_gn=HOOK2;refGene_tx=XM_023245857.1;refGene_gn=HOOK2;refGene_tx=XM_02
3245861.1;refGene_gn=HOOK2;refGene_tx=XM_023245877.1;refGene_gn=HOOK2;
refGene_tx=XM_023245882.1
A2 9106720 9107764 refGene_gn=JUNB;refGene_tx=XM_003981915.4
A2 9110660 9110746 refGene_gn=PRDX2;refGene_tx=XM_003981916.5

A2	9111306	9111437	refGene_gn=PRDX2;refGene_tx=XM_003981916.5
A2	9111580	9111703	refGene_gn=PRDX2;refGene_tx=XM_003981916.5
A2	9112317	9112471	refGene_gn=PRDX2;refGene_tx=XM_003981916.5
A2	9112559	9112662	refGene_gn=PRDX2;refGene_tx=XM_003981916.5
A2	9115982	9116109
refGene_gn=RNASEH2A;refGene_tx=XM_003981917.5;refGene_gn=RNASEH2A;refGene_tx=XM_011289361.3;refGene_gn=RNASEH2A;refGene_tx=XM_011289362.3
A2	9116298	9116370
refGene_gn=RNASEH2A;refGene_tx=XM_003981917.5;refGene_gn=RNASEH2A;refGene_tx=XM_011289361.3;refGene_gn=RNASEH2A;refGene_tx=XM_011289362.3
A2	9116495	9116622
refGene_gn=RNASEH2A;refGene_tx=XM_003981917.5;refGene_gn=RNASEH2A;refGene_tx=XM_011289361.3;refGene_gn=RNASEH2A;refGene_tx=XM_011289362.3
A2	9116714	9116802
refGene_gn=RNASEH2A;refGene_tx=XM_003981917.5;refGene_gn=RNASEH2A;refGene_tx=XM_011289361.3;refGene_gn=RNASEH2A;refGene_tx=XM_011289362.3
A2	9119311	9119449
refGene_gn=RNASEH2A;refGene_tx=XM_003981917.5;refGene_gn=RNASEH2A;refGene_tx=XM_011289361.3;refGene_gn=RNASEH2A;refGene_tx=XM_011289362.3
A2	9119556	9119644
refGene_gn=RNASEH2A;refGene_tx=XM_003981917.5;refGene_gn=RNASEH2A;refGene_tx=XM_011289361.3;refGene_gn=RNASEH2A;refGene_tx=XM_011289362.3
A2	9120546	9120670
refGene_gn=RNASEH2A;refGene_tx=XM_003981917.5;refGene_gn=RNASEH2A;refGene_tx=XM_011289361.3;refGene_gn=RNASEH2A;refGene_tx=XM_011289362.3
A2	9120761	9120897
refGene_gn=RNASEH2A;refGene_tx=XM_003981917.5;refGene_gn=RNASEH2A;refGene_tx=XM_011289361.3;refGene_gn=RNASEH2A;refGene_tx=XM_011289362.3
A2	9128272	9128500
refGene_gn=RTBDN;refGene_tx=XM_003981918.4;refGene_gn=RTBDN;refGene_tx=XM_019818148.2;refGene_gn=RTBDN;refGene_tx=XM_019818153.1;refGene_gn=RTBDN;refGene_tx=XM_019818160.2;refGene_gn=RTBDN;refGene_tx=XM_023245984.1;refGene_gn=RTBDN;refGene_tx=XM_023245992.1;refGene_gn=RTBDN;refGene_tx=XM_023246009.1
A2	9129039	9129136
refGene_gn=RTBDN;refGene_tx=XM_003981918.4;refGene_gn=RTBDN;refGene_tx=XM_019818148.2;refGene_gn=RTBDN;refGene_tx=XM_019818153.1;refGene_gn=RTBDN;refGene_tx=XM_019818160.2;refGene_gn=RTBDN;refGene_tx=XM_023245984.1;refGene_gn=RTBDN;refGene_tx=XM_023245992.1;refGene_gn=RTBDN;refGene_tx=XM_023246009.1
A2	9129645	9129756
refGene_gn=RTBDN;refGene_tx=XM_003981918.4;refGene_gn=RTBDN;refGene_tx=XM_019818148.2;refGene_gn=RTBDN;refGene_tx=XM_019818153.1;refGene_gn=RTBDN;refGene_tx=XM_019818160.2;refGene_gn=RTBDN;refGene_tx=XM_023245984.1;refGene_gn=RTBDN;refGene_tx=XM_023245992.1;refGene_gn=RTBDN;refGene_tx=XM_023246009.1
A2	9129852	9129919
refGene_gn=RTBDN;refGene_tx=XM_003981918.4;refGene_gn=RTBDN;refGene_tx=XM_019818148.2;refGene_gn=RTBDN;refGene_tx=XM_019818153.1;refGene_gn=RTBDN;refGene_tx=XM_019818160.2;refGene_gn=RTBDN;refGene_tx=XM_0232459

84.1;refGene_gn=RTBDN;refGene_tx=XM_023245992.1;refGene_gn=RTBDN;refGene_tx=XM_023246009.1
A2 9130497 9130672 refGene_gn=RTBDN;refGene_tx=XM_003981918.4;refGene_gn=RTBDN;refGene_tx=XM_019818148.2;refGene_gn=RTBDN;refGene_tx=XM_019818153.1;refGene_gn=RTBDN;refGene_tx=XM_019818160.2;refGene_gn=RTBDN;refGene_tx=XM_023245984.1;refGene_gn=RTBDN;refGene_tx=XM_023245992.1;refGene_gn=RTBDN;refGene_tx=XM_023246009.1
A2 9132880 9132895 refGene_gn=RTBDN;refGene_tx=XM_019818148.2;refGene_gn=RTBDN;refGene_tx=XM_023245984.1;refGene_gn=RTBDN;refGene_tx=XM_023245992.1
A2 9134493 9134532 refGene_gn=MAST1;refGene_tx=XM_023245943.1;refGene_gn=MAST1;refGene_tx=XM_023245950.1
A2 9136373 9136379 refGene_gn=MAST1;refGene_tx=XM_023245958.1
A2 9138179 9138211 refGene_gn=MAST1;refGene_tx=XM_023245943.1;refGene_gn=MAST1;refGene_tx=XM_023245950.1;refGene_gn=MAST1;refGene_tx=XM_023245958.1
A2 9138466 9138549 refGene_gn=MAST1;refGene_tx=XM_023245932.1;refGene_gn=MAST1;refGene_tx=XM_023245939.1
A2 9140154 9140243 refGene_gn=MAST1;refGene_tx=XM_023245932.1;refGene_gn=MAST1;refGene_tx=XM_023245939.1;refGene_gn=MAST1;refGene_tx=XM_023245943.1;refGene_gn=MAST1;refGene_tx=XM_023245950.1;refGene_gn=MAST1;refGene_tx=XM_023245958.1
A2 9140713 9140789 refGene_gn=MAST1;refGene_tx=XM_023245932.1;refGene_gn=MAST1;refGene_tx=XM_023245939.1;refGene_gn=MAST1;refGene_tx=XM_023245943.1;refGene_gn=MAST1;refGene_tx=XM_023245950.1;refGene_gn=MAST1;refGene_tx=XM_023245958.1
A2 9142671 9142750 refGene_gn=MAST1;refGene_tx=XM_023245932.1;refGene_gn=MAST1;refGene_tx=XM_023245939.1;refGene_gn=MAST1;refGene_tx=XM_023245943.1;refGene_gn=MAST1;refGene_tx=XM_023245950.1;refGene_gn=MAST1;refGene_tx=XM_023245958.1
A2 9144614 9144775 refGene_gn=MAST1;refGene_tx=XM_023245932.1;refGene_gn=MAST1;refGene_tx=XM_023245939.1;refGene_gn=MAST1;refGene_tx=XM_023245943.1;refGene_gn=MAST1;refGene_tx=XM_023245950.1;refGene_gn=MAST1;refGene_tx=XM_023245958.1
A2 9144970 9145046 refGene_gn=MAST1;refGene_tx=XM_023245932.1;refGene_gn=MAST1;refGene_tx=XM_023245939.1;refGene_gn=MAST1;refGene_tx=XM_023245943.1;refGene_gn=MAST1;refGene_tx=XM_023245950.1;refGene_gn=MAST1;refGene_tx=XM_023245958.1
A2 9145215 9145425 refGene_gn=MAST1;refGene_tx=XM_023245932.1;refGene_gn=MAST1;refGene_tx=XM_023245939.1;refGene_gn=MAST1;refGene_tx=XM_023245943.1;refGene_gn=MAST1;refGene_tx=XM_023245950.1;refGene_gn=MAST1;refGene_tx=XM_0232459

58.1
A2	9147884	9147986
refGene_gn=MAST1;refGene_tx=XM_023245932.1;refGene_gn=MAST1;refGene_tx=XM_023245939.1;refGene_gn=MAST1;refGene_tx=XM_023245943.1;refGene_gn=MAST1;refGene_tx=XM_023245950.1;refGene_gn=MAST1;refGene_tx=XM_023245958.1
A2	9148075	9148208
refGene_gn=MAST1;refGene_tx=XM_023245932.1;refGene_gn=MAST1;refGene_tx=XM_023245939.1;refGene_gn=MAST1;refGene_tx=XM_023245943.1;refGene_gn=MAST1;refGene_tx=XM_023245950.1;refGene_gn=MAST1;refGene_tx=XM_023245958.1
A2	9148286	9148354
refGene_gn=MAST1;refGene_tx=XM_023245932.1;refGene_gn=MAST1;refGene_tx=XM_023245939.1;refGene_gn=MAST1;refGene_tx=XM_023245943.1;refGene_gn=MAST1;refGene_tx=XM_023245950.1;refGene_gn=MAST1;refGene_tx=XM_023245958.1
A2	9151282	9151368
refGene_gn=MAST1;refGene_tx=XM_023245932.1;refGene_gn=MAST1;refGene_tx=XM_023245939.1;refGene_gn=MAST1;refGene_tx=XM_023245943.1;refGene_gn=MAST1;refGene_tx=XM_023245950.1;refGene_gn=MAST1;refGene_tx=XM_023245958.1
A2	9151457	9151666
refGene_gn=MAST1;refGene_tx=XM_023245932.1;refGene_gn=MAST1;refGene_tx=XM_023245939.1;refGene_gn=MAST1;refGene_tx=XM_023245943.1;refGene_gn=MAST1;refGene_tx=XM_023245950.1;refGene_gn=MAST1;refGene_tx=XM_023245958.1
A2	9154371	9154510
refGene_gn=MAST1;refGene_tx=XM_023245932.1;refGene_gn=MAST1;refGene_tx=XM_023245939.1;refGene_gn=MAST1;refGene_tx=XM_023245943.1;refGene_gn=MAST1;refGene_tx=XM_023245950.1;refGene_gn=MAST1;refGene_tx=XM_023245958.1
A2	9154629	9154762
refGene_gn=MAST1;refGene_tx=XM_023245932.1;refGene_gn=MAST1;refGene_tx=XM_023245939.1;refGene_gn=MAST1;refGene_tx=XM_023245943.1;refGene_gn=MAST1;refGene_tx=XM_023245950.1;refGene_gn=MAST1;refGene_tx=XM_023245958.1
A2	9154898	9155064
refGene_gn=MAST1;refGene_tx=XM_023245932.1;refGene_gn=MAST1;refGene_tx=XM_023245939.1;refGene_gn=MAST1;refGene_tx=XM_023245943.1;refGene_gn=MAST1;refGene_tx=XM_023245950.1;refGene_gn=MAST1;refGene_tx=XM_023245958.1
A2	9155165	9155267
refGene_gn=MAST1;refGene_tx=XM_023245932.1;refGene_gn=MAST1;refGene_tx=XM_023245939.1;refGene_gn=MAST1;refGene_tx=XM_023245943.1;refGene_gn=MAST1;refGene_tx=XM_023245950.1;refGene_gn=MAST1;refGene_tx=XM_023245958.1
A2	9155419	9155542
refGene_gn=MAST1;refGene_tx=XM_023245932.1;refGene_gn=MAST1;refGene_tx=XM_023245939.1;refGene_gn=MAST1;refGene_tx=XM_023245943.1;refGene_gn=MAST1;refGene_tx=XM_023245950.1;refGene_gn=MAST1;refGene_tx=XM_0232459

58.1
A2	9156021	9156131
refGene_gn=MAST1;refGene_tx=XM_023245932.1;refGene_gn=MAST1;refGene_tx
=XM_023245939.1;refGene_gn=MAST1;refGene_tx=XM_023245943.1;refGene_gn=
MAST1;refGene_tx=XM_023245950.1;refGene_gn=MAST1;refGene_tx=XM_0232459
58.1
A2	9156749	9156928
refGene_gn=MAST1;refGene_tx=XM_023245932.1;refGene_gn=MAST1;refGene_tx
=XM_023245939.1;refGene_gn=MAST1;refGene_tx=XM_023245943.1;refGene_gn=
MAST1;refGene_tx=XM_023245950.1;refGene_gn=MAST1;refGene_tx=XM_0232459
58.1
A2	9157011	9157262
refGene_gn=MAST1;refGene_tx=XM_023245932.1;refGene_gn=MAST1;refGene_tx
=XM_023245939.1;refGene_gn=MAST1;refGene_tx=XM_023245943.1;refGene_gn=
MAST1;refGene_tx=XM_023245950.1;refGene_gn=MAST1;refGene_tx=XM_0232459
58.1
A2	9157593	9157800
refGene_gn=MAST1;refGene_tx=XM_023245932.1;refGene_gn=MAST1;refGene_tx
=XM_023245939.1;refGene_gn=MAST1;refGene_tx=XM_023245943.1;refGene_gn=
MAST1;refGene_tx=XM_023245950.1;refGene_gn=MAST1;refGene_tx=XM_0232459
58.1
A2	9158015	9158245
refGene_gn=MAST1;refGene_tx=XM_023245932.1;refGene_gn=MAST1;refGene_tx
=XM_023245939.1;refGene_gn=MAST1;refGene_tx=XM_023245943.1;refGene_gn=
MAST1;refGene_tx=XM_023245950.1;refGene_gn=MAST1;refGene_tx=XM_0232459
58.1
A2	9158452	9158575
refGene_gn=MAST1;refGene_tx=XM_023245932.1;refGene_gn=MAST1;refGene_tx
=XM_023245939.1;refGene_gn=MAST1;refGene_tx=XM_023245943.1;refGene_gn=
MAST1;refGene_tx=XM_023245950.1;refGene_gn=MAST1;refGene_tx=XM_0232459
58.1
A2	9158693	9158830
refGene_gn=MAST1;refGene_tx=XM_023245932.1;refGene_gn=MAST1;refGene_tx
=XM_023245939.1;refGene_gn=MAST1;refGene_tx=XM_023245943.1;refGene_gn=
MAST1;refGene_tx=XM_023245950.1;refGene_gn=MAST1;refGene_tx=XM_0232459
58.1
A2	9159694	9159882
refGene_gn=MAST1;refGene_tx=XM_023245932.1;refGene_gn=MAST1;refGene_tx
=XM_023245939.1;refGene_gn=MAST1;refGene_tx=XM_023245943.1;refGene_gn=
MAST1;refGene_tx=XM_023245950.1;refGene_gn=MAST1;refGene_tx=XM_0232459
58.1
A2	9159985	9161244
refGene_gn=MAST1;refGene_tx=XM_023245932.1;refGene_gn=MAST1;refGene_tx
=XM_023245939.1;refGene_gn=MAST1;refGene_tx=XM_023245943.1;refGene_gn=
MAST1;refGene_tx=XM_023245950.1;refGene_gn=MAST1;refGene_tx=XM_0232459
58.1
A2	9161754	9161790	refGene_gn=DNASE2;refGene_tx=XM_003981919.4
A2	9161792	9162139	refGene_gn=DNASE2;refGene_tx=XM_003981919.4
A2	9162543	9162741	refGene_gn=DNASE2;refGene_tx=XM_003981919.4
A2	9162830	9162995	refGene_gn=DNASE2;refGene_tx=XM_003981919.4

A2	9163321	9163400	refGene_gn=DNASE2;refGene_tx=XM_003981919.4
A2	9163489	9163667	refGene_gn=DNASE2;refGene_tx=XM_003981919.4
A2	9163793	9163888	refGene_gn=DNASE2;refGene_tx=XM_003981919.4
A2	9165288	9165464	refGene_gn=KLF1;refGene_tx=XM_023245968.1
A2	9166679	9167553	refGene_gn=KLF1;refGene_tx=XM_023245968.1
A2	9168334	9168421	refGene_gn=KLF1;refGene_tx=XM_023245968.1
A2	9173154	9173245	refGene_gn=GCDH;refGene_tx=XM_003981920.5
A2	9173334	9173370	refGene_gn=GCDH;refGene_tx=XM_003981920.5
A2	9173662	9173806	refGene_gn=GCDH;refGene_tx=XM_003981920.5
A2	9173956	9174019	refGene_gn=GCDH;refGene_tx=XM_003981920.5
A2	9174288	9174459	refGene_gn=GCDH;refGene_tx=XM_003981920.5
A2	9175163	9175293	refGene_gn=GCDH;refGene_tx=XM_003981920.5
A2	9175374	9175591	refGene_gn=GCDH;refGene_tx=XM_003981920.5
A2	9176129	9176233	refGene_gn=GCDH;refGene_tx=XM_003981920.5
A2	9176493	9176619	refGene_gn=GCDH;refGene_tx=XM_003981920.5
A2	9176959	9177120	refGene_gn=GCDH;refGene_tx=XM_003981920.5
A2	9178432	9178480
refGene_gn=SYCE2;refGene_tx=XM_023246018.1;refGene_gn=SYCE2;refGene_tx
=XM_023246023.1;refGene_gn=SYCE2;refGene_tx=XM_023246026.1;refGene_gn=
SYCE2;refGene_tx=XM_023246032.1;refGene_gn=SYCE2;refGene_tx=XM_0232460
33.1
A2	9178569	9178643	refGene_gn=GCDH;refGene_tx=XM_003981920.5
A2	9179606	9179795
refGene_gn=SYCE2;refGene_tx=XM_023246018.1;refGene_gn=SYCE2;refGene_tx
=XM_023246023.1;refGene_gn=SYCE2;refGene_tx=XM_023246026.1;refGene_gn=
SYCE2;refGene_tx=XM_023246032.1;refGene_gn=SYCE2;refGene_tx=XM_0232460
33.1
A2	9181480	9181691
refGene_gn=SYCE2;refGene_tx=XM_023246018.1;refGene_gn=SYCE2;refGene_tx
=XM_023246023.1;refGene_gn=SYCE2;refGene_tx=XM_023246026.1;refGene_gn=
SYCE2;refGene_tx=XM_023246032.1;refGene_gn=SYCE2;refGene_tx=XM_0232460
33.1
A2	9192237	9192353
refGene_gn=SYCE2;refGene_tx=XM_023246018.1;refGene_gn=SYCE2;refGene_tx
=XM_023246023.1;refGene_gn=SYCE2;refGene_tx=XM_023246026.1;refGene_gn=
SYCE2;refGene_tx=XM_023246032.1;refGene_gn=SYCE2;refGene_tx=XM_0232460
33.1
A2	9192654	9192777
refGene_gn=SYCE2;refGene_tx=XM_023246018.1;refGene_gn=SYCE2;refGene_tx
=XM_023246023.1;refGene_gn=SYCE2;refGene_tx=XM_023246026.1
A2	9192998	9193013
refGene_gn=SYCE2;refGene_tx=XM_023246032.1;refGene_gn=SYCE2;refGene_tx
=XM_023246033.1
A2	9197327	9197466
refGene_gn=FARSA;refGene_tx=XM_003981921.4;refGene_gn=FARSA;refGene_tx
=XM_019818179.2
A2	9198135	9198250
refGene_gn=FARSA;refGene_tx=XM_003981921.4;refGene_gn=FARSA;refGene_tx
=XM_019818179.2
A2	9198403	9198481

refGene_gn=FARSA;refGene_tx=XM_003981921.4;refGene_gn=FARSA;refGene_tx=XM_019818179.2
A2	9198583	9198752	refGene_gn=FARSA;refGene_tx=XM_003981921.4;refGene_gn=FARSA;refGene_tx=XM_019818179.2
A2	9198843	9198943	refGene_gn=FARSA;refGene_tx=XM_003981921.4;refGene_gn=FARSA;refGene_tx=XM_019818179.2
A2	9199038	9199144	refGene_gn=FARSA;refGene_tx=XM_003981921.4;refGene_gn=FARSA;refGene_tx=XM_019818179.2
A2	9200573	9200689	refGene_gn=FARSA;refGene_tx=XM_003981921.4;refGene_gn=FARSA;refGene_tx=XM_019818179.2
A2	9200765	9200894	refGene_gn=FARSA;refGene_tx=XM_003981921.4;refGene_gn=FARSA;refGene_tx=XM_019818179.2
A2	9200978	9201071	refGene_gn=FARSA;refGene_tx=XM_003981921.4;refGene_gn=FARSA;refGene_tx=XM_019818179.2
A2	9202004	9202123	refGene_gn=FARSA;refGene_tx=XM_003981921.4;refGene_gn=FARSA;refGene_tx=XM_019818179.2
A2	9202206	9202305	refGene_gn=FARSA;refGene_tx=XM_003981921.4;refGene_gn=FARSA;refGene_tx=XM_019818179.2
A2	9202387	9202525	refGene_gn=FARSA;refGene_tx=XM_003981921.4;refGene_gn=FARSA;refGene_tx=XM_019818179.2
A2	9205044	9205191	refGene_gn=FARSA;refGene_tx=XM_003981921.4;refGene_gn=FARSA;refGene_tx=XM_019818179.2
A2	9208736	9208827	refGene_gn=CALR;refGene_tx=XM_003981922.5
A2	9209195	9209297	refGene_gn=CALR;refGene_tx=XM_003981922.5
A2	9209498	9209702	refGene_gn=CALR;refGene_tx=XM_003981922.5
A2	9210136	9210231	refGene_gn=CALR;refGene_tx=XM_003981922.5
A2	9210320	9210530	refGene_gn=CALR;refGene_tx=XM_003981922.5
A2	9210609	9210723	refGene_gn=CALR;refGene_tx=XM_003981922.5
A2	9210809	9210953	refGene_gn=CALR;refGene_tx=XM_003981922.5
A2	9211607	9211700	refGene_gn=CALR;refGene_tx=XM_003981922.5
A2	9211790	9211991	refGene_gn=CALR;refGene_tx=XM_003981922.5
A2	9214544	9214616	refGene_gn=RAD23A;refGene_tx=XM_003981923.4
A2	9216003	9216165	refGene_gn=RAD23A;refGene_tx=XM_003981923.4
A2	9216338	9216520	refGene_gn=RAD23A;refGene_tx=XM_003981923.4
A2	9216658	9216714	refGene_gn=RAD23A;refGene_tx=XM_003981923.4
A2	9216847	9216975	refGene_gn=RAD23A;refGene_tx=XM_003981923.4
A2	9217249	9217328	refGene_gn=RAD23A;refGene_tx=XM_003981923.4
A2	9217445	9217576	refGene_gn=RAD23A;refGene_tx=XM_003981923.4
A2	9219648	9219813	refGene_gn=RAD23A;refGene_tx=XM_003981923.4
A2	9219892	9220006	refGene_gn=RAD23A;refGene_tx=XM_003981923.4

A2 9221216 9221535
refGene_gn=GADD45GIP1;refGene_tx=XM_023246055.1
A2 9223004 9223354
refGene_gn=GADD45GIP1;refGene_tx=XM_023246055.1
A2 9234830 9235157 refGene_gn=DAND5;refGene_tx=XM_004001085.4
A2 9237877 9238123 refGene_gn=DAND5;refGene_tx=XM_004001085.4
A2 9249851 9249878 refGene_gn=NFIX;refGene_tx=XM_023246090.1
A2 9276941 9276944 refGene_gn=NFIX;refGene_tx=XM_023246091.1
A2 9277338 9277604
refGene_gn=NFIX;refGene_tx=XM_003981925.5;refGene_gn=NFIX;refGene_tx=XM_019818198.2;refGene_gn=NFIX;refGene_tx=XM_019818203.2
A2 9277948 9278504
refGene_gn=NFIX;refGene_tx=XM_003981925.5;refGene_gn=NFIX;refGene_tx=XM_019818198.2;refGene_gn=NFIX;refGene_tx=XM_019818203.2;refGene_gn=NFIX;refGene_tx=XM_019818211.2;refGene_gn=NFIX;refGene_tx=XM_023246090.1;refGene_gn=NFIX;refGene_tx=XM_023246091.1
A2 9325263 9325326
refGene_gn=NFIX;refGene_tx=XM_003981925.5;refGene_gn=NFIX;refGene_tx=XM_019818198.2;refGene_gn=NFIX;refGene_tx=XM_019818203.2;refGene_gn=NFIX;refGene_tx=XM_019818211.2;refGene_gn=NFIX;refGene_tx=XM_023246090.1;refGene_gn=NFIX;refGene_tx=XM_023246091.1
A2 9325688 9325763
refGene_gn=NFIX;refGene_tx=XM_003981925.5;refGene_gn=NFIX;refGene_tx=XM_019818198.2;refGene_gn=NFIX;refGene_tx=XM_019818203.2;refGene_gn=NFIX;refGene_tx=XM_019818211.2;refGene_gn=NFIX;refGene_tx=XM_023246090.1;refGene_gn=NFIX;refGene_tx=XM_023246091.1
A2 9326127 9326248
refGene_gn=NFIX;refGene_tx=XM_003981925.5;refGene_gn=NFIX;refGene_tx=XM_019818198.2;refGene_gn=NFIX;refGene_tx=XM_019818203.2;refGene_gn=NFIX;refGene_tx=XM_019818211.2;refGene_gn=NFIX;refGene_tx=XM_023246090.1;refGene_gn=NFIX;refGene_tx=XM_023246091.1
A2 9327369 9327506
refGene_gn=NFIX;refGene_tx=XM_003981925.5;refGene_gn=NFIX;refGene_tx=XM_019818198.2;refGene_gn=NFIX;refGene_tx=XM_019818203.2;refGene_gn=NFIX;refGene_tx=XM_019818211.2;refGene_gn=NFIX;refGene_tx=XM_023246090.1;refGene_gn=NFIX;refGene_tx=XM_023246091.1
A2 9330347 9330470
refGene_gn=NFIX;refGene_tx=XM_003981925.5;refGene_gn=NFIX;refGene_tx=XM_019818203.2;refGene_gn=NFIX;refGene_tx=XM_019818211.2;refGene_gn=NFIX;refGene_tx=XM_023246090.1;refGene_gn=NFIX;refGene_tx=XM_023246091.1
A2 9332681 9332857
refGene_gn=NFIX;refGene_tx=XM_003981925.5;refGene_gn=NFIX;refGene_tx=XM_019818198.2;refGene_gn=NFIX;refGene_tx=XM_019818203.2;refGene_gn=NFIX;refGene_tx=XM_019818211.2;refGene_gn=NFIX;refGene_tx=XM_023246090.1;refGene_gn=NFIX;refGene_tx=XM_023246091.1
A2 9338704 9338852
refGene_gn=NFIX;refGene_tx=XM_003981925.5;refGene_gn=NFIX;refGene_tx=XM_019818198.2;refGene_gn=NFIX;refGene_tx=XM_023246090.1;refGene_gn=NFIX;refGene_tx=XM_023246091.1
A2 9340975 9341067

refGene_gn=NFIX;refGene_tx=XM_003981925.5;refGene_gn=NFIX;refGene_tx=XM_019818198.2;refGene_gn=NFIX;refGene_tx=XM_019818203.2;refGene_gn=NFIX;refGene_tx=XM_019818211.2;refGene_gn=NFIX;refGene_tx=XM_023246090.1;refGene_gn=NFIX;refGene_tx=XM_023246091.1
A2 9345430 9345445
refGene_gn=NFIX;refGene_tx=XM_003981925.5;refGene_gn=NFIX;refGene_tx=XM_019818198.2;refGene_gn=NFIX;refGene_tx=XM_023246090.1;refGene_gn=NFIX;refGene_tx=XM_023246091.1
A2 9350311 9350782 refGene_gn=LYL1;refGene_tx=XM_023241021.1
A2 9351022 9351139 refGene_gn=LYL1;refGene_tx=XM_023241021.1
A2 9351226 9351344 refGene_gn=LYL1;refGene_tx=XM_023241021.1
A2 9351704 9351901 refGene_gn=LYL1;refGene_tx=XM_023241021.1
A2 9355112 9355262
refGene_gn=TRMT1;refGene_tx=XM_003981951.5;refGene_gn=TRMT1;refGene_tx=XM_006928473.4;refGene_gn=TRMT1;refGene_tx=XM_023246062.1
A2 9355434 9355564
refGene_gn=TRMT1;refGene_tx=XM_003981951.5;refGene_gn=TRMT1;refGene_tx=XM_006928473.4;refGene_gn=TRMT1;refGene_tx=XM_023246062.1
A2 9355640 9355760
refGene_gn=TRMT1;refGene_tx=XM_003981951.5;refGene_gn=TRMT1;refGene_tx=XM_006928473.4;refGene_gn=TRMT1;refGene_tx=XM_023246062.1
A2 9356627 9356704
refGene_gn=TRMT1;refGene_tx=XM_003981951.5;refGene_gn=TRMT1;refGene_tx=XM_006928473.4;refGene_gn=TRMT1;refGene_tx=XM_023246062.1
A2 9356790 9356899
refGene_gn=TRMT1;refGene_tx=XM_003981951.5;refGene_gn=TRMT1;refGene_tx=XM_006928473.4;refGene_gn=TRMT1;refGene_tx=XM_023246062.1
A2 9357216 9357302
refGene_gn=TRMT1;refGene_tx=XM_003981951.5;refGene_gn=TRMT1;refGene_tx=XM_006928473.4;refGene_gn=TRMT1;refGene_tx=XM_023246062.1
A2 9357393 9357528
refGene_gn=TRMT1;refGene_tx=XM_003981951.5;refGene_gn=TRMT1;refGene_tx=XM_006928473.4;refGene_gn=TRMT1;refGene_tx=XM_023246062.1
A2 9357627 9357697
refGene_gn=TRMT1;refGene_tx=XM_003981951.5;refGene_gn=TRMT1;refGene_tx=XM_006928473.4;refGene_gn=TRMT1;refGene_tx=XM_023246062.1
A2 9357777 9357864
refGene_gn=TRMT1;refGene_tx=XM_003981951.5;refGene_gn=TRMT1;refGene_tx=XM_006928473.4;refGene_gn=TRMT1;refGene_tx=XM_023246062.1
A2 9358021 9358170
refGene_gn=TRMT1;refGene_tx=XM_003981951.5;refGene_gn=TRMT1;refGene_tx=XM_006928473.4;refGene_gn=TRMT1;refGene_tx=XM_023246062.1
A2 9358951 9359064
refGene_gn=TRMT1;refGene_tx=XM_003981951.5;refGene_gn=TRMT1;refGene_tx=XM_006928473.4;refGene_gn=TRMT1;refGene_tx=XM_023246062.1
A2 9359142 9359258
refGene_gn=TRMT1;refGene_tx=XM_003981951.5;refGene_gn=TRMT1;refGene_tx=XM_006928473.4;refGene_gn=TRMT1;refGene_tx=XM_023246062.1
A2 9363563 9363751
refGene_gn=TRMT1;refGene_tx=XM_003981951.5;refGene_gn=TRMT1;refGene_tx

=XM_006928473.4
A2 9363916 9364059
refGene_gn=TRMT1;refGene_tx=XM_003981951.5;refGene_gn=TRMT1;refGene_tx
=XM_006928473.4
A2 9364311 9364367
refGene_gn=TRMT1;refGene_tx=XM_003981951.5;refGene_gn=TRMT1;refGene_tx
=XM_006928473.4;refGene_gn=TRMT1;refGene_tx=XM_023246062.1
A2 9364460 9364720
refGene_gn=TRMT1;refGene_tx=XM_003981951.5;refGene_gn=TRMT1;refGene_tx
=XM_006928473.4
A2 9379165 9380138
refGene_gn=NACC1;refGene_tx=XM_011289451.3;refGene_gn=NACC1;refGene_tx
=XM_023246066.1;refGene_gn=NACC1;refGene_tx=XM_023246074.1
A2 9380225 9380399
refGene_gn=NACC1;refGene_tx=XM_011289451.3;refGene_gn=NACC1;refGene_tx
=XM_023246066.1;refGene_gn=NACC1;refGene_tx=XM_023246074.1
A2 9380985 9381091
refGene_gn=NACC1;refGene_tx=XM_011289451.3;refGene_gn=NACC1;refGene_tx
=XM_023246066.1;refGene_gn=NACC1;refGene_tx=XM_023246074.1
A2 9381189 9381287
refGene_gn=NACC1;refGene_tx=XM_011289451.3;refGene_gn=NACC1;refGene_tx
=XM_023246066.1;refGene_gn=NACC1;refGene_tx=XM_023246074.1
A2 9381858 9382118
refGene_gn=NACC1;refGene_tx=XM_011289451.3;refGene_gn=NACC1;refGene_tx
=XM_023246066.1;refGene_gn=NACC1;refGene_tx=XM_023246074.1
A2 9384648 9384745
refGene_gn=NACC1;refGene_tx=XM_023246066.1;refGene_gn=NACC1;refGene_tx
=XM_023246074.1
A2 9386975 9387052
refGene_gn=STX10;refGene_tx=XM_003981927.4;refGene_gn=STX10;refGene_tx
=XM_023246093.1
A2 9387149 9387244
refGene_gn=STX10;refGene_tx=XM_003981927.4;refGene_gn=STX10;refGene_tx
=XM_011289462.3;refGene_gn=STX10;refGene_tx=XM_023246093.1
A2 9387341 9387448
refGene_gn=STX10;refGene_tx=XM_003981927.4;refGene_gn=STX10;refGene_tx
=XM_011289462.3;refGene_gn=STX10;refGene_tx=XM_023246093.1
A2 9387542 9387650
refGene_gn=STX10;refGene_tx=XM_003981927.4;refGene_gn=STX10;refGene_tx
=XM_011289462.3;refGene_gn=STX10;refGene_tx=XM_023246093.1
A2 9388537 9388600
refGene_gn=STX10;refGene_tx=XM_003981927.4;refGene_gn=STX10;refGene_tx
=XM_011289462.3;refGene_gn=STX10;refGene_tx=XM_023246093.1
A2 9388677 9388772
refGene_gn=STX10;refGene_tx=XM_003981927.4;refGene_gn=STX10;refGene_tx
=XM_011289462.3;refGene_gn=STX10;refGene_tx=XM_023246093.1
A2 9388914 9389084
refGene_gn=STX10;refGene_tx=XM_003981927.4;refGene_gn=STX10;refGene_tx
=XM_011289462.3;refGene_gn=STX10;refGene_tx=XM_023246093.1
A2 9389342 9389377

refGene_gn=STX10;refGene_tx=XM_003981927.4;refGene_gn=STX10;refGene_tx
=XM_011289462.3

| | | | |
|---|---|---|---|
| A2 | 9389506 | 9389574 | refGene_gn=STX10;refGene_tx=XM_023246093.1 |
| A2 | 9392631 | 9393303 | refGene_gn=IER2;refGene_tx=XM_003981952.5 |
| A2 | 9424210 | 9424906 | refGene_gn=CACNA1A;refGene_tx=XM_023246121.1 |
| A2 | 9425755 | 9426006 | refGene_gn=CACNA1A;refGene_tx=XM_023246121.1 |
| A2 | 9426310 | 9426497 | refGene_gn=CACNA1A;refGene_tx=XM_023246121.1 |
| A2 | 9427556 | 9427592 | refGene_gn=CACNA1A;refGene_tx=XM_023246121.1 |
| A2 | 9428950 | 9429064 | refGene_gn=CACNA1A;refGene_tx=XM_023246121.1 |
| A2 | 9429197 | 9429336 | refGene_gn=CACNA1A;refGene_tx=XM_023246121.1 |
| A2 | 9429450 | 9429560 | refGene_gn=CACNA1A;refGene_tx=XM_023246121.1 |
| A2 | 9430104 | 9430205 | refGene_gn=CACNA1A;refGene_tx=XM_023246121.1 |
| A2 | 9430351 | 9430459 | refGene_gn=CACNA1A;refGene_tx=XM_023246121.1 |
| A2 | 9438970 | 9439076 | refGene_gn=CACNA1A;refGene_tx=XM_023246121.1 |
| A2 | 9441539 | 9441636 | refGene_gn=CACNA1A;refGene_tx=XM_023246121.1 |
| A2 | 9443842 | 9443970 | refGene_gn=CACNA1A;refGene_tx=XM_023246121.1 |
| A2 | 9446854 | 9447005 | refGene_gn=CACNA1A;refGene_tx=XM_023246121.1 |
| A2 | 9449112 | 9449228 | refGene_gn=CACNA1A;refGene_tx=XM_023246121.1 |
| A2 | 9449399 | 9449465 | refGene_gn=CACNA1A;refGene_tx=XM_023246121.1 |
| A2 | 9449817 | 9449934 | refGene_gn=CACNA1A;refGene_tx=XM_023246121.1 |
| A2 | 9454109 | 9454115 | refGene_gn=CACNA1A;refGene_tx=XM_023246121.1 |
| A2 | 9457145 | 9457229 | refGene_gn=CACNA1A;refGene_tx=XM_023246121.1 |
| A2 | 9460869 | 9460980 | refGene_gn=CACNA1A;refGene_tx=XM_023246121.1 |
| A2 | 9464402 | 9464567 | refGene_gn=CACNA1A;refGene_tx=XM_023246121.1 |
| A2 | 9465592 | 9465794 | refGene_gn=CACNA1A;refGene_tx=XM_023246121.1 |
| A2 | 9467054 | 9467192 | refGene_gn=CACNA1A;refGene_tx=XM_023246121.1 |
| A2 | 9469182 | 9469343 | refGene_gn=CACNA1A;refGene_tx=XM_023246121.1 |
| A2 | 9471299 | 9471399 | refGene_gn=CACNA1A;refGene_tx=XM_023246121.1 |
| A2 | 9478843 | 9478950 | refGene_gn=CACNA1A;refGene_tx=XM_023246121.1 |
| A2 | 9479643 | 9479703 | refGene_gn=CACNA1A;refGene_tx=XM_023246121.1 |
| A2 | 9484120 | 9484250 | refGene_gn=CACNA1A;refGene_tx=XM_023246121.1 |
| A2 | 9486036 | 9486175 | refGene_gn=CACNA1A;refGene_tx=XM_023246121.1 |
| A2 | 9487628 | 9488092 | refGene_gn=CACNA1A;refGene_tx=XM_023246121.1 |
| A2 | 9495769 | 9496221 | refGene_gn=CACNA1A;refGene_tx=XM_023246121.1 |
| A2 | 9496222 | 9496586 | refGene_gn=CACNA1A;refGene_tx=XM_023246121.1 |
| A2 | 9498443 | 9498550 | refGene_gn=CACNA1A;refGene_tx=XM_023246121.1 |
| A2 | 9500547 | 9500615 | refGene_gn=CACNA1A;refGene_tx=XM_023246121.1 |
| A2 | 9500778 | 9500896 | refGene_gn=CACNA1A;refGene_tx=XM_023246121.1 |
| A2 | 9503391 | 9503464 | refGene_gn=CACNA1A;refGene_tx=XM_023246121.1 |
| A2 | 9503760 | 9503892 | refGene_gn=CACNA1A;refGene_tx=XM_023246121.1 |
| A2 | 9504030 | 9504143 | refGene_gn=CACNA1A;refGene_tx=XM_023246121.1 |
| A2 | 9505091 | 9505204 | refGene_gn=CACNA1A;refGene_tx=XM_023246121.1 |
| A2 | 9509158 | 9509368 | refGene_gn=CACNA1A;refGene_tx=XM_023246121.1 |
| A2 | 9516508 | 9516598 | refGene_gn=CACNA1A;refGene_tx=XM_023246121.1 |
| A2 | 9517702 | 9517759 | refGene_gn=CACNA1A;refGene_tx=XM_023246121.1 |
| A2 | 9518833 | 9518949 | refGene_gn=CACNA1A;refGene_tx=XM_023246121.1 |
| A2 | 9520082 | 9520186 | refGene_gn=CACNA1A;refGene_tx=XM_023246121.1 |
| A2 | 9534446 | 9534640 | refGene_gn=CACNA1A;refGene_tx=XM_023246121.1 |
| A2 | 9538067 | 9538220 | refGene_gn=CACNA1A;refGene_tx=XM_023246121.1 |
| A2 | 9540677 | 9540769 | refGene_gn=CACNA1A;refGene_tx=XM_023246121.1 |

| | | | |
|---|---|---|---|
| A2 | 9593439 | 9593579 | refGene_gn=CACNA1A;refGene_tx=XM_023246121.1 |
| A2 | 9596252 | 9596358 | refGene_gn=CACNA1A;refGene_tx=XM_023246121.1 |
| A2 | 9634393 | 9634687 | refGene_gn=CACNA1A;refGene_tx=XM_023246121.1 |
| A2 | 9693534 | 9693794 | refGene_gn=CACNA1A;refGene_tx=XM_023246121.1 |
| A2 | 9778676 | 9778679 | refGene_gn=CCDC130;refGene_tx=XM_023246129.1 |
| A2 | 9779947 | 9780001 | refGene_gn=CCDC130;refGene_tx=XM_023246129.1 |
| A2 | 9781720 | 9781803 | refGene_gn=CCDC130;refGene_tx=XM_023246129.1 |
| A2 | 9783020 | 9783076 | refGene_gn=CCDC130;refGene_tx=XM_023246129.1 |
| A2 | 9784444 | 9784505 | refGene_gn=CCDC130;refGene_tx=XM_023246129.1 |
| A2 | 9785333 | 9785476 | refGene_gn=CCDC130;refGene_tx=XM_023246129.1 |
| A2 | 9785568 | 9785741 | refGene_gn=CCDC130;refGene_tx=XM_023246129.1 |
| A2 | 9786525 | 9786607 | refGene_gn=CCDC130;refGene_tx=XM_023246129.1 |
| A2 | 9786609 | 9786663 | refGene_gn=CCDC130;refGene_tx=XM_023246129.1 |
| A2 | 9786796 | 9786870 | refGene_gn=CCDC130;refGene_tx=XM_023246129.1 |
| A2 | 9786871 | 9786927 | refGene_gn=CCDC130;refGene_tx=XM_023246129.1 |
| A2 | 9786928 | 9787271 | refGene_gn=CCDC130;refGene_tx=XM_023246129.1 |
| A2 | 9788569 | 9788701 | refGene_gn=MRI1;refGene_tx=XM_023246141.1 |
| A2 | 9788777 | 9788943 | refGene_gn=MRI1;refGene_tx=XM_023246141.1 |
| A2 | 9788945 | 9789015 | refGene_gn=MRI1;refGene_tx=XM_023246141.1 |
| A2 | 9789820 | 9789997 | refGene_gn=MRI1;refGene_tx=XM_023246141.1 |
| A2 | 9791297 | 9791362 | refGene_gn=MRI1;refGene_tx=XM_023246141.1 |
| A2 | 9791364 | 9791481 | refGene_gn=MRI1;refGene_tx=XM_023246141.1 |
| A2 | 9791559 | 9791783 | refGene_gn=MRI1;refGene_tx=XM_023246141.1 |
| A2 | 9792982 | 9792983 | refGene_gn=MRI1;refGene_tx=XM_023246141.1 |
| A2 | 9793154 | 9793217 | refGene_gn=MRI1;refGene_tx=XM_023246141.1 |
| A2 | 9793219 | 9793276 | refGene_gn=MRI1;refGene_tx=XM_023246141.1 |
| A2 | 9793278 | 9793313 | refGene_gn=MRI1;refGene_tx=XM_023246141.1 |
| A2 | 9794107 | 9794204 | refGene_gn=CA2H19orf53;refGene_tx=XM_023246135.1 |
| A2 | 9794281 | 9794337 | refGene_gn=CA2H19orf53;refGene_tx=XM_023246135.1 |
| A2 | 9796366 | 9796513 | refGene_gn=CA2H19orf53;refGene_tx=XM_023246135.1 |
| A2 | 9804405 | 9804570 | refGene_gn=ZSWIM4;refGene_tx=XM_023246142.1 |
| A2 | 9806565 | 9806767 | refGene_gn=ZSWIM4;refGene_tx=XM_023246142.1 |
| A2 | 9808402 | 9808759 | refGene_gn=ZSWIM4;refGene_tx=XM_023246142.1 |
| A2 | 9809374 | 9809523 | refGene_gn=ZSWIM4;refGene_tx=XM_023246142.1 |
| A2 | 9809614 | 9809765 | refGene_gn=ZSWIM4;refGene_tx=XM_023246142.1 |
| A2 | 9810881 | 9811049 | refGene_gn=ZSWIM4;refGene_tx=XM_023246142.1 |
| A2 | 9811606 | 9811963 | refGene_gn=ZSWIM4;refGene_tx=XM_023246142.1 |
| A2 | 9813405 | 9813543 | refGene_gn=ZSWIM4;refGene_tx=XM_023246142.1 |
| A2 | 9813861 | 9814116 | refGene_gn=ZSWIM4;refGene_tx=XM_023246142.1 |
| A2 | 9814669 | 9814805 | refGene_gn=ZSWIM4;refGene_tx=XM_023246142.1 |
| A2 | 9816865 | 9817020 | refGene_gn=ZSWIM4;refGene_tx=XM_023246142.1 |
| A2 | 9818254 | 9818418 | refGene_gn=ZSWIM4;refGene_tx=XM_023246142.1 |
| A2 | 9819905 | 9819987 | refGene_gn=ZSWIM4;refGene_tx=XM_023246142.1 |
| A2 | 9820391 | 9821251 | refGene_gn=ZSWIM4;refGene_tx=XM_023246142.1 |
| A2 | 9853572 | 9854238 | refGene_gn=NANOS3;refGene_tx=XM_023246151.1 |
| A2 | 9854239 | 9854411 | refGene_gn=NANOS3;refGene_tx=XM_023246151.1 |
| A2 | 9856351 | 9856419 | refGene_gn=NANOS3;refGene_tx=XM_023246151.1 |

A2	9859089	9859240
refGene_gn=CA2H19orf57;refGene_tx=XM_006928481.4;refGene_gn=CA2H19orf57;refGene_tx=XM_006928482.4;refGene_gn=CA2H19orf57;refGene_tx=XM_006928483.4;refGene_gn=CA2H19orf57;refGene_tx=XM_019818350.2
A2	9859482	9859575
refGene_gn=CA2H19orf57;refGene_tx=XM_006928481.4;refGene_gn=CA2H19orf57;refGene_tx=XM_006928482.4;refGene_gn=CA2H19orf57;refGene_tx=XM_006928483.4;refGene_gn=CA2H19orf57;refGene_tx=XM_019818350.2
A2	9861603	9861692
refGene_gn=CA2H19orf57;refGene_tx=XM_006928481.4;refGene_gn=CA2H19orf57;refGene_tx=XM_006928482.4;refGene_gn=CA2H19orf57;refGene_tx=XM_006928483.4;refGene_gn=CA2H19orf57;refGene_tx=XM_019818350.2
A2	9861934	9862045
refGene_gn=CA2H19orf57;refGene_tx=XM_006928481.4;refGene_gn=CA2H19orf57;refGene_tx=XM_006928482.4;refGene_gn=CA2H19orf57;refGene_tx=XM_019818350.2
A2	9865154	9866459
refGene_gn=CA2H19orf57;refGene_tx=XM_006928481.4;refGene_gn=CA2H19orf57;refGene_tx=XM_006928482.4;refGene_gn=CA2H19orf57;refGene_tx=XM_006928483.4
A2	9870344	9870455
refGene_gn=CA2H19orf57;refGene_tx=XM_006928481.4;refGene_gn=CA2H19orf57;refGene_tx=XM_006928483.4;refGene_gn=CA2H19orf57;refGene_tx=XM_019818350.2
A2	9870689	9870771
refGene_gn=CA2H19orf57;refGene_tx=XM_006928481.4;refGene_gn=CA2H19orf57;refGene_tx=XM_006928482.4;refGene_gn=CA2H19orf57;refGene_tx=XM_006928483.4;refGene_gn=CA2H19orf57;refGene_tx=XM_019818350.2
A2	9873753	9873931
refGene_gn=CA2H19orf57;refGene_tx=XM_006928481.4;refGene_gn=CA2H19orf57;refGene_tx=XM_006928482.4;refGene_gn=CA2H19orf57;refGene_tx=XM_006928483.4;refGene_gn=CA2H19orf57;refGene_tx=XM_019818350.2
A2	9878029	9878081
refGene_gn=CA2H19orf57;refGene_tx=XM_006928481.4;refGene_gn=CA2H19orf57;refGene_tx=XM_006928482.4;refGene_gn=CA2H19orf57;refGene_tx=XM_006928483.4;refGene_gn=CA2H19orf57;refGene_tx=XM_019818350.2
A2	9878521	9878585
refGene_gn=CA2H19orf57;refGene_tx=XM_006928481.4;refGene_gn=CA2H19orf57;refGene_tx=XM_006928482.4;refGene_gn=CA2H19orf57;refGene_tx=XM_006928483.4;refGene_gn=CA2H19orf57;refGene_tx=XM_019818350.2
A2	9878880	9878912
refGene_gn=CA2H19orf57;refGene_tx=XM_006928481.4;refGene_gn=CA2H19orf57;refGene_tx=XM_006928482.4;refGene_gn=CA2H19orf57;refGene_tx=XM_006928483.4;refGene_gn=CA2H19orf57;refGene_tx=XM_019818350.2
A2	9879292	9879352
refGene_gn=CC2D1A;refGene_tx=XM_003981932.4;refGene_gn=CC2D1A;refGene_tx=XM_006928486.3;refGene_gn=CC2D1A;refGene_tx=XM_006928487.3
A2	9880919	9881055
refGene_gn=CC2D1A;refGene_tx=XM_003981932.4;refGene_gn=CC2D1A;refGene_tx=XM_006928486.3;refGene_gn=CC2D1A;refGene_tx=XM_006928487.3

A2	9882917	9883030
refGene_gn=CC2D1A;refGene_tx=XM_003981932.4;refGene_gn=CC2D1A;refGene_tx=XM_006928486.3;refGene_gn=CC2D1A;refGene_tx=XM_006928487.3
A2	9883124	9883190
refGene_gn=CC2D1A;refGene_tx=XM_003981932.4;refGene_gn=CC2D1A;refGene_tx=XM_006928486.3;refGene_gn=CC2D1A;refGene_tx=XM_006928487.3
A2	9883474	9883609
refGene_gn=CC2D1A;refGene_tx=XM_003981932.4;refGene_gn=CC2D1A;refGene_tx=XM_006928486.3;refGene_gn=CC2D1A;refGene_tx=XM_006928487.3
A2	9883712	9883950
refGene_gn=CC2D1A;refGene_tx=XM_003981932.4;refGene_gn=CC2D1A;refGene_tx=XM_006928486.3;refGene_gn=CC2D1A;refGene_tx=XM_006928487.3
A2	9887527	9887652
refGene_gn=CC2D1A;refGene_tx=XM_003981932.4;refGene_gn=CC2D1A;refGene_tx=XM_006928486.3;refGene_gn=CC2D1A;refGene_tx=XM_006928487.3
A2	9887934	9888007
refGene_gn=CC2D1A;refGene_tx=XM_003981932.4;refGene_gn=CC2D1A;refGene_tx=XM_006928486.3;refGene_gn=CC2D1A;refGene_tx=XM_006928487.3
A2	9888152	9888224
refGene_gn=CC2D1A;refGene_tx=XM_003981932.4;refGene_gn=CC2D1A;refGene_tx=XM_006928486.3;refGene_gn=CC2D1A;refGene_tx=XM_006928487.3
A2	9888332	9888463
refGene_gn=CC2D1A;refGene_tx=XM_003981932.4;refGene_gn=CC2D1A;refGene_tx=XM_006928486.3;refGene_gn=CC2D1A;refGene_tx=XM_006928487.3
A2	9888541	9888614
refGene_gn=CC2D1A;refGene_tx=XM_003981932.4;refGene_gn=CC2D1A;refGene_tx=XM_006928486.3;refGene_gn=CC2D1A;refGene_tx=XM_006928487.3
A2	9889301	9889435
refGene_gn=CC2D1A;refGene_tx=XM_003981932.4;refGene_gn=CC2D1A;refGene_tx=XM_006928486.3;refGene_gn=CC2D1A;refGene_tx=XM_006928487.3
A2	9889773	9889888
refGene_gn=CC2D1A;refGene_tx=XM_003981932.4;refGene_gn=CC2D1A;refGene_tx=XM_006928486.3;refGene_gn=CC2D1A;refGene_tx=XM_006928487.3
A2	9889969	9890142
refGene_gn=CC2D1A;refGene_tx=XM_003981932.4;refGene_gn=CC2D1A;refGene_tx=XM_006928486.3;refGene_gn=CC2D1A;refGene_tx=XM_006928487.3
A2	9890802	9890925
refGene_gn=CC2D1A;refGene_tx=XM_003981932.4;refGene_gn=CC2D1A;refGene_tx=XM_006928486.3;refGene_gn=CC2D1A;refGene_tx=XM_006928487.3
A2	9891007	9891066
refGene_gn=CC2D1A;refGene_tx=XM_003981932.4;refGene_gn=CC2D1A;refGene_tx=XM_006928486.3;refGene_gn=CC2D1A;refGene_tx=XM_006928487.3
A2	9891157	9891274
refGene_gn=CC2D1A;refGene_tx=XM_003981932.4;refGene_gn=CC2D1A;refGene_tx=XM_006928486.3;refGene_gn=CC2D1A;refGene_tx=XM_006928487.3
A2	9892592	9892666
refGene_gn=CC2D1A;refGene_tx=XM_003981932.4;refGene_gn=CC2D1A;refGene_tx=XM_006928486.3;refGene_gn=CC2D1A;refGene_tx=XM_006928487.3
A2	9892740	9892799
refGene_gn=CC2D1A;refGene_tx=XM_003981932.4;refGene_gn=CC2D1A;refGene_

tx=XM_006928486.3;refGene_gn=CC2D1A;refGene_tx=XM_006928487.3
A2	9892923	9892975
refGene_gn=CC2D1A;refGene_tx=XM_003981932.4;refGene_gn=CC2D1A;refGene_
tx=XM_006928486.3;refGene_gn=CC2D1A;refGene_tx=XM_006928487.3
A2	9893094	9893194
refGene_gn=CC2D1A;refGene_tx=XM_003981932.4;refGene_gn=CC2D1A;refGene_
tx=XM_006928486.3;refGene_gn=CC2D1A;refGene_tx=XM_006928487.3
A2	9893280	9893371
refGene_gn=CC2D1A;refGene_tx=XM_003981932.4;refGene_gn=CC2D1A;refGene_
tx=XM_006928486.3;refGene_gn=CC2D1A;refGene_tx=XM_006928487.3
A2	9893685	9893823
refGene_gn=CC2D1A;refGene_tx=XM_003981932.4;refGene_gn=CC2D1A;refGene_
tx=XM_006928486.3;refGene_gn=CC2D1A;refGene_tx=XM_006928487.3
A2	9893910	9893975
refGene_gn=CC2D1A;refGene_tx=XM_003981932.4;refGene_gn=CC2D1A;refGene_
tx=XM_006928486.3;refGene_gn=CC2D1A;refGene_tx=XM_006928487.3
A2	9894559	9894623
refGene_gn=CC2D1A;refGene_tx=XM_003981932.4;refGene_gn=CC2D1A;refGene_
tx=XM_006928486.3;refGene_gn=CC2D1A;refGene_tx=XM_006928487.3
A2	9894718	9894845
refGene_gn=CC2D1A;refGene_tx=XM_003981932.4;refGene_gn=CC2D1A;refGene_
tx=XM_006928486.3;refGene_gn=CC2D1A;refGene_tx=XM_006928487.3
A2	9894923	9895000
refGene_gn=CC2D1A;refGene_tx=XM_003981932.4;refGene_gn=CC2D1A;refGene_
tx=XM_006928486.3;refGene_gn=CC2D1A;refGene_tx=XM_006928487.3
A2	9895079	9895127
refGene_gn=CC2D1A;refGene_tx=XM_003981932.4;refGene_gn=CC2D1A;refGene_
tx=XM_006928486.3;refGene_gn=CC2D1A;refGene_tx=XM_006928487.3
A2	9895211	9895232
refGene_gn=CC2D1A;refGene_tx=XM_003981932.4;refGene_gn=CC2D1A;refGene_
tx=XM_006928486.3;refGene_gn=CC2D1A;refGene_tx=XM_006928487.3
A2	9896379	9896554	refGene_gn=PODNL1;refGene_tx=XM_023241031.1
A2	9896710	9896859	refGene_gn=PODNL1;refGene_tx=XM_023241031.1
A2	9897514	9898286	refGene_gn=PODNL1;refGene_tx=XM_023241031.1
A2	9898776	9898892	refGene_gn=PODNL1;refGene_tx=XM_023241031.1
A2	9899146	9899303	refGene_gn=PODNL1;refGene_tx=XM_023241031.1
A2	9899632	9899742	refGene_gn=PODNL1;refGene_tx=XM_023241031.1
A2	9899876	9899941	refGene_gn=PODNL1;refGene_tx=XM_023241031.1
A2	9900229	9900323	refGene_gn=PODNL1;refGene_tx=XM_023241031.1
A2	9900888	9901113	refGene_gn=PODNL1;refGene_tx=XM_023241031.1
A2	9901269	9901272	refGene_gn=PODNL1;refGene_tx=XM_023241031.1
A2	9914035	9914167	refGene_gn=DCAF15;refGene_tx=XM_023246201.1
A2	9915867	9915965	refGene_gn=DCAF15;refGene_tx=XM_023246201.1
A2	9916057	9916193	refGene_gn=DCAF15;refGene_tx=XM_023246201.1
A2	9916955	9917062	refGene_gn=DCAF15;refGene_tx=XM_023246201.1
A2	9917137	9917277	refGene_gn=DCAF15;refGene_tx=XM_023246201.1
A2	9917373	9917544	refGene_gn=DCAF15;refGene_tx=XM_023246201.1
A2	9919478	9919913	refGene_gn=DCAF15;refGene_tx=XM_023246201.1
A2	9920006	9920098	refGene_gn=DCAF15;refGene_tx=XM_023246201.1
A2	9920194	9920323	refGene_gn=DCAF15;refGene_tx=XM_023246201.1

A2	9920437	9920523	refGene_gn=DCAF15;refGene_tx=XM_023246201.1
A2	9920738	9920843	refGene_gn=DCAF15;refGene_tx=XM_023246201.1
A2	9920945	9921061	refGene_gn=DCAF15;refGene_tx=XM_023246201.1
A2	9921466	9921522	refGene_gn=DCAF15;refGene_tx=XM_023246201.1
A2	9923345	9923515
refGene_gn=RFX1;refGene_tx=XM_023246171.1;refGene_gn=RFX1;refGene_tx=XM_023246176.1;refGene_gn=RFX1;refGene_tx=XM_023246181.1;refGene_gn=RFX1;refGene_tx=XM_023246185.1;refGene_gn=RFX1;refGene_tx=XM_023246189.1;refGene_gn=RFX1;refGene_tx=XM_023246191.1;refGene_gn=RFX1;refGene_tx=XM_023246192.1;refGene_gn=RFX1;refGene_tx=XM_023246193.1;refGene_gn=RFX1;refGene_tx=XM_023246194.1;refGene_gn=RFX1;refGene_tx=XM_023246195.1;refGene_gn=RFX1;refGene_tx=XM_023246196.1;refGene_gn=RFX1;refGene_tx=XM_023246197.1;refGene_gn=RFX1;refGene_tx=XM_023246198.1;refGene_gn=RFX1;refGene_tx=XM_023246199.1
A2	9923652	9923698
refGene_gn=RFX1;refGene_tx=XM_023246171.1;refGene_gn=RFX1;refGene_tx=XM_023246176.1;refGene_gn=RFX1;refGene_tx=XM_023246181.1;refGene_gn=RFX1;refGene_tx=XM_023246185.1;refGene_gn=RFX1;refGene_tx=XM_023246189.1;refGene_gn=RFX1;refGene_tx=XM_023246191.1;refGene_gn=RFX1;refGene_tx=XM_023246192.1;refGene_gn=RFX1;refGene_tx=XM_023246193.1;refGene_gn=RFX1;refGene_tx=XM_023246194.1;refGene_gn=RFX1;refGene_tx=XM_023246195.1;refGene_gn=RFX1;refGene_tx=XM_023246196.1;refGene_gn=RFX1;refGene_tx=XM_023246197.1;refGene_gn=RFX1;refGene_tx=XM_023246198.1;refGene_gn=RFX1;refGene_tx=XM_023246199.1
A2	9923785	9923939
refGene_gn=RFX1;refGene_tx=XM_023246171.1;refGene_gn=RFX1;refGene_tx=XM_023246176.1;refGene_gn=RFX1;refGene_tx=XM_023246181.1;refGene_gn=RFX1;refGene_tx=XM_023246185.1;refGene_gn=RFX1;refGene_tx=XM_023246189.1;refGene_gn=RFX1;refGene_tx=XM_023246191.1;refGene_gn=RFX1;refGene_tx=XM_023246192.1;refGene_gn=RFX1;refGene_tx=XM_023246193.1;refGene_gn=RFX1;refGene_tx=XM_023246194.1;refGene_gn=RFX1;refGene_tx=XM_023246195.1;refGene_gn=RFX1;refGene_tx=XM_023246196.1;refGene_gn=RFX1;refGene_tx=XM_023246197.1;refGene_gn=RFX1;refGene_tx=XM_023246198.1;refGene_gn=RFX1;refGene_tx=XM_023246199.1
A2	9924364	9924573
refGene_gn=RFX1;refGene_tx=XM_023246171.1;refGene_gn=RFX1;refGene_tx=XM_023246176.1;refGene_gn=RFX1;refGene_tx=XM_023246181.1;refGene_gn=RFX1;refGene_tx=XM_023246185.1;refGene_gn=RFX1;refGene_tx=XM_023246189.1;refGene_gn=RFX1;refGene_tx=XM_023246191.1;refGene_gn=RFX1;refGene_tx=XM_023246192.1;refGene_gn=RFX1;refGene_tx=XM_023246193.1;refGene_gn=RFX1;refGene_tx=XM_023246194.1;refGene_gn=RFX1;refGene_tx=XM_023246195.1;refGene_gn=RFX1;refGene_tx=XM_023246196.1;refGene_gn=RFX1;refGene_tx=XM_023246197.1;refGene_gn=RFX1;refGene_tx=XM_023246198.1;refGene_gn=RFX1;refGene_tx=XM_023246199.1
A2	9924658	9924808
refGene_gn=RFX1;refGene_tx=XM_023246171.1;refGene_gn=RFX1;refGene_tx=XM_023246176.1;refGene_gn=RFX1;refGene_tx=XM_023246181.1;refGene_gn=RFX1;refGene_tx=XM_023246185.1;refGene_gn=RFX1;refGene_tx=XM_023246189.1;refGene_gn=RFX1;refGene_tx=XM_023246191.1;refGene_gn=RFX1;refGene_tx=XM_023246192.1;refGene_gn=RFX1;refGene_tx=XM_023246193.1;refGene_gn=RFX

1;refGene_tx=XM_023246194.1;refGene_gn=RFX1;refGene_tx=XM_023246195.1;
refGene_gn=RFX1;refGene_tx=XM_023246196.1;refGene_gn=RFX1;refGene_tx=X
M_023246197.1;refGene_gn=RFX1;refGene_tx=XM_023246198.1;refGene_gn=RFX
1;refGene_tx=XM_023246199.1
A2 9926066 9926164
refGene_gn=RFX1;refGene_tx=XM_023246171.1;refGene_gn=RFX1;refGene_tx=X
M_023246176.1;refGene_gn=RFX1;refGene_tx=XM_023246181.1;refGene_gn=RFX
1;refGene_tx=XM_023246185.1;refGene_gn=RFX1;refGene_tx=XM_023246189.1;
refGene_gn=RFX1;refGene_tx=XM_023246191.1;refGene_gn=RFX1;refGene_tx=X
M_023246192.1;refGene_gn=RFX1;refGene_tx=XM_023246193.1;refGene_gn=RFX
1;refGene_tx=XM_023246194.1;refGene_gn=RFX1;refGene_tx=XM_023246195.1;
refGene_gn=RFX1;refGene_tx=XM_023246196.1;refGene_gn=RFX1;refGene_tx=X
M_023246197.1;refGene_gn=RFX1;refGene_tx=XM_023246198.1;refGene_gn=RFX
1;refGene_tx=XM_023246199.1
A2 9926256 9926408
refGene_gn=RFX1;refGene_tx=XM_023246171.1;refGene_gn=RFX1;refGene_tx=X
M_023246176.1;refGene_gn=RFX1;refGene_tx=XM_023246181.1;refGene_gn=RFX
1;refGene_tx=XM_023246185.1;refGene_gn=RFX1;refGene_tx=XM_023246189.1;
refGene_gn=RFX1;refGene_tx=XM_023246191.1;refGene_gn=RFX1;refGene_tx=X
M_023246192.1;refGene_gn=RFX1;refGene_tx=XM_023246193.1;refGene_gn=RFX
1;refGene_tx=XM_023246194.1;refGene_gn=RFX1;refGene_tx=XM_023246195.1;
refGene_gn=RFX1;refGene_tx=XM_023246196.1;refGene_gn=RFX1;refGene_tx=X
M_023246197.1;refGene_gn=RFX1;refGene_tx=XM_023246198.1;refGene_gn=RFX
1;refGene_tx=XM_023246199.1
A2 9926697 9926807
refGene_gn=RFX1;refGene_tx=XM_023246171.1;refGene_gn=RFX1;refGene_tx=X
M_023246176.1;refGene_gn=RFX1;refGene_tx=XM_023246181.1;refGene_gn=RFX
1;refGene_tx=XM_023246185.1;refGene_gn=RFX1;refGene_tx=XM_023246189.1;
refGene_gn=RFX1;refGene_tx=XM_023246191.1;refGene_gn=RFX1;refGene_tx=X
M_023246192.1;refGene_gn=RFX1;refGene_tx=XM_023246193.1;refGene_gn=RFX
1;refGene_tx=XM_023246194.1;refGene_gn=RFX1;refGene_tx=XM_023246195.1;
refGene_gn=RFX1;refGene_tx=XM_023246196.1;refGene_gn=RFX1;refGene_tx=X
M_023246197.1;refGene_gn=RFX1;refGene_tx=XM_023246198.1;refGene_gn=RFX
1;refGene_tx=XM_023246199.1
A2 9926896 9927015
refGene_gn=RFX1;refGene_tx=XM_023246171.1;refGene_gn=RFX1;refGene_tx=X
M_023246176.1;refGene_gn=RFX1;refGene_tx=XM_023246181.1;refGene_gn=RFX
1;refGene_tx=XM_023246185.1;refGene_gn=RFX1;refGene_tx=XM_023246189.1;
refGene_gn=RFX1;refGene_tx=XM_023246191.1;refGene_gn=RFX1;refGene_tx=X
M_023246192.1;refGene_gn=RFX1;refGene_tx=XM_023246193.1;refGene_gn=RFX
1;refGene_tx=XM_023246194.1;refGene_gn=RFX1;refGene_tx=XM_023246195.1;
refGene_gn=RFX1;refGene_tx=XM_023246196.1;refGene_gn=RFX1;refGene_tx=X
M_023246197.1;refGene_gn=RFX1;refGene_tx=XM_023246198.1;refGene_gn=RFX
1;refGene_tx=XM_023246199.1
A2 9927267 9927383
refGene_gn=RFX1;refGene_tx=XM_023246171.1;refGene_gn=RFX1;refGene_tx=X
M_023246176.1;refGene_gn=RFX1;refGene_tx=XM_023246181.1;refGene_gn=RFX
1;refGene_tx=XM_023246185.1;refGene_gn=RFX1;refGene_tx=XM_023246189.1;
refGene_gn=RFX1;refGene_tx=XM_023246191.1;refGene_gn=RFX1;refGene_tx=X
M_023246192.1;refGene_gn=RFX1;refGene_tx=XM_023246193.1;refGene_gn=RFX

1;refGene_tx=XM_023246194.1;refGene_gn=RFX1;refGene_tx=XM_023246195.1;
refGene_gn=RFX1;refGene_tx=XM_023246196.1;refGene_gn=RFX1;refGene_tx=X
M_023246197.1;refGene_gn=RFX1;refGene_tx=XM_023246198.1;refGene_gn=RFX
1;refGene_tx=XM_023246199.1
A2 9927472 9927592
refGene_gn=RFX1;refGene_tx=XM_023246171.1;refGene_gn=RFX1;refGene_tx=X
M_023246176.1;refGene_gn=RFX1;refGene_tx=XM_023246181.1;refGene_gn=RFX
1;refGene_tx=XM_023246185.1;refGene_gn=RFX1;refGene_tx=XM_023246189.1;
refGene_gn=RFX1;refGene_tx=XM_023246191.1;refGene_gn=RFX1;refGene_tx=X
M_023246192.1;refGene_gn=RFX1;refGene_tx=XM_023246193.1;refGene_gn=RFX
1;refGene_tx=XM_023246194.1;refGene_gn=RFX1;refGene_tx=XM_023246195.1;
refGene_gn=RFX1;refGene_tx=XM_023246196.1;refGene_gn=RFX1;refGene_tx=X
M_023246197.1;refGene_gn=RFX1;refGene_tx=XM_023246198.1;refGene_gn=RFX
1;refGene_tx=XM_023246199.1
A2 9928586 9928768
refGene_gn=RFX1;refGene_tx=XM_023246171.1;refGene_gn=RFX1;refGene_tx=X
M_023246176.1;refGene_gn=RFX1;refGene_tx=XM_023246181.1;refGene_gn=RFX
1;refGene_tx=XM_023246185.1;refGene_gn=RFX1;refGene_tx=XM_023246189.1;
refGene_gn=RFX1;refGene_tx=XM_023246191.1;refGene_gn=RFX1;refGene_tx=X
M_023246192.1;refGene_gn=RFX1;refGene_tx=XM_023246193.1;refGene_gn=RFX
1;refGene_tx=XM_023246194.1;refGene_gn=RFX1;refGene_tx=XM_023246195.1;
refGene_gn=RFX1;refGene_tx=XM_023246196.1;refGene_gn=RFX1;refGene_tx=X
M_023246197.1;refGene_gn=RFX1;refGene_tx=XM_023246198.1;refGene_gn=RFX
1;refGene_tx=XM_023246199.1
A2 9930947 9931332
refGene_gn=RFX1;refGene_tx=XM_023246171.1;refGene_gn=RFX1;refGene_tx=X
M_023246176.1;refGene_gn=RFX1;refGene_tx=XM_023246181.1;refGene_gn=RFX
1;refGene_tx=XM_023246185.1;refGene_gn=RFX1;refGene_tx=XM_023246189.1;
refGene_gn=RFX1;refGene_tx=XM_023246191.1;refGene_gn=RFX1;refGene_tx=X
M_023246192.1;refGene_gn=RFX1;refGene_tx=XM_023246193.1;refGene_gn=RFX
1;refGene_tx=XM_023246194.1;refGene_gn=RFX1;refGene_tx=XM_023246195.1;
refGene_gn=RFX1;refGene_tx=XM_023246196.1;refGene_gn=RFX1;refGene_tx=X
M_023246197.1;refGene_gn=RFX1;refGene_tx=XM_023246198.1;refGene_gn=RFX
1;refGene_tx=XM_023246199.1
A2 9934171 9934266
refGene_gn=RFX1;refGene_tx=XM_023246171.1;refGene_gn=RFX1;refGene_tx=X
M_023246176.1;refGene_gn=RFX1;refGene_tx=XM_023246181.1;refGene_gn=RFX
1;refGene_tx=XM_023246185.1;refGene_gn=RFX1;refGene_tx=XM_023246189.1;
refGene_gn=RFX1;refGene_tx=XM_023246191.1;refGene_gn=RFX1;refGene_tx=X
M_023246192.1;refGene_gn=RFX1;refGene_tx=XM_023246193.1;refGene_gn=RFX
1;refGene_tx=XM_023246194.1;refGene_gn=RFX1;refGene_tx=XM_023246195.1;
refGene_gn=RFX1;refGene_tx=XM_023246196.1;refGene_gn=RFX1;refGene_tx=X
M_023246197.1;refGene_gn=RFX1;refGene_tx=XM_023246198.1;refGene_gn=RFX
1;refGene_tx=XM_023246199.1
A2 9935074 9935458
refGene_gn=RFX1;refGene_tx=XM_023246171.1;refGene_gn=RFX1;refGene_tx=X
M_023246176.1;refGene_gn=RFX1;refGene_tx=XM_023246181.1;refGene_gn=RFX
1;refGene_tx=XM_023246185.1;refGene_gn=RFX1;refGene_tx=XM_023246189.1;
refGene_gn=RFX1;refGene_tx=XM_023246191.1;refGene_gn=RFX1;refGene_tx=X
M_023246192.1;refGene_gn=RFX1;refGene_tx=XM_023246193.1;refGene_gn=RFX

1;refGene_tx=XM_023246194.1;refGene_gn=RFX1;refGene_tx=XM_023246195.1;
refGene_gn=RFX1;refGene_tx=XM_023246196.1
A2 9935854 9935950
refGene_gn=RFX1;refGene_tx=XM_023246171.1;refGene_gn=RFX1;refGene_tx=X
M_023246176.1;refGene_gn=RFX1;refGene_tx=XM_023246181.1;refGene_gn=RFX
1;refGene_tx=XM_023246185.1;refGene_gn=RFX1;refGene_tx=XM_023246189.1;
refGene_gn=RFX1;refGene_tx=XM_023246191.1;refGene_gn=RFX1;refGene_tx=X
M_023246192.1;refGene_gn=RFX1;refGene_tx=XM_023246193.1;refGene_gn=RFX
1;refGene_tx=XM_023246194.1;refGene_gn=RFX1;refGene_tx=XM_023246195.1;
refGene_gn=RFX1;refGene_tx=XM_023246196.1;refGene_gn=RFX1;refGene_tx=X
M_023246197.1;refGene_gn=RFX1;refGene_tx=XM_023246198.1;refGene_gn=RFX
1;refGene_tx=XM_023246199.1
A2 9937026 9937143
refGene_gn=RFX1;refGene_tx=XM_023246171.1;refGene_gn=RFX1;refGene_tx=X
M_023246176.1;refGene_gn=RFX1;refGene_tx=XM_023246181.1;refGene_gn=RFX
1;refGene_tx=XM_023246185.1;refGene_gn=RFX1;refGene_tx=XM_023246189.1;
refGene_gn=RFX1;refGene_tx=XM_023246191.1;refGene_gn=RFX1;refGene_tx=X
M_023246192.1;refGene_gn=RFX1;refGene_tx=XM_023246194.1;refGene_gn=RFX
1;refGene_tx=XM_023246195.1;refGene_gn=RFX1;refGene_tx=XM_023246196.1;
refGene_gn=RFX1;refGene_tx=XM_023246197.1;refGene_gn=RFX1;refGene_tx=X
M_023246198.1
A2 9938110 9938218
refGene_gn=RFX1;refGene_tx=XM_023246171.1;refGene_gn=RFX1;refGene_tx=X
M_023246176.1;refGene_gn=RFX1;refGene_tx=XM_023246181.1;refGene_gn=RFX
1;refGene_tx=XM_023246185.1;refGene_gn=RFX1;refGene_tx=XM_023246189.1;
refGene_gn=RFX1;refGene_tx=XM_023246191.1;refGene_gn=RFX1;refGene_tx=X
M_023246192.1;refGene_gn=RFX1;refGene_tx=XM_023246193.1;refGene_gn=RFX
1;refGene_tx=XM_023246194.1;refGene_gn=RFX1;refGene_tx=XM_023246195.1;
refGene_gn=RFX1;refGene_tx=XM_023246196.1;refGene_gn=RFX1;refGene_tx=X
M_023246197.1;refGene_gn=RFX1;refGene_tx=XM_023246198.1;refGene_gn=RFX
1;refGene_tx=XM_023246199.1
A2 9938887 9938971
refGene_gn=RFX1;refGene_tx=XM_023246171.1;refGene_gn=RFX1;refGene_tx=X
M_023246176.1;refGene_gn=RFX1;refGene_tx=XM_023246181.1;refGene_gn=RFX
1;refGene_tx=XM_023246185.1;refGene_gn=RFX1;refGene_tx=XM_023246189.1;
refGene_gn=RFX1;refGene_tx=XM_023246191.1;refGene_gn=RFX1;refGene_tx=X
M_023246193.1;refGene_gn=RFX1;refGene_tx=XM_023246194.1;refGene_gn=RFX
1;refGene_tx=XM_023246195.1;refGene_gn=RFX1;refGene_tx=XM_023246196.1;
refGene_gn=RFX1;refGene_tx=XM_023246197.1;refGene_gn=RFX1;refGene_tx=X
M_023246198.1;refGene_gn=RFX1;refGene_tx=XM_023246199.1
A2 9939187 9939297
refGene_gn=RFX1;refGene_tx=XM_023246171.1;refGene_gn=RFX1;refGene_tx=X
M_023246176.1;refGene_gn=RFX1;refGene_tx=XM_023246181.1;refGene_gn=RFX
1;refGene_tx=XM_023246185.1;refGene_gn=RFX1;refGene_tx=XM_023246189.1;
refGene_gn=RFX1;refGene_tx=XM_023246191.1;refGene_gn=RFX1;refGene_tx=X
M_023246192.1;refGene_gn=RFX1;refGene_tx=XM_023246193.1;refGene_gn=RFX
1;refGene_tx=XM_023246194.1;refGene_gn=RFX1;refGene_tx=XM_023246195.1;
refGene_gn=RFX1;refGene_tx=XM_023246196.1;refGene_gn=RFX1;refGene_tx=X
M_023246197.1;refGene_gn=RFX1;refGene_tx=XM_023246198.1;refGene_gn=RFX
1;refGene_tx=XM_023246199.1

A2 9945991 9946298
refGene_gn=RFX1;refGene_tx=XM_023246171.1;refGene_gn=RFX1;refGene_tx=XM_023246176.1;refGene_gn=RFX1;refGene_tx=XM_023246181.1;refGene_gn=RFX1;refGene_tx=XM_023246185.1;refGene_gn=RFX1;refGene_tx=XM_023246189.1;refGene_gn=RFX1;refGene_tx=XM_023246191.1;refGene_gn=RFX1;refGene_tx=XM_023246192.1;refGene_gn=RFX1;refGene_tx=XM_023246193.1;refGene_gn=RFX1;refGene_tx=XM_023246197.1;refGene_gn=RFX1;refGene_tx=XM_023246198.1;refGene_gn=RFX1;refGene_tx=XM_023246199.1
A2 9970399 9970589 refGene_gn=RLN3;refGene_tx=XM_019818403.2
A2 9971992 9972207 refGene_gn=RLN3;refGene_tx=XM_019818403.2
A2 9973096 9973190 refGene_gn=IL27RA;refGene_tx=XM_006928488.4
A2 9973616 9973734 refGene_gn=IL27RA;refGene_tx=XM_006928488.4
A2 9977129 9977287 refGene_gn=IL27RA;refGene_tx=XM_006928488.4
A2 9977380 9977538 refGene_gn=IL27RA;refGene_tx=XM_006928488.4
A2 9979101 9979261 refGene_gn=IL27RA;refGene_tx=XM_006928488.4
A2 9979370 9979435 refGene_gn=IL27RA;refGene_tx=XM_006928488.4
A2 9982824 9983008 refGene_gn=IL27RA;refGene_tx=XM_006928488.4
A2 9983118 9983307 refGene_gn=IL27RA;refGene_tx=XM_006928488.4
A2 9985168 9985270 refGene_gn=IL27RA;refGene_tx=XM_006928488.4
A2 9985345 9985504 refGene_gn=IL27RA;refGene_tx=XM_006928488.4
A2 9988395 9988521 refGene_gn=IL27RA;refGene_tx=XM_006928488.4
A2 9988889 9988983 refGene_gn=IL27RA;refGene_tx=XM_006928488.4
A2 9989170 9989264 refGene_gn=IL27RA;refGene_tx=XM_006928488.4
A2 9989390 9989552 refGene_gn=IL27RA;refGene_tx=XM_006928488.4
A2 9989554 9989578 refGene_gn=IL27RA;refGene_tx=XM_006928488.4
A2 9990470 9990608 refGene_gn=PALM3;refGene_tx=XM_019818408.2
A2 9990609 9990935 refGene_gn=PALM3;refGene_tx=XM_019818408.2
A2 9991124 9992270 refGene_gn=PALM3;refGene_tx=XM_019818408.2
A2 9992386 9992432 refGene_gn=PALM3;refGene_tx=XM_019818408.2
A2 9993909 9993994 refGene_gn=PALM3;refGene_tx=XM_019818408.2
A2 9994134 9994277 refGene_gn=PALM3;refGene_tx=XM_019818408.2
A2 9994839 9994920 refGene_gn=PALM3;refGene_tx=XM_019818408.2
A2 9995943 9995992 refGene_gn=PALM3;refGene_tx=XM_019818408.2
A2 9998461 9998502 refGene_gn=PALM3;refGene_tx=XM_019818408.2
A2 10004747 10005315 refGene_gn=MISP3;refGene_tx=XM_023246229.1
A2 10005834 10005908 refGene_gn=MISP3;refGene_tx=XM_023246229.1
A2 10006184 10006202 refGene_gn=MISP3;refGene_tx=XM_023246229.1
A2 10011249 10011424
refGene_gn=CA2H19orf67;refGene_tx=XM_006928489.3;refGene_gn=CA2H19orf67;refGene_tx=XM_019818425.2
A2 10011906 10012041
refGene_gn=CA2H19orf67;refGene_tx=XM_006928489.3;refGene_gn=CA2H19orf67;refGene_tx=XM_019818425.2
A2 10012393 10012579
refGene_gn=CA2H19orf67;refGene_tx=XM_006928489.3;refGene_gn=CA2H19orf67;refGene_tx=XM_019818425.2
A2 10012665 10012766
refGene_gn=CA2H19orf67;refGene_tx=XM_006928489.3;refGene_gn=CA2H19orf67;refGene_tx=XM_019818425.2
A2 10012888 10013033

refGene_gn=CA2H19orf67;refGene_tx=XM_006928489.3;refGene_gn=CA2H19orf67;refGene_tx=XM_019818425.2
A2 10013693 10014028
refGene_gn=CA2H19orf67;refGene_tx=XM_006928489.3;refGene_gn=CA2H19orf67;refGene_tx=XM_019818425.2
A2 10015837 10015996 refGene_gn=SAMD1;refGene_tx=XM_023246248.1
A2 10016094 10016206 refGene_gn=SAMD1;refGene_tx=XM_023246248.1
A2 10016305 10016402 refGene_gn=SAMD1;refGene_tx=XM_023246248.1
A2 10016493 10016788 refGene_gn=SAMD1;refGene_tx=XM_023246248.1
A2 10017225 10017636 refGene_gn=SAMD1;refGene_tx=XM_023246248.1
A2 10017637 10018186 refGene_gn=SAMD1;refGene_tx=XM_023246248.1
A2 10021012 10021138
refGene_gn=PRKACA;refGene_tx=XM_003981935.5;refGene_gn=PRKACA;refGene_tx=XM_006928490.4;refGene_gn=PRKACA;refGene_tx=XM_023246261.1
A2 10021556 10021721
refGene_gn=PRKACA;refGene_tx=XM_003981935.5;refGene_gn=PRKACA;refGene_tx=XM_006928490.4;refGene_gn=PRKACA;refGene_tx=XM_023246261.1
A2 10023404 10023527
refGene_gn=PRKACA;refGene_tx=XM_003981935.5;refGene_gn=PRKACA;refGene_tx=XM_006928490.4;refGene_gn=PRKACA;refGene_tx=XM_023246261.1
A2 10023614 10023710
refGene_gn=PRKACA;refGene_tx=XM_003981935.5;refGene_gn=PRKACA;refGene_tx=XM_006928490.4;refGene_gn=PRKACA;refGene_tx=XM_023246261.1
A2 10023810 10023937
refGene_gn=PRKACA;refGene_tx=XM_003981935.5;refGene_gn=PRKACA;refGene_tx=XM_006928490.4;refGene_gn=PRKACA;refGene_tx=XM_023246261.1
A2 10024740 10024823
refGene_gn=PRKACA;refGene_tx=XM_003981935.5;refGene_gn=PRKACA;refGene_tx=XM_006928490.4;refGene_gn=PRKACA;refGene_tx=XM_023246261.1
A2 10025769 10025868
refGene_gn=PRKACA;refGene_tx=XM_003981935.5;refGene_gn=PRKACA;refGene_tx=XM_006928490.4;refGene_gn=PRKACA;refGene_tx=XM_023246261.1
A2 10027797 10027926
refGene_gn=PRKACA;refGene_tx=XM_003981935.5;refGene_gn=PRKACA;refGene_tx=XM_006928490.4;refGene_gn=PRKACA;refGene_tx=XM_023246261.1
A2 10028411 10028473
refGene_gn=PRKACA;refGene_tx=XM_003981935.5;refGene_gn=PRKACA;refGene_tx=XM_006928490.4;refGene_gn=PRKACA;refGene_tx=XM_023246261.1
A2 10031371 10031444 refGene_gn=PRKACA;refGene_tx=XM_006928490.4
A2 10033268 10033293 refGene_gn=PRKACA;refGene_tx=XM_003981935.5
A2 10036693 10036739 refGene_gn=PRKACA;refGene_tx=XM_023246261.1
A2 10039661 10039868 refGene_gn=ASF1B;refGene_tx=XM_003981937.5
A2 10040252 10040429 refGene_gn=ASF1B;refGene_tx=XM_003981937.5
A2 10042979 10043095 refGene_gn=ASF1B;refGene_tx=XM_003981937.5
A2 10047270 10047379 refGene_gn=ASF1B;refGene_tx=XM_003981937.5
A2 10063317 10064060 refGene_gn=ADGRL1;refGene_tx=XM_023246270.1
A2 10064464 10064482 refGene_gn=ADGRL1;refGene_tx=XM_023246270.1
A2 10064655 10064784 refGene_gn=ADGRL1;refGene_tx=XM_023246270.1
A2 10064876 10064973 refGene_gn=ADGRL1;refGene_tx=XM_023246270.1
A2 10065158 10065287 refGene_gn=ADGRL1;refGene_tx=XM_023246270.1

A2	10066604 10066773 refGene_gn=ADGRL1;refGene_tx=XM_023246270.1
A2	10067323 10067415 refGene_gn=ADGRL1;refGene_tx=XM_023246270.1
A2	10067851 10067918 refGene_gn=ADGRL1;refGene_tx=XM_023246270.1
A2	10068133 10068354 refGene_gn=ADGRL1;refGene_tx=XM_023246270.1
A2	10068448 10068658 refGene_gn=ADGRL1;refGene_tx=XM_023246270.1
A2	10069107 10069278 refGene_gn=ADGRL1;refGene_tx=XM_023246270.1
A2	10069573 10069788 refGene_gn=ADGRL1;refGene_tx=XM_023246270.1
A2	10070312 10070435 refGene_gn=ADGRL1;refGene_tx=XM_023246270.1
A2	10070587 10070771 refGene_gn=ADGRL1;refGene_tx=XM_023246270.1
A2	10070963 10071002 refGene_gn=ADGRL1;refGene_tx=XM_023246270.1
A2	10071321 10071507 refGene_gn=ADGRL1;refGene_tx=XM_023246270.1
A2	10071820 10071924 refGene_gn=ADGRL1;refGene_tx=XM_023246270.1
A2	10072622 10072937 refGene_gn=ADGRL1;refGene_tx=XM_023246270.1
A2	10073747 10074225 refGene_gn=ADGRL1;refGene_tx=XM_023246270.1
A2	10074226 10074343 refGene_gn=ADGRL1;refGene_tx=XM_023246270.1
A2	10074344 10074550 refGene_gn=ADGRL1;refGene_tx=XM_023246270.1
A2	10078087 10078102 refGene_gn=ADGRL1;refGene_tx=XM_023246270.1
A2	10081820 10081930 refGene_gn=ADGRL1;refGene_tx=XM_023246270.1
A2	10085061 10085275 refGene_gn=ADGRL1;refGene_tx=XM_023246270.1
A2	10088216 10088286 refGene_gn=ADGRL1;refGene_tx=XM_023246270.1
A2	10220500 10220540 refGene_gn=ADGRE5;refGene_tx=XM_023246285.1
A2	10223708 10223759 refGene_gn=ADGRE5;refGene_tx=XM_023246285.1
A2	10223974 10224094 refGene_gn=ADGRE5;refGene_tx=XM_023246285.1
A2	10224635 10224791 refGene_gn=ADGRE5;refGene_tx=XM_023246285.1
A2	10227665 10227797 refGene_gn=ADGRE5;refGene_tx=XM_023246285.1
A2	10228374 10228521 refGene_gn=ADGRE5;refGene_tx=XM_023246285.1
A2	10228981 10229128 refGene_gn=ADGRE5;refGene_tx=XM_023246285.1
A2	10229216 10229263 refGene_gn=ADGRE5;refGene_tx=XM_023246285.1
A2	10229407 10229485 refGene_gn=ADGRE5;refGene_tx=XM_023246285.1
A2	10230499 10230677 refGene_gn=ADGRE5;refGene_tx=XM_023246285.1
A2	10230747 10230855 refGene_gn=ADGRE5;refGene_tx=XM_023246285.1
A2	10231078 10231302 refGene_gn=ADGRE5;refGene_tx=XM_023246285.1
A2	10231864 10232044 refGene_gn=ADGRE5;refGene_tx=XM_023246285.1
A2	10232752 10232944 refGene_gn=ADGRE5;refGene_tx=XM_023246285.1
A2	10233186 10233345 refGene_gn=ADGRE5;refGene_tx=XM_023246285.1
A2	10233346 10233414 refGene_gn=ADGRE5;refGene_tx=XM_023246285.1
A2	10233515 10233582 refGene_gn=ADGRE5;refGene_tx=XM_023246285.1
A2	10233734 10233826 refGene_gn=ADGRE5;refGene_tx=XM_023246285.1
A2	10233914 10234083 refGene_gn=ADGRE5;refGene_tx=XM_023246285.1
A2	10234181 10234268 refGene_gn=ADGRE5;refGene_tx=XM_023246285.1
A2	10234352 10234382 refGene_gn=ADGRE5;refGene_tx=XM_023246285.1
A2	10235214 10235231
refGene_gn=DDX39A;refGene_tx=XM_003981942.5;refGene_gn=DDX39A;refGene_tx=XM_006928497.4
A2	10235318 10235466
refGene_gn=DDX39A;refGene_tx=XM_003981942.5;refGene_gn=DDX39A;refGene_tx=XM_006928497.4
A2	10235621 10235766
refGene_gn=DDX39A;refGene_tx=XM_003981942.5;refGene_gn=DDX39A;refGene_tx=XM_006928497.4

A2 10235875 10235985
refGene_gn=DDX39A;refGene_tx=XM_003981942.5;refGene_gn=DDX39A;refGene_tx=XM_006928497.4
A2 10236083 10236215
refGene_gn=DDX39A;refGene_tx=XM_003981942.5;refGene_gn=DDX39A;refGene_tx=XM_006928497.4
A2 10236562 10236681
refGene_gn=DDX39A;refGene_tx=XM_003981942.5;refGene_gn=DDX39A;refGene_tx=XM_006928497.4
A2 10237370 10237554
refGene_gn=DDX39A;refGene_tx=XM_003981942.5;refGene_gn=DDX39A;refGene_tx=XM_006928497.4
A2 10237921 10238014
refGene_gn=DDX39A;refGene_tx=XM_003981942.5;refGene_gn=DDX39A;refGene_tx=XM_006928497.4
A2 10239075 10239203
refGene_gn=DDX39A;refGene_tx=XM_003981942.5;refGene_gn=DDX39A;refGene_tx=XM_006928497.4
A2 10239335 10239543
refGene_gn=DDX39A;refGene_tx=XM_003981942.5;refGene_gn=DDX39A;refGene_tx=XM_006928497.4
A2 10261336 10261357 refGene_gn=PKN1;refGene_tx=XM_023246301.1
A2 10262361 10262418 refGene_gn=PKN1;refGene_tx=XM_023246299.1
A2 10264675 10264714 refGene_gn=PKN1;refGene_tx=XM_023246300.1
A2 10265685 10265989
refGene_gn=PKN1;refGene_tx=XM_023246299.1;refGene_gn=PKN1;refGene_tx=XM_023246300.1;refGene_gn=PKN1;refGene_tx=XM_023246301.1
A2 10267909 10268061
refGene_gn=PKN1;refGene_tx=XM_023246299.1;refGene_gn=PKN1;refGene_tx=XM_023246300.1;refGene_gn=PKN1;refGene_tx=XM_023246301.1
A2 10269184 10269311
refGene_gn=PKN1;refGene_tx=XM_023246299.1;refGene_gn=PKN1;refGene_tx=XM_023246300.1;refGene_gn=PKN1;refGene_tx=XM_023246301.1
A2 10272440 10272586
refGene_gn=PKN1;refGene_tx=XM_023246299.1;refGene_gn=PKN1;refGene_tx=XM_023246300.1;refGene_gn=PKN1;refGene_tx=XM_023246301.1
A2 10272849 10273063
refGene_gn=PKN1;refGene_tx=XM_023246299.1;refGene_gn=PKN1;refGene_tx=XM_023246300.1;refGene_gn=PKN1;refGene_tx=XM_023246301.1
A2 10273820 10274015
refGene_gn=PKN1;refGene_tx=XM_023246299.1;refGene_gn=PKN1;refGene_tx=XM_023246300.1;refGene_gn=PKN1;refGene_tx=XM_023246301.1
A2 10275844 10275954
refGene_gn=PKN1;refGene_tx=XM_023246299.1;refGene_gn=PKN1;refGene_tx=XM_023246300.1;refGene_gn=PKN1;refGene_tx=XM_023246301.1
A2 10276049 10276193
refGene_gn=PKN1;refGene_tx=XM_023246299.1;refGene_gn=PKN1;refGene_tx=XM_023246300.1;refGene_gn=PKN1;refGene_tx=XM_023246301.1
A2 10278005 10278081
refGene_gn=PKN1;refGene_tx=XM_023246299.1;refGene_gn=PKN1;refGene_tx=X

M_023246300.1;refGene_gn=PKN1;refGene_tx=XM_023246301.1
A2 10278165 10278307
refGene_gn=PKN1;refGene_tx=XM_023246299.1;refGene_gn=PKN1;refGene_tx=XM_023246300.1;refGene_gn=PKN1;refGene_tx=XM_023246301.1
A2 10278397 10278491
refGene_gn=PKN1;refGene_tx=XM_023246299.1;refGene_gn=PKN1;refGene_tx=XM_023246300.1;refGene_gn=PKN1;refGene_tx=XM_023246301.1
A2 10280367 10280447
refGene_gn=PKN1;refGene_tx=XM_023246299.1;refGene_gn=PKN1;refGene_tx=XM_023246300.1;refGene_gn=PKN1;refGene_tx=XM_023246301.1
A2 10280523 10280599
refGene_gn=PKN1;refGene_tx=XM_023246299.1;refGene_gn=PKN1;refGene_tx=XM_023246300.1;refGene_gn=PKN1;refGene_tx=XM_023246301.1
A2 10280681 10280773
refGene_gn=PKN1;refGene_tx=XM_023246299.1;refGene_gn=PKN1;refGene_tx=XM_023246300.1;refGene_gn=PKN1;refGene_tx=XM_023246301.1
A2 10282434 10282611
refGene_gn=PKN1;refGene_tx=XM_023246299.1;refGene_gn=PKN1;refGene_tx=XM_023246300.1;refGene_gn=PKN1;refGene_tx=XM_023246301.1
A2 10282815 10282878
refGene_gn=PKN1;refGene_tx=XM_023246299.1;refGene_gn=PKN1;refGene_tx=XM_023246300.1;refGene_gn=PKN1;refGene_tx=XM_023246301.1
A2 10282997 10283074
refGene_gn=PKN1;refGene_tx=XM_023246299.1;refGene_gn=PKN1;refGene_tx=XM_023246300.1;refGene_gn=PKN1;refGene_tx=XM_023246301.1
A2 10283315 10283458
refGene_gn=PKN1;refGene_tx=XM_023246299.1;refGene_gn=PKN1;refGene_tx=XM_023246300.1;refGene_gn=PKN1;refGene_tx=XM_023246301.1
A2 10283812 10283920
refGene_gn=PKN1;refGene_tx=XM_023246299.1;refGene_gn=PKN1;refGene_tx=XM_023246300.1;refGene_gn=PKN1;refGene_tx=XM_023246301.1
A2 10283998 10284079
refGene_gn=PKN1;refGene_tx=XM_023246299.1;refGene_gn=PKN1;refGene_tx=XM_023246300.1;refGene_gn=PKN1;refGene_tx=XM_023246301.1
A2 10284478 10284682
refGene_gn=PKN1;refGene_tx=XM_023246299.1;refGene_gn=PKN1;refGene_tx=XM_023246300.1;refGene_gn=PKN1;refGene_tx=XM_023246301.1
A2 10285489 10285756 refGene_gn=PTGER1;refGene_tx=XM_011290063.3
A2 10286319 10287042 refGene_gn=PTGER1;refGene_tx=XM_011290063.3
A2 10287043 10287262 refGene_gn=PTGER1;refGene_tx=XM_011290063.3
A2 10291219 10291371
refGene_gn=GIPC1;refGene_tx=XM_019818512.2;refGene_gn=GIPC1;refGene_tx=XM_023246319.1;refGene_gn=GIPC1;refGene_tx=XM_023246323.1;refGene_gn=GIPC1;refGene_tx=XM_023246326.1;refGene_gn=GIPC1;refGene_tx=XM_023246329.1;refGene_gn=GIPC1;refGene_tx=XM_023246334.1
A2 10291501 10291583
refGene_gn=GIPC1;refGene_tx=XM_019818512.2;refGene_gn=GIPC1;refGene_tx=XM_023246319.1;refGene_gn=GIPC1;refGene_tx=XM_023246323.1;refGene_gn=GIPC1;refGene_tx=XM_023246326.1;refGene_gn=GIPC1;refGene_tx=XM_023246329.1;refGene_gn=GIPC1;refGene_tx=XM_023246334.1

A2	10291902	10292015
refGene_gn=GIPC1;refGene_tx=XM_019818512.2;refGene_gn=GIPC1;refGene_tx
=XM_023246319.1;refGene_gn=GIPC1;refGene_tx=XM_023246323.1;refGene_gn=
GIPC1;refGene_tx=XM_023246326.1;refGene_gn=GIPC1;refGene_tx=XM_0232463
29.1;refGene_gn=GIPC1;refGene_tx=XM_023246334.1
A2	10292762	10292943
refGene_gn=GIPC1;refGene_tx=XM_019818512.2;refGene_gn=GIPC1;refGene_tx
=XM_023246319.1;refGene_gn=GIPC1;refGene_tx=XM_023246323.1;refGene_gn=
GIPC1;refGene_tx=XM_023246326.1;refGene_gn=GIPC1;refGene_tx=XM_0232463
29.1;refGene_gn=GIPC1;refGene_tx=XM_023246334.1
A2	10293042	10293228
refGene_gn=GIPC1;refGene_tx=XM_019818512.2;refGene_gn=GIPC1;refGene_tx
=XM_023246319.1;refGene_gn=GIPC1;refGene_tx=XM_023246323.1;refGene_gn=
GIPC1;refGene_tx=XM_023246326.1;refGene_gn=GIPC1;refGene_tx=XM_0232463
29.1;refGene_gn=GIPC1;refGene_tx=XM_023246334.1
A2	10293992	10294280
refGene_gn=GIPC1;refGene_tx=XM_019818512.2;refGene_gn=GIPC1;refGene_tx
=XM_023246319.1;refGene_gn=GIPC1;refGene_tx=XM_023246323.1;refGene_gn=
GIPC1;refGene_tx=XM_023246326.1;refGene_gn=GIPC1;refGene_tx=XM_0232463
29.1;refGene_gn=GIPC1;refGene_tx=XM_023246334.1
A2	10314898	10315129
refGene_gn=DNAJB1;refGene_tx=XM_003981944.5;refGene_gn=DNAJB1;refGene_
tx=XM_011289899.3;refGene_gn=DNAJB1;refGene_tx=XM_019818617.2
A2	10315372	10315953
refGene_gn=DNAJB1;refGene_tx=XM_003981944.5;refGene_gn=DNAJB1;refGene_
tx=XM_011289899.3;refGene_gn=DNAJB1;refGene_tx=XM_019818617.2
A2	10317143	10317354	refGene_gn=DNAJB1;refGene_tx=XM_003981944.5
A2	10324379	10324394	refGene_gn=TECR;refGene_tx=XM_003981940.4
A2	10328070	10328091	refGene_gn=TECR;refGene_tx=XM_019818561.2
A2	10338052	10338502
refGene_gn=TECR;refGene_tx=XM_019818545.1;refGene_gn=TECR;refGene_tx=X
M_023246345.1
A2	10345500	10345689
refGene_gn=TECR;refGene_tx=XM_003981940.4;refGene_gn=TECR;refGene_tx=X
M_006928499.3;refGene_gn=TECR;refGene_tx=XM_019818545.1;refGene_gn=TEC
R;refGene_tx=XM_019818561.2;refGene_gn=TECR;refGene_tx=XM_023246345.1;
refGene_gn=TECR;refGene_tx=XM_023246353.1
A2	10346317	10346369
refGene_gn=TECR;refGene_tx=XM_003981940.4;refGene_gn=TECR;refGene_tx=X
M_006928499.3;refGene_gn=TECR;refGene_tx=XM_019818545.1;refGene_gn=TEC
R;refGene_tx=XM_019818561.2;refGene_gn=TECR;refGene_tx=XM_023246345.1;
refGene_gn=TECR;refGene_tx=XM_023246353.1
A2	10346703	10346748
refGene_gn=TECR;refGene_tx=XM_003981940.4;refGene_gn=TECR;refGene_tx=X
M_006928499.3;refGene_gn=TECR;refGene_tx=XM_019818545.1;refGene_gn=TEC
R;refGene_tx=XM_019818561.2
A2	10346843	10346947
refGene_gn=TECR;refGene_tx=XM_003981940.4;refGene_gn=TECR;refGene_tx=X
M_006928499.3;refGene_gn=TECR;refGene_tx=XM_019818545.1;refGene_gn=TEC
R;refGene_tx=XM_019818561.2;refGene_gn=TECR;refGene_tx=XM_023246345.1;

refGene_gn=TECR;refGene_tx=XM_023246353.1
A2 10347024 10347140
refGene_gn=TECR;refGene_tx=XM_003981940.4;refGene_gn=TECR;refGene_tx=XM_006928499.3;refGene_gn=TECR;refGene_tx=XM_019818545.1;refGene_gn=TECR;refGene_tx=XM_019818561.2;refGene_gn=TECR;refGene_tx=XM_023246345.1;refGene_gn=TECR;refGene_tx=XM_023246353.1
A2 10347229 10347335
refGene_gn=TECR;refGene_tx=XM_003981940.4;refGene_gn=TECR;refGene_tx=XM_006928499.3;refGene_gn=TECR;refGene_tx=XM_019818545.1;refGene_gn=TECR;refGene_tx=XM_019818561.2;refGene_gn=TECR;refGene_tx=XM_023246345.1;refGene_gn=TECR;refGene_tx=XM_023246353.1
A2 10347462 10347535
refGene_gn=TECR;refGene_tx=XM_003981940.4;refGene_gn=TECR;refGene_tx=XM_006928499.3;refGene_gn=TECR;refGene_tx=XM_019818545.1;refGene_gn=TECR;refGene_tx=XM_019818561.2;refGene_gn=TECR;refGene_tx=XM_023246345.1;refGene_gn=TECR;refGene_tx=XM_023246353.1
A2 10347631 10347675
refGene_gn=TECR;refGene_tx=XM_003981940.4;refGene_gn=TECR;refGene_tx=XM_006928499.3;refGene_gn=TECR;refGene_tx=XM_019818545.1;refGene_gn=TECR;refGene_tx=XM_019818561.2;refGene_gn=TECR;refGene_tx=XM_023246345.1;refGene_gn=TECR;refGene_tx=XM_023246353.1
A2 10347748 10347806
refGene_gn=TECR;refGene_tx=XM_003981940.4;refGene_gn=TECR;refGene_tx=XM_006928499.3;refGene_gn=TECR;refGene_tx=XM_019818545.1;refGene_gn=TECR;refGene_tx=XM_019818561.2;refGene_gn=TECR;refGene_tx=XM_023246345.1;refGene_gn=TECR;refGene_tx=XM_023246353.1
A2 10347888 10347977
refGene_gn=TECR;refGene_tx=XM_003981940.4;refGene_gn=TECR;refGene_tx=XM_006928499.3;refGene_gn=TECR;refGene_tx=XM_019818545.1;refGene_gn=TECR;refGene_tx=XM_019818561.2;refGene_gn=TECR;refGene_tx=XM_023246345.1;refGene_gn=TECR;refGene_tx=XM_023246353.1
A2 10348320 10348366
refGene_gn=TECR;refGene_tx=XM_003981940.4;refGene_gn=TECR;refGene_tx=XM_006928499.3;refGene_gn=TECR;refGene_tx=XM_019818545.1;refGene_gn=TECR;refGene_tx=XM_019818561.2;refGene_gn=TECR;refGene_tx=XM_023246345.1;refGene_gn=TECR;refGene_tx=XM_023246353.1
A2 10348454 10348582
refGene_gn=TECR;refGene_tx=XM_003981940.4;refGene_gn=TECR;refGene_tx=XM_006928499.3;refGene_gn=TECR;refGene_tx=XM_019818545.1;refGene_gn=TECR;refGene_tx=XM_019818561.2;refGene_gn=TECR;refGene_tx=XM_023246345.1;refGene_gn=TECR;refGene_tx=XM_023246353.1
A2 10348854 10348987 refGene_gn=NDUFB7;refGene_tx=XM_003981941.3
A2 10349414 10349583 refGene_gn=NDUFB7;refGene_tx=XM_003981941.3
A2 10352986 10353098 refGene_gn=NDUFB7;refGene_tx=XM_003981941.3
A2 10364240 10364280
refGene_gn=CLEC17A;refGene_tx=XM_019818590.2;refGene_gn=CLEC17A;refGene_tx=XM_019818597.2;refGene_gn=CLEC17A;refGene_tx=XM_023246415.1
A2 10364427 10364505
refGene_gn=CLEC17A;refGene_tx=XM_019818590.2;refGene_gn=CLEC17A;refGene_tx=XM_019818597.2;refGene_gn=CLEC17A;refGene_tx=XM_023246415.1

A2 10365817 10365850
refGene_gn=CLEC17A;refGene_tx=XM_019818590.2;refGene_gn=CLEC17A;refGene_tx=XM_019818597.2;refGene_gn=CLEC17A;refGene_tx=XM_023246415.1
A2 10365953 10366004
refGene_gn=CLEC17A;refGene_tx=XM_019818590.2;refGene_gn=CLEC17A;refGene_tx=XM_019818597.2;refGene_gn=CLEC17A;refGene_tx=XM_023246415.1
A2 10366087 10366129
refGene_gn=CLEC17A;refGene_tx=XM_019818590.2;refGene_gn=CLEC17A;refGene_tx=XM_019818597.2;refGene_gn=CLEC17A;refGene_tx=XM_023246415.1
A2 10366288 10366351
refGene_gn=CLEC17A;refGene_tx=XM_019818590.2;refGene_gn=CLEC17A;refGene_tx=XM_019818597.2;refGene_gn=CLEC17A;refGene_tx=XM_023246415.1
A2 10367123 10367264
refGene_gn=CLEC17A;refGene_tx=XM_019818590.2;refGene_gn=CLEC17A;refGene_tx=XM_019818597.2;refGene_gn=CLEC17A;refGene_tx=XM_023246415.1
A2 10367349 10367412
refGene_gn=CLEC17A;refGene_tx=XM_019818590.2;refGene_gn=CLEC17A;refGene_tx=XM_019818597.2;refGene_gn=CLEC17A;refGene_tx=XM_023246415.1
A2 10369113 10369209
refGene_gn=CLEC17A;refGene_tx=XM_019818590.2;refGene_gn=CLEC17A;refGene_tx=XM_019818597.2;refGene_gn=CLEC17A;refGene_tx=XM_023246415.1
A2 10369420 10369572
refGene_gn=CLEC17A;refGene_tx=XM_019818590.2;refGene_gn=CLEC17A;refGene_tx=XM_019818597.2;refGene_gn=CLEC17A;refGene_tx=XM_023246415.1
A2 10371143 10371253
refGene_gn=CLEC17A;refGene_tx=XM_019818590.2;refGene_gn=CLEC17A;refGene_tx=XM_023246415.1
A2 10374461 10374594
refGene_gn=CLEC17A;refGene_tx=XM_019818590.2;refGene_gn=CLEC17A;refGene_tx=XM_019818597.2;refGene_gn=CLEC17A;refGene_tx=XM_023246415.1
A2 10381934 10381961
refGene_gn=ADGRE3;refGene_tx=XM_023246420.1;refGene_gn=ADGRE3;refGene_tx=XM_023246424.1;refGene_gn=ADGRE3;refGene_tx=XM_023246427.1;refGene_gn=ADGRE3;refGene_tx=XM_023246431.1;refGene_gn=ADGRE3;refGene_tx=XM_023246435.1
A2 10385616 10385712
refGene_gn=ADGRE3;refGene_tx=XM_023246420.1;refGene_gn=ADGRE3;refGene_tx=XM_023246424.1;refGene_gn=ADGRE3;refGene_tx=XM_023246427.1;refGene_gn=ADGRE3;refGene_tx=XM_023246431.1;refGene_gn=ADGRE3;refGene_tx=XM_023246435.1
A2 10386989 10387158
refGene_gn=ADGRE3;refGene_tx=XM_023246420.1;refGene_gn=ADGRE3;refGene_tx=XM_023246424.1;refGene_gn=ADGRE3;refGene_tx=XM_023246427.1;refGene_gn=ADGRE3;refGene_tx=XM_023246431.1;refGene_gn=ADGRE3;refGene_tx=XM_023246435.1
A2 10388002 10388094
refGene_gn=ADGRE3;refGene_tx=XM_023246420.1;refGene_gn=ADGRE3;refGene_tx=XM_023246424.1;refGene_gn=ADGRE3;refGene_tx=XM_023246427.1;refGene_gn=ADGRE3;refGene_tx=XM_023246431.1;refGene_gn=ADGRE3;refGene_tx=XM_023246435.1

A2 10388509 10388576
refGene_gn=ADGRE3;refGene_tx=XM_023246420.1;refGene_gn=ADGRE3;refGene_tx=XM_023246424.1;refGene_gn=ADGRE3;refGene_tx=XM_023246427.1;refGene_gn=ADGRE3;refGene_tx=XM_023246431.1;refGene_gn=ADGRE3;refGene_tx=XM_023246435.1
A2 10388868 10389104
refGene_gn=ADGRE3;refGene_tx=XM_023246420.1;refGene_gn=ADGRE3;refGene_tx=XM_023246424.1;refGene_gn=ADGRE3;refGene_tx=XM_023246427.1;refGene_gn=ADGRE3;refGene_tx=XM_023246431.1;refGene_gn=ADGRE3;refGene_tx=XM_023246435.1
A2 10389390 10389588
refGene_gn=ADGRE3;refGene_tx=XM_023246420.1;refGene_gn=ADGRE3;refGene_tx=XM_023246424.1;refGene_gn=ADGRE3;refGene_tx=XM_023246427.1;refGene_gn=ADGRE3;refGene_tx=XM_023246431.1;refGene_gn=ADGRE3;refGene_tx=XM_023246435.1
A2 10390620 10390788
refGene_gn=ADGRE3;refGene_tx=XM_023246420.1;refGene_gn=ADGRE3;refGene_tx=XM_023246424.1;refGene_gn=ADGRE3;refGene_tx=XM_023246427.1;refGene_gn=ADGRE3;refGene_tx=XM_023246431.1;refGene_gn=ADGRE3;refGene_tx=XM_023246435.1
A2 10392447 10392632
refGene_gn=ADGRE3;refGene_tx=XM_023246420.1;refGene_gn=ADGRE3;refGene_tx=XM_023246424.1;refGene_gn=ADGRE3;refGene_tx=XM_023246427.1;refGene_gn=ADGRE3;refGene_tx=XM_023246431.1;refGene_gn=ADGRE3;refGene_tx=XM_023246435.1
A2 10398734 10398854
refGene_gn=ADGRE3;refGene_tx=XM_023246420.1;refGene_gn=ADGRE3;refGene_tx=XM_023246424.1;refGene_gn=ADGRE3;refGene_tx=XM_023246427.1;refGene_gn=ADGRE3;refGene_tx=XM_023246431.1;refGene_gn=ADGRE3;refGene_tx=XM_023246435.1
A2 10401574 10401758
refGene_gn=ADGRE3;refGene_tx=XM_023246420.1;refGene_gn=ADGRE3;refGene_tx=XM_023246424.1;refGene_gn=ADGRE3;refGene_tx=XM_023246427.1;refGene_gn=ADGRE3;refGene_tx=XM_023246431.1
A2 10408301 10408339
refGene_gn=ADGRE3;refGene_tx=XM_023246420.1;refGene_gn=ADGRE3;refGene_tx=XM_023246424.1;refGene_gn=ADGRE3;refGene_tx=XM_023246427.1;refGene_gn=ADGRE3;refGene_tx=XM_023246431.1
A2 10412918 10413102
refGene_gn=ADGRE3;refGene_tx=XM_023246420.1;refGene_gn=ADGRE3;refGene_tx=XM_023246424.1;refGene_gn=ADGRE3;refGene_tx=XM_023246427.1;refGene_gn=ADGRE3;refGene_tx=XM_023246431.1
A2 10416976 10417102
refGene_gn=ADGRE3;refGene_tx=XM_023246420.1;refGene_gn=ADGRE3;refGene_tx=XM_023246424.1
A2 10418708 10418759
refGene_gn=ADGRE3;refGene_tx=XM_023246420.1;refGene_gn=ADGRE3;refGene_tx=XM_023246424.1
A2 10425113 10425138
refGene_gn=ADGRE3;refGene_tx=XM_023246420.1;refGene_gn=ADGRE3;refGene_

tx=XM_023246424.1
A2 10439765 10439768
refGene_gn=ZNF333;refGene_tx=XM_023246441.1;refGene_gn=ZNF333;refGene_tx=XM_023246448.1;refGene_gn=ZNF333;refGene_tx=XM_023246457.1
A2 10441041 10441165
refGene_gn=ZNF333;refGene_tx=XM_023246441.1;refGene_gn=ZNF333;refGene_tx=XM_023246448.1;refGene_gn=ZNF333;refGene_tx=XM_023246457.1
A2 10443705 10443788
refGene_gn=ZNF333;refGene_tx=XM_023246441.1;refGene_gn=ZNF333;refGene_tx=XM_023246448.1;refGene_gn=ZNF333;refGene_tx=XM_023246453.1;refGene_gn=ZNF333;refGene_tx=XM_023246454.1;refGene_gn=ZNF333;refGene_tx=XM_023246457.1
A2 10446406 10446523
refGene_gn=ZNF333;refGene_tx=XM_011289918.3;refGene_gn=ZNF333;refGene_tx=XM_023246451.1;refGene_gn=ZNF333;refGene_tx=XM_023246457.1
A2 10447714 10447802
refGene_gn=ZNF333;refGene_tx=XM_011289918.3;refGene_gn=ZNF333;refGene_tx=XM_019818672.2;refGene_gn=ZNF333;refGene_tx=XM_023246441.1;refGene_gn=ZNF333;refGene_tx=XM_023246448.1;refGene_gn=ZNF333;refGene_tx=XM_023246451.1;refGene_gn=ZNF333;refGene_tx=XM_023246453.1;refGene_gn=ZNF333;refGene_tx=XM_023246454.1;refGene_gn=ZNF333;refGene_tx=XM_023246457.1
A2 10455956 10456045
refGene_gn=ZNF333;refGene_tx=XM_011289918.3;refGene_gn=ZNF333;refGene_tx=XM_019818672.2;refGene_gn=ZNF333;refGene_tx=XM_023246441.1;refGene_gn=ZNF333;refGene_tx=XM_023246448.1;refGene_gn=ZNF333;refGene_tx=XM_023246451.1;refGene_gn=ZNF333;refGene_tx=XM_023246453.1;refGene_gn=ZNF333;refGene_tx=XM_023246454.1;refGene_gn=ZNF333;refGene_tx=XM_023246457.1
A2 10456658 10456785
refGene_gn=ZNF333;refGene_tx=XM_011289918.3;refGene_gn=ZNF333;refGene_tx=XM_019818672.2;refGene_gn=ZNF333;refGene_tx=XM_023246441.1;refGene_gn=ZNF333;refGene_tx=XM_023246448.1;refGene_gn=ZNF333;refGene_tx=XM_023246451.1;refGene_gn=ZNF333;refGene_tx=XM_023246453.1;refGene_gn=ZNF333;refGene_tx=XM_023246454.1;refGene_gn=ZNF333;refGene_tx=XM_023246457.1
A2 10457223 10457319
refGene_gn=ZNF333;refGene_tx=XM_011289918.3;refGene_gn=ZNF333;refGene_tx=XM_019818672.2;refGene_gn=ZNF333;refGene_tx=XM_023246441.1;refGene_gn=ZNF333;refGene_tx=XM_023246451.1;refGene_gn=ZNF333;refGene_tx=XM_023246453.1;refGene_gn=ZNF333;refGene_tx=XM_023246454.1;refGene_gn=ZNF333;refGene_tx=XM_023246457.1
A2 10457821 10457895
refGene_gn=ZNF333;refGene_tx=XM_011289918.3;refGene_gn=ZNF333;refGene_tx=XM_019818672.2;refGene_gn=ZNF333;refGene_tx=XM_023246441.1;refGene_gn=ZNF333;refGene_tx=XM_023246448.1;refGene_gn=ZNF333;refGene_tx=XM_023246451.1;refGene_gn=ZNF333;refGene_tx=XM_023246453.1;refGene_gn=ZNF333;refGene_tx=XM_023246454.1;refGene_gn=ZNF333;refGene_tx=XM_023246457.1
A2 10459025 10460123

refGene_gn=ZNF333;refGene_tx=XM_011289918.3;refGene_gn=ZNF333;refGene_tx=XM_019818672.2;refGene_gn=ZNF333;refGene_tx=XM_023246441.1;refGene_gn=ZNF333;refGene_tx=XM_023246448.1;refGene_gn=ZNF333;refGene_tx=XM_023246451.1;refGene_gn=ZNF333;refGene_tx=XM_023246453.1;refGene_gn=ZNF333;refGene_tx=XM_023246454.1;refGene_gn=ZNF333;refGene_tx=XM_023246457.1
A2 10474304 10474335 refGene_gn=ADGRE2;refGene_tx=XM_023241032.1
A2 10474457 10474532 refGene_gn=ADGRE2;refGene_tx=XM_023241032.1
A2 10478278 10478332 refGene_gn=ADGRE2;refGene_tx=XM_023241032.1
A2 10478639 10478759 refGene_gn=ADGRE2;refGene_tx=XM_023241032.1
A2 10480963 10481119 refGene_gn=ADGRE2;refGene_tx=XM_023241032.1
A2 10483978 10484110 refGene_gn=ADGRE2;refGene_tx=XM_023241032.1
A2 10484685 10484832 refGene_gn=ADGRE2;refGene_tx=XM_023241032.1
A2 10485292 10485439 refGene_gn=ADGRE2;refGene_tx=XM_023241032.1
A2 10485527 10485574 refGene_gn=ADGRE2;refGene_tx=XM_023241032.1
A2 10485718 10485796 refGene_gn=ADGRE2;refGene_tx=XM_023241032.1
A2 10486184 10486362 refGene_gn=ADGRE2;refGene_tx=XM_023241032.1
A2 10489202 10489310 refGene_gn=ADGRE2;refGene_tx=XM_023241032.1
A2 10489474 10489695 refGene_gn=ADGRE2;refGene_tx=XM_023241032.1
A2 10489885 10490065 refGene_gn=ADGRE2;refGene_tx=XM_023241032.1
A2 10490622 10490820 refGene_gn=ADGRE2;refGene_tx=XM_023241032.1
A2 10491183 10491419 refGene_gn=ADGRE2;refGene_tx=XM_023241032.1
A2 10499811 10499878 refGene_gn=ADGRE2;refGene_tx=XM_023241032.1
A2 10500158 10500250 refGene_gn=ADGRE2;refGene_tx=XM_023241032.1
A2 10501276 10501445 refGene_gn=ADGRE2;refGene_tx=XM_023241032.1
A2 10501530 10501641 refGene_gn=ADGRE2;refGene_tx=XM_023241032.1
A2 10505962 10505971 refGene_gn=ADGRE2;refGene_tx=XM_023241032.1
A2 10526227 10526269
refGene_gn=LOC101083497;refGene_tx=XM_023246468.1;refGene_gn=LOC101083497;refGene_tx=XM_023246473.1;refGene_gn=LOC101083497;refGene_tx=XM_023246475.1;refGene_gn=LOC101083497;refGene_tx=XM_023246483.1;refGene_gn=LOC101083497;refGene_tx=XM_023246489.1;refGene_gn=LOC101083497;refGene_tx=XM_023246495.1
A2 10526985 10527096
refGene_gn=LOC101083497;refGene_tx=XM_023246468.1;refGene_gn=LOC101083497;refGene_tx=XM_023246473.1;refGene_gn=LOC101083497;refGene_tx=XM_023246475.1;refGene_gn=LOC101083497;refGene_tx=XM_023246483.1;refGene_gn=LOC101083497;refGene_tx=XM_023246489.1;refGene_gn=LOC101083497;refGene_tx=XM_023246495.1
A2 10527181 10527350
refGene_gn=LOC101083497;refGene_tx=XM_023246468.1;refGene_gn=LOC101083497;refGene_tx=XM_023246473.1;refGene_gn=LOC101083497;refGene_tx=XM_023246475.1;refGene_gn=LOC101083497;refGene_tx=XM_023246483.1;refGene_gn=LOC101083497;refGene_tx=XM_023246489.1;refGene_gn=LOC101083497;refGene_tx=XM_023246495.1
A2 10528194 10528286
refGene_gn=LOC101083497;refGene_tx=XM_023246468.1;refGene_gn=LOC101083497;refGene_tx=XM_023246473.1;refGene_gn=LOC101083497;refGene_tx=XM_023246475.1;refGene_gn=LOC101083497;refGene_tx=XM_023246483.1;refGene_gn=LOC101083497;refGene_tx=XM_023246489.1;refGene_gn=LOC101083497;refGen

e_tx=XM_023246495.1
A2	10528708	10528775
refGene_gn=LOC101083497;refGene_tx=XM_023246468.1;refGene_gn=LOC101083497;refGene_tx=XM_023246473.1;refGene_gn=LOC101083497;refGene_tx=XM_023246475.1;refGene_gn=LOC101083497;refGene_tx=XM_023246483.1;refGene_gn=LOC101083497;refGene_tx=XM_023246489.1;refGene_gn=LOC101083497;refGene_tx=XM_023246495.1
A2	10529067	10529303
refGene_gn=LOC101083497;refGene_tx=XM_023246468.1;refGene_gn=LOC101083497;refGene_tx=XM_023246473.1;refGene_gn=LOC101083497;refGene_tx=XM_023246475.1;refGene_gn=LOC101083497;refGene_tx=XM_023246483.1;refGene_gn=LOC101083497;refGene_tx=XM_023246489.1;refGene_gn=LOC101083497;refGene_tx=XM_023246495.1
A2	10529591	10529789
refGene_gn=LOC101083497;refGene_tx=XM_023246468.1;refGene_gn=LOC101083497;refGene_tx=XM_023246473.1;refGene_gn=LOC101083497;refGene_tx=XM_023246475.1;refGene_gn=LOC101083497;refGene_tx=XM_023246483.1;refGene_gn=LOC101083497;refGene_tx=XM_023246489.1;refGene_gn=LOC101083497;refGene_tx=XM_023246495.1
A2	10530346	10530526
refGene_gn=LOC101083497;refGene_tx=XM_023246468.1;refGene_gn=LOC101083497;refGene_tx=XM_023246473.1;refGene_gn=LOC101083497;refGene_tx=XM_023246475.1;refGene_gn=LOC101083497;refGene_tx=XM_023246483.1;refGene_gn=LOC101083497;refGene_tx=XM_023246489.1;refGene_gn=LOC101083497;refGene_tx=XM_023246495.1
A2	10530716	10530937
refGene_gn=LOC101083497;refGene_tx=XM_023246468.1;refGene_gn=LOC101083497;refGene_tx=XM_023246473.1;refGene_gn=LOC101083497;refGene_tx=XM_023246475.1;refGene_gn=LOC101083497;refGene_tx=XM_023246483.1;refGene_gn=LOC101083497;refGene_tx=XM_023246489.1;refGene_gn=LOC101083497;refGene_tx=XM_023246495.1
A2	10531101	10531209
refGene_gn=LOC101083497;refGene_tx=XM_023246468.1;refGene_gn=LOC101083497;refGene_tx=XM_023246473.1;refGene_gn=LOC101083497;refGene_tx=XM_023246475.1;refGene_gn=LOC101083497;refGene_tx=XM_023246483.1;refGene_gn=LOC101083497;refGene_tx=XM_023246489.1;refGene_gn=LOC101083497;refGene_tx=XM_023246495.1
A2	10534000	10534178
refGene_gn=LOC101083497;refGene_tx=XM_023246468.1;refGene_gn=LOC101083497;refGene_tx=XM_023246473.1;refGene_gn=LOC101083497;refGene_tx=XM_023246475.1;refGene_gn=LOC101083497;refGene_tx=XM_023246483.1;refGene_gn=LOC101083497;refGene_tx=XM_023246489.1;refGene_gn=LOC101083497;refGene_tx=XM_023246495.1
A2	10534283	10534361
refGene_gn=LOC101083497;refGene_tx=XM_023246468.1;refGene_gn=LOC101083497;refGene_tx=XM_023246473.1;refGene_gn=LOC101083497;refGene_tx=XM_023246475.1;refGene_gn=LOC101083497;refGene_tx=XM_023246483.1;refGene_gn=LOC101083497;refGene_tx=XM_023246489.1;refGene_gn=LOC101083497;refGene_tx=XM_023246495.1
A2	10534503	10534550

refGene_gn=LOC101083497;refGene_tx=XM_023246468.1;refGene_gn=LOC101083
497;refGene_tx=XM_023246473.1;refGene_gn=LOC101083497;refGene_tx=XM_02
3246475.1;refGene_gn=LOC101083497;refGene_tx=XM_023246483.1;refGene_gn
=LOC101083497;refGene_tx=XM_023246489.1;refGene_gn=LOC101083497;refGen
e_tx=XM_023246495.1
A2 10535054 10535231
refGene_gn=LOC101083497;refGene_tx=XM_023246468.1;refGene_gn=LOC101083
497;refGene_tx=XM_023246473.1;refGene_gn=LOC101083497;refGene_tx=XM_02
3246475.1;refGene_gn=LOC101083497;refGene_tx=XM_023246483.1;refGene_gn
=LOC101083497;refGene_tx=XM_023246489.1;refGene_gn=LOC101083497;refGen
e_tx=XM_023246495.1
A2 10535795 10535951
refGene_gn=LOC101083497;refGene_tx=XM_023246468.1;refGene_gn=LOC101083
497;refGene_tx=XM_023246473.1;refGene_gn=LOC101083497;refGene_tx=XM_02
3246475.1;refGene_gn=LOC101083497;refGene_tx=XM_023246483.1;refGene_gn
=LOC101083497;refGene_tx=XM_023246489.1;refGene_gn=LOC101083497;refGen
e_tx=XM_023246495.1
A2 10539438 10539573
refGene_gn=LOC101083497;refGene_tx=XM_023246468.1;refGene_gn=LOC101083
497;refGene_tx=XM_023246473.1;refGene_gn=LOC101083497;refGene_tx=XM_02
3246475.1;refGene_gn=LOC101083497;refGene_tx=XM_023246483.1;refGene_gn
=LOC101083497;refGene_tx=XM_023246489.1;refGene_gn=LOC101083497;refGen
e_tx=XM_023246495.1
A2 10539869 10539920
refGene_gn=LOC101083497;refGene_tx=XM_023246468.1;refGene_gn=LOC101083
497;refGene_tx=XM_023246473.1;refGene_gn=LOC101083497;refGene_tx=XM_02
3246475.1;refGene_gn=LOC101083497;refGene_tx=XM_023246483.1;refGene_gn
=LOC101083497;refGene_tx=XM_023246489.1;refGene_gn=LOC101083497;refGen
e_tx=XM_023246495.1
A2 10541846 10541996
refGene_gn=LOC101083497;refGene_tx=XM_023246468.1;refGene_gn=LOC101083
497;refGene_tx=XM_023246473.1;refGene_gn=LOC101083497;refGene_tx=XM_02
3246475.1;refGene_gn=LOC101083497;refGene_tx=XM_023246483.1;refGene_gn
=LOC101083497;refGene_tx=XM_023246489.1;refGene_gn=LOC101083497;refGen
e_tx=XM_023246495.1
A2 10546946 10547147 refGene_gn=BBIP1;refGene_tx=XM_011290032.3
A2 10550649 10550703
refGene_gn=LOC101083497;refGene_tx=XM_023246468.1;refGene_gn=LOC101083
497;refGene_tx=XM_023246473.1;refGene_gn=LOC101083497;refGene_tx=XM_02
3246475.1;refGene_gn=LOC101083497;refGene_tx=XM_023246483.1;refGene_gn
=LOC101083497;refGene_tx=XM_023246489.1;refGene_gn=LOC101083497;refGen
e_tx=XM_023246495.1
A2 10553168 10553196
refGene_gn=LOC101083497;refGene_tx=XM_023246468.1;refGene_gn=LOC101083
497;refGene_tx=XM_023246473.1;refGene_gn=LOC101083497;refGene_tx=XM_02
3246475.1;refGene_gn=LOC101083497;refGene_tx=XM_023246483.1;refGene_gn
=LOC101083497;refGene_tx=XM_023246489.1;refGene_gn=LOC101083497;refGen
e_tx=XM_023246495.1
A2 10563614 10563701
refGene_gn=LOC101100284;refGene_tx=XM_023246508.1;refGene_gn=LOC101100

284;refGene_tx=XM_023246512.1;refGene_gn=LOC101100284;refGene_tx=XM_023246523.1
A2 10566294 10566405
refGene_gn=LOC101100284;refGene_tx=XM_023246508.1;refGene_gn=LOC101100284;refGene_tx=XM_023246512.1;refGene_gn=LOC101100284;refGene_tx=XM_023246523.1
A2 10566490 10566659
refGene_gn=LOC101100284;refGene_tx=XM_023246508.1;refGene_gn=LOC101100284;refGene_tx=XM_023246512.1;refGene_gn=LOC101100284;refGene_tx=XM_023246523.1
A2 10567503 10567595
refGene_gn=LOC101100284;refGene_tx=XM_023246508.1;refGene_gn=LOC101100284;refGene_tx=XM_023246512.1;refGene_gn=LOC101100284;refGene_tx=XM_023246523.1
A2 10568013 10568080
refGene_gn=LOC101100284;refGene_tx=XM_023246508.1;refGene_gn=LOC101100284;refGene_tx=XM_023246512.1;refGene_gn=LOC101100284;refGene_tx=XM_023246523.1
A2 10568372 10568608
refGene_gn=LOC101100284;refGene_tx=XM_023246508.1;refGene_gn=LOC101100284;refGene_tx=XM_023246512.1;refGene_gn=LOC101100284;refGene_tx=XM_023246523.1
A2 10568942 10569137
refGene_gn=LOC101100284;refGene_tx=XM_023246508.1;refGene_gn=LOC101100284;refGene_tx=XM_023246512.1;refGene_gn=LOC101100284;refGene_tx=XM_023246523.1
A2 10572867 10573047
refGene_gn=LOC101100284;refGene_tx=XM_023246508.1;refGene_gn=LOC101100284;refGene_tx=XM_023246512.1;refGene_gn=LOC101100284;refGene_tx=XM_023246523.1
A2 10573237 10573458
refGene_gn=LOC101100284;refGene_tx=XM_023246508.1;refGene_gn=LOC101100284;refGene_tx=XM_023246512.1;refGene_gn=LOC101100284;refGene_tx=XM_023246523.1
A2 10573645 10573753
refGene_gn=LOC101100284;refGene_tx=XM_023246508.1;refGene_gn=LOC101100284;refGene_tx=XM_023246512.1
A2 10577451 10577629
refGene_gn=LOC101100284;refGene_tx=XM_023246508.1;refGene_gn=LOC101100284;refGene_tx=XM_023246512.1;refGene_gn=LOC101100284;refGene_tx=XM_023246523.1
A2 10578040 10578118
refGene_gn=LOC101100284;refGene_tx=XM_023246508.1;refGene_gn=LOC101100284;refGene_tx=XM_023246523.1
A2 10578240 10578287
refGene_gn=LOC101100284;refGene_tx=XM_023246508.1;refGene_gn=LOC101100284;refGene_tx=XM_023246512.1;refGene_gn=LOC101100284;refGene_tx=XM_023246523.1
A2 10578482 10578629
refGene_gn=LOC101100284;refGene_tx=XM_023246508.1;refGene_gn=LOC101100

284;refGene_tx=XM_023246512.1;refGene_gn=LOC101100284;refGene_tx=XM_023246523.1
A2 10583030 10583186
refGene_gn=LOC101100284;refGene_tx=XM_023246508.1;refGene_gn=LOC101100284;refGene_tx=XM_023246512.1;refGene_gn=LOC101100284;refGene_tx=XM_023246523.1
A2 10585194 10585317
refGene_gn=LOC101100284;refGene_tx=XM_023246508.1;refGene_gn=LOC101100284;refGene_tx=XM_023246512.1;refGene_gn=LOC101100284;refGene_tx=XM_023246523.1
A2 10585581 10585632
refGene_gn=LOC101100284;refGene_tx=XM_023246508.1;refGene_gn=LOC101100284;refGene_tx=XM_023246512.1;refGene_gn=LOC101100284;refGene_tx=XM_023246523.1
A2 10586635 10586666
refGene_gn=LOC101100284;refGene_tx=XM_023246508.1;refGene_gn=LOC101100284;refGene_tx=XM_023246512.1;refGene_gn=LOC101100284;refGene_tx=XM_023246523.1
A2 10709681 10710650
refGene_gn=LOC101101292;refGene_tx=XM_019826093.2
A2 10721546 10722506
refGene_gn=LOC101101549;refGene_tx=XM_019825000.2
A2 10741519 10742488
refGene_gn=LOC101080364;refGene_tx=XM_023241067.1
A2 10761312 10762281
refGene_gn=LOC105260290;refGene_tx=XM_011280078.2
A2 10770757 10771696
refGene_gn=LOC101080874;refGene_tx=XM_011290201.2
A2 10781980 10782949
refGene_gn=LOC101081122;refGene_tx=XM_019826092.2
A2 10802116 10802947
refGene_gn=LOC101081366;refGene_tx=XM_011290016.3
A2 10840187 10841120
refGene_gn=LOC101085613;refGene_tx=XM_004001250.1
A2 10848984 10849336
refGene_gn=LOC101083405;refGene_tx=XM_023241097.1
A2 10849621 10849814
refGene_gn=LOC101083405;refGene_tx=XM_023241097.1
A2 10856550 10856848
refGene_gn=LOC101083405;refGene_tx=XM_023241097.1
A2 10858246 10858504
refGene_gn=LOC111557133;refGene_tx=XM_023241108.1
A2 10858789 10859137
refGene_gn=LOC111557133;refGene_tx=XM_023241108.1
A2 10867731 10867940
refGene_gn=LOC101083158;refGene_tx=XM_023241117.1
A2 10868225 10868580
refGene_gn=LOC101083158;refGene_tx=XM_023241117.1
A2 10886373 10887300
refGene_gn=LOC101084693;refGene_tx=XM_004001272.3

A2 10895883 10896238
refGene_gn=LOC111557136;refGene_tx=XM_023241139.1
A2 10896523 10896713
refGene_gn=LOC111557136;refGene_tx=XM_023241139.1
A2 10901106 10901218
refGene_gn=LOC111557136;refGene_tx=XM_023241139.1
A2 10905430 10906399
refGene_gn=LOC101082634;refGene_tx=XM_011290174.1
A2 10914833 10915070
refGene_gn=LOC101085871;refGene_tx=XM_023241154.1
A2 10915355 10915703
refGene_gn=LOC101085871;refGene_tx=XM_023241154.1
A2 10939107 10939729
refGene_gn=LOC101082131;refGene_tx=XM_011290160.2
A2 10949716 10950099
refGene_gn=LOC101082131;refGene_tx=XM_011290160.2
A2 10960987 10961188
refGene_gn=LOC101083150;refGene_tx=XM_023241165.1
A2 10961473 10961821
refGene_gn=LOC101083150;refGene_tx=XM_023241165.1
A2 10976296 10976441
refGene_gn=LOC111557140;refGene_tx=XM_023241170.1
A2 10981218 10981411
refGene_gn=LOC111557140;refGene_tx=XM_023241170.1
A2 10981696 10982048
refGene_gn=LOC111557140;refGene_tx=XM_023241170.1
A2 11010361 11011315
refGene_gn=LOC101084174;refGene_tx=XM_003981988.1
A2 11032548 11032748
refGene_gn=LOC111558239;refGene_tx=XM_023246540.1
A2 11040322 11040671
refGene_gn=LOC111558239;refGene_tx=XM_023246540.1
A2 11050430 11051396
refGene_gn=LOC101084939;refGene_tx=XM_003981990.2
A2 11064711 11065165
refGene_gn=LOC101085186;refGene_tx=XM_023241189.1
A2 11065450 11065644
refGene_gn=LOC101085186;refGene_tx=XM_023241189.1
A2 11077770 11078668
refGene_gn=LOC101093132;refGene_tx=XM_011290123.2
A2 11082273 11082337
refGene_gn=LOC101093132;refGene_tx=XM_011290123.2
A2 11082528 11082583
refGene_gn=LOC101093132;refGene_tx=XM_011290123.2
A2 11095690 11095735
refGene_gn=LOC101093132;refGene_tx=XM_011290123.2
A2 11095817 11096172
refGene_gn=LOC101085443;refGene_tx=XM_023241190.1
A2 11096457 11096687
refGene_gn=LOC101085443;refGene_tx=XM_023241190.1

A2 11107331 11108264
refGene_gn=LOC101085956;refGene_tx=XM_006928537.2
A2 11121352 11122369
refGene_gn=LOC101086209;refGene_tx=XM_011290109.2
A2 11173333 11174287
refGene_gn=LOC101086704;refGene_tx=XM_003981995.3
A2 11183113 11184043
refGene_gn=LOC101086955;refGene_tx=XM_003981996.1
A2 11202294 11203245
refGene_gn=LOC101087209;refGene_tx=XM_003981997.1
A2 11217396 11218521
refGene_gn=LOC101087457;refGene_tx=XM_003981998.3
A2 11220833 11221029
refGene_gn=SLC1A6;refGene_tx=XM_003981968.4;refGene_gn=SLC1A6;refGene_tx=XM_023246561.1;refGene_gn=SLC1A6;refGene_tx=XM_023246578.1
A2 11221880 11222015
refGene_gn=SLC1A6;refGene_tx=XM_003981968.4;refGene_gn=SLC1A6;refGene_tx=XM_023246561.1;refGene_gn=SLC1A6;refGene_tx=XM_023246578.1
A2 11222123 11222136 refGene_gn=SLC1A6;refGene_tx=XM_023246584.1
A2 11222787 11222982
refGene_gn=SLC1A6;refGene_tx=XM_003981968.4;refGene_gn=SLC1A6;refGene_tx=XM_023246561.1;refGene_gn=SLC1A6;refGene_tx=XM_023246578.1;refGene_gn=SLC1A6;refGene_tx=XM_023246584.1
A2 11224168 11224402
refGene_gn=SLC1A6;refGene_tx=XM_003981968.4;refGene_gn=SLC1A6;refGene_tx=XM_023246561.1;refGene_gn=SLC1A6;refGene_tx=XM_023246578.1;refGene_gn=SLC1A6;refGene_tx=XM_023246584.1
A2 11226898 11227242
refGene_gn=SLC1A6;refGene_tx=XM_003981968.4;refGene_gn=SLC1A6;refGene_tx=XM_023246561.1;refGene_gn=SLC1A6;refGene_tx=XM_023246578.1;refGene_gn=SLC1A6;refGene_tx=XM_023246584.1
A2 11227748 11227791
refGene_gn=SLC1A6;refGene_tx=XM_003981968.4;refGene_gn=SLC1A6;refGene_tx=XM_023246561.1;refGene_gn=SLC1A6;refGene_tx=XM_023246578.1;refGene_gn=SLC1A6;refGene_tx=XM_023246584.1
A2 11228535 11228740
refGene_gn=SLC1A6;refGene_tx=XM_003981968.4;refGene_gn=SLC1A6;refGene_tx=XM_023246561.1;refGene_gn=SLC1A6;refGene_tx=XM_023246578.1;refGene_gn=SLC1A6;refGene_tx=XM_023246584.1
A2 11230229 11230367
refGene_gn=SLC1A6;refGene_tx=XM_003981968.4;refGene_gn=SLC1A6;refGene_tx=XM_023246561.1;refGene_gn=SLC1A6;refGene_tx=XM_023246578.1;refGene_gn=SLC1A6;refGene_tx=XM_023246584.1
A2 11230572 11230777
refGene_gn=SLC1A6;refGene_tx=XM_003981968.4;refGene_gn=SLC1A6;refGene_tx=XM_023246561.1;refGene_gn=SLC1A6;refGene_tx=XM_023246578.1;refGene_gn=SLC1A6;refGene_tx=XM_023246584.1
A2 11256447 11257017 refGene_gn=CCDC105;refGene_tx=XM_023246592.1
A2 11257902 11258007 refGene_gn=CCDC105;refGene_tx=XM_023246592.1
A2 11263284 11263483 refGene_gn=CCDC105;refGene_tx=XM_023246592.1

A2	11263874 11264041 refGene_gn=CCDC105;refGene_tx=XM_023246592.1
A2	11264193 11264289 refGene_gn=CCDC105;refGene_tx=XM_023246592.1
A2	11264383 11264497 refGene_gn=CCDC105;refGene_tx=XM_023246592.1
A2	11265312 11265564 refGene_gn=CCDC105;refGene_tx=XM_023246592.1
A2	11294400 11294427 refGene_gn=CASP14;refGene_tx=XM_003981969.3
A2	11295452 11295602 refGene_gn=CASP14;refGene_tx=XM_003981969.3
A2	11295693 11295919 refGene_gn=CASP14;refGene_tx=XM_003981969.3
A2	11296060 11296177 refGene_gn=CASP14;refGene_tx=XM_003981969.3
A2	11296372 11296476 refGene_gn=CASP14;refGene_tx=XM_003981969.3
A2	11296974 11297079 refGene_gn=CASP14;refGene_tx=XM_003981969.3
A2	11320968 11321059
refGene_gn=LOC101099278;refGene_tx=XM_011290088.3
A2	11321061 11321088
refGene_gn=LOC101099278;refGene_tx=XM_011290088.3
A2	11321090 11321201
refGene_gn=LOC101099278;refGene_tx=XM_011290088.3
A2	11321203 11321415
refGene_gn=LOC101099278;refGene_tx=XM_011290088.3
A2	11321417 11321620
refGene_gn=LOC101099278;refGene_tx=XM_011290088.3
A2	11321621 11321739
refGene_gn=LOC101099278;refGene_tx=XM_011290088.3
A2	11321740 11321981
refGene_gn=LOC101099278;refGene_tx=XM_011290088.3
A2	11321983 11322027
refGene_gn=LOC101099278;refGene_tx=XM_011290088.3
A2	11330281 11330369 refGene_gn=SYDE1;refGene_tx=XM_023241193.1
A2	11332014 11332407 refGene_gn=SYDE1;refGene_tx=XM_023241193.1
A2	11333148 11333364 refGene_gn=SYDE1;refGene_tx=XM_023241193.1
A2	11333518 11333733 refGene_gn=SYDE1;refGene_tx=XM_023241193.1
A2	11334331 11334458 refGene_gn=SYDE1;refGene_tx=XM_023241193.1
A2	11334669 11334830 refGene_gn=SYDE1;refGene_tx=XM_023241193.1
A2	11335412 11335638 refGene_gn=SYDE1;refGene_tx=XM_023241193.1
A2	11336020 11336424 refGene_gn=SYDE1;refGene_tx=XM_023241193.1
A2	11337746 11337922
refGene_gn=ILVBL;refGene_tx=XM_019818809.2;refGene_gn=ILVBL;refGene_tx
=XM_023246606.1
A2	11338056 11338162
refGene_gn=ILVBL;refGene_tx=XM_019818809.2;refGene_gn=ILVBL;refGene_tx
=XM_023246606.1
A2	11338325 11338397
refGene_gn=ILVBL;refGene_tx=XM_019818809.2;refGene_gn=ILVBL;refGene_tx
=XM_023246606.1
A2	11338476 11338547
refGene_gn=ILVBL;refGene_tx=XM_019818809.2;refGene_gn=ILVBL;refGene_tx
=XM_023246606.1
A2	11338623 11338793
refGene_gn=ILVBL;refGene_tx=XM_019818809.2;refGene_gn=ILVBL;refGene_tx
=XM_023246606.1
A2	11338896 11339011

refGene_gn=ILVBL;refGene_tx=XM_019818809.2;refGene_gn=ILVBL;refGene_tx=XM_023246606.1
A2 11340038 11340173 refGene_gn=ILVBL;refGene_tx=XM_019818809.2;refGene_gn=ILVBL;refGene_tx=XM_023246606.1
A2 11341973 11342110 refGene_gn=ILVBL;refGene_tx=XM_019818809.2;refGene_gn=ILVBL;refGene_tx=XM_023246606.1
A2 11342227 11342325 refGene_gn=ILVBL;refGene_tx=XM_019818809.2;refGene_gn=ILVBL;refGene_tx=XM_023246606.1
A2 11342466 11342566 refGene_gn=ILVBL;refGene_tx=XM_019818809.2;refGene_gn=ILVBL;refGene_tx=XM_023246606.1
A2 11345230 11345342 refGene_gn=ILVBL;refGene_tx=XM_019818809.2;refGene_gn=ILVBL;refGene_tx=XM_023246606.1
A2 11345436 11345575 refGene_gn=ILVBL;refGene_tx=XM_019818809.2;refGene_gn=ILVBL;refGene_tx=XM_023246606.1
A2 11345661 11345789 refGene_gn=ILVBL;refGene_tx=XM_019818809.2;refGene_gn=ILVBL;refGene_tx=XM_023246606.1
A2 11345911 11346113 refGene_gn=ILVBL;refGene_tx=XM_019818809.2;refGene_gn=ILVBL;refGene_tx=XM_023246606.1
A2 11347267 11347412 refGene_gn=ILVBL;refGene_tx=XM_019818809.2;refGene_gn=ILVBL;refGene_tx=XM_023246606.1
A2 11347947 11347988 refGene_gn=ILVBL;refGene_tx=XM_019818809.2
A2 11359761 11359907 refGene_gn=LOC111557226;refGene_tx=XM_023241194.1
A2 11359993 11360324 refGene_gn=LOC111557226;refGene_tx=XM_023241194.1
A2 11360497 11360668 refGene_gn=LOC111557226;refGene_tx=XM_023241194.1
A2 11362934 11362946 refGene_gn=LOC111557226;refGene_tx=XM_023241194.1
A2 11377452 11378505 refGene_gn=NOTCH3;refGene_tx=XM_023246608.1
A2 11379555 11379653 refGene_gn=NOTCH3;refGene_tx=XM_023246608.1
A2 11383157 11383305 refGene_gn=NOTCH3;refGene_tx=XM_023246608.1
A2 11383580 11383885 refGene_gn=NOTCH3;refGene_tx=XM_023246608.1
A2 11385496 11385659 refGene_gn=NOTCH3;refGene_tx=XM_023246608.1
A2 11388058 11388143 refGene_gn=NOTCH3;refGene_tx=XM_023246608.1
A2 11389152 11389375 refGene_gn=NOTCH3;refGene_tx=XM_023246608.1
A2 11389464 11389619 refGene_gn=NOTCH3;refGene_tx=XM_023246608.1
A2 11391605 11391938 refGene_gn=NOTCH3;refGene_tx=XM_023246608.1
A2 11393631 11394197 refGene_gn=NOTCH3;refGene_tx=XM_023246608.1
A2 11394537 11394656 refGene_gn=NOTCH3;refGene_tx=XM_023246608.1
A2 11394738 11394996 refGene_gn=NOTCH3;refGene_tx=XM_023246608.1

A2 11395084 11395217 refGene_gn=NOTCH3;refGene_tx=XM_023246608.1
A2 11395788 11395973 refGene_gn=NOTCH3;refGene_tx=XM_023246608.1
A2 11396405 11396559 refGene_gn=NOTCH3;refGene_tx=XM_023246608.1
A2 11396705 11396907 refGene_gn=NOTCH3;refGene_tx=XM_023246608.1
A2 11397333 11397559 refGene_gn=NOTCH3;refGene_tx=XM_023246608.1
A2 11398743 11398899 refGene_gn=NOTCH3;refGene_tx=XM_023246608.1
A2 11399400 11399514 refGene_gn=NOTCH3;refGene_tx=XM_023246608.1
A2 11399737 11399889 refGene_gn=NOTCH3;refGene_tx=XM_023246608.1
A2 11399965 11400158 refGene_gn=NOTCH3;refGene_tx=XM_023246608.1
A2 11400872 11400983 refGene_gn=NOTCH3;refGene_tx=XM_023246608.1
A2 11401095 11401329 refGene_gn=NOTCH3;refGene_tx=XM_023246608.1
A2 11401909 11402023 refGene_gn=NOTCH3;refGene_tx=XM_023246608.1
A2 11402529 11402643 refGene_gn=NOTCH3;refGene_tx=XM_023246608.1
A2 11403309 11403495 refGene_gn=NOTCH3;refGene_tx=XM_023246608.1
A2 11403585 11403741 refGene_gn=NOTCH3;refGene_tx=XM_023246608.1
A2 11404344 11404578 refGene_gn=NOTCH3;refGene_tx=XM_023246608.1
A2 11404666 11404789 refGene_gn=NOTCH3;refGene_tx=XM_023246608.1
A2 11404882 11405221 refGene_gn=NOTCH3;refGene_tx=XM_023246608.1
A2 11405303 11405446 refGene_gn=NOTCH3;refGene_tx=XM_023246608.1
A2 11409294 11409373 refGene_gn=NOTCH3;refGene_tx=XM_023246608.1
A2 11411921 11412039 refGene_gn=NOTCH3;refGene_tx=XM_023246608.1
A2 11421938 11422164 refGene_gn=EPHX3;refGene_tx=XM_003981971.5
A2 11422251 11422388 refGene_gn=EPHX3;refGene_tx=XM_003981971.5
A2 11422482 11422586 refGene_gn=EPHX3;refGene_tx=XM_003981971.5
A2 11423410 11423539 refGene_gn=EPHX3;refGene_tx=XM_003981971.5
A2 11423632 11423790 refGene_gn=EPHX3;refGene_tx=XM_003981971.5
A2 11424133 11424219 refGene_gn=EPHX3;refGene_tx=XM_003981971.5
A2 11424441 11424684 refGene_gn=EPHX3;refGene_tx=XM_003981971.5
A2 11429639 11429708
refGene_gn=BRD4;refGene_tx=XM_023246610.1;refGene_gn=BRD4;refGene_tx=XM_023246612.1;refGene_gn=BRD4;refGene_tx=XM_023246615.1;refGene_gn=BRD4;refGene_tx=XM_023246621.1;refGene_gn=BRD4;refGene_tx=XM_023246623.1;refGene_gn=BRD4;refGene_tx=XM_023246624.1
A2 11430100 11430338
refGene_gn=BRD4;refGene_tx=XM_023246610.1;refGene_gn=BRD4;refGene_tx=XM_023246612.1;refGene_gn=BRD4;refGene_tx=XM_023246615.1;refGene_gn=BRD4;refGene_tx=XM_023246621.1;refGene_gn=BRD4;refGene_tx=XM_023246623.1;refGene_gn=BRD4;refGene_tx=XM_023246624.1
A2 11430447 11430653
refGene_gn=BRD4;refGene_tx=XM_023246610.1;refGene_gn=BRD4;refGene_tx=XM_023246612.1;refGene_gn=BRD4;refGene_tx=XM_023246615.1;refGene_gn=BRD4;refGene_tx=XM_023246621.1;refGene_gn=BRD4;refGene_tx=XM_023246623.1;refGene_gn=BRD4;refGene_tx=XM_023246624.1
A2 11430777 11430908
refGene_gn=BRD4;refGene_tx=XM_023246610.1;refGene_gn=BRD4;refGene_tx=XM_023246612.1;refGene_gn=BRD4;refGene_tx=XM_023246615.1;refGene_gn=BRD4;refGene_tx=XM_023246621.1;refGene_gn=BRD4;refGene_tx=XM_023246623.1;refGene_gn=BRD4;refGene_tx=XM_023246624.1
A2 11431039 11431414
refGene_gn=BRD4;refGene_tx=XM_023246610.1;refGene_gn=BRD4;refGene_tx=X

M_023246612.1;refGene_gn=BRD4;refGene_tx=XM_023246615.1;refGene_gn=BRD4;refGene_tx=XM_023246621.1;refGene_gn=BRD4;refGene_tx=XM_023246623.1;refGene_gn=BRD4;refGene_tx=XM_023246624.1
A2 11433987 11434575
refGene_gn=BRD4;refGene_tx=XM_023246610.1;refGene_gn=BRD4;refGene_tx=XM_023246612.1;refGene_gn=BRD4;refGene_tx=XM_023246615.1;refGene_gn=BRD4;refGene_tx=XM_023246621.1;refGene_gn=BRD4;refGene_tx=XM_023246623.1;refGene_gn=BRD4;refGene_tx=XM_023246624.1
A2 11435210 11435577
refGene_gn=BRD4;refGene_tx=XM_023246610.1;refGene_gn=BRD4;refGene_tx=XM_023246612.1;refGene_gn=BRD4;refGene_tx=XM_023246615.1;refGene_gn=BRD4;refGene_tx=XM_023246621.1;refGene_gn=BRD4;refGene_tx=XM_023246623.1;refGene_gn=BRD4;refGene_tx=XM_023246624.1
A2 11435662 11435745
refGene_gn=BRD4;refGene_tx=XM_023246610.1;refGene_gn=BRD4;refGene_tx=XM_023246612.1;refGene_gn=BRD4;refGene_tx=XM_023246615.1;refGene_gn=BRD4;refGene_tx=XM_023246621.1;refGene_gn=BRD4;refGene_tx=XM_023246623.1;refGene_gn=BRD4;refGene_tx=XM_023246624.1
A2 11445197 11445308
refGene_gn=BRD4;refGene_tx=XM_023246610.1;refGene_gn=BRD4;refGene_tx=XM_023246612.1;refGene_gn=BRD4;refGene_tx=XM_023246615.1;refGene_gn=BRD4;refGene_tx=XM_023246621.1;refGene_gn=BRD4;refGene_tx=XM_023246623.1;refGene_gn=BRD4;refGene_tx=XM_023246624.1
A2 11446335 11446631
refGene_gn=BRD4;refGene_tx=XM_023246610.1;refGene_gn=BRD4;refGene_tx=XM_023246612.1;refGene_gn=BRD4;refGene_tx=XM_023246615.1;refGene_gn=BRD4;refGene_tx=XM_023246621.1;refGene_gn=BRD4;refGene_tx=XM_023246623.1;refGene_gn=BRD4;refGene_tx=XM_023246624.1
A2 11446954 11447154
refGene_gn=BRD4;refGene_tx=XM_023246610.1;refGene_gn=BRD4;refGene_tx=XM_023246612.1;refGene_gn=BRD4;refGene_tx=XM_023246615.1;refGene_gn=BRD4;refGene_tx=XM_023246621.1;refGene_gn=BRD4;refGene_tx=XM_023246623.1;refGene_gn=BRD4;refGene_tx=XM_023246624.1
A2 11447991 11448201
refGene_gn=BRD4;refGene_tx=XM_023246610.1;refGene_gn=BRD4;refGene_tx=XM_023246612.1;refGene_gn=BRD4;refGene_tx=XM_023246615.1;refGene_gn=BRD4;refGene_tx=XM_023246621.1;refGene_gn=BRD4;refGene_tx=XM_023246623.1;refGene_gn=BRD4;refGene_tx=XM_023246624.1
A2 11454845 11454974
refGene_gn=BRD4;refGene_tx=XM_023246610.1;refGene_gn=BRD4;refGene_tx=XM_023246612.1;refGene_gn=BRD4;refGene_tx=XM_023246615.1;refGene_gn=BRD4;refGene_tx=XM_023246621.1;refGene_gn=BRD4;refGene_tx=XM_023246623.1;refGene_gn=BRD4;refGene_tx=XM_023246624.1
A2 11455869 11456232
refGene_gn=BRD4;refGene_tx=XM_023246610.1;refGene_gn=BRD4;refGene_tx=XM_023246612.1;refGene_gn=BRD4;refGene_tx=XM_023246615.1;refGene_gn=BRD4;refGene_tx=XM_023246621.1;refGene_gn=BRD4;refGene_tx=XM_023246623.1;refGene_gn=BRD4;refGene_tx=XM_023246624.1
A2 11456930 11457220
refGene_gn=BRD4;refGene_tx=XM_023246610.1;refGene_gn=BRD4;refGene_tx=X

M_023246612.1;refGene_gn=BRD4;refGene_tx=XM_023246615.1;refGene_gn=BRD4;refGene_tx=XM_023246621.1;refGene_gn=BRD4;refGene_tx=XM_023246623.1;refGene_gn=BRD4;refGene_tx=XM_023246624.1
A2	11458951	11459087	refGene_gn=BRD4;refGene_tx=XM_023246610.1;refGene_gn=BRD4;refGene_tx=XM_023246612.1;refGene_gn=BRD4;refGene_tx=XM_023246615.1;refGene_gn=BRD4;refGene_tx=XM_023246621.1;refGene_gn=BRD4;refGene_tx=XM_023246623.1;refGene_gn=BRD4;refGene_tx=XM_023246624.1
A2	11460410	11460548	refGene_gn=BRD4;refGene_tx=XM_023246610.1;refGene_gn=BRD4;refGene_tx=XM_023246612.1;refGene_gn=BRD4;refGene_tx=XM_023246615.1;refGene_gn=BRD4;refGene_tx=XM_023246621.1;refGene_gn=BRD4;refGene_tx=XM_023246623.1
A2	11464182	11464467	refGene_gn=BRD4;refGene_tx=XM_023246610.1;refGene_gn=BRD4;refGene_tx=XM_023246612.1;refGene_gn=BRD4;refGene_tx=XM_023246615.1;refGene_gn=BRD4;refGene_tx=XM_023246621.1;refGene_gn=BRD4;refGene_tx=XM_023246623.1
A2	11527302	11527839	refGene_gn=AKAP8;refGene_tx=XM_011289980.3
A2	11529704	11529800	refGene_gn=AKAP8;refGene_tx=XM_011289980.3
A2	11530918	11531049	refGene_gn=AKAP8;refGene_tx=XM_011289980.3
A2	11531480	11531574	refGene_gn=AKAP8;refGene_tx=XM_011289980.3
A2	11531904	11532046	refGene_gn=AKAP8;refGene_tx=XM_011289980.3
A2	11534478	11534566	refGene_gn=AKAP8;refGene_tx=XM_011289980.3
A2	11536530	11536564	refGene_gn=AKAP8;refGene_tx=XM_011289980.3
A2	11537918	11537965	refGene_gn=AKAP8;refGene_tx=XM_011289980.3
A2	11538216	11538346	refGene_gn=AKAP8;refGene_tx=XM_011289980.3
A2	11538851	11539353	refGene_gn=AKAP8;refGene_tx=XM_011289980.3
A2	11539796	11540073	refGene_gn=AKAP8;refGene_tx=XM_011289980.3
A2	11540581	11540614	refGene_gn=AKAP8;refGene_tx=XM_011289980.3
A2	11542091	11542130	refGene_gn=AKAP8;refGene_tx=XM_011289980.3
A2	11544618	11544637	refGene_gn=AKAP8;refGene_tx=XM_011289980.3
A2	11545083	11545404	refGene_gn=AKAP8L;refGene_tx=XM_019818776.2;refGene_gn=AKAP8L;refGene_tx=XM_023246649.1
A2	11545485	11545581	refGene_gn=AKAP8L;refGene_tx=XM_019818776.2;refGene_gn=AKAP8L;refGene_tx=XM_023246649.1
A2	11555119	11555250	refGene_gn=AKAP8L;refGene_tx=XM_019818776.2;refGene_gn=AKAP8L;refGene_tx=XM_023246649.1
A2	11555482	11555588	refGene_gn=AKAP8L;refGene_tx=XM_019818776.2;refGene_gn=AKAP8L;refGene_tx=XM_023246649.1
A2	11555686	11555828	refGene_gn=AKAP8L;refGene_tx=XM_019818776.2;refGene_gn=AKAP8L;refGene_tx=XM_023246649.1
A2	11557278	11557387	refGene_gn=AKAP8L;refGene_tx=XM_019818776.2;refGene_gn=AKAP8L;refGene_tx=XM_023246649.1
A2	11558316	11558380	refGene_gn=AKAP8L;refGene_tx=XM_019818776.2;refGene_gn=AKAP8L;refGene_

tx=XM_023246649.1
A2 11558732 11558803
refGene_gn=AKAP8L;refGene_tx=XM_019818776.2;refGene_gn=AKAP8L;refGene_tx=XM_023246649.1
A2 11558885 11558982
refGene_gn=AKAP8L;refGene_tx=XM_019818776.2;refGene_gn=AKAP8L;refGene_tx=XM_023246649.1
A2 11559077 11559531
refGene_gn=AKAP8L;refGene_tx=XM_019818776.2;refGene_gn=AKAP8L;refGene_tx=XM_023246649.1
A2 11561378 11561619
refGene_gn=AKAP8L;refGene_tx=XM_019818776.2;refGene_gn=AKAP8L;refGene_tx=XM_023246649.1
A2 11561917 11561950
refGene_gn=AKAP8L;refGene_tx=XM_019818776.2;refGene_gn=AKAP8L;refGene_tx=XM_023246649.1
A2 11566418 11566493
refGene_gn=AKAP8L;refGene_tx=XM_019818776.2;refGene_gn=AKAP8L;refGene_tx=XM_023246649.1
A2 11572338 11572351
refGene_gn=AKAP8L;refGene_tx=XM_019818776.2;refGene_gn=AKAP8L;refGene_tx=XM_023246649.1
A2 11576582 11576742
refGene_gn=WIZ;refGene_tx=XM_023246657.1;refGene_gn=WIZ;refGene_tx=XM_023246663.1;refGene_gn=WIZ;refGene_tx=XM_023246666.1;refGene_gn=WIZ;refGene_tx=XM_023246668.1;refGene_gn=WIZ;refGene_tx=XM_023246674.1;refGene_gn=WIZ;refGene_tx=XM_023246679.1;refGene_gn=WIZ;refGene_tx=XM_023246688.1
A2 11577770 11577966
refGene_gn=WIZ;refGene_tx=XM_023246657.1;refGene_gn=WIZ;refGene_tx=XM_023246663.1;refGene_gn=WIZ;refGene_tx=XM_023246666.1;refGene_gn=WIZ;refGene_tx=XM_023246668.1;refGene_gn=WIZ;refGene_tx=XM_023246674.1;refGene_gn=WIZ;refGene_tx=XM_023246679.1;refGene_gn=WIZ;refGene_tx=XM_023246688.1
A2 11578246 11578666
refGene_gn=WIZ;refGene_tx=XM_023246657.1;refGene_gn=WIZ;refGene_tx=XM_023246663.1;refGene_gn=WIZ;refGene_tx=XM_023246666.1;refGene_gn=WIZ;refGene_tx=XM_023246668.1;refGene_gn=WIZ;refGene_tx=XM_023246674.1;refGene_gn=WIZ;refGene_tx=XM_023246679.1;refGene_gn=WIZ;refGene_tx=XM_023246688.1
A2 11578868 11579396
refGene_gn=WIZ;refGene_tx=XM_023246657.1;refGene_gn=WIZ;refGene_tx=XM_023246663.1;refGene_gn=WIZ;refGene_tx=XM_023246666.1;refGene_gn=WIZ;refGene_tx=XM_023246668.1;refGene_gn=WIZ;refGene_tx=XM_023246674.1;refGene_gn=WIZ;refGene_tx=XM_023246679.1;refGene_gn=WIZ;refGene_tx=XM_023246688.1
A2 11580488 11581040
refGene_gn=WIZ;refGene_tx=XM_023246657.1;refGene_gn=WIZ;refGene_tx=XM_023246663.1;refGene_gn=WIZ;refGene_tx=XM_023246666.1;refGene_gn=WIZ;refGene_tx=XM_023246668.1;refGene_gn=WIZ;refGene_tx=XM_023246674.1;refGe

ne_gn=WIZ;refGene_tx=XM_023246679.1;refGene_gn=WIZ;refGene_tx=XM_02324
6688.1
A2 11581629 11582028
refGene_gn=WIZ;refGene_tx=XM_023246657.1;refGene_gn=WIZ;refGene_tx=XM_
023246663.1;refGene_gn=WIZ;refGene_tx=XM_023246666.1;refGene_gn=WIZ;re
fGene_tx=XM_023246679.1
A2 11583102 11583606
refGene_gn=WIZ;refGene_tx=XM_023246657.1;refGene_gn=WIZ;refGene_tx=XM_
023246663.1;refGene_gn=WIZ;refGene_tx=XM_023246666.1;refGene_gn=WIZ;re
fGene_tx=XM_023246668.1;refGene_gn=WIZ;refGene_tx=XM_023246674.1;refGe
ne_gn=WIZ;refGene_tx=XM_023246679.1;refGene_gn=WIZ;refGene_tx=XM_02324
6688.1
A2 11584512 11584683
refGene_gn=WIZ;refGene_tx=XM_023246657.1;refGene_gn=WIZ;refGene_tx=XM_
023246663.1;refGene_gn=WIZ;refGene_tx=XM_023246668.1;refGene_gn=WIZ;re
fGene_tx=XM_023246679.1
A2 11588886 11589210
refGene_gn=WIZ;refGene_tx=XM_023246657.1;refGene_gn=WIZ;refGene_tx=XM_
023246663.1;refGene_gn=WIZ;refGene_tx=XM_023246666.1;refGene_gn=WIZ;re
fGene_tx=XM_023246668.1;refGene_gn=WIZ;refGene_tx=XM_023246674.1;refGe
ne_gn=WIZ;refGene_tx=XM_023246679.1;refGene_gn=WIZ;refGene_tx=XM_02324
6688.1
A2 11590413 11592551
refGene_gn=WIZ;refGene_tx=XM_023246657.1;refGene_gn=WIZ;refGene_tx=XM_
023246663.1;refGene_gn=WIZ;refGene_tx=XM_023246666.1;refGene_gn=WIZ;re
fGene_tx=XM_023246668.1;refGene_gn=WIZ;refGene_tx=XM_023246674.1
A2 11594374 11594447
refGene_gn=WIZ;refGene_tx=XM_023246657.1;refGene_gn=WIZ;refGene_tx=XM_
023246663.1;refGene_gn=WIZ;refGene_tx=XM_023246666.1;refGene_gn=WIZ;re
fGene_tx=XM_023246668.1;refGene_gn=WIZ;refGene_tx=XM_023246674.1
A2 11599381 11599586
refGene_gn=WIZ;refGene_tx=XM_023246657.1;refGene_gn=WIZ;refGene_tx=XM_
023246663.1;refGene_gn=WIZ;refGene_tx=XM_023246666.1;refGene_gn=WIZ;re
fGene_tx=XM_023246668.1;refGene_gn=WIZ;refGene_tx=XM_023246674.1;refGe
ne_gn=WIZ;refGene_tx=XM_023246679.1;refGene_gn=WIZ;refGene_tx=XM_02324
6688.1
A2 11603125 11603269 refGene_gn=RASAL3;refGene_tx=XM_019818691.2
A2 11603373 11603437 refGene_gn=RASAL3;refGene_tx=XM_019818691.2
A2 11603968 11604126 refGene_gn=RASAL3;refGene_tx=XM_019818691.2
A2 11604412 11604803 refGene_gn=RASAL3;refGene_tx=XM_019818691.2
A2 11605717 11605836 refGene_gn=RASAL3;refGene_tx=XM_019818691.2
A2 11605934 11606136 refGene_gn=RASAL3;refGene_tx=XM_019818691.2
A2 11606269 11606506 refGene_gn=RASAL3;refGene_tx=XM_019818691.2
A2 11607371 11607516 refGene_gn=RASAL3;refGene_tx=XM_019818691.2
A2 11607770 11607915 refGene_gn=RASAL3;refGene_tx=XM_019818691.2
A2 11608288 11608831 refGene_gn=RASAL3;refGene_tx=XM_019818691.2
A2 11609290 11609389 refGene_gn=RASAL3;refGene_tx=XM_019818691.2
A2 11609470 11609597 refGene_gn=RASAL3;refGene_tx=XM_019818691.2
A2 11610223 11610294 refGene_gn=RASAL3;refGene_tx=XM_019818691.2
A2 11610667 11610729 refGene_gn=RASAL3;refGene_tx=XM_019818691.2

A2 11610839 11610918 refGene_gn=RASAL3;refGene_tx=XM_019818691.2
A2 11611054 11611185 refGene_gn=RASAL3;refGene_tx=XM_019818691.2
A2 11612954 11613276 refGene_gn=RASAL3;refGene_tx=XM_019818691.2
A2 11615878 11616033
refGene_gn=PGLYRP2;refGene_tx=XM_003982008.3;refGene_gn=PGLYRP2;refGene_tx=XM_019818713.2
A2 11616855 11617153
refGene_gn=PGLYRP2;refGene_tx=XM_003982008.3;refGene_gn=PGLYRP2;refGene_tx=XM_019818713.2
A2 11617746 11617957
refGene_gn=PGLYRP2;refGene_tx=XM_003982008.3;refGene_gn=PGLYRP2;refGene_tx=XM_019818713.2
A2 11618989 11620117
refGene_gn=PGLYRP2;refGene_tx=XM_003982008.3;refGene_gn=PGLYRP2;refGene_tx=XM_019818713.2
A2 11621485 11621576
refGene_gn=PGLYRP2;refGene_tx=XM_003982008.3;refGene_gn=PGLYRP2;refGene_tx=XM_019818713.2
A2 11642294 11642650
refGene_gn=LOC101089989;refGene_tx=XM_011289937.3
A2 11652456 11652679
refGene_gn=LOC101089989;refGene_tx=XM_011289937.3;refGene_gn=LOC101089989;refGene_tx=XM_019818723.2;refGene_gn=LOC101089989;refGene_tx=XM_019818729.2;refGene_gn=LOC101089989;refGene_tx=XM_023246696.1
A2 11656270 11656415
refGene_gn=LOC101089989;refGene_tx=XM_011289937.3;refGene_gn=LOC101089989;refGene_tx=XM_019818723.2;refGene_gn=LOC101089989;refGene_tx=XM_019818729.2;refGene_gn=LOC101089989;refGene_tx=XM_023246696.1
A2 11661115 11661169
refGene_gn=LOC101089989;refGene_tx=XM_011289937.3;refGene_gn=LOC101089989;refGene_tx=XM_019818723.2;refGene_gn=LOC101089989;refGene_tx=XM_019818729.2;refGene_gn=LOC101089989;refGene_tx=XM_023246696.1
A2 11661288 11661416
refGene_gn=LOC101089989;refGene_tx=XM_011289937.3;refGene_gn=LOC101089989;refGene_tx=XM_019818723.2;refGene_gn=LOC101089989;refGene_tx=XM_019818729.2;refGene_gn=LOC101089989;refGene_tx=XM_023246696.1
A2 11661626 11661748
refGene_gn=LOC101089989;refGene_tx=XM_011289937.3;refGene_gn=LOC101089989;refGene_tx=XM_019818723.2;refGene_gn=LOC101089989;refGene_tx=XM_019818729.2;refGene_gn=LOC101089989;refGene_tx=XM_023246696.1
A2 11664516 11664784
refGene_gn=LOC101089989;refGene_tx=XM_011289937.3;refGene_gn=LOC101089989;refGene_tx=XM_019818723.2;refGene_gn=LOC101089989;refGene_tx=XM_019818729.2;refGene_gn=LOC101089989;refGene_tx=XM_023246696.1
A2 11666117 11666184
refGene_gn=LOC101089989;refGene_tx=XM_011289937.3;refGene_gn=LOC101089989;refGene_tx=XM_019818723.2;refGene_gn=LOC101089989;refGene_tx=XM_019818729.2;refGene_gn=LOC101089989;refGene_tx=XM_023246696.1
A2 11666285 11666415
refGene_gn=LOC101089989;refGene_tx=XM_011289937.3;refGene_gn=LOC101089

989;refGene_tx=XM_019818723.2;refGene_gn=LOC101089989;refGene_tx=XM_019818729.2;refGene_gn=LOC101089989;refGene_tx=XM_023246696.1
A2 11667865 11667999
refGene_gn=LOC101089989;refGene_tx=XM_011289937.3;refGene_gn=LOC101089989;refGene_tx=XM_019818723.2;refGene_gn=LOC101089989;refGene_tx=XM_019818729.2;refGene_gn=LOC101089989;refGene_tx=XM_023246696.1
A2 11668902 11668967
refGene_gn=LOC101089989;refGene_tx=XM_011289937.3;refGene_gn=LOC101089989;refGene_tx=XM_019818723.2;refGene_gn=LOC101089989;refGene_tx=XM_019818729.2;refGene_gn=LOC101089989;refGene_tx=XM_023246696.1
A2 11669628 11669711
refGene_gn=LOC101089989;refGene_tx=XM_011289937.3;refGene_gn=LOC101089989;refGene_tx=XM_019818723.2;refGene_gn=LOC101089989;refGene_tx=XM_019818729.2;refGene_gn=LOC101089989;refGene_tx=XM_023246696.1
A2 11670103 11670281
refGene_gn=LOC101089989;refGene_tx=XM_011289937.3;refGene_gn=LOC101089989;refGene_tx=XM_019818723.2;refGene_gn=LOC101089989;refGene_tx=XM_019818729.2;refGene_gn=LOC101089989;refGene_tx=XM_023246696.1
A2 11680735 11680744
refGene_gn=LOC105259722;refGene_tx=XM_023241195.1
A2 11680964 11681004
refGene_gn=LOC105259722;refGene_tx=XM_023241195.1
A2 11681405 11681470
refGene_gn=LOC105259722;refGene_tx=XM_023241195.1
A2 11687409 11687724
refGene_gn=LOC105259722;refGene_tx=XM_023241195.1
A2 11691327 11691472
refGene_gn=LOC105259722;refGene_tx=XM_023241195.1
A2 11693085 11693139
refGene_gn=LOC105259722;refGene_tx=XM_023241195.1
A2 11693232 11693360
refGene_gn=LOC105259722;refGene_tx=XM_023241195.1
A2 11694810 11694932
refGene_gn=LOC105259722;refGene_tx=XM_023241195.1
A2 11695630 11695901
refGene_gn=LOC105259722;refGene_tx=XM_023241195.1
A2 11696232 11696299
refGene_gn=LOC105259722;refGene_tx=XM_023241195.1
A2 11696489 11696619
refGene_gn=LOC105259722;refGene_tx=XM_023241195.1
A2 11697107 11697413
refGene_gn=LOC105259722;refGene_tx=XM_023241195.1
A2 11702856 11703015
refGene_gn=LOC105259722;refGene_tx=XM_023241195.1
A2 11703388 11703566
refGene_gn=LOC105259722;refGene_tx=XM_023241195.1
A2 11729702 11730653
refGene_gn=LOC101091513;refGene_tx=XM_003982014.1
A2 11765071 11766033
refGene_gn=LOC105261280;refGene_tx=XM_023241202.1

A2 11781148 11781269
refGene_gn=LOC105261280;refGene_tx=XM_023241202.1
A2 11782741 11783692
refGene_gn=LOC101091766;refGene_tx=XM_003982015.2
A2 11791181 11792141
refGene_gn=LOC105261287;refGene_tx=XM_023241386.1
A2 11803033 11803172
refGene_gn=LINC01835;refGene_tx=XM_023246720.1
A2 11804815 11804919
refGene_gn=LINC01835;refGene_tx=XM_023246720.1
A2 11805794 11805946
refGene_gn=LINC01835;refGene_tx=XM_023246720.1
A2 11808221 11808311
refGene_gn=LINC01835;refGene_tx=XM_023246720.1
A2 11808407 11808461
refGene_gn=LINC01835;refGene_tx=XM_023246720.1
A2 11808962 11809070
refGene_gn=LINC01835;refGene_tx=XM_023246720.1
A2 11809404 11809486
refGene_gn=LINC01835;refGene_tx=XM_023246720.1
A2 11820389 11821334
refGene_gn=LOC101099942;refGene_tx=XM_003982048.2
A2 11860710 11860824
refGene_gn=TPM4;refGene_tx=XM_003982017.5;refGene_gn=TPM4;refGene_tx=XM_003982018.5
A2 11866224 11866350
refGene_gn=TPM4;refGene_tx=XM_003982017.5;refGene_gn=TPM4;refGene_tx=XM_003982018.5
A2 11866801 11866933
refGene_gn=TPM4;refGene_tx=XM_023246735.1;refGene_gn=TPM4;refGene_tx=XM_023246737.1
A2 11871435 11871569
refGene_gn=TPM4;refGene_tx=XM_003982017.5;refGene_gn=TPM4;refGene_tx=XM_003982018.5;refGene_gn=TPM4;refGene_tx=XM_023246735.1;refGene_gn=TPM4;refGene_tx=XM_023246737.1
A2 11873910 11874028
refGene_gn=TPM4;refGene_tx=XM_003982017.5;refGene_gn=TPM4;refGene_tx=XM_003982018.5;refGene_gn=TPM4;refGene_tx=XM_023246735.1;refGene_gn=TPM4;refGene_tx=XM_023246737.1
A2 11874464 11874535
refGene_gn=TPM4;refGene_tx=XM_003982017.5;refGene_gn=TPM4;refGene_tx=XM_003982018.5;refGene_gn=TPM4;refGene_tx=XM_023246735.1;refGene_gn=TPM4;refGene_tx=XM_023246737.1
A2 11875446 11875522
refGene_gn=TPM4;refGene_tx=XM_003982017.5;refGene_gn=TPM4;refGene_tx=XM_003982018.5;refGene_gn=TPM4;refGene_tx=XM_023246735.1;refGene_gn=TPM4;refGene_tx=XM_023246737.1
A2 11877245 11877308
refGene_gn=TPM4;refGene_tx=XM_003982017.5;refGene_gn=TPM4;refGene_tx=XM_003982018.5;refGene_gn=TPM4;refGene_tx=XM_023246735.1;refGene_gn=TPM

4;refGene_tx=XM_023246737.1
A2 11877393 11877463
refGene_gn=TPM4;refGene_tx=XM_003982017.5;refGene_gn=TPM4;refGene_tx=XM_003982018.5;refGene_gn=TPM4;refGene_tx=XM_023246735.1;refGene_gn=TPM4;refGene_tx=XM_023246737.1
A2 11878489 11878572
refGene_gn=TPM4;refGene_tx=XM_003982018.5;refGene_gn=TPM4;refGene_tx=XM_023246737.1
A2 11881493 11881576
refGene_gn=TPM4;refGene_tx=XM_003982017.5;refGene_gn=TPM4;refGene_tx=XM_023246735.1
A2 11886100 11886224 refGene_gn=RAB8A;refGene_tx=XM_003982019.5
A2 11890572 11890633 refGene_gn=RAB8A;refGene_tx=XM_003982019.5
A2 11892506 11892567 refGene_gn=RAB8A;refGene_tx=XM_003982019.5
A2 11895950 11896028 refGene_gn=RAB8A;refGene_tx=XM_003982019.5
A2 11897552 11897642 refGene_gn=RAB8A;refGene_tx=XM_003982019.5
A2 11898208 11898274 refGene_gn=RAB8A;refGene_tx=XM_003982019.5
A2 11899585 11899636 refGene_gn=RAB8A;refGene_tx=XM_003982019.5
A2 11902958 11903051 refGene_gn=RAB8A;refGene_tx=XM_003982019.5
A2 11910080 11910205
refGene_gn=HSH2D;refGene_tx=XM_003982050.5;refGene_gn=HSH2D;refGene_tx=XM_019818910.2
A2 11911628 11911718
refGene_gn=HSH2D;refGene_tx=XM_003982050.5;refGene_gn=HSH2D;refGene_tx=XM_019818910.2
A2 11912712 11912878
refGene_gn=HSH2D;refGene_tx=XM_003982050.5;refGene_gn=HSH2D;refGene_tx=XM_019818910.2
A2 11913315 11913450
refGene_gn=HSH2D;refGene_tx=XM_003982050.5;refGene_gn=HSH2D;refGene_tx=XM_019818910.2
A2 11915693 11916260
refGene_gn=HSH2D;refGene_tx=XM_003982050.5;refGene_gn=HSH2D;refGene_tx=XM_019818910.2
A2 11917938 11917960 refGene_gn=CIB3;refGene_tx=XM_023241207.1
A2 11919157 11919353 refGene_gn=CIB3;refGene_tx=XM_023241207.1
A2 11920251 11920399 refGene_gn=CIB3;refGene_tx=XM_023241207.1
A2 11921811 11922009 refGene_gn=CIB3;refGene_tx=XM_023241207.1
A2 11922496 11922553 refGene_gn=CIB3;refGene_tx=XM_023241207.1
A2 11934138 11934248 refGene_gn=FAM32A;refGene_tx=XM_003982021.4
A2 11934370 11934512 refGene_gn=FAM32A;refGene_tx=XM_003982021.4
A2 11938928 11938982 refGene_gn=FAM32A;refGene_tx=XM_003982021.4
A2 11939281 11939350 refGene_gn=FAM32A;refGene_tx=XM_003982021.4
A2 11942634 11942676 refGene_gn=AP1M1;refGene_tx=XM_003982023.5
A2 11948345 11948502 refGene_gn=AP1M1;refGene_tx=XM_003982023.5
A2 11950423 11950491 refGene_gn=AP1M1;refGene_tx=XM_003982023.5
A2 11952039 11952170 refGene_gn=AP1M1;refGene_tx=XM_003982023.5
A2 11952883 11953031 refGene_gn=AP1M1;refGene_tx=XM_003982023.5
A2 11962621 11962748 refGene_gn=AP1M1;refGene_tx=XM_003982023.5
A2 11963866 11964009 refGene_gn=AP1M1;refGene_tx=XM_003982023.5

A2 11964426 11964498 refGene_gn=AP1M1;refGene_tx=XM_003982023.5
A2 11964925 11965084 refGene_gn=AP1M1;refGene_tx=XM_003982023.5
A2 11970320 11970446 refGene_gn=AP1M1;refGene_tx=XM_003982023.5
A2 11971030 11971106 refGene_gn=AP1M1;refGene_tx=XM_003982023.5
A2 11971270 11971293 refGene_gn=AP1M1;refGene_tx=XM_003982023.5
A2 12042557 12042632 refGene_gn=KLF2;refGene_tx=XM_023246767.1
A2 12042875 12043310 refGene_gn=KLF2;refGene_tx=XM_023246767.1
A2 12043311 12043714 refGene_gn=KLF2;refGene_tx=XM_023246767.1
A2 12044322 12044498 refGene_gn=KLF2;refGene_tx=XM_023246767.1
A2 12066712 12066859
refGene_gn=EPS15L1;refGene_tx=XM_023246774.1;refGene_gn=EPS15L1;refGene_tx=XM_023246779.1;refGene_gn=EPS15L1;refGene_tx=XM_023246782.1;refGene_gn=EPS15L1;refGene_tx=XM_023246793.1;refGene_gn=EPS15L1;refGene_tx=XM_023246796.1;refGene_gn=EPS15L1;refGene_tx=XM_023246805.1;refGene_gn=EPS15L1;refGene_tx=XM_023246812.1;refGene_gn=EPS15L1;refGene_tx=XM_023246815.1;refGene_gn=EPS15L1;refGene_tx=XM_023246818.1
A2 12071459 12071492 refGene_gn=EPS15L1;refGene_tx=XM_023246786.1
A2 12071800 12072006
refGene_gn=EPS15L1;refGene_tx=XM_023246774.1;refGene_gn=EPS15L1;refGene_tx=XM_023246779.1;refGene_gn=EPS15L1;refGene_tx=XM_023246782.1;refGene_gn=EPS15L1;refGene_tx=XM_023246786.1;refGene_gn=EPS15L1;refGene_tx=XM_023246793.1;refGene_gn=EPS15L1;refGene_tx=XM_023246796.1;refGene_gn=EPS15L1;refGene_tx=XM_023246816.1
A2 12084188 12084236
refGene_gn=EPS15L1;refGene_tx=XM_023246774.1;refGene_gn=EPS15L1;refGene_tx=XM_023246779.1;refGene_gn=EPS15L1;refGene_tx=XM_023246786.1;refGene_gn=EPS15L1;refGene_tx=XM_023246793.1;refGene_gn=EPS15L1;refGene_tx=XM_023246796.1;refGene_gn=EPS15L1;refGene_tx=XM_023246805.1
A2 12086375 12086508
refGene_gn=EPS15L1;refGene_tx=XM_023246774.1;refGene_gn=EPS15L1;refGene_tx=XM_023246779.1;refGene_gn=EPS15L1;refGene_tx=XM_023246782.1;refGene_gn=EPS15L1;refGene_tx=XM_023246786.1;refGene_gn=EPS15L1;refGene_tx=XM_023246793.1;refGene_gn=EPS15L1;refGene_tx=XM_023246796.1;refGene_gn=EPS15L1;refGene_tx=XM_023246805.1;refGene_gn=EPS15L1;refGene_tx=XM_023246815.1
A2 12092747 12092830
refGene_gn=EPS15L1;refGene_tx=XM_023246774.1;refGene_gn=EPS15L1;refGene_tx=XM_023246779.1;refGene_gn=EPS15L1;refGene_tx=XM_023246782.1;refGene_gn=EPS15L1;refGene_tx=XM_023246786.1;refGene_gn=EPS15L1;refGene_tx=XM_023246796.1;refGene_gn=EPS15L1;refGene_tx=XM_023246805.1;refGene_gn=EPS15L1;refGene_tx=XM_023246812.1;refGene_gn=EPS15L1;refGene_tx=XM_023246815.1;refGene_gn=EPS15L1;refGene_tx=XM_023246816.1
A2 12093749 12093810
refGene_gn=EPS15L1;refGene_tx=XM_023246774.1;refGene_gn=EPS15L1;refGene_tx=XM_023246779.1;refGene_gn=EPS15L1;refGene_tx=XM_023246782.1;refGene_gn=EPS15L1;refGene_tx=XM_023246786.1;refGene_gn=EPS15L1;refGene_tx=XM_023246796.1;refGene_gn=EPS15L1;refGene_tx=XM_023246805.1;refGene_gn=EPS15L1;refGene_tx=XM_023246812.1;refGene_gn=EPS15L1;refGene_tx=XM_023246815.1;refGene_gn=EPS15L1;refGene_tx=XM_023246816.1
A2 12097795 12097932

refGene_gn=EPS15L1;refGene_tx=XM_023246774.1;refGene_gn=EPS15L1;refGene_tx=XM_023246779.1;refGene_gn=EPS15L1;refGene_tx=XM_023246782.1;refGene_gn=EPS15L1;refGene_tx=XM_023246786.1;refGene_gn=EPS15L1;refGene_tx=XM_023246793.1;refGene_gn=EPS15L1;refGene_tx=XM_023246796.1;refGene_gn=EPS15L1;refGene_tx=XM_023246805.1;refGene_gn=EPS15L1;refGene_tx=XM_023246812.1;refGene_gn=EPS15L1;refGene_tx=XM_023246815.1;refGene_gn=EPS15L1;refGene_tx=XM_023246816.1;refGene_gn=EPS15L1;refGene_tx=XM_023246818.1
A2 12098584 12098635
refGene_gn=EPS15L1;refGene_tx=XM_023246774.1;refGene_gn=EPS15L1;refGene_tx=XM_023246779.1;refGene_gn=EPS15L1;refGene_tx=XM_023246782.1;refGene_gn=EPS15L1;refGene_tx=XM_023246786.1;refGene_gn=EPS15L1;refGene_tx=XM_023246793.1;refGene_gn=EPS15L1;refGene_tx=XM_023246796.1;refGene_gn=EPS15L1;refGene_tx=XM_023246805.1;refGene_gn=EPS15L1;refGene_tx=XM_023246812.1;refGene_gn=EPS15L1;refGene_tx=XM_023246815.1;refGene_gn=EPS15L1;refGene_tx=XM_023246816.1;refGene_gn=EPS15L1;refGene_tx=XM_023246818.1
A2 12099595 12099719
refGene_gn=EPS15L1;refGene_tx=XM_023246774.1;refGene_gn=EPS15L1;refGene_tx=XM_023246779.1;refGene_gn=EPS15L1;refGene_tx=XM_023246782.1;refGene_gn=EPS15L1;refGene_tx=XM_023246786.1;refGene_gn=EPS15L1;refGene_tx=XM_023246793.1;refGene_gn=EPS15L1;refGene_tx=XM_023246796.1;refGene_gn=EPS15L1;refGene_tx=XM_023246805.1;refGene_gn=EPS15L1;refGene_tx=XM_023246812.1;refGene_gn=EPS15L1;refGene_tx=XM_023246815.1;refGene_gn=EPS15L1;refGene_tx=XM_023246816.1;refGene_gn=EPS15L1;refGene_tx=XM_023246818.1
A2 12104607 12104772
refGene_gn=EPS15L1;refGene_tx=XM_023246774.1;refGene_gn=EPS15L1;refGene_tx=XM_023246779.1;refGene_gn=EPS15L1;refGene_tx=XM_023246782.1;refGene_gn=EPS15L1;refGene_tx=XM_023246786.1;refGene_gn=EPS15L1;refGene_tx=XM_023246793.1;refGene_gn=EPS15L1;refGene_tx=XM_023246796.1;refGene_gn=EPS15L1;refGene_tx=XM_023246805.1;refGene_gn=EPS15L1;refGene_tx=XM_023246812.1;refGene_gn=EPS15L1;refGene_tx=XM_023246815.1;refGene_gn=EPS15L1;refGene_tx=XM_023246816.1;refGene_gn=EPS15L1;refGene_tx=XM_023246818.1
A2 12106544 12106742
refGene_gn=EPS15L1;refGene_tx=XM_023246774.1;refGene_gn=EPS15L1;refGene_tx=XM_023246779.1;refGene_gn=EPS15L1;refGene_tx=XM_023246782.1;refGene_gn=EPS15L1;refGene_tx=XM_023246786.1;refGene_gn=EPS15L1;refGene_tx=XM_023246793.1;refGene_gn=EPS15L1;refGene_tx=XM_023246796.1;refGene_gn=EPS15L1;refGene_tx=XM_023246805.1;refGene_gn=EPS15L1;refGene_tx=XM_023246812.1;refGene_gn=EPS15L1;refGene_tx=XM_023246815.1;refGene_gn=EPS15L1;refGene_tx=XM_023246816.1;refGene_gn=EPS15L1;refGene_tx=XM_023246818.1
A2 12107364 12107526
refGene_gn=EPS15L1;refGene_tx=XM_023246774.1;refGene_gn=EPS15L1;refGene_tx=XM_023246779.1;refGene_gn=EPS15L1;refGene_tx=XM_023246782.1;refGene_gn=EPS15L1;refGene_tx=XM_023246786.1;refGene_gn=EPS15L1;refGene_tx=XM_023246793.1;refGene_gn=EPS15L1;refGene_tx=XM_023246796.1;refGene_gn=EPS15L1;refGene_tx=XM_023246805.1;refGene_gn=EPS15L1;refGene_tx=XM_02

3246812.1;refGene_gn=EPS15L1;refGene_tx=XM_023246815.1;refGene_gn=EPS15L1;refGene_tx=XM_023246816.1;refGene_gn=EPS15L1;refGene_tx=XM_023246818.1
A2 12113261 12113334
refGene_gn=EPS15L1;refGene_tx=XM_023246774.1;refGene_gn=EPS15L1;refGene_tx=XM_023246779.1;refGene_gn=EPS15L1;refGene_tx=XM_023246782.1;refGene_gn=EPS15L1;refGene_tx=XM_023246786.1;refGene_gn=EPS15L1;refGene_tx=XM_023246793.1;refGene_gn=EPS15L1;refGene_tx=XM_023246796.1;refGene_gn=EPS15L1;refGene_tx=XM_023246805.1;refGene_gn=EPS15L1;refGene_tx=XM_023246812.1;refGene_gn=EPS15L1;refGene_tx=XM_023246815.1;refGene_gn=EPS15L1;refGene_tx=XM_023246816.1;refGene_gn=EPS15L1;refGene_tx=XM_023246818.1
A2 12114192 12114278
refGene_gn=EPS15L1;refGene_tx=XM_023246774.1;refGene_gn=EPS15L1;refGene_tx=XM_023246779.1;refGene_gn=EPS15L1;refGene_tx=XM_023246782.1;refGene_gn=EPS15L1;refGene_tx=XM_023246786.1;refGene_gn=EPS15L1;refGene_tx=XM_023246793.1;refGene_gn=EPS15L1;refGene_tx=XM_023246796.1;refGene_gn=EPS15L1;refGene_tx=XM_023246805.1;refGene_gn=EPS15L1;refGene_tx=XM_023246812.1;refGene_gn=EPS15L1;refGene_tx=XM_023246815.1;refGene_gn=EPS15L1;refGene_tx=XM_023246816.1;refGene_gn=EPS15L1;refGene_tx=XM_023246818.1
A2 12114940 12115097
refGene_gn=EPS15L1;refGene_tx=XM_023246774.1;refGene_gn=EPS15L1;refGene_tx=XM_023246779.1;refGene_gn=EPS15L1;refGene_tx=XM_023246782.1;refGene_gn=EPS15L1;refGene_tx=XM_023246786.1;refGene_gn=EPS15L1;refGene_tx=XM_023246793.1;refGene_gn=EPS15L1;refGene_tx=XM_023246796.1;refGene_gn=EPS15L1;refGene_tx=XM_023246805.1;refGene_gn=EPS15L1;refGene_tx=XM_023246812.1;refGene_gn=EPS15L1;refGene_tx=XM_023246815.1;refGene_gn=EPS15L1;refGene_tx=XM_023246816.1;refGene_gn=EPS15L1;refGene_tx=XM_023246818.1
A2 12119731 12119889
refGene_gn=EPS15L1;refGene_tx=XM_023246774.1;refGene_gn=EPS15L1;refGene_tx=XM_023246779.1;refGene_gn=EPS15L1;refGene_tx=XM_023246782.1;refGene_gn=EPS15L1;refGene_tx=XM_023246786.1;refGene_gn=EPS15L1;refGene_tx=XM_023246793.1;refGene_gn=EPS15L1;refGene_tx=XM_023246796.1;refGene_gn=EPS15L1;refGene_tx=XM_023246805.1;refGene_gn=EPS15L1;refGene_tx=XM_023246812.1;refGene_gn=EPS15L1;refGene_tx=XM_023246815.1;refGene_gn=EPS15L1;refGene_tx=XM_023246816.1;refGene_gn=EPS15L1;refGene_tx=XM_023246818.1
A2 12124700 12124934
refGene_gn=EPS15L1;refGene_tx=XM_023246774.1;refGene_gn=EPS15L1;refGene_tx=XM_023246779.1;refGene_gn=EPS15L1;refGene_tx=XM_023246782.1;refGene_gn=EPS15L1;refGene_tx=XM_023246786.1;refGene_gn=EPS15L1;refGene_tx=XM_023246793.1;refGene_gn=EPS15L1;refGene_tx=XM_023246796.1;refGene_gn=EPS15L1;refGene_tx=XM_023246805.1;refGene_gn=EPS15L1;refGene_tx=XM_023246812.1;refGene_gn=EPS15L1;refGene_tx=XM_023246815.1;refGene_gn=EPS15L1;refGene_tx=XM_023246816.1;refGene_gn=EPS15L1;refGene_tx=XM_023246818.1
A2 12129281 12129341
refGene_gn=EPS15L1;refGene_tx=XM_023246774.1;refGene_gn=EPS15L1;refGen

e_tx=XM_023246779.1;refGene_gn=EPS15L1;refGene_tx=XM_023246782.1;refGene_gn=EPS15L1;refGene_tx=XM_023246786.1;refGene_gn=EPS15L1;refGene_tx=XM_023246793.1;refGene_gn=EPS15L1;refGene_tx=XM_023246796.1;refGene_gn=EPS15L1;refGene_tx=XM_023246805.1;refGene_gn=EPS15L1;refGene_tx=XM_023246812.1;refGene_gn=EPS15L1;refGene_tx=XM_023246815.1;refGene_gn=EPS15L1;refGene_tx=XM_023246816.1;refGene_gn=EPS15L1;refGene_tx=XM_023246818.1
A2 12133505 12133631
refGene_gn=EPS15L1;refGene_tx=XM_023246774.1;refGene_gn=EPS15L1;refGene_tx=XM_023246779.1;refGene_gn=EPS15L1;refGene_tx=XM_023246782.1;refGene_gn=EPS15L1;refGene_tx=XM_023246786.1;refGene_gn=EPS15L1;refGene_tx=XM_023246793.1;refGene_gn=EPS15L1;refGene_tx=XM_023246796.1;refGene_gn=EPS15L1;refGene_tx=XM_023246805.1;refGene_gn=EPS15L1;refGene_tx=XM_023246812.1;refGene_gn=EPS15L1;refGene_tx=XM_023246815.1;refGene_gn=EPS15L1;refGene_tx=XM_023246816.1;refGene_gn=EPS15L1;refGene_tx=XM_023246818.1
A2 12136465 12136528
refGene_gn=EPS15L1;refGene_tx=XM_023246774.1;refGene_gn=EPS15L1;refGene_tx=XM_023246779.1;refGene_gn=EPS15L1;refGene_tx=XM_023246782.1;refGene_gn=EPS15L1;refGene_tx=XM_023246786.1;refGene_gn=EPS15L1;refGene_tx=XM_023246793.1;refGene_gn=EPS15L1;refGene_tx=XM_023246796.1;refGene_gn=EPS15L1;refGene_tx=XM_023246805.1;refGene_gn=EPS15L1;refGene_tx=XM_023246812.1;refGene_gn=EPS15L1;refGene_tx=XM_023246815.1;refGene_gn=EPS15L1;refGene_tx=XM_023246816.1;refGene_gn=EPS15L1;refGene_tx=XM_023246818.1
A2 12137169 12137265
refGene_gn=EPS15L1;refGene_tx=XM_023246774.1;refGene_gn=EPS15L1;refGene_tx=XM_023246779.1;refGene_gn=EPS15L1;refGene_tx=XM_023246782.1;refGene_gn=EPS15L1;refGene_tx=XM_023246786.1;refGene_gn=EPS15L1;refGene_tx=XM_023246793.1;refGene_gn=EPS15L1;refGene_tx=XM_023246796.1;refGene_gn=EPS15L1;refGene_tx=XM_023246805.1;refGene_gn=EPS15L1;refGene_tx=XM_023246812.1;refGene_gn=EPS15L1;refGene_tx=XM_023246815.1;refGene_gn=EPS15L1;refGene_tx=XM_023246816.1;refGene_gn=EPS15L1;refGene_tx=XM_023246818.1
A2 12139767 12139815
refGene_gn=EPS15L1;refGene_tx=XM_023246774.1;refGene_gn=EPS15L1;refGene_tx=XM_023246779.1;refGene_gn=EPS15L1;refGene_tx=XM_023246782.1;refGene_gn=EPS15L1;refGene_tx=XM_023246786.1;refGene_gn=EPS15L1;refGene_tx=XM_023246793.1;refGene_gn=EPS15L1;refGene_tx=XM_023246805.1;refGene_gn=EPS15L1;refGene_tx=XM_023246812.1;refGene_gn=EPS15L1;refGene_tx=XM_023246815.1;refGene_gn=EPS15L1;refGene_tx=XM_023246816.1;refGene_gn=EPS15L1;refGene_tx=XM_023246818.1
A2 12140282 12140372
refGene_gn=EPS15L1;refGene_tx=XM_023246774.1;refGene_gn=EPS15L1;refGene_tx=XM_023246779.1;refGene_gn=EPS15L1;refGene_tx=XM_023246782.1;refGene_gn=EPS15L1;refGene_tx=XM_023246786.1;refGene_gn=EPS15L1;refGene_tx=XM_023246793.1;refGene_gn=EPS15L1;refGene_tx=XM_023246805.1;refGene_gn=EPS15L1;refGene_tx=XM_023246812.1;refGene_gn=EPS15L1;refGene_tx=XM_023246815.1;refGene_gn=EPS15L1;refGene_tx=XM_023246816.1;refGene_gn=EPS15L1;refGene_tx=XM_023246818.1

A2	12140571	12140613
refGene_gn=EPS15L1;refGene_tx=XM_023246774.1;refGene_gn=EPS15L1;refGene_tx=XM_023246782.1;refGene_gn=EPS15L1;refGene_tx=XM_023246786.1;refGene_gn=EPS15L1;refGene_tx=XM_023246793.1;refGene_gn=EPS15L1;refGene_tx=XM_023246805.1;refGene_gn=EPS15L1;refGene_tx=XM_023246812.1;refGene_gn=EPS15L1;refGene_tx=XM_023246815.1;refGene_gn=EPS15L1;refGene_tx=XM_023246816.1;refGene_gn=EPS15L1;refGene_tx=XM_023246818.1
A2	12165749	12165782
refGene_gn=EPS15L1;refGene_tx=XM_023246774.1;refGene_gn=EPS15L1;refGene_tx=XM_023246779.1;refGene_gn=EPS15L1;refGene_tx=XM_023246782.1;refGene_gn=EPS15L1;refGene_tx=XM_023246786.1;refGene_gn=EPS15L1;refGene_tx=XM_023246793.1;refGene_gn=EPS15L1;refGene_tx=XM_023246805.1;refGene_gn=EPS15L1;refGene_tx=XM_023246812.1;refGene_gn=EPS15L1;refGene_tx=XM_023246815.1;refGene_gn=EPS15L1;refGene_tx=XM_023246816.1;refGene_gn=EPS15L1;refGene_tx=XM_023246818.1
A2	12170647	12170791	refGene_gn=CALR3;refGene_tx=XM_011290407.3
A2	12175041	12175134	refGene_gn=CALR3;refGene_tx=XM_011290407.3
A2	12176156	12176288	refGene_gn=CALR3;refGene_tx=XM_011290407.3
A2	12176383	12176491	refGene_gn=CALR3;refGene_tx=XM_011290407.3
A2	12180588	12180777	refGene_gn=CALR3;refGene_tx=XM_011290407.3
A2	12183773	12183868	refGene_gn=CALR3;refGene_tx=XM_011290407.3
A2	12190410	12190614	refGene_gn=CALR3;refGene_tx=XM_011290407.3
A2	12193564	12193666	refGene_gn=CALR3;refGene_tx=XM_011290407.3
A2	12193937	12194028	refGene_gn=CALR3;refGene_tx=XM_011290407.3
A2	12199549	12200311
refGene_gn=CA2H19orf44;refGene_tx=XM_019819011.2;refGene_gn=CA2H19orf44;refGene_tx=XM_019819016.2;refGene_gn=CA2H19orf44;refGene_tx=XM_019819024.2;refGene_gn=CA2H19orf44;refGene_tx=XM_019819031.2;refGene_gn=CA2H19orf44;refGene_tx=XM_019819036.2;refGene_gn=CA2H19orf44;refGene_tx=XM_019819038.2;refGene_gn=CA2H19orf44;refGene_tx=XM_019819047.2;refGene_gn=CA2H19orf44;refGene_tx=XM_019819056.2;refGene_gn=CA2H19orf44;refGene_tx=XM_019819066.2;refGene_gn=CA2H19orf44;refGene_tx=XM_019819070.2;refGene_gn=CA2H19orf44;refGene_tx=XM_019819078.2
A2	12202384	12202697
refGene_gn=CA2H19orf44;refGene_tx=XM_019819011.2;refGene_gn=CA2H19orf44;refGene_tx=XM_019819016.2;refGene_gn=CA2H19orf44;refGene_tx=XM_019819024.2;refGene_gn=CA2H19orf44;refGene_tx=XM_019819031.2;refGene_gn=CA2H19orf44;refGene_tx=XM_019819036.2;refGene_gn=CA2H19orf44;refGene_tx=XM_019819038.2;refGene_gn=CA2H19orf44;refGene_tx=XM_019819047.2;refGene_gn=CA2H19orf44;refGene_tx=XM_019819056.2;refGene_gn=CA2H19orf44;refGene_tx=XM_019819066.2;refGene_gn=CA2H19orf44;refGene_tx=XM_019819070.2;refGene_gn=CA2H19orf44;refGene_tx=XM_019819078.2
A2	12209588	12209662
refGene_gn=CA2H19orf44;refGene_tx=XM_019819011.2;refGene_gn=CA2H19orf44;refGene_tx=XM_019819016.2;refGene_gn=CA2H19orf44;refGene_tx=XM_019819024.2;refGene_gn=CA2H19orf44;refGene_tx=XM_019819031.2;refGene_gn=CA2H19orf44;refGene_tx=XM_019819036.2;refGene_gn=CA2H19orf44;refGene_tx=XM_019819038.2;refGene_gn=CA2H19orf44;refGene_tx=XM_019819047.2;refGene_gn=CA2H19orf44;refGene_tx=XM_019819056.2;refGene_gn=CA2H19orf44;refGene_tx=XM_019819066.2;refGene_gn=CA2H19orf44;refGene_tx=XM_019819070.2;

refGene_gn=CA2H19orf44;refGene_tx=XM_019819078.2
A2 12210598 12211013
refGene_gn=CA2H19orf44;refGene_tx=XM_019819011.2;refGene_gn=CA2H19orf44;refGene_tx=XM_019819016.2;refGene_gn=CA2H19orf44;refGene_tx=XM_019819024.2;refGene_gn=CA2H19orf44;refGene_tx=XM_019819031.2;refGene_gn=CA2H19orf44;refGene_tx=XM_019819036.2;refGene_gn=CA2H19orf44;refGene_tx=XM_019819038.2;refGene_gn=CA2H19orf44;refGene_tx=XM_019819047.2;refGene_gn=CA2H19orf44;refGene_tx=XM_019819056.2;refGene_gn=CA2H19orf44;refGene_tx=XM_019819066.2;refGene_gn=CA2H19orf44;refGene_tx=XM_019819070.2;refGene_gn=CA2H19orf44;refGene_tx=XM_019819078.2
A2 12212371 12212467
refGene_gn=CA2H19orf44;refGene_tx=XM_019819011.2;refGene_gn=CA2H19orf44;refGene_tx=XM_019819016.2;refGene_gn=CA2H19orf44;refGene_tx=XM_019819024.2;refGene_gn=CA2H19orf44;refGene_tx=XM_019819031.2;refGene_gn=CA2H19orf44;refGene_tx=XM_019819036.2;refGene_gn=CA2H19orf44;refGene_tx=XM_019819038.2;refGene_gn=CA2H19orf44;refGene_tx=XM_019819047.2;refGene_gn=CA2H19orf44;refGene_tx=XM_019819056.2;refGene_gn=CA2H19orf44;refGene_tx=XM_019819066.2;refGene_gn=CA2H19orf44;refGene_tx=XM_019819070.2;refGene_gn=CA2H19orf44;refGene_tx=XM_019819078.2
A2 12214290 12214796
refGene_gn=CA2H19orf44;refGene_tx=XM_019819011.2;refGene_gn=CA2H19orf44;refGene_tx=XM_019819016.2;refGene_gn=CA2H19orf44;refGene_tx=XM_019819024.2;refGene_gn=CA2H19orf44;refGene_tx=XM_019819031.2;refGene_gn=CA2H19orf44;refGene_tx=XM_019819036.2;refGene_gn=CA2H19orf44;refGene_tx=XM_019819038.2;refGene_gn=CA2H19orf44;refGene_tx=XM_019819047.2;refGene_gn=CA2H19orf44;refGene_tx=XM_019819056.2;refGene_gn=CA2H19orf44;refGene_tx=XM_019819066.2;refGene_gn=CA2H19orf44;refGene_tx=XM_019819070.2;refGene_gn=CA2H19orf44;refGene_tx=XM_019819078.2
A2 12215494 12215590
refGene_gn=CA2H19orf44;refGene_tx=XM_019819078.2
A2 12215851 12216072
refGene_gn=CA2H19orf44;refGene_tx=XM_019819011.2;refGene_gn=CA2H19orf44;refGene_tx=XM_019819016.2;refGene_gn=CA2H19orf44;refGene_tx=XM_019819024.2;refGene_gn=CA2H19orf44;refGene_tx=XM_019819031.2;refGene_gn=CA2H19orf44;refGene_tx=XM_019819036.2;refGene_gn=CA2H19orf44;refGene_tx=XM_019819038.2;refGene_gn=CA2H19orf44;refGene_tx=XM_019819047.2;refGene_gn=CA2H19orf44;refGene_tx=XM_019819056.2;refGene_gn=CA2H19orf44;refGene_tx=XM_019819070.2
A2 12216215 12216455
refGene_gn=CA2H19orf44;refGene_tx=XM_019819011.2;refGene_gn=CA2H19orf44;refGene_tx=XM_019819016.2;refGene_gn=CA2H19orf44;refGene_tx=XM_019819024.2;refGene_gn=CA2H19orf44;refGene_tx=XM_019819031.2;refGene_gn=CA2H19orf44;refGene_tx=XM_019819036.2;refGene_gn=CA2H19orf44;refGene_tx=XM_019819038.2;refGene_gn=CA2H19orf44;refGene_tx=XM_019819047.2;refGene_gn=CA2H19orf44;refGene_tx=XM_019819056.2;refGene_gn=CA2H19orf44;refGene_tx=XM_019819070.2
A2 12217886 12218080
refGene_gn=CHERP;refGene_tx=XM_011290401.3;refGene_gn=CHERP;refGene_tx=XM_011290402.3
A2 12218349 12218444

refGene_gn=CHERP;refGene_tx=XM_011290401.3;refGene_gn=CHERP;refGene_tx=XM_011290402.3
A2 12218927 12219044
refGene_gn=CHERP;refGene_tx=XM_011290401.3;refGene_gn=CHERP;refGene_tx=XM_011290402.3
A2 12219134 12219278
refGene_gn=CHERP;refGene_tx=XM_011290401.3;refGene_gn=CHERP;refGene_tx=XM_011290402.3
A2 12219555 12219642
refGene_gn=CHERP;refGene_tx=XM_011290401.3;refGene_gn=CHERP;refGene_tx=XM_011290402.3
A2 12220242 12220376
refGene_gn=CHERP;refGene_tx=XM_011290401.3;refGene_gn=CHERP;refGene_tx=XM_011290402.3
A2 12221376 12221615
refGene_gn=CHERP;refGene_tx=XM_011290401.3;refGene_gn=CHERP;refGene_tx=XM_011290402.3
A2 12222828 12223264
refGene_gn=CHERP;refGene_tx=XM_011290401.3;refGene_gn=CHERP;refGene_tx=XM_011290402.3
A2 12224984 12225160
refGene_gn=CHERP;refGene_tx=XM_011290401.3;refGene_gn=CHERP;refGene_tx=XM_011290402.3
A2 12225946 12226205
refGene_gn=CHERP;refGene_tx=XM_011290401.3;refGene_gn=CHERP;refGene_tx=XM_011290402.3
A2 12227777 12227867
refGene_gn=CHERP;refGene_tx=XM_011290401.3;refGene_gn=CHERP;refGene_tx=XM_011290402.3
A2 12227981 12228126
refGene_gn=CHERP;refGene_tx=XM_011290401.3;refGene_gn=CHERP;refGene_tx=XM_011290402.3
A2 12229757 12229909
refGene_gn=CHERP;refGene_tx=XM_011290401.3;refGene_gn=CHERP;refGene_tx=XM_011290402.3
A2 12230226 12230364
refGene_gn=CHERP;refGene_tx=XM_011290401.3;refGene_gn=CHERP;refGene_tx=XM_011290402.3
A2 12231498 12231683
refGene_gn=CHERP;refGene_tx=XM_011290401.3;refGene_gn=CHERP;refGene_tx=XM_011290402.3
A2 12234893 12235067
refGene_gn=CHERP;refGene_tx=XM_011290401.3;refGene_gn=CHERP;refGene_tx=XM_011290402.3
A2 12235376 12235401
refGene_gn=CHERP;refGene_tx=XM_011290401.3;refGene_gn=CHERP;refGene_tx=XM_011290402.3
A2 12244297 12244528 refGene_gn=SLC35E1;refGene_tx=XM_023246863.1
A2 12245021 12245267 refGene_gn=SLC35E1;refGene_tx=XM_023246863.1
A2 12257139 12257265 refGene_gn=SLC35E1;refGene_tx=XM_023246863.1

A2 12258655 12258793 refGene_gn=SLC35E1;refGene_tx=XM_023246863.1
A2 12260366 12260437 refGene_gn=SLC35E1;refGene_tx=XM_023246863.1
A2 12260840 12261264 refGene_gn=SLC35E1;refGene_tx=XM_023246863.1
A2 12264933 12266586
refGene_gn=MED26;refGene_tx=XM_023246873.1;refGene_gn=MED26;refGene_tx
=XM_023246877.1;refGene_gn=MED26;refGene_tx=XM_023246880.1
A2 12267220 12267295
refGene_gn=MED26;refGene_tx=XM_023246873.1;refGene_gn=MED26;refGene_tx
=XM_023246877.1;refGene_gn=MED26;refGene_tx=XM_023246880.1
A2 12316328 12316400 refGene_gn=MED26;refGene_tx=XM_023246873.1
A2 12334869 12334885 refGene_gn=SMIM7;refGene_tx=XM_003982029.5
A2 12340085 12340176 refGene_gn=SMIM7;refGene_tx=XM_003982029.5
A2 12344813 12344866 refGene_gn=SMIM7;refGene_tx=XM_003982029.5
A2 12345346 12345388 refGene_gn=SMIM7;refGene_tx=XM_003982029.5
A2 12345470 12345496 refGene_gn=SMIM7;refGene_tx=XM_003982029.5
A2 12346638 12346762 refGene_gn=TMEM38A;refGene_tx=XM_003982055.4
A2 12359472 12359629 refGene_gn=TMEM38A;refGene_tx=XM_003982055.4
A2 12359892 12360077 refGene_gn=TMEM38A;refGene_tx=XM_003982055.4
A2 12362325 12362413 refGene_gn=TMEM38A;refGene_tx=XM_003982055.4
A2 12363983 12364101 refGene_gn=TMEM38A;refGene_tx=XM_003982055.4
A2 12364922 12365162 refGene_gn=TMEM38A;refGene_tx=XM_003982055.4
A2 12385472 12385570 refGene_gn=NWD1;refGene_tx=XM_019819120.2
A2 12385708 12385743 refGene_gn=NWD1;refGene_tx=XM_019819120.2
A2 12389952 12390038 refGene_gn=NWD1;refGene_tx=XM_019819120.2
A2 12391412 12391529 refGene_gn=NWD1;refGene_tx=XM_019819120.2
A2 12396449 12396747 refGene_gn=NWD1;refGene_tx=XM_019819120.2
A2 12402574 12403847 refGene_gn=NWD1;refGene_tx=XM_019819120.2
A2 12412292 12412496 refGene_gn=NWD1;refGene_tx=XM_019819120.2
A2 12413876 12414036 refGene_gn=NWD1;refGene_tx=XM_019819120.2
A2 12414662 12414780 refGene_gn=NWD1;refGene_tx=XM_019819120.2
A2 12415452 12415611 refGene_gn=NWD1;refGene_tx=XM_019819120.2
A2 12419491 12419689 refGene_gn=NWD1;refGene_tx=XM_019819120.2
A2 12422782 12422905 refGene_gn=NWD1;refGene_tx=XM_019819120.2
A2 12430069 12430278 refGene_gn=NWD1;refGene_tx=XM_019819120.2
A2 12433645 12433918 refGene_gn=NWD1;refGene_tx=XM_019819120.2
A2 12435679 12435770 refGene_gn=NWD1;refGene_tx=XM_019819120.2
A2 12441570 12441725 refGene_gn=NWD1;refGene_tx=XM_019819120.2
A2 12443262 12443539 refGene_gn=NWD1;refGene_tx=XM_019819120.2
A2 12445897 12446448 refGene_gn=NWD1;refGene_tx=XM_019819120.2
A2 12449672 12449778 refGene_gn=NWD1;refGene_tx=XM_019819120.2
A2 12451325 12451399 refGene_gn=NWD1;refGene_tx=XM_019819120.2
A2 12451400 12451628 refGene_gn=NWD1;refGene_tx=XM_019819120.2
A2 12464619 12464733 refGene_gn=SIN3B;refGene_tx=XM_023246897.1
A2 12464978 12465085 refGene_gn=SIN3B;refGene_tx=XM_023246897.1
A2 12466363 12466517 refGene_gn=SIN3B;refGene_tx=XM_023246897.1
A2 12473925 12474126 refGene_gn=SIN3B;refGene_tx=XM_023246897.1
A2 12476399 12476543 refGene_gn=SIN3B;refGene_tx=XM_023246897.1
A2 12482312 12482435 refGene_gn=SIN3B;refGene_tx=XM_023246897.1
A2 12483776 12483845 refGene_gn=SIN3B;refGene_tx=XM_023246897.1
A2 12483846 12483867 refGene_gn=SIN3B;refGene_tx=XM_023246897.1

A2	12484336	12484455	refGene_gn=SIN3B;refGene_tx=XM_023246897.1
A2	12488718	12488926	refGene_gn=SIN3B;refGene_tx=XM_023246897.1
A2	12489719	12489836	refGene_gn=SIN3B;refGene_tx=XM_023246897.1
A2	12490998	12491237	refGene_gn=SIN3B;refGene_tx=XM_023246897.1
A2	12491983	12492167	refGene_gn=SIN3B;refGene_tx=XM_023246897.1
A2	12494462	12495066	refGene_gn=SIN3B;refGene_tx=XM_023246897.1
A2	12495270	12495440	refGene_gn=SIN3B;refGene_tx=XM_023246897.1
A2	12498232	12498406	refGene_gn=SIN3B;refGene_tx=XM_023246897.1
A2	12498650	12498743	refGene_gn=SIN3B;refGene_tx=XM_023246897.1
A2	12499740	12499835	refGene_gn=SIN3B;refGene_tx=XM_023246897.1
A2	12500068	12500273	refGene_gn=SIN3B;refGene_tx=XM_023246897.1
A2	12500385	12500616	refGene_gn=SIN3B;refGene_tx=XM_023246897.1
A2	12506385	12506644
refGene_gn=F2RL3;refGene_tx=XM_023246905.1;refGene_gn=F2RL3;refGene_tx
=XM_023246910.1
A2	12506922	12507992
refGene_gn=F2RL3;refGene_tx=XM_023246905.1;refGene_gn=F2RL3;refGene_tx
=XM_023246910.1
A2	12508363	12508489	refGene_gn=F2RL3;refGene_tx=XM_023246905.1
A2	12510648	12510670	refGene_gn=CPAMD8;refGene_tx=XM_023246922.1
A2	12513272	12513396	refGene_gn=CPAMD8;refGene_tx=XM_023246922.1
A2	12514000	12514210	refGene_gn=CPAMD8;refGene_tx=XM_023246922.1
A2	12515560	12515671	refGene_gn=CPAMD8;refGene_tx=XM_023246922.1
A2	12515758	12515864	refGene_gn=CPAMD8;refGene_tx=XM_023246922.1
A2	12516420	12516495	refGene_gn=CPAMD8;refGene_tx=XM_023246922.1
A2	12517362	12517450	refGene_gn=CPAMD8;refGene_tx=XM_023246922.1
A2	12519392	12519577	refGene_gn=CPAMD8;refGene_tx=XM_023246922.1
A2	12520347	12520410	refGene_gn=CPAMD8;refGene_tx=XM_023246922.1
A2	12520521	12520677	refGene_gn=CPAMD8;refGene_tx=XM_023246922.1
A2	12521111	12521248	refGene_gn=CPAMD8;refGene_tx=XM_023246922.1
A2	12521325	12521412	refGene_gn=CPAMD8;refGene_tx=XM_023246922.1
A2	12522402	12522568	refGene_gn=CPAMD8;refGene_tx=XM_023246922.1
A2	12525496	12525571	refGene_gn=CPAMD8;refGene_tx=XM_023246922.1
A2	12525705	12525862	refGene_gn=CPAMD8;refGene_tx=XM_023246922.1
A2	12528669	12528751	refGene_gn=CPAMD8;refGene_tx=XM_023246922.1
A2	12533057	12533234	refGene_gn=CPAMD8;refGene_tx=XM_023246922.1
A2	12534781	12535007	refGene_gn=CPAMD8;refGene_tx=XM_023246922.1
A2	12535636	12535935	refGene_gn=CPAMD8;refGene_tx=XM_023246922.1
A2	12541307	12541359	refGene_gn=CPAMD8;refGene_tx=XM_023246922.1
A2	12543204	12543335	refGene_gn=CPAMD8;refGene_tx=XM_023246922.1
A2	12543580	12543734	refGene_gn=CPAMD8;refGene_tx=XM_023246922.1
A2	12549683	12549915	refGene_gn=CPAMD8;refGene_tx=XM_023246922.1
A2	12556254	12556317	refGene_gn=CPAMD8;refGene_tx=XM_023246922.1
A2	12559194	12559337	refGene_gn=CPAMD8;refGene_tx=XM_023246922.1
A2	12560781	12560940	refGene_gn=CPAMD8;refGene_tx=XM_023246922.1
A2	12561296	12561446	refGene_gn=CPAMD8;refGene_tx=XM_023246922.1
A2	12562007	12562180	refGene_gn=CPAMD8;refGene_tx=XM_023246922.1
A2	12564722	12564906	refGene_gn=CPAMD8;refGene_tx=XM_023246922.1
A2	12571556	12571685	refGene_gn=CPAMD8;refGene_tx=XM_023246922.1
A2	12575185	12575356	refGene_gn=CPAMD8;refGene_tx=XM_023246922.1

A2 12578256 12578484 refGene_gn=CPAMD8;refGene_tx=XM_023246922.1
A2 12580826 12580935 refGene_gn=CPAMD8;refGene_tx=XM_023246922.1
A2 12582250 12582335 refGene_gn=CPAMD8;refGene_tx=XM_023246922.1
A2 12583607 12583721 refGene_gn=CPAMD8;refGene_tx=XM_023246922.1
A2 12585946 12586001 refGene_gn=CPAMD8;refGene_tx=XM_023246922.1
A2 12586412 12586430 refGene_gn=CPAMD8;refGene_tx=XM_023246922.1
A2 12586777 12586830 refGene_gn=CPAMD8;refGene_tx=XM_023246922.1
A2 12586907 12587073 refGene_gn=CPAMD8;refGene_tx=XM_023246922.1
A2 12588802 12588825 refGene_gn=CPAMD8;refGene_tx=XM_023246922.1
A2 12590467 12590619 refGene_gn=CPAMD8;refGene_tx=XM_023246922.1
A2 12594348 12594578 refGene_gn=CPAMD8;refGene_tx=XM_023246922.1
A2 12631128 12631327
refGene_gn=HAUS8;refGene_tx=XM_006928568.4;refGene_gn=HAUS8;refGene_tx=XM_019819160.2;refGene_gn=HAUS8;refGene_tx=XM_019819165.2;refGene_gn=HAUS8;refGene_tx=XM_019819169.2
A2 12632280 12632422
refGene_gn=HAUS8;refGene_tx=XM_006928568.4;refGene_gn=HAUS8;refGene_tx=XM_019819160.2;refGene_gn=HAUS8;refGene_tx=XM_019819165.2;refGene_gn=HAUS8;refGene_tx=XM_019819169.2
A2 12634275 12634417
refGene_gn=HAUS8;refGene_tx=XM_006928568.4;refGene_gn=HAUS8;refGene_tx=XM_019819160.2;refGene_gn=HAUS8;refGene_tx=XM_019819165.2;refGene_gn=HAUS8;refGene_tx=XM_019819169.2
A2 12635566 12635725
refGene_gn=HAUS8;refGene_tx=XM_006928568.4;refGene_gn=HAUS8;refGene_tx=XM_019819160.2;refGene_gn=HAUS8;refGene_tx=XM_019819165.2;refGene_gn=HAUS8;refGene_tx=XM_019819169.2
A2 12635829 12635895
refGene_gn=HAUS8;refGene_tx=XM_006928568.4;refGene_gn=HAUS8;refGene_tx=XM_019819160.2;refGene_gn=HAUS8;refGene_tx=XM_019819165.2;refGene_gn=HAUS8;refGene_tx=XM_019819169.2
A2 12636508 12636603
refGene_gn=HAUS8;refGene_tx=XM_006928568.4;refGene_gn=HAUS8;refGene_tx=XM_019819160.2;refGene_gn=HAUS8;refGene_tx=XM_019819165.2;refGene_gn=HAUS8;refGene_tx=XM_019819169.2
A2 12636705 12636798
refGene_gn=HAUS8;refGene_tx=XM_006928568.4;refGene_gn=HAUS8;refGene_tx=XM_019819160.2;refGene_gn=HAUS8;refGene_tx=XM_019819165.2;refGene_gn=HAUS8;refGene_tx=XM_019819169.2
A2 12639354 12639436
refGene_gn=HAUS8;refGene_tx=XM_006928568.4;refGene_gn=HAUS8;refGene_tx=XM_019819160.2;refGene_gn=HAUS8;refGene_tx=XM_019819165.2;refGene_gn=HAUS8;refGene_tx=XM_019819169.2
A2 12643642 12643698
refGene_gn=HAUS8;refGene_tx=XM_006928568.4;refGene_gn=HAUS8;refGene_tx=XM_019819160.2;refGene_gn=HAUS8;refGene_tx=XM_019819165.2
A2 12646558 12646623
refGene_gn=HAUS8;refGene_tx=XM_006928568.4;refGene_gn=HAUS8;refGene_tx=XM_019819160.2;refGene_gn=HAUS8;refGene_tx=XM_019819165.2
A2 12649107 12649136 refGene_gn=HAUS8;refGene_tx=XM_006928568.4

A2 12650767 12650868 refGene_gn=HAUS8;refGene_tx=XM_019819160.2
A2 12669260 12670100
refGene_gn=MYO9B;refGene_tx=XM_006928569.3;refGene_gn=MYO9B;refGene_tx=XM_023246941.1;refGene_gn=MYO9B;refGene_tx=XM_023246946.1;refGene_gn=MYO9B;refGene_tx=XM_023246950.1;refGene_gn=MYO9B;refGene_tx=XM_023246954.1;refGene_gn=MYO9B;refGene_tx=XM_023246957.1;refGene_gn=MYO9B;refGene_tx=XM_023246958.1;refGene_gn=MYO9B;refGene_tx=XM_023246959.1;refGene_gn=MYO9B;refGene_tx=XM_023246962.1
A2 12696329 12696424
refGene_gn=MYO9B;refGene_tx=XM_006928569.3;refGene_gn=MYO9B;refGene_tx=XM_023246941.1;refGene_gn=MYO9B;refGene_tx=XM_023246946.1;refGene_gn=MYO9B;refGene_tx=XM_023246950.1;refGene_gn=MYO9B;refGene_tx=XM_023246954.1;refGene_gn=MYO9B;refGene_tx=XM_023246957.1;refGene_gn=MYO9B;refGene_tx=XM_023246958.1;refGene_gn=MYO9B;refGene_tx=XM_023246959.1;refGene_gn=MYO9B;refGene_tx=XM_023246962.1
A2 12702519 12702582
refGene_gn=MYO9B;refGene_tx=XM_006928569.3;refGene_gn=MYO9B;refGene_tx=XM_023246941.1;refGene_gn=MYO9B;refGene_tx=XM_023246946.1;refGene_gn=MYO9B;refGene_tx=XM_023246950.1;refGene_gn=MYO9B;refGene_tx=XM_023246954.1;refGene_gn=MYO9B;refGene_tx=XM_023246957.1;refGene_gn=MYO9B;refGene_tx=XM_023246958.1;refGene_gn=MYO9B;refGene_tx=XM_023246959.1;refGene_gn=MYO9B;refGene_tx=XM_023246962.1
A2 12702995 12703095
refGene_gn=MYO9B;refGene_tx=XM_006928569.3;refGene_gn=MYO9B;refGene_tx=XM_023246941.1;refGene_gn=MYO9B;refGene_tx=XM_023246946.1;refGene_gn=MYO9B;refGene_tx=XM_023246950.1;refGene_gn=MYO9B;refGene_tx=XM_023246954.1;refGene_gn=MYO9B;refGene_tx=XM_023246957.1;refGene_gn=MYO9B;refGene_tx=XM_023246958.1;refGene_gn=MYO9B;refGene_tx=XM_023246959.1;refGene_gn=MYO9B;refGene_tx=XM_023246962.1
A2 12703359 12703460
refGene_gn=MYO9B;refGene_tx=XM_006928569.3;refGene_gn=MYO9B;refGene_tx=XM_023246941.1;refGene_gn=MYO9B;refGene_tx=XM_023246946.1;refGene_gn=MYO9B;refGene_tx=XM_023246950.1;refGene_gn=MYO9B;refGene_tx=XM_023246954.1;refGene_gn=MYO9B;refGene_tx=XM_023246957.1;refGene_gn=MYO9B;refGene_tx=XM_023246958.1;refGene_gn=MYO9B;refGene_tx=XM_023246959.1;refGene_gn=MYO9B;refGene_tx=XM_023246962.1
A2 12705091 12705221
refGene_gn=MYO9B;refGene_tx=XM_006928569.3;refGene_gn=MYO9B;refGene_tx=XM_023246941.1;refGene_gn=MYO9B;refGene_tx=XM_023246946.1;refGene_gn=MYO9B;refGene_tx=XM_023246950.1;refGene_gn=MYO9B;refGene_tx=XM_023246954.1;refGene_gn=MYO9B;refGene_tx=XM_023246957.1;refGene_gn=MYO9B;refGene_tx=XM_023246958.1;refGene_gn=MYO9B;refGene_tx=XM_023246959.1;refGene_gn=MYO9B;refGene_tx=XM_023246962.1
A2 12706083 12706173
refGene_gn=MYO9B;refGene_tx=XM_006928569.3;refGene_gn=MYO9B;refGene_tx=XM_023246941.1;refGene_gn=MYO9B;refGene_tx=XM_023246946.1;refGene_gn=MYO9B;refGene_tx=XM_023246950.1;refGene_gn=MYO9B;refGene_tx=XM_023246954.1;refGene_gn=MYO9B;refGene_tx=XM_023246957.1;refGene_gn=MYO9B;refGene_tx=XM_023246958.1;refGene_gn=MYO9B;refGene_tx=XM_023246959.1;refGene_gn=MYO9B;refGene_tx=XM_023246962.1

A2 12708219 12708336
refGene_gn=MYO9B;refGene_tx=XM_006928569.3;refGene_gn=MYO9B;refGene_tx=XM_023246941.1;refGene_gn=MYO9B;refGene_tx=XM_023246946.1;refGene_gn=MYO9B;refGene_tx=XM_023246950.1;refGene_gn=MYO9B;refGene_tx=XM_023246954.1;refGene_gn=MYO9B;refGene_tx=XM_023246957.1;refGene_gn=MYO9B;refGene_tx=XM_023246958.1;refGene_gn=MYO9B;refGene_tx=XM_023246959.1;refGene_gn=MYO9B;refGene_tx=XM_023246962.1
A2 12708677 12708812
refGene_gn=MYO9B;refGene_tx=XM_006928569.3;refGene_gn=MYO9B;refGene_tx=XM_023246941.1;refGene_gn=MYO9B;refGene_tx=XM_023246946.1;refGene_gn=MYO9B;refGene_tx=XM_023246950.1;refGene_gn=MYO9B;refGene_tx=XM_023246954.1;refGene_gn=MYO9B;refGene_tx=XM_023246957.1;refGene_gn=MYO9B;refGene_tx=XM_023246958.1;refGene_gn=MYO9B;refGene_tx=XM_023246959.1;refGene_gn=MYO9B;refGene_tx=XM_023246962.1
A2 12710221 12710343
refGene_gn=MYO9B;refGene_tx=XM_006928569.3;refGene_gn=MYO9B;refGene_tx=XM_023246941.1;refGene_gn=MYO9B;refGene_tx=XM_023246946.1;refGene_gn=MYO9B;refGene_tx=XM_023246950.1;refGene_gn=MYO9B;refGene_tx=XM_023246954.1;refGene_gn=MYO9B;refGene_tx=XM_023246957.1;refGene_gn=MYO9B;refGene_tx=XM_023246958.1;refGene_gn=MYO9B;refGene_tx=XM_023246959.1;refGene_gn=MYO9B;refGene_tx=XM_023246962.1
A2 12712740 12712882
refGene_gn=MYO9B;refGene_tx=XM_006928569.3;refGene_gn=MYO9B;refGene_tx=XM_023246941.1;refGene_gn=MYO9B;refGene_tx=XM_023246946.1;refGene_gn=MYO9B;refGene_tx=XM_023246950.1;refGene_gn=MYO9B;refGene_tx=XM_023246954.1;refGene_gn=MYO9B;refGene_tx=XM_023246957.1;refGene_gn=MYO9B;refGene_tx=XM_023246958.1;refGene_gn=MYO9B;refGene_tx=XM_023246959.1;refGene_gn=MYO9B;refGene_tx=XM_023246962.1
A2 12713184 12713389
refGene_gn=MYO9B;refGene_tx=XM_006928569.3;refGene_gn=MYO9B;refGene_tx=XM_023246941.1;refGene_gn=MYO9B;refGene_tx=XM_023246946.1;refGene_gn=MYO9B;refGene_tx=XM_023246950.1;refGene_gn=MYO9B;refGene_tx=XM_023246954.1;refGene_gn=MYO9B;refGene_tx=XM_023246957.1;refGene_gn=MYO9B;refGene_tx=XM_023246958.1;refGene_gn=MYO9B;refGene_tx=XM_023246959.1;refGene_gn=MYO9B;refGene_tx=XM_023246962.1
A2 12714254 12714336
refGene_gn=MYO9B;refGene_tx=XM_006928569.3;refGene_gn=MYO9B;refGene_tx=XM_023246941.1;refGene_gn=MYO9B;refGene_tx=XM_023246946.1;refGene_gn=MYO9B;refGene_tx=XM_023246950.1;refGene_gn=MYO9B;refGene_tx=XM_023246954.1;refGene_gn=MYO9B;refGene_tx=XM_023246957.1;refGene_gn=MYO9B;refGene_tx=XM_023246958.1;refGene_gn=MYO9B;refGene_tx=XM_023246959.1;refGene_gn=MYO9B;refGene_tx=XM_023246962.1
A2 12715795 12715810
refGene_gn=MYO9B;refGene_tx=XM_023246957.1;refGene_gn=MYO9B;refGene_tx=XM_023246958.1
A2 12716906 12717020
refGene_gn=MYO9B;refGene_tx=XM_006928569.3;refGene_gn=MYO9B;refGene_tx=XM_023246941.1;refGene_gn=MYO9B;refGene_tx=XM_023246946.1;refGene_gn=MYO9B;refGene_tx=XM_023246950.1;refGene_gn=MYO9B;refGene_tx=XM_023246954.1

A2 12718464 12718504
refGene_gn=MYO9B;refGene_tx=XM_006928569.3;refGene_gn=MYO9B;refGene_tx=XM_023246941.1;refGene_gn=MYO9B;refGene_tx=XM_023246946.1;refGene_gn=MYO9B;refGene_tx=XM_023246950.1;refGene_gn=MYO9B;refGene_tx=XM_023246954.1;refGene_gn=MYO9B;refGene_tx=XM_023246957.1;refGene_gn=MYO9B;refGene_tx=XM_023246958.1;refGene_gn=MYO9B;refGene_tx=XM_023246959.1;refGene_gn=MYO9B;refGene_tx=XM_023246962.1
A2 12719609 12719732
refGene_gn=MYO9B;refGene_tx=XM_006928569.3;refGene_gn=MYO9B;refGene_tx=XM_023246941.1;refGene_gn=MYO9B;refGene_tx=XM_023246946.1;refGene_gn=MYO9B;refGene_tx=XM_023246950.1;refGene_gn=MYO9B;refGene_tx=XM_023246954.1;refGene_gn=MYO9B;refGene_tx=XM_023246957.1;refGene_gn=MYO9B;refGene_tx=XM_023246958.1;refGene_gn=MYO9B;refGene_tx=XM_023246959.1;refGene_gn=MYO9B;refGene_tx=XM_023246962.1
A2 12720585 12720666
refGene_gn=MYO9B;refGene_tx=XM_006928569.3;refGene_gn=MYO9B;refGene_tx=XM_023246941.1;refGene_gn=MYO9B;refGene_tx=XM_023246946.1;refGene_gn=MYO9B;refGene_tx=XM_023246950.1;refGene_gn=MYO9B;refGene_tx=XM_023246954.1;refGene_gn=MYO9B;refGene_tx=XM_023246957.1;refGene_gn=MYO9B;refGene_tx=XM_023246958.1;refGene_gn=MYO9B;refGene_tx=XM_023246959.1;refGene_gn=MYO9B;refGene_tx=XM_023246962.1
A2 12721832 12721943
refGene_gn=MYO9B;refGene_tx=XM_006928569.3;refGene_gn=MYO9B;refGene_tx=XM_023246941.1;refGene_gn=MYO9B;refGene_tx=XM_023246946.1;refGene_gn=MYO9B;refGene_tx=XM_023246950.1;refGene_gn=MYO9B;refGene_tx=XM_023246954.1;refGene_gn=MYO9B;refGene_tx=XM_023246957.1;refGene_gn=MYO9B;refGene_tx=XM_023246958.1;refGene_gn=MYO9B;refGene_tx=XM_023246959.1;refGene_gn=MYO9B;refGene_tx=XM_023246962.1
A2 12724021 12724144
refGene_gn=MYO9B;refGene_tx=XM_006928569.3;refGene_gn=MYO9B;refGene_tx=XM_023246941.1;refGene_gn=MYO9B;refGene_tx=XM_023246946.1;refGene_gn=MYO9B;refGene_tx=XM_023246950.1;refGene_gn=MYO9B;refGene_tx=XM_023246954.1;refGene_gn=MYO9B;refGene_tx=XM_023246957.1;refGene_gn=MYO9B;refGene_tx=XM_023246958.1;refGene_gn=MYO9B;refGene_tx=XM_023246959.1;refGene_gn=MYO9B;refGene_tx=XM_023246962.1
A2 12725272 12725422
refGene_gn=MYO9B;refGene_tx=XM_006928569.3;refGene_gn=MYO9B;refGene_tx=XM_023246941.1;refGene_gn=MYO9B;refGene_tx=XM_023246946.1;refGene_gn=MYO9B;refGene_tx=XM_023246950.1;refGene_gn=MYO9B;refGene_tx=XM_023246954.1;refGene_gn=MYO9B;refGene_tx=XM_023246957.1;refGene_gn=MYO9B;refGene_tx=XM_023246958.1;refGene_gn=MYO9B;refGene_tx=XM_023246959.1;refGene_gn=MYO9B;refGene_tx=XM_023246962.1
A2 12725515 12725682
refGene_gn=MYO9B;refGene_tx=XM_006928569.3;refGene_gn=MYO9B;refGene_tx=XM_023246941.1;refGene_gn=MYO9B;refGene_tx=XM_023246946.1;refGene_gn=MYO9B;refGene_tx=XM_023246950.1;refGene_gn=MYO9B;refGene_tx=XM_023246954.1;refGene_gn=MYO9B;refGene_tx=XM_023246957.1;refGene_gn=MYO9B;refGene_tx=XM_023246958.1;refGene_gn=MYO9B;refGene_tx=XM_023246959.1;refGene_gn=MYO9B;refGene_tx=XM_023246962.1
A2 12726391 12727306

refGene_gn=MYO9B;refGene_tx=XM_006928569.3;refGene_gn=MYO9B;refGene_tx=XM_023246941.1;refGene_gn=MYO9B;refGene_tx=XM_023246946.1;refGene_gn=MYO9B;refGene_tx=XM_023246950.1;refGene_gn=MYO9B;refGene_tx=XM_023246954.1;refGene_gn=MYO9B;refGene_tx=XM_023246957.1;refGene_gn=MYO9B;refGene_tx=XM_023246958.1;refGene_gn=MYO9B;refGene_tx=XM_023246959.1;refGene_gn=MYO9B;refGene_tx=XM_023246962.1
A2 12727997 12728070
refGene_gn=MYO9B;refGene_tx=XM_006928569.3;refGene_gn=MYO9B;refGene_tx=XM_023246941.1;refGene_gn=MYO9B;refGene_tx=XM_023246946.1;refGene_gn=MYO9B;refGene_tx=XM_023246950.1;refGene_gn=MYO9B;refGene_tx=XM_023246954.1;refGene_gn=MYO9B;refGene_tx=XM_023246957.1;refGene_gn=MYO9B;refGene_tx=XM_023246958.1;refGene_gn=MYO9B;refGene_tx=XM_023246959.1;refGene_gn=MYO9B;refGene_tx=XM_023246962.1
A2 12728263 12728388
refGene_gn=MYO9B;refGene_tx=XM_006928569.3;refGene_gn=MYO9B;refGene_tx=XM_023246941.1;refGene_gn=MYO9B;refGene_tx=XM_023246946.1;refGene_gn=MYO9B;refGene_tx=XM_023246950.1;refGene_gn=MYO9B;refGene_tx=XM_023246954.1;refGene_gn=MYO9B;refGene_tx=XM_023246957.1;refGene_gn=MYO9B;refGene_tx=XM_023246958.1;refGene_gn=MYO9B;refGene_tx=XM_023246959.1;refGene_gn=MYO9B;refGene_tx=XM_023246962.1
A2 12730430 12730564
refGene_gn=MYO9B;refGene_tx=XM_006928569.3;refGene_gn=MYO9B;refGene_tx=XM_023246941.1;refGene_gn=MYO9B;refGene_tx=XM_023246946.1;refGene_gn=MYO9B;refGene_tx=XM_023246950.1;refGene_gn=MYO9B;refGene_tx=XM_023246954.1;refGene_gn=MYO9B;refGene_tx=XM_023246957.1;refGene_gn=MYO9B;refGene_tx=XM_023246958.1;refGene_gn=MYO9B;refGene_tx=XM_023246959.1;refGene_gn=MYO9B;refGene_tx=XM_023246962.1
A2 12730795 12730986
refGene_gn=MYO9B;refGene_tx=XM_006928569.3;refGene_gn=MYO9B;refGene_tx=XM_023246941.1;refGene_gn=MYO9B;refGene_tx=XM_023246946.1;refGene_gn=MYO9B;refGene_tx=XM_023246950.1;refGene_gn=MYO9B;refGene_tx=XM_023246954.1;refGene_gn=MYO9B;refGene_tx=XM_023246957.1;refGene_gn=MYO9B;refGene_tx=XM_023246958.1;refGene_gn=MYO9B;refGene_tx=XM_023246959.1;refGene_gn=MYO9B;refGene_tx=XM_023246962.1
A2 12732106 12732205
refGene_gn=MYO9B;refGene_tx=XM_006928569.3;refGene_gn=MYO9B;refGene_tx=XM_023246941.1;refGene_gn=MYO9B;refGene_tx=XM_023246946.1;refGene_gn=MYO9B;refGene_tx=XM_023246950.1;refGene_gn=MYO9B;refGene_tx=XM_023246954.1;refGene_gn=MYO9B;refGene_tx=XM_023246957.1;refGene_gn=MYO9B;refGene_tx=XM_023246958.1;refGene_gn=MYO9B;refGene_tx=XM_023246959.1;refGene_gn=MYO9B;refGene_tx=XM_023246962.1
A2 12732316 12732490
refGene_gn=MYO9B;refGene_tx=XM_006928569.3;refGene_gn=MYO9B;refGene_tx=XM_023246941.1;refGene_gn=MYO9B;refGene_tx=XM_023246946.1;refGene_gn=MYO9B;refGene_tx=XM_023246950.1;refGene_gn=MYO9B;refGene_tx=XM_023246954.1;refGene_gn=MYO9B;refGene_tx=XM_023246957.1;refGene_gn=MYO9B;refGene_tx=XM_023246958.1;refGene_gn=MYO9B;refGene_tx=XM_023246959.1;refGene_gn=MYO9B;refGene_tx=XM_023246962.1
A2 12732573 12732615
refGene_gn=MYO9B;refGene_tx=XM_006928569.3;refGene_gn=MYO9B;refGene_tx

=XM_023246941.1;refGene_gn=MYO9B;refGene_tx=XM_023246946.1;refGene_gn=MYO9B;refGene_tx=XM_023246950.1;refGene_gn=MYO9B;refGene_tx=XM_023246954.1;refGene_gn=MYO9B;refGene_tx=XM_023246957.1;refGene_gn=MYO9B;refGene_tx=XM_023246958.1;refGene_gn=MYO9B;refGene_tx=XM_023246959.1;refGene_gn=MYO9B;refGene_tx=XM_023246962.1
A2 12732810 12732922
refGene_gn=MYO9B;refGene_tx=XM_006928569.3;refGene_gn=MYO9B;refGene_tx=XM_023246941.1;refGene_gn=MYO9B;refGene_tx=XM_023246946.1;refGene_gn=MYO9B;refGene_tx=XM_023246950.1;refGene_gn=MYO9B;refGene_tx=XM_023246954.1;refGene_gn=MYO9B;refGene_tx=XM_023246957.1;refGene_gn=MYO9B;refGene_tx=XM_023246958.1;refGene_gn=MYO9B;refGene_tx=XM_023246959.1;refGene_gn=MYO9B;refGene_tx=XM_023246962.1
A2 12734089 12734163
refGene_gn=MYO9B;refGene_tx=XM_006928569.3;refGene_gn=MYO9B;refGene_tx=XM_023246941.1;refGene_gn=MYO9B;refGene_tx=XM_023246946.1;refGene_gn=MYO9B;refGene_tx=XM_023246950.1;refGene_gn=MYO9B;refGene_tx=XM_023246954.1;refGene_gn=MYO9B;refGene_tx=XM_023246957.1;refGene_gn=MYO9B;refGene_tx=XM_023246958.1;refGene_gn=MYO9B;refGene_tx=XM_023246959.1;refGene_gn=MYO9B;refGene_tx=XM_023246962.1
A2 12734619 12734812
refGene_gn=MYO9B;refGene_tx=XM_006928569.3;refGene_gn=MYO9B;refGene_tx=XM_023246941.1;refGene_gn=MYO9B;refGene_tx=XM_023246946.1;refGene_gn=MYO9B;refGene_tx=XM_023246950.1;refGene_gn=MYO9B;refGene_tx=XM_023246954.1;refGene_gn=MYO9B;refGene_tx=XM_023246957.1;refGene_gn=MYO9B;refGene_tx=XM_023246958.1;refGene_gn=MYO9B;refGene_tx=XM_023246959.1;refGene_gn=MYO9B;refGene_tx=XM_023246962.1
A2 12734900 12735029
refGene_gn=MYO9B;refGene_tx=XM_006928569.3;refGene_gn=MYO9B;refGene_tx=XM_023246941.1;refGene_gn=MYO9B;refGene_tx=XM_023246946.1;refGene_gn=MYO9B;refGene_tx=XM_023246950.1;refGene_gn=MYO9B;refGene_tx=XM_023246954.1;refGene_gn=MYO9B;refGene_tx=XM_023246957.1;refGene_gn=MYO9B;refGene_tx=XM_023246958.1;refGene_gn=MYO9B;refGene_tx=XM_023246959.1;refGene_gn=MYO9B;refGene_tx=XM_023246962.1
A2 12735294 12735400
refGene_gn=MYO9B;refGene_tx=XM_006928569.3;refGene_gn=MYO9B;refGene_tx=XM_023246941.1;refGene_gn=MYO9B;refGene_tx=XM_023246946.1;refGene_gn=MYO9B;refGene_tx=XM_023246950.1;refGene_gn=MYO9B;refGene_tx=XM_023246954.1;refGene_gn=MYO9B;refGene_tx=XM_023246957.1;refGene_gn=MYO9B;refGene_tx=XM_023246958.1;refGene_gn=MYO9B;refGene_tx=XM_023246959.1;refGene_gn=MYO9B;refGene_tx=XM_023246962.1
A2 12735694 12735826
refGene_gn=MYO9B;refGene_tx=XM_006928569.3;refGene_gn=MYO9B;refGene_tx=XM_023246941.1;refGene_gn=MYO9B;refGene_tx=XM_023246946.1;refGene_gn=MYO9B;refGene_tx=XM_023246950.1;refGene_gn=MYO9B;refGene_tx=XM_023246954.1;refGene_gn=MYO9B;refGene_tx=XM_023246957.1;refGene_gn=MYO9B;refGene_tx=XM_023246958.1;refGene_gn=MYO9B;refGene_tx=XM_023246959.1;refGene_gn=MYO9B;refGene_tx=XM_023246962.1
A2 12736784 12736908
refGene_gn=MYO9B;refGene_tx=XM_006928569.3;refGene_gn=MYO9B;refGene_tx=XM_023246941.1;refGene_gn=MYO9B;refGene_tx=XM_023246946.1;refGene_gn=

MYO9B;refGene_tx=XM_023246950.1;refGene_gn=MYO9B;refGene_tx=XM_0232469
54.1;refGene_gn=MYO9B;refGene_tx=XM_023246957.1;refGene_gn=MYO9B;refGe
ne_tx=XM_023246958.1;refGene_gn=MYO9B;refGene_tx=XM_023246959.1;refGen
e_gn=MYO9B;refGene_tx=XM_023246962.1
A2 12737359 12737407
refGene_gn=MYO9B;refGene_tx=XM_006928569.3;refGene_gn=MYO9B;refGene_tx
=XM_023246941.1;refGene_gn=MYO9B;refGene_tx=XM_023246946.1;refGene_gn=
MYO9B;refGene_tx=XM_023246957.1;refGene_gn=MYO9B;refGene_tx=XM_0232469
58.1;refGene_gn=MYO9B;refGene_tx=XM_023246959.1;refGene_gn=MYO9B;refGe
ne_tx=XM_023246962.1
A2 12737752 12737895
refGene_gn=MYO9B;refGene_tx=XM_006928569.3;refGene_gn=MYO9B;refGene_tx
=XM_023246941.1;refGene_gn=MYO9B;refGene_tx=XM_023246946.1;refGene_gn=
MYO9B;refGene_tx=XM_023246950.1;refGene_gn=MYO9B;refGene_tx=XM_0232469
54.1;refGene_gn=MYO9B;refGene_tx=XM_023246957.1;refGene_gn=MYO9B;refGe
ne_tx=XM_023246958.1;refGene_gn=MYO9B;refGene_tx=XM_023246959.1;refGen
e_gn=MYO9B;refGene_tx=XM_023246962.1
A2 12738566 12738694
refGene_gn=MYO9B;refGene_tx=XM_006928569.3;refGene_gn=MYO9B;refGene_tx
=XM_023246941.1;refGene_gn=MYO9B;refGene_tx=XM_023246946.1;refGene_gn=
MYO9B;refGene_tx=XM_023246950.1;refGene_gn=MYO9B;refGene_tx=XM_0232469
54.1;refGene_gn=MYO9B;refGene_tx=XM_023246957.1;refGene_gn=MYO9B;refGe
ne_tx=XM_023246958.1;refGene_gn=MYO9B;refGene_tx=XM_023246959.1;refGen
e_gn=MYO9B;refGene_tx=XM_023246962.1
A2 12738814 12739230
refGene_gn=MYO9B;refGene_tx=XM_006928569.3;refGene_gn=MYO9B;refGene_tx
=XM_023246941.1;refGene_gn=MYO9B;refGene_tx=XM_023246946.1;refGene_gn=
MYO9B;refGene_tx=XM_023246950.1;refGene_gn=MYO9B;refGene_tx=XM_0232469
54.1;refGene_gn=MYO9B;refGene_tx=XM_023246957.1;refGene_gn=MYO9B;refGe
ne_tx=XM_023246958.1;refGene_gn=MYO9B;refGene_tx=XM_023246959.1;refGen
e_gn=MYO9B;refGene_tx=XM_023246962.1
A2 12741723 12741864
refGene_gn=USE1;refGene_tx=XM_003982032.5;refGene_gn=USE1;refGene_tx=X
M_006928574.4
A2 12742191 12742241
refGene_gn=USE1;refGene_tx=XM_003982032.5;refGene_gn=USE1;refGene_tx=X
M_006928574.4
A2 12742401 12742480
refGene_gn=USE1;refGene_tx=XM_003982032.5;refGene_gn=USE1;refGene_tx=X
M_006928574.4
A2 12742570 12742723
refGene_gn=USE1;refGene_tx=XM_003982032.5;refGene_gn=USE1;refGene_tx=X
M_006928574.4
A2 12743237 12743250
refGene_gn=USE1;refGene_tx=XM_003982032.5;refGene_gn=USE1;refGene_tx=X
M_006928574.4
A2 12743345 12743453 refGene_gn=USE1;refGene_tx=XM_006928574.4
A2 12743689 12743717
refGene_gn=USE1;refGene_tx=XM_003982032.5;refGene_gn=USE1;refGene_tx=X
M_006928574.4

A2 12744006 12744181
refGene_gn=USE1;refGene_tx=XM_003982032.5;refGene_gn=USE1;refGene_tx=XM_006928574.4
A2 12744422 12744605
refGene_gn=USE1;refGene_tx=XM_003982032.5;refGene_gn=USE1;refGene_tx=XM_006928574.4
A2 12746326 12746395 refGene_gn=OCEL1;refGene_tx=XM_011290569.3
A2 12746752 12746929 refGene_gn=OCEL1;refGene_tx=XM_011290569.3
A2 12747057 12747257 refGene_gn=OCEL1;refGene_tx=XM_011290569.3
A2 12747479 12747645 refGene_gn=OCEL1;refGene_tx=XM_011290569.3
A2 12747903 12747957 refGene_gn=OCEL1;refGene_tx=XM_011290569.3
A2 12748140 12748263 refGene_gn=OCEL1;refGene_tx=XM_011290569.3
A2 12751203 12751478 refGene_gn=NR2F6;refGene_tx=XM_023247004.1
A2 12752840 12753428 refGene_gn=NR2F6;refGene_tx=XM_023247004.1
A2 12756422 12756517 refGene_gn=NR2F6;refGene_tx=XM_023247004.1
A2 12760150 12760437 refGene_gn=NR2F6;refGene_tx=XM_023247004.1
A2 12764577 12764767 refGene_gn=USHBP1;refGene_tx=XM_003982033.5
A2 12765044 12765167 refGene_gn=USHBP1;refGene_tx=XM_003982033.5
A2 12765367 12765474 refGene_gn=USHBP1;refGene_tx=XM_003982033.5
A2 12766443 12766665 refGene_gn=USHBP1;refGene_tx=XM_003982033.5
A2 12767211 12767461 refGene_gn=USHBP1;refGene_tx=XM_003982033.5
A2 12768156 12768330 refGene_gn=USHBP1;refGene_tx=XM_003982033.5
A2 12768585 12768726 refGene_gn=USHBP1;refGene_tx=XM_003982033.5
A2 12768892 12769029 refGene_gn=USHBP1;refGene_tx=XM_003982033.5
A2 12769200 12769326 refGene_gn=USHBP1;refGene_tx=XM_003982033.5
A2 12772767 12773206 refGene_gn=USHBP1;refGene_tx=XM_003982033.5
A2 12773421 12773570 refGene_gn=USHBP1;refGene_tx=XM_003982033.5
A2 12773661 12773715 refGene_gn=USHBP1;refGene_tx=XM_003982033.5
A2 12778319 12778619 refGene_gn=BABAM1;refGene_tx=XM_006928576.3
A2 12779324 12779383 refGene_gn=BABAM1;refGene_tx=XM_006928576.3
A2 12780056 12780177 refGene_gn=BABAM1;refGene_tx=XM_006928576.3
A2 12780260 12780339 refGene_gn=BABAM1;refGene_tx=XM_006928576.3
A2 12781529 12781554 refGene_gn=BABAM1;refGene_tx=XM_006928576.3
A2 12782793 12782923 refGene_gn=BABAM1;refGene_tx=XM_006928576.3
A2 12783111 12783198 refGene_gn=BABAM1;refGene_tx=XM_006928576.3
A2 12783935 12783971 refGene_gn=BABAM1;refGene_tx=XM_006928576.3
A2 12783972 12784140 refGene_gn=BABAM1;refGene_tx=XM_006928576.3
A2 12785644 12785868 refGene_gn=ANKLE1;refGene_tx=XM_006928575.4
A2 12785940 12786093 refGene_gn=ANKLE1;refGene_tx=XM_006928575.4
A2 12786543 12786649 refGene_gn=ANKLE1;refGene_tx=XM_006928575.4
A2 12786748 12786893 refGene_gn=ANKLE1;refGene_tx=XM_006928575.4
A2 12787120 12787894 refGene_gn=ANKLE1;refGene_tx=XM_006928575.4
A2 12788055 12788233 refGene_gn=ANKLE1;refGene_tx=XM_006928575.4
A2 12788985 12789145 refGene_gn=ANKLE1;refGene_tx=XM_006928575.4
A2 12789250 12789380 refGene_gn=ANKLE1;refGene_tx=XM_006928575.4
A2 12789638 12789826 refGene_gn=ANKLE1;refGene_tx=XM_006928575.4
A2 12792931 12793102
refGene_gn=ABHD8;refGene_tx=XM_003982035.4;refGene_gn=ABHD8;refGene_tx=XM_011290588.3;refGene_gn=ABHD8;refGene_tx=XM_019819246.2
A2 12794357 12794574

refGene_gn=ABHD8;refGene_tx=XM_003982035.4;refGene_gn=ABHD8;refGene_tx=XM_011290588.3;refGene_gn=ABHD8;refGene_tx=XM_019819246.2
A2 12794719 12794890
refGene_gn=ABHD8;refGene_tx=XM_003982035.4;refGene_gn=ABHD8;refGene_tx=XM_011290588.3;refGene_gn=ABHD8;refGene_tx=XM_019819246.2
A2 12796417 12797157
refGene_gn=ABHD8;refGene_tx=XM_003982035.4;refGene_gn=ABHD8;refGene_tx=XM_011290588.3;refGene_gn=ABHD8;refGene_tx=XM_019819246.2
A2 12800526 12800573 refGene_gn=MRPL34;refGene_tx=XM_003982036.5
A2 12800653 12800712 refGene_gn=MRPL34;refGene_tx=XM_006928579.1
A2 12800811 12801025
refGene_gn=MRPL34;refGene_tx=XM_003982036.5;refGene_gn=MRPL34;refGene_tx=XM_006928579.1
A2 12802699 12802702 refGene_gn=DDA1;refGene_tx=XM_003982037.4
A2 12803272 12803278 refGene_gn=DDA1;refGene_tx=XM_019819262.2
A2 12807280 12807361
refGene_gn=DDA1;refGene_tx=XM_003982037.4;refGene_gn=DDA1;refGene_tx=XM_019819262.2
A2 12807587 12807639
refGene_gn=DDA1;refGene_tx=XM_003982037.4;refGene_gn=DDA1;refGene_tx=XM_019819262.2
A2 12808448 12808510
refGene_gn=DDA1;refGene_tx=XM_003982037.4;refGene_gn=DDA1;refGene_tx=XM_019819262.2
A2 12809699 12809810
refGene_gn=DDA1;refGene_tx=XM_003982037.4;refGene_gn=DDA1;refGene_tx=XM_019819262.2
A2 12812753 12813133
refGene_gn=ANO8;refGene_tx=XM_023247024.1;refGene_gn=ANO8;refGene_tx=XM_023247026.1;refGene_gn=ANO8;refGene_tx=XM_023247030.1
A2 12814003 12814670
refGene_gn=ANO8;refGene_tx=XM_023247024.1;refGene_gn=ANO8;refGene_tx=XM_023247026.1;refGene_gn=ANO8;refGene_tx=XM_023247030.1
A2 12815888 12815999
refGene_gn=ANO8;refGene_tx=XM_023247024.1;refGene_gn=ANO8;refGene_tx=XM_023247026.1;refGene_gn=ANO8;refGene_tx=XM_023247030.1
A2 12816103 12816235
refGene_gn=ANO8;refGene_tx=XM_023247024.1;refGene_gn=ANO8;refGene_tx=XM_023247026.1;refGene_gn=ANO8;refGene_tx=XM_023247030.1
A2 12816373 12816565
refGene_gn=ANO8;refGene_tx=XM_023247024.1;refGene_gn=ANO8;refGene_tx=XM_023247026.1;refGene_gn=ANO8;refGene_tx=XM_023247030.1
A2 12816821 12817631
refGene_gn=ANO8;refGene_tx=XM_023247024.1;refGene_gn=ANO8;refGene_tx=XM_023247026.1;refGene_gn=ANO8;refGene_tx=XM_023247030.1
A2 12818104 12818236
refGene_gn=ANO8;refGene_tx=XM_023247024.1;refGene_gn=ANO8;refGene_tx=XM_023247030.1
A2 12818439 12818514
refGene_gn=ANO8;refGene_tx=XM_023247024.1;refGene_gn=ANO8;refGene_tx=X

M_023247026.1;refGene_gn=ANO8;refGene_tx=XM_023247030.1
A2 12818611 12818667
refGene_gn=ANO8;refGene_tx=XM_023247024.1;refGene_gn=ANO8;refGene_tx=XM_023247026.1;refGene_gn=ANO8;refGene_tx=XM_023247030.1
A2 12818786 12818913
refGene_gn=ANO8;refGene_tx=XM_023247024.1;refGene_gn=ANO8;refGene_tx=XM_023247026.1;refGene_gn=ANO8;refGene_tx=XM_023247030.1
A2 12819049 12819202
refGene_gn=ANO8;refGene_tx=XM_023247024.1;refGene_gn=ANO8;refGene_tx=XM_023247026.1;refGene_gn=ANO8;refGene_tx=XM_023247030.1
A2 12819526 12819688
refGene_gn=ANO8;refGene_tx=XM_023247024.1;refGene_gn=ANO8;refGene_tx=XM_023247026.1
A2 12819782 12819910
refGene_gn=ANO8;refGene_tx=XM_023247024.1;refGene_gn=ANO8;refGene_tx=XM_023247026.1;refGene_gn=ANO8;refGene_tx=XM_023247030.1
A2 12820014 12820131
refGene_gn=ANO8;refGene_tx=XM_023247024.1;refGene_gn=ANO8;refGene_tx=XM_023247026.1;refGene_gn=ANO8;refGene_tx=XM_023247030.1
A2 12820775 12820872
refGene_gn=ANO8;refGene_tx=XM_023247024.1;refGene_gn=ANO8;refGene_tx=XM_023247026.1;refGene_gn=ANO8;refGene_tx=XM_023247030.1
A2 12820952 12821091
refGene_gn=ANO8;refGene_tx=XM_023247024.1;refGene_gn=ANO8;refGene_tx=XM_023247026.1;refGene_gn=ANO8;refGene_tx=XM_023247030.1
A2 12821271 12821404
refGene_gn=ANO8;refGene_tx=XM_023247024.1;refGene_gn=ANO8;refGene_tx=XM_023247026.1;refGene_gn=ANO8;refGene_tx=XM_023247030.1
A2 12821518 12821629
refGene_gn=ANO8;refGene_tx=XM_023247024.1;refGene_gn=ANO8;refGene_tx=XM_023247026.1;refGene_gn=ANO8;refGene_tx=XM_023247030.1
A2 12822397 12822503
refGene_gn=ANO8;refGene_tx=XM_023247024.1;refGene_gn=ANO8;refGene_tx=XM_023247026.1;refGene_gn=ANO8;refGene_tx=XM_023247030.1
A2 12824489 12824542
refGene_gn=GTPBP3;refGene_tx=XM_006928581.4;refGene_gn=GTPBP3;refGene_tx=XM_006928582.4
A2 12824961 12825215
refGene_gn=GTPBP3;refGene_tx=XM_006928581.4;refGene_gn=GTPBP3;refGene_tx=XM_006928582.4
A2 12825321 12825408
refGene_gn=GTPBP3;refGene_tx=XM_006928581.4;refGene_gn=GTPBP3;refGene_tx=XM_006928582.4
A2 12825506 12825709
refGene_gn=GTPBP3;refGene_tx=XM_006928581.4;refGene_gn=GTPBP3;refGene_tx=XM_006928582.4
A2 12825877 12826181
refGene_gn=GTPBP3;refGene_tx=XM_006928581.4;refGene_gn=GTPBP3;refGene_tx=XM_006928582.4
A2 12826374 12826540

refGene_gn=GTPBP3;refGene_tx=XM_006928581.4;refGene_gn=GTPBP3;refGene_tx=XM_006928582.4
A2 12826770 12827049
refGene_gn=GTPBP3;refGene_tx=XM_006928581.4;refGene_gn=GTPBP3;refGene_tx=XM_006928582.4
A2 12827249 12827475
refGene_gn=GTPBP3;refGene_tx=XM_006928581.4;refGene_gn=GTPBP3;refGene_tx=XM_006928582.4
A2 12833910 12833917 refGene_gn=PLVAP;refGene_tx=XM_006928583.4
A2 12838399 12838478 refGene_gn=PLVAP;refGene_tx=XM_006928583.4
A2 12838615 12838673 refGene_gn=PLVAP;refGene_tx=XM_006928583.4
A2 12840140 12840853 refGene_gn=PLVAP;refGene_tx=XM_006928583.4
A2 12840975 12841072 refGene_gn=PLVAP;refGene_tx=XM_006928583.4
A2 12845562 12845928 refGene_gn=PLVAP;refGene_tx=XM_006928583.4
A2 12855561 12855574
refGene_gn=LOC109494756;refGene_tx=XM_019819303.2
A2 12855790 12855937
refGene_gn=LOC109494756;refGene_tx=XM_019819303.2
A2 12856489 12856622
refGene_gn=LOC109494756;refGene_tx=XM_019819303.2
A2 12857029 12857126
refGene_gn=LOC109494756;refGene_tx=XM_019819303.2
A2 12858207 12858531
refGene_gn=LOC109494756;refGene_tx=XM_019819303.2
A2 12866724 12866845 refGene_gn=BST2;refGene_tx=NM_001243085.1
A2 12867084 12867145 refGene_gn=BST2;refGene_tx=NM_001243085.1
A2 12867422 12867498 refGene_gn=BST2;refGene_tx=NM_001243085.1
A2 12868267 12868568 refGene_gn=BST2;refGene_tx=NM_001243085.1
A2 12882898 12883163
refGene_gn=MVB12A;refGene_tx=XM_003982040.4;refGene_gn=MVB12A;refGene_tx=XM_006928584.3
A2 12883271 12883370
refGene_gn=MVB12A;refGene_tx=XM_003982040.4;refGene_gn=MVB12A;refGene_tx=XM_006928584.3
A2 12883500 12883597 refGene_gn=MVB12A;refGene_tx=XM_003982040.4
A2 12884900 12885027
refGene_gn=MVB12A;refGene_tx=XM_003982040.4;refGene_gn=MVB12A;refGene_tx=XM_006928584.3
A2 12885582 12885702
refGene_gn=MVB12A;refGene_tx=XM_003982040.4;refGene_gn=MVB12A;refGene_tx=XM_006928584.3
A2 12885777 12885884
refGene_gn=MVB12A;refGene_tx=XM_003982040.4;refGene_gn=MVB12A;refGene_tx=XM_006928584.3
A2 12886122 12886184
refGene_gn=MVB12A;refGene_tx=XM_003982040.4;refGene_gn=MVB12A;refGene_tx=XM_006928584.3
A2 12886484 12886541
refGene_gn=MVB12A;refGene_tx=XM_003982040.4;refGene_gn=MVB12A;refGene_tx=XM_006928584.3

A2 12886716 12886779
refGene_gn=MVB12A;refGene_tx=XM_003982040.4;refGene_gn=MVB12A;refGene_tx=XM_006928584.3
A2 12894768 12895238 refGene_gn=TMEM221;refGene_tx=XM_003982041.4
A2 12898270 12898356 refGene_gn=TMEM221;refGene_tx=XM_003982041.4
A2 12900409 12900729 refGene_gn=TMEM221;refGene_tx=XM_003982041.4
A2 12902592 12902923 refGene_gn=NXNL1;refGene_tx=XM_011280048.3
A2 12905843 12906169 refGene_gn=NXNL1;refGene_tx=XM_011280048.3
A2 12914605 12914772
refGene_gn=SLC27A1;refGene_tx=XM_011290625.2;refGene_gn=SLC27A1;refGene_tx=XM_023247108.1;refGene_gn=SLC27A1;refGene_tx=XM_023247114.1
A2 12934237 12934632
refGene_gn=SLC27A1;refGene_tx=XM_011290625.2;refGene_gn=SLC27A1;refGene_tx=XM_023247108.1;refGene_gn=SLC27A1;refGene_tx=XM_023247114.1
A2 12934737 12934899
refGene_gn=SLC27A1;refGene_tx=XM_011290625.2;refGene_gn=SLC27A1;refGene_tx=XM_023247108.1;refGene_gn=SLC27A1;refGene_tx=XM_023247114.1
A2 12934996 12935066
refGene_gn=SLC27A1;refGene_tx=XM_011290625.2;refGene_gn=SLC27A1;refGene_tx=XM_023247108.1;refGene_gn=SLC27A1;refGene_tx=XM_023247114.1
A2 12935766 12935858
refGene_gn=SLC27A1;refGene_tx=XM_011290625.2;refGene_gn=SLC27A1;refGene_tx=XM_023247108.1;refGene_gn=SLC27A1;refGene_tx=XM_023247114.1
A2 12935927 12936037
refGene_gn=SLC27A1;refGene_tx=XM_011290625.2;refGene_gn=SLC27A1;refGene_tx=XM_023247108.1;refGene_gn=SLC27A1;refGene_tx=XM_023247114.1
A2 12941176 12941386
refGene_gn=SLC27A1;refGene_tx=XM_011290625.2;refGene_gn=SLC27A1;refGene_tx=XM_023247108.1;refGene_gn=SLC27A1;refGene_tx=XM_023247114.1
A2 12942554 12942681
refGene_gn=SLC27A1;refGene_tx=XM_011290625.2;refGene_gn=SLC27A1;refGene_tx=XM_023247108.1;refGene_gn=SLC27A1;refGene_tx=XM_023247114.1
A2 12942769 12942907
refGene_gn=SLC27A1;refGene_tx=XM_011290625.2;refGene_gn=SLC27A1;refGene_tx=XM_023247108.1;refGene_gn=SLC27A1;refGene_tx=XM_023247114.1
A2 12942979 12943144
refGene_gn=SLC27A1;refGene_tx=XM_011290625.2;refGene_gn=SLC27A1;refGene_tx=XM_023247108.1
A2 12943535 12943682 refGene_gn=SLC27A1;refGene_tx=XM_011290625.2
A2 12945302 12945460
refGene_gn=SLC27A1;refGene_tx=XM_011290625.2;refGene_gn=SLC27A1;refGene_tx=XM_023247108.1;refGene_gn=SLC27A1;refGene_tx=XM_023247114.1
A2 12953176 12953464 refGene_gn=PGLS;refGene_tx=XM_023247120.1
A2 12956870 12956978 refGene_gn=PGLS;refGene_tx=XM_023247120.1
A2 12957410 12957512 refGene_gn=PGLS;refGene_tx=XM_023247120.1
A2 12957798 12957939 refGene_gn=PGLS;refGene_tx=XM_023247120.1
A2 12959962 12960100 refGene_gn=PGLS;refGene_tx=XM_023247120.1
A2 12964340 12964395
refGene_gn=FAM129C;refGene_tx=XM_023247127.1;refGene_gn=FAM129C;refGene_tx=XM_023247130.1

A2 12965915 12966046
refGene_gn=FAM129C;refGene_tx=XM_023247127.1;refGene_gn=FAM129C;refGen
e_tx=XM_023247130.1
A2 12968623 12968749
refGene_gn=FAM129C;refGene_tx=XM_023247127.1;refGene_gn=FAM129C;refGen
e_tx=XM_023247130.1
A2 12969322 12969437
refGene_gn=FAM129C;refGene_tx=XM_023247127.1;refGene_gn=FAM129C;refGen
e_tx=XM_023247130.1
A2 12971076 12971273
refGene_gn=FAM129C;refGene_tx=XM_023247127.1;refGene_gn=FAM129C;refGen
e_tx=XM_023247130.1;refGene_gn=FAM129C;refGene_tx=XM_023247137.1;refGe
ne_gn=FAM129C;refGene_tx=XM_023247142.1
A2 12971846 12971962
refGene_gn=FAM129C;refGene_tx=XM_023247127.1;refGene_gn=FAM129C;refGen
e_tx=XM_023247130.1;refGene_gn=FAM129C;refGene_tx=XM_023247137.1;refGe
ne_gn=FAM129C;refGene_tx=XM_023247142.1
A2 12972036 12972501
refGene_gn=FAM129C;refGene_tx=XM_023247127.1;refGene_gn=FAM129C;refGen
e_tx=XM_023247130.1;refGene_gn=FAM129C;refGene_tx=XM_023247137.1;refGe
ne_gn=FAM129C;refGene_tx=XM_023247142.1
A2 12973079 12973270
refGene_gn=FAM129C;refGene_tx=XM_023247127.1;refGene_gn=FAM129C;refGen
e_tx=XM_023247130.1;refGene_gn=FAM129C;refGene_tx=XM_023247137.1;refGe
ne_gn=FAM129C;refGene_tx=XM_023247142.1
A2 12974165 12974324
refGene_gn=FAM129C;refGene_tx=XM_023247127.1;refGene_gn=FAM129C;refGen
e_tx=XM_023247130.1;refGene_gn=FAM129C;refGene_tx=XM_023247137.1;refGe
ne_gn=FAM129C;refGene_tx=XM_023247142.1
A2 12975284 12975401
refGene_gn=FAM129C;refGene_tx=XM_023247127.1;refGene_gn=FAM129C;refGen
e_tx=XM_023247130.1;refGene_gn=FAM129C;refGene_tx=XM_023247137.1;refGe
ne_gn=FAM129C;refGene_tx=XM_023247142.1
A2 12975491 12975599
refGene_gn=FAM129C;refGene_tx=XM_023247127.1;refGene_gn=FAM129C;refGen
e_tx=XM_023247130.1;refGene_gn=FAM129C;refGene_tx=XM_023247137.1;refGe
ne_gn=FAM129C;refGene_tx=XM_023247142.1
A2 12976563 12976675
refGene_gn=FAM129C;refGene_tx=XM_023247127.1;refGene_gn=FAM129C;refGen
e_tx=XM_023247130.1;refGene_gn=FAM129C;refGene_tx=XM_023247137.1;refGe
ne_gn=FAM129C;refGene_tx=XM_023247142.1
A2 12977814 12977898
refGene_gn=FAM129C;refGene_tx=XM_023247127.1;refGene_gn=FAM129C;refGen
e_tx=XM_023247130.1;refGene_gn=FAM129C;refGene_tx=XM_023247137.1;refGe
ne_gn=FAM129C;refGene_tx=XM_023247142.1
A2 12979397 12979543
refGene_gn=FAM129C;refGene_tx=XM_023247127.1;refGene_gn=FAM129C;refGen
e_tx=XM_023247130.1;refGene_gn=FAM129C;refGene_tx=XM_023247137.1;refGe
ne_gn=FAM129C;refGene_tx=XM_023247142.1
A2 12982420 12982683 refGene_gn=COLGALT1;refGene_tx=XM_023247144.1

A2 12984423 12984534 refGene_gn=COLGALT1;refGene_tx=XM_023247144.1
A2 12985208 12985326 refGene_gn=COLGALT1;refGene_tx=XM_023247144.1
A2 12989265 12989400 refGene_gn=COLGALT1;refGene_tx=XM_023247144.1
A2 12991240 12991445 refGene_gn=COLGALT1;refGene_tx=XM_023247144.1
A2 12993812 12993932 refGene_gn=COLGALT1;refGene_tx=XM_023247144.1
A2 12998060 12998137 refGene_gn=COLGALT1;refGene_tx=XM_023247144.1
A2 12998221 12998328 refGene_gn=COLGALT1;refGene_tx=XM_023247144.1
A2 12998465 12998598 refGene_gn=COLGALT1;refGene_tx=XM_023247144.1
A2 12999134 12999262 refGene_gn=COLGALT1;refGene_tx=XM_023247144.1
A2 13000151 13000358 refGene_gn=COLGALT1;refGene_tx=XM_023247144.1
A2 13000604 13000872 refGene_gn=COLGALT1;refGene_tx=XM_023247144.1
A2 13013798 13014099
refGene_gn=UNC13A;refGene_tx=XM_023247146.1;refGene_gn=UNC13A;refGene_
tx=XM_023247152.1;refGene_gn=UNC13A;refGene_tx=XM_023247153.1;refGene_
gn=UNC13A;refGene_tx=XM_023247156.1;refGene_gn=UNC13A;refGene_tx=XM_02
3247158.1;refGene_gn=UNC13A;refGene_tx=XM_023247159.1;refGene_gn=UNC13
A;refGene_tx=XM_023247161.1
A2 13015631 13015791
refGene_gn=UNC13A;refGene_tx=XM_023247146.1;refGene_gn=UNC13A;refGene_
tx=XM_023247152.1;refGene_gn=UNC13A;refGene_tx=XM_023247153.1;refGene_
gn=UNC13A;refGene_tx=XM_023247156.1;refGene_gn=UNC13A;refGene_tx=XM_02
3247158.1;refGene_gn=UNC13A;refGene_tx=XM_023247159.1;refGene_gn=UNC13
A;refGene_tx=XM_023247161.1
A2 13016794 13016887
refGene_gn=UNC13A;refGene_tx=XM_023247146.1;refGene_gn=UNC13A;refGene_
tx=XM_023247152.1;refGene_gn=UNC13A;refGene_tx=XM_023247153.1;refGene_
gn=UNC13A;refGene_tx=XM_023247156.1;refGene_gn=UNC13A;refGene_tx=XM_02
3247158.1;refGene_gn=UNC13A;refGene_tx=XM_023247159.1;refGene_gn=UNC13
A;refGene_tx=XM_023247161.1
A2 13019508 13019565
refGene_gn=UNC13A;refGene_tx=XM_023247146.1;refGene_gn=UNC13A;refGene_
tx=XM_023247152.1;refGene_gn=UNC13A;refGene_tx=XM_023247153.1;refGene_
gn=UNC13A;refGene_tx=XM_023247158.1;refGene_gn=UNC13A;refGene_tx=XM_02
3247159.1
A2 13020615 13020763
refGene_gn=UNC13A;refGene_tx=XM_023247146.1;refGene_gn=UNC13A;refGene_
tx=XM_023247152.1;refGene_gn=UNC13A;refGene_tx=XM_023247153.1;refGene_
gn=UNC13A;refGene_tx=XM_023247156.1;refGene_gn=UNC13A;refGene_tx=XM_02
3247158.1;refGene_gn=UNC13A;refGene_tx=XM_023247159.1;refGene_gn=UNC13
A;refGene_tx=XM_023247161.1
A2 13021630 13021711
refGene_gn=UNC13A;refGene_tx=XM_023247146.1;refGene_gn=UNC13A;refGene_
tx=XM_023247152.1;refGene_gn=UNC13A;refGene_tx=XM_023247153.1;refGene_
gn=UNC13A;refGene_tx=XM_023247156.1;refGene_gn=UNC13A;refGene_tx=XM_02
3247158.1;refGene_gn=UNC13A;refGene_tx=XM_023247159.1;refGene_gn=UNC13
A;refGene_tx=XM_023247161.1
A2 13022081 13022138
refGene_gn=UNC13A;refGene_tx=XM_023247146.1;refGene_gn=UNC13A;refGene_
tx=XM_023247152.1;refGene_gn=UNC13A;refGene_tx=XM_023247153.1;refGene_
gn=UNC13A;refGene_tx=XM_023247156.1;refGene_gn=UNC13A;refGene_tx=XM_02

3247158.1;refGene_gn=UNC13A;refGene_tx=XM_023247159.1;refGene_gn=UNC13A;refGene_tx=XM_023247161.1
A2 13023536 13023566
refGene_gn=UNC13A;refGene_tx=XM_023247146.1;refGene_gn=UNC13A;refGene_tx=XM_023247152.1;refGene_gn=UNC13A;refGene_tx=XM_023247153.1;refGene_gn=UNC13A;refGene_tx=XM_023247156.1
A2 13024770 13024809
refGene_gn=UNC13A;refGene_tx=XM_023247146.1;refGene_gn=UNC13A;refGene_tx=XM_023247152.1;refGene_gn=UNC13A;refGene_tx=XM_023247153.1;refGene_gn=UNC13A;refGene_tx=XM_023247156.1
A2 13026530 13026536
refGene_gn=UNC13A;refGene_tx=XM_023247146.1;refGene_gn=UNC13A;refGene_tx=XM_023247152.1;refGene_gn=UNC13A;refGene_tx=XM_023247153.1;refGene_gn=UNC13A;refGene_tx=XM_023247156.1;refGene_gn=UNC13A;refGene_tx=XM_023247158.1
A2 13027451 13027575
refGene_gn=UNC13A;refGene_tx=XM_023247146.1;refGene_gn=UNC13A;refGene_tx=XM_023247152.1;refGene_gn=UNC13A;refGene_tx=XM_023247153.1;refGene_gn=UNC13A;refGene_tx=XM_023247156.1;refGene_gn=UNC13A;refGene_tx=XM_023247158.1;refGene_gn=UNC13A;refGene_tx=XM_023247159.1;refGene_gn=UNC13A;refGene_tx=XM_023247161.1
A2 13028780 13028933
refGene_gn=UNC13A;refGene_tx=XM_023247146.1;refGene_gn=UNC13A;refGene_tx=XM_023247152.1;refGene_gn=UNC13A;refGene_tx=XM_023247153.1;refGene_gn=UNC13A;refGene_tx=XM_023247156.1;refGene_gn=UNC13A;refGene_tx=XM_023247158.1;refGene_gn=UNC13A;refGene_tx=XM_023247159.1;refGene_gn=UNC13A;refGene_tx=XM_023247161.1
A2 13029390 13029479
refGene_gn=UNC13A;refGene_tx=XM_023247146.1;refGene_gn=UNC13A;refGene_tx=XM_023247152.1;refGene_gn=UNC13A;refGene_tx=XM_023247153.1;refGene_gn=UNC13A;refGene_tx=XM_023247156.1;refGene_gn=UNC13A;refGene_tx=XM_023247158.1;refGene_gn=UNC13A;refGene_tx=XM_023247159.1;refGene_gn=UNC13A;refGene_tx=XM_023247161.1
A2 13029701 13029779
refGene_gn=UNC13A;refGene_tx=XM_023247146.1;refGene_gn=UNC13A;refGene_tx=XM_023247152.1;refGene_gn=UNC13A;refGene_tx=XM_023247153.1;refGene_gn=UNC13A;refGene_tx=XM_023247156.1;refGene_gn=UNC13A;refGene_tx=XM_023247158.1;refGene_gn=UNC13A;refGene_tx=XM_023247159.1;refGene_gn=UNC13A;refGene_tx=XM_023247161.1
A2 13030324 13030408
refGene_gn=UNC13A;refGene_tx=XM_023247146.1;refGene_gn=UNC13A;refGene_tx=XM_023247152.1;refGene_gn=UNC13A;refGene_tx=XM_023247153.1;refGene_gn=UNC13A;refGene_tx=XM_023247156.1;refGene_gn=UNC13A;refGene_tx=XM_023247158.1;refGene_gn=UNC13A;refGene_tx=XM_023247159.1;refGene_gn=UNC13A;refGene_tx=XM_023247161.1
A2 13031183 13031327
refGene_gn=UNC13A;refGene_tx=XM_023247146.1;refGene_gn=UNC13A;refGene_tx=XM_023247152.1;refGene_gn=UNC13A;refGene_tx=XM_023247153.1;refGene_gn=UNC13A;refGene_tx=XM_023247156.1;refGene_gn=UNC13A;refGene_tx=XM_023247158.1;refGene_gn=UNC13A;refGene_tx=XM_023247159.1;refGene_gn=UNC13

A;refGene_tx=XM_023247161.1
A2 13031707 13031804
refGene_gn=UNC13A;refGene_tx=XM_023247146.1;refGene_gn=UNC13A;refGene_tx=XM_023247152.1;refGene_gn=UNC13A;refGene_tx=XM_023247153.1;refGene_gn=UNC13A;refGene_tx=XM_023247156.1;refGene_gn=UNC13A;refGene_tx=XM_023247158.1;refGene_gn=UNC13A;refGene_tx=XM_023247159.1;refGene_gn=UNC13A;refGene_tx=XM_023247161.1
A2 13032567 13032694
refGene_gn=UNC13A;refGene_tx=XM_023247146.1;refGene_gn=UNC13A;refGene_tx=XM_023247152.1;refGene_gn=UNC13A;refGene_tx=XM_023247153.1;refGene_gn=UNC13A;refGene_tx=XM_023247156.1;refGene_gn=UNC13A;refGene_tx=XM_023247158.1;refGene_gn=UNC13A;refGene_tx=XM_023247159.1;refGene_gn=UNC13A;refGene_tx=XM_023247161.1
A2 13032896 13032982
refGene_gn=UNC13A;refGene_tx=XM_023247146.1;refGene_gn=UNC13A;refGene_tx=XM_023247152.1;refGene_gn=UNC13A;refGene_tx=XM_023247153.1;refGene_gn=UNC13A;refGene_tx=XM_023247156.1;refGene_gn=UNC13A;refGene_tx=XM_023247158.1;refGene_gn=UNC13A;refGene_tx=XM_023247159.1;refGene_gn=UNC13A;refGene_tx=XM_023247161.1
A2 13034525 13034659
refGene_gn=UNC13A;refGene_tx=XM_023247146.1;refGene_gn=UNC13A;refGene_tx=XM_023247152.1;refGene_gn=UNC13A;refGene_tx=XM_023247153.1;refGene_gn=UNC13A;refGene_tx=XM_023247156.1;refGene_gn=UNC13A;refGene_tx=XM_023247158.1;refGene_gn=UNC13A;refGene_tx=XM_023247159.1;refGene_gn=UNC13A;refGene_tx=XM_023247161.1
A2 13036224 13036359
refGene_gn=UNC13A;refGene_tx=XM_023247146.1;refGene_gn=UNC13A;refGene_tx=XM_023247152.1;refGene_gn=UNC13A;refGene_tx=XM_023247153.1;refGene_gn=UNC13A;refGene_tx=XM_023247156.1;refGene_gn=UNC13A;refGene_tx=XM_023247158.1;refGene_gn=UNC13A;refGene_tx=XM_023247159.1;refGene_gn=UNC13A;refGene_tx=XM_023247161.1
A2 13036577 13036667
refGene_gn=UNC13A;refGene_tx=XM_023247146.1;refGene_gn=UNC13A;refGene_tx=XM_023247152.1;refGene_gn=UNC13A;refGene_tx=XM_023247153.1;refGene_gn=UNC13A;refGene_tx=XM_023247156.1;refGene_gn=UNC13A;refGene_tx=XM_023247158.1;refGene_gn=UNC13A;refGene_tx=XM_023247159.1;refGene_gn=UNC13A;refGene_tx=XM_023247161.1
A2 13036983 13037052
refGene_gn=UNC13A;refGene_tx=XM_023247146.1;refGene_gn=UNC13A;refGene_tx=XM_023247152.1;refGene_gn=UNC13A;refGene_tx=XM_023247153.1;refGene_gn=UNC13A;refGene_tx=XM_023247156.1;refGene_gn=UNC13A;refGene_tx=XM_023247158.1;refGene_gn=UNC13A;refGene_tx=XM_023247159.1;refGene_gn=UNC13A;refGene_tx=XM_023247161.1
A2 13037368 13037519
refGene_gn=UNC13A;refGene_tx=XM_023247146.1;refGene_gn=UNC13A;refGene_tx=XM_023247152.1;refGene_gn=UNC13A;refGene_tx=XM_023247153.1;refGene_gn=UNC13A;refGene_tx=XM_023247156.1;refGene_gn=UNC13A;refGene_tx=XM_023247158.1;refGene_gn=UNC13A;refGene_tx=XM_023247159.1;refGene_gn=UNC13A;refGene_tx=XM_023247161.1
A2 13037915 13038079

refGene_gn=UNC13A;refGene_tx=XM_023247146.1;refGene_gn=UNC13A;refGene_tx=XM_023247152.1;refGene_gn=UNC13A;refGene_tx=XM_023247153.1;refGene_gn=UNC13A;refGene_tx=XM_023247156.1;refGene_gn=UNC13A;refGene_tx=XM_023247158.1;refGene_gn=UNC13A;refGene_tx=XM_023247159.1;refGene_gn=UNC13A;refGene_tx=XM_023247161.1
A2 13039153 13039269
refGene_gn=UNC13A;refGene_tx=XM_023247146.1;refGene_gn=UNC13A;refGene_tx=XM_023247152.1;refGene_gn=UNC13A;refGene_tx=XM_023247153.1;refGene_gn=UNC13A;refGene_tx=XM_023247156.1;refGene_gn=UNC13A;refGene_tx=XM_023247158.1;refGene_gn=UNC13A;refGene_tx=XM_023247159.1;refGene_gn=UNC13A;refGene_tx=XM_023247161.1
A2 13040148 13040318
refGene_gn=UNC13A;refGene_tx=XM_023247146.1;refGene_gn=UNC13A;refGene_tx=XM_023247152.1;refGene_gn=UNC13A;refGene_tx=XM_023247153.1;refGene_gn=UNC13A;refGene_tx=XM_023247156.1;refGene_gn=UNC13A;refGene_tx=XM_023247158.1;refGene_gn=UNC13A;refGene_tx=XM_023247159.1;refGene_gn=UNC13A;refGene_tx=XM_023247161.1
A2 13040433 13040575
refGene_gn=UNC13A;refGene_tx=XM_023247146.1;refGene_gn=UNC13A;refGene_tx=XM_023247152.1;refGene_gn=UNC13A;refGene_tx=XM_023247153.1;refGene_gn=UNC13A;refGene_tx=XM_023247156.1;refGene_gn=UNC13A;refGene_tx=XM_023247158.1;refGene_gn=UNC13A;refGene_tx=XM_023247159.1;refGene_gn=UNC13A;refGene_tx=XM_023247161.1
A2 13041452 13041680
refGene_gn=UNC13A;refGene_tx=XM_023247146.1;refGene_gn=UNC13A;refGene_tx=XM_023247152.1;refGene_gn=UNC13A;refGene_tx=XM_023247153.1;refGene_gn=UNC13A;refGene_tx=XM_023247156.1;refGene_gn=UNC13A;refGene_tx=XM_023247158.1;refGene_gn=UNC13A;refGene_tx=XM_023247159.1;refGene_gn=UNC13A;refGene_tx=XM_023247161.1
A2 13042315 13042535
refGene_gn=UNC13A;refGene_tx=XM_023247146.1;refGene_gn=UNC13A;refGene_tx=XM_023247152.1;refGene_gn=UNC13A;refGene_tx=XM_023247153.1;refGene_gn=UNC13A;refGene_tx=XM_023247156.1;refGene_gn=UNC13A;refGene_tx=XM_023247158.1;refGene_gn=UNC13A;refGene_tx=XM_023247159.1;refGene_gn=UNC13A;refGene_tx=XM_023247161.1
A2 13042812 13042884
refGene_gn=UNC13A;refGene_tx=XM_023247146.1;refGene_gn=UNC13A;refGene_tx=XM_023247152.1;refGene_gn=UNC13A;refGene_tx=XM_023247153.1;refGene_gn=UNC13A;refGene_tx=XM_023247156.1;refGene_gn=UNC13A;refGene_tx=XM_023247158.1;refGene_gn=UNC13A;refGene_tx=XM_023247159.1;refGene_gn=UNC13A;refGene_tx=XM_023247161.1
A2 13043353 13043359
refGene_gn=UNC13A;refGene_tx=XM_023247146.1;refGene_gn=UNC13A;refGene_tx=XM_023247152.1;refGene_gn=UNC13A;refGene_tx=XM_023247153.1;refGene_gn=UNC13A;refGene_tx=XM_023247156.1;refGene_gn=UNC13A;refGene_tx=XM_023247158.1;refGene_gn=UNC13A;refGene_tx=XM_023247159.1;refGene_gn=UNC13A;refGene_tx=XM_023247161.1
A2 13043522 13043601
refGene_gn=UNC13A;refGene_tx=XM_023247146.1;refGene_gn=UNC13A;refGene_tx=XM_023247152.1;refGene_gn=UNC13A;refGene_tx=XM_023247153.1;refGene_

gn=UNC13A;refGene_tx=XM_023247156.1;refGene_gn=UNC13A;refGene_tx=XM_02
3247158.1;refGene_gn=UNC13A;refGene_tx=XM_023247159.1;refGene_gn=UNC13
A;refGene_tx=XM_023247161.1
A2 13046096 13046143
refGene_gn=UNC13A;refGene_tx=XM_023247146.1;refGene_gn=UNC13A;refGene_
tx=XM_023247152.1;refGene_gn=UNC13A;refGene_tx=XM_023247153.1;refGene_
gn=UNC13A;refGene_tx=XM_023247156.1;refGene_gn=UNC13A;refGene_tx=XM_02
3247158.1;refGene_gn=UNC13A;refGene_tx=XM_023247159.1;refGene_gn=UNC13
A;refGene_tx=XM_023247161.1
A2 13047891 13048000
refGene_gn=UNC13A;refGene_tx=XM_023247146.1;refGene_gn=UNC13A;refGene_
tx=XM_023247152.1;refGene_gn=UNC13A;refGene_tx=XM_023247153.1;refGene_
gn=UNC13A;refGene_tx=XM_023247156.1;refGene_gn=UNC13A;refGene_tx=XM_02
3247158.1;refGene_gn=UNC13A;refGene_tx=XM_023247159.1;refGene_gn=UNC13
A;refGene_tx=XM_023247161.1
A2 13048243 13048804
refGene_gn=UNC13A;refGene_tx=XM_023247146.1;refGene_gn=UNC13A;refGene_
tx=XM_023247152.1;refGene_gn=UNC13A;refGene_tx=XM_023247153.1;refGene_
gn=UNC13A;refGene_tx=XM_023247156.1;refGene_gn=UNC13A;refGene_tx=XM_02
3247158.1;refGene_gn=UNC13A;refGene_tx=XM_023247159.1;refGene_gn=UNC13
A;refGene_tx=XM_023247161.1
A2 13051583 13051791
refGene_gn=UNC13A;refGene_tx=XM_023247146.1;refGene_gn=UNC13A;refGene_
tx=XM_023247152.1;refGene_gn=UNC13A;refGene_tx=XM_023247153.1;refGene_
gn=UNC13A;refGene_tx=XM_023247156.1;refGene_gn=UNC13A;refGene_tx=XM_02
3247158.1;refGene_gn=UNC13A;refGene_tx=XM_023247159.1;refGene_gn=UNC13
A;refGene_tx=XM_023247161.1
A2 13055186 13055222
refGene_gn=UNC13A;refGene_tx=XM_023247146.1;refGene_gn=UNC13A;refGene_
tx=XM_023247152.1;refGene_gn=UNC13A;refGene_tx=XM_023247156.1;refGene_
gn=UNC13A;refGene_tx=XM_023247158.1;refGene_gn=UNC13A;refGene_tx=XM_02
3247159.1;refGene_gn=UNC13A;refGene_tx=XM_023247161.1
A2 13057130 13057185
refGene_gn=UNC13A;refGene_tx=XM_023247146.1;refGene_gn=UNC13A;refGene_
tx=XM_023247152.1;refGene_gn=UNC13A;refGene_tx=XM_023247153.1;refGene_
gn=UNC13A;refGene_tx=XM_023247156.1;refGene_gn=UNC13A;refGene_tx=XM_02
3247158.1;refGene_gn=UNC13A;refGene_tx=XM_023247159.1;refGene_gn=UNC13
A;refGene_tx=XM_023247161.1
A2 13058649 13058723
refGene_gn=UNC13A;refGene_tx=XM_023247146.1;refGene_gn=UNC13A;refGene_
tx=XM_023247152.1;refGene_gn=UNC13A;refGene_tx=XM_023247153.1;refGene_
gn=UNC13A;refGene_tx=XM_023247156.1;refGene_gn=UNC13A;refGene_tx=XM_02
3247158.1;refGene_gn=UNC13A;refGene_tx=XM_023247159.1;refGene_gn=UNC13
A;refGene_tx=XM_023247161.1
A2 13059325 13059449
refGene_gn=UNC13A;refGene_tx=XM_023247146.1;refGene_gn=UNC13A;refGene_
tx=XM_023247152.1;refGene_gn=UNC13A;refGene_tx=XM_023247153.1;refGene_
gn=UNC13A;refGene_tx=XM_023247156.1;refGene_gn=UNC13A;refGene_tx=XM_02
3247158.1;refGene_gn=UNC13A;refGene_tx=XM_023247159.1;refGene_gn=UNC13
A;refGene_tx=XM_023247161.1

A2 13060961 13061079
refGene_gn=UNC13A;refGene_tx=XM_023247146.1;refGene_gn=UNC13A;refGene_tx=XM_023247152.1;refGene_gn=UNC13A;refGene_tx=XM_023247153.1;refGene_gn=UNC13A;refGene_tx=XM_023247156.1;refGene_gn=UNC13A;refGene_tx=XM_023247158.1;refGene_gn=UNC13A;refGene_tx=XM_023247159.1;refGene_gn=UNC13A;refGene_tx=XM_023247161.1
A2 13062931 13063031
refGene_gn=UNC13A;refGene_tx=XM_023247146.1;refGene_gn=UNC13A;refGene_tx=XM_023247152.1;refGene_gn=UNC13A;refGene_tx=XM_023247153.1;refGene_gn=UNC13A;refGene_tx=XM_023247156.1;refGene_gn=UNC13A;refGene_tx=XM_023247158.1;refGene_gn=UNC13A;refGene_tx=XM_023247159.1;refGene_gn=UNC13A;refGene_tx=XM_023247161.1
A2 13063901 13063931
refGene_gn=UNC13A;refGene_tx=XM_023247146.1;refGene_gn=UNC13A;refGene_tx=XM_023247152.1;refGene_gn=UNC13A;refGene_tx=XM_023247153.1;refGene_gn=UNC13A;refGene_tx=XM_023247156.1;refGene_gn=UNC13A;refGene_tx=XM_023247158.1;refGene_gn=UNC13A;refGene_tx=XM_023247159.1;refGene_gn=UNC13A;refGene_tx=XM_023247161.1
A2 13073118 13073140
refGene_gn=UNC13A;refGene_tx=XM_023247146.1;refGene_gn=UNC13A;refGene_tx=XM_023247152.1;refGene_gn=UNC13A;refGene_tx=XM_023247153.1;refGene_gn=UNC13A;refGene_tx=XM_023247156.1;refGene_gn=UNC13A;refGene_tx=XM_023247158.1;refGene_gn=UNC13A;refGene_tx=XM_023247159.1;refGene_gn=UNC13A;refGene_tx=XM_023247161.1
A2 13087954 13088075 refGene_gn=MAP1S;refGene_tx=XM_023247163.1
A2 13089372 13089474 refGene_gn=MAP1S;refGene_tx=XM_023247163.1
A2 13090938 13091021 refGene_gn=MAP1S;refGene_tx=XM_023247163.1
A2 13092027 13092076 refGene_gn=MAP1S;refGene_tx=XM_023247163.1
A2 13092078 13092167 refGene_gn=MAP1S;refGene_tx=XM_023247163.1
A2 13092826 13092905 refGene_gn=MAP1S;refGene_tx=XM_023247163.1
A2 13092907 13092949 refGene_gn=MAP1S;refGene_tx=XM_023247163.1
A2 13092951 13093041 refGene_gn=MAP1S;refGene_tx=XM_023247163.1
A2 13093043 13093688 refGene_gn=MAP1S;refGene_tx=XM_023247163.1
A2 13093690 13093724 refGene_gn=MAP1S;refGene_tx=XM_023247163.1
A2 13093725 13094323 refGene_gn=MAP1S;refGene_tx=XM_023247163.1
A2 13094325 13094408 refGene_gn=MAP1S;refGene_tx=XM_023247163.1
A2 13094409 13094494 refGene_gn=MAP1S;refGene_tx=XM_023247163.1
A2 13094495 13094540 refGene_gn=MAP1S;refGene_tx=XM_023247163.1
A2 13094542 13094590 refGene_gn=MAP1S;refGene_tx=XM_023247163.1
A2 13094591 13094630 refGene_gn=MAP1S;refGene_tx=XM_023247163.1
A2 13094631 13094691 refGene_gn=MAP1S;refGene_tx=XM_023247163.1
A2 13094692 13094903 refGene_gn=MAP1S;refGene_tx=XM_023247163.1
A2 13094905 13094938 refGene_gn=MAP1S;refGene_tx=XM_023247163.1
A2 13094940 13095096 refGene_gn=MAP1S;refGene_tx=XM_023247163.1
A2 13095098 13095185 refGene_gn=MAP1S;refGene_tx=XM_023247163.1
A2 13103911 13104147 refGene_gn=MAP1S;refGene_tx=XM_023247163.1
A2 13104973 13105129 refGene_gn=MAP1S;refGene_tx=XM_023247163.1
A2 13124342 13124369
refGene_gn=FCHO1;refGene_tx=XM_006928598.4;refGene_gn=FCHO1;refGene_tx=XM_019819447.2;refGene_gn=FCHO1;refGene_tx=XM_019819454.1;refGene_gn=

FCHO1;refGene_tx=XM_019819460.2;refGene_gn=FCHO1;refGene_tx=XM_0198194
62.2
A2 13127474 13127566
refGene_gn=FCHO1;refGene_tx=XM_006928598.4;refGene_gn=FCHO1;refGene_tx
=XM_019819447.2;refGene_gn=FCHO1;refGene_tx=XM_019819454.1;refGene_gn=
FCHO1;refGene_tx=XM_019819460.2;refGene_gn=FCHO1;refGene_tx=XM_0198194
62.2
A2 13128597 13128672
refGene_gn=FCHO1;refGene_tx=XM_006928598.4;refGene_gn=FCHO1;refGene_tx
=XM_019819447.2;refGene_gn=FCHO1;refGene_tx=XM_019819454.1;refGene_gn=
FCHO1;refGene_tx=XM_019819460.2;refGene_gn=FCHO1;refGene_tx=XM_0198194
62.2
A2 13129724 13129866
refGene_gn=FCHO1;refGene_tx=XM_006928598.4;refGene_gn=FCHO1;refGene_tx
=XM_019819447.2;refGene_gn=FCHO1;refGene_tx=XM_019819454.1;refGene_gn=
FCHO1;refGene_tx=XM_019819460.2;refGene_gn=FCHO1;refGene_tx=XM_0198194
62.2
A2 13131497 13131650
refGene_gn=FCHO1;refGene_tx=XM_006928598.4;refGene_gn=FCHO1;refGene_tx
=XM_019819447.2;refGene_gn=FCHO1;refGene_tx=XM_019819454.1;refGene_gn=
FCHO1;refGene_tx=XM_019819460.2;refGene_gn=FCHO1;refGene_tx=XM_0198194
62.2
A2 13131749 13131854
refGene_gn=FCHO1;refGene_tx=XM_006928598.4;refGene_gn=FCHO1;refGene_tx
=XM_019819447.2;refGene_gn=FCHO1;refGene_tx=XM_019819454.1;refGene_gn=
FCHO1;refGene_tx=XM_019819460.2;refGene_gn=FCHO1;refGene_tx=XM_0198194
62.2
A2 13133195 13133294
refGene_gn=FCHO1;refGene_tx=XM_006928598.4;refGene_gn=FCHO1;refGene_tx
=XM_019819447.2;refGene_gn=FCHO1;refGene_tx=XM_019819454.1;refGene_gn=
FCHO1;refGene_tx=XM_019819460.2;refGene_gn=FCHO1;refGene_tx=XM_0198194
62.2
A2 13133387 13133484
refGene_gn=FCHO1;refGene_tx=XM_006928598.4;refGene_gn=FCHO1;refGene_tx
=XM_019819447.2;refGene_gn=FCHO1;refGene_tx=XM_019819454.1;refGene_gn=
FCHO1;refGene_tx=XM_019819460.2;refGene_gn=FCHO1;refGene_tx=XM_0198194
62.2
A2 13134408 13134453
refGene_gn=FCHO1;refGene_tx=XM_006928598.4;refGene_gn=FCHO1;refGene_tx
=XM_019819447.2;refGene_gn=FCHO1;refGene_tx=XM_019819454.1;refGene_gn=
FCHO1;refGene_tx=XM_019819460.2;refGene_gn=FCHO1;refGene_tx=XM_0198194
62.2
A2 13134570 13134649
refGene_gn=FCHO1;refGene_tx=XM_006928598.4;refGene_gn=FCHO1;refGene_tx
=XM_019819447.2;refGene_gn=FCHO1;refGene_tx=XM_019819454.1;refGene_gn=
FCHO1;refGene_tx=XM_019819460.2;refGene_gn=FCHO1;refGene_tx=XM_0198194
62.2
A2 13135150 13135175
refGene_gn=FCHO1;refGene_tx=XM_006928598.4;refGene_gn=FCHO1;refGene_tx
=XM_019819447.2;refGene_gn=FCHO1;refGene_tx=XM_019819454.1;refGene_gn=

FCHO1;refGene_tx=XM_019819460.2;refGene_gn=FCHO1;refGene_tx=XM_0198194
62.2
A2 13135500 13135558
refGene_gn=FCHO1;refGene_tx=XM_006928598.4;refGene_gn=FCHO1;refGene_tx
=XM_019819447.2;refGene_gn=FCHO1;refGene_tx=XM_019819454.1;refGene_gn=
FCHO1;refGene_tx=XM_019819460.2;refGene_gn=FCHO1;refGene_tx=XM_0198194
62.2
A2 13135989 13136168
refGene_gn=FCHO1;refGene_tx=XM_006928598.4;refGene_gn=FCHO1;refGene_tx
=XM_019819447.2;refGene_gn=FCHO1;refGene_tx=XM_019819454.1;refGene_gn=
FCHO1;refGene_tx=XM_019819460.2;refGene_gn=FCHO1;refGene_tx=XM_0198194
62.2
A2 13136253 13136278
refGene_gn=FCHO1;refGene_tx=XM_006928598.4;refGene_gn=FCHO1;refGene_tx
=XM_019819447.2;refGene_gn=FCHO1;refGene_tx=XM_019819454.1;refGene_gn=
FCHO1;refGene_tx=XM_019819460.2;refGene_gn=FCHO1;refGene_tx=XM_0198194
62.2
A2 13136615 13136667
refGene_gn=FCHO1;refGene_tx=XM_006928598.4;refGene_gn=FCHO1;refGene_tx
=XM_019819447.2;refGene_gn=FCHO1;refGene_tx=XM_019819454.1;refGene_gn=
FCHO1;refGene_tx=XM_019819460.2;refGene_gn=FCHO1;refGene_tx=XM_0198194
62.2
A2 13137457 13137549
refGene_gn=FCHO1;refGene_tx=XM_006928598.4;refGene_gn=FCHO1;refGene_tx
=XM_019819447.2;refGene_gn=FCHO1;refGene_tx=XM_019819454.1;refGene_gn=
FCHO1;refGene_tx=XM_019819460.2;refGene_gn=FCHO1;refGene_tx=XM_0198194
62.2
A2 13137907 13138183
refGene_gn=FCHO1;refGene_tx=XM_006928598.4;refGene_gn=FCHO1;refGene_tx
=XM_019819447.2;refGene_gn=FCHO1;refGene_tx=XM_019819454.1;refGene_gn=
FCHO1;refGene_tx=XM_019819460.2
A2 13140010 13140123
refGene_gn=FCHO1;refGene_tx=XM_006928598.4;refGene_gn=FCHO1;refGene_tx
=XM_019819447.2;refGene_gn=FCHO1;refGene_tx=XM_019819454.1;refGene_gn=
FCHO1;refGene_tx=XM_019819460.2;refGene_gn=FCHO1;refGene_tx=XM_0198194
62.2
A2 13140238 13140326
refGene_gn=FCHO1;refGene_tx=XM_006928598.4;refGene_gn=FCHO1;refGene_tx
=XM_019819447.2;refGene_gn=FCHO1;refGene_tx=XM_019819454.1;refGene_gn=
FCHO1;refGene_tx=XM_019819460.2;refGene_gn=FCHO1;refGene_tx=XM_0198194
62.2
A2 13140462 13140571
refGene_gn=FCHO1;refGene_tx=XM_006928598.4;refGene_gn=FCHO1;refGene_tx
=XM_019819447.2;refGene_gn=FCHO1;refGene_tx=XM_019819454.1;refGene_gn=
FCHO1;refGene_tx=XM_019819460.2;refGene_gn=FCHO1;refGene_tx=XM_0198194
62.2
A2 13141269 13141425
refGene_gn=FCHO1;refGene_tx=XM_006928598.4;refGene_gn=FCHO1;refGene_tx
=XM_019819447.2;refGene_gn=FCHO1;refGene_tx=XM_019819454.1;refGene_gn=
FCHO1;refGene_tx=XM_019819460.2;refGene_gn=FCHO1;refGene_tx=XM_0198194

62.2
A2	13141747 13141880
refGene_gn=FCHO1;refGene_tx=XM_006928598.4;refGene_gn=FCHO1;refGene_tx=XM_019819447.2;refGene_gn=FCHO1;refGene_tx=XM_019819454.1;refGene_gn=FCHO1;refGene_tx=XM_019819460.2;refGene_gn=FCHO1;refGene_tx=XM_019819462.2
A2	13142356 13142556
refGene_gn=FCHO1;refGene_tx=XM_006928598.4;refGene_gn=FCHO1;refGene_tx=XM_019819447.2;refGene_gn=FCHO1;refGene_tx=XM_019819454.1;refGene_gn=FCHO1;refGene_tx=XM_019819460.2;refGene_gn=FCHO1;refGene_tx=XM_019819462.2
A2	13143447 13143503
refGene_gn=FCHO1;refGene_tx=XM_006928598.4;refGene_gn=FCHO1;refGene_tx=XM_019819447.2;refGene_gn=FCHO1;refGene_tx=XM_019819454.1;refGene_gn=FCHO1;refGene_tx=XM_019819460.2;refGene_gn=FCHO1;refGene_tx=XM_019819462.2
A2	13143874 13144039
refGene_gn=FCHO1;refGene_tx=XM_006928598.4;refGene_gn=FCHO1;refGene_tx=XM_019819447.2;refGene_gn=FCHO1;refGene_tx=XM_019819454.1;refGene_gn=FCHO1;refGene_tx=XM_019819460.2;refGene_gn=FCHO1;refGene_tx=XM_019819462.2
A2	13144447 13144470
refGene_gn=FCHO1;refGene_tx=XM_006928598.4;refGene_gn=FCHO1;refGene_tx=XM_019819447.2;refGene_gn=FCHO1;refGene_tx=XM_019819454.1;refGene_gn=FCHO1;refGene_tx=XM_019819460.2;refGene_gn=FCHO1;refGene_tx=XM_019819462.2
A2	13166135 13166190 refGene_gn=B3GNT3;refGene_tx=XM_003982099.5
A2	13167438 13168037
refGene_gn=B3GNT3;refGene_tx=XM_003982099.5;refGene_gn=B3GNT3;refGene_tx=XM_006928599.4;refGene_gn=B3GNT3;refGene_tx=XM_006928600.4;refGene_gn=B3GNT3;refGene_tx=XM_006928601.4;refGene_gn=B3GNT3;refGene_tx=XM_006928603.4;refGene_gn=B3GNT3;refGene_tx=XM_006928604.3;refGene_gn=B3GNT3;refGene_tx=XM_019819477.2
A2	13169653 13170205
refGene_gn=B3GNT3;refGene_tx=XM_003982099.5;refGene_gn=B3GNT3;refGene_tx=XM_006928599.4;refGene_gn=B3GNT3;refGene_tx=XM_006928600.4;refGene_gn=B3GNT3;refGene_tx=XM_006928601.4;refGene_gn=B3GNT3;refGene_tx=XM_006928603.4;refGene_gn=B3GNT3;refGene_tx=XM_006928604.3;refGene_gn=B3GNT3;refGene_tx=XM_019819477.2
A2	13174208 13174411
refGene_gn=INSL3;refGene_tx=XM_003982069.5;refGene_gn=INSL3;refGene_tx=XM_019819501.2
A2	13174564 13174651 refGene_gn=INSL3;refGene_tx=XM_019819501.2
A2	13175780 13175973
refGene_gn=INSL3;refGene_tx=XM_003982069.5;refGene_gn=INSL3;refGene_tx=XM_019819501.2
A2	13180078 13180192
refGene_gn=JAK3;refGene_tx=XM_023247201.1;refGene_gn=JAK3;refGene_tx=XM_023247208.1
A2	13181092 13181203

refGene_gn=JAK3;refGene_tx=XM_023247201.1;refGene_gn=JAK3;refGene_tx=XM_023247208.1
A2 13181530 13181648
refGene_gn=JAK3;refGene_tx=XM_023247201.1;refGene_gn=JAK3;refGene_tx=XM_023247208.1
A2 13182259 13182432
refGene_gn=JAK3;refGene_tx=XM_023247201.1;refGene_gn=JAK3;refGene_tx=XM_023247208.1
A2 13182697 13182822
refGene_gn=JAK3;refGene_tx=XM_023247201.1;refGene_gn=JAK3;refGene_tx=XM_023247208.1
A2 13183095 13183285
refGene_gn=JAK3;refGene_tx=XM_023247201.1;refGene_gn=JAK3;refGene_tx=XM_023247208.1
A2 13183361 13183501
refGene_gn=JAK3;refGene_tx=XM_023247201.1;refGene_gn=JAK3;refGene_tx=XM_023247208.1
A2 13184097 13184248
refGene_gn=JAK3;refGene_tx=XM_023247201.1;refGene_gn=JAK3;refGene_tx=XM_023247208.1
A2 13184367 13184519
refGene_gn=JAK3;refGene_tx=XM_023247201.1;refGene_gn=JAK3;refGene_tx=XM_023247208.1
A2 13184594 13184727
refGene_gn=JAK3;refGene_tx=XM_023247201.1;refGene_gn=JAK3;refGene_tx=XM_023247208.1
A2 13185376 13185504
refGene_gn=JAK3;refGene_tx=XM_023247201.1;refGene_gn=JAK3;refGene_tx=XM_023247208.1
A2 13185939 13186024
refGene_gn=JAK3;refGene_tx=XM_023247201.1;refGene_gn=JAK3;refGene_tx=XM_023247208.1
A2 13186727 13186859
refGene_gn=JAK3;refGene_tx=XM_023247201.1;refGene_gn=JAK3;refGene_tx=XM_023247208.1
A2 13187026 13187154
refGene_gn=JAK3;refGene_tx=XM_023247201.1;refGene_gn=JAK3;refGene_tx=XM_023247208.1
A2 13187902 13188089
refGene_gn=JAK3;refGene_tx=XM_023247201.1;refGene_gn=JAK3;refGene_tx=XM_023247208.1
A2 13188349 13188461
refGene_gn=JAK3;refGene_tx=XM_023247201.1;refGene_gn=JAK3;refGene_tx=XM_023247208.1
A2 13188912 13189070
refGene_gn=JAK3;refGene_tx=XM_023247201.1;refGene_gn=JAK3;refGene_tx=XM_023247208.1
A2 13189181 13189304
refGene_gn=JAK3;refGene_tx=XM_023247201.1;refGene_gn=JAK3;refGene_tx=XM_023247208.1

A2 13189878 13190173
refGene_gn=JAK3;refGene_tx=XM_023247201.1;refGene_gn=JAK3;refGene_tx=XM_023247208.1
A2 13190507 13190653
refGene_gn=JAK3;refGene_tx=XM_023247201.1;refGene_gn=JAK3;refGene_tx=XM_023247208.1
A2 13190844 13190956
refGene_gn=JAK3;refGene_tx=XM_023247201.1;refGene_gn=JAK3;refGene_tx=XM_023247208.1
A2 13191764 13191888
refGene_gn=JAK3;refGene_tx=XM_023247201.1;refGene_gn=JAK3;refGene_tx=XM_023247208.1
A2 13192338 13192522
refGene_gn=JAK3;refGene_tx=XM_023247201.1;refGene_gn=JAK3;refGene_tx=XM_023247208.1
A2 13207582 13207600 refGene_gn=RPL18A;refGene_tx=XM_003982070.3
A2 13208901 13209081 refGene_gn=RPL18A;refGene_tx=XM_003982070.3
A2 13209713 13209843 refGene_gn=RPL18A;refGene_tx=XM_003982070.3
A2 13210434 13210544 refGene_gn=RPL18A;refGene_tx=XM_003982070.3
A2 13210763 13210856 refGene_gn=RPL18A;refGene_tx=XM_003982070.3
A2 13216615 13216972
refGene_gn=SLC5A5;refGene_tx=XM_011280067.3;refGene_gn=SLC5A5;refGene_tx=XM_023247222.1;refGene_gn=SLC5A5;refGene_tx=XM_023247227.1;refGene_gn=SLC5A5;refGene_tx=XM_023247234.1;refGene_gn=SLC5A5;refGene_tx=XM_023247235.1
A2 13217777 13217843
refGene_gn=SLC5A5;refGene_tx=XM_011280067.3;refGene_gn=SLC5A5;refGene_tx=XM_023247222.1;refGene_gn=SLC5A5;refGene_tx=XM_023247227.1;refGene_gn=SLC5A5;refGene_tx=XM_023247235.1
A2 13218219 13218271
refGene_gn=SLC5A5;refGene_tx=XM_011280067.3;refGene_gn=SLC5A5;refGene_tx=XM_023247222.1;refGene_gn=SLC5A5;refGene_tx=XM_023247227.1;refGene_gn=SLC5A5;refGene_tx=XM_023247234.1;refGene_gn=SLC5A5;refGene_tx=XM_023247235.1
A2 13218462 13218530
refGene_gn=SLC5A5;refGene_tx=XM_011280067.3;refGene_gn=SLC5A5;refGene_tx=XM_023247222.1;refGene_gn=SLC5A5;refGene_tx=XM_023247227.1;refGene_gn=SLC5A5;refGene_tx=XM_023247234.1;refGene_gn=SLC5A5;refGene_tx=XM_023247235.1
A2 13218828 13218983
refGene_gn=SLC5A5;refGene_tx=XM_011280067.3;refGene_gn=SLC5A5;refGene_tx=XM_023247222.1;refGene_gn=SLC5A5;refGene_tx=XM_023247227.1;refGene_gn=SLC5A5;refGene_tx=XM_023247234.1;refGene_gn=SLC5A5;refGene_tx=XM_023247235.1
A2 13219391 13219532
refGene_gn=SLC5A5;refGene_tx=XM_011280067.3;refGene_gn=SLC5A5;refGene_tx=XM_023247222.1;refGene_gn=SLC5A5;refGene_tx=XM_023247227.1;refGene_gn=SLC5A5;refGene_tx=XM_023247234.1;refGene_gn=SLC5A5;refGene_tx=XM_023247235.1
A2 13219613 13219743

refGene_gn=SLC5A5;refGene_tx=XM_011280067.3;refGene_gn=SLC5A5;refGene_tx=XM_023247222.1;refGene_gn=SLC5A5;refGene_tx=XM_023247227.1;refGene_gn=SLC5A5;refGene_tx=XM_023247234.1;refGene_gn=SLC5A5;refGene_tx=XM_023247235.1
A2 13220624 13220713
refGene_gn=SLC5A5;refGene_tx=XM_011280067.3;refGene_gn=SLC5A5;refGene_tx=XM_023247222.1;refGene_gn=SLC5A5;refGene_tx=XM_023247227.1;refGene_gn=SLC5A5;refGene_tx=XM_023247234.1;refGene_gn=SLC5A5;refGene_tx=XM_023247235.1
A2 13221329 13221442
refGene_gn=SLC5A5;refGene_tx=XM_011280067.3;refGene_gn=SLC5A5;refGene_tx=XM_023247222.1;refGene_gn=SLC5A5;refGene_tx=XM_023247227.1;refGene_gn=SLC5A5;refGene_tx=XM_023247234.1;refGene_gn=SLC5A5;refGene_tx=XM_023247235.1
A2 13221538 13221609
refGene_gn=SLC5A5;refGene_tx=XM_011280067.3;refGene_gn=SLC5A5;refGene_tx=XM_023247222.1;refGene_gn=SLC5A5;refGene_tx=XM_023247227.1;refGene_gn=SLC5A5;refGene_tx=XM_023247234.1;refGene_gn=SLC5A5;refGene_tx=XM_023247235.1
A2 13222039 13222126
refGene_gn=SLC5A5;refGene_tx=XM_011280067.3;refGene_gn=SLC5A5;refGene_tx=XM_023247222.1;refGene_gn=SLC5A5;refGene_tx=XM_023247227.1;refGene_gn=SLC5A5;refGene_tx=XM_023247234.1;refGene_gn=SLC5A5;refGene_tx=XM_023247235.1
A2 13222202 13222396
refGene_gn=SLC5A5;refGene_tx=XM_011280067.3;refGene_gn=SLC5A5;refGene_tx=XM_023247222.1;refGene_gn=SLC5A5;refGene_tx=XM_023247227.1;refGene_gn=SLC5A5;refGene_tx=XM_023247234.1;refGene_gn=SLC5A5;refGene_tx=XM_023247235.1
A2 13225718 13225843
refGene_gn=SLC5A5;refGene_tx=XM_011280067.3;refGene_gn=SLC5A5;refGene_tx=XM_023247222.1;refGene_gn=SLC5A5;refGene_tx=XM_023247227.1;refGene_gn=SLC5A5;refGene_tx=XM_023247234.1;refGene_gn=SLC5A5;refGene_tx=XM_023247235.1
A2 13226579 13226695
refGene_gn=SLC5A5;refGene_tx=XM_011280067.3;refGene_gn=SLC5A5;refGene_tx=XM_023247222.1;refGene_gn=SLC5A5;refGene_tx=XM_023247227.1;refGene_gn=SLC5A5;refGene_tx=XM_023247234.1
A2 13227726 13227891
refGene_gn=SLC5A5;refGene_tx=XM_011280067.3;refGene_gn=SLC5A5;refGene_tx=XM_023247222.1;refGene_gn=SLC5A5;refGene_tx=XM_023247227.1;refGene_gn=SLC5A5;refGene_tx=XM_023247234.1;refGene_gn=SLC5A5;refGene_tx=XM_023247235.1
A2 13248273 13248380 refGene_gn=CCDC124;refGene_tx=XM_003982071.3
A2 13253472 13253644 refGene_gn=CCDC124;refGene_tx=XM_003982071.3
A2 13258143 13258333 refGene_gn=CCDC124;refGene_tx=XM_003982071.3
A2 13258840 13258955 refGene_gn=CCDC124;refGene_tx=XM_003982071.3
A2 13259091 13259299 refGene_gn=CCDC124;refGene_tx=XM_003982071.3
A2 13278348 13278750
refGene_gn=KCNN1;refGene_tx=XM_023247240.1;refGene_gn=KCNN1;refGene_tx

=XM_023247244.1;refGene_gn=KCNN1;refGene_tx=XM_023247245.1;refGene_gn=KCNN1;refGene_tx=XM_023247247.1;refGene_gn=KCNN1;refGene_tx=XM_023247255.1;refGene_gn=KCNN1;refGene_tx=XM_023247256.1;refGene_gn=KCNN1;refGene_tx=XM_023247258.1;refGene_gn=KCNN1;refGene_tx=XM_023247265.1;refGene_gn=KCNN1;refGene_tx=XM_023247277.1
A2 13279506 13279602
refGene_gn=KCNN1;refGene_tx=XM_023247240.1;refGene_gn=KCNN1;refGene_tx=XM_023247244.1;refGene_gn=KCNN1;refGene_tx=XM_023247245.1;refGene_gn=KCNN1;refGene_tx=XM_023247247.1;refGene_gn=KCNN1;refGene_tx=XM_023247255.1;refGene_gn=KCNN1;refGene_tx=XM_023247256.1;refGene_gn=KCNN1;refGene_tx=XM_023247258.1;refGene_gn=KCNN1;refGene_tx=XM_023247265.1;refGene_gn=KCNN1;refGene_tx=XM_023247277.1
A2 13282080 13282499
refGene_gn=KCNN1;refGene_tx=XM_023247240.1;refGene_gn=KCNN1;refGene_tx=XM_023247244.1;refGene_gn=KCNN1;refGene_tx=XM_023247245.1;refGene_gn=KCNN1;refGene_tx=XM_023247247.1;refGene_gn=KCNN1;refGene_tx=XM_023247255.1;refGene_gn=KCNN1;refGene_tx=XM_023247256.1;refGene_gn=KCNN1;refGene_tx=XM_023247258.1;refGene_gn=KCNN1;refGene_tx=XM_023247265.1;refGene_gn=KCNN1;refGene_tx=XM_023247277.1
A2 13282915 13282924
refGene_gn=KCNN1;refGene_tx=XM_023247240.1;refGene_gn=KCNN1;refGene_tx=XM_023247245.1;refGene_gn=KCNN1;refGene_tx=XM_023247247.1;refGene_gn=KCNN1;refGene_tx=XM_023247255.1;refGene_gn=KCNN1;refGene_tx=XM_023247256.1;refGene_gn=KCNN1;refGene_tx=XM_023247258.1;refGene_gn=KCNN1;refGene_tx=XM_023247265.1
A2 13285301 13285443
refGene_gn=KCNN1;refGene_tx=XM_023247240.1;refGene_gn=KCNN1;refGene_tx=XM_023247244.1;refGene_gn=KCNN1;refGene_tx=XM_023247245.1;refGene_gn=KCNN1;refGene_tx=XM_023247247.1;refGene_gn=KCNN1;refGene_tx=XM_023247255.1;refGene_gn=KCNN1;refGene_tx=XM_023247256.1;refGene_gn=KCNN1;refGene_tx=XM_023247258.1;refGene_gn=KCNN1;refGene_tx=XM_023247265.1;refGene_gn=KCNN1;refGene_tx=XM_023247277.1
A2 13288728 13288840
refGene_gn=KCNN1;refGene_tx=XM_023247240.1;refGene_gn=KCNN1;refGene_tx=XM_023247244.1;refGene_gn=KCNN1;refGene_tx=XM_023247245.1;refGene_gn=KCNN1;refGene_tx=XM_023247247.1;refGene_gn=KCNN1;refGene_tx=XM_023247255.1;refGene_gn=KCNN1;refGene_tx=XM_023247256.1;refGene_gn=KCNN1;refGene_tx=XM_023247258.1;refGene_gn=KCNN1;refGene_tx=XM_023247265.1;refGene_gn=KCNN1;refGene_tx=XM_023247277.1
A2 13289440 13289567
refGene_gn=KCNN1;refGene_tx=XM_023247240.1;refGene_gn=KCNN1;refGene_tx=XM_023247244.1;refGene_gn=KCNN1;refGene_tx=XM_023247245.1;refGene_gn=KCNN1;refGene_tx=XM_023247247.1;refGene_gn=KCNN1;refGene_tx=XM_023247255.1;refGene_gn=KCNN1;refGene_tx=XM_023247256.1;refGene_gn=KCNN1;refGene_tx=XM_023247258.1;refGene_gn=KCNN1;refGene_tx=XM_023247265.1;refGene_gn=KCNN1;refGene_tx=XM_023247277.1
A2 13293728 13293737
refGene_gn=KCNN1;refGene_tx=XM_023247240.1;refGene_gn=KCNN1;refGene_tx=XM_023247244.1;refGene_gn=KCNN1;refGene_tx=XM_023247247.1;refGene_gn=KCNN1;refGene_tx=XM_023247256.1;refGene_gn=KCNN1;refGene_tx=XM_0232472

58.1;refGene_gn=KCNN1;refGene_tx=XM_023247265.1
A2 13294153 13294223
refGene_gn=KCNN1;refGene_tx=XM_023247240.1;refGene_gn=KCNN1;refGene_tx=XM_023247244.1;refGene_gn=KCNN1;refGene_tx=XM_023247245.1;refGene_gn=KCNN1;refGene_tx=XM_023247247.1;refGene_gn=KCNN1;refGene_tx=XM_023247255.1;refGene_gn=KCNN1;refGene_tx=XM_023247256.1;refGene_gn=KCNN1;refGene_tx=XM_023247258.1;refGene_gn=KCNN1;refGene_tx=XM_023247265.1;refGene_gn=KCNN1;refGene_tx=XM_023247277.1
A2 13294785 13294980
refGene_gn=KCNN1;refGene_tx=XM_023247247.1;refGene_gn=KCNN1;refGene_tx=XM_023247255.1
A2 13296613 13296868
refGene_gn=KCNN1;refGene_tx=XM_023247240.1;refGene_gn=KCNN1;refGene_tx=XM_023247244.1;refGene_gn=KCNN1;refGene_tx=XM_023247245.1;refGene_gn=KCNN1;refGene_tx=XM_023247256.1;refGene_gn=KCNN1;refGene_tx=XM_023247258.1;refGene_gn=KCNN1;refGene_tx=XM_023247265.1;refGene_gn=KCNN1;refGene_tx=XM_023247277.1
A2 13298977 13299236 refGene_gn=ARRDC2;refGene_tx=XM_023247295.1
A2 13301867 13302141
refGene_gn=ARRDC2;refGene_tx=XM_006928631.3;refGene_gn=ARRDC2;refGene_tx=XM_023247292.1
A2 13302234 13302301
refGene_gn=ARRDC2;refGene_tx=XM_006928631.3;refGene_gn=ARRDC2;refGene_tx=XM_023247292.1;refGene_gn=ARRDC2;refGene_tx=XM_023247295.1
A2 13302542 13302690
refGene_gn=ARRDC2;refGene_tx=XM_006928631.3;refGene_gn=ARRDC2;refGene_tx=XM_023247292.1;refGene_gn=ARRDC2;refGene_tx=XM_023247295.1
A2 13303210 13303313
refGene_gn=ARRDC2;refGene_tx=XM_006928631.3;refGene_gn=ARRDC2;refGene_tx=XM_023247292.1;refGene_gn=ARRDC2;refGene_tx=XM_023247295.1
A2 13303412 13303669
refGene_gn=ARRDC2;refGene_tx=XM_006928631.3;refGene_gn=ARRDC2;refGene_tx=XM_023247292.1;refGene_gn=ARRDC2;refGene_tx=XM_023247295.1
A2 13303782 13303945
refGene_gn=ARRDC2;refGene_tx=XM_006928631.3;refGene_gn=ARRDC2;refGene_tx=XM_023247292.1;refGene_gn=ARRDC2;refGene_tx=XM_023247295.1
A2 13304218 13304376
refGene_gn=ARRDC2;refGene_tx=XM_006928631.3;refGene_gn=ARRDC2;refGene_tx=XM_023247292.1;refGene_gn=ARRDC2;refGene_tx=XM_023247295.1
A2 13304619 13304673
refGene_gn=ARRDC2;refGene_tx=XM_006928631.3;refGene_gn=ARRDC2;refGene_tx=XM_023247292.1;refGene_gn=ARRDC2;refGene_tx=XM_023247295.1
A2 13332581 13332695 refGene_gn=IL12RB1;refGene_tx=XM_023241208.1
A2 13332696 13332885 refGene_gn=IL12RB1;refGene_tx=XM_023241208.1
A2 13333660 13333742 refGene_gn=IL12RB1;refGene_tx=XM_023241208.1
A2 13334032 13334135 refGene_gn=IL12RB1;refGene_tx=XM_023241208.1
A2 13335084 13335219 refGene_gn=IL12RB1;refGene_tx=XM_023241208.1
A2 13336752 13336908 refGene_gn=IL12RB1;refGene_tx=XM_023241208.1
A2 13337273 13337411 refGene_gn=IL12RB1;refGene_tx=XM_023241208.1
A2 13338111 13338273 refGene_gn=IL12RB1;refGene_tx=XM_023241208.1

A2 13340832 13341076 refGene_gn=IL12RB1;refGene_tx=XM_023241208.1
A2 13341939 13342022 refGene_gn=IL12RB1;refGene_tx=XM_023241208.1
A2 13342990 13343116 refGene_gn=IL12RB1;refGene_tx=XM_023241208.1
A2 13344018 13344052 refGene_gn=IL12RB1;refGene_tx=XM_023241208.1
A2 13345404 13345421 refGene_gn=IL12RB1;refGene_tx=XM_023241208.1
A2 13348515 13348748 refGene_gn=IL12RB1;refGene_tx=XM_023241208.1
A2 13349277 13349392 refGene_gn=IL12RB1;refGene_tx=XM_023241208.1
A2 13350240 13350303 refGene_gn=IL12RB1;refGene_tx=XM_023241208.1
A2 13351330 13351400 refGene_gn=IL12RB1;refGene_tx=XM_023241208.1
A2 13357842 13357881
refGene_gn=MAST3;refGene_tx=XM_019819687.2;refGene_gn=MAST3;refGene_tx=XM_019819706.2;refGene_gn=MAST3;refGene_tx=XM_023247329.1;refGene_gn=MAST3;refGene_tx=XM_023247333.1;refGene_gn=MAST3;refGene_tx=XM_023247337.1;refGene_gn=MAST3;refGene_tx=XM_023247346.1;refGene_gn=MAST3;refGene_tx=XM_023247348.1;refGene_gn=MAST3;refGene_tx=XM_023247352.1;refGene_gn=MAST3;refGene_tx=XM_023247353.1;refGene_gn=MAST3;refGene_tx=XM_023247357.1;refGene_gn=MAST3;refGene_tx=XM_023247363.1;refGene_gn=MAST3;refGene_tx=XM_023247370.1
A2 13363874 13363906
refGene_gn=MAST3;refGene_tx=XM_019819687.2;refGene_gn=MAST3;refGene_tx=XM_019819706.2;refGene_gn=MAST3;refGene_tx=XM_023247329.1;refGene_gn=MAST3;refGene_tx=XM_023247333.1;refGene_gn=MAST3;refGene_tx=XM_023247337.1;refGene_gn=MAST3;refGene_tx=XM_023247346.1;refGene_gn=MAST3;refGene_tx=XM_023247348.1;refGene_gn=MAST3;refGene_tx=XM_023247352.1;refGene_gn=MAST3;refGene_tx=XM_023247353.1;refGene_gn=MAST3;refGene_tx=XM_023247357.1;refGene_gn=MAST3;refGene_tx=XM_023247363.1;refGene_gn=MAST3;refGene_tx=XM_023247370.1
A2 13366049 13366096
refGene_gn=MAST3;refGene_tx=XM_006928610.4;refGene_gn=MAST3;refGene_tx=XM_019819694.2;refGene_gn=MAST3;refGene_tx=XM_023247385.1
A2 13366412 13366502
refGene_gn=MAST3;refGene_tx=XM_006928610.4;refGene_gn=MAST3;refGene_tx=XM_019819687.2;refGene_gn=MAST3;refGene_tx=XM_019819694.2;refGene_gn=MAST3;refGene_tx=XM_023247329.1;refGene_gn=MAST3;refGene_tx=XM_023247333.1;refGene_gn=MAST3;refGene_tx=XM_023247337.1;refGene_gn=MAST3;refGene_tx=XM_023247346.1;refGene_gn=MAST3;refGene_tx=XM_023247348.1;refGene_gn=MAST3;refGene_tx=XM_023247352.1;refGene_gn=MAST3;refGene_tx=XM_023247353.1;refGene_gn=MAST3;refGene_tx=XM_023247357.1;refGene_gn=MAST3;refGene_tx=XM_023247363.1;refGene_gn=MAST3;refGene_tx=XM_023247385.1
A2 13371446 13371535
refGene_gn=MAST3;refGene_tx=XM_023247372.1;refGene_gn=MAST3;refGene_tx=XM_023247374.1;refGene_gn=MAST3;refGene_tx=XM_023247390.1
A2 13374043 13374132
refGene_gn=MAST3;refGene_tx=XM_006928610.4;refGene_gn=MAST3;refGene_tx=XM_019819687.2;refGene_gn=MAST3;refGene_tx=XM_019819694.2;refGene_gn=MAST3;refGene_tx=XM_019819706.2;refGene_gn=MAST3;refGene_tx=XM_023247329.1;refGene_gn=MAST3;refGene_tx=XM_023247333.1;refGene_gn=MAST3;refGene_tx=XM_023247337.1;refGene_gn=MAST3;refGene_tx=XM_023247346.1;refGene_gn=MAST3;refGene_tx=XM_023247348.1;refGene_gn=MAST3;refGene_tx=XM_023247352.1;refGene_gn=MAST3;refGene_tx=XM_023247353.1;refGene_gn=MAST3;

refGene_tx=XM_023247357.1;refGene_gn=MAST3;refGene_tx=XM_023247363.1;refGene_gn=MAST3;refGene_tx=XM_023247370.1;refGene_gn=MAST3;refGene_tx=XM_023247372.1;refGene_gn=MAST3;refGene_tx=XM_023247374.1;refGene_gn=MAST3;refGene_tx=XM_023247385.1;refGene_gn=MAST3;refGene_tx=XM_023247390.1
A2 13374215 13374288
refGene_gn=MAST3;refGene_tx=XM_006928610.4;refGene_gn=MAST3;refGene_tx=XM_019819687.2;refGene_gn=MAST3;refGene_tx=XM_019819694.2;refGene_gn=MAST3;refGene_tx=XM_019819706.2;refGene_gn=MAST3;refGene_tx=XM_023247329.1;refGene_gn=MAST3;refGene_tx=XM_023247333.1;refGene_gn=MAST3;refGene_tx=XM_023247337.1;refGene_gn=MAST3;refGene_tx=XM_023247346.1;refGene_gn=MAST3;refGene_tx=XM_023247348.1;refGene_gn=MAST3;refGene_tx=XM_023247352.1;refGene_gn=MAST3;refGene_tx=XM_023247353.1;refGene_gn=MAST3;refGene_tx=XM_023247357.1;refGene_gn=MAST3;refGene_tx=XM_023247363.1;refGene_gn=MAST3;refGene_tx=XM_023247370.1;refGene_gn=MAST3;refGene_tx=XM_023247372.1;refGene_gn=MAST3;refGene_tx=XM_023247374.1;refGene_gn=MAST3;refGene_tx=XM_023247385.1;refGene_gn=MAST3;refGene_tx=XM_023247390.1
A2 13374466 13374487
refGene_gn=MAST3;refGene_tx=XM_019819687.2;refGene_gn=MAST3;refGene_tx=XM_019819694.2;refGene_gn=MAST3;refGene_tx=XM_019819706.2;refGene_gn=MAST3;refGene_tx=XM_019819713.2;refGene_gn=MAST3;refGene_tx=XM_023247329.1;refGene_gn=MAST3;refGene_tx=XM_023247333.1;refGene_gn=MAST3;refGene_tx=XM_023247348.1;refGene_gn=MAST3;refGene_tx=XM_023247352.1;refGene_gn=MAST3;refGene_tx=XM_023247370.1
A2 13375001 13375116
refGene_gn=MAST3;refGene_tx=XM_006928610.4;refGene_gn=MAST3;refGene_tx=XM_019819687.2;refGene_gn=MAST3;refGene_tx=XM_019819694.2;refGene_gn=MAST3;refGene_tx=XM_019819706.2;refGene_gn=MAST3;refGene_tx=XM_019819713.2;refGene_gn=MAST3;refGene_tx=XM_023247329.1;refGene_gn=MAST3;refGene_tx=XM_023247333.1;refGene_gn=MAST3;refGene_tx=XM_023247337.1;refGene_gn=MAST3;refGene_tx=XM_023247346.1;refGene_gn=MAST3;refGene_tx=XM_023247348.1;refGene_gn=MAST3;refGene_tx=XM_023247352.1;refGene_gn=MAST3;refGene_tx=XM_023247353.1;refGene_gn=MAST3;refGene_tx=XM_023247357.1;refGene_gn=MAST3;refGene_tx=XM_023247363.1;refGene_gn=MAST3;refGene_tx=XM_023247370.1;refGene_gn=MAST3;refGene_tx=XM_023247372.1;refGene_gn=MAST3;refGene_tx=XM_023247374.1;refGene_gn=MAST3;refGene_tx=XM_023247385.1;refGene_gn=MAST3;refGene_tx=XM_023247390.1
A2 13375526 13375687
refGene_gn=MAST3;refGene_tx=XM_006928610.4;refGene_gn=MAST3;refGene_tx=XM_019819687.2;refGene_gn=MAST3;refGene_tx=XM_019819694.2;refGene_gn=MAST3;refGene_tx=XM_019819706.2;refGene_gn=MAST3;refGene_tx=XM_019819713.2;refGene_gn=MAST3;refGene_tx=XM_023247329.1;refGene_gn=MAST3;refGene_tx=XM_023247333.1;refGene_gn=MAST3;refGene_tx=XM_023247337.1;refGene_gn=MAST3;refGene_tx=XM_023247346.1;refGene_gn=MAST3;refGene_tx=XM_023247348.1;refGene_gn=MAST3;refGene_tx=XM_023247352.1;refGene_gn=MAST3;refGene_tx=XM_023247353.1;refGene_gn=MAST3;refGene_tx=XM_023247357.1;refGene_gn=MAST3;refGene_tx=XM_023247363.1;refGene_gn=MAST3;refGene_tx=XM_023247370.1;refGene_gn=MAST3;refGene_tx=XM_023247372.1;refGene_gn=MAST3;refGene_tx=XM_023247374.1;refGene_gn=MAST3;refGene_tx=XM_02324738

5.1;refGene_gn=MAST3;refGene_tx=XM_023247390.1
A2 13375872 13375948
refGene_gn=MAST3;refGene_tx=XM_006928610.4;refGene_gn=MAST3;refGene_tx=XM_019819687.2;refGene_gn=MAST3;refGene_tx=XM_019819694.2;refGene_gn=MAST3;refGene_tx=XM_019819706.2;refGene_gn=MAST3;refGene_tx=XM_019819713.2;refGene_gn=MAST3;refGene_tx=XM_023247329.1;refGene_gn=MAST3;refGene_tx=XM_023247333.1;refGene_gn=MAST3;refGene_tx=XM_023247337.1;refGene_gn=MAST3;refGene_tx=XM_023247346.1;refGene_gn=MAST3;refGene_tx=XM_023247348.1;refGene_gn=MAST3;refGene_tx=XM_023247352.1;refGene_gn=MAST3;refGene_tx=XM_023247353.1;refGene_gn=MAST3;refGene_tx=XM_023247357.1;refGene_gn=MAST3;refGene_tx=XM_023247363.1;refGene_gn=MAST3;refGene_tx=XM_023247370.1;refGene_gn=MAST3;refGene_tx=XM_023247372.1;refGene_gn=MAST3;refGene_tx=XM_023247374.1;refGene_gn=MAST3;refGene_tx=XM_023247385.1;refGene_gn=MAST3;refGene_tx=XM_023247390.1
A2 13376255 13376465
refGene_gn=MAST3;refGene_tx=XM_006928610.4;refGene_gn=MAST3;refGene_tx=XM_019819687.2;refGene_gn=MAST3;refGene_tx=XM_019819694.2;refGene_gn=MAST3;refGene_tx=XM_019819706.2;refGene_gn=MAST3;refGene_tx=XM_019819713.2;refGene_gn=MAST3;refGene_tx=XM_023247329.1;refGene_gn=MAST3;refGene_tx=XM_023247333.1;refGene_gn=MAST3;refGene_tx=XM_023247337.1;refGene_gn=MAST3;refGene_tx=XM_023247346.1;refGene_gn=MAST3;refGene_tx=XM_023247348.1;refGene_gn=MAST3;refGene_tx=XM_023247352.1;refGene_gn=MAST3;refGene_tx=XM_023247353.1;refGene_gn=MAST3;refGene_tx=XM_023247357.1;refGene_gn=MAST3;refGene_tx=XM_023247363.1;refGene_gn=MAST3;refGene_tx=XM_023247370.1;refGene_gn=MAST3;refGene_tx=XM_023247372.1;refGene_gn=MAST3;refGene_tx=XM_023247374.1;refGene_gn=MAST3;refGene_tx=XM_023247385.1;refGene_gn=MAST3;refGene_tx=XM_023247390.1
A2 13376557 13376659
refGene_gn=MAST3;refGene_tx=XM_006928610.4;refGene_gn=MAST3;refGene_tx=XM_019819687.2;refGene_gn=MAST3;refGene_tx=XM_019819694.2;refGene_gn=MAST3;refGene_tx=XM_019819706.2;refGene_gn=MAST3;refGene_tx=XM_019819713.2;refGene_gn=MAST3;refGene_tx=XM_023247329.1;refGene_gn=MAST3;refGene_tx=XM_023247333.1;refGene_gn=MAST3;refGene_tx=XM_023247337.1;refGene_gn=MAST3;refGene_tx=XM_023247346.1;refGene_gn=MAST3;refGene_tx=XM_023247348.1;refGene_gn=MAST3;refGene_tx=XM_023247352.1;refGene_gn=MAST3;refGene_tx=XM_023247353.1;refGene_gn=MAST3;refGene_tx=XM_023247357.1;refGene_gn=MAST3;refGene_tx=XM_023247363.1;refGene_gn=MAST3;refGene_tx=XM_023247370.1;refGene_gn=MAST3;refGene_tx=XM_023247372.1;refGene_gn=MAST3;refGene_tx=XM_023247374.1;refGene_gn=MAST3;refGene_tx=XM_023247385.1;refGene_gn=MAST3;refGene_tx=XM_023247390.1
A2 13376942 13377075
refGene_gn=MAST3;refGene_tx=XM_006928610.4;refGene_gn=MAST3;refGene_tx=XM_019819687.2;refGene_gn=MAST3;refGene_tx=XM_019819694.2;refGene_gn=MAST3;refGene_tx=XM_019819706.2;refGene_gn=MAST3;refGene_tx=XM_019819713.2;refGene_gn=MAST3;refGene_tx=XM_023247329.1;refGene_gn=MAST3;refGene_tx=XM_023247333.1;refGene_gn=MAST3;refGene_tx=XM_023247337.1;refGene_gn=MAST3;refGene_tx=XM_023247346.1;refGene_gn=MAST3;refGene_tx=XM_023247348.1;refGene_gn=MAST3;refGene_tx=XM_023247352.1;refGene_gn=MAST3;refGene_tx=XM_023247353.1;refGene_gn=MAST3;refGene_tx=XM_023247357.1;refGene_gn=MAST3;refGene_tx=XM_023247363.1;refGene_gn=MAST3;refGene_tx=

XM_023247370.1;refGene_gn=MAST3;refGene_tx=XM_023247372.1;refGene_gn=MAST3;refGene_tx=XM_023247374.1;refGene_gn=MAST3;refGene_tx=XM_023247385.1;refGene_gn=MAST3;refGene_tx=XM_023247390.1
A2 13378540 13378602
refGene_gn=MAST3;refGene_tx=XM_006928610.4;refGene_gn=MAST3;refGene_tx=XM_019819687.2;refGene_gn=MAST3;refGene_tx=XM_019819694.2;refGene_gn=MAST3;refGene_tx=XM_019819706.2;refGene_gn=MAST3;refGene_tx=XM_019819713.2;refGene_gn=MAST3;refGene_tx=XM_023247329.1;refGene_gn=MAST3;refGene_tx=XM_023247333.1;refGene_gn=MAST3;refGene_tx=XM_023247337.1;refGene_gn=MAST3;refGene_tx=XM_023247346.1;refGene_gn=MAST3;refGene_tx=XM_023247348.1;refGene_gn=MAST3;refGene_tx=XM_023247352.1;refGene_gn=MAST3;refGene_tx=XM_023247353.1;refGene_gn=MAST3;refGene_tx=XM_023247357.1;refGene_gn=MAST3;refGene_tx=XM_023247363.1;refGene_gn=MAST3;refGene_tx=XM_023247370.1;refGene_gn=MAST3;refGene_tx=XM_023247372.1;refGene_gn=MAST3;refGene_tx=XM_023247374.1;refGene_gn=MAST3;refGene_tx=XM_023247385.1;refGene_gn=MAST3;refGene_tx=XM_023247390.1
A2 13379002 13379088
refGene_gn=MAST3;refGene_tx=XM_006928610.4;refGene_gn=MAST3;refGene_tx=XM_019819687.2;refGene_gn=MAST3;refGene_tx=XM_019819694.2;refGene_gn=MAST3;refGene_tx=XM_019819706.2;refGene_gn=MAST3;refGene_tx=XM_019819713.2;refGene_gn=MAST3;refGene_tx=XM_023247329.1;refGene_gn=MAST3;refGene_tx=XM_023247333.1;refGene_gn=MAST3;refGene_tx=XM_023247337.1;refGene_gn=MAST3;refGene_tx=XM_023247346.1;refGene_gn=MAST3;refGene_tx=XM_023247348.1;refGene_gn=MAST3;refGene_tx=XM_023247352.1;refGene_gn=MAST3;refGene_tx=XM_023247353.1;refGene_gn=MAST3;refGene_tx=XM_023247357.1;refGene_gn=MAST3;refGene_tx=XM_023247363.1;refGene_gn=MAST3;refGene_tx=XM_023247370.1;refGene_gn=MAST3;refGene_tx=XM_023247372.1;refGene_gn=MAST3;refGene_tx=XM_023247374.1;refGene_gn=MAST3;refGene_tx=XM_023247385.1;refGene_gn=MAST3;refGene_tx=XM_023247390.1
A2 13379902 13380111
refGene_gn=MAST3;refGene_tx=XM_006928610.4;refGene_gn=MAST3;refGene_tx=XM_019819687.2;refGene_gn=MAST3;refGene_tx=XM_019819694.2;refGene_gn=MAST3;refGene_tx=XM_019819706.2;refGene_gn=MAST3;refGene_tx=XM_019819713.2;refGene_gn=MAST3;refGene_tx=XM_023247329.1;refGene_gn=MAST3;refGene_tx=XM_023247333.1;refGene_gn=MAST3;refGene_tx=XM_023247337.1;refGene_gn=MAST3;refGene_tx=XM_023247346.1;refGene_gn=MAST3;refGene_tx=XM_023247348.1;refGene_gn=MAST3;refGene_tx=XM_023247352.1;refGene_gn=MAST3;refGene_tx=XM_023247353.1;refGene_gn=MAST3;refGene_tx=XM_023247357.1;refGene_gn=MAST3;refGene_tx=XM_023247363.1;refGene_gn=MAST3;refGene_tx=XM_023247370.1;refGene_gn=MAST3;refGene_tx=XM_023247372.1;refGene_gn=MAST3;refGene_tx=XM_023247374.1;refGene_gn=MAST3;refGene_tx=XM_023247385.1;refGene_gn=MAST3;refGene_tx=XM_023247390.1
A2 13380792 13380931
refGene_gn=MAST3;refGene_tx=XM_006928610.4;refGene_gn=MAST3;refGene_tx=XM_019819687.2;refGene_gn=MAST3;refGene_tx=XM_019819694.2;refGene_gn=MAST3;refGene_tx=XM_019819706.2;refGene_gn=MAST3;refGene_tx=XM_019819713.2;refGene_gn=MAST3;refGene_tx=XM_023247329.1;refGene_gn=MAST3;refGene_tx=XM_023247333.1;refGene_gn=MAST3;refGene_tx=XM_023247337.1;refGene_gn=MAST3;refGene_tx=XM_023247346.1;refGene_gn=MAST3;refGene_tx=XM_023247348.1;refGene_gn=MAST3;refGene_tx=XM_023247352.1;refGene_gn=MAST3;

refGene_tx=XM_023247353.1;refGene_gn=MAST3;refGene_tx=XM_023247357.1;refGene_gn=MAST3;refGene_tx=XM_023247363.1;refGene_gn=MAST3;refGene_tx=XM_023247370.1;refGene_gn=MAST3;refGene_tx=XM_023247372.1;refGene_gn=MAST3;refGene_tx=XM_023247374.1;refGene_gn=MAST3;refGene_tx=XM_023247385.1;refGene_gn=MAST3;refGene_tx=XM_023247390.1
A2 13383592 13383725
refGene_gn=MAST3;refGene_tx=XM_006928610.4;refGene_gn=MAST3;refGene_tx=XM_019819687.2;refGene_gn=MAST3;refGene_tx=XM_019819694.2;refGene_gn=MAST3;refGene_tx=XM_019819706.2;refGene_gn=MAST3;refGene_tx=XM_019819713.2;refGene_gn=MAST3;refGene_tx=XM_023247329.1;refGene_gn=MAST3;refGene_tx=XM_023247333.1;refGene_gn=MAST3;refGene_tx=XM_023247337.1;refGene_gn=MAST3;refGene_tx=XM_023247346.1;refGene_gn=MAST3;refGene_tx=XM_023247348.1;refGene_gn=MAST3;refGene_tx=XM_023247352.1;refGene_gn=MAST3;refGene_tx=XM_023247353.1;refGene_gn=MAST3;refGene_tx=XM_023247357.1;refGene_gn=MAST3;refGene_tx=XM_023247363.1;refGene_gn=MAST3;refGene_tx=XM_023247370.1;refGene_gn=MAST3;refGene_tx=XM_023247372.1;refGene_gn=MAST3;refGene_tx=XM_023247374.1;refGene_gn=MAST3;refGene_tx=XM_023247385.1;refGene_gn=MAST3;refGene_tx=XM_023247390.1
A2 13383963 13384129
refGene_gn=MAST3;refGene_tx=XM_006928610.4;refGene_gn=MAST3;refGene_tx=XM_019819687.2;refGene_gn=MAST3;refGene_tx=XM_019819694.2;refGene_gn=MAST3;refGene_tx=XM_019819706.2;refGene_gn=MAST3;refGene_tx=XM_019819713.2;refGene_gn=MAST3;refGene_tx=XM_023247329.1;refGene_gn=MAST3;refGene_tx=XM_023247333.1;refGene_gn=MAST3;refGene_tx=XM_023247337.1;refGene_gn=MAST3;refGene_tx=XM_023247346.1;refGene_gn=MAST3;refGene_tx=XM_023247348.1;refGene_gn=MAST3;refGene_tx=XM_023247352.1;refGene_gn=MAST3;refGene_tx=XM_023247353.1;refGene_gn=MAST3;refGene_tx=XM_023247357.1;refGene_gn=MAST3;refGene_tx=XM_023247363.1;refGene_gn=MAST3;refGene_tx=XM_023247370.1;refGene_gn=MAST3;refGene_tx=XM_023247372.1;refGene_gn=MAST3;refGene_tx=XM_023247374.1;refGene_gn=MAST3;refGene_tx=XM_023247385.1;refGene_gn=MAST3;refGene_tx=XM_023247390.1
A2 13384733 13384835
refGene_gn=MAST3;refGene_tx=XM_006928610.4;refGene_gn=MAST3;refGene_tx=XM_019819687.2;refGene_gn=MAST3;refGene_tx=XM_019819694.2;refGene_gn=MAST3;refGene_tx=XM_019819706.2;refGene_gn=MAST3;refGene_tx=XM_019819713.2;refGene_gn=MAST3;refGene_tx=XM_023247329.1;refGene_gn=MAST3;refGene_tx=XM_023247333.1;refGene_gn=MAST3;refGene_tx=XM_023247337.1;refGene_gn=MAST3;refGene_tx=XM_023247346.1;refGene_gn=MAST3;refGene_tx=XM_023247348.1;refGene_gn=MAST3;refGene_tx=XM_023247352.1;refGene_gn=MAST3;refGene_tx=XM_023247353.1;refGene_gn=MAST3;refGene_tx=XM_023247357.1;refGene_gn=MAST3;refGene_tx=XM_023247363.1;refGene_gn=MAST3;refGene_tx=XM_023247370.1;refGene_gn=MAST3;refGene_tx=XM_023247372.1;refGene_gn=MAST3;refGene_tx=XM_023247374.1;refGene_gn=MAST3;refGene_tx=XM_023247385.1;refGene_gn=MAST3;refGene_tx=XM_023247390.1
A2 13385468 13385591
refGene_gn=MAST3;refGene_tx=XM_006928610.4;refGene_gn=MAST3;refGene_tx=XM_019819687.2;refGene_gn=MAST3;refGene_tx=XM_019819694.2;refGene_gn=MAST3;refGene_tx=XM_019819706.2;refGene_gn=MAST3;refGene_tx=XM_019819713.2;refGene_gn=MAST3;refGene_tx=XM_023247329.1;refGene_gn=MAST3;refGene_tx=XM_023247333.1;refGene_gn=MAST3;refGene_tx=XM_023247337.1;refGen

e_gn=MAST3;refGene_tx=XM_023247346.1;refGene_gn=MAST3;refGene_tx=XM_023247348.1;refGene_gn=MAST3;refGene_tx=XM_023247352.1;refGene_gn=MAST3;refGene_tx=XM_023247353.1;refGene_gn=MAST3;refGene_tx=XM_023247357.1;refGene_gn=MAST3;refGene_tx=XM_023247363.1;refGene_gn=MAST3;refGene_tx=XM_023247370.1;refGene_gn=MAST3;refGene_tx=XM_023247372.1;refGene_gn=MAST3;refGene_tx=XM_023247374.1;refGene_gn=MAST3;refGene_tx=XM_023247385.1;refGene_gn=MAST3;refGene_tx=XM_023247390.1
A2 13386472 13386582
refGene_gn=MAST3;refGene_tx=XM_006928610.4;refGene_gn=MAST3;refGene_tx=XM_019819687.2;refGene_gn=MAST3;refGene_tx=XM_019819694.2;refGene_gn=MAST3;refGene_tx=XM_019819706.2;refGene_gn=MAST3;refGene_tx=XM_019819713.2;refGene_gn=MAST3;refGene_tx=XM_023247329.1;refGene_gn=MAST3;refGene_tx=XM_023247333.1;refGene_gn=MAST3;refGene_tx=XM_023247337.1;refGene_gn=MAST3;refGene_tx=XM_023247346.1;refGene_gn=MAST3;refGene_tx=XM_023247348.1;refGene_gn=MAST3;refGene_tx=XM_023247352.1;refGene_gn=MAST3;refGene_tx=XM_023247353.1;refGene_gn=MAST3;refGene_tx=XM_023247357.1;refGene_gn=MAST3;refGene_tx=XM_023247363.1;refGene_gn=MAST3;refGene_tx=XM_023247370.1;refGene_gn=MAST3;refGene_tx=XM_023247372.1;refGene_gn=MAST3;refGene_tx=XM_023247374.1;refGene_gn=MAST3;refGene_tx=XM_023247385.1;refGene_gn=MAST3;refGene_tx=XM_023247390.1
A2 13387908 13388045
refGene_gn=MAST3;refGene_tx=XM_006928610.4;refGene_gn=MAST3;refGene_tx=XM_019819687.2;refGene_gn=MAST3;refGene_tx=XM_019819694.2;refGene_gn=MAST3;refGene_tx=XM_019819706.2;refGene_gn=MAST3;refGene_tx=XM_019819713.2;refGene_gn=MAST3;refGene_tx=XM_023247329.1;refGene_gn=MAST3;refGene_tx=XM_023247333.1;refGene_gn=MAST3;refGene_tx=XM_023247337.1;refGene_gn=MAST3;refGene_tx=XM_023247346.1;refGene_gn=MAST3;refGene_tx=XM_023247348.1;refGene_gn=MAST3;refGene_tx=XM_023247352.1;refGene_gn=MAST3;refGene_tx=XM_023247353.1;refGene_gn=MAST3;refGene_tx=XM_023247357.1;refGene_gn=MAST3;refGene_tx=XM_023247363.1;refGene_gn=MAST3;refGene_tx=XM_023247370.1;refGene_gn=MAST3;refGene_tx=XM_023247372.1;refGene_gn=MAST3;refGene_tx=XM_023247374.1;refGene_gn=MAST3;refGene_tx=XM_023247385.1;refGene_gn=MAST3;refGene_tx=XM_023247390.1
A2 13388569 13388814
refGene_gn=MAST3;refGene_tx=XM_006928610.4;refGene_gn=MAST3;refGene_tx=XM_019819687.2;refGene_gn=MAST3;refGene_tx=XM_019819694.2;refGene_gn=MAST3;refGene_tx=XM_019819706.2;refGene_gn=MAST3;refGene_tx=XM_019819713.2;refGene_gn=MAST3;refGene_tx=XM_023247329.1;refGene_gn=MAST3;refGene_tx=XM_023247333.1;refGene_gn=MAST3;refGene_tx=XM_023247337.1;refGene_gn=MAST3;refGene_tx=XM_023247346.1;refGene_gn=MAST3;refGene_tx=XM_023247348.1;refGene_gn=MAST3;refGene_tx=XM_023247352.1;refGene_gn=MAST3;refGene_tx=XM_023247353.1;refGene_gn=MAST3;refGene_tx=XM_023247357.1;refGene_gn=MAST3;refGene_tx=XM_023247363.1;refGene_gn=MAST3;refGene_tx=XM_023247370.1;refGene_gn=MAST3;refGene_tx=XM_023247372.1;refGene_gn=MAST3;refGene_tx=XM_023247374.1;refGene_gn=MAST3;refGene_tx=XM_023247385.1;refGene_gn=MAST3;refGene_tx=XM_023247390.1
A2 13389306 13389513
refGene_gn=MAST3;refGene_tx=XM_006928610.4;refGene_gn=MAST3;refGene_tx=XM_019819687.2;refGene_gn=MAST3;refGene_tx=XM_019819694.2;refGene_gn=MAST3;refGene_tx=XM_019819706.2;refGene_gn=MAST3;refGene_tx=XM_0198197

13.2;refGene_gn=MAST3;refGene_tx=XM_023247329.1;refGene_gn=MAST3;refGene_tx=XM_023247333.1;refGene_gn=MAST3;refGene_tx=XM_023247337.1;refGene_gn=MAST3;refGene_tx=XM_023247346.1;refGene_gn=MAST3;refGene_tx=XM_023247348.1;refGene_gn=MAST3;refGene_tx=XM_023247352.1;refGene_gn=MAST3;refGene_tx=XM_023247353.1;refGene_gn=MAST3;refGene_tx=XM_023247357.1;refGene_gn=MAST3;refGene_tx=XM_023247363.1;refGene_gn=MAST3;refGene_tx=XM_023247370.1;refGene_gn=MAST3;refGene_tx=XM_023247372.1;refGene_gn=MAST3;refGene_tx=XM_023247374.1;refGene_gn=MAST3;refGene_tx=XM_023247385.1;refGene_gn=MAST3;refGene_tx=XM_023247390.1
A2 13389807 13390034
refGene_gn=MAST3;refGene_tx=XM_006928610.4;refGene_gn=MAST3;refGene_tx=XM_019819687.2;refGene_gn=MAST3;refGene_tx=XM_019819694.2;refGene_gn=MAST3;refGene_tx=XM_019819706.2;refGene_gn=MAST3;refGene_tx=XM_019819713.2;refGene_gn=MAST3;refGene_tx=XM_023247329.1;refGene_gn=MAST3;refGene_tx=XM_023247333.1;refGene_gn=MAST3;refGene_tx=XM_023247337.1;refGene_gn=MAST3;refGene_tx=XM_023247346.1;refGene_gn=MAST3;refGene_tx=XM_023247348.1;refGene_gn=MAST3;refGene_tx=XM_023247352.1;refGene_gn=MAST3;refGene_tx=XM_023247353.1;refGene_gn=MAST3;refGene_tx=XM_023247357.1;refGene_gn=MAST3;refGene_tx=XM_023247363.1;refGene_gn=MAST3;refGene_tx=XM_023247370.1;refGene_gn=MAST3;refGene_tx=XM_023247372.1;refGene_gn=MAST3;refGene_tx=XM_023247374.1;refGene_gn=MAST3;refGene_tx=XM_023247385.1;refGene_gn=MAST3;refGene_tx=XM_023247390.1
A2 13390489 13390612
refGene_gn=MAST3;refGene_tx=XM_006928610.4;refGene_gn=MAST3;refGene_tx=XM_019819687.2;refGene_gn=MAST3;refGene_tx=XM_019819694.2;refGene_gn=MAST3;refGene_tx=XM_019819706.2;refGene_gn=MAST3;refGene_tx=XM_019819713.2;refGene_gn=MAST3;refGene_tx=XM_023247329.1;refGene_gn=MAST3;refGene_tx=XM_023247333.1;refGene_gn=MAST3;refGene_tx=XM_023247337.1;refGene_gn=MAST3;refGene_tx=XM_023247346.1;refGene_gn=MAST3;refGene_tx=XM_023247348.1;refGene_gn=MAST3;refGene_tx=XM_023247352.1;refGene_gn=MAST3;refGene_tx=XM_023247353.1;refGene_gn=MAST3;refGene_tx=XM_023247357.1;refGene_gn=MAST3;refGene_tx=XM_023247363.1;refGene_gn=MAST3;refGene_tx=XM_023247370.1;refGene_gn=MAST3;refGene_tx=XM_023247372.1;refGene_gn=MAST3;refGene_tx=XM_023247374.1;refGene_gn=MAST3;refGene_tx=XM_023247385.1;refGene_gn=MAST3;refGene_tx=XM_023247390.1
A2 13391218 13391355
refGene_gn=MAST3;refGene_tx=XM_006928610.4;refGene_gn=MAST3;refGene_tx=XM_019819687.2;refGene_gn=MAST3;refGene_tx=XM_019819694.2;refGene_gn=MAST3;refGene_tx=XM_019819706.2;refGene_gn=MAST3;refGene_tx=XM_019819713.2;refGene_gn=MAST3;refGene_tx=XM_023247329.1;refGene_gn=MAST3;refGene_tx=XM_023247333.1;refGene_gn=MAST3;refGene_tx=XM_023247337.1;refGene_gn=MAST3;refGene_tx=XM_023247346.1;refGene_gn=MAST3;refGene_tx=XM_023247348.1;refGene_gn=MAST3;refGene_tx=XM_023247352.1;refGene_gn=MAST3;refGene_tx=XM_023247353.1;refGene_gn=MAST3;refGene_tx=XM_023247357.1;refGene_gn=MAST3;refGene_tx=XM_023247363.1;refGene_gn=MAST3;refGene_tx=XM_023247370.1;refGene_gn=MAST3;refGene_tx=XM_023247372.1;refGene_gn=MAST3;refGene_tx=XM_023247374.1;refGene_gn=MAST3;refGene_tx=XM_023247385.1;refGene_gn=MAST3;refGene_tx=XM_023247390.1
A2 13391460 13391645
refGene_gn=MAST3;refGene_tx=XM_006928610.4;refGene_gn=MAST3;refGene_tx

=XM_019819687.2;refGene_gn=MAST3;refGene_tx=XM_019819694.2;refGene_gn=MAST3;refGene_tx=XM_019819706.2;refGene_gn=MAST3;refGene_tx=XM_019819713.2;refGene_gn=MAST3;refGene_tx=XM_023247329.1;refGene_gn=MAST3;refGene_tx=XM_023247333.1;refGene_gn=MAST3;refGene_tx=XM_023247337.1;refGene_gn=MAST3;refGene_tx=XM_023247346.1;refGene_gn=MAST3;refGene_tx=XM_023247348.1;refGene_gn=MAST3;refGene_tx=XM_023247352.1;refGene_gn=MAST3;refGene_tx=XM_023247353.1;refGene_gn=MAST3;refGene_tx=XM_023247357.1;refGene_gn=MAST3;refGene_tx=XM_023247363.1;refGene_gn=MAST3;refGene_tx=XM_023247370.1;refGene_gn=MAST3;refGene_tx=XM_023247372.1;refGene_gn=MAST3;refGene_tx=XM_023247374.1;refGene_gn=MAST3;refGene_tx=XM_023247385.1;refGene_gn=MAST3;refGene_tx=XM_023247390.1
A2 13392613 13393146
refGene_gn=MAST3;refGene_tx=XM_006928610.4;refGene_gn=MAST3;refGene_tx=XM_019819687.2;refGene_gn=MAST3;refGene_tx=XM_019819694.2;refGene_gn=MAST3;refGene_tx=XM_019819706.2;refGene_gn=MAST3;refGene_tx=XM_019819713.2;refGene_gn=MAST3;refGene_tx=XM_023247329.1;refGene_gn=MAST3;refGene_tx=XM_023247333.1;refGene_gn=MAST3;refGene_tx=XM_023247337.1;refGene_gn=MAST3;refGene_tx=XM_023247346.1;refGene_gn=MAST3;refGene_tx=XM_023247348.1;refGene_gn=MAST3;refGene_tx=XM_023247352.1;refGene_gn=MAST3;refGene_tx=XM_023247353.1;refGene_gn=MAST3;refGene_tx=XM_023247357.1;refGene_gn=MAST3;refGene_tx=XM_023247363.1;refGene_gn=MAST3;refGene_tx=XM_023247370.1;refGene_gn=MAST3;refGene_tx=XM_023247372.1;refGene_gn=MAST3;refGene_tx=XM_023247374.1;refGene_gn=MAST3;refGene_tx=XM_023247385.1;refGene_gn=MAST3;refGene_tx=XM_023247390.1
A2 13398020 13398342 refGene_gn=PIK3R2;refGene_tx=XM_023247400.1
A2 13401138 13401231 refGene_gn=PIK3R2;refGene_tx=XM_023247400.1
A2 13401596 13401647 refGene_gn=PIK3R2;refGene_tx=XM_023247400.1
A2 13401736 13401868 refGene_gn=PIK3R2;refGene_tx=XM_023247400.1
A2 13401957 13402168 refGene_gn=PIK3R2;refGene_tx=XM_023247400.1
A2 13402660 13402746 refGene_gn=PIK3R2;refGene_tx=XM_023247400.1
A2 13402904 13403013 refGene_gn=PIK3R2;refGene_tx=XM_023247400.1
A2 13403100 13403199 refGene_gn=PIK3R2;refGene_tx=XM_023247400.1
A2 13403357 13403538 refGene_gn=PIK3R2;refGene_tx=XM_023247400.1
A2 13403622 13403748 refGene_gn=PIK3R2;refGene_tx=XM_023247400.1
A2 13405238 13405381 refGene_gn=PIK3R2;refGene_tx=XM_023247400.1
A2 13405614 13405791 refGene_gn=PIK3R2;refGene_tx=XM_023247400.1
A2 13406724 13406796 refGene_gn=PIK3R2;refGene_tx=XM_023247400.1
A2 13406969 13407140 refGene_gn=PIK3R2;refGene_tx=XM_023247400.1
A2 13407508 13407716 refGene_gn=PIK3R2;refGene_tx=XM_023247400.1
A2 13411290 13411419 refGene_gn=IFI30;refGene_tx=XM_023247408.1
A2 13412418 13412601 refGene_gn=IFI30;refGene_tx=XM_023247408.1
A2 13412683 13412758 refGene_gn=IFI30;refGene_tx=XM_023247408.1
A2 13413006 13413099 refGene_gn=IFI30;refGene_tx=XM_023247408.1
A2 13413926 13414013 refGene_gn=IFI30;refGene_tx=XM_023247408.1
A2 13414014 13414080 refGene_gn=IFI30;refGene_tx=XM_023247408.1
A2 13414264 13414318 refGene_gn=IFI30;refGene_tx=XM_023247408.1
A2 13414402 13414465 refGene_gn=IFI30;refGene_tx=XM_023247408.1
A2 13425537 13425724
refGene_gn=MPV17L2;refGene_tx=XM_011290730.3;refGene_gn=MPV17L2;refGene_tx=XM_011290735.3;refGene_gn=MPV17L2;refGene_tx=XM_011290739.3

A2 13426380 13426551
refGene_gn=MPV17L2;refGene_tx=XM_011290730.3;refGene_gn=MPV17L2;refGene_tx=XM_011290735.3;refGene_gn=MPV17L2;refGene_tx=XM_011290739.3
A2 13427265 13427342
refGene_gn=MPV17L2;refGene_tx=XM_011290730.3;refGene_gn=MPV17L2;refGene_tx=XM_011290735.3;refGene_gn=MPV17L2;refGene_tx=XM_011290739.3
A2 13427434 13427620
refGene_gn=MPV17L2;refGene_tx=XM_011290730.3;refGene_gn=MPV17L2;refGene_tx=XM_011290735.3;refGene_gn=MPV17L2;refGene_tx=XM_011290739.3
A2 13427766 13427796 refGene_gn=MPV17L2;refGene_tx=XM_011290739.3
A2 13428756 13428947 refGene_gn=RAB3A;refGene_tx=XM_003982073.5
A2 13429818 13429943 refGene_gn=RAB3A;refGene_tx=XM_003982073.5
A2 13430524 13430643 refGene_gn=RAB3A;refGene_tx=XM_003982073.5
A2 13431775 13432003 refGene_gn=RAB3A;refGene_tx=XM_003982073.5
A2 13438750 13439149
refGene_gn=PDE4C;refGene_tx=XM_006928616.3;refGene_gn=PDE4C;refGene_tx=XM_006928617.3;refGene_gn=PDE4C;refGene_tx=XM_019819750.2;refGene_gn=PDE4C;refGene_tx=XM_019819753.2
A2 13439717 13439900
refGene_gn=PDE4C;refGene_tx=XM_006928616.3;refGene_gn=PDE4C;refGene_tx=XM_006928617.3;refGene_gn=PDE4C;refGene_tx=XM_019819750.2;refGene_gn=PDE4C;refGene_tx=XM_019819753.2
A2 13440158 13440281
refGene_gn=PDE4C;refGene_tx=XM_006928616.3;refGene_gn=PDE4C;refGene_tx=XM_006928617.3;refGene_gn=PDE4C;refGene_tx=XM_019819750.2;refGene_gn=PDE4C;refGene_tx=XM_019819753.2
A2 13441755 13441910
refGene_gn=PDE4C;refGene_tx=XM_006928616.3;refGene_gn=PDE4C;refGene_tx=XM_006928617.3;refGene_gn=PDE4C;refGene_tx=XM_019819750.2;refGene_gn=PDE4C;refGene_tx=XM_019819753.2
A2 13442518 13442618
refGene_gn=PDE4C;refGene_tx=XM_006928616.3;refGene_gn=PDE4C;refGene_tx=XM_006928617.3;refGene_gn=PDE4C;refGene_tx=XM_019819750.2;refGene_gn=PDE4C;refGene_tx=XM_019819753.2
A2 13442704 13442869
refGene_gn=PDE4C;refGene_tx=XM_006928616.3;refGene_gn=PDE4C;refGene_tx=XM_006928617.3;refGene_gn=PDE4C;refGene_tx=XM_019819750.2;refGene_gn=PDE4C;refGene_tx=XM_019819753.2
A2 13443314 13443413
refGene_gn=PDE4C;refGene_tx=XM_006928616.3;refGene_gn=PDE4C;refGene_tx=XM_006928617.3;refGene_gn=PDE4C;refGene_tx=XM_019819750.2;refGene_gn=PDE4C;refGene_tx=XM_019819753.2
A2 13443612 13443776
refGene_gn=PDE4C;refGene_tx=XM_006928616.3;refGene_gn=PDE4C;refGene_tx=XM_006928617.3;refGene_gn=PDE4C;refGene_tx=XM_019819750.2;refGene_gn=PDE4C;refGene_tx=XM_019819753.2
A2 13444305 13444399
refGene_gn=PDE4C;refGene_tx=XM_006928616.3;refGene_gn=PDE4C;refGene_tx=XM_006928617.3;refGene_gn=PDE4C;refGene_tx=XM_019819750.2;refGene_gn=PDE4C;refGene_tx=XM_019819753.2

A2 13444479 13444592
refGene_gn=PDE4C;refGene_tx=XM_006928616.3;refGene_gn=PDE4C;refGene_tx=XM_006928617.3;refGene_gn=PDE4C;refGene_tx=XM_019819750.2;refGene_gn=PDE4C;refGene_tx=XM_019819753.2
A2 13444966 13445019
refGene_gn=PDE4C;refGene_tx=XM_006928616.3;refGene_gn=PDE4C;refGene_tx=XM_006928617.3;refGene_gn=PDE4C;refGene_tx=XM_019819750.2;refGene_gn=PDE4C;refGene_tx=XM_019819753.2
A2 13445308 13445382
refGene_gn=PDE4C;refGene_tx=XM_006928616.3;refGene_gn=PDE4C;refGene_tx=XM_006928617.3;refGene_gn=PDE4C;refGene_tx=XM_019819750.2;refGene_gn=PDE4C;refGene_tx=XM_019819753.2
A2 13445472 13445509
refGene_gn=PDE4C;refGene_tx=XM_006928616.3;refGene_gn=PDE4C;refGene_tx=XM_006928617.3;refGene_gn=PDE4C;refGene_tx=XM_019819750.2;refGene_gn=PDE4C;refGene_tx=XM_019819753.2
A2 13446403 13446595
refGene_gn=PDE4C;refGene_tx=XM_006928616.3;refGene_gn=PDE4C;refGene_tx=XM_006928617.3;refGene_gn=PDE4C;refGene_tx=XM_019819750.2;refGene_gn=PDE4C;refGene_tx=XM_019819753.2
A2 13449652 13449798 refGene_gn=PDE4C;refGene_tx=XM_006928617.3
A2 13454643 13454885 refGene_gn=PDE4C;refGene_tx=XM_019819753.2
A2 13457293 13457499 refGene_gn=PDE4C;refGene_tx=XM_019819750.2
A2 13462068 13462110 refGene_gn=PDE4C;refGene_tx=XM_019819750.2
A2 13464430 13464519
refGene_gn=LOC109494881;refGene_tx=XM_023247464.1
A2 13464637 13464777
refGene_gn=LOC109494881;refGene_tx=XM_023247461.1
A2 13466835 13467132
refGene_gn=LOC109494881;refGene_tx=XM_023247461.1;refGene_gn=LOC109494881;refGene_tx=XM_023247464.1
A2 13470871 13471133
refGene_gn=LOC109494881;refGene_tx=XM_023247461.1;refGene_gn=LOC109494881;refGene_tx=XM_023247464.1
A2 13471919 13472846
refGene_gn=IQCN;refGene_tx=XM_011290750.3;refGene_gn=IQCN;refGene_tx=XM_023247439.1;refGene_gn=IQCN;refGene_tx=XM_023247441.1
A2 13480084 13483474
refGene_gn=IQCN;refGene_tx=XM_011290750.3;refGene_gn=IQCN;refGene_tx=XM_023247439.1;refGene_gn=IQCN;refGene_tx=XM_023247441.1
A2 13484596 13484680
refGene_gn=IQCN;refGene_tx=XM_011290750.3;refGene_gn=IQCN;refGene_tx=XM_023247439.1
A2 13492170 13493214 refGene_gn=JUND;refGene_tx=XM_011279661.2
A2 13510342 13510434
refGene_gn=LSM4;refGene_tx=XM_003982075.5;refGene_gn=LSM4;refGene_tx=XM_023247504.1
A2 13512250 13512434
refGene_gn=LSM4;refGene_tx=XM_003982075.5;refGene_gn=LSM4;refGene_tx=XM_023247504.1

A2 13515434 13515533
refGene_gn=LSM4;refGene_tx=XM_003982075.5;refGene_gn=LSM4;refGene_tx=XM_023247504.1
A2 13519083 13519125
refGene_gn=LSM4;refGene_tx=XM_003982075.5;refGene_gn=LSM4;refGene_tx=XM_023247504.1
A2 13523718 13523721 refGene_gn=LSM4;refGene_tx=XM_003982075.5
A2 13533158 13533192
refGene_gn=PGPEP1;refGene_tx=XM_023247483.1;refGene_gn=PGPEP1;refGene_tx=XM_023247491.1
A2 13533794 13533798 refGene_gn=PGPEP1;refGene_tx=XM_006928618.4
A2 13535179 13535232
refGene_gn=PGPEP1;refGene_tx=XM_006928618.4;refGene_gn=PGPEP1;refGene_tx=XM_023247483.1;refGene_gn=PGPEP1;refGene_tx=XM_023247491.1
A2 13549837 13549954
refGene_gn=PGPEP1;refGene_tx=XM_006928618.4;refGene_gn=PGPEP1;refGene_tx=XM_023247483.1;refGene_gn=PGPEP1;refGene_tx=XM_023247491.1
A2 13552963 13553196
refGene_gn=PGPEP1;refGene_tx=XM_006928618.4;refGene_gn=PGPEP1;refGene_tx=XM_023247483.1;refGene_gn=PGPEP1;refGene_tx=XM_023247494.1
A2 13556926 13557130
refGene_gn=PGPEP1;refGene_tx=XM_006928618.4;refGene_gn=PGPEP1;refGene_tx=XM_023247483.1;refGene_gn=PGPEP1;refGene_tx=XM_023247491.1;refGene_gn=PGPEP1;refGene_tx=XM_023247494.1
A2 13589481 13589764 refGene_gn=GDF15;refGene_tx=XM_023247515.1
A2 13591035 13591205 refGene_gn=GDF15;refGene_tx=XM_023247515.1
A2 13591206 13591683 refGene_gn=GDF15;refGene_tx=XM_023247515.1
A2 13595487 13595626
refGene_gn=LRRC25;refGene_tx=XM_003982077.5;refGene_gn=LRRC25;refGene_tx=XM_011290761.3
A2 13597902 13598687
refGene_gn=LRRC25;refGene_tx=XM_003982077.5;refGene_gn=LRRC25;refGene_tx=XM_011290761.3
A2 13611345 13611354 refGene_gn=SSBP4;refGene_tx=XM_023241209.1
A2 13611400 13611555 refGene_gn=SSBP4;refGene_tx=XM_023241209.1
A2 13618132 13618205 refGene_gn=SSBP4;refGene_tx=XM_023241209.1
A2 13618563 13618625 refGene_gn=SSBP4;refGene_tx=XM_023241209.1
A2 13618720 13618805 refGene_gn=SSBP4;refGene_tx=XM_023241209.1
A2 13622417 13622507 refGene_gn=SSBP4;refGene_tx=XM_023241209.1
A2 13622995 13623061 refGene_gn=SSBP4;refGene_tx=XM_023241209.1
A2 13623285 13623345 refGene_gn=SSBP4;refGene_tx=XM_023241209.1
A2 13623431 13623501 refGene_gn=SSBP4;refGene_tx=XM_023241209.1
A2 13623667 13623735 refGene_gn=SSBP4;refGene_tx=XM_023241209.1
A2 13623818 13623886 refGene_gn=SSBP4;refGene_tx=XM_023241209.1
A2 13624195 13624241 refGene_gn=SSBP4;refGene_tx=XM_023241209.1
A2 13624331 13624367 refGene_gn=SSBP4;refGene_tx=XM_023241209.1
A2 13624490 13624545 refGene_gn=SSBP4;refGene_tx=XM_023241209.1
A2 13624631 13624702 refGene_gn=SSBP4;refGene_tx=XM_023241209.1
A2 13624797 13624876 refGene_gn=SSBP4;refGene_tx=XM_023241209.1
A2 13625246 13625275 refGene_gn=SSBP4;refGene_tx=XM_023241209.1

A2 13625436 13625544 refGene_gn=SSBP4;refGene_tx=XM_023241209.1
A2 13625918 13625948 refGene_gn=SSBP4;refGene_tx=XM_023241209.1
A2 13626628 13626830
refGene_gn=ISYNA1;refGene_tx=XM_003982078.5;refGene_gn=ISYNA1;refGene_tx=XM_023247525.1
A2 13626955 13627173
refGene_gn=ISYNA1;refGene_tx=XM_003982078.5;refGene_gn=ISYNA1;refGene_tx=XM_023247525.1
A2 13627249 13627363
refGene_gn=ISYNA1;refGene_tx=XM_003982078.5;refGene_gn=ISYNA1;refGene_tx=XM_023247525.1
A2 13627466 13627631
refGene_gn=ISYNA1;refGene_tx=XM_003982078.5;refGene_gn=ISYNA1;refGene_tx=XM_023247525.1
A2 13627741 13627957
refGene_gn=ISYNA1;refGene_tx=XM_003982078.5;refGene_gn=ISYNA1;refGene_tx=XM_023247525.1
A2 13628042 13628192
refGene_gn=ISYNA1;refGene_tx=XM_003982078.5;refGene_gn=ISYNA1;refGene_tx=XM_023247525.1
A2 13628385 13628579
refGene_gn=ISYNA1;refGene_tx=XM_003982078.5;refGene_gn=ISYNA1;refGene_tx=XM_023247525.1
A2 13628690 13628823
refGene_gn=ISYNA1;refGene_tx=XM_003982078.5;refGene_gn=ISYNA1;refGene_tx=XM_023247525.1
A2 13629101 13629263 refGene_gn=ISYNA1;refGene_tx=XM_003982078.5
A2 13629406 13629526
refGene_gn=ISYNA1;refGene_tx=XM_003982078.5;refGene_gn=ISYNA1;refGene_tx=XM_023247525.1
A2 13635014 13635131 refGene_gn=ELL;refGene_tx=XM_003982080.5
A2 13635819 13635864 refGene_gn=ELL;refGene_tx=XM_003982080.5
A2 13637149 13637321 refGene_gn=ELL;refGene_tx=XM_003982080.5
A2 13637568 13637635 refGene_gn=ELL;refGene_tx=XM_003982080.5
A2 13640150 13640625 refGene_gn=ELL;refGene_tx=XM_003982080.5
A2 13641777 13641874 refGene_gn=ELL;refGene_tx=XM_003982080.5
A2 13649380 13649505 refGene_gn=ELL;refGene_tx=XM_003982080.5
A2 13652846 13653121 refGene_gn=ELL;refGene_tx=XM_003982080.5
A2 13655224 13655388 refGene_gn=ELL;refGene_tx=XM_003982080.5
A2 13655805 13655927 refGene_gn=ELL;refGene_tx=XM_003982080.5
A2 13662862 13662910 refGene_gn=ELL;refGene_tx=XM_003982080.5
A2 13709720 13709855 refGene_gn=ELL;refGene_tx=XM_003982080.5
A2 13720334 13720421
refGene_gn=FKBP8;refGene_tx=XM_003982081.5;refGene_gn=FKBP8;refGene_tx=XM_023247536.1;refGene_gn=FKBP8;refGene_tx=XM_023247545.1
A2 13720842 13720974
refGene_gn=FKBP8;refGene_tx=XM_003982081.5;refGene_gn=FKBP8;refGene_tx=XM_023247536.1;refGene_gn=FKBP8;refGene_tx=XM_023247545.1
A2 13721481 13721559
refGene_gn=FKBP8;refGene_tx=XM_003982081.5;refGene_gn=FKBP8;refGene_tx

=XM_023247536.1;refGene_gn=FKBP8;refGene_tx=XM_023247545.1
A2	13723605	13723778
refGene_gn=FKBP8;refGene_tx=XM_003982081.5;refGene_gn=FKBP8;refGene_tx=XM_023247536.1;refGene_gn=FKBP8;refGene_tx=XM_023247545.1
A2	13724431	13724652
refGene_gn=FKBP8;refGene_tx=XM_003982081.5;refGene_gn=FKBP8;refGene_tx=XM_023247536.1;refGene_gn=FKBP8;refGene_tx=XM_023247545.1
A2	13724928	13725017
refGene_gn=FKBP8;refGene_tx=XM_003982081.5;refGene_gn=FKBP8;refGene_tx=XM_023247536.1;refGene_gn=FKBP8;refGene_tx=XM_023247545.1
A2	13725207	13725377
refGene_gn=FKBP8;refGene_tx=XM_003982081.5;refGene_gn=FKBP8;refGene_tx=XM_023247536.1;refGene_gn=FKBP8;refGene_tx=XM_023247545.1
A2	13727519	13727820
refGene_gn=FKBP8;refGene_tx=XM_003982081.5;refGene_gn=FKBP8;refGene_tx=XM_023247536.1;refGene_gn=FKBP8;refGene_tx=XM_023247545.1
A2	13738772	13738873	refGene_gn=KXD1;refGene_tx=XM_003982082.5
A2	13739840	13739993	refGene_gn=KXD1;refGene_tx=XM_003982082.5
A2	13741580	13741627	refGene_gn=KXD1;refGene_tx=XM_003982082.5
A2	13742570	13742800	refGene_gn=KXD1;refGene_tx=XM_003982082.5
A2	13745785	13745888
refGene_gn=UBA52;refGene_tx=NM_001123354.1;refGene_gn=UBA52;refGene_tx=XM_006928596.2;refGene_gn=UBA52;refGene_tx=XM_006928597.3;refGene_gn=UBA52;refGene_tx=XM_011290464.2
A2	13746159	13746246
refGene_gn=UBA52;refGene_tx=NM_001123354.1;refGene_gn=UBA52;refGene_tx=XM_006928596.2;refGene_gn=UBA52;refGene_tx=XM_006928597.3;refGene_gn=UBA52;refGene_tx=XM_011290464.2
A2	13747435	13747538
refGene_gn=UBA52;refGene_tx=NM_001123354.1;refGene_gn=UBA52;refGene_tx=XM_006928596.2;refGene_gn=UBA52;refGene_tx=XM_006928597.3;refGene_gn=UBA52;refGene_tx=XM_011290464.2
A2	13747634	13747728
refGene_gn=UBA52;refGene_tx=NM_001123354.1;refGene_gn=UBA52;refGene_tx=XM_006928596.2;refGene_gn=UBA52;refGene_tx=XM_006928597.3;refGene_gn=UBA52;refGene_tx=XM_011290464.2
A2	13749331	13749703
refGene_gn=REX1BD;refGene_tx=XM_023247586.1;refGene_gn=REX1BD;refGene_tx=XM_023247590.1
A2	13749797	13749880
refGene_gn=REX1BD;refGene_tx=XM_023247586.1;refGene_gn=REX1BD;refGene_tx=XM_023247590.1
A2	13750127	13750132	refGene_gn=CRLF1;refGene_tx=XM_023247582.1
A2	13750239	13750468
refGene_gn=REX1BD;refGene_tx=XM_023247586.1;refGene_gn=REX1BD;refGene_tx=XM_023247590.1
A2	13751369	13751449
refGene_gn=REX1BD;refGene_tx=XM_023247586.1;refGene_gn=REX1BD;refGene_tx=XM_023247590.1
A2	13752077	13752150

refGene_gn=REX1BD;refGene_tx=XM_023247586.1;refGene_gn=REX1BD;refGene_tx=XM_023247590.1
A2 13752925 13752939 refGene_gn=CRLF1;refGene_tx=XM_023247571.1
A2 13753483 13753526
refGene_gn=CRLF1;refGene_tx=XM_023247571.1;refGene_gn=CRLF1;refGene_tx=XM_023247582.1
A2 13753639 13753827
refGene_gn=CRLF1;refGene_tx=XM_023247571.1;refGene_gn=CRLF1;refGene_tx=XM_023247582.1
A2 13755961 13756130
refGene_gn=CRLF1;refGene_tx=XM_023247571.1;refGene_gn=CRLF1;refGene_tx=XM_023247582.1
A2 13756210 13756368
refGene_gn=CRLF1;refGene_tx=XM_023247571.1;refGene_gn=CRLF1;refGene_tx=XM_023247582.1
A2 13757409 13757579
refGene_gn=CRLF1;refGene_tx=XM_023247571.1;refGene_gn=CRLF1;refGene_tx=XM_023247582.1
A2 13757753 13757883
refGene_gn=CRLF1;refGene_tx=XM_023247571.1;refGene_gn=CRLF1;refGene_tx=XM_023247582.1
A2 13758180 13758462
refGene_gn=CRLF1;refGene_tx=XM_023247571.1;refGene_gn=CRLF1;refGene_tx=XM_023247582.1
A2 13763066 13763178
refGene_gn=CRLF1;refGene_tx=XM_023247571.1;refGene_gn=CRLF1;refGene_tx=XM_023247582.1
A2 13769338 13769506 refGene_gn=TMEM59L;refGene_tx=XM_003982083.4
A2 13770469 13770626 refGene_gn=TMEM59L;refGene_tx=XM_003982083.4
A2 13770727 13770837 refGene_gn=TMEM59L;refGene_tx=XM_003982083.4
A2 13771984 13772137 refGene_gn=TMEM59L;refGene_tx=XM_003982083.4
A2 13772723 13772826 refGene_gn=TMEM59L;refGene_tx=XM_003982083.4
A2 13773127 13773245 refGene_gn=TMEM59L;refGene_tx=XM_003982083.4
A2 13773347 13773465 refGene_gn=TMEM59L;refGene_tx=XM_003982083.4
A2 13773715 13773844 refGene_gn=TMEM59L;refGene_tx=XM_003982083.4
A2 13787187 13787264 refGene_gn=KLHL26;refGene_tx=XM_023247604.1
A2 13788975 13789007 refGene_gn=KLHL26;refGene_tx=XM_023247608.1
A2 13810019 13810202
refGene_gn=KLHL26;refGene_tx=XM_023247604.1;refGene_gn=KLHL26;refGene_tx=XM_023247608.1
A2 13812374 13813956
refGene_gn=KLHL26;refGene_tx=XM_023247604.1;refGene_gn=KLHL26;refGene_tx=XM_023247608.1
A2 13824232 13824358
refGene_gn=CRTC1;refGene_tx=XM_023247614.1;refGene_gn=CRTC1;refGene_tx=XM_023247619.1;refGene_gn=CRTC1;refGene_tx=XM_023247627.1;refGene_gn=CRTC1;refGene_tx=XM_023247631.1;refGene_gn=CRTC1;refGene_tx=XM_023247633.1;refGene_gn=CRTC1;refGene_tx=XM_023247639.1;refGene_gn=CRTC1;refGene_tx=XM_023247643.1;refGene_gn=CRTC1;refGene_tx=XM_023247644.1;refGene_gn=CRTC1;refGene_tx=XM_023247651.1;refGene_gn=CRTC1;refGene_tx=XM_02

3247653.1
A2 13866019 13866136
refGene_gn=CRTC1;refGene_tx=XM_023247614.1;refGene_gn=CRTC1;refGene_tx=XM_023247619.1;refGene_gn=CRTC1;refGene_tx=XM_023247627.1;refGene_gn=CRTC1;refGene_tx=XM_023247631.1;refGene_gn=CRTC1;refGene_tx=XM_023247633.1;refGene_gn=CRTC1;refGene_tx=XM_023247639.1;refGene_gn=CRTC1;refGene_tx=XM_023247643.1;refGene_gn=CRTC1;refGene_tx=XM_023247644.1;refGene_gn=CRTC1;refGene_tx=XM_023247651.1;refGene_gn=CRTC1;refGene_tx=XM_023247653.1
A2 13867334 13867382
refGene_gn=CRTC1;refGene_tx=XM_023247614.1;refGene_gn=CRTC1;refGene_tx=XM_023247627.1;refGene_gn=CRTC1;refGene_tx=XM_023247631.1;refGene_gn=CRTC1;refGene_tx=XM_023247633.1;refGene_gn=CRTC1;refGene_tx=XM_023247639.1;refGene_gn=CRTC1;refGene_tx=XM_023247643.1;refGene_gn=CRTC1;refGene_tx=XM_023247644.1
A2 13868908 13869046
refGene_gn=CRTC1;refGene_tx=XM_023247614.1;refGene_gn=CRTC1;refGene_tx=XM_023247619.1;refGene_gn=CRTC1;refGene_tx=XM_023247627.1;refGene_gn=CRTC1;refGene_tx=XM_023247631.1;refGene_gn=CRTC1;refGene_tx=XM_023247633.1;refGene_gn=CRTC1;refGene_tx=XM_023247639.1;refGene_gn=CRTC1;refGene_tx=XM_023247643.1;refGene_gn=CRTC1;refGene_tx=XM_023247644.1;refGene_gn=CRTC1;refGene_tx=XM_023247651.1;refGene_gn=CRTC1;refGene_tx=XM_023247653.1
A2 13870049 13870111
refGene_gn=CRTC1;refGene_tx=XM_023247614.1;refGene_gn=CRTC1;refGene_tx=XM_023247619.1;refGene_gn=CRTC1;refGene_tx=XM_023247627.1;refGene_gn=CRTC1;refGene_tx=XM_023247631.1;refGene_gn=CRTC1;refGene_tx=XM_023247633.1;refGene_gn=CRTC1;refGene_tx=XM_023247639.1;refGene_gn=CRTC1;refGene_tx=XM_023247643.1;refGene_gn=CRTC1;refGene_tx=XM_023247644.1;refGene_gn=CRTC1;refGene_tx=XM_023247651.1;refGene_gn=CRTC1;refGene_tx=XM_023247653.1
A2 13871942 13872037
refGene_gn=CRTC1;refGene_tx=XM_023247614.1;refGene_gn=CRTC1;refGene_tx=XM_023247619.1;refGene_gn=CRTC1;refGene_tx=XM_023247627.1;refGene_gn=CRTC1;refGene_tx=XM_023247631.1;refGene_gn=CRTC1;refGene_tx=XM_023247633.1;refGene_gn=CRTC1;refGene_tx=XM_023247639.1;refGene_gn=CRTC1;refGene_tx=XM_023247643.1;refGene_gn=CRTC1;refGene_tx=XM_023247644.1;refGene_gn=CRTC1;refGene_tx=XM_023247651.1;refGene_gn=CRTC1;refGene_tx=XM_023247653.1
A2 13874025 13874111
refGene_gn=CRTC1;refGene_tx=XM_023247614.1;refGene_gn=CRTC1;refGene_tx=XM_023247619.1;refGene_gn=CRTC1;refGene_tx=XM_023247627.1;refGene_gn=CRTC1;refGene_tx=XM_023247631.1;refGene_gn=CRTC1;refGene_tx=XM_023247633.1;refGene_gn=CRTC1;refGene_tx=XM_023247639.1;refGene_gn=CRTC1;refGene_tx=XM_023247643.1;refGene_gn=CRTC1;refGene_tx=XM_023247644.1;refGene_gn=CRTC1;refGene_tx=XM_023247651.1;refGene_gn=CRTC1;refGene_tx=XM_023247653.1
A2 13878048 13878089
refGene_gn=CRTC1;refGene_tx=XM_023247614.1;refGene_gn=CRTC1;refGene_tx=XM_023247619.1;refGene_gn=CRTC1;refGene_tx=XM_023247627.1;refGene_gn=

CRTC1;refGene_tx=XM_023247631.1;refGene_gn=CRTC1;refGene_tx=XM_023247633.1;refGene_gn=CRTC1;refGene_tx=XM_023247639.1;refGene_gn=CRTC1;refGene_tx=XM_023247643.1;refGene_gn=CRTC1;refGene_tx=XM_023247644.1;refGene_gn=CRTC1;refGene_tx=XM_023247651.1;refGene_gn=CRTC1;refGene_tx=XM_023247653.1
A2	13878575	13878796
refGene_gn=CRTC1;refGene_tx=XM_023247614.1;refGene_gn=CRTC1;refGene_tx=XM_023247619.1;refGene_gn=CRTC1;refGene_tx=XM_023247627.1;refGene_gn=CRTC1;refGene_tx=XM_023247631.1;refGene_gn=CRTC1;refGene_tx=XM_023247633.1;refGene_gn=CRTC1;refGene_tx=XM_023247639.1;refGene_gn=CRTC1;refGene_tx=XM_023247643.1;refGene_gn=CRTC1;refGene_tx=XM_023247644.1;refGene_gn=CRTC1;refGene_tx=XM_023247651.1;refGene_gn=CRTC1;refGene_tx=XM_023247653.1
A2	13883523	13883657
refGene_gn=CRTC1;refGene_tx=XM_023247614.1;refGene_gn=CRTC1;refGene_tx=XM_023247619.1;refGene_gn=CRTC1;refGene_tx=XM_023247627.1;refGene_gn=CRTC1;refGene_tx=XM_023247631.1;refGene_gn=CRTC1;refGene_tx=XM_023247633.1;refGene_gn=CRTC1;refGene_tx=XM_023247639.1;refGene_gn=CRTC1;refGene_tx=XM_023247643.1;refGene_gn=CRTC1;refGene_tx=XM_023247644.1;refGene_gn=CRTC1;refGene_tx=XM_023247651.1;refGene_gn=CRTC1;refGene_tx=XM_023247653.1
A2	13886864	13887176
refGene_gn=CRTC1;refGene_tx=XM_023247614.1;refGene_gn=CRTC1;refGene_tx=XM_023247619.1;refGene_gn=CRTC1;refGene_tx=XM_023247627.1;refGene_gn=CRTC1;refGene_tx=XM_023247631.1;refGene_gn=CRTC1;refGene_tx=XM_023247633.1;refGene_gn=CRTC1;refGene_tx=XM_023247639.1;refGene_gn=CRTC1;refGene_tx=XM_023247643.1;refGene_gn=CRTC1;refGene_tx=XM_023247644.1;refGene_gn=CRTC1;refGene_tx=XM_023247651.1;refGene_gn=CRTC1;refGene_tx=XM_023247653.1
A2	13889347	13889452
refGene_gn=CRTC1;refGene_tx=XM_023247614.1;refGene_gn=CRTC1;refGene_tx=XM_023247619.1;refGene_gn=CRTC1;refGene_tx=XM_023247627.1;refGene_gn=CRTC1;refGene_tx=XM_023247633.1;refGene_gn=CRTC1;refGene_tx=XM_023247651.1;refGene_gn=CRTC1;refGene_tx=XM_023247653.1
A2	13892487	13892574
refGene_gn=CRTC1;refGene_tx=XM_023247614.1;refGene_gn=CRTC1;refGene_tx=XM_023247619.1;refGene_gn=CRTC1;refGene_tx=XM_023247631.1;refGene_gn=CRTC1;refGene_tx=XM_023247633.1;refGene_gn=CRTC1;refGene_tx=XM_023247643.1;refGene_gn=CRTC1;refGene_tx=XM_023247651.1
A2	13893196	13893377
refGene_gn=CRTC1;refGene_tx=XM_023247614.1;refGene_gn=CRTC1;refGene_tx=XM_023247619.1;refGene_gn=CRTC1;refGene_tx=XM_023247627.1;refGene_gn=CRTC1;refGene_tx=XM_023247631.1;refGene_gn=CRTC1;refGene_tx=XM_023247633.1;refGene_gn=CRTC1;refGene_tx=XM_023247639.1;refGene_gn=CRTC1;refGene_tx=XM_023247643.1;refGene_gn=CRTC1;refGene_tx=XM_023247644.1;refGene_gn=CRTC1;refGene_tx=XM_023247651.1;refGene_gn=CRTC1;refGene_tx=XM_023247653.1
A2	13894756	13894968
refGene_gn=CRTC1;refGene_tx=XM_023247614.1;refGene_gn=CRTC1;refGene_tx=XM_023247619.1;refGene_gn=CRTC1;refGene_tx=XM_023247627.1;refGene_gn=

CRTC1;refGene_tx=XM_023247631.1;refGene_gn=CRTC1;refGene_tx=XM_023247633.1;refGene_gn=CRTC1;refGene_tx=XM_023247639.1;refGene_gn=CRTC1;refGene_tx=XM_023247643.1;refGene_gn=CRTC1;refGene_tx=XM_023247644.1;refGene_gn=CRTC1;refGene_tx=XM_023247651.1;refGene_gn=CRTC1;refGene_tx=XM_023247653.1

| | | | |
|---|---|---|---|
| A2 | 13901080 | 13901127 | refGene_gn=COMP;refGene_tx=XM_003982085.4 |
| A2 | 13901216 | 13901356 | refGene_gn=COMP;refGene_tx=XM_003982085.4 |
| A2 | 13901783 | 13901956 | refGene_gn=COMP;refGene_tx=XM_003982085.4 |
| A2 | 13902580 | 13902777 | refGene_gn=COMP;refGene_tx=XM_003982085.4 |
| A2 | 13903095 | 13903144 | refGene_gn=COMP;refGene_tx=XM_003982085.4 |
| A2 | 13903254 | 13903433 | refGene_gn=COMP;refGene_tx=XM_003982085.4 |
| A2 | 13903596 | 13903778 | refGene_gn=COMP;refGene_tx=XM_003982085.4 |
| A2 | 13903881 | 13903934 | refGene_gn=COMP;refGene_tx=XM_003982085.4 |
| A2 | 13904123 | 13904242 | refGene_gn=COMP;refGene_tx=XM_003982085.4 |
| A2 | 13904813 | 13904973 | refGene_gn=COMP;refGene_tx=XM_003982085.4 |
| A2 | 13905197 | 13905305 | refGene_gn=COMP;refGene_tx=XM_003982085.4 |
| A2 | 13905392 | 13905497 | refGene_gn=COMP;refGene_tx=XM_003982085.4 |
| A2 | 13905594 | 13905753 | refGene_gn=COMP;refGene_tx=XM_003982085.4 |
| A2 | 13905849 | 13905924 | refGene_gn=COMP;refGene_tx=XM_003982085.4 |
| A2 | 13906191 | 13906329 | refGene_gn=COMP;refGene_tx=XM_003982085.4 |
| A2 | 13907020 | 13907190 | refGene_gn=COMP;refGene_tx=XM_003982085.4 |
| A2 | 13907648 | 13907700 | refGene_gn=COMP;refGene_tx=XM_003982085.4 |
| A2 | 13908108 | 13908194 | refGene_gn=COMP;refGene_tx=XM_003982085.4 |
| A2 | 13908437 | 13908516 | refGene_gn=COMP;refGene_tx=XM_003982085.4 |
| A2 | 13938560 | 13938791 | refGene_gn=UPF1;refGene_tx=XM_023247663.1 |
| A2 | 13952833 | 13952976 | refGene_gn=UPF1;refGene_tx=XM_023247663.1 |
| A2 | 13954587 | 13954677 | refGene_gn=UPF1;refGene_tx=XM_023247663.1 |
| A2 | 13957169 | 13957337 | refGene_gn=UPF1;refGene_tx=XM_023247663.1 |
| A2 | 13957688 | 13957869 | refGene_gn=UPF1;refGene_tx=XM_023247663.1 |
| A2 | 13959559 | 13959721 | refGene_gn=UPF1;refGene_tx=XM_023247663.1 |
| A2 | 13960371 | 13960489 | refGene_gn=UPF1;refGene_tx=XM_023247663.1 |
| A2 | 13960644 | 13960743 | refGene_gn=UPF1;refGene_tx=XM_023247663.1 |
| A2 | 13961214 | 13961323 | refGene_gn=UPF1;refGene_tx=XM_023247663.1 |
| A2 | 13961492 | 13961652 | refGene_gn=UPF1;refGene_tx=XM_023247663.1 |
| A2 | 13961768 | 13961887 | refGene_gn=UPF1;refGene_tx=XM_023247663.1 |
| A2 | 13962347 | 13962512 | refGene_gn=UPF1;refGene_tx=XM_023247663.1 |
| A2 | 13962613 | 13962728 | refGene_gn=UPF1;refGene_tx=XM_023247663.1 |
| A2 | 13963216 | 13963360 | refGene_gn=UPF1;refGene_tx=XM_023247663.1 |
| A2 | 13964029 | 13964243 | refGene_gn=UPF1;refGene_tx=XM_023247663.1 |
| A2 | 13966964 | 13967082 | refGene_gn=UPF1;refGene_tx=XM_023247663.1 |
| A2 | 13968322 | 13968479 | refGene_gn=UPF1;refGene_tx=XM_023247663.1 |
| A2 | 13969138 | 13969281 | refGene_gn=UPF1;refGene_tx=XM_023247663.1 |
| A2 | 13970401 | 13970576 | refGene_gn=UPF1;refGene_tx=XM_023247663.1 |
| A2 | 13971108 | 13971190 | refGene_gn=UPF1;refGene_tx=XM_023247663.1 |
| A2 | 13972611 | 13972773 | refGene_gn=UPF1;refGene_tx=XM_023247663.1 |
| A2 | 13972908 | 13973039 | refGene_gn=UPF1;refGene_tx=XM_023247663.1 |
| A2 | 13973040 | 13973127 | refGene_gn=UPF1;refGene_tx=XM_023247663.1 |
| A2 | 13973473 | 13973593 | refGene_gn=UPF1;refGene_tx=XM_023247663.1 |
| A2 | 13976432 | 13977220 | refGene_gn=GDF1;refGene_tx=XM_023247668.1 |
| A2 | 13977878 | 13978203 | refGene_gn=GDF1;refGene_tx=XM_023247668.1 |

A2 13978563 13978606 refGene_gn=CERS1;refGene_tx=XM_023247676.1
A2 13985278 13985388 refGene_gn=CERS1;refGene_tx=XM_023247676.1
A2 13985643 13985791 refGene_gn=CERS1;refGene_tx=XM_023247676.1
A2 13986212 13986374 refGene_gn=CERS1;refGene_tx=XM_023247676.1
A2 13988042 13988223 refGene_gn=CERS1;refGene_tx=XM_023247676.1
A2 13993017 13993177 refGene_gn=CERS1;refGene_tx=XM_023247676.1
A2 13995514 13995763 refGene_gn=CERS1;refGene_tx=XM_023247676.1
A2 13998968 13999016 refGene_gn=COPE;refGene_tx=XM_003982087.5
A2 13999148 13999223 refGene_gn=COPE;refGene_tx=XM_003982087.5
A2 14000272 14000341 refGene_gn=COPE;refGene_tx=XM_003982087.5
A2 14002854 14003010 refGene_gn=COPE;refGene_tx=XM_003982087.5
A2 14003906 14003988 refGene_gn=COPE;refGene_tx=XM_003982087.5
A2 14004425 14004479 refGene_gn=COPE;refGene_tx=XM_003982087.5
A2 14005756 14005909 refGene_gn=COPE;refGene_tx=XM_003982087.5
A2 14008253 14008354 refGene_gn=COPE;refGene_tx=XM_003982087.5
A2 14009503 14009566 refGene_gn=COPE;refGene_tx=XM_003982087.5
A2 14014689 14014815 refGene_gn=COPE;refGene_tx=XM_003982087.5
A2 14015201 14015316 refGene_gn=DDX49;refGene_tx=XM_003982090.5
A2 14016145 14016269 refGene_gn=DDX49;refGene_tx=XM_003982090.5
A2 14017259 14017345 refGene_gn=DDX49;refGene_tx=XM_003982090.5
A2 14017434 14017556 refGene_gn=DDX49;refGene_tx=XM_003982090.5
A2 14017885 14018073 refGene_gn=DDX49;refGene_tx=XM_003982090.5
A2 14018216 14018357 refGene_gn=DDX49;refGene_tx=XM_003982090.5
A2 14019092 14019168 refGene_gn=DDX49;refGene_tx=XM_003982090.5
A2 14019438 14019515 refGene_gn=DDX49;refGene_tx=XM_003982090.5
A2 14019675 14019773 refGene_gn=DDX49;refGene_tx=XM_003982090.5
A2 14020411 14020486 refGene_gn=DDX49;refGene_tx=XM_003982090.5
A2 14021217 14021306 refGene_gn=DDX49;refGene_tx=XM_003982090.5
A2 14021387 14021459 refGene_gn=DDX49;refGene_tx=XM_003982090.5
A2 14021546 14021735 refGene_gn=DDX49;refGene_tx=XM_003982090.5
A2 14023245 14023437
refGene_gn=HOMER3;refGene_tx=XM_011290871.3;refGene_gn=HOMER3;refGene_tx=XM_011290873.3;refGene_gn=HOMER3;refGene_tx=XM_011290875.3
A2 14024540 14024627
refGene_gn=HOMER3;refGene_tx=XM_011290871.3;refGene_gn=HOMER3;refGene_tx=XM_011290873.3;refGene_gn=HOMER3;refGene_tx=XM_011290875.3
A2 14024722 14024833
refGene_gn=HOMER3;refGene_tx=XM_011290871.3;refGene_gn=HOMER3;refGene_tx=XM_011290873.3;refGene_gn=HOMER3;refGene_tx=XM_011290875.3
A2 14025150 14025307
refGene_gn=HOMER3;refGene_tx=XM_011290871.3;refGene_gn=HOMER3;refGene_tx=XM_011290873.3;refGene_gn=HOMER3;refGene_tx=XM_011290875.3
A2 14025830 14025952
refGene_gn=HOMER3;refGene_tx=XM_011290871.3;refGene_gn=HOMER3;refGene_tx=XM_011290873.3;refGene_gn=HOMER3;refGene_tx=XM_011290875.3
A2 14027026 14027134
refGene_gn=HOMER3;refGene_tx=XM_011290871.3;refGene_gn=HOMER3;refGene_tx=XM_011290873.3;refGene_gn=HOMER3;refGene_tx=XM_011290875.3
A2 14028316 14028448
refGene_gn=HOMER3;refGene_tx=XM_011290871.3;refGene_gn=HOMER3;refGene_

tx=XM_011290873.3;refGene_gn=HOMER3;refGene_tx=XM_011290875.3
A2	14028695	14028852
refGene_gn=HOMER3;refGene_tx=XM_011290871.3;refGene_gn=HOMER3;refGene_tx=XM_011290873.3;refGene_gn=HOMER3;refGene_tx=XM_011290875.3
A2	14028932	14028946
refGene_gn=HOMER3;refGene_tx=XM_011290871.3;refGene_gn=HOMER3;refGene_tx=XM_011290873.3;refGene_gn=HOMER3;refGene_tx=XM_011290875.3
A2	14071223	14071344
refGene_gn=SUGP2;refGene_tx=XM_003982092.5;refGene_gn=SUGP2;refGene_tx=XM_011290885.2;refGene_gn=SUGP2;refGene_tx=XM_023247721.1;refGene_gn=SUGP2;refGene_tx=XM_023247727.1;refGene_gn=SUGP2;refGene_tx=XM_023247729.1;refGene_gn=SUGP2;refGene_tx=XM_023247730.1;refGene_gn=SUGP2;refGene_tx=XM_023247731.1;refGene_gn=SUGP2;refGene_tx=XM_023247733.1;refGene_gn=SUGP2;refGene_tx=XM_023247735.1
A2	14072042	14072179
refGene_gn=SUGP2;refGene_tx=XM_003982092.5;refGene_gn=SUGP2;refGene_tx=XM_011290885.2;refGene_gn=SUGP2;refGene_tx=XM_023247721.1;refGene_gn=SUGP2;refGene_tx=XM_023247727.1;refGene_gn=SUGP2;refGene_tx=XM_023247729.1;refGene_gn=SUGP2;refGene_tx=XM_023247730.1;refGene_gn=SUGP2;refGene_tx=XM_023247731.1;refGene_gn=SUGP2;refGene_tx=XM_023247733.1;refGene_gn=SUGP2;refGene_tx=XM_023247735.1
A2	14074901	14074963
refGene_gn=SUGP2;refGene_tx=XM_003982092.5;refGene_gn=SUGP2;refGene_tx=XM_011290885.2;refGene_gn=SUGP2;refGene_tx=XM_023247721.1;refGene_gn=SUGP2;refGene_tx=XM_023247727.1;refGene_gn=SUGP2;refGene_tx=XM_023247729.1;refGene_gn=SUGP2;refGene_tx=XM_023247730.1;refGene_gn=SUGP2;refGene_tx=XM_023247731.1;refGene_gn=SUGP2;refGene_tx=XM_023247733.1;refGene_gn=SUGP2;refGene_tx=XM_023247735.1
A2	14076838	14077371
refGene_gn=SUGP2;refGene_tx=XM_003982092.5;refGene_gn=SUGP2;refGene_tx=XM_011290885.2;refGene_gn=SUGP2;refGene_tx=XM_023247721.1;refGene_gn=SUGP2;refGene_tx=XM_023247727.1;refGene_gn=SUGP2;refGene_tx=XM_023247729.1;refGene_gn=SUGP2;refGene_tx=XM_023247730.1;refGene_gn=SUGP2;refGene_tx=XM_023247731.1;refGene_gn=SUGP2;refGene_tx=XM_023247733.1;refGene_gn=SUGP2;refGene_tx=XM_023247735.1
A2	14083845	14083957
refGene_gn=SUGP2;refGene_tx=XM_003982092.5;refGene_gn=SUGP2;refGene_tx=XM_011290885.2;refGene_gn=SUGP2;refGene_tx=XM_023247721.1;refGene_gn=SUGP2;refGene_tx=XM_023247727.1;refGene_gn=SUGP2;refGene_tx=XM_023247729.1;refGene_gn=SUGP2;refGene_tx=XM_023247730.1;refGene_gn=SUGP2;refGene_tx=XM_023247731.1;refGene_gn=SUGP2;refGene_tx=XM_023247733.1;refGene_gn=SUGP2;refGene_tx=XM_023247735.1
A2	14084995	14085483
refGene_gn=SUGP2;refGene_tx=XM_003982092.5;refGene_gn=SUGP2;refGene_tx=XM_011290885.2;refGene_gn=SUGP2;refGene_tx=XM_023247721.1;refGene_gn=SUGP2;refGene_tx=XM_023247727.1;refGene_gn=SUGP2;refGene_tx=XM_023247729.1;refGene_gn=SUGP2;refGene_tx=XM_023247730.1;refGene_gn=SUGP2;refGene_tx=XM_023247731.1;refGene_gn=SUGP2;refGene_tx=XM_023247733.1;refGene_gn=SUGP2;refGene_tx=XM_023247735.1
A2	14090774	14090895

refGene_gn=SUGP2;refGene_tx=XM_003982092.5;refGene_gn=SUGP2;refGene_tx=XM_011290885.2;refGene_gn=SUGP2;refGene_tx=XM_023247721.1;refGene_gn=SUGP2;refGene_tx=XM_023247727.1;refGene_gn=SUGP2;refGene_tx=XM_023247729.1;refGene_gn=SUGP2;refGene_tx=XM_023247730.1;refGene_gn=SUGP2;refGene_tx=XM_023247731.1;refGene_gn=SUGP2;refGene_tx=XM_023247733.1;refGene_gn=SUGP2;refGene_tx=XM_023247735.1
A2 14094128 14095733
refGene_gn=SUGP2;refGene_tx=XM_003982092.5;refGene_gn=SUGP2;refGene_tx=XM_011290885.2;refGene_gn=SUGP2;refGene_tx=XM_023247721.1;refGene_gn=SUGP2;refGene_tx=XM_023247727.1;refGene_gn=SUGP2;refGene_tx=XM_023247729.1;refGene_gn=SUGP2;refGene_tx=XM_023247730.1;refGene_gn=SUGP2;refGene_tx=XM_023247731.1;refGene_gn=SUGP2;refGene_tx=XM_023247733.1;refGene_gn=SUGP2;refGene_tx=XM_023247735.1
A2 14099130 14099262
refGene_gn=SUGP2;refGene_tx=XM_003982092.5;refGene_gn=SUGP2;refGene_tx=XM_011290885.2;refGene_gn=SUGP2;refGene_tx=XM_023247721.1;refGene_gn=SUGP2;refGene_tx=XM_023247727.1;refGene_gn=SUGP2;refGene_tx=XM_023247729.1;refGene_gn=SUGP2;refGene_tx=XM_023247730.1;refGene_gn=SUGP2;refGene_tx=XM_023247731.1;refGene_gn=SUGP2;refGene_tx=XM_023247733.1
A2 14100176 14100204 refGene_gn=SUGP2;refGene_tx=XM_023247721.1
A2 14100613 14100653
refGene_gn=SUGP2;refGene_tx=XM_003982092.5;refGene_gn=SUGP2;refGene_tx=XM_011290885.2;refGene_gn=SUGP2;refGene_tx=XM_023247733.1
A2 14108294 14108421 refGene_gn=ARMC6;refGene_tx=XM_011290882.2
A2 14109491 14109574 refGene_gn=ARMC6;refGene_tx=XM_011290882.2
A2 14114159 14114733 refGene_gn=ARMC6;refGene_tx=XM_011290882.2
A2 14116347 14116493 refGene_gn=ARMC6;refGene_tx=XM_011290882.2
A2 14117810 14117942 refGene_gn=ARMC6;refGene_tx=XM_011290882.2
A2 14118242 14118380 refGene_gn=ARMC6;refGene_tx=XM_011290882.2
A2 14119451 14119664 refGene_gn=ARMC6;refGene_tx=XM_011290882.2
A2 14144345 14144426
refGene_gn=SLC25A42;refGene_tx=XM_003982095.4;refGene_gn=SLC25A42;refGene_tx=XM_019820106.2;refGene_gn=SLC25A42;refGene_tx=XM_019820110.2
A2 14147854 14147960
refGene_gn=SLC25A42;refGene_tx=XM_003982095.4;refGene_gn=SLC25A42;refGene_tx=XM_019820106.2;refGene_gn=SLC25A42;refGene_tx=XM_019820110.2
A2 14149674 14149700
refGene_gn=SLC25A42;refGene_tx=XM_003982095.4;refGene_gn=SLC25A42;refGene_tx=XM_019820106.2;refGene_gn=SLC25A42;refGene_tx=XM_019820110.2
A2 14150332 14150499
refGene_gn=SLC25A42;refGene_tx=XM_003982095.4;refGene_gn=SLC25A42;refGene_tx=XM_019820106.2;refGene_gn=SLC25A42;refGene_tx=XM_019820110.2
A2 14150881 14150998
refGene_gn=SLC25A42;refGene_tx=XM_003982095.4;refGene_gn=SLC25A42;refGene_tx=XM_019820106.2;refGene_gn=SLC25A42;refGene_tx=XM_019820110.2
A2 14151625 14151777
refGene_gn=SLC25A42;refGene_tx=XM_003982095.4;refGene_gn=SLC25A42;refGene_tx=XM_019820106.2;refGene_gn=SLC25A42;refGene_tx=XM_019820110.2
A2 14153618 14153926
refGene_gn=SLC25A42;refGene_tx=XM_003982095.4;refGene_gn=SLC25A42;refG

ene_tx=XM_019820106.2
A2 14154473 14154616 refGene_gn=SLC25A42;refGene_tx=XM_019820110.2
A2 14160791 14161045
refGene_gn=TMEM161A;refGene_tx=XM_023247758.1;refGene_gn=TMEM161A;refGene_tx=XM_023247765.1;refGene_gn=TMEM161A;refGene_tx=XM_023247766.1
A2 14161382 14161479
refGene_gn=TMEM161A;refGene_tx=XM_023247758.1;refGene_gn=TMEM161A;refGene_tx=XM_023247765.1;refGene_gn=TMEM161A;refGene_tx=XM_023247766.1
A2 14161618 14161793
refGene_gn=TMEM161A;refGene_tx=XM_023247758.1;refGene_gn=TMEM161A;refGene_tx=XM_023247765.1;refGene_gn=TMEM161A;refGene_tx=XM_023247766.1
A2 14161940 14162054
refGene_gn=TMEM161A;refGene_tx=XM_023247758.1;refGene_gn=TMEM161A;refGene_tx=XM_023247765.1;refGene_gn=TMEM161A;refGene_tx=XM_023247766.1
A2 14162156 14162300
refGene_gn=TMEM161A;refGene_tx=XM_023247758.1;refGene_gn=TMEM161A;refGene_tx=XM_023247765.1;refGene_gn=TMEM161A;refGene_tx=XM_023247766.1
A2 14162398 14162459
refGene_gn=TMEM161A;refGene_tx=XM_023247758.1;refGene_gn=TMEM161A;refGene_tx=XM_023247765.1;refGene_gn=TMEM161A;refGene_tx=XM_023247766.1
A2 14171276 14171428
refGene_gn=TMEM161A;refGene_tx=XM_023247758.1;refGene_gn=TMEM161A;refGene_tx=XM_023247765.1;refGene_gn=TMEM161A;refGene_tx=XM_023247766.1
A2 14172412 14172569
refGene_gn=TMEM161A;refGene_tx=XM_023247758.1;refGene_gn=TMEM161A;refGene_tx=XM_023247765.1;refGene_gn=TMEM161A;refGene_tx=XM_023247766.1
A2 14172686 14172784
refGene_gn=TMEM161A;refGene_tx=XM_023247758.1;refGene_gn=TMEM161A;refGene_tx=XM_023247765.1;refGene_gn=TMEM161A;refGene_tx=XM_023247766.1
A2 14173148 14173229
refGene_gn=TMEM161A;refGene_tx=XM_023247758.1;refGene_gn=TMEM161A;refGene_tx=XM_023247765.1;refGene_gn=TMEM161A;refGene_tx=XM_023247766.1
A2 14174169 14174273
refGene_gn=TMEM161A;refGene_tx=XM_023247758.1;refGene_gn=TMEM161A;refGene_tx=XM_023247765.1;refGene_gn=TMEM161A;refGene_tx=XM_023247766.1
A2 14177415 14177418
refGene_gn=TMEM161A;refGene_tx=XM_023247758.1;refGene_gn=TMEM161A;refGene_tx=XM_023247766.1
A2 14181655 14181930
refGene_gn=MEF2B;refGene_tx=XM_003982170.5;refGene_gn=MEF2B;refGene_tx=XM_019820130.2;refGene_gn=MEF2B;refGene_tx=XM_019820131.2;refGene_gn=MEF2B;refGene_tx=XM_023247773.1
A2 14182241 14182353
refGene_gn=MEF2B;refGene_tx=XM_003982170.5;refGene_gn=MEF2B;refGene_tx=XM_019820130.2;refGene_gn=MEF2B;refGene_tx=XM_019820131.2;refGene_gn=MEF2B;refGene_tx=XM_023247773.1
A2 14182525 14182619
refGene_gn=MEF2B;refGene_tx=XM_003982170.5;refGene_gn=MEF2B;refGene_tx=XM_019820130.2;refGene_gn=MEF2B;refGene_tx=XM_019820131.2
A2 14182713 14182847

refGene_gn=MEF2B;refGene_tx=XM_003982170.5;refGene_gn=MEF2B;refGene_tx=XM_019820130.2;refGene_gn=MEF2B;refGene_tx=XM_019820131.2;refGene_gn=MEF2B;refGene_tx=XM_023247773.1
A2 14183017 14183165
refGene_gn=MEF2B;refGene_tx=XM_003982170.5;refGene_gn=MEF2B;refGene_tx=XM_019820130.2;refGene_gn=MEF2B;refGene_tx=XM_019820131.2;refGene_gn=MEF2B;refGene_tx=XM_023247773.1
A2 14183876 14184011
refGene_gn=MEF2B;refGene_tx=XM_003982170.5;refGene_gn=MEF2B;refGene_tx=XM_019820130.2;refGene_gn=MEF2B;refGene_tx=XM_019820131.2;refGene_gn=MEF2B;refGene_tx=XM_023247773.1
A2 14184977 14185181
refGene_gn=MEF2B;refGene_tx=XM_003982170.5;refGene_gn=MEF2B;refGene_tx=XM_019820130.2;refGene_gn=MEF2B;refGene_tx=XM_019820131.2;refGene_gn=MEF2B;refGene_tx=XM_023247773.1
A2 14185969 14186023
refGene_gn=MEF2B;refGene_tx=XM_003982170.5;refGene_gn=MEF2B;refGene_tx=XM_019820130.2;refGene_gn=MEF2B;refGene_tx=XM_019820131.2;refGene_gn=MEF2B;refGene_tx=XM_023247773.1
A2 14205093 14206302 refGene_gn=BORCS8;refGene_tx=XM_023247793.1
A2 14206687 14206721
refGene_gn=BORCS8;refGene_tx=XM_011290914.3;refGene_gn=BORCS8;refGene_tx=XM_011290917.3;refGene_gn=BORCS8;refGene_tx=XM_011290921.3;refGene_gn=BORCS8;refGene_tx=XM_011290925.3
A2 14208547 14208662
refGene_gn=BORCS8;refGene_tx=XM_011290914.3;refGene_gn=BORCS8;refGene_tx=XM_011290917.3;refGene_gn=BORCS8;refGene_tx=XM_011290921.3;refGene_gn=BORCS8;refGene_tx=XM_011290925.3;refGene_gn=BORCS8;refGene_tx=XM_023247816.1;refGene_gn=BORCS8;refGene_tx=XM_023247819.1
A2 14212155 14212220
refGene_gn=BORCS8;refGene_tx=XM_011290914.3;refGene_gn=BORCS8;refGene_tx=XM_011290917.3;refGene_gn=BORCS8;refGene_tx=XM_011290921.3;refGene_gn=BORCS8;refGene_tx=XM_011290925.3;refGene_gn=BORCS8;refGene_tx=XM_023247816.1;refGene_gn=BORCS8;refGene_tx=XM_023247819.1
A2 14212695 14212808
refGene_gn=BORCS8;refGene_tx=XM_011290914.3;refGene_gn=BORCS8;refGene_tx=XM_011290917.3;refGene_gn=BORCS8;refGene_tx=XM_011290921.3;refGene_gn=BORCS8;refGene_tx=XM_011290925.3;refGene_gn=BORCS8;refGene_tx=XM_023247816.1;refGene_gn=BORCS8;refGene_tx=XM_023247819.1
A2 14213524 14213662
refGene_gn=BORCS8;refGene_tx=XM_011290914.3;refGene_gn=BORCS8;refGene_tx=XM_011290917.3;refGene_gn=BORCS8;refGene_tx=XM_011290921.3;refGene_gn=BORCS8;refGene_tx=XM_023247816.1
A2 14215400 14215437
refGene_gn=BORCS8;refGene_tx=XM_011290914.3;refGene_gn=BORCS8;refGene_tx=XM_011290917.3;refGene_gn=BORCS8;refGene_tx=XM_011290921.3;refGene_gn=BORCS8;refGene_tx=XM_011290925.3;refGene_gn=BORCS8;refGene_tx=XM_023247816.1;refGene_gn=BORCS8;refGene_tx=XM_023247819.1
A2 14216959 14217146
refGene_gn=RFXANK;refGene_tx=XM_006928657.4;refGene_gn=RFXANK;refGene_

tx=XM_019820169.2
A2 14219205 14219289
refGene_gn=RFXANK;refGene_tx=XM_006928657.4;refGene_gn=RFXANK;refGene_tx=XM_019820169.2
A2 14219436 14219502
refGene_gn=RFXANK;refGene_tx=XM_006928657.4;refGene_gn=RFXANK;refGene_tx=XM_019820169.2
A2 14219727 14219828
refGene_gn=RFXANK;refGene_tx=XM_006928657.4;refGene_gn=RFXANK;refGene_tx=XM_019820169.2
A2 14219997 14220123
refGene_gn=RFXANK;refGene_tx=XM_006928657.4;refGene_gn=RFXANK;refGene_tx=XM_019820169.2
A2 14220509 14220576
refGene_gn=RFXANK;refGene_tx=XM_006928657.4;refGene_gn=RFXANK;refGene_tx=XM_019820169.2
A2 14220983 14221064
refGene_gn=RFXANK;refGene_tx=XM_006928657.4;refGene_gn=RFXANK;refGene_tx=XM_019820169.2
A2 14221482 14221547 refGene_gn=RFXANK;refGene_tx=XM_019820169.2
A2 14221984 14222055 refGene_gn=RFXANK;refGene_tx=XM_006928657.4
A2 14222267 14222384 refGene_gn=NR2C2AP;refGene_tx=XM_003982171.5
A2 14222649 14222717 refGene_gn=NR2C2AP;refGene_tx=XM_003982171.5
A2 14222794 14222900 refGene_gn=NR2C2AP;refGene_tx=XM_003982171.5
A2 14223139 14223230 refGene_gn=NR2C2AP;refGene_tx=XM_003982171.5
A2 14223397 14223435 refGene_gn=NR2C2AP;refGene_tx=XM_003982171.5
A2 14233668 14233741 refGene_gn=NCAN;refGene_tx=XM_023247827.1
A2 14234611 14235025 refGene_gn=NCAN;refGene_tx=XM_023247827.1
A2 14237275 14237450 refGene_gn=NCAN;refGene_tx=XM_023247827.1
A2 14237560 14237688 refGene_gn=NCAN;refGene_tx=XM_023247827.1
A2 14238234 14238528 refGene_gn=NCAN;refGene_tx=XM_023247827.1
A2 14239415 14240003 refGene_gn=NCAN;refGene_tx=XM_023247827.1
A2 14240202 14241672 refGene_gn=NCAN;refGene_tx=XM_023247827.1
A2 14245555 14245672 refGene_gn=NCAN;refGene_tx=XM_023247827.1
A2 14246716 14246830 refGene_gn=NCAN;refGene_tx=XM_023247827.1
A2 14248375 14248534 refGene_gn=NCAN;refGene_tx=XM_023247827.1
A2 14249544 14249627 refGene_gn=NCAN;refGene_tx=XM_023247827.1
A2 14251892 14252037 refGene_gn=NCAN;refGene_tx=XM_023247827.1
A2 14252755 14252938 refGene_gn=NCAN;refGene_tx=XM_023247827.1
A2 14253455 14253601 refGene_gn=NCAN;refGene_tx=XM_023247827.1
A2 14261272 14261664 refGene_gn=HAPLN4;refGene_tx=XM_023247830.1
A2 14262022 14262358 refGene_gn=HAPLN4;refGene_tx=XM_023247830.1
A2 14263916 14264279 refGene_gn=HAPLN4;refGene_tx=XM_023247830.1
A2 14264568 14264686 refGene_gn=HAPLN4;refGene_tx=XM_023247830.1
A2 14265605 14265608 refGene_gn=HAPLN4;refGene_tx=XM_023247830.1
A2 14267240 14267522
refGene_gn=TM6SF2;refGene_tx=XM_023247832.1;refGene_gn=TM6SF2;refGene_tx=XM_023247834.1;refGene_gn=TM6SF2;refGene_tx=XM_023247835.1;refGene_gn=TM6SF2;refGene_tx=XM_023247837.1;refGene_gn=TM6SF2;refGene_tx=XM_023247838.1

A2	14268371 14268491
refGene_gn=TM6SF2;refGene_tx=XM_023247832.1;refGene_gn=TM6SF2;refGene_tx=XM_023247834.1;refGene_gn=TM6SF2;refGene_tx=XM_023247835.1;refGene_gn=TM6SF2;refGene_tx=XM_023247837.1;refGene_gn=TM6SF2;refGene_tx=XM_023247838.1
A2	14269395 14269488
refGene_gn=TM6SF2;refGene_tx=XM_023247832.1;refGene_gn=TM6SF2;refGene_tx=XM_023247834.1;refGene_gn=TM6SF2;refGene_tx=XM_023247835.1;refGene_gn=TM6SF2;refGene_tx=XM_023247837.1;refGene_gn=TM6SF2;refGene_tx=XM_023247838.1
A2	14269797 14269899
refGene_gn=TM6SF2;refGene_tx=XM_023247832.1;refGene_gn=TM6SF2;refGene_tx=XM_023247834.1;refGene_gn=TM6SF2;refGene_tx=XM_023247835.1;refGene_gn=TM6SF2;refGene_tx=XM_023247837.1;refGene_gn=TM6SF2;refGene_tx=XM_023247838.1
A2	14270084 14270213
refGene_gn=TM6SF2;refGene_tx=XM_023247832.1;refGene_gn=TM6SF2;refGene_tx=XM_023247834.1;refGene_gn=TM6SF2;refGene_tx=XM_023247835.1;refGene_gn=TM6SF2;refGene_tx=XM_023247837.1;refGene_gn=TM6SF2;refGene_tx=XM_023247838.1
A2	14270807 14270890
refGene_gn=TM6SF2;refGene_tx=XM_023247832.1;refGene_gn=TM6SF2;refGene_tx=XM_023247834.1;refGene_gn=TM6SF2;refGene_tx=XM_023247835.1;refGene_gn=TM6SF2;refGene_tx=XM_023247837.1;refGene_gn=TM6SF2;refGene_tx=XM_023247838.1
A2	14271283 14271387
refGene_gn=TM6SF2;refGene_tx=XM_023247832.1;refGene_gn=TM6SF2;refGene_tx=XM_023247834.1;refGene_gn=TM6SF2;refGene_tx=XM_023247835.1;refGene_gn=TM6SF2;refGene_tx=XM_023247837.1;refGene_gn=TM6SF2;refGene_tx=XM_023247838.1
A2	14271456 14271554
refGene_gn=TM6SF2;refGene_tx=XM_023247832.1;refGene_gn=TM6SF2;refGene_tx=XM_023247834.1;refGene_gn=TM6SF2;refGene_tx=XM_023247835.1;refGene_gn=TM6SF2;refGene_tx=XM_023247837.1;refGene_gn=TM6SF2;refGene_tx=XM_023247838.1
A2	14271993 14272097
refGene_gn=TM6SF2;refGene_tx=XM_023247832.1;refGene_gn=TM6SF2;refGene_tx=XM_023247834.1;refGene_gn=TM6SF2;refGene_tx=XM_023247835.1;refGene_gn=TM6SF2;refGene_tx=XM_023247837.1;refGene_gn=TM6SF2;refGene_tx=XM_023247838.1
A2	14273885 14273980
refGene_gn=TM6SF2;refGene_tx=XM_023247832.1;refGene_gn=TM6SF2;refGene_tx=XM_023247834.1;refGene_gn=TM6SF2;refGene_tx=XM_023247835.1;refGene_gn=TM6SF2;refGene_tx=XM_023247837.1;refGene_gn=TM6SF2;refGene_tx=XM_023247838.1
A2	14277005 14277032
refGene_gn=SUGP1;refGene_tx=XM_003982120.5;refGene_gn=SUGP1;refGene_tx=XM_006928659.4
A2	14277301 14277431
refGene_gn=SUGP1;refGene_tx=XM_003982120.5;refGene_gn=SUGP1;refGene_tx

=XM_006928659.4
A2 14277512 14277658
refGene_gn=SUGP1;refGene_tx=XM_003982120.5;refGene_gn=SUGP1;refGene_tx
=XM_006928659.4
A2 14278504 14278611
refGene_gn=SUGP1;refGene_tx=XM_003982120.5;refGene_gn=SUGP1;refGene_tx
=XM_006928659.4
A2 14279037 14279215
refGene_gn=SUGP1;refGene_tx=XM_003982120.5;refGene_gn=SUGP1;refGene_tx
=XM_006928659.4
A2 14280103 14280210
refGene_gn=SUGP1;refGene_tx=XM_003982120.5;refGene_gn=SUGP1;refGene_tx
=XM_006928659.4
A2 14287996 14288352
refGene_gn=SUGP1;refGene_tx=XM_003982120.5;refGene_gn=SUGP1;refGene_tx
=XM_006928659.4
A2 14297961 14298085
refGene_gn=SUGP1;refGene_tx=XM_003982120.5;refGene_gn=SUGP1;refGene_tx
=XM_006928659.4
A2 14298953 14299054
refGene_gn=SUGP1;refGene_tx=XM_003982120.5;refGene_gn=SUGP1;refGene_tx
=XM_006928659.4
A2 14299382 14299506
refGene_gn=SUGP1;refGene_tx=XM_003982120.5;refGene_gn=SUGP1;refGene_tx
=XM_006928659.4
A2 14301607 14301808
refGene_gn=SUGP1;refGene_tx=XM_003982120.5;refGene_gn=SUGP1;refGene_tx
=XM_006928659.4
A2 14303232 14303333
refGene_gn=SUGP1;refGene_tx=XM_003982120.5;refGene_gn=SUGP1;refGene_tx
=XM_006928659.4
A2 14307448 14307623
refGene_gn=SUGP1;refGene_tx=XM_003982120.5;refGene_gn=SUGP1;refGene_tx
=XM_006928659.4
A2 14309736 14309770
refGene_gn=SUGP1;refGene_tx=XM_003982120.5;refGene_gn=SUGP1;refGene_tx
=XM_006928659.4
A2 14310125 14310401
refGene_gn=MAU2;refGene_tx=XM_003982121.5;refGene_gn=MAU2;refGene_tx=X
M_006928660.4;refGene_gn=MAU2;refGene_tx=XM_006928661.4
A2 14318784 14318802
refGene_gn=MAU2;refGene_tx=XM_003982121.5;refGene_gn=MAU2;refGene_tx=X
M_006928660.4;refGene_gn=MAU2;refGene_tx=XM_006928661.4
A2 14319203 14319269
refGene_gn=MAU2;refGene_tx=XM_003982121.5;refGene_gn=MAU2;refGene_tx=X
M_006928660.4;refGene_gn=MAU2;refGene_tx=XM_006928661.4
A2 14319931 14320027
refGene_gn=MAU2;refGene_tx=XM_003982121.5;refGene_gn=MAU2;refGene_tx=X
M_006928660.4;refGene_gn=MAU2;refGene_tx=XM_006928661.4
A2 14321697 14321792

refGene_gn=MAU2;refGene_tx=XM_003982121.5;refGene_gn=MAU2;refGene_tx=XM_006928660.4;refGene_gn=MAU2;refGene_tx=XM_006928661.4
A2 14324121 14324149
refGene_gn=MAU2;refGene_tx=XM_003982121.5;refGene_gn=MAU2;refGene_tx=XM_006928660.4;refGene_gn=MAU2;refGene_tx=XM_006928661.4
A2 14324453 14324609
refGene_gn=MAU2;refGene_tx=XM_003982121.5;refGene_gn=MAU2;refGene_tx=XM_006928660.4;refGene_gn=MAU2;refGene_tx=XM_006928661.4
A2 14325828 14325975
refGene_gn=MAU2;refGene_tx=XM_003982121.5;refGene_gn=MAU2;refGene_tx=XM_006928660.4;refGene_gn=MAU2;refGene_tx=XM_006928661.4
A2 14326063 14326154
refGene_gn=MAU2;refGene_tx=XM_003982121.5;refGene_gn=MAU2;refGene_tx=XM_006928660.4;refGene_gn=MAU2;refGene_tx=XM_006928661.4
A2 14327307 14327411
refGene_gn=MAU2;refGene_tx=XM_003982121.5;refGene_gn=MAU2;refGene_tx=XM_006928660.4;refGene_gn=MAU2;refGene_tx=XM_006928661.4
A2 14328352 14328430
refGene_gn=MAU2;refGene_tx=XM_003982121.5;refGene_gn=MAU2;refGene_tx=XM_006928660.4;refGene_gn=MAU2;refGene_tx=XM_006928661.4
A2 14328710 14328776
refGene_gn=MAU2;refGene_tx=XM_003982121.5;refGene_gn=MAU2;refGene_tx=XM_006928660.4;refGene_gn=MAU2;refGene_tx=XM_006928661.4
A2 14330847 14330934
refGene_gn=MAU2;refGene_tx=XM_003982121.5;refGene_gn=MAU2;refGene_tx=XM_006928660.4;refGene_gn=MAU2;refGene_tx=XM_006928661.4
A2 14332482 14332532
refGene_gn=MAU2;refGene_tx=XM_003982121.5;refGene_gn=MAU2;refGene_tx=XM_006928660.4;refGene_gn=MAU2;refGene_tx=XM_006928661.4
A2 14332729 14332807
refGene_gn=MAU2;refGene_tx=XM_003982121.5;refGene_gn=MAU2;refGene_tx=XM_006928660.4;refGene_gn=MAU2;refGene_tx=XM_006928661.4
A2 14332984 14333096
refGene_gn=MAU2;refGene_tx=XM_003982121.5;refGene_gn=MAU2;refGene_tx=XM_006928660.4;refGene_gn=MAU2;refGene_tx=XM_006928661.4
A2 14336014 14336105
refGene_gn=MAU2;refGene_tx=XM_003982121.5;refGene_gn=MAU2;refGene_tx=XM_006928660.4;refGene_gn=MAU2;refGene_tx=XM_006928661.4
A2 14336980 14337108
refGene_gn=MAU2;refGene_tx=XM_003982121.5;refGene_gn=MAU2;refGene_tx=XM_006928660.4;refGene_gn=MAU2;refGene_tx=XM_006928661.4
A2 14337476 14337551
refGene_gn=MAU2;refGene_tx=XM_003982121.5;refGene_gn=MAU2;refGene_tx=XM_006928660.4;refGene_gn=MAU2;refGene_tx=XM_006928661.4
A2 14399164 14399179
refGene_gn=GATAD2A;refGene_tx=XM_006928662.4;refGene_gn=GATAD2A;refGene_tx=XM_023247857.1;refGene_gn=GATAD2A;refGene_tx=XM_023247858.1
A2 14422426 14422695
refGene_gn=GATAD2A;refGene_tx=XM_006928662.4;refGene_gn=GATAD2A;refGene_tx=XM_019820232.2;refGene_gn=GATAD2A;refGene_tx=XM_023247853.1;refGe

ne_gn=GATAD2A;refGene_tx=XM_023247857.1;refGene_gn=GATAD2A;refGene_tx=XM_023247858.1
A2 14446006 14446139
refGene_gn=GATAD2A;refGene_tx=XM_006928662.4;refGene_gn=GATAD2A;refGene_tx=XM_019820232.2;refGene_gn=GATAD2A;refGene_tx=XM_023247853.1;refGene_gn=GATAD2A;refGene_tx=XM_023247857.1;refGene_gn=GATAD2A;refGene_tx=XM_023247858.1
A2 14446265 14446397
refGene_gn=GATAD2A;refGene_tx=XM_006928662.4;refGene_gn=GATAD2A;refGene_tx=XM_019820232.2;refGene_gn=GATAD2A;refGene_tx=XM_023247853.1;refGene_gn=GATAD2A;refGene_tx=XM_023247857.1;refGene_gn=GATAD2A;refGene_tx=XM_023247858.1
A2 14448147 14448237
refGene_gn=GATAD2A;refGene_tx=XM_006928662.4;refGene_gn=GATAD2A;refGene_tx=XM_019820232.2;refGene_gn=GATAD2A;refGene_tx=XM_023247853.1;refGene_gn=GATAD2A;refGene_tx=XM_023247857.1;refGene_gn=GATAD2A;refGene_tx=XM_023247858.1
A2 14448851 14448983
refGene_gn=GATAD2A;refGene_tx=XM_006928662.4;refGene_gn=GATAD2A;refGene_tx=XM_019820232.2;refGene_gn=GATAD2A;refGene_tx=XM_023247853.1;refGene_gn=GATAD2A;refGene_tx=XM_023247857.1;refGene_gn=GATAD2A;refGene_tx=XM_023247858.1
A2 14449150 14449318
refGene_gn=GATAD2A;refGene_tx=XM_006928662.4;refGene_gn=GATAD2A;refGene_tx=XM_019820232.2;refGene_gn=GATAD2A;refGene_tx=XM_023247853.1;refGene_gn=GATAD2A;refGene_tx=XM_023247857.1;refGene_gn=GATAD2A;refGene_tx=XM_023247858.1
A2 14450923 14451203
refGene_gn=GATAD2A;refGene_tx=XM_006928662.4;refGene_gn=GATAD2A;refGene_tx=XM_019820232.2;refGene_gn=GATAD2A;refGene_tx=XM_023247853.1;refGene_gn=GATAD2A;refGene_tx=XM_023247857.1;refGene_gn=GATAD2A;refGene_tx=XM_023247858.1
A2 14454099 14454398
refGene_gn=GATAD2A;refGene_tx=XM_006928662.4;refGene_gn=GATAD2A;refGene_tx=XM_019820232.2;refGene_gn=GATAD2A;refGene_tx=XM_023247853.1;refGene_gn=GATAD2A;refGene_tx=XM_023247857.1;refGene_gn=GATAD2A;refGene_tx=XM_023247858.1
A2 14455077 14455152
refGene_gn=GATAD2A;refGene_tx=XM_006928662.4;refGene_gn=GATAD2A;refGene_tx=XM_019820232.2;refGene_gn=GATAD2A;refGene_tx=XM_023247857.1;refGene_gn=GATAD2A;refGene_tx=XM_023247858.1
A2 14455436 14455632
refGene_gn=GATAD2A;refGene_tx=XM_006928662.4;refGene_gn=GATAD2A;refGene_tx=XM_019820232.2;refGene_gn=GATAD2A;refGene_tx=XM_023247853.1;refGene_gn=GATAD2A;refGene_tx=XM_023247857.1;refGene_gn=GATAD2A;refGene_tx=XM_023247858.1
A2 14458007 14458138
refGene_gn=GATAD2A;refGene_tx=XM_006928662.4;refGene_gn=GATAD2A;refGene_tx=XM_019820232.2;refGene_gn=GATAD2A;refGene_tx=XM_023247853.1;refGene_gn=GATAD2A;refGene_tx=XM_023247857.1;refGene_gn=GATAD2A;refGene_tx=

XM_023247858.1
A2	14464780 14465602 refGene_gn=TSSK6;refGene_tx=XM_003982123.5
A2	14466437 14466531 refGene_gn=NDUFA13;refGene_tx=XM_003982124.5
A2	14472896 14472975 refGene_gn=NDUFA13;refGene_tx=XM_003982124.5
A2	14474179 14474251 refGene_gn=NDUFA13;refGene_tx=XM_003982124.5
A2	14474593 14474663 refGene_gn=NDUFA13;refGene_tx=XM_003982124.5
A2	14474886 14475006 refGene_gn=NDUFA13;refGene_tx=XM_003982124.5
A2	14475713 14475772
refGene_gn=YJEFN3;refGene_tx=XM_003982125.5;refGene_gn=YJEFN3;refGene_tx=XM_019820260.2
A2	14478997 14479106
refGene_gn=YJEFN3;refGene_tx=XM_003982125.5;refGene_gn=YJEFN3;refGene_tx=XM_019820260.2
A2	14480947 14481058
refGene_gn=YJEFN3;refGene_tx=XM_003982125.5;refGene_gn=YJEFN3;refGene_tx=XM_019820260.2
A2	14481283 14481397
refGene_gn=YJEFN3;refGene_tx=XM_003982125.5;refGene_gn=YJEFN3;refGene_tx=XM_019820260.2
A2	14481477 14481628
refGene_gn=YJEFN3;refGene_tx=XM_003982125.5;refGene_gn=YJEFN3;refGene_tx=XM_019820260.2
A2	14482977 14483226
refGene_gn=YJEFN3;refGene_tx=XM_003982125.5;refGene_gn=YJEFN3;refGene_tx=XM_019820260.2
A2	14483899 14483963 refGene_gn=CILP2;refGene_tx=XM_003982126.4
A2	14484990 14485089 refGene_gn=CILP2;refGene_tx=XM_003982126.4
A2	14486061 14486334 refGene_gn=CILP2;refGene_tx=XM_003982126.4
A2	14486986 14487142 refGene_gn=CILP2;refGene_tx=XM_003982126.4
A2	14488209 14488488 refGene_gn=CILP2;refGene_tx=XM_003982126.4
A2	14488891 14489000 refGene_gn=CILP2;refGene_tx=XM_003982126.4
A2	14489194 14489352 refGene_gn=CILP2;refGene_tx=XM_003982126.4
A2	14489591 14491927 refGene_gn=CILP2;refGene_tx=XM_003982126.4
A2	14505438 14505531
refGene_gn=PBX4;refGene_tx=XM_003982174.4;refGene_gn=PBX4;refGene_tx=XM_023247888.1
A2	14507877 14507984
refGene_gn=PBX4;refGene_tx=XM_003982174.4;refGene_gn=PBX4;refGene_tx=XM_023247888.1
A2	14508195 14508337
refGene_gn=PBX4;refGene_tx=XM_003982174.4;refGene_gn=PBX4;refGene_tx=XM_023247888.1
A2	14511235 14511371
refGene_gn=PBX4;refGene_tx=XM_003982174.4;refGene_gn=PBX4;refGene_tx=XM_023247888.1
A2	14511639 14511830
refGene_gn=PBX4;refGene_tx=XM_003982174.4;refGene_gn=PBX4;refGene_tx=XM_023247888.1
A2	14512108 14512356
refGene_gn=PBX4;refGene_tx=XM_003982174.4;refGene_gn=PBX4;refGene_tx=X

M_023247888.1
A2	14529258	14529332
refGene_gn=PBX4;refGene_tx=XM_003982174.4;refGene_gn=PBX4;refGene_tx=X
M_023247888.1
A2	14547861	14548013	refGene_gn=PBX4;refGene_tx=XM_003982174.4
A2	14552971	14553285	refGene_gn=LPAR2;refGene_tx=XM_023247902.1
A2	14554013	14554755	refGene_gn=LPAR2;refGene_tx=XM_023247902.1
A2	14557271	14557645
refGene_gn=GMIP;refGene_tx=XM_003982128.4;refGene_gn=GMIP;refGene_tx=X
M_019820297.2;refGene_gn=GMIP;refGene_tx=XM_019820303.2;refGene_gn=GMI
P;refGene_tx=XM_019820304.2;refGene_gn=GMIP;refGene_tx=XM_019820305.2
A2	14557774	14557838
refGene_gn=GMIP;refGene_tx=XM_003982128.4;refGene_gn=GMIP;refGene_tx=X
M_019820297.2;refGene_gn=GMIP;refGene_tx=XM_019820303.2;refGene_gn=GMI
P;refGene_tx=XM_019820304.2;refGene_gn=GMIP;refGene_tx=XM_019820305.2
A2	14558273	14558324
refGene_gn=GMIP;refGene_tx=XM_019820297.2;refGene_gn=GMIP;refGene_tx=X
M_019820303.2;refGene_gn=GMIP;refGene_tx=XM_019820305.2
A2	14559227	14559615
refGene_gn=GMIP;refGene_tx=XM_003982128.4;refGene_gn=GMIP;refGene_tx=X
M_019820297.2;refGene_gn=GMIP;refGene_tx=XM_019820303.2;refGene_gn=GMI
P;refGene_tx=XM_019820304.2;refGene_gn=GMIP;refGene_tx=XM_019820305.2
A2	14559954	14560151
refGene_gn=GMIP;refGene_tx=XM_003982128.4;refGene_gn=GMIP;refGene_tx=X
M_019820297.2;refGene_gn=GMIP;refGene_tx=XM_019820303.2;refGene_gn=GMI
P;refGene_tx=XM_019820304.2;refGene_gn=GMIP;refGene_tx=XM_019820305.2
A2	14560252	14560390
refGene_gn=GMIP;refGene_tx=XM_003982128.4;refGene_gn=GMIP;refGene_tx=X
M_019820297.2;refGene_gn=GMIP;refGene_tx=XM_019820303.2;refGene_gn=GMI
P;refGene_tx=XM_019820304.2;refGene_gn=GMIP;refGene_tx=XM_019820305.2
A2	14560485	14560674
refGene_gn=GMIP;refGene_tx=XM_003982128.4;refGene_gn=GMIP;refGene_tx=X
M_019820297.2;refGene_gn=GMIP;refGene_tx=XM_019820303.2;refGene_gn=GMI
P;refGene_tx=XM_019820304.2;refGene_gn=GMIP;refGene_tx=XM_019820305.2
A2	14560890	14561045
refGene_gn=GMIP;refGene_tx=XM_003982128.4;refGene_gn=GMIP;refGene_tx=X
M_019820297.2;refGene_gn=GMIP;refGene_tx=XM_019820303.2;refGene_gn=GMI
P;refGene_tx=XM_019820304.2;refGene_gn=GMIP;refGene_tx=XM_019820305.2
A2	14561113	14561191
refGene_gn=GMIP;refGene_tx=XM_003982128.4;refGene_gn=GMIP;refGene_tx=X
M_019820297.2;refGene_gn=GMIP;refGene_tx=XM_019820305.2
A2	14561550	14561640
refGene_gn=GMIP;refGene_tx=XM_003982128.4;refGene_gn=GMIP;refGene_tx=X
M_019820297.2;refGene_gn=GMIP;refGene_tx=XM_019820303.2;refGene_gn=GMI
P;refGene_tx=XM_019820304.2;refGene_gn=GMIP;refGene_tx=XM_019820305.2
A2	14561713	14561820
refGene_gn=GMIP;refGene_tx=XM_003982128.4;refGene_gn=GMIP;refGene_tx=X
M_019820297.2;refGene_gn=GMIP;refGene_tx=XM_019820303.2;refGene_gn=GMI
P;refGene_tx=XM_019820304.2;refGene_gn=GMIP;refGene_tx=XM_019820305.2
A2	14562139	14562336

refGene_gn=GMIP;refGene_tx=XM_003982128.4;refGene_gn=GMIP;refGene_tx=XM_019820297.2;refGene_gn=GMIP;refGene_tx=XM_019820303.2;refGene_gn=GMIP;refGene_tx=XM_019820304.2;refGene_gn=GMIP;refGene_tx=XM_019820305.2
A2 14562620 14562758
refGene_gn=GMIP;refGene_tx=XM_003982128.4;refGene_gn=GMIP;refGene_tx=XM_019820297.2;refGene_gn=GMIP;refGene_tx=XM_019820303.2;refGene_gn=GMIP;refGene_tx=XM_019820304.2;refGene_gn=GMIP;refGene_tx=XM_019820305.2
A2 14562851 14563022
refGene_gn=GMIP;refGene_tx=XM_003982128.4;refGene_gn=GMIP;refGene_tx=XM_019820297.2;refGene_gn=GMIP;refGene_tx=XM_019820303.2;refGene_gn=GMIP;refGene_tx=XM_019820304.2;refGene_gn=GMIP;refGene_tx=XM_019820305.2
A2 14563124 14563205
refGene_gn=GMIP;refGene_tx=XM_003982128.4;refGene_gn=GMIP;refGene_tx=XM_019820297.2;refGene_gn=GMIP;refGene_tx=XM_019820303.2;refGene_gn=GMIP;refGene_tx=XM_019820304.2;refGene_gn=GMIP;refGene_tx=XM_019820305.2
A2 14563614 14563722
refGene_gn=GMIP;refGene_tx=XM_003982128.4;refGene_gn=GMIP;refGene_tx=XM_019820297.2;refGene_gn=GMIP;refGene_tx=XM_019820303.2;refGene_gn=GMIP;refGene_tx=XM_019820304.2;refGene_gn=GMIP;refGene_tx=XM_019820305.2
A2 14563797 14563862
refGene_gn=GMIP;refGene_tx=XM_003982128.4;refGene_gn=GMIP;refGene_tx=XM_019820297.2;refGene_gn=GMIP;refGene_tx=XM_019820303.2;refGene_gn=GMIP;refGene_tx=XM_019820304.2;refGene_gn=GMIP;refGene_tx=XM_019820305.2
A2 14563946 14564072
refGene_gn=GMIP;refGene_tx=XM_003982128.4;refGene_gn=GMIP;refGene_tx=XM_019820297.2;refGene_gn=GMIP;refGene_tx=XM_019820303.2;refGene_gn=GMIP;refGene_tx=XM_019820304.2
A2 14564381 14564439
refGene_gn=GMIP;refGene_tx=XM_003982128.4;refGene_gn=GMIP;refGene_tx=XM_019820297.2;refGene_gn=GMIP;refGene_tx=XM_019820303.2;refGene_gn=GMIP;refGene_tx=XM_019820304.2
A2 14564551 14564627
refGene_gn=GMIP;refGene_tx=XM_003982128.4;refGene_gn=GMIP;refGene_tx=XM_019820297.2;refGene_gn=GMIP;refGene_tx=XM_019820303.2;refGene_gn=GMIP;refGene_tx=XM_019820304.2
A2 14565088 14565173
refGene_gn=GMIP;refGene_tx=XM_003982128.4;refGene_gn=GMIP;refGene_tx=XM_019820297.2;refGene_gn=GMIP;refGene_tx=XM_019820303.2;refGene_gn=GMIP;refGene_tx=XM_019820304.2
A2 14566091 14566110
refGene_gn=GMIP;refGene_tx=XM_003982128.4;refGene_gn=GMIP;refGene_tx=XM_019820297.2;refGene_gn=GMIP;refGene_tx=XM_019820303.2;refGene_gn=GMIP;refGene_tx=XM_019820304.2
A2 14567970 14568081 refGene_gn=ATP13A1;refGene_tx=XM_023247905.1
A2 14568168 14568312 refGene_gn=ATP13A1;refGene_tx=XM_023247905.1
A2 14568398 14568510 refGene_gn=ATP13A1;refGene_tx=XM_023247905.1
A2 14568708 14568851 refGene_gn=ATP13A1;refGene_tx=XM_023247905.1
A2 14569649 14569846 refGene_gn=ATP13A1;refGene_tx=XM_023247905.1
A2 14569938 14570053 refGene_gn=ATP13A1;refGene_tx=XM_023247905.1
A2 14570122 14570283 refGene_gn=ATP13A1;refGene_tx=XM_023247905.1

A2 14571262 14571359 refGene_gn=ATP13A1;refGene_tx=XM_023247905.1
A2 14571441 14571641 refGene_gn=ATP13A1;refGene_tx=XM_023247905.1
A2 14573062 14573171 refGene_gn=ATP13A1;refGene_tx=XM_023247905.1
A2 14574013 14574139 refGene_gn=ATP13A1;refGene_tx=XM_023247905.1
A2 14575142 14575253 refGene_gn=ATP13A1;refGene_tx=XM_023247905.1
A2 14575324 14575500 refGene_gn=ATP13A1;refGene_tx=XM_023247905.1
A2 14575888 14576046 refGene_gn=ATP13A1;refGene_tx=XM_023247905.1
A2 14576409 14576530 refGene_gn=ATP13A1;refGene_tx=XM_023247905.1
A2 14576610 14576748 refGene_gn=ATP13A1;refGene_tx=XM_023247905.1
A2 14576825 14576952 refGene_gn=ATP13A1;refGene_tx=XM_023247905.1
A2 14577539 14577595 refGene_gn=ATP13A1;refGene_tx=XM_023247905.1
A2 14577721 14577851 refGene_gn=ATP13A1;refGene_tx=XM_023247905.1
A2 14578339 14578446 refGene_gn=ATP13A1;refGene_tx=XM_023247905.1
A2 14578523 14578593 refGene_gn=ATP13A1;refGene_tx=XM_023247905.1
A2 14578753 14578909 refGene_gn=ATP13A1;refGene_tx=XM_023247905.1
A2 14579129 14579202 refGene_gn=ATP13A1;refGene_tx=XM_023247905.1
A2 14581542 14581733 refGene_gn=ATP13A1;refGene_tx=XM_023247905.1
A2 14581859 14581949 refGene_gn=ATP13A1;refGene_tx=XM_023247905.1
A2 14584225 14584621 refGene_gn=ATP13A1;refGene_tx=XM_023247905.1
A2 14636432 14636487
refGene_gn=ZNF502;refGene_tx=XM_003982131.5;refGene_gn=ZNF502;refGene_tx=XM_019820328.2;refGene_gn=ZNF502;refGene_tx=XM_019820332.2
A2 14636969 14638543
refGene_gn=ZNF502;refGene_tx=XM_003982131.5;refGene_gn=ZNF502;refGene_tx=XM_019820328.2;refGene_gn=ZNF502;refGene_tx=XM_019820332.2
A2 14645948 14646023 refGene_gn=ZNF502;refGene_tx=XM_019820332.2
A2 14648405 14648443 refGene_gn=ZNF502;refGene_tx=XM_019820332.2
A2 14653201 14653422 refGene_gn=KIAA1143;refGene_tx=XM_003982132.4
A2 14654099 14654244 refGene_gn=KIAA1143;refGene_tx=XM_003982132.4
A2 14658874 14658982 refGene_gn=KIAA1143;refGene_tx=XM_003982132.4
A2 14659243 14659262
refGene_gn=KIF15;refGene_tx=XM_023247936.1;refGene_gn=KIF15;refGene_tx=XM_023247945.1;refGene_gn=KIF15;refGene_tx=XM_023247950.1;refGene_gn=KIF15;refGene_tx=XM_023247951.1
A2 14667883 14667926
refGene_gn=KIF15;refGene_tx=XM_023247936.1;refGene_gn=KIF15;refGene_tx=XM_023247945.1;refGene_gn=KIF15;refGene_tx=XM_023247950.1;refGene_gn=KIF15;refGene_tx=XM_023247951.1
A2 14668711 14668895
refGene_gn=KIF15;refGene_tx=XM_023247936.1;refGene_gn=KIF15;refGene_tx=XM_023247945.1;refGene_gn=KIF15;refGene_tx=XM_023247950.1;refGene_gn=KIF15;refGene_tx=XM_023247951.1
A2 14671160 14671237
refGene_gn=KIF15;refGene_tx=XM_023247936.1;refGene_gn=KIF15;refGene_tx=XM_023247945.1;refGene_gn=KIF15;refGene_tx=XM_023247950.1;refGene_gn=KIF15;refGene_tx=XM_023247951.1
A2 14676995 14677033
refGene_gn=KIF15;refGene_tx=XM_023247936.1;refGene_gn=KIF15;refGene_tx=XM_023247945.1;refGene_gn=KIF15;refGene_tx=XM_023247950.1;refGene_gn=KIF15;refGene_tx=XM_023247951.1

A2 14679038 14679136
refGene_gn=KIF15;refGene_tx=XM_023247936.1;refGene_gn=KIF15;refGene_tx=XM_023247945.1;refGene_gn=KIF15;refGene_tx=XM_023247950.1;refGene_gn=KIF15;refGene_tx=XM_023247951.1
A2 14680448 14680628
refGene_gn=KIF15;refGene_tx=XM_023247936.1;refGene_gn=KIF15;refGene_tx=XM_023247945.1;refGene_gn=KIF15;refGene_tx=XM_023247950.1;refGene_gn=KIF15;refGene_tx=XM_023247951.1
A2 14688137 14688347
refGene_gn=KIF15;refGene_tx=XM_023247936.1;refGene_gn=KIF15;refGene_tx=XM_023247945.1;refGene_gn=KIF15;refGene_tx=XM_023247950.1;refGene_gn=KIF15;refGene_tx=XM_023247951.1
A2 14691552 14691678
refGene_gn=KIF15;refGene_tx=XM_023247936.1;refGene_gn=KIF15;refGene_tx=XM_023247945.1;refGene_gn=KIF15;refGene_tx=XM_023247950.1;refGene_gn=KIF15;refGene_tx=XM_023247951.1
A2 14691838 14691961
refGene_gn=KIF15;refGene_tx=XM_023247936.1;refGene_gn=KIF15;refGene_tx=XM_023247945.1;refGene_gn=KIF15;refGene_tx=XM_023247950.1;refGene_gn=KIF15;refGene_tx=XM_023247951.1
A2 14695426 14695550
refGene_gn=KIF15;refGene_tx=XM_023247936.1;refGene_gn=KIF15;refGene_tx=XM_023247945.1;refGene_gn=KIF15;refGene_tx=XM_023247950.1;refGene_gn=KIF15;refGene_tx=XM_023247951.1
A2 14699106 14699183
refGene_gn=KIF15;refGene_tx=XM_023247936.1;refGene_gn=KIF15;refGene_tx=XM_023247945.1;refGene_gn=KIF15;refGene_tx=XM_023247950.1;refGene_gn=KIF15;refGene_tx=XM_023247951.1
A2 14699415 14699625
refGene_gn=KIF15;refGene_tx=XM_023247945.1;refGene_gn=KIF15;refGene_tx=XM_023247950.1;refGene_gn=KIF15;refGene_tx=XM_023247951.1
A2 14700724 14700906
refGene_gn=KIF15;refGene_tx=XM_023247936.1;refGene_gn=KIF15;refGene_tx=XM_023247945.1;refGene_gn=KIF15;refGene_tx=XM_023247950.1;refGene_gn=KIF15;refGene_tx=XM_023247951.1
A2 14705500 14705638
refGene_gn=KIF15;refGene_tx=XM_023247936.1;refGene_gn=KIF15;refGene_tx=XM_023247945.1;refGene_gn=KIF15;refGene_tx=XM_023247950.1;refGene_gn=KIF15;refGene_tx=XM_023247951.1
A2 14706749 14706891
refGene_gn=KIF15;refGene_tx=XM_023247936.1;refGene_gn=KIF15;refGene_tx=XM_023247945.1;refGene_gn=KIF15;refGene_tx=XM_023247950.1;refGene_gn=KIF15;refGene_tx=XM_023247951.1
A2 14714317 14714515
refGene_gn=KIF15;refGene_tx=XM_023247936.1;refGene_gn=KIF15;refGene_tx=XM_023247945.1;refGene_gn=KIF15;refGene_tx=XM_023247950.1;refGene_gn=KIF15;refGene_tx=XM_023247951.1
A2 14715574 14715682
refGene_gn=KIF15;refGene_tx=XM_023247936.1;refGene_gn=KIF15;refGene_tx=XM_023247945.1;refGene_gn=KIF15;refGene_tx=XM_023247950.1;refGene_gn=

KIF15;refGene_tx=XM_023247951.1
A2 14716108 14716214
refGene_gn=KIF15;refGene_tx=XM_023247936.1;refGene_gn=KIF15;refGene_tx=XM_023247945.1;refGene_gn=KIF15;refGene_tx=XM_023247950.1;refGene_gn=KIF15;refGene_tx=XM_023247951.1
A2 14717161 14717327
refGene_gn=KIF15;refGene_tx=XM_023247936.1;refGene_gn=KIF15;refGene_tx=XM_023247945.1;refGene_gn=KIF15;refGene_tx=XM_023247950.1;refGene_gn=KIF15;refGene_tx=XM_023247951.1
A2 14720351 14720502
refGene_gn=KIF15;refGene_tx=XM_023247936.1;refGene_gn=KIF15;refGene_tx=XM_023247950.1;refGene_gn=KIF15;refGene_tx=XM_023247951.1
A2 14720710 14720796
refGene_gn=KIF15;refGene_tx=XM_023247936.1;refGene_gn=KIF15;refGene_tx=XM_023247950.1;refGene_gn=KIF15;refGene_tx=XM_023247951.1
A2 14721789 14721859
refGene_gn=KIF15;refGene_tx=XM_023247936.1;refGene_gn=KIF15;refGene_tx=XM_023247945.1;refGene_gn=KIF15;refGene_tx=XM_023247950.1;refGene_gn=KIF15;refGene_tx=XM_023247951.1
A2 14722248 14722335
refGene_gn=KIF15;refGene_tx=XM_023247936.1;refGene_gn=KIF15;refGene_tx=XM_023247945.1;refGene_gn=KIF15;refGene_tx=XM_023247950.1;refGene_gn=KIF15;refGene_tx=XM_023247951.1
A2 14723135 14723240
refGene_gn=KIF15;refGene_tx=XM_023247936.1;refGene_gn=KIF15;refGene_tx=XM_023247945.1;refGene_gn=KIF15;refGene_tx=XM_023247950.1;refGene_gn=KIF15;refGene_tx=XM_023247951.1
A2 14724183 14724306
refGene_gn=KIF15;refGene_tx=XM_023247936.1;refGene_gn=KIF15;refGene_tx=XM_023247945.1;refGene_gn=KIF15;refGene_tx=XM_023247950.1;refGene_gn=KIF15;refGene_tx=XM_023247951.1
A2 14729263 14729410
refGene_gn=KIF15;refGene_tx=XM_023247936.1;refGene_gn=KIF15;refGene_tx=XM_023247945.1;refGene_gn=KIF15;refGene_tx=XM_023247950.1;refGene_gn=KIF15;refGene_tx=XM_023247951.1
A2 14730443 14730545
refGene_gn=KIF15;refGene_tx=XM_023247936.1;refGene_gn=KIF15;refGene_tx=XM_023247945.1;refGene_gn=KIF15;refGene_tx=XM_023247950.1;refGene_gn=KIF15;refGene_tx=XM_023247951.1
A2 14730814 14730979
refGene_gn=KIF15;refGene_tx=XM_023247936.1;refGene_gn=KIF15;refGene_tx=XM_023247945.1;refGene_gn=KIF15;refGene_tx=XM_023247950.1;refGene_gn=KIF15;refGene_tx=XM_023247951.1
A2 14732716 14732826
refGene_gn=KIF15;refGene_tx=XM_023247936.1;refGene_gn=KIF15;refGene_tx=XM_023247945.1;refGene_gn=KIF15;refGene_tx=XM_023247950.1;refGene_gn=KIF15;refGene_tx=XM_023247951.1
A2 14735555 14735628
refGene_gn=KIF15;refGene_tx=XM_023247936.1;refGene_gn=KIF15;refGene_tx=XM_023247945.1;refGene_gn=KIF15;refGene_tx=XM_023247950.1;refGene_gn=

KIF15;refGene_tx=XM_023247951.1
A2 14736105 14736143
refGene_gn=KIF15;refGene_tx=XM_023247936.1;refGene_gn=KIF15;refGene_tx=XM_023247945.1;refGene_gn=KIF15;refGene_tx=XM_023247951.1
A2 14736199 14736226 refGene_gn=KIF15;refGene_tx=XM_023247950.1
A2 14740303 14740469
refGene_gn=KIF15;refGene_tx=XM_023247936.1;refGene_gn=KIF15;refGene_tx=XM_023247945.1;refGene_gn=KIF15;refGene_tx=XM_023247950.1;refGene_gn=KIF15;refGene_tx=XM_023247951.1
A2 14740691 14740823
refGene_gn=KIF15;refGene_tx=XM_023247936.1;refGene_gn=KIF15;refGene_tx=XM_023247945.1;refGene_gn=KIF15;refGene_tx=XM_023247951.1
A2 14741151 14741214
refGene_gn=KIF15;refGene_tx=XM_023247936.1;refGene_gn=KIF15;refGene_tx=XM_023247945.1;refGene_gn=KIF15;refGene_tx=XM_023247951.1
A2 14744506 14744698
refGene_gn=TMEM42;refGene_tx=XM_003982134.5;refGene_gn=TMEM42;refGene_tx=XM_019820401.2
A2 14745103 14745244 refGene_gn=TMEM42;refGene_tx=XM_019820410.2
A2 14746746 14746893
refGene_gn=TMEM42;refGene_tx=XM_003982134.5;refGene_gn=TMEM42;refGene_tx=XM_019820401.2;refGene_gn=TMEM42;refGene_tx=XM_019820410.2
A2 14747250 14747484
refGene_gn=TMEM42;refGene_tx=XM_019820401.2;refGene_gn=TMEM42;refGene_tx=XM_019820410.2
A2 14747640 14747781 refGene_gn=TMEM42;refGene_tx=XM_003982134.5
A2 14763223 14763481
refGene_gn=ZDHHC3;refGene_tx=XM_019820386.2;refGene_gn=ZDHHC3;refGene_tx=XM_023247979.1
A2 14772248 14772407
refGene_gn=ZDHHC3;refGene_tx=XM_011291062.3;refGene_gn=ZDHHC3;refGene_tx=XM_019820393.2
A2 14774017 14774281
refGene_gn=ZDHHC3;refGene_tx=XM_019820367.2;refGene_gn=ZDHHC3;refGene_tx=XM_019820369.2;refGene_gn=ZDHHC3;refGene_tx=XM_019820377.2;refGene_gn=ZDHHC3;refGene_tx=XM_019820390.2
A2 14774895 14775026
refGene_gn=ZDHHC3;refGene_tx=XM_011291062.3;refGene_gn=ZDHHC3;refGene_tx=XM_019820367.2;refGene_gn=ZDHHC3;refGene_tx=XM_019820369.2;refGene_gn=ZDHHC3;refGene_tx=XM_019820377.2;refGene_gn=ZDHHC3;refGene_tx=XM_019820386.2;refGene_gn=ZDHHC3;refGene_tx=XM_019820390.2;refGene_gn=ZDHHC3;refGene_tx=XM_019820393.2;refGene_gn=ZDHHC3;refGene_tx=XM_023247979.1
A2 14778715 14778808
refGene_gn=ZDHHC3;refGene_tx=XM_019820367.2;refGene_gn=ZDHHC3;refGene_tx=XM_019820369.2;refGene_gn=ZDHHC3;refGene_tx=XM_019820377.2;refGene_gn=ZDHHC3;refGene_tx=XM_019820386.2;refGene_gn=ZDHHC3;refGene_tx=XM_019820393.2
A2 14778918 14779000
refGene_gn=ZDHHC3;refGene_tx=XM_011291062.3;refGene_gn=ZDHHC3;refGene_

tx=XM_019820367.2;refGene_gn=ZDHHC3;refGene_tx=XM_019820369.2;refGene_gn=ZDHHC3;refGene_tx=XM_019820377.2;refGene_gn=ZDHHC3;refGene_tx=XM_019820386.2;refGene_gn=ZDHHC3;refGene_tx=XM_019820390.2;refGene_gn=ZDHHC3;refGene_tx=XM_019820393.2;refGene_gn=ZDHHC3;refGene_tx=XM_023247979.1
A2 14779656 14779753
refGene_gn=ZDHHC3;refGene_tx=XM_011291062.3;refGene_gn=ZDHHC3;refGene_tx=XM_019820367.2;refGene_gn=ZDHHC3;refGene_tx=XM_019820369.2;refGene_gn=ZDHHC3;refGene_tx=XM_019820377.2;refGene_gn=ZDHHC3;refGene_tx=XM_019820386.2;refGene_gn=ZDHHC3;refGene_tx=XM_019820390.2;refGene_gn=ZDHHC3;refGene_tx=XM_019820393.2;refGene_gn=ZDHHC3;refGene_tx=XM_023247979.1
A2 14786451 14786576
refGene_gn=ZDHHC3;refGene_tx=XM_011291062.3;refGene_gn=ZDHHC3;refGene_tx=XM_019820367.2;refGene_gn=ZDHHC3;refGene_tx=XM_019820369.2;refGene_gn=ZDHHC3;refGene_tx=XM_019820377.2;refGene_gn=ZDHHC3;refGene_tx=XM_019820386.2;refGene_gn=ZDHHC3;refGene_tx=XM_019820390.2;refGene_gn=ZDHHC3;refGene_tx=XM_019820393.2;refGene_gn=ZDHHC3;refGene_tx=XM_023247979.1
A2 14803822 14804128
refGene_gn=ZDHHC3;refGene_tx=XM_011291062.3;refGene_gn=ZDHHC3;refGene_tx=XM_019820367.2;refGene_gn=ZDHHC3;refGene_tx=XM_019820369.2;refGene_gn=ZDHHC3;refGene_tx=XM_019820377.2;refGene_gn=ZDHHC3;refGene_tx=XM_019820386.2;refGene_gn=ZDHHC3;refGene_tx=XM_019820390.2;refGene_gn=ZDHHC3;refGene_tx=XM_019820393.2;refGene_gn=ZDHHC3;refGene_tx=XM_023247979.1
A2 14822591 14822648 refGene_gn=EXOSC7;refGene_tx=XM_003982137.5
A2 14834923 14835025 refGene_gn=EXOSC7;refGene_tx=XM_003982137.5
A2 14835314 14835409 refGene_gn=EXOSC7;refGene_tx=XM_003982137.5
A2 14843932 14844098 refGene_gn=EXOSC7;refGene_tx=XM_003982137.5
A2 14849307 14849378 refGene_gn=EXOSC7;refGene_tx=XM_003982137.5
A2 14854655 14854779 refGene_gn=EXOSC7;refGene_tx=XM_003982137.5
A2 14855850 14856006 refGene_gn=EXOSC7;refGene_tx=XM_003982137.5
A2 14859654 14859759 refGene_gn=EXOSC7;refGene_tx=XM_003982137.5
A2 14871820 14871929 refGene_gn=CLEC3B;refGene_tx=XM_003982138.5
A2 14875380 14875479 refGene_gn=CLEC3B;refGene_tx=XM_003982138.5
A2 14878514 14878915 refGene_gn=CLEC3B;refGene_tx=XM_003982138.5
A2 14907039 14907049 refGene_gn=CDCP1;refGene_tx=XM_023248005.1
A2 14911408 14911823 refGene_gn=CDCP1;refGene_tx=XM_023248002.1
A2 14914276 14914364
refGene_gn=CDCP1;refGene_tx=XM_023248002.1;refGene_gn=CDCP1;refGene_tx=XM_023248005.1
A2 14916543 14916909
refGene_gn=CDCP1;refGene_tx=XM_023248002.1;refGene_gn=CDCP1;refGene_tx=XM_023248005.1
A2 14919273 14919654
refGene_gn=CDCP1;refGene_tx=XM_023248002.1;refGene_gn=CDCP1;refGene_tx=XM_023248005.1
A2 14921308 14921530
refGene_gn=CDCP1;refGene_tx=XM_023248002.1;refGene_gn=CDCP1;refGene_tx

=XM_023248005.1
A2 14930807 14931176
refGene_gn=CDCP1;refGene_tx=XM_023248002.1;refGene_gn=CDCP1;refGene_tx
=XM_023248005.1
A2 14932742 14933105
refGene_gn=CDCP1;refGene_tx=XM_023248002.1;refGene_gn=CDCP1;refGene_tx
=XM_023248005.1
A2 14939387 14939597
refGene_gn=CDCP1;refGene_tx=XM_023248002.1;refGene_gn=CDCP1;refGene_tx
=XM_023248005.1
A2 14968984 14969060
refGene_gn=CDCP1;refGene_tx=XM_023248002.1;refGene_gn=CDCP1;refGene_tx
=XM_023248005.1
A2 15034660 15035029 refGene_gn=TMEM158;refGene_tx=XM_023248010.1
A2 15035030 15035573 refGene_gn=TMEM158;refGene_tx=XM_023248010.1
A2 15161294 15161528 refGene_gn=LARS2;refGene_tx=XM_003982139.5
A2 15168360 15168489 refGene_gn=LARS2;refGene_tx=XM_003982139.5
A2 15183438 15183530 refGene_gn=LARS2;refGene_tx=XM_003982139.5
A2 15185984 15186045 refGene_gn=LARS2;refGene_tx=XM_003982139.5
A2 15221054 15221144 refGene_gn=LARS2;refGene_tx=XM_003982139.5
A2 15237279 15237423 refGene_gn=LARS2;refGene_tx=XM_003982139.5
A2 15259658 15259766 refGene_gn=LARS2;refGene_tx=XM_003982139.5
A2 15262130 15262290 refGene_gn=LARS2;refGene_tx=XM_003982139.5
A2 15271492 15271597 refGene_gn=LARS2;refGene_tx=XM_003982139.5
A2 15274548 15274664 refGene_gn=LARS2;refGene_tx=XM_003982139.5
A2 15278075 15278359 refGene_gn=LARS2;refGene_tx=XM_003982139.5
A2 15281789 15281888 refGene_gn=LARS2;refGene_tx=XM_003982139.5
A2 15285354 15285492 refGene_gn=LARS2;refGene_tx=XM_003982139.5
A2 15303009 15303110 refGene_gn=LARS2;refGene_tx=XM_003982139.5
A2 15305746 15305929 refGene_gn=LARS2;refGene_tx=XM_003982139.5
A2 15307364 15307375 refGene_gn=LARS2;refGene_tx=XM_003982139.5
A2 15307376 15307535 refGene_gn=LARS2;refGene_tx=XM_003982139.5
A2 15309406 15309484 refGene_gn=LARS2;refGene_tx=XM_003982139.5
A2 15312869 15312981 refGene_gn=LARS2;refGene_tx=XM_003982139.5
A2 15325090 15325218 refGene_gn=LARS2;refGene_tx=XM_003982139.5
A2 15330973 15331153 refGene_gn=LARS2;refGene_tx=XM_003982139.5
A2 15358029 15359428 refGene_gn=LIMD1;refGene_tx=XM_003982176.4
A2 15367807 15368270 refGene_gn=LIMD1;refGene_tx=XM_006928675.4
A2 15389858 15389960
refGene_gn=LIMD1;refGene_tx=XM_003982176.4;refGene_gn=LIMD1;refGene_tx
=XM_006928675.4
A2 15414291 15414359
refGene_gn=LIMD1;refGene_tx=XM_003982176.4;refGene_gn=LIMD1;refGene_tx
=XM_006928675.4
A2 15416061 15416124
refGene_gn=LIMD1;refGene_tx=XM_003982176.4;refGene_gn=LIMD1;refGene_tx
=XM_006928675.4
A2 15421658 15421789
refGene_gn=LIMD1;refGene_tx=XM_003982176.4;refGene_gn=LIMD1;refGene_tx
=XM_006928675.4

A2 15422373 15422425
refGene_gn=LIMD1;refGene_tx=XM_003982176.4;refGene_gn=LIMD1;refGene_tx
=XM_006928675.4
A2 15423418 15423487
refGene_gn=LIMD1;refGene_tx=XM_003982176.4;refGene_gn=LIMD1;refGene_tx
=XM_006928675.4
A2 15424325 15424463
refGene_gn=LIMD1;refGene_tx=XM_003982176.4;refGene_gn=LIMD1;refGene_tx
=XM_006928675.4
A2 15434178 15434210 refGene_gn=SACM1L;refGene_tx=XM_003982140.4
A2 15450528 15450626 refGene_gn=SACM1L;refGene_tx=XM_003982140.4
A2 15453305 15453380 refGene_gn=SACM1L;refGene_tx=XM_003982140.4
A2 15454634 15454762 refGene_gn=SACM1L;refGene_tx=XM_003982140.4
A2 15457420 15457570 refGene_gn=SACM1L;refGene_tx=XM_003982140.4
A2 15460505 15460565 refGene_gn=SACM1L;refGene_tx=XM_003982140.4
A2 15461466 15461500 refGene_gn=SACM1L;refGene_tx=XM_003982140.4
A2 15466913 15467015 refGene_gn=SACM1L;refGene_tx=XM_003982140.4
A2 15469057 15469143 refGene_gn=SACM1L;refGene_tx=XM_003982140.4
A2 15469957 15470044 refGene_gn=SACM1L;refGene_tx=XM_003982140.4
A2 15470538 15470607 refGene_gn=SACM1L;refGene_tx=XM_003982140.4
A2 15477789 15477869 refGene_gn=SACM1L;refGene_tx=XM_003982140.4
A2 15482459 15482558 refGene_gn=SACM1L;refGene_tx=XM_003982140.4
A2 15484461 15484600 refGene_gn=SACM1L;refGene_tx=XM_003982140.4
A2 15486680 15486750 refGene_gn=SACM1L;refGene_tx=XM_003982140.4
A2 15486871 15486944 refGene_gn=SACM1L;refGene_tx=XM_003982140.4
A2 15487968 15488062 refGene_gn=SACM1L;refGene_tx=XM_003982140.4
A2 15488179 15488272 refGene_gn=SACM1L;refGene_tx=XM_003982140.4
A2 15488901 15488959 refGene_gn=SACM1L;refGene_tx=XM_003982140.4
A2 15492201 15492338 refGene_gn=SACM1L;refGene_tx=XM_003982140.4
A2 15500501 15500651 refGene_gn=SLC6A20;refGene_tx=XM_023248037.1
A2 15502252 15502418 refGene_gn=SLC6A20;refGene_tx=XM_023248037.1
A2 15504804 15504964 refGene_gn=SLC6A20;refGene_tx=XM_023248037.1
A2 15506425 15506630 refGene_gn=SLC6A20;refGene_tx=XM_023248037.1
A2 15509719 15509781 refGene_gn=SLC6A20;refGene_tx=XM_023248037.1
A2 15509783 15509881 refGene_gn=SLC6A20;refGene_tx=XM_023248037.1
A2 15511220 15511462 refGene_gn=SLC6A20;refGene_tx=XM_023248037.1
A2 15512243 15512354 refGene_gn=SLC6A20;refGene_tx=XM_023248037.1
A2 15515481 15515709 refGene_gn=SLC6A20;refGene_tx=XM_023248037.1
A2 15519620 15519712 refGene_gn=SLC6A20;refGene_tx=XM_023248037.1
A2 15521178 15521319 refGene_gn=SLC6A20;refGene_tx=XM_023248037.1
A2 15537084 15537645
refGene_gn=LOC109495144;refGene_tx=XM_023248038.1;refGene_gn=SLC6A20;r
efGene_tx=XM_023248037.1
A2 15543265 15543415
refGene_gn=LOC109495144;refGene_tx=XM_023248038.1
A2 15556243 15556262
refGene_gn=LZTFL1;refGene_tx=XM_006928676.3;refGene_gn=LZTFL1;refGene_
tx=XM_011291098.3;refGene_gn=LZTFL1;refGene_tx=XM_011291101.3
A2 15557284 15557388
refGene_gn=LZTFL1;refGene_tx=XM_006928676.3;refGene_gn=LZTFL1;refGene_

tx=XM_011291098.3;refGene_gn=LZTFL1;refGene_tx=XM_011291101.3
A2 15558415 15558592
refGene_gn=LZTFL1;refGene_tx=XM_006928676.3;refGene_gn=LZTFL1;refGene_tx=XM_011291098.3;refGene_gn=LZTFL1;refGene_tx=XM_011291101.3
A2 15560866 15560944
refGene_gn=LZTFL1;refGene_tx=XM_006928676.3;refGene_gn=LZTFL1;refGene_tx=XM_011291098.3;refGene_gn=LZTFL1;refGene_tx=XM_011291101.3
A2 15561030 15561096
refGene_gn=LZTFL1;refGene_tx=XM_006928676.3;refGene_gn=LZTFL1;refGene_tx=XM_011291098.3;refGene_gn=LZTFL1;refGene_tx=XM_011291101.3
A2 15562498 15562567
refGene_gn=LZTFL1;refGene_tx=XM_006928676.3;refGene_gn=LZTFL1;refGene_tx=XM_011291098.3;refGene_gn=LZTFL1;refGene_tx=XM_011291101.3
A2 15563897 15563958
refGene_gn=LZTFL1;refGene_tx=XM_006928676.3;refGene_gn=LZTFL1;refGene_tx=XM_011291098.3;refGene_gn=LZTFL1;refGene_tx=XM_011291101.3
A2 15566050 15566245
refGene_gn=LZTFL1;refGene_tx=XM_006928676.3;refGene_gn=LZTFL1;refGene_tx=XM_011291098.3;refGene_gn=LZTFL1;refGene_tx=XM_011291101.3
A2 15568736 15568861
refGene_gn=LZTFL1;refGene_tx=XM_006928676.3;refGene_gn=LZTFL1;refGene_tx=XM_011291098.3;refGene_gn=LZTFL1;refGene_tx=XM_011291101.3
A2 15571540 15571543
refGene_gn=LZTFL1;refGene_tx=XM_006928676.3;refGene_gn=LZTFL1;refGene_tx=XM_011291098.3;refGene_gn=LZTFL1;refGene_tx=XM_011291101.3
A2 15616289 15616310
refGene_gn=CCR9;refGene_tx=XM_023248069.1;refGene_gn=CCR9;refGene_tx=XM_023248071.1
A2 15618249 15618270
refGene_gn=CCR9;refGene_tx=XM_003982142.3;refGene_gn=CCR9;refGene_tx=XM_019820560.1
A2 15625665 15626748
refGene_gn=CCR9;refGene_tx=XM_003982142.3;refGene_gn=CCR9;refGene_tx=XM_019820560.1;refGene_gn=CCR9;refGene_tx=XM_023248069.1;refGene_gn=CCR9;refGene_tx=XM_023248071.1
A2 15642520 15642596
refGene_gn=FYCO1;refGene_tx=XM_023248072.1;refGene_gn=FYCO1;refGene_tx=XM_023248080.1
A2 15644532 15644642
refGene_gn=FYCO1;refGene_tx=XM_023248072.1;refGene_gn=FYCO1;refGene_tx=XM_023248080.1
A2 15650344 15650555
refGene_gn=FYCO1;refGene_tx=XM_023248072.1;refGene_gn=FYCO1;refGene_tx=XM_023248080.1
A2 15653275 15653371
refGene_gn=FYCO1;refGene_tx=XM_023248072.1;refGene_gn=FYCO1;refGene_tx=XM_023248080.1
A2 15664165 15665191 refGene_gn=CXCR6;refGene_tx=XM_011291106.3
A2 15671356 15671498
refGene_gn=FYCO1;refGene_tx=XM_023248072.1;refGene_gn=FYCO1;refGene_tx

=XM_023248080.1
A2 15675477 15675695
refGene_gn=FYCO1;refGene_tx=XM_023248072.1;refGene_gn=FYCO1;refGene_tx
=XM_023248080.1
A2 15676382 15676532
refGene_gn=FYCO1;refGene_tx=XM_023248072.1;refGene_gn=FYCO1;refGene_tx
=XM_023248080.1
A2 15679431 15679599
refGene_gn=FYCO1;refGene_tx=XM_023248072.1;refGene_gn=FYCO1;refGene_tx
=XM_023248080.1
A2 15681881 15682000
refGene_gn=FYCO1;refGene_tx=XM_023248072.1;refGene_gn=FYCO1;refGene_tx
=XM_023248080.1
A2 15682630 15682723
refGene_gn=FYCO1;refGene_tx=XM_023248072.1;refGene_gn=FYCO1;refGene_tx
=XM_023248080.1
A2 15683777 15686072
refGene_gn=FYCO1;refGene_tx=XM_023248072.1;refGene_gn=FYCO1;refGene_tx
=XM_023248080.1
A2 15687236 15687327
refGene_gn=FYCO1;refGene_tx=XM_023248072.1;refGene_gn=FYCO1;refGene_tx
=XM_023248080.1
A2 15690775 15690919
refGene_gn=FYCO1;refGene_tx=XM_023248072.1;refGene_gn=FYCO1;refGene_tx
=XM_023248080.1
A2 15692868 15692975
refGene_gn=FYCO1;refGene_tx=XM_023248072.1;refGene_gn=FYCO1;refGene_tx
=XM_023248080.1
A2 15696591 15696717
refGene_gn=FYCO1;refGene_tx=XM_023248072.1;refGene_gn=FYCO1;refGene_tx
=XM_023248080.1
A2 15698362 15698469
refGene_gn=FYCO1;refGene_tx=XM_023248072.1;refGene_gn=FYCO1;refGene_tx
=XM_023248080.1
A2 15702264 15702319
refGene_gn=FYCO1;refGene_tx=XM_023248072.1;refGene_gn=FYCO1;refGene_tx
=XM_023248080.1
A2 15721199 15722195 refGene_gn=XCR1;refGene_tx=XM_023248091.1
A2 15803648 15804731
refGene_gn=CCR1;refGene_tx=NM_001135969.1;refGene_gn=CCR1;refGene_tx=X
M_006928640.4;refGene_gn=CCR1;refGene_tx=XM_019814626.2;refGene_gn=CCR
1;refGene_tx=XM_019814628.2
A2 15832648 15832685
refGene_gn=CCR3;refGene_tx=XM_006928641.3;refGene_gn=CCR3;refGene_tx=X
M_019814642.2;refGene_gn=CCR3;refGene_tx=XM_019814655.2
A2 15842449 15843540
refGene_gn=CCR3;refGene_tx=NM_001135970.1;refGene_gn=CCR3;refGene_tx=X
M_006928641.3;refGene_gn=CCR3;refGene_tx=XM_019814642.2;refGene_gn=CCR
3;refGene_tx=XM_019814655.2
A2 15923669 15924791

refGene_gn=CCR2;refGene_tx=NM_001097580.1;refGene_gn=CCR2;refGene_tx=XM_019814564.2
A2 15935967 15937026
refGene_gn=CCR5;refGene_tx=NM_001009248.1;refGene_gn=CCR5;refGene_tx=XM_006928652.3;refGene_gn=CCR5;refGene_tx=XM_019814950.2
A2 15961479 15962529
refGene_gn=CCRL2;refGene_tx=XM_003982179.4;refGene_gn=CCRL2;refGene_tx=XM_006928678.4
A2 15964997 15965032 refGene_gn=LTF;refGene_tx=XM_011291127.2
A2 15966442 15966632 refGene_gn=LTF;refGene_tx=XM_011291127.2
A2 15967797 15967982 refGene_gn=LTF;refGene_tx=XM_011291127.2
A2 15970722 15970790 refGene_gn=LTF;refGene_tx=XM_011291127.2
A2 15976180 15976322 refGene_gn=LTF;refGene_tx=XM_011291127.2
A2 15978021 15978177 refGene_gn=LTF;refGene_tx=XM_011291127.2
A2 15979294 15979342 refGene_gn=LTF;refGene_tx=XM_011291127.2
A2 15980382 15980473 refGene_gn=LTF;refGene_tx=XM_011291127.2
A2 15982252 15982407 refGene_gn=LTF;refGene_tx=XM_011291127.2
A2 15983218 15983393 refGene_gn=LTF;refGene_tx=XM_011291127.2
A2 15983861 15984040 refGene_gn=LTF;refGene_tx=XM_011291127.2
A2 15987043 15987099 refGene_gn=LTF;refGene_tx=XM_011291127.2
A2 15988004 15988152 refGene_gn=LTF;refGene_tx=XM_011291127.2
A2 15988493 15988676 refGene_gn=LTF;refGene_tx=XM_011291127.2
A2 15988986 15989095 refGene_gn=LTF;refGene_tx=XM_011291127.2
A2 15990618 15990782 refGene_gn=LTF;refGene_tx=XM_011291127.2
A2 15996854 15996897 refGene_gn=LTF;refGene_tx=XM_011291127.2
A2 16032082 16032237 refGene_gn=RTP3;refGene_tx=XM_011291135.3
A2 16035110 16035310 refGene_gn=RTP3;refGene_tx=XM_011291135.3
A2 16035312 16035537 refGene_gn=RTP3;refGene_tx=XM_011291135.3
A2 16035539 16035832 refGene_gn=RTP3;refGene_tx=XM_011291135.3
A2 16045173 16045279
refGene_gn=LRRC2;refGene_tx=XM_003982145.5;refGene_gn=LRRC2;refGene_tx=XM_006928680.3
A2 16049287 16049443
refGene_gn=LRRC2;refGene_tx=XM_003982145.5;refGene_gn=LRRC2;refGene_tx=XM_006928680.3
A2 16051223 16051369
refGene_gn=LRRC2;refGene_tx=XM_003982145.5;refGene_gn=LRRC2;refGene_tx=XM_006928680.3
A2 16056039 16056176
refGene_gn=LRRC2;refGene_tx=XM_003982145.5;refGene_gn=LRRC2;refGene_tx=XM_006928680.3
A2 16056911 16057068
refGene_gn=LRRC2;refGene_tx=XM_003982145.5;refGene_gn=LRRC2;refGene_tx=XM_006928680.3
A2 16058999 16059207
refGene_gn=LRRC2;refGene_tx=XM_003982145.5;refGene_gn=LRRC2;refGene_tx=XM_006928680.3
A2 16066412 16066537
refGene_gn=LRRC2;refGene_tx=XM_003982145.5;refGene_gn=LRRC2;refGene_tx=XM_006928680.3

| | | | |
|---|---|---|---|
| A2 | 16091958 | 16092032 | refGene_gn=TDGF1;refGene_tx=XM_019826095.2 |
| A2 | 16092797 | 16092856 | refGene_gn=TDGF1;refGene_tx=XM_019826095.2 |
| A2 | 16092937 | 16093075 | refGene_gn=TDGF1;refGene_tx=XM_019826095.2 |
| A2 | 16093707 | 16093822 | refGene_gn=TDGF1;refGene_tx=XM_019826095.2 |
| A2 | 16093920 | 16094030 | refGene_gn=TDGF1;refGene_tx=XM_019826095.2 |
| A2 | 16095149 | 16095265 | refGene_gn=TDGF1;refGene_tx=XM_019826095.2 |
| A2 | 16123743 | 16123886 | refGene_gn=FAM240A;refGene_tx=XM_023248118.1 |
| A2 | 16128245 | 16128378 | refGene_gn=FAM240A;refGene_tx=XM_023248118.1 |
| A2 | 16150430 | 16150511 | refGene_gn=ALS2CL;refGene_tx=XM_011291142.3 |
| A2 | 16150907 | 16151004 | refGene_gn=ALS2CL;refGene_tx=XM_011291142.3 |
| A2 | 16151311 | 16151461 | refGene_gn=ALS2CL;refGene_tx=XM_011291142.3 |
| A2 | 16151569 | 16151631 | refGene_gn=ALS2CL;refGene_tx=XM_011291142.3 |
| A2 | 16152061 | 16152104 | refGene_gn=ALS2CL;refGene_tx=XM_011291142.3 |
| A2 | 16153311 | 16153485 | refGene_gn=ALS2CL;refGene_tx=XM_011291142.3 |
| A2 | 16154358 | 16154427 | refGene_gn=ALS2CL;refGene_tx=XM_011291142.3 |
| A2 | 16154946 | 16155104 | refGene_gn=ALS2CL;refGene_tx=XM_011291142.3 |
| A2 | 16155306 | 16155403 | refGene_gn=ALS2CL;refGene_tx=XM_011291142.3 |
| A2 | 16155575 | 16155749 | refGene_gn=ALS2CL;refGene_tx=XM_011291142.3 |
| A2 | 16156953 | 16157084 | refGene_gn=ALS2CL;refGene_tx=XM_011291142.3 |
| A2 | 16157941 | 16158019 | refGene_gn=ALS2CL;refGene_tx=XM_011291142.3 |
| A2 | 16159108 | 16159220 | refGene_gn=ALS2CL;refGene_tx=XM_011291142.3 |
| A2 | 16159961 | 16160123 | refGene_gn=ALS2CL;refGene_tx=XM_011291142.3 |
| A2 | 16160225 | 16160324 | refGene_gn=ALS2CL;refGene_tx=XM_011291142.3 |
| A2 | 16160749 | 16160815 | refGene_gn=ALS2CL;refGene_tx=XM_011291142.3 |
| A2 | 16161856 | 16162053 | refGene_gn=ALS2CL;refGene_tx=XM_011291142.3 |
| A2 | 16162487 | 16162554 | refGene_gn=ALS2CL;refGene_tx=XM_011291142.3 |
| A2 | 16162692 | 16162751 | refGene_gn=ALS2CL;refGene_tx=XM_011291142.3 |
| A2 | 16163941 | 16164061 | refGene_gn=ALS2CL;refGene_tx=XM_011291142.3 |
| A2 | 16164620 | 16164752 | refGene_gn=ALS2CL;refGene_tx=XM_011291142.3 |
| A2 | 16165292 | 16165458 | refGene_gn=ALS2CL;refGene_tx=XM_011291142.3 |
| A2 | 16165921 | 16165987 | refGene_gn=ALS2CL;refGene_tx=XM_011291142.3 |
| A2 | 16166287 | 16166480 | refGene_gn=ALS2CL;refGene_tx=XM_011291142.3 |
| A2 | 16167214 | 16167317 | refGene_gn=ALS2CL;refGene_tx=XM_011291142.3 |
| A2 | 16176200 | 16176446 | refGene_gn=TMIE;refGene_tx=XM_003982146.5 |
| A2 | 16179931 | 16180049 | refGene_gn=TMIE;refGene_tx=XM_003982146.5 |
| A2 | 16183329 | 16183479 | refGene_gn=TMIE;refGene_tx=XM_003982146.5 |
| A2 | 16183785 | 16183886 | refGene_gn=TMIE;refGene_tx=XM_003982146.5 |
| A2 | 16186343 | 16186574 | refGene_gn=PRSS50;refGene_tx=XM_023241215.1 |
| A2 | 16186930 | 16187097 | refGene_gn=PRSS50;refGene_tx=XM_023241215.1 |
| A2 | 16188213 | 16188497 | refGene_gn=PRSS50;refGene_tx=XM_023241215.1 |
| A2 | 16189543 | 16189706 | refGene_gn=PRSS50;refGene_tx=XM_023241215.1 |
| A2 | 16190963 | 16191155 | refGene_gn=PRSS50;refGene_tx=XM_023241215.1 |
| A2 | 16191233 | 16191417 | refGene_gn=PRSS50;refGene_tx=XM_023241215.1 |
| A2 | 16191524 | 16191680 | refGene_gn=PRSS50;refGene_tx=XM_023241215.1 |
| A2 | 16193051 | 16193300 | refGene_gn=PRSS46P;refGene_tx=XM_006928682.4 |
| A2 | 16212028 | 16212192 | refGene_gn=PRSS46P;refGene_tx=XM_006928682.4 |
| A2 | 16212997 | 16213257 | refGene_gn=PRSS46P;refGene_tx=XM_006928682.4 |
| A2 | 16213437 | 16213615 | refGene_gn=PRSS46P;refGene_tx=XM_006928682.4 |
| A2 | 16215790 | 16215872 | refGene_gn=PRSS46P;refGene_tx=XM_006928682.4 |
| A2 | 16224742 | 16224958 | refGene_gn=PRSS45;refGene_tx=XM_023241221.1 |

A2	16225294	16225458	refGene_gn=PRSS45;refGene_tx=XM_023241221.1
A2	16226295	16226570	refGene_gn=PRSS45;refGene_tx=XM_023241221.1
A2	16227124	16227272	refGene_gn=PRSS45;refGene_tx=XM_023241221.1
A2	16230263	16230338	refGene_gn=PRSS45;refGene_tx=XM_023241221.1
A2	16232142	16232788	refGene_gn=PRSS45;refGene_tx=XM_023241221.1
A2	16234386	16234567	refGene_gn=PRSS45;refGene_tx=XM_023241221.1
A2	16236426	16236582	refGene_gn=PRSS45;refGene_tx=XM_023241221.1
A2	16236972	16237236	refGene_gn=PRSS45;refGene_tx=XM_023241221.1
A2	16237594	16237734	refGene_gn=PRSS45;refGene_tx=XM_023241221.1
A2	16241974	16242007
refGene_gn=LOC101095475;refGene_tx=XM_023248126.1
A2	16245502	16245736
refGene_gn=LOC101095475;refGene_tx=XM_023248119.1
A2	16246285	16246449
refGene_gn=LOC101095475;refGene_tx=XM_023248119.1;refGene_gn=LOC101095
475;refGene_tx=XM_023248126.1
A2	16247445	16247711
refGene_gn=LOC101095475;refGene_tx=XM_023248119.1;refGene_gn=LOC101095
475;refGene_tx=XM_023248126.1
A2	16248381	16248538
refGene_gn=LOC101095475;refGene_tx=XM_023248119.1;refGene_gn=LOC101095
475;refGene_tx=XM_023248126.1
A2	16248860	16249155
refGene_gn=LOC101095475;refGene_tx=XM_023248119.1;refGene_gn=LOC101095
475;refGene_tx=XM_023248126.1
A2	16262698	16262905	refGene_gn=PRSS42;refGene_tx=XM_019820703.2
A2	16262906	16262936	refGene_gn=PRSS42;refGene_tx=XM_019820703.2
A2	16264283	16264444	refGene_gn=PRSS42;refGene_tx=XM_019820703.2
A2	16265366	16265629	refGene_gn=PRSS42;refGene_tx=XM_019820703.2
A2	16266273	16266430	refGene_gn=PRSS42;refGene_tx=XM_019820703.2
A2	16266767	16266996	refGene_gn=PRSS42;refGene_tx=XM_019820703.2
A2	16290627	16290656
refGene_gn=MYL3;refGene_tx=XM_003982147.5;refGene_gn=MYL3;refGene_tx=X
M_006928689.4
A2	16290774	16290852
refGene_gn=MYL3;refGene_tx=XM_003982147.5;refGene_gn=MYL3;refGene_tx=X
M_006928689.4
A2	16291936	16292110
refGene_gn=MYL3;refGene_tx=XM_003982147.5;refGene_gn=MYL3;refGene_tx=X
M_006928689.4
A2	16292892	16293042
refGene_gn=MYL3;refGene_tx=XM_003982147.5;refGene_gn=MYL3;refGene_tx=X
M_006928689.4
A2	16293174	16293202
refGene_gn=MYL3;refGene_tx=XM_003982147.5;refGene_gn=MYL3;refGene_tx=X
M_006928689.4
A2	16295450	16295582
refGene_gn=MYL3;refGene_tx=XM_003982147.5;refGene_gn=MYL3;refGene_tx=X
M_006928689.4
A2	16314189	16314264	refGene_gn=PTH1R;refGene_tx=XM_023248158.1

A2 16324695 16324798 refGene_gn=PTH1R;refGene_tx=XM_023248158.1
A2 16326751 16326886 refGene_gn=PTH1R;refGene_tx=XM_023248158.1
A2 16328493 16328604 refGene_gn=PTH1R;refGene_tx=XM_023248158.1
A2 16328707 16328826 refGene_gn=PTH1R;refGene_tx=XM_023248158.1
A2 16328981 16329076 refGene_gn=PTH1R;refGene_tx=XM_023248158.1
A2 16329263 16329435 refGene_gn=PTH1R;refGene_tx=XM_023248158.1
A2 16329436 16329460 refGene_gn=PTH1R;refGene_tx=XM_023248158.1
A2 16329990 16330145 refGene_gn=PTH1R;refGene_tx=XM_023248158.1
A2 16331680 16331740 refGene_gn=PTH1R;refGene_tx=XM_023248158.1
A2 16332080 16332147 refGene_gn=PTH1R;refGene_tx=XM_023248158.1
A2 16332422 16332517 refGene_gn=PTH1R;refGene_tx=XM_023248158.1
A2 16333128 16333270 refGene_gn=PTH1R;refGene_tx=XM_023248158.1
A2 16333387 16333429 refGene_gn=PTH1R;refGene_tx=XM_023248158.1
A2 16333934 16334312 refGene_gn=PTH1R;refGene_tx=XM_023248158.1
A2 16351160 16351243 refGene_gn=CCDC12;refGene_tx=XM_006928690.3
A2 16351338 16351415 refGene_gn=CCDC12;refGene_tx=XM_006928690.3
A2 16351926 16352086 refGene_gn=CCDC12;refGene_tx=XM_023248173.1
A2 16352488 16352523
refGene_gn=CCDC12;refGene_tx=XM_006928690.3;refGene_gn=CCDC12;refGene_tx=XM_023248173.1
A2 16352743 16352805
refGene_gn=CCDC12;refGene_tx=XM_006928690.3;refGene_gn=CCDC12;refGene_tx=XM_023248173.1
A2 16354422 16354502
refGene_gn=CCDC12;refGene_tx=XM_006928690.3;refGene_gn=CCDC12;refGene_tx=XM_023248173.1
A2 16371639 16371707
refGene_gn=CCDC12;refGene_tx=XM_006928690.3;refGene_gn=CCDC12;refGene_tx=XM_023248173.1
A2 16404883 16404979
refGene_gn=CCDC12;refGene_tx=XM_006928690.3;refGene_gn=CCDC12;refGene_tx=XM_023248173.1
A2 16407671 16407722
refGene_gn=NBEAL2;refGene_tx=XM_023248189.1;refGene_gn=NBEAL2;refGene_tx=XM_023248199.1;refGene_gn=NBEAL2;refGene_tx=XM_023248202.1;refGene_gn=NBEAL2;refGene_tx=XM_023248209.1
A2 16415149 16415179 refGene_gn=NBEAL2;refGene_tx=XM_003982191.5
A2 16415783 16415872
refGene_gn=NBEAL2;refGene_tx=XM_003982191.5;refGene_gn=NBEAL2;refGene_tx=XM_023248189.1;refGene_gn=NBEAL2;refGene_tx=XM_023248199.1;refGene_gn=NBEAL2;refGene_tx=XM_023248202.1;refGene_gn=NBEAL2;refGene_tx=XM_023248209.1
A2 16415954 16416083
refGene_gn=NBEAL2;refGene_tx=XM_003982191.5;refGene_gn=NBEAL2;refGene_tx=XM_023248189.1;refGene_gn=NBEAL2;refGene_tx=XM_023248199.1;refGene_gn=NBEAL2;refGene_tx=XM_023248202.1;refGene_gn=NBEAL2;refGene_tx=XM_023248209.1
A2 16416186 16416268
refGene_gn=NBEAL2;refGene_tx=XM_003982191.5;refGene_gn=NBEAL2;refGene_tx=XM_023248189.1;refGene_gn=NBEAL2;refGene_tx=XM_023248199.1;refGene_

gn=NBEAL2;refGene_tx=XM_023248202.1;refGene_gn=NBEAL2;refGene_tx=XM_023248209.1
A2 16416359 16416484
refGene_gn=NBEAL2;refGene_tx=XM_003982191.5;refGene_gn=NBEAL2;refGene_tx=XM_023248189.1;refGene_gn=NBEAL2;refGene_tx=XM_023248199.1;refGene_gn=NBEAL2;refGene_tx=XM_023248202.1;refGene_gn=NBEAL2;refGene_tx=XM_023248209.1
A2 16416694 16416777
refGene_gn=NBEAL2;refGene_tx=XM_003982191.5;refGene_gn=NBEAL2;refGene_tx=XM_023248189.1;refGene_gn=NBEAL2;refGene_tx=XM_023248199.1;refGene_gn=NBEAL2;refGene_tx=XM_023248202.1;refGene_gn=NBEAL2;refGene_tx=XM_023248209.1
A2 16418395 16418481
refGene_gn=NBEAL2;refGene_tx=XM_003982191.5;refGene_gn=NBEAL2;refGene_tx=XM_023248189.1;refGene_gn=NBEAL2;refGene_tx=XM_023248199.1;refGene_gn=NBEAL2;refGene_tx=XM_023248202.1;refGene_gn=NBEAL2;refGene_tx=XM_023248209.1
A2 16418577 16418860
refGene_gn=NBEAL2;refGene_tx=XM_003982191.5;refGene_gn=NBEAL2;refGene_tx=XM_023248189.1;refGene_gn=NBEAL2;refGene_tx=XM_023248199.1;refGene_gn=NBEAL2;refGene_tx=XM_023248202.1;refGene_gn=NBEAL2;refGene_tx=XM_023248209.1
A2 16419021 16419128
refGene_gn=NBEAL2;refGene_tx=XM_003982191.5;refGene_gn=NBEAL2;refGene_tx=XM_023248189.1;refGene_gn=NBEAL2;refGene_tx=XM_023248199.1;refGene_gn=NBEAL2;refGene_tx=XM_023248202.1;refGene_gn=NBEAL2;refGene_tx=XM_023248209.1
A2 16419652 16419733
refGene_gn=NBEAL2;refGene_tx=XM_003982191.5;refGene_gn=NBEAL2;refGene_tx=XM_023248189.1;refGene_gn=NBEAL2;refGene_tx=XM_023248199.1;refGene_gn=NBEAL2;refGene_tx=XM_023248209.1
A2 16421134 16421218
refGene_gn=NBEAL2;refGene_tx=XM_003982191.5;refGene_gn=NBEAL2;refGene_tx=XM_023248189.1;refGene_gn=NBEAL2;refGene_tx=XM_023248199.1;refGene_gn=NBEAL2;refGene_tx=XM_023248202.1;refGene_gn=NBEAL2;refGene_tx=XM_023248209.1
A2 16421646 16421745
refGene_gn=NBEAL2;refGene_tx=XM_003982191.5;refGene_gn=NBEAL2;refGene_tx=XM_023248189.1;refGene_gn=NBEAL2;refGene_tx=XM_023248199.1;refGene_gn=NBEAL2;refGene_tx=XM_023248202.1;refGene_gn=NBEAL2;refGene_tx=XM_023248209.1
A2 16422184 16422786
refGene_gn=NBEAL2;refGene_tx=XM_003982191.5;refGene_gn=NBEAL2;refGene_tx=XM_023248189.1;refGene_gn=NBEAL2;refGene_tx=XM_023248199.1;refGene_gn=NBEAL2;refGene_tx=XM_023248202.1;refGene_gn=NBEAL2;refGene_tx=XM_023248209.1
A2 16422864 16422997
refGene_gn=NBEAL2;refGene_tx=XM_003982191.5;refGene_gn=NBEAL2;refGene_tx=XM_023248189.1;refGene_gn=NBEAL2;refGene_tx=XM_023248199.1;refGene_gn=NBEAL2;refGene_tx=XM_023248202.1;refGene_gn=NBEAL2;refGene_tx=XM_02

3248209.1
A2 16423088 16423208
refGene_gn=NBEAL2;refGene_tx=XM_003982191.5;refGene_gn=NBEAL2;refGene_tx=XM_023248189.1;refGene_gn=NBEAL2;refGene_tx=XM_023248199.1;refGene_gn=NBEAL2;refGene_tx=XM_023248202.1;refGene_gn=NBEAL2;refGene_tx=XM_023248209.1
A2 16423425 16423747
refGene_gn=NBEAL2;refGene_tx=XM_003982191.5;refGene_gn=NBEAL2;refGene_tx=XM_023248189.1;refGene_gn=NBEAL2;refGene_tx=XM_023248199.1;refGene_gn=NBEAL2;refGene_tx=XM_023248202.1;refGene_gn=NBEAL2;refGene_tx=XM_023248209.1
A2 16423962 16424045
refGene_gn=NBEAL2;refGene_tx=XM_003982191.5;refGene_gn=NBEAL2;refGene_tx=XM_023248189.1;refGene_gn=NBEAL2;refGene_tx=XM_023248199.1;refGene_gn=NBEAL2;refGene_tx=XM_023248202.1;refGene_gn=NBEAL2;refGene_tx=XM_023248209.1
A2 16424151 16424244
refGene_gn=NBEAL2;refGene_tx=XM_003982191.5;refGene_gn=NBEAL2;refGene_tx=XM_023248189.1;refGene_gn=NBEAL2;refGene_tx=XM_023248199.1;refGene_gn=NBEAL2;refGene_tx=XM_023248202.1;refGene_gn=NBEAL2;refGene_tx=XM_023248209.1
A2 16424450 16424625
refGene_gn=NBEAL2;refGene_tx=XM_003982191.5;refGene_gn=NBEAL2;refGene_tx=XM_023248189.1;refGene_gn=NBEAL2;refGene_tx=XM_023248199.1;refGene_gn=NBEAL2;refGene_tx=XM_023248202.1;refGene_gn=NBEAL2;refGene_tx=XM_023248209.1
A2 16424746 16424880
refGene_gn=NBEAL2;refGene_tx=XM_003982191.5;refGene_gn=NBEAL2;refGene_tx=XM_023248189.1;refGene_gn=NBEAL2;refGene_tx=XM_023248199.1;refGene_gn=NBEAL2;refGene_tx=XM_023248202.1;refGene_gn=NBEAL2;refGene_tx=XM_023248209.1
A2 16425241 16425401
refGene_gn=NBEAL2;refGene_tx=XM_003982191.5;refGene_gn=NBEAL2;refGene_tx=XM_023248189.1;refGene_gn=NBEAL2;refGene_tx=XM_023248199.1;refGene_gn=NBEAL2;refGene_tx=XM_023248202.1;refGene_gn=NBEAL2;refGene_tx=XM_023248209.1
A2 16425604 16425707
refGene_gn=NBEAL2;refGene_tx=XM_003982191.5;refGene_gn=NBEAL2;refGene_tx=XM_023248189.1;refGene_gn=NBEAL2;refGene_tx=XM_023248199.1;refGene_gn=NBEAL2;refGene_tx=XM_023248202.1;refGene_gn=NBEAL2;refGene_tx=XM_023248209.1
A2 16425872 16426026
refGene_gn=NBEAL2;refGene_tx=XM_003982191.5;refGene_gn=NBEAL2;refGene_tx=XM_023248189.1;refGene_gn=NBEAL2;refGene_tx=XM_023248199.1;refGene_gn=NBEAL2;refGene_tx=XM_023248202.1;refGene_gn=NBEAL2;refGene_tx=XM_023248209.1
A2 16426137 16426296
refGene_gn=NBEAL2;refGene_tx=XM_003982191.5;refGene_gn=NBEAL2;refGene_tx=XM_023248189.1;refGene_gn=NBEAL2;refGene_tx=XM_023248199.1;refGene_gn=NBEAL2;refGene_tx=XM_023248202.1;refGene_gn=NBEAL2;refGene_tx=XM_02

3248209.1
A2 16426443 16426603
refGene_gn=NBEAL2;refGene_tx=XM_003982191.5;refGene_gn=NBEAL2;refGene_tx=XM_023248189.1;refGene_gn=NBEAL2;refGene_tx=XM_023248199.1;refGene_gn=NBEAL2;refGene_tx=XM_023248202.1;refGene_gn=NBEAL2;refGene_tx=XM_023248209.1
A2 16426764 16426850
refGene_gn=NBEAL2;refGene_tx=XM_003982191.5;refGene_gn=NBEAL2;refGene_tx=XM_023248189.1;refGene_gn=NBEAL2;refGene_tx=XM_023248199.1;refGene_gn=NBEAL2;refGene_tx=XM_023248202.1;refGene_gn=NBEAL2;refGene_tx=XM_023248209.1
A2 16427027 16427540
refGene_gn=NBEAL2;refGene_tx=XM_003982191.5;refGene_gn=NBEAL2;refGene_tx=XM_023248189.1;refGene_gn=NBEAL2;refGene_tx=XM_023248199.1;refGene_gn=NBEAL2;refGene_tx=XM_023248202.1;refGene_gn=NBEAL2;refGene_tx=XM_023248209.1
A2 16428194 16428373
refGene_gn=NBEAL2;refGene_tx=XM_003982191.5;refGene_gn=NBEAL2;refGene_tx=XM_023248189.1;refGene_gn=NBEAL2;refGene_tx=XM_023248199.1;refGene_gn=NBEAL2;refGene_tx=XM_023248202.1;refGene_gn=NBEAL2;refGene_tx=XM_023248209.1
A2 16428468 16428628
refGene_gn=NBEAL2;refGene_tx=XM_003982191.5;refGene_gn=NBEAL2;refGene_tx=XM_023248189.1;refGene_gn=NBEAL2;refGene_tx=XM_023248199.1;refGene_gn=NBEAL2;refGene_tx=XM_023248202.1;refGene_gn=NBEAL2;refGene_tx=XM_023248209.1
A2 16428860 16428998
refGene_gn=NBEAL2;refGene_tx=XM_003982191.5;refGene_gn=NBEAL2;refGene_tx=XM_023248189.1;refGene_gn=NBEAL2;refGene_tx=XM_023248199.1;refGene_gn=NBEAL2;refGene_tx=XM_023248202.1;refGene_gn=NBEAL2;refGene_tx=XM_023248209.1
A2 16429089 16429449
refGene_gn=NBEAL2;refGene_tx=XM_003982191.5;refGene_gn=NBEAL2;refGene_tx=XM_023248189.1;refGene_gn=NBEAL2;refGene_tx=XM_023248199.1;refGene_gn=NBEAL2;refGene_tx=XM_023248202.1;refGene_gn=NBEAL2;refGene_tx=XM_023248209.1
A2 16429537 16429687
refGene_gn=NBEAL2;refGene_tx=XM_003982191.5;refGene_gn=NBEAL2;refGene_tx=XM_023248189.1;refGene_gn=NBEAL2;refGene_tx=XM_023248199.1;refGene_gn=NBEAL2;refGene_tx=XM_023248202.1;refGene_gn=NBEAL2;refGene_tx=XM_023248209.1
A2 16429824 16429982
refGene_gn=NBEAL2;refGene_tx=XM_003982191.5;refGene_gn=NBEAL2;refGene_tx=XM_023248189.1;refGene_gn=NBEAL2;refGene_tx=XM_023248199.1;refGene_gn=NBEAL2;refGene_tx=XM_023248202.1;refGene_gn=NBEAL2;refGene_tx=XM_023248209.1
A2 16430136 16430261
refGene_gn=NBEAL2;refGene_tx=XM_003982191.5;refGene_gn=NBEAL2;refGene_tx=XM_023248189.1;refGene_gn=NBEAL2;refGene_tx=XM_023248199.1;refGene_gn=NBEAL2;refGene_tx=XM_023248202.1;refGene_gn=NBEAL2;refGene_tx=XM_02

3248209.1
A2 16430350 16430486
refGene_gn=NBEAL2;refGene_tx=XM_003982191.5;refGene_gn=NBEAL2;refGene_tx=XM_023248189.1;refGene_gn=NBEAL2;refGene_tx=XM_023248199.1;refGene_gn=NBEAL2;refGene_tx=XM_023248202.1;refGene_gn=NBEAL2;refGene_tx=XM_023248209.1
A2 16430923 16431084
refGene_gn=NBEAL2;refGene_tx=XM_003982191.5;refGene_gn=NBEAL2;refGene_tx=XM_023248189.1;refGene_gn=NBEAL2;refGene_tx=XM_023248199.1;refGene_gn=NBEAL2;refGene_tx=XM_023248202.1;refGene_gn=NBEAL2;refGene_tx=XM_023248209.1
A2 16431183 16431500
refGene_gn=NBEAL2;refGene_tx=XM_003982191.5;refGene_gn=NBEAL2;refGene_tx=XM_023248189.1;refGene_gn=NBEAL2;refGene_tx=XM_023248199.1;refGene_gn=NBEAL2;refGene_tx=XM_023248202.1;refGene_gn=NBEAL2;refGene_tx=XM_023248209.1
A2 16431605 16431701
refGene_gn=NBEAL2;refGene_tx=XM_003982191.5;refGene_gn=NBEAL2;refGene_tx=XM_023248189.1;refGene_gn=NBEAL2;refGene_tx=XM_023248199.1;refGene_gn=NBEAL2;refGene_tx=XM_023248202.1;refGene_gn=NBEAL2;refGene_tx=XM_023248209.1
A2 16432060 16432185
refGene_gn=NBEAL2;refGene_tx=XM_003982191.5;refGene_gn=NBEAL2;refGene_tx=XM_023248189.1;refGene_gn=NBEAL2;refGene_tx=XM_023248199.1;refGene_gn=NBEAL2;refGene_tx=XM_023248202.1;refGene_gn=NBEAL2;refGene_tx=XM_023248209.1
A2 16432271 16432412
refGene_gn=NBEAL2;refGene_tx=XM_003982191.5;refGene_gn=NBEAL2;refGene_tx=XM_023248189.1;refGene_gn=NBEAL2;refGene_tx=XM_023248199.1;refGene_gn=NBEAL2;refGene_tx=XM_023248202.1;refGene_gn=NBEAL2;refGene_tx=XM_023248209.1
A2 16432592 16432723
refGene_gn=NBEAL2;refGene_tx=XM_003982191.5;refGene_gn=NBEAL2;refGene_tx=XM_023248189.1;refGene_gn=NBEAL2;refGene_tx=XM_023248199.1;refGene_gn=NBEAL2;refGene_tx=XM_023248202.1;refGene_gn=NBEAL2;refGene_tx=XM_023248209.1
A2 16432822 16432932
refGene_gn=NBEAL2;refGene_tx=XM_003982191.5;refGene_gn=NBEAL2;refGene_tx=XM_023248189.1;refGene_gn=NBEAL2;refGene_tx=XM_023248199.1;refGene_gn=NBEAL2;refGene_tx=XM_023248202.1;refGene_gn=NBEAL2;refGene_tx=XM_023248209.1
A2 16433031 16433149
refGene_gn=NBEAL2;refGene_tx=XM_003982191.5;refGene_gn=NBEAL2;refGene_tx=XM_023248189.1;refGene_gn=NBEAL2;refGene_tx=XM_023248199.1;refGene_gn=NBEAL2;refGene_tx=XM_023248202.1;refGene_gn=NBEAL2;refGene_tx=XM_023248209.1
A2 16433250 16433348
refGene_gn=NBEAL2;refGene_tx=XM_003982191.5;refGene_gn=NBEAL2;refGene_tx=XM_023248189.1;refGene_gn=NBEAL2;refGene_tx=XM_023248199.1;refGene_gn=NBEAL2;refGene_tx=XM_023248202.1;refGene_gn=NBEAL2;refGene_tx=XM_02

3248209.1
A2 16433856 16433973
refGene_gn=NBEAL2;refGene_tx=XM_003982191.5;refGene_gn=NBEAL2;refGene_tx=XM_023248189.1;refGene_gn=NBEAL2;refGene_tx=XM_023248199.1;refGene_gn=NBEAL2;refGene_tx=XM_023248202.1;refGene_gn=NBEAL2;refGene_tx=XM_023248209.1
A2 16434611 16434701
refGene_gn=NBEAL2;refGene_tx=XM_003982191.5;refGene_gn=NBEAL2;refGene_tx=XM_023248189.1;refGene_gn=NBEAL2;refGene_tx=XM_023248199.1;refGene_gn=NBEAL2;refGene_tx=XM_023248202.1;refGene_gn=NBEAL2;refGene_tx=XM_023248209.1
A2 16434785 16434895
refGene_gn=NBEAL2;refGene_tx=XM_003982191.5;refGene_gn=NBEAL2;refGene_tx=XM_023248189.1;refGene_gn=NBEAL2;refGene_tx=XM_023248199.1;refGene_gn=NBEAL2;refGene_tx=XM_023248202.1;refGene_gn=NBEAL2;refGene_tx=XM_023248209.1
A2 16435059 16435232
refGene_gn=NBEAL2;refGene_tx=XM_003982191.5;refGene_gn=NBEAL2;refGene_tx=XM_023248189.1;refGene_gn=NBEAL2;refGene_tx=XM_023248199.1;refGene_gn=NBEAL2;refGene_tx=XM_023248202.1;refGene_gn=NBEAL2;refGene_tx=XM_023248209.1
A2 16435331 16435426
refGene_gn=NBEAL2;refGene_tx=XM_003982191.5;refGene_gn=NBEAL2;refGene_tx=XM_023248189.1;refGene_gn=NBEAL2;refGene_tx=XM_023248199.1;refGene_gn=NBEAL2;refGene_tx=XM_023248202.1;refGene_gn=NBEAL2;refGene_tx=XM_023248209.1
A2 16435630 16435747
refGene_gn=NBEAL2;refGene_tx=XM_003982191.5;refGene_gn=NBEAL2;refGene_tx=XM_023248189.1;refGene_gn=NBEAL2;refGene_tx=XM_023248199.1;refGene_gn=NBEAL2;refGene_tx=XM_023248202.1;refGene_gn=NBEAL2;refGene_tx=XM_023248209.1
A2 16435845 16436004
refGene_gn=NBEAL2;refGene_tx=XM_003982191.5;refGene_gn=NBEAL2;refGene_tx=XM_023248189.1;refGene_gn=NBEAL2;refGene_tx=XM_023248199.1;refGene_gn=NBEAL2;refGene_tx=XM_023248202.1;refGene_gn=NBEAL2;refGene_tx=XM_023248209.1
A2 16436080 16436229
refGene_gn=NBEAL2;refGene_tx=XM_003982191.5;refGene_gn=NBEAL2;refGene_tx=XM_023248189.1;refGene_gn=NBEAL2;refGene_tx=XM_023248199.1;refGene_gn=NBEAL2;refGene_tx=XM_023248202.1;refGene_gn=NBEAL2;refGene_tx=XM_023248209.1
A2 16436395 16436423 refGene_gn=NBEAL2;refGene_tx=XM_023248209.1
A2 16436474 16436610
refGene_gn=NBEAL2;refGene_tx=XM_003982191.5;refGene_gn=NBEAL2;refGene_tx=XM_023248189.1;refGene_gn=NBEAL2;refGene_tx=XM_023248199.1;refGene_gn=NBEAL2;refGene_tx=XM_023248202.1
A2 16436694 16436796
refGene_gn=NBEAL2;refGene_tx=XM_003982191.5;refGene_gn=NBEAL2;refGene_tx=XM_023248189.1;refGene_gn=NBEAL2;refGene_tx=XM_023248199.1;refGene_gn=NBEAL2;refGene_tx=XM_023248202.1

A2 16438957 16439048
refGene_gn=LOC101096707;refGene_tx=XM_006928694.4
A2 16439718 16439902
refGene_gn=LOC101096707;refGene_tx=XM_006928694.4
A2 16440017 16440160
refGene_gn=LOC101096707;refGene_tx=XM_006928694.4
A2 16440258 16440578
refGene_gn=LOC101096707;refGene_tx=XM_006928694.4
A2 16443973 16444135 refGene_gn=SETD2;refGene_tx=XM_023248221.1
A2 16444478 16444580 refGene_gn=SETD2;refGene_tx=XM_023248221.1
A2 16446477 16446558 refGene_gn=SETD2;refGene_tx=XM_023248221.1
A2 16462957 16463069 refGene_gn=SETD2;refGene_tx=XM_023248221.1
A2 16465428 16465568 refGene_gn=SETD2;refGene_tx=XM_023248221.1
A2 16468082 16468217 refGene_gn=SETD2;refGene_tx=XM_023248221.1
A2 16474478 16475148 refGene_gn=SETD2;refGene_tx=XM_023248221.1
A2 16481450 16481634 refGene_gn=SETD2;refGene_tx=XM_023248221.1
A2 16484447 16484496 refGene_gn=SETD2;refGene_tx=XM_023248221.1
A2 16508181 16508844 refGene_gn=SETD2;refGene_tx=XM_023248221.1
A2 16510420 16510540 refGene_gn=SETD2;refGene_tx=XM_023248221.1
A2 16510792 16510927 refGene_gn=SETD2;refGene_tx=XM_023248221.1
A2 16518689 16518816 refGene_gn=SETD2;refGene_tx=XM_023248221.1
A2 16521596 16521694 refGene_gn=SETD2;refGene_tx=XM_023248221.1
A2 16523658 16523736 refGene_gn=SETD2;refGene_tx=XM_023248221.1
A2 16524701 16524825 refGene_gn=SETD2;refGene_tx=XM_023248221.1
A2 16527080 16527209 refGene_gn=SETD2;refGene_tx=XM_023248221.1
A2 16530808 16530940 refGene_gn=SETD2;refGene_tx=XM_023248221.1
A2 16534367 16538731 refGene_gn=SETD2;refGene_tx=XM_023248221.1
A2 16540811 16540827 refGene_gn=SETD2;refGene_tx=XM_023248221.1
A2 16570812 16570883 refGene_gn=SETD2;refGene_tx=XM_023248221.1
A2 16621235 16621286
refGene_gn=KIF9;refGene_tx=XM_023248228.1;refGene_gn=KIF9;refGene_tx=XM_023248232.1;refGene_gn=KIF9;refGene_tx=XM_023248233.1;refGene_gn=KIF9;refGene_tx=XM_023248236.1;refGene_gn=KIF9;refGene_tx=XM_023248240.1;refGene_gn=KIF9;refGene_tx=XM_023248242.1;refGene_gn=KIF9;refGene_tx=XM_023248245.1;refGene_gn=KIF9;refGene_tx=XM_023248246.1;refGene_gn=KIF9;refGene_tx=XM_023248250.1;refGene_gn=KIF9;refGene_tx=XM_023248255.1;refGene_gn=KIF9;refGene_tx=XM_023248258.1
A2 16632764 16632869
refGene_gn=KIF9;refGene_tx=XM_023248228.1;refGene_gn=KIF9;refGene_tx=XM_023248232.1;refGene_gn=KIF9;refGene_tx=XM_023248233.1;refGene_gn=KIF9;refGene_tx=XM_023248236.1;refGene_gn=KIF9;refGene_tx=XM_023248240.1;refGene_gn=KIF9;refGene_tx=XM_023248242.1;refGene_gn=KIF9;refGene_tx=XM_023248245.1;refGene_gn=KIF9;refGene_tx=XM_023248246.1;refGene_gn=KIF9;refGene_tx=XM_023248250.1;refGene_gn=KIF9;refGene_tx=XM_023248255.1;refGene_gn=KIF9;refGene_tx=XM_023248258.1
A2 16633278 16633394
refGene_gn=KIF9;refGene_tx=XM_023248228.1;refGene_gn=KIF9;refGene_tx=XM_023248232.1;refGene_gn=KIF9;refGene_tx=XM_023248233.1;refGene_gn=KIF9;refGene_tx=XM_023248236.1;refGene_gn=KIF9;refGene_tx=XM_023248240.1;refGene_gn=KIF9;refGene_tx=XM_023248242.1;refGene_gn=KIF9;refGene_tx=X

M_023248245.1;refGene_gn=KIF9;refGene_tx=XM_023248246.1;refGene_gn=KIF9;refGene_tx=XM_023248250.1;refGene_gn=KIF9;refGene_tx=XM_023248255.1;refGene_gn=KIF9;refGene_tx=XM_023248258.1
A2 16633843 16634020
refGene_gn=KIF9;refGene_tx=XM_023248228.1;refGene_gn=KIF9;refGene_tx=XM_023248232.1;refGene_gn=KIF9;refGene_tx=XM_023248233.1;refGene_gn=KIF9;refGene_tx=XM_023248236.1;refGene_gn=KIF9;refGene_tx=XM_023248240.1;refGene_gn=KIF9;refGene_tx=XM_023248242.1;refGene_gn=KIF9;refGene_tx=XM_023248245.1;refGene_gn=KIF9;refGene_tx=XM_023248246.1;refGene_gn=KIF9;refGene_tx=XM_023248250.1;refGene_gn=KIF9;refGene_tx=XM_023248255.1;refGene_gn=KIF9;refGene_tx=XM_023248258.1
A2 16637847 16638062
refGene_gn=KIF9;refGene_tx=XM_023248228.1;refGene_gn=KIF9;refGene_tx=XM_023248232.1;refGene_gn=KIF9;refGene_tx=XM_023248233.1;refGene_gn=KIF9;refGene_tx=XM_023248236.1;refGene_gn=KIF9;refGene_tx=XM_023248240.1;refGene_gn=KIF9;refGene_tx=XM_023248242.1;refGene_gn=KIF9;refGene_tx=XM_023248245.1;refGene_gn=KIF9;refGene_tx=XM_023248246.1;refGene_gn=KIF9;refGene_tx=XM_023248250.1;refGene_gn=KIF9;refGene_tx=XM_023248255.1;refGene_gn=KIF9;refGene_tx=XM_023248258.1
A2 16640081 16640273
refGene_gn=KIF9;refGene_tx=XM_023248228.1;refGene_gn=KIF9;refGene_tx=XM_023248232.1;refGene_gn=KIF9;refGene_tx=XM_023248233.1;refGene_gn=KIF9;refGene_tx=XM_023248236.1;refGene_gn=KIF9;refGene_tx=XM_023248240.1;refGene_gn=KIF9;refGene_tx=XM_023248242.1;refGene_gn=KIF9;refGene_tx=XM_023248245.1;refGene_gn=KIF9;refGene_tx=XM_023248246.1;refGene_gn=KIF9;refGene_tx=XM_023248250.1;refGene_gn=KIF9;refGene_tx=XM_023248255.1;refGene_gn=KIF9;refGene_tx=XM_023248258.1
A2 16641603 16641737
refGene_gn=KIF9;refGene_tx=XM_023248228.1;refGene_gn=KIF9;refGene_tx=XM_023248232.1;refGene_gn=KIF9;refGene_tx=XM_023248233.1;refGene_gn=KIF9;refGene_tx=XM_023248236.1;refGene_gn=KIF9;refGene_tx=XM_023248240.1;refGene_gn=KIF9;refGene_tx=XM_023248242.1;refGene_gn=KIF9;refGene_tx=XM_023248245.1;refGene_gn=KIF9;refGene_tx=XM_023248246.1;refGene_gn=KIF9;refGene_tx=XM_023248250.1;refGene_gn=KIF9;refGene_tx=XM_023248255.1;refGene_gn=KIF9;refGene_tx=XM_023248258.1
A2 16642193 16642284
refGene_gn=KIF9;refGene_tx=XM_023248228.1;refGene_gn=KIF9;refGene_tx=XM_023248232.1;refGene_gn=KIF9;refGene_tx=XM_023248233.1;refGene_gn=KIF9;refGene_tx=XM_023248236.1;refGene_gn=KIF9;refGene_tx=XM_023248240.1;refGene_gn=KIF9;refGene_tx=XM_023248242.1;refGene_gn=KIF9;refGene_tx=XM_023248245.1;refGene_gn=KIF9;refGene_tx=XM_023248246.1;refGene_gn=KIF9;refGene_tx=XM_023248250.1;refGene_gn=KIF9;refGene_tx=XM_023248255.1;refGene_gn=KIF9;refGene_tx=XM_023248258.1
A2 16643294 16643350
refGene_gn=KIF9;refGene_tx=XM_023248228.1;refGene_gn=KIF9;refGene_tx=XM_023248232.1;refGene_gn=KIF9;refGene_tx=XM_023248233.1;refGene_gn=KIF9;refGene_tx=XM_023248236.1;refGene_gn=KIF9;refGene_tx=XM_023248240.1;refGene_gn=KIF9;refGene_tx=XM_023248242.1;refGene_gn=KIF9;refGene_tx=XM_023248245.1;refGene_gn=KIF9;refGene_tx=XM_023248246.1;refGene_gn=KIF9;refGene_tx=XM_023248250.1;refGene_gn=KIF9;refGene_tx=XM_023248255.1;

refGene_gn=KIF9;refGene_tx=XM_023248258.1
A2 16644510 16644615
refGene_gn=KIF9;refGene_tx=XM_023248228.1;refGene_gn=KIF9;refGene_tx=XM_023248232.1;refGene_gn=KIF9;refGene_tx=XM_023248233.1;refGene_gn=KIF9;refGene_tx=XM_023248236.1;refGene_gn=KIF9;refGene_tx=XM_023248240.1;refGene_gn=KIF9;refGene_tx=XM_023248242.1;refGene_gn=KIF9;refGene_tx=XM_023248245.1;refGene_gn=KIF9;refGene_tx=XM_023248246.1;refGene_gn=KIF9;refGene_tx=XM_023248250.1;refGene_gn=KIF9;refGene_tx=XM_023248255.1;refGene_gn=KIF9;refGene_tx=XM_023248258.1
A2 16645361 16645430
refGene_gn=KIF9;refGene_tx=XM_023248228.1;refGene_gn=KIF9;refGene_tx=XM_023248232.1;refGene_gn=KIF9;refGene_tx=XM_023248233.1;refGene_gn=KIF9;refGene_tx=XM_023248236.1;refGene_gn=KIF9;refGene_tx=XM_023248240.1;refGene_gn=KIF9;refGene_tx=XM_023248242.1;refGene_gn=KIF9;refGene_tx=XM_023248245.1;refGene_gn=KIF9;refGene_tx=XM_023248246.1;refGene_gn=KIF9;refGene_tx=XM_023248250.1;refGene_gn=KIF9;refGene_tx=XM_023248255.1;refGene_gn=KIF9;refGene_tx=XM_023248258.1
A2 16652783 16652861
refGene_gn=KIF9;refGene_tx=XM_023248228.1;refGene_gn=KIF9;refGene_tx=XM_023248232.1;refGene_gn=KIF9;refGene_tx=XM_023248233.1;refGene_gn=KIF9;refGene_tx=XM_023248236.1;refGene_gn=KIF9;refGene_tx=XM_023248240.1;refGene_gn=KIF9;refGene_tx=XM_023248242.1;refGene_gn=KIF9;refGene_tx=XM_023248245.1;refGene_gn=KIF9;refGene_tx=XM_023248250.1;refGene_gn=KIF9;refGene_tx=XM_023248255.1;refGene_gn=KIF9;refGene_tx=XM_023248258.1
A2 16655442 16655507
refGene_gn=KIF9;refGene_tx=XM_023248228.1;refGene_gn=KIF9;refGene_tx=XM_023248232.1;refGene_gn=KIF9;refGene_tx=XM_023248233.1;refGene_gn=KIF9;refGene_tx=XM_023248236.1;refGene_gn=KIF9;refGene_tx=XM_023248240.1;refGene_gn=KIF9;refGene_tx=XM_023248242.1;refGene_gn=KIF9;refGene_tx=XM_023248245.1;refGene_gn=KIF9;refGene_tx=XM_023248246.1;refGene_gn=KIF9;refGene_tx=XM_023248250.1;refGene_gn=KIF9;refGene_tx=XM_023248255.1;refGene_gn=KIF9;refGene_tx=XM_023248258.1
A2 16656577 16656725
refGene_gn=KIF9;refGene_tx=XM_023248228.1;refGene_gn=KIF9;refGene_tx=XM_023248232.1;refGene_gn=KIF9;refGene_tx=XM_023248233.1;refGene_gn=KIF9;refGene_tx=XM_023248236.1;refGene_gn=KIF9;refGene_tx=XM_023248240.1;refGene_gn=KIF9;refGene_tx=XM_023248242.1;refGene_gn=KIF9;refGene_tx=XM_023248245.1;refGene_gn=KIF9;refGene_tx=XM_023248246.1;refGene_gn=KIF9;refGene_tx=XM_023248250.1;refGene_gn=KIF9;refGene_tx=XM_023248255.1;refGene_gn=KIF9;refGene_tx=XM_023248258.1
A2 16657496 16657589
refGene_gn=KIF9;refGene_tx=XM_023248228.1;refGene_gn=KIF9;refGene_tx=XM_023248232.1;refGene_gn=KIF9;refGene_tx=XM_023248233.1;refGene_gn=KIF9;refGene_tx=XM_023248236.1;refGene_gn=KIF9;refGene_tx=XM_023248240.1;refGene_gn=KIF9;refGene_tx=XM_023248242.1;refGene_gn=KIF9;refGene_tx=XM_023248245.1;refGene_gn=KIF9;refGene_tx=XM_023248246.1;refGene_gn=KIF9;refGene_tx=XM_023248250.1;refGene_gn=KIF9;refGene_tx=XM_023248255.1;refGene_gn=KIF9;refGene_tx=XM_023248258.1
A2 16657719 16657803
refGene_gn=KIF9;refGene_tx=XM_023248228.1;refGene_gn=KIF9;refGene_tx=X

M_023248232.1;refGene_gn=KIF9;refGene_tx=XM_023248233.1;refGene_gn=KIF9;refGene_tx=XM_023248236.1;refGene_gn=KIF9;refGene_tx=XM_023248240.1;refGene_gn=KIF9;refGene_tx=XM_023248242.1;refGene_gn=KIF9;refGene_tx=XM_023248245.1;refGene_gn=KIF9;refGene_tx=XM_023248246.1;refGene_gn=KIF9;refGene_tx=XM_023248250.1;refGene_gn=KIF9;refGene_tx=XM_023248255.1;refGene_gn=KIF9;refGene_tx=XM_023248258.1
A2 16664252 16664477
refGene_gn=KIF9;refGene_tx=XM_023248228.1;refGene_gn=KIF9;refGene_tx=XM_023248232.1;refGene_gn=KIF9;refGene_tx=XM_023248233.1;refGene_gn=KIF9;refGene_tx=XM_023248236.1;refGene_gn=KIF9;refGene_tx=XM_023248240.1;refGene_gn=KIF9;refGene_tx=XM_023248242.1;refGene_gn=KIF9;refGene_tx=XM_023248245.1;refGene_gn=KIF9;refGene_tx=XM_023248246.1;refGene_gn=KIF9;refGene_tx=XM_023248250.1;refGene_gn=KIF9;refGene_tx=XM_023248255.1;refGene_gn=KIF9;refGene_tx=XM_023248258.1
A2 16667477 16667584
refGene_gn=KIF9;refGene_tx=XM_023248228.1;refGene_gn=KIF9;refGene_tx=XM_023248232.1;refGene_gn=KIF9;refGene_tx=XM_023248233.1;refGene_gn=KIF9;refGene_tx=XM_023248236.1;refGene_gn=KIF9;refGene_tx=XM_023248240.1;refGene_gn=KIF9;refGene_tx=XM_023248242.1;refGene_gn=KIF9;refGene_tx=XM_023248245.1;refGene_gn=KIF9;refGene_tx=XM_023248246.1;refGene_gn=KIF9;refGene_tx=XM_023248250.1;refGene_gn=KIF9;refGene_tx=XM_023248255.1;refGene_gn=KIF9;refGene_tx=XM_023248258.1
A2 16669355 16669521
refGene_gn=KIF9;refGene_tx=XM_023248228.1;refGene_gn=KIF9;refGene_tx=XM_023248232.1;refGene_gn=KIF9;refGene_tx=XM_023248233.1;refGene_gn=KIF9;refGene_tx=XM_023248236.1;refGene_gn=KIF9;refGene_tx=XM_023248240.1;refGene_gn=KIF9;refGene_tx=XM_023248242.1;refGene_gn=KIF9;refGene_tx=XM_023248245.1;refGene_gn=KIF9;refGene_tx=XM_023248246.1;refGene_gn=KIF9;refGene_tx=XM_023248250.1;refGene_gn=KIF9;refGene_tx=XM_023248255.1;refGene_gn=KIF9;refGene_tx=XM_023248258.1
A2 16671268 16671366
refGene_gn=KIF9;refGene_tx=XM_023248228.1;refGene_gn=KIF9;refGene_tx=XM_023248232.1;refGene_gn=KIF9;refGene_tx=XM_023248233.1;refGene_gn=KIF9;refGene_tx=XM_023248236.1;refGene_gn=KIF9;refGene_tx=XM_023248240.1;refGene_gn=KIF9;refGene_tx=XM_023248242.1;refGene_gn=KIF9;refGene_tx=XM_023248245.1;refGene_gn=KIF9;refGene_tx=XM_023248246.1;refGene_gn=KIF9;refGene_tx=XM_023248250.1;refGene_gn=KIF9;refGene_tx=XM_023248255.1
A2 16674767 16674804
refGene_gn=KIF9;refGene_tx=XM_023248232.1;refGene_gn=KIF9;refGene_tx=XM_023248233.1;refGene_gn=KIF9;refGene_tx=XM_023248236.1;refGene_gn=KIF9;refGene_tx=XM_023248240.1;refGene_gn=KIF9;refGene_tx=XM_023248242.1;refGene_gn=KIF9;refGene_tx=XM_023248245.1;refGene_gn=KIF9;refGene_tx=XM_023248258.1
A2 16675574 16675653
refGene_gn=KIF9;refGene_tx=XM_023248228.1;refGene_gn=KIF9;refGene_tx=XM_023248246.1
A2 16676473 16676602
refGene_gn=KLHL18;refGene_tx=XM_003982150.5;refGene_gn=KLHL18;refGene_tx=XM_006928698.4
A2 16709638 16709769

refGene_gn=KLHL18;refGene_tx=XM_003982150.5;refGene_gn=KLHL18;refGene_tx=XM_006928698.4;refGene_gn=KLHL18;refGene_tx=XM_006928699.4
A2	16712197	16712338	refGene_gn=KLHL18;refGene_tx=XM_003982150.5;refGene_gn=KLHL18;refGene_tx=XM_006928698.4;refGene_gn=KLHL18;refGene_tx=XM_006928699.4
A2	16716399	16716598	refGene_gn=KLHL18;refGene_tx=XM_003982150.5;refGene_gn=KLHL18;refGene_tx=XM_006928698.4;refGene_gn=KLHL18;refGene_tx=XM_006928699.4
A2	16719403	16719564	refGene_gn=KLHL18;refGene_tx=XM_003982150.5;refGene_gn=KLHL18;refGene_tx=XM_006928698.4;refGene_gn=KLHL18;refGene_tx=XM_006928699.4
A2	16720778	16720915	refGene_gn=KLHL18;refGene_tx=XM_003982150.5;refGene_gn=KLHL18;refGene_tx=XM_006928698.4;refGene_gn=KLHL18;refGene_tx=XM_006928699.4
A2	16722270	16722508	refGene_gn=KLHL18;refGene_tx=XM_003982150.5;refGene_gn=KLHL18;refGene_tx=XM_006928698.4;refGene_gn=KLHL18;refGene_tx=XM_006928699.4
A2	16725667	16725772	refGene_gn=KLHL18;refGene_tx=XM_003982150.5;refGene_gn=KLHL18;refGene_tx=XM_006928698.4;refGene_gn=KLHL18;refGene_tx=XM_006928699.4
A2	16727777	16727889	refGene_gn=KLHL18;refGene_tx=XM_003982150.5;refGene_gn=KLHL18;refGene_tx=XM_006928698.4;refGene_gn=KLHL18;refGene_tx=XM_006928699.4
A2	16728626	16729013	refGene_gn=KLHL18;refGene_tx=XM_003982150.5;refGene_gn=KLHL18;refGene_tx=XM_006928698.4;refGene_gn=KLHL18;refGene_tx=XM_006928699.4
A2	16759357	16759441	refGene_gn=PTPN23;refGene_tx=XM_003982194.5;refGene_gn=PTPN23;refGene_tx=XM_023248297.1
A2	16767772	16767847	refGene_gn=PTPN23;refGene_tx=XM_003982194.5
A2	16785910	16786038	refGene_gn=PTPN23;refGene_tx=XM_003982194.5;refGene_gn=PTPN23;refGene_tx=XM_023248297.1
A2	16786290	16786367	refGene_gn=PTPN23;refGene_tx=XM_003982194.5;refGene_gn=PTPN23;refGene_tx=XM_023248297.1
A2	16787018	16787068	refGene_gn=PTPN23;refGene_tx=XM_003982194.5;refGene_gn=PTPN23;refGene_tx=XM_023248297.1
A2	16787225	16787357	refGene_gn=PTPN23;refGene_tx=XM_003982194.5;refGene_gn=PTPN23;refGene_tx=XM_023248297.1
A2	16787623	16787704	refGene_gn=PTPN23;refGene_tx=XM_003982194.5;refGene_gn=PTPN23;refGene_tx=XM_023248297.1
A2	16787798	16787930	refGene_gn=PTPN23;refGene_tx=XM_003982194.5;refGene_gn=PTPN23;refGene_tx=XM_023248297.1
A2	16788033	16788081	refGene_gn=PTPN23;refGene_tx=XM_003982194.5;refGene_gn=PTPN23;refGene_

tx=XM_023248297.1
A2 16788435 16788492
refGene_gn=PTPN23;refGene_tx=XM_003982194.5;refGene_gn=PTPN23;refGene_tx=XM_023248297.1
A2 16788630 16788689
refGene_gn=PTPN23;refGene_tx=XM_003982194.5;refGene_gn=PTPN23;refGene_tx=XM_023248297.1
A2 16788848 16788928
refGene_gn=PTPN23;refGene_tx=XM_003982194.5;refGene_gn=PTPN23;refGene_tx=XM_023248297.1
A2 16789042 16789157
refGene_gn=PTPN23;refGene_tx=XM_003982194.5;refGene_gn=PTPN23;refGene_tx=XM_023248297.1
A2 16789250 16789316
refGene_gn=PTPN23;refGene_tx=XM_003982194.5;refGene_gn=PTPN23;refGene_tx=XM_023248297.1
A2 16789687 16789833
refGene_gn=PTPN23;refGene_tx=XM_003982194.5;refGene_gn=PTPN23;refGene_tx=XM_023248297.1
A2 16790121 16790442
refGene_gn=PTPN23;refGene_tx=XM_003982194.5;refGene_gn=PTPN23;refGene_tx=XM_023248297.1
A2 16790516 16790671
refGene_gn=PTPN23;refGene_tx=XM_003982194.5;refGene_gn=PTPN23;refGene_tx=XM_023248297.1
A2 16790758 16790910
refGene_gn=PTPN23;refGene_tx=XM_003982194.5;refGene_gn=PTPN23;refGene_tx=XM_023248297.1
A2 16791003 16791183
refGene_gn=PTPN23;refGene_tx=XM_003982194.5;refGene_gn=PTPN23;refGene_tx=XM_023248297.1
A2 16791256 16793072
refGene_gn=PTPN23;refGene_tx=XM_003982194.5;refGene_gn=PTPN23;refGene_tx=XM_023248297.1
A2 16793171 16793356
refGene_gn=PTPN23;refGene_tx=XM_003982194.5;refGene_gn=PTPN23;refGene_tx=XM_023248297.1
A2 16793442 16793547
refGene_gn=PTPN23;refGene_tx=XM_003982194.5;refGene_gn=PTPN23;refGene_tx=XM_023248297.1
A2 16793625 16793764
refGene_gn=PTPN23;refGene_tx=XM_003982194.5;refGene_gn=PTPN23;refGene_tx=XM_023248297.1
A2 16793862 16793976
refGene_gn=PTPN23;refGene_tx=XM_003982194.5;refGene_gn=PTPN23;refGene_tx=XM_023248297.1
A2 16794054 16794534
refGene_gn=PTPN23;refGene_tx=XM_003982194.5;refGene_gn=PTPN23;refGene_tx=XM_023248297.1
A2 16795209 16795455 refGene_gn=SCAP;refGene_tx=XM_023248298.1

A2 16795536 16795743 refGene_gn=SCAP;refGene_tx=XM_023248298.1
A2 16795925 16796006 refGene_gn=SCAP;refGene_tx=XM_023248298.1
A2 16796152 16796310 refGene_gn=SCAP;refGene_tx=XM_023248298.1
A2 16796420 16796503 refGene_gn=SCAP;refGene_tx=XM_023248298.1
A2 16798383 16798469 refGene_gn=SCAP;refGene_tx=XM_023248298.1
A2 16798564 16799087 refGene_gn=SCAP;refGene_tx=XM_023248298.1
A2 16799194 16799310 refGene_gn=SCAP;refGene_tx=XM_023248298.1
A2 16799388 16799590 refGene_gn=SCAP;refGene_tx=XM_023248298.1
A2 16799668 16799857 refGene_gn=SCAP;refGene_tx=XM_023248298.1
A2 16800343 16800720 refGene_gn=SCAP;refGene_tx=XM_023248298.1
A2 16801557 16801776 refGene_gn=SCAP;refGene_tx=XM_023248298.1
A2 16801924 16802023 refGene_gn=SCAP;refGene_tx=XM_023248298.1
A2 16803236 16803331 refGene_gn=SCAP;refGene_tx=XM_023248298.1
A2 16804348 16804461 refGene_gn=SCAP;refGene_tx=XM_023248298.1
A2 16806002 16806129 refGene_gn=SCAP;refGene_tx=XM_023248298.1
A2 16806525 16806698 refGene_gn=SCAP;refGene_tx=XM_023248298.1
A2 16807338 16807444 refGene_gn=SCAP;refGene_tx=XM_023248298.1
A2 16807623 16807844 refGene_gn=SCAP;refGene_tx=XM_023248298.1
A2 16808630 16808788 refGene_gn=SCAP;refGene_tx=XM_023248298.1
A2 16817996 16818126 refGene_gn=SCAP;refGene_tx=XM_023248298.1
A2 16831386 16831508 refGene_gn=SCAP;refGene_tx=XM_023248298.1
A2 16879070 16879199
refGene_gn=ELP6;refGene_tx=XM_003982151.4;refGene_gn=ELP6;refGene_tx=XM_019820917.2;refGene_gn=ELP6;refGene_tx=XM_019820922.1;refGene_gn=ELP6;refGene_tx=XM_019820928.2
A2 16881291 16881438
refGene_gn=ELP6;refGene_tx=XM_003982151.4;refGene_gn=ELP6;refGene_tx=XM_019820917.2;refGene_gn=ELP6;refGene_tx=XM_019820922.1;refGene_gn=ELP6;refGene_tx=XM_019820928.2
A2 16883488 16883690
refGene_gn=ELP6;refGene_tx=XM_003982151.4;refGene_gn=ELP6;refGene_tx=XM_019820917.2;refGene_gn=ELP6;refGene_tx=XM_019820922.1;refGene_gn=ELP6;refGene_tx=XM_019820928.2
A2 16885066 16885224
refGene_gn=ELP6;refGene_tx=XM_003982151.4;refGene_gn=ELP6;refGene_tx=XM_019820917.2;refGene_gn=ELP6;refGene_tx=XM_019820922.1;refGene_gn=ELP6;refGene_tx=XM_019820928.2
A2 16888068 16888139
refGene_gn=ELP6;refGene_tx=XM_003982151.4;refGene_gn=ELP6;refGene_tx=XM_019820917.2
A2 16889368 16889447
refGene_gn=ELP6;refGene_tx=XM_003982151.4;refGene_gn=ELP6;refGene_tx=XM_019820917.2;refGene_gn=ELP6;refGene_tx=XM_019820922.1;refGene_gn=ELP6;refGene_tx=XM_019820928.2
A2 16891737 16891791
refGene_gn=ELP6;refGene_tx=XM_003982151.4;refGene_gn=ELP6;refGene_tx=XM_019820917.2
A2 16899327 16899459
refGene_gn=CSPG5;refGene_tx=XM_023248321.1;refGene_gn=CSPG5;refGene_tx=XM_023248323.1

A2	16932969	16933064	refGene_gn=CSPG5;refGene_tx=XM_023248321.1
A2	16938432	16938594	refGene_gn=CSPG5;refGene_tx=XM_023248324.1
A2	16944907	16944983	refGene_gn=CSPG5;refGene_tx=XM_023248321.1;refGene_gn=CSPG5;refGene_tx=XM_023248323.1;refGene_gn=CSPG5;refGene_tx=XM_023248324.1
A2	16946807	16946996	refGene_gn=CSPG5;refGene_tx=XM_023248321.1;refGene_gn=CSPG5;refGene_tx=XM_023248323.1;refGene_gn=CSPG5;refGene_tx=XM_023248324.1
A2	16950222	16951333	refGene_gn=CSPG5;refGene_tx=XM_023248321.1;refGene_gn=CSPG5;refGene_tx=XM_023248323.1;refGene_gn=CSPG5;refGene_tx=XM_023248324.1
A2	16951934	16952031	refGene_gn=CSPG5;refGene_tx=XM_023248324.1
A2	16960231	16960329	refGene_gn=SMARCC1;refGene_tx=XM_023248317.1
A2	16962560	16962724	refGene_gn=SMARCC1;refGene_tx=XM_023248317.1
A2	16974496	16974506	refGene_gn=SMARCC1;refGene_tx=XM_023248317.1
A2	16986387	16986649	refGene_gn=SMARCC1;refGene_tx=XM_023248317.1
A2	16997028	16997163	refGene_gn=SMARCC1;refGene_tx=XM_023248317.1
A2	17011624	17011779	refGene_gn=SMARCC1;refGene_tx=XM_023248317.1
A2	17012853	17012968	refGene_gn=SMARCC1;refGene_tx=XM_023248317.1
A2	17016839	17016895	refGene_gn=SMARCC1;refGene_tx=XM_023248317.1
A2	17041680	17041842	refGene_gn=SMARCC1;refGene_tx=XM_023248317.1
A2	17042553	17042812	refGene_gn=SMARCC1;refGene_tx=XM_023248317.1
A2	17051117	17051177	refGene_gn=SMARCC1;refGene_tx=XM_023248317.1
A2	17058007	17058121	refGene_gn=SMARCC1;refGene_tx=XM_023248317.1
A2	17058613	17058767	refGene_gn=SMARCC1;refGene_tx=XM_023248317.1
A2	17059394	17059508	refGene_gn=SMARCC1;refGene_tx=XM_023248317.1
A2	17062256	17062328	refGene_gn=SMARCC1;refGene_tx=XM_023248317.1
A2	17066213	17066335	refGene_gn=SMARCC1;refGene_tx=XM_023248317.1
A2	17069694	17069732	refGene_gn=SMARCC1;refGene_tx=XM_023248317.1
A2	17074318	17074378	refGene_gn=SMARCC1;refGene_tx=XM_023248317.1
A2	17079460	17079585	refGene_gn=SMARCC1;refGene_tx=XM_023248317.1
A2	17082951	17083073	refGene_gn=SMARCC1;refGene_tx=XM_023248317.1
A2	17086279	17086405	refGene_gn=SMARCC1;refGene_tx=XM_023248317.1
A2	17091086	17091162	refGene_gn=SMARCC1;refGene_tx=XM_023248317.1
A2	17094958	17095028	refGene_gn=SMARCC1;refGene_tx=XM_023248317.1
A2	17100096	17100166	refGene_gn=SMARCC1;refGene_tx=XM_023248317.1
A2	17102671	17102764	refGene_gn=SMARCC1;refGene_tx=XM_023248317.1
A2	17103888	17103970	refGene_gn=SMARCC1;refGene_tx=XM_023248317.1
A2	17113192	17113278	refGene_gn=SMARCC1;refGene_tx=XM_023248317.1
A2	17129287	17129407	refGene_gn=SMARCC1;refGene_tx=XM_023248317.1
A2	17140584	17140779	refGene_gn=SMARCC1;refGene_tx=XM_023248317.1
A2	17184016	17184041	refGene_gn=DHX30;refGene_tx=XM_023248328.1;refGene_gn=DHX30;refGene_tx=XM_023248330.1
A2	17185159	17185214	refGene_gn=DHX30;refGene_tx=XM_023248328.1;refGene_gn=DHX30;refGene_tx=XM_023248330.1;refGene_gn=DHX30;refGene_tx=XM_023248331.1;refGene_gn=DHX30;refGene_tx=XM_023248334.1;refGene_gn=DHX30;refGene_tx=XM_023248338.1;refGene_gn=DHX30;refGene_tx=XM_023248340.1;refGene_gn=DHX30;refGene_tx=XM_023248345.1

A2 17190086 17190182
refGene_gn=DHX30;refGene_tx=XM_023248328.1;refGene_gn=DHX30;refGene_tx=XM_023248330.1;refGene_gn=DHX30;refGene_tx=XM_023248331.1;refGene_gn=DHX30;refGene_tx=XM_023248334.1;refGene_gn=DHX30;refGene_tx=XM_023248338.1
A2 17191297 17191326
refGene_gn=DHX30;refGene_tx=XM_023248328.1;refGene_gn=DHX30;refGene_tx=XM_023248330.1
A2 17193707 17193767
refGene_gn=DHX30;refGene_tx=XM_023248328.1;refGene_gn=DHX30;refGene_tx=XM_023248331.1;refGene_gn=DHX30;refGene_tx=XM_023248334.1
A2 17195406 17195614 refGene_gn=DHX30;refGene_tx=XM_003982154.5
A2 17197378 17197509
refGene_gn=DHX30;refGene_tx=XM_003982154.5;refGene_gn=DHX30;refGene_tx=XM_023248328.1;refGene_gn=DHX30;refGene_tx=XM_023248330.1;refGene_gn=DHX30;refGene_tx=XM_023248331.1;refGene_gn=DHX30;refGene_tx=XM_023248334.1;refGene_gn=DHX30;refGene_tx=XM_023248338.1;refGene_gn=DHX30;refGene_tx=XM_023248340.1;refGene_gn=DHX30;refGene_tx=XM_023248345.1
A2 17198148 17198259
refGene_gn=DHX30;refGene_tx=XM_003982154.5;refGene_gn=DHX30;refGene_tx=XM_023248328.1;refGene_gn=DHX30;refGene_tx=XM_023248330.1;refGene_gn=DHX30;refGene_tx=XM_023248331.1;refGene_gn=DHX30;refGene_tx=XM_023248334.1;refGene_gn=DHX30;refGene_tx=XM_023248338.1;refGene_gn=DHX30;refGene_tx=XM_023248340.1;refGene_gn=DHX30;refGene_tx=XM_023248345.1
A2 17202487 17202789
refGene_gn=DHX30;refGene_tx=XM_003982154.5;refGene_gn=DHX30;refGene_tx=XM_023248328.1;refGene_gn=DHX30;refGene_tx=XM_023248330.1;refGene_gn=DHX30;refGene_tx=XM_023248331.1;refGene_gn=DHX30;refGene_tx=XM_023248334.1;refGene_gn=DHX30;refGene_tx=XM_023248338.1;refGene_gn=DHX30;refGene_tx=XM_023248340.1;refGene_gn=DHX30;refGene_tx=XM_023248345.1
A2 17203286 17203407
refGene_gn=DHX30;refGene_tx=XM_003982154.5;refGene_gn=DHX30;refGene_tx=XM_023248328.1;refGene_gn=DHX30;refGene_tx=XM_023248330.1;refGene_gn=DHX30;refGene_tx=XM_023248331.1;refGene_gn=DHX30;refGene_tx=XM_023248334.1;refGene_gn=DHX30;refGene_tx=XM_023248338.1;refGene_gn=DHX30;refGene_tx=XM_023248340.1;refGene_gn=DHX30;refGene_tx=XM_023248345.1
A2 17204136 17204286
refGene_gn=DHX30;refGene_tx=XM_003982154.5;refGene_gn=DHX30;refGene_tx=XM_023248328.1;refGene_gn=DHX30;refGene_tx=XM_023248330.1;refGene_gn=DHX30;refGene_tx=XM_023248331.1;refGene_gn=DHX30;refGene_tx=XM_023248334.1;refGene_gn=DHX30;refGene_tx=XM_023248338.1;refGene_gn=DHX30;refGene_tx=XM_023248340.1;refGene_gn=DHX30;refGene_tx=XM_023248345.1
A2 17206217 17206370
refGene_gn=DHX30;refGene_tx=XM_003982154.5;refGene_gn=DHX30;refGene_tx=XM_023248328.1;refGene_gn=DHX30;refGene_tx=XM_023248330.1;refGene_gn=DHX30;refGene_tx=XM_023248331.1;refGene_gn=DHX30;refGene_tx=XM_023248334.1;refGene_gn=DHX30;refGene_tx=XM_023248338.1;refGene_gn=DHX30;refGene_tx=XM_023248340.1;refGene_gn=DHX30;refGene_tx=XM_023248345.1
A2 17206667 17207504
refGene_gn=DHX30;refGene_tx=XM_003982154.5;refGene_gn=DHX30;refGene_tx

=XM_023248328.1;refGene_gn=DHX30;refGene_tx=XM_023248330.1;refGene_gn=DHX30;refGene_tx=XM_023248331.1;refGene_gn=DHX30;refGene_tx=XM_023248334.1;refGene_gn=DHX30;refGene_tx=XM_023248338.1;refGene_gn=DHX30;refGene_tx=XM_023248340.1;refGene_gn=DHX30;refGene_tx=XM_023248345.1
A2 17207786 17207862
refGene_gn=DHX30;refGene_tx=XM_003982154.5;refGene_gn=DHX30;refGene_tx=XM_023248328.1;refGene_gn=DHX30;refGene_tx=XM_023248330.1;refGene_gn=DHX30;refGene_tx=XM_023248331.1;refGene_gn=DHX30;refGene_tx=XM_023248334.1;refGene_gn=DHX30;refGene_tx=XM_023248338.1;refGene_gn=DHX30;refGene_tx=XM_023248340.1;refGene_gn=DHX30;refGene_tx=XM_023248345.1
A2 17207948 17208053
refGene_gn=DHX30;refGene_tx=XM_003982154.5;refGene_gn=DHX30;refGene_tx=XM_023248328.1;refGene_gn=DHX30;refGene_tx=XM_023248330.1;refGene_gn=DHX30;refGene_tx=XM_023248331.1;refGene_gn=DHX30;refGene_tx=XM_023248334.1;refGene_gn=DHX30;refGene_tx=XM_023248338.1;refGene_gn=DHX30;refGene_tx=XM_023248340.1;refGene_gn=DHX30;refGene_tx=XM_023248345.1
A2 17208311 17208487
refGene_gn=DHX30;refGene_tx=XM_003982154.5;refGene_gn=DHX30;refGene_tx=XM_023248328.1;refGene_gn=DHX30;refGene_tx=XM_023248330.1;refGene_gn=DHX30;refGene_tx=XM_023248331.1;refGene_gn=DHX30;refGene_tx=XM_023248334.1;refGene_gn=DHX30;refGene_tx=XM_023248338.1;refGene_gn=DHX30;refGene_tx=XM_023248340.1;refGene_gn=DHX30;refGene_tx=XM_023248345.1
A2 17208731 17208938
refGene_gn=DHX30;refGene_tx=XM_003982154.5;refGene_gn=DHX30;refGene_tx=XM_023248328.1;refGene_gn=DHX30;refGene_tx=XM_023248330.1;refGene_gn=DHX30;refGene_tx=XM_023248331.1;refGene_gn=DHX30;refGene_tx=XM_023248334.1;refGene_gn=DHX30;refGene_tx=XM_023248338.1;refGene_gn=DHX30;refGene_tx=XM_023248340.1;refGene_gn=DHX30;refGene_tx=XM_023248345.1
A2 17209031 17209113
refGene_gn=DHX30;refGene_tx=XM_003982154.5;refGene_gn=DHX30;refGene_tx=XM_023248328.1;refGene_gn=DHX30;refGene_tx=XM_023248330.1;refGene_gn=DHX30;refGene_tx=XM_023248331.1;refGene_gn=DHX30;refGene_tx=XM_023248334.1;refGene_gn=DHX30;refGene_tx=XM_023248338.1;refGene_gn=DHX30;refGene_tx=XM_023248340.1;refGene_gn=DHX30;refGene_tx=XM_023248345.1
A2 17209189 17209383
refGene_gn=DHX30;refGene_tx=XM_003982154.5;refGene_gn=DHX30;refGene_tx=XM_023248328.1;refGene_gn=DHX30;refGene_tx=XM_023248330.1;refGene_gn=DHX30;refGene_tx=XM_023248331.1;refGene_gn=DHX30;refGene_tx=XM_023248334.1;refGene_gn=DHX30;refGene_tx=XM_023248338.1;refGene_gn=DHX30;refGene_tx=XM_023248340.1;refGene_gn=DHX30;refGene_tx=XM_023248345.1
A2 17209508 17209668
refGene_gn=DHX30;refGene_tx=XM_003982154.5;refGene_gn=DHX30;refGene_tx=XM_023248328.1;refGene_gn=DHX30;refGene_tx=XM_023248330.1;refGene_gn=DHX30;refGene_tx=XM_023248331.1;refGene_gn=DHX30;refGene_tx=XM_023248334.1;refGene_gn=DHX30;refGene_tx=XM_023248338.1;refGene_gn=DHX30;refGene_tx=XM_023248340.1;refGene_gn=DHX30;refGene_tx=XM_023248345.1
A2 17209797 17209955
refGene_gn=DHX30;refGene_tx=XM_003982154.5;refGene_gn=DHX30;refGene_tx=XM_023248328.1;refGene_gn=DHX30;refGene_tx=XM_023248330.1;refGene_gn=DHX30;refGene_tx=XM_023248331.1;refGene_gn=DHX30;refGene_tx=XM_0232483

34.1;refGene_gn=DHX30;refGene_tx=XM_023248338.1;refGene_gn=DHX30;refGene_tx=XM_023248340.1;refGene_gn=DHX30;refGene_tx=XM_023248345.1
A2 17210050 17210154
refGene_gn=DHX30;refGene_tx=XM_003982154.5;refGene_gn=DHX30;refGene_tx=XM_023248328.1;refGene_gn=DHX30;refGene_tx=XM_023248330.1;refGene_gn=DHX30;refGene_tx=XM_023248331.1;refGene_gn=DHX30;refGene_tx=XM_023248334.1;refGene_gn=DHX30;refGene_tx=XM_023248338.1;refGene_gn=DHX30;refGene_tx=XM_023248340.1;refGene_gn=DHX30;refGene_tx=XM_023248345.1
A2 17210234 17210374
refGene_gn=DHX30;refGene_tx=XM_003982154.5;refGene_gn=DHX30;refGene_tx=XM_023248328.1;refGene_gn=DHX30;refGene_tx=XM_023248330.1;refGene_gn=DHX30;refGene_tx=XM_023248331.1;refGene_gn=DHX30;refGene_tx=XM_023248334.1;refGene_gn=DHX30;refGene_tx=XM_023248338.1;refGene_gn=DHX30;refGene_tx=XM_023248340.1;refGene_gn=DHX30;refGene_tx=XM_023248345.1
A2 17210482 17210736
refGene_gn=DHX30;refGene_tx=XM_003982154.5;refGene_gn=DHX30;refGene_tx=XM_023248328.1;refGene_gn=DHX30;refGene_tx=XM_023248330.1;refGene_gn=DHX30;refGene_tx=XM_023248331.1;refGene_gn=DHX30;refGene_tx=XM_023248334.1;refGene_gn=DHX30;refGene_tx=XM_023248338.1;refGene_gn=DHX30;refGene_tx=XM_023248340.1;refGene_gn=DHX30;refGene_tx=XM_023248345.1
A2 17213683 17213691 refGene_gn=MAP4;refGene_tx=NM_001164033.2
A2 17213825 17214015 refGene_gn=MAP4;refGene_tx=NM_001164033.2
A2 17215906 17216019 refGene_gn=MAP4;refGene_tx=NM_001164033.2
A2 17217618 17217700 refGene_gn=MAP4;refGene_tx=NM_001164033.2
A2 17223588 17223681 refGene_gn=MAP4;refGene_tx=NM_001164033.2
A2 17225660 17225774 refGene_gn=MAP4;refGene_tx=NM_001164033.2
A2 17227106 17227399 refGene_gn=MAP4;refGene_tx=NM_001164033.2
A2 17227515 17227575 refGene_gn=MAP4;refGene_tx=NM_001164033.2
A2 17228126 17228310 refGene_gn=MAP4;refGene_tx=NM_001164033.2
A2 17231986 17232202 refGene_gn=MAP4;refGene_tx=NM_001164033.2
A2 17233020 17233127 refGene_gn=MAP4;refGene_tx=NM_001164033.2
A2 17266248 17266371 refGene_gn=MAP4;refGene_tx=NM_001164033.2
A2 17267053 17267634 refGene_gn=MAP4;refGene_tx=NM_001164033.2
A2 17267635 17268215 refGene_gn=MAP4;refGene_tx=NM_001164033.2
A2 17269004 17269139 refGene_gn=MAP4;refGene_tx=NM_001164033.2
A2 17271674 17271788 refGene_gn=MAP4;refGene_tx=NM_001164033.2
A2 17278231 17278354 refGene_gn=MAP4;refGene_tx=NM_001164033.2
A2 17319536 17319605 refGene_gn=MAP4;refGene_tx=NM_001164033.2
A2 17341561 17341784 refGene_gn=MAP4;refGene_tx=NM_001164033.2
A2 17448836 17448977
refGene_gn=CDC25A;refGene_tx=XM_011291281.3;refGene_gn=CDC25A;refGene_tx=XM_011291284.3;refGene_gn=CDC25A;refGene_tx=XM_011291287.3;refGene_gn=CDC25A;refGene_tx=XM_011291290.3;refGene_gn=CDC25A;refGene_tx=XM_023248391.1
A2 17449183 17449295
refGene_gn=CDC25A;refGene_tx=XM_011291281.3;refGene_gn=CDC25A;refGene_tx=XM_011291284.3;refGene_gn=CDC25A;refGene_tx=XM_011291287.3;refGene_gn=CDC25A;refGene_tx=XM_011291290.3;refGene_gn=CDC25A;refGene_tx=XM_023248391.1
A2 17450823 17450954

refGene_gn=CDC25A;refGene_tx=XM_011291281.3;refGene_gn=CDC25A;refGene_tx=XM_011291284.3;refGene_gn=CDC25A;refGene_tx=XM_011291287.3;refGene_gn=CDC25A;refGene_tx=XM_011291290.3;refGene_gn=CDC25A;refGene_tx=XM_023248391.1
A2 17451884 17451983
refGene_gn=CDC25A;refGene_tx=XM_011291281.3;refGene_gn=CDC25A;refGene_tx=XM_011291284.3;refGene_gn=CDC25A;refGene_tx=XM_011291287.3;refGene_gn=CDC25A;refGene_tx=XM_011291290.3;refGene_gn=CDC25A;refGene_tx=XM_023248391.1
A2 17452078 17452141
refGene_gn=CDC25A;refGene_tx=XM_011291281.3;refGene_gn=CDC25A;refGene_tx=XM_011291284.3;refGene_gn=CDC25A;refGene_tx=XM_011291287.3;refGene_gn=CDC25A;refGene_tx=XM_011291290.3;refGene_gn=CDC25A;refGene_tx=XM_023248391.1
A2 17453658 17453757
refGene_gn=CDC25A;refGene_tx=XM_011291281.3;refGene_gn=CDC25A;refGene_tx=XM_011291284.3;refGene_gn=CDC25A;refGene_tx=XM_011291287.3;refGene_gn=CDC25A;refGene_tx=XM_011291290.3;refGene_gn=CDC25A;refGene_tx=XM_023248391.1
A2 17457346 17457520
refGene_gn=CDC25A;refGene_tx=XM_011291281.3;refGene_gn=CDC25A;refGene_tx=XM_011291284.3;refGene_gn=CDC25A;refGene_tx=XM_011291287.3;refGene_gn=CDC25A;refGene_tx=XM_011291290.3
A2 17462756 17462831
refGene_gn=CDC25A;refGene_tx=XM_011291281.3;refGene_gn=CDC25A;refGene_tx=XM_011291284.3;refGene_gn=CDC25A;refGene_tx=XM_011291287.3;refGene_gn=CDC25A;refGene_tx=XM_011291290.3;refGene_gn=CDC25A;refGene_tx=XM_023248391.1
A2 17463330 17463465
refGene_gn=CDC25A;refGene_tx=XM_011291281.3;refGene_gn=CDC25A;refGene_tx=XM_011291284.3;refGene_gn=CDC25A;refGene_tx=XM_011291287.3;refGene_gn=CDC25A;refGene_tx=XM_011291290.3;refGene_gn=CDC25A;refGene_tx=XM_023248391.1
A2 17465868 17465988
refGene_gn=CDC25A;refGene_tx=XM_011291281.3;refGene_gn=CDC25A;refGene_tx=XM_011291284.3;refGene_gn=CDC25A;refGene_tx=XM_023248391.1
A2 17467567 17467669
refGene_gn=CDC25A;refGene_tx=XM_011291281.3;refGene_gn=CDC25A;refGene_tx=XM_011291284.3;refGene_gn=CDC25A;refGene_tx=XM_011291287.3;refGene_gn=CDC25A;refGene_tx=XM_011291290.3;refGene_gn=CDC25A;refGene_tx=XM_023248391.1
A2 17468150 17468187
refGene_gn=CDC25A;refGene_tx=XM_011291281.3;refGene_gn=CDC25A;refGene_tx=XM_011291284.3;refGene_gn=CDC25A;refGene_tx=XM_011291287.3;refGene_gn=CDC25A;refGene_tx=XM_011291290.3;refGene_gn=CDC25A;refGene_tx=XM_023248391.1
A2 17468697 17468740
refGene_gn=CDC25A;refGene_tx=XM_011291281.3;refGene_gn=CDC25A;refGene_tx=XM_011291284.3;refGene_gn=CDC25A;refGene_tx=XM_011291287.3;refGene_gn=CDC25A;refGene_tx=XM_011291290.3;refGene_gn=CDC25A;refGene_tx=XM_02

3248391.1
A2	17470088	17470162
refGene_gn=CDC25A;refGene_tx=XM_011291281.3;refGene_gn=CDC25A;refGene_tx=XM_011291284.3;refGene_gn=CDC25A;refGene_tx=XM_011291287.3;refGene_gn=CDC25A;refGene_tx=XM_011291290.3;refGene_gn=CDC25A;refGene_tx=XM_023248391.1
A2	17471162	17471332
refGene_gn=CDC25A;refGene_tx=XM_011291281.3;refGene_gn=CDC25A;refGene_tx=XM_011291284.3;refGene_gn=CDC25A;refGene_tx=XM_011291287.3;refGene_gn=CDC25A;refGene_tx=XM_011291290.3;refGene_gn=CDC25A;refGene_tx=XM_023248391.1
A2	17505992	17506190	refGene_gn=CAMP;refGene_tx=NM_001204778.1
A2	17507066	17507174	refGene_gn=CAMP;refGene_tx=NM_001204778.1
A2	17507311	17507383	refGene_gn=CAMP;refGene_tx=NM_001204778.1
A2	17507950	17508088	refGene_gn=CAMP;refGene_tx=NM_001204778.1
A2	17541797	17541985
refGene_gn=NME6;refGene_tx=XM_003982155.5;refGene_gn=NME6;refGene_tx=XM_006928709.4;refGene_gn=NME6;refGene_tx=XM_006928710.4;refGene_gn=NME6;refGene_tx=XM_019821003.2;refGene_gn=NME6;refGene_tx=XM_019821006.2
A2	17542261	17542422
refGene_gn=NME6;refGene_tx=XM_003982155.5;refGene_gn=NME6;refGene_tx=XM_006928709.4;refGene_gn=NME6;refGene_tx=XM_006928710.4;refGene_gn=NME6;refGene_tx=XM_019821003.2;refGene_gn=NME6;refGene_tx=XM_019821006.2
A2	17542846	17542886
refGene_gn=NME6;refGene_tx=XM_003982155.5;refGene_gn=NME6;refGene_tx=XM_006928709.4;refGene_gn=NME6;refGene_tx=XM_006928710.4;refGene_gn=NME6;refGene_tx=XM_019821003.2;refGene_gn=NME6;refGene_tx=XM_019821006.2
A2	17543430	17543533
refGene_gn=NME6;refGene_tx=XM_003982155.5;refGene_gn=NME6;refGene_tx=XM_006928709.4;refGene_gn=NME6;refGene_tx=XM_006928710.4;refGene_gn=NME6;refGene_tx=XM_019821003.2;refGene_gn=NME6;refGene_tx=XM_019821006.2
A2	17545583	17545673
refGene_gn=NME6;refGene_tx=XM_003982155.5;refGene_gn=NME6;refGene_tx=XM_006928709.4;refGene_gn=NME6;refGene_tx=XM_006928710.4;refGene_gn=NME6;refGene_tx=XM_019821003.2
A2	17565949	17566025	refGene_gn=SPINK8;refGene_tx=XM_023241835.1
A2	17573028	17573153	refGene_gn=SPINK8;refGene_tx=XM_023241835.1
A2	17574063	17574069	refGene_gn=SPINK8;refGene_tx=XM_023241835.1
A2	17584308	17584329	refGene_gn=FBXW12;refGene_tx=XM_023248416.1
A2	17602790	17602828	refGene_gn=FBXW12;refGene_tx=XM_023248416.1
A2	17603041	17603199	refGene_gn=FBXW12;refGene_tx=XM_023248416.1
A2	17604013	17604132	refGene_gn=FBXW12;refGene_tx=XM_023248416.1
A2	17610690	17610831	refGene_gn=FBXW12;refGene_tx=XM_023248416.1
A2	17610832	17610901	refGene_gn=FBXW12;refGene_tx=XM_023248416.1
A2	17611702	17611861	refGene_gn=FBXW12;refGene_tx=XM_023248416.1
A2	17612821	17613032	refGene_gn=FBXW12;refGene_tx=XM_023248416.1
A2	17613987	17614166	refGene_gn=FBXW12;refGene_tx=XM_023248416.1
A2	17614247	17614381	refGene_gn=FBXW12;refGene_tx=XM_023248416.1
A2	17623646	17623746	refGene_gn=FBXW12;refGene_tx=XM_023248416.1
A2	17635080	17635185

refGene_gn=PLXNB1;refGene_tx=XM_003982157.5;refGene_gn=PLXNB1;refGene_tx=XM_023248458.1;refGene_gn=PLXNB1;refGene_tx=XM_023248461.1;refGene_gn=PLXNB1;refGene_tx=XM_023248463.1;refGene_gn=PLXNB1;refGene_tx=XM_023248469.1
A2 17636253 17636328
refGene_gn=PLXNB1;refGene_tx=XM_003982157.5;refGene_gn=PLXNB1;refGene_tx=XM_023248458.1;refGene_gn=PLXNB1;refGene_tx=XM_023248461.1;refGene_gn=PLXNB1;refGene_tx=XM_023248463.1;refGene_gn=PLXNB1;refGene_tx=XM_023248469.1
A2 17637232 17637308
refGene_gn=PLXNB1;refGene_tx=XM_003982157.5;refGene_gn=PLXNB1;refGene_tx=XM_023248458.1;refGene_gn=PLXNB1;refGene_tx=XM_023248461.1;refGene_gn=PLXNB1;refGene_tx=XM_023248463.1;refGene_gn=PLXNB1;refGene_tx=XM_023248469.1
A2 17637444 17637509
refGene_gn=PLXNB1;refGene_tx=XM_003982157.5;refGene_gn=PLXNB1;refGene_tx=XM_023248458.1;refGene_gn=PLXNB1;refGene_tx=XM_023248461.1;refGene_gn=PLXNB1;refGene_tx=XM_023248463.1;refGene_gn=PLXNB1;refGene_tx=XM_023248469.1
A2 17639351 17639499
refGene_gn=PLXNB1;refGene_tx=XM_003982157.5;refGene_gn=PLXNB1;refGene_tx=XM_023248458.1;refGene_gn=PLXNB1;refGene_tx=XM_023248461.1;refGene_gn=PLXNB1;refGene_tx=XM_023248463.1;refGene_gn=PLXNB1;refGene_tx=XM_023248469.1
A2 17639596 17639757
refGene_gn=PLXNB1;refGene_tx=XM_003982157.5;refGene_gn=PLXNB1;refGene_tx=XM_023248458.1;refGene_gn=PLXNB1;refGene_tx=XM_023248461.1;refGene_gn=PLXNB1;refGene_tx=XM_023248463.1;refGene_gn=PLXNB1;refGene_tx=XM_023248469.1
A2 17639965 17640138
refGene_gn=PLXNB1;refGene_tx=XM_003982157.5;refGene_gn=PLXNB1;refGene_tx=XM_023248458.1;refGene_gn=PLXNB1;refGene_tx=XM_023248461.1;refGene_gn=PLXNB1;refGene_tx=XM_023248463.1;refGene_gn=PLXNB1;refGene_tx=XM_023248469.1
A2 17640373 17640458
refGene_gn=PLXNB1;refGene_tx=XM_003982157.5;refGene_gn=PLXNB1;refGene_tx=XM_023248458.1;refGene_gn=PLXNB1;refGene_tx=XM_023248461.1;refGene_gn=PLXNB1;refGene_tx=XM_023248463.1;refGene_gn=PLXNB1;refGene_tx=XM_023248469.1
A2 17640532 17640636
refGene_gn=PLXNB1;refGene_tx=XM_003982157.5;refGene_gn=PLXNB1;refGene_tx=XM_023248458.1;refGene_gn=PLXNB1;refGene_tx=XM_023248461.1;refGene_gn=PLXNB1;refGene_tx=XM_023248463.1;refGene_gn=PLXNB1;refGene_tx=XM_023248469.1
A2 17640900 17641069
refGene_gn=PLXNB1;refGene_tx=XM_003982157.5;refGene_gn=PLXNB1;refGene_tx=XM_023248458.1;refGene_gn=PLXNB1;refGene_tx=XM_023248461.1;refGene_gn=PLXNB1;refGene_tx=XM_023248463.1;refGene_gn=PLXNB1;refGene_tx=XM_023248469.1
A2 17641989 17642136

refGene_gn=PLXNB1;refGene_tx=XM_003982157.5;refGene_gn=PLXNB1;refGene_tx=XM_023248458.1;refGene_gn=PLXNB1;refGene_tx=XM_023248461.1;refGene_gn=PLXNB1;refGene_tx=XM_023248463.1;refGene_gn=PLXNB1;refGene_tx=XM_023248469.1
A2 17642366 17642433
refGene_gn=PLXNB1;refGene_tx=XM_003982157.5;refGene_gn=PLXNB1;refGene_tx=XM_023248458.1;refGene_gn=PLXNB1;refGene_tx=XM_023248461.1;refGene_gn=PLXNB1;refGene_tx=XM_023248463.1;refGene_gn=PLXNB1;refGene_tx=XM_023248469.1
A2 17642586 17642765
refGene_gn=PLXNB1;refGene_tx=XM_003982157.5;refGene_gn=PLXNB1;refGene_tx=XM_023248458.1;refGene_gn=PLXNB1;refGene_tx=XM_023248461.1;refGene_gn=PLXNB1;refGene_tx=XM_023248463.1;refGene_gn=PLXNB1;refGene_tx=XM_023248469.1
A2 17642905 17643123
refGene_gn=PLXNB1;refGene_tx=XM_003982157.5;refGene_gn=PLXNB1;refGene_tx=XM_023248458.1;refGene_gn=PLXNB1;refGene_tx=XM_023248461.1;refGene_gn=PLXNB1;refGene_tx=XM_023248463.1;refGene_gn=PLXNB1;refGene_tx=XM_023248469.1
A2 17643227 17643328
refGene_gn=PLXNB1;refGene_tx=XM_003982157.5;refGene_gn=PLXNB1;refGene_tx=XM_023248458.1;refGene_gn=PLXNB1;refGene_tx=XM_023248461.1;refGene_gn=PLXNB1;refGene_tx=XM_023248463.1;refGene_gn=PLXNB1;refGene_tx=XM_023248469.1
A2 17643838 17643987
refGene_gn=PLXNB1;refGene_tx=XM_003982157.5;refGene_gn=PLXNB1;refGene_tx=XM_023248458.1;refGene_gn=PLXNB1;refGene_tx=XM_023248461.1;refGene_gn=PLXNB1;refGene_tx=XM_023248463.1;refGene_gn=PLXNB1;refGene_tx=XM_023248469.1
A2 17644064 17644241
refGene_gn=PLXNB1;refGene_tx=XM_003982157.5;refGene_gn=PLXNB1;refGene_tx=XM_023248458.1;refGene_gn=PLXNB1;refGene_tx=XM_023248461.1;refGene_gn=PLXNB1;refGene_tx=XM_023248463.1;refGene_gn=PLXNB1;refGene_tx=XM_023248469.1
A2 17644944 17645187
refGene_gn=PLXNB1;refGene_tx=XM_003982157.5;refGene_gn=PLXNB1;refGene_tx=XM_023248458.1;refGene_gn=PLXNB1;refGene_tx=XM_023248461.1;refGene_gn=PLXNB1;refGene_tx=XM_023248463.1;refGene_gn=PLXNB1;refGene_tx=XM_023248469.1
A2 17645367 17645539
refGene_gn=PLXNB1;refGene_tx=XM_003982157.5;refGene_gn=PLXNB1;refGene_tx=XM_023248458.1;refGene_gn=PLXNB1;refGene_tx=XM_023248461.1;refGene_gn=PLXNB1;refGene_tx=XM_023248463.1;refGene_gn=PLXNB1;refGene_tx=XM_023248469.1
A2 17645731 17645908
refGene_gn=PLXNB1;refGene_tx=XM_003982157.5;refGene_gn=PLXNB1;refGene_tx=XM_023248458.1;refGene_gn=PLXNB1;refGene_tx=XM_023248461.1;refGene_gn=PLXNB1;refGene_tx=XM_023248463.1;refGene_gn=PLXNB1;refGene_tx=XM_023248469.1
A2 17646198 17646335

refGene_gn=PLXNB1;refGene_tx=XM_003982157.5;refGene_gn=PLXNB1;refGene_tx=XM_023248458.1;refGene_gn=PLXNB1;refGene_tx=XM_023248461.1;refGene_gn=PLXNB1;refGene_tx=XM_023248463.1;refGene_gn=PLXNB1;refGene_tx=XM_023248469.1
A2 17646478 17646584
refGene_gn=PLXNB1;refGene_tx=XM_003982157.5;refGene_gn=PLXNB1;refGene_tx=XM_023248458.1;refGene_gn=PLXNB1;refGene_tx=XM_023248461.1;refGene_gn=PLXNB1;refGene_tx=XM_023248463.1;refGene_gn=PLXNB1;refGene_tx=XM_023248469.1
A2 17648308 17648460
refGene_gn=PLXNB1;refGene_tx=XM_003982157.5;refGene_gn=PLXNB1;refGene_tx=XM_023248458.1;refGene_gn=PLXNB1;refGene_tx=XM_023248461.1;refGene_gn=PLXNB1;refGene_tx=XM_023248463.1;refGene_gn=PLXNB1;refGene_tx=XM_023248469.1
A2 17648621 17648794
refGene_gn=PLXNB1;refGene_tx=XM_003982157.5;refGene_gn=PLXNB1;refGene_tx=XM_023248458.1;refGene_gn=PLXNB1;refGene_tx=XM_023248461.1;refGene_gn=PLXNB1;refGene_tx=XM_023248463.1;refGene_gn=PLXNB1;refGene_tx=XM_023248469.1
A2 17648877 17648971
refGene_gn=PLXNB1;refGene_tx=XM_003982157.5;refGene_gn=PLXNB1;refGene_tx=XM_023248458.1;refGene_gn=PLXNB1;refGene_tx=XM_023248461.1;refGene_gn=PLXNB1;refGene_tx=XM_023248463.1;refGene_gn=PLXNB1;refGene_tx=XM_023248469.1
A2 17649272 17649395
refGene_gn=PLXNB1;refGene_tx=XM_003982157.5;refGene_gn=PLXNB1;refGene_tx=XM_023248458.1;refGene_gn=PLXNB1;refGene_tx=XM_023248461.1;refGene_gn=PLXNB1;refGene_tx=XM_023248463.1;refGene_gn=PLXNB1;refGene_tx=XM_023248469.1
A2 17649585 17649708
refGene_gn=PLXNB1;refGene_tx=XM_003982157.5;refGene_gn=PLXNB1;refGene_tx=XM_023248458.1;refGene_gn=PLXNB1;refGene_tx=XM_023248461.1;refGene_gn=PLXNB1;refGene_tx=XM_023248463.1;refGene_gn=PLXNB1;refGene_tx=XM_023248469.1
A2 17649888 17650569
refGene_gn=PLXNB1;refGene_tx=XM_003982157.5;refGene_gn=PLXNB1;refGene_tx=XM_023248458.1;refGene_gn=PLXNB1;refGene_tx=XM_023248461.1;refGene_gn=PLXNB1;refGene_tx=XM_023248463.1;refGene_gn=PLXNB1;refGene_tx=XM_023248469.1
A2 17650980 17651089
refGene_gn=PLXNB1;refGene_tx=XM_003982157.5;refGene_gn=PLXNB1;refGene_tx=XM_023248458.1;refGene_gn=PLXNB1;refGene_tx=XM_023248461.1;refGene_gn=PLXNB1;refGene_tx=XM_023248463.1;refGene_gn=PLXNB1;refGene_tx=XM_023248469.1
A2 17651165 17651274
refGene_gn=PLXNB1;refGene_tx=XM_003982157.5;refGene_gn=PLXNB1;refGene_tx=XM_023248458.1;refGene_gn=PLXNB1;refGene_tx=XM_023248461.1;refGene_gn=PLXNB1;refGene_tx=XM_023248463.1;refGene_gn=PLXNB1;refGene_tx=XM_023248469.1
A2 17651519 17651676

refGene_gn=PLXNB1;refGene_tx=XM_003982157.5;refGene_gn=PLXNB1;refGene_tx=XM_023248458.1;refGene_gn=PLXNB1;refGene_tx=XM_023248461.1;refGene_gn=PLXNB1;refGene_tx=XM_023248463.1;refGene_gn=PLXNB1;refGene_tx=XM_023248469.1
A2 17651957 17652090
refGene_gn=PLXNB1;refGene_tx=XM_003982157.5;refGene_gn=PLXNB1;refGene_tx=XM_023248458.1;refGene_gn=PLXNB1;refGene_tx=XM_023248461.1;refGene_gn=PLXNB1;refGene_tx=XM_023248463.1;refGene_gn=PLXNB1;refGene_tx=XM_023248469.1
A2 17652380 17652481
refGene_gn=PLXNB1;refGene_tx=XM_003982157.5;refGene_gn=PLXNB1;refGene_tx=XM_023248458.1;refGene_gn=PLXNB1;refGene_tx=XM_023248461.1;refGene_gn=PLXNB1;refGene_tx=XM_023248463.1;refGene_gn=PLXNB1;refGene_tx=XM_023248469.1
A2 17652574 17652703
refGene_gn=PLXNB1;refGene_tx=XM_003982157.5;refGene_gn=PLXNB1;refGene_tx=XM_023248458.1;refGene_gn=PLXNB1;refGene_tx=XM_023248461.1;refGene_gn=PLXNB1;refGene_tx=XM_023248463.1;refGene_gn=PLXNB1;refGene_tx=XM_023248469.1
A2 17653024 17653207
refGene_gn=PLXNB1;refGene_tx=XM_003982157.5;refGene_gn=PLXNB1;refGene_tx=XM_023248458.1;refGene_gn=PLXNB1;refGene_tx=XM_023248461.1;refGene_gn=PLXNB1;refGene_tx=XM_023248463.1;refGene_gn=PLXNB1;refGene_tx=XM_023248469.1
A2 17653990 17655097
refGene_gn=PLXNB1;refGene_tx=XM_003982157.5;refGene_gn=PLXNB1;refGene_tx=XM_023248458.1;refGene_gn=PLXNB1;refGene_tx=XM_023248461.1;refGene_gn=PLXNB1;refGene_tx=XM_023248463.1;refGene_gn=PLXNB1;refGene_tx=XM_023248469.1
A2 17662890 17663646
refGene_gn=CCDC51;refGene_tx=XM_003982158.4;refGene_gn=CCDC51;refGene_tx=XM_006928712.4;refGene_gn=CCDC51;refGene_tx=XM_006928716.4;refGene_gn=CCDC51;refGene_tx=XM_006928717.4;refGene_gn=CCDC51;refGene_tx=XM_011291332.2;refGene_gn=CCDC51;refGene_tx=XM_019821081.1;refGene_gn=CCDC51;refGene_tx=XM_023248485.1;refGene_gn=CCDC51;refGene_tx=XM_023248487.1;refGene_gn=CCDC51;refGene_tx=XM_023248494.1
A2 17664193 17664358
refGene_gn=CCDC51;refGene_tx=XM_003982158.4;refGene_gn=CCDC51;refGene_tx=XM_006928712.4;refGene_gn=CCDC51;refGene_tx=XM_006928716.4;refGene_gn=CCDC51;refGene_tx=XM_006928717.4;refGene_gn=CCDC51;refGene_tx=XM_011291332.2;refGene_gn=CCDC51;refGene_tx=XM_019821081.1;refGene_gn=CCDC51;refGene_tx=XM_023248485.1;refGene_gn=CCDC51;refGene_tx=XM_023248487.1;refGene_gn=CCDC51;refGene_tx=XM_023248494.1
A2 17665367 17665684
refGene_gn=CCDC51;refGene_tx=XM_003982158.4;refGene_gn=CCDC51;refGene_tx=XM_006928712.4;refGene_gn=CCDC51;refGene_tx=XM_011291332.2;refGene_gn=CCDC51;refGene_tx=XM_023248485.1;refGene_gn=CCDC51;refGene_tx=XM_023248487.1
A2 17670994 17671031
refGene_gn=CCDC51;refGene_tx=XM_023248485.1;refGene_gn=CCDC51;refGene_

tx=XM_023248487.1;refGene_gn=CCDC51;refGene_tx=XM_023248494.1
A2 17676287 17676534
refGene_gn=ATRIP;refGene_tx=XM_006928718.4;refGene_gn=ATRIP;refGene_tx=XM_006928719.4;refGene_gn=ATRIP;refGene_tx=XM_019821036.2;refGene_gn=ATRIP;refGene_tx=XM_019821041.2
A2 17678426 17678557
refGene_gn=ATRIP;refGene_tx=XM_006928718.4;refGene_gn=ATRIP;refGene_tx=XM_006928719.4;refGene_gn=ATRIP;refGene_tx=XM_019821036.2;refGene_gn=ATRIP;refGene_tx=XM_019821041.2
A2 17679678 17679849
refGene_gn=ATRIP;refGene_tx=XM_006928718.4;refGene_gn=ATRIP;refGene_tx=XM_006928719.4;refGene_gn=ATRIP;refGene_tx=XM_019821036.2;refGene_gn=ATRIP;refGene_tx=XM_019821041.2;refGene_gn=ATRIP;refGene_tx=XM_019821052.1
A2 17681678 17681797
refGene_gn=ATRIP;refGene_tx=XM_006928718.4;refGene_gn=ATRIP;refGene_tx=XM_006928719.4;refGene_gn=ATRIP;refGene_tx=XM_019821036.2;refGene_gn=ATRIP;refGene_tx=XM_019821041.2;refGene_gn=ATRIP;refGene_tx=XM_019821052.1
A2 17682990 17683148
refGene_gn=ATRIP;refGene_tx=XM_006928718.4;refGene_gn=ATRIP;refGene_tx=XM_006928719.4;refGene_gn=ATRIP;refGene_tx=XM_019821036.2;refGene_gn=ATRIP;refGene_tx=XM_019821041.2;refGene_gn=ATRIP;refGene_tx=XM_019821052.1
A2 17684513 17684615
refGene_gn=ATRIP;refGene_tx=XM_006928718.4;refGene_gn=ATRIP;refGene_tx=XM_006928719.4;refGene_gn=ATRIP;refGene_tx=XM_019821036.2;refGene_gn=ATRIP;refGene_tx=XM_019821041.2;refGene_gn=ATRIP;refGene_tx=XM_019821052.1
A2 17684929 17685059
refGene_gn=ATRIP;refGene_tx=XM_006928718.4;refGene_gn=ATRIP;refGene_tx=XM_006928719.4;refGene_gn=ATRIP;refGene_tx=XM_019821036.2;refGene_gn=ATRIP;refGene_tx=XM_019821041.2;refGene_gn=ATRIP;refGene_tx=XM_019821052.1
A2 17685242 17685932
refGene_gn=ATRIP;refGene_tx=XM_006928718.4;refGene_gn=ATRIP;refGene_tx=XM_006928719.4;refGene_gn=ATRIP;refGene_tx=XM_019821036.2;refGene_gn=ATRIP;refGene_tx=XM_019821041.2;refGene_gn=ATRIP;refGene_tx=XM_019821052.1
A2 17688499 17688636
refGene_gn=ATRIP;refGene_tx=XM_006928718.4;refGene_gn=ATRIP;refGene_tx=XM_006928719.4;refGene_gn=ATRIP;refGene_tx=XM_019821036.2;refGene_gn=ATRIP;refGene_tx=XM_019821041.2;refGene_gn=ATRIP;refGene_tx=XM_019821052.1
A2 17688798 17688890
refGene_gn=ATRIP;refGene_tx=XM_006928718.4;refGene_gn=ATRIP;refGene_tx=XM_006928719.4;refGene_gn=ATRIP;refGene_tx=XM_019821036.2;refGene_gn=ATRIP;refGene_tx=XM_019821041.2;refGene_gn=ATRIP;refGene_tx=XM_019821052.1
A2 17689322 17689403

refGene_gn=ATRIP;refGene_tx=XM_006928718.4;refGene_gn=ATRIP;refGene_tx=XM_006928719.4;refGene_gn=ATRIP;refGene_tx=XM_019821036.2;refGene_gn=ATRIP;refGene_tx=XM_019821041.2;refGene_gn=ATRIP;refGene_tx=XM_019821052.1
A2 17689609 17689862
refGene_gn=ATRIP;refGene_tx=XM_006928718.4;refGene_gn=ATRIP;refGene_tx=XM_006928719.4;refGene_gn=ATRIP;refGene_tx=XM_019821036.2;refGene_gn=ATRIP;refGene_tx=XM_019821041.2;refGene_gn=ATRIP;refGene_tx=XM_019821052.1
A2 17690134 17690171 refGene_gn=ATRIP;refGene_tx=XM_006928719.4
A2 17690267 17690335
refGene_gn=ATRIP;refGene_tx=XM_006928718.4;refGene_gn=ATRIP;refGene_tx=XM_006928719.4;refGene_gn=ATRIP;refGene_tx=XM_019821036.2;refGene_gn=ATRIP;refGene_tx=XM_019821041.2;refGene_gn=ATRIP;refGene_tx=XM_019821052.1
A2 17691410 17692355 refGene_gn=TREX1;refGene_tx=XM_006928720.4
A2 17693791 17693871
refGene_gn=SHISA5;refGene_tx=XM_006928721.4;refGene_gn=SHISA5;refGene_tx=XM_006928722.4;refGene_gn=SHISA5;refGene_tx=XM_006928723.3;refGene_gn=SHISA5;refGene_tx=XM_019821106.2
A2 17693987 17694200
refGene_gn=SHISA5;refGene_tx=XM_006928721.4;refGene_gn=SHISA5;refGene_tx=XM_006928722.4;refGene_gn=SHISA5;refGene_tx=XM_006928723.3;refGene_gn=SHISA5;refGene_tx=XM_019821106.2
A2 17694348 17694464
refGene_gn=SHISA5;refGene_tx=XM_006928721.4;refGene_gn=SHISA5;refGene_tx=XM_006928722.4;refGene_gn=SHISA5;refGene_tx=XM_006928723.3;refGene_gn=SHISA5;refGene_tx=XM_019821106.2
A2 17696076 17696081 refGene_gn=SHISA5;refGene_tx=XM_006928723.3
A2 17701739 17701823
refGene_gn=SHISA5;refGene_tx=XM_006928721.4;refGene_gn=SHISA5;refGene_tx=XM_006928722.4;refGene_gn=SHISA5;refGene_tx=XM_019821106.2
A2 17710577 17710734
refGene_gn=SHISA5;refGene_tx=XM_006928721.4;refGene_gn=SHISA5;refGene_tx=XM_006928722.4
A2 17713079 17713143
refGene_gn=SHISA5;refGene_tx=XM_006928721.4;refGene_gn=SHISA5;refGene_tx=XM_006928722.4
A2 17728820 17728880
refGene_gn=PFKFB4;refGene_tx=XM_003982162.3;refGene_gn=PFKFB4;refGene_tx=XM_003982163.3;refGene_gn=PFKFB4;refGene_tx=XM_019821149.1;refGene_gn=PFKFB4;refGene_tx=XM_019821154.1;refGene_gn=PFKFB4;refGene_tx=XM_019821162.2;refGene_gn=PFKFB4;refGene_tx=XM_019821166.1
A2 17730977 17731042
refGene_gn=PFKFB4;refGene_tx=XM_003982162.3;refGene_gn=PFKFB4;refGene_tx=XM_003982163.3;refGene_gn=PFKFB4;refGene_tx=XM_019821149.1;refGene_gn=PFKFB4;refGene_tx=XM_019821154.1;refGene_gn=PFKFB4;refGene_tx=XM_019821162.2;refGene_gn=PFKFB4;refGene_tx=XM_019821166.1
A2 17732049 17732112
refGene_gn=PFKFB4;refGene_tx=XM_003982162.3;refGene_gn=PFKFB4;refGene_

tx=XM_003982163.3;refGene_gn=PFKFB4;refGene_tx=XM_019821149.1;refGene_gn=PFKFB4;refGene_tx=XM_019821162.2;refGene_gn=PFKFB4;refGene_tx=XM_019821166.1
A2 17732219 17732349
refGene_gn=PFKFB4;refGene_tx=XM_003982162.3;refGene_gn=PFKFB4;refGene_tx=XM_003982163.3;refGene_gn=PFKFB4;refGene_tx=XM_019821149.1;refGene_gn=PFKFB4;refGene_tx=XM_019821154.1;refGene_gn=PFKFB4;refGene_tx=XM_019821162.2;refGene_gn=PFKFB4;refGene_tx=XM_019821166.1
A2 17733854 17733959
refGene_gn=PFKFB4;refGene_tx=XM_003982162.3;refGene_gn=PFKFB4;refGene_tx=XM_003982163.3;refGene_gn=PFKFB4;refGene_tx=XM_019821149.1;refGene_gn=PFKFB4;refGene_tx=XM_019821154.1;refGene_gn=PFKFB4;refGene_tx=XM_019821162.2;refGene_gn=PFKFB4;refGene_tx=XM_019821166.1
A2 17741147 17741294
refGene_gn=PFKFB4;refGene_tx=XM_003982162.3;refGene_gn=PFKFB4;refGene_tx=XM_003982163.3;refGene_gn=PFKFB4;refGene_tx=XM_019821149.1;refGene_gn=PFKFB4;refGene_tx=XM_019821154.1;refGene_gn=PFKFB4;refGene_tx=XM_019821162.2;refGene_gn=PFKFB4;refGene_tx=XM_019821166.1
A2 17741850 17742058
refGene_gn=PFKFB4;refGene_tx=XM_003982162.3;refGene_gn=PFKFB4;refGene_tx=XM_003982163.3;refGene_gn=PFKFB4;refGene_tx=XM_019821149.1;refGene_gn=PFKFB4;refGene_tx=XM_019821154.1;refGene_gn=PFKFB4;refGene_tx=XM_019821162.2;refGene_gn=PFKFB4;refGene_tx=XM_019821166.1
A2 17744261 17744383
refGene_gn=PFKFB4;refGene_tx=XM_003982162.3;refGene_gn=PFKFB4;refGene_tx=XM_003982163.3;refGene_gn=PFKFB4;refGene_tx=XM_019821149.1;refGene_gn=PFKFB4;refGene_tx=XM_019821154.1;refGene_gn=PFKFB4;refGene_tx=XM_019821162.2;refGene_gn=PFKFB4;refGene_tx=XM_019821166.1
A2 17745043 17745100
refGene_gn=PFKFB4;refGene_tx=XM_003982162.3;refGene_gn=PFKFB4;refGene_tx=XM_003982163.3;refGene_gn=PFKFB4;refGene_tx=XM_019821154.1;refGene_gn=PFKFB4;refGene_tx=XM_019821162.2
A2 17745447 17745522
refGene_gn=PFKFB4;refGene_tx=XM_003982162.3;refGene_gn=PFKFB4;refGene_tx=XM_003982163.3;refGene_gn=PFKFB4;refGene_tx=XM_019821149.1;refGene_gn=PFKFB4;refGene_tx=XM_019821154.1;refGene_gn=PFKFB4;refGene_tx=XM_019821162.2
A2 17748049 17748116
refGene_gn=PFKFB4;refGene_tx=XM_003982162.3;refGene_gn=PFKFB4;refGene_tx=XM_003982163.3;refGene_gn=PFKFB4;refGene_tx=XM_019821149.1;refGene_gn=PFKFB4;refGene_tx=XM_019821154.1;refGene_gn=PFKFB4;refGene_tx=XM_019821162.2
A2 17751676 17751773
refGene_gn=PFKFB4;refGene_tx=XM_003982162.3;refGene_gn=PFKFB4;refGene_tx=XM_003982163.3;refGene_gn=PFKFB4;refGene_tx=XM_019821149.1;refGene_gn=PFKFB4;refGene_tx=XM_019821154.1;refGene_gn=PFKFB4;refGene_tx=XM_019821162.2
A2 17751916 17752033
refGene_gn=PFKFB4;refGene_tx=XM_003982162.3;refGene_gn=PFKFB4;refGene_tx=XM_003982163.3;refGene_gn=PFKFB4;refGene_tx=XM_019821149.1;refGene_

gn=PFKFB4;refGene_tx=XM_019821154.1;refGene_gn=PFKFB4;refGene_tx=XM_01
9821162.2
A2 17757657 17757754
refGene_gn=PFKFB4;refGene_tx=XM_003982162.3;refGene_gn=PFKFB4;refGene_
tx=XM_019821149.1;refGene_gn=PFKFB4;refGene_tx=XM_019821154.1
A2 17761354 17761418 refGene_gn=PFKFB4;refGene_tx=XM_003982163.3
A2 17762959 17763310 refGene_gn=UCN2;refGene_tx=XM_006928725.2
A2 17763934 17764039 refGene_gn=UCN2;refGene_tx=XM_006928725.2
A2 17764623 17764640
refGene_gn=COL7A1;refGene_tx=XM_003982165.5;refGene_gn=COL7A1;refGene_
tx=XM_019821172.1
A2 17764970 17765168
refGene_gn=COL7A1;refGene_tx=XM_003982165.5;refGene_gn=COL7A1;refGene_
tx=XM_019821172.1
A2 17765294 17765384
refGene_gn=COL7A1;refGene_tx=XM_003982165.5;refGene_gn=COL7A1;refGene_
tx=XM_019821172.1
A2 17765617 17765704 refGene_gn=COL7A1;refGene_tx=XM_003982165.5
A2 17765738 17765767 refGene_gn=COL7A1;refGene_tx=XM_019821176.2
A2 17765852 17765885
refGene_gn=COL7A1;refGene_tx=XM_003982165.5;refGene_gn=COL7A1;refGene_
tx=XM_019821172.1;refGene_gn=COL7A1;refGene_tx=XM_019821176.2
A2 17766533 17766582
refGene_gn=COL7A1;refGene_tx=XM_003982165.5;refGene_gn=COL7A1;refGene_
tx=XM_019821172.1;refGene_gn=COL7A1;refGene_tx=XM_019821176.2
A2 17766711 17766765
refGene_gn=COL7A1;refGene_tx=XM_003982165.5;refGene_gn=COL7A1;refGene_
tx=XM_019821172.1;refGene_gn=COL7A1;refGene_tx=XM_019821176.2
A2 17766862 17766940
refGene_gn=COL7A1;refGene_tx=XM_003982165.5;refGene_gn=COL7A1;refGene_
tx=XM_019821172.1;refGene_gn=COL7A1;refGene_tx=XM_019821176.2
A2 17767110 17767227
refGene_gn=COL7A1;refGene_tx=XM_003982165.5;refGene_gn=COL7A1;refGene_
tx=XM_019821172.1;refGene_gn=COL7A1;refGene_tx=XM_019821176.2
A2 17767326 17767389
refGene_gn=COL7A1;refGene_tx=XM_003982165.5;refGene_gn=COL7A1;refGene_
tx=XM_019821172.1;refGene_gn=COL7A1;refGene_tx=XM_019821176.2
A2 17767843 17767906
refGene_gn=COL7A1;refGene_tx=XM_003982165.5;refGene_gn=COL7A1;refGene_
tx=XM_019821172.1;refGene_gn=COL7A1;refGene_tx=XM_019821176.2
A2 17767978 17768032
refGene_gn=COL7A1;refGene_tx=XM_003982165.5;refGene_gn=COL7A1;refGene_
tx=XM_019821172.1;refGene_gn=COL7A1;refGene_tx=XM_019821176.2
A2 17768109 17768163
refGene_gn=COL7A1;refGene_tx=XM_003982165.5;refGene_gn=COL7A1;refGene_
tx=XM_019821172.1;refGene_gn=COL7A1;refGene_tx=XM_019821176.2
A2 17768412 17768493
refGene_gn=COL7A1;refGene_tx=XM_003982165.5;refGene_gn=COL7A1;refGene_
tx=XM_019821172.1;refGene_gn=COL7A1;refGene_tx=XM_019821176.2
A2 17768809 17768845

refGene_gn=COL7A1;refGene_tx=XM_003982165.5;refGene_gn=COL7A1;refGene_tx=XM_019821172.1;refGene_gn=COL7A1;refGene_tx=XM_019821176.2
A2 17769058 17769130
refGene_gn=COL7A1;refGene_tx=XM_003982165.5;refGene_gn=COL7A1;refGene_tx=XM_019821172.1;refGene_gn=COL7A1;refGene_tx=XM_019821176.2
A2 17769611 17769683
refGene_gn=COL7A1;refGene_tx=XM_003982165.5;refGene_gn=COL7A1;refGene_tx=XM_019821172.1;refGene_gn=COL7A1;refGene_tx=XM_019821176.2
A2 17769832 17769889
refGene_gn=COL7A1;refGene_tx=XM_003982165.5;refGene_gn=COL7A1;refGene_tx=XM_019821172.1;refGene_gn=COL7A1;refGene_tx=XM_019821176.2
A2 17769968 17770004
refGene_gn=COL7A1;refGene_tx=XM_003982165.5;refGene_gn=COL7A1;refGene_tx=XM_019821172.1;refGene_gn=COL7A1;refGene_tx=XM_019821176.2
A2 17770114 17770150
refGene_gn=COL7A1;refGene_tx=XM_003982165.5;refGene_gn=COL7A1;refGene_tx=XM_019821172.1;refGene_gn=COL7A1;refGene_tx=XM_019821176.2
A2 17770371 17770416
refGene_gn=COL7A1;refGene_tx=XM_003982165.5;refGene_gn=COL7A1;refGene_tx=XM_019821172.1;refGene_gn=COL7A1;refGene_tx=XM_019821176.2
A2 17770513 17770573
refGene_gn=COL7A1;refGene_tx=XM_003982165.5;refGene_gn=COL7A1;refGene_tx=XM_019821172.1;refGene_gn=COL7A1;refGene_tx=XM_019821176.2
A2 17770698 17770734
refGene_gn=COL7A1;refGene_tx=XM_003982165.5;refGene_gn=COL7A1;refGene_tx=XM_019821172.1;refGene_gn=COL7A1;refGene_tx=XM_019821176.2
A2 17770875 17770947
refGene_gn=COL7A1;refGene_tx=XM_003982165.5;refGene_gn=COL7A1;refGene_tx=XM_019821172.1;refGene_gn=COL7A1;refGene_tx=XM_019821176.2
A2 17771123 17771231
refGene_gn=COL7A1;refGene_tx=XM_003982165.5;refGene_gn=COL7A1;refGene_tx=XM_019821172.1;refGene_gn=COL7A1;refGene_tx=XM_019821176.2
A2 17771365 17771425
refGene_gn=COL7A1;refGene_tx=XM_003982165.5;refGene_gn=COL7A1;refGene_tx=XM_019821172.1;refGene_gn=COL7A1;refGene_tx=XM_019821176.2
A2 17771507 17771543
refGene_gn=COL7A1;refGene_tx=XM_003982165.5;refGene_gn=COL7A1;refGene_tx=XM_019821172.1;refGene_gn=COL7A1;refGene_tx=XM_019821176.2
A2 17772154 17772199
refGene_gn=COL7A1;refGene_tx=XM_003982165.5;refGene_gn=COL7A1;refGene_tx=XM_019821172.1;refGene_gn=COL7A1;refGene_tx=XM_019821176.2
A2 17772288 17772333
refGene_gn=COL7A1;refGene_tx=XM_003982165.5;refGene_gn=COL7A1;refGene_tx=XM_019821172.1;refGene_gn=COL7A1;refGene_tx=XM_019821176.2
A2 17772486 17772528
refGene_gn=COL7A1;refGene_tx=XM_003982165.5;refGene_gn=COL7A1;refGene_tx=XM_019821172.1;refGene_gn=COL7A1;refGene_tx=XM_019821176.2
A2 17772602 17772638
refGene_gn=COL7A1;refGene_tx=XM_003982165.5;refGene_gn=COL7A1;refGene_tx=XM_019821172.1;refGene_gn=COL7A1;refGene_tx=XM_019821176.2

A2 17772757 17772826
refGene_gn=COL7A1;refGene_tx=XM_003982165.5;refGene_gn=COL7A1;refGene_tx=XM_019821172.1;refGene_gn=COL7A1;refGene_tx=XM_019821176.2
A2 17772965 17773046
refGene_gn=COL7A1;refGene_tx=XM_003982165.5;refGene_gn=COL7A1;refGene_tx=XM_019821172.1;refGene_gn=COL7A1;refGene_tx=XM_019821176.2
A2 17773126 17773162
refGene_gn=COL7A1;refGene_tx=XM_003982165.5;refGene_gn=COL7A1;refGene_tx=XM_019821172.1;refGene_gn=COL7A1;refGene_tx=XM_019821176.2
A2 17773252 17773315
refGene_gn=COL7A1;refGene_tx=XM_003982165.5;refGene_gn=COL7A1;refGene_tx=XM_019821172.1;refGene_gn=COL7A1;refGene_tx=XM_019821176.2
A2 17773394 17773427
refGene_gn=COL7A1;refGene_tx=XM_003982165.5;refGene_gn=COL7A1;refGene_tx=XM_019821172.1;refGene_gn=COL7A1;refGene_tx=XM_019821176.2
A2 17773647 17773692
refGene_gn=COL7A1;refGene_tx=XM_003982165.5;refGene_gn=COL7A1;refGene_tx=XM_019821172.1;refGene_gn=COL7A1;refGene_tx=XM_019821176.2
A2 17773833 17773869
refGene_gn=COL7A1;refGene_tx=XM_003982165.5;refGene_gn=COL7A1;refGene_tx=XM_019821172.1;refGene_gn=COL7A1;refGene_tx=XM_019821176.2
A2 17773996 17774032
refGene_gn=COL7A1;refGene_tx=XM_003982165.5;refGene_gn=COL7A1;refGene_tx=XM_019821172.1;refGene_gn=COL7A1;refGene_tx=XM_019821176.2
A2 17774363 17774408
refGene_gn=COL7A1;refGene_tx=XM_003982165.5;refGene_gn=COL7A1;refGene_tx=XM_019821172.1;refGene_gn=COL7A1;refGene_tx=XM_019821176.2
A2 17774556 17774619
refGene_gn=COL7A1;refGene_tx=XM_003982165.5;refGene_gn=COL7A1;refGene_tx=XM_019821172.1;refGene_gn=COL7A1;refGene_tx=XM_019821176.2
A2 17774735 17774780
refGene_gn=COL7A1;refGene_tx=XM_003982165.5;refGene_gn=COL7A1;refGene_tx=XM_019821172.1;refGene_gn=COL7A1;refGene_tx=XM_019821176.2
A2 17774857 17774926
refGene_gn=COL7A1;refGene_tx=XM_003982165.5;refGene_gn=COL7A1;refGene_tx=XM_019821172.1;refGene_gn=COL7A1;refGene_tx=XM_019821176.2
A2 17775108 17775162
refGene_gn=COL7A1;refGene_tx=XM_003982165.5;refGene_gn=COL7A1;refGene_tx=XM_019821172.1;refGene_gn=COL7A1;refGene_tx=XM_019821176.2
A2 17775243 17775279
refGene_gn=COL7A1;refGene_tx=XM_003982165.5;refGene_gn=COL7A1;refGene_tx=XM_019821172.1;refGene_gn=COL7A1;refGene_tx=XM_019821176.2
A2 17775373 17775574
refGene_gn=COL7A1;refGene_tx=XM_003982165.5;refGene_gn=COL7A1;refGene_tx=XM_019821172.1;refGene_gn=COL7A1;refGene_tx=XM_019821176.2
A2 17775655 17775778
refGene_gn=COL7A1;refGene_tx=XM_003982165.5;refGene_gn=COL7A1;refGene_tx=XM_019821172.1;refGene_gn=COL7A1;refGene_tx=XM_019821176.2
A2 17775910 17775946
refGene_gn=COL7A1;refGene_tx=XM_003982165.5;refGene_gn=COL7A1;refGene_

tx=XM_019821172.1;refGene_gn=COL7A1;refGene_tx=XM_019821176.2
A2 17776267 17776315
refGene_gn=COL7A1;refGene_tx=XM_003982165.5;refGene_gn=COL7A1;refGene_tx=XM_019821172.1;refGene_gn=COL7A1;refGene_tx=XM_019821176.2
A2 17776423 17776459
refGene_gn=COL7A1;refGene_tx=XM_003982165.5;refGene_gn=COL7A1;refGene_tx=XM_019821172.1;refGene_gn=COL7A1;refGene_tx=XM_019821176.2
A2 17776555 17776591
refGene_gn=COL7A1;refGene_tx=XM_003982165.5;refGene_gn=COL7A1;refGene_tx=XM_019821172.1;refGene_gn=COL7A1;refGene_tx=XM_019821176.2
A2 17776712 17776808
refGene_gn=COL7A1;refGene_tx=XM_003982165.5;refGene_gn=COL7A1;refGene_tx=XM_019821172.1;refGene_gn=COL7A1;refGene_tx=XM_019821176.2
A2 17776920 17776956
refGene_gn=COL7A1;refGene_tx=XM_003982165.5;refGene_gn=COL7A1;refGene_tx=XM_019821172.1;refGene_gn=COL7A1;refGene_tx=XM_019821176.2
A2 17777029 17777065
refGene_gn=COL7A1;refGene_tx=XM_003982165.5;refGene_gn=COL7A1;refGene_tx=XM_019821172.1;refGene_gn=COL7A1;refGene_tx=XM_019821176.2
A2 17778052 17778097
refGene_gn=COL7A1;refGene_tx=XM_003982165.5;refGene_gn=COL7A1;refGene_tx=XM_019821172.1;refGene_gn=COL7A1;refGene_tx=XM_019821176.2
A2 17778333 17778396
refGene_gn=COL7A1;refGene_tx=XM_003982165.5;refGene_gn=COL7A1;refGene_tx=XM_019821172.1;refGene_gn=COL7A1;refGene_tx=XM_019821176.2
A2 17779047 17779083
refGene_gn=COL7A1;refGene_tx=XM_003982165.5;refGene_gn=COL7A1;refGene_tx=XM_019821172.1;refGene_gn=COL7A1;refGene_tx=XM_019821176.2
A2 17779338 17779419
refGene_gn=COL7A1;refGene_tx=XM_003982165.5;refGene_gn=COL7A1;refGene_tx=XM_019821172.1;refGene_gn=COL7A1;refGene_tx=XM_019821176.2
A2 17779508 17779544
refGene_gn=COL7A1;refGene_tx=XM_003982165.5;refGene_gn=COL7A1;refGene_tx=XM_019821172.1;refGene_gn=COL7A1;refGene_tx=XM_019821176.2
A2 17779620 17779656
refGene_gn=COL7A1;refGene_tx=XM_003982165.5;refGene_gn=COL7A1;refGene_tx=XM_019821172.1;refGene_gn=COL7A1;refGene_tx=XM_019821176.2
A2 17779732 17779813
refGene_gn=COL7A1;refGene_tx=XM_003982165.5;refGene_gn=COL7A1;refGene_tx=XM_019821172.1;refGene_gn=COL7A1;refGene_tx=XM_019821176.2
A2 17779931 17779961
refGene_gn=COL7A1;refGene_tx=XM_003982165.5;refGene_gn=COL7A1;refGene_tx=XM_019821172.1;refGene_gn=COL7A1;refGene_tx=XM_019821176.2
A2 17780157 17780184
refGene_gn=COL7A1;refGene_tx=XM_003982165.5;refGene_gn=COL7A1;refGene_tx=XM_019821172.1;refGene_gn=COL7A1;refGene_tx=XM_019821176.2
A2 17780431 17780476
refGene_gn=COL7A1;refGene_tx=XM_003982165.5;refGene_gn=COL7A1;refGene_tx=XM_019821172.1;refGene_gn=COL7A1;refGene_tx=XM_019821176.2
A2 17780693 17780765

refGene_gn=COL7A1;refGene_tx=XM_003982165.5;refGene_gn=COL7A1;refGene_tx=XM_019821172.1;refGene_gn=COL7A1;refGene_tx=XM_019821176.2
A2 17780918 17780963
refGene_gn=COL7A1;refGene_tx=XM_003982165.5;refGene_gn=COL7A1;refGene_tx=XM_019821172.1;refGene_gn=COL7A1;refGene_tx=XM_019821176.2
A2 17781160 17781196
refGene_gn=COL7A1;refGene_tx=XM_003982165.5;refGene_gn=COL7A1;refGene_tx=XM_019821172.1;refGene_gn=COL7A1;refGene_tx=XM_019821176.2
A2 17781298 17781379
refGene_gn=COL7A1;refGene_tx=XM_003982165.5;refGene_gn=COL7A1;refGene_tx=XM_019821172.1;refGene_gn=COL7A1;refGene_tx=XM_019821176.2
A2 17781486 17781522
refGene_gn=COL7A1;refGene_tx=XM_003982165.5;refGene_gn=COL7A1;refGene_tx=XM_019821172.1;refGene_gn=COL7A1;refGene_tx=XM_019821176.2
A2 17781614 17781674
refGene_gn=COL7A1;refGene_tx=XM_003982165.5;refGene_gn=COL7A1;refGene_tx=XM_019821172.1;refGene_gn=COL7A1;refGene_tx=XM_019821176.2
A2 17781751 17781805
refGene_gn=COL7A1;refGene_tx=XM_003982165.5;refGene_gn=COL7A1;refGene_tx=XM_019821172.1;refGene_gn=COL7A1;refGene_tx=XM_019821176.2
A2 17781983 17782016
refGene_gn=COL7A1;refGene_tx=XM_003982165.5;refGene_gn=COL7A1;refGene_tx=XM_019821172.1;refGene_gn=COL7A1;refGene_tx=XM_019821176.2
A2 17782378 17782411
refGene_gn=COL7A1;refGene_tx=XM_003982165.5;refGene_gn=COL7A1;refGene_tx=XM_019821172.1;refGene_gn=COL7A1;refGene_tx=XM_019821176.2
A2 17782538 17782574
refGene_gn=COL7A1;refGene_tx=XM_003982165.5;refGene_gn=COL7A1;refGene_tx=XM_019821172.1;refGene_gn=COL7A1;refGene_tx=XM_019821176.2
A2 17782657 17782702
refGene_gn=COL7A1;refGene_tx=XM_003982165.5;refGene_gn=COL7A1;refGene_tx=XM_019821172.1;refGene_gn=COL7A1;refGene_tx=XM_019821176.2
A2 17783057 17783093
refGene_gn=COL7A1;refGene_tx=XM_003982165.5;refGene_gn=COL7A1;refGene_tx=XM_019821172.1;refGene_gn=COL7A1;refGene_tx=XM_019821176.2
A2 17783172 17783217
refGene_gn=COL7A1;refGene_tx=XM_003982165.5;refGene_gn=COL7A1;refGene_tx=XM_019821172.1;refGene_gn=COL7A1;refGene_tx=XM_019821176.2
A2 17783414 17783450
refGene_gn=COL7A1;refGene_tx=XM_003982165.5;refGene_gn=COL7A1;refGene_tx=XM_019821172.1;refGene_gn=COL7A1;refGene_tx=XM_019821176.2
A2 17783554 17783614
refGene_gn=COL7A1;refGene_tx=XM_003982165.5;refGene_gn=COL7A1;refGene_tx=XM_019821172.1;refGene_gn=COL7A1;refGene_tx=XM_019821176.2
A2 17783701 17783764
refGene_gn=COL7A1;refGene_tx=XM_003982165.5;refGene_gn=COL7A1;refGene_tx=XM_019821172.1;refGene_gn=COL7A1;refGene_tx=XM_019821176.2
A2 17783906 17783960
refGene_gn=COL7A1;refGene_tx=XM_003982165.5;refGene_gn=COL7A1;refGene_tx=XM_019821172.1;refGene_gn=COL7A1;refGene_tx=XM_019821176.2

A2	17784042 17784069
refGene_gn=COL7A1;refGene_tx=XM_003982165.5;refGene_gn=COL7A1;refGene_tx=XM_019821172.1;refGene_gn=COL7A1;refGene_tx=XM_019821176.2
A2	17784313 17784391
refGene_gn=COL7A1;refGene_tx=XM_003982165.5;refGene_gn=COL7A1;refGene_tx=XM_019821172.1;refGene_gn=COL7A1;refGene_tx=XM_019821176.2
A2	17784502 17784574
refGene_gn=COL7A1;refGene_tx=XM_003982165.5;refGene_gn=COL7A1;refGene_tx=XM_019821172.1;refGene_gn=COL7A1;refGene_tx=XM_019821176.2
A2	17784739 17784775
refGene_gn=COL7A1;refGene_tx=XM_003982165.5;refGene_gn=COL7A1;refGene_tx=XM_019821172.1;refGene_gn=COL7A1;refGene_tx=XM_019821176.2
A2	17784908 17784944
refGene_gn=COL7A1;refGene_tx=XM_003982165.5;refGene_gn=COL7A1;refGene_tx=XM_019821172.1;refGene_gn=COL7A1;refGene_tx=XM_019821176.2
A2	17785039 17785111
refGene_gn=COL7A1;refGene_tx=XM_003982165.5;refGene_gn=COL7A1;refGene_tx=XM_019821172.1;refGene_gn=COL7A1;refGene_tx=XM_019821176.2
A2	17785548 17785611
refGene_gn=COL7A1;refGene_tx=XM_003982165.5;refGene_gn=COL7A1;refGene_tx=XM_019821172.1;refGene_gn=COL7A1;refGene_tx=XM_019821176.2
A2	17785684 17785729
refGene_gn=COL7A1;refGene_tx=XM_003982165.5;refGene_gn=COL7A1;refGene_tx=XM_019821172.1;refGene_gn=COL7A1;refGene_tx=XM_019821176.2
A2	17785821 17785848
refGene_gn=COL7A1;refGene_tx=XM_003982165.5;refGene_gn=COL7A1;refGene_tx=XM_019821172.1;refGene_gn=COL7A1;refGene_tx=XM_019821176.2
A2	17785932 17785968
refGene_gn=COL7A1;refGene_tx=XM_003982165.5;refGene_gn=COL7A1;refGene_tx=XM_019821172.1;refGene_gn=COL7A1;refGene_tx=XM_019821176.2
A2	17786068 17786241
refGene_gn=COL7A1;refGene_tx=XM_003982165.5;refGene_gn=COL7A1;refGene_tx=XM_019821172.1;refGene_gn=COL7A1;refGene_tx=XM_019821176.2
A2	17786328 17786475
refGene_gn=COL7A1;refGene_tx=XM_003982165.5;refGene_gn=COL7A1;refGene_tx=XM_019821172.1;refGene_gn=COL7A1;refGene_tx=XM_019821176.2
A2	17786559 17786686
refGene_gn=COL7A1;refGene_tx=XM_003982165.5;refGene_gn=COL7A1;refGene_tx=XM_019821172.1;refGene_gn=COL7A1;refGene_tx=XM_019821176.2
A2	17786983 17787120
refGene_gn=COL7A1;refGene_tx=XM_003982165.5;refGene_gn=COL7A1;refGene_tx=XM_019821172.1;refGene_gn=COL7A1;refGene_tx=XM_019821176.2
A2	17787201 17787348
refGene_gn=COL7A1;refGene_tx=XM_003982165.5;refGene_gn=COL7A1;refGene_tx=XM_019821172.1;refGene_gn=COL7A1;refGene_tx=XM_019821176.2
A2	17787433 17787568
refGene_gn=COL7A1;refGene_tx=XM_003982165.5;refGene_gn=COL7A1;refGene_tx=XM_019821172.1;refGene_gn=COL7A1;refGene_tx=XM_019821176.2
A2	17787824 17787971
refGene_gn=COL7A1;refGene_tx=XM_003982165.5;refGene_gn=COL7A1;refGene_

tx=XM_019821172.1;refGene_gn=COL7A1;refGene_tx=XM_019821176.2
A2 17788268 17788391
refGene_gn=COL7A1;refGene_tx=XM_003982165.5;refGene_gn=COL7A1;refGene_tx=XM_019821172.1;refGene_gn=COL7A1;refGene_tx=XM_019821176.2
A2 17788576 17788723
refGene_gn=COL7A1;refGene_tx=XM_003982165.5;refGene_gn=COL7A1;refGene_tx=XM_019821172.1;refGene_gn=COL7A1;refGene_tx=XM_019821176.2
A2 17788822 17788948
refGene_gn=COL7A1;refGene_tx=XM_003982165.5;refGene_gn=COL7A1;refGene_tx=XM_019821172.1;refGene_gn=COL7A1;refGene_tx=XM_019821176.2
A2 17789256 17789400
refGene_gn=COL7A1;refGene_tx=XM_003982165.5;refGene_gn=COL7A1;refGene_tx=XM_019821172.1;refGene_gn=COL7A1;refGene_tx=XM_019821176.2
A2 17789559 17789679
refGene_gn=COL7A1;refGene_tx=XM_003982165.5;refGene_gn=COL7A1;refGene_tx=XM_019821172.1;refGene_gn=COL7A1;refGene_tx=XM_019821176.2
A2 17790276 17790420
refGene_gn=COL7A1;refGene_tx=XM_003982165.5;refGene_gn=COL7A1;refGene_tx=XM_019821172.1;refGene_gn=COL7A1;refGene_tx=XM_019821176.2
A2 17790522 17790648
refGene_gn=COL7A1;refGene_tx=XM_003982165.5;refGene_gn=COL7A1;refGene_tx=XM_019821172.1;refGene_gn=COL7A1;refGene_tx=XM_019821176.2
A2 17790736 17790880
refGene_gn=COL7A1;refGene_tx=XM_003982165.5;refGene_gn=COL7A1;refGene_tx=XM_019821172.1;refGene_gn=COL7A1;refGene_tx=XM_019821176.2
A2 17791460 17791589
refGene_gn=COL7A1;refGene_tx=XM_003982165.5;refGene_gn=COL7A1;refGene_tx=XM_019821172.1;refGene_gn=COL7A1;refGene_tx=XM_019821176.2
A2 17791671 17791821
refGene_gn=COL7A1;refGene_tx=XM_003982165.5;refGene_gn=COL7A1;refGene_tx=XM_019821172.1;refGene_gn=COL7A1;refGene_tx=XM_019821176.2
A2 17791895 17792012
refGene_gn=COL7A1;refGene_tx=XM_003982165.5;refGene_gn=COL7A1;refGene_tx=XM_019821172.1;refGene_gn=COL7A1;refGene_tx=XM_019821176.2
A2 17792097 17792244
refGene_gn=COL7A1;refGene_tx=XM_003982165.5;refGene_gn=COL7A1;refGene_tx=XM_019821172.1;refGene_gn=COL7A1;refGene_tx=XM_019821176.2
A2 17792360 17792477
refGene_gn=COL7A1;refGene_tx=XM_003982165.5;refGene_gn=COL7A1;refGene_tx=XM_019821172.1;refGene_gn=COL7A1;refGene_tx=XM_019821176.2
A2 17792574 17792704
refGene_gn=COL7A1;refGene_tx=XM_003982165.5;refGene_gn=COL7A1;refGene_tx=XM_019821172.1;refGene_gn=COL7A1;refGene_tx=XM_019821176.2
A2 17792781 17792945
refGene_gn=COL7A1;refGene_tx=XM_003982165.5;refGene_gn=COL7A1;refGene_tx=XM_019821172.1;refGene_gn=COL7A1;refGene_tx=XM_019821176.2
A2 17793196 17793358
refGene_gn=COL7A1;refGene_tx=XM_003982165.5;refGene_gn=COL7A1;refGene_tx=XM_019821172.1;refGene_gn=COL7A1;refGene_tx=XM_019821176.2
A2 17793449 17793543

refGene_gn=COL7A1;refGene_tx=XM_003982165.5;refGene_gn=COL7A1;refGene_tx=XM_019821172.1;refGene_gn=COL7A1;refGene_tx=XM_019821176.2
A2 17793622 17793788
refGene_gn=COL7A1;refGene_tx=XM_003982165.5;refGene_gn=COL7A1;refGene_tx=XM_019821172.1;refGene_gn=COL7A1;refGene_tx=XM_019821176.2
A2 17794429 17794610
refGene_gn=COL7A1;refGene_tx=XM_003982165.5;refGene_gn=COL7A1;refGene_tx=XM_019821172.1;refGene_gn=COL7A1;refGene_tx=XM_019821176.2
A2 17795115 17795200
refGene_gn=COL7A1;refGene_tx=XM_003982165.5;refGene_gn=COL7A1;refGene_tx=XM_019821172.1;refGene_gn=COL7A1;refGene_tx=XM_019821176.2
A2 17799258 17799323
refGene_gn=LOC101089499;refGene_tx=XM_003982166.4;refGene_gn=LOC101089499;refGene_tx=XM_011291380.2
A2 17799764 17799840
refGene_gn=LOC101089499;refGene_tx=XM_003982166.4;refGene_gn=LOC101089499;refGene_tx=XM_011291380.2
A2 17800192 17800281
refGene_gn=LOC101089499;refGene_tx=XM_003982166.4;refGene_gn=LOC101089499;refGene_tx=XM_011291380.2
A2 17800615 17800701
refGene_gn=LOC101089499;refGene_tx=XM_003982166.4;refGene_gn=LOC101089499;refGene_tx=XM_011291380.2
A2 17800811 17800972
refGene_gn=LOC101089499;refGene_tx=XM_003982166.4;refGene_gn=LOC101089499;refGene_tx=XM_011291380.2
A2 17801103 17801247
refGene_gn=LOC101089499;refGene_tx=XM_003982166.4;refGene_gn=LOC101089499;refGene_tx=XM_011291380.2
A2 17801486 17801602
refGene_gn=LOC101089499;refGene_tx=XM_003982166.4;refGene_gn=LOC101089499;refGene_tx=XM_011291380.2
A2 17803020 17803100
refGene_gn=LOC101089499;refGene_tx=XM_003982166.4;refGene_gn=LOC101089499;refGene_tx=XM_011291380.2
A2 17803637 17803836
refGene_gn=LOC101089499;refGene_tx=XM_003982166.4;refGene_gn=LOC101089499;refGene_tx=XM_011291380.2
A2 17804055 17804185
refGene_gn=LOC101089499;refGene_tx=XM_003982166.4;refGene_gn=LOC101089499;refGene_tx=XM_011291380.2
A2 17805254 17805341
refGene_gn=LOC101089499;refGene_tx=XM_003982166.4;refGene_gn=LOC101089499;refGene_tx=XM_011291380.2
A2 17807825 17807969
refGene_gn=LOC101089499;refGene_tx=XM_003982166.4;refGene_gn=LOC101089499;refGene_tx=XM_011291380.2
A2 17808220 17808289
refGene_gn=LOC101089499;refGene_tx=XM_003982166.4;refGene_gn=LOC101089499;refGene_tx=XM_011291380.2

| | | | |
|---|---|---|---|
| A2 | 17817388 | 17817574 | refGene_gn=TMEM89;refGene_tx=XM_003982201.5 |
| A2 | 17817966 | 17818527 | refGene_gn=TMEM89;refGene_tx=XM_003982201.5 |
| A2 | 17820233 | 17820248 | refGene_gn=CELSR3;refGene_tx=XM_006928732.4 |
| A2 | 17820485 | 17820622 | refGene_gn=CELSR3;refGene_tx=XM_006928732.4 |
| A2 | 17820871 | 17820926 | refGene_gn=CELSR3;refGene_tx=XM_006928732.4 |
| A2 | 17821116 | 17821296 | refGene_gn=CELSR3;refGene_tx=XM_006928732.4 |
| A2 | 17821931 | 17822102 | refGene_gn=CELSR3;refGene_tx=XM_006928732.4 |
| A2 | 17822473 | 17822569 | refGene_gn=CELSR3;refGene_tx=XM_006928732.4 |
| A2 | 17822654 | 17822747 | refGene_gn=CELSR3;refGene_tx=XM_006928732.4 |
| A2 | 17823674 | 17823744 | refGene_gn=CELSR3;refGene_tx=XM_006928732.4 |
| A2 | 17823890 | 17823997 | refGene_gn=CELSR3;refGene_tx=XM_006928732.4 |
| A2 | 17824086 | 17824182 | refGene_gn=CELSR3;refGene_tx=XM_006928732.4 |
| A2 | 17824483 | 17824561 | refGene_gn=CELSR3;refGene_tx=XM_006928732.4 |
| A2 | 17824680 | 17824794 | refGene_gn=CELSR3;refGene_tx=XM_006928732.4 |
| A2 | 17825288 | 17825436 | refGene_gn=CELSR3;refGene_tx=XM_006928732.4 |
| A2 | 17825520 | 17825603 | refGene_gn=CELSR3;refGene_tx=XM_006928732.4 |
| A2 | 17825882 | 17826035 | refGene_gn=CELSR3;refGene_tx=XM_006928732.4 |
| A2 | 17826114 | 17826279 | refGene_gn=CELSR3;refGene_tx=XM_006928732.4 |
| A2 | 17826489 | 17826653 | refGene_gn=CELSR3;refGene_tx=XM_006928732.4 |
| A2 | 17827210 | 17827292 | refGene_gn=CELSR3;refGene_tx=XM_006928732.4 |
| A2 | 17827293 | 17827322 | refGene_gn=CELSR3;refGene_tx=XM_006928732.4 |
| A2 | 17827553 | 17827693 | refGene_gn=CELSR3;refGene_tx=XM_006928732.4 |
| A2 | 17827773 | 17827932 | refGene_gn=CELSR3;refGene_tx=XM_006928732.4 |
| A2 | 17833925 | 17834814 | refGene_gn=CELSR3;refGene_tx=XM_006928732.4 |
| A2 | 17835195 | 17835396 | refGene_gn=CELSR3;refGene_tx=XM_006928732.4 |
| A2 | 17835966 | 17836125 | refGene_gn=CELSR3;refGene_tx=XM_006928732.4 |
| A2 | 17836437 | 17836553 | refGene_gn=CELSR3;refGene_tx=XM_006928732.4 |
| A2 | 17836828 | 17836977 | refGene_gn=CELSR3;refGene_tx=XM_006928732.4 |
| A2 | 17837084 | 17837201 | refGene_gn=CELSR3;refGene_tx=XM_006928732.4 |
| A2 | 17837638 | 17837765 | refGene_gn=CELSR3;refGene_tx=XM_006928732.4 |
| A2 | 17838227 | 17838307 | refGene_gn=CELSR3;refGene_tx=XM_006928732.4 |
| A2 | 17838752 | 17838865 | refGene_gn=CELSR3;refGene_tx=XM_006928732.4 |
| A2 | 17839094 | 17839269 | refGene_gn=CELSR3;refGene_tx=XM_006928732.4 |
| A2 | 17839475 | 17839682 | refGene_gn=CELSR3;refGene_tx=XM_006928732.4 |
| A2 | 17839770 | 17839897 | refGene_gn=CELSR3;refGene_tx=XM_006928732.4 |
| A2 | 17840120 | 17840288 | refGene_gn=CELSR3;refGene_tx=XM_006928732.4 |
| A2 | 17840930 | 17841105 | refGene_gn=CELSR3;refGene_tx=XM_006928732.4 |
| A2 | 17841503 | 17841659 | refGene_gn=CELSR3;refGene_tx=XM_006928732.4 |
| A2 | 17841930 | 17842126 | refGene_gn=CELSR3;refGene_tx=XM_006928732.4 |
| A2 | 17842351 | 17842561 | refGene_gn=CELSR3;refGene_tx=XM_006928732.4 |
| A2 | 17842785 | 17842880 | refGene_gn=CELSR3;refGene_tx=XM_006928732.4 |
| A2 | 17844286 | 17844388 | refGene_gn=CELSR3;refGene_tx=XM_006928732.4 |
| A2 | 17844703 | 17844887 | refGene_gn=CELSR3;refGene_tx=XM_006928732.4 |
| A2 | 17845205 | 17845326 | refGene_gn=CELSR3;refGene_tx=XM_006928732.4 |
| A2 | 17845434 | 17845576 | refGene_gn=CELSR3;refGene_tx=XM_006928732.4 |
| A2 | 17845717 | 17845889 | refGene_gn=CELSR3;refGene_tx=XM_006928732.4 |
| A2 | 17846258 | 17846375 | refGene_gn=CELSR3;refGene_tx=XM_006928732.4 |
| A2 | 17846925 | 17847111 | refGene_gn=CELSR3;refGene_tx=XM_006928732.4 |
| A2 | 17847526 | 17847696 | refGene_gn=CELSR3;refGene_tx=XM_006928732.4 |
| A2 | 17847786 | 17847912 | refGene_gn=CELSR3;refGene_tx=XM_006928732.4 |

A2 17848173 17848337 refGene_gn=CELSR3;refGene_tx=XM_006928732.4
A2 17848870 17849028 refGene_gn=CELSR3;refGene_tx=XM_006928732.4
A2 17849136 17849225 refGene_gn=CELSR3;refGene_tx=XM_006928732.4
A2 17849594 17849710 refGene_gn=CELSR3;refGene_tx=XM_006928732.4
A2 17850038 17850264 refGene_gn=CELSR3;refGene_tx=XM_006928732.4
A2 17850624 17851275 refGene_gn=CELSR3;refGene_tx=XM_006928732.4
A2 17852786 17856534 refGene_gn=CELSR3;refGene_tx=XM_006928732.4
A2 17865400 17865604
refGene_gn=NCKIPSD;refGene_tx=XM_019821282.2;refGene_gn=NCKIPSD;refGene_tx=XM_023248586.1
A2 17867845 17868018
refGene_gn=NCKIPSD;refGene_tx=XM_019821282.2;refGene_gn=NCKIPSD;refGene_tx=XM_023248586.1
A2 17868152 17868245
refGene_gn=NCKIPSD;refGene_tx=XM_019821282.2;refGene_gn=NCKIPSD;refGene_tx=XM_023248586.1
A2 17868339 17868468
refGene_gn=NCKIPSD;refGene_tx=XM_019821282.2;refGene_gn=NCKIPSD;refGene_tx=XM_023248586.1
A2 17868618 17868699
refGene_gn=NCKIPSD;refGene_tx=XM_019821282.2;refGene_gn=NCKIPSD;refGene_tx=XM_023248586.1
A2 17868779 17868918
refGene_gn=NCKIPSD;refGene_tx=XM_019821282.2;refGene_gn=NCKIPSD;refGene_tx=XM_023248586.1
A2 17869006 17869093
refGene_gn=NCKIPSD;refGene_tx=XM_019821282.2;refGene_gn=NCKIPSD;refGene_tx=XM_023248586.1
A2 17869242 17869413
refGene_gn=NCKIPSD;refGene_tx=XM_019821282.2;refGene_gn=NCKIPSD;refGene_tx=XM_023248586.1
A2 17870698 17871189
refGene_gn=NCKIPSD;refGene_tx=XM_019821282.2;refGene_gn=NCKIPSD;refGene_tx=XM_023248586.1
A2 17871394 17871485
refGene_gn=NCKIPSD;refGene_tx=XM_019821282.2;refGene_gn=NCKIPSD;refGene_tx=XM_023248586.1
A2 17871665 17871870
refGene_gn=NCKIPSD;refGene_tx=XM_019821282.2;refGene_gn=NCKIPSD;refGene_tx=XM_023248586.1
A2 17872118 17872228
refGene_gn=NCKIPSD;refGene_tx=XM_019821282.2;refGene_gn=NCKIPSD;refGene_tx=XM_023248586.1
A2 17874741 17874912 refGene_gn=NCKIPSD;refGene_tx=XM_023248586.1
A2 17877050 17877551
refGene_gn=IP6K2;refGene_tx=XM_011291409.3;refGene_gn=IP6K2;refGene_tx=XM_019821285.2;refGene_gn=IP6K2;refGene_tx=XM_023248590.1
A2 17878006 17878182
refGene_gn=IP6K2;refGene_tx=XM_011291409.3;refGene_gn=IP6K2;refGene_tx=XM_019821285.2;refGene_gn=IP6K2;refGene_tx=XM_023248590.1

A2	17882058	17882234
refGene_gn=IP6K2;refGene_tx=XM_011291409.3;refGene_gn=IP6K2;refGene_tx=XM_019821285.2;refGene_gn=IP6K2;refGene_tx=XM_023248590.1
A2	17883131	17883354
refGene_gn=IP6K2;refGene_tx=XM_011291409.3;refGene_gn=IP6K2;refGene_tx=XM_019821285.2;refGene_gn=IP6K2;refGene_tx=XM_023248590.1
A2	17885292	17885494
refGene_gn=IP6K2;refGene_tx=XM_011291409.3;refGene_gn=IP6K2;refGene_tx=XM_019821285.2;refGene_gn=IP6K2;refGene_tx=XM_023248590.1
A2	17923962	17924096	refGene_gn=PRKAR2A;refGene_tx=XM_011291413.3
A2	17924784	17924926	refGene_gn=PRKAR2A;refGene_tx=XM_011291413.3
A2	17927947	17928013	refGene_gn=PRKAR2A;refGene_tx=XM_011291413.3
A2	17940258	17940333	refGene_gn=PRKAR2A;refGene_tx=XM_011291413.3
A2	17940501	17940603	refGene_gn=PRKAR2A;refGene_tx=XM_011291413.3
A2	17949980	17950134	refGene_gn=PRKAR2A;refGene_tx=XM_011291413.3
A2	17952056	17952163	refGene_gn=PRKAR2A;refGene_tx=XM_011291413.3
A2	17958458	17958542	refGene_gn=PRKAR2A;refGene_tx=XM_011291413.3
A2	17961212	17961265	refGene_gn=PRKAR2A;refGene_tx=XM_011291413.3
A2	17973093	17973129	refGene_gn=PRKAR2A;refGene_tx=XM_011291413.3
A2	18033722	18033975	refGene_gn=PRKAR2A;refGene_tx=XM_011291413.3
A2	18040638	18040701	refGene_gn=SLC25A20;refGene_tx=XM_003982205.5
A2	18041133	18041258	refGene_gn=SLC25A20;refGene_tx=XM_003982205.5
A2	18041743	18041853	refGene_gn=SLC25A20;refGene_tx=XM_003982205.5
A2	18042230	18042303	refGene_gn=SLC25A20;refGene_tx=XM_003982205.5
A2	18043027	18043145	refGene_gn=SLC25A20;refGene_tx=XM_003982205.5
A2	18057040	18057131	refGene_gn=SLC25A20;refGene_tx=XM_003982205.5
A2	18060149	18060277	refGene_gn=SLC25A20;refGene_tx=XM_003982205.5
A2	18064632	18064725	refGene_gn=SLC25A20;refGene_tx=XM_003982205.5
A2	18069317	18069422	refGene_gn=SLC25A20;refGene_tx=XM_003982205.5
A2	18084226	18084424	refGene_gn=ARIH2OS;refGene_tx=XM_023241842.1
A2	18084425	18084459	refGene_gn=ARIH2OS;refGene_tx=XM_023241842.1
A2	18084460	18084568	refGene_gn=ARIH2OS;refGene_tx=XM_023241842.1
A2	18084624	18084809	refGene_gn=ARIH2OS;refGene_tx=XM_023241842.1
A2	18084810	18084914	refGene_gn=ARIH2OS;refGene_tx=XM_023241842.1
A2	18084916	18084968	refGene_gn=ARIH2OS;refGene_tx=XM_023241842.1
A2	18084970	18085077	refGene_gn=ARIH2OS;refGene_tx=XM_023241842.1
A2	18085362	18085411	refGene_gn=ARIH2OS;refGene_tx=XM_023241842.1
A2	18089870	18090119
refGene_gn=ARIH2;refGene_tx=XM_003982206.5;refGene_gn=ARIH2;refGene_tx=XM_006928747.4;refGene_gn=ARIH2;refGene_tx=XM_019821311.2
A2	18104313	18104352	refGene_gn=ARIH2;refGene_tx=XM_006928748.2
A2	18115655	18115723
refGene_gn=ARIH2;refGene_tx=XM_003982206.5;refGene_gn=ARIH2;refGene_tx=XM_006928747.4;refGene_gn=ARIH2;refGene_tx=XM_006928748.2;refGene_gn=ARIH2;refGene_tx=XM_019821311.2
A2	18120706	18120770
refGene_gn=ARIH2;refGene_tx=XM_003982206.5;refGene_gn=ARIH2;refGene_tx=XM_006928747.4;refGene_gn=ARIH2;refGene_tx=XM_006928748.2;refGene_gn=ARIH2;refGene_tx=XM_019821311.2
A2	18122319	18122470

refGene_gn=ARIH2;refGene_tx=XM_003982206.5;refGene_gn=ARIH2;refGene_tx=XM_006928747.4;refGene_gn=ARIH2;refGene_tx=XM_006928748.2;refGene_gn=ARIH2;refGene_tx=XM_019821311.2
A2 18123794 18123916 refGene_gn=ARIH2;refGene_tx=XM_003982206.5;refGene_gn=ARIH2;refGene_tx=XM_006928747.4;refGene_gn=ARIH2;refGene_tx=XM_006928748.2;refGene_gn=ARIH2;refGene_tx=XM_019821311.2
A2 18125372 18125482 refGene_gn=ARIH2;refGene_tx=XM_003982206.5;refGene_gn=ARIH2;refGene_tx=XM_006928747.4;refGene_gn=ARIH2;refGene_tx=XM_006928748.2;refGene_gn=ARIH2;refGene_tx=XM_019821311.2
A2 18126928 18127046 refGene_gn=ARIH2;refGene_tx=XM_003982206.5;refGene_gn=ARIH2;refGene_tx=XM_006928747.4;refGene_gn=ARIH2;refGene_tx=XM_006928748.2;refGene_gn=ARIH2;refGene_tx=XM_019821311.2
A2 18128045 18128096 refGene_gn=ARIH2;refGene_tx=XM_003982206.5;refGene_gn=ARIH2;refGene_tx=XM_006928747.4;refGene_gn=ARIH2;refGene_tx=XM_006928748.2;refGene_gn=ARIH2;refGene_tx=XM_019821311.2
A2 18128186 18128208 refGene_gn=ARIH2;refGene_tx=XM_003982206.5;refGene_gn=ARIH2;refGene_tx=XM_006928747.4;refGene_gn=ARIH2;refGene_tx=XM_006928748.2;refGene_gn=ARIH2;refGene_tx=XM_019821311.2
A2 18130063 18130215 refGene_gn=ARIH2;refGene_tx=XM_003982206.5;refGene_gn=ARIH2;refGene_tx=XM_006928747.4;refGene_gn=ARIH2;refGene_tx=XM_006928748.2;refGene_gn=ARIH2;refGene_tx=XM_019821311.2
A2 18130797 18130941 refGene_gn=ARIH2;refGene_tx=XM_003982206.5;refGene_gn=ARIH2;refGene_tx=XM_006928747.4;refGene_gn=ARIH2;refGene_tx=XM_006928748.2;refGene_gn=ARIH2;refGene_tx=XM_019821311.2
A2 18131575 18131644 refGene_gn=ARIH2;refGene_tx=XM_003982206.5;refGene_gn=ARIH2;refGene_tx=XM_006928747.4;refGene_gn=ARIH2;refGene_tx=XM_006928748.2
A2 18132927 18133011 refGene_gn=ARIH2;refGene_tx=XM_003982206.5;refGene_gn=ARIH2;refGene_tx=XM_006928747.4;refGene_gn=ARIH2;refGene_tx=XM_006928748.2;refGene_gn=ARIH2;refGene_tx=XM_019821311.2
A2 18133230 18133302 refGene_gn=ARIH2;refGene_tx=XM_003982206.5;refGene_gn=ARIH2;refGene_tx=XM_006928747.4;refGene_gn=ARIH2;refGene_tx=XM_006928748.2;refGene_gn=ARIH2;refGene_tx=XM_019821311.2
A2 18139407 18139737 refGene_gn=P4HTM;refGene_tx=XM_023248612.1
A2 18139939 18140021 refGene_gn=P4HTM;refGene_tx=XM_023248612.1
A2 18143959 18144276 refGene_gn=LOC102902069;refGene_tx=XM_019821343.2
A2 18144611 18144698 refGene_gn=LOC102902069;refGene_tx=XM_019821343.2
A2 18144942 18145112 refGene_gn=LOC102902069;refGene_tx=XM_019821343.2

A2 18146103 18146387
refGene_gn=LOC102902069;refGene_tx=XM_019821343.2
A2 18148037 18148228 refGene_gn=P4HTM;refGene_tx=XM_023248612.1
A2 18149084 18149181 refGene_gn=P4HTM;refGene_tx=XM_023248612.1
A2 18150591 18150754 refGene_gn=P4HTM;refGene_tx=XM_023248612.1
A2 18151326 18151512 refGene_gn=P4HTM;refGene_tx=XM_023248612.1
A2 18152236 18152327 refGene_gn=P4HTM;refGene_tx=XM_023248612.1
A2 18152528 18152652 refGene_gn=P4HTM;refGene_tx=XM_023248612.1
A2 18153170 18153391 refGene_gn=P4HTM;refGene_tx=XM_023248612.1
A2 18153920 18154020 refGene_gn=WDR6;refGene_tx=XM_006928752.3
A2 18157419 18159892 refGene_gn=WDR6;refGene_tx=XM_006928752.3
A2 18159993 18160077 refGene_gn=WDR6;refGene_tx=XM_006928752.3
A2 18160172 18160289 refGene_gn=WDR6;refGene_tx=XM_006928752.3
A2 18160387 18160506 refGene_gn=WDR6;refGene_tx=XM_006928752.3
A2 18160605 18161066 refGene_gn=WDR6;refGene_tx=XM_006928752.3
A2 18161383 18161503
refGene_gn=DALRD3;refGene_tx=XM_003982208.4;refGene_gn=DALRD3;refGene_tx=XM_011291433.3;refGene_gn=DALRD3;refGene_tx=XM_019821328.2
A2 18161589 18161658
refGene_gn=DALRD3;refGene_tx=XM_003982208.4;refGene_gn=DALRD3;refGene_tx=XM_011291433.3;refGene_gn=DALRD3;refGene_tx=XM_019821328.2
A2 18161759 18161873
refGene_gn=DALRD3;refGene_tx=XM_003982208.4;refGene_gn=DALRD3;refGene_tx=XM_011291433.3
A2 18161949 18162132
refGene_gn=DALRD3;refGene_tx=XM_003982208.4;refGene_gn=DALRD3;refGene_tx=XM_011291433.3;refGene_gn=DALRD3;refGene_tx=XM_019821328.2
A2 18162207 18162290
refGene_gn=DALRD3;refGene_tx=XM_003982208.4;refGene_gn=DALRD3;refGene_tx=XM_011291433.3;refGene_gn=DALRD3;refGene_tx=XM_019821328.2
A2 18162384 18162446
refGene_gn=DALRD3;refGene_tx=XM_003982208.4;refGene_gn=DALRD3;refGene_tx=XM_011291433.3;refGene_gn=DALRD3;refGene_tx=XM_019821328.2
A2 18162546 18162617
refGene_gn=DALRD3;refGene_tx=XM_003982208.4;refGene_gn=DALRD3;refGene_tx=XM_011291433.3;refGene_gn=DALRD3;refGene_tx=XM_019821328.2
A2 18162952 18163081
refGene_gn=DALRD3;refGene_tx=XM_003982208.4;refGene_gn=DALRD3;refGene_tx=XM_011291433.3;refGene_gn=DALRD3;refGene_tx=XM_019821328.2
A2 18163168 18163248
refGene_gn=DALRD3;refGene_tx=XM_003982208.4;refGene_gn=DALRD3;refGene_tx=XM_011291433.3;refGene_gn=DALRD3;refGene_tx=XM_019821328.2
A2 18163333 18163590
refGene_gn=DALRD3;refGene_tx=XM_003982208.4;refGene_gn=DALRD3;refGene_tx=XM_011291433.3;refGene_gn=DALRD3;refGene_tx=XM_019821328.2
A2 18163711 18164007
refGene_gn=DALRD3;refGene_tx=XM_003982208.4;refGene_gn=DALRD3;refGene_tx=XM_011291433.3;refGene_gn=DALRD3;refGene_tx=XM_019821328.2
A2 18164083 18164465
refGene_gn=DALRD3;refGene_tx=XM_003982208.4;refGene_gn=DALRD3;refGene_

tx=XM_011291433.3;refGene_gn=DALRD3;refGene_tx=XM_019821328.2
A2 18167462 18167539
refGene_gn=NDUFAF3;refGene_tx=XM_003982209.5;refGene_gn=NDUFAF3;refGene_tx=XM_006928753.4
A2 18167642 18167856
refGene_gn=NDUFAF3;refGene_tx=XM_003982209.5;refGene_gn=NDUFAF3;refGene_tx=XM_006928753.4
A2 18167992 18168059
refGene_gn=NDUFAF3;refGene_tx=XM_003982209.5;refGene_gn=NDUFAF3;refGene_tx=XM_006928753.4
A2 18168164 18168265
refGene_gn=NDUFAF3;refGene_tx=XM_003982209.5;refGene_gn=NDUFAF3;refGene_tx=XM_006928753.4
A2 18168344 18168461
refGene_gn=NDUFAF3;refGene_tx=XM_003982209.5;refGene_gn=NDUFAF3;refGene_tx=XM_006928753.4
A2 18169227 18169249
refGene_gn=IMPDH2;refGene_tx=XM_003982210.4;refGene_gn=IMPDH2;refGene_tx=XM_011291449.2
A2 18169351 18169435
refGene_gn=IMPDH2;refGene_tx=XM_003982210.4;refGene_gn=IMPDH2;refGene_tx=XM_011291449.2
A2 18169514 18169658
refGene_gn=IMPDH2;refGene_tx=XM_003982210.4;refGene_gn=IMPDH2;refGene_tx=XM_011291449.2
A2 18169756 18169901
refGene_gn=IMPDH2;refGene_tx=XM_003982210.4;refGene_gn=IMPDH2;refGene_tx=XM_011291449.2
A2 18169987 18170131
refGene_gn=IMPDH2;refGene_tx=XM_003982210.4;refGene_gn=IMPDH2;refGene_tx=XM_011291449.2
A2 18170785 18170857 refGene_gn=IMPDH2;refGene_tx=XM_011291449.2
A2 18171131 18171227
refGene_gn=IMPDH2;refGene_tx=XM_003982210.4;refGene_gn=IMPDH2;refGene_tx=XM_011291449.2
A2 18171319 18171410
refGene_gn=IMPDH2;refGene_tx=XM_003982210.4;refGene_gn=IMPDH2;refGene_tx=XM_011291449.2
A2 18171489 18171689
refGene_gn=IMPDH2;refGene_tx=XM_003982210.4;refGene_gn=IMPDH2;refGene_tx=XM_011291449.2
A2 18171762 18171850
refGene_gn=IMPDH2;refGene_tx=XM_003982210.4;refGene_gn=IMPDH2;refGene_tx=XM_011291449.2
A2 18172352 18172559
refGene_gn=IMPDH2;refGene_tx=XM_003982210.4;refGene_gn=IMPDH2;refGene_tx=XM_011291449.2
A2 18172874 18172949
refGene_gn=IMPDH2;refGene_tx=XM_003982210.4;refGene_gn=IMPDH2;refGene_tx=XM_011291449.2

A2 18173273 18173375
refGene_gn=IMPDH2;refGene_tx=XM_003982210.4;refGene_gn=IMPDH2;refGene_tx=XM_011291449.2
A2 18173593 18173642
refGene_gn=IMPDH2;refGene_tx=XM_003982210.4;refGene_gn=IMPDH2;refGene_tx=XM_011291449.2
A2 18174085 18174183
refGene_gn=IMPDH2;refGene_tx=XM_003982210.4;refGene_gn=IMPDH2;refGene_tx=XM_011291449.2
A2 18175279 18175472
refGene_gn=QRICH1;refGene_tx=XM_006928754.4;refGene_gn=QRICH1;refGene_tx=XM_011291451.3;refGene_gn=QRICH1;refGene_tx=XM_019821354.2;refGene_gn=QRICH1;refGene_tx=XM_019821359.2;refGene_gn=QRICH1;refGene_tx=XM_023248672.1;refGene_gn=QRICH1;refGene_tx=XM_023248678.1;refGene_gn=QRICH1;refGene_tx=XM_023248679.1;refGene_gn=QRICH1;refGene_tx=XM_023248682.1
A2 18177138 18177229
refGene_gn=QRICH1;refGene_tx=XM_006928754.4;refGene_gn=QRICH1;refGene_tx=XM_011291451.3;refGene_gn=QRICH1;refGene_tx=XM_019821354.2;refGene_gn=QRICH1;refGene_tx=XM_019821359.2;refGene_gn=QRICH1;refGene_tx=XM_023248672.1;refGene_gn=QRICH1;refGene_tx=XM_023248678.1;refGene_gn=QRICH1;refGene_tx=XM_023248679.1;refGene_gn=QRICH1;refGene_tx=XM_023248682.1
A2 18177550 18177702
refGene_gn=QRICH1;refGene_tx=XM_006928754.4;refGene_gn=QRICH1;refGene_tx=XM_011291451.3;refGene_gn=QRICH1;refGene_tx=XM_019821354.2;refGene_gn=QRICH1;refGene_tx=XM_019821359.2;refGene_gn=QRICH1;refGene_tx=XM_023248672.1;refGene_gn=QRICH1;refGene_tx=XM_023248678.1;refGene_gn=QRICH1;refGene_tx=XM_023248679.1;refGene_gn=QRICH1;refGene_tx=XM_023248682.1
A2 18178058 18178167
refGene_gn=QRICH1;refGene_tx=XM_006928754.4;refGene_gn=QRICH1;refGene_tx=XM_011291451.3;refGene_gn=QRICH1;refGene_tx=XM_019821354.2;refGene_gn=QRICH1;refGene_tx=XM_019821359.2;refGene_gn=QRICH1;refGene_tx=XM_023248672.1;refGene_gn=QRICH1;refGene_tx=XM_023248678.1;refGene_gn=QRICH1;refGene_tx=XM_023248679.1;refGene_gn=QRICH1;refGene_tx=XM_023248682.1
A2 18184517 18184632
refGene_gn=QRICH1;refGene_tx=XM_006928754.4;refGene_gn=QRICH1;refGene_tx=XM_011291451.3;refGene_gn=QRICH1;refGene_tx=XM_019821354.2;refGene_gn=QRICH1;refGene_tx=XM_019821359.2;refGene_gn=QRICH1;refGene_tx=XM_023248672.1;refGene_gn=QRICH1;refGene_tx=XM_023248678.1;refGene_gn=QRICH1;refGene_tx=XM_023248679.1;refGene_gn=QRICH1;refGene_tx=XM_023248682.1
A2 18185712 18185867
refGene_gn=QRICH1;refGene_tx=XM_006928754.4;refGene_gn=QRICH1;refGene_tx=XM_011291451.3;refGene_gn=QRICH1;refGene_tx=XM_019821354.2;refGene_gn=QRICH1;refGene_tx=XM_019821359.2;refGene_gn=QRICH1;refGene_tx=XM_023248672.1;refGene_gn=QRICH1;refGene_tx=XM_023248678.1;refGene_gn=QRICH1;refGene_tx=XM_023248679.1;refGene_gn=QRICH1;refGene_tx=XM_023248682.

1
A2 18186501 18186679
refGene_gn=QRICH1;refGene_tx=XM_006928754.4;refGene_gn=QRICH1;refGene_tx=XM_011291451.3;refGene_gn=QRICH1;refGene_tx=XM_019821354.2;refGene_gn=QRICH1;refGene_tx=XM_019821359.2;refGene_gn=QRICH1;refGene_tx=XM_023248672.1;refGene_gn=QRICH1;refGene_tx=XM_023248678.1;refGene_gn=QRICH1;refGene_tx=XM_023248679.1;refGene_gn=QRICH1;refGene_tx=XM_023248682.1
A2 18193044 18194082
refGene_gn=QRICH1;refGene_tx=XM_006928754.4;refGene_gn=QRICH1;refGene_tx=XM_011291451.3;refGene_gn=QRICH1;refGene_tx=XM_019821354.2;refGene_gn=QRICH1;refGene_tx=XM_019821359.2;refGene_gn=QRICH1;refGene_tx=XM_023248672.1;refGene_gn=QRICH1;refGene_tx=XM_023248678.1;refGene_gn=QRICH1;refGene_tx=XM_023248679.1;refGene_gn=QRICH1;refGene_tx=XM_023248682.1
A2 18215912 18216221
refGene_gn=QRICH1;refGene_tx=XM_006928754.4;refGene_gn=QRICH1;refGene_tx=XM_011291451.3;refGene_gn=QRICH1;refGene_tx=XM_019821354.2;refGene_gn=QRICH1;refGene_tx=XM_019821359.2;refGene_gn=QRICH1;refGene_tx=XM_023248672.1;refGene_gn=QRICH1;refGene_tx=XM_023248678.1;refGene_gn=QRICH1;refGene_tx=XM_023248679.1;refGene_gn=QRICH1;refGene_tx=XM_023248682.1
A2 18227328 18227379 refGene_gn=QARS;refGene_tx=XM_003982213.4
A2 18227772 18227898 refGene_gn=QARS;refGene_tx=XM_003982213.4
A2 18227972 18228039 refGene_gn=QARS;refGene_tx=XM_003982213.4
A2 18228132 18228260 refGene_gn=QARS;refGene_tx=XM_003982213.4
A2 18228378 18228471 refGene_gn=QARS;refGene_tx=XM_003982213.4
A2 18228660 18228765 refGene_gn=QARS;refGene_tx=XM_003982213.4
A2 18228890 18229034 refGene_gn=QARS;refGene_tx=XM_003982213.4
A2 18229119 18229207 refGene_gn=QARS;refGene_tx=XM_003982213.4
A2 18229294 18229432 refGene_gn=QARS;refGene_tx=XM_003982213.4
A2 18229543 18229636 refGene_gn=QARS;refGene_tx=XM_003982213.4
A2 18229743 18229874 refGene_gn=QARS;refGene_tx=XM_003982213.4
A2 18229979 18230088 refGene_gn=QARS;refGene_tx=XM_003982213.4
A2 18230169 18230248 refGene_gn=QARS;refGene_tx=XM_003982213.4
A2 18230359 18230459 refGene_gn=QARS;refGene_tx=XM_003982213.4
A2 18230870 18230957 refGene_gn=QARS;refGene_tx=XM_003982213.4
A2 18231134 18231220 refGene_gn=QARS;refGene_tx=XM_003982213.4
A2 18231325 18231397 refGene_gn=QARS;refGene_tx=XM_003982213.4
A2 18231762 18231823 refGene_gn=QARS;refGene_tx=XM_003982213.4
A2 18231959 18232013 refGene_gn=QARS;refGene_tx=XM_003982213.4
A2 18232613 18232678 refGene_gn=QARS;refGene_tx=XM_003982213.4
A2 18233086 18233162 refGene_gn=QARS;refGene_tx=XM_003982213.4
A2 18233320 18233430 refGene_gn=QARS;refGene_tx=XM_003982213.4
A2 18233790 18233938 refGene_gn=QARS;refGene_tx=XM_003982213.4
A2 18234089 18234206 refGene_gn=QARS;refGene_tx=XM_003982213.4
A2 18237768 18237885
refGene_gn=USP19;refGene_tx=XM_006928763.4;refGene_gn=USP19;refGene_tx=XM_006928764.3;refGene_gn=USP19;refGene_tx=XM_006928765.4;refGene_gn=USP19;refGene_tx=XM_006928766.4;refGene_gn=USP19;refGene_tx=XM_0069287

68.2;refGene_gn=USP19;refGene_tx=XM_023248652.1;refGene_gn=USP19;refGene_tx=XM_023248657.1
A2 18238311 18238539
refGene_gn=USP19;refGene_tx=XM_006928756.3;refGene_gn=USP19;refGene_tx=XM_006928757.4;refGene_gn=USP19;refGene_tx=XM_006928758.3;refGene_gn=USP19;refGene_tx=XM_006928759.4;refGene_gn=USP19;refGene_tx=XM_006928760.4;refGene_gn=USP19;refGene_tx=XM_006928761.4;refGene_gn=USP19;refGene_tx=XM_006928762.4;refGene_gn=USP19;refGene_tx=XM_006928767.2;refGene_gn=USP19;refGene_tx=XM_019821381.2
A2 18239538 18239717
refGene_gn=USP19;refGene_tx=XM_006928756.3;refGene_gn=USP19;refGene_tx=XM_006928757.4;refGene_gn=USP19;refGene_tx=XM_006928758.3;refGene_gn=USP19;refGene_tx=XM_006928759.4;refGene_gn=USP19;refGene_tx=XM_006928760.4;refGene_gn=USP19;refGene_tx=XM_006928761.4;refGene_gn=USP19;refGene_tx=XM_006928762.4;refGene_gn=USP19;refGene_tx=XM_006928763.4;refGene_gn=USP19;refGene_tx=XM_006928764.3;refGene_gn=USP19;refGene_tx=XM_006928765.4;refGene_gn=USP19;refGene_tx=XM_006928766.4;refGene_gn=USP19;refGene_tx=XM_006928767.2;refGene_gn=USP19;refGene_tx=XM_006928768.2;refGene_gn=USP19;refGene_tx=XM_019821381.2;refGene_gn=USP19;refGene_tx=XM_023248652.1;refGene_gn=USP19;refGene_tx=XM_023248657.1
A2 18239798 18239959
refGene_gn=USP19;refGene_tx=XM_006928756.3;refGene_gn=USP19;refGene_tx=XM_006928757.4;refGene_gn=USP19;refGene_tx=XM_006928758.3;refGene_gn=USP19;refGene_tx=XM_006928759.4;refGene_gn=USP19;refGene_tx=XM_006928760.4;refGene_gn=USP19;refGene_tx=XM_006928761.4;refGene_gn=USP19;refGene_tx=XM_006928762.4;refGene_gn=USP19;refGene_tx=XM_006928763.4;refGene_gn=USP19;refGene_tx=XM_006928764.3;refGene_gn=USP19;refGene_tx=XM_006928765.4;refGene_gn=USP19;refGene_tx=XM_006928766.4;refGene_gn=USP19;refGene_tx=XM_006928767.2;refGene_gn=USP19;refGene_tx=XM_006928768.2;refGene_gn=USP19;refGene_tx=XM_019821381.2;refGene_gn=USP19;refGene_tx=XM_023248652.1;refGene_gn=USP19;refGene_tx=XM_023248657.1
A2 18240054 18240207
refGene_gn=USP19;refGene_tx=XM_006928756.3;refGene_gn=USP19;refGene_tx=XM_006928757.4;refGene_gn=USP19;refGene_tx=XM_006928758.3;refGene_gn=USP19;refGene_tx=XM_006928759.4;refGene_gn=USP19;refGene_tx=XM_006928760.4;refGene_gn=USP19;refGene_tx=XM_006928761.4;refGene_gn=USP19;refGene_tx=XM_006928762.4;refGene_gn=USP19;refGene_tx=XM_006928763.4;refGene_gn=USP19;refGene_tx=XM_006928764.3;refGene_gn=USP19;refGene_tx=XM_006928765.4;refGene_gn=USP19;refGene_tx=XM_006928766.4;refGene_gn=USP19;refGene_tx=XM_006928767.2;refGene_gn=USP19;refGene_tx=XM_006928768.2;refGene_gn=USP19;refGene_tx=XM_019821381.2;refGene_gn=USP19;refGene_tx=XM_023248652.1;refGene_gn=USP19;refGene_tx=XM_023248657.1
A2 18240290 18240507
refGene_gn=USP19;refGene_tx=XM_006928756.3;refGene_gn=USP19;refGene_tx=XM_006928757.4;refGene_gn=USP19;refGene_tx=XM_006928758.3;refGene_gn=USP19;refGene_tx=XM_006928759.4;refGene_gn=USP19;refGene_tx=XM_006928760.4;refGene_gn=USP19;refGene_tx=XM_006928761.4;refGene_gn=USP19;refGene_tx=XM_006928762.4;refGene_gn=USP19;refGene_tx=XM_006928763.4;refGene_gn=USP19;refGene_tx=XM_006928764.3;refGene_gn=USP19;refGene_tx=XM_006928765.4;refGene_gn=USP19;refGene_tx=XM_006928766.4;refGene_gn=USP19;

refGene_tx=XM_006928767.2;refGene_gn=USP19;refGene_tx=XM_006928768.2;refGene_gn=USP19;refGene_tx=XM_019821381.2;refGene_gn=USP19;refGene_tx=XM_023248652.1;refGene_gn=USP19;refGene_tx=XM_023248657.1
A2 18240594 18240705
refGene_gn=USP19;refGene_tx=XM_006928756.3;refGene_gn=USP19;refGene_tx=XM_006928757.4;refGene_gn=USP19;refGene_tx=XM_006928758.3;refGene_gn=USP19;refGene_tx=XM_006928759.4;refGene_gn=USP19;refGene_tx=XM_006928760.4;refGene_gn=USP19;refGene_tx=XM_006928761.4;refGene_gn=USP19;refGene_tx=XM_006928762.4;refGene_gn=USP19;refGene_tx=XM_006928763.4;refGene_gn=USP19;refGene_tx=XM_006928764.3;refGene_gn=USP19;refGene_tx=XM_006928765.4;refGene_gn=USP19;refGene_tx=XM_006928766.4;refGene_gn=USP19;refGene_tx=XM_006928767.2;refGene_gn=USP19;refGene_tx=XM_006928768.2;refGene_gn=USP19;refGene_tx=XM_019821381.2;refGene_gn=USP19;refGene_tx=XM_023248652.1;refGene_gn=USP19;refGene_tx=XM_023248657.1
A2 18240808 18241122
refGene_gn=USP19;refGene_tx=XM_006928756.3;refGene_gn=USP19;refGene_tx=XM_006928757.4;refGene_gn=USP19;refGene_tx=XM_006928758.3;refGene_gn=USP19;refGene_tx=XM_006928759.4;refGene_gn=USP19;refGene_tx=XM_006928760.4;refGene_gn=USP19;refGene_tx=XM_006928761.4;refGene_gn=USP19;refGene_tx=XM_006928762.4;refGene_gn=USP19;refGene_tx=XM_006928763.4;refGene_gn=USP19;refGene_tx=XM_006928764.3;refGene_gn=USP19;refGene_tx=XM_006928765.4;refGene_gn=USP19;refGene_tx=XM_006928766.4;refGene_gn=USP19;refGene_tx=XM_006928767.2;refGene_gn=USP19;refGene_tx=XM_006928768.2;refGene_gn=USP19;refGene_tx=XM_019821381.2;refGene_gn=USP19;refGene_tx=XM_023248652.1;refGene_gn=USP19;refGene_tx=XM_023248657.1
A2 18241221 18241359
refGene_gn=USP19;refGene_tx=XM_006928756.3;refGene_gn=USP19;refGene_tx=XM_006928757.4;refGene_gn=USP19;refGene_tx=XM_006928758.3;refGene_gn=USP19;refGene_tx=XM_006928759.4;refGene_gn=USP19;refGene_tx=XM_006928760.4;refGene_gn=USP19;refGene_tx=XM_006928761.4;refGene_gn=USP19;refGene_tx=XM_006928762.4;refGene_gn=USP19;refGene_tx=XM_006928763.4;refGene_gn=USP19;refGene_tx=XM_006928764.3;refGene_gn=USP19;refGene_tx=XM_006928765.4;refGene_gn=USP19;refGene_tx=XM_006928766.4;refGene_gn=USP19;refGene_tx=XM_006928767.2;refGene_gn=USP19;refGene_tx=XM_006928768.2;refGene_gn=USP19;refGene_tx=XM_019821381.2;refGene_gn=USP19;refGene_tx=XM_023248652.1;refGene_gn=USP19;refGene_tx=XM_023248657.1
A2 18241595 18241714
refGene_gn=USP19;refGene_tx=XM_006928756.3;refGene_gn=USP19;refGene_tx=XM_006928757.4;refGene_gn=USP19;refGene_tx=XM_006928758.3;refGene_gn=USP19;refGene_tx=XM_006928759.4;refGene_gn=USP19;refGene_tx=XM_006928760.4;refGene_gn=USP19;refGene_tx=XM_006928761.4;refGene_gn=USP19;refGene_tx=XM_006928762.4;refGene_gn=USP19;refGene_tx=XM_006928763.4;refGene_gn=USP19;refGene_tx=XM_006928764.3;refGene_gn=USP19;refGene_tx=XM_006928765.4;refGene_gn=USP19;refGene_tx=XM_006928766.4;refGene_gn=USP19;refGene_tx=XM_006928767.2;refGene_gn=USP19;refGene_tx=XM_006928768.2;refGene_gn=USP19;refGene_tx=XM_019821381.2;refGene_gn=USP19;refGene_tx=XM_023248652.1;refGene_gn=USP19;refGene_tx=XM_023248657.1
A2 18241805 18241946
refGene_gn=USP19;refGene_tx=XM_006928756.3;refGene_gn=USP19;refGene_tx=XM_006928757.4;refGene_gn=USP19;refGene_tx=XM_006928758.3;refGene_gn=

USP19;refGene_tx=XM_006928759.4;refGene_gn=USP19;refGene_tx=XM_0069287
60.4;refGene_gn=USP19;refGene_tx=XM_006928761.4;refGene_gn=USP19;refGe
ne_tx=XM_006928762.4;refGene_gn=USP19;refGene_tx=XM_006928763.4;refGen
e_gn=USP19;refGene_tx=XM_006928764.3;refGene_gn=USP19;refGene_tx=XM_00
6928765.4;refGene_gn=USP19;refGene_tx=XM_006928766.4;refGene_gn=USP19;
refGene_tx=XM_006928767.2;refGene_gn=USP19;refGene_tx=XM_006928768.2;r
efGene_gn=USP19;refGene_tx=XM_019821381.2;refGene_gn=USP19;refGene_tx=
XM_023248652.1;refGene_gn=USP19;refGene_tx=XM_023248657.1
A2 18242722 18242824
refGene_gn=USP19;refGene_tx=XM_006928756.3;refGene_gn=USP19;refGene_tx
=XM_006928757.4;refGene_gn=USP19;refGene_tx=XM_006928758.3;refGene_gn=
USP19;refGene_tx=XM_006928759.4;refGene_gn=USP19;refGene_tx=XM_0069287
60.4;refGene_gn=USP19;refGene_tx=XM_006928761.4;refGene_gn=USP19;refGe
ne_tx=XM_006928762.4;refGene_gn=USP19;refGene_tx=XM_006928763.4;refGen
e_gn=USP19;refGene_tx=XM_006928764.3;refGene_gn=USP19;refGene_tx=XM_00
6928765.4;refGene_gn=USP19;refGene_tx=XM_006928766.4;refGene_gn=USP19;
refGene_tx=XM_006928767.2;refGene_gn=USP19;refGene_tx=XM_006928768.2;r
efGene_gn=USP19;refGene_tx=XM_019821381.2;refGene_gn=USP19;refGene_tx=
XM_023248652.1;refGene_gn=USP19;refGene_tx=XM_023248657.1
A2 18242905 18243016
refGene_gn=USP19;refGene_tx=XM_006928756.3;refGene_gn=USP19;refGene_tx
=XM_006928757.4;refGene_gn=USP19;refGene_tx=XM_006928758.3;refGene_gn=
USP19;refGene_tx=XM_006928759.4;refGene_gn=USP19;refGene_tx=XM_0069287
60.4;refGene_gn=USP19;refGene_tx=XM_006928761.4;refGene_gn=USP19;refGe
ne_tx=XM_006928762.4;refGene_gn=USP19;refGene_tx=XM_006928763.4;refGen
e_gn=USP19;refGene_tx=XM_006928764.3;refGene_gn=USP19;refGene_tx=XM_00
6928765.4;refGene_gn=USP19;refGene_tx=XM_006928766.4;refGene_gn=USP19;
refGene_tx=XM_006928767.2;refGene_gn=USP19;refGene_tx=XM_006928768.2;r
efGene_gn=USP19;refGene_tx=XM_019821381.2;refGene_gn=USP19;refGene_tx=
XM_023248652.1;refGene_gn=USP19;refGene_tx=XM_023248657.1
A2 18243478 18243589
refGene_gn=USP19;refGene_tx=XM_006928756.3;refGene_gn=USP19;refGene_tx
=XM_006928757.4;refGene_gn=USP19;refGene_tx=XM_006928758.3;refGene_gn=
USP19;refGene_tx=XM_006928759.4;refGene_gn=USP19;refGene_tx=XM_0069287
60.4;refGene_gn=USP19;refGene_tx=XM_006928761.4;refGene_gn=USP19;refGe
ne_tx=XM_006928762.4;refGene_gn=USP19;refGene_tx=XM_006928763.4;refGen
e_gn=USP19;refGene_tx=XM_006928764.3;refGene_gn=USP19;refGene_tx=XM_00
6928765.4;refGene_gn=USP19;refGene_tx=XM_006928766.4;refGene_gn=USP19;
refGene_tx=XM_006928767.2;refGene_gn=USP19;refGene_tx=XM_006928768.2;r
efGene_gn=USP19;refGene_tx=XM_019821381.2;refGene_gn=USP19;refGene_tx=
XM_023248652.1;refGene_gn=USP19;refGene_tx=XM_023248657.1
A2 18243677 18243836
refGene_gn=USP19;refGene_tx=XM_006928756.3;refGene_gn=USP19;refGene_tx
=XM_006928757.4;refGene_gn=USP19;refGene_tx=XM_006928758.3;refGene_gn=
USP19;refGene_tx=XM_006928759.4;refGene_gn=USP19;refGene_tx=XM_0069287
60.4;refGene_gn=USP19;refGene_tx=XM_006928761.4;refGene_gn=USP19;refGe
ne_tx=XM_006928762.4;refGene_gn=USP19;refGene_tx=XM_006928763.4;refGen
e_gn=USP19;refGene_tx=XM_006928764.3;refGene_gn=USP19;refGene_tx=XM_00
6928765.4;refGene_gn=USP19;refGene_tx=XM_006928766.4;refGene_gn=USP19;
refGene_tx=XM_006928767.2;refGene_gn=USP19;refGene_tx=XM_006928768.2;r

efGene_gn=USP19;refGene_tx=XM_019821381.2;refGene_gn=USP19;refGene_tx=XM_023248652.1;refGene_gn=USP19;refGene_tx=XM_023248657.1
A2 18243945 18244079
refGene_gn=USP19;refGene_tx=XM_006928756.3;refGene_gn=USP19;refGene_tx=XM_006928757.4;refGene_gn=USP19;refGene_tx=XM_006928758.3;refGene_gn=USP19;refGene_tx=XM_006928759.4;refGene_gn=USP19;refGene_tx=XM_006928760.4;refGene_gn=USP19;refGene_tx=XM_006928761.4;refGene_gn=USP19;refGene_tx=XM_006928762.4;refGene_gn=USP19;refGene_tx=XM_006928763.4;refGene_gn=USP19;refGene_tx=XM_006928764.3;refGene_gn=USP19;refGene_tx=XM_006928765.4;refGene_gn=USP19;refGene_tx=XM_006928766.4;refGene_gn=USP19;refGene_tx=XM_006928767.2;refGene_gn=USP19;refGene_tx=XM_006928768.2;refGene_gn=USP19;refGene_tx=XM_019821381.2;refGene_gn=USP19;refGene_tx=XM_023248652.1;refGene_gn=USP19;refGene_tx=XM_023248657.1
A2 18244162 18244358
refGene_gn=USP19;refGene_tx=XM_006928756.3;refGene_gn=USP19;refGene_tx=XM_006928757.4;refGene_gn=USP19;refGene_tx=XM_006928758.3;refGene_gn=USP19;refGene_tx=XM_006928759.4;refGene_gn=USP19;refGene_tx=XM_006928760.4;refGene_gn=USP19;refGene_tx=XM_006928761.4;refGene_gn=USP19;refGene_tx=XM_006928762.4;refGene_gn=USP19;refGene_tx=XM_006928763.4;refGene_gn=USP19;refGene_tx=XM_006928764.3;refGene_gn=USP19;refGene_tx=XM_006928765.4;refGene_gn=USP19;refGene_tx=XM_006928766.4;refGene_gn=USP19;refGene_tx=XM_006928767.2;refGene_gn=USP19;refGene_tx=XM_006928768.2;refGene_gn=USP19;refGene_tx=XM_019821381.2;refGene_gn=USP19;refGene_tx=XM_023248652.1;refGene_gn=USP19;refGene_tx=XM_023248657.1
A2 18244442 18244663
refGene_gn=USP19;refGene_tx=XM_006928756.3;refGene_gn=USP19;refGene_tx=XM_006928757.4;refGene_gn=USP19;refGene_tx=XM_006928758.3;refGene_gn=USP19;refGene_tx=XM_006928759.4;refGene_gn=USP19;refGene_tx=XM_006928760.4;refGene_gn=USP19;refGene_tx=XM_006928761.4;refGene_gn=USP19;refGene_tx=XM_006928762.4;refGene_gn=USP19;refGene_tx=XM_006928763.4;refGene_gn=USP19;refGene_tx=XM_006928764.3;refGene_gn=USP19;refGene_tx=XM_006928765.4;refGene_gn=USP19;refGene_tx=XM_006928766.4;refGene_gn=USP19;refGene_tx=XM_006928767.2;refGene_gn=USP19;refGene_tx=XM_006928768.2;refGene_gn=USP19;refGene_tx=XM_019821381.2;refGene_gn=USP19;refGene_tx=XM_023248652.1;refGene_gn=USP19;refGene_tx=XM_023248657.1
A2 18244764 18244880
refGene_gn=USP19;refGene_tx=XM_006928756.3;refGene_gn=USP19;refGene_tx=XM_006928757.4;refGene_gn=USP19;refGene_tx=XM_006928758.3;refGene_gn=USP19;refGene_tx=XM_006928759.4;refGene_gn=USP19;refGene_tx=XM_006928760.4;refGene_gn=USP19;refGene_tx=XM_006928761.4;refGene_gn=USP19;refGene_tx=XM_006928762.4;refGene_gn=USP19;refGene_tx=XM_006928763.4;refGene_gn=USP19;refGene_tx=XM_006928764.3;refGene_gn=USP19;refGene_tx=XM_006928765.4;refGene_gn=USP19;refGene_tx=XM_006928766.4;refGene_gn=USP19;refGene_tx=XM_006928767.2;refGene_gn=USP19;refGene_tx=XM_006928768.2;refGene_gn=USP19;refGene_tx=XM_019821381.2;refGene_gn=USP19;refGene_tx=XM_023248652.1;refGene_gn=USP19;refGene_tx=XM_023248657.1
A2 18244995 18245067
refGene_gn=USP19;refGene_tx=XM_006928756.3;refGene_gn=USP19;refGene_tx=XM_006928757.4;refGene_gn=USP19;refGene_tx=XM_006928758.3;refGene_gn=USP19;refGene_tx=XM_006928759.4;refGene_gn=USP19;refGene_tx=XM_0069287

60.4;refGene_gn=USP19;refGene_tx=XM_006928761.4;refGene_gn=USP19;refGene_tx=XM_006928762.4;refGene_gn=USP19;refGene_tx=XM_006928763.4;refGene_gn=USP19;refGene_tx=XM_006928764.3;refGene_gn=USP19;refGene_tx=XM_006928765.4;refGene_gn=USP19;refGene_tx=XM_006928766.4;refGene_gn=USP19;refGene_tx=XM_006928767.2;refGene_gn=USP19;refGene_tx=XM_006928768.2;refGene_gn=USP19;refGene_tx=XM_019821381.2;refGene_gn=USP19;refGene_tx=XM_023248652.1;refGene_gn=USP19;refGene_tx=XM_023248657.1
A2 18245178 18245335
refGene_gn=USP19;refGene_tx=XM_006928756.3;refGene_gn=USP19;refGene_tx=XM_006928757.4;refGene_gn=USP19;refGene_tx=XM_006928758.3;refGene_gn=USP19;refGene_tx=XM_006928759.4;refGene_gn=USP19;refGene_tx=XM_006928760.4;refGene_gn=USP19;refGene_tx=XM_006928761.4;refGene_gn=USP19;refGene_tx=XM_006928762.4;refGene_gn=USP19;refGene_tx=XM_006928763.4;refGene_gn=USP19;refGene_tx=XM_006928764.3;refGene_gn=USP19;refGene_tx=XM_006928765.4;refGene_gn=USP19;refGene_tx=XM_006928766.4;refGene_gn=USP19;refGene_tx=XM_006928767.2;refGene_gn=USP19;refGene_tx=XM_006928768.2;refGene_gn=USP19;refGene_tx=XM_019821381.2;refGene_gn=USP19;refGene_tx=XM_023248652.1;refGene_gn=USP19;refGene_tx=XM_023248657.1
A2 18245458 18245681
refGene_gn=USP19;refGene_tx=XM_006928756.3;refGene_gn=USP19;refGene_tx=XM_006928757.4;refGene_gn=USP19;refGene_tx=XM_006928758.3;refGene_gn=USP19;refGene_tx=XM_006928759.4;refGene_gn=USP19;refGene_tx=XM_006928760.4;refGene_gn=USP19;refGene_tx=XM_006928761.4;refGene_gn=USP19;refGene_tx=XM_006928762.4;refGene_gn=USP19;refGene_tx=XM_006928763.4;refGene_gn=USP19;refGene_tx=XM_006928764.3;refGene_gn=USP19;refGene_tx=XM_006928765.4;refGene_gn=USP19;refGene_tx=XM_006928766.4;refGene_gn=USP19;refGene_tx=XM_006928767.2;refGene_gn=USP19;refGene_tx=XM_006928768.2;refGene_gn=USP19;refGene_tx=XM_019821381.2;refGene_gn=USP19;refGene_tx=XM_023248652.1;refGene_gn=USP19;refGene_tx=XM_023248657.1
A2 18245789 18246092
refGene_gn=USP19;refGene_tx=XM_006928756.3;refGene_gn=USP19;refGene_tx=XM_006928757.4;refGene_gn=USP19;refGene_tx=XM_006928758.3;refGene_gn=USP19;refGene_tx=XM_006928759.4;refGene_gn=USP19;refGene_tx=XM_006928760.4;refGene_gn=USP19;refGene_tx=XM_006928761.4;refGene_gn=USP19;refGene_tx=XM_006928762.4;refGene_gn=USP19;refGene_tx=XM_006928763.4;refGene_gn=USP19;refGene_tx=XM_006928764.3;refGene_gn=USP19;refGene_tx=XM_006928765.4;refGene_gn=USP19;refGene_tx=XM_006928766.4;refGene_gn=USP19;refGene_tx=XM_006928767.2;refGene_gn=USP19;refGene_tx=XM_006928768.2;refGene_gn=USP19;refGene_tx=XM_019821381.2;refGene_gn=USP19;refGene_tx=XM_023248652.1;refGene_gn=USP19;refGene_tx=XM_023248657.1
A2 18246165 18246299
refGene_gn=USP19;refGene_tx=XM_006928756.3;refGene_gn=USP19;refGene_tx=XM_006928757.4;refGene_gn=USP19;refGene_tx=XM_006928758.3;refGene_gn=USP19;refGene_tx=XM_006928759.4;refGene_gn=USP19;refGene_tx=XM_006928760.4;refGene_gn=USP19;refGene_tx=XM_006928761.4;refGene_gn=USP19;refGene_tx=XM_006928762.4;refGene_gn=USP19;refGene_tx=XM_006928763.4;refGene_gn=USP19;refGene_tx=XM_006928764.3;refGene_gn=USP19;refGene_tx=XM_006928765.4;refGene_gn=USP19;refGene_tx=XM_006928766.4;refGene_gn=USP19;refGene_tx=XM_006928767.2;refGene_gn=USP19;refGene_tx=XM_006928768.2;refGene_gn=USP19;refGene_tx=XM_019821381.2;refGene_gn=USP19;refGene_tx=

XM_023248652.1;refGene_gn=USP19;refGene_tx=XM_023248657.1
A2 18246385 18246514
refGene_gn=USP19;refGene_tx=XM_006928756.3;refGene_gn=USP19;refGene_tx=XM_006928757.4;refGene_gn=USP19;refGene_tx=XM_006928758.3;refGene_gn=USP19;refGene_tx=XM_006928759.4;refGene_gn=USP19;refGene_tx=XM_006928760.4;refGene_gn=USP19;refGene_tx=XM_006928761.4;refGene_gn=USP19;refGene_tx=XM_006928762.4;refGene_gn=USP19;refGene_tx=XM_006928763.4;refGene_gn=USP19;refGene_tx=XM_006928764.3;refGene_gn=USP19;refGene_tx=XM_006928765.4;refGene_gn=USP19;refGene_tx=XM_006928766.4;refGene_gn=USP19;refGene_tx=XM_006928767.2;refGene_gn=USP19;refGene_tx=XM_006928768.2;refGene_gn=USP19;refGene_tx=XM_019821381.2;refGene_gn=USP19;refGene_tx=XM_023248652.1;refGene_gn=USP19;refGene_tx=XM_023248657.1
A2 18246685 18246721
refGene_gn=USP19;refGene_tx=XM_006928756.3;refGene_gn=USP19;refGene_tx=XM_006928757.4;refGene_gn=USP19;refGene_tx=XM_006928758.3;refGene_gn=USP19;refGene_tx=XM_006928759.4;refGene_gn=USP19;refGene_tx=XM_006928760.4;refGene_gn=USP19;refGene_tx=XM_006928761.4;refGene_gn=USP19;refGene_tx=XM_006928762.4;refGene_gn=USP19;refGene_tx=XM_006928763.4;refGene_gn=USP19;refGene_tx=XM_006928764.3;refGene_gn=USP19;refGene_tx=XM_006928765.4;refGene_gn=USP19;refGene_tx=XM_006928766.4;refGene_gn=USP19;refGene_tx=XM_006928767.2;refGene_gn=USP19;refGene_tx=XM_006928768.2;refGene_gn=USP19;refGene_tx=XM_019821381.2;refGene_gn=USP19;refGene_tx=XM_023248652.1;refGene_gn=USP19;refGene_tx=XM_023248657.1
A2 18246975 18247099
refGene_gn=USP19;refGene_tx=XM_006928756.3;refGene_gn=USP19;refGene_tx=XM_006928757.4;refGene_gn=USP19;refGene_tx=XM_006928758.3;refGene_gn=USP19;refGene_tx=XM_006928759.4;refGene_gn=USP19;refGene_tx=XM_006928760.4;refGene_gn=USP19;refGene_tx=XM_006928761.4;refGene_gn=USP19;refGene_tx=XM_006928762.4;refGene_gn=USP19;refGene_tx=XM_006928763.4;refGene_gn=USP19;refGene_tx=XM_006928764.3;refGene_gn=USP19;refGene_tx=XM_006928765.4;refGene_gn=USP19;refGene_tx=XM_006928766.4;refGene_gn=USP19;refGene_tx=XM_006928767.2;refGene_gn=USP19;refGene_tx=XM_006928768.2;refGene_gn=USP19;refGene_tx=XM_019821381.2;refGene_gn=USP19;refGene_tx=XM_023248652.1;refGene_gn=USP19;refGene_tx=XM_023248657.1
A2 18249174 18249311
refGene_gn=LAMB2;refGene_tx=XM_003982216.3;refGene_gn=LAMB2;refGene_tx=XM_019821375.1
A2 18249390 18249550
refGene_gn=LAMB2;refGene_tx=XM_003982216.3;refGene_gn=LAMB2;refGene_tx=XM_019821375.1
A2 18249640 18249817
refGene_gn=LAMB2;refGene_tx=XM_003982216.3;refGene_gn=LAMB2;refGene_tx=XM_019821375.1
A2 18249900 18250042
refGene_gn=LAMB2;refGene_tx=XM_003982216.3;refGene_gn=LAMB2;refGene_tx=XM_019821375.1
A2 18250119 18250327
refGene_gn=LAMB2;refGene_tx=XM_003982216.3;refGene_gn=LAMB2;refGene_tx=XM_019821375.1
A2 18250656 18251005

refGene_gn=LAMB2;refGene_tx=XM_003982216.3;refGene_gn=LAMB2;refGene_tx=XM_019821375.1
A2 18251084 18251326
refGene_gn=LAMB2;refGene_tx=XM_003982216.3;refGene_gn=LAMB2;refGene_tx=XM_019821375.1
A2 18251403 18251588
refGene_gn=LAMB2;refGene_tx=XM_003982216.3;refGene_gn=LAMB2;refGene_tx=XM_019821375.1
A2 18251676 18252049
refGene_gn=LAMB2;refGene_tx=XM_003982216.3;refGene_gn=LAMB2;refGene_tx=XM_019821375.1
A2 18252136 18252233
refGene_gn=LAMB2;refGene_tx=XM_003982216.3;refGene_gn=LAMB2;refGene_tx=XM_019821375.1
A2 18252344 18252562
refGene_gn=LAMB2;refGene_tx=XM_003982216.3;refGene_gn=LAMB2;refGene_tx=XM_019821375.1
A2 18252640 18252865
refGene_gn=LAMB2;refGene_tx=XM_003982216.3;refGene_gn=LAMB2;refGene_tx=XM_019821375.1
A2 18252940 18253104
refGene_gn=LAMB2;refGene_tx=XM_003982216.3;refGene_gn=LAMB2;refGene_tx=XM_019821375.1
A2 18253187 18253419
refGene_gn=LAMB2;refGene_tx=XM_003982216.3;refGene_gn=LAMB2;refGene_tx=XM_019821375.1
A2 18253643 18253787
refGene_gn=LAMB2;refGene_tx=XM_003982216.3;refGene_gn=LAMB2;refGene_tx=XM_019821375.1
A2 18253866 18254059
refGene_gn=LAMB2;refGene_tx=XM_003982216.3;refGene_gn=LAMB2;refGene_tx=XM_019821375.1
A2 18254244 18254377
refGene_gn=LAMB2;refGene_tx=XM_003982216.3;refGene_gn=LAMB2;refGene_tx=XM_019821375.1
A2 18256054 18256182
refGene_gn=LAMB2;refGene_tx=XM_003982216.3;refGene_gn=LAMB2;refGene_tx=XM_019821375.1
A2 18256259 18256418
refGene_gn=LAMB2;refGene_tx=XM_003982216.3;refGene_gn=LAMB2;refGene_tx=XM_019821375.1
A2 18256593 18256726
refGene_gn=LAMB2;refGene_tx=XM_003982216.3;refGene_gn=LAMB2;refGene_tx=XM_019821375.1
A2 18256810 18256890
refGene_gn=LAMB2;refGene_tx=XM_003982216.3;refGene_gn=LAMB2;refGene_tx=XM_019821375.1
A2 18257142 18257255
refGene_gn=LAMB2;refGene_tx=XM_003982216.3;refGene_gn=LAMB2;refGene_tx=XM_019821375.1

A2 18257374 18257554
refGene_gn=LAMB2;refGene_tx=XM_003982216.3;refGene_gn=LAMB2;refGene_tx=XM_019821375.1
A2 18257765 18257954
refGene_gn=LAMB2;refGene_tx=XM_003982216.3;refGene_gn=LAMB2;refGene_tx=XM_019821375.1
A2 18258285 18258406
refGene_gn=LAMB2;refGene_tx=XM_003982216.3;refGene_gn=LAMB2;refGene_tx=XM_019821375.1
A2 18258494 18258697
refGene_gn=LAMB2;refGene_tx=XM_003982216.3;refGene_gn=LAMB2;refGene_tx=XM_019821375.1
A2 18258931 18258995
refGene_gn=LAMB2;refGene_tx=XM_003982216.3;refGene_gn=LAMB2;refGene_tx=XM_019821375.1
A2 18259087 18259276 refGene_gn=LAMB2;refGene_tx=XM_003982216.3
A2 18259642 18259716 refGene_gn=LAMB2;refGene_tx=XM_003982216.3
A2 18259797 18259933
refGene_gn=LAMB2;refGene_tx=XM_003982216.3;refGene_gn=LAMB2;refGene_tx=XM_019821375.1
A2 18260019 18260192 refGene_gn=LAMB2;refGene_tx=XM_003982216.3
A2 18260309 18260394 refGene_gn=LAMB2;refGene_tx=XM_003982216.3
A2 18271980 18273399 refGene_gn=CCDC71;refGene_tx=XM_023248702.1
A2 18281090 18281466
refGene_gn=KLHDC8B;refGene_tx=XM_003982218.4;refGene_gn=KLHDC8B;refGene_tx=XM_019821410.2;refGene_gn=KLHDC8B;refGene_tx=XM_019821415.2
A2 18282539 18282704
refGene_gn=KLHDC8B;refGene_tx=XM_003982218.4;refGene_gn=KLHDC8B;refGene_tx=XM_019821410.2
A2 18283057 18283282
refGene_gn=KLHDC8B;refGene_tx=XM_003982218.4;refGene_gn=KLHDC8B;refGene_tx=XM_019821410.2;refGene_gn=KLHDC8B;refGene_tx=XM_019821415.2
A2 18283380 18283482
refGene_gn=KLHDC8B;refGene_tx=XM_003982218.4;refGene_gn=KLHDC8B;refGene_tx=XM_019821410.2;refGene_gn=KLHDC8B;refGene_tx=XM_019821415.2
A2 18283873 18284070
refGene_gn=KLHDC8B;refGene_tx=XM_003982218.4;refGene_gn=KLHDC8B;refGene_tx=XM_019821410.2;refGene_gn=KLHDC8B;refGene_tx=XM_019821415.2
A2 18285447 18285858
refGene_gn=CA2H3orf84;refGene_tx=XM_003982237.4;refGene_gn=CA2H3orf84;refGene_tx=XM_019821421.2;refGene_gn=CA2H3orf84;refGene_tx=XM_019821428.2;refGene_gn=CA2H3orf84;refGene_tx=XM_019821433.2;refGene_gn=CA2H3orf84;refGene_tx=XM_019821442.2;refGene_gn=CA2H3orf84;refGene_tx=XM_019821450.2;refGene_gn=CA2H3orf84;refGene_tx=XM_019821452.2;refGene_gn=CA2H3orf84;refGene_tx=XM_019821455.2;refGene_gn=CA2H3orf84;refGene_tx=XM_019821473.2;refGene_gn=CA2H3orf84;refGene_tx=XM_019821476.2;refGene_gn=CA2H3orf84;refGene_tx=XM_019821481.2;refGene_gn=CA2H3orf84;refGene_tx=XM_019821483.2;refGene_gn=CA2H3orf84;refGene_tx=XM_023248749.1
A2 18286227 18286284
refGene_gn=CA2H3orf84;refGene_tx=XM_003982237.4;refGene_gn=CA2H3orf84;

refGene_tx=XM_019821421.2;refGene_gn=CA2H3orf84;refGene_tx=XM_01982142
8.2;refGene_gn=CA2H3orf84;refGene_tx=XM_019821433.2;refGene_gn=CA2H3or
f84;refGene_tx=XM_019821442.2;refGene_gn=CA2H3orf84;refGene_tx=XM_0198
21450.2;refGene_gn=CA2H3orf84;refGene_tx=XM_019821452.2;refGene_gn=CA2
H3orf84;refGene_tx=XM_019821455.2;refGene_gn=CA2H3orf84;refGene_tx=XM_
019821473.2;refGene_gn=CA2H3orf84;refGene_tx=XM_019821476.2;refGene_gn
=CA2H3orf84;refGene_tx=XM_019821481.2;refGene_gn=CA2H3orf84;refGene_tx
=XM_019821483.2;refGene_gn=CA2H3orf84;refGene_tx=XM_023248749.1
A2 18292428 18292460
refGene_gn=CA2H3orf84;refGene_tx=XM_019821433.2;refGene_gn=CA2H3orf84;
refGene_tx=XM_019821442.2;refGene_gn=CA2H3orf84;refGene_tx=XM_01982145
5.2;refGene_gn=CA2H3orf84;refGene_tx=XM_019821473.2;refGene_gn=CA2H3or
f84;refGene_tx=XM_023248749.1
A2 18292703 18292745
refGene_gn=CA2H3orf84;refGene_tx=XM_019821433.2;refGene_gn=CA2H3orf84;
refGene_tx=XM_019821442.2;refGene_gn=CA2H3orf84;refGene_tx=XM_01982145
0.2;refGene_gn=CA2H3orf84;refGene_tx=XM_019821452.2;refGene_gn=CA2H3or
f84;refGene_tx=XM_019821483.2;refGene_gn=CA2H3orf84;refGene_tx=XM_0232
48749.1
A2 18293361 18293475
refGene_gn=CA2H3orf84;refGene_tx=XM_003982237.4;refGene_gn=CA2H3orf84;
refGene_tx=XM_019821421.2;refGene_gn=CA2H3orf84;refGene_tx=XM_01982142
8.2;refGene_gn=CA2H3orf84;refGene_tx=XM_019821455.2;refGene_gn=CA2H3or
f84;refGene_tx=XM_019821473.2
A2 18295458 18295485
refGene_gn=CA2H3orf84;refGene_tx=XM_003982237.4;refGene_gn=CA2H3orf84;
refGene_tx=XM_019821421.2;refGene_gn=CA2H3orf84;refGene_tx=XM_01982142
8.2;refGene_gn=CA2H3orf84;refGene_tx=XM_019821476.2;refGene_gn=CA2H3or
f84;refGene_tx=XM_019821481.2
A2 18308524 18308580
refGene_gn=CCDC36;refGene_tx=XM_003982219.4;refGene_gn=CCDC36;refGene_
tx=XM_023248722.1
A2 18312634 18312809
refGene_gn=CCDC36;refGene_tx=XM_003982219.4;refGene_gn=CCDC36;refGene_
tx=XM_023248722.1
A2 18314965 18315129
refGene_gn=CCDC36;refGene_tx=XM_003982219.4;refGene_gn=CCDC36;refGene_
tx=XM_023248722.1
A2 18327530 18327579
refGene_gn=CCDC36;refGene_tx=XM_003982219.4;refGene_gn=CCDC36;refGene_
tx=XM_023248722.1
A2 18327769 18327857
refGene_gn=CCDC36;refGene_tx=XM_003982219.4;refGene_gn=CCDC36;refGene_
tx=XM_023248722.1
A2 18334796 18334900
refGene_gn=CCDC36;refGene_tx=XM_003982219.4;refGene_gn=CCDC36;refGene_
tx=XM_023248722.1
A2 18335362 18336502
refGene_gn=CCDC36;refGene_tx=XM_003982219.4;refGene_gn=CCDC36;refGene_
tx=XM_023248722.1

A2 18342663 18342719
refGene_gn=CA2H3orf62;refGene_tx=XM_023248782.1
A2 18344623 18344889
refGene_gn=CA2H3orf62;refGene_tx=XM_003982220.5
A2 18346018 18346110
refGene_gn=CA2H3orf62;refGene_tx=XM_003982220.5;refGene_gn=CA2H3orf62;
refGene_tx=XM_023248782.1
A2 18347047 18347781
refGene_gn=CA2H3orf62;refGene_tx=XM_003982220.5;refGene_gn=CA2H3orf62;
refGene_tx=XM_023248782.1
A2 18348940 18349102
refGene_gn=USP4;refGene_tx=XM_003982221.5;refGene_gn=USP4;refGene_tx=X
M_003982222.5
A2 18349463 18349552
refGene_gn=USP4;refGene_tx=XM_003982221.5;refGene_gn=USP4;refGene_tx=X
M_003982222.5
A2 18352483 18352587
refGene_gn=USP4;refGene_tx=XM_003982221.5;refGene_gn=USP4;refGene_tx=X
M_003982222.5
A2 18353625 18353775
refGene_gn=USP4;refGene_tx=XM_003982221.5;refGene_gn=USP4;refGene_tx=X
M_003982222.5
A2 18354164 18354283
refGene_gn=USP4;refGene_tx=XM_003982221.5;refGene_gn=USP4;refGene_tx=X
M_003982222.5
A2 18354545 18354616
refGene_gn=USP4;refGene_tx=XM_003982221.5;refGene_gn=USP4;refGene_tx=X
M_003982222.5
A2 18356115 18356343
refGene_gn=USP4;refGene_tx=XM_003982221.5;refGene_gn=USP4;refGene_tx=X
M_003982222.5
A2 18361141 18361230
refGene_gn=USP4;refGene_tx=XM_003982221.5;refGene_gn=USP4;refGene_tx=X
M_003982222.5
A2 18362089 18362281
refGene_gn=USP4;refGene_tx=XM_003982221.5;refGene_gn=USP4;refGene_tx=X
M_003982222.5
A2 18362917 18362987 refGene_gn=USP4;refGene_tx=XM_019821522.2
A2 18364529 18364624
refGene_gn=USP4;refGene_tx=XM_003982221.5;refGene_gn=USP4;refGene_tx=X
M_003982222.5;refGene_gn=USP4;refGene_tx=XM_019821522.2
A2 18365539 18365623
refGene_gn=USP4;refGene_tx=XM_003982221.5;refGene_gn=USP4;refGene_tx=X
M_003982222.5;refGene_gn=USP4;refGene_tx=XM_019821522.2
A2 18366356 18366581
refGene_gn=USP4;refGene_tx=XM_003982221.5;refGene_gn=USP4;refGene_tx=X
M_003982222.5;refGene_gn=USP4;refGene_tx=XM_019821522.2
A2 18367450 18367609
refGene_gn=USP4;refGene_tx=XM_003982221.5;refGene_gn=USP4;refGene_tx=X
M_003982222.5;refGene_gn=USP4;refGene_tx=XM_019821522.2

A2 18371171 18371345
refGene_gn=USP4;refGene_tx=XM_003982221.5;refGene_gn=USP4;refGene_tx=XM_003982222.5;refGene_gn=USP4;refGene_tx=XM_019821522.2
A2 18375162 18375280
refGene_gn=USP4;refGene_tx=XM_003982221.5;refGene_gn=USP4;refGene_tx=XM_003982222.5;refGene_gn=USP4;refGene_tx=XM_019821522.2
A2 18376108 18376249
refGene_gn=USP4;refGene_tx=XM_003982221.5;refGene_gn=USP4;refGene_tx=XM_019821522.2
A2 18386218 18386280
refGene_gn=USP4;refGene_tx=XM_003982221.5;refGene_gn=USP4;refGene_tx=XM_003982222.5;refGene_gn=USP4;refGene_tx=XM_019821522.2
A2 18386404 18386550
refGene_gn=USP4;refGene_tx=XM_003982221.5;refGene_gn=USP4;refGene_tx=XM_003982222.5;refGene_gn=USP4;refGene_tx=XM_019821522.2
A2 18387162 18387289
refGene_gn=USP4;refGene_tx=XM_003982221.5;refGene_gn=USP4;refGene_tx=XM_003982222.5;refGene_gn=USP4;refGene_tx=XM_019821522.2
A2 18388427 18388558
refGene_gn=USP4;refGene_tx=XM_003982221.5;refGene_gn=USP4;refGene_tx=XM_003982222.5;refGene_gn=USP4;refGene_tx=XM_019821522.2
A2 18394151 18394279
refGene_gn=USP4;refGene_tx=XM_003982221.5;refGene_gn=USP4;refGene_tx=XM_003982222.5;refGene_gn=USP4;refGene_tx=XM_019821522.2
A2 18397335 18397436
refGene_gn=USP4;refGene_tx=XM_003982221.5;refGene_gn=USP4;refGene_tx=XM_003982222.5;refGene_gn=USP4;refGene_tx=XM_019821522.2
A2 18408111 18408471 refGene_gn=GPX1;refGene_tx=XM_004001361.5
A2 18408707 18408968 refGene_gn=GPX1;refGene_tx=XM_004001361.5
A2 18411126 18411300
refGene_gn=RHOA;refGene_tx=XM_023248787.1;refGene_gn=RHOA;refGene_tx=XM_023248788.1
A2 18412240 18412371
refGene_gn=RHOA;refGene_tx=XM_023248787.1;refGene_gn=RHOA;refGene_tx=XM_023248788.1
A2 18418024 18418145
refGene_gn=RHOA;refGene_tx=XM_023248787.1;refGene_gn=RHOA;refGene_tx=XM_023248788.1
A2 18421478 18421634
refGene_gn=RHOA;refGene_tx=XM_023248787.1;refGene_gn=RHOA;refGene_tx=XM_023248788.1
A2 18474422 18474645
refGene_gn=TCTA;refGene_tx=XM_019821557.2;refGene_gn=TCTA;refGene_tx=XM_019821566.2;refGene_gn=TCTA;refGene_tx=XM_019821570.2
A2 18474925 18475028
refGene_gn=TCTA;refGene_tx=XM_019821557.2;refGene_gn=TCTA;refGene_tx=XM_019821566.2;refGene_gn=TCTA;refGene_tx=XM_019821570.2
A2 18476156 18476199
refGene_gn=TCTA;refGene_tx=XM_019821557.2;refGene_gn=TCTA;refGene_tx=XM_019821566.2;refGene_gn=TCTA;refGene_tx=XM_019821570.2

A2 18478831 18479010 refGene_gn=AMT;refGene_tx=XM_003982225.5
A2 18479109 18479265 refGene_gn=AMT;refGene_tx=XM_003982225.5
A2 18479572 18479753 refGene_gn=AMT;refGene_tx=XM_003982225.5
A2 18479837 18479983 refGene_gn=AMT;refGene_tx=XM_003982225.5
A2 18480288 18480367 refGene_gn=AMT;refGene_tx=XM_003982225.5
A2 18480765 18480897 refGene_gn=AMT;refGene_tx=XM_003982225.5
A2 18481770 18481851 refGene_gn=AMT;refGene_tx=XM_003982225.5
A2 18482382 18482550 refGene_gn=AMT;refGene_tx=XM_003982225.5
A2 18482651 18482741 refGene_gn=AMT;refGene_tx=XM_003982225.5
A2 18484611 18484653
refGene_gn=NICN1;refGene_tx=XM_003982228.5;refGene_gn=NICN1;refGene_tx
=XM_023248802.1
A2 18484726 18484831
refGene_gn=NICN1;refGene_tx=XM_003982228.5;refGene_gn=NICN1;refGene_tx
=XM_023248802.1
A2 18485171 18485243
refGene_gn=NICN1;refGene_tx=XM_003982228.5;refGene_gn=NICN1;refGene_tx
=XM_023248802.1
A2 18485693 18485807
refGene_gn=NICN1;refGene_tx=XM_003982228.5;refGene_gn=NICN1;refGene_tx
=XM_023248802.1
A2 18486062 18486239
refGene_gn=NICN1;refGene_tx=XM_003982228.5;refGene_gn=NICN1;refGene_tx
=XM_023248802.1
A2 18487418 18487550
refGene_gn=NICN1;refGene_tx=XM_003982228.5;refGene_gn=NICN1;refGene_tx
=XM_023248802.1
A2 18564526 18564802
refGene_gn=DAG1;refGene_tx=NM_001130839.1;refGene_gn=DAG1;refGene_tx=X
M_006928746.4;refGene_gn=DAG1;refGene_tx=XM_023240650.1;refGene_gn=DAG
1;refGene_tx=XM_023240658.1;refGene_gn=DAG1;refGene_tx=XM_023240664.1
A2 18580142 18582545
refGene_gn=DAG1;refGene_tx=NM_001130839.1;refGene_gn=DAG1;refGene_tx=X
M_006928746.4;refGene_gn=DAG1;refGene_tx=XM_023240650.1;refGene_gn=DAG
1;refGene_tx=XM_023240658.1;refGene_gn=DAG1;refGene_tx=XM_023240664.1
A2 18594216 18594464 refGene_gn=BSN;refGene_tx=XM_003982229.5
A2 18648435 18648844 refGene_gn=BSN;refGene_tx=XM_003982229.5
A2 18660252 18661137 refGene_gn=BSN;refGene_tx=XM_003982229.5
A2 18668360 18668807 refGene_gn=BSN;refGene_tx=XM_003982229.5
A2 18669208 18675856 refGene_gn=BSN;refGene_tx=XM_003982229.5
A2 18678140 18680217 refGene_gn=BSN;refGene_tx=XM_003982229.5
A2 18680505 18681302 refGene_gn=BSN;refGene_tx=XM_003982229.5
A2 18681435 18681535 refGene_gn=BSN;refGene_tx=XM_003982229.5
A2 18682066 18682198 refGene_gn=BSN;refGene_tx=XM_003982229.5
A2 18682454 18682495 refGene_gn=BSN;refGene_tx=XM_003982229.5
A2 18691356 18691368 refGene_gn=APEH;refGene_tx=XM_003982230.5
A2 18691443 18691576 refGene_gn=APEH;refGene_tx=XM_003982230.5
A2 18692144 18692271 refGene_gn=APEH;refGene_tx=XM_003982230.5
A2 18692677 18692771 refGene_gn=APEH;refGene_tx=XM_003982230.5
A2 18692877 18692953 refGene_gn=APEH;refGene_tx=XM_003982230.5

A2 18693042 18693206 refGene_gn=APEH;refGene_tx=XM_003982230.5
A2 18693375 18693513 refGene_gn=APEH;refGene_tx=XM_003982230.5
A2 18693610 18693702 refGene_gn=APEH;refGene_tx=XM_003982230.5
A2 18693779 18693820 refGene_gn=APEH;refGene_tx=XM_003982230.5
A2 18693904 18694026 refGene_gn=APEH;refGene_tx=XM_003982230.5
A2 18694561 18694622 refGene_gn=APEH;refGene_tx=XM_003982230.5
A2 18695816 18695914 refGene_gn=APEH;refGene_tx=XM_003982230.5
A2 18696547 18696599 refGene_gn=APEH;refGene_tx=XM_003982230.5
A2 18697426 18697515 refGene_gn=APEH;refGene_tx=XM_003982230.5
A2 18697966 18698105 refGene_gn=APEH;refGene_tx=XM_003982230.5
A2 18698596 18698680 refGene_gn=APEH;refGene_tx=XM_003982230.5
A2 18698760 18698841 refGene_gn=APEH;refGene_tx=XM_003982230.5
A2 18699184 18699273 refGene_gn=APEH;refGene_tx=XM_003982230.5
A2 18699362 18699553 refGene_gn=APEH;refGene_tx=XM_003982230.5
A2 18699661 18699764 refGene_gn=APEH;refGene_tx=XM_003982230.5
A2 18699849 18699956 refGene_gn=APEH;refGene_tx=XM_003982230.5
A2 18700045 18700151 refGene_gn=APEH;refGene_tx=XM_003982230.5
A2 18700794 18700956
refGene_gn=MST1;refGene_tx=XM_003982231.5;refGene_gn=MST1;refGene_tx=XM_011291572.3;refGene_gn=MST1;refGene_tx=XM_019821637.2;refGene_gn=MST1;refGene_tx=XM_019821640.2
A2 18701083 18701223
refGene_gn=MST1;refGene_tx=XM_003982231.5;refGene_gn=MST1;refGene_tx=XM_011291572.3;refGene_gn=MST1;refGene_tx=XM_019821637.2;refGene_gn=MST1;refGene_tx=XM_019821640.2
A2 18701325 18701432
refGene_gn=MST1;refGene_tx=XM_003982231.5;refGene_gn=MST1;refGene_tx=XM_019821637.2;refGene_gn=MST1;refGene_tx=XM_019821640.2
A2 18701513 18701660
refGene_gn=MST1;refGene_tx=XM_003982231.5;refGene_gn=MST1;refGene_tx=XM_019821637.2;refGene_gn=MST1;refGene_tx=XM_019821640.2
A2 18701792 18701870
refGene_gn=MST1;refGene_tx=XM_003982231.5;refGene_gn=MST1;refGene_tx=XM_011291572.3;refGene_gn=MST1;refGene_tx=XM_019821637.2;refGene_gn=MST1;refGene_tx=XM_019821640.2
A2 18702036 18702157
refGene_gn=MST1;refGene_tx=XM_003982231.5;refGene_gn=MST1;refGene_tx=XM_011291572.3;refGene_gn=MST1;refGene_tx=XM_019821637.2;refGene_gn=MST1;refGene_tx=XM_019821640.2
A2 18702243 18702279
refGene_gn=MST1;refGene_tx=XM_003982231.5;refGene_gn=MST1;refGene_tx=XM_011291572.3;refGene_gn=MST1;refGene_tx=XM_019821637.2;refGene_gn=MST1;refGene_tx=XM_019821640.2
A2 18702364 18702501
refGene_gn=MST1;refGene_tx=XM_003982231.5;refGene_gn=MST1;refGene_tx=XM_011291572.3;refGene_gn=MST1;refGene_tx=XM_019821637.2;refGene_gn=MST1;refGene_tx=XM_019821640.2
A2 18702627 18702749
refGene_gn=MST1;refGene_tx=XM_003982231.5;refGene_gn=MST1;refGene_tx=XM_011291572.3;refGene_gn=MST1;refGene_tx=XM_019821637.2;refGene_gn=MST

1;refGene_tx=XM_019821640.2
A2 18702812 18702943
refGene_gn=MST1;refGene_tx=XM_003982231.5;refGene_gn=MST1;refGene_tx=XM_011291572.3;refGene_gn=MST1;refGene_tx=XM_019821637.2;refGene_gn=MST1;refGene_tx=XM_019821640.2
A2 18703114 18703283
refGene_gn=MST1;refGene_tx=XM_003982231.5;refGene_gn=MST1;refGene_tx=XM_011291572.3;refGene_gn=MST1;refGene_tx=XM_019821637.2
A2 18703463 18703582
refGene_gn=MST1;refGene_tx=XM_003982231.5;refGene_gn=MST1;refGene_tx=XM_011291572.3;refGene_gn=MST1;refGene_tx=XM_019821637.2;refGene_gn=MST1;refGene_tx=XM_019821640.2
A2 18703721 18703842
refGene_gn=MST1;refGene_tx=XM_003982231.5;refGene_gn=MST1;refGene_tx=XM_011291572.3;refGene_gn=MST1;refGene_tx=XM_019821637.2;refGene_gn=MST1;refGene_tx=XM_019821640.2
A2 18703920 18704057
refGene_gn=MST1;refGene_tx=XM_003982231.5;refGene_gn=MST1;refGene_tx=XM_011291572.3;refGene_gn=MST1;refGene_tx=XM_019821637.2;refGene_gn=MST1;refGene_tx=XM_019821640.2
A2 18704138 18704253
refGene_gn=MST1;refGene_tx=XM_003982231.5;refGene_gn=MST1;refGene_tx=XM_011291572.3;refGene_gn=MST1;refGene_tx=XM_019821637.2;refGene_gn=MST1;refGene_tx=XM_019821640.2
A2 18704336 18704449
refGene_gn=MST1;refGene_tx=XM_003982231.5;refGene_gn=MST1;refGene_tx=XM_011291572.3;refGene_gn=MST1;refGene_tx=XM_019821637.2;refGene_gn=MST1;refGene_tx=XM_019821640.2
A2 18704521 18704669
refGene_gn=MST1;refGene_tx=XM_003982231.5;refGene_gn=MST1;refGene_tx=XM_011291572.3;refGene_gn=MST1;refGene_tx=XM_019821637.2;refGene_gn=MST1;refGene_tx=XM_019821640.2
A2 18705303 18705397
refGene_gn=MST1;refGene_tx=XM_003982231.5;refGene_gn=MST1;refGene_tx=XM_011291572.3;refGene_gn=MST1;refGene_tx=XM_019821637.2;refGene_gn=MST1;refGene_tx=XM_019821640.2
A2 18707813 18707895
refGene_gn=RNF123;refGene_tx=XM_003982232.3;refGene_gn=RNF123;refGene_tx=XM_006928771.4;refGene_gn=RNF123;refGene_tx=XM_006928772.2;refGene_gn=RNF123;refGene_tx=XM_019821601.1;refGene_gn=RNF123;refGene_tx=XM_019821605.2;refGene_gn=RNF123;refGene_tx=XM_019821607.2;refGene_gn=RNF123;refGene_tx=XM_019821616.1
A2 18708176 18708261
refGene_gn=RNF123;refGene_tx=XM_003982232.3;refGene_gn=RNF123;refGene_tx=XM_006928771.4;refGene_gn=RNF123;refGene_tx=XM_006928772.2;refGene_gn=RNF123;refGene_tx=XM_019821601.1;refGene_gn=RNF123;refGene_tx=XM_019821605.2;refGene_gn=RNF123;refGene_tx=XM_019821607.2;refGene_gn=RNF123;refGene_tx=XM_019821616.1
A2 18710876 18710956
refGene_gn=RNF123;refGene_tx=XM_003982232.3;refGene_gn=RNF123;refGene_

tx=XM_006928771.4;refGene_gn=RNF123;refGene_tx=XM_006928772.2;refGene_gn=RNF123;refGene_tx=XM_019821601.1;refGene_gn=RNF123;refGene_tx=XM_019821605.2;refGene_gn=RNF123;refGene_tx=XM_019821607.2;refGene_gn=RNF123;refGene_tx=XM_019821616.1
A2 18711101 18711196
refGene_gn=RNF123;refGene_tx=XM_003982232.3;refGene_gn=RNF123;refGene_tx=XM_006928771.4;refGene_gn=RNF123;refGene_tx=XM_006928772.2;refGene_gn=RNF123;refGene_tx=XM_019821601.1;refGene_gn=RNF123;refGene_tx=XM_019821605.2;refGene_gn=RNF123;refGene_tx=XM_019821607.2;refGene_gn=RNF123;refGene_tx=XM_019821616.1
A2 18711585 18711640
refGene_gn=RNF123;refGene_tx=XM_003982232.3;refGene_gn=RNF123;refGene_tx=XM_006928771.4;refGene_gn=RNF123;refGene_tx=XM_006928772.2;refGene_gn=RNF123;refGene_tx=XM_019821601.1;refGene_gn=RNF123;refGene_tx=XM_019821605.2;refGene_gn=RNF123;refGene_tx=XM_019821607.2;refGene_gn=RNF123;refGene_tx=XM_019821616.1
A2 18711761 18711847
refGene_gn=RNF123;refGene_tx=XM_003982232.3;refGene_gn=RNF123;refGene_tx=XM_006928771.4;refGene_gn=RNF123;refGene_tx=XM_006928772.2;refGene_gn=RNF123;refGene_tx=XM_019821601.1;refGene_gn=RNF123;refGene_tx=XM_019821605.2;refGene_gn=RNF123;refGene_tx=XM_019821607.2;refGene_gn=RNF123;refGene_tx=XM_019821616.1
A2 18712118 18712205
refGene_gn=RNF123;refGene_tx=XM_003982232.3;refGene_gn=RNF123;refGene_tx=XM_006928771.4;refGene_gn=RNF123;refGene_tx=XM_006928772.2;refGene_gn=RNF123;refGene_tx=XM_019821601.1;refGene_gn=RNF123;refGene_tx=XM_019821605.2;refGene_gn=RNF123;refGene_tx=XM_019821607.2;refGene_gn=RNF123;refGene_tx=XM_019821616.1
A2 18712432 18712500
refGene_gn=RNF123;refGene_tx=XM_003982232.3;refGene_gn=RNF123;refGene_tx=XM_006928771.4;refGene_gn=RNF123;refGene_tx=XM_006928772.2;refGene_gn=RNF123;refGene_tx=XM_019821601.1;refGene_gn=RNF123;refGene_tx=XM_019821605.2;refGene_gn=RNF123;refGene_tx=XM_019821607.2;refGene_gn=RNF123;refGene_tx=XM_019821616.1
A2 18712696 18712822
refGene_gn=RNF123;refGene_tx=XM_003982232.3;refGene_gn=RNF123;refGene_tx=XM_006928771.4;refGene_gn=RNF123;refGene_tx=XM_006928772.2;refGene_gn=RNF123;refGene_tx=XM_019821601.1;refGene_gn=RNF123;refGene_tx=XM_019821605.2;refGene_gn=RNF123;refGene_tx=XM_019821607.2;refGene_gn=RNF123;refGene_tx=XM_019821616.1
A2 18713204 18713319
refGene_gn=RNF123;refGene_tx=XM_003982232.3;refGene_gn=RNF123;refGene_tx=XM_006928771.4;refGene_gn=RNF123;refGene_tx=XM_006928772.2;refGene_gn=RNF123;refGene_tx=XM_019821601.1;refGene_gn=RNF123;refGene_tx=XM_019821605.2;refGene_gn=RNF123;refGene_tx=XM_019821607.2;refGene_gn=RNF123;refGene_tx=XM_019821616.1
A2 18713397 18713502
refGene_gn=RNF123;refGene_tx=XM_003982232.3;refGene_gn=RNF123;refGene_tx=XM_006928771.4;refGene_gn=RNF123;refGene_tx=XM_006928772.2;refGene_gn=RNF123;refGene_tx=XM_019821601.1;refGene_gn=RNF123;refGene_tx=XM_01

9821605.2;refGene_gn=RNF123;refGene_tx=XM_019821607.2;refGene_gn=RNF123;refGene_tx=XM_019821616.1
A2 18713920 18714046
refGene_gn=RNF123;refGene_tx=XM_003982232.3;refGene_gn=RNF123;refGene_tx=XM_006928771.4;refGene_gn=RNF123;refGene_tx=XM_006928772.2;refGene_gn=RNF123;refGene_tx=XM_019821601.1;refGene_gn=RNF123;refGene_tx=XM_019821605.2;refGene_gn=RNF123;refGene_tx=XM_019821607.2;refGene_gn=RNF123;refGene_tx=XM_019821616.1
A2 18714197 18714290
refGene_gn=RNF123;refGene_tx=XM_003982232.3;refGene_gn=RNF123;refGene_tx=XM_006928771.4;refGene_gn=RNF123;refGene_tx=XM_006928772.2;refGene_gn=RNF123;refGene_tx=XM_019821601.1;refGene_gn=RNF123;refGene_tx=XM_019821605.2;refGene_gn=RNF123;refGene_tx=XM_019821607.2;refGene_gn=RNF123;refGene_tx=XM_019821616.1
A2 18714362 18714436
refGene_gn=RNF123;refGene_tx=XM_003982232.3;refGene_gn=RNF123;refGene_tx=XM_006928771.4;refGene_gn=RNF123;refGene_tx=XM_006928772.2;refGene_gn=RNF123;refGene_tx=XM_019821601.1;refGene_gn=RNF123;refGene_tx=XM_019821605.2;refGene_gn=RNF123;refGene_tx=XM_019821607.2;refGene_gn=RNF123;refGene_tx=XM_019821616.1
A2 18715234 18715352
refGene_gn=RNF123;refGene_tx=XM_003982232.3;refGene_gn=RNF123;refGene_tx=XM_006928771.4;refGene_gn=RNF123;refGene_tx=XM_006928772.2;refGene_gn=RNF123;refGene_tx=XM_019821601.1;refGene_gn=RNF123;refGene_tx=XM_019821605.2;refGene_gn=RNF123;refGene_tx=XM_019821607.2;refGene_gn=RNF123;refGene_tx=XM_019821616.1
A2 18715545 18715645
refGene_gn=RNF123;refGene_tx=XM_003982232.3;refGene_gn=RNF123;refGene_tx=XM_006928771.4;refGene_gn=RNF123;refGene_tx=XM_006928772.2;refGene_gn=RNF123;refGene_tx=XM_019821601.1;refGene_gn=RNF123;refGene_tx=XM_019821605.2;refGene_gn=RNF123;refGene_tx=XM_019821607.2;refGene_gn=RNF123;refGene_tx=XM_019821616.1
A2 18715888 18715950
refGene_gn=RNF123;refGene_tx=XM_003982232.3;refGene_gn=RNF123;refGene_tx=XM_006928771.4;refGene_gn=RNF123;refGene_tx=XM_006928772.2;refGene_gn=RNF123;refGene_tx=XM_019821601.1;refGene_gn=RNF123;refGene_tx=XM_019821605.2;refGene_gn=RNF123;refGene_tx=XM_019821607.2;refGene_gn=RNF123;refGene_tx=XM_019821616.1
A2 18716133 18716205
refGene_gn=RNF123;refGene_tx=XM_003982232.3;refGene_gn=RNF123;refGene_tx=XM_006928771.4;refGene_gn=RNF123;refGene_tx=XM_006928772.2;refGene_gn=RNF123;refGene_tx=XM_019821601.1;refGene_gn=RNF123;refGene_tx=XM_019821605.2;refGene_gn=RNF123;refGene_tx=XM_019821607.2;refGene_gn=RNF123;refGene_tx=XM_019821616.1
A2 18716423 18716544
refGene_gn=RNF123;refGene_tx=XM_003982232.3;refGene_gn=RNF123;refGene_tx=XM_006928771.4;refGene_gn=RNF123;refGene_tx=XM_006928772.2;refGene_gn=RNF123;refGene_tx=XM_019821601.1;refGene_gn=RNF123;refGene_tx=XM_019821605.2;refGene_gn=RNF123;refGene_tx=XM_019821607.2;refGene_gn=RNF123;refGene_tx=XM_019821616.1

A2 18717175 18717277
refGene_gn=RNF123;refGene_tx=XM_003982232.3;refGene_gn=RNF123;refGene_tx=XM_006928771.4;refGene_gn=RNF123;refGene_tx=XM_006928772.2;refGene_gn=RNF123;refGene_tx=XM_019821601.1;refGene_gn=RNF123;refGene_tx=XM_019821605.2;refGene_gn=RNF123;refGene_tx=XM_019821607.2;refGene_gn=RNF123;refGene_tx=XM_019821616.1
A2 18718496 18718611
refGene_gn=RNF123;refGene_tx=XM_003982232.3;refGene_gn=RNF123;refGene_tx=XM_006928771.4;refGene_gn=RNF123;refGene_tx=XM_006928772.2;refGene_gn=RNF123;refGene_tx=XM_019821601.1;refGene_gn=RNF123;refGene_tx=XM_019821605.2;refGene_gn=RNF123;refGene_tx=XM_019821607.2;refGene_gn=RNF123;refGene_tx=XM_019821616.1
A2 18718847 18719046
refGene_gn=RNF123;refGene_tx=XM_003982232.3;refGene_gn=RNF123;refGene_tx=XM_006928771.4;refGene_gn=RNF123;refGene_tx=XM_006928772.2;refGene_gn=RNF123;refGene_tx=XM_019821601.1;refGene_gn=RNF123;refGene_tx=XM_019821605.2;refGene_gn=RNF123;refGene_tx=XM_019821607.2;refGene_gn=RNF123;refGene_tx=XM_019821616.1
A2 18719382 18719528
refGene_gn=RNF123;refGene_tx=XM_003982232.3;refGene_gn=RNF123;refGene_tx=XM_006928771.4;refGene_gn=RNF123;refGene_tx=XM_006928772.2;refGene_gn=RNF123;refGene_tx=XM_019821601.1;refGene_gn=RNF123;refGene_tx=XM_019821605.2;refGene_gn=RNF123;refGene_tx=XM_019821607.2;refGene_gn=RNF123;refGene_tx=XM_019821616.1
A2 18719818 18719902
refGene_gn=RNF123;refGene_tx=XM_003982232.3;refGene_gn=RNF123;refGene_tx=XM_006928771.4;refGene_gn=RNF123;refGene_tx=XM_006928772.2;refGene_gn=RNF123;refGene_tx=XM_019821601.1;refGene_gn=RNF123;refGene_tx=XM_019821605.2;refGene_gn=RNF123;refGene_tx=XM_019821607.2;refGene_gn=RNF123;refGene_tx=XM_019821616.1
A2 18720692 18720800
refGene_gn=RNF123;refGene_tx=XM_003982232.3;refGene_gn=RNF123;refGene_tx=XM_006928771.4;refGene_gn=RNF123;refGene_tx=XM_006928772.2;refGene_gn=RNF123;refGene_tx=XM_019821601.1;refGene_gn=RNF123;refGene_tx=XM_019821605.2;refGene_gn=RNF123;refGene_tx=XM_019821607.2;refGene_gn=RNF123;refGene_tx=XM_019821616.1
A2 18728970 18729148
refGene_gn=RNF123;refGene_tx=XM_003982232.3;refGene_gn=RNF123;refGene_tx=XM_006928771.4;refGene_gn=RNF123;refGene_tx=XM_006928772.2;refGene_gn=RNF123;refGene_tx=XM_019821601.1;refGene_gn=RNF123;refGene_tx=XM_019821605.2;refGene_gn=RNF123;refGene_tx=XM_019821607.2;refGene_gn=RNF123;refGene_tx=XM_019821616.1
A2 18730019 18730094
refGene_gn=RNF123;refGene_tx=XM_003982232.3;refGene_gn=RNF123;refGene_tx=XM_006928771.4;refGene_gn=RNF123;refGene_tx=XM_006928772.2;refGene_gn=RNF123;refGene_tx=XM_019821601.1;refGene_gn=RNF123;refGene_tx=XM_019821605.2;refGene_gn=RNF123;refGene_tx=XM_019821607.2;refGene_gn=RNF123;refGene_tx=XM_019821616.1
A2 18730243 18730331
refGene_gn=RNF123;refGene_tx=XM_003982232.3;refGene_gn=RNF123;refGene_

tx=XM_006928771.4;refGene_gn=RNF123;refGene_tx=XM_006928772.2;refGene_gn=RNF123;refGene_tx=XM_019821601.1;refGene_gn=RNF123;refGene_tx=XM_019821605.2;refGene_gn=RNF123;refGene_tx=XM_019821607.2;refGene_gn=RNF123;refGene_tx=XM_019821616.1
A2 18730424 18730512 refGene_gn=RNF123;refGene_tx=XM_003982232.3;refGene_gn=RNF123;refGene_tx=XM_006928771.4;refGene_gn=RNF123;refGene_tx=XM_006928772.2;refGene_gn=RNF123;refGene_tx=XM_019821601.1;refGene_gn=RNF123;refGene_tx=XM_019821605.2;refGene_gn=RNF123;refGene_tx=XM_019821607.2;refGene_gn=RNF123;refGene_tx=XM_019821616.1
A2 18730599 18730684 refGene_gn=RNF123;refGene_tx=XM_003982232.3;refGene_gn=RNF123;refGene_tx=XM_006928771.4;refGene_gn=RNF123;refGene_tx=XM_006928772.2;refGene_gn=RNF123;refGene_tx=XM_019821601.1;refGene_gn=RNF123;refGene_tx=XM_019821605.2;refGene_gn=RNF123;refGene_tx=XM_019821607.2;refGene_gn=RNF123;refGene_tx=XM_019821616.1
A2 18732013 18732153 refGene_gn=RNF123;refGene_tx=XM_003982232.3;refGene_gn=RNF123;refGene_tx=XM_006928771.4;refGene_gn=RNF123;refGene_tx=XM_006928772.2;refGene_gn=RNF123;refGene_tx=XM_019821601.1;refGene_gn=RNF123;refGene_tx=XM_019821605.2;refGene_gn=RNF123;refGene_tx=XM_019821607.2;refGene_gn=RNF123;refGene_tx=XM_019821616.1
A2 18732252 18732441 refGene_gn=RNF123;refGene_tx=XM_003982232.3;refGene_gn=RNF123;refGene_tx=XM_006928771.4;refGene_gn=RNF123;refGene_tx=XM_006928772.2;refGene_gn=RNF123;refGene_tx=XM_019821601.1;refGene_gn=RNF123;refGene_tx=XM_019821605.2;refGene_gn=RNF123;refGene_tx=XM_019821607.2;refGene_gn=RNF123;refGene_tx=XM_019821616.1
A2 18732533 18732609 refGene_gn=RNF123;refGene_tx=XM_003982232.3;refGene_gn=RNF123;refGene_tx=XM_006928771.4;refGene_gn=RNF123;refGene_tx=XM_006928772.2;refGene_gn=RNF123;refGene_tx=XM_019821601.1;refGene_gn=RNF123;refGene_tx=XM_019821605.2;refGene_gn=RNF123;refGene_tx=XM_019821607.2;refGene_gn=RNF123;refGene_tx=XM_019821616.1
A2 18732820 18732905 refGene_gn=RNF123;refGene_tx=XM_003982232.3;refGene_gn=RNF123;refGene_tx=XM_006928771.4;refGene_gn=RNF123;refGene_tx=XM_006928772.2;refGene_gn=RNF123;refGene_tx=XM_019821601.1;refGene_gn=RNF123;refGene_tx=XM_019821605.2;refGene_gn=RNF123;refGene_tx=XM_019821607.2;refGene_gn=RNF123;refGene_tx=XM_019821616.1
A2 18734248 18735772 refGene_gn=AMIGO3;refGene_tx=XM_006928770.3
A2 18736483 18736626 refGene_gn=RNF123;refGene_tx=XM_003982232.3;refGene_gn=RNF123;refGene_tx=XM_006928771.4;refGene_gn=RNF123;refGene_tx=XM_006928772.2;refGene_gn=RNF123;refGene_tx=XM_019821601.1;refGene_gn=RNF123;refGene_tx=XM_019821605.2;refGene_gn=RNF123;refGene_tx=XM_019821607.2;refGene_gn=RNF123;refGene_tx=XM_019821616.1
A2 18736738 18736755 refGene_gn=RNF123;refGene_tx=XM_019821616.1
A2 18736775 18736870 refGene_gn=RNF123;refGene_tx=XM_003982232.3;refGene_gn=RNF123;refGene_

tx=XM_006928771.4;refGene_gn=RNF123;refGene_tx=XM_006928772.2;refGene_gn=RNF123;refGene_tx=XM_019821601.1;refGene_gn=RNF123;refGene_tx=XM_019821605.2;refGene_gn=RNF123;refGene_tx=XM_019821607.2
A2 18737005 18737091
refGene_gn=RNF123;refGene_tx=XM_003982232.3;refGene_gn=RNF123;refGene_tx=XM_006928771.4;refGene_gn=RNF123;refGene_tx=XM_006928772.2;refGene_gn=RNF123;refGene_tx=XM_019821601.1;refGene_gn=RNF123;refGene_tx=XM_019821605.2;refGene_gn=RNF123;refGene_tx=XM_019821607.2
A2 18737167 18737288
refGene_gn=RNF123;refGene_tx=XM_003982232.3;refGene_gn=RNF123;refGene_tx=XM_006928771.4;refGene_gn=RNF123;refGene_tx=XM_006928772.2;refGene_gn=RNF123;refGene_tx=XM_019821601.1;refGene_gn=RNF123;refGene_tx=XM_019821605.2;refGene_gn=RNF123;refGene_tx=XM_019821607.2
A2 18737723 18737855 refGene_gn=GMPPB;refGene_tx=XM_003982233.5
A2 18737937 18738120 refGene_gn=GMPPB;refGene_tx=XM_003982233.5
A2 18738249 18738377 refGene_gn=GMPPB;refGene_tx=XM_003982233.5
A2 18738451 18738530 refGene_gn=GMPPB;refGene_tx=XM_003982233.5
A2 18738616 18738775 refGene_gn=GMPPB;refGene_tx=XM_003982233.5
A2 18738989 18739132 refGene_gn=GMPPB;refGene_tx=XM_003982233.5
A2 18739267 18739316 refGene_gn=GMPPB;refGene_tx=XM_003982233.5
A2 18739407 18739488 refGene_gn=GMPPB;refGene_tx=XM_003982233.5
A2 18739602 18739731 refGene_gn=GMPPB;refGene_tx=XM_003982233.5
A2 18743259 18743793 refGene_gn=IP6K1;refGene_tx=XM_011291603.3
A2 18744148 18744324 refGene_gn=IP6K1;refGene_tx=XM_011291603.3
A2 18747317 18747499 refGene_gn=IP6K1;refGene_tx=XM_011291603.3
A2 18752221 18752432 refGene_gn=IP6K1;refGene_tx=XM_011291603.3
A2 18762710 18762933 refGene_gn=IP6K1;refGene_tx=XM_011291603.3
A2 18801135 18801191
refGene_gn=CDHR4;refGene_tx=XM_003982238.5;refGene_gn=CDHR4;refGene_tx=XM_006928777.4
A2 18801577 18801718
refGene_gn=CDHR4;refGene_tx=XM_003982238.5;refGene_gn=CDHR4;refGene_tx=XM_006928776.4;refGene_gn=CDHR4;refGene_tx=XM_006928777.4;refGene_gn=CDHR4;refGene_tx=XM_019821670.2;refGene_gn=CDHR4;refGene_tx=XM_019821671.2;refGene_gn=CDHR4;refGene_tx=XM_019821685.2
A2 18801942 18802033
refGene_gn=CDHR4;refGene_tx=XM_003982238.5;refGene_gn=CDHR4;refGene_tx=XM_006928776.4;refGene_gn=CDHR4;refGene_tx=XM_006928777.4;refGene_gn=CDHR4;refGene_tx=XM_019821670.2;refGene_gn=CDHR4;refGene_tx=XM_019821671.2;refGene_gn=CDHR4;refGene_tx=XM_019821685.2
A2 18802134 18802191
refGene_gn=CDHR4;refGene_tx=XM_003982238.5;refGene_gn=CDHR4;refGene_tx=XM_006928776.4;refGene_gn=CDHR4;refGene_tx=XM_006928777.4;refGene_gn=CDHR4;refGene_tx=XM_019821670.2;refGene_gn=CDHR4;refGene_tx=XM_019821671.2;refGene_gn=CDHR4;refGene_tx=XM_019821685.2
A2 18802689 18802852
refGene_gn=CDHR4;refGene_tx=XM_003982238.5;refGene_gn=CDHR4;refGene_tx=XM_006928776.4;refGene_gn=CDHR4;refGene_tx=XM_006928777.4;refGene_gn=CDHR4;refGene_tx=XM_019821670.2;refGene_gn=CDHR4;refGene_tx=XM_019821671.2;refGene_gn=CDHR4;refGene_tx=XM_019821685.2;refGene_gn=CDHR4;refGe

ne_tx=XM_019821701.2
A2	18803062	18803283
refGene_gn=CDHR4;refGene_tx=XM_003982238.5;refGene_gn=CDHR4;refGene_tx=XM_006928776.4;refGene_gn=CDHR4;refGene_tx=XM_006928777.4;refGene_gn=CDHR4;refGene_tx=XM_019821670.2;refGene_gn=CDHR4;refGene_tx=XM_019821671.2;refGene_gn=CDHR4;refGene_tx=XM_019821685.2;refGene_gn=CDHR4;refGene_tx=XM_019821701.2
A2	18803362	18803513
refGene_gn=CDHR4;refGene_tx=XM_003982238.5;refGene_gn=CDHR4;refGene_tx=XM_006928776.4;refGene_gn=CDHR4;refGene_tx=XM_006928777.4;refGene_gn=CDHR4;refGene_tx=XM_019821670.2;refGene_gn=CDHR4;refGene_tx=XM_019821671.2;refGene_gn=CDHR4;refGene_tx=XM_019821685.2;refGene_gn=CDHR4;refGene_tx=XM_019821701.2
A2	18803734	18803874
refGene_gn=CDHR4;refGene_tx=XM_003982238.5;refGene_gn=CDHR4;refGene_tx=XM_006928776.4;refGene_gn=CDHR4;refGene_tx=XM_006928777.4;refGene_gn=CDHR4;refGene_tx=XM_019821670.2;refGene_gn=CDHR4;refGene_tx=XM_019821671.2;refGene_gn=CDHR4;refGene_tx=XM_019821685.2;refGene_gn=CDHR4;refGene_tx=XM_019821701.2
A2	18803954	18804158
refGene_gn=CDHR4;refGene_tx=XM_003982238.5;refGene_gn=CDHR4;refGene_tx=XM_006928776.4;refGene_gn=CDHR4;refGene_tx=XM_006928777.4;refGene_gn=CDHR4;refGene_tx=XM_019821670.2;refGene_gn=CDHR4;refGene_tx=XM_019821671.2;refGene_gn=CDHR4;refGene_tx=XM_019821685.2;refGene_gn=CDHR4;refGene_tx=XM_019821701.2
A2	18804462	18804556
refGene_gn=CDHR4;refGene_tx=XM_003982238.5;refGene_gn=CDHR4;refGene_tx=XM_006928776.4;refGene_gn=CDHR4;refGene_tx=XM_006928777.4;refGene_gn=CDHR4;refGene_tx=XM_019821670.2;refGene_gn=CDHR4;refGene_tx=XM_019821671.2;refGene_gn=CDHR4;refGene_tx=XM_019821685.2;refGene_gn=CDHR4;refGene_tx=XM_019821701.2
A2	18804812	18804966
refGene_gn=CDHR4;refGene_tx=XM_003982238.5;refGene_gn=CDHR4;refGene_tx=XM_006928776.4;refGene_gn=CDHR4;refGene_tx=XM_006928777.4;refGene_gn=CDHR4;refGene_tx=XM_019821670.2;refGene_gn=CDHR4;refGene_tx=XM_019821671.2;refGene_gn=CDHR4;refGene_tx=XM_019821685.2;refGene_gn=CDHR4;refGene_tx=XM_019821701.2
A2	18805064	18805248
refGene_gn=CDHR4;refGene_tx=XM_003982238.5;refGene_gn=CDHR4;refGene_tx=XM_006928776.4;refGene_gn=CDHR4;refGene_tx=XM_006928777.4;refGene_gn=CDHR4;refGene_tx=XM_019821670.2;refGene_gn=CDHR4;refGene_tx=XM_019821671.2;refGene_gn=CDHR4;refGene_tx=XM_019821685.2;refGene_gn=CDHR4;refGene_tx=XM_019821701.2
A2	18805531	18805668
refGene_gn=CDHR4;refGene_tx=XM_003982238.5;refGene_gn=CDHR4;refGene_tx=XM_006928776.4;refGene_gn=CDHR4;refGene_tx=XM_006928777.4;refGene_gn=CDHR4;refGene_tx=XM_019821670.2;refGene_gn=CDHR4;refGene_tx=XM_019821671.2;refGene_gn=CDHR4;refGene_tx=XM_019821685.2;refGene_gn=CDHR4;refGene_tx=XM_019821701.2
A2	18805862	18805966

refGene_gn=CDHR4;refGene_tx=XM_003982238.5;refGene_gn=CDHR4;refGene_tx=XM_006928776.4;refGene_gn=CDHR4;refGene_tx=XM_006928777.4;refGene_gn=CDHR4;refGene_tx=XM_019821670.2;refGene_gn=CDHR4;refGene_tx=XM_019821671.2;refGene_gn=CDHR4;refGene_tx=XM_019821685.2;refGene_gn=CDHR4;refGene_tx=XM_019821701.2
A2 18807313 18807424
refGene_gn=CDHR4;refGene_tx=XM_003982238.5;refGene_gn=CDHR4;refGene_tx=XM_006928776.4;refGene_gn=CDHR4;refGene_tx=XM_006928777.4;refGene_gn=CDHR4;refGene_tx=XM_019821670.2;refGene_gn=CDHR4;refGene_tx=XM_019821671.2;refGene_gn=CDHR4;refGene_tx=XM_019821685.2;refGene_gn=CDHR4;refGene_tx=XM_019821701.2
A2 18807958 18808050
refGene_gn=CDHR4;refGene_tx=XM_003982238.5;refGene_gn=CDHR4;refGene_tx=XM_006928776.4;refGene_gn=CDHR4;refGene_tx=XM_006928777.4;refGene_gn=CDHR4;refGene_tx=XM_019821670.2;refGene_gn=CDHR4;refGene_tx=XM_019821671.2;refGene_gn=CDHR4;refGene_tx=XM_019821685.2;refGene_gn=CDHR4;refGene_tx=XM_019821701.2
A2 18808126 18808289
refGene_gn=CDHR4;refGene_tx=XM_003982238.5;refGene_gn=CDHR4;refGene_tx=XM_006928776.4;refGene_gn=CDHR4;refGene_tx=XM_006928777.4;refGene_gn=CDHR4;refGene_tx=XM_019821670.2;refGene_gn=CDHR4;refGene_tx=XM_019821671.2;refGene_gn=CDHR4;refGene_tx=XM_019821685.2;refGene_gn=CDHR4;refGene_tx=XM_019821701.2
A2 18808413 18808598
refGene_gn=CDHR4;refGene_tx=XM_003982238.5;refGene_gn=CDHR4;refGene_tx=XM_006928776.4;refGene_gn=CDHR4;refGene_tx=XM_006928777.4;refGene_gn=CDHR4;refGene_tx=XM_019821670.2;refGene_gn=CDHR4;refGene_tx=XM_019821671.2;refGene_gn=CDHR4;refGene_tx=XM_019821685.2;refGene_gn=CDHR4;refGene_tx=XM_019821701.2
A2 18808945 18808994
refGene_gn=CDHR4;refGene_tx=XM_003982238.5;refGene_gn=CDHR4;refGene_tx=XM_006928776.4;refGene_gn=CDHR4;refGene_tx=XM_006928777.4;refGene_gn=CDHR4;refGene_tx=XM_019821670.2;refGene_gn=CDHR4;refGene_tx=XM_019821671.2;refGene_gn=CDHR4;refGene_tx=XM_019821685.2;refGene_gn=CDHR4;refGene_tx=XM_019821701.2
A2 18812564 18812777 refGene_gn=FAM212A;refGene_tx=XM_003982239.5
A2 18813525 18814317 refGene_gn=FAM212A;refGene_tx=XM_003982239.5
A2 18814621 18814751
refGene_gn=UBA7;refGene_tx=NM_001278841.1;refGene_gn=UBA7;refGene_tx=XM_011290546.3
A2 18815243 18815344
refGene_gn=UBA7;refGene_tx=NM_001278841.1;refGene_gn=UBA7;refGene_tx=XM_011290546.3
A2 18815421 18815520
refGene_gn=UBA7;refGene_tx=NM_001278841.1;refGene_gn=UBA7;refGene_tx=XM_011290546.3
A2 18815745 18815949 refGene_gn=UBA7;refGene_tx=XM_006928773.4
A2 18817109 18817301
refGene_gn=UBA7;refGene_tx=NM_001278841.1;refGene_gn=UBA7;refGene_tx=XM_006928773.4;refGene_gn=UBA7;refGene_tx=XM_011290546.3

A2	18817393	18817486
refGene_gn=UBA7;refGene_tx=NM_001278841.1;refGene_gn=UBA7;refGene_tx=XM_006928773.4;refGene_gn=UBA7;refGene_tx=XM_011290546.3
A2	18817768	18817851
refGene_gn=UBA7;refGene_tx=NM_001278841.1;refGene_gn=UBA7;refGene_tx=XM_006928773.4;refGene_gn=UBA7;refGene_tx=XM_011290546.3
A2	18818322	18818506
refGene_gn=UBA7;refGene_tx=NM_001278841.1;refGene_gn=UBA7;refGene_tx=XM_006928773.4;refGene_gn=UBA7;refGene_tx=XM_011290546.3
A2	18818752	18818827
refGene_gn=UBA7;refGene_tx=NM_001278841.1;refGene_gn=UBA7;refGene_tx=XM_006928773.4;refGene_gn=UBA7;refGene_tx=XM_011290546.3
A2	18818900	18819084
refGene_gn=UBA7;refGene_tx=NM_001278841.1;refGene_gn=UBA7;refGene_tx=XM_006928773.4;refGene_gn=UBA7;refGene_tx=XM_011290546.3
A2	18819433	18819498
refGene_gn=UBA7;refGene_tx=NM_001278841.1;refGene_gn=UBA7;refGene_tx=XM_006928773.4;refGene_gn=UBA7;refGene_tx=XM_011290546.3
A2	18819596	18819802
refGene_gn=UBA7;refGene_tx=NM_001278841.1;refGene_gn=UBA7;refGene_tx=XM_006928773.4;refGene_gn=UBA7;refGene_tx=XM_011290546.3
A2	18819882	18820048
refGene_gn=UBA7;refGene_tx=NM_001278841.1;refGene_gn=UBA7;refGene_tx=XM_006928773.4;refGene_gn=UBA7;refGene_tx=XM_011290546.3
A2	18820140	18820296
refGene_gn=UBA7;refGene_tx=NM_001278841.1;refGene_gn=UBA7;refGene_tx=XM_006928773.4;refGene_gn=UBA7;refGene_tx=XM_011290546.3
A2	18820376	18820457
refGene_gn=UBA7;refGene_tx=NM_001278841.1;refGene_gn=UBA7;refGene_tx=XM_006928773.4;refGene_gn=UBA7;refGene_tx=XM_011290546.3
A2	18820614	18820722
refGene_gn=UBA7;refGene_tx=NM_001278841.1;refGene_gn=UBA7;refGene_tx=XM_006928773.4;refGene_gn=UBA7;refGene_tx=XM_011290546.3
A2	18820900	18821083
refGene_gn=UBA7;refGene_tx=NM_001278841.1;refGene_gn=UBA7;refGene_tx=XM_006928773.4
A2	18821514	18821661
refGene_gn=UBA7;refGene_tx=NM_001278841.1;refGene_gn=UBA7;refGene_tx=XM_006928773.4;refGene_gn=UBA7;refGene_tx=XM_011290546.3
A2	18821746	18821844
refGene_gn=UBA7;refGene_tx=NM_001278841.1;refGene_gn=UBA7;refGene_tx=XM_006928773.4;refGene_gn=UBA7;refGene_tx=XM_011290546.3
A2	18822047	18822177
refGene_gn=UBA7;refGene_tx=NM_001278841.1;refGene_gn=UBA7;refGene_tx=XM_006928773.4;refGene_gn=UBA7;refGene_tx=XM_011290546.3
A2	18822250	18822341
refGene_gn=UBA7;refGene_tx=NM_001278841.1;refGene_gn=UBA7;refGene_tx=XM_006928773.4;refGene_gn=UBA7;refGene_tx=XM_011290546.3
A2	18822677	18822784
refGene_gn=UBA7;refGene_tx=NM_001278841.1;refGene_gn=UBA7;refGene_tx=X

M_006928773.4;refGene_gn=UBA7;refGene_tx=XM_011290546.3
A2	18822882	18823017
refGene_gn=UBA7;refGene_tx=NM_001278841.1;refGene_gn=UBA7;refGene_tx=XM_006928773.4;refGene_gn=UBA7;refGene_tx=XM_011290546.3
A2	18823119	18823288
refGene_gn=UBA7;refGene_tx=NM_001278841.1;refGene_gn=UBA7;refGene_tx=XM_006928773.4;refGene_gn=UBA7;refGene_tx=XM_011290546.3
A2	18823372	18823428
refGene_gn=UBA7;refGene_tx=NM_001278841.1;refGene_gn=UBA7;refGene_tx=XM_006928773.4;refGene_gn=UBA7;refGene_tx=XM_011290546.3
A2	18830119	18830239
refGene_gn=TRAIP;refGene_tx=XM_003982241.5;refGene_gn=TRAIP;refGene_tx=XM_019821719.2
A2	18830449	18830500
refGene_gn=TRAIP;refGene_tx=XM_003982241.5;refGene_gn=TRAIP;refGene_tx=XM_019821719.2
A2	18830621	18830771
refGene_gn=TRAIP;refGene_tx=XM_003982241.5;refGene_gn=TRAIP;refGene_tx=XM_019821719.2
A2	18831049	18831098
refGene_gn=TRAIP;refGene_tx=XM_003982241.5;refGene_gn=TRAIP;refGene_tx=XM_019821719.2
A2	18832944	18833100
refGene_gn=TRAIP;refGene_tx=XM_003982241.5;refGene_gn=TRAIP;refGene_tx=XM_019821719.2
A2	18836961	18837050
refGene_gn=TRAIP;refGene_tx=XM_003982241.5;refGene_gn=TRAIP;refGene_tx=XM_019821719.2
A2	18837447	18837537
refGene_gn=TRAIP;refGene_tx=XM_003982241.5;refGene_gn=TRAIP;refGene_tx=XM_019821719.2
A2	18838159	18838247
refGene_gn=TRAIP;refGene_tx=XM_003982241.5;refGene_gn=TRAIP;refGene_tx=XM_019821719.2
A2	18839236	18839350
refGene_gn=TRAIP;refGene_tx=XM_003982241.5;refGene_gn=TRAIP;refGene_tx=XM_019821719.2
A2	18839841	18839936
refGene_gn=TRAIP;refGene_tx=XM_003982241.5;refGene_gn=TRAIP;refGene_tx=XM_019821719.2
A2	18841160	18841288	refGene_gn=TRAIP;refGene_tx=XM_003982241.5
A2	18841879	18841919	refGene_gn=TRAIP;refGene_tx=XM_003982241.5
A2	18844199	18844283
refGene_gn=TRAIP;refGene_tx=XM_003982241.5;refGene_gn=TRAIP;refGene_tx=XM_019821719.2
A2	18844800	18844858
refGene_gn=TRAIP;refGene_tx=XM_003982241.5;refGene_gn=TRAIP;refGene_tx=XM_019821719.2
A2	18851597	18851893
refGene_gn=TRAIP;refGene_tx=XM_003982241.5;refGene_gn=TRAIP;refGene_tx

=XM_019821719.2
A2	18854549	18855131
refGene_gn=CAMKV;refGene_tx=XM_023248856.1;refGene_gn=CAMKV;refGene_tx=XM_023248857.1;refGene_gn=CAMKV;refGene_tx=XM_023248859.1
A2	18855429	18855517
refGene_gn=CAMKV;refGene_tx=XM_023248856.1;refGene_gn=CAMKV;refGene_tx=XM_023248857.1;refGene_gn=CAMKV;refGene_tx=XM_023248859.1
A2	18855751	18855830
refGene_gn=CAMKV;refGene_tx=XM_023248856.1;refGene_gn=CAMKV;refGene_tx=XM_023248857.1;refGene_gn=CAMKV;refGene_tx=XM_023248859.1
A2	18855990	18856127
refGene_gn=CAMKV;refGene_tx=XM_023248856.1;refGene_gn=CAMKV;refGene_tx=XM_023248857.1;refGene_gn=CAMKV;refGene_tx=XM_023248859.1
A2	18856213	18856289
refGene_gn=CAMKV;refGene_tx=XM_023248856.1;refGene_gn=CAMKV;refGene_tx=XM_023248857.1;refGene_gn=CAMKV;refGene_tx=XM_023248859.1
A2	18856445	18856566
refGene_gn=CAMKV;refGene_tx=XM_023248856.1;refGene_gn=CAMKV;refGene_tx=XM_023248857.1;refGene_gn=CAMKV;refGene_tx=XM_023248859.1
A2	18856701	18856840
refGene_gn=CAMKV;refGene_tx=XM_023248856.1;refGene_gn=CAMKV;refGene_tx=XM_023248857.1;refGene_gn=CAMKV;refGene_tx=XM_023248859.1
A2	18857053	18857128
refGene_gn=CAMKV;refGene_tx=XM_023248856.1;refGene_gn=CAMKV;refGene_tx=XM_023248857.1;refGene_gn=CAMKV;refGene_tx=XM_023248859.1
A2	18857308	18857440
refGene_gn=CAMKV;refGene_tx=XM_023248856.1;refGene_gn=CAMKV;refGene_tx=XM_023248857.1
A2	18857550	18857645
refGene_gn=CAMKV;refGene_tx=XM_023248856.1;refGene_gn=CAMKV;refGene_tx=XM_023248857.1;refGene_gn=CAMKV;refGene_tx=XM_023248859.1
A2	18881319	18881575
refGene_gn=MST1R;refGene_tx=XM_003982244.5;refGene_gn=MST1R;refGene_tx=XM_006928778.4;refGene_gn=MST1R;refGene_tx=XM_006928780.4;refGene_gn=MST1R;refGene_tx=XM_006928781.4;refGene_gn=MST1R;refGene_tx=XM_006928782.4;refGene_gn=MST1R;refGene_tx=XM_006928783.4;refGene_gn=MST1R;refGene_tx=XM_019821741.2;refGene_gn=MST1R;refGene_tx=XM_019821760.2;refGene_gn=MST1R;refGene_tx=XM_019821762.2;refGene_gn=MST1R;refGene_tx=XM_019821771.2;refGene_gn=MST1R;refGene_tx=XM_019821775.2;refGene_gn=MST1R;refGene_tx=XM_019821784.2
A2	18883771	18883908
refGene_gn=MST1R;refGene_tx=XM_003982244.5;refGene_gn=MST1R;refGene_tx=XM_006928778.4;refGene_gn=MST1R;refGene_tx=XM_006928780.4;refGene_gn=MST1R;refGene_tx=XM_006928781.4;refGene_gn=MST1R;refGene_tx=XM_006928782.4;refGene_gn=MST1R;refGene_tx=XM_006928783.4;refGene_gn=MST1R;refGene_tx=XM_019821741.2;refGene_gn=MST1R;refGene_tx=XM_019821760.2;refGene_gn=MST1R;refGene_tx=XM_019821762.2;refGene_gn=MST1R;refGene_tx=XM_019821771.2;refGene_gn=MST1R;refGene_tx=XM_019821775.2;refGene_gn=MST1R;refGene_tx=XM_019821784.2
A2	18884315	18884481

refGene_gn=MST1R;refGene_tx=XM_003982244.5;refGene_gn=MST1R;refGene_tx
=XM_006928778.4;refGene_gn=MST1R;refGene_tx=XM_006928780.4;refGene_gn=
MST1R;refGene_tx=XM_006928781.4;refGene_gn=MST1R;refGene_tx=XM_0069287
82.4;refGene_gn=MST1R;refGene_tx=XM_006928783.4;refGene_gn=MST1R;refGe
ne_tx=XM_019821741.2;refGene_gn=MST1R;refGene_tx=XM_019821760.2;refGen
e_gn=MST1R;refGene_tx=XM_019821762.2;refGene_gn=MST1R;refGene_tx=XM_01
9821771.2;refGene_gn=MST1R;refGene_tx=XM_019821775.2;refGene_gn=MST1R;
refGene_tx=XM_019821784.2
A2 18884745 18884855
refGene_gn=MST1R;refGene_tx=XM_003982244.5;refGene_gn=MST1R;refGene_tx
=XM_006928778.4;refGene_gn=MST1R;refGene_tx=XM_006928780.4;refGene_gn=
MST1R;refGene_tx=XM_006928781.4;refGene_gn=MST1R;refGene_tx=XM_0069287
82.4;refGene_gn=MST1R;refGene_tx=XM_006928783.4;refGene_gn=MST1R;refGe
ne_tx=XM_019821741.2;refGene_gn=MST1R;refGene_tx=XM_019821760.2;refGen
e_gn=MST1R;refGene_tx=XM_019821762.2;refGene_gn=MST1R;refGene_tx=XM_01
9821771.2;refGene_gn=MST1R;refGene_tx=XM_019821775.2;refGene_gn=MST1R;
refGene_tx=XM_019821784.2
A2 18884938 18885120
refGene_gn=MST1R;refGene_tx=XM_003982244.5;refGene_gn=MST1R;refGene_tx
=XM_006928778.4;refGene_gn=MST1R;refGene_tx=XM_006928780.4;refGene_gn=
MST1R;refGene_tx=XM_006928781.4;refGene_gn=MST1R;refGene_tx=XM_0069287
82.4;refGene_gn=MST1R;refGene_tx=XM_006928783.4;refGene_gn=MST1R;refGe
ne_tx=XM_019821741.2;refGene_gn=MST1R;refGene_tx=XM_019821760.2;refGen
e_gn=MST1R;refGene_tx=XM_019821762.2;refGene_gn=MST1R;refGene_tx=XM_01
9821771.2;refGene_gn=MST1R;refGene_tx=XM_019821775.2;refGene_gn=MST1R;
refGene_tx=XM_019821784.2
A2 18885271 18885352
refGene_gn=MST1R;refGene_tx=XM_003982244.5;refGene_gn=MST1R;refGene_tx
=XM_006928778.4;refGene_gn=MST1R;refGene_tx=XM_006928780.4;refGene_gn=
MST1R;refGene_tx=XM_006928781.4;refGene_gn=MST1R;refGene_tx=XM_0069287
82.4;refGene_gn=MST1R;refGene_tx=XM_006928783.4;refGene_gn=MST1R;refGe
ne_tx=XM_019821741.2;refGene_gn=MST1R;refGene_tx=XM_019821760.2;refGen
e_gn=MST1R;refGene_tx=XM_019821762.2;refGene_gn=MST1R;refGene_tx=XM_01
9821771.2;refGene_gn=MST1R;refGene_tx=XM_019821775.2;refGene_gn=MST1R;
refGene_tx=XM_019821784.2
A2 18885714 18886098
refGene_gn=MST1R;refGene_tx=XM_003982244.5;refGene_gn=MST1R;refGene_tx
=XM_006928778.4;refGene_gn=MST1R;refGene_tx=XM_006928780.4;refGene_gn=
MST1R;refGene_tx=XM_006928781.4;refGene_gn=MST1R;refGene_tx=XM_0069287
82.4;refGene_gn=MST1R;refGene_tx=XM_006928783.4;refGene_gn=MST1R;refGe
ne_tx=XM_019821741.2;refGene_gn=MST1R;refGene_tx=XM_019821760.2;refGen
e_gn=MST1R;refGene_tx=XM_019821762.2;refGene_gn=MST1R;refGene_tx=XM_01
9821771.2;refGene_gn=MST1R;refGene_tx=XM_019821775.2;refGene_gn=MST1R;
refGene_tx=XM_019821784.2
A2 18886244 18886407
refGene_gn=MST1R;refGene_tx=XM_003982244.5;refGene_gn=MST1R;refGene_tx
=XM_006928778.4;refGene_gn=MST1R;refGene_tx=XM_006928780.4;refGene_gn=
MST1R;refGene_tx=XM_006928781.4;refGene_gn=MST1R;refGene_tx=XM_0069287
82.4;refGene_gn=MST1R;refGene_tx=XM_006928783.4;refGene_gn=MST1R;refGe
ne_tx=XM_019821741.2;refGene_gn=MST1R;refGene_tx=XM_019821760.2;refGen

e_gn=MST1R;refGene_tx=XM_019821762.2;refGene_gn=MST1R;refGene_tx=XM_01
9821771.2;refGene_gn=MST1R;refGene_tx=XM_019821775.2;refGene_gn=MST1R;
refGene_tx=XM_019821784.2
A2 18886486 18886633
refGene_gn=MST1R;refGene_tx=XM_003982244.5;refGene_gn=MST1R;refGene_tx
=XM_006928778.4;refGene_gn=MST1R;refGene_tx=XM_006928780.4;refGene_gn=
MST1R;refGene_tx=XM_006928781.4;refGene_gn=MST1R;refGene_tx=XM_0069287
82.4;refGene_gn=MST1R;refGene_tx=XM_006928783.4;refGene_gn=MST1R;refGe
ne_tx=XM_019821741.2;refGene_gn=MST1R;refGene_tx=XM_019821760.2;refGen
e_gn=MST1R;refGene_tx=XM_019821762.2;refGene_gn=MST1R;refGene_tx=XM_01
9821771.2;refGene_gn=MST1R;refGene_tx=XM_019821775.2;refGene_gn=MST1R;
refGene_tx=XM_019821784.2
A2 18886733 18886946
refGene_gn=MST1R;refGene_tx=XM_003982244.5;refGene_gn=MST1R;refGene_tx
=XM_006928778.4;refGene_gn=MST1R;refGene_tx=XM_006928780.4;refGene_gn=
MST1R;refGene_tx=XM_006928781.4;refGene_gn=MST1R;refGene_tx=XM_0069287
82.4;refGene_gn=MST1R;refGene_tx=XM_006928783.4;refGene_gn=MST1R;refGe
ne_tx=XM_019821741.2;refGene_gn=MST1R;refGene_tx=XM_019821760.2;refGen
e_gn=MST1R;refGene_tx=XM_019821762.2;refGene_gn=MST1R;refGene_tx=XM_01
9821771.2;refGene_gn=MST1R;refGene_tx=XM_019821775.2;refGene_gn=MST1R;
refGene_tx=XM_019821784.2
A2 18887101 18887195
refGene_gn=MST1R;refGene_tx=XM_003982244.5;refGene_gn=MST1R;refGene_tx
=XM_006928778.4;refGene_gn=MST1R;refGene_tx=XM_006928780.4;refGene_gn=
MST1R;refGene_tx=XM_006928781.4;refGene_gn=MST1R;refGene_tx=XM_0069287
82.4;refGene_gn=MST1R;refGene_tx=XM_006928783.4;refGene_gn=MST1R;refGe
ne_tx=XM_019821741.2;refGene_gn=MST1R;refGene_tx=XM_019821760.2;refGen
e_gn=MST1R;refGene_tx=XM_019821762.2;refGene_gn=MST1R;refGene_tx=XM_01
9821771.2;refGene_gn=MST1R;refGene_tx=XM_019821775.2;refGene_gn=MST1R;
refGene_tx=XM_019821784.2
A2 18887291 18887453
refGene_gn=MST1R;refGene_tx=XM_003982244.5;refGene_gn=MST1R;refGene_tx
=XM_006928778.4;refGene_gn=MST1R;refGene_tx=XM_006928780.4;refGene_gn=
MST1R;refGene_tx=XM_006928781.4;refGene_gn=MST1R;refGene_tx=XM_0069287
82.4;refGene_gn=MST1R;refGene_tx=XM_006928783.4;refGene_gn=MST1R;refGe
ne_tx=XM_019821741.2;refGene_gn=MST1R;refGene_tx=XM_019821760.2;refGen
e_gn=MST1R;refGene_tx=XM_019821762.2;refGene_gn=MST1R;refGene_tx=XM_01
9821771.2;refGene_gn=MST1R;refGene_tx=XM_019821775.2;refGene_gn=MST1R;
refGene_tx=XM_019821784.2
A2 18887883 18888020
refGene_gn=MST1R;refGene_tx=XM_003982244.5;refGene_gn=MST1R;refGene_tx
=XM_006928778.4;refGene_gn=MST1R;refGene_tx=XM_006928780.4;refGene_gn=
MST1R;refGene_tx=XM_006928781.4;refGene_gn=MST1R;refGene_tx=XM_0069287
82.4;refGene_gn=MST1R;refGene_tx=XM_006928783.4;refGene_gn=MST1R;refGe
ne_tx=XM_019821741.2;refGene_gn=MST1R;refGene_tx=XM_019821760.2;refGen
e_gn=MST1R;refGene_tx=XM_019821762.2;refGene_gn=MST1R;refGene_tx=XM_01
9821771.2;refGene_gn=MST1R;refGene_tx=XM_019821775.2;refGene_gn=MST1R;
refGene_tx=XM_019821784.2
A2 18888133 18888299
refGene_gn=MST1R;refGene_tx=XM_003982244.5;refGene_gn=MST1R;refGene_tx

=XM_006928778.4;refGene_gn=MST1R;refGene_tx=XM_006928780.4;refGene_gn=MST1R;refGene_tx=XM_006928781.4;refGene_gn=MST1R;refGene_tx=XM_006928782.4;refGene_gn=MST1R;refGene_tx=XM_006928783.4;refGene_gn=MST1R;refGene_tx=XM_019821741.2;refGene_gn=MST1R;refGene_tx=XM_019821760.2;refGene_gn=MST1R;refGene_tx=XM_019821762.2;refGene_gn=MST1R;refGene_tx=XM_019821771.2;refGene_gn=MST1R;refGene_tx=XM_019821775.2;refGene_gn=MST1R;refGene_tx=XM_019821784.2
A2 18888662 18888823
refGene_gn=MST1R;refGene_tx=XM_003982244.5;refGene_gn=MST1R;refGene_tx=XM_006928778.4;refGene_gn=MST1R;refGene_tx=XM_006928780.4;refGene_gn=MST1R;refGene_tx=XM_006928781.4;refGene_gn=MST1R;refGene_tx=XM_006928782.4;refGene_gn=MST1R;refGene_tx=XM_006928783.4;refGene_gn=MST1R;refGene_tx=XM_019821741.2;refGene_gn=MST1R;refGene_tx=XM_019821760.2;refGene_gn=MST1R;refGene_tx=XM_019821762.2;refGene_gn=MST1R;refGene_tx=XM_019821771.2;refGene_gn=MST1R;refGene_tx=XM_019821775.2;refGene_gn=MST1R;refGene_tx=XM_019821784.2
A2 18889127 18889298
refGene_gn=MST1R;refGene_tx=XM_003982244.5;refGene_gn=MST1R;refGene_tx=XM_006928778.4;refGene_gn=MST1R;refGene_tx=XM_006928780.4;refGene_gn=MST1R;refGene_tx=XM_006928781.4;refGene_gn=MST1R;refGene_tx=XM_006928782.4;refGene_gn=MST1R;refGene_tx=XM_006928783.4;refGene_gn=MST1R;refGene_tx=XM_019821741.2;refGene_gn=MST1R;refGene_tx=XM_019821760.2;refGene_gn=MST1R;refGene_tx=XM_019821762.2;refGene_gn=MST1R;refGene_tx=XM_019821771.2;refGene_gn=MST1R;refGene_tx=XM_019821775.2;refGene_gn=MST1R;refGene_tx=XM_019821784.2
A2 18889475 18889604
refGene_gn=MST1R;refGene_tx=XM_003982244.5;refGene_gn=MST1R;refGene_tx=XM_006928778.4;refGene_gn=MST1R;refGene_tx=XM_006928780.4;refGene_gn=MST1R;refGene_tx=XM_006928781.4;refGene_gn=MST1R;refGene_tx=XM_006928782.4;refGene_gn=MST1R;refGene_tx=XM_006928783.4;refGene_gn=MST1R;refGene_tx=XM_019821741.2;refGene_gn=MST1R;refGene_tx=XM_019821760.2;refGene_gn=MST1R;refGene_tx=XM_019821762.2;refGene_gn=MST1R;refGene_tx=XM_019821771.2;refGene_gn=MST1R;refGene_tx=XM_019821775.2;refGene_gn=MST1R;refGene_tx=XM_019821784.2
A2 18889684 18889873
refGene_gn=MST1R;refGene_tx=XM_003982244.5;refGene_gn=MST1R;refGene_tx=XM_006928778.4;refGene_gn=MST1R;refGene_tx=XM_006928780.4;refGene_gn=MST1R;refGene_tx=XM_006928781.4;refGene_gn=MST1R;refGene_tx=XM_006928782.4;refGene_gn=MST1R;refGene_tx=XM_006928783.4;refGene_gn=MST1R;refGene_tx=XM_019821741.2;refGene_gn=MST1R;refGene_tx=XM_019821760.2;refGene_gn=MST1R;refGene_tx=XM_019821762.2;refGene_gn=MST1R;refGene_tx=XM_019821771.2;refGene_gn=MST1R;refGene_tx=XM_019821775.2;refGene_gn=MST1R;refGene_tx=XM_019821784.2
A2 18892736 18893969
refGene_gn=MST1R;refGene_tx=XM_003982244.5;refGene_gn=MST1R;refGene_tx=XM_006928778.4;refGene_gn=MST1R;refGene_tx=XM_006928780.4;refGene_gn=MST1R;refGene_tx=XM_006928781.4;refGene_gn=MST1R;refGene_tx=XM_006928782.4;refGene_gn=MST1R;refGene_tx=XM_006928783.4;refGene_gn=MST1R;refGene_tx=XM_019821741.2;refGene_gn=MST1R;refGene_tx=XM_019821760.2;refGene_gn=MST1R;refGene_tx=XM_019821762.2;refGene_gn=MST1R;refGene_tx=XM_01

9821771.2;refGene_gn=MST1R;refGene_tx=XM_019821775.2;refGene_gn=MST1R;
refGene_tx=XM_019821784.2
A2 18898337 18898478
refGene_gn=MON1A;refGene_tx=XM_019821814.2;refGene_gn=MON1A;refGene_tx
=XM_023248929.1
A2 18898577 18898725
refGene_gn=MON1A;refGene_tx=XM_019821814.2;refGene_gn=MON1A;refGene_tx
=XM_023248929.1
A2 18899433 18900199
refGene_gn=MON1A;refGene_tx=XM_019821814.2;refGene_gn=MON1A;refGene_tx
=XM_023248929.1
A2 18900749 18901235
refGene_gn=MON1A;refGene_tx=XM_019821814.2;refGene_gn=MON1A;refGene_tx
=XM_023248929.1
A2 18902519 18902659
refGene_gn=MON1A;refGene_tx=XM_019821814.2;refGene_gn=MON1A;refGene_tx
=XM_023248929.1
A2 18909949 18910130 refGene_gn=MON1A;refGene_tx=XM_023248929.1
A2 18910327 18910352 refGene_gn=MON1A;refGene_tx=XM_023248929.1
A2 18938812 18938825
refGene_gn=RBM6;refGene_tx=XM_006928786.3;refGene_gn=RBM6;refGene_tx=X
M_019821830.2
A2 18940059 18940168
refGene_gn=RBM6;refGene_tx=XM_003982247.4;refGene_gn=RBM6;refGene_tx=X
M_006928786.3;refGene_gn=RBM6;refGene_tx=XM_006928787.3;refGene_gn=RBM
6;refGene_tx=XM_019821830.2;refGene_gn=RBM6;refGene_tx=XM_023248940.1
A2 18944212 18945491
refGene_gn=RBM6;refGene_tx=XM_003982247.4;refGene_gn=RBM6;refGene_tx=X
M_006928786.3;refGene_gn=RBM6;refGene_tx=XM_006928787.3;refGene_gn=RBM
6;refGene_tx=XM_019821830.2;refGene_gn=RBM6;refGene_tx=XM_019821842.1;
refGene_gn=RBM6;refGene_tx=XM_023248940.1
A2 18946925 18947015
refGene_gn=RBM6;refGene_tx=XM_003982247.4;refGene_gn=RBM6;refGene_tx=X
M_006928786.3;refGene_gn=RBM6;refGene_tx=XM_006928787.3;refGene_gn=RBM
6;refGene_tx=XM_019821830.2;refGene_gn=RBM6;refGene_tx=XM_019821842.1;
refGene_gn=RBM6;refGene_tx=XM_023248940.1
A2 18948707 18948777
refGene_gn=RBM6;refGene_tx=XM_003982247.4;refGene_gn=RBM6;refGene_tx=X
M_006928786.3;refGene_gn=RBM6;refGene_tx=XM_006928787.3;refGene_gn=RBM
6;refGene_tx=XM_019821830.2;refGene_gn=RBM6;refGene_tx=XM_019821842.1;
refGene_gn=RBM6;refGene_tx=XM_023248940.1
A2 18968226 18968300
refGene_gn=RBM6;refGene_tx=XM_003982247.4;refGene_gn=RBM6;refGene_tx=X
M_006928786.3;refGene_gn=RBM6;refGene_tx=XM_006928787.3;refGene_gn=RBM
6;refGene_tx=XM_019821830.2;refGene_gn=RBM6;refGene_tx=XM_019821842.1;
refGene_gn=RBM6;refGene_tx=XM_023248940.1
A2 19004702 19004777
refGene_gn=RBM6;refGene_tx=XM_003982247.4;refGene_gn=RBM6;refGene_tx=X
M_006928786.3;refGene_gn=RBM6;refGene_tx=XM_006928787.3;refGene_gn=RBM
6;refGene_tx=XM_006928788.4;refGene_gn=RBM6;refGene_tx=XM_006928789.2;

refGene_gn=RBM6;refGene_tx=XM_006928790.4;refGene_gn=RBM6;refGene_tx=XM_019821830.2;refGene_gn=RBM6;refGene_tx=XM_019821842.1;refGene_gn=RBM6;refGene_tx=XM_023248940.1
A2 19011086 19011147
refGene_gn=RBM6;refGene_tx=XM_003982247.4;refGene_gn=RBM6;refGene_tx=XM_006928786.3;refGene_gn=RBM6;refGene_tx=XM_006928787.3;refGene_gn=RBM6;refGene_tx=XM_006928788.4;refGene_gn=RBM6;refGene_tx=XM_006928789.2;refGene_gn=RBM6;refGene_tx=XM_006928790.4;refGene_gn=RBM6;refGene_tx=XM_019821830.2;refGene_gn=RBM6;refGene_tx=XM_019821842.1;refGene_gn=RBM6;refGene_tx=XM_023248940.1
A2 19012950 19013226
refGene_gn=RBM6;refGene_tx=XM_003982247.4;refGene_gn=RBM6;refGene_tx=XM_006928786.3;refGene_gn=RBM6;refGene_tx=XM_006928787.3;refGene_gn=RBM6;refGene_tx=XM_006928788.4;refGene_gn=RBM6;refGene_tx=XM_006928789.2;refGene_gn=RBM6;refGene_tx=XM_006928790.4;refGene_gn=RBM6;refGene_tx=XM_019821830.2;refGene_gn=RBM6;refGene_tx=XM_019821842.1;refGene_gn=RBM6;refGene_tx=XM_023248940.1
A2 19014048 19014209
refGene_gn=RBM6;refGene_tx=XM_003982247.4;refGene_gn=RBM6;refGene_tx=XM_006928786.3;refGene_gn=RBM6;refGene_tx=XM_006928787.3;refGene_gn=RBM6;refGene_tx=XM_006928788.4;refGene_gn=RBM6;refGene_tx=XM_006928789.2;refGene_gn=RBM6;refGene_tx=XM_006928790.4;refGene_gn=RBM6;refGene_tx=XM_019821830.2;refGene_gn=RBM6;refGene_tx=XM_019821842.1;refGene_gn=RBM6;refGene_tx=XM_023248940.1
A2 19015226 19015324
refGene_gn=RBM6;refGene_tx=XM_003982247.4;refGene_gn=RBM6;refGene_tx=XM_006928786.3;refGene_gn=RBM6;refGene_tx=XM_006928787.3;refGene_gn=RBM6;refGene_tx=XM_006928788.4;refGene_gn=RBM6;refGene_tx=XM_006928789.2;refGene_gn=RBM6;refGene_tx=XM_006928790.4;refGene_gn=RBM6;refGene_tx=XM_019821830.2;refGene_gn=RBM6;refGene_tx=XM_019821842.1;refGene_gn=RBM6;refGene_tx=XM_023248940.1
A2 19015997 19016040
refGene_gn=RBM6;refGene_tx=XM_003982247.4;refGene_gn=RBM6;refGene_tx=XM_006928786.3;refGene_gn=RBM6;refGene_tx=XM_006928787.3;refGene_gn=RBM6;refGene_tx=XM_006928788.4;refGene_gn=RBM6;refGene_tx=XM_006928789.2;refGene_gn=RBM6;refGene_tx=XM_006928790.4;refGene_gn=RBM6;refGene_tx=XM_019821830.2;refGene_gn=RBM6;refGene_tx=XM_019821842.1;refGene_gn=RBM6;refGene_tx=XM_023248940.1
A2 19016115 19016197
refGene_gn=RBM6;refGene_tx=XM_003982247.4;refGene_gn=RBM6;refGene_tx=XM_006928786.3;refGene_gn=RBM6;refGene_tx=XM_006928787.3;refGene_gn=RBM6;refGene_tx=XM_006928788.4;refGene_gn=RBM6;refGene_tx=XM_006928789.2;refGene_gn=RBM6;refGene_tx=XM_006928790.4;refGene_gn=RBM6;refGene_tx=XM_019821830.2;refGene_gn=RBM6;refGene_tx=XM_019821842.1;refGene_gn=RBM6;refGene_tx=XM_023248940.1
A2 19016443 19016529
refGene_gn=RBM6;refGene_tx=XM_003982247.4;refGene_gn=RBM6;refGene_tx=XM_006928786.3;refGene_gn=RBM6;refGene_tx=XM_006928787.3;refGene_gn=RBM6;refGene_tx=XM_006928788.4;refGene_gn=RBM6;refGene_tx=XM_006928789.2;refGene_gn=RBM6;refGene_tx=XM_006928790.4;refGene_gn=RBM6;refGene_tx=X

M_019821830.2;refGene_gn=RBM6;refGene_tx=XM_019821842.1;refGene_gn=RBM
6;refGene_tx=XM_023248940.1
A2 19017075 19017222
refGene_gn=RBM6;refGene_tx=XM_003982247.4;refGene_gn=RBM6;refGene_tx=X
M_006928786.3;refGene_gn=RBM6;refGene_tx=XM_006928787.3;refGene_gn=RBM
6;refGene_tx=XM_006928788.4;refGene_gn=RBM6;refGene_tx=XM_006928789.2;
refGene_gn=RBM6;refGene_tx=XM_006928790.4;refGene_gn=RBM6;refGene_tx=X
M_019821830.2;refGene_gn=RBM6;refGene_tx=XM_019821842.1;refGene_gn=RBM
6;refGene_tx=XM_023248940.1
A2 19018295 19018391
refGene_gn=RBM6;refGene_tx=XM_003982247.4;refGene_gn=RBM6;refGene_tx=X
M_006928786.3;refGene_gn=RBM6;refGene_tx=XM_006928787.3;refGene_gn=RBM
6;refGene_tx=XM_006928788.4;refGene_gn=RBM6;refGene_tx=XM_006928789.2;
refGene_gn=RBM6;refGene_tx=XM_006928790.4;refGene_gn=RBM6;refGene_tx=X
M_019821830.2;refGene_gn=RBM6;refGene_tx=XM_019821842.1;refGene_gn=RBM
6;refGene_tx=XM_023248940.1
A2 19019544 19019805
refGene_gn=RBM6;refGene_tx=XM_003982247.4;refGene_gn=RBM6;refGene_tx=X
M_006928786.3;refGene_gn=RBM6;refGene_tx=XM_006928787.3;refGene_gn=RBM
6;refGene_tx=XM_006928788.4;refGene_gn=RBM6;refGene_tx=XM_006928789.2;
refGene_gn=RBM6;refGene_tx=XM_006928790.4;refGene_gn=RBM6;refGene_tx=X
M_019821830.2;refGene_gn=RBM6;refGene_tx=XM_019821842.1;refGene_gn=RBM
6;refGene_tx=XM_023248940.1
A2 19021520 19021595
refGene_gn=RBM6;refGene_tx=XM_003982247.4;refGene_gn=RBM6;refGene_tx=X
M_006928786.3;refGene_gn=RBM6;refGene_tx=XM_006928787.3;refGene_gn=RBM
6;refGene_tx=XM_006928788.4;refGene_gn=RBM6;refGene_tx=XM_006928789.2;
refGene_gn=RBM6;refGene_tx=XM_006928790.4;refGene_gn=RBM6;refGene_tx=X
M_019821830.2;refGene_gn=RBM6;refGene_tx=XM_019821842.1;refGene_gn=RBM
6;refGene_tx=XM_023248940.1
A2 19022809 19022907
refGene_gn=RBM6;refGene_tx=XM_003982247.4;refGene_gn=RBM6;refGene_tx=X
M_006928786.3;refGene_gn=RBM6;refGene_tx=XM_006928787.3;refGene_gn=RBM
6;refGene_tx=XM_006928788.4;refGene_gn=RBM6;refGene_tx=XM_006928789.2;
refGene_gn=RBM6;refGene_tx=XM_006928790.4;refGene_gn=RBM6;refGene_tx=X
M_019821830.2;refGene_gn=RBM6;refGene_tx=XM_019821842.1;refGene_gn=RBM
6;refGene_tx=XM_023248940.1
A2 19027115 19027245
refGene_gn=RBM6;refGene_tx=XM_003982247.4;refGene_gn=RBM6;refGene_tx=X
M_006928786.3;refGene_gn=RBM6;refGene_tx=XM_006928787.3;refGene_gn=RBM
6;refGene_tx=XM_006928788.4;refGene_gn=RBM6;refGene_tx=XM_006928789.2;
refGene_gn=RBM6;refGene_tx=XM_006928790.4;refGene_gn=RBM6;refGene_tx=X
M_019821830.2;refGene_gn=RBM6;refGene_tx=XM_019821842.1;refGene_gn=RBM
6;refGene_tx=XM_023248940.1
A2 19027990 19028116
refGene_gn=RBM6;refGene_tx=XM_003982247.4;refGene_gn=RBM6;refGene_tx=X
M_006928786.3;refGene_gn=RBM6;refGene_tx=XM_006928787.3;refGene_gn=RBM
6;refGene_tx=XM_006928788.4;refGene_gn=RBM6;refGene_tx=XM_006928789.2;
refGene_gn=RBM6;refGene_tx=XM_006928790.4;refGene_gn=RBM6;refGene_tx=X
M_019821830.2;refGene_gn=RBM6;refGene_tx=XM_019821842.1;refGene_gn=RBM

6;refGene_tx=XM_023248940.1
A2 19035348 19035365 refGene_gn=RBM5;refGene_tx=XM_003982250.4
A2 19037011 19037177 refGene_gn=RBM5;refGene_tx=XM_003982250.4
A2 19037943 19038099
refGene_gn=RBM5;refGene_tx=XM_003982250.4;refGene_gn=RBM5;refGene_tx=XM_019821852.1
A2 19042870 19042940
refGene_gn=RBM5;refGene_tx=XM_003982250.4;refGene_gn=RBM5;refGene_tx=XM_019821852.1
A2 19043430 19043504
refGene_gn=RBM5;refGene_tx=XM_003982250.4;refGene_gn=RBM5;refGene_tx=XM_019821852.1
A2 19045974 19046058
refGene_gn=RBM5;refGene_tx=XM_003982250.4;refGene_gn=RBM5;refGene_tx=XM_019821852.1
A2 19047439 19047500
refGene_gn=RBM5;refGene_tx=XM_003982250.4;refGene_gn=RBM5;refGene_tx=XM_019821852.1
A2 19048139 19048205
refGene_gn=RBM5;refGene_tx=XM_003982250.4;refGene_gn=RBM5;refGene_tx=XM_019821852.1
A2 19048464 19048625
refGene_gn=RBM5;refGene_tx=XM_003982250.4;refGene_gn=RBM5;refGene_tx=XM_019821852.1
A2 19049217 19049315
refGene_gn=RBM5;refGene_tx=XM_003982250.4;refGene_gn=RBM5;refGene_tx=XM_019821852.1
A2 19049918 19050006
refGene_gn=RBM5;refGene_tx=XM_003982250.4;refGene_gn=RBM5;refGene_tx=XM_019821852.1
A2 19050166 19050244
refGene_gn=RBM5;refGene_tx=XM_003982250.4;refGene_gn=RBM5;refGene_tx=XM_019821852.1
A2 19050325 19050398
refGene_gn=RBM5;refGene_tx=XM_003982250.4;refGene_gn=RBM5;refGene_tx=XM_019821852.1
A2 19051712 19051798
refGene_gn=RBM5;refGene_tx=XM_003982250.4;refGene_gn=RBM5;refGene_tx=XM_019821852.1
A2 19052197 19052282
refGene_gn=RBM5;refGene_tx=XM_003982250.4;refGene_gn=RBM5;refGene_tx=XM_019821852.1
A2 19052497 19052589
refGene_gn=RBM5;refGene_tx=XM_003982250.4;refGene_gn=RBM5;refGene_tx=XM_019821852.1
A2 19055208 19055370
refGene_gn=RBM5;refGene_tx=XM_003982250.4;refGene_gn=RBM5;refGene_tx=XM_019821852.1
A2 19055883 19056033
refGene_gn=RBM5;refGene_tx=XM_003982250.4;refGene_gn=RBM5;refGene_tx=X

M_019821852.1
A2	19056116 19056188
refGene_gn=RBM5;refGene_tx=XM_003982250.4;refGene_gn=RBM5;refGene_tx=X
M_019821852.1
A2	19056829 19057009
refGene_gn=RBM5;refGene_tx=XM_003982250.4;refGene_gn=RBM5;refGene_tx=X
M_019821852.1
A2	19057294 19057369
refGene_gn=RBM5;refGene_tx=XM_003982250.4;refGene_gn=RBM5;refGene_tx=X
M_019821852.1
A2	19058828 19058926
refGene_gn=RBM5;refGene_tx=XM_003982250.4;refGene_gn=RBM5;refGene_tx=X
M_019821852.1
A2	19059004 19059134
refGene_gn=RBM5;refGene_tx=XM_003982250.4;refGene_gn=RBM5;refGene_tx=X
M_019821852.1
A2	19060073 19060199
refGene_gn=RBM5;refGene_tx=XM_003982250.4;refGene_gn=RBM5;refGene_tx=X
M_019821852.1
A2	19095026 19095138
refGene_gn=SEMA3F;refGene_tx=XM_003982251.4;refGene_gn=SEMA3F;refGene_
tx=XM_006928792.3;refGene_gn=SEMA3F;refGene_tx=XM_019821873.2
A2	19108996 19109157
refGene_gn=SEMA3F;refGene_tx=XM_003982251.4;refGene_gn=SEMA3F;refGene_
tx=XM_006928792.3;refGene_gn=SEMA3F;refGene_tx=XM_019821873.2
A2	19109241 19109304
refGene_gn=SEMA3F;refGene_tx=XM_003982251.4;refGene_gn=SEMA3F;refGene_
tx=XM_006928792.3;refGene_gn=SEMA3F;refGene_tx=XM_019821873.2
A2	19109420 19109540
refGene_gn=SEMA3F;refGene_tx=XM_003982251.4;refGene_gn=SEMA3F;refGene_
tx=XM_006928792.3;refGene_gn=SEMA3F;refGene_tx=XM_019821873.2
A2	19110247 19110340
refGene_gn=SEMA3F;refGene_tx=XM_003982251.4;refGene_gn=SEMA3F;refGene_
tx=XM_019821873.2
A2	19111885 19111979
refGene_gn=SEMA3F;refGene_tx=XM_003982251.4;refGene_gn=SEMA3F;refGene_
tx=XM_006928792.3;refGene_gn=SEMA3F;refGene_tx=XM_019821873.2
A2	19113781 19113901
refGene_gn=SEMA3F;refGene_tx=XM_003982251.4;refGene_gn=SEMA3F;refGene_
tx=XM_006928792.3;refGene_gn=SEMA3F;refGene_tx=XM_019821873.2
A2	19114105 19114245
refGene_gn=SEMA3F;refGene_tx=XM_003982251.4;refGene_gn=SEMA3F;refGene_
tx=XM_006928792.3;refGene_gn=SEMA3F;refGene_tx=XM_019821873.2
A2	19114358 19114473
refGene_gn=SEMA3F;refGene_tx=XM_003982251.4;refGene_gn=SEMA3F;refGene_
tx=XM_006928792.3;refGene_gn=SEMA3F;refGene_tx=XM_019821873.2
A2	19114636 19114706
refGene_gn=SEMA3F;refGene_tx=XM_003982251.4;refGene_gn=SEMA3F;refGene_
tx=XM_006928792.3;refGene_gn=SEMA3F;refGene_tx=XM_019821873.2
A2	19114875 19115020

refGene_gn=SEMA3F;refGene_tx=XM_003982251.4;refGene_gn=SEMA3F;refGene_tx=XM_006928792.3;refGene_gn=SEMA3F;refGene_tx=XM_019821873.2
A2 19116131 19116354
refGene_gn=SEMA3F;refGene_tx=XM_003982251.4;refGene_gn=SEMA3F;refGene_tx=XM_006928792.3;refGene_gn=SEMA3F;refGene_tx=XM_019821873.2
A2 19116957 19117046
refGene_gn=SEMA3F;refGene_tx=XM_003982251.4;refGene_gn=SEMA3F;refGene_tx=XM_006928792.3;refGene_gn=SEMA3F;refGene_tx=XM_019821873.2
A2 19117182 19117224
refGene_gn=SEMA3F;refGene_tx=XM_003982251.4;refGene_gn=SEMA3F;refGene_tx=XM_006928792.3;refGene_gn=SEMA3F;refGene_tx=XM_019821873.2
A2 19117376 19117534
refGene_gn=SEMA3F;refGene_tx=XM_003982251.4;refGene_gn=SEMA3F;refGene_tx=XM_006928792.3;refGene_gn=SEMA3F;refGene_tx=XM_019821873.2
A2 19117725 19117793
refGene_gn=SEMA3F;refGene_tx=XM_003982251.4;refGene_gn=SEMA3F;refGene_tx=XM_006928792.3;refGene_gn=SEMA3F;refGene_tx=XM_019821873.2
A2 19118086 19118220
refGene_gn=SEMA3F;refGene_tx=XM_003982251.4;refGene_gn=SEMA3F;refGene_tx=XM_006928792.3;refGene_gn=SEMA3F;refGene_tx=XM_019821873.2
A2 19118868 19119279
refGene_gn=SEMA3F;refGene_tx=XM_003982251.4;refGene_gn=SEMA3F;refGene_tx=XM_006928792.3;refGene_gn=SEMA3F;refGene_tx=XM_019821873.2
A2 19122941 19123047 refGene_gn=GNAT1;refGene_tx=XM_003982253.4
A2 19124216 19124259
refGene_gn=GNAT1;refGene_tx=XM_003982253.4;refGene_gn=GNAT1;refGene_tx=XM_019821885.2
A2 19124365 19124507
refGene_gn=GNAT1;refGene_tx=XM_003982253.4;refGene_gn=GNAT1;refGene_tx=XM_019821885.2
A2 19124629 19124787
refGene_gn=GNAT1;refGene_tx=XM_003982253.4;refGene_gn=GNAT1;refGene_tx=XM_019821885.2
A2 19124876 19125005
refGene_gn=GNAT1;refGene_tx=XM_003982253.4;refGene_gn=GNAT1;refGene_tx=XM_019821885.2
A2 19125272 19125402
refGene_gn=GNAT1;refGene_tx=XM_003982253.4;refGene_gn=GNAT1;refGene_tx=XM_019821885.2
A2 19125668 19125822
refGene_gn=GNAT1;refGene_tx=XM_003982253.4;refGene_gn=GNAT1;refGene_tx=XM_019821885.2
A2 19125946 19126137
refGene_gn=GNAT1;refGene_tx=XM_003982253.4;refGene_gn=GNAT1;refGene_tx=XM_019821885.2
A2 19141100 19141201
refGene_gn=SLC38A3;refGene_tx=XM_003982254.5;refGene_gn=SLC38A3;refGene_tx=XM_019821897.2
A2 19141291 19141373
refGene_gn=SLC38A3;refGene_tx=XM_003982254.5;refGene_gn=SLC38A3;refGen

e_tx=XM_019821897.2
A2 19141540 19141656
refGene_gn=SLC38A3;refGene_tx=XM_003982254.5;refGene_gn=SLC38A3;refGene_tx=XM_019821897.2
A2 19142197 19142271
refGene_gn=SLC38A3;refGene_tx=XM_003982254.5;refGene_gn=SLC38A3;refGene_tx=XM_019821897.2
A2 19142353 19142446
refGene_gn=SLC38A3;refGene_tx=XM_003982254.5;refGene_gn=SLC38A3;refGene_tx=XM_019821897.2
A2 19142548 19142630
refGene_gn=SLC38A3;refGene_tx=XM_003982254.5;refGene_gn=SLC38A3;refGene_tx=XM_019821897.2
A2 19143738 19143821
refGene_gn=SLC38A3;refGene_tx=XM_003982254.5;refGene_gn=SLC38A3;refGene_tx=XM_019821897.2
A2 19143913 19143972
refGene_gn=SLC38A3;refGene_tx=XM_003982254.5;refGene_gn=SLC38A3;refGene_tx=XM_019821897.2
A2 19144169 19144334
refGene_gn=SLC38A3;refGene_tx=XM_003982254.5;refGene_gn=SLC38A3;refGene_tx=XM_019821897.2
A2 19144413 19144493
refGene_gn=SLC38A3;refGene_tx=XM_003982254.5;refGene_gn=SLC38A3;refGene_tx=XM_019821897.2
A2 19144773 19144874
refGene_gn=SLC38A3;refGene_tx=XM_003982254.5;refGene_gn=SLC38A3;refGene_tx=XM_019821897.2
A2 19145108 19145233
refGene_gn=SLC38A3;refGene_tx=XM_003982254.5;refGene_gn=SLC38A3;refGene_tx=XM_019821897.2
A2 19145317 19145462
refGene_gn=SLC38A3;refGene_tx=XM_003982254.5;refGene_gn=SLC38A3;refGene_tx=XM_019821897.2
A2 19146393 19146497
refGene_gn=SLC38A3;refGene_tx=XM_003982254.5;refGene_gn=SLC38A3;refGene_tx=XM_019821897.2
A2 19146587 19146692
refGene_gn=SLC38A3;refGene_tx=XM_003982254.5;refGene_gn=SLC38A3;refGene_tx=XM_019821897.2
A2 19159109 19159227 refGene_gn=GNAI2;refGene_tx=XM_011291675.3
A2 19174199 19174242 refGene_gn=GNAI2;refGene_tx=XM_011291675.3
A2 19174528 19174670 refGene_gn=GNAI2;refGene_tx=XM_011291675.3
A2 19174835 19174996 refGene_gn=GNAI2;refGene_tx=XM_011291675.3
A2 19177803 19177932 refGene_gn=GNAI2;refGene_tx=XM_011291675.3
A2 19178348 19178478 refGene_gn=GNAI2;refGene_tx=XM_011291675.3
A2 19178558 19178712 refGene_gn=GNAI2;refGene_tx=XM_011291675.3
A2 19179063 19179254 refGene_gn=GNAI2;refGene_tx=XM_011291675.3
A2 19189644 19189753
refGene_gn=SEMA3B;refGene_tx=XM_003982255.5;refGene_gn=SEMA3B;refGene_

tx=XM_006928797.4;refGene_gn=SEMA3B;refGene_tx=XM_006928798.4;refGene_gn=SEMA3B;refGene_tx=XM_006928799.4;refGene_gn=SEMA3B;refGene_tx=XM_006928800.4;refGene_gn=SEMA3B;refGene_tx=XM_023248969.1;refGene_gn=SEMA3B;refGene_tx=XM_023248978.1
A2 19190427 19190585
refGene_gn=SEMA3B;refGene_tx=XM_003982255.5;refGene_gn=SEMA3B;refGene_tx=XM_006928797.4;refGene_gn=SEMA3B;refGene_tx=XM_006928798.4;refGene_gn=SEMA3B;refGene_tx=XM_006928799.4;refGene_gn=SEMA3B;refGene_tx=XM_006928800.4;refGene_gn=SEMA3B;refGene_tx=XM_023248969.1;refGene_gn=SEMA3B;refGene_tx=XM_023248978.1
A2 19190705 19190768
refGene_gn=SEMA3B;refGene_tx=XM_003982255.5;refGene_gn=SEMA3B;refGene_tx=XM_006928797.4;refGene_gn=SEMA3B;refGene_tx=XM_006928798.4;refGene_gn=SEMA3B;refGene_tx=XM_006928799.4;refGene_gn=SEMA3B;refGene_tx=XM_006928800.4;refGene_gn=SEMA3B;refGene_tx=XM_023248969.1;refGene_gn=SEMA3B;refGene_tx=XM_023248978.1
A2 19191160 19191280
refGene_gn=SEMA3B;refGene_tx=XM_003982255.5;refGene_gn=SEMA3B;refGene_tx=XM_006928797.4;refGene_gn=SEMA3B;refGene_tx=XM_006928798.4;refGene_gn=SEMA3B;refGene_tx=XM_006928799.4;refGene_gn=SEMA3B;refGene_tx=XM_006928800.4;refGene_gn=SEMA3B;refGene_tx=XM_023248969.1;refGene_gn=SEMA3B;refGene_tx=XM_023248978.1
A2 19191359 19191453
refGene_gn=SEMA3B;refGene_tx=XM_003982255.5;refGene_gn=SEMA3B;refGene_tx=XM_006928797.4;refGene_gn=SEMA3B;refGene_tx=XM_006928798.4;refGene_gn=SEMA3B;refGene_tx=XM_006928799.4;refGene_gn=SEMA3B;refGene_tx=XM_006928800.4;refGene_gn=SEMA3B;refGene_tx=XM_023248969.1;refGene_gn=SEMA3B;refGene_tx=XM_023248978.1
A2 19191607 19191727
refGene_gn=SEMA3B;refGene_tx=XM_003982255.5;refGene_gn=SEMA3B;refGene_tx=XM_006928797.4;refGene_gn=SEMA3B;refGene_tx=XM_006928798.4;refGene_gn=SEMA3B;refGene_tx=XM_006928799.4;refGene_gn=SEMA3B;refGene_tx=XM_006928800.4;refGene_gn=SEMA3B;refGene_tx=XM_019821949.2;refGene_gn=SEMA3B;refGene_tx=XM_023248969.1;refGene_gn=SEMA3B;refGene_tx=XM_023248978.1
A2 19192604 19192750
refGene_gn=SEMA3B;refGene_tx=XM_003982255.5;refGene_gn=SEMA3B;refGene_tx=XM_006928797.4;refGene_gn=SEMA3B;refGene_tx=XM_006928798.4;refGene_gn=SEMA3B;refGene_tx=XM_006928799.4;refGene_gn=SEMA3B;refGene_tx=XM_006928800.4;refGene_gn=SEMA3B;refGene_tx=XM_019821949.2;refGene_gn=SEMA3B;refGene_tx=XM_023248969.1;refGene_gn=SEMA3B;refGene_tx=XM_023248978.1
A2 19192841 19192953
refGene_gn=SEMA3B;refGene_tx=XM_003982255.5;refGene_gn=SEMA3B;refGene_tx=XM_006928797.4;refGene_gn=SEMA3B;refGene_tx=XM_006928798.4;refGene_gn=SEMA3B;refGene_tx=XM_006928799.4;refGene_gn=SEMA3B;refGene_tx=XM_006928800.4;refGene_gn=SEMA3B;refGene_tx=XM_019821949.2;refGene_gn=SEMA3B;refGene_tx=XM_023248969.1;refGene_gn=SEMA3B;refGene_tx=XM_023248978.1
A2 19193069 19193139

refGene_gn=SEMA3B;refGene_tx=XM_003982255.5;refGene_gn=SEMA3B;refGene_tx=XM_006928797.4;refGene_gn=SEMA3B;refGene_tx=XM_006928798.4;refGene_gn=SEMA3B;refGene_tx=XM_006928799.4;refGene_gn=SEMA3B;refGene_tx=XM_006928800.4;refGene_gn=SEMA3B;refGene_tx=XM_019821949.2;refGene_gn=SEMA3B;refGene_tx=XM_023248969.1;refGene_gn=SEMA3B;refGene_tx=XM_023248978.1
A2 19193216 19193361
refGene_gn=SEMA3B;refGene_tx=XM_003982255.5;refGene_gn=SEMA3B;refGene_tx=XM_006928797.4;refGene_gn=SEMA3B;refGene_tx=XM_006928798.4;refGene_gn=SEMA3B;refGene_tx=XM_006928799.4;refGene_gn=SEMA3B;refGene_tx=XM_006928800.4;refGene_gn=SEMA3B;refGene_tx=XM_019821949.2;refGene_gn=SEMA3B;refGene_tx=XM_023248969.1;refGene_gn=SEMA3B;refGene_tx=XM_023248978.1
A2 19193683 19193903
refGene_gn=SEMA3B;refGene_tx=XM_003982255.5;refGene_gn=SEMA3B;refGene_tx=XM_006928797.4;refGene_gn=SEMA3B;refGene_tx=XM_006928798.4;refGene_gn=SEMA3B;refGene_tx=XM_006928799.4;refGene_gn=SEMA3B;refGene_tx=XM_006928800.4;refGene_gn=SEMA3B;refGene_tx=XM_019821949.2;refGene_gn=SEMA3B;refGene_tx=XM_023248969.1;refGene_gn=SEMA3B;refGene_tx=XM_023248978.1
A2 19194163 19194255
refGene_gn=SEMA3B;refGene_tx=XM_003982255.5;refGene_gn=SEMA3B;refGene_tx=XM_006928797.4;refGene_gn=SEMA3B;refGene_tx=XM_006928798.4;refGene_gn=SEMA3B;refGene_tx=XM_006928799.4;refGene_gn=SEMA3B;refGene_tx=XM_006928800.4;refGene_gn=SEMA3B;refGene_tx=XM_019821949.2;refGene_gn=SEMA3B;refGene_tx=XM_023248969.1;refGene_gn=SEMA3B;refGene_tx=XM_023248978.1
A2 19194334 19194376
refGene_gn=SEMA3B;refGene_tx=XM_003982255.5;refGene_gn=SEMA3B;refGene_tx=XM_006928797.4;refGene_gn=SEMA3B;refGene_tx=XM_006928798.4;refGene_gn=SEMA3B;refGene_tx=XM_006928799.4;refGene_gn=SEMA3B;refGene_tx=XM_006928800.4;refGene_gn=SEMA3B;refGene_tx=XM_019821949.2;refGene_gn=SEMA3B;refGene_tx=XM_023248969.1;refGene_gn=SEMA3B;refGene_tx=XM_023248978.1
A2 19194609 19194767
refGene_gn=SEMA3B;refGene_tx=XM_003982255.5;refGene_gn=SEMA3B;refGene_tx=XM_006928797.4;refGene_gn=SEMA3B;refGene_tx=XM_006928798.4;refGene_gn=SEMA3B;refGene_tx=XM_006928799.4;refGene_gn=SEMA3B;refGene_tx=XM_006928800.4;refGene_gn=SEMA3B;refGene_tx=XM_019821949.2;refGene_gn=SEMA3B;refGene_tx=XM_023248969.1;refGene_gn=SEMA3B;refGene_tx=XM_023248978.1
A2 19194873 19194929
refGene_gn=SEMA3B;refGene_tx=XM_003982255.5;refGene_gn=SEMA3B;refGene_tx=XM_006928797.4;refGene_gn=SEMA3B;refGene_tx=XM_006928798.4;refGene_gn=SEMA3B;refGene_tx=XM_006928799.4;refGene_gn=SEMA3B;refGene_tx=XM_006928800.4;refGene_gn=SEMA3B;refGene_tx=XM_019821949.2;refGene_gn=SEMA3B;refGene_tx=XM_023248969.1;refGene_gn=SEMA3B;refGene_tx=XM_023248978.1
A2 19195016 19195156
refGene_gn=SEMA3B;refGene_tx=XM_003982255.5;refGene_gn=SEMA3B;refGene_

tx=XM_006928797.4;refGene_gn=SEMA3B;refGene_tx=XM_006928798.4;refGene_gn=SEMA3B;refGene_tx=XM_006928799.4;refGene_gn=SEMA3B;refGene_tx=XM_006928800.4;refGene_gn=SEMA3B;refGene_tx=XM_019821949.2;refGene_gn=SEMA3B;refGene_tx=XM_023248969.1;refGene_gn=SEMA3B;refGene_tx=XM_023248978.1
A2 19195677 19196079
refGene_gn=SEMA3B;refGene_tx=XM_003982255.5;refGene_gn=SEMA3B;refGene_tx=XM_006928797.4;refGene_gn=SEMA3B;refGene_tx=XM_006928798.4;refGene_gn=SEMA3B;refGene_tx=XM_006928799.4;refGene_gn=SEMA3B;refGene_tx=XM_006928800.4;refGene_gn=SEMA3B;refGene_tx=XM_019821949.2;refGene_gn=SEMA3B;refGene_tx=XM_023248969.1;refGene_gn=SEMA3B;refGene_tx=XM_023248978.1
A2 19197866 19197888
refGene_gn=LSMEM2;refGene_tx=XM_006928796.4;refGene_gn=LSMEM2;refGene_tx=XM_019821981.1
A2 19200227 19200341
refGene_gn=LSMEM2;refGene_tx=XM_006928796.4;refGene_gn=LSMEM2;refGene_tx=XM_019821981.1
A2 19200433 19200616
refGene_gn=LSMEM2;refGene_tx=XM_006928796.4;refGene_gn=LSMEM2;refGene_tx=XM_019821981.1
A2 19200784 19200918
refGene_gn=LSMEM2;refGene_tx=XM_006928796.4;refGene_gn=LSMEM2;refGene_tx=XM_019821981.1
A2 19201868 19201949 refGene_gn=IFRD2;refGene_tx=XM_003982256.5
A2 19202086 19202182
refGene_gn=IFRD2;refGene_tx=XM_003982256.5;refGene_gn=IFRD2;refGene_tx=XM_006928795.4
A2 19202261 19202390
refGene_gn=IFRD2;refGene_tx=XM_003982256.5;refGene_gn=IFRD2;refGene_tx=XM_006928795.4
A2 19202476 19202614
refGene_gn=IFRD2;refGene_tx=XM_003982256.5;refGene_gn=IFRD2;refGene_tx=XM_006928795.4
A2 19202937 19203043 refGene_gn=IFRD2;refGene_tx=XM_003982256.5
A2 19203127 19203306
refGene_gn=IFRD2;refGene_tx=XM_003982256.5;refGene_gn=IFRD2;refGene_tx=XM_006928795.4
A2 19203392 19203443
refGene_gn=IFRD2;refGene_tx=XM_003982256.5;refGene_gn=IFRD2;refGene_tx=XM_006928795.4
A2 19203605 19203763
refGene_gn=IFRD2;refGene_tx=XM_003982256.5;refGene_gn=IFRD2;refGene_tx=XM_006928795.4
A2 19203847 19203972
refGene_gn=IFRD2;refGene_tx=XM_003982256.5;refGene_gn=IFRD2;refGene_tx=XM_006928795.4
A2 19204078 19204163
refGene_gn=IFRD2;refGene_tx=XM_003982256.5;refGene_gn=IFRD2;refGene_tx=XM_006928795.4

A2 19204247 19204367
refGene_gn=IFRD2;refGene_tx=XM_003982256.5;refGene_gn=IFRD2;refGene_tx
=XM_006928795.4
A2 19206552 19206610
refGene_gn=IFRD2;refGene_tx=XM_003982256.5;refGene_gn=IFRD2;refGene_tx
=XM_006928795.4
A2 19207641 19207917 refGene_gn=HYAL3;refGene_tx=XM_003982257.5
A2 19208034 19208124 refGene_gn=HYAL3;refGene_tx=XM_003982257.5
A2 19208520 19209414 refGene_gn=HYAL3;refGene_tx=XM_003982257.5
A2 19210373 19211237 refGene_gn=NAT6;refGene_tx=XM_023248993.1
A2 19211238 19211268 refGene_gn=NAT6;refGene_tx=XM_023248993.1
A2 19215020 19215338
refGene_gn=HYAL2;refGene_tx=XM_006928802.3;refGene_gn=HYAL2;refGene_tx
=XM_006928803.3;refGene_gn=HYAL2;refGene_tx=XM_006928804.3;refGene_gn=
HYAL2;refGene_tx=XM_019821914.2
A2 19215538 19215628
refGene_gn=HYAL2;refGene_tx=XM_006928802.3;refGene_gn=HYAL2;refGene_tx
=XM_006928803.3;refGene_gn=HYAL2;refGene_tx=XM_006928804.3
A2 19215972 19216896
refGene_gn=HYAL2;refGene_tx=XM_006928802.3;refGene_gn=HYAL2;refGene_tx
=XM_006928803.3;refGene_gn=HYAL2;refGene_tx=XM_006928804.3;refGene_gn=
HYAL2;refGene_tx=XM_019821914.2
A2 19217356 19217377 refGene_gn=HYAL2;refGene_tx=XM_006928804.3
A2 19221357 19221768 refGene_gn=HYAL2;refGene_tx=XM_003982261.4
A2 19222096 19222186
refGene_gn=HYAL2;refGene_tx=XM_003982261.4;refGene_gn=HYAL2;refGene_tx
=XM_006928802.3;refGene_gn=HYAL2;refGene_tx=XM_006928803.3;refGene_gn=
HYAL2;refGene_tx=XM_019821914.2
A2 19222687 19223608
refGene_gn=HYAL2;refGene_tx=XM_003982261.4;refGene_gn=HYAL2;refGene_tx
=XM_006928802.3;refGene_gn=HYAL2;refGene_tx=XM_006928803.3;refGene_gn=
HYAL2;refGene_tx=XM_019821914.2
A2 19229228 19229294 refGene_gn=TUSC2;refGene_tx=XM_003982263.3
A2 19229424 19229545 refGene_gn=TUSC2;refGene_tx=XM_003982263.3
A2 19230628 19230774 refGene_gn=TUSC2;refGene_tx=XM_003982263.3
A2 19232580 19232727
refGene_gn=RASSF1;refGene_tx=XM_003982264.4;refGene_gn=RASSF1;refGene_
tx=XM_023249048.1;refGene_gn=RASSF1;refGene_tx=XM_023249050.1
A2 19233367 19233483
refGene_gn=RASSF1;refGene_tx=XM_003982264.4;refGene_gn=RASSF1;refGene_
tx=XM_023249048.1;refGene_gn=RASSF1;refGene_tx=XM_023249050.1
A2 19233600 19233898
refGene_gn=RASSF1;refGene_tx=XM_003982264.4;refGene_gn=RASSF1;refGene_
tx=XM_023249048.1;refGene_gn=RASSF1;refGene_tx=XM_023249050.1
A2 19234103 19234208
refGene_gn=RASSF1;refGene_tx=XM_003982264.4;refGene_gn=RASSF1;refGene_
tx=XM_023249048.1;refGene_gn=RASSF1;refGene_tx=XM_023249050.1
A2 19237589 19237742 refGene_gn=RASSF1;refGene_tx=XM_003982264.4
A2 19238297 19238404
refGene_gn=RASSF1;refGene_tx=XM_023249048.1;refGene_gn=RASSF1;refGene_

tx=XM_023249050.1
A2 19240450 19240955
refGene_gn=RASSF1;refGene_tx=XM_023249048.1;refGene_gn=RASSF1;refGene_tx=XM_023249050.1
A2 19241329 19241405
refGene_gn=ZMYND10;refGene_tx=XM_003982265.5;refGene_gn=ZMYND10;refGene_tx=XM_006928808.4;refGene_gn=ZMYND10;refGene_tx=XM_011291731.3;refGene_gn=ZMYND10;refGene_tx=XM_011291736.3
A2 19241493 19241619
refGene_gn=ZMYND10;refGene_tx=XM_003982265.5;refGene_gn=ZMYND10;refGene_tx=XM_006928808.4;refGene_gn=ZMYND10;refGene_tx=XM_011291731.3;refGene_gn=ZMYND10;refGene_tx=XM_011291736.3
A2 19241739 19241861
refGene_gn=ZMYND10;refGene_tx=XM_003982265.5;refGene_gn=ZMYND10;refGene_tx=XM_006928808.4;refGene_gn=ZMYND10;refGene_tx=XM_011291731.3;refGene_gn=ZMYND10;refGene_tx=XM_011291736.3
A2 19241943 19242069
refGene_gn=ZMYND10;refGene_tx=XM_003982265.5;refGene_gn=ZMYND10;refGene_tx=XM_006928808.4;refGene_gn=ZMYND10;refGene_tx=XM_011291731.3
A2 19242294 19242467
refGene_gn=ZMYND10;refGene_tx=XM_003982265.5;refGene_gn=ZMYND10;refGene_tx=XM_006928808.4;refGene_gn=ZMYND10;refGene_tx=XM_011291731.3;refGene_gn=ZMYND10;refGene_tx=XM_011291736.3
A2 19242797 19242898
refGene_gn=ZMYND10;refGene_tx=XM_003982265.5;refGene_gn=ZMYND10;refGene_tx=XM_006928808.4;refGene_gn=ZMYND10;refGene_tx=XM_011291731.3;refGene_gn=ZMYND10;refGene_tx=XM_011291736.3
A2 19242990 19243103
refGene_gn=ZMYND10;refGene_tx=XM_003982265.5;refGene_gn=ZMYND10;refGene_tx=XM_006928808.4;refGene_gn=ZMYND10;refGene_tx=XM_011291731.3;refGene_gn=ZMYND10;refGene_tx=XM_011291736.3
A2 19243179 19243317
refGene_gn=ZMYND10;refGene_tx=XM_003982265.5;refGene_gn=ZMYND10;refGene_tx=XM_011291731.3;refGene_gn=ZMYND10;refGene_tx=XM_011291736.3
A2 19243437 19243491
refGene_gn=ZMYND10;refGene_tx=XM_003982265.5;refGene_gn=ZMYND10;refGene_tx=XM_006928808.4;refGene_gn=ZMYND10;refGene_tx=XM_011291731.3;refGene_gn=ZMYND10;refGene_tx=XM_011291736.3
A2 19243612 19243729
refGene_gn=ZMYND10;refGene_tx=XM_003982265.5;refGene_gn=ZMYND10;refGene_tx=XM_006928808.4;refGene_gn=ZMYND10;refGene_tx=XM_011291731.3;refGene_gn=ZMYND10;refGene_tx=XM_011291736.3
A2 19244649 19244758
refGene_gn=ZMYND10;refGene_tx=XM_003982265.5;refGene_gn=ZMYND10;refGene_tx=XM_006928808.4;refGene_gn=ZMYND10;refGene_tx=XM_011291731.3;refGene_gn=ZMYND10;refGene_tx=XM_011291736.3
A2 19245024 19245116
refGene_gn=ZMYND10;refGene_tx=XM_003982265.5;refGene_gn=ZMYND10;refGene_tx=XM_006928808.4;refGene_gn=ZMYND10;refGene_tx=XM_011291731.3;refGene_gn=ZMYND10;refGene_tx=XM_011291736.3

A2	19246548 19246616 refGene_gn=NPRL2;refGene_tx=XM_006928811.3
A2	19246702 19246845 refGene_gn=NPRL2;refGene_tx=XM_006928811.3
A2	19247118 19247236 refGene_gn=NPRL2;refGene_tx=XM_006928811.3
A2	19247311 19247405 refGene_gn=NPRL2;refGene_tx=XM_006928811.3
A2	19247523 19247560 refGene_gn=NPRL2;refGene_tx=XM_006928811.3
A2	19247681 19247779 refGene_gn=NPRL2;refGene_tx=XM_006928811.3
A2	19247870 19248007 refGene_gn=NPRL2;refGene_tx=XM_006928811.3
A2	19248479 19248588 refGene_gn=NPRL2;refGene_tx=XM_006928811.3
A2	19248727 19248896 refGene_gn=NPRL2;refGene_tx=XM_006928811.3
A2	19248997 19249089 refGene_gn=NPRL2;refGene_tx=XM_006928811.3
A2	19249631 19249868
refGene_gn=LOC101095404;refGene_tx=XM_019822078.1;refGene_gn=NPRL2;refGene_tx=XM_006928811.3
A2	19250415 19250567
refGene_gn=LOC101095404;refGene_tx=XM_006928809.4;refGene_gn=LOC101095404;refGene_tx=XM_006928810.4;refGene_gn=LOC101095404;refGene_tx=XM_019822078.1;refGene_gn=LOC101095404;refGene_tx=XM_023249076.1
A2	19251032 19251070
refGene_gn=LOC101095404;refGene_tx=XM_006928809.4;refGene_gn=LOC101095404;refGene_tx=XM_006928810.4;refGene_gn=LOC101095404;refGene_tx=XM_019822078.1;refGene_gn=LOC101095404;refGene_tx=XM_023249076.1
A2	19251755 19252259
refGene_gn=LOC101095404;refGene_tx=XM_006928809.4;refGene_gn=LOC101095404;refGene_tx=XM_006928810.4;refGene_gn=LOC101095404;refGene_tx=XM_019822078.1;refGene_gn=LOC101095404;refGene_tx=XM_023249076.1
A2	19253809 19254014 refGene_gn=TMEM115;refGene_tx=XM_003982268.4
A2	19256634 19257485 refGene_gn=TMEM115;refGene_tx=XM_003982268.4
A2	19262183 19262391
refGene_gn=CACNA2D2;refGene_tx=XM_023249003.1;refGene_gn=CACNA2D2;refGene_tx=XM_023249011.1;refGene_gn=CACNA2D2;refGene_tx=XM_023249014.1;refGene_gn=CACNA2D2;refGene_tx=XM_023249017.1;refGene_gn=CACNA2D2;refGene_tx=XM_023249020.1;refGene_gn=CACNA2D2;refGene_tx=XM_023249025.1;refGene_gn=CACNA2D2;refGene_tx=XM_023249027.1;refGene_gn=CACNA2D2;refGene_tx=XM_023249031.1;refGene_gn=CACNA2D2;refGene_tx=XM_023249036.1
A2	19262478 19262567
refGene_gn=CACNA2D2;refGene_tx=XM_023249003.1;refGene_gn=CACNA2D2;refGene_tx=XM_023249011.1;refGene_gn=CACNA2D2;refGene_tx=XM_023249014.1;refGene_gn=CACNA2D2;refGene_tx=XM_023249017.1;refGene_gn=CACNA2D2;refGene_tx=XM_023249020.1;refGene_gn=CACNA2D2;refGene_tx=XM_023249025.1;refGene_gn=CACNA2D2;refGene_tx=XM_023249027.1;refGene_gn=CACNA2D2;refGene_tx=XM_023249031.1;refGene_gn=CACNA2D2;refGene_tx=XM_023249036.1
A2	19262658 19262768
refGene_gn=CACNA2D2;refGene_tx=XM_023249003.1;refGene_gn=CACNA2D2;refGene_tx=XM_023249011.1;refGene_gn=CACNA2D2;refGene_tx=XM_023249014.1;refGene_gn=CACNA2D2;refGene_tx=XM_023249017.1;refGene_gn=CACNA2D2;refGene_tx=XM_023249020.1;refGene_gn=CACNA2D2;refGene_tx=XM_023249025.1;refGene_gn=CACNA2D2;refGene_tx=XM_023249027.1;refGene_gn=CACNA2D2;refGene_tx=XM_023249031.1;refGene_gn=CACNA2D2;refGene_tx=XM_023249036.1
A2	19262921 19263048
refGene_gn=CACNA2D2;refGene_tx=XM_023249003.1;refGene_gn=CACNA2D2;refG

ene_tx=XM_023249011.1;refGene_gn=CACNA2D2;refGene_tx=XM_023249014.1;refGene_gn=CACNA2D2;refGene_tx=XM_023249017.1;refGene_gn=CACNA2D2;refGene_tx=XM_023249020.1;refGene_gn=CACNA2D2;refGene_tx=XM_023249025.1;refGene_gn=CACNA2D2;refGene_tx=XM_023249027.1;refGene_gn=CACNA2D2;refGene_tx=XM_023249031.1;refGene_gn=CACNA2D2;refGene_tx=XM_023249036.1
A2 19263204 19263260
refGene_gn=CACNA2D2;refGene_tx=XM_023249003.1;refGene_gn=CACNA2D2;refGene_tx=XM_023249011.1;refGene_gn=CACNA2D2;refGene_tx=XM_023249014.1;refGene_gn=CACNA2D2;refGene_tx=XM_023249017.1;refGene_gn=CACNA2D2;refGene_tx=XM_023249020.1;refGene_gn=CACNA2D2;refGene_tx=XM_023249025.1;refGene_gn=CACNA2D2;refGene_tx=XM_023249027.1;refGene_gn=CACNA2D2;refGene_tx=XM_023249031.1;refGene_gn=CACNA2D2;refGene_tx=XM_023249036.1
A2 19263382 19263435
refGene_gn=CACNA2D2;refGene_tx=XM_023249003.1;refGene_gn=CACNA2D2;refGene_tx=XM_023249011.1;refGene_gn=CACNA2D2;refGene_tx=XM_023249014.1;refGene_gn=CACNA2D2;refGene_tx=XM_023249017.1;refGene_gn=CACNA2D2;refGene_tx=XM_023249020.1;refGene_gn=CACNA2D2;refGene_tx=XM_023249025.1;refGene_gn=CACNA2D2;refGene_tx=XM_023249027.1;refGene_gn=CACNA2D2;refGene_tx=XM_023249031.1;refGene_gn=CACNA2D2;refGene_tx=XM_023249036.1
A2 19263605 19263758
refGene_gn=CACNA2D2;refGene_tx=XM_023249003.1;refGene_gn=CACNA2D2;refGene_tx=XM_023249011.1;refGene_gn=CACNA2D2;refGene_tx=XM_023249014.1;refGene_gn=CACNA2D2;refGene_tx=XM_023249017.1;refGene_gn=CACNA2D2;refGene_tx=XM_023249020.1;refGene_gn=CACNA2D2;refGene_tx=XM_023249025.1;refGene_gn=CACNA2D2;refGene_tx=XM_023249027.1;refGene_gn=CACNA2D2;refGene_tx=XM_023249031.1;refGene_gn=CACNA2D2;refGene_tx=XM_023249036.1
A2 19263861 19263933
refGene_gn=CACNA2D2;refGene_tx=XM_023249003.1;refGene_gn=CACNA2D2;refGene_tx=XM_023249011.1;refGene_gn=CACNA2D2;refGene_tx=XM_023249014.1;refGene_gn=CACNA2D2;refGene_tx=XM_023249017.1;refGene_gn=CACNA2D2;refGene_tx=XM_023249020.1;refGene_gn=CACNA2D2;refGene_tx=XM_023249025.1;refGene_gn=CACNA2D2;refGene_tx=XM_023249027.1;refGene_gn=CACNA2D2;refGene_tx=XM_023249031.1;refGene_gn=CACNA2D2;refGene_tx=XM_023249036.1
A2 19264161 19264212
refGene_gn=CACNA2D2;refGene_tx=XM_023249003.1;refGene_gn=CACNA2D2;refGene_tx=XM_023249011.1;refGene_gn=CACNA2D2;refGene_tx=XM_023249014.1;refGene_gn=CACNA2D2;refGene_tx=XM_023249017.1;refGene_gn=CACNA2D2;refGene_tx=XM_023249020.1;refGene_gn=CACNA2D2;refGene_tx=XM_023249025.1;refGene_gn=CACNA2D2;refGene_tx=XM_023249027.1;refGene_gn=CACNA2D2;refGene_tx=XM_023249031.1;refGene_gn=CACNA2D2;refGene_tx=XM_023249036.1
A2 19264409 19264498
refGene_gn=CACNA2D2;refGene_tx=XM_023249003.1;refGene_gn=CACNA2D2;refGene_tx=XM_023249011.1;refGene_gn=CACNA2D2;refGene_tx=XM_023249014.1;refGene_gn=CACNA2D2;refGene_tx=XM_023249017.1;refGene_gn=CACNA2D2;refGene_tx=XM_023249020.1;refGene_gn=CACNA2D2;refGene_tx=XM_023249025.1;refGene_gn=CACNA2D2;refGene_tx=XM_023249027.1;refGene_gn=CACNA2D2;refGene_tx=XM_023249031.1;refGene_gn=CACNA2D2;refGene_tx=XM_023249036.1
A2 19264590 19264689
refGene_gn=CACNA2D2;refGene_tx=XM_023249003.1;refGene_gn=CACNA2D2;refGene_tx=XM_023249011.1;refGene_gn=CACNA2D2;refGene_tx=XM_023249014.1;re

fGene_gn=CACNA2D2;refGene_tx=XM_023249017.1;refGene_gn=CACNA2D2;refGene_tx=XM_023249020.1;refGene_gn=CACNA2D2;refGene_tx=XM_023249025.1;refGene_gn=CACNA2D2;refGene_tx=XM_023249027.1;refGene_gn=CACNA2D2;refGene_tx=XM_023249031.1;refGene_gn=CACNA2D2;refGene_tx=XM_023249036.1
A2 19264954 19265058
refGene_gn=CACNA2D2;refGene_tx=XM_023249003.1;refGene_gn=CACNA2D2;refGene_tx=XM_023249011.1;refGene_gn=CACNA2D2;refGene_tx=XM_023249014.1;refGene_gn=CACNA2D2;refGene_tx=XM_023249017.1;refGene_gn=CACNA2D2;refGene_tx=XM_023249020.1;refGene_gn=CACNA2D2;refGene_tx=XM_023249025.1;refGene_gn=CACNA2D2;refGene_tx=XM_023249027.1;refGene_gn=CACNA2D2;refGene_tx=XM_023249031.1;refGene_gn=CACNA2D2;refGene_tx=XM_023249036.1
A2 19265207 19265270
refGene_gn=CACNA2D2;refGene_tx=XM_023249003.1;refGene_gn=CACNA2D2;refGene_tx=XM_023249011.1;refGene_gn=CACNA2D2;refGene_tx=XM_023249014.1;refGene_gn=CACNA2D2;refGene_tx=XM_023249017.1;refGene_gn=CACNA2D2;refGene_tx=XM_023249020.1;refGene_gn=CACNA2D2;refGene_tx=XM_023249025.1;refGene_gn=CACNA2D2;refGene_tx=XM_023249027.1;refGene_gn=CACNA2D2;refGene_tx=XM_023249031.1;refGene_gn=CACNA2D2;refGene_tx=XM_023249036.1
A2 19265377 19265468
refGene_gn=CACNA2D2;refGene_tx=XM_023249003.1;refGene_gn=CACNA2D2;refGene_tx=XM_023249011.1;refGene_gn=CACNA2D2;refGene_tx=XM_023249014.1;refGene_gn=CACNA2D2;refGene_tx=XM_023249017.1;refGene_gn=CACNA2D2;refGene_tx=XM_023249020.1;refGene_gn=CACNA2D2;refGene_tx=XM_023249025.1;refGene_gn=CACNA2D2;refGene_tx=XM_023249027.1;refGene_gn=CACNA2D2;refGene_tx=XM_023249031.1;refGene_gn=CACNA2D2;refGene_tx=XM_023249036.1
A2 19265689 19265787
refGene_gn=CACNA2D2;refGene_tx=XM_023249003.1;refGene_gn=CACNA2D2;refGene_tx=XM_023249011.1;refGene_gn=CACNA2D2;refGene_tx=XM_023249014.1;refGene_gn=CACNA2D2;refGene_tx=XM_023249017.1;refGene_gn=CACNA2D2;refGene_tx=XM_023249020.1;refGene_gn=CACNA2D2;refGene_tx=XM_023249025.1;refGene_gn=CACNA2D2;refGene_tx=XM_023249027.1;refGene_gn=CACNA2D2;refGene_tx=XM_023249031.1;refGene_gn=CACNA2D2;refGene_tx=XM_023249036.1
A2 19267822 19267883
refGene_gn=CACNA2D2;refGene_tx=XM_023249003.1;refGene_gn=CACNA2D2;refGene_tx=XM_023249011.1;refGene_gn=CACNA2D2;refGene_tx=XM_023249014.1;refGene_gn=CACNA2D2;refGene_tx=XM_023249017.1;refGene_gn=CACNA2D2;refGene_tx=XM_023249020.1;refGene_gn=CACNA2D2;refGene_tx=XM_023249025.1;refGene_gn=CACNA2D2;refGene_tx=XM_023249027.1;refGene_gn=CACNA2D2;refGene_tx=XM_023249031.1;refGene_gn=CACNA2D2;refGene_tx=XM_023249036.1
A2 19270598 19270619
refGene_gn=CACNA2D2;refGene_tx=XM_023249003.1;refGene_gn=CACNA2D2;refGene_tx=XM_023249017.1;refGene_gn=CACNA2D2;refGene_tx=XM_023249020.1;refGene_gn=CACNA2D2;refGene_tx=XM_023249025.1;refGene_gn=CACNA2D2;refGene_tx=XM_023249036.1
A2 19271818 19271895
refGene_gn=CACNA2D2;refGene_tx=XM_023249003.1;refGene_gn=CACNA2D2;refGene_tx=XM_023249011.1;refGene_gn=CACNA2D2;refGene_tx=XM_023249014.1;refGene_gn=CACNA2D2;refGene_tx=XM_023249017.1;refGene_gn=CACNA2D2;refGene_tx=XM_023249020.1;refGene_gn=CACNA2D2;refGene_tx=XM_023249025.1;refGene_gn=CACNA2D2;refGene_tx=XM_023249027.1;refGene_gn=CACNA2D2;refGene_

tx=XM_023249031.1;refGene_gn=CACNA2D2;refGene_tx=XM_023249036.1
A2 19272774 19272836
refGene_gn=CACNA2D2;refGene_tx=XM_023249003.1;refGene_gn=CACNA2D2;refGene_tx=XM_023249011.1;refGene_gn=CACNA2D2;refGene_tx=XM_023249014.1;refGene_gn=CACNA2D2;refGene_tx=XM_023249017.1;refGene_gn=CACNA2D2;refGene_tx=XM_023249020.1;refGene_gn=CACNA2D2;refGene_tx=XM_023249025.1;refGene_gn=CACNA2D2;refGene_tx=XM_023249027.1;refGene_gn=CACNA2D2;refGene_tx=XM_023249031.1;refGene_gn=CACNA2D2;refGene_tx=XM_023249036.1
A2 19272940 19273012
refGene_gn=CACNA2D2;refGene_tx=XM_023249003.1;refGene_gn=CACNA2D2;refGene_tx=XM_023249011.1;refGene_gn=CACNA2D2;refGene_tx=XM_023249014.1;refGene_gn=CACNA2D2;refGene_tx=XM_023249017.1;refGene_gn=CACNA2D2;refGene_tx=XM_023249020.1;refGene_gn=CACNA2D2;refGene_tx=XM_023249025.1;refGene_gn=CACNA2D2;refGene_tx=XM_023249027.1;refGene_gn=CACNA2D2;refGene_tx=XM_023249031.1;refGene_gn=CACNA2D2;refGene_tx=XM_023249036.1
A2 19273098 19273170
refGene_gn=CACNA2D2;refGene_tx=XM_023249003.1;refGene_gn=CACNA2D2;refGene_tx=XM_023249011.1;refGene_gn=CACNA2D2;refGene_tx=XM_023249014.1;refGene_gn=CACNA2D2;refGene_tx=XM_023249017.1;refGene_gn=CACNA2D2;refGene_tx=XM_023249020.1;refGene_gn=CACNA2D2;refGene_tx=XM_023249025.1;refGene_gn=CACNA2D2;refGene_tx=XM_023249027.1;refGene_gn=CACNA2D2;refGene_tx=XM_023249031.1;refGene_gn=CACNA2D2;refGene_tx=XM_023249036.1
A2 19273248 19273323
refGene_gn=CACNA2D2;refGene_tx=XM_023249003.1;refGene_gn=CACNA2D2;refGene_tx=XM_023249011.1;refGene_gn=CACNA2D2;refGene_tx=XM_023249014.1;refGene_gn=CACNA2D2;refGene_tx=XM_023249017.1;refGene_gn=CACNA2D2;refGene_tx=XM_023249020.1;refGene_gn=CACNA2D2;refGene_tx=XM_023249025.1;refGene_gn=CACNA2D2;refGene_tx=XM_023249027.1;refGene_gn=CACNA2D2;refGene_tx=XM_023249031.1;refGene_gn=CACNA2D2;refGene_tx=XM_023249036.1
A2 19274544 19274619
refGene_gn=CACNA2D2;refGene_tx=XM_023249003.1;refGene_gn=CACNA2D2;refGene_tx=XM_023249011.1;refGene_gn=CACNA2D2;refGene_tx=XM_023249014.1;refGene_gn=CACNA2D2;refGene_tx=XM_023249017.1;refGene_gn=CACNA2D2;refGene_tx=XM_023249020.1;refGene_gn=CACNA2D2;refGene_tx=XM_023249025.1;refGene_gn=CACNA2D2;refGene_tx=XM_023249027.1;refGene_gn=CACNA2D2;refGene_tx=XM_023249031.1;refGene_gn=CACNA2D2;refGene_tx=XM_023249036.1
A2 19274808 19274880
refGene_gn=CACNA2D2;refGene_tx=XM_023249003.1;refGene_gn=CACNA2D2;refGene_tx=XM_023249011.1;refGene_gn=CACNA2D2;refGene_tx=XM_023249014.1;refGene_gn=CACNA2D2;refGene_tx=XM_023249017.1;refGene_gn=CACNA2D2;refGene_tx=XM_023249020.1;refGene_gn=CACNA2D2;refGene_tx=XM_023249025.1;refGene_gn=CACNA2D2;refGene_tx=XM_023249027.1;refGene_gn=CACNA2D2;refGene_tx=XM_023249031.1;refGene_gn=CACNA2D2;refGene_tx=XM_023249036.1
A2 19275058 19275148
refGene_gn=CACNA2D2;refGene_tx=XM_023249003.1;refGene_gn=CACNA2D2;refGene_tx=XM_023249011.1;refGene_gn=CACNA2D2;refGene_tx=XM_023249014.1;refGene_gn=CACNA2D2;refGene_tx=XM_023249017.1;refGene_gn=CACNA2D2;refGene_tx=XM_023249020.1;refGene_gn=CACNA2D2;refGene_tx=XM_023249025.1;refGene_gn=CACNA2D2;refGene_tx=XM_023249027.1;refGene_gn=CACNA2D2;refGene_tx=XM_023249031.1;refGene_gn=CACNA2D2;refGene_tx=XM_023249036.1

A2 19275329 19275379
refGene_gn=CACNA2D2;refGene_tx=XM_023249003.1;refGene_gn=CACNA2D2;refGene_tx=XM_023249011.1;refGene_gn=CACNA2D2;refGene_tx=XM_023249014.1;refGene_gn=CACNA2D2;refGene_tx=XM_023249017.1;refGene_gn=CACNA2D2;refGene_tx=XM_023249020.1;refGene_gn=CACNA2D2;refGene_tx=XM_023249025.1;refGene_gn=CACNA2D2;refGene_tx=XM_023249027.1;refGene_gn=CACNA2D2;refGene_tx=XM_023249031.1;refGene_gn=CACNA2D2;refGene_tx=XM_023249036.1
A2 19275954 19276033
refGene_gn=CACNA2D2;refGene_tx=XM_023249003.1;refGene_gn=CACNA2D2;refGene_tx=XM_023249011.1;refGene_gn=CACNA2D2;refGene_tx=XM_023249014.1;refGene_gn=CACNA2D2;refGene_tx=XM_023249017.1;refGene_gn=CACNA2D2;refGene_tx=XM_023249020.1;refGene_gn=CACNA2D2;refGene_tx=XM_023249025.1;refGene_gn=CACNA2D2;refGene_tx=XM_023249027.1;refGene_gn=CACNA2D2;refGene_tx=XM_023249031.1;refGene_gn=CACNA2D2;refGene_tx=XM_023249036.1
A2 19276131 19276239
refGene_gn=CACNA2D2;refGene_tx=XM_023249003.1;refGene_gn=CACNA2D2;refGene_tx=XM_023249011.1;refGene_gn=CACNA2D2;refGene_tx=XM_023249014.1;refGene_gn=CACNA2D2;refGene_tx=XM_023249017.1;refGene_gn=CACNA2D2;refGene_tx=XM_023249020.1;refGene_gn=CACNA2D2;refGene_tx=XM_023249025.1;refGene_gn=CACNA2D2;refGene_tx=XM_023249027.1;refGene_gn=CACNA2D2;refGene_tx=XM_023249031.1;refGene_gn=CACNA2D2;refGene_tx=XM_023249036.1
A2 19276474 19276633
refGene_gn=CACNA2D2;refGene_tx=XM_023249003.1;refGene_gn=CACNA2D2;refGene_tx=XM_023249011.1;refGene_gn=CACNA2D2;refGene_tx=XM_023249014.1;refGene_gn=CACNA2D2;refGene_tx=XM_023249017.1;refGene_gn=CACNA2D2;refGene_tx=XM_023249020.1;refGene_gn=CACNA2D2;refGene_tx=XM_023249025.1;refGene_gn=CACNA2D2;refGene_tx=XM_023249027.1;refGene_gn=CACNA2D2;refGene_tx=XM_023249031.1
A2 19276753 19276853
refGene_gn=CACNA2D2;refGene_tx=XM_023249003.1;refGene_gn=CACNA2D2;refGene_tx=XM_023249011.1;refGene_gn=CACNA2D2;refGene_tx=XM_023249014.1;refGene_gn=CACNA2D2;refGene_tx=XM_023249017.1;refGene_gn=CACNA2D2;refGene_tx=XM_023249020.1;refGene_gn=CACNA2D2;refGene_tx=XM_023249025.1;refGene_gn=CACNA2D2;refGene_tx=XM_023249027.1;refGene_gn=CACNA2D2;refGene_tx=XM_023249031.1;refGene_gn=CACNA2D2;refGene_tx=XM_023249036.1
A2 19276981 19277032
refGene_gn=CACNA2D2;refGene_tx=XM_023249003.1;refGene_gn=CACNA2D2;refGene_tx=XM_023249011.1;refGene_gn=CACNA2D2;refGene_tx=XM_023249014.1;refGene_gn=CACNA2D2;refGene_tx=XM_023249017.1;refGene_gn=CACNA2D2;refGene_tx=XM_023249020.1;refGene_gn=CACNA2D2;refGene_tx=XM_023249025.1;refGene_gn=CACNA2D2;refGene_tx=XM_023249027.1;refGene_gn=CACNA2D2;refGene_tx=XM_023249031.1;refGene_gn=CACNA2D2;refGene_tx=XM_023249036.1
A2 19277691 19277749
refGene_gn=CACNA2D2;refGene_tx=XM_023249003.1;refGene_gn=CACNA2D2;refGene_tx=XM_023249011.1;refGene_gn=CACNA2D2;refGene_tx=XM_023249014.1;refGene_gn=CACNA2D2;refGene_tx=XM_023249017.1;refGene_gn=CACNA2D2;refGene_tx=XM_023249020.1;refGene_gn=CACNA2D2;refGene_tx=XM_023249025.1;refGene_gn=CACNA2D2;refGene_tx=XM_023249027.1;refGene_gn=CACNA2D2;refGene_tx=XM_023249031.1;refGene_gn=CACNA2D2;refGene_tx=XM_023249036.1
A2 19277937 19278069

refGene_gn=CACNA2D2;refGene_tx=XM_023249003.1;refGene_gn=CACNA2D2;refGene_tx=XM_023249011.1;refGene_gn=CACNA2D2;refGene_tx=XM_023249014.1;refGene_gn=CACNA2D2;refGene_tx=XM_023249017.1;refGene_gn=CACNA2D2;refGene_tx=XM_023249020.1;refGene_gn=CACNA2D2;refGene_tx=XM_023249025.1;refGene_gn=CACNA2D2;refGene_tx=XM_023249027.1;refGene_gn=CACNA2D2;refGene_tx=XM_023249031.1;refGene_gn=CACNA2D2;refGene_tx=XM_023249036.1
A2 19281361 19281503
refGene_gn=CACNA2D2;refGene_tx=XM_023249003.1;refGene_gn=CACNA2D2;refGene_tx=XM_023249011.1;refGene_gn=CACNA2D2;refGene_tx=XM_023249014.1;refGene_gn=CACNA2D2;refGene_tx=XM_023249017.1;refGene_gn=CACNA2D2;refGene_tx=XM_023249020.1;refGene_gn=CACNA2D2;refGene_tx=XM_023249025.1;refGene_gn=CACNA2D2;refGene_tx=XM_023249027.1;refGene_gn=CACNA2D2;refGene_tx=XM_023249031.1;refGene_gn=CACNA2D2;refGene_tx=XM_023249036.1
A2 19284734 19284779
refGene_gn=CACNA2D2;refGene_tx=XM_023249003.1;refGene_gn=CACNA2D2;refGene_tx=XM_023249011.1;refGene_gn=CACNA2D2;refGene_tx=XM_023249014.1;refGene_gn=CACNA2D2;refGene_tx=XM_023249017.1;refGene_gn=CACNA2D2;refGene_tx=XM_023249020.1;refGene_gn=CACNA2D2;refGene_tx=XM_023249025.1;refGene_gn=CACNA2D2;refGene_tx=XM_023249027.1;refGene_gn=CACNA2D2;refGene_tx=XM_023249031.1;refGene_gn=CACNA2D2;refGene_tx=XM_023249036.1
A2 19292562 19292622
refGene_gn=CACNA2D2;refGene_tx=XM_023249003.1;refGene_gn=CACNA2D2;refGene_tx=XM_023249011.1;refGene_gn=CACNA2D2;refGene_tx=XM_023249014.1;refGene_gn=CACNA2D2;refGene_tx=XM_023249017.1;refGene_gn=CACNA2D2;refGene_tx=XM_023249020.1;refGene_gn=CACNA2D2;refGene_tx=XM_023249025.1;refGene_gn=CACNA2D2;refGene_tx=XM_023249027.1;refGene_gn=CACNA2D2;refGene_tx=XM_023249031.1;refGene_gn=CACNA2D2;refGene_tx=XM_023249036.1
A2 19331893 19332010
refGene_gn=CACNA2D2;refGene_tx=XM_023249003.1;refGene_gn=CACNA2D2;refGene_tx=XM_023249011.1;refGene_gn=CACNA2D2;refGene_tx=XM_023249014.1;refGene_gn=CACNA2D2;refGene_tx=XM_023249017.1;refGene_gn=CACNA2D2;refGene_tx=XM_023249020.1;refGene_gn=CACNA2D2;refGene_tx=XM_023249025.1;refGene_gn=CACNA2D2;refGene_tx=XM_023249027.1;refGene_gn=CACNA2D2;refGene_tx=XM_023249031.1;refGene_gn=CACNA2D2;refGene_tx=XM_023249036.1
A2 19377468 19377550
refGene_gn=CACNA2D2;refGene_tx=XM_023249003.1;refGene_gn=CACNA2D2;refGene_tx=XM_023249011.1;refGene_gn=CACNA2D2;refGene_tx=XM_023249014.1;refGene_gn=CACNA2D2;refGene_tx=XM_023249017.1;refGene_gn=CACNA2D2;refGene_tx=XM_023249020.1;refGene_gn=CACNA2D2;refGene_tx=XM_023249025.1;refGene_gn=CACNA2D2;refGene_tx=XM_023249027.1;refGene_gn=CACNA2D2;refGene_tx=XM_023249031.1;refGene_gn=CACNA2D2;refGene_tx=XM_023249036.1
A2 19404169 19404384
refGene_gn=CACNA2D2;refGene_tx=XM_023249003.1;refGene_gn=CACNA2D2;refGene_tx=XM_023249011.1;refGene_gn=CACNA2D2;refGene_tx=XM_023249014.1;refGene_gn=CACNA2D2;refGene_tx=XM_023249017.1;refGene_gn=CACNA2D2;refGene_tx=XM_023249020.1;refGene_gn=CACNA2D2;refGene_tx=XM_023249025.1;refGene_gn=CACNA2D2;refGene_tx=XM_023249027.1;refGene_gn=CACNA2D2;refGene_tx=XM_023249031.1;refGene_gn=CACNA2D2;refGene_tx=XM_023249036.1
A2 19455745 19455826
refGene_gn=CA2H3orf18;refGene_tx=XM_006928813.3;refGene_gn=CA2H3orf18;

refGene_tx=XM_006928814.3;refGene_gn=CA2H3orf18;refGene_tx=XM_01982213
3.2
A2 19457075 19457223
refGene_gn=CA2H3orf18;refGene_tx=XM_006928813.3;refGene_gn=CA2H3orf18;
refGene_tx=XM_006928814.3;refGene_gn=CA2H3orf18;refGene_tx=XM_01982213
3.2
A2 19457752 19457778
refGene_gn=CA2H3orf18;refGene_tx=XM_006928813.3;refGene_gn=CA2H3orf18;
refGene_tx=XM_006928814.3;refGene_gn=CA2H3orf18;refGene_tx=XM_01982213
3.2
A2 19459346 19459484
refGene_gn=CA2H3orf18;refGene_tx=XM_019822133.2
A2 19460437 19460671
refGene_gn=CA2H3orf18;refGene_tx=XM_006928813.3;refGene_gn=CA2H3orf18;
refGene_tx=XM_006928814.3
A2 19465355 19465583
refGene_gn=HEMK1;refGene_tx=XM_006928815.4;refGene_gn=HEMK1;refGene_tx
=XM_019822143.1;refGene_gn=HEMK1;refGene_tx=XM_019822165.1
A2 19465881 19465973
refGene_gn=HEMK1;refGene_tx=XM_006928815.4;refGene_gn=HEMK1;refGene_tx
=XM_019822143.1
A2 19466285 19466379
refGene_gn=HEMK1;refGene_tx=XM_006928815.4;refGene_gn=HEMK1;refGene_tx
=XM_019822143.1
A2 19470348 19470483
refGene_gn=HEMK1;refGene_tx=XM_006928815.4;refGene_gn=HEMK1;refGene_tx
=XM_019822143.1;refGene_gn=HEMK1;refGene_tx=XM_019822165.1
A2 19470811 19470876
refGene_gn=HEMK1;refGene_tx=XM_006928815.4;refGene_gn=HEMK1;refGene_tx
=XM_019822143.1;refGene_gn=HEMK1;refGene_tx=XM_019822165.1
A2 19471153 19471203
refGene_gn=HEMK1;refGene_tx=XM_006928815.4;refGene_gn=HEMK1;refGene_tx
=XM_019822143.1;refGene_gn=HEMK1;refGene_tx=XM_019822165.1
A2 19472137 19472246
refGene_gn=HEMK1;refGene_tx=XM_006928815.4;refGene_gn=HEMK1;refGene_tx
=XM_019822143.1;refGene_gn=HEMK1;refGene_tx=XM_019822165.1
A2 19473132 19473228
refGene_gn=HEMK1;refGene_tx=XM_006928815.4;refGene_gn=HEMK1;refGene_tx
=XM_019822165.1
A2 19473379 19473493
refGene_gn=HEMK1;refGene_tx=XM_006928815.4;refGene_gn=HEMK1;refGene_tx
=XM_019822143.1;refGene_gn=HEMK1;refGene_tx=XM_019822165.1
A2 19473656 19473717
refGene_gn=HEMK1;refGene_tx=XM_006928815.4;refGene_gn=HEMK1;refGene_tx
=XM_019822143.1;refGene_gn=HEMK1;refGene_tx=XM_019822165.1
A2 19497000 19497533 refGene_gn=CISH;refGene_tx=XM_003982272.5
A2 19497773 19497994 refGene_gn=CISH;refGene_tx=XM_003982272.5
A2 19501058 19501078 refGene_gn=CISH;refGene_tx=XM_003982272.5
A2 19506004 19506229
refGene_gn=MAPKAPK3;refGene_tx=XM_023249100.1;refGene_gn=MAPKAPK3;refG

ene_tx=XM_023249102.1
A2 19525211 19525351
refGene_gn=MAPKAPK3;refGene_tx=XM_019822168.2;refGene_gn=MAPKAPK3;refGene_tx=XM_023249100.1;refGene_gn=MAPKAPK3;refGene_tx=XM_023249102.1
A2 19526535 19526600
refGene_gn=MAPKAPK3;refGene_tx=XM_019822168.2;refGene_gn=MAPKAPK3;refGene_tx=XM_023249100.1;refGene_gn=MAPKAPK3;refGene_tx=XM_023249102.1
A2 19527069 19527149
refGene_gn=MAPKAPK3;refGene_tx=XM_019822168.2;refGene_gn=MAPKAPK3;refGene_tx=XM_023249100.1;refGene_gn=MAPKAPK3;refGene_tx=XM_023249102.1
A2 19528937 19529061
refGene_gn=MAPKAPK3;refGene_tx=XM_019822168.2;refGene_gn=MAPKAPK3;refGene_tx=XM_023249100.1;refGene_gn=MAPKAPK3;refGene_tx=XM_023249102.1
A2 19529639 19529769 refGene_gn=MAPKAPK3;refGene_tx=XM_019822168.2
A2 19530234 19530310
refGene_gn=MAPKAPK3;refGene_tx=XM_019822168.2;refGene_gn=MAPKAPK3;refGene_tx=XM_023249100.1;refGene_gn=MAPKAPK3;refGene_tx=XM_023249102.1
A2 19530656 19530781
refGene_gn=MAPKAPK3;refGene_tx=XM_019822168.2;refGene_gn=MAPKAPK3;refGene_tx=XM_023249100.1;refGene_gn=MAPKAPK3;refGene_tx=XM_023249102.1
A2 19531206 19531292
refGene_gn=MAPKAPK3;refGene_tx=XM_019822168.2;refGene_gn=MAPKAPK3;refGene_tx=XM_023249100.1;refGene_gn=MAPKAPK3;refGene_tx=XM_023249102.1
A2 19531565 19531646
refGene_gn=MAPKAPK3;refGene_tx=XM_019822168.2;refGene_gn=MAPKAPK3;refGene_tx=XM_023249100.1;refGene_gn=MAPKAPK3;refGene_tx=XM_023249102.1
A2 19532333 19532486
refGene_gn=MAPKAPK3;refGene_tx=XM_019822168.2;refGene_gn=MAPKAPK3;refGene_tx=XM_023249100.1;refGene_gn=MAPKAPK3;refGene_tx=XM_023249102.1
A2 19558320 19558357
refGene_gn=DOCK3;refGene_tx=XM_023249110.1;refGene_gn=DOCK3;refGene_tx=XM_023249114.1;refGene_gn=DOCK3;refGene_tx=XM_023249115.1;refGene_gn=DOCK3;refGene_tx=XM_023249117.1;refGene_gn=DOCK3;refGene_tx=XM_023249119.1;refGene_gn=DOCK3;refGene_tx=XM_023249123.1;refGene_gn=DOCK3;refGene_tx=XM_023249126.1;refGene_gn=DOCK3;refGene_tx=XM_023249134.1;refGene_gn=DOCK3;refGene_tx=XM_023249135.1;refGene_gn=DOCK3;refGene_tx=XM_023249140.1
A2 19659892 19659976
refGene_gn=DOCK3;refGene_tx=XM_023249110.1;refGene_gn=DOCK3;refGene_tx=XM_023249114.1;refGene_gn=DOCK3;refGene_tx=XM_023249115.1;refGene_gn=DOCK3;refGene_tx=XM_023249117.1;refGene_gn=DOCK3;refGene_tx=XM_023249119.1;refGene_gn=DOCK3;refGene_tx=XM_023249123.1;refGene_gn=DOCK3;refGene_tx=XM_023249126.1;refGene_gn=DOCK3;refGene_tx=XM_023249134.1;refGene_gn=DOCK3;refGene_tx=XM_023249135.1;refGene_gn=DOCK3;refGene_tx=XM_023249140.1
A2 19705237 19705278
refGene_gn=DOCK3;refGene_tx=XM_023249110.1;refGene_gn=DOCK3;refGene_tx=XM_023249114.1;refGene_gn=DOCK3;refGene_tx=XM_023249115.1;refGene_gn=DOCK3;refGene_tx=XM_023249117.1;refGene_gn=DOCK3;refGene_tx=XM_023249119.1;refGene_gn=DOCK3;refGene_tx=XM_023249123.1;refGene_gn=DOCK3;refGe

ne_tx=XM_023249126.1;refGene_gn=DOCK3;refGene_tx=XM_023249134.1;refGene_gn=DOCK3;refGene_tx=XM_023249135.1;refGene_gn=DOCK3;refGene_tx=XM_023249140.1
A2 19709291 19709312 refGene_gn=DOCK3;refGene_tx=XM_023249130.1
A2 19747645 19747701
refGene_gn=DOCK3;refGene_tx=XM_023249110.1;refGene_gn=DOCK3;refGene_tx=XM_023249114.1;refGene_gn=DOCK3;refGene_tx=XM_023249115.1;refGene_gn=DOCK3;refGene_tx=XM_023249117.1;refGene_gn=DOCK3;refGene_tx=XM_023249119.1;refGene_gn=DOCK3;refGene_tx=XM_023249123.1;refGene_gn=DOCK3;refGene_tx=XM_023249126.1;refGene_gn=DOCK3;refGene_tx=XM_023249130.1;refGene_gn=DOCK3;refGene_tx=XM_023249134.1;refGene_gn=DOCK3;refGene_tx=XM_023249135.1;refGene_gn=DOCK3;refGene_tx=XM_023249140.1
A2 19778615 19778712
refGene_gn=DOCK3;refGene_tx=XM_023249110.1;refGene_gn=DOCK3;refGene_tx=XM_023249114.1;refGene_gn=DOCK3;refGene_tx=XM_023249115.1;refGene_gn=DOCK3;refGene_tx=XM_023249117.1;refGene_gn=DOCK3;refGene_tx=XM_023249119.1;refGene_gn=DOCK3;refGene_tx=XM_023249123.1;refGene_gn=DOCK3;refGene_tx=XM_023249126.1;refGene_gn=DOCK3;refGene_tx=XM_023249130.1;refGene_gn=DOCK3;refGene_tx=XM_023249134.1;refGene_gn=DOCK3;refGene_tx=XM_023249135.1;refGene_gn=DOCK3;refGene_tx=XM_023249140.1
A2 19927440 19927589
refGene_gn=DOCK3;refGene_tx=XM_023249110.1;refGene_gn=DOCK3;refGene_tx=XM_023249114.1;refGene_gn=DOCK3;refGene_tx=XM_023249115.1;refGene_gn=DOCK3;refGene_tx=XM_023249117.1;refGene_gn=DOCK3;refGene_tx=XM_023249119.1;refGene_gn=DOCK3;refGene_tx=XM_023249123.1;refGene_gn=DOCK3;refGene_tx=XM_023249126.1;refGene_gn=DOCK3;refGene_tx=XM_023249130.1;refGene_gn=DOCK3;refGene_tx=XM_023249134.1;refGene_gn=DOCK3;refGene_tx=XM_023249135.1;refGene_gn=DOCK3;refGene_tx=XM_023249137.1;refGene_gn=DOCK3;refGene_tx=XM_023249140.1
A2 19943416 19943501
refGene_gn=DOCK3;refGene_tx=XM_023249110.1;refGene_gn=DOCK3;refGene_tx=XM_023249114.1;refGene_gn=DOCK3;refGene_tx=XM_023249115.1;refGene_gn=DOCK3;refGene_tx=XM_023249117.1;refGene_gn=DOCK3;refGene_tx=XM_023249119.1;refGene_gn=DOCK3;refGene_tx=XM_023249123.1;refGene_gn=DOCK3;refGene_tx=XM_023249126.1;refGene_gn=DOCK3;refGene_tx=XM_023249130.1;refGene_gn=DOCK3;refGene_tx=XM_023249134.1;refGene_gn=DOCK3;refGene_tx=XM_023249135.1;refGene_gn=DOCK3;refGene_tx=XM_023249137.1;refGene_gn=DOCK3;refGene_tx=XM_023249140.1
A2 19961803 19961845
refGene_gn=DOCK3;refGene_tx=XM_023249110.1;refGene_gn=DOCK3;refGene_tx=XM_023249114.1;refGene_gn=DOCK3;refGene_tx=XM_023249115.1;refGene_gn=DOCK3;refGene_tx=XM_023249117.1;refGene_gn=DOCK3;refGene_tx=XM_023249119.1;refGene_gn=DOCK3;refGene_tx=XM_023249123.1;refGene_gn=DOCK3;refGene_tx=XM_023249126.1;refGene_gn=DOCK3;refGene_tx=XM_023249130.1;refGene_gn=DOCK3;refGene_tx=XM_023249134.1;refGene_gn=DOCK3;refGene_tx=XM_023249135.1;refGene_gn=DOCK3;refGene_tx=XM_023249137.1;refGene_gn=DOCK3;refGene_tx=XM_023249140.1
A2 19962486 19962641
refGene_gn=DOCK3;refGene_tx=XM_023249110.1;refGene_gn=DOCK3;refGene_tx=XM_023249114.1;refGene_gn=DOCK3;refGene_tx=XM_023249115.1;refGene_gn=

DOCK3;refGene_tx=XM_023249117.1;refGene_gn=DOCK3;refGene_tx=XM_0232491
19.1;refGene_gn=DOCK3;refGene_tx=XM_023249123.1;refGene_gn=DOCK3;refGe
ne_tx=XM_023249126.1;refGene_gn=DOCK3;refGene_tx=XM_023249130.1;refGen
e_gn=DOCK3;refGene_tx=XM_023249134.1;refGene_gn=DOCK3;refGene_tx=XM_02
3249135.1;refGene_gn=DOCK3;refGene_tx=XM_023249137.1;refGene_gn=DOCK3;
refGene_tx=XM_023249140.1
A2 20021069 20021151
refGene_gn=DOCK3;refGene_tx=XM_023249110.1;refGene_gn=DOCK3;refGene_tx
=XM_023249114.1;refGene_gn=DOCK3;refGene_tx=XM_023249115.1;refGene_gn=
DOCK3;refGene_tx=XM_023249117.1;refGene_gn=DOCK3;refGene_tx=XM_0232491
19.1;refGene_gn=DOCK3;refGene_tx=XM_023249123.1;refGene_gn=DOCK3;refGe
ne_tx=XM_023249126.1;refGene_gn=DOCK3;refGene_tx=XM_023249130.1;refGen
e_gn=DOCK3;refGene_tx=XM_023249134.1;refGene_gn=DOCK3;refGene_tx=XM_02
3249135.1;refGene_gn=DOCK3;refGene_tx=XM_023249137.1;refGene_gn=DOCK3;
refGene_tx=XM_023249140.1
A2 20026089 20026150
refGene_gn=DOCK3;refGene_tx=XM_023249110.1;refGene_gn=DOCK3;refGene_tx
=XM_023249114.1;refGene_gn=DOCK3;refGene_tx=XM_023249115.1;refGene_gn=
DOCK3;refGene_tx=XM_023249117.1;refGene_gn=DOCK3;refGene_tx=XM_0232491
19.1;refGene_gn=DOCK3;refGene_tx=XM_023249123.1;refGene_gn=DOCK3;refGe
ne_tx=XM_023249126.1;refGene_gn=DOCK3;refGene_tx=XM_023249130.1;refGen
e_gn=DOCK3;refGene_tx=XM_023249134.1;refGene_gn=DOCK3;refGene_tx=XM_02
3249135.1;refGene_gn=DOCK3;refGene_tx=XM_023249137.1;refGene_gn=DOCK3;
refGene_tx=XM_023249140.1
A2 20027436 20027584
refGene_gn=DOCK3;refGene_tx=XM_023249110.1;refGene_gn=DOCK3;refGene_tx
=XM_023249114.1;refGene_gn=DOCK3;refGene_tx=XM_023249115.1;refGene_gn=
DOCK3;refGene_tx=XM_023249117.1;refGene_gn=DOCK3;refGene_tx=XM_0232491
19.1;refGene_gn=DOCK3;refGene_tx=XM_023249123.1;refGene_gn=DOCK3;refGe
ne_tx=XM_023249126.1;refGene_gn=DOCK3;refGene_tx=XM_023249130.1;refGen
e_gn=DOCK3;refGene_tx=XM_023249134.1;refGene_gn=DOCK3;refGene_tx=XM_02
3249135.1;refGene_gn=DOCK3;refGene_tx=XM_023249137.1;refGene_gn=DOCK3;
refGene_tx=XM_023249140.1
A2 20044194 20044283
refGene_gn=DOCK3;refGene_tx=XM_023249110.1;refGene_gn=DOCK3;refGene_tx
=XM_023249114.1;refGene_gn=DOCK3;refGene_tx=XM_023249115.1;refGene_gn=
DOCK3;refGene_tx=XM_023249117.1;refGene_gn=DOCK3;refGene_tx=XM_0232491
19.1;refGene_gn=DOCK3;refGene_tx=XM_023249123.1;refGene_gn=DOCK3;refGe
ne_tx=XM_023249126.1;refGene_gn=DOCK3;refGene_tx=XM_023249130.1;refGen
e_gn=DOCK3;refGene_tx=XM_023249134.1;refGene_gn=DOCK3;refGene_tx=XM_02
3249135.1;refGene_gn=DOCK3;refGene_tx=XM_023249137.1;refGene_gn=DOCK3;
refGene_tx=XM_023249140.1
A2 20049646 20049772
refGene_gn=DOCK3;refGene_tx=XM_023249110.1;refGene_gn=DOCK3;refGene_tx
=XM_023249114.1;refGene_gn=DOCK3;refGene_tx=XM_023249115.1;refGene_gn=
DOCK3;refGene_tx=XM_023249117.1;refGene_gn=DOCK3;refGene_tx=XM_0232491
19.1;refGene_gn=DOCK3;refGene_tx=XM_023249123.1;refGene_gn=DOCK3;refGe
ne_tx=XM_023249130.1;refGene_gn=DOCK3;refGene_tx=XM_023249134.1;refGen
e_gn=DOCK3;refGene_tx=XM_023249135.1;refGene_gn=DOCK3;refGene_tx=XM_02
3249137.1;refGene_gn=DOCK3;refGene_tx=XM_023249140.1

A2 20061564 20061689
refGene_gn=DOCK3;refGene_tx=XM_023249110.1;refGene_gn=DOCK3;refGene_tx=XM_023249114.1;refGene_gn=DOCK3;refGene_tx=XM_023249115.1;refGene_gn=DOCK3;refGene_tx=XM_023249117.1;refGene_gn=DOCK3;refGene_tx=XM_023249119.1;refGene_gn=DOCK3;refGene_tx=XM_023249123.1;refGene_gn=DOCK3;refGene_tx=XM_023249126.1;refGene_gn=DOCK3;refGene_tx=XM_023249130.1;refGene_gn=DOCK3;refGene_tx=XM_023249134.1;refGene_gn=DOCK3;refGene_tx=XM_023249135.1;refGene_gn=DOCK3;refGene_tx=XM_023249137.1;refGene_gn=DOCK3;refGene_tx=XM_023249140.1
A2 20063206 20063369
refGene_gn=DOCK3;refGene_tx=XM_023249110.1;refGene_gn=DOCK3;refGene_tx=XM_023249114.1;refGene_gn=DOCK3;refGene_tx=XM_023249115.1;refGene_gn=DOCK3;refGene_tx=XM_023249117.1;refGene_gn=DOCK3;refGene_tx=XM_023249119.1;refGene_gn=DOCK3;refGene_tx=XM_023249123.1;refGene_gn=DOCK3;refGene_tx=XM_023249126.1;refGene_gn=DOCK3;refGene_tx=XM_023249130.1;refGene_gn=DOCK3;refGene_tx=XM_023249134.1;refGene_gn=DOCK3;refGene_tx=XM_023249135.1;refGene_gn=DOCK3;refGene_tx=XM_023249137.1;refGene_gn=DOCK3;refGene_tx=XM_023249140.1
A2 20063906 20064013
refGene_gn=DOCK3;refGene_tx=XM_023249110.1;refGene_gn=DOCK3;refGene_tx=XM_023249114.1;refGene_gn=DOCK3;refGene_tx=XM_023249115.1;refGene_gn=DOCK3;refGene_tx=XM_023249117.1;refGene_gn=DOCK3;refGene_tx=XM_023249119.1;refGene_gn=DOCK3;refGene_tx=XM_023249123.1;refGene_gn=DOCK3;refGene_tx=XM_023249126.1;refGene_gn=DOCK3;refGene_tx=XM_023249130.1;refGene_gn=DOCK3;refGene_tx=XM_023249134.1;refGene_gn=DOCK3;refGene_tx=XM_023249135.1;refGene_gn=DOCK3;refGene_tx=XM_023249137.1;refGene_gn=DOCK3;refGene_tx=XM_023249140.1
A2 20064578 20064750
refGene_gn=DOCK3;refGene_tx=XM_023249110.1;refGene_gn=DOCK3;refGene_tx=XM_023249114.1;refGene_gn=DOCK3;refGene_tx=XM_023249115.1;refGene_gn=DOCK3;refGene_tx=XM_023249117.1;refGene_gn=DOCK3;refGene_tx=XM_023249119.1;refGene_gn=DOCK3;refGene_tx=XM_023249123.1;refGene_gn=DOCK3;refGene_tx=XM_023249126.1;refGene_gn=DOCK3;refGene_tx=XM_023249130.1;refGene_gn=DOCK3;refGene_tx=XM_023249134.1;refGene_gn=DOCK3;refGene_tx=XM_023249135.1;refGene_gn=DOCK3;refGene_tx=XM_023249137.1;refGene_gn=DOCK3;refGene_tx=XM_023249140.1
A2 20065122 20065220
refGene_gn=DOCK3;refGene_tx=XM_023249110.1;refGene_gn=DOCK3;refGene_tx=XM_023249114.1;refGene_gn=DOCK3;refGene_tx=XM_023249115.1;refGene_gn=DOCK3;refGene_tx=XM_023249117.1;refGene_gn=DOCK3;refGene_tx=XM_023249119.1;refGene_gn=DOCK3;refGene_tx=XM_023249123.1;refGene_gn=DOCK3;refGene_tx=XM_023249126.1;refGene_gn=DOCK3;refGene_tx=XM_023249130.1;refGene_gn=DOCK3;refGene_tx=XM_023249134.1;refGene_gn=DOCK3;refGene_tx=XM_023249135.1;refGene_gn=DOCK3;refGene_tx=XM_023249137.1;refGene_gn=DOCK3;refGene_tx=XM_023249140.1
A2 20067998 20068082
refGene_gn=DOCK3;refGene_tx=XM_023249110.1;refGene_gn=DOCK3;refGene_tx=XM_023249114.1;refGene_gn=DOCK3;refGene_tx=XM_023249115.1;refGene_gn=DOCK3;refGene_tx=XM_023249117.1;refGene_gn=DOCK3;refGene_tx=XM_023249119.1;refGene_gn=DOCK3;refGene_tx=XM_023249123.1;refGene_gn=DOCK3;refGe

ne_tx=XM_023249126.1;refGene_gn=DOCK3;refGene_tx=XM_023249130.1;refGene_gn=DOCK3;refGene_tx=XM_023249134.1;refGene_gn=DOCK3;refGene_tx=XM_023249135.1;refGene_gn=DOCK3;refGene_tx=XM_023249137.1;refGene_gn=DOCK3;refGene_tx=XM_023249140.1
A2 20069169 20069270
refGene_gn=DOCK3;refGene_tx=XM_023249110.1;refGene_gn=DOCK3;refGene_tx=XM_023249114.1;refGene_gn=DOCK3;refGene_tx=XM_023249115.1;refGene_gn=DOCK3;refGene_tx=XM_023249117.1;refGene_gn=DOCK3;refGene_tx=XM_023249119.1;refGene_gn=DOCK3;refGene_tx=XM_023249123.1;refGene_gn=DOCK3;refGene_tx=XM_023249126.1;refGene_gn=DOCK3;refGene_tx=XM_023249130.1;refGene_gn=DOCK3;refGene_tx=XM_023249134.1;refGene_gn=DOCK3;refGene_tx=XM_023249135.1;refGene_gn=DOCK3;refGene_tx=XM_023249137.1;refGene_gn=DOCK3;refGene_tx=XM_023249140.1
A2 20072086 20072168
refGene_gn=DOCK3;refGene_tx=XM_023249110.1;refGene_gn=DOCK3;refGene_tx=XM_023249114.1;refGene_gn=DOCK3;refGene_tx=XM_023249115.1;refGene_gn=DOCK3;refGene_tx=XM_023249117.1;refGene_gn=DOCK3;refGene_tx=XM_023249119.1;refGene_gn=DOCK3;refGene_tx=XM_023249123.1;refGene_gn=DOCK3;refGene_tx=XM_023249126.1;refGene_gn=DOCK3;refGene_tx=XM_023249130.1;refGene_gn=DOCK3;refGene_tx=XM_023249134.1;refGene_gn=DOCK3;refGene_tx=XM_023249135.1;refGene_gn=DOCK3;refGene_tx=XM_023249137.1;refGene_gn=DOCK3;refGene_tx=XM_023249140.1
A2 20076346 20076517
refGene_gn=DOCK3;refGene_tx=XM_023249110.1;refGene_gn=DOCK3;refGene_tx=XM_023249114.1;refGene_gn=DOCK3;refGene_tx=XM_023249115.1;refGene_gn=DOCK3;refGene_tx=XM_023249117.1;refGene_gn=DOCK3;refGene_tx=XM_023249119.1;refGene_gn=DOCK3;refGene_tx=XM_023249123.1;refGene_gn=DOCK3;refGene_tx=XM_023249126.1;refGene_gn=DOCK3;refGene_tx=XM_023249130.1;refGene_gn=DOCK3;refGene_tx=XM_023249134.1;refGene_gn=DOCK3;refGene_tx=XM_023249135.1;refGene_gn=DOCK3;refGene_tx=XM_023249137.1;refGene_gn=DOCK3;refGene_tx=XM_023249140.1
A2 20080169 20080362
refGene_gn=DOCK3;refGene_tx=XM_023249110.1;refGene_gn=DOCK3;refGene_tx=XM_023249114.1;refGene_gn=DOCK3;refGene_tx=XM_023249115.1;refGene_gn=DOCK3;refGene_tx=XM_023249117.1;refGene_gn=DOCK3;refGene_tx=XM_023249119.1;refGene_gn=DOCK3;refGene_tx=XM_023249123.1;refGene_gn=DOCK3;refGene_tx=XM_023249126.1;refGene_gn=DOCK3;refGene_tx=XM_023249130.1;refGene_gn=DOCK3;refGene_tx=XM_023249134.1;refGene_gn=DOCK3;refGene_tx=XM_023249135.1;refGene_gn=DOCK3;refGene_tx=XM_023249137.1;refGene_gn=DOCK3;refGene_tx=XM_023249140.1
A2 20083459 20083587
refGene_gn=DOCK3;refGene_tx=XM_023249110.1;refGene_gn=DOCK3;refGene_tx=XM_023249114.1;refGene_gn=DOCK3;refGene_tx=XM_023249115.1;refGene_gn=DOCK3;refGene_tx=XM_023249117.1;refGene_gn=DOCK3;refGene_tx=XM_023249119.1;refGene_gn=DOCK3;refGene_tx=XM_023249123.1;refGene_gn=DOCK3;refGene_tx=XM_023249126.1;refGene_gn=DOCK3;refGene_tx=XM_023249130.1;refGene_gn=DOCK3;refGene_tx=XM_023249134.1;refGene_gn=DOCK3;refGene_tx=XM_023249135.1;refGene_gn=DOCK3;refGene_tx=XM_023249137.1;refGene_gn=DOCK3;refGene_tx=XM_023249140.1
A2 20086215 20086362

refGene_gn=DOCK3;refGene_tx=XM_023249110.1;refGene_gn=DOCK3;refGene_tx=XM_023249114.1;refGene_gn=DOCK3;refGene_tx=XM_023249115.1;refGene_gn=DOCK3;refGene_tx=XM_023249117.1;refGene_gn=DOCK3;refGene_tx=XM_023249119.1;refGene_gn=DOCK3;refGene_tx=XM_023249123.1;refGene_gn=DOCK3;refGene_tx=XM_023249126.1;refGene_gn=DOCK3;refGene_tx=XM_023249130.1;refGene_gn=DOCK3;refGene_tx=XM_023249135.1;refGene_gn=DOCK3;refGene_tx=XM_023249137.1;refGene_gn=DOCK3;refGene_tx=XM_023249140.1
A2 20088291 20088390
refGene_gn=DOCK3;refGene_tx=XM_023249110.1;refGene_gn=DOCK3;refGene_tx=XM_023249114.1;refGene_gn=DOCK3;refGene_tx=XM_023249115.1;refGene_gn=DOCK3;refGene_tx=XM_023249117.1;refGene_gn=DOCK3;refGene_tx=XM_023249119.1;refGene_gn=DOCK3;refGene_tx=XM_023249123.1;refGene_gn=DOCK3;refGene_tx=XM_023249126.1;refGene_gn=DOCK3;refGene_tx=XM_023249130.1;refGene_gn=DOCK3;refGene_tx=XM_023249134.1;refGene_gn=DOCK3;refGene_tx=XM_023249135.1;refGene_gn=DOCK3;refGene_tx=XM_023249137.1;refGene_gn=DOCK3;refGene_tx=XM_023249140.1
A2 20099261 20099356
refGene_gn=DOCK3;refGene_tx=XM_023249110.1;refGene_gn=DOCK3;refGene_tx=XM_023249114.1;refGene_gn=DOCK3;refGene_tx=XM_023249115.1;refGene_gn=DOCK3;refGene_tx=XM_023249117.1;refGene_gn=DOCK3;refGene_tx=XM_023249119.1;refGene_gn=DOCK3;refGene_tx=XM_023249123.1;refGene_gn=DOCK3;refGene_tx=XM_023249126.1;refGene_gn=DOCK3;refGene_tx=XM_023249130.1;refGene_gn=DOCK3;refGene_tx=XM_023249134.1;refGene_gn=DOCK3;refGene_tx=XM_023249135.1;refGene_gn=DOCK3;refGene_tx=XM_023249137.1;refGene_gn=DOCK3;refGene_tx=XM_023249140.1
A2 20101122 20101198
refGene_gn=DOCK3;refGene_tx=XM_023249110.1;refGene_gn=DOCK3;refGene_tx=XM_023249114.1;refGene_gn=DOCK3;refGene_tx=XM_023249115.1;refGene_gn=DOCK3;refGene_tx=XM_023249117.1;refGene_gn=DOCK3;refGene_tx=XM_023249119.1;refGene_gn=DOCK3;refGene_tx=XM_023249123.1;refGene_gn=DOCK3;refGene_tx=XM_023249126.1;refGene_gn=DOCK3;refGene_tx=XM_023249130.1;refGene_gn=DOCK3;refGene_tx=XM_023249134.1;refGene_gn=DOCK3;refGene_tx=XM_023249135.1;refGene_gn=DOCK3;refGene_tx=XM_023249137.1;refGene_gn=DOCK3;refGene_tx=XM_023249140.1
A2 20101719 20101820
refGene_gn=DOCK3;refGene_tx=XM_023249110.1;refGene_gn=DOCK3;refGene_tx=XM_023249114.1;refGene_gn=DOCK3;refGene_tx=XM_023249115.1;refGene_gn=DOCK3;refGene_tx=XM_023249117.1;refGene_gn=DOCK3;refGene_tx=XM_023249119.1;refGene_gn=DOCK3;refGene_tx=XM_023249123.1;refGene_gn=DOCK3;refGene_tx=XM_023249126.1;refGene_gn=DOCK3;refGene_tx=XM_023249130.1;refGene_gn=DOCK3;refGene_tx=XM_023249134.1;refGene_gn=DOCK3;refGene_tx=XM_023249135.1;refGene_gn=DOCK3;refGene_tx=XM_023249137.1;refGene_gn=DOCK3;refGene_tx=XM_023249140.1
A2 20102082 20102141
refGene_gn=DOCK3;refGene_tx=XM_023249110.1;refGene_gn=DOCK3;refGene_tx=XM_023249114.1;refGene_gn=DOCK3;refGene_tx=XM_023249115.1;refGene_gn=DOCK3;refGene_tx=XM_023249117.1;refGene_gn=DOCK3;refGene_tx=XM_023249119.1;refGene_gn=DOCK3;refGene_tx=XM_023249123.1;refGene_gn=DOCK3;refGene_tx=XM_023249126.1;refGene_gn=DOCK3;refGene_tx=XM_023249130.1;refGene_gn=DOCK3;refGene_tx=XM_023249134.1;refGene_gn=DOCK3;refGene_tx=XM_02

3249135.1;refGene_gn=DOCK3;refGene_tx=XM_023249137.1;refGene_gn=DOCK3;
refGene_tx=XM_023249140.1
A2 20104629 20104778
refGene_gn=DOCK3;refGene_tx=XM_023249110.1;refGene_gn=DOCK3;refGene_tx
=XM_023249114.1;refGene_gn=DOCK3;refGene_tx=XM_023249115.1;refGene_gn=
DOCK3;refGene_tx=XM_023249117.1;refGene_gn=DOCK3;refGene_tx=XM_0232491
19.1;refGene_gn=DOCK3;refGene_tx=XM_023249123.1;refGene_gn=DOCK3;refGe
ne_tx=XM_023249126.1;refGene_gn=DOCK3;refGene_tx=XM_023249130.1;refGen
e_gn=DOCK3;refGene_tx=XM_023249134.1;refGene_gn=DOCK3;refGene_tx=XM_02
3249135.1;refGene_gn=DOCK3;refGene_tx=XM_023249137.1;refGene_gn=DOCK3;
refGene_tx=XM_023249140.1
A2 20119600 20119686
refGene_gn=DOCK3;refGene_tx=XM_023249110.1;refGene_gn=DOCK3;refGene_tx
=XM_023249114.1;refGene_gn=DOCK3;refGene_tx=XM_023249115.1;refGene_gn=
DOCK3;refGene_tx=XM_023249117.1;refGene_gn=DOCK3;refGene_tx=XM_0232491
19.1;refGene_gn=DOCK3;refGene_tx=XM_023249123.1;refGene_gn=DOCK3;refGe
ne_tx=XM_023249126.1;refGene_gn=DOCK3;refGene_tx=XM_023249130.1;refGen
e_gn=DOCK3;refGene_tx=XM_023249134.1;refGene_gn=DOCK3;refGene_tx=XM_02
3249135.1;refGene_gn=DOCK3;refGene_tx=XM_023249137.1;refGene_gn=DOCK3;
refGene_tx=XM_023249140.1
A2 20122286 20122313
refGene_gn=DOCK3;refGene_tx=XM_023249110.1;refGene_gn=DOCK3;refGene_tx
=XM_023249115.1;refGene_gn=DOCK3;refGene_tx=XM_023249117.1;refGene_gn=
DOCK3;refGene_tx=XM_023249119.1;refGene_gn=DOCK3;refGene_tx=XM_0232491
23.1;refGene_gn=DOCK3;refGene_tx=XM_023249126.1;refGene_gn=DOCK3;refGe
ne_tx=XM_023249130.1;refGene_gn=DOCK3;refGene_tx=XM_023249134.1;refGen
e_gn=DOCK3;refGene_tx=XM_023249135.1;refGene_gn=DOCK3;refGene_tx=XM_02
3249137.1;refGene_gn=DOCK3;refGene_tx=XM_023249140.1
A2 20122450 20122546
refGene_gn=DOCK3;refGene_tx=XM_023249110.1;refGene_gn=DOCK3;refGene_tx
=XM_023249114.1;refGene_gn=DOCK3;refGene_tx=XM_023249115.1;refGene_gn=
DOCK3;refGene_tx=XM_023249117.1;refGene_gn=DOCK3;refGene_tx=XM_0232491
19.1;refGene_gn=DOCK3;refGene_tx=XM_023249123.1;refGene_gn=DOCK3;refGe
ne_tx=XM_023249126.1;refGene_gn=DOCK3;refGene_tx=XM_023249130.1;refGen
e_gn=DOCK3;refGene_tx=XM_023249134.1;refGene_gn=DOCK3;refGene_tx=XM_02
3249135.1;refGene_gn=DOCK3;refGene_tx=XM_023249137.1;refGene_gn=DOCK3;
refGene_tx=XM_023249140.1
A2 20126917 20126978
refGene_gn=DOCK3;refGene_tx=XM_023249110.1;refGene_gn=DOCK3;refGene_tx
=XM_023249114.1;refGene_gn=DOCK3;refGene_tx=XM_023249115.1;refGene_gn=
DOCK3;refGene_tx=XM_023249117.1;refGene_gn=DOCK3;refGene_tx=XM_0232491
19.1;refGene_gn=DOCK3;refGene_tx=XM_023249123.1;refGene_gn=DOCK3;refGe
ne_tx=XM_023249126.1;refGene_gn=DOCK3;refGene_tx=XM_023249130.1;refGen
e_gn=DOCK3;refGene_tx=XM_023249134.1;refGene_gn=DOCK3;refGene_tx=XM_02
3249135.1;refGene_gn=DOCK3;refGene_tx=XM_023249137.1;refGene_gn=DOCK3;
refGene_tx=XM_023249140.1
A2 20127484 20127578
refGene_gn=DOCK3;refGene_tx=XM_023249110.1;refGene_gn=DOCK3;refGene_tx
=XM_023249114.1;refGene_gn=DOCK3;refGene_tx=XM_023249115.1;refGene_gn=
DOCK3;refGene_tx=XM_023249117.1;refGene_gn=DOCK3;refGene_tx=XM_0232491

19.1;refGene_gn=DOCK3;refGene_tx=XM_023249123.1;refGene_gn=DOCK3;refGene_tx=XM_023249126.1;refGene_gn=DOCK3;refGene_tx=XM_023249130.1;refGene_gn=DOCK3;refGene_tx=XM_023249134.1;refGene_gn=DOCK3;refGene_tx=XM_023249135.1;refGene_gn=DOCK3;refGene_tx=XM_023249137.1;refGene_gn=DOCK3;refGene_tx=XM_023249140.1
A2 20129871 20130020
refGene_gn=DOCK3;refGene_tx=XM_023249110.1;refGene_gn=DOCK3;refGene_tx=XM_023249114.1;refGene_gn=DOCK3;refGene_tx=XM_023249115.1;refGene_gn=DOCK3;refGene_tx=XM_023249117.1;refGene_gn=DOCK3;refGene_tx=XM_023249119.1;refGene_gn=DOCK3;refGene_tx=XM_023249123.1;refGene_gn=DOCK3;refGene_tx=XM_023249126.1;refGene_gn=DOCK3;refGene_tx=XM_023249130.1;refGene_gn=DOCK3;refGene_tx=XM_023249134.1;refGene_gn=DOCK3;refGene_tx=XM_023249135.1;refGene_gn=DOCK3;refGene_tx=XM_023249137.1;refGene_gn=DOCK3;refGene_tx=XM_023249140.1
A2 20135810 20135897
refGene_gn=DOCK3;refGene_tx=XM_023249110.1;refGene_gn=DOCK3;refGene_tx=XM_023249114.1;refGene_gn=DOCK3;refGene_tx=XM_023249115.1;refGene_gn=DOCK3;refGene_tx=XM_023249117.1;refGene_gn=DOCK3;refGene_tx=XM_023249119.1;refGene_gn=DOCK3;refGene_tx=XM_023249123.1;refGene_gn=DOCK3;refGene_tx=XM_023249126.1;refGene_gn=DOCK3;refGene_tx=XM_023249130.1;refGene_gn=DOCK3;refGene_tx=XM_023249134.1;refGene_gn=DOCK3;refGene_tx=XM_023249135.1;refGene_gn=DOCK3;refGene_tx=XM_023249137.1;refGene_gn=DOCK3;refGene_tx=XM_023249140.1
A2 20137277 20137382
refGene_gn=DOCK3;refGene_tx=XM_023249110.1;refGene_gn=DOCK3;refGene_tx=XM_023249114.1;refGene_gn=DOCK3;refGene_tx=XM_023249115.1;refGene_gn=DOCK3;refGene_tx=XM_023249117.1;refGene_gn=DOCK3;refGene_tx=XM_023249119.1;refGene_gn=DOCK3;refGene_tx=XM_023249123.1;refGene_gn=DOCK3;refGene_tx=XM_023249126.1;refGene_gn=DOCK3;refGene_tx=XM_023249130.1;refGene_gn=DOCK3;refGene_tx=XM_023249134.1;refGene_gn=DOCK3;refGene_tx=XM_023249135.1;refGene_gn=DOCK3;refGene_tx=XM_023249137.1;refGene_gn=DOCK3;refGene_tx=XM_023249140.1
A2 20141245 20141387
refGene_gn=DOCK3;refGene_tx=XM_023249110.1;refGene_gn=DOCK3;refGene_tx=XM_023249114.1;refGene_gn=DOCK3;refGene_tx=XM_023249115.1;refGene_gn=DOCK3;refGene_tx=XM_023249117.1;refGene_gn=DOCK3;refGene_tx=XM_023249119.1;refGene_gn=DOCK3;refGene_tx=XM_023249123.1;refGene_gn=DOCK3;refGene_tx=XM_023249126.1;refGene_gn=DOCK3;refGene_tx=XM_023249130.1;refGene_gn=DOCK3;refGene_tx=XM_023249134.1;refGene_gn=DOCK3;refGene_tx=XM_023249135.1;refGene_gn=DOCK3;refGene_tx=XM_023249137.1;refGene_gn=DOCK3;refGene_tx=XM_023249140.1
A2 20142421 20142588
refGene_gn=DOCK3;refGene_tx=XM_023249110.1;refGene_gn=DOCK3;refGene_tx=XM_023249114.1;refGene_gn=DOCK3;refGene_tx=XM_023249115.1;refGene_gn=DOCK3;refGene_tx=XM_023249117.1;refGene_gn=DOCK3;refGene_tx=XM_023249119.1;refGene_gn=DOCK3;refGene_tx=XM_023249123.1;refGene_gn=DOCK3;refGene_tx=XM_023249126.1;refGene_gn=DOCK3;refGene_tx=XM_023249130.1;refGene_gn=DOCK3;refGene_tx=XM_023249134.1;refGene_gn=DOCK3;refGene_tx=XM_023249135.1;refGene_gn=DOCK3;refGene_tx=XM_023249137.1;refGene_gn=DOCK3;refGene_tx=XM_023249140.1

A2 20142731 20142818
refGene_gn=DOCK3;refGene_tx=XM_023249110.1;refGene_gn=DOCK3;refGene_tx=XM_023249114.1;refGene_gn=DOCK3;refGene_tx=XM_023249115.1;refGene_gn=DOCK3;refGene_tx=XM_023249117.1;refGene_gn=DOCK3;refGene_tx=XM_023249119.1;refGene_gn=DOCK3;refGene_tx=XM_023249123.1;refGene_gn=DOCK3;refGene_tx=XM_023249126.1;refGene_gn=DOCK3;refGene_tx=XM_023249130.1;refGene_gn=DOCK3;refGene_tx=XM_023249134.1;refGene_gn=DOCK3;refGene_tx=XM_023249135.1;refGene_gn=DOCK3;refGene_tx=XM_023249137.1;refGene_gn=DOCK3;refGene_tx=XM_023249140.1
A2 20143296 20143476
refGene_gn=DOCK3;refGene_tx=XM_023249110.1;refGene_gn=DOCK3;refGene_tx=XM_023249114.1;refGene_gn=DOCK3;refGene_tx=XM_023249115.1;refGene_gn=DOCK3;refGene_tx=XM_023249117.1;refGene_gn=DOCK3;refGene_tx=XM_023249119.1;refGene_gn=DOCK3;refGene_tx=XM_023249123.1;refGene_gn=DOCK3;refGene_tx=XM_023249126.1;refGene_gn=DOCK3;refGene_tx=XM_023249130.1;refGene_gn=DOCK3;refGene_tx=XM_023249134.1;refGene_gn=DOCK3;refGene_tx=XM_023249135.1;refGene_gn=DOCK3;refGene_tx=XM_023249137.1;refGene_gn=DOCK3;refGene_tx=XM_023249140.1
A2 20144110 20144194
refGene_gn=DOCK3;refGene_tx=XM_023249110.1;refGene_gn=DOCK3;refGene_tx=XM_023249114.1;refGene_gn=DOCK3;refGene_tx=XM_023249115.1;refGene_gn=DOCK3;refGene_tx=XM_023249117.1;refGene_gn=DOCK3;refGene_tx=XM_023249119.1;refGene_gn=DOCK3;refGene_tx=XM_023249123.1;refGene_gn=DOCK3;refGene_tx=XM_023249126.1;refGene_gn=DOCK3;refGene_tx=XM_023249130.1;refGene_gn=DOCK3;refGene_tx=XM_023249134.1;refGene_gn=DOCK3;refGene_tx=XM_023249135.1;refGene_gn=DOCK3;refGene_tx=XM_023249137.1;refGene_gn=DOCK3;refGene_tx=XM_023249140.1
A2 20144316 20144433
refGene_gn=DOCK3;refGene_tx=XM_023249110.1;refGene_gn=DOCK3;refGene_tx=XM_023249114.1;refGene_gn=DOCK3;refGene_tx=XM_023249115.1;refGene_gn=DOCK3;refGene_tx=XM_023249117.1;refGene_gn=DOCK3;refGene_tx=XM_023249119.1;refGene_gn=DOCK3;refGene_tx=XM_023249123.1;refGene_gn=DOCK3;refGene_tx=XM_023249126.1;refGene_gn=DOCK3;refGene_tx=XM_023249130.1;refGene_gn=DOCK3;refGene_tx=XM_023249134.1;refGene_gn=DOCK3;refGene_tx=XM_023249135.1;refGene_gn=DOCK3;refGene_tx=XM_023249137.1;refGene_gn=DOCK3;refGene_tx=XM_023249140.1
A2 20146381 20146503
refGene_gn=DOCK3;refGene_tx=XM_023249110.1;refGene_gn=DOCK3;refGene_tx=XM_023249114.1;refGene_gn=DOCK3;refGene_tx=XM_023249115.1;refGene_gn=DOCK3;refGene_tx=XM_023249117.1;refGene_gn=DOCK3;refGene_tx=XM_023249119.1;refGene_gn=DOCK3;refGene_tx=XM_023249123.1;refGene_gn=DOCK3;refGene_tx=XM_023249126.1;refGene_gn=DOCK3;refGene_tx=XM_023249130.1;refGene_gn=DOCK3;refGene_tx=XM_023249134.1;refGene_gn=DOCK3;refGene_tx=XM_023249135.1;refGene_gn=DOCK3;refGene_tx=XM_023249137.1;refGene_gn=DOCK3;refGene_tx=XM_023249140.1
A2 20147776 20147915
refGene_gn=DOCK3;refGene_tx=XM_023249110.1;refGene_gn=DOCK3;refGene_tx=XM_023249114.1;refGene_gn=DOCK3;refGene_tx=XM_023249115.1;refGene_gn=DOCK3;refGene_tx=XM_023249117.1;refGene_gn=DOCK3;refGene_tx=XM_023249119.1;refGene_gn=DOCK3;refGene_tx=XM_023249123.1;refGene_gn=DOCK3;refGe

ne_tx=XM_023249126.1;refGene_gn=DOCK3;refGene_tx=XM_023249130.1;refGene_gn=DOCK3;refGene_tx=XM_023249134.1;refGene_gn=DOCK3;refGene_tx=XM_023249135.1;refGene_gn=DOCK3;refGene_tx=XM_023249137.1;refGene_gn=DOCK3;refGene_tx=XM_023249140.1
A2	20148437	20148585
refGene_gn=DOCK3;refGene_tx=XM_023249110.1;refGene_gn=DOCK3;refGene_tx=XM_023249114.1;refGene_gn=DOCK3;refGene_tx=XM_023249115.1;refGene_gn=DOCK3;refGene_tx=XM_023249117.1;refGene_gn=DOCK3;refGene_tx=XM_023249119.1;refGene_gn=DOCK3;refGene_tx=XM_023249123.1;refGene_gn=DOCK3;refGene_tx=XM_023249126.1;refGene_gn=DOCK3;refGene_tx=XM_023249130.1;refGene_gn=DOCK3;refGene_tx=XM_023249134.1;refGene_gn=DOCK3;refGene_tx=XM_023249135.1;refGene_gn=DOCK3;refGene_tx=XM_023249137.1;refGene_gn=DOCK3;refGene_tx=XM_023249140.1
A2	20152364	20152483
refGene_gn=DOCK3;refGene_tx=XM_023249110.1;refGene_gn=DOCK3;refGene_tx=XM_023249114.1;refGene_gn=DOCK3;refGene_tx=XM_023249115.1;refGene_gn=DOCK3;refGene_tx=XM_023249117.1;refGene_gn=DOCK3;refGene_tx=XM_023249119.1;refGene_gn=DOCK3;refGene_tx=XM_023249123.1;refGene_gn=DOCK3;refGene_tx=XM_023249126.1;refGene_gn=DOCK3;refGene_tx=XM_023249130.1;refGene_gn=DOCK3;refGene_tx=XM_023249134.1;refGene_gn=DOCK3;refGene_tx=XM_023249135.1;refGene_gn=DOCK3;refGene_tx=XM_023249137.1;refGene_gn=DOCK3;refGene_tx=XM_023249140.1
A2	20153614	20153702
refGene_gn=DOCK3;refGene_tx=XM_023249110.1;refGene_gn=DOCK3;refGene_tx=XM_023249114.1;refGene_gn=DOCK3;refGene_tx=XM_023249115.1;refGene_gn=DOCK3;refGene_tx=XM_023249117.1;refGene_gn=DOCK3;refGene_tx=XM_023249119.1;refGene_gn=DOCK3;refGene_tx=XM_023249123.1;refGene_gn=DOCK3;refGene_tx=XM_023249126.1;refGene_gn=DOCK3;refGene_tx=XM_023249130.1;refGene_gn=DOCK3;refGene_tx=XM_023249134.1;refGene_gn=DOCK3;refGene_tx=XM_023249135.1;refGene_gn=DOCK3;refGene_tx=XM_023249137.1
A2	20157490	20157613
refGene_gn=DOCK3;refGene_tx=XM_023249110.1;refGene_gn=DOCK3;refGene_tx=XM_023249114.1;refGene_gn=DOCK3;refGene_tx=XM_023249115.1;refGene_gn=DOCK3;refGene_tx=XM_023249117.1;refGene_gn=DOCK3;refGene_tx=XM_023249119.1;refGene_gn=DOCK3;refGene_tx=XM_023249126.1;refGene_gn=DOCK3;refGene_tx=XM_023249130.1;refGene_gn=DOCK3;refGene_tx=XM_023249134.1;refGene_gn=DOCK3;refGene_tx=XM_023249135.1;refGene_gn=DOCK3;refGene_tx=XM_023249137.1;refGene_gn=DOCK3;refGene_tx=XM_023249140.1
A2	20158140	20158223
refGene_gn=DOCK3;refGene_tx=XM_023249110.1;refGene_gn=DOCK3;refGene_tx=XM_023249114.1;refGene_gn=DOCK3;refGene_tx=XM_023249115.1;refGene_gn=DOCK3;refGene_tx=XM_023249117.1;refGene_gn=DOCK3;refGene_tx=XM_023249119.1;refGene_gn=DOCK3;refGene_tx=XM_023249123.1;refGene_gn=DOCK3;refGene_tx=XM_023249126.1;refGene_gn=DOCK3;refGene_tx=XM_023249130.1;refGene_gn=DOCK3;refGene_tx=XM_023249134.1;refGene_gn=DOCK3;refGene_tx=XM_023249135.1;refGene_gn=DOCK3;refGene_tx=XM_023249137.1;refGene_gn=DOCK3;refGene_tx=XM_023249140.1
A2	20158999	20159509
refGene_gn=DOCK3;refGene_tx=XM_023249110.1;refGene_gn=DOCK3;refGene_tx=XM_023249114.1;refGene_gn=DOCK3;refGene_tx=XM_023249115.1;refGene_gn=

DOCK3;refGene_tx=XM_023249117.1;refGene_gn=DOCK3;refGene_tx=XM_0232491
19.1;refGene_gn=DOCK3;refGene_tx=XM_023249123.1;refGene_gn=DOCK3;refGe
ne_tx=XM_023249126.1;refGene_gn=DOCK3;refGene_tx=XM_023249130.1;refGen
e_gn=DOCK3;refGene_tx=XM_023249134.1;refGene_gn=DOCK3;refGene_tx=XM_02
3249135.1;refGene_gn=DOCK3;refGene_tx=XM_023249137.1;refGene_gn=DOCK3;
refGene_tx=XM_023249140.1
A2 20162866 20162944 refGene_gn=MANF;refGene_tx=XM_006928821.4
A2 20163038 20163258 refGene_gn=MANF;refGene_tx=XM_006928821.4
A2 20164000 20164128 refGene_gn=MANF;refGene_tx=XM_006928821.4
A2 20165124 20165266 refGene_gn=MANF;refGene_tx=XM_006928821.4
A2 20166461 20166646 refGene_gn=MANF;refGene_tx=XM_006928821.4
A2 20169218 20171918 refGene_gn=RBM15B;refGene_tx=XM_023249173.1
A2 20176669 20176728
refGene_gn=DCAF1;refGene_tx=XM_006928823.4;refGene_gn=DCAF1;refGene_tx
=XM_006928824.4;refGene_gn=DCAF1;refGene_tx=XM_006928825.4;refGene_gn=
DCAF1;refGene_tx=XM_006928826.4;refGene_gn=DCAF1;refGene_tx=XM_0198222
06.2;refGene_gn=DCAF1;refGene_tx=XM_019822218.2;refGene_gn=DCAF1;refGe
ne_tx=XM_019822224.2
A2 20180985 20181238
refGene_gn=DCAF1;refGene_tx=XM_006928823.4;refGene_gn=DCAF1;refGene_tx
=XM_006928824.4;refGene_gn=DCAF1;refGene_tx=XM_006928825.4;refGene_gn=
DCAF1;refGene_tx=XM_006928826.4;refGene_gn=DCAF1;refGene_tx=XM_0198222
06.2;refGene_gn=DCAF1;refGene_tx=XM_019822224.2
A2 20186631 20186640
refGene_gn=DCAF1;refGene_tx=XM_006928823.4;refGene_gn=DCAF1;refGene_tx
=XM_006928825.4;refGene_gn=DCAF1;refGene_tx=XM_006928826.4;refGene_gn=
DCAF1;refGene_tx=XM_019822206.2;refGene_gn=DCAF1;refGene_tx=XM_0198222
24.2
A2 20200703 20200805
refGene_gn=DCAF1;refGene_tx=XM_006928823.4;refGene_gn=DCAF1;refGene_tx
=XM_006928824.4;refGene_gn=DCAF1;refGene_tx=XM_006928825.4;refGene_gn=
DCAF1;refGene_tx=XM_006928826.4;refGene_gn=DCAF1;refGene_tx=XM_0198222
06.2;refGene_gn=DCAF1;refGene_tx=XM_019822218.2;refGene_gn=DCAF1;refGe
ne_tx=XM_019822224.2
A2 20201338 20201412
refGene_gn=DCAF1;refGene_tx=XM_006928823.4;refGene_gn=DCAF1;refGene_tx
=XM_006928824.4;refGene_gn=DCAF1;refGene_tx=XM_006928825.4;refGene_gn=
DCAF1;refGene_tx=XM_006928826.4;refGene_gn=DCAF1;refGene_tx=XM_0198222
06.2;refGene_gn=DCAF1;refGene_tx=XM_019822218.2;refGene_gn=DCAF1;refGe
ne_tx=XM_019822224.2
A2 20201620 20201725
refGene_gn=DCAF1;refGene_tx=XM_006928823.4;refGene_gn=DCAF1;refGene_tx
=XM_006928824.4;refGene_gn=DCAF1;refGene_tx=XM_006928825.4;refGene_gn=
DCAF1;refGene_tx=XM_006928826.4;refGene_gn=DCAF1;refGene_tx=XM_0198222
06.2;refGene_gn=DCAF1;refGene_tx=XM_019822218.2;refGene_gn=DCAF1;refGe
ne_tx=XM_019822224.2
A2 20202240 20202334
refGene_gn=DCAF1;refGene_tx=XM_006928823.4;refGene_gn=DCAF1;refGene_tx
=XM_006928824.4;refGene_gn=DCAF1;refGene_tx=XM_006928825.4;refGene_gn=
DCAF1;refGene_tx=XM_006928826.4;refGene_gn=DCAF1;refGene_tx=XM_0198222

06.2;refGene_gn=DCAF1;refGene_tx=XM_019822218.2;refGene_gn=DCAF1;refGene_tx=XM_019822224.2
A2 20203166 20203400
refGene_gn=DCAF1;refGene_tx=XM_006928823.4;refGene_gn=DCAF1;refGene_tx=XM_006928824.4;refGene_gn=DCAF1;refGene_tx=XM_006928825.4;refGene_gn=DCAF1;refGene_tx=XM_006928826.4;refGene_gn=DCAF1;refGene_tx=XM_019822206.2;refGene_gn=DCAF1;refGene_tx=XM_019822218.2;refGene_gn=DCAF1;refGene_tx=XM_019822224.2;refGene_gn=DCAF1;refGene_tx=XM_019822231.2
A2 20206396 20206481
refGene_gn=DCAF1;refGene_tx=XM_006928823.4;refGene_gn=DCAF1;refGene_tx=XM_006928824.4;refGene_gn=DCAF1;refGene_tx=XM_006928825.4;refGene_gn=DCAF1;refGene_tx=XM_006928826.4;refGene_gn=DCAF1;refGene_tx=XM_019822206.2;refGene_gn=DCAF1;refGene_tx=XM_019822218.2;refGene_gn=DCAF1;refGene_tx=XM_019822224.2;refGene_gn=DCAF1;refGene_tx=XM_019822231.2
A2 20207053 20207136
refGene_gn=DCAF1;refGene_tx=XM_006928823.4;refGene_gn=DCAF1;refGene_tx=XM_006928824.4;refGene_gn=DCAF1;refGene_tx=XM_006928825.4;refGene_gn=DCAF1;refGene_tx=XM_006928826.4;refGene_gn=DCAF1;refGene_tx=XM_019822206.2;refGene_gn=DCAF1;refGene_tx=XM_019822218.2;refGene_gn=DCAF1;refGene_tx=XM_019822224.2;refGene_gn=DCAF1;refGene_tx=XM_019822231.2
A2 20208309 20208508
refGene_gn=DCAF1;refGene_tx=XM_006928823.4;refGene_gn=DCAF1;refGene_tx=XM_006928824.4;refGene_gn=DCAF1;refGene_tx=XM_006928825.4;refGene_gn=DCAF1;refGene_tx=XM_006928826.4;refGene_gn=DCAF1;refGene_tx=XM_019822206.2;refGene_gn=DCAF1;refGene_tx=XM_019822218.2;refGene_gn=DCAF1;refGene_tx=XM_019822224.2;refGene_gn=DCAF1;refGene_tx=XM_019822231.2
A2 20209135 20210399
refGene_gn=DCAF1;refGene_tx=XM_006928823.4;refGene_gn=DCAF1;refGene_tx=XM_006928824.4;refGene_gn=DCAF1;refGene_tx=XM_006928825.4;refGene_gn=DCAF1;refGene_tx=XM_006928826.4;refGene_gn=DCAF1;refGene_tx=XM_019822206.2;refGene_gn=DCAF1;refGene_tx=XM_019822218.2;refGene_gn=DCAF1;refGene_tx=XM_019822224.2;refGene_gn=DCAF1;refGene_tx=XM_019822231.2
A2 20211547 20211672
refGene_gn=DCAF1;refGene_tx=XM_006928823.4;refGene_gn=DCAF1;refGene_tx=XM_006928824.4;refGene_gn=DCAF1;refGene_tx=XM_006928825.4;refGene_gn=DCAF1;refGene_tx=XM_006928826.4;refGene_gn=DCAF1;refGene_tx=XM_019822206.2;refGene_gn=DCAF1;refGene_tx=XM_019822218.2;refGene_gn=DCAF1;refGene_tx=XM_019822224.2;refGene_gn=DCAF1;refGene_tx=XM_019822231.2
A2 20215468 20215638
refGene_gn=DCAF1;refGene_tx=XM_006928823.4;refGene_gn=DCAF1;refGene_tx=XM_006928824.4;refGene_gn=DCAF1;refGene_tx=XM_006928825.4;refGene_gn=DCAF1;refGene_tx=XM_006928826.4;refGene_gn=DCAF1;refGene_tx=XM_019822206.2;refGene_gn=DCAF1;refGene_tx=XM_019822218.2;refGene_gn=DCAF1;refGene_tx=XM_019822224.2;refGene_gn=DCAF1;refGene_tx=XM_019822231.2
A2 20216293 20216503
refGene_gn=DCAF1;refGene_tx=XM_006928823.4;refGene_gn=DCAF1;refGene_tx=XM_006928824.4;refGene_gn=DCAF1;refGene_tx=XM_006928825.4;refGene_gn=DCAF1;refGene_tx=XM_006928826.4;refGene_gn=DCAF1;refGene_tx=XM_019822206.2;refGene_gn=DCAF1;refGene_tx=XM_019822218.2;refGene_gn=DCAF1;refGene_tx=XM_019822224.2;refGene_gn=DCAF1;refGene_tx=XM_019822231.2

A2 20217112 20217292
refGene_gn=DCAF1;refGene_tx=XM_006928823.4;refGene_gn=DCAF1;refGene_tx=XM_006928824.4;refGene_gn=DCAF1;refGene_tx=XM_006928825.4;refGene_gn=DCAF1;refGene_tx=XM_006928826.4;refGene_gn=DCAF1;refGene_tx=XM_019822206.2;refGene_gn=DCAF1;refGene_tx=XM_019822218.2;refGene_gn=DCAF1;refGene_tx=XM_019822224.2;refGene_gn=DCAF1;refGene_tx=XM_019822231.2
A2 20219018 20219177
refGene_gn=DCAF1;refGene_tx=XM_006928823.4;refGene_gn=DCAF1;refGene_tx=XM_006928824.4;refGene_gn=DCAF1;refGene_tx=XM_006928825.4;refGene_gn=DCAF1;refGene_tx=XM_006928826.4;refGene_gn=DCAF1;refGene_tx=XM_019822206.2;refGene_gn=DCAF1;refGene_tx=XM_019822218.2;refGene_gn=DCAF1;refGene_tx=XM_019822224.2;refGene_gn=DCAF1;refGene_tx=XM_019822231.2
A2 20222918 20223020
refGene_gn=DCAF1;refGene_tx=XM_006928823.4;refGene_gn=DCAF1;refGene_tx=XM_006928824.4;refGene_gn=DCAF1;refGene_tx=XM_006928825.4;refGene_gn=DCAF1;refGene_tx=XM_006928826.4;refGene_gn=DCAF1;refGene_tx=XM_019822206.2;refGene_gn=DCAF1;refGene_tx=XM_019822218.2;refGene_gn=DCAF1;refGene_tx=XM_019822224.2;refGene_gn=DCAF1;refGene_tx=XM_019822231.2
A2 20223337 20223850
refGene_gn=DCAF1;refGene_tx=XM_006928823.4;refGene_gn=DCAF1;refGene_tx=XM_006928824.4;refGene_gn=DCAF1;refGene_tx=XM_006928825.4;refGene_gn=DCAF1;refGene_tx=XM_006928826.4;refGene_gn=DCAF1;refGene_tx=XM_019822206.2;refGene_gn=DCAF1;refGene_tx=XM_019822218.2;refGene_gn=DCAF1;refGene_tx=XM_019822224.2;refGene_gn=DCAF1;refGene_tx=XM_019822231.2
A2 20225687 20225825
refGene_gn=DCAF1;refGene_tx=XM_006928823.4;refGene_gn=DCAF1;refGene_tx=XM_006928824.4;refGene_gn=DCAF1;refGene_tx=XM_006928825.4;refGene_gn=DCAF1;refGene_tx=XM_006928826.4;refGene_gn=DCAF1;refGene_tx=XM_019822206.2;refGene_gn=DCAF1;refGene_tx=XM_019822218.2;refGene_gn=DCAF1;refGene_tx=XM_019822224.2;refGene_gn=DCAF1;refGene_tx=XM_019822231.2
A2 20237634 20237748
refGene_gn=DCAF1;refGene_tx=XM_006928823.4;refGene_gn=DCAF1;refGene_tx=XM_006928824.4;refGene_gn=DCAF1;refGene_tx=XM_006928825.4;refGene_gn=DCAF1;refGene_tx=XM_006928826.4;refGene_gn=DCAF1;refGene_tx=XM_019822206.2;refGene_gn=DCAF1;refGene_tx=XM_019822218.2;refGene_gn=DCAF1;refGene_tx=XM_019822224.2;refGene_gn=DCAF1;refGene_tx=XM_019822231.2
A2 20240175 20240249
refGene_gn=DCAF1;refGene_tx=XM_006928823.4;refGene_gn=DCAF1;refGene_tx=XM_006928824.4;refGene_gn=DCAF1;refGene_tx=XM_006928825.4;refGene_gn=DCAF1;refGene_tx=XM_006928826.4;refGene_gn=DCAF1;refGene_tx=XM_019822206.2;refGene_gn=DCAF1;refGene_tx=XM_019822218.2;refGene_gn=DCAF1;refGene_tx=XM_019822231.2
A2 20242372 20242449
refGene_gn=DCAF1;refGene_tx=XM_006928823.4;refGene_gn=DCAF1;refGene_tx=XM_006928824.4;refGene_gn=DCAF1;refGene_tx=XM_006928825.4;refGene_gn=DCAF1;refGene_tx=XM_006928826.4;refGene_gn=DCAF1;refGene_tx=XM_019822206.2;refGene_gn=DCAF1;refGene_tx=XM_019822218.2;refGene_gn=DCAF1;refGene_tx=XM_019822231.2
A2 20247335 20247453
refGene_gn=DCAF1;refGene_tx=XM_006928823.4;refGene_gn=DCAF1;refGene_tx

=XM_006928824.4;refGene_gn=DCAF1;refGene_tx=XM_006928825.4;refGene_gn=DCAF1;refGene_tx=XM_006928826.4;refGene_gn=DCAF1;refGene_tx=XM_019822206.2;refGene_gn=DCAF1;refGene_tx=XM_019822218.2;refGene_gn=DCAF1;refGene_tx=XM_019822231.2
A2 20260191 20260288
refGene_gn=DCAF1;refGene_tx=XM_006928823.4;refGene_gn=DCAF1;refGene_tx=XM_006928824.4;refGene_gn=DCAF1;refGene_tx=XM_019822206.2;refGene_gn=DCAF1;refGene_tx=XM_019822218.2;refGene_gn=DCAF1;refGene_tx=XM_019822231.2
A2 20339072 20339211
refGene_gn=RAD54L2;refGene_tx=XM_006928828.3;refGene_gn=RAD54L2;refGene_tx=XM_023249176.1;refGene_gn=RAD54L2;refGene_tx=XM_023249178.1
A2 20374662 20374864
refGene_gn=RAD54L2;refGene_tx=XM_006928828.3;refGene_gn=RAD54L2;refGene_tx=XM_023249176.1;refGene_gn=RAD54L2;refGene_tx=XM_023249178.1
A2 20375848 20375988
refGene_gn=RAD54L2;refGene_tx=XM_006928828.3;refGene_gn=RAD54L2;refGene_tx=XM_023249176.1;refGene_gn=RAD54L2;refGene_tx=XM_023249178.1
A2 20376759 20376876
refGene_gn=RAD54L2;refGene_tx=XM_006928828.3;refGene_gn=RAD54L2;refGene_tx=XM_023249176.1;refGene_gn=RAD54L2;refGene_tx=XM_023249178.1
A2 20377197 20377424
refGene_gn=RAD54L2;refGene_tx=XM_006928828.3;refGene_gn=RAD54L2;refGene_tx=XM_023249176.1;refGene_gn=RAD54L2;refGene_tx=XM_023249178.1
A2 20379687 20379870
refGene_gn=RAD54L2;refGene_tx=XM_006928828.3;refGene_gn=RAD54L2;refGene_tx=XM_023249176.1;refGene_gn=RAD54L2;refGene_tx=XM_023249178.1
A2 20379995 20380129
refGene_gn=RAD54L2;refGene_tx=XM_006928828.3;refGene_gn=RAD54L2;refGene_tx=XM_023249176.1;refGene_gn=RAD54L2;refGene_tx=XM_023249178.1
A2 20381331 20381528
refGene_gn=RAD54L2;refGene_tx=XM_006928828.3;refGene_gn=RAD54L2;refGene_tx=XM_019822251.1;refGene_gn=RAD54L2;refGene_tx=XM_023249176.1;refGene_gn=RAD54L2;refGene_tx=XM_023249178.1
A2 20382818 20383240
refGene_gn=RAD54L2;refGene_tx=XM_006928828.3;refGene_gn=RAD54L2;refGene_tx=XM_019822251.1;refGene_gn=RAD54L2;refGene_tx=XM_023249176.1;refGene_gn=RAD54L2;refGene_tx=XM_023249178.1
A2 20383926 20384104
refGene_gn=RAD54L2;refGene_tx=XM_006928828.3;refGene_gn=RAD54L2;refGene_tx=XM_019822251.1;refGene_gn=RAD54L2;refGene_tx=XM_023249176.1;refGene_gn=RAD54L2;refGene_tx=XM_023249178.1
A2 20385348 20385600
refGene_gn=RAD54L2;refGene_tx=XM_006928828.3;refGene_gn=RAD54L2;refGene_tx=XM_019822251.1;refGene_gn=RAD54L2;refGene_tx=XM_023249176.1;refGene_gn=RAD54L2;refGene_tx=XM_023249178.1
A2 20385827 20385946
refGene_gn=RAD54L2;refGene_tx=XM_006928828.3;refGene_gn=RAD54L2;refGene_tx=XM_019822251.1;refGene_gn=RAD54L2;refGene_tx=XM_023249176.1;refGene_gn=RAD54L2;refGene_tx=XM_023249178.1

A2 20387285 20387404
refGene_gn=RAD54L2;refGene_tx=XM_006928828.3;refGene_gn=RAD54L2;refGene_tx=XM_019822251.1;refGene_gn=RAD54L2;refGene_tx=XM_023249176.1;refGene_gn=RAD54L2;refGene_tx=XM_023249178.1
A2 20390522 20390622
refGene_gn=RAD54L2;refGene_tx=XM_006928828.3;refGene_gn=RAD54L2;refGene_tx=XM_019822251.1;refGene_gn=RAD54L2;refGene_tx=XM_023249176.1;refGene_gn=RAD54L2;refGene_tx=XM_023249178.1
A2 20391373 20391579
refGene_gn=RAD54L2;refGene_tx=XM_006928828.3;refGene_gn=RAD54L2;refGene_tx=XM_019822251.1;refGene_gn=RAD54L2;refGene_tx=XM_023249176.1;refGene_gn=RAD54L2;refGene_tx=XM_023249178.1
A2 20391983 20392156
refGene_gn=RAD54L2;refGene_tx=XM_006928828.3;refGene_gn=RAD54L2;refGene_tx=XM_019822251.1;refGene_gn=RAD54L2;refGene_tx=XM_023249176.1;refGene_gn=RAD54L2;refGene_tx=XM_023249178.1
A2 20392895 20393092
refGene_gn=RAD54L2;refGene_tx=XM_006928828.3;refGene_gn=RAD54L2;refGene_tx=XM_019822251.1;refGene_gn=RAD54L2;refGene_tx=XM_023249176.1;refGene_gn=RAD54L2;refGene_tx=XM_023249178.1
A2 20396116 20396316
refGene_gn=RAD54L2;refGene_tx=XM_006928828.3;refGene_gn=RAD54L2;refGene_tx=XM_019822251.1;refGene_gn=RAD54L2;refGene_tx=XM_023249176.1;refGene_gn=RAD54L2;refGene_tx=XM_023249178.1
A2 20397390 20397480
refGene_gn=RAD54L2;refGene_tx=XM_006928828.3;refGene_gn=RAD54L2;refGene_tx=XM_019822251.1;refGene_gn=RAD54L2;refGene_tx=XM_023249176.1;refGene_gn=RAD54L2;refGene_tx=XM_023249178.1
A2 20399689 20399782
refGene_gn=RAD54L2;refGene_tx=XM_006928828.3;refGene_gn=RAD54L2;refGene_tx=XM_019822251.1;refGene_gn=RAD54L2;refGene_tx=XM_023249176.1;refGene_gn=RAD54L2;refGene_tx=XM_023249178.1
A2 20401674 20402669
refGene_gn=RAD54L2;refGene_tx=XM_006928828.3;refGene_gn=RAD54L2;refGene_tx=XM_019822251.1;refGene_gn=RAD54L2;refGene_tx=XM_023249176.1;refGene_gn=RAD54L2;refGene_tx=XM_023249178.1
A2 20408691 20408699 refGene_gn=TEX264;refGene_tx=XM_011291827.3
A2 20409942 20410124 refGene_gn=TEX264;refGene_tx=XM_023249227.1
A2 20410543 20410566 refGene_gn=TEX264;refGene_tx=XM_011291824.3
A2 20410890 20410925 refGene_gn=TEX264;refGene_tx=XM_011291818.3
A2 20411059 20411135
refGene_gn=TEX264;refGene_tx=XM_011291818.3;refGene_gn=TEX264;refGene_tx=XM_011291824.3;refGene_gn=TEX264;refGene_tx=XM_011291827.3;refGene_gn=TEX264;refGene_tx=XM_023249227.1
A2 20413682 20413940
refGene_gn=TEX264;refGene_tx=XM_006928830.4;refGene_gn=TEX264;refGene_tx=XM_019822284.2;refGene_gn=TEX264;refGene_tx=XM_023249205.1;refGene_gn=TEX264;refGene_tx=XM_023249219.1;refGene_gn=TEX264;refGene_tx=XM_023249245.1;refGene_gn=TEX264;refGene_tx=XM_023249248.1
A2 20423453 20423675

refGene_gn=TEX264;refGene_tx=XM_006928830.4;refGene_gn=TEX264;refGene_tx=XM_011291818.3;refGene_gn=TEX264;refGene_tx=XM_011291824.3;refGene_gn=TEX264;refGene_tx=XM_011291827.3;refGene_gn=TEX264;refGene_tx=XM_019822284.2;refGene_gn=TEX264;refGene_tx=XM_023249205.1;refGene_gn=TEX264;refGene_tx=XM_023249219.1;refGene_gn=TEX264;refGene_tx=XM_023249227.1;refGene_gn=TEX264;refGene_tx=XM_023249245.1;refGene_gn=TEX264;refGene_tx=XM_023249248.1
A2 20436842 20437011 refGene_gn=TEX264;refGene_tx=XM_006928830.4;refGene_gn=TEX264;refGene_tx=XM_011291818.3;refGene_gn=TEX264;refGene_tx=XM_011291824.3;refGene_gn=TEX264;refGene_tx=XM_011291827.3;refGene_gn=TEX264;refGene_tx=XM_019822284.2;refGene_gn=TEX264;refGene_tx=XM_023249205.1;refGene_gn=TEX264;refGene_tx=XM_023249219.1;refGene_gn=TEX264;refGene_tx=XM_023249227.1;refGene_gn=TEX264;refGene_tx=XM_023249245.1;refGene_gn=TEX264;refGene_tx=XM_023249248.1
A2 20437181 20437654 refGene_gn=TEX264;refGene_tx=XM_006928830.4;refGene_gn=TEX264;refGene_tx=XM_011291818.3;refGene_gn=TEX264;refGene_tx=XM_011291824.3;refGene_gn=TEX264;refGene_tx=XM_011291827.3;refGene_gn=TEX264;refGene_tx=XM_019822284.2;refGene_gn=TEX264;refGene_tx=XM_023249205.1;refGene_gn=TEX264;refGene_tx=XM_023249219.1;refGene_gn=TEX264;refGene_tx=XM_023249227.1
A2 20440453 20440746 refGene_gn=TEX264;refGene_tx=XM_023249245.1
A2 20442877 20443032 refGene_gn=TEX264;refGene_tx=XM_023249248.1
A2 20445769 20446219 refGene_gn=GRM2;refGene_tx=XM_006928832.4;refGene_gn=GRM2;refGene_tx=XM_019822261.2;refGene_gn=GRM2;refGene_tx=XM_023249255.1
A2 20448597 20449435 refGene_gn=GRM2;refGene_tx=XM_006928832.4;refGene_gn=GRM2;refGene_tx=XM_019822261.2;refGene_gn=GRM2;refGene_tx=XM_019822266.2;refGene_gn=GRM2;refGene_tx=XM_019822269.2;refGene_gn=GRM2;refGene_tx=XM_023249255.1
A2 20450935 20452011 refGene_gn=GRM2;refGene_tx=XM_006928832.4;refGene_gn=GRM2;refGene_tx=XM_019822261.2;refGene_gn=GRM2;refGene_tx=XM_019822266.2;refGene_gn=GRM2;refGene_tx=XM_019822269.2;refGene_gn=GRM2;refGene_tx=XM_023249255.1
A2 20453927 20454108 refGene_gn=GRM2;refGene_tx=XM_006928832.4;refGene_gn=GRM2;refGene_tx=XM_019822261.2;refGene_gn=GRM2;refGene_tx=XM_019822266.2;refGene_gn=GRM2;refGene_tx=XM_019822269.2;refGene_gn=GRM2;refGene_tx=XM_023249255.1
A2 20454337 20454411 refGene_gn=GRM2;refGene_tx=XM_006928832.4;refGene_gn=GRM2;refGene_tx=XM_019822261.2;refGene_gn=GRM2;refGene_tx=XM_019822266.2;refGene_gn=GRM2;refGene_tx=XM_019822269.2;refGene_gn=GRM2;refGene_tx=XM_023249255.1
A2 20482772 20483141 refGene_gn=IQCF6;refGene_tx=XM_003982281.4
A2 20483352 20483367 refGene_gn=IQCF6;refGene_tx=XM_003982281.4
A2 20510843 20510909 refGene_gn=LOC111558745;refGene_tx=XM_023249270.1
A2 20511979 20512414 refGene_gn=LOC111558745;refGene_tx=XM_023249270.1
A2 20522492 20522510 refGene_gn=IQCF3;refGene_tx=XM_003982282.4

```
A2	20522914	20522959	refGene_gn=IQCF3;refGene_tx=XM_003982282.4
A2	20523707	20524103	refGene_gn=IQCF3;refGene_tx=XM_003982282.4
A2	20536756	20536774	refGene_gn=IQCF2;refGene_tx=XM_003982340.4
A2	20536950	20537034	refGene_gn=IQCF2;refGene_tx=XM_003982340.4
A2	20537794	20538178	refGene_gn=IQCF2;refGene_tx=XM_003982340.4
A2	20550535	20550975
refGene_gn=IQCF5;refGene_tx=XM_003982283.5;refGene_gn=IQCF5;refGene_tx
=XM_019822361.2
A2	20552369	20552373	refGene_gn=IQCF5;refGene_tx=XM_003982283.5
A2	20553557	20553567	refGene_gn=IQCF5;refGene_tx=XM_019822361.2
A2	20563075	20563525
refGene_gn=IQCF1;refGene_tx=XM_011279385.3;refGene_gn=IQCF1;refGene_tx
=XM_019822346.2;refGene_gn=IQCF1;refGene_tx=XM_019822352.2
A2	20564295	20564358
refGene_gn=IQCF1;refGene_tx=XM_011279385.3;refGene_gn=IQCF1;refGene_tx
=XM_019822346.2;refGene_gn=IQCF1;refGene_tx=XM_019822352.2
A2	20564492	20564605
refGene_gn=IQCF1;refGene_tx=XM_019822346.2;refGene_gn=IQCF1;refGene_tx
=XM_019822352.2
A2	20564777	20564784
refGene_gn=IQCF1;refGene_tx=XM_011279385.3;refGene_gn=IQCF1;refGene_tx
=XM_019822346.2;refGene_gn=IQCF1;refGene_tx=XM_019822352.2
A2	20576013	20576107
refGene_gn=RRP9;refGene_tx=XM_006928835.4;refGene_gn=RRP9;refGene_tx=X
M_019822427.2
A2	20576220	20576294
refGene_gn=RRP9;refGene_tx=XM_006928835.4;refGene_gn=RRP9;refGene_tx=X
M_019822427.2
A2	20577054	20577134
refGene_gn=RRP9;refGene_tx=XM_006928835.4;refGene_gn=RRP9;refGene_tx=X
M_019822427.2
A2	20577216	20577362
refGene_gn=RRP9;refGene_tx=XM_006928835.4;refGene_gn=RRP9;refGene_tx=X
M_019822427.2
A2	20577512	20577575
refGene_gn=RRP9;refGene_tx=XM_006928835.4;refGene_gn=RRP9;refGene_tx=X
M_019822427.2
A2	20577651	20577786
refGene_gn=RRP9;refGene_tx=XM_006928835.4;refGene_gn=RRP9;refGene_tx=X
M_019822427.2
A2	20577865	20577966
refGene_gn=RRP9;refGene_tx=XM_006928835.4;refGene_gn=RRP9;refGene_tx=X
M_019822427.2
A2	20578295	20578388
refGene_gn=RRP9;refGene_tx=XM_006928835.4;refGene_gn=RRP9;refGene_tx=X
M_019822427.2
A2	20578470	20578595
refGene_gn=RRP9;refGene_tx=XM_006928835.4;refGene_gn=RRP9;refGene_tx=X
M_019822427.2
A2	20579429	20579556
```

refGene_gn=RRP9;refGene_tx=XM_006928835.4;refGene_gn=RRP9;refGene_tx=XM_019822427.2
A2 20579794 20579836
refGene_gn=RRP9;refGene_tx=XM_006928835.4;refGene_gn=RRP9;refGene_tx=XM_019822427.2
A2 20579917 20579985
refGene_gn=RRP9;refGene_tx=XM_006928835.4;refGene_gn=RRP9;refGene_tx=XM_019822427.2
A2 20580351 20580461
refGene_gn=RRP9;refGene_tx=XM_006928835.4;refGene_gn=RRP9;refGene_tx=XM_019822427.2
A2 20582780 20582847 refGene_gn=RRP9;refGene_tx=XM_019822427.2
A2 20583047 20583136
refGene_gn=RRP9;refGene_tx=XM_006928835.4;refGene_gn=RRP9;refGene_tx=XM_019822427.2
A2 20583415 20583502 refGene_gn=RRP9;refGene_tx=XM_006928835.4
A2 20585086 20585269
refGene_gn=PARP3;refGene_tx=XM_003982285.5;refGene_gn=PARP3;refGene_tx=XM_006928836.4;refGene_gn=PARP3;refGene_tx=XM_019822398.2;refGene_gn=PARP3;refGene_tx=XM_019822406.2;refGene_gn=PARP3;refGene_tx=XM_019822410.2;refGene_gn=PARP3;refGene_tx=XM_019822416.2;refGene_gn=PARP3;refGene_tx=XM_019822420.2
A2 20585916 20586042
refGene_gn=PARP3;refGene_tx=XM_003982285.5;refGene_gn=PARP3;refGene_tx=XM_006928836.4;refGene_gn=PARP3;refGene_tx=XM_019822398.2;refGene_gn=PARP3;refGene_tx=XM_019822406.2;refGene_gn=PARP3;refGene_tx=XM_019822410.2;refGene_gn=PARP3;refGene_tx=XM_019822416.2;refGene_gn=PARP3;refGene_tx=XM_019822420.2
A2 20586258 20586447
refGene_gn=PARP3;refGene_tx=XM_003982285.5;refGene_gn=PARP3;refGene_tx=XM_006928836.4;refGene_gn=PARP3;refGene_tx=XM_019822398.2;refGene_gn=PARP3;refGene_tx=XM_019822406.2;refGene_gn=PARP3;refGene_tx=XM_019822410.2;refGene_gn=PARP3;refGene_tx=XM_019822420.2
A2 20586658 20586791
refGene_gn=PARP3;refGene_tx=XM_003982285.5;refGene_gn=PARP3;refGene_tx=XM_006928836.4;refGene_gn=PARP3;refGene_tx=XM_019822398.2;refGene_gn=PARP3;refGene_tx=XM_019822406.2;refGene_gn=PARP3;refGene_tx=XM_019822410.2;refGene_gn=PARP3;refGene_tx=XM_019822416.2;refGene_gn=PARP3;refGene_tx=XM_019822420.2
A2 20586880 20587107
refGene_gn=PARP3;refGene_tx=XM_003982285.5;refGene_gn=PARP3;refGene_tx=XM_006928836.4;refGene_gn=PARP3;refGene_tx=XM_019822398.2;refGene_gn=PARP3;refGene_tx=XM_019822406.2;refGene_gn=PARP3;refGene_tx=XM_019822410.2;refGene_gn=PARP3;refGene_tx=XM_019822416.2;refGene_gn=PARP3;refGene_tx=XM_019822420.2
A2 20587350 20587497
refGene_gn=PARP3;refGene_tx=XM_003982285.5;refGene_gn=PARP3;refGene_tx=XM_006928836.4;refGene_gn=PARP3;refGene_tx=XM_019822398.2;refGene_gn=PARP3;refGene_tx=XM_019822406.2;refGene_gn=PARP3;refGene_tx=XM_019822410.2;refGene_gn=PARP3;refGene_tx=XM_019822416.2;refGene_gn=PARP3;refGe

ne_tx=XM_019822420.2
A2 20587704 20587791
refGene_gn=PARP3;refGene_tx=XM_003982285.5;refGene_gn=PARP3;refGene_tx=XM_006928836.4;refGene_gn=PARP3;refGene_tx=XM_019822398.2;refGene_gn=PARP3;refGene_tx=XM_019822406.2;refGene_gn=PARP3;refGene_tx=XM_019822410.2;refGene_gn=PARP3;refGene_tx=XM_019822416.2;refGene_gn=PARP3;refGene_tx=XM_019822420.2
A2 20588047 20588225
refGene_gn=PARP3;refGene_tx=XM_003982285.5;refGene_gn=PARP3;refGene_tx=XM_006928836.4;refGene_gn=PARP3;refGene_tx=XM_019822398.2;refGene_gn=PARP3;refGene_tx=XM_019822406.2;refGene_gn=PARP3;refGene_tx=XM_019822410.2;refGene_gn=PARP3;refGene_tx=XM_019822416.2;refGene_gn=PARP3;refGene_tx=XM_019822420.2
A2 20588881 20588907 refGene_gn=PARP3;refGene_tx=XM_019822420.2
A2 20589431 20589587
refGene_gn=PARP3;refGene_tx=XM_003982285.5;refGene_gn=PARP3;refGene_tx=XM_006928836.4;refGene_gn=PARP3;refGene_tx=XM_019822398.2;refGene_gn=PARP3;refGene_tx=XM_019822406.2;refGene_gn=PARP3;refGene_tx=XM_019822410.2;refGene_gn=PARP3;refGene_tx=XM_019822416.2
A2 20590037 20590207
refGene_gn=PARP3;refGene_tx=XM_003982285.5;refGene_gn=PARP3;refGene_tx=XM_006928836.4;refGene_gn=PARP3;refGene_tx=XM_019822398.2;refGene_gn=PARP3;refGene_tx=XM_019822406.2;refGene_gn=PARP3;refGene_tx=XM_019822410.2;refGene_gn=PARP3;refGene_tx=XM_019822416.2
A2 20595667 20596759 refGene_gn=GPR62;refGene_tx=XM_011279407.3
A2 20598085 20598374
refGene_gn=PCBP4;refGene_tx=XM_019822437.2;refGene_gn=PCBP4;refGene_tx=XM_023249366.1;refGene_gn=PCBP4;refGene_tx=XM_023249370.1
A2 20598795 20598965
refGene_gn=PCBP4;refGene_tx=XM_019822437.2;refGene_gn=PCBP4;refGene_tx=XM_023249366.1;refGene_gn=PCBP4;refGene_tx=XM_023249370.1
A2 20599054 20599107
refGene_gn=PCBP4;refGene_tx=XM_019822437.2;refGene_gn=PCBP4;refGene_tx=XM_023249366.1;refGene_gn=PCBP4;refGene_tx=XM_023249370.1
A2 20599230 20599294
refGene_gn=PCBP4;refGene_tx=XM_019822437.2;refGene_gn=PCBP4;refGene_tx=XM_023249366.1;refGene_gn=PCBP4;refGene_tx=XM_023249370.1
A2 20599370 20599415
refGene_gn=PCBP4;refGene_tx=XM_019822437.2;refGene_gn=PCBP4;refGene_tx=XM_023249366.1;refGene_gn=PCBP4;refGene_tx=XM_023249370.1
A2 20599579 20599654
refGene_gn=PCBP4;refGene_tx=XM_019822437.2;refGene_gn=PCBP4;refGene_tx=XM_023249366.1;refGene_gn=PCBP4;refGene_tx=XM_023249370.1
A2 20599900 20600029
refGene_gn=PCBP4;refGene_tx=XM_019822437.2;refGene_gn=PCBP4;refGene_tx=XM_023249366.1;refGene_gn=PCBP4;refGene_tx=XM_023249370.1
A2 20600189 20600321
refGene_gn=PCBP4;refGene_tx=XM_019822437.2;refGene_gn=PCBP4;refGene_tx=XM_023249366.1;refGene_gn=PCBP4;refGene_tx=XM_023249370.1
A2 20600542 20600659

refGene_gn=PCBP4;refGene_tx=XM_019822437.2;refGene_gn=PCBP4;refGene_tx=XM_023249366.1;refGene_gn=PCBP4;refGene_tx=XM_023249370.1
A2 20600884 20600917
refGene_gn=PCBP4;refGene_tx=XM_023249366.1;refGene_gn=PCBP4;refGene_tx=XM_023249370.1
A2 20601023 20601047
refGene_gn=PCBP4;refGene_tx=XM_023249366.1;refGene_gn=PCBP4;refGene_tx=XM_023249370.1
A2 20601191 20601272
refGene_gn=PCBP4;refGene_tx=XM_023249366.1;refGene_gn=PCBP4;refGene_tx=XM_023249370.1
A2 20609532 20609712 refGene_gn=ABHD14B;refGene_tx=XM_006928843.4
A2 20610093 20610335 refGene_gn=ABHD14B;refGene_tx=XM_006928843.4
A2 20611534 20611745 refGene_gn=ABHD14B;refGene_tx=XM_006928843.4
A2 20612342 20612552
refGene_gn=LOC101101118;refGene_tx=XM_019822446.2;refGene_gn=LOC101101118;refGene_tx=XM_019822453.2
A2 20613858 20613942
refGene_gn=LOC101101118;refGene_tx=XM_003982288.3;refGene_gn=LOC101101118;refGene_tx=XM_019822447.2
A2 20616242 20616454
refGene_gn=LOC101101118;refGene_tx=XM_003982288.3;refGene_gn=LOC101101118;refGene_tx=XM_019822446.2;refGene_gn=LOC101101118;refGene_tx=XM_019822447.2;refGene_gn=LOC101101118;refGene_tx=XM_019822453.2
A2 20616630 20616746
refGene_gn=LOC101101118;refGene_tx=XM_003982288.3;refGene_gn=LOC101101118;refGene_tx=XM_019822446.2;refGene_gn=LOC101101118;refGene_tx=XM_019822447.2;refGene_gn=LOC101101118;refGene_tx=XM_019822453.2
A2 20617877 20618113
refGene_gn=LOC101101118;refGene_tx=XM_003982288.3;refGene_gn=LOC101101118;refGene_tx=XM_019822446.2;refGene_gn=LOC101101118;refGene_tx=XM_019822447.2;refGene_gn=LOC101101118;refGene_tx=XM_019822453.2
A2 20618319 20618502
refGene_gn=LOC101101118;refGene_tx=XM_003982288.3
A2 20620823 20620935
refGene_gn=LOC101101118;refGene_tx=XM_011279419.3;refGene_gn=LOC101101118;refGene_tx=XM_011279424.3;refGene_gn=LOC101101118;refGene_tx=XM_019822446.2;refGene_gn=LOC101101118;refGene_tx=XM_019822447.2;refGene_gn=LOC101101118;refGene_tx=XM_019822453.2;refGene_gn=LOC101101118;refGene_tx=XM_019822469.2
A2 20621632 20621697
refGene_gn=LOC101101118;refGene_tx=XM_011279419.3;refGene_gn=LOC101101118;refGene_tx=XM_011279424.3;refGene_gn=LOC101101118;refGene_tx=XM_019822446.2;refGene_gn=LOC101101118;refGene_tx=XM_019822447.2;refGene_gn=LOC101101118;refGene_tx=XM_019822453.2;refGene_gn=LOC101101118;refGene_tx=XM_019822469.2
A2 20621768 20621873
refGene_gn=LOC101101118;refGene_tx=XM_011279419.3;refGene_gn=LOC101101118;refGene_tx=XM_011279424.3;refGene_gn=LOC101101118;refGene_tx=XM_019822446.2;refGene_gn=LOC101101118;refGene_tx=XM_019822447.2;refGene_gn

=LOC101101118;refGene_tx=XM_019822453.2;refGene_gn=LOC101101118;refGene_tx=XM_019822469.2
A2 20622234 20622329
refGene_gn=LOC101101118;refGene_tx=XM_011279419.3;refGene_gn=LOC101101118;refGene_tx=XM_011279424.3;refGene_gn=LOC101101118;refGene_tx=XM_019822446.2;refGene_gn=LOC101101118;refGene_tx=XM_019822447.2;refGene_gn=LOC101101118;refGene_tx=XM_019822453.2;refGene_gn=LOC101101118;refGene_tx=XM_019822469.2
A2 20622617 20622694
refGene_gn=LOC101101118;refGene_tx=XM_011279419.3;refGene_gn=LOC101101118;refGene_tx=XM_011279424.3;refGene_gn=LOC101101118;refGene_tx=XM_019822446.2;refGene_gn=LOC101101118;refGene_tx=XM_019822447.2;refGene_gn=LOC101101118;refGene_tx=XM_019822453.2;refGene_gn=LOC101101118;refGene_tx=XM_019822469.2
A2 20622785 20622875
refGene_gn=LOC101101118;refGene_tx=XM_011279419.3;refGene_gn=LOC101101118;refGene_tx=XM_011279424.3;refGene_gn=LOC101101118;refGene_tx=XM_019822446.2;refGene_gn=LOC101101118;refGene_tx=XM_019822447.2;refGene_gn=LOC101101118;refGene_tx=XM_019822453.2;refGene_gn=LOC101101118;refGene_tx=XM_019822469.2
A2 20622975 20623032
refGene_gn=LOC101101118;refGene_tx=XM_011279419.3;refGene_gn=LOC101101118;refGene_tx=XM_011279424.3;refGene_gn=LOC101101118;refGene_tx=XM_019822446.2;refGene_gn=LOC101101118;refGene_tx=XM_019822447.2;refGene_gn=LOC101101118;refGene_tx=XM_019822453.2;refGene_gn=LOC101101118;refGene_tx=XM_019822469.2
A2 20623370 20623444
refGene_gn=LOC101101118;refGene_tx=XM_011279419.3;refGene_gn=LOC101101118;refGene_tx=XM_011279424.3;refGene_gn=LOC101101118;refGene_tx=XM_019822446.2;refGene_gn=LOC101101118;refGene_tx=XM_019822447.2;refGene_gn=LOC101101118;refGene_tx=XM_019822453.2;refGene_gn=LOC101101118;refGene_tx=XM_019822469.2
A2 20623529 20623579
refGene_gn=LOC101101118;refGene_tx=XM_011279419.3;refGene_gn=LOC101101118;refGene_tx=XM_011279424.3;refGene_gn=LOC101101118;refGene_tx=XM_019822446.2;refGene_gn=LOC101101118;refGene_tx=XM_019822447.2;refGene_gn=LOC101101118;refGene_tx=XM_019822453.2;refGene_gn=LOC101101118;refGene_tx=XM_019822469.2
A2 20623679 20623824
refGene_gn=LOC101101118;refGene_tx=XM_011279419.3;refGene_gn=LOC101101118;refGene_tx=XM_011279424.3;refGene_gn=LOC101101118;refGene_tx=XM_019822446.2;refGene_gn=LOC101101118;refGene_tx=XM_019822447.2;refGene_gn=LOC101101118;refGene_tx=XM_019822453.2;refGene_gn=LOC101101118;refGene_tx=XM_019822469.2
A2 20623925 20623994
refGene_gn=LOC101101118;refGene_tx=XM_011279419.3;refGene_gn=LOC101101118;refGene_tx=XM_011279424.3;refGene_gn=LOC101101118;refGene_tx=XM_019822446.2;refGene_gn=LOC101101118;refGene_tx=XM_019822447.2;refGene_gn=LOC101101118;refGene_tx=XM_019822453.2;refGene_gn=LOC101101118;refGene_tx=XM_019822469.2

A2 20624624 20624704
refGene_gn=LOC101101118;refGene_tx=XM_011279419.3;refGene_gn=LOC101101118;refGene_tx=XM_011279424.3;refGene_gn=LOC101101118;refGene_tx=XM_019822446.2;refGene_gn=LOC101101118;refGene_tx=XM_019822447.2;refGene_gn=LOC101101118;refGene_tx=XM_019822453.2;refGene_gn=LOC101101118;refGene_tx=XM_019822469.2
A2 20624837 20624898
refGene_gn=LOC101101118;refGene_tx=XM_011279419.3;refGene_gn=LOC101101118;refGene_tx=XM_011279424.3;refGene_gn=LOC101101118;refGene_tx=XM_019822446.2;refGene_gn=LOC101101118;refGene_tx=XM_019822447.2;refGene_gn=LOC101101118;refGene_tx=XM_019822453.2;refGene_gn=LOC101101118;refGene_tx=XM_019822469.2
A2 20624979 20625144
refGene_gn=LOC101101118;refGene_tx=XM_011279419.3;refGene_gn=LOC101101118;refGene_tx=XM_011279424.3;refGene_gn=LOC101101118;refGene_tx=XM_019822446.2;refGene_gn=LOC101101118;refGene_tx=XM_019822447.2;refGene_gn=LOC101101118;refGene_tx=XM_019822453.2;refGene_gn=LOC101101118;refGene_tx=XM_019822469.2
A2 20627533 20627917 refGene_gn=RPL29;refGene_tx=XM_003982291.4
A2 20628945 20629010 refGene_gn=RPL29;refGene_tx=XM_003982291.4
A2 20629340 20629377 refGene_gn=RPL29;refGene_tx=XM_003982291.4
A2 20676304 20676612 refGene_gn=DUSP7;refGene_tx=XM_003982292.5
A2 20679336 20679771 refGene_gn=DUSP7;refGene_tx=XM_003982292.5
A2 20681248 20681765 refGene_gn=DUSP7;refGene_tx=XM_003982292.5
A2 20699519 20699618
refGene_gn=POC1A;refGene_tx=XM_003982293.4;refGene_gn=POC1A;refGene_tx=XM_019822527.2;refGene_gn=POC1A;refGene_tx=XM_019822531.2;refGene_gn=POC1A;refGene_tx=XM_019822537.2;refGene_gn=POC1A;refGene_tx=XM_019822556.2;refGene_gn=POC1A;refGene_tx=XM_023249399.1;refGene_gn=POC1A;refGene_tx=XM_023249402.1;refGene_gn=POC1A;refGene_tx=XM_023249403.1
A2 20719208 20719352
refGene_gn=POC1A;refGene_tx=XM_003982293.4;refGene_gn=POC1A;refGene_tx=XM_019822527.2;refGene_gn=POC1A;refGene_tx=XM_019822531.2;refGene_gn=POC1A;refGene_tx=XM_019822537.2;refGene_gn=POC1A;refGene_tx=XM_019822556.2;refGene_gn=POC1A;refGene_tx=XM_023249399.1;refGene_gn=POC1A;refGene_tx=XM_023249402.1;refGene_gn=POC1A;refGene_tx=XM_023249403.1
A2 20734639 20734729
refGene_gn=POC1A;refGene_tx=XM_019822527.2;refGene_gn=POC1A;refGene_tx=XM_019822531.2;refGene_gn=POC1A;refGene_tx=XM_019822556.2;refGene_gn=POC1A;refGene_tx=XM_023249399.1;refGene_gn=POC1A;refGene_tx=XM_023249402.1;refGene_gn=POC1A;refGene_tx=XM_023249403.1
A2 20744706 20744805
refGene_gn=POC1A;refGene_tx=XM_003982293.4;refGene_gn=POC1A;refGene_tx=XM_019822527.2;refGene_gn=POC1A;refGene_tx=XM_019822531.2;refGene_gn=POC1A;refGene_tx=XM_019822537.2;refGene_gn=POC1A;refGene_tx=XM_019822556.2;refGene_gn=POC1A;refGene_tx=XM_023249399.1;refGene_gn=POC1A;refGene_tx=XM_023249402.1;refGene_gn=POC1A;refGene_tx=XM_023249403.1
A2 20747727 20747796
refGene_gn=POC1A;refGene_tx=XM_003982293.4;refGene_gn=POC1A;refGene_tx=XM_019822527.2;refGene_gn=POC1A;refGene_tx=XM_019822531.2;refGene_gn=

POC1A;refGene_tx=XM_019822537.2;refGene_gn=POC1A;refGene_tx=XM_0198225
56.2;refGene_gn=POC1A;refGene_tx=XM_023249399.1;refGene_gn=POC1A;refGe
ne_tx=XM_023249402.1;refGene_gn=POC1A;refGene_tx=XM_023249403.1
A2 20758241 20758375
refGene_gn=POC1A;refGene_tx=XM_003982293.4;refGene_gn=POC1A;refGene_tx
=XM_019822527.2;refGene_gn=POC1A;refGene_tx=XM_019822531.2;refGene_gn=
POC1A;refGene_tx=XM_019822537.2;refGene_gn=POC1A;refGene_tx=XM_0198225
56.2;refGene_gn=POC1A;refGene_tx=XM_023249399.1;refGene_gn=POC1A;refGe
ne_tx=XM_023249402.1;refGene_gn=POC1A;refGene_tx=XM_023249403.1
A2 20766497 20766613
refGene_gn=POC1A;refGene_tx=XM_003982293.4;refGene_gn=POC1A;refGene_tx
=XM_019822527.2;refGene_gn=POC1A;refGene_tx=XM_019822531.2;refGene_gn=
POC1A;refGene_tx=XM_019822537.2;refGene_gn=POC1A;refGene_tx=XM_0198225
56.2;refGene_gn=POC1A;refGene_tx=XM_023249399.1;refGene_gn=POC1A;refGe
ne_tx=XM_023249402.1;refGene_gn=POC1A;refGene_tx=XM_023249403.1
A2 20767815 20767923
refGene_gn=POC1A;refGene_tx=XM_003982293.4;refGene_gn=POC1A;refGene_tx
=XM_019822527.2;refGene_gn=POC1A;refGene_tx=XM_019822531.2;refGene_gn=
POC1A;refGene_tx=XM_019822537.2;refGene_gn=POC1A;refGene_tx=XM_0198225
56.2;refGene_gn=POC1A;refGene_tx=XM_023249399.1;refGene_gn=POC1A;refGe
ne_tx=XM_023249402.1;refGene_gn=POC1A;refGene_tx=XM_023249403.1
A2 20768828 20769008
refGene_gn=POC1A;refGene_tx=XM_003982293.4;refGene_gn=POC1A;refGene_tx
=XM_019822527.2;refGene_gn=POC1A;refGene_tx=XM_019822531.2;refGene_gn=
POC1A;refGene_tx=XM_019822537.2;refGene_gn=POC1A;refGene_tx=XM_0198225
56.2;refGene_gn=POC1A;refGene_tx=XM_023249399.1;refGene_gn=POC1A;refGe
ne_tx=XM_023249402.1;refGene_gn=POC1A;refGene_tx=XM_023249403.1
A2 20769383 20769555
refGene_gn=POC1A;refGene_tx=XM_003982293.4;refGene_gn=POC1A;refGene_tx
=XM_019822527.2;refGene_gn=POC1A;refGene_tx=XM_019822531.2;refGene_gn=
POC1A;refGene_tx=XM_019822537.2;refGene_gn=POC1A;refGene_tx=XM_0198225
56.2;refGene_gn=POC1A;refGene_tx=XM_023249399.1;refGene_gn=POC1A;refGe
ne_tx=XM_023249402.1;refGene_gn=POC1A;refGene_tx=XM_023249403.1
A2 20770522 20770607
refGene_gn=POC1A;refGene_tx=XM_003982293.4;refGene_gn=POC1A;refGene_tx
=XM_019822527.2;refGene_gn=POC1A;refGene_tx=XM_019822531.2;refGene_gn=
POC1A;refGene_tx=XM_023249399.1;refGene_gn=POC1A;refGene_tx=XM_0232494
02.1;refGene_gn=POC1A;refGene_tx=XM_023249403.1
A2 20773895 20773913
refGene_gn=POC1A;refGene_tx=XM_003982293.4;refGene_gn=POC1A;refGene_tx
=XM_019822531.2
A2 20793285 20793359
refGene_gn=POC1A;refGene_tx=XM_019822527.2;refGene_gn=POC1A;refGene_tx
=XM_023249399.1;refGene_gn=POC1A;refGene_tx=XM_023249402.1;refGene_gn=
POC1A;refGene_tx=XM_023249403.1
A2 20799092 20799102
refGene_gn=POC1A;refGene_tx=XM_019822527.2;refGene_gn=POC1A;refGene_tx
=XM_023249399.1;refGene_gn=POC1A;refGene_tx=XM_023249402.1;refGene_gn=
POC1A;refGene_tx=XM_023249403.1
A2 20801383 20801582 refGene_gn=ALAS1;refGene_tx=XM_003982295.5

A2	20804284	20804512	refGene_gn=ALAS1;refGene_tx=XM_003982295.5
A2	20805312	20805462	refGene_gn=ALAS1;refGene_tx=XM_003982295.5
A2	20805878	20806101	refGene_gn=ALAS1;refGene_tx=XM_003982295.5
A2	20807265	20807450	refGene_gn=ALAS1;refGene_tx=XM_003982295.5
A2	20808005	20808185	refGene_gn=ALAS1;refGene_tx=XM_003982295.5
A2	20808535	20808700	refGene_gn=ALAS1;refGene_tx=XM_003982295.5
A2	20811208	20811477	refGene_gn=ALAS1;refGene_tx=XM_003982295.5
A2	20812154	20812317	refGene_gn=ALAS1;refGene_tx=XM_003982295.5
A2	20813904	20814065	refGene_gn=ALAS1;refGene_tx=XM_003982295.5
A2	20820465	20823558
refGene_gn=TLR9;refGene_tx=NM_001009285.1;refGene_gn=TLR9;refGene_tx=XM_019814986.2
A2	20824455	20824558
refGene_gn=TLR9;refGene_tx=NM_001009285.1;refGene_gn=TLR9;refGene_tx=XM_019814986.2
A2	20828488	20828610	refGene_gn=TLR9;refGene_tx=XM_019814986.2
A2	20828744	20828810	refGene_gn=TLR9;refGene_tx=XM_019814986.2
A2	20846808	20847032	refGene_gn=PPM1M;refGene_tx=XM_023249427.1
A2	20847704	20847828	refGene_gn=PPM1M;refGene_tx=XM_023249427.1
A2	20847964	20848219	refGene_gn=PPM1M;refGene_tx=XM_023249427.1
A2	20848685	20848798	refGene_gn=PPM1M;refGene_tx=XM_023249427.1
A2	20849125	20849210	refGene_gn=PPM1M;refGene_tx=XM_023249427.1
A2	20849316	20849435	refGene_gn=PPM1M;refGene_tx=XM_023249427.1
A2	20849596	20849660	refGene_gn=PPM1M;refGene_tx=XM_023249427.1
A2	20849889	20849997	refGene_gn=PPM1M;refGene_tx=XM_023249427.1
A2	20850106	20850255	refGene_gn=PPM1M;refGene_tx=XM_023249427.1
A2	20850625	20850770	refGene_gn=PPM1M;refGene_tx=XM_023249427.1
A2	20856472	20856502	refGene_gn=WDR82;refGene_tx=XM_003982298.5
A2	20857492	20857635	refGene_gn=WDR82;refGene_tx=XM_003982298.5
A2	20858658	20858728	refGene_gn=WDR82;refGene_tx=XM_003982298.5
A2	20859166	20859322	refGene_gn=WDR82;refGene_tx=XM_003982298.5
A2	20859555	20859672	refGene_gn=WDR82;refGene_tx=XM_003982298.5
A2	20860158	20860258	refGene_gn=WDR82;refGene_tx=XM_003982298.5
A2	20864810	20864877	refGene_gn=WDR82;refGene_tx=XM_003982298.5
A2	20867889	20867987	refGene_gn=WDR82;refGene_tx=XM_003982298.5
A2	20872031	20872192	refGene_gn=WDR82;refGene_tx=XM_003982298.5
A2	20894191	20894568
refGene_gn=GLYCTK;refGene_tx=XM_003982299.5;refGene_gn=GLYCTK;refGene_tx=XM_006928845.4;refGene_gn=GLYCTK;refGene_tx=XM_006928846.4;refGene_gn=GLYCTK;refGene_tx=XM_019822576.2
A2	20894791	20894943
refGene_gn=GLYCTK;refGene_tx=XM_003982299.5;refGene_gn=GLYCTK;refGene_tx=XM_006928845.4;refGene_gn=GLYCTK;refGene_tx=XM_006928846.4;refGene_gn=GLYCTK;refGene_tx=XM_019822576.2
A2	20895580	20895756
refGene_gn=GLYCTK;refGene_tx=XM_003982299.5;refGene_gn=GLYCTK;refGene_tx=XM_006928845.4;refGene_gn=GLYCTK;refGene_tx=XM_006928846.4;refGene_gn=GLYCTK;refGene_tx=XM_019822576.2
A2	20896092	20896959
refGene_gn=GLYCTK;refGene_tx=XM_003982299.5;refGene_gn=GLYCTK;refGene_

tx=XM_006928845.4;refGene_gn=GLYCTK;refGene_tx=XM_006928846.4;refGene_gn=GLYCTK;refGene_tx=XM_019822576.2
A2 20923919 20924255
refGene_gn=DNAH1;refGene_tx=XM_011279464.3;refGene_gn=DNAH1;refGene_tx=XM_019822581.2;refGene_gn=DNAH1;refGene_tx=XM_019822582.2;refGene_gn=DNAH1;refGene_tx=XM_019822587.2;refGene_gn=DNAH1;refGene_tx=XM_019822590.2;refGene_gn=DNAH1;refGene_tx=XM_019822591.2
A2 20925285 20925358
refGene_gn=DNAH1;refGene_tx=XM_011279464.3;refGene_gn=DNAH1;refGene_tx=XM_019822581.2;refGene_gn=DNAH1;refGene_tx=XM_019822582.2;refGene_gn=DNAH1;refGene_tx=XM_019822587.2;refGene_gn=DNAH1;refGene_tx=XM_019822590.2;refGene_gn=DNAH1;refGene_tx=XM_019822591.2
A2 20926529 20926704
refGene_gn=DNAH1;refGene_tx=XM_011279464.3;refGene_gn=DNAH1;refGene_tx=XM_019822581.2;refGene_gn=DNAH1;refGene_tx=XM_019822582.2;refGene_gn=DNAH1;refGene_tx=XM_019822587.2;refGene_gn=DNAH1;refGene_tx=XM_019822590.2;refGene_gn=DNAH1;refGene_tx=XM_019822591.2
A2 20927328 20927485
refGene_gn=DNAH1;refGene_tx=XM_011279464.3;refGene_gn=DNAH1;refGene_tx=XM_019822581.2;refGene_gn=DNAH1;refGene_tx=XM_019822582.2;refGene_gn=DNAH1;refGene_tx=XM_019822587.2;refGene_gn=DNAH1;refGene_tx=XM_019822590.2;refGene_gn=DNAH1;refGene_tx=XM_019822591.2
A2 20928433 20928566
refGene_gn=DNAH1;refGene_tx=XM_011279464.3;refGene_gn=DNAH1;refGene_tx=XM_019822581.2;refGene_gn=DNAH1;refGene_tx=XM_019822582.2;refGene_gn=DNAH1;refGene_tx=XM_019822587.2;refGene_gn=DNAH1;refGene_tx=XM_019822590.2;refGene_gn=DNAH1;refGene_tx=XM_019822591.2;refGene_gn=DNAH1;refGene_tx=XM_023249453.1
A2 20930860 20931056
refGene_gn=DNAH1;refGene_tx=XM_011279464.3;refGene_gn=DNAH1;refGene_tx=XM_019822581.2;refGene_gn=DNAH1;refGene_tx=XM_019822582.2;refGene_gn=DNAH1;refGene_tx=XM_019822587.2;refGene_gn=DNAH1;refGene_tx=XM_019822590.2;refGene_gn=DNAH1;refGene_tx=XM_019822591.2;refGene_gn=DNAH1;refGene_tx=XM_023249453.1
A2 20931643 20931896
refGene_gn=DNAH1;refGene_tx=XM_011279464.3;refGene_gn=DNAH1;refGene_tx=XM_019822581.2;refGene_gn=DNAH1;refGene_tx=XM_019822582.2;refGene_gn=DNAH1;refGene_tx=XM_019822587.2;refGene_gn=DNAH1;refGene_tx=XM_019822590.2;refGene_gn=DNAH1;refGene_tx=XM_019822591.2;refGene_gn=DNAH1;refGene_tx=XM_023249453.1
A2 20946047 20946205
refGene_gn=DNAH1;refGene_tx=XM_011279464.3;refGene_gn=DNAH1;refGene_tx=XM_019822581.2;refGene_gn=DNAH1;refGene_tx=XM_019822582.2;refGene_gn=DNAH1;refGene_tx=XM_019822587.2;refGene_gn=DNAH1;refGene_tx=XM_019822590.2;refGene_gn=DNAH1;refGene_tx=XM_019822591.2;refGene_gn=DNAH1;refGene_tx=XM_023249453.1
A2 20946968 20947180
refGene_gn=DNAH1;refGene_tx=XM_011279464.3;refGene_gn=DNAH1;refGene_tx=XM_019822581.2;refGene_gn=DNAH1;refGene_tx=XM_019822582.2;refGene_gn=DNAH1;refGene_tx=XM_019822587.2;refGene_gn=DNAH1;refGene_tx=XM_0198225

90.2;refGene_gn=DNAH1;refGene_tx=XM_019822591.2;refGene_gn=DNAH1;refGene_tx=XM_023249453.1
A2 20948022 20948321
refGene_gn=DNAH1;refGene_tx=XM_011279464.3;refGene_gn=DNAH1;refGene_tx=XM_019822581.2;refGene_gn=DNAH1;refGene_tx=XM_019822582.2;refGene_gn=DNAH1;refGene_tx=XM_019822587.2;refGene_gn=DNAH1;refGene_tx=XM_019822590.2;refGene_gn=DNAH1;refGene_tx=XM_019822591.2;refGene_gn=DNAH1;refGene_tx=XM_023249453.1
A2 20949704 20949855
refGene_gn=DNAH1;refGene_tx=XM_011279464.3;refGene_gn=DNAH1;refGene_tx=XM_019822581.2;refGene_gn=DNAH1;refGene_tx=XM_019822582.2;refGene_gn=DNAH1;refGene_tx=XM_019822587.2;refGene_gn=DNAH1;refGene_tx=XM_019822590.2;refGene_gn=DNAH1;refGene_tx=XM_019822591.2;refGene_gn=DNAH1;refGene_tx=XM_023249453.1
A2 20950642 20950836
refGene_gn=DNAH1;refGene_tx=XM_011279464.3;refGene_gn=DNAH1;refGene_tx=XM_019822581.2;refGene_gn=DNAH1;refGene_tx=XM_019822582.2;refGene_gn=DNAH1;refGene_tx=XM_019822587.2;refGene_gn=DNAH1;refGene_tx=XM_019822590.2;refGene_gn=DNAH1;refGene_tx=XM_019822591.2;refGene_gn=DNAH1;refGene_tx=XM_023249453.1
A2 20950931 20951157
refGene_gn=DNAH1;refGene_tx=XM_011279464.3;refGene_gn=DNAH1;refGene_tx=XM_019822581.2;refGene_gn=DNAH1;refGene_tx=XM_019822582.2;refGene_gn=DNAH1;refGene_tx=XM_019822587.2;refGene_gn=DNAH1;refGene_tx=XM_019822590.2;refGene_gn=DNAH1;refGene_tx=XM_019822591.2;refGene_gn=DNAH1;refGene_tx=XM_023249453.1
A2 20951649 20951769
refGene_gn=DNAH1;refGene_tx=XM_011279464.3;refGene_gn=DNAH1;refGene_tx=XM_019822581.2;refGene_gn=DNAH1;refGene_tx=XM_019822582.2;refGene_gn=DNAH1;refGene_tx=XM_019822587.2;refGene_gn=DNAH1;refGene_tx=XM_019822590.2;refGene_gn=DNAH1;refGene_tx=XM_019822591.2;refGene_gn=DNAH1;refGene_tx=XM_023249453.1
A2 20952067 20952150
refGene_gn=DNAH1;refGene_tx=XM_011279464.3;refGene_gn=DNAH1;refGene_tx=XM_019822581.2;refGene_gn=DNAH1;refGene_tx=XM_019822582.2;refGene_gn=DNAH1;refGene_tx=XM_019822587.2;refGene_gn=DNAH1;refGene_tx=XM_019822590.2;refGene_gn=DNAH1;refGene_tx=XM_019822591.2;refGene_gn=DNAH1;refGene_tx=XM_023249453.1
A2 20953283 20953425
refGene_gn=DNAH1;refGene_tx=XM_011279464.3;refGene_gn=DNAH1;refGene_tx=XM_019822581.2;refGene_gn=DNAH1;refGene_tx=XM_019822582.2;refGene_gn=DNAH1;refGene_tx=XM_019822587.2;refGene_gn=DNAH1;refGene_tx=XM_019822590.2;refGene_gn=DNAH1;refGene_tx=XM_019822591.2;refGene_gn=DNAH1;refGene_tx=XM_023249453.1
A2 20953907 20954063
refGene_gn=DNAH1;refGene_tx=XM_011279464.3;refGene_gn=DNAH1;refGene_tx=XM_019822581.2;refGene_gn=DNAH1;refGene_tx=XM_019822582.2;refGene_gn=DNAH1;refGene_tx=XM_019822587.2;refGene_gn=DNAH1;refGene_tx=XM_019822590.2;refGene_gn=DNAH1;refGene_tx=XM_019822591.2;refGene_gn=DNAH1;refGene_tx=XM_023249453.1

A2	20954446	20954645
refGene_gn=DNAH1;refGene_tx=XM_011279464.3;refGene_gn=DNAH1;refGene_tx=XM_019822581.2;refGene_gn=DNAH1;refGene_tx=XM_019822582.2;refGene_gn=DNAH1;refGene_tx=XM_019822587.2;refGene_gn=DNAH1;refGene_tx=XM_019822590.2;refGene_gn=DNAH1;refGene_tx=XM_019822591.2;refGene_gn=DNAH1;refGene_tx=XM_023249453.1
A2	20954724	20954978
refGene_gn=DNAH1;refGene_tx=XM_011279464.3;refGene_gn=DNAH1;refGene_tx=XM_019822581.2;refGene_gn=DNAH1;refGene_tx=XM_019822582.2;refGene_gn=DNAH1;refGene_tx=XM_019822587.2;refGene_gn=DNAH1;refGene_tx=XM_019822590.2;refGene_gn=DNAH1;refGene_tx=XM_019822591.2;refGene_gn=DNAH1;refGene_tx=XM_023249453.1
A2	20955484	20955697
refGene_gn=DNAH1;refGene_tx=XM_011279464.3;refGene_gn=DNAH1;refGene_tx=XM_019822581.2;refGene_gn=DNAH1;refGene_tx=XM_019822582.2;refGene_gn=DNAH1;refGene_tx=XM_019822587.2;refGene_gn=DNAH1;refGene_tx=XM_019822590.2;refGene_gn=DNAH1;refGene_tx=XM_019822591.2;refGene_gn=DNAH1;refGene_tx=XM_023249453.1
A2	20957195	20957360
refGene_gn=DNAH1;refGene_tx=XM_011279464.3;refGene_gn=DNAH1;refGene_tx=XM_019822581.2;refGene_gn=DNAH1;refGene_tx=XM_019822582.2;refGene_gn=DNAH1;refGene_tx=XM_019822587.2;refGene_gn=DNAH1;refGene_tx=XM_019822590.2;refGene_gn=DNAH1;refGene_tx=XM_019822591.2;refGene_gn=DNAH1;refGene_tx=XM_023249453.1
A2	20958069	20958191
refGene_gn=DNAH1;refGene_tx=XM_011279464.3;refGene_gn=DNAH1;refGene_tx=XM_019822581.2;refGene_gn=DNAH1;refGene_tx=XM_019822582.2;refGene_gn=DNAH1;refGene_tx=XM_019822587.2;refGene_gn=DNAH1;refGene_tx=XM_019822590.2;refGene_gn=DNAH1;refGene_tx=XM_019822591.2;refGene_gn=DNAH1;refGene_tx=XM_023249453.1
A2	20958291	20958397
refGene_gn=DNAH1;refGene_tx=XM_011279464.3;refGene_gn=DNAH1;refGene_tx=XM_019822581.2;refGene_gn=DNAH1;refGene_tx=XM_019822582.2;refGene_gn=DNAH1;refGene_tx=XM_019822587.2;refGene_gn=DNAH1;refGene_tx=XM_019822590.2;refGene_gn=DNAH1;refGene_tx=XM_019822591.2;refGene_gn=DNAH1;refGene_tx=XM_023249453.1
A2	20958855	20959035
refGene_gn=DNAH1;refGene_tx=XM_011279464.3;refGene_gn=DNAH1;refGene_tx=XM_019822581.2;refGene_gn=DNAH1;refGene_tx=XM_019822582.2;refGene_gn=DNAH1;refGene_tx=XM_019822587.2;refGene_gn=DNAH1;refGene_tx=XM_019822590.2;refGene_gn=DNAH1;refGene_tx=XM_019822591.2;refGene_gn=DNAH1;refGene_tx=XM_023249453.1
A2	20959682	20959823
refGene_gn=DNAH1;refGene_tx=XM_011279464.3;refGene_gn=DNAH1;refGene_tx=XM_019822581.2;refGene_gn=DNAH1;refGene_tx=XM_019822582.2;refGene_gn=DNAH1;refGene_tx=XM_019822587.2;refGene_gn=DNAH1;refGene_tx=XM_019822590.2;refGene_gn=DNAH1;refGene_tx=XM_019822591.2;refGene_gn=DNAH1;refGene_tx=XM_023249453.1
A2	20960448	20960612
refGene_gn=DNAH1;refGene_tx=XM_011279464.3;refGene_gn=DNAH1;refGene_tx

=XM_019822581.2;refGene_gn=DNAH1;refGene_tx=XM_019822582.2;refGene_gn=DNAH1;refGene_tx=XM_019822587.2;refGene_gn=DNAH1;refGene_tx=XM_019822590.2;refGene_gn=DNAH1;refGene_tx=XM_019822591.2;refGene_gn=DNAH1;refGene_tx=XM_023249453.1
A2 20960801 20960915
refGene_gn=DNAH1;refGene_tx=XM_011279464.3;refGene_gn=DNAH1;refGene_tx=XM_019822581.2;refGene_gn=DNAH1;refGene_tx=XM_019822582.2;refGene_gn=DNAH1;refGene_tx=XM_019822587.2;refGene_gn=DNAH1;refGene_tx=XM_019822590.2;refGene_gn=DNAH1;refGene_tx=XM_019822591.2;refGene_gn=DNAH1;refGene_tx=XM_023249453.1
A2 20961662 20961851
refGene_gn=DNAH1;refGene_tx=XM_011279464.3;refGene_gn=DNAH1;refGene_tx=XM_019822581.2;refGene_gn=DNAH1;refGene_tx=XM_019822582.2;refGene_gn=DNAH1;refGene_tx=XM_019822587.2;refGene_gn=DNAH1;refGene_tx=XM_019822590.2;refGene_gn=DNAH1;refGene_tx=XM_019822591.2;refGene_gn=DNAH1;refGene_tx=XM_023249453.1
A2 20962144 20962250
refGene_gn=DNAH1;refGene_tx=XM_011279464.3;refGene_gn=DNAH1;refGene_tx=XM_019822581.2;refGene_gn=DNAH1;refGene_tx=XM_019822582.2;refGene_gn=DNAH1;refGene_tx=XM_019822587.2;refGene_gn=DNAH1;refGene_tx=XM_019822590.2;refGene_gn=DNAH1;refGene_tx=XM_019822591.2;refGene_gn=DNAH1;refGene_tx=XM_023249453.1
A2 20962961 20963075
refGene_gn=DNAH1;refGene_tx=XM_011279464.3;refGene_gn=DNAH1;refGene_tx=XM_019822581.2;refGene_gn=DNAH1;refGene_tx=XM_019822582.2;refGene_gn=DNAH1;refGene_tx=XM_019822587.2;refGene_gn=DNAH1;refGene_tx=XM_019822590.2;refGene_gn=DNAH1;refGene_tx=XM_019822591.2;refGene_gn=DNAH1;refGene_tx=XM_023249453.1
A2 20963492 20963642
refGene_gn=DNAH1;refGene_tx=XM_011279464.3;refGene_gn=DNAH1;refGene_tx=XM_019822581.2;refGene_gn=DNAH1;refGene_tx=XM_019822582.2;refGene_gn=DNAH1;refGene_tx=XM_019822587.2;refGene_gn=DNAH1;refGene_tx=XM_019822590.2;refGene_gn=DNAH1;refGene_tx=XM_019822591.2;refGene_gn=DNAH1;refGene_tx=XM_023249453.1
A2 20965261 20965348
refGene_gn=DNAH1;refGene_tx=XM_011279464.3;refGene_gn=DNAH1;refGene_tx=XM_019822581.2;refGene_gn=DNAH1;refGene_tx=XM_019822582.2;refGene_gn=DNAH1;refGene_tx=XM_019822587.2;refGene_gn=DNAH1;refGene_tx=XM_019822590.2;refGene_gn=DNAH1;refGene_tx=XM_019822591.2;refGene_gn=DNAH1;refGene_tx=XM_023249453.1
A2 20965457 20965644
refGene_gn=DNAH1;refGene_tx=XM_011279464.3;refGene_gn=DNAH1;refGene_tx=XM_019822581.2;refGene_gn=DNAH1;refGene_tx=XM_019822582.2;refGene_gn=DNAH1;refGene_tx=XM_019822587.2;refGene_gn=DNAH1;refGene_tx=XM_019822590.2;refGene_gn=DNAH1;refGene_tx=XM_019822591.2;refGene_gn=DNAH1;refGene_tx=XM_023249453.1
A2 20967956 20968048
refGene_gn=DNAH1;refGene_tx=XM_011279464.3;refGene_gn=DNAH1;refGene_tx=XM_019822581.2;refGene_gn=DNAH1;refGene_tx=XM_019822582.2;refGene_gn=DNAH1;refGene_tx=XM_019822587.2;refGene_gn=DNAH1;refGene_tx=XM_0198225

90.2;refGene_gn=DNAH1;refGene_tx=XM_019822591.2;refGene_gn=DNAH1;refGene_tx=XM_023249453.1
A2 20968243 20968398
refGene_gn=DNAH1;refGene_tx=XM_011279464.3;refGene_gn=DNAH1;refGene_tx=XM_019822581.2;refGene_gn=DNAH1;refGene_tx=XM_019822582.2;refGene_gn=DNAH1;refGene_tx=XM_019822587.2;refGene_gn=DNAH1;refGene_tx=XM_019822590.2;refGene_gn=DNAH1;refGene_tx=XM_019822591.2;refGene_gn=DNAH1;refGene_tx=XM_023249453.1
A2 20969526 20969704
refGene_gn=DNAH1;refGene_tx=XM_011279464.3;refGene_gn=DNAH1;refGene_tx=XM_019822581.2;refGene_gn=DNAH1;refGene_tx=XM_019822582.2;refGene_gn=DNAH1;refGene_tx=XM_019822587.2;refGene_gn=DNAH1;refGene_tx=XM_019822590.2;refGene_gn=DNAH1;refGene_tx=XM_019822591.2;refGene_gn=DNAH1;refGene_tx=XM_023249453.1
A2 20970507 20970702
refGene_gn=DNAH1;refGene_tx=XM_011279464.3;refGene_gn=DNAH1;refGene_tx=XM_019822581.2;refGene_gn=DNAH1;refGene_tx=XM_019822582.2;refGene_gn=DNAH1;refGene_tx=XM_019822587.2;refGene_gn=DNAH1;refGene_tx=XM_019822590.2;refGene_gn=DNAH1;refGene_tx=XM_019822591.2;refGene_gn=DNAH1;refGene_tx=XM_023249453.1
A2 20970791 20970911
refGene_gn=DNAH1;refGene_tx=XM_011279464.3;refGene_gn=DNAH1;refGene_tx=XM_019822581.2;refGene_gn=DNAH1;refGene_tx=XM_019822582.2;refGene_gn=DNAH1;refGene_tx=XM_019822587.2;refGene_gn=DNAH1;refGene_tx=XM_019822590.2;refGene_gn=DNAH1;refGene_tx=XM_019822591.2;refGene_gn=DNAH1;refGene_tx=XM_023249453.1
A2 20971178 20971337
refGene_gn=DNAH1;refGene_tx=XM_011279464.3;refGene_gn=DNAH1;refGene_tx=XM_019822581.2;refGene_gn=DNAH1;refGene_tx=XM_019822582.2;refGene_gn=DNAH1;refGene_tx=XM_019822587.2;refGene_gn=DNAH1;refGene_tx=XM_019822590.2;refGene_gn=DNAH1;refGene_tx=XM_019822591.2;refGene_gn=DNAH1;refGene_tx=XM_023249453.1
A2 20971515 20971623
refGene_gn=DNAH1;refGene_tx=XM_011279464.3;refGene_gn=DNAH1;refGene_tx=XM_019822581.2;refGene_gn=DNAH1;refGene_tx=XM_019822582.2;refGene_gn=DNAH1;refGene_tx=XM_019822587.2;refGene_gn=DNAH1;refGene_tx=XM_019822590.2;refGene_gn=DNAH1;refGene_tx=XM_019822591.2;refGene_gn=DNAH1;refGene_tx=XM_023249453.1
A2 20972692 20972833
refGene_gn=DNAH1;refGene_tx=XM_011279464.3;refGene_gn=DNAH1;refGene_tx=XM_019822581.2;refGene_gn=DNAH1;refGene_tx=XM_019822582.2;refGene_gn=DNAH1;refGene_tx=XM_019822587.2;refGene_gn=DNAH1;refGene_tx=XM_019822590.2;refGene_gn=DNAH1;refGene_tx=XM_019822591.2;refGene_gn=DNAH1;refGene_tx=XM_023249453.1
A2 20972976 20973137
refGene_gn=DNAH1;refGene_tx=XM_011279464.3;refGene_gn=DNAH1;refGene_tx=XM_019822581.2;refGene_gn=DNAH1;refGene_tx=XM_019822582.2;refGene_gn=DNAH1;refGene_tx=XM_019822587.2;refGene_gn=DNAH1;refGene_tx=XM_019822590.2;refGene_gn=DNAH1;refGene_tx=XM_019822591.2;refGene_gn=DNAH1;refGene_tx=XM_023249453.1

A2 20973571 20973729
refGene_gn=DNAH1;refGene_tx=XM_011279464.3;refGene_gn=DNAH1;refGene_tx
=XM_019822581.2;refGene_gn=DNAH1;refGene_tx=XM_019822582.2;refGene_gn=
DNAH1;refGene_tx=XM_019822587.2;refGene_gn=DNAH1;refGene_tx=XM_0198225
90.2;refGene_gn=DNAH1;refGene_tx=XM_019822591.2;refGene_gn=DNAH1;refGe
ne_tx=XM_023249453.1
A2 20974287 20974461
refGene_gn=DNAH1;refGene_tx=XM_011279464.3;refGene_gn=DNAH1;refGene_tx
=XM_019822581.2;refGene_gn=DNAH1;refGene_tx=XM_019822582.2;refGene_gn=
DNAH1;refGene_tx=XM_019822587.2;refGene_gn=DNAH1;refGene_tx=XM_0198225
90.2;refGene_gn=DNAH1;refGene_tx=XM_019822591.2;refGene_gn=DNAH1;refGe
ne_tx=XM_023249453.1
A2 20975265 20975304
refGene_gn=DNAH1;refGene_tx=XM_011279464.3;refGene_gn=DNAH1;refGene_tx
=XM_019822581.2;refGene_gn=DNAH1;refGene_tx=XM_019822582.2;refGene_gn=
DNAH1;refGene_tx=XM_019822587.2;refGene_gn=DNAH1;refGene_tx=XM_0198225
90.2;refGene_gn=DNAH1;refGene_tx=XM_019822591.2;refGene_gn=DNAH1;refGe
ne_tx=XM_023249453.1
A2 20977743 20977922
refGene_gn=DNAH1;refGene_tx=XM_011279464.3;refGene_gn=DNAH1;refGene_tx
=XM_019822581.2;refGene_gn=DNAH1;refGene_tx=XM_019822582.2;refGene_gn=
DNAH1;refGene_tx=XM_019822587.2;refGene_gn=DNAH1;refGene_tx=XM_0198225
90.2;refGene_gn=DNAH1;refGene_tx=XM_019822591.2;refGene_gn=DNAH1;refGe
ne_tx=XM_023249453.1
A2 20978886 20979117
refGene_gn=DNAH1;refGene_tx=XM_011279464.3;refGene_gn=DNAH1;refGene_tx
=XM_019822581.2;refGene_gn=DNAH1;refGene_tx=XM_019822582.2;refGene_gn=
DNAH1;refGene_tx=XM_019822587.2;refGene_gn=DNAH1;refGene_tx=XM_0198225
90.2;refGene_gn=DNAH1;refGene_tx=XM_019822591.2;refGene_gn=DNAH1;refGe
ne_tx=XM_023249453.1
A2 20979788 20979985
refGene_gn=DNAH1;refGene_tx=XM_011279464.3;refGene_gn=DNAH1;refGene_tx
=XM_019822581.2;refGene_gn=DNAH1;refGene_tx=XM_019822582.2;refGene_gn=
DNAH1;refGene_tx=XM_019822587.2;refGene_gn=DNAH1;refGene_tx=XM_0198225
90.2;refGene_gn=DNAH1;refGene_tx=XM_019822591.2;refGene_gn=DNAH1;refGe
ne_tx=XM_023249453.1
A2 20980462 20980598
refGene_gn=DNAH1;refGene_tx=XM_011279464.3;refGene_gn=DNAH1;refGene_tx
=XM_019822581.2;refGene_gn=DNAH1;refGene_tx=XM_019822582.2;refGene_gn=
DNAH1;refGene_tx=XM_019822587.2;refGene_gn=DNAH1;refGene_tx=XM_0198225
90.2;refGene_gn=DNAH1;refGene_tx=XM_019822591.2;refGene_gn=DNAH1;refGe
ne_tx=XM_023249453.1
A2 20981512 20981721
refGene_gn=DNAH1;refGene_tx=XM_011279464.3;refGene_gn=DNAH1;refGene_tx
=XM_019822581.2;refGene_gn=DNAH1;refGene_tx=XM_019822582.2;refGene_gn=
DNAH1;refGene_tx=XM_019822587.2;refGene_gn=DNAH1;refGene_tx=XM_0198225
90.2;refGene_gn=DNAH1;refGene_tx=XM_019822591.2;refGene_gn=DNAH1;refGe
ne_tx=XM_023249453.1
A2 20981811 20981983
refGene_gn=DNAH1;refGene_tx=XM_011279464.3;refGene_gn=DNAH1;refGene_tx

=XM_019822581.2;refGene_gn=DNAH1;refGene_tx=XM_019822582.2;refGene_gn=DNAH1;refGene_tx=XM_019822587.2;refGene_gn=DNAH1;refGene_tx=XM_019822590.2;refGene_gn=DNAH1;refGene_tx=XM_019822591.2;refGene_gn=DNAH1;refGene_tx=XM_023249453.1
A2 20982722 20982914
refGene_gn=DNAH1;refGene_tx=XM_011279464.3;refGene_gn=DNAH1;refGene_tx=XM_019822581.2;refGene_gn=DNAH1;refGene_tx=XM_019822582.2;refGene_gn=DNAH1;refGene_tx=XM_019822590.2;refGene_gn=DNAH1;refGene_tx=XM_019822591.2;refGene_gn=DNAH1;refGene_tx=XM_023249453.1
A2 20983451 20983562
refGene_gn=DNAH1;refGene_tx=XM_011279464.3;refGene_gn=DNAH1;refGene_tx=XM_019822581.2;refGene_gn=DNAH1;refGene_tx=XM_019822582.2;refGene_gn=DNAH1;refGene_tx=XM_019822587.2;refGene_gn=DNAH1;refGene_tx=XM_019822590.2;refGene_gn=DNAH1;refGene_tx=XM_019822591.2;refGene_gn=DNAH1;refGene_tx=XM_023249453.1
A2 20984356 20984542
refGene_gn=DNAH1;refGene_tx=XM_011279464.3;refGene_gn=DNAH1;refGene_tx=XM_019822581.2;refGene_gn=DNAH1;refGene_tx=XM_019822582.2;refGene_gn=DNAH1;refGene_tx=XM_019822587.2;refGene_gn=DNAH1;refGene_tx=XM_019822590.2;refGene_gn=DNAH1;refGene_tx=XM_019822591.2;refGene_gn=DNAH1;refGene_tx=XM_023249453.1
A2 20984958 20985150
refGene_gn=DNAH1;refGene_tx=XM_011279464.3;refGene_gn=DNAH1;refGene_tx=XM_019822581.2;refGene_gn=DNAH1;refGene_tx=XM_019822582.2;refGene_gn=DNAH1;refGene_tx=XM_019822587.2;refGene_gn=DNAH1;refGene_tx=XM_019822590.2;refGene_gn=DNAH1;refGene_tx=XM_019822591.2;refGene_gn=DNAH1;refGene_tx=XM_023249453.1
A2 20986334 20986502
refGene_gn=DNAH1;refGene_tx=XM_011279464.3;refGene_gn=DNAH1;refGene_tx=XM_019822581.2;refGene_gn=DNAH1;refGene_tx=XM_019822582.2;refGene_gn=DNAH1;refGene_tx=XM_019822587.2;refGene_gn=DNAH1;refGene_tx=XM_019822590.2;refGene_gn=DNAH1;refGene_tx=XM_019822591.2;refGene_gn=DNAH1;refGene_tx=XM_023249453.1
A2 20986594 20986786
refGene_gn=DNAH1;refGene_tx=XM_011279464.3;refGene_gn=DNAH1;refGene_tx=XM_019822581.2;refGene_gn=DNAH1;refGene_tx=XM_019822582.2;refGene_gn=DNAH1;refGene_tx=XM_019822587.2;refGene_gn=DNAH1;refGene_tx=XM_019822590.2;refGene_gn=DNAH1;refGene_tx=XM_019822591.2;refGene_gn=DNAH1;refGene_tx=XM_023249453.1
A2 20986911 20987043
refGene_gn=DNAH1;refGene_tx=XM_011279464.3;refGene_gn=DNAH1;refGene_tx=XM_019822581.2;refGene_gn=DNAH1;refGene_tx=XM_019822582.2;refGene_gn=DNAH1;refGene_tx=XM_019822587.2;refGene_gn=DNAH1;refGene_tx=XM_019822590.2;refGene_gn=DNAH1;refGene_tx=XM_019822591.2;refGene_gn=DNAH1;refGene_tx=XM_023249453.1
A2 20987557 20987683
refGene_gn=DNAH1;refGene_tx=XM_011279464.3;refGene_gn=DNAH1;refGene_tx=XM_019822581.2;refGene_gn=DNAH1;refGene_tx=XM_019822582.2;refGene_gn=DNAH1;refGene_tx=XM_019822587.2;refGene_gn=DNAH1;refGene_tx=XM_019822590.2;refGene_gn=DNAH1;refGene_tx=XM_019822591.2;refGene_gn=DNAH1;refGe

ne_tx=XM_023249453.1
A2 20988317 20988437
refGene_gn=DNAH1;refGene_tx=XM_011279464.3;refGene_gn=DNAH1;refGene_tx=XM_019822581.2;refGene_gn=DNAH1;refGene_tx=XM_019822582.2;refGene_gn=DNAH1;refGene_tx=XM_019822587.2;refGene_gn=DNAH1;refGene_tx=XM_019822590.2;refGene_gn=DNAH1;refGene_tx=XM_019822591.2;refGene_gn=DNAH1;refGene_tx=XM_023249453.1
A2 20988574 20988724
refGene_gn=DNAH1;refGene_tx=XM_011279464.3;refGene_gn=DNAH1;refGene_tx=XM_019822581.2;refGene_gn=DNAH1;refGene_tx=XM_019822582.2;refGene_gn=DNAH1;refGene_tx=XM_019822587.2;refGene_gn=DNAH1;refGene_tx=XM_019822590.2;refGene_gn=DNAH1;refGene_tx=XM_019822591.2;refGene_gn=DNAH1;refGene_tx=XM_023249453.1
A2 20988851 20989012
refGene_gn=DNAH1;refGene_tx=XM_011279464.3;refGene_gn=DNAH1;refGene_tx=XM_019822581.2;refGene_gn=DNAH1;refGene_tx=XM_019822582.2;refGene_gn=DNAH1;refGene_tx=XM_019822587.2;refGene_gn=DNAH1;refGene_tx=XM_019822590.2;refGene_gn=DNAH1;refGene_tx=XM_019822591.2;refGene_gn=DNAH1;refGene_tx=XM_023249453.1
A2 20989824 20990050
refGene_gn=DNAH1;refGene_tx=XM_011279464.3;refGene_gn=DNAH1;refGene_tx=XM_019822581.2;refGene_gn=DNAH1;refGene_tx=XM_019822582.2;refGene_gn=DNAH1;refGene_tx=XM_019822587.2;refGene_gn=DNAH1;refGene_tx=XM_019822590.2;refGene_gn=DNAH1;refGene_tx=XM_019822591.2;refGene_gn=DNAH1;refGene_tx=XM_023249453.1
A2 20990185 20990381
refGene_gn=DNAH1;refGene_tx=XM_011279464.3;refGene_gn=DNAH1;refGene_tx=XM_019822581.2;refGene_gn=DNAH1;refGene_tx=XM_019822582.2;refGene_gn=DNAH1;refGene_tx=XM_019822587.2;refGene_gn=DNAH1;refGene_tx=XM_019822590.2;refGene_gn=DNAH1;refGene_tx=XM_019822591.2;refGene_gn=DNAH1;refGene_tx=XM_023249453.1
A2 20990714 20990866
refGene_gn=DNAH1;refGene_tx=XM_011279464.3;refGene_gn=DNAH1;refGene_tx=XM_019822581.2;refGene_gn=DNAH1;refGene_tx=XM_019822582.2;refGene_gn=DNAH1;refGene_tx=XM_019822587.2;refGene_gn=DNAH1;refGene_tx=XM_019822590.2;refGene_gn=DNAH1;refGene_tx=XM_019822591.2;refGene_gn=DNAH1;refGene_tx=XM_023249453.1
A2 20991535 20991732
refGene_gn=DNAH1;refGene_tx=XM_011279464.3;refGene_gn=DNAH1;refGene_tx=XM_019822581.2;refGene_gn=DNAH1;refGene_tx=XM_019822582.2;refGene_gn=DNAH1;refGene_tx=XM_019822587.2;refGene_gn=DNAH1;refGene_tx=XM_019822590.2;refGene_gn=DNAH1;refGene_tx=XM_019822591.2;refGene_gn=DNAH1;refGene_tx=XM_023249453.1
A2 20991979 20992124
refGene_gn=DNAH1;refGene_tx=XM_011279464.3;refGene_gn=DNAH1;refGene_tx=XM_019822581.2;refGene_gn=DNAH1;refGene_tx=XM_019822582.2;refGene_gn=DNAH1;refGene_tx=XM_019822587.2;refGene_gn=DNAH1;refGene_tx=XM_019822590.2;refGene_gn=DNAH1;refGene_tx=XM_019822591.2;refGene_gn=DNAH1;refGene_tx=XM_023249453.1
A2 20992380 20992539

refGene_gn=DNAH1;refGene_tx=XM_011279464.3;refGene_gn=DNAH1;refGene_tx
=XM_019822581.2;refGene_gn=DNAH1;refGene_tx=XM_019822582.2;refGene_gn=
DNAH1;refGene_tx=XM_019822587.2;refGene_gn=DNAH1;refGene_tx=XM_0198225
90.2;refGene_gn=DNAH1;refGene_tx=XM_019822591.2;refGene_gn=DNAH1;refGe
ne_tx=XM_023249453.1
A2 20992619 20992751
refGene_gn=DNAH1;refGene_tx=XM_011279464.3;refGene_gn=DNAH1;refGene_tx
=XM_019822581.2;refGene_gn=DNAH1;refGene_tx=XM_019822587.2;refGene_gn=
DNAH1;refGene_tx=XM_023249453.1
A2 20993134 20993305
refGene_gn=DNAH1;refGene_tx=XM_011279464.3;refGene_gn=DNAH1;refGene_tx
=XM_019822581.2;refGene_gn=DNAH1;refGene_tx=XM_019822582.2;refGene_gn=
DNAH1;refGene_tx=XM_019822587.2;refGene_gn=DNAH1;refGene_tx=XM_0232494
53.1
A2 20993390 20993570
refGene_gn=DNAH1;refGene_tx=XM_011279464.3;refGene_gn=DNAH1;refGene_tx
=XM_019822581.2;refGene_gn=DNAH1;refGene_tx=XM_019822582.2;refGene_gn=
DNAH1;refGene_tx=XM_019822587.2;refGene_gn=DNAH1;refGene_tx=XM_0232494
53.1
A2 20993646 20993823
refGene_gn=DNAH1;refGene_tx=XM_011279464.3;refGene_gn=DNAH1;refGene_tx
=XM_019822581.2;refGene_gn=DNAH1;refGene_tx=XM_019822582.2;refGene_gn=
DNAH1;refGene_tx=XM_019822587.2;refGene_gn=DNAH1;refGene_tx=XM_0232494
53.1
A2 20994402 20994573
refGene_gn=DNAH1;refGene_tx=XM_011279464.3;refGene_gn=DNAH1;refGene_tx
=XM_019822581.2;refGene_gn=DNAH1;refGene_tx=XM_019822582.2;refGene_gn=
DNAH1;refGene_tx=XM_019822587.2;refGene_gn=DNAH1;refGene_tx=XM_0232494
53.1
A2 20994730 20994861
refGene_gn=DNAH1;refGene_tx=XM_011279464.3;refGene_gn=DNAH1;refGene_tx
=XM_019822581.2;refGene_gn=DNAH1;refGene_tx=XM_019822582.2;refGene_gn=
DNAH1;refGene_tx=XM_019822587.2;refGene_gn=DNAH1;refGene_tx=XM_0232494
53.1
A2 20995212 20995564
refGene_gn=DNAH1;refGene_tx=XM_011279464.3;refGene_gn=DNAH1;refGene_tx
=XM_019822581.2;refGene_gn=DNAH1;refGene_tx=XM_019822582.2;refGene_gn=
DNAH1;refGene_tx=XM_019822587.2;refGene_gn=DNAH1;refGene_tx=XM_0232494
53.1
A2 20995855 20996090
refGene_gn=DNAH1;refGene_tx=XM_011279464.3;refGene_gn=DNAH1;refGene_tx
=XM_019822581.2;refGene_gn=DNAH1;refGene_tx=XM_019822582.2;refGene_gn=
DNAH1;refGene_tx=XM_019822587.2;refGene_gn=DNAH1;refGene_tx=XM_0232494
53.1
A2 20996603 20996725
refGene_gn=DNAH1;refGene_tx=XM_011279464.3;refGene_gn=DNAH1;refGene_tx
=XM_019822581.2;refGene_gn=DNAH1;refGene_tx=XM_019822582.2;refGene_gn=
DNAH1;refGene_tx=XM_019822587.2;refGene_gn=DNAH1;refGene_tx=XM_0232494
53.1
A2 20998520 20998654

refGene_gn=BAP1;refGene_tx=XM_023249464.1;refGene_gn=BAP1;refGene_tx=XM_023249465.1
A2 20998840 20998913
refGene_gn=BAP1;refGene_tx=XM_023249464.1;refGene_gn=BAP1;refGene_tx=XM_023249465.1
A2 20999036 20999129
refGene_gn=BAP1;refGene_tx=XM_023249464.1;refGene_gn=BAP1;refGene_tx=XM_023249465.1
A2 20999395 20999556
refGene_gn=BAP1;refGene_tx=XM_023249464.1;refGene_gn=BAP1;refGene_tx=XM_023249465.1
A2 20999663 21000142
refGene_gn=BAP1;refGene_tx=XM_023249464.1;refGene_gn=BAP1;refGene_tx=XM_023249465.1
A2 21000661 21000795
refGene_gn=BAP1;refGene_tx=XM_023249464.1;refGene_gn=BAP1;refGene_tx=XM_023249465.1
A2 21001274 21001459
refGene_gn=BAP1;refGene_tx=XM_023249464.1;refGene_gn=BAP1;refGene_tx=XM_023249465.1
A2 21001961 21002109
refGene_gn=BAP1;refGene_tx=XM_023249464.1;refGene_gn=BAP1;refGene_tx=XM_023249465.1
A2 21002389 21002513
refGene_gn=BAP1;refGene_tx=XM_023249464.1;refGene_gn=BAP1;refGene_tx=XM_023249465.1
A2 21002971 21003050
refGene_gn=BAP1;refGene_tx=XM_023249464.1;refGene_gn=BAP1;refGene_tx=XM_023249465.1
A2 21003306 21003449
refGene_gn=BAP1;refGene_tx=XM_023249464.1;refGene_gn=BAP1;refGene_tx=XM_023249465.1
A2 21003532 21003594
refGene_gn=BAP1;refGene_tx=XM_023249464.1;refGene_gn=BAP1;refGene_tx=XM_023249465.1
A2 21004039 21004159
refGene_gn=BAP1;refGene_tx=XM_023249464.1;refGene_gn=BAP1;refGene_tx=XM_023249465.1
A2 21004535 21004668 refGene_gn=BAP1;refGene_tx=XM_023249464.1
A2 21005647 21005702
refGene_gn=BAP1;refGene_tx=XM_023249464.1;refGene_gn=BAP1;refGene_tx=XM_023249465.1
A2 21005826 21005856 refGene_gn=BAP1;refGene_tx=XM_023249464.1
A2 21005955 21005992 refGene_gn=BAP1;refGene_tx=XM_023249464.1
A2 21008377 21008421
refGene_gn=PHF7;refGene_tx=XM_003982301.5;refGene_gn=PHF7;refGene_tx=XM_006928848.4;refGene_gn=PHF7;refGene_tx=XM_011279468.3;refGene_gn=PHF7;refGene_tx=XM_019822612.2;refGene_gn=PHF7;refGene_tx=XM_019822615.2;refGene_gn=PHF7;refGene_tx=XM_019822621.2
A2 21009137 21009190

refGene_gn=PHF7;refGene_tx=XM_003982301.5;refGene_gn=PHF7;refGene_tx=XM_006928848.4;refGene_gn=PHF7;refGene_tx=XM_011279468.3;refGene_gn=PHF7;refGene_tx=XM_019822612.2;refGene_gn=PHF7;refGene_tx=XM_019822615.2;refGene_gn=PHF7;refGene_tx=XM_019822621.2
A2 21009564 21009656
refGene_gn=PHF7;refGene_tx=XM_003982301.5;refGene_gn=PHF7;refGene_tx=XM_006928848.4;refGene_gn=PHF7;refGene_tx=XM_011279468.3;refGene_gn=PHF7;refGene_tx=XM_019822612.2;refGene_gn=PHF7;refGene_tx=XM_019822615.2;refGene_gn=PHF7;refGene_tx=XM_019822621.2
A2 21015348 21015450
refGene_gn=PHF7;refGene_tx=XM_003982301.5;refGene_gn=PHF7;refGene_tx=XM_006928848.4;refGene_gn=PHF7;refGene_tx=XM_011279468.3;refGene_gn=PHF7;refGene_tx=XM_019822612.2;refGene_gn=PHF7;refGene_tx=XM_019822615.2;refGene_gn=PHF7;refGene_tx=XM_019822621.2
A2 21015835 21015960
refGene_gn=PHF7;refGene_tx=XM_003982301.5;refGene_gn=PHF7;refGene_tx=XM_006928848.4;refGene_gn=PHF7;refGene_tx=XM_011279468.3;refGene_gn=PHF7;refGene_tx=XM_019822612.2;refGene_gn=PHF7;refGene_tx=XM_019822615.2;refGene_gn=PHF7;refGene_tx=XM_019822621.2
A2 21016421 21016581
refGene_gn=PHF7;refGene_tx=XM_003982301.5;refGene_gn=PHF7;refGene_tx=XM_006928848.4;refGene_gn=PHF7;refGene_tx=XM_011279468.3;refGene_gn=PHF7;refGene_tx=XM_019822612.2;refGene_gn=PHF7;refGene_tx=XM_019822615.2;refGene_gn=PHF7;refGene_tx=XM_019822621.2
A2 21017144 21017251
refGene_gn=PHF7;refGene_tx=XM_003982301.5;refGene_gn=PHF7;refGene_tx=XM_006928848.4;refGene_gn=PHF7;refGene_tx=XM_011279468.3;refGene_gn=PHF7;refGene_tx=XM_019822612.2;refGene_gn=PHF7;refGene_tx=XM_019822615.2;refGene_gn=PHF7;refGene_tx=XM_019822621.2
A2 21017734 21017851
refGene_gn=PHF7;refGene_tx=XM_003982301.5;refGene_gn=PHF7;refGene_tx=XM_006928848.4;refGene_gn=PHF7;refGene_tx=XM_011279468.3;refGene_gn=PHF7;refGene_tx=XM_019822612.2;refGene_gn=PHF7;refGene_tx=XM_019822615.2
A2 21018268 21018393
refGene_gn=PHF7;refGene_tx=XM_003982301.5;refGene_gn=PHF7;refGene_tx=XM_006928848.4;refGene_gn=PHF7;refGene_tx=XM_011279468.3;refGene_gn=PHF7;refGene_tx=XM_019822612.2;refGene_gn=PHF7;refGene_tx=XM_019822615.2;refGene_gn=PHF7;refGene_tx=XM_019822621.2
A2 21018602 21018829
refGene_gn=PHF7;refGene_tx=XM_003982301.5;refGene_gn=PHF7;refGene_tx=XM_011279468.3;refGene_gn=PHF7;refGene_tx=XM_019822612.2;refGene_gn=PHF7;refGene_tx=XM_019822615.2;refGene_gn=PHF7;refGene_tx=XM_019822621.2
A2 21021793 21021858 refGene_gn=PHF7;refGene_tx=XM_006928848.4
A2 21029953 21030394
refGene_gn=SEMA3G;refGene_tx=XM_003982303.4;refGene_gn=SEMA3G;refGene_tx=XM_006928849.4;refGene_gn=SEMA3G;refGene_tx=XM_019822598.2
A2 21031766 21031906
refGene_gn=SEMA3G;refGene_tx=XM_003982303.4;refGene_gn=SEMA3G;refGene_tx=XM_006928849.4;refGene_gn=SEMA3G;refGene_tx=XM_019822598.2
A2 21032171 21032400

refGene_gn=SEMA3G;refGene_tx=XM_003982303.4;refGene_gn=SEMA3G;refGene_tx=XM_006928849.4;refGene_gn=SEMA3G;refGene_tx=XM_019822598.2
A2 21033413 21033455
refGene_gn=SEMA3G;refGene_tx=XM_003982303.4;refGene_gn=SEMA3G;refGene_tx=XM_006928849.4;refGene_gn=SEMA3G;refGene_tx=XM_019822598.2
A2 21034125 21034217
refGene_gn=SEMA3G;refGene_tx=XM_003982303.4;refGene_gn=SEMA3G;refGene_tx=XM_006928849.4;refGene_gn=SEMA3G;refGene_tx=XM_019822598.2
A2 21034324 21034556
refGene_gn=SEMA3G;refGene_tx=XM_003982303.4;refGene_gn=SEMA3G;refGene_tx=XM_006928849.4;refGene_gn=SEMA3G;refGene_tx=XM_019822598.2
A2 21034831 21034976
refGene_gn=SEMA3G;refGene_tx=XM_003982303.4;refGene_gn=SEMA3G;refGene_tx=XM_006928849.4;refGene_gn=SEMA3G;refGene_tx=XM_019822598.2
A2 21035280 21035350
refGene_gn=SEMA3G;refGene_tx=XM_003982303.4;refGene_gn=SEMA3G;refGene_tx=XM_006928849.4;refGene_gn=SEMA3G;refGene_tx=XM_019822598.2
A2 21035443 21035558
refGene_gn=SEMA3G;refGene_tx=XM_003982303.4;refGene_gn=SEMA3G;refGene_tx=XM_006928849.4;refGene_gn=SEMA3G;refGene_tx=XM_019822598.2
A2 21035758 21035904
refGene_gn=SEMA3G;refGene_tx=XM_003982303.4;refGene_gn=SEMA3G;refGene_tx=XM_006928849.4;refGene_gn=SEMA3G;refGene_tx=XM_019822598.2
A2 21036137 21036254
refGene_gn=SEMA3G;refGene_tx=XM_003982303.4;refGene_gn=SEMA3G;refGene_tx=XM_006928849.4;refGene_gn=SEMA3G;refGene_tx=XM_019822598.2
A2 21036379 21036470
refGene_gn=SEMA3G;refGene_tx=XM_003982303.4;refGene_gn=SEMA3G;refGene_tx=XM_019822598.2
A2 21036744 21036864
refGene_gn=SEMA3G;refGene_tx=XM_003982303.4;refGene_gn=SEMA3G;refGene_tx=XM_006928849.4
A2 21037125 21037188
refGene_gn=SEMA3G;refGene_tx=XM_003982303.4;refGene_gn=SEMA3G;refGene_tx=XM_006928849.4;refGene_gn=SEMA3G;refGene_tx=XM_019822598.2
A2 21037288 21037449
refGene_gn=SEMA3G;refGene_tx=XM_003982303.4;refGene_gn=SEMA3G;refGene_tx=XM_006928849.4;refGene_gn=SEMA3G;refGene_tx=XM_019822598.2
A2 21039246 21039355
refGene_gn=SEMA3G;refGene_tx=XM_003982303.4;refGene_gn=SEMA3G;refGene_tx=XM_019822598.2
A2 21044755 21044787 refGene_gn=TNNC1;refGene_tx=XM_003982304.4
A2 21044872 21045009 refGene_gn=TNNC1;refGene_tx=XM_003982304.4
A2 21045252 21045367 refGene_gn=TNNC1;refGene_tx=XM_003982304.4
A2 21045643 21045790 refGene_gn=TNNC1;refGene_tx=XM_003982304.4
A2 21045995 21046026 refGene_gn=TNNC1;refGene_tx=XM_003982304.4
A2 21047548 21047572 refGene_gn=TNNC1;refGene_tx=XM_003982304.4
A2 21049373 21049469
refGene_gn=NISCH;refGene_tx=XM_003982305.5;refGene_gn=NISCH;refGene_tx=XM_006928851.3;refGene_gn=NISCH;refGene_tx=XM_023249492.1

A2	21051617	21051701
refGene_gn=NISCH;refGene_tx=XM_003982305.5;refGene_gn=NISCH;refGene_tx
=XM_006928851.3;refGene_gn=NISCH;refGene_tx=XM_023249492.1
A2	21052293	21052476
refGene_gn=NISCH;refGene_tx=XM_003982305.5;refGene_gn=NISCH;refGene_tx
=XM_006928851.3;refGene_gn=NISCH;refGene_tx=XM_023249492.1
A2	21061667	21061716
refGene_gn=NISCH;refGene_tx=XM_003982305.5;refGene_gn=NISCH;refGene_tx
=XM_006928851.3;refGene_gn=NISCH;refGene_tx=XM_023249492.1
A2	21062691	21062855
refGene_gn=NISCH;refGene_tx=XM_003982305.5;refGene_gn=NISCH;refGene_tx
=XM_006928851.3;refGene_gn=NISCH;refGene_tx=XM_023249492.1
A2	21063220	21063316
refGene_gn=NISCH;refGene_tx=XM_003982305.5;refGene_gn=NISCH;refGene_tx
=XM_006928851.3;refGene_gn=NISCH;refGene_tx=XM_023249492.1
A2	21064672	21064768
refGene_gn=NISCH;refGene_tx=XM_003982305.5;refGene_gn=NISCH;refGene_tx
=XM_006928851.3;refGene_gn=NISCH;refGene_tx=XM_023249492.1;refGene_gn=
NISCH;refGene_tx=XM_023249494.1
A2	21066193	21066346
refGene_gn=NISCH;refGene_tx=XM_003982305.5;refGene_gn=NISCH;refGene_tx
=XM_006928851.3;refGene_gn=NISCH;refGene_tx=XM_023249492.1;refGene_gn=
NISCH;refGene_tx=XM_023249494.1
A2	21066994	21067063
refGene_gn=NISCH;refGene_tx=XM_003982305.5;refGene_gn=NISCH;refGene_tx
=XM_006928851.3;refGene_gn=NISCH;refGene_tx=XM_023249492.1;refGene_gn=
NISCH;refGene_tx=XM_023249494.1
A2	21067566	21067752
refGene_gn=NISCH;refGene_tx=XM_003982305.5;refGene_gn=NISCH;refGene_tx
=XM_006928851.3;refGene_gn=NISCH;refGene_tx=XM_023249492.1;refGene_gn=
NISCH;refGene_tx=XM_023249494.1
A2	21068161	21068290
refGene_gn=NISCH;refGene_tx=XM_003982305.5;refGene_gn=NISCH;refGene_tx
=XM_006928851.3;refGene_gn=NISCH;refGene_tx=XM_023249492.1;refGene_gn=
NISCH;refGene_tx=XM_023249494.1
A2	21069405	21069519
refGene_gn=NISCH;refGene_tx=XM_003982305.5;refGene_gn=NISCH;refGene_tx
=XM_006928851.3;refGene_gn=NISCH;refGene_tx=XM_023249492.1;refGene_gn=
NISCH;refGene_tx=XM_023249494.1
A2	21069873	21069985
refGene_gn=NISCH;refGene_tx=XM_003982305.5;refGene_gn=NISCH;refGene_tx
=XM_006928851.3;refGene_gn=NISCH;refGene_tx=XM_023249492.1;refGene_gn=
NISCH;refGene_tx=XM_023249494.1
A2	21070576	21070596	refGene_gn=NISCH;refGene_tx=XM_006928851.3
A2	21074191	21074316
refGene_gn=NISCH;refGene_tx=XM_003982305.5;refGene_gn=NISCH;refGene_tx
=XM_023249492.1;refGene_gn=NISCH;refGene_tx=XM_023249494.1
A2	21075434	21075484
refGene_gn=NISCH;refGene_tx=XM_003982305.5;refGene_gn=NISCH;refGene_tx
=XM_023249492.1;refGene_gn=NISCH;refGene_tx=XM_023249494.1

A2	21076358	21077756
refGene_gn=NISCH;refGene_tx=XM_003982305.5;refGene_gn=NISCH;refGene_tx=XM_023249492.1;refGene_gn=NISCH;refGene_tx=XM_023249494.1
A2	21079489	21079907
refGene_gn=NISCH;refGene_tx=XM_003982305.5;refGene_gn=NISCH;refGene_tx=XM_023249492.1;refGene_gn=NISCH;refGene_tx=XM_023249494.1
A2	21080425	21080582
refGene_gn=NISCH;refGene_tx=XM_003982305.5;refGene_gn=NISCH;refGene_tx=XM_023249492.1;refGene_gn=NISCH;refGene_tx=XM_023249494.1
A2	21081095	21081224
refGene_gn=NISCH;refGene_tx=XM_003982305.5;refGene_gn=NISCH;refGene_tx=XM_023249492.1;refGene_gn=NISCH;refGene_tx=XM_023249494.1
A2	21081703	21081865
refGene_gn=NISCH;refGene_tx=XM_003982305.5;refGene_gn=NISCH;refGene_tx=XM_023249494.1
A2	21082281	21082892
refGene_gn=NISCH;refGene_tx=XM_003982305.5;refGene_gn=NISCH;refGene_tx=XM_023249492.1;refGene_gn=NISCH;refGene_tx=XM_023249494.1
A2	21085712	21085793
refGene_gn=STAB1;refGene_tx=XM_011279480.2;refGene_gn=STAB1;refGene_tx=XM_011279482.2;refGene_gn=STAB1;refGene_tx=XM_011279484.2;refGene_gn=STAB1;refGene_tx=XM_011279487.2;refGene_gn=STAB1;refGene_tx=XM_019822624.1;refGene_gn=STAB1;refGene_tx=XM_019822625.1;refGene_gn=STAB1;refGene_tx=XM_019822633.2
A2	21089739	21089876
refGene_gn=STAB1;refGene_tx=XM_011279480.2;refGene_gn=STAB1;refGene_tx=XM_011279482.2;refGene_gn=STAB1;refGene_tx=XM_011279484.2;refGene_gn=STAB1;refGene_tx=XM_011279487.2;refGene_gn=STAB1;refGene_tx=XM_019822624.1;refGene_gn=STAB1;refGene_tx=XM_019822625.1;refGene_gn=STAB1;refGene_tx=XM_019822633.2
A2	21090221	21090337
refGene_gn=STAB1;refGene_tx=XM_011279480.2;refGene_gn=STAB1;refGene_tx=XM_011279482.2;refGene_gn=STAB1;refGene_tx=XM_011279484.2;refGene_gn=STAB1;refGene_tx=XM_011279487.2;refGene_gn=STAB1;refGene_tx=XM_019822624.1;refGene_gn=STAB1;refGene_tx=XM_019822625.1;refGene_gn=STAB1;refGene_tx=XM_019822633.2
A2	21090573	21090659
refGene_gn=STAB1;refGene_tx=XM_011279480.2;refGene_gn=STAB1;refGene_tx=XM_011279482.2;refGene_gn=STAB1;refGene_tx=XM_011279484.2;refGene_gn=STAB1;refGene_tx=XM_011279487.2;refGene_gn=STAB1;refGene_tx=XM_019822624.1;refGene_gn=STAB1;refGene_tx=XM_019822625.1;refGene_gn=STAB1;refGene_tx=XM_019822633.2
A2	21090728	21090798
refGene_gn=STAB1;refGene_tx=XM_011279480.2;refGene_gn=STAB1;refGene_tx=XM_011279482.2;refGene_gn=STAB1;refGene_tx=XM_011279484.2;refGene_gn=STAB1;refGene_tx=XM_011279487.2;refGene_gn=STAB1;refGene_tx=XM_019822624.1;refGene_gn=STAB1;refGene_tx=XM_019822625.1;refGene_gn=STAB1;refGene_tx=XM_019822633.2
A2	21091157	21091253
refGene_gn=STAB1;refGene_tx=XM_011279480.2;refGene_gn=STAB1;refGene_tx

=XM_011279482.2;refGene_gn=STAB1;refGene_tx=XM_011279484.2;refGene_gn=STAB1;refGene_tx=XM_011279487.2;refGene_gn=STAB1;refGene_tx=XM_019822624.1;refGene_gn=STAB1;refGene_tx=XM_019822625.1;refGene_gn=STAB1;refGene_tx=XM_019822633.2
A2 21091581 21091692
refGene_gn=STAB1;refGene_tx=XM_011279480.2;refGene_gn=STAB1;refGene_tx=XM_011279482.2;refGene_gn=STAB1;refGene_tx=XM_011279484.2;refGene_gn=STAB1;refGene_tx=XM_011279487.2;refGene_gn=STAB1;refGene_tx=XM_019822624.1;refGene_gn=STAB1;refGene_tx=XM_019822625.1;refGene_gn=STAB1;refGene_tx=XM_019822633.2
A2 21092179 21092376
refGene_gn=STAB1;refGene_tx=XM_011279480.2;refGene_gn=STAB1;refGene_tx=XM_011279482.2;refGene_gn=STAB1;refGene_tx=XM_011279484.2;refGene_gn=STAB1;refGene_tx=XM_011279487.2;refGene_gn=STAB1;refGene_tx=XM_019822624.1;refGene_gn=STAB1;refGene_tx=XM_019822625.1;refGene_gn=STAB1;refGene_tx=XM_019822633.2
A2 21092624 21092755
refGene_gn=STAB1;refGene_tx=XM_011279480.2;refGene_gn=STAB1;refGene_tx=XM_011279482.2;refGene_gn=STAB1;refGene_tx=XM_011279484.2;refGene_gn=STAB1;refGene_tx=XM_011279487.2;refGene_gn=STAB1;refGene_tx=XM_019822624.1;refGene_gn=STAB1;refGene_tx=XM_019822625.1;refGene_gn=STAB1;refGene_tx=XM_019822633.2
A2 21092935 21093063
refGene_gn=STAB1;refGene_tx=XM_011279480.2;refGene_gn=STAB1;refGene_tx=XM_011279482.2;refGene_gn=STAB1;refGene_tx=XM_011279484.2;refGene_gn=STAB1;refGene_tx=XM_011279487.2;refGene_gn=STAB1;refGene_tx=XM_019822624.1;refGene_gn=STAB1;refGene_tx=XM_019822625.1;refGene_gn=STAB1;refGene_tx=XM_019822633.2
A2 21093355 21093444
refGene_gn=STAB1;refGene_tx=XM_011279480.2;refGene_gn=STAB1;refGene_tx=XM_011279482.2;refGene_gn=STAB1;refGene_tx=XM_011279484.2;refGene_gn=STAB1;refGene_tx=XM_011279487.2;refGene_gn=STAB1;refGene_tx=XM_019822624.1;refGene_gn=STAB1;refGene_tx=XM_019822625.1;refGene_gn=STAB1;refGene_tx=XM_019822633.2
A2 21093642 21093780
refGene_gn=STAB1;refGene_tx=XM_011279480.2;refGene_gn=STAB1;refGene_tx=XM_011279482.2;refGene_gn=STAB1;refGene_tx=XM_011279484.2;refGene_gn=STAB1;refGene_tx=XM_011279487.2;refGene_gn=STAB1;refGene_tx=XM_019822624.1;refGene_gn=STAB1;refGene_tx=XM_019822625.1;refGene_gn=STAB1;refGene_tx=XM_019822633.2
A2 21093919 21094063
refGene_gn=STAB1;refGene_tx=XM_011279480.2;refGene_gn=STAB1;refGene_tx=XM_011279482.2;refGene_gn=STAB1;refGene_tx=XM_011279484.2;refGene_gn=STAB1;refGene_tx=XM_011279487.2;refGene_gn=STAB1;refGene_tx=XM_019822624.1;refGene_gn=STAB1;refGene_tx=XM_019822625.1;refGene_gn=STAB1;refGene_tx=XM_019822633.2
A2 21094242 21094305
refGene_gn=STAB1;refGene_tx=XM_011279480.2;refGene_gn=STAB1;refGene_tx=XM_011279482.2;refGene_gn=STAB1;refGene_tx=XM_011279484.2;refGene_gn=STAB1;refGene_tx=XM_011279487.2;refGene_gn=STAB1;refGene_tx=XM_0198226

24.1;refGene_gn=STAB1;refGene_tx=XM_019822625.1;refGene_gn=STAB1;refGene_tx=XM_019822633.2
A2 21094585 21094699
refGene_gn=STAB1;refGene_tx=XM_011279480.2;refGene_gn=STAB1;refGene_tx=XM_011279482.2;refGene_gn=STAB1;refGene_tx=XM_011279484.2;refGene_gn=STAB1;refGene_tx=XM_011279487.2;refGene_gn=STAB1;refGene_tx=XM_019822624.1;refGene_gn=STAB1;refGene_tx=XM_019822625.1;refGene_gn=STAB1;refGene_tx=XM_019822633.2
A2 21094783 21094837
refGene_gn=STAB1;refGene_tx=XM_011279480.2;refGene_gn=STAB1;refGene_tx=XM_011279482.2;refGene_gn=STAB1;refGene_tx=XM_011279484.2;refGene_gn=STAB1;refGene_tx=XM_011279487.2;refGene_gn=STAB1;refGene_tx=XM_019822624.1;refGene_gn=STAB1;refGene_tx=XM_019822625.1;refGene_gn=STAB1;refGene_tx=XM_019822633.2
A2 21095078 21095159
refGene_gn=STAB1;refGene_tx=XM_011279480.2;refGene_gn=STAB1;refGene_tx=XM_011279482.2;refGene_gn=STAB1;refGene_tx=XM_011279484.2;refGene_gn=STAB1;refGene_tx=XM_011279487.2;refGene_gn=STAB1;refGene_tx=XM_019822624.1;refGene_gn=STAB1;refGene_tx=XM_019822625.1;refGene_gn=STAB1;refGene_tx=XM_019822633.2
A2 21095606 21095765
refGene_gn=STAB1;refGene_tx=XM_011279480.2;refGene_gn=STAB1;refGene_tx=XM_011279482.2;refGene_gn=STAB1;refGene_tx=XM_011279484.2;refGene_gn=STAB1;refGene_tx=XM_011279487.2;refGene_gn=STAB1;refGene_tx=XM_019822624.1;refGene_gn=STAB1;refGene_tx=XM_019822625.1;refGene_gn=STAB1;refGene_tx=XM_019822633.2
A2 21096131 21096194
refGene_gn=STAB1;refGene_tx=XM_011279480.2;refGene_gn=STAB1;refGene_tx=XM_011279482.2;refGene_gn=STAB1;refGene_tx=XM_011279484.2;refGene_gn=STAB1;refGene_tx=XM_011279487.2;refGene_gn=STAB1;refGene_tx=XM_019822624.1;refGene_gn=STAB1;refGene_tx=XM_019822625.1;refGene_gn=STAB1;refGene_tx=XM_019822633.2
A2 21096426 21096522
refGene_gn=STAB1;refGene_tx=XM_011279480.2;refGene_gn=STAB1;refGene_tx=XM_011279482.2;refGene_gn=STAB1;refGene_tx=XM_011279484.2;refGene_gn=STAB1;refGene_tx=XM_011279487.2;refGene_gn=STAB1;refGene_tx=XM_019822624.1;refGene_gn=STAB1;refGene_tx=XM_019822625.1;refGene_gn=STAB1;refGene_tx=XM_019822633.2
A2 21096711 21096798
refGene_gn=STAB1;refGene_tx=XM_011279480.2;refGene_gn=STAB1;refGene_tx=XM_011279482.2;refGene_gn=STAB1;refGene_tx=XM_011279484.2;refGene_gn=STAB1;refGene_tx=XM_011279487.2;refGene_gn=STAB1;refGene_tx=XM_019822624.1;refGene_gn=STAB1;refGene_tx=XM_019822625.1;refGene_gn=STAB1;refGene_tx=XM_019822633.2
A2 21097287 21097399
refGene_gn=STAB1;refGene_tx=XM_011279480.2;refGene_gn=STAB1;refGene_tx=XM_011279482.2;refGene_gn=STAB1;refGene_tx=XM_011279484.2;refGene_gn=STAB1;refGene_tx=XM_011279487.2;refGene_gn=STAB1;refGene_tx=XM_019822624.1;refGene_gn=STAB1;refGene_tx=XM_019822625.1;refGene_gn=STAB1;refGene_tx=XM_019822633.2

A2	21098096 21098283
refGene_gn=STAB1;refGene_tx=XM_011279480.2;refGene_gn=STAB1;refGene_tx=XM_011279482.2;refGene_gn=STAB1;refGene_tx=XM_011279484.2;refGene_gn=STAB1;refGene_tx=XM_011279487.2;refGene_gn=STAB1;refGene_tx=XM_019822624.1;refGene_gn=STAB1;refGene_tx=XM_019822625.1;refGene_gn=STAB1;refGene_tx=XM_019822633.2
A2	21098393 21098487
refGene_gn=STAB1;refGene_tx=XM_011279480.2;refGene_gn=STAB1;refGene_tx=XM_011279482.2;refGene_gn=STAB1;refGene_tx=XM_011279484.2;refGene_gn=STAB1;refGene_tx=XM_011279487.2;refGene_gn=STAB1;refGene_tx=XM_019822624.1;refGene_gn=STAB1;refGene_tx=XM_019822625.1;refGene_gn=STAB1;refGene_tx=XM_019822633.2
A2	21098585 21098744
refGene_gn=STAB1;refGene_tx=XM_011279480.2;refGene_gn=STAB1;refGene_tx=XM_011279482.2;refGene_gn=STAB1;refGene_tx=XM_011279484.2;refGene_gn=STAB1;refGene_tx=XM_011279487.2;refGene_gn=STAB1;refGene_tx=XM_019822624.1;refGene_gn=STAB1;refGene_tx=XM_019822625.1;refGene_gn=STAB1;refGene_tx=XM_019822633.2
A2	21100027 21100123
refGene_gn=STAB1;refGene_tx=XM_011279480.2;refGene_gn=STAB1;refGene_tx=XM_011279482.2;refGene_gn=STAB1;refGene_tx=XM_011279484.2;refGene_gn=STAB1;refGene_tx=XM_011279487.2;refGene_gn=STAB1;refGene_tx=XM_019822624.1;refGene_gn=STAB1;refGene_tx=XM_019822625.1;refGene_gn=STAB1;refGene_tx=XM_019822633.2
A2	21100735 21100831
refGene_gn=STAB1;refGene_tx=XM_011279480.2;refGene_gn=STAB1;refGene_tx=XM_011279482.2;refGene_gn=STAB1;refGene_tx=XM_011279484.2;refGene_gn=STAB1;refGene_tx=XM_011279487.2;refGene_gn=STAB1;refGene_tx=XM_019822624.1;refGene_gn=STAB1;refGene_tx=XM_019822625.1;refGene_gn=STAB1;refGene_tx=XM_019822633.2
A2	21100987 21101035
refGene_gn=STAB1;refGene_tx=XM_011279480.2;refGene_gn=STAB1;refGene_tx=XM_011279482.2;refGene_gn=STAB1;refGene_tx=XM_011279484.2;refGene_gn=STAB1;refGene_tx=XM_011279487.2;refGene_gn=STAB1;refGene_tx=XM_019822624.1;refGene_gn=STAB1;refGene_tx=XM_019822625.1;refGene_gn=STAB1;refGene_tx=XM_019822633.2
A2	21101213 21101344
refGene_gn=STAB1;refGene_tx=XM_011279480.2;refGene_gn=STAB1;refGene_tx=XM_011279482.2;refGene_gn=STAB1;refGene_tx=XM_011279484.2;refGene_gn=STAB1;refGene_tx=XM_011279487.2;refGene_gn=STAB1;refGene_tx=XM_019822624.1;refGene_gn=STAB1;refGene_tx=XM_019822625.1;refGene_gn=STAB1;refGene_tx=XM_019822633.2
A2	21101500 21101612
refGene_gn=STAB1;refGene_tx=XM_011279480.2;refGene_gn=STAB1;refGene_tx=XM_011279482.2;refGene_gn=STAB1;refGene_tx=XM_011279484.2;refGene_gn=STAB1;refGene_tx=XM_011279487.2;refGene_gn=STAB1;refGene_tx=XM_019822624.1;refGene_gn=STAB1;refGene_tx=XM_019822625.1;refGene_gn=STAB1;refGene_tx=XM_019822633.2
A2	21102003 21102081
refGene_gn=STAB1;refGene_tx=XM_011279480.2;refGene_gn=STAB1;refGene_tx

=XM_011279482.2;refGene_gn=STAB1;refGene_tx=XM_011279484.2;refGene_gn=STAB1;refGene_tx=XM_011279487.2;refGene_gn=STAB1;refGene_tx=XM_019822624.1;refGene_gn=STAB1;refGene_tx=XM_019822625.1;refGene_gn=STAB1;refGene_tx=XM_019822633.2
A2 21102163 21102262
refGene_gn=STAB1;refGene_tx=XM_011279480.2;refGene_gn=STAB1;refGene_tx=XM_011279482.2;refGene_gn=STAB1;refGene_tx=XM_011279484.2;refGene_gn=STAB1;refGene_tx=XM_011279487.2;refGene_gn=STAB1;refGene_tx=XM_019822624.1;refGene_gn=STAB1;refGene_tx=XM_019822625.1;refGene_gn=STAB1;refGene_tx=XM_019822633.2
A2 21102394 21102493
refGene_gn=STAB1;refGene_tx=XM_011279480.2;refGene_gn=STAB1;refGene_tx=XM_011279482.2;refGene_gn=STAB1;refGene_tx=XM_011279484.2;refGene_gn=STAB1;refGene_tx=XM_011279487.2;refGene_gn=STAB1;refGene_tx=XM_019822624.1;refGene_gn=STAB1;refGene_tx=XM_019822625.1;refGene_gn=STAB1;refGene_tx=XM_019822633.2
A2 21102638 21102770
refGene_gn=STAB1;refGene_tx=XM_011279480.2;refGene_gn=STAB1;refGene_tx=XM_011279482.2;refGene_gn=STAB1;refGene_tx=XM_011279484.2;refGene_gn=STAB1;refGene_tx=XM_011279487.2;refGene_gn=STAB1;refGene_tx=XM_019822624.1;refGene_gn=STAB1;refGene_tx=XM_019822625.1;refGene_gn=STAB1;refGene_tx=XM_019822633.2
A2 21102916 21103045
refGene_gn=STAB1;refGene_tx=XM_011279480.2;refGene_gn=STAB1;refGene_tx=XM_011279482.2;refGene_gn=STAB1;refGene_tx=XM_011279484.2;refGene_gn=STAB1;refGene_tx=XM_011279487.2;refGene_gn=STAB1;refGene_tx=XM_019822624.1;refGene_gn=STAB1;refGene_tx=XM_019822625.1;refGene_gn=STAB1;refGene_tx=XM_019822633.2
A2 21103217 21103274
refGene_gn=STAB1;refGene_tx=XM_011279480.2;refGene_gn=STAB1;refGene_tx=XM_011279482.2;refGene_gn=STAB1;refGene_tx=XM_011279484.2;refGene_gn=STAB1;refGene_tx=XM_011279487.2;refGene_gn=STAB1;refGene_tx=XM_019822624.1;refGene_gn=STAB1;refGene_tx=XM_019822625.1;refGene_gn=STAB1;refGene_tx=XM_019822633.2
A2 21103651 21103735
refGene_gn=STAB1;refGene_tx=XM_011279480.2;refGene_gn=STAB1;refGene_tx=XM_011279482.2;refGene_gn=STAB1;refGene_tx=XM_011279484.2;refGene_gn=STAB1;refGene_tx=XM_011279487.2;refGene_gn=STAB1;refGene_tx=XM_019822624.1;refGene_gn=STAB1;refGene_tx=XM_019822625.1;refGene_gn=STAB1;refGene_tx=XM_019822633.2
A2 21104249 21104445
refGene_gn=STAB1;refGene_tx=XM_011279480.2;refGene_gn=STAB1;refGene_tx=XM_011279482.2;refGene_gn=STAB1;refGene_tx=XM_011279484.2;refGene_gn=STAB1;refGene_tx=XM_011279487.2;refGene_gn=STAB1;refGene_tx=XM_019822624.1;refGene_gn=STAB1;refGene_tx=XM_019822625.1;refGene_gn=STAB1;refGene_tx=XM_019822633.2
A2 21104544 21104640
refGene_gn=STAB1;refGene_tx=XM_011279480.2;refGene_gn=STAB1;refGene_tx=XM_011279482.2;refGene_gn=STAB1;refGene_tx=XM_011279484.2;refGene_gn=STAB1;refGene_tx=XM_011279487.2;refGene_gn=STAB1;refGene_tx=XM_0198226

24.1;refGene_gn=STAB1;refGene_tx=XM_019822625.1;refGene_gn=STAB1;refGene_tx=XM_019822633.2
A2 21104729 21104775
refGene_gn=STAB1;refGene_tx=XM_011279480.2;refGene_gn=STAB1;refGene_tx=XM_011279482.2;refGene_gn=STAB1;refGene_tx=XM_011279484.2;refGene_gn=STAB1;refGene_tx=XM_011279487.2;refGene_gn=STAB1;refGene_tx=XM_019822624.1;refGene_gn=STAB1;refGene_tx=XM_019822625.1;refGene_gn=STAB1;refGene_tx=XM_019822633.2
A2 21104876 21104953
refGene_gn=STAB1;refGene_tx=XM_011279480.2;refGene_gn=STAB1;refGene_tx=XM_011279482.2;refGene_gn=STAB1;refGene_tx=XM_011279484.2;refGene_gn=STAB1;refGene_tx=XM_011279487.2;refGene_gn=STAB1;refGene_tx=XM_019822624.1;refGene_gn=STAB1;refGene_tx=XM_019822625.1;refGene_gn=STAB1;refGene_tx=XM_019822633.2
A2 21105173 21105299
refGene_gn=STAB1;refGene_tx=XM_011279480.2;refGene_gn=STAB1;refGene_tx=XM_011279482.2;refGene_gn=STAB1;refGene_tx=XM_011279484.2;refGene_gn=STAB1;refGene_tx=XM_011279487.2;refGene_gn=STAB1;refGene_tx=XM_019822624.1;refGene_gn=STAB1;refGene_tx=XM_019822625.1;refGene_gn=STAB1;refGene_tx=XM_019822633.2
A2 21105481 21105555
refGene_gn=STAB1;refGene_tx=XM_011279480.2;refGene_gn=STAB1;refGene_tx=XM_011279482.2;refGene_gn=STAB1;refGene_tx=XM_011279484.2;refGene_gn=STAB1;refGene_tx=XM_011279487.2;refGene_gn=STAB1;refGene_tx=XM_019822624.1;refGene_gn=STAB1;refGene_tx=XM_019822625.1;refGene_gn=STAB1;refGene_tx=XM_019822633.2
A2 21105700 21105775
refGene_gn=STAB1;refGene_tx=XM_011279480.2;refGene_gn=STAB1;refGene_tx=XM_011279482.2;refGene_gn=STAB1;refGene_tx=XM_011279484.2;refGene_gn=STAB1;refGene_tx=XM_011279487.2;refGene_gn=STAB1;refGene_tx=XM_019822624.1;refGene_gn=STAB1;refGene_tx=XM_019822625.1;refGene_gn=STAB1;refGene_tx=XM_019822633.2
A2 21106023 21106146
refGene_gn=STAB1;refGene_tx=XM_011279480.2;refGene_gn=STAB1;refGene_tx=XM_011279482.2;refGene_gn=STAB1;refGene_tx=XM_011279484.2;refGene_gn=STAB1;refGene_tx=XM_011279487.2;refGene_gn=STAB1;refGene_tx=XM_019822624.1;refGene_gn=STAB1;refGene_tx=XM_019822625.1;refGene_gn=STAB1;refGene_tx=XM_019822633.2
A2 21106448 21106496
refGene_gn=STAB1;refGene_tx=XM_011279480.2;refGene_gn=STAB1;refGene_tx=XM_011279482.2;refGene_gn=STAB1;refGene_tx=XM_011279484.2;refGene_gn=STAB1;refGene_tx=XM_011279487.2;refGene_gn=STAB1;refGene_tx=XM_019822624.1;refGene_gn=STAB1;refGene_tx=XM_019822625.1;refGene_gn=STAB1;refGene_tx=XM_019822633.2
A2 21106710 21106788
refGene_gn=STAB1;refGene_tx=XM_011279480.2;refGene_gn=STAB1;refGene_tx=XM_011279482.2;refGene_gn=STAB1;refGene_tx=XM_011279484.2;refGene_gn=STAB1;refGene_tx=XM_011279487.2;refGene_gn=STAB1;refGene_tx=XM_019822624.1;refGene_gn=STAB1;refGene_tx=XM_019822625.1;refGene_gn=STAB1;refGene_tx=XM_019822633.2

A2 21106905 21107052
refGene_gn=STAB1;refGene_tx=XM_011279480.2;refGene_gn=STAB1;refGene_tx=XM_011279482.2;refGene_gn=STAB1;refGene_tx=XM_011279484.2;refGene_gn=STAB1;refGene_tx=XM_011279487.2;refGene_gn=STAB1;refGene_tx=XM_019822624.1;refGene_gn=STAB1;refGene_tx=XM_019822625.1;refGene_gn=STAB1;refGene_tx=XM_019822633.2
A2 21107404 21107545
refGene_gn=STAB1;refGene_tx=XM_011279480.2;refGene_gn=STAB1;refGene_tx=XM_011279482.2;refGene_gn=STAB1;refGene_tx=XM_011279484.2;refGene_gn=STAB1;refGene_tx=XM_011279487.2;refGene_gn=STAB1;refGene_tx=XM_019822624.1;refGene_gn=STAB1;refGene_tx=XM_019822625.1;refGene_gn=STAB1;refGene_tx=XM_019822633.2
A2 21107648 21107708
refGene_gn=STAB1;refGene_tx=XM_011279480.2;refGene_gn=STAB1;refGene_tx=XM_011279482.2;refGene_gn=STAB1;refGene_tx=XM_011279484.2;refGene_gn=STAB1;refGene_tx=XM_011279487.2;refGene_gn=STAB1;refGene_tx=XM_019822624.1;refGene_gn=STAB1;refGene_tx=XM_019822625.1;refGene_gn=STAB1;refGene_tx=XM_019822633.2
A2 21108108 21108285
refGene_gn=STAB1;refGene_tx=XM_011279480.2;refGene_gn=STAB1;refGene_tx=XM_011279482.2;refGene_gn=STAB1;refGene_tx=XM_011279484.2;refGene_gn=STAB1;refGene_tx=XM_011279487.2;refGene_gn=STAB1;refGene_tx=XM_019822624.1;refGene_gn=STAB1;refGene_tx=XM_019822625.1;refGene_gn=STAB1;refGene_tx=XM_019822633.2
A2 21108368 21108455
refGene_gn=STAB1;refGene_tx=XM_011279480.2;refGene_gn=STAB1;refGene_tx=XM_011279482.2;refGene_gn=STAB1;refGene_tx=XM_011279484.2;refGene_gn=STAB1;refGene_tx=XM_011279487.2;refGene_gn=STAB1;refGene_tx=XM_019822624.1;refGene_gn=STAB1;refGene_tx=XM_019822625.1;refGene_gn=STAB1;refGene_tx=XM_019822633.2
A2 21108564 21108714
refGene_gn=STAB1;refGene_tx=XM_011279480.2;refGene_gn=STAB1;refGene_tx=XM_011279482.2;refGene_gn=STAB1;refGene_tx=XM_011279484.2;refGene_gn=STAB1;refGene_tx=XM_011279487.2;refGene_gn=STAB1;refGene_tx=XM_019822624.1;refGene_gn=STAB1;refGene_tx=XM_019822625.1;refGene_gn=STAB1;refGene_tx=XM_019822633.2
A2 21108819 21108876
refGene_gn=STAB1;refGene_tx=XM_011279480.2;refGene_gn=STAB1;refGene_tx=XM_011279482.2;refGene_gn=STAB1;refGene_tx=XM_011279484.2;refGene_gn=STAB1;refGene_tx=XM_011279487.2;refGene_gn=STAB1;refGene_tx=XM_019822624.1;refGene_gn=STAB1;refGene_tx=XM_019822625.1;refGene_gn=STAB1;refGene_tx=XM_019822633.2
A2 21108952 21109154
refGene_gn=STAB1;refGene_tx=XM_011279480.2;refGene_gn=STAB1;refGene_tx=XM_011279482.2;refGene_gn=STAB1;refGene_tx=XM_011279484.2;refGene_gn=STAB1;refGene_tx=XM_011279487.2;refGene_gn=STAB1;refGene_tx=XM_019822624.1;refGene_gn=STAB1;refGene_tx=XM_019822625.1;refGene_gn=STAB1;refGene_tx=XM_019822633.2
A2 21109477 21109627
refGene_gn=STAB1;refGene_tx=XM_011279480.2;refGene_gn=STAB1;refGene_tx

=XM_011279482.2;refGene_gn=STAB1;refGene_tx=XM_011279484.2;refGene_gn=STAB1;refGene_tx=XM_011279487.2;refGene_gn=STAB1;refGene_tx=XM_019822624.1;refGene_gn=STAB1;refGene_tx=XM_019822625.1;refGene_gn=STAB1;refGene_tx=XM_019822633.2
A2 21109718 21109918
refGene_gn=STAB1;refGene_tx=XM_011279480.2;refGene_gn=STAB1;refGene_tx=XM_011279482.2;refGene_gn=STAB1;refGene_tx=XM_011279484.2;refGene_gn=STAB1;refGene_tx=XM_011279487.2;refGene_gn=STAB1;refGene_tx=XM_019822624.1;refGene_gn=STAB1;refGene_tx=XM_019822625.1;refGene_gn=STAB1;refGene_tx=XM_019822633.2
A2 21109996 21110104
refGene_gn=STAB1;refGene_tx=XM_011279480.2;refGene_gn=STAB1;refGene_tx=XM_011279482.2;refGene_gn=STAB1;refGene_tx=XM_011279484.2;refGene_gn=STAB1;refGene_tx=XM_011279487.2;refGene_gn=STAB1;refGene_tx=XM_019822624.1;refGene_gn=STAB1;refGene_tx=XM_019822625.1
A2 21110184 21110378
refGene_gn=STAB1;refGene_tx=XM_011279480.2;refGene_gn=STAB1;refGene_tx=XM_011279482.2;refGene_gn=STAB1;refGene_tx=XM_011279484.2;refGene_gn=STAB1;refGene_tx=XM_011279487.2;refGene_gn=STAB1;refGene_tx=XM_019822624.1;refGene_gn=STAB1;refGene_tx=XM_019822625.1
A2 21110489 21110634
refGene_gn=STAB1;refGene_tx=XM_011279480.2;refGene_gn=STAB1;refGene_tx=XM_011279482.2;refGene_gn=STAB1;refGene_tx=XM_011279484.2;refGene_gn=STAB1;refGene_tx=XM_011279487.2;refGene_gn=STAB1;refGene_tx=XM_019822624.1;refGene_gn=STAB1;refGene_tx=XM_019822625.1
A2 21110713 21110847
refGene_gn=STAB1;refGene_tx=XM_011279480.2;refGene_gn=STAB1;refGene_tx=XM_011279482.2;refGene_gn=STAB1;refGene_tx=XM_011279484.2;refGene_gn=STAB1;refGene_tx=XM_011279487.2;refGene_gn=STAB1;refGene_tx=XM_019822624.1;refGene_gn=STAB1;refGene_tx=XM_019822625.1
A2 21110936 21111102
refGene_gn=STAB1;refGene_tx=XM_011279480.2;refGene_gn=STAB1;refGene_tx=XM_011279482.2;refGene_gn=STAB1;refGene_tx=XM_011279484.2;refGene_gn=STAB1;refGene_tx=XM_011279487.2;refGene_gn=STAB1;refGene_tx=XM_019822624.1;refGene_gn=STAB1;refGene_tx=XM_019822625.1
A2 21111181 21111291
refGene_gn=STAB1;refGene_tx=XM_011279480.2;refGene_gn=STAB1;refGene_tx=XM_011279482.2;refGene_gn=STAB1;refGene_tx=XM_011279484.2;refGene_gn=STAB1;refGene_tx=XM_011279487.2;refGene_gn=STAB1;refGene_tx=XM_019822624.1;refGene_gn=STAB1;refGene_tx=XM_019822625.1
A2 21111382 21111502
refGene_gn=STAB1;refGene_tx=XM_011279480.2;refGene_gn=STAB1;refGene_tx=XM_011279482.2;refGene_gn=STAB1;refGene_tx=XM_011279484.2;refGene_gn=STAB1;refGene_tx=XM_011279487.2;refGene_gn=STAB1;refGene_tx=XM_019822624.1;refGene_gn=STAB1;refGene_tx=XM_019822625.1
A2 21111588 21111738
refGene_gn=STAB1;refGene_tx=XM_011279480.2;refGene_gn=STAB1;refGene_tx=XM_011279482.2;refGene_gn=STAB1;refGene_tx=XM_011279484.2;refGene_gn=STAB1;refGene_tx=XM_011279487.2;refGene_gn=STAB1;refGene_tx=XM_019822624.1;refGene_gn=STAB1;refGene_tx=XM_019822625.1

A2 21111816 21111921
refGene_gn=STAB1;refGene_tx=XM_011279480.2;refGene_gn=STAB1;refGene_tx=XM_011279482.2;refGene_gn=STAB1;refGene_tx=XM_011279484.2;refGene_gn=STAB1;refGene_tx=XM_011279487.2;refGene_gn=STAB1;refGene_tx=XM_019822624.1;refGene_gn=STAB1;refGene_tx=XM_019822625.1
A2 21112009 21112156
refGene_gn=STAB1;refGene_tx=XM_011279480.2;refGene_gn=STAB1;refGene_tx=XM_011279482.2;refGene_gn=STAB1;refGene_tx=XM_011279484.2;refGene_gn=STAB1;refGene_tx=XM_011279487.2;refGene_gn=STAB1;refGene_tx=XM_019822624.1;refGene_gn=STAB1;refGene_tx=XM_019822625.1
A2 21112251 21112368
refGene_gn=STAB1;refGene_tx=XM_011279480.2;refGene_gn=STAB1;refGene_tx=XM_011279482.2;refGene_gn=STAB1;refGene_tx=XM_011279484.2;refGene_gn=STAB1;refGene_tx=XM_011279487.2;refGene_gn=STAB1;refGene_tx=XM_019822624.1;refGene_gn=STAB1;refGene_tx=XM_019822625.1
A2 21112433 21112626
refGene_gn=STAB1;refGene_tx=XM_011279480.2;refGene_gn=STAB1;refGene_tx=XM_011279482.2;refGene_gn=STAB1;refGene_tx=XM_011279484.2;refGene_gn=STAB1;refGene_tx=XM_011279487.2;refGene_gn=STAB1;refGene_tx=XM_019822624.1;refGene_gn=STAB1;refGene_tx=XM_019822625.1
A2 21112634 21112896
refGene_gn=NT5DC2;refGene_tx=XM_019822635.2;refGene_gn=NT5DC2;refGene_tx=XM_023249524.1
A2 21112974 21113039 refGene_gn=NT5DC2;refGene_tx=XM_019822635.2
A2 21113120 21113261
refGene_gn=NT5DC2;refGene_tx=XM_019822635.2;refGene_gn=NT5DC2;refGene_tx=XM_023249524.1
A2 21113334 21113421
refGene_gn=NT5DC2;refGene_tx=XM_019822635.2;refGene_gn=NT5DC2;refGene_tx=XM_023249524.1
A2 21115154 21115236
refGene_gn=NT5DC2;refGene_tx=XM_019822635.2;refGene_gn=NT5DC2;refGene_tx=XM_023249524.1
A2 21115431 21115533
refGene_gn=NT5DC2;refGene_tx=XM_019822635.2;refGene_gn=NT5DC2;refGene_tx=XM_023249524.1
A2 21115643 21115746
refGene_gn=NT5DC2;refGene_tx=XM_019822635.2;refGene_gn=NT5DC2;refGene_tx=XM_023249524.1
A2 21115825 21115886
refGene_gn=NT5DC2;refGene_tx=XM_019822635.2;refGene_gn=NT5DC2;refGene_tx=XM_023249524.1
A2 21115996 21116125
refGene_gn=NT5DC2;refGene_tx=XM_019822635.2;refGene_gn=NT5DC2;refGene_tx=XM_023249524.1
A2 21116317 21116411
refGene_gn=NT5DC2;refGene_tx=XM_019822635.2;refGene_gn=NT5DC2;refGene_tx=XM_023249524.1
A2 21116528 21116584
refGene_gn=NT5DC2;refGene_tx=XM_019822635.2;refGene_gn=NT5DC2;refGene_

tx=XM_023249524.1
A2	21116823	21116898
refGene_gn=NT5DC2;refGene_tx=XM_019822635.2;refGene_gn=NT5DC2;refGene_tx=XM_023249524.1
A2	21117034	21117219
refGene_gn=NT5DC2;refGene_tx=XM_019822635.2;refGene_gn=NT5DC2;refGene_tx=XM_023249524.1
A2	21121335	21121567
refGene_gn=NT5DC2;refGene_tx=XM_019822635.2;refGene_gn=NT5DC2;refGene_tx=XM_023249524.1
A2	21124410	21124561	refGene_gn=SMIM4;refGene_tx=XM_003982308.5
A2	21127813	21127905	refGene_gn=SMIM4;refGene_tx=XM_003982308.5
A2	21134072	21134245
refGene_gn=PBRM1;refGene_tx=XM_003982314.5;refGene_gn=PBRM1;refGene_tx=XM_019822655.2;refGene_gn=PBRM1;refGene_tx=XM_019822656.2;refGene_gn=PBRM1;refGene_tx=XM_019822657.2;refGene_gn=PBRM1;refGene_tx=XM_019822659.2;refGene_gn=PBRM1;refGene_tx=XM_019822665.2;refGene_gn=PBRM1;refGene_tx=XM_019822680.2;refGene_gn=PBRM1;refGene_tx=XM_019822682.2;refGene_gn=PBRM1;refGene_tx=XM_019822686.2;refGene_gn=PBRM1;refGene_tx=XM_019822693.2;refGene_gn=PBRM1;refGene_tx=XM_019822696.2;refGene_gn=PBRM1;refGene_tx=XM_019822697.2;refGene_gn=PBRM1;refGene_tx=XM_019822700.2;refGene_gn=PBRM1;refGene_tx=XM_019822704.2;refGene_gn=PBRM1;refGene_tx=XM_019822708.2;refGene_gn=PBRM1;refGene_tx=XM_019822711.2;refGene_gn=PBRM1;refGene_tx=XM_019822713.2;refGene_gn=PBRM1;refGene_tx=XM_019822716.2;refGene_gn=PBRM1;refGene_tx=XM_019822718.2;refGene_gn=PBRM1;refGene_tx=XM_019822719.2;refGene_gn=PBRM1;refGene_tx=XM_019822722.2;refGene_gn=PBRM1;refGene_tx=XM_019822723.2;refGene_gn=PBRM1;refGene_tx=XM_019822726.2;refGene_gn=PBRM1;refGene_tx=XM_019822731.1;refGene_gn=PBRM1;refGene_tx=XM_023249537.1;refGene_gn=PBRM1;refGene_tx=XM_023249539.1;refGene_gn=PBRM1;refGene_tx=XM_023249543.1;refGene_gn=PBRM1;refGene_tx=XM_023249544.1;refGene_gn=PBRM1;refGene_tx=XM_023249570.1;refGene_gn=PBRM1;refGene_tx=XM_023249576.1;refGene_gn=PBRM1;refGene_tx=XM_023249582.1
A2	21135422	21135639
refGene_gn=PBRM1;refGene_tx=XM_003982314.5;refGene_gn=PBRM1;refGene_tx=XM_019822655.2;refGene_gn=PBRM1;refGene_tx=XM_019822656.2;refGene_gn=PBRM1;refGene_tx=XM_019822657.2;refGene_gn=PBRM1;refGene_tx=XM_019822659.2;refGene_gn=PBRM1;refGene_tx=XM_019822665.2;refGene_gn=PBRM1;refGene_tx=XM_019822680.2;refGene_gn=PBRM1;refGene_tx=XM_019822682.2;refGene_gn=PBRM1;refGene_tx=XM_019822686.2;refGene_gn=PBRM1;refGene_tx=XM_019822693.2;refGene_gn=PBRM1;refGene_tx=XM_019822696.2;refGene_gn=PBRM1;refGene_tx=XM_019822697.2;refGene_gn=PBRM1;refGene_tx=XM_019822700.2;refGene_gn=PBRM1;refGene_tx=XM_019822704.2;refGene_gn=PBRM1;refGene_tx=XM_019822708.2;refGene_gn=PBRM1;refGene_tx=XM_019822711.2;refGene_gn=PBRM1;refGene_tx=XM_019822713.2;refGene_gn=PBRM1;refGene_tx=XM_019822716.2;refGene_gn=PBRM1;refGene_tx=XM_019822718.2;refGene_gn=PBRM1;refGene_tx=XM_019822719.2;refGene_gn=PBRM1;refGene_tx=XM_019822722.2;refGene_gn=PBRM1;refGene_tx=XM_019822723.2;refGene_gn=PBRM1;refGene_tx=XM_019822726.2;refGene_gn=PBRM1;refGene_tx=XM_019822731.1;refGene_gn=PBRM1;refGene_tx=XM_023249537.1;refGene_gn=PBRM1;refGene_tx=XM_023249539.1;re

fGene_gn=PBRM1;refGene_tx=XM_023249543.1;refGene_gn=PBRM1;refGene_tx=XM_023249544.1;refGene_gn=PBRM1;refGene_tx=XM_023249570.1;refGene_gn=PBRM1;refGene_tx=XM_023249576.1;refGene_gn=PBRM1;refGene_tx=XM_023249582.1
A2 21135749 21135820
refGene_gn=PBRM1;refGene_tx=XM_003982314.5;refGene_gn=PBRM1;refGene_tx=XM_019822655.2;refGene_gn=PBRM1;refGene_tx=XM_019822656.2;refGene_gn=PBRM1;refGene_tx=XM_019822657.2;refGene_gn=PBRM1;refGene_tx=XM_019822659.2;refGene_gn=PBRM1;refGene_tx=XM_019822665.2;refGene_gn=PBRM1;refGene_tx=XM_019822680.2;refGene_gn=PBRM1;refGene_tx=XM_019822682.2;refGene_gn=PBRM1;refGene_tx=XM_019822686.2;refGene_gn=PBRM1;refGene_tx=XM_019822693.2;refGene_gn=PBRM1;refGene_tx=XM_019822696.2;refGene_gn=PBRM1;refGene_tx=XM_019822697.2;refGene_gn=PBRM1;refGene_tx=XM_019822700.2;refGene_gn=PBRM1;refGene_tx=XM_019822704.2;refGene_gn=PBRM1;refGene_tx=XM_019822708.2;refGene_gn=PBRM1;refGene_tx=XM_019822711.2;refGene_gn=PBRM1;refGene_tx=XM_019822713.2;refGene_gn=PBRM1;refGene_tx=XM_019822716.2;refGene_gn=PBRM1;refGene_tx=XM_019822718.2;refGene_gn=PBRM1;refGene_tx=XM_019822719.2;refGene_gn=PBRM1;refGene_tx=XM_019822722.2;refGene_gn=PBRM1;refGene_tx=XM_019822723.2;refGene_gn=PBRM1;refGene_tx=XM_019822726.2;refGene_gn=PBRM1;refGene_tx=XM_019822731.1;refGene_gn=PBRM1;refGene_tx=XM_023249537.1;refGene_gn=PBRM1;refGene_tx=XM_023249539.1;refGene_gn=PBRM1;refGene_tx=XM_023249543.1;refGene_gn=PBRM1;refGene_tx=XM_023249544.1;refGene_gn=PBRM1;refGene_tx=XM_023249570.1;refGene_gn=PBRM1;refGene_tx=XM_023249576.1;refGene_gn=PBRM1;refGene_tx=XM_023249582.1
A2 21138880 21139036
refGene_gn=PBRM1;refGene_tx=XM_003982314.5;refGene_gn=PBRM1;refGene_tx=XM_019822655.2;refGene_gn=PBRM1;refGene_tx=XM_019822656.2;refGene_gn=PBRM1;refGene_tx=XM_019822657.2;refGene_gn=PBRM1;refGene_tx=XM_019822659.2;refGene_gn=PBRM1;refGene_tx=XM_019822665.2;refGene_gn=PBRM1;refGene_tx=XM_019822680.2;refGene_gn=PBRM1;refGene_tx=XM_019822682.2;refGene_gn=PBRM1;refGene_tx=XM_019822686.2;refGene_gn=PBRM1;refGene_tx=XM_019822693.2;refGene_gn=PBRM1;refGene_tx=XM_019822696.2;refGene_gn=PBRM1;refGene_tx=XM_019822697.2;refGene_gn=PBRM1;refGene_tx=XM_019822700.2;refGene_gn=PBRM1;refGene_tx=XM_019822704.2;refGene_gn=PBRM1;refGene_tx=XM_019822711.2;refGene_gn=PBRM1;refGene_tx=XM_019822713.2;refGene_gn=PBRM1;refGene_tx=XM_019822718.2;refGene_gn=PBRM1;refGene_tx=XM_019822722.2;refGene_gn=PBRM1;refGene_tx=XM_019822723.2;refGene_gn=PBRM1;refGene_tx=XM_023249537.1;refGene_gn=PBRM1;refGene_tx=XM_023249539.1;refGene_gn=PBRM1;refGene_tx=XM_023249543.1;refGene_gn=PBRM1;refGene_tx=XM_023249544.1
A2 21142417 21142582
refGene_gn=PBRM1;refGene_tx=XM_003982314.5;refGene_gn=PBRM1;refGene_tx=XM_019822655.2;refGene_gn=PBRM1;refGene_tx=XM_019822656.2;refGene_gn=PBRM1;refGene_tx=XM_019822657.2;refGene_gn=PBRM1;refGene_tx=XM_019822659.2;refGene_gn=PBRM1;refGene_tx=XM_019822665.2;refGene_gn=PBRM1;refGene_tx=XM_019822680.2;refGene_gn=PBRM1;refGene_tx=XM_019822682.2;refGene_gn=PBRM1;refGene_tx=XM_019822686.2;refGene_gn=PBRM1;refGene_tx=XM_019822693.2;refGene_gn=PBRM1;refGene_tx=XM_019822696.2;refGene_gn=PBRM1;refGene_tx=XM_019822697.2;refGene_gn=PBRM1;refGene_tx=XM_019822700.2;r

efGene_gn=PBRM1;refGene_tx=XM_019822704.2;refGene_gn=PBRM1;refGene_tx=XM_019822708.2;refGene_gn=PBRM1;refGene_tx=XM_019822711.2;refGene_gn=PBRM1;refGene_tx=XM_019822716.2;refGene_gn=PBRM1;refGene_tx=XM_023249537.1;refGene_gn=PBRM1;refGene_tx=XM_023249539.1;refGene_gn=PBRM1;refGene_tx=XM_023249543.1;refGene_gn=PBRM1;refGene_tx=XM_023249544.1
A2 21145982 21146184
refGene_gn=PBRM1;refGene_tx=XM_003982314.5;refGene_gn=PBRM1;refGene_tx=XM_019822655.2;refGene_gn=PBRM1;refGene_tx=XM_019822656.2;refGene_gn=PBRM1;refGene_tx=XM_019822657.2;refGene_gn=PBRM1;refGene_tx=XM_019822659.2;refGene_gn=PBRM1;refGene_tx=XM_019822665.2;refGene_gn=PBRM1;refGene_tx=XM_019822680.2;refGene_gn=PBRM1;refGene_tx=XM_019822682.2;refGene_gn=PBRM1;refGene_tx=XM_019822686.2;refGene_gn=PBRM1;refGene_tx=XM_019822693.2;refGene_gn=PBRM1;refGene_tx=XM_019822696.2;refGene_gn=PBRM1;refGene_tx=XM_019822697.2;refGene_gn=PBRM1;refGene_tx=XM_019822700.2;refGene_gn=PBRM1;refGene_tx=XM_019822704.2;refGene_gn=PBRM1;refGene_tx=XM_019822708.2;refGene_gn=PBRM1;refGene_tx=XM_019822711.2;refGene_gn=PBRM1;refGene_tx=XM_019822713.2;refGene_gn=PBRM1;refGene_tx=XM_019822716.2;refGene_gn=PBRM1;refGene_tx=XM_019822718.2;refGene_gn=PBRM1;refGene_tx=XM_019822719.2;refGene_gn=PBRM1;refGene_tx=XM_019822722.2;refGene_gn=PBRM1;refGene_tx=XM_019822723.2;refGene_gn=PBRM1;refGene_tx=XM_019822726.2;refGene_gn=PBRM1;refGene_tx=XM_019822731.1;refGene_gn=PBRM1;refGene_tx=XM_023249537.1;refGene_gn=PBRM1;refGene_tx=XM_023249539.1;refGene_gn=PBRM1;refGene_tx=XM_023249543.1;refGene_gn=PBRM1;refGene_tx=XM_023249544.1;refGene_gn=PBRM1;refGene_tx=XM_023249570.1;refGene_gn=PBRM1;refGene_tx=XM_023249576.1;refGene_gn=PBRM1;refGene_tx=XM_023249582.1
A2 21146762 21146973
refGene_gn=PBRM1;refGene_tx=XM_003982314.5;refGene_gn=PBRM1;refGene_tx=XM_019822655.2;refGene_gn=PBRM1;refGene_tx=XM_019822656.2;refGene_gn=PBRM1;refGene_tx=XM_019822657.2;refGene_gn=PBRM1;refGene_tx=XM_019822659.2;refGene_gn=PBRM1;refGene_tx=XM_019822665.2;refGene_gn=PBRM1;refGene_tx=XM_019822680.2;refGene_gn=PBRM1;refGene_tx=XM_019822682.2;refGene_gn=PBRM1;refGene_tx=XM_019822686.2;refGene_gn=PBRM1;refGene_tx=XM_019822693.2;refGene_gn=PBRM1;refGene_tx=XM_019822696.2;refGene_gn=PBRM1;refGene_tx=XM_019822697.2;refGene_gn=PBRM1;refGene_tx=XM_019822700.2;refGene_gn=PBRM1;refGene_tx=XM_019822704.2;refGene_gn=PBRM1;refGene_tx=XM_019822708.2;refGene_gn=PBRM1;refGene_tx=XM_019822711.2;refGene_gn=PBRM1;refGene_tx=XM_019822713.2;refGene_gn=PBRM1;refGene_tx=XM_019822716.2;refGene_gn=PBRM1;refGene_tx=XM_019822718.2;refGene_gn=PBRM1;refGene_tx=XM_019822719.2;refGene_gn=PBRM1;refGene_tx=XM_019822722.2;refGene_gn=PBRM1;refGene_tx=XM_019822723.2;refGene_gn=PBRM1;refGene_tx=XM_019822726.2;refGene_gn=PBRM1;refGene_tx=XM_019822731.1;refGene_gn=PBRM1;refGene_tx=XM_023249537.1;refGene_gn=PBRM1;refGene_tx=XM_023249539.1;refGene_gn=PBRM1;refGene_tx=XM_023249543.1;refGene_gn=PBRM1;refGene_tx=XM_023249544.1;refGene_gn=PBRM1;refGene_tx=XM_023249570.1;refGene_gn=PBRM1;refGene_tx=XM_023249576.1;refGene_gn=PBRM1;refGene_tx=XM_023249582.1
A2 21147534 21147718
refGene_gn=PBRM1;refGene_tx=XM_003982314.5;refGene_gn=PBRM1;refGene_tx=XM_019822655.2;refGene_gn=PBRM1;refGene_tx=XM_019822656.2;refGene_gn=

PBRM1;refGene_tx=XM_019822657.2;refGene_gn=PBRM1;refGene_tx=XM_0198226
59.2;refGene_gn=PBRM1;refGene_tx=XM_019822665.2;refGene_gn=PBRM1;refGe
ne_tx=XM_019822680.2;refGene_gn=PBRM1;refGene_tx=XM_019822682.2;refGen
e_gn=PBRM1;refGene_tx=XM_019822686.2;refGene_gn=PBRM1;refGene_tx=XM_01
9822693.2;refGene_gn=PBRM1;refGene_tx=XM_019822696.2;refGene_gn=PBRM1;
refGene_tx=XM_019822697.2;refGene_gn=PBRM1;refGene_tx=XM_019822700.2;r
efGene_gn=PBRM1;refGene_tx=XM_019822704.2;refGene_gn=PBRM1;refGene_tx=
XM_019822708.2;refGene_gn=PBRM1;refGene_tx=XM_019822711.2;refGene_gn=P
BRM1;refGene_tx=XM_019822713.2;refGene_gn=PBRM1;refGene_tx=XM_01982271
6.2;refGene_gn=PBRM1;refGene_tx=XM_019822718.2;refGene_gn=PBRM1;refGen
e_tx=XM_019822719.2;refGene_gn=PBRM1;refGene_tx=XM_019822722.2;refGene
_gn=PBRM1;refGene_tx=XM_019822723.2;refGene_gn=PBRM1;refGene_tx=XM_019
822726.2;refGene_gn=PBRM1;refGene_tx=XM_019822731.1;refGene_gn=PBRM1;r
efGene_tx=XM_023249537.1;refGene_gn=PBRM1;refGene_tx=XM_023249539.1;re
fGene_gn=PBRM1;refGene_tx=XM_023249543.1;refGene_gn=PBRM1;refGene_tx=X
M_023249544.1;refGene_gn=PBRM1;refGene_tx=XM_023249570.1;refGene_gn=PB
RM1;refGene_tx=XM_023249576.1;refGene_gn=PBRM1;refGene_tx=XM_023249582
.1
A2 21157964 21158122
refGene_gn=PBRM1;refGene_tx=XM_003982314.5;refGene_gn=PBRM1;refGene_tx
=XM_019822655.2;refGene_gn=PBRM1;refGene_tx=XM_019822656.2;refGene_gn=
PBRM1;refGene_tx=XM_019822657.2;refGene_gn=PBRM1;refGene_tx=XM_0198226
59.2;refGene_gn=PBRM1;refGene_tx=XM_019822665.2;refGene_gn=PBRM1;refGe
ne_tx=XM_019822680.2;refGene_gn=PBRM1;refGene_tx=XM_019822682.2;refGen
e_gn=PBRM1;refGene_tx=XM_019822686.2;refGene_gn=PBRM1;refGene_tx=XM_01
9822693.2;refGene_gn=PBRM1;refGene_tx=XM_019822696.2;refGene_gn=PBRM1;
refGene_tx=XM_019822697.2;refGene_gn=PBRM1;refGene_tx=XM_019822700.2;r
efGene_gn=PBRM1;refGene_tx=XM_019822704.2;refGene_gn=PBRM1;refGene_tx=
XM_019822708.2;refGene_gn=PBRM1;refGene_tx=XM_019822711.2;refGene_gn=P
BRM1;refGene_tx=XM_019822713.2;refGene_gn=PBRM1;refGene_tx=XM_01982271
6.2;refGene_gn=PBRM1;refGene_tx=XM_019822718.2;refGene_gn=PBRM1;refGen
e_tx=XM_019822719.2;refGene_gn=PBRM1;refGene_tx=XM_019822722.2;refGene
_gn=PBRM1;refGene_tx=XM_019822723.2;refGene_gn=PBRM1;refGene_tx=XM_019
822726.2;refGene_gn=PBRM1;refGene_tx=XM_019822731.1;refGene_gn=PBRM1;r
efGene_tx=XM_023249537.1;refGene_gn=PBRM1;refGene_tx=XM_023249539.1;re
fGene_gn=PBRM1;refGene_tx=XM_023249543.1;refGene_gn=PBRM1;refGene_tx=X
M_023249544.1;refGene_gn=PBRM1;refGene_tx=XM_023249570.1;refGene_gn=PB
RM1;refGene_tx=XM_023249576.1;refGene_gn=PBRM1;refGene_tx=XM_023249582
.1
A2 21160731 21160877
refGene_gn=PBRM1;refGene_tx=XM_003982314.5;refGene_gn=PBRM1;refGene_tx
=XM_019822655.2;refGene_gn=PBRM1;refGene_tx=XM_019822656.2;refGene_gn=
PBRM1;refGene_tx=XM_019822657.2;refGene_gn=PBRM1;refGene_tx=XM_0198226
59.2;refGene_gn=PBRM1;refGene_tx=XM_019822665.2;refGene_gn=PBRM1;refGe
ne_tx=XM_019822680.2;refGene_gn=PBRM1;refGene_tx=XM_019822682.2;refGen
e_gn=PBRM1;refGene_tx=XM_019822686.2;refGene_gn=PBRM1;refGene_tx=XM_01
9822693.2;refGene_gn=PBRM1;refGene_tx=XM_019822696.2;refGene_gn=PBRM1;
refGene_tx=XM_019822697.2;refGene_gn=PBRM1;refGene_tx=XM_019822700.2;r
efGene_gn=PBRM1;refGene_tx=XM_019822704.2;refGene_gn=PBRM1;refGene_tx=
XM_019822708.2;refGene_gn=PBRM1;refGene_tx=XM_019822711.2;refGene_gn=P

BRM1;refGene_tx=XM_019822713.2;refGene_gn=PBRM1;refGene_tx=XM_01982271
6.2;refGene_gn=PBRM1;refGene_tx=XM_019822718.2;refGene_gn=PBRM1;refGen
e_tx=XM_019822719.2;refGene_gn=PBRM1;refGene_tx=XM_019822722.2;refGene
_gn=PBRM1;refGene_tx=XM_019822723.2;refGene_gn=PBRM1;refGene_tx=XM_019
822726.2;refGene_gn=PBRM1;refGene_tx=XM_019822731.1;refGene_gn=PBRM1;r
efGene_tx=XM_023249537.1;refGene_gn=PBRM1;refGene_tx=XM_023249539.1;re
fGene_gn=PBRM1;refGene_tx=XM_023249543.1;refGene_gn=PBRM1;refGene_tx=X
M_023249544.1;refGene_gn=PBRM1;refGene_tx=XM_023249570.1;refGene_gn=PB
RM1;refGene_tx=XM_023249576.1;refGene_gn=PBRM1;refGene_tx=XM_023249582
.1
A2 21165572 21165836
refGene_gn=PBRM1;refGene_tx=XM_003982314.5;refGene_gn=PBRM1;refGene_tx
=XM_019822655.2;refGene_gn=PBRM1;refGene_tx=XM_019822656.2;refGene_gn=
PBRM1;refGene_tx=XM_019822657.2;refGene_gn=PBRM1;refGene_tx=XM_0198226
59.2;refGene_gn=PBRM1;refGene_tx=XM_019822665.2;refGene_gn=PBRM1;refGe
ne_tx=XM_019822680.2;refGene_gn=PBRM1;refGene_tx=XM_019822682.2;refGen
e_gn=PBRM1;refGene_tx=XM_019822686.2;refGene_gn=PBRM1;refGene_tx=XM_01
9822693.2;refGene_gn=PBRM1;refGene_tx=XM_019822696.2;refGene_gn=PBRM1;
refGene_tx=XM_019822697.2;refGene_gn=PBRM1;refGene_tx=XM_019822700.2;r
efGene_gn=PBRM1;refGene_tx=XM_019822704.2;refGene_gn=PBRM1;refGene_tx=
XM_019822708.2;refGene_gn=PBRM1;refGene_tx=XM_019822711.2;refGene_gn=P
BRM1;refGene_tx=XM_019822713.2;refGene_gn=PBRM1;refGene_tx=XM_01982271
6.2;refGene_gn=PBRM1;refGene_tx=XM_019822718.2;refGene_gn=PBRM1;refGen
e_tx=XM_019822719.2;refGene_gn=PBRM1;refGene_tx=XM_019822722.2;refGene
_gn=PBRM1;refGene_tx=XM_019822723.2;refGene_gn=PBRM1;refGene_tx=XM_019
822726.2;refGene_gn=PBRM1;refGene_tx=XM_019822731.1;refGene_gn=PBRM1;r
efGene_tx=XM_023249537.1;refGene_gn=PBRM1;refGene_tx=XM_023249539.1;re
fGene_gn=PBRM1;refGene_tx=XM_023249543.1;refGene_gn=PBRM1;refGene_tx=X
M_023249544.1;refGene_gn=PBRM1;refGene_tx=XM_023249570.1;refGene_gn=PB
RM1;refGene_tx=XM_023249576.1;refGene_gn=PBRM1;refGene_tx=XM_023249582
.1
A2 21166725 21166883
refGene_gn=PBRM1;refGene_tx=XM_003982314.5;refGene_gn=PBRM1;refGene_tx
=XM_019822655.2;refGene_gn=PBRM1;refGene_tx=XM_019822656.2;refGene_gn=
PBRM1;refGene_tx=XM_019822657.2;refGene_gn=PBRM1;refGene_tx=XM_0198226
59.2;refGene_gn=PBRM1;refGene_tx=XM_019822665.2;refGene_gn=PBRM1;refGe
ne_tx=XM_019822680.2;refGene_gn=PBRM1;refGene_tx=XM_019822682.2;refGen
e_gn=PBRM1;refGene_tx=XM_019822686.2;refGene_gn=PBRM1;refGene_tx=XM_01
9822693.2;refGene_gn=PBRM1;refGene_tx=XM_019822696.2;refGene_gn=PBRM1;
refGene_tx=XM_019822697.2;refGene_gn=PBRM1;refGene_tx=XM_019822700.2;r
efGene_gn=PBRM1;refGene_tx=XM_019822704.2;refGene_gn=PBRM1;refGene_tx=
XM_019822708.2;refGene_gn=PBRM1;refGene_tx=XM_019822711.2;refGene_gn=P
BRM1;refGene_tx=XM_019822713.2;refGene_gn=PBRM1;refGene_tx=XM_01982271
6.2;refGene_gn=PBRM1;refGene_tx=XM_019822718.2;refGene_gn=PBRM1;refGen
e_tx=XM_019822719.2;refGene_gn=PBRM1;refGene_tx=XM_019822722.2;refGene
_gn=PBRM1;refGene_tx=XM_019822723.2;refGene_gn=PBRM1;refGene_tx=XM_019
822726.2;refGene_gn=PBRM1;refGene_tx=XM_019822731.1;refGene_gn=PBRM1;r
efGene_tx=XM_023249537.1;refGene_gn=PBRM1;refGene_tx=XM_023249539.1;re
fGene_gn=PBRM1;refGene_tx=XM_023249543.1;refGene_gn=PBRM1;refGene_tx=X
M_023249544.1;refGene_gn=PBRM1;refGene_tx=XM_023249570.1;refGene_gn=PB

RM1;refGene_tx=XM_023249576.1;refGene_gn=PBRM1;refGene_tx=XM_023249582
.1
A2 21167780 21167966
refGene_gn=PBRM1;refGene_tx=XM_003982314.5;refGene_gn=PBRM1;refGene_tx
=XM_019822655.2;refGene_gn=PBRM1;refGene_tx=XM_019822656.2;refGene_gn=
PBRM1;refGene_tx=XM_019822657.2;refGene_gn=PBRM1;refGene_tx=XM_0198226
59.2;refGene_gn=PBRM1;refGene_tx=XM_019822665.2;refGene_gn=PBRM1;refGe
ne_tx=XM_019822680.2;refGene_gn=PBRM1;refGene_tx=XM_019822682.2;refGen
e_gn=PBRM1;refGene_tx=XM_019822686.2;refGene_gn=PBRM1;refGene_tx=XM_01
9822693.2;refGene_gn=PBRM1;refGene_tx=XM_019822696.2;refGene_gn=PBRM1;
refGene_tx=XM_019822697.2;refGene_gn=PBRM1;refGene_tx=XM_019822700.2;r
efGene_gn=PBRM1;refGene_tx=XM_019822704.2;refGene_gn=PBRM1;refGene_tx=
XM_019822708.2;refGene_gn=PBRM1;refGene_tx=XM_019822711.2;refGene_gn=P
BRM1;refGene_tx=XM_019822713.2;refGene_gn=PBRM1;refGene_tx=XM_01982271
6.2;refGene_gn=PBRM1;refGene_tx=XM_019822718.2;refGene_gn=PBRM1;refGen
e_tx=XM_019822719.2;refGene_gn=PBRM1;refGene_tx=XM_019822722.2;refGene
_gn=PBRM1;refGene_tx=XM_019822723.2;refGene_gn=PBRM1;refGene_tx=XM_019
822726.2;refGene_gn=PBRM1;refGene_tx=XM_019822731.1;refGene_gn=PBRM1;r
efGene_tx=XM_023249537.1;refGene_gn=PBRM1;refGene_tx=XM_023249539.1;re
fGene_gn=PBRM1;refGene_tx=XM_023249543.1;refGene_gn=PBRM1;refGene_tx=X
M_023249544.1;refGene_gn=PBRM1;refGene_tx=XM_023249570.1;refGene_gn=PB
RM1;refGene_tx=XM_023249576.1;refGene_gn=PBRM1;refGene_tx=XM_023249582
.1
A2 21179811 21180023
refGene_gn=PBRM1;refGene_tx=XM_003982314.5;refGene_gn=PBRM1;refGene_tx
=XM_019822655.2;refGene_gn=PBRM1;refGene_tx=XM_019822656.2;refGene_gn=
PBRM1;refGene_tx=XM_019822657.2;refGene_gn=PBRM1;refGene_tx=XM_0198226
59.2;refGene_gn=PBRM1;refGene_tx=XM_019822665.2;refGene_gn=PBRM1;refGe
ne_tx=XM_019822680.2;refGene_gn=PBRM1;refGene_tx=XM_019822682.2;refGen
e_gn=PBRM1;refGene_tx=XM_019822686.2;refGene_gn=PBRM1;refGene_tx=XM_01
9822693.2;refGene_gn=PBRM1;refGene_tx=XM_019822696.2;refGene_gn=PBRM1;
refGene_tx=XM_019822697.2;refGene_gn=PBRM1;refGene_tx=XM_019822700.2;r
efGene_gn=PBRM1;refGene_tx=XM_019822704.2;refGene_gn=PBRM1;refGene_tx=
XM_019822708.2;refGene_gn=PBRM1;refGene_tx=XM_019822711.2;refGene_gn=P
BRM1;refGene_tx=XM_019822713.2;refGene_gn=PBRM1;refGene_tx=XM_01982271
6.2;refGene_gn=PBRM1;refGene_tx=XM_019822718.2;refGene_gn=PBRM1;refGen
e_tx=XM_019822719.2;refGene_gn=PBRM1;refGene_tx=XM_019822722.2;refGene
_gn=PBRM1;refGene_tx=XM_019822723.2;refGene_gn=PBRM1;refGene_tx=XM_019
822726.2;refGene_gn=PBRM1;refGene_tx=XM_019822731.1;refGene_gn=PBRM1;r
efGene_tx=XM_023249537.1;refGene_gn=PBRM1;refGene_tx=XM_023249539.1;re
fGene_gn=PBRM1;refGene_tx=XM_023249543.1;refGene_gn=PBRM1;refGene_tx=X
M_023249544.1;refGene_gn=PBRM1;refGene_tx=XM_023249570.1;refGene_gn=PB
RM1;refGene_tx=XM_023249576.1;refGene_gn=PBRM1;refGene_tx=XM_023249582
.1
A2 21184194 21184837
refGene_gn=PBRM1;refGene_tx=XM_003982314.5;refGene_gn=PBRM1;refGene_tx
=XM_019822655.2;refGene_gn=PBRM1;refGene_tx=XM_019822656.2;refGene_gn=
PBRM1;refGene_tx=XM_019822657.2;refGene_gn=PBRM1;refGene_tx=XM_0198226
59.2;refGene_gn=PBRM1;refGene_tx=XM_019822665.2;refGene_gn=PBRM1;refGe
ne_tx=XM_019822680.2;refGene_gn=PBRM1;refGene_tx=XM_019822682.2;refGen

e_gn=PBRM1;refGene_tx=XM_019822686.2;refGene_gn=PBRM1;refGene_tx=XM_01
9822693.2;refGene_gn=PBRM1;refGene_tx=XM_019822696.2;refGene_gn=PBRM1;
refGene_tx=XM_019822697.2;refGene_gn=PBRM1;refGene_tx=XM_019822700.2;r
efGene_gn=PBRM1;refGene_tx=XM_019822704.2;refGene_gn=PBRM1;refGene_tx=
XM_019822708.2;refGene_gn=PBRM1;refGene_tx=XM_019822711.2;refGene_gn=P
BRM1;refGene_tx=XM_019822713.2;refGene_gn=PBRM1;refGene_tx=XM_01982271
6.2;refGene_gn=PBRM1;refGene_tx=XM_019822718.2;refGene_gn=PBRM1;refGen
e_tx=XM_019822719.2;refGene_gn=PBRM1;refGene_tx=XM_019822722.2;refGene
_gn=PBRM1;refGene_tx=XM_019822723.2;refGene_gn=PBRM1;refGene_tx=XM_019
822726.2;refGene_gn=PBRM1;refGene_tx=XM_019822731.1;refGene_gn=PBRM1;r
efGene_tx=XM_023249537.1;refGene_gn=PBRM1;refGene_tx=XM_023249539.1;re
fGene_gn=PBRM1;refGene_tx=XM_023249543.1;refGene_gn=PBRM1;refGene_tx=X
M_023249544.1;refGene_gn=PBRM1;refGene_tx=XM_023249570.1;refGene_gn=PB
RM1;refGene_tx=XM_023249576.1;refGene_gn=PBRM1;refGene_tx=XM_023249582
.1
A2 21188921 21189027
refGene_gn=PBRM1;refGene_tx=XM_003982314.5;refGene_gn=PBRM1;refGene_tx
=XM_019822655.2;refGene_gn=PBRM1;refGene_tx=XM_019822656.2;refGene_gn=
PBRM1;refGene_tx=XM_019822657.2;refGene_gn=PBRM1;refGene_tx=XM_0198226
59.2;refGene_gn=PBRM1;refGene_tx=XM_019822665.2;refGene_gn=PBRM1;refGe
ne_tx=XM_019822680.2;refGene_gn=PBRM1;refGene_tx=XM_019822682.2;refGen
e_gn=PBRM1;refGene_tx=XM_019822686.2;refGene_gn=PBRM1;refGene_tx=XM_01
9822693.2;refGene_gn=PBRM1;refGene_tx=XM_019822696.2;refGene_gn=PBRM1;
refGene_tx=XM_019822697.2;refGene_gn=PBRM1;refGene_tx=XM_019822700.2;r
efGene_gn=PBRM1;refGene_tx=XM_019822704.2;refGene_gn=PBRM1;refGene_tx=
XM_019822708.2;refGene_gn=PBRM1;refGene_tx=XM_019822711.2;refGene_gn=P
BRM1;refGene_tx=XM_019822713.2;refGene_gn=PBRM1;refGene_tx=XM_01982271
6.2;refGene_gn=PBRM1;refGene_tx=XM_019822718.2;refGene_gn=PBRM1;refGen
e_tx=XM_019822719.2;refGene_gn=PBRM1;refGene_tx=XM_019822722.2;refGene
_gn=PBRM1;refGene_tx=XM_019822723.2;refGene_gn=PBRM1;refGene_tx=XM_019
822726.2;refGene_gn=PBRM1;refGene_tx=XM_019822731.1;refGene_gn=PBRM1;r
efGene_tx=XM_023249537.1;refGene_gn=PBRM1;refGene_tx=XM_023249539.1;re
fGene_gn=PBRM1;refGene_tx=XM_023249543.1;refGene_gn=PBRM1;refGene_tx=X
M_023249544.1;refGene_gn=PBRM1;refGene_tx=XM_023249570.1;refGene_gn=PB
RM1;refGene_tx=XM_023249576.1;refGene_gn=PBRM1;refGene_tx=XM_023249582
.1
A2 21191164 21191441
refGene_gn=PBRM1;refGene_tx=XM_003982314.5;refGene_gn=PBRM1;refGene_tx
=XM_019822655.2;refGene_gn=PBRM1;refGene_tx=XM_019822656.2;refGene_gn=
PBRM1;refGene_tx=XM_019822657.2;refGene_gn=PBRM1;refGene_tx=XM_0198226
59.2;refGene_gn=PBRM1;refGene_tx=XM_019822665.2;refGene_gn=PBRM1;refGe
ne_tx=XM_019822680.2;refGene_gn=PBRM1;refGene_tx=XM_019822682.2;refGen
e_gn=PBRM1;refGene_tx=XM_019822686.2;refGene_gn=PBRM1;refGene_tx=XM_01
9822693.2;refGene_gn=PBRM1;refGene_tx=XM_019822696.2;refGene_gn=PBRM1;
refGene_tx=XM_019822697.2;refGene_gn=PBRM1;refGene_tx=XM_019822700.2;r
efGene_gn=PBRM1;refGene_tx=XM_019822704.2;refGene_gn=PBRM1;refGene_tx=
XM_019822708.2;refGene_gn=PBRM1;refGene_tx=XM_019822711.2;refGene_gn=P
BRM1;refGene_tx=XM_019822713.2;refGene_gn=PBRM1;refGene_tx=XM_01982271
6.2;refGene_gn=PBRM1;refGene_tx=XM_019822718.2;refGene_gn=PBRM1;refGen
e_tx=XM_019822719.2;refGene_gn=PBRM1;refGene_tx=XM_019822722.2;refGene

_gn=PBRM1;refGene_tx=XM_019822723.2;refGene_gn=PBRM1;refGene_tx=XM_019822726.2;refGene_gn=PBRM1;refGene_tx=XM_019822731.1;refGene_gn=PBRM1;refGene_tx=XM_023249537.1;refGene_gn=PBRM1;refGene_tx=XM_023249539.1;refGene_gn=PBRM1;refGene_tx=XM_023249543.1;refGene_gn=PBRM1;refGene_tx=XM_023249544.1;refGene_gn=PBRM1;refGene_tx=XM_023249570.1;refGene_gn=PBRM1;refGene_tx=XM_023249576.1;refGene_gn=PBRM1;refGene_tx=XM_023249582.1
A2 21198672 21198717
refGene_gn=PBRM1;refGene_tx=XM_003982314.5;refGene_gn=PBRM1;refGene_tx=XM_019822655.2;refGene_gn=PBRM1;refGene_tx=XM_019822656.2;refGene_gn=PBRM1;refGene_tx=XM_019822657.2;refGene_gn=PBRM1;refGene_tx=XM_019822659.2;refGene_gn=PBRM1;refGene_tx=XM_019822665.2;refGene_gn=PBRM1;refGene_tx=XM_019822682.2;refGene_gn=PBRM1;refGene_tx=XM_019822686.2;refGene_gn=PBRM1;refGene_tx=XM_019822693.2;refGene_gn=PBRM1;refGene_tx=XM_019822696.2;refGene_gn=PBRM1;refGene_tx=XM_019822697.2;refGene_gn=PBRM1;refGene_tx=XM_019822704.2;refGene_gn=PBRM1;refGene_tx=XM_019822708.2;refGene_gn=PBRM1;refGene_tx=XM_019822713.2;refGene_gn=PBRM1;refGene_tx=XM_019822716.2;refGene_gn=PBRM1;refGene_tx=XM_019822719.2;refGene_gn=PBRM1;refGene_tx=XM_019822726.2;refGene_gn=PBRM1;refGene_tx=XM_023249537.1;refGene_gn=PBRM1;refGene_tx=XM_023249539.1;refGene_gn=PBRM1;refGene_tx=XM_023249543.1;refGene_gn=PBRM1;refGene_tx=XM_023249544.1;refGene_gn=PBRM1;refGene_tx=XM_023249570.1;refGene_gn=PBRM1;refGene_tx=XM_023249576.1
A2 21200848 21200946
refGene_gn=PBRM1;refGene_tx=XM_003982314.5;refGene_gn=PBRM1;refGene_tx=XM_019822655.2;refGene_gn=PBRM1;refGene_tx=XM_019822656.2;refGene_gn=PBRM1;refGene_tx=XM_019822657.2;refGene_gn=PBRM1;refGene_tx=XM_019822659.2;refGene_gn=PBRM1;refGene_tx=XM_019822665.2;refGene_gn=PBRM1;refGene_tx=XM_019822680.2;refGene_gn=PBRM1;refGene_tx=XM_019822682.2;refGene_gn=PBRM1;refGene_tx=XM_019822686.2;refGene_gn=PBRM1;refGene_tx=XM_019822693.2;refGene_gn=PBRM1;refGene_tx=XM_019822696.2;refGene_gn=PBRM1;refGene_tx=XM_019822697.2;refGene_gn=PBRM1;refGene_tx=XM_019822700.2;refGene_gn=PBRM1;refGene_tx=XM_019822704.2;refGene_gn=PBRM1;refGene_tx=XM_019822708.2;refGene_gn=PBRM1;refGene_tx=XM_019822711.2;refGene_gn=PBRM1;refGene_tx=XM_019822713.2;refGene_gn=PBRM1;refGene_tx=XM_019822716.2;refGene_gn=PBRM1;refGene_tx=XM_019822718.2;refGene_gn=PBRM1;refGene_tx=XM_019822719.2;refGene_gn=PBRM1;refGene_tx=XM_019822722.2;refGene_gn=PBRM1;refGene_tx=XM_019822723.2;refGene_gn=PBRM1;refGene_tx=XM_019822726.2;refGene_gn=PBRM1;refGene_tx=XM_019822731.1;refGene_gn=PBRM1;refGene_tx=XM_023249537.1;refGene_gn=PBRM1;refGene_tx=XM_023249539.1;refGene_gn=PBRM1;refGene_tx=XM_023249543.1;refGene_gn=PBRM1;refGene_tx=XM_023249544.1;refGene_gn=PBRM1;refGene_tx=XM_023249570.1;refGene_gn=PBRM1;refGene_tx=XM_023249576.1;refGene_gn=PBRM1;refGene_tx=XM_023249582.1
A2 21202059 21202201
refGene_gn=PBRM1;refGene_tx=XM_003982314.5;refGene_gn=PBRM1;refGene_tx=XM_019822655.2;refGene_gn=PBRM1;refGene_tx=XM_019822656.2;refGene_gn=PBRM1;refGene_tx=XM_019822657.2;refGene_gn=PBRM1;refGene_tx=XM_019822659.2;refGene_gn=PBRM1;refGene_tx=XM_019822665.2;refGene_gn=PBRM1;refGene_tx=XM_019822680.2;refGene_gn=PBRM1;refGene_tx=XM_019822682.2;refGen

e_gn=PBRM1;refGene_tx=XM_019822686.2;refGene_gn=PBRM1;refGene_tx=XM_01
9822693.2;refGene_gn=PBRM1;refGene_tx=XM_019822696.2;refGene_gn=PBRM1;
refGene_tx=XM_019822697.2;refGene_gn=PBRM1;refGene_tx=XM_019822700.2;r
efGene_gn=PBRM1;refGene_tx=XM_019822704.2;refGene_gn=PBRM1;refGene_tx=
XM_019822708.2;refGene_gn=PBRM1;refGene_tx=XM_019822711.2;refGene_gn=P
BRM1;refGene_tx=XM_019822713.2;refGene_gn=PBRM1;refGene_tx=XM_01982271
6.2;refGene_gn=PBRM1;refGene_tx=XM_019822718.2;refGene_gn=PBRM1;refGen
e_tx=XM_019822719.2;refGene_gn=PBRM1;refGene_tx=XM_019822722.2;refGene
_gn=PBRM1;refGene_tx=XM_019822723.2;refGene_gn=PBRM1;refGene_tx=XM_019
822726.2;refGene_gn=PBRM1;refGene_tx=XM_019822731.1;refGene_gn=PBRM1;r
efGene_tx=XM_023249537.1;refGene_gn=PBRM1;refGene_tx=XM_023249539.1;re
fGene_gn=PBRM1;refGene_tx=XM_023249543.1;refGene_gn=PBRM1;refGene_tx=X
M_023249544.1;refGene_gn=PBRM1;refGene_tx=XM_023249570.1;refGene_gn=PB
RM1;refGene_tx=XM_023249576.1;refGene_gn=PBRM1;refGene_tx=XM_023249582
.1
A2 21205592 21205806
refGene_gn=PBRM1;refGene_tx=XM_003982314.5;refGene_gn=PBRM1;refGene_tx
=XM_019822655.2;refGene_gn=PBRM1;refGene_tx=XM_019822656.2;refGene_gn=
PBRM1;refGene_tx=XM_019822657.2;refGene_gn=PBRM1;refGene_tx=XM_0198226
59.2;refGene_gn=PBRM1;refGene_tx=XM_019822665.2;refGene_gn=PBRM1;refGe
ne_tx=XM_019822680.2;refGene_gn=PBRM1;refGene_tx=XM_019822682.2;refGen
e_gn=PBRM1;refGene_tx=XM_019822686.2;refGene_gn=PBRM1;refGene_tx=XM_01
9822693.2;refGene_gn=PBRM1;refGene_tx=XM_019822696.2;refGene_gn=PBRM1;
refGene_tx=XM_019822697.2;refGene_gn=PBRM1;refGene_tx=XM_019822700.2;r
efGene_gn=PBRM1;refGene_tx=XM_019822704.2;refGene_gn=PBRM1;refGene_tx=
XM_019822708.2;refGene_gn=PBRM1;refGene_tx=XM_019822711.2;refGene_gn=P
BRM1;refGene_tx=XM_019822713.2;refGene_gn=PBRM1;refGene_tx=XM_01982271
6.2;refGene_gn=PBRM1;refGene_tx=XM_019822718.2;refGene_gn=PBRM1;refGen
e_tx=XM_019822719.2;refGene_gn=PBRM1;refGene_tx=XM_019822722.2;refGene
_gn=PBRM1;refGene_tx=XM_019822723.2;refGene_gn=PBRM1;refGene_tx=XM_019
822726.2;refGene_gn=PBRM1;refGene_tx=XM_019822731.1;refGene_gn=PBRM1;r
efGene_tx=XM_023249537.1;refGene_gn=PBRM1;refGene_tx=XM_023249539.1;re
fGene_gn=PBRM1;refGene_tx=XM_023249543.1;refGene_gn=PBRM1;refGene_tx=X
M_023249544.1;refGene_gn=PBRM1;refGene_tx=XM_023249570.1;refGene_gn=PB
RM1;refGene_tx=XM_023249576.1;refGene_gn=PBRM1;refGene_tx=XM_023249582
.1
A2 21209337 21209426
refGene_gn=PBRM1;refGene_tx=XM_003982314.5;refGene_gn=PBRM1;refGene_tx
=XM_019822655.2;refGene_gn=PBRM1;refGene_tx=XM_019822656.2;refGene_gn=
PBRM1;refGene_tx=XM_019822657.2;refGene_gn=PBRM1;refGene_tx=XM_0198226
59.2;refGene_gn=PBRM1;refGene_tx=XM_019822665.2;refGene_gn=PBRM1;refGe
ne_tx=XM_019822680.2;refGene_gn=PBRM1;refGene_tx=XM_019822682.2;refGen
e_gn=PBRM1;refGene_tx=XM_019822686.2;refGene_gn=PBRM1;refGene_tx=XM_01
9822693.2;refGene_gn=PBRM1;refGene_tx=XM_019822696.2;refGene_gn=PBRM1;
refGene_tx=XM_019822697.2;refGene_gn=PBRM1;refGene_tx=XM_019822700.2;r
efGene_gn=PBRM1;refGene_tx=XM_019822704.2;refGene_gn=PBRM1;refGene_tx=
XM_019822708.2;refGene_gn=PBRM1;refGene_tx=XM_019822711.2;refGene_gn=P
BRM1;refGene_tx=XM_019822713.2;refGene_gn=PBRM1;refGene_tx=XM_01982271
6.2;refGene_gn=PBRM1;refGene_tx=XM_019822718.2;refGene_gn=PBRM1;refGen
e_tx=XM_019822719.2;refGene_gn=PBRM1;refGene_tx=XM_019822722.2;refGene

_gn=PBRM1;refGene_tx=XM_019822723.2;refGene_gn=PBRM1;refGene_tx=XM_019822726.2;refGene_gn=PBRM1;refGene_tx=XM_019822731.1;refGene_gn=PBRM1;refGene_tx=XM_023249537.1;refGene_gn=PBRM1;refGene_tx=XM_023249539.1;refGene_gn=PBRM1;refGene_tx=XM_023249543.1;refGene_gn=PBRM1;refGene_tx=XM_023249544.1;refGene_gn=PBRM1;refGene_tx=XM_023249570.1;refGene_gn=PBRM1;refGene_tx=XM_023249576.1;refGene_gn=PBRM1;refGene_tx=XM_023249582.1
A2 21210055 21210151
refGene_gn=PBRM1;refGene_tx=XM_003982314.5;refGene_gn=PBRM1;refGene_tx=XM_019822655.2;refGene_gn=PBRM1;refGene_tx=XM_019822656.2;refGene_gn=PBRM1;refGene_tx=XM_019822657.2;refGene_gn=PBRM1;refGene_tx=XM_019822659.2;refGene_gn=PBRM1;refGene_tx=XM_019822665.2;refGene_gn=PBRM1;refGene_tx=XM_019822680.2;refGene_gn=PBRM1;refGene_tx=XM_019822682.2;refGene_gn=PBRM1;refGene_tx=XM_019822686.2;refGene_gn=PBRM1;refGene_tx=XM_019822693.2;refGene_gn=PBRM1;refGene_tx=XM_019822696.2;refGene_gn=PBRM1;refGene_tx=XM_019822697.2;refGene_gn=PBRM1;refGene_tx=XM_019822700.2;refGene_gn=PBRM1;refGene_tx=XM_019822708.2;refGene_gn=PBRM1;refGene_tx=XM_019822711.2;refGene_gn=PBRM1;refGene_tx=XM_019822713.2;refGene_gn=PBRM1;refGene_tx=XM_019822716.2;refGene_gn=PBRM1;refGene_tx=XM_019822718.2;refGene_gn=PBRM1;refGene_tx=XM_019822719.2;refGene_gn=PBRM1;refGene_tx=XM_019822722.2;refGene_gn=PBRM1;refGene_tx=XM_019822726.2;refGene_gn=PBRM1;refGene_tx=XM_019822731.1;refGene_gn=PBRM1;refGene_tx=XM_023249537.1;refGene_gn=PBRM1;refGene_tx=XM_023249539.1;refGene_gn=PBRM1;refGene_tx=XM_023249543.1;refGene_gn=PBRM1;refGene_tx=XM_023249544.1;refGene_gn=PBRM1;refGene_tx=XM_023249570.1;refGene_gn=PBRM1;refGene_tx=XM_023249576.1;refGene_gn=PBRM1;refGene_tx=XM_023249582.1
A2 21210903 21210989
refGene_gn=PBRM1;refGene_tx=XM_003982314.5;refGene_gn=PBRM1;refGene_tx=XM_019822655.2;refGene_gn=PBRM1;refGene_tx=XM_019822656.2;refGene_gn=PBRM1;refGene_tx=XM_019822657.2;refGene_gn=PBRM1;refGene_tx=XM_019822659.2;refGene_gn=PBRM1;refGene_tx=XM_019822665.2;refGene_gn=PBRM1;refGene_tx=XM_019822680.2;refGene_gn=PBRM1;refGene_tx=XM_019822682.2;refGene_gn=PBRM1;refGene_tx=XM_019822686.2;refGene_gn=PBRM1;refGene_tx=XM_019822693.2;refGene_gn=PBRM1;refGene_tx=XM_019822696.2;refGene_gn=PBRM1;refGene_tx=XM_019822697.2;refGene_gn=PBRM1;refGene_tx=XM_019822700.2;refGene_gn=PBRM1;refGene_tx=XM_019822704.2;refGene_gn=PBRM1;refGene_tx=XM_019822708.2;refGene_gn=PBRM1;refGene_tx=XM_019822711.2;refGene_gn=PBRM1;refGene_tx=XM_019822713.2;refGene_gn=PBRM1;refGene_tx=XM_019822716.2;refGene_gn=PBRM1;refGene_tx=XM_019822718.2;refGene_gn=PBRM1;refGene_tx=XM_019822719.2;refGene_gn=PBRM1;refGene_tx=XM_019822722.2;refGene_gn=PBRM1;refGene_tx=XM_019822723.2;refGene_gn=PBRM1;refGene_tx=XM_019822726.2;refGene_gn=PBRM1;refGene_tx=XM_019822731.1;refGene_gn=PBRM1;refGene_tx=XM_023249537.1;refGene_gn=PBRM1;refGene_tx=XM_023249539.1;refGene_gn=PBRM1;refGene_tx=XM_023249543.1;refGene_gn=PBRM1;refGene_tx=XM_023249544.1;refGene_gn=PBRM1;refGene_tx=XM_023249570.1;refGene_gn=PBRM1;refGene_tx=XM_023249576.1;refGene_gn=PBRM1;refGene_tx=XM_023249582.1
A2 21213707 21213806
refGene_gn=PBRM1;refGene_tx=XM_003982314.5;refGene_gn=PBRM1;refGene_tx=XM_019822655.2;refGene_gn=PBRM1;refGene_tx=XM_019822656.2;refGene_gn=

PBRM1;refGene_tx=XM_019822657.2;refGene_gn=PBRM1;refGene_tx=XM_0198226
59.2;refGene_gn=PBRM1;refGene_tx=XM_019822665.2;refGene_gn=PBRM1;refGe
ne_tx=XM_019822680.2;refGene_gn=PBRM1;refGene_tx=XM_019822682.2;refGen
e_gn=PBRM1;refGene_tx=XM_019822686.2;refGene_gn=PBRM1;refGene_tx=XM_01
9822693.2;refGene_gn=PBRM1;refGene_tx=XM_019822696.2;refGene_gn=PBRM1;
refGene_tx=XM_019822697.2;refGene_gn=PBRM1;refGene_tx=XM_019822700.2;r
efGene_gn=PBRM1;refGene_tx=XM_019822704.2;refGene_gn=PBRM1;refGene_tx=
XM_019822708.2;refGene_gn=PBRM1;refGene_tx=XM_019822711.2;refGene_gn=P
BRM1;refGene_tx=XM_019822713.2;refGene_gn=PBRM1;refGene_tx=XM_01982271
6.2;refGene_gn=PBRM1;refGene_tx=XM_019822718.2;refGene_gn=PBRM1;refGen
e_tx=XM_019822719.2;refGene_gn=PBRM1;refGene_tx=XM_019822722.2;refGene
_gn=PBRM1;refGene_tx=XM_019822723.2;refGene_gn=PBRM1;refGene_tx=XM_019
822726.2;refGene_gn=PBRM1;refGene_tx=XM_019822731.1;refGene_gn=PBRM1;r
efGene_tx=XM_023249537.1;refGene_gn=PBRM1;refGene_tx=XM_023249539.1;re
fGene_gn=PBRM1;refGene_tx=XM_023249543.1;refGene_gn=PBRM1;refGene_tx=X
M_023249544.1;refGene_gn=PBRM1;refGene_tx=XM_023249570.1;refGene_gn=PB
RM1;refGene_tx=XM_023249576.1;refGene_gn=PBRM1;refGene_tx=XM_023249582
.1
A2 21217345 21217414
refGene_gn=PBRM1;refGene_tx=XM_003982314.5;refGene_gn=PBRM1;refGene_tx
=XM_019822655.2;refGene_gn=PBRM1;refGene_tx=XM_019822656.2;refGene_gn=
PBRM1;refGene_tx=XM_019822657.2;refGene_gn=PBRM1;refGene_tx=XM_0198226
59.2;refGene_gn=PBRM1;refGene_tx=XM_019822665.2;refGene_gn=PBRM1;refGe
ne_tx=XM_019822680.2;refGene_gn=PBRM1;refGene_tx=XM_019822682.2;refGen
e_gn=PBRM1;refGene_tx=XM_019822686.2;refGene_gn=PBRM1;refGene_tx=XM_01
9822693.2;refGene_gn=PBRM1;refGene_tx=XM_019822696.2;refGene_gn=PBRM1;
refGene_tx=XM_019822697.2;refGene_gn=PBRM1;refGene_tx=XM_019822700.2;r
efGene_gn=PBRM1;refGene_tx=XM_019822704.2;refGene_gn=PBRM1;refGene_tx=
XM_019822708.2;refGene_gn=PBRM1;refGene_tx=XM_019822711.2;refGene_gn=P
BRM1;refGene_tx=XM_019822713.2;refGene_gn=PBRM1;refGene_tx=XM_01982271
6.2;refGene_gn=PBRM1;refGene_tx=XM_019822718.2;refGene_gn=PBRM1;refGen
e_tx=XM_019822719.2;refGene_gn=PBRM1;refGene_tx=XM_019822722.2;refGene
_gn=PBRM1;refGene_tx=XM_019822723.2;refGene_gn=PBRM1;refGene_tx=XM_019
822726.2;refGene_gn=PBRM1;refGene_tx=XM_019822731.1;refGene_gn=PBRM1;r
efGene_tx=XM_023249537.1;refGene_gn=PBRM1;refGene_tx=XM_023249539.1;re
fGene_gn=PBRM1;refGene_tx=XM_023249543.1;refGene_gn=PBRM1;refGene_tx=X
M_023249544.1;refGene_gn=PBRM1;refGene_tx=XM_023249570.1;refGene_gn=PB
RM1;refGene_tx=XM_023249576.1;refGene_gn=PBRM1;refGene_tx=XM_023249582
.1
A2 21221185 21221344
refGene_gn=PBRM1;refGene_tx=XM_003982314.5;refGene_gn=PBRM1;refGene_tx
=XM_019822655.2;refGene_gn=PBRM1;refGene_tx=XM_019822656.2;refGene_gn=
PBRM1;refGene_tx=XM_019822657.2;refGene_gn=PBRM1;refGene_tx=XM_0198226
59.2;refGene_gn=PBRM1;refGene_tx=XM_019822665.2;refGene_gn=PBRM1;refGe
ne_tx=XM_019822680.2;refGene_gn=PBRM1;refGene_tx=XM_019822682.2;refGen
e_gn=PBRM1;refGene_tx=XM_019822686.2;refGene_gn=PBRM1;refGene_tx=XM_01
9822693.2;refGene_gn=PBRM1;refGene_tx=XM_019822696.2;refGene_gn=PBRM1;
refGene_tx=XM_019822697.2;refGene_gn=PBRM1;refGene_tx=XM_019822700.2;r
efGene_gn=PBRM1;refGene_tx=XM_019822704.2;refGene_gn=PBRM1;refGene_tx=
XM_019822708.2;refGene_gn=PBRM1;refGene_tx=XM_019822711.2;refGene_gn=P

BRM1;refGene_tx=XM_019822713.2;refGene_gn=PBRM1;refGene_tx=XM_01982271
6.2;refGene_gn=PBRM1;refGene_tx=XM_019822718.2;refGene_gn=PBRM1;refGen
e_tx=XM_019822719.2;refGene_gn=PBRM1;refGene_tx=XM_019822722.2;refGene
_gn=PBRM1;refGene_tx=XM_019822723.2;refGene_gn=PBRM1;refGene_tx=XM_019
822726.2;refGene_gn=PBRM1;refGene_tx=XM_019822731.1;refGene_gn=PBRM1;r
efGene_tx=XM_023249537.1;refGene_gn=PBRM1;refGene_tx=XM_023249539.1;re
fGene_gn=PBRM1;refGene_tx=XM_023249543.1;refGene_gn=PBRM1;refGene_tx=X
M_023249544.1;refGene_gn=PBRM1;refGene_tx=XM_023249570.1;refGene_gn=PB
RM1;refGene_tx=XM_023249576.1;refGene_gn=PBRM1;refGene_tx=XM_023249582
.1
A2 21224509 21224653
refGene_gn=PBRM1;refGene_tx=XM_003982314.5;refGene_gn=PBRM1;refGene_tx
=XM_019822655.2;refGene_gn=PBRM1;refGene_tx=XM_019822656.2;refGene_gn=
PBRM1;refGene_tx=XM_019822657.2;refGene_gn=PBRM1;refGene_tx=XM_0198226
59.2;refGene_gn=PBRM1;refGene_tx=XM_019822665.2;refGene_gn=PBRM1;refGe
ne_tx=XM_019822680.2;refGene_gn=PBRM1;refGene_tx=XM_019822682.2;refGen
e_gn=PBRM1;refGene_tx=XM_019822686.2;refGene_gn=PBRM1;refGene_tx=XM_01
9822693.2;refGene_gn=PBRM1;refGene_tx=XM_019822696.2;refGene_gn=PBRM1;
refGene_tx=XM_019822697.2;refGene_gn=PBRM1;refGene_tx=XM_019822700.2;r
efGene_gn=PBRM1;refGene_tx=XM_019822704.2;refGene_gn=PBRM1;refGene_tx=
XM_019822708.2;refGene_gn=PBRM1;refGene_tx=XM_019822711.2;refGene_gn=P
BRM1;refGene_tx=XM_019822713.2;refGene_gn=PBRM1;refGene_tx=XM_01982271
6.2;refGene_gn=PBRM1;refGene_tx=XM_019822718.2;refGene_gn=PBRM1;refGen
e_tx=XM_019822719.2;refGene_gn=PBRM1;refGene_tx=XM_019822722.2;refGene
_gn=PBRM1;refGene_tx=XM_019822723.2;refGene_gn=PBRM1;refGene_tx=XM_019
822726.2;refGene_gn=PBRM1;refGene_tx=XM_019822731.1;refGene_gn=PBRM1;r
efGene_tx=XM_023249537.1;refGene_gn=PBRM1;refGene_tx=XM_023249539.1;re
fGene_gn=PBRM1;refGene_tx=XM_023249543.1;refGene_gn=PBRM1;refGene_tx=X
M_023249544.1;refGene_gn=PBRM1;refGene_tx=XM_023249570.1;refGene_gn=PB
RM1;refGene_tx=XM_023249576.1;refGene_gn=PBRM1;refGene_tx=XM_023249582
.1
A2 21232100 21232248
refGene_gn=PBRM1;refGene_tx=XM_003982314.5;refGene_gn=PBRM1;refGene_tx
=XM_019822655.2;refGene_gn=PBRM1;refGene_tx=XM_019822656.2;refGene_gn=
PBRM1;refGene_tx=XM_019822657.2;refGene_gn=PBRM1;refGene_tx=XM_0198226
59.2;refGene_gn=PBRM1;refGene_tx=XM_019822665.2;refGene_gn=PBRM1;refGe
ne_tx=XM_019822680.2;refGene_gn=PBRM1;refGene_tx=XM_019822682.2;refGen
e_gn=PBRM1;refGene_tx=XM_019822686.2;refGene_gn=PBRM1;refGene_tx=XM_01
9822693.2;refGene_gn=PBRM1;refGene_tx=XM_019822696.2;refGene_gn=PBRM1;
refGene_tx=XM_019822697.2;refGene_gn=PBRM1;refGene_tx=XM_019822700.2;r
efGene_gn=PBRM1;refGene_tx=XM_019822704.2;refGene_gn=PBRM1;refGene_tx=
XM_019822708.2;refGene_gn=PBRM1;refGene_tx=XM_019822711.2;refGene_gn=P
BRM1;refGene_tx=XM_019822713.2;refGene_gn=PBRM1;refGene_tx=XM_01982271
6.2;refGene_gn=PBRM1;refGene_tx=XM_019822718.2;refGene_gn=PBRM1;refGen
e_tx=XM_019822719.2;refGene_gn=PBRM1;refGene_tx=XM_019822722.2;refGene
_gn=PBRM1;refGene_tx=XM_019822723.2;refGene_gn=PBRM1;refGene_tx=XM_019
822726.2;refGene_gn=PBRM1;refGene_tx=XM_019822731.1;refGene_gn=PBRM1;r
efGene_tx=XM_023249537.1;refGene_gn=PBRM1;refGene_tx=XM_023249539.1;re
fGene_gn=PBRM1;refGene_tx=XM_023249543.1;refGene_gn=PBRM1;refGene_tx=X
M_023249544.1;refGene_gn=PBRM1;refGene_tx=XM_023249570.1;refGene_gn=PB

RM1;refGene_tx=XM_023249576.1;refGene_gn=PBRM1;refGene_tx=XM_023249582
.1
A2 21241526 21241624
refGene_gn=PBRM1;refGene_tx=XM_003982314.5;refGene_gn=PBRM1;refGene_tx
=XM_019822655.2;refGene_gn=PBRM1;refGene_tx=XM_019822656.2;refGene_gn=
PBRM1;refGene_tx=XM_019822657.2;refGene_gn=PBRM1;refGene_tx=XM_0198226
59.2;refGene_gn=PBRM1;refGene_tx=XM_019822665.2;refGene_gn=PBRM1;refGe
ne_tx=XM_019822680.2;refGene_gn=PBRM1;refGene_tx=XM_019822682.2;refGen
e_gn=PBRM1;refGene_tx=XM_019822686.2;refGene_gn=PBRM1;refGene_tx=XM_01
9822693.2;refGene_gn=PBRM1;refGene_tx=XM_019822696.2;refGene_gn=PBRM1;
refGene_tx=XM_019822697.2;refGene_gn=PBRM1;refGene_tx=XM_019822700.2;r
efGene_gn=PBRM1;refGene_tx=XM_019822704.2;refGene_gn=PBRM1;refGene_tx=
XM_019822708.2;refGene_gn=PBRM1;refGene_tx=XM_019822711.2;refGene_gn=P
BRM1;refGene_tx=XM_019822713.2;refGene_gn=PBRM1;refGene_tx=XM_01982271
6.2;refGene_gn=PBRM1;refGene_tx=XM_019822718.2;refGene_gn=PBRM1;refGen
e_tx=XM_019822719.2;refGene_gn=PBRM1;refGene_tx=XM_019822722.2;refGene
_gn=PBRM1;refGene_tx=XM_019822723.2;refGene_gn=PBRM1;refGene_tx=XM_019
822726.2;refGene_gn=PBRM1;refGene_tx=XM_019822731.1;refGene_gn=PBRM1;r
efGene_tx=XM_023249537.1;refGene_gn=PBRM1;refGene_tx=XM_023249539.1;re
fGene_gn=PBRM1;refGene_tx=XM_023249543.1;refGene_gn=PBRM1;refGene_tx=X
M_023249544.1;refGene_gn=PBRM1;refGene_tx=XM_023249570.1;refGene_gn=PB
RM1;refGene_tx=XM_023249576.1;refGene_gn=PBRM1;refGene_tx=XM_023249582
.1
A2 21243817 21243970
refGene_gn=PBRM1;refGene_tx=XM_003982314.5;refGene_gn=PBRM1;refGene_tx
=XM_019822655.2;refGene_gn=PBRM1;refGene_tx=XM_019822656.2;refGene_gn=
PBRM1;refGene_tx=XM_019822657.2;refGene_gn=PBRM1;refGene_tx=XM_0198226
59.2;refGene_gn=PBRM1;refGene_tx=XM_019822665.2;refGene_gn=PBRM1;refGe
ne_tx=XM_019822680.2;refGene_gn=PBRM1;refGene_tx=XM_019822682.2;refGen
e_gn=PBRM1;refGene_tx=XM_019822686.2;refGene_gn=PBRM1;refGene_tx=XM_01
9822693.2;refGene_gn=PBRM1;refGene_tx=XM_019822696.2;refGene_gn=PBRM1;
refGene_tx=XM_019822697.2;refGene_gn=PBRM1;refGene_tx=XM_019822700.2;r
efGene_gn=PBRM1;refGene_tx=XM_019822704.2;refGene_gn=PBRM1;refGene_tx=
XM_019822708.2;refGene_gn=PBRM1;refGene_tx=XM_019822711.2;refGene_gn=P
BRM1;refGene_tx=XM_019822713.2;refGene_gn=PBRM1;refGene_tx=XM_01982271
6.2;refGene_gn=PBRM1;refGene_tx=XM_019822718.2;refGene_gn=PBRM1;refGen
e_tx=XM_019822719.2;refGene_gn=PBRM1;refGene_tx=XM_019822722.2;refGene
_gn=PBRM1;refGene_tx=XM_019822723.2;refGene_gn=PBRM1;refGene_tx=XM_019
822726.2;refGene_gn=PBRM1;refGene_tx=XM_019822731.1;refGene_gn=PBRM1;r
efGene_tx=XM_023249537.1;refGene_gn=PBRM1;refGene_tx=XM_023249539.1;re
fGene_gn=PBRM1;refGene_tx=XM_023249543.1;refGene_gn=PBRM1;refGene_tx=X
M_023249544.1;refGene_gn=PBRM1;refGene_tx=XM_023249570.1;refGene_gn=PB
RM1;refGene_tx=XM_023249576.1;refGene_gn=PBRM1;refGene_tx=XM_023249582
.1
A2 21245620 21245668
refGene_gn=PBRM1;refGene_tx=XM_003982314.5;refGene_gn=PBRM1;refGene_tx
=XM_019822655.2;refGene_gn=PBRM1;refGene_tx=XM_019822656.2;refGene_gn=
PBRM1;refGene_tx=XM_019822657.2;refGene_gn=PBRM1;refGene_tx=XM_0198226
59.2;refGene_gn=PBRM1;refGene_tx=XM_019822665.2;refGene_gn=PBRM1;refGe
ne_tx=XM_019822680.2;refGene_gn=PBRM1;refGene_tx=XM_019822697.2;refGen

e_gn=PBRM1;refGene_tx=XM_019822700.2;refGene_gn=PBRM1;refGene_tx=XM_01
9822704.2;refGene_gn=PBRM1;refGene_tx=XM_019822708.2;refGene_gn=PBRM1;
refGene_tx=XM_019822711.2;refGene_gn=PBRM1;refGene_tx=XM_019822713.2;r
efGene_gn=PBRM1;refGene_tx=XM_019822716.2;refGene_gn=PBRM1;refGene_tx=
XM_019822718.2;refGene_gn=PBRM1;refGene_tx=XM_019822719.2;refGene_gn=P
BRM1;refGene_tx=XM_019822722.2;refGene_gn=PBRM1;refGene_tx=XM_01982272
3.2;refGene_gn=PBRM1;refGene_tx=XM_019822726.2;refGene_gn=PBRM1;refGen
e_tx=XM_019822731.1;refGene_gn=PBRM1;refGene_tx=XM_023249537.1;refGene
_gn=PBRM1;refGene_tx=XM_023249539.1;refGene_gn=PBRM1;refGene_tx=XM_023
249570.1;refGene_gn=PBRM1;refGene_tx=XM_023249576.1;refGene_gn=PBRM1;r
efGene_tx=XM_023249582.1
A2 21248868 21248881
refGene_gn=GNL3;refGene_tx=XM_003982345.5;refGene_gn=GNL3;refGene_tx=X
M_006928852.4
A2 21249510 21249569
refGene_gn=GNL3;refGene_tx=XM_003982345.5;refGene_gn=GNL3;refGene_tx=X
M_006928852.4
A2 21249923 21250061
refGene_gn=GNL3;refGene_tx=XM_003982345.5;refGene_gn=GNL3;refGene_tx=X
M_006928852.4
A2 21250177 21250291
refGene_gn=GNL3;refGene_tx=XM_003982345.5;refGene_gn=GNL3;refGene_tx=X
M_006928852.4
A2 21250512 21250596
refGene_gn=GNL3;refGene_tx=XM_003982345.5;refGene_gn=GNL3;refGene_tx=X
M_006928852.4
A2 21251810 21251943
refGene_gn=GNL3;refGene_tx=XM_003982345.5;refGene_gn=GNL3;refGene_tx=X
M_006928852.4
A2 21252684 21252797
refGene_gn=GNL3;refGene_tx=XM_003982345.5;refGene_gn=GNL3;refGene_tx=X
M_006928852.4
A2 21253035 21253159
refGene_gn=GNL3;refGene_tx=XM_003982345.5;refGene_gn=GNL3;refGene_tx=X
M_006928852.4
A2 21253580 21253668
refGene_gn=GNL3;refGene_tx=XM_003982345.5;refGene_gn=GNL3;refGene_tx=X
M_006928852.4
A2 21254117 21254292
refGene_gn=GNL3;refGene_tx=XM_003982345.5;refGene_gn=GNL3;refGene_tx=X
M_006928852.4
A2 21254429 21254572
refGene_gn=GNL3;refGene_tx=XM_003982345.5;refGene_gn=GNL3;refGene_tx=X
M_006928852.4
A2 21254655 21254792
refGene_gn=GNL3;refGene_tx=XM_003982345.5;refGene_gn=GNL3;refGene_tx=X
M_006928852.4
A2 21254877 21255059
refGene_gn=GNL3;refGene_tx=XM_003982345.5;refGene_gn=GNL3;refGene_tx=X
M_006928852.4

A2 21255205 21255408
refGene_gn=GLT8D1;refGene_tx=XM_019822778.2;refGene_gn=GNL3;refGene_tx=XM_003982345.5;refGene_gn=GNL3;refGene_tx=XM_006928852.4
A2 21255489 21255572 refGene_gn=GNL3;refGene_tx=XM_003982345.5
A2 21256178 21256369
refGene_gn=GLT8D1;refGene_tx=XM_003982317.5;refGene_gn=GLT8D1;refGene_tx=XM_023249606.1
A2 21256537 21256650
refGene_gn=GLT8D1;refGene_tx=XM_003982317.5;refGene_gn=GLT8D1;refGene_tx=XM_019822778.2;refGene_gn=GLT8D1;refGene_tx=XM_023249606.1
A2 21256770 21256937
refGene_gn=GLT8D1;refGene_tx=XM_003982317.5;refGene_gn=GLT8D1;refGene_tx=XM_019822778.2;refGene_gn=GLT8D1;refGene_tx=XM_023249606.1
A2 21257494 21257607
refGene_gn=GLT8D1;refGene_tx=XM_003982317.5;refGene_gn=GLT8D1;refGene_tx=XM_019822778.2;refGene_gn=GLT8D1;refGene_tx=XM_023249606.1
A2 21257795 21257880
refGene_gn=GLT8D1;refGene_tx=XM_003982317.5;refGene_gn=GLT8D1;refGene_tx=XM_019822778.2;refGene_gn=GLT8D1;refGene_tx=XM_023249606.1
A2 21258372 21258490
refGene_gn=GLT8D1;refGene_tx=XM_003982317.5;refGene_gn=GLT8D1;refGene_tx=XM_019822778.2;refGene_gn=GLT8D1;refGene_tx=XM_023249606.1
A2 21260311 21260325 refGene_gn=GNL3;refGene_tx=XM_006928852.4
A2 21260484 21260698
refGene_gn=GLT8D1;refGene_tx=XM_003982317.5;refGene_gn=GLT8D1;refGene_tx=XM_019822778.2;refGene_gn=GLT8D1;refGene_tx=XM_023249606.1
A2 21262556 21262655
refGene_gn=GLT8D1;refGene_tx=XM_003982317.5;refGene_gn=GLT8D1;refGene_tx=XM_019822778.2;refGene_gn=GLT8D1;refGene_tx=XM_023249606.1
A2 21262740 21262846
refGene_gn=GLT8D1;refGene_tx=XM_003982317.5;refGene_gn=GLT8D1;refGene_tx=XM_019822778.2;refGene_gn=GLT8D1;refGene_tx=XM_023249606.1
A2 21267848 21268064 refGene_gn=SPCS1;refGene_tx=XM_003982318.5
A2 21268375 21268435 refGene_gn=SPCS1;refGene_tx=XM_003982318.5
A2 21268541 21268628 refGene_gn=SPCS1;refGene_tx=XM_003982318.5
A2 21269137 21269263 refGene_gn=SPCS1;refGene_tx=XM_003982318.5
A2 21269752 21269785 refGene_gn=NEK4;refGene_tx=XM_011279511.3
A2 21273475 21273568
refGene_gn=NEK4;refGene_tx=XM_011279507.3;refGene_gn=NEK4;refGene_tx=XM_019822755.2;refGene_gn=NEK4;refGene_tx=XM_023249596.1
A2 21279871 21279898
refGene_gn=NEK4;refGene_tx=XM_011279513.3;refGene_gn=NEK4;refGene_tx=XM_011279516.3
A2 21296457 21296544 refGene_gn=NEK4;refGene_tx=XM_011279510.3
A2 21297757 21297891
refGene_gn=NEK4;refGene_tx=XM_011279507.3;refGene_gn=NEK4;refGene_tx=XM_011279510.3;refGene_gn=NEK4;refGene_tx=XM_011279511.3;refGene_gn=NEK4;refGene_tx=XM_011279513.3;refGene_gn=NEK4;refGene_tx=XM_011279516.3;refGene_gn=NEK4;refGene_tx=XM_019822755.2;refGene_gn=NEK4;refGene_tx=XM_023249596.1

A2	21298348 21298554
refGene_gn=NEK4;refGene_tx=XM_011279507.3;refGene_gn=NEK4;refGene_tx=XM_011279510.3;refGene_gn=NEK4;refGene_tx=XM_011279511.3;refGene_gn=NEK4;refGene_tx=XM_011279513.3;refGene_gn=NEK4;refGene_tx=XM_011279516.3;refGene_gn=NEK4;refGene_tx=XM_019822755.2;refGene_gn=NEK4;refGene_tx=XM_023249596.1
A2	21300031 21300120
refGene_gn=NEK4;refGene_tx=XM_011279507.3;refGene_gn=NEK4;refGene_tx=XM_011279510.3;refGene_gn=NEK4;refGene_tx=XM_011279511.3;refGene_gn=NEK4;refGene_tx=XM_011279513.3;refGene_gn=NEK4;refGene_tx=XM_011279516.3;refGene_gn=NEK4;refGene_tx=XM_019822755.2;refGene_gn=NEK4;refGene_tx=XM_023249596.1
A2	21301448 21301558
refGene_gn=NEK4;refGene_tx=XM_011279507.3;refGene_gn=NEK4;refGene_tx=XM_011279510.3;refGene_gn=NEK4;refGene_tx=XM_011279511.3;refGene_gn=NEK4;refGene_tx=XM_011279513.3;refGene_gn=NEK4;refGene_tx=XM_011279516.3;refGene_gn=NEK4;refGene_tx=XM_019822755.2;refGene_gn=NEK4;refGene_tx=XM_023249596.1
A2	21302340 21302407
refGene_gn=NEK4;refGene_tx=XM_011279507.3;refGene_gn=NEK4;refGene_tx=XM_011279510.3;refGene_gn=NEK4;refGene_tx=XM_011279511.3;refGene_gn=NEK4;refGene_tx=XM_011279513.3;refGene_gn=NEK4;refGene_tx=XM_011279516.3;refGene_gn=NEK4;refGene_tx=XM_019822755.2;refGene_gn=NEK4;refGene_tx=XM_023249596.1
A2	21302872 21303025
refGene_gn=NEK4;refGene_tx=XM_011279507.3;refGene_gn=NEK4;refGene_tx=XM_011279510.3;refGene_gn=NEK4;refGene_tx=XM_011279511.3;refGene_gn=NEK4;refGene_tx=XM_011279513.3;refGene_gn=NEK4;refGene_tx=XM_011279516.3;refGene_gn=NEK4;refGene_tx=XM_019822755.2;refGene_gn=NEK4;refGene_tx=XM_023249596.1
A2	21303533 21303704
refGene_gn=NEK4;refGene_tx=XM_011279507.3;refGene_gn=NEK4;refGene_tx=XM_011279510.3;refGene_gn=NEK4;refGene_tx=XM_011279511.3;refGene_gn=NEK4;refGene_tx=XM_011279513.3;refGene_gn=NEK4;refGene_tx=XM_011279516.3;refGene_gn=NEK4;refGene_tx=XM_019822755.2;refGene_gn=NEK4;refGene_tx=XM_023249596.1
A2	21309979 21310357
refGene_gn=NEK4;refGene_tx=XM_011279507.3;refGene_gn=NEK4;refGene_tx=XM_011279510.3;refGene_gn=NEK4;refGene_tx=XM_011279511.3;refGene_gn=NEK4;refGene_tx=XM_011279513.3;refGene_gn=NEK4;refGene_tx=XM_011279516.3;refGene_gn=NEK4;refGene_tx=XM_019822755.2;refGene_gn=NEK4;refGene_tx=XM_023249596.1
A2	21312522 21312664
refGene_gn=NEK4;refGene_tx=XM_011279507.3;refGene_gn=NEK4;refGene_tx=XM_011279510.3;refGene_gn=NEK4;refGene_tx=XM_011279511.3;refGene_gn=NEK4;refGene_tx=XM_011279513.3;refGene_gn=NEK4;refGene_tx=XM_011279516.3;refGene_gn=NEK4;refGene_tx=XM_019822755.2;refGene_gn=NEK4;refGene_tx=XM_023249596.1
A2	21314483 21314638
refGene_gn=NEK4;refGene_tx=XM_011279507.3;refGene_gn=NEK4;refGene_tx=X

M_011279510.3;refGene_gn=NEK4;refGene_tx=XM_011279511.3;refGene_gn=NEK4;refGene_tx=XM_011279513.3;refGene_gn=NEK4;refGene_tx=XM_011279516.3;refGene_gn=NEK4;refGene_tx=XM_019822755.2;refGene_gn=NEK4;refGene_tx=XM_023249596.1
A2 21316305 21316413 refGene_gn=NEK4;refGene_tx=XM_011279507.3;refGene_gn=NEK4;refGene_tx=XM_011279510.3;refGene_gn=NEK4;refGene_tx=XM_011279511.3;refGene_gn=NEK4;refGene_tx=XM_011279513.3;refGene_gn=NEK4;refGene_tx=XM_011279516.3;refGene_gn=NEK4;refGene_tx=XM_019822755.2;refGene_gn=NEK4;refGene_tx=XM_023249596.1
A2 21316604 21316802 refGene_gn=NEK4;refGene_tx=XM_011279507.3;refGene_gn=NEK4;refGene_tx=XM_011279510.3;refGene_gn=NEK4;refGene_tx=XM_011279511.3;refGene_gn=NEK4;refGene_tx=XM_011279513.3;refGene_gn=NEK4;refGene_tx=XM_011279516.3;refGene_gn=NEK4;refGene_tx=XM_023249596.1
A2 21317717 21317984 refGene_gn=NEK4;refGene_tx=XM_011279507.3;refGene_gn=NEK4;refGene_tx=XM_011279510.3;refGene_gn=NEK4;refGene_tx=XM_011279511.3;refGene_gn=NEK4;refGene_tx=XM_011279513.3;refGene_gn=NEK4;refGene_tx=XM_011279516.3;refGene_gn=NEK4;refGene_tx=XM_023249596.1
A2 21319625 21319718 refGene_gn=NEK4;refGene_tx=XM_011279507.3;refGene_gn=NEK4;refGene_tx=XM_011279510.3;refGene_gn=NEK4;refGene_tx=XM_011279511.3;refGene_gn=NEK4;refGene_tx=XM_011279513.3;refGene_gn=NEK4;refGene_tx=XM_011279516.3;refGene_gn=NEK4;refGene_tx=XM_019822755.2
A2 21325789 21325903 refGene_gn=ITIH1;refGene_tx=XM_003982319.3
A2 21326164 21326185 refGene_gn=ITIH1;refGene_tx=XM_003982319.3
A2 21326484 21326651 refGene_gn=ITIH1;refGene_tx=XM_003982319.3
A2 21327103 21327208 refGene_gn=ITIH1;refGene_tx=XM_003982319.3
A2 21327561 21327724 refGene_gn=ITIH1;refGene_tx=XM_003982319.3
A2 21328275 21328389 refGene_gn=ITIH1;refGene_tx=XM_003982319.3
A2 21329748 21329874 refGene_gn=ITIH1;refGene_tx=XM_003982319.3
A2 21329959 21330076 refGene_gn=ITIH1;refGene_tx=XM_003982319.3
A2 21330777 21330946 refGene_gn=ITIH1;refGene_tx=XM_003982319.3
A2 21331034 21331160 refGene_gn=ITIH1;refGene_tx=XM_003982319.3
A2 21332123 21332305 refGene_gn=ITIH1;refGene_tx=XM_003982319.3
A2 21332591 21332777 refGene_gn=ITIH1;refGene_tx=XM_003982319.3
A2 21333853 21333993 refGene_gn=ITIH1;refGene_tx=XM_003982319.3
A2 21334578 21334733 refGene_gn=ITIH1;refGene_tx=XM_003982319.3
A2 21334832 21334847 refGene_gn=ITIH1;refGene_tx=XM_003982319.3
A2 21335192 21335213 refGene_gn=ITIH1;refGene_tx=XM_003982319.3
A2 21335597 21335678 refGene_gn=ITIH1;refGene_tx=XM_003982319.3
A2 21335782 21335896 refGene_gn=ITIH1;refGene_tx=XM_003982319.3
A2 21336862 21337064 refGene_gn=ITIH1;refGene_tx=XM_003982319.3
A2 21337910 21338083 refGene_gn=ITIH1;refGene_tx=XM_003982319.3
A2 21338678 21338790 refGene_gn=ITIH1;refGene_tx=XM_003982319.3
A2 21338945 21339075 refGene_gn=ITIH1;refGene_tx=XM_003982319.3
A2 21342899 21342992 refGene_gn=ITIH3;refGene_tx=XM_006928854.2
A2 21343619 21343640 refGene_gn=ITIH3;refGene_tx=XM_006928854.2
A2 21344804 21344971 refGene_gn=ITIH3;refGene_tx=XM_006928854.2

A2 21345052 21345157 refGene_gn=ITIH3;refGene_tx=XM_006928854.2
A2 21345435 21345598 refGene_gn=ITIH3;refGene_tx=XM_006928854.2
A2 21345984 21346098 refGene_gn=ITIH3;refGene_tx=XM_006928854.2
A2 21347069 21347195 refGene_gn=ITIH3;refGene_tx=XM_006928854.2
A2 21347482 21347599 refGene_gn=ITIH3;refGene_tx=XM_006928854.2
A2 21347842 21348011 refGene_gn=ITIH3;refGene_tx=XM_006928854.2
A2 21348603 21348729 refGene_gn=ITIH3;refGene_tx=XM_006928854.2
A2 21348984 21349166 refGene_gn=ITIH3;refGene_tx=XM_006928854.2
A2 21350411 21350597 refGene_gn=ITIH3;refGene_tx=XM_006928854.2
A2 21350751 21350891 refGene_gn=ITIH3;refGene_tx=XM_006928854.2
A2 21351639 21351794 refGene_gn=ITIH3;refGene_tx=XM_006928854.2
A2 21352478 21352496 refGene_gn=ITIH3;refGene_tx=XM_006928854.2
A2 21353563 21353587 refGene_gn=ITIH3;refGene_tx=XM_006928854.2
A2 21353852 21353888 refGene_gn=ITIH3;refGene_tx=XM_006928854.2
A2 21354051 21354165 refGene_gn=ITIH3;refGene_tx=XM_006928854.2
A2 21354577 21354782 refGene_gn=ITIH3;refGene_tx=XM_006928854.2
A2 21355686 21355859 refGene_gn=ITIH3;refGene_tx=XM_006928854.2
A2 21356067 21356179 refGene_gn=ITIH3;refGene_tx=XM_006928854.2
A2 21356526 21356656 refGene_gn=ITIH3;refGene_tx=XM_006928854.2
A2 21360334 21360404
refGene_gn=ITIH4;refGene_tx=XM_006928856.3;refGene_gn=ITIH4;refGene_tx=XM_006928857.3;refGene_gn=ITIH4;refGene_tx=XM_011279531.2;refGene_gn=ITIH4;refGene_tx=XM_011279532.2;refGene_gn=ITIH4;refGene_tx=XM_019822794.1;refGene_gn=ITIH4;refGene_tx=XM_019822800.1
A2 21360862 21360959
refGene_gn=ITIH4;refGene_tx=XM_006928856.3;refGene_gn=ITIH4;refGene_tx=XM_006928857.3;refGene_gn=ITIH4;refGene_tx=XM_011279531.2;refGene_gn=ITIH4;refGene_tx=XM_011279532.2;refGene_gn=ITIH4;refGene_tx=XM_019822794.1;refGene_gn=ITIH4;refGene_tx=XM_019822800.1
A2 21361086 21361241
refGene_gn=ITIH4;refGene_tx=XM_006928856.3;refGene_gn=ITIH4;refGene_tx=XM_006928857.3;refGene_gn=ITIH4;refGene_tx=XM_011279531.2;refGene_gn=ITIH4;refGene_tx=XM_011279532.2;refGene_gn=ITIH4;refGene_tx=XM_019822794.1;refGene_gn=ITIH4;refGene_tx=XM_019822800.1
A2 21362674 21362849
refGene_gn=ITIH4;refGene_tx=XM_006928856.3;refGene_gn=ITIH4;refGene_tx=XM_006928857.3;refGene_gn=ITIH4;refGene_tx=XM_011279531.2;refGene_gn=ITIH4;refGene_tx=XM_011279532.2;refGene_gn=ITIH4;refGene_tx=XM_019822794.1;refGene_gn=ITIH4;refGene_tx=XM_019822800.1
A2 21363936 21364053
refGene_gn=ITIH4;refGene_tx=XM_006928856.3;refGene_gn=ITIH4;refGene_tx=XM_006928857.3;refGene_gn=ITIH4;refGene_tx=XM_011279531.2;refGene_gn=ITIH4;refGene_tx=XM_011279532.2;refGene_gn=ITIH4;refGene_tx=XM_019822794.1;refGene_gn=ITIH4;refGene_tx=XM_019822800.1
A2 21364141 21364168
refGene_gn=ITIH4;refGene_tx=XM_006928856.3;refGene_gn=ITIH4;refGene_tx=XM_006928857.3;refGene_gn=ITIH4;refGene_tx=XM_011279531.2;refGene_gn=ITIH4;refGene_tx=XM_011279532.2;refGene_gn=ITIH4;refGene_tx=XM_019822794.1;refGene_gn=ITIH4;refGene_tx=XM_019822800.1
A2 21364292 21364415

refGene_gn=ITIH4;refGene_tx=XM_006928856.3;refGene_gn=ITIH4;refGene_tx
=XM_006928857.3;refGene_gn=ITIH4;refGene_tx=XM_011279531.2;refGene_gn=
ITIH4;refGene_tx=XM_011279532.2;refGene_gn=ITIH4;refGene_tx=XM_0198227
94.1;refGene_gn=ITIH4;refGene_tx=XM_019822800.1
A2 21364993 21365119
refGene_gn=ITIH4;refGene_tx=XM_006928856.3;refGene_gn=ITIH4;refGene_tx
=XM_006928857.3;refGene_gn=ITIH4;refGene_tx=XM_011279531.2;refGene_gn=
ITIH4;refGene_tx=XM_011279532.2;refGene_gn=ITIH4;refGene_tx=XM_0198227
94.1;refGene_gn=ITIH4;refGene_tx=XM_019822800.1
A2 21365376 21365430
refGene_gn=ITIH4;refGene_tx=XM_006928856.3;refGene_gn=ITIH4;refGene_tx
=XM_011279531.2;refGene_gn=ITIH4;refGene_tx=XM_019822794.1;refGene_gn=
ITIH4;refGene_tx=XM_019822800.1
A2 21365585 21365639
refGene_gn=ITIH4;refGene_tx=XM_011279531.2;refGene_gn=ITIH4;refGene_tx
=XM_011279532.2;refGene_gn=ITIH4;refGene_tx=XM_019822794.1
A2 21365917 21365944
refGene_gn=ITIH4;refGene_tx=XM_006928856.3;refGene_gn=ITIH4;refGene_tx
=XM_006928857.3;refGene_gn=ITIH4;refGene_tx=XM_011279531.2;refGene_gn=
ITIH4;refGene_tx=XM_011279532.2;refGene_gn=ITIH4;refGene_tx=XM_0198227
94.1;refGene_gn=ITIH4;refGene_tx=XM_019822800.1
A2 21366243 21366398
refGene_gn=ITIH4;refGene_tx=XM_006928856.3;refGene_gn=ITIH4;refGene_tx
=XM_006928857.3;refGene_gn=ITIH4;refGene_tx=XM_011279531.2;refGene_gn=
ITIH4;refGene_tx=XM_011279532.2;refGene_gn=ITIH4;refGene_tx=XM_0198227
94.1;refGene_gn=ITIH4;refGene_tx=XM_019822800.1
A2 21366662 21366802
refGene_gn=ITIH4;refGene_tx=XM_006928856.3;refGene_gn=ITIH4;refGene_tx
=XM_006928857.3;refGene_gn=ITIH4;refGene_tx=XM_011279531.2;refGene_gn=
ITIH4;refGene_tx=XM_011279532.2;refGene_gn=ITIH4;refGene_tx=XM_0198227
94.1;refGene_gn=ITIH4;refGene_tx=XM_019822800.1
A2 21368903 21369089
refGene_gn=ITIH4;refGene_tx=XM_006928856.3;refGene_gn=ITIH4;refGene_tx
=XM_006928857.3;refGene_gn=ITIH4;refGene_tx=XM_011279531.2;refGene_gn=
ITIH4;refGene_tx=XM_011279532.2;refGene_gn=ITIH4;refGene_tx=XM_0198227
94.1;refGene_gn=ITIH4;refGene_tx=XM_019822800.1
A2 21369183 21369365
refGene_gn=ITIH4;refGene_tx=XM_006928856.3;refGene_gn=ITIH4;refGene_tx
=XM_006928857.3;refGene_gn=ITIH4;refGene_tx=XM_011279531.2;refGene_gn=
ITIH4;refGene_tx=XM_011279532.2;refGene_gn=ITIH4;refGene_tx=XM_0198227
94.1;refGene_gn=ITIH4;refGene_tx=XM_019822800.1
A2 21369546 21369672
refGene_gn=ITIH4;refGene_tx=XM_006928856.3;refGene_gn=ITIH4;refGene_tx
=XM_006928857.3;refGene_gn=ITIH4;refGene_tx=XM_011279531.2;refGene_gn=
ITIH4;refGene_tx=XM_011279532.2;refGene_gn=ITIH4;refGene_tx=XM_0198227
94.1;refGene_gn=ITIH4;refGene_tx=XM_019822800.1
A2 21369773 21369942
refGene_gn=ITIH4;refGene_tx=XM_006928856.3;refGene_gn=ITIH4;refGene_tx
=XM_006928857.3;refGene_gn=ITIH4;refGene_tx=XM_011279531.2;refGene_gn=
ITIH4;refGene_tx=XM_011279532.2;refGene_gn=ITIH4;refGene_tx=XM_0198227

94.1;refGene_gn=ITIH4;refGene_tx=XM_019822800.1
A2 21370204 21370321
refGene_gn=ITIH4;refGene_tx=XM_006928856.3;refGene_gn=ITIH4;refGene_tx=XM_006928857.3;refGene_gn=ITIH4;refGene_tx=XM_011279531.2;refGene_gn=ITIH4;refGene_tx=XM_011279532.2;refGene_gn=ITIH4;refGene_tx=XM_019822794.1;refGene_gn=ITIH4;refGene_tx=XM_019822800.1
A2 21371212 21371338
refGene_gn=ITIH4;refGene_tx=XM_006928856.3;refGene_gn=ITIH4;refGene_tx=XM_006928857.3;refGene_gn=ITIH4;refGene_tx=XM_011279531.2;refGene_gn=ITIH4;refGene_tx=XM_011279532.2;refGene_gn=ITIH4;refGene_tx=XM_019822794.1;refGene_gn=ITIH4;refGene_tx=XM_019822800.1
A2 21371830 21371941
refGene_gn=ITIH4;refGene_tx=XM_006928856.3;refGene_gn=ITIH4;refGene_tx=XM_006928857.3;refGene_gn=ITIH4;refGene_tx=XM_011279531.2;refGene_gn=ITIH4;refGene_tx=XM_011279532.2;refGene_gn=ITIH4;refGene_tx=XM_019822794.1;refGene_gn=ITIH4;refGene_tx=XM_019822800.1
A2 21372079 21372242
refGene_gn=ITIH4;refGene_tx=XM_006928856.3;refGene_gn=ITIH4;refGene_tx=XM_006928857.3;refGene_gn=ITIH4;refGene_tx=XM_011279531.2;refGene_gn=ITIH4;refGene_tx=XM_011279532.2;refGene_gn=ITIH4;refGene_tx=XM_019822794.1;refGene_gn=ITIH4;refGene_tx=XM_019822800.1
A2 21372388 21372493
refGene_gn=ITIH4;refGene_tx=XM_006928856.3;refGene_gn=ITIH4;refGene_tx=XM_006928857.3;refGene_gn=ITIH4;refGene_tx=XM_011279531.2;refGene_gn=ITIH4;refGene_tx=XM_011279532.2;refGene_gn=ITIH4;refGene_tx=XM_019822794.1;refGene_gn=ITIH4;refGene_tx=XM_019822800.1
A2 21374494 21374655
refGene_gn=ITIH4;refGene_tx=XM_006928856.3;refGene_gn=ITIH4;refGene_tx=XM_006928857.3;refGene_gn=ITIH4;refGene_tx=XM_011279531.2;refGene_gn=ITIH4;refGene_tx=XM_011279532.2;refGene_gn=ITIH4;refGene_tx=XM_019822794.1;refGene_gn=ITIH4;refGene_tx=XM_019822800.1
A2 21376198 21376288
refGene_gn=ITIH4;refGene_tx=XM_006928856.3;refGene_gn=ITIH4;refGene_tx=XM_006928857.3;refGene_gn=ITIH4;refGene_tx=XM_011279531.2;refGene_gn=ITIH4;refGene_tx=XM_011279532.2;refGene_gn=ITIH4;refGene_tx=XM_019822794.1;refGene_gn=ITIH4;refGene_tx=XM_019822800.1
A2 21378844 21378951 refGene_gn=MUSTN1;refGene_tx=XM_006928855.3
A2 21379114 21379247 refGene_gn=MUSTN1;refGene_tx=XM_006928855.3
A2 21380358 21380367 refGene_gn=MUSTN1;refGene_tx=XM_006928855.3
A2 21385660 21385777 refGene_gn=STIMATE;refGene_tx=XM_006928858.4
A2 21387643 21387793 refGene_gn=STIMATE;refGene_tx=XM_006928858.4
A2 21388571 21388649 refGene_gn=STIMATE;refGene_tx=XM_006928858.4
A2 21389686 21389799 refGene_gn=STIMATE;refGene_tx=XM_006928858.4
A2 21394384 21394506 refGene_gn=STIMATE;refGene_tx=XM_006928858.4
A2 21397804 21397900 refGene_gn=STIMATE;refGene_tx=XM_006928858.4
A2 21399726 21399775 refGene_gn=STIMATE;refGene_tx=XM_006928858.4
A2 21435025 21435185 refGene_gn=STIMATE;refGene_tx=XM_006928858.4
A2 21443112 21443253
refGene_gn=SFMBT1;refGene_tx=XM_011279552.3;refGene_gn=SFMBT1;refGene_tx=XM_023249657.1;refGene_gn=SFMBT1;refGene_tx=XM_023249661.1;refGene_

gn=SFMBT1;refGene_tx=XM_023249669.1;refGene_gn=SFMBT1;refGene_tx=XM_02
3249676.1;refGene_gn=SFMBT1;refGene_tx=XM_023249681.1;refGene_gn=SFMBT
1;refGene_tx=XM_023249685.1;refGene_gn=SFMBT1;refGene_tx=XM_023249688.
1;refGene_gn=SFMBT1;refGene_tx=XM_023249693.1
A2 21444131 21444260
refGene_gn=SFMBT1;refGene_tx=XM_011279552.3;refGene_gn=SFMBT1;refGene_
tx=XM_023249657.1;refGene_gn=SFMBT1;refGene_tx=XM_023249661.1;refGene_
gn=SFMBT1;refGene_tx=XM_023249669.1;refGene_gn=SFMBT1;refGene_tx=XM_02
3249676.1;refGene_gn=SFMBT1;refGene_tx=XM_023249681.1;refGene_gn=SFMBT
1;refGene_tx=XM_023249685.1;refGene_gn=SFMBT1;refGene_tx=XM_023249688.
1;refGene_gn=SFMBT1;refGene_tx=XM_023249693.1
A2 21445498 21445744
refGene_gn=SFMBT1;refGene_tx=XM_011279552.3;refGene_gn=SFMBT1;refGene_
tx=XM_023249657.1;refGene_gn=SFMBT1;refGene_tx=XM_023249661.1;refGene_
gn=SFMBT1;refGene_tx=XM_023249669.1;refGene_gn=SFMBT1;refGene_tx=XM_02
3249676.1;refGene_gn=SFMBT1;refGene_tx=XM_023249681.1;refGene_gn=SFMBT
1;refGene_tx=XM_023249685.1;refGene_gn=SFMBT1;refGene_tx=XM_023249688.
1;refGene_gn=SFMBT1;refGene_tx=XM_023249693.1
A2 21445927 21446106
refGene_gn=SFMBT1;refGene_tx=XM_011279552.3;refGene_gn=SFMBT1;refGene_
tx=XM_023249657.1;refGene_gn=SFMBT1;refGene_tx=XM_023249661.1;refGene_
gn=SFMBT1;refGene_tx=XM_023249669.1;refGene_gn=SFMBT1;refGene_tx=XM_02
3249676.1;refGene_gn=SFMBT1;refGene_tx=XM_023249681.1;refGene_gn=SFMBT
1;refGene_tx=XM_023249685.1;refGene_gn=SFMBT1;refGene_tx=XM_023249688.
1;refGene_gn=SFMBT1;refGene_tx=XM_023249693.1
A2 21449451 21449627
refGene_gn=SFMBT1;refGene_tx=XM_011279552.3;refGene_gn=SFMBT1;refGene_
tx=XM_023249657.1;refGene_gn=SFMBT1;refGene_tx=XM_023249661.1;refGene_
gn=SFMBT1;refGene_tx=XM_023249669.1;refGene_gn=SFMBT1;refGene_tx=XM_02
3249676.1;refGene_gn=SFMBT1;refGene_tx=XM_023249681.1;refGene_gn=SFMBT
1;refGene_tx=XM_023249685.1;refGene_gn=SFMBT1;refGene_tx=XM_023249688.
1;refGene_gn=SFMBT1;refGene_tx=XM_023249693.1
A2 21450342 21450452
refGene_gn=SFMBT1;refGene_tx=XM_011279552.3;refGene_gn=SFMBT1;refGene_
tx=XM_023249657.1;refGene_gn=SFMBT1;refGene_tx=XM_023249661.1;refGene_
gn=SFMBT1;refGene_tx=XM_023249669.1;refGene_gn=SFMBT1;refGene_tx=XM_02
3249676.1;refGene_gn=SFMBT1;refGene_tx=XM_023249681.1;refGene_gn=SFMBT
1;refGene_tx=XM_023249685.1;refGene_gn=SFMBT1;refGene_tx=XM_023249688.
1;refGene_gn=SFMBT1;refGene_tx=XM_023249693.1
A2 21451115 21451255
refGene_gn=SFMBT1;refGene_tx=XM_011279552.3;refGene_gn=SFMBT1;refGene_
tx=XM_023249657.1;refGene_gn=SFMBT1;refGene_tx=XM_023249661.1;refGene_
gn=SFMBT1;refGene_tx=XM_023249669.1;refGene_gn=SFMBT1;refGene_tx=XM_02
3249676.1;refGene_gn=SFMBT1;refGene_tx=XM_023249681.1;refGene_gn=SFMBT
1;refGene_tx=XM_023249685.1;refGene_gn=SFMBT1;refGene_tx=XM_023249688.
1;refGene_gn=SFMBT1;refGene_tx=XM_023249693.1
A2 21456551 21456616
refGene_gn=SFMBT1;refGene_tx=XM_011279552.3;refGene_gn=SFMBT1;refGene_
tx=XM_023249657.1;refGene_gn=SFMBT1;refGene_tx=XM_023249661.1;refGene_
gn=SFMBT1;refGene_tx=XM_023249669.1;refGene_gn=SFMBT1;refGene_tx=XM_02

3249676.1;refGene_gn=SFMBT1;refGene_tx=XM_023249681.1;refGene_gn=SFMBT1;refGene_tx=XM_023249685.1;refGene_gn=SFMBT1;refGene_tx=XM_023249688.1;refGene_gn=SFMBT1;refGene_tx=XM_023249693.1
A2 21458139 21458182
refGene_gn=SFMBT1;refGene_tx=XM_011279552.3;refGene_gn=SFMBT1;refGene_tx=XM_023249657.1;refGene_gn=SFMBT1;refGene_tx=XM_023249661.1;refGene_gn=SFMBT1;refGene_tx=XM_023249669.1;refGene_gn=SFMBT1;refGene_tx=XM_023249676.1;refGene_gn=SFMBT1;refGene_tx=XM_023249681.1;refGene_gn=SFMBT1;refGene_tx=XM_023249685.1;refGene_gn=SFMBT1;refGene_tx=XM_023249688.1;refGene_gn=SFMBT1;refGene_tx=XM_023249693.1
A2 21460749 21460863
refGene_gn=SFMBT1;refGene_tx=XM_011279552.3;refGene_gn=SFMBT1;refGene_tx=XM_023249657.1;refGene_gn=SFMBT1;refGene_tx=XM_023249661.1;refGene_gn=SFMBT1;refGene_tx=XM_023249669.1;refGene_gn=SFMBT1;refGene_tx=XM_023249676.1;refGene_gn=SFMBT1;refGene_tx=XM_023249681.1;refGene_gn=SFMBT1;refGene_tx=XM_023249685.1;refGene_gn=SFMBT1;refGene_tx=XM_023249688.1;refGene_gn=SFMBT1;refGene_tx=XM_023249693.1
A2 21462110 21462237
refGene_gn=SFMBT1;refGene_tx=XM_011279552.3;refGene_gn=SFMBT1;refGene_tx=XM_023249657.1;refGene_gn=SFMBT1;refGene_tx=XM_023249661.1;refGene_gn=SFMBT1;refGene_tx=XM_023249669.1;refGene_gn=SFMBT1;refGene_tx=XM_023249676.1;refGene_gn=SFMBT1;refGene_tx=XM_023249681.1;refGene_gn=SFMBT1;refGene_tx=XM_023249685.1;refGene_gn=SFMBT1;refGene_tx=XM_023249688.1;refGene_gn=SFMBT1;refGene_tx=XM_023249693.1
A2 21466735 21466818
refGene_gn=SFMBT1;refGene_tx=XM_011279552.3;refGene_gn=SFMBT1;refGene_tx=XM_023249657.1;refGene_gn=SFMBT1;refGene_tx=XM_023249661.1;refGene_gn=SFMBT1;refGene_tx=XM_023249669.1;refGene_gn=SFMBT1;refGene_tx=XM_023249676.1;refGene_gn=SFMBT1;refGene_tx=XM_023249681.1;refGene_gn=SFMBT1;refGene_tx=XM_023249685.1;refGene_gn=SFMBT1;refGene_tx=XM_023249688.1;refGene_gn=SFMBT1;refGene_tx=XM_023249693.1
A2 21469119 21469270
refGene_gn=SFMBT1;refGene_tx=XM_011279552.3;refGene_gn=SFMBT1;refGene_tx=XM_023249657.1;refGene_gn=SFMBT1;refGene_tx=XM_023249661.1;refGene_gn=SFMBT1;refGene_tx=XM_023249669.1;refGene_gn=SFMBT1;refGene_tx=XM_023249676.1;refGene_gn=SFMBT1;refGene_tx=XM_023249681.1;refGene_gn=SFMBT1;refGene_tx=XM_023249685.1;refGene_gn=SFMBT1;refGene_tx=XM_023249688.1;refGene_gn=SFMBT1;refGene_tx=XM_023249693.1
A2 21470738 21470840
refGene_gn=SFMBT1;refGene_tx=XM_011279552.3;refGene_gn=SFMBT1;refGene_tx=XM_023249657.1;refGene_gn=SFMBT1;refGene_tx=XM_023249661.1;refGene_gn=SFMBT1;refGene_tx=XM_023249669.1;refGene_gn=SFMBT1;refGene_tx=XM_023249676.1;refGene_gn=SFMBT1;refGene_tx=XM_023249681.1;refGene_gn=SFMBT1;refGene_tx=XM_023249685.1;refGene_gn=SFMBT1;refGene_tx=XM_023249688.1;refGene_gn=SFMBT1;refGene_tx=XM_023249693.1
A2 21471355 21471450
refGene_gn=SFMBT1;refGene_tx=XM_011279552.3;refGene_gn=SFMBT1;refGene_tx=XM_023249657.1;refGene_gn=SFMBT1;refGene_tx=XM_023249661.1;refGene_gn=SFMBT1;refGene_tx=XM_023249669.1;refGene_gn=SFMBT1;refGene_tx=XM_023249676.1;refGene_gn=SFMBT1;refGene_tx=XM_023249681.1;refGene_gn=SFMBT

1;refGene_tx=XM_023249685.1;refGene_gn=SFMBT1;refGene_tx=XM_023249688.
1;refGene_gn=SFMBT1;refGene_tx=XM_023249693.1
A2 21472113 21472360
refGene_gn=SFMBT1;refGene_tx=XM_011279552.3;refGene_gn=SFMBT1;refGene_
tx=XM_023249657.1;refGene_gn=SFMBT1;refGene_tx=XM_023249661.1;refGene_
gn=SFMBT1;refGene_tx=XM_023249669.1;refGene_gn=SFMBT1;refGene_tx=XM_02
3249676.1;refGene_gn=SFMBT1;refGene_tx=XM_023249681.1;refGene_gn=SFMBT
1;refGene_tx=XM_023249685.1;refGene_gn=SFMBT1;refGene_tx=XM_023249688.
1;refGene_gn=SFMBT1;refGene_tx=XM_023249693.1
A2 21474471 21474560
refGene_gn=SFMBT1;refGene_tx=XM_011279552.3;refGene_gn=SFMBT1;refGene_
tx=XM_023249657.1;refGene_gn=SFMBT1;refGene_tx=XM_023249661.1;refGene_
gn=SFMBT1;refGene_tx=XM_023249669.1;refGene_gn=SFMBT1;refGene_tx=XM_02
3249676.1;refGene_gn=SFMBT1;refGene_tx=XM_023249681.1;refGene_gn=SFMBT
1;refGene_tx=XM_023249685.1;refGene_gn=SFMBT1;refGene_tx=XM_023249688.
1;refGene_gn=SFMBT1;refGene_tx=XM_023249693.1
A2 21482617 21482858
refGene_gn=SFMBT1;refGene_tx=XM_011279552.3;refGene_gn=SFMBT1;refGene_
tx=XM_023249657.1;refGene_gn=SFMBT1;refGene_tx=XM_023249661.1;refGene_
gn=SFMBT1;refGene_tx=XM_023249669.1;refGene_gn=SFMBT1;refGene_tx=XM_02
3249676.1;refGene_gn=SFMBT1;refGene_tx=XM_023249681.1;refGene_gn=SFMBT
1;refGene_tx=XM_023249685.1;refGene_gn=SFMBT1;refGene_tx=XM_023249688.
1;refGene_gn=SFMBT1;refGene_tx=XM_023249693.1
A2 21494461 21494556
refGene_gn=SFMBT1;refGene_tx=XM_011279552.3;refGene_gn=SFMBT1;refGene_
tx=XM_023249657.1;refGene_gn=SFMBT1;refGene_tx=XM_023249661.1;refGene_
gn=SFMBT1;refGene_tx=XM_023249669.1;refGene_gn=SFMBT1;refGene_tx=XM_02
3249676.1;refGene_gn=SFMBT1;refGene_tx=XM_023249681.1;refGene_gn=SFMBT
1;refGene_tx=XM_023249685.1;refGene_gn=SFMBT1;refGene_tx=XM_023249688.
1;refGene_gn=SFMBT1;refGene_tx=XM_023249693.1
A2 21504606 21504634
refGene_gn=SFMBT1;refGene_tx=XM_011279552.3;refGene_gn=SFMBT1;refGene_
tx=XM_023249657.1;refGene_gn=SFMBT1;refGene_tx=XM_023249661.1;refGene_
gn=SFMBT1;refGene_tx=XM_023249669.1;refGene_gn=SFMBT1;refGene_tx=XM_02
3249676.1;refGene_gn=SFMBT1;refGene_tx=XM_023249681.1;refGene_gn=SFMBT
1;refGene_tx=XM_023249685.1;refGene_gn=SFMBT1;refGene_tx=XM_023249688.
1;refGene_gn=SFMBT1;refGene_tx=XM_023249693.1
A2 21571359 21571410
refGene_gn=LOC111557313;refGene_tx=XM_023241847.1
A2 21572158 21573080
refGene_gn=LOC111557313;refGene_tx=XM_023241847.1
A2 21573165 21573701
refGene_gn=LOC111557313;refGene_tx=XM_023241847.1
A2 21605901 21606069 refGene_gn=RFT1;refGene_tx=XM_011279564.3
A2 21606630 21606880 refGene_gn=RFT1;refGene_tx=XM_011279564.3
A2 21611696 21611802 refGene_gn=RFT1;refGene_tx=XM_011279564.3
A2 21615409 21615554 refGene_gn=RFT1;refGene_tx=XM_011279564.3
A2 21617133 21617264 refGene_gn=RFT1;refGene_tx=XM_011279564.3
A2 21618184 21618235 refGene_gn=RFT1;refGene_tx=XM_011279564.3
A2 21621109 21621188 refGene_gn=RFT1;refGene_tx=XM_011279564.3

A2 21627740 21627878 refGene_gn=RFT1;refGene_tx=XM_011279564.3
A2 21632258 21632360 refGene_gn=RFT1;refGene_tx=XM_011279564.3
A2 21633365 21633555 refGene_gn=RFT1;refGene_tx=XM_011279564.3
A2 21634659 21634776 refGene_gn=RFT1;refGene_tx=XM_011279564.3
A2 21637097 21637183 refGene_gn=RFT1;refGene_tx=XM_011279564.3
A2 21640663 21640726 refGene_gn=RFT1;refGene_tx=XM_011279564.3
A2 21686312 21686427 refGene_gn=PRKCD;refGene_tx=XM_003982325.5
A2 21687537 21687737 refGene_gn=PRKCD;refGene_tx=XM_003982325.5
A2 21689070 21689131 refGene_gn=PRKCD;refGene_tx=XM_003982325.5
A2 21689302 21689465 refGene_gn=PRKCD;refGene_tx=XM_003982325.5
A2 21689555 21689587 refGene_gn=PRKCD;refGene_tx=XM_003982325.5
A2 21691010 21691096 refGene_gn=PRKCD;refGene_tx=XM_003982325.5
A2 21691387 21691517 refGene_gn=PRKCD;refGene_tx=XM_003982325.5
A2 21692198 21692299 refGene_gn=PRKCD;refGene_tx=XM_003982325.5
A2 21693123 21693211 refGene_gn=PRKCD;refGene_tx=XM_003982325.5
A2 21693430 21693528 refGene_gn=PRKCD;refGene_tx=XM_003982325.5
A2 21693698 21693872 refGene_gn=PRKCD;refGene_tx=XM_003982325.5
A2 21694061 21694153 refGene_gn=PRKCD;refGene_tx=XM_003982325.5
A2 21694734 21694797 refGene_gn=PRKCD;refGene_tx=XM_003982325.5
A2 21695812 21695951 refGene_gn=PRKCD;refGene_tx=XM_003982325.5
A2 21696114 21696303 refGene_gn=PRKCD;refGene_tx=XM_003982325.5
A2 21696906 21697035 refGene_gn=PRKCD;refGene_tx=XM_003982325.5
A2 21699493 21699652 refGene_gn=PRKCD;refGene_tx=XM_003982325.5
A2 21720143 21720319
refGene_gn=TKT;refGene_tx=XM_019822871.1;refGene_gn=TKT;refGene_tx=XM_023249713.1;refGene_gn=TKT;refGene_tx=XM_023249714.1
A2 21721038 21721161
refGene_gn=TKT;refGene_tx=XM_019822871.1;refGene_gn=TKT;refGene_tx=XM_023249713.1;refGene_gn=TKT;refGene_tx=XM_023249714.1
A2 21722370 21722464
refGene_gn=TKT;refGene_tx=XM_019822871.1;refGene_gn=TKT;refGene_tx=XM_023249713.1;refGene_gn=TKT;refGene_tx=XM_023249714.1
A2 21722570 21722654
refGene_gn=TKT;refGene_tx=XM_019822871.1;refGene_gn=TKT;refGene_tx=XM_023249713.1;refGene_gn=TKT;refGene_tx=XM_023249714.1
A2 21723549 21723680
refGene_gn=TKT;refGene_tx=XM_019822871.1;refGene_gn=TKT;refGene_tx=XM_023249713.1;refGene_gn=TKT;refGene_tx=XM_023249714.1
A2 21723810 21723967
refGene_gn=TKT;refGene_tx=XM_019822871.1;refGene_gn=TKT;refGene_tx=XM_023249713.1;refGene_gn=TKT;refGene_tx=XM_023249714.1
A2 21725057 21725222
refGene_gn=TKT;refGene_tx=XM_019822871.1;refGene_gn=TKT;refGene_tx=XM_023249713.1;refGene_gn=TKT;refGene_tx=XM_023249714.1
A2 21725992 21726186
refGene_gn=TKT;refGene_tx=XM_019822871.1;refGene_gn=TKT;refGene_tx=XM_023249713.1;refGene_gn=TKT;refGene_tx=XM_023249714.1
A2 21727645 21727764
refGene_gn=TKT;refGene_tx=XM_019822871.1;refGene_gn=TKT;refGene_tx=XM_023249713.1;refGene_gn=TKT;refGene_tx=XM_023249714.1

A2 21729739 21729931
refGene_gn=TKT;refGene_tx=XM_019822871.1;refGene_gn=TKT;refGene_tx=XM_023249713.1;refGene_gn=TKT;refGene_tx=XM_023249714.1
A2 21733753 21733851
refGene_gn=TKT;refGene_tx=XM_019822871.1;refGene_gn=TKT;refGene_tx=XM_023249713.1;refGene_gn=TKT;refGene_tx=XM_023249714.1
A2 21734607 21734721
refGene_gn=TKT;refGene_tx=XM_019822871.1;refGene_gn=TKT;refGene_tx=XM_023249713.1;refGene_gn=TKT;refGene_tx=XM_023249714.1
A2 21735580 21735698
refGene_gn=TKT;refGene_tx=XM_019822871.1;refGene_gn=TKT;refGene_tx=XM_023249713.1;refGene_gn=TKT;refGene_tx=XM_023249714.1
A2 21737049 21737095
refGene_gn=TKT;refGene_tx=XM_019822871.1;refGene_gn=TKT;refGene_tx=XM_023249714.1
A2 21737638 21737720 refGene_gn=TKT;refGene_tx=XM_019822871.1
A2 21745792 21745899 refGene_gn=TKT;refGene_tx=XM_023249713.1
A2 21772570 21772651
refGene_gn=DCP1A;refGene_tx=XM_003982326.5;refGene_gn=DCP1A;refGene_tx=XM_019822878.2
A2 21772961 21773180
refGene_gn=DCP1A;refGene_tx=XM_003982326.5;refGene_gn=DCP1A;refGene_tx=XM_019822878.2
A2 21774700 21774766
refGene_gn=DCP1A;refGene_tx=XM_003982326.5;refGene_gn=DCP1A;refGene_tx=XM_019822878.2
A2 21776703 21777456
refGene_gn=DCP1A;refGene_tx=XM_003982326.5;refGene_gn=DCP1A;refGene_tx=XM_019822878.2
A2 21787531 21787645
refGene_gn=DCP1A;refGene_tx=XM_003982326.5;refGene_gn=DCP1A;refGene_tx=XM_019822878.2
A2 21792649 21792788
refGene_gn=DCP1A;refGene_tx=XM_003982326.5;refGene_gn=DCP1A;refGene_tx=XM_019822878.2
A2 21798270 21798337
refGene_gn=DCP1A;refGene_tx=XM_003982326.5;refGene_gn=DCP1A;refGene_tx=XM_019822878.2
A2 21822372 21822500 refGene_gn=DCP1A;refGene_tx=XM_003982326.5
A2 21824620 21824661 refGene_gn=DCP1A;refGene_tx=XM_003982326.5
A2 21826672 21826807 refGene_gn=DCP1A;refGene_tx=XM_003982326.5
A2 21937439 21937511
refGene_gn=LOC111557316;refGene_tx=XM_023241849.1
A2 21938821 21939357
refGene_gn=LOC111557316;refGene_tx=XM_023241849.1
A2 21939413 21939648
refGene_gn=LOC111557316;refGene_tx=XM_023241849.1
A2 21939998 21940065
refGene_gn=CACNA1D;refGene_tx=XM_003982328.4;refGene_gn=CACNA1D;refGene_tx=XM_003982329.4;refGene_gn=CACNA1D;refGene_tx=XM_003982331.4;refGe

ne_gn=CACNA1D;refGene_tx=XM_019822887.2;refGene_gn=CACNA1D;refGene_tx=XM_019822894.2;refGene_gn=CACNA1D;refGene_tx=XM_019822897.2;refGene_gn=CACNA1D;refGene_tx=XM_019822906.2;refGene_gn=CACNA1D;refGene_tx=XM_019822918.1;refGene_gn=CACNA1D;refGene_tx=XM_019822924.2;refGene_gn=CACNA1D;refGene_tx=XM_019822928.2;refGene_gn=CACNA1D;refGene_tx=XM_019822930.2;refGene_gn=CACNA1D;refGene_tx=XM_019822934.2;refGene_gn=CACNA1D;refGene_tx=XM_023249752.1;refGene_gn=CACNA1D;refGene_tx=XM_023249761.1;refGene_gn=CACNA1D;refGene_tx=XM_023249772.1;refGene_gn=CACNA1D;refGene_tx=XM_023249799.1;refGene_gn=CACNA1D;refGene_tx=XM_023249806.1;refGene_gn=CACNA1D;refGene_tx=XM_023249808.1;refGene_gn=CACNA1D;refGene_tx=XM_023249814.1;refGene_gn=CACNA1D;refGene_tx=XM_023249815.1
A2 21942011 21942321
refGene_gn=CACNA1D;refGene_tx=XM_003982328.4;refGene_gn=CACNA1D;refGene_tx=XM_003982329.4;refGene_gn=CACNA1D;refGene_tx=XM_003982331.4;refGene_gn=CACNA1D;refGene_tx=XM_019822887.2;refGene_gn=CACNA1D;refGene_tx=XM_019822894.2;refGene_gn=CACNA1D;refGene_tx=XM_019822897.2;refGene_gn=CACNA1D;refGene_tx=XM_019822906.2;refGene_gn=CACNA1D;refGene_tx=XM_019822918.1;refGene_gn=CACNA1D;refGene_tx=XM_019822924.2;refGene_gn=CACNA1D;refGene_tx=XM_019822928.2;refGene_gn=CACNA1D;refGene_tx=XM_019822930.2;refGene_gn=CACNA1D;refGene_tx=XM_019822934.2;refGene_gn=CACNA1D;refGene_tx=XM_023249752.1;refGene_gn=CACNA1D;refGene_tx=XM_023249761.1;refGene_gn=CACNA1D;refGene_tx=XM_023249772.1;refGene_gn=CACNA1D;refGene_tx=XM_023249799.1;refGene_gn=CACNA1D;refGene_tx=XM_023249806.1;refGene_gn=CACNA1D;refGene_tx=XM_023249808.1;refGene_gn=CACNA1D;refGene_tx=XM_023249814.1;refGene_gn=CACNA1D;refGene_tx=XM_023249815.1
A2 21946029 21946135
refGene_gn=CACNA1D;refGene_tx=XM_003982328.4;refGene_gn=CACNA1D;refGene_tx=XM_003982329.4;refGene_gn=CACNA1D;refGene_tx=XM_003982331.4;refGene_gn=CACNA1D;refGene_tx=XM_019822887.2;refGene_gn=CACNA1D;refGene_tx=XM_019822894.2;refGene_gn=CACNA1D;refGene_tx=XM_019822897.2;refGene_gn=CACNA1D;refGene_tx=XM_019822906.2;refGene_gn=CACNA1D;refGene_tx=XM_019822918.1;refGene_gn=CACNA1D;refGene_tx=XM_019822924.2;refGene_gn=CACNA1D;refGene_tx=XM_019822928.2;refGene_gn=CACNA1D;refGene_tx=XM_019822930.2;refGene_gn=CACNA1D;refGene_tx=XM_019822934.2;refGene_gn=CACNA1D;refGene_tx=XM_023249752.1;refGene_gn=CACNA1D;refGene_tx=XM_023249761.1;refGene_gn=CACNA1D;refGene_tx=XM_023249772.1;refGene_gn=CACNA1D;refGene_tx=XM_023249799.1;refGene_gn=CACNA1D;refGene_tx=XM_023249806.1;refGene_gn=CACNA1D;refGene_tx=XM_023249808.1;refGene_gn=CACNA1D;refGene_tx=XM_023249814.1;refGene_gn=CACNA1D;refGene_tx=XM_023249815.1
A2 22086604 22086744
refGene_gn=CACNA1D;refGene_tx=XM_003982328.4;refGene_gn=CACNA1D;refGene_tx=XM_003982329.4;refGene_gn=CACNA1D;refGene_tx=XM_003982331.4;refGene_gn=CACNA1D;refGene_tx=XM_019822887.2;refGene_gn=CACNA1D;refGene_tx=XM_019822894.2;refGene_gn=CACNA1D;refGene_tx=XM_019822897.2;refGene_gn=CACNA1D;refGene_tx=XM_019822906.2;refGene_gn=CACNA1D;refGene_tx=XM_019822918.1;refGene_gn=CACNA1D;refGene_tx=XM_019822924.2;refGene_gn=CACNA1D;refGene_tx=XM_019822928.2;refGene_gn=CACNA1D;refGene_tx=XM_019822930.2;refGene_gn=CACNA1D;refGene_tx=XM_019822934.2;refGene_gn=CACNA1D;refGene_tx=XM_023249752.1;refGene_gn=CACNA1D;refGene_tx=XM_023249761.1;refGene_gn=CACNA1D;refGene_tx=XM_023249772.1;refGene_gn=CACNA1D;refGen

e_tx=XM_023249799.1;refGene_gn=CACNA1D;refGene_tx=XM_023249806.1;refGene_gn=CACNA1D;refGene_tx=XM_023249808.1;refGene_gn=CACNA1D;refGene_tx=XM_023249814.1;refGene_gn=CACNA1D;refGene_tx=XM_023249815.1
A2 22095533 22095676
refGene_gn=CACNA1D;refGene_tx=XM_003982328.4;refGene_gn=CACNA1D;refGene_tx=XM_003982329.4;refGene_gn=CACNA1D;refGene_tx=XM_003982331.4;refGene_gn=CACNA1D;refGene_tx=XM_019822887.2;refGene_gn=CACNA1D;refGene_tx=XM_019822894.2;refGene_gn=CACNA1D;refGene_tx=XM_019822897.2;refGene_gn=CACNA1D;refGene_tx=XM_019822906.2;refGene_gn=CACNA1D;refGene_tx=XM_019822918.1;refGene_gn=CACNA1D;refGene_tx=XM_019822924.2;refGene_gn=CACNA1D;refGene_tx=XM_019822928.2;refGene_gn=CACNA1D;refGene_tx=XM_019822930.2;refGene_gn=CACNA1D;refGene_tx=XM_019822934.2;refGene_gn=CACNA1D;refGene_tx=XM_023249752.1;refGene_gn=CACNA1D;refGene_tx=XM_023249761.1;refGene_gn=CACNA1D;refGene_tx=XM_023249772.1;refGene_gn=CACNA1D;refGene_tx=XM_023249799.1;refGene_gn=CACNA1D;refGene_tx=XM_023249806.1;refGene_gn=CACNA1D;refGene_tx=XM_023249808.1;refGene_gn=CACNA1D;refGene_tx=XM_023249814.1;refGene_gn=CACNA1D;refGene_tx=XM_023249815.1
A2 22100025 22100178
refGene_gn=CACNA1D;refGene_tx=XM_003982328.4;refGene_gn=CACNA1D;refGene_tx=XM_003982329.4;refGene_gn=CACNA1D;refGene_tx=XM_003982331.4;refGene_gn=CACNA1D;refGene_tx=XM_019822887.2;refGene_gn=CACNA1D;refGene_tx=XM_019822894.2;refGene_gn=CACNA1D;refGene_tx=XM_019822897.2;refGene_gn=CACNA1D;refGene_tx=XM_019822906.2;refGene_gn=CACNA1D;refGene_tx=XM_019822918.1;refGene_gn=CACNA1D;refGene_tx=XM_019822924.2;refGene_gn=CACNA1D;refGene_tx=XM_019822928.2;refGene_gn=CACNA1D;refGene_tx=XM_019822930.2;refGene_gn=CACNA1D;refGene_tx=XM_019822934.2;refGene_gn=CACNA1D;refGene_tx=XM_023249752.1;refGene_gn=CACNA1D;refGene_tx=XM_023249761.1;refGene_gn=CACNA1D;refGene_tx=XM_023249772.1;refGene_gn=CACNA1D;refGene_tx=XM_023249799.1;refGene_gn=CACNA1D;refGene_tx=XM_023249806.1;refGene_gn=CACNA1D;refGene_tx=XM_023249808.1;refGene_gn=CACNA1D;refGene_tx=XM_023249814.1;refGene_gn=CACNA1D;refGene_tx=XM_023249815.1
A2 22100699 22100896
refGene_gn=CACNA1D;refGene_tx=XM_003982328.4;refGene_gn=CACNA1D;refGene_tx=XM_003982329.4;refGene_gn=CACNA1D;refGene_tx=XM_003982331.4;refGene_gn=CACNA1D;refGene_tx=XM_019822887.2;refGene_gn=CACNA1D;refGene_tx=XM_019822894.2;refGene_gn=CACNA1D;refGene_tx=XM_019822897.2;refGene_gn=CACNA1D;refGene_tx=XM_019822906.2;refGene_gn=CACNA1D;refGene_tx=XM_019822918.1;refGene_gn=CACNA1D;refGene_tx=XM_019822924.2;refGene_gn=CACNA1D;refGene_tx=XM_019822928.2;refGene_gn=CACNA1D;refGene_tx=XM_019822930.2;refGene_gn=CACNA1D;refGene_tx=XM_019822934.2;refGene_gn=CACNA1D;refGene_tx=XM_023249752.1;refGene_gn=CACNA1D;refGene_tx=XM_023249761.1;refGene_gn=CACNA1D;refGene_tx=XM_023249772.1;refGene_gn=CACNA1D;refGene_tx=XM_023249799.1;refGene_gn=CACNA1D;refGene_tx=XM_023249806.1;refGene_gn=CACNA1D;refGene_tx=XM_023249808.1;refGene_gn=CACNA1D;refGene_tx=XM_023249814.1;refGene_gn=CACNA1D;refGene_tx=XM_023249815.1
A2 22106959 22107063
refGene_gn=CACNA1D;refGene_tx=XM_003982328.4;refGene_gn=CACNA1D;refGene_tx=XM_003982331.4;refGene_gn=CACNA1D;refGene_tx=XM_019822887.2;refGene_gn=CACNA1D;refGene_tx=XM_019822897.2;refGene_gn=CACNA1D;refGene_tx=XM_019822934.2;refGene_gn=CACNA1D;refGene_tx=XM_023249752.1;refGene_gn

=CACNA1D;refGene_tx=XM_023249761.1;refGene_gn=CACNA1D;refGene_tx=XM_023249772.1;refGene_gn=CACNA1D;refGene_tx=XM_023249799.1;refGene_gn=CACNA1D;refGene_tx=XM_023249806.1;refGene_gn=CACNA1D;refGene_tx=XM_023249808.1;refGene_gn=CACNA1D;refGene_tx=XM_023249814.1;refGene_gn=CACNA1D;refGene_tx=XM_023249815.1
A2 22107608 22107712
refGene_gn=CACNA1D;refGene_tx=XM_003982329.4;refGene_gn=CACNA1D;refGene_tx=XM_019822894.2;refGene_gn=CACNA1D;refGene_tx=XM_019822906.2;refGene_gn=CACNA1D;refGene_tx=XM_019822918.1;refGene_gn=CACNA1D;refGene_tx=XM_019822924.2;refGene_gn=CACNA1D;refGene_tx=XM_019822928.2;refGene_gn=CACNA1D;refGene_tx=XM_019822930.2
A2 22138874 22139044
refGene_gn=CACNA1D;refGene_tx=XM_003982328.4;refGene_gn=CACNA1D;refGene_tx=XM_003982329.4;refGene_gn=CACNA1D;refGene_tx=XM_003982331.4;refGene_gn=CACNA1D;refGene_tx=XM_019822887.2;refGene_gn=CACNA1D;refGene_tx=XM_019822894.2;refGene_gn=CACNA1D;refGene_tx=XM_019822897.2;refGene_gn=CACNA1D;refGene_tx=XM_019822906.2;refGene_gn=CACNA1D;refGene_tx=XM_019822918.1;refGene_gn=CACNA1D;refGene_tx=XM_019822924.2;refGene_gn=CACNA1D;refGene_tx=XM_019822928.2;refGene_gn=CACNA1D;refGene_tx=XM_019822930.2;refGene_gn=CACNA1D;refGene_tx=XM_019822934.2;refGene_gn=CACNA1D;refGene_tx=XM_023249752.1;refGene_gn=CACNA1D;refGene_tx=XM_023249761.1;refGene_gn=CACNA1D;refGene_tx=XM_023249772.1;refGene_gn=CACNA1D;refGene_tx=XM_023249799.1;refGene_gn=CACNA1D;refGene_tx=XM_023249806.1;refGene_gn=CACNA1D;refGene_tx=XM_023249808.1;refGene_gn=CACNA1D;refGene_tx=XM_023249814.1;refGene_gn=CACNA1D;refGene_tx=XM_023249815.1
A2 22154940 22155028
refGene_gn=CACNA1D;refGene_tx=XM_003982328.4;refGene_gn=CACNA1D;refGene_tx=XM_003982329.4;refGene_gn=CACNA1D;refGene_tx=XM_003982331.4;refGene_gn=CACNA1D;refGene_tx=XM_019822887.2;refGene_gn=CACNA1D;refGene_tx=XM_019822894.2;refGene_gn=CACNA1D;refGene_tx=XM_019822897.2;refGene_gn=CACNA1D;refGene_tx=XM_019822906.2;refGene_gn=CACNA1D;refGene_tx=XM_019822918.1;refGene_gn=CACNA1D;refGene_tx=XM_019822924.2;refGene_gn=CACNA1D;refGene_tx=XM_019822928.2;refGene_gn=CACNA1D;refGene_tx=XM_019822930.2;refGene_gn=CACNA1D;refGene_tx=XM_019822934.2;refGene_gn=CACNA1D;refGene_tx=XM_023249752.1;refGene_gn=CACNA1D;refGene_tx=XM_023249761.1;refGene_gn=CACNA1D;refGene_tx=XM_023249772.1;refGene_gn=CACNA1D;refGene_tx=XM_023249799.1;refGene_gn=CACNA1D;refGene_tx=XM_023249806.1;refGene_gn=CACNA1D;refGene_tx=XM_023249808.1;refGene_gn=CACNA1D;refGene_tx=XM_023249814.1;refGene_gn=CACNA1D;refGene_tx=XM_023249815.1
A2 22155307 22155367
refGene_gn=CACNA1D;refGene_tx=XM_003982329.4;refGene_gn=CACNA1D;refGene_tx=XM_019822887.2;refGene_gn=CACNA1D;refGene_tx=XM_019822894.2;refGene_gn=CACNA1D;refGene_tx=XM_019822924.2;refGene_gn=CACNA1D;refGene_tx=XM_023249752.1;refGene_gn=CACNA1D;refGene_tx=XM_023249761.1;refGene_gn=CACNA1D;refGene_tx=XM_023249772.1;refGene_gn=CACNA1D;refGene_tx=XM_023249799.1;refGene_gn=CACNA1D;refGene_tx=XM_023249806.1;refGene_gn=CACNA1D;refGene_tx=XM_023249808.1;refGene_gn=CACNA1D;refGene_tx=XM_023249814.1;refGene_gn=CACNA1D;refGene_tx=XM_023249815.1
A2 22156379 22156406
refGene_gn=CACNA1D;refGene_tx=XM_003982328.4;refGene_gn=CACNA1D;refGen

e_tx=XM_003982329.4;refGene_gn=CACNA1D;refGene_tx=XM_003982331.4;refGene_gn=CACNA1D;refGene_tx=XM_019822887.2;refGene_gn=CACNA1D;refGene_tx=XM_019822894.2;refGene_gn=CACNA1D;refGene_tx=XM_019822897.2;refGene_gn=CACNA1D;refGene_tx=XM_019822906.2;refGene_gn=CACNA1D;refGene_tx=XM_019822918.1;refGene_gn=CACNA1D;refGene_tx=XM_019822924.2;refGene_gn=CACNA1D;refGene_tx=XM_019822928.2;refGene_gn=CACNA1D;refGene_tx=XM_019822930.2;refGene_gn=CACNA1D;refGene_tx=XM_019822934.2;refGene_gn=CACNA1D;refGene_tx=XM_023249752.1;refGene_gn=CACNA1D;refGene_tx=XM_023249761.1;refGene_gn=CACNA1D;refGene_tx=XM_023249772.1;refGene_gn=CACNA1D;refGene_tx=XM_023249799.1;refGene_gn=CACNA1D;refGene_tx=XM_023249806.1;refGene_gn=CACNA1D;refGene_tx=XM_023249808.1;refGene_gn=CACNA1D;refGene_tx=XM_023249814.1;refGene_gn=CACNA1D;refGene_tx=XM_023249815.1
A2 22159764 22159925
refGene_gn=CACNA1D;refGene_tx=XM_003982328.4;refGene_gn=CACNA1D;refGene_tx=XM_003982329.4;refGene_gn=CACNA1D;refGene_tx=XM_003982331.4;refGene_gn=CACNA1D;refGene_tx=XM_019822887.2;refGene_gn=CACNA1D;refGene_tx=XM_019822894.2;refGene_gn=CACNA1D;refGene_tx=XM_019822897.2;refGene_gn=CACNA1D;refGene_tx=XM_019822906.2;refGene_gn=CACNA1D;refGene_tx=XM_019822918.1;refGene_gn=CACNA1D;refGene_tx=XM_019822924.2;refGene_gn=CACNA1D;refGene_tx=XM_019822928.2;refGene_gn=CACNA1D;refGene_tx=XM_019822930.2;refGene_gn=CACNA1D;refGene_tx=XM_019822934.2;refGene_gn=CACNA1D;refGene_tx=XM_023249752.1;refGene_gn=CACNA1D;refGene_tx=XM_023249761.1;refGene_gn=CACNA1D;refGene_tx=XM_023249772.1;refGene_gn=CACNA1D;refGene_tx=XM_023249799.1;refGene_gn=CACNA1D;refGene_tx=XM_023249806.1;refGene_gn=CACNA1D;refGene_tx=XM_023249808.1;refGene_gn=CACNA1D;refGene_tx=XM_023249814.1;refGene_gn=CACNA1D;refGene_tx=XM_023249815.1
A2 22160706 22160932
refGene_gn=CACNA1D;refGene_tx=XM_003982328.4;refGene_gn=CACNA1D;refGene_tx=XM_003982329.4;refGene_gn=CACNA1D;refGene_tx=XM_003982331.4;refGene_gn=CACNA1D;refGene_tx=XM_019822887.2;refGene_gn=CACNA1D;refGene_tx=XM_019822894.2;refGene_gn=CACNA1D;refGene_tx=XM_019822897.2;refGene_gn=CACNA1D;refGene_tx=XM_019822906.2;refGene_gn=CACNA1D;refGene_tx=XM_019822918.1;refGene_gn=CACNA1D;refGene_tx=XM_019822924.2;refGene_gn=CACNA1D;refGene_tx=XM_019822928.2;refGene_gn=CACNA1D;refGene_tx=XM_019822930.2;refGene_gn=CACNA1D;refGene_tx=XM_019822934.2;refGene_gn=CACNA1D;refGene_tx=XM_023249752.1;refGene_gn=CACNA1D;refGene_tx=XM_023249761.1;refGene_gn=CACNA1D;refGene_tx=XM_023249772.1;refGene_gn=CACNA1D;refGene_tx=XM_023249799.1;refGene_gn=CACNA1D;refGene_tx=XM_023249806.1;refGene_gn=CACNA1D;refGene_tx=XM_023249808.1;refGene_gn=CACNA1D;refGene_tx=XM_023249814.1;refGene_gn=CACNA1D;refGene_tx=XM_023249815.1
A2 22161059 22161267
refGene_gn=CACNA1D;refGene_tx=XM_003982328.4;refGene_gn=CACNA1D;refGene_tx=XM_003982329.4;refGene_gn=CACNA1D;refGene_tx=XM_003982331.4;refGene_gn=CACNA1D;refGene_tx=XM_019822887.2;refGene_gn=CACNA1D;refGene_tx=XM_019822894.2;refGene_gn=CACNA1D;refGene_tx=XM_019822897.2;refGene_gn=CACNA1D;refGene_tx=XM_019822906.2;refGene_gn=CACNA1D;refGene_tx=XM_019822918.1;refGene_gn=CACNA1D;refGene_tx=XM_019822924.2;refGene_gn=CACNA1D;refGene_tx=XM_019822928.2;refGene_gn=CACNA1D;refGene_tx=XM_019822930.2;refGene_gn=CACNA1D;refGene_tx=XM_019822934.2;refGene_gn=CACNA1D;refGene_tx=XM_023249752.1;refGene_gn=CACNA1D;refGene_tx=XM_023249761.1;

refGene_gn=CACNA1D;refGene_tx=XM_023249772.1;refGene_gn=CACNA1D;refGene_tx=XM_023249799.1;refGene_gn=CACNA1D;refGene_tx=XM_023249806.1;refGene_gn=CACNA1D;refGene_tx=XM_023249808.1;refGene_gn=CACNA1D;refGene_tx=XM_023249814.1;refGene_gn=CACNA1D;refGene_tx=XM_023249815.1
A2 22164204 22164325
refGene_gn=CACNA1D;refGene_tx=XM_003982328.4;refGene_gn=CACNA1D;refGene_tx=XM_003982329.4;refGene_gn=CACNA1D;refGene_tx=XM_003982331.4;refGene_gn=CACNA1D;refGene_tx=XM_019822887.2;refGene_gn=CACNA1D;refGene_tx=XM_019822894.2;refGene_gn=CACNA1D;refGene_tx=XM_019822897.2;refGene_gn=CACNA1D;refGene_tx=XM_019822906.2;refGene_gn=CACNA1D;refGene_tx=XM_019822918.1;refGene_gn=CACNA1D;refGene_tx=XM_019822924.2;refGene_gn=CACNA1D;refGene_tx=XM_019822928.2;refGene_gn=CACNA1D;refGene_tx=XM_019822930.2;refGene_gn=CACNA1D;refGene_tx=XM_019822934.2;refGene_gn=CACNA1D;refGene_tx=XM_023249752.1;refGene_gn=CACNA1D;refGene_tx=XM_023249761.1;refGene_gn=CACNA1D;refGene_tx=XM_023249772.1;refGene_gn=CACNA1D;refGene_tx=XM_023249799.1;refGene_gn=CACNA1D;refGene_tx=XM_023249806.1;refGene_gn=CACNA1D;refGene_tx=XM_023249808.1;refGene_gn=CACNA1D;refGene_tx=XM_023249814.1;refGene_gn=CACNA1D;refGene_tx=XM_023249815.1
A2 22167846 22167961
refGene_gn=CACNA1D;refGene_tx=XM_003982328.4;refGene_gn=CACNA1D;refGene_tx=XM_003982329.4;refGene_gn=CACNA1D;refGene_tx=XM_003982331.4;refGene_gn=CACNA1D;refGene_tx=XM_019822887.2;refGene_gn=CACNA1D;refGene_tx=XM_019822894.2;refGene_gn=CACNA1D;refGene_tx=XM_019822897.2;refGene_gn=CACNA1D;refGene_tx=XM_019822906.2;refGene_gn=CACNA1D;refGene_tx=XM_019822918.1;refGene_gn=CACNA1D;refGene_tx=XM_019822924.2;refGene_gn=CACNA1D;refGene_tx=XM_019822928.2;refGene_gn=CACNA1D;refGene_tx=XM_019822930.2;refGene_gn=CACNA1D;refGene_tx=XM_019822934.2;refGene_gn=CACNA1D;refGene_tx=XM_023249752.1;refGene_gn=CACNA1D;refGene_tx=XM_023249761.1;refGene_gn=CACNA1D;refGene_tx=XM_023249772.1;refGene_gn=CACNA1D;refGene_tx=XM_023249799.1;refGene_gn=CACNA1D;refGene_tx=XM_023249806.1;refGene_gn=CACNA1D;refGene_tx=XM_023249808.1;refGene_gn=CACNA1D;refGene_tx=XM_023249814.1;refGene_gn=CACNA1D;refGene_tx=XM_023249815.1
A2 22168342 22168412
refGene_gn=CACNA1D;refGene_tx=XM_003982328.4;refGene_gn=CACNA1D;refGene_tx=XM_003982329.4;refGene_gn=CACNA1D;refGene_tx=XM_003982331.4;refGene_gn=CACNA1D;refGene_tx=XM_019822887.2;refGene_gn=CACNA1D;refGene_tx=XM_019822894.2;refGene_gn=CACNA1D;refGene_tx=XM_019822897.2;refGene_gn=CACNA1D;refGene_tx=XM_019822906.2;refGene_gn=CACNA1D;refGene_tx=XM_019822918.1;refGene_gn=CACNA1D;refGene_tx=XM_019822924.2;refGene_gn=CACNA1D;refGene_tx=XM_019822928.2;refGene_gn=CACNA1D;refGene_tx=XM_019822930.2;refGene_gn=CACNA1D;refGene_tx=XM_019822934.2;refGene_gn=CACNA1D;refGene_tx=XM_023249752.1;refGene_gn=CACNA1D;refGene_tx=XM_023249761.1;refGene_gn=CACNA1D;refGene_tx=XM_023249772.1;refGene_gn=CACNA1D;refGene_tx=XM_023249799.1;refGene_gn=CACNA1D;refGene_tx=XM_023249806.1;refGene_gn=CACNA1D;refGene_tx=XM_023249808.1;refGene_gn=CACNA1D;refGene_tx=XM_023249814.1;refGene_gn=CACNA1D;refGene_tx=XM_023249815.1
A2 22169275 22169342
refGene_gn=CACNA1D;refGene_tx=XM_003982328.4;refGene_gn=CACNA1D;refGene_tx=XM_003982329.4;refGene_gn=CACNA1D;refGene_tx=XM_003982331.4;refGene_gn=CACNA1D;refGene_tx=XM_019822887.2;refGene_gn=CACNA1D;refGene_tx=

XM_019822894.2;refGene_gn=CACNA1D;refGene_tx=XM_019822897.2;refGene_gn=CACNA1D;refGene_tx=XM_019822906.2;refGene_gn=CACNA1D;refGene_tx=XM_019822918.1;refGene_gn=CACNA1D;refGene_tx=XM_019822924.2;refGene_gn=CACNA1D;refGene_tx=XM_019822928.2;refGene_gn=CACNA1D;refGene_tx=XM_019822930.2;refGene_gn=CACNA1D;refGene_tx=XM_019822934.2;refGene_gn=CACNA1D;refGene_tx=XM_023249752.1;refGene_gn=CACNA1D;refGene_tx=XM_023249761.1;refGene_gn=CACNA1D;refGene_tx=XM_023249772.1;refGene_gn=CACNA1D;refGene_tx=XM_023249799.1;refGene_gn=CACNA1D;refGene_tx=XM_023249806.1;refGene_gn=CACNA1D;refGene_tx=XM_023249808.1;refGene_gn=CACNA1D;refGene_tx=XM_023249814.1;refGene_gn=CACNA1D;refGene_tx=XM_023249815.1

A2 22170091 22170239

refGene_gn=CACNA1D;refGene_tx=XM_003982328.4;refGene_gn=CACNA1D;refGene_tx=XM_003982329.4;refGene_gn=CACNA1D;refGene_tx=XM_003982331.4;refGene_gn=CACNA1D;refGene_tx=XM_019822887.2;refGene_gn=CACNA1D;refGene_tx=XM_019822894.2;refGene_gn=CACNA1D;refGene_tx=XM_019822897.2;refGene_gn=CACNA1D;refGene_tx=XM_019822906.2;refGene_gn=CACNA1D;refGene_tx=XM_019822918.1;refGene_gn=CACNA1D;refGene_tx=XM_019822924.2;refGene_gn=CACNA1D;refGene_tx=XM_019822928.2;refGene_gn=CACNA1D;refGene_tx=XM_019822930.2;refGene_gn=CACNA1D;refGene_tx=XM_019822934.2;refGene_gn=CACNA1D;refGene_tx=XM_023249752.1;refGene_gn=CACNA1D;refGene_tx=XM_023249761.1;refGene_gn=CACNA1D;refGene_tx=XM_023249772.1;refGene_gn=CACNA1D;refGene_tx=XM_023249799.1;refGene_gn=CACNA1D;refGene_tx=XM_023249806.1;refGene_gn=CACNA1D;refGene_tx=XM_023249808.1;refGene_gn=CACNA1D;refGene_tx=XM_023249814.1;refGene_gn=CACNA1D;refGene_tx=XM_023249815.1

A2 22172854 22172984

refGene_gn=CACNA1D;refGene_tx=XM_003982328.4;refGene_gn=CACNA1D;refGene_tx=XM_003982329.4;refGene_gn=CACNA1D;refGene_tx=XM_003982331.4;refGene_gn=CACNA1D;refGene_tx=XM_019822887.2;refGene_gn=CACNA1D;refGene_tx=XM_019822894.2;refGene_gn=CACNA1D;refGene_tx=XM_019822897.2;refGene_gn=CACNA1D;refGene_tx=XM_019822906.2;refGene_gn=CACNA1D;refGene_tx=XM_019822918.1;refGene_gn=CACNA1D;refGene_tx=XM_019822924.2;refGene_gn=CACNA1D;refGene_tx=XM_019822928.2;refGene_gn=CACNA1D;refGene_tx=XM_019822930.2;refGene_gn=CACNA1D;refGene_tx=XM_019822934.2;refGene_gn=CACNA1D;refGene_tx=XM_023249752.1;refGene_gn=CACNA1D;refGene_tx=XM_023249761.1;refGene_gn=CACNA1D;refGene_tx=XM_023249772.1;refGene_gn=CACNA1D;refGene_tx=XM_023249799.1;refGene_gn=CACNA1D;refGene_tx=XM_023249806.1;refGene_gn=CACNA1D;refGene_tx=XM_023249808.1;refGene_gn=CACNA1D;refGene_tx=XM_023249814.1;refGene_gn=CACNA1D;refGene_tx=XM_023249815.1

A2 22176257 22176317

refGene_gn=CACNA1D;refGene_tx=XM_003982328.4;refGene_gn=CACNA1D;refGene_tx=XM_003982329.4;refGene_gn=CACNA1D;refGene_tx=XM_003982331.4;refGene_gn=CACNA1D;refGene_tx=XM_019822887.2;refGene_gn=CACNA1D;refGene_tx=XM_019822894.2;refGene_gn=CACNA1D;refGene_tx=XM_019822897.2;refGene_gn=CACNA1D;refGene_tx=XM_019822906.2;refGene_gn=CACNA1D;refGene_tx=XM_019822918.1;refGene_gn=CACNA1D;refGene_tx=XM_019822924.2;refGene_gn=CACNA1D;refGene_tx=XM_019822928.2;refGene_gn=CACNA1D;refGene_tx=XM_019822930.2;refGene_gn=CACNA1D;refGene_tx=XM_019822934.2;refGene_gn=CACNA1D;refGene_tx=XM_023249752.1;refGene_gn=CACNA1D;refGene_tx=XM_023249761.1;refGene_gn=CACNA1D;refGene_tx=XM_023249772.1;refGene_gn=CACNA1D;refGene_tx=XM_023249799.1;refGene_gn=CACNA1D;refGene_tx=XM_023249806.1;refGe

ne_gn=CACNA1D;refGene_tx=XM_023249808.1;refGene_gn=CACNA1D;refGene_tx=XM_023249814.1;refGene_gn=CACNA1D;refGene_tx=XM_023249815.1
A2 22179639 22179746
refGene_gn=CACNA1D;refGene_tx=XM_003982328.4;refGene_gn=CACNA1D;refGene_tx=XM_003982329.4;refGene_gn=CACNA1D;refGene_tx=XM_003982331.4;refGene_gn=CACNA1D;refGene_tx=XM_019822887.2;refGene_gn=CACNA1D;refGene_tx=XM_019822894.2;refGene_gn=CACNA1D;refGene_tx=XM_019822897.2;refGene_gn=CACNA1D;refGene_tx=XM_019822906.2;refGene_gn=CACNA1D;refGene_tx=XM_019822918.1;refGene_gn=CACNA1D;refGene_tx=XM_019822924.2;refGene_gn=CACNA1D;refGene_tx=XM_019822928.2;refGene_gn=CACNA1D;refGene_tx=XM_019822930.2;refGene_gn=CACNA1D;refGene_tx=XM_019822934.2;refGene_gn=CACNA1D;refGene_tx=XM_023249752.1;refGene_gn=CACNA1D;refGene_tx=XM_023249761.1;refGene_gn=CACNA1D;refGene_tx=XM_023249772.1;refGene_gn=CACNA1D;refGene_tx=XM_023249799.1;refGene_gn=CACNA1D;refGene_tx=XM_023249806.1;refGene_gn=CACNA1D;refGene_tx=XM_023249808.1;refGene_gn=CACNA1D;refGene_tx=XM_023249814.1;refGene_gn=CACNA1D;refGene_tx=XM_023249815.1
A2 22181344 22181432
refGene_gn=CACNA1D;refGene_tx=XM_003982328.4;refGene_gn=CACNA1D;refGene_tx=XM_003982329.4;refGene_gn=CACNA1D;refGene_tx=XM_003982331.4;refGene_gn=CACNA1D;refGene_tx=XM_019822887.2;refGene_gn=CACNA1D;refGene_tx=XM_019822894.2;refGene_gn=CACNA1D;refGene_tx=XM_019822897.2;refGene_gn=CACNA1D;refGene_tx=XM_019822906.2;refGene_gn=CACNA1D;refGene_tx=XM_019822918.1;refGene_gn=CACNA1D;refGene_tx=XM_019822924.2;refGene_gn=CACNA1D;refGene_tx=XM_019822928.2;refGene_gn=CACNA1D;refGene_tx=XM_019822930.2;refGene_gn=CACNA1D;refGene_tx=XM_019822934.2;refGene_gn=CACNA1D;refGene_tx=XM_023249752.1;refGene_gn=CACNA1D;refGene_tx=XM_023249761.1;refGene_gn=CACNA1D;refGene_tx=XM_023249772.1;refGene_gn=CACNA1D;refGene_tx=XM_023249799.1;refGene_gn=CACNA1D;refGene_tx=XM_023249806.1;refGene_gn=CACNA1D;refGene_tx=XM_023249808.1;refGene_gn=CACNA1D;refGene_tx=XM_023249814.1;refGene_gn=CACNA1D;refGene_tx=XM_023249815.1
A2 22182448 22182556
refGene_gn=CACNA1D;refGene_tx=XM_003982328.4;refGene_gn=CACNA1D;refGene_tx=XM_003982329.4;refGene_gn=CACNA1D;refGene_tx=XM_003982331.4;refGene_gn=CACNA1D;refGene_tx=XM_019822887.2;refGene_gn=CACNA1D;refGene_tx=XM_019822894.2;refGene_gn=CACNA1D;refGene_tx=XM_019822897.2;refGene_gn=CACNA1D;refGene_tx=XM_019822906.2;refGene_gn=CACNA1D;refGene_tx=XM_019822918.1;refGene_gn=CACNA1D;refGene_tx=XM_019822924.2;refGene_gn=CACNA1D;refGene_tx=XM_019822928.2;refGene_gn=CACNA1D;refGene_tx=XM_019822930.2;refGene_gn=CACNA1D;refGene_tx=XM_019822934.2;refGene_gn=CACNA1D;refGene_tx=XM_023249752.1;refGene_gn=CACNA1D;refGene_tx=XM_023249761.1;refGene_gn=CACNA1D;refGene_tx=XM_023249772.1;refGene_gn=CACNA1D;refGene_tx=XM_023249799.1;refGene_gn=CACNA1D;refGene_tx=XM_023249806.1;refGene_gn=CACNA1D;refGene_tx=XM_023249808.1;refGene_gn=CACNA1D;refGene_tx=XM_023249814.1;refGene_gn=CACNA1D;refGene_tx=XM_023249815.1
A2 22182642 22182695
refGene_gn=CACNA1D;refGene_tx=XM_003982328.4;refGene_gn=CACNA1D;refGene_tx=XM_003982329.4;refGene_gn=CACNA1D;refGene_tx=XM_003982331.4;refGene_gn=CACNA1D;refGene_tx=XM_019822887.2;refGene_gn=CACNA1D;refGene_tx=XM_019822894.2;refGene_gn=CACNA1D;refGene_tx=XM_019822897.2;refGene_gn=CACNA1D;refGene_tx=XM_019822906.2;refGene_gn=CACNA1D;refGene_tx=XM_01

9822918.1;refGene_gn=CACNA1D;refGene_tx=XM_019822924.2;refGene_gn=CACNA1D;refGene_tx=XM_019822928.2;refGene_gn=CACNA1D;refGene_tx=XM_019822930.2;refGene_gn=CACNA1D;refGene_tx=XM_019822934.2;refGene_gn=CACNA1D;refGene_tx=XM_023249752.1;refGene_gn=CACNA1D;refGene_tx=XM_023249761.1;refGene_gn=CACNA1D;refGene_tx=XM_023249772.1;refGene_gn=CACNA1D;refGene_tx=XM_023249799.1;refGene_gn=CACNA1D;refGene_tx=XM_023249806.1;refGene_gn=CACNA1D;refGene_tx=XM_023249808.1;refGene_gn=CACNA1D;refGene_tx=XM_023249814.1;refGene_gn=CACNA1D;refGene_tx=XM_023249815.1
A2 22184164 22184311
refGene_gn=CACNA1D;refGene_tx=XM_003982328.4;refGene_gn=CACNA1D;refGene_tx=XM_003982329.4;refGene_gn=CACNA1D;refGene_tx=XM_003982331.4;refGene_gn=CACNA1D;refGene_tx=XM_019822887.2;refGene_gn=CACNA1D;refGene_tx=XM_019822894.2;refGene_gn=CACNA1D;refGene_tx=XM_019822897.2;refGene_gn=CACNA1D;refGene_tx=XM_019822906.2;refGene_gn=CACNA1D;refGene_tx=XM_019822918.1;refGene_gn=CACNA1D;refGene_tx=XM_019822924.2;refGene_gn=CACNA1D;refGene_tx=XM_019822928.2;refGene_gn=CACNA1D;refGene_tx=XM_019822930.2;refGene_gn=CACNA1D;refGene_tx=XM_019822934.2;refGene_gn=CACNA1D;refGene_tx=XM_023249752.1;refGene_gn=CACNA1D;refGene_tx=XM_023249761.1;refGene_gn=CACNA1D;refGene_tx=XM_023249772.1;refGene_gn=CACNA1D;refGene_tx=XM_023249799.1;refGene_gn=CACNA1D;refGene_tx=XM_023249806.1;refGene_gn=CACNA1D;refGene_tx=XM_023249808.1;refGene_gn=CACNA1D;refGene_tx=XM_023249814.1;refGene_gn=CACNA1D;refGene_tx=XM_023249815.1
A2 22185754 22185956
refGene_gn=CACNA1D;refGene_tx=XM_003982328.4;refGene_gn=CACNA1D;refGene_tx=XM_003982329.4;refGene_gn=CACNA1D;refGene_tx=XM_003982331.4;refGene_gn=CACNA1D;refGene_tx=XM_019822887.2;refGene_gn=CACNA1D;refGene_tx=XM_019822894.2;refGene_gn=CACNA1D;refGene_tx=XM_019822897.2;refGene_gn=CACNA1D;refGene_tx=XM_019822906.2;refGene_gn=CACNA1D;refGene_tx=XM_019822918.1;refGene_gn=CACNA1D;refGene_tx=XM_019822924.2;refGene_gn=CACNA1D;refGene_tx=XM_019822928.2;refGene_gn=CACNA1D;refGene_tx=XM_019822930.2;refGene_gn=CACNA1D;refGene_tx=XM_019822934.2;refGene_gn=CACNA1D;refGene_tx=XM_023249752.1;refGene_gn=CACNA1D;refGene_tx=XM_023249761.1;refGene_gn=CACNA1D;refGene_tx=XM_023249772.1;refGene_gn=CACNA1D;refGene_tx=XM_023249799.1;refGene_gn=CACNA1D;refGene_tx=XM_023249806.1;refGene_gn=CACNA1D;refGene_tx=XM_023249808.1;refGene_gn=CACNA1D;refGene_tx=XM_023249814.1;refGene_gn=CACNA1D;refGene_tx=XM_023249815.1
A2 22188026 22188185
refGene_gn=CACNA1D;refGene_tx=XM_003982328.4;refGene_gn=CACNA1D;refGene_tx=XM_003982329.4;refGene_gn=CACNA1D;refGene_tx=XM_003982331.4;refGene_gn=CACNA1D;refGene_tx=XM_019822887.2;refGene_gn=CACNA1D;refGene_tx=XM_019822894.2;refGene_gn=CACNA1D;refGene_tx=XM_019822897.2;refGene_gn=CACNA1D;refGene_tx=XM_019822906.2;refGene_gn=CACNA1D;refGene_tx=XM_019822918.1;refGene_gn=CACNA1D;refGene_tx=XM_019822924.2;refGene_gn=CACNA1D;refGene_tx=XM_019822928.2;refGene_gn=CACNA1D;refGene_tx=XM_019822930.2;refGene_gn=CACNA1D;refGene_tx=XM_019822934.2;refGene_gn=CACNA1D;refGene_tx=XM_023249752.1;refGene_gn=CACNA1D;refGene_tx=XM_023249761.1;refGene_gn=CACNA1D;refGene_tx=XM_023249772.1;refGene_gn=CACNA1D;refGene_tx=XM_023249799.1;refGene_gn=CACNA1D;refGene_tx=XM_023249806.1;refGene_gn=CACNA1D;refGene_tx=XM_023249808.1;refGene_gn=CACNA1D;refGene_tx=XM_023249814.1;refGene_gn=CACNA1D;refGene_tx=XM_023249815.1

A2 22189509 22189620
refGene_gn=CACNA1D;refGene_tx=XM_003982328.4;refGene_gn=CACNA1D;refGene_tx=XM_003982329.4;refGene_gn=CACNA1D;refGene_tx=XM_003982331.4;refGene_gn=CACNA1D;refGene_tx=XM_019822887.2;refGene_gn=CACNA1D;refGene_tx=XM_019822894.2;refGene_gn=CACNA1D;refGene_tx=XM_019822897.2;refGene_gn=CACNA1D;refGene_tx=XM_019822906.2;refGene_gn=CACNA1D;refGene_tx=XM_019822918.1;refGene_gn=CACNA1D;refGene_tx=XM_019822924.2;refGene_gn=CACNA1D;refGene_tx=XM_019822928.2;refGene_gn=CACNA1D;refGene_tx=XM_019822930.2;refGene_gn=CACNA1D;refGene_tx=XM_019822934.2;refGene_gn=CACNA1D;refGene_tx=XM_023249752.1;refGene_gn=CACNA1D;refGene_tx=XM_023249761.1;refGene_gn=CACNA1D;refGene_tx=XM_023249772.1;refGene_gn=CACNA1D;refGene_tx=XM_023249799.1;refGene_gn=CACNA1D;refGene_tx=XM_023249806.1;refGene_gn=CACNA1D;refGene_tx=XM_023249808.1;refGene_gn=CACNA1D;refGene_tx=XM_023249814.1;refGene_gn=CACNA1D;refGene_tx=XM_023249815.1
A2 22195987 22196071
refGene_gn=CACNA1D;refGene_tx=XM_003982328.4;refGene_gn=CACNA1D;refGene_tx=XM_003982329.4;refGene_gn=CACNA1D;refGene_tx=XM_003982331.4;refGene_gn=CACNA1D;refGene_tx=XM_019822887.2;refGene_gn=CACNA1D;refGene_tx=XM_019822894.2;refGene_gn=CACNA1D;refGene_tx=XM_019822897.2;refGene_gn=CACNA1D;refGene_tx=XM_019822906.2;refGene_gn=CACNA1D;refGene_tx=XM_019822918.1;refGene_gn=CACNA1D;refGene_tx=XM_019822924.2;refGene_gn=CACNA1D;refGene_tx=XM_019822928.2;refGene_gn=CACNA1D;refGene_tx=XM_023249752.1;refGene_gn=CACNA1D;refGene_tx=XM_023249761.1;refGene_gn=CACNA1D;refGene_tx=XM_023249772.1;refGene_gn=CACNA1D;refGene_tx=XM_023249806.1;refGene_gn=CACNA1D;refGene_tx=XM_023249814.1;refGene_gn=CACNA1D;refGene_tx=XM_023249815.1
A2 22196534 22196618
refGene_gn=CACNA1D;refGene_tx=XM_019822887.2;refGene_gn=CACNA1D;refGene_tx=XM_019822894.2;refGene_gn=CACNA1D;refGene_tx=XM_019822897.2;refGene_gn=CACNA1D;refGene_tx=XM_019822906.2;refGene_gn=CACNA1D;refGene_tx=XM_019822928.2;refGene_gn=CACNA1D;refGene_tx=XM_019822930.2;refGene_gn=CACNA1D;refGene_tx=XM_019822934.2;refGene_gn=CACNA1D;refGene_tx=XM_023249752.1;refGene_gn=CACNA1D;refGene_tx=XM_023249761.1;refGene_gn=CACNA1D;refGene_tx=XM_023249772.1;refGene_gn=CACNA1D;refGene_tx=XM_023249799.1;refGene_gn=CACNA1D;refGene_tx=XM_023249808.1;refGene_gn=CACNA1D;refGene_tx=XM_023249814.1;refGene_gn=CACNA1D;refGene_tx=XM_023249815.1
A2 22203838 22203883
refGene_gn=CACNA1D;refGene_tx=XM_003982328.4;refGene_gn=CACNA1D;refGene_tx=XM_003982329.4;refGene_gn=CACNA1D;refGene_tx=XM_019822887.2;refGene_gn=CACNA1D;refGene_tx=XM_019822894.2;refGene_gn=CACNA1D;refGene_tx=XM_019822897.2;refGene_gn=CACNA1D;refGene_tx=XM_019822906.2;refGene_gn=CACNA1D;refGene_tx=XM_023249752.1;refGene_gn=CACNA1D;refGene_tx=XM_023249761.1;refGene_gn=CACNA1D;refGene_tx=XM_023249799.1;refGene_gn=CACNA1D;refGene_tx=XM_023249814.1;refGene_gn=CACNA1D;refGene_tx=XM_023249815.1
A2 22204285 22204414
refGene_gn=CACNA1D;refGene_tx=XM_003982328.4;refGene_gn=CACNA1D;refGene_tx=XM_003982329.4;refGene_gn=CACNA1D;refGene_tx=XM_003982331.4;refGene_gn=CACNA1D;refGene_tx=XM_019822887.2;refGene_gn=CACNA1D;refGene_tx=XM_019822894.2;refGene_gn=CACNA1D;refGene_tx=XM_019822897.2;refGene_gn

=CACNA1D;refGene_tx=XM_019822906.2;refGene_gn=CACNA1D;refGene_tx=XM_019822918.1;refGene_gn=CACNA1D;refGene_tx=XM_019822924.2;refGene_gn=CACNA1D;refGene_tx=XM_019822928.2;refGene_gn=CACNA1D;refGene_tx=XM_019822930.2;refGene_gn=CACNA1D;refGene_tx=XM_019822934.2;refGene_gn=CACNA1D;refGene_tx=XM_023249752.1;refGene_gn=CACNA1D;refGene_tx=XM_023249761.1;refGene_gn=CACNA1D;refGene_tx=XM_023249772.1;refGene_gn=CACNA1D;refGene_tx=XM_023249799.1;refGene_gn=CACNA1D;refGene_tx=XM_023249806.1;refGene_gn=CACNA1D;refGene_tx=XM_023249808.1;refGene_gn=CACNA1D;refGene_tx=XM_023249814.1;refGene_gn=CACNA1D;refGene_tx=XM_023249815.1
A2 22206652 22206718
refGene_gn=CACNA1D;refGene_tx=XM_003982328.4;refGene_gn=CACNA1D;refGene_tx=XM_003982329.4;refGene_gn=CACNA1D;refGene_tx=XM_003982331.4;refGene_gn=CACNA1D;refGene_tx=XM_019822887.2;refGene_gn=CACNA1D;refGene_tx=XM_019822894.2;refGene_gn=CACNA1D;refGene_tx=XM_019822897.2;refGene_gn=CACNA1D;refGene_tx=XM_019822906.2;refGene_gn=CACNA1D;refGene_tx=XM_019822918.1;refGene_gn=CACNA1D;refGene_tx=XM_019822924.2;refGene_gn=CACNA1D;refGene_tx=XM_019822928.2;refGene_gn=CACNA1D;refGene_tx=XM_019822930.2;refGene_gn=CACNA1D;refGene_tx=XM_019822934.2;refGene_gn=CACNA1D;refGene_tx=XM_023249752.1;refGene_gn=CACNA1D;refGene_tx=XM_023249761.1;refGene_gn=CACNA1D;refGene_tx=XM_023249772.1;refGene_gn=CACNA1D;refGene_tx=XM_023249799.1;refGene_gn=CACNA1D;refGene_tx=XM_023249806.1;refGene_gn=CACNA1D;refGene_tx=XM_023249808.1;refGene_gn=CACNA1D;refGene_tx=XM_023249814.1;refGene_gn=CACNA1D;refGene_tx=XM_023249815.1
A2 22208415 22208507
refGene_gn=CACNA1D;refGene_tx=XM_003982328.4;refGene_gn=CACNA1D;refGene_tx=XM_003982329.4;refGene_gn=CACNA1D;refGene_tx=XM_003982331.4;refGene_gn=CACNA1D;refGene_tx=XM_019822887.2;refGene_gn=CACNA1D;refGene_tx=XM_019822894.2;refGene_gn=CACNA1D;refGene_tx=XM_019822897.2;refGene_gn=CACNA1D;refGene_tx=XM_019822906.2;refGene_gn=CACNA1D;refGene_tx=XM_019822918.1;refGene_gn=CACNA1D;refGene_tx=XM_019822924.2;refGene_gn=CACNA1D;refGene_tx=XM_019822928.2;refGene_gn=CACNA1D;refGene_tx=XM_019822930.2;refGene_gn=CACNA1D;refGene_tx=XM_019822934.2;refGene_gn=CACNA1D;refGene_tx=XM_023249752.1;refGene_gn=CACNA1D;refGene_tx=XM_023249761.1;refGene_gn=CACNA1D;refGene_tx=XM_023249772.1;refGene_gn=CACNA1D;refGene_tx=XM_023249799.1;refGene_gn=CACNA1D;refGene_tx=XM_023249806.1;refGene_gn=CACNA1D;refGene_tx=XM_023249808.1;refGene_gn=CACNA1D;refGene_tx=XM_023249814.1;refGene_gn=CACNA1D;refGene_tx=XM_023249815.1
A2 22209637 22209797
refGene_gn=CACNA1D;refGene_tx=XM_003982328.4;refGene_gn=CACNA1D;refGene_tx=XM_003982329.4;refGene_gn=CACNA1D;refGene_tx=XM_003982331.4;refGene_gn=CACNA1D;refGene_tx=XM_019822887.2;refGene_gn=CACNA1D;refGene_tx=XM_019822894.2;refGene_gn=CACNA1D;refGene_tx=XM_019822897.2;refGene_gn=CACNA1D;refGene_tx=XM_019822906.2;refGene_gn=CACNA1D;refGene_tx=XM_019822918.1;refGene_gn=CACNA1D;refGene_tx=XM_019822924.2;refGene_gn=CACNA1D;refGene_tx=XM_019822928.2;refGene_gn=CACNA1D;refGene_tx=XM_019822930.2;refGene_gn=CACNA1D;refGene_tx=XM_019822934.2;refGene_gn=CACNA1D;refGene_tx=XM_023249752.1;refGene_gn=CACNA1D;refGene_tx=XM_023249761.1;refGene_gn=CACNA1D;refGene_tx=XM_023249772.1;refGene_gn=CACNA1D;refGene_tx=XM_023249799.1;refGene_gn=CACNA1D;refGene_tx=XM_023249806.1;refGene_gn=CACNA1D;refGene_tx=XM_023249808.1;refGene_gn=CACNA1D;refGene_tx=

XM_023249814.1;refGene_gn=CACNA1D;refGene_tx=XM_023249815.1
A2 22210604 22210732
refGene_gn=CACNA1D;refGene_tx=XM_003982328.4;refGene_gn=CACNA1D;refGene_tx=XM_003982329.4;refGene_gn=CACNA1D;refGene_tx=XM_003982331.4;refGene_gn=CACNA1D;refGene_tx=XM_019822887.2;refGene_gn=CACNA1D;refGene_tx=XM_019822894.2;refGene_gn=CACNA1D;refGene_tx=XM_019822897.2;refGene_gn=CACNA1D;refGene_tx=XM_019822906.2;refGene_gn=CACNA1D;refGene_tx=XM_019822918.1;refGene_gn=CACNA1D;refGene_tx=XM_019822924.2;refGene_gn=CACNA1D;refGene_tx=XM_019822928.2;refGene_gn=CACNA1D;refGene_tx=XM_019822930.2;refGene_gn=CACNA1D;refGene_tx=XM_019822934.2;refGene_gn=CACNA1D;refGene_tx=XM_023249752.1;refGene_gn=CACNA1D;refGene_tx=XM_023249761.1;refGene_gn=CACNA1D;refGene_tx=XM_023249772.1;refGene_gn=CACNA1D;refGene_tx=XM_023249799.1;refGene_gn=CACNA1D;refGene_tx=XM_023249806.1;refGene_gn=CACNA1D;refGene_tx=XM_023249808.1;refGene_gn=CACNA1D;refGene_tx=XM_023249814.1;refGene_gn=CACNA1D;refGene_tx=XM_023249815.1
A2 22210870 22210967
refGene_gn=CACNA1D;refGene_tx=XM_003982328.4;refGene_gn=CACNA1D;refGene_tx=XM_003982329.4;refGene_gn=CACNA1D;refGene_tx=XM_003982331.4;refGene_gn=CACNA1D;refGene_tx=XM_019822887.2;refGene_gn=CACNA1D;refGene_tx=XM_019822894.2;refGene_gn=CACNA1D;refGene_tx=XM_019822897.2;refGene_gn=CACNA1D;refGene_tx=XM_019822906.2;refGene_gn=CACNA1D;refGene_tx=XM_019822918.1;refGene_gn=CACNA1D;refGene_tx=XM_019822924.2;refGene_gn=CACNA1D;refGene_tx=XM_019822928.2;refGene_gn=CACNA1D;refGene_tx=XM_019822930.2;refGene_gn=CACNA1D;refGene_tx=XM_019822934.2;refGene_gn=CACNA1D;refGene_tx=XM_023249752.1;refGene_gn=CACNA1D;refGene_tx=XM_023249761.1;refGene_gn=CACNA1D;refGene_tx=XM_023249772.1;refGene_gn=CACNA1D;refGene_tx=XM_023249799.1;refGene_gn=CACNA1D;refGene_tx=XM_023249806.1;refGene_gn=CACNA1D;refGene_tx=XM_023249808.1;refGene_gn=CACNA1D;refGene_tx=XM_023249814.1;refGene_gn=CACNA1D;refGene_tx=XM_023249815.1
A2 22213347 22213450
refGene_gn=CACNA1D;refGene_tx=XM_003982328.4;refGene_gn=CACNA1D;refGene_tx=XM_003982329.4;refGene_gn=CACNA1D;refGene_tx=XM_003982331.4;refGene_gn=CACNA1D;refGene_tx=XM_019822887.2;refGene_gn=CACNA1D;refGene_tx=XM_019822894.2;refGene_gn=CACNA1D;refGene_tx=XM_019822897.2;refGene_gn=CACNA1D;refGene_tx=XM_019822906.2;refGene_gn=CACNA1D;refGene_tx=XM_019822918.1;refGene_gn=CACNA1D;refGene_tx=XM_019822924.2;refGene_gn=CACNA1D;refGene_tx=XM_019822928.2;refGene_gn=CACNA1D;refGene_tx=XM_019822930.2;refGene_gn=CACNA1D;refGene_tx=XM_019822934.2;refGene_gn=CACNA1D;refGene_tx=XM_023249752.1;refGene_gn=CACNA1D;refGene_tx=XM_023249761.1;refGene_gn=CACNA1D;refGene_tx=XM_023249772.1;refGene_gn=CACNA1D;refGene_tx=XM_023249799.1;refGene_gn=CACNA1D;refGene_tx=XM_023249806.1;refGene_gn=CACNA1D;refGene_tx=XM_023249808.1;refGene_gn=CACNA1D;refGene_tx=XM_023249814.1;refGene_gn=CACNA1D;refGene_tx=XM_023249815.1
A2 22214780 22214882
refGene_gn=CACNA1D;refGene_tx=XM_003982328.4;refGene_gn=CACNA1D;refGene_tx=XM_003982329.4;refGene_gn=CACNA1D;refGene_tx=XM_003982331.4;refGene_gn=CACNA1D;refGene_tx=XM_019822887.2;refGene_gn=CACNA1D;refGene_tx=XM_019822894.2;refGene_gn=CACNA1D;refGene_tx=XM_019822897.2;refGene_gn=CACNA1D;refGene_tx=XM_019822906.2;refGene_gn=CACNA1D;refGene_tx=XM_019822918.1;refGene_gn=CACNA1D;refGene_tx=XM_019822924.2;refGene_gn=CACN

A1D;refGene_tx=XM_019822928.2;refGene_gn=CACNA1D;refGene_tx=XM_0198229
30.2;refGene_gn=CACNA1D;refGene_tx=XM_019822934.2;refGene_gn=CACNA1D;r
efGene_tx=XM_023249752.1;refGene_gn=CACNA1D;refGene_tx=XM_023249761.1;
refGene_gn=CACNA1D;refGene_tx=XM_023249772.1;refGene_gn=CACNA1D;refGen
e_tx=XM_023249799.1;refGene_gn=CACNA1D;refGene_tx=XM_023249806.1;refGe
ne_gn=CACNA1D;refGene_tx=XM_023249808.1;refGene_gn=CACNA1D;refGene_tx=
XM_023249814.1;refGene_gn=CACNA1D;refGene_tx=XM_023249815.1
A2 22219814 22219945
refGene_gn=CACNA1D;refGene_tx=XM_003982328.4;refGene_gn=CACNA1D;refGen
e_tx=XM_003982329.4;refGene_gn=CACNA1D;refGene_tx=XM_003982331.4;refGe
ne_gn=CACNA1D;refGene_tx=XM_019822887.2;refGene_gn=CACNA1D;refGene_tx=
XM_019822894.2;refGene_gn=CACNA1D;refGene_tx=XM_019822897.2;refGene_gn
=CACNA1D;refGene_tx=XM_019822906.2;refGene_gn=CACNA1D;refGene_tx=XM_01
9822918.1;refGene_gn=CACNA1D;refGene_tx=XM_019822924.2;refGene_gn=CACN
A1D;refGene_tx=XM_019822928.2;refGene_gn=CACNA1D;refGene_tx=XM_0198229
30.2;refGene_gn=CACNA1D;refGene_tx=XM_019822934.2;refGene_gn=CACNA1D;r
efGene_tx=XM_023249752.1;refGene_gn=CACNA1D;refGene_tx=XM_023249761.1;
refGene_gn=CACNA1D;refGene_tx=XM_023249772.1;refGene_gn=CACNA1D;refGen
e_tx=XM_023249799.1;refGene_gn=CACNA1D;refGene_tx=XM_023249806.1;refGe
ne_gn=CACNA1D;refGene_tx=XM_023249808.1;refGene_gn=CACNA1D;refGene_tx=
XM_023249814.1;refGene_gn=CACNA1D;refGene_tx=XM_023249815.1
A2 22223853 22223874 refGene_gn=CACNA1D;refGene_tx=XM_023249815.1
A2 22232839 22232956
refGene_gn=CACNA1D;refGene_tx=XM_003982328.4;refGene_gn=CACNA1D;refGen
e_tx=XM_003982329.4;refGene_gn=CACNA1D;refGene_tx=XM_003982331.4;refGe
ne_gn=CACNA1D;refGene_tx=XM_019822887.2;refGene_gn=CACNA1D;refGene_tx=
XM_019822894.2;refGene_gn=CACNA1D;refGene_tx=XM_019822897.2;refGene_gn
=CACNA1D;refGene_tx=XM_019822906.2;refGene_gn=CACNA1D;refGene_tx=XM_01
9822918.1;refGene_gn=CACNA1D;refGene_tx=XM_019822924.2;refGene_gn=CACN
A1D;refGene_tx=XM_019822928.2;refGene_gn=CACNA1D;refGene_tx=XM_0198229
30.2;refGene_gn=CACNA1D;refGene_tx=XM_019822934.2;refGene_gn=CACNA1D;r
efGene_tx=XM_023249752.1;refGene_gn=CACNA1D;refGene_tx=XM_023249761.1;
refGene_gn=CACNA1D;refGene_tx=XM_023249772.1;refGene_gn=CACNA1D;refGen
e_tx=XM_023249799.1;refGene_gn=CACNA1D;refGene_tx=XM_023249806.1;refGe
ne_gn=CACNA1D;refGene_tx=XM_023249808.1;refGene_gn=CACNA1D;refGene_tx=
XM_023249814.1
A2 22233643 22234011
refGene_gn=CACNA1D;refGene_tx=XM_003982328.4;refGene_gn=CACNA1D;refGen
e_tx=XM_003982329.4;refGene_gn=CACNA1D;refGene_tx=XM_003982331.4;refGe
ne_gn=CACNA1D;refGene_tx=XM_019822887.2;refGene_gn=CACNA1D;refGene_tx=
XM_019822894.2;refGene_gn=CACNA1D;refGene_tx=XM_019822897.2;refGene_gn
=CACNA1D;refGene_tx=XM_019822906.2;refGene_gn=CACNA1D;refGene_tx=XM_01
9822918.1;refGene_gn=CACNA1D;refGene_tx=XM_019822924.2;refGene_gn=CACN
A1D;refGene_tx=XM_019822928.2;refGene_gn=CACNA1D;refGene_tx=XM_0198229
30.2;refGene_gn=CACNA1D;refGene_tx=XM_019822934.2;refGene_gn=CACNA1D;r
efGene_tx=XM_023249752.1;refGene_gn=CACNA1D;refGene_tx=XM_023249761.1;
refGene_gn=CACNA1D;refGene_tx=XM_023249772.1;refGene_gn=CACNA1D;refGen
e_tx=XM_023249799.1;refGene_gn=CACNA1D;refGene_tx=XM_023249806.1;refGe
ne_gn=CACNA1D;refGene_tx=XM_023249808.1;refGene_gn=CACNA1D;refGene_tx=
XM_023249814.1

A2 22234712 22234739
refGene_gn=CACNA1D;refGene_tx=XM_003982328.4;refGene_gn=CACNA1D;refGene_tx=XM_003982329.4;refGene_gn=CACNA1D;refGene_tx=XM_019822887.2;refGene_gn=CACNA1D;refGene_tx=XM_019822894.2;refGene_gn=CACNA1D;refGene_tx=XM_019822897.2;refGene_gn=CACNA1D;refGene_tx=XM_019822906.2;refGene_gn=CACNA1D;refGene_tx=XM_019822918.1;refGene_gn=CACNA1D;refGene_tx=XM_019822924.2;refGene_gn=CACNA1D;refGene_tx=XM_019822928.2;refGene_gn=CACNA1D;refGene_tx=XM_019822930.2;refGene_gn=CACNA1D;refGene_tx=XM_019822934.2;refGene_gn=CACNA1D;refGene_tx=XM_023249752.1;refGene_gn=CACNA1D;refGene_tx=XM_023249772.1;refGene_gn=CACNA1D;refGene_tx=XM_023249799.1;refGene_gn=CACNA1D;refGene_tx=XM_023249806.1;refGene_gn=CACNA1D;refGene_tx=XM_023249808.1
A2 22235896 22236046
refGene_gn=CACNA1D;refGene_tx=XM_003982328.4;refGene_gn=CACNA1D;refGene_tx=XM_003982329.4;refGene_gn=CACNA1D;refGene_tx=XM_003982331.4;refGene_gn=CACNA1D;refGene_tx=XM_019822887.2;refGene_gn=CACNA1D;refGene_tx=XM_019822894.2;refGene_gn=CACNA1D;refGene_tx=XM_019822897.2;refGene_gn=CACNA1D;refGene_tx=XM_019822906.2;refGene_gn=CACNA1D;refGene_tx=XM_019822918.1;refGene_gn=CACNA1D;refGene_tx=XM_019822924.2;refGene_gn=CACNA1D;refGene_tx=XM_019822928.2;refGene_gn=CACNA1D;refGene_tx=XM_019822930.2;refGene_gn=CACNA1D;refGene_tx=XM_019822934.2;refGene_gn=CACNA1D;refGene_tx=XM_023249752.1;refGene_gn=CACNA1D;refGene_tx=XM_023249761.1;refGene_gn=CACNA1D;refGene_tx=XM_023249772.1;refGene_gn=CACNA1D;refGene_tx=XM_023249799.1;refGene_gn=CACNA1D;refGene_tx=XM_023249806.1;refGene_gn=CACNA1D;refGene_tx=XM_023249808.1
A2 22236409 22236573
refGene_gn=CACNA1D;refGene_tx=XM_003982328.4;refGene_gn=CACNA1D;refGene_tx=XM_003982329.4;refGene_gn=CACNA1D;refGene_tx=XM_003982331.4;refGene_gn=CACNA1D;refGene_tx=XM_019822887.2;refGene_gn=CACNA1D;refGene_tx=XM_019822894.2;refGene_gn=CACNA1D;refGene_tx=XM_019822897.2;refGene_gn=CACNA1D;refGene_tx=XM_019822906.2;refGene_gn=CACNA1D;refGene_tx=XM_019822918.1;refGene_gn=CACNA1D;refGene_tx=XM_019822924.2;refGene_gn=CACNA1D;refGene_tx=XM_019822928.2;refGene_gn=CACNA1D;refGene_tx=XM_019822930.2;refGene_gn=CACNA1D;refGene_tx=XM_019822934.2;refGene_gn=CACNA1D;refGene_tx=XM_023249752.1;refGene_gn=CACNA1D;refGene_tx=XM_023249761.1;refGene_gn=CACNA1D;refGene_tx=XM_023249772.1;refGene_gn=CACNA1D;refGene_tx=XM_023249799.1;refGene_gn=CACNA1D;refGene_tx=XM_023249806.1;refGene_gn=CACNA1D;refGene_tx=XM_023249808.1
A2 22239227 22239349
refGene_gn=CACNA1D;refGene_tx=XM_003982328.4;refGene_gn=CACNA1D;refGene_tx=XM_003982329.4;refGene_gn=CACNA1D;refGene_tx=XM_003982331.4;refGene_gn=CACNA1D;refGene_tx=XM_019822887.2;refGene_gn=CACNA1D;refGene_tx=XM_019822894.2;refGene_gn=CACNA1D;refGene_tx=XM_019822897.2;refGene_gn=CACNA1D;refGene_tx=XM_019822906.2;refGene_gn=CACNA1D;refGene_tx=XM_019822918.1;refGene_gn=CACNA1D;refGene_tx=XM_019822924.2;refGene_gn=CACNA1D;refGene_tx=XM_019822928.2;refGene_gn=CACNA1D;refGene_tx=XM_019822930.2;refGene_gn=CACNA1D;refGene_tx=XM_019822934.2;refGene_gn=CACNA1D;refGene_tx=XM_023249752.1;refGene_gn=CACNA1D;refGene_tx=XM_023249761.1;refGene_gn=CACNA1D;refGene_tx=XM_023249772.1;refGene_gn=CACNA1D;refGene_tx=XM_023249799.1;refGene_gn=CACNA1D;refGene_tx=XM_023249806.1;refGe

ne_gn=CACNA1D;refGene_tx=XM_023249808.1
A2 22240898 22241219
refGene_gn=CACNA1D;refGene_tx=XM_003982328.4;refGene_gn=CACNA1D;refGene_tx=XM_003982329.4;refGene_gn=CACNA1D;refGene_tx=XM_003982331.4;refGene_gn=CACNA1D;refGene_tx=XM_019822887.2;refGene_gn=CACNA1D;refGene_tx=XM_019822894.2;refGene_gn=CACNA1D;refGene_tx=XM_019822897.2;refGene_gn=CACNA1D;refGene_tx=XM_019822906.2;refGene_gn=CACNA1D;refGene_tx=XM_019822918.1;refGene_gn=CACNA1D;refGene_tx=XM_019822924.2;refGene_gn=CACNA1D;refGene_tx=XM_019822928.2;refGene_gn=CACNA1D;refGene_tx=XM_019822930.2;refGene_gn=CACNA1D;refGene_tx=XM_019822934.2;refGene_gn=CACNA1D;refGene_tx=XM_023249752.1;refGene_gn=CACNA1D;refGene_tx=XM_023249761.1;refGene_gn=CACNA1D;refGene_tx=XM_023249772.1;refGene_gn=CACNA1D;refGene_tx=XM_023249799.1;refGene_gn=CACNA1D;refGene_tx=XM_023249806.1;refGene_gn=CACNA1D;refGene_tx=XM_023249808.1
A2 22241705 22241999
refGene_gn=CACNA1D;refGene_tx=XM_003982328.4;refGene_gn=CACNA1D;refGene_tx=XM_003982329.4;refGene_gn=CACNA1D;refGene_tx=XM_003982331.4;refGene_gn=CACNA1D;refGene_tx=XM_019822887.2;refGene_gn=CACNA1D;refGene_tx=XM_019822894.2;refGene_gn=CACNA1D;refGene_tx=XM_019822897.2;refGene_gn=CACNA1D;refGene_tx=XM_019822906.2;refGene_gn=CACNA1D;refGene_tx=XM_019822918.1;refGene_gn=CACNA1D;refGene_tx=XM_019822924.2;refGene_gn=CACNA1D;refGene_tx=XM_019822928.2;refGene_gn=CACNA1D;refGene_tx=XM_019822930.2;refGene_gn=CACNA1D;refGene_tx=XM_019822934.2;refGene_gn=CACNA1D;refGene_tx=XM_023249752.1;refGene_gn=CACNA1D;refGene_tx=XM_023249761.1;refGene_gn=CACNA1D;refGene_tx=XM_023249772.1;refGene_gn=CACNA1D;refGene_tx=XM_023249799.1;refGene_gn=CACNA1D;refGene_tx=XM_023249806.1;refGene_gn=CACNA1D;refGene_tx=XM_023249808.1
A2 22247630 22248049
refGene_gn=CHDH;refGene_tx=XM_011279616.3;refGene_gn=CHDH;refGene_tx=XM_019822965.1;refGene_gn=CHDH;refGene_tx=XM_019822966.2;refGene_gn=CHDH;refGene_tx=XM_023249823.1
A2 22248774 22248877
refGene_gn=CHDH;refGene_tx=XM_011279616.3;refGene_gn=CHDH;refGene_tx=XM_019822965.1;refGene_gn=CHDH;refGene_tx=XM_019822966.2;refGene_gn=CHDH;refGene_tx=XM_023249823.1
A2 22249370 22249513
refGene_gn=CHDH;refGene_tx=XM_011279616.3;refGene_gn=CHDH;refGene_tx=XM_019822965.1;refGene_gn=CHDH;refGene_tx=XM_019822966.2;refGene_gn=CHDH;refGene_tx=XM_023249823.1
A2 22250733 22250868
refGene_gn=CHDH;refGene_tx=XM_011279616.3;refGene_gn=CHDH;refGene_tx=XM_019822965.1;refGene_gn=CHDH;refGene_tx=XM_019822966.2;refGene_gn=CHDH;refGene_tx=XM_023249823.1
A2 22251837 22251967
refGene_gn=CHDH;refGene_tx=XM_011279616.3;refGene_gn=CHDH;refGene_tx=XM_019822965.1;refGene_gn=CHDH;refGene_tx=XM_019822966.2;refGene_gn=CHDH;refGene_tx=XM_023249823.1
A2 22252832 22252984
refGene_gn=CHDH;refGene_tx=XM_011279616.3;refGene_gn=CHDH;refGene_tx=XM_019822965.1;refGene_gn=CHDH;refGene_tx=XM_019822966.2;refGene_gn=CHD

H;refGene_tx=XM_023249823.1
A2	22253583	22254286
refGene_gn=CHDH;refGene_tx=XM_011279616.3;refGene_gn=CHDH;refGene_tx=XM_019822965.1;refGene_gn=CHDH;refGene_tx=XM_019822966.2;refGene_gn=CHDH;refGene_tx=XM_023249823.1
A2	22285526	22285586	refGene_gn=IL17RB;refGene_tx=XM_006928869.3
A2	22287595	22287620	refGene_gn=IL17RB;refGene_tx=XM_006928869.3
A2	22288363	22288501
refGene_gn=IL17RB;refGene_tx=XM_006928869.3;refGene_gn=IL17RB;refGene_tx=XM_006928870.3
A2	22290367	22290495
refGene_gn=IL17RB;refGene_tx=XM_006928869.3;refGene_gn=IL17RB;refGene_tx=XM_006928870.3
A2	22290947	22291074
refGene_gn=IL17RB;refGene_tx=XM_006928869.3;refGene_gn=IL17RB;refGene_tx=XM_006928870.3
A2	22292962	22293010
refGene_gn=IL17RB;refGene_tx=XM_006928869.3;refGene_gn=IL17RB;refGene_tx=XM_006928870.3
A2	22294385	22294528
refGene_gn=IL17RB;refGene_tx=XM_006928869.3;refGene_gn=IL17RB;refGene_tx=XM_006928870.3
A2	22294906	22294975
refGene_gn=IL17RB;refGene_tx=XM_006928869.3;refGene_gn=IL17RB;refGene_tx=XM_006928870.3
A2	22296011	22296111
refGene_gn=IL17RB;refGene_tx=XM_006928869.3;refGene_gn=IL17RB;refGene_tx=XM_006928870.3
A2	22297500	22297599
refGene_gn=IL17RB;refGene_tx=XM_006928869.3;refGene_gn=IL17RB;refGene_tx=XM_006928870.3
A2	22298754	22299302
refGene_gn=IL17RB;refGene_tx=XM_006928869.3;refGene_gn=IL17RB;refGene_tx=XM_006928870.3
A2	22302145	22302289
refGene_gn=ACTR8;refGene_tx=XM_011279636.2;refGene_gn=ACTR8;refGene_tx=XM_019822943.1;refGene_gn=ACTR8;refGene_tx=XM_019822947.2;refGene_gn=ACTR8;refGene_tx=XM_019822949.2;refGene_gn=ACTR8;refGene_tx=XM_019822953.2
A2	22303091	22303255
refGene_gn=ACTR8;refGene_tx=XM_011279636.2;refGene_gn=ACTR8;refGene_tx=XM_019822947.2;refGene_gn=ACTR8;refGene_tx=XM_019822949.2;refGene_gn=ACTR8;refGene_tx=XM_019822953.2
A2	22303973	22304238
refGene_gn=ACTR8;refGene_tx=XM_011279636.2;refGene_gn=ACTR8;refGene_tx=XM_019822943.1;refGene_gn=ACTR8;refGene_tx=XM_019822947.2;refGene_gn=ACTR8;refGene_tx=XM_019822949.2;refGene_gn=ACTR8;refGene_tx=XM_019822953.2
A2	22305178	22305319
refGene_gn=ACTR8;refGene_tx=XM_011279636.2;refGene_gn=ACTR8;refGene_tx

=XM_019822943.1;refGene_gn=ACTR8;refGene_tx=XM_019822947.2;refGene_gn=ACTR8;refGene_tx=XM_019822949.2;refGene_gn=ACTR8;refGene_tx=XM_019822953.2
A2 22306307 22306403
refGene_gn=ACTR8;refGene_tx=XM_011279636.2;refGene_gn=ACTR8;refGene_tx=XM_019822943.1;refGene_gn=ACTR8;refGene_tx=XM_019822947.2;refGene_gn=ACTR8;refGene_tx=XM_019822949.2;refGene_gn=ACTR8;refGene_tx=XM_019822953.2
A2 22307402 22307556
refGene_gn=ACTR8;refGene_tx=XM_011279636.2;refGene_gn=ACTR8;refGene_tx=XM_019822943.1;refGene_gn=ACTR8;refGene_tx=XM_019822947.2;refGene_gn=ACTR8;refGene_tx=XM_019822949.2;refGene_gn=ACTR8;refGene_tx=XM_019822953.2
A2 22309245 22309378
refGene_gn=ACTR8;refGene_tx=XM_011279636.2;refGene_gn=ACTR8;refGene_tx=XM_019822943.1;refGene_gn=ACTR8;refGene_tx=XM_019822947.2;refGene_gn=ACTR8;refGene_tx=XM_019822949.2;refGene_gn=ACTR8;refGene_tx=XM_019822953.2
A2 22309594 22309688
refGene_gn=ACTR8;refGene_tx=XM_011279636.2;refGene_gn=ACTR8;refGene_tx=XM_019822943.1;refGene_gn=ACTR8;refGene_tx=XM_019822947.2;refGene_gn=ACTR8;refGene_tx=XM_019822949.2;refGene_gn=ACTR8;refGene_tx=XM_019822953.2
A2 22311873 22312047
refGene_gn=ACTR8;refGene_tx=XM_011279636.2;refGene_gn=ACTR8;refGene_tx=XM_019822943.1;refGene_gn=ACTR8;refGene_tx=XM_019822947.2;refGene_gn=ACTR8;refGene_tx=XM_019822949.2;refGene_gn=ACTR8;refGene_tx=XM_019822953.2
A2 22312594 22312699
refGene_gn=ACTR8;refGene_tx=XM_011279636.2;refGene_gn=ACTR8;refGene_tx=XM_019822943.1;refGene_gn=ACTR8;refGene_tx=XM_019822947.2;refGene_gn=ACTR8;refGene_tx=XM_019822949.2;refGene_gn=ACTR8;refGene_tx=XM_019822953.2
A2 22313358 22313469
refGene_gn=ACTR8;refGene_tx=XM_011279636.2;refGene_gn=ACTR8;refGene_tx=XM_019822943.1;refGene_gn=ACTR8;refGene_tx=XM_019822947.2;refGene_gn=ACTR8;refGene_tx=XM_019822949.2;refGene_gn=ACTR8;refGene_tx=XM_019822953.2
A2 22314711 22314882
refGene_gn=ACTR8;refGene_tx=XM_011279636.2;refGene_gn=ACTR8;refGene_tx=XM_019822943.1
A2 22317844 22317967
refGene_gn=ACTR8;refGene_tx=XM_011279636.2;refGene_gn=ACTR8;refGene_tx=XM_019822943.1
A2 22320853 22320857 refGene_gn=SELENOK;refGene_tx=XM_003982351.5
A2 22321119 22321206 refGene_gn=SELENOK;refGene_tx=XM_003982351.5
A2 22321774 22321858 refGene_gn=SELENOK;refGene_tx=XM_003982351.5
A2 22323581 22323672 refGene_gn=SELENOK;refGene_tx=XM_003982351.5
A2 22327149 22327168 refGene_gn=SELENOK;refGene_tx=XM_003982351.5
A2 22510683 22510805

refGene_gn=CACNA2D3;refGene_tx=XM_023249840.1;refGene_gn=CACNA2D3;refGene_tx=XM_023249841.1;refGene_gn=CACNA2D3;refGene_tx=XM_023249842.1
A2 22511472 22511554
refGene_gn=CACNA2D3;refGene_tx=XM_023249840.1;refGene_gn=CACNA2D3;refGene_tx=XM_023249841.1;refGene_gn=CACNA2D3;refGene_tx=XM_023249842.1
A2 22688716 22688833
refGene_gn=CACNA2D3;refGene_tx=XM_019823012.1;refGene_gn=CACNA2D3;refGene_tx=XM_023249840.1;refGene_gn=CACNA2D3;refGene_tx=XM_023249841.1;refGene_gn=CACNA2D3;refGene_tx=XM_023249842.1
A2 22761591 22761651
refGene_gn=CACNA2D3;refGene_tx=XM_019823012.1;refGene_gn=CACNA2D3;refGene_tx=XM_023249840.1;refGene_gn=CACNA2D3;refGene_tx=XM_023249841.1;refGene_gn=CACNA2D3;refGene_tx=XM_023249842.1
A2 22840595 22840758
refGene_gn=CACNA2D3;refGene_tx=XM_019823012.1;refGene_gn=CACNA2D3;refGene_tx=XM_023249840.1;refGene_gn=CACNA2D3;refGene_tx=XM_023249841.1;refGene_gn=CACNA2D3;refGene_tx=XM_023249842.1
A2 22897893 22898025
refGene_gn=CACNA2D3;refGene_tx=XM_019823012.1;refGene_gn=CACNA2D3;refGene_tx=XM_023249840.1;refGene_gn=CACNA2D3;refGene_tx=XM_023249841.1;refGene_gn=CACNA2D3;refGene_tx=XM_023249842.1
A2 22906252 22906313
refGene_gn=CACNA2D3;refGene_tx=XM_019823012.1;refGene_gn=CACNA2D3;refGene_tx=XM_023249840.1;refGene_gn=CACNA2D3;refGene_tx=XM_023249841.1;refGene_gn=CACNA2D3;refGene_tx=XM_023249842.1
A2 22906405 22906556
refGene_gn=CACNA2D3;refGene_tx=XM_019823012.1;refGene_gn=CACNA2D3;refGene_tx=XM_023249840.1;refGene_gn=CACNA2D3;refGene_tx=XM_023249841.1;refGene_gn=CACNA2D3;refGene_tx=XM_023249842.1
A2 22918399 22918474
refGene_gn=CACNA2D3;refGene_tx=XM_019823012.1;refGene_gn=CACNA2D3;refGene_tx=XM_023249840.1;refGene_gn=CACNA2D3;refGene_tx=XM_023249841.1;refGene_gn=CACNA2D3;refGene_tx=XM_023249842.1
A2 22968187 22968277
refGene_gn=CACNA2D3;refGene_tx=XM_019823012.1;refGene_gn=CACNA2D3;refGene_tx=XM_023249840.1;refGene_gn=CACNA2D3;refGene_tx=XM_023249841.1;refGene_gn=CACNA2D3;refGene_tx=XM_023249842.1
A2 22977009 22977123
refGene_gn=CACNA2D3;refGene_tx=XM_019823012.1;refGene_gn=CACNA2D3;refGene_tx=XM_023249840.1;refGene_gn=CACNA2D3;refGene_tx=XM_023249841.1;refGene_gn=CACNA2D3;refGene_tx=XM_023249842.1
A2 23073646 23073725
refGene_gn=CACNA2D3;refGene_tx=XM_019823012.1;refGene_gn=CACNA2D3;refGene_tx=XM_023249840.1;refGene_gn=CACNA2D3;refGene_tx=XM_023249841.1;refGene_gn=CACNA2D3;refGene_tx=XM_023249842.1
A2 23084281 23084415
refGene_gn=CACNA2D3;refGene_tx=XM_019823012.1;refGene_gn=CACNA2D3;refGene_tx=XM_023249840.1;refGene_gn=CACNA2D3;refGene_tx=XM_023249841.1;refGene_gn=CACNA2D3;refGene_tx=XM_023249842.1
A2 23131560 23131578

refGene_gn=CACNA2D3;refGene_tx=XM_019823012.1;refGene_gn=CACNA2D3;refGene_tx=XM_023249840.1
A2 23150973 23151045
refGene_gn=CACNA2D3;refGene_tx=XM_019823012.1;refGene_gn=CACNA2D3;refGene_tx=XM_023249840.1;refGene_gn=CACNA2D3;refGene_tx=XM_023249841.1
A2 23152286 23152367
refGene_gn=CACNA2D3;refGene_tx=XM_019823012.1;refGene_gn=CACNA2D3;refGene_tx=XM_023249840.1;refGene_gn=CACNA2D3;refGene_tx=XM_023249841.1
A2 23161754 23161829
refGene_gn=CACNA2D3;refGene_tx=XM_019823012.1;refGene_gn=CACNA2D3;refGene_tx=XM_023249840.1;refGene_gn=CACNA2D3;refGene_tx=XM_023249841.1
A2 23186416 23186500
refGene_gn=CACNA2D3;refGene_tx=XM_019823012.1;refGene_gn=CACNA2D3;refGene_tx=XM_023249840.1;refGene_gn=CACNA2D3;refGene_tx=XM_023249841.1;refGene_gn=CACNA2D3;refGene_tx=XM_023249842.1
A2 23194771 23194843
refGene_gn=CACNA2D3;refGene_tx=XM_019823012.1;refGene_gn=CACNA2D3;refGene_tx=XM_023249840.1;refGene_gn=CACNA2D3;refGene_tx=XM_023249841.1;refGene_gn=CACNA2D3;refGene_tx=XM_023249842.1
A2 23195100 23195162
refGene_gn=CACNA2D3;refGene_tx=XM_019823012.1;refGene_gn=CACNA2D3;refGene_tx=XM_023249840.1;refGene_gn=CACNA2D3;refGene_tx=XM_023249841.1;refGene_gn=CACNA2D3;refGene_tx=XM_023249842.1
A2 23196523 23196591
refGene_gn=CACNA2D3;refGene_tx=XM_019823012.1;refGene_gn=CACNA2D3;refGene_tx=XM_023249840.1;refGene_gn=CACNA2D3;refGene_tx=XM_023249841.1;refGene_gn=CACNA2D3;refGene_tx=XM_023249842.1
A2 23200850 23200896
refGene_gn=CACNA2D3;refGene_tx=XM_019823012.1;refGene_gn=CACNA2D3;refGene_tx=XM_023249840.1;refGene_gn=CACNA2D3;refGene_tx=XM_023249841.1;refGene_gn=CACNA2D3;refGene_tx=XM_023249842.1
A2 23201054 23201152
refGene_gn=CACNA2D3;refGene_tx=XM_019823012.1;refGene_gn=CACNA2D3;refGene_tx=XM_023249840.1;refGene_gn=CACNA2D3;refGene_tx=XM_023249841.1;refGene_gn=CACNA2D3;refGene_tx=XM_023249842.1
A2 23203244 23203338
refGene_gn=CACNA2D3;refGene_tx=XM_019823012.1;refGene_gn=CACNA2D3;refGene_tx=XM_023249840.1;refGene_gn=CACNA2D3;refGene_tx=XM_023249841.1;refGene_gn=CACNA2D3;refGene_tx=XM_023249842.1
A2 23206278 23206374
refGene_gn=CACNA2D3;refGene_tx=XM_019823012.1;refGene_gn=CACNA2D3;refGene_tx=XM_023249840.1;refGene_gn=CACNA2D3;refGene_tx=XM_023249841.1;refGene_gn=CACNA2D3;refGene_tx=XM_023249842.1
A2 23212401 23212523
refGene_gn=CACNA2D3;refGene_tx=XM_019823012.1;refGene_gn=CACNA2D3;refGene_tx=XM_023249840.1;refGene_gn=CACNA2D3;refGene_tx=XM_023249841.1;refGene_gn=CACNA2D3;refGene_tx=XM_023249842.1
A2 23215130 23215211
refGene_gn=CACNA2D3;refGene_tx=XM_019823012.1;refGene_gn=CACNA2D3;refGene_tx=XM_023249840.1;refGene_gn=CACNA2D3;refGene_tx=XM_023249841.1;re

fGene_gn=CACNA2D3;refGene_tx=XM_023249842.1
A2	23233518 23233952 refGene_gn=LRTM1;refGene_tx=XM_019823025.1
A2	23239587 23240184 refGene_gn=LRTM1;refGene_tx=XM_019823025.1
A2	23242967 23242974 refGene_gn=LRTM1;refGene_tx=XM_019823025.1
A2	23281003 23281065
refGene_gn=CACNA2D3;refGene_tx=XM_019823012.1;refGene_gn=CACNA2D3;refGene_tx=XM_023249840.1;refGene_gn=CACNA2D3;refGene_tx=XM_023249841.1;refGene_gn=CACNA2D3;refGene_tx=XM_023249842.1
A2	23282292 23282337
refGene_gn=CACNA2D3;refGene_tx=XM_019823012.1;refGene_gn=CACNA2D3;refGene_tx=XM_023249840.1;refGene_gn=CACNA2D3;refGene_tx=XM_023249841.1;refGene_gn=CACNA2D3;refGene_tx=XM_023249842.1
A2	23289857 23289920
refGene_gn=CACNA2D3;refGene_tx=XM_019823012.1;refGene_gn=CACNA2D3;refGene_tx=XM_023249840.1;refGene_gn=CACNA2D3;refGene_tx=XM_023249841.1;refGene_gn=CACNA2D3;refGene_tx=XM_023249842.1
A2	23293075 23293146
refGene_gn=CACNA2D3;refGene_tx=XM_019823012.1;refGene_gn=CACNA2D3;refGene_tx=XM_023249840.1;refGene_gn=CACNA2D3;refGene_tx=XM_023249841.1;refGene_gn=CACNA2D3;refGene_tx=XM_023249842.1
A2	23306567 23306643
refGene_gn=CACNA2D3;refGene_tx=XM_019823012.1;refGene_gn=CACNA2D3;refGene_tx=XM_023249840.1;refGene_gn=CACNA2D3;refGene_tx=XM_023249841.1;refGene_gn=CACNA2D3;refGene_tx=XM_023249842.1
A2	23309274 23309327
refGene_gn=CACNA2D3;refGene_tx=XM_019823012.1;refGene_gn=CACNA2D3;refGene_tx=XM_023249840.1;refGene_gn=CACNA2D3;refGene_tx=XM_023249841.1;refGene_gn=CACNA2D3;refGene_tx=XM_023249842.1
A2	23310850 23310906
refGene_gn=CACNA2D3;refGene_tx=XM_019823012.1;refGene_gn=CACNA2D3;refGene_tx=XM_023249840.1;refGene_gn=CACNA2D3;refGene_tx=XM_023249841.1;refGene_gn=CACNA2D3;refGene_tx=XM_023249842.1
A2	23319879 23319991
refGene_gn=CACNA2D3;refGene_tx=XM_019823012.1;refGene_gn=CACNA2D3;refGene_tx=XM_023249840.1;refGene_gn=CACNA2D3;refGene_tx=XM_023249841.1;refGene_gn=CACNA2D3;refGene_tx=XM_023249842.1
A2	23367944 23368057
refGene_gn=CACNA2D3;refGene_tx=XM_019823012.1;refGene_gn=CACNA2D3;refGene_tx=XM_023249840.1;refGene_gn=CACNA2D3;refGene_tx=XM_023249841.1;refGene_gn=CACNA2D3;refGene_tx=XM_023249842.1
A2	23368276 23368359
refGene_gn=CACNA2D3;refGene_tx=XM_019823012.1;refGene_gn=CACNA2D3;refGene_tx=XM_023249840.1;refGene_gn=CACNA2D3;refGene_tx=XM_023249841.1;refGene_gn=CACNA2D3;refGene_tx=XM_023249842.1
A2	23370598 23370691
refGene_gn=CACNA2D3;refGene_tx=XM_019823012.1;refGene_gn=CACNA2D3;refGene_tx=XM_023249840.1;refGene_gn=CACNA2D3;refGene_tx=XM_023249841.1;refGene_gn=CACNA2D3;refGene_tx=XM_023249842.1
A2	23686033 23686492
refGene_gn=WNT5A;refGene_tx=XM_023249844.1;refGene_gn=WNT5A;refGene_tx

=XM_023249845.1;refGene_gn=WNT5A;refGene_tx=XM_023249847.1
A2	23690736	23691029
refGene_gn=WNT5A;refGene_tx=XM_023249844.1;refGene_gn=WNT5A;refGene_tx=XM_023249845.1;refGene_gn=WNT5A;refGene_tx=XM_023249847.1
A2	23695391	23695642
refGene_gn=WNT5A;refGene_tx=XM_023249844.1;refGene_gn=WNT5A;refGene_tx=XM_023249845.1;refGene_gn=WNT5A;refGene_tx=XM_023249847.1
A2	23696867	23697001
refGene_gn=WNT5A;refGene_tx=XM_023249844.1;refGene_gn=WNT5A;refGene_tx=XM_023249845.1;refGene_gn=WNT5A;refGene_tx=XM_023249847.1
A2	23702990	23702996	refGene_gn=WNT5A;refGene_tx=XM_023249845.1
A2	23727513	23727756	refGene_gn=ERC2;refGene_tx=XM_023249849.1
A2	23909352	23909487	refGene_gn=ERC2;refGene_tx=XM_023249849.1
A2	23943564	23943712	refGene_gn=ERC2;refGene_tx=XM_023249849.1
A2	24087350	24087511	refGene_gn=ERC2;refGene_tx=XM_023249849.1
A2	24149036	24149172	refGene_gn=ERC2;refGene_tx=XM_023249849.1
A2	24183668	24183680	refGene_gn=ERC2;refGene_tx=XM_023249849.1
A2	24189514	24189708	refGene_gn=ERC2;refGene_tx=XM_023249849.1
A2	24204084	24204225	refGene_gn=ERC2;refGene_tx=XM_023249849.1
A2	24207357	24207498	refGene_gn=ERC2;refGene_tx=XM_023249849.1
A2	24208450	24208459	refGene_gn=ERC2;refGene_tx=XM_023249849.1
A2	24215978	24216107	refGene_gn=ERC2;refGene_tx=XM_023249849.1
A2	24266584	24266752	refGene_gn=ERC2;refGene_tx=XM_023249849.1
A2	24325271	24325439	refGene_gn=ERC2;refGene_tx=XM_023249849.1
A2	24340219	24340375	refGene_gn=ERC2;refGene_tx=XM_023249849.1
A2	24363440	24363515	refGene_gn=ERC2;refGene_tx=XM_023249849.1
A2	24479407	24479824	refGene_gn=ERC2;refGene_tx=XM_023249849.1
A2	24618518	24619175	refGene_gn=ERC2;refGene_tx=XM_023249849.1
A2	24736673	24736684
refGene_gn=CCDC66;refGene_tx=XM_006928872.2;refGene_gn=CCDC66;refGene_tx=XM_006928873.2;refGene_gn=CCDC66;refGene_tx=XM_006928875.2;refGene_gn=CCDC66;refGene_tx=XM_019823123.1;refGene_gn=CCDC66;refGene_tx=XM_019823139.1;refGene_gn=CCDC66;refGene_tx=XM_019823149.1
A2	24738314	24738379
refGene_gn=CCDC66;refGene_tx=XM_006928872.2;refGene_gn=CCDC66;refGene_tx=XM_006928873.2;refGene_gn=CCDC66;refGene_tx=XM_006928875.2;refGene_gn=CCDC66;refGene_tx=XM_019823123.1;refGene_gn=CCDC66;refGene_tx=XM_019823128.2;refGene_gn=CCDC66;refGene_tx=XM_019823139.1;refGene_gn=CCDC66;refGene_tx=XM_019823149.1
A2	24739018	24739044
refGene_gn=CCDC66;refGene_tx=XM_006928872.2;refGene_gn=CCDC66;refGene_tx=XM_006928873.2;refGene_gn=CCDC66;refGene_tx=XM_006928875.2;refGene_gn=CCDC66;refGene_tx=XM_019823123.1;refGene_gn=CCDC66;refGene_tx=XM_019823139.1;refGene_gn=CCDC66;refGene_tx=XM_019823149.1
A2	24748162	24748607
refGene_gn=CCDC66;refGene_tx=XM_006928872.2;refGene_gn=CCDC66;refGene_tx=XM_006928873.2;refGene_gn=CCDC66;refGene_tx=XM_006928874.3;refGene_gn=CCDC66;refGene_tx=XM_006928875.2;refGene_gn=CCDC66;refGene_tx=XM_019823123.1;refGene_gn=CCDC66;refGene_tx=XM_019823128.2;refGene_gn=CCDC66;refGene_tx=XM_019823131.2;refGene_gn=CCDC66;refGene_tx=XM_019823139.

1;refGene_gn=CCDC66;refGene_tx=XM_019823149.1;refGene_gn=CCDC66;refGene_tx=XM_019823158.1
A2 24750042 24750208
refGene_gn=CCDC66;refGene_tx=XM_006928872.2;refGene_gn=CCDC66;refGene_tx=XM_006928873.2;refGene_gn=CCDC66;refGene_tx=XM_006928874.3;refGene_gn=CCDC66;refGene_tx=XM_006928875.2;refGene_gn=CCDC66;refGene_tx=XM_019823123.1;refGene_gn=CCDC66;refGene_tx=XM_019823128.2;refGene_gn=CCDC66;refGene_tx=XM_019823131.2;refGene_gn=CCDC66;refGene_tx=XM_019823139.1;refGene_gn=CCDC66;refGene_tx=XM_019823149.1;refGene_gn=CCDC66;refGene_tx=XM_019823158.1
A2 24750387 24750495
refGene_gn=CCDC66;refGene_tx=XM_006928872.2;refGene_gn=CCDC66;refGene_tx=XM_006928873.2;refGene_gn=CCDC66;refGene_tx=XM_006928874.3;refGene_gn=CCDC66;refGene_tx=XM_006928875.2;refGene_gn=CCDC66;refGene_tx=XM_019823123.1;refGene_gn=CCDC66;refGene_tx=XM_019823128.2;refGene_gn=CCDC66;refGene_tx=XM_019823131.2;refGene_gn=CCDC66;refGene_tx=XM_019823139.1;refGene_gn=CCDC66;refGene_tx=XM_019823149.1;refGene_gn=CCDC66;refGene_tx=XM_019823158.1
A2 24752240 24752376
refGene_gn=CCDC66;refGene_tx=XM_006928872.2;refGene_gn=CCDC66;refGene_tx=XM_006928873.2;refGene_gn=CCDC66;refGene_tx=XM_006928874.3;refGene_gn=CCDC66;refGene_tx=XM_006928875.2;refGene_gn=CCDC66;refGene_tx=XM_019823123.1;refGene_gn=CCDC66;refGene_tx=XM_019823128.2;refGene_gn=CCDC66;refGene_tx=XM_019823131.2;refGene_gn=CCDC66;refGene_tx=XM_019823139.1;refGene_gn=CCDC66;refGene_tx=XM_019823149.1;refGene_gn=CCDC66;refGene_tx=XM_019823158.1
A2 24753193 24753295
refGene_gn=CCDC66;refGene_tx=XM_006928872.2;refGene_gn=CCDC66;refGene_tx=XM_006928873.2;refGene_gn=CCDC66;refGene_tx=XM_006928874.3;refGene_gn=CCDC66;refGene_tx=XM_006928875.2;refGene_gn=CCDC66;refGene_tx=XM_019823123.1;refGene_gn=CCDC66;refGene_tx=XM_019823128.2;refGene_gn=CCDC66;refGene_tx=XM_019823131.2;refGene_gn=CCDC66;refGene_tx=XM_019823158.1
A2 24754083 24754092 refGene_gn=CCDC66;refGene_tx=XM_023249853.1
A2 24761576 24761624 refGene_gn=CCDC66;refGene_tx=XM_023249852.1
A2 24764775 24764907
refGene_gn=CCDC66;refGene_tx=XM_006928872.2;refGene_gn=CCDC66;refGene_tx=XM_006928873.2;refGene_gn=CCDC66;refGene_tx=XM_006928874.3;refGene_gn=CCDC66;refGene_tx=XM_006928875.2;refGene_gn=CCDC66;refGene_tx=XM_019823123.1;refGene_gn=CCDC66;refGene_tx=XM_019823128.2;refGene_gn=CCDC66;refGene_tx=XM_019823131.2;refGene_gn=CCDC66;refGene_tx=XM_019823139.1;refGene_gn=CCDC66;refGene_tx=XM_019823149.1;refGene_gn=CCDC66;refGene_tx=XM_019823158.1;refGene_gn=CCDC66;refGene_tx=XM_023249852.1;refGene_gn=CCDC66;refGene_tx=XM_023249853.1
A2 24765507 24765758
refGene_gn=CCDC66;refGene_tx=XM_006928872.2;refGene_gn=CCDC66;refGene_tx=XM_006928873.2;refGene_gn=CCDC66;refGene_tx=XM_006928874.3;refGene_gn=CCDC66;refGene_tx=XM_006928875.2;refGene_gn=CCDC66;refGene_tx=XM_019823123.1;refGene_gn=CCDC66;refGene_tx=XM_019823128.2;refGene_gn=CCDC66;refGene_tx=XM_019823131.2;refGene_gn=CCDC66;refGene_tx=XM_019823139.

1;refGene_gn=CCDC66;refGene_tx=XM_019823149.1;refGene_gn=CCDC66;refGene_tx=XM_019823158.1;refGene_gn=CCDC66;refGene_tx=XM_023249852.1;refGene_gn=CCDC66;refGene_tx=XM_023249853.1
A2 24765976 24766061
refGene_gn=CCDC66;refGene_tx=XM_006928872.2;refGene_gn=CCDC66;refGene_tx=XM_006928873.2;refGene_gn=CCDC66;refGene_tx=XM_006928874.3;refGene_gn=CCDC66;refGene_tx=XM_006928875.2;refGene_gn=CCDC66;refGene_tx=XM_019823123.1;refGene_gn=CCDC66;refGene_tx=XM_019823128.2;refGene_gn=CCDC66;refGene_tx=XM_019823131.2;refGene_gn=CCDC66;refGene_tx=XM_019823139.1;refGene_gn=CCDC66;refGene_tx=XM_019823149.1;refGene_gn=CCDC66;refGene_tx=XM_019823158.1;refGene_gn=CCDC66;refGene_tx=XM_023249852.1;refGene_gn=CCDC66;refGene_tx=XM_023249853.1
A2 24767134 24767296
refGene_gn=CCDC66;refGene_tx=XM_006928872.2;refGene_gn=CCDC66;refGene_tx=XM_006928873.2;refGene_gn=CCDC66;refGene_tx=XM_006928874.3;refGene_gn=CCDC66;refGene_tx=XM_006928875.2;refGene_gn=CCDC66;refGene_tx=XM_019823123.1;refGene_gn=CCDC66;refGene_tx=XM_019823128.2;refGene_gn=CCDC66;refGene_tx=XM_019823131.2;refGene_gn=CCDC66;refGene_tx=XM_019823139.1;refGene_gn=CCDC66;refGene_tx=XM_019823149.1;refGene_gn=CCDC66;refGene_tx=XM_019823158.1;refGene_gn=CCDC66;refGene_tx=XM_023249852.1;refGene_gn=CCDC66;refGene_tx=XM_023249853.1
A2 24770166 24770311
refGene_gn=CCDC66;refGene_tx=XM_006928872.2;refGene_gn=CCDC66;refGene_tx=XM_006928873.2;refGene_gn=CCDC66;refGene_tx=XM_006928874.3;refGene_gn=CCDC66;refGene_tx=XM_006928875.2;refGene_gn=CCDC66;refGene_tx=XM_019823123.1;refGene_gn=CCDC66;refGene_tx=XM_019823128.2;refGene_gn=CCDC66;refGene_tx=XM_019823131.2;refGene_gn=CCDC66;refGene_tx=XM_019823139.1;refGene_gn=CCDC66;refGene_tx=XM_019823149.1;refGene_gn=CCDC66;refGene_tx=XM_019823158.1;refGene_gn=CCDC66;refGene_tx=XM_023249852.1;refGene_gn=CCDC66;refGene_tx=XM_023249853.1
A2 24771053 24771140
refGene_gn=CCDC66;refGene_tx=XM_006928872.2;refGene_gn=CCDC66;refGene_tx=XM_006928873.2;refGene_gn=CCDC66;refGene_tx=XM_006928874.3;refGene_gn=CCDC66;refGene_tx=XM_006928875.2;refGene_gn=CCDC66;refGene_tx=XM_019823123.1;refGene_gn=CCDC66;refGene_tx=XM_019823128.2;refGene_gn=CCDC66;refGene_tx=XM_019823131.2;refGene_gn=CCDC66;refGene_tx=XM_019823139.1;refGene_gn=CCDC66;refGene_tx=XM_019823149.1;refGene_gn=CCDC66;refGene_tx=XM_019823158.1;refGene_gn=CCDC66;refGene_tx=XM_023249852.1;refGene_gn=CCDC66;refGene_tx=XM_023249853.1
A2 24782238 24782726
refGene_gn=CCDC66;refGene_tx=XM_006928872.2;refGene_gn=CCDC66;refGene_tx=XM_006928873.2;refGene_gn=CCDC66;refGene_tx=XM_006928874.3;refGene_gn=CCDC66;refGene_tx=XM_006928875.2;refGene_gn=CCDC66;refGene_tx=XM_019823123.1;refGene_gn=CCDC66;refGene_tx=XM_019823128.2;refGene_gn=CCDC66;refGene_tx=XM_019823131.2;refGene_gn=CCDC66;refGene_tx=XM_019823139.1;refGene_gn=CCDC66;refGene_tx=XM_019823149.1;refGene_gn=CCDC66;refGene_tx=XM_019823158.1;refGene_gn=CCDC66;refGene_tx=XM_023249852.1;refGene_gn=CCDC66;refGene_tx=XM_023249853.1
A2 24783308 24783349
refGene_gn=CCDC66;refGene_tx=XM_006928872.2;refGene_gn=CCDC66;refGene_

tx=XM_006928873.2;refGene_gn=CCDC66;refGene_tx=XM_006928874.3;refGene_gn=CCDC66;refGene_tx=XM_006928875.2;refGene_gn=CCDC66;refGene_tx=XM_019823123.1;refGene_gn=CCDC66;refGene_tx=XM_019823128.2;refGene_gn=CCDC66;refGene_tx=XM_019823131.2;refGene_gn=CCDC66;refGene_tx=XM_019823139.1;refGene_gn=CCDC66;refGene_tx=XM_019823149.1;refGene_gn=CCDC66;refGene_tx=XM_019823158.1;refGene_gn=CCDC66;refGene_tx=XM_023249852.1;refGene_gn=CCDC66;refGene_tx=XM_023249853.1
A2 24784117 24784359
refGene_gn=CCDC66;refGene_tx=XM_006928872.2;refGene_gn=CCDC66;refGene_tx=XM_006928873.2;refGene_gn=CCDC66;refGene_tx=XM_006928874.3;refGene_gn=CCDC66;refGene_tx=XM_006928875.2;refGene_gn=CCDC66;refGene_tx=XM_019823123.1;refGene_gn=CCDC66;refGene_tx=XM_019823128.2;refGene_gn=CCDC66;refGene_tx=XM_019823131.2;refGene_gn=CCDC66;refGene_tx=XM_019823139.1;refGene_gn=CCDC66;refGene_tx=XM_019823149.1;refGene_gn=CCDC66;refGene_tx=XM_019823158.1;refGene_gn=CCDC66;refGene_tx=XM_023249852.1;refGene_gn=CCDC66;refGene_tx=XM_023249853.1
A2 24784547 24784672
refGene_gn=CCDC66;refGene_tx=XM_006928872.2;refGene_gn=CCDC66;refGene_tx=XM_006928873.2;refGene_gn=CCDC66;refGene_tx=XM_006928874.3;refGene_gn=CCDC66;refGene_tx=XM_006928875.2;refGene_gn=CCDC66;refGene_tx=XM_019823123.1;refGene_gn=CCDC66;refGene_tx=XM_019823128.2;refGene_gn=CCDC66;refGene_tx=XM_019823131.2;refGene_gn=CCDC66;refGene_tx=XM_019823139.1;refGene_gn=CCDC66;refGene_tx=XM_019823149.1;refGene_gn=CCDC66;refGene_tx=XM_019823158.1;refGene_gn=CCDC66;refGene_tx=XM_023249852.1;refGene_gn=CCDC66;refGene_tx=XM_023249853.1
A2 24786055 24786142
refGene_gn=CCDC66;refGene_tx=XM_006928872.2;refGene_gn=CCDC66;refGene_tx=XM_006928873.2;refGene_gn=CCDC66;refGene_tx=XM_006928874.3;refGene_gn=CCDC66;refGene_tx=XM_006928875.2;refGene_gn=CCDC66;refGene_tx=XM_019823123.1;refGene_gn=CCDC66;refGene_tx=XM_019823128.2;refGene_gn=CCDC66;refGene_tx=XM_019823131.2;refGene_gn=CCDC66;refGene_tx=XM_019823139.1;refGene_gn=CCDC66;refGene_tx=XM_019823149.1;refGene_gn=CCDC66;refGene_tx=XM_019823158.1;refGene_gn=CCDC66;refGene_tx=XM_023249852.1;refGene_gn=CCDC66;refGene_tx=XM_023249853.1
A2 24787510 24788043
refGene_gn=FAM208A;refGene_tx=XM_006928876.3;refGene_gn=FAM208A;refGene_tx=XM_006928878.3;refGene_gn=FAM208A;refGene_tx=XM_019823105.2;refGene_gn=FAM208A;refGene_tx=XM_019823108.2;refGene_gn=FAM208A;refGene_tx=XM_019823116.2
A2 24788264 24788320
refGene_gn=FAM208A;refGene_tx=XM_003982354.5;refGene_gn=FAM208A;refGene_tx=XM_006928879.4
A2 24788923 24789088
refGene_gn=FAM208A;refGene_tx=XM_003982354.5;refGene_gn=FAM208A;refGene_tx=XM_006928876.3;refGene_gn=FAM208A;refGene_tx=XM_006928878.3;refGene_gn=FAM208A;refGene_tx=XM_006928879.4;refGene_gn=FAM208A;refGene_tx=XM_019823105.2;refGene_gn=FAM208A;refGene_tx=XM_019823108.2;refGene_gn=FAM208A;refGene_tx=XM_019823116.2
A2 24789254 24789433
refGene_gn=FAM208A;refGene_tx=XM_003982354.5;refGene_gn=FAM208A;refGen

e_tx=XM_006928876.3;refGene_gn=FAM208A;refGene_tx=XM_006928878.3;refGene_gn=FAM208A;refGene_tx=XM_006928879.4;refGene_gn=FAM208A;refGene_tx=XM_019823105.2;refGene_gn=FAM208A;refGene_tx=XM_019823108.2;refGene_gn=FAM208A;refGene_tx=XM_019823116.2
A2 24790914 24791023
refGene_gn=FAM208A;refGene_tx=XM_003982354.5;refGene_gn=FAM208A;refGene_tx=XM_006928876.3;refGene_gn=FAM208A;refGene_tx=XM_006928878.3;refGene_gn=FAM208A;refGene_tx=XM_006928879.4;refGene_gn=FAM208A;refGene_tx=XM_019823105.2;refGene_gn=FAM208A;refGene_tx=XM_019823108.2;refGene_gn=FAM208A;refGene_tx=XM_019823116.2
A2 24791803 24791963
refGene_gn=FAM208A;refGene_tx=XM_003982354.5;refGene_gn=FAM208A;refGene_tx=XM_006928876.3;refGene_gn=FAM208A;refGene_tx=XM_006928878.3;refGene_gn=FAM208A;refGene_tx=XM_006928879.4;refGene_gn=FAM208A;refGene_tx=XM_019823105.2;refGene_gn=FAM208A;refGene_tx=XM_019823108.2;refGene_gn=FAM208A;refGene_tx=XM_019823116.2
A2 24792740 24792863
refGene_gn=FAM208A;refGene_tx=XM_003982354.5;refGene_gn=FAM208A;refGene_tx=XM_006928876.3;refGene_gn=FAM208A;refGene_tx=XM_006928878.3;refGene_gn=FAM208A;refGene_tx=XM_006928879.4;refGene_gn=FAM208A;refGene_tx=XM_019823105.2;refGene_gn=FAM208A;refGene_tx=XM_019823108.2;refGene_gn=FAM208A;refGene_tx=XM_019823116.2
A2 24794957 24795892
refGene_gn=FAM208A;refGene_tx=XM_003982354.5;refGene_gn=FAM208A;refGene_tx=XM_006928876.3;refGene_gn=FAM208A;refGene_tx=XM_006928878.3;refGene_gn=FAM208A;refGene_tx=XM_006928879.4;refGene_gn=FAM208A;refGene_tx=XM_019823105.2;refGene_gn=FAM208A;refGene_tx=XM_019823108.2;refGene_gn=FAM208A;refGene_tx=XM_019823116.2
A2 24799224 24799284
refGene_gn=FAM208A;refGene_tx=XM_003982354.5;refGene_gn=FAM208A;refGene_tx=XM_006928876.3;refGene_gn=FAM208A;refGene_tx=XM_006928878.3;refGene_gn=FAM208A;refGene_tx=XM_006928879.4;refGene_gn=FAM208A;refGene_tx=XM_019823108.2
A2 24800547 24800689
refGene_gn=FAM208A;refGene_tx=XM_003982354.5;refGene_gn=FAM208A;refGene_tx=XM_006928876.3;refGene_gn=FAM208A;refGene_tx=XM_006928878.3;refGene_gn=FAM208A;refGene_tx=XM_006928879.4;refGene_gn=FAM208A;refGene_tx=XM_019823105.2;refGene_gn=FAM208A;refGene_tx=XM_019823108.2;refGene_gn=FAM208A;refGene_tx=XM_019823116.2
A2 24801965 24802372
refGene_gn=FAM208A;refGene_tx=XM_003982354.5;refGene_gn=FAM208A;refGene_tx=XM_006928876.3;refGene_gn=FAM208A;refGene_tx=XM_006928878.3;refGene_gn=FAM208A;refGene_tx=XM_006928879.4;refGene_gn=FAM208A;refGene_tx=XM_019823105.2;refGene_gn=FAM208A;refGene_tx=XM_019823108.2;refGene_gn=FAM208A;refGene_tx=XM_019823116.2
A2 24805303 24806005
refGene_gn=FAM208A;refGene_tx=XM_003982354.5;refGene_gn=FAM208A;refGene_tx=XM_006928876.3;refGene_gn=FAM208A;refGene_tx=XM_006928878.3;refGene_gn=FAM208A;refGene_tx=XM_006928879.4;refGene_gn=FAM208A;refGene_tx=XM_019823105.2;refGene_gn=FAM208A;refGene_tx=XM_019823108.2;refGene_gn

=FAM208A;refGene_tx=XM_019823116.2
A2 24807440 24807512
refGene_gn=FAM208A;refGene_tx=XM_003982354.5;refGene_gn=FAM208A;refGene_tx=XM_006928876.3;refGene_gn=FAM208A;refGene_tx=XM_006928878.3;refGene_gn=FAM208A;refGene_tx=XM_006928879.4;refGene_gn=FAM208A;refGene_tx=XM_019823105.2;refGene_gn=FAM208A;refGene_tx=XM_019823108.2;refGene_gn=FAM208A;refGene_tx=XM_019823116.2
A2 24807614 24807687
refGene_gn=FAM208A;refGene_tx=XM_003982354.5;refGene_gn=FAM208A;refGene_tx=XM_006928876.3;refGene_gn=FAM208A;refGene_tx=XM_006928878.3;refGene_gn=FAM208A;refGene_tx=XM_006928879.4;refGene_gn=FAM208A;refGene_tx=XM_019823105.2;refGene_gn=FAM208A;refGene_tx=XM_019823108.2;refGene_gn=FAM208A;refGene_tx=XM_019823116.2
A2 24819757 24819861
refGene_gn=FAM208A;refGene_tx=XM_003982354.5;refGene_gn=FAM208A;refGene_tx=XM_006928876.3;refGene_gn=FAM208A;refGene_tx=XM_006928878.3;refGene_gn=FAM208A;refGene_tx=XM_006928879.4;refGene_gn=FAM208A;refGene_tx=XM_019823105.2;refGene_gn=FAM208A;refGene_tx=XM_019823108.2;refGene_gn=FAM208A;refGene_tx=XM_019823116.2
A2 24819940 24820044
refGene_gn=FAM208A;refGene_tx=XM_003982354.5;refGene_gn=FAM208A;refGene_tx=XM_006928876.3;refGene_gn=FAM208A;refGene_tx=XM_006928878.3;refGene_gn=FAM208A;refGene_tx=XM_006928879.4;refGene_gn=FAM208A;refGene_tx=XM_019823105.2;refGene_gn=FAM208A;refGene_tx=XM_019823108.2;refGene_gn=FAM208A;refGene_tx=XM_019823116.2
A2 24820812 24820918
refGene_gn=FAM208A;refGene_tx=XM_003982354.5;refGene_gn=FAM208A;refGene_tx=XM_006928876.3;refGene_gn=FAM208A;refGene_tx=XM_006928878.3;refGene_gn=FAM208A;refGene_tx=XM_006928879.4;refGene_gn=FAM208A;refGene_tx=XM_019823105.2;refGene_gn=FAM208A;refGene_tx=XM_019823108.2;refGene_gn=FAM208A;refGene_tx=XM_019823116.2
A2 24821652 24821684
refGene_gn=FAM208A;refGene_tx=XM_003982354.5;refGene_gn=FAM208A;refGene_tx=XM_006928876.3;refGene_gn=FAM208A;refGene_tx=XM_006928878.3;refGene_gn=FAM208A;refGene_tx=XM_006928879.4;refGene_gn=FAM208A;refGene_tx=XM_019823105.2;refGene_gn=FAM208A;refGene_tx=XM_019823108.2;refGene_gn=FAM208A;refGene_tx=XM_019823116.2
A2 24824046 24824171
refGene_gn=FAM208A;refGene_tx=XM_003982354.5;refGene_gn=FAM208A;refGene_tx=XM_006928876.3;refGene_gn=FAM208A;refGene_tx=XM_006928878.3;refGene_gn=FAM208A;refGene_tx=XM_006928879.4;refGene_gn=FAM208A;refGene_tx=XM_019823105.2;refGene_gn=FAM208A;refGene_tx=XM_019823108.2;refGene_gn=FAM208A;refGene_tx=XM_019823116.2
A2 24826858 24826993
refGene_gn=FAM208A;refGene_tx=XM_003982354.5;refGene_gn=FAM208A;refGene_tx=XM_006928876.3;refGene_gn=FAM208A;refGene_tx=XM_006928878.3;refGene_gn=FAM208A;refGene_tx=XM_006928879.4;refGene_gn=FAM208A;refGene_tx=XM_019823105.2;refGene_gn=FAM208A;refGene_tx=XM_019823108.2;refGene_gn=FAM208A;refGene_tx=XM_019823116.2
A2 24827678 24827770

refGene_gn=FAM208A;refGene_tx=XM_003982354.5;refGene_gn=FAM208A;refGene_tx=XM_006928876.3;refGene_gn=FAM208A;refGene_tx=XM_006928878.3;refGene_gn=FAM208A;refGene_tx=XM_006928879.4;refGene_gn=FAM208A;refGene_tx=XM_019823105.2;refGene_gn=FAM208A;refGene_tx=XM_019823108.2;refGene_gn=FAM208A;refGene_tx=XM_019823116.2
A2 24828214 24828287
refGene_gn=FAM208A;refGene_tx=XM_003982354.5;refGene_gn=FAM208A;refGene_tx=XM_006928876.3;refGene_gn=FAM208A;refGene_tx=XM_006928878.3;refGene_gn=FAM208A;refGene_tx=XM_006928879.4;refGene_gn=FAM208A;refGene_tx=XM_019823105.2;refGene_gn=FAM208A;refGene_tx=XM_019823108.2;refGene_gn=FAM208A;refGene_tx=XM_019823116.2
A2 24828955 24829048
refGene_gn=FAM208A;refGene_tx=XM_003982354.5;refGene_gn=FAM208A;refGene_tx=XM_006928876.3;refGene_gn=FAM208A;refGene_tx=XM_006928878.3;refGene_gn=FAM208A;refGene_tx=XM_006928879.4;refGene_gn=FAM208A;refGene_tx=XM_019823105.2;refGene_gn=FAM208A;refGene_tx=XM_019823108.2;refGene_gn=FAM208A;refGene_tx=XM_019823116.2
A2 24830214 24830360
refGene_gn=FAM208A;refGene_tx=XM_003982354.5;refGene_gn=FAM208A;refGene_tx=XM_006928876.3;refGene_gn=FAM208A;refGene_tx=XM_006928878.3;refGene_gn=FAM208A;refGene_tx=XM_006928879.4;refGene_gn=FAM208A;refGene_tx=XM_019823105.2;refGene_gn=FAM208A;refGene_tx=XM_019823108.2;refGene_gn=FAM208A;refGene_tx=XM_019823116.2
A2 24838336 24838664
refGene_gn=FAM208A;refGene_tx=XM_003982354.5;refGene_gn=FAM208A;refGene_tx=XM_006928876.3;refGene_gn=FAM208A;refGene_tx=XM_006928878.3;refGene_gn=FAM208A;refGene_tx=XM_006928879.4;refGene_gn=FAM208A;refGene_tx=XM_019823105.2;refGene_gn=FAM208A;refGene_tx=XM_019823108.2;refGene_gn=FAM208A;refGene_tx=XM_019823116.2
A2 24874625 24874978
refGene_gn=ARHGEF3;refGene_tx=XM_003982357.5;refGene_gn=ARHGEF3;refGene_tx=XM_006928881.4;refGene_gn=ARHGEF3;refGene_tx=XM_019823177.2;refGene_gn=ARHGEF3;refGene_tx=XM_023249854.1;refGene_gn=ARHGEF3;refGene_tx=XM_023249855.1;refGene_gn=ARHGEF3;refGene_tx=XM_023249856.1;refGene_gn=ARHGEF3;refGene_tx=XM_023249857.1
A2 24876743 24876930
refGene_gn=ARHGEF3;refGene_tx=XM_003982357.5;refGene_gn=ARHGEF3;refGene_tx=XM_006928881.4;refGene_gn=ARHGEF3;refGene_tx=XM_019823177.2;refGene_gn=ARHGEF3;refGene_tx=XM_023249854.1;refGene_gn=ARHGEF3;refGene_tx=XM_023249855.1;refGene_gn=ARHGEF3;refGene_tx=XM_023249856.1;refGene_gn=ARHGEF3;refGene_tx=XM_023249857.1
A2 24880209 24880380
refGene_gn=ARHGEF3;refGene_tx=XM_003982357.5;refGene_gn=ARHGEF3;refGene_tx=XM_006928881.4;refGene_gn=ARHGEF3;refGene_tx=XM_019823177.2;refGene_gn=ARHGEF3;refGene_tx=XM_023249854.1;refGene_gn=ARHGEF3;refGene_tx=XM_023249855.1;refGene_gn=ARHGEF3;refGene_tx=XM_023249856.1;refGene_gn=ARHGEF3;refGene_tx=XM_023249857.1
A2 24889061 24889319
refGene_gn=ARHGEF3;refGene_tx=XM_003982357.5;refGene_gn=ARHGEF3;refGene_tx=XM_006928881.4;refGene_gn=ARHGEF3;refGene_tx=XM_019823177.2;refGe

ne_gn=ARHGEF3;refGene_tx=XM_023249854.1;refGene_gn=ARHGEF3;refGene_tx=XM_023249855.1;refGene_gn=ARHGEF3;refGene_tx=XM_023249856.1;refGene_gn=ARHGEF3;refGene_tx=XM_023249857.1
A2 24895076 24895153 refGene_gn=ARHGEF3;refGene_tx=XM_003982357.5;refGene_gn=ARHGEF3;refGene_tx=XM_006928881.4;refGene_gn=ARHGEF3;refGene_tx=XM_019823177.2;refGene_gn=ARHGEF3;refGene_tx=XM_023249854.1;refGene_gn=ARHGEF3;refGene_tx=XM_023249855.1;refGene_gn=ARHGEF3;refGene_tx=XM_023249856.1;refGene_gn=ARHGEF3;refGene_tx=XM_023249857.1
A2 24895325 24895422 refGene_gn=ARHGEF3;refGene_tx=XM_003982357.5;refGene_gn=ARHGEF3;refGene_tx=XM_006928881.4;refGene_gn=ARHGEF3;refGene_tx=XM_019823177.2;refGene_gn=ARHGEF3;refGene_tx=XM_023249854.1;refGene_gn=ARHGEF3;refGene_tx=XM_023249855.1;refGene_gn=ARHGEF3;refGene_tx=XM_023249856.1;refGene_gn=ARHGEF3;refGene_tx=XM_023249857.1
A2 24897519 24897582 refGene_gn=ARHGEF3;refGene_tx=XM_003982357.5;refGene_gn=ARHGEF3;refGene_tx=XM_006928881.4;refGene_gn=ARHGEF3;refGene_tx=XM_019823177.2;refGene_gn=ARHGEF3;refGene_tx=XM_023249854.1;refGene_gn=ARHGEF3;refGene_tx=XM_023249855.1;refGene_gn=ARHGEF3;refGene_tx=XM_023249856.1;refGene_gn=ARHGEF3;refGene_tx=XM_023249857.1
A2 24900816 24900987 refGene_gn=ARHGEF3;refGene_tx=XM_003982357.5;refGene_gn=ARHGEF3;refGene_tx=XM_006928881.4;refGene_gn=ARHGEF3;refGene_tx=XM_019823177.2;refGene_gn=ARHGEF3;refGene_tx=XM_023249854.1;refGene_gn=ARHGEF3;refGene_tx=XM_023249855.1;refGene_gn=ARHGEF3;refGene_tx=XM_023249856.1;refGene_gn=ARHGEF3;refGene_tx=XM_023249857.1
A2 24916153 24916261 refGene_gn=ARHGEF3;refGene_tx=XM_003982357.5;refGene_gn=ARHGEF3;refGene_tx=XM_006928881.4;refGene_gn=ARHGEF3;refGene_tx=XM_019823177.2;refGene_gn=ARHGEF3;refGene_tx=XM_023249854.1;refGene_gn=ARHGEF3;refGene_tx=XM_023249855.1;refGene_gn=ARHGEF3;refGene_tx=XM_023249856.1;refGene_gn=ARHGEF3;refGene_tx=XM_023249857.1
A2 24917140 24917281 refGene_gn=ARHGEF3;refGene_tx=XM_019823177.2
A2 24942560 24942656 refGene_gn=ARHGEF3;refGene_tx=XM_003982357.5
A2 25016382 25016445 refGene_gn=ARHGEF3;refGene_tx=XM_006928881.4;refGene_gn=ARHGEF3;refGene_tx=XM_023249854.1;refGene_gn=ARHGEF3;refGene_tx=XM_023249855.1;refGene_gn=ARHGEF3;refGene_tx=XM_023249856.1;refGene_gn=ARHGEF3;refGene_tx=XM_023249857.1
A2 25049449 25049512 refGene_gn=ARHGEF3;refGene_tx=XM_023249856.1
A2 25052171 25052222 refGene_gn=ARHGEF3;refGene_tx=XM_006928881.4
A2 25083478 25083545 refGene_gn=ARHGEF3;refGene_tx=XM_023249855.1;refGene_gn=ARHGEF3;refGene_tx=XM_023249857.1
A2 25141656 25141749 refGene_gn=ARHGEF3;refGene_tx=XM_023249854.1;refGene_gn=ARHGEF3;refGene_tx=XM_023249855.1
A2 25174218 25174334 refGene_gn=SPATA12;refGene_tx=XM_019825021.1
A2 25174335 25174490 refGene_gn=SPATA12;refGene_tx=XM_019825021.1

A2 25174492 25174580 refGene_gn=SPATA12;refGene_tx=XM_019825021.1
A2 25174581 25174804 refGene_gn=SPATA12;refGene_tx=XM_019825021.1
A2 25179665 25179722 refGene_gn=ARHGEF3;refGene_tx=XM_023249854.1
A2 25195028 25195141
refGene_gn=IL17RD;refGene_tx=XM_019823182.2;refGene_gn=IL17RD;refGene_tx=XM_019823191.1
A2 25196155 25197089
refGene_gn=IL17RD;refGene_tx=XM_019823182.2;refGene_gn=IL17RD;refGene_tx=XM_019823191.1
A2 25200939 25201124
refGene_gn=IL17RD;refGene_tx=XM_019823182.2;refGene_gn=IL17RD;refGene_tx=XM_019823191.1
A2 25201919 25202030
refGene_gn=IL17RD;refGene_tx=XM_019823182.2;refGene_gn=IL17RD;refGene_tx=XM_019823191.1
A2 25202797 25202852
refGene_gn=IL17RD;refGene_tx=XM_019823182.2;refGene_gn=IL17RD;refGene_tx=XM_019823191.1
A2 25204842 25204908
refGene_gn=IL17RD;refGene_tx=XM_019823182.2;refGene_gn=IL17RD;refGene_tx=XM_019823191.1
A2 25206225 25206377
refGene_gn=IL17RD;refGene_tx=XM_019823182.2;refGene_gn=IL17RD;refGene_tx=XM_019823191.1
A2 25206469 25206514
refGene_gn=IL17RD;refGene_tx=XM_019823182.2;refGene_gn=IL17RD;refGene_tx=XM_019823191.1
A2 25208916 25209037
refGene_gn=IL17RD;refGene_tx=XM_019823182.2;refGene_gn=IL17RD;refGene_tx=XM_019823191.1
A2 25209534 25209653
refGene_gn=IL17RD;refGene_tx=XM_019823182.2;refGene_gn=IL17RD;refGene_tx=XM_019823191.1
A2 25214221 25214347
refGene_gn=IL17RD;refGene_tx=XM_019823182.2;refGene_gn=IL17RD;refGene_tx=XM_019823191.1
A2 25218734 25218792
refGene_gn=IL17RD;refGene_tx=XM_019823182.2;refGene_gn=IL17RD;refGene_tx=XM_019823191.1
A2 25260340 25260466
refGene_gn=IL17RD;refGene_tx=XM_019823182.2;refGene_gn=IL17RD;refGene_tx=XM_019823191.1
A2 25303587 25303686 refGene_gn=HESX1;refGene_tx=XM_003982360.4
A2 25303782 25303884 refGene_gn=HESX1;refGene_tx=XM_003982360.4
A2 25304197 25304397 refGene_gn=HESX1;refGene_tx=XM_003982360.4
A2 25304911 25305068 refGene_gn=HESX1;refGene_tx=XM_003982360.4
A2 25332332 25332386
refGene_gn=APPL1;refGene_tx=XM_011279730.2;refGene_gn=APPL1;refGene_tx=XM_019823206.2
A2 25338891 25338990

refGene_gn=APPL1;refGene_tx=XM_011279730.2;refGene_gn=APPL1;refGene_tx=XM_019823206.2
A2	25340224	25340284	refGene_gn=APPL1;refGene_tx=XM_011279730.2;refGene_gn=APPL1;refGene_tx=XM_019823206.2
A2	25341829	25341901	refGene_gn=APPL1;refGene_tx=XM_011279730.2;refGene_gn=APPL1;refGene_tx=XM_019823206.2
A2	25344523	25344611	refGene_gn=APPL1;refGene_tx=XM_011279730.2;refGene_gn=APPL1;refGene_tx=XM_019823206.2
A2	25346250	25346292	refGene_gn=APPL1;refGene_tx=XM_011279730.2;refGene_gn=APPL1;refGene_tx=XM_019823206.2
A2	25346860	25346919	refGene_gn=APPL1;refGene_tx=XM_011279730.2;refGene_gn=APPL1;refGene_tx=XM_019823206.2
A2	25349933	25350080	refGene_gn=APPL1;refGene_tx=XM_011279730.2;refGene_gn=APPL1;refGene_tx=XM_019823206.2
A2	25350940	25351023	refGene_gn=APPL1;refGene_tx=XM_011279730.2;refGene_gn=APPL1;refGene_tx=XM_019823206.2
A2	25351720	25351879	refGene_gn=APPL1;refGene_tx=XM_011279730.2
A2	25352789	25352978	refGene_gn=APPL1;refGene_tx=XM_011279730.2;refGene_gn=APPL1;refGene_tx=XM_019823206.2
A2	25355171	25355214	refGene_gn=APPL1;refGene_tx=XM_011279730.2;refGene_gn=APPL1;refGene_tx=XM_019823206.2
A2	25356451	25356508	refGene_gn=APPL1;refGene_tx=XM_011279730.2;refGene_gn=APPL1;refGene_tx=XM_019823206.2
A2	25359033	25359125	refGene_gn=APPL1;refGene_tx=XM_011279730.2;refGene_gn=APPL1;refGene_tx=XM_019823206.2
A2	25359980	25360163	refGene_gn=APPL1;refGene_tx=XM_011279730.2;refGene_gn=APPL1;refGene_tx=XM_019823206.2
A2	25362100	25362153	refGene_gn=APPL1;refGene_tx=XM_011279730.2;refGene_gn=APPL1;refGene_tx=XM_019823206.2
A2	25362725	25362900	refGene_gn=APPL1;refGene_tx=XM_011279730.2;refGene_gn=APPL1;refGene_tx=XM_019823206.2
A2	25363217	25363254	refGene_gn=APPL1;refGene_tx=XM_011279730.2;refGene_gn=APPL1;refGene_tx=XM_019823206.2
A2	25363868	25364015	refGene_gn=APPL1;refGene_tx=XM_011279730.2;refGene_gn=APPL1;refGene_tx

=XM_019823206.2
A2	25364591	25364642
refGene_gn=APPL1;refGene_tx=XM_011279730.2;refGene_gn=APPL1;refGene_tx=XM_019823206.2
A2	25364931	25365042
refGene_gn=APPL1;refGene_tx=XM_011279730.2;refGene_gn=APPL1;refGene_tx=XM_019823206.2
A2	25366031	25366178	refGene_gn=APPL1;refGene_tx=XM_011279730.2
A2	25375549	25375728
refGene_gn=ASB14;refGene_tx=XM_006928884.3;refGene_gn=ASB14;refGene_tx=XM_006928887.3;refGene_gn=ASB14;refGene_tx=XM_019823213.1;refGene_gn=ASB14;refGene_tx=XM_019823223.1;refGene_gn=ASB14;refGene_tx=XM_019823225.1;refGene_gn=ASB14;refGene_tx=XM_019823231.1
A2	25376483	25376637
refGene_gn=ASB14;refGene_tx=XM_006928884.3;refGene_gn=ASB14;refGene_tx=XM_006928887.3;refGene_gn=ASB14;refGene_tx=XM_019823213.1;refGene_gn=ASB14;refGene_tx=XM_019823223.1;refGene_gn=ASB14;refGene_tx=XM_019823225.1;refGene_gn=ASB14;refGene_tx=XM_019823231.1
A2	25378698	25379242
refGene_gn=ASB14;refGene_tx=XM_006928884.3;refGene_gn=ASB14;refGene_tx=XM_006928887.3;refGene_gn=ASB14;refGene_tx=XM_019823213.1;refGene_gn=ASB14;refGene_tx=XM_019823223.1;refGene_gn=ASB14;refGene_tx=XM_019823225.1;refGene_gn=ASB14;refGene_tx=XM_019823231.1
A2	25379927	25380099
refGene_gn=ASB14;refGene_tx=XM_006928884.3;refGene_gn=ASB14;refGene_tx=XM_006928887.3;refGene_gn=ASB14;refGene_tx=XM_019823213.1;refGene_gn=ASB14;refGene_tx=XM_019823223.1;refGene_gn=ASB14;refGene_tx=XM_019823225.1;refGene_gn=ASB14;refGene_tx=XM_019823231.1
A2	25382517	25382763
refGene_gn=ASB14;refGene_tx=XM_006928884.3;refGene_gn=ASB14;refGene_tx=XM_006928887.3;refGene_gn=ASB14;refGene_tx=XM_019823213.1;refGene_gn=ASB14;refGene_tx=XM_019823223.1;refGene_gn=ASB14;refGene_tx=XM_019823231.1
A2	25385272	25385431
refGene_gn=ASB14;refGene_tx=XM_006928884.3;refGene_gn=ASB14;refGene_tx=XM_006928887.3;refGene_gn=ASB14;refGene_tx=XM_019823213.1;refGene_gn=ASB14;refGene_tx=XM_019823225.1
A2	25385527	25385659
refGene_gn=ASB14;refGene_tx=XM_006928884.3;refGene_gn=ASB14;refGene_tx=XM_006928887.3;refGene_gn=ASB14;refGene_tx=XM_019823213.1;refGene_gn=ASB14;refGene_tx=XM_019823223.1;refGene_gn=ASB14;refGene_tx=XM_019823225.1
A2	25386106	25386162
refGene_gn=ASB14;refGene_tx=XM_006928884.3;refGene_gn=ASB14;refGene_tx=XM_006928887.3;refGene_gn=ASB14;refGene_tx=XM_019823213.1;refGene_gn=ASB14;refGene_tx=XM_019823223.1;refGene_gn=ASB14;refGene_tx=XM_019823225.1
A2	25387854	25387976
refGene_gn=ASB14;refGene_tx=XM_006928884.3;refGene_gn=ASB14;refGene_tx=XM_006928887.3;refGene_gn=ASB14;refGene_tx=XM_019823213.1;refGene_gn=

ASB14;refGene_tx=XM_019823223.1;refGene_gn=ASB14;refGene_tx=XM_0198232
25.1

| | | | |
|---|---|---|---|
| A2 | 25390226 | 25390417 | refGene_gn=DNAH12;refGene_tx=XM_023249860.1 |
| A2 | 25392249 | 25392317 | refGene_gn=DNAH12;refGene_tx=XM_023249860.1 |
| A2 | 25393941 | 25394033 | refGene_gn=DNAH12;refGene_tx=XM_023249860.1 |
| A2 | 25394474 | 25394612 | refGene_gn=DNAH12;refGene_tx=XM_023249860.1 |
| A2 | 25400292 | 25400497 | refGene_gn=DNAH12;refGene_tx=XM_023249860.1 |
| A2 | 25402087 | 25402191 | refGene_gn=DNAH12;refGene_tx=XM_023249860.1 |
| A2 | 25404952 | 25405141 | refGene_gn=DNAH12;refGene_tx=XM_023249860.1 |
| A2 | 25405425 | 25405659 | refGene_gn=DNAH12;refGene_tx=XM_023249860.1 |
| A2 | 25409121 | 25409259 | refGene_gn=DNAH12;refGene_tx=XM_023249860.1 |
| A2 | 25412203 | 25412344 | refGene_gn=DNAH12;refGene_tx=XM_023249860.1 |
| A2 | 25413710 | 25413964 | refGene_gn=DNAH12;refGene_tx=XM_023249860.1 |
| A2 | 25414174 | 25414325 | refGene_gn=DNAH12;refGene_tx=XM_023249860.1 |
| A2 | 25422407 | 25422552 | refGene_gn=DNAH12;refGene_tx=XM_023249860.1 |
| A2 | 25422675 | 25422834 | refGene_gn=DNAH12;refGene_tx=XM_023249860.1 |
| A2 | 25435786 | 25435927 | refGene_gn=DNAH12;refGene_tx=XM_023249860.1 |
| A2 | 25440333 | 25440506 | refGene_gn=DNAH12;refGene_tx=XM_023249860.1 |
| A2 | 25442046 | 25442239 | refGene_gn=DNAH12;refGene_tx=XM_023249860.1 |
| A2 | 25444066 | 25444212 | refGene_gn=DNAH12;refGene_tx=XM_023249860.1 |
| A2 | 25444213 | 25444260 | refGene_gn=DNAH12;refGene_tx=XM_023249860.1 |
| A2 | 25445000 | 25445215 | refGene_gn=DNAH12;refGene_tx=XM_023249860.1 |
| A2 | 25452990 | 25453136 | refGene_gn=DNAH12;refGene_tx=XM_023249860.1 |
| A2 | 25453679 | 25453827 | refGene_gn=DNAH12;refGene_tx=XM_023249860.1 |
| A2 | 25454870 | 25455112 | refGene_gn=DNAH12;refGene_tx=XM_023249860.1 |
| A2 | 25457020 | 25457161 | refGene_gn=DNAH12;refGene_tx=XM_023249860.1 |
| A2 | 25457755 | 25457845 | refGene_gn=DNAH12;refGene_tx=XM_023249860.1 |
| A2 | 25459928 | 25460060 | refGene_gn=DNAH12;refGene_tx=XM_023249860.1 |
| A2 | 25461313 | 25461490 | refGene_gn=DNAH12;refGene_tx=XM_023249860.1 |
| A2 | 25462084 | 25462165 | refGene_gn=DNAH12;refGene_tx=XM_023249860.1 |
| A2 | 25462920 | 25463083 | refGene_gn=DNAH12;refGene_tx=XM_023249860.1 |
| A2 | 25464883 | 25465017 | refGene_gn=DNAH12;refGene_tx=XM_023249860.1 |
| A2 | 25471643 | 25471838 | refGene_gn=DNAH12;refGene_tx=XM_023249860.1 |
| A2 | 25478006 | 25478168 | refGene_gn=DNAH12;refGene_tx=XM_023249860.1 |
| A2 | 25485968 | 25486161 | refGene_gn=DNAH12;refGene_tx=XM_023249860.1 |
| A2 | 25487609 | 25487788 | refGene_gn=DNAH12;refGene_tx=XM_023249860.1 |
| A2 | 25488572 | 25488872 | refGene_gn=DNAH12;refGene_tx=XM_023249860.1 |
| A2 | 25492967 | 25493223 | refGene_gn=DNAH12;refGene_tx=XM_023249860.1 |
| A2 | 25496078 | 25496245 | refGene_gn=DNAH12;refGene_tx=XM_023249860.1 |
| A2 | 25497846 | 25497985 | refGene_gn=DNAH12;refGene_tx=XM_023249860.1 |
| A2 | 25502443 | 25502595 | refGene_gn=DNAH12;refGene_tx=XM_023249860.1 |
| A2 | 25504221 | 25504410 | refGene_gn=DNAH12;refGene_tx=XM_023249860.1 |
| A2 | 25506652 | 25506772 | refGene_gn=DNAH12;refGene_tx=XM_023249860.1 |
| A2 | 25523722 | 25523911 | refGene_gn=DNAH12;refGene_tx=XM_023249860.1 |
| A2 | 25524415 | 25524499 | refGene_gn=DNAH12;refGene_tx=XM_023249860.1 |
| A2 | 25529770 | 25529911 | refGene_gn=DNAH12;refGene_tx=XM_023249860.1 |
| A2 | 25530036 | 25530220 | refGene_gn=DNAH12;refGene_tx=XM_023249860.1 |
| A2 | 25532918 | 25533028 | refGene_gn=DNAH12;refGene_tx=XM_023249860.1 |
| A2 | 25540084 | 25540204 | refGene_gn=DNAH12;refGene_tx=XM_023249860.1 |
| A2 | 25540772 | 25541018 | refGene_gn=DNAH12;refGene_tx=XM_023249860.1 |

A2	25541360 25541600 refGene_gn=DNAH12;refGene_tx=XM_023249860.1
A2	25542116 25542269 refGene_gn=DNAH12;refGene_tx=XM_023249860.1
A2	25546009 25546182 refGene_gn=DNAH12;refGene_tx=XM_023249860.1
A2	25546411 25546568 refGene_gn=DNAH12;refGene_tx=XM_023249860.1
A2	25547119 25547239 refGene_gn=DNAH12;refGene_tx=XM_023249860.1
A2	25549735 25550018 refGene_gn=DNAH12;refGene_tx=XM_023249860.1
A2	25550117 25550239 refGene_gn=DNAH12;refGene_tx=XM_023249860.1
A2	25553145 25553340 refGene_gn=DNAH12;refGene_tx=XM_023249860.1
A2	25554126 25554327 refGene_gn=DNAH12;refGene_tx=XM_023249860.1
A2	25555439 25555625 refGene_gn=DNAH12;refGene_tx=XM_023249860.1
A2	25571951 25572195 refGene_gn=DNAH12;refGene_tx=XM_023249860.1
A2	25572930 25573124 refGene_gn=DNAH12;refGene_tx=XM_023249860.1
A2	25573472 25573607 refGene_gn=DNAH12;refGene_tx=XM_023249860.1
A2	25575211 25575337 refGene_gn=DNAH12;refGene_tx=XM_023249860.1
A2	25578283 25578419 refGene_gn=DNAH12;refGene_tx=XM_023249860.1
A2	25583501 25583680 refGene_gn=DNAH12;refGene_tx=XM_023249860.1
A2	25584572 25584664 refGene_gn=DNAH12;refGene_tx=XM_023249860.1
A2	25585303 25585460 refGene_gn=DNAH12;refGene_tx=XM_023249860.1
A2	25586241 25586430 refGene_gn=DNAH12;refGene_tx=XM_023249860.1
A2	25589903 25590099 refGene_gn=DNAH12;refGene_tx=XM_023249860.1
A2	25590303 25590462 refGene_gn=DNAH12;refGene_tx=XM_023249860.1
A2	25591495 25591568 refGene_gn=DNAH12;refGene_tx=XM_023249860.1
A2	25592404 25592594 refGene_gn=DNAH12;refGene_tx=XM_023249860.1
A2	25595759 25595786 refGene_gn=DNAH12;refGene_tx=XM_023249860.1
A2	25595958 25596040 refGene_gn=DNAH12;refGene_tx=XM_023249860.1
A2	25605380 25605550 refGene_gn=DNAH12;refGene_tx=XM_023249860.1
A2	25614924 25616229 refGene_gn=PDE12;refGene_tx=XM_003982364.5
A2	25617644 25617723 refGene_gn=PDE12;refGene_tx=XM_003982364.5
A2	25618219 25618662 refGene_gn=PDE12;refGene_tx=XM_003982364.5
A2	25631466 25631553 refGene_gn=ARF4;refGene_tx=XM_003982365.5
A2	25633633 25633759 refGene_gn=ARF4;refGene_tx=XM_003982365.5
A2	25636987 25637059 refGene_gn=ARF4;refGene_tx=XM_003982365.5
A2	25643024 25643134 refGene_gn=ARF4;refGene_tx=XM_003982365.5
A2	25643622 25643703 refGene_gn=ARF4;refGene_tx=XM_003982365.5
A2	25652605 25652672 refGene_gn=ARF4;refGene_tx=XM_003982365.5
A2	25675741 25675873
refGene_gn=DENND6A;refGene_tx=XM_006928909.4;refGene_gn=DENND6A;refGene_tx=XM_011279762.3;refGene_gn=DENND6A;refGene_tx=XM_019823348.2
A2	25676572 25676647
refGene_gn=DENND6A;refGene_tx=XM_006928909.4;refGene_gn=DENND6A;refGene_tx=XM_011279762.3;refGene_gn=DENND6A;refGene_tx=XM_019823348.2
A2	25677814 25677917
refGene_gn=DENND6A;refGene_tx=XM_006928909.4;refGene_gn=DENND6A;refGene_tx=XM_011279762.3;refGene_gn=DENND6A;refGene_tx=XM_019823348.2
A2	25678574 25678684
refGene_gn=DENND6A;refGene_tx=XM_006928909.4;refGene_gn=DENND6A;refGene_tx=XM_011279762.3;refGene_gn=DENND6A;refGene_tx=XM_019823348.2
A2	25678788 25678842
refGene_gn=DENND6A;refGene_tx=XM_006928909.4;refGene_gn=DENND6A;refGene_tx=XM_011279762.3;refGene_gn=DENND6A;refGene_tx=XM_019823348.2

A2 25680045 25680135
refGene_gn=DENND6A;refGene_tx=XM_006928909.4;refGene_gn=DENND6A;refGene_tx=XM_011279762.3;refGene_gn=DENND6A;refGene_tx=XM_019823348.2
A2 25680716 25680781
refGene_gn=DENND6A;refGene_tx=XM_006928909.4;refGene_gn=DENND6A;refGene_tx=XM_011279762.3;refGene_gn=DENND6A;refGene_tx=XM_019823348.2
A2 25680879 25680945
refGene_gn=DENND6A;refGene_tx=XM_006928909.4;refGene_gn=DENND6A;refGene_tx=XM_011279762.3;refGene_gn=DENND6A;refGene_tx=XM_019823348.2
A2 25694945 25695040
refGene_gn=DENND6A;refGene_tx=XM_006928909.4;refGene_gn=DENND6A;refGene_tx=XM_011279762.3;refGene_gn=DENND6A;refGene_tx=XM_019823348.2
A2 25697152 25697248
refGene_gn=DENND6A;refGene_tx=XM_006928909.4;refGene_gn=DENND6A;refGene_tx=XM_011279762.3;refGene_gn=DENND6A;refGene_tx=XM_019823348.2
A2 25697774 25697897
refGene_gn=DENND6A;refGene_tx=XM_006928909.4;refGene_gn=DENND6A;refGene_tx=XM_011279762.3;refGene_gn=DENND6A;refGene_tx=XM_019823348.2
A2 25706218 25706274
refGene_gn=DENND6A;refGene_tx=XM_006928909.4;refGene_gn=DENND6A;refGene_tx=XM_011279762.3;refGene_gn=DENND6A;refGene_tx=XM_019823348.2
A2 25707259 25707322
refGene_gn=DENND6A;refGene_tx=XM_006928909.4;refGene_gn=DENND6A;refGene_tx=XM_011279762.3;refGene_gn=DENND6A;refGene_tx=XM_019823348.2
A2 25708375 25708455
refGene_gn=DENND6A;refGene_tx=XM_006928909.4;refGene_gn=DENND6A;refGene_tx=XM_011279762.3;refGene_gn=DENND6A;refGene_tx=XM_019823348.2
A2 25709671 25709777
refGene_gn=DENND6A;refGene_tx=XM_006928909.4;refGene_gn=DENND6A;refGene_tx=XM_011279762.3;refGene_gn=DENND6A;refGene_tx=XM_019823348.2
A2 25711361 25711442
refGene_gn=DENND6A;refGene_tx=XM_006928909.4;refGene_gn=DENND6A;refGene_tx=XM_011279762.3;refGene_gn=DENND6A;refGene_tx=XM_019823348.2
A2 25713443 25713556
refGene_gn=DENND6A;refGene_tx=XM_006928909.4;refGene_gn=DENND6A;refGene_tx=XM_011279762.3;refGene_gn=DENND6A;refGene_tx=XM_019823348.2
A2 25725132 25725175
refGene_gn=DENND6A;refGene_tx=XM_006928909.4;refGene_gn=DENND6A;refGene_tx=XM_011279762.3;refGene_gn=DENND6A;refGene_tx=XM_019823348.2
A2 25725275 25725314
refGene_gn=DENND6A;refGene_tx=XM_006928909.4;refGene_gn=DENND6A;refGene_tx=XM_011279762.3;refGene_gn=DENND6A;refGene_tx=XM_019823348.2
A2 25728758 25728836 refGene_gn=DENND6A;refGene_tx=XM_019823348.2
A2 25739893 25740130
refGene_gn=DENND6A;refGene_tx=XM_006928909.4;refGene_gn=DENND6A;refGene_tx=XM_011279762.3
A2 25786169 25786367
refGene_gn=SLMAP;refGene_tx=XM_003982366.4;refGene_gn=SLMAP;refGene_tx=XM_011279767.2;refGene_gn=SLMAP;refGene_tx=XM_011279778.2;refGene_gn=SLMAP;refGene_tx=XM_011279797.2;refGene_gn=SLMAP;refGene_tx=XM_0198233

52.1;refGene_gn=SLMAP;refGene_tx=XM_019823363.1;refGene_gn=SLMAP;refGene_tx=XM_019823372.1;refGene_gn=SLMAP;refGene_tx=XM_019823379.1;refGene_gn=SLMAP;refGene_tx=XM_019823389.1;refGene_gn=SLMAP;refGene_tx=XM_019823395.1;refGene_gn=SLMAP;refGene_tx=XM_019823398.1;refGene_gn=SLMAP;refGene_tx=XM_019823399.1;refGene_gn=SLMAP;refGene_tx=XM_019823400.1;refGene_gn=SLMAP;refGene_tx=XM_019823404.1;refGene_gn=SLMAP;refGene_tx=XM_019823408.1;refGene_gn=SLMAP;refGene_tx=XM_019823412.1
A2 25852337 25852491
refGene_gn=SLMAP;refGene_tx=XM_003982366.4;refGene_gn=SLMAP;refGene_tx=XM_011279767.2;refGene_gn=SLMAP;refGene_tx=XM_011279778.2;refGene_gn=SLMAP;refGene_tx=XM_011279797.2;refGene_gn=SLMAP;refGene_tx=XM_019823352.1;refGene_gn=SLMAP;refGene_tx=XM_019823363.1;refGene_gn=SLMAP;refGene_tx=XM_019823372.1;refGene_gn=SLMAP;refGene_tx=XM_019823379.1;refGene_gn=SLMAP;refGene_tx=XM_019823389.1;refGene_gn=SLMAP;refGene_tx=XM_019823395.1;refGene_gn=SLMAP;refGene_tx=XM_019823398.1;refGene_gn=SLMAP;refGene_tx=XM_019823399.1;refGene_gn=SLMAP;refGene_tx=XM_019823400.1;refGene_gn=SLMAP;refGene_tx=XM_019823404.1;refGene_gn=SLMAP;refGene_tx=XM_019823408.1;refGene_gn=SLMAP;refGene_tx=XM_019823412.1
A2 25867918 25867991
refGene_gn=SLMAP;refGene_tx=XM_003982366.4;refGene_gn=SLMAP;refGene_tx=XM_011279767.2;refGene_gn=SLMAP;refGene_tx=XM_011279778.2;refGene_gn=SLMAP;refGene_tx=XM_011279797.2;refGene_gn=SLMAP;refGene_tx=XM_019823352.1;refGene_gn=SLMAP;refGene_tx=XM_019823363.1;refGene_gn=SLMAP;refGene_tx=XM_019823372.1;refGene_gn=SLMAP;refGene_tx=XM_019823379.1;refGene_gn=SLMAP;refGene_tx=XM_019823389.1;refGene_gn=SLMAP;refGene_tx=XM_019823395.1;refGene_gn=SLMAP;refGene_tx=XM_019823398.1;refGene_gn=SLMAP;refGene_tx=XM_019823399.1;refGene_gn=SLMAP;refGene_tx=XM_019823400.1;refGene_gn=SLMAP;refGene_tx=XM_019823404.1;refGene_gn=SLMAP;refGene_tx=XM_019823408.1;refGene_gn=SLMAP;refGene_tx=XM_019823412.1;refGene_gn=SLMAP;refGene_tx=XM_023249863.1
A2 25874028 25874065
refGene_gn=SLMAP;refGene_tx=XM_003982366.4;refGene_gn=SLMAP;refGene_tx=XM_011279767.2;refGene_gn=SLMAP;refGene_tx=XM_011279778.2;refGene_gn=SLMAP;refGene_tx=XM_011279797.2;refGene_gn=SLMAP;refGene_tx=XM_019823352.1;refGene_gn=SLMAP;refGene_tx=XM_019823363.1;refGene_gn=SLMAP;refGene_tx=XM_019823372.1;refGene_gn=SLMAP;refGene_tx=XM_019823379.1;refGene_gn=SLMAP;refGene_tx=XM_019823389.1;refGene_gn=SLMAP;refGene_tx=XM_019823395.1;refGene_gn=SLMAP;refGene_tx=XM_019823398.1;refGene_gn=SLMAP;refGene_tx=XM_019823399.1;refGene_gn=SLMAP;refGene_tx=XM_019823400.1;refGene_gn=SLMAP;refGene_tx=XM_019823404.1;refGene_gn=SLMAP;refGene_tx=XM_019823408.1;refGene_gn=SLMAP;refGene_tx=XM_019823412.1;refGene_gn=SLMAP;refGene_tx=XM_023249863.1
A2 25876008 25876071
refGene_gn=SLMAP;refGene_tx=XM_003982366.4;refGene_gn=SLMAP;refGene_tx=XM_011279767.2;refGene_gn=SLMAP;refGene_tx=XM_011279778.2;refGene_gn=SLMAP;refGene_tx=XM_011279797.2;refGene_gn=SLMAP;refGene_tx=XM_019823352.1;refGene_gn=SLMAP;refGene_tx=XM_019823363.1;refGene_gn=SLMAP;refGene_tx=XM_019823372.1;refGene_gn=SLMAP;refGene_tx=XM_019823379.1;refGene_gn=SLMAP;refGene_tx=XM_019823389.1;refGene_gn=SLMAP;refGene_tx=XM_019823395.1;refGene_gn=SLMAP;refGene_tx=XM_019823398.1;refGene_gn=SLMAP;

refGene_tx=XM_019823399.1;refGene_gn=SLMAP;refGene_tx=XM_019823400.1;refGene_gn=SLMAP;refGene_tx=XM_019823404.1;refGene_gn=SLMAP;refGene_tx=XM_019823408.1;refGene_gn=SLMAP;refGene_tx=XM_019823412.1;refGene_gn=SLMAP;refGene_tx=XM_023249863.1
A2 25879513 25879609
refGene_gn=SLMAP;refGene_tx=XM_003982366.4;refGene_gn=SLMAP;refGene_tx=XM_011279767.2;refGene_gn=SLMAP;refGene_tx=XM_011279778.2;refGene_gn=SLMAP;refGene_tx=XM_011279797.2;refGene_gn=SLMAP;refGene_tx=XM_019823352.1;refGene_gn=SLMAP;refGene_tx=XM_019823363.1;refGene_gn=SLMAP;refGene_tx=XM_019823372.1;refGene_gn=SLMAP;refGene_tx=XM_019823379.1;refGene_gn=SLMAP;refGene_tx=XM_019823389.1;refGene_gn=SLMAP;refGene_tx=XM_019823395.1;refGene_gn=SLMAP;refGene_tx=XM_019823398.1;refGene_gn=SLMAP;refGene_tx=XM_019823399.1;refGene_gn=SLMAP;refGene_tx=XM_019823400.1;refGene_gn=SLMAP;refGene_tx=XM_019823404.1;refGene_gn=SLMAP;refGene_tx=XM_019823408.1;refGene_gn=SLMAP;refGene_tx=XM_019823412.1;refGene_gn=SLMAP;refGene_tx=XM_023249863.1
A2 25879853 25879925
refGene_gn=SLMAP;refGene_tx=XM_003982366.4;refGene_gn=SLMAP;refGene_tx=XM_011279767.2;refGene_gn=SLMAP;refGene_tx=XM_011279778.2;refGene_gn=SLMAP;refGene_tx=XM_011279797.2;refGene_gn=SLMAP;refGene_tx=XM_019823352.1;refGene_gn=SLMAP;refGene_tx=XM_019823363.1;refGene_gn=SLMAP;refGene_tx=XM_019823372.1;refGene_gn=SLMAP;refGene_tx=XM_019823379.1;refGene_gn=SLMAP;refGene_tx=XM_019823389.1;refGene_gn=SLMAP;refGene_tx=XM_019823395.1;refGene_gn=SLMAP;refGene_tx=XM_019823398.1;refGene_gn=SLMAP;refGene_tx=XM_019823399.1;refGene_gn=SLMAP;refGene_tx=XM_019823400.1;refGene_gn=SLMAP;refGene_tx=XM_019823404.1;refGene_gn=SLMAP;refGene_tx=XM_019823408.1;refGene_gn=SLMAP;refGene_tx=XM_019823412.1;refGene_gn=SLMAP;refGene_tx=XM_023249863.1
A2 25881279 25881420
refGene_gn=SLMAP;refGene_tx=XM_003982366.4;refGene_gn=SLMAP;refGene_tx=XM_011279767.2;refGene_gn=SLMAP;refGene_tx=XM_011279778.2;refGene_gn=SLMAP;refGene_tx=XM_011279797.2;refGene_gn=SLMAP;refGene_tx=XM_019823352.1;refGene_gn=SLMAP;refGene_tx=XM_019823363.1;refGene_gn=SLMAP;refGene_tx=XM_019823372.1;refGene_gn=SLMAP;refGene_tx=XM_019823379.1;refGene_gn=SLMAP;refGene_tx=XM_019823389.1;refGene_gn=SLMAP;refGene_tx=XM_019823395.1;refGene_gn=SLMAP;refGene_tx=XM_019823398.1;refGene_gn=SLMAP;refGene_tx=XM_019823399.1;refGene_gn=SLMAP;refGene_tx=XM_019823400.1;refGene_gn=SLMAP;refGene_tx=XM_019823404.1;refGene_gn=SLMAP;refGene_tx=XM_019823408.1;refGene_gn=SLMAP;refGene_tx=XM_019823412.1;refGene_gn=SLMAP;refGene_tx=XM_023249863.1
A2 25882744 25882882
refGene_gn=SLMAP;refGene_tx=XM_003982366.4;refGene_gn=SLMAP;refGene_tx=XM_011279767.2;refGene_gn=SLMAP;refGene_tx=XM_011279778.2;refGene_gn=SLMAP;refGene_tx=XM_011279797.2;refGene_gn=SLMAP;refGene_tx=XM_019823352.1;refGene_gn=SLMAP;refGene_tx=XM_019823363.1;refGene_gn=SLMAP;refGene_tx=XM_019823372.1;refGene_gn=SLMAP;refGene_tx=XM_019823379.1;refGene_gn=SLMAP;refGene_tx=XM_019823389.1;refGene_gn=SLMAP;refGene_tx=XM_019823395.1;refGene_gn=SLMAP;refGene_tx=XM_019823398.1;refGene_gn=SLMAP;refGene_tx=XM_019823399.1;refGene_gn=SLMAP;refGene_tx=XM_019823400.1;refGene_gn=SLMAP;refGene_tx=XM_019823404.1;refGene_gn=SLMAP;refGene_tx=

XM_019823408.1;refGene_gn=SLMAP;refGene_tx=XM_019823412.1;refGene_gn=SLMAP;refGene_tx=XM_023249863.1
A2 25883951 25884120
refGene_gn=SLMAP;refGene_tx=XM_003982366.4;refGene_gn=SLMAP;refGene_tx=XM_011279767.2;refGene_gn=SLMAP;refGene_tx=XM_011279778.2;refGene_gn=SLMAP;refGene_tx=XM_011279797.2;refGene_gn=SLMAP;refGene_tx=XM_019823352.1;refGene_gn=SLMAP;refGene_tx=XM_019823363.1;refGene_gn=SLMAP;refGene_tx=XM_019823372.1;refGene_gn=SLMAP;refGene_tx=XM_019823379.1;refGene_gn=SLMAP;refGene_tx=XM_019823389.1;refGene_gn=SLMAP;refGene_tx=XM_019823395.1;refGene_gn=SLMAP;refGene_tx=XM_019823398.1;refGene_gn=SLMAP;refGene_tx=XM_019823399.1;refGene_gn=SLMAP;refGene_tx=XM_019823400.1;refGene_gn=SLMAP;refGene_tx=XM_019823404.1;refGene_gn=SLMAP;refGene_tx=XM_019823408.1;refGene_gn=SLMAP;refGene_tx=XM_019823412.1;refGene_gn=SLMAP;refGene_tx=XM_023249863.1
A2 25884207 25884258
refGene_gn=SLMAP;refGene_tx=XM_011279767.2;refGene_gn=SLMAP;refGene_tx=XM_011279778.2;refGene_gn=SLMAP;refGene_tx=XM_019823352.1;refGene_gn=SLMAP;refGene_tx=XM_019823363.1;refGene_gn=SLMAP;refGene_tx=XM_019823372.1;refGene_gn=SLMAP;refGene_tx=XM_019823389.1;refGene_gn=SLMAP;refGene_tx=XM_019823395.1;refGene_gn=SLMAP;refGene_tx=XM_019823399.1;refGene_gn=SLMAP;refGene_tx=XM_019823404.1;refGene_gn=SLMAP;refGene_tx=XM_023249863.1
A2 25884719 25884770
refGene_gn=SLMAP;refGene_tx=XM_011279767.2;refGene_gn=SLMAP;refGene_tx=XM_019823352.1;refGene_gn=SLMAP;refGene_tx=XM_019823363.1;refGene_gn=SLMAP;refGene_tx=XM_019823379.1;refGene_gn=SLMAP;refGene_tx=XM_019823389.1;refGene_gn=SLMAP;refGene_tx=XM_019823395.1;refGene_gn=SLMAP;refGene_tx=XM_019823400.1;refGene_gn=SLMAP;refGene_tx=XM_019823404.1;refGene_gn=SLMAP;refGene_tx=XM_023249863.1
A2 25889509 25889572
refGene_gn=SLMAP;refGene_tx=XM_003982366.4;refGene_gn=SLMAP;refGene_tx=XM_011279767.2;refGene_gn=SLMAP;refGene_tx=XM_011279778.2;refGene_gn=SLMAP;refGene_tx=XM_019823352.1;refGene_gn=SLMAP;refGene_tx=XM_019823363.1;refGene_gn=SLMAP;refGene_tx=XM_019823372.1;refGene_gn=SLMAP;refGene_tx=XM_019823379.1;refGene_gn=SLMAP;refGene_tx=XM_019823398.1;refGene_gn=SLMAP;refGene_tx=XM_019823404.1;refGene_gn=SLMAP;refGene_tx=XM_019823412.1;refGene_gn=SLMAP;refGene_tx=XM_023249863.1
A2 25901696 25901756
refGene_gn=SLMAP;refGene_tx=XM_003982366.4;refGene_gn=SLMAP;refGene_tx=XM_011279767.2;refGene_gn=SLMAP;refGene_tx=XM_011279778.2;refGene_gn=SLMAP;refGene_tx=XM_011279797.2;refGene_gn=SLMAP;refGene_tx=XM_019823352.1;refGene_gn=SLMAP;refGene_tx=XM_019823363.1;refGene_gn=SLMAP;refGene_tx=XM_019823372.1;refGene_gn=SLMAP;refGene_tx=XM_019823379.1;refGene_gn=SLMAP;refGene_tx=XM_019823389.1;refGene_gn=SLMAP;refGene_tx=XM_019823395.1;refGene_gn=SLMAP;refGene_tx=XM_019823398.1;refGene_gn=SLMAP;refGene_tx=XM_019823399.1;refGene_gn=SLMAP;refGene_tx=XM_019823400.1;refGene_gn=SLMAP;refGene_tx=XM_019823404.1;refGene_gn=SLMAP;refGene_tx=XM_019823408.1;refGene_gn=SLMAP;refGene_tx=XM_019823412.1;refGene_gn=SLMAP;refGene_tx=XM_023249863.1
A2 25906945 25907026

refGene_gn=SLMAP;refGene_tx=XM_003982366.4;refGene_gn=SLMAP;refGene_tx=XM_011279767.2;refGene_gn=SLMAP;refGene_tx=XM_011279778.2;refGene_gn=SLMAP;refGene_tx=XM_011279797.2;refGene_gn=SLMAP;refGene_tx=XM_019823352.1;refGene_gn=SLMAP;refGene_tx=XM_019823363.1;refGene_gn=SLMAP;refGene_tx=XM_019823372.1;refGene_gn=SLMAP;refGene_tx=XM_019823379.1;refGene_gn=SLMAP;refGene_tx=XM_019823389.1;refGene_gn=SLMAP;refGene_tx=XM_019823395.1;refGene_gn=SLMAP;refGene_tx=XM_019823398.1;refGene_gn=SLMAP;refGene_tx=XM_019823399.1;refGene_gn=SLMAP;refGene_tx=XM_019823400.1;refGene_gn=SLMAP;refGene_tx=XM_019823404.1;refGene_gn=SLMAP;refGene_tx=XM_019823408.1;refGene_gn=SLMAP;refGene_tx=XM_019823412.1;refGene_gn=SLMAP;refGene_tx=XM_023249863.1
A2 25907332 25907392
refGene_gn=SLMAP;refGene_tx=XM_003982366.4;refGene_gn=SLMAP;refGene_tx=XM_011279767.2;refGene_gn=SLMAP;refGene_tx=XM_011279778.2;refGene_gn=SLMAP;refGene_tx=XM_011279797.2;refGene_gn=SLMAP;refGene_tx=XM_019823352.1;refGene_gn=SLMAP;refGene_tx=XM_019823363.1;refGene_gn=SLMAP;refGene_tx=XM_019823372.1;refGene_gn=SLMAP;refGene_tx=XM_019823379.1;refGene_gn=SLMAP;refGene_tx=XM_019823389.1;refGene_gn=SLMAP;refGene_tx=XM_019823395.1;refGene_gn=SLMAP;refGene_tx=XM_019823398.1;refGene_gn=SLMAP;refGene_tx=XM_019823399.1;refGene_gn=SLMAP;refGene_tx=XM_019823400.1;refGene_gn=SLMAP;refGene_tx=XM_019823404.1;refGene_gn=SLMAP;refGene_tx=XM_019823408.1;refGene_gn=SLMAP;refGene_tx=XM_019823412.1;refGene_gn=SLMAP;refGene_tx=XM_023249863.1;refGene_gn=SLMAP;refGene_tx=XM_023249864.1
A2 25920067 25920190
refGene_gn=SLMAP;refGene_tx=XM_003982366.4;refGene_gn=SLMAP;refGene_tx=XM_011279767.2;refGene_gn=SLMAP;refGene_tx=XM_011279778.2;refGene_gn=SLMAP;refGene_tx=XM_011279797.2;refGene_gn=SLMAP;refGene_tx=XM_019823352.1;refGene_gn=SLMAP;refGene_tx=XM_019823363.1;refGene_gn=SLMAP;refGene_tx=XM_019823372.1;refGene_gn=SLMAP;refGene_tx=XM_019823379.1;refGene_gn=SLMAP;refGene_tx=XM_019823389.1;refGene_gn=SLMAP;refGene_tx=XM_019823395.1;refGene_gn=SLMAP;refGene_tx=XM_019823398.1;refGene_gn=SLMAP;refGene_tx=XM_019823399.1;refGene_gn=SLMAP;refGene_tx=XM_019823400.1;refGene_gn=SLMAP;refGene_tx=XM_019823408.1;refGene_gn=SLMAP;refGene_tx=XM_023249863.1;refGene_gn=SLMAP;refGene_tx=XM_023249864.1
A2 25921243 25921318
refGene_gn=SLMAP;refGene_tx=XM_003982366.4;refGene_gn=SLMAP;refGene_tx=XM_011279767.2;refGene_gn=SLMAP;refGene_tx=XM_011279778.2;refGene_gn=SLMAP;refGene_tx=XM_011279797.2;refGene_gn=SLMAP;refGene_tx=XM_019823352.1;refGene_gn=SLMAP;refGene_tx=XM_019823363.1;refGene_gn=SLMAP;refGene_tx=XM_019823372.1;refGene_gn=SLMAP;refGene_tx=XM_019823379.1;refGene_gn=SLMAP;refGene_tx=XM_019823389.1;refGene_gn=SLMAP;refGene_tx=XM_019823395.1;refGene_gn=SLMAP;refGene_tx=XM_019823398.1;refGene_gn=SLMAP;refGene_tx=XM_019823399.1;refGene_gn=SLMAP;refGene_tx=XM_019823400.1;refGene_gn=SLMAP;refGene_tx=XM_019823404.1;refGene_gn=SLMAP;refGene_tx=XM_019823408.1;refGene_gn=SLMAP;refGene_tx=XM_019823412.1;refGene_gn=SLMAP;refGene_tx=XM_023249863.1;refGene_gn=SLMAP;refGene_tx=XM_023249864.1
A2 25923493 25923814
refGene_gn=SLMAP;refGene_tx=XM_003982366.4;refGene_gn=SLMAP;refGene_tx

=XM_011279767.2;refGene_gn=SLMAP;refGene_tx=XM_011279778.2;refGene_gn=SLMAP;refGene_tx=XM_011279797.2;refGene_gn=SLMAP;refGene_tx=XM_019823352.1;refGene_gn=SLMAP;refGene_tx=XM_019823363.1;refGene_gn=SLMAP;refGene_tx=XM_019823372.1;refGene_gn=SLMAP;refGene_tx=XM_019823379.1;refGene_gn=SLMAP;refGene_tx=XM_019823389.1;refGene_gn=SLMAP;refGene_tx=XM_019823395.1;refGene_gn=SLMAP;refGene_tx=XM_019823398.1;refGene_gn=SLMAP;refGene_tx=XM_019823399.1;refGene_gn=SLMAP;refGene_tx=XM_019823400.1;refGene_gn=SLMAP;refGene_tx=XM_019823404.1;refGene_gn=SLMAP;refGene_tx=XM_019823408.1;refGene_gn=SLMAP;refGene_tx=XM_019823412.1;refGene_gn=SLMAP;refGene_tx=XM_023249863.1;refGene_gn=SLMAP;refGene_tx=XM_023249864.1
A2 25924245 25924363
refGene_gn=SLMAP;refGene_tx=XM_003982366.4;refGene_gn=SLMAP;refGene_tx=XM_011279767.2;refGene_gn=SLMAP;refGene_tx=XM_011279778.2;refGene_gn=SLMAP;refGene_tx=XM_011279797.2;refGene_gn=SLMAP;refGene_tx=XM_019823352.1;refGene_gn=SLMAP;refGene_tx=XM_019823363.1;refGene_gn=SLMAP;refGene_tx=XM_019823372.1;refGene_gn=SLMAP;refGene_tx=XM_019823379.1;refGene_gn=SLMAP;refGene_tx=XM_019823389.1;refGene_gn=SLMAP;refGene_tx=XM_019823395.1;refGene_gn=SLMAP;refGene_tx=XM_019823398.1;refGene_gn=SLMAP;refGene_tx=XM_019823399.1;refGene_gn=SLMAP;refGene_tx=XM_019823400.1;refGene_gn=SLMAP;refGene_tx=XM_019823404.1;refGene_gn=SLMAP;refGene_tx=XM_019823408.1;refGene_gn=SLMAP;refGene_tx=XM_019823412.1;refGene_gn=SLMAP;refGene_tx=XM_023249863.1;refGene_gn=SLMAP;refGene_tx=XM_023249864.1
A2 25928047 25928219
refGene_gn=SLMAP;refGene_tx=XM_003982366.4;refGene_gn=SLMAP;refGene_tx=XM_011279767.2;refGene_gn=SLMAP;refGene_tx=XM_011279778.2;refGene_gn=SLMAP;refGene_tx=XM_011279797.2;refGene_gn=SLMAP;refGene_tx=XM_019823352.1;refGene_gn=SLMAP;refGene_tx=XM_019823363.1;refGene_gn=SLMAP;refGene_tx=XM_019823372.1;refGene_gn=SLMAP;refGene_tx=XM_019823379.1;refGene_gn=SLMAP;refGene_tx=XM_019823389.1;refGene_gn=SLMAP;refGene_tx=XM_019823395.1;refGene_gn=SLMAP;refGene_tx=XM_019823398.1;refGene_gn=SLMAP;refGene_tx=XM_019823399.1;refGene_gn=SLMAP;refGene_tx=XM_019823400.1;refGene_gn=SLMAP;refGene_tx=XM_019823404.1;refGene_gn=SLMAP;refGene_tx=XM_019823408.1;refGene_gn=SLMAP;refGene_tx=XM_019823412.1;refGene_gn=SLMAP;refGene_tx=XM_023249863.1;refGene_gn=SLMAP;refGene_tx=XM_023249864.1
A2 25932306 25932441
refGene_gn=SLMAP;refGene_tx=XM_003982366.4;refGene_gn=SLMAP;refGene_tx=XM_011279767.2;refGene_gn=SLMAP;refGene_tx=XM_011279778.2;refGene_gn=SLMAP;refGene_tx=XM_011279797.2;refGene_gn=SLMAP;refGene_tx=XM_019823352.1;refGene_gn=SLMAP;refGene_tx=XM_019823363.1;refGene_gn=SLMAP;refGene_tx=XM_019823372.1;refGene_gn=SLMAP;refGene_tx=XM_019823379.1;refGene_gn=SLMAP;refGene_tx=XM_019823389.1;refGene_gn=SLMAP;refGene_tx=XM_019823395.1;refGene_gn=SLMAP;refGene_tx=XM_019823398.1;refGene_gn=SLMAP;refGene_tx=XM_019823399.1;refGene_gn=SLMAP;refGene_tx=XM_019823400.1;refGene_gn=SLMAP;refGene_tx=XM_019823404.1;refGene_gn=SLMAP;refGene_tx=XM_019823408.1;refGene_gn=SLMAP;refGene_tx=XM_019823412.1;refGene_gn=SLMAP;refGene_tx=XM_023249863.1;refGene_gn=SLMAP;refGene_tx=XM_023249864.1

A2 25934433 25934517
refGene_gn=SLMAP;refGene_tx=XM_019823363.1;refGene_gn=SLMAP;refGene_tx=XM_019823395.1
A2 25935934 25936027
refGene_gn=SLMAP;refGene_tx=XM_003982366.4;refGene_gn=SLMAP;refGene_tx=XM_011279767.2;refGene_gn=SLMAP;refGene_tx=XM_011279778.2;refGene_gn=SLMAP;refGene_tx=XM_011279797.2;refGene_gn=SLMAP;refGene_tx=XM_019823352.1;refGene_gn=SLMAP;refGene_tx=XM_019823372.1;refGene_gn=SLMAP;refGene_tx=XM_019823379.1;refGene_gn=SLMAP;refGene_tx=XM_019823389.1;refGene_gn=SLMAP;refGene_tx=XM_019823398.1;refGene_gn=SLMAP;refGene_tx=XM_019823399.1;refGene_gn=SLMAP;refGene_tx=XM_019823400.1;refGene_gn=SLMAP;refGene_tx=XM_019823404.1;refGene_gn=SLMAP;refGene_tx=XM_019823408.1;refGene_gn=SLMAP;refGene_tx=XM_019823412.1;refGene_gn=SLMAP;refGene_tx=XM_023249863.1;refGene_gn=SLMAP;refGene_tx=XM_023249864.1
A2 25996118 25996410
refGene_gn=FLNB;refGene_tx=XM_003982367.4;refGene_gn=FLNB;refGene_tx=XM_003982370.4;refGene_gn=FLNB;refGene_tx=XM_019823423.2;refGene_gn=FLNB;refGene_tx=XM_019823424.2;refGene_gn=FLNB;refGene_tx=XM_019823426.2;refGene_gn=FLNB;refGene_tx=XM_019823428.2
A2 26054596 26054845
refGene_gn=FLNB;refGene_tx=XM_003982367.4;refGene_gn=FLNB;refGene_tx=XM_003982370.4;refGene_gn=FLNB;refGene_tx=XM_019823423.2;refGene_gn=FLNB;refGene_tx=XM_019823424.2;refGene_gn=FLNB;refGene_tx=XM_019823426.2;refGene_gn=FLNB;refGene_tx=XM_019823428.2
A2 26057451 26057549
refGene_gn=FLNB;refGene_tx=XM_003982367.4;refGene_gn=FLNB;refGene_tx=XM_003982370.4;refGene_gn=FLNB;refGene_tx=XM_019823423.2;refGene_gn=FLNB;refGene_tx=XM_019823424.2;refGene_gn=FLNB;refGene_tx=XM_019823426.2;refGene_gn=FLNB;refGene_tx=XM_019823428.2
A2 26060030 26060178
refGene_gn=FLNB;refGene_tx=XM_003982367.4;refGene_gn=FLNB;refGene_tx=XM_003982370.4;refGene_gn=FLNB;refGene_tx=XM_019823423.2;refGene_gn=FLNB;refGene_tx=XM_019823424.2;refGene_gn=FLNB;refGene_tx=XM_019823426.2;refGene_gn=FLNB;refGene_tx=XM_019823428.2
A2 26067828 26067947
refGene_gn=FLNB;refGene_tx=XM_003982367.4;refGene_gn=FLNB;refGene_tx=XM_003982370.4;refGene_gn=FLNB;refGene_tx=XM_019823423.2;refGene_gn=FLNB;refGene_tx=XM_019823424.2;refGene_gn=FLNB;refGene_tx=XM_019823426.2;refGene_gn=FLNB;refGene_tx=XM_019823428.2
A2 26068985 26069063
refGene_gn=FLNB;refGene_tx=XM_003982367.4;refGene_gn=FLNB;refGene_tx=XM_003982370.4;refGene_gn=FLNB;refGene_tx=XM_019823423.2;refGene_gn=FLNB;refGene_tx=XM_019823424.2;refGene_gn=FLNB;refGene_tx=XM_019823426.2;refGene_gn=FLNB;refGene_tx=XM_019823428.2
A2 26070498 26070661
refGene_gn=FLNB;refGene_tx=XM_003982367.4;refGene_gn=FLNB;refGene_tx=XM_003982370.4;refGene_gn=FLNB;refGene_tx=XM_019823423.2;refGene_gn=FLNB;refGene_tx=XM_019823424.2;refGene_gn=FLNB;refGene_tx=XM_019823426.2;refGene_gn=FLNB;refGene_tx=XM_019823428.2
A2 26071488 26071686

refGene_gn=FLNB;refGene_tx=XM_003982367.4;refGene_gn=FLNB;refGene_tx=XM_003982370.4;refGene_gn=FLNB;refGene_tx=XM_019823423.2;refGene_gn=FLNB;refGene_tx=XM_019823424.2;refGene_gn=FLNB;refGene_tx=XM_019823426.2;refGene_gn=FLNB;refGene_tx=XM_019823428.2
A2 26074286 26074424
refGene_gn=FLNB;refGene_tx=XM_003982367.4;refGene_gn=FLNB;refGene_tx=XM_003982370.4;refGene_gn=FLNB;refGene_tx=XM_019823423.2;refGene_gn=FLNB;refGene_tx=XM_019823424.2;refGene_gn=FLNB;refGene_tx=XM_019823426.2;refGene_gn=FLNB;refGene_tx=XM_019823428.2
A2 26075951 26076078
refGene_gn=FLNB;refGene_tx=XM_003982367.4;refGene_gn=FLNB;refGene_tx=XM_003982370.4;refGene_gn=FLNB;refGene_tx=XM_019823423.2;refGene_gn=FLNB;refGene_tx=XM_019823424.2;refGene_gn=FLNB;refGene_tx=XM_019823426.2;refGene_gn=FLNB;refGene_tx=XM_019823428.2
A2 26077156 26077293
refGene_gn=FLNB;refGene_tx=XM_003982367.4;refGene_gn=FLNB;refGene_tx=XM_003982370.4;refGene_gn=FLNB;refGene_tx=XM_019823423.2;refGene_gn=FLNB;refGene_tx=XM_019823424.2;refGene_gn=FLNB;refGene_tx=XM_019823426.2;refGene_gn=FLNB;refGene_tx=XM_019823428.2
A2 26079285 26079479
refGene_gn=FLNB;refGene_tx=XM_003982367.4;refGene_gn=FLNB;refGene_tx=XM_003982370.4;refGene_gn=FLNB;refGene_tx=XM_019823423.2;refGene_gn=FLNB;refGene_tx=XM_019823424.2;refGene_gn=FLNB;refGene_tx=XM_019823426.2;refGene_gn=FLNB;refGene_tx=XM_019823428.2
A2 26080707 26080821
refGene_gn=FLNB;refGene_tx=XM_003982367.4;refGene_gn=FLNB;refGene_tx=XM_003982370.4;refGene_gn=FLNB;refGene_tx=XM_019823423.2;refGene_gn=FLNB;refGene_tx=XM_019823424.2;refGene_gn=FLNB;refGene_tx=XM_019823426.2;refGene_gn=FLNB;refGene_tx=XM_019823428.2
A2 26081421 26081565
refGene_gn=FLNB;refGene_tx=XM_003982367.4;refGene_gn=FLNB;refGene_tx=XM_003982370.4;refGene_gn=FLNB;refGene_tx=XM_019823423.2;refGene_gn=FLNB;refGene_tx=XM_019823424.2;refGene_gn=FLNB;refGene_tx=XM_019823426.2;refGene_gn=FLNB;refGene_tx=XM_019823428.2
A2 26081807 26081931
refGene_gn=FLNB;refGene_tx=XM_003982367.4;refGene_gn=FLNB;refGene_tx=XM_003982370.4;refGene_gn=FLNB;refGene_tx=XM_019823423.2;refGene_gn=FLNB;refGene_tx=XM_019823424.2;refGene_gn=FLNB;refGene_tx=XM_019823426.2;refGene_gn=FLNB;refGene_tx=XM_019823428.2
A2 26082269 26082430
refGene_gn=FLNB;refGene_tx=XM_003982367.4;refGene_gn=FLNB;refGene_tx=XM_003982370.4;refGene_gn=FLNB;refGene_tx=XM_019823423.2;refGene_gn=FLNB;refGene_tx=XM_019823424.2;refGene_gn=FLNB;refGene_tx=XM_019823426.2;refGene_gn=FLNB;refGene_tx=XM_019823428.2
A2 26083666 26083757
refGene_gn=FLNB;refGene_tx=XM_003982367.4;refGene_gn=FLNB;refGene_tx=XM_003982370.4;refGene_gn=FLNB;refGene_tx=XM_019823423.2;refGene_gn=FLNB;refGene_tx=XM_019823424.2;refGene_gn=FLNB;refGene_tx=XM_019823426.2;refGene_gn=FLNB;refGene_tx=XM_019823428.2
A2 26084017 26084187

refGene_gn=FLNB;refGene_tx=XM_003982367.4;refGene_gn=FLNB;refGene_tx=XM_003982370.4;refGene_gn=FLNB;refGene_tx=XM_019823423.2;refGene_gn=FLNB;refGene_tx=XM_019823424.2;refGene_gn=FLNB;refGene_tx=XM_019823426.2;refGene_gn=FLNB;refGene_tx=XM_019823428.2
A2 26089331 26089449
refGene_gn=FLNB;refGene_tx=XM_003982367.4;refGene_gn=FLNB;refGene_tx=XM_003982370.4;refGene_gn=FLNB;refGene_tx=XM_019823423.2;refGene_gn=FLNB;refGene_tx=XM_019823424.2;refGene_gn=FLNB;refGene_tx=XM_019823426.2;refGene_gn=FLNB;refGene_tx=XM_019823428.2
A2 26091596 26091859
refGene_gn=FLNB;refGene_tx=XM_003982367.4;refGene_gn=FLNB;refGene_tx=XM_003982370.4;refGene_gn=FLNB;refGene_tx=XM_019823423.2;refGene_gn=FLNB;refGene_tx=XM_019823424.2;refGene_gn=FLNB;refGene_tx=XM_019823426.2;refGene_gn=FLNB;refGene_tx=XM_019823428.2
A2 26093237 26093835
refGene_gn=FLNB;refGene_tx=XM_003982367.4;refGene_gn=FLNB;refGene_tx=XM_003982370.4;refGene_gn=FLNB;refGene_tx=XM_019823423.2;refGene_gn=FLNB;refGene_tx=XM_019823424.2;refGene_gn=FLNB;refGene_tx=XM_019823426.2;refGene_gn=FLNB;refGene_tx=XM_019823428.2
A2 26094488 26094662
refGene_gn=FLNB;refGene_tx=XM_003982367.4;refGene_gn=FLNB;refGene_tx=XM_003982370.4;refGene_gn=FLNB;refGene_tx=XM_019823423.2;refGene_gn=FLNB;refGene_tx=XM_019823424.2;refGene_gn=FLNB;refGene_tx=XM_019823426.2;refGene_gn=FLNB;refGene_tx=XM_019823428.2
A2 26095651 26095814
refGene_gn=FLNB;refGene_tx=XM_003982367.4;refGene_gn=FLNB;refGene_tx=XM_003982370.4;refGene_gn=FLNB;refGene_tx=XM_019823423.2;refGene_gn=FLNB;refGene_tx=XM_019823424.2;refGene_gn=FLNB;refGene_tx=XM_019823426.2;refGene_gn=FLNB;refGene_tx=XM_019823428.2
A2 26096497 26096658
refGene_gn=FLNB;refGene_tx=XM_003982367.4;refGene_gn=FLNB;refGene_tx=XM_003982370.4;refGene_gn=FLNB;refGene_tx=XM_019823423.2;refGene_gn=FLNB;refGene_tx=XM_019823424.2;refGene_gn=FLNB;refGene_tx=XM_019823426.2;refGene_gn=FLNB;refGene_tx=XM_019823428.2
A2 26100335 26100503
refGene_gn=FLNB;refGene_tx=XM_003982367.4;refGene_gn=FLNB;refGene_tx=XM_003982370.4;refGene_gn=FLNB;refGene_tx=XM_019823423.2;refGene_gn=FLNB;refGene_tx=XM_019823424.2;refGene_gn=FLNB;refGene_tx=XM_019823426.2;refGene_gn=FLNB;refGene_tx=XM_019823428.2
A2 26101830 26101923
refGene_gn=FLNB;refGene_tx=XM_019823423.2;refGene_gn=FLNB;refGene_tx=XM_019823424.2;refGene_gn=FLNB;refGene_tx=XM_019823426.2
A2 26102714 26102838
refGene_gn=FLNB;refGene_tx=XM_003982367.4;refGene_gn=FLNB;refGene_tx=XM_003982370.4;refGene_gn=FLNB;refGene_tx=XM_019823423.2;refGene_gn=FLNB;refGene_tx=XM_019823424.2;refGene_gn=FLNB;refGene_tx=XM_019823426.2;refGene_gn=FLNB;refGene_tx=XM_019823428.2
A2 26104308 26104465
refGene_gn=FLNB;refGene_tx=XM_003982367.4;refGene_gn=FLNB;refGene_tx=XM_003982370.4;refGene_gn=FLNB;refGene_tx=XM_019823423.2;refGene_gn=FLN

B;refGene_tx=XM_019823424.2;refGene_gn=FLNB;refGene_tx=XM_019823426.2;
refGene_gn=FLNB;refGene_tx=XM_019823428.2
A2 26105513 26105703
refGene_gn=FLNB;refGene_tx=XM_003982367.4;refGene_gn=FLNB;refGene_tx=X
M_003982370.4;refGene_gn=FLNB;refGene_tx=XM_019823423.2;refGene_gn=FLN
B;refGene_tx=XM_019823424.2;refGene_gn=FLNB;refGene_tx=XM_019823426.2;
refGene_gn=FLNB;refGene_tx=XM_019823428.2
A2 26107402 26107650
refGene_gn=FLNB;refGene_tx=XM_003982367.4;refGene_gn=FLNB;refGene_tx=X
M_003982370.4;refGene_gn=FLNB;refGene_tx=XM_019823423.2;refGene_gn=FLN
B;refGene_tx=XM_019823424.2;refGene_gn=FLNB;refGene_tx=XM_019823426.2;
refGene_gn=FLNB;refGene_tx=XM_019823428.2
A2 26110078 26110150
refGene_gn=FLNB;refGene_tx=XM_003982367.4;refGene_gn=FLNB;refGene_tx=X
M_019823423.2;refGene_gn=FLNB;refGene_tx=XM_019823424.2;refGene_gn=FLN
B;refGene_tx=XM_019823428.2
A2 26110866 26110969
refGene_gn=FLNB;refGene_tx=XM_003982367.4;refGene_gn=FLNB;refGene_tx=X
M_003982370.4;refGene_gn=FLNB;refGene_tx=XM_019823423.2;refGene_gn=FLN
B;refGene_tx=XM_019823424.2;refGene_gn=FLNB;refGene_tx=XM_019823426.2;
refGene_gn=FLNB;refGene_tx=XM_019823428.2
A2 26111559 26111700
refGene_gn=FLNB;refGene_tx=XM_003982367.4;refGene_gn=FLNB;refGene_tx=X
M_003982370.4;refGene_gn=FLNB;refGene_tx=XM_019823423.2;refGene_gn=FLN
B;refGene_tx=XM_019823424.2;refGene_gn=FLNB;refGene_tx=XM_019823426.2;
refGene_gn=FLNB;refGene_tx=XM_019823428.2
A2 26115134 26115263
refGene_gn=FLNB;refGene_tx=XM_003982367.4;refGene_gn=FLNB;refGene_tx=X
M_003982370.4;refGene_gn=FLNB;refGene_tx=XM_019823423.2;refGene_gn=FLN
B;refGene_tx=XM_019823424.2;refGene_gn=FLNB;refGene_tx=XM_019823426.2;
refGene_gn=FLNB;refGene_tx=XM_019823428.2
A2 26116336 26116510
refGene_gn=FLNB;refGene_tx=XM_003982367.4;refGene_gn=FLNB;refGene_tx=X
M_003982370.4;refGene_gn=FLNB;refGene_tx=XM_019823423.2;refGene_gn=FLN
B;refGene_tx=XM_019823424.2;refGene_gn=FLNB;refGene_tx=XM_019823426.2;
refGene_gn=FLNB;refGene_tx=XM_019823428.2
A2 26117694 26117853
refGene_gn=FLNB;refGene_tx=XM_003982367.4;refGene_gn=FLNB;refGene_tx=X
M_003982370.4;refGene_gn=FLNB;refGene_tx=XM_019823423.2;refGene_gn=FLN
B;refGene_tx=XM_019823424.2;refGene_gn=FLNB;refGene_tx=XM_019823426.2;
refGene_gn=FLNB;refGene_tx=XM_019823428.2
A2 26118140 26118344
refGene_gn=FLNB;refGene_tx=XM_003982367.4;refGene_gn=FLNB;refGene_tx=X
M_003982370.4;refGene_gn=FLNB;refGene_tx=XM_019823423.2;refGene_gn=FLN
B;refGene_tx=XM_019823424.2;refGene_gn=FLNB;refGene_tx=XM_019823426.2;
refGene_gn=FLNB;refGene_tx=XM_019823428.2
A2 26119655 26119808
refGene_gn=FLNB;refGene_tx=XM_003982367.4;refGene_gn=FLNB;refGene_tx=X
M_003982370.4;refGene_gn=FLNB;refGene_tx=XM_019823423.2;refGene_gn=FLN
B;refGene_tx=XM_019823424.2;refGene_gn=FLNB;refGene_tx=XM_019823426.2;

refGene_gn=FLNB;refGene_tx=XM_019823428.2
A2 26119914 26120037
refGene_gn=FLNB;refGene_tx=XM_003982367.4;refGene_gn=FLNB;refGene_tx=XM_003982370.4;refGene_gn=FLNB;refGene_tx=XM_019823423.2;refGene_gn=FLNB;refGene_tx=XM_019823424.2;refGene_gn=FLNB;refGene_tx=XM_019823426.2;refGene_gn=FLNB;refGene_tx=XM_019823428.2
A2 26122590 26122857
refGene_gn=FLNB;refGene_tx=XM_003982367.4;refGene_gn=FLNB;refGene_tx=XM_003982370.4;refGene_gn=FLNB;refGene_tx=XM_019823423.2;refGene_gn=FLNB;refGene_tx=XM_019823424.2;refGene_gn=FLNB;refGene_tx=XM_019823426.2;refGene_gn=FLNB;refGene_tx=XM_019823428.2
A2 26123705 26123843
refGene_gn=FLNB;refGene_tx=XM_003982367.4;refGene_gn=FLNB;refGene_tx=XM_003982370.4;refGene_gn=FLNB;refGene_tx=XM_019823423.2;refGene_gn=FLNB;refGene_tx=XM_019823424.2;refGene_gn=FLNB;refGene_tx=XM_019823426.2;refGene_gn=FLNB;refGene_tx=XM_019823428.2
A2 26124871 26124987
refGene_gn=FLNB;refGene_tx=XM_003982367.4;refGene_gn=FLNB;refGene_tx=XM_003982370.4;refGene_gn=FLNB;refGene_tx=XM_019823423.2;refGene_gn=FLNB;refGene_tx=XM_019823424.2;refGene_gn=FLNB;refGene_tx=XM_019823426.2;refGene_gn=FLNB;refGene_tx=XM_019823428.2
A2 26129153 26129286
refGene_gn=FLNB;refGene_tx=XM_003982367.4;refGene_gn=FLNB;refGene_tx=XM_003982370.4;refGene_gn=FLNB;refGene_tx=XM_019823423.2;refGene_gn=FLNB;refGene_tx=XM_019823424.2;refGene_gn=FLNB;refGene_tx=XM_019823426.2;refGene_gn=FLNB;refGene_tx=XM_019823428.2
A2 26130705 26130882
refGene_gn=FLNB;refGene_tx=XM_003982367.4;refGene_gn=FLNB;refGene_tx=XM_003982370.4;refGene_gn=FLNB;refGene_tx=XM_019823423.2;refGene_gn=FLNB;refGene_tx=XM_019823424.2;refGene_gn=FLNB;refGene_tx=XM_019823426.2;refGene_gn=FLNB;refGene_tx=XM_019823428.2
A2 26135596 26135815
refGene_gn=FLNB;refGene_tx=XM_003982367.4;refGene_gn=FLNB;refGene_tx=XM_003982370.4;refGene_gn=FLNB;refGene_tx=XM_019823423.2;refGene_gn=FLNB;refGene_tx=XM_019823424.2;refGene_gn=FLNB;refGene_tx=XM_019823426.2;refGene_gn=FLNB;refGene_tx=XM_019823428.2
A2 26136973 26137177
refGene_gn=FLNB;refGene_tx=XM_003982367.4;refGene_gn=FLNB;refGene_tx=XM_003982370.4;refGene_gn=FLNB;refGene_tx=XM_019823423.2;refGene_gn=FLNB;refGene_tx=XM_019823424.2;refGene_gn=FLNB;refGene_tx=XM_019823426.2;refGene_gn=FLNB;refGene_tx=XM_019823428.2
A2 26138704 26138892
refGene_gn=FLNB;refGene_tx=XM_003982367.4;refGene_gn=FLNB;refGene_tx=XM_003982370.4;refGene_gn=FLNB;refGene_tx=XM_019823423.2;refGene_gn=FLNB;refGene_tx=XM_019823424.2;refGene_gn=FLNB;refGene_tx=XM_019823426.2;refGene_gn=FLNB;refGene_tx=XM_019823428.2
A2 26147661 26148169
refGene_gn=LOC109496337;refGene_tx=XM_023249865.1
A2 26150784 26150909
refGene_gn=LOC109496337;refGene_tx=XM_023249865.1

A2 26159606 26159720 refGene_gn=DNASE1L3;refGene_tx=XM_006928890.3
A2 26160022 26160119 refGene_gn=DNASE1L3;refGene_tx=XM_006928890.3
A2 26163324 26163482 refGene_gn=DNASE1L3;refGene_tx=XM_006928890.3
A2 26166111 26166224 refGene_gn=DNASE1L3;refGene_tx=XM_006928890.3
A2 26168958 26169071 refGene_gn=DNASE1L3;refGene_tx=XM_006928890.3
A2 26169657 26169765 refGene_gn=DNASE1L3;refGene_tx=XM_006928890.3
A2 26173178 26173267 refGene_gn=DNASE1L3;refGene_tx=XM_006928890.3
A2 26180851 26180992 refGene_gn=DNASE1L3;refGene_tx=XM_006928890.3
A2 26216755 26216874
refGene_gn=ABHD6;refGene_tx=XM_006928891.4;refGene_gn=ABHD6;refGene_tx
=XM_019823435.2;refGene_gn=ABHD6;refGene_tx=XM_019823438.2;refGene_gn=
ABHD6;refGene_tx=XM_023249866.1
A2 26230353 26230510
refGene_gn=ABHD6;refGene_tx=XM_006928891.4;refGene_gn=ABHD6;refGene_tx
=XM_019823435.2;refGene_gn=ABHD6;refGene_tx=XM_019823438.2;refGene_gn=
ABHD6;refGene_tx=XM_023249866.1
A2 26232936 26233050
refGene_gn=ABHD6;refGene_tx=XM_006928891.4;refGene_gn=ABHD6;refGene_tx
=XM_019823435.2;refGene_gn=ABHD6;refGene_tx=XM_019823438.2;refGene_gn=
ABHD6;refGene_tx=XM_023249866.1
A2 26234491 26234624
refGene_gn=ABHD6;refGene_tx=XM_006928891.4;refGene_gn=ABHD6;refGene_tx
=XM_019823435.2;refGene_gn=ABHD6;refGene_tx=XM_019823438.2;refGene_gn=
ABHD6;refGene_tx=XM_023249866.1
A2 26238262 26238420
refGene_gn=ABHD6;refGene_tx=XM_006928891.4;refGene_gn=ABHD6;refGene_tx
=XM_019823435.2;refGene_gn=ABHD6;refGene_tx=XM_019823438.2;refGene_gn=
ABHD6;refGene_tx=XM_023249866.1
A2 26241817 26241872
refGene_gn=ABHD6;refGene_tx=XM_006928891.4;refGene_gn=ABHD6;refGene_tx
=XM_019823435.2;refGene_gn=ABHD6;refGene_tx=XM_019823438.2;refGene_gn=
ABHD6;refGene_tx=XM_023249866.1
A2 26242106 26242207
refGene_gn=ABHD6;refGene_tx=XM_006928891.4;refGene_gn=ABHD6;refGene_tx
=XM_019823435.2;refGene_gn=ABHD6;refGene_tx=XM_019823438.2;refGene_gn=
ABHD6;refGene_tx=XM_023249866.1
A2 26249489 26249666
refGene_gn=ABHD6;refGene_tx=XM_006928891.4;refGene_gn=ABHD6;refGene_tx
=XM_019823435.2;refGene_gn=ABHD6;refGene_tx=XM_019823438.2;refGene_gn=
ABHD6;refGene_tx=XM_023249866.1
A2 26263131 26263229
refGene_gn=RPP14;refGene_tx=XM_003982373.5;refGene_gn=RPP14;refGene_tx
=XM_006928893.3;refGene_gn=RPP14;refGene_tx=XM_023249869.1
A2 26263321 26263406
refGene_gn=RPP14;refGene_tx=XM_003982373.5;refGene_gn=RPP14;refGene_tx
=XM_006928893.3;refGene_gn=RPP14;refGene_tx=XM_023249869.1
A2 26266781 26266858
refGene_gn=RPP14;refGene_tx=XM_003982373.5;refGene_gn=RPP14;refGene_tx
=XM_006928893.3;refGene_gn=RPP14;refGene_tx=XM_023249869.1
A2 26267412 26267491

refGene_gn=RPP14;refGene_tx=XM_003982373.5;refGene_gn=RPP14;refGene_tx=XM_006928893.3;refGene_gn=RPP14;refGene_tx=XM_023249869.1
A2 26267947 26268004
refGene_gn=RPP14;refGene_tx=XM_003982373.5;refGene_gn=RPP14;refGene_tx=XM_006928893.3;refGene_gn=RPP14;refGene_tx=XM_023249869.1
A2 26268124 26268628
refGene_gn=LOC111559427;refGene_tx=XM_023249867.1
A2 26277715 26277817
refGene_gn=PXK;refGene_tx=XM_023249870.1;refGene_gn=PXK;refGene_tx=XM_023249871.1;refGene_gn=PXK;refGene_tx=XM_023249872.1;refGene_gn=PXK;refGene_tx=XM_023249873.1;refGene_gn=PXK;refGene_tx=XM_023249874.1;refGene_gn=PXK;refGene_tx=XM_023249875.1;refGene_gn=PXK;refGene_tx=XM_023249876.1;refGene_gn=PXK;refGene_tx=XM_023249877.1;refGene_gn=PXK;refGene_tx=XM_023249878.1;refGene_gn=PXK;refGene_tx=XM_023249879.1;refGene_gn=PXK;refGene_tx=XM_023249880.1;refGene_gn=PXK;refGene_tx=XM_023249881.1;refGene_gn=PXK;refGene_tx=XM_023249882.1;refGene_gn=PXK;refGene_tx=XM_023249883.1;refGene_gn=PXK;refGene_tx=XM_023249884.1;refGene_gn=PXK;refGene_tx=XM_023249885.1
A2 26302199 26302250
refGene_gn=PXK;refGene_tx=XM_023249870.1;refGene_gn=PXK;refGene_tx=XM_023249871.1;refGene_gn=PXK;refGene_tx=XM_023249872.1;refGene_gn=PXK;refGene_tx=XM_023249873.1;refGene_gn=PXK;refGene_tx=XM_023249874.1;refGene_gn=PXK;refGene_tx=XM_023249875.1;refGene_gn=PXK;refGene_tx=XM_023249876.1;refGene_gn=PXK;refGene_tx=XM_023249877.1;refGene_gn=PXK;refGene_tx=XM_023249878.1;refGene_gn=PXK;refGene_tx=XM_023249879.1;refGene_gn=PXK;refGene_tx=XM_023249880.1;refGene_gn=PXK;refGene_tx=XM_023249881.1;refGene_gn=PXK;refGene_tx=XM_023249882.1;refGene_gn=PXK;refGene_tx=XM_023249883.1;refGene_gn=PXK;refGene_tx=XM_023249884.1;refGene_gn=PXK;refGene_tx=XM_023249885.1
A2 26304155 26304203
refGene_gn=PXK;refGene_tx=XM_023249870.1;refGene_gn=PXK;refGene_tx=XM_023249871.1;refGene_gn=PXK;refGene_tx=XM_023249872.1;refGene_gn=PXK;refGene_tx=XM_023249873.1;refGene_gn=PXK;refGene_tx=XM_023249874.1;refGene_gn=PXK;refGene_tx=XM_023249875.1;refGene_gn=PXK;refGene_tx=XM_023249876.1;refGene_gn=PXK;refGene_tx=XM_023249877.1;refGene_gn=PXK;refGene_tx=XM_023249878.1;refGene_gn=PXK;refGene_tx=XM_023249879.1;refGene_gn=PXK;refGene_tx=XM_023249880.1;refGene_gn=PXK;refGene_tx=XM_023249881.1;refGene_gn=PXK;refGene_tx=XM_023249882.1;refGene_gn=PXK;refGene_tx=XM_023249883.1;refGene_gn=PXK;refGene_tx=XM_023249884.1;refGene_gn=PXK;refGene_tx=XM_023249885.1
A2 26312561 26312748
refGene_gn=PXK;refGene_tx=XM_023249870.1;refGene_gn=PXK;refGene_tx=XM_023249871.1;refGene_gn=PXK;refGene_tx=XM_023249872.1;refGene_gn=PXK;refGene_tx=XM_023249873.1;refGene_gn=PXK;refGene_tx=XM_023249874.1;refGene_gn=PXK;refGene_tx=XM_023249875.1;refGene_gn=PXK;refGene_tx=XM_023249876.1;refGene_gn=PXK;refGene_tx=XM_023249877.1;refGene_gn=PXK;refGene_tx=XM_023249878.1;refGene_gn=PXK;refGene_tx=XM_023249879.1;refGene_gn=PXK;refGene_tx=XM_023249880.1;refGene_gn=PXK;refGene_tx=XM_023249881.1;refGene_gn=PXK;refGene_tx=XM_023249882.1;refGene_gn=PXK;refGene_tx=XM_023249883.1;refGene_gn=PXK;refGene_tx=XM_023249884.1;refGene_gn=PXK;

refGene_tx=XM_023249885.1
A2 26320070 26320148
refGene_gn=PXK;refGene_tx=XM_019823480.2;refGene_gn=PXK;refGene_tx=XM_023249870.1;refGene_gn=PXK;refGene_tx=XM_023249871.1;refGene_gn=PXK;refGene_tx=XM_023249872.1;refGene_gn=PXK;refGene_tx=XM_023249873.1;refGene_gn=PXK;refGene_tx=XM_023249874.1;refGene_gn=PXK;refGene_tx=XM_023249875.1;refGene_gn=PXK;refGene_tx=XM_023249876.1;refGene_gn=PXK;refGene_tx=XM_023249877.1;refGene_gn=PXK;refGene_tx=XM_023249878.1;refGene_gn=PXK;refGene_tx=XM_023249879.1;refGene_gn=PXK;refGene_tx=XM_023249880.1;refGene_gn=PXK;refGene_tx=XM_023249881.1;refGene_gn=PXK;refGene_tx=XM_023249882.1;refGene_gn=PXK;refGene_tx=XM_023249883.1;refGene_gn=PXK;refGene_tx=XM_023249884.1;refGene_gn=PXK;refGene_tx=XM_023249885.1
A2 26320680 26320754
refGene_gn=PXK;refGene_tx=XM_019823480.2;refGene_gn=PXK;refGene_tx=XM_023249870.1;refGene_gn=PXK;refGene_tx=XM_023249871.1;refGene_gn=PXK;refGene_tx=XM_023249872.1;refGene_gn=PXK;refGene_tx=XM_023249873.1;refGene_gn=PXK;refGene_tx=XM_023249874.1;refGene_gn=PXK;refGene_tx=XM_023249875.1;refGene_gn=PXK;refGene_tx=XM_023249876.1;refGene_gn=PXK;refGene_tx=XM_023249877.1;refGene_gn=PXK;refGene_tx=XM_023249878.1;refGene_gn=PXK;refGene_tx=XM_023249879.1;refGene_gn=PXK;refGene_tx=XM_023249880.1;refGene_gn=PXK;refGene_tx=XM_023249881.1;refGene_gn=PXK;refGene_tx=XM_023249882.1;refGene_gn=PXK;refGene_tx=XM_023249883.1;refGene_gn=PXK;refGene_tx=XM_023249884.1;refGene_gn=PXK;refGene_tx=XM_023249885.1
A2 26321316 26321391
refGene_gn=PXK;refGene_tx=XM_019823480.2;refGene_gn=PXK;refGene_tx=XM_023249870.1;refGene_gn=PXK;refGene_tx=XM_023249871.1;refGene_gn=PXK;refGene_tx=XM_023249872.1;refGene_gn=PXK;refGene_tx=XM_023249873.1;refGene_gn=PXK;refGene_tx=XM_023249874.1;refGene_gn=PXK;refGene_tx=XM_023249875.1;refGene_gn=PXK;refGene_tx=XM_023249876.1;refGene_gn=PXK;refGene_tx=XM_023249877.1;refGene_gn=PXK;refGene_tx=XM_023249878.1;refGene_gn=PXK;refGene_tx=XM_023249879.1;refGene_gn=PXK;refGene_tx=XM_023249880.1;refGene_gn=PXK;refGene_tx=XM_023249881.1;refGene_gn=PXK;refGene_tx=XM_023249882.1;refGene_gn=PXK;refGene_tx=XM_023249883.1;refGene_gn=PXK;refGene_tx=XM_023249884.1;refGene_gn=PXK;refGene_tx=XM_023249885.1
A2 26324360 26324465
refGene_gn=PXK;refGene_tx=XM_019823480.2;refGene_gn=PXK;refGene_tx=XM_023249870.1;refGene_gn=PXK;refGene_tx=XM_023249871.1;refGene_gn=PXK;refGene_tx=XM_023249872.1;refGene_gn=PXK;refGene_tx=XM_023249873.1;refGene_gn=PXK;refGene_tx=XM_023249874.1;refGene_gn=PXK;refGene_tx=XM_023249875.1;refGene_gn=PXK;refGene_tx=XM_023249876.1;refGene_gn=PXK;refGene_tx=XM_023249877.1;refGene_gn=PXK;refGene_tx=XM_023249878.1;refGene_gn=PXK;refGene_tx=XM_023249879.1;refGene_gn=PXK;refGene_tx=XM_023249880.1;refGene_gn=PXK;refGene_tx=XM_023249881.1;refGene_gn=PXK;refGene_tx=XM_023249882.1;refGene_gn=PXK;refGene_tx=XM_023249883.1;refGene_gn=PXK;refGene_tx=XM_023249884.1;refGene_gn=PXK;refGene_tx=XM_023249885.1
A2 26325024 26325126
refGene_gn=PXK;refGene_tx=XM_019823480.2;refGene_gn=PXK;refGene_tx=XM_023249870.1;refGene_gn=PXK;refGene_tx=XM_023249871.1;refGene_gn=PXK;refGene_tx=XM_023249872.1;refGene_gn=PXK;refGene_tx=XM_023249873.1;refGene_gn=PXK;refGene_tx=XM_023249874.1;refGene_gn=PXK;refGene_tx=XM_02324

9875.1;refGene_gn=PXK;refGene_tx=XM_023249876.1;refGene_gn=PXK;refGene_tx=XM_023249877.1;refGene_gn=PXK;refGene_tx=XM_023249878.1;refGene_gn=PXK;refGene_tx=XM_023249879.1;refGene_gn=PXK;refGene_tx=XM_023249880.1;refGene_gn=PXK;refGene_tx=XM_023249881.1;refGene_gn=PXK;refGene_tx=XM_023249882.1;refGene_gn=PXK;refGene_tx=XM_023249883.1;refGene_gn=PXK;refGene_tx=XM_023249884.1;refGene_gn=PXK;refGene_tx=XM_023249885.1
A2 26326578 26326740
refGene_gn=PXK;refGene_tx=XM_019823480.2;refGene_gn=PXK;refGene_tx=XM_023249870.1;refGene_gn=PXK;refGene_tx=XM_023249871.1;refGene_gn=PXK;refGene_tx=XM_023249873.1;refGene_gn=PXK;refGene_tx=XM_023249874.1;refGene_gn=PXK;refGene_tx=XM_023249875.1;refGene_gn=PXK;refGene_tx=XM_023249876.1;refGene_gn=PXK;refGene_tx=XM_023249877.1;refGene_gn=PXK;refGene_tx=XM_023249878.1;refGene_gn=PXK;refGene_tx=XM_023249879.1;refGene_gn=PXK;refGene_tx=XM_023249880.1;refGene_gn=PXK;refGene_tx=XM_023249881.1;refGene_gn=PXK;refGene_tx=XM_023249882.1;refGene_gn=PXK;refGene_tx=XM_023249883.1;refGene_gn=PXK;refGene_tx=XM_023249884.1;refGene_gn=PXK;refGene_tx=XM_023249885.1
A2 26327173 26327291
refGene_gn=PXK;refGene_tx=XM_019823480.2;refGene_gn=PXK;refGene_tx=XM_023249870.1;refGene_gn=PXK;refGene_tx=XM_023249871.1;refGene_gn=PXK;refGene_tx=XM_023249872.1;refGene_gn=PXK;refGene_tx=XM_023249873.1;refGene_gn=PXK;refGene_tx=XM_023249874.1;refGene_gn=PXK;refGene_tx=XM_023249875.1;refGene_gn=PXK;refGene_tx=XM_023249876.1;refGene_gn=PXK;refGene_tx=XM_023249877.1;refGene_gn=PXK;refGene_tx=XM_023249878.1;refGene_gn=PXK;refGene_tx=XM_023249879.1;refGene_gn=PXK;refGene_tx=XM_023249880.1;refGene_gn=PXK;refGene_tx=XM_023249881.1;refGene_gn=PXK;refGene_tx=XM_023249882.1;refGene_gn=PXK;refGene_tx=XM_023249883.1;refGene_gn=PXK;refGene_tx=XM_023249884.1;refGene_gn=PXK;refGene_tx=XM_023249885.1
A2 26328545 26328624
refGene_gn=PXK;refGene_tx=XM_019823480.2;refGene_gn=PXK;refGene_tx=XM_023249870.1;refGene_gn=PXK;refGene_tx=XM_023249871.1;refGene_gn=PXK;refGene_tx=XM_023249872.1;refGene_gn=PXK;refGene_tx=XM_023249873.1;refGene_gn=PXK;refGene_tx=XM_023249874.1;refGene_gn=PXK;refGene_tx=XM_023249875.1;refGene_gn=PXK;refGene_tx=XM_023249876.1;refGene_gn=PXK;refGene_tx=XM_023249877.1;refGene_gn=PXK;refGene_tx=XM_023249878.1;refGene_gn=PXK;refGene_tx=XM_023249879.1;refGene_gn=PXK;refGene_tx=XM_023249880.1;refGene_gn=PXK;refGene_tx=XM_023249881.1;refGene_gn=PXK;refGene_tx=XM_023249882.1;refGene_gn=PXK;refGene_tx=XM_023249883.1;refGene_gn=PXK;refGene_tx=XM_023249884.1;refGene_gn=PXK;refGene_tx=XM_023249885.1
A2 26331239 26331288
refGene_gn=PXK;refGene_tx=XM_019823480.2;refGene_gn=PXK;refGene_tx=XM_023249870.1;refGene_gn=PXK;refGene_tx=XM_023249871.1;refGene_gn=PXK;refGene_tx=XM_023249872.1;refGene_gn=PXK;refGene_tx=XM_023249873.1;refGene_gn=PXK;refGene_tx=XM_023249874.1;refGene_gn=PXK;refGene_tx=XM_023249875.1;refGene_gn=PXK;refGene_tx=XM_023249876.1;refGene_gn=PXK;refGene_tx=XM_023249877.1;refGene_gn=PXK;refGene_tx=XM_023249878.1;refGene_gn=PXK;refGene_tx=XM_023249879.1;refGene_gn=PXK;refGene_tx=XM_023249880.1;refGene_gn=PXK;refGene_tx=XM_023249881.1;refGene_gn=PXK;refGene_tx=XM_023249882.1;refGene_gn=PXK;refGene_tx=XM_023249883.1;refGene_gn=PXK;refGene_tx=XM_023249884.1;refGene_gn=PXK;refGene_tx=XM_023249885.1

A2	26336819 26336897
refGene_gn=PXK;refGene_tx=XM_019823480.2;refGene_gn=PXK;refGene_tx=XM_023249870.1;refGene_gn=PXK;refGene_tx=XM_023249871.1;refGene_gn=PXK;refGene_tx=XM_023249872.1;refGene_gn=PXK;refGene_tx=XM_023249873.1;refGene_gn=PXK;refGene_tx=XM_023249874.1;refGene_gn=PXK;refGene_tx=XM_023249875.1;refGene_gn=PXK;refGene_tx=XM_023249876.1;refGene_gn=PXK;refGene_tx=XM_023249877.1;refGene_gn=PXK;refGene_tx=XM_023249878.1;refGene_gn=PXK;refGene_tx=XM_023249879.1;refGene_gn=PXK;refGene_tx=XM_023249880.1;refGene_gn=PXK;refGene_tx=XM_023249881.1;refGene_gn=PXK;refGene_tx=XM_023249882.1;refGene_gn=PXK;refGene_tx=XM_023249883.1;refGene_gn=PXK;refGene_tx=XM_023249884.1;refGene_gn=PXK;refGene_tx=XM_023249885.1
A2	26337430 26337517
refGene_gn=PXK;refGene_tx=XM_019823480.2;refGene_gn=PXK;refGene_tx=XM_023249870.1;refGene_gn=PXK;refGene_tx=XM_023249871.1;refGene_gn=PXK;refGene_tx=XM_023249872.1;refGene_gn=PXK;refGene_tx=XM_023249873.1;refGene_gn=PXK;refGene_tx=XM_023249874.1;refGene_gn=PXK;refGene_tx=XM_023249875.1;refGene_gn=PXK;refGene_tx=XM_023249876.1;refGene_gn=PXK;refGene_tx=XM_023249877.1;refGene_gn=PXK;refGene_tx=XM_023249878.1;refGene_gn=PXK;refGene_tx=XM_023249879.1;refGene_gn=PXK;refGene_tx=XM_023249880.1;refGene_gn=PXK;refGene_tx=XM_023249881.1;refGene_gn=PXK;refGene_tx=XM_023249882.1;refGene_gn=PXK;refGene_tx=XM_023249883.1;refGene_gn=PXK;refGene_tx=XM_023249884.1;refGene_gn=PXK;refGene_tx=XM_023249885.1
A2	26337971 26338041
refGene_gn=PXK;refGene_tx=XM_019823480.2;refGene_gn=PXK;refGene_tx=XM_023249870.1;refGene_gn=PXK;refGene_tx=XM_023249871.1;refGene_gn=PXK;refGene_tx=XM_023249872.1;refGene_gn=PXK;refGene_tx=XM_023249873.1;refGene_gn=PXK;refGene_tx=XM_023249874.1;refGene_gn=PXK;refGene_tx=XM_023249875.1;refGene_gn=PXK;refGene_tx=XM_023249876.1;refGene_gn=PXK;refGene_tx=XM_023249877.1;refGene_gn=PXK;refGene_tx=XM_023249878.1;refGene_gn=PXK;refGene_tx=XM_023249879.1;refGene_gn=PXK;refGene_tx=XM_023249880.1;refGene_gn=PXK;refGene_tx=XM_023249881.1;refGene_gn=PXK;refGene_tx=XM_023249882.1;refGene_gn=PXK;refGene_tx=XM_023249883.1;refGene_gn=PXK;refGene_tx=XM_023249884.1;refGene_gn=PXK;refGene_tx=XM_023249885.1
A2	26340689 26340752
refGene_gn=PXK;refGene_tx=XM_019823480.2;refGene_gn=PXK;refGene_tx=XM_023249870.1;refGene_gn=PXK;refGene_tx=XM_023249871.1;refGene_gn=PXK;refGene_tx=XM_023249872.1;refGene_gn=PXK;refGene_tx=XM_023249873.1;refGene_gn=PXK;refGene_tx=XM_023249874.1;refGene_gn=PXK;refGene_tx=XM_023249875.1;refGene_gn=PXK;refGene_tx=XM_023249876.1;refGene_gn=PXK;refGene_tx=XM_023249877.1;refGene_gn=PXK;refGene_tx=XM_023249878.1;refGene_gn=PXK;refGene_tx=XM_023249879.1;refGene_gn=PXK;refGene_tx=XM_023249880.1;refGene_gn=PXK;refGene_tx=XM_023249881.1;refGene_gn=PXK;refGene_tx=XM_023249882.1;refGene_gn=PXK;refGene_tx=XM_023249883.1;refGene_gn=PXK;refGene_tx=XM_023249884.1;refGene_gn=PXK;refGene_tx=XM_023249885.1
A2	26346623 26346643
refGene_gn=PXK;refGene_tx=XM_023249873.1;refGene_gn=PXK;refGene_tx=XM_023249874.1;refGene_gn=PXK;refGene_tx=XM_023249875.1;refGene_gn=PXK;refGene_tx=XM_023249876.1;refGene_gn=PXK;refGene_tx=XM_023249877.1;refGene_gn=PXK;refGene_tx=XM_023249878.1;refGene_gn=PXK;refGene_tx=XM_023249879.1;refGene_gn=PXK;refGene_tx=XM_023249880.1;refGene_gn=PXK;refGene

_tx=XM_023249881.1;refGene_gn=PXK;refGene_tx=XM_023249882.1;refGene_gn=PXK;refGene_tx=XM_023249883.1
A2 26349210 26349247
refGene_gn=PXK;refGene_tx=XM_023249884.1;refGene_gn=PXK;refGene_tx=XM_023249885.1
A2 26350510 26350716
refGene_gn=PXK;refGene_tx=XM_019823480.2;refGene_gn=PXK;refGene_tx=XM_023249870.1;refGene_gn=PXK;refGene_tx=XM_023249871.1;refGene_gn=PXK;refGene_tx=XM_023249872.1;refGene_gn=PXK;refGene_tx=XM_023249884.1;refGene_gn=PXK;refGene_tx=XM_023249885.1
A2 26352660 26352806 refGene_gn=PDHB;refGene_tx=XM_003982375.4
A2 26353106 26353248 refGene_gn=PDHB;refGene_tx=XM_003982375.4
A2 26354145 26354237 refGene_gn=PDHB;refGene_tx=XM_003982375.4
A2 26354576 26354687 refGene_gn=PDHB;refGene_tx=XM_003982375.4
A2 26355527 26355813 refGene_gn=PDHB;refGene_tx=XM_003982375.4
A2 26355955 26355991 refGene_gn=PDHB;refGene_tx=XM_003982375.4
A2 26356094 26356157 refGene_gn=PDHB;refGene_tx=XM_003982375.4
A2 26356247 26356355 refGene_gn=PDHB;refGene_tx=XM_003982375.4
A2 26357798 26357852 refGene_gn=PDHB;refGene_tx=XM_003982375.4
A2 26357931 26357973 refGene_gn=PDHB;refGene_tx=XM_003982375.4
A2 26394480 26394507
refGene_gn=KCTD6;refGene_tx=XM_006928902.4;refGene_gn=KCTD6;refGene_tx=XM_006928903.4;refGene_gn=KCTD6;refGene_tx=XM_019823494.2;refGene_gn=KCTD6;refGene_tx=XM_023249886.1
A2 26396020 26396707
refGene_gn=KCTD6;refGene_tx=XM_006928902.4;refGene_gn=KCTD6;refGene_tx=XM_006928903.4;refGene_gn=KCTD6;refGene_tx=XM_019823494.2;refGene_gn=KCTD6;refGene_tx=XM_023249886.1
A2 26399984 26400047
refGene_gn=ACOX2;refGene_tx=XM_003982378.3;refGene_gn=ACOX2;refGene_tx=XM_019823487.2;refGene_gn=ACOX2;refGene_tx=XM_019823488.1
A2 26403175 26403308
refGene_gn=ACOX2;refGene_tx=XM_003982378.3;refGene_gn=ACOX2;refGene_tx=XM_019823487.2;refGene_gn=ACOX2;refGene_tx=XM_019823488.1
A2 26410443 26410661
refGene_gn=ACOX2;refGene_tx=XM_003982378.3;refGene_gn=ACOX2;refGene_tx=XM_019823487.2;refGene_gn=ACOX2;refGene_tx=XM_019823488.1
A2 26415270 26415376
refGene_gn=ACOX2;refGene_tx=XM_003982378.3;refGene_gn=ACOX2;refGene_tx=XM_019823487.2;refGene_gn=ACOX2;refGene_tx=XM_019823488.1
A2 26416850 26417030
refGene_gn=ACOX2;refGene_tx=XM_003982378.3;refGene_gn=ACOX2;refGene_tx=XM_019823487.2;refGene_gn=ACOX2;refGene_tx=XM_019823488.1
A2 26418473 26418664
refGene_gn=ACOX2;refGene_tx=XM_003982378.3;refGene_gn=ACOX2;refGene_tx=XM_019823487.2;refGene_gn=ACOX2;refGene_tx=XM_019823488.1
A2 26419820 26419983
refGene_gn=ACOX2;refGene_tx=XM_003982378.3;refGene_gn=ACOX2;refGene_tx=XM_019823487.2;refGene_gn=ACOX2;refGene_tx=XM_019823488.1
A2 26422207 26422380

refGene_gn=ACOX2;refGene_tx=XM_003982378.3;refGene_gn=ACOX2;refGene_tx
=XM_019823487.2;refGene_gn=ACOX2;refGene_tx=XM_019823488.1
A2	26423049	26423165
refGene_gn=ACOX2;refGene_tx=XM_003982378.3;refGene_gn=ACOX2;refGene_tx
=XM_019823487.2;refGene_gn=ACOX2;refGene_tx=XM_019823488.1
A2	26423479	26423599
refGene_gn=ACOX2;refGene_tx=XM_003982378.3;refGene_gn=ACOX2;refGene_tx
=XM_019823487.2;refGene_gn=ACOX2;refGene_tx=XM_019823488.1
A2	26425630	26425738
refGene_gn=ACOX2;refGene_tx=XM_003982378.3;refGene_gn=ACOX2;refGene_tx
=XM_019823487.2;refGene_gn=ACOX2;refGene_tx=XM_019823488.1
A2	26426152	26426304
refGene_gn=ACOX2;refGene_tx=XM_003982378.3;refGene_gn=ACOX2;refGene_tx
=XM_019823487.2;refGene_gn=ACOX2;refGene_tx=XM_019823488.1
A2	26426518	26426681
refGene_gn=ACOX2;refGene_tx=XM_003982378.3;refGene_gn=ACOX2;refGene_tx
=XM_019823487.2;refGene_gn=ACOX2;refGene_tx=XM_019823488.1
A2	26427080	26427240
refGene_gn=ACOX2;refGene_tx=XM_003982378.3;refGene_gn=ACOX2;refGene_tx
=XM_019823487.2;refGene_gn=ACOX2;refGene_tx=XM_019823488.1
A2	26443091	26443199
refGene_gn=FAM107A;refGene_tx=XM_003982380.5;refGene_gn=FAM107A;refGen
e_tx=XM_003982381.5;refGene_gn=FAM107A;refGene_tx=XM_006928901.4
A2	26443718	26443875
refGene_gn=FAM107A;refGene_tx=XM_003982380.5;refGene_gn=FAM107A;refGen
e_tx=XM_003982381.5;refGene_gn=FAM107A;refGene_tx=XM_006928901.4
A2	26445447	26445622
refGene_gn=FAM107A;refGene_tx=XM_003982380.5;refGene_gn=FAM107A;refGen
e_tx=XM_003982381.5;refGene_gn=FAM107A;refGene_tx=XM_006928901.4
A2	26462148	26462227	refGene_gn=FAM107A;refGene_tx=XM_003982381.5
A2	26464206	26464270	refGene_gn=FAM107A;refGene_tx=XM_006928901.4
A2	26494097	26494121	refGene_gn=FAM107A;refGene_tx=XM_006928901.4
A2	26503874	26503964
refGene_gn=FAM3D;refGene_tx=XM_006928898.3;refGene_gn=FAM3D;refGene_tx
=XM_019823495.1;refGene_gn=FAM3D;refGene_tx=XM_019823506.2;refGene_gn=
FAM3D;refGene_tx=XM_019823510.2;refGene_gn=FAM3D;refGene_tx=XM_0198235
18.2;refGene_gn=FAM3D;refGene_tx=XM_023249887.1;refGene_gn=FAM3D;refGe
ne_tx=XM_023249888.1
A2	26506718	26506845
refGene_gn=FAM3D;refGene_tx=XM_006928898.3;refGene_gn=FAM3D;refGene_tx
=XM_019823495.1;refGene_gn=FAM3D;refGene_tx=XM_019823506.2;refGene_gn=
FAM3D;refGene_tx=XM_019823510.2;refGene_gn=FAM3D;refGene_tx=XM_0198235
18.2;refGene_gn=FAM3D;refGene_tx=XM_023249887.1;refGene_gn=FAM3D;refGe
ne_tx=XM_023249888.1
A2	26507538	26507623
refGene_gn=FAM3D;refGene_tx=XM_006928898.3;refGene_gn=FAM3D;refGene_tx
=XM_019823495.1;refGene_gn=FAM3D;refGene_tx=XM_019823506.2;refGene_gn=
FAM3D;refGene_tx=XM_019823510.2;refGene_gn=FAM3D;refGene_tx=XM_0198235
18.2;refGene_gn=FAM3D;refGene_tx=XM_023249887.1;refGene_gn=FAM3D;refGe
ne_tx=XM_023249888.1

A2	26511124	26511175
refGene_gn=FAM3D;refGene_tx=XM_006928898.3;refGene_gn=FAM3D;refGene_tx=XM_019823495.1;refGene_gn=FAM3D;refGene_tx=XM_019823506.2;refGene_gn=FAM3D;refGene_tx=XM_019823510.2;refGene_gn=FAM3D;refGene_tx=XM_019823518.2;refGene_gn=FAM3D;refGene_tx=XM_023249887.1;refGene_gn=FAM3D;refGene_tx=XM_023249888.1
A2	26514186	26514245
refGene_gn=FAM3D;refGene_tx=XM_006928898.3;refGene_gn=FAM3D;refGene_tx=XM_019823495.1;refGene_gn=FAM3D;refGene_tx=XM_019823506.2;refGene_gn=FAM3D;refGene_tx=XM_019823510.2;refGene_gn=FAM3D;refGene_tx=XM_019823518.2;refGene_gn=FAM3D;refGene_tx=XM_023249887.1;refGene_gn=FAM3D;refGene_tx=XM_023249888.1
A2	26515656	26515774
refGene_gn=FAM3D;refGene_tx=XM_006928898.3;refGene_gn=FAM3D;refGene_tx=XM_019823495.1;refGene_gn=FAM3D;refGene_tx=XM_019823506.2;refGene_gn=FAM3D;refGene_tx=XM_019823510.2;refGene_gn=FAM3D;refGene_tx=XM_019823518.2;refGene_gn=FAM3D;refGene_tx=XM_023249887.1;refGene_gn=FAM3D;refGene_tx=XM_023249888.1
A2	26519690	26519711
refGene_gn=FAM3D;refGene_tx=XM_006928898.3;refGene_gn=FAM3D;refGene_tx=XM_019823495.1;refGene_gn=FAM3D;refGene_tx=XM_019823506.2;refGene_gn=FAM3D;refGene_tx=XM_019823510.2;refGene_gn=FAM3D;refGene_tx=XM_019823518.2;refGene_gn=FAM3D;refGene_tx=XM_023249887.1;refGene_gn=FAM3D;refGene_tx=XM_023249888.1
A2	26522057	26522165
refGene_gn=FAM3D;refGene_tx=XM_006928898.3;refGene_gn=FAM3D;refGene_tx=XM_019823495.1;refGene_gn=FAM3D;refGene_tx=XM_019823506.2;refGene_gn=FAM3D;refGene_tx=XM_019823510.2;refGene_gn=FAM3D;refGene_tx=XM_019823518.2;refGene_gn=FAM3D;refGene_tx=XM_023249887.1;refGene_gn=FAM3D;refGene_tx=XM_023249888.1
A2	26523961	26523974
refGene_gn=FAM3D;refGene_tx=XM_006928898.3;refGene_gn=FAM3D;refGene_tx=XM_019823495.1;refGene_gn=FAM3D;refGene_tx=XM_019823506.2;refGene_gn=FAM3D;refGene_tx=XM_019823510.2;refGene_gn=FAM3D;refGene_tx=XM_019823518.2;refGene_gn=FAM3D;refGene_tx=XM_023249887.1;refGene_gn=FAM3D;refGene_tx=XM_023249888.1
A2	26598083	26598202
refGene_gn=CA2H3orf67;refGene_tx=XM_011279961.3;refGene_gn=CA2H3orf67;refGene_tx=XM_011279970.3;refGene_gn=CA2H3orf67;refGene_tx=XM_023249889.1
A2	26610494	26610589
refGene_gn=CA2H3orf67;refGene_tx=XM_011279961.3;refGene_gn=CA2H3orf67;refGene_tx=XM_011279970.3;refGene_gn=CA2H3orf67;refGene_tx=XM_023249889.1
A2	26638196	26638533
refGene_gn=CA2H3orf67;refGene_tx=XM_019823544.2;refGene_gn=CA2H3orf67;refGene_tx=XM_019823545.2;refGene_gn=CA2H3orf67;refGene_tx=XM_019823553.2;refGene_gn=CA2H3orf67;refGene_tx=XM_019823566.2;refGene_gn=CA2H3orf67;refGene_tx=XM_019823568.2;refGene_gn=CA2H3orf67;refGene_tx=XM_019823570.2;refGene_gn=CA2H3orf67;refGene_tx=XM_023249890.1;refGene_gn=CA2

H3orf67;refGene_tx=XM_023249891.1;refGene_gn=CA2H3orf67;refGene_tx=XM_023249892.1
A2 26645093 26645155
refGene_gn=CA2H3orf67;refGene_tx=XM_011279961.3;refGene_gn=CA2H3orf67;refGene_tx=XM_011279970.3;refGene_gn=CA2H3orf67;refGene_tx=XM_019823544.2;refGene_gn=CA2H3orf67;refGene_tx=XM_019823545.2;refGene_gn=CA2H3orf67;refGene_tx=XM_019823553.2;refGene_gn=CA2H3orf67;refGene_tx=XM_019823566.2;refGene_gn=CA2H3orf67;refGene_tx=XM_019823568.2;refGene_gn=CA2H3orf67;refGene_tx=XM_019823570.2;refGene_gn=CA2H3orf67;refGene_tx=XM_023249889.1;refGene_gn=CA2H3orf67;refGene_tx=XM_023249890.1;refGene_gn=CA2H3orf67;refGene_tx=XM_023249891.1;refGene_gn=CA2H3orf67;refGene_tx=XM_023249892.1
A2 26656193 26656397
refGene_gn=CA2H3orf67;refGene_tx=XM_011279961.3;refGene_gn=CA2H3orf67;refGene_tx=XM_011279970.3;refGene_gn=CA2H3orf67;refGene_tx=XM_019823544.2;refGene_gn=CA2H3orf67;refGene_tx=XM_019823545.2;refGene_gn=CA2H3orf67;refGene_tx=XM_019823553.2;refGene_gn=CA2H3orf67;refGene_tx=XM_019823566.2;refGene_gn=CA2H3orf67;refGene_tx=XM_019823568.2;refGene_gn=CA2H3orf67;refGene_tx=XM_019823570.2;refGene_gn=CA2H3orf67;refGene_tx=XM_023249889.1;refGene_gn=CA2H3orf67;refGene_tx=XM_023249890.1;refGene_gn=CA2H3orf67;refGene_tx=XM_023249891.1;refGene_gn=CA2H3orf67;refGene_tx=XM_023249892.1
A2 26657637 26657686
refGene_gn=CA2H3orf67;refGene_tx=XM_011279961.3;refGene_gn=CA2H3orf67;refGene_tx=XM_019823544.2;refGene_gn=CA2H3orf67;refGene_tx=XM_019823545.2;refGene_gn=CA2H3orf67;refGene_tx=XM_019823566.2;refGene_gn=CA2H3orf67;refGene_tx=XM_019823568.2;refGene_gn=CA2H3orf67;refGene_tx=XM_019823578.2;refGene_gn=CA2H3orf67;refGene_tx=XM_023249890.1;refGene_gn=CA2H3orf67;refGene_tx=XM_023249891.1;refGene_gn=CA2H3orf67;refGene_tx=XM_023249892.1
A2 26674633 26675008
refGene_gn=CA2H3orf67;refGene_tx=XM_011279961.3;refGene_gn=CA2H3orf67;refGene_tx=XM_019823544.2;refGene_gn=CA2H3orf67;refGene_tx=XM_019823545.2;refGene_gn=CA2H3orf67;refGene_tx=XM_019823553.2;refGene_gn=CA2H3orf67;refGene_tx=XM_019823566.2;refGene_gn=CA2H3orf67;refGene_tx=XM_019823568.2;refGene_gn=CA2H3orf67;refGene_tx=XM_019823578.2;refGene_gn=CA2H3orf67;refGene_tx=XM_023249889.1;refGene_gn=CA2H3orf67;refGene_tx=XM_023249890.1;refGene_gn=CA2H3orf67;refGene_tx=XM_023249891.1;refGene_gn=CA2H3orf67;refGene_tx=XM_023249892.1
A2 26684405 26684737
refGene_gn=CA2H3orf67;refGene_tx=XM_011279961.3;refGene_gn=CA2H3orf67;refGene_tx=XM_011279970.3;refGene_gn=CA2H3orf67;refGene_tx=XM_019823544.2;refGene_gn=CA2H3orf67;refGene_tx=XM_019823545.2;refGene_gn=CA2H3orf67;refGene_tx=XM_019823553.2;refGene_gn=CA2H3orf67;refGene_tx=XM_019823566.2;refGene_gn=CA2H3orf67;refGene_tx=XM_019823568.2;refGene_gn=CA2H3orf67;refGene_tx=XM_019823570.2;refGene_gn=CA2H3orf67;refGene_tx=XM_019823578.2;refGene_gn=CA2H3orf67;refGene_tx=XM_023249889.1;refGene_gn=CA2H3orf67;refGene_tx=XM_023249890.1;refGene_gn=CA2H3orf67;refGene_tx=XM_023249891.1;refGene_gn=CA2H3orf67;refGene_tx=XM_023249892.1
A2 26687382 26687502

refGene_gn=CA2H3orf67;refGene_tx=XM_011279961.3;refGene_gn=CA2H3orf67;refGene_tx=XM_011279970.3;refGene_gn=CA2H3orf67;refGene_tx=XM_019823544.2;refGene_gn=CA2H3orf67;refGene_tx=XM_019823545.2;refGene_gn=CA2H3orf67;refGene_tx=XM_019823553.2;refGene_gn=CA2H3orf67;refGene_tx=XM_019823566.2;refGene_gn=CA2H3orf67;refGene_tx=XM_019823568.2;refGene_gn=CA2H3orf67;refGene_tx=XM_019823570.2;refGene_gn=CA2H3orf67;refGene_tx=XM_019823578.2;refGene_gn=CA2H3orf67;refGene_tx=XM_023249889.1;refGene_gn=CA2H3orf67;refGene_tx=XM_023249890.1;refGene_gn=CA2H3orf67;refGene_tx=XM_023249891.1;refGene_gn=CA2H3orf67;refGene_tx=XM_023249892.1
A2 26688651 26688768
refGene_gn=CA2H3orf67;refGene_tx=XM_011279961.3;refGene_gn=CA2H3orf67;refGene_tx=XM_011279970.3;refGene_gn=CA2H3orf67;refGene_tx=XM_019823544.2;refGene_gn=CA2H3orf67;refGene_tx=XM_019823545.2;refGene_gn=CA2H3orf67;refGene_tx=XM_019823553.2;refGene_gn=CA2H3orf67;refGene_tx=XM_019823566.2;refGene_gn=CA2H3orf67;refGene_tx=XM_019823568.2;refGene_gn=CA2H3orf67;refGene_tx=XM_019823570.2;refGene_gn=CA2H3orf67;refGene_tx=XM_019823578.2;refGene_gn=CA2H3orf67;refGene_tx=XM_023249889.1;refGene_gn=CA2H3orf67;refGene_tx=XM_023249890.1;refGene_gn=CA2H3orf67;refGene_tx=XM_023249891.1;refGene_gn=CA2H3orf67;refGene_tx=XM_023249892.1
A2 26690399 26690562
refGene_gn=CA2H3orf67;refGene_tx=XM_011279961.3;refGene_gn=CA2H3orf67;refGene_tx=XM_011279970.3;refGene_gn=CA2H3orf67;refGene_tx=XM_019823544.2;refGene_gn=CA2H3orf67;refGene_tx=XM_019823545.2;refGene_gn=CA2H3orf67;refGene_tx=XM_019823553.2;refGene_gn=CA2H3orf67;refGene_tx=XM_019823566.2;refGene_gn=CA2H3orf67;refGene_tx=XM_019823568.2;refGene_gn=CA2H3orf67;refGene_tx=XM_019823570.2;refGene_gn=CA2H3orf67;refGene_tx=XM_019823578.2;refGene_gn=CA2H3orf67;refGene_tx=XM_023249889.1;refGene_gn=CA2H3orf67;refGene_tx=XM_023249890.1;refGene_gn=CA2H3orf67;refGene_tx=XM_023249891.1;refGene_gn=CA2H3orf67;refGene_tx=XM_023249892.1
A2 26691246 26691383
refGene_gn=CA2H3orf67;refGene_tx=XM_011279961.3;refGene_gn=CA2H3orf67;refGene_tx=XM_011279970.3;refGene_gn=CA2H3orf67;refGene_tx=XM_019823544.2;refGene_gn=CA2H3orf67;refGene_tx=XM_019823545.2;refGene_gn=CA2H3orf67;refGene_tx=XM_019823553.2;refGene_gn=CA2H3orf67;refGene_tx=XM_019823566.2;refGene_gn=CA2H3orf67;refGene_tx=XM_019823568.2;refGene_gn=CA2H3orf67;refGene_tx=XM_019823570.2;refGene_gn=CA2H3orf67;refGene_tx=XM_019823578.2;refGene_gn=CA2H3orf67;refGene_tx=XM_023249889.1;refGene_gn=CA2H3orf67;refGene_tx=XM_023249890.1;refGene_gn=CA2H3orf67;refGene_tx=XM_023249891.1;refGene_gn=CA2H3orf67;refGene_tx=XM_023249892.1
A2 26705363 26705528
refGene_gn=CA2H3orf67;refGene_tx=XM_011279961.3;refGene_gn=CA2H3orf67;refGene_tx=XM_011279970.3;refGene_gn=CA2H3orf67;refGene_tx=XM_019823544.2;refGene_gn=CA2H3orf67;refGene_tx=XM_019823545.2;refGene_gn=CA2H3orf67;refGene_tx=XM_019823553.2;refGene_gn=CA2H3orf67;refGene_tx=XM_019823566.2;refGene_gn=CA2H3orf67;refGene_tx=XM_019823568.2;refGene_gn=CA2H3orf67;refGene_tx=XM_019823570.2;refGene_gn=CA2H3orf67;refGene_tx=XM_019823578.2;refGene_gn=CA2H3orf67;refGene_tx=XM_023249889.1;refGene_gn=CA2H3orf67;refGene_tx=XM_023249890.1;refGene_gn=CA2H3orf67;refGene_tx=XM_023249891.1;refGene_gn=CA2H3orf67;refGene_tx=XM_023249892.1
A2 26722168 26722325

refGene_gn=CA2H3orf67;refGene_tx=XM_011279961.3;refGene_gn=CA2H3orf67;refGene_tx=XM_011279970.3;refGene_gn=CA2H3orf67;refGene_tx=XM_019823544.2;refGene_gn=CA2H3orf67;refGene_tx=XM_019823545.2;refGene_gn=CA2H3orf67;refGene_tx=XM_019823553.2;refGene_gn=CA2H3orf67;refGene_tx=XM_019823566.2;refGene_gn=CA2H3orf67;refGene_tx=XM_019823568.2;refGene_gn=CA2H3orf67;refGene_tx=XM_019823570.2;refGene_gn=CA2H3orf67;refGene_tx=XM_019823578.2;refGene_gn=CA2H3orf67;refGene_tx=XM_023249889.1;refGene_gn=CA2H3orf67;refGene_tx=XM_023249890.1;refGene_gn=CA2H3orf67;refGene_tx=XM_023249891.1;refGene_gn=CA2H3orf67;refGene_tx=XM_023249892.1
A2 26747958 26748073
refGene_gn=CA2H3orf67;refGene_tx=XM_011279961.3;refGene_gn=CA2H3orf67;refGene_tx=XM_011279970.3;refGene_gn=CA2H3orf67;refGene_tx=XM_019823544.2;refGene_gn=CA2H3orf67;refGene_tx=XM_019823545.2;refGene_gn=CA2H3orf67;refGene_tx=XM_019823553.2;refGene_gn=CA2H3orf67;refGene_tx=XM_019823566.2;refGene_gn=CA2H3orf67;refGene_tx=XM_019823568.2;refGene_gn=CA2H3orf67;refGene_tx=XM_019823570.2;refGene_gn=CA2H3orf67;refGene_tx=XM_019823578.2;refGene_gn=CA2H3orf67;refGene_tx=XM_023249889.1;refGene_gn=CA2H3orf67;refGene_tx=XM_023249890.1;refGene_gn=CA2H3orf67;refGene_tx=XM_023249891.1;refGene_gn=CA2H3orf67;refGene_tx=XM_023249892.1
A2 26817093 26817166
refGene_gn=CA2H3orf67;refGene_tx=XM_011279961.3;refGene_gn=CA2H3orf67;refGene_tx=XM_011279970.3;refGene_gn=CA2H3orf67;refGene_tx=XM_019823544.2;refGene_gn=CA2H3orf67;refGene_tx=XM_019823545.2;refGene_gn=CA2H3orf67;refGene_tx=XM_019823553.2;refGene_gn=CA2H3orf67;refGene_tx=XM_019823570.2;refGene_gn=CA2H3orf67;refGene_tx=XM_019823578.2;refGene_gn=CA2H3orf67;refGene_tx=XM_023249889.1
A2 26835336 26835430
refGene_gn=CA2H3orf67;refGene_tx=XM_011279961.3;refGene_gn=CA2H3orf67;refGene_tx=XM_011279970.3;refGene_gn=CA2H3orf67;refGene_tx=XM_019823544.2;refGene_gn=CA2H3orf67;refGene_tx=XM_019823545.2;refGene_gn=CA2H3orf67;refGene_tx=XM_019823553.2;refGene_gn=CA2H3orf67;refGene_tx=XM_019823570.2;refGene_gn=CA2H3orf67;refGene_tx=XM_019823578.2;refGene_gn=CA2H3orf67;refGene_tx=XM_023249889.1
A2 26838335 26838425
refGene_gn=CA2H3orf67;refGene_tx=XM_011279961.3;refGene_gn=CA2H3orf67;refGene_tx=XM_011279970.3;refGene_gn=CA2H3orf67;refGene_tx=XM_019823544.2;refGene_gn=CA2H3orf67;refGene_tx=XM_019823545.2;refGene_gn=CA2H3orf67;refGene_tx=XM_019823553.2;refGene_gn=CA2H3orf67;refGene_tx=XM_019823570.2;refGene_gn=CA2H3orf67;refGene_tx=XM_019823578.2;refGene_gn=CA2H3orf67;refGene_tx=XM_023249889.1
A2 26840153 26840174
refGene_gn=CA2H3orf67;refGene_tx=XM_011279961.3;refGene_gn=CA2H3orf67;refGene_tx=XM_011279970.3;refGene_gn=CA2H3orf67;refGene_tx=XM_019823544.2;refGene_gn=CA2H3orf67;refGene_tx=XM_019823545.2;refGene_gn=CA2H3orf67;refGene_tx=XM_019823553.2;refGene_gn=CA2H3orf67;refGene_tx=XM_019823570.2;refGene_gn=CA2H3orf67;refGene_tx=XM_019823578.2;refGene_gn=CA2H3orf67;refGene_tx=XM_023249889.1
A2 27488357 27488459
refGene_gn=FHIT;refGene_tx=XM_006928913.4;refGene_gn=FHIT;refGene_tx=XM_019823698.2;refGene_gn=FHIT;refGene_tx=XM_019823727.2;refGene_gn=FHI

T;refGene_tx=XM_023249894.1;refGene_gn=FHIT;refGene_tx=XM_023249895.1;
refGene_gn=FHIT;refGene_tx=XM_023249896.1;refGene_gn=FHIT;refGene_tx=X
M_023249898.1
A2 27640418 27640487
refGene_gn=FHIT;refGene_tx=XM_006928913.4;refGene_gn=FHIT;refGene_tx=X
M_019823698.2;refGene_gn=FHIT;refGene_tx=XM_019823727.2;refGene_gn=FHI
T;refGene_tx=XM_023249894.1;refGene_gn=FHIT;refGene_tx=XM_023249896.1;
refGene_gn=FHIT;refGene_tx=XM_023249898.1
A2 27725934 27725964
refGene_gn=FHIT;refGene_tx=XM_006928913.4;refGene_gn=FHIT;refGene_tx=X
M_019823698.2;refGene_gn=FHIT;refGene_tx=XM_019823727.2;refGene_gn=FHI
T;refGene_tx=XM_023249894.1;refGene_gn=FHIT;refGene_tx=XM_023249895.1;
refGene_gn=FHIT;refGene_tx=XM_023249896.1;refGene_gn=FHIT;refGene_tx=X
M_023249898.1
A2 27728249 27728395
refGene_gn=FHIT;refGene_tx=XM_006928913.4;refGene_gn=FHIT;refGene_tx=X
M_019823698.2;refGene_gn=FHIT;refGene_tx=XM_019823727.2;refGene_gn=FHI
T;refGene_tx=XM_023249894.1;refGene_gn=FHIT;refGene_tx=XM_023249895.1;
refGene_gn=FHIT;refGene_tx=XM_023249896.1;refGene_gn=FHIT;refGene_tx=X
M_023249898.1
A2 27753070 27753129
refGene_gn=LOC109496477;refGene_tx=XM_019823729.2;refGene_gn=LOC109496
477;refGene_tx=XM_023249899.1
A2 27753264 27753398
refGene_gn=LOC109496477;refGene_tx=XM_019823729.2;refGene_gn=LOC109496
477;refGene_tx=XM_023249899.1
A2 27763644 27763787
refGene_gn=LOC109496477;refGene_tx=XM_019823729.2;refGene_gn=LOC109496
477;refGene_tx=XM_023249899.1
A2 28242497 28242617
refGene_gn=FHIT;refGene_tx=XM_006928913.4;refGene_gn=FHIT;refGene_tx=X
M_019823698.2;refGene_gn=FHIT;refGene_tx=XM_019823727.2;refGene_gn=FHI
T;refGene_tx=XM_023249894.1;refGene_gn=FHIT;refGene_tx=XM_023249895.1;
refGene_gn=FHIT;refGene_tx=XM_023249896.1;refGene_gn=FHIT;refGene_tx=X
M_023249898.1
A2 28248309 28248376
refGene_gn=FHIT;refGene_tx=XM_019823698.2;refGene_gn=FHIT;refGene_tx=X
M_023249895.1
A2 28329048 28329094 refGene_gn=FHIT;refGene_tx=XM_023249894.1
A2 29176203 29176288 refGene_gn=PTPRG;refGene_tx=XM_011280147.2
A2 29353036 29353141 refGene_gn=PTPRG;refGene_tx=XM_011280147.2
A2 29589369 29589549 refGene_gn=PTPRG;refGene_tx=XM_011280147.2
A2 29602802 29602951 refGene_gn=PTPRG;refGene_tx=XM_011280147.2
A2 29672732 29672828 refGene_gn=PTPRG;refGene_tx=XM_011280147.2
A2 29723528 29723595 refGene_gn=PTPRG;refGene_tx=XM_011280147.2
A2 29745478 29745636 refGene_gn=PTPRG;refGene_tx=XM_011280147.2
A2 29757329 29757522 refGene_gn=PTPRG;refGene_tx=XM_011280147.2
A2 29781194 29781379 refGene_gn=PTPRG;refGene_tx=XM_011280147.2
A2 29784839 29784948 refGene_gn=PTPRG;refGene_tx=XM_011280147.2
A2 29790983 29791033 refGene_gn=PTPRG;refGene_tx=XM_011280147.2

A2 29792776 29793467 refGene_gn=PTPRG;refGene_tx=XM_011280147.2
A2 29793469 29793553 refGene_gn=PTPRG;refGene_tx=XM_011280147.2
A2 29808392 29808525 refGene_gn=PTPRG;refGene_tx=XM_011280147.2
A2 29821679 29821766 refGene_gn=PTPRG;refGene_tx=XM_011280147.2
A2 29835013 29835105 refGene_gn=PTPRG;refGene_tx=XM_011280147.2
A2 29848239 29848331 refGene_gn=PTPRG;refGene_tx=XM_011280147.2
A2 29855760 29855857 refGene_gn=PTPRG;refGene_tx=XM_011280147.2
A2 29860316 29860399 refGene_gn=PTPRG;refGene_tx=XM_011280147.2
A2 29860587 29860722 refGene_gn=PTPRG;refGene_tx=XM_011280147.2
A2 29861724 29861859 refGene_gn=PTPRG;refGene_tx=XM_011280147.2
A2 29863850 29864023 refGene_gn=PTPRG;refGene_tx=XM_011280147.2
A2 29865845 29865981 refGene_gn=PTPRG;refGene_tx=XM_011280147.2
A2 29866551 29866698 refGene_gn=PTPRG;refGene_tx=XM_011280147.2
A2 29869367 29869461 refGene_gn=PTPRG;refGene_tx=XM_011280147.2
A2 29870354 29870431 refGene_gn=PTPRG;refGene_tx=XM_011280147.2
A2 29870924 29871053 refGene_gn=PTPRG;refGene_tx=XM_011280147.2
A2 29874215 29874362 refGene_gn=PTPRG;refGene_tx=XM_011280147.2
A2 29875104 29875247 refGene_gn=PTPRG;refGene_tx=XM_011280147.2
A2 29888037 29888173 refGene_gn=PTPRG;refGene_tx=XM_011280147.2
A2 29888764 29888911 refGene_gn=PTPRG;refGene_tx=XM_011280147.2
A2 29974492 29974752 refGene_gn=FEZF2;refGene_tx=XM_023249900.1
A2 29975666 29975799 refGene_gn=FEZF2;refGene_tx=XM_023249900.1
A2 29975990 29976125 refGene_gn=FEZF2;refGene_tx=XM_023249900.1
A2 29976477 29976879 refGene_gn=FEZF2;refGene_tx=XM_023249900.1
A2 29976880 29977345 refGene_gn=FEZF2;refGene_tx=XM_023249900.1
A2 30003880 30004060
refGene_gn=CADPS;refGene_tx=XM_019823806.2;refGene_gn=CADPS;refGene_tx=XM_019823812.2;refGene_gn=CADPS;refGene_tx=XM_019823818.2;refGene_gn=CADPS;refGene_tx=XM_019823827.2;refGene_gn=CADPS;refGene_tx=XM_019823832.2;refGene_gn=CADPS;refGene_tx=XM_019823837.2;refGene_gn=CADPS;refGene_tx=XM_019823841.2;refGene_gn=CADPS;refGene_tx=XM_019823845.2;refGene_gn=CADPS;refGene_tx=XM_019823856.2;refGene_gn=CADPS;refGene_tx=XM_019823857.2;refGene_gn=CADPS;refGene_tx=XM_019823865.2;refGene_gn=CADPS;refGene_tx=XM_019823869.2;refGene_gn=CADPS;refGene_tx=XM_019823872.2;refGene_gn=CADPS;refGene_tx=XM_019823877.2;refGene_gn=CADPS;refGene_tx=XM_019823879.2;refGene_gn=CADPS;refGene_tx=XM_023249901.1;refGene_gn=CADPS;refGene_tx=XM_023249902.1;refGene_gn=CADPS;refGene_tx=XM_023249903.1;refGene_gn=CADPS;refGene_tx=XM_023249904.1;refGene_gn=CADPS;refGene_tx=XM_023249905.1;refGene_gn=CADPS;refGene_tx=XM_023249906.1
A2 30007338 30007443
refGene_gn=CADPS;refGene_tx=XM_019823806.2;refGene_gn=CADPS;refGene_tx=XM_019823812.2;refGene_gn=CADPS;refGene_tx=XM_019823818.2;refGene_gn=CADPS;refGene_tx=XM_019823827.2;refGene_gn=CADPS;refGene_tx=XM_019823832.2;refGene_gn=CADPS;refGene_tx=XM_019823837.2;refGene_gn=CADPS;refGene_tx=XM_019823841.2;refGene_gn=CADPS;refGene_tx=XM_019823845.2;refGene_gn=CADPS;refGene_tx=XM_019823856.2;refGene_gn=CADPS;refGene_tx=XM_019823857.2;refGene_gn=CADPS;refGene_tx=XM_019823865.2;refGene_gn=CADPS;refGene_tx=XM_019823869.2;refGene_gn=CADPS;refGene_tx=XM_019823872.2;refGene_gn=CADPS;refGene_tx=XM_019823877.2;refGene_gn=CADPS;refGene_tx=XM_019823879.2;refGene_gn=CADPS;refGene_tx=XM_023249901.1;refGene_gn=C

ADPS;refGene_tx=XM_023249902.1;refGene_gn=CADPS;refGene_tx=XM_02324990
3.1;refGene_gn=CADPS;refGene_tx=XM_023249904.1;refGene_gn=CADPS;refGen
e_tx=XM_023249905.1;refGene_gn=CADPS;refGene_tx=XM_023249906.1
A2 30042593 30042701
refGene_gn=CADPS;refGene_tx=XM_019823806.2;refGene_gn=CADPS;refGene_tx
=XM_019823812.2;refGene_gn=CADPS;refGene_tx=XM_019823818.2;refGene_gn=
CADPS;refGene_tx=XM_019823827.2;refGene_gn=CADPS;refGene_tx=XM_0198238
32.2;refGene_gn=CADPS;refGene_tx=XM_019823837.2;refGene_gn=CADPS;refGe
ne_tx=XM_019823841.2;refGene_gn=CADPS;refGene_tx=XM_019823845.2;refGen
e_gn=CADPS;refGene_tx=XM_019823856.2;refGene_gn=CADPS;refGene_tx=XM_01
9823857.2;refGene_gn=CADPS;refGene_tx=XM_019823865.2;refGene_gn=CADPS;
refGene_tx=XM_019823869.2;refGene_gn=CADPS;refGene_tx=XM_019823872.2;r
efGene_gn=CADPS;refGene_tx=XM_019823877.2;refGene_gn=CADPS;refGene_tx=
XM_019823879.2;refGene_gn=CADPS;refGene_tx=XM_023249901.1;refGene_gn=C
ADPS;refGene_tx=XM_023249902.1;refGene_gn=CADPS;refGene_tx=XM_02324990
3.1;refGene_gn=CADPS;refGene_tx=XM_023249904.1;refGene_gn=CADPS;refGen
e_tx=XM_023249905.1;refGene_gn=CADPS;refGene_tx=XM_023249906.1
A2 30049552 30049585
refGene_gn=CADPS;refGene_tx=XM_019823806.2;refGene_gn=CADPS;refGene_tx
=XM_019823812.2;refGene_gn=CADPS;refGene_tx=XM_019823818.2;refGene_gn=
CADPS;refGene_tx=XM_019823827.2;refGene_gn=CADPS;refGene_tx=XM_0198238
32.2;refGene_gn=CADPS;refGene_tx=XM_019823837.2;refGene_gn=CADPS;refGe
ne_tx=XM_019823841.2;refGene_gn=CADPS;refGene_tx=XM_019823845.2;refGen
e_gn=CADPS;refGene_tx=XM_019823856.2;refGene_gn=CADPS;refGene_tx=XM_01
9823857.2;refGene_gn=CADPS;refGene_tx=XM_019823865.2;refGene_gn=CADPS;
refGene_tx=XM_019823869.2;refGene_gn=CADPS;refGene_tx=XM_019823872.2;r
efGene_gn=CADPS;refGene_tx=XM_019823877.2;refGene_gn=CADPS;refGene_tx=
XM_019823879.2;refGene_gn=CADPS;refGene_tx=XM_023249901.1;refGene_gn=C
ADPS;refGene_tx=XM_023249902.1;refGene_gn=CADPS;refGene_tx=XM_02324990
3.1;refGene_gn=CADPS;refGene_tx=XM_023249904.1;refGene_gn=CADPS;refGen
e_tx=XM_023249905.1;refGene_gn=CADPS;refGene_tx=XM_023249906.1
A2 30073284 30073368
refGene_gn=CADPS;refGene_tx=XM_019823806.2;refGene_gn=CADPS;refGene_tx
=XM_019823812.2;refGene_gn=CADPS;refGene_tx=XM_019823818.2;refGene_gn=
CADPS;refGene_tx=XM_019823827.2;refGene_gn=CADPS;refGene_tx=XM_0198238
32.2;refGene_gn=CADPS;refGene_tx=XM_019823837.2;refGene_gn=CADPS;refGe
ne_tx=XM_019823841.2;refGene_gn=CADPS;refGene_tx=XM_019823845.2;refGen
e_gn=CADPS;refGene_tx=XM_019823856.2;refGene_gn=CADPS;refGene_tx=XM_01
9823857.2;refGene_gn=CADPS;refGene_tx=XM_019823865.2;refGene_gn=CADPS;
refGene_tx=XM_019823869.2;refGene_gn=CADPS;refGene_tx=XM_019823872.2;r
efGene_gn=CADPS;refGene_tx=XM_019823877.2;refGene_gn=CADPS;refGene_tx=
XM_019823879.2;refGene_gn=CADPS;refGene_tx=XM_023249901.1;refGene_gn=C
ADPS;refGene_tx=XM_023249902.1;refGene_gn=CADPS;refGene_tx=XM_02324990
3.1;refGene_gn=CADPS;refGene_tx=XM_023249904.1;refGene_gn=CADPS;refGen
e_tx=XM_023249905.1;refGene_gn=CADPS;refGene_tx=XM_023249906.1
A2 30074256 30074331
refGene_gn=CADPS;refGene_tx=XM_019823806.2;refGene_gn=CADPS;refGene_tx
=XM_019823812.2;refGene_gn=CADPS;refGene_tx=XM_019823818.2;refGene_gn=
CADPS;refGene_tx=XM_019823827.2;refGene_gn=CADPS;refGene_tx=XM_0198238
32.2;refGene_gn=CADPS;refGene_tx=XM_019823837.2;refGene_gn=CADPS;refGe

ne_tx=XM_019823841.2;refGene_gn=CADPS;refGene_tx=XM_019823845.2;refGene_gn=CADPS;refGene_tx=XM_019823856.2;refGene_gn=CADPS;refGene_tx=XM_019823857.2;refGene_gn=CADPS;refGene_tx=XM_019823865.2;refGene_gn=CADPS;refGene_tx=XM_019823869.2;refGene_gn=CADPS;refGene_tx=XM_019823872.2;refGene_gn=CADPS;refGene_tx=XM_019823877.2;refGene_gn=CADPS;refGene_tx=XM_019823879.2;refGene_gn=CADPS;refGene_tx=XM_023249901.1;refGene_gn=CADPS;refGene_tx=XM_023249902.1;refGene_gn=CADPS;refGene_tx=XM_023249903.1;refGene_gn=CADPS;refGene_tx=XM_023249904.1;refGene_gn=CADPS;refGene_tx=XM_023249905.1;refGene_gn=CADPS;refGene_tx=XM_023249906.1
A2 30075230 30075245
refGene_gn=CADPS;refGene_tx=XM_019823806.2;refGene_gn=CADPS;refGene_tx=XM_019823812.2;refGene_gn=CADPS;refGene_tx=XM_019823818.2;refGene_gn=CADPS;refGene_tx=XM_019823827.2;refGene_gn=CADPS;refGene_tx=XM_019823832.2;refGene_gn=CADPS;refGene_tx=XM_019823845.2;refGene_gn=CADPS;refGene_tx=XM_019823872.2;refGene_gn=CADPS;refGene_tx=XM_023249901.1;refGene_gn=CADPS;refGene_tx=XM_023249903.1;refGene_gn=CADPS;refGene_tx=XM_023249904.1;refGene_gn=CADPS;refGene_tx=XM_023249905.1;refGene_gn=CADPS;refGene_tx=XM_023249906.1
A2 30081967 30082115
refGene_gn=CADPS;refGene_tx=XM_019823806.2;refGene_gn=CADPS;refGene_tx=XM_019823812.2;refGene_gn=CADPS;refGene_tx=XM_019823818.2;refGene_gn=CADPS;refGene_tx=XM_019823827.2;refGene_gn=CADPS;refGene_tx=XM_019823832.2;refGene_gn=CADPS;refGene_tx=XM_019823837.2;refGene_gn=CADPS;refGene_tx=XM_019823841.2;refGene_gn=CADPS;refGene_tx=XM_019823845.2;refGene_gn=CADPS;refGene_tx=XM_019823856.2;refGene_gn=CADPS;refGene_tx=XM_019823857.2;refGene_gn=CADPS;refGene_tx=XM_019823865.2;refGene_gn=CADPS;refGene_tx=XM_019823869.2;refGene_gn=CADPS;refGene_tx=XM_019823872.2;refGene_gn=CADPS;refGene_tx=XM_019823877.2;refGene_gn=CADPS;refGene_tx=XM_019823879.2;refGene_gn=CADPS;refGene_tx=XM_023249901.1;refGene_gn=CADPS;refGene_tx=XM_023249902.1;refGene_gn=CADPS;refGene_tx=XM_023249903.1;refGene_gn=CADPS;refGene_tx=XM_023249904.1;refGene_gn=CADPS;refGene_tx=XM_023249905.1;refGene_gn=CADPS;refGene_tx=XM_023249906.1
A2 30085922 30086078
refGene_gn=CADPS;refGene_tx=XM_019823806.2;refGene_gn=CADPS;refGene_tx=XM_019823812.2;refGene_gn=CADPS;refGene_tx=XM_019823818.2;refGene_gn=CADPS;refGene_tx=XM_019823827.2;refGene_gn=CADPS;refGene_tx=XM_019823832.2;refGene_gn=CADPS;refGene_tx=XM_019823837.2;refGene_gn=CADPS;refGene_tx=XM_019823841.2;refGene_gn=CADPS;refGene_tx=XM_019823845.2;refGene_gn=CADPS;refGene_tx=XM_019823856.2;refGene_gn=CADPS;refGene_tx=XM_019823857.2;refGene_gn=CADPS;refGene_tx=XM_019823865.2;refGene_gn=CADPS;refGene_tx=XM_019823869.2;refGene_gn=CADPS;refGene_tx=XM_019823872.2;refGene_gn=CADPS;refGene_tx=XM_019823877.2;refGene_gn=CADPS;refGene_tx=XM_019823879.2;refGene_gn=CADPS;refGene_tx=XM_023249901.1;refGene_gn=CADPS;refGene_tx=XM_023249902.1;refGene_gn=CADPS;refGene_tx=XM_023249903.1;refGene_gn=CADPS;refGene_tx=XM_023249904.1;refGene_gn=CADPS;refGene_tx=XM_023249905.1;refGene_gn=CADPS;refGene_tx=XM_023249906.1
A2 30089463 30089610
refGene_gn=CADPS;refGene_tx=XM_019823806.2;refGene_gn=CADPS;refGene_tx=XM_019823812.2;refGene_gn=CADPS;refGene_tx=XM_019823818.2;refGene_gn=CADPS;refGene_tx=XM_019823827.2;refGene_gn=CADPS;refGene_tx=XM_0198238

32.2;refGene_gn=CADPS;refGene_tx=XM_019823837.2;refGene_gn=CADPS;refGene_tx=XM_019823841.2;refGene_gn=CADPS;refGene_tx=XM_019823845.2;refGene_gn=CADPS;refGene_tx=XM_019823856.2;refGene_gn=CADPS;refGene_tx=XM_019823857.2;refGene_gn=CADPS;refGene_tx=XM_019823879.2;refGene_gn=CADPS;refGene_tx=XM_023249901.1;refGene_gn=CADPS;refGene_tx=XM_023249902.1;refGene_gn=CADPS;refGene_tx=XM_023249903.1;refGene_gn=CADPS;refGene_tx=XM_023249904.1;refGene_gn=CADPS;refGene_tx=XM_023249906.1
A2 30097658 30097800
refGene_gn=CADPS;refGene_tx=XM_019823806.2;refGene_gn=CADPS;refGene_tx=XM_019823812.2;refGene_gn=CADPS;refGene_tx=XM_019823818.2;refGene_gn=CADPS;refGene_tx=XM_019823827.2;refGene_gn=CADPS;refGene_tx=XM_019823832.2;refGene_gn=CADPS;refGene_tx=XM_019823837.2;refGene_gn=CADPS;refGene_tx=XM_019823841.2;refGene_gn=CADPS;refGene_tx=XM_019823845.2;refGene_gn=CADPS;refGene_tx=XM_019823856.2;refGene_gn=CADPS;refGene_tx=XM_019823857.2;refGene_gn=CADPS;refGene_tx=XM_019823865.2;refGene_gn=CADPS;refGene_tx=XM_019823869.2;refGene_gn=CADPS;refGene_tx=XM_019823872.2;refGene_gn=CADPS;refGene_tx=XM_019823877.2;refGene_gn=CADPS;refGene_tx=XM_019823879.2;refGene_gn=CADPS;refGene_tx=XM_023249901.1;refGene_gn=CADPS;refGene_tx=XM_023249902.1;refGene_gn=CADPS;refGene_tx=XM_023249903.1;refGene_gn=CADPS;refGene_tx=XM_023249904.1;refGene_gn=CADPS;refGene_tx=XM_023249905.1;refGene_gn=CADPS;refGene_tx=XM_023249906.1
A2 30098876 30099033
refGene_gn=CADPS;refGene_tx=XM_019823806.2;refGene_gn=CADPS;refGene_tx=XM_019823812.2;refGene_gn=CADPS;refGene_tx=XM_019823818.2;refGene_gn=CADPS;refGene_tx=XM_019823827.2;refGene_gn=CADPS;refGene_tx=XM_019823832.2;refGene_gn=CADPS;refGene_tx=XM_019823837.2;refGene_gn=CADPS;refGene_tx=XM_019823841.2;refGene_gn=CADPS;refGene_tx=XM_019823845.2;refGene_gn=CADPS;refGene_tx=XM_019823856.2;refGene_gn=CADPS;refGene_tx=XM_019823857.2;refGene_gn=CADPS;refGene_tx=XM_019823865.2;refGene_gn=CADPS;refGene_tx=XM_019823869.2;refGene_gn=CADPS;refGene_tx=XM_019823872.2;refGene_gn=CADPS;refGene_tx=XM_019823877.2;refGene_gn=CADPS;refGene_tx=XM_019823879.2;refGene_gn=CADPS;refGene_tx=XM_023249901.1;refGene_gn=CADPS;refGene_tx=XM_023249902.1;refGene_gn=CADPS;refGene_tx=XM_023249903.1;refGene_gn=CADPS;refGene_tx=XM_023249904.1;refGene_gn=CADPS;refGene_tx=XM_023249905.1;refGene_gn=CADPS;refGene_tx=XM_023249906.1
A2 30100236 30100257
refGene_gn=CADPS;refGene_tx=XM_019823806.2;refGene_gn=CADPS;refGene_tx=XM_019823812.2;refGene_gn=CADPS;refGene_tx=XM_019823818.2;refGene_gn=CADPS;refGene_tx=XM_019823827.2;refGene_gn=CADPS;refGene_tx=XM_019823832.2;refGene_gn=CADPS;refGene_tx=XM_019823841.2;refGene_gn=CADPS;refGene_tx=XM_019823845.2;refGene_gn=CADPS;refGene_tx=XM_019823856.2;refGene_gn=CADPS;refGene_tx=XM_019823879.2;refGene_gn=CADPS;refGene_tx=XM_023249901.1;refGene_gn=CADPS;refGene_tx=XM_023249902.1;refGene_gn=CADPS;refGene_tx=XM_023249905.1;refGene_gn=CADPS;refGene_tx=XM_023249906.1
A2 30104486 30104495
refGene_gn=CADPS;refGene_tx=XM_019823806.2;refGene_gn=CADPS;refGene_tx=XM_019823812.2;refGene_gn=CADPS;refGene_tx=XM_019823818.2;refGene_gn=CADPS;refGene_tx=XM_019823827.2;refGene_gn=CADPS;refGene_tx=XM_019823832.2;refGene_gn=CADPS;refGene_tx=XM_019823845.2;refGene_gn=CADPS;refGene_tx=XM_019823857.2;refGene_gn=CADPS;refGene_tx=XM_019823879.2;refGen

e_gn=CADPS;refGene_tx=XM_023249902.1;refGene_gn=CADPS;refGene_tx=XM_02
3249903.1;refGene_gn=CADPS;refGene_tx=XM_023249905.1;refGene_gn=CADPS;
refGene_tx=XM_023249906.1
A2 30105531 30105638
refGene_gn=CADPS;refGene_tx=XM_019823806.2;refGene_gn=CADPS;refGene_tx
=XM_019823812.2;refGene_gn=CADPS;refGene_tx=XM_019823818.2;refGene_gn=
CADPS;refGene_tx=XM_019823827.2;refGene_gn=CADPS;refGene_tx=XM_0198238
32.2;refGene_gn=CADPS;refGene_tx=XM_019823837.2;refGene_gn=CADPS;refGe
ne_tx=XM_019823841.2;refGene_gn=CADPS;refGene_tx=XM_019823845.2;refGen
e_gn=CADPS;refGene_tx=XM_019823856.2;refGene_gn=CADPS;refGene_tx=XM_01
9823857.2;refGene_gn=CADPS;refGene_tx=XM_019823865.2;refGene_gn=CADPS;
refGene_tx=XM_019823869.2;refGene_gn=CADPS;refGene_tx=XM_019823872.2;r
efGene_gn=CADPS;refGene_tx=XM_019823877.2;refGene_gn=CADPS;refGene_tx=
XM_019823879.2;refGene_gn=CADPS;refGene_tx=XM_023249901.1;refGene_gn=C
ADPS;refGene_tx=XM_023249902.1;refGene_gn=CADPS;refGene_tx=XM_02324990
3.1;refGene_gn=CADPS;refGene_tx=XM_023249904.1;refGene_gn=CADPS;refGen
e_tx=XM_023249905.1;refGene_gn=CADPS;refGene_tx=XM_023249906.1
A2 30108869 30108962
refGene_gn=CADPS;refGene_tx=XM_019823806.2;refGene_gn=CADPS;refGene_tx
=XM_023249901.1;refGene_gn=CADPS;refGene_tx=XM_023249902.1;refGene_gn=
CADPS;refGene_tx=XM_023249903.1;refGene_gn=CADPS;refGene_tx=XM_0232499
04.1;refGene_gn=CADPS;refGene_tx=XM_023249905.1;refGene_gn=CADPS;refGe
ne_tx=XM_023249906.1
A2 30118002 30118020
refGene_gn=CADPS;refGene_tx=XM_019823812.2;refGene_gn=CADPS;refGene_tx
=XM_019823827.2;refGene_gn=CADPS;refGene_tx=XM_019823841.2;refGene_gn=
CADPS;refGene_tx=XM_019823856.2;refGene_gn=CADPS;refGene_tx=XM_0198238
57.2
A2 30118891 30118960
refGene_gn=CADPS;refGene_tx=XM_019823812.2;refGene_gn=CADPS;refGene_tx
=XM_019823818.2;refGene_gn=CADPS;refGene_tx=XM_019823837.2;refGene_gn=
CADPS;refGene_tx=XM_019823869.2
A2 30121293 30121417
refGene_gn=CADPS;refGene_tx=XM_019823806.2;refGene_gn=CADPS;refGene_tx
=XM_019823812.2;refGene_gn=CADPS;refGene_tx=XM_019823818.2;refGene_gn=
CADPS;refGene_tx=XM_019823827.2;refGene_gn=CADPS;refGene_tx=XM_0198238
32.2;refGene_gn=CADPS;refGene_tx=XM_019823837.2;refGene_gn=CADPS;refGe
ne_tx=XM_019823841.2;refGene_gn=CADPS;refGene_tx=XM_019823845.2;refGen
e_gn=CADPS;refGene_tx=XM_019823856.2;refGene_gn=CADPS;refGene_tx=XM_01
9823857.2;refGene_gn=CADPS;refGene_tx=XM_019823865.2;refGene_gn=CADPS;
refGene_tx=XM_019823869.2;refGene_gn=CADPS;refGene_tx=XM_019823872.2;r
efGene_gn=CADPS;refGene_tx=XM_019823877.2;refGene_gn=CADPS;refGene_tx=
XM_019823879.2;refGene_gn=CADPS;refGene_tx=XM_023249901.1;refGene_gn=C
ADPS;refGene_tx=XM_023249902.1;refGene_gn=CADPS;refGene_tx=XM_02324990
3.1;refGene_gn=CADPS;refGene_tx=XM_023249904.1;refGene_gn=CADPS;refGen
e_tx=XM_023249905.1;refGene_gn=CADPS;refGene_tx=XM_023249906.1
A2 30121832 30121896
refGene_gn=CADPS;refGene_tx=XM_019823806.2;refGene_gn=CADPS;refGene_tx
=XM_019823812.2;refGene_gn=CADPS;refGene_tx=XM_019823818.2;refGene_gn=
CADPS;refGene_tx=XM_019823827.2;refGene_gn=CADPS;refGene_tx=XM_0198238

32.2;refGene_gn=CADPS;refGene_tx=XM_019823837.2;refGene_gn=CADPS;refGene_tx=XM_019823841.2;refGene_gn=CADPS;refGene_tx=XM_019823845.2;refGene_gn=CADPS;refGene_tx=XM_019823856.2;refGene_gn=CADPS;refGene_tx=XM_019823857.2;refGene_gn=CADPS;refGene_tx=XM_019823865.2;refGene_gn=CADPS;refGene_tx=XM_019823869.2;refGene_gn=CADPS;refGene_tx=XM_019823872.2;refGene_gn=CADPS;refGene_tx=XM_019823877.2;refGene_gn=CADPS;refGene_tx=XM_019823879.2;refGene_gn=CADPS;refGene_tx=XM_023249901.1;refGene_gn=CADPS;refGene_tx=XM_023249902.1;refGene_gn=CADPS;refGene_tx=XM_023249903.1;refGene_gn=CADPS;refGene_tx=XM_023249904.1;refGene_gn=CADPS;refGene_tx=XM_023249905.1;refGene_gn=CADPS;refGene_tx=XM_023249906.1
A2 30123878 30123980
refGene_gn=CADPS;refGene_tx=XM_019823806.2;refGene_gn=CADPS;refGene_tx=XM_019823812.2;refGene_gn=CADPS;refGene_tx=XM_019823818.2;refGene_gn=CADPS;refGene_tx=XM_019823827.2;refGene_gn=CADPS;refGene_tx=XM_019823832.2;refGene_gn=CADPS;refGene_tx=XM_019823837.2;refGene_gn=CADPS;refGene_tx=XM_019823841.2;refGene_gn=CADPS;refGene_tx=XM_019823845.2;refGene_gn=CADPS;refGene_tx=XM_019823856.2;refGene_gn=CADPS;refGene_tx=XM_019823857.2;refGene_gn=CADPS;refGene_tx=XM_019823865.2;refGene_gn=CADPS;refGene_tx=XM_019823869.2;refGene_gn=CADPS;refGene_tx=XM_019823872.2;refGene_gn=CADPS;refGene_tx=XM_019823877.2;refGene_gn=CADPS;refGene_tx=XM_019823879.2;refGene_gn=CADPS;refGene_tx=XM_023249901.1;refGene_gn=CADPS;refGene_tx=XM_023249902.1;refGene_gn=CADPS;refGene_tx=XM_023249903.1;refGene_gn=CADPS;refGene_tx=XM_023249904.1;refGene_gn=CADPS;refGene_tx=XM_023249905.1;refGene_gn=CADPS;refGene_tx=XM_023249906.1
A2 30139748 30139907
refGene_gn=CADPS;refGene_tx=XM_019823806.2;refGene_gn=CADPS;refGene_tx=XM_019823812.2;refGene_gn=CADPS;refGene_tx=XM_019823818.2;refGene_gn=CADPS;refGene_tx=XM_019823827.2;refGene_gn=CADPS;refGene_tx=XM_019823832.2;refGene_gn=CADPS;refGene_tx=XM_019823865.2;refGene_gn=CADPS;refGene_tx=XM_023249901.1;refGene_gn=CADPS;refGene_tx=XM_023249902.1;refGene_gn=CADPS;refGene_tx=XM_023249903.1;refGene_gn=CADPS;refGene_tx=XM_023249904.1;refGene_gn=CADPS;refGene_tx=XM_023249905.1
A2 30141942 30142130
refGene_gn=CADPS;refGene_tx=XM_019823806.2;refGene_gn=CADPS;refGene_tx=XM_019823812.2;refGene_gn=CADPS;refGene_tx=XM_019823818.2;refGene_gn=CADPS;refGene_tx=XM_019823827.2;refGene_gn=CADPS;refGene_tx=XM_019823832.2;refGene_gn=CADPS;refGene_tx=XM_019823837.2;refGene_gn=CADPS;refGene_tx=XM_019823841.2;refGene_gn=CADPS;refGene_tx=XM_019823845.2;refGene_gn=CADPS;refGene_tx=XM_019823856.2;refGene_gn=CADPS;refGene_tx=XM_019823857.2;refGene_gn=CADPS;refGene_tx=XM_019823865.2;refGene_gn=CADPS;refGene_tx=XM_019823869.2;refGene_gn=CADPS;refGene_tx=XM_019823872.2;refGene_gn=CADPS;refGene_tx=XM_019823877.2;refGene_gn=CADPS;refGene_tx=XM_019823879.2;refGene_gn=CADPS;refGene_tx=XM_023249901.1;refGene_gn=CADPS;refGene_tx=XM_023249902.1;refGene_gn=CADPS;refGene_tx=XM_023249903.1;refGene_gn=CADPS;refGene_tx=XM_023249904.1;refGene_gn=CADPS;refGene_tx=XM_023249905.1;refGene_gn=CADPS;refGene_tx=XM_023249906.1
A2 30145849 30145986
refGene_gn=CADPS;refGene_tx=XM_019823806.2;refGene_gn=CADPS;refGene_tx=XM_019823812.2;refGene_gn=CADPS;refGene_tx=XM_019823818.2;refGene_gn=CADPS;refGene_tx=XM_019823827.2;refGene_gn=CADPS;refGene_tx=XM_0198238

32.2;refGene_gn=CADPS;refGene_tx=XM_019823837.2;refGene_gn=CADPS;refGene_tx=XM_019823841.2;refGene_gn=CADPS;refGene_tx=XM_019823845.2;refGene_gn=CADPS;refGene_tx=XM_019823856.2;refGene_gn=CADPS;refGene_tx=XM_019823857.2;refGene_gn=CADPS;refGene_tx=XM_019823865.2;refGene_gn=CADPS;refGene_tx=XM_019823869.2;refGene_gn=CADPS;refGene_tx=XM_019823872.2;refGene_gn=CADPS;refGene_tx=XM_019823877.2;refGene_gn=CADPS;refGene_tx=XM_019823879.2;refGene_gn=CADPS;refGene_tx=XM_023249901.1;refGene_gn=CADPS;refGene_tx=XM_023249902.1;refGene_gn=CADPS;refGene_tx=XM_023249903.1;refGene_gn=CADPS;refGene_tx=XM_023249904.1;refGene_gn=CADPS;refGene_tx=XM_023249905.1;refGene_gn=CADPS;refGene_tx=XM_023249906.1
A2 30154208 30154220 refGene_gn=CADPS;refGene_tx=XM_019823841.2
A2 30158977 30159190
refGene_gn=CADPS;refGene_tx=XM_019823806.2;refGene_gn=CADPS;refGene_tx=XM_019823812.2;refGene_gn=CADPS;refGene_tx=XM_019823818.2;refGene_gn=CADPS;refGene_tx=XM_019823827.2;refGene_gn=CADPS;refGene_tx=XM_019823832.2;refGene_gn=CADPS;refGene_tx=XM_019823837.2;refGene_gn=CADPS;refGene_tx=XM_019823841.2;refGene_gn=CADPS;refGene_tx=XM_019823845.2;refGene_gn=CADPS;refGene_tx=XM_019823856.2;refGene_gn=CADPS;refGene_tx=XM_019823857.2;refGene_gn=CADPS;refGene_tx=XM_019823865.2;refGene_gn=CADPS;refGene_tx=XM_019823869.2;refGene_gn=CADPS;refGene_tx=XM_019823872.2;refGene_gn=CADPS;refGene_tx=XM_019823877.2;refGene_gn=CADPS;refGene_tx=XM_019823879.2;refGene_gn=CADPS;refGene_tx=XM_023249901.1;refGene_gn=CADPS;refGene_tx=XM_023249902.1;refGene_gn=CADPS;refGene_tx=XM_023249903.1;refGene_gn=CADPS;refGene_tx=XM_023249904.1;refGene_gn=CADPS;refGene_tx=XM_023249905.1;refGene_gn=CADPS;refGene_tx=XM_023249906.1
A2 30166578 30166687
refGene_gn=CADPS;refGene_tx=XM_019823806.2;refGene_gn=CADPS;refGene_tx=XM_019823812.2;refGene_gn=CADPS;refGene_tx=XM_019823818.2;refGene_gn=CADPS;refGene_tx=XM_019823827.2;refGene_gn=CADPS;refGene_tx=XM_019823832.2;refGene_gn=CADPS;refGene_tx=XM_019823837.2;refGene_gn=CADPS;refGene_tx=XM_019823841.2;refGene_gn=CADPS;refGene_tx=XM_019823845.2;refGene_gn=CADPS;refGene_tx=XM_019823856.2;refGene_gn=CADPS;refGene_tx=XM_019823857.2;refGene_gn=CADPS;refGene_tx=XM_019823865.2;refGene_gn=CADPS;refGene_tx=XM_019823869.2;refGene_gn=CADPS;refGene_tx=XM_019823872.2;refGene_gn=CADPS;refGene_tx=XM_019823877.2;refGene_gn=CADPS;refGene_tx=XM_019823879.2;refGene_gn=CADPS;refGene_tx=XM_023249901.1;refGene_gn=CADPS;refGene_tx=XM_023249902.1;refGene_gn=CADPS;refGene_tx=XM_023249903.1;refGene_gn=CADPS;refGene_tx=XM_023249904.1;refGene_gn=CADPS;refGene_tx=XM_023249905.1;refGene_gn=CADPS;refGene_tx=XM_023249906.1
A2 30177782 30177849
refGene_gn=CADPS;refGene_tx=XM_019823806.2;refGene_gn=CADPS;refGene_tx=XM_019823812.2;refGene_gn=CADPS;refGene_tx=XM_019823818.2;refGene_gn=CADPS;refGene_tx=XM_019823827.2;refGene_gn=CADPS;refGene_tx=XM_019823832.2;refGene_gn=CADPS;refGene_tx=XM_019823837.2;refGene_gn=CADPS;refGene_tx=XM_019823841.2;refGene_gn=CADPS;refGene_tx=XM_019823845.2;refGene_gn=CADPS;refGene_tx=XM_019823856.2;refGene_gn=CADPS;refGene_tx=XM_019823857.2;refGene_gn=CADPS;refGene_tx=XM_019823865.2;refGene_gn=CADPS;refGene_tx=XM_019823869.2;refGene_gn=CADPS;refGene_tx=XM_019823872.2;refGene_gn=CADPS;refGene_tx=XM_019823877.2;refGene_gn=CADPS;refGene_tx=XM_019823879.2;refGene_gn=CADPS;refGene_tx=XM_023249901.1;refGene_gn=C

ADPS;refGene_tx=XM_023249902.1;refGene_gn=CADPS;refGene_tx=XM_02324990
3.1;refGene_gn=CADPS;refGene_tx=XM_023249904.1;refGene_gn=CADPS;refGen
e_tx=XM_023249905.1;refGene_gn=CADPS;refGene_tx=XM_023249906.1
A2 30189326 30189466
refGene_gn=CADPS;refGene_tx=XM_019823806.2;refGene_gn=CADPS;refGene_tx
=XM_019823812.2;refGene_gn=CADPS;refGene_tx=XM_019823818.2;refGene_gn=
CADPS;refGene_tx=XM_019823827.2;refGene_gn=CADPS;refGene_tx=XM_0198238
32.2;refGene_gn=CADPS;refGene_tx=XM_019823837.2;refGene_gn=CADPS;refGe
ne_tx=XM_019823841.2;refGene_gn=CADPS;refGene_tx=XM_019823845.2;refGen
e_gn=CADPS;refGene_tx=XM_019823856.2;refGene_gn=CADPS;refGene_tx=XM_01
9823857.2;refGene_gn=CADPS;refGene_tx=XM_019823865.2;refGene_gn=CADPS;
refGene_tx=XM_019823869.2;refGene_gn=CADPS;refGene_tx=XM_019823872.2;r
efGene_gn=CADPS;refGene_tx=XM_019823877.2;refGene_gn=CADPS;refGene_tx=
XM_019823879.2;refGene_gn=CADPS;refGene_tx=XM_023249901.1;refGene_gn=C
ADPS;refGene_tx=XM_023249902.1;refGene_gn=CADPS;refGene_tx=XM_02324990
3.1;refGene_gn=CADPS;refGene_tx=XM_023249904.1;refGene_gn=CADPS;refGen
e_tx=XM_023249905.1;refGene_gn=CADPS;refGene_tx=XM_023249906.1
A2 30197912 30198024
refGene_gn=CADPS;refGene_tx=XM_019823806.2;refGene_gn=CADPS;refGene_tx
=XM_019823812.2;refGene_gn=CADPS;refGene_tx=XM_019823818.2;refGene_gn=
CADPS;refGene_tx=XM_019823827.2;refGene_gn=CADPS;refGene_tx=XM_0198238
32.2;refGene_gn=CADPS;refGene_tx=XM_019823837.2;refGene_gn=CADPS;refGe
ne_tx=XM_019823841.2;refGene_gn=CADPS;refGene_tx=XM_019823845.2;refGen
e_gn=CADPS;refGene_tx=XM_019823856.2;refGene_gn=CADPS;refGene_tx=XM_01
9823857.2;refGene_gn=CADPS;refGene_tx=XM_019823865.2;refGene_gn=CADPS;
refGene_tx=XM_019823869.2;refGene_gn=CADPS;refGene_tx=XM_019823872.2;r
efGene_gn=CADPS;refGene_tx=XM_019823877.2;refGene_gn=CADPS;refGene_tx=
XM_019823879.2;refGene_gn=CADPS;refGene_tx=XM_023249901.1;refGene_gn=C
ADPS;refGene_tx=XM_023249902.1;refGene_gn=CADPS;refGene_tx=XM_02324990
3.1;refGene_gn=CADPS;refGene_tx=XM_023249904.1;refGene_gn=CADPS;refGen
e_tx=XM_023249905.1;refGene_gn=CADPS;refGene_tx=XM_023249906.1
A2 30244707 30244829
refGene_gn=CADPS;refGene_tx=XM_019823806.2;refGene_gn=CADPS;refGene_tx
=XM_019823812.2;refGene_gn=CADPS;refGene_tx=XM_019823818.2;refGene_gn=
CADPS;refGene_tx=XM_019823827.2;refGene_gn=CADPS;refGene_tx=XM_0198238
32.2;refGene_gn=CADPS;refGene_tx=XM_019823837.2;refGene_gn=CADPS;refGe
ne_tx=XM_019823841.2;refGene_gn=CADPS;refGene_tx=XM_019823845.2;refGen
e_gn=CADPS;refGene_tx=XM_019823856.2;refGene_gn=CADPS;refGene_tx=XM_01
9823857.2;refGene_gn=CADPS;refGene_tx=XM_019823865.2;refGene_gn=CADPS;
refGene_tx=XM_019823869.2;refGene_gn=CADPS;refGene_tx=XM_019823872.2;r
efGene_gn=CADPS;refGene_tx=XM_019823877.2;refGene_gn=CADPS;refGene_tx=
XM_019823879.2;refGene_gn=CADPS;refGene_tx=XM_023249901.1;refGene_gn=C
ADPS;refGene_tx=XM_023249902.1;refGene_gn=CADPS;refGene_tx=XM_02324990
3.1;refGene_gn=CADPS;refGene_tx=XM_023249904.1;refGene_gn=CADPS;refGen
e_tx=XM_023249905.1;refGene_gn=CADPS;refGene_tx=XM_023249906.1
A2 30249142 30249376
refGene_gn=CADPS;refGene_tx=XM_019823806.2;refGene_gn=CADPS;refGene_tx
=XM_019823812.2;refGene_gn=CADPS;refGene_tx=XM_019823818.2;refGene_gn=
CADPS;refGene_tx=XM_019823827.2;refGene_gn=CADPS;refGene_tx=XM_0198238
32.2;refGene_gn=CADPS;refGene_tx=XM_019823837.2;refGene_gn=CADPS;refGe

ne_tx=XM_019823841.2;refGene_gn=CADPS;refGene_tx=XM_019823845.2;refGene_gn=CADPS;refGene_tx=XM_019823856.2;refGene_gn=CADPS;refGene_tx=XM_019823857.2;refGene_gn=CADPS;refGene_tx=XM_019823865.2;refGene_gn=CADPS;refGene_tx=XM_019823869.2;refGene_gn=CADPS;refGene_tx=XM_019823872.2;refGene_gn=CADPS;refGene_tx=XM_019823877.2;refGene_gn=CADPS;refGene_tx=XM_019823879.2;refGene_gn=CADPS;refGene_tx=XM_023249901.1;refGene_gn=CADPS;refGene_tx=XM_023249902.1;refGene_gn=CADPS;refGene_tx=XM_023249903.1;refGene_gn=CADPS;refGene_tx=XM_023249904.1;refGene_gn=CADPS;refGene_tx=XM_023249905.1;refGene_gn=CADPS;refGene_tx=XM_023249906.1
A2 30266390 30266471
refGene_gn=CADPS;refGene_tx=XM_019823806.2;refGene_gn=CADPS;refGene_tx=XM_019823812.2;refGene_gn=CADPS;refGene_tx=XM_019823818.2;refGene_gn=CADPS;refGene_tx=XM_019823827.2;refGene_gn=CADPS;refGene_tx=XM_019823832.2;refGene_gn=CADPS;refGene_tx=XM_019823837.2;refGene_gn=CADPS;refGene_tx=XM_019823841.2;refGene_gn=CADPS;refGene_tx=XM_019823845.2;refGene_gn=CADPS;refGene_tx=XM_019823856.2;refGene_gn=CADPS;refGene_tx=XM_019823857.2;refGene_gn=CADPS;refGene_tx=XM_019823865.2;refGene_gn=CADPS;refGene_tx=XM_019823869.2;refGene_gn=CADPS;refGene_tx=XM_019823872.2;refGene_gn=CADPS;refGene_tx=XM_019823877.2;refGene_gn=CADPS;refGene_tx=XM_019823879.2;refGene_gn=CADPS;refGene_tx=XM_023249901.1;refGene_gn=CADPS;refGene_tx=XM_023249902.1;refGene_gn=CADPS;refGene_tx=XM_023249903.1;refGene_gn=CADPS;refGene_tx=XM_023249904.1;refGene_gn=CADPS;refGene_tx=XM_023249905.1;refGene_gn=CADPS;refGene_tx=XM_023249906.1
A2 30359705 30360038
refGene_gn=CADPS;refGene_tx=XM_019823806.2;refGene_gn=CADPS;refGene_tx=XM_019823812.2;refGene_gn=CADPS;refGene_tx=XM_019823818.2;refGene_gn=CADPS;refGene_tx=XM_019823827.2;refGene_gn=CADPS;refGene_tx=XM_019823832.2;refGene_gn=CADPS;refGene_tx=XM_019823837.2;refGene_gn=CADPS;refGene_tx=XM_019823841.2;refGene_gn=CADPS;refGene_tx=XM_019823845.2;refGene_gn=CADPS;refGene_tx=XM_019823856.2;refGene_gn=CADPS;refGene_tx=XM_019823857.2;refGene_gn=CADPS;refGene_tx=XM_019823865.2;refGene_gn=CADPS;refGene_tx=XM_019823869.2;refGene_gn=CADPS;refGene_tx=XM_019823872.2;refGene_gn=CADPS;refGene_tx=XM_019823877.2;refGene_gn=CADPS;refGene_tx=XM_019823879.2;refGene_gn=CADPS;refGene_tx=XM_023249901.1;refGene_gn=CADPS;refGene_tx=XM_023249902.1;refGene_gn=CADPS;refGene_tx=XM_023249903.1;refGene_gn=CADPS;refGene_tx=XM_023249904.1;refGene_gn=CADPS;refGene_tx=XM_023249905.1;refGene_gn=CADPS;refGene_tx=XM_023249906.1
A2 30373314 30373428
refGene_gn=CADPS;refGene_tx=XM_019823806.2;refGene_gn=CADPS;refGene_tx=XM_019823812.2;refGene_gn=CADPS;refGene_tx=XM_019823818.2;refGene_gn=CADPS;refGene_tx=XM_019823827.2;refGene_gn=CADPS;refGene_tx=XM_019823832.2;refGene_gn=CADPS;refGene_tx=XM_019823837.2;refGene_gn=CADPS;refGene_tx=XM_019823841.2;refGene_gn=CADPS;refGene_tx=XM_019823845.2;refGene_gn=CADPS;refGene_tx=XM_019823856.2;refGene_gn=CADPS;refGene_tx=XM_019823857.2;refGene_gn=CADPS;refGene_tx=XM_019823865.2;refGene_gn=CADPS;refGene_tx=XM_019823869.2;refGene_gn=CADPS;refGene_tx=XM_019823872.2;refGene_gn=CADPS;refGene_tx=XM_019823877.2;refGene_gn=CADPS;refGene_tx=XM_019823879.2;refGene_gn=CADPS;refGene_tx=XM_023249901.1;refGene_gn=CADPS;refGene_tx=XM_023249902.1;refGene_gn=CADPS;refGene_tx=XM_023249903.1;refGene_gn=CADPS;refGene_tx=XM_023249904.1;refGene_gn=CADPS;refGen

e_tx=XM_023249905.1;refGene_gn=CADPS;refGene_tx=XM_023249906.1
A2 30478655 30479105
refGene_gn=CADPS;refGene_tx=XM_019823806.2;refGene_gn=CADPS;refGene_tx=XM_019823812.2;refGene_gn=CADPS;refGene_tx=XM_019823818.2;refGene_gn=CADPS;refGene_tx=XM_019823827.2;refGene_gn=CADPS;refGene_tx=XM_019823832.2;refGene_gn=CADPS;refGene_tx=XM_019823837.2;refGene_gn=CADPS;refGene_tx=XM_019823841.2;refGene_gn=CADPS;refGene_tx=XM_019823845.2;refGene_gn=CADPS;refGene_tx=XM_019823856.2;refGene_gn=CADPS;refGene_tx=XM_019823857.2;refGene_gn=CADPS;refGene_tx=XM_019823865.2;refGene_gn=CADPS;refGene_tx=XM_019823869.2;refGene_gn=CADPS;refGene_tx=XM_019823872.2;refGene_gn=CADPS;refGene_tx=XM_019823877.2;refGene_gn=CADPS;refGene_tx=XM_019823879.2;refGene_gn=CADPS;refGene_tx=XM_023249901.1;refGene_gn=CADPS;refGene_tx=XM_023249902.1;refGene_gn=CADPS;refGene_tx=XM_023249903.1;refGene_gn=CADPS;refGene_tx=XM_023249904.1;refGene_gn=CADPS;refGene_tx=XM_023249905.1;refGene_gn=CADPS;refGene_tx=XM_023249906.1
A2 30851440 30851458
refGene_gn=SYNPR;refGene_tx=XM_023249907.1;refGene_gn=SYNPR;refGene_tx=XM_023249908.1
A2 30851565 30851631
refGene_gn=SYNPR;refGene_tx=XM_023249907.1;refGene_gn=SYNPR;refGene_tx=XM_023249908.1
A2 31007555 31007579 refGene_gn=SYNPR;refGene_tx=XM_011280148.2
A2 31043847 31043972
refGene_gn=SYNPR;refGene_tx=XM_011280148.2;refGene_gn=SYNPR;refGene_tx=XM_023249907.1;refGene_gn=SYNPR;refGene_tx=XM_023249908.1
A2 31115267 31115466
refGene_gn=SYNPR;refGene_tx=XM_011280148.2;refGene_gn=SYNPR;refGene_tx=XM_023249907.1;refGene_gn=SYNPR;refGene_tx=XM_023249908.1
A2 31156278 31156470
refGene_gn=SYNPR;refGene_tx=XM_011280148.2;refGene_gn=SYNPR;refGene_tx=XM_023249907.1;refGene_gn=SYNPR;refGene_tx=XM_023249908.1
A2 31163427 31163685
refGene_gn=SYNPR;refGene_tx=XM_011280148.2;refGene_gn=SYNPR;refGene_tx=XM_023249907.1;refGene_gn=SYNPR;refGene_tx=XM_023249908.1
A2 31191759 31191869 refGene_gn=SNTN;refGene_tx=XM_003982395.1
A2 31193849 31193884 refGene_gn=SNTN;refGene_tx=XM_003982395.1
A2 31201236 31201376 refGene_gn=SNTN;refGene_tx=XM_003982395.1
A2 31206238 31206397 refGene_gn=SNTN;refGene_tx=XM_003982395.1
A2 31349864 31349932 refGene_gn=THOC7;refGene_tx=XM_023249910.1
A2 31351375 31351445 refGene_gn=THOC7;refGene_tx=XM_023249910.1
A2 31351527 31351594 refGene_gn=THOC7;refGene_tx=XM_023249910.1
A2 31353185 31353243 refGene_gn=THOC7;refGene_tx=XM_023249910.1
A2 31355577 31355664 refGene_gn=THOC7;refGene_tx=XM_023249910.1
A2 31356213 31356341 refGene_gn=THOC7;refGene_tx=XM_023249910.1
A2 31357270 31357388 refGene_gn=THOC7;refGene_tx=XM_023249910.1
A2 31373307 31373326 refGene_gn=THOC7;refGene_tx=XM_023249910.1
A2 31422006 31422262
refGene_gn=ATXN7;refGene_tx=XM_023249911.1;refGene_gn=ATXN7;refGene_tx=XM_023249912.1;refGene_gn=ATXN7;refGene_tx=XM_023249913.1;refGene_gn=ATXN7;refGene_tx=XM_023249916.1

A2 31422657 31422996
refGene_gn=ATXN7;refGene_tx=XM_023249911.1;refGene_gn=ATXN7;refGene_tx=XM_023249912.1;refGene_gn=ATXN7;refGene_tx=XM_023249913.1;refGene_gn=ATXN7;refGene_tx=XM_023249914.1;refGene_gn=ATXN7;refGene_tx=XM_023249915.1;refGene_gn=ATXN7;refGene_tx=XM_023249916.1
A2 31423225 31423294
refGene_gn=ATXN7;refGene_tx=XM_023249911.1;refGene_gn=ATXN7;refGene_tx=XM_023249912.1;refGene_gn=ATXN7;refGene_tx=XM_023249913.1;refGene_gn=ATXN7;refGene_tx=XM_023249914.1;refGene_gn=ATXN7;refGene_tx=XM_023249915.1;refGene_gn=ATXN7;refGene_tx=XM_023249916.1
A2 31465180 31465285
refGene_gn=ATXN7;refGene_tx=XM_023249911.1;refGene_gn=ATXN7;refGene_tx=XM_023249912.1;refGene_gn=ATXN7;refGene_tx=XM_023249913.1;refGene_gn=ATXN7;refGene_tx=XM_023249914.1;refGene_gn=ATXN7;refGene_tx=XM_023249915.1;refGene_gn=ATXN7;refGene_tx=XM_023249916.1
A2 31495071 31495324
refGene_gn=ATXN7;refGene_tx=XM_023249911.1;refGene_gn=ATXN7;refGene_tx=XM_023249912.1;refGene_gn=ATXN7;refGene_tx=XM_023249913.1;refGene_gn=ATXN7;refGene_tx=XM_023249914.1;refGene_gn=ATXN7;refGene_tx=XM_023249915.1;refGene_gn=ATXN7;refGene_tx=XM_023249916.1
A2 31497692 31497952
refGene_gn=ATXN7;refGene_tx=XM_023249911.1;refGene_gn=ATXN7;refGene_tx=XM_023249912.1;refGene_gn=ATXN7;refGene_tx=XM_023249913.1;refGene_gn=ATXN7;refGene_tx=XM_023249914.1;refGene_gn=ATXN7;refGene_tx=XM_023249915.1;refGene_gn=ATXN7;refGene_tx=XM_023249916.1
A2 31498426 31498509
refGene_gn=ATXN7;refGene_tx=XM_023249911.1;refGene_gn=ATXN7;refGene_tx=XM_023249912.1;refGene_gn=ATXN7;refGene_tx=XM_023249913.1;refGene_gn=ATXN7;refGene_tx=XM_023249914.1;refGene_gn=ATXN7;refGene_tx=XM_023249915.1;refGene_gn=ATXN7;refGene_tx=XM_023249916.1
A2 31503436 31503702
refGene_gn=ATXN7;refGene_tx=XM_023249911.1;refGene_gn=ATXN7;refGene_tx=XM_023249912.1;refGene_gn=ATXN7;refGene_tx=XM_023249913.1;refGene_gn=ATXN7;refGene_tx=XM_023249914.1;refGene_gn=ATXN7;refGene_tx=XM_023249915.1;refGene_gn=ATXN7;refGene_tx=XM_023249916.1
A2 31505151 31505350
refGene_gn=ATXN7;refGene_tx=XM_023249911.1;refGene_gn=ATXN7;refGene_tx=XM_023249912.1;refGene_gn=ATXN7;refGene_tx=XM_023249913.1;refGene_gn=ATXN7;refGene_tx=XM_023249914.1;refGene_gn=ATXN7;refGene_tx=XM_023249915.1;refGene_gn=ATXN7;refGene_tx=XM_023249916.1
A2 31505723 31505845
refGene_gn=ATXN7;refGene_tx=XM_023249911.1;refGene_gn=ATXN7;refGene_tx=XM_023249912.1;refGene_gn=ATXN7;refGene_tx=XM_023249913.1;refGene_gn=ATXN7;refGene_tx=XM_023249914.1;refGene_gn=ATXN7;refGene_tx=XM_023249915.1;refGene_gn=ATXN7;refGene_tx=XM_023249916.1
A2 31510169 31511186
refGene_gn=ATXN7;refGene_tx=XM_023249911.1;refGene_gn=ATXN7;refGene_tx=XM_023249912.1;refGene_gn=ATXN7;refGene_tx=XM_023249913.1;refGene_gn=ATXN7;refGene_tx=XM_023249914.1;refGene_gn=ATXN7;refGene_tx=XM_023249915.1;refGene_gn=ATXN7;refGene_tx=XM_023249916.1

A2 31511985 31512052 refGene_gn=ATXN7;refGene_tx=XM_023249916.1
A2 31513723 31513833 refGene_gn=ATXN7;refGene_tx=XM_023249916.1
A2 31526283 31526380 refGene_gn=PSMD6;refGene_tx=XM_003982397.5
A2 31526493 31526571 refGene_gn=PSMD6;refGene_tx=XM_003982397.5
A2 31531128 31531297 refGene_gn=PSMD6;refGene_tx=XM_003982397.5
A2 31536364 31536473 refGene_gn=PSMD6;refGene_tx=XM_003982397.5
A2 31536579 31536799 refGene_gn=PSMD6;refGene_tx=XM_003982397.5
A2 31537050 31537196 refGene_gn=PSMD6;refGene_tx=XM_003982397.5
A2 31540692 31540898 refGene_gn=PSMD6;refGene_tx=XM_003982397.5
A2 31541622 31541767 refGene_gn=PSMD6;refGene_tx=XM_003982397.5
A2 31629099 31629974
refGene_gn=PRICKLE2;refGene_tx=XM_011280150.3;refGene_gn=PRICKLE2;refGene_tx=XM_011280151.3;refGene_gn=PRICKLE2;refGene_tx=XM_019823948.2
A2 31667564 31668440
refGene_gn=PRICKLE2;refGene_tx=XM_011280150.3;refGene_gn=PRICKLE2;refGene_tx=XM_011280151.3;refGene_gn=PRICKLE2;refGene_tx=XM_019823948.2
A2 31673461 31673648
refGene_gn=PRICKLE2;refGene_tx=XM_011280150.3;refGene_gn=PRICKLE2;refGene_tx=XM_011280151.3;refGene_gn=PRICKLE2;refGene_tx=XM_019823948.2
A2 31676692 31676896
refGene_gn=PRICKLE2;refGene_tx=XM_011280150.3;refGene_gn=PRICKLE2;refGene_tx=XM_011280151.3;refGene_gn=PRICKLE2;refGene_tx=XM_019823948.2
A2 31680941 31681079
refGene_gn=PRICKLE2;refGene_tx=XM_011280150.3;refGene_gn=PRICKLE2;refGene_tx=XM_011280151.3;refGene_gn=PRICKLE2;refGene_tx=XM_019823948.2
A2 31684000 31684114
refGene_gn=PRICKLE2;refGene_tx=XM_011280150.3;refGene_gn=PRICKLE2;refGene_tx=XM_011280151.3;refGene_gn=PRICKLE2;refGene_tx=XM_019823948.2
A2 31715687 31715871
refGene_gn=PRICKLE2;refGene_tx=XM_011280150.3;refGene_gn=PRICKLE2;refGene_tx=XM_011280151.3;refGene_gn=PRICKLE2;refGene_tx=XM_019823948.2
A2 31954256 31954342 refGene_gn=PRICKLE2;refGene_tx=XM_019823948.2
A2 31955114 31955156 refGene_gn=PRICKLE2;refGene_tx=XM_019823948.2
A2 32024840 32024930
refGene_gn=ADAMTS9;refGene_tx=XM_003982398.5;refGene_gn=ADAMTS9;refGene_tx=XM_006928919.4;refGene_gn=ADAMTS9;refGene_tx=XM_011280154.2
A2 32035471 32035576
refGene_gn=ADAMTS9;refGene_tx=XM_003982398.5;refGene_gn=ADAMTS9;refGene_tx=XM_006928919.4;refGene_gn=ADAMTS9;refGene_tx=XM_011280154.2
A2 32040382 32040474
refGene_gn=ADAMTS9;refGene_tx=XM_003982398.5;refGene_gn=ADAMTS9;refGene_tx=XM_006928919.4;refGene_gn=ADAMTS9;refGene_tx=XM_011280154.2
A2 32042272 32042406
refGene_gn=ADAMTS9;refGene_tx=XM_003982398.5;refGene_gn=ADAMTS9;refGene_tx=XM_006928919.4;refGene_gn=ADAMTS9;refGene_tx=XM_011280154.2
A2 32042498 32042593
refGene_gn=ADAMTS9;refGene_tx=XM_003982398.5;refGene_gn=ADAMTS9;refGene_tx=XM_006928919.4;refGene_gn=ADAMTS9;refGene_tx=XM_011280154.2
A2 32042708 32042803
refGene_gn=ADAMTS9;refGene_tx=XM_003982398.5;refGene_gn=ADAMTS9;refGen

e_tx=XM_006928919.4;refGene_gn=ADAMTS9;refGene_tx=XM_011280154.2
A2 32043343 32043476
refGene_gn=ADAMTS9;refGene_tx=XM_003982398.5;refGene_gn=ADAMTS9;refGene_tx=XM_006928919.4;refGene_gn=ADAMTS9;refGene_tx=XM_011280154.2
A2 32044535 32044730
refGene_gn=ADAMTS9;refGene_tx=XM_003982398.5;refGene_gn=ADAMTS9;refGene_tx=XM_006928919.4;refGene_gn=ADAMTS9;refGene_tx=XM_011280154.2
A2 32048628 32048802
refGene_gn=ADAMTS9;refGene_tx=XM_003982398.5;refGene_gn=ADAMTS9;refGene_tx=XM_006928919.4;refGene_gn=ADAMTS9;refGene_tx=XM_011280154.2
A2 32060757 32060931
refGene_gn=ADAMTS9;refGene_tx=XM_003982398.5;refGene_gn=ADAMTS9;refGene_tx=XM_006928919.4;refGene_gn=ADAMTS9;refGene_tx=XM_011280154.2
A2 32067870 32068038
refGene_gn=ADAMTS9;refGene_tx=XM_003982398.5;refGene_gn=ADAMTS9;refGene_tx=XM_006928919.4
A2 32093410 32093587
refGene_gn=ADAMTS9;refGene_tx=XM_003982398.5;refGene_gn=ADAMTS9;refGene_tx=XM_006928919.4;refGene_gn=ADAMTS9;refGene_tx=XM_011280154.2
A2 32095687 32095849
refGene_gn=ADAMTS9;refGene_tx=XM_003982398.5;refGene_gn=ADAMTS9;refGene_tx=XM_006928919.4;refGene_gn=ADAMTS9;refGene_tx=XM_011280154.2
A2 32101706 32101976
refGene_gn=ADAMTS9;refGene_tx=XM_003982398.5;refGene_gn=ADAMTS9;refGene_tx=XM_006928919.4;refGene_gn=ADAMTS9;refGene_tx=XM_011280154.2
A2 32103780 32103948
refGene_gn=ADAMTS9;refGene_tx=XM_003982398.5;refGene_gn=ADAMTS9;refGene_tx=XM_006928919.4;refGene_gn=ADAMTS9;refGene_tx=XM_011280154.2
A2 32104085 32104190
refGene_gn=ADAMTS9;refGene_tx=XM_003982398.5;refGene_gn=ADAMTS9;refGene_tx=XM_006928919.4;refGene_gn=ADAMTS9;refGene_tx=XM_011280154.2
A2 32107075 32107195
refGene_gn=ADAMTS9;refGene_tx=XM_003982398.5;refGene_gn=ADAMTS9;refGene_tx=XM_006928919.4;refGene_gn=ADAMTS9;refGene_tx=XM_011280154.2
A2 32110227 32110392
refGene_gn=ADAMTS9;refGene_tx=XM_003982398.5;refGene_gn=ADAMTS9;refGene_tx=XM_006928919.4;refGene_gn=ADAMTS9;refGene_tx=XM_011280154.2
A2 32112661 32112826
refGene_gn=ADAMTS9;refGene_tx=XM_003982398.5;refGene_gn=ADAMTS9;refGene_tx=XM_006928919.4;refGene_gn=ADAMTS9;refGene_tx=XM_011280154.2
A2 32113318 32113529
refGene_gn=ADAMTS9;refGene_tx=XM_003982398.5;refGene_gn=ADAMTS9;refGene_tx=XM_006928919.4;refGene_gn=ADAMTS9;refGene_tx=XM_011280154.2
A2 32119309 32119436
refGene_gn=ADAMTS9;refGene_tx=XM_003982398.5;refGene_gn=ADAMTS9;refGene_tx=XM_006928919.4;refGene_gn=ADAMTS9;refGene_tx=XM_011280154.2
A2 32120049 32120179
refGene_gn=ADAMTS9;refGene_tx=XM_003982398.5;refGene_gn=ADAMTS9;refGene_tx=XM_006928919.4;refGene_gn=ADAMTS9;refGene_tx=XM_011280154.2
A2 32120270 32120437

refGene_gn=ADAMTS9;refGene_tx=XM_003982398.5;refGene_gn=ADAMTS9;refGene_tx=XM_006928919.4;refGene_gn=ADAMTS9;refGene_tx=XM_011280154.2
A2 32128715 32128811
refGene_gn=ADAMTS9;refGene_tx=XM_003982398.5;refGene_gn=ADAMTS9;refGene_tx=XM_006928919.4;refGene_gn=ADAMTS9;refGene_tx=XM_011280154.2
A2 32129061 32129179
refGene_gn=ADAMTS9;refGene_tx=XM_003982398.5;refGene_gn=ADAMTS9;refGene_tx=XM_006928919.4;refGene_gn=ADAMTS9;refGene_tx=XM_011280154.2
A2 32130721 32130858
refGene_gn=ADAMTS9;refGene_tx=XM_003982398.5;refGene_gn=ADAMTS9;refGene_tx=XM_006928919.4;refGene_gn=ADAMTS9;refGene_tx=XM_011280154.2
A2 32130952 32131134
refGene_gn=ADAMTS9;refGene_tx=XM_003982398.5;refGene_gn=ADAMTS9;refGene_tx=XM_006928919.4;refGene_gn=ADAMTS9;refGene_tx=XM_011280154.2
A2 32139644 32139790
refGene_gn=ADAMTS9;refGene_tx=XM_003982398.5;refGene_gn=ADAMTS9;refGene_tx=XM_006928919.4;refGene_gn=ADAMTS9;refGene_tx=XM_011280154.2
A2 32145395 32145500
refGene_gn=ADAMTS9;refGene_tx=XM_003982398.5;refGene_gn=ADAMTS9;refGene_tx=XM_006928919.4;refGene_gn=ADAMTS9;refGene_tx=XM_011280154.2
A2 32146705 32146847
refGene_gn=ADAMTS9;refGene_tx=XM_003982398.5;refGene_gn=ADAMTS9;refGene_tx=XM_006928919.4;refGene_gn=ADAMTS9;refGene_tx=XM_011280154.2
A2 32148108 32148255
refGene_gn=ADAMTS9;refGene_tx=XM_003982398.5;refGene_gn=ADAMTS9;refGene_tx=XM_006928919.4;refGene_gn=ADAMTS9;refGene_tx=XM_011280154.2
A2 32153237 32153343
refGene_gn=ADAMTS9;refGene_tx=XM_003982398.5;refGene_gn=ADAMTS9;refGene_tx=XM_006928919.4;refGene_gn=ADAMTS9;refGene_tx=XM_011280154.2
A2 32153472 32153513
refGene_gn=ADAMTS9;refGene_tx=XM_003982398.5;refGene_gn=ADAMTS9;refGene_tx=XM_006928919.4;refGene_gn=ADAMTS9;refGene_tx=XM_011280154.2
A2 32154218 32154331
refGene_gn=ADAMTS9;refGene_tx=XM_003982398.5;refGene_gn=ADAMTS9;refGene_tx=XM_006928919.4;refGene_gn=ADAMTS9;refGene_tx=XM_011280154.2
A2 32154432 32154516
refGene_gn=ADAMTS9;refGene_tx=XM_003982398.5;refGene_gn=ADAMTS9;refGene_tx=XM_011280154.2
A2 32156986 32157276
refGene_gn=ADAMTS9;refGene_tx=XM_003982398.5;refGene_gn=ADAMTS9;refGene_tx=XM_006928919.4;refGene_gn=ADAMTS9;refGene_tx=XM_011280154.2
A2 32178862 32179025
refGene_gn=ADAMTS9;refGene_tx=XM_003982398.5;refGene_gn=ADAMTS9;refGene_tx=XM_006928919.4;refGene_gn=ADAMTS9;refGene_tx=XM_011280154.2
A2 32182310 32182679
refGene_gn=LOC102899165;refGene_tx=XM_019823988.2;refGene_gn=LOC102899165;refGene_tx=XM_019823992.2;refGene_gn=LOC102899165;refGene_tx=XM_019823997.2;refGene_gn=LOC102899165;refGene_tx=XM_019823998.2;refGene_gn=LOC102899165;refGene_tx=XM_019824011.2;refGene_gn=LOC102899165;refGene_tx=XM_019824027.2;refGene_gn=LOC102899165;refGene_tx=XM_019824033.2

A2 32183903 32184316
refGene_gn=ADAMTS9;refGene_tx=XM_003982398.5;refGene_gn=ADAMTS9;refGene_tx=XM_006928919.4;refGene_gn=ADAMTS9;refGene_tx=XM_011280154.2
A2 32184927 32185042
refGene_gn=ADAMTS9;refGene_tx=XM_003982398.5;refGene_gn=ADAMTS9;refGene_tx=XM_006928919.4;refGene_gn=ADAMTS9;refGene_tx=XM_011280154.2
A2 32285772 32285864
refGene_gn=LOC102899165;refGene_tx=XM_019824033.2
A2 32349899 32350002
refGene_gn=LOC102899165;refGene_tx=XM_019823988.2;refGene_gn=LOC102899165;refGene_tx=XM_019823992.2;refGene_gn=LOC102899165;refGene_tx=XM_019823997.2;refGene_gn=LOC102899165;refGene_tx=XM_019823998.2;refGene_gn=LOC102899165;refGene_tx=XM_019824011.2;refGene_gn=LOC102899165;refGene_tx=XM_019824027.2;refGene_gn=LOC102899165;refGene_tx=XM_019824033.2
A2 32353931 32354002
refGene_gn=LOC102899165;refGene_tx=XM_019823998.2
A2 32354616 32354633
refGene_gn=LOC102899165;refGene_tx=XM_019824027.2
A2 32394283 32394399
refGene_gn=LOC102899165;refGene_tx=XM_019823992.2
A2 32407237 32407342
refGene_gn=LOC102899165;refGene_tx=XM_019823988.2;refGene_gn=LOC102899165;refGene_tx=XM_019824033.2
A2 32407703 32407810
refGene_gn=LOC102899165;refGene_tx=XM_019823988.2;refGene_gn=LOC102899165;refGene_tx=XM_019823997.2;refGene_gn=LOC102899165;refGene_tx=XM_019824033.2
A2 32496336 32496368
refGene_gn=LOC102899165;refGene_tx=XM_019824011.2
A2 32829961 32830695
refGene_gn=MAGI1;refGene_tx=XM_023249917.1;refGene_gn=MAGI1;refGene_tx=XM_023249918.1;refGene_gn=MAGI1;refGene_tx=XM_023249919.1;refGene_gn=MAGI1;refGene_tx=XM_023249920.1;refGene_gn=MAGI1;refGene_tx=XM_023249921.1;refGene_gn=MAGI1;refGene_tx=XM_023249922.1;refGene_gn=MAGI1;refGene_tx=XM_023249923.1;refGene_gn=MAGI1;refGene_tx=XM_023249924.1;refGene_gn=MAGI1;refGene_tx=XM_023249925.1;refGene_gn=MAGI1;refGene_tx=XM_023249926.1;refGene_gn=MAGI1;refGene_tx=XM_023249927.1;refGene_gn=MAGI1;refGene_tx=XM_023249928.1;refGene_gn=MAGI1;refGene_tx=XM_023249929.1;refGene_gn=MAGI1;refGene_tx=XM_023249931.1;refGene_gn=MAGI1;refGene_tx=XM_023249932.1;refGene_gn=MAGI1;refGene_tx=XM_023249933.1
A2 32832593 32832682
refGene_gn=MAGI1;refGene_tx=XM_023249927.1;refGene_gn=MAGI1;refGene_tx=XM_023249928.1;refGene_gn=MAGI1;refGene_tx=XM_023249929.1;refGene_gn=MAGI1;refGene_tx=XM_023249932.1
A2 32834698 32834887
refGene_gn=MAGI1;refGene_tx=XM_023249917.1;refGene_gn=MAGI1;refGene_tx=XM_023249918.1;refGene_gn=MAGI1;refGene_tx=XM_023249919.1;refGene_gn=MAGI1;refGene_tx=XM_023249920.1;refGene_gn=MAGI1;refGene_tx=XM_023249921.1;refGene_gn=MAGI1;refGene_tx=XM_023249922.1;refGene_gn=MAGI1;refGene_tx=XM_023249923.1;refGene_gn=MAGI1;refGene_tx=XM_023249924.1;refGen

e_gn=MAGI1;refGene_tx=XM_023249925.1;refGene_gn=MAGI1;refGene_tx=XM_02
3249926.1;refGene_gn=MAGI1;refGene_tx=XM_023249927.1;refGene_gn=MAGI1;
refGene_tx=XM_023249928.1;refGene_gn=MAGI1;refGene_tx=XM_023249929.1;r
efGene_gn=MAGI1;refGene_tx=XM_023249930.1;refGene_gn=MAGI1;refGene_tx=
XM_023249931.1;refGene_gn=MAGI1;refGene_tx=XM_023249932.1;refGene_gn=M
AGI1;refGene_tx=XM_023249933.1
A2 32837334 32837478
refGene_gn=MAGI1;refGene_tx=XM_023249917.1;refGene_gn=MAGI1;refGene_tx
=XM_023249918.1;refGene_gn=MAGI1;refGene_tx=XM_023249919.1;refGene_gn=
MAGI1;refGene_tx=XM_023249920.1;refGene_gn=MAGI1;refGene_tx=XM_0232499
21.1;refGene_gn=MAGI1;refGene_tx=XM_023249922.1;refGene_gn=MAGI1;refGe
ne_tx=XM_023249923.1;refGene_gn=MAGI1;refGene_tx=XM_023249924.1;refGen
e_gn=MAGI1;refGene_tx=XM_023249925.1;refGene_gn=MAGI1;refGene_tx=XM_02
3249926.1;refGene_gn=MAGI1;refGene_tx=XM_023249927.1;refGene_gn=MAGI1;
refGene_tx=XM_023249928.1;refGene_gn=MAGI1;refGene_tx=XM_023249929.1;r
efGene_gn=MAGI1;refGene_tx=XM_023249930.1;refGene_gn=MAGI1;refGene_tx=
XM_023249931.1;refGene_gn=MAGI1;refGene_tx=XM_023249932.1;refGene_gn=M
AGI1;refGene_tx=XM_023249933.1
A2 32838369 32838430
refGene_gn=MAGI1;refGene_tx=XM_023249917.1;refGene_gn=MAGI1;refGene_tx
=XM_023249918.1;refGene_gn=MAGI1;refGene_tx=XM_023249919.1;refGene_gn=
MAGI1;refGene_tx=XM_023249920.1;refGene_gn=MAGI1;refGene_tx=XM_0232499
21.1;refGene_gn=MAGI1;refGene_tx=XM_023249922.1;refGene_gn=MAGI1;refGe
ne_tx=XM_023249923.1;refGene_gn=MAGI1;refGene_tx=XM_023249924.1;refGen
e_gn=MAGI1;refGene_tx=XM_023249925.1;refGene_gn=MAGI1;refGene_tx=XM_02
3249926.1;refGene_gn=MAGI1;refGene_tx=XM_023249927.1;refGene_gn=MAGI1;
refGene_tx=XM_023249928.1;refGene_gn=MAGI1;refGene_tx=XM_023249929.1;r
efGene_gn=MAGI1;refGene_tx=XM_023249930.1;refGene_gn=MAGI1;refGene_tx=
XM_023249931.1;refGene_gn=MAGI1;refGene_tx=XM_023249932.1;refGene_gn=M
AGI1;refGene_tx=XM_023249933.1
A2 32838575 32838669
refGene_gn=MAGI1;refGene_tx=XM_023249917.1;refGene_gn=MAGI1;refGene_tx
=XM_023249918.1;refGene_gn=MAGI1;refGene_tx=XM_023249919.1;refGene_gn=
MAGI1;refGene_tx=XM_023249920.1;refGene_gn=MAGI1;refGene_tx=XM_0232499
21.1;refGene_gn=MAGI1;refGene_tx=XM_023249922.1;refGene_gn=MAGI1;refGe
ne_tx=XM_023249923.1;refGene_gn=MAGI1;refGene_tx=XM_023249924.1;refGen
e_gn=MAGI1;refGene_tx=XM_023249925.1;refGene_gn=MAGI1;refGene_tx=XM_02
3249926.1;refGene_gn=MAGI1;refGene_tx=XM_023249927.1;refGene_gn=MAGI1;
refGene_tx=XM_023249928.1;refGene_gn=MAGI1;refGene_tx=XM_023249929.1;r
efGene_gn=MAGI1;refGene_tx=XM_023249930.1;refGene_gn=MAGI1;refGene_tx=
XM_023249931.1;refGene_gn=MAGI1;refGene_tx=XM_023249932.1;refGene_gn=M
AGI1;refGene_tx=XM_023249933.1
A2 32846578 32846782
refGene_gn=MAGI1;refGene_tx=XM_023249917.1;refGene_gn=MAGI1;refGene_tx
=XM_023249918.1;refGene_gn=MAGI1;refGene_tx=XM_023249919.1;refGene_gn=
MAGI1;refGene_tx=XM_023249920.1;refGene_gn=MAGI1;refGene_tx=XM_0232499
21.1;refGene_gn=MAGI1;refGene_tx=XM_023249922.1;refGene_gn=MAGI1;refGe
ne_tx=XM_023249923.1;refGene_gn=MAGI1;refGene_tx=XM_023249924.1;refGen
e_gn=MAGI1;refGene_tx=XM_023249925.1;refGene_gn=MAGI1;refGene_tx=XM_02
3249927.1;refGene_gn=MAGI1;refGene_tx=XM_023249928.1;refGene_gn=MAGI1;

refGene_tx=XM_023249929.1;refGene_gn=MAGI1;refGene_tx=XM_023249930.1;refGene_gn=MAGI1;refGene_tx=XM_023249931.1;refGene_gn=MAGI1;refGene_tx=XM_023249932.1;refGene_gn=MAGI1;refGene_tx=XM_023249933.1
A2 32849756 32850050
refGene_gn=MAGI1;refGene_tx=XM_023249917.1;refGene_gn=MAGI1;refGene_tx=XM_023249918.1;refGene_gn=MAGI1;refGene_tx=XM_023249919.1;refGene_gn=MAGI1;refGene_tx=XM_023249920.1;refGene_gn=MAGI1;refGene_tx=XM_023249921.1;refGene_gn=MAGI1;refGene_tx=XM_023249922.1;refGene_gn=MAGI1;refGene_tx=XM_023249923.1;refGene_gn=MAGI1;refGene_tx=XM_023249924.1;refGene_gn=MAGI1;refGene_tx=XM_023249925.1;refGene_gn=MAGI1;refGene_tx=XM_023249926.1;refGene_gn=MAGI1;refGene_tx=XM_023249927.1;refGene_gn=MAGI1;refGene_tx=XM_023249928.1;refGene_gn=MAGI1;refGene_tx=XM_023249929.1;refGene_gn=MAGI1;refGene_tx=XM_023249930.1;refGene_gn=MAGI1;refGene_tx=XM_023249931.1;refGene_gn=MAGI1;refGene_tx=XM_023249932.1;refGene_gn=MAGI1;refGene_tx=XM_023249933.1
A2 32852430 32852623
refGene_gn=MAGI1;refGene_tx=XM_023249917.1;refGene_gn=MAGI1;refGene_tx=XM_023249918.1;refGene_gn=MAGI1;refGene_tx=XM_023249919.1;refGene_gn=MAGI1;refGene_tx=XM_023249920.1;refGene_gn=MAGI1;refGene_tx=XM_023249921.1;refGene_gn=MAGI1;refGene_tx=XM_023249922.1;refGene_gn=MAGI1;refGene_tx=XM_023249923.1;refGene_gn=MAGI1;refGene_tx=XM_023249924.1;refGene_gn=MAGI1;refGene_tx=XM_023249925.1;refGene_gn=MAGI1;refGene_tx=XM_023249926.1;refGene_gn=MAGI1;refGene_tx=XM_023249927.1;refGene_gn=MAGI1;refGene_tx=XM_023249928.1;refGene_gn=MAGI1;refGene_tx=XM_023249929.1;refGene_gn=MAGI1;refGene_tx=XM_023249930.1;refGene_gn=MAGI1;refGene_tx=XM_023249931.1;refGene_gn=MAGI1;refGene_tx=XM_023249932.1;refGene_gn=MAGI1;refGene_tx=XM_023249933.1
A2 32854070 32854162
refGene_gn=MAGI1;refGene_tx=XM_023249917.1;refGene_gn=MAGI1;refGene_tx=XM_023249918.1;refGene_gn=MAGI1;refGene_tx=XM_023249919.1;refGene_gn=MAGI1;refGene_tx=XM_023249920.1;refGene_gn=MAGI1;refGene_tx=XM_023249921.1;refGene_gn=MAGI1;refGene_tx=XM_023249922.1;refGene_gn=MAGI1;refGene_tx=XM_023249923.1;refGene_gn=MAGI1;refGene_tx=XM_023249924.1;refGene_gn=MAGI1;refGene_tx=XM_023249925.1;refGene_gn=MAGI1;refGene_tx=XM_023249926.1;refGene_gn=MAGI1;refGene_tx=XM_023249927.1;refGene_gn=MAGI1;refGene_tx=XM_023249928.1;refGene_gn=MAGI1;refGene_tx=XM_023249929.1;refGene_gn=MAGI1;refGene_tx=XM_023249930.1;refGene_gn=MAGI1;refGene_tx=XM_023249931.1;refGene_gn=MAGI1;refGene_tx=XM_023249932.1;refGene_gn=MAGI1;refGene_tx=XM_023249933.1
A2 32856998 32857082
refGene_gn=MAGI1;refGene_tx=XM_023249917.1;refGene_gn=MAGI1;refGene_tx=XM_023249918.1;refGene_gn=MAGI1;refGene_tx=XM_023249919.1;refGene_gn=MAGI1;refGene_tx=XM_023249920.1;refGene_gn=MAGI1;refGene_tx=XM_023249921.1;refGene_gn=MAGI1;refGene_tx=XM_023249922.1;refGene_gn=MAGI1;refGene_tx=XM_023249923.1;refGene_gn=MAGI1;refGene_tx=XM_023249926.1;refGene_gn=MAGI1;refGene_tx=XM_023249927.1;refGene_gn=MAGI1;refGene_tx=XM_023249928.1;refGene_gn=MAGI1;refGene_tx=XM_023249929.1;refGene_gn=MAGI1;refGene_tx=XM_023249930.1;refGene_gn=MAGI1;refGene_tx=XM_023249932.1
A2 32860765 32860982
refGene_gn=MAGI1;refGene_tx=XM_023249917.1;refGene_gn=MAGI1;refGene_tx

=XM_023249918.1;refGene_gn=MAGI1;refGene_tx=XM_023249919.1;refGene_gn=MAGI1;refGene_tx=XM_023249920.1;refGene_gn=MAGI1;refGene_tx=XM_023249921.1;refGene_gn=MAGI1;refGene_tx=XM_023249922.1;refGene_gn=MAGI1;refGene_tx=XM_023249923.1;refGene_gn=MAGI1;refGene_tx=XM_023249924.1;refGene_gn=MAGI1;refGene_tx=XM_023249925.1;refGene_gn=MAGI1;refGene_tx=XM_023249926.1;refGene_gn=MAGI1;refGene_tx=XM_023249927.1;refGene_gn=MAGI1;refGene_tx=XM_023249928.1;refGene_gn=MAGI1;refGene_tx=XM_023249929.1;refGene_gn=MAGI1;refGene_tx=XM_023249930.1;refGene_gn=MAGI1;refGene_tx=XM_023249931.1;refGene_gn=MAGI1;refGene_tx=XM_023249932.1;refGene_gn=MAGI1;refGene_tx=XM_023249933.1

A2 32871461 32871493

refGene_gn=MAGI1;refGene_tx=XM_023249917.1;refGene_gn=MAGI1;refGene_tx=XM_023249918.1;refGene_gn=MAGI1;refGene_tx=XM_023249919.1;refGene_gn=MAGI1;refGene_tx=XM_023249920.1;refGene_gn=MAGI1;refGene_tx=XM_023249921.1;refGene_gn=MAGI1;refGene_tx=XM_023249922.1;refGene_gn=MAGI1;refGene_tx=XM_023249923.1;refGene_gn=MAGI1;refGene_tx=XM_023249924.1;refGene_gn=MAGI1;refGene_tx=XM_023249925.1;refGene_gn=MAGI1;refGene_tx=XM_023249926.1;refGene_gn=MAGI1;refGene_tx=XM_023249927.1;refGene_gn=MAGI1;refGene_tx=XM_023249928.1;refGene_gn=MAGI1;refGene_tx=XM_023249929.1;refGene_gn=MAGI1;refGene_tx=XM_023249930.1;refGene_gn=MAGI1;refGene_tx=XM_023249931.1;refGene_gn=MAGI1;refGene_tx=XM_023249932.1;refGene_gn=MAGI1;refGene_tx=XM_023249933.1

A2 32902364 32902985

refGene_gn=MAGI1;refGene_tx=XM_023249917.1;refGene_gn=MAGI1;refGene_tx=XM_023249918.1;refGene_gn=MAGI1;refGene_tx=XM_023249919.1;refGene_gn=MAGI1;refGene_tx=XM_023249920.1;refGene_gn=MAGI1;refGene_tx=XM_023249921.1;refGene_gn=MAGI1;refGene_tx=XM_023249922.1;refGene_gn=MAGI1;refGene_tx=XM_023249923.1;refGene_gn=MAGI1;refGene_tx=XM_023249924.1;refGene_gn=MAGI1;refGene_tx=XM_023249925.1;refGene_gn=MAGI1;refGene_tx=XM_023249926.1;refGene_gn=MAGI1;refGene_tx=XM_023249927.1;refGene_gn=MAGI1;refGene_tx=XM_023249928.1;refGene_gn=MAGI1;refGene_tx=XM_023249929.1;refGene_gn=MAGI1;refGene_tx=XM_023249930.1;refGene_gn=MAGI1;refGene_tx=XM_023249931.1;refGene_gn=MAGI1;refGene_tx=XM_023249932.1;refGene_gn=MAGI1;refGene_tx=XM_023249933.1

A2 32903766 32903949

refGene_gn=MAGI1;refGene_tx=XM_023249917.1;refGene_gn=MAGI1;refGene_tx=XM_023249918.1;refGene_gn=MAGI1;refGene_tx=XM_023249919.1;refGene_gn=MAGI1;refGene_tx=XM_023249920.1;refGene_gn=MAGI1;refGene_tx=XM_023249921.1;refGene_gn=MAGI1;refGene_tx=XM_023249922.1;refGene_gn=MAGI1;refGene_tx=XM_023249923.1;refGene_gn=MAGI1;refGene_tx=XM_023249924.1;refGene_gn=MAGI1;refGene_tx=XM_023249925.1;refGene_gn=MAGI1;refGene_tx=XM_023249926.1;refGene_gn=MAGI1;refGene_tx=XM_023249927.1;refGene_gn=MAGI1;refGene_tx=XM_023249928.1;refGene_gn=MAGI1;refGene_tx=XM_023249929.1;refGene_gn=MAGI1;refGene_tx=XM_023249930.1;refGene_gn=MAGI1;refGene_tx=XM_023249931.1;refGene_gn=MAGI1;refGene_tx=XM_023249932.1;refGene_gn=MAGI1;refGene_tx=XM_023249933.1

A2 32909156 32909249

refGene_gn=MAGI1;refGene_tx=XM_023249917.1;refGene_gn=MAGI1;refGene_tx=XM_023249918.1;refGene_gn=MAGI1;refGene_tx=XM_023249919.1;refGene_gn=MAGI1;refGene_tx=XM_023249920.1;refGene_gn=MAGI1;refGene_tx=XM_0232499

21.1;refGene_gn=MAGI1;refGene_tx=XM_023249922.1;refGene_gn=MAGI1;refGene_tx=XM_023249923.1;refGene_gn=MAGI1;refGene_tx=XM_023249924.1;refGene_gn=MAGI1;refGene_tx=XM_023249925.1;refGene_gn=MAGI1;refGene_tx=XM_023249926.1;refGene_gn=MAGI1;refGene_tx=XM_023249927.1;refGene_gn=MAGI1;refGene_tx=XM_023249928.1;refGene_gn=MAGI1;refGene_tx=XM_023249929.1;refGene_gn=MAGI1;refGene_tx=XM_023249930.1;refGene_gn=MAGI1;refGene_tx=XM_023249931.1;refGene_gn=MAGI1;refGene_tx=XM_023249932.1;refGene_gn=MAGI1;refGene_tx=XM_023249933.1
A2 32911991 32912098
refGene_gn=MAGI1;refGene_tx=XM_023249917.1;refGene_gn=MAGI1;refGene_tx=XM_023249918.1;refGene_gn=MAGI1;refGene_tx=XM_023249919.1;refGene_gn=MAGI1;refGene_tx=XM_023249920.1;refGene_gn=MAGI1;refGene_tx=XM_023249921.1;refGene_gn=MAGI1;refGene_tx=XM_023249922.1;refGene_gn=MAGI1;refGene_tx=XM_023249923.1;refGene_gn=MAGI1;refGene_tx=XM_023249924.1;refGene_gn=MAGI1;refGene_tx=XM_023249925.1;refGene_gn=MAGI1;refGene_tx=XM_023249926.1;refGene_gn=MAGI1;refGene_tx=XM_023249927.1;refGene_gn=MAGI1;refGene_tx=XM_023249928.1;refGene_gn=MAGI1;refGene_tx=XM_023249929.1;refGene_gn=MAGI1;refGene_tx=XM_023249930.1;refGene_gn=MAGI1;refGene_tx=XM_023249931.1;refGene_gn=MAGI1;refGene_tx=XM_023249932.1;refGene_gn=MAGI1;refGene_tx=XM_023249933.1
A2 32914212 32914270
refGene_gn=MAGI1;refGene_tx=XM_023249917.1;refGene_gn=MAGI1;refGene_tx=XM_023249918.1;refGene_gn=MAGI1;refGene_tx=XM_023249919.1;refGene_gn=MAGI1;refGene_tx=XM_023249920.1;refGene_gn=MAGI1;refGene_tx=XM_023249921.1;refGene_gn=MAGI1;refGene_tx=XM_023249922.1;refGene_gn=MAGI1;refGene_tx=XM_023249923.1;refGene_gn=MAGI1;refGene_tx=XM_023249924.1;refGene_gn=MAGI1;refGene_tx=XM_023249925.1;refGene_gn=MAGI1;refGene_tx=XM_023249926.1;refGene_gn=MAGI1;refGene_tx=XM_023249927.1;refGene_gn=MAGI1;refGene_tx=XM_023249928.1;refGene_gn=MAGI1;refGene_tx=XM_023249929.1;refGene_gn=MAGI1;refGene_tx=XM_023249930.1;refGene_gn=MAGI1;refGene_tx=XM_023249931.1;refGene_gn=MAGI1;refGene_tx=XM_023249932.1;refGene_gn=MAGI1;refGene_tx=XM_023249933.1
A2 32919710 32919749
refGene_gn=MAGI1;refGene_tx=XM_023249917.1;refGene_gn=MAGI1;refGene_tx=XM_023249918.1;refGene_gn=MAGI1;refGene_tx=XM_023249919.1;refGene_gn=MAGI1;refGene_tx=XM_023249920.1;refGene_gn=MAGI1;refGene_tx=XM_023249921.1;refGene_gn=MAGI1;refGene_tx=XM_023249924.1;refGene_gn=MAGI1;refGene_tx=XM_023249925.1;refGene_gn=MAGI1;refGene_tx=XM_023249926.1;refGene_gn=MAGI1;refGene_tx=XM_023249927.1;refGene_gn=MAGI1;refGene_tx=XM_023249928.1;refGene_gn=MAGI1;refGene_tx=XM_023249929.1;refGene_gn=MAGI1;refGene_tx=XM_023249930.1;refGene_gn=MAGI1;refGene_tx=XM_023249931.1;refGene_gn=MAGI1;refGene_tx=XM_023249932.1
A2 32924291 32924374
refGene_gn=MAGI1;refGene_tx=XM_023249917.1;refGene_gn=MAGI1;refGene_tx=XM_023249918.1;refGene_gn=MAGI1;refGene_tx=XM_023249919.1;refGene_gn=MAGI1;refGene_tx=XM_023249920.1;refGene_gn=MAGI1;refGene_tx=XM_023249921.1;refGene_gn=MAGI1;refGene_tx=XM_023249922.1;refGene_gn=MAGI1;refGene_tx=XM_023249923.1;refGene_gn=MAGI1;refGene_tx=XM_023249924.1;refGene_gn=MAGI1;refGene_tx=XM_023249925.1;refGene_gn=MAGI1;refGene_tx=XM_023249926.1;refGene_gn=MAGI1;refGene_tx=XM_023249927.1;refGene_gn=MAGI1;

refGene_tx=XM_023249928.1;refGene_gn=MAGI1;refGene_tx=XM_023249929.1;refGene_gn=MAGI1;refGene_tx=XM_023249930.1;refGene_gn=MAGI1;refGene_tx=XM_023249931.1;refGene_gn=MAGI1;refGene_tx=XM_023249932.1;refGene_gn=MAGI1;refGene_tx=XM_023249933.1
A2 32933342 32933544
refGene_gn=MAGI1;refGene_tx=XM_023249917.1;refGene_gn=MAGI1;refGene_tx=XM_023249918.1;refGene_gn=MAGI1;refGene_tx=XM_023249919.1;refGene_gn=MAGI1;refGene_tx=XM_023249920.1;refGene_gn=MAGI1;refGene_tx=XM_023249921.1;refGene_gn=MAGI1;refGene_tx=XM_023249922.1;refGene_gn=MAGI1;refGene_tx=XM_023249923.1;refGene_gn=MAGI1;refGene_tx=XM_023249924.1;refGene_gn=MAGI1;refGene_tx=XM_023249925.1;refGene_gn=MAGI1;refGene_tx=XM_023249926.1;refGene_gn=MAGI1;refGene_tx=XM_023249927.1;refGene_gn=MAGI1;refGene_tx=XM_023249928.1;refGene_gn=MAGI1;refGene_tx=XM_023249929.1;refGene_gn=MAGI1;refGene_tx=XM_023249930.1;refGene_gn=MAGI1;refGene_tx=XM_023249931.1;refGene_gn=MAGI1;refGene_tx=XM_023249932.1;refGene_gn=MAGI1;refGene_tx=XM_023249933.1
A2 32941946 32942153
refGene_gn=MAGI1;refGene_tx=XM_023249917.1;refGene_gn=MAGI1;refGene_tx=XM_023249918.1;refGene_gn=MAGI1;refGene_tx=XM_023249919.1;refGene_gn=MAGI1;refGene_tx=XM_023249920.1;refGene_gn=MAGI1;refGene_tx=XM_023249921.1;refGene_gn=MAGI1;refGene_tx=XM_023249922.1;refGene_gn=MAGI1;refGene_tx=XM_023249923.1;refGene_gn=MAGI1;refGene_tx=XM_023249924.1;refGene_gn=MAGI1;refGene_tx=XM_023249925.1;refGene_gn=MAGI1;refGene_tx=XM_023249926.1;refGene_gn=MAGI1;refGene_tx=XM_023249927.1;refGene_gn=MAGI1;refGene_tx=XM_023249928.1;refGene_gn=MAGI1;refGene_tx=XM_023249929.1;refGene_gn=MAGI1;refGene_tx=XM_023249930.1;refGene_gn=MAGI1;refGene_tx=XM_023249931.1;refGene_gn=MAGI1;refGene_tx=XM_023249932.1;refGene_gn=MAGI1;refGene_tx=XM_023249933.1
A2 32956824 32956944
refGene_gn=MAGI1;refGene_tx=XM_023249917.1;refGene_gn=MAGI1;refGene_tx=XM_023249918.1;refGene_gn=MAGI1;refGene_tx=XM_023249919.1;refGene_gn=MAGI1;refGene_tx=XM_023249920.1;refGene_gn=MAGI1;refGene_tx=XM_023249921.1;refGene_gn=MAGI1;refGene_tx=XM_023249922.1;refGene_gn=MAGI1;refGene_tx=XM_023249923.1;refGene_gn=MAGI1;refGene_tx=XM_023249924.1;refGene_gn=MAGI1;refGene_tx=XM_023249925.1;refGene_gn=MAGI1;refGene_tx=XM_023249926.1;refGene_gn=MAGI1;refGene_tx=XM_023249927.1;refGene_gn=MAGI1;refGene_tx=XM_023249928.1;refGene_gn=MAGI1;refGene_tx=XM_023249929.1;refGene_gn=MAGI1;refGene_tx=XM_023249930.1;refGene_gn=MAGI1;refGene_tx=XM_023249931.1;refGene_gn=MAGI1;refGene_tx=XM_023249932.1;refGene_gn=MAGI1;refGene_tx=XM_023249933.1
A2 33082330 33082447
refGene_gn=MAGI1;refGene_tx=XM_023249917.1;refGene_gn=MAGI1;refGene_tx=XM_023249918.1;refGene_gn=MAGI1;refGene_tx=XM_023249919.1;refGene_gn=MAGI1;refGene_tx=XM_023249920.1;refGene_gn=MAGI1;refGene_tx=XM_023249921.1;refGene_gn=MAGI1;refGene_tx=XM_023249922.1;refGene_gn=MAGI1;refGene_tx=XM_023249923.1;refGene_gn=MAGI1;refGene_tx=XM_023249924.1;refGene_gn=MAGI1;refGene_tx=XM_023249925.1;refGene_gn=MAGI1;refGene_tx=XM_023249926.1;refGene_gn=MAGI1;refGene_tx=XM_023249927.1;refGene_gn=MAGI1;refGene_tx=XM_023249928.1;refGene_gn=MAGI1;refGene_tx=XM_023249929.1;refGene_gn=MAGI1;refGene_tx=XM_023249930.1;refGene_gn=MAGI1;refGene_tx=

XM_023249931.1;refGene_gn=MAGI1;refGene_tx=XM_023249932.1;refGene_gn=MAGI1;refGene_tx=XM_023249933.1
A2 33465602 33465915
refGene_gn=MAGI1;refGene_tx=XM_023249917.1;refGene_gn=MAGI1;refGene_tx=XM_023249918.1;refGene_gn=MAGI1;refGene_tx=XM_023249919.1;refGene_gn=MAGI1;refGene_tx=XM_023249920.1;refGene_gn=MAGI1;refGene_tx=XM_023249921.1;refGene_gn=MAGI1;refGene_tx=XM_023249922.1;refGene_gn=MAGI1;refGene_tx=XM_023249923.1;refGene_gn=MAGI1;refGene_tx=XM_023249924.1;refGene_gn=MAGI1;refGene_tx=XM_023249925.1;refGene_gn=MAGI1;refGene_tx=XM_023249926.1;refGene_gn=MAGI1;refGene_tx=XM_023249927.1;refGene_gn=MAGI1;refGene_tx=XM_023249928.1;refGene_gn=MAGI1;refGene_tx=XM_023249929.1;refGene_gn=MAGI1;refGene_tx=XM_023249930.1;refGene_gn=MAGI1;refGene_tx=XM_023249931.1;refGene_gn=MAGI1;refGene_tx=XM_023249932.1;refGene_gn=MAGI1;refGene_tx=XM_023249933.1
A2 33627072 33627171
refGene_gn=SLC25A26;refGene_tx=XM_011280156.3;refGene_gn=SLC25A26;refGene_tx=XM_019824115.2
A2 33641627 33641784
refGene_gn=SLC25A26;refGene_tx=XM_011280156.3;refGene_gn=SLC25A26;refGene_tx=XM_019824115.2
A2 33645983 33646093
refGene_gn=SLC25A26;refGene_tx=XM_011280156.3;refGene_gn=SLC25A26;refGene_tx=XM_019824115.2
A2 33667129 33667234
refGene_gn=SLC25A26;refGene_tx=XM_011280156.3;refGene_gn=SLC25A26;refGene_tx=XM_019824115.2
A2 33668779 33668827
refGene_gn=SLC25A26;refGene_tx=XM_011280156.3;refGene_gn=SLC25A26;refGene_tx=XM_019824115.2
A2 33735005 33735050 refGene_gn=SLC25A26;refGene_tx=XM_011280156.3
A2 33755832 33755902
refGene_gn=SLC25A26;refGene_tx=XM_011280156.3;refGene_gn=SLC25A26;refGene_tx=XM_019824115.2
A2 33762097 33762162
refGene_gn=SLC25A26;refGene_tx=XM_011280156.3;refGene_gn=SLC25A26;refGene_tx=XM_019824115.2
A2 33763168 33763242
refGene_gn=SLC25A26;refGene_tx=XM_011280156.3;refGene_gn=SLC25A26;refGene_tx=XM_019824115.2
A2 33767656 33767774
refGene_gn=SLC25A26;refGene_tx=XM_011280156.3;refGene_gn=SLC25A26;refGene_tx=XM_019824115.2
A2 33770224 33770484 refGene_gn=LRIG1;refGene_tx=XM_011280158.3
A2 33770582 33770867 refGene_gn=LRIG1;refGene_tx=XM_011280158.3
A2 33771429 33771576 refGene_gn=LRIG1;refGene_tx=XM_011280158.3
A2 33772186 33772312 refGene_gn=LRIG1;refGene_tx=XM_011280158.3
A2 33772678 33773098 refGene_gn=LRIG1;refGene_tx=XM_011280158.3
A2 33773654 33773936 refGene_gn=LRIG1;refGene_tx=XM_011280158.3
A2 33775228 33775549 refGene_gn=LRIG1;refGene_tx=XM_011280158.3
A2 33781518 33781682 refGene_gn=LRIG1;refGene_tx=XM_011280158.3

A2	33786033 33786105 refGene_gn=LRIG1;refGene_tx=XM_011280158.3
A2	33786944 33787016 refGene_gn=LRIG1;refGene_tx=XM_011280158.3
A2	33792431 33792512 refGene_gn=LRIG1;refGene_tx=XM_011280158.3
A2	33794443 33794587 refGene_gn=LRIG1;refGene_tx=XM_011280158.3
A2	33796519 33796663 refGene_gn=LRIG1;refGene_tx=XM_011280158.3
A2	33798894 33799038 refGene_gn=LRIG1;refGene_tx=XM_011280158.3
A2	33800827 33800971 refGene_gn=LRIG1;refGene_tx=XM_011280158.3
A2	33803332 33803470 refGene_gn=LRIG1;refGene_tx=XM_011280158.3
A2	33834326 33834401 refGene_gn=LRIG1;refGene_tx=XM_011280158.3
A2	33844073 33844145 refGene_gn=LRIG1;refGene_tx=XM_011280158.3
A2	33882109 33882330 refGene_gn=LRIG1;refGene_tx=XM_011280158.3
A2	34349372 34349388 refGene_gn=KBTBD8;refGene_tx=XM_006928926.3
A2	34349965 34350176 refGene_gn=KBTBD8;refGene_tx=XM_006928926.3
A2	34354238 34355353
refGene_gn=KBTBD8;refGene_tx=XM_006928926.3;refGene_gn=KBTBD8;refGene_tx=XM_011280157.2
A2	34358584 34359048
refGene_gn=KBTBD8;refGene_tx=XM_006928926.3;refGene_gn=KBTBD8;refGene_tx=XM_011280157.2
A2	34620981 34621053 refGene_gn=SUCLG2;refGene_tx=XM_023249936.1
A2	34622641 34622826
refGene_gn=SUCLG2;refGene_tx=XM_023249935.1;refGene_gn=SUCLG2;refGene_tx=XM_023249936.1;refGene_gn=SUCLG2;refGene_tx=XM_023249939.1
A2	34677416 34677532 refGene_gn=SUCLG2;refGene_tx=XM_023249938.1
A2	34688013 34688153 refGene_gn=SUCLG2;refGene_tx=XM_023249937.1
A2	34704361 34704396 refGene_gn=SUCLG2;refGene_tx=XM_023249940.1
A2	34704472 34704593
refGene_gn=SUCLG2;refGene_tx=XM_023249935.1;refGene_gn=SUCLG2;refGene_tx=XM_023249936.1;refGene_gn=SUCLG2;refGene_tx=XM_023249937.1;refGene_gn=SUCLG2;refGene_tx=XM_023249938.1;refGene_gn=SUCLG2;refGene_tx=XM_023249939.1;refGene_gn=SUCLG2;refGene_tx=XM_023249940.1
A2	34789210 34789353
refGene_gn=SUCLG2;refGene_tx=XM_023249935.1;refGene_gn=SUCLG2;refGene_tx=XM_023249936.1;refGene_gn=SUCLG2;refGene_tx=XM_023249937.1;refGene_gn=SUCLG2;refGene_tx=XM_023249938.1;refGene_gn=SUCLG2;refGene_tx=XM_023249939.1;refGene_gn=SUCLG2;refGene_tx=XM_023249940.1
A2	34791593 34791755
refGene_gn=SUCLG2;refGene_tx=XM_023249935.1;refGene_gn=SUCLG2;refGene_tx=XM_023249936.1;refGene_gn=SUCLG2;refGene_tx=XM_023249937.1;refGene_gn=SUCLG2;refGene_tx=XM_023249938.1;refGene_gn=SUCLG2;refGene_tx=XM_023249939.1;refGene_gn=SUCLG2;refGene_tx=XM_023249940.1
A2	34805313 34805410
refGene_gn=SUCLG2;refGene_tx=XM_023249935.1;refGene_gn=SUCLG2;refGene_tx=XM_023249936.1;refGene_gn=SUCLG2;refGene_tx=XM_023249937.1;refGene_gn=SUCLG2;refGene_tx=XM_023249938.1;refGene_gn=SUCLG2;refGene_tx=XM_023249939.1;refGene_gn=SUCLG2;refGene_tx=XM_023249940.1
A2	34813920 34814010
refGene_gn=SUCLG2;refGene_tx=XM_023249935.1;refGene_gn=SUCLG2;refGene_tx=XM_023249936.1;refGene_gn=SUCLG2;refGene_tx=XM_023249937.1;refGene_gn=SUCLG2;refGene_tx=XM_023249938.1;refGene_gn=SUCLG2;refGene_tx=XM_02

3249939.1;refGene_gn=SUCLG2;refGene_tx=XM_023249940.1
A2 34815529 34815682
refGene_gn=SUCLG2;refGene_tx=XM_023249935.1;refGene_gn=SUCLG2;refGene_tx=XM_023249936.1;refGene_gn=SUCLG2;refGene_tx=XM_023249937.1;refGene_gn=SUCLG2;refGene_tx=XM_023249938.1;refGene_gn=SUCLG2;refGene_tx=XM_023249939.1;refGene_gn=SUCLG2;refGene_tx=XM_023249940.1
A2 34820040 34820131
refGene_gn=SUCLG2;refGene_tx=XM_023249935.1;refGene_gn=SUCLG2;refGene_tx=XM_023249936.1;refGene_gn=SUCLG2;refGene_tx=XM_023249937.1;refGene_gn=SUCLG2;refGene_tx=XM_023249938.1;refGene_gn=SUCLG2;refGene_tx=XM_023249939.1;refGene_gn=SUCLG2;refGene_tx=XM_023249940.1
A2 34821147 34821247
refGene_gn=SUCLG2;refGene_tx=XM_023249935.1;refGene_gn=SUCLG2;refGene_tx=XM_023249936.1;refGene_gn=SUCLG2;refGene_tx=XM_023249937.1;refGene_gn=SUCLG2;refGene_tx=XM_023249938.1;refGene_gn=SUCLG2;refGene_tx=XM_023249939.1;refGene_gn=SUCLG2;refGene_tx=XM_023249940.1
A2 34900454 34900596
refGene_gn=SUCLG2;refGene_tx=XM_023249935.1;refGene_gn=SUCLG2;refGene_tx=XM_023249936.1;refGene_gn=SUCLG2;refGene_tx=XM_023249937.1;refGene_gn=SUCLG2;refGene_tx=XM_023249938.1;refGene_gn=SUCLG2;refGene_tx=XM_023249939.1;refGene_gn=SUCLG2;refGene_tx=XM_023249940.1
A2 34952740 34952824
refGene_gn=SUCLG2;refGene_tx=XM_023249935.1;refGene_gn=SUCLG2;refGene_tx=XM_023249936.1;refGene_gn=SUCLG2;refGene_tx=XM_023249937.1;refGene_gn=SUCLG2;refGene_tx=XM_023249938.1;refGene_gn=SUCLG2;refGene_tx=XM_023249940.1
A2 35295616 35295734
refGene_gn=FAM19A1;refGene_tx=XM_019824172.2;refGene_gn=FAM19A1;refGene_tx=XM_019824181.2;refGene_gn=FAM19A1;refGene_tx=XM_019824189.2
A2 35684520 35684661
refGene_gn=FAM19A1;refGene_tx=XM_019824172.2;refGene_gn=FAM19A1;refGene_tx=XM_019824181.2;refGene_gn=FAM19A1;refGene_tx=XM_019824189.2
A2 35802218 35802343
refGene_gn=FAM19A1;refGene_tx=XM_019824172.2;refGene_gn=FAM19A1;refGene_tx=XM_019824181.2;refGene_gn=FAM19A1;refGene_tx=XM_019824189.2
A2 35808152 35808170
refGene_gn=FAM19A1;refGene_tx=XM_019824172.2;refGene_gn=FAM19A1;refGene_tx=XM_019824181.2;refGene_gn=FAM19A1;refGene_tx=XM_019824189.2
A2 35972089 35972277
refGene_gn=FAM19A4;refGene_tx=XM_019824220.2;refGene_gn=FAM19A4;refGene_tx=XM_019824228.2
A2 35979856 35979981
refGene_gn=FAM19A4;refGene_tx=XM_019824220.2;refGene_gn=FAM19A4;refGene_tx=XM_019824228.2
A2 35998034 35998190
refGene_gn=FAM19A4;refGene_tx=XM_019824220.2;refGene_gn=FAM19A4;refGene_tx=XM_019824228.2
A2 36096170 36096286
refGene_gn=FAM19A4;refGene_tx=XM_019824220.2;refGene_gn=FAM19A4;refGene_tx=XM_019824228.2

A2 36102798 36102812
refGene_gn=FAM19A4;refGene_tx=XM_019824220.2;refGene_gn=FAM19A4;refGene_tx=XM_019824228.2
A2 36189392 36189539 refGene_gn=EOGT;refGene_tx=XM_023249942.1
A2 36190111 36190214 refGene_gn=EOGT;refGene_tx=XM_023249942.1
A2 36192388 36192508 refGene_gn=EOGT;refGene_tx=XM_023249942.1
A2 36195383 36195445 refGene_gn=EOGT;refGene_tx=XM_023249942.1
A2 36200103 36200172 refGene_gn=EOGT;refGene_tx=XM_023249942.1
A2 36200692 36200779 refGene_gn=EOGT;refGene_tx=XM_023249942.1
A2 36200925 36200997 refGene_gn=EOGT;refGene_tx=XM_023249942.1
A2 36201347 36201440 refGene_gn=EOGT;refGene_tx=XM_023249942.1
A2 36207190 36207294 refGene_gn=EOGT;refGene_tx=XM_023249942.1
A2 36211006 36211113 refGene_gn=EOGT;refGene_tx=XM_023249942.1
A2 36215393 36215498 refGene_gn=EOGT;refGene_tx=XM_023249942.1
A2 36215839 36215934 refGene_gn=EOGT;refGene_tx=XM_023249942.1
A2 36217935 36218044 refGene_gn=EOGT;refGene_tx=XM_023249942.1
A2 36220533 36220634 refGene_gn=EOGT;refGene_tx=XM_023249942.1
A2 36221779 36222003 refGene_gn=EOGT;refGene_tx=XM_023249942.1
A2 36224567 36224577 refGene_gn=EOGT;refGene_tx=XM_023249942.1
A2 36234454 36234598 refGene_gn=TMF1;refGene_tx=XM_003982410.5
A2 36235342 36235468 refGene_gn=TMF1;refGene_tx=XM_003982410.5
A2 36236609 36236762 refGene_gn=TMF1;refGene_tx=XM_003982410.5
A2 36237278 36237380 refGene_gn=TMF1;refGene_tx=XM_003982410.5
A2 36238746 36238839 refGene_gn=TMF1;refGene_tx=XM_003982410.5
A2 36239111 36239181 refGene_gn=TMF1;refGene_tx=XM_003982410.5
A2 36239969 36240162 refGene_gn=TMF1;refGene_tx=XM_003982410.5
A2 36243464 36243621 refGene_gn=TMF1;refGene_tx=XM_003982410.5
A2 36244302 36244395 refGene_gn=TMF1;refGene_tx=XM_003982410.5
A2 36247639 36247796 refGene_gn=TMF1;refGene_tx=XM_003982410.5
A2 36248083 36248250 refGene_gn=TMF1;refGene_tx=XM_003982410.5
A2 36248488 36248631 refGene_gn=TMF1;refGene_tx=XM_003982410.5
A2 36257416 36257522 refGene_gn=TMF1;refGene_tx=XM_003982410.5
A2 36260348 36260475 refGene_gn=TMF1;refGene_tx=XM_003982410.5
A2 36261076 36261180 refGene_gn=TMF1;refGene_tx=XM_003982410.5
A2 36262933 36264129 refGene_gn=TMF1;refGene_tx=XM_003982410.5
A2 36267195 36267337 refGene_gn=TMF1;refGene_tx=XM_003982410.5
A2 36270439 36270528
refGene_gn=UBA3;refGene_tx=XM_003982411.4;refGene_gn=UBA3;refGene_tx=XM_023249944.1;refGene_gn=UBA3;refGene_tx=XM_023249945.1
A2 36271074 36271129
refGene_gn=UBA3;refGene_tx=XM_003982411.4;refGene_gn=UBA3;refGene_tx=XM_023249944.1;refGene_gn=UBA3;refGene_tx=XM_023249945.1
A2 36271270 36271334
refGene_gn=UBA3;refGene_tx=XM_003982411.4;refGene_gn=UBA3;refGene_tx=XM_023249944.1;refGene_gn=UBA3;refGene_tx=XM_023249945.1
A2 36271466 36271567
refGene_gn=UBA3;refGene_tx=XM_003982411.4;refGene_gn=UBA3;refGene_tx=XM_023249944.1;refGene_gn=UBA3;refGene_tx=XM_023249945.1
A2 36271905 36271987
refGene_gn=UBA3;refGene_tx=XM_003982411.4;refGene_gn=UBA3;refGene_tx=X

M_023249944.1;refGene_gn=UBA3;refGene_tx=XM_023249945.1
A2 36272073 36272110
refGene_gn=UBA3;refGene_tx=XM_003982411.4;refGene_gn=UBA3;refGene_tx=XM_023249944.1;refGene_gn=UBA3;refGene_tx=XM_023249945.1
A2 36273108 36273162
refGene_gn=UBA3;refGene_tx=XM_003982411.4;refGene_gn=UBA3;refGene_tx=XM_023249944.1;refGene_gn=UBA3;refGene_tx=XM_023249945.1
A2 36278295 36278409
refGene_gn=UBA3;refGene_tx=XM_003982411.4;refGene_gn=UBA3;refGene_tx=XM_023249944.1;refGene_gn=UBA3;refGene_tx=XM_023249945.1
A2 36278583 36278686
refGene_gn=UBA3;refGene_tx=XM_003982411.4;refGene_gn=UBA3;refGene_tx=XM_023249944.1;refGene_gn=UBA3;refGene_tx=XM_023249945.1
A2 36279645 36279801
refGene_gn=UBA3;refGene_tx=XM_003982411.4;refGene_gn=UBA3;refGene_tx=XM_023249944.1;refGene_gn=UBA3;refGene_tx=XM_023249945.1
A2 36280103 36280168
refGene_gn=UBA3;refGene_tx=XM_003982411.4;refGene_gn=UBA3;refGene_tx=XM_023249944.1;refGene_gn=UBA3;refGene_tx=XM_023249945.1
A2 36280513 36280557
refGene_gn=UBA3;refGene_tx=XM_003982411.4;refGene_gn=UBA3;refGene_tx=XM_023249944.1;refGene_gn=UBA3;refGene_tx=XM_023249945.1
A2 36284645 36284726
refGene_gn=UBA3;refGene_tx=XM_003982411.4;refGene_gn=UBA3;refGene_tx=XM_023249944.1;refGene_gn=UBA3;refGene_tx=XM_023249945.1
A2 36287543 36287626
refGene_gn=UBA3;refGene_tx=XM_003982411.4;refGene_gn=UBA3;refGene_tx=XM_023249944.1;refGene_gn=UBA3;refGene_tx=XM_023249945.1
A2 36291318 36291399
refGene_gn=UBA3;refGene_tx=XM_003982411.4;refGene_gn=UBA3;refGene_tx=XM_023249944.1;refGene_gn=UBA3;refGene_tx=XM_023249945.1
A2 36292689 36292810
refGene_gn=UBA3;refGene_tx=XM_003982411.4;refGene_gn=UBA3;refGene_tx=XM_023249944.1;refGene_gn=UBA3;refGene_tx=XM_023249945.1
A2 36294204 36294224 refGene_gn=UBA3;refGene_tx=XM_003982411.4
A2 36294648 36294690 refGene_gn=UBA3;refGene_tx=XM_023249944.1
A2 36294876 36294896
refGene_gn=UBA3;refGene_tx=XM_023249944.1;refGene_gn=UBA3;refGene_tx=XM_023249945.1
A2 36299525 36299701 refGene_gn=ARL6IP5;refGene_tx=XM_003982412.5
A2 36314996 36315214 refGene_gn=ARL6IP5;refGene_tx=XM_003982412.5
A2 36317558 36317731 refGene_gn=ARL6IP5;refGene_tx=XM_003982412.5
A2 36322775 36322802 refGene_gn=LMOD3;refGene_tx=XM_006928927.4
A2 36331668 36333081 refGene_gn=LMOD3;refGene_tx=XM_006928927.4
A2 36335903 36336197 refGene_gn=LMOD3;refGene_tx=XM_006928927.4
A2 36380142 36380263
refGene_gn=FRMD4B;refGene_tx=XM_003982414.5;refGene_gn=FRMD4B;refGene_tx=XM_019824270.2;refGene_gn=FRMD4B;refGene_tx=XM_019824283.2;refGene_gn=FRMD4B;refGene_tx=XM_019824294.2
A2 36381319 36381411

refGene_gn=FRMD4B;refGene_tx=XM_019824270.2;refGene_gn=FRMD4B;refGene_tx=XM_019824283.2;refGene_gn=FRMD4B;refGene_tx=XM_019824294.2
A2 36384913 36385046
refGene_gn=FRMD4B;refGene_tx=XM_003982414.5;refGene_gn=FRMD4B;refGene_tx=XM_019824270.2;refGene_gn=FRMD4B;refGene_tx=XM_019824283.2;refGene_gn=FRMD4B;refGene_tx=XM_019824294.2
A2 36388151 36388963
refGene_gn=FRMD4B;refGene_tx=XM_003982414.5;refGene_gn=FRMD4B;refGene_tx=XM_019824270.2;refGene_gn=FRMD4B;refGene_tx=XM_019824283.2;refGene_gn=FRMD4B;refGene_tx=XM_019824294.2
A2 36390136 36390256
refGene_gn=FRMD4B;refGene_tx=XM_003982414.5;refGene_gn=FRMD4B;refGene_tx=XM_019824270.2;refGene_gn=FRMD4B;refGene_tx=XM_019824283.2;refGene_gn=FRMD4B;refGene_tx=XM_019824294.2
A2 36393237 36393382
refGene_gn=FRMD4B;refGene_tx=XM_003982414.5;refGene_gn=FRMD4B;refGene_tx=XM_019824270.2;refGene_gn=FRMD4B;refGene_tx=XM_019824283.2;refGene_gn=FRMD4B;refGene_tx=XM_019824294.2
A2 36395635 36395692
refGene_gn=FRMD4B;refGene_tx=XM_003982414.5;refGene_gn=FRMD4B;refGene_tx=XM_019824270.2;refGene_gn=FRMD4B;refGene_tx=XM_019824283.2;refGene_gn=FRMD4B;refGene_tx=XM_019824294.2
A2 36398281 36398510
refGene_gn=FRMD4B;refGene_tx=XM_003982414.5;refGene_gn=FRMD4B;refGene_tx=XM_019824270.2;refGene_gn=FRMD4B;refGene_tx=XM_019824283.2;refGene_gn=FRMD4B;refGene_tx=XM_019824294.2
A2 36399646 36399766
refGene_gn=FRMD4B;refGene_tx=XM_003982414.5;refGene_gn=FRMD4B;refGene_tx=XM_019824270.2;refGene_gn=FRMD4B;refGene_tx=XM_019824283.2;refGene_gn=FRMD4B;refGene_tx=XM_019824294.2
A2 36399863 36399997
refGene_gn=FRMD4B;refGene_tx=XM_003982414.5;refGene_gn=FRMD4B;refGene_tx=XM_019824270.2;refGene_gn=FRMD4B;refGene_tx=XM_019824283.2;refGene_gn=FRMD4B;refGene_tx=XM_019824294.2
A2 36400868 36401010
refGene_gn=FRMD4B;refGene_tx=XM_003982414.5;refGene_gn=FRMD4B;refGene_tx=XM_019824270.2;refGene_gn=FRMD4B;refGene_tx=XM_019824283.2;refGene_gn=FRMD4B;refGene_tx=XM_019824294.2
A2 36401711 36401850
refGene_gn=FRMD4B;refGene_tx=XM_003982414.5;refGene_gn=FRMD4B;refGene_tx=XM_019824270.2;refGene_gn=FRMD4B;refGene_tx=XM_019824283.2;refGene_gn=FRMD4B;refGene_tx=XM_019824294.2
A2 36403138 36403215
refGene_gn=FRMD4B;refGene_tx=XM_003982414.5;refGene_gn=FRMD4B;refGene_tx=XM_019824270.2;refGene_gn=FRMD4B;refGene_tx=XM_019824283.2;refGene_gn=FRMD4B;refGene_tx=XM_019824294.2
A2 36414638 36414725
refGene_gn=FRMD4B;refGene_tx=XM_003982414.5;refGene_gn=FRMD4B;refGene_tx=XM_019824270.2;refGene_gn=FRMD4B;refGene_tx=XM_019824283.2;refGene_gn=FRMD4B;refGene_tx=XM_019824294.2

A2 36417031 36417089
refGene_gn=FRMD4B;refGene_tx=XM_003982414.5;refGene_gn=FRMD4B;refGene_tx=XM_019824270.2;refGene_gn=FRMD4B;refGene_tx=XM_019824283.2;refGene_gn=FRMD4B;refGene_tx=XM_019824294.2
A2 36419815 36419881
refGene_gn=FRMD4B;refGene_tx=XM_003982414.5;refGene_gn=FRMD4B;refGene_tx=XM_019824270.2;refGene_gn=FRMD4B;refGene_tx=XM_019824283.2;refGene_gn=FRMD4B;refGene_tx=XM_019824294.2
A2 36422008 36422092
refGene_gn=FRMD4B;refGene_tx=XM_003982414.5;refGene_gn=FRMD4B;refGene_tx=XM_019824270.2;refGene_gn=FRMD4B;refGene_tx=XM_019824283.2;refGene_gn=FRMD4B;refGene_tx=XM_019824294.2
A2 36448613 36448636
refGene_gn=FRMD4B;refGene_tx=XM_003982414.5;refGene_gn=FRMD4B;refGene_tx=XM_019824270.2;refGene_gn=FRMD4B;refGene_tx=XM_019824283.2;refGene_gn=FRMD4B;refGene_tx=XM_019824294.2
A2 36449430 36449487
refGene_gn=FRMD4B;refGene_tx=XM_003982414.5;refGene_gn=FRMD4B;refGene_tx=XM_019824270.2;refGene_gn=FRMD4B;refGene_tx=XM_019824283.2;refGene_gn=FRMD4B;refGene_tx=XM_019824294.2
A2 36480218 36480303
refGene_gn=FRMD4B;refGene_tx=XM_003982414.5;refGene_gn=FRMD4B;refGene_tx=XM_019824270.2;refGene_gn=FRMD4B;refGene_tx=XM_019824283.2;refGene_gn=FRMD4B;refGene_tx=XM_019824294.2
A2 36491781 36491874
refGene_gn=FRMD4B;refGene_tx=XM_003982414.5;refGene_gn=FRMD4B;refGene_tx=XM_019824270.2;refGene_gn=FRMD4B;refGene_tx=XM_019824283.2;refGene_gn=FRMD4B;refGene_tx=XM_019824294.2
A2 36499057 36499152
refGene_gn=FRMD4B;refGene_tx=XM_003982414.5;refGene_gn=FRMD4B;refGene_tx=XM_019824270.2;refGene_gn=FRMD4B;refGene_tx=XM_019824283.2;refGene_gn=FRMD4B;refGene_tx=XM_019824294.2
A2 36501448 36501514
refGene_gn=FRMD4B;refGene_tx=XM_003982414.5;refGene_gn=FRMD4B;refGene_tx=XM_019824270.2;refGene_gn=FRMD4B;refGene_tx=XM_019824283.2;refGene_gn=FRMD4B;refGene_tx=XM_019824294.2
A2 36525635 36525659 refGene_gn=FRMD4B;refGene_tx=XM_019824283.2
A2 36549686 36549848
refGene_gn=FRMD4B;refGene_tx=XM_003982414.5;refGene_gn=FRMD4B;refGene_tx=XM_019824270.2
A2 36825125 36825268
refGene_gn=LOC111559439;refGene_tx=XM_023249948.1;refGene_gn=LOC111559439;refGene_tx=XM_023249949.1
A2 36825431 36825498
refGene_gn=LOC111559439;refGene_tx=XM_023249948.1;refGene_gn=LOC111559439;refGene_tx=XM_023249949.1
A2 36826237 36826300
refGene_gn=LOC111559439;refGene_tx=XM_023249948.1
A2 36826554 36826643
refGene_gn=LOC111559439;refGene_tx=XM_023249948.1;refGene_gn=LOC111559

439;refGene_tx=XM_023249949.1
A2	36827586	36827608
refGene_gn=LOC111559439;refGene_tx=XM_023249948.1;refGene_gn=LOC111559
439;refGene_tx=XM_023249949.1
A2	36836474	36836540
refGene_gn=LOC111559439;refGene_tx=XM_023249948.1;refGene_gn=LOC111559
439;refGene_tx=XM_023249949.1
A2	36890042	36890103
refGene_gn=LOC111556285;refGene_tx=XM_023249946.1;refGene_gn=LOC111556
285;refGene_tx=XM_023249947.1
A2	36890209	36890245
refGene_gn=LOC111556285;refGene_tx=XM_023249946.1;refGene_gn=LOC111556
285;refGene_tx=XM_023249947.1
A2	36890327	36890395
refGene_gn=LOC111556285;refGene_tx=XM_023249946.1;refGene_gn=LOC111556
285;refGene_tx=XM_023249947.1
A2	36890519	36890613
refGene_gn=LOC111556285;refGene_tx=XM_023249946.1;refGene_gn=LOC111556
285;refGene_tx=XM_023249947.1
A2	36892493	36892724
refGene_gn=LOC111556285;refGene_tx=XM_023249946.1;refGene_gn=LOC111556
285;refGene_tx=XM_023249947.1
A2	37078195	37078251	refGene_gn=MITF;refGene_tx=XM_006928932.3
A2	37089949	37090202	refGene_gn=MITF;refGene_tx=XM_006928932.3
A2	37149852	37150080	refGene_gn=MITF;refGene_tx=XM_006928932.3
A2	37151144	37151228	refGene_gn=MITF;refGene_tx=XM_006928932.3
A2	37153430	37153490	refGene_gn=MITF;refGene_tx=XM_006928932.3
A2	37153491	37153527	refGene_gn=MITF;refGene_tx=XM_006928932.3
A2	37162301	37162419	refGene_gn=MITF;refGene_tx=XM_006928932.3
A2	37165399	37165456	refGene_gn=MITF;refGene_tx=XM_006928932.3
A2	37171394	37171470	refGene_gn=MITF;refGene_tx=XM_006928932.3
A2	37174821	37174969	refGene_gn=MITF;refGene_tx=XM_006928932.3
A2	37180679	37181081	refGene_gn=MITF;refGene_tx=XM_006928932.3
A2	38053533	38053678
refGene_gn=FOXP1;refGene_tx=XM_019824396.2;refGene_gn=FOXP1;refGene_tx
=XM_019824399.2;refGene_gn=FOXP1;refGene_tx=XM_019824410.2;refGene_gn=
FOXP1;refGene_tx=XM_023249950.1;refGene_gn=FOXP1;refGene_tx=XM_0232499
51.1;refGene_gn=FOXP1;refGene_tx=XM_023249952.1;refGene_gn=FOXP1;refGe
ne_tx=XM_023249953.1;refGene_gn=FOXP1;refGene_tx=XM_023249954.1;refGen
e_gn=FOXP1;refGene_tx=XM_023249956.1
A2	38059301	38059468
refGene_gn=FOXP1;refGene_tx=XM_019824396.2;refGene_gn=FOXP1;refGene_tx
=XM_019824399.2;refGene_gn=FOXP1;refGene_tx=XM_019824410.2;refGene_gn=
FOXP1;refGene_tx=XM_023249950.1;refGene_gn=FOXP1;refGene_tx=XM_0232499
51.1;refGene_gn=FOXP1;refGene_tx=XM_023249952.1;refGene_gn=FOXP1;refGe
ne_tx=XM_023249953.1;refGene_gn=FOXP1;refGene_tx=XM_023249954.1;refGen
e_gn=FOXP1;refGene_tx=XM_023249956.1
A2	38063813	38063883
refGene_gn=FOXP1;refGene_tx=XM_019824396.2;refGene_gn=FOXP1;refGene_tx
=XM_019824399.2;refGene_gn=FOXP1;refGene_tx=XM_019824410.2;refGene_gn=

FOXP1;refGene_tx=XM_023249950.1;refGene_gn=FOXP1;refGene_tx=XM_0232499
51.1;refGene_gn=FOXP1;refGene_tx=XM_023249952.1;refGene_gn=FOXP1;refGe
ne_tx=XM_023249953.1;refGene_gn=FOXP1;refGene_tx=XM_023249954.1;refGen
e_gn=FOXP1;refGene_tx=XM_023249955.1;refGene_gn=FOXP1;refGene_tx=XM_02
3249956.1
A2 38065016 38065186 refGene_gn=FOXP1;refGene_tx=XM_023249956.1
A2 38065563 38065685
refGene_gn=FOXP1;refGene_tx=XM_019824396.2;refGene_gn=FOXP1;refGene_tx
=XM_019824399.2;refGene_gn=FOXP1;refGene_tx=XM_019824410.2;refGene_gn=
FOXP1;refGene_tx=XM_023249950.1;refGene_gn=FOXP1;refGene_tx=XM_0232499
51.1;refGene_gn=FOXP1;refGene_tx=XM_023249952.1;refGene_gn=FOXP1;refGe
ne_tx=XM_023249953.1;refGene_gn=FOXP1;refGene_tx=XM_023249954.1
A2 38069425 38069527
refGene_gn=FOXP1;refGene_tx=XM_019824396.2;refGene_gn=FOXP1;refGene_tx
=XM_019824399.2;refGene_gn=FOXP1;refGene_tx=XM_019824410.2;refGene_gn=
FOXP1;refGene_tx=XM_023249950.1;refGene_gn=FOXP1;refGene_tx=XM_0232499
51.1;refGene_gn=FOXP1;refGene_tx=XM_023249952.1;refGene_gn=FOXP1;refGe
ne_tx=XM_023249953.1;refGene_gn=FOXP1;refGene_tx=XM_023249954.1;refGen
e_gn=FOXP1;refGene_tx=XM_023249955.1;refGene_gn=FOXP1;refGene_tx=XM_02
3249956.1
A2 38070131 38070211
refGene_gn=FOXP1;refGene_tx=XM_019824396.2;refGene_gn=FOXP1;refGene_tx
=XM_019824399.2;refGene_gn=FOXP1;refGene_tx=XM_019824410.2;refGene_gn=
FOXP1;refGene_tx=XM_023249950.1;refGene_gn=FOXP1;refGene_tx=XM_0232499
51.1;refGene_gn=FOXP1;refGene_tx=XM_023249952.1;refGene_gn=FOXP1;refGe
ne_tx=XM_023249953.1;refGene_gn=FOXP1;refGene_tx=XM_023249954.1;refGen
e_gn=FOXP1;refGene_tx=XM_023249955.1;refGene_gn=FOXP1;refGene_tx=XM_02
3249956.1
A2 38070315 38070517
refGene_gn=FOXP1;refGene_tx=XM_019824396.2;refGene_gn=FOXP1;refGene_tx
=XM_019824399.2;refGene_gn=FOXP1;refGene_tx=XM_019824410.2;refGene_gn=
FOXP1;refGene_tx=XM_023249950.1;refGene_gn=FOXP1;refGene_tx=XM_0232499
51.1;refGene_gn=FOXP1;refGene_tx=XM_023249952.1;refGene_gn=FOXP1;refGe
ne_tx=XM_023249953.1;refGene_gn=FOXP1;refGene_tx=XM_023249954.1;refGen
e_gn=FOXP1;refGene_tx=XM_023249955.1;refGene_gn=FOXP1;refGene_tx=XM_02
3249956.1
A2 38080247 38080331
refGene_gn=FOXP1;refGene_tx=XM_019824396.2;refGene_gn=FOXP1;refGene_tx
=XM_019824399.2;refGene_gn=FOXP1;refGene_tx=XM_019824410.2;refGene_gn=
FOXP1;refGene_tx=XM_023249950.1;refGene_gn=FOXP1;refGene_tx=XM_0232499
51.1;refGene_gn=FOXP1;refGene_tx=XM_023249952.1;refGene_gn=FOXP1;refGe
ne_tx=XM_023249953.1;refGene_gn=FOXP1;refGene_tx=XM_023249954.1;refGen
e_gn=FOXP1;refGene_tx=XM_023249955.1;refGene_gn=FOXP1;refGene_tx=XM_02
3249956.1
A2 38093159 38093247
refGene_gn=FOXP1;refGene_tx=XM_019824396.2;refGene_gn=FOXP1;refGene_tx
=XM_019824399.2;refGene_gn=FOXP1;refGene_tx=XM_019824410.2;refGene_gn=
FOXP1;refGene_tx=XM_023249950.1;refGene_gn=FOXP1;refGene_tx=XM_0232499
51.1;refGene_gn=FOXP1;refGene_tx=XM_023249952.1;refGene_gn=FOXP1;refGe
ne_tx=XM_023249953.1;refGene_gn=FOXP1;refGene_tx=XM_023249954.1;refGen

e_gn=FOXP1;refGene_tx=XM_023249955.1;refGene_gn=FOXP1;refGene_tx=XM_023249956.1
A2 38106152 38106257
refGene_gn=FOXP1;refGene_tx=XM_019824396.2;refGene_gn=FOXP1;refGene_tx=XM_019824399.2;refGene_gn=FOXP1;refGene_tx=XM_019824410.2;refGene_gn=FOXP1;refGene_tx=XM_023249950.1;refGene_gn=FOXP1;refGene_tx=XM_023249951.1;refGene_gn=FOXP1;refGene_tx=XM_023249952.1;refGene_gn=FOXP1;refGene_tx=XM_023249953.1;refGene_gn=FOXP1;refGene_tx=XM_023249954.1;refGene_gn=FOXP1;refGene_tx=XM_023249955.1;refGene_gn=FOXP1;refGene_tx=XM_023249956.1
A2 38133100 38133305
refGene_gn=FOXP1;refGene_tx=XM_019824396.2;refGene_gn=FOXP1;refGene_tx=XM_019824399.2;refGene_gn=FOXP1;refGene_tx=XM_019824410.2;refGene_gn=FOXP1;refGene_tx=XM_023249950.1;refGene_gn=FOXP1;refGene_tx=XM_023249951.1;refGene_gn=FOXP1;refGene_tx=XM_023249952.1;refGene_gn=FOXP1;refGene_tx=XM_023249953.1;refGene_gn=FOXP1;refGene_tx=XM_023249954.1;refGene_gn=FOXP1;refGene_tx=XM_023249955.1;refGene_gn=FOXP1;refGene_tx=XM_023249956.1
A2 38138930 38139084
refGene_gn=FOXP1;refGene_tx=XM_019824396.2;refGene_gn=FOXP1;refGene_tx=XM_019824399.2;refGene_gn=FOXP1;refGene_tx=XM_019824410.2;refGene_gn=FOXP1;refGene_tx=XM_023249950.1;refGene_gn=FOXP1;refGene_tx=XM_023249951.1;refGene_gn=FOXP1;refGene_tx=XM_023249952.1;refGene_gn=FOXP1;refGene_tx=XM_023249953.1;refGene_gn=FOXP1;refGene_tx=XM_023249954.1;refGene_gn=FOXP1;refGene_tx=XM_023249955.1;refGene_gn=FOXP1;refGene_tx=XM_023249956.1
A2 38144746 38144836
refGene_gn=FOXP1;refGene_tx=XM_019824396.2;refGene_gn=FOXP1;refGene_tx=XM_019824399.2;refGene_gn=FOXP1;refGene_tx=XM_019824410.2;refGene_gn=FOXP1;refGene_tx=XM_023249950.1;refGene_gn=FOXP1;refGene_tx=XM_023249951.1;refGene_gn=FOXP1;refGene_tx=XM_023249952.1;refGene_gn=FOXP1;refGene_tx=XM_023249953.1;refGene_gn=FOXP1;refGene_tx=XM_023249954.1;refGene_gn=FOXP1;refGene_tx=XM_023249955.1;refGene_gn=FOXP1;refGene_tx=XM_023249956.1
A2 38145869 38146007
refGene_gn=FOXP1;refGene_tx=XM_019824396.2;refGene_gn=FOXP1;refGene_tx=XM_019824399.2;refGene_gn=FOXP1;refGene_tx=XM_019824410.2;refGene_gn=FOXP1;refGene_tx=XM_023249950.1;refGene_gn=FOXP1;refGene_tx=XM_023249951.1;refGene_gn=FOXP1;refGene_tx=XM_023249952.1;refGene_gn=FOXP1;refGene_tx=XM_023249953.1;refGene_gn=FOXP1;refGene_tx=XM_023249954.1;refGene_gn=FOXP1;refGene_tx=XM_023249955.1;refGene_gn=FOXP1;refGene_tx=XM_023249956.1
A2 38204391 38204493
refGene_gn=FOXP1;refGene_tx=XM_019824396.2;refGene_gn=FOXP1;refGene_tx=XM_019824399.2;refGene_gn=FOXP1;refGene_tx=XM_019824410.2;refGene_gn=FOXP1;refGene_tx=XM_023249950.1;refGene_gn=FOXP1;refGene_tx=XM_023249951.1;refGene_gn=FOXP1;refGene_tx=XM_023249952.1;refGene_gn=FOXP1;refGene_tx=XM_023249953.1;refGene_gn=FOXP1;refGene_tx=XM_023249955.1;refGene_gn=FOXP1;refGene_tx=XM_023249956.1
A2 38223769 38223955

refGene_gn=FOXP1;refGene_tx=XM_019824396.2;refGene_gn=FOXP1;refGene_tx=XM_019824399.2
A2 38225379 38225415 refGene_gn=FOXP1;refGene_tx=XM_019824410.2
A2 38287246 38287432
refGene_gn=FOXP1;refGene_tx=XM_023249950.1;refGene_gn=FOXP1;refGene_tx=XM_023249951.1;refGene_gn=FOXP1;refGene_tx=XM_023249952.1;refGene_gn=FOXP1;refGene_tx=XM_023249953.1;refGene_gn=FOXP1;refGene_tx=XM_023249955.1;refGene_gn=FOXP1;refGene_tx=XM_023249956.1
A2 38643067 38643412
refGene_gn=LOC102900768;refGene_tx=XM_023241872.1
A2 38643720 38644431
refGene_gn=LOC102900768;refGene_tx=XM_023241872.1
A2 38644808 38644901
refGene_gn=LOC102900768;refGene_tx=XM_023241872.1
A2 38744519 38744566
refGene_gn=EIF4E3;refGene_tx=XM_023249958.1;refGene_gn=EIF4E3;refGene_tx=XM_023249959.1
A2 38748942 38749098
refGene_gn=EIF4E3;refGene_tx=XM_023249958.1;refGene_gn=EIF4E3;refGene_tx=XM_023249959.1
A2 38752491 38752558
refGene_gn=EIF4E3;refGene_tx=XM_023249958.1;refGene_gn=EIF4E3;refGene_tx=XM_023249959.1
A2 38755107 38755168
refGene_gn=EIF4E3;refGene_tx=XM_023249958.1;refGene_gn=EIF4E3;refGene_tx=XM_023249959.1
A2 38756805 38756900
refGene_gn=EIF4E3;refGene_tx=XM_023249958.1;refGene_gn=EIF4E3;refGene_tx=XM_023249959.1
A2 38767672 38767745 refGene_gn=EIF4E3;refGene_tx=XM_023249958.1
A2 38782404 38782577 refGene_gn=EIF4E3;refGene_tx=XM_023249958.1
A2 38809563 38810694 refGene_gn=GPR27;refGene_tx=XM_023249957.1
A2 38825274 38825379
refGene_gn=PROK2;refGene_tx=XM_023249960.1;refGene_gn=PROK2;refGene_tx=XM_023249961.1
A2 38826710 38826773 refGene_gn=PROK2;refGene_tx=XM_023249960.1
A2 38835452 38835578
refGene_gn=PROK2;refGene_tx=XM_023249960.1;refGene_gn=PROK2;refGene_tx=XM_023249961.1
A2 38838973 38839057
refGene_gn=PROK2;refGene_tx=XM_023249960.1;refGene_gn=PROK2;refGene_tx=XM_023249961.1
A2 39373581 39373834 refGene_gn=RYBP;refGene_tx=XM_023249963.1
A2 39374267 39374366 refGene_gn=RYBP;refGene_tx=XM_023249963.1
A2 39374462 39374639 refGene_gn=RYBP;refGene_tx=XM_023249963.1
A2 39453334 39453460 refGene_gn=RYBP;refGene_tx=XM_023249963.1
A2 39454216 39454248 refGene_gn=RYBP;refGene_tx=XM_023249963.1
A2 39717525 39718054 refGene_gn=SHQ1;refGene_tx=XM_003982422.5
A2 39759346 39759467 refGene_gn=SHQ1;refGene_tx=XM_003982422.5
A2 39778258 39778382 refGene_gn=SHQ1;refGene_tx=XM_003982422.5

A2 39781205 39781259 refGene_gn=SHQ1;refGene_tx=XM_003982422.5
A2 39783066 39783221 refGene_gn=SHQ1;refGene_tx=XM_003982422.5
A2 39792319 39792447 refGene_gn=SHQ1;refGene_tx=XM_003982422.5
A2 39799259 39799372 refGene_gn=SHQ1;refGene_tx=XM_003982422.5
A2 39808084 39808245 refGene_gn=SHQ1;refGene_tx=XM_003982422.5
A2 39809469 39809592 refGene_gn=SHQ1;refGene_tx=XM_003982422.5
A2 39811798 39811863 refGene_gn=SHQ1;refGene_tx=XM_003982422.5
A2 39816320 39816463 refGene_gn=SHQ1;refGene_tx=XM_003982422.5
A2 39849354 39849629 refGene_gn=GXYLT2;refGene_tx=XM_019825029.2
A2 39871953 39872146 refGene_gn=GXYLT2;refGene_tx=XM_019825029.2
A2 39880184 39880316 refGene_gn=GXYLT2;refGene_tx=XM_019825029.2
A2 39924210 39924462 refGene_gn=GXYLT2;refGene_tx=XM_019825029.2
A2 39925584 39925708 refGene_gn=GXYLT2;refGene_tx=XM_019825029.2
A2 39936269 39936442 refGene_gn=GXYLT2;refGene_tx=XM_019825029.2
A2 39941794 39941977 refGene_gn=GXYLT2;refGene_tx=XM_019825029.2
A2 39962621 39962655 refGene_gn=PPP4R2;refGene_tx=XM_023249965.1
A2 39963667 39963749 refGene_gn=PPP4R2;refGene_tx=XM_023249965.1
A2 39997968 39998139 refGene_gn=PPP4R2;refGene_tx=XM_023249965.1
A2 40007390 40007484 refGene_gn=PPP4R2;refGene_tx=XM_023249965.1
A2 40009300 40009338 refGene_gn=PPP4R2;refGene_tx=XM_023249965.1
A2 40010219 40010294 refGene_gn=PPP4R2;refGene_tx=XM_023249965.1
A2 40010530 40010674 refGene_gn=PPP4R2;refGene_tx=XM_023249965.1
A2 40011379 40011669 refGene_gn=PPP4R2;refGene_tx=XM_023249965.1
A2 40011923 40012243 refGene_gn=PPP4R2;refGene_tx=XM_023249965.1
A2 40287982 40289539
refGene_gn=PDZRN3;refGene_tx=XM_011280160.2;refGene_gn=PDZRN3;refGene_tx=XM_019824509.2;refGene_gn=PDZRN3;refGene_tx=XM_019824520.2
A2 40290234 40290351
refGene_gn=PDZRN3;refGene_tx=XM_011280160.2;refGene_gn=PDZRN3;refGene_tx=XM_019824509.2;refGene_gn=PDZRN3;refGene_tx=XM_019824520.2
A2 40291940 40292042
refGene_gn=PDZRN3;refGene_tx=XM_011280160.2;refGene_gn=PDZRN3;refGene_tx=XM_019824509.2;refGene_gn=PDZRN3;refGene_tx=XM_019824520.2
A2 40293842 40293905
refGene_gn=PDZRN3;refGene_tx=XM_011280160.2;refGene_gn=PDZRN3;refGene_tx=XM_019824509.2;refGene_gn=PDZRN3;refGene_tx=XM_019824520.2
A2 40294978 40295077
refGene_gn=PDZRN3;refGene_tx=XM_011280160.2;refGene_gn=PDZRN3;refGene_tx=XM_019824509.2;refGene_gn=PDZRN3;refGene_tx=XM_019824520.2
A2 40305586 40305674
refGene_gn=PDZRN3;refGene_tx=XM_011280160.2;refGene_gn=PDZRN3;refGene_tx=XM_019824509.2;refGene_gn=PDZRN3;refGene_tx=XM_019824520.2
A2 40308804 40309052
refGene_gn=PDZRN3;refGene_tx=XM_011280160.2;refGene_gn=PDZRN3;refGene_tx=XM_019824509.2;refGene_gn=PDZRN3;refGene_tx=XM_019824520.2
A2 40336350 40336419 refGene_gn=PDZRN3;refGene_tx=XM_019824509.2
A2 40356936 40356954 refGene_gn=PDZRN3;refGene_tx=XM_019824520.2
A2 40506079 40506187 refGene_gn=PDZRN3;refGene_tx=XM_011280160.2
A2 40511723 40511810 refGene_gn=PDZRN3;refGene_tx=XM_011280160.2
A2 40527659 40528382 refGene_gn=PDZRN3;refGene_tx=XM_011280160.2

A2 41161565 41161612 refGene_gn=CNTN3;refGene_tx=XM_023249968.1
A2 41162016 41162123
refGene_gn=CNTN3;refGene_tx=XM_023249967.1;refGene_gn=CNTN3;refGene_tx=XM_023249968.1;refGene_gn=CNTN3;refGene_tx=XM_023249969.1
A2 41163897 41164066
refGene_gn=CNTN3;refGene_tx=XM_023249967.1;refGene_gn=CNTN3;refGene_tx=XM_023249968.1;refGene_gn=CNTN3;refGene_tx=XM_023249969.1
A2 41164700 41164813
refGene_gn=CNTN3;refGene_tx=XM_023249967.1;refGene_gn=CNTN3;refGene_tx=XM_023249968.1;refGene_gn=CNTN3;refGene_tx=XM_023249969.1
A2 41180374 41180558
refGene_gn=CNTN3;refGene_tx=XM_023249967.1;refGene_gn=CNTN3;refGene_tx=XM_023249968.1;refGene_gn=CNTN3;refGene_tx=XM_023249969.1
A2 41191520 41191636
refGene_gn=CNTN3;refGene_tx=XM_023249967.1;refGene_gn=CNTN3;refGene_tx=XM_023249968.1;refGene_gn=CNTN3;refGene_tx=XM_023249969.1
A2 41194109 41194344
refGene_gn=CNTN3;refGene_tx=XM_023249967.1;refGene_gn=CNTN3;refGene_tx=XM_023249968.1;refGene_gn=CNTN3;refGene_tx=XM_023249969.1
A2 41195358 41195429
refGene_gn=CNTN3;refGene_tx=XM_023249967.1;refGene_gn=CNTN3;refGene_tx=XM_023249968.1;refGene_gn=CNTN3;refGene_tx=XM_023249969.1
A2 41197362 41197512
refGene_gn=CNTN3;refGene_tx=XM_023249967.1;refGene_gn=CNTN3;refGene_tx=XM_023249968.1
A2 41197611 41197770
refGene_gn=CNTN3;refGene_tx=XM_023249967.1;refGene_gn=CNTN3;refGene_tx=XM_023249968.1;refGene_gn=CNTN3;refGene_tx=XM_023249969.1
A2 41198629 41198747
refGene_gn=CNTN3;refGene_tx=XM_023249967.1;refGene_gn=CNTN3;refGene_tx=XM_023249968.1;refGene_gn=CNTN3;refGene_tx=XM_023249969.1
A2 41223883 41224059
refGene_gn=CNTN3;refGene_tx=XM_023249967.1;refGene_gn=CNTN3;refGene_tx=XM_023249968.1;refGene_gn=CNTN3;refGene_tx=XM_023249969.1
A2 41226853 41226981
refGene_gn=CNTN3;refGene_tx=XM_023249967.1;refGene_gn=CNTN3;refGene_tx=XM_023249968.1;refGene_gn=CNTN3;refGene_tx=XM_023249969.1
A2 41251002 41251153
refGene_gn=CNTN3;refGene_tx=XM_023249967.1;refGene_gn=CNTN3;refGene_tx=XM_023249968.1;refGene_gn=CNTN3;refGene_tx=XM_023249969.1
A2 41253852 41253982
refGene_gn=CNTN3;refGene_tx=XM_023249967.1;refGene_gn=CNTN3;refGene_tx=XM_023249968.1;refGene_gn=CNTN3;refGene_tx=XM_023249969.1
A2 41254941 41255078
refGene_gn=CNTN3;refGene_tx=XM_023249967.1;refGene_gn=CNTN3;refGene_tx=XM_023249968.1;refGene_gn=CNTN3;refGene_tx=XM_023249969.1
A2 41258723 41258908
refGene_gn=CNTN3;refGene_tx=XM_023249967.1;refGene_gn=CNTN3;refGene_tx=XM_023249968.1;refGene_gn=CNTN3;refGene_tx=XM_023249969.1
A2 41259668 41259771

refGene_gn=CNTN3;refGene_tx=XM_023249967.1;refGene_gn=CNTN3;refGene_tx
=XM_023249968.1;refGene_gn=CNTN3;refGene_tx=XM_023249969.1
A2 41261226 41261430
refGene_gn=CNTN3;refGene_tx=XM_023249967.1;refGene_gn=CNTN3;refGene_tx
=XM_023249968.1;refGene_gn=CNTN3;refGene_tx=XM_023249969.1
A2 41313237 41313333
refGene_gn=CNTN3;refGene_tx=XM_023249967.1;refGene_gn=CNTN3;refGene_tx
=XM_023249968.1;refGene_gn=CNTN3;refGene_tx=XM_023249969.1
A2 41371957 41372133
refGene_gn=CNTN3;refGene_tx=XM_023249967.1;refGene_gn=CNTN3;refGene_tx
=XM_023249968.1;refGene_gn=CNTN3;refGene_tx=XM_023249969.1
A2 41383258 41383385
refGene_gn=CNTN3;refGene_tx=XM_023249967.1;refGene_gn=CNTN3;refGene_tx
=XM_023249968.1;refGene_gn=CNTN3;refGene_tx=XM_023249969.1
A2 41411677 41411732
refGene_gn=CNTN3;refGene_tx=XM_023249967.1;refGene_gn=CNTN3;refGene_tx
=XM_023249968.1;refGene_gn=CNTN3;refGene_tx=XM_023249969.1
A2 42348786 42348871
refGene_gn=CHL1;refGene_tx=XM_006928940.4;refGene_gn=CHL1;refGene_tx=X
M_023249970.1;refGene_gn=CHL1;refGene_tx=XM_023249971.1;refGene_gn=CHL
1;refGene_tx=XM_023249972.1;refGene_gn=CHL1;refGene_tx=XM_023249973.1;
refGene_gn=CHL1;refGene_tx=XM_023249974.1;refGene_gn=CHL1;refGene_tx=X
M_023249975.1;refGene_gn=CHL1;refGene_tx=XM_023249976.1;refGene_gn=CHL
1;refGene_tx=XM_023249977.1
A2 42355372 42355478
refGene_gn=CHL1;refGene_tx=XM_006928940.4;refGene_gn=CHL1;refGene_tx=X
M_023249970.1;refGene_gn=CHL1;refGene_tx=XM_023249971.1;refGene_gn=CHL
1;refGene_tx=XM_023249972.1;refGene_gn=CHL1;refGene_tx=XM_023249973.1;
refGene_gn=CHL1;refGene_tx=XM_023249974.1;refGene_gn=CHL1;refGene_tx=X
M_023249975.1;refGene_gn=CHL1;refGene_tx=XM_023249976.1;refGene_gn=CHL
1;refGene_tx=XM_023249977.1
A2 42358622 42358810
refGene_gn=CHL1;refGene_tx=XM_006928940.4;refGene_gn=CHL1;refGene_tx=X
M_023249970.1;refGene_gn=CHL1;refGene_tx=XM_023249971.1;refGene_gn=CHL
1;refGene_tx=XM_023249972.1;refGene_gn=CHL1;refGene_tx=XM_023249973.1;
refGene_gn=CHL1;refGene_tx=XM_023249974.1;refGene_gn=CHL1;refGene_tx=X
M_023249975.1;refGene_gn=CHL1;refGene_tx=XM_023249976.1;refGene_gn=CHL
1;refGene_tx=XM_023249977.1
A2 42369909 42370032
refGene_gn=CHL1;refGene_tx=XM_006928940.4;refGene_gn=CHL1;refGene_tx=X
M_023249970.1;refGene_gn=CHL1;refGene_tx=XM_023249971.1;refGene_gn=CHL
1;refGene_tx=XM_023249972.1;refGene_gn=CHL1;refGene_tx=XM_023249973.1;
refGene_gn=CHL1;refGene_tx=XM_023249974.1;refGene_gn=CHL1;refGene_tx=X
M_023249975.1;refGene_gn=CHL1;refGene_tx=XM_023249976.1;refGene_gn=CHL
1;refGene_tx=XM_023249977.1
A2 42371148 42371319
refGene_gn=CHL1;refGene_tx=XM_006928940.4;refGene_gn=CHL1;refGene_tx=X
M_023249970.1;refGene_gn=CHL1;refGene_tx=XM_023249971.1;refGene_gn=CHL
1;refGene_tx=XM_023249972.1;refGene_gn=CHL1;refGene_tx=XM_023249973.1;
refGene_gn=CHL1;refGene_tx=XM_023249974.1;refGene_gn=CHL1;refGene_tx=X

M_023249975.1;refGene_gn=CHL1;refGene_tx=XM_023249976.1;refGene_gn=CHL1;refGene_tx=XM_023249977.1
A2 42372280 42372328
refGene_gn=CHL1;refGene_tx=XM_006928940.4;refGene_gn=CHL1;refGene_tx=XM_023249970.1;refGene_gn=CHL1;refGene_tx=XM_023249971.1;refGene_gn=CHL1;refGene_tx=XM_023249972.1;refGene_gn=CHL1;refGene_tx=XM_023249973.1;refGene_gn=CHL1;refGene_tx=XM_023249974.1;refGene_gn=CHL1;refGene_tx=XM_023249975.1;refGene_gn=CHL1;refGene_tx=XM_023249977.1
A2 42374674 42374795
refGene_gn=CHL1;refGene_tx=XM_006928940.4;refGene_gn=CHL1;refGene_tx=XM_023249970.1;refGene_gn=CHL1;refGene_tx=XM_023249971.1;refGene_gn=CHL1;refGene_tx=XM_023249972.1;refGene_gn=CHL1;refGene_tx=XM_023249973.1;refGene_gn=CHL1;refGene_tx=XM_023249974.1;refGene_gn=CHL1;refGene_tx=XM_023249975.1;refGene_gn=CHL1;refGene_tx=XM_023249976.1;refGene_gn=CHL1;refGene_tx=XM_023249977.1
A2 42379058 42379243
refGene_gn=CHL1;refGene_tx=XM_006928940.4;refGene_gn=CHL1;refGene_tx=XM_023249970.1;refGene_gn=CHL1;refGene_tx=XM_023249971.1;refGene_gn=CHL1;refGene_tx=XM_023249972.1;refGene_gn=CHL1;refGene_tx=XM_023249973.1;refGene_gn=CHL1;refGene_tx=XM_023249974.1;refGene_gn=CHL1;refGene_tx=XM_023249975.1;refGene_gn=CHL1;refGene_tx=XM_023249976.1;refGene_gn=CHL1;refGene_tx=XM_023249977.1
A2 42385201 42385333
refGene_gn=CHL1;refGene_tx=XM_006928940.4;refGene_gn=CHL1;refGene_tx=XM_023249970.1;refGene_gn=CHL1;refGene_tx=XM_023249971.1;refGene_gn=CHL1;refGene_tx=XM_023249972.1;refGene_gn=CHL1;refGene_tx=XM_023249973.1;refGene_gn=CHL1;refGene_tx=XM_023249974.1;refGene_gn=CHL1;refGene_tx=XM_023249975.1;refGene_gn=CHL1;refGene_tx=XM_023249976.1;refGene_gn=CHL1;refGene_tx=XM_023249977.1
A2 42390907 42391048
refGene_gn=CHL1;refGene_tx=XM_006928940.4;refGene_gn=CHL1;refGene_tx=XM_023249970.1;refGene_gn=CHL1;refGene_tx=XM_023249971.1;refGene_gn=CHL1;refGene_tx=XM_023249972.1;refGene_gn=CHL1;refGene_tx=XM_023249973.1;refGene_gn=CHL1;refGene_tx=XM_023249974.1;refGene_gn=CHL1;refGene_tx=XM_023249975.1;refGene_gn=CHL1;refGene_tx=XM_023249976.1;refGene_gn=CHL1;refGene_tx=XM_023249977.1
A2 42391375 42391487
refGene_gn=CHL1;refGene_tx=XM_006928940.4;refGene_gn=CHL1;refGene_tx=XM_023249970.1;refGene_gn=CHL1;refGene_tx=XM_023249971.1;refGene_gn=CHL1;refGene_tx=XM_023249972.1;refGene_gn=CHL1;refGene_tx=XM_023249973.1;refGene_gn=CHL1;refGene_tx=XM_023249974.1;refGene_gn=CHL1;refGene_tx=XM_023249975.1;refGene_gn=CHL1;refGene_tx=XM_023249976.1;refGene_gn=CHL1;refGene_tx=XM_023249977.1
A2 42393422 42393589
refGene_gn=CHL1;refGene_tx=XM_006928940.4;refGene_gn=CHL1;refGene_tx=XM_023249970.1;refGene_gn=CHL1;refGene_tx=XM_023249971.1;refGene_gn=CHL1;refGene_tx=XM_023249972.1;refGene_gn=CHL1;refGene_tx=XM_023249973.1;refGene_gn=CHL1;refGene_tx=XM_023249974.1;refGene_gn=CHL1;refGene_tx=XM_023249975.1;refGene_gn=CHL1;refGene_tx=XM_023249976.1;refGene_gn=CHL1;refGene_tx=XM_023249977.1

A2 42396630 42396796
refGene_gn=CHL1;refGene_tx=XM_006928940.4;refGene_gn=CHL1;refGene_tx=XM_023249970.1;refGene_gn=CHL1;refGene_tx=XM_023249971.1;refGene_gn=CHL1;refGene_tx=XM_023249972.1;refGene_gn=CHL1;refGene_tx=XM_023249973.1;refGene_gn=CHL1;refGene_tx=XM_023249974.1;refGene_gn=CHL1;refGene_tx=XM_023249975.1;refGene_gn=CHL1;refGene_tx=XM_023249976.1;refGene_gn=CHL1;refGene_tx=XM_023249977.1
A2 42400045 42400170
refGene_gn=CHL1;refGene_tx=XM_006928940.4;refGene_gn=CHL1;refGene_tx=XM_023249970.1;refGene_gn=CHL1;refGene_tx=XM_023249971.1;refGene_gn=CHL1;refGene_tx=XM_023249972.1;refGene_gn=CHL1;refGene_tx=XM_023249973.1;refGene_gn=CHL1;refGene_tx=XM_023249974.1;refGene_gn=CHL1;refGene_tx=XM_023249975.1;refGene_gn=CHL1;refGene_tx=XM_023249976.1;refGene_gn=CHL1;refGene_tx=XM_023249977.1
A2 42404770 42404872
refGene_gn=CHL1;refGene_tx=XM_006928940.4;refGene_gn=CHL1;refGene_tx=XM_023249970.1;refGene_gn=CHL1;refGene_tx=XM_023249971.1;refGene_gn=CHL1;refGene_tx=XM_023249972.1;refGene_gn=CHL1;refGene_tx=XM_023249973.1;refGene_gn=CHL1;refGene_tx=XM_023249974.1;refGene_gn=CHL1;refGene_tx=XM_023249975.1;refGene_gn=CHL1;refGene_tx=XM_023249976.1;refGene_gn=CHL1;refGene_tx=XM_023249977.1
A2 42405321 42405519
refGene_gn=CHL1;refGene_tx=XM_006928940.4;refGene_gn=CHL1;refGene_tx=XM_023249970.1;refGene_gn=CHL1;refGene_tx=XM_023249971.1;refGene_gn=CHL1;refGene_tx=XM_023249972.1;refGene_gn=CHL1;refGene_tx=XM_023249973.1;refGene_gn=CHL1;refGene_tx=XM_023249974.1;refGene_gn=CHL1;refGene_tx=XM_023249975.1;refGene_gn=CHL1;refGene_tx=XM_023249976.1;refGene_gn=CHL1;refGene_tx=XM_023249977.1
A2 42407520 42407591
refGene_gn=CHL1;refGene_tx=XM_006928940.4;refGene_gn=CHL1;refGene_tx=XM_023249970.1;refGene_gn=CHL1;refGene_tx=XM_023249971.1;refGene_gn=CHL1;refGene_tx=XM_023249972.1;refGene_gn=CHL1;refGene_tx=XM_023249973.1;refGene_gn=CHL1;refGene_tx=XM_023249974.1;refGene_gn=CHL1;refGene_tx=XM_023249975.1;refGene_gn=CHL1;refGene_tx=XM_023249976.1;refGene_gn=CHL1;refGene_tx=XM_023249977.1
A2 42410590 42410813
refGene_gn=CHL1;refGene_tx=XM_006928940.4;refGene_gn=CHL1;refGene_tx=XM_023249970.1;refGene_gn=CHL1;refGene_tx=XM_023249971.1;refGene_gn=CHL1;refGene_tx=XM_023249972.1;refGene_gn=CHL1;refGene_tx=XM_023249973.1;refGene_gn=CHL1;refGene_tx=XM_023249974.1;refGene_gn=CHL1;refGene_tx=XM_023249975.1;refGene_gn=CHL1;refGene_tx=XM_023249976.1;refGene_gn=CHL1;refGene_tx=XM_023249977.1
A2 42411989 42412105
refGene_gn=CHL1;refGene_tx=XM_006928940.4;refGene_gn=CHL1;refGene_tx=XM_023249970.1;refGene_gn=CHL1;refGene_tx=XM_023249971.1;refGene_gn=CHL1;refGene_tx=XM_023249972.1;refGene_gn=CHL1;refGene_tx=XM_023249973.1;refGene_gn=CHL1;refGene_tx=XM_023249974.1;refGene_gn=CHL1;refGene_tx=XM_023249975.1;refGene_gn=CHL1;refGene_tx=XM_023249976.1;refGene_gn=CHL1;refGene_tx=XM_023249977.1
A2 42412237 42412442

refGene_gn=CHL1;refGene_tx=XM_006928940.4;refGene_gn=CHL1;refGene_tx=XM_023249970.1;refGene_gn=CHL1;refGene_tx=XM_023249971.1;refGene_gn=CHL1;refGene_tx=XM_023249972.1;refGene_gn=CHL1;refGene_tx=XM_023249973.1;refGene_gn=CHL1;refGene_tx=XM_023249974.1;refGene_gn=CHL1;refGene_tx=XM_023249975.1;refGene_gn=CHL1;refGene_tx=XM_023249976.1;refGene_gn=CHL1;refGene_tx=XM_023249977.1
A2 42412649 42412772
refGene_gn=CHL1;refGene_tx=XM_006928940.4;refGene_gn=CHL1;refGene_tx=XM_023249970.1;refGene_gn=CHL1;refGene_tx=XM_023249971.1;refGene_gn=CHL1;refGene_tx=XM_023249972.1;refGene_gn=CHL1;refGene_tx=XM_023249973.1;refGene_gn=CHL1;refGene_tx=XM_023249974.1;refGene_gn=CHL1;refGene_tx=XM_023249975.1;refGene_gn=CHL1;refGene_tx=XM_023249976.1;refGene_gn=CHL1;refGene_tx=XM_023249977.1
A2 42415890 42416070
refGene_gn=CHL1;refGene_tx=XM_006928940.4;refGene_gn=CHL1;refGene_tx=XM_023249970.1;refGene_gn=CHL1;refGene_tx=XM_023249971.1;refGene_gn=CHL1;refGene_tx=XM_023249972.1;refGene_gn=CHL1;refGene_tx=XM_023249973.1;refGene_gn=CHL1;refGene_tx=XM_023249974.1;refGene_gn=CHL1;refGene_tx=XM_023249975.1;refGene_gn=CHL1;refGene_tx=XM_023249976.1;refGene_gn=CHL1;refGene_tx=XM_023249977.1
A2 42419689 42419851
refGene_gn=CHL1;refGene_tx=XM_006928940.4;refGene_gn=CHL1;refGene_tx=XM_023249970.1;refGene_gn=CHL1;refGene_tx=XM_023249971.1;refGene_gn=CHL1;refGene_tx=XM_023249972.1;refGene_gn=CHL1;refGene_tx=XM_023249973.1;refGene_gn=CHL1;refGene_tx=XM_023249974.1;refGene_gn=CHL1;refGene_tx=XM_023249975.1;refGene_gn=CHL1;refGene_tx=XM_023249976.1
A2 42420494 42420626
refGene_gn=CHL1;refGene_tx=XM_006928940.4;refGene_gn=CHL1;refGene_tx=XM_023249970.1;refGene_gn=CHL1;refGene_tx=XM_023249971.1;refGene_gn=CHL1;refGene_tx=XM_023249972.1;refGene_gn=CHL1;refGene_tx=XM_023249973.1;refGene_gn=CHL1;refGene_tx=XM_023249974.1;refGene_gn=CHL1;refGene_tx=XM_023249975.1;refGene_gn=CHL1;refGene_tx=XM_023249976.1;refGene_gn=CHL1;refGene_tx=XM_023249977.1
A2 42422546 42422619
refGene_gn=CHL1;refGene_tx=XM_006928940.4;refGene_gn=CHL1;refGene_tx=XM_023249970.1;refGene_gn=CHL1;refGene_tx=XM_023249971.1;refGene_gn=CHL1;refGene_tx=XM_023249972.1;refGene_gn=CHL1;refGene_tx=XM_023249973.1;refGene_gn=CHL1;refGene_tx=XM_023249974.1;refGene_gn=CHL1;refGene_tx=XM_023249975.1;refGene_gn=CHL1;refGene_tx=XM_023249976.1;refGene_gn=CHL1;refGene_tx=XM_023249977.1
A2 42425820 42426037
refGene_gn=CHL1;refGene_tx=XM_006928940.4;refGene_gn=CHL1;refGene_tx=XM_023249970.1;refGene_gn=CHL1;refGene_tx=XM_023249971.1;refGene_gn=CHL1;refGene_tx=XM_023249972.1;refGene_gn=CHL1;refGene_tx=XM_023249973.1;refGene_gn=CHL1;refGene_tx=XM_023249974.1;refGene_gn=CHL1;refGene_tx=XM_023249975.1;refGene_gn=CHL1;refGene_tx=XM_023249976.1;refGene_gn=CHL1;refGene_tx=XM_023249977.1
A2 43135349 43135404 refGene_gn=CNTN6;refGene_tx=XM_023249978.1
A2 43159563 43159659 refGene_gn=CNTN6;refGene_tx=XM_023249978.1
A2 43194034 43194161 refGene_gn=CNTN6;refGene_tx=XM_023249978.1

A2 43201183 43201359 refGene_gn=CNTN6;refGene_tx=XM_023249978.1
A2 43243287 43243383 refGene_gn=CNTN6;refGene_tx=XM_023249978.1
A2 43257993 43258197 refGene_gn=CNTN6;refGene_tx=XM_023249978.1
A2 43260459 43260562 refGene_gn=CNTN6;refGene_tx=XM_023249978.1
A2 43281416 43281601 refGene_gn=CNTN6;refGene_tx=XM_023249978.1
A2 43285369 43285506 refGene_gn=CNTN6;refGene_tx=XM_023249978.1
A2 43286986 43287116 refGene_gn=CNTN6;refGene_tx=XM_023249978.1
A2 43289087 43289238 refGene_gn=CNTN6;refGene_tx=XM_023249978.1
A2 43316044 43316172 refGene_gn=CNTN6;refGene_tx=XM_023249978.1
A2 43337590 43337766 refGene_gn=CNTN6;refGene_tx=XM_023249978.1
A2 43338110 43338228 refGene_gn=CNTN6;refGene_tx=XM_023249978.1
A2 43338881 43339040 refGene_gn=CNTN6;refGene_tx=XM_023249978.1
A2 43339153 43339303 refGene_gn=CNTN6;refGene_tx=XM_023249978.1
A2 43343234 43343305 refGene_gn=CNTN6;refGene_tx=XM_023249978.1
A2 43350772 43351007 refGene_gn=CNTN6;refGene_tx=XM_023249978.1
A2 43351120 43351236 refGene_gn=CNTN6;refGene_tx=XM_023249978.1
A2 43352599 43352786 refGene_gn=CNTN6;refGene_tx=XM_023249978.1
A2 43362326 43362439 refGene_gn=CNTN6;refGene_tx=XM_023249978.1
A2 43363217 43363386 refGene_gn=CNTN6;refGene_tx=XM_023249978.1
A2 43364235 43364296 refGene_gn=CNTN6;refGene_tx=XM_023249978.1
A2 43364297 43364340 refGene_gn=CNTN6;refGene_tx=XM_023249978.1
A2 44420530 44420585
refGene_gn=CNTN4;refGene_tx=XM_011280163.3;refGene_gn=CNTN4;refGene_tx
=XM_019824593.2;refGene_gn=CNTN4;refGene_tx=XM_019824595.2
A2 44572793 44572920
refGene_gn=CNTN4;refGene_tx=XM_011280163.3;refGene_gn=CNTN4;refGene_tx
=XM_019824593.2;refGene_gn=CNTN4;refGene_tx=XM_019824595.2
A2 44581033 44581209
refGene_gn=CNTN4;refGene_tx=XM_011280163.3;refGene_gn=CNTN4;refGene_tx
=XM_019824593.2;refGene_gn=CNTN4;refGene_tx=XM_019824595.2
A2 44649360 44649456
refGene_gn=CNTN4;refGene_tx=XM_011280163.3;refGene_gn=CNTN4;refGene_tx
=XM_019824593.2;refGene_gn=CNTN4;refGene_tx=XM_019824595.2
A2 44681234 44681250 refGene_gn=CNTN4;refGene_tx=XM_023249979.1
A2 44695862 44696060
refGene_gn=CNTN4;refGene_tx=XM_011280163.3;refGene_gn=CNTN4;refGene_tx
=XM_019824593.2;refGene_gn=CNTN4;refGene_tx=XM_019824595.2;refGene_gn=
CNTN4;refGene_tx=XM_023249979.1
A2 44712002 44712105
refGene_gn=CNTN4;refGene_tx=XM_011280163.3;refGene_gn=CNTN4;refGene_tx
=XM_019824593.2;refGene_gn=CNTN4;refGene_tx=XM_019824595.2;refGene_gn=
CNTN4;refGene_tx=XM_023249979.1
A2 44714748 44714933
refGene_gn=CNTN4;refGene_tx=XM_011280163.3;refGene_gn=CNTN4;refGene_tx
=XM_019824593.2;refGene_gn=CNTN4;refGene_tx=XM_019824595.2;refGene_gn=
CNTN4;refGene_tx=XM_023249979.1
A2 44727863 44728000
refGene_gn=CNTN4;refGene_tx=XM_011280163.3;refGene_gn=CNTN4;refGene_tx
=XM_019824593.2;refGene_gn=CNTN4;refGene_tx=XM_019824595.2;refGene_gn=
CNTN4;refGene_tx=XM_023249979.1

A2 44732041 44732171
refGene_gn=CNTN4;refGene_tx=XM_011280163.3;refGene_gn=CNTN4;refGene_tx=XM_019824593.2;refGene_gn=CNTN4;refGene_tx=XM_019824595.2;refGene_gn=CNTN4;refGene_tx=XM_023249979.1
A2 44750094 44750245
refGene_gn=CNTN4;refGene_tx=XM_011280163.3;refGene_gn=CNTN4;refGene_tx=XM_019824593.2;refGene_gn=CNTN4;refGene_tx=XM_019824595.2;refGene_gn=CNTN4;refGene_tx=XM_023249979.1
A2 44811486 44811614
refGene_gn=CNTN4;refGene_tx=XM_011280163.3;refGene_gn=CNTN4;refGene_tx=XM_019824593.2;refGene_gn=CNTN4;refGene_tx=XM_019824595.2;refGene_gn=CNTN4;refGene_tx=XM_023249979.1
A2 44850865 44851041
refGene_gn=CNTN4;refGene_tx=XM_011280163.3;refGene_gn=CNTN4;refGene_tx=XM_019824593.2;refGene_gn=CNTN4;refGene_tx=XM_019824595.2;refGene_gn=CNTN4;refGene_tx=XM_023249979.1
A2 44855176 44855297
refGene_gn=CNTN4;refGene_tx=XM_011280163.3;refGene_gn=CNTN4;refGene_tx=XM_019824593.2;refGene_gn=CNTN4;refGene_tx=XM_019824595.2;refGene_gn=CNTN4;refGene_tx=XM_023249979.1
A2 44859417 44859576
refGene_gn=CNTN4;refGene_tx=XM_011280163.3;refGene_gn=CNTN4;refGene_tx=XM_019824593.2;refGene_gn=CNTN4;refGene_tx=XM_019824595.2;refGene_gn=CNTN4;refGene_tx=XM_023249979.1
A2 44862243 44862393
refGene_gn=CNTN4;refGene_tx=XM_011280163.3;refGene_gn=CNTN4;refGene_tx=XM_019824593.2;refGene_gn=CNTN4;refGene_tx=XM_019824595.2;refGene_gn=CNTN4;refGene_tx=XM_023249979.1
A2 44863924 44863995
refGene_gn=CNTN4;refGene_tx=XM_011280163.3;refGene_gn=CNTN4;refGene_tx=XM_019824593.2;refGene_gn=CNTN4;refGene_tx=XM_019824595.2;refGene_gn=CNTN4;refGene_tx=XM_023249979.1
A2 44865156 44865391
refGene_gn=CNTN4;refGene_tx=XM_011280163.3;refGene_gn=CNTN4;refGene_tx=XM_019824593.2;refGene_gn=CNTN4;refGene_tx=XM_019824595.2;refGene_gn=CNTN4;refGene_tx=XM_023249979.1
A2 44866338 44866451
refGene_gn=CNTN4;refGene_tx=XM_011280163.3;refGene_gn=CNTN4;refGene_tx=XM_019824593.2;refGene_gn=CNTN4;refGene_tx=XM_019824595.2;refGene_gn=CNTN4;refGene_tx=XM_023249979.1
A2 44867230 44867417
refGene_gn=CNTN4;refGene_tx=XM_011280163.3;refGene_gn=CNTN4;refGene_tx=XM_019824593.2;refGene_gn=CNTN4;refGene_tx=XM_019824595.2;refGene_gn=CNTN4;refGene_tx=XM_023249979.1
A2 44867849 44867962
refGene_gn=CNTN4;refGene_tx=XM_011280163.3;refGene_gn=CNTN4;refGene_tx=XM_019824593.2;refGene_gn=CNTN4;refGene_tx=XM_019824595.2;refGene_gn=CNTN4;refGene_tx=XM_023249979.1
A2 44873375 44873544
refGene_gn=CNTN4;refGene_tx=XM_011280163.3;refGene_gn=CNTN4;refGene_tx

=XM_019824593.2;refGene_gn=CNTN4;refGene_tx=XM_019824595.2;refGene_gn=CNTN4;refGene_tx=XM_023249979.1
A2 44876487 44876588
refGene_gn=CNTN4;refGene_tx=XM_011280163.3;refGene_gn=CNTN4;refGene_tx=XM_019824593.2;refGene_gn=CNTN4;refGene_tx=XM_019824595.2;refGene_gn=CNTN4;refGene_tx=XM_023249979.1
A2 44889940 44890027
refGene_gn=IL5RA;refGene_tx=XM_011280164.3;refGene_gn=IL5RA;refGene_tx=XM_019824621.2
A2 44893195 44893280
refGene_gn=IL5RA;refGene_tx=XM_011280164.3;refGene_gn=IL5RA;refGene_tx=XM_019824621.2
A2 44894795 44894892
refGene_gn=IL5RA;refGene_tx=XM_011280164.3;refGene_gn=IL5RA;refGene_tx=XM_019824621.2;refGene_gn=IL5RA;refGene_tx=XM_023249982.1
A2 44907066 44907205
refGene_gn=IL5RA;refGene_tx=XM_011280164.3;refGene_gn=IL5RA;refGene_tx=XM_019824621.2;refGene_gn=IL5RA;refGene_tx=XM_023249982.1
A2 44910536 44910682
refGene_gn=IL5RA;refGene_tx=XM_011280164.3;refGene_gn=IL5RA;refGene_tx=XM_019824621.2;refGene_gn=IL5RA;refGene_tx=XM_023249982.1
A2 44912720 44912908
refGene_gn=IL5RA;refGene_tx=XM_011280164.3;refGene_gn=IL5RA;refGene_tx=XM_019824621.2;refGene_gn=IL5RA;refGene_tx=XM_023249982.1
A2 44912982 44913136
refGene_gn=IL5RA;refGene_tx=XM_011280164.3;refGene_gn=IL5RA;refGene_tx=XM_019824621.2;refGene_gn=IL5RA;refGene_tx=XM_023249982.1
A2 44916142 44916281
refGene_gn=IL5RA;refGene_tx=XM_011280164.3;refGene_gn=IL5RA;refGene_tx=XM_019824621.2;refGene_gn=IL5RA;refGene_tx=XM_023249982.1
A2 44917361 44917507
refGene_gn=IL5RA;refGene_tx=XM_011280164.3;refGene_gn=IL5RA;refGene_tx=XM_019824621.2;refGene_gn=IL5RA;refGene_tx=XM_023249982.1
A2 44919003 44919076
refGene_gn=IL5RA;refGene_tx=XM_011280164.3;refGene_gn=IL5RA;refGene_tx=XM_019824621.2;refGene_gn=IL5RA;refGene_tx=XM_023249982.1
A2 44937684 44937832
refGene_gn=TRNT1;refGene_tx=XM_006928945.4;refGene_gn=TRNT1;refGene_tx=XM_006928946.4;refGene_gn=TRNT1;refGene_tx=XM_023249981.1
A2 44939732 44939742 refGene_gn=TRNT1;refGene_tx=XM_006928948.4
A2 44943912 44944106
refGene_gn=TRNT1;refGene_tx=XM_006928945.4;refGene_gn=TRNT1;refGene_tx=XM_006928946.4;refGene_gn=TRNT1;refGene_tx=XM_006928948.4;refGene_gn=TRNT1;refGene_tx=XM_023249981.1
A2 44947894 44948033
refGene_gn=TRNT1;refGene_tx=XM_006928945.4;refGene_gn=TRNT1;refGene_tx=XM_006928946.4;refGene_gn=TRNT1;refGene_tx=XM_006928948.4;refGene_gn=TRNT1;refGene_tx=XM_023249981.1
A2 44952124 44952251
refGene_gn=TRNT1;refGene_tx=XM_006928945.4;refGene_gn=TRNT1;refGene_tx

=XM_006928946.4;refGene_gn=TRNT1;refGene_tx=XM_006928948.4;refGene_gn=TRNT1;refGene_tx=XM_023249981.1
A2 44954304 44954498 refGene_gn=TRNT1;refGene_tx=XM_006928945.4;refGene_gn=TRNT1;refGene_tx=XM_006928946.4;refGene_gn=TRNT1;refGene_tx=XM_006928948.4;refGene_gn=TRNT1;refGene_tx=XM_023249981.1
A2 44955285 44955539 refGene_gn=TRNT1;refGene_tx=XM_006928945.4;refGene_gn=TRNT1;refGene_tx=XM_006928946.4;refGene_gn=TRNT1;refGene_tx=XM_006928948.4;refGene_gn=TRNT1;refGene_tx=XM_023249981.1
A2 44955715 44955970 refGene_gn=TRNT1;refGene_tx=XM_006928945.4;refGene_gn=TRNT1;refGene_tx=XM_006928946.4;refGene_gn=TRNT1;refGene_tx=XM_006928948.4;refGene_gn=TRNT1;refGene_tx=XM_023249981.1
A2 44958336 44958517 refGene_gn=CRBN;refGene_tx=XM_003982431.4;refGene_gn=CRBN;refGene_tx=XM_023249980.1
A2 44959449 44959581 refGene_gn=CRBN;refGene_tx=XM_003982431.4;refGene_gn=CRBN;refGene_tx=XM_023249980.1
A2 44960230 44960295 refGene_gn=CRBN;refGene_tx=XM_003982431.4;refGene_gn=CRBN;refGene_tx=XM_023249980.1
A2 44960931 44961047 refGene_gn=CRBN;refGene_tx=XM_003982431.4;refGene_gn=CRBN;refGene_tx=XM_023249980.1
A2 44961647 44961732 refGene_gn=CRBN;refGene_tx=XM_003982431.4;refGene_gn=CRBN;refGene_tx=XM_023249980.1
A2 44962955 44963018 refGene_gn=CRBN;refGene_tx=XM_003982431.4;refGene_gn=CRBN;refGene_tx=XM_023249980.1
A2 44971323 44971483 refGene_gn=CRBN;refGene_tx=XM_003982431.4;refGene_gn=CRBN;refGene_tx=XM_023249980.1
A2 44975401 44975551 refGene_gn=CRBN;refGene_tx=XM_003982431.4;refGene_gn=CRBN;refGene_tx=XM_023249980.1
A2 44976387 44976590 refGene_gn=CRBN;refGene_tx=XM_003982431.4;refGene_gn=CRBN;refGene_tx=XM_023249980.1
A2 44977196 44977309 refGene_gn=CRBN;refGene_tx=XM_003982431.4;refGene_gn=CRBN;refGene_tx=XM_023249980.1
A2 44981456 44981523 refGene_gn=CRBN;refGene_tx=XM_003982431.4
A2 45362840 45363143 refGene_gn=LOC102902070;refGene_tx=XM_019825030.2
A2 45363258 45364077 refGene_gn=LOC102902070;refGene_tx=XM_019825030.2
A2 45364078 45366202

refGene_gn=LOC102902070;refGene_tx=XM_019825030.2
A2 45600935 45601451 refGene_gn=LRRN1;refGene_tx=XM_011280165.2
A2 45601452 45601926 refGene_gn=LRRN1;refGene_tx=XM_011280165.2
A2 45601927 45603088 refGene_gn=LRRN1;refGene_tx=XM_011280165.2
A2 46021495 46021612
refGene_gn=LOC101088560;refGene_tx=XM_011280168.3
A2 46030979 46031765
refGene_gn=LOC101088560;refGene_tx=XM_011280168.3
A2 46054637 46054748
refGene_gn=SUMF1;refGene_tx=XM_011280169.2;refGene_gn=SUMF1;refGene_tx=XM_019824640.1
A2 46065417 46065477
refGene_gn=SUMF1;refGene_tx=XM_011280169.2;refGene_gn=SUMF1;refGene_tx=XM_019824640.1
A2 46094175 46094289 refGene_gn=SUMF1;refGene_tx=XM_011280169.2
A2 46099578 46099693
refGene_gn=SUMF1;refGene_tx=XM_011280169.2;refGene_gn=SUMF1;refGene_tx=XM_019824640.1
A2 46100445 46100568
refGene_gn=SUMF1;refGene_tx=XM_011280169.2;refGene_gn=SUMF1;refGene_tx=XM_019824640.1
A2 46102456 46102539
refGene_gn=SUMF1;refGene_tx=XM_011280169.2;refGene_gn=SUMF1;refGene_tx=XM_019824640.1
A2 46127085 46127160
refGene_gn=SUMF1;refGene_tx=XM_011280169.2;refGene_gn=SUMF1;refGene_tx=XM_019824640.1
A2 46131077 46131251
refGene_gn=SUMF1;refGene_tx=XM_011280169.2;refGene_gn=SUMF1;refGene_tx=XM_019824640.1
A2 46149177 46149453
refGene_gn=SUMF1;refGene_tx=XM_011280169.2;refGene_gn=SUMF1;refGene_tx=XM_019824640.1
A2 46203746 46203838
refGene_gn=ITPR1;refGene_tx=XM_011280170.2;refGene_gn=ITPR1;refGene_tx=XM_019824643.1;refGene_gn=ITPR1;refGene_tx=XM_019824645.1;refGene_gn=ITPR1;refGene_tx=XM_019824647.1;refGene_gn=ITPR1;refGene_tx=XM_019824649.1;refGene_gn=ITPR1;refGene_tx=XM_019824650.1;refGene_gn=ITPR1;refGene_tx=XM_023249983.1
A2 46208728 46208799
refGene_gn=ITPR1;refGene_tx=XM_011280170.2;refGene_gn=ITPR1;refGene_tx=XM_019824643.1;refGene_gn=ITPR1;refGene_tx=XM_019824645.1;refGene_gn=ITPR1;refGene_tx=XM_019824647.1;refGene_gn=ITPR1;refGene_tx=XM_019824649.1;refGene_gn=ITPR1;refGene_tx=XM_019824650.1;refGene_gn=ITPR1;refGene_tx=XM_023249983.1
A2 46299300 46299416
refGene_gn=ITPR1;refGene_tx=XM_011280170.2;refGene_gn=ITPR1;refGene_tx=XM_019824643.1;refGene_gn=ITPR1;refGene_tx=XM_019824645.1;refGene_gn=ITPR1;refGene_tx=XM_019824647.1;refGene_gn=ITPR1;refGene_tx=XM_019824649.1;refGene_gn=ITPR1;refGene_tx=XM_019824650.1;refGene_gn=ITPR1;refGe

ne_tx=XM_023249983.1
A2	46308750	46308837
refGene_gn=ITPR1;refGene_tx=XM_011280170.2;refGene_gn=ITPR1;refGene_tx=XM_019824643.1;refGene_gn=ITPR1;refGene_tx=XM_019824645.1;refGene_gn=ITPR1;refGene_tx=XM_019824647.1;refGene_gn=ITPR1;refGene_tx=XM_019824649.1;refGene_gn=ITPR1;refGene_tx=XM_019824650.1;refGene_gn=ITPR1;refGene_tx=XM_023249983.1
A2	46311077	46311236
refGene_gn=ITPR1;refGene_tx=XM_011280170.2;refGene_gn=ITPR1;refGene_tx=XM_019824643.1;refGene_gn=ITPR1;refGene_tx=XM_019824645.1;refGene_gn=ITPR1;refGene_tx=XM_019824647.1;refGene_gn=ITPR1;refGene_tx=XM_019824649.1;refGene_gn=ITPR1;refGene_tx=XM_019824650.1;refGene_gn=ITPR1;refGene_tx=XM_023249983.1
A2	46313136	46313235
refGene_gn=ITPR1;refGene_tx=XM_011280170.2;refGene_gn=ITPR1;refGene_tx=XM_019824643.1;refGene_gn=ITPR1;refGene_tx=XM_019824645.1;refGene_gn=ITPR1;refGene_tx=XM_019824647.1;refGene_gn=ITPR1;refGene_tx=XM_019824649.1;refGene_gn=ITPR1;refGene_tx=XM_019824650.1;refGene_gn=ITPR1;refGene_tx=XM_023249983.1
A2	46314254	46314338
refGene_gn=ITPR1;refGene_tx=XM_011280170.2;refGene_gn=ITPR1;refGene_tx=XM_019824643.1;refGene_gn=ITPR1;refGene_tx=XM_019824645.1;refGene_gn=ITPR1;refGene_tx=XM_019824647.1;refGene_gn=ITPR1;refGene_tx=XM_019824649.1;refGene_gn=ITPR1;refGene_tx=XM_019824650.1;refGene_gn=ITPR1;refGene_tx=XM_023249983.1
A2	46314446	46314593
refGene_gn=ITPR1;refGene_tx=XM_011280170.2;refGene_gn=ITPR1;refGene_tx=XM_019824643.1;refGene_gn=ITPR1;refGene_tx=XM_019824645.1;refGene_gn=ITPR1;refGene_tx=XM_019824647.1;refGene_gn=ITPR1;refGene_tx=XM_019824649.1;refGene_gn=ITPR1;refGene_tx=XM_019824650.1;refGene_gn=ITPR1;refGene_tx=XM_023249983.1
A2	46318577	46318673
refGene_gn=ITPR1;refGene_tx=XM_011280170.2;refGene_gn=ITPR1;refGene_tx=XM_019824643.1;refGene_gn=ITPR1;refGene_tx=XM_019824645.1;refGene_gn=ITPR1;refGene_tx=XM_019824647.1;refGene_gn=ITPR1;refGene_tx=XM_019824649.1;refGene_gn=ITPR1;refGene_tx=XM_019824650.1;refGene_gn=ITPR1;refGene_tx=XM_023249983.1
A2	46320166	46320211
refGene_gn=ITPR1;refGene_tx=XM_011280170.2;refGene_gn=ITPR1;refGene_tx=XM_019824643.1;refGene_gn=ITPR1;refGene_tx=XM_019824647.1;refGene_gn=ITPR1;refGene_tx=XM_023249983.1
A2	46324110	46324265
refGene_gn=ITPR1;refGene_tx=XM_011280170.2;refGene_gn=ITPR1;refGene_tx=XM_019824643.1;refGene_gn=ITPR1;refGene_tx=XM_019824645.1;refGene_gn=ITPR1;refGene_tx=XM_019824647.1;refGene_gn=ITPR1;refGene_tx=XM_019824649.1;refGene_gn=ITPR1;refGene_tx=XM_019824650.1;refGene_gn=ITPR1;refGene_tx=XM_023249983.1
A2	46332028	46332128
refGene_gn=ITPR1;refGene_tx=XM_011280170.2;refGene_gn=ITPR1;refGene_tx=XM_019824643.1;refGene_gn=ITPR1;refGene_tx=XM_019824645.1;refGene_gn=

ITPR1;refGene_tx=XM_019824647.1;refGene_gn=ITPR1;refGene_tx=XM_019824649.1;refGene_gn=ITPR1;refGene_tx=XM_019824650.1;refGene_gn=ITPR1;refGene_tx=XM_023249983.1
A2 46333544 46333705
refGene_gn=ITPR1;refGene_tx=XM_011280170.2;refGene_gn=ITPR1;refGene_tx=XM_019824643.1;refGene_gn=ITPR1;refGene_tx=XM_019824645.1;refGene_gn=ITPR1;refGene_tx=XM_019824647.1;refGene_gn=ITPR1;refGene_tx=XM_019824649.1;refGene_gn=ITPR1;refGene_tx=XM_019824650.1;refGene_gn=ITPR1;refGene_tx=XM_023249983.1
A2 46334203 46334345
refGene_gn=ITPR1;refGene_tx=XM_011280170.2;refGene_gn=ITPR1;refGene_tx=XM_019824643.1;refGene_gn=ITPR1;refGene_tx=XM_019824645.1;refGene_gn=ITPR1;refGene_tx=XM_019824647.1;refGene_gn=ITPR1;refGene_tx=XM_019824649.1;refGene_gn=ITPR1;refGene_tx=XM_019824650.1;refGene_gn=ITPR1;refGene_tx=XM_023249983.1
A2 46336221 46336380
refGene_gn=ITPR1;refGene_tx=XM_011280170.2;refGene_gn=ITPR1;refGene_tx=XM_019824643.1;refGene_gn=ITPR1;refGene_tx=XM_019824645.1;refGene_gn=ITPR1;refGene_tx=XM_019824647.1;refGene_gn=ITPR1;refGene_tx=XM_019824649.1;refGene_gn=ITPR1;refGene_tx=XM_019824650.1;refGene_gn=ITPR1;refGene_tx=XM_023249983.1
A2 46338495 46338668
refGene_gn=ITPR1;refGene_tx=XM_011280170.2;refGene_gn=ITPR1;refGene_tx=XM_019824643.1;refGene_gn=ITPR1;refGene_tx=XM_019824645.1;refGene_gn=ITPR1;refGene_tx=XM_019824647.1;refGene_gn=ITPR1;refGene_tx=XM_019824649.1;refGene_gn=ITPR1;refGene_tx=XM_019824650.1;refGene_gn=ITPR1;refGene_tx=XM_023249983.1
A2 46339873 46339993
refGene_gn=ITPR1;refGene_tx=XM_011280170.2;refGene_gn=ITPR1;refGene_tx=XM_019824643.1;refGene_gn=ITPR1;refGene_tx=XM_019824645.1;refGene_gn=ITPR1;refGene_tx=XM_019824647.1;refGene_gn=ITPR1;refGene_tx=XM_019824649.1;refGene_gn=ITPR1;refGene_tx=XM_019824650.1;refGene_gn=ITPR1;refGene_tx=XM_023249983.1
A2 46341515 46341710
refGene_gn=ITPR1;refGene_tx=XM_011280170.2;refGene_gn=ITPR1;refGene_tx=XM_019824643.1;refGene_gn=ITPR1;refGene_tx=XM_019824645.1;refGene_gn=ITPR1;refGene_tx=XM_019824647.1;refGene_gn=ITPR1;refGene_tx=XM_019824649.1;refGene_gn=ITPR1;refGene_tx=XM_019824650.1;refGene_gn=ITPR1;refGene_tx=XM_023249983.1
A2 46344292 46344544 refGene_gn=ITPR1;refGene_tx=XM_023249983.1
A2 46345472 46345724
refGene_gn=ITPR1;refGene_tx=XM_011280170.2;refGene_gn=ITPR1;refGene_tx=XM_019824643.1;refGene_gn=ITPR1;refGene_tx=XM_019824645.1;refGene_gn=ITPR1;refGene_tx=XM_019824647.1;refGene_gn=ITPR1;refGene_tx=XM_019824649.1;refGene_gn=ITPR1;refGene_tx=XM_019824650.1
A2 46346626 46346768
refGene_gn=ITPR1;refGene_tx=XM_011280170.2;refGene_gn=ITPR1;refGene_tx=XM_019824643.1;refGene_gn=ITPR1;refGene_tx=XM_019824645.1;refGene_gn=ITPR1;refGene_tx=XM_019824647.1;refGene_gn=ITPR1;refGene_tx=XM_019824649.1;refGene_gn=ITPR1;refGene_tx=XM_019824650.1;refGene_gn=ITPR1;refGe

ne_tx=XM_023249983.1
A2	46347928 46348082
refGene_gn=ITPR1;refGene_tx=XM_011280170.2;refGene_gn=ITPR1;refGene_tx=XM_019824643.1;refGene_gn=ITPR1;refGene_tx=XM_019824645.1;refGene_gn=ITPR1;refGene_tx=XM_019824647.1;refGene_gn=ITPR1;refGene_tx=XM_019824649.1;refGene_gn=ITPR1;refGene_tx=XM_019824650.1;refGene_gn=ITPR1;refGene_tx=XM_023249983.1
A2	46349082 46349270
refGene_gn=ITPR1;refGene_tx=XM_011280170.2;refGene_gn=ITPR1;refGene_tx=XM_019824643.1;refGene_gn=ITPR1;refGene_tx=XM_019824645.1;refGene_gn=ITPR1;refGene_tx=XM_019824647.1;refGene_gn=ITPR1;refGene_tx=XM_019824649.1;refGene_gn=ITPR1;refGene_tx=XM_019824650.1;refGene_gn=ITPR1;refGene_tx=XM_023249983.1
A2	46352373 46352512
refGene_gn=ITPR1;refGene_tx=XM_011280170.2;refGene_gn=ITPR1;refGene_tx=XM_019824643.1;refGene_gn=ITPR1;refGene_tx=XM_019824645.1;refGene_gn=ITPR1;refGene_tx=XM_019824647.1;refGene_gn=ITPR1;refGene_tx=XM_019824649.1;refGene_gn=ITPR1;refGene_tx=XM_019824650.1;refGene_gn=ITPR1;refGene_tx=XM_023249983.1
A2	46353184 46353239
refGene_gn=ITPR1;refGene_tx=XM_011280170.2;refGene_gn=ITPR1;refGene_tx=XM_019824643.1;refGene_gn=ITPR1;refGene_tx=XM_019824645.1;refGene_gn=ITPR1;refGene_tx=XM_019824647.1;refGene_gn=ITPR1;refGene_tx=XM_019824649.1;refGene_gn=ITPR1;refGene_tx=XM_019824650.1;refGene_gn=ITPR1;refGene_tx=XM_023249983.1
A2	46355354 46355520
refGene_gn=ITPR1;refGene_tx=XM_011280170.2;refGene_gn=ITPR1;refGene_tx=XM_019824643.1;refGene_gn=ITPR1;refGene_tx=XM_019824645.1;refGene_gn=ITPR1;refGene_tx=XM_019824647.1;refGene_gn=ITPR1;refGene_tx=XM_019824649.1;refGene_gn=ITPR1;refGene_tx=XM_019824650.1;refGene_gn=ITPR1;refGene_tx=XM_023249983.1
A2	46355610 46355781
refGene_gn=ITPR1;refGene_tx=XM_011280170.2;refGene_gn=ITPR1;refGene_tx=XM_019824643.1;refGene_gn=ITPR1;refGene_tx=XM_019824645.1;refGene_gn=ITPR1;refGene_tx=XM_019824647.1;refGene_gn=ITPR1;refGene_tx=XM_019824649.1;refGene_gn=ITPR1;refGene_tx=XM_019824650.1;refGene_gn=ITPR1;refGene_tx=XM_023249983.1
A2	46356280 46356346
refGene_gn=ITPR1;refGene_tx=XM_011280170.2;refGene_gn=ITPR1;refGene_tx=XM_019824643.1;refGene_gn=ITPR1;refGene_tx=XM_019824645.1;refGene_gn=ITPR1;refGene_tx=XM_019824647.1;refGene_gn=ITPR1;refGene_tx=XM_019824649.1;refGene_gn=ITPR1;refGene_tx=XM_019824650.1;refGene_gn=ITPR1;refGene_tx=XM_023249983.1
A2	46357856 46357994
refGene_gn=ITPR1;refGene_tx=XM_011280170.2;refGene_gn=ITPR1;refGene_tx=XM_019824643.1;refGene_gn=ITPR1;refGene_tx=XM_019824645.1;refGene_gn=ITPR1;refGene_tx=XM_019824647.1;refGene_gn=ITPR1;refGene_tx=XM_019824649.1;refGene_gn=ITPR1;refGene_tx=XM_019824650.1;refGene_gn=ITPR1;refGene_tx=XM_023249983.1
A2	46361501 46361627

refGene_gn=ITPR1;refGene_tx=XM_011280170.2;refGene_gn=ITPR1;refGene_tx=XM_019824643.1;refGene_gn=ITPR1;refGene_tx=XM_019824645.1;refGene_gn=ITPR1;refGene_tx=XM_019824647.1;refGene_gn=ITPR1;refGene_tx=XM_019824649.1;refGene_gn=ITPR1;refGene_tx=XM_019824650.1;refGene_gn=ITPR1;refGene_tx=XM_023249983.1
A2 46363993 46364194
refGene_gn=ITPR1;refGene_tx=XM_011280170.2;refGene_gn=ITPR1;refGene_tx=XM_019824643.1;refGene_gn=ITPR1;refGene_tx=XM_019824645.1;refGene_gn=ITPR1;refGene_tx=XM_019824647.1;refGene_gn=ITPR1;refGene_tx=XM_019824649.1;refGene_gn=ITPR1;refGene_tx=XM_019824650.1;refGene_gn=ITPR1;refGene_tx=XM_023249983.1
A2 46366267 46366519
refGene_gn=ITPR1;refGene_tx=XM_011280170.2;refGene_gn=ITPR1;refGene_tx=XM_019824643.1;refGene_gn=ITPR1;refGene_tx=XM_019824645.1;refGene_gn=ITPR1;refGene_tx=XM_019824647.1;refGene_gn=ITPR1;refGene_tx=XM_019824649.1;refGene_gn=ITPR1;refGene_tx=XM_019824650.1;refGene_gn=ITPR1;refGene_tx=XM_023249983.1
A2 46367895 46368021
refGene_gn=ITPR1;refGene_tx=XM_011280170.2;refGene_gn=ITPR1;refGene_tx=XM_019824643.1;refGene_gn=ITPR1;refGene_tx=XM_019824645.1;refGene_gn=ITPR1;refGene_tx=XM_019824647.1;refGene_gn=ITPR1;refGene_tx=XM_019824649.1;refGene_gn=ITPR1;refGene_tx=XM_019824650.1;refGene_gn=ITPR1;refGene_tx=XM_023249983.1
A2 46370250 46370379
refGene_gn=ITPR1;refGene_tx=XM_011280170.2;refGene_gn=ITPR1;refGene_tx=XM_019824643.1;refGene_gn=ITPR1;refGene_tx=XM_019824645.1;refGene_gn=ITPR1;refGene_tx=XM_019824647.1;refGene_gn=ITPR1;refGene_tx=XM_019824649.1;refGene_gn=ITPR1;refGene_tx=XM_019824650.1;refGene_gn=ITPR1;refGene_tx=XM_023249983.1
A2 46374623 46374744
refGene_gn=ITPR1;refGene_tx=XM_011280170.2;refGene_gn=ITPR1;refGene_tx=XM_019824643.1;refGene_gn=ITPR1;refGene_tx=XM_019824645.1;refGene_gn=ITPR1;refGene_tx=XM_019824647.1;refGene_gn=ITPR1;refGene_tx=XM_019824649.1;refGene_gn=ITPR1;refGene_tx=XM_019824650.1;refGene_gn=ITPR1;refGene_tx=XM_023249983.1
A2 46377207 46377392
refGene_gn=ITPR1;refGene_tx=XM_011280170.2;refGene_gn=ITPR1;refGene_tx=XM_019824643.1;refGene_gn=ITPR1;refGene_tx=XM_019824645.1;refGene_gn=ITPR1;refGene_tx=XM_019824647.1;refGene_gn=ITPR1;refGene_tx=XM_019824649.1;refGene_gn=ITPR1;refGene_tx=XM_019824650.1;refGene_gn=ITPR1;refGene_tx=XM_023249983.1
A2 46381404 46381553
refGene_gn=ITPR1;refGene_tx=XM_011280170.2;refGene_gn=ITPR1;refGene_tx=XM_019824643.1;refGene_gn=ITPR1;refGene_tx=XM_019824645.1;refGene_gn=ITPR1;refGene_tx=XM_019824647.1;refGene_gn=ITPR1;refGene_tx=XM_019824649.1;refGene_gn=ITPR1;refGene_tx=XM_019824650.1;refGene_gn=ITPR1;refGene_tx=XM_023249983.1
A2 46382510 46382622
refGene_gn=ITPR1;refGene_tx=XM_011280170.2;refGene_gn=ITPR1;refGene_tx=XM_019824643.1;refGene_gn=ITPR1;refGene_tx=XM_019824645.1;refGene_gn=

ITPR1;refGene_tx=XM_019824647.1;refGene_gn=ITPR1;refGene_tx=XM_019824649.1;refGene_gn=ITPR1;refGene_tx=XM_019824650.1;refGene_gn=ITPR1;refGene_tx=XM_023249983.1
A2 46388267 46388300
refGene_gn=ITPR1;refGene_tx=XM_011280170.2;refGene_gn=ITPR1;refGene_tx=XM_019824643.1;refGene_gn=ITPR1;refGene_tx=XM_019824645.1;refGene_gn=ITPR1;refGene_tx=XM_019824649.1;refGene_gn=ITPR1;refGene_tx=XM_023249983.1
A2 46396056 46396092
refGene_gn=ITPR1;refGene_tx=XM_011280170.2;refGene_gn=ITPR1;refGene_tx=XM_019824643.1;refGene_gn=ITPR1;refGene_tx=XM_019824645.1;refGene_gn=ITPR1;refGene_tx=XM_019824649.1;refGene_gn=ITPR1;refGene_tx=XM_023249983.1
A2 46397231 46397282
refGene_gn=ITPR1;refGene_tx=XM_011280170.2;refGene_gn=ITPR1;refGene_tx=XM_019824643.1;refGene_gn=ITPR1;refGene_tx=XM_019824645.1;refGene_gn=ITPR1;refGene_tx=XM_019824649.1;refGene_gn=ITPR1;refGene_tx=XM_023249983.1
A2 46402885 46403018
refGene_gn=ITPR1;refGene_tx=XM_011280170.2;refGene_gn=ITPR1;refGene_tx=XM_019824643.1;refGene_gn=ITPR1;refGene_tx=XM_019824645.1;refGene_gn=ITPR1;refGene_tx=XM_019824647.1;refGene_gn=ITPR1;refGene_tx=XM_019824649.1;refGene_gn=ITPR1;refGene_tx=XM_019824650.1;refGene_gn=ITPR1;refGene_tx=XM_023249983.1
A2 46405031 46405222
refGene_gn=ITPR1;refGene_tx=XM_011280170.2;refGene_gn=ITPR1;refGene_tx=XM_019824643.1;refGene_gn=ITPR1;refGene_tx=XM_019824645.1;refGene_gn=ITPR1;refGene_tx=XM_019824647.1;refGene_gn=ITPR1;refGene_tx=XM_019824649.1;refGene_gn=ITPR1;refGene_tx=XM_019824650.1;refGene_gn=ITPR1;refGene_tx=XM_023249983.1
A2 46434859 46435040
refGene_gn=ITPR1;refGene_tx=XM_011280170.2;refGene_gn=ITPR1;refGene_tx=XM_019824643.1;refGene_gn=ITPR1;refGene_tx=XM_019824645.1;refGene_gn=ITPR1;refGene_tx=XM_019824647.1;refGene_gn=ITPR1;refGene_tx=XM_019824649.1;refGene_gn=ITPR1;refGene_tx=XM_019824650.1;refGene_gn=ITPR1;refGene_tx=XM_023249983.1
A2 46436752 46437006
refGene_gn=ITPR1;refGene_tx=XM_011280170.2;refGene_gn=ITPR1;refGene_tx=XM_019824643.1;refGene_gn=ITPR1;refGene_tx=XM_019824645.1;refGene_gn=ITPR1;refGene_tx=XM_019824647.1;refGene_gn=ITPR1;refGene_tx=XM_019824649.1;refGene_gn=ITPR1;refGene_tx=XM_019824650.1;refGene_gn=ITPR1;refGene_tx=XM_023249983.1
A2 46442940 46443141
refGene_gn=ITPR1;refGene_tx=XM_011280170.2;refGene_gn=ITPR1;refGene_tx=XM_019824643.1;refGene_gn=ITPR1;refGene_tx=XM_019824645.1;refGene_gn=ITPR1;refGene_tx=XM_019824647.1;refGene_gn=ITPR1;refGene_tx=XM_019824649.1;refGene_gn=ITPR1;refGene_tx=XM_019824650.1;refGene_gn=ITPR1;refGene_tx=XM_023249983.1
A2 46444680 46444791
refGene_gn=ITPR1;refGene_tx=XM_011280170.2;refGene_gn=ITPR1;refGene_tx

=XM_019824643.1;refGene_gn=ITPR1;refGene_tx=XM_019824645.1;refGene_gn=ITPR1;refGene_tx=XM_019824647.1;refGene_gn=ITPR1;refGene_tx=XM_019824649.1;refGene_gn=ITPR1;refGene_tx=XM_019824650.1;refGene_gn=ITPR1;refGene_tx=XM_023249983.1
A2 46446934 46447030
refGene_gn=ITPR1;refGene_tx=XM_011280170.2;refGene_gn=ITPR1;refGene_tx=XM_019824643.1;refGene_gn=ITPR1;refGene_tx=XM_019824645.1;refGene_gn=ITPR1;refGene_tx=XM_019824647.1;refGene_gn=ITPR1;refGene_tx=XM_019824649.1;refGene_gn=ITPR1;refGene_tx=XM_019824650.1;refGene_gn=ITPR1;refGene_tx=XM_023249983.1
A2 46449543 46449666
refGene_gn=ITPR1;refGene_tx=XM_011280170.2;refGene_gn=ITPR1;refGene_tx=XM_019824643.1;refGene_gn=ITPR1;refGene_tx=XM_019824645.1;refGene_gn=ITPR1;refGene_tx=XM_019824647.1;refGene_gn=ITPR1;refGene_tx=XM_019824649.1;refGene_gn=ITPR1;refGene_tx=XM_019824650.1;refGene_gn=ITPR1;refGene_tx=XM_023249983.1
A2 46450751 46450856
refGene_gn=ITPR1;refGene_tx=XM_011280170.2;refGene_gn=ITPR1;refGene_tx=XM_019824643.1;refGene_gn=ITPR1;refGene_tx=XM_019824645.1;refGene_gn=ITPR1;refGene_tx=XM_019824647.1;refGene_gn=ITPR1;refGene_tx=XM_019824649.1;refGene_gn=ITPR1;refGene_tx=XM_019824650.1;refGene_gn=ITPR1;refGene_tx=XM_023249983.1
A2 46453711 46453904
refGene_gn=ITPR1;refGene_tx=XM_011280170.2;refGene_gn=ITPR1;refGene_tx=XM_019824643.1;refGene_gn=ITPR1;refGene_tx=XM_019824645.1;refGene_gn=ITPR1;refGene_tx=XM_019824647.1;refGene_gn=ITPR1;refGene_tx=XM_019824649.1;refGene_gn=ITPR1;refGene_tx=XM_019824650.1;refGene_gn=ITPR1;refGene_tx=XM_023249983.1
A2 46460462 46460585
refGene_gn=ITPR1;refGene_tx=XM_011280170.2;refGene_gn=ITPR1;refGene_tx=XM_019824643.1;refGene_gn=ITPR1;refGene_tx=XM_019824645.1;refGene_gn=ITPR1;refGene_tx=XM_019824647.1;refGene_gn=ITPR1;refGene_tx=XM_019824649.1;refGene_gn=ITPR1;refGene_tx=XM_019824650.1;refGene_gn=ITPR1;refGene_tx=XM_023249983.1
A2 46466473 46466649
refGene_gn=ITPR1;refGene_tx=XM_011280170.2;refGene_gn=ITPR1;refGene_tx=XM_019824643.1;refGene_gn=ITPR1;refGene_tx=XM_019824645.1;refGene_gn=ITPR1;refGene_tx=XM_019824647.1;refGene_gn=ITPR1;refGene_tx=XM_019824649.1;refGene_gn=ITPR1;refGene_tx=XM_019824650.1;refGene_gn=ITPR1;refGene_tx=XM_023249983.1
A2 46470808 46470973
refGene_gn=ITPR1;refGene_tx=XM_011280170.2;refGene_gn=ITPR1;refGene_tx=XM_019824643.1;refGene_gn=ITPR1;refGene_tx=XM_019824645.1;refGene_gn=ITPR1;refGene_tx=XM_019824647.1;refGene_gn=ITPR1;refGene_tx=XM_019824649.1;refGene_gn=ITPR1;refGene_tx=XM_019824650.1;refGene_gn=ITPR1;refGene_tx=XM_023249983.1
A2 46475633 46475829
refGene_gn=ITPR1;refGene_tx=XM_011280170.2;refGene_gn=ITPR1;refGene_tx=XM_019824643.1;refGene_gn=ITPR1;refGene_tx=XM_019824645.1;refGene_gn=ITPR1;refGene_tx=XM_019824647.1;refGene_gn=ITPR1;refGene_tx=XM_0198246

49.1;refGene_gn=ITPR1;refGene_tx=XM_019824650.1;refGene_gn=ITPR1;refGene_tx=XM_023249983.1
A2 46477178 46477271
refGene_gn=ITPR1;refGene_tx=XM_011280170.2;refGene_gn=ITPR1;refGene_tx=XM_019824643.1;refGene_gn=ITPR1;refGene_tx=XM_019824645.1;refGene_gn=ITPR1;refGene_tx=XM_019824647.1;refGene_gn=ITPR1;refGene_tx=XM_019824649.1;refGene_gn=ITPR1;refGene_tx=XM_019824650.1;refGene_gn=ITPR1;refGene_tx=XM_023249983.1
A2 46478479 46478619
refGene_gn=ITPR1;refGene_tx=XM_011280170.2;refGene_gn=ITPR1;refGene_tx=XM_019824643.1;refGene_gn=ITPR1;refGene_tx=XM_019824645.1;refGene_gn=ITPR1;refGene_tx=XM_019824647.1;refGene_gn=ITPR1;refGene_tx=XM_019824649.1;refGene_gn=ITPR1;refGene_tx=XM_019824650.1;refGene_gn=ITPR1;refGene_tx=XM_023249983.1
A2 46479180 46479346
refGene_gn=ITPR1;refGene_tx=XM_011280170.2;refGene_gn=ITPR1;refGene_tx=XM_019824643.1;refGene_gn=ITPR1;refGene_tx=XM_019824645.1;refGene_gn=ITPR1;refGene_tx=XM_019824647.1;refGene_gn=ITPR1;refGene_tx=XM_019824649.1;refGene_gn=ITPR1;refGene_tx=XM_019824650.1;refGene_gn=ITPR1;refGene_tx=XM_023249983.1
A2 46482252 46482413
refGene_gn=ITPR1;refGene_tx=XM_011280170.2;refGene_gn=ITPR1;refGene_tx=XM_019824643.1;refGene_gn=ITPR1;refGene_tx=XM_019824645.1;refGene_gn=ITPR1;refGene_tx=XM_019824647.1;refGene_gn=ITPR1;refGene_tx=XM_019824649.1;refGene_gn=ITPR1;refGene_tx=XM_019824650.1;refGene_gn=ITPR1;refGene_tx=XM_023249983.1
A2 46500186 46500348
refGene_gn=ITPR1;refGene_tx=XM_011280170.2;refGene_gn=ITPR1;refGene_tx=XM_019824643.1;refGene_gn=ITPR1;refGene_tx=XM_019824645.1;refGene_gn=ITPR1;refGene_tx=XM_019824647.1;refGene_gn=ITPR1;refGene_tx=XM_019824649.1;refGene_gn=ITPR1;refGene_tx=XM_019824650.1;refGene_gn=ITPR1;refGene_tx=XM_023249983.1
A2 46507488 46507575
refGene_gn=ITPR1;refGene_tx=XM_011280170.2;refGene_gn=ITPR1;refGene_tx=XM_019824643.1;refGene_gn=ITPR1;refGene_tx=XM_019824645.1;refGene_gn=ITPR1;refGene_tx=XM_019824647.1;refGene_gn=ITPR1;refGene_tx=XM_019824649.1;refGene_gn=ITPR1;refGene_tx=XM_019824650.1;refGene_gn=ITPR1;refGene_tx=XM_023249983.1
A2 46609610 46609690 refGene_gn=BHLHE40;refGene_tx=XM_003982442.4
A2 46609857 46609927 refGene_gn=BHLHE40;refGene_tx=XM_003982442.4
A2 46610185 46610293 refGene_gn=BHLHE40;refGene_tx=XM_003982442.4
A2 46611379 46611503 refGene_gn=BHLHE40;refGene_tx=XM_003982442.4
A2 46612786 46613643 refGene_gn=BHLHE40;refGene_tx=XM_003982442.4
A2 46735375 46735498 refGene_gn=ARL8B;refGene_tx=XM_003982443.4
A2 46770471 46770552 refGene_gn=ARL8B;refGene_tx=XM_003982443.4
A2 46771430 46771504 refGene_gn=ARL8B;refGene_tx=XM_003982443.4
A2 46771906 46772000 refGene_gn=ARL8B;refGene_tx=XM_003982443.4
A2 46775509 46775577 refGene_gn=ARL8B;refGene_tx=XM_003982443.4
A2 46776413 46776484 refGene_gn=ARL8B;refGene_tx=XM_003982443.4
A2 46782572 46782622 refGene_gn=ARL8B;refGene_tx=XM_003982443.4

A2 46792725 46793222 refGene_gn=EDEM1;refGene_tx=XM_003982475.4
A2 46799481 46799554 refGene_gn=EDEM1;refGene_tx=XM_003982475.4
A2 46802494 46802598 refGene_gn=EDEM1;refGene_tx=XM_003982475.4
A2 46803836 46804008 refGene_gn=EDEM1;refGene_tx=XM_003982475.4
A2 46805219 46805403 refGene_gn=EDEM1;refGene_tx=XM_003982475.4
A2 46807993 46808168 refGene_gn=EDEM1;refGene_tx=XM_003982475.4
A2 46809743 46809864 refGene_gn=EDEM1;refGene_tx=XM_003982475.4
A2 46810360 46810531 refGene_gn=EDEM1;refGene_tx=XM_003982475.4
A2 46812382 46812456 refGene_gn=EDEM1;refGene_tx=XM_003982475.4
A2 46813303 46813400 refGene_gn=EDEM1;refGene_tx=XM_003982475.4
A2 46815515 46815719 refGene_gn=EDEM1;refGene_tx=XM_003982475.4
A2 46818232 46818322 refGene_gn=EDEM1;refGene_tx=XM_003982475.4
A2 48335025 48335544
refGene_gn=GRM7;refGene_tx=XM_011280176.3;refGene_gn=GRM7;refGene_tx=XM_019824714.2;refGene_gn=GRM7;refGene_tx=XM_019824718.2
A2 48612038 48612255
refGene_gn=GRM7;refGene_tx=XM_011280176.3;refGene_gn=GRM7;refGene_tx=XM_019824714.2;refGene_gn=GRM7;refGene_tx=XM_019824718.2
A2 48779865 48780007
refGene_gn=GRM7;refGene_tx=XM_011280176.3;refGene_gn=GRM7;refGene_tx=XM_019824714.2;refGene_gn=GRM7;refGene_tx=XM_019824718.2
A2 48788172 48788327
refGene_gn=GRM7;refGene_tx=XM_011280176.3;refGene_gn=GRM7;refGene_tx=XM_019824714.2;refGene_gn=GRM7;refGene_tx=XM_019824718.2
A2 48882365 48882506
refGene_gn=GRM7;refGene_tx=XM_011280176.3;refGene_gn=GRM7;refGene_tx=XM_019824714.2;refGene_gn=GRM7;refGene_tx=XM_019824718.2
A2 48930054 48930255
refGene_gn=GRM7;refGene_tx=XM_011280176.3;refGene_gn=GRM7;refGene_tx=XM_019824714.2;refGene_gn=GRM7;refGene_tx=XM_019824718.2
A2 48937695 48937835
refGene_gn=GRM7;refGene_tx=XM_011280176.3;refGene_gn=GRM7;refGene_tx=XM_019824714.2;refGene_gn=GRM7;refGene_tx=XM_019824718.2
A2 49050368 49051304
refGene_gn=GRM7;refGene_tx=XM_011280176.3;refGene_gn=GRM7;refGene_tx=XM_019824714.2;refGene_gn=GRM7;refGene_tx=XM_019824718.2
A2 49154522 49154793
refGene_gn=GRM7;refGene_tx=XM_011280176.3;refGene_gn=GRM7;refGene_tx=XM_019824714.2;refGene_gn=GRM7;refGene_tx=XM_019824718.2
A2 49160312 49160383 refGene_gn=GRM7;refGene_tx=XM_011280176.3
A2 49210839 49210889 refGene_gn=GRM7;refGene_tx=XM_019824714.2
A2 49913544 49913586 refGene_gn=LMCD1;refGene_tx=XM_003982444.4
A2 49943750 49943839 refGene_gn=LMCD1;refGene_tx=XM_003982444.4
A2 49949633 49949889 refGene_gn=LMCD1;refGene_tx=XM_003982444.4
A2 49956638 49956968 refGene_gn=LMCD1;refGene_tx=XM_003982444.4
A2 49970256 49970472 refGene_gn=LMCD1;refGene_tx=XM_003982444.4
A2 49971185 49971344 refGene_gn=LMCD1;refGene_tx=XM_003982444.4
A2 50017117 50017264 refGene_gn=SSUH2;refGene_tx=XM_023249984.1
A2 50021736 50021844 refGene_gn=SSUH2;refGene_tx=XM_023249984.1
A2 50024180 50024286 refGene_gn=SSUH2;refGene_tx=XM_023249984.1

A2	50024828	50024921	refGene_gn=SSUH2;refGene_tx=XM_023249984.1
A2	50026079	50026165	refGene_gn=SSUH2;refGene_tx=XM_023249984.1
A2	50027859	50027922	refGene_gn=SSUH2;refGene_tx=XM_023249984.1
A2	50029436	50029561	refGene_gn=SSUH2;refGene_tx=XM_023249984.1
A2	50030565	50030626	refGene_gn=SSUH2;refGene_tx=XM_023249984.1
A2	50031881	50032011	refGene_gn=SSUH2;refGene_tx=XM_023249984.1
A2	50033605	50033687	refGene_gn=SSUH2;refGene_tx=XM_023249984.1
A2	50034068	50034167	refGene_gn=SSUH2;refGene_tx=XM_023249984.1
A2	50047846	50047874	refGene_gn=SSUH2;refGene_tx=XM_023249984.1
A2	50099529	50099643	refGene_gn=CAV3;refGene_tx=XM_003982445.4
A2	50111383	50111725	refGene_gn=CAV3;refGene_tx=XM_003982445.4
A2	50119629	50119877
refGene_gn=OXTR;refGene_tx=XM_003982446.3;refGene_gn=OXTR;refGene_tx=XM_023249985.1
A2	50136571	50137493
refGene_gn=OXTR;refGene_tx=XM_003982446.3;refGene_gn=OXTR;refGene_tx=XM_023249985.1
A2	50242429	50242532
refGene_gn=RAD18;refGene_tx=XM_003982447.4;refGene_gn=RAD18;refGene_tx=XM_019824935.1
A2	50251142	50251209
refGene_gn=RAD18;refGene_tx=XM_003982447.4;refGene_gn=RAD18;refGene_tx=XM_019824936.1
A2	50283646	50283821
refGene_gn=RAD18;refGene_tx=XM_003982447.4;refGene_gn=RAD18;refGene_tx=XM_019824935.1;refGene_gn=RAD18;refGene_tx=XM_019824936.1
A2	50288513	50288651
refGene_gn=RAD18;refGene_tx=XM_003982447.4;refGene_gn=RAD18;refGene_tx=XM_019824935.1;refGene_gn=RAD18;refGene_tx=XM_019824936.1
A2	50298782	50298843
refGene_gn=RAD18;refGene_tx=XM_003982447.4;refGene_gn=RAD18;refGene_tx=XM_019824935.1;refGene_gn=RAD18;refGene_tx=XM_019824936.1
A2	50300113	50300190
refGene_gn=RAD18;refGene_tx=XM_003982447.4;refGene_gn=RAD18;refGene_tx=XM_019824935.1;refGene_gn=RAD18;refGene_tx=XM_019824936.1
A2	50313334	50313519
refGene_gn=RAD18;refGene_tx=XM_003982447.4;refGene_gn=RAD18;refGene_tx=XM_019824935.1;refGene_gn=RAD18;refGene_tx=XM_019824936.1
A2	50316720	50316820
refGene_gn=RAD18;refGene_tx=XM_003982447.4;refGene_gn=RAD18;refGene_tx=XM_019824935.1;refGene_gn=RAD18;refGene_tx=XM_019824936.1
A2	50318140	50318478
refGene_gn=RAD18;refGene_tx=XM_003982447.4;refGene_gn=RAD18;refGene_tx=XM_019824935.1;refGene_gn=RAD18;refGene_tx=XM_019824936.1
A2	50330909	50330980
refGene_gn=RAD18;refGene_tx=XM_003982447.4;refGene_gn=RAD18;refGene_tx=XM_019824935.1;refGene_gn=RAD18;refGene_tx=XM_019824936.1
A2	50333449	50333511
refGene_gn=RAD18;refGene_tx=XM_003982447.4;refGene_gn=RAD18;refGene_tx=XM_019824935.1;refGene_gn=RAD18;refGene_tx=XM_019824936.1

A2	50347384 50347466
refGene_gn=RAD18;refGene_tx=XM_003982447.4;refGene_gn=RAD18;refGene_tx=XM_019824935.1;refGene_gn=RAD18;refGene_tx=XM_019824936.1
A2	50352919 50352970
refGene_gn=RAD18;refGene_tx=XM_003982447.4;refGene_gn=RAD18;refGene_tx=XM_019824935.1;refGene_gn=RAD18;refGene_tx=XM_019824936.1
A2	50380696 50381110
refGene_gn=SRGAP3;refGene_tx=XM_003982448.5;refGene_gn=SRGAP3;refGene_tx=XM_011280178.3;refGene_gn=SRGAP3;refGene_tx=XM_019824937.2;refGene_gn=SRGAP3;refGene_tx=XM_019824938.2;refGene_gn=SRGAP3;refGene_tx=XM_019824939.2;refGene_gn=SRGAP3;refGene_tx=XM_019824940.2
A2	50383934 50384262
refGene_gn=SRGAP3;refGene_tx=XM_003982448.5;refGene_gn=SRGAP3;refGene_tx=XM_011280178.3;refGene_gn=SRGAP3;refGene_tx=XM_019824937.2;refGene_gn=SRGAP3;refGene_tx=XM_019824938.2;refGene_gn=SRGAP3;refGene_tx=XM_019824939.2
A2	50386335 50386485
refGene_gn=SRGAP3;refGene_tx=XM_003982448.5;refGene_gn=SRGAP3;refGene_tx=XM_011280178.3;refGene_gn=SRGAP3;refGene_tx=XM_019824937.2;refGene_gn=SRGAP3;refGene_tx=XM_019824938.2;refGene_gn=SRGAP3;refGene_tx=XM_019824939.2;refGene_gn=SRGAP3;refGene_tx=XM_019824940.2
A2	50387823 50388004
refGene_gn=SRGAP3;refGene_tx=XM_003982448.5;refGene_gn=SRGAP3;refGene_tx=XM_011280178.3;refGene_gn=SRGAP3;refGene_tx=XM_019824937.2;refGene_gn=SRGAP3;refGene_tx=XM_019824938.2;refGene_gn=SRGAP3;refGene_tx=XM_019824939.2;refGene_gn=SRGAP3;refGene_tx=XM_019824940.2
A2	50399251 50399331
refGene_gn=SRGAP3;refGene_tx=XM_003982448.5;refGene_gn=SRGAP3;refGene_tx=XM_011280178.3;refGene_gn=SRGAP3;refGene_tx=XM_019824937.2;refGene_gn=SRGAP3;refGene_tx=XM_019824938.2;refGene_gn=SRGAP3;refGene_tx=XM_019824939.2;refGene_gn=SRGAP3;refGene_tx=XM_019824940.2
A2	50402136 50402364
refGene_gn=SRGAP3;refGene_tx=XM_003982448.5;refGene_gn=SRGAP3;refGene_tx=XM_011280178.3;refGene_gn=SRGAP3;refGene_tx=XM_019824937.2;refGene_gn=SRGAP3;refGene_tx=XM_019824938.2;refGene_gn=SRGAP3;refGene_tx=XM_019824939.2;refGene_gn=SRGAP3;refGene_tx=XM_019824940.2
A2	50402577 50402683
refGene_gn=SRGAP3;refGene_tx=XM_003982448.5;refGene_gn=SRGAP3;refGene_tx=XM_011280178.3;refGene_gn=SRGAP3;refGene_tx=XM_019824937.2;refGene_gn=SRGAP3;refGene_tx=XM_019824938.2;refGene_gn=SRGAP3;refGene_tx=XM_019824939.2;refGene_gn=SRGAP3;refGene_tx=XM_019824940.2
A2	50405772 50405907
refGene_gn=SRGAP3;refGene_tx=XM_003982448.5;refGene_gn=SRGAP3;refGene_tx=XM_011280178.3;refGene_gn=SRGAP3;refGene_tx=XM_019824937.2;refGene_gn=SRGAP3;refGene_tx=XM_019824938.2;refGene_gn=SRGAP3;refGene_tx=XM_019824939.2;refGene_gn=SRGAP3;refGene_tx=XM_019824940.2
A2	50415570 50415648
refGene_gn=SRGAP3;refGene_tx=XM_003982448.5;refGene_gn=SRGAP3;refGene_tx=XM_011280178.3;refGene_gn=SRGAP3;refGene_tx=XM_019824937.2;refGene_gn=SRGAP3;refGene_tx=XM_019824938.2;refGene_gn=SRGAP3;refGene_tx=XM_01

9824939.2;refGene_gn=SRGAP3;refGene_tx=XM_019824940.2
A2 50417370 50417431
refGene_gn=SRGAP3;refGene_tx=XM_003982448.5;refGene_gn=SRGAP3;refGene_tx=XM_011280178.3;refGene_gn=SRGAP3;refGene_tx=XM_019824937.2;refGene_gn=SRGAP3;refGene_tx=XM_019824938.2;refGene_gn=SRGAP3;refGene_tx=XM_019824939.2;refGene_gn=SRGAP3;refGene_tx=XM_019824940.2;refGene_gn=SRGAP3;refGene_tx=XM_019824942.2
A2 50418549 50418595
refGene_gn=SRGAP3;refGene_tx=XM_011280178.3;refGene_gn=SRGAP3;refGene_tx=XM_019824937.2;refGene_gn=SRGAP3;refGene_tx=XM_019824938.2;refGene_gn=SRGAP3;refGene_tx=XM_019824939.2;refGene_gn=SRGAP3;refGene_tx=XM_019824940.2
A2 50422646 50422749 refGene_gn=SRGAP3;refGene_tx=XM_003982448.5
A2 50425166 50425253
refGene_gn=SRGAP3;refGene_tx=XM_019824937.2;refGene_gn=SRGAP3;refGene_tx=XM_019824938.2;refGene_gn=SRGAP3;refGene_tx=XM_019824940.2
A2 50427701 50427729
refGene_gn=SRGAP3;refGene_tx=XM_003982448.5;refGene_gn=SRGAP3;refGene_tx=XM_011280178.3;refGene_gn=SRGAP3;refGene_tx=XM_019824937.2;refGene_gn=SRGAP3;refGene_tx=XM_019824938.2;refGene_gn=SRGAP3;refGene_tx=XM_019824939.2;refGene_gn=SRGAP3;refGene_tx=XM_019824940.2;refGene_gn=SRGAP3;refGene_tx=XM_019824942.2
A2 50435641 50435726
refGene_gn=SRGAP3;refGene_tx=XM_003982448.5;refGene_gn=SRGAP3;refGene_tx=XM_011280178.3;refGene_gn=SRGAP3;refGene_tx=XM_019824937.2;refGene_gn=SRGAP3;refGene_tx=XM_019824938.2;refGene_gn=SRGAP3;refGene_tx=XM_019824939.2;refGene_gn=SRGAP3;refGene_tx=XM_019824940.2;refGene_gn=SRGAP3;refGene_tx=XM_019824942.2
A2 50442846 50443044
refGene_gn=SRGAP3;refGene_tx=XM_003982448.5;refGene_gn=SRGAP3;refGene_tx=XM_011280178.3;refGene_gn=SRGAP3;refGene_tx=XM_019824937.2;refGene_gn=SRGAP3;refGene_tx=XM_019824938.2;refGene_gn=SRGAP3;refGene_tx=XM_019824939.2;refGene_gn=SRGAP3;refGene_tx=XM_019824940.2;refGene_gn=SRGAP3;refGene_tx=XM_019824942.2
A2 50445203 50445305
refGene_gn=SRGAP3;refGene_tx=XM_003982448.5;refGene_gn=SRGAP3;refGene_tx=XM_011280178.3;refGene_gn=SRGAP3;refGene_tx=XM_019824937.2;refGene_gn=SRGAP3;refGene_tx=XM_019824938.2;refGene_gn=SRGAP3;refGene_tx=XM_019824939.2;refGene_gn=SRGAP3;refGene_tx=XM_019824940.2;refGene_gn=SRGAP3;refGene_tx=XM_019824942.2
A2 50446778 50447000
refGene_gn=SRGAP3;refGene_tx=XM_003982448.5;refGene_gn=SRGAP3;refGene_tx=XM_011280178.3;refGene_gn=SRGAP3;refGene_tx=XM_019824937.2;refGene_gn=SRGAP3;refGene_tx=XM_019824938.2;refGene_gn=SRGAP3;refGene_tx=XM_019824939.2;refGene_gn=SRGAP3;refGene_tx=XM_019824940.2;refGene_gn=SRGAP3;refGene_tx=XM_019824942.2
A2 50448359 50448488
refGene_gn=SRGAP3;refGene_tx=XM_003982448.5;refGene_gn=SRGAP3;refGene_tx=XM_011280178.3;refGene_gn=SRGAP3;refGene_tx=XM_019824937.2;refGene_gn=SRGAP3;refGene_tx=XM_019824938.2;refGene_gn=SRGAP3;refGene_tx=XM_01

9824939.2;refGene_gn=SRGAP3;refGene_tx=XM_019824940.2;refGene_gn=SRGAP3;refGene_tx=XM_019824942.2
A2 50453118 50453304
refGene_gn=SRGAP3;refGene_tx=XM_003982448.5;refGene_gn=SRGAP3;refGene_tx=XM_011280178.3;refGene_gn=SRGAP3;refGene_tx=XM_019824937.2;refGene_gn=SRGAP3;refGene_tx=XM_019824938.2;refGene_gn=SRGAP3;refGene_tx=XM_019824939.2;refGene_gn=SRGAP3;refGene_tx=XM_019824940.2;refGene_gn=SRGAP3;refGene_tx=XM_019824942.2
A2 50467381 50467444
refGene_gn=SRGAP3;refGene_tx=XM_003982448.5;refGene_gn=SRGAP3;refGene_tx=XM_011280178.3;refGene_gn=SRGAP3;refGene_tx=XM_019824937.2;refGene_gn=SRGAP3;refGene_tx=XM_019824938.2;refGene_gn=SRGAP3;refGene_tx=XM_019824939.2;refGene_gn=SRGAP3;refGene_tx=XM_019824940.2;refGene_gn=SRGAP3;refGene_tx=XM_019824942.2
A2 50490568 50490731
refGene_gn=SRGAP3;refGene_tx=XM_003982448.5;refGene_gn=SRGAP3;refGene_tx=XM_011280178.3;refGene_gn=SRGAP3;refGene_tx=XM_019824937.2;refGene_gn=SRGAP3;refGene_tx=XM_019824938.2;refGene_gn=SRGAP3;refGene_tx=XM_019824939.2;refGene_gn=SRGAP3;refGene_tx=XM_019824940.2;refGene_gn=SRGAP3;refGene_tx=XM_019824942.2
A2 50512012 50512205
refGene_gn=SRGAP3;refGene_tx=XM_003982448.5;refGene_gn=SRGAP3;refGene_tx=XM_011280178.3;refGene_gn=SRGAP3;refGene_tx=XM_019824937.2;refGene_gn=SRGAP3;refGene_tx=XM_019824938.2;refGene_gn=SRGAP3;refGene_tx=XM_019824939.2;refGene_gn=SRGAP3;refGene_tx=XM_019824940.2;refGene_gn=SRGAP3;refGene_tx=XM_019824942.2
A2 50638537 50638604
refGene_gn=SRGAP3;refGene_tx=XM_003982448.5;refGene_gn=SRGAP3;refGene_tx=XM_011280178.3;refGene_gn=SRGAP3;refGene_tx=XM_019824937.2;refGene_gn=SRGAP3;refGene_tx=XM_019824938.2;refGene_gn=SRGAP3;refGene_tx=XM_019824939.2;refGene_gn=SRGAP3;refGene_tx=XM_019824940.2;refGene_gn=SRGAP3;refGene_tx=XM_019824942.2
A2 50736502 50736757
refGene_gn=THUMPD3;refGene_tx=XM_004001397.3;refGene_gn=THUMPD3;refGene_tx=XM_011280179.3;refGene_gn=THUMPD3;refGene_tx=XM_011280180.3
A2 50737541 50737616 refGene_gn=THUMPD3;refGene_tx=XM_019824943.2
A2 50738095 50738173
refGene_gn=THUMPD3;refGene_tx=XM_004001397.3;refGene_gn=THUMPD3;refGene_tx=XM_011280179.3;refGene_gn=THUMPD3;refGene_tx=XM_011280180.3;refGene_gn=THUMPD3;refGene_tx=XM_019824943.2
A2 50745700 50746174
refGene_gn=THUMPD3;refGene_tx=XM_004001397.3;refGene_gn=THUMPD3;refGene_tx=XM_011280179.3;refGene_gn=THUMPD3;refGene_tx=XM_011280180.3;refGene_gn=THUMPD3;refGene_tx=XM_019824943.2
A2 50749220 50749351
refGene_gn=THUMPD3;refGene_tx=XM_004001397.3;refGene_gn=THUMPD3;refGene_tx=XM_011280179.3;refGene_gn=THUMPD3;refGene_tx=XM_011280180.3;refGene_gn=THUMPD3;refGene_tx=XM_019824943.2
A2 50750176 50750246
refGene_gn=THUMPD3;refGene_tx=XM_004001397.3;refGene_gn=THUMPD3;refGen

e_tx=XM_011280179.3;refGene_gn=THUMPD3;refGene_tx=XM_011280180.3;refGene_gn=THUMPD3;refGene_tx=XM_019824943.2
A2 50752539 50752655
refGene_gn=THUMPD3;refGene_tx=XM_004001397.3;refGene_gn=THUMPD3;refGene_tx=XM_011280179.3;refGene_gn=THUMPD3;refGene_tx=XM_011280180.3;refGene_gn=THUMPD3;refGene_tx=XM_019824943.2
A2 50755935 50756046
refGene_gn=THUMPD3;refGene_tx=XM_004001397.3;refGene_gn=THUMPD3;refGene_tx=XM_011280179.3;refGene_gn=THUMPD3;refGene_tx=XM_011280180.3;refGene_gn=THUMPD3;refGene_tx=XM_019824943.2
A2 50756617 50756741
refGene_gn=THUMPD3;refGene_tx=XM_004001397.3;refGene_gn=THUMPD3;refGene_tx=XM_011280179.3;refGene_gn=THUMPD3;refGene_tx=XM_011280180.3;refGene_gn=THUMPD3;refGene_tx=XM_019824943.2
A2 50756925 50757075
refGene_gn=THUMPD3;refGene_tx=XM_004001397.3;refGene_gn=THUMPD3;refGene_tx=XM_011280179.3;refGene_gn=THUMPD3;refGene_tx=XM_011280180.3;refGene_gn=THUMPD3;refGene_tx=XM_019824943.2
A2 50803609 50803680
refGene_gn=SETD5;refGene_tx=XM_023249990.1;refGene_gn=SETD5;refGene_tx=XM_023249994.1;refGene_gn=SETD5;refGene_tx=XM_023249997.1
A2 50808204 50808296
refGene_gn=SETD5;refGene_tx=XM_023249986.1;refGene_gn=SETD5;refGene_tx=XM_023249987.1;refGene_gn=SETD5;refGene_tx=XM_023249988.1;refGene_gn=SETD5;refGene_tx=XM_023249989.1;refGene_gn=SETD5;refGene_tx=XM_023249991.1;refGene_gn=SETD5;refGene_tx=XM_023249992.1;refGene_gn=SETD5;refGene_tx=XM_023249993.1;refGene_gn=SETD5;refGene_tx=XM_023249996.1
A2 50808576 50808682
refGene_gn=SETD5;refGene_tx=XM_023249986.1;refGene_gn=SETD5;refGene_tx=XM_023249987.1;refGene_gn=SETD5;refGene_tx=XM_023249988.1;refGene_gn=SETD5;refGene_tx=XM_023249989.1;refGene_gn=SETD5;refGene_tx=XM_023249990.1;refGene_gn=SETD5;refGene_tx=XM_023249991.1;refGene_gn=SETD5;refGene_tx=XM_023249992.1;refGene_gn=SETD5;refGene_tx=XM_023249993.1;refGene_gn=SETD5;refGene_tx=XM_023249994.1;refGene_gn=SETD5;refGene_tx=XM_023249996.1;refGene_gn=SETD5;refGene_tx=XM_023249997.1
A2 50808850 50808913 refGene_gn=SETD5;refGene_tx=XM_023249995.1
A2 50809008 50809217
refGene_gn=SETD5;refGene_tx=XM_023249986.1;refGene_gn=SETD5;refGene_tx=XM_023249987.1;refGene_gn=SETD5;refGene_tx=XM_023249988.1;refGene_gn=SETD5;refGene_tx=XM_023249989.1;refGene_gn=SETD5;refGene_tx=XM_023249990.1;refGene_gn=SETD5;refGene_tx=XM_023249991.1;refGene_gn=SETD5;refGene_tx=XM_023249992.1;refGene_gn=SETD5;refGene_tx=XM_023249993.1;refGene_gn=SETD5;refGene_tx=XM_023249994.1;refGene_gn=SETD5;refGene_tx=XM_023249995.1;refGene_gn=SETD5;refGene_tx=XM_023249996.1;refGene_gn=SETD5;refGene_tx=XM_023249997.1
A2 50809555 50809614
refGene_gn=SETD5;refGene_tx=XM_023249986.1;refGene_gn=SETD5;refGene_tx=XM_023249987.1;refGene_gn=SETD5;refGene_tx=XM_023249988.1;refGene_gn=SETD5;refGene_tx=XM_023249989.1;refGene_gn=SETD5;refGene_tx=XM_023249990.1;refGene_gn=SETD5;refGene_tx=XM_023249991.1;refGene_gn=SETD5;refGe

ne_tx=XM_023249992.1;refGene_gn=SETD5;refGene_tx=XM_023249993.1;refGene_gn=SETD5;refGene_tx=XM_023249994.1;refGene_gn=SETD5;refGene_tx=XM_023249995.1;refGene_gn=SETD5;refGene_tx=XM_023249996.1;refGene_gn=SETD5;refGene_tx=XM_023249997.1;refGene_gn=SETD5;refGene_tx=XM_023249998.1
A2 50810125 50810304
refGene_gn=SETD5;refGene_tx=XM_023249986.1;refGene_gn=SETD5;refGene_tx=XM_023249987.1;refGene_gn=SETD5;refGene_tx=XM_023249988.1;refGene_gn=SETD5;refGene_tx=XM_023249989.1;refGene_gn=SETD5;refGene_tx=XM_023249990.1;refGene_gn=SETD5;refGene_tx=XM_023249991.1;refGene_gn=SETD5;refGene_tx=XM_023249992.1;refGene_gn=SETD5;refGene_tx=XM_023249993.1;refGene_gn=SETD5;refGene_tx=XM_023249994.1;refGene_gn=SETD5;refGene_tx=XM_023249995.1;refGene_gn=SETD5;refGene_tx=XM_023249996.1;refGene_gn=SETD5;refGene_tx=XM_023249997.1;refGene_gn=SETD5;refGene_tx=XM_023249998.1
A2 50811237 50811276
refGene_gn=SETD5;refGene_tx=XM_023249986.1;refGene_gn=SETD5;refGene_tx=XM_023249987.1;refGene_gn=SETD5;refGene_tx=XM_023249988.1;refGene_gn=SETD5;refGene_tx=XM_023249989.1;refGene_gn=SETD5;refGene_tx=XM_023249990.1;refGene_gn=SETD5;refGene_tx=XM_023249992.1;refGene_gn=SETD5;refGene_tx=XM_023249993.1;refGene_gn=SETD5;refGene_tx=XM_023249995.1;refGene_gn=SETD5;refGene_tx=XM_023249998.1
A2 50814421 50814664
refGene_gn=SETD5;refGene_tx=XM_023249986.1;refGene_gn=SETD5;refGene_tx=XM_023249987.1;refGene_gn=SETD5;refGene_tx=XM_023249988.1;refGene_gn=SETD5;refGene_tx=XM_023249989.1;refGene_gn=SETD5;refGene_tx=XM_023249990.1;refGene_gn=SETD5;refGene_tx=XM_023249991.1;refGene_gn=SETD5;refGene_tx=XM_023249992.1;refGene_gn=SETD5;refGene_tx=XM_023249993.1;refGene_gn=SETD5;refGene_tx=XM_023249994.1;refGene_gn=SETD5;refGene_tx=XM_023249995.1;refGene_gn=SETD5;refGene_tx=XM_023249996.1;refGene_gn=SETD5;refGene_tx=XM_023249997.1;refGene_gn=SETD5;refGene_tx=XM_023249998.1
A2 50815225 50815374
refGene_gn=SETD5;refGene_tx=XM_023249986.1;refGene_gn=SETD5;refGene_tx=XM_023249987.1;refGene_gn=SETD5;refGene_tx=XM_023249988.1;refGene_gn=SETD5;refGene_tx=XM_023249989.1;refGene_gn=SETD5;refGene_tx=XM_023249990.1;refGene_gn=SETD5;refGene_tx=XM_023249991.1;refGene_gn=SETD5;refGene_tx=XM_023249992.1;refGene_gn=SETD5;refGene_tx=XM_023249993.1;refGene_gn=SETD5;refGene_tx=XM_023249994.1;refGene_gn=SETD5;refGene_tx=XM_023249995.1;refGene_gn=SETD5;refGene_tx=XM_023249996.1;refGene_gn=SETD5;refGene_tx=XM_023249997.1;refGene_gn=SETD5;refGene_tx=XM_023249998.1
A2 50815797 50815915
refGene_gn=SETD5;refGene_tx=XM_023249986.1;refGene_gn=SETD5;refGene_tx=XM_023249987.1;refGene_gn=SETD5;refGene_tx=XM_023249988.1;refGene_gn=SETD5;refGene_tx=XM_023249989.1;refGene_gn=SETD5;refGene_tx=XM_023249990.1;refGene_gn=SETD5;refGene_tx=XM_023249991.1;refGene_gn=SETD5;refGene_tx=XM_023249992.1;refGene_gn=SETD5;refGene_tx=XM_023249993.1;refGene_gn=SETD5;refGene_tx=XM_023249994.1;refGene_gn=SETD5;refGene_tx=XM_023249995.1;refGene_gn=SETD5;refGene_tx=XM_023249996.1;refGene_gn=SETD5;refGene_tx=XM_023249997.1;refGene_gn=SETD5;refGene_tx=XM_023249998.1
A2 50816600 50816710
refGene_gn=SETD5;refGene_tx=XM_023249986.1;refGene_gn=SETD5;refGene_tx=XM_023249987.1;refGene_gn=SETD5;refGene_tx=XM_023249988.1;refGene_gn=

SETD5;refGene_tx=XM_023249989.1;refGene_gn=SETD5;refGene_tx=XM_0232499
90.1;refGene_gn=SETD5;refGene_tx=XM_023249991.1;refGene_gn=SETD5;refGe
ne_tx=XM_023249992.1;refGene_gn=SETD5;refGene_tx=XM_023249993.1;refGen
e_gn=SETD5;refGene_tx=XM_023249994.1;refGene_gn=SETD5;refGene_tx=XM_02
3249995.1;refGene_gn=SETD5;refGene_tx=XM_023249996.1;refGene_gn=SETD5;
refGene_tx=XM_023249997.1;refGene_gn=SETD5;refGene_tx=XM_023249998.1
A2 50817649 50817899
refGene_gn=SETD5;refGene_tx=XM_023249986.1;refGene_gn=SETD5;refGene_tx
=XM_023249987.1;refGene_gn=SETD5;refGene_tx=XM_023249988.1;refGene_gn=
SETD5;refGene_tx=XM_023249989.1;refGene_gn=SETD5;refGene_tx=XM_0232499
90.1;refGene_gn=SETD5;refGene_tx=XM_023249991.1;refGene_gn=SETD5;refGe
ne_tx=XM_023249992.1;refGene_gn=SETD5;refGene_tx=XM_023249993.1;refGen
e_gn=SETD5;refGene_tx=XM_023249994.1;refGene_gn=SETD5;refGene_tx=XM_02
3249995.1;refGene_gn=SETD5;refGene_tx=XM_023249996.1;refGene_gn=SETD5;
refGene_tx=XM_023249997.1;refGene_gn=SETD5;refGene_tx=XM_023249998.1
A2 50818297 50818381
refGene_gn=SETD5;refGene_tx=XM_023249986.1;refGene_gn=SETD5;refGene_tx
=XM_023249987.1;refGene_gn=SETD5;refGene_tx=XM_023249988.1;refGene_gn=
SETD5;refGene_tx=XM_023249989.1;refGene_gn=SETD5;refGene_tx=XM_0232499
90.1;refGene_gn=SETD5;refGene_tx=XM_023249991.1;refGene_gn=SETD5;refGe
ne_tx=XM_023249992.1;refGene_gn=SETD5;refGene_tx=XM_023249993.1;refGen
e_gn=SETD5;refGene_tx=XM_023249994.1;refGene_gn=SETD5;refGene_tx=XM_02
3249995.1;refGene_gn=SETD5;refGene_tx=XM_023249996.1;refGene_gn=SETD5;
refGene_tx=XM_023249997.1;refGene_gn=SETD5;refGene_tx=XM_023249998.1
A2 50819647 50819905
refGene_gn=SETD5;refGene_tx=XM_023249986.1;refGene_gn=SETD5;refGene_tx
=XM_023249987.1;refGene_gn=SETD5;refGene_tx=XM_023249988.1;refGene_gn=
SETD5;refGene_tx=XM_023249989.1;refGene_gn=SETD5;refGene_tx=XM_0232499
90.1;refGene_gn=SETD5;refGene_tx=XM_023249991.1;refGene_gn=SETD5;refGe
ne_tx=XM_023249992.1;refGene_gn=SETD5;refGene_tx=XM_023249993.1;refGen
e_gn=SETD5;refGene_tx=XM_023249994.1;refGene_gn=SETD5;refGene_tx=XM_02
3249995.1;refGene_gn=SETD5;refGene_tx=XM_023249996.1;refGene_gn=SETD5;
refGene_tx=XM_023249997.1;refGene_gn=SETD5;refGene_tx=XM_023249998.1
A2 50820852 50821173
refGene_gn=SETD5;refGene_tx=XM_023249986.1;refGene_gn=SETD5;refGene_tx
=XM_023249987.1;refGene_gn=SETD5;refGene_tx=XM_023249988.1;refGene_gn=
SETD5;refGene_tx=XM_023249989.1;refGene_gn=SETD5;refGene_tx=XM_0232499
90.1;refGene_gn=SETD5;refGene_tx=XM_023249991.1;refGene_gn=SETD5;refGe
ne_tx=XM_023249992.1;refGene_gn=SETD5;refGene_tx=XM_023249993.1;refGen
e_gn=SETD5;refGene_tx=XM_023249994.1;refGene_gn=SETD5;refGene_tx=XM_02
3249995.1;refGene_gn=SETD5;refGene_tx=XM_023249996.1;refGene_gn=SETD5;
refGene_tx=XM_023249997.1;refGene_gn=SETD5;refGene_tx=XM_023249998.1
A2 50821583 50821826
refGene_gn=SETD5;refGene_tx=XM_023249986.1;refGene_gn=SETD5;refGene_tx
=XM_023249987.1;refGene_gn=SETD5;refGene_tx=XM_023249988.1;refGene_gn=
SETD5;refGene_tx=XM_023249989.1;refGene_gn=SETD5;refGene_tx=XM_0232499
90.1;refGene_gn=SETD5;refGene_tx=XM_023249991.1;refGene_gn=SETD5;refGe
ne_tx=XM_023249992.1;refGene_gn=SETD5;refGene_tx=XM_023249993.1;refGen
e_gn=SETD5;refGene_tx=XM_023249994.1;refGene_gn=SETD5;refGene_tx=XM_02
3249995.1;refGene_gn=SETD5;refGene_tx=XM_023249996.1;refGene_gn=SETD5;

refGene_tx=XM_023249997.1;refGene_gn=SETD5;refGene_tx=XM_023249998.1
A2 50827039 50827169
refGene_gn=SETD5;refGene_tx=XM_023249986.1;refGene_gn=SETD5;refGene_tx=XM_023249987.1;refGene_gn=SETD5;refGene_tx=XM_023249988.1;refGene_gn=SETD5;refGene_tx=XM_023249989.1;refGene_gn=SETD5;refGene_tx=XM_023249990.1;refGene_gn=SETD5;refGene_tx=XM_023249991.1;refGene_gn=SETD5;refGene_tx=XM_023249992.1;refGene_gn=SETD5;refGene_tx=XM_023249993.1;refGene_gn=SETD5;refGene_tx=XM_023249994.1;refGene_gn=SETD5;refGene_tx=XM_023249995.1;refGene_gn=SETD5;refGene_tx=XM_023249996.1;refGene_gn=SETD5;refGene_tx=XM_023249997.1;refGene_gn=SETD5;refGene_tx=XM_023249998.1
A2 50837038 50837283
refGene_gn=SETD5;refGene_tx=XM_023249986.1;refGene_gn=SETD5;refGene_tx=XM_023249987.1;refGene_gn=SETD5;refGene_tx=XM_023249988.1;refGene_gn=SETD5;refGene_tx=XM_023249989.1;refGene_gn=SETD5;refGene_tx=XM_023249990.1;refGene_gn=SETD5;refGene_tx=XM_023249991.1;refGene_gn=SETD5;refGene_tx=XM_023249992.1;refGene_gn=SETD5;refGene_tx=XM_023249993.1;refGene_gn=SETD5;refGene_tx=XM_023249994.1;refGene_gn=SETD5;refGene_tx=XM_023249995.1;refGene_gn=SETD5;refGene_tx=XM_023249996.1;refGene_gn=SETD5;refGene_tx=XM_023249997.1;refGene_gn=SETD5;refGene_tx=XM_023249998.1
A2 50840472 50840529
refGene_gn=SETD5;refGene_tx=XM_023249986.1;refGene_gn=SETD5;refGene_tx=XM_023249987.1;refGene_gn=SETD5;refGene_tx=XM_023249988.1;refGene_gn=SETD5;refGene_tx=XM_023249989.1;refGene_gn=SETD5;refGene_tx=XM_023249990.1;refGene_gn=SETD5;refGene_tx=XM_023249991.1;refGene_gn=SETD5;refGene_tx=XM_023249993.1;refGene_gn=SETD5;refGene_tx=XM_023249995.1;refGene_gn=SETD5;refGene_tx=XM_023249998.1
A2 50842442 50842913
refGene_gn=SETD5;refGene_tx=XM_023249986.1;refGene_gn=SETD5;refGene_tx=XM_023249987.1;refGene_gn=SETD5;refGene_tx=XM_023249988.1;refGene_gn=SETD5;refGene_tx=XM_023249989.1;refGene_gn=SETD5;refGene_tx=XM_023249990.1;refGene_gn=SETD5;refGene_tx=XM_023249991.1;refGene_gn=SETD5;refGene_tx=XM_023249992.1;refGene_gn=SETD5;refGene_tx=XM_023249993.1;refGene_gn=SETD5;refGene_tx=XM_023249994.1;refGene_gn=SETD5;refGene_tx=XM_023249995.1;refGene_gn=SETD5;refGene_tx=XM_023249996.1;refGene_gn=SETD5;refGene_tx=XM_023249997.1;refGene_gn=SETD5;refGene_tx=XM_023249998.1
A2 50845567 50845866
refGene_gn=SETD5;refGene_tx=XM_023249986.1;refGene_gn=SETD5;refGene_tx=XM_023249987.1;refGene_gn=SETD5;refGene_tx=XM_023249988.1;refGene_gn=SETD5;refGene_tx=XM_023249989.1;refGene_gn=SETD5;refGene_tx=XM_023249990.1;refGene_gn=SETD5;refGene_tx=XM_023249991.1;refGene_gn=SETD5;refGene_tx=XM_023249992.1;refGene_gn=SETD5;refGene_tx=XM_023249993.1;refGene_gn=SETD5;refGene_tx=XM_023249994.1;refGene_gn=SETD5;refGene_tx=XM_023249995.1;refGene_gn=SETD5;refGene_tx=XM_023249996.1;refGene_gn=SETD5;refGene_tx=XM_023249997.1;refGene_gn=SETD5;refGene_tx=XM_023249998.1
A2 50846723 50846857
refGene_gn=SETD5;refGene_tx=XM_023249986.1;refGene_gn=SETD5;refGene_tx=XM_023249987.1;refGene_gn=SETD5;refGene_tx=XM_023249988.1;refGene_gn=SETD5;refGene_tx=XM_023249989.1;refGene_gn=SETD5;refGene_tx=XM_023249990.1;refGene_gn=SETD5;refGene_tx=XM_023249991.1;refGene_gn=SETD5;refGene_tx=XM_023249992.1;refGene_gn=SETD5;refGene_tx=XM_023249993.1;refGen

e_gn=SETD5;refGene_tx=XM_023249994.1;refGene_gn=SETD5;refGene_tx=XM_023249995.1;refGene_gn=SETD5;refGene_tx=XM_023249996.1;refGene_gn=SETD5;refGene_tx=XM_023249997.1;refGene_gn=SETD5;refGene_tx=XM_023249998.1
A2 50847397 50847486
refGene_gn=SETD5;refGene_tx=XM_023249986.1;refGene_gn=SETD5;refGene_tx=XM_023249987.1;refGene_gn=SETD5;refGene_tx=XM_023249988.1;refGene_gn=SETD5;refGene_tx=XM_023249989.1;refGene_gn=SETD5;refGene_tx=XM_023249990.1;refGene_gn=SETD5;refGene_tx=XM_023249991.1;refGene_gn=SETD5;refGene_tx=XM_023249992.1;refGene_gn=SETD5;refGene_tx=XM_023249993.1;refGene_gn=SETD5;refGene_tx=XM_023249994.1;refGene_gn=SETD5;refGene_tx=XM_023249995.1;refGene_gn=SETD5;refGene_tx=XM_023249996.1;refGene_gn=SETD5;refGene_tx=XM_023249997.1;refGene_gn=SETD5;refGene_tx=XM_023249998.1
A2 50847861 50848467
refGene_gn=SETD5;refGene_tx=XM_023249986.1;refGene_gn=SETD5;refGene_tx=XM_023249987.1;refGene_gn=SETD5;refGene_tx=XM_023249988.1;refGene_gn=SETD5;refGene_tx=XM_023249989.1;refGene_gn=SETD5;refGene_tx=XM_023249990.1;refGene_gn=SETD5;refGene_tx=XM_023249991.1;refGene_gn=SETD5;refGene_tx=XM_023249992.1;refGene_gn=SETD5;refGene_tx=XM_023249993.1;refGene_gn=SETD5;refGene_tx=XM_023249994.1;refGene_gn=SETD5;refGene_tx=XM_023249995.1;refGene_gn=SETD5;refGene_tx=XM_023249996.1;refGene_gn=SETD5;refGene_tx=XM_023249997.1;refGene_gn=SETD5;refGene_tx=XM_023249998.1
A2 50873408 50873509 refGene_gn=LHFPL4;refGene_tx=XM_011280191.3
A2 50876015 50876252 refGene_gn=LHFPL4;refGene_tx=XM_011280191.3
A2 50900656 50901062 refGene_gn=LHFPL4;refGene_tx=XM_011280191.3
A2 50943823 50943979
refGene_gn=MTMR14;refGene_tx=XM_003982451.4;refGene_gn=MTMR14;refGene_tx=XM_003982452.4;refGene_gn=MTMR14;refGene_tx=XM_003982453.4
A2 50946112 50946261
refGene_gn=MTMR14;refGene_tx=XM_003982451.4;refGene_gn=MTMR14;refGene_tx=XM_003982452.4;refGene_gn=MTMR14;refGene_tx=XM_003982453.4
A2 50951272 50951381
refGene_gn=MTMR14;refGene_tx=XM_003982451.4;refGene_gn=MTMR14;refGene_tx=XM_003982452.4;refGene_gn=MTMR14;refGene_tx=XM_003982453.4
A2 50957842 50957918
refGene_gn=MTMR14;refGene_tx=XM_003982451.4;refGene_gn=MTMR14;refGene_tx=XM_003982452.4;refGene_gn=MTMR14;refGene_tx=XM_003982453.4
A2 50958196 50958257
refGene_gn=MTMR14;refGene_tx=XM_003982451.4;refGene_gn=MTMR14;refGene_tx=XM_003982452.4;refGene_gn=MTMR14;refGene_tx=XM_003982453.4
A2 50959764 50959887
refGene_gn=MTMR14;refGene_tx=XM_003982451.4;refGene_gn=MTMR14;refGene_tx=XM_003982452.4;refGene_gn=MTMR14;refGene_tx=XM_003982453.4
A2 50961178 50961252
refGene_gn=MTMR14;refGene_tx=XM_003982451.4;refGene_gn=MTMR14;refGene_tx=XM_003982452.4;refGene_gn=MTMR14;refGene_tx=XM_003982453.4
A2 50965313 50965384
refGene_gn=MTMR14;refGene_tx=XM_003982451.4;refGene_gn=MTMR14;refGene_tx=XM_003982452.4;refGene_gn=MTMR14;refGene_tx=XM_003982453.4
A2 50966425 50966500
refGene_gn=MTMR14;refGene_tx=XM_003982451.4;refGene_gn=MTMR14;refGene_

tx=XM_003982452.4;refGene_gn=MTMR14;refGene_tx=XM_003982453.4
A2 50971159 50971226
refGene_gn=MTMR14;refGene_tx=XM_003982451.4;refGene_gn=MTMR14;refGene_tx=XM_003982452.4;refGene_gn=MTMR14;refGene_tx=XM_003982453.4
A2 50972491 50972577
refGene_gn=MTMR14;refGene_tx=XM_003982451.4;refGene_gn=MTMR14;refGene_tx=XM_003982452.4;refGene_gn=MTMR14;refGene_tx=XM_003982453.4
A2 50972862 50972939
refGene_gn=MTMR14;refGene_tx=XM_003982451.4;refGene_gn=MTMR14;refGene_tx=XM_003982452.4;refGene_gn=MTMR14;refGene_tx=XM_003982453.4
A2 50973201 50973238
refGene_gn=MTMR14;refGene_tx=XM_003982451.4;refGene_gn=MTMR14;refGene_tx=XM_003982452.4;refGene_gn=MTMR14;refGene_tx=XM_003982453.4
A2 50975938 50976009
refGene_gn=MTMR14;refGene_tx=XM_003982451.4;refGene_gn=MTMR14;refGene_tx=XM_003982452.4;refGene_gn=MTMR14;refGene_tx=XM_003982453.4
A2 50976765 50976824
refGene_gn=MTMR14;refGene_tx=XM_003982451.4;refGene_gn=MTMR14;refGene_tx=XM_003982452.4;refGene_gn=MTMR14;refGene_tx=XM_003982453.4
A2 50976997 50977136
refGene_gn=MTMR14;refGene_tx=XM_003982451.4;refGene_gn=MTMR14;refGene_tx=XM_003982452.4;refGene_gn=MTMR14;refGene_tx=XM_003982453.4
A2 50977885 50978062
refGene_gn=MTMR14;refGene_tx=XM_003982451.4;refGene_gn=MTMR14;refGene_tx=XM_003982452.4
A2 50985196 50985352 refGene_gn=MTMR14;refGene_tx=XM_003982451.4
A2 50988261 50988445
refGene_gn=MTMR14;refGene_tx=XM_003982451.4;refGene_gn=MTMR14;refGene_tx=XM_003982452.4;refGene_gn=MTMR14;refGene_tx=XM_003982453.4
A2 50990161 50990229
refGene_gn=CPNE9;refGene_tx=XM_011280193.2;refGene_gn=CPNE9;refGene_tx=XM_019824958.2;refGene_gn=CPNE9;refGene_tx=XM_019824959.1
A2 50990759 50990800
refGene_gn=CPNE9;refGene_tx=XM_011280193.2;refGene_gn=CPNE9;refGene_tx=XM_019824958.2;refGene_gn=CPNE9;refGene_tx=XM_019824959.1
A2 50990910 50990957
refGene_gn=CPNE9;refGene_tx=XM_011280193.2;refGene_gn=CPNE9;refGene_tx=XM_019824958.2;refGene_gn=CPNE9;refGene_tx=XM_019824959.1
A2 50991055 50991159
refGene_gn=CPNE9;refGene_tx=XM_011280193.2;refGene_gn=CPNE9;refGene_tx=XM_019824958.2;refGene_gn=CPNE9;refGene_tx=XM_019824959.1;refGene_gn=CPNE9;refGene_tx=XM_019824960.1
A2 50991639 50991676
refGene_gn=CPNE9;refGene_tx=XM_011280193.2;refGene_gn=CPNE9;refGene_tx=XM_019824958.2;refGene_gn=CPNE9;refGene_tx=XM_019824959.1;refGene_gn=CPNE9;refGene_tx=XM_019824960.1
A2 50991901 50991904
refGene_gn=CPNE9;refGene_tx=XM_011280193.2;refGene_gn=CPNE9;refGene_tx=XM_019824958.2;refGene_gn=CPNE9;refGene_tx=XM_019824959.1;refGene_gn=CPNE9;refGene_tx=XM_019824960.1

A2	50992178	50992255
refGene_gn=CPNE9;refGene_tx=XM_011280193.2;refGene_gn=CPNE9;refGene_tx=XM_019824958.2;refGene_gn=CPNE9;refGene_tx=XM_019824959.1
A2	50995720	50995784
refGene_gn=CPNE9;refGene_tx=XM_011280193.2;refGene_gn=CPNE9;refGene_tx=XM_019824958.2
A2	50995900	50996004
refGene_gn=CPNE9;refGene_tx=XM_011280193.2;refGene_gn=CPNE9;refGene_tx=XM_019824958.2
A2	50996252	50996357
refGene_gn=CPNE9;refGene_tx=XM_011280193.2;refGene_gn=CPNE9;refGene_tx=XM_019824958.2
A2	50998017	50998059
refGene_gn=CPNE9;refGene_tx=XM_011280193.2;refGene_gn=CPNE9;refGene_tx=XM_019824958.2;refGene_gn=CPNE9;refGene_tx=XM_019824959.1;refGene_gn=CPNE9;refGene_tx=XM_019824960.1
A2	50998360	50998436
refGene_gn=CPNE9;refGene_tx=XM_011280193.2;refGene_gn=CPNE9;refGene_tx=XM_019824958.2;refGene_gn=CPNE9;refGene_tx=XM_019824959.1;refGene_gn=CPNE9;refGene_tx=XM_019824960.1
A2	50998544	50998598
refGene_gn=CPNE9;refGene_tx=XM_011280193.2;refGene_gn=CPNE9;refGene_tx=XM_019824958.2;refGene_gn=CPNE9;refGene_tx=XM_019824959.1;refGene_gn=CPNE9;refGene_tx=XM_019824960.1
A2	50998996	50999058
refGene_gn=CPNE9;refGene_tx=XM_011280193.2;refGene_gn=CPNE9;refGene_tx=XM_019824958.2;refGene_gn=CPNE9;refGene_tx=XM_019824959.1;refGene_gn=CPNE9;refGene_tx=XM_019824960.1
A2	50999775	50999822
refGene_gn=CPNE9;refGene_tx=XM_011280193.2;refGene_gn=CPNE9;refGene_tx=XM_019824958.2;refGene_gn=CPNE9;refGene_tx=XM_019824959.1;refGene_gn=CPNE9;refGene_tx=XM_019824960.1
A2	51000677	51000859
refGene_gn=CPNE9;refGene_tx=XM_011280193.2;refGene_gn=CPNE9;refGene_tx=XM_019824958.2;refGene_gn=CPNE9;refGene_tx=XM_019824959.1;refGene_gn=CPNE9;refGene_tx=XM_019824960.1
A2	51001122	51001250
refGene_gn=CPNE9;refGene_tx=XM_011280193.2;refGene_gn=CPNE9;refGene_tx=XM_019824958.2;refGene_gn=CPNE9;refGene_tx=XM_019824959.1;refGene_gn=CPNE9;refGene_tx=XM_019824960.1
A2	51005842	51005945
refGene_gn=CPNE9;refGene_tx=XM_011280193.2;refGene_gn=CPNE9;refGene_tx=XM_019824958.2;refGene_gn=CPNE9;refGene_tx=XM_019824959.1;refGene_gn=CPNE9;refGene_tx=XM_019824960.1
A2	51006540	51006598
refGene_gn=CPNE9;refGene_tx=XM_011280193.2;refGene_gn=CPNE9;refGene_tx=XM_019824958.2;refGene_gn=CPNE9;refGene_tx=XM_019824959.1;refGene_gn=CPNE9;refGene_tx=XM_019824960.1
A2	51006979	51007053
refGene_gn=CPNE9;refGene_tx=XM_011280193.2;refGene_gn=CPNE9;refGene_tx

=XM_019824958.2;refGene_gn=CPNE9;refGene_tx=XM_019824959.1;refGene_gn=CPNE9;refGene_tx=XM_019824960.1
A2 51008628 51008814
refGene_gn=CPNE9;refGene_tx=XM_011280193.2;refGene_gn=CPNE9;refGene_tx=XM_019824958.2;refGene_gn=CPNE9;refGene_tx=XM_019824959.1;refGene_gn=CPNE9;refGene_tx=XM_019824960.1
A2 51013186 51013785
refGene_gn=BRPF1;refGene_tx=XM_003982454.5;refGene_gn=BRPF1;refGene_tx=XM_003982455.5;refGene_gn=BRPF1;refGene_tx=XM_006928959.4;refGene_gn=BRPF1;refGene_tx=XM_006928961.4;refGene_gn=BRPF1;refGene_tx=XM_006928962.4;refGene_gn=BRPF1;refGene_tx=XM_006928963.4;refGene_gn=BRPF1;refGene_tx=XM_006928964.4;refGene_gn=BRPF1;refGene_tx=XM_019824961.2;refGene_gn=BRPF1;refGene_tx=XM_019824962.2;refGene_gn=BRPF1;refGene_tx=XM_023249999.1
A2 51017279 51018239
refGene_gn=BRPF1;refGene_tx=XM_003982454.5;refGene_gn=BRPF1;refGene_tx=XM_003982455.5;refGene_gn=BRPF1;refGene_tx=XM_006928959.4;refGene_gn=BRPF1;refGene_tx=XM_006928961.4;refGene_gn=BRPF1;refGene_tx=XM_006928962.4;refGene_gn=BRPF1;refGene_tx=XM_006928963.4;refGene_gn=BRPF1;refGene_tx=XM_006928964.4;refGene_gn=BRPF1;refGene_tx=XM_019824961.2;refGene_gn=BRPF1;refGene_tx=XM_019824962.2;refGene_gn=BRPF1;refGene_tx=XM_023249999.1
A2 51019043 51019206
refGene_gn=BRPF1;refGene_tx=XM_003982454.5;refGene_gn=BRPF1;refGene_tx=XM_003982455.5;refGene_gn=BRPF1;refGene_tx=XM_006928959.4;refGene_gn=BRPF1;refGene_tx=XM_006928961.4;refGene_gn=BRPF1;refGene_tx=XM_006928962.4;refGene_gn=BRPF1;refGene_tx=XM_006928963.4;refGene_gn=BRPF1;refGene_tx=XM_006928964.4;refGene_gn=BRPF1;refGene_tx=XM_019824961.2;refGene_gn=BRPF1;refGene_tx=XM_019824962.2;refGene_gn=BRPF1;refGene_tx=XM_023249999.1
A2 51019624 51019756
refGene_gn=BRPF1;refGene_tx=XM_003982454.5;refGene_gn=BRPF1;refGene_tx=XM_003982455.5;refGene_gn=BRPF1;refGene_tx=XM_006928959.4;refGene_gn=BRPF1;refGene_tx=XM_006928961.4;refGene_gn=BRPF1;refGene_tx=XM_006928962.4;refGene_gn=BRPF1;refGene_tx=XM_006928963.4;refGene_gn=BRPF1;refGene_tx=XM_006928964.4;refGene_gn=BRPF1;refGene_tx=XM_019824961.2;refGene_gn=BRPF1;refGene_tx=XM_019824962.2;refGene_gn=BRPF1;refGene_tx=XM_023249999.1
A2 51020163 51020310
refGene_gn=BRPF1;refGene_tx=XM_003982454.5;refGene_gn=BRPF1;refGene_tx=XM_003982455.5;refGene_gn=BRPF1;refGene_tx=XM_006928959.4;refGene_gn=BRPF1;refGene_tx=XM_006928961.4;refGene_gn=BRPF1;refGene_tx=XM_006928962.4;refGene_gn=BRPF1;refGene_tx=XM_006928963.4;refGene_gn=BRPF1;refGene_tx=XM_006928964.4;refGene_gn=BRPF1;refGene_tx=XM_019824961.2;refGene_gn=BRPF1;refGene_tx=XM_019824962.2;refGene_gn=BRPF1;refGene_tx=XM_023249999.1
A2 51021088 51021398
refGene_gn=BRPF1;refGene_tx=XM_003982454.5;refGene_gn=BRPF1;refGene_tx=XM_003982455.5;refGene_gn=BRPF1;refGene_tx=XM_006928959.4;refGene_gn=BRPF1;refGene_tx=XM_006928961.4;refGene_gn=BRPF1;refGene_tx=XM_0069289

62.4;refGene_gn=BRPF1;refGene_tx=XM_006928963.4;refGene_gn=BRPF1;refGene_tx=XM_006928964.4;refGene_gn=BRPF1;refGene_tx=XM_019824961.2;refGene_gn=BRPF1;refGene_tx=XM_019824962.2;refGene_gn=BRPF1;refGene_tx=XM_023249999.1
A2 51021726 51022050
refGene_gn=BRPF1;refGene_tx=XM_003982454.5;refGene_gn=BRPF1;refGene_tx=XM_003982455.5;refGene_gn=BRPF1;refGene_tx=XM_006928959.4;refGene_gn=BRPF1;refGene_tx=XM_006928961.4;refGene_gn=BRPF1;refGene_tx=XM_006928962.4;refGene_gn=BRPF1;refGene_tx=XM_006928963.4;refGene_gn=BRPF1;refGene_tx=XM_006928964.4;refGene_gn=BRPF1;refGene_tx=XM_019824961.2;refGene_gn=BRPF1;refGene_tx=XM_019824962.2;refGene_gn=BRPF1;refGene_tx=XM_023249999.1
A2 51022369 51022654
refGene_gn=BRPF1;refGene_tx=XM_003982454.5;refGene_gn=BRPF1;refGene_tx=XM_003982455.5;refGene_gn=BRPF1;refGene_tx=XM_006928959.4;refGene_gn=BRPF1;refGene_tx=XM_006928961.4;refGene_gn=BRPF1;refGene_tx=XM_006928962.4;refGene_gn=BRPF1;refGene_tx=XM_006928963.4;refGene_gn=BRPF1;refGene_tx=XM_019824961.2;refGene_gn=BRPF1;refGene_tx=XM_023249999.1
A2 51023151 51023299
refGene_gn=BRPF1;refGene_tx=XM_003982454.5;refGene_gn=BRPF1;refGene_tx=XM_003982455.5;refGene_gn=BRPF1;refGene_tx=XM_006928959.4;refGene_gn=BRPF1;refGene_tx=XM_006928961.4;refGene_gn=BRPF1;refGene_tx=XM_006928962.4;refGene_gn=BRPF1;refGene_tx=XM_006928963.4;refGene_gn=BRPF1;refGene_tx=XM_006928964.4;refGene_gn=BRPF1;refGene_tx=XM_019824961.2;refGene_gn=BRPF1;refGene_tx=XM_019824962.2;refGene_gn=BRPF1;refGene_tx=XM_023249999.1
A2 51023858 51024216
refGene_gn=BRPF1;refGene_tx=XM_003982454.5;refGene_gn=BRPF1;refGene_tx=XM_003982455.5;refGene_gn=BRPF1;refGene_tx=XM_006928959.4;refGene_gn=BRPF1;refGene_tx=XM_006928961.4;refGene_gn=BRPF1;refGene_tx=XM_006928962.4;refGene_gn=BRPF1;refGene_tx=XM_006928963.4;refGene_gn=BRPF1;refGene_tx=XM_006928964.4;refGene_gn=BRPF1;refGene_tx=XM_019824961.2;refGene_gn=BRPF1;refGene_tx=XM_019824962.2;refGene_gn=BRPF1;refGene_tx=XM_023249999.1
A2 51024561 51024716
refGene_gn=BRPF1;refGene_tx=XM_003982454.5;refGene_gn=BRPF1;refGene_tx=XM_003982455.5;refGene_gn=BRPF1;refGene_tx=XM_006928959.4;refGene_gn=BRPF1;refGene_tx=XM_006928961.4;refGene_gn=BRPF1;refGene_tx=XM_006928962.4;refGene_gn=BRPF1;refGene_tx=XM_006928963.4;refGene_gn=BRPF1;refGene_tx=XM_006928964.4;refGene_gn=BRPF1;refGene_tx=XM_019824961.2;refGene_gn=BRPF1;refGene_tx=XM_019824962.2;refGene_gn=BRPF1;refGene_tx=XM_023249999.1
A2 51025453 51025637
refGene_gn=BRPF1;refGene_tx=XM_003982454.5;refGene_gn=BRPF1;refGene_tx=XM_003982455.5;refGene_gn=BRPF1;refGene_tx=XM_006928959.4;refGene_gn=BRPF1;refGene_tx=XM_006928961.4;refGene_gn=BRPF1;refGene_tx=XM_006928962.4;refGene_gn=BRPF1;refGene_tx=XM_006928963.4;refGene_gn=BRPF1;refGene_tx=XM_006928964.4;refGene_gn=BRPF1;refGene_tx=XM_019824961.2;refGene_gn=BRPF1;refGene_tx=XM_019824962.2;refGene_gn=BRPF1;refGene_tx=XM_023249999.1

A2	51028355	51028462
refGene_gn=OGG1;refGene_tx=XM_006928965.4;refGene_gn=OGG1;refGene_tx=XM_006928966.3;refGene_gn=OGG1;refGene_tx=XM_019824971.2
A2	51028941	51029189
refGene_gn=OGG1;refGene_tx=XM_006928965.4;refGene_gn=OGG1;refGene_tx=XM_006928966.3;refGene_gn=OGG1;refGene_tx=XM_019824971.2
A2	51029742	51029930
refGene_gn=OGG1;refGene_tx=XM_006928965.4;refGene_gn=OGG1;refGene_tx=XM_006928966.3;refGene_gn=OGG1;refGene_tx=XM_019824971.2
A2	51031370	51031552
refGene_gn=OGG1;refGene_tx=XM_006928965.4;refGene_gn=OGG1;refGene_tx=XM_019824971.2
A2	51032795	51032946
refGene_gn=OGG1;refGene_tx=XM_006928965.4;refGene_gn=OGG1;refGene_tx=XM_019824971.2
A2	51033086	51033136
refGene_gn=OGG1;refGene_tx=XM_006928965.4;refGene_gn=OGG1;refGene_tx=XM_019824971.2
A2	51033373	51033482
refGene_gn=OGG1;refGene_tx=XM_006928965.4;refGene_gn=OGG1;refGene_tx=XM_019824971.2
A2	51033848	51033931
refGene_gn=CAMK1;refGene_tx=XM_003982456.5;refGene_gn=CAMK1;refGene_tx=XM_011280194.3
A2	51034034	51034152
refGene_gn=CAMK1;refGene_tx=XM_003982456.5;refGene_gn=CAMK1;refGene_tx=XM_011280194.3
A2	51035753	51035841
refGene_gn=CAMK1;refGene_tx=XM_003982456.5;refGene_gn=CAMK1;refGene_tx=XM_011280194.3
A2	51035937	51036016
refGene_gn=CAMK1;refGene_tx=XM_003982456.5;refGene_gn=CAMK1;refGene_tx=XM_011280194.3
A2	51036566	51036679
refGene_gn=CAMK1;refGene_tx=XM_003982456.5;refGene_gn=CAMK1;refGene_tx=XM_011280194.3
A2	51037350	51037426
refGene_gn=CAMK1;refGene_tx=XM_003982456.5;refGene_gn=CAMK1;refGene_tx=XM_011280194.3
A2	51037520	51037722
refGene_gn=CAMK1;refGene_tx=XM_003982456.5;refGene_gn=CAMK1;refGene_tx=XM_011280194.3
A2	51038489	51038628
refGene_gn=CAMK1;refGene_tx=XM_003982456.5;refGene_gn=CAMK1;refGene_tx=XM_011280194.3
A2	51038713	51038788
refGene_gn=CAMK1;refGene_tx=XM_003982456.5;refGene_gn=CAMK1;refGene_tx=XM_011280194.3
A2	51040504	51040636
refGene_gn=CAMK1;refGene_tx=XM_003982456.5;refGene_gn=CAMK1;refGene_tx

=XM_011280194.3
A2 51042396 51042479
refGene_gn=CAMK1;refGene_tx=XM_003982456.5;refGene_gn=CAMK1;refGene_tx
=XM_011280194.3
A2 51056440 51056633
refGene_gn=TADA3;refGene_tx=XM_003982457.5;refGene_gn=TADA3;refGene_tx
=XM_006928968.4;refGene_gn=TADA3;refGene_tx=XM_023250006.1
A2 51058981 51059167
refGene_gn=TADA3;refGene_tx=XM_003982457.5;refGene_gn=TADA3;refGene_tx
=XM_006928968.4
A2 51061509 51061619
refGene_gn=TADA3;refGene_tx=XM_003982457.5;refGene_gn=TADA3;refGene_tx
=XM_006928968.4;refGene_gn=TADA3;refGene_tx=XM_023250006.1
A2 51062836 51062940
refGene_gn=TADA3;refGene_tx=XM_003982457.5;refGene_gn=TADA3;refGene_tx
=XM_006928968.4;refGene_gn=TADA3;refGene_tx=XM_023250006.1
A2 51063033 51063175
refGene_gn=TADA3;refGene_tx=XM_003982457.5;refGene_gn=TADA3;refGene_tx
=XM_006928968.4;refGene_gn=TADA3;refGene_tx=XM_023250006.1
A2 51064694 51064800
refGene_gn=TADA3;refGene_tx=XM_003982457.5;refGene_gn=TADA3;refGene_tx
=XM_006928968.4;refGene_gn=TADA3;refGene_tx=XM_023250006.1
A2 51064919 51065170
refGene_gn=TADA3;refGene_tx=XM_003982457.5;refGene_gn=TADA3;refGene_tx
=XM_006928968.4;refGene_gn=TADA3;refGene_tx=XM_023250006.1
A2 51066500 51066707
refGene_gn=TADA3;refGene_tx=XM_003982457.5;refGene_gn=TADA3;refGene_tx
=XM_006928968.4;refGene_gn=TADA3;refGene_tx=XM_023250006.1
A2 51068597 51068600 refGene_gn=ARPC4;refGene_tx=XM_003982458.5
A2 51071558 51071677 refGene_gn=ARPC4;refGene_tx=XM_003982458.5
A2 51072891 51073003 refGene_gn=ARPC4;refGene_tx=XM_003982458.5
A2 51074199 51074295 refGene_gn=ARPC4;refGene_tx=XM_003982458.5
A2 51075259 51075430 refGene_gn=ARPC4;refGene_tx=XM_003982458.5
A2 51077588 51077594 refGene_gn=ARPC4;refGene_tx=XM_003982458.5
A2 51081276 51081742 refGene_gn=TTLL3;refGene_tx=XM_019824972.2
A2 51081887 51081979 refGene_gn=TTLL3;refGene_tx=XM_019824972.2
A2 51083740 51083909 refGene_gn=TTLL3;refGene_tx=XM_019824972.2
A2 51084050 51084148 refGene_gn=TTLL3;refGene_tx=XM_019824972.2
A2 51085256 51085385 refGene_gn=TTLL3;refGene_tx=XM_019824972.2
A2 51086509 51086624 refGene_gn=TTLL3;refGene_tx=XM_019824972.2
A2 51087364 51087463 refGene_gn=TTLL3;refGene_tx=XM_019824972.2
A2 51088714 51088910 refGene_gn=TTLL3;refGene_tx=XM_019824972.2
A2 51092145 51092294 refGene_gn=TTLL3;refGene_tx=XM_019824972.2
A2 51095869 51096113 refGene_gn=TTLL3;refGene_tx=XM_019824972.2
A2 51097263 51097699 refGene_gn=TTLL3;refGene_tx=XM_019824972.2
A2 51104102 51104247 refGene_gn=TTLL3;refGene_tx=XM_019824972.2
A2 51105238 51105476 refGene_gn=TTLL3;refGene_tx=XM_019824972.2
A2 51105667 51105779 refGene_gn=TTLL3;refGene_tx=XM_019824972.2
A2 51105781 51105815 refGene_gn=TTLL3;refGene_tx=XM_019824972.2
A2 51105817 51105901 refGene_gn=TTLL3;refGene_tx=XM_019824972.2

A2 51105903 51105995 refGene_gn=TTLL3;refGene_tx=XM_019824972.2
A2 51108202 51108394
refGene_gn=RPUSD3;refGene_tx=XM_003982482.4;refGene_gn=RPUSD3;refGene_tx=XM_006928972.3;refGene_gn=RPUSD3;refGene_tx=XM_023250007.1
A2 51109211 51109351
refGene_gn=RPUSD3;refGene_tx=XM_003982482.4;refGene_gn=RPUSD3;refGene_tx=XM_006928972.3
A2 51111700 51111824
refGene_gn=RPUSD3;refGene_tx=XM_003982482.4;refGene_gn=RPUSD3;refGene_tx=XM_006928972.3;refGene_gn=RPUSD3;refGene_tx=XM_023250007.1
A2 51112049 51112134
refGene_gn=RPUSD3;refGene_tx=XM_003982482.4;refGene_gn=RPUSD3;refGene_tx=XM_006928972.3;refGene_gn=RPUSD3;refGene_tx=XM_023250007.1
A2 51112215 51112323
refGene_gn=RPUSD3;refGene_tx=XM_003982482.4;refGene_gn=RPUSD3;refGene_tx=XM_006928972.3;refGene_gn=RPUSD3;refGene_tx=XM_023250007.1
A2 51113042 51113142
refGene_gn=RPUSD3;refGene_tx=XM_003982482.4;refGene_gn=RPUSD3;refGene_tx=XM_006928972.3;refGene_gn=RPUSD3;refGene_tx=XM_023250007.1
A2 51113283 51113328
refGene_gn=RPUSD3;refGene_tx=XM_003982482.4;refGene_gn=RPUSD3;refGene_tx=XM_023250007.1
A2 51114188 51114328
refGene_gn=RPUSD3;refGene_tx=XM_003982482.4;refGene_gn=RPUSD3;refGene_tx=XM_006928972.3;refGene_gn=RPUSD3;refGene_tx=XM_023250007.1
A2 51114651 51114776
refGene_gn=RPUSD3;refGene_tx=XM_003982482.4;refGene_gn=RPUSD3;refGene_tx=XM_006928972.3;refGene_gn=RPUSD3;refGene_tx=XM_023250007.1
A2 51116386 51116409
refGene_gn=IL17RE;refGene_tx=XM_023250000.1;refGene_gn=IL17RE;refGene_tx=XM_023250001.1;refGene_gn=IL17RE;refGene_tx=XM_023250002.1;refGene_gn=IL17RE;refGene_tx=XM_023250003.1
A2 51119803 51119966 refGene_gn=CIDEC;refGene_tx=XM_006928974.3
A2 51122989 51123177 refGene_gn=CIDEC;refGene_tx=XM_006928974.3
A2 51123282 51123441 refGene_gn=CIDEC;refGene_tx=XM_006928974.3
A2 51128552 51128712 refGene_gn=CIDEC;refGene_tx=XM_006928974.3
A2 51129610 51129663 refGene_gn=CIDEC;refGene_tx=XM_006928974.3
A2 51142109 51142198 refGene_gn=JAGN1;refGene_tx=XM_003982463.3
A2 51143823 51144286 refGene_gn=JAGN1;refGene_tx=XM_003982463.3
A2 51150711 51150883
refGene_gn=IL17RE;refGene_tx=XM_023250000.1;refGene_gn=IL17RE;refGene_tx=XM_023250001.1;refGene_gn=IL17RE;refGene_tx=XM_023250002.1;refGene_gn=IL17RE;refGene_tx=XM_023250003.1;refGene_gn=IL17RE;refGene_tx=XM_023250004.1;refGene_gn=IL17RE;refGene_tx=XM_023250005.1
A2 51151226 51151242
refGene_gn=IL17RE;refGene_tx=XM_023250000.1;refGene_gn=IL17RE;refGene_tx=XM_023250001.1;refGene_gn=IL17RE;refGene_tx=XM_023250002.1;refGene_gn=IL17RE;refGene_tx=XM_023250003.1;refGene_gn=IL17RE;refGene_tx=XM_023250004.1;refGene_gn=IL17RE;refGene_tx=XM_023250005.1
A2 51151814 51151937

refGene_gn=IL17RE;refGene_tx=XM_023250000.1;refGene_gn=IL17RE;refGene_tx=XM_023250001.1;refGene_gn=IL17RE;refGene_tx=XM_023250002.1;refGene_gn=IL17RE;refGene_tx=XM_023250003.1;refGene_gn=IL17RE;refGene_tx=XM_023250004.1;refGene_gn=IL17RE;refGene_tx=XM_023250005.1
A2 51152569 51152667
refGene_gn=IL17RE;refGene_tx=XM_023250000.1;refGene_gn=IL17RE;refGene_tx=XM_023250001.1;refGene_gn=IL17RE;refGene_tx=XM_023250002.1;refGene_gn=IL17RE;refGene_tx=XM_023250003.1;refGene_gn=IL17RE;refGene_tx=XM_023250004.1;refGene_gn=IL17RE;refGene_tx=XM_023250005.1
A2 51152886 51153070
refGene_gn=IL17RE;refGene_tx=XM_023250000.1;refGene_gn=IL17RE;refGene_tx=XM_023250001.1;refGene_gn=IL17RE;refGene_tx=XM_023250002.1;refGene_gn=IL17RE;refGene_tx=XM_023250004.1;refGene_gn=IL17RE;refGene_tx=XM_023250005.1
A2 51153164 51153304
refGene_gn=IL17RE;refGene_tx=XM_023250000.1;refGene_gn=IL17RE;refGene_tx=XM_023250001.1;refGene_gn=IL17RE;refGene_tx=XM_023250002.1;refGene_gn=IL17RE;refGene_tx=XM_023250004.1;refGene_gn=IL17RE;refGene_tx=XM_023250005.1
A2 51154177 51154246
refGene_gn=IL17RE;refGene_tx=XM_023250000.1;refGene_gn=IL17RE;refGene_tx=XM_023250001.1;refGene_gn=IL17RE;refGene_tx=XM_023250002.1;refGene_gn=IL17RE;refGene_tx=XM_023250003.1;refGene_gn=IL17RE;refGene_tx=XM_023250004.1;refGene_gn=IL17RE;refGene_tx=XM_023250005.1
A2 51155146 51155213
refGene_gn=IL17RE;refGene_tx=XM_023250000.1;refGene_gn=IL17RE;refGene_tx=XM_023250001.1;refGene_gn=IL17RE;refGene_tx=XM_023250002.1;refGene_gn=IL17RE;refGene_tx=XM_023250003.1;refGene_gn=IL17RE;refGene_tx=XM_023250004.1;refGene_gn=IL17RE;refGene_tx=XM_023250005.1
A2 51156439 51156615
refGene_gn=IL17RE;refGene_tx=XM_023250000.1;refGene_gn=IL17RE;refGene_tx=XM_023250001.1;refGene_gn=IL17RE;refGene_tx=XM_023250003.1;refGene_gn=IL17RE;refGene_tx=XM_023250004.1;refGene_gn=IL17RE;refGene_tx=XM_023250005.1
A2 51156700 51156748
refGene_gn=IL17RE;refGene_tx=XM_023250000.1;refGene_gn=IL17RE;refGene_tx=XM_023250001.1;refGene_gn=IL17RE;refGene_tx=XM_023250003.1;refGene_gn=IL17RE;refGene_tx=XM_023250004.1;refGene_gn=IL17RE;refGene_tx=XM_023250005.1
A2 51156830 51156876
refGene_gn=IL17RE;refGene_tx=XM_023250000.1;refGene_gn=IL17RE;refGene_tx=XM_023250001.1;refGene_gn=IL17RE;refGene_tx=XM_023250003.1;refGene_gn=IL17RE;refGene_tx=XM_023250004.1;refGene_gn=IL17RE;refGene_tx=XM_023250005.1
A2 51157046 51157195
refGene_gn=IL17RE;refGene_tx=XM_023250000.1;refGene_gn=IL17RE;refGene_tx=XM_023250001.1;refGene_gn=IL17RE;refGene_tx=XM_023250002.1;refGene_gn=IL17RE;refGene_tx=XM_023250003.1;refGene_gn=IL17RE;refGene_tx=XM_023250004.1;refGene_gn=IL17RE;refGene_tx=XM_023250005.1
A2 51158933 51159002

refGene_gn=IL17RE;refGene_tx=XM_023250000.1;refGene_gn=IL17RE;refGene_tx=XM_023250001.1;refGene_gn=IL17RE;refGene_tx=XM_023250002.1;refGene_gn=IL17RE;refGene_tx=XM_023250003.1;refGene_gn=IL17RE;refGene_tx=XM_023250004.1;refGene_gn=IL17RE;refGene_tx=XM_023250005.1
A2 51159498 51159550
refGene_gn=IL17RE;refGene_tx=XM_023250000.1;refGene_gn=IL17RE;refGene_tx=XM_023250001.1;refGene_gn=IL17RE;refGene_tx=XM_023250002.1;refGene_gn=IL17RE;refGene_tx=XM_023250003.1;refGene_gn=IL17RE;refGene_tx=XM_023250004.1;refGene_gn=IL17RE;refGene_tx=XM_023250005.1
A2 51159633 51159732
refGene_gn=IL17RE;refGene_tx=XM_023250000.1;refGene_gn=IL17RE;refGene_tx=XM_023250001.1;refGene_gn=IL17RE;refGene_tx=XM_023250002.1;refGene_gn=IL17RE;refGene_tx=XM_023250003.1;refGene_gn=IL17RE;refGene_tx=XM_023250004.1;refGene_gn=IL17RE;refGene_tx=XM_023250005.1
A2 51160246 51160842
refGene_gn=IL17RE;refGene_tx=XM_023250000.1;refGene_gn=IL17RE;refGene_tx=XM_023250001.1;refGene_gn=IL17RE;refGene_tx=XM_023250002.1;refGene_gn=IL17RE;refGene_tx=XM_023250003.1;refGene_gn=IL17RE;refGene_tx=XM_023250004.1;refGene_gn=IL17RE;refGene_tx=XM_023250005.1
A2 51162313 51162418
refGene_gn=IL17RC;refGene_tx=XM_003982484.3;refGene_gn=IL17RC;refGene_tx=XM_011280197.2;refGene_gn=IL17RC;refGene_tx=XM_019824964.1;refGene_gn=IL17RC;refGene_tx=XM_019824965.1;refGene_gn=IL17RC;refGene_tx=XM_019824966.2;refGene_gn=IL17RC;refGene_tx=XM_019824967.1;refGene_gn=IL17RC;refGene_tx=XM_019824968.2;refGene_gn=IL17RC;refGene_tx=XM_019824969.2
A2 51162706 51162728
refGene_gn=IL17RC;refGene_tx=XM_003982484.3;refGene_gn=IL17RC;refGene_tx=XM_011280197.2;refGene_gn=IL17RC;refGene_tx=XM_019824964.1;refGene_gn=IL17RC;refGene_tx=XM_019824965.1;refGene_gn=IL17RC;refGene_tx=XM_019824966.2;refGene_gn=IL17RC;refGene_tx=XM_019824967.1;refGene_gn=IL17RC;refGene_tx=XM_019824968.2;refGene_gn=IL17RC;refGene_tx=XM_019824969.2
A2 51162919 51163072
refGene_gn=IL17RC;refGene_tx=XM_003982484.3;refGene_gn=IL17RC;refGene_tx=XM_011280197.2;refGene_gn=IL17RC;refGene_tx=XM_019824964.1;refGene_gn=IL17RC;refGene_tx=XM_019824965.1;refGene_gn=IL17RC;refGene_tx=XM_019824966.2;refGene_gn=IL17RC;refGene_tx=XM_019824967.1;refGene_gn=IL17RC;refGene_tx=XM_019824968.2;refGene_gn=IL17RC;refGene_tx=XM_019824969.2;refGene_gn=IL17RC;refGene_tx=XM_019824970.1
A2 51163366 51163441
refGene_gn=IL17RC;refGene_tx=XM_003982484.3;refGene_gn=IL17RC;refGene_tx=XM_011280197.2;refGene_gn=IL17RC;refGene_tx=XM_019824964.1;refGene_gn=IL17RC;refGene_tx=XM_019824965.1;refGene_gn=IL17RC;refGene_tx=XM_019824966.2;refGene_gn=IL17RC;refGene_tx=XM_019824967.1;refGene_gn=IL17RC;refGene_tx=XM_019824968.2;refGene_gn=IL17RC;refGene_tx=XM_019824969.2;refGene_gn=IL17RC;refGene_tx=XM_019824970.1
A2 51163532 51163642
refGene_gn=IL17RC;refGene_tx=XM_003982484.3;refGene_gn=IL17RC;refGene_tx=XM_011280197.2;refGene_gn=IL17RC;refGene_tx=XM_019824964.1;refGene_

gn=IL17RC;refGene_tx=XM_019824965.1;refGene_gn=IL17RC;refGene_tx=XM_019824966.2;refGene_gn=IL17RC;refGene_tx=XM_019824967.1;refGene_gn=IL17RC;refGene_tx=XM_019824968.2;refGene_gn=IL17RC;refGene_tx=XM_019824969.2;refGene_gn=IL17RC;refGene_tx=XM_019824970.1
A2 51164845 51164957
refGene_gn=IL17RC;refGene_tx=XM_003982484.3;refGene_gn=IL17RC;refGene_tx=XM_011280197.2;refGene_gn=IL17RC;refGene_tx=XM_019824964.1;refGene_gn=IL17RC;refGene_tx=XM_019824965.1;refGene_gn=IL17RC;refGene_tx=XM_019824966.2;refGene_gn=IL17RC;refGene_tx=XM_019824967.1;refGene_gn=IL17RC;refGene_tx=XM_019824968.2;refGene_gn=IL17RC;refGene_tx=XM_019824969.2;refGene_gn=IL17RC;refGene_tx=XM_019824970.1
A2 51165279 51165324
refGene_gn=IL17RC;refGene_tx=XM_003982484.3;refGene_gn=IL17RC;refGene_tx=XM_011280197.2;refGene_gn=IL17RC;refGene_tx=XM_019824964.1;refGene_gn=IL17RC;refGene_tx=XM_019824965.1;refGene_gn=IL17RC;refGene_tx=XM_019824966.2;refGene_gn=IL17RC;refGene_tx=XM_019824967.1;refGene_gn=IL17RC;refGene_tx=XM_019824968.2;refGene_gn=IL17RC;refGene_tx=XM_019824969.2;refGene_gn=IL17RC;refGene_tx=XM_019824970.1
A2 51166836 51166976
refGene_gn=IL17RC;refGene_tx=XM_003982484.3;refGene_gn=IL17RC;refGene_tx=XM_011280197.2;refGene_gn=IL17RC;refGene_tx=XM_019824964.1;refGene_gn=IL17RC;refGene_tx=XM_019824965.1;refGene_gn=IL17RC;refGene_tx=XM_019824966.2;refGene_gn=IL17RC;refGene_tx=XM_019824967.1;refGene_gn=IL17RC;refGene_tx=XM_019824968.2;refGene_gn=IL17RC;refGene_tx=XM_019824969.2;refGene_gn=IL17RC;refGene_tx=XM_019824970.1
A2 51167199 51167259
refGene_gn=IL17RC;refGene_tx=XM_003982484.3;refGene_gn=IL17RC;refGene_tx=XM_011280197.2;refGene_gn=IL17RC;refGene_tx=XM_019824964.1;refGene_gn=IL17RC;refGene_tx=XM_019824965.1;refGene_gn=IL17RC;refGene_tx=XM_019824966.2;refGene_gn=IL17RC;refGene_tx=XM_019824967.1;refGene_gn=IL17RC;refGene_tx=XM_019824968.2;refGene_gn=IL17RC;refGene_tx=XM_019824970.1
A2 51169485 51169552
refGene_gn=IL17RC;refGene_tx=XM_003982484.3;refGene_gn=IL17RC;refGene_tx=XM_011280197.2;refGene_gn=IL17RC;refGene_tx=XM_019824964.1;refGene_gn=IL17RC;refGene_tx=XM_019824965.1;refGene_gn=IL17RC;refGene_tx=XM_019824966.2;refGene_gn=IL17RC;refGene_tx=XM_019824967.1;refGene_gn=IL17RC;refGene_tx=XM_019824968.2;refGene_gn=IL17RC;refGene_tx=XM_019824969.2;refGene_gn=IL17RC;refGene_tx=XM_019824970.1
A2 51169638 51169820
refGene_gn=IL17RC;refGene_tx=XM_003982484.3;refGene_gn=IL17RC;refGene_tx=XM_011280197.2;refGene_gn=IL17RC;refGene_tx=XM_019824964.1;refGene_gn=IL17RC;refGene_tx=XM_019824965.1;refGene_gn=IL17RC;refGene_tx=XM_019824966.2;refGene_gn=IL17RC;refGene_tx=XM_019824967.1;refGene_gn=IL17RC;refGene_tx=XM_019824968.2;refGene_gn=IL17RC;refGene_tx=XM_019824969.2;refGene_gn=IL17RC;refGene_tx=XM_019824970.1
A2 51169908 51169959
refGene_gn=IL17RC;refGene_tx=XM_003982484.3;refGene_gn=IL17RC;refGene_tx=XM_011280197.2;refGene_gn=IL17RC;refGene_tx=XM_019824964.1;refGene_gn=IL17RC;refGene_tx=XM_019824965.1;refGene_gn=IL17RC;refGene_tx=XM_01

9824966.2;refGene_gn=IL17RC;refGene_tx=XM_019824967.1;refGene_gn=IL17RC;refGene_tx=XM_019824968.2;refGene_gn=IL17RC;refGene_tx=XM_019824969.2;refGene_gn=IL17RC;refGene_tx=XM_019824970.1
A2 51170643 51170692
refGene_gn=IL17RC;refGene_tx=XM_003982484.3;refGene_gn=IL17RC;refGene_tx=XM_011280197.2;refGene_gn=IL17RC;refGene_tx=XM_019824964.1;refGene_gn=IL17RC;refGene_tx=XM_019824965.1;refGene_gn=IL17RC;refGene_tx=XM_019824966.2;refGene_gn=IL17RC;refGene_tx=XM_019824967.1;refGene_gn=IL17RC;refGene_tx=XM_019824968.2;refGene_gn=IL17RC;refGene_tx=XM_019824969.2;refGene_gn=IL17RC;refGene_tx=XM_019824970.1
A2 51170865 51170987
refGene_gn=IL17RC;refGene_tx=XM_003982484.3;refGene_gn=IL17RC;refGene_tx=XM_011280197.2;refGene_gn=IL17RC;refGene_tx=XM_019824964.1;refGene_gn=IL17RC;refGene_tx=XM_019824965.1;refGene_gn=IL17RC;refGene_tx=XM_019824966.2;refGene_gn=IL17RC;refGene_tx=XM_019824967.1;refGene_gn=IL17RC;refGene_tx=XM_019824968.2;refGene_gn=IL17RC;refGene_tx=XM_019824969.2;refGene_gn=IL17RC;refGene_tx=XM_019824970.1
A2 51171252 51171312
refGene_gn=IL17RC;refGene_tx=XM_003982484.3;refGene_gn=IL17RC;refGene_tx=XM_011280197.2;refGene_gn=IL17RC;refGene_tx=XM_019824964.1;refGene_gn=IL17RC;refGene_tx=XM_019824965.1;refGene_gn=IL17RC;refGene_tx=XM_019824966.2;refGene_gn=IL17RC;refGene_tx=XM_019824967.1;refGene_gn=IL17RC;refGene_tx=XM_019824968.2;refGene_gn=IL17RC;refGene_tx=XM_019824969.2;refGene_gn=IL17RC;refGene_tx=XM_019824970.1
A2 51171874 51171923
refGene_gn=IL17RC;refGene_tx=XM_003982484.3;refGene_gn=IL17RC;refGene_tx=XM_011280197.2;refGene_gn=IL17RC;refGene_tx=XM_019824964.1;refGene_gn=IL17RC;refGene_tx=XM_019824965.1;refGene_gn=IL17RC;refGene_tx=XM_019824966.2;refGene_gn=IL17RC;refGene_tx=XM_019824967.1;refGene_gn=IL17RC;refGene_tx=XM_019824968.2;refGene_gn=IL17RC;refGene_tx=XM_019824969.2;refGene_gn=IL17RC;refGene_tx=XM_019824970.1
A2 51172987 51173083
refGene_gn=IL17RC;refGene_tx=XM_003982484.3;refGene_gn=IL17RC;refGene_tx=XM_011280197.2;refGene_gn=IL17RC;refGene_tx=XM_019824964.1;refGene_gn=IL17RC;refGene_tx=XM_019824965.1;refGene_gn=IL17RC;refGene_tx=XM_019824966.2;refGene_gn=IL17RC;refGene_tx=XM_019824967.1;refGene_gn=IL17RC;refGene_tx=XM_019824968.2;refGene_gn=IL17RC;refGene_tx=XM_019824969.2;refGene_gn=IL17RC;refGene_tx=XM_019824970.1
A2 51173187 51173232
refGene_gn=IL17RC;refGene_tx=XM_003982484.3;refGene_gn=IL17RC;refGene_tx=XM_019824964.1;refGene_gn=IL17RC;refGene_tx=XM_019824965.1;refGene_gn=IL17RC;refGene_tx=XM_019824966.2;refGene_gn=IL17RC;refGene_tx=XM_019824967.1;refGene_gn=IL17RC;refGene_tx=XM_019824969.2;refGene_gn=IL17RC;refGene_tx=XM_019824970.1
A2 51173319 51173939
refGene_gn=IL17RC;refGene_tx=XM_003982484.3;refGene_gn=IL17RC;refGene_tx=XM_011280197.2;refGene_gn=IL17RC;refGene_tx=XM_019824964.1;refGene_gn=IL17RC;refGene_tx=XM_019824965.1;refGene_gn=IL17RC;refGene_tx=XM_019824966.2;refGene_gn=IL17RC;refGene_tx=XM_019824967.1;refGene_gn=IL17RC;refGene_tx=XM_019824968.2;refGene_gn=IL17RC;refGene_tx=XM_019824969.

2;refGene_gn=IL17RC;refGene_tx=XM_019824970.1
A2 51174168 51174241
refGene_gn=CRELD1;refGene_tx=XM_003982460.4;refGene_gn=CRELD1;refGene_tx=XM_006928976.3
A2 51174769 51174963
refGene_gn=CRELD1;refGene_tx=XM_003982460.4;refGene_gn=CRELD1;refGene_tx=XM_006928976.3
A2 51175195 51175278
refGene_gn=CRELD1;refGene_tx=XM_003982460.4;refGene_gn=CRELD1;refGene_tx=XM_006928976.3
A2 51178609 51178720
refGene_gn=CRELD1;refGene_tx=XM_003982460.4;refGene_gn=CRELD1;refGene_tx=XM_006928976.3
A2 51179050 51179142
refGene_gn=CRELD1;refGene_tx=XM_003982460.4;refGene_gn=CRELD1;refGene_tx=XM_006928976.3
A2 51179852 51180029
refGene_gn=CRELD1;refGene_tx=XM_003982460.4;refGene_gn=CRELD1;refGene_tx=XM_006928976.3
A2 51180113 51180209
refGene_gn=CRELD1;refGene_tx=XM_003982460.4;refGene_gn=CRELD1;refGene_tx=XM_006928976.3
A2 51181135 51181219
refGene_gn=CRELD1;refGene_tx=XM_003982460.4;refGene_gn=CRELD1;refGene_tx=XM_006928976.3
A2 51181419 51181515
refGene_gn=CRELD1;refGene_tx=XM_003982460.4;refGene_gn=CRELD1;refGene_tx=XM_006928976.3
A2 51181729 51181864
refGene_gn=CRELD1;refGene_tx=XM_003982460.4;refGene_gn=CRELD1;refGene_tx=XM_006928976.3
A2 51182324 51182356 refGene_gn=CRELD1;refGene_tx=XM_006928976.3
A2 51182671 51182886 refGene_gn=CRELD1;refGene_tx=XM_003982460.4
A2 51184223 51185989
refGene_gn=PRRT3;refGene_tx=XM_003982461.5;refGene_gn=PRRT3;refGene_tx=XM_019824963.1
A2 51186570 51186726
refGene_gn=PRRT3;refGene_tx=XM_003982461.5;refGene_gn=PRRT3;refGene_tx=XM_019824963.1
A2 51186937 51187964
refGene_gn=PRRT3;refGene_tx=XM_003982461.5;refGene_gn=PRRT3;refGene_tx=XM_019824963.1
A2 51202030 51202159
refGene_gn=EMC3;refGene_tx=XM_003982464.4;refGene_gn=EMC3;refGene_tx=XM_019824974.2
A2 51206349 51206432
refGene_gn=EMC3;refGene_tx=XM_003982464.4;refGene_gn=EMC3;refGene_tx=XM_019824974.2
A2 51207340 51207420
refGene_gn=EMC3;refGene_tx=XM_003982464.4;refGene_gn=EMC3;refGene_tx=X

M_019824974.2
A2	51207702	51207784
refGene_gn=EMC3;refGene_tx=XM_003982464.4;refGene_gn=EMC3;refGene_tx=XM_019824974.2
A2	51208361	51208466
refGene_gn=EMC3;refGene_tx=XM_003982464.4;refGene_gn=EMC3;refGene_tx=XM_019824974.2
A2	51209403	51209605	refGene_gn=EMC3;refGene_tx=XM_019824974.2
A2	51209994	51210088	refGene_gn=EMC3;refGene_tx=XM_003982464.4
A2	51210413	51210471	refGene_gn=EMC3;refGene_tx=XM_003982464.4
A2	51219751	51219906	refGene_gn=EMC3;refGene_tx=XM_003982464.4
A2	51222398	51222462
refGene_gn=FANCD2;refGene_tx=XM_011280201.3;refGene_gn=FANCD2;refGene_tx=XM_011280202.3;refGene_gn=FANCD2;refGene_tx=XM_019824973.2
A2	51224378	51224519
refGene_gn=FANCD2;refGene_tx=XM_011280201.3;refGene_gn=FANCD2;refGene_tx=XM_011280202.3;refGene_gn=FANCD2;refGene_tx=XM_019824973.2
A2	51225091	51225159
refGene_gn=FANCD2;refGene_tx=XM_011280201.3;refGene_gn=FANCD2;refGene_tx=XM_011280202.3;refGene_gn=FANCD2;refGene_tx=XM_019824973.2
A2	51225564	51225668
refGene_gn=FANCD2;refGene_tx=XM_011280201.3;refGene_gn=FANCD2;refGene_tx=XM_011280202.3;refGene_gn=FANCD2;refGene_tx=XM_019824973.2
A2	51226274	51226335
refGene_gn=FANCD2;refGene_tx=XM_011280201.3;refGene_gn=FANCD2;refGene_tx=XM_011280202.3;refGene_gn=FANCD2;refGene_tx=XM_019824973.2
A2	51227270	51227323
refGene_gn=FANCD2;refGene_tx=XM_011280201.3;refGene_gn=FANCD2;refGene_tx=XM_011280202.3;refGene_gn=FANCD2;refGene_tx=XM_019824973.2
A2	51230037	51230116
refGene_gn=FANCD2;refGene_tx=XM_011280201.3;refGene_gn=FANCD2;refGene_tx=XM_011280202.3;refGene_gn=FANCD2;refGene_tx=XM_019824973.2
A2	51231145	51231270
refGene_gn=FANCD2;refGene_tx=XM_011280201.3;refGene_gn=FANCD2;refGene_tx=XM_011280202.3;refGene_gn=FANCD2;refGene_tx=XM_019824973.2
A2	51232619	51232707
refGene_gn=FANCD2;refGene_tx=XM_011280201.3;refGene_gn=FANCD2;refGene_tx=XM_011280202.3;refGene_gn=FANCD2;refGene_tx=XM_019824973.2
A2	51233964	51234069
refGene_gn=FANCD2;refGene_tx=XM_011280201.3;refGene_gn=FANCD2;refGene_tx=XM_011280202.3;refGene_gn=FANCD2;refGene_tx=XM_019824973.2
A2	51234570	51234671
refGene_gn=FANCD2;refGene_tx=XM_011280201.3;refGene_gn=FANCD2;refGene_tx=XM_011280202.3;refGene_gn=FANCD2;refGene_tx=XM_019824973.2
A2	51235075	51235184
refGene_gn=FANCD2;refGene_tx=XM_011280201.3;refGene_gn=FANCD2;refGene_tx=XM_011280202.3;refGene_gn=FANCD2;refGene_tx=XM_019824973.2
A2	51235416	51235452
refGene_gn=FANCD2;refGene_tx=XM_011280201.3;refGene_gn=FANCD2;refGene_tx=XM_011280202.3;refGene_gn=FANCD2;refGene_tx=XM_019824973.2

A2 51240066 51240210
refGene_gn=FANCD2;refGene_tx=XM_011280201.3;refGene_gn=FANCD2;refGene_tx=XM_011280202.3;refGene_gn=FANCD2;refGene_tx=XM_019824973.2
A2 51242349 51242484
refGene_gn=FANCD2;refGene_tx=XM_011280201.3;refGene_gn=FANCD2;refGene_tx=XM_011280202.3;refGene_gn=FANCD2;refGene_tx=XM_019824973.2
A2 51244263 51244395
refGene_gn=FANCD2;refGene_tx=XM_011280201.3;refGene_gn=FANCD2;refGene_tx=XM_011280202.3;refGene_gn=FANCD2;refGene_tx=XM_019824973.2
A2 51245761 51245872
refGene_gn=FANCD2;refGene_tx=XM_011280201.3;refGene_gn=FANCD2;refGene_tx=XM_011280202.3;refGene_gn=FANCD2;refGene_tx=XM_019824973.2
A2 51249678 51249788
refGene_gn=FANCD2;refGene_tx=XM_011280201.3;refGene_gn=FANCD2;refGene_tx=XM_011280202.3;refGene_gn=FANCD2;refGene_tx=XM_019824973.2
A2 51250735 51250796
refGene_gn=FANCD2;refGene_tx=XM_011280201.3;refGene_gn=FANCD2;refGene_tx=XM_011280202.3;refGene_gn=FANCD2;refGene_tx=XM_019824973.2
A2 51251454 51251574
refGene_gn=FANCD2;refGene_tx=XM_011280201.3;refGene_gn=FANCD2;refGene_tx=XM_011280202.3;refGene_gn=FANCD2;refGene_tx=XM_019824973.2
A2 51252226 51252300
refGene_gn=FANCD2;refGene_tx=XM_011280201.3;refGene_gn=FANCD2;refGene_tx=XM_011280202.3;refGene_gn=FANCD2;refGene_tx=XM_019824973.2
A2 51252597 51252744
refGene_gn=FANCD2;refGene_tx=XM_011280201.3;refGene_gn=FANCD2;refGene_tx=XM_011280202.3;refGene_gn=FANCD2;refGene_tx=XM_019824973.2
A2 51253252 51253353
refGene_gn=FANCD2;refGene_tx=XM_011280201.3;refGene_gn=FANCD2;refGene_tx=XM_011280202.3;refGene_gn=FANCD2;refGene_tx=XM_019824973.2
A2 51253768 51253884
refGene_gn=FANCD2;refGene_tx=XM_011280201.3;refGene_gn=FANCD2;refGene_tx=XM_011280202.3;refGene_gn=FANCD2;refGene_tx=XM_019824973.2
A2 51255717 51255826
refGene_gn=FANCD2;refGene_tx=XM_011280201.3;refGene_gn=FANCD2;refGene_tx=XM_011280202.3;refGene_gn=FANCD2;refGene_tx=XM_019824973.2
A2 51258678 51258789
refGene_gn=FANCD2;refGene_tx=XM_011280201.3;refGene_gn=FANCD2;refGene_tx=XM_011280202.3;refGene_gn=FANCD2;refGene_tx=XM_019824973.2
A2 51259039 51259152
refGene_gn=FANCD2;refGene_tx=XM_011280201.3;refGene_gn=FANCD2;refGene_tx=XM_011280202.3;refGene_gn=FANCD2;refGene_tx=XM_019824973.2
A2 51260129 51260273
refGene_gn=FANCD2;refGene_tx=XM_011280201.3;refGene_gn=FANCD2;refGene_tx=XM_011280202.3;refGene_gn=FANCD2;refGene_tx=XM_019824973.2
A2 51262458 51262575
refGene_gn=FANCD2;refGene_tx=XM_011280201.3;refGene_gn=FANCD2;refGene_tx=XM_011280202.3;refGene_gn=FANCD2;refGene_tx=XM_019824973.2
A2 51263327 51263456
refGene_gn=FANCD2;refGene_tx=XM_011280201.3;refGene_gn=FANCD2;refGene_

tx=XM_011280202.3;refGene_gn=FANCD2;refGene_tx=XM_019824973.2
A2 51263594 51263713
refGene_gn=FANCD2;refGene_tx=XM_011280201.3;refGene_gn=FANCD2;refGene_tx=XM_011280202.3;refGene_gn=FANCD2;refGene_tx=XM_019824973.2
A2 51269692 51269803
refGene_gn=FANCD2;refGene_tx=XM_011280201.3;refGene_gn=FANCD2;refGene_tx=XM_011280202.3;refGene_gn=FANCD2;refGene_tx=XM_019824973.2
A2 51271391 51271519
refGene_gn=FANCD2;refGene_tx=XM_011280201.3;refGene_gn=FANCD2;refGene_tx=XM_011280202.3;refGene_gn=FANCD2;refGene_tx=XM_019824973.2
A2 51272587 51272681
refGene_gn=FANCD2;refGene_tx=XM_011280201.3;refGene_gn=FANCD2;refGene_tx=XM_011280202.3;refGene_gn=FANCD2;refGene_tx=XM_019824973.2
A2 51273698 51273821
refGene_gn=FANCD2;refGene_tx=XM_011280201.3;refGene_gn=FANCD2;refGene_tx=XM_011280202.3;refGene_gn=FANCD2;refGene_tx=XM_019824973.2
A2 51274621 51274715
refGene_gn=FANCD2;refGene_tx=XM_011280201.3;refGene_gn=FANCD2;refGene_tx=XM_011280202.3;refGene_gn=FANCD2;refGene_tx=XM_019824973.2
A2 51275394 51275466
refGene_gn=FANCD2;refGene_tx=XM_011280201.3;refGene_gn=FANCD2;refGene_tx=XM_011280202.3;refGene_gn=FANCD2;refGene_tx=XM_019824973.2
A2 51276424 51276463
refGene_gn=FANCD2;refGene_tx=XM_011280201.3;refGene_gn=FANCD2;refGene_tx=XM_019824973.2
A2 51277454 51277529
refGene_gn=FANCD2;refGene_tx=XM_011280201.3;refGene_gn=FANCD2;refGene_tx=XM_011280202.3;refGene_gn=FANCD2;refGene_tx=XM_019824973.2
A2 51278278 51278353
refGene_gn=FANCD2;refGene_tx=XM_011280201.3;refGene_gn=FANCD2;refGene_tx=XM_011280202.3;refGene_gn=FANCD2;refGene_tx=XM_019824973.2
A2 51279300 51279447
refGene_gn=FANCD2;refGene_tx=XM_011280201.3;refGene_gn=FANCD2;refGene_tx=XM_011280202.3;refGene_gn=FANCD2;refGene_tx=XM_019824973.2
A2 51280977 51281073
refGene_gn=FANCD2;refGene_tx=XM_011280201.3;refGene_gn=FANCD2;refGene_tx=XM_011280202.3;refGene_gn=FANCD2;refGene_tx=XM_019824973.2
A2 51284363 51284435
refGene_gn=FANCD2;refGene_tx=XM_011280201.3;refGene_gn=FANCD2;refGene_tx=XM_011280202.3;refGene_gn=FANCD2;refGene_tx=XM_019824973.2
A2 51285849 51286386
refGene_gn=FANCD2OS;refGene_tx=XM_003982465.4;refGene_gn=FANCD2OS;refGene_tx=XM_006928978.3;refGene_gn=FANCD2OS;refGene_tx=XM_006928979.3;refGene_gn=FANCD2OS;refGene_tx=XM_006928980.3;refGene_gn=FANCD2OS;refGene_tx=XM_019824975.2
A2 51295512 51295630
refGene_gn=BRK1;refGene_tx=XM_003982466.5;refGene_gn=BRK1;refGene_tx=XM_019824977.2
A2 51303308 51303391
refGene_gn=BRK1;refGene_tx=XM_003982466.5;refGene_gn=BRK1;refGene_tx=X

M_019824977.2
A2	51303651	51303678
refGene_gn=BRK1;refGene_tx=XM_003982466.5;refGene_gn=BRK1;refGene_tx=XM_019824977.2
A2	51310457	51310585	refGene_gn=BRK1;refGene_tx=XM_019824977.2
A2	51313639	51313979	refGene_gn=VHL;refGene_tx=XM_003982486.5
A2	51315968	51316091	refGene_gn=VHL;refGene_tx=XM_003982486.5
A2	51319646	51319840	refGene_gn=VHL;refGene_tx=XM_003982486.5
A2	51331786	51331880
refGene_gn=IRAK2;refGene_tx=XM_003982467.5;refGene_gn=IRAK2;refGene_tx=XM_019824978.2;refGene_gn=IRAK2;refGene_tx=XM_019824979.2
A2	51339980	51340163
refGene_gn=IRAK2;refGene_tx=XM_003982467.5;refGene_gn=IRAK2;refGene_tx=XM_019824978.2;refGene_gn=IRAK2;refGene_tx=XM_019824979.2
A2	51359672	51359810
refGene_gn=IRAK2;refGene_tx=XM_003982467.5;refGene_gn=IRAK2;refGene_tx=XM_019824978.2
A2	51366136	51366228
refGene_gn=IRAK2;refGene_tx=XM_003982467.5;refGene_gn=IRAK2;refGene_tx=XM_019824978.2;refGene_gn=IRAK2;refGene_tx=XM_019824979.2
A2	51368356	51368551
refGene_gn=IRAK2;refGene_tx=XM_003982467.5;refGene_gn=IRAK2;refGene_tx=XM_019824978.2;refGene_gn=IRAK2;refGene_tx=XM_019824979.2
A2	51368634	51368699
refGene_gn=IRAK2;refGene_tx=XM_003982467.5;refGene_gn=IRAK2;refGene_tx=XM_019824978.2;refGene_gn=IRAK2;refGene_tx=XM_019824979.2
A2	51371181	51371296
refGene_gn=IRAK2;refGene_tx=XM_003982467.5;refGene_gn=IRAK2;refGene_tx=XM_019824978.2;refGene_gn=IRAK2;refGene_tx=XM_019824979.2
A2	51373866	51373976
refGene_gn=IRAK2;refGene_tx=XM_003982467.5;refGene_gn=IRAK2;refGene_tx=XM_019824978.2;refGene_gn=IRAK2;refGene_tx=XM_019824979.2
A2	51376071	51376267
refGene_gn=IRAK2;refGene_tx=XM_003982467.5;refGene_gn=IRAK2;refGene_tx=XM_019824978.2;refGene_gn=IRAK2;refGene_tx=XM_019824979.2
A2	51380149	51380212
refGene_gn=IRAK2;refGene_tx=XM_003982467.5;refGene_gn=IRAK2;refGene_tx=XM_019824979.2
A2	51388258	51388459
refGene_gn=IRAK2;refGene_tx=XM_003982467.5;refGene_gn=IRAK2;refGene_tx=XM_019824978.2;refGene_gn=IRAK2;refGene_tx=XM_019824979.2
A2	51391673	51391965
refGene_gn=IRAK2;refGene_tx=XM_003982467.5;refGene_gn=IRAK2;refGene_tx=XM_019824978.2;refGene_gn=IRAK2;refGene_tx=XM_019824979.2
A2	51393045	51393158
refGene_gn=IRAK2;refGene_tx=XM_003982467.5;refGene_gn=IRAK2;refGene_tx=XM_019824978.2;refGene_gn=IRAK2;refGene_tx=XM_019824979.2
A2	51397344	51397770	refGene_gn=TATDN2;refGene_tx=XM_003982468.5
A2	51404933	51405428	refGene_gn=TATDN2;refGene_tx=XM_003982468.5
A2	51412745	51413624	refGene_gn=TATDN2;refGene_tx=XM_003982468.5

A2 51418243 51418371 refGene_gn=TATDN2;refGene_tx=XM_003982468.5
A2 51419906 51420090 refGene_gn=TATDN2;refGene_tx=XM_003982468.5
A2 51420506 51420647 refGene_gn=TATDN2;refGene_tx=XM_003982468.5
A2 51428103 51428123 refGene_gn=GHRL;refGene_tx=NM_001009853.1
A2 51428912 51429021 refGene_gn=GHRL;refGene_tx=NM_001009853.1
A2 51431980 51432094 refGene_gn=GHRL;refGene_tx=NM_001009853.1
A2 51432336 51432444 refGene_gn=GHRL;refGene_tx=NM_001009853.1
A2 51433054 51433219
refGene_gn=SEC13;refGene_tx=XM_019824980.2;refGene_gn=SEC13;refGene_tx=XM_019824981.2
A2 51453415 51453529 refGene_gn=SEC13;refGene_tx=XM_003982469.4
A2 51463391 51463538
refGene_gn=SEC13;refGene_tx=XM_003982469.4;refGene_gn=SEC13;refGene_tx=XM_019824980.2;refGene_gn=SEC13;refGene_tx=XM_019824981.2
A2 51464671 51464795
refGene_gn=SEC13;refGene_tx=XM_003982469.4;refGene_gn=SEC13;refGene_tx=XM_019824980.2;refGene_gn=SEC13;refGene_tx=XM_019824981.2
A2 51465218 51465352
refGene_gn=SEC13;refGene_tx=XM_003982469.4;refGene_gn=SEC13;refGene_tx=XM_019824980.2;refGene_gn=SEC13;refGene_tx=XM_019824981.2
A2 51479219 51479353
refGene_gn=SEC13;refGene_tx=XM_003982469.4;refGene_gn=SEC13;refGene_tx=XM_019824980.2;refGene_gn=SEC13;refGene_tx=XM_019824981.2
A2 51479895 51480047
refGene_gn=SEC13;refGene_tx=XM_003982469.4;refGene_gn=SEC13;refGene_tx=XM_019824980.2;refGene_gn=SEC13;refGene_tx=XM_019824981.2
A2 51483024 51483140
refGene_gn=SEC13;refGene_tx=XM_003982469.4;refGene_gn=SEC13;refGene_tx=XM_019824980.2;refGene_gn=SEC13;refGene_tx=XM_019824981.2
A2 51485363 51485408
refGene_gn=SEC13;refGene_tx=XM_003982469.4;refGene_gn=SEC13;refGene_tx=XM_019824980.2;refGene_gn=SEC13;refGene_tx=XM_019824981.2
A2 51488727 51488730
refGene_gn=SEC13;refGene_tx=XM_003982469.4;refGene_gn=SEC13;refGene_tx=XM_019824981.2
A2 51495816 51496128
refGene_gn=ATP2B2;refGene_tx=XM_023250008.1;refGene_gn=ATP2B2;refGene_tx=XM_023250009.1;refGene_gn=ATP2B2;refGene_tx=XM_023250010.1;refGene_gn=ATP2B2;refGene_tx=XM_023250011.1;refGene_gn=ATP2B2;refGene_tx=XM_023250012.1;refGene_gn=ATP2B2;refGene_tx=XM_023250013.1;refGene_gn=ATP2B2;refGene_tx=XM_023250014.1;refGene_gn=ATP2B2;refGene_tx=XM_023250015.1;refGene_gn=ATP2B2;refGene_tx=XM_023250017.1;refGene_gn=ATP2B2;refGene_tx=XM_023250018.1;refGene_gn=ATP2B2;refGene_tx=XM_023250019.1
A2 51498974 51498982 refGene_gn=ATP2B2;refGene_tx=XM_023250016.1
A2 51502881 51503053
refGene_gn=ATP2B2;refGene_tx=XM_023250015.1;refGene_gn=ATP2B2;refGene_tx=XM_023250016.1;refGene_gn=ATP2B2;refGene_tx=XM_023250018.1;refGene_gn=ATP2B2;refGene_tx=XM_023250019.1
A2 51504299 51504482
refGene_gn=ATP2B2;refGene_tx=XM_023250008.1;refGene_gn=ATP2B2;refGene_

tx=XM_023250009.1;refGene_gn=ATP2B2;refGene_tx=XM_023250010.1;refGene_gn=ATP2B2;refGene_tx=XM_023250011.1;refGene_gn=ATP2B2;refGene_tx=XM_023250012.1;refGene_gn=ATP2B2;refGene_tx=XM_023250013.1;refGene_gn=ATP2B2;refGene_tx=XM_023250014.1;refGene_gn=ATP2B2;refGene_tx=XM_023250015.1;refGene_gn=ATP2B2;refGene_tx=XM_023250016.1;refGene_gn=ATP2B2;refGene_tx=XM_023250017.1;refGene_gn=ATP2B2;refGene_tx=XM_023250018.1;refGene_gn=ATP2B2;refGene_tx=XM_023250019.1
A2 51506313 51506421
refGene_gn=ATP2B2;refGene_tx=XM_023250008.1;refGene_gn=ATP2B2;refGene_tx=XM_023250009.1;refGene_gn=ATP2B2;refGene_tx=XM_023250010.1;refGene_gn=ATP2B2;refGene_tx=XM_023250011.1;refGene_gn=ATP2B2;refGene_tx=XM_023250012.1;refGene_gn=ATP2B2;refGene_tx=XM_023250013.1;refGene_gn=ATP2B2;refGene_tx=XM_023250014.1;refGene_gn=ATP2B2;refGene_tx=XM_023250015.1;refGene_gn=ATP2B2;refGene_tx=XM_023250016.1;refGene_gn=ATP2B2;refGene_tx=XM_023250017.1;refGene_gn=ATP2B2;refGene_tx=XM_023250018.1;refGene_gn=ATP2B2;refGene_tx=XM_023250019.1
A2 51506595 51506807
refGene_gn=ATP2B2;refGene_tx=XM_023250008.1;refGene_gn=ATP2B2;refGene_tx=XM_023250009.1;refGene_gn=ATP2B2;refGene_tx=XM_023250010.1;refGene_gn=ATP2B2;refGene_tx=XM_023250011.1;refGene_gn=ATP2B2;refGene_tx=XM_023250012.1;refGene_gn=ATP2B2;refGene_tx=XM_023250013.1;refGene_gn=ATP2B2;refGene_tx=XM_023250014.1;refGene_gn=ATP2B2;refGene_tx=XM_023250015.1;refGene_gn=ATP2B2;refGene_tx=XM_023250016.1;refGene_gn=ATP2B2;refGene_tx=XM_023250017.1;refGene_gn=ATP2B2;refGene_tx=XM_023250018.1;refGene_gn=ATP2B2;refGene_tx=XM_023250019.1
A2 51508515 51508729
refGene_gn=ATP2B2;refGene_tx=XM_023250008.1;refGene_gn=ATP2B2;refGene_tx=XM_023250009.1;refGene_gn=ATP2B2;refGene_tx=XM_023250010.1;refGene_gn=ATP2B2;refGene_tx=XM_023250011.1;refGene_gn=ATP2B2;refGene_tx=XM_023250012.1;refGene_gn=ATP2B2;refGene_tx=XM_023250013.1;refGene_gn=ATP2B2;refGene_tx=XM_023250014.1;refGene_gn=ATP2B2;refGene_tx=XM_023250015.1;refGene_gn=ATP2B2;refGene_tx=XM_023250016.1;refGene_gn=ATP2B2;refGene_tx=XM_023250017.1;refGene_gn=ATP2B2;refGene_tx=XM_023250018.1;refGene_gn=ATP2B2;refGene_tx=XM_023250019.1
A2 51511548 51511740
refGene_gn=ATP2B2;refGene_tx=XM_023250008.1;refGene_gn=ATP2B2;refGene_tx=XM_023250009.1;refGene_gn=ATP2B2;refGene_tx=XM_023250010.1;refGene_gn=ATP2B2;refGene_tx=XM_023250011.1;refGene_gn=ATP2B2;refGene_tx=XM_023250012.1;refGene_gn=ATP2B2;refGene_tx=XM_023250013.1;refGene_gn=ATP2B2;refGene_tx=XM_023250014.1;refGene_gn=ATP2B2;refGene_tx=XM_023250015.1;refGene_gn=ATP2B2;refGene_tx=XM_023250016.1;refGene_gn=ATP2B2;refGene_tx=XM_023250017.1;refGene_gn=ATP2B2;refGene_tx=XM_023250018.1;refGene_gn=ATP2B2;refGene_tx=XM_023250019.1
A2 51512209 51512316
refGene_gn=ATP2B2;refGene_tx=XM_023250008.1;refGene_gn=ATP2B2;refGene_tx=XM_023250009.1;refGene_gn=ATP2B2;refGene_tx=XM_023250010.1;refGene_gn=ATP2B2;refGene_tx=XM_023250011.1;refGene_gn=ATP2B2;refGene_tx=XM_023250012.1;refGene_gn=ATP2B2;refGene_tx=XM_023250013.1;refGene_gn=ATP2B2;refGene_tx=XM_023250014.1;refGene_gn=ATP2B2;refGene_tx=XM_023250015.1;refGene_gn=ATP2B2;refGene_tx=XM_023250016.1;refGene_gn=ATP2B2;refGen

e_tx=XM_023250017.1;refGene_gn=ATP2B2;refGene_tx=XM_023250018.1;refGene_gn=ATP2B2;refGene_tx=XM_023250019.1
A2 51516395 51516483
refGene_gn=ATP2B2;refGene_tx=XM_023250008.1;refGene_gn=ATP2B2;refGene_tx=XM_023250009.1;refGene_gn=ATP2B2;refGene_tx=XM_023250010.1;refGene_gn=ATP2B2;refGene_tx=XM_023250011.1;refGene_gn=ATP2B2;refGene_tx=XM_023250012.1;refGene_gn=ATP2B2;refGene_tx=XM_023250013.1;refGene_gn=ATP2B2;refGene_tx=XM_023250014.1;refGene_gn=ATP2B2;refGene_tx=XM_023250015.1;refGene_gn=ATP2B2;refGene_tx=XM_023250016.1;refGene_gn=ATP2B2;refGene_tx=XM_023250017.1;refGene_gn=ATP2B2;refGene_tx=XM_023250018.1;refGene_gn=ATP2B2;refGene_tx=XM_023250019.1
A2 51516664 51516844
refGene_gn=ATP2B2;refGene_tx=XM_023250008.1;refGene_gn=ATP2B2;refGene_tx=XM_023250009.1;refGene_gn=ATP2B2;refGene_tx=XM_023250010.1;refGene_gn=ATP2B2;refGene_tx=XM_023250011.1;refGene_gn=ATP2B2;refGene_tx=XM_023250012.1;refGene_gn=ATP2B2;refGene_tx=XM_023250013.1;refGene_gn=ATP2B2;refGene_tx=XM_023250014.1;refGene_gn=ATP2B2;refGene_tx=XM_023250015.1;refGene_gn=ATP2B2;refGene_tx=XM_023250016.1;refGene_gn=ATP2B2;refGene_tx=XM_023250017.1;refGene_gn=ATP2B2;refGene_tx=XM_023250018.1;refGene_gn=ATP2B2;refGene_tx=XM_023250019.1
A2 51525303 51525538
refGene_gn=ATP2B2;refGene_tx=XM_023250008.1;refGene_gn=ATP2B2;refGene_tx=XM_023250009.1;refGene_gn=ATP2B2;refGene_tx=XM_023250010.1;refGene_gn=ATP2B2;refGene_tx=XM_023250011.1;refGene_gn=ATP2B2;refGene_tx=XM_023250012.1;refGene_gn=ATP2B2;refGene_tx=XM_023250013.1;refGene_gn=ATP2B2;refGene_tx=XM_023250014.1;refGene_gn=ATP2B2;refGene_tx=XM_023250015.1;refGene_gn=ATP2B2;refGene_tx=XM_023250016.1;refGene_gn=ATP2B2;refGene_tx=XM_023250017.1;refGene_gn=ATP2B2;refGene_tx=XM_023250018.1;refGene_gn=ATP2B2;refGene_tx=XM_023250019.1
A2 51526666 51526908
refGene_gn=ATP2B2;refGene_tx=XM_023250008.1;refGene_gn=ATP2B2;refGene_tx=XM_023250009.1;refGene_gn=ATP2B2;refGene_tx=XM_023250010.1;refGene_gn=ATP2B2;refGene_tx=XM_023250011.1;refGene_gn=ATP2B2;refGene_tx=XM_023250012.1;refGene_gn=ATP2B2;refGene_tx=XM_023250013.1;refGene_gn=ATP2B2;refGene_tx=XM_023250014.1;refGene_gn=ATP2B2;refGene_tx=XM_023250015.1;refGene_gn=ATP2B2;refGene_tx=XM_023250016.1;refGene_gn=ATP2B2;refGene_tx=XM_023250017.1;refGene_gn=ATP2B2;refGene_tx=XM_023250018.1;refGene_gn=ATP2B2;refGene_tx=XM_023250019.1
A2 51538917 51539160
refGene_gn=ATP2B2;refGene_tx=XM_023250008.1;refGene_gn=ATP2B2;refGene_tx=XM_023250009.1;refGene_gn=ATP2B2;refGene_tx=XM_023250010.1;refGene_gn=ATP2B2;refGene_tx=XM_023250011.1;refGene_gn=ATP2B2;refGene_tx=XM_023250012.1;refGene_gn=ATP2B2;refGene_tx=XM_023250013.1;refGene_gn=ATP2B2;refGene_tx=XM_023250014.1;refGene_gn=ATP2B2;refGene_tx=XM_023250015.1;refGene_gn=ATP2B2;refGene_tx=XM_023250016.1;refGene_gn=ATP2B2;refGene_tx=XM_023250017.1;refGene_gn=ATP2B2;refGene_tx=XM_023250018.1;refGene_gn=ATP2B2;refGene_tx=XM_023250019.1
A2 51542844 51543059
refGene_gn=ATP2B2;refGene_tx=XM_023250008.1;refGene_gn=ATP2B2;refGene_tx=XM_023250009.1;refGene_gn=ATP2B2;refGene_tx=XM_023250010.1;refGene_

gn=ATP2B2;refGene_tx=XM_023250011.1;refGene_gn=ATP2B2;refGene_tx=XM_023250012.1;refGene_gn=ATP2B2;refGene_tx=XM_023250013.1;refGene_gn=ATP2B2;refGene_tx=XM_023250014.1;refGene_gn=ATP2B2;refGene_tx=XM_023250015.1;refGene_gn=ATP2B2;refGene_tx=XM_023250016.1;refGene_gn=ATP2B2;refGene_tx=XM_023250017.1;refGene_gn=ATP2B2;refGene_tx=XM_023250018.1;refGene_gn=ATP2B2;refGene_tx=XM_023250019.1
A2 51545854 51546013
refGene_gn=ATP2B2;refGene_tx=XM_023250008.1;refGene_gn=ATP2B2;refGene_tx=XM_023250009.1;refGene_gn=ATP2B2;refGene_tx=XM_023250010.1;refGene_gn=ATP2B2;refGene_tx=XM_023250011.1;refGene_gn=ATP2B2;refGene_tx=XM_023250012.1;refGene_gn=ATP2B2;refGene_tx=XM_023250013.1;refGene_gn=ATP2B2;refGene_tx=XM_023250014.1;refGene_gn=ATP2B2;refGene_tx=XM_023250015.1;refGene_gn=ATP2B2;refGene_tx=XM_023250016.1;refGene_gn=ATP2B2;refGene_tx=XM_023250017.1;refGene_gn=ATP2B2;refGene_tx=XM_023250018.1;refGene_gn=ATP2B2;refGene_tx=XM_023250019.1
A2 51546846 51546888
refGene_gn=ATP2B2;refGene_tx=XM_023250008.1;refGene_gn=ATP2B2;refGene_tx=XM_023250009.1;refGene_gn=ATP2B2;refGene_tx=XM_023250010.1;refGene_gn=ATP2B2;refGene_tx=XM_023250011.1;refGene_gn=ATP2B2;refGene_tx=XM_023250012.1;refGene_gn=ATP2B2;refGene_tx=XM_023250013.1;refGene_gn=ATP2B2;refGene_tx=XM_023250014.1;refGene_gn=ATP2B2;refGene_tx=XM_023250015.1;refGene_gn=ATP2B2;refGene_tx=XM_023250016.1;refGene_gn=ATP2B2;refGene_tx=XM_023250018.1
A2 51552033 51552093
refGene_gn=ATP2B2;refGene_tx=XM_023250008.1;refGene_gn=ATP2B2;refGene_tx=XM_023250009.1;refGene_gn=ATP2B2;refGene_tx=XM_023250010.1;refGene_gn=ATP2B2;refGene_tx=XM_023250011.1;refGene_gn=ATP2B2;refGene_tx=XM_023250012.1;refGene_gn=ATP2B2;refGene_tx=XM_023250015.1;refGene_gn=ATP2B2;refGene_tx=XM_023250016.1
A2 51553227 51553260
refGene_gn=ATP2B2;refGene_tx=XM_023250008.1;refGene_gn=ATP2B2;refGene_tx=XM_023250009.1;refGene_gn=ATP2B2;refGene_tx=XM_023250010.1;refGene_gn=ATP2B2;refGene_tx=XM_023250011.1;refGene_gn=ATP2B2;refGene_tx=XM_023250013.1;refGene_gn=ATP2B2;refGene_tx=XM_023250015.1;refGene_gn=ATP2B2;refGene_tx=XM_023250016.1
A2 51554955 51555081
refGene_gn=ATP2B2;refGene_tx=XM_023250008.1;refGene_gn=ATP2B2;refGene_tx=XM_023250009.1;refGene_gn=ATP2B2;refGene_tx=XM_023250010.1;refGene_gn=ATP2B2;refGene_tx=XM_023250011.1;refGene_gn=ATP2B2;refGene_tx=XM_023250012.1;refGene_gn=ATP2B2;refGene_tx=XM_023250013.1;refGene_gn=ATP2B2;refGene_tx=XM_023250014.1;refGene_gn=ATP2B2;refGene_tx=XM_023250015.1;refGene_gn=ATP2B2;refGene_tx=XM_023250016.1;refGene_gn=ATP2B2;refGene_tx=XM_023250017.1;refGene_gn=ATP2B2;refGene_tx=XM_023250018.1;refGene_gn=ATP2B2;refGene_tx=XM_023250019.1
A2 51568101 51568227
refGene_gn=ATP2B2;refGene_tx=XM_023250008.1;refGene_gn=ATP2B2;refGene_tx=XM_023250009.1;refGene_gn=ATP2B2;refGene_tx=XM_023250010.1;refGene_gn=ATP2B2;refGene_tx=XM_023250011.1;refGene_gn=ATP2B2;refGene_tx=XM_023250012.1;refGene_gn=ATP2B2;refGene_tx=XM_023250013.1;refGene_gn=ATP2B2;refGene_tx=XM_023250014.1;refGene_gn=ATP2B2;refGene_tx=XM_023250015.

1;refGene_gn=ATP2B2;refGene_tx=XM_023250016.1;refGene_gn=ATP2B2;refGene_tx=XM_023250017.1;refGene_gn=ATP2B2;refGene_tx=XM_023250018.1;refGene_gn=ATP2B2;refGene_tx=XM_023250019.1
A2 51569229 51569487
refGene_gn=ATP2B2;refGene_tx=XM_023250008.1;refGene_gn=ATP2B2;refGene_tx=XM_023250009.1;refGene_gn=ATP2B2;refGene_tx=XM_023250010.1;refGene_gn=ATP2B2;refGene_tx=XM_023250011.1;refGene_gn=ATP2B2;refGene_tx=XM_023250012.1;refGene_gn=ATP2B2;refGene_tx=XM_023250013.1;refGene_gn=ATP2B2;refGene_tx=XM_023250014.1;refGene_gn=ATP2B2;refGene_tx=XM_023250015.1;refGene_gn=ATP2B2;refGene_tx=XM_023250016.1;refGene_gn=ATP2B2;refGene_tx=XM_023250017.1;refGene_gn=ATP2B2;refGene_tx=XM_023250018.1;refGene_gn=ATP2B2;refGene_tx=XM_023250019.1
A2 51575666 51575864
refGene_gn=ATP2B2;refGene_tx=XM_023250008.1;refGene_gn=ATP2B2;refGene_tx=XM_023250009.1;refGene_gn=ATP2B2;refGene_tx=XM_023250010.1;refGene_gn=ATP2B2;refGene_tx=XM_023250011.1;refGene_gn=ATP2B2;refGene_tx=XM_023250012.1;refGene_gn=ATP2B2;refGene_tx=XM_023250013.1;refGene_gn=ATP2B2;refGene_tx=XM_023250014.1;refGene_gn=ATP2B2;refGene_tx=XM_023250015.1;refGene_gn=ATP2B2;refGene_tx=XM_023250016.1;refGene_gn=ATP2B2;refGene_tx=XM_023250017.1;refGene_gn=ATP2B2;refGene_tx=XM_023250018.1;refGene_gn=ATP2B2;refGene_tx=XM_023250019.1
A2 51614623 51614822
refGene_gn=ATP2B2;refGene_tx=XM_023250008.1;refGene_gn=ATP2B2;refGene_tx=XM_023250009.1;refGene_gn=ATP2B2;refGene_tx=XM_023250010.1;refGene_gn=ATP2B2;refGene_tx=XM_023250011.1;refGene_gn=ATP2B2;refGene_tx=XM_023250012.1;refGene_gn=ATP2B2;refGene_tx=XM_023250013.1;refGene_gn=ATP2B2;refGene_tx=XM_023250014.1;refGene_gn=ATP2B2;refGene_tx=XM_023250015.1;refGene_gn=ATP2B2;refGene_tx=XM_023250016.1;refGene_gn=ATP2B2;refGene_tx=XM_023250017.1;refGene_gn=ATP2B2;refGene_tx=XM_023250018.1;refGene_gn=ATP2B2;refGene_tx=XM_023250019.1
A2 51987744 51988003
refGene_gn=SLC6A11;refGene_tx=XM_003982487.5;refGene_gn=SLC6A11;refGene_tx=XM_023250020.1
A2 51990970 51991105
refGene_gn=SLC6A11;refGene_tx=XM_003982487.5;refGene_gn=SLC6A11;refGene_tx=XM_023250020.1
A2 51991221 51991362
refGene_gn=SLC6A11;refGene_tx=XM_003982487.5;refGene_gn=SLC6A11;refGene_tx=XM_023250020.1
A2 51994649 51994740
refGene_gn=SLC6A11;refGene_tx=XM_003982487.5;refGene_gn=SLC6A11;refGene_tx=XM_023250020.1
A2 52016164 52016297
refGene_gn=SLC6A11;refGene_tx=XM_003982487.5;refGene_gn=SLC6A11;refGene_tx=XM_023250020.1
A2 52050579 52050714 refGene_gn=SLC6A11;refGene_tx=XM_003982487.5
A2 52087807 52087911
refGene_gn=SLC6A11;refGene_tx=XM_003982487.5;refGene_gn=SLC6A11;refGene_tx=XM_023250020.1
A2 52093112 52093237

refGene_gn=SLC6A11;refGene_tx=XM_003982487.5;refGene_gn=SLC6A11;refGene_tx=XM_023250020.1
A2 52106529 52106642
refGene_gn=SLC6A11;refGene_tx=XM_003982487.5;refGene_gn=SLC6A11;refGene_tx=XM_023250020.1
A2 52109699 52109837
refGene_gn=SLC6A11;refGene_tx=XM_003982487.5;refGene_gn=SLC6A11;refGene_tx=XM_023250020.1
A2 52113854 52113957
refGene_gn=SLC6A11;refGene_tx=XM_003982487.5;refGene_gn=SLC6A11;refGene_tx=XM_023250020.1
A2 52114818 52114919
refGene_gn=SLC6A11;refGene_tx=XM_003982487.5;refGene_gn=SLC6A11;refGene_tx=XM_023250020.1
A2 52115698 52115869
refGene_gn=SLC6A11;refGene_tx=XM_003982487.5;refGene_gn=SLC6A11;refGene_tx=XM_023250020.1
A2 52119831 52119984
refGene_gn=SLC6A11;refGene_tx=XM_003982487.5;refGene_gn=SLC6A11;refGene_tx=XM_023250020.1
A2 52194019 52194257
refGene_gn=SLC6A1;refGene_tx=XM_019824992.2;refGene_gn=SLC6A1;refGene_tx=XM_023250021.1
A2 52194663 52194795
refGene_gn=SLC6A1;refGene_tx=XM_019824992.2;refGene_gn=SLC6A1;refGene_tx=XM_023250021.1
A2 52195600 52195701
refGene_gn=SLC6A1;refGene_tx=XM_019824992.2;refGene_gn=SLC6A1;refGene_tx=XM_023250021.1
A2 52196582 52196692
refGene_gn=SLC6A1;refGene_tx=XM_019824992.2;refGene_gn=SLC6A1;refGene_tx=XM_023250021.1
A2 52198391 52198524
refGene_gn=SLC6A1;refGene_tx=XM_019824992.2;refGene_gn=SLC6A1;refGene_tx=XM_023250021.1
A2 52201340 52201475
refGene_gn=SLC6A1;refGene_tx=XM_019824992.2;refGene_gn=SLC6A1;refGene_tx=XM_023250021.1
A2 52201693 52201797
refGene_gn=SLC6A1;refGene_tx=XM_019824992.2;refGene_gn=SLC6A1;refGene_tx=XM_023250021.1
A2 52202118 52202243
refGene_gn=SLC6A1;refGene_tx=XM_019824992.2;refGene_gn=SLC6A1;refGene_tx=XM_023250021.1
A2 52203613 52203726
refGene_gn=SLC6A1;refGene_tx=XM_019824992.2;refGene_gn=SLC6A1;refGene_tx=XM_023250021.1
A2 52204141 52204273
refGene_gn=SLC6A1;refGene_tx=XM_019824992.2;refGene_gn=SLC6A1;refGene_tx=XM_023250021.1

A2	52205772	52205875
refGene_gn=SLC6A1;refGene_tx=XM_019824992.2;refGene_gn=SLC6A1;refGene_tx=XM_023250021.1
A2	52207468	52207569
refGene_gn=SLC6A1;refGene_tx=XM_019824992.2;refGene_gn=SLC6A1;refGene_tx=XM_023250021.1
A2	52208406	52208574
refGene_gn=SLC6A1;refGene_tx=XM_019824992.2;refGene_gn=SLC6A1;refGene_tx=XM_023250021.1
A2	52210470	52210575
refGene_gn=SLC6A1;refGene_tx=XM_019824992.2;refGene_gn=SLC6A1;refGene_tx=XM_023250021.1
A2	52366354	52366400	refGene_gn=HRH1;refGene_tx=XM_023250022.1
A2	52387715	52389214
refGene_gn=HRH1;refGene_tx=XM_003982473.4;refGene_gn=HRH1;refGene_tx=XM_023250022.1;refGene_gn=HRH1;refGene_tx=XM_023250023.1
A2	52414276	52414436
refGene_gn=ATG7;refGene_tx=XM_011280206.3;refGene_gn=ATG7;refGene_tx=XM_011280207.2;refGene_gn=ATG7;refGene_tx=XM_019824993.2
A2	52415181	52415236
refGene_gn=ATG7;refGene_tx=XM_011280206.3;refGene_gn=ATG7;refGene_tx=XM_011280207.2;refGene_gn=ATG7;refGene_tx=XM_019824993.2
A2	52423085	52423203
refGene_gn=ATG7;refGene_tx=XM_011280206.3;refGene_gn=ATG7;refGene_tx=XM_011280207.2;refGene_gn=ATG7;refGene_tx=XM_019824993.2
A2	52425078	52425156
refGene_gn=ATG7;refGene_tx=XM_011280206.3;refGene_gn=ATG7;refGene_tx=XM_011280207.2;refGene_gn=ATG7;refGene_tx=XM_019824993.2
A2	52428776	52428893
refGene_gn=ATG7;refGene_tx=XM_011280206.3;refGene_gn=ATG7;refGene_tx=XM_011280207.2;refGene_gn=ATG7;refGene_tx=XM_019824993.2
A2	52428913	52428919	refGene_gn=ATG7;refGene_tx=XM_023250024.1
A2	52430275	52430425
refGene_gn=ATG7;refGene_tx=XM_011280206.3;refGene_gn=ATG7;refGene_tx=XM_011280207.2;refGene_gn=ATG7;refGene_tx=XM_019824993.2;refGene_gn=ATG7;refGene_tx=XM_023250024.1
A2	52440049	52440138
refGene_gn=ATG7;refGene_tx=XM_011280206.3;refGene_gn=ATG7;refGene_tx=XM_011280207.2;refGene_gn=ATG7;refGene_tx=XM_019824993.2;refGene_gn=ATG7;refGene_tx=XM_023250024.1
A2	52441758	52441880
refGene_gn=ATG7;refGene_tx=XM_011280206.3;refGene_gn=ATG7;refGene_tx=XM_011280207.2;refGene_gn=ATG7;refGene_tx=XM_019824993.2;refGene_gn=ATG7;refGene_tx=XM_023250024.1
A2	52445613	52445704
refGene_gn=ATG7;refGene_tx=XM_011280206.3;refGene_gn=ATG7;refGene_tx=XM_011280207.2;refGene_gn=ATG7;refGene_tx=XM_019824993.2;refGene_gn=ATG7;refGene_tx=XM_023250024.1
A2	52446411	52446556
refGene_gn=ATG7;refGene_tx=XM_011280206.3;refGene_gn=ATG7;refGene_tx=X

M_011280207.2;refGene_gn=ATG7;refGene_tx=XM_019824993.2;refGene_gn=ATG7;refGene_tx=XM_023250024.1
A2 52449827 52449986
refGene_gn=ATG7;refGene_tx=XM_011280206.3;refGene_gn=ATG7;refGene_tx=XM_011280207.2;refGene_gn=ATG7;refGene_tx=XM_019824993.2;refGene_gn=ATG7;refGene_tx=XM_023250024.1
A2 52456594 52456789
refGene_gn=ATG7;refGene_tx=XM_011280206.3;refGene_gn=ATG7;refGene_tx=XM_011280207.2;refGene_gn=ATG7;refGene_tx=XM_019824993.2;refGene_gn=ATG7;refGene_tx=XM_023250024.1
A2 52464118 52464322
refGene_gn=ATG7;refGene_tx=XM_011280206.3;refGene_gn=ATG7;refGene_tx=XM_011280207.2;refGene_gn=ATG7;refGene_tx=XM_019824993.2;refGene_gn=ATG7;refGene_tx=XM_023250024.1
A2 52465994 52466110
refGene_gn=ATG7;refGene_tx=XM_011280206.3;refGene_gn=ATG7;refGene_tx=XM_011280207.2;refGene_gn=ATG7;refGene_tx=XM_019824993.2;refGene_gn=ATG7;refGene_tx=XM_023250024.1
A2 52470925 52471001
refGene_gn=ATG7;refGene_tx=XM_011280206.3;refGene_gn=ATG7;refGene_tx=XM_011280207.2;refGene_gn=ATG7;refGene_tx=XM_019824993.2;refGene_gn=ATG7;refGene_tx=XM_023250024.1
A2 52484530 52484611
refGene_gn=ATG7;refGene_tx=XM_011280206.3;refGene_gn=ATG7;refGene_tx=XM_011280207.2;refGene_gn=ATG7;refGene_tx=XM_019824993.2;refGene_gn=ATG7;refGene_tx=XM_023250024.1
A2 52522327 52522450
refGene_gn=ATG7;refGene_tx=XM_011280206.3;refGene_gn=ATG7;refGene_tx=XM_011280207.2;refGene_gn=ATG7;refGene_tx=XM_019824993.2;refGene_gn=ATG7;refGene_tx=XM_023250024.1
A2 52650335 52650368
refGene_gn=ATG7;refGene_tx=XM_011280206.3;refGene_gn=ATG7;refGene_tx=XM_011280207.2;refGene_gn=ATG7;refGene_tx=XM_019824993.2;refGene_gn=ATG7;refGene_tx=XM_023250024.1
A2 52653894 52654166
refGene_gn=VGLL4;refGene_tx=XM_003982489.5;refGene_gn=VGLL4;refGene_tx=XM_003982490.5;refGene_gn=VGLL4;refGene_tx=XM_019824994.2;refGene_gn=VGLL4;refGene_tx=XM_019824995.2;refGene_gn=VGLL4;refGene_tx=XM_019824996.2;refGene_gn=VGLL4;refGene_tx=XM_023250025.1
A2 52654584 52654708
refGene_gn=VGLL4;refGene_tx=XM_003982489.5;refGene_gn=VGLL4;refGene_tx=XM_003982490.5;refGene_gn=VGLL4;refGene_tx=XM_019824994.2;refGene_gn=VGLL4;refGene_tx=XM_019824995.2;refGene_gn=VGLL4;refGene_tx=XM_019824996.2;refGene_gn=VGLL4;refGene_tx=XM_023250025.1
A2 52659714 52659937
refGene_gn=VGLL4;refGene_tx=XM_003982489.5;refGene_gn=VGLL4;refGene_tx=XM_003982490.5;refGene_gn=VGLL4;refGene_tx=XM_019824994.2;refGene_gn=VGLL4;refGene_tx=XM_019824995.2;refGene_gn=VGLL4;refGene_tx=XM_019824996.2;refGene_gn=VGLL4;refGene_tx=XM_023250025.1
A2 52663579 52663659 refGene_gn=VGLL4;refGene_tx=XM_019824994.2

A2 52666172 52666174 refGene_gn=VGLL4;refGene_tx=XM_019824996.2
A2 52676437 52676451 refGene_gn=VGLL4;refGene_tx=XM_019824995.2
A2 52693048 52693238
refGene_gn=VGLL4;refGene_tx=XM_003982489.5;refGene_gn=VGLL4;refGene_tx=XM_003982490.5;refGene_gn=VGLL4;refGene_tx=XM_023250025.1
A2 52732243 52732325 refGene_gn=VGLL4;refGene_tx=XM_003982490.5
A2 52803624 52803688
refGene_gn=VGLL4;refGene_tx=XM_003982489.5;refGene_gn=VGLL4;refGene_tx=XM_023250025.1
A2 52869067 52869144
refGene_gn=TAMM41;refGene_tx=XM_003982492.5;refGene_gn=TAMM41;refGene_tx=XM_011280210.3;refGene_gn=TAMM41;refGene_tx=XM_011280212.3;refGene_gn=TAMM41;refGene_tx=XM_019825040.2;refGene_gn=TAMM41;refGene_tx=XM_019825041.1;refGene_gn=TAMM41;refGene_tx=XM_019825042.1
A2 52880164 52880227
refGene_gn=TAMM41;refGene_tx=XM_003982492.5;refGene_gn=TAMM41;refGene_tx=XM_011280210.3;refGene_gn=TAMM41;refGene_tx=XM_011280212.3;refGene_gn=TAMM41;refGene_tx=XM_019825040.2;refGene_gn=TAMM41;refGene_tx=XM_019825041.1;refGene_gn=TAMM41;refGene_tx=XM_019825042.1
A2 52881484 52881650
refGene_gn=TAMM41;refGene_tx=XM_003982492.5;refGene_gn=TAMM41;refGene_tx=XM_011280210.3;refGene_gn=TAMM41;refGene_tx=XM_011280212.3;refGene_gn=TAMM41;refGene_tx=XM_019825040.2;refGene_gn=TAMM41;refGene_tx=XM_019825041.1;refGene_gn=TAMM41;refGene_tx=XM_019825042.1
A2 52891207 52891353
refGene_gn=TAMM41;refGene_tx=XM_003982492.5;refGene_gn=TAMM41;refGene_tx=XM_011280210.3;refGene_gn=TAMM41;refGene_tx=XM_019825040.2;refGene_gn=TAMM41;refGene_tx=XM_019825041.1;refGene_gn=TAMM41;refGene_tx=XM_019825042.1
A2 52905048 52905199
refGene_gn=TAMM41;refGene_tx=XM_003982492.5;refGene_gn=TAMM41;refGene_tx=XM_011280210.3;refGene_gn=TAMM41;refGene_tx=XM_011280212.3;refGene_gn=TAMM41;refGene_tx=XM_019825040.2;refGene_gn=TAMM41;refGene_tx=XM_019825041.1;refGene_gn=TAMM41;refGene_tx=XM_019825042.1
A2 52905585 52905621
refGene_gn=TAMM41;refGene_tx=XM_019825040.2;refGene_gn=TAMM41;refGene_tx=XM_019825041.1
A2 52912627 52912720
refGene_gn=TAMM41;refGene_tx=XM_003982492.5;refGene_gn=TAMM41;refGene_tx=XM_011280210.3;refGene_gn=TAMM41;refGene_tx=XM_011280212.3
A2 52915770 52915953
refGene_gn=TAMM41;refGene_tx=XM_003982492.5;refGene_gn=TAMM41;refGene_tx=XM_011280210.3;refGene_gn=TAMM41;refGene_tx=XM_011280212.3
A2 52918448 52918637
refGene_gn=TAMM41;refGene_tx=XM_003982492.5;refGene_gn=TAMM41;refGene_tx=XM_011280210.3;refGene_gn=TAMM41;refGene_tx=XM_011280212.3
A2 53002454 53002846 refGene_gn=SYN2;refGene_tx=XM_023250026.1
A2 53148006 53148064 refGene_gn=SYN2;refGene_tx=XM_023250026.1
A2 53149379 53149471 refGene_gn=SYN2;refGene_tx=XM_023250026.1
A2 53151668 53151825 refGene_gn=SYN2;refGene_tx=XM_023250026.1

A2	53156106	53156196	refGene_gn=SYN2;refGene_tx=XM_023250026.1
A2	53158431	53158629	refGene_gn=TIMP4;refGene_tx=XM_003982494.5
A2	53159505	53159630	refGene_gn=TIMP4;refGene_tx=XM_003982494.5
A2	53162249	53162364	refGene_gn=TIMP4;refGene_tx=XM_003982494.5
A2	53162759	53162857	refGene_gn=TIMP4;refGene_tx=XM_003982494.5
A2	53164041	53164180	refGene_gn=TIMP4;refGene_tx=XM_003982494.5
A2	53167226	53167289	refGene_gn=SYN2;refGene_tx=XM_023250026.1
A2	53167673	53167816	refGene_gn=SYN2;refGene_tx=XM_023250026.1
A2	53174855	53174930	refGene_gn=SYN2;refGene_tx=XM_023250026.1
A2	53175394	53175497	refGene_gn=SYN2;refGene_tx=XM_023250026.1
A2	53176809	53176959	refGene_gn=SYN2;refGene_tx=XM_023250026.1
A2	53191620	53191681	refGene_gn=SYN2;refGene_tx=XM_023250026.1
A2	53195757	53195977	refGene_gn=SYN2;refGene_tx=XM_023250026.1
A2	53198487	53198623	refGene_gn=SYN2;refGene_tx=XM_023250026.1
A2	53353069	53353151	refGene_gn=PPARG;refGene_tx=XM_003982496.5
A2	53376906	53377134
refGene_gn=PPARG;refGene_tx=XM_003982496.5;refGene_gn=PPARG;refGene_tx=XM_006928991.3;refGene_gn=PPARG;refGene_tx=XM_011280215.3;refGene_gn=PPARG;refGene_tx=XM_011280216.2;refGene_gn=PPARG;refGene_tx=XM_019825091.2
A2	53378709	53378879
refGene_gn=PPARG;refGene_tx=XM_003982496.5;refGene_gn=PPARG;refGene_tx=XM_006928991.3;refGene_gn=PPARG;refGene_tx=XM_011280215.3;refGene_gn=PPARG;refGene_tx=XM_011280216.2;refGene_gn=PPARG;refGene_tx=XM_019825091.2
A2	53387935	53388074
refGene_gn=PPARG;refGene_tx=XM_003982496.5;refGene_gn=PPARG;refGene_tx=XM_006928991.3;refGene_gn=PPARG;refGene_tx=XM_011280215.3;refGene_gn=PPARG;refGene_tx=XM_011280216.2;refGene_gn=PPARG;refGene_tx=XM_019825091.2
A2	53400359	53400559
refGene_gn=PPARG;refGene_tx=XM_003982496.5;refGene_gn=PPARG;refGene_tx=XM_006928991.3;refGene_gn=PPARG;refGene_tx=XM_011280215.3;refGene_gn=PPARG;refGene_tx=XM_011280216.2;refGene_gn=PPARG;refGene_tx=XM_019825091.2
A2	53409729	53410180
refGene_gn=PPARG;refGene_tx=XM_003982496.5;refGene_gn=PPARG;refGene_tx=XM_006928991.3;refGene_gn=PPARG;refGene_tx=XM_011280215.3;refGene_gn=PPARG;refGene_tx=XM_011280216.2;refGene_gn=PPARG;refGene_tx=XM_019825091.2
A2	53427202	53427450
refGene_gn=PPARG;refGene_tx=XM_003982496.5;refGene_gn=PPARG;refGene_tx=XM_006928991.3;refGene_gn=PPARG;refGene_tx=XM_011280215.3;refGene_gn=PPARG;refGene_tx=XM_011280216.2;refGene_gn=PPARG;refGene_tx=XM_019825091.2
A2	53478561	53478750
refGene_gn=TSEN2;refGene_tx=XM_003982498.5;refGene_gn=TSEN2;refGene_tx=XM_011280217.2;refGene_gn=TSEN2;refGene_tx=XM_019825092.1
A2	53479895	53479977
refGene_gn=TSEN2;refGene_tx=XM_003982498.5;refGene_gn=TSEN2;refGene_tx

=XM_011280217.2;refGene_gn=TSEN2;refGene_tx=XM_019825092.1
A2 53487164 53487201
refGene_gn=TSEN2;refGene_tx=XM_003982498.5;refGene_gn=TSEN2;refGene_tx
=XM_011280217.2;refGene_gn=TSEN2;refGene_tx=XM_019825092.1;refGene_gn=
TSEN2;refGene_tx=XM_019825094.2
A2 53494615 53495147
refGene_gn=TSEN2;refGene_tx=XM_003982498.5;refGene_gn=TSEN2;refGene_tx
=XM_011280217.2;refGene_gn=TSEN2;refGene_tx=XM_019825092.1;refGene_gn=
TSEN2;refGene_tx=XM_019825094.2
A2 53496318 53496396
refGene_gn=TSEN2;refGene_tx=XM_003982498.5;refGene_gn=TSEN2;refGene_tx
=XM_011280217.2;refGene_gn=TSEN2;refGene_tx=XM_019825092.1;refGene_gn=
TSEN2;refGene_tx=XM_019825094.2
A2 53505674 53505725
refGene_gn=TSEN2;refGene_tx=XM_003982498.5;refGene_gn=TSEN2;refGene_tx
=XM_011280217.2;refGene_gn=TSEN2;refGene_tx=XM_019825092.1;refGene_gn=
TSEN2;refGene_tx=XM_019825094.2
A2 53509573 53509712
refGene_gn=TSEN2;refGene_tx=XM_003982498.5;refGene_gn=TSEN2;refGene_tx
=XM_011280217.2;refGene_gn=TSEN2;refGene_tx=XM_019825092.1;refGene_gn=
TSEN2;refGene_tx=XM_019825094.2
A2 53517646 53517683
refGene_gn=TSEN2;refGene_tx=XM_003982498.5;refGene_gn=TSEN2;refGene_tx
=XM_011280217.2;refGene_gn=TSEN2;refGene_tx=XM_019825092.1;refGene_gn=
TSEN2;refGene_tx=XM_019825094.2
A2 53518411 53518523
refGene_gn=TSEN2;refGene_tx=XM_003982498.5;refGene_gn=TSEN2;refGene_tx
=XM_011280217.2;refGene_gn=TSEN2;refGene_tx=XM_019825092.1;refGene_gn=
TSEN2;refGene_tx=XM_019825094.2
A2 53519545 53519635
refGene_gn=TSEN2;refGene_tx=XM_003982498.5;refGene_gn=TSEN2;refGene_tx
=XM_011280217.2;refGene_gn=TSEN2;refGene_tx=XM_019825092.1;refGene_gn=
TSEN2;refGene_tx=XM_019825094.2
A2 53520571 53520631
refGene_gn=TSEN2;refGene_tx=XM_003982498.5;refGene_gn=TSEN2;refGene_tx
=XM_011280217.2;refGene_gn=TSEN2;refGene_tx=XM_019825092.1;refGene_gn=
TSEN2;refGene_tx=XM_019825094.2
A2 53530222 53530463 refGene_gn=MKRN2OS;refGene_tx=XM_003982499.3
A2 53531734 53531897 refGene_gn=MKRN2OS;refGene_tx=XM_003982499.3
A2 53533128 53533178 refGene_gn=MKRN2OS;refGene_tx=XM_003982499.3
A2 53536588 53536806 refGene_gn=MKRN2OS;refGene_tx=XM_003982499.3
A2 53568389 53568415 refGene_gn=MKRN2;refGene_tx=XM_003982501.5
A2 53577875 53577898 refGene_gn=MKRN2;refGene_tx=XM_019825095.1
A2 53581830 53581959
refGene_gn=MKRN2;refGene_tx=XM_003982501.5;refGene_gn=MKRN2;refGene_tx
=XM_019825095.1;refGene_gn=MKRN2;refGene_tx=XM_019825097.2
A2 53583287 53583469
refGene_gn=MKRN2;refGene_tx=XM_003982501.5;refGene_gn=MKRN2;refGene_tx
=XM_019825095.1;refGene_gn=MKRN2;refGene_tx=XM_019825097.2
A2 53584617 53584922

refGene_gn=MKRN2;refGene_tx=XM_003982501.5;refGene_gn=MKRN2;refGene_tx=XM_019825095.1;refGene_gn=MKRN2;refGene_tx=XM_019825097.2
A2 53586203 53586418
refGene_gn=MKRN2;refGene_tx=XM_003982501.5;refGene_gn=MKRN2;refGene_tx=XM_019825095.1;refGene_gn=MKRN2;refGene_tx=XM_019825097.2
A2 53588470 53588581
refGene_gn=MKRN2;refGene_tx=XM_003982501.5;refGene_gn=MKRN2;refGene_tx=XM_019825095.1;refGene_gn=MKRN2;refGene_tx=XM_019825097.2
A2 53594354 53594499
refGene_gn=MKRN2;refGene_tx=XM_003982501.5;refGene_gn=MKRN2;refGene_tx=XM_019825095.1;refGene_gn=MKRN2;refGene_tx=XM_019825097.2
A2 53594671 53594809
refGene_gn=MKRN2;refGene_tx=XM_003982501.5;refGene_gn=MKRN2;refGene_tx=XM_019825095.1;refGene_gn=MKRN2;refGene_tx=XM_019825097.2
A2 53596946 53597090
refGene_gn=RAF1;refGene_tx=XM_003982500.4;refGene_gn=RAF1;refGene_tx=XM_006928993.3;refGene_gn=RAF1;refGene_tx=XM_023250029.1;refGene_gn=RAF1;refGene_tx=XM_023250030.1
A2 53597309 53597444
refGene_gn=RAF1;refGene_tx=XM_003982500.4;refGene_gn=RAF1;refGene_tx=XM_006928993.3;refGene_gn=RAF1;refGene_tx=XM_023250029.1;refGene_gn=RAF1;refGene_tx=XM_023250030.1
A2 53597587 53597719
refGene_gn=RAF1;refGene_tx=XM_003982500.4;refGene_gn=RAF1;refGene_tx=XM_006928993.3;refGene_gn=RAF1;refGene_tx=XM_023250029.1;refGene_gn=RAF1;refGene_tx=XM_023250030.1
A2 53598077 53598196
refGene_gn=RAF1;refGene_tx=XM_003982500.4;refGene_gn=RAF1;refGene_tx=XM_006928993.3;refGene_gn=RAF1;refGene_tx=XM_023250029.1;refGene_gn=RAF1;refGene_tx=XM_023250030.1
A2 53599648 53599695
refGene_gn=RAF1;refGene_tx=XM_003982500.4;refGene_gn=RAF1;refGene_tx=XM_006928993.3;refGene_gn=RAF1;refGene_tx=XM_023250029.1;refGene_gn=RAF1;refGene_tx=XM_023250030.1
A2 53600921 53601098
refGene_gn=RAF1;refGene_tx=XM_003982500.4;refGene_gn=RAF1;refGene_tx=XM_006928993.3;refGene_gn=RAF1;refGene_tx=XM_023250029.1;refGene_gn=RAF1;refGene_tx=XM_023250030.1
A2 53601846 53601931
refGene_gn=RAF1;refGene_tx=XM_003982500.4;refGene_gn=RAF1;refGene_tx=XM_006928993.3;refGene_gn=RAF1;refGene_tx=XM_023250029.1;refGene_gn=RAF1;refGene_tx=XM_023250030.1
A2 53610751 53610869
refGene_gn=RAF1;refGene_tx=XM_003982500.4;refGene_gn=RAF1;refGene_tx=XM_006928993.3;refGene_gn=RAF1;refGene_tx=XM_023250029.1;refGene_gn=RAF1;refGene_tx=XM_023250030.1
A2 53611248 53611376
refGene_gn=RAF1;refGene_tx=XM_003982500.4;refGene_gn=RAF1;refGene_tx=XM_006928993.3;refGene_gn=RAF1;refGene_tx=XM_023250029.1;refGene_gn=RAF1;refGene_tx=XM_023250030.1

A2 53611484 53611512
refGene_gn=RAF1;refGene_tx=XM_003982500.4;refGene_gn=RAF1;refGene_tx=XM_006928993.3;refGene_gn=RAF1;refGene_tx=XM_023250029.1;refGene_gn=RAF1;refGene_tx=XM_023250030.1
A2 53614182 53614242
refGene_gn=RAF1;refGene_tx=XM_006928993.3;refGene_gn=RAF1;refGene_tx=XM_023250029.1
A2 53614851 53615005
refGene_gn=RAF1;refGene_tx=XM_003982500.4;refGene_gn=RAF1;refGene_tx=XM_006928993.3;refGene_gn=RAF1;refGene_tx=XM_023250029.1;refGene_gn=RAF1;refGene_tx=XM_023250030.1
A2 53616599 53616698
refGene_gn=RAF1;refGene_tx=XM_003982500.4;refGene_gn=RAF1;refGene_tx=XM_006928993.3;refGene_gn=RAF1;refGene_tx=XM_023250029.1;refGene_gn=RAF1;refGene_tx=XM_023250030.1
A2 53618701 53618859
refGene_gn=RAF1;refGene_tx=XM_003982500.4;refGene_gn=RAF1;refGene_tx=XM_006928993.3;refGene_gn=RAF1;refGene_tx=XM_023250029.1;refGene_gn=RAF1;refGene_tx=XM_023250030.1
A2 53619176 53619279
refGene_gn=RAF1;refGene_tx=XM_003982500.4;refGene_gn=RAF1;refGene_tx=XM_006928993.3;refGene_gn=RAF1;refGene_tx=XM_023250029.1;refGene_gn=RAF1;refGene_tx=XM_023250030.1
A2 53622346 53622459
refGene_gn=RAF1;refGene_tx=XM_003982500.4;refGene_gn=RAF1;refGene_tx=XM_006928993.3
A2 53627133 53627340
refGene_gn=RAF1;refGene_tx=XM_003982500.4;refGene_gn=RAF1;refGene_tx=XM_006928993.3;refGene_gn=RAF1;refGene_tx=XM_023250029.1;refGene_gn=RAF1;refGene_tx=XM_023250030.1
A2 53731266 53731322
refGene_gn=TMEM40;refGene_tx=XM_019825099.2;refGene_gn=TMEM40;refGene_tx=XM_019825100.1;refGene_gn=TMEM40;refGene_tx=XM_019825101.1;refGene_gn=TMEM40;refGene_tx=XM_019825102.1;refGene_gn=TMEM40;refGene_tx=XM_019825103.2;refGene_gn=TMEM40;refGene_tx=XM_019825105.1;refGene_gn=TMEM40;refGene_tx=XM_019825106.1
A2 53732253 53732273 refGene_gn=TMEM40;refGene_tx=XM_006928995.3
A2 53733059 53733122
refGene_gn=TMEM40;refGene_tx=XM_006928995.3;refGene_gn=TMEM40;refGene_tx=XM_019825099.2;refGene_gn=TMEM40;refGene_tx=XM_019825100.1;refGene_gn=TMEM40;refGene_tx=XM_019825101.1;refGene_gn=TMEM40;refGene_tx=XM_019825102.1;refGene_gn=TMEM40;refGene_tx=XM_019825103.2;refGene_gn=TMEM40;refGene_tx=XM_019825105.1;refGene_gn=TMEM40;refGene_tx=XM_019825106.1
A2 53733620 53733695
refGene_gn=TMEM40;refGene_tx=XM_006928995.3;refGene_gn=TMEM40;refGene_tx=XM_019825099.2;refGene_gn=TMEM40;refGene_tx=XM_019825100.1;refGene_gn=TMEM40;refGene_tx=XM_019825101.1;refGene_gn=TMEM40;refGene_tx=XM_019825102.1;refGene_gn=TMEM40;refGene_tx=XM_019825103.2;refGene_gn=TMEM40;refGene_tx=XM_019825105.1;refGene_gn=TMEM40;refGene_tx=XM_019825106.

1
A2 53733783 53733855
refGene_gn=TMEM40;refGene_tx=XM_006928995.3;refGene_gn=TMEM40;refGene_tx=XM_019825099.2;refGene_gn=TMEM40;refGene_tx=XM_019825100.1;refGene_gn=TMEM40;refGene_tx=XM_019825101.1;refGene_gn=TMEM40;refGene_tx=XM_019825102.1;refGene_gn=TMEM40;refGene_tx=XM_019825103.2;refGene_gn=TMEM40;refGene_tx=XM_019825105.1;refGene_gn=TMEM40;refGene_tx=XM_019825106.1
A2 53737296 53737344
refGene_gn=TMEM40;refGene_tx=XM_006928995.3;refGene_gn=TMEM40;refGene_tx=XM_019825099.2;refGene_gn=TMEM40;refGene_tx=XM_019825100.1;refGene_gn=TMEM40;refGene_tx=XM_019825101.1;refGene_gn=TMEM40;refGene_tx=XM_019825102.1;refGene_gn=TMEM40;refGene_tx=XM_019825103.2;refGene_gn=TMEM40;refGene_tx=XM_019825105.1;refGene_gn=TMEM40;refGene_tx=XM_019825106.1
A2 53737732 53737765
refGene_gn=TMEM40;refGene_tx=XM_006928995.3;refGene_gn=TMEM40;refGene_tx=XM_019825099.2;refGene_gn=TMEM40;refGene_tx=XM_019825100.1;refGene_gn=TMEM40;refGene_tx=XM_019825101.1;refGene_gn=TMEM40;refGene_tx=XM_019825102.1;refGene_gn=TMEM40;refGene_tx=XM_019825103.2;refGene_gn=TMEM40;refGene_tx=XM_019825105.1;refGene_gn=TMEM40;refGene_tx=XM_019825106.1
A2 53738131 53738167
refGene_gn=TMEM40;refGene_tx=XM_006928995.3;refGene_gn=TMEM40;refGene_tx=XM_019825099.2;refGene_gn=TMEM40;refGene_tx=XM_019825100.1;refGene_gn=TMEM40;refGene_tx=XM_019825101.1;refGene_gn=TMEM40;refGene_tx=XM_019825102.1;refGene_gn=TMEM40;refGene_tx=XM_019825103.2;refGene_gn=TMEM40;refGene_tx=XM_019825105.1;refGene_gn=TMEM40;refGene_tx=XM_019825106.1
A2 53740595 53740649
refGene_gn=TMEM40;refGene_tx=XM_006928995.3;refGene_gn=TMEM40;refGene_tx=XM_019825099.2;refGene_gn=TMEM40;refGene_tx=XM_019825100.1;refGene_gn=TMEM40;refGene_tx=XM_019825101.1;refGene_gn=TMEM40;refGene_tx=XM_019825102.1;refGene_gn=TMEM40;refGene_tx=XM_019825103.2;refGene_gn=TMEM40;refGene_tx=XM_019825105.1;refGene_gn=TMEM40;refGene_tx=XM_019825106.1
A2 53742256 53742346
refGene_gn=TMEM40;refGene_tx=XM_006928995.3;refGene_gn=TMEM40;refGene_tx=XM_019825099.2;refGene_gn=TMEM40;refGene_tx=XM_019825100.1;refGene_gn=TMEM40;refGene_tx=XM_019825102.1;refGene_gn=TMEM40;refGene_tx=XM_019825105.1;refGene_gn=TMEM40;refGene_tx=XM_019825106.1
A2 53745637 53745781
refGene_gn=TMEM40;refGene_tx=XM_006928995.3;refGene_gn=TMEM40;refGene_tx=XM_019825099.2;refGene_gn=TMEM40;refGene_tx=XM_019825100.1;refGene_gn=TMEM40;refGene_tx=XM_019825101.1;refGene_gn=TMEM40;refGene_tx=XM_019825105.1;refGene_gn=TMEM40;refGene_tx=XM_019825106.1
A2 53746445 53746526
refGene_gn=TMEM40;refGene_tx=XM_006928995.3;refGene_gn=TMEM40;refGene_tx=XM_019825099.2;refGene_gn=TMEM40;refGene_tx=XM_019825100.1;refGene_gn=TMEM40;refGene_tx=XM_019825101.1;refGene_gn=TMEM40;refGene_tx=XM_01

9825102.1;refGene_gn=TMEM40;refGene_tx=XM_019825103.2;refGene_gn=TMEM4
0;refGene_tx=XM_019825105.1;refGene_gn=TMEM40;refGene_tx=XM_019825106.
1
A2 53796939 53797100
refGene_gn=TMEM40;refGene_tx=XM_006928995.3;refGene_gn=TMEM40;refGene_
tx=XM_019825099.2;refGene_gn=TMEM40;refGene_tx=XM_019825100.1;refGene_
gn=TMEM40;refGene_tx=XM_019825101.1;refGene_gn=TMEM40;refGene_tx=XM_01
9825102.1;refGene_gn=TMEM40;refGene_tx=XM_019825103.2
A2 53797238 53797480
refGene_gn=TMEM40;refGene_tx=XM_006928995.3;refGene_gn=TMEM40;refGene_
tx=XM_019825099.2;refGene_gn=TMEM40;refGene_tx=XM_019825100.1;refGene_
gn=TMEM40;refGene_tx=XM_019825101.1;refGene_gn=TMEM40;refGene_tx=XM_01
9825102.1;refGene_gn=TMEM40;refGene_tx=XM_019825103.2
A2 53797519 53797587
refGene_gn=CAND2;refGene_tx=XM_023250032.1;refGene_gn=CAND2;refGene_tx
=XM_023250033.1
A2 53802560 53802704
refGene_gn=CAND2;refGene_tx=XM_023250031.1;refGene_gn=CAND2;refGene_tx
=XM_023250032.1;refGene_gn=CAND2;refGene_tx=XM_023250033.1
A2 53805545 53805700
refGene_gn=CAND2;refGene_tx=XM_023250031.1;refGene_gn=CAND2;refGene_tx
=XM_023250032.1;refGene_gn=CAND2;refGene_tx=XM_023250033.1
A2 53806384 53806508
refGene_gn=CAND2;refGene_tx=XM_023250031.1;refGene_gn=CAND2;refGene_tx
=XM_023250032.1;refGene_gn=CAND2;refGene_tx=XM_023250033.1
A2 53809869 53810135
refGene_gn=CAND2;refGene_tx=XM_023250031.1;refGene_gn=CAND2;refGene_tx
=XM_023250032.1;refGene_gn=CAND2;refGene_tx=XM_023250033.1
A2 53811527 53811633
refGene_gn=CAND2;refGene_tx=XM_023250031.1;refGene_gn=CAND2;refGene_tx
=XM_023250032.1;refGene_gn=CAND2;refGene_tx=XM_023250033.1
A2 53811810 53811953
refGene_gn=CAND2;refGene_tx=XM_023250031.1;refGene_gn=CAND2;refGene_tx
=XM_023250032.1;refGene_gn=CAND2;refGene_tx=XM_023250033.1
A2 53814006 53814299
refGene_gn=CAND2;refGene_tx=XM_023250031.1;refGene_gn=CAND2;refGene_tx
=XM_023250032.1;refGene_gn=CAND2;refGene_tx=XM_023250033.1
A2 53815387 53815529
refGene_gn=CAND2;refGene_tx=XM_023250031.1;refGene_gn=CAND2;refGene_tx
=XM_023250032.1;refGene_gn=CAND2;refGene_tx=XM_023250033.1
A2 53815856 53817359
refGene_gn=CAND2;refGene_tx=XM_023250031.1;refGene_gn=CAND2;refGene_tx
=XM_023250032.1;refGene_gn=CAND2;refGene_tx=XM_023250033.1
A2 53819745 53819841
refGene_gn=CAND2;refGene_tx=XM_023250031.1;refGene_gn=CAND2;refGene_tx
=XM_023250032.1;refGene_gn=CAND2;refGene_tx=XM_023250033.1
A2 53824033 53824203
refGene_gn=CAND2;refGene_tx=XM_023250031.1;refGene_gn=CAND2;refGene_tx
=XM_023250032.1;refGene_gn=CAND2;refGene_tx=XM_023250033.1
A2 53825479 53825644

refGene_gn=CAND2;refGene_tx=XM_023250031.1;refGene_gn=CAND2;refGene_tx=XM_023250032.1;refGene_gn=CAND2;refGene_tx=XM_023250033.1
A2 53827369 53827477
refGene_gn=CAND2;refGene_tx=XM_023250031.1;refGene_gn=CAND2;refGene_tx=XM_023250032.1;refGene_gn=CAND2;refGene_tx=XM_023250033.1
A2 53829813 53830041
refGene_gn=CAND2;refGene_tx=XM_023250031.1;refGene_gn=CAND2;refGene_tx=XM_023250033.1
A2 53832719 53832849 refGene_gn=RPL32;refGene_tx=XM_003982504.4
A2 53834623 53834752 refGene_gn=CAND2;refGene_tx=XM_023250032.1
A2 53835032 53835214 refGene_gn=RPL32;refGene_tx=XM_003982504.4
A2 53835811 53835907 refGene_gn=RPL32;refGene_tx=XM_003982504.4
A2 53839106 53839365
refGene_gn=EFCAB12;refGene_tx=XM_023250036.1;refGene_gn=EFCAB12;refGene_tx=XM_023250037.1
A2 53840264 53840321
refGene_gn=EFCAB12;refGene_tx=XM_023250036.1;refGene_gn=EFCAB12;refGene_tx=XM_023250037.1
A2 53841955 53842115
refGene_gn=EFCAB12;refGene_tx=XM_023250036.1;refGene_gn=EFCAB12;refGene_tx=XM_023250037.1
A2 53846633 53846847
refGene_gn=EFCAB12;refGene_tx=XM_023250036.1;refGene_gn=EFCAB12;refGene_tx=XM_023250037.1
A2 53848406 53848600
refGene_gn=EFCAB12;refGene_tx=XM_023250036.1;refGene_gn=EFCAB12;refGene_tx=XM_023250037.1
A2 53851242 53851399
refGene_gn=EFCAB12;refGene_tx=XM_023250036.1;refGene_gn=EFCAB12;refGene_tx=XM_023250037.1
A2 53853412 53853595
refGene_gn=EFCAB12;refGene_tx=XM_023250036.1;refGene_gn=EFCAB12;refGene_tx=XM_023250037.1
A2 53856711 53857160
refGene_gn=EFCAB12;refGene_tx=XM_023250036.1;refGene_gn=EFCAB12;refGene_tx=XM_023250037.1
A2 53861323 53861405 refGene_gn=EFCAB12;refGene_tx=XM_023250036.1
A2 53862604 53862653 refGene_gn=EFCAB12;refGene_tx=XM_023250037.1
A2 53865445 53865523
refGene_gn=MBD4;refGene_tx=XM_019825112.2;refGene_gn=MBD4;refGene_tx=XM_023250043.1
A2 53866430 53866534
refGene_gn=MBD4;refGene_tx=XM_019825112.2;refGene_gn=MBD4;refGene_tx=XM_023250043.1
A2 53867034 53867184
refGene_gn=MBD4;refGene_tx=XM_019825112.2;refGene_gn=MBD4;refGene_tx=XM_023250043.1
A2 53868291 53868426
refGene_gn=MBD4;refGene_tx=XM_019825112.2;refGene_gn=MBD4;refGene_tx=XM_023250043.1

A2	53868503	53868578
refGene_gn=MBD4;refGene_tx=XM_019825112.2;refGene_gn=MBD4;refGene_tx=XM_023250043.1
A2	53871767	53872631
refGene_gn=MBD4;refGene_tx=XM_023250042.1;refGene_gn=MBD4;refGene_tx=XM_023250043.1
A2	53873234	53873465
refGene_gn=MBD4;refGene_tx=XM_019825112.2;refGene_gn=MBD4;refGene_tx=XM_023250042.1
A2	53874485	53874574
refGene_gn=MBD4;refGene_tx=XM_019825112.2;refGene_gn=MBD4;refGene_tx=XM_023250042.1
A2	53875061	53875102
refGene_gn=IFT122;refGene_tx=XM_003982509.4;refGene_gn=IFT122;refGene_tx=XM_003982510.4
A2	53878374	53878388
refGene_gn=IFT122;refGene_tx=XM_023250039.1;refGene_gn=IFT122;refGene_tx=XM_023250040.1
A2	53879677	53879836
refGene_gn=IFT122;refGene_tx=XM_019825107.2;refGene_gn=IFT122;refGene_tx=XM_023250038.1;refGene_gn=IFT122;refGene_tx=XM_023250039.1;refGene_gn=IFT122;refGene_tx=XM_023250040.1
A2	53882973	53883040
refGene_gn=IFT122;refGene_tx=XM_003982509.4;refGene_gn=IFT122;refGene_tx=XM_003982510.4;refGene_gn=IFT122;refGene_tx=XM_019825107.2;refGene_gn=IFT122;refGene_tx=XM_023250038.1;refGene_gn=IFT122;refGene_tx=XM_023250039.1;refGene_gn=IFT122;refGene_tx=XM_023250040.1
A2	53884391	53884476
refGene_gn=IFT122;refGene_tx=XM_003982509.4;refGene_gn=IFT122;refGene_tx=XM_003982510.4;refGene_gn=IFT122;refGene_tx=XM_019825107.2;refGene_gn=IFT122;refGene_tx=XM_023250038.1;refGene_gn=IFT122;refGene_tx=XM_023250039.1;refGene_gn=IFT122;refGene_tx=XM_023250040.1
A2	53889895	53889974
refGene_gn=IFT122;refGene_tx=XM_003982509.4;refGene_gn=IFT122;refGene_tx=XM_003982510.4;refGene_gn=IFT122;refGene_tx=XM_019825107.2;refGene_gn=IFT122;refGene_tx=XM_023250038.1;refGene_gn=IFT122;refGene_tx=XM_023250039.1;refGene_gn=IFT122;refGene_tx=XM_023250040.1
A2	53891678	53891755
refGene_gn=IFT122;refGene_tx=XM_003982509.4;refGene_gn=IFT122;refGene_tx=XM_003982510.4;refGene_gn=IFT122;refGene_tx=XM_019825107.2;refGene_gn=IFT122;refGene_tx=XM_023250038.1;refGene_gn=IFT122;refGene_tx=XM_023250039.1;refGene_gn=IFT122;refGene_tx=XM_023250040.1
A2	53894120	53894187
refGene_gn=IFT122;refGene_tx=XM_003982509.4;refGene_gn=IFT122;refGene_tx=XM_003982510.4;refGene_gn=IFT122;refGene_tx=XM_019825107.2;refGene_gn=IFT122;refGene_tx=XM_023250038.1;refGene_gn=IFT122;refGene_tx=XM_023250039.1;refGene_gn=IFT122;refGene_tx=XM_023250040.1
A2	53895132	53895279
refGene_gn=IFT122;refGene_tx=XM_003982509.4;refGene_gn=IFT122;refGene_tx=XM_003982510.4;refGene_gn=IFT122;refGene_tx=XM_019825107.2;refGene_

gn=IFT122;refGene_tx=XM_019825108.2;refGene_gn=IFT122;refGene_tx=XM_023250038.1;refGene_gn=IFT122;refGene_tx=XM_023250039.1;refGene_gn=IFT122;refGene_tx=XM_023250040.1
A2 53896343 53896520
refGene_gn=IFT122;refGene_tx=XM_003982509.4;refGene_gn=IFT122;refGene_tx=XM_019825107.2;refGene_gn=IFT122;refGene_tx=XM_019825108.2;refGene_gn=IFT122;refGene_tx=XM_023250038.1;refGene_gn=IFT122;refGene_tx=XM_023250039.1;refGene_gn=IFT122;refGene_tx=XM_023250040.1
A2 53898470 53898546
refGene_gn=IFT122;refGene_tx=XM_003982509.4;refGene_gn=IFT122;refGene_tx=XM_003982510.4;refGene_gn=IFT122;refGene_tx=XM_019825107.2;refGene_gn=IFT122;refGene_tx=XM_019825108.2;refGene_gn=IFT122;refGene_tx=XM_023250038.1;refGene_gn=IFT122;refGene_tx=XM_023250039.1;refGene_gn=IFT122;refGene_tx=XM_023250040.1
A2 53904786 53904978
refGene_gn=IFT122;refGene_tx=XM_003982509.4;refGene_gn=IFT122;refGene_tx=XM_003982510.4;refGene_gn=IFT122;refGene_tx=XM_019825107.2;refGene_gn=IFT122;refGene_tx=XM_019825108.2;refGene_gn=IFT122;refGene_tx=XM_023250038.1;refGene_gn=IFT122;refGene_tx=XM_023250039.1;refGene_gn=IFT122;refGene_tx=XM_023250040.1
A2 53905134 53905273
refGene_gn=IFT122;refGene_tx=XM_003982509.4;refGene_gn=IFT122;refGene_tx=XM_003982510.4;refGene_gn=IFT122;refGene_tx=XM_019825107.2;refGene_gn=IFT122;refGene_tx=XM_019825108.2;refGene_gn=IFT122;refGene_tx=XM_023250038.1;refGene_gn=IFT122;refGene_tx=XM_023250039.1;refGene_gn=IFT122;refGene_tx=XM_023250040.1
A2 53906455 53906658
refGene_gn=IFT122;refGene_tx=XM_003982509.4;refGene_gn=IFT122;refGene_tx=XM_003982510.4;refGene_gn=IFT122;refGene_tx=XM_019825107.2;refGene_gn=IFT122;refGene_tx=XM_019825108.2;refGene_gn=IFT122;refGene_tx=XM_023250038.1;refGene_gn=IFT122;refGene_tx=XM_023250039.1;refGene_gn=IFT122;refGene_tx=XM_023250040.1
A2 53907877 53908015
refGene_gn=IFT122;refGene_tx=XM_003982509.4;refGene_gn=IFT122;refGene_tx=XM_003982510.4;refGene_gn=IFT122;refGene_tx=XM_019825107.2;refGene_gn=IFT122;refGene_tx=XM_019825108.2;refGene_gn=IFT122;refGene_tx=XM_023250038.1;refGene_gn=IFT122;refGene_tx=XM_023250039.1;refGene_gn=IFT122;refGene_tx=XM_023250040.1
A2 53909594 53909759
refGene_gn=IFT122;refGene_tx=XM_003982509.4;refGene_gn=IFT122;refGene_tx=XM_003982510.4;refGene_gn=IFT122;refGene_tx=XM_019825107.2;refGene_gn=IFT122;refGene_tx=XM_019825108.2;refGene_gn=IFT122;refGene_tx=XM_023250038.1;refGene_gn=IFT122;refGene_tx=XM_023250039.1;refGene_gn=IFT122;refGene_tx=XM_023250040.1
A2 53912340 53912538
refGene_gn=IFT122;refGene_tx=XM_003982509.4;refGene_gn=IFT122;refGene_tx=XM_003982510.4;refGene_gn=IFT122;refGene_tx=XM_019825107.2;refGene_gn=IFT122;refGene_tx=XM_019825108.2;refGene_gn=IFT122;refGene_tx=XM_023250038.1;refGene_gn=IFT122;refGene_tx=XM_023250039.1;refGene_gn=IFT122;refGene_tx=XM_023250040.1

A2 53917366 53917507
refGene_gn=IFT122;refGene_tx=XM_003982509.4;refGene_gn=IFT122;refGene_tx=XM_003982510.4;refGene_gn=IFT122;refGene_tx=XM_019825107.2;refGene_gn=IFT122;refGene_tx=XM_019825108.2;refGene_gn=IFT122;refGene_tx=XM_023250038.1;refGene_gn=IFT122;refGene_tx=XM_023250039.1;refGene_gn=IFT122;refGene_tx=XM_023250040.1
A2 53921455 53921509
refGene_gn=IFT122;refGene_tx=XM_003982509.4;refGene_gn=IFT122;refGene_tx=XM_003982510.4;refGene_gn=IFT122;refGene_tx=XM_019825107.2;refGene_gn=IFT122;refGene_tx=XM_019825108.2;refGene_gn=IFT122;refGene_tx=XM_023250039.1;refGene_gn=IFT122;refGene_tx=XM_023250040.1
A2 53924885 53925047
refGene_gn=IFT122;refGene_tx=XM_003982509.4;refGene_gn=IFT122;refGene_tx=XM_003982510.4;refGene_gn=IFT122;refGene_tx=XM_019825107.2;refGene_gn=IFT122;refGene_tx=XM_019825108.2;refGene_gn=IFT122;refGene_tx=XM_023250038.1;refGene_gn=IFT122;refGene_tx=XM_023250039.1;refGene_gn=IFT122;refGene_tx=XM_023250040.1
A2 53929079 53929246
refGene_gn=IFT122;refGene_tx=XM_003982509.4;refGene_gn=IFT122;refGene_tx=XM_003982510.4;refGene_gn=IFT122;refGene_tx=XM_019825107.2;refGene_gn=IFT122;refGene_tx=XM_019825108.2;refGene_gn=IFT122;refGene_tx=XM_023250038.1;refGene_gn=IFT122;refGene_tx=XM_023250039.1;refGene_gn=IFT122;refGene_tx=XM_023250040.1
A2 53932014 53932186
refGene_gn=IFT122;refGene_tx=XM_003982509.4;refGene_gn=IFT122;refGene_tx=XM_003982510.4;refGene_gn=IFT122;refGene_tx=XM_019825107.2;refGene_gn=IFT122;refGene_tx=XM_019825108.2;refGene_gn=IFT122;refGene_tx=XM_023250038.1;refGene_gn=IFT122;refGene_tx=XM_023250039.1;refGene_gn=IFT122;refGene_tx=XM_023250040.1
A2 53934373 53934476
refGene_gn=IFT122;refGene_tx=XM_003982509.4;refGene_gn=IFT122;refGene_tx=XM_003982510.4;refGene_gn=IFT122;refGene_tx=XM_019825107.2;refGene_gn=IFT122;refGene_tx=XM_019825108.2;refGene_gn=IFT122;refGene_tx=XM_023250038.1;refGene_gn=IFT122;refGene_tx=XM_023250039.1;refGene_gn=IFT122;refGene_tx=XM_023250040.1
A2 53936380 53936521
refGene_gn=IFT122;refGene_tx=XM_003982509.4;refGene_gn=IFT122;refGene_tx=XM_003982510.4;refGene_gn=IFT122;refGene_tx=XM_019825107.2;refGene_gn=IFT122;refGene_tx=XM_019825108.2;refGene_gn=IFT122;refGene_tx=XM_023250038.1;refGene_gn=IFT122;refGene_tx=XM_023250039.1;refGene_gn=IFT122;refGene_tx=XM_023250040.1
A2 53937666 53937761
refGene_gn=IFT122;refGene_tx=XM_003982509.4;refGene_gn=IFT122;refGene_tx=XM_003982510.4;refGene_gn=IFT122;refGene_tx=XM_019825107.2;refGene_gn=IFT122;refGene_tx=XM_019825108.2;refGene_gn=IFT122;refGene_tx=XM_023250038.1;refGene_gn=IFT122;refGene_tx=XM_023250039.1;refGene_gn=IFT122;refGene_tx=XM_023250040.1
A2 53940981 53941082
refGene_gn=IFT122;refGene_tx=XM_003982509.4;refGene_gn=IFT122;refGene_tx=XM_003982510.4;refGene_gn=IFT122;refGene_tx=XM_019825107.2;refGene_

gn=IFT122;refGene_tx=XM_019825108.2;refGene_gn=IFT122;refGene_tx=XM_023250038.1;refGene_gn=IFT122;refGene_tx=XM_023250039.1;refGene_gn=IFT122;refGene_tx=XM_023250040.1
A2 53943121 53943287
refGene_gn=IFT122;refGene_tx=XM_003982509.4;refGene_gn=IFT122;refGene_tx=XM_003982510.4;refGene_gn=IFT122;refGene_tx=XM_019825107.2;refGene_gn=IFT122;refGene_tx=XM_019825108.2;refGene_gn=IFT122;refGene_tx=XM_023250038.1;refGene_gn=IFT122;refGene_tx=XM_023250039.1;refGene_gn=IFT122;refGene_tx=XM_023250040.1
A2 53944273 53944385
refGene_gn=IFT122;refGene_tx=XM_003982509.4;refGene_gn=IFT122;refGene_tx=XM_003982510.4;refGene_gn=IFT122;refGene_tx=XM_019825107.2;refGene_gn=IFT122;refGene_tx=XM_019825108.2;refGene_gn=IFT122;refGene_tx=XM_023250038.1;refGene_gn=IFT122;refGene_tx=XM_023250039.1;refGene_gn=IFT122;refGene_tx=XM_023250040.1
A2 53944902 53944967 refGene_gn=IFT122;refGene_tx=XM_023250039.1
A2 53945045 53945171
refGene_gn=IFT122;refGene_tx=XM_003982509.4;refGene_gn=IFT122;refGene_tx=XM_003982510.4;refGene_gn=IFT122;refGene_tx=XM_019825107.2;refGene_gn=IFT122;refGene_tx=XM_019825108.2;refGene_gn=IFT122;refGene_tx=XM_023250038.1;refGene_gn=IFT122;refGene_tx=XM_023250040.1
A2 53946843 53946923
refGene_gn=IFT122;refGene_tx=XM_003982509.4;refGene_gn=IFT122;refGene_tx=XM_003982510.4;refGene_gn=IFT122;refGene_tx=XM_019825107.2;refGene_gn=IFT122;refGene_tx=XM_019825108.2;refGene_gn=IFT122;refGene_tx=XM_023250038.1
A2 53947258 53947423
refGene_gn=IFT122;refGene_tx=XM_003982509.4;refGene_gn=IFT122;refGene_tx=XM_003982510.4;refGene_gn=IFT122;refGene_tx=XM_019825108.2;refGene_gn=IFT122;refGene_tx=XM_023250038.1
A2 53947821 53947911
refGene_gn=IFT122;refGene_tx=XM_003982509.4;refGene_gn=IFT122;refGene_tx=XM_003982510.4;refGene_gn=IFT122;refGene_tx=XM_019825107.2;refGene_gn=IFT122;refGene_tx=XM_019825108.2;refGene_gn=IFT122;refGene_tx=XM_023250038.1
A2 53955073 53955434 refGene_gn=RHO;refGene_tx=NM_001009242.1
A2 53957908 53958077 refGene_gn=RHO;refGene_tx=NM_001009242.1
A2 53959350 53959516 refGene_gn=RHO;refGene_tx=NM_001009242.1
A2 53959622 53959862 refGene_gn=RHO;refGene_tx=NM_001009242.1
A2 53960726 53960837 refGene_gn=RHO;refGene_tx=NM_001009242.1
A2 53968305 53968678
refGene_gn=LOC101098613;refGene_tx=XM_019825113.2
A2 53976099 53976359
refGene_gn=LOC101098613;refGene_tx=XM_019825113.2
A2 53978098 53978462
refGene_gn=LOC101098613;refGene_tx=XM_019825113.2
A2 53979564 53979626
refGene_gn=LOC101098613;refGene_tx=XM_019825113.2
A2 53979943 53980192
refGene_gn=LOC101098613;refGene_tx=XM_019825113.2

A2 53985464 53985581
refGene_gn=PLXND1;refGene_tx=XM_011280229.3;refGene_gn=PLXND1;refGene_tx=XM_023250044.1
A2 53985838 53985913
refGene_gn=PLXND1;refGene_tx=XM_011280229.3;refGene_gn=PLXND1;refGene_tx=XM_023250044.1
A2 53986073 53986214
refGene_gn=PLXND1;refGene_tx=XM_011280229.3;refGene_gn=PLXND1;refGene_tx=XM_023250044.1
A2 53987329 53987477
refGene_gn=PLXND1;refGene_tx=XM_011280229.3;refGene_gn=PLXND1;refGene_tx=XM_023250044.1
A2 53988644 53988808
refGene_gn=PLXND1;refGene_tx=XM_011280229.3;refGene_gn=PLXND1;refGene_tx=XM_023250044.1
A2 53989312 53989417
refGene_gn=PLXND1;refGene_tx=XM_011280229.3;refGene_gn=PLXND1;refGene_tx=XM_023250044.1
A2 53989688 53989723
refGene_gn=PLXND1;refGene_tx=XM_011280229.3;refGene_gn=PLXND1;refGene_tx=XM_023250044.1
A2 53990438 53990502
refGene_gn=PLXND1;refGene_tx=XM_011280229.3;refGene_gn=PLXND1;refGene_tx=XM_023250044.1
A2 53990585 53990689
refGene_gn=PLXND1;refGene_tx=XM_011280229.3;refGene_gn=PLXND1;refGene_tx=XM_023250044.1
A2 53991503 53991660
refGene_gn=PLXND1;refGene_tx=XM_011280229.3;refGene_gn=PLXND1;refGene_tx=XM_023250044.1
A2 53991859 53992006
refGene_gn=PLXND1;refGene_tx=XM_011280229.3;refGene_gn=PLXND1;refGene_tx=XM_023250044.1
A2 53993075 53993274
refGene_gn=PLXND1;refGene_tx=XM_011280229.3;refGene_gn=PLXND1;refGene_tx=XM_023250044.1
A2 53993656 53993787
refGene_gn=PLXND1;refGene_tx=XM_011280229.3;refGene_gn=PLXND1;refGene_tx=XM_023250044.1
A2 53994306 53994411
refGene_gn=PLXND1;refGene_tx=XM_011280229.3;refGene_gn=PLXND1;refGene_tx=XM_023250044.1
A2 53995266 53995379
refGene_gn=PLXND1;refGene_tx=XM_011280229.3;refGene_gn=PLXND1;refGene_tx=XM_023250044.1
A2 53995474 53995582
refGene_gn=PLXND1;refGene_tx=XM_011280229.3;refGene_gn=PLXND1;refGene_tx=XM_023250044.1
A2 54003929 54004044
refGene_gn=PLXND1;refGene_tx=XM_011280229.3;refGene_gn=PLXND1;refGene_

tx=XM_023250044.1
A2 54004870 54005020
refGene_gn=PLXND1;refGene_tx=XM_011280229.3;refGene_gn=PLXND1;refGene_tx=XM_023250044.1
A2 54005107 54005371
refGene_gn=PLXND1;refGene_tx=XM_011280229.3;refGene_gn=PLXND1;refGene_tx=XM_023250044.1
A2 54005568 54005659
refGene_gn=PLXND1;refGene_tx=XM_011280229.3;refGene_gn=PLXND1;refGene_tx=XM_023250044.1
A2 54005741 54005909
refGene_gn=PLXND1;refGene_tx=XM_011280229.3;refGene_gn=PLXND1;refGene_tx=XM_023250044.1
A2 54006701 54006841
refGene_gn=PLXND1;refGene_tx=XM_011280229.3;refGene_gn=PLXND1;refGene_tx=XM_023250044.1
A2 54006934 54007034
refGene_gn=PLXND1;refGene_tx=XM_011280229.3;refGene_gn=PLXND1;refGene_tx=XM_023250044.1
A2 54007964 54008116
refGene_gn=PLXND1;refGene_tx=XM_011280229.3;refGene_gn=PLXND1;refGene_tx=XM_023250044.1
A2 54009150 54009305
refGene_gn=PLXND1;refGene_tx=XM_011280229.3;refGene_gn=PLXND1;refGene_tx=XM_023250044.1
A2 54010541 54010635
refGene_gn=PLXND1;refGene_tx=XM_011280229.3;refGene_gn=PLXND1;refGene_tx=XM_023250044.1
A2 54010853 54010943
refGene_gn=PLXND1;refGene_tx=XM_011280229.3;refGene_gn=PLXND1;refGene_tx=XM_023250044.1
A2 54013409 54013511
refGene_gn=PLXND1;refGene_tx=XM_011280229.3;refGene_gn=PLXND1;refGene_tx=XM_023250044.1
A2 54019661 54019764
refGene_gn=PLXND1;refGene_tx=XM_011280229.3;refGene_gn=PLXND1;refGene_tx=XM_023250044.1
A2 54020213 54020322
refGene_gn=PLXND1;refGene_tx=XM_011280229.3;refGene_gn=PLXND1;refGene_tx=XM_023250044.1
A2 54020468 54020646
refGene_gn=PLXND1;refGene_tx=XM_011280229.3;refGene_gn=PLXND1;refGene_tx=XM_023250044.1
A2 54022123 54022253
refGene_gn=PLXND1;refGene_tx=XM_011280229.3;refGene_gn=PLXND1;refGene_tx=XM_023250044.1
A2 54022343 54022444
refGene_gn=PLXND1;refGene_tx=XM_011280229.3;refGene_gn=PLXND1;refGene_tx=XM_023250044.1
A2 54022702 54022834

refGene_gn=PLXND1;refGene_tx=XM_011280229.3;refGene_gn=PLXND1;refGene_tx=XM_023250044.1
A2 54025764 54025941
refGene_gn=PLXND1;refGene_tx=XM_011280229.3;refGene_gn=PLXND1;refGene_tx=XM_023250044.1
A2 54044263 54045583
refGene_gn=PLXND1;refGene_tx=XM_011280229.3;refGene_gn=PLXND1;refGene_tx=XM_023250044.1
A2 54085128 54085443
refGene_gn=TMCC1;refGene_tx=XM_006929007.4;refGene_gn=TMCC1;refGene_tx=XM_019825115.1;refGene_gn=TMCC1;refGene_tx=XM_023250045.1;refGene_gn=TMCC1;refGene_tx=XM_023250046.1
A2 54088278 54088414
refGene_gn=TMCC1;refGene_tx=XM_006929007.4;refGene_gn=TMCC1;refGene_tx=XM_019825115.1;refGene_gn=TMCC1;refGene_tx=XM_023250045.1;refGene_gn=TMCC1;refGene_tx=XM_023250046.1
A2 54097285 54097303
refGene_gn=TMCC1;refGene_tx=XM_006929007.4;refGene_gn=TMCC1;refGene_tx=XM_023250045.1
A2 54106043 54106978
refGene_gn=TMCC1;refGene_tx=XM_006929007.4;refGene_gn=TMCC1;refGene_tx=XM_019825115.1;refGene_gn=TMCC1;refGene_tx=XM_023250045.1;refGene_gn=TMCC1;refGene_tx=XM_023250046.1
A2 54120752 54120815 refGene_gn=TMCC1;refGene_tx=XM_023250046.1
A2 54272476 54273043
refGene_gn=TMCC1;refGene_tx=XM_006929007.4;refGene_gn=TMCC1;refGene_tx=XM_019825115.1;refGene_gn=TMCC1;refGene_tx=XM_023250045.1
A2 54398517 54398593 refGene_gn=TRH;refGene_tx=XM_023250047.1
A2 54399235 54399448
refGene_gn=TRH;refGene_tx=XM_003982516.4;refGene_gn=TRH;refGene_tx=XM_023250047.1
A2 54400100 54400609
refGene_gn=TRH;refGene_tx=XM_003982516.4;refGene_gn=TRH;refGene_tx=XM_023250047.1
A2 54438842 54438990
refGene_gn=RBSN;refGene_tx=XM_003982517.5;refGene_gn=RBSN;refGene_tx=XM_011280220.3;refGene_gn=RBSN;refGene_tx=XM_011280221.3
A2 54443194 54443335
refGene_gn=RBSN;refGene_tx=XM_003982517.5;refGene_gn=RBSN;refGene_tx=XM_011280220.3;refGene_gn=RBSN;refGene_tx=XM_011280221.3
A2 54445925 54446026
refGene_gn=RBSN;refGene_tx=XM_003982517.5;refGene_gn=RBSN;refGene_tx=XM_011280220.3;refGene_gn=RBSN;refGene_tx=XM_011280221.3
A2 54447197 54447243
refGene_gn=RBSN;refGene_tx=XM_003982517.5;refGene_gn=RBSN;refGene_tx=XM_011280220.3;refGene_gn=RBSN;refGene_tx=XM_011280221.3
A2 54447355 54447517
refGene_gn=RBSN;refGene_tx=XM_003982517.5;refGene_gn=RBSN;refGene_tx=XM_011280220.3;refGene_gn=RBSN;refGene_tx=XM_011280221.3
A2 54449587 54449829

refGene_gn=RBSN;refGene_tx=XM_003982517.5;refGene_gn=RBSN;refGene_tx=XM_011280220.3;refGene_gn=RBSN;refGene_tx=XM_011280221.3
A2 54451912 54451983
refGene_gn=RBSN;refGene_tx=XM_003982517.5;refGene_gn=RBSN;refGene_tx=XM_011280220.3;refGene_gn=RBSN;refGene_tx=XM_011280221.3
A2 54455983 54456070
refGene_gn=RBSN;refGene_tx=XM_003982517.5;refGene_gn=RBSN;refGene_tx=XM_011280220.3;refGene_gn=RBSN;refGene_tx=XM_011280221.3
A2 54456942 54457045
refGene_gn=RBSN;refGene_tx=XM_003982517.5;refGene_gn=RBSN;refGene_tx=XM_011280220.3;refGene_gn=RBSN;refGene_tx=XM_011280221.3
A2 54458320 54458425
refGene_gn=RBSN;refGene_tx=XM_003982517.5;refGene_gn=RBSN;refGene_tx=XM_011280220.3;refGene_gn=RBSN;refGene_tx=XM_011280221.3
A2 54459101 54460247
refGene_gn=RBSN;refGene_tx=XM_003982517.5;refGene_gn=RBSN;refGene_tx=XM_011280220.3;refGene_gn=RBSN;refGene_tx=XM_011280221.3
A2 54467410 54467544 refGene_gn=MRPS25;refGene_tx=XM_003982518.4
A2 54472802 54472909 refGene_gn=MRPS25;refGene_tx=XM_003982518.4
A2 54478673 54478761 refGene_gn=MRPS25;refGene_tx=XM_003982518.4
A2 54479426 54479619 refGene_gn=MRPS25;refGene_tx=XM_003982518.4
A2 54487925 54488100
refGene_gn=NR2C2;refGene_tx=XM_023250051.1;refGene_gn=NR2C2;refGene_tx=XM_023250052.1
A2 54490924 54491030
refGene_gn=NR2C2;refGene_tx=XM_023250051.1;refGene_gn=NR2C2;refGene_tx=XM_023250052.1
A2 54492005 54492143
refGene_gn=NR2C2;refGene_tx=XM_023250051.1;refGene_gn=NR2C2;refGene_tx=XM_023250052.1
A2 54494661 54494801
refGene_gn=NR2C2;refGene_tx=XM_023250051.1;refGene_gn=NR2C2;refGene_tx=XM_023250052.1
A2 54495608 54495730
refGene_gn=NR2C2;refGene_tx=XM_023250051.1;refGene_gn=NR2C2;refGene_tx=XM_023250052.1
A2 54496851 54497029
refGene_gn=NR2C2;refGene_tx=XM_023250051.1;refGene_gn=NR2C2;refGene_tx=XM_023250052.1
A2 54497970 54498104
refGene_gn=NR2C2;refGene_tx=XM_023250051.1;refGene_gn=NR2C2;refGene_tx=XM_023250052.1
A2 54501641 54501735
refGene_gn=NR2C2;refGene_tx=XM_023250051.1;refGene_gn=NR2C2;refGene_tx=XM_023250052.1
A2 54502499 54502647
refGene_gn=NR2C2;refGene_tx=XM_023250051.1;refGene_gn=NR2C2;refGene_tx=XM_023250052.1
A2 54504016 54504196
refGene_gn=NR2C2;refGene_tx=XM_023250051.1;refGene_gn=NR2C2;refGene_tx

=XM_023250052.1
A2 54506804 54506907
refGene_gn=NR2C2;refGene_tx=XM_023250051.1;refGene_gn=NR2C2;refGene_tx
=XM_023250052.1
A2 54509526 54509727
refGene_gn=NR2C2;refGene_tx=XM_023250051.1;refGene_gn=NR2C2;refGene_tx
=XM_023250052.1
A2 54530717 54530828
refGene_gn=NR2C2;refGene_tx=XM_023250051.1;refGene_gn=NR2C2;refGene_tx
=XM_023250052.1
A2 54571187 54571247 refGene_gn=NR2C2;refGene_tx=XM_023250051.1
A2 54598624 54598661
refGene_gn=FGD5;refGene_tx=XM_023250048.1;refGene_gn=FGD5;refGene_tx=X
M_023250049.1
A2 54599018 54599173
refGene_gn=FGD5;refGene_tx=XM_023250048.1;refGene_gn=FGD5;refGene_tx=X
M_023250049.1
A2 54604048 54604177
refGene_gn=FGD5;refGene_tx=XM_023250048.1;refGene_gn=FGD5;refGene_tx=X
M_023250049.1
A2 54609310 54609441
refGene_gn=FGD5;refGene_tx=XM_023250048.1;refGene_gn=FGD5;refGene_tx=X
M_023250049.1
A2 54610266 54610396
refGene_gn=FGD5;refGene_tx=XM_023250048.1;refGene_gn=FGD5;refGene_tx=X
M_023250049.1
A2 54610870 54611008
refGene_gn=FGD5;refGene_tx=XM_023250048.1;refGene_gn=FGD5;refGene_tx=X
M_023250049.1
A2 54611348 54611448
refGene_gn=FGD5;refGene_tx=XM_023250048.1;refGene_gn=FGD5;refGene_tx=X
M_023250049.1
A2 54613068 54613148
refGene_gn=FGD5;refGene_tx=XM_023250048.1;refGene_gn=FGD5;refGene_tx=X
M_023250049.1
A2 54614241 54614325
refGene_gn=FGD5;refGene_tx=XM_023250048.1;refGene_gn=FGD5;refGene_tx=X
M_023250049.1
A2 54618252 54618321
refGene_gn=FGD5;refGene_tx=XM_023250048.1;refGene_gn=FGD5;refGene_tx=X
M_023250049.1
A2 54619338 54619410
refGene_gn=FGD5;refGene_tx=XM_023250048.1;refGene_gn=FGD5;refGene_tx=X
M_023250049.1
A2 54627289 54627348
refGene_gn=FGD5;refGene_tx=XM_023250048.1;refGene_gn=FGD5;refGene_tx=X
M_023250049.1
A2 54627956 54628007
refGene_gn=FGD5;refGene_tx=XM_023250048.1;refGene_gn=FGD5;refGene_tx=X
M_023250049.1

A2 54629750 54629838
refGene_gn=FGD5;refGene_tx=XM_023250048.1;refGene_gn=FGD5;refGene_tx=XM_023250049.1
A2 54630480 54630637
refGene_gn=FGD5;refGene_tx=XM_023250048.1;refGene_gn=FGD5;refGene_tx=XM_023250049.1
A2 54630900 54631061
refGene_gn=FGD5;refGene_tx=XM_023250048.1;refGene_gn=FGD5;refGene_tx=XM_023250049.1
A2 54649910 54649941
refGene_gn=FGD5;refGene_tx=XM_023250048.1;refGene_gn=FGD5;refGene_tx=XM_023250049.1
A2 54650053 54650112
refGene_gn=FGD5;refGene_tx=XM_023250048.1;refGene_gn=FGD5;refGene_tx=XM_023250049.1
A2 54666913 54667046
refGene_gn=FGD5;refGene_tx=XM_023250048.1;refGene_gn=FGD5;refGene_tx=XM_023250049.1
A2 54708948 54711589
refGene_gn=FGD5;refGene_tx=XM_023250048.1;refGene_gn=FGD5;refGene_tx=XM_023250049.1
A2 54720625 54720638
refGene_gn=FGD5;refGene_tx=XM_023250048.1;refGene_gn=FGD5;refGene_tx=XM_023250049.1
A2 54772490 54772587
refGene_gn=CA2H3orf20;refGene_tx=XM_019825118.2
A2 54773209 54773341
refGene_gn=CA2H3orf20;refGene_tx=XM_019825118.2
A2 54779520 54779663
refGene_gn=CA2H3orf20;refGene_tx=XM_019825118.2
A2 54780995 54781058
refGene_gn=CA2H3orf20;refGene_tx=XM_019825118.2
A2 54781059 54781092
refGene_gn=CA2H3orf20;refGene_tx=XM_019825118.2
A2 54781093 54781105
refGene_gn=CA2H3orf20;refGene_tx=XM_019825118.2
A2 54783665 54783969
refGene_gn=CA2H3orf20;refGene_tx=XM_019825118.2
A2 54819227 54819480
refGene_gn=CA2H3orf20;refGene_tx=XM_019825118.2
A2 54820826 54820941
refGene_gn=CA2H3orf20;refGene_tx=XM_019825118.2
A2 54825835 54825967
refGene_gn=CA2H3orf20;refGene_tx=XM_019825118.2
A2 54831917 54832038
refGene_gn=CA2H3orf20;refGene_tx=XM_019825118.2
A2 54832811 54832964
refGene_gn=CA2H3orf20;refGene_tx=XM_019825118.2
A2 54847037 54847310
refGene_gn=CA2H3orf20;refGene_tx=XM_019825118.2

A2	54848372	54848514
refGene_gn=CA2H3orf20;refGene_tx=XM_019825118.2
A2	54855405	54855525
refGene_gn=CA2H3orf20;refGene_tx=XM_019825118.2
A2	54861480	54861621
refGene_gn=CA2H3orf20;refGene_tx=XM_019825118.2
A2	54862377	54862849
refGene_gn=CA2H3orf20;refGene_tx=XM_019825118.2
A2	54874927	54875226	refGene_gn=CCDC174;refGene_tx=XM_003982522.4
A2	54876314	54876461	refGene_gn=CCDC174;refGene_tx=XM_003982522.4
A2	54878199	54878332	refGene_gn=CCDC174;refGene_tx=XM_003982522.4
A2	54881399	54881494	refGene_gn=CCDC174;refGene_tx=XM_003982522.4
A2	54881854	54881997	refGene_gn=CCDC174;refGene_tx=XM_003982522.4
A2	54882974	54883070	refGene_gn=CCDC174;refGene_tx=XM_003982522.4
A2	54886608	54886786	refGene_gn=CCDC174;refGene_tx=XM_003982522.4
A2	54888857	54888916	refGene_gn=CCDC174;refGene_tx=XM_003982522.4
A2	54893335	54893436	refGene_gn=CCDC174;refGene_tx=XM_003982522.4
A2	54894120	54894225	refGene_gn=CCDC174;refGene_tx=XM_003982522.4
A2	54897541	54897655	refGene_gn=CCDC174;refGene_tx=XM_003982522.4
A2	54943351	54943406
refGene_gn=GRIP2;refGene_tx=XM_023250053.1;refGene_gn=GRIP2;refGene_tx=XM_023250054.1;refGene_gn=GRIP2;refGene_tx=XM_023250055.1
A2	54980917	54980972	refGene_gn=GRIP2;refGene_tx=XM_006929013.3
A2	54995606	54995646	refGene_gn=GRIP2;refGene_tx=XM_006929014.3
A2	55008160	55008241
refGene_gn=GRIP2;refGene_tx=XM_006929013.3;refGene_gn=GRIP2;refGene_tx=XM_006929014.3;refGene_gn=GRIP2;refGene_tx=XM_023250053.1;refGene_gn=GRIP2;refGene_tx=XM_023250054.1;refGene_gn=GRIP2;refGene_tx=XM_023250055.1
A2	55008520	55008656
refGene_gn=GRIP2;refGene_tx=XM_006929013.3;refGene_gn=GRIP2;refGene_tx=XM_006929014.3;refGene_gn=GRIP2;refGene_tx=XM_023250053.1;refGene_gn=GRIP2;refGene_tx=XM_023250054.1;refGene_gn=GRIP2;refGene_tx=XM_023250055.1
A2	55009548	55009694
refGene_gn=GRIP2;refGene_tx=XM_006929013.3;refGene_gn=GRIP2;refGene_tx=XM_006929014.3;refGene_gn=GRIP2;refGene_tx=XM_023250053.1;refGene_gn=GRIP2;refGene_tx=XM_023250054.1;refGene_gn=GRIP2;refGene_tx=XM_023250055.1
A2	55010367	55010454
refGene_gn=GRIP2;refGene_tx=XM_006929013.3;refGene_gn=GRIP2;refGene_tx=XM_006929014.3;refGene_gn=GRIP2;refGene_tx=XM_023250053.1;refGene_gn=GRIP2;refGene_tx=XM_023250054.1;refGene_gn=GRIP2;refGene_tx=XM_023250055.1
A2	55010965	55011041
refGene_gn=GRIP2;refGene_tx=XM_006929013.3;refGene_gn=GRIP2;refGene_tx=XM_006929014.3;refGene_gn=GRIP2;refGene_tx=XM_023250053.1;refGene_gn=GRIP2;refGene_tx=XM_023250054.1;refGene_gn=GRIP2;refGene_tx=XM_023250055.1
A2	55012248	55012394

refGene_gn=GRIP2;refGene_tx=XM_006929013.3;refGene_gn=GRIP2;refGene_tx
=XM_006929014.3;refGene_gn=GRIP2;refGene_tx=XM_023250053.1;refGene_gn=
GRIP2;refGene_tx=XM_023250054.1;refGene_gn=GRIP2;refGene_tx=XM_0232500
55.1
A2 55013250 55013398
refGene_gn=GRIP2;refGene_tx=XM_006929013.3;refGene_gn=GRIP2;refGene_tx
=XM_006929014.3;refGene_gn=GRIP2;refGene_tx=XM_023250053.1;refGene_gn=
GRIP2;refGene_tx=XM_023250054.1;refGene_gn=GRIP2;refGene_tx=XM_0232500
55.1
A2 55013510 55013680
refGene_gn=GRIP2;refGene_tx=XM_006929013.3;refGene_gn=GRIP2;refGene_tx
=XM_006929014.3;refGene_gn=GRIP2;refGene_tx=XM_023250053.1;refGene_gn=
GRIP2;refGene_tx=XM_023250054.1;refGene_gn=GRIP2;refGene_tx=XM_0232500
55.1
A2 55015769 55015895
refGene_gn=GRIP2;refGene_tx=XM_006929013.3;refGene_gn=GRIP2;refGene_tx
=XM_006929014.3;refGene_gn=GRIP2;refGene_tx=XM_023250053.1;refGene_gn=
GRIP2;refGene_tx=XM_023250055.1
A2 55016223 55016373
refGene_gn=GRIP2;refGene_tx=XM_006929013.3;refGene_gn=GRIP2;refGene_tx
=XM_006929014.3;refGene_gn=GRIP2;refGene_tx=XM_023250053.1;refGene_gn=
GRIP2;refGene_tx=XM_023250054.1;refGene_gn=GRIP2;refGene_tx=XM_0232500
55.1
A2 55019240 55019427
refGene_gn=GRIP2;refGene_tx=XM_006929013.3;refGene_gn=GRIP2;refGene_tx
=XM_006929014.3;refGene_gn=GRIP2;refGene_tx=XM_023250053.1;refGene_gn=
GRIP2;refGene_tx=XM_023250054.1;refGene_gn=GRIP2;refGene_tx=XM_0232500
55.1
A2 55019920 55020066
refGene_gn=GRIP2;refGene_tx=XM_006929013.3;refGene_gn=GRIP2;refGene_tx
=XM_006929014.3;refGene_gn=GRIP2;refGene_tx=XM_023250053.1;refGene_gn=
GRIP2;refGene_tx=XM_023250054.1;refGene_gn=GRIP2;refGene_tx=XM_0232500
55.1
A2 55020827 55020908
refGene_gn=GRIP2;refGene_tx=XM_006929013.3;refGene_gn=GRIP2;refGene_tx
=XM_006929014.3;refGene_gn=GRIP2;refGene_tx=XM_023250053.1;refGene_gn=
GRIP2;refGene_tx=XM_023250054.1;refGene_gn=GRIP2;refGene_tx=XM_0232500
55.1
A2 55023688 55023755
refGene_gn=GRIP2;refGene_tx=XM_006929013.3;refGene_gn=GRIP2;refGene_tx
=XM_006929014.3;refGene_gn=GRIP2;refGene_tx=XM_023250053.1;refGene_gn=
GRIP2;refGene_tx=XM_023250054.1;refGene_gn=GRIP2;refGene_tx=XM_0232500
55.1
A2 55023847 55023993
refGene_gn=GRIP2;refGene_tx=XM_006929013.3;refGene_gn=GRIP2;refGene_tx
=XM_006929014.3;refGene_gn=GRIP2;refGene_tx=XM_023250053.1;refGene_gn=
GRIP2;refGene_tx=XM_023250054.1;refGene_gn=GRIP2;refGene_tx=XM_0232500
55.1
A2 55024805 55024950
refGene_gn=GRIP2;refGene_tx=XM_006929013.3;refGene_gn=GRIP2;refGene_tx

=XM_006929014.3;refGene_gn=GRIP2;refGene_tx=XM_023250053.1;refGene_gn=GRIP2;refGene_tx=XM_023250054.1;refGene_gn=GRIP2;refGene_tx=XM_023250055.1
A2 55026846 55026986
refGene_gn=GRIP2;refGene_tx=XM_006929013.3;refGene_gn=GRIP2;refGene_tx=XM_006929014.3;refGene_gn=GRIP2;refGene_tx=XM_023250053.1;refGene_gn=GRIP2;refGene_tx=XM_023250054.1;refGene_gn=GRIP2;refGene_tx=XM_023250055.1
A2 55027610 55027790
refGene_gn=GRIP2;refGene_tx=XM_006929013.3;refGene_gn=GRIP2;refGene_tx=XM_006929014.3;refGene_gn=GRIP2;refGene_tx=XM_023250053.1;refGene_gn=GRIP2;refGene_tx=XM_023250054.1;refGene_gn=GRIP2;refGene_tx=XM_023250055.1
A2 55028813 55029006
refGene_gn=GRIP2;refGene_tx=XM_006929013.3;refGene_gn=GRIP2;refGene_tx=XM_006929014.3;refGene_gn=GRIP2;refGene_tx=XM_023250053.1;refGene_gn=GRIP2;refGene_tx=XM_023250054.1;refGene_gn=GRIP2;refGene_tx=XM_023250055.1
A2 55030318 55030367 refGene_gn=GRIP2;refGene_tx=XM_023250055.1
A2 55030620 55030726
refGene_gn=GRIP2;refGene_tx=XM_006929013.3;refGene_gn=GRIP2;refGene_tx=XM_006929014.3;refGene_gn=GRIP2;refGene_tx=XM_023250053.1;refGene_gn=GRIP2;refGene_tx=XM_023250054.1
A2 55034754 55034898
refGene_gn=GRIP2;refGene_tx=XM_006929013.3;refGene_gn=GRIP2;refGene_tx=XM_006929014.3;refGene_gn=GRIP2;refGene_tx=XM_023250053.1;refGene_gn=GRIP2;refGene_tx=XM_023250054.1
A2 55035649 55035796
refGene_gn=GRIP2;refGene_tx=XM_006929013.3;refGene_gn=GRIP2;refGene_tx=XM_006929014.3;refGene_gn=GRIP2;refGene_tx=XM_023250053.1;refGene_gn=GRIP2;refGene_tx=XM_023250054.1
A2 55036830 55036989
refGene_gn=GRIP2;refGene_tx=XM_006929013.3;refGene_gn=GRIP2;refGene_tx=XM_006929014.3;refGene_gn=GRIP2;refGene_tx=XM_023250053.1;refGene_gn=GRIP2;refGene_tx=XM_023250054.1
A2 55046596 55046737
refGene_gn=SLC6A6;refGene_tx=XM_003982523.5;refGene_gn=SLC6A6;refGene_tx=XM_019825127.2;refGene_gn=SLC6A6;refGene_tx=XM_023250056.1
A2 55050047 55050218
refGene_gn=SLC6A6;refGene_tx=XM_003982523.5;refGene_gn=SLC6A6;refGene_tx=XM_019825127.2;refGene_gn=SLC6A6;refGene_tx=XM_023250056.1
A2 55052699 55052800
refGene_gn=SLC6A6;refGene_tx=XM_003982523.5;refGene_gn=SLC6A6;refGene_tx=XM_019825127.2;refGene_gn=SLC6A6;refGene_tx=XM_023250056.1
A2 55053374 55053477
refGene_gn=SLC6A6;refGene_tx=XM_003982523.5;refGene_gn=SLC6A6;refGene_tx=XM_019825127.2;refGene_gn=SLC6A6;refGene_tx=XM_023250056.1
A2 55054807 55054945
refGene_gn=SLC6A6;refGene_tx=XM_003982523.5;refGene_gn=SLC6A6;refGene_tx=XM_019825127.2;refGene_gn=SLC6A6;refGene_tx=XM_023250056.1

A2 55059439 55059552
refGene_gn=SLC6A6;refGene_tx=XM_003982523.5;refGene_gn=SLC6A6;refGene_tx=XM_019825127.2;refGene_gn=SLC6A6;refGene_tx=XM_023250056.1
A2 55063523 55063648
refGene_gn=SLC6A6;refGene_tx=XM_003982523.5;refGene_gn=SLC6A6;refGene_tx=XM_019825127.2;refGene_gn=SLC6A6;refGene_tx=XM_023250056.1
A2 55063779 55063883
refGene_gn=SLC6A6;refGene_tx=XM_003982523.5;refGene_gn=SLC6A6;refGene_tx=XM_019825127.2;refGene_gn=SLC6A6;refGene_tx=XM_023250056.1
A2 55065093 55065228
refGene_gn=SLC6A6;refGene_tx=XM_003982523.5;refGene_gn=SLC6A6;refGene_tx=XM_019825127.2;refGene_gn=SLC6A6;refGene_tx=XM_023250056.1
A2 55072998 55073131
refGene_gn=SLC6A6;refGene_tx=XM_003982523.5;refGene_gn=SLC6A6;refGene_tx=XM_019825127.2;refGene_gn=SLC6A6;refGene_tx=XM_023250056.1
A2 55083348 55083587
refGene_gn=SLC6A6;refGene_tx=XM_003982523.5;refGene_gn=SLC6A6;refGene_tx=XM_019825127.2;refGene_gn=SLC6A6;refGene_tx=XM_019825128.2;refGene_gn=SLC6A6;refGene_tx=XM_023250056.1
A2 55084960 55085095
refGene_gn=SLC6A6;refGene_tx=XM_003982523.5;refGene_gn=SLC6A6;refGene_tx=XM_019825127.2;refGene_gn=SLC6A6;refGene_tx=XM_019825128.2;refGene_gn=SLC6A6;refGene_tx=XM_023250056.1
A2 55086977 55087206
refGene_gn=SLC6A6;refGene_tx=XM_003982523.5;refGene_gn=SLC6A6;refGene_tx=XM_019825127.2;refGene_gn=SLC6A6;refGene_tx=XM_019825128.2;refGene_gn=SLC6A6;refGene_tx=XM_023250056.1
A2 55329890 55329971 refGene_gn=LSM3;refGene_tx=XM_003982524.4
A2 55335615 55335711 refGene_gn=LSM3;refGene_tx=XM_003982524.4
A2 55338760 55338871 refGene_gn=LSM3;refGene_tx=XM_003982524.4
A2 55341060 55341081 refGene_gn=LSM3;refGene_tx=XM_003982524.4
A2 55341394 55341497
refGene_gn=XPC;refGene_tx=XM_011280222.3;refGene_gn=XPC;refGene_tx=XM_011280223.3
A2 55346999 55347192
refGene_gn=XPC;refGene_tx=XM_011280222.3;refGene_gn=XPC;refGene_tx=XM_011280223.3
A2 55349542 55349661
refGene_gn=XPC;refGene_tx=XM_011280222.3;refGene_gn=XPC;refGene_tx=XM_011280223.3
A2 55352372 55352496
refGene_gn=XPC;refGene_tx=XM_011280222.3;refGene_gn=XPC;refGene_tx=XM_011280223.3
A2 55353511 55353596
refGene_gn=XPC;refGene_tx=XM_011280222.3;refGene_gn=XPC;refGene_tx=XM_011280223.3
A2 55354734 55354892
refGene_gn=XPC;refGene_tx=XM_011280222.3;refGene_gn=XPC;refGene_tx=XM_011280223.3
A2 55355385 55355506

refGene_gn=XPC;refGene_tx=XM_011280222.3;refGene_gn=XPC;refGene_tx=XM_011280223.3
A2 55358550 55358640 refGene_gn=XPC;refGene_tx=XM_011280222.3;refGene_gn=XPC;refGene_tx=XM_011280223.3
A2 55359842 55360736 refGene_gn=XPC;refGene_tx=XM_011280222.3;refGene_gn=XPC;refGene_tx=XM_011280223.3
A2 55362204 55362365 refGene_gn=XPC;refGene_tx=XM_011280222.3;refGene_gn=XPC;refGene_tx=XM_011280223.3
A2 55365174 55365256 refGene_gn=XPC;refGene_tx=XM_011280222.3;refGene_gn=XPC;refGene_tx=XM_011280223.3
A2 55366186 55366321 refGene_gn=XPC;refGene_tx=XM_011280222.3;refGene_gn=XPC;refGene_tx=XM_011280223.3
A2 55366406 55366576 refGene_gn=XPC;refGene_tx=XM_011280222.3;refGene_gn=XPC;refGene_tx=XM_011280223.3
A2 55367096 55367190 refGene_gn=XPC;refGene_tx=XM_011280222.3;refGene_gn=XPC;refGene_tx=XM_011280223.3
A2 55367427 55367529 refGene_gn=XPC;refGene_tx=XM_011280223.3
A2 55367955 55368045 refGene_gn=XPC;refGene_tx=XM_011280222.3
A2 55369160 55369376 refGene_gn=XPC;refGene_tx=XM_011280222.3
A2 55373197 55373400 refGene_gn=TMEM43;refGene_tx=XM_003982527.5;refGene_gn=TMEM43;refGene_tx=XM_006929015.4;refGene_gn=TMEM43;refGene_tx=XM_006929016.3
A2 55375568 55375686 refGene_gn=TMEM43;refGene_tx=XM_003982527.5;refGene_gn=TMEM43;refGene_tx=XM_006929015.4;refGene_gn=TMEM43;refGene_tx=XM_006929016.3
A2 55378164 55378266 refGene_gn=TMEM43;refGene_tx=XM_003982527.5;refGene_gn=TMEM43;refGene_tx=XM_006929015.4;refGene_gn=TMEM43;refGene_tx=XM_006929016.3
A2 55378882 55378957 refGene_gn=TMEM43;refGene_tx=XM_003982527.5;refGene_gn=TMEM43;refGene_tx=XM_006929015.4;refGene_gn=TMEM43;refGene_tx=XM_006929016.3
A2 55379233 55379355 refGene_gn=TMEM43;refGene_tx=XM_003982527.5;refGene_gn=TMEM43;refGene_tx=XM_006929015.4;refGene_gn=TMEM43;refGene_tx=XM_006929016.3
A2 55380240 55380311 refGene_gn=TMEM43;refGene_tx=XM_003982527.5;refGene_gn=TMEM43;refGene_tx=XM_006929015.4;refGene_gn=TMEM43;refGene_tx=XM_006929016.3
A2 55381118 55381188 refGene_gn=TMEM43;refGene_tx=XM_003982527.5;refGene_gn=TMEM43;refGene_tx=XM_006929015.4;refGene_gn=TMEM43;refGene_tx=XM_006929016.3
A2 55381450 55381500 refGene_gn=TMEM43;refGene_tx=XM_003982527.5;refGene_gn=TMEM43;refGene_tx=XM_006929015.4;refGene_gn=TMEM43;refGene_tx=XM_006929016.3

A2 55382529 55382624
refGene_gn=TMEM43;refGene_tx=XM_003982527.5;refGene_gn=TMEM43;refGene_tx=XM_006929015.4;refGene_gn=TMEM43;refGene_tx=XM_006929016.3
A2 55383274 55383409
refGene_gn=TMEM43;refGene_tx=XM_003982527.5;refGene_gn=TMEM43;refGene_tx=XM_006929015.4;refGene_gn=TMEM43;refGene_tx=XM_006929016.3
A2 55384282 55384432
refGene_gn=TMEM43;refGene_tx=XM_003982527.5;refGene_gn=TMEM43;refGene_tx=XM_006929015.4;refGene_gn=TMEM43;refGene_tx=XM_006929016.3
A2 55387466 55387487 refGene_gn=TMEM43;refGene_tx=XM_006929015.4
A2 55388645 55388657 refGene_gn=TMEM43;refGene_tx=XM_003982527.5
A2 55389193 55389215 refGene_gn=CHCHD4;refGene_tx=XM_003982528.4
A2 55389414 55389460 refGene_gn=CHCHD4;refGene_tx=XM_019825130.2
A2 55396614 55396713
refGene_gn=CHCHD4;refGene_tx=XM_003982528.4;refGene_gn=CHCHD4;refGene_tx=XM_019825130.2
A2 55400161 55400466
refGene_gn=CHCHD4;refGene_tx=XM_003982528.4;refGene_gn=CHCHD4;refGene_tx=XM_019825130.2
A2 55461221 55461292 refGene_gn=WNT7A;refGene_tx=XM_011280231.3
A2 55466316 55466543 refGene_gn=WNT7A;refGene_tx=XM_011280231.3
A2 55485055 55485327 refGene_gn=WNT7A;refGene_tx=XM_011280231.3
A2 55522619 55523099 refGene_gn=WNT7A;refGene_tx=XM_011280231.3
A2 55669350 55669708
refGene_gn=FBLN2;refGene_tx=XM_011280224.2;refGene_gn=FBLN2;refGene_tx=XM_011280225.2;refGene_gn=FBLN2;refGene_tx=XM_023250057.1
A2 55672275 55672403
refGene_gn=FBLN2;refGene_tx=XM_011280224.2;refGene_gn=FBLN2;refGene_tx=XM_011280225.2;refGene_gn=FBLN2;refGene_tx=XM_023250057.1;refGene_gn=FBLN2;refGene_tx=XM_023250058.1
A2 55675838 55675967
refGene_gn=FBLN2;refGene_tx=XM_011280224.2;refGene_gn=FBLN2;refGene_tx=XM_011280225.2;refGene_gn=FBLN2;refGene_tx=XM_023250057.1;refGene_gn=FBLN2;refGene_tx=XM_023250058.1
A2 55676488 55676605
refGene_gn=FBLN2;refGene_tx=XM_011280224.2;refGene_gn=FBLN2;refGene_tx=XM_011280225.2;refGene_gn=FBLN2;refGene_tx=XM_023250057.1;refGene_gn=FBLN2;refGene_tx=XM_023250058.1
A2 55678727 55678853
refGene_gn=FBLN2;refGene_tx=XM_011280224.2;refGene_gn=FBLN2;refGene_tx=XM_011280225.2;refGene_gn=FBLN2;refGene_tx=XM_023250057.1;refGene_gn=FBLN2;refGene_tx=XM_023250058.1
A2 55679312 55679441
refGene_gn=FBLN2;refGene_tx=XM_011280224.2;refGene_gn=FBLN2;refGene_tx=XM_011280225.2;refGene_gn=FBLN2;refGene_tx=XM_023250057.1;refGene_gn=FBLN2;refGene_tx=XM_023250058.1
A2 55679568 55679712
refGene_gn=FBLN2;refGene_tx=XM_011280224.2;refGene_gn=FBLN2;refGene_tx=XM_011280225.2;refGene_gn=FBLN2;refGene_tx=XM_023250057.1;refGene_gn=FBLN2;refGene_tx=XM_023250058.1

A2	55681622	55681760
refGene_gn=FBLN2;refGene_tx=XM_011280224.2;refGene_gn=FBLN2;refGene_tx=XM_011280225.2;refGene_gn=FBLN2;refGene_tx=XM_023250057.1;refGene_gn=FBLN2;refGene_tx=XM_023250058.1
A2	55683658	55683793
refGene_gn=FBLN2;refGene_tx=XM_011280224.2;refGene_gn=FBLN2;refGene_tx=XM_011280225.2;refGene_gn=FBLN2;refGene_tx=XM_023250057.1;refGene_gn=FBLN2;refGene_tx=XM_023250058.1
A2	55687950	55688091
refGene_gn=FBLN2;refGene_tx=XM_011280224.2;refGene_gn=FBLN2;refGene_tx=XM_023250057.1
A2	55690304	55690406
refGene_gn=FBLN2;refGene_tx=XM_011280224.2;refGene_gn=FBLN2;refGene_tx=XM_011280225.2;refGene_gn=FBLN2;refGene_tx=XM_023250057.1;refGene_gn=FBLN2;refGene_tx=XM_023250058.1
A2	55691010	55691124
refGene_gn=FBLN2;refGene_tx=XM_011280224.2;refGene_gn=FBLN2;refGene_tx=XM_011280225.2;refGene_gn=FBLN2;refGene_tx=XM_023250057.1;refGene_gn=FBLN2;refGene_tx=XM_023250058.1
A2	55691739	55691949
refGene_gn=FBLN2;refGene_tx=XM_011280224.2;refGene_gn=FBLN2;refGene_tx=XM_011280225.2;refGene_gn=FBLN2;refGene_tx=XM_023250057.1;refGene_gn=FBLN2;refGene_tx=XM_023250058.1
A2	55694550	55694731
refGene_gn=FBLN2;refGene_tx=XM_011280224.2;refGene_gn=FBLN2;refGene_tx=XM_011280225.2;refGene_gn=FBLN2;refGene_tx=XM_023250057.1;refGene_gn=FBLN2;refGene_tx=XM_023250058.1
A2	55696913	55697043
refGene_gn=FBLN2;refGene_tx=XM_011280224.2;refGene_gn=FBLN2;refGene_tx=XM_011280225.2;refGene_gn=FBLN2;refGene_tx=XM_023250057.1;refGene_gn=FBLN2;refGene_tx=XM_023250058.1
A2	55698473	55698585
refGene_gn=FBLN2;refGene_tx=XM_011280224.2;refGene_gn=FBLN2;refGene_tx=XM_011280225.2;refGene_gn=FBLN2;refGene_tx=XM_023250057.1;refGene_gn=FBLN2;refGene_tx=XM_023250058.1
A2	55728218	55729500
refGene_gn=FBLN2;refGene_tx=XM_011280224.2;refGene_gn=FBLN2;refGene_tx=XM_011280225.2;refGene_gn=FBLN2;refGene_tx=XM_023250057.1;refGene_gn=FBLN2;refGene_tx=XM_023250058.1
A2	55793708	55793924	refGene_gn=HDAC11;refGene_tx=XM_023250059.1
A2	55794069	55794248	refGene_gn=HDAC11;refGene_tx=XM_023250059.1
A2	55795142	55795239	refGene_gn=HDAC11;refGene_tx=XM_023250059.1
A2	55796242	55796305	refGene_gn=HDAC11;refGene_tx=XM_023250059.1
A2	55797361	55797438	refGene_gn=HDAC11;refGene_tx=XM_023250059.1
A2	55799185	55799228	refGene_gn=HDAC11;refGene_tx=XM_023250059.1
A2	55800488	55800605	refGene_gn=HDAC11;refGene_tx=XM_023250059.1
A2	55811274	55811375	refGene_gn=HDAC11;refGene_tx=XM_023250059.1
A2	55813562	55813711	refGene_gn=HDAC11;refGene_tx=XM_023250059.1
A2	55814659	55814661	refGene_gn=HDAC11;refGene_tx=XM_023250059.1
A2	55864838	55865011

refGene_gn=NUP210;refGene_tx=XM_023250060.1;refGene_gn=NUP210;refGene_tx=XM_023250061.1;refGene_gn=NUP210;refGene_tx=XM_023250062.1
A2 55883627 55883764
refGene_gn=NUP210;refGene_tx=XM_023250060.1;refGene_gn=NUP210;refGene_tx=XM_023250061.1;refGene_gn=NUP210;refGene_tx=XM_023250062.1
A2 55885652 55885784
refGene_gn=NUP210;refGene_tx=XM_023250060.1;refGene_gn=NUP210;refGene_tx=XM_023250061.1;refGene_gn=NUP210;refGene_tx=XM_023250062.1
A2 55889037 55889134
refGene_gn=NUP210;refGene_tx=XM_023250060.1;refGene_gn=NUP210;refGene_tx=XM_023250061.1;refGene_gn=NUP210;refGene_tx=XM_023250062.1
A2 55892687 55892838
refGene_gn=NUP210;refGene_tx=XM_023250060.1;refGene_gn=NUP210;refGene_tx=XM_023250061.1;refGene_gn=NUP210;refGene_tx=XM_023250062.1
A2 55894858 55894991
refGene_gn=NUP210;refGene_tx=XM_023250060.1;refGene_gn=NUP210;refGene_tx=XM_023250061.1;refGene_gn=NUP210;refGene_tx=XM_023250062.1
A2 55900043 55900202
refGene_gn=NUP210;refGene_tx=XM_023250060.1;refGene_gn=NUP210;refGene_tx=XM_023250061.1;refGene_gn=NUP210;refGene_tx=XM_023250062.1
A2 55900781 55900850
refGene_gn=NUP210;refGene_tx=XM_023250060.1;refGene_gn=NUP210;refGene_tx=XM_023250061.1;refGene_gn=NUP210;refGene_tx=XM_023250062.1
A2 55904193 55904300
refGene_gn=NUP210;refGene_tx=XM_023250060.1;refGene_gn=NUP210;refGene_tx=XM_023250061.1;refGene_gn=NUP210;refGene_tx=XM_023250062.1
A2 55905285 55905426
refGene_gn=NUP210;refGene_tx=XM_023250060.1;refGene_gn=NUP210;refGene_tx=XM_023250061.1;refGene_gn=NUP210;refGene_tx=XM_023250062.1
A2 55906308 55906446
refGene_gn=NUP210;refGene_tx=XM_023250060.1;refGene_gn=NUP210;refGene_tx=XM_023250061.1;refGene_gn=NUP210;refGene_tx=XM_023250062.1
A2 55907447 55907603
refGene_gn=NUP210;refGene_tx=XM_023250060.1;refGene_gn=NUP210;refGene_tx=XM_023250061.1;refGene_gn=NUP210;refGene_tx=XM_023250062.1
A2 55909090 55909289
refGene_gn=NUP210;refGene_tx=XM_023250060.1;refGene_gn=NUP210;refGene_tx=XM_023250061.1;refGene_gn=NUP210;refGene_tx=XM_023250062.1
A2 55914383 55914529
refGene_gn=NUP210;refGene_tx=XM_023250060.1;refGene_gn=NUP210;refGene_tx=XM_023250061.1;refGene_gn=NUP210;refGene_tx=XM_023250062.1
A2 55924208 55924430
refGene_gn=NUP210;refGene_tx=XM_023250060.1;refGene_gn=NUP210;refGene_tx=XM_023250061.1;refGene_gn=NUP210;refGene_tx=XM_023250062.1
A2 55926521 55926695
refGene_gn=NUP210;refGene_tx=XM_023250060.1;refGene_gn=NUP210;refGene_tx=XM_023250061.1;refGene_gn=NUP210;refGene_tx=XM_023250062.1
A2 55932251 55932444
refGene_gn=NUP210;refGene_tx=XM_023250060.1;refGene_gn=NUP210;refGene_tx=XM_023250061.1;refGene_gn=NUP210;refGene_tx=XM_023250062.1

A2 55932957 55933064
refGene_gn=NUP210;refGene_tx=XM_023250060.1;refGene_gn=NUP210;refGene_tx=XM_023250061.1;refGene_gn=NUP210;refGene_tx=XM_023250062.1
A2 55934810 55934915
refGene_gn=NUP210;refGene_tx=XM_023250060.1;refGene_gn=NUP210;refGene_tx=XM_023250061.1;refGene_gn=NUP210;refGene_tx=XM_023250062.1
A2 55934997 55935099
refGene_gn=NUP210;refGene_tx=XM_023250060.1;refGene_gn=NUP210;refGene_tx=XM_023250061.1;refGene_gn=NUP210;refGene_tx=XM_023250062.1
A2 55943422 55943551
refGene_gn=NUP210;refGene_tx=XM_023250060.1;refGene_gn=NUP210;refGene_tx=XM_023250061.1;refGene_gn=NUP210;refGene_tx=XM_023250062.1
A2 55944633 55944761
refGene_gn=NUP210;refGene_tx=XM_023250060.1;refGene_gn=NUP210;refGene_tx=XM_023250061.1;refGene_gn=NUP210;refGene_tx=XM_023250062.1
A2 55944850 55944986
refGene_gn=NUP210;refGene_tx=XM_023250060.1;refGene_gn=NUP210;refGene_tx=XM_023250061.1;refGene_gn=NUP210;refGene_tx=XM_023250062.1
A2 55946582 55946645
refGene_gn=NUP210;refGene_tx=XM_023250060.1;refGene_gn=NUP210;refGene_tx=XM_023250061.1;refGene_gn=NUP210;refGene_tx=XM_023250062.1
A2 55946812 55946992
refGene_gn=NUP210;refGene_tx=XM_023250060.1;refGene_gn=NUP210;refGene_tx=XM_023250061.1;refGene_gn=NUP210;refGene_tx=XM_023250062.1
A2 55950057 55950138
refGene_gn=NUP210;refGene_tx=XM_023250060.1;refGene_gn=NUP210;refGene_tx=XM_023250061.1;refGene_gn=NUP210;refGene_tx=XM_023250062.1
A2 55951200 55951332
refGene_gn=NUP210;refGene_tx=XM_023250060.1;refGene_gn=NUP210;refGene_tx=XM_023250061.1;refGene_gn=NUP210;refGene_tx=XM_023250062.1
A2 55952427 55952586
refGene_gn=NUP210;refGene_tx=XM_023250060.1;refGene_gn=NUP210;refGene_tx=XM_023250061.1;refGene_gn=NUP210;refGene_tx=XM_023250062.1
A2 55955677 55955769
refGene_gn=NUP210;refGene_tx=XM_023250060.1;refGene_gn=NUP210;refGene_tx=XM_023250061.1;refGene_gn=NUP210;refGene_tx=XM_023250062.1
A2 55957350 55957525
refGene_gn=NUP210;refGene_tx=XM_023250060.1;refGene_gn=NUP210;refGene_tx=XM_023250061.1;refGene_gn=NUP210;refGene_tx=XM_023250062.1
A2 55958951 55959127
refGene_gn=NUP210;refGene_tx=XM_023250060.1;refGene_gn=NUP210;refGene_tx=XM_023250061.1;refGene_gn=NUP210;refGene_tx=XM_023250062.1
A2 55961049 55961270
refGene_gn=NUP210;refGene_tx=XM_023250060.1;refGene_gn=NUP210;refGene_tx=XM_023250061.1;refGene_gn=NUP210;refGene_tx=XM_023250062.1
A2 55962486 55962623
refGene_gn=NUP210;refGene_tx=XM_023250060.1;refGene_gn=NUP210;refGene_tx=XM_023250061.1;refGene_gn=NUP210;refGene_tx=XM_023250062.1
A2 55965415 55965533
refGene_gn=NUP210;refGene_tx=XM_023250060.1;refGene_gn=NUP210;refGene_

tx=XM_023250061.1;refGene_gn=NUP210;refGene_tx=XM_023250062.1
A2 55965672 55965701 refGene_gn=NUP210;refGene_tx=XM_023250062.1
A2 55966551 55966698
refGene_gn=NUP210;refGene_tx=XM_023250060.1;refGene_gn=NUP210;refGene_
tx=XM_023250061.1
A2 55968260 55968511
refGene_gn=NUP210;refGene_tx=XM_023250060.1;refGene_gn=NUP210;refGene_
tx=XM_023250061.1
A2 55969498 55969715
refGene_gn=NUP210;refGene_tx=XM_023250060.1;refGene_gn=NUP210;refGene_
tx=XM_023250061.1
A2 55971448 55971544
refGene_gn=NUP210;refGene_tx=XM_023250060.1;refGene_gn=NUP210;refGene_
tx=XM_023250061.1
A2 55971622 55971706
refGene_gn=NUP210;refGene_tx=XM_023250060.1;refGene_gn=NUP210;refGene_
tx=XM_023250061.1
A2 55972905 55973006
refGene_gn=NUP210;refGene_tx=XM_023250060.1;refGene_gn=NUP210;refGene_
tx=XM_023250061.1
A2 56000018 56000290 refGene_gn=IQSEC1;refGene_tx=XM_023250063.1
A2 56133277 56133307 refGene_gn=IQSEC1;refGene_tx=XM_023250063.1
A2 56499274 56499566 refGene_gn=IQSEC1;refGene_tx=XM_023250063.1
A2 56504121 56505371 refGene_gn=IQSEC1;refGene_tx=XM_023250063.1
A2 56517812 56517974 refGene_gn=IQSEC1;refGene_tx=XM_023250063.1
A2 56520743 56520866 refGene_gn=IQSEC1;refGene_tx=XM_023250063.1
A2 56522398 56522565 refGene_gn=IQSEC1;refGene_tx=XM_023250063.1
A2 56527474 56527614 refGene_gn=IQSEC1;refGene_tx=XM_023250063.1
A2 56528341 56528371 refGene_gn=IQSEC1;refGene_tx=XM_023250063.1
A2 56529858 56529984 refGene_gn=IQSEC1;refGene_tx=XM_023250063.1
A2 56531200 56531287 refGene_gn=IQSEC1;refGene_tx=XM_023250063.1
A2 56531288 56531301 refGene_gn=IQSEC1;refGene_tx=XM_023250063.1
A2 56533818 56533980 refGene_gn=IQSEC1;refGene_tx=XM_023250063.1
A2 56534817 56534889 refGene_gn=IQSEC1;refGene_tx=XM_023250063.1
A2 56534891 56534936 refGene_gn=IQSEC1;refGene_tx=XM_023250063.1
A2 56534938 56534992 refGene_gn=IQSEC1;refGene_tx=XM_023250063.1
A2 56540410 56540460 refGene_gn=IQSEC1;refGene_tx=XM_023250063.1
A2 56541742 56541920 refGene_gn=IQSEC1;refGene_tx=XM_023250063.1
A2 56541922 56542080 refGene_gn=IQSEC1;refGene_tx=XM_023250063.1
A2 56542082 56542191 refGene_gn=IQSEC1;refGene_tx=XM_023250063.1
A2 56542193 56542279 refGene_gn=IQSEC1;refGene_tx=XM_023250063.1
A2 56575213 56575363 refGene_gn=ACAD9;refGene_tx=XM_006929031.4
A2 56581219 56581313 refGene_gn=ACAD9;refGene_tx=XM_006929031.4
A2 56598882 56598984 refGene_gn=ACAD9;refGene_tx=XM_006929031.4
A2 56602986 56603093 refGene_gn=ACAD9;refGene_tx=XM_006929031.4
A2 56604041 56604142 refGene_gn=ACAD9;refGene_tx=XM_006929031.4
A2 56605213 56605292 refGene_gn=ACAD9;refGene_tx=XM_006929031.4
A2 56606269 56606302 refGene_gn=ACAD9;refGene_tx=XM_006929031.4
A2 56606303 56606445 refGene_gn=ACAD9;refGene_tx=XM_006929031.4
A2 56609064 56609138 refGene_gn=ACAD9;refGene_tx=XM_006929031.4

A2 56612757 56612833 refGene_gn=ACAD9;refGene_tx=XM_006929031.4
A2 56614394 56614465 refGene_gn=ACAD9;refGene_tx=XM_006929031.4
A2 56614723 56614843 refGene_gn=ACAD9;refGene_tx=XM_006929031.4
A2 56615977 56616106 refGene_gn=ACAD9;refGene_tx=XM_006929031.4
A2 56618018 56618098 refGene_gn=ACAD9;refGene_tx=XM_006929031.4
A2 56618827 56618954 refGene_gn=ACAD9;refGene_tx=XM_006929031.4
A2 56619169 56619247 refGene_gn=ACAD9;refGene_tx=XM_006929031.4
A2 56619711 56619840 refGene_gn=ACAD9;refGene_tx=XM_006929031.4
A2 56620030 56620093
refGene_gn=KIAA1257;refGene_tx=XM_011280234.3;refGene_gn=KIAA1257;refGene_tx=XM_019825137.2
A2 56620476 56620549 refGene_gn=ACAD9;refGene_tx=XM_006929031.4
A2 56621400 56621501 refGene_gn=ACAD9;refGene_tx=XM_006929031.4
A2 56623635 56623890
refGene_gn=KIAA1257;refGene_tx=XM_011280234.3;refGene_gn=KIAA1257;refGene_tx=XM_019825135.2;refGene_gn=KIAA1257;refGene_tx=XM_019825136.2;refGene_gn=KIAA1257;refGene_tx=XM_019825137.2;refGene_gn=KIAA1257;refGene_tx=XM_023250066.1
A2 56623977 56624190
refGene_gn=KIAA1257;refGene_tx=XM_011280234.3;refGene_gn=KIAA1257;refGene_tx=XM_019825135.2;refGene_gn=KIAA1257;refGene_tx=XM_019825136.2;refGene_gn=KIAA1257;refGene_tx=XM_019825137.2;refGene_gn=KIAA1257;refGene_tx=XM_023250066.1
A2 56624743 56624908
refGene_gn=KIAA1257;refGene_tx=XM_011280234.3;refGene_gn=KIAA1257;refGene_tx=XM_019825135.2;refGene_gn=KIAA1257;refGene_tx=XM_019825136.2;refGene_gn=KIAA1257;refGene_tx=XM_019825137.2;refGene_gn=KIAA1257;refGene_tx=XM_023250066.1
A2 56625519 56625537 refGene_gn=KIAA1257;refGene_tx=XM_023250067.1
A2 56645254 56645552
refGene_gn=KIAA1257;refGene_tx=XM_011280234.3;refGene_gn=KIAA1257;refGene_tx=XM_019825135.2;refGene_gn=KIAA1257;refGene_tx=XM_019825136.2;refGene_gn=KIAA1257;refGene_tx=XM_019825137.2;refGene_gn=KIAA1257;refGene_tx=XM_023250066.1;refGene_gn=KIAA1257;refGene_tx=XM_023250067.1
A2 56647487 56647682
refGene_gn=KIAA1257;refGene_tx=XM_011280234.3;refGene_gn=KIAA1257;refGene_tx=XM_019825135.2;refGene_gn=KIAA1257;refGene_tx=XM_019825136.2;refGene_gn=KIAA1257;refGene_tx=XM_019825137.2;refGene_gn=KIAA1257;refGene_tx=XM_023250066.1;refGene_gn=KIAA1257;refGene_tx=XM_023250067.1
A2 56653009 56653914
refGene_gn=KIAA1257;refGene_tx=XM_011280234.3;refGene_gn=KIAA1257;refGene_tx=XM_019825135.2;refGene_gn=KIAA1257;refGene_tx=XM_019825136.2;refGene_gn=KIAA1257;refGene_tx=XM_019825137.2;refGene_gn=KIAA1257;refGene_tx=XM_023250066.1;refGene_gn=KIAA1257;refGene_tx=XM_023250067.1;refGene_gn=KIAA1257;refGene_tx=XM_023250068.1
A2 56676561 56676746
refGene_gn=KIAA1257;refGene_tx=XM_011280234.3;refGene_gn=KIAA1257;refGene_tx=XM_019825135.2;refGene_gn=KIAA1257;refGene_tx=XM_019825136.2;refGene_gn=KIAA1257;refGene_tx=XM_019825137.2;refGene_gn=KIAA1257;refGene_tx=XM_023250066.1;refGene_gn=KIAA1257;refGene_tx=XM_023250067.1

A2 56680453 56680597
refGene_gn=KIAA1257;refGene_tx=XM_011280234.3;refGene_gn=KIAA1257;refGene_tx=XM_019825135.2;refGene_gn=KIAA1257;refGene_tx=XM_019825136.2;refGene_gn=KIAA1257;refGene_tx=XM_019825137.2;refGene_gn=KIAA1257;refGene_tx=XM_023250066.1;refGene_gn=KIAA1257;refGene_tx=XM_023250067.1;refGene_gn=KIAA1257;refGene_tx=XM_023250068.1
A2 56684577 56684699
refGene_gn=KIAA1257;refGene_tx=XM_011280234.3;refGene_gn=KIAA1257;refGene_tx=XM_019825135.2;refGene_gn=KIAA1257;refGene_tx=XM_019825136.2;refGene_gn=KIAA1257;refGene_tx=XM_019825137.2;refGene_gn=KIAA1257;refGene_tx=XM_023250066.1;refGene_gn=KIAA1257;refGene_tx=XM_023250067.1;refGene_gn=KIAA1257;refGene_tx=XM_023250068.1
A2 56686028 56686104
refGene_gn=KIAA1257;refGene_tx=XM_011280234.3;refGene_gn=KIAA1257;refGene_tx=XM_019825135.2;refGene_gn=KIAA1257;refGene_tx=XM_019825136.2;refGene_gn=KIAA1257;refGene_tx=XM_019825137.2;refGene_gn=KIAA1257;refGene_tx=XM_023250066.1;refGene_gn=KIAA1257;refGene_tx=XM_023250067.1;refGene_gn=KIAA1257;refGene_tx=XM_023250068.1
A2 56687037 56687169
refGene_gn=KIAA1257;refGene_tx=XM_011280234.3;refGene_gn=KIAA1257;refGene_tx=XM_019825135.2;refGene_gn=KIAA1257;refGene_tx=XM_019825136.2;refGene_gn=KIAA1257;refGene_tx=XM_019825137.2;refGene_gn=KIAA1257;refGene_tx=XM_023250066.1;refGene_gn=KIAA1257;refGene_tx=XM_023250067.1;refGene_gn=KIAA1257;refGene_tx=XM_023250068.1
A2 56689254 56689459
refGene_gn=KIAA1257;refGene_tx=XM_011280234.3;refGene_gn=KIAA1257;refGene_tx=XM_019825135.2;refGene_gn=KIAA1257;refGene_tx=XM_019825136.2;refGene_gn=KIAA1257;refGene_tx=XM_019825137.2;refGene_gn=KIAA1257;refGene_tx=XM_023250066.1;refGene_gn=KIAA1257;refGene_tx=XM_023250067.1;refGene_gn=KIAA1257;refGene_tx=XM_023250068.1
A2 56690046 56690237
refGene_gn=KIAA1257;refGene_tx=XM_011280234.3;refGene_gn=KIAA1257;refGene_tx=XM_019825135.2;refGene_gn=KIAA1257;refGene_tx=XM_019825136.2;refGene_gn=KIAA1257;refGene_tx=XM_019825137.2;refGene_gn=KIAA1257;refGene_tx=XM_023250066.1;refGene_gn=KIAA1257;refGene_tx=XM_023250067.1;refGene_gn=KIAA1257;refGene_tx=XM_023250068.1
A2 56691551 56691810
refGene_gn=KIAA1257;refGene_tx=XM_011280234.3;refGene_gn=KIAA1257;refGene_tx=XM_019825135.2;refGene_gn=KIAA1257;refGene_tx=XM_019825136.2;refGene_gn=KIAA1257;refGene_tx=XM_019825137.2;refGene_gn=KIAA1257;refGene_tx=XM_023250066.1;refGene_gn=KIAA1257;refGene_tx=XM_023250067.1;refGene_gn=KIAA1257;refGene_tx=XM_023250068.1
A2 56703230 56703926 refGene_gn=EFCC1;refGene_tx=XM_023250069.1
A2 56705163 56705447 refGene_gn=EFCC1;refGene_tx=XM_023250069.1
A2 56733899 56734057 refGene_gn=EFCC1;refGene_tx=XM_023250069.1
A2 56735822 56735964 refGene_gn=EFCC1;refGene_tx=XM_023250069.1
A2 56737159 56737325 refGene_gn=EFCC1;refGene_tx=XM_023250069.1
A2 56739154 56739295 refGene_gn=EFCC1;refGene_tx=XM_023250069.1
A2 56741028 56741098 refGene_gn=EFCC1;refGene_tx=XM_023250069.1
A2 56741921 56742058 refGene_gn=EFCC1;refGene_tx=XM_023250069.1

A2 56766618 56767152
refGene_gn=GP9;refGene_tx=XM_019825142.2;refGene_gn=GP9;refGene_tx=XM_019825143.2
A2 56780957 56781208 refGene_gn=RAB43;refGene_tx=XM_011280237.3
A2 56785166 56785350 refGene_gn=RAB43;refGene_tx=XM_011280237.3
A2 56810923 56811013 refGene_gn=RAB43;refGene_tx=XM_011280237.3
A2 56811014 56811128 refGene_gn=RAB43;refGene_tx=XM_011280237.3
A2 56817124 56817232 refGene_gn=ISY1;refGene_tx=XM_003982538.5
A2 56817618 56817705 refGene_gn=ISY1;refGene_tx=XM_003982538.5
A2 56820956 56821078 refGene_gn=ISY1;refGene_tx=XM_003982538.5
A2 56821873 56821996 refGene_gn=ISY1;refGene_tx=XM_003982538.5
A2 56825596 56825714 refGene_gn=ISY1;refGene_tx=XM_003982538.5
A2 56831277 56831390 refGene_gn=ISY1;refGene_tx=XM_003982538.5
A2 56836044 56836087 refGene_gn=ISY1;refGene_tx=XM_003982538.5
A2 56837090 56837156 refGene_gn=ISY1;refGene_tx=XM_003982538.5
A2 56838621 56838673 refGene_gn=ISY1;refGene_tx=XM_003982538.5
A2 56839278 56839301 refGene_gn=ISY1;refGene_tx=XM_003982538.5
A2 56841366 56841369 refGene_gn=ISY1;refGene_tx=XM_003982538.5
A2 56859644 56859762
refGene_gn=CNBP;refGene_tx=XM_003982539.5;refGene_gn=CNBP;refGene_tx=XM_003982540.5;refGene_gn=CNBP;refGene_tx=XM_011280238.3;refGene_gn=CNBP;refGene_tx=XM_019825145.2
A2 56861064 56861266
refGene_gn=CNBP;refGene_tx=XM_003982539.5;refGene_gn=CNBP;refGene_tx=XM_003982540.5;refGene_gn=CNBP;refGene_tx=XM_011280238.3;refGene_gn=CNBP;refGene_tx=XM_019825145.2
A2 56861446 56861539
refGene_gn=CNBP;refGene_tx=XM_003982539.5;refGene_gn=CNBP;refGene_tx=XM_003982540.5;refGene_gn=CNBP;refGene_tx=XM_011280238.3;refGene_gn=CNBP;refGene_tx=XM_019825145.2
A2 56861637 56861761
refGene_gn=CNBP;refGene_tx=XM_003982539.5;refGene_gn=CNBP;refGene_tx=XM_003982540.5;refGene_gn=CNBP;refGene_tx=XM_011280238.3;refGene_gn=CNBP;refGene_tx=XM_019825145.2
A2 56931983 56932020 refGene_gn=COPG1;refGene_tx=XM_003982543.4
A2 56932936 56932989 refGene_gn=COPG1;refGene_tx=XM_003982543.4
A2 56933417 56933498 refGene_gn=COPG1;refGene_tx=XM_003982543.4
A2 56933759 56933831
refGene_gn=COPG1;refGene_tx=XM_003982543.4;refGene_gn=COPG1;refGene_tx=XM_019825146.2
A2 56933939 56934019
refGene_gn=COPG1;refGene_tx=XM_003982543.4;refGene_gn=COPG1;refGene_tx=XM_019825146.2
A2 56935531 56935607
refGene_gn=COPG1;refGene_tx=XM_003982543.4;refGene_gn=COPG1;refGene_tx=XM_019825146.2
A2 56935881 56935974
refGene_gn=COPG1;refGene_tx=XM_003982543.4;refGene_gn=COPG1;refGene_tx=XM_019825146.2
A2 56936380 56936467

refGene_gn=COPG1;refGene_tx=XM_003982543.4;refGene_gn=COPG1;refGene_tx=XM_019825146.2
A2 56937752 56937910
refGene_gn=COPG1;refGene_tx=XM_003982543.4;refGene_gn=COPG1;refGene_tx=XM_019825146.2
A2 56938001 56938135
refGene_gn=COPG1;refGene_tx=XM_003982543.4;refGene_gn=COPG1;refGene_tx=XM_019825146.2
A2 56939236 56939304
refGene_gn=COPG1;refGene_tx=XM_003982543.4;refGene_gn=COPG1;refGene_tx=XM_019825146.2
A2 56939543 56939732
refGene_gn=COPG1;refGene_tx=XM_003982543.4;refGene_gn=COPG1;refGene_tx=XM_019825146.2
A2 56940666 56940762
refGene_gn=COPG1;refGene_tx=XM_003982543.4;refGene_gn=COPG1;refGene_tx=XM_019825146.2
A2 56941857 56942101
refGene_gn=COPG1;refGene_tx=XM_003982543.4;refGene_gn=COPG1;refGene_tx=XM_019825146.2
A2 56942920 56942996
refGene_gn=COPG1;refGene_tx=XM_003982543.4;refGene_gn=COPG1;refGene_tx=XM_019825146.2
A2 56945366 56945470
refGene_gn=COPG1;refGene_tx=XM_003982543.4;refGene_gn=COPG1;refGene_tx=XM_019825146.2
A2 56945938 56946064
refGene_gn=COPG1;refGene_tx=XM_003982543.4;refGene_gn=COPG1;refGene_tx=XM_019825146.2
A2 56946376 56946445
refGene_gn=COPG1;refGene_tx=XM_003982543.4;refGene_gn=COPG1;refGene_tx=XM_019825146.2
A2 56949389 56949532
refGene_gn=COPG1;refGene_tx=XM_003982543.4;refGene_gn=COPG1;refGene_tx=XM_019825146.2
A2 56949841 56950013
refGene_gn=COPG1;refGene_tx=XM_003982543.4;refGene_gn=COPG1;refGene_tx=XM_019825146.2
A2 56950719 56950817
refGene_gn=COPG1;refGene_tx=XM_003982543.4;refGene_gn=COPG1;refGene_tx=XM_019825146.2
A2 56951780 56951919
refGene_gn=COPG1;refGene_tx=XM_003982543.4;refGene_gn=COPG1;refGene_tx=XM_019825146.2
A2 56952507 56952606
refGene_gn=COPG1;refGene_tx=XM_003982543.4;refGene_gn=COPG1;refGene_tx=XM_019825146.2
A2 56955034 56955165
refGene_gn=COPG1;refGene_tx=XM_003982543.4;refGene_gn=COPG1;refGene_tx=XM_019825146.2

A2 56957176 56957359
refGene_gn=HMCES;refGene_tx=XM_023250070.1;refGene_gn=HMCES;refGene_tx
=XM_023250071.1
A2 56963822 56963966
refGene_gn=HMCES;refGene_tx=XM_023250070.1;refGene_gn=HMCES;refGene_tx
=XM_023250071.1
A2 56964475 56964601
refGene_gn=HMCES;refGene_tx=XM_023250070.1;refGene_gn=HMCES;refGene_tx
=XM_023250071.1
A2 56970913 56971095
refGene_gn=HMCES;refGene_tx=XM_023250070.1;refGene_gn=HMCES;refGene_tx
=XM_023250071.1
A2 56972373 56972566
refGene_gn=HMCES;refGene_tx=XM_023250070.1;refGene_gn=HMCES;refGene_tx
=XM_023250071.1
A2 56974417 56974645
refGene_gn=HMCES;refGene_tx=XM_023250070.1;refGene_gn=HMCES;refGene_tx
=XM_023250071.1
A2 56985387 56986029
refGene_gn=LOC111559457;refGene_tx=XM_023250072.1
A2 57014307 57014403 refGene_gn=RAB7A;refGene_tx=XM_011280239.3
A2 57019319 57019448 refGene_gn=RAB7A;refGene_tx=XM_011280239.3
A2 57020996 57021215 refGene_gn=RAB7A;refGene_tx=XM_011280239.3
A2 57031394 57031521 refGene_gn=RAB7A;refGene_tx=XM_011280239.3
A2 57033041 57033094 refGene_gn=RAB7A;refGene_tx=XM_011280239.3
A2 57556266 57556527 refGene_gn=RPN1;refGene_tx=XM_023250073.1
A2 57559867 57559932 refGene_gn=RPN1;refGene_tx=XM_023250073.1
A2 57564407 57564714 refGene_gn=RPN1;refGene_tx=XM_023250073.1
A2 57572590 57572800 refGene_gn=RPN1;refGene_tx=XM_023250073.1
A2 57574705 57574898 refGene_gn=RPN1;refGene_tx=XM_023250073.1
A2 57579233 57579333 refGene_gn=RPN1;refGene_tx=XM_023250073.1
A2 57581222 57581361 refGene_gn=RPN1;refGene_tx=XM_023250073.1
A2 57581594 57581714 refGene_gn=RPN1;refGene_tx=XM_023250073.1
A2 57584831 57585077 refGene_gn=RPN1;refGene_tx=XM_023250073.1
A2 57586287 57586470 refGene_gn=RPN1;refGene_tx=XM_023250073.1
A2 57704512 57704595
refGene_gn=LOC102901658;refGene_tx=XM_019825149.2
A2 57709816 57709926
refGene_gn=LOC102901658;refGene_tx=XM_023250075.1
A2 57710803 57711094
refGene_gn=LOC102901658;refGene_tx=XM_019825149.2;refGene_gn=LOC102901
658;refGene_tx=XM_023250075.1
A2 57711455 57711663
refGene_gn=LOC102901658;refGene_tx=XM_019825149.2;refGene_gn=LOC102901
658;refGene_tx=XM_023250075.1
A2 57713910 57714139
refGene_gn=GATA2;refGene_tx=XM_011280258.3;refGene_gn=GATA2;refGene_tx
=XM_019825148.2
A2 57714573 57715215
refGene_gn=GATA2;refGene_tx=XM_011280258.3;refGene_gn=GATA2;refGene_tx

=XM_019825148.2
A2 57716996 57717142
refGene_gn=GATA2;refGene_tx=XM_011280258.3;refGene_gn=GATA2;refGene_tx
=XM_019825148.2
A2 57718966 57719092
refGene_gn=GATA2;refGene_tx=XM_011280258.3;refGene_gn=GATA2;refGene_tx
=XM_019825148.2
A2 57719591 57719891
refGene_gn=GATA2;refGene_tx=XM_011280258.3;refGene_gn=GATA2;refGene_tx
=XM_019825148.2
A2 57739148 57739841 refGene_gn=DNAJB8;refGene_tx=XM_003982554.4
A2 57789466 57789657
refGene_gn=EEFSEC;refGene_tx=XM_023250076.1;refGene_gn=EEFSEC;refGene_
tx=XM_023250078.1;refGene_gn=EEFSEC;refGene_tx=XM_023250079.1
A2 57837157 57837314
refGene_gn=EEFSEC;refGene_tx=XM_023250076.1;refGene_gn=EEFSEC;refGene_
tx=XM_023250078.1;refGene_gn=EEFSEC;refGene_tx=XM_023250079.1
A2 57854699 57855356
refGene_gn=EEFSEC;refGene_tx=XM_023250076.1;refGene_gn=EEFSEC;refGene_
tx=XM_023250078.1;refGene_gn=EEFSEC;refGene_tx=XM_023250079.1
A2 57925121 57925286
refGene_gn=EEFSEC;refGene_tx=XM_023250076.1;refGene_gn=EEFSEC;refGene_
tx=XM_023250079.1
A2 57927528 57927625
refGene_gn=EEFSEC;refGene_tx=XM_023250076.1;refGene_gn=EEFSEC;refGene_
tx=XM_023250078.1;refGene_gn=EEFSEC;refGene_tx=XM_023250079.1
A2 57952349 57952557
refGene_gn=EEFSEC;refGene_tx=XM_023250076.1;refGene_gn=EEFSEC;refGene_
tx=XM_023250078.1;refGene_gn=EEFSEC;refGene_tx=XM_023250079.1
A2 58038639 58038961
refGene_gn=EEFSEC;refGene_tx=XM_023250076.1;refGene_gn=EEFSEC;refGene_
tx=XM_023250078.1
A2 58052486 58052627
refGene_gn=RUVBL1;refGene_tx=XM_003982556.3;refGene_gn=RUVBL1;refGene_
tx=XM_019825151.1
A2 58056326 58056413
refGene_gn=RUVBL1;refGene_tx=XM_003982556.3;refGene_gn=RUVBL1;refGene_
tx=XM_019825151.1
A2 58064745 58064878
refGene_gn=RUVBL1;refGene_tx=XM_003982556.3;refGene_gn=RUVBL1;refGene_
tx=XM_019825151.1
A2 58073663 58073815
refGene_gn=RUVBL1;refGene_tx=XM_003982556.3;refGene_gn=RUVBL1;refGene_
tx=XM_019825151.1
A2 58077600 58077690 refGene_gn=RUVBL1;refGene_tx=XM_003982556.3
A2 58078479 58078629 refGene_gn=RUVBL1;refGene_tx=XM_003982556.3
A2 58080340 58080404
refGene_gn=RUVBL1;refGene_tx=XM_003982556.3;refGene_gn=RUVBL1;refGene_
tx=XM_019825151.1
A2 58081831 58082030

refGene_gn=RUVBL1;refGene_tx=XM_003982556.3;refGene_gn=RUVBL1;refGene_tx=XM_019825151.1
A2 58090722 58090825
refGene_gn=RUVBL1;refGene_tx=XM_003982556.3;refGene_gn=RUVBL1;refGene_tx=XM_019825151.1
A2 58095093 58095185
refGene_gn=RUVBL1;refGene_tx=XM_003982556.3;refGene_gn=RUVBL1;refGene_tx=XM_019825151.1
A2 58096198 58096358
refGene_gn=RUVBL1;refGene_tx=XM_003982556.3;refGene_gn=RUVBL1;refGene_tx=XM_019825151.1
A2 58107462 58107649 refGene_gn=SEC61A1;refGene_tx=XM_003982557.5
A2 58109328 58109405 refGene_gn=SEC61A1;refGene_tx=XM_003982557.5
A2 58109794 58109986 refGene_gn=SEC61A1;refGene_tx=XM_003982557.5
A2 58110258 58110456 refGene_gn=SEC61A1;refGene_tx=XM_003982557.5
A2 58111687 58111848 refGene_gn=SEC61A1;refGene_tx=XM_003982557.5
A2 58115447 58115601 refGene_gn=SEC61A1;refGene_tx=XM_003982557.5
A2 58115897 58116007 refGene_gn=SEC61A1;refGene_tx=XM_003982557.5
A2 58117937 58118069 refGene_gn=SEC61A1;refGene_tx=XM_003982557.5
A2 58119033 58119112 refGene_gn=SEC61A1;refGene_tx=XM_003982557.5
A2 58119201 58119267 refGene_gn=SEC61A1;refGene_tx=XM_003982557.5
A2 58122457 58122525 refGene_gn=SEC61A1;refGene_tx=XM_003982557.5
A2 58122785 58122792 refGene_gn=SEC61A1;refGene_tx=XM_003982557.5
A2 58153878 58154060 refGene_gn=KBTBD12;refGene_tx=XM_011280253.3
A2 58176056 58176254 refGene_gn=KBTBD12;refGene_tx=XM_011280253.3
A2 58213664 58213815 refGene_gn=KBTBD12;refGene_tx=XM_011280253.3
A2 58215908 58216179 refGene_gn=KBTBD12;refGene_tx=XM_011280253.3
A2 58219662 58220423 refGene_gn=KBTBD12;refGene_tx=XM_011280253.3
A2 58220424 58220730 refGene_gn=KBTBD12;refGene_tx=XM_011280253.3
A2 58265249 58265289
refGene_gn=MGLL;refGene_tx=XM_023250083.1;refGene_gn=MGLL;refGene_tx=XM_023250085.1
A2 58308183 58308292
refGene_gn=MGLL;refGene_tx=XM_023250081.1;refGene_gn=MGLL;refGene_tx=XM_023250082.1
A2 58308771 58308916
refGene_gn=MGLL;refGene_tx=XM_023250081.1;refGene_gn=MGLL;refGene_tx=XM_023250082.1;refGene_gn=MGLL;refGene_tx=XM_023250083.1;refGene_gn=MGLL;refGene_tx=XM_023250084.1;refGene_gn=MGLL;refGene_tx=XM_023250085.1
A2 58350513 58350620
refGene_gn=MGLL;refGene_tx=XM_023250081.1;refGene_gn=MGLL;refGene_tx=XM_023250082.1;refGene_gn=MGLL;refGene_tx=XM_023250083.1;refGene_gn=MGLL;refGene_tx=XM_023250084.1
A2 58393767 58393904
refGene_gn=MGLL;refGene_tx=XM_023250081.1;refGene_gn=MGLL;refGene_tx=XM_023250082.1;refGene_gn=MGLL;refGene_tx=XM_023250083.1;refGene_gn=MGLL;refGene_tx=XM_023250084.1;refGene_gn=MGLL;refGene_tx=XM_023250085.1
A2 58395227 58395338
refGene_gn=MGLL;refGene_tx=XM_023250081.1;refGene_gn=MGLL;refGene_tx=XM_023250082.1;refGene_gn=MGLL;refGene_tx=XM_023250083.1;refGene_gn=MGL

L;refGene_tx=XM_023250084.1;refGene_gn=MGLL;refGene_tx=XM_023250085.1
A2 58400962 58401031
refGene_gn=MGLL;refGene_tx=XM_023250081.1;refGene_gn=MGLL;refGene_tx=XM_023250083.1;refGene_gn=MGLL;refGene_tx=XM_023250084.1;refGene_gn=MGLL;refGene_tx=XM_023250085.1
A2 58406156 58406246
refGene_gn=MGLL;refGene_tx=XM_023250081.1;refGene_gn=MGLL;refGene_tx=XM_023250082.1;refGene_gn=MGLL;refGene_tx=XM_023250083.1;refGene_gn=MGLL;refGene_tx=XM_023250084.1;refGene_gn=MGLL;refGene_tx=XM_023250085.1
A2 58420839 58421055
refGene_gn=MGLL;refGene_tx=XM_023250081.1;refGene_gn=MGLL;refGene_tx=XM_023250082.1;refGene_gn=MGLL;refGene_tx=XM_023250083.1;refGene_gn=MGLL;refGene_tx=XM_023250084.1;refGene_gn=MGLL;refGene_tx=XM_023250085.1
A2 58424526 58424652
refGene_gn=MGLL;refGene_tx=XM_023250081.1;refGene_gn=MGLL;refGene_tx=XM_023250082.1;refGene_gn=MGLL;refGene_tx=XM_023250083.1;refGene_gn=MGLL;refGene_tx=XM_023250084.1;refGene_gn=MGLL;refGene_tx=XM_023250085.1
A2 58433639 58433846
refGene_gn=ABTB1;refGene_tx=XM_011280251.3;refGene_gn=ABTB1;refGene_tx=XM_011280252.3
A2 58433925 58434126
refGene_gn=ABTB1;refGene_tx=XM_011280251.3;refGene_gn=ABTB1;refGene_tx=XM_011280252.3
A2 58436060 58436228
refGene_gn=ABTB1;refGene_tx=XM_011280251.3;refGene_gn=ABTB1;refGene_tx=XM_011280252.3
A2 58436334 58436433
refGene_gn=ABTB1;refGene_tx=XM_011280251.3;refGene_gn=ABTB1;refGene_tx=XM_011280252.3
A2 58436697 58436816
refGene_gn=ABTB1;refGene_tx=XM_011280251.3;refGene_gn=ABTB1;refGene_tx=XM_011280252.3
A2 58436901 58437018
refGene_gn=ABTB1;refGene_tx=XM_011280251.3;refGene_gn=ABTB1;refGene_tx=XM_011280252.3
A2 58437364 58437410
refGene_gn=ABTB1;refGene_tx=XM_011280251.3;refGene_gn=ABTB1;refGene_tx=XM_011280252.3
A2 58437506 58437666
refGene_gn=ABTB1;refGene_tx=XM_011280251.3;refGene_gn=ABTB1;refGene_tx=XM_011280252.3
A2 58437812 58437957
refGene_gn=ABTB1;refGene_tx=XM_011280251.3;refGene_gn=ABTB1;refGene_tx=XM_011280252.3
A2 58438994 58439050
refGene_gn=ABTB1;refGene_tx=XM_011280251.3;refGene_gn=ABTB1;refGene_tx=XM_011280252.3
A2 58439139 58439202
refGene_gn=ABTB1;refGene_tx=XM_011280251.3;refGene_gn=ABTB1;refGene_tx=XM_011280252.3

A2 58440425 58440481
refGene_gn=ABTB1;refGene_tx=XM_011280251.3;refGene_gn=ABTB1;refGene_tx
=XM_011280252.3
A2 58441026 58441239 refGene_gn=PODXL2;refGene_tx=XM_023250087.1
A2 58441710 58441890 refGene_gn=PODXL2;refGene_tx=XM_023250087.1
A2 58444186 58444248 refGene_gn=PODXL2;refGene_tx=XM_023250087.1
A2 58444805 58444962 refGene_gn=PODXL2;refGene_tx=XM_023250087.1
A2 58456288 58456363 refGene_gn=PODXL2;refGene_tx=XM_023250087.1
A2 58457054 58457651 refGene_gn=PODXL2;refGene_tx=XM_023250087.1
A2 58457652 58457852 refGene_gn=PODXL2;refGene_tx=XM_023250087.1
A2 58474652 58474931 refGene_gn=PODXL2;refGene_tx=XM_023250087.1
A2 58480570 58480640 refGene_gn=PODXL2;refGene_tx=XM_023250087.1
A2 58491583 58491694
refGene_gn=MCM2;refGene_tx=XM_003982560.4;refGene_gn=MCM2;refGene_tx=X
M_019825158.1
A2 58492147 58492303
refGene_gn=MCM2;refGene_tx=XM_003982560.4;refGene_gn=MCM2;refGene_tx=X
M_019825158.1
A2 58492547 58492730
refGene_gn=MCM2;refGene_tx=XM_003982560.4;refGene_gn=MCM2;refGene_tx=X
M_019825158.1
A2 58493477 58493726
refGene_gn=MCM2;refGene_tx=XM_003982560.4;refGene_gn=MCM2;refGene_tx=X
M_019825158.1
A2 58494632 58494745
refGene_gn=MCM2;refGene_tx=XM_003982560.4;refGene_gn=MCM2;refGene_tx=X
M_019825158.1
A2 58495347 58495474
refGene_gn=MCM2;refGene_tx=XM_003982560.4;refGene_gn=MCM2;refGene_tx=X
M_019825158.1
A2 58495594 58495845
refGene_gn=MCM2;refGene_tx=XM_003982560.4;refGene_gn=MCM2;refGene_tx=X
M_019825158.1
A2 58497107 58497201
refGene_gn=MCM2;refGene_tx=XM_003982560.4;refGene_gn=MCM2;refGene_tx=X
M_019825158.1
A2 58503370 58503562
refGene_gn=MCM2;refGene_tx=XM_003982560.4;refGene_gn=MCM2;refGene_tx=X
M_019825158.1
A2 58503879 58504014
refGene_gn=MCM2;refGene_tx=XM_003982560.4;refGene_gn=MCM2;refGene_tx=X
M_019825158.1
A2 58505546 58505754
refGene_gn=MCM2;refGene_tx=XM_003982560.4;refGene_gn=MCM2;refGene_tx=X
M_019825158.1
A2 58506020 58506240
refGene_gn=MCM2;refGene_tx=XM_003982560.4;refGene_gn=MCM2;refGene_tx=X
M_019825158.1
A2 58507229 58507490
refGene_gn=MCM2;refGene_tx=XM_003982560.4;refGene_gn=MCM2;refGene_tx=X

M_019825158.1
A2	58507612	58507788
refGene_gn=MCM2;refGene_tx=XM_003982560.4;refGene_gn=MCM2;refGene_tx=XM_019825158.1
A2	58512038	58512268
refGene_gn=MCM2;refGene_tx=XM_003982560.4;refGene_gn=MCM2;refGene_tx=XM_019825158.1
A2	58512732	58512816	refGene_gn=MCM2;refGene_tx=XM_019825158.1
A2	58513174	58513180	refGene_gn=MCM2;refGene_tx=XM_003982560.4
A2	58526088	58526213
refGene_gn=TPRA1;refGene_tx=XM_011280249.3;refGene_gn=TPRA1;refGene_tx=XM_011280250.3
A2	58526367	58526500
refGene_gn=TPRA1;refGene_tx=XM_011280249.3;refGene_gn=TPRA1;refGene_tx=XM_011280250.3
A2	58529173	58529260
refGene_gn=TPRA1;refGene_tx=XM_011280249.3;refGene_gn=TPRA1;refGene_tx=XM_011280250.3
A2	58529359	58529432
refGene_gn=TPRA1;refGene_tx=XM_011280249.3;refGene_gn=TPRA1;refGene_tx=XM_011280250.3
A2	58529541	58529621
refGene_gn=TPRA1;refGene_tx=XM_011280249.3;refGene_gn=TPRA1;refGene_tx=XM_011280250.3
A2	58530207	58530318
refGene_gn=TPRA1;refGene_tx=XM_011280249.3;refGene_gn=TPRA1;refGene_tx=XM_011280250.3
A2	58530448	58530509
refGene_gn=TPRA1;refGene_tx=XM_011280249.3;refGene_gn=TPRA1;refGene_tx=XM_011280250.3
A2	58530695	58530798
refGene_gn=TPRA1;refGene_tx=XM_011280249.3;refGene_gn=TPRA1;refGene_tx=XM_011280250.3
A2	58530929	58531010
refGene_gn=TPRA1;refGene_tx=XM_011280249.3;refGene_gn=TPRA1;refGene_tx=XM_011280250.3
A2	58532279	58532523
refGene_gn=TPRA1;refGene_tx=XM_011280249.3;refGene_gn=TPRA1;refGene_tx=XM_011280250.3
A2	58765101	58765155
refGene_gn=LOC102899298;refGene_tx=XM_011280264.3
A2	58766679	58766722
refGene_gn=LOC102899298;refGene_tx=XM_011280264.3
A2	58772436	58772984
refGene_gn=LOC102899298;refGene_tx=XM_011280264.3
A2	58830913	58830943
refGene_gn=CA2H3orf56;refGene_tx=XM_011280263.2
A2	58831038	58831389
refGene_gn=CA2H3orf56;refGene_tx=XM_011280263.2
A2	58831391	58831547

refGene_gn=CA2H3orf56;refGene_tx=XM_011280263.2
A2 58998438 58998534
refGene_gn=PLXNA1;refGene_tx=XM_003982562.5;refGene_gn=PLXNA1;refGene_tx=XM_023250088.1
A2 59000322 59000473
refGene_gn=PLXNA1;refGene_tx=XM_003982562.5;refGene_gn=PLXNA1;refGene_tx=XM_023250088.1
A2 59000552 59000765
refGene_gn=PLXNA1;refGene_tx=XM_003982562.5;refGene_gn=PLXNA1;refGene_tx=XM_023250088.1
A2 59002324 59002494
refGene_gn=PLXNA1;refGene_tx=XM_003982562.5;refGene_gn=PLXNA1;refGene_tx=XM_023250088.1
A2 59002661 59002852
refGene_gn=PLXNA1;refGene_tx=XM_003982562.5;refGene_gn=PLXNA1;refGene_tx=XM_023250088.1
A2 59003186 59003283
refGene_gn=PLXNA1;refGene_tx=XM_003982562.5;refGene_gn=PLXNA1;refGene_tx=XM_023250088.1
A2 59003535 59003639
refGene_gn=PLXNA1;refGene_tx=XM_003982562.5;refGene_gn=PLXNA1;refGene_tx=XM_023250088.1
A2 59004428 59004588
refGene_gn=PLXNA1;refGene_tx=XM_003982562.5;refGene_gn=PLXNA1;refGene_tx=XM_023250088.1
A2 59004696 59004843
refGene_gn=PLXNA1;refGene_tx=XM_003982562.5;refGene_gn=PLXNA1;refGene_tx=XM_023250088.1
A2 59007367 59007434
refGene_gn=PLXNA1;refGene_tx=XM_003982562.5;refGene_gn=PLXNA1;refGene_tx=XM_023250088.1
A2 59007705 59007962
refGene_gn=PLXNA1;refGene_tx=XM_003982562.5;refGene_gn=PLXNA1;refGene_tx=XM_023250088.1
A2 59009463 59009606
refGene_gn=PLXNA1;refGene_tx=XM_003982562.5;refGene_gn=PLXNA1;refGene_tx=XM_023250088.1
A2 59012059 59012294
refGene_gn=PLXNA1;refGene_tx=XM_003982562.5;refGene_gn=PLXNA1;refGene_tx=XM_023250088.1
A2 59012596 59012740
refGene_gn=PLXNA1;refGene_tx=XM_003982562.5;refGene_gn=PLXNA1;refGene_tx=XM_023250088.1
A2 59012824 59013064
refGene_gn=PLXNA1;refGene_tx=XM_003982562.5;refGene_gn=PLXNA1;refGene_tx=XM_023250088.1
A2 59013263 59013357
refGene_gn=PLXNA1;refGene_tx=XM_003982562.5;refGene_gn=PLXNA1;refGene_tx=XM_023250088.1
A2 59013843 59014011

refGene_gn=PLXNA1;refGene_tx=XM_003982562.5;refGene_gn=PLXNA1;refGene_tx=XM_023250088.1
A2 59014965 59015102
refGene_gn=PLXNA1;refGene_tx=XM_003982562.5;refGene_gn=PLXNA1;refGene_tx=XM_023250088.1
A2 59015410 59015531
refGene_gn=PLXNA1;refGene_tx=XM_003982562.5;refGene_gn=PLXNA1;refGene_tx=XM_023250088.1
A2 59015629 59015781
refGene_gn=PLXNA1;refGene_tx=XM_003982562.5;refGene_gn=PLXNA1;refGene_tx=XM_023250088.1
A2 59015905 59016099
refGene_gn=PLXNA1;refGene_tx=XM_003982562.5;refGene_gn=PLXNA1;refGene_tx=XM_023250088.1
A2 59016169 59016266
refGene_gn=PLXNA1;refGene_tx=XM_003982562.5;refGene_gn=PLXNA1;refGene_tx=XM_023250088.1
A2 59018209 59018410
refGene_gn=PLXNA1;refGene_tx=XM_003982562.5;refGene_gn=PLXNA1;refGene_tx=XM_023250088.1
A2 59020770 59020885
refGene_gn=PLXNA1;refGene_tx=XM_003982562.5;refGene_gn=PLXNA1;refGene_tx=XM_023250088.1
A2 59022038 59022138
refGene_gn=PLXNA1;refGene_tx=XM_003982562.5;refGene_gn=PLXNA1;refGene_tx=XM_023250088.1
A2 59022468 59022622
refGene_gn=PLXNA1;refGene_tx=XM_003982562.5;refGene_gn=PLXNA1;refGene_tx=XM_023250088.1
A2 59022723 59022847
refGene_gn=PLXNA1;refGene_tx=XM_003982562.5;refGene_gn=PLXNA1;refGene_tx=XM_023250088.1
A2 59023090 59023191
refGene_gn=PLXNA1;refGene_tx=XM_003982562.5;refGene_gn=PLXNA1;refGene_tx=XM_023250088.1
A2 59024020 59024161
refGene_gn=PLXNA1;refGene_tx=XM_003982562.5;refGene_gn=PLXNA1;refGene_tx=XM_023250088.1
A2 59035487 59035670
refGene_gn=PLXNA1;refGene_tx=XM_003982562.5;refGene_gn=PLXNA1;refGene_tx=XM_023250088.1
A2 59037454 59038813
refGene_gn=PLXNA1;refGene_tx=XM_003982562.5;refGene_gn=PLXNA1;refGene_tx=XM_023250088.1
A2 59062146 59062152
refGene_gn=CHCHD6;refGene_tx=XM_006929038.3;refGene_gn=CHCHD6;refGene_tx=XM_019825159.1
A2 59064806 59064942
refGene_gn=CHCHD6;refGene_tx=XM_006929038.3;refGene_gn=CHCHD6;refGene_tx=XM_019825159.1

A2	59109530	59109601
refGene_gn=CHCHD6;refGene_tx=XM_006929038.3;refGene_gn=CHCHD6;refGene_tx=XM_019825159.1
A2	59158830	59158944	refGene_gn=CHCHD6;refGene_tx=XM_019825160.2
A2	59178385	59178469
refGene_gn=CHCHD6;refGene_tx=XM_006929038.3;refGene_gn=CHCHD6;refGene_tx=XM_019825159.1;refGene_gn=CHCHD6;refGene_tx=XM_019825160.2
A2	59287299	59287444
refGene_gn=CHCHD6;refGene_tx=XM_006929038.3;refGene_gn=CHCHD6;refGene_tx=XM_019825159.1;refGene_gn=CHCHD6;refGene_tx=XM_019825160.2
A2	59290198	59290271
refGene_gn=CHCHD6;refGene_tx=XM_006929038.3;refGene_gn=CHCHD6;refGene_tx=XM_019825159.1;refGene_gn=CHCHD6;refGene_tx=XM_019825160.2
A2	59293974	59294095
refGene_gn=CHCHD6;refGene_tx=XM_006929038.3;refGene_gn=CHCHD6;refGene_tx=XM_019825159.1;refGene_gn=CHCHD6;refGene_tx=XM_019825160.2
A2	59302027	59302042	refGene_gn=CHCHD6;refGene_tx=XM_019825159.1
A2	59313677	59313764
refGene_gn=CHCHD6;refGene_tx=XM_006929038.3;refGene_gn=CHCHD6;refGene_tx=XM_019825160.2
A2	59348663	59348822	refGene_gn=TXNRD3;refGene_tx=XM_023250090.1
A2	59363793	59363854	refGene_gn=TXNRD3;refGene_tx=XM_023250090.1
A2	59365129	59365239
refGene_gn=TXNRD3;refGene_tx=XM_011280248.3;refGene_gn=TXNRD3;refGene_tx=XM_023250090.1
A2	59367867	59367972
refGene_gn=TXNRD3;refGene_tx=XM_011280248.3;refGene_gn=TXNRD3;refGene_tx=XM_023250090.1
A2	59368420	59368493
refGene_gn=TXNRD3;refGene_tx=XM_011280248.3;refGene_gn=TXNRD3;refGene_tx=XM_023250090.1
A2	59370888	59371008
refGene_gn=TXNRD3;refGene_tx=XM_011280248.3;refGene_gn=TXNRD3;refGene_tx=XM_023250090.1
A2	59378474	59378617
refGene_gn=TXNRD3;refGene_tx=XM_011280248.3;refGene_gn=TXNRD3;refGene_tx=XM_023250090.1
A2	59380599	59380715
refGene_gn=TXNRD3;refGene_tx=XM_011280248.3;refGene_gn=TXNRD3;refGene_tx=XM_023250090.1
A2	59381986	59382212
refGene_gn=TXNRD3;refGene_tx=XM_011280248.3;refGene_gn=TXNRD3;refGene_tx=XM_023250090.1
A2	59383658	59383751
refGene_gn=TXNRD3;refGene_tx=XM_011280248.3;refGene_gn=TXNRD3;refGene_tx=XM_023250090.1
A2	59390453	59390530
refGene_gn=TXNRD3;refGene_tx=XM_011280248.3;refGene_gn=TXNRD3;refGene_tx=XM_023250090.1
A2	59391581	59391738

refGene_gn=TXNRD3;refGene_tx=XM_011280248.3;refGene_gn=TXNRD3;refGene_tx=XM_023250090.1
A2 59395667 59395775
refGene_gn=TXNRD3;refGene_tx=XM_011280248.3;refGene_gn=TXNRD3;refGene_tx=XM_023250090.1
A2 59413109 59413205
refGene_gn=TXNRD3;refGene_tx=XM_011280248.3;refGene_gn=TXNRD3;refGene_tx=XM_023250090.1
A2 59415533 59415668
refGene_gn=TXNRD3;refGene_tx=XM_011280248.3;refGene_gn=TXNRD3;refGene_tx=XM_023250090.1
A2 59415888 59415951
refGene_gn=TXNRD3;refGene_tx=XM_011280248.3;refGene_gn=TXNRD3;refGene_tx=XM_023250090.1
A2 59453527 59453571
refGene_gn=CA2H3orf22;refGene_tx=XM_003982571.4
A2 59454483 59454686
refGene_gn=CA2H3orf22;refGene_tx=XM_023250091.1
A2 59456944 59457034
refGene_gn=CA2H3orf22;refGene_tx=XM_003982571.4;refGene_gn=CA2H3orf22;refGene_tx=XM_011280246.3;refGene_gn=CA2H3orf22;refGene_tx=XM_011280247.3;refGene_gn=CA2H3orf22;refGene_tx=XM_019825166.2;refGene_gn=CA2H3orf22;refGene_tx=XM_023250091.1
A2 59457739 59457871
refGene_gn=CA2H3orf22;refGene_tx=XM_003982571.4;refGene_gn=CA2H3orf22;refGene_tx=XM_011280246.3;refGene_gn=CA2H3orf22;refGene_tx=XM_011280247.3;refGene_gn=CA2H3orf22;refGene_tx=XM_019825166.2;refGene_gn=CA2H3orf22;refGene_tx=XM_023250091.1
A2 59459871 59460043
refGene_gn=CA2H3orf22;refGene_tx=XM_003982571.4;refGene_gn=CA2H3orf22;refGene_tx=XM_011280246.3;refGene_gn=CA2H3orf22;refGene_tx=XM_011280247.3;refGene_gn=CA2H3orf22;refGene_tx=XM_019825166.2;refGene_gn=CA2H3orf22;refGene_tx=XM_023250091.1
A2 59466555 59467401
refGene_gn=CHST13;refGene_tx=XM_019825163.2;refGene_gn=CHST13;refGene_tx=XM_023250093.1
A2 59472841 59472921
refGene_gn=CHST13;refGene_tx=XM_019825163.2;refGene_gn=CHST13;refGene_tx=XM_023250093.1
A2 59492143 59492240 refGene_gn=CHST13;refGene_tx=XM_023250093.1
A2 59499391 59499517 refGene_gn=UROC1;refGene_tx=XM_003982564.3
A2 59509311 59509442 refGene_gn=UROC1;refGene_tx=XM_003982564.3
A2 59510525 59510619 refGene_gn=UROC1;refGene_tx=XM_003982564.3
A2 59511606 59511666 refGene_gn=UROC1;refGene_tx=XM_003982564.3
A2 59511989 59512118 refGene_gn=UROC1;refGene_tx=XM_003982564.3
A2 59512257 59512319 refGene_gn=UROC1;refGene_tx=XM_003982564.3
A2 59516105 59516172 refGene_gn=UROC1;refGene_tx=XM_003982564.3
A2 59516257 59516401 refGene_gn=UROC1;refGene_tx=XM_003982564.3
A2 59518289 59518378 refGene_gn=UROC1;refGene_tx=XM_003982564.3
A2 59519548 59519611 refGene_gn=UROC1;refGene_tx=XM_003982564.3

A2 59519904 59520084 refGene_gn=UROC1;refGene_tx=XM_003982564.3
A2 59520366 59520464 refGene_gn=UROC1;refGene_tx=XM_003982564.3
A2 59521261 59521334 refGene_gn=UROC1;refGene_tx=XM_003982564.3
A2 59522195 59522317 refGene_gn=UROC1;refGene_tx=XM_003982564.3
A2 59524216 59524287 refGene_gn=UROC1;refGene_tx=XM_003982564.3
A2 59527430 59527529 refGene_gn=UROC1;refGene_tx=XM_003982564.3
A2 59530115 59530215 refGene_gn=UROC1;refGene_tx=XM_003982564.3
A2 59533536 59533618 refGene_gn=UROC1;refGene_tx=XM_003982564.3
A2 59537956 59538056 refGene_gn=UROC1;refGene_tx=XM_003982564.3
A2 59539191 59539332 refGene_gn=UROC1;refGene_tx=XM_003982564.3
A2 59555919 59556889
refGene_gn=ZXDC;refGene_tx=XM_006929041.4;refGene_gn=ZXDC;refGene_tx=XM_023250094.1;refGene_gn=ZXDC;refGene_tx=XM_023250095.1;refGene_gn=ZXDC;refGene_tx=XM_023250097.1
A2 59559411 59559564
refGene_gn=ZXDC;refGene_tx=XM_006929041.4;refGene_gn=ZXDC;refGene_tx=XM_023250094.1;refGene_gn=ZXDC;refGene_tx=XM_023250095.1;refGene_gn=ZXDC;refGene_tx=XM_023250097.1
A2 59559662 59559741
refGene_gn=ZXDC;refGene_tx=XM_006929041.4;refGene_gn=ZXDC;refGene_tx=XM_023250094.1;refGene_gn=ZXDC;refGene_tx=XM_023250095.1;refGene_gn=ZXDC;refGene_tx=XM_023250097.1
A2 59560662 59560793
refGene_gn=ZXDC;refGene_tx=XM_006929041.4;refGene_gn=ZXDC;refGene_tx=XM_023250094.1;refGene_gn=ZXDC;refGene_tx=XM_023250095.1;refGene_gn=ZXDC;refGene_tx=XM_023250097.1
A2 59565906 59566077
refGene_gn=ZXDC;refGene_tx=XM_006929041.4;refGene_gn=ZXDC;refGene_tx=XM_023250094.1;refGene_gn=ZXDC;refGene_tx=XM_023250095.1;refGene_gn=ZXDC;refGene_tx=XM_023250097.1
A2 59569703 59570392
refGene_gn=ZXDC;refGene_tx=XM_006929041.4;refGene_gn=ZXDC;refGene_tx=XM_023250094.1;refGene_gn=ZXDC;refGene_tx=XM_023250095.1;refGene_gn=ZXDC;refGene_tx=XM_023250097.1
A2 59571555 59571618
refGene_gn=ZXDC;refGene_tx=XM_006929041.4;refGene_gn=ZXDC;refGene_tx=XM_023250095.1
A2 59572276 59572361
refGene_gn=ZXDC;refGene_tx=XM_006929041.4;refGene_gn=ZXDC;refGene_tx=XM_023250094.1;refGene_gn=ZXDC;refGene_tx=XM_023250095.1;refGene_gn=ZXDC;refGene_tx=XM_023250097.1
A2 59589380 59589550
refGene_gn=ZXDC;refGene_tx=XM_006929041.4;refGene_gn=ZXDC;refGene_tx=XM_023250094.1;refGene_gn=ZXDC;refGene_tx=XM_023250095.1;refGene_gn=ZXDC;refGene_tx=XM_023250097.1
A2 59590898 59591000
refGene_gn=ZXDC;refGene_tx=XM_006929041.4;refGene_gn=ZXDC;refGene_tx=XM_023250094.1;refGene_gn=ZXDC;refGene_tx=XM_023250095.1;refGene_gn=ZXDC;refGene_tx=XM_023250097.1
A2 59592172 59592259

refGene_gn=ZXDC;refGene_tx=XM_006929041.4;refGene_gn=ZXDC;refGene_tx=XM_023250094.1;refGene_gn=ZXDC;refGene_tx=XM_023250095.1;refGene_gn=ZXDC;refGene_tx=XM_023250097.1
A2 59594379 59594493
refGene_gn=CFAP100;refGene_tx=XM_011280245.3;refGene_gn=CFAP100;refGene_tx=XM_019825167.2;refGene_gn=CFAP100;refGene_tx=XM_019825169.2
A2 59595437 59595531
refGene_gn=CFAP100;refGene_tx=XM_011280245.3;refGene_gn=CFAP100;refGene_tx=XM_019825167.2;refGene_gn=CFAP100;refGene_tx=XM_019825169.2
A2 59596488 59596694
refGene_gn=CFAP100;refGene_tx=XM_011280245.3;refGene_gn=CFAP100;refGene_tx=XM_019825167.2;refGene_gn=CFAP100;refGene_tx=XM_019825169.2
A2 59597826 59597962
refGene_gn=CFAP100;refGene_tx=XM_011280245.3;refGene_gn=CFAP100;refGene_tx=XM_019825167.2;refGene_gn=CFAP100;refGene_tx=XM_019825169.2
A2 59615727 59615992 refGene_gn=CFAP100;refGene_tx=XM_019825170.2
A2 59616333 59616485
refGene_gn=CFAP100;refGene_tx=XM_011280245.3;refGene_gn=CFAP100;refGene_tx=XM_019825167.2;refGene_gn=CFAP100;refGene_tx=XM_019825169.2;refGene_gn=CFAP100;refGene_tx=XM_019825170.2
A2 59616570 59616619
refGene_gn=CFAP100;refGene_tx=XM_011280245.3;refGene_gn=CFAP100;refGene_tx=XM_019825167.2;refGene_gn=CFAP100;refGene_tx=XM_019825169.2;refGene_gn=CFAP100;refGene_tx=XM_019825170.2
A2 59620639 59620766
refGene_gn=CFAP100;refGene_tx=XM_011280245.3;refGene_gn=CFAP100;refGene_tx=XM_019825167.2;refGene_gn=CFAP100;refGene_tx=XM_019825169.2;refGene_gn=CFAP100;refGene_tx=XM_019825170.2
A2 59620842 59620875
refGene_gn=CFAP100;refGene_tx=XM_011280245.3;refGene_gn=CFAP100;refGene_tx=XM_019825167.2;refGene_gn=CFAP100;refGene_tx=XM_019825170.2
A2 59621254 59621436
refGene_gn=CFAP100;refGene_tx=XM_011280245.3;refGene_gn=CFAP100;refGene_tx=XM_019825167.2;refGene_gn=CFAP100;refGene_tx=XM_019825169.2;refGene_gn=CFAP100;refGene_tx=XM_019825170.2
A2 59621831 59621912
refGene_gn=CFAP100;refGene_tx=XM_011280245.3;refGene_gn=CFAP100;refGene_tx=XM_019825167.2;refGene_gn=CFAP100;refGene_tx=XM_019825169.2;refGene_gn=CFAP100;refGene_tx=XM_019825170.2
A2 59622122 59622286
refGene_gn=CFAP100;refGene_tx=XM_011280245.3;refGene_gn=CFAP100;refGene_tx=XM_019825167.2;refGene_gn=CFAP100;refGene_tx=XM_019825169.2;refGene_gn=CFAP100;refGene_tx=XM_019825170.2
A2 59622378 59622446
refGene_gn=CFAP100;refGene_tx=XM_011280245.3;refGene_gn=CFAP100;refGene_tx=XM_019825167.2;refGene_gn=CFAP100;refGene_tx=XM_019825169.2;refGene_gn=CFAP100;refGene_tx=XM_019825170.2
A2 59624614 59624807
refGene_gn=CFAP100;refGene_tx=XM_011280245.3;refGene_gn=CFAP100;refGene_tx=XM_019825167.2;refGene_gn=CFAP100;refGene_tx=XM_019825169.2;refGe

ne_gn=CFAP100;refGene_tx=XM_019825170.2
A2 59627369 59627470
refGene_gn=CFAP100;refGene_tx=XM_011280245.3;refGene_gn=CFAP100;refGene_tx=XM_019825167.2;refGene_gn=CFAP100;refGene_tx=XM_019825169.2;refGene_gn=CFAP100;refGene_tx=XM_019825170.2
A2 59634558 59634633
refGene_gn=CFAP100;refGene_tx=XM_011280245.3;refGene_gn=CFAP100;refGene_tx=XM_019825167.2;refGene_gn=CFAP100;refGene_tx=XM_019825169.2;refGene_gn=CFAP100;refGene_tx=XM_019825170.2
A2 59644435 59644484
refGene_gn=CFAP100;refGene_tx=XM_011280245.3;refGene_gn=CFAP100;refGene_tx=XM_019825167.2;refGene_gn=CFAP100;refGene_tx=XM_019825169.2;refGene_gn=CFAP100;refGene_tx=XM_019825170.2
A2 59686648 59686752 refGene_gn=KLF15;refGene_tx=XM_011280244.3
A2 59688040 59689141
refGene_gn=KLF15;refGene_tx=XM_011280244.3;refGene_gn=KLF15;refGene_tx=XM_023250098.1
A2 59696822 59696991
refGene_gn=KLF15;refGene_tx=XM_011280244.3;refGene_gn=KLF15;refGene_tx=XM_023250098.1
A2 59881175 59881302
refGene_gn=ALDH1L1;refGene_tx=XM_023250099.1;refGene_gn=ALDH1L1;refGene_tx=XM_023250100.1;refGene_gn=ALDH1L1;refGene_tx=XM_023250101.1
A2 59883386 59883621
refGene_gn=ALDH1L1;refGene_tx=XM_023250099.1;refGene_gn=ALDH1L1;refGene_tx=XM_023250100.1;refGene_gn=ALDH1L1;refGene_tx=XM_023250101.1
A2 59884388 59884554
refGene_gn=ALDH1L1;refGene_tx=XM_023250099.1;refGene_gn=ALDH1L1;refGene_tx=XM_023250100.1;refGene_gn=ALDH1L1;refGene_tx=XM_023250101.1
A2 59886337 59886439
refGene_gn=ALDH1L1;refGene_tx=XM_023250099.1;refGene_gn=ALDH1L1;refGene_tx=XM_023250100.1;refGene_gn=ALDH1L1;refGene_tx=XM_023250101.1
A2 59887274 59887364
refGene_gn=ALDH1L1;refGene_tx=XM_023250099.1;refGene_gn=ALDH1L1;refGene_tx=XM_023250100.1;refGene_gn=ALDH1L1;refGene_tx=XM_023250101.1
A2 59888307 59888445
refGene_gn=ALDH1L1;refGene_tx=XM_023250099.1;refGene_gn=ALDH1L1;refGene_tx=XM_023250100.1;refGene_gn=ALDH1L1;refGene_tx=XM_023250101.1
A2 59891152 59891278
refGene_gn=ALDH1L1;refGene_tx=XM_023250099.1;refGene_gn=ALDH1L1;refGene_tx=XM_023250100.1;refGene_gn=ALDH1L1;refGene_tx=XM_023250101.1
A2 59894458 59894550
refGene_gn=ALDH1L1;refGene_tx=XM_023250099.1;refGene_gn=ALDH1L1;refGene_tx=XM_023250100.1;refGene_gn=ALDH1L1;refGene_tx=XM_023250101.1
A2 59928905 59929053
refGene_gn=ALDH1L1;refGene_tx=XM_023250099.1;refGene_gn=ALDH1L1;refGene_tx=XM_023250100.1;refGene_gn=ALDH1L1;refGene_tx=XM_023250101.1
A2 59930672 59930792
refGene_gn=ALDH1L1;refGene_tx=XM_023250099.1;refGene_gn=ALDH1L1;refGene_tx=XM_023250100.1;refGene_gn=ALDH1L1;refGene_tx=XM_023250101.1

A2 59932062 59932190
refGene_gn=ALDH1L1;refGene_tx=XM_023250099.1;refGene_gn=ALDH1L1;refGene_tx=XM_023250100.1;refGene_gn=ALDH1L1;refGene_tx=XM_023250101.1
A2 59936077 59936228
refGene_gn=ALDH1L1;refGene_tx=XM_023250099.1;refGene_gn=ALDH1L1;refGene_tx=XM_023250100.1;refGene_gn=ALDH1L1;refGene_tx=XM_023250101.1
A2 59937387 59937458
refGene_gn=ALDH1L1;refGene_tx=XM_023250099.1;refGene_gn=ALDH1L1;refGene_tx=XM_023250100.1;refGene_gn=ALDH1L1;refGene_tx=XM_023250101.1
A2 59940565 59940671
refGene_gn=ALDH1L1;refGene_tx=XM_023250099.1;refGene_gn=ALDH1L1;refGene_tx=XM_023250100.1;refGene_gn=ALDH1L1;refGene_tx=XM_023250101.1
A2 59942125 59942213
refGene_gn=ALDH1L1;refGene_tx=XM_023250099.1;refGene_gn=ALDH1L1;refGene_tx=XM_023250100.1;refGene_gn=ALDH1L1;refGene_tx=XM_023250101.1
A2 59963696 59963790
refGene_gn=ALDH1L1;refGene_tx=XM_023250099.1;refGene_gn=ALDH1L1;refGene_tx=XM_023250100.1;refGene_gn=ALDH1L1;refGene_tx=XM_023250101.1
A2 59966211 59966311
refGene_gn=ALDH1L1;refGene_tx=XM_023250099.1;refGene_gn=ALDH1L1;refGene_tx=XM_023250100.1;refGene_gn=ALDH1L1;refGene_tx=XM_023250101.1
A2 59967591 59967690
refGene_gn=ALDH1L1;refGene_tx=XM_023250099.1;refGene_gn=ALDH1L1;refGene_tx=XM_023250100.1;refGene_gn=ALDH1L1;refGene_tx=XM_023250101.1
A2 59970639 59970805
refGene_gn=ALDH1L1;refGene_tx=XM_023250099.1;refGene_gn=ALDH1L1;refGene_tx=XM_023250100.1;refGene_gn=ALDH1L1;refGene_tx=XM_023250101.1
A2 59973833 59973939
refGene_gn=ALDH1L1;refGene_tx=XM_023250099.1;refGene_gn=ALDH1L1;refGene_tx=XM_023250100.1;refGene_gn=ALDH1L1;refGene_tx=XM_023250101.1
A2 59975113 59975313
refGene_gn=ALDH1L1;refGene_tx=XM_023250099.1;refGene_gn=ALDH1L1;refGene_tx=XM_023250100.1;refGene_gn=ALDH1L1;refGene_tx=XM_023250101.1
A2 59977116 59977172
refGene_gn=ALDH1L1;refGene_tx=XM_023250099.1;refGene_gn=ALDH1L1;refGene_tx=XM_023250100.1;refGene_gn=ALDH1L1;refGene_tx=XM_023250101.1
A2 62255363 62255647
refGene_gn=MRPS24;refGene_tx=XM_003982575.4;refGene_gn=MRPS24;refGene_tx=XM_019825178.2
A2 62263408 62263520
refGene_gn=MRPS24;refGene_tx=XM_003982575.4;refGene_gn=MRPS24;refGene_tx=XM_019825178.2
A2 62263605 62263704
refGene_gn=MRPS24;refGene_tx=XM_003982575.4;refGene_gn=MRPS24;refGene_tx=XM_019825178.2
A2 62263754 62263793
refGene_gn=MRPS24;refGene_tx=XM_003982575.4;refGene_gn=MRPS24;refGene_tx=XM_019825178.2
A2 62272325 62274919
refGene_gn=LOC101097628;refGene_tx=XM_011280280.3;refGene_gn=LOC101097

628;refGene_tx=XM_019825173.2;refGene_gn=LOC101097628;refGene_tx=XM_019825174.2;refGene_gn=LOC101097628;refGene_tx=XM_019825175.1;refGene_gn=LOC101097628;refGene_tx=XM_019825176.2;refGene_gn=LOC101097628;refGene_tx=XM_023250102.1;refGene_gn=LOC101097628;refGene_tx=XM_023250103.1;refGene_gn=LOC101097628;refGene_tx=XM_023250104.1;refGene_gn=LOC101097628;refGene_tx=XM_023250105.1
A2 62276689 62276728
refGene_gn=LOC101097628;refGene_tx=XM_011280280.3;refGene_gn=LOC101097628;refGene_tx=XM_019825173.2;refGene_gn=LOC101097628;refGene_tx=XM_019825174.2;refGene_gn=LOC101097628;refGene_tx=XM_019825175.1;refGene_gn=LOC101097628;refGene_tx=XM_019825176.2;refGene_gn=LOC101097628;refGene_tx=XM_023250102.1;refGene_gn=LOC101097628;refGene_tx=XM_023250103.1;refGene_gn=LOC101097628;refGene_tx=XM_023250104.1;refGene_gn=LOC101097628;refGene_tx=XM_023250105.1
A2 62276951 62276984
refGene_gn=LOC101097628;refGene_tx=XM_011280280.3;refGene_gn=LOC101097628;refGene_tx=XM_019825173.2;refGene_gn=LOC101097628;refGene_tx=XM_019825174.2;refGene_gn=LOC101097628;refGene_tx=XM_019825175.1;refGene_gn=LOC101097628;refGene_tx=XM_019825176.2;refGene_gn=LOC101097628;refGene_tx=XM_023250102.1;refGene_gn=LOC101097628;refGene_tx=XM_023250103.1;refGene_gn=LOC101097628;refGene_tx=XM_023250104.1;refGene_gn=LOC101097628;refGene_tx=XM_023250105.1
A2 62286406 62286477
refGene_gn=LOC101097628;refGene_tx=XM_023250102.1;refGene_gn=LOC101097628;refGene_tx=XM_023250103.1;refGene_gn=LOC101097628;refGene_tx=XM_023250104.1
A2 62286786 62286813
refGene_gn=LOC101097628;refGene_tx=XM_023250102.1;refGene_gn=LOC101097628;refGene_tx=XM_023250103.1
A2 62289040 62289183
refGene_gn=LOC101097628;refGene_tx=XM_023250102.1
A2 62300081 62300095
refGene_gn=LOC101097628;refGene_tx=XM_023250103.1;refGene_gn=LOC101097628;refGene_tx=XM_023250104.1
A2 62340529 62340612
refGene_gn=DBNL;refGene_tx=XM_023250106.1;refGene_gn=DBNL;refGene_tx=XM_023250107.1
A2 62340909 62340989 refGene_gn=DBNL;refGene_tx=XM_019825179.2
A2 62345099 62345155
refGene_gn=DBNL;refGene_tx=XM_019825179.2;refGene_gn=DBNL;refGene_tx=XM_023250106.1;refGene_gn=DBNL;refGene_tx=XM_023250107.1
A2 62346383 62346496
refGene_gn=DBNL;refGene_tx=XM_019825179.2;refGene_gn=DBNL;refGene_tx=XM_023250106.1;refGene_gn=DBNL;refGene_tx=XM_023250107.1;refGene_gn=DBNL;refGene_tx=XM_023250108.1
A2 62347045 62347120
refGene_gn=DBNL;refGene_tx=XM_019825179.2;refGene_gn=DBNL;refGene_tx=XM_023250106.1;refGene_gn=DBNL;refGene_tx=XM_023250107.1;refGene_gn=DBNL;refGene_tx=XM_023250108.1
A2 62348118 62348265

refGene_gn=DBNL;refGene_tx=XM_019825179.2;refGene_gn=DBNL;refGene_tx=XM_023250106.1;refGene_gn=DBNL;refGene_tx=XM_023250107.1;refGene_gn=DBNL;refGene_tx=XM_023250108.1
A2 62349015 62349093
refGene_gn=DBNL;refGene_tx=XM_019825179.2;refGene_gn=DBNL;refGene_tx=XM_023250106.1;refGene_gn=DBNL;refGene_tx=XM_023250107.1;refGene_gn=DBNL;refGene_tx=XM_023250108.1
A2 62349385 62349540
refGene_gn=DBNL;refGene_tx=XM_019825179.2;refGene_gn=DBNL;refGene_tx=XM_023250106.1;refGene_gn=DBNL;refGene_tx=XM_023250107.1;refGene_gn=DBNL;refGene_tx=XM_023250108.1
A2 62349618 62349667
refGene_gn=DBNL;refGene_tx=XM_019825179.2;refGene_gn=DBNL;refGene_tx=XM_023250106.1;refGene_gn=DBNL;refGene_tx=XM_023250107.1;refGene_gn=DBNL;refGene_tx=XM_023250108.1
A2 62349893 62349987
refGene_gn=DBNL;refGene_tx=XM_019825179.2;refGene_gn=DBNL;refGene_tx=XM_023250106.1;refGene_gn=DBNL;refGene_tx=XM_023250107.1;refGene_gn=DBNL;refGene_tx=XM_023250108.1
A2 62350524 62350620
refGene_gn=DBNL;refGene_tx=XM_019825179.2;refGene_gn=DBNL;refGene_tx=XM_023250106.1;refGene_gn=DBNL;refGene_tx=XM_023250107.1;refGene_gn=DBNL;refGene_tx=XM_023250108.1
A2 62350727 62350843
refGene_gn=DBNL;refGene_tx=XM_019825179.2;refGene_gn=DBNL;refGene_tx=XM_023250106.1;refGene_gn=DBNL;refGene_tx=XM_023250107.1;refGene_gn=DBNL;refGene_tx=XM_023250108.1
A2 62351114 62351211
refGene_gn=DBNL;refGene_tx=XM_019825179.2;refGene_gn=DBNL;refGene_tx=XM_023250106.1;refGene_gn=DBNL;refGene_tx=XM_023250107.1;refGene_gn=DBNL;refGene_tx=XM_023250108.1
A2 62351562 62351702
refGene_gn=DBNL;refGene_tx=XM_019825179.2;refGene_gn=DBNL;refGene_tx=XM_023250106.1;refGene_gn=DBNL;refGene_tx=XM_023250107.1;refGene_gn=DBNL;refGene_tx=XM_023250108.1
A2 62353356 62353523 refGene_gn=PGAM2;refGene_tx=XM_003982576.2
A2 62355037 62355218 refGene_gn=PGAM2;refGene_tx=XM_003982576.2
A2 62355330 62355744 refGene_gn=PGAM2;refGene_tx=XM_003982576.2
A2 62367386 62367473
refGene_gn=POLM;refGene_tx=XM_011280274.2;refGene_gn=POLM;refGene_tx=XM_011280275.2;refGene_gn=POLM;refGene_tx=XM_019825180.1;refGene_gn=POLM;refGene_tx=XM_019825181.1;refGene_gn=POLM;refGene_tx=XM_019825183.1
A2 62367558 62367642
refGene_gn=POLM;refGene_tx=XM_011280274.2;refGene_gn=POLM;refGene_tx=XM_011280275.2;refGene_gn=POLM;refGene_tx=XM_019825180.1;refGene_gn=POLM;refGene_tx=XM_019825181.1;refGene_gn=POLM;refGene_tx=XM_019825183.1
A2 62367713 62367941
refGene_gn=POLM;refGene_tx=XM_011280274.2;refGene_gn=POLM;refGene_tx=XM_011280275.2;refGene_gn=POLM;refGene_tx=XM_019825180.1;refGene_gn=POLM;refGene_tx=XM_019825181.1;refGene_gn=POLM;refGene_tx=XM_019825183.1

A2 62368038 62368156
refGene_gn=POLM;refGene_tx=XM_011280274.2;refGene_gn=POLM;refGene_tx=XM_011280275.2;refGene_gn=POLM;refGene_tx=XM_019825180.1;refGene_gn=POLM;refGene_tx=XM_019825181.1;refGene_gn=POLM;refGene_tx=XM_019825183.1
A2 62368289 62368422
refGene_gn=POLM;refGene_tx=XM_011280274.2;refGene_gn=POLM;refGene_tx=XM_011280275.2;refGene_gn=POLM;refGene_tx=XM_019825180.1;refGene_gn=POLM;refGene_tx=XM_019825181.1;refGene_gn=POLM;refGene_tx=XM_019825183.1
A2 62371334 62371455
refGene_gn=POLM;refGene_tx=XM_011280274.2;refGene_gn=POLM;refGene_tx=XM_011280275.2;refGene_gn=POLM;refGene_tx=XM_019825180.1;refGene_gn=POLM;refGene_tx=XM_019825181.1;refGene_gn=POLM;refGene_tx=XM_019825183.1
A2 62373110 62373182
refGene_gn=POLM;refGene_tx=XM_011280274.2;refGene_gn=POLM;refGene_tx=XM_011280275.2;refGene_gn=POLM;refGene_tx=XM_019825180.1;refGene_gn=POLM;refGene_tx=XM_019825181.1;refGene_gn=POLM;refGene_tx=XM_019825183.1
A2 62374227 62374398
refGene_gn=POLM;refGene_tx=XM_011280274.2;refGene_gn=POLM;refGene_tx=XM_011280275.2;refGene_gn=POLM;refGene_tx=XM_019825180.1;refGene_gn=POLM;refGene_tx=XM_019825181.1;refGene_gn=POLM;refGene_tx=XM_019825183.1
A2 62374542 62374641
refGene_gn=POLM;refGene_tx=XM_011280274.2;refGene_gn=POLM;refGene_tx=XM_011280275.2;refGene_gn=POLM;refGene_tx=XM_019825180.1;refGene_gn=POLM;refGene_tx=XM_019825181.1
A2 62375249 62375436
refGene_gn=POLM;refGene_tx=XM_011280274.2;refGene_gn=POLM;refGene_tx=XM_011280275.2;refGene_gn=POLM;refGene_tx=XM_019825180.1;refGene_gn=POLM;refGene_tx=XM_019825181.1;refGene_gn=POLM;refGene_tx=XM_019825183.1
A2 62376806 62377183
refGene_gn=POLM;refGene_tx=XM_011280274.2;refGene_gn=POLM;refGene_tx=XM_011280275.2;refGene_gn=POLM;refGene_tx=XM_019825180.1;refGene_gn=POLM;refGene_tx=XM_019825181.1;refGene_gn=POLM;refGene_tx=XM_019825183.1
A2 62395082 62395335
refGene_gn=AEBP1;refGene_tx=XM_023250109.1;refGene_gn=AEBP1;refGene_tx=XM_023250110.1
A2 62396941 62397319
refGene_gn=AEBP1;refGene_tx=XM_023250109.1;refGene_gn=AEBP1;refGene_tx=XM_023250110.1
A2 62397833 62397893
refGene_gn=AEBP1;refGene_tx=XM_023250109.1;refGene_gn=AEBP1;refGene_tx=XM_023250110.1
A2 62398019 62398091
refGene_gn=AEBP1;refGene_tx=XM_023250109.1;refGene_gn=AEBP1;refGene_tx=XM_023250110.1
A2 62398253 62398388
refGene_gn=AEBP1;refGene_tx=XM_023250109.1;refGene_gn=AEBP1;refGene_tx=XM_023250110.1
A2 62398476 62398554
refGene_gn=AEBP1;refGene_tx=XM_023250109.1;refGene_gn=AEBP1;refGene_tx=XM_023250110.1

A2 62399177 62399255
refGene_gn=AEBP1;refGene_tx=XM_023250109.1;refGene_gn=AEBP1;refGene_tx
=XM_023250110.1
A2 62399617 62399701 refGene_gn=AEBP1;refGene_tx=XM_023250109.1
A2 62399810 62399864
refGene_gn=AEBP1;refGene_tx=XM_023250109.1;refGene_gn=AEBP1;refGene_tx
=XM_023250110.1
A2 62400539 62400649
refGene_gn=AEBP1;refGene_tx=XM_023250109.1;refGene_gn=AEBP1;refGene_tx
=XM_023250110.1
A2 62400728 62400868
refGene_gn=AEBP1;refGene_tx=XM_023250109.1;refGene_gn=AEBP1;refGene_tx
=XM_023250110.1
A2 62401099 62401184
refGene_gn=AEBP1;refGene_tx=XM_023250109.1;refGene_gn=AEBP1;refGene_tx
=XM_023250110.1
A2 62401281 62401426
refGene_gn=AEBP1;refGene_tx=XM_023250109.1;refGene_gn=AEBP1;refGene_tx
=XM_023250110.1
A2 62401529 62401615
refGene_gn=AEBP1;refGene_tx=XM_023250109.1;refGene_gn=AEBP1;refGene_tx
=XM_023250110.1
A2 62401897 62402021
refGene_gn=AEBP1;refGene_tx=XM_023250109.1;refGene_gn=AEBP1;refGene_tx
=XM_023250110.1
A2 62402263 62402460
refGene_gn=AEBP1;refGene_tx=XM_023250109.1;refGene_gn=AEBP1;refGene_tx
=XM_023250110.1
A2 62402582 62402762
refGene_gn=AEBP1;refGene_tx=XM_023250109.1;refGene_gn=AEBP1;refGene_tx
=XM_023250110.1
A2 62402931 62403283
refGene_gn=AEBP1;refGene_tx=XM_023250109.1;refGene_gn=AEBP1;refGene_tx
=XM_023250110.1
A2 62403366 62403506
refGene_gn=AEBP1;refGene_tx=XM_023250109.1;refGene_gn=AEBP1;refGene_tx
=XM_023250110.1
A2 62403640 62403740
refGene_gn=AEBP1;refGene_tx=XM_023250109.1;refGene_gn=AEBP1;refGene_tx
=XM_023250110.1
A2 62403956 62404597
refGene_gn=AEBP1;refGene_tx=XM_023250109.1;refGene_gn=AEBP1;refGene_tx
=XM_023250110.1
A2 62404977 62405138
refGene_gn=POLD2;refGene_tx=XM_003982577.5;refGene_gn=POLD2;refGene_tx
=XM_006929050.4;refGene_gn=POLD2;refGene_tx=XM_006929051.4;refGene_gn=
POLD2;refGene_tx=XM_023250111.1
A2 62405498 62405600
refGene_gn=POLD2;refGene_tx=XM_003982577.5;refGene_gn=POLD2;refGene_tx
=XM_006929050.4;refGene_gn=POLD2;refGene_tx=XM_006929051.4;refGene_gn=

POLD2;refGene_tx=XM_023250111.1
A2 62406026 62406154
refGene_gn=POLD2;refGene_tx=XM_003982577.5;refGene_gn=POLD2;refGene_tx=XM_006929050.4;refGene_gn=POLD2;refGene_tx=XM_006929051.4;refGene_gn=POLD2;refGene_tx=XM_023250111.1
A2 62406373 62406531
refGene_gn=POLD2;refGene_tx=XM_003982577.5;refGene_gn=POLD2;refGene_tx=XM_006929050.4;refGene_gn=POLD2;refGene_tx=XM_006929051.4;refGene_gn=POLD2;refGene_tx=XM_023250111.1
A2 62406693 62406774
refGene_gn=POLD2;refGene_tx=XM_003982577.5;refGene_gn=POLD2;refGene_tx=XM_006929050.4;refGene_gn=POLD2;refGene_tx=XM_006929051.4;refGene_gn=POLD2;refGene_tx=XM_023250111.1
A2 62406991 62407190
refGene_gn=POLD2;refGene_tx=XM_003982577.5;refGene_gn=POLD2;refGene_tx=XM_006929050.4;refGene_gn=POLD2;refGene_tx=XM_006929051.4;refGene_gn=POLD2;refGene_tx=XM_023250111.1
A2 62407347 62407462
refGene_gn=POLD2;refGene_tx=XM_003982577.5;refGene_gn=POLD2;refGene_tx=XM_006929050.4;refGene_gn=POLD2;refGene_tx=XM_006929051.4;refGene_gn=POLD2;refGene_tx=XM_023250111.1
A2 62407589 62407713
refGene_gn=POLD2;refGene_tx=XM_003982577.5;refGene_gn=POLD2;refGene_tx=XM_006929050.4;refGene_gn=POLD2;refGene_tx=XM_006929051.4;refGene_gn=POLD2;refGene_tx=XM_023250111.1
A2 62407889 62408011
refGene_gn=POLD2;refGene_tx=XM_003982577.5;refGene_gn=POLD2;refGene_tx=XM_006929050.4;refGene_gn=POLD2;refGene_tx=XM_006929051.4;refGene_gn=POLD2;refGene_tx=XM_023250111.1
A2 62410596 62410816
refGene_gn=POLD2;refGene_tx=XM_003982577.5;refGene_gn=POLD2;refGene_tx=XM_006929050.4;refGene_gn=POLD2;refGene_tx=XM_006929051.4;refGene_gn=POLD2;refGene_tx=XM_023250111.1
A2 62431210 62431312 refGene_gn=MYL7;refGene_tx=XM_019826096.2
A2 62431815 62431864 refGene_gn=MYL7;refGene_tx=XM_019826096.2
A2 62432017 62432096 refGene_gn=MYL7;refGene_tx=XM_019826096.2
A2 62432469 62432574 refGene_gn=MYL7;refGene_tx=XM_019826096.2
A2 62432798 62432874 refGene_gn=MYL7;refGene_tx=XM_019826096.2
A2 62433013 62433127 refGene_gn=MYL7;refGene_tx=XM_019826096.2
A2 62433339 62433342 refGene_gn=MYL7;refGene_tx=XM_019826096.2
A2 62436426 62436571 refGene_gn=GCK;refGene_tx=NM_001083954.1
A2 62436845 62437079 refGene_gn=GCK;refGene_tx=NM_001083954.1
A2 62437600 62437756 refGene_gn=GCK;refGene_tx=NM_001083954.1
A2 62438656 62438840 refGene_gn=GCK;refGene_tx=NM_001083954.1
A2 62440776 62440876 refGene_gn=GCK;refGene_tx=NM_001083954.1
A2 62441008 62441104 refGene_gn=GCK;refGene_tx=NM_001083954.1
A2 62442064 62442184 refGene_gn=GCK;refGene_tx=NM_001083954.1
A2 62443136 62443291 refGene_gn=GCK;refGene_tx=NM_001083954.1
A2 62443806 62443969 refGene_gn=GCK;refGene_tx=NM_001083954.1
A2 62476104 62476149 refGene_gn=GCK;refGene_tx=NM_001083954.1

A2 62488366 62488470 refGene_gn=YKT6;refGene_tx=XM_003982579.5
A2 62490619 62490702 refGene_gn=YKT6;refGene_tx=XM_003982579.5
A2 62492787 62492888 refGene_gn=YKT6;refGene_tx=XM_003982579.5
A2 62493838 62493943 refGene_gn=YKT6;refGene_tx=XM_003982579.5
A2 62494229 62494295 refGene_gn=YKT6;refGene_tx=XM_003982579.5
A2 62496386 62496488 refGene_gn=YKT6;refGene_tx=XM_003982579.5
A2 62497122 62497158 refGene_gn=YKT6;refGene_tx=XM_003982579.5
A2 62502161 62502394
refGene_gn=CAMK2B;refGene_tx=XM_023250112.1;refGene_gn=CAMK2B;refGene_tx=XM_023250113.1;refGene_gn=CAMK2B;refGene_tx=XM_023250114.1;refGene_gn=CAMK2B;refGene_tx=XM_023250115.1;refGene_gn=CAMK2B;refGene_tx=XM_023250116.1;refGene_gn=CAMK2B;refGene_tx=XM_023250117.1;refGene_gn=CAMK2B;refGene_tx=XM_023250118.1;refGene_gn=CAMK2B;refGene_tx=XM_023250119.1;refGene_gn=CAMK2B;refGene_tx=XM_023250120.1
A2 62502591 62502686
refGene_gn=CAMK2B;refGene_tx=XM_023250112.1;refGene_gn=CAMK2B;refGene_tx=XM_023250113.1;refGene_gn=CAMK2B;refGene_tx=XM_023250114.1;refGene_gn=CAMK2B;refGene_tx=XM_023250115.1;refGene_gn=CAMK2B;refGene_tx=XM_023250116.1;refGene_gn=CAMK2B;refGene_tx=XM_023250117.1;refGene_gn=CAMK2B;refGene_tx=XM_023250118.1;refGene_gn=CAMK2B;refGene_tx=XM_023250119.1;refGene_gn=CAMK2B;refGene_tx=XM_023250120.1
A2 62502791 62502867
refGene_gn=CAMK2B;refGene_tx=XM_023250112.1;refGene_gn=CAMK2B;refGene_tx=XM_023250113.1;refGene_gn=CAMK2B;refGene_tx=XM_023250114.1;refGene_gn=CAMK2B;refGene_tx=XM_023250115.1;refGene_gn=CAMK2B;refGene_tx=XM_023250116.1;refGene_gn=CAMK2B;refGene_tx=XM_023250117.1;refGene_gn=CAMK2B;refGene_tx=XM_023250118.1;refGene_gn=CAMK2B;refGene_tx=XM_023250119.1;refGene_gn=CAMK2B;refGene_tx=XM_023250120.1
A2 62506922 62507036
refGene_gn=CAMK2B;refGene_tx=XM_023250112.1;refGene_gn=CAMK2B;refGene_tx=XM_023250113.1;refGene_gn=CAMK2B;refGene_tx=XM_023250114.1;refGene_gn=CAMK2B;refGene_tx=XM_023250115.1;refGene_gn=CAMK2B;refGene_tx=XM_023250116.1;refGene_gn=CAMK2B;refGene_tx=XM_023250118.1
A2 62507825 62507906
refGene_gn=CAMK2B;refGene_tx=XM_023250112.1;refGene_gn=CAMK2B;refGene_tx=XM_023250113.1;refGene_gn=CAMK2B;refGene_tx=XM_023250114.1
A2 62509483 62509532
refGene_gn=CAMK2B;refGene_tx=XM_023250112.1;refGene_gn=CAMK2B;refGene_tx=XM_023250113.1;refGene_gn=CAMK2B;refGene_tx=XM_023250114.1;refGene_gn=CAMK2B;refGene_tx=XM_023250115.1;refGene_gn=CAMK2B;refGene_tx=XM_023250116.1;refGene_gn=CAMK2B;refGene_tx=XM_023250117.1;refGene_gn=CAMK2B;refGene_tx=XM_023250118.1;refGene_gn=CAMK2B;refGene_tx=XM_023250119.1;refGene_gn=CAMK2B;refGene_tx=XM_023250120.1
A2 62510900 62510945
refGene_gn=CAMK2B;refGene_tx=XM_023250112.1;refGene_gn=CAMK2B;refGene_tx=XM_023250114.1;refGene_gn=CAMK2B;refGene_tx=XM_023250115.1;refGene_gn=CAMK2B;refGene_tx=XM_023250117.1;refGene_gn=CAMK2B;refGene_tx=XM_023250120.1
A2 62512163 62512235
refGene_gn=CAMK2B;refGene_tx=XM_023250112.1;refGene_gn=CAMK2B;refGene_

tx=XM_023250113.1;refGene_gn=CAMK2B;refGene_tx=XM_023250114.1;refGene_gn=CAMK2B;refGene_tx=XM_023250115.1;refGene_gn=CAMK2B;refGene_tx=XM_023250116.1;refGene_gn=CAMK2B;refGene_tx=XM_023250117.1;refGene_gn=CAMK2B;refGene_tx=XM_023250118.1;refGene_gn=CAMK2B;refGene_tx=XM_023250119.1;refGene_gn=CAMK2B;refGene_tx=XM_023250120.1
A2 62512364 62512402
refGene_gn=CAMK2B;refGene_tx=XM_023250112.1;refGene_gn=CAMK2B;refGene_tx=XM_023250113.1;refGene_gn=CAMK2B;refGene_tx=XM_023250114.1;refGene_gn=CAMK2B;refGene_tx=XM_023250115.1;refGene_gn=CAMK2B;refGene_tx=XM_023250116.1;refGene_gn=CAMK2B;refGene_tx=XM_023250117.1;refGene_gn=CAMK2B;refGene_tx=XM_023250118.1;refGene_gn=CAMK2B;refGene_tx=XM_023250119.1;refGene_gn=CAMK2B;refGene_tx=XM_023250120.1
A2 62516548 62516623
refGene_gn=CAMK2B;refGene_tx=XM_023250112.1;refGene_gn=CAMK2B;refGene_tx=XM_023250113.1;refGene_gn=CAMK2B;refGene_tx=XM_023250115.1;refGene_gn=CAMK2B;refGene_tx=XM_023250116.1;refGene_gn=CAMK2B;refGene_tx=XM_023250117.1;refGene_gn=CAMK2B;refGene_tx=XM_023250119.1
A2 62517558 62517601
refGene_gn=CAMK2B;refGene_tx=XM_023250112.1;refGene_gn=CAMK2B;refGene_tx=XM_023250113.1;refGene_gn=CAMK2B;refGene_tx=XM_023250114.1;refGene_gn=CAMK2B;refGene_tx=XM_023250115.1;refGene_gn=CAMK2B;refGene_tx=XM_023250116.1;refGene_gn=CAMK2B;refGene_tx=XM_023250117.1;refGene_gn=CAMK2B;refGene_tx=XM_023250118.1;refGene_gn=CAMK2B;refGene_tx=XM_023250119.1;refGene_gn=CAMK2B;refGene_tx=XM_023250120.1
A2 62518202 62518286
refGene_gn=CAMK2B;refGene_tx=XM_023250112.1;refGene_gn=CAMK2B;refGene_tx=XM_023250113.1;refGene_gn=CAMK2B;refGene_tx=XM_023250114.1;refGene_gn=CAMK2B;refGene_tx=XM_023250115.1;refGene_gn=CAMK2B;refGene_tx=XM_023250116.1;refGene_gn=CAMK2B;refGene_tx=XM_023250117.1;refGene_gn=CAMK2B;refGene_tx=XM_023250118.1;refGene_gn=CAMK2B;refGene_tx=XM_023250119.1;refGene_gn=CAMK2B;refGene_tx=XM_023250120.1
A2 62518670 62518793
refGene_gn=CAMK2B;refGene_tx=XM_023250112.1;refGene_gn=CAMK2B;refGene_tx=XM_023250113.1;refGene_gn=CAMK2B;refGene_tx=XM_023250114.1;refGene_gn=CAMK2B;refGene_tx=XM_023250115.1;refGene_gn=CAMK2B;refGene_tx=XM_023250116.1;refGene_gn=CAMK2B;refGene_tx=XM_023250117.1;refGene_gn=CAMK2B;refGene_tx=XM_023250118.1;refGene_gn=CAMK2B;refGene_tx=XM_023250119.1;refGene_gn=CAMK2B;refGene_tx=XM_023250120.1
A2 62518950 62519045
refGene_gn=CAMK2B;refGene_tx=XM_023250112.1;refGene_gn=CAMK2B;refGene_tx=XM_023250113.1;refGene_gn=CAMK2B;refGene_tx=XM_023250114.1;refGene_gn=CAMK2B;refGene_tx=XM_023250115.1;refGene_gn=CAMK2B;refGene_tx=XM_023250116.1;refGene_gn=CAMK2B;refGene_tx=XM_023250117.1;refGene_gn=CAMK2B;refGene_tx=XM_023250118.1;refGene_gn=CAMK2B;refGene_tx=XM_023250119.1;refGene_gn=CAMK2B;refGene_tx=XM_023250120.1
A2 62520424 62520508
refGene_gn=CAMK2B;refGene_tx=XM_023250112.1;refGene_gn=CAMK2B;refGene_tx=XM_023250113.1;refGene_gn=CAMK2B;refGene_tx=XM_023250114.1;refGene_gn=CAMK2B;refGene_tx=XM_023250115.1;refGene_gn=CAMK2B;refGene_tx=XM_023250116.1;refGene_gn=CAMK2B;refGene_tx=XM_023250117.1;refGene_gn=CAMK2

B;refGene_tx=XM_023250118.1;refGene_gn=CAMK2B;refGene_tx=XM_023250119.1;refGene_gn=CAMK2B;refGene_tx=XM_023250120.1
A2 62520598 62520701
refGene_gn=CAMK2B;refGene_tx=XM_023250112.1;refGene_gn=CAMK2B;refGene_tx=XM_023250113.1;refGene_gn=CAMK2B;refGene_tx=XM_023250114.1;refGene_gn=CAMK2B;refGene_tx=XM_023250115.1;refGene_gn=CAMK2B;refGene_tx=XM_023250116.1;refGene_gn=CAMK2B;refGene_tx=XM_023250117.1;refGene_gn=CAMK2B;refGene_tx=XM_023250118.1;refGene_gn=CAMK2B;refGene_tx=XM_023250119.1;refGene_gn=CAMK2B;refGene_tx=XM_023250120.1
A2 62523315 62523388
refGene_gn=CAMK2B;refGene_tx=XM_023250112.1;refGene_gn=CAMK2B;refGene_tx=XM_023250113.1;refGene_gn=CAMK2B;refGene_tx=XM_023250114.1;refGene_gn=CAMK2B;refGene_tx=XM_023250115.1;refGene_gn=CAMK2B;refGene_tx=XM_023250116.1;refGene_gn=CAMK2B;refGene_tx=XM_023250117.1;refGene_gn=CAMK2B;refGene_tx=XM_023250118.1;refGene_gn=CAMK2B;refGene_tx=XM_023250119.1;refGene_gn=CAMK2B;refGene_tx=XM_023250120.1
A2 62529032 62529098
refGene_gn=CAMK2B;refGene_tx=XM_023250112.1;refGene_gn=CAMK2B;refGene_tx=XM_023250113.1;refGene_gn=CAMK2B;refGene_tx=XM_023250114.1;refGene_gn=CAMK2B;refGene_tx=XM_023250115.1;refGene_gn=CAMK2B;refGene_tx=XM_023250116.1;refGene_gn=CAMK2B;refGene_tx=XM_023250117.1;refGene_gn=CAMK2B;refGene_tx=XM_023250118.1;refGene_gn=CAMK2B;refGene_tx=XM_023250119.1;refGene_gn=CAMK2B;refGene_tx=XM_023250120.1
A2 62532201 62532256
refGene_gn=CAMK2B;refGene_tx=XM_023250112.1;refGene_gn=CAMK2B;refGene_tx=XM_023250113.1;refGene_gn=CAMK2B;refGene_tx=XM_023250114.1;refGene_gn=CAMK2B;refGene_tx=XM_023250115.1;refGene_gn=CAMK2B;refGene_tx=XM_023250116.1;refGene_gn=CAMK2B;refGene_tx=XM_023250117.1;refGene_gn=CAMK2B;refGene_tx=XM_023250118.1;refGene_gn=CAMK2B;refGene_tx=XM_023250119.1;refGene_gn=CAMK2B;refGene_tx=XM_023250120.1
A2 62536979 62537039
refGene_gn=CAMK2B;refGene_tx=XM_023250112.1;refGene_gn=CAMK2B;refGene_tx=XM_023250113.1;refGene_gn=CAMK2B;refGene_tx=XM_023250114.1;refGene_gn=CAMK2B;refGene_tx=XM_023250115.1;refGene_gn=CAMK2B;refGene_tx=XM_023250116.1;refGene_gn=CAMK2B;refGene_tx=XM_023250117.1;refGene_gn=CAMK2B;refGene_tx=XM_023250118.1;refGene_gn=CAMK2B;refGene_tx=XM_023250119.1;refGene_gn=CAMK2B;refGene_tx=XM_023250120.1
A2 62552798 62552893
refGene_gn=CAMK2B;refGene_tx=XM_023250112.1;refGene_gn=CAMK2B;refGene_tx=XM_023250113.1;refGene_gn=CAMK2B;refGene_tx=XM_023250114.1;refGene_gn=CAMK2B;refGene_tx=XM_023250115.1;refGene_gn=CAMK2B;refGene_tx=XM_023250116.1;refGene_gn=CAMK2B;refGene_tx=XM_023250117.1;refGene_gn=CAMK2B;refGene_tx=XM_023250118.1;refGene_gn=CAMK2B;refGene_tx=XM_023250119.1;refGene_gn=CAMK2B;refGene_tx=XM_023250120.1
A2 62582560 62582625
refGene_gn=CAMK2B;refGene_tx=XM_023250112.1;refGene_gn=CAMK2B;refGene_tx=XM_023250113.1;refGene_gn=CAMK2B;refGene_tx=XM_023250114.1;refGene_gn=CAMK2B;refGene_tx=XM_023250115.1;refGene_gn=CAMK2B;refGene_tx=XM_023250116.1;refGene_gn=CAMK2B;refGene_tx=XM_023250117.1;refGene_gn=CAMK2B;refGene_tx=XM_023250118.1;refGene_gn=CAMK2B;refGene_tx=XM_023250119.

1;refGene_gn=CAMK2B;refGene_tx=XM_023250120.1
A2 62614952 62614979 refGene_gn=NUDCD3;refGene_tx=XM_023250122.1
A2 62616096 62616207 refGene_gn=NUDCD3;refGene_tx=XM_023250121.1
A2 62618271 62618460
refGene_gn=NUDCD3;refGene_tx=XM_023250121.1;refGene_gn=NUDCD3;refGene_tx=XM_023250122.1
A2 62625745 62625889
refGene_gn=NUDCD3;refGene_tx=XM_023250121.1;refGene_gn=NUDCD3;refGene_tx=XM_023250122.1
A2 62638625 62638758
refGene_gn=NUDCD3;refGene_tx=XM_023250121.1;refGene_gn=NUDCD3;refGene_tx=XM_023250122.1
A2 62671611 62671910
refGene_gn=NUDCD3;refGene_tx=XM_023250121.1;refGene_gn=NUDCD3;refGene_tx=XM_023250122.1
A2 62676912 62677104
refGene_gn=NUDCD3;refGene_tx=XM_023250121.1;refGene_gn=NUDCD3;refGene_tx=XM_023250122.1
A2 62682026 62682223
refGene_gn=NPC1L1;refGene_tx=XM_003982593.4;refGene_gn=NPC1L1;refGene_tx=XM_019825187.1
A2 62682548 62682711
refGene_gn=NPC1L1;refGene_tx=XM_003982593.4;refGene_gn=NPC1L1;refGene_tx=XM_019825187.1
A2 62682793 62682907
refGene_gn=NPC1L1;refGene_tx=XM_003982593.4;refGene_gn=NPC1L1;refGene_tx=XM_019825187.1
A2 62682993 62683225
refGene_gn=NPC1L1;refGene_tx=XM_003982593.4;refGene_gn=NPC1L1;refGene_tx=XM_019825187.1
A2 62683335 62683486
refGene_gn=NPC1L1;refGene_tx=XM_003982593.4;refGene_gn=NPC1L1;refGene_tx=XM_019825187.1
A2 62684228 62684284
refGene_gn=NPC1L1;refGene_tx=XM_003982593.4;refGene_gn=NPC1L1;refGene_tx=XM_019825187.1
A2 62684369 62684496
refGene_gn=NPC1L1;refGene_tx=XM_003982593.4;refGene_gn=NPC1L1;refGene_tx=XM_019825187.1
A2 62684990 62685115
refGene_gn=NPC1L1;refGene_tx=XM_003982593.4;refGene_gn=NPC1L1;refGene_tx=XM_019825187.1
A2 62685229 62685420
refGene_gn=NPC1L1;refGene_tx=XM_003982593.4;refGene_gn=NPC1L1;refGene_tx=XM_019825187.1
A2 62689172 62689262 refGene_gn=NPC1L1;refGene_tx=XM_003982593.4
A2 62689382 62689520
refGene_gn=NPC1L1;refGene_tx=XM_003982593.4;refGene_gn=NPC1L1;refGene_tx=XM_019825187.1
A2 62689820 62689948

refGene_gn=NPC1L1;refGene_tx=XM_003982593.4;refGene_gn=NPC1L1;refGene_tx=XM_019825187.1
A2 62690054 62690169
refGene_gn=NPC1L1;refGene_tx=XM_003982593.4;refGene_gn=NPC1L1;refGene_tx=XM_019825187.1
A2 62690616 62690799
refGene_gn=NPC1L1;refGene_tx=XM_003982593.4;refGene_gn=NPC1L1;refGene_tx=XM_019825187.1
A2 62691082 62691211
refGene_gn=NPC1L1;refGene_tx=XM_003982593.4;refGene_gn=NPC1L1;refGene_tx=XM_019825187.1
A2 62691407 62691580
refGene_gn=NPC1L1;refGene_tx=XM_003982593.4;refGene_gn=NPC1L1;refGene_tx=XM_019825187.1
A2 62692157 62692258
refGene_gn=NPC1L1;refGene_tx=XM_003982593.4;refGene_gn=NPC1L1;refGene_tx=XM_019825187.1
A2 62694142 62695671
refGene_gn=NPC1L1;refGene_tx=XM_003982593.4;refGene_gn=NPC1L1;refGene_tx=XM_019825187.1
A2 62696493 62696547
refGene_gn=NPC1L1;refGene_tx=XM_003982593.4;refGene_gn=NPC1L1;refGene_tx=XM_019825187.1
A2 62712677 62712755 refGene_gn=DDX56;refGene_tx=XM_003982594.4
A2 62713064 62713141 refGene_gn=DDX56;refGene_tx=XM_003982594.4
A2 62715208 62715314 refGene_gn=DDX56;refGene_tx=XM_003982594.4
A2 62715456 62715546 refGene_gn=DDX56;refGene_tx=XM_003982594.4
A2 62715678 62715752 refGene_gn=DDX56;refGene_tx=XM_003982594.4
A2 62716860 62716955 refGene_gn=DDX56;refGene_tx=XM_003982594.4
A2 62717031 62717145 refGene_gn=DDX56;refGene_tx=XM_003982594.4
A2 62717636 62717756 refGene_gn=DDX56;refGene_tx=XM_003982594.4
A2 62718039 62718284 refGene_gn=DDX56;refGene_tx=XM_003982594.4
A2 62718798 62718889 refGene_gn=DDX56;refGene_tx=XM_003982594.4
A2 62719006 62719177 refGene_gn=DDX56;refGene_tx=XM_003982594.4
A2 62719323 62719484 refGene_gn=DDX56;refGene_tx=XM_003982594.4
A2 62719935 62720097 refGene_gn=DDX56;refGene_tx=XM_003982594.4
A2 62720185 62720245 refGene_gn=DDX56;refGene_tx=XM_003982594.4
A2 62723602 62723752 refGene_gn=TMED4;refGene_tx=XM_019825201.2
A2 62724125 62724272 refGene_gn=TMED4;refGene_tx=XM_019825201.2
A2 62724526 62724652 refGene_gn=TMED4;refGene_tx=XM_019825201.2
A2 62724815 62724916 refGene_gn=TMED4;refGene_tx=XM_019825201.2
A2 62725035 62725162 refGene_gn=TMED4;refGene_tx=XM_019825201.2
A2 62749922 62750144
refGene_gn=OGDH;refGene_tx=XM_003982581.3;refGene_gn=OGDH;refGene_tx=XM_019825202.1;refGene_gn=OGDH;refGene_tx=XM_019825203.1;refGene_gn=OGDH;refGene_tx=XM_019825204.1
A2 62760761 62760953
refGene_gn=OGDH;refGene_tx=XM_003982581.3;refGene_gn=OGDH;refGene_tx=XM_019825202.1;refGene_gn=OGDH;refGene_tx=XM_019825203.1;refGene_gn=OGDH;refGene_tx=XM_019825204.1

A2 62762266 62762357
refGene_gn=OGDH;refGene_tx=XM_019825203.1;refGene_gn=OGDH;refGene_tx=XM_019825204.1
A2 62762473 62762576
refGene_gn=OGDH;refGene_tx=XM_003982581.3;refGene_gn=OGDH;refGene_tx=XM_019825202.1
A2 62768140 62768185
refGene_gn=OGDH;refGene_tx=XM_019825202.1;refGene_gn=OGDH;refGene_tx=XM_019825203.1
A2 62776086 62776202
refGene_gn=OGDH;refGene_tx=XM_003982581.3;refGene_gn=OGDH;refGene_tx=XM_019825202.1;refGene_gn=OGDH;refGene_tx=XM_019825203.1;refGene_gn=OGDH;refGene_tx=XM_019825204.1
A2 62782497 62782652
refGene_gn=OGDH;refGene_tx=XM_003982581.3;refGene_gn=OGDH;refGene_tx=XM_019825202.1;refGene_gn=OGDH;refGene_tx=XM_019825203.1;refGene_gn=OGDH;refGene_tx=XM_019825204.1
A2 62782764 62782911
refGene_gn=OGDH;refGene_tx=XM_003982581.3;refGene_gn=OGDH;refGene_tx=XM_019825202.1;refGene_gn=OGDH;refGene_tx=XM_019825203.1;refGene_gn=OGDH;refGene_tx=XM_019825204.1
A2 62783097 62783188
refGene_gn=OGDH;refGene_tx=XM_003982581.3;refGene_gn=OGDH;refGene_tx=XM_019825202.1;refGene_gn=OGDH;refGene_tx=XM_019825203.1;refGene_gn=OGDH;refGene_tx=XM_019825204.1
A2 62783539 62783719
refGene_gn=OGDH;refGene_tx=XM_003982581.3;refGene_gn=OGDH;refGene_tx=XM_019825202.1;refGene_gn=OGDH;refGene_tx=XM_019825203.1;refGene_gn=OGDH;refGene_tx=XM_019825204.1
A2 62786475 62786604
refGene_gn=OGDH;refGene_tx=XM_003982581.3;refGene_gn=OGDH;refGene_tx=XM_019825202.1;refGene_gn=OGDH;refGene_tx=XM_019825203.1;refGene_gn=OGDH;refGene_tx=XM_019825204.1
A2 62795592 62795772
refGene_gn=OGDH;refGene_tx=XM_003982581.3;refGene_gn=OGDH;refGene_tx=XM_019825202.1;refGene_gn=OGDH;refGene_tx=XM_019825203.1;refGene_gn=OGDH;refGene_tx=XM_019825204.1
A2 62796076 62796229
refGene_gn=OGDH;refGene_tx=XM_003982581.3;refGene_gn=OGDH;refGene_tx=XM_019825202.1;refGene_gn=OGDH;refGene_tx=XM_019825203.1;refGene_gn=OGDH;refGene_tx=XM_019825204.1
A2 62796728 62796831
refGene_gn=OGDH;refGene_tx=XM_003982581.3;refGene_gn=OGDH;refGene_tx=XM_019825202.1;refGene_gn=OGDH;refGene_tx=XM_019825203.1;refGene_gn=OGDH;refGene_tx=XM_019825204.1
A2 62796946 62797075
refGene_gn=OGDH;refGene_tx=XM_003982581.3;refGene_gn=OGDH;refGene_tx=XM_019825202.1;refGene_gn=OGDH;refGene_tx=XM_019825203.1;refGene_gn=OGDH;refGene_tx=XM_019825204.1
A2 62797196 62797347

refGene_gn=OGDH;refGene_tx=XM_003982581.3;refGene_gn=OGDH;refGene_tx=XM_019825202.1;refGene_gn=OGDH;refGene_tx=XM_019825203.1;refGene_gn=OGDH;refGene_tx=XM_019825204.1
A2 62797587 62797715
refGene_gn=OGDH;refGene_tx=XM_003982581.3;refGene_gn=OGDH;refGene_tx=XM_019825202.1;refGene_gn=OGDH;refGene_tx=XM_019825203.1;refGene_gn=OGDH;refGene_tx=XM_019825204.1
A2 62797819 62797998
refGene_gn=OGDH;refGene_tx=XM_003982581.3;refGene_gn=OGDH;refGene_tx=XM_019825202.1;refGene_gn=OGDH;refGene_tx=XM_019825203.1;refGene_gn=OGDH;refGene_tx=XM_019825204.1
A2 62798367 62798439
refGene_gn=OGDH;refGene_tx=XM_003982581.3;refGene_gn=OGDH;refGene_tx=XM_019825202.1;refGene_gn=OGDH;refGene_tx=XM_019825203.1;refGene_gn=OGDH;refGene_tx=XM_019825204.1
A2 62798747 62798876
refGene_gn=OGDH;refGene_tx=XM_003982581.3;refGene_gn=OGDH;refGene_tx=XM_019825202.1;refGene_gn=OGDH;refGene_tx=XM_019825203.1;refGene_gn=OGDH;refGene_tx=XM_019825204.1
A2 62799222 62799295
refGene_gn=OGDH;refGene_tx=XM_003982581.3;refGene_gn=OGDH;refGene_tx=XM_019825202.1;refGene_gn=OGDH;refGene_tx=XM_019825203.1;refGene_gn=OGDH;refGene_tx=XM_019825204.1
A2 62801160 62801324
refGene_gn=OGDH;refGene_tx=XM_003982581.3;refGene_gn=OGDH;refGene_tx=XM_019825202.1;refGene_gn=OGDH;refGene_tx=XM_019825203.1;refGene_gn=OGDH;refGene_tx=XM_019825204.1
A2 62801411 62801566
refGene_gn=OGDH;refGene_tx=XM_003982581.3;refGene_gn=OGDH;refGene_tx=XM_019825202.1;refGene_gn=OGDH;refGene_tx=XM_019825203.1;refGene_gn=OGDH;refGene_tx=XM_019825204.1
A2 62801701 62801822
refGene_gn=OGDH;refGene_tx=XM_003982581.3;refGene_gn=OGDH;refGene_tx=XM_019825202.1;refGene_gn=OGDH;refGene_tx=XM_019825203.1;refGene_gn=OGDH;refGene_tx=XM_019825204.1
A2 62822350 62822400
refGene_gn=ZMIZ2;refGene_tx=XM_003982585.5;refGene_gn=ZMIZ2;refGene_tx=XM_003982586.5;refGene_gn=ZMIZ2;refGene_tx=XM_019825205.2;refGene_gn=ZMIZ2;refGene_tx=XM_019825206.2;refGene_gn=ZMIZ2;refGene_tx=XM_023250123.1;refGene_gn=ZMIZ2;refGene_tx=XM_023250124.1
A2 62822559 62822674
refGene_gn=ZMIZ2;refGene_tx=XM_003982585.5;refGene_gn=ZMIZ2;refGene_tx=XM_003982586.5;refGene_gn=ZMIZ2;refGene_tx=XM_019825205.2;refGene_gn=ZMIZ2;refGene_tx=XM_019825206.2;refGene_gn=ZMIZ2;refGene_tx=XM_023250123.1;refGene_gn=ZMIZ2;refGene_tx=XM_023250124.1
A2 62822973 62823176
refGene_gn=ZMIZ2;refGene_tx=XM_003982585.5;refGene_gn=ZMIZ2;refGene_tx=XM_003982586.5;refGene_gn=ZMIZ2;refGene_tx=XM_019825205.2;refGene_gn=ZMIZ2;refGene_tx=XM_019825206.2;refGene_gn=ZMIZ2;refGene_tx=XM_023250123.1;refGene_gn=ZMIZ2;refGene_tx=XM_023250124.1

A2 62823356 62823612
refGene_gn=ZMIZ2;refGene_tx=XM_003982585.5;refGene_gn=ZMIZ2;refGene_tx=XM_003982586.5;refGene_gn=ZMIZ2;refGene_tx=XM_019825205.2;refGene_gn=ZMIZ2;refGene_tx=XM_019825206.2;refGene_gn=ZMIZ2;refGene_tx=XM_019825208.2;refGene_gn=ZMIZ2;refGene_tx=XM_023250123.1;refGene_gn=ZMIZ2;refGene_tx=XM_023250124.1
A2 62823802 62824066
refGene_gn=ZMIZ2;refGene_tx=XM_003982585.5;refGene_gn=ZMIZ2;refGene_tx=XM_003982586.5;refGene_gn=ZMIZ2;refGene_tx=XM_019825205.2;refGene_gn=ZMIZ2;refGene_tx=XM_019825206.2;refGene_gn=ZMIZ2;refGene_tx=XM_019825208.2;refGene_gn=ZMIZ2;refGene_tx=XM_023250123.1;refGene_gn=ZMIZ2;refGene_tx=XM_023250124.1
A2 62824227 62824404
refGene_gn=ZMIZ2;refGene_tx=XM_003982585.5;refGene_gn=ZMIZ2;refGene_tx=XM_003982586.5;refGene_gn=ZMIZ2;refGene_tx=XM_019825205.2;refGene_gn=ZMIZ2;refGene_tx=XM_019825206.2;refGene_gn=ZMIZ2;refGene_tx=XM_019825208.2;refGene_gn=ZMIZ2;refGene_tx=XM_023250123.1;refGene_gn=ZMIZ2;refGene_tx=XM_023250124.1
A2 62824930 62825008
refGene_gn=ZMIZ2;refGene_tx=XM_003982585.5;refGene_gn=ZMIZ2;refGene_tx=XM_019825205.2;refGene_gn=ZMIZ2;refGene_tx=XM_019825208.2;refGene_gn=ZMIZ2;refGene_tx=XM_023250123.1;refGene_gn=ZMIZ2;refGene_tx=XM_023250124.1
A2 62825193 62825362
refGene_gn=ZMIZ2;refGene_tx=XM_003982585.5;refGene_gn=ZMIZ2;refGene_tx=XM_003982586.5;refGene_gn=ZMIZ2;refGene_tx=XM_019825205.2;refGene_gn=ZMIZ2;refGene_tx=XM_019825206.2;refGene_gn=ZMIZ2;refGene_tx=XM_019825208.2;refGene_gn=ZMIZ2;refGene_tx=XM_023250123.1;refGene_gn=ZMIZ2;refGene_tx=XM_023250124.1
A2 62825986 62826131
refGene_gn=ZMIZ2;refGene_tx=XM_003982585.5;refGene_gn=ZMIZ2;refGene_tx=XM_003982586.5;refGene_gn=ZMIZ2;refGene_tx=XM_019825205.2;refGene_gn=ZMIZ2;refGene_tx=XM_019825206.2;refGene_gn=ZMIZ2;refGene_tx=XM_019825208.2;refGene_gn=ZMIZ2;refGene_tx=XM_023250123.1;refGene_gn=ZMIZ2;refGene_tx=XM_023250124.1
A2 62826582 62826793
refGene_gn=ZMIZ2;refGene_tx=XM_003982585.5;refGene_gn=ZMIZ2;refGene_tx=XM_003982586.5;refGene_gn=ZMIZ2;refGene_tx=XM_019825205.2;refGene_gn=ZMIZ2;refGene_tx=XM_019825206.2;refGene_gn=ZMIZ2;refGene_tx=XM_019825208.2;refGene_gn=ZMIZ2;refGene_tx=XM_023250123.1;refGene_gn=ZMIZ2;refGene_tx=XM_023250124.1
A2 62827142 62827248
refGene_gn=ZMIZ2;refGene_tx=XM_003982585.5;refGene_gn=ZMIZ2;refGene_tx=XM_003982586.5;refGene_gn=ZMIZ2;refGene_tx=XM_019825205.2;refGene_gn=ZMIZ2;refGene_tx=XM_019825206.2;refGene_gn=ZMIZ2;refGene_tx=XM_019825208.2;refGene_gn=ZMIZ2;refGene_tx=XM_023250123.1;refGene_gn=ZMIZ2;refGene_tx=XM_023250124.1
A2 62827382 62827540
refGene_gn=ZMIZ2;refGene_tx=XM_003982585.5;refGene_gn=ZMIZ2;refGene_tx=XM_003982586.5;refGene_gn=ZMIZ2;refGene_tx=XM_019825205.2;refGene_gn=

ZMIZ2;refGene_tx=XM_019825206.2;refGene_gn=ZMIZ2;refGene_tx=XM_019825208.2;refGene_gn=ZMIZ2;refGene_tx=XM_023250123.1;refGene_gn=ZMIZ2;refGene_tx=XM_023250124.1
A2 62828090 62828158
refGene_gn=ZMIZ2;refGene_tx=XM_003982585.5;refGene_gn=ZMIZ2;refGene_tx=XM_003982586.5;refGene_gn=ZMIZ2;refGene_tx=XM_019825205.2;refGene_gn=ZMIZ2;refGene_tx=XM_019825206.2;refGene_gn=ZMIZ2;refGene_tx=XM_019825208.2;refGene_gn=ZMIZ2;refGene_tx=XM_023250123.1;refGene_gn=ZMIZ2;refGene_tx=XM_023250124.1
A2 62828469 62828538
refGene_gn=ZMIZ2;refGene_tx=XM_003982585.5;refGene_gn=ZMIZ2;refGene_tx=XM_003982586.5;refGene_gn=ZMIZ2;refGene_tx=XM_019825205.2;refGene_gn=ZMIZ2;refGene_tx=XM_019825206.2;refGene_gn=ZMIZ2;refGene_tx=XM_019825208.2;refGene_gn=ZMIZ2;refGene_tx=XM_023250123.1;refGene_gn=ZMIZ2;refGene_tx=XM_023250124.1
A2 62828770 62829021
refGene_gn=ZMIZ2;refGene_tx=XM_003982585.5;refGene_gn=ZMIZ2;refGene_tx=XM_003982586.5;refGene_gn=ZMIZ2;refGene_tx=XM_019825205.2;refGene_gn=ZMIZ2;refGene_tx=XM_019825206.2;refGene_gn=ZMIZ2;refGene_tx=XM_019825208.2;refGene_gn=ZMIZ2;refGene_tx=XM_023250123.1;refGene_gn=ZMIZ2;refGene_tx=XM_023250124.1
A2 62829345 62829557
refGene_gn=ZMIZ2;refGene_tx=XM_003982585.5;refGene_gn=ZMIZ2;refGene_tx=XM_003982586.5;refGene_gn=ZMIZ2;refGene_tx=XM_019825205.2;refGene_gn=ZMIZ2;refGene_tx=XM_019825206.2;refGene_gn=ZMIZ2;refGene_tx=XM_019825208.2;refGene_gn=ZMIZ2;refGene_tx=XM_023250123.1;refGene_gn=ZMIZ2;refGene_tx=XM_023250124.1
A2 62829636 62829876
refGene_gn=ZMIZ2;refGene_tx=XM_003982585.5;refGene_gn=ZMIZ2;refGene_tx=XM_003982586.5;refGene_gn=ZMIZ2;refGene_tx=XM_019825205.2;refGene_gn=ZMIZ2;refGene_tx=XM_019825206.2;refGene_gn=ZMIZ2;refGene_tx=XM_019825208.2;refGene_gn=ZMIZ2;refGene_tx=XM_023250123.1;refGene_gn=ZMIZ2;refGene_tx=XM_023250124.1
A2 62830545 62830653
refGene_gn=ZMIZ2;refGene_tx=XM_003982585.5;refGene_gn=ZMIZ2;refGene_tx=XM_003982586.5;refGene_gn=ZMIZ2;refGene_tx=XM_019825205.2;refGene_gn=ZMIZ2;refGene_tx=XM_019825206.2;refGene_gn=ZMIZ2;refGene_tx=XM_019825208.2;refGene_gn=ZMIZ2;refGene_tx=XM_023250123.1;refGene_gn=ZMIZ2;refGene_tx=XM_023250124.1
A2 62846740 62846809 refGene_gn=PPIA;refGene_tx=NM_001009370.1
A2 62847869 62847900 refGene_gn=PPIA;refGene_tx=NM_001009370.1
A2 62847979 62848068 refGene_gn=PPIA;refGene_tx=NM_001009370.1
A2 62848283 62848456 refGene_gn=PPIA;refGene_tx=NM_001009370.1
A2 62849033 62849166 refGene_gn=PPIA;refGene_tx=NM_001009370.1
A2 62855859 62855921
refGene_gn=LOC101099622;refGene_tx=XM_023250125.1;refGene_gn=LOC101099622;refGene_tx=XM_023250126.1
A2 62856547 62856677
refGene_gn=LOC101099622;refGene_tx=XM_023250125.1;refGene_gn=LOC101099622;refGene_tx=XM_023250126.1

A2 62858983 62859097
refGene_gn=LOC101099622;refGene_tx=XM_023250125.1
A2 62861350 62861428
refGene_gn=LOC101099622;refGene_tx=XM_023250125.1;refGene_gn=LOC101099
622;refGene_tx=XM_023250126.1
A2 62865752 62865755
refGene_gn=LOC101099622;refGene_tx=XM_023250125.1;refGene_gn=LOC101099
622;refGene_tx=XM_023250126.1
A2 62880606 62881518 refGene_gn=PURB;refGene_tx=XM_023250127.1
A2 62901071 62901228
refGene_gn=MYO1G;refGene_tx=XM_023250128.1;refGene_gn=MYO1G;refGene_tx
=XM_023250129.1
A2 62901553 62901708
refGene_gn=MYO1G;refGene_tx=XM_023250128.1;refGene_gn=MYO1G;refGene_tx
=XM_023250129.1
A2 62902232 62902346
refGene_gn=MYO1G;refGene_tx=XM_023250128.1;refGene_gn=MYO1G;refGene_tx
=XM_023250129.1
A2 62902495 62902600
refGene_gn=MYO1G;refGene_tx=XM_023250128.1;refGene_gn=MYO1G;refGene_tx
=XM_023250129.1
A2 62903184 62903329
refGene_gn=MYO1G;refGene_tx=XM_023250128.1;refGene_gn=MYO1G;refGene_tx
=XM_023250129.1
A2 62903779 62904003
refGene_gn=MYO1G;refGene_tx=XM_023250128.1;refGene_gn=MYO1G;refGene_tx
=XM_023250129.1
A2 62904969 62905119 refGene_gn=MYO1G;refGene_tx=XM_023250130.1
A2 62905555 62905763
refGene_gn=MYO1G;refGene_tx=XM_023250128.1;refGene_gn=MYO1G;refGene_tx
=XM_023250129.1;refGene_gn=MYO1G;refGene_tx=XM_023250130.1
A2 62906054 62906221
refGene_gn=MYO1G;refGene_tx=XM_023250128.1;refGene_gn=MYO1G;refGene_tx
=XM_023250129.1;refGene_gn=MYO1G;refGene_tx=XM_023250130.1
A2 62906780 62906913
refGene_gn=MYO1G;refGene_tx=XM_023250128.1;refGene_gn=MYO1G;refGene_tx
=XM_023250129.1;refGene_gn=MYO1G;refGene_tx=XM_023250130.1
A2 62907105 62907180
refGene_gn=MYO1G;refGene_tx=XM_023250128.1;refGene_gn=MYO1G;refGene_tx
=XM_023250129.1;refGene_gn=MYO1G;refGene_tx=XM_023250130.1
A2 62908283 62908354
refGene_gn=MYO1G;refGene_tx=XM_023250128.1;refGene_gn=MYO1G;refGene_tx
=XM_023250129.1;refGene_gn=MYO1G;refGene_tx=XM_023250130.1
A2 62908535 62908706
refGene_gn=MYO1G;refGene_tx=XM_023250128.1;refGene_gn=MYO1G;refGene_tx
=XM_023250129.1;refGene_gn=MYO1G;refGene_tx=XM_023250130.1
A2 62908819 62908934
refGene_gn=MYO1G;refGene_tx=XM_023250128.1;refGene_gn=MYO1G;refGene_tx
=XM_023250129.1;refGene_gn=MYO1G;refGene_tx=XM_023250130.1
A2 62909367 62909513

refGene_gn=MYO1G;refGene_tx=XM_023250128.1;refGene_gn=MYO1G;refGene_tx=XM_023250129.1;refGene_gn=MYO1G;refGene_tx=XM_023250130.1
A2 62909623 62909848
refGene_gn=MYO1G;refGene_tx=XM_023250128.1;refGene_gn=MYO1G;refGene_tx=XM_023250129.1;refGene_gn=MYO1G;refGene_tx=XM_023250130.1
A2 62910413 62910533
refGene_gn=MYO1G;refGene_tx=XM_023250128.1;refGene_gn=MYO1G;refGene_tx=XM_023250129.1;refGene_gn=MYO1G;refGene_tx=XM_023250130.1
A2 62910770 62910866
refGene_gn=MYO1G;refGene_tx=XM_023250128.1;refGene_gn=MYO1G;refGene_tx=XM_023250129.1;refGene_gn=MYO1G;refGene_tx=XM_023250130.1
A2 62911260 62911314
refGene_gn=MYO1G;refGene_tx=XM_023250128.1;refGene_gn=MYO1G;refGene_tx=XM_023250130.1
A2 62911488 62911654
refGene_gn=MYO1G;refGene_tx=XM_023250128.1;refGene_gn=MYO1G;refGene_tx=XM_023250129.1;refGene_gn=MYO1G;refGene_tx=XM_023250130.1
A2 62912293 62912387
refGene_gn=MYO1G;refGene_tx=XM_023250128.1;refGene_gn=MYO1G;refGene_tx=XM_023250129.1;refGene_gn=MYO1G;refGene_tx=XM_023250130.1
A2 62912511 62912720
refGene_gn=MYO1G;refGene_tx=XM_023250128.1;refGene_gn=MYO1G;refGene_tx=XM_023250129.1;refGene_gn=MYO1G;refGene_tx=XM_023250130.1
A2 62914189 62914281
refGene_gn=MYO1G;refGene_tx=XM_023250128.1;refGene_gn=MYO1G;refGene_tx=XM_023250129.1;refGene_gn=MYO1G;refGene_tx=XM_023250130.1
A2 62917627 62917666
refGene_gn=LOC109497364;refGene_tx=XM_023250131.1;refGene_gn=LOC109497364;refGene_tx=XM_023250132.1;refGene_gn=LOC109497364;refGene_tx=XM_023250133.1
A2 62917897 62918102
refGene_gn=LOC109497364;refGene_tx=XM_023250131.1;refGene_gn=LOC109497364;refGene_tx=XM_023250132.1;refGene_gn=LOC109497364;refGene_tx=XM_023250133.1
A2 62918249 62918287
refGene_gn=LOC109497364;refGene_tx=XM_023250131.1;refGene_gn=LOC109497364;refGene_tx=XM_023250132.1;refGene_gn=LOC109497364;refGene_tx=XM_023250133.1
A2 62919216 62919585
refGene_gn=LOC109497364;refGene_tx=XM_023250131.1;refGene_gn=LOC109497364;refGene_tx=XM_023250132.1;refGene_gn=LOC109497364;refGene_tx=XM_023250133.1
A2 62926194 62926224
refGene_gn=CCM2;refGene_tx=XM_023250134.1;refGene_gn=CCM2;refGene_tx=XM_023250135.1;refGene_gn=CCM2;refGene_tx=XM_023250136.1;refGene_gn=CCM2;refGene_tx=XM_023250137.1
A2 62944010 62944022 refGene_gn=CCM2;refGene_tx=XM_006929057.2
A2 62948451 62948625
refGene_gn=CCM2;refGene_tx=XM_006929057.2;refGene_gn=CCM2;refGene_tx=XM_023250134.1;refGene_gn=CCM2;refGene_tx=XM_023250135.1;refGene_gn=CCM

2;refGene_tx=XM_023250136.1
A2	62951142 62951217 refGene_gn=CCM2;refGene_tx=XM_019825214.2
A2	62958741 62958825
refGene_gn=CCM2;refGene_tx=XM_006929057.2;refGene_gn=CCM2;refGene_tx=XM_019825214.2;refGene_gn=CCM2;refGene_tx=XM_023250134.1;refGene_gn=CCM2;refGene_tx=XM_023250135.1;refGene_gn=CCM2;refGene_tx=XM_023250137.1
A2	62959240 62959424
refGene_gn=CCM2;refGene_tx=XM_006929057.2;refGene_gn=CCM2;refGene_tx=XM_019825214.2;refGene_gn=CCM2;refGene_tx=XM_023250134.1;refGene_gn=CCM2;refGene_tx=XM_023250135.1;refGene_gn=CCM2;refGene_tx=XM_023250136.1;refGene_gn=CCM2;refGene_tx=XM_023250137.1
A2	62963012 62963149
refGene_gn=CCM2;refGene_tx=XM_006929057.2;refGene_gn=CCM2;refGene_tx=XM_019825214.2;refGene_gn=CCM2;refGene_tx=XM_023250134.1;refGene_gn=CCM2;refGene_tx=XM_023250135.1;refGene_gn=CCM2;refGene_tx=XM_023250136.1;refGene_gn=CCM2;refGene_tx=XM_023250137.1
A2	62964351 62964487
refGene_gn=CCM2;refGene_tx=XM_006929057.2;refGene_gn=CCM2;refGene_tx=XM_019825214.2;refGene_gn=CCM2;refGene_tx=XM_023250134.1;refGene_gn=CCM2;refGene_tx=XM_023250135.1;refGene_gn=CCM2;refGene_tx=XM_023250136.1;refGene_gn=CCM2;refGene_tx=XM_023250137.1
A2	62965511 62965569
refGene_gn=CCM2;refGene_tx=XM_006929057.2;refGene_gn=CCM2;refGene_tx=XM_019825214.2;refGene_gn=CCM2;refGene_tx=XM_023250134.1;refGene_gn=CCM2;refGene_tx=XM_023250135.1;refGene_gn=CCM2;refGene_tx=XM_023250136.1;refGene_gn=CCM2;refGene_tx=XM_023250137.1
A2	62965711 62965850
refGene_gn=CCM2;refGene_tx=XM_006929057.2;refGene_gn=CCM2;refGene_tx=XM_019825214.2;refGene_gn=CCM2;refGene_tx=XM_023250134.1;refGene_gn=CCM2;refGene_tx=XM_023250135.1;refGene_gn=CCM2;refGene_tx=XM_023250136.1;refGene_gn=CCM2;refGene_tx=XM_023250137.1
A2	62966715 62966854
refGene_gn=CCM2;refGene_tx=XM_006929057.2;refGene_gn=CCM2;refGene_tx=XM_019825214.2;refGene_gn=CCM2;refGene_tx=XM_023250134.1;refGene_gn=CCM2;refGene_tx=XM_023250135.1;refGene_gn=CCM2;refGene_tx=XM_023250136.1;refGene_gn=CCM2;refGene_tx=XM_023250137.1
A2	62967631 62967912
refGene_gn=CCM2;refGene_tx=XM_006929057.2;refGene_gn=CCM2;refGene_tx=XM_019825214.2;refGene_gn=CCM2;refGene_tx=XM_023250134.1;refGene_gn=CCM2;refGene_tx=XM_023250135.1;refGene_gn=CCM2;refGene_tx=XM_023250136.1;refGene_gn=CCM2;refGene_tx=XM_023250137.1
A2	62969784 62969799 refGene_gn=NACAD;refGene_tx=XM_023241978.1
A2	62969886 62970009 refGene_gn=NACAD;refGene_tx=XM_023241978.1
A2	62970110 62970260 refGene_gn=NACAD;refGene_tx=XM_023241978.1
A2	62970332 62970479 refGene_gn=NACAD;refGene_tx=XM_023241978.1
A2	62970735 62970810 refGene_gn=NACAD;refGene_tx=XM_023241978.1
A2	62970885 62971013 refGene_gn=NACAD;refGene_tx=XM_023241978.1
A2	62971141 62972497 refGene_gn=NACAD;refGene_tx=XM_023241978.1
A2	62972797 62973412 refGene_gn=NACAD;refGene_tx=XM_023241978.1
A2	62975098 62978311 refGene_gn=NACAD;refGene_tx=XM_023241978.1

A2 62980125 62980192 refGene_gn=NACAD;refGene_tx=XM_023241978.1
A2 62994965 62995067
refGene_gn=TBRG4;refGene_tx=XM_011280270.3;refGene_gn=TBRG4;refGene_tx=XM_019825215.2;refGene_gn=TBRG4;refGene_tx=XM_023250138.1;refGene_gn=TBRG4;refGene_tx=XM_023250139.1;refGene_gn=TBRG4;refGene_tx=XM_023250140.1;refGene_gn=TBRG4;refGene_tx=XM_023250141.1
A2 62995516 62995631
refGene_gn=TBRG4;refGene_tx=XM_011280270.3;refGene_gn=TBRG4;refGene_tx=XM_019825215.2;refGene_gn=TBRG4;refGene_tx=XM_023250138.1;refGene_gn=TBRG4;refGene_tx=XM_023250139.1;refGene_gn=TBRG4;refGene_tx=XM_023250140.1;refGene_gn=TBRG4;refGene_tx=XM_023250141.1
A2 62995737 62995849
refGene_gn=TBRG4;refGene_tx=XM_011280270.3;refGene_gn=TBRG4;refGene_tx=XM_019825215.2;refGene_gn=TBRG4;refGene_tx=XM_023250138.1;refGene_gn=TBRG4;refGene_tx=XM_023250139.1;refGene_gn=TBRG4;refGene_tx=XM_023250140.1;refGene_gn=TBRG4;refGene_tx=XM_023250141.1
A2 62995962 62996208
refGene_gn=TBRG4;refGene_tx=XM_011280270.3;refGene_gn=TBRG4;refGene_tx=XM_019825215.2;refGene_gn=TBRG4;refGene_tx=XM_023250138.1;refGene_gn=TBRG4;refGene_tx=XM_023250139.1;refGene_gn=TBRG4;refGene_tx=XM_023250140.1;refGene_gn=TBRG4;refGene_tx=XM_023250141.1
A2 62996503 62996648
refGene_gn=TBRG4;refGene_tx=XM_011280270.3;refGene_gn=TBRG4;refGene_tx=XM_019825215.2;refGene_gn=TBRG4;refGene_tx=XM_023250138.1;refGene_gn=TBRG4;refGene_tx=XM_023250139.1;refGene_gn=TBRG4;refGene_tx=XM_023250140.1;refGene_gn=TBRG4;refGene_tx=XM_023250141.1
A2 62997327 62997438
refGene_gn=TBRG4;refGene_tx=XM_011280270.3;refGene_gn=TBRG4;refGene_tx=XM_019825215.2;refGene_gn=TBRG4;refGene_tx=XM_023250138.1;refGene_gn=TBRG4;refGene_tx=XM_023250139.1;refGene_gn=TBRG4;refGene_tx=XM_023250140.1;refGene_gn=TBRG4;refGene_tx=XM_023250141.1
A2 62997922 62998080
refGene_gn=TBRG4;refGene_tx=XM_011280270.3;refGene_gn=TBRG4;refGene_tx=XM_019825215.2;refGene_gn=TBRG4;refGene_tx=XM_023250138.1;refGene_gn=TBRG4;refGene_tx=XM_023250139.1;refGene_gn=TBRG4;refGene_tx=XM_023250140.1;refGene_gn=TBRG4;refGene_tx=XM_023250141.1
A2 62998357 62998529
refGene_gn=TBRG4;refGene_tx=XM_011280270.3;refGene_gn=TBRG4;refGene_tx=XM_019825215.2;refGene_gn=TBRG4;refGene_tx=XM_023250138.1;refGene_gn=TBRG4;refGene_tx=XM_023250139.1;refGene_gn=TBRG4;refGene_tx=XM_023250140.1;refGene_gn=TBRG4;refGene_tx=XM_023250141.1
A2 62999019 62999337
refGene_gn=TBRG4;refGene_tx=XM_011280270.3;refGene_gn=TBRG4;refGene_tx=XM_019825215.2;refGene_gn=TBRG4;refGene_tx=XM_023250138.1;refGene_gn=TBRG4;refGene_tx=XM_023250139.1;refGene_gn=TBRG4;refGene_tx=XM_023250140.1;refGene_gn=TBRG4;refGene_tx=XM_023250141.1
A2 63001393 63001807
refGene_gn=TBRG4;refGene_tx=XM_011280270.3;refGene_gn=TBRG4;refGene_tx=XM_019825215.2;refGene_gn=TBRG4;refGene_tx=XM_023250138.1;refGene_gn=TBRG4;refGene_tx=XM_023250139.1;refGene_gn=TBRG4;refGene_tx=XM_0232501

40.1;refGene_gn=TBRG4;refGene_tx=XM_023250141.1
A2	63011427	63011444
refGene_gn=LOC101100876;refGene_tx=XM_019825217.2
A2	63011771	63011930
refGene_gn=LOC101100876;refGene_tx=XM_019825217.2
A2	63012204	63012337
refGene_gn=LOC101100876;refGene_tx=XM_019825217.2
A2	63039318	63039382	refGene_gn=RAMP3;refGene_tx=XM_011280269.3
A2	63045940	63046073	refGene_gn=RAMP3;refGene_tx=XM_011280269.3
A2	63050246	63050502	refGene_gn=RAMP3;refGene_tx=XM_011280269.3
A2	63189490	63189604	refGene_gn=ADCY1;refGene_tx=XM_006929062.4
A2	63207312	63207431	refGene_gn=ADCY1;refGene_tx=XM_006929062.4
A2	63211783	63211895	refGene_gn=ADCY1;refGene_tx=XM_006929062.4
A2	63224199	63224327	refGene_gn=ADCY1;refGene_tx=XM_006929062.4
A2	63230462	63230621	refGene_gn=ADCY1;refGene_tx=XM_006929062.4
A2	63232878	63233020	refGene_gn=ADCY1;refGene_tx=XM_006929062.4
A2	63234014	63234170	refGene_gn=ADCY1;refGene_tx=XM_006929062.4
A2	63241747	63241942	refGene_gn=ADCY1;refGene_tx=XM_006929062.4
A2	63242113	63242211	refGene_gn=ADCY1;refGene_tx=XM_006929062.4
A2	63243230	63243315	refGene_gn=ADCY1;refGene_tx=XM_006929062.4
A2	63248902	63248992	refGene_gn=ADCY1;refGene_tx=XM_006929062.4
A2	63249408	63249665	refGene_gn=ADCY1;refGene_tx=XM_006929062.4
A2	63249922	63250049	refGene_gn=ADCY1;refGene_tx=XM_006929062.4
A2	63267669	63267786	refGene_gn=ADCY1;refGene_tx=XM_006929062.4
A2	63268245	63268392	refGene_gn=ADCY1;refGene_tx=XM_006929062.4
A2	63268899	63268998	refGene_gn=ADCY1;refGene_tx=XM_006929062.4
A2	63274896	63275011	refGene_gn=ADCY1;refGene_tx=XM_006929062.4
A2	63276062	63276187	refGene_gn=ADCY1;refGene_tx=XM_006929062.4
A2	63277889	63278216	refGene_gn=ADCY1;refGene_tx=XM_006929062.4
A2	63289766	63289931
refGene_gn=LOC102900793;refGene_tx=XM_006929064.4
A2	63292295	63292721
refGene_gn=LOC102900793;refGene_tx=XM_006929064.4
A2	63298225	63298243
refGene_gn=LOC102900793;refGene_tx=XM_006929064.4
A2	63311061	63311401	refGene_gn=IGFBP1;refGene_tx=XM_023250142.1
A2	63312860	63313012	refGene_gn=IGFBP1;refGene_tx=XM_023250142.1
A2	63313913	63314039	refGene_gn=IGFBP1;refGene_tx=XM_023250142.1
A2	63314932	63315058	refGene_gn=IGFBP1;refGene_tx=XM_023250142.1
A2	63327536	63327662	refGene_gn=IGFBP3;refGene_tx=XM_023250143.1
A2	63328438	63328558	refGene_gn=IGFBP3;refGene_tx=XM_023250143.1
A2	63329714	63329768	refGene_gn=IGFBP3;refGene_tx=XM_023250143.1
A2	63329770	63329946	refGene_gn=IGFBP3;refGene_tx=XM_023250143.1
A2	63333007	63333416	refGene_gn=IGFBP3;refGene_tx=XM_023250143.1
A2	63973284	63973429	refGene_gn=TNS3;refGene_tx=XM_023250144.1
A2	63974901	63974928	refGene_gn=TNS3;refGene_tx=XM_023250144.1
A2	63975030	63975099	refGene_gn=TNS3;refGene_tx=XM_023250144.1
A2	63976591	63976760	refGene_gn=TNS3;refGene_tx=XM_023250144.1
A2	63985133	63985211	refGene_gn=TNS3;refGene_tx=XM_023250144.1
A2	63986041	63986119	refGene_gn=TNS3;refGene_tx=XM_023250144.1

| | | | |
|---|---|---|---|
| A2 | 63987005 | 63987101 | refGene_gn=TNS3;refGene_tx=XM_023250144.1 |
| A2 | 63990919 | 63991051 | refGene_gn=TNS3;refGene_tx=XM_023250144.1 |
| A2 | 63994484 | 63994571 | refGene_gn=TNS3;refGene_tx=XM_023250144.1 |
| A2 | 63995320 | 63995946 | refGene_gn=TNS3;refGene_tx=XM_023250144.1 |
| A2 | 63997330 | 63997502 | refGene_gn=TNS3;refGene_tx=XM_023250144.1 |
| A2 | 64033170 | 64033254 | refGene_gn=TNS3;refGene_tx=XM_023250144.1 |
| A2 | 64033341 | 64033468 | refGene_gn=TNS3;refGene_tx=XM_023250144.1 |
| A2 | 64034289 | 64034462 | refGene_gn=TNS3;refGene_tx=XM_023250144.1 |
| A2 | 64043517 | 64044756 | refGene_gn=TNS3;refGene_tx=XM_023250144.1 |
| A2 | 64062121 | 64062226 | refGene_gn=TNS3;refGene_tx=XM_023250144.1 |
| A2 | 64063586 | 64063652 | refGene_gn=TNS3;refGene_tx=XM_023250144.1 |
| A2 | 64063933 | 64064063 | refGene_gn=TNS3;refGene_tx=XM_023250144.1 |
| A2 | 64075479 | 64075555 | refGene_gn=TNS3;refGene_tx=XM_023250144.1 |
| A2 | 64076623 | 64076684 | refGene_gn=TNS3;refGene_tx=XM_023250144.1 |
| A2 | 64076971 | 64077084 | refGene_gn=TNS3;refGene_tx=XM_023250144.1 |
| A2 | 64085231 | 64085315 | refGene_gn=TNS3;refGene_tx=XM_023250144.1 |
| A2 | 64087779 | 64087844 | refGene_gn=TNS3;refGene_tx=XM_023250144.1 |
| A2 | 64091167 | 64091290 | refGene_gn=TNS3;refGene_tx=XM_023250144.1 |
| A2 | 64093132 | 64093183 | refGene_gn=TNS3;refGene_tx=XM_023250144.1 |
| A2 | 64094905 | 64095077 | refGene_gn=TNS3;refGene_tx=XM_023250144.1 |
| A2 | 64097079 | 64097132 | refGene_gn=TNS3;refGene_tx=XM_023250144.1 |
| A2 | 64138887 | 64138925 | refGene_gn=TNS3;refGene_tx=XM_023250144.1 |
| A2 | 64145555 | 64145664 | refGene_gn=TNS3;refGene_tx=XM_023250144.1 |
| A2 | 64155372 | 64155450 | refGene_gn=TNS3;refGene_tx=XM_023250144.1 |
| A2 | 64296690 | 64296837 | refGene_gn=PKD1L1;refGene_tx=XM_023250145.1 |
| A2 | 64300042 | 64300252 | refGene_gn=PKD1L1;refGene_tx=XM_023250145.1 |
| A2 | 64302367 | 64302502 | refGene_gn=PKD1L1;refGene_tx=XM_023250145.1 |
| A2 | 64304007 | 64304148 | refGene_gn=PKD1L1;refGene_tx=XM_023250145.1 |
| A2 | 64304422 | 64304527 | refGene_gn=PKD1L1;refGene_tx=XM_023250145.1 |
| A2 | 64305251 | 64305486 | refGene_gn=PKD1L1;refGene_tx=XM_023250145.1 |
| A2 | 64306517 | 64306690 | refGene_gn=PKD1L1;refGene_tx=XM_023250145.1 |
| A2 | 64307605 | 64307689 | refGene_gn=PKD1L1;refGene_tx=XM_023250145.1 |
| A2 | 64309386 | 64309510 | refGene_gn=PKD1L1;refGene_tx=XM_023250145.1 |
| A2 | 64313533 | 64313644 | refGene_gn=PKD1L1;refGene_tx=XM_023250145.1 |
| A2 | 64319262 | 64319381 | refGene_gn=PKD1L1;refGene_tx=XM_023250145.1 |
| A2 | 64321134 | 64321287 | refGene_gn=PKD1L1;refGene_tx=XM_023250145.1 |
| A2 | 64321575 | 64321660 | refGene_gn=PKD1L1;refGene_tx=XM_023250145.1 |
| A2 | 64322622 | 64322761 | refGene_gn=PKD1L1;refGene_tx=XM_023250145.1 |
| A2 | 64325618 | 64325754 | refGene_gn=PKD1L1;refGene_tx=XM_023250145.1 |
| A2 | 64326616 | 64326663 | refGene_gn=PKD1L1;refGene_tx=XM_023250145.1 |
| A2 | 64327056 | 64327126 | refGene_gn=PKD1L1;refGene_tx=XM_023250145.1 |
| A2 | 64327213 | 64327333 | refGene_gn=PKD1L1;refGene_tx=XM_023250145.1 |
| A2 | 64328469 | 64328643 | refGene_gn=PKD1L1;refGene_tx=XM_023250145.1 |
| A2 | 64329876 | 64330099 | refGene_gn=PKD1L1;refGene_tx=XM_023250145.1 |
| A2 | 64330847 | 64330954 | refGene_gn=PKD1L1;refGene_tx=XM_023250145.1 |
| A2 | 64332142 | 64332350 | refGene_gn=PKD1L1;refGene_tx=XM_023250145.1 |
| A2 | 64334436 | 64334520 | refGene_gn=PKD1L1;refGene_tx=XM_023250145.1 |
| A2 | 64336157 | 64336350 | refGene_gn=PKD1L1;refGene_tx=XM_023250145.1 |
| A2 | 64339549 | 64339650 | refGene_gn=PKD1L1;refGene_tx=XM_023250145.1 |
| A2 | 64341044 | 64341202 | refGene_gn=PKD1L1;refGene_tx=XM_023250145.1 |

A2 64341313 64341424 refGene_gn=PKD1L1;refGene_tx=XM_023250145.1
A2 64342855 64343098 refGene_gn=PKD1L1;refGene_tx=XM_023250145.1
A2 64343936 64344149 refGene_gn=PKD1L1;refGene_tx=XM_023250145.1
A2 64348990 64349047 refGene_gn=PKD1L1;refGene_tx=XM_023250145.1
A2 64349770 64349966 refGene_gn=PKD1L1;refGene_tx=XM_023250145.1
A2 64352052 64352164 refGene_gn=PKD1L1;refGene_tx=XM_023250145.1
A2 64355093 64355214 refGene_gn=PKD1L1;refGene_tx=XM_023250145.1
A2 64356270 64356413 refGene_gn=PKD1L1;refGene_tx=XM_023250145.1
A2 64358506 64358584 refGene_gn=PKD1L1;refGene_tx=XM_023250145.1
A2 64359753 64359930 refGene_gn=PKD1L1;refGene_tx=XM_023250145.1
A2 64361573 64361639 refGene_gn=PKD1L1;refGene_tx=XM_023250145.1
A2 64361851 64362184 refGene_gn=PKD1L1;refGene_tx=XM_023250145.1
A2 64363421 64363582 refGene_gn=PKD1L1;refGene_tx=XM_023250145.1
A2 64365560 64365782 refGene_gn=PKD1L1;refGene_tx=XM_023250145.1
A2 64368095 64368277 refGene_gn=PKD1L1;refGene_tx=XM_023250145.1
A2 64371818 64372025 refGene_gn=PKD1L1;refGene_tx=XM_023250145.1
A2 64374120 64374253 refGene_gn=PKD1L1;refGene_tx=XM_023250145.1
A2 64374811 64375051 refGene_gn=PKD1L1;refGene_tx=XM_023250145.1
A2 64375780 64375949 refGene_gn=PKD1L1;refGene_tx=XM_023250145.1
A2 64376371 64376491 refGene_gn=PKD1L1;refGene_tx=XM_023250145.1
A2 64378408 64378582 refGene_gn=PKD1L1;refGene_tx=XM_023250145.1
A2 64383921 64384220 refGene_gn=PKD1L1;refGene_tx=XM_023250145.1
A2 64384586 64384798 refGene_gn=PKD1L1;refGene_tx=XM_023250145.1
A2 64385344 64385465 refGene_gn=PKD1L1;refGene_tx=XM_023250145.1
A2 64387232 64387321 refGene_gn=PKD1L1;refGene_tx=XM_023250145.1
A2 64394015 64394149 refGene_gn=PKD1L1;refGene_tx=XM_023250145.1
A2 64405555 64405679 refGene_gn=PKD1L1;refGene_tx=XM_023250145.1
A2 64412086 64412149 refGene_gn=PKD1L1;refGene_tx=XM_023250145.1
A2 64412841 64412924
refGene_gn=HUS1;refGene_tx=XM_006929065.4;refGene_gn=HUS1;refGene_tx=XM_011280267.2
A2 64414915 64415035
refGene_gn=HUS1;refGene_tx=XM_006929065.4;refGene_gn=HUS1;refGene_tx=XM_011280267.2
A2 64415990 64416090
refGene_gn=HUS1;refGene_tx=XM_006929065.4;refGene_gn=HUS1;refGene_tx=XM_011280267.2
A2 64418605 64418680
refGene_gn=HUS1;refGene_tx=XM_006929065.4;refGene_gn=HUS1;refGene_tx=XM_011280267.2
A2 64420426 64420534
refGene_gn=HUS1;refGene_tx=XM_006929065.4;refGene_gn=HUS1;refGene_tx=XM_011280267.2
A2 64422659 64422836
refGene_gn=HUS1;refGene_tx=XM_006929065.4;refGene_gn=HUS1;refGene_tx=XM_011280267.2
A2 64422920 64423048
refGene_gn=HUS1;refGene_tx=XM_006929065.4;refGene_gn=HUS1;refGene_tx=XM_011280267.2
A2 64423194 64423255 refGene_gn=HUS1;refGene_tx=XM_006929065.4

A2	64424091 64424143 refGene_gn=HUS1;refGene_tx=XM_011280267.2
A2	64433964 64434228
refGene_gn=SUN3;refGene_tx=XM_006929066.3;refGene_gn=SUN3;refGene_tx=XM_006929067.3
A2	64435385 64435478
refGene_gn=SUN3;refGene_tx=XM_006929066.3;refGene_gn=SUN3;refGene_tx=XM_006929067.3
A2	64442129 64442297
refGene_gn=SUN3;refGene_tx=XM_006929066.3;refGene_gn=SUN3;refGene_tx=XM_006929067.3
A2	64443241 64443357
refGene_gn=SUN3;refGene_tx=XM_006929066.3;refGene_gn=SUN3;refGene_tx=XM_006929067.3
A2	64447696 64447781
refGene_gn=SUN3;refGene_tx=XM_006929066.3;refGene_gn=SUN3;refGene_tx=XM_006929067.3
A2	64449309 64449475
refGene_gn=SUN3;refGene_tx=XM_006929066.3;refGene_gn=SUN3;refGene_tx=XM_006929067.3
A2	64450809 64450850
refGene_gn=SUN3;refGene_tx=XM_006929066.3;refGene_gn=SUN3;refGene_tx=XM_006929067.3
A2	64456095 64456199
refGene_gn=SUN3;refGene_tx=XM_006929066.3;refGene_gn=SUN3;refGene_tx=XM_006929067.3
A2	64462749 64462811
refGene_gn=SUN3;refGene_tx=XM_006929066.3;refGene_gn=SUN3;refGene_tx=XM_006929067.3
A2	64465087 64465209
refGene_gn=SUN3;refGene_tx=XM_006929066.3;refGene_gn=SUN3;refGene_tx=XM_006929067.3
A2	64469108 64469163
refGene_gn=CA2H7orf57;refGene_tx=XM_006929068.3
A2	64473425 64473611
refGene_gn=CA2H7orf57;refGene_tx=XM_006929068.3
A2	64475266 64475378
refGene_gn=CA2H7orf57;refGene_tx=XM_006929068.3
A2	64476445 64476599
refGene_gn=CA2H7orf57;refGene_tx=XM_006929068.3
A2	64478006 64478104
refGene_gn=CA2H7orf57;refGene_tx=XM_006929068.3
A2	64479747 64479965
refGene_gn=CA2H7orf57;refGene_tx=XM_006929068.3
A2	64481964 64481979
refGene_gn=CA2H7orf57;refGene_tx=XM_006929068.3
A2	64483979 64484026
refGene_gn=CA2H7orf57;refGene_tx=XM_006929068.3
A2	64514518 64514596
refGene_gn=UPP1;refGene_tx=XM_011280266.3;refGene_gn=UPP1;refGene_tx=XM_019825231.2;refGene_gn=UPP1;refGene_tx=XM_019825232.2;refGene_gn=UPP

1;refGene_tx=XM_019825233.2
A2 64517578 64517737
refGene_gn=UPP1;refGene_tx=XM_011280266.3;refGene_gn=UPP1;refGene_tx=XM_019825231.2;refGene_gn=UPP1;refGene_tx=XM_019825232.2;refGene_gn=UPP1;refGene_tx=XM_019825233.2;refGene_gn=UPP1;refGene_tx=XM_023250146.1;refGene_gn=UPP1;refGene_tx=XM_023250147.1
A2 64518717 64518832
refGene_gn=UPP1;refGene_tx=XM_011280266.3;refGene_gn=UPP1;refGene_tx=XM_019825231.2;refGene_gn=UPP1;refGene_tx=XM_019825232.2;refGene_gn=UPP1;refGene_tx=XM_019825233.2;refGene_gn=UPP1;refGene_tx=XM_023250146.1;refGene_gn=UPP1;refGene_tx=XM_023250147.1
A2 64519599 64519621 refGene_gn=UPP1;refGene_tx=XM_019825235.2
A2 64521052 64521262
refGene_gn=UPP1;refGene_tx=XM_011280266.3;refGene_gn=UPP1;refGene_tx=XM_019825231.2;refGene_gn=UPP1;refGene_tx=XM_019825232.2;refGene_gn=UPP1;refGene_tx=XM_019825233.2;refGene_gn=UPP1;refGene_tx=XM_019825235.2;refGene_gn=UPP1;refGene_tx=XM_023250146.1;refGene_gn=UPP1;refGene_tx=XM_023250147.1
A2 64521547 64521694
refGene_gn=UPP1;refGene_tx=XM_011280266.3;refGene_gn=UPP1;refGene_tx=XM_019825231.2;refGene_gn=UPP1;refGene_tx=XM_019825232.2;refGene_gn=UPP1;refGene_tx=XM_019825233.2;refGene_gn=UPP1;refGene_tx=XM_019825235.2;refGene_gn=UPP1;refGene_tx=XM_023250146.1;refGene_gn=UPP1;refGene_tx=XM_023250147.1
A2 64522207 64522365
refGene_gn=UPP1;refGene_tx=XM_011280266.3;refGene_gn=UPP1;refGene_tx=XM_019825231.2;refGene_gn=UPP1;refGene_tx=XM_019825232.2;refGene_gn=UPP1;refGene_tx=XM_019825233.2;refGene_gn=UPP1;refGene_tx=XM_019825235.2;refGene_gn=UPP1;refGene_tx=XM_023250146.1;refGene_gn=UPP1;refGene_tx=XM_023250147.1
A2 64542042 64542111 refGene_gn=ABCA13;refGene_tx=XM_023241981.1
A2 64565928 64566022 refGene_gn=ABCA13;refGene_tx=XM_023241981.1
A2 64569629 64569753 refGene_gn=ABCA13;refGene_tx=XM_023241981.1
A2 64587826 64587981 refGene_gn=ABCA13;refGene_tx=XM_023241981.1
A2 64588176 64588205 refGene_gn=ABCA13;refGene_tx=XM_023241981.1
A2 64591606 64591770 refGene_gn=ABCA13;refGene_tx=XM_023241981.1
A2 64593528 64593659 refGene_gn=ABCA13;refGene_tx=XM_023241981.1
A2 64597816 64597950 refGene_gn=ABCA13;refGene_tx=XM_023241981.1
A2 64601444 64601609 refGene_gn=ABCA13;refGene_tx=XM_023241981.1
A2 64603296 64603496 refGene_gn=ABCA13;refGene_tx=XM_023241981.1
A2 64607016 64607144 refGene_gn=ABCA13;refGene_tx=XM_023241981.1
A2 64607815 64607916 refGene_gn=ABCA13;refGene_tx=XM_023241981.1
A2 64608761 64608929 refGene_gn=ABCA13;refGene_tx=XM_023241981.1
A2 64610261 64610467 refGene_gn=ABCA13;refGene_tx=XM_023241981.1
A2 64611873 64612013 refGene_gn=ABCA13;refGene_tx=XM_023241981.1
A2 64627099 64627214 refGene_gn=ABCA13;refGene_tx=XM_023241981.1
A2 64630554 64635336 refGene_gn=ABCA13;refGene_tx=XM_023241981.1
A2 64637326 64639138 refGene_gn=ABCA13;refGene_tx=XM_023241981.1
A2 64643521 64643628 refGene_gn=ABCA13;refGene_tx=XM_023241981.1
A2 64649230 64649349 refGene_gn=ABCA13;refGene_tx=XM_023241981.1

| | | | |
|---|---|---|---|
| A2 | 64654408 | 64654572 | refGene_gn=ABCA13;refGene_tx=XM_023241981.1 |
| A2 | 64656378 | 64656458 | refGene_gn=ABCA13;refGene_tx=XM_023241981.1 |
| A2 | 64657152 | 64657274 | refGene_gn=ABCA13;refGene_tx=XM_023241981.1 |
| A2 | 64665512 | 64665707 | refGene_gn=ABCA13;refGene_tx=XM_023241981.1 |
| A2 | 64667710 | 64667875 | refGene_gn=ABCA13;refGene_tx=XM_023241981.1 |
| A2 | 64668813 | 64668991 | refGene_gn=ABCA13;refGene_tx=XM_023241981.1 |
| A2 | 64673243 | 64673383 | refGene_gn=ABCA13;refGene_tx=XM_023241981.1 |
| A2 | 64687845 | 64687959 | refGene_gn=ABCA13;refGene_tx=XM_023241981.1 |
| A2 | 64691063 | 64691154 | refGene_gn=ABCA13;refGene_tx=XM_023241981.1 |
| A2 | 64694615 | 64694792 | refGene_gn=ABCA13;refGene_tx=XM_023241981.1 |
| A2 | 64696786 | 64697093 | refGene_gn=ABCA13;refGene_tx=XM_023241981.1 |
| A2 | 64714842 | 64714957 | refGene_gn=ABCA13;refGene_tx=XM_023241981.1 |
| A2 | 64716336 | 64716666 | refGene_gn=ABCA13;refGene_tx=XM_023241981.1 |
| A2 | 64719004 | 64719074 | refGene_gn=ABCA13;refGene_tx=XM_023241981.1 |
| A2 | 64719783 | 64719915 | refGene_gn=ABCA13;refGene_tx=XM_023241981.1 |
| A2 | 64729676 | 64729814 | refGene_gn=ABCA13;refGene_tx=XM_023241981.1 |
| A2 | 64731040 | 64731218 | refGene_gn=ABCA13;refGene_tx=XM_023241981.1 |
| A2 | 64732841 | 64733060 | refGene_gn=ABCA13;refGene_tx=XM_023241981.1 |
| A2 | 64738133 | 64738330 | refGene_gn=ABCA13;refGene_tx=XM_023241981.1 |
| A2 | 64742732 | 64742890 | refGene_gn=ABCA13;refGene_tx=XM_023241981.1 |
| A2 | 64744515 | 64744746 | refGene_gn=ABCA13;refGene_tx=XM_023241981.1 |
| A2 | 64761138 | 64761411 | refGene_gn=ABCA13;refGene_tx=XM_023241981.1 |
| A2 | 64761597 | 64761722 | refGene_gn=ABCA13;refGene_tx=XM_023241981.1 |
| A2 | 64761807 | 64761911 | refGene_gn=ABCA13;refGene_tx=XM_023241981.1 |
| A2 | 64781598 | 64781839 | refGene_gn=ABCA13;refGene_tx=XM_023241981.1 |
| A2 | 64796119 | 64796221 | refGene_gn=ABCA13;refGene_tx=XM_023241981.1 |
| A2 | 64801278 | 64801348 | refGene_gn=ABCA13;refGene_tx=XM_023241981.1 |
| A2 | 64805190 | 64805309 | refGene_gn=ABCA13;refGene_tx=XM_023241981.1 |
| A2 | 64808226 | 64808314 | refGene_gn=ABCA13;refGene_tx=XM_023241981.1 |
| A2 | 64813082 | 64813191 | refGene_gn=ABCA13;refGene_tx=XM_023241981.1 |
| A2 | 64828394 | 64828449 | refGene_gn=ABCA13;refGene_tx=XM_023241981.1 |
| A2 | 64830217 | 64830395 | refGene_gn=ABCA13;refGene_tx=XM_023241981.1 |
| A2 | 64832050 | 64832166 | refGene_gn=ABCA13;refGene_tx=XM_023241981.1 |
| A2 | 64838065 | 64838222 | refGene_gn=ABCA13;refGene_tx=XM_023241981.1 |
| A2 | 64843636 | 64843890 | refGene_gn=ABCA13;refGene_tx=XM_023241981.1 |
| A2 | 64850104 | 64850297 | refGene_gn=ABCA13;refGene_tx=XM_023241981.1 |
| A2 | 64852946 | 64853056 | refGene_gn=ABCA13;refGene_tx=XM_023241981.1 |
| A2 | 64893805 | 64893956 | refGene_gn=ABCA13;refGene_tx=XM_023241981.1 |
| A2 | 64899697 | 64899832 | refGene_gn=ABCA13;refGene_tx=XM_023241981.1 |
| A2 | 64902396 | 64902500 | refGene_gn=ABCA13;refGene_tx=XM_023241981.1 |
| A2 | 64910902 | 64910995 | refGene_gn=ABCA13;refGene_tx=XM_023241981.1 |
| A2 | 64936878 | 64936984 | refGene_gn=ABCA13;refGene_tx=XM_023241981.1 |
| A2 | 64938447 | 64938585 | refGene_gn=ABCA13;refGene_tx=XM_023241981.1 |
| A2 | 64940121 | 64940217 | refGene_gn=ABCA13;refGene_tx=XM_023241981.1 |
| A2 | 65982126 | 65982846 | refGene_gn=VWC2;refGene_tx=XM_023250155.1 |
| A2 | 66006170 | 66006300 | refGene_gn=VWC2;refGene_tx=XM_023250155.1 |
| A2 | 66111415 | 66111567 | refGene_gn=VWC2;refGene_tx=XM_023250155.1 |
| A2 | 66142676 | 66142771 | refGene_gn=ZPBP;refGene_tx=XM_006929088.4;refGene_gn=ZPBP;refGene_tx=XM_023250153.1;refGene_gn=ZPBP;refGene_tx=XM_023250154.1 |

A2 66180067 66180245
refGene_gn=ZPBP;refGene_tx=XM_006929088.4;refGene_gn=ZPBP;refGene_tx=XM_023250153.1;refGene_gn=ZPBP;refGene_tx=XM_023250154.1
A2 66202925 66203002
refGene_gn=ZPBP;refGene_tx=XM_006929088.4;refGene_gn=ZPBP;refGene_tx=XM_023250153.1;refGene_gn=ZPBP;refGene_tx=XM_023250154.1
A2 66222533 66222752
refGene_gn=ZPBP;refGene_tx=XM_006929088.4;refGene_gn=ZPBP;refGene_tx=XM_023250153.1
A2 66235346 66235496
refGene_gn=ZPBP;refGene_tx=XM_006929088.4;refGene_gn=ZPBP;refGene_tx=XM_023250153.1;refGene_gn=ZPBP;refGene_tx=XM_023250154.1
A2 66244412 66244538
refGene_gn=ZPBP;refGene_tx=XM_006929088.4;refGene_gn=ZPBP;refGene_tx=XM_023250154.1
A2 66253233 66253314
refGene_gn=ZPBP;refGene_tx=XM_006929088.4;refGene_gn=ZPBP;refGene_tx=XM_023250153.1;refGene_gn=ZPBP;refGene_tx=XM_023250154.1
A2 66255027 66255199
refGene_gn=ZPBP;refGene_tx=XM_006929088.4;refGene_gn=ZPBP;refGene_tx=XM_023250153.1;refGene_gn=ZPBP;refGene_tx=XM_023250154.1
A2 66258540 66259093
refGene_gn=SPATA48;refGene_tx=XM_023250148.1;refGene_gn=SPATA48;refGene_tx=XM_023250149.1;refGene_gn=SPATA48;refGene_tx=XM_023250150.1;refGene_gn=SPATA48;refGene_tx=XM_023250151.1
A2 66265868 66265966
refGene_gn=SPATA48;refGene_tx=XM_023250148.1;refGene_gn=SPATA48;refGene_tx=XM_023250149.1;refGene_gn=SPATA48;refGene_tx=XM_023250150.1;refGene_gn=SPATA48;refGene_tx=XM_023250151.1
A2 66284244 66284321
refGene_gn=SPATA48;refGene_tx=XM_023250148.1;refGene_gn=SPATA48;refGene_tx=XM_023250149.1;refGene_gn=SPATA48;refGene_tx=XM_023250150.1;refGene_gn=SPATA48;refGene_tx=XM_023250151.1
A2 66290049 66290188
refGene_gn=SPATA48;refGene_tx=XM_023250148.1;refGene_gn=SPATA48;refGene_tx=XM_023250149.1;refGene_gn=SPATA48;refGene_tx=XM_023250150.1;refGene_gn=SPATA48;refGene_tx=XM_023250151.1
A2 66292809 66292877
refGene_gn=SPATA48;refGene_tx=XM_023250148.1;refGene_gn=SPATA48;refGene_tx=XM_023250149.1;refGene_gn=SPATA48;refGene_tx=XM_023250150.1;refGene_gn=SPATA48;refGene_tx=XM_023250151.1
A2 66296683 66296725
refGene_gn=SPATA48;refGene_tx=XM_023250148.1;refGene_gn=SPATA48;refGene_tx=XM_023250149.1;refGene_gn=SPATA48;refGene_tx=XM_023250150.1;refGene_gn=SPATA48;refGene_tx=XM_023250151.1
A2 66297951 66298059
refGene_gn=SPATA48;refGene_tx=XM_023250148.1;refGene_gn=SPATA48;refGene_tx=XM_023250149.1;refGene_gn=SPATA48;refGene_tx=XM_023250150.1;refGene_gn=SPATA48;refGene_tx=XM_023250151.1
A2 66302170 66302174 refGene_gn=SPATA48;refGene_tx=XM_023250151.1

A2 66307081 66307189
refGene_gn=SPATA48;refGene_tx=XM_023250148.1;refGene_gn=SPATA48;refGene_tx=XM_023250149.1;refGene_gn=SPATA48;refGene_tx=XM_023250150.1
A2 66307307 66307314 refGene_gn=SPATA48;refGene_tx=XM_023250150.1
A2 66307426 66307577 refGene_gn=SPATA48;refGene_tx=XM_023250148.1
A2 66312480 66312604 refGene_gn=SPATA48;refGene_tx=XM_023250149.1
A2 66481322 66481362
refGene_gn=IKZF1;refGene_tx=XM_023250156.1;refGene_gn=IKZF1;refGene_tx=XM_023250157.1;refGene_gn=IKZF1;refGene_tx=XM_023250158.1;refGene_gn=IKZF1;refGene_tx=XM_023250159.1;refGene_gn=IKZF1;refGene_tx=XM_023250160.1;refGene_gn=IKZF1;refGene_tx=XM_023250161.1;refGene_gn=IKZF1;refGene_tx=XM_023250162.1;refGene_gn=IKZF1;refGene_tx=XM_023250163.1;refGene_gn=IKZF1;refGene_tx=XM_023250164.1;refGene_gn=IKZF1;refGene_tx=XM_023250165.1;refGene_gn=IKZF1;refGene_tx=XM_023250166.1;refGene_gn=IKZF1;refGene_tx=XM_023250168.1;refGene_gn=IKZF1;refGene_tx=XM_023250169.1;refGene_gn=IKZF1;refGene_tx=XM_023250170.1
A2 66489451 66489571
refGene_gn=IKZF1;refGene_tx=XM_023250156.1;refGene_gn=IKZF1;refGene_tx=XM_023250157.1;refGene_gn=IKZF1;refGene_tx=XM_023250158.1;refGene_gn=IKZF1;refGene_tx=XM_023250159.1;refGene_gn=IKZF1;refGene_tx=XM_023250160.1;refGene_gn=IKZF1;refGene_tx=XM_023250161.1;refGene_gn=IKZF1;refGene_tx=XM_023250162.1;refGene_gn=IKZF1;refGene_tx=XM_023250163.1;refGene_gn=IKZF1;refGene_tx=XM_023250164.1;refGene_gn=IKZF1;refGene_tx=XM_023250165.1;refGene_gn=IKZF1;refGene_tx=XM_023250166.1;refGene_gn=IKZF1;refGene_tx=XM_023250168.1;refGene_gn=IKZF1;refGene_tx=XM_023250169.1;refGene_gn=IKZF1;refGene_tx=XM_023250170.1
A2 66530610 66530670
refGene_gn=IKZF1;refGene_tx=XM_023250156.1;refGene_gn=IKZF1;refGene_tx=XM_023250157.1;refGene_gn=IKZF1;refGene_tx=XM_023250158.1;refGene_gn=IKZF1;refGene_tx=XM_023250159.1;refGene_gn=IKZF1;refGene_tx=XM_023250160.1;refGene_gn=IKZF1;refGene_tx=XM_023250161.1;refGene_gn=IKZF1;refGene_tx=XM_023250163.1;refGene_gn=IKZF1;refGene_tx=XM_023250164.1
A2 66538680 66538941
refGene_gn=IKZF1;refGene_tx=XM_023250156.1;refGene_gn=IKZF1;refGene_tx=XM_023250157.1;refGene_gn=IKZF1;refGene_tx=XM_023250158.1;refGene_gn=IKZF1;refGene_tx=XM_023250159.1;refGene_gn=IKZF1;refGene_tx=XM_023250160.1;refGene_gn=IKZF1;refGene_tx=XM_023250161.1;refGene_gn=IKZF1;refGene_tx=XM_023250162.1;refGene_gn=IKZF1;refGene_tx=XM_023250163.1;refGene_gn=IKZF1;refGene_tx=XM_023250164.1;refGene_gn=IKZF1;refGene_tx=XM_023250165.1;refGene_gn=IKZF1;refGene_tx=XM_023250167.1
A2 66545354 66545522
refGene_gn=IKZF1;refGene_tx=XM_023250156.1;refGene_gn=IKZF1;refGene_tx=XM_023250157.1;refGene_gn=IKZF1;refGene_tx=XM_023250158.1;refGene_gn=IKZF1;refGene_tx=XM_023250159.1;refGene_gn=IKZF1;refGene_tx=XM_023250160.1;refGene_gn=IKZF1;refGene_tx=XM_023250161.1;refGene_gn=IKZF1;refGene_tx=XM_023250162.1;refGene_gn=IKZF1;refGene_tx=XM_023250163.1;refGene_gn=IKZF1;refGene_tx=XM_023250164.1;refGene_gn=IKZF1;refGene_tx=XM_023250165.1;refGene_gn=IKZF1;refGene_tx=XM_023250166.1;refGene_gn=IKZF1;refGene_tx=XM_023250167.1;refGene_gn=IKZF1;refGene_tx=XM_023250168.1;refGene_gn=IKZF1;refGene_tx=XM_023250169.1;refGene_gn=IKZF1;refGene_tx=

XM_023250170.1
A2 66549205 66549331
refGene_gn=IKZF1;refGene_tx=XM_023250156.1;refGene_gn=IKZF1;refGene_tx=XM_023250157.1;refGene_gn=IKZF1;refGene_tx=XM_023250158.1;refGene_gn=IKZF1;refGene_tx=XM_023250159.1;refGene_gn=IKZF1;refGene_tx=XM_023250160.1;refGene_gn=IKZF1;refGene_tx=XM_023250161.1;refGene_gn=IKZF1;refGene_tx=XM_023250162.1;refGene_gn=IKZF1;refGene_tx=XM_023250164.1;refGene_gn=IKZF1;refGene_tx=XM_023250166.1;refGene_gn=IKZF1;refGene_tx=XM_023250167.1;refGene_gn=IKZF1;refGene_tx=XM_023250168.1;refGene_gn=IKZF1;refGene_tx=XM_023250171.1
A2 66553852 66553987
refGene_gn=IKZF1;refGene_tx=XM_023250156.1;refGene_gn=IKZF1;refGene_tx=XM_023250157.1;refGene_gn=IKZF1;refGene_tx=XM_023250158.1;refGene_gn=IKZF1;refGene_tx=XM_023250159.1;refGene_gn=IKZF1;refGene_tx=XM_023250160.1;refGene_gn=IKZF1;refGene_tx=XM_023250161.1;refGene_gn=IKZF1;refGene_tx=XM_023250162.1;refGene_gn=IKZF1;refGene_tx=XM_023250163.1;refGene_gn=IKZF1;refGene_tx=XM_023250165.1;refGene_gn=IKZF1;refGene_tx=XM_023250166.1;refGene_gn=IKZF1;refGene_tx=XM_023250167.1;refGene_gn=IKZF1;refGene_tx=XM_023250168.1;refGene_gn=IKZF1;refGene_tx=XM_023250169.1;refGene_gn=IKZF1;refGene_tx=XM_023250170.1;refGene_gn=IKZF1;refGene_tx=XM_023250171.1
A2 66562689 66563399
refGene_gn=IKZF1;refGene_tx=XM_023250156.1;refGene_gn=IKZF1;refGene_tx=XM_023250157.1;refGene_gn=IKZF1;refGene_tx=XM_023250158.1;refGene_gn=IKZF1;refGene_tx=XM_023250159.1;refGene_gn=IKZF1;refGene_tx=XM_023250160.1;refGene_gn=IKZF1;refGene_tx=XM_023250161.1;refGene_gn=IKZF1;refGene_tx=XM_023250162.1;refGene_gn=IKZF1;refGene_tx=XM_023250163.1;refGene_gn=IKZF1;refGene_tx=XM_023250164.1;refGene_gn=IKZF1;refGene_tx=XM_023250165.1;refGene_gn=IKZF1;refGene_tx=XM_023250166.1;refGene_gn=IKZF1;refGene_tx=XM_023250167.1;refGene_gn=IKZF1;refGene_tx=XM_023250168.1;refGene_gn=IKZF1;refGene_tx=XM_023250169.1;refGene_gn=IKZF1;refGene_tx=XM_023250170.1;refGene_gn=IKZF1;refGene_tx=XM_023250171.1
A2 66602687 66604728
refGene_gn=FIGNL1;refGene_tx=XM_003982612.4;refGene_gn=FIGNL1;refGene_tx=XM_019825240.2;refGene_gn=FIGNL1;refGene_tx=XM_023250172.1;refGene_gn=FIGNL1;refGene_tx=XM_023250173.1
A2 66606582 66606617 refGene_gn=FIGNL1;refGene_tx=XM_019825240.2
A2 66616776 66616977
refGene_gn=DDC;refGene_tx=XM_003982613.4;refGene_gn=DDC;refGene_tx=XM_019825243.2;refGene_gn=DDC;refGene_tx=XM_023250174.1
A2 66620063 66620165
refGene_gn=DDC;refGene_tx=XM_003982613.4;refGene_gn=DDC;refGene_tx=XM_019825243.2;refGene_gn=DDC;refGene_tx=XM_023250174.1
A2 66623504 66623603
refGene_gn=DDC;refGene_tx=XM_003982613.4;refGene_gn=DDC;refGene_tx=XM_019825243.2;refGene_gn=DDC;refGene_tx=XM_023250174.1;refGene_gn=DDC;refGene_tx=XM_023250175.1
A2 66627472 66627492
refGene_gn=DDC;refGene_tx=XM_003982613.4;refGene_gn=DDC;refGene_tx=XM_019825243.2;refGene_gn=DDC;refGene_tx=XM_023250174.1

A2 66632551 66632628
refGene_gn=DDC;refGene_tx=XM_003982613.4;refGene_gn=DDC;refGene_tx=XM_019825243.2;refGene_gn=DDC;refGene_tx=XM_023250174.1;refGene_gn=DDC;refGene_tx=XM_023250175.1
A2 66642091 66642159
refGene_gn=DDC;refGene_tx=XM_003982613.4;refGene_gn=DDC;refGene_tx=XM_019825243.2;refGene_gn=DDC;refGene_tx=XM_023250174.1;refGene_gn=DDC;refGene_tx=XM_023250175.1
A2 66647719 66647814
refGene_gn=DDC;refGene_tx=XM_003982613.4;refGene_gn=DDC;refGene_tx=XM_019825243.2;refGene_gn=DDC;refGene_tx=XM_023250174.1;refGene_gn=DDC;refGene_tx=XM_023250175.1
A2 66652762 66652829
refGene_gn=DDC;refGene_tx=XM_003982613.4;refGene_gn=DDC;refGene_tx=XM_019825243.2;refGene_gn=DDC;refGene_tx=XM_023250174.1;refGene_gn=DDC;refGene_tx=XM_023250175.1
A2 66668680 66668824
refGene_gn=DDC;refGene_tx=XM_003982613.4;refGene_gn=DDC;refGene_tx=XM_019825243.2;refGene_gn=DDC;refGene_tx=XM_023250174.1;refGene_gn=DDC;refGene_tx=XM_023250175.1
A2 66669523 66669658
refGene_gn=DDC;refGene_tx=XM_003982613.4;refGene_gn=DDC;refGene_tx=XM_019825243.2;refGene_gn=DDC;refGene_tx=XM_023250174.1;refGene_gn=DDC;refGene_tx=XM_023250175.1
A2 66675218 66675338
refGene_gn=DDC;refGene_tx=XM_003982613.4;refGene_gn=DDC;refGene_tx=XM_019825243.2;refGene_gn=DDC;refGene_tx=XM_023250174.1;refGene_gn=DDC;refGene_tx=XM_023250175.1
A2 66677881 66677995
refGene_gn=DDC;refGene_tx=XM_003982613.4;refGene_gn=DDC;refGene_tx=XM_019825243.2;refGene_gn=DDC;refGene_tx=XM_023250174.1;refGene_gn=DDC;refGene_tx=XM_023250175.1
A2 66680731 66680960
refGene_gn=DDC;refGene_tx=XM_003982613.4;refGene_gn=DDC;refGene_tx=XM_019825243.2;refGene_gn=DDC;refGene_tx=XM_023250174.1;refGene_gn=DDC;refGene_tx=XM_023250175.1
A2 66691942 66692004 refGene_gn=DDC;refGene_tx=XM_023250175.1
A2 66714800 66714947
refGene_gn=GRB10;refGene_tx=XM_003982618.5;refGene_gn=GRB10;refGene_tx=XM_006929085.4;refGene_gn=GRB10;refGene_tx=XM_023250176.1;refGene_gn=GRB10;refGene_tx=XM_023250177.1;refGene_gn=GRB10;refGene_tx=XM_023250178.1;refGene_gn=GRB10;refGene_tx=XM_023250179.1
A2 66717195 66717289
refGene_gn=GRB10;refGene_tx=XM_003982618.5;refGene_gn=GRB10;refGene_tx=XM_006929085.4;refGene_gn=GRB10;refGene_tx=XM_023250176.1;refGene_gn=GRB10;refGene_tx=XM_023250177.1;refGene_gn=GRB10;refGene_tx=XM_023250178.1;refGene_gn=GRB10;refGene_tx=XM_023250179.1
A2 66723248 66723336
refGene_gn=GRB10;refGene_tx=XM_003982618.5;refGene_gn=GRB10;refGene_tx=XM_006929085.4;refGene_gn=GRB10;refGene_tx=XM_023250176.1;refGene_gn=

GRB10;refGene_tx=XM_023250177.1;refGene_gn=GRB10;refGene_tx=XM_023250178.1;refGene_gn=GRB10;refGene_tx=XM_023250179.1
A2 66723555 66723622
refGene_gn=GRB10;refGene_tx=XM_003982618.5;refGene_gn=GRB10;refGene_tx=XM_006929085.4;refGene_gn=GRB10;refGene_tx=XM_023250176.1;refGene_gn=GRB10;refGene_tx=XM_023250177.1;refGene_gn=GRB10;refGene_tx=XM_023250178.1;refGene_gn=GRB10;refGene_tx=XM_023250179.1
A2 66724579 66724696
refGene_gn=GRB10;refGene_tx=XM_003982618.5;refGene_gn=GRB10;refGene_tx=XM_006929085.4;refGene_gn=GRB10;refGene_tx=XM_023250176.1;refGene_gn=GRB10;refGene_tx=XM_023250177.1;refGene_gn=GRB10;refGene_tx=XM_023250178.1;refGene_gn=GRB10;refGene_tx=XM_023250179.1
A2 66725543 66725621
refGene_gn=GRB10;refGene_tx=XM_003982618.5;refGene_gn=GRB10;refGene_tx=XM_006929085.4;refGene_gn=GRB10;refGene_tx=XM_023250176.1;refGene_gn=GRB10;refGene_tx=XM_023250177.1;refGene_gn=GRB10;refGene_tx=XM_023250178.1;refGene_gn=GRB10;refGene_tx=XM_023250179.1
A2 66727767 66727866
refGene_gn=GRB10;refGene_tx=XM_003982618.5;refGene_gn=GRB10;refGene_tx=XM_006929085.4;refGene_gn=GRB10;refGene_tx=XM_023250176.1;refGene_gn=GRB10;refGene_tx=XM_023250177.1;refGene_gn=GRB10;refGene_tx=XM_023250178.1;refGene_gn=GRB10;refGene_tx=XM_023250179.1
A2 66729923 66730034
refGene_gn=GRB10;refGene_tx=XM_003982618.5;refGene_gn=GRB10;refGene_tx=XM_006929085.4;refGene_gn=GRB10;refGene_tx=XM_023250176.1;refGene_gn=GRB10;refGene_tx=XM_023250177.1;refGene_gn=GRB10;refGene_tx=XM_023250178.1;refGene_gn=GRB10;refGene_tx=XM_023250179.1
A2 66730872 66731010
refGene_gn=GRB10;refGene_tx=XM_003982618.5;refGene_gn=GRB10;refGene_tx=XM_006929085.4;refGene_gn=GRB10;refGene_tx=XM_023250176.1;refGene_gn=GRB10;refGene_tx=XM_023250177.1;refGene_gn=GRB10;refGene_tx=XM_023250178.1;refGene_gn=GRB10;refGene_tx=XM_023250179.1
A2 66731904 66731973
refGene_gn=GRB10;refGene_tx=XM_003982618.5;refGene_gn=GRB10;refGene_tx=XM_006929085.4;refGene_gn=GRB10;refGene_tx=XM_023250176.1;refGene_gn=GRB10;refGene_tx=XM_023250177.1;refGene_gn=GRB10;refGene_tx=XM_023250178.1;refGene_gn=GRB10;refGene_tx=XM_023250179.1
A2 66733411 66733527
refGene_gn=GRB10;refGene_tx=XM_003982618.5;refGene_gn=GRB10;refGene_tx=XM_006929085.4;refGene_gn=GRB10;refGene_tx=XM_023250176.1;refGene_gn=GRB10;refGene_tx=XM_023250177.1;refGene_gn=GRB10;refGene_tx=XM_023250178.1;refGene_gn=GRB10;refGene_tx=XM_023250179.1
A2 66740800 66740957
refGene_gn=GRB10;refGene_tx=XM_003982618.5;refGene_gn=GRB10;refGene_tx=XM_006929085.4;refGene_gn=GRB10;refGene_tx=XM_023250176.1;refGene_gn=GRB10;refGene_tx=XM_023250177.1;refGene_gn=GRB10;refGene_tx=XM_023250178.1;refGene_gn=GRB10;refGene_tx=XM_023250179.1
A2 66772940 66773082
refGene_gn=GRB10;refGene_tx=XM_003982618.5;refGene_gn=GRB10;refGene_tx=XM_006929085.4;refGene_gn=GRB10;refGene_tx=XM_023250176.1;refGene_gn=

GRB10;refGene_tx=XM_023250177.1;refGene_gn=GRB10;refGene_tx=XM_0232501
78.1;refGene_gn=GRB10;refGene_tx=XM_023250179.1
A2 66779287 66779507
refGene_gn=GRB10;refGene_tx=XM_003982618.5;refGene_gn=GRB10;refGene_tx
=XM_006929085.4;refGene_gn=GRB10;refGene_tx=XM_023250176.1;refGene_gn=
GRB10;refGene_tx=XM_023250177.1;refGene_gn=GRB10;refGene_tx=XM_0232501
78.1;refGene_gn=GRB10;refGene_tx=XM_023250179.1
A2 66808077 66808168
refGene_gn=GRB10;refGene_tx=XM_003982618.5;refGene_gn=GRB10;refGene_tx
=XM_023250176.1;refGene_gn=GRB10;refGene_tx=XM_023250177.1
A2 66835507 66835558 refGene_gn=GRB10;refGene_tx=XM_023250177.1
A2 66835560 66835611
refGene_gn=GRB10;refGene_tx=XM_003982618.5;refGene_gn=GRB10;refGene_tx
=XM_023250176.1
A2 67058418 67058553 refGene_gn=COBL;refGene_tx=XM_023250180.1
A2 67064444 67064699 refGene_gn=COBL;refGene_tx=XM_023250180.1
A2 67065305 67065425 refGene_gn=COBL;refGene_tx=XM_023250180.1
A2 67066300 67068096 refGene_gn=COBL;refGene_tx=XM_023250180.1
A2 67069511 67069609 refGene_gn=COBL;refGene_tx=XM_023250180.1
A2 67083428 67083738 refGene_gn=COBL;refGene_tx=XM_023250180.1
A2 67119686 67119860 refGene_gn=COBL;refGene_tx=XM_023250180.1
A2 67123862 67124001 refGene_gn=COBL;refGene_tx=XM_023250180.1
A2 67171157 67171340 refGene_gn=COBL;refGene_tx=XM_023250180.1
A2 67207823 67207921 refGene_gn=COBL;refGene_tx=XM_023250180.1
A2 67213564 67213793 refGene_gn=COBL;refGene_tx=XM_023250180.1
A2 67215434 67215559 refGene_gn=COBL;refGene_tx=XM_023250180.1
A2 67215561 67215644 refGene_gn=COBL;refGene_tx=XM_023250180.1
A2 67230708 67230909 refGene_gn=COBL;refGene_tx=XM_023250180.1
A2 67331172 67331213 refGene_gn=COBL;refGene_tx=XM_023250180.1
A2 68900720 68901443
refGene_gn=POM121L12;refGene_tx=XM_006929091.4
A2 70376235 70376314
refGene_gn=VSTM2A;refGene_tx=XM_011280292.2;refGene_gn=VSTM2A;refGene_
tx=XM_011280293.2;refGene_gn=VSTM2A;refGene_tx=XM_011280294.2;refGene_
gn=VSTM2A;refGene_tx=XM_019825254.1
A2 70378112 70378279
refGene_gn=VSTM2A;refGene_tx=XM_011280292.2;refGene_gn=VSTM2A;refGene_
tx=XM_011280293.2;refGene_gn=VSTM2A;refGene_tx=XM_011280294.2;refGene_
gn=VSTM2A;refGene_tx=XM_019825254.1
A2 70379414 70379465
refGene_gn=VSTM2A;refGene_tx=XM_011280292.2;refGene_gn=VSTM2A;refGene_
tx=XM_011280293.2;refGene_gn=VSTM2A;refGene_tx=XM_011280294.2;refGene_
gn=VSTM2A;refGene_tx=XM_019825254.1
A2 70382330 70382667
refGene_gn=VSTM2A;refGene_tx=XM_011280292.2;refGene_gn=VSTM2A;refGene_
tx=XM_011280293.2;refGene_gn=VSTM2A;refGene_tx=XM_011280294.2;refGene_
gn=VSTM2A;refGene_tx=XM_019825254.1
A2 70401811 70401858 refGene_gn=VSTM2A;refGene_tx=XM_011280294.2
A2 70401949 70402029 refGene_gn=VSTM2A;refGene_tx=XM_011280293.2
A2 70402074 70402163

refGene_gn=VSTM2A;refGene_tx=XM_011280292.2;refGene_gn=VSTM2A;refGene_tx=XM_019825254.1
A2 70641492 70641502 refGene_gn=SEC61G;refGene_tx=XM_003982620.4
A2 70645759 70645862 refGene_gn=SEC61G;refGene_tx=XM_003982620.4
A2 70647366 70647460 refGene_gn=SEC61G;refGene_tx=XM_003982620.4
A2 70930686 70930774 refGene_gn=EGFR;refGene_tx=XM_006929086.3
A2 71076058 71076210 refGene_gn=EGFR;refGene_tx=XM_006929086.3
A2 71077170 71077185 refGene_gn=EGFR;refGene_tx=XM_019825256.1
A2 71077277 71077461
refGene_gn=EGFR;refGene_tx=XM_006929086.3;refGene_gn=EGFR;refGene_tx=XM_019825256.1
A2 71080810 71080945
refGene_gn=EGFR;refGene_tx=XM_006929086.3;refGene_gn=EGFR;refGene_tx=XM_019825256.1
A2 71084383 71084452
refGene_gn=EGFR;refGene_tx=XM_006929086.3;refGene_gn=EGFR;refGene_tx=XM_019825256.1
A2 71085305 71085424
refGene_gn=EGFR;refGene_tx=XM_006929086.3;refGene_gn=EGFR;refGene_tx=XM_019825256.1
A2 71087025 71087167
refGene_gn=EGFR;refGene_tx=XM_006929086.3;refGene_gn=EGFR;refGene_tx=XM_019825256.1
A2 71089448 71089565
refGene_gn=EGFR;refGene_tx=XM_006929086.3;refGene_gn=EGFR;refGene_tx=XM_019825256.1
A2 71093000 71093127
refGene_gn=EGFR;refGene_tx=XM_006929086.3;refGene_gn=EGFR;refGene_tx=XM_019825256.1
A2 71093226 71093300
refGene_gn=EGFR;refGene_tx=XM_006929086.3;refGene_gn=EGFR;refGene_tx=XM_019825256.1
A2 71094141 71094232
refGene_gn=EGFR;refGene_tx=XM_006929086.3;refGene_gn=EGFR;refGene_tx=XM_019825256.1
A2 71097021 71097221
refGene_gn=EGFR;refGene_tx=XM_006929086.3;refGene_gn=EGFR;refGene_tx=XM_019825256.1
A2 71098155 71098288
refGene_gn=EGFR;refGene_tx=XM_006929086.3;refGene_gn=EGFR;refGene_tx=XM_019825256.1
A2 71100300 71100391
refGene_gn=EGFR;refGene_tx=XM_006929086.3;refGene_gn=EGFR;refGene_tx=XM_019825256.1
A2 71102279 71102437
refGene_gn=EGFR;refGene_tx=XM_006929086.3;refGene_gn=EGFR;refGene_tx=XM_019825256.1
A2 71104498 71104537
refGene_gn=EGFR;refGene_tx=XM_006929086.3;refGene_gn=EGFR;refGene_tx=XM_019825256.1

A2	71107493	71107635
refGene_gn=EGFR;refGene_tx=XM_006929086.3;refGene_gn=EGFR;refGene_tx=XM_019825256.1
A2	71108506	71108629
refGene_gn=EGFR;refGene_tx=XM_006929086.3;refGene_gn=EGFR;refGene_tx=XM_019825256.1
A2	71109299	71109398
refGene_gn=EGFR;refGene_tx=XM_006929086.3;refGene_gn=EGFR;refGene_tx=XM_019825256.1
A2	71116143	71116329
refGene_gn=EGFR;refGene_tx=XM_006929086.3;refGene_gn=EGFR;refGene_tx=XM_019825256.1
A2	71127586	71127742
refGene_gn=EGFR;refGene_tx=XM_006929086.3;refGene_gn=EGFR;refGene_tx=XM_019825256.1
A2	71128657	71128733
refGene_gn=EGFR;refGene_tx=XM_006929086.3;refGene_gn=EGFR;refGene_tx=XM_019825256.1
A2	71131924	71132071
refGene_gn=EGFR;refGene_tx=XM_006929086.3;refGene_gn=EGFR;refGene_tx=XM_019825256.1
A2	71133522	71133620
refGene_gn=EGFR;refGene_tx=XM_006929086.3;refGene_gn=EGFR;refGene_tx=XM_019825256.1
A2	71135031	71135199
refGene_gn=EGFR;refGene_tx=XM_006929086.3;refGene_gn=EGFR;refGene_tx=XM_019825256.1
A2	71135611	71135659
refGene_gn=EGFR;refGene_tx=XM_006929086.3;refGene_gn=EGFR;refGene_tx=XM_019825256.1
A2	71136373	71136476
refGene_gn=EGFR;refGene_tx=XM_006929086.3;refGene_gn=EGFR;refGene_tx=XM_019825256.1
A2	71137152	71137511
refGene_gn=EGFR;refGene_tx=XM_006929086.3;refGene_gn=EGFR;refGene_tx=XM_019825256.1
A2	71286793	71286997	refGene_gn=LANCL2;refGene_tx=XM_003982621.3
A2	71311782	71311900	refGene_gn=LANCL2;refGene_tx=XM_003982621.3
A2	71325579	71325787	refGene_gn=LANCL2;refGene_tx=XM_003982621.3
A2	71327059	71327207	refGene_gn=LANCL2;refGene_tx=XM_003982621.3
A2	71328225	71328372	refGene_gn=LANCL2;refGene_tx=XM_003982621.3
A2	71331917	71332100	refGene_gn=LANCL2;refGene_tx=XM_003982621.3
A2	71340538	71340715	refGene_gn=LANCL2;refGene_tx=XM_003982621.3
A2	71340993	71341066	refGene_gn=LANCL2;refGene_tx=XM_003982621.3
A2	71344445	71344552	refGene_gn=LANCL2;refGene_tx=XM_003982621.3
A2	71390594	71390785
refGene_gn=VOPP1;refGene_tx=XM_023250182.1;refGene_gn=VOPP1;refGene_tx=XM_023250183.1
A2	71414464	71414591
refGene_gn=VOPP1;refGene_tx=XM_023250182.1;refGene_gn=VOPP1;refGene_tx

=XM_023250183.1
A2	72111799	72111823
refGene_gn=LOC105259735;refGene_tx=XM_023242000.1
A2	72113562	72113670
refGene_gn=LOC105259735;refGene_tx=XM_023242000.1
A2	72114620	72114814
refGene_gn=LOC105259735;refGene_tx=XM_023242000.1
A2	72115549	72115609
refGene_gn=LOC105259735;refGene_tx=XM_023242000.1
A2	72116551	72116718
refGene_gn=LOC105259735;refGene_tx=XM_023242000.1
A2	72116988	72117431
refGene_gn=LOC105259735;refGene_tx=XM_023242000.1
A2	72179037	72179296
refGene_gn=BLVRA;refGene_tx=XM_023250187.1;refGene_gn=BLVRA;refGene_tx
=XM_023250188.1;refGene_gn=BLVRA;refGene_tx=XM_023250189.1;refGene_gn=
BLVRA;refGene_tx=XM_023250190.1;refGene_gn=BLVRA;refGene_tx=XM_0232501
91.1
A2	72183582	72183754
refGene_gn=BLVRA;refGene_tx=XM_023250187.1;refGene_gn=BLVRA;refGene_tx
=XM_023250188.1;refGene_gn=BLVRA;refGene_tx=XM_023250189.1;refGene_gn=
BLVRA;refGene_tx=XM_023250190.1;refGene_gn=BLVRA;refGene_tx=XM_0232501
91.1
A2	72186369	72186477
refGene_gn=BLVRA;refGene_tx=XM_023250187.1;refGene_gn=BLVRA;refGene_tx
=XM_023250188.1;refGene_gn=BLVRA;refGene_tx=XM_023250189.1;refGene_gn=
BLVRA;refGene_tx=XM_023250190.1;refGene_gn=BLVRA;refGene_tx=XM_0232501
91.1
A2	72204446	72204544
refGene_gn=BLVRA;refGene_tx=XM_023250187.1;refGene_gn=BLVRA;refGene_tx
=XM_023250188.1;refGene_gn=BLVRA;refGene_tx=XM_023250189.1;refGene_gn=
BLVRA;refGene_tx=XM_023250190.1;refGene_gn=BLVRA;refGene_tx=XM_0232501
91.1
A2	72206278	72206398
refGene_gn=BLVRA;refGene_tx=XM_023250187.1;refGene_gn=BLVRA;refGene_tx
=XM_023250188.1;refGene_gn=BLVRA;refGene_tx=XM_023250189.1;refGene_gn=
BLVRA;refGene_tx=XM_023250190.1;refGene_gn=BLVRA;refGene_tx=XM_0232501
91.1
A2	72208355	72208477
refGene_gn=BLVRA;refGene_tx=XM_023250187.1;refGene_gn=BLVRA;refGene_tx
=XM_023250188.1;refGene_gn=BLVRA;refGene_tx=XM_023250189.1;refGene_gn=
BLVRA;refGene_tx=XM_023250190.1
A2	72208920	72208938	refGene_gn=BLVRA;refGene_tx=XM_023250187.1
A2	72228647	72228659
refGene_gn=BLVRA;refGene_tx=XM_023250188.1;refGene_gn=BLVRA;refGene_tx
=XM_023250189.1;refGene_gn=BLVRA;refGene_tx=XM_023250190.1;refGene_gn=
BLVRA;refGene_tx=XM_023250191.1
A2	72355818	72355903
refGene_gn=LOC101088997;refGene_tx=XM_023250192.1
A2	72366086	72366234

refGene_gn=LOC101088997;refGene_tx=XM_023250192.1
A2	72369919	72370079
refGene_gn=LOC101088997;refGene_tx=XM_023250192.1
A2	72390360	72390685
refGene_gn=STK17A;refGene_tx=XM_023250193.1;refGene_gn=STK17A;refGene_tx=XM_023250194.1
A2	72391522	72391702
refGene_gn=STK17A;refGene_tx=XM_023250193.1;refGene_gn=STK17A;refGene_tx=XM_023250194.1
A2	72391790	72391839
refGene_gn=STK17A;refGene_tx=XM_023250193.1;refGene_gn=STK17A;refGene_tx=XM_023250194.1
A2	72395551	72395678
refGene_gn=STK17A;refGene_tx=XM_023250193.1;refGene_gn=STK17A;refGene_tx=XM_023250194.1
A2	72406987	72407132
refGene_gn=STK17A;refGene_tx=XM_023250193.1;refGene_gn=STK17A;refGene_tx=XM_023250194.1
A2	72417906	72418119
refGene_gn=STK17A;refGene_tx=XM_023250193.1;refGene_gn=STK17A;refGene_tx=XM_023250194.1
A2	72429715	72429912
refGene_gn=STK17A;refGene_tx=XM_023250193.1;refGene_gn=STK17A;refGene_tx=XM_023250194.1
A2	72448702	72448814
refGene_gn=HECW1;refGene_tx=XM_006929092.4;refGene_gn=HECW1;refGene_tx=XM_019825270.2;refGene_gn=HECW1;refGene_tx=XM_023250195.1
A2	72457767	72457966
refGene_gn=HECW1;refGene_tx=XM_006929092.4;refGene_gn=HECW1;refGene_tx=XM_019825270.2;refGene_gn=HECW1;refGene_tx=XM_023250195.1
A2	72459901	72460016
refGene_gn=HECW1;refGene_tx=XM_006929092.4;refGene_gn=HECW1;refGene_tx=XM_019825270.2;refGene_gn=HECW1;refGene_tx=XM_023250195.1
A2	72461398	72461545
refGene_gn=HECW1;refGene_tx=XM_006929092.4;refGene_gn=HECW1;refGene_tx=XM_019825270.2;refGene_gn=HECW1;refGene_tx=XM_023250195.1
A2	72466521	72466651
refGene_gn=HECW1;refGene_tx=XM_006929092.4;refGene_gn=HECW1;refGene_tx=XM_019825270.2;refGene_gn=HECW1;refGene_tx=XM_023250195.1
A2	72467197	72467296
refGene_gn=HECW1;refGene_tx=XM_006929092.4;refGene_gn=HECW1;refGene_tx=XM_019825270.2;refGene_gn=HECW1;refGene_tx=XM_023250195.1
A2	72496554	72496707
refGene_gn=HECW1;refGene_tx=XM_006929092.4;refGene_gn=HECW1;refGene_tx=XM_019825270.2;refGene_gn=HECW1;refGene_tx=XM_023250195.1
A2	72497528	72497642
refGene_gn=HECW1;refGene_tx=XM_006929092.4;refGene_gn=HECW1;refGene_tx=XM_019825270.2;refGene_gn=HECW1;refGene_tx=XM_023250195.1
A2	72498354	72498475
refGene_gn=HECW1;refGene_tx=XM_006929092.4;refGene_gn=HECW1;refGene_tx

=XM_019825270.2;refGene_gn=HECW1;refGene_tx=XM_023250195.1
A2 72503976 72504086
refGene_gn=HECW1;refGene_tx=XM_006929092.4;refGene_gn=HECW1;refGene_tx=XM_019825270.2;refGene_gn=HECW1;refGene_tx=XM_023250195.1
A2 72504964 72505048
refGene_gn=HECW1;refGene_tx=XM_006929092.4;refGene_gn=HECW1;refGene_tx=XM_019825270.2;refGene_gn=HECW1;refGene_tx=XM_023250195.1
A2 72510247 72510344
refGene_gn=HECW1;refGene_tx=XM_006929092.4;refGene_gn=HECW1;refGene_tx=XM_019825270.2;refGene_gn=HECW1;refGene_tx=XM_023250195.1
A2 72511263 72511369
refGene_gn=HECW1;refGene_tx=XM_006929092.4;refGene_gn=HECW1;refGene_tx=XM_019825270.2;refGene_gn=HECW1;refGene_tx=XM_023250195.1
A2 72519994 72520129
refGene_gn=HECW1;refGene_tx=XM_006929092.4;refGene_gn=HECW1;refGene_tx=XM_019825270.2;refGene_gn=HECW1;refGene_tx=XM_023250195.1
A2 72530875 72531061
refGene_gn=HECW1;refGene_tx=XM_006929092.4;refGene_gn=HECW1;refGene_tx=XM_019825270.2;refGene_gn=HECW1;refGene_tx=XM_023250195.1
A2 72533830 72533952
refGene_gn=HECW1;refGene_tx=XM_006929092.4;refGene_gn=HECW1;refGene_tx=XM_019825270.2;refGene_gn=HECW1;refGene_tx=XM_023250195.1
A2 72535709 72535849
refGene_gn=HECW1;refGene_tx=XM_006929092.4;refGene_gn=HECW1;refGene_tx=XM_019825270.2;refGene_gn=HECW1;refGene_tx=XM_023250195.1
A2 72543567 72543718
refGene_gn=HECW1;refGene_tx=XM_006929092.4;refGene_gn=HECW1;refGene_tx=XM_019825270.2;refGene_gn=HECW1;refGene_tx=XM_023250195.1
A2 72548237 72548339
refGene_gn=HECW1;refGene_tx=XM_006929092.4;refGene_gn=HECW1;refGene_tx=XM_023250195.1
A2 72553408 72554746
refGene_gn=HECW1;refGene_tx=XM_006929092.4;refGene_gn=HECW1;refGene_tx=XM_019825270.2;refGene_gn=HECW1;refGene_tx=XM_023250195.1
A2 72556285 72556386
refGene_gn=HECW1;refGene_tx=XM_006929092.4;refGene_gn=HECW1;refGene_tx=XM_019825270.2;refGene_gn=HECW1;refGene_tx=XM_023250195.1
A2 72558378 72558521
refGene_gn=HECW1;refGene_tx=XM_006929092.4;refGene_gn=HECW1;refGene_tx=XM_019825270.2;refGene_gn=HECW1;refGene_tx=XM_023250195.1
A2 72569725 72569824 refGene_gn=HECW1;refGene_tx=XM_023250195.1
A2 72593057 72593227
refGene_gn=HECW1;refGene_tx=XM_006929092.4;refGene_gn=HECW1;refGene_tx=XM_019825270.2;refGene_gn=HECW1;refGene_tx=XM_023250195.1
A2 72598906 72598982
refGene_gn=HECW1;refGene_tx=XM_006929092.4;refGene_gn=HECW1;refGene_tx=XM_019825270.2;refGene_gn=HECW1;refGene_tx=XM_023250195.1
A2 72643630 72643725
refGene_gn=HECW1;refGene_tx=XM_006929092.4;refGene_gn=HECW1;refGene_tx=XM_019825270.2;refGene_gn=HECW1;refGene_tx=XM_023250195.1

A2 72678533 72678641
refGene_gn=HECW1;refGene_tx=XM_006929092.4;refGene_gn=HECW1;refGene_tx=XM_019825270.2;refGene_gn=HECW1;refGene_tx=XM_023250195.1
A2 72686138 72686463
refGene_gn=HECW1;refGene_tx=XM_006929092.4;refGene_gn=HECW1;refGene_tx=XM_019825270.2;refGene_gn=HECW1;refGene_tx=XM_023250195.1
A2 72866254 72866377
refGene_gn=HECW1;refGene_tx=XM_006929092.4;refGene_gn=HECW1;refGene_tx=XM_019825270.2
A2 72866610 72867004
refGene_gn=LOC109497386;refGene_tx=XM_019825274.2
A2 72901575 72901634
refGene_gn=LOC109497386;refGene_tx=XM_019825274.2
A2 73006582 73006837 refGene_gn=MRPL32;refGene_tx=XM_003982630.4
A2 73008921 73009103 refGene_gn=MRPL32;refGene_tx=XM_003982630.4
A2 73010478 73010608 refGene_gn=MRPL32;refGene_tx=XM_003982630.4
A2 73010848 73010889 refGene_gn=PSMA2;refGene_tx=XM_003982631.3
A2 73014555 73014632 refGene_gn=PSMA2;refGene_tx=XM_003982631.3
A2 73015602 73015735 refGene_gn=PSMA2;refGene_tx=XM_003982631.3
A2 73017205 73017328 refGene_gn=PSMA2;refGene_tx=XM_003982631.3
A2 73018082 73018164 refGene_gn=PSMA2;refGene_tx=XM_003982631.3
A2 73019847 73019921 refGene_gn=PSMA2;refGene_tx=XM_003982631.3
A2 73020955 73021013 refGene_gn=PSMA2;refGene_tx=XM_003982631.3
A2 73021100 73021217 refGene_gn=PSMA2;refGene_tx=XM_003982631.3
A2 73027448 73028714
refGene_gn=CA2H7orf25;refGene_tx=XM_006929094.4;refGene_gn=CA2H7orf25;refGene_tx=XM_023250196.1
A2 73456296 73456474
refGene_gn=LOC101089756;refGene_tx=XM_011280458.2
A2 73458174 73458260
refGene_gn=LOC101089756;refGene_tx=XM_011280458.2
A2 73458734 73458904
refGene_gn=LOC101089756;refGene_tx=XM_011280458.2
A2 73461950 73462041
refGene_gn=LOC101089756;refGene_tx=XM_011280458.2
A2 73495623 73495686
refGene_gn=LOC101089756;refGene_tx=XM_011280458.2
A2 73498202 73498407
refGene_gn=LOC101089756;refGene_tx=XM_011280458.2
A2 73498409 73498636
refGene_gn=LOC101089756;refGene_tx=XM_011280458.2
A2 73498638 73498965
refGene_gn=LOC101089756;refGene_tx=XM_011280458.2
A2 73662762 73662886
refGene_gn=GLI3;refGene_tx=XM_023250197.1;refGene_gn=GLI3;refGene_tx=XM_023250198.1
A2 73739045 73739288
refGene_gn=GLI3;refGene_tx=XM_011280298.3;refGene_gn=GLI3;refGene_tx=XM_019825278.2;refGene_gn=GLI3;refGene_tx=XM_023250197.1;refGene_gn=GLI3;refGene_tx=XM_023250198.1

A2 73789022 73789146 refGene_gn=GLI3;refGene_tx=XM_019825279.2
A2 73803688 73803794
refGene_gn=GLI3;refGene_tx=XM_011280298.3;refGene_gn=GLI3;refGene_tx=XM_019825278.2;refGene_gn=GLI3;refGene_tx=XM_019825279.2;refGene_gn=GLI3;refGene_tx=XM_023250197.1;refGene_gn=GLI3;refGene_tx=XM_023250198.1
A2 73832873 73833076
refGene_gn=GLI3;refGene_tx=XM_011280298.3;refGene_gn=GLI3;refGene_tx=XM_019825278.2;refGene_gn=GLI3;refGene_tx=XM_019825279.2;refGene_gn=GLI3;refGene_tx=XM_019825280.2;refGene_gn=GLI3;refGene_tx=XM_023250197.1;refGene_gn=GLI3;refGene_tx=XM_023250198.1
A2 73836241 73836388
refGene_gn=GLI3;refGene_tx=XM_011280298.3;refGene_gn=GLI3;refGene_tx=XM_019825278.2;refGene_gn=GLI3;refGene_tx=XM_019825279.2;refGene_gn=GLI3;refGene_tx=XM_019825280.2;refGene_gn=GLI3;refGene_tx=XM_023250197.1;refGene_gn=GLI3;refGene_tx=XM_023250198.1
A2 73841291 73841493
refGene_gn=GLI3;refGene_tx=XM_011280298.3;refGene_gn=GLI3;refGene_tx=XM_019825278.2;refGene_gn=GLI3;refGene_tx=XM_019825279.2;refGene_gn=GLI3;refGene_tx=XM_019825280.2;refGene_gn=GLI3;refGene_tx=XM_023250197.1;refGene_gn=GLI3;refGene_tx=XM_023250198.1
A2 73854693 73854907
refGene_gn=GLI3;refGene_tx=XM_011280298.3;refGene_gn=GLI3;refGene_tx=XM_019825278.2;refGene_gn=GLI3;refGene_tx=XM_019825279.2;refGene_gn=GLI3;refGene_tx=XM_019825280.2;refGene_gn=GLI3;refGene_tx=XM_023250197.1;refGene_gn=GLI3;refGene_tx=XM_023250198.1
A2 73855693 73855807
refGene_gn=GLI3;refGene_tx=XM_011280298.3;refGene_gn=GLI3;refGene_tx=XM_019825278.2;refGene_gn=GLI3;refGene_tx=XM_019825279.2;refGene_gn=GLI3;refGene_tx=XM_019825280.2;refGene_gn=GLI3;refGene_tx=XM_023250197.1;refGene_gn=GLI3;refGene_tx=XM_023250198.1
A2 73857684 73857825
refGene_gn=GLI3;refGene_tx=XM_011280298.3;refGene_gn=GLI3;refGene_tx=XM_019825278.2;refGene_gn=GLI3;refGene_tx=XM_019825279.2;refGene_gn=GLI3;refGene_tx=XM_019825280.2;refGene_gn=GLI3;refGene_tx=XM_023250197.1;refGene_gn=GLI3;refGene_tx=XM_023250198.1
A2 73909596 73909746
refGene_gn=GLI3;refGene_tx=XM_011280298.3;refGene_gn=GLI3;refGene_tx=XM_019825278.2;refGene_gn=GLI3;refGene_tx=XM_019825279.2;refGene_gn=GLI3;refGene_tx=XM_019825280.2;refGene_gn=GLI3;refGene_tx=XM_023250197.1;refGene_gn=GLI3;refGene_tx=XM_023250198.1
A2 73911106 73911271
refGene_gn=GLI3;refGene_tx=XM_011280298.3;refGene_gn=GLI3;refGene_tx=XM_019825278.2;refGene_gn=GLI3;refGene_tx=XM_019825279.2;refGene_gn=GLI3;refGene_tx=XM_019825280.2;refGene_gn=GLI3;refGene_tx=XM_023250197.1;refGene_gn=GLI3;refGene_tx=XM_023250198.1
A2 73916855 73917146
refGene_gn=GLI3;refGene_tx=XM_011280298.3;refGene_gn=GLI3;refGene_tx=XM_019825278.2;refGene_gn=GLI3;refGene_tx=XM_019825279.2;refGene_gn=GLI3;refGene_tx=XM_019825280.2;refGene_gn=GLI3;refGene_tx=XM_023250197.1;refGene_gn=GLI3;refGene_tx=XM_023250198.1

A2 73921557 73921888
refGene_gn=GLI3;refGene_tx=XM_011280298.3;refGene_gn=GLI3;refGene_tx=XM_019825278.2;refGene_gn=GLI3;refGene_tx=XM_019825279.2;refGene_gn=GLI3;refGene_tx=XM_019825280.2;refGene_gn=GLI3;refGene_tx=XM_023250197.1;refGene_gn=GLI3;refGene_tx=XM_023250198.1
A2 73922890 73925196
refGene_gn=GLI3;refGene_tx=XM_011280298.3;refGene_gn=GLI3;refGene_tx=XM_019825278.2;refGene_gn=GLI3;refGene_tx=XM_019825279.2;refGene_gn=GLI3;refGene_tx=XM_019825280.2;refGene_gn=GLI3;refGene_tx=XM_023250197.1;refGene_gn=GLI3;refGene_tx=XM_023250198.1
A2 74139844 74140232
refGene_gn=INHBA;refGene_tx=NM_001009856.1;refGene_gn=INHBA;refGene_tx=XM_023240895.1;refGene_gn=INHBA;refGene_tx=XM_023240897.1
A2 74149625 74150512
refGene_gn=INHBA;refGene_tx=NM_001009856.1;refGene_gn=INHBA;refGene_tx=XM_023240895.1;refGene_gn=INHBA;refGene_tx=XM_023240897.1
A2 74954640 74954804 refGene_gn=SUGCT;refGene_tx=XM_011280299.3
A2 75057987 75058051 refGene_gn=SUGCT;refGene_tx=XM_011280299.3
A2 75332312 75332415 refGene_gn=SUGCT;refGene_tx=XM_011280299.3
A2 75380101 75380199 refGene_gn=SUGCT;refGene_tx=XM_011280299.3
A2 75388274 75388346 refGene_gn=SUGCT;refGene_tx=XM_011280299.3
A2 75531161 75531257 refGene_gn=SUGCT;refGene_tx=XM_011280299.3
A2 75579128 75579272 refGene_gn=SUGCT;refGene_tx=XM_011280299.3
A2 75616155 75616247 refGene_gn=SUGCT;refGene_tx=XM_011280299.3
A2 75672671 75672792 refGene_gn=SUGCT;refGene_tx=XM_011280299.3
A2 75680168 75680219 refGene_gn=SUGCT;refGene_tx=XM_011280299.3
A2 75681499 75681585 refGene_gn=SUGCT;refGene_tx=XM_011280299.3
A2 75690129 75690203 refGene_gn=SUGCT;refGene_tx=XM_011280299.3
A2 75690853 75690905 refGene_gn=SUGCT;refGene_tx=XM_011280299.3
A2 75719995 75720095 refGene_gn=SUGCT;refGene_tx=XM_011280299.3
A2 75720460 75720925 refGene_gn=MPLKIP;refGene_tx=XM_003982636.5
A2 75721988 75722189 refGene_gn=MPLKIP;refGene_tx=XM_003982636.5
A2 75755845 75756699
refGene_gn=CDK13;refGene_tx=XM_019825281.2;refGene_gn=CDK13;refGene_tx=XM_023250200.1;refGene_gn=CDK13;refGene_tx=XM_023250201.1;refGene_gn=CDK13;refGene_tx=XM_023250202.1
A2 75757312 75757765
refGene_gn=CDK13;refGene_tx=XM_019825281.2;refGene_gn=CDK13;refGene_tx=XM_023250200.1;refGene_gn=CDK13;refGene_tx=XM_023250201.1;refGene_gn=CDK13;refGene_tx=XM_023250202.1
A2 75759948 75760154
refGene_gn=CDK13;refGene_tx=XM_019825281.2;refGene_gn=CDK13;refGene_tx=XM_023250200.1;refGene_gn=CDK13;refGene_tx=XM_023250201.1;refGene_gn=CDK13;refGene_tx=XM_023250202.1
A2 75761894 75761942
refGene_gn=CDK13;refGene_tx=XM_019825281.2;refGene_gn=CDK13;refGene_tx=XM_023250201.1
A2 75766167 75766299
refGene_gn=CDK13;refGene_tx=XM_019825281.2;refGene_gn=CDK13;refGene_tx=XM_023250200.1;refGene_gn=CDK13;refGene_tx=XM_023250201.1;refGene_gn=

CDK13;refGene_tx=XM_023250202.1
A2 75766850 75766967
refGene_gn=CDK13;refGene_tx=XM_019825281.2;refGene_gn=CDK13;refGene_tx=XM_023250200.1;refGene_gn=CDK13;refGene_tx=XM_023250201.1;refGene_gn=CDK13;refGene_tx=XM_023250202.1
A2 75776178 75776256
refGene_gn=CDK13;refGene_tx=XM_019825281.2;refGene_gn=CDK13;refGene_tx=XM_023250200.1;refGene_gn=CDK13;refGene_tx=XM_023250201.1;refGene_gn=CDK13;refGene_tx=XM_023250202.1
A2 75776351 75776453
refGene_gn=CDK13;refGene_tx=XM_019825281.2;refGene_gn=CDK13;refGene_tx=XM_023250200.1;refGene_gn=CDK13;refGene_tx=XM_023250201.1;refGene_gn=CDK13;refGene_tx=XM_023250202.1
A2 75795944 75796001
refGene_gn=CDK13;refGene_tx=XM_019825281.2;refGene_gn=CDK13;refGene_tx=XM_023250200.1;refGene_gn=CDK13;refGene_tx=XM_023250201.1;refGene_gn=CDK13;refGene_tx=XM_023250202.1
A2 75797267 75797457
refGene_gn=CDK13;refGene_tx=XM_019825281.2;refGene_gn=CDK13;refGene_tx=XM_023250200.1;refGene_gn=CDK13;refGene_tx=XM_023250201.1;refGene_gn=CDK13;refGene_tx=XM_023250202.1
A2 75838764 75838935
refGene_gn=CDK13;refGene_tx=XM_019825281.2;refGene_gn=CDK13;refGene_tx=XM_023250200.1;refGene_gn=CDK13;refGene_tx=XM_023250201.1;refGene_gn=CDK13;refGene_tx=XM_023250202.1
A2 75840746 75840886
refGene_gn=CDK13;refGene_tx=XM_019825281.2;refGene_gn=CDK13;refGene_tx=XM_023250200.1;refGene_gn=CDK13;refGene_tx=XM_023250201.1;refGene_gn=CDK13;refGene_tx=XM_023250202.1
A2 75842226 75842397
refGene_gn=CDK13;refGene_tx=XM_019825281.2;refGene_gn=CDK13;refGene_tx=XM_023250200.1;refGene_gn=CDK13;refGene_tx=XM_023250201.1;refGene_gn=CDK13;refGene_tx=XM_023250202.1
A2 75850486 75851146
refGene_gn=CDK13;refGene_tx=XM_019825281.2;refGene_gn=CDK13;refGene_tx=XM_023250200.1;refGene_gn=CDK13;refGene_tx=XM_023250201.1;refGene_gn=CDK13;refGene_tx=XM_023250202.1
A2 75875707 75876921
refGene_gn=CDK13;refGene_tx=XM_019825281.2;refGene_gn=CDK13;refGene_tx=XM_023250200.1;refGene_gn=CDK13;refGene_tx=XM_023250201.1;refGene_gn=CDK13;refGene_tx=XM_023250202.1
A2 75974806 75974929 refGene_gn=RALA;refGene_tx=XM_023250203.1
A2 75983082 75983257 refGene_gn=RALA;refGene_tx=XM_023250203.1
A2 75988103 75988312 refGene_gn=RALA;refGene_tx=XM_023250203.1
A2 75990749 75990863 refGene_gn=RALA;refGene_tx=XM_023250203.1
A2 76085785 76086215 refGene_gn=YAE1D1;refGene_tx=XM_003982638.5
A2 76088239 76088361 refGene_gn=YAE1D1;refGene_tx=XM_003982638.5
A2 76089889 76090018 refGene_gn=YAE1D1;refGene_tx=XM_003982638.5
A2 76190804 76191309
refGene_gn=POU6F2;refGene_tx=XM_023250204.1;refGene_gn=POU6F2;refGene_

tx=XM_023250205.1;refGene_gn=POU6F2;refGene_tx=XM_023250206.1
A2	76194846	76195015
refGene_gn=POU6F2;refGene_tx=XM_023250204.1;refGene_gn=POU6F2;refGene_tx=XM_023250205.1;refGene_gn=POU6F2;refGene_tx=XM_023250206.1
A2	76204316	76204485
refGene_gn=POU6F2;refGene_tx=XM_023250204.1;refGene_gn=POU6F2;refGene_tx=XM_023250205.1;refGene_gn=POU6F2;refGene_tx=XM_023250206.1
A2	76217230	76217437
refGene_gn=POU6F2;refGene_tx=XM_023250204.1;refGene_gn=POU6F2;refGene_tx=XM_023250205.1;refGene_gn=POU6F2;refGene_tx=XM_023250206.1
A2	76244682	76244823
refGene_gn=POU6F2;refGene_tx=XM_023250204.1;refGene_gn=POU6F2;refGene_tx=XM_023250205.1;refGene_gn=POU6F2;refGene_tx=XM_023250206.1
A2	76313895	76314284
refGene_gn=POU6F2;refGene_tx=XM_023250204.1;refGene_gn=POU6F2;refGene_tx=XM_023250205.1;refGene_gn=POU6F2;refGene_tx=XM_023250206.1
A2	76449674	76449903
refGene_gn=POU6F2;refGene_tx=XM_023250204.1;refGene_gn=POU6F2;refGene_tx=XM_023250205.1;refGene_gn=POU6F2;refGene_tx=XM_023250206.1
A2	76453450	76453542
refGene_gn=POU6F2;refGene_tx=XM_023250204.1;refGene_gn=POU6F2;refGene_tx=XM_023250205.1;refGene_gn=POU6F2;refGene_tx=XM_023250206.1
A2	76576039	76576211
refGene_gn=POU6F2;refGene_tx=XM_023250204.1;refGene_gn=POU6F2;refGene_tx=XM_023250205.1;refGene_gn=POU6F2;refGene_tx=XM_023250206.1
A2	76657162	76657180
refGene_gn=POU6F2;refGene_tx=XM_023250204.1;refGene_gn=POU6F2;refGene_tx=XM_023250206.1
A2	76760287	76760308	refGene_gn=VPS41;refGene_tx=XM_006929098.3
A2	76770057	76770096	refGene_gn=VPS41;refGene_tx=XM_006929098.3
A2	76801277	76801385	refGene_gn=VPS41;refGene_tx=XM_006929098.3
A2	76807687	76807765	refGene_gn=VPS41;refGene_tx=XM_006929098.3
A2	76839238	76839313	refGene_gn=VPS41;refGene_tx=XM_006929098.3
A2	76847452	76847515	refGene_gn=VPS41;refGene_tx=XM_006929098.3
A2	76849476	76849542	refGene_gn=VPS41;refGene_tx=XM_006929098.3
A2	76868539	76868659
refGene_gn=VPS41;refGene_tx=XM_006929098.3;refGene_gn=VPS41;refGene_tx=XM_006929099.4
A2	76870272	76870419
refGene_gn=VPS41;refGene_tx=XM_006929098.3;refGene_gn=VPS41;refGene_tx=XM_006929099.4
A2	76877953	76878020
refGene_gn=VPS41;refGene_tx=XM_006929098.3;refGene_gn=VPS41;refGene_tx=XM_006929099.4
A2	76891119	76891217
refGene_gn=VPS41;refGene_tx=XM_006929098.3;refGene_gn=VPS41;refGene_tx=XM_006929099.4
A2	76894635	76894765
refGene_gn=VPS41;refGene_tx=XM_006929098.3;refGene_gn=VPS41;refGene_tx=XM_006929099.4

A2 76896404 76896520
refGene_gn=VPS41;refGene_tx=XM_006929098.3;refGene_gn=VPS41;refGene_tx
=XM_006929099.4
A2 76897830 76897887
refGene_gn=VPS41;refGene_tx=XM_006929098.3;refGene_gn=VPS41;refGene_tx
=XM_006929099.4
A2 76899195 76899257
refGene_gn=VPS41;refGene_tx=XM_006929098.3;refGene_gn=VPS41;refGene_tx
=XM_006929099.4
A2 76900916 76900998
refGene_gn=VPS41;refGene_tx=XM_006929098.3;refGene_gn=VPS41;refGene_tx
=XM_006929099.4
A2 76904845 76904938
refGene_gn=VPS41;refGene_tx=XM_006929098.3;refGene_gn=VPS41;refGene_tx
=XM_006929099.4
A2 76909825 76909953
refGene_gn=VPS41;refGene_tx=XM_006929098.3;refGene_gn=VPS41;refGene_tx
=XM_006929099.4
A2 76911717 76911862
refGene_gn=VPS41;refGene_tx=XM_006929098.3;refGene_gn=VPS41;refGene_tx
=XM_006929099.4
A2 76913979 76914021
refGene_gn=VPS41;refGene_tx=XM_006929098.3;refGene_gn=VPS41;refGene_tx
=XM_006929099.4
A2 76914168 76914219
refGene_gn=VPS41;refGene_tx=XM_006929098.3;refGene_gn=VPS41;refGene_tx
=XM_006929099.4
A2 76918227 76918365
refGene_gn=VPS41;refGene_tx=XM_006929098.3;refGene_gn=VPS41;refGene_tx
=XM_006929099.4
A2 76926516 76926571
refGene_gn=VPS41;refGene_tx=XM_006929098.3;refGene_gn=VPS41;refGene_tx
=XM_006929099.4
A2 76929690 76929831
refGene_gn=VPS41;refGene_tx=XM_006929098.3;refGene_gn=VPS41;refGene_tx
=XM_006929099.4
A2 76931364 76931501
refGene_gn=VPS41;refGene_tx=XM_006929098.3;refGene_gn=VPS41;refGene_tx
=XM_006929099.4
A2 76943876 76943976
refGene_gn=VPS41;refGene_tx=XM_006929098.3;refGene_gn=VPS41;refGene_tx
=XM_006929099.4
A2 76944082 76944127
refGene_gn=VPS41;refGene_tx=XM_006929098.3;refGene_gn=VPS41;refGene_tx
=XM_006929099.4
A2 76946030 76946110
refGene_gn=VPS41;refGene_tx=XM_006929098.3;refGene_gn=VPS41;refGene_tx
=XM_006929099.4
A2 76947138 76947219
refGene_gn=VPS41;refGene_tx=XM_006929098.3;refGene_gn=VPS41;refGene_tx

=XM_006929099.4
A2	77051298	77051367
refGene_gn=AMPH;refGene_tx=XM_023250207.1;refGene_gn=AMPH;refGene_tx=XM_023250208.1;refGene_gn=AMPH;refGene_tx=XM_023250209.1;refGene_gn=AMPH;refGene_tx=XM_023250210.1;refGene_gn=AMPH;refGene_tx=XM_023250211.1;refGene_gn=AMPH;refGene_tx=XM_023250212.1;refGene_gn=AMPH;refGene_tx=XM_023250213.1
A2	77158887	77158968
refGene_gn=AMPH;refGene_tx=XM_023250207.1;refGene_gn=AMPH;refGene_tx=XM_023250208.1;refGene_gn=AMPH;refGene_tx=XM_023250209.1;refGene_gn=AMPH;refGene_tx=XM_023250210.1;refGene_gn=AMPH;refGene_tx=XM_023250211.1;refGene_gn=AMPH;refGene_tx=XM_023250212.1;refGene_gn=AMPH;refGene_tx=XM_023250213.1
A2	77189970	77190025
refGene_gn=AMPH;refGene_tx=XM_023250207.1;refGene_gn=AMPH;refGene_tx=XM_023250208.1;refGene_gn=AMPH;refGene_tx=XM_023250209.1;refGene_gn=AMPH;refGene_tx=XM_023250210.1;refGene_gn=AMPH;refGene_tx=XM_023250211.1;refGene_gn=AMPH;refGene_tx=XM_023250212.1;refGene_gn=AMPH;refGene_tx=XM_023250213.1
A2	77198859	77198954
refGene_gn=AMPH;refGene_tx=XM_023250207.1;refGene_gn=AMPH;refGene_tx=XM_023250208.1;refGene_gn=AMPH;refGene_tx=XM_023250209.1;refGene_gn=AMPH;refGene_tx=XM_023250210.1;refGene_gn=AMPH;refGene_tx=XM_023250211.1;refGene_gn=AMPH;refGene_tx=XM_023250212.1;refGene_gn=AMPH;refGene_tx=XM_023250213.1
A2	77201111	77201207
refGene_gn=AMPH;refGene_tx=XM_023250207.1;refGene_gn=AMPH;refGene_tx=XM_023250208.1;refGene_gn=AMPH;refGene_tx=XM_023250209.1;refGene_gn=AMPH;refGene_tx=XM_023250210.1;refGene_gn=AMPH;refGene_tx=XM_023250211.1;refGene_gn=AMPH;refGene_tx=XM_023250212.1;refGene_gn=AMPH;refGene_tx=XM_023250213.1
A2	77210026	77210134
refGene_gn=AMPH;refGene_tx=XM_023250207.1;refGene_gn=AMPH;refGene_tx=XM_023250208.1;refGene_gn=AMPH;refGene_tx=XM_023250209.1;refGene_gn=AMPH;refGene_tx=XM_023250210.1;refGene_gn=AMPH;refGene_tx=XM_023250211.1;refGene_gn=AMPH;refGene_tx=XM_023250212.1;refGene_gn=AMPH;refGene_tx=XM_023250213.1
A2	77212302	77212388
refGene_gn=AMPH;refGene_tx=XM_023250207.1;refGene_gn=AMPH;refGene_tx=XM_023250208.1;refGene_gn=AMPH;refGene_tx=XM_023250209.1;refGene_gn=AMPH;refGene_tx=XM_023250210.1;refGene_gn=AMPH;refGene_tx=XM_023250211.1;refGene_gn=AMPH;refGene_tx=XM_023250212.1;refGene_gn=AMPH;refGene_tx=XM_023250213.1
A2	77221853	77221929
refGene_gn=AMPH;refGene_tx=XM_023250207.1;refGene_gn=AMPH;refGene_tx=XM_023250208.1;refGene_gn=AMPH;refGene_tx=XM_023250209.1;refGene_gn=AMPH;refGene_tx=XM_023250210.1;refGene_gn=AMPH;refGene_tx=XM_023250211.1;refGene_gn=AMPH;refGene_tx=XM_023250212.1;refGene_gn=AMPH;refGene_tx=XM_023250213.1
A2	77222815	77222898

refGene_gn=AMPH;refGene_tx=XM_023250207.1;refGene_gn=AMPH;refGene_tx=XM_023250208.1;refGene_gn=AMPH;refGene_tx=XM_023250209.1;refGene_gn=AMPH;refGene_tx=XM_023250210.1;refGene_gn=AMPH;refGene_tx=XM_023250211.1;refGene_gn=AMPH;refGene_tx=XM_023250212.1;refGene_gn=AMPH;refGene_tx=XM_023250213.1
A2 77224349 77224358
refGene_gn=AMPH;refGene_tx=XM_023250207.1;refGene_gn=AMPH;refGene_tx=XM_023250209.1;refGene_gn=AMPH;refGene_tx=XM_023250210.1;refGene_gn=AMPH;refGene_tx=XM_023250212.1
A2 77225319 77225458
refGene_gn=AMPH;refGene_tx=XM_023250207.1;refGene_gn=AMPH;refGene_tx=XM_023250208.1;refGene_gn=AMPH;refGene_tx=XM_023250209.1;refGene_gn=AMPH;refGene_tx=XM_023250210.1;refGene_gn=AMPH;refGene_tx=XM_023250211.1;refGene_gn=AMPH;refGene_tx=XM_023250212.1;refGene_gn=AMPH;refGene_tx=XM_023250213.1
A2 77226527 77226656
refGene_gn=AMPH;refGene_tx=XM_023250207.1;refGene_gn=AMPH;refGene_tx=XM_023250208.1;refGene_gn=AMPH;refGene_tx=XM_023250209.1;refGene_gn=AMPH;refGene_tx=XM_023250210.1;refGene_gn=AMPH;refGene_tx=XM_023250211.1;refGene_gn=AMPH;refGene_tx=XM_023250212.1;refGene_gn=AMPH;refGene_tx=XM_023250213.1
A2 77249761 77249878
refGene_gn=AMPH;refGene_tx=XM_023250207.1;refGene_gn=AMPH;refGene_tx=XM_023250208.1;refGene_gn=AMPH;refGene_tx=XM_023250209.1;refGene_gn=AMPH;refGene_tx=XM_023250210.1;refGene_gn=AMPH;refGene_tx=XM_023250211.1;refGene_gn=AMPH;refGene_tx=XM_023250212.1;refGene_gn=AMPH;refGene_tx=XM_023250213.1
A2 77253704 77253728
refGene_gn=AMPH;refGene_tx=XM_023250207.1;refGene_gn=AMPH;refGene_tx=XM_023250208.1;refGene_gn=AMPH;refGene_tx=XM_023250209.1;refGene_gn=AMPH;refGene_tx=XM_023250210.1;refGene_gn=AMPH;refGene_tx=XM_023250211.1;refGene_gn=AMPH;refGene_tx=XM_023250212.1;refGene_gn=AMPH;refGene_tx=XM_023250213.1
A2 77256445 77256469
refGene_gn=AMPH;refGene_tx=XM_023250207.1;refGene_gn=AMPH;refGene_tx=XM_023250208.1;refGene_gn=AMPH;refGene_tx=XM_023250209.1;refGene_gn=AMPH;refGene_tx=XM_023250210.1;refGene_gn=AMPH;refGene_tx=XM_023250211.1;refGene_gn=AMPH;refGene_tx=XM_023250212.1;refGene_gn=AMPH;refGene_tx=XM_023250213.1
A2 77256754 77257276
refGene_gn=AMPH;refGene_tx=XM_023250207.1;refGene_gn=AMPH;refGene_tx=XM_023250208.1;refGene_gn=AMPH;refGene_tx=XM_023250209.1
A2 77259228 77259261
refGene_gn=AMPH;refGene_tx=XM_023250207.1;refGene_gn=AMPH;refGene_tx=XM_023250208.1;refGene_gn=AMPH;refGene_tx=XM_023250209.1;refGene_gn=AMPH;refGene_tx=XM_023250210.1;refGene_gn=AMPH;refGene_tx=XM_023250211.1;refGene_gn=AMPH;refGene_tx=XM_023250212.1;refGene_gn=AMPH;refGene_tx=XM_023250213.1
A2 77264894 77264951
refGene_gn=AMPH;refGene_tx=XM_023250207.1;refGene_gn=AMPH;refGene_tx=X

M_023250208.1;refGene_gn=AMPH;refGene_tx=XM_023250209.1;refGene_gn=AMPH;refGene_tx=XM_023250210.1;refGene_gn=AMPH;refGene_tx=XM_023250211.1;refGene_gn=AMPH;refGene_tx=XM_023250212.1;refGene_gn=AMPH;refGene_tx=XM_023250213.1
A2 77270270 77270396
refGene_gn=AMPH;refGene_tx=XM_023250207.1;refGene_gn=AMPH;refGene_tx=XM_023250208.1;refGene_gn=AMPH;refGene_tx=XM_023250210.1;refGene_gn=AMPH;refGene_tx=XM_023250211.1
A2 77288551 77288761
refGene_gn=AMPH;refGene_tx=XM_023250207.1;refGene_gn=AMPH;refGene_tx=XM_023250208.1;refGene_gn=AMPH;refGene_tx=XM_023250209.1;refGene_gn=AMPH;refGene_tx=XM_023250210.1;refGene_gn=AMPH;refGene_tx=XM_023250211.1;refGene_gn=AMPH;refGene_tx=XM_023250212.1;refGene_gn=AMPH;refGene_tx=XM_023250213.1
A2 77290852 77291083
refGene_gn=AMPH;refGene_tx=XM_023250207.1;refGene_gn=AMPH;refGene_tx=XM_023250208.1;refGene_gn=AMPH;refGene_tx=XM_023250209.1;refGene_gn=AMPH;refGene_tx=XM_023250210.1;refGene_gn=AMPH;refGene_tx=XM_023250211.1;refGene_gn=AMPH;refGene_tx=XM_023250212.1;refGene_gn=AMPH;refGene_tx=XM_023250213.1
A2 77294202 77294304
refGene_gn=AMPH;refGene_tx=XM_023250207.1;refGene_gn=AMPH;refGene_tx=XM_023250208.1;refGene_gn=AMPH;refGene_tx=XM_023250209.1;refGene_gn=AMPH;refGene_tx=XM_023250210.1;refGene_gn=AMPH;refGene_tx=XM_023250211.1;refGene_gn=AMPH;refGene_tx=XM_023250212.1;refGene_gn=AMPH;refGene_tx=XM_023250213.1
A2 77300066 77300174
refGene_gn=AMPH;refGene_tx=XM_023250207.1;refGene_gn=AMPH;refGene_tx=XM_023250208.1;refGene_gn=AMPH;refGene_tx=XM_023250209.1;refGene_gn=AMPH;refGene_tx=XM_023250210.1;refGene_gn=AMPH;refGene_tx=XM_023250211.1;refGene_gn=AMPH;refGene_tx=XM_023250212.1;refGene_gn=AMPH;refGene_tx=XM_023250213.1
A2 77314087 77314130 refGene_gn=TRG;refGene_tx=NM_001305026.1
A2 77362711 77363054
refGene_gn=LOC101091772;refGene_tx=XM_023250215.1
A2 77363149 77363467
refGene_gn=LOC101091772;refGene_tx=XM_023250215.1
A2 77374989 77375040
refGene_gn=LOC101091772;refGene_tx=XM_023250215.1
A2 77383384 77383714
refGene_gn=LOC101091772;refGene_tx=XM_023250215.1
A2 77385505 77385631
refGene_gn=LOC101091772;refGene_tx=XM_023250215.1
A2 77391950 77391998
refGene_gn=LOC101091772;refGene_tx=XM_023250215.1
A2 77393597 77393740
refGene_gn=LOC101091772;refGene_tx=XM_023250215.1
A2 77403737 77404046 refGene_gn=TRG;refGene_tx=NM_001305026.1
A2 77424054 77424108 refGene_gn=TRG;refGene_tx=NM_001305026.1
A2 77429166 77429208 refGene_gn=TRG;refGene_tx=NM_001305026.1

A2 77452914 77453057 refGene_gn=TARP;refGene_tx=XM_019825293.2
A2 77455135 77455183 refGene_gn=TARP;refGene_tx=XM_019825293.2
A2 77465093 77465219 refGene_gn=TARP;refGene_tx=XM_019825293.2
A2 77466726 77467056 refGene_gn=TARP;refGene_tx=XM_019825293.2
A2 77474867 77474928 refGene_gn=TARP;refGene_tx=XM_019825293.2
A2 77511607 77511750
refGene_gn=LOC101092254;refGene_tx=XM_023250216.1;refGene_gn=LOC101092
254;refGene_tx=XM_023250217.1
A2 77513745 77513793
refGene_gn=LOC101092254;refGene_tx=XM_023250216.1;refGene_gn=LOC101092
254;refGene_tx=XM_023250217.1
A2 77516913 77517232
refGene_gn=LOC101092254;refGene_tx=XM_023250216.1;refGene_gn=LOC101092
254;refGene_tx=XM_023250217.1
A2 77526654 77526776
refGene_gn=LOC111557335;refGene_tx=XM_023242015.1
A2 77528665 77529133
refGene_gn=LOC111557335;refGene_tx=XM_023242015.1
A2 77529391 77529538
refGene_gn=LOC111557335;refGene_tx=XM_023242015.1
A2 77536377 77536513
refGene_gn=LOC111557335;refGene_tx=XM_023242015.1
A2 77558560 77589945 refGene_gn=LOC101092505
A2 77595971 77596030
refGene_gn=STARD3NL;refGene_tx=XM_006929102.4;refGene_gn=STARD3NL;refG
ene_tx=XM_019825306.2;refGene_gn=STARD3NL;refGene_tx=XM_019825307.2
A2 77608095 77608191
refGene_gn=STARD3NL;refGene_tx=XM_006929102.4;refGene_gn=STARD3NL;refG
ene_tx=XM_019825306.2;refGene_gn=STARD3NL;refGene_tx=XM_019825307.2
A2 77610147 77610265
refGene_gn=STARD3NL;refGene_tx=XM_006929102.4;refGene_gn=STARD3NL;refG
ene_tx=XM_019825306.2;refGene_gn=STARD3NL;refGene_tx=XM_019825307.2
A2 77610379 77610433
refGene_gn=STARD3NL;refGene_tx=XM_006929102.4;refGene_gn=STARD3NL;refG
ene_tx=XM_019825306.2;refGene_gn=STARD3NL;refGene_tx=XM_019825307.2
A2 77612611 77612689
refGene_gn=STARD3NL;refGene_tx=XM_006929102.4;refGene_gn=STARD3NL;refG
ene_tx=XM_019825306.2;refGene_gn=STARD3NL;refGene_tx=XM_019825307.2
A2 77614949 77615027
refGene_gn=STARD3NL;refGene_tx=XM_006929102.4;refGene_gn=STARD3NL;refG
ene_tx=XM_019825306.2;refGene_gn=STARD3NL;refGene_tx=XM_019825307.2
A2 77621850 77622075
refGene_gn=STARD3NL;refGene_tx=XM_006929102.4;refGene_gn=STARD3NL;refG
ene_tx=XM_019825306.2;refGene_gn=STARD3NL;refGene_tx=XM_019825307.2
A2 77860490 77860687
refGene_gn=EPDR1;refGene_tx=XM_019825308.2;refGene_gn=EPDR1;refGene_tx
=XM_023250223.1
A2 77862086 77862295 refGene_gn=EPDR1;refGene_tx=XM_019825308.2
A2 77886369 77886677
refGene_gn=EPDR1;refGene_tx=XM_019825308.2;refGene_gn=EPDR1;refGene_tx

=XM_023250223.1
A2	77890984	77891429
refGene_gn=SFRP4;refGene_tx=XM_011280303.2;refGene_gn=SFRP4;refGene_tx
=XM_011280304.3
A2	77892790	77892871
refGene_gn=SFRP4;refGene_tx=XM_011280303.2;refGene_gn=SFRP4;refGene_tx
=XM_011280304.3
A2	77892980	77893046
refGene_gn=SFRP4;refGene_tx=XM_011280303.2;refGene_gn=SFRP4;refGene_tx
=XM_011280304.3
A2	77895179	77895378
refGene_gn=SFRP4;refGene_tx=XM_011280303.2;refGene_gn=SFRP4;refGene_tx
=XM_011280304.3
A2	77896125	77896189
refGene_gn=SFRP4;refGene_tx=XM_011280303.2;refGene_gn=SFRP4;refGene_tx
=XM_011280304.3
A2	77896978	77896999 refGene_gn=SFRP4;refGene_tx=XM_011280304.3
A2	77898573	77898759 refGene_gn=SFRP4;refGene_tx=XM_011280303.2
A2	77919681	77919919
refGene_gn=NME8;refGene_tx=XM_003982666.5;refGene_gn=NME8;refGene_tx=X
M_019825295.2;refGene_gn=NME8;refGene_tx=XM_019825298.2;refGene_gn=NME
8;refGene_tx=XM_019825299.2;refGene_gn=NME8;refGene_tx=XM_023250219.1;
refGene_gn=NME8;refGene_tx=XM_023250220.1;refGene_gn=NME8;refGene_tx=X
M_023250222.1
A2	77921199	77921344
refGene_gn=NME8;refGene_tx=XM_003982666.5;refGene_gn=NME8;refGene_tx=X
M_019825295.2;refGene_gn=NME8;refGene_tx=XM_019825298.2;refGene_gn=NME
8;refGene_tx=XM_019825299.2;refGene_gn=NME8;refGene_tx=XM_023250219.1;
refGene_gn=NME8;refGene_tx=XM_023250220.1;refGene_gn=NME8;refGene_tx=X
M_023250222.1
A2	77923385	77923537
refGene_gn=NME8;refGene_tx=XM_003982666.5;refGene_gn=NME8;refGene_tx=X
M_019825295.2;refGene_gn=NME8;refGene_tx=XM_019825298.2;refGene_gn=NME
8;refGene_tx=XM_019825299.2;refGene_gn=NME8;refGene_tx=XM_023250219.1;
refGene_gn=NME8;refGene_tx=XM_023250220.1;refGene_gn=NME8;refGene_tx=X
M_023250222.1
A2	77925720	77925828
refGene_gn=NME8;refGene_tx=XM_003982666.5;refGene_gn=NME8;refGene_tx=X
M_019825295.2;refGene_gn=NME8;refGene_tx=XM_019825298.2;refGene_gn=NME
8;refGene_tx=XM_019825299.2;refGene_gn=NME8;refGene_tx=XM_023250219.1;
refGene_gn=NME8;refGene_tx=XM_023250220.1;refGene_gn=NME8;refGene_tx=X
M_023250222.1
A2	77926712	77926857
refGene_gn=NME8;refGene_tx=XM_003982666.5;refGene_gn=NME8;refGene_tx=X
M_019825295.2;refGene_gn=NME8;refGene_tx=XM_019825298.2;refGene_gn=NME
8;refGene_tx=XM_019825299.2;refGene_gn=NME8;refGene_tx=XM_023250219.1;
refGene_gn=NME8;refGene_tx=XM_023250220.1;refGene_gn=NME8;refGene_tx=X
M_023250222.1
A2	77934955	77935131
refGene_gn=NME8;refGene_tx=XM_003982666.5;refGene_gn=NME8;refGene_tx=X

M_019825295.2;refGene_gn=NME8;refGene_tx=XM_019825298.2;refGene_gn=NME
8;refGene_tx=XM_019825299.2;refGene_gn=NME8;refGene_tx=XM_023250219.1;
refGene_gn=NME8;refGene_tx=XM_023250220.1;refGene_gn=NME8;refGene_tx=X
M_023250222.1
A2 77942635 77942823
refGene_gn=NME8;refGene_tx=XM_003982666.5;refGene_gn=NME8;refGene_tx=X
M_019825295.2;refGene_gn=NME8;refGene_tx=XM_019825298.2;refGene_gn=NME
8;refGene_tx=XM_019825299.2;refGene_gn=NME8;refGene_tx=XM_023250219.1;
refGene_gn=NME8;refGene_tx=XM_023250220.1;refGene_gn=NME8;refGene_tx=X
M_023250222.1
A2 77944187 77944280
refGene_gn=NME8;refGene_tx=XM_003982666.5;refGene_gn=NME8;refGene_tx=X
M_019825295.2;refGene_gn=NME8;refGene_tx=XM_019825298.2;refGene_gn=NME
8;refGene_tx=XM_019825299.2;refGene_gn=NME8;refGene_tx=XM_023250219.1;
refGene_gn=NME8;refGene_tx=XM_023250220.1;refGene_gn=NME8;refGene_tx=X
M_023250222.1
A2 77948173 77948247
refGene_gn=NME8;refGene_tx=XM_003982666.5;refGene_gn=NME8;refGene_tx=X
M_019825295.2;refGene_gn=NME8;refGene_tx=XM_019825298.2;refGene_gn=NME
8;refGene_tx=XM_019825299.2;refGene_gn=NME8;refGene_tx=XM_023250219.1;
refGene_gn=NME8;refGene_tx=XM_023250220.1;refGene_gn=NME8;refGene_tx=X
M_023250222.1
A2 77949338 77949405
refGene_gn=NME8;refGene_tx=XM_003982666.5;refGene_gn=NME8;refGene_tx=X
M_019825295.2;refGene_gn=NME8;refGene_tx=XM_019825298.2;refGene_gn=NME
8;refGene_tx=XM_019825299.2;refGene_gn=NME8;refGene_tx=XM_023250219.1;
refGene_gn=NME8;refGene_tx=XM_023250220.1;refGene_gn=NME8;refGene_tx=X
M_023250222.1
A2 77952423 77952540
refGene_gn=NME8;refGene_tx=XM_003982666.5;refGene_gn=NME8;refGene_tx=X
M_019825295.2;refGene_gn=NME8;refGene_tx=XM_019825298.2;refGene_gn=NME
8;refGene_tx=XM_019825299.2;refGene_gn=NME8;refGene_tx=XM_023250219.1;
refGene_gn=NME8;refGene_tx=XM_023250220.1;refGene_gn=NME8;refGene_tx=X
M_023250222.1
A2 77957290 77957362
refGene_gn=NME8;refGene_tx=XM_003982666.5;refGene_gn=NME8;refGene_tx=X
M_019825295.2;refGene_gn=NME8;refGene_tx=XM_019825298.2;refGene_gn=NME
8;refGene_tx=XM_019825299.2;refGene_gn=NME8;refGene_tx=XM_023250219.1;
refGene_gn=NME8;refGene_tx=XM_023250220.1;refGene_gn=NME8;refGene_tx=X
M_023250222.1
A2 77963721 77963828
refGene_gn=NME8;refGene_tx=XM_003982666.5;refGene_gn=NME8;refGene_tx=X
M_019825295.2;refGene_gn=NME8;refGene_tx=XM_019825298.2;refGene_gn=NME
8;refGene_tx=XM_019825299.2;refGene_gn=NME8;refGene_tx=XM_023250219.1;
refGene_gn=NME8;refGene_tx=XM_023250220.1;refGene_gn=NME8;refGene_tx=X
M_023250222.1
A2 77964027 77964085
refGene_gn=NME8;refGene_tx=XM_003982666.5;refGene_gn=NME8;refGene_tx=X
M_019825295.2;refGene_gn=NME8;refGene_tx=XM_019825298.2;refGene_gn=NME
8;refGene_tx=XM_019825299.2;refGene_gn=NME8;refGene_tx=XM_023250219.1;

refGene_gn=NME8;refGene_tx=XM_023250220.1;refGene_gn=NME8;refGene_tx=XM_023250222.1
A2 77964164 77964197
refGene_gn=NME8;refGene_tx=XM_003982666.5;refGene_gn=NME8;refGene_tx=XM_019825295.2;refGene_gn=NME8;refGene_tx=XM_019825298.2;refGene_gn=NME8;refGene_tx=XM_019825299.2;refGene_gn=NME8;refGene_tx=XM_023250219.1;refGene_gn=NME8;refGene_tx=XM_023250220.1;refGene_gn=NME8;refGene_tx=XM_023250222.1
A2 77977892 77978780
refGene_gn=LOC109497404;refGene_tx=XM_019825309.1;refGene_gn=LOC109497404;refGene_tx=XM_019825310.1
A2 78012129 78013182
refGene_gn=PNPLA8;refGene_tx=XM_011280305.2;refGene_gn=PNPLA8;refGene_tx=XM_011280306.2
A2 78013332 78013482
refGene_gn=PNPLA8;refGene_tx=XM_011280305.2;refGene_gn=PNPLA8;refGene_tx=XM_011280306.2
A2 78020258 78020410
refGene_gn=PNPLA8;refGene_tx=XM_011280305.2;refGene_gn=PNPLA8;refGene_tx=XM_011280306.2
A2 78024587 78024682
refGene_gn=PNPLA8;refGene_tx=XM_011280305.2;refGene_gn=PNPLA8;refGene_tx=XM_011280306.2
A2 78027606 78027778
refGene_gn=PNPLA8;refGene_tx=XM_011280305.2;refGene_gn=PNPLA8;refGene_tx=XM_011280306.2
A2 78030586 78030644
refGene_gn=PNPLA8;refGene_tx=XM_011280305.2;refGene_gn=PNPLA8;refGene_tx=XM_011280306.2
A2 78033226 78033421
refGene_gn=PNPLA8;refGene_tx=XM_011280305.2;refGene_gn=PNPLA8;refGene_tx=XM_011280306.2
A2 78042742 78042938
refGene_gn=PNPLA8;refGene_tx=XM_011280305.2;refGene_gn=PNPLA8;refGene_tx=XM_011280306.2
A2 78044115 78044207 refGene_gn=PNPLA8;refGene_tx=XM_011280306.2
A2 78049842 78050117 refGene_gn=PNPLA8;refGene_tx=XM_011280305.2
A2 78289948 78290054
refGene_gn=NRCAM;refGene_tx=XM_023250225.1;refGene_gn=NRCAM;refGene_tx=XM_023250226.1;refGene_gn=NRCAM;refGene_tx=XM_023250227.1;refGene_gn=NRCAM;refGene_tx=XM_023250228.1;refGene_gn=NRCAM;refGene_tx=XM_023250230.1;refGene_gn=NRCAM;refGene_tx=XM_023250231.1;refGene_gn=NRCAM;refGene_tx=XM_023250232.1;refGene_gn=NRCAM;refGene_tx=XM_023250233.1;refGene_gn=NRCAM;refGene_tx=XM_023250234.1;refGene_gn=NRCAM;refGene_tx=XM_023250235.1;refGene_gn=NRCAM;refGene_tx=XM_023250236.1;refGene_gn=NRCAM;refGene_tx=XM_023250237.1;refGene_gn=NRCAM;refGene_tx=XM_023250238.1;refGene_gn=NRCAM;refGene_tx=XM_023250239.1;refGene_gn=NRCAM;refGene_tx=XM_023250240.1;refGene_gn=NRCAM;refGene_tx=XM_023250241.1
A2 78292556 78292574
refGene_gn=NRCAM;refGene_tx=XM_023250225.1;refGene_gn=NRCAM;refGene_tx

=XM_023250226.1;refGene_gn=NRCAM;refGene_tx=XM_023250227.1;refGene_gn=NRCAM;refGene_tx=XM_023250228.1;refGene_gn=NRCAM;refGene_tx=XM_023250231.1;refGene_gn=NRCAM;refGene_tx=XM_023250232.1;refGene_gn=NRCAM;refGene_tx=XM_023250233.1;refGene_gn=NRCAM;refGene_tx=XM_023250234.1;refGene_gn=NRCAM;refGene_tx=XM_023250235.1;refGene_gn=NRCAM;refGene_tx=XM_023250236.1;refGene_gn=NRCAM;refGene_tx=XM_023250237.1;refGene_gn=NRCAM;refGene_tx=XM_023250238.1;refGene_gn=NRCAM;refGene_tx=XM_023250239.1
A2 78295487 78295593
refGene_gn=NRCAM;refGene_tx=XM_023250225.1;refGene_gn=NRCAM;refGene_tx=XM_023250226.1;refGene_gn=NRCAM;refGene_tx=XM_023250227.1;refGene_gn=NRCAM;refGene_tx=XM_023250228.1;refGene_gn=NRCAM;refGene_tx=XM_023250230.1;refGene_gn=NRCAM;refGene_tx=XM_023250231.1;refGene_gn=NRCAM;refGene_tx=XM_023250232.1;refGene_gn=NRCAM;refGene_tx=XM_023250233.1;refGene_gn=NRCAM;refGene_tx=XM_023250234.1;refGene_gn=NRCAM;refGene_tx=XM_023250235.1;refGene_gn=NRCAM;refGene_tx=XM_023250236.1;refGene_gn=NRCAM;refGene_tx=XM_023250237.1;refGene_gn=NRCAM;refGene_tx=XM_023250238.1;refGene_gn=NRCAM;refGene_tx=XM_023250239.1;refGene_gn=NRCAM;refGene_tx=XM_023250240.1;refGene_gn=NRCAM;refGene_tx=XM_023250241.1
A2 78297720 78297917
refGene_gn=NRCAM;refGene_tx=XM_023250225.1;refGene_gn=NRCAM;refGene_tx=XM_023250226.1;refGene_gn=NRCAM;refGene_tx=XM_023250227.1;refGene_gn=NRCAM;refGene_tx=XM_023250228.1;refGene_gn=NRCAM;refGene_tx=XM_023250230.1;refGene_gn=NRCAM;refGene_tx=XM_023250231.1;refGene_gn=NRCAM;refGene_tx=XM_023250232.1;refGene_gn=NRCAM;refGene_tx=XM_023250233.1;refGene_gn=NRCAM;refGene_tx=XM_023250234.1;refGene_gn=NRCAM;refGene_tx=XM_023250235.1;refGene_gn=NRCAM;refGene_tx=XM_023250236.1;refGene_gn=NRCAM;refGene_tx=XM_023250237.1;refGene_gn=NRCAM;refGene_tx=XM_023250238.1;refGene_gn=NRCAM;refGene_tx=XM_023250239.1;refGene_gn=NRCAM;refGene_tx=XM_023250240.1;refGene_gn=NRCAM;refGene_tx=XM_023250241.1
A2 78299353 78299476
refGene_gn=NRCAM;refGene_tx=XM_023250225.1;refGene_gn=NRCAM;refGene_tx=XM_023250226.1;refGene_gn=NRCAM;refGene_tx=XM_023250227.1;refGene_gn=NRCAM;refGene_tx=XM_023250228.1;refGene_gn=NRCAM;refGene_tx=XM_023250230.1;refGene_gn=NRCAM;refGene_tx=XM_023250231.1;refGene_gn=NRCAM;refGene_tx=XM_023250232.1;refGene_gn=NRCAM;refGene_tx=XM_023250233.1;refGene_gn=NRCAM;refGene_tx=XM_023250234.1;refGene_gn=NRCAM;refGene_tx=XM_023250235.1;refGene_gn=NRCAM;refGene_tx=XM_023250236.1;refGene_gn=NRCAM;refGene_tx=XM_023250237.1;refGene_gn=NRCAM;refGene_tx=XM_023250238.1;refGene_gn=NRCAM;refGene_tx=XM_023250239.1;refGene_gn=NRCAM;refGene_tx=XM_023250240.1;refGene_gn=NRCAM;refGene_tx=XM_023250241.1
A2 78305053 78305217
refGene_gn=NRCAM;refGene_tx=XM_023250225.1;refGene_gn=NRCAM;refGene_tx=XM_023250226.1;refGene_gn=NRCAM;refGene_tx=XM_023250227.1;refGene_gn=NRCAM;refGene_tx=XM_023250228.1;refGene_gn=NRCAM;refGene_tx=XM_023250230.1;refGene_gn=NRCAM;refGene_tx=XM_023250231.1;refGene_gn=NRCAM;refGene_tx=XM_023250232.1;refGene_gn=NRCAM;refGene_tx=XM_023250233.1;refGene_gn=NRCAM;refGene_tx=XM_023250234.1;refGene_gn=NRCAM;refGene_tx=XM_023250235.1;refGene_gn=NRCAM;refGene_tx=XM_023250236.1;refGene_gn=NRCAM;refGene_tx=XM_023250237.1;refGene_gn=NRCAM;refGene_tx=XM_023250238.1;refGene_gn=NRCAM;refGene_tx=XM_023250239.1;refGene_gn=NRCAM;refGene_tx=

XM_023250240.1;refGene_gn=NRCAM;refGene_tx=XM_023250241.1
A2 78305954 78306018
refGene_gn=NRCAM;refGene_tx=XM_023250225.1;refGene_gn=NRCAM;refGene_tx=XM_023250226.1;refGene_gn=NRCAM;refGene_tx=XM_023250227.1;refGene_gn=NRCAM;refGene_tx=XM_023250228.1;refGene_gn=NRCAM;refGene_tx=XM_023250230.1;refGene_gn=NRCAM;refGene_tx=XM_023250231.1;refGene_gn=NRCAM;refGene_tx=XM_023250232.1;refGene_gn=NRCAM;refGene_tx=XM_023250233.1;refGene_gn=NRCAM;refGene_tx=XM_023250234.1;refGene_gn=NRCAM;refGene_tx=XM_023250235.1;refGene_gn=NRCAM;refGene_tx=XM_023250236.1;refGene_gn=NRCAM;refGene_tx=XM_023250237.1;refGene_gn=NRCAM;refGene_tx=XM_023250238.1;refGene_gn=NRCAM;refGene_tx=XM_023250239.1;refGene_gn=NRCAM;refGene_tx=XM_023250240.1;refGene_gn=NRCAM;refGene_tx=XM_023250241.1
A2 78308079 78308191
refGene_gn=NRCAM;refGene_tx=XM_023250225.1;refGene_gn=NRCAM;refGene_tx=XM_023250226.1;refGene_gn=NRCAM;refGene_tx=XM_023250227.1;refGene_gn=NRCAM;refGene_tx=XM_023250228.1;refGene_gn=NRCAM;refGene_tx=XM_023250230.1;refGene_gn=NRCAM;refGene_tx=XM_023250231.1;refGene_gn=NRCAM;refGene_tx=XM_023250232.1;refGene_gn=NRCAM;refGene_tx=XM_023250233.1;refGene_gn=NRCAM;refGene_tx=XM_023250234.1;refGene_gn=NRCAM;refGene_tx=XM_023250235.1;refGene_gn=NRCAM;refGene_tx=XM_023250236.1;refGene_gn=NRCAM;refGene_tx=XM_023250237.1;refGene_gn=NRCAM;refGene_tx=XM_023250238.1;refGene_gn=NRCAM;refGene_tx=XM_023250239.1;refGene_gn=NRCAM;refGene_tx=XM_023250240.1;refGene_gn=NRCAM;refGene_tx=XM_023250241.1
A2 78311158 78311343
refGene_gn=NRCAM;refGene_tx=XM_023250225.1;refGene_gn=NRCAM;refGene_tx=XM_023250226.1;refGene_gn=NRCAM;refGene_tx=XM_023250227.1;refGene_gn=NRCAM;refGene_tx=XM_023250228.1;refGene_gn=NRCAM;refGene_tx=XM_023250230.1;refGene_gn=NRCAM;refGene_tx=XM_023250231.1;refGene_gn=NRCAM;refGene_tx=XM_023250232.1;refGene_gn=NRCAM;refGene_tx=XM_023250233.1;refGene_gn=NRCAM;refGene_tx=XM_023250234.1;refGene_gn=NRCAM;refGene_tx=XM_023250235.1;refGene_gn=NRCAM;refGene_tx=XM_023250236.1;refGene_gn=NRCAM;refGene_tx=XM_023250237.1;refGene_gn=NRCAM;refGene_tx=XM_023250238.1;refGene_gn=NRCAM;refGene_tx=XM_023250239.1;refGene_gn=NRCAM;refGene_tx=XM_023250240.1;refGene_gn=NRCAM;refGene_tx=XM_023250241.1
A2 78312810 78312942
refGene_gn=NRCAM;refGene_tx=XM_023250225.1;refGene_gn=NRCAM;refGene_tx=XM_023250226.1;refGene_gn=NRCAM;refGene_tx=XM_023250227.1;refGene_gn=NRCAM;refGene_tx=XM_023250228.1;refGene_gn=NRCAM;refGene_tx=XM_023250230.1;refGene_gn=NRCAM;refGene_tx=XM_023250231.1;refGene_gn=NRCAM;refGene_tx=XM_023250232.1;refGene_gn=NRCAM;refGene_tx=XM_023250233.1;refGene_gn=NRCAM;refGene_tx=XM_023250234.1;refGene_gn=NRCAM;refGene_tx=XM_023250235.1;refGene_gn=NRCAM;refGene_tx=XM_023250236.1;refGene_gn=NRCAM;refGene_tx=XM_023250237.1;refGene_gn=NRCAM;refGene_tx=XM_023250238.1;refGene_gn=NRCAM;refGene_tx=XM_023250239.1;refGene_gn=NRCAM;refGene_tx=XM_023250240.1;refGene_gn=NRCAM;refGene_tx=XM_023250241.1
A2 78320956 78321100
refGene_gn=NRCAM;refGene_tx=XM_023250225.1;refGene_gn=NRCAM;refGene_tx=XM_023250226.1;refGene_gn=NRCAM;refGene_tx=XM_023250227.1;refGene_gn=NRCAM;refGene_tx=XM_023250228.1;refGene_gn=NRCAM;refGene_tx=XM_023250230.1;refGene_gn=NRCAM;refGene_tx=XM_023250231.1;refGene_gn=NRCAM;refGe

ne_tx=XM_023250232.1;refGene_gn=NRCAM;refGene_tx=XM_023250233.1;refGen
e_gn=NRCAM;refGene_tx=XM_023250234.1;refGene_gn=NRCAM;refGene_tx=XM_02
3250235.1;refGene_gn=NRCAM;refGene_tx=XM_023250236.1;refGene_gn=NRCAM;
refGene_tx=XM_023250237.1;refGene_gn=NRCAM;refGene_tx=XM_023250238.1;r
efGene_gn=NRCAM;refGene_tx=XM_023250239.1;refGene_gn=NRCAM;refGene_tx=
XM_023250240.1;refGene_gn=NRCAM;refGene_tx=XM_023250241.1
A2 78324147 78324259
refGene_gn=NRCAM;refGene_tx=XM_023250225.1;refGene_gn=NRCAM;refGene_tx
=XM_023250226.1;refGene_gn=NRCAM;refGene_tx=XM_023250227.1;refGene_gn=
NRCAM;refGene_tx=XM_023250228.1;refGene_gn=NRCAM;refGene_tx=XM_0232502
30.1;refGene_gn=NRCAM;refGene_tx=XM_023250231.1;refGene_gn=NRCAM;refGe
ne_tx=XM_023250232.1;refGene_gn=NRCAM;refGene_tx=XM_023250233.1;refGen
e_gn=NRCAM;refGene_tx=XM_023250234.1;refGene_gn=NRCAM;refGene_tx=XM_02
3250235.1;refGene_gn=NRCAM;refGene_tx=XM_023250236.1;refGene_gn=NRCAM;
refGene_tx=XM_023250237.1;refGene_gn=NRCAM;refGene_tx=XM_023250238.1;r
efGene_gn=NRCAM;refGene_tx=XM_023250239.1;refGene_gn=NRCAM;refGene_tx=
XM_023250240.1;refGene_gn=NRCAM;refGene_tx=XM_023250241.1
A2 78325769 78325936
refGene_gn=NRCAM;refGene_tx=XM_023250225.1;refGene_gn=NRCAM;refGene_tx
=XM_023250226.1;refGene_gn=NRCAM;refGene_tx=XM_023250227.1;refGene_gn=
NRCAM;refGene_tx=XM_023250228.1;refGene_gn=NRCAM;refGene_tx=XM_0232502
30.1;refGene_gn=NRCAM;refGene_tx=XM_023250231.1;refGene_gn=NRCAM;refGe
ne_tx=XM_023250232.1;refGene_gn=NRCAM;refGene_tx=XM_023250233.1;refGen
e_gn=NRCAM;refGene_tx=XM_023250234.1;refGene_gn=NRCAM;refGene_tx=XM_02
3250235.1;refGene_gn=NRCAM;refGene_tx=XM_023250236.1;refGene_gn=NRCAM;
refGene_tx=XM_023250237.1;refGene_gn=NRCAM;refGene_tx=XM_023250238.1;r
efGene_gn=NRCAM;refGene_tx=XM_023250239.1;refGene_gn=NRCAM;refGene_tx=
XM_023250240.1;refGene_gn=NRCAM;refGene_tx=XM_023250241.1
A2 78326028 78326176
refGene_gn=NRCAM;refGene_tx=XM_023250225.1;refGene_gn=NRCAM;refGene_tx
=XM_023250226.1;refGene_gn=NRCAM;refGene_tx=XM_023250227.1;refGene_gn=
NRCAM;refGene_tx=XM_023250228.1;refGene_gn=NRCAM;refGene_tx=XM_0232502
30.1;refGene_gn=NRCAM;refGene_tx=XM_023250231.1;refGene_gn=NRCAM;refGe
ne_tx=XM_023250232.1;refGene_gn=NRCAM;refGene_tx=XM_023250233.1;refGen
e_gn=NRCAM;refGene_tx=XM_023250234.1;refGene_gn=NRCAM;refGene_tx=XM_02
3250235.1;refGene_gn=NRCAM;refGene_tx=XM_023250236.1;refGene_gn=NRCAM;
refGene_tx=XM_023250237.1;refGene_gn=NRCAM;refGene_tx=XM_023250238.1;r
efGene_gn=NRCAM;refGene_tx=XM_023250239.1;refGene_gn=NRCAM;refGene_tx=
XM_023250240.1;refGene_gn=NRCAM;refGene_tx=XM_023250241.1
A2 78328413 78328538
refGene_gn=NRCAM;refGene_tx=XM_023250225.1;refGene_gn=NRCAM;refGene_tx
=XM_023250226.1;refGene_gn=NRCAM;refGene_tx=XM_023250227.1;refGene_gn=
NRCAM;refGene_tx=XM_023250228.1;refGene_gn=NRCAM;refGene_tx=XM_0232502
30.1;refGene_gn=NRCAM;refGene_tx=XM_023250231.1;refGene_gn=NRCAM;refGe
ne_tx=XM_023250232.1;refGene_gn=NRCAM;refGene_tx=XM_023250233.1;refGen
e_gn=NRCAM;refGene_tx=XM_023250234.1;refGene_gn=NRCAM;refGene_tx=XM_02
3250235.1;refGene_gn=NRCAM;refGene_tx=XM_023250236.1;refGene_gn=NRCAM;
refGene_tx=XM_023250237.1;refGene_gn=NRCAM;refGene_tx=XM_023250238.1;r
efGene_gn=NRCAM;refGene_tx=XM_023250239.1;refGene_gn=NRCAM;refGene_tx=
XM_023250240.1;refGene_gn=NRCAM;refGene_tx=XM_023250241.1

A2 78328989 78329019
refGene_gn=NRCAM;refGene_tx=XM_023250225.1;refGene_gn=NRCAM;refGene_tx=XM_023250226.1;refGene_gn=NRCAM;refGene_tx=XM_023250227.1;refGene_gn=NRCAM;refGene_tx=XM_023250228.1;refGene_gn=NRCAM;refGene_tx=XM_023250230.1;refGene_gn=NRCAM;refGene_tx=XM_023250231.1;refGene_gn=NRCAM;refGene_tx=XM_023250233.1;refGene_gn=NRCAM;refGene_tx=XM_023250234.1;refGene_gn=NRCAM;refGene_tx=XM_023250235.1;refGene_gn=NRCAM;refGene_tx=XM_023250236.1;refGene_gn=NRCAM;refGene_tx=XM_023250239.1
A2 78330829 78330931
refGene_gn=NRCAM;refGene_tx=XM_023250225.1;refGene_gn=NRCAM;refGene_tx=XM_023250226.1;refGene_gn=NRCAM;refGene_tx=XM_023250227.1;refGene_gn=NRCAM;refGene_tx=XM_023250228.1;refGene_gn=NRCAM;refGene_tx=XM_023250230.1;refGene_gn=NRCAM;refGene_tx=XM_023250231.1;refGene_gn=NRCAM;refGene_tx=XM_023250232.1;refGene_gn=NRCAM;refGene_tx=XM_023250233.1;refGene_gn=NRCAM;refGene_tx=XM_023250234.1;refGene_gn=NRCAM;refGene_tx=XM_023250235.1;refGene_gn=NRCAM;refGene_tx=XM_023250236.1;refGene_gn=NRCAM;refGene_tx=XM_023250237.1;refGene_gn=NRCAM;refGene_tx=XM_023250238.1;refGene_gn=NRCAM;refGene_tx=XM_023250239.1;refGene_gn=NRCAM;refGene_tx=XM_023250240.1;refGene_gn=NRCAM;refGene_tx=XM_023250241.1
A2 78335214 78335412
refGene_gn=NRCAM;refGene_tx=XM_023250225.1;refGene_gn=NRCAM;refGene_tx=XM_023250226.1;refGene_gn=NRCAM;refGene_tx=XM_023250227.1;refGene_gn=NRCAM;refGene_tx=XM_023250228.1;refGene_gn=NRCAM;refGene_tx=XM_023250230.1;refGene_gn=NRCAM;refGene_tx=XM_023250231.1;refGene_gn=NRCAM;refGene_tx=XM_023250232.1;refGene_gn=NRCAM;refGene_tx=XM_023250233.1;refGene_gn=NRCAM;refGene_tx=XM_023250234.1;refGene_gn=NRCAM;refGene_tx=XM_023250235.1;refGene_gn=NRCAM;refGene_tx=XM_023250236.1;refGene_gn=NRCAM;refGene_tx=XM_023250237.1;refGene_gn=NRCAM;refGene_tx=XM_023250238.1;refGene_gn=NRCAM;refGene_tx=XM_023250239.1;refGene_gn=NRCAM;refGene_tx=XM_023250240.1;refGene_gn=NRCAM;refGene_tx=XM_023250241.1
A2 78335532 78335603
refGene_gn=NRCAM;refGene_tx=XM_023250225.1;refGene_gn=NRCAM;refGene_tx=XM_023250226.1;refGene_gn=NRCAM;refGene_tx=XM_023250227.1;refGene_gn=NRCAM;refGene_tx=XM_023250228.1;refGene_gn=NRCAM;refGene_tx=XM_023250230.1;refGene_gn=NRCAM;refGene_tx=XM_023250231.1;refGene_gn=NRCAM;refGene_tx=XM_023250232.1;refGene_gn=NRCAM;refGene_tx=XM_023250233.1;refGene_gn=NRCAM;refGene_tx=XM_023250234.1;refGene_gn=NRCAM;refGene_tx=XM_023250235.1;refGene_gn=NRCAM;refGene_tx=XM_023250236.1;refGene_gn=NRCAM;refGene_tx=XM_023250237.1;refGene_gn=NRCAM;refGene_tx=XM_023250238.1;refGene_gn=NRCAM;refGene_tx=XM_023250239.1;refGene_gn=NRCAM;refGene_tx=XM_023250240.1;refGene_gn=NRCAM;refGene_tx=XM_023250241.1
A2 78336974 78337200
refGene_gn=NRCAM;refGene_tx=XM_023250225.1;refGene_gn=NRCAM;refGene_tx=XM_023250226.1;refGene_gn=NRCAM;refGene_tx=XM_023250227.1;refGene_gn=NRCAM;refGene_tx=XM_023250228.1;refGene_gn=NRCAM;refGene_tx=XM_023250230.1;refGene_gn=NRCAM;refGene_tx=XM_023250231.1;refGene_gn=NRCAM;refGene_tx=XM_023250232.1;refGene_gn=NRCAM;refGene_tx=XM_023250233.1;refGene_gn=NRCAM;refGene_tx=XM_023250234.1;refGene_gn=NRCAM;refGene_tx=XM_023250235.1;refGene_gn=NRCAM;refGene_tx=XM_023250236.1;refGene_gn=NRCAM;refGene_tx=XM_023250237.1;refGene_gn=NRCAM;refGene_tx=XM_023250238.1;r

efGene_gn=NRCAM;refGene_tx=XM_023250239.1;refGene_gn=NRCAM;refGene_tx=XM_023250240.1;refGene_gn=NRCAM;refGene_tx=XM_023250241.1
A2 78337572 78337688
refGene_gn=NRCAM;refGene_tx=XM_023250225.1;refGene_gn=NRCAM;refGene_tx=XM_023250226.1;refGene_gn=NRCAM;refGene_tx=XM_023250227.1;refGene_gn=NRCAM;refGene_tx=XM_023250228.1;refGene_gn=NRCAM;refGene_tx=XM_023250230.1;refGene_gn=NRCAM;refGene_tx=XM_023250231.1;refGene_gn=NRCAM;refGene_tx=XM_023250232.1;refGene_gn=NRCAM;refGene_tx=XM_023250233.1;refGene_gn=NRCAM;refGene_tx=XM_023250234.1;refGene_gn=NRCAM;refGene_tx=XM_023250235.1;refGene_gn=NRCAM;refGene_tx=XM_023250236.1;refGene_gn=NRCAM;refGene_tx=XM_023250237.1;refGene_gn=NRCAM;refGene_tx=XM_023250238.1;refGene_gn=NRCAM;refGene_tx=XM_023250239.1;refGene_gn=NRCAM;refGene_tx=XM_023250240.1;refGene_gn=NRCAM;refGene_tx=XM_023250241.1
A2 78339667 78339875
refGene_gn=NRCAM;refGene_tx=XM_023250225.1;refGene_gn=NRCAM;refGene_tx=XM_023250226.1;refGene_gn=NRCAM;refGene_tx=XM_023250227.1;refGene_gn=NRCAM;refGene_tx=XM_023250228.1;refGene_gn=NRCAM;refGene_tx=XM_023250230.1;refGene_gn=NRCAM;refGene_tx=XM_023250231.1;refGene_gn=NRCAM;refGene_tx=XM_023250232.1;refGene_gn=NRCAM;refGene_tx=XM_023250233.1;refGene_gn=NRCAM;refGene_tx=XM_023250234.1;refGene_gn=NRCAM;refGene_tx=XM_023250235.1;refGene_gn=NRCAM;refGene_tx=XM_023250236.1;refGene_gn=NRCAM;refGene_tx=XM_023250237.1;refGene_gn=NRCAM;refGene_tx=XM_023250238.1;refGene_gn=NRCAM;refGene_tx=XM_023250239.1;refGene_gn=NRCAM;refGene_tx=XM_023250240.1;refGene_gn=NRCAM;refGene_tx=XM_023250241.1
A2 78342308 78342431
refGene_gn=NRCAM;refGene_tx=XM_023250225.1;refGene_gn=NRCAM;refGene_tx=XM_023250226.1;refGene_gn=NRCAM;refGene_tx=XM_023250227.1;refGene_gn=NRCAM;refGene_tx=XM_023250228.1;refGene_gn=NRCAM;refGene_tx=XM_023250230.1;refGene_gn=NRCAM;refGene_tx=XM_023250231.1;refGene_gn=NRCAM;refGene_tx=XM_023250232.1;refGene_gn=NRCAM;refGene_tx=XM_023250233.1;refGene_gn=NRCAM;refGene_tx=XM_023250234.1;refGene_gn=NRCAM;refGene_tx=XM_023250235.1;refGene_gn=NRCAM;refGene_tx=XM_023250236.1;refGene_gn=NRCAM;refGene_tx=XM_023250237.1;refGene_gn=NRCAM;refGene_tx=XM_023250238.1;refGene_gn=NRCAM;refGene_tx=XM_023250239.1;refGene_gn=NRCAM;refGene_tx=XM_023250240.1;refGene_gn=NRCAM;refGene_tx=XM_023250241.1
A2 78343307 78343493
refGene_gn=NRCAM;refGene_tx=XM_023250225.1;refGene_gn=NRCAM;refGene_tx=XM_023250226.1;refGene_gn=NRCAM;refGene_tx=XM_023250227.1;refGene_gn=NRCAM;refGene_tx=XM_023250228.1;refGene_gn=NRCAM;refGene_tx=XM_023250230.1;refGene_gn=NRCAM;refGene_tx=XM_023250231.1;refGene_gn=NRCAM;refGene_tx=XM_023250232.1;refGene_gn=NRCAM;refGene_tx=XM_023250233.1;refGene_gn=NRCAM;refGene_tx=XM_023250234.1;refGene_gn=NRCAM;refGene_tx=XM_023250235.1;refGene_gn=NRCAM;refGene_tx=XM_023250236.1;refGene_gn=NRCAM;refGene_tx=XM_023250237.1;refGene_gn=NRCAM;refGene_tx=XM_023250238.1;refGene_gn=NRCAM;refGene_tx=XM_023250239.1;refGene_gn=NRCAM;refGene_tx=XM_023250240.1;refGene_gn=NRCAM;refGene_tx=XM_023250241.1
A2 78344534 78344570
refGene_gn=NRCAM;refGene_tx=XM_023250225.1;refGene_gn=NRCAM;refGene_tx=XM_023250226.1;refGene_gn=NRCAM;refGene_tx=XM_023250227.1;refGene_gn=NRCAM;refGene_tx=XM_023250228.1;refGene_gn=NRCAM;refGene_tx=XM_0232502

30.1;refGene_gn=NRCAM;refGene_tx=XM_023250231.1;refGene_gn=NRCAM;refGene_tx=XM_023250232.1;refGene_gn=NRCAM;refGene_tx=XM_023250239.1
A2 78357184 78357316
refGene_gn=NRCAM;refGene_tx=XM_023250225.1;refGene_gn=NRCAM;refGene_tx=XM_023250226.1;refGene_gn=NRCAM;refGene_tx=XM_023250227.1;refGene_gn=NRCAM;refGene_tx=XM_023250228.1;refGene_gn=NRCAM;refGene_tx=XM_023250230.1;refGene_gn=NRCAM;refGene_tx=XM_023250231.1;refGene_gn=NRCAM;refGene_tx=XM_023250232.1;refGene_gn=NRCAM;refGene_tx=XM_023250233.1;refGene_gn=NRCAM;refGene_tx=XM_023250234.1;refGene_gn=NRCAM;refGene_tx=XM_023250235.1;refGene_gn=NRCAM;refGene_tx=XM_023250236.1;refGene_gn=NRCAM;refGene_tx=XM_023250237.1;refGene_gn=NRCAM;refGene_tx=XM_023250238.1;refGene_gn=NRCAM;refGene_tx=XM_023250239.1;refGene_gn=NRCAM;refGene_tx=XM_023250240.1;refGene_gn=NRCAM;refGene_tx=XM_023250241.1
A2 78358135 78358214
refGene_gn=NRCAM;refGene_tx=XM_023250225.1;refGene_gn=NRCAM;refGene_tx=XM_023250226.1;refGene_gn=NRCAM;refGene_tx=XM_023250227.1;refGene_gn=NRCAM;refGene_tx=XM_023250228.1;refGene_gn=NRCAM;refGene_tx=XM_023250230.1;refGene_gn=NRCAM;refGene_tx=XM_023250231.1;refGene_gn=NRCAM;refGene_tx=XM_023250232.1;refGene_gn=NRCAM;refGene_tx=XM_023250233.1;refGene_gn=NRCAM;refGene_tx=XM_023250234.1;refGene_gn=NRCAM;refGene_tx=XM_023250235.1;refGene_gn=NRCAM;refGene_tx=XM_023250236.1;refGene_gn=NRCAM;refGene_tx=XM_023250237.1;refGene_gn=NRCAM;refGene_tx=XM_023250238.1;refGene_gn=NRCAM;refGene_tx=XM_023250239.1;refGene_gn=NRCAM;refGene_tx=XM_023250240.1;refGene_gn=NRCAM;refGene_tx=XM_023250241.1
A2 78369004 78369016
refGene_gn=NRCAM;refGene_tx=XM_023250225.1;refGene_gn=NRCAM;refGene_tx=XM_023250226.1;refGene_gn=NRCAM;refGene_tx=XM_023250227.1;refGene_gn=NRCAM;refGene_tx=XM_023250230.1;refGene_gn=NRCAM;refGene_tx=XM_023250232.1;refGene_gn=NRCAM;refGene_tx=XM_023250233.1;refGene_gn=NRCAM;refGene_tx=XM_023250234.1
A2 78369326 78369456 refGene_gn=NRCAM;refGene_tx=XM_023250239.1
A2 78369583 78369821
refGene_gn=NRCAM;refGene_tx=XM_023250225.1;refGene_gn=NRCAM;refGene_tx=XM_023250226.1;refGene_gn=NRCAM;refGene_tx=XM_023250227.1;refGene_gn=NRCAM;refGene_tx=XM_023250228.1;refGene_gn=NRCAM;refGene_tx=XM_023250230.1;refGene_gn=NRCAM;refGene_tx=XM_023250231.1;refGene_gn=NRCAM;refGene_tx=XM_023250232.1;refGene_gn=NRCAM;refGene_tx=XM_023250233.1;refGene_gn=NRCAM;refGene_tx=XM_023250234.1;refGene_gn=NRCAM;refGene_tx=XM_023250235.1;refGene_gn=NRCAM;refGene_tx=XM_023250236.1;refGene_gn=NRCAM;refGene_tx=XM_023250237.1;refGene_gn=NRCAM;refGene_tx=XM_023250238.1;refGene_gn=NRCAM;refGene_tx=XM_023250240.1;refGene_gn=NRCAM;refGene_tx=XM_023250241.1
A2 78413185 78413219
refGene_gn=LAMB4;refGene_tx=XM_023250242.1;refGene_gn=LAMB4;refGene_tx=XM_023250243.1;refGene_gn=LAMB4;refGene_tx=XM_023250244.1;refGene_gn=LAMB4;refGene_tx=XM_023250245.1;refGene_gn=LAMB4;refGene_tx=XM_023250246.1
A2 78418134 78418292
refGene_gn=LAMB4;refGene_tx=XM_023250242.1;refGene_gn=LAMB4;refGene_tx=XM_023250243.1;refGene_gn=LAMB4;refGene_tx=XM_023250244.1;refGene_gn=

LAMB4;refGene_tx=XM_023250245.1;refGene_gn=LAMB4;refGene_tx=XM_0232502
46.1
A2 78421701 78421837
refGene_gn=LAMB4;refGene_tx=XM_023250242.1;refGene_gn=LAMB4;refGene_tx
=XM_023250243.1;refGene_gn=LAMB4;refGene_tx=XM_023250244.1;refGene_gn=
LAMB4;refGene_tx=XM_023250245.1;refGene_gn=LAMB4;refGene_tx=XM_0232502
46.1
A2 78424069 78424143
refGene_gn=LAMB4;refGene_tx=XM_023250242.1;refGene_gn=LAMB4;refGene_tx
=XM_023250243.1;refGene_gn=LAMB4;refGene_tx=XM_023250244.1;refGene_gn=
LAMB4;refGene_tx=XM_023250245.1;refGene_gn=LAMB4;refGene_tx=XM_0232502
46.1
A2 78425255 78425444
refGene_gn=LAMB4;refGene_tx=XM_023250242.1;refGene_gn=LAMB4;refGene_tx
=XM_023250243.1;refGene_gn=LAMB4;refGene_tx=XM_023250244.1;refGene_gn=
LAMB4;refGene_tx=XM_023250245.1;refGene_gn=LAMB4;refGene_tx=XM_0232502
46.1
A2 78426890 78426954
refGene_gn=LAMB4;refGene_tx=XM_023250242.1;refGene_gn=LAMB4;refGene_tx
=XM_023250243.1;refGene_gn=LAMB4;refGene_tx=XM_023250244.1;refGene_gn=
LAMB4;refGene_tx=XM_023250245.1;refGene_gn=LAMB4;refGene_tx=XM_0232502
46.1
A2 78427078 78427293
refGene_gn=LAMB4;refGene_tx=XM_023250242.1;refGene_gn=LAMB4;refGene_tx
=XM_023250243.1;refGene_gn=LAMB4;refGene_tx=XM_023250244.1;refGene_gn=
LAMB4;refGene_tx=XM_023250245.1;refGene_gn=LAMB4;refGene_tx=XM_0232502
46.1
A2 78428315 78428436
refGene_gn=LAMB4;refGene_tx=XM_023250242.1;refGene_gn=LAMB4;refGene_tx
=XM_023250243.1;refGene_gn=LAMB4;refGene_tx=XM_023250244.1;refGene_gn=
LAMB4;refGene_tx=XM_023250245.1;refGene_gn=LAMB4;refGene_tx=XM_0232502
46.1
A2 78429658 78429847
refGene_gn=LAMB4;refGene_tx=XM_023250242.1;refGene_gn=LAMB4;refGene_tx
=XM_023250243.1;refGene_gn=LAMB4;refGene_tx=XM_023250244.1;refGene_gn=
LAMB4;refGene_tx=XM_023250245.1;refGene_gn=LAMB4;refGene_tx=XM_0232502
46.1
A2 78435590 78435770
refGene_gn=LAMB4;refGene_tx=XM_023250242.1;refGene_gn=LAMB4;refGene_tx
=XM_023250243.1;refGene_gn=LAMB4;refGene_tx=XM_023250244.1;refGene_gn=
LAMB4;refGene_tx=XM_023250245.1;refGene_gn=LAMB4;refGene_tx=XM_0232502
46.1
A2 78437964 78438074
refGene_gn=LAMB4;refGene_tx=XM_023250242.1;refGene_gn=LAMB4;refGene_tx
=XM_023250243.1;refGene_gn=LAMB4;refGene_tx=XM_023250244.1;refGene_gn=
LAMB4;refGene_tx=XM_023250245.1;refGene_gn=LAMB4;refGene_tx=XM_0232502
46.1
A2 78440902 78440982
refGene_gn=LAMB4;refGene_tx=XM_023250242.1;refGene_gn=LAMB4;refGene_tx
=XM_023250243.1;refGene_gn=LAMB4;refGene_tx=XM_023250244.1;refGene_gn=

LAMB4;refGene_tx=XM_023250245.1;refGene_gn=LAMB4;refGene_tx=XM_023250246.1
A2 78441537 78441688
refGene_gn=LAMB4;refGene_tx=XM_023250242.1;refGene_gn=LAMB4;refGene_tx=XM_023250243.1;refGene_gn=LAMB4;refGene_tx=XM_023250244.1;refGene_gn=LAMB4;refGene_tx=XM_023250245.1;refGene_gn=LAMB4;refGene_tx=XM_023250246.1
A2 78447173 78447305
refGene_gn=LAMB4;refGene_tx=XM_023250242.1;refGene_gn=LAMB4;refGene_tx=XM_023250243.1;refGene_gn=LAMB4;refGene_tx=XM_023250244.1;refGene_gn=LAMB4;refGene_tx=XM_023250245.1;refGene_gn=LAMB4;refGene_tx=XM_023250246.1
A2 78455130 78455316
refGene_gn=LAMB4;refGene_tx=XM_023250242.1;refGene_gn=LAMB4;refGene_tx=XM_023250243.1;refGene_gn=LAMB4;refGene_tx=XM_023250244.1;refGene_gn=LAMB4;refGene_tx=XM_023250245.1;refGene_gn=LAMB4;refGene_tx=XM_023250246.1
A2 78457503 78457619
refGene_gn=LAMB4;refGene_tx=XM_023250242.1;refGene_gn=LAMB4;refGene_tx=XM_023250243.1;refGene_gn=LAMB4;refGene_tx=XM_023250244.1;refGene_gn=LAMB4;refGene_tx=XM_023250245.1;refGene_gn=LAMB4;refGene_tx=XM_023250246.1
A2 78458626 78458747
refGene_gn=LAMB4;refGene_tx=XM_023250242.1;refGene_gn=LAMB4;refGene_tx=XM_023250243.1;refGene_gn=LAMB4;refGene_tx=XM_023250244.1;refGene_gn=LAMB4;refGene_tx=XM_023250245.1;refGene_gn=LAMB4;refGene_tx=XM_023250246.1
A2 78466813 78466991
refGene_gn=LAMB4;refGene_tx=XM_023250242.1;refGene_gn=LAMB4;refGene_tx=XM_023250243.1;refGene_gn=LAMB4;refGene_tx=XM_023250244.1;refGene_gn=LAMB4;refGene_tx=XM_023250245.1;refGene_gn=LAMB4;refGene_tx=XM_023250246.1
A2 78468381 78468525
refGene_gn=LAMB4;refGene_tx=XM_023250242.1;refGene_gn=LAMB4;refGene_tx=XM_023250243.1;refGene_gn=LAMB4;refGene_tx=XM_023250244.1;refGene_gn=LAMB4;refGene_tx=XM_023250245.1;refGene_gn=LAMB4;refGene_tx=XM_023250246.1
A2 78469941 78470173
refGene_gn=LAMB4;refGene_tx=XM_023250242.1;refGene_gn=LAMB4;refGene_tx=XM_023250243.1;refGene_gn=LAMB4;refGene_tx=XM_023250244.1;refGene_gn=LAMB4;refGene_tx=XM_023250245.1;refGene_gn=LAMB4;refGene_tx=XM_023250246.1
A2 78470913 78471091
refGene_gn=LAMB4;refGene_tx=XM_023250242.1;refGene_gn=LAMB4;refGene_tx=XM_023250243.1;refGene_gn=LAMB4;refGene_tx=XM_023250244.1;refGene_gn=LAMB4;refGene_tx=XM_023250245.1;refGene_gn=LAMB4;refGene_tx=XM_023250246.1
A2 78473035 78473260
refGene_gn=LAMB4;refGene_tx=XM_023250242.1;refGene_gn=LAMB4;refGene_tx=XM_023250243.1;refGene_gn=LAMB4;refGene_tx=XM_023250244.1;refGene_gn=

LAMB4;refGene_tx=XM_023250245.1
A2	78474374 78474595
refGene_gn=LAMB4;refGene_tx=XM_023250242.1;refGene_gn=LAMB4;refGene_tx
=XM_023250243.1;refGene_gn=LAMB4;refGene_tx=XM_023250244.1;refGene_gn=
LAMB4;refGene_tx=XM_023250245.1
A2	78480510 78480607
refGene_gn=LAMB4;refGene_tx=XM_023250242.1;refGene_gn=LAMB4;refGene_tx
=XM_023250243.1;refGene_gn=LAMB4;refGene_tx=XM_023250244.1;refGene_gn=
LAMB4;refGene_tx=XM_023250245.1
A2	78484763 78485139
refGene_gn=LAMB4;refGene_tx=XM_023250242.1;refGene_gn=LAMB4;refGene_tx
=XM_023250243.1;refGene_gn=LAMB4;refGene_tx=XM_023250244.1;refGene_gn=
LAMB4;refGene_tx=XM_023250245.1
A2	78488704 78488865
refGene_gn=LAMB4;refGene_tx=XM_023250242.1;refGene_gn=LAMB4;refGene_tx
=XM_023250243.1;refGene_gn=LAMB4;refGene_tx=XM_023250244.1;refGene_gn=
LAMB4;refGene_tx=XM_023250245.1
A2	78491838 78492044
refGene_gn=LAMB4;refGene_tx=XM_023250242.1;refGene_gn=LAMB4;refGene_tx
=XM_023250243.1;refGene_gn=LAMB4;refGene_tx=XM_023250244.1;refGene_gn=
LAMB4;refGene_tx=XM_023250245.1
A2	78492659 78492821
refGene_gn=LAMB4;refGene_tx=XM_023250242.1;refGene_gn=LAMB4;refGene_tx
=XM_023250243.1;refGene_gn=LAMB4;refGene_tx=XM_023250244.1;refGene_gn=
LAMB4;refGene_tx=XM_023250245.1
A2	78493139 78493208
refGene_gn=LAMB4;refGene_tx=XM_023250244.1;refGene_gn=LAMB4;refGene_tx
=XM_023250245.1
A2	78496827 78496972
refGene_gn=LAMB4;refGene_tx=XM_023250242.1;refGene_gn=LAMB4;refGene_tx
=XM_023250243.1;refGene_gn=LAMB4;refGene_tx=XM_023250244.1
A2	78500225 78500433
refGene_gn=LAMB4;refGene_tx=XM_023250242.1;refGene_gn=LAMB4;refGene_tx
=XM_023250243.1
A2	78503724 78503863
refGene_gn=LAMB4;refGene_tx=XM_023250242.1;refGene_gn=LAMB4;refGene_tx
=XM_023250243.1
A2	78507134 78507308
refGene_gn=LAMB4;refGene_tx=XM_023250242.1;refGene_gn=LAMB4;refGene_tx
=XM_023250243.1
A2	78508605 78508759
refGene_gn=LAMB4;refGene_tx=XM_023250242.1;refGene_gn=LAMB4;refGene_tx
=XM_023250243.1
A2	78513347 78513487
refGene_gn=LAMB4;refGene_tx=XM_023250242.1;refGene_gn=LAMB4;refGene_tx
=XM_023250243.1
A2	78534943 78534980 refGene_gn=LAMB1;refGene_tx=XM_006929110.2
A2	78536060 78536236 refGene_gn=LAMB1;refGene_tx=XM_006929110.2
A2	78538596 78538732 refGene_gn=LAMB1;refGene_tx=XM_006929110.2
A2	78541880 78541954 refGene_gn=LAMB1;refGene_tx=XM_006929110.2

| | | | |
|---|---|---|---|
| A2 | 78549324 | 78549513 | refGene_gn=LAMB1;refGene_tx=XM_006929110.2 |
| A2 | 78549601 | 78549665 | refGene_gn=LAMB1;refGene_tx=XM_006929110.2 |
| A2 | 78553372 | 78553575 | refGene_gn=LAMB1;refGene_tx=XM_006929110.2 |
| A2 | 78555642 | 78555763 | refGene_gn=LAMB1;refGene_tx=XM_006929110.2 |
| A2 | 78557224 | 78557413 | refGene_gn=LAMB1;refGene_tx=XM_006929110.2 |
| A2 | 78557899 | 78558079 | refGene_gn=LAMB1;refGene_tx=XM_006929110.2 |
| A2 | 78558213 | 78558326 | refGene_gn=LAMB1;refGene_tx=XM_006929110.2 |
| A2 | 78559813 | 78559893 | refGene_gn=LAMB1;refGene_tx=XM_006929110.2 |
| A2 | 78569104 | 78569240 | refGene_gn=LAMB1;refGene_tx=XM_006929110.2 |
| A2 | 78570725 | 78570884 | refGene_gn=LAMB1;refGene_tx=XM_006929110.2 |
| A2 | 78571728 | 78571856 | refGene_gn=LAMB1;refGene_tx=XM_006929110.2 |
| A2 | 78572090 | 78572214 | refGene_gn=LAMB1;refGene_tx=XM_006929110.2 |
| A2 | 78572908 | 78573113 | refGene_gn=LAMB1;refGene_tx=XM_006929110.2 |
| A2 | 78574265 | 78574409 | refGene_gn=LAMB1;refGene_tx=XM_006929110.2 |
| A2 | 78574620 | 78574852 | refGene_gn=LAMB1;refGene_tx=XM_006929110.2 |
| A2 | 78579615 | 78579779 | refGene_gn=LAMB1;refGene_tx=XM_006929110.2 |
| A2 | 78580804 | 78581029 | refGene_gn=LAMB1;refGene_tx=XM_006929110.2 |
| A2 | 78582064 | 78582279 | refGene_gn=LAMB1;refGene_tx=XM_006929110.2 |
| A2 | 78582978 | 78583075 | refGene_gn=LAMB1;refGene_tx=XM_006929110.2 |
| A2 | 78593358 | 78593728 | refGene_gn=LAMB1;refGene_tx=XM_006929110.2 |
| A2 | 78596614 | 78596799 | refGene_gn=LAMB1;refGene_tx=XM_006929110.2 |
| A2 | 78597664 | 78597906 | refGene_gn=LAMB1;refGene_tx=XM_006929110.2 |
| A2 | 78601274 | 78601478 | refGene_gn=LAMB1;refGene_tx=XM_006929110.2 |
| A2 | 78602116 | 78602261 | refGene_gn=LAMB1;refGene_tx=XM_006929110.2 |
| A2 | 78604276 | 78604484 | refGene_gn=LAMB1;refGene_tx=XM_006929110.2 |
| A2 | 78604694 | 78604836 | refGene_gn=LAMB1;refGene_tx=XM_006929110.2 |
| A2 | 78607134 | 78607311 | refGene_gn=LAMB1;refGene_tx=XM_006929110.2 |
| A2 | 78608695 | 78608855 | refGene_gn=LAMB1;refGene_tx=XM_006929110.2 |
| A2 | 78609008 | 78609145 | refGene_gn=LAMB1;refGene_tx=XM_006929110.2 |
| A2 | 78612553 | 78612619 | refGene_gn=DLD;refGene_tx=XM_003982645.4 |
| A2 | 78612713 | 78612803 | refGene_gn=DLD;refGene_tx=XM_003982645.4 |
| A2 | 78613697 | 78613835 | refGene_gn=DLD;refGene_tx=XM_003982645.4 |
| A2 | 78614477 | 78614667 | refGene_gn=DLD;refGene_tx=XM_003982645.4 |
| A2 | 78614990 | 78615161 | refGene_gn=DLD;refGene_tx=XM_003982645.4 |
| A2 | 78615604 | 78615795 | refGene_gn=DLD;refGene_tx=XM_003982645.4 |
| A2 | 78633159 | 78633261 | refGene_gn=DLD;refGene_tx=XM_003982645.4 |
| A2 | 78633785 | 78633929 | refGene_gn=DLD;refGene_tx=XM_003982645.4 |
| A2 | 78634729 | 78634830 | refGene_gn=DLD;refGene_tx=XM_003982645.4 |
| A2 | 78636467 | 78636537 | refGene_gn=DLD;refGene_tx=XM_003982645.4 |
| A2 | 78637103 | 78637172 | refGene_gn=DLD;refGene_tx=XM_003982645.4 |
| A2 | 78637654 | 78637734 | refGene_gn=DLD;refGene_tx=XM_003982645.4 |
| A2 | 78645512 | 78645591 | refGene_gn=DLD;refGene_tx=XM_003982645.4 |
| A2 | 78647086 | 78647125 | refGene_gn=DLD;refGene_tx=XM_003982645.4 |
| A2 | 78730275 | 78730406 | refGene_gn=SLC26A3;refGene_tx=XM_006929111.3 |
| A2 | 78731061 | 78731201 | refGene_gn=SLC26A3;refGene_tx=XM_006929111.3 |
| A2 | 78732924 | 78733035 | refGene_gn=SLC26A3;refGene_tx=XM_006929111.3 |
| A2 | 78733290 | 78733463 | refGene_gn=SLC26A3;refGene_tx=XM_006929111.3 |
| A2 | 78734767 | 78734932 | refGene_gn=SLC26A3;refGene_tx=XM_006929111.3 |
| A2 | 78737219 | 78737372 | refGene_gn=SLC26A3;refGene_tx=XM_006929111.3 |
| A2 | 78737782 | 78737865 | refGene_gn=SLC26A3;refGene_tx=XM_006929111.3 |

A2 78742084 78742232 refGene_gn=SLC26A3;refGene_tx=XM_006929111.3
A2 78742330 78742444 refGene_gn=SLC26A3;refGene_tx=XM_006929111.3
A2 78742550 78742628 refGene_gn=SLC26A3;refGene_tx=XM_006929111.3
A2 78745994 78746090 refGene_gn=SLC26A3;refGene_tx=XM_006929111.3
A2 78746991 78747098 refGene_gn=SLC26A3;refGene_tx=XM_006929111.3
A2 78748198 78748268 refGene_gn=SLC26A3;refGene_tx=XM_006929111.3
A2 78748359 78748452 refGene_gn=SLC26A3;refGene_tx=XM_006929111.3
A2 78749997 78750093 refGene_gn=SLC26A3;refGene_tx=XM_006929111.3
A2 78750647 78750881 refGene_gn=SLC26A3;refGene_tx=XM_006929111.3
A2 78752699 78752754 refGene_gn=SLC26A3;refGene_tx=XM_006929111.3
A2 78754707 78754850 refGene_gn=SLC26A3;refGene_tx=XM_006929111.3
A2 78754974 78755040 refGene_gn=SLC26A3;refGene_tx=XM_006929111.3
A2 78757475 78757499 refGene_gn=SLC26A3;refGene_tx=XM_006929111.3
A2 78765360 78766396
refGene_gn=CBLL1;refGene_tx=XM_004001604.5;refGene_gn=CBLL1;refGene_tx=XM_011280323.3;refGene_gn=CBLL1;refGene_tx=XM_019825337.2;refGene_gn=CBLL1;refGene_tx=XM_019825338.2
A2 78768907 78768981
refGene_gn=CBLL1;refGene_tx=XM_004001604.5;refGene_gn=CBLL1;refGene_tx=XM_011280323.3;refGene_gn=CBLL1;refGene_tx=XM_019825337.2;refGene_gn=CBLL1;refGene_tx=XM_019825338.2
A2 78771645 78771729
refGene_gn=CBLL1;refGene_tx=XM_004001604.5;refGene_gn=CBLL1;refGene_tx=XM_011280323.3;refGene_gn=CBLL1;refGene_tx=XM_019825337.2;refGene_gn=CBLL1;refGene_tx=XM_019825338.2
A2 78772388 78772489
refGene_gn=CBLL1;refGene_tx=XM_004001604.5;refGene_gn=CBLL1;refGene_tx=XM_011280323.3;refGene_gn=CBLL1;refGene_tx=XM_019825337.2;refGene_gn=CBLL1;refGene_tx=XM_019825338.2
A2 78775751 78775919
refGene_gn=CBLL1;refGene_tx=XM_004001604.5;refGene_gn=CBLL1;refGene_tx=XM_011280323.3;refGene_gn=CBLL1;refGene_tx=XM_019825337.2;refGene_gn=CBLL1;refGene_tx=XM_019825338.2
A2 78779903 78780171 refGene_gn=CBLL1;refGene_tx=XM_019825337.2
A2 78780377 78780528 refGene_gn=CBLL1;refGene_tx=XM_019825338.2
A2 78780562 78780575 refGene_gn=CBLL1;refGene_tx=XM_004001604.5
A2 78781594 78781643 refGene_gn=CBLL1;refGene_tx=XM_011280323.3
A2 78808597 78808621
refGene_gn=SLC26A4;refGene_tx=XM_003982649.5;refGene_gn=SLC26A4;refGene_tx=XM_019825339.2;refGene_gn=SLC26A4;refGene_tx=XM_019825340.2
A2 78810845 78810929
refGene_gn=SLC26A4;refGene_tx=XM_003982649.5;refGene_gn=SLC26A4;refGene_tx=XM_019825339.2;refGene_gn=SLC26A4;refGene_tx=XM_019825340.2
A2 78813390 78813536
refGene_gn=SLC26A4;refGene_tx=XM_003982649.5;refGene_gn=SLC26A4;refGene_tx=XM_019825339.2;refGene_gn=SLC26A4;refGene_tx=XM_019825340.2
A2 78814633 78814688
refGene_gn=SLC26A4;refGene_tx=XM_003982649.5;refGene_gn=SLC26A4;refGene_tx=XM_019825339.2;refGene_gn=SLC26A4;refGene_tx=XM_019825340.2
A2 78818558 78818789

refGene_gn=SLC26A4;refGene_tx=XM_003982649.5;refGene_gn=SLC26A4;refGene_tx=XM_019825339.2;refGene_gn=SLC26A4;refGene_tx=XM_019825340.2
A2 78819400 78819496
refGene_gn=SLC26A4;refGene_tx=XM_003982649.5;refGene_gn=SLC26A4;refGene_tx=XM_019825339.2;refGene_gn=SLC26A4;refGene_tx=XM_019825340.2
A2 78820384 78820477
refGene_gn=SLC26A4;refGene_tx=XM_003982649.5;refGene_gn=SLC26A4;refGene_tx=XM_019825339.2;refGene_gn=SLC26A4;refGene_tx=XM_019825340.2
A2 78822224 78822294
refGene_gn=SLC26A4;refGene_tx=XM_003982649.5;refGene_gn=SLC26A4;refGene_tx=XM_019825339.2;refGene_gn=SLC26A4;refGene_tx=XM_019825340.2
A2 78825027 78825134
refGene_gn=SLC26A4;refGene_tx=XM_003982649.5;refGene_gn=SLC26A4;refGene_tx=XM_019825339.2;refGene_gn=SLC26A4;refGene_tx=XM_019825340.2
A2 78826517 78826613
refGene_gn=SLC26A4;refGene_tx=XM_003982649.5;refGene_gn=SLC26A4;refGene_tx=XM_019825339.2;refGene_gn=SLC26A4;refGene_tx=XM_019825340.2
A2 78826754 78826832
refGene_gn=SLC26A4;refGene_tx=XM_003982649.5;refGene_gn=SLC26A4;refGene_tx=XM_019825339.2;refGene_gn=SLC26A4;refGene_tx=XM_019825340.2
A2 78830651 78830765
refGene_gn=SLC26A4;refGene_tx=XM_003982649.5;refGene_gn=SLC26A4;refGene_tx=XM_019825339.2;refGene_gn=SLC26A4;refGene_tx=XM_019825340.2
A2 78831705 78831853
refGene_gn=SLC26A4;refGene_tx=XM_003982649.5;refGene_gn=SLC26A4;refGene_tx=XM_019825339.2;refGene_gn=SLC26A4;refGene_tx=XM_019825340.2
A2 78836314 78836397
refGene_gn=SLC26A4;refGene_tx=XM_003982649.5;refGene_gn=SLC26A4;refGene_tx=XM_019825339.2;refGene_gn=SLC26A4;refGene_tx=XM_019825340.2
A2 78837092 78837245
refGene_gn=SLC26A4;refGene_tx=XM_003982649.5;refGene_gn=SLC26A4;refGene_tx=XM_019825339.2;refGene_gn=SLC26A4;refGene_tx=XM_019825340.2
A2 78845251 78845416
refGene_gn=SLC26A4;refGene_tx=XM_003982649.5;refGene_gn=SLC26A4;refGene_tx=XM_019825339.2;refGene_gn=SLC26A4;refGene_tx=XM_019825340.2
A2 78846438 78846623
refGene_gn=SLC26A4;refGene_tx=XM_003982649.5;refGene_gn=SLC26A4;refGene_tx=XM_019825339.2;refGene_gn=SLC26A4;refGene_tx=XM_019825340.2
A2 78848235 78848346
refGene_gn=SLC26A4;refGene_tx=XM_003982649.5;refGene_gn=SLC26A4;refGene_tx=XM_019825339.2;refGene_gn=SLC26A4;refGene_tx=XM_019825340.2
A2 78856965 78857105
refGene_gn=SLC26A4;refGene_tx=XM_003982649.5;refGene_gn=SLC26A4;refGene_tx=XM_019825339.2;refGene_gn=SLC26A4;refGene_tx=XM_019825340.2
A2 78859096 78859260
refGene_gn=SLC26A4;refGene_tx=XM_003982649.5;refGene_gn=SLC26A4;refGene_tx=XM_019825339.2;refGene_gn=SLC26A4;refGene_tx=XM_019825340.2
A2 78886275 78886311
refGene_gn=BCAP29;refGene_tx=XM_006929112.4;refGene_gn=BCAP29;refGene_tx=XM_023250248.1;refGene_gn=BCAP29;refGene_tx=XM_023250249.1;refGene_

gn=BCAP29;refGene_tx=XM_023250250.1
A2 78892824 78892925
refGene_gn=BCAP29;refGene_tx=XM_006929112.4;refGene_gn=BCAP29;refGene_tx=XM_023250248.1;refGene_gn=BCAP29;refGene_tx=XM_023250249.1;refGene_gn=BCAP29;refGene_tx=XM_023250250.1
A2 78924093 78924202
refGene_gn=BCAP29;refGene_tx=XM_006929112.4;refGene_gn=BCAP29;refGene_tx=XM_023250248.1;refGene_gn=BCAP29;refGene_tx=XM_023250249.1;refGene_gn=BCAP29;refGene_tx=XM_023250250.1
A2 78930106 78930242
refGene_gn=BCAP29;refGene_tx=XM_006929112.4;refGene_gn=BCAP29;refGene_tx=XM_023250248.1;refGene_gn=BCAP29;refGene_tx=XM_023250249.1;refGene_gn=BCAP29;refGene_tx=XM_023250250.1
A2 78931638 78931789
refGene_gn=BCAP29;refGene_tx=XM_006929112.4;refGene_gn=BCAP29;refGene_tx=XM_023250248.1;refGene_gn=BCAP29;refGene_tx=XM_023250249.1;refGene_gn=BCAP29;refGene_tx=XM_023250250.1
A2 78936449 78936550
refGene_gn=BCAP29;refGene_tx=XM_006929112.4;refGene_gn=BCAP29;refGene_tx=XM_023250248.1;refGene_gn=BCAP29;refGene_tx=XM_023250249.1;refGene_gn=BCAP29;refGene_tx=XM_023250250.1
A2 78940110 78940202
refGene_gn=BCAP29;refGene_tx=XM_006929112.4;refGene_gn=BCAP29;refGene_tx=XM_023250248.1;refGene_gn=BCAP29;refGene_tx=XM_023250249.1;refGene_gn=BCAP29;refGene_tx=XM_023250250.1
A2 78943732 78943980 refGene_gn=DUS4L;refGene_tx=XM_003982650.4
A2 78944774 78945001 refGene_gn=DUS4L;refGene_tx=XM_003982650.4
A2 78945775 78945898 refGene_gn=DUS4L;refGene_tx=XM_003982650.4
A2 78946950 78947068 refGene_gn=DUS4L;refGene_tx=XM_003982650.4
A2 78950428 78950550 refGene_gn=DUS4L;refGene_tx=XM_003982650.4
A2 78954802 78954918 refGene_gn=DUS4L;refGene_tx=XM_003982650.4
A2 78959729 78959804 refGene_gn=COG5;refGene_tx=XM_006929113.3
A2 78959875 78960089 refGene_gn=COG5;refGene_tx=XM_006929113.3
A2 78964824 78964964 refGene_gn=COG5;refGene_tx=XM_006929113.3
A2 78969089 78969147 refGene_gn=COG5;refGene_tx=XM_006929113.3
A2 78974435 78974490 refGene_gn=COG5;refGene_tx=XM_006929113.3
A2 78974591 78974661 refGene_gn=COG5;refGene_tx=XM_006929113.3
A2 78993143 78993444
refGene_gn=LOC102901356;refGene_tx=XM_019825346.1
A2 78993542 78993625
refGene_gn=LOC102901356;refGene_tx=XM_019825346.1
A2 78997900 78998021 refGene_gn=COG5;refGene_tx=XM_006929113.3
A2 79037400 79038702
refGene_gn=GPR22;refGene_tx=XM_003982652.4;refGene_gn=GPR22;refGene_tx=XM_019825345.1
A2 79097909 79098040 refGene_gn=COG5;refGene_tx=XM_006929113.3
A2 79131662 79131828 refGene_gn=COG5;refGene_tx=XM_006929113.3
A2 79140133 79140246 refGene_gn=COG5;refGene_tx=XM_006929113.3
A2 79140501 79140579 refGene_gn=COG5;refGene_tx=XM_006929113.3
A2 79178383 79178465 refGene_gn=COG5;refGene_tx=XM_006929113.3

A2 79193903 79194108 refGene_gn=COG5;refGene_tx=XM_006929113.3
A2 79204651 79204813 refGene_gn=COG5;refGene_tx=XM_006929113.3
A2 79207000 79207100 refGene_gn=COG5;refGene_tx=XM_006929113.3
A2 79230176 79230287 refGene_gn=COG5;refGene_tx=XM_006929113.3
A2 79232034 79232097 refGene_gn=COG5;refGene_tx=XM_006929113.3
A2 79241242 79241346 refGene_gn=COG5;refGene_tx=XM_006929113.3
A2 79245720 79245958 refGene_gn=COG5;refGene_tx=XM_006929113.3
A2 79252013 79252090 refGene_gn=COG5;refGene_tx=XM_006929113.3
A2 79261180 79261307 refGene_gn=COG5;refGene_tx=XM_006929113.3
A2 79261989 79262069 refGene_gn=COG5;refGene_tx=XM_006929113.3
A2 79268816 79268934 refGene_gn=COG5;refGene_tx=XM_006929113.3
A2 79270944 79270962
refGene_gn=HBP1;refGene_tx=XM_011280326.3;refGene_gn=HBP1;refGene_tx=XM_023250251.1
A2 79272015 79272157
refGene_gn=HBP1;refGene_tx=XM_011280326.3;refGene_gn=HBP1;refGene_tx=XM_023250251.1
A2 79274277 79274595
refGene_gn=HBP1;refGene_tx=XM_011280326.3;refGene_gn=HBP1;refGene_tx=XM_023250251.1
A2 79281449 79281594
refGene_gn=HBP1;refGene_tx=XM_011280326.3;refGene_gn=HBP1;refGene_tx=XM_023250251.1
A2 79282268 79282425
refGene_gn=HBP1;refGene_tx=XM_011280326.3;refGene_gn=HBP1;refGene_tx=XM_023250251.1
A2 79284508 79284621
refGene_gn=HBP1;refGene_tx=XM_011280326.3;refGene_gn=HBP1;refGene_tx=XM_023250251.1
A2 79284707 79284819
refGene_gn=HBP1;refGene_tx=XM_011280326.3;refGene_gn=HBP1;refGene_tx=XM_023250251.1
A2 79285186 79285325
refGene_gn=HBP1;refGene_tx=XM_011280326.3;refGene_gn=HBP1;refGene_tx=XM_023250251.1
A2 79288975 79289204
refGene_gn=HBP1;refGene_tx=XM_011280326.3;refGene_gn=HBP1;refGene_tx=XM_023250251.1
A2 79291182 79291366
refGene_gn=HBP1;refGene_tx=XM_011280326.3;refGene_gn=HBP1;refGene_tx=XM_023250251.1
A2 79299358 79299706
refGene_gn=LOC111559479;refGene_tx=XM_023250254.1;refGene_gn=LOC111559479;refGene_tx=XM_023250255.1;refGene_gn=LOC111559479;refGene_tx=XM_023250256.1;refGene_gn=LOC111559479;refGene_tx=XM_023250257.1
A2 79299899 79300031 refGene_gn=HBP1;refGene_tx=XM_023250251.1
A2 79300057 79300072 refGene_gn=HBP1;refGene_tx=XM_011280326.3
A2 79300263 79300282
refGene_gn=LOC111559479;refGene_tx=XM_023250256.1
A2 79306590 79306677

refGene_gn=LOC111559479;refGene_tx=XM_023250254.1
A2	79310030	79310164	refGene_gn=PRKAR2B;refGene_tx=XM_023250252.1
A2	79311258	79311360
refGene_gn=LOC111559479;refGene_tx=XM_023250254.1;refGene_gn=LOC111559479;refGene_tx=XM_023250255.1
A2	79313273	79313412	refGene_gn=PRKAR2B;refGene_tx=XM_023250252.1
A2	79313545	79313611	refGene_gn=PRKAR2B;refGene_tx=XM_023250252.1
A2	79316738	79316813	refGene_gn=PRKAR2B;refGene_tx=XM_023250252.1
A2	79317005	79317015
refGene_gn=LOC111559479;refGene_tx=XM_023250257.1
A2	79318555	79318657	refGene_gn=PRKAR2B;refGene_tx=XM_023250252.1
A2	79323250	79323404	refGene_gn=PRKAR2B;refGene_tx=XM_023250252.1
A2	79326398	79326505	refGene_gn=PRKAR2B;refGene_tx=XM_023250252.1
A2	79337047	79337131	refGene_gn=PRKAR2B;refGene_tx=XM_023250252.1
A2	79344904	79344957	refGene_gn=PRKAR2B;refGene_tx=XM_023250252.1
A2	79385916	79385952	refGene_gn=PRKAR2B;refGene_tx=XM_023250252.1
A2	79414561	79414871	refGene_gn=PRKAR2B;refGene_tx=XM_023250252.1
A2	79537238	79537517
refGene_gn=PIK3CG;refGene_tx=XM_019825341.2;refGene_gn=PIK3CG;refGene_tx=XM_019825342.2;refGene_gn=PIK3CG;refGene_tx=XM_019825343.2;refGene_gn=PIK3CG;refGene_tx=XM_019825344.2
A2	79549239	79549397
refGene_gn=PIK3CG;refGene_tx=XM_019825341.2;refGene_gn=PIK3CG;refGene_tx=XM_019825342.2;refGene_gn=PIK3CG;refGene_tx=XM_019825343.2;refGene_gn=PIK3CG;refGene_tx=XM_019825344.2
A2	79550891	79551003
refGene_gn=PIK3CG;refGene_tx=XM_019825341.2;refGene_gn=PIK3CG;refGene_tx=XM_019825342.2;refGene_gn=PIK3CG;refGene_tx=XM_019825343.2;refGene_gn=PIK3CG;refGene_tx=XM_019825344.2
A2	79551989	79552252
refGene_gn=PIK3CG;refGene_tx=XM_019825341.2;refGene_gn=PIK3CG;refGene_tx=XM_019825342.2;refGene_gn=PIK3CG;refGene_tx=XM_019825343.2;refGene_gn=PIK3CG;refGene_tx=XM_019825344.2
A2	79553376	79553467
refGene_gn=PIK3CG;refGene_tx=XM_019825341.2;refGene_gn=PIK3CG;refGene_tx=XM_019825342.2;refGene_gn=PIK3CG;refGene_tx=XM_019825343.2;refGene_gn=PIK3CG;refGene_tx=XM_019825344.2
A2	79555019	79555166
refGene_gn=PIK3CG;refGene_tx=XM_019825341.2;refGene_gn=PIK3CG;refGene_tx=XM_019825342.2;refGene_gn=PIK3CG;refGene_tx=XM_019825343.2;refGene_gn=PIK3CG;refGene_tx=XM_019825344.2
A2	79559409	79559513
refGene_gn=PIK3CG;refGene_tx=XM_019825341.2;refGene_gn=PIK3CG;refGene_tx=XM_019825342.2;refGene_gn=PIK3CG;refGene_tx=XM_019825343.2;refGene_gn=PIK3CG;refGene_tx=XM_019825344.2
A2	79561277	79561503
refGene_gn=PIK3CG;refGene_tx=XM_019825341.2;refGene_gn=PIK3CG;refGene_tx=XM_019825342.2;refGene_gn=PIK3CG;refGene_tx=XM_019825343.2;refGene_gn=PIK3CG;refGene_tx=XM_019825344.2
A2	79561608	79561674

refGene_gn=PIK3CG;refGene_tx=XM_019825341.2;refGene_gn=PIK3CG;refGene_tx=XM_019825342.2;refGene_gn=PIK3CG;refGene_tx=XM_019825343.2;refGene_gn=PIK3CG;refGene_tx=XM_019825344.2
A2 79565316 79567323
refGene_gn=PIK3CG;refGene_tx=XM_019825341.2;refGene_gn=PIK3CG;refGene_tx=XM_019825342.2;refGene_gn=PIK3CG;refGene_tx=XM_019825343.2;refGene_gn=PIK3CG;refGene_tx=XM_019825344.2
A2 79569268 79569340
refGene_gn=PIK3CG;refGene_tx=XM_019825341.2;refGene_gn=PIK3CG;refGene_tx=XM_019825344.2
A2 79620762 79621622
refGene_gn=LOC101096798;refGene_tx=XM_023250258.1
A2 79621624 79621667
refGene_gn=LOC101096798;refGene_tx=XM_023250258.1
A2 79736913 79737630 refGene_gn=CCDC71L;refGene_tx=XM_023250259.1
A2 80130827 80130884 refGene_gn=NAMPT;refGene_tx=XM_011280328.3
A2 80146228 80146385 refGene_gn=NAMPT;refGene_tx=XM_011280328.3
A2 80147743 80147847 refGene_gn=NAMPT;refGene_tx=XM_011280328.3
A2 80149779 80149908 refGene_gn=NAMPT;refGene_tx=XM_011280328.3
A2 80153830 80153989 refGene_gn=NAMPT;refGene_tx=XM_011280328.3
A2 80154760 80154897 refGene_gn=NAMPT;refGene_tx=XM_011280328.3
A2 80159756 80159982 refGene_gn=NAMPT;refGene_tx=XM_011280328.3
A2 80160780 80160900 refGene_gn=NAMPT;refGene_tx=XM_011280328.3
A2 80163629 80163770 refGene_gn=NAMPT;refGene_tx=XM_011280328.3
A2 80164849 80164984 refGene_gn=NAMPT;refGene_tx=XM_011280328.3
A2 80166961 80167072 refGene_gn=NAMPT;refGene_tx=XM_011280328.3
A2 80335404 80335455 refGene_gn=SYPL1;refGene_tx=XM_023250260.1
A2 80335707 80335779 refGene_gn=SYPL1;refGene_tx=XM_023250260.1
A2 80348954 80349076 refGene_gn=SYPL1;refGene_tx=XM_023250260.1
A2 80350299 80350507 refGene_gn=SYPL1;refGene_tx=XM_023250260.1
A2 80355714 80355900 refGene_gn=SYPL1;refGene_tx=XM_023250260.1
A2 80357533 80357668 refGene_gn=SYPL1;refGene_tx=XM_023250260.1
A2 80419738 80420040 refGene_gn=CDHR3;refGene_tx=XM_023242030.1
A2 80421494 80421543 refGene_gn=CDHR3;refGene_tx=XM_023242030.1
A2 80422054 80422277 refGene_gn=CDHR3;refGene_tx=XM_023242030.1
A2 80428321 80428503 refGene_gn=CDHR3;refGene_tx=XM_023242030.1
A2 80430645 80430896 refGene_gn=CDHR3;refGene_tx=XM_023242030.1
A2 80432432 80432604 refGene_gn=CDHR3;refGene_tx=XM_023242030.1
A2 80434542 80434769 refGene_gn=CDHR3;refGene_tx=XM_023242030.1
A2 80436662 80436761 refGene_gn=CDHR3;refGene_tx=XM_023242030.1
A2 80437645 80437748 refGene_gn=CDHR3;refGene_tx=XM_023242030.1
A2 80439269 80439441 refGene_gn=CDHR3;refGene_tx=XM_023242030.1
A2 80446672 80446862 refGene_gn=CDHR3;refGene_tx=XM_023242030.1
A2 80449070 80449219 refGene_gn=CDHR3;refGene_tx=XM_023242030.1
A2 80454539 80454644 refGene_gn=CDHR3;refGene_tx=XM_023242030.1
A2 80455438 80455533 refGene_gn=CDHR3;refGene_tx=XM_023242030.1
A2 80467694 80467792 refGene_gn=CDHR3;refGene_tx=XM_023242030.1
A2 80470837 80471003 refGene_gn=CDHR3;refGene_tx=XM_023242030.1
A2 80476946 80477149 refGene_gn=CDHR3;refGene_tx=XM_023242030.1
A2 80486058 80486065 refGene_gn=CDHR3;refGene_tx=XM_023242030.1

A2	80566078 80566259
refGene_gn=ATXN7L1;refGene_tx=XM_023250261.1;refGene_gn=ATXN7L1;refGene_tx=XM_023250262.1;refGene_gn=ATXN7L1;refGene_tx=XM_023250263.1;refGene_gn=ATXN7L1;refGene_tx=XM_023250264.1;refGene_gn=ATXN7L1;refGene_tx=XM_023250265.1;refGene_gn=ATXN7L1;refGene_tx=XM_023250266.1
A2	80566782 80566851
refGene_gn=ATXN7L1;refGene_tx=XM_023250261.1;refGene_gn=ATXN7L1;refGene_tx=XM_023250262.1;refGene_gn=ATXN7L1;refGene_tx=XM_023250263.1;refGene_gn=ATXN7L1;refGene_tx=XM_023250264.1;refGene_gn=ATXN7L1;refGene_tx=XM_023250265.1;refGene_gn=ATXN7L1;refGene_tx=XM_023250266.1
A2	80652131 80652236
refGene_gn=ATXN7L1;refGene_tx=XM_023250261.1;refGene_gn=ATXN7L1;refGene_tx=XM_023250262.1;refGene_gn=ATXN7L1;refGene_tx=XM_023250264.1;refGene_gn=ATXN7L1;refGene_tx=XM_023250265.1;refGene_gn=ATXN7L1;refGene_tx=XM_023250266.1
A2	80676484 80676570 refGene_gn=ATXN7L1;refGene_tx=XM_023250266.1
A2	80756957 80757180
refGene_gn=ATXN7L1;refGene_tx=XM_006929126.4;refGene_gn=ATXN7L1;refGene_tx=XM_006929127.4;refGene_gn=ATXN7L1;refGene_tx=XM_019825347.2;refGene_gn=ATXN7L1;refGene_tx=XM_019825348.2;refGene_gn=ATXN7L1;refGene_tx=XM_019825349.2;refGene_gn=ATXN7L1;refGene_tx=XM_023250261.1;refGene_gn=ATXN7L1;refGene_tx=XM_023250262.1;refGene_gn=ATXN7L1;refGene_tx=XM_023250263.1;refGene_gn=ATXN7L1;refGene_tx=XM_023250264.1;refGene_gn=ATXN7L1;refGene_tx=XM_023250265.1
A2	80780274 80780558
refGene_gn=ATXN7L1;refGene_tx=XM_006929126.4;refGene_gn=ATXN7L1;refGene_tx=XM_006929127.4;refGene_gn=ATXN7L1;refGene_tx=XM_019825347.2;refGene_gn=ATXN7L1;refGene_tx=XM_019825348.2;refGene_gn=ATXN7L1;refGene_tx=XM_019825349.2;refGene_gn=ATXN7L1;refGene_tx=XM_019825351.2;refGene_gn=ATXN7L1;refGene_tx=XM_023250261.1;refGene_gn=ATXN7L1;refGene_tx=XM_023250262.1;refGene_gn=ATXN7L1;refGene_tx=XM_023250263.1;refGene_gn=ATXN7L1;refGene_tx=XM_023250264.1;refGene_gn=ATXN7L1;refGene_tx=XM_023250265.1
A2	80783724 80783807
refGene_gn=ATXN7L1;refGene_tx=XM_006929126.4;refGene_gn=ATXN7L1;refGene_tx=XM_006929127.4;refGene_gn=ATXN7L1;refGene_tx=XM_019825347.2;refGene_gn=ATXN7L1;refGene_tx=XM_019825348.2;refGene_gn=ATXN7L1;refGene_tx=XM_019825349.2;refGene_gn=ATXN7L1;refGene_tx=XM_019825351.2;refGene_gn=ATXN7L1;refGene_tx=XM_023250261.1;refGene_gn=ATXN7L1;refGene_tx=XM_023250262.1;refGene_gn=ATXN7L1;refGene_tx=XM_023250263.1;refGene_gn=ATXN7L1;refGene_tx=XM_023250264.1;refGene_gn=ATXN7L1;refGene_tx=XM_023250265.1
A2	80784670 80784927
refGene_gn=ATXN7L1;refGene_tx=XM_006929126.4;refGene_gn=ATXN7L1;refGene_tx=XM_006929127.4;refGene_gn=ATXN7L1;refGene_tx=XM_019825347.2;refGene_gn=ATXN7L1;refGene_tx=XM_019825348.2;refGene_gn=ATXN7L1;refGene_tx=XM_019825349.2;refGene_gn=ATXN7L1;refGene_tx=XM_019825351.2;refGene_gn=ATXN7L1;refGene_tx=XM_023250261.1;refGene_gn=ATXN7L1;refGene_tx=XM_023250262.1;refGene_gn=ATXN7L1;refGene_tx=XM_023250263.1;refGene_gn=ATXN7L1;refGene_tx=XM_023250264.1;refGene_gn=ATXN7L1;refGene_tx=XM_0232502

65.1
A2 80797208 80797401
refGene_gn=ATXN7L1;refGene_tx=XM_006929126.4;refGene_gn=ATXN7L1;refGene_tx=XM_006929127.4;refGene_gn=ATXN7L1;refGene_tx=XM_019825347.2;refGene_gn=ATXN7L1;refGene_tx=XM_019825348.2;refGene_gn=ATXN7L1;refGene_tx=XM_019825349.2;refGene_gn=ATXN7L1;refGene_tx=XM_019825351.2;refGene_gn=ATXN7L1;refGene_tx=XM_023250261.1;refGene_gn=ATXN7L1;refGene_tx=XM_023250262.1;refGene_gn=ATXN7L1;refGene_tx=XM_023250263.1;refGene_gn=ATXN7L1;refGene_tx=XM_023250264.1;refGene_gn=ATXN7L1;refGene_tx=XM_023250265.1
A2 80801798 80801920
refGene_gn=ATXN7L1;refGene_tx=XM_006929126.4;refGene_gn=ATXN7L1;refGene_tx=XM_006929127.4;refGene_gn=ATXN7L1;refGene_tx=XM_019825347.2;refGene_gn=ATXN7L1;refGene_tx=XM_019825348.2;refGene_gn=ATXN7L1;refGene_tx=XM_019825349.2;refGene_gn=ATXN7L1;refGene_tx=XM_019825351.2;refGene_gn=ATXN7L1;refGene_tx=XM_023250261.1;refGene_gn=ATXN7L1;refGene_tx=XM_023250262.1;refGene_gn=ATXN7L1;refGene_tx=XM_023250263.1;refGene_gn=ATXN7L1;refGene_tx=XM_023250264.1;refGene_gn=ATXN7L1;refGene_tx=XM_023250265.1
A2 80803355 80803661
refGene_gn=ATXN7L1;refGene_tx=XM_006929126.4;refGene_gn=ATXN7L1;refGene_tx=XM_006929127.4;refGene_gn=ATXN7L1;refGene_tx=XM_019825347.2;refGene_gn=ATXN7L1;refGene_tx=XM_019825348.2;refGene_gn=ATXN7L1;refGene_tx=XM_019825349.2;refGene_gn=ATXN7L1;refGene_tx=XM_019825351.2;refGene_gn=ATXN7L1;refGene_tx=XM_023250261.1;refGene_gn=ATXN7L1;refGene_tx=XM_023250262.1;refGene_gn=ATXN7L1;refGene_tx=XM_023250263.1
A2 80806211 80807151
refGene_gn=ATXN7L1;refGene_tx=XM_006929126.4;refGene_gn=ATXN7L1;refGene_tx=XM_006929127.4;refGene_gn=ATXN7L1;refGene_tx=XM_019825347.2;refGene_gn=ATXN7L1;refGene_tx=XM_019825348.2;refGene_gn=ATXN7L1;refGene_tx=XM_019825349.2;refGene_gn=ATXN7L1;refGene_tx=XM_019825351.2;refGene_gn=ATXN7L1;refGene_tx=XM_023250261.1;refGene_gn=ATXN7L1;refGene_tx=XM_023250262.1;refGene_gn=ATXN7L1;refGene_tx=XM_023250263.1;refGene_gn=ATXN7L1;refGene_tx=XM_023250264.1;refGene_gn=ATXN7L1;refGene_tx=XM_023250265.1
A2 80810370 80810445
refGene_gn=ATXN7L1;refGene_tx=XM_006929126.4;refGene_gn=ATXN7L1;refGene_tx=XM_006929127.4;refGene_gn=ATXN7L1;refGene_tx=XM_019825347.2;refGene_gn=ATXN7L1;refGene_tx=XM_019825348.2;refGene_gn=ATXN7L1;refGene_tx=XM_019825349.2;refGene_gn=ATXN7L1;refGene_tx=XM_019825351.2;refGene_gn=ATXN7L1;refGene_tx=XM_023250261.1;refGene_gn=ATXN7L1;refGene_tx=XM_023250262.1;refGene_gn=ATXN7L1;refGene_tx=XM_023250263.1;refGene_gn=ATXN7L1;refGene_tx=XM_023250264.1;refGene_gn=ATXN7L1;refGene_tx=XM_023250265.1
A2 80812016 80812058
refGene_gn=ATXN7L1;refGene_tx=XM_006929126.4;refGene_gn=ATXN7L1;refGene_tx=XM_006929127.4;refGene_gn=ATXN7L1;refGene_tx=XM_019825347.2;refGene_gn=ATXN7L1;refGene_tx=XM_019825348.2;refGene_gn=ATXN7L1;refGene_tx=XM_019825349.2;refGene_gn=ATXN7L1;refGene_tx=XM_019825351.2;refGene_gn=ATXN7L1;refGene_tx=XM_023250261.1;refGene_gn=ATXN7L1;refGene_tx=XM_02

3250262.1;refGene_gn=ATXN7L1;refGene_tx=XM_023250263.1;refGene_gn=ATXN7L1;refGene_tx=XM_023250264.1;refGene_gn=ATXN7L1;refGene_tx=XM_023250265.1
A2 80830804 80830913 refGene_gn=LOC102902529;refGene_tx=XM_006929129.4;refGene_gn=LOC102902529;refGene_tx=XM_023250267.1;refGene_gn=LOC102902529;refGene_tx=XM_023250268.1;refGene_gn=LOC102902529;refGene_tx=XM_023250269.1
A2 80836962 80837112 refGene_gn=LOC102902529;refGene_tx=XM_023250267.1
A2 80839039 80839204 refGene_gn=LOC102902529;refGene_tx=XM_006929129.4;refGene_gn=LOC102902529;refGene_tx=XM_023250267.1;refGene_gn=LOC102902529;refGene_tx=XM_023250268.1
A2 80839313 80839401 refGene_gn=LOC102902529;refGene_tx=XM_006929129.4;refGene_gn=LOC102902529;refGene_tx=XM_023250267.1;refGene_gn=LOC102902529;refGene_tx=XM_023250268.1;refGene_gn=LOC102902529;refGene_tx=XM_023250269.1
A2 80840997 80841034 refGene_gn=LOC102902529;refGene_tx=XM_006929129.4;refGene_gn=LOC102902529;refGene_tx=XM_023250269.1
A2 80841179 80841372 refGene_gn=RINT1;refGene_tx=XM_011280336.3;refGene_gn=RINT1;refGene_tx=XM_019825352.2
A2 80842222 80842262 refGene_gn=LOC102902529;refGene_tx=XM_023250267.1;refGene_gn=LOC102902529;refGene_tx=XM_023250268.1
A2 80842304 80842423 refGene_gn=RINT1;refGene_tx=XM_011280336.3;refGene_gn=RINT1;refGene_tx=XM_019825352.2
A2 80842497 80842678 refGene_gn=RINT1;refGene_tx=XM_011280336.3;refGene_gn=RINT1;refGene_tx=XM_019825352.2
A2 80846429 80846478 refGene_gn=RINT1;refGene_tx=XM_019825354.2
A2 80846563 80846778 refGene_gn=RINT1;refGene_tx=XM_011280336.3;refGene_gn=RINT1;refGene_tx=XM_019825352.2;refGene_gn=RINT1;refGene_tx=XM_019825354.2
A2 80851492 80851692 refGene_gn=RINT1;refGene_tx=XM_011280336.3;refGene_gn=RINT1;refGene_tx=XM_019825352.2;refGene_gn=RINT1;refGene_tx=XM_019825354.2
A2 80853337 80853475 refGene_gn=RINT1;refGene_tx=XM_011280336.3;refGene_gn=RINT1;refGene_tx=XM_019825352.2;refGene_gn=RINT1;refGene_tx=XM_019825354.2
A2 80853995 80854221 refGene_gn=RINT1;refGene_tx=XM_011280336.3;refGene_gn=RINT1;refGene_tx=XM_019825352.2;refGene_gn=RINT1;refGene_tx=XM_019825354.2
A2 80854320 80854431 refGene_gn=RINT1;refGene_tx=XM_011280336.3;refGene_gn=RINT1;refGene_tx=XM_019825352.2;refGene_gn=RINT1;refGene_tx=XM_019825354.2
A2 80856958 80857115 refGene_gn=RINT1;refGene_tx=XM_011280336.3;refGene_gn=RINT1;refGene_tx

=XM_019825352.2;refGene_gn=RINT1;refGene_tx=XM_019825354.2
A2	80857700 80857850
refGene_gn=RINT1;refGene_tx=XM_011280336.3;refGene_gn=RINT1;refGene_tx=XM_019825352.2;refGene_gn=RINT1;refGene_tx=XM_019825354.2
A2	80857949 80858123
refGene_gn=RINT1;refGene_tx=XM_011280336.3;refGene_gn=RINT1;refGene_tx=XM_019825352.2;refGene_gn=RINT1;refGene_tx=XM_019825354.2
A2	80861568 80861810
refGene_gn=RINT1;refGene_tx=XM_011280336.3;refGene_gn=RINT1;refGene_tx=XM_019825352.2;refGene_gn=RINT1;refGene_tx=XM_019825354.2
A2	80866181 80866366
refGene_gn=RINT1;refGene_tx=XM_011280336.3;refGene_gn=RINT1;refGene_tx=XM_019825352.2;refGene_gn=RINT1;refGene_tx=XM_019825354.2
A2	80870578 80870624
refGene_gn=RINT1;refGene_tx=XM_011280336.3;refGene_gn=RINT1;refGene_tx=XM_019825352.2;refGene_gn=RINT1;refGene_tx=XM_019825354.2
A2	80871073 80871115
refGene_gn=RINT1;refGene_tx=XM_011280336.3;refGene_gn=RINT1;refGene_tx=XM_019825352.2;refGene_gn=RINT1;refGene_tx=XM_019825354.2
A2	80876475 80876602
refGene_gn=PUS7;refGene_tx=XM_023250270.1;refGene_gn=PUS7;refGene_tx=XM_023250271.1;refGene_gn=PUS7;refGene_tx=XM_023250272.1;refGene_gn=PUS7;refGene_tx=XM_023250273.1
A2	80887757 80888190
refGene_gn=PUS7;refGene_tx=XM_023250270.1;refGene_gn=PUS7;refGene_tx=XM_023250271.1;refGene_gn=PUS7;refGene_tx=XM_023250272.1;refGene_gn=PUS7;refGene_tx=XM_023250273.1
A2	80890316 80890401
refGene_gn=PUS7;refGene_tx=XM_023250270.1;refGene_gn=PUS7;refGene_tx=XM_023250271.1;refGene_gn=PUS7;refGene_tx=XM_023250272.1;refGene_gn=PUS7;refGene_tx=XM_023250273.1
A2	80890574 80890676
refGene_gn=PUS7;refGene_tx=XM_023250270.1;refGene_gn=PUS7;refGene_tx=XM_023250271.1;refGene_gn=PUS7;refGene_tx=XM_023250272.1;refGene_gn=PUS7;refGene_tx=XM_023250273.1
A2	80892840 80892985
refGene_gn=PUS7;refGene_tx=XM_023250270.1;refGene_gn=PUS7;refGene_tx=XM_023250271.1;refGene_gn=PUS7;refGene_tx=XM_023250272.1;refGene_gn=PUS7;refGene_tx=XM_023250273.1
A2	80899020 80899038
refGene_gn=PUS7;refGene_tx=XM_023250270.1;refGene_gn=PUS7;refGene_tx=XM_023250272.1;refGene_gn=PUS7;refGene_tx=XM_023250273.1
A2	80900503 80900615
refGene_gn=PUS7;refGene_tx=XM_023250270.1;refGene_gn=PUS7;refGene_tx=XM_023250271.1;refGene_gn=PUS7;refGene_tx=XM_023250272.1;refGene_gn=PUS7;refGene_tx=XM_023250273.1
A2	80902158 80902236
refGene_gn=PUS7;refGene_tx=XM_023250270.1;refGene_gn=PUS7;refGene_tx=XM_023250271.1;refGene_gn=PUS7;refGene_tx=XM_023250272.1;refGene_gn=PUS7;refGene_tx=XM_023250273.1

A2 80913482 80913611
refGene_gn=PUS7;refGene_tx=XM_023250270.1;refGene_gn=PUS7;refGene_tx=XM_023250271.1
A2 80914501 80914627
refGene_gn=PUS7;refGene_tx=XM_023250270.1;refGene_gn=PUS7;refGene_tx=XM_023250271.1;refGene_gn=PUS7;refGene_tx=XM_023250272.1
A2 80923399 80923461
refGene_gn=PUS7;refGene_tx=XM_023250270.1;refGene_gn=PUS7;refGene_tx=XM_023250271.1;refGene_gn=PUS7;refGene_tx=XM_023250272.1;refGene_gn=PUS7;refGene_tx=XM_023250273.1
A2 80924558 80924719
refGene_gn=PUS7;refGene_tx=XM_023250270.1;refGene_gn=PUS7;refGene_tx=XM_023250271.1;refGene_gn=PUS7;refGene_tx=XM_023250272.1;refGene_gn=PUS7;refGene_tx=XM_023250273.1
A2 80925759 80925886
refGene_gn=PUS7;refGene_tx=XM_023250270.1;refGene_gn=PUS7;refGene_tx=XM_023250271.1;refGene_gn=PUS7;refGene_tx=XM_023250272.1;refGene_gn=PUS7;refGene_tx=XM_023250273.1
A2 80927139 80927241
refGene_gn=PUS7;refGene_tx=XM_023250270.1;refGene_gn=PUS7;refGene_tx=XM_023250271.1;refGene_gn=PUS7;refGene_tx=XM_023250272.1;refGene_gn=PUS7;refGene_tx=XM_023250273.1
A2 80929735 80929865
refGene_gn=PUS7;refGene_tx=XM_023250270.1;refGene_gn=PUS7;refGene_tx=XM_023250271.1;refGene_gn=PUS7;refGene_tx=XM_023250272.1;refGene_gn=PUS7;refGene_tx=XM_023250273.1
A2 80931153 80931245
refGene_gn=PUS7;refGene_tx=XM_023250270.1;refGene_gn=PUS7;refGene_tx=XM_023250271.1;refGene_gn=PUS7;refGene_tx=XM_023250272.1;refGene_gn=PUS7;refGene_tx=XM_023250273.1
A2 80931789 80931926
refGene_gn=PUS7;refGene_tx=XM_023250270.1;refGene_gn=PUS7;refGene_tx=XM_023250271.1;refGene_gn=PUS7;refGene_tx=XM_023250272.1;refGene_gn=PUS7;refGene_tx=XM_023250273.1
A2 80974708 80974779 refGene_gn=SRPK2;refGene_tx=XM_019825360.1
A2 80982913 80983040
refGene_gn=SRPK2;refGene_tx=XM_023250274.1;refGene_gn=SRPK2;refGene_tx=XM_023250275.1;refGene_gn=SRPK2;refGene_tx=XM_023250276.1;refGene_gn=SRPK2;refGene_tx=XM_023250277.1;refGene_gn=SRPK2;refGene_tx=XM_023250278.1;refGene_gn=SRPK2;refGene_tx=XM_023250279.1;refGene_gn=SRPK2;refGene_tx=XM_023250281.1
A2 80983506 80983522
refGene_gn=SRPK2;refGene_tx=XM_023250280.1;refGene_gn=SRPK2;refGene_tx=XM_023250284.1
A2 80983612 80983667
refGene_gn=SRPK2;refGene_tx=XM_023250274.1;refGene_gn=SRPK2;refGene_tx=XM_023250275.1;refGene_gn=SRPK2;refGene_tx=XM_023250276.1;refGene_gn=SRPK2;refGene_tx=XM_023250277.1;refGene_gn=SRPK2;refGene_tx=XM_023250278.1;refGene_gn=SRPK2;refGene_tx=XM_023250279.1;refGene_gn=SRPK2;refGene_tx=XM_023250280.1;refGene_gn=SRPK2;refGene_tx=XM_023250281.1;refGen

e_gn=SRPK2;refGene_tx=XM_023250284.1
A2 81074404 81074468
refGene_gn=SRPK2;refGene_tx=XM_023250276.1;refGene_gn=SRPK2;refGene_tx=XM_023250281.1;refGene_gn=SRPK2;refGene_tx=XM_023250282.1;refGene_gn=SRPK2;refGene_tx=XM_023250283.1
A2 81075032 81075049 refGene_gn=SRPK2;refGene_tx=XM_019825362.2
A2 81155935 81156087
refGene_gn=SRPK2;refGene_tx=XM_019825360.1;refGene_gn=SRPK2;refGene_tx=XM_019825362.2;refGene_gn=SRPK2;refGene_tx=XM_023250274.1;refGene_gn=SRPK2;refGene_tx=XM_023250275.1;refGene_gn=SRPK2;refGene_tx=XM_023250276.1;refGene_gn=SRPK2;refGene_tx=XM_023250277.1;refGene_gn=SRPK2;refGene_tx=XM_023250278.1;refGene_gn=SRPK2;refGene_tx=XM_023250279.1;refGene_gn=SRPK2;refGene_tx=XM_023250280.1;refGene_gn=SRPK2;refGene_tx=XM_023250281.1;refGene_gn=SRPK2;refGene_tx=XM_023250282.1;refGene_gn=SRPK2;refGene_tx=XM_023250283.1;refGene_gn=SRPK2;refGene_tx=XM_023250284.1
A2 81188977 81189086
refGene_gn=SRPK2;refGene_tx=XM_019825360.1;refGene_gn=SRPK2;refGene_tx=XM_019825362.2;refGene_gn=SRPK2;refGene_tx=XM_023250274.1;refGene_gn=SRPK2;refGene_tx=XM_023250275.1;refGene_gn=SRPK2;refGene_tx=XM_023250276.1;refGene_gn=SRPK2;refGene_tx=XM_023250277.1;refGene_gn=SRPK2;refGene_tx=XM_023250278.1;refGene_gn=SRPK2;refGene_tx=XM_023250279.1;refGene_gn=SRPK2;refGene_tx=XM_023250280.1;refGene_gn=SRPK2;refGene_tx=XM_023250281.1;refGene_gn=SRPK2;refGene_tx=XM_023250282.1;refGene_gn=SRPK2;refGene_tx=XM_023250283.1;refGene_gn=SRPK2;refGene_tx=XM_023250284.1
A2 81189748 81189836
refGene_gn=SRPK2;refGene_tx=XM_019825360.1;refGene_gn=SRPK2;refGene_tx=XM_019825362.2;refGene_gn=SRPK2;refGene_tx=XM_023250274.1;refGene_gn=SRPK2;refGene_tx=XM_023250275.1;refGene_gn=SRPK2;refGene_tx=XM_023250276.1;refGene_gn=SRPK2;refGene_tx=XM_023250277.1;refGene_gn=SRPK2;refGene_tx=XM_023250278.1;refGene_gn=SRPK2;refGene_tx=XM_023250279.1;refGene_gn=SRPK2;refGene_tx=XM_023250280.1;refGene_gn=SRPK2;refGene_tx=XM_023250281.1;refGene_gn=SRPK2;refGene_tx=XM_023250282.1;refGene_gn=SRPK2;refGene_tx=XM_023250283.1;refGene_gn=SRPK2;refGene_tx=XM_023250284.1
A2 81190058 81190146
refGene_gn=SRPK2;refGene_tx=XM_019825360.1;refGene_gn=SRPK2;refGene_tx=XM_019825362.2;refGene_gn=SRPK2;refGene_tx=XM_023250274.1;refGene_gn=SRPK2;refGene_tx=XM_023250275.1;refGene_gn=SRPK2;refGene_tx=XM_023250276.1;refGene_gn=SRPK2;refGene_tx=XM_023250277.1;refGene_gn=SRPK2;refGene_tx=XM_023250278.1;refGene_gn=SRPK2;refGene_tx=XM_023250279.1;refGene_gn=SRPK2;refGene_tx=XM_023250280.1;refGene_gn=SRPK2;refGene_tx=XM_023250281.1;refGene_gn=SRPK2;refGene_tx=XM_023250282.1;refGene_gn=SRPK2;refGene_tx=XM_023250283.1;refGene_gn=SRPK2;refGene_tx=XM_023250284.1
A2 81192421 81192528
refGene_gn=SRPK2;refGene_tx=XM_019825360.1;refGene_gn=SRPK2;refGene_tx=XM_019825362.2;refGene_gn=SRPK2;refGene_tx=XM_023250274.1;refGene_gn=SRPK2;refGene_tx=XM_023250275.1;refGene_gn=SRPK2;refGene_tx=XM_023250276.1;refGene_gn=SRPK2;refGene_tx=XM_023250277.1;refGene_gn=SRPK2;refGene_tx=XM_023250278.1;refGene_gn=SRPK2;refGene_tx=XM_023250279.1;refGene_gn=SRPK2;refGene_tx=XM_023250280.1;refGene_gn=SRPK2;refGene_tx=XM_023250281.1;refGene_gn=SRPK2;refGene_tx=XM_023250282.1;refGene_gn=SRPK2;

refGene_tx=XM_023250283.1;refGene_gn=SRPK2;refGene_tx=XM_023250284.1
A2 81204038 81204204
refGene_gn=SRPK2;refGene_tx=XM_019825360.1;refGene_gn=SRPK2;refGene_tx=XM_019825362.2;refGene_gn=SRPK2;refGene_tx=XM_023250274.1;refGene_gn=SRPK2;refGene_tx=XM_023250275.1;refGene_gn=SRPK2;refGene_tx=XM_023250276.1;refGene_gn=SRPK2;refGene_tx=XM_023250277.1;refGene_gn=SRPK2;refGene_tx=XM_023250278.1;refGene_gn=SRPK2;refGene_tx=XM_023250279.1;refGene_gn=SRPK2;refGene_tx=XM_023250280.1;refGene_gn=SRPK2;refGene_tx=XM_023250281.1;refGene_gn=SRPK2;refGene_tx=XM_023250282.1;refGene_gn=SRPK2;refGene_tx=XM_023250283.1;refGene_gn=SRPK2;refGene_tx=XM_023250284.1
A2 81205368 81205394
refGene_gn=SRPK2;refGene_tx=XM_019825360.1;refGene_gn=SRPK2;refGene_tx=XM_019825362.2;refGene_gn=SRPK2;refGene_tx=XM_023250274.1;refGene_gn=SRPK2;refGene_tx=XM_023250275.1;refGene_gn=SRPK2;refGene_tx=XM_023250276.1;refGene_gn=SRPK2;refGene_tx=XM_023250277.1;refGene_gn=SRPK2;refGene_tx=XM_023250278.1;refGene_gn=SRPK2;refGene_tx=XM_023250279.1;refGene_gn=SRPK2;refGene_tx=XM_023250280.1;refGene_gn=SRPK2;refGene_tx=XM_023250281.1;refGene_gn=SRPK2;refGene_tx=XM_023250282.1;refGene_gn=SRPK2;refGene_tx=XM_023250283.1;refGene_gn=SRPK2;refGene_tx=XM_023250284.1
A2 81206871 81207118
refGene_gn=SRPK2;refGene_tx=XM_019825360.1;refGene_gn=SRPK2;refGene_tx=XM_019825362.2;refGene_gn=SRPK2;refGene_tx=XM_023250274.1;refGene_gn=SRPK2;refGene_tx=XM_023250275.1;refGene_gn=SRPK2;refGene_tx=XM_023250276.1;refGene_gn=SRPK2;refGene_tx=XM_023250277.1;refGene_gn=SRPK2;refGene_tx=XM_023250278.1;refGene_gn=SRPK2;refGene_tx=XM_023250279.1;refGene_gn=SRPK2;refGene_tx=XM_023250280.1;refGene_gn=SRPK2;refGene_tx=XM_023250281.1;refGene_gn=SRPK2;refGene_tx=XM_023250282.1;refGene_gn=SRPK2;refGene_tx=XM_023250283.1;refGene_gn=SRPK2;refGene_tx=XM_023250284.1
A2 81208017 81208500
refGene_gn=SRPK2;refGene_tx=XM_019825360.1;refGene_gn=SRPK2;refGene_tx=XM_019825362.2;refGene_gn=SRPK2;refGene_tx=XM_023250274.1;refGene_gn=SRPK2;refGene_tx=XM_023250275.1;refGene_gn=SRPK2;refGene_tx=XM_023250276.1;refGene_gn=SRPK2;refGene_tx=XM_023250277.1;refGene_gn=SRPK2;refGene_tx=XM_023250278.1;refGene_gn=SRPK2;refGene_tx=XM_023250279.1;refGene_gn=SRPK2;refGene_tx=XM_023250280.1;refGene_gn=SRPK2;refGene_tx=XM_023250281.1;refGene_gn=SRPK2;refGene_tx=XM_023250282.1;refGene_gn=SRPK2;refGene_tx=XM_023250283.1;refGene_gn=SRPK2;refGene_tx=XM_023250284.1
A2 81214755 81214856
refGene_gn=SRPK2;refGene_tx=XM_019825360.1;refGene_gn=SRPK2;refGene_tx=XM_019825362.2;refGene_gn=SRPK2;refGene_tx=XM_023250274.1;refGene_gn=SRPK2;refGene_tx=XM_023250275.1;refGene_gn=SRPK2;refGene_tx=XM_023250276.1;refGene_gn=SRPK2;refGene_tx=XM_023250277.1;refGene_gn=SRPK2;refGene_tx=XM_023250278.1;refGene_gn=SRPK2;refGene_tx=XM_023250279.1;refGene_gn=SRPK2;refGene_tx=XM_023250280.1;refGene_gn=SRPK2;refGene_tx=XM_023250281.1;refGene_gn=SRPK2;refGene_tx=XM_023250282.1;refGene_gn=SRPK2;refGene_tx=XM_023250283.1;refGene_gn=SRPK2;refGene_tx=XM_023250284.1
A2 81214976 81215084
refGene_gn=SRPK2;refGene_tx=XM_019825360.1;refGene_gn=SRPK2;refGene_tx=XM_019825362.2;refGene_gn=SRPK2;refGene_tx=XM_023250274.1;refGene_gn=SRPK2;refGene_tx=XM_023250275.1;refGene_gn=SRPK2;refGene_tx=XM_0232502

76.1;refGene_gn=SRPK2;refGene_tx=XM_023250277.1;refGene_gn=SRPK2;refGene_tx=XM_023250278.1;refGene_gn=SRPK2;refGene_tx=XM_023250279.1;refGene_gn=SRPK2;refGene_tx=XM_023250280.1;refGene_gn=SRPK2;refGene_tx=XM_023250281.1;refGene_gn=SRPK2;refGene_tx=XM_023250282.1;refGene_gn=SRPK2;refGene_tx=XM_023250283.1;refGene_gn=SRPK2;refGene_tx=XM_023250284.1
A2 81219873 81219943
refGene_gn=SRPK2;refGene_tx=XM_019825360.1;refGene_gn=SRPK2;refGene_tx=XM_019825362.2;refGene_gn=SRPK2;refGene_tx=XM_023250274.1;refGene_gn=SRPK2;refGene_tx=XM_023250275.1;refGene_gn=SRPK2;refGene_tx=XM_023250276.1;refGene_gn=SRPK2;refGene_tx=XM_023250277.1;refGene_gn=SRPK2;refGene_tx=XM_023250278.1;refGene_gn=SRPK2;refGene_tx=XM_023250279.1;refGene_gn=SRPK2;refGene_tx=XM_023250280.1;refGene_gn=SRPK2;refGene_tx=XM_023250281.1;refGene_gn=SRPK2;refGene_tx=XM_023250282.1;refGene_gn=SRPK2;refGene_tx=XM_023250283.1;refGene_gn=SRPK2;refGene_tx=XM_023250284.1
A2 81220561 81220654
refGene_gn=SRPK2;refGene_tx=XM_019825360.1;refGene_gn=SRPK2;refGene_tx=XM_019825362.2;refGene_gn=SRPK2;refGene_tx=XM_023250274.1;refGene_gn=SRPK2;refGene_tx=XM_023250275.1;refGene_gn=SRPK2;refGene_tx=XM_023250276.1;refGene_gn=SRPK2;refGene_tx=XM_023250277.1;refGene_gn=SRPK2;refGene_tx=XM_023250280.1;refGene_gn=SRPK2;refGene_tx=XM_023250281.1;refGene_gn=SRPK2;refGene_tx=XM_023250282.1;refGene_gn=SRPK2;refGene_tx=XM_023250283.1;refGene_gn=SRPK2;refGene_tx=XM_023250284.1
A2 81220982 81221075
refGene_gn=SRPK2;refGene_tx=XM_019825360.1;refGene_gn=SRPK2;refGene_tx=XM_019825362.2;refGene_gn=SRPK2;refGene_tx=XM_023250274.1;refGene_gn=SRPK2;refGene_tx=XM_023250275.1;refGene_gn=SRPK2;refGene_tx=XM_023250276.1;refGene_gn=SRPK2;refGene_tx=XM_023250278.1;refGene_gn=SRPK2;refGene_tx=XM_023250279.1;refGene_gn=SRPK2;refGene_tx=XM_023250280.1;refGene_gn=SRPK2;refGene_tx=XM_023250282.1;refGene_gn=SRPK2;refGene_tx=XM_023250283.1
A2 81230963 81231145
refGene_gn=SRPK2;refGene_tx=XM_023250275.1;refGene_gn=SRPK2;refGene_tx=XM_023250279.1
A2 81231168 81231353
refGene_gn=SRPK2;refGene_tx=XM_019825360.1;refGene_gn=SRPK2;refGene_tx=XM_019825362.2;refGene_gn=SRPK2;refGene_tx=XM_023250274.1;refGene_gn=SRPK2;refGene_tx=XM_023250276.1;refGene_gn=SRPK2;refGene_tx=XM_023250277.1;refGene_gn=SRPK2;refGene_tx=XM_023250278.1;refGene_gn=SRPK2;refGene_tx=XM_023250280.1;refGene_gn=SRPK2;refGene_tx=XM_023250281.1;refGene_gn=SRPK2;refGene_tx=XM_023250282.1;refGene_gn=SRPK2;refGene_tx=XM_023250283.1;refGene_gn=SRPK2;refGene_tx=XM_023250284.1
A2 81235990 81237502
refGene_gn=KMT2E;refGene_tx=XM_003982682.5;refGene_gn=KMT2E;refGene_tx=XM_006929135.4;refGene_gn=KMT2E;refGene_tx=XM_019825358.2;refGene_gn=KMT2E;refGene_tx=XM_019825359.2
A2 81238256 81238354
refGene_gn=KMT2E;refGene_tx=XM_003982682.5;refGene_gn=KMT2E;refGene_tx=XM_006929135.4;refGene_gn=KMT2E;refGene_tx=XM_019825358.2;refGene_gn=KMT2E;refGene_tx=XM_019825359.2
A2 81238536 81238662

refGene_gn=KMT2E;refGene_tx=XM_003982682.5;refGene_gn=KMT2E;refGene_tx=XM_006929135.4;refGene_gn=KMT2E;refGene_tx=XM_019825359.2
A2 81238765 81238854
refGene_gn=KMT2E;refGene_tx=XM_003982682.5;refGene_gn=KMT2E;refGene_tx=XM_006929135.4;refGene_gn=KMT2E;refGene_tx=XM_019825358.2;refGene_gn=KMT2E;refGene_tx=XM_019825359.2
A2 81239720 81240007
refGene_gn=KMT2E;refGene_tx=XM_003982682.5;refGene_gn=KMT2E;refGene_tx=XM_006929135.4;refGene_gn=KMT2E;refGene_tx=XM_019825358.2;refGene_gn=KMT2E;refGene_tx=XM_019825359.2
A2 81240982 81241546
refGene_gn=KMT2E;refGene_tx=XM_003982682.5;refGene_gn=KMT2E;refGene_tx=XM_006929135.4;refGene_gn=KMT2E;refGene_tx=XM_019825358.2;refGene_gn=KMT2E;refGene_tx=XM_019825359.2
A2 81241685 81241742
refGene_gn=KMT2E;refGene_tx=XM_003982682.5;refGene_gn=KMT2E;refGene_tx=XM_006929135.4;refGene_gn=KMT2E;refGene_tx=XM_019825358.2;refGene_gn=KMT2E;refGene_tx=XM_019825359.2
A2 81242100 81242351
refGene_gn=KMT2E;refGene_tx=XM_003982682.5;refGene_gn=KMT2E;refGene_tx=XM_006929135.4;refGene_gn=KMT2E;refGene_tx=XM_019825358.2;refGene_gn=KMT2E;refGene_tx=XM_019825359.2
A2 81243041 81243186
refGene_gn=KMT2E;refGene_tx=XM_003982682.5;refGene_gn=KMT2E;refGene_tx=XM_006929135.4;refGene_gn=KMT2E;refGene_tx=XM_019825358.2;refGene_gn=KMT2E;refGene_tx=XM_019825359.2
A2 81243345 81243600
refGene_gn=KMT2E;refGene_tx=XM_003982682.5;refGene_gn=KMT2E;refGene_tx=XM_006929135.4;refGene_gn=KMT2E;refGene_tx=XM_019825358.2;refGene_gn=KMT2E;refGene_tx=XM_019825359.2
A2 81246753 81247062
refGene_gn=KMT2E;refGene_tx=XM_003982682.5;refGene_gn=KMT2E;refGene_tx=XM_006929135.4;refGene_gn=KMT2E;refGene_tx=XM_019825358.2;refGene_gn=KMT2E;refGene_tx=XM_019825359.2
A2 81247368 81247533
refGene_gn=KMT2E;refGene_tx=XM_003982682.5;refGene_gn=KMT2E;refGene_tx=XM_006929135.4;refGene_gn=KMT2E;refGene_tx=XM_019825358.2;refGene_gn=KMT2E;refGene_tx=XM_019825359.2
A2 81255587 81255686
refGene_gn=KMT2E;refGene_tx=XM_003982682.5;refGene_gn=KMT2E;refGene_tx=XM_006929135.4;refGene_gn=KMT2E;refGene_tx=XM_019825358.2;refGene_gn=KMT2E;refGene_tx=XM_019825359.2
A2 81256619 81256884
refGene_gn=KMT2E;refGene_tx=XM_003982682.5;refGene_gn=KMT2E;refGene_tx=XM_006929135.4;refGene_gn=KMT2E;refGene_tx=XM_019825358.2;refGene_gn=KMT2E;refGene_tx=XM_019825359.2
A2 81259941 81260051
refGene_gn=KMT2E;refGene_tx=XM_003982682.5;refGene_gn=KMT2E;refGene_tx=XM_006929135.4;refGene_gn=KMT2E;refGene_tx=XM_019825358.2;refGene_gn=KMT2E;refGene_tx=XM_019825359.2

A2 81261021 81261139
refGene_gn=KMT2E;refGene_tx=XM_003982682.5;refGene_gn=KMT2E;refGene_tx=XM_006929135.4;refGene_gn=KMT2E;refGene_tx=XM_019825358.2;refGene_gn=KMT2E;refGene_tx=XM_019825359.2
A2 81262125 81262256
refGene_gn=KMT2E;refGene_tx=XM_003982682.5;refGene_gn=KMT2E;refGene_tx=XM_006929135.4;refGene_gn=KMT2E;refGene_tx=XM_019825358.2;refGene_gn=KMT2E;refGene_tx=XM_019825359.2
A2 81262374 81262605
refGene_gn=KMT2E;refGene_tx=XM_003982682.5;refGene_gn=KMT2E;refGene_tx=XM_006929135.4;refGene_gn=KMT2E;refGene_tx=XM_019825358.2;refGene_gn=KMT2E;refGene_tx=XM_019825359.2
A2 81265441 81265480
refGene_gn=KMT2E;refGene_tx=XM_003982682.5;refGene_gn=KMT2E;refGene_tx=XM_006929135.4;refGene_gn=KMT2E;refGene_tx=XM_019825358.2;refGene_gn=KMT2E;refGene_tx=XM_019825359.2
A2 81268700 81268873
refGene_gn=KMT2E;refGene_tx=XM_003982682.5;refGene_gn=KMT2E;refGene_tx=XM_006929135.4;refGene_gn=KMT2E;refGene_tx=XM_019825358.2;refGene_gn=KMT2E;refGene_tx=XM_019825359.2
A2 81269823 81269882
refGene_gn=KMT2E;refGene_tx=XM_003982682.5;refGene_gn=KMT2E;refGene_tx=XM_006929135.4;refGene_gn=KMT2E;refGene_tx=XM_019825358.2;refGene_gn=KMT2E;refGene_tx=XM_019825359.2
A2 81276706 81276787
refGene_gn=KMT2E;refGene_tx=XM_003982682.5;refGene_gn=KMT2E;refGene_tx=XM_006929135.4;refGene_gn=KMT2E;refGene_tx=XM_019825358.2;refGene_gn=KMT2E;refGene_tx=XM_019825359.2
A2 81279615 81279845
refGene_gn=KMT2E;refGene_tx=XM_003982682.5;refGene_gn=KMT2E;refGene_tx=XM_006929135.4;refGene_gn=KMT2E;refGene_tx=XM_019825358.2
A2 81281117 81281232
refGene_gn=KMT2E;refGene_tx=XM_003982682.5;refGene_gn=KMT2E;refGene_tx=XM_006929135.4;refGene_gn=KMT2E;refGene_tx=XM_019825358.2
A2 81307154 81307225
refGene_gn=KMT2E;refGene_tx=XM_003982682.5;refGene_gn=KMT2E;refGene_tx=XM_006929135.4;refGene_gn=KMT2E;refGene_tx=XM_019825358.2
A2 81431433 81431462 refGene_gn=LHFPL3;refGene_tx=XM_011280338.3
A2 81487917 81487934
refGene_gn=LHFPL3;refGene_tx=XM_023250285.1;refGene_gn=LHFPL3;refGene_tx=XM_023250286.1
A2 81604890 81605127
refGene_gn=LHFPL3;refGene_tx=XM_011280338.3;refGene_gn=LHFPL3;refGene_tx=XM_023250285.1;refGene_gn=LHFPL3;refGene_tx=XM_023250286.1
A2 82008247 82008689
refGene_gn=LHFPL3;refGene_tx=XM_011280338.3;refGene_gn=LHFPL3;refGene_tx=XM_023250285.1;refGene_gn=LHFPL3;refGene_tx=XM_023250286.1
A2 82112675 82112747 refGene_gn=ORC5;refGene_tx=XM_003982684.5
A2 82116915 82116990 refGene_gn=ORC5;refGene_tx=XM_003982684.5
A2 82116991 82117009 refGene_gn=ORC5;refGene_tx=XM_003982684.5

A2 82118126 82118327 refGene_gn=ORC5;refGene_tx=XM_003982684.5
A2 82128038 82128113 refGene_gn=ORC5;refGene_tx=XM_003982684.5
A2 82131738 82131850 refGene_gn=ORC5;refGene_tx=XM_003982684.5
A2 82137279 82137410 refGene_gn=ORC5;refGene_tx=XM_003982684.5
A2 82146018 82146067 refGene_gn=ORC5;refGene_tx=XM_003982684.5
A2 82146155 82146246 refGene_gn=ORC5;refGene_tx=XM_003982684.5
A2 82150431 82150484 refGene_gn=ORC5;refGene_tx=XM_003982684.5
A2 82151840 82151953 refGene_gn=ORC5;refGene_tx=XM_003982684.5
A2 82153072 82153120 refGene_gn=ORC5;refGene_tx=XM_003982684.5
A2 82159907 82160018 refGene_gn=ORC5;refGene_tx=XM_003982684.5
A2 82184218 82184331 refGene_gn=ORC5;refGene_tx=XM_003982684.5
A2 82193413 82193459 refGene_gn=ORC5;refGene_tx=XM_003982684.5
A2 82324174 82324400
refGene_gn=RELN;refGene_tx=XM_011280339.3;refGene_gn=RELN;refGene_tx=XM_019825367.2;refGene_gn=RELN;refGene_tx=XM_019825368.2
A2 82400461 82400572
refGene_gn=RELN;refGene_tx=XM_011280339.3;refGene_gn=RELN;refGene_tx=XM_019825367.2;refGene_gn=RELN;refGene_tx=XM_019825368.2
A2 82478111 82478247
refGene_gn=RELN;refGene_tx=XM_011280339.3;refGene_gn=RELN;refGene_tx=XM_019825367.2;refGene_gn=RELN;refGene_tx=XM_019825368.2
A2 82548302 82548373
refGene_gn=RELN;refGene_tx=XM_011280339.3;refGene_gn=RELN;refGene_tx=XM_019825367.2;refGene_gn=RELN;refGene_tx=XM_019825368.2
A2 82569701 82569734
refGene_gn=RELN;refGene_tx=XM_011280339.3;refGene_gn=RELN;refGene_tx=XM_019825367.2;refGene_gn=RELN;refGene_tx=XM_019825368.2
A2 82572725 82572804
refGene_gn=RELN;refGene_tx=XM_011280339.3;refGene_gn=RELN;refGene_tx=XM_019825367.2;refGene_gn=RELN;refGene_tx=XM_019825368.2
A2 82591671 82591768
refGene_gn=RELN;refGene_tx=XM_011280339.3;refGene_gn=RELN;refGene_tx=XM_019825367.2;refGene_gn=RELN;refGene_tx=XM_019825368.2
A2 82597082 82597134
refGene_gn=RELN;refGene_tx=XM_011280339.3;refGene_gn=RELN;refGene_tx=XM_019825367.2;refGene_gn=RELN;refGene_tx=XM_019825368.2
A2 82621175 82621272
refGene_gn=RELN;refGene_tx=XM_011280339.3;refGene_gn=RELN;refGene_tx=XM_019825367.2;refGene_gn=RELN;refGene_tx=XM_019825368.2
A2 82624429 82624670
refGene_gn=RELN;refGene_tx=XM_011280339.3;refGene_gn=RELN;refGene_tx=XM_019825367.2;refGene_gn=RELN;refGene_tx=XM_019825368.2
A2 82642670 82642816
refGene_gn=RELN;refGene_tx=XM_011280339.3;refGene_gn=RELN;refGene_tx=XM_019825367.2;refGene_gn=RELN;refGene_tx=XM_019825368.2
A2 82667912 82668064
refGene_gn=RELN;refGene_tx=XM_011280339.3;refGene_gn=RELN;refGene_tx=XM_019825367.2;refGene_gn=RELN;refGene_tx=XM_019825368.2
A2 82682732 82682845
refGene_gn=RELN;refGene_tx=XM_011280339.3;refGene_gn=RELN;refGene_tx=X

M_019825367.2;refGene_gn=RELN;refGene_tx=XM_019825368.2
A2 82684520 82684729
refGene_gn=RELN;refGene_tx=XM_011280339.3;refGene_gn=RELN;refGene_tx=XM_019825367.2;refGene_gn=RELN;refGene_tx=XM_019825368.2
A2 82685860 82685989
refGene_gn=RELN;refGene_tx=XM_011280339.3;refGene_gn=RELN;refGene_tx=XM_019825367.2;refGene_gn=RELN;refGene_tx=XM_019825368.2
A2 82687009 82687119
refGene_gn=RELN;refGene_tx=XM_011280339.3;refGene_gn=RELN;refGene_tx=XM_019825367.2;refGene_gn=RELN;refGene_tx=XM_019825368.2
A2 82691881 82691948
refGene_gn=RELN;refGene_tx=XM_011280339.3;refGene_gn=RELN;refGene_tx=XM_019825367.2;refGene_gn=RELN;refGene_tx=XM_019825368.2
A2 82696567 82696801
refGene_gn=RELN;refGene_tx=XM_011280339.3;refGene_gn=RELN;refGene_tx=XM_019825367.2;refGene_gn=RELN;refGene_tx=XM_019825368.2
A2 82697464 82697626
refGene_gn=RELN;refGene_tx=XM_011280339.3;refGene_gn=RELN;refGene_tx=XM_019825367.2;refGene_gn=RELN;refGene_tx=XM_019825368.2
A2 82702212 82702449
refGene_gn=RELN;refGene_tx=XM_011280339.3;refGene_gn=RELN;refGene_tx=XM_019825367.2;refGene_gn=RELN;refGene_tx=XM_019825368.2
A2 82721763 82721956
refGene_gn=RELN;refGene_tx=XM_011280339.3;refGene_gn=RELN;refGene_tx=XM_019825367.2;refGene_gn=RELN;refGene_tx=XM_019825368.2
A2 82722750 82722863
refGene_gn=RELN;refGene_tx=XM_011280339.3;refGene_gn=RELN;refGene_tx=XM_019825367.2;refGene_gn=RELN;refGene_tx=XM_019825368.2
A2 82729979 82730117
refGene_gn=RELN;refGene_tx=XM_011280339.3;refGene_gn=RELN;refGene_tx=XM_019825367.2;refGene_gn=RELN;refGene_tx=XM_019825368.2
A2 82731449 82731636
refGene_gn=RELN;refGene_tx=XM_011280339.3;refGene_gn=RELN;refGene_tx=XM_019825367.2;refGene_gn=RELN;refGene_tx=XM_019825368.2
A2 82739037 82739243
refGene_gn=RELN;refGene_tx=XM_011280339.3;refGene_gn=RELN;refGene_tx=XM_019825367.2;refGene_gn=RELN;refGene_tx=XM_019825368.2
A2 82743902 82744074
refGene_gn=RELN;refGene_tx=XM_011280339.3;refGene_gn=RELN;refGene_tx=XM_019825367.2;refGene_gn=RELN;refGene_tx=XM_019825368.2
A2 82744555 82744756
refGene_gn=RELN;refGene_tx=XM_011280339.3;refGene_gn=RELN;refGene_tx=XM_019825367.2;refGene_gn=RELN;refGene_tx=XM_019825368.2
A2 82750877 82751110
refGene_gn=RELN;refGene_tx=XM_011280339.3;refGene_gn=RELN;refGene_tx=XM_019825367.2;refGene_gn=RELN;refGene_tx=XM_019825368.2
A2 82760474 82760632
refGene_gn=RELN;refGene_tx=XM_011280339.3;refGene_gn=RELN;refGene_tx=XM_019825367.2;refGene_gn=RELN;refGene_tx=XM_019825368.2
A2 82761888 82762096

refGene_gn=RELN;refGene_tx=XM_011280339.3;refGene_gn=RELN;refGene_tx=XM_019825367.2;refGene_gn=RELN;refGene_tx=XM_019825368.2
A2 82763423 82763500
refGene_gn=RELN;refGene_tx=XM_011280339.3;refGene_gn=RELN;refGene_tx=XM_019825367.2;refGene_gn=RELN;refGene_tx=XM_019825368.2
A2 82770172 82770331
refGene_gn=RELN;refGene_tx=XM_011280339.3;refGene_gn=RELN;refGene_tx=XM_019825367.2;refGene_gn=RELN;refGene_tx=XM_019825368.2
A2 82770517 82770706
refGene_gn=RELN;refGene_tx=XM_011280339.3;refGene_gn=RELN;refGene_tx=XM_019825367.2;refGene_gn=RELN;refGene_tx=XM_019825368.2
A2 82771215 82771489
refGene_gn=RELN;refGene_tx=XM_011280339.3;refGene_gn=RELN;refGene_tx=XM_019825367.2;refGene_gn=RELN;refGene_tx=XM_019825368.2
A2 82773974 82774115
refGene_gn=RELN;refGene_tx=XM_011280339.3;refGene_gn=RELN;refGene_tx=XM_019825367.2;refGene_gn=RELN;refGene_tx=XM_019825368.2
A2 82774220 82774398
refGene_gn=RELN;refGene_tx=XM_011280339.3;refGene_gn=RELN;refGene_tx=XM_019825367.2;refGene_gn=RELN;refGene_tx=XM_019825368.2
A2 82777754 82777839
refGene_gn=RELN;refGene_tx=XM_011280339.3;refGene_gn=RELN;refGene_tx=XM_019825367.2;refGene_gn=RELN;refGene_tx=XM_019825368.2
A2 82778667 82778850
refGene_gn=RELN;refGene_tx=XM_011280339.3;refGene_gn=RELN;refGene_tx=XM_019825367.2;refGene_gn=RELN;refGene_tx=XM_019825368.2
A2 82781747 82781919
refGene_gn=RELN;refGene_tx=XM_011280339.3;refGene_gn=RELN;refGene_tx=XM_019825367.2;refGene_gn=RELN;refGene_tx=XM_019825368.2
A2 82782017 82782120
refGene_gn=RELN;refGene_tx=XM_011280339.3;refGene_gn=RELN;refGene_tx=XM_019825367.2;refGene_gn=RELN;refGene_tx=XM_019825368.2
A2 82784760 82784990
refGene_gn=RELN;refGene_tx=XM_011280339.3;refGene_gn=RELN;refGene_tx=XM_019825367.2;refGene_gn=RELN;refGene_tx=XM_019825368.2
A2 82790842 82791063
refGene_gn=RELN;refGene_tx=XM_011280339.3;refGene_gn=RELN;refGene_tx=XM_019825367.2;refGene_gn=RELN;refGene_tx=XM_019825368.2
A2 82793436 82793584
refGene_gn=RELN;refGene_tx=XM_011280339.3;refGene_gn=RELN;refGene_tx=XM_019825367.2;refGene_gn=RELN;refGene_tx=XM_019825368.2
A2 82795852 82796111
refGene_gn=RELN;refGene_tx=XM_011280339.3;refGene_gn=RELN;refGene_tx=XM_019825367.2;refGene_gn=RELN;refGene_tx=XM_019825368.2
A2 82796970 82797220
refGene_gn=RELN;refGene_tx=XM_011280339.3;refGene_gn=RELN;refGene_tx=XM_019825367.2;refGene_gn=RELN;refGene_tx=XM_019825368.2
A2 82801219 82801388
refGene_gn=RELN;refGene_tx=XM_011280339.3;refGene_gn=RELN;refGene_tx=XM_019825367.2;refGene_gn=RELN;refGene_tx=XM_019825368.2

A2 82814565 82814706
refGene_gn=RELN;refGene_tx=XM_011280339.3;refGene_gn=RELN;refGene_tx=XM_019825367.2;refGene_gn=RELN;refGene_tx=XM_019825368.2
A2 82816089 82816267
refGene_gn=RELN;refGene_tx=XM_011280339.3;refGene_gn=RELN;refGene_tx=XM_019825367.2;refGene_gn=RELN;refGene_tx=XM_019825368.2
A2 82819450 82819644
refGene_gn=RELN;refGene_tx=XM_011280339.3;refGene_gn=RELN;refGene_tx=XM_019825367.2;refGene_gn=RELN;refGene_tx=XM_019825368.2
A2 82823916 82824173
refGene_gn=RELN;refGene_tx=XM_011280339.3;refGene_gn=RELN;refGene_tx=XM_019825367.2;refGene_gn=RELN;refGene_tx=XM_019825368.2
A2 82827470 82827625
refGene_gn=RELN;refGene_tx=XM_011280339.3;refGene_gn=RELN;refGene_tx=XM_019825367.2;refGene_gn=RELN;refGene_tx=XM_019825368.2
A2 82828713 82828928
refGene_gn=RELN;refGene_tx=XM_011280339.3;refGene_gn=RELN;refGene_tx=XM_019825367.2;refGene_gn=RELN;refGene_tx=XM_019825368.2
A2 82831056 82831234
refGene_gn=RELN;refGene_tx=XM_011280339.3;refGene_gn=RELN;refGene_tx=XM_019825367.2;refGene_gn=RELN;refGene_tx=XM_019825368.2
A2 82834087 82834263
refGene_gn=RELN;refGene_tx=XM_011280339.3;refGene_gn=RELN;refGene_tx=XM_019825367.2;refGene_gn=RELN;refGene_tx=XM_019825368.2
A2 82834411 82834518
refGene_gn=RELN;refGene_tx=XM_011280339.3;refGene_gn=RELN;refGene_tx=XM_019825367.2;refGene_gn=RELN;refGene_tx=XM_019825368.2
A2 82835619 82835862
refGene_gn=RELN;refGene_tx=XM_011280339.3;refGene_gn=RELN;refGene_tx=XM_019825367.2;refGene_gn=RELN;refGene_tx=XM_019825368.2
A2 82836438 82836614
refGene_gn=RELN;refGene_tx=XM_011280339.3;refGene_gn=RELN;refGene_tx=XM_019825367.2;refGene_gn=RELN;refGene_tx=XM_019825368.2
A2 82842692 82842766
refGene_gn=RELN;refGene_tx=XM_011280339.3;refGene_gn=RELN;refGene_tx=XM_019825367.2;refGene_gn=RELN;refGene_tx=XM_019825368.2
A2 82844043 82844205
refGene_gn=RELN;refGene_tx=XM_011280339.3;refGene_gn=RELN;refGene_tx=XM_019825367.2;refGene_gn=RELN;refGene_tx=XM_019825368.2
A2 82844958 82845116
refGene_gn=RELN;refGene_tx=XM_011280339.3;refGene_gn=RELN;refGene_tx=XM_019825367.2;refGene_gn=RELN;refGene_tx=XM_019825368.2
A2 82847358 82847578
refGene_gn=RELN;refGene_tx=XM_011280339.3;refGene_gn=RELN;refGene_tx=XM_019825367.2;refGene_gn=RELN;refGene_tx=XM_019825368.2
A2 82849410 82849608
refGene_gn=RELN;refGene_tx=XM_011280339.3;refGene_gn=RELN;refGene_tx=XM_019825367.2;refGene_gn=RELN;refGene_tx=XM_019825368.2
A2 82850238 82850337
refGene_gn=RELN;refGene_tx=XM_011280339.3;refGene_gn=RELN;refGene_tx=X

M_019825367.2;refGene_gn=RELN;refGene_tx=XM_019825368.2
A2 82851043 82851133 refGene_gn=RELN;refGene_tx=XM_019825367.2
A2 82855248 82855254
refGene_gn=RELN;refGene_tx=XM_019825367.2;refGene_gn=RELN;refGene_tx=XM_019825368.2
A2 82860346 82860443
refGene_gn=RELN;refGene_tx=XM_011280339.3;refGene_gn=RELN;refGene_tx=XM_019825367.2;refGene_gn=RELN;refGene_tx=XM_019825368.2
A2 82900394 82900534 refGene_gn=SLC26A5;refGene_tx=NM_001137656.1
A2 82906903 82907014 refGene_gn=SLC26A5;refGene_tx=NM_001137656.1
A2 82908096 82908263 refGene_gn=SLC26A5;refGene_tx=NM_001137656.1
A2 82908816 82908981 refGene_gn=SLC26A5;refGene_tx=NM_001137656.1
A2 82910921 82911074 refGene_gn=SLC26A5;refGene_tx=NM_001137656.1
A2 82912400 82912483 refGene_gn=SLC26A5;refGene_tx=NM_001137656.1
A2 82916880 82917028 refGene_gn=SLC26A5;refGene_tx=NM_001137656.1
A2 82918052 82918166 refGene_gn=SLC26A5;refGene_tx=NM_001137656.1
A2 82919521 82919599 refGene_gn=SLC26A5;refGene_tx=NM_001137656.1
A2 82920286 82920382 refGene_gn=SLC26A5;refGene_tx=NM_001137656.1
A2 82920590 82920697 refGene_gn=SLC26A5;refGene_tx=NM_001137656.1
A2 82921781 82921851 refGene_gn=SLC26A5;refGene_tx=NM_001137656.1
A2 82922365 82922458 refGene_gn=SLC26A5;refGene_tx=NM_001137656.1
A2 82923142 82923250 refGene_gn=SLC26A5;refGene_tx=NM_001137656.1
A2 82923908 82924109 refGene_gn=SLC26A5;refGene_tx=NM_001137656.1
A2 82924719 82924774 refGene_gn=SLC26A5;refGene_tx=NM_001137656.1
A2 82926121 82926306 refGene_gn=SLC26A5;refGene_tx=NM_001137656.1
A2 82931682 82931840 refGene_gn=PSMC2;refGene_tx=XM_003982686.3
A2 82931928 82932025 refGene_gn=PSMC2;refGene_tx=XM_003982686.3
A2 82932114 82932317 refGene_gn=PSMC2;refGene_tx=XM_003982686.3
A2 82934153 82934241 refGene_gn=PSMC2;refGene_tx=XM_003982686.3
A2 82935768 82935933 refGene_gn=PSMC2;refGene_tx=XM_003982686.3
A2 82936509 82936605 refGene_gn=PSMC2;refGene_tx=XM_003982686.3
A2 82936992 82937065 refGene_gn=PSMC2;refGene_tx=XM_003982686.3
A2 82937466 82937598 refGene_gn=PSMC2;refGene_tx=XM_003982686.3
A2 82941190 82941290 refGene_gn=PSMC2;refGene_tx=XM_003982686.3
A2 82942006 82942088 refGene_gn=PSMC2;refGene_tx=XM_003982686.3
A2 82942649 82942687 refGene_gn=PSMC2;refGene_tx=XM_003982686.3
A2 82944515 82944585 refGene_gn=PSMC2;refGene_tx=XM_003982686.3
A2 82947975 82948039 refGene_gn=DNAJC2;refGene_tx=XM_004001540.5
A2 82950250 82950441
refGene_gn=DNAJC2;refGene_tx=XM_004001540.5;refGene_gn=DNAJC2;refGene_tx=XM_011280341.2
A2 82954728 82954804
refGene_gn=DNAJC2;refGene_tx=XM_004001540.5;refGene_gn=DNAJC2;refGene_tx=XM_011280341.2
A2 82970682 82970781
refGene_gn=DNAJC2;refGene_tx=XM_004001540.5;refGene_gn=DNAJC2;refGene_tx=XM_011280341.2;refGene_gn=DNAJC2;refGene_tx=XM_011280342.3
A2 82972004 82972146
refGene_gn=DNAJC2;refGene_tx=XM_004001540.5;refGene_gn=DNAJC2;refGene_tx=XM_011280341.2;refGene_gn=DNAJC2;refGene_tx=XM_011280342.3

A2 82974136 82974217
refGene_gn=DNAJC2;refGene_tx=XM_004001540.5;refGene_gn=DNAJC2;refGene_tx=XM_011280341.2;refGene_gn=DNAJC2;refGene_tx=XM_011280342.3
A2 82974789 82974855
refGene_gn=DNAJC2;refGene_tx=XM_004001540.5;refGene_gn=DNAJC2;refGene_tx=XM_011280341.2;refGene_gn=DNAJC2;refGene_tx=XM_011280342.3
A2 82975297 82975389
refGene_gn=DNAJC2;refGene_tx=XM_004001540.5;refGene_gn=DNAJC2;refGene_tx=XM_011280341.2;refGene_gn=DNAJC2;refGene_tx=XM_011280342.3
A2 82975674 82975796
refGene_gn=DNAJC2;refGene_tx=XM_004001540.5;refGene_gn=DNAJC2;refGene_tx=XM_011280341.2;refGene_gn=DNAJC2;refGene_tx=XM_011280342.3
A2 82976446 82976596
refGene_gn=DNAJC2;refGene_tx=XM_004001540.5;refGene_gn=DNAJC2;refGene_tx=XM_011280341.2;refGene_gn=DNAJC2;refGene_tx=XM_011280342.3
A2 82979104 82979193
refGene_gn=DNAJC2;refGene_tx=XM_004001540.5;refGene_gn=DNAJC2;refGene_tx=XM_011280341.2;refGene_gn=DNAJC2;refGene_tx=XM_011280342.3
A2 82979266 82979336
refGene_gn=DNAJC2;refGene_tx=XM_004001540.5;refGene_gn=DNAJC2;refGene_tx=XM_011280341.2;refGene_gn=DNAJC2;refGene_tx=XM_011280342.3
A2 82981011 82981196
refGene_gn=DNAJC2;refGene_tx=XM_004001540.5;refGene_gn=DNAJC2;refGene_tx=XM_011280341.2;refGene_gn=DNAJC2;refGene_tx=XM_011280342.3
A2 82981968 82982069
refGene_gn=DNAJC2;refGene_tx=XM_004001540.5;refGene_gn=DNAJC2;refGene_tx=XM_011280341.2;refGene_gn=DNAJC2;refGene_tx=XM_011280342.3
A2 82982183 82982291
refGene_gn=DNAJC2;refGene_tx=XM_004001540.5;refGene_gn=DNAJC2;refGene_tx=XM_011280341.2;refGene_gn=DNAJC2;refGene_tx=XM_011280342.3
A2 82984185 82984340
refGene_gn=DNAJC2;refGene_tx=XM_004001540.5;refGene_gn=DNAJC2;refGene_tx=XM_011280341.2;refGene_gn=DNAJC2;refGene_tx=XM_011280342.3
A2 82984797 82984872
refGene_gn=DNAJC2;refGene_tx=XM_004001540.5;refGene_gn=DNAJC2;refGene_tx=XM_011280341.2;refGene_gn=DNAJC2;refGene_tx=XM_011280342.3
A2 82985142 82985207 refGene_gn=PMPCB;refGene_tx=XM_003982687.4
A2 82985301 82985377 refGene_gn=PMPCB;refGene_tx=XM_003982687.4
A2 82985537 82985626 refGene_gn=PMPCB;refGene_tx=XM_003982687.4
A2 82985707 82985793 refGene_gn=PMPCB;refGene_tx=XM_003982687.4
A2 82987195 82987356 refGene_gn=PMPCB;refGene_tx=XM_003982687.4
A2 82988163 82988307 refGene_gn=PMPCB;refGene_tx=XM_003982687.4
A2 82989545 82989658 refGene_gn=PMPCB;refGene_tx=XM_003982687.4
A2 82992215 82992295 refGene_gn=PMPCB;refGene_tx=XM_003982687.4
A2 82992667 82992866 refGene_gn=PMPCB;refGene_tx=XM_003982687.4
A2 82997354 82997484 refGene_gn=PMPCB;refGene_tx=XM_003982687.4
A2 82997812 82997899 refGene_gn=PMPCB;refGene_tx=XM_003982687.4
A2 82998796 82998937 refGene_gn=PMPCB;refGene_tx=XM_003982687.4
A2 82999891 82999990 refGene_gn=PMPCB;refGene_tx=XM_003982687.4
A2 83068714 83069008

refGene_gn=NAPEPLD;refGene_tx=XM_003982699.5;refGene_gn=NAPEPLD;refGene_tx=XM_023250287.1;refGene_gn=NAPEPLD;refGene_tx=XM_023250288.1;refGene_gn=NAPEPLD;refGene_tx=XM_023250289.1;refGene_gn=NAPEPLD;refGene_tx=XM_023250290.1
A2 83084577 83085224
refGene_gn=NAPEPLD;refGene_tx=XM_003982699.5;refGene_gn=NAPEPLD;refGene_tx=XM_023250287.1;refGene_gn=NAPEPLD;refGene_tx=XM_023250288.1;refGene_gn=NAPEPLD;refGene_tx=XM_023250289.1;refGene_gn=NAPEPLD;refGene_tx=XM_023250290.1
A2 83088501 83088616
refGene_gn=NAPEPLD;refGene_tx=XM_003982699.5;refGene_gn=NAPEPLD;refGene_tx=XM_023250287.1;refGene_gn=NAPEPLD;refGene_tx=XM_023250288.1;refGene_gn=NAPEPLD;refGene_tx=XM_023250289.1;refGene_gn=NAPEPLD;refGene_tx=XM_023250290.1
A2 83091811 83091940
refGene_gn=NAPEPLD;refGene_tx=XM_003982699.5;refGene_gn=NAPEPLD;refGene_tx=XM_023250287.1;refGene_gn=NAPEPLD;refGene_tx=XM_023250288.1;refGene_gn=NAPEPLD;refGene_tx=XM_023250289.1;refGene_gn=NAPEPLD;refGene_tx=XM_023250290.1
A2 83095887 83096139 refGene_gn=ARMC10;refGene_tx=XM_003982688.5
A2 83097035 83097107 refGene_gn=ARMC10;refGene_tx=XM_003982688.5
A2 83101690 83101867 refGene_gn=ARMC10;refGene_tx=XM_003982688.5
A2 83104903 83105038 refGene_gn=ARMC10;refGene_tx=XM_003982688.5
A2 83107849 83107998 refGene_gn=ARMC10;refGene_tx=XM_003982688.5
A2 83113186 83113325 refGene_gn=ARMC10;refGene_tx=XM_003982688.5
A2 83113839 83113914 refGene_gn=FBXL13;refGene_tx=XM_011280343.3
A2 83129961 83130065 refGene_gn=FBXL13;refGene_tx=XM_011280343.3
A2 83130752 83130843 refGene_gn=FBXL13;refGene_tx=XM_011280343.3
A2 83149275 83149343 refGene_gn=FBXL13;refGene_tx=XM_011280343.3
A2 83150397 83150552 refGene_gn=FBXL13;refGene_tx=XM_011280343.3
A2 83151911 83152021 refGene_gn=FBXL13;refGene_tx=XM_011280343.3
A2 83154743 83154911 refGene_gn=FBXL13;refGene_tx=XM_011280343.3
A2 83199893 83199989 refGene_gn=FBXL13;refGene_tx=XM_011280343.3
A2 83202384 83202517 refGene_gn=FBXL13;refGene_tx=XM_011280343.3
A2 83213846 83214244 refGene_gn=LRRC17;refGene_tx=XM_003982691.4
A2 83218476 83218632 refGene_gn=LRRC17;refGene_tx=XM_003982691.4
A2 83227656 83228428 refGene_gn=LRRC17;refGene_tx=XM_003982691.4
A2 83231563 83231616 refGene_gn=FBXL13;refGene_tx=XM_011280343.3
A2 83235518 83235619 refGene_gn=FBXL13;refGene_tx=XM_011280343.3
A2 83244724 83244854 refGene_gn=FBXL13;refGene_tx=XM_011280343.3
A2 83264188 83264287 refGene_gn=FBXL13;refGene_tx=XM_011280343.3
A2 83264825 83264943 refGene_gn=FBXL13;refGene_tx=XM_011280343.3
A2 83265038 83265201 refGene_gn=FBXL13;refGene_tx=XM_011280343.3
A2 83270885 83271005 refGene_gn=FBXL13;refGene_tx=XM_011280343.3
A2 83272020 83272147 refGene_gn=FBXL13;refGene_tx=XM_011280343.3
A2 83294720 83294804 refGene_gn=FBXL13;refGene_tx=XM_011280343.3
A2 83308926 83309061 refGene_gn=FBXL13;refGene_tx=XM_011280343.3
A2 83314228 83314392 refGene_gn=FBXL13;refGene_tx=XM_011280343.3
A2 83324045 83324232 refGene_gn=FBXL13;refGene_tx=XM_011280343.3
A2 83422257 83422416

refGene_gn=CCDC146;refGene_tx=XM_003982693.5;refGene_gn=CCDC146;refGene_tx=XM_019825375.2;refGene_gn=CCDC146;refGene_tx=XM_019825376.2;refGene_gn=CCDC146;refGene_tx=XM_019825377.2
A2 83443923 83444630 refGene_gn=FGL2;refGene_tx=XM_003982694.5
A2 83447107 83447723 refGene_gn=FGL2;refGene_tx=XM_003982694.5
A2 83479791 83479874
refGene_gn=CCDC146;refGene_tx=XM_003982693.5;refGene_gn=CCDC146;refGene_tx=XM_019825375.2;refGene_gn=CCDC146;refGene_tx=XM_019825376.2;refGene_gn=CCDC146;refGene_tx=XM_019825377.2
A2 83483077 83483287
refGene_gn=CCDC146;refGene_tx=XM_003982693.5;refGene_gn=CCDC146;refGene_tx=XM_019825375.2;refGene_gn=CCDC146;refGene_tx=XM_019825376.2;refGene_gn=CCDC146;refGene_tx=XM_019825377.2
A2 83488781 83488839
refGene_gn=CCDC146;refGene_tx=XM_003982693.5;refGene_gn=CCDC146;refGene_tx=XM_019825375.2;refGene_gn=CCDC146;refGene_tx=XM_019825376.2;refGene_gn=CCDC146;refGene_tx=XM_019825377.2
A2 83491069 83491246
refGene_gn=CCDC146;refGene_tx=XM_003982693.5;refGene_gn=CCDC146;refGene_tx=XM_019825375.2;refGene_gn=CCDC146;refGene_tx=XM_019825376.2;refGene_gn=CCDC146;refGene_tx=XM_019825377.2
A2 83493125 83493199
refGene_gn=CCDC146;refGene_tx=XM_003982693.5;refGene_gn=CCDC146;refGene_tx=XM_019825375.2;refGene_gn=CCDC146;refGene_tx=XM_019825376.2;refGene_gn=CCDC146;refGene_tx=XM_019825377.2
A2 83494128 83494356
refGene_gn=CCDC146;refGene_tx=XM_003982693.5;refGene_gn=CCDC146;refGene_tx=XM_019825375.2;refGene_gn=CCDC146;refGene_tx=XM_019825376.2;refGene_gn=CCDC146;refGene_tx=XM_019825377.2
A2 83495206 83495393
refGene_gn=CCDC146;refGene_tx=XM_003982693.5;refGene_gn=CCDC146;refGene_tx=XM_019825375.2;refGene_gn=CCDC146;refGene_tx=XM_019825376.2;refGene_gn=CCDC146;refGene_tx=XM_019825377.2
A2 83501594 83501690
refGene_gn=CCDC146;refGene_tx=XM_003982693.5;refGene_gn=CCDC146;refGene_tx=XM_019825375.2;refGene_gn=CCDC146;refGene_tx=XM_019825376.2;refGene_gn=CCDC146;refGene_tx=XM_019825377.2
A2 83502301 83502472
refGene_gn=CCDC146;refGene_tx=XM_003982693.5;refGene_gn=CCDC146;refGene_tx=XM_019825375.2;refGene_gn=CCDC146;refGene_tx=XM_019825376.2;refGene_gn=CCDC146;refGene_tx=XM_019825377.2
A2 83503687 83503776
refGene_gn=CCDC146;refGene_tx=XM_003982693.5;refGene_gn=CCDC146;refGene_tx=XM_019825375.2;refGene_gn=CCDC146;refGene_tx=XM_019825376.2;refGene_gn=CCDC146;refGene_tx=XM_019825377.2
A2 83503865 83504030
refGene_gn=CCDC146;refGene_tx=XM_003982693.5;refGene_gn=CCDC146;refGene_tx=XM_019825375.2;refGene_gn=CCDC146;refGene_tx=XM_019825376.2;refGene_gn=CCDC146;refGene_tx=XM_019825377.2
A2 83506137 83506362

refGene_gn=CCDC146;refGene_tx=XM_003982693.5;refGene_gn=CCDC146;refGene_tx=XM_019825375.2;refGene_gn=CCDC146;refGene_tx=XM_019825376.2;refGene_gn=CCDC146;refGene_tx=XM_019825377.2
A2 83508156 83508385
refGene_gn=CCDC146;refGene_tx=XM_003982693.5;refGene_gn=CCDC146;refGene_tx=XM_019825375.2;refGene_gn=CCDC146;refGene_tx=XM_019825376.2;refGene_gn=CCDC146;refGene_tx=XM_019825377.2
A2 83513620 83513749
refGene_gn=CCDC146;refGene_tx=XM_003982693.5;refGene_gn=CCDC146;refGene_tx=XM_019825375.2;refGene_gn=CCDC146;refGene_tx=XM_019825376.2;refGene_gn=CCDC146;refGene_tx=XM_019825377.2
A2 83514527 83514665
refGene_gn=CCDC146;refGene_tx=XM_003982693.5;refGene_gn=CCDC146;refGene_tx=XM_019825375.2;refGene_gn=CCDC146;refGene_tx=XM_019825376.2;refGene_gn=CCDC146;refGene_tx=XM_019825377.2
A2 83517639 83517888
refGene_gn=CCDC146;refGene_tx=XM_003982693.5;refGene_gn=CCDC146;refGene_tx=XM_019825375.2;refGene_gn=CCDC146;refGene_tx=XM_019825376.2;refGene_gn=CCDC146;refGene_tx=XM_019825377.2
A2 83519493 83519697
refGene_gn=CCDC146;refGene_tx=XM_003982693.5;refGene_gn=CCDC146;refGene_tx=XM_019825375.2;refGene_gn=CCDC146;refGene_tx=XM_019825376.2;refGene_gn=CCDC146;refGene_tx=XM_019825377.2
A2 83522822 83522914
refGene_gn=GSAP;refGene_tx=XM_023250292.1;refGene_gn=GSAP;refGene_tx=XM_023250293.1;refGene_gn=GSAP;refGene_tx=XM_023250295.1;refGene_gn=GSAP;refGene_tx=XM_023250296.1
A2 83523342 83523442
refGene_gn=GSAP;refGene_tx=XM_023250292.1;refGene_gn=GSAP;refGene_tx=XM_023250293.1;refGene_gn=GSAP;refGene_tx=XM_023250295.1;refGene_gn=GSAP;refGene_tx=XM_023250296.1
A2 83523602 83523704
refGene_gn=GSAP;refGene_tx=XM_023250292.1;refGene_gn=GSAP;refGene_tx=XM_023250293.1;refGene_gn=GSAP;refGene_tx=XM_023250295.1;refGene_gn=GSAP;refGene_tx=XM_023250296.1;refGene_gn=GSAP;refGene_tx=XM_023250297.1
A2 83525996 83526058
refGene_gn=GSAP;refGene_tx=XM_023250292.1;refGene_gn=GSAP;refGene_tx=XM_023250293.1;refGene_gn=GSAP;refGene_tx=XM_023250295.1;refGene_gn=GSAP;refGene_tx=XM_023250296.1
A2 83526908 83527028
refGene_gn=GSAP;refGene_tx=XM_023250292.1;refGene_gn=GSAP;refGene_tx=XM_023250293.1;refGene_gn=GSAP;refGene_tx=XM_023250295.1
A2 83527114 83527134 refGene_gn=GSAP;refGene_tx=XM_023250298.1
A2 83532039 83532134
refGene_gn=GSAP;refGene_tx=XM_023250292.1;refGene_gn=GSAP;refGene_tx=XM_023250293.1;refGene_gn=GSAP;refGene_tx=XM_023250295.1;refGene_gn=GSAP;refGene_tx=XM_023250296.1;refGene_gn=GSAP;refGene_tx=XM_023250297.1;refGene_gn=GSAP;refGene_tx=XM_023250298.1
A2 83532822 83532893
refGene_gn=GSAP;refGene_tx=XM_023250292.1;refGene_gn=GSAP;refGene_tx=X

M_023250293.1;refGene_gn=GSAP;refGene_tx=XM_023250295.1;refGene_gn=GSAP;refGene_tx=XM_023250296.1;refGene_gn=GSAP;refGene_tx=XM_023250297.1;refGene_gn=GSAP;refGene_tx=XM_023250298.1
A2 83536598 83536694
refGene_gn=GSAP;refGene_tx=XM_023250292.1;refGene_gn=GSAP;refGene_tx=XM_023250293.1;refGene_gn=GSAP;refGene_tx=XM_023250295.1;refGene_gn=GSAP;refGene_tx=XM_023250296.1;refGene_gn=GSAP;refGene_tx=XM_023250297.1;refGene_gn=GSAP;refGene_tx=XM_023250298.1
A2 83540165 83540227
refGene_gn=GSAP;refGene_tx=XM_023250292.1;refGene_gn=GSAP;refGene_tx=XM_023250293.1;refGene_gn=GSAP;refGene_tx=XM_023250295.1;refGene_gn=GSAP;refGene_tx=XM_023250296.1;refGene_gn=GSAP;refGene_tx=XM_023250297.1;refGene_gn=GSAP;refGene_tx=XM_023250298.1
A2 83542537 83542569
refGene_gn=GSAP;refGene_tx=XM_023250292.1;refGene_gn=GSAP;refGene_tx=XM_023250293.1;refGene_gn=GSAP;refGene_tx=XM_023250295.1;refGene_gn=GSAP;refGene_tx=XM_023250296.1;refGene_gn=GSAP;refGene_tx=XM_023250297.1;refGene_gn=GSAP;refGene_tx=XM_023250298.1
A2 83543226 83543285
refGene_gn=GSAP;refGene_tx=XM_023250292.1;refGene_gn=GSAP;refGene_tx=XM_023250293.1;refGene_gn=GSAP;refGene_tx=XM_023250295.1;refGene_gn=GSAP;refGene_tx=XM_023250296.1;refGene_gn=GSAP;refGene_tx=XM_023250297.1;refGene_gn=GSAP;refGene_tx=XM_023250298.1
A2 83544140 83544269
refGene_gn=GSAP;refGene_tx=XM_023250292.1;refGene_gn=GSAP;refGene_tx=XM_023250293.1;refGene_gn=GSAP;refGene_tx=XM_023250295.1;refGene_gn=GSAP;refGene_tx=XM_023250296.1;refGene_gn=GSAP;refGene_tx=XM_023250297.1;refGene_gn=GSAP;refGene_tx=XM_023250298.1
A2 83554894 83554948
refGene_gn=GSAP;refGene_tx=XM_023250292.1;refGene_gn=GSAP;refGene_tx=XM_023250293.1;refGene_gn=GSAP;refGene_tx=XM_023250295.1;refGene_gn=GSAP;refGene_tx=XM_023250296.1;refGene_gn=GSAP;refGene_tx=XM_023250297.1;refGene_gn=GSAP;refGene_tx=XM_023250298.1
A2 83558171 83558254
refGene_gn=GSAP;refGene_tx=XM_023250292.1;refGene_gn=GSAP;refGene_tx=XM_023250293.1;refGene_gn=GSAP;refGene_tx=XM_023250295.1;refGene_gn=GSAP;refGene_tx=XM_023250296.1;refGene_gn=GSAP;refGene_tx=XM_023250297.1;refGene_gn=GSAP;refGene_tx=XM_023250298.1
A2 83558799 83558869
refGene_gn=GSAP;refGene_tx=XM_023250292.1;refGene_gn=GSAP;refGene_tx=XM_023250293.1;refGene_gn=GSAP;refGene_tx=XM_023250295.1;refGene_gn=GSAP;refGene_tx=XM_023250296.1;refGene_gn=GSAP;refGene_tx=XM_023250297.1;refGene_gn=GSAP;refGene_tx=XM_023250298.1
A2 83561180 83561398
refGene_gn=GSAP;refGene_tx=XM_023250292.1;refGene_gn=GSAP;refGene_tx=XM_023250293.1;refGene_gn=GSAP;refGene_tx=XM_023250295.1;refGene_gn=GSAP;refGene_tx=XM_023250296.1;refGene_gn=GSAP;refGene_tx=XM_023250297.1;refGene_gn=GSAP;refGene_tx=XM_023250298.1
A2 83561531 83561624
refGene_gn=GSAP;refGene_tx=XM_023250292.1;refGene_gn=GSAP;refGene_tx=X

M_023250293.1;refGene_gn=GSAP;refGene_tx=XM_023250295.1;refGene_gn=GSAP;refGene_tx=XM_023250296.1;refGene_gn=GSAP;refGene_tx=XM_023250297.1;refGene_gn=GSAP;refGene_tx=XM_023250298.1
A2 83565133 83565211
refGene_gn=GSAP;refGene_tx=XM_023250292.1;refGene_gn=GSAP;refGene_tx=XM_023250293.1;refGene_gn=GSAP;refGene_tx=XM_023250295.1;refGene_gn=GSAP;refGene_tx=XM_023250296.1;refGene_gn=GSAP;refGene_tx=XM_023250297.1;refGene_gn=GSAP;refGene_tx=XM_023250298.1
A2 83565881 83565986
refGene_gn=GSAP;refGene_tx=XM_023250292.1;refGene_gn=GSAP;refGene_tx=XM_023250293.1;refGene_gn=GSAP;refGene_tx=XM_023250296.1;refGene_gn=GSAP;refGene_tx=XM_023250297.1;refGene_gn=GSAP;refGene_tx=XM_023250298.1
A2 83573434 83573520
refGene_gn=GSAP;refGene_tx=XM_023250292.1;refGene_gn=GSAP;refGene_tx=XM_023250293.1;refGene_gn=GSAP;refGene_tx=XM_023250295.1;refGene_gn=GSAP;refGene_tx=XM_023250296.1;refGene_gn=GSAP;refGene_tx=XM_023250297.1;refGene_gn=GSAP;refGene_tx=XM_023250298.1
A2 83575981 83576025
refGene_gn=GSAP;refGene_tx=XM_023250292.1;refGene_gn=GSAP;refGene_tx=XM_023250293.1;refGene_gn=GSAP;refGene_tx=XM_023250295.1;refGene_gn=GSAP;refGene_tx=XM_023250296.1;refGene_gn=GSAP;refGene_tx=XM_023250297.1;refGene_gn=GSAP;refGene_tx=XM_023250298.1
A2 83576977 83577037
refGene_gn=GSAP;refGene_tx=XM_023250292.1;refGene_gn=GSAP;refGene_tx=XM_023250293.1;refGene_gn=GSAP;refGene_tx=XM_023250295.1;refGene_gn=GSAP;refGene_tx=XM_023250296.1;refGene_gn=GSAP;refGene_tx=XM_023250297.1;refGene_gn=GSAP;refGene_tx=XM_023250298.1
A2 83577286 83577391
refGene_gn=GSAP;refGene_tx=XM_023250292.1;refGene_gn=GSAP;refGene_tx=XM_023250293.1;refGene_gn=GSAP;refGene_tx=XM_023250295.1;refGene_gn=GSAP;refGene_tx=XM_023250296.1;refGene_gn=GSAP;refGene_tx=XM_023250297.1;refGene_gn=GSAP;refGene_tx=XM_023250298.1
A2 83584055 83584105
refGene_gn=GSAP;refGene_tx=XM_023250292.1;refGene_gn=GSAP;refGene_tx=XM_023250293.1;refGene_gn=GSAP;refGene_tx=XM_023250295.1;refGene_gn=GSAP;refGene_tx=XM_023250296.1;refGene_gn=GSAP;refGene_tx=XM_023250297.1;refGene_gn=GSAP;refGene_tx=XM_023250298.1
A2 83585294 83585364
refGene_gn=GSAP;refGene_tx=XM_023250292.1;refGene_gn=GSAP;refGene_tx=XM_023250293.1;refGene_gn=GSAP;refGene_tx=XM_023250295.1;refGene_gn=GSAP;refGene_tx=XM_023250296.1;refGene_gn=GSAP;refGene_tx=XM_023250297.1;refGene_gn=GSAP;refGene_tx=XM_023250298.1
A2 83589420 83589509
refGene_gn=GSAP;refGene_tx=XM_023250292.1;refGene_gn=GSAP;refGene_tx=XM_023250293.1;refGene_gn=GSAP;refGene_tx=XM_023250295.1;refGene_gn=GSAP;refGene_tx=XM_023250296.1;refGene_gn=GSAP;refGene_tx=XM_023250297.1;refGene_gn=GSAP;refGene_tx=XM_023250298.1
A2 83596181 83596235
refGene_gn=GSAP;refGene_tx=XM_023250292.1;refGene_gn=GSAP;refGene_tx=XM_023250293.1;refGene_gn=GSAP;refGene_tx=XM_023250295.1;refGene_gn=GSA

P;refGene_tx=XM_023250296.1;refGene_gn=GSAP;refGene_tx=XM_023250297.1;
refGene_gn=GSAP;refGene_tx=XM_023250298.1
A2 83596557 83596627
refGene_gn=GSAP;refGene_tx=XM_023250292.1;refGene_gn=GSAP;refGene_tx=X
M_023250293.1;refGene_gn=GSAP;refGene_tx=XM_023250295.1;refGene_gn=GSA
P;refGene_tx=XM_023250296.1;refGene_gn=GSAP;refGene_tx=XM_023250297.1;
refGene_gn=GSAP;refGene_tx=XM_023250298.1
A2 83598800 83598857
refGene_gn=GSAP;refGene_tx=XM_023250292.1;refGene_gn=GSAP;refGene_tx=X
M_023250293.1;refGene_gn=GSAP;refGene_tx=XM_023250295.1;refGene_gn=GSA
P;refGene_tx=XM_023250296.1;refGene_gn=GSAP;refGene_tx=XM_023250297.1;
refGene_gn=GSAP;refGene_tx=XM_023250298.1
A2 83599957 83600034
refGene_gn=GSAP;refGene_tx=XM_023250292.1;refGene_gn=GSAP;refGene_tx=X
M_023250293.1;refGene_gn=GSAP;refGene_tx=XM_023250295.1;refGene_gn=GSA
P;refGene_tx=XM_023250296.1;refGene_gn=GSAP;refGene_tx=XM_023250297.1;
refGene_gn=GSAP;refGene_tx=XM_023250298.1
A2 83610635 83610744
refGene_gn=GSAP;refGene_tx=XM_023250292.1;refGene_gn=GSAP;refGene_tx=X
M_023250293.1;refGene_gn=GSAP;refGene_tx=XM_023250295.1;refGene_gn=GSA
P;refGene_tx=XM_023250296.1;refGene_gn=GSAP;refGene_tx=XM_023250297.1;
refGene_gn=GSAP;refGene_tx=XM_023250298.1
A2 83727935 83728034
refGene_gn=PTPN12;refGene_tx=XM_011280344.3;refGene_gn=PTPN12;refGene_
tx=XM_019825379.2
A2 83768806 83768915
refGene_gn=PTPN12;refGene_tx=XM_011280344.3;refGene_gn=PTPN12;refGene_
tx=XM_019825379.2
A2 83777947 83777960 refGene_gn=PTPN12;refGene_tx=XM_006929139.4
A2 83781658 83781735
refGene_gn=PTPN12;refGene_tx=XM_006929139.4;refGene_gn=PTPN12;refGene_
tx=XM_011280344.3;refGene_gn=PTPN12;refGene_tx=XM_019825379.2
A2 83784490 83784586
refGene_gn=PTPN12;refGene_tx=XM_006929139.4;refGene_gn=PTPN12;refGene_
tx=XM_011280344.3;refGene_gn=PTPN12;refGene_tx=XM_019825379.2
A2 83785895 83785934
refGene_gn=PTPN12;refGene_tx=XM_006929139.4;refGene_gn=PTPN12;refGene_
tx=XM_011280344.3;refGene_gn=PTPN12;refGene_tx=XM_019825379.2
A2 83786792 83786864
refGene_gn=PTPN12;refGene_tx=XM_006929139.4;refGene_gn=PTPN12;refGene_
tx=XM_011280344.3;refGene_gn=PTPN12;refGene_tx=XM_019825379.2
A2 83790389 83790449
refGene_gn=PTPN12;refGene_tx=XM_006929139.4;refGene_gn=PTPN12;refGene_
tx=XM_011280344.3;refGene_gn=PTPN12;refGene_tx=XM_019825379.2
A2 83791842 83791985
refGene_gn=PTPN12;refGene_tx=XM_006929139.4;refGene_gn=PTPN12;refGene_
tx=XM_011280344.3;refGene_gn=PTPN12;refGene_tx=XM_019825379.2
A2 83795196 83795263
refGene_gn=PTPN12;refGene_tx=XM_006929139.4;refGene_gn=PTPN12;refGene_
tx=XM_011280344.3;refGene_gn=PTPN12;refGene_tx=XM_019825379.2

A2 83798300 83798378
refGene_gn=PTPN12;refGene_tx=XM_006929139.4;refGene_gn=PTPN12;refGene_tx=XM_011280344.3;refGene_gn=PTPN12;refGene_tx=XM_019825379.2
A2 83798487 83798586
refGene_gn=PTPN12;refGene_tx=XM_006929139.4;refGene_gn=PTPN12;refGene_tx=XM_011280344.3;refGene_gn=PTPN12;refGene_tx=XM_019825379.2
A2 83812933 83813019
refGene_gn=PTPN12;refGene_tx=XM_006929139.4;refGene_gn=PTPN12;refGene_tx=XM_011280344.3;refGene_gn=PTPN12;refGene_tx=XM_019825379.2
A2 83820834 83821811
refGene_gn=PTPN12;refGene_tx=XM_006929139.4;refGene_gn=PTPN12;refGene_tx=XM_011280344.3;refGene_gn=PTPN12;refGene_tx=XM_019825379.2
A2 83826127 83826205
refGene_gn=PTPN12;refGene_tx=XM_006929139.4;refGene_gn=PTPN12;refGene_tx=XM_011280344.3;refGene_gn=PTPN12;refGene_tx=XM_019825379.2
A2 83827048 83827116
refGene_gn=PTPN12;refGene_tx=XM_006929139.4;refGene_gn=PTPN12;refGene_tx=XM_011280344.3
A2 83828046 83828077
refGene_gn=PTPN12;refGene_tx=XM_006929139.4;refGene_gn=PTPN12;refGene_tx=XM_011280344.3
A2 83830953 83831061
refGene_gn=PTPN12;refGene_tx=XM_006929139.4;refGene_gn=PTPN12;refGene_tx=XM_011280344.3
A2 83831288 83831350
refGene_gn=PTPN12;refGene_tx=XM_006929139.4;refGene_gn=PTPN12;refGene_tx=XM_011280344.3
A2 83895217 83895782 refGene_gn=RSBN1L;refGene_tx=XM_003982695.5
A2 83927571 83927682 refGene_gn=RSBN1L;refGene_tx=XM_003982695.5
A2 83935900 83936541 refGene_gn=RSBN1L;refGene_tx=XM_003982695.5
A2 83952779 83952917 refGene_gn=RSBN1L;refGene_tx=XM_003982695.5
A2 83954676 83954819 refGene_gn=RSBN1L;refGene_tx=XM_003982695.5
A2 83957600 83957768 refGene_gn=RSBN1L;refGene_tx=XM_003982695.5
A2 83958885 83958994 refGene_gn=RSBN1L;refGene_tx=XM_003982695.5
A2 83959074 83959713 refGene_gn=RSBN1L;refGene_tx=XM_003982695.5
A2 83970574 83970976 refGene_gn=TMEM60;refGene_tx=XM_004001387.5
A2 84007809 84007854
refGene_gn=PHTF2;refGene_tx=XM_006929140.4;refGene_gn=PHTF2;refGene_tx=XM_019825381.2;refGene_gn=PHTF2;refGene_tx=XM_023250299.1;refGene_gn=PHTF2;refGene_tx=XM_023250300.1;refGene_gn=PHTF2;refGene_tx=XM_023250301.1
A2 84045655 84045712
refGene_gn=PHTF2;refGene_tx=XM_006929140.4;refGene_gn=PHTF2;refGene_tx=XM_019825381.2;refGene_gn=PHTF2;refGene_tx=XM_023250299.1;refGene_gn=PHTF2;refGene_tx=XM_023250300.1;refGene_gn=PHTF2;refGene_tx=XM_023250301.1
A2 84046065 84046077
refGene_gn=PHTF2;refGene_tx=XM_006929140.4;refGene_gn=PHTF2;refGene_tx=XM_019825381.2;refGene_gn=PHTF2;refGene_tx=XM_023250299.1;refGene_gn=PHTF2;refGene_tx=XM_023250301.1

A2 84053110 84053180
refGene_gn=PHTF2;refGene_tx=XM_006929140.4;refGene_gn=PHTF2;refGene_tx=XM_019825381.2;refGene_gn=PHTF2;refGene_tx=XM_023250299.1;refGene_gn=PHTF2;refGene_tx=XM_023250300.1;refGene_gn=PHTF2;refGene_tx=XM_023250301.1
A2 84053745 84053904
refGene_gn=PHTF2;refGene_tx=XM_006929140.4;refGene_gn=PHTF2;refGene_tx=XM_019825381.2;refGene_gn=PHTF2;refGene_tx=XM_023250299.1;refGene_gn=PHTF2;refGene_tx=XM_023250300.1;refGene_gn=PHTF2;refGene_tx=XM_023250301.1
A2 84061127 84061293
refGene_gn=PHTF2;refGene_tx=XM_006929140.4;refGene_gn=PHTF2;refGene_tx=XM_019825381.2;refGene_gn=PHTF2;refGene_tx=XM_023250299.1;refGene_gn=PHTF2;refGene_tx=XM_023250300.1;refGene_gn=PHTF2;refGene_tx=XM_023250301.1
A2 84062225 84062390
refGene_gn=PHTF2;refGene_tx=XM_006929140.4;refGene_gn=PHTF2;refGene_tx=XM_019825381.2;refGene_gn=PHTF2;refGene_tx=XM_023250299.1;refGene_gn=PHTF2;refGene_tx=XM_023250300.1;refGene_gn=PHTF2;refGene_tx=XM_023250301.1
A2 84072665 84072852
refGene_gn=PHTF2;refGene_tx=XM_006929140.4;refGene_gn=PHTF2;refGene_tx=XM_019825381.2;refGene_gn=PHTF2;refGene_tx=XM_023250299.1;refGene_gn=PHTF2;refGene_tx=XM_023250300.1;refGene_gn=PHTF2;refGene_tx=XM_023250301.1
A2 84074600 84074756
refGene_gn=PHTF2;refGene_tx=XM_006929140.4;refGene_gn=PHTF2;refGene_tx=XM_019825381.2;refGene_gn=PHTF2;refGene_tx=XM_023250299.1;refGene_gn=PHTF2;refGene_tx=XM_023250300.1;refGene_gn=PHTF2;refGene_tx=XM_023250301.1
A2 84081851 84082070
refGene_gn=PHTF2;refGene_tx=XM_006929140.4;refGene_gn=PHTF2;refGene_tx=XM_019825381.2;refGene_gn=PHTF2;refGene_tx=XM_023250299.1;refGene_gn=PHTF2;refGene_tx=XM_023250300.1
A2 84088494 84088623
refGene_gn=PHTF2;refGene_tx=XM_019825381.2;refGene_gn=PHTF2;refGene_tx=XM_023250299.1;refGene_gn=PHTF2;refGene_tx=XM_023250300.1;refGene_gn=PHTF2;refGene_tx=XM_023250301.1
A2 84090280 84090553
refGene_gn=PHTF2;refGene_tx=XM_019825381.2;refGene_gn=PHTF2;refGene_tx=XM_023250299.1;refGene_gn=PHTF2;refGene_tx=XM_023250300.1;refGene_gn=PHTF2;refGene_tx=XM_023250301.1
A2 84090782 84090914
refGene_gn=PHTF2;refGene_tx=XM_019825381.2;refGene_gn=PHTF2;refGene_tx=XM_023250299.1;refGene_gn=PHTF2;refGene_tx=XM_023250300.1;refGene_gn=PHTF2;refGene_tx=XM_023250301.1
A2 84093803 84093890
refGene_gn=PHTF2;refGene_tx=XM_019825381.2;refGene_gn=PHTF2;refGene_tx=XM_023250299.1;refGene_gn=PHTF2;refGene_tx=XM_023250300.1;refGene_gn=PHTF2;refGene_tx=XM_023250301.1

A2 84095521 84095539 refGene_gn=PHTF2;refGene_tx=XM_019825381.2
A2 84097244 84097400
refGene_gn=PHTF2;refGene_tx=XM_023250299.1;refGene_gn=PHTF2;refGene_tx=XM_023250300.1;refGene_gn=PHTF2;refGene_tx=XM_023250301.1
A2 84098284 84098380
refGene_gn=PHTF2;refGene_tx=XM_023250299.1;refGene_gn=PHTF2;refGene_tx=XM_023250300.1;refGene_gn=PHTF2;refGene_tx=XM_023250301.1
A2 84100524 84100650
refGene_gn=PHTF2;refGene_tx=XM_023250299.1;refGene_gn=PHTF2;refGene_tx=XM_023250300.1;refGene_gn=PHTF2;refGene_tx=XM_023250301.1
A2 84101706 84101727
refGene_gn=PHTF2;refGene_tx=XM_023250299.1;refGene_gn=PHTF2;refGene_tx=XM_023250300.1;refGene_gn=PHTF2;refGene_tx=XM_023250301.1
A2 84180727 84181401
refGene_gn=MAGI2;refGene_tx=XM_019825382.2;refGene_gn=MAGI2;refGene_tx=XM_019825384.2;refGene_gn=MAGI2;refGene_tx=XM_023250302.1;refGene_gn=MAGI2;refGene_tx=XM_023250303.1
A2 84241138 84241249
refGene_gn=MAGI2;refGene_tx=XM_019825382.2;refGene_gn=MAGI2;refGene_tx=XM_019825383.2;refGene_gn=MAGI2;refGene_tx=XM_019825384.2
A2 84251031 84251170
refGene_gn=MAGI2;refGene_tx=XM_019825382.2;refGene_gn=MAGI2;refGene_tx=XM_019825383.2;refGene_gn=MAGI2;refGene_tx=XM_019825384.2;refGene_gn=MAGI2;refGene_tx=XM_023250302.1;refGene_gn=MAGI2;refGene_tx=XM_023250303.1
A2 84295485 84295629
refGene_gn=MAGI2;refGene_tx=XM_019825382.2;refGene_gn=MAGI2;refGene_tx=XM_019825383.2;refGene_gn=MAGI2;refGene_tx=XM_019825384.2;refGene_gn=MAGI2;refGene_tx=XM_023250302.1;refGene_gn=MAGI2;refGene_tx=XM_023250303.1
A2 84296668 84296888
refGene_gn=MAGI2;refGene_tx=XM_019825382.2;refGene_gn=MAGI2;refGene_tx=XM_019825383.2;refGene_gn=MAGI2;refGene_tx=XM_019825384.2;refGene_gn=MAGI2;refGene_tx=XM_023250302.1;refGene_gn=MAGI2;refGene_tx=XM_023250303.1
A2 84302012 84302166
refGene_gn=MAGI2;refGene_tx=XM_019825382.2;refGene_gn=MAGI2;refGene_tx=XM_019825383.2;refGene_gn=MAGI2;refGene_tx=XM_019825384.2;refGene_gn=MAGI2;refGene_tx=XM_023250302.1;refGene_gn=MAGI2;refGene_tx=XM_023250303.1
A2 84304106 84304292
refGene_gn=MAGI2;refGene_tx=XM_019825382.2;refGene_gn=MAGI2;refGene_tx=XM_019825383.2;refGene_gn=MAGI2;refGene_tx=XM_019825384.2;refGene_gn=MAGI2;refGene_tx=XM_023250302.1;refGene_gn=MAGI2;refGene_tx=XM_023250303.1
A2 84329291 84329540
refGene_gn=MAGI2;refGene_tx=XM_019825382.2;refGene_gn=MAGI2;refGene_tx=XM_019825383.2;refGene_gn=MAGI2;refGene_tx=XM_019825384.2;refGene_gn=MAGI2;refGene_tx=XM_023250302.1;refGene_gn=MAGI2;refGene_tx=XM_023250303.1

A2 84335909 84336102
refGene_gn=MAGI2;refGene_tx=XM_019825382.2;refGene_gn=MAGI2;refGene_tx=XM_019825383.2;refGene_gn=MAGI2;refGene_tx=XM_019825384.2;refGene_gn=MAGI2;refGene_tx=XM_023250302.1;refGene_gn=MAGI2;refGene_tx=XM_023250303.1
A2 84345050 84345142
refGene_gn=MAGI2;refGene_tx=XM_019825382.2;refGene_gn=MAGI2;refGene_tx=XM_019825383.2;refGene_gn=MAGI2;refGene_tx=XM_019825384.2;refGene_gn=MAGI2;refGene_tx=XM_023250302.1;refGene_gn=MAGI2;refGene_tx=XM_023250303.1
A2 84352176 84352218
refGene_gn=MAGI2;refGene_tx=XM_019825382.2;refGene_gn=MAGI2;refGene_tx=XM_019825383.2;refGene_gn=MAGI2;refGene_tx=XM_019825384.2;refGene_gn=MAGI2;refGene_tx=XM_023250302.1
A2 84361032 84361222
refGene_gn=MAGI2;refGene_tx=XM_019825382.2;refGene_gn=MAGI2;refGene_tx=XM_019825383.2;refGene_gn=MAGI2;refGene_tx=XM_019825384.2;refGene_gn=MAGI2;refGene_tx=XM_023250302.1;refGene_gn=MAGI2;refGene_tx=XM_023250303.1
A2 84367549 84367581
refGene_gn=MAGI2;refGene_tx=XM_019825382.2;refGene_gn=MAGI2;refGene_tx=XM_019825383.2;refGene_gn=MAGI2;refGene_tx=XM_019825384.2;refGene_gn=MAGI2;refGene_tx=XM_023250302.1;refGene_gn=MAGI2;refGene_tx=XM_023250303.1
A2 84421441 84422080
refGene_gn=MAGI2;refGene_tx=XM_019825382.2;refGene_gn=MAGI2;refGene_tx=XM_019825383.2;refGene_gn=MAGI2;refGene_tx=XM_019825384.2;refGene_gn=MAGI2;refGene_tx=XM_023250302.1;refGene_gn=MAGI2;refGene_tx=XM_023250303.1
A2 84488893 84489076
refGene_gn=MAGI2;refGene_tx=XM_019825382.2;refGene_gn=MAGI2;refGene_tx=XM_019825383.2;refGene_gn=MAGI2;refGene_tx=XM_019825384.2;refGene_gn=MAGI2;refGene_tx=XM_023250302.1;refGene_gn=MAGI2;refGene_tx=XM_023250303.1
A2 84490855 84490984
refGene_gn=MAGI2;refGene_tx=XM_019825382.2;refGene_gn=MAGI2;refGene_tx=XM_019825383.2
A2 84491332 84491454
refGene_gn=MAGI2;refGene_tx=XM_019825382.2;refGene_gn=MAGI2;refGene_tx=XM_019825383.2;refGene_gn=MAGI2;refGene_tx=XM_019825384.2;refGene_gn=MAGI2;refGene_tx=XM_023250302.1;refGene_gn=MAGI2;refGene_tx=XM_023250303.1
A2 84515064 84515122
refGene_gn=MAGI2;refGene_tx=XM_019825382.2;refGene_gn=MAGI2;refGene_tx=XM_019825383.2;refGene_gn=MAGI2;refGene_tx=XM_019825384.2;refGene_gn=MAGI2;refGene_tx=XM_023250302.1;refGene_gn=MAGI2;refGene_tx=XM_023250303.1
A2 84645164 84645244
refGene_gn=MAGI2;refGene_tx=XM_019825382.2;refGene_gn=MAGI2;refGene_tx=XM_019825383.2;refGene_gn=MAGI2;refGene_tx=XM_019825384.2;refGene_gn=

MAGI2;refGene_tx=XM_023250302.1;refGene_gn=MAGI2;refGene_tx=XM_0232503
03.1
A2	84657971	84658182
refGene_gn=MAGI2;refGene_tx=XM_019825382.2;refGene_gn=MAGI2;refGene_tx
=XM_019825383.2;refGene_gn=MAGI2;refGene_tx=XM_019825384.2;refGene_gn=
MAGI2;refGene_tx=XM_023250302.1;refGene_gn=MAGI2;refGene_tx=XM_0232503
03.1
A2	84677736	84677949
refGene_gn=MAGI2;refGene_tx=XM_019825382.2;refGene_gn=MAGI2;refGene_tx
=XM_019825383.2;refGene_gn=MAGI2;refGene_tx=XM_019825384.2;refGene_gn=
MAGI2;refGene_tx=XM_023250302.1;refGene_gn=MAGI2;refGene_tx=XM_0232503
03.1
A2	84776807	84776927
refGene_gn=MAGI2;refGene_tx=XM_019825382.2;refGene_gn=MAGI2;refGene_tx
=XM_019825383.2;refGene_gn=MAGI2;refGene_tx=XM_019825384.2;refGene_gn=
MAGI2;refGene_tx=XM_023250302.1;refGene_gn=MAGI2;refGene_tx=XM_0232503
03.1
A2	85143179	85143296
refGene_gn=MAGI2;refGene_tx=XM_019825382.2;refGene_gn=MAGI2;refGene_tx
=XM_019825383.2;refGene_gn=MAGI2;refGene_tx=XM_019825384.2;refGene_gn=
MAGI2;refGene_tx=XM_023250302.1;refGene_gn=MAGI2;refGene_tx=XM_0232503
03.1
A2	85536978	85537279
refGene_gn=MAGI2;refGene_tx=XM_019825382.2;refGene_gn=MAGI2;refGene_tx
=XM_019825383.2;refGene_gn=MAGI2;refGene_tx=XM_019825384.2;refGene_gn=
MAGI2;refGene_tx=XM_023250302.1;refGene_gn=MAGI2;refGene_tx=XM_0232503
03.1
A2	85937150	85937586
refGene_gn=LOC111557341;refGene_tx=XM_023242045.1
A2	85937587	85938204
refGene_gn=LOC111557341;refGene_tx=XM_023242045.1
A2	85938417	85938567
refGene_gn=LOC111557341;refGene_tx=XM_023242045.1
A2	85938700	85938952
refGene_gn=LOC111557341;refGene_tx=XM_023242045.1
A2	86170510	86170628
refGene_gn=GNAI1;refGene_tx=XM_011280345.2;refGene_gn=GNAI1;refGene_tx
=XM_023250305.1
A2	86226974	86227017
refGene_gn=GNAI1;refGene_tx=XM_011280345.2;refGene_gn=GNAI1;refGene_tx
=XM_019825386.1;refGene_gn=GNAI1;refGene_tx=XM_023250305.1
A2	86227117	86227259
refGene_gn=GNAI1;refGene_tx=XM_011280345.2;refGene_gn=GNAI1;refGene_tx
=XM_019825386.1;refGene_gn=GNAI1;refGene_tx=XM_023250305.1
A2	86236740	86236898
refGene_gn=GNAI1;refGene_tx=XM_011280345.2;refGene_gn=GNAI1;refGene_tx
=XM_019825386.1;refGene_gn=GNAI1;refGene_tx=XM_023250305.1
A2	86240641	86240770
refGene_gn=GNAI1;refGene_tx=XM_011280345.2;refGene_gn=GNAI1;refGene_tx
=XM_019825386.1;refGene_gn=GNAI1;refGene_tx=XM_023250305.1

A2 86247175 86247305
refGene_gn=GNAI1;refGene_tx=XM_011280345.2;refGene_gn=GNAI1;refGene_tx=XM_019825386.1;refGene_gn=GNAI1;refGene_tx=XM_023250305.1
A2 86248725 86248879
refGene_gn=GNAI1;refGene_tx=XM_011280345.2;refGene_gn=GNAI1;refGene_tx=XM_019825386.1;refGene_gn=GNAI1;refGene_tx=XM_023250305.1
A2 86254779 86254970
refGene_gn=GNAI1;refGene_tx=XM_011280345.2;refGene_gn=GNAI1;refGene_tx=XM_019825386.1;refGene_gn=GNAI1;refGene_tx=XM_023250305.1
A2 86513703 86513894 refGene_gn=GNAT3;refGene_tx=XM_003982705.4
A2 86516947 86517101 refGene_gn=GNAT3;refGene_tx=XM_003982705.4
A2 86517318 86517448 refGene_gn=GNAT3;refGene_tx=XM_003982705.4
A2 86529424 86529553 refGene_gn=GNAT3;refGene_tx=XM_003982705.4
A2 86535147 86535305 refGene_gn=GNAT3;refGene_tx=XM_003982705.4
A2 86543250 86543392 refGene_gn=GNAT3;refGene_tx=XM_003982705.4
A2 86549424 86549467 refGene_gn=GNAT3;refGene_tx=XM_003982705.4
A2 86573734 86573852 refGene_gn=GNAT3;refGene_tx=XM_003982705.4
A2 86698587 86698707
refGene_gn=CD36;refGene_tx=XM_003982706.4;refGene_gn=CD36;refGene_tx=XM_011280346.3;refGene_gn=CD36;refGene_tx=XM_011280347.3;refGene_gn=CD36;refGene_tx=XM_019825387.2;refGene_gn=CD36;refGene_tx=XM_019825388.2
A2 86708205 86708366
refGene_gn=CD36;refGene_tx=XM_003982706.4;refGene_gn=CD36;refGene_tx=XM_011280346.3;refGene_gn=CD36;refGene_tx=XM_011280347.3;refGene_gn=CD36;refGene_tx=XM_019825387.2;refGene_gn=CD36;refGene_tx=XM_019825388.2
A2 86712957 86713105
refGene_gn=CD36;refGene_tx=XM_003982706.4;refGene_gn=CD36;refGene_tx=XM_011280346.3;refGene_gn=CD36;refGene_tx=XM_011280347.3;refGene_gn=CD36;refGene_tx=XM_019825387.2;refGene_gn=CD36;refGene_tx=XM_019825388.2
A2 86713544 86713724
refGene_gn=CD36;refGene_tx=XM_003982706.4;refGene_gn=CD36;refGene_tx=XM_011280346.3;refGene_gn=CD36;refGene_tx=XM_011280347.3;refGene_gn=CD36;refGene_tx=XM_019825387.2;refGene_gn=CD36;refGene_tx=XM_019825388.2
A2 86715035 86715127
refGene_gn=CD36;refGene_tx=XM_003982706.4;refGene_gn=CD36;refGene_tx=XM_011280346.3;refGene_gn=CD36;refGene_tx=XM_011280347.3;refGene_gn=CD36;refGene_tx=XM_019825387.2;refGene_gn=CD36;refGene_tx=XM_019825388.2
A2 86717439 86717486
refGene_gn=CD36;refGene_tx=XM_003982706.4;refGene_gn=CD36;refGene_tx=XM_011280346.3;refGene_gn=CD36;refGene_tx=XM_011280347.3;refGene_gn=CD36;refGene_tx=XM_019825387.2;refGene_gn=CD36;refGene_tx=XM_019825388.2
A2 86720113 86720183
refGene_gn=CD36;refGene_tx=XM_003982706.4;refGene_gn=CD36;refGene_tx=XM_011280346.3;refGene_gn=CD36;refGene_tx=XM_011280347.3;refGene_gn=CD36;refGene_tx=XM_019825387.2;refGene_gn=CD36;refGene_tx=XM_019825388.2
A2 86721549 86721737
refGene_gn=CD36;refGene_tx=XM_003982706.4;refGene_gn=CD36;refGene_tx=XM_011280346.3;refGene_gn=CD36;refGene_tx=XM_011280347.3;refGene_gn=CD36;refGene_tx=XM_019825387.2;refGene_gn=CD36;refGene_tx=XM_019825388.2
A2 86724259 86724378

refGene_gn=CD36;refGene_tx=XM_003982706.4;refGene_gn=CD36;refGene_tx=XM_011280346.3;refGene_gn=CD36;refGene_tx=XM_011280347.3;refGene_gn=CD36;refGene_tx=XM_019825387.2;refGene_gn=CD36;refGene_tx=XM_019825388.2
A2 86725394 86725468
refGene_gn=CD36;refGene_tx=XM_003982706.4;refGene_gn=CD36;refGene_tx=XM_011280346.3;refGene_gn=CD36;refGene_tx=XM_011280347.3;refGene_gn=CD36;refGene_tx=XM_019825387.2;refGene_gn=CD36;refGene_tx=XM_019825388.2
A2 86726055 86726110
refGene_gn=CD36;refGene_tx=XM_003982706.4;refGene_gn=CD36;refGene_tx=XM_011280346.3;refGene_gn=CD36;refGene_tx=XM_011280347.3;refGene_gn=CD36;refGene_tx=XM_019825387.2;refGene_gn=CD36;refGene_tx=XM_019825388.2
A2 86726907 86727072
refGene_gn=CD36;refGene_tx=XM_003982706.4;refGene_gn=CD36;refGene_tx=XM_011280346.3;refGene_gn=CD36;refGene_tx=XM_011280347.3;refGene_gn=CD36;refGene_tx=XM_019825387.2;refGene_gn=CD36;refGene_tx=XM_019825388.2
A2 86820058 86820472 refGene_gn=SEMA3C;refGene_tx=XM_023250307.1
A2 86824827 86824958 refGene_gn=SEMA3C;refGene_tx=XM_023250307.1
A2 86827123 86827191 refGene_gn=SEMA3C;refGene_tx=XM_023250307.1
A2 86835761 86835919 refGene_gn=SEMA3C;refGene_tx=XM_023250307.1
A2 86839059 86839101 refGene_gn=SEMA3C;refGene_tx=XM_023250307.1
A2 86842510 86842599 refGene_gn=SEMA3C;refGene_tx=XM_023250307.1
A2 86869343 86869566 refGene_gn=SEMA3C;refGene_tx=XM_023250307.1
A2 86878753 86878898 refGene_gn=SEMA3C;refGene_tx=XM_023250307.1
A2 86880664 86880734 refGene_gn=SEMA3C;refGene_tx=XM_023250307.1
A2 86888720 86888835 refGene_gn=SEMA3C;refGene_tx=XM_023250307.1
A2 86890364 86890507 refGene_gn=SEMA3C;refGene_tx=XM_023250307.1
A2 86892320 86892440 refGene_gn=SEMA3C;refGene_tx=XM_023250307.1
A2 86896371 86896454 refGene_gn=SEMA3C;refGene_tx=XM_023250307.1
A2 86898986 86898995 refGene_gn=SEMA3C;refGene_tx=XM_023250307.1
A2 86902894 86902965 refGene_gn=SEMA3C;refGene_tx=XM_023250307.1
A2 86902966 86903014 refGene_gn=SEMA3C;refGene_tx=XM_023250307.1
A2 86912561 86912624 refGene_gn=SEMA3C;refGene_tx=XM_023250307.1
A2 86913981 86914142 refGene_gn=SEMA3C;refGene_tx=XM_023250307.1
A2 86998686 86998789 refGene_gn=SEMA3C;refGene_tx=XM_023250307.1
A2 87723934 87724111
refGene_gn=HGF;refGene_tx=NM_001009830.1;refGene_gn=HGF;refGene_tx=XM_006929143.3;refGene_gn=HGF;refGene_tx=XM_019814912.1
A2 87731712 87731858
refGene_gn=HGF;refGene_tx=NM_001009830.1;refGene_gn=HGF;refGene_tx=XM_006929143.3;refGene_gn=HGF;refGene_tx=XM_019814912.1
A2 87731978 87732085
refGene_gn=HGF;refGene_tx=NM_001009830.1;refGene_gn=HGF;refGene_tx=XM_006929143.3;refGene_gn=HGF;refGene_tx=XM_019814912.1
A2 87732590 87732737
refGene_gn=HGF;refGene_tx=NM_001009830.1;refGene_gn=HGF;refGene_tx=XM_006929143.3;refGene_gn=HGF;refGene_tx=XM_019814912.1
A2 87733588 87733663
refGene_gn=HGF;refGene_tx=NM_001009830.1;refGene_gn=HGF;refGene_tx=XM_006929143.3;refGene_gn=HGF;refGene_tx=XM_019814912.1
A2 87736441 87736538

refGene_gn=HGF;refGene_tx=NM_001009830.1;refGene_gn=HGF;refGene_tx=XM_006929143.3;refGene_gn=HGF;refGene_tx=XM_019814912.1
A2	87737748	87737787
refGene_gn=HGF;refGene_tx=NM_001009830.1;refGene_gn=HGF;refGene_tx=XM_006929143.3;refGene_gn=HGF;refGene_tx=XM_019814912.1
A2	87743532	87743666
refGene_gn=HGF;refGene_tx=NM_001009830.1;refGene_gn=HGF;refGene_tx=XM_006929143.3;refGene_gn=HGF;refGene_tx=XM_019814912.1
A2	87746702	87746805
refGene_gn=HGF;refGene_tx=NM_001009830.1;refGene_gn=HGF;refGene_tx=XM_006929143.3;refGene_gn=HGF;refGene_tx=XM_019814912.1
A2	87755188	87755316
refGene_gn=HGF;refGene_tx=NM_001009830.1;refGene_gn=HGF;refGene_tx=XM_006929143.3;refGene_gn=HGF;refGene_tx=XM_019814912.1
A2	87757903	87758078
refGene_gn=HGF;refGene_tx=NM_001009830.1;refGene_gn=HGF;refGene_tx=XM_006929143.3;refGene_gn=HGF;refGene_tx=XM_019814912.1
A2	87774132	87774251
refGene_gn=HGF;refGene_tx=NM_001009830.1;refGene_gn=HGF;refGene_tx=XM_006929143.3;refGene_gn=HGF;refGene_tx=XM_019814912.1
A2	87775618	87775739
refGene_gn=HGF;refGene_tx=NM_001009830.1;refGene_gn=HGF;refGene_tx=XM_006929143.3;refGene_gn=HGF;refGene_tx=XM_019814912.1
A2	87781465	87781608
refGene_gn=HGF;refGene_tx=NM_001009830.1;refGene_gn=HGF;refGene_tx=XM_006929143.3;refGene_gn=HGF;refGene_tx=XM_019814912.1
A2	87786465	87786580
refGene_gn=HGF;refGene_tx=NM_001009830.1;refGene_gn=HGF;refGene_tx=XM_006929143.3;refGene_gn=HGF;refGene_tx=XM_019814912.1
A2	87788058	87788171
refGene_gn=HGF;refGene_tx=NM_001009830.1;refGene_gn=HGF;refGene_tx=XM_006929143.3;refGene_gn=HGF;refGene_tx=XM_019814912.1
A2	87792583	87792749
refGene_gn=HGF;refGene_tx=NM_001009830.1;refGene_gn=HGF;refGene_tx=XM_006929143.3;refGene_gn=HGF;refGene_tx=XM_019814912.1
A2	87799396	87799478
refGene_gn=HGF;refGene_tx=NM_001009830.1;refGene_gn=HGF;refGene_tx=XM_006929143.3;refGene_gn=HGF;refGene_tx=XM_019814912.1
A2	87967591	87967708
refGene_gn=CACNA2D1;refGene_tx=XM_011280348.3;refGene_gn=CACNA2D1;refGene_tx=XM_011280349.3;refGene_gn=CACNA2D1;refGene_tx=XM_011280350.3;refGene_gn=CACNA2D1;refGene_tx=XM_011280351.3;refGene_gn=CACNA2D1;refGene_tx=XM_011280352.3;refGene_gn=CACNA2D1;refGene_tx=XM_011280353.3;refGene_gn=CACNA2D1;refGene_tx=XM_019825390.2;refGene_gn=CACNA2D1;refGene_tx=XM_019825391.2
A2	87985795	87985878
refGene_gn=CACNA2D1;refGene_tx=XM_011280348.3;refGene_gn=CACNA2D1;refGene_tx=XM_011280349.3;refGene_gn=CACNA2D1;refGene_tx=XM_011280350.3;refGene_gn=CACNA2D1;refGene_tx=XM_011280351.3;refGene_gn=CACNA2D1;refGene_tx=XM_011280352.3;refGene_gn=CACNA2D1;refGene_tx=XM_011280353.3;refG

ene_gn=CACNA2D1;refGene_tx=XM_019825390.2;refGene_gn=CACNA2D1;refGene_tx=XM_019825391.2
A2 87986912 87987022
refGene_gn=CACNA2D1;refGene_tx=XM_011280348.3;refGene_gn=CACNA2D1;refGene_tx=XM_011280349.3;refGene_gn=CACNA2D1;refGene_tx=XM_011280350.3;refGene_gn=CACNA2D1;refGene_tx=XM_011280351.3;refGene_gn=CACNA2D1;refGene_tx=XM_011280352.3;refGene_gn=CACNA2D1;refGene_tx=XM_011280353.3;refGene_gn=CACNA2D1;refGene_tx=XM_019825390.2;refGene_gn=CACNA2D1;refGene_tx=XM_019825391.2
A2 87988966 87989096
refGene_gn=CACNA2D1;refGene_tx=XM_011280348.3;refGene_gn=CACNA2D1;refGene_tx=XM_011280349.3;refGene_gn=CACNA2D1;refGene_tx=XM_011280350.3;refGene_gn=CACNA2D1;refGene_tx=XM_011280351.3;refGene_gn=CACNA2D1;refGene_tx=XM_011280352.3;refGene_gn=CACNA2D1;refGene_tx=XM_011280353.3;refGene_gn=CACNA2D1;refGene_tx=XM_019825390.2;refGene_gn=CACNA2D1;refGene_tx=XM_019825391.2
A2 87989401 87989457
refGene_gn=CACNA2D1;refGene_tx=XM_011280348.3;refGene_gn=CACNA2D1;refGene_tx=XM_011280349.3;refGene_gn=CACNA2D1;refGene_tx=XM_011280350.3;refGene_gn=CACNA2D1;refGene_tx=XM_011280351.3;refGene_gn=CACNA2D1;refGene_tx=XM_011280352.3;refGene_gn=CACNA2D1;refGene_tx=XM_011280353.3;refGene_gn=CACNA2D1;refGene_tx=XM_019825390.2;refGene_gn=CACNA2D1;refGene_tx=XM_019825391.2
A2 87990990 87991043
refGene_gn=CACNA2D1;refGene_tx=XM_011280348.3;refGene_gn=CACNA2D1;refGene_tx=XM_011280349.3;refGene_gn=CACNA2D1;refGene_tx=XM_011280350.3;refGene_gn=CACNA2D1;refGene_tx=XM_011280351.3;refGene_gn=CACNA2D1;refGene_tx=XM_011280352.3;refGene_gn=CACNA2D1;refGene_tx=XM_011280353.3;refGene_gn=CACNA2D1;refGene_tx=XM_019825390.2;refGene_gn=CACNA2D1;refGene_tx=XM_019825391.2
A2 87991126 87991279
refGene_gn=CACNA2D1;refGene_tx=XM_011280348.3;refGene_gn=CACNA2D1;refGene_tx=XM_011280349.3;refGene_gn=CACNA2D1;refGene_tx=XM_011280350.3;refGene_gn=CACNA2D1;refGene_tx=XM_011280351.3;refGene_gn=CACNA2D1;refGene_tx=XM_011280352.3;refGene_gn=CACNA2D1;refGene_tx=XM_011280353.3;refGene_gn=CACNA2D1;refGene_tx=XM_019825390.2;refGene_gn=CACNA2D1;refGene_tx=XM_019825391.2
A2 87992332 87992404
refGene_gn=CACNA2D1;refGene_tx=XM_011280348.3;refGene_gn=CACNA2D1;refGene_tx=XM_011280349.3;refGene_gn=CACNA2D1;refGene_tx=XM_011280350.3;refGene_gn=CACNA2D1;refGene_tx=XM_011280351.3;refGene_gn=CACNA2D1;refGene_tx=XM_011280352.3;refGene_gn=CACNA2D1;refGene_tx=XM_011280353.3;refGene_gn=CACNA2D1;refGene_tx=XM_019825390.2;refGene_gn=CACNA2D1;refGene_tx=XM_019825391.2
A2 87994541 87994580
refGene_gn=CACNA2D1;refGene_tx=XM_011280348.3;refGene_gn=CACNA2D1;refGene_tx=XM_011280349.3;refGene_gn=CACNA2D1;refGene_tx=XM_011280350.3;refGene_gn=CACNA2D1;refGene_tx=XM_011280351.3;refGene_gn=CACNA2D1;refGene_tx=XM_011280352.3;refGene_gn=CACNA2D1;refGene_tx=XM_011280353.3;refGene_gn=CACNA2D1;refGene_tx=XM_019825390.2;refGene_gn=CACNA2D1;refGene_

tx=XM_019825391.2
A2 87994962 87995030
refGene_gn=CACNA2D1;refGene_tx=XM_011280348.3;refGene_gn=CACNA2D1;refGene_tx=XM_011280349.3;refGene_gn=CACNA2D1;refGene_tx=XM_011280350.3;refGene_gn=CACNA2D1;refGene_tx=XM_011280351.3;refGene_gn=CACNA2D1;refGene_tx=XM_011280352.3;refGene_gn=CACNA2D1;refGene_tx=XM_011280353.3;refGene_gn=CACNA2D1;refGene_tx=XM_019825390.2;refGene_gn=CACNA2D1;refGene_tx=XM_019825391.2
A2 87996248 87996335
refGene_gn=CACNA2D1;refGene_tx=XM_011280348.3;refGene_gn=CACNA2D1;refGene_tx=XM_011280349.3;refGene_gn=CACNA2D1;refGene_tx=XM_011280350.3;refGene_gn=CACNA2D1;refGene_tx=XM_011280351.3;refGene_gn=CACNA2D1;refGene_tx=XM_011280352.3;refGene_gn=CACNA2D1;refGene_tx=XM_011280353.3;refGene_gn=CACNA2D1;refGene_tx=XM_019825390.2;refGene_gn=CACNA2D1;refGene_tx=XM_019825391.2
A2 87997186 87997290
refGene_gn=CACNA2D1;refGene_tx=XM_011280348.3;refGene_gn=CACNA2D1;refGene_tx=XM_011280349.3;refGene_gn=CACNA2D1;refGene_tx=XM_011280350.3;refGene_gn=CACNA2D1;refGene_tx=XM_011280351.3;refGene_gn=CACNA2D1;refGene_tx=XM_011280352.3;refGene_gn=CACNA2D1;refGene_tx=XM_011280353.3;refGene_gn=CACNA2D1;refGene_tx=XM_019825390.2;refGene_gn=CACNA2D1;refGene_tx=XM_019825391.2
A2 87997890 87997953
refGene_gn=CACNA2D1;refGene_tx=XM_011280348.3;refGene_gn=CACNA2D1;refGene_tx=XM_011280349.3;refGene_gn=CACNA2D1;refGene_tx=XM_011280350.3;refGene_gn=CACNA2D1;refGene_tx=XM_011280351.3;refGene_gn=CACNA2D1;refGene_tx=XM_011280352.3;refGene_gn=CACNA2D1;refGene_tx=XM_011280353.3;refGene_gn=CACNA2D1;refGene_tx=XM_019825390.2;refGene_gn=CACNA2D1;refGene_tx=XM_019825391.2
A2 87999161 87999249
refGene_gn=CACNA2D1;refGene_tx=XM_011280348.3;refGene_gn=CACNA2D1;refGene_tx=XM_011280349.3;refGene_gn=CACNA2D1;refGene_tx=XM_011280350.3;refGene_gn=CACNA2D1;refGene_tx=XM_011280351.3;refGene_gn=CACNA2D1;refGene_tx=XM_011280352.3;refGene_gn=CACNA2D1;refGene_tx=XM_011280353.3;refGene_gn=CACNA2D1;refGene_tx=XM_019825390.2;refGene_gn=CACNA2D1;refGene_tx=XM_019825391.2
A2 88002264 88002362
refGene_gn=CACNA2D1;refGene_tx=XM_011280348.3;refGene_gn=CACNA2D1;refGene_tx=XM_011280349.3;refGene_gn=CACNA2D1;refGene_tx=XM_011280350.3;refGene_gn=CACNA2D1;refGene_tx=XM_011280351.3;refGene_gn=CACNA2D1;refGene_tx=XM_011280352.3;refGene_gn=CACNA2D1;refGene_tx=XM_011280353.3;refGene_gn=CACNA2D1;refGene_tx=XM_019825390.2;refGene_gn=CACNA2D1;refGene_tx=XM_019825391.2
A2 88008852 88008913
refGene_gn=CACNA2D1;refGene_tx=XM_011280348.3;refGene_gn=CACNA2D1;refGene_tx=XM_011280349.3;refGene_gn=CACNA2D1;refGene_tx=XM_011280350.3;refGene_gn=CACNA2D1;refGene_tx=XM_011280351.3;refGene_gn=CACNA2D1;refGene_tx=XM_011280352.3;refGene_gn=CACNA2D1;refGene_tx=XM_011280353.3;refGene_gn=CACNA2D1;refGene_tx=XM_019825390.2;refGene_gn=CACNA2D1;refGene_tx=XM_019825391.2;refGene_gn=CACNA2D1;refGene_tx=XM_019825392.2

A2 88009591 88009612
refGene_gn=CACNA2D1;refGene_tx=XM_011280348.3;refGene_gn=CACNA2D1;refGene_tx=XM_011280350.3;refGene_gn=CACNA2D1;refGene_tx=XM_011280351.3;refGene_gn=CACNA2D1;refGene_tx=XM_019825390.2;refGene_gn=CACNA2D1;refGene_tx=XM_019825391.2
A2 88010871 88010948
refGene_gn=CACNA2D1;refGene_tx=XM_011280348.3;refGene_gn=CACNA2D1;refGene_tx=XM_011280349.3;refGene_gn=CACNA2D1;refGene_tx=XM_011280350.3;refGene_gn=CACNA2D1;refGene_tx=XM_011280351.3;refGene_gn=CACNA2D1;refGene_tx=XM_011280352.3;refGene_gn=CACNA2D1;refGene_tx=XM_011280353.3;refGene_gn=CACNA2D1;refGene_tx=XM_019825390.2;refGene_gn=CACNA2D1;refGene_tx=XM_019825391.2
A2 88014846 88014908
refGene_gn=CACNA2D1;refGene_tx=XM_011280348.3;refGene_gn=CACNA2D1;refGene_tx=XM_011280349.3;refGene_gn=CACNA2D1;refGene_tx=XM_011280350.3;refGene_gn=CACNA2D1;refGene_tx=XM_011280351.3;refGene_gn=CACNA2D1;refGene_tx=XM_011280352.3;refGene_gn=CACNA2D1;refGene_tx=XM_011280353.3;refGene_gn=CACNA2D1;refGene_tx=XM_019825390.2;refGene_gn=CACNA2D1;refGene_tx=XM_019825391.2;refGene_gn=CACNA2D1;refGene_tx=XM_019825392.2
A2 88017114 88017186
refGene_gn=CACNA2D1;refGene_tx=XM_011280348.3;refGene_gn=CACNA2D1;refGene_tx=XM_011280349.3;refGene_gn=CACNA2D1;refGene_tx=XM_011280350.3;refGene_gn=CACNA2D1;refGene_tx=XM_011280351.3;refGene_gn=CACNA2D1;refGene_tx=XM_011280352.3;refGene_gn=CACNA2D1;refGene_tx=XM_011280353.3;refGene_gn=CACNA2D1;refGene_tx=XM_019825390.2;refGene_gn=CACNA2D1;refGene_tx=XM_019825391.2;refGene_gn=CACNA2D1;refGene_tx=XM_019825392.2
A2 88018529 88018601
refGene_gn=CACNA2D1;refGene_tx=XM_011280348.3;refGene_gn=CACNA2D1;refGene_tx=XM_011280349.3;refGene_gn=CACNA2D1;refGene_tx=XM_011280350.3;refGene_gn=CACNA2D1;refGene_tx=XM_011280351.3;refGene_gn=CACNA2D1;refGene_tx=XM_011280352.3;refGene_gn=CACNA2D1;refGene_tx=XM_011280353.3;refGene_gn=CACNA2D1;refGene_tx=XM_019825390.2;refGene_gn=CACNA2D1;refGene_tx=XM_019825391.2;refGene_gn=CACNA2D1;refGene_tx=XM_019825392.2
A2 88022608 88022665
refGene_gn=CACNA2D1;refGene_tx=XM_011280348.3;refGene_gn=CACNA2D1;refGene_tx=XM_011280349.3;refGene_gn=CACNA2D1;refGene_tx=XM_019825390.2;refGene_gn=CACNA2D1;refGene_tx=XM_019825391.2;refGene_gn=CACNA2D1;refGene_tx=XM_019825392.2
A2 88025725 88025800
refGene_gn=CACNA2D1;refGene_tx=XM_011280348.3;refGene_gn=CACNA2D1;refGene_tx=XM_011280349.3;refGene_gn=CACNA2D1;refGene_tx=XM_011280350.3;refGene_gn=CACNA2D1;refGene_tx=XM_011280351.3;refGene_gn=CACNA2D1;refGene_tx=XM_011280352.3;refGene_gn=CACNA2D1;refGene_tx=XM_011280353.3;refGene_gn=CACNA2D1;refGene_tx=XM_019825390.2;refGene_gn=CACNA2D1;refGene_tx=XM_019825391.2;refGene_gn=CACNA2D1;refGene_tx=XM_019825392.2
A2 88026055 88026130
refGene_gn=CACNA2D1;refGene_tx=XM_011280348.3;refGene_gn=CACNA2D1;refGene_tx=XM_011280349.3;refGene_gn=CACNA2D1;refGene_tx=XM_011280350.3;refGene_gn=CACNA2D1;refGene_tx=XM_011280351.3;refGene_gn=CACNA2D1;refGene_tx=XM_011280352.3;refGene_gn=CACNA2D1;refGene_tx=XM_011280353.3;refG

ene_gn=CACNA2D1;refGene_tx=XM_019825390.2;refGene_gn=CACNA2D1;refGene_tx=XM_019825391.2;refGene_gn=CACNA2D1;refGene_tx=XM_019825392.2
A2 88028155 88028233
refGene_gn=CACNA2D1;refGene_tx=XM_011280348.3;refGene_gn=CACNA2D1;refGene_tx=XM_011280349.3;refGene_gn=CACNA2D1;refGene_tx=XM_011280350.3;refGene_gn=CACNA2D1;refGene_tx=XM_011280351.3;refGene_gn=CACNA2D1;refGene_tx=XM_011280352.3;refGene_gn=CACNA2D1;refGene_tx=XM_011280353.3;refGene_gn=CACNA2D1;refGene_tx=XM_019825390.2;refGene_gn=CACNA2D1;refGene_tx=XM_019825391.2;refGene_gn=CACNA2D1;refGene_tx=XM_019825392.2
A2 88032128 88032218
refGene_gn=CACNA2D1;refGene_tx=XM_011280348.3;refGene_gn=CACNA2D1;refGene_tx=XM_011280349.3;refGene_gn=CACNA2D1;refGene_tx=XM_011280350.3;refGene_gn=CACNA2D1;refGene_tx=XM_011280351.3;refGene_gn=CACNA2D1;refGene_tx=XM_011280352.3;refGene_gn=CACNA2D1;refGene_tx=XM_011280353.3;refGene_gn=CACNA2D1;refGene_tx=XM_019825390.2;refGene_gn=CACNA2D1;refGene_tx=XM_019825391.2;refGene_gn=CACNA2D1;refGene_tx=XM_019825392.2
A2 88033635 88033685
refGene_gn=CACNA2D1;refGene_tx=XM_011280348.3;refGene_gn=CACNA2D1;refGene_tx=XM_011280349.3;refGene_gn=CACNA2D1;refGene_tx=XM_011280350.3;refGene_gn=CACNA2D1;refGene_tx=XM_011280351.3;refGene_gn=CACNA2D1;refGene_tx=XM_011280352.3;refGene_gn=CACNA2D1;refGene_tx=XM_011280353.3;refGene_gn=CACNA2D1;refGene_tx=XM_019825390.2;refGene_gn=CACNA2D1;refGene_tx=XM_019825391.2;refGene_gn=CACNA2D1;refGene_tx=XM_019825392.2
A2 88034564 88034643
refGene_gn=CACNA2D1;refGene_tx=XM_011280348.3;refGene_gn=CACNA2D1;refGene_tx=XM_011280349.3;refGene_gn=CACNA2D1;refGene_tx=XM_011280350.3;refGene_gn=CACNA2D1;refGene_tx=XM_011280351.3;refGene_gn=CACNA2D1;refGene_tx=XM_011280352.3;refGene_gn=CACNA2D1;refGene_tx=XM_011280353.3;refGene_gn=CACNA2D1;refGene_tx=XM_019825390.2;refGene_gn=CACNA2D1;refGene_tx=XM_019825391.2;refGene_gn=CACNA2D1;refGene_tx=XM_019825392.2
A2 88050712 88050817
refGene_gn=CACNA2D1;refGene_tx=XM_011280348.3;refGene_gn=CACNA2D1;refGene_tx=XM_011280349.3;refGene_gn=CACNA2D1;refGene_tx=XM_011280350.3;refGene_gn=CACNA2D1;refGene_tx=XM_011280351.3;refGene_gn=CACNA2D1;refGene_tx=XM_011280352.3;refGene_gn=CACNA2D1;refGene_tx=XM_011280353.3;refGene_gn=CACNA2D1;refGene_tx=XM_019825390.2;refGene_gn=CACNA2D1;refGene_tx=XM_019825391.2;refGene_gn=CACNA2D1;refGene_tx=XM_019825392.2
A2 88055744 88055903
refGene_gn=CACNA2D1;refGene_tx=XM_011280348.3;refGene_gn=CACNA2D1;refGene_tx=XM_011280349.3;refGene_gn=CACNA2D1;refGene_tx=XM_011280350.3;refGene_gn=CACNA2D1;refGene_tx=XM_011280351.3;refGene_gn=CACNA2D1;refGene_tx=XM_011280352.3;refGene_gn=CACNA2D1;refGene_tx=XM_011280353.3;refGene_gn=CACNA2D1;refGene_tx=XM_019825390.2;refGene_gn=CACNA2D1;refGene_tx=XM_019825391.2;refGene_gn=CACNA2D1;refGene_tx=XM_019825392.2
A2 88080078 88080178
refGene_gn=CACNA2D1;refGene_tx=XM_011280348.3;refGene_gn=CACNA2D1;refGene_tx=XM_011280349.3;refGene_gn=CACNA2D1;refGene_tx=XM_011280350.3;refGene_gn=CACNA2D1;refGene_tx=XM_011280351.3;refGene_gn=CACNA2D1;refGene_tx=XM_011280352.3;refGene_gn=CACNA2D1;refGene_tx=XM_011280353.3;refGene_gn=CACNA2D1;refGene_tx=XM_019825390.2;refGene_gn=CACNA2D1;refGene_

tx=XM_019825391.2;refGene_gn=CACNA2D1;refGene_tx=XM_019825392.2
A2 88082770 88082821
refGene_gn=CACNA2D1;refGene_tx=XM_011280348.3;refGene_gn=CACNA2D1;refGene_tx=XM_011280349.3;refGene_gn=CACNA2D1;refGene_tx=XM_011280350.3;refGene_gn=CACNA2D1;refGene_tx=XM_011280351.3;refGene_gn=CACNA2D1;refGene_tx=XM_011280352.3;refGene_gn=CACNA2D1;refGene_tx=XM_011280353.3;refGene_gn=CACNA2D1;refGene_tx=XM_019825390.2;refGene_gn=CACNA2D1;refGene_tx=XM_019825391.2;refGene_gn=CACNA2D1;refGene_tx=XM_019825392.2
A2 88084955 88085025
refGene_gn=CACNA2D1;refGene_tx=XM_011280348.3;refGene_gn=CACNA2D1;refGene_tx=XM_011280349.3;refGene_gn=CACNA2D1;refGene_tx=XM_011280350.3;refGene_gn=CACNA2D1;refGene_tx=XM_011280351.3;refGene_gn=CACNA2D1;refGene_tx=XM_011280352.3;refGene_gn=CACNA2D1;refGene_tx=XM_011280353.3;refGene_gn=CACNA2D1;refGene_tx=XM_019825390.2;refGene_gn=CACNA2D1;refGene_tx=XM_019825391.2;refGene_gn=CACNA2D1;refGene_tx=XM_019825392.2
A2 88104026 88104158
refGene_gn=CACNA2D1;refGene_tx=XM_011280348.3;refGene_gn=CACNA2D1;refGene_tx=XM_011280349.3;refGene_gn=CACNA2D1;refGene_tx=XM_011280350.3;refGene_gn=CACNA2D1;refGene_tx=XM_011280351.3;refGene_gn=CACNA2D1;refGene_tx=XM_011280352.3;refGene_gn=CACNA2D1;refGene_tx=XM_011280353.3;refGene_gn=CACNA2D1;refGene_tx=XM_019825390.2;refGene_gn=CACNA2D1;refGene_tx=XM_019825391.2;refGene_gn=CACNA2D1;refGene_tx=XM_019825392.2
A2 88140355 88140485
refGene_gn=CACNA2D1;refGene_tx=XM_011280348.3;refGene_gn=CACNA2D1;refGene_tx=XM_011280349.3;refGene_gn=CACNA2D1;refGene_tx=XM_011280350.3;refGene_gn=CACNA2D1;refGene_tx=XM_011280351.3;refGene_gn=CACNA2D1;refGene_tx=XM_011280352.3;refGene_gn=CACNA2D1;refGene_tx=XM_011280353.3;refGene_gn=CACNA2D1;refGene_tx=XM_019825390.2;refGene_gn=CACNA2D1;refGene_tx=XM_019825391.2;refGene_gn=CACNA2D1;refGene_tx=XM_019825392.2
A2 88159884 88159926
refGene_gn=CACNA2D1;refGene_tx=XM_011280348.3;refGene_gn=CACNA2D1;refGene_tx=XM_011280349.3;refGene_gn=CACNA2D1;refGene_tx=XM_011280350.3;refGene_gn=CACNA2D1;refGene_tx=XM_011280351.3;refGene_gn=CACNA2D1;refGene_tx=XM_011280352.3;refGene_gn=CACNA2D1;refGene_tx=XM_011280353.3;refGene_gn=CACNA2D1;refGene_tx=XM_019825390.2;refGene_gn=CACNA2D1;refGene_tx=XM_019825391.2;refGene_gn=CACNA2D1;refGene_tx=XM_019825392.2
A2 88197901 88197961
refGene_gn=CACNA2D1;refGene_tx=XM_011280348.3;refGene_gn=CACNA2D1;refGene_tx=XM_011280349.3;refGene_gn=CACNA2D1;refGene_tx=XM_011280350.3;refGene_gn=CACNA2D1;refGene_tx=XM_011280351.3;refGene_gn=CACNA2D1;refGene_tx=XM_011280352.3;refGene_gn=CACNA2D1;refGene_tx=XM_011280353.3;refGene_gn=CACNA2D1;refGene_tx=XM_019825391.2;refGene_gn=CACNA2D1;refGene_tx=XM_019825392.2
A2 88362315 88362432
refGene_gn=CACNA2D1;refGene_tx=XM_011280348.3;refGene_gn=CACNA2D1;refGene_tx=XM_011280349.3;refGene_gn=CACNA2D1;refGene_tx=XM_011280350.3;refGene_gn=CACNA2D1;refGene_tx=XM_011280351.3;refGene_gn=CACNA2D1;refGene_tx=XM_011280352.3;refGene_gn=CACNA2D1;refGene_tx=XM_011280353.3;refGene_gn=CACNA2D1;refGene_tx=XM_019825390.2;refGene_gn=CACNA2D1;refGene_tx=XM_019825391.2;refGene_gn=CACNA2D1;refGene_tx=XM_019825392.2

A2 88378414 88378496
refGene_gn=CACNA2D1;refGene_tx=XM_011280348.3;refGene_gn=CACNA2D1;refGene_tx=XM_011280349.3;refGene_gn=CACNA2D1;refGene_tx=XM_011280350.3;refGene_gn=CACNA2D1;refGene_tx=XM_011280351.3;refGene_gn=CACNA2D1;refGene_tx=XM_011280352.3;refGene_gn=CACNA2D1;refGene_tx=XM_011280353.3;refGene_gn=CACNA2D1;refGene_tx=XM_019825390.2;refGene_gn=CACNA2D1;refGene_tx=XM_019825391.2;refGene_gn=CACNA2D1;refGene_tx=XM_019825392.2
A2 88468765 88468860
refGene_gn=CACNA2D1;refGene_tx=XM_011280348.3;refGene_gn=CACNA2D1;refGene_tx=XM_011280349.3;refGene_gn=CACNA2D1;refGene_tx=XM_011280350.3;refGene_gn=CACNA2D1;refGene_tx=XM_011280351.3;refGene_gn=CACNA2D1;refGene_tx=XM_011280352.3;refGene_gn=CACNA2D1;refGene_tx=XM_011280353.3;refGene_gn=CACNA2D1;refGene_tx=XM_019825390.2;refGene_gn=CACNA2D1;refGene_tx=XM_019825392.2
A2 88774075 88774216 refGene_gn=PCLO;refGene_tx=XM_023250308.1
A2 88776345 88776491 refGene_gn=PCLO;refGene_tx=XM_023250308.1
A2 88776910 88777045 refGene_gn=PCLO;refGene_tx=XM_023250308.1
A2 88817889 88817963 refGene_gn=PCLO;refGene_tx=XM_023250308.1
A2 88822595 88822737 refGene_gn=PCLO;refGene_tx=XM_023250308.1
A2 88839219 88839414 refGene_gn=PCLO;refGene_tx=XM_023250308.1
A2 88841030 88841211 refGene_gn=PCLO;refGene_tx=XM_023250308.1
A2 88843205 88843277 refGene_gn=PCLO;refGene_tx=XM_023250308.1
A2 88844902 88844996 refGene_gn=PCLO;refGene_tx=XM_023250308.1
A2 88852752 88852779 refGene_gn=PCLO;refGene_tx=XM_023250308.1
A2 88855257 88855382 refGene_gn=PCLO;refGene_tx=XM_023250308.1
A2 88858759 88858810 refGene_gn=PCLO;refGene_tx=XM_023250308.1
A2 88863857 88864072 refGene_gn=PCLO;refGene_tx=XM_023250308.1
A2 88864848 88864916 refGene_gn=PCLO;refGene_tx=XM_023250308.1
A2 88865463 88865572 refGene_gn=PCLO;refGene_tx=XM_023250308.1
A2 88892080 88892206 refGene_gn=PCLO;refGene_tx=XM_023250308.1
A2 88911189 88911280 refGene_gn=PCLO;refGene_tx=XM_023250308.1
A2 88917545 88917682 refGene_gn=PCLO;refGene_tx=XM_023250308.1
A2 88922772 88924960 refGene_gn=PCLO;refGene_tx=XM_023250308.1
A2 88969538 88971553 refGene_gn=PCLO;refGene_tx=XM_023250308.1
A2 88971916 88977011 refGene_gn=PCLO;refGene_tx=XM_023250308.1
A2 88987677 88988337 refGene_gn=PCLO;refGene_tx=XM_023250308.1
A2 89160207 89161617 refGene_gn=PCLO;refGene_tx=XM_023250308.1
A2 89181428 89183136 refGene_gn=PCLO;refGene_tx=XM_023250308.1
A2 89189137 89189391 refGene_gn=PCLO;refGene_tx=XM_023250308.1
A2 89362138 89362591 refGene_gn=SEMA3E;refGene_tx=XM_011280354.3
A2 89380265 89380405 refGene_gn=SEMA3E;refGene_tx=XM_011280354.3
A2 89381963 89382031 refGene_gn=SEMA3E;refGene_tx=XM_011280354.3
A2 89387056 89387223 refGene_gn=SEMA3E;refGene_tx=XM_011280354.3
A2 89389108 89389150 refGene_gn=SEMA3E;refGene_tx=XM_011280354.3
A2 89391374 89391466 refGene_gn=SEMA3E;refGene_tx=XM_011280354.3
A2 89393962 89394185 refGene_gn=SEMA3E;refGene_tx=XM_011280354.3
A2 89396349 89396494 refGene_gn=SEMA3E;refGene_tx=XM_011280354.3
A2 89399449 89399519 refGene_gn=SEMA3E;refGene_tx=XM_011280354.3
A2 89399927 89400042 refGene_gn=SEMA3E;refGene_tx=XM_011280354.3
A2 89401290 89401433 refGene_gn=SEMA3E;refGene_tx=XM_011280354.3

A2 89402544 89402664 refGene_gn=SEMA3E;refGene_tx=XM_011280354.3
A2 89412979 89413073 refGene_gn=SEMA3E;refGene_tx=XM_011280354.3
A2 89444931 89445051 refGene_gn=SEMA3E;refGene_tx=XM_011280354.3
A2 89447721 89447781 refGene_gn=SEMA3E;refGene_tx=XM_011280354.3
A2 89468886 89469047 refGene_gn=SEMA3E;refGene_tx=XM_011280354.3
A2 89597634 89597749 refGene_gn=SEMA3E;refGene_tx=XM_011280354.3
A2 89899196 89899652
refGene_gn=SEMA3A;refGene_tx=XM_003982709.5;refGene_gn=SEMA3A;refGene_tx=XM_011280355.3;refGene_gn=SEMA3A;refGene_tx=XM_019825395.2;refGene_gn=SEMA3A;refGene_tx=XM_023250310.1
A2 89901034 89901177
refGene_gn=SEMA3A;refGene_tx=XM_003982709.5;refGene_gn=SEMA3A;refGene_tx=XM_011280355.3;refGene_gn=SEMA3A;refGene_tx=XM_019825395.2;refGene_gn=SEMA3A;refGene_tx=XM_023250310.1
A2 89914147 89914215
refGene_gn=SEMA3A;refGene_tx=XM_003982709.5;refGene_gn=SEMA3A;refGene_tx=XM_011280355.3;refGene_gn=SEMA3A;refGene_tx=XM_019825395.2;refGene_gn=SEMA3A;refGene_tx=XM_023250310.1
A2 89917828 89917986
refGene_gn=SEMA3A;refGene_tx=XM_003982709.5;refGene_gn=SEMA3A;refGene_tx=XM_011280355.3;refGene_gn=SEMA3A;refGene_tx=XM_019825395.2;refGene_gn=SEMA3A;refGene_tx=XM_023250310.1
A2 89922711 89922753
refGene_gn=SEMA3A;refGene_tx=XM_003982709.5;refGene_gn=SEMA3A;refGene_tx=XM_011280355.3;refGene_gn=SEMA3A;refGene_tx=XM_019825395.2;refGene_gn=SEMA3A;refGene_tx=XM_023250310.1
A2 89932035 89932127
refGene_gn=SEMA3A;refGene_tx=XM_003982709.5;refGene_gn=SEMA3A;refGene_tx=XM_011280355.3;refGene_gn=SEMA3A;refGene_tx=XM_019825395.2;refGene_gn=SEMA3A;refGene_tx=XM_023250310.1
A2 89935308 89935528
refGene_gn=SEMA3A;refGene_tx=XM_003982709.5;refGene_gn=SEMA3A;refGene_tx=XM_011280355.3;refGene_gn=SEMA3A;refGene_tx=XM_019825395.2;refGene_gn=SEMA3A;refGene_tx=XM_023250310.1
A2 89938801 89938946
refGene_gn=SEMA3A;refGene_tx=XM_003982709.5;refGene_gn=SEMA3A;refGene_tx=XM_011280355.3;refGene_gn=SEMA3A;refGene_tx=XM_019825395.2;refGene_gn=SEMA3A;refGene_tx=XM_023250310.1
A2 89940708 89940778
refGene_gn=SEMA3A;refGene_tx=XM_003982709.5;refGene_gn=SEMA3A;refGene_tx=XM_011280355.3;refGene_gn=SEMA3A;refGene_tx=XM_019825395.2;refGene_gn=SEMA3A;refGene_tx=XM_023250310.1
A2 89940870 89940985
refGene_gn=SEMA3A;refGene_tx=XM_003982709.5;refGene_gn=SEMA3A;refGene_tx=XM_011280355.3;refGene_gn=SEMA3A;refGene_tx=XM_019825395.2;refGene_gn=SEMA3A;refGene_tx=XM_023250310.1
A2 89942801 89942944
refGene_gn=SEMA3A;refGene_tx=XM_003982709.5;refGene_gn=SEMA3A;refGene_tx=XM_011280355.3;refGene_gn=SEMA3A;refGene_tx=XM_019825395.2;refGene_gn=SEMA3A;refGene_tx=XM_023250310.1

A2 89978018 89978138
refGene_gn=SEMA3A;refGene_tx=XM_003982709.5;refGene_gn=SEMA3A;refGene_tx=XM_011280355.3;refGene_gn=SEMA3A;refGene_tx=XM_019825395.2;refGene_gn=SEMA3A;refGene_tx=XM_023250310.1
A2 89986752 89986846
refGene_gn=SEMA3A;refGene_tx=XM_003982709.5;refGene_gn=SEMA3A;refGene_tx=XM_011280355.3;refGene_gn=SEMA3A;refGene_tx=XM_019825395.2;refGene_gn=SEMA3A;refGene_tx=XM_023250310.1
A2 90025496 90025616
refGene_gn=SEMA3A;refGene_tx=XM_003982709.5;refGene_gn=SEMA3A;refGene_tx=XM_011280355.3;refGene_gn=SEMA3A;refGene_tx=XM_019825395.2;refGene_gn=SEMA3A;refGene_tx=XM_023250310.1
A2 90043685 90043748
refGene_gn=SEMA3A;refGene_tx=XM_003982709.5;refGene_gn=SEMA3A;refGene_tx=XM_011280355.3;refGene_gn=SEMA3A;refGene_tx=XM_019825395.2;refGene_gn=SEMA3A;refGene_tx=XM_023250310.1
A2 90048707 90048865
refGene_gn=SEMA3A;refGene_tx=XM_003982709.5;refGene_gn=SEMA3A;refGene_tx=XM_011280355.3;refGene_gn=SEMA3A;refGene_tx=XM_019825395.2;refGene_gn=SEMA3A;refGene_tx=XM_023250310.1
A2 90063275 90063315 refGene_gn=SEMA3A;refGene_tx=XM_011280355.3
A2 90101079 90101191
refGene_gn=SEMA3A;refGene_tx=XM_003982709.5;refGene_gn=SEMA3A;refGene_tx=XM_019825395.2
A2 90760008 90760434
refGene_gn=SEMA3D;refGene_tx=XM_011280356.2;refGene_gn=SEMA3D;refGene_tx=XM_019825397.2;refGene_gn=SEMA3D;refGene_tx=XM_019825398.2;refGene_gn=SEMA3D;refGene_tx=XM_019825399.2
A2 90768163 90768303
refGene_gn=SEMA3D;refGene_tx=XM_011280356.2;refGene_gn=SEMA3D;refGene_tx=XM_019825397.2;refGene_gn=SEMA3D;refGene_tx=XM_019825398.2;refGene_gn=SEMA3D;refGene_tx=XM_019825399.2
A2 90775109 90775174
refGene_gn=SEMA3D;refGene_tx=XM_011280356.2;refGene_gn=SEMA3D;refGene_tx=XM_019825397.2;refGene_gn=SEMA3D;refGene_tx=XM_019825398.2;refGene_gn=SEMA3D;refGene_tx=XM_019825399.2
A2 90777396 90777554
refGene_gn=SEMA3D;refGene_tx=XM_011280356.2;refGene_gn=SEMA3D;refGene_tx=XM_019825397.2;refGene_gn=SEMA3D;refGene_tx=XM_019825398.2;refGene_gn=SEMA3D;refGene_tx=XM_019825399.2
A2 90780741 90780783
refGene_gn=SEMA3D;refGene_tx=XM_011280356.2;refGene_gn=SEMA3D;refGene_tx=XM_019825397.2;refGene_gn=SEMA3D;refGene_tx=XM_019825398.2;refGene_gn=SEMA3D;refGene_tx=XM_019825399.2
A2 90782615 90782704
refGene_gn=SEMA3D;refGene_tx=XM_011280356.2;refGene_gn=SEMA3D;refGene_tx=XM_019825397.2;refGene_gn=SEMA3D;refGene_tx=XM_019825398.2;refGene_gn=SEMA3D;refGene_tx=XM_019825399.2
A2 90785272 90785495
refGene_gn=SEMA3D;refGene_tx=XM_011280356.2;refGene_gn=SEMA3D;refGene_

tx=XM_019825397.2;refGene_gn=SEMA3D;refGene_tx=XM_019825398.2;refGene_gn=SEMA3D;refGene_tx=XM_019825399.2
A2 90796316 90796461
refGene_gn=SEMA3D;refGene_tx=XM_011280356.2;refGene_gn=SEMA3D;refGene_tx=XM_019825397.2;refGene_gn=SEMA3D;refGene_tx=XM_019825398.2;refGene_gn=SEMA3D;refGene_tx=XM_019825399.2
A2 90799362 90799432
refGene_gn=SEMA3D;refGene_tx=XM_011280356.2;refGene_gn=SEMA3D;refGene_tx=XM_019825397.2;refGene_gn=SEMA3D;refGene_tx=XM_019825398.2;refGene_gn=SEMA3D;refGene_tx=XM_019825399.2
A2 90800662 90800777
refGene_gn=SEMA3D;refGene_tx=XM_011280356.2;refGene_gn=SEMA3D;refGene_tx=XM_019825397.2;refGene_gn=SEMA3D;refGene_tx=XM_019825398.2;refGene_gn=SEMA3D;refGene_tx=XM_019825399.2
A2 90813613 90813756
refGene_gn=SEMA3D;refGene_tx=XM_011280356.2;refGene_gn=SEMA3D;refGene_tx=XM_019825397.2;refGene_gn=SEMA3D;refGene_tx=XM_019825398.2;refGene_gn=SEMA3D;refGene_tx=XM_019825399.2
A2 90825172 90825301
refGene_gn=SEMA3D;refGene_tx=XM_011280356.2;refGene_gn=SEMA3D;refGene_tx=XM_019825397.2;refGene_gn=SEMA3D;refGene_tx=XM_019825398.2;refGene_gn=SEMA3D;refGene_tx=XM_019825399.2
A2 90828076 90828170
refGene_gn=SEMA3D;refGene_tx=XM_011280356.2;refGene_gn=SEMA3D;refGene_tx=XM_019825397.2;refGene_gn=SEMA3D;refGene_tx=XM_019825398.2;refGene_gn=SEMA3D;refGene_tx=XM_019825399.2
A2 90833010 90833130
refGene_gn=SEMA3D;refGene_tx=XM_011280356.2;refGene_gn=SEMA3D;refGene_tx=XM_019825397.2;refGene_gn=SEMA3D;refGene_tx=XM_019825398.2;refGene_gn=SEMA3D;refGene_tx=XM_019825399.2
A2 90842494 90842557
refGene_gn=SEMA3D;refGene_tx=XM_011280356.2;refGene_gn=SEMA3D;refGene_tx=XM_019825397.2;refGene_gn=SEMA3D;refGene_tx=XM_019825398.2;refGene_gn=SEMA3D;refGene_tx=XM_019825399.2
A2 90862158 90862319
refGene_gn=SEMA3D;refGene_tx=XM_011280356.2;refGene_gn=SEMA3D;refGene_tx=XM_019825397.2;refGene_gn=SEMA3D;refGene_tx=XM_019825398.2;refGene_gn=SEMA3D;refGene_tx=XM_019825399.2
A2 90880822 90880973
refGene_gn=SEMA3D;refGene_tx=XM_011280356.2;refGene_gn=SEMA3D;refGene_tx=XM_019825397.2;refGene_gn=SEMA3D;refGene_tx=XM_019825398.2;refGene_gn=SEMA3D;refGene_tx=XM_019825399.2
A2 92428022 92428490
refGene_gn=GRM3;refGene_tx=XM_006929146.4;refGene_gn=GRM3;refGene_tx=XM_019825400.1
A2 92449366 92450222
refGene_gn=GRM3;refGene_tx=XM_006929146.4;refGene_gn=GRM3;refGene_tx=XM_019825400.1;refGene_gn=GRM3;refGene_tx=XM_019825401.2
A2 92501271 92502338
refGene_gn=GRM3;refGene_tx=XM_006929146.4;refGene_gn=GRM3;refGene_tx=X

M_019825400.1;refGene_gn=GRM3;refGene_tx=XM_019825401.2
A2 92512864 92513039
refGene_gn=GRM3;refGene_tx=XM_006929146.4;refGene_gn=GRM3;refGene_tx=XM_019825400.1;refGene_gn=GRM3;refGene_tx=XM_019825401.2
A2 92513870 92513878 refGene_gn=GRM3;refGene_tx=XM_019825400.1
A2 92526489 92526563
refGene_gn=GRM3;refGene_tx=XM_006929146.4;refGene_gn=GRM3;refGene_tx=XM_019825401.2
A2 92534695 92534755
refGene_gn=KIAA1324L;refGene_tx=XM_023250311.1;refGene_gn=KIAA1324L;refGene_tx=XM_023250312.1
A2 92546587 92546753
refGene_gn=KIAA1324L;refGene_tx=XM_023250311.1;refGene_gn=KIAA1324L;refGene_tx=XM_023250312.1
A2 92547696 92547875
refGene_gn=KIAA1324L;refGene_tx=XM_023250311.1;refGene_gn=KIAA1324L;refGene_tx=XM_023250312.1
A2 92552945 92553072
refGene_gn=KIAA1324L;refGene_tx=XM_023250311.1;refGene_gn=KIAA1324L;refGene_tx=XM_023250312.1
A2 92568110 92568212
refGene_gn=KIAA1324L;refGene_tx=XM_023250311.1;refGene_gn=KIAA1324L;refGene_tx=XM_023250312.1
A2 92568787 92568884
refGene_gn=KIAA1324L;refGene_tx=XM_023250311.1;refGene_gn=KIAA1324L;refGene_tx=XM_023250312.1
A2 92570349 92570539
refGene_gn=KIAA1324L;refGene_tx=XM_023250311.1;refGene_gn=KIAA1324L;refGene_tx=XM_023250312.1
A2 92572571 92572745
refGene_gn=KIAA1324L;refGene_tx=XM_023250311.1;refGene_gn=KIAA1324L;refGene_tx=XM_023250312.1
A2 92573446 92573710
refGene_gn=KIAA1324L;refGene_tx=XM_023250311.1;refGene_gn=KIAA1324L;refGene_tx=XM_023250312.1
A2 92575201 92575339
refGene_gn=KIAA1324L;refGene_tx=XM_023250311.1;refGene_gn=KIAA1324L;refGene_tx=XM_023250312.1
A2 92578531 92578634
refGene_gn=KIAA1324L;refGene_tx=XM_023250311.1;refGene_gn=KIAA1324L;refGene_tx=XM_023250312.1
A2 92579310 92579401
refGene_gn=KIAA1324L;refGene_tx=XM_023250311.1;refGene_gn=KIAA1324L;refGene_tx=XM_023250312.1
A2 92584186 92584315
refGene_gn=KIAA1324L;refGene_tx=XM_023250311.1;refGene_gn=KIAA1324L;refGene_tx=XM_023250312.1
A2 92586194 92586375
refGene_gn=KIAA1324L;refGene_tx=XM_023250311.1;refGene_gn=KIAA1324L;refGene_tx=XM_023250312.1

A2 92596226 92596315
refGene_gn=KIAA1324L;refGene_tx=XM_023250311.1;refGene_gn=KIAA1324L;re
fGene_tx=XM_023250312.1
A2 92596919 92597072
refGene_gn=KIAA1324L;refGene_tx=XM_023250311.1;refGene_gn=KIAA1324L;re
fGene_tx=XM_023250312.1
A2 92598046 92598152
refGene_gn=KIAA1324L;refGene_tx=XM_023250311.1;refGene_gn=KIAA1324L;re
fGene_tx=XM_023250312.1
A2 92600234 92600321
refGene_gn=KIAA1324L;refGene_tx=XM_023250311.1;refGene_gn=KIAA1324L;re
fGene_tx=XM_023250312.1
A2 92603558 92603706
refGene_gn=KIAA1324L;refGene_tx=XM_023250311.1;refGene_gn=KIAA1324L;re
fGene_tx=XM_023250312.1
A2 92609835 92610031
refGene_gn=KIAA1324L;refGene_tx=XM_023250311.1;refGene_gn=KIAA1324L;re
fGene_tx=XM_023250312.1
A2 92614672 92614682
refGene_gn=KIAA1324L;refGene_tx=XM_023250312.1
A2 92821849 92821958
refGene_gn=DMTF1;refGene_tx=XM_003982713.5;refGene_gn=DMTF1;refGene_tx
=XM_019825402.1;refGene_gn=DMTF1;refGene_tx=XM_019825403.2;refGene_gn=
DMTF1;refGene_tx=XM_019825404.2;refGene_gn=DMTF1;refGene_tx=XM_0198254
05.2;refGene_gn=DMTF1;refGene_tx=XM_023250313.1;refGene_gn=DMTF1;refGe
ne_tx=XM_023250314.1
A2 92823622 92823745
refGene_gn=DMTF1;refGene_tx=XM_003982713.5;refGene_gn=DMTF1;refGene_tx
=XM_019825402.1;refGene_gn=DMTF1;refGene_tx=XM_019825403.2;refGene_gn=
DMTF1;refGene_tx=XM_019825404.2;refGene_gn=DMTF1;refGene_tx=XM_0232503
13.1
A2 92825355 92825389 refGene_gn=DMTF1;refGene_tx=XM_019825406.2
A2 92828106 92828201
refGene_gn=DMTF1;refGene_tx=XM_003982713.5;refGene_gn=DMTF1;refGene_tx
=XM_019825402.1;refGene_gn=DMTF1;refGene_tx=XM_019825403.2;refGene_gn=
DMTF1;refGene_tx=XM_019825404.2;refGene_gn=DMTF1;refGene_tx=XM_0198254
05.2;refGene_gn=DMTF1;refGene_tx=XM_019825406.2;refGene_gn=DMTF1;refGe
ne_tx=XM_023250313.1;refGene_gn=DMTF1;refGene_tx=XM_023250314.1
A2 92833718 92833833
refGene_gn=DMTF1;refGene_tx=XM_003982713.5;refGene_gn=DMTF1;refGene_tx
=XM_019825402.1;refGene_gn=DMTF1;refGene_tx=XM_019825403.2;refGene_gn=
DMTF1;refGene_tx=XM_019825404.2;refGene_gn=DMTF1;refGene_tx=XM_0198254
05.2;refGene_gn=DMTF1;refGene_tx=XM_019825406.2;refGene_gn=DMTF1;refGe
ne_tx=XM_023250313.1;refGene_gn=DMTF1;refGene_tx=XM_023250314.1
A2 92834711 92834788
refGene_gn=DMTF1;refGene_tx=XM_003982713.5;refGene_gn=DMTF1;refGene_tx
=XM_019825402.1;refGene_gn=DMTF1;refGene_tx=XM_019825403.2;refGene_gn=
DMTF1;refGene_tx=XM_019825404.2;refGene_gn=DMTF1;refGene_tx=XM_0198254
05.2;refGene_gn=DMTF1;refGene_tx=XM_019825406.2;refGene_gn=DMTF1;refGe
ne_tx=XM_019825407.1;refGene_gn=DMTF1;refGene_tx=XM_023250313.1;refGen

e_gn=DMTF1;refGene_tx=XM_023250314.1
A2 92839476 92839634
refGene_gn=DMTF1;refGene_tx=XM_003982713.5;refGene_gn=DMTF1;refGene_tx=XM_019825402.1;refGene_gn=DMTF1;refGene_tx=XM_019825403.2;refGene_gn=DMTF1;refGene_tx=XM_019825404.2;refGene_gn=DMTF1;refGene_tx=XM_019825405.2;refGene_gn=DMTF1;refGene_tx=XM_019825406.2;refGene_gn=DMTF1;refGene_tx=XM_019825407.1;refGene_gn=DMTF1;refGene_tx=XM_023250313.1;refGene_gn=DMTF1;refGene_tx=XM_023250314.1
A2 92840767 92840800
refGene_gn=DMTF1;refGene_tx=XM_003982713.5;refGene_gn=DMTF1;refGene_tx=XM_019825402.1;refGene_gn=DMTF1;refGene_tx=XM_019825403.2;refGene_gn=DMTF1;refGene_tx=XM_019825404.2;refGene_gn=DMTF1;refGene_tx=XM_019825405.2;refGene_gn=DMTF1;refGene_tx=XM_019825406.2;refGene_gn=DMTF1;refGene_tx=XM_019825407.1;refGene_gn=DMTF1;refGene_tx=XM_023250313.1;refGene_gn=DMTF1;refGene_tx=XM_023250314.1
A2 92841696 92841806
refGene_gn=DMTF1;refGene_tx=XM_003982713.5;refGene_gn=DMTF1;refGene_tx=XM_019825402.1;refGene_gn=DMTF1;refGene_tx=XM_019825403.2;refGene_gn=DMTF1;refGene_tx=XM_019825404.2;refGene_gn=DMTF1;refGene_tx=XM_019825405.2;refGene_gn=DMTF1;refGene_tx=XM_019825406.2;refGene_gn=DMTF1;refGene_tx=XM_019825407.1;refGene_gn=DMTF1;refGene_tx=XM_023250313.1;refGene_gn=DMTF1;refGene_tx=XM_023250314.1
A2 92842533 92842710
refGene_gn=DMTF1;refGene_tx=XM_019825402.1;refGene_gn=DMTF1;refGene_tx=XM_019825403.2;refGene_gn=DMTF1;refGene_tx=XM_019825404.2;refGene_gn=DMTF1;refGene_tx=XM_019825405.2;refGene_gn=DMTF1;refGene_tx=XM_019825406.2;refGene_gn=DMTF1;refGene_tx=XM_019825407.1;refGene_gn=DMTF1;refGene_tx=XM_023250313.1
A2 92845516 92845745
refGene_gn=DMTF1;refGene_tx=XM_003982713.5;refGene_gn=DMTF1;refGene_tx=XM_019825402.1;refGene_gn=DMTF1;refGene_tx=XM_019825403.2;refGene_gn=DMTF1;refGene_tx=XM_019825404.2;refGene_gn=DMTF1;refGene_tx=XM_019825405.2;refGene_gn=DMTF1;refGene_tx=XM_019825406.2;refGene_gn=DMTF1;refGene_tx=XM_019825407.1;refGene_gn=DMTF1;refGene_tx=XM_023250313.1;refGene_gn=DMTF1;refGene_tx=XM_023250314.1
A2 92846965 92847117
refGene_gn=DMTF1;refGene_tx=XM_003982713.5;refGene_gn=DMTF1;refGene_tx=XM_019825402.1;refGene_gn=DMTF1;refGene_tx=XM_019825403.2;refGene_gn=DMTF1;refGene_tx=XM_019825404.2;refGene_gn=DMTF1;refGene_tx=XM_019825405.2;refGene_gn=DMTF1;refGene_tx=XM_019825406.2;refGene_gn=DMTF1;refGene_tx=XM_019825407.1;refGene_gn=DMTF1;refGene_tx=XM_023250313.1;refGene_gn=DMTF1;refGene_tx=XM_023250314.1
A2 92848417 92848627
refGene_gn=DMTF1;refGene_tx=XM_003982713.5;refGene_gn=DMTF1;refGene_tx=XM_019825402.1;refGene_gn=DMTF1;refGene_tx=XM_019825403.2;refGene_gn=DMTF1;refGene_tx=XM_019825404.2;refGene_gn=DMTF1;refGene_tx=XM_019825405.2;refGene_gn=DMTF1;refGene_tx=XM_019825406.2;refGene_gn=DMTF1;refGene_tx=XM_019825407.1;refGene_gn=DMTF1;refGene_tx=XM_023250313.1;refGene_gn=DMTF1;refGene_tx=XM_023250314.1
A2 92851178 92851264

refGene_gn=DMTF1;refGene_tx=XM_003982713.5;refGene_gn=DMTF1;refGene_tx=XM_019825402.1;refGene_gn=DMTF1;refGene_tx=XM_019825403.2;refGene_gn=DMTF1;refGene_tx=XM_019825404.2;refGene_gn=DMTF1;refGene_tx=XM_019825405.2;refGene_gn=DMTF1;refGene_tx=XM_019825406.2;refGene_gn=DMTF1;refGene_tx=XM_019825407.1;refGene_gn=DMTF1;refGene_tx=XM_023250313.1;refGene_gn=DMTF1;refGene_tx=XM_023250314.1
A2 92853846 92854002
refGene_gn=DMTF1;refGene_tx=XM_003982713.5;refGene_gn=DMTF1;refGene_tx=XM_019825402.1;refGene_gn=DMTF1;refGene_tx=XM_019825403.2;refGene_gn=DMTF1;refGene_tx=XM_019825404.2;refGene_gn=DMTF1;refGene_tx=XM_019825405.2;refGene_gn=DMTF1;refGene_tx=XM_019825406.2;refGene_gn=DMTF1;refGene_tx=XM_019825407.1;refGene_gn=DMTF1;refGene_tx=XM_023250313.1;refGene_gn=DMTF1;refGene_tx=XM_023250314.1
A2 92854508 92854883
refGene_gn=DMTF1;refGene_tx=XM_003982713.5;refGene_gn=DMTF1;refGene_tx=XM_019825402.1;refGene_gn=DMTF1;refGene_tx=XM_019825403.2;refGene_gn=DMTF1;refGene_tx=XM_019825404.2;refGene_gn=DMTF1;refGene_tx=XM_019825405.2;refGene_gn=DMTF1;refGene_tx=XM_019825406.2;refGene_gn=DMTF1;refGene_tx=XM_019825407.1;refGene_gn=DMTF1;refGene_tx=XM_023250313.1;refGene_gn=DMTF1;refGene_tx=XM_023250314.1
A2 92855473 92855621
refGene_gn=DMTF1;refGene_tx=XM_003982713.5;refGene_gn=DMTF1;refGene_tx=XM_019825402.1;refGene_gn=DMTF1;refGene_tx=XM_019825403.2;refGene_gn=DMTF1;refGene_tx=XM_019825404.2;refGene_gn=DMTF1;refGene_tx=XM_019825405.2;refGene_gn=DMTF1;refGene_tx=XM_019825406.2;refGene_gn=DMTF1;refGene_tx=XM_019825407.1;refGene_gn=DMTF1;refGene_tx=XM_023250313.1;refGene_gn=DMTF1;refGene_tx=XM_023250314.1
A2 92855811 92855921
refGene_gn=DMTF1;refGene_tx=XM_003982713.5;refGene_gn=DMTF1;refGene_tx=XM_019825402.1;refGene_gn=DMTF1;refGene_tx=XM_019825403.2;refGene_gn=DMTF1;refGene_tx=XM_019825404.2;refGene_gn=DMTF1;refGene_tx=XM_019825405.2;refGene_gn=DMTF1;refGene_tx=XM_019825406.2;refGene_gn=DMTF1;refGene_tx=XM_019825407.1;refGene_gn=DMTF1;refGene_tx=XM_023250313.1;refGene_gn=DMTF1;refGene_tx=XM_023250314.1
A2 92857483 92857606
refGene_gn=TMEM243;refGene_tx=XM_011280357.3;refGene_gn=TMEM243;refGene_tx=XM_011280358.3;refGene_gn=TMEM243;refGene_tx=XM_011280359.2;refGene_gn=TMEM243;refGene_tx=XM_011280361.3
A2 92858963 92859068
refGene_gn=TMEM243;refGene_tx=XM_011280357.3;refGene_gn=TMEM243;refGene_tx=XM_011280358.3;refGene_gn=TMEM243;refGene_tx=XM_011280359.2;refGene_gn=TMEM243;refGene_tx=XM_011280361.3
A2 92860029 92860080
refGene_gn=TMEM243;refGene_tx=XM_011280357.3;refGene_gn=TMEM243;refGene_tx=XM_011280358.3;refGene_gn=TMEM243;refGene_tx=XM_011280359.2;refGene_gn=TMEM243;refGene_tx=XM_011280361.3
A2 92877774 92877852
refGene_gn=TMEM243;refGene_tx=XM_011280357.3;refGene_gn=TMEM243;refGene_tx=XM_011280358.3;refGene_gn=TMEM243;refGene_tx=XM_011280359.2;refGene_gn=TMEM243;refGene_tx=XM_011280361.3

A2 92961184 92961299
refGene_gn=CROT;refGene_tx=XM_011280362.2;refGene_gn=CROT;refGene_tx=XM_011280363.2
A2 92964527 92964652
refGene_gn=CROT;refGene_tx=XM_011280362.2;refGene_gn=CROT;refGene_tx=XM_011280363.2
A2 92966627 92966809
refGene_gn=CROT;refGene_tx=XM_011280362.2;refGene_gn=CROT;refGene_tx=XM_011280363.2
A2 92966978 92967103
refGene_gn=CROT;refGene_tx=XM_011280362.2;refGene_gn=CROT;refGene_tx=XM_011280363.2
A2 92977272 92977381
refGene_gn=CROT;refGene_tx=XM_011280362.2;refGene_gn=CROT;refGene_tx=XM_011280363.2
A2 92984680 92984774
refGene_gn=CROT;refGene_tx=XM_011280362.2;refGene_gn=CROT;refGene_tx=XM_011280363.2
A2 92985550 92985676
refGene_gn=CROT;refGene_tx=XM_011280362.2;refGene_gn=CROT;refGene_tx=XM_011280363.2
A2 92987240 92987342
refGene_gn=CROT;refGene_tx=XM_011280362.2;refGene_gn=CROT;refGene_tx=XM_011280363.2
A2 92991141 92991225
refGene_gn=CROT;refGene_tx=XM_011280362.2;refGene_gn=CROT;refGene_tx=XM_011280363.2
A2 92991305 92991413
refGene_gn=CROT;refGene_tx=XM_011280362.2;refGene_gn=CROT;refGene_tx=XM_011280363.2
A2 92991640 92991771
refGene_gn=CROT;refGene_tx=XM_011280362.2;refGene_gn=CROT;refGene_tx=XM_011280363.2
A2 92997557 92997681
refGene_gn=CROT;refGene_tx=XM_011280362.2;refGene_gn=CROT;refGene_tx=XM_011280363.2
A2 92999536 92999615
refGene_gn=CROT;refGene_tx=XM_011280362.2;refGene_gn=CROT;refGene_tx=XM_011280363.2
A2 92999702 92999796
refGene_gn=CROT;refGene_tx=XM_011280362.2;refGene_gn=CROT;refGene_tx=XM_011280363.2
A2 93000142 93000262
refGene_gn=CROT;refGene_tx=XM_011280362.2;refGene_gn=CROT;refGene_tx=XM_011280363.2
A2 93003074 93003195
refGene_gn=CROT;refGene_tx=XM_011280362.2;refGene_gn=CROT;refGene_tx=XM_011280363.2
A2 93005933 93006140
refGene_gn=ABCB4;refGene_tx=XM_023250316.1;refGene_gn=ABCB4;refGene_tx

=XM_023250319.1;refGene_gn=ABCB4;refGene_tx=XM_023250320.1
A2 93006586 93006733
refGene_gn=ABCB4;refGene_tx=XM_023250316.1;refGene_gn=ABCB4;refGene_tx=XM_023250319.1;refGene_gn=ABCB4;refGene_tx=XM_023250320.1
A2 93009508 93009715
refGene_gn=ABCB4;refGene_tx=XM_023250316.1;refGene_gn=ABCB4;refGene_tx=XM_023250319.1;refGene_gn=ABCB4;refGene_tx=XM_023250320.1
A2 93009736 93009748 refGene_gn=ABCB4;refGene_tx=XM_023250318.1
A2 93009939 93009984 refGene_gn=ABCB4;refGene_tx=XM_023250317.1
A2 93010642 93010840
refGene_gn=ABCB4;refGene_tx=XM_023250316.1;refGene_gn=ABCB4;refGene_tx=XM_023250317.1;refGene_gn=ABCB4;refGene_tx=XM_023250318.1;refGene_gn=ABCB4;refGene_tx=XM_023250319.1;refGene_gn=ABCB4;refGene_tx=XM_023250320.1
A2 93012324 93012481
refGene_gn=ABCB4;refGene_tx=XM_023250316.1;refGene_gn=ABCB4;refGene_tx=XM_023250317.1;refGene_gn=ABCB4;refGene_tx=XM_023250318.1;refGene_gn=ABCB4;refGene_tx=XM_023250319.1;refGene_gn=ABCB4;refGene_tx=XM_023250320.1
A2 93015647 93015788
refGene_gn=ABCB4;refGene_tx=XM_023250316.1;refGene_gn=ABCB4;refGene_tx=XM_023250317.1;refGene_gn=ABCB4;refGene_tx=XM_023250318.1;refGene_gn=ABCB4;refGene_tx=XM_023250319.1;refGene_gn=ABCB4;refGene_tx=XM_023250320.1
A2 93017359 93017460
refGene_gn=ABCB4;refGene_tx=XM_023250316.1;refGene_gn=ABCB4;refGene_tx=XM_023250317.1;refGene_gn=ABCB4;refGene_tx=XM_023250318.1;refGene_gn=ABCB4;refGene_tx=XM_023250319.1;refGene_gn=ABCB4;refGene_tx=XM_023250320.1
A2 93021661 93021865
refGene_gn=ABCB4;refGene_tx=XM_023250316.1;refGene_gn=ABCB4;refGene_tx=XM_023250317.1;refGene_gn=ABCB4;refGene_tx=XM_023250318.1;refGene_gn=ABCB4;refGene_tx=XM_023250319.1;refGene_gn=ABCB4;refGene_tx=XM_023250320.1
A2 93022973 93023057
refGene_gn=ABCB4;refGene_tx=XM_023250316.1;refGene_gn=ABCB4;refGene_tx=XM_023250317.1;refGene_gn=ABCB4;refGene_tx=XM_023250318.1;refGene_gn=ABCB4;refGene_tx=XM_023250319.1;refGene_gn=ABCB4;refGene_tx=XM_023250320.1
A2 93024402 93024480
refGene_gn=ABCB4;refGene_tx=XM_023250316.1;refGene_gn=ABCB4;refGene_tx=XM_023250317.1;refGene_gn=ABCB4;refGene_tx=XM_023250318.1;refGene_gn=ABCB4;refGene_tx=XM_023250319.1;refGene_gn=ABCB4;refGene_tx=XM_023250320.1
A2 93026338 93026443
refGene_gn=ABCB4;refGene_tx=XM_023250316.1;refGene_gn=ABCB4;refGene_tx=XM_023250317.1;refGene_gn=ABCB4;refGene_tx=XM_023250318.1;refGene_gn=ABCB4;refGene_tx=XM_023250319.1;refGene_gn=ABCB4;refGene_tx=XM_023250320.1
A2 93028331 93028478

refGene_gn=ABCB4;refGene_tx=XM_023250316.1;refGene_gn=ABCB4;refGene_tx=XM_023250317.1;refGene_gn=ABCB4;refGene_tx=XM_023250318.1;refGene_gn=ABCB4;refGene_tx=XM_023250319.1;refGene_gn=ABCB4;refGene_tx=XM_023250320.1
A2 93031355 93031532
refGene_gn=ABCB4;refGene_tx=XM_023250316.1;refGene_gn=ABCB4;refGene_tx=XM_023250317.1;refGene_gn=ABCB4;refGene_tx=XM_023250318.1;refGene_gn=ABCB4;refGene_tx=XM_023250319.1;refGene_gn=ABCB4;refGene_tx=XM_023250320.1
A2 93038012 93038174
refGene_gn=ABCB4;refGene_tx=XM_023250316.1;refGene_gn=ABCB4;refGene_tx=XM_023250317.1;refGene_gn=ABCB4;refGene_tx=XM_023250318.1;refGene_gn=ABCB4;refGene_tx=XM_023250319.1;refGene_gn=ABCB4;refGene_tx=XM_023250320.1
A2 93046242 93046413
refGene_gn=ABCB4;refGene_tx=XM_023250316.1;refGene_gn=ABCB4;refGene_tx=XM_023250317.1;refGene_gn=ABCB4;refGene_tx=XM_023250318.1;refGene_gn=ABCB4;refGene_tx=XM_023250319.1;refGene_gn=ABCB4;refGene_tx=XM_023250320.1
A2 93046768 93046972
refGene_gn=ABCB4;refGene_tx=XM_023250316.1;refGene_gn=ABCB4;refGene_tx=XM_023250317.1;refGene_gn=ABCB4;refGene_tx=XM_023250318.1;refGene_gn=ABCB4;refGene_tx=XM_023250319.1;refGene_gn=ABCB4;refGene_tx=XM_023250320.1
A2 93050039 93050165
refGene_gn=ABCB4;refGene_tx=XM_023250316.1;refGene_gn=ABCB4;refGene_tx=XM_023250317.1;refGene_gn=ABCB4;refGene_tx=XM_023250318.1;refGene_gn=ABCB4;refGene_tx=XM_023250319.1;refGene_gn=ABCB4;refGene_tx=XM_023250320.1
A2 93050381 93050492
refGene_gn=ABCB4;refGene_tx=XM_023250316.1;refGene_gn=ABCB4;refGene_tx=XM_023250317.1;refGene_gn=ABCB4;refGene_tx=XM_023250318.1;refGene_gn=ABCB4;refGene_tx=XM_023250319.1
A2 93051583 93051697
refGene_gn=ABCB4;refGene_tx=XM_023250316.1;refGene_gn=ABCB4;refGene_tx=XM_023250317.1;refGene_gn=ABCB4;refGene_tx=XM_023250318.1;refGene_gn=ABCB4;refGene_tx=XM_023250319.1
A2 93054410 93054582
refGene_gn=ABCB4;refGene_tx=XM_023250316.1;refGene_gn=ABCB4;refGene_tx=XM_023250317.1;refGene_gn=ABCB4;refGene_tx=XM_023250318.1;refGene_gn=ABCB4;refGene_tx=XM_023250319.1
A2 93057251 93057376
refGene_gn=ABCB4;refGene_tx=XM_023250316.1;refGene_gn=ABCB4;refGene_tx=XM_023250317.1;refGene_gn=ABCB4;refGene_tx=XM_023250318.1;refGene_gn=ABCB4;refGene_tx=XM_023250319.1
A2 93058636 93058808
refGene_gn=ABCB4;refGene_tx=XM_023250316.1;refGene_gn=ABCB4;refGene_tx=XM_023250317.1;refGene_gn=ABCB4;refGene_tx=XM_023250318.1;refGene_gn=ABCB4;refGene_tx=XM_023250319.1
A2 93059896 93060088

refGene_gn=ABCB4;refGene_tx=XM_023250316.1;refGene_gn=ABCB4;refGene_tx=XM_023250317.1;refGene_gn=ABCB4;refGene_tx=XM_023250318.1
A2 93061615 93061673
refGene_gn=ABCB4;refGene_tx=XM_023250316.1;refGene_gn=ABCB4;refGene_tx=XM_023250317.1;refGene_gn=ABCB4;refGene_tx=XM_023250318.1
A2 93068858 93069009
refGene_gn=ABCB4;refGene_tx=XM_023250316.1;refGene_gn=ABCB4;refGene_tx=XM_023250317.1;refGene_gn=ABCB4;refGene_tx=XM_023250318.1
A2 93072400 93072455
refGene_gn=ABCB4;refGene_tx=XM_023250316.1;refGene_gn=ABCB4;refGene_tx=XM_023250317.1;refGene_gn=ABCB4;refGene_tx=XM_023250318.1
A2 93075689 93075769
refGene_gn=ABCB4;refGene_tx=XM_023250316.1;refGene_gn=ABCB4;refGene_tx=XM_023250317.1;refGene_gn=ABCB4;refGene_tx=XM_023250318.1
A2 93100013 93100217
refGene_gn=ABCB1;refGene_tx=NM_001171064.2;refGene_gn=ABCB1;refGene_tx=XM_019814933.2;refGene_gn=ABCB1;refGene_tx=XM_019814935.2;refGene_gn=ABCB1;refGene_tx=XM_019814938.1
A2 93101383 93101530
refGene_gn=ABCB1;refGene_tx=NM_001171064.2;refGene_gn=ABCB1;refGene_tx=XM_019814933.2;refGene_gn=ABCB1;refGene_tx=XM_019814935.2;refGene_gn=ABCB1;refGene_tx=XM_019814938.1
A2 93104621 93104828
refGene_gn=ABCB1;refGene_tx=NM_001171064.2;refGene_gn=ABCB1;refGene_tx=XM_019814933.2;refGene_gn=ABCB1;refGene_tx=XM_019814935.2;refGene_gn=ABCB1;refGene_tx=XM_019814938.1
A2 93116924 93117122
refGene_gn=ABCB1;refGene_tx=NM_001171064.2;refGene_gn=ABCB1;refGene_tx=XM_019814933.2;refGene_gn=ABCB1;refGene_tx=XM_019814935.2;refGene_gn=ABCB1;refGene_tx=XM_019814938.1
A2 93117620 93117777
refGene_gn=ABCB1;refGene_tx=NM_001171064.2;refGene_gn=ABCB1;refGene_tx=XM_019814933.2;refGene_gn=ABCB1;refGene_tx=XM_019814935.2;refGene_gn=ABCB1;refGene_tx=XM_019814938.1
A2 93121017 93121158
refGene_gn=ABCB1;refGene_tx=NM_001171064.2;refGene_gn=ABCB1;refGene_tx=XM_019814933.2;refGene_gn=ABCB1;refGene_tx=XM_019814935.2;refGene_gn=ABCB1;refGene_tx=XM_019814938.1
A2 93122400 93122501
refGene_gn=ABCB1;refGene_tx=NM_001171064.2;refGene_gn=ABCB1;refGene_tx=XM_019814933.2;refGene_gn=ABCB1;refGene_tx=XM_019814935.2;refGene_gn=ABCB1;refGene_tx=XM_019814938.1
A2 93126613 93126817
refGene_gn=ABCB1;refGene_tx=NM_001171064.2;refGene_gn=ABCB1;refGene_tx=XM_019814933.2;refGene_gn=ABCB1;refGene_tx=XM_019814938.1
A2 93133084 93133168
refGene_gn=ABCB1;refGene_tx=NM_001171064.2;refGene_gn=ABCB1;refGene_tx=XM_019814933.2;refGene_gn=ABCB1;refGene_tx=XM_019814935.2;refGene_gn=ABCB1;refGene_tx=XM_019814938.1
A2 93136064 93136142

refGene_gn=ABCB1;refGene_tx=NM_001171064.2;refGene_gn=ABCB1;refGene_tx=XM_019814933.2;refGene_gn=ABCB1;refGene_tx=XM_019814935.2;refGene_gn=ABCB1;refGene_tx=XM_019814938.1
A2 93141175 93141283
refGene_gn=ABCB1;refGene_tx=NM_001171064.2;refGene_gn=ABCB1;refGene_tx=XM_019814933.2;refGene_gn=ABCB1;refGene_tx=XM_019814935.2;refGene_gn=ABCB1;refGene_tx=XM_019814938.1
A2 93143528 93143675
refGene_gn=ABCB1;refGene_tx=NM_001171064.2;refGene_gn=ABCB1;refGene_tx=XM_019814933.2;refGene_gn=ABCB1;refGene_tx=XM_019814935.2;refGene_gn=ABCB1;refGene_tx=XM_019814938.1
A2 93144221 93144398
refGene_gn=ABCB1;refGene_tx=NM_001171064.2;refGene_gn=ABCB1;refGene_tx=XM_019814933.2;refGene_gn=ABCB1;refGene_tx=XM_019814935.2;refGene_gn=ABCB1;refGene_tx=XM_019814938.1
A2 93145480 93145642
refGene_gn=ABCB1;refGene_tx=NM_001171064.2;refGene_gn=ABCB1;refGene_tx=XM_019814933.2;refGene_gn=ABCB1;refGene_tx=XM_019814935.2;refGene_gn=ABCB1;refGene_tx=XM_019814938.1
A2 93154510 93154681
refGene_gn=ABCB1;refGene_tx=NM_001171064.2;refGene_gn=ABCB1;refGene_tx=XM_019814933.2;refGene_gn=ABCB1;refGene_tx=XM_019814935.2;refGene_gn=ABCB1;refGene_tx=XM_019814938.1
A2 93155002 93155206
refGene_gn=ABCB1;refGene_tx=NM_001171064.2;refGene_gn=ABCB1;refGene_tx=XM_019814933.2;refGene_gn=ABCB1;refGene_tx=XM_019814935.2;refGene_gn=ABCB1;refGene_tx=XM_019814938.1
A2 93155315 93155441
refGene_gn=ABCB1;refGene_tx=NM_001171064.2;refGene_gn=ABCB1;refGene_tx=XM_019814933.2;refGene_gn=ABCB1;refGene_tx=XM_019814935.2;refGene_gn=ABCB1;refGene_tx=XM_019814938.1
A2 93155620 93155731
refGene_gn=ABCB1;refGene_tx=NM_001171064.2;refGene_gn=ABCB1;refGene_tx=XM_019814933.2;refGene_gn=ABCB1;refGene_tx=XM_019814935.2;refGene_gn=ABCB1;refGene_tx=XM_019814938.1
A2 93155848 93155962
refGene_gn=ABCB1;refGene_tx=NM_001171064.2;refGene_gn=ABCB1;refGene_tx=XM_019814933.2;refGene_gn=ABCB1;refGene_tx=XM_019814935.2;refGene_gn=ABCB1;refGene_tx=XM_019814938.1
A2 93158272 93158444
refGene_gn=ABCB1;refGene_tx=NM_001171064.2;refGene_gn=ABCB1;refGene_tx=XM_019814933.2;refGene_gn=ABCB1;refGene_tx=XM_019814935.2;refGene_gn=ABCB1;refGene_tx=XM_019814938.1
A2 93166814 93166939
refGene_gn=ABCB1;refGene_tx=NM_001171064.2;refGene_gn=ABCB1;refGene_tx=XM_019814933.2;refGene_gn=ABCB1;refGene_tx=XM_019814935.2;refGene_gn=ABCB1;refGene_tx=XM_019814938.1
A2 93179703 93179875
refGene_gn=ABCB1;refGene_tx=NM_001171064.2;refGene_gn=ABCB1;refGene_tx=XM_019814933.2;refGene_gn=ABCB1;refGene_tx=XM_019814935.2;refGene_gn=

ABCB1;refGene_tx=XM_019814938.1
A2 93180287 93180479
refGene_gn=ABCB1;refGene_tx=NM_001171064.2;refGene_gn=ABCB1;refGene_tx=XM_019814933.2;refGene_gn=ABCB1;refGene_tx=XM_019814935.2;refGene_gn=ABCB1;refGene_tx=XM_019814938.1
A2 93183003 93183055
refGene_gn=ABCB1;refGene_tx=NM_001171064.2;refGene_gn=ABCB1;refGene_tx=XM_019814933.2;refGene_gn=ABCB1;refGene_tx=XM_019814935.2;refGene_gn=ABCB1;refGene_tx=XM_019814938.1
A2 93195454 93195626
refGene_gn=ABCB1;refGene_tx=NM_001171064.2;refGene_gn=ABCB1;refGene_tx=XM_019814933.2;refGene_gn=ABCB1;refGene_tx=XM_019814935.2
A2 93207780 93207829
refGene_gn=ABCB1;refGene_tx=NM_001171064.2;refGene_gn=ABCB1;refGene_tx=XM_019814933.2;refGene_gn=ABCB1;refGene_tx=XM_019814935.2
A2 93212956 93213021
refGene_gn=ABCB1;refGene_tx=NM_001171064.2;refGene_gn=ABCB1;refGene_tx=XM_019814933.2;refGene_gn=ABCB1;refGene_tx=XM_019814935.2
A2 93263881 93264003
refGene_gn=RUNDC3B;refGene_tx=XM_003982716.4;refGene_gn=RUNDC3B;refGene_tx=XM_003982717.5;refGene_gn=RUNDC3B;refGene_tx=XM_003982718.4;refGene_gn=RUNDC3B;refGene_tx=XM_006929153.3
A2 93282643 93282759
refGene_gn=RUNDC3B;refGene_tx=XM_003982716.4;refGene_gn=RUNDC3B;refGene_tx=XM_003982717.5;refGene_gn=RUNDC3B;refGene_tx=XM_003982718.4;refGene_gn=RUNDC3B;refGene_tx=XM_006929153.3
A2 93315498 93315549
refGene_gn=RUNDC3B;refGene_tx=XM_003982716.4;refGene_gn=RUNDC3B;refGene_tx=XM_006929153.3
A2 93318323 93318498 refGene_gn=RUNDC3B;refGene_tx=XM_019825408.2
A2 93321274 93321408
refGene_gn=RUNDC3B;refGene_tx=XM_003982716.4;refGene_gn=RUNDC3B;refGene_tx=XM_003982717.5;refGene_gn=RUNDC3B;refGene_tx=XM_003982718.4;refGene_gn=RUNDC3B;refGene_tx=XM_006929153.3;refGene_gn=RUNDC3B;refGene_tx=XM_019825408.2
A2 93332224 93332310
refGene_gn=RUNDC3B;refGene_tx=XM_003982716.4;refGene_gn=RUNDC3B;refGene_tx=XM_003982717.5;refGene_gn=RUNDC3B;refGene_tx=XM_003982718.4;refGene_gn=RUNDC3B;refGene_tx=XM_006929153.3;refGene_gn=RUNDC3B;refGene_tx=XM_019825408.2
A2 93342734 93342824
refGene_gn=RUNDC3B;refGene_tx=XM_003982716.4;refGene_gn=RUNDC3B;refGene_tx=XM_003982717.5;refGene_gn=RUNDC3B;refGene_tx=XM_003982718.4;refGene_gn=RUNDC3B;refGene_tx=XM_006929153.3;refGene_gn=RUNDC3B;refGene_tx=XM_019825408.2
A2 93344902 93344983
refGene_gn=RUNDC3B;refGene_tx=XM_003982716.4;refGene_gn=RUNDC3B;refGene_tx=XM_003982717.5;refGene_gn=RUNDC3B;refGene_tx=XM_003982718.4;refGene_gn=RUNDC3B;refGene_tx=XM_006929153.3;refGene_gn=RUNDC3B;refGene_tx=XM_019825408.2

A2	93357212 93357381
refGene_gn=RUNDC3B;refGene_tx=XM_003982716.4;refGene_gn=RUNDC3B;refGene_tx=XM_003982717.5;refGene_gn=RUNDC3B;refGene_tx=XM_003982718.4;refGene_gn=RUNDC3B;refGene_tx=XM_006929153.3;refGene_gn=RUNDC3B;refGene_tx=XM_019825408.2
A2	93358251 93358409
refGene_gn=RUNDC3B;refGene_tx=XM_003982716.4;refGene_gn=RUNDC3B;refGene_tx=XM_003982717.5;refGene_gn=RUNDC3B;refGene_tx=XM_003982718.4;refGene_gn=RUNDC3B;refGene_tx=XM_006929153.3;refGene_gn=RUNDC3B;refGene_tx=XM_019825408.2
A2	93387205 93387352
refGene_gn=RUNDC3B;refGene_tx=XM_003982716.4;refGene_gn=RUNDC3B;refGene_tx=XM_003982718.4;refGene_gn=RUNDC3B;refGene_tx=XM_019825408.2
A2	93393939 93394061
refGene_gn=RUNDC3B;refGene_tx=XM_003982716.4;refGene_gn=RUNDC3B;refGene_tx=XM_003982717.5;refGene_gn=RUNDC3B;refGene_tx=XM_003982718.4;refGene_gn=RUNDC3B;refGene_tx=XM_006929153.3;refGene_gn=RUNDC3B;refGene_tx=XM_019825408.2
A2	93409139 93409285
refGene_gn=RUNDC3B;refGene_tx=XM_003982716.4;refGene_gn=RUNDC3B;refGene_tx=XM_003982717.5;refGene_gn=RUNDC3B;refGene_tx=XM_003982718.4;refGene_gn=RUNDC3B;refGene_tx=XM_006929153.3;refGene_gn=RUNDC3B;refGene_tx=XM_019825408.2
A2	93415251 93415367
refGene_gn=SLC25A40;refGene_tx=XM_011280365.3;refGene_gn=SLC25A40;refGene_tx=XM_019825410.2
A2	93415718 93415799
refGene_gn=SLC25A40;refGene_tx=XM_011280365.3;refGene_gn=SLC25A40;refGene_tx=XM_019825410.2
A2	93419969 93420051
refGene_gn=SLC25A40;refGene_tx=XM_011280365.3;refGene_gn=SLC25A40;refGene_tx=XM_019825410.2
A2	93422376 93422486
refGene_gn=SLC25A40;refGene_tx=XM_011280365.3;refGene_gn=SLC25A40;refGene_tx=XM_019825410.2
A2	93432502 93432676
refGene_gn=SLC25A40;refGene_tx=XM_011280365.3;refGene_gn=SLC25A40;refGene_tx=XM_019825410.2
A2	93433546 93433671
refGene_gn=SLC25A40;refGene_tx=XM_011280365.3;refGene_gn=SLC25A40;refGene_tx=XM_019825410.2
A2	93436004 93436072
refGene_gn=SLC25A40;refGene_tx=XM_011280365.3;refGene_gn=SLC25A40;refGene_tx=XM_019825410.2
A2	93438277 93438376 refGene_gn=SLC25A40;refGene_tx=XM_019825410.2
A2	93447461 93447568 refGene_gn=SLC25A40;refGene_tx=XM_011280365.3
A2	93449379 93449439 refGene_gn=SLC25A40;refGene_tx=XM_011280365.3
A2	93450659 93450756 refGene_gn=SLC25A40;refGene_tx=XM_011280365.3
A2	93467544 93467590
refGene_gn=DBF4;refGene_tx=XM_011280366.3;refGene_gn=DBF4;refGene_tx=X

M_019825411.2
A2 93469082 93469255
refGene_gn=DBF4;refGene_tx=XM_011280366.3;refGene_gn=DBF4;refGene_tx=XM_019825411.2
A2 93475412 93475592
refGene_gn=DBF4;refGene_tx=XM_011280366.3;refGene_gn=DBF4;refGene_tx=XM_019825411.2
A2 93477049 93477100
refGene_gn=DBF4;refGene_tx=XM_011280366.3;refGene_gn=DBF4;refGene_tx=XM_019825411.2
A2 93477584 93477654
refGene_gn=DBF4;refGene_tx=XM_011280366.3;refGene_gn=DBF4;refGene_tx=XM_019825411.2
A2 93477858 93477935
refGene_gn=DBF4;refGene_tx=XM_011280366.3;refGene_gn=DBF4;refGene_tx=XM_019825411.2
A2 93486232 93486269
refGene_gn=DBF4;refGene_tx=XM_011280366.3;refGene_gn=DBF4;refGene_tx=XM_019825411.2
A2 93487529 93487575
refGene_gn=DBF4;refGene_tx=XM_011280366.3;refGene_gn=DBF4;refGene_tx=XM_019825411.2;refGene_gn=DBF4;refGene_tx=XM_019825412.1
A2 93490554 93490683
refGene_gn=DBF4;refGene_tx=XM_011280366.3;refGene_gn=DBF4;refGene_tx=XM_019825411.2;refGene_gn=DBF4;refGene_tx=XM_019825412.1
A2 93491142 93491254
refGene_gn=DBF4;refGene_tx=XM_011280366.3;refGene_gn=DBF4;refGene_tx=XM_019825411.2;refGene_gn=DBF4;refGene_tx=XM_019825412.1
A2 93494392 93494517
refGene_gn=DBF4;refGene_tx=XM_011280366.3;refGene_gn=DBF4;refGene_tx=XM_019825411.2;refGene_gn=DBF4;refGene_tx=XM_019825412.1
A2 93496751 93497736
refGene_gn=DBF4;refGene_tx=XM_011280366.3;refGene_gn=DBF4;refGene_tx=XM_019825411.2;refGene_gn=DBF4;refGene_tx=XM_019825412.1
A2 93513573 93513658
refGene_gn=ADAM22;refGene_tx=XM_011280369.3;refGene_gn=ADAM22;refGene_tx=XM_011280370.3;refGene_gn=ADAM22;refGene_tx=XM_011280371.3;refGene_gn=ADAM22;refGene_tx=XM_019825413.2;refGene_gn=ADAM22;refGene_tx=XM_019825415.2;refGene_gn=ADAM22;refGene_tx=XM_019825416.2;refGene_gn=ADAM22;refGene_tx=XM_019825417.2;refGene_gn=ADAM22;refGene_tx=XM_019825418.2;refGene_gn=ADAM22;refGene_tx=XM_019825419.2;refGene_gn=ADAM22;refGene_tx=XM_019825420.2;refGene_gn=ADAM22;refGene_tx=XM_019825421.2;refGene_gn=ADAM22;refGene_tx=XM_019825422.2;refGene_gn=ADAM22;refGene_tx=XM_019825423.2;refGene_gn=ADAM22;refGene_tx=XM_019825424.2;refGene_gn=ADAM22;refGene_tx=XM_019825425.2;refGene_gn=ADAM22;refGene_tx=XM_019825426.2;refGene_gn=ADAM22;refGene_tx=XM_019825428.2;refGene_gn=ADAM22;refGene_tx=XM_019825431.2;refGene_gn=ADAM22;refGene_tx=XM_019825433.2;refGene_gn=ADAM22;refGene_tx=XM_019825434.2;refGene_gn=ADAM22;refGene_tx=XM_019825436.2;refGene_gn=ADAM22;refGene_tx=XM_019825437.2
A2 93514121 93514282

refGene_gn=ADAM22;refGene_tx=XM_011280369.3;refGene_gn=ADAM22;refGene_tx=XM_011280370.3;refGene_gn=ADAM22;refGene_tx=XM_011280371.3;refGene_gn=ADAM22;refGene_tx=XM_019825413.2;refGene_gn=ADAM22;refGene_tx=XM_019825415.2;refGene_gn=ADAM22;refGene_tx=XM_019825416.2;refGene_gn=ADAM22;refGene_tx=XM_019825417.2;refGene_gn=ADAM22;refGene_tx=XM_019825418.2;refGene_gn=ADAM22;refGene_tx=XM_019825419.2;refGene_gn=ADAM22;refGene_tx=XM_019825420.2;refGene_gn=ADAM22;refGene_tx=XM_019825421.2;refGene_gn=ADAM22;refGene_tx=XM_019825422.2;refGene_gn=ADAM22;refGene_tx=XM_019825423.2;refGene_gn=ADAM22;refGene_tx=XM_019825424.2;refGene_gn=ADAM22;refGene_tx=XM_019825425.2;refGene_gn=ADAM22;refGene_tx=XM_019825426.2;refGene_gn=ADAM22;refGene_tx=XM_019825428.2;refGene_gn=ADAM22;refGene_tx=XM_019825431.2;refGene_gn=ADAM22;refGene_tx=XM_019825433.2;refGene_gn=ADAM22;refGene_tx=XM_019825434.2;refGene_gn=ADAM22;refGene_tx=XM_019825436.2;refGene_gn=ADAM22;refGene_tx=XM_019825437.2
A2 93555722 93555799
refGene_gn=ADAM22;refGene_tx=XM_011280369.3;refGene_gn=ADAM22;refGene_tx=XM_011280370.3;refGene_gn=ADAM22;refGene_tx=XM_011280371.3;refGene_gn=ADAM22;refGene_tx=XM_019825413.2;refGene_gn=ADAM22;refGene_tx=XM_019825415.2;refGene_gn=ADAM22;refGene_tx=XM_019825416.2;refGene_gn=ADAM22;refGene_tx=XM_019825417.2;refGene_gn=ADAM22;refGene_tx=XM_019825418.2;refGene_gn=ADAM22;refGene_tx=XM_019825419.2;refGene_gn=ADAM22;refGene_tx=XM_019825420.2;refGene_gn=ADAM22;refGene_tx=XM_019825421.2;refGene_gn=ADAM22;refGene_tx=XM_019825422.2;refGene_gn=ADAM22;refGene_tx=XM_019825423.2;refGene_gn=ADAM22;refGene_tx=XM_019825424.2;refGene_gn=ADAM22;refGene_tx=XM_019825425.2;refGene_gn=ADAM22;refGene_tx=XM_019825426.2;refGene_gn=ADAM22;refGene_tx=XM_019825428.2;refGene_gn=ADAM22;refGene_tx=XM_019825431.2;refGene_gn=ADAM22;refGene_tx=XM_019825433.2;refGene_gn=ADAM22;refGene_tx=XM_019825434.2;refGene_gn=ADAM22;refGene_tx=XM_019825436.2;refGene_gn=ADAM22;refGene_tx=XM_019825437.2
A2 93638504 93638571
refGene_gn=ADAM22;refGene_tx=XM_011280369.3;refGene_gn=ADAM22;refGene_tx=XM_011280370.3;refGene_gn=ADAM22;refGene_tx=XM_011280371.3;refGene_gn=ADAM22;refGene_tx=XM_019825413.2;refGene_gn=ADAM22;refGene_tx=XM_019825415.2;refGene_gn=ADAM22;refGene_tx=XM_019825416.2;refGene_gn=ADAM22;refGene_tx=XM_019825417.2;refGene_gn=ADAM22;refGene_tx=XM_019825418.2;refGene_gn=ADAM22;refGene_tx=XM_019825419.2;refGene_gn=ADAM22;refGene_tx=XM_019825420.2;refGene_gn=ADAM22;refGene_tx=XM_019825421.2;refGene_gn=ADAM22;refGene_tx=XM_019825422.2;refGene_gn=ADAM22;refGene_tx=XM_019825423.2;refGene_gn=ADAM22;refGene_tx=XM_019825424.2;refGene_gn=ADAM22;refGene_tx=XM_019825425.2;refGene_gn=ADAM22;refGene_tx=XM_019825426.2;refGene_gn=ADAM22;refGene_tx=XM_019825428.2;refGene_gn=ADAM22;refGene_tx=XM_019825431.2;refGene_gn=ADAM22;refGene_tx=XM_019825433.2;refGene_gn=ADAM22;refGene_tx=XM_019825434.2;refGene_gn=ADAM22;refGene_tx=XM_019825436.2;refGene_gn=ADAM22;refGene_tx=XM_019825437.2
A2 93666655 93666738
refGene_gn=ADAM22;refGene_tx=XM_011280369.3;refGene_gn=ADAM22;refGene_tx=XM_011280370.3;refGene_gn=ADAM22;refGene_tx=XM_011280371.3;refGene_gn=ADAM22;refGene_tx=XM_019825413.2;refGene_gn=ADAM22;refGene_tx=XM_019825415.2;refGene_gn=ADAM22;refGene_tx=XM_019825416.2;refGene_gn=ADAM22;refGene_tx=XM_019825417.2;refGene_gn=ADAM22;refGene_tx=XM_019825418.

2;refGene_gn=ADAM22;refGene_tx=XM_019825419.2;refGene_gn=ADAM22;refGene_tx=XM_019825420.2;refGene_gn=ADAM22;refGene_tx=XM_019825421.2;refGene_gn=ADAM22;refGene_tx=XM_019825422.2;refGene_gn=ADAM22;refGene_tx=XM_019825423.2;refGene_gn=ADAM22;refGene_tx=XM_019825424.2;refGene_gn=ADAM22;refGene_tx=XM_019825425.2;refGene_gn=ADAM22;refGene_tx=XM_019825426.2;refGene_gn=ADAM22;refGene_tx=XM_019825428.2;refGene_gn=ADAM22;refGene_tx=XM_019825431.2;refGene_gn=ADAM22;refGene_tx=XM_019825433.2;refGene_gn=ADAM22;refGene_tx=XM_019825434.2;refGene_gn=ADAM22;refGene_tx=XM_019825436.2;refGene_gn=ADAM22;refGene_tx=XM_019825437.2
A2 93672647 93672711
refGene_gn=ADAM22;refGene_tx=XM_011280369.3;refGene_gn=ADAM22;refGene_tx=XM_011280370.3;refGene_gn=ADAM22;refGene_tx=XM_011280371.3;refGene_gn=ADAM22;refGene_tx=XM_019825413.2;refGene_gn=ADAM22;refGene_tx=XM_019825415.2;refGene_gn=ADAM22;refGene_tx=XM_019825416.2;refGene_gn=ADAM22;refGene_tx=XM_019825417.2;refGene_gn=ADAM22;refGene_tx=XM_019825418.2;refGene_gn=ADAM22;refGene_tx=XM_019825419.2;refGene_gn=ADAM22;refGene_tx=XM_019825420.2;refGene_gn=ADAM22;refGene_tx=XM_019825421.2;refGene_gn=ADAM22;refGene_tx=XM_019825422.2;refGene_gn=ADAM22;refGene_tx=XM_019825423.2;refGene_gn=ADAM22;refGene_tx=XM_019825424.2;refGene_gn=ADAM22;refGene_tx=XM_019825425.2;refGene_gn=ADAM22;refGene_tx=XM_019825426.2;refGene_gn=ADAM22;refGene_tx=XM_019825428.2;refGene_gn=ADAM22;refGene_tx=XM_019825431.2;refGene_gn=ADAM22;refGene_tx=XM_019825433.2;refGene_gn=ADAM22;refGene_tx=XM_019825434.2;refGene_gn=ADAM22;refGene_tx=XM_019825436.2;refGene_gn=ADAM22;refGene_tx=XM_019825437.2
A2 93674363 93674433
refGene_gn=ADAM22;refGene_tx=XM_011280369.3;refGene_gn=ADAM22;refGene_tx=XM_011280370.3;refGene_gn=ADAM22;refGene_tx=XM_011280371.3;refGene_gn=ADAM22;refGene_tx=XM_019825413.2;refGene_gn=ADAM22;refGene_tx=XM_019825415.2;refGene_gn=ADAM22;refGene_tx=XM_019825416.2;refGene_gn=ADAM22;refGene_tx=XM_019825417.2;refGene_gn=ADAM22;refGene_tx=XM_019825418.2;refGene_gn=ADAM22;refGene_tx=XM_019825419.2;refGene_gn=ADAM22;refGene_tx=XM_019825420.2;refGene_gn=ADAM22;refGene_tx=XM_019825421.2;refGene_gn=ADAM22;refGene_tx=XM_019825422.2;refGene_gn=ADAM22;refGene_tx=XM_019825423.2;refGene_gn=ADAM22;refGene_tx=XM_019825424.2;refGene_gn=ADAM22;refGene_tx=XM_019825425.2;refGene_gn=ADAM22;refGene_tx=XM_019825426.2;refGene_gn=ADAM22;refGene_tx=XM_019825428.2;refGene_gn=ADAM22;refGene_tx=XM_019825431.2;refGene_gn=ADAM22;refGene_tx=XM_019825433.2;refGene_gn=ADAM22;refGene_tx=XM_019825434.2;refGene_gn=ADAM22;refGene_tx=XM_019825436.2;refGene_gn=ADAM22;refGene_tx=XM_019825437.2
A2 93679588 93679659
refGene_gn=ADAM22;refGene_tx=XM_011280369.3;refGene_gn=ADAM22;refGene_tx=XM_011280370.3;refGene_gn=ADAM22;refGene_tx=XM_011280371.3;refGene_gn=ADAM22;refGene_tx=XM_019825413.2;refGene_gn=ADAM22;refGene_tx=XM_019825415.2;refGene_gn=ADAM22;refGene_tx=XM_019825416.2;refGene_gn=ADAM22;refGene_tx=XM_019825417.2;refGene_gn=ADAM22;refGene_tx=XM_019825418.2;refGene_gn=ADAM22;refGene_tx=XM_019825419.2;refGene_gn=ADAM22;refGene_tx=XM_019825420.2;refGene_gn=ADAM22;refGene_tx=XM_019825421.2;refGene_gn=ADAM22;refGene_tx=XM_019825422.2;refGene_gn=ADAM22;refGene_tx=XM_019825423.2;refGene_gn=ADAM22;refGene_tx=XM_019825424.2;refGene_gn=ADAM22;refGene_tx=XM_019825425.2;refGene_gn=ADAM22;refGene_tx=XM_01982542

6.2;refGene_gn=ADAM22;refGene_tx=XM_019825428.2;refGene_gn=ADAM22;refGene_tx=XM_019825431.2;refGene_gn=ADAM22;refGene_tx=XM_019825433.2;refGene_gn=ADAM22;refGene_tx=XM_019825434.2;refGene_gn=ADAM22;refGene_tx=XM_019825436.2;refGene_gn=ADAM22;refGene_tx=XM_019825437.2
A2 93683653 93683728
refGene_gn=ADAM22;refGene_tx=XM_011280369.3;refGene_gn=ADAM22;refGene_tx=XM_011280370.3;refGene_gn=ADAM22;refGene_tx=XM_011280371.3;refGene_gn=ADAM22;refGene_tx=XM_019825413.2;refGene_gn=ADAM22;refGene_tx=XM_019825415.2;refGene_gn=ADAM22;refGene_tx=XM_019825416.2;refGene_gn=ADAM22;refGene_tx=XM_019825417.2;refGene_gn=ADAM22;refGene_tx=XM_019825418.2;refGene_gn=ADAM22;refGene_tx=XM_019825419.2;refGene_gn=ADAM22;refGene_tx=XM_019825420.2;refGene_gn=ADAM22;refGene_tx=XM_019825421.2;refGene_gn=ADAM22;refGene_tx=XM_019825422.2;refGene_gn=ADAM22;refGene_tx=XM_019825423.2;refGene_gn=ADAM22;refGene_tx=XM_019825424.2;refGene_gn=ADAM22;refGene_tx=XM_019825425.2;refGene_gn=ADAM22;refGene_tx=XM_019825426.2;refGene_gn=ADAM22;refGene_tx=XM_019825428.2;refGene_gn=ADAM22;refGene_tx=XM_019825431.2;refGene_gn=ADAM22;refGene_tx=XM_019825433.2;refGene_gn=ADAM22;refGene_tx=XM_019825434.2;refGene_gn=ADAM22;refGene_tx=XM_019825436.2;refGene_gn=ADAM22;refGene_tx=XM_019825437.2
A2 93685334 93685406
refGene_gn=ADAM22;refGene_tx=XM_011280369.3;refGene_gn=ADAM22;refGene_tx=XM_011280370.3;refGene_gn=ADAM22;refGene_tx=XM_011280371.3;refGene_gn=ADAM22;refGene_tx=XM_019825413.2;refGene_gn=ADAM22;refGene_tx=XM_019825415.2;refGene_gn=ADAM22;refGene_tx=XM_019825416.2;refGene_gn=ADAM22;refGene_tx=XM_019825417.2;refGene_gn=ADAM22;refGene_tx=XM_019825418.2;refGene_gn=ADAM22;refGene_tx=XM_019825419.2;refGene_gn=ADAM22;refGene_tx=XM_019825420.2;refGene_gn=ADAM22;refGene_tx=XM_019825421.2;refGene_gn=ADAM22;refGene_tx=XM_019825422.2;refGene_gn=ADAM22;refGene_tx=XM_019825423.2;refGene_gn=ADAM22;refGene_tx=XM_019825424.2;refGene_gn=ADAM22;refGene_tx=XM_019825425.2;refGene_gn=ADAM22;refGene_tx=XM_019825426.2;refGene_gn=ADAM22;refGene_tx=XM_019825428.2;refGene_gn=ADAM22;refGene_tx=XM_019825431.2;refGene_gn=ADAM22;refGene_tx=XM_019825433.2;refGene_gn=ADAM22;refGene_tx=XM_019825434.2;refGene_gn=ADAM22;refGene_tx=XM_019825436.2;refGene_gn=ADAM22;refGene_tx=XM_019825437.2
A2 93686220 93686387
refGene_gn=ADAM22;refGene_tx=XM_011280369.3;refGene_gn=ADAM22;refGene_tx=XM_011280370.3;refGene_gn=ADAM22;refGene_tx=XM_011280371.3;refGene_gn=ADAM22;refGene_tx=XM_019825413.2;refGene_gn=ADAM22;refGene_tx=XM_019825415.2;refGene_gn=ADAM22;refGene_tx=XM_019825416.2;refGene_gn=ADAM22;refGene_tx=XM_019825417.2;refGene_gn=ADAM22;refGene_tx=XM_019825418.2;refGene_gn=ADAM22;refGene_tx=XM_019825419.2;refGene_gn=ADAM22;refGene_tx=XM_019825420.2;refGene_gn=ADAM22;refGene_tx=XM_019825421.2;refGene_gn=ADAM22;refGene_tx=XM_019825422.2;refGene_gn=ADAM22;refGene_tx=XM_019825423.2;refGene_gn=ADAM22;refGene_tx=XM_019825424.2;refGene_gn=ADAM22;refGene_tx=XM_019825425.2;refGene_gn=ADAM22;refGene_tx=XM_019825426.2;refGene_gn=ADAM22;refGene_tx=XM_019825428.2;refGene_gn=ADAM22;refGene_tx=XM_019825431.2;refGene_gn=ADAM22;refGene_tx=XM_019825433.2;refGene_gn=ADAM22;refGene_tx=XM_019825434.2;refGene_gn=ADAM22;refGene_tx=XM_019825436.2;refGene_gn=ADAM22;refGene_tx=XM_019825437.2
A2 93688083 93688168

refGene_gn=ADAM22;refGene_tx=XM_011280369.3;refGene_gn=ADAM22;refGene_tx=XM_011280370.3;refGene_gn=ADAM22;refGene_tx=XM_011280371.3;refGene_gn=ADAM22;refGene_tx=XM_019825413.2;refGene_gn=ADAM22;refGene_tx=XM_019825415.2;refGene_gn=ADAM22;refGene_tx=XM_019825416.2;refGene_gn=ADAM22;refGene_tx=XM_019825417.2;refGene_gn=ADAM22;refGene_tx=XM_019825418.2;refGene_gn=ADAM22;refGene_tx=XM_019825419.2;refGene_gn=ADAM22;refGene_tx=XM_019825420.2;refGene_gn=ADAM22;refGene_tx=XM_019825421.2;refGene_gn=ADAM22;refGene_tx=XM_019825422.2;refGene_gn=ADAM22;refGene_tx=XM_019825423.2;refGene_gn=ADAM22;refGene_tx=XM_019825424.2;refGene_gn=ADAM22;refGene_tx=XM_019825425.2;refGene_gn=ADAM22;refGene_tx=XM_019825426.2;refGene_gn=ADAM22;refGene_tx=XM_019825428.2;refGene_gn=ADAM22;refGene_tx=XM_019825431.2;refGene_gn=ADAM22;refGene_tx=XM_019825433.2;refGene_gn=ADAM22;refGene_tx=XM_019825434.2;refGene_gn=ADAM22;refGene_tx=XM_019825436.2;refGene_gn=ADAM22;refGene_tx=XM_019825437.2
A2 93689187 93689278
refGene_gn=ADAM22;refGene_tx=XM_011280369.3;refGene_gn=ADAM22;refGene_tx=XM_011280370.3;refGene_gn=ADAM22;refGene_tx=XM_011280371.3;refGene_gn=ADAM22;refGene_tx=XM_019825413.2;refGene_gn=ADAM22;refGene_tx=XM_019825415.2;refGene_gn=ADAM22;refGene_tx=XM_019825416.2;refGene_gn=ADAM22;refGene_tx=XM_019825417.2;refGene_gn=ADAM22;refGene_tx=XM_019825418.2;refGene_gn=ADAM22;refGene_tx=XM_019825419.2;refGene_gn=ADAM22;refGene_tx=XM_019825420.2;refGene_gn=ADAM22;refGene_tx=XM_019825421.2;refGene_gn=ADAM22;refGene_tx=XM_019825422.2;refGene_gn=ADAM22;refGene_tx=XM_019825423.2;refGene_gn=ADAM22;refGene_tx=XM_019825424.2;refGene_gn=ADAM22;refGene_tx=XM_019825425.2;refGene_gn=ADAM22;refGene_tx=XM_019825426.2;refGene_gn=ADAM22;refGene_tx=XM_019825428.2;refGene_gn=ADAM22;refGene_tx=XM_019825431.2;refGene_gn=ADAM22;refGene_tx=XM_019825433.2;refGene_gn=ADAM22;refGene_tx=XM_019825434.2;refGene_gn=ADAM22;refGene_tx=XM_019825436.2;refGene_gn=ADAM22;refGene_tx=XM_019825437.2
A2 93690703 93690755
refGene_gn=ADAM22;refGene_tx=XM_011280369.3;refGene_gn=ADAM22;refGene_tx=XM_011280370.3;refGene_gn=ADAM22;refGene_tx=XM_011280371.3;refGene_gn=ADAM22;refGene_tx=XM_019825413.2;refGene_gn=ADAM22;refGene_tx=XM_019825415.2;refGene_gn=ADAM22;refGene_tx=XM_019825416.2;refGene_gn=ADAM22;refGene_tx=XM_019825417.2;refGene_gn=ADAM22;refGene_tx=XM_019825418.2;refGene_gn=ADAM22;refGene_tx=XM_019825419.2;refGene_gn=ADAM22;refGene_tx=XM_019825420.2;refGene_gn=ADAM22;refGene_tx=XM_019825421.2;refGene_gn=ADAM22;refGene_tx=XM_019825422.2;refGene_gn=ADAM22;refGene_tx=XM_019825423.2;refGene_gn=ADAM22;refGene_tx=XM_019825424.2;refGene_gn=ADAM22;refGene_tx=XM_019825425.2;refGene_gn=ADAM22;refGene_tx=XM_019825426.2;refGene_gn=ADAM22;refGene_tx=XM_019825428.2;refGene_gn=ADAM22;refGene_tx=XM_019825431.2;refGene_gn=ADAM22;refGene_tx=XM_019825433.2;refGene_gn=ADAM22;refGene_tx=XM_019825434.2;refGene_gn=ADAM22;refGene_tx=XM_019825436.2;refGene_gn=ADAM22;refGene_tx=XM_019825437.2
A2 93695487 93695587
refGene_gn=ADAM22;refGene_tx=XM_011280369.3;refGene_gn=ADAM22;refGene_tx=XM_011280370.3;refGene_gn=ADAM22;refGene_tx=XM_011280371.3;refGene_gn=ADAM22;refGene_tx=XM_019825413.2;refGene_gn=ADAM22;refGene_tx=XM_019825415.2;refGene_gn=ADAM22;refGene_tx=XM_019825416.2;refGene_gn=ADAM22;refGene_tx=XM_019825417.2;refGene_gn=ADAM22;refGene_tx=XM_019825418.

2;refGene_gn=ADAM22;refGene_tx=XM_019825419.2;refGene_gn=ADAM22;refGene_tx=XM_019825420.2;refGene_gn=ADAM22;refGene_tx=XM_019825421.2;refGene_gn=ADAM22;refGene_tx=XM_019825422.2;refGene_gn=ADAM22;refGene_tx=XM_019825423.2;refGene_gn=ADAM22;refGene_tx=XM_019825424.2;refGene_gn=ADAM22;refGene_tx=XM_019825425.2;refGene_gn=ADAM22;refGene_tx=XM_019825426.2;refGene_gn=ADAM22;refGene_tx=XM_019825428.2;refGene_gn=ADAM22;refGene_tx=XM_019825431.2;refGene_gn=ADAM22;refGene_tx=XM_019825433.2;refGene_gn=ADAM22;refGene_tx=XM_019825434.2;refGene_gn=ADAM22;refGene_tx=XM_019825436.2;refGene_gn=ADAM22;refGene_tx=XM_019825437.2
A2 93698030 93698102
refGene_gn=ADAM22;refGene_tx=XM_011280369.3;refGene_gn=ADAM22;refGene_tx=XM_011280370.3;refGene_gn=ADAM22;refGene_tx=XM_011280371.3;refGene_gn=ADAM22;refGene_tx=XM_019825413.2;refGene_gn=ADAM22;refGene_tx=XM_019825415.2;refGene_gn=ADAM22;refGene_tx=XM_019825416.2;refGene_gn=ADAM22;refGene_tx=XM_019825417.2;refGene_gn=ADAM22;refGene_tx=XM_019825418.2;refGene_gn=ADAM22;refGene_tx=XM_019825419.2;refGene_gn=ADAM22;refGene_tx=XM_019825420.2;refGene_gn=ADAM22;refGene_tx=XM_019825421.2;refGene_gn=ADAM22;refGene_tx=XM_019825422.2;refGene_gn=ADAM22;refGene_tx=XM_019825423.2;refGene_gn=ADAM22;refGene_tx=XM_019825424.2;refGene_gn=ADAM22;refGene_tx=XM_019825425.2;refGene_gn=ADAM22;refGene_tx=XM_019825426.2;refGene_gn=ADAM22;refGene_tx=XM_019825428.2;refGene_gn=ADAM22;refGene_tx=XM_019825431.2;refGene_gn=ADAM22;refGene_tx=XM_019825433.2;refGene_gn=ADAM22;refGene_tx=XM_019825434.2;refGene_gn=ADAM22;refGene_tx=XM_019825436.2;refGene_gn=ADAM22;refGene_tx=XM_019825437.2
A2 93698324 93698417
refGene_gn=ADAM22;refGene_tx=XM_011280369.3;refGene_gn=ADAM22;refGene_tx=XM_011280370.3;refGene_gn=ADAM22;refGene_tx=XM_011280371.3;refGene_gn=ADAM22;refGene_tx=XM_019825413.2;refGene_gn=ADAM22;refGene_tx=XM_019825415.2;refGene_gn=ADAM22;refGene_tx=XM_019825416.2;refGene_gn=ADAM22;refGene_tx=XM_019825417.2;refGene_gn=ADAM22;refGene_tx=XM_019825418.2;refGene_gn=ADAM22;refGene_tx=XM_019825419.2;refGene_gn=ADAM22;refGene_tx=XM_019825420.2;refGene_gn=ADAM22;refGene_tx=XM_019825421.2;refGene_gn=ADAM22;refGene_tx=XM_019825422.2;refGene_gn=ADAM22;refGene_tx=XM_019825423.2;refGene_gn=ADAM22;refGene_tx=XM_019825424.2;refGene_gn=ADAM22;refGene_tx=XM_019825425.2;refGene_gn=ADAM22;refGene_tx=XM_019825426.2;refGene_gn=ADAM22;refGene_tx=XM_019825428.2;refGene_gn=ADAM22;refGene_tx=XM_019825431.2;refGene_gn=ADAM22;refGene_tx=XM_019825433.2;refGene_gn=ADAM22;refGene_tx=XM_019825434.2;refGene_gn=ADAM22;refGene_tx=XM_019825436.2;refGene_gn=ADAM22;refGene_tx=XM_019825437.2
A2 93702258 93702339
refGene_gn=ADAM22;refGene_tx=XM_011280369.3;refGene_gn=ADAM22;refGene_tx=XM_011280370.3;refGene_gn=ADAM22;refGene_tx=XM_011280371.3;refGene_gn=ADAM22;refGene_tx=XM_019825413.2;refGene_gn=ADAM22;refGene_tx=XM_019825415.2;refGene_gn=ADAM22;refGene_tx=XM_019825416.2;refGene_gn=ADAM22;refGene_tx=XM_019825417.2;refGene_gn=ADAM22;refGene_tx=XM_019825418.2;refGene_gn=ADAM22;refGene_tx=XM_019825419.2;refGene_gn=ADAM22;refGene_tx=XM_019825420.2;refGene_gn=ADAM22;refGene_tx=XM_019825421.2;refGene_gn=ADAM22;refGene_tx=XM_019825422.2;refGene_gn=ADAM22;refGene_tx=XM_019825423.2;refGene_gn=ADAM22;refGene_tx=XM_019825424.2;refGene_gn=ADAM22;refGene_tx=XM_019825425.2;refGene_gn=ADAM22;refGene_tx=XM_01982542

6.2;refGene_gn=ADAM22;refGene_tx=XM_019825428.2;refGene_gn=ADAM22;refGene_tx=XM_019825431.2;refGene_gn=ADAM22;refGene_tx=XM_019825433.2;refGene_gn=ADAM22;refGene_tx=XM_019825434.2;refGene_gn=ADAM22;refGene_tx=XM_019825436.2;refGene_gn=ADAM22;refGene_tx=XM_019825437.2
A2 93705365 93705416
refGene_gn=ADAM22;refGene_tx=XM_011280369.3;refGene_gn=ADAM22;refGene_tx=XM_011280370.3;refGene_gn=ADAM22;refGene_tx=XM_011280371.3;refGene_gn=ADAM22;refGene_tx=XM_019825413.2;refGene_gn=ADAM22;refGene_tx=XM_019825415.2;refGene_gn=ADAM22;refGene_tx=XM_019825417.2;refGene_gn=ADAM22;refGene_tx=XM_019825418.2;refGene_gn=ADAM22;refGene_tx=XM_019825419.2;refGene_gn=ADAM22;refGene_tx=XM_019825420.2;refGene_gn=ADAM22;refGene_tx=XM_019825421.2;refGene_gn=ADAM22;refGene_tx=XM_019825422.2;refGene_gn=ADAM22;refGene_tx=XM_019825423.2;refGene_gn=ADAM22;refGene_tx=XM_019825424.2;refGene_gn=ADAM22;refGene_tx=XM_019825425.2;refGene_gn=ADAM22;refGene_tx=XM_019825426.2;refGene_gn=ADAM22;refGene_tx=XM_019825428.2;refGene_gn=ADAM22;refGene_tx=XM_019825431.2;refGene_gn=ADAM22;refGene_tx=XM_019825433.2;refGene_gn=ADAM22;refGene_tx=XM_019825434.2;refGene_gn=ADAM22;refGene_tx=XM_019825436.2;refGene_gn=ADAM22;refGene_tx=XM_019825437.2
A2 93705640 93705704
refGene_gn=ADAM22;refGene_tx=XM_011280369.3;refGene_gn=ADAM22;refGene_tx=XM_011280370.3;refGene_gn=ADAM22;refGene_tx=XM_011280371.3;refGene_gn=ADAM22;refGene_tx=XM_019825413.2;refGene_gn=ADAM22;refGene_tx=XM_019825415.2;refGene_gn=ADAM22;refGene_tx=XM_019825416.2;refGene_gn=ADAM22;refGene_tx=XM_019825417.2;refGene_gn=ADAM22;refGene_tx=XM_019825418.2;refGene_gn=ADAM22;refGene_tx=XM_019825419.2;refGene_gn=ADAM22;refGene_tx=XM_019825420.2;refGene_gn=ADAM22;refGene_tx=XM_019825421.2;refGene_gn=ADAM22;refGene_tx=XM_019825422.2;refGene_gn=ADAM22;refGene_tx=XM_019825423.2;refGene_gn=ADAM22;refGene_tx=XM_019825424.2;refGene_gn=ADAM22;refGene_tx=XM_019825425.2;refGene_gn=ADAM22;refGene_tx=XM_019825426.2;refGene_gn=ADAM22;refGene_tx=XM_019825428.2;refGene_gn=ADAM22;refGene_tx=XM_019825431.2;refGene_gn=ADAM22;refGene_tx=XM_019825433.2;refGene_gn=ADAM22;refGene_tx=XM_019825434.2;refGene_gn=ADAM22;refGene_tx=XM_019825436.2;refGene_gn=ADAM22;refGene_tx=XM_019825437.2
A2 93707643 93707749
refGene_gn=ADAM22;refGene_tx=XM_011280369.3;refGene_gn=ADAM22;refGene_tx=XM_011280370.3;refGene_gn=ADAM22;refGene_tx=XM_011280371.3;refGene_gn=ADAM22;refGene_tx=XM_019825413.2;refGene_gn=ADAM22;refGene_tx=XM_019825415.2;refGene_gn=ADAM22;refGene_tx=XM_019825416.2;refGene_gn=ADAM22;refGene_tx=XM_019825417.2;refGene_gn=ADAM22;refGene_tx=XM_019825418.2;refGene_gn=ADAM22;refGene_tx=XM_019825419.2;refGene_gn=ADAM22;refGene_tx=XM_019825420.2;refGene_gn=ADAM22;refGene_tx=XM_019825421.2;refGene_gn=ADAM22;refGene_tx=XM_019825422.2;refGene_gn=ADAM22;refGene_tx=XM_019825423.2;refGene_gn=ADAM22;refGene_tx=XM_019825424.2;refGene_gn=ADAM22;refGene_tx=XM_019825425.2;refGene_gn=ADAM22;refGene_tx=XM_019825426.2;refGene_gn=ADAM22;refGene_tx=XM_019825428.2;refGene_gn=ADAM22;refGene_tx=XM_019825431.2;refGene_gn=ADAM22;refGene_tx=XM_019825433.2;refGene_gn=ADAM22;refGene_tx=XM_019825434.2;refGene_gn=ADAM22;refGene_tx=XM_019825436.2;refGene_gn=ADAM22;refGene_tx=XM_019825437.2
A2 93709860 93709980

refGene_gn=ADAM22;refGene_tx=XM_011280369.3;refGene_gn=ADAM22;refGene_tx=XM_011280370.3;refGene_gn=ADAM22;refGene_tx=XM_011280371.3;refGene_gn=ADAM22;refGene_tx=XM_019825413.2;refGene_gn=ADAM22;refGene_tx=XM_019825415.2;refGene_gn=ADAM22;refGene_tx=XM_019825416.2;refGene_gn=ADAM22;refGene_tx=XM_019825417.2;refGene_gn=ADAM22;refGene_tx=XM_019825418.2;refGene_gn=ADAM22;refGene_tx=XM_019825419.2;refGene_gn=ADAM22;refGene_tx=XM_019825420.2;refGene_gn=ADAM22;refGene_tx=XM_019825421.2;refGene_gn=ADAM22;refGene_tx=XM_019825422.2;refGene_gn=ADAM22;refGene_tx=XM_019825423.2;refGene_gn=ADAM22;refGene_tx=XM_019825424.2;refGene_gn=ADAM22;refGene_tx=XM_019825425.2;refGene_gn=ADAM22;refGene_tx=XM_019825426.2;refGene_gn=ADAM22;refGene_tx=XM_019825428.2;refGene_gn=ADAM22;refGene_tx=XM_019825431.2;refGene_gn=ADAM22;refGene_tx=XM_019825433.2;refGene_gn=ADAM22;refGene_tx=XM_019825434.2;refGene_gn=ADAM22;refGene_tx=XM_019825436.2;refGene_gn=ADAM22;refGene_tx=XM_019825437.2

A2 93713386 93713555

refGene_gn=ADAM22;refGene_tx=XM_011280369.3;refGene_gn=ADAM22;refGene_tx=XM_011280370.3;refGene_gn=ADAM22;refGene_tx=XM_011280371.3;refGene_gn=ADAM22;refGene_tx=XM_019825413.2;refGene_gn=ADAM22;refGene_tx=XM_019825415.2;refGene_gn=ADAM22;refGene_tx=XM_019825416.2;refGene_gn=ADAM22;refGene_tx=XM_019825417.2;refGene_gn=ADAM22;refGene_tx=XM_019825418.2;refGene_gn=ADAM22;refGene_tx=XM_019825419.2;refGene_gn=ADAM22;refGene_tx=XM_019825420.2;refGene_gn=ADAM22;refGene_tx=XM_019825421.2;refGene_gn=ADAM22;refGene_tx=XM_019825422.2;refGene_gn=ADAM22;refGene_tx=XM_019825423.2;refGene_gn=ADAM22;refGene_tx=XM_019825424.2;refGene_gn=ADAM22;refGene_tx=XM_019825425.2;refGene_gn=ADAM22;refGene_tx=XM_019825426.2;refGene_gn=ADAM22;refGene_tx=XM_019825428.2;refGene_gn=ADAM22;refGene_tx=XM_019825431.2;refGene_gn=ADAM22;refGene_tx=XM_019825433.2;refGene_gn=ADAM22;refGene_tx=XM_019825434.2;refGene_gn=ADAM22;refGene_tx=XM_019825436.2;refGene_gn=ADAM22;refGene_tx=XM_019825437.2

A2 93715069 93715184

refGene_gn=ADAM22;refGene_tx=XM_011280369.3;refGene_gn=ADAM22;refGene_tx=XM_011280370.3;refGene_gn=ADAM22;refGene_tx=XM_011280371.3;refGene_gn=ADAM22;refGene_tx=XM_019825413.2;refGene_gn=ADAM22;refGene_tx=XM_019825415.2;refGene_gn=ADAM22;refGene_tx=XM_019825416.2;refGene_gn=ADAM22;refGene_tx=XM_019825417.2;refGene_gn=ADAM22;refGene_tx=XM_019825418.2;refGene_gn=ADAM22;refGene_tx=XM_019825419.2;refGene_gn=ADAM22;refGene_tx=XM_019825420.2;refGene_gn=ADAM22;refGene_tx=XM_019825421.2;refGene_gn=ADAM22;refGene_tx=XM_019825422.2;refGene_gn=ADAM22;refGene_tx=XM_019825423.2;refGene_gn=ADAM22;refGene_tx=XM_019825424.2;refGene_gn=ADAM22;refGene_tx=XM_019825425.2;refGene_gn=ADAM22;refGene_tx=XM_019825426.2;refGene_gn=ADAM22;refGene_tx=XM_019825428.2;refGene_gn=ADAM22;refGene_tx=XM_019825431.2;refGene_gn=ADAM22;refGene_tx=XM_019825433.2;refGene_gn=ADAM22;refGene_tx=XM_019825434.2;refGene_gn=ADAM22;refGene_tx=XM_019825436.2;refGene_gn=ADAM22;refGene_tx=XM_019825437.2

A2 93718037 93718128

refGene_gn=ADAM22;refGene_tx=XM_011280369.3;refGene_gn=ADAM22;refGene_tx=XM_011280370.3;refGene_gn=ADAM22;refGene_tx=XM_011280371.3;refGene_gn=ADAM22;refGene_tx=XM_019825413.2;refGene_gn=ADAM22;refGene_tx=XM_019825415.2;refGene_gn=ADAM22;refGene_tx=XM_019825416.2;refGene_gn=ADAM22;refGene_tx=XM_019825417.2;refGene_gn=ADAM22;refGene_tx=XM_019825418.

2;refGene_gn=ADAM22;refGene_tx=XM_019825419.2;refGene_gn=ADAM22;refGene_tx=XM_019825420.2;refGene_gn=ADAM22;refGene_tx=XM_019825421.2;refGene_gn=ADAM22;refGene_tx=XM_019825422.2;refGene_gn=ADAM22;refGene_tx=XM_019825423.2;refGene_gn=ADAM22;refGene_tx=XM_019825424.2;refGene_gn=ADAM22;refGene_tx=XM_019825425.2;refGene_gn=ADAM22;refGene_tx=XM_019825426.2;refGene_gn=ADAM22;refGene_tx=XM_019825428.2
A2 93722198 93722216
refGene_gn=ADAM22;refGene_tx=XM_011280369.3;refGene_gn=ADAM22;refGene_tx=XM_011280370.3;refGene_gn=ADAM22;refGene_tx=XM_011280371.3;refGene_gn=ADAM22;refGene_tx=XM_019825413.2;refGene_gn=ADAM22;refGene_tx=XM_019825415.2;refGene_gn=ADAM22;refGene_tx=XM_019825416.2;refGene_gn=ADAM22;refGene_tx=XM_019825417.2;refGene_gn=ADAM22;refGene_tx=XM_019825418.2;refGene_gn=ADAM22;refGene_tx=XM_019825419.2;refGene_gn=ADAM22;refGene_tx=XM_019825420.2;refGene_gn=ADAM22;refGene_tx=XM_019825421.2;refGene_gn=ADAM22;refGene_tx=XM_019825422.2;refGene_gn=ADAM22;refGene_tx=XM_019825423.2;refGene_gn=ADAM22;refGene_tx=XM_019825425.2;refGene_gn=ADAM22;refGene_tx=XM_019825426.2;refGene_gn=ADAM22;refGene_tx=XM_019825428.2;refGene_gn=ADAM22;refGene_tx=XM_019825431.2;refGene_gn=ADAM22;refGene_tx=XM_019825433.2;refGene_gn=ADAM22;refGene_tx=XM_019825436.2;refGene_gn=ADAM22;refGene_tx=XM_019825437.2
A2 93729536 93729731
refGene_gn=ADAM22;refGene_tx=XM_011280369.3;refGene_gn=ADAM22;refGene_tx=XM_019825413.2;refGene_gn=ADAM22;refGene_tx=XM_019825415.2;refGene_gn=ADAM22;refGene_tx=XM_019825416.2;refGene_gn=ADAM22;refGene_tx=XM_019825417.2;refGene_gn=ADAM22;refGene_tx=XM_019825418.2;refGene_gn=ADAM22;refGene_tx=XM_019825419.2;refGene_gn=ADAM22;refGene_tx=XM_019825420.2;refGene_gn=ADAM22;refGene_tx=XM_019825421.2;refGene_gn=ADAM22;refGene_tx=XM_019825422.2;refGene_gn=ADAM22;refGene_tx=XM_019825425.2;refGene_gn=ADAM22;refGene_tx=XM_019825426.2;refGene_gn=ADAM22;refGene_tx=XM_019825428.2
A2 93732043 93732144
refGene_gn=ADAM22;refGene_tx=XM_011280369.3;refGene_gn=ADAM22;refGene_tx=XM_011280370.3;refGene_gn=ADAM22;refGene_tx=XM_011280371.3;refGene_gn=ADAM22;refGene_tx=XM_019825413.2;refGene_gn=ADAM22;refGene_tx=XM_019825415.2;refGene_gn=ADAM22;refGene_tx=XM_019825416.2;refGene_gn=ADAM22;refGene_tx=XM_019825417.2;refGene_gn=ADAM22;refGene_tx=XM_019825418.2;refGene_gn=ADAM22;refGene_tx=XM_019825419.2;refGene_gn=ADAM22;refGene_tx=XM_019825420.2;refGene_gn=ADAM22;refGene_tx=XM_019825421.2;refGene_gn=ADAM22;refGene_tx=XM_019825422.2;refGene_gn=ADAM22;refGene_tx=XM_019825423.2;refGene_gn=ADAM22;refGene_tx=XM_019825424.2;refGene_gn=ADAM22;refGene_tx=XM_019825425.2;refGene_gn=ADAM22;refGene_tx=XM_019825426.2;refGene_gn=ADAM22;refGene_tx=XM_019825428.2;refGene_gn=ADAM22;refGene_tx=XM_019825431.2;refGene_gn=ADAM22;refGene_tx=XM_019825433.2;refGene_gn=ADAM22;refGene_tx=XM_019825434.2;refGene_gn=ADAM22;refGene_tx=XM_019825436.2;refGene_gn=ADAM22;refGene_tx=XM_019825437.2
A2 93732491 93732558
refGene_gn=ADAM22;refGene_tx=XM_011280369.3;refGene_gn=ADAM22;refGene_tx=XM_011280370.3;refGene_gn=ADAM22;refGene_tx=XM_011280371.3;refGene_gn=ADAM22;refGene_tx=XM_019825413.2;refGene_gn=ADAM22;refGene_tx=XM_019825415.2;refGene_gn=ADAM22;refGene_tx=XM_019825416.2;refGene_gn=ADAM2

2;refGene_tx=XM_019825417.2;refGene_gn=ADAM22;refGene_tx=XM_019825418.2;refGene_gn=ADAM22;refGene_tx=XM_019825419.2;refGene_gn=ADAM22;refGene_tx=XM_019825420.2;refGene_gn=ADAM22;refGene_tx=XM_019825421.2;refGene_gn=ADAM22;refGene_tx=XM_019825422.2;refGene_gn=ADAM22;refGene_tx=XM_019825423.2;refGene_gn=ADAM22;refGene_tx=XM_019825424.2;refGene_gn=ADAM22;refGene_tx=XM_019825425.2;refGene_gn=ADAM22;refGene_tx=XM_019825426.2;refGene_gn=ADAM22;refGene_tx=XM_019825428.2;refGene_gn=ADAM22;refGene_tx=XM_019825431.2;refGene_gn=ADAM22;refGene_tx=XM_019825433.2;refGene_gn=ADAM22;refGene_tx=XM_019825434.2;refGene_gn=ADAM22;refGene_tx=XM_019825436.2;refGene_gn=ADAM22;refGene_tx=XM_019825437.2
A2 93734802 93734913
refGene_gn=ADAM22;refGene_tx=XM_019825413.2;refGene_gn=ADAM22;refGene_tx=XM_019825415.2;refGene_gn=ADAM22;refGene_tx=XM_019825416.2;refGene_gn=ADAM22;refGene_tx=XM_019825417.2;refGene_gn=ADAM22;refGene_tx=XM_019825418.2;refGene_gn=ADAM22;refGene_tx=XM_019825419.2;refGene_gn=ADAM22;refGene_tx=XM_019825421.2;refGene_gn=ADAM22;refGene_tx=XM_019825423.2;refGene_gn=ADAM22;refGene_tx=XM_019825424.2;refGene_gn=ADAM22;refGene_tx=XM_019825436.2;refGene_gn=ADAM22;refGene_tx=XM_019825437.2
A2 93737129 93737216
refGene_gn=ADAM22;refGene_tx=XM_011280370.3;refGene_gn=ADAM22;refGene_tx=XM_019825413.2;refGene_gn=ADAM22;refGene_tx=XM_019825415.2;refGene_gn=ADAM22;refGene_tx=XM_019825416.2;refGene_gn=ADAM22;refGene_tx=XM_019825417.2;refGene_gn=ADAM22;refGene_tx=XM_019825419.2;refGene_gn=ADAM22;refGene_tx=XM_019825420.2;refGene_gn=ADAM22;refGene_tx=XM_019825422.2;refGene_gn=ADAM22;refGene_tx=XM_019825423.2;refGene_gn=ADAM22;refGene_tx=XM_019825424.2;refGene_gn=ADAM22;refGene_tx=XM_019825426.2
A2 93738542 93738572
refGene_gn=ADAM22;refGene_tx=XM_019825413.2;refGene_gn=ADAM22;refGene_tx=XM_019825416.2;refGene_gn=ADAM22;refGene_tx=XM_019825417.2;refGene_gn=ADAM22;refGene_tx=XM_019825418.2;refGene_gn=ADAM22;refGene_tx=XM_019825419.2;refGene_gn=ADAM22;refGene_tx=XM_019825420.2;refGene_gn=ADAM22;refGene_tx=XM_019825423.2;refGene_gn=ADAM22;refGene_tx=XM_019825424.2
A2 93742957 93743081
refGene_gn=ADAM22;refGene_tx=XM_011280369.3;refGene_gn=ADAM22;refGene_tx=XM_011280370.3;refGene_gn=ADAM22;refGene_tx=XM_011280371.3;refGene_gn=ADAM22;refGene_tx=XM_019825413.2;refGene_gn=ADAM22;refGene_tx=XM_019825415.2;refGene_gn=ADAM22;refGene_tx=XM_019825416.2;refGene_gn=ADAM22;refGene_tx=XM_019825417.2;refGene_gn=ADAM22;refGene_tx=XM_019825418.2;refGene_gn=ADAM22;refGene_tx=XM_019825419.2;refGene_gn=ADAM22;refGene_tx=XM_019825420.2;refGene_gn=ADAM22;refGene_tx=XM_019825421.2;refGene_gn=ADAM22;refGene_tx=XM_019825422.2;refGene_gn=ADAM22;refGene_tx=XM_019825423.2;refGene_gn=ADAM22;refGene_tx=XM_019825424.2;refGene_gn=ADAM22;refGene_tx=XM_019825425.2;refGene_gn=ADAM22;refGene_tx=XM_019825426.2;refGene_gn=ADAM22;refGene_tx=XM_019825428.2;refGene_gn=ADAM22;refGene_tx=XM_019825431.2;refGene_gn=ADAM22;refGene_tx=XM_019825433.2;refGene_gn=ADAM22;refGene_tx=XM_019825434.2
A2 93747265 93747286
refGene_gn=ADAM22;refGene_tx=XM_011280369.3;refGene_gn=ADAM22;refGene_tx=XM_011280370.3;refGene_gn=ADAM22;refGene_tx=XM_011280371.3;refGene_

gn=ADAM22;refGene_tx=XM_019825419.2;refGene_gn=ADAM22;refGene_tx=XM_01
9825426.2;refGene_gn=ADAM22;refGene_tx=XM_019825428.2
A2 93747338 93747467
refGene_gn=ADAM22;refGene_tx=XM_019825413.2;refGene_gn=ADAM22;refGene_
tx=XM_019825415.2;refGene_gn=ADAM22;refGene_tx=XM_019825416.2;refGene_
gn=ADAM22;refGene_tx=XM_019825417.2;refGene_gn=ADAM22;refGene_tx=XM_01
9825418.2;refGene_gn=ADAM22;refGene_tx=XM_019825420.2;refGene_gn=ADAM2
2;refGene_tx=XM_019825421.2;refGene_gn=ADAM22;refGene_tx=XM_019825422.
2;refGene_gn=ADAM22;refGene_tx=XM_019825423.2;refGene_gn=ADAM22;refGen
e_tx=XM_019825424.2;refGene_gn=ADAM22;refGene_tx=XM_019825425.2
A2 93757286 93757313 refGene_gn=SRI;refGene_tx=XM_023250323.1
A2 93758908 93758967 refGene_gn=SRI;refGene_tx=XM_023250323.1
A2 93760420 93760534 refGene_gn=SRI;refGene_tx=XM_023250323.1
A2 93760799 93760947 refGene_gn=SRI;refGene_tx=XM_023250323.1
A2 93761748 93761792 refGene_gn=SRI;refGene_tx=XM_023250323.1
A2 93768906 93768976 refGene_gn=SRI;refGene_tx=XM_023250323.1
A2 93771080 93771164 refGene_gn=SRI;refGene_tx=XM_023250323.1
A2 93772136 93772187 refGene_gn=SRI;refGene_tx=XM_023250323.1
A2 93819263 93819551 refGene_gn=STEAP4;refGene_tx=XM_003982729.4
A2 93821161 93821326 refGene_gn=STEAP4;refGene_tx=XM_003982729.4
A2 93822847 93823375 refGene_gn=STEAP4;refGene_tx=XM_003982729.4
A2 93824011 93824467 refGene_gn=STEAP4;refGene_tx=XM_003982729.4
A2 94255847 94255955 refGene_gn=ZNF804B;refGene_tx=XM_019825438.2
A2 94285527 94286289 refGene_gn=TEX47;refGene_tx=XM_023250325.1
A2 94668571 94668715 refGene_gn=ZNF804B;refGene_tx=XM_019825438.2
A2 94762880 94763011 refGene_gn=ZNF804B;refGene_tx=XM_019825438.2
A2 94768794 94772461 refGene_gn=ZNF804B;refGene_tx=XM_019825438.2
A2 95577566 95577694 refGene_gn=STEAP1;refGene_tx=XM_003982731.3
A2 95581465 95581577
refGene_gn=STEAP1;refGene_tx=XM_003982731.3;refGene_gn=STEAP1;refGene_
tx=XM_006929163.2
A2 95583806 95584319
refGene_gn=STEAP1;refGene_tx=XM_003982731.3;refGene_gn=STEAP1;refGene_
tx=XM_006929163.2
A2 95584907 95585072
refGene_gn=STEAP1;refGene_tx=XM_003982731.3;refGene_gn=STEAP1;refGene_
tx=XM_006929163.2
A2 95587193 95587451
refGene_gn=STEAP1;refGene_tx=XM_003982731.3;refGene_gn=STEAP1;refGene_
tx=XM_006929163.2
A2 95627851 95628343
refGene_gn=STEAP2;refGene_tx=XM_003982732.5;refGene_gn=STEAP2;refGene_
tx=XM_019826145.2;refGene_gn=STEAP2;refGene_tx=XM_019826146.2;refGene_
gn=STEAP2;refGene_tx=XM_019826147.1
A2 95631365 95631893
refGene_gn=STEAP2;refGene_tx=XM_003982732.5;refGene_gn=STEAP2;refGene_
tx=XM_019826145.2;refGene_gn=STEAP2;refGene_tx=XM_019826146.2;refGene_
gn=STEAP2;refGene_tx=XM_019826147.1
A2 95634276 95634441
refGene_gn=STEAP2;refGene_tx=XM_003982732.5;refGene_gn=STEAP2;refGene_

tx=XM_019826145.2;refGene_gn=STEAP2;refGene_tx=XM_019826146.2;refGene_gn=STEAP2;refGene_tx=XM_019826147.1
A2 95636398 95636686
refGene_gn=STEAP2;refGene_tx=XM_003982732.5;refGene_gn=STEAP2;refGene_tx=XM_019826145.2;refGene_gn=STEAP2;refGene_tx=XM_019826146.2;refGene_gn=STEAP2;refGene_tx=XM_019826147.1
A2 95649518 95649638
refGene_gn=CFAP69;refGene_tx=XM_023250327.1;refGene_gn=CFAP69;refGene_tx=XM_023250328.1;refGene_gn=CFAP69;refGene_tx=XM_023250329.1
A2 95660238 95660298
refGene_gn=CFAP69;refGene_tx=XM_023250327.1;refGene_gn=CFAP69;refGene_tx=XM_023250328.1;refGene_gn=CFAP69;refGene_tx=XM_023250329.1;refGene_gn=CFAP69;refGene_tx=XM_023250330.1
A2 95662107 95662173
refGene_gn=CFAP69;refGene_tx=XM_023250327.1;refGene_gn=CFAP69;refGene_tx=XM_023250328.1;refGene_gn=CFAP69;refGene_tx=XM_023250329.1;refGene_gn=CFAP69;refGene_tx=XM_023250330.1
A2 95664530 95664640
refGene_gn=CFAP69;refGene_tx=XM_023250327.1;refGene_gn=CFAP69;refGene_tx=XM_023250328.1;refGene_gn=CFAP69;refGene_tx=XM_023250329.1;refGene_gn=CFAP69;refGene_tx=XM_023250330.1
A2 95667532 95667609
refGene_gn=CFAP69;refGene_tx=XM_023250327.1;refGene_gn=CFAP69;refGene_tx=XM_023250328.1;refGene_gn=CFAP69;refGene_tx=XM_023250329.1;refGene_gn=CFAP69;refGene_tx=XM_023250330.1
A2 95669959 95670058
refGene_gn=CFAP69;refGene_tx=XM_023250327.1;refGene_gn=CFAP69;refGene_tx=XM_023250328.1;refGene_gn=CFAP69;refGene_tx=XM_023250329.1;refGene_gn=CFAP69;refGene_tx=XM_023250330.1;refGene_gn=CFAP69;refGene_tx=XM_023250331.1
A2 95673435 95673585
refGene_gn=CFAP69;refGene_tx=XM_023250327.1;refGene_gn=CFAP69;refGene_tx=XM_023250328.1;refGene_gn=CFAP69;refGene_tx=XM_023250329.1;refGene_gn=CFAP69;refGene_tx=XM_023250330.1;refGene_gn=CFAP69;refGene_tx=XM_023250331.1
A2 95673689 95673867
refGene_gn=CFAP69;refGene_tx=XM_023250327.1;refGene_gn=CFAP69;refGene_tx=XM_023250328.1;refGene_gn=CFAP69;refGene_tx=XM_023250329.1;refGene_gn=CFAP69;refGene_tx=XM_023250330.1;refGene_gn=CFAP69;refGene_tx=XM_023250331.1
A2 95675913 95676037
refGene_gn=CFAP69;refGene_tx=XM_023250327.1;refGene_gn=CFAP69;refGene_tx=XM_023250328.1;refGene_gn=CFAP69;refGene_tx=XM_023250329.1;refGene_gn=CFAP69;refGene_tx=XM_023250330.1;refGene_gn=CFAP69;refGene_tx=XM_023250331.1
A2 95678839 95678888
refGene_gn=CFAP69;refGene_tx=XM_023250327.1;refGene_gn=CFAP69;refGene_tx=XM_023250328.1;refGene_gn=CFAP69;refGene_tx=XM_023250329.1;refGene_gn=CFAP69;refGene_tx=XM_023250330.1;refGene_gn=CFAP69;refGene_tx=XM_023250331.1

A2 95678978 95679100
refGene_gn=CFAP69;refGene_tx=XM_023250327.1;refGene_gn=CFAP69;refGene_tx=XM_023250328.1;refGene_gn=CFAP69;refGene_tx=XM_023250329.1;refGene_gn=CFAP69;refGene_tx=XM_023250330.1;refGene_gn=CFAP69;refGene_tx=XM_023250331.1
A2 95682032 95682249
refGene_gn=CFAP69;refGene_tx=XM_023250327.1;refGene_gn=CFAP69;refGene_tx=XM_023250328.1;refGene_gn=CFAP69;refGene_tx=XM_023250329.1;refGene_gn=CFAP69;refGene_tx=XM_023250330.1;refGene_gn=CFAP69;refGene_tx=XM_023250331.1
A2 95685324 95685489
refGene_gn=CFAP69;refGene_tx=XM_023250327.1;refGene_gn=CFAP69;refGene_tx=XM_023250328.1;refGene_gn=CFAP69;refGene_tx=XM_023250329.1;refGene_gn=CFAP69;refGene_tx=XM_023250330.1;refGene_gn=CFAP69;refGene_tx=XM_023250331.1
A2 95688818 95688934
refGene_gn=CFAP69;refGene_tx=XM_023250327.1;refGene_gn=CFAP69;refGene_tx=XM_023250328.1;refGene_gn=CFAP69;refGene_tx=XM_023250329.1;refGene_gn=CFAP69;refGene_tx=XM_023250330.1;refGene_gn=CFAP69;refGene_tx=XM_023250331.1
A2 95694091 95694210
refGene_gn=CFAP69;refGene_tx=XM_023250327.1;refGene_gn=CFAP69;refGene_tx=XM_023250328.1;refGene_gn=CFAP69;refGene_tx=XM_023250329.1;refGene_gn=CFAP69;refGene_tx=XM_023250330.1;refGene_gn=CFAP69;refGene_tx=XM_023250331.1
A2 95697932 95698014
refGene_gn=CFAP69;refGene_tx=XM_023250327.1;refGene_gn=CFAP69;refGene_tx=XM_023250328.1;refGene_gn=CFAP69;refGene_tx=XM_023250329.1;refGene_gn=CFAP69;refGene_tx=XM_023250330.1;refGene_gn=CFAP69;refGene_tx=XM_023250331.1
A2 95700938 95701131
refGene_gn=CFAP69;refGene_tx=XM_023250327.1;refGene_gn=CFAP69;refGene_tx=XM_023250328.1;refGene_gn=CFAP69;refGene_tx=XM_023250329.1;refGene_gn=CFAP69;refGene_tx=XM_023250330.1;refGene_gn=CFAP69;refGene_tx=XM_023250331.1
A2 95704381 95704519
refGene_gn=CFAP69;refGene_tx=XM_023250327.1;refGene_gn=CFAP69;refGene_tx=XM_023250328.1;refGene_gn=CFAP69;refGene_tx=XM_023250329.1;refGene_gn=CFAP69;refGene_tx=XM_023250330.1;refGene_gn=CFAP69;refGene_tx=XM_023250331.1
A2 95705150 95705227
refGene_gn=CFAP69;refGene_tx=XM_023250327.1;refGene_gn=CFAP69;refGene_tx=XM_023250328.1;refGene_gn=CFAP69;refGene_tx=XM_023250329.1;refGene_gn=CFAP69;refGene_tx=XM_023250330.1;refGene_gn=CFAP69;refGene_tx=XM_023250331.1
A2 95706557 95706755
refGene_gn=CFAP69;refGene_tx=XM_023250327.1;refGene_gn=CFAP69;refGene_tx=XM_023250328.1;refGene_gn=CFAP69;refGene_tx=XM_023250329.1;refGene_gn=CFAP69;refGene_tx=XM_023250330.1;refGene_gn=CFAP69;refGene_tx=XM_023250331.1

A2 95708452 95708539
refGene_gn=CFAP69;refGene_tx=XM_023250327.1;refGene_gn=CFAP69;refGene_tx=XM_023250328.1;refGene_gn=CFAP69;refGene_tx=XM_023250329.1;refGene_gn=CFAP69;refGene_tx=XM_023250330.1;refGene_gn=CFAP69;refGene_tx=XM_023250331.1
A2 95710411 95710516
refGene_gn=CFAP69;refGene_tx=XM_023250327.1;refGene_gn=CFAP69;refGene_tx=XM_023250328.1;refGene_gn=CFAP69;refGene_tx=XM_023250329.1;refGene_gn=CFAP69;refGene_tx=XM_023250330.1;refGene_gn=CFAP69;refGene_tx=XM_023250331.1
A2 95710828 95710999
refGene_gn=CFAP69;refGene_tx=XM_023250327.1;refGene_gn=CFAP69;refGene_tx=XM_023250328.1;refGene_gn=CFAP69;refGene_tx=XM_023250329.1;refGene_gn=CFAP69;refGene_tx=XM_023250330.1;refGene_gn=CFAP69;refGene_tx=XM_023250331.1
A2 95744847 95744880 refGene_gn=GTPBP10;refGene_tx=XM_004001474.5
A2 95747468 95747662
refGene_gn=GTPBP10;refGene_tx=XM_004001474.5;refGene_gn=GTPBP10;refGene_tx=XM_011280373.3;refGene_gn=GTPBP10;refGene_tx=XM_019826158.2
A2 95754071 95754163
refGene_gn=GTPBP10;refGene_tx=XM_004001474.5;refGene_gn=GTPBP10;refGene_tx=XM_011280373.3;refGene_gn=GTPBP10;refGene_tx=XM_019826158.2
A2 95754934 95755079
refGene_gn=GTPBP10;refGene_tx=XM_004001474.5;refGene_gn=GTPBP10;refGene_tx=XM_011280373.3;refGene_gn=GTPBP10;refGene_tx=XM_019826158.2
A2 95759440 95759514
refGene_gn=GTPBP10;refGene_tx=XM_004001474.5;refGene_gn=GTPBP10;refGene_tx=XM_011280373.3;refGene_gn=GTPBP10;refGene_tx=XM_019826158.2
A2 95760855 95760908
refGene_gn=GTPBP10;refGene_tx=XM_004001474.5;refGene_gn=GTPBP10;refGene_tx=XM_011280373.3;refGene_gn=GTPBP10;refGene_tx=XM_019826158.2
A2 95763290 95763398
refGene_gn=GTPBP10;refGene_tx=XM_004001474.5;refGene_gn=GTPBP10;refGene_tx=XM_011280373.3;refGene_gn=GTPBP10;refGene_tx=XM_019826158.2
A2 95763963 95764041
refGene_gn=GTPBP10;refGene_tx=XM_004001474.5;refGene_gn=GTPBP10;refGene_tx=XM_011280373.3;refGene_gn=GTPBP10;refGene_tx=XM_019826158.2
A2 95766766 95766890
refGene_gn=GTPBP10;refGene_tx=XM_004001474.5;refGene_gn=GTPBP10;refGene_tx=XM_011280373.3;refGene_gn=GTPBP10;refGene_tx=XM_019826158.2
A2 95768689 95768940
refGene_gn=GTPBP10;refGene_tx=XM_004001474.5;refGene_gn=GTPBP10;refGene_tx=XM_011280373.3;refGene_gn=GTPBP10;refGene_tx=XM_019826158.2
A2 95787853 95788588 refGene_gn=CLDN12;refGene_tx=XM_003982734.5
A2 95943316 95943407 refGene_gn=CDK14;refGene_tx=XM_023250334.1
A2 95950459 95950491 refGene_gn=CDK14;refGene_tx=XM_023250334.1
A2 96075297 96075543 refGene_gn=CDK14;refGene_tx=XM_023250334.1
A2 96089113 96089208 refGene_gn=CDK14;refGene_tx=XM_023250334.1
A2 96124864 96124944 refGene_gn=CDK14;refGene_tx=XM_023250334.1
A2 96188720 96188815 refGene_gn=CDK14;refGene_tx=XM_023250334.1

A2 96220749 96220812 refGene_gn=CDK14;refGene_tx=XM_023250334.1
A2 96245325 96245449 refGene_gn=CDK14;refGene_tx=XM_023250334.1
A2 96281426 96281547 refGene_gn=CDK14;refGene_tx=XM_023250334.1
A2 96305956 96306050 refGene_gn=CDK14;refGene_tx=XM_023250334.1
A2 96364259 96364323 refGene_gn=CDK14;refGene_tx=XM_023250334.1
A2 96389420 96389469 refGene_gn=CDK14;refGene_tx=XM_023250334.1
A2 96423989 96424075 refGene_gn=CDK14;refGene_tx=XM_023250334.1
A2 96424077 96424128 refGene_gn=CDK14;refGene_tx=XM_023250334.1
A2 96430139 96430255 refGene_gn=CDK14;refGene_tx=XM_023250334.1
A2 96564579 96566538 refGene_gn=FZD1;refGene_tx=XM_003982740.5
A2 97028317 97029506
refGene_gn=MTERF1;refGene_tx=XM_003982749.4;refGene_gn=MTERF1;refGene_tx=XM_006929171.3
A2 97035717 97035749
refGene_gn=MTERF1;refGene_tx=XM_003982749.4;refGene_gn=MTERF1;refGene_tx=XM_006929171.3
A2 97073289 97073475
refGene_gn=LOC109497451;refGene_tx=XM_023242073.1
A2 97073618 97073668
refGene_gn=LOC109497451;refGene_tx=XM_023242073.1
A2 97078269 97078597
refGene_gn=LOC109497451;refGene_tx=XM_023242073.1
A2 97078690 97078781
refGene_gn=AKAP9;refGene_tx=XM_011280375.3;refGene_gn=LOC109497451;refGene_tx=XM_023242073.1
A2 97101289 97101517 refGene_gn=AKAP9;refGene_tx=XM_011280375.3
A2 97105070 97105115 refGene_gn=AKAP9;refGene_tx=XM_011280375.3
A2 97114797 97114851 refGene_gn=AKAP9;refGene_tx=XM_011280375.3
A2 97115216 97115387 refGene_gn=AKAP9;refGene_tx=XM_011280375.3
A2 97116605 97116761 refGene_gn=AKAP9;refGene_tx=XM_011280375.3
A2 97117600 97117798 refGene_gn=AKAP9;refGene_tx=XM_011280375.3
A2 97123018 97123106 refGene_gn=AKAP9;refGene_tx=XM_011280375.3
A2 97123108 97125402 refGene_gn=AKAP9;refGene_tx=XM_011280375.3
A2 97130518 97130732 refGene_gn=AKAP9;refGene_tx=XM_011280375.3
A2 97132818 97132898 refGene_gn=AKAP9;refGene_tx=XM_011280375.3
A2 97135612 97135751 refGene_gn=AKAP9;refGene_tx=XM_011280375.3
A2 97136176 97136262 refGene_gn=AKAP9;refGene_tx=XM_011280375.3
A2 97140202 97140314 refGene_gn=AKAP9;refGene_tx=XM_011280375.3
A2 97140981 97141180 refGene_gn=AKAP9;refGene_tx=XM_011280375.3
A2 97145407 97145504 refGene_gn=AKAP9;refGene_tx=XM_011280375.3
A2 97146699 97146792 refGene_gn=AKAP9;refGene_tx=XM_011280375.3
A2 97154250 97154592 refGene_gn=AKAP9;refGene_tx=XM_011280375.3
A2 97155845 97156070 refGene_gn=AKAP9;refGene_tx=XM_011280375.3
A2 97157146 97157287 refGene_gn=AKAP9;refGene_tx=XM_011280375.3
A2 97157774 97157878 refGene_gn=AKAP9;refGene_tx=XM_011280375.3
A2 97160275 97160481 refGene_gn=AKAP9;refGene_tx=XM_011280375.3
A2 97167443 97167676 refGene_gn=AKAP9;refGene_tx=XM_011280375.3
A2 97184821 97184984 refGene_gn=AKAP9;refGene_tx=XM_011280375.3
A2 97185901 97186114 refGene_gn=AKAP9;refGene_tx=XM_011280375.3
A2 97188500 97188733 refGene_gn=AKAP9;refGene_tx=XM_011280375.3

A2 97189663 97189783 refGene_gn=AKAP9;refGene_tx=XM_011280375.3
A2 97193702 97193879 refGene_gn=AKAP9;refGene_tx=XM_011280375.3
A2 97194535 97194640 refGene_gn=AKAP9;refGene_tx=XM_011280375.3
A2 97199753 97199906 refGene_gn=AKAP9;refGene_tx=XM_011280375.3
A2 97200260 97200440 refGene_gn=AKAP9;refGene_tx=XM_011280375.3
A2 97201284 97202358 refGene_gn=AKAP9;refGene_tx=XM_011280375.3
A2 97203690 97203831 refGene_gn=AKAP9;refGene_tx=XM_011280375.3
A2 97204308 97204797 refGene_gn=AKAP9;refGene_tx=XM_011280375.3
A2 97205235 97205299 refGene_gn=AKAP9;refGene_tx=XM_011280375.3
A2 97205617 97205739 refGene_gn=AKAP9;refGene_tx=XM_011280375.3
A2 97206565 97206757 refGene_gn=AKAP9;refGene_tx=XM_011280375.3
A2 97207021 97207210 refGene_gn=AKAP9;refGene_tx=XM_011280375.3
A2 97212059 97212204 refGene_gn=AKAP9;refGene_tx=XM_011280375.3
A2 97213528 97213748 refGene_gn=AKAP9;refGene_tx=XM_011280375.3
A2 97214456 97214607 refGene_gn=AKAP9;refGene_tx=XM_011280375.3
A2 97216207 97216876 refGene_gn=AKAP9;refGene_tx=XM_011280375.3
A2 97217322 97217531 refGene_gn=AKAP9;refGene_tx=XM_011280375.3
A2 97218275 97218381 refGene_gn=AKAP9;refGene_tx=XM_011280375.3
A2 97219891 97220074 refGene_gn=AKAP9;refGene_tx=XM_011280375.3
A2 97221288 97221489 refGene_gn=AKAP9;refGene_tx=XM_011280375.3
A2 97222344 97222577 refGene_gn=AKAP9;refGene_tx=XM_011280375.3
A2 97224481 97224567 refGene_gn=AKAP9;refGene_tx=XM_011280375.3
A2 97225733 97225863 refGene_gn=AKAP9;refGene_tx=XM_011280375.3
A2 97227483 97227623 refGene_gn=AKAP9;refGene_tx=XM_011280375.3
A2 97231814 97231849 refGene_gn=AKAP9;refGene_tx=XM_011280375.3
A2 97234207 97234386
refGene_gn=LOC101083499;refGene_tx=XM_003982742.5;refGene_gn=LOC101083499;refGene_tx=XM_006929172.4;refGene_gn=LOC101083499;refGene_tx=XM_006929174.3;refGene_gn=LOC101083499;refGene_tx=XM_019825448.1;refGene_gn=LOC101083499;refGene_tx=XM_019825449.1;refGene_gn=LOC101083499;refGene_tx=XM_019825450.2;refGene_gn=LOC101083499;refGene_tx=XM_019825452.2;refGene_gn=LOC101083499;refGene_tx=XM_023250337.1;refGene_gn=LOC101083499;refGene_tx=XM_023250338.1
A2 97240710 97240879
refGene_gn=LOC101083499;refGene_tx=XM_003982742.5;refGene_gn=LOC101083499;refGene_tx=XM_006929172.4;refGene_gn=LOC101083499;refGene_tx=XM_006929174.3;refGene_gn=LOC101083499;refGene_tx=XM_019825448.1;refGene_gn=LOC101083499;refGene_tx=XM_019825449.1;refGene_gn=LOC101083499;refGene_tx=XM_019825450.2;refGene_gn=LOC101083499;refGene_tx=XM_019825452.2;refGene_gn=LOC101083499;refGene_tx=XM_023250337.1;refGene_gn=LOC101083499;refGene_tx=XM_023250338.1
A2 97241462 97241558
refGene_gn=LOC101083499;refGene_tx=XM_003982742.5;refGene_gn=LOC101083499;refGene_tx=XM_006929172.4;refGene_gn=LOC101083499;refGene_tx=XM_006929174.3;refGene_gn=LOC101083499;refGene_tx=XM_019825448.1;refGene_gn=LOC101083499;refGene_tx=XM_019825449.1;refGene_gn=LOC101083499;refGene_tx=XM_019825450.2;refGene_gn=LOC101083499;refGene_tx=XM_019825452.2;refGene_gn=LOC101083499;refGene_tx=XM_023250337.1;refGene_gn=LOC101083499;refGene_tx=XM_023250338.1
A2 97243148 97243344

refGene_gn=LOC101083499;refGene_tx=XM_003982742.5;refGene_gn=LOC101083499;refGene_tx=XM_006929172.4;refGene_gn=LOC101083499;refGene_tx=XM_006929174.3;refGene_gn=LOC101083499;refGene_tx=XM_019825448.1;refGene_gn=LOC101083499;refGene_tx=XM_019825449.1;refGene_gn=LOC101083499;refGene_tx=XM_019825450.2;refGene_gn=LOC101083499;refGene_tx=XM_019825452.2;refGene_gn=LOC101083499;refGene_tx=XM_023250337.1;refGene_gn=LOC101083499;refGene_tx=XM_023250338.1
A2 97243993 97244113
refGene_gn=LOC101083499;refGene_tx=XM_003982742.5;refGene_gn=LOC101083499;refGene_tx=XM_006929172.4;refGene_gn=LOC101083499;refGene_tx=XM_006929174.3;refGene_gn=LOC101083499;refGene_tx=XM_019825448.1;refGene_gn=LOC101083499;refGene_tx=XM_019825449.1;refGene_gn=LOC101083499;refGene_tx=XM_019825450.2;refGene_gn=LOC101083499;refGene_tx=XM_019825452.2;refGene_gn=LOC101083499;refGene_tx=XM_023250337.1;refGene_gn=LOC101083499;refGene_tx=XM_023250338.1
A2 97246098 97246273
refGene_gn=LOC101083499;refGene_tx=XM_003982742.5;refGene_gn=LOC101083499;refGene_tx=XM_006929172.4;refGene_gn=LOC101083499;refGene_tx=XM_006929174.3;refGene_gn=LOC101083499;refGene_tx=XM_019825448.1;refGene_gn=LOC101083499;refGene_tx=XM_019825449.1;refGene_gn=LOC101083499;refGene_tx=XM_019825450.2;refGene_gn=LOC101083499;refGene_tx=XM_019825452.2;refGene_gn=LOC101083499;refGene_tx=XM_023250337.1;refGene_gn=LOC101083499;refGene_tx=XM_023250338.1
A2 97247757 97247884
refGene_gn=LOC101083499;refGene_tx=XM_003982742.5;refGene_gn=LOC101083499;refGene_tx=XM_006929172.4;refGene_gn=LOC101083499;refGene_tx=XM_006929174.3;refGene_gn=LOC101083499;refGene_tx=XM_019825448.1;refGene_gn=LOC101083499;refGene_tx=XM_019825449.1;refGene_gn=LOC101083499;refGene_tx=XM_019825450.2;refGene_gn=LOC101083499;refGene_tx=XM_019825452.2;refGene_gn=LOC101083499;refGene_tx=XM_023250337.1;refGene_gn=LOC101083499;refGene_tx=XM_023250338.1
A2 97249124 97249301
refGene_gn=LOC101083499;refGene_tx=XM_003982742.5;refGene_gn=LOC101083499;refGene_tx=XM_006929172.4;refGene_gn=LOC101083499;refGene_tx=XM_006929174.3;refGene_gn=LOC101083499;refGene_tx=XM_019825448.1;refGene_gn=LOC101083499;refGene_tx=XM_019825449.1;refGene_gn=LOC101083499;refGene_tx=XM_019825450.2;refGene_gn=LOC101083499;refGene_tx=XM_019825452.2;refGene_gn=LOC101083499;refGene_tx=XM_023250337.1;refGene_gn=LOC101083499;refGene_tx=XM_023250338.1
A2 97253026 97253125
refGene_gn=LOC101083499;refGene_tx=XM_003982742.5;refGene_gn=LOC101083499;refGene_tx=XM_006929172.4;refGene_gn=LOC101083499;refGene_tx=XM_006929174.3;refGene_gn=LOC101083499;refGene_tx=XM_019825448.1;refGene_gn=LOC101083499;refGene_tx=XM_019825449.1;refGene_gn=LOC101083499;refGene_tx=XM_019825450.2
A2 97254275 97254449
refGene_gn=LOC101083499;refGene_tx=XM_003982742.5
A2 97254625 97254724
refGene_gn=LOC101083499;refGene_tx=XM_006929172.4;refGene_gn=LOC101083499;refGene_tx=XM_019825448.1;refGene_gn=LOC101083499;refGene_tx=XM_01

9825449.1;refGene_gn=LOC101083499;refGene_tx=XM_019825450.2
A2 97255787 97255829
refGene_gn=LOC101083499;refGene_tx=XM_006929172.4;refGene_gn=LOC101083499;refGene_tx=XM_006929174.3;refGene_gn=LOC101083499;refGene_tx=XM_019825448.1;refGene_gn=LOC101083499;refGene_tx=XM_019825449.1;refGene_gn=LOC101083499;refGene_tx=XM_019825450.2
A2 97274862 97275049 refGene_gn=LRRD1;refGene_tx=XM_011280376.2
A2 97275691 97275809 refGene_gn=LRRD1;refGene_tx=XM_011280376.2
A2 97278124 97278286 refGene_gn=LRRD1;refGene_tx=XM_011280376.2
A2 97292002 97292201 refGene_gn=LRRD1;refGene_tx=XM_011280376.2
A2 97295983 97296468 refGene_gn=LRRD1;refGene_tx=XM_011280376.2
A2 97296470 97296740 refGene_gn=LRRD1;refGene_tx=XM_011280376.2
A2 97296742 97297509 refGene_gn=LRRD1;refGene_tx=XM_011280376.2
A2 97297510 97297917 refGene_gn=LRRD1;refGene_tx=XM_011280376.2
A2 97318469 97318538
refGene_gn=KRIT1;refGene_tx=XM_003982743.5;refGene_gn=KRIT1;refGene_tx=XM_011280377.3;refGene_gn=KRIT1;refGene_tx=XM_011280378.3;refGene_gn=KRIT1;refGene_tx=XM_011280380.3;refGene_gn=KRIT1;refGene_tx=XM_019825447.1
A2 97319025 97319142
refGene_gn=KRIT1;refGene_tx=XM_003982743.5;refGene_gn=KRIT1;refGene_tx=XM_011280377.3;refGene_gn=KRIT1;refGene_tx=XM_011280378.3;refGene_gn=KRIT1;refGene_tx=XM_011280380.3;refGene_gn=KRIT1;refGene_tx=XM_019825447.1
A2 97325665 97325872
refGene_gn=KRIT1;refGene_tx=XM_003982743.5;refGene_gn=KRIT1;refGene_tx=XM_011280377.3;refGene_gn=KRIT1;refGene_tx=XM_011280378.3;refGene_gn=KRIT1;refGene_tx=XM_011280380.3;refGene_gn=KRIT1;refGene_tx=XM_019825447.1
A2 97326468 97326556
refGene_gn=KRIT1;refGene_tx=XM_003982743.5;refGene_gn=KRIT1;refGene_tx=XM_011280377.3;refGene_gn=KRIT1;refGene_tx=XM_011280378.3;refGene_gn=KRIT1;refGene_tx=XM_011280380.3;refGene_gn=KRIT1;refGene_tx=XM_019825447.1
A2 97327289 97327456
refGene_gn=KRIT1;refGene_tx=XM_003982743.5;refGene_gn=KRIT1;refGene_tx=XM_011280377.3;refGene_gn=KRIT1;refGene_tx=XM_011280378.3;refGene_gn=KRIT1;refGene_tx=XM_011280380.3;refGene_gn=KRIT1;refGene_tx=XM_019825447.1
A2 97332607 97332759
refGene_gn=KRIT1;refGene_tx=XM_003982743.5;refGene_gn=KRIT1;refGene_tx=XM_011280377.3;refGene_gn=KRIT1;refGene_tx=XM_011280378.3;refGene_gn=KRIT1;refGene_tx=XM_011280380.3;refGene_gn=KRIT1;refGene_tx=XM_019825447.1
A2 97334789 97334946
refGene_gn=KRIT1;refGene_tx=XM_003982743.5;refGene_gn=KRIT1;refGene_tx=XM_011280377.3;refGene_gn=KRIT1;refGene_tx=XM_011280378.3;refGene_gn=KRIT1;refGene_tx=XM_011280380.3;refGene_gn=KRIT1;refGene_tx=XM_019825447.1
A2 97337275 97337383

refGene_gn=KRIT1;refGene_tx=XM_003982743.5;refGene_gn=KRIT1;refGene_tx=XM_011280377.3;refGene_gn=KRIT1;refGene_tx=XM_011280378.3;refGene_gn=KRIT1;refGene_tx=XM_011280380.3;refGene_gn=KRIT1;refGene_tx=XM_019825447.1
A2 97338289 97338446
refGene_gn=KRIT1;refGene_tx=XM_003982743.5;refGene_gn=KRIT1;refGene_tx=XM_011280377.3;refGene_gn=KRIT1;refGene_tx=XM_011280378.3;refGene_gn=KRIT1;refGene_tx=XM_011280380.3;refGene_gn=KRIT1;refGene_tx=XM_019825447.1
A2 97343752 97343896
refGene_gn=KRIT1;refGene_tx=XM_003982743.5;refGene_gn=KRIT1;refGene_tx=XM_011280377.3;refGene_gn=KRIT1;refGene_tx=XM_011280378.3;refGene_gn=KRIT1;refGene_tx=XM_011280380.3;refGene_gn=KRIT1;refGene_tx=XM_019825447.1
A2 97344126 97344242
refGene_gn=KRIT1;refGene_tx=XM_003982743.5;refGene_gn=KRIT1;refGene_tx=XM_011280377.3;refGene_gn=KRIT1;refGene_tx=XM_011280378.3;refGene_gn=KRIT1;refGene_tx=XM_011280380.3;refGene_gn=KRIT1;refGene_tx=XM_019825447.1
A2 97344436 97344680
refGene_gn=KRIT1;refGene_tx=XM_003982743.5;refGene_gn=KRIT1;refGene_tx=XM_011280377.3;refGene_gn=KRIT1;refGene_tx=XM_011280378.3;refGene_gn=KRIT1;refGene_tx=XM_011280380.3;refGene_gn=KRIT1;refGene_tx=XM_019825447.1
A2 97346367 97346497
refGene_gn=KRIT1;refGene_tx=XM_003982743.5;refGene_gn=KRIT1;refGene_tx=XM_011280377.3;refGene_gn=KRIT1;refGene_tx=XM_011280378.3;refGene_gn=KRIT1;refGene_tx=XM_011280380.3;refGene_gn=KRIT1;refGene_tx=XM_019825447.1
A2 97347840 97347933
refGene_gn=KRIT1;refGene_tx=XM_003982743.5;refGene_gn=KRIT1;refGene_tx=XM_011280377.3;refGene_gn=KRIT1;refGene_tx=XM_011280378.3;refGene_gn=KRIT1;refGene_tx=XM_011280380.3;refGene_gn=KRIT1;refGene_tx=XM_019825447.1
A2 97352130 97352290
refGene_gn=KRIT1;refGene_tx=XM_003982743.5;refGene_gn=KRIT1;refGene_tx=XM_011280377.3;refGene_gn=KRIT1;refGene_tx=XM_011280378.3;refGene_gn=KRIT1;refGene_tx=XM_011280380.3
A2 97353223 97353325
refGene_gn=KRIT1;refGene_tx=XM_003982743.5;refGene_gn=KRIT1;refGene_tx=XM_011280377.3;refGene_gn=KRIT1;refGene_tx=XM_011280378.3;refGene_gn=KRIT1;refGene_tx=XM_011280380.3
A2 97413261 97413449
refGene_gn=ANKIB1;refGene_tx=XM_011280381.3;refGene_gn=ANKIB1;refGene_tx=XM_011280382.3;refGene_gn=ANKIB1;refGene_tx=XM_019825453.1
A2 97423242 97423543
refGene_gn=ANKIB1;refGene_tx=XM_011280381.3;refGene_gn=ANKIB1;refGene_tx=XM_011280382.3;refGene_gn=ANKIB1;refGene_tx=XM_019825453.1
A2 97436262 97436445
refGene_gn=ANKIB1;refGene_tx=XM_011280381.3;refGene_gn=ANKIB1;refGene_

tx=XM_011280382.3;refGene_gn=ANKIB1;refGene_tx=XM_019825453.1
A2 97446336 97446454
refGene_gn=ANKIB1;refGene_tx=XM_011280381.3;refGene_gn=ANKIB1;refGene_
tx=XM_011280382.3;refGene_gn=ANKIB1;refGene_tx=XM_019825453.1
A2 97453934 97454143
refGene_gn=ANKIB1;refGene_tx=XM_011280381.3;refGene_gn=ANKIB1;refGene_
tx=XM_011280382.3;refGene_gn=ANKIB1;refGene_tx=XM_019825453.1
A2 97455334 97455423
refGene_gn=ANKIB1;refGene_tx=XM_011280381.3;refGene_gn=ANKIB1;refGene_
tx=XM_011280382.3;refGene_gn=ANKIB1;refGene_tx=XM_019825453.1
A2 97462280 97462425
refGene_gn=ANKIB1;refGene_tx=XM_011280381.3;refGene_gn=ANKIB1;refGene_
tx=XM_011280382.3;refGene_gn=ANKIB1;refGene_tx=XM_019825453.1
A2 97463916 97464083
refGene_gn=ANKIB1;refGene_tx=XM_011280381.3;refGene_gn=ANKIB1;refGene_
tx=XM_011280382.3;refGene_gn=ANKIB1;refGene_tx=XM_019825453.1
A2 97473952 97474041
refGene_gn=ANKIB1;refGene_tx=XM_011280381.3;refGene_gn=ANKIB1;refGene_
tx=XM_011280382.3;refGene_gn=ANKIB1;refGene_tx=XM_019825453.1
A2 97482367 97482498
refGene_gn=ANKIB1;refGene_tx=XM_011280381.3;refGene_gn=ANKIB1;refGene_
tx=XM_011280382.3;refGene_gn=ANKIB1;refGene_tx=XM_019825453.1
A2 97490291 97490426
refGene_gn=ANKIB1;refGene_tx=XM_011280381.3;refGene_gn=ANKIB1;refGene_
tx=XM_011280382.3;refGene_gn=ANKIB1;refGene_tx=XM_019825453.1
A2 97491352 97491439
refGene_gn=ANKIB1;refGene_tx=XM_011280381.3;refGene_gn=ANKIB1;refGene_
tx=XM_011280382.3;refGene_gn=ANKIB1;refGene_tx=XM_019825453.1
A2 97491525 97491592
refGene_gn=ANKIB1;refGene_tx=XM_011280381.3;refGene_gn=ANKIB1;refGene_
tx=XM_011280382.3;refGene_gn=ANKIB1;refGene_tx=XM_019825453.1
A2 97493951 97494097
refGene_gn=ANKIB1;refGene_tx=XM_011280381.3;refGene_gn=ANKIB1;refGene_
tx=XM_011280382.3;refGene_gn=ANKIB1;refGene_tx=XM_019825453.1
A2 97495112 97495291
refGene_gn=ANKIB1;refGene_tx=XM_011280381.3;refGene_gn=ANKIB1;refGene_
tx=XM_011280382.3;refGene_gn=ANKIB1;refGene_tx=XM_019825453.1
A2 97496097 97496149
refGene_gn=ANKIB1;refGene_tx=XM_011280381.3;refGene_gn=ANKIB1;refGene_
tx=XM_011280382.3;refGene_gn=ANKIB1;refGene_tx=XM_019825453.1
A2 97500214 97500329
refGene_gn=ANKIB1;refGene_tx=XM_011280381.3;refGene_gn=ANKIB1;refGene_
tx=XM_011280382.3;refGene_gn=ANKIB1;refGene_tx=XM_019825453.1
A2 97502176 97502310
refGene_gn=ANKIB1;refGene_tx=XM_011280381.3;refGene_gn=ANKIB1;refGene_
tx=XM_011280382.3;refGene_gn=ANKIB1;refGene_tx=XM_019825453.1
A2 97502650 97503391
refGene_gn=ANKIB1;refGene_tx=XM_011280381.3;refGene_gn=ANKIB1;refGene_
tx=XM_011280382.3;refGene_gn=ANKIB1;refGene_tx=XM_019825453.1
A2 97520805 97520958

refGene_gn=TMBIM7P;refGene_tx=XM_006929177.3;refGene_gn=TMBIM7P;refGene_tx=XM_011280383.3;refGene_gn=TMBIM7P;refGene_tx=XM_023250339.1
A2 97525020 97525068
refGene_gn=TMBIM7P;refGene_tx=XM_006929177.3;refGene_gn=TMBIM7P;refGene_tx=XM_011280383.3;refGene_gn=TMBIM7P;refGene_tx=XM_023250339.1
A2 97527641 97527731
refGene_gn=TMBIM7P;refGene_tx=XM_006929177.3;refGene_gn=TMBIM7P;refGene_tx=XM_011280383.3;refGene_gn=TMBIM7P;refGene_tx=XM_023250339.1
A2 97529461 97529549
refGene_gn=TMBIM7P;refGene_tx=XM_006929177.3;refGene_gn=TMBIM7P;refGene_tx=XM_011280383.3;refGene_gn=TMBIM7P;refGene_tx=XM_023250339.1
A2 97529632 97529670
refGene_gn=TMBIM7P;refGene_tx=XM_006929177.3;refGene_gn=TMBIM7P;refGene_tx=XM_011280383.3;refGene_gn=TMBIM7P;refGene_tx=XM_023250339.1
A2 97534914 97535005
refGene_gn=TMBIM7P;refGene_tx=XM_006929177.3;refGene_gn=TMBIM7P;refGene_tx=XM_011280383.3;refGene_gn=TMBIM7P;refGene_tx=XM_023250339.1
A2 97535921 97535975
refGene_gn=TMBIM7P;refGene_tx=XM_006929177.3;refGene_gn=TMBIM7P;refGene_tx=XM_011280383.3;refGene_gn=TMBIM7P;refGene_tx=XM_023250339.1
A2 97536688 97536767
refGene_gn=TMBIM7P;refGene_tx=XM_006929177.3;refGene_gn=TMBIM7P;refGene_tx=XM_011280383.3;refGene_gn=TMBIM7P;refGene_tx=XM_023250339.1
A2 97547812 97548081
refGene_gn=TMBIM7P;refGene_tx=XM_006929177.3;refGene_gn=TMBIM7P;refGene_tx=XM_011280383.3;refGene_gn=TMBIM7P;refGene_tx=XM_023250339.1
A2 97551291 97551293
refGene_gn=TMBIM7P;refGene_tx=XM_006929177.3;refGene_gn=TMBIM7P;refGene_tx=XM_011280383.3;refGene_gn=TMBIM7P;refGene_tx=XM_023250339.1
A2 97562014 97562266 refGene_gn=GATAD1;refGene_tx=XM_023250340.1
A2 97563053 97563146 refGene_gn=GATAD1;refGene_tx=XM_023250340.1
A2 97563147 97563180 refGene_gn=GATAD1;refGene_tx=XM_023250340.1
A2 97564825 97564885 refGene_gn=GATAD1;refGene_tx=XM_023250340.1
A2 97568401 97568585 refGene_gn=GATAD1;refGene_tx=XM_023250340.1
A2 97570915 97571106 refGene_gn=GATAD1;refGene_tx=XM_023250340.1
A2 97584787 97584872 refGene_gn=PEX1;refGene_tx=XM_023250341.1
A2 97586594 97586725
refGene_gn=PEX1;refGene_tx=XM_023250341.1;refGene_gn=PEX1;refGene_tx=XM_023250342.1
A2 97587010 97587208
refGene_gn=PEX1;refGene_tx=XM_023250341.1;refGene_gn=PEX1;refGene_tx=XM_023250342.1
A2 97588361 97588592
refGene_gn=PEX1;refGene_tx=XM_023250341.1;refGene_gn=PEX1;refGene_tx=XM_023250342.1;refGene_gn=PEX1;refGene_tx=XM_023250343.1
A2 97589984 97590161
refGene_gn=PEX1;refGene_tx=XM_023250341.1;refGene_gn=PEX1;refGene_tx=XM_023250342.1;refGene_gn=PEX1;refGene_tx=XM_023250343.1
A2 97590957 97591061
refGene_gn=PEX1;refGene_tx=XM_023250341.1;refGene_gn=PEX1;refGene_tx=X

M_023250342.1;refGene_gn=PEX1;refGene_tx=XM_023250343.1
A2	97591139 97591282
refGene_gn=PEX1;refGene_tx=XM_023250341.1;refGene_gn=PEX1;refGene_tx=XM_023250342.1;refGene_gn=PEX1;refGene_tx=XM_023250343.1
A2	97594416 97594481
refGene_gn=PEX1;refGene_tx=XM_023250341.1;refGene_gn=PEX1;refGene_tx=XM_023250342.1;refGene_gn=PEX1;refGene_tx=XM_023250343.1
A2	97597520 97597655
refGene_gn=PEX1;refGene_tx=XM_023250341.1;refGene_gn=PEX1;refGene_tx=XM_023250342.1;refGene_gn=PEX1;refGene_tx=XM_023250343.1
A2	97599090 97599257
refGene_gn=PEX1;refGene_tx=XM_023250341.1;refGene_gn=PEX1;refGene_tx=XM_023250342.1;refGene_gn=PEX1;refGene_tx=XM_023250343.1
A2	97599460 97599650
refGene_gn=PEX1;refGene_tx=XM_023250341.1;refGene_gn=PEX1;refGene_tx=XM_023250342.1;refGene_gn=PEX1;refGene_tx=XM_023250343.1
A2	97604802 97604957
refGene_gn=PEX1;refGene_tx=XM_023250341.1;refGene_gn=PEX1;refGene_tx=XM_023250342.1;refGene_gn=PEX1;refGene_tx=XM_023250343.1
A2	97606382 97606553
refGene_gn=PEX1;refGene_tx=XM_023250341.1;refGene_gn=PEX1;refGene_tx=XM_023250342.1;refGene_gn=PEX1;refGene_tx=XM_023250343.1
A2	97608515 97608612
refGene_gn=PEX1;refGene_tx=XM_023250341.1;refGene_gn=PEX1;refGene_tx=XM_023250342.1;refGene_gn=PEX1;refGene_tx=XM_023250343.1
A2	97614241 97614374
refGene_gn=PEX1;refGene_tx=XM_023250341.1;refGene_gn=PEX1;refGene_tx=XM_023250342.1;refGene_gn=PEX1;refGene_tx=XM_023250343.1
A2	97615490 97615573
refGene_gn=PEX1;refGene_tx=XM_023250341.1;refGene_gn=PEX1;refGene_tx=XM_023250342.1;refGene_gn=PEX1;refGene_tx=XM_023250343.1
A2	97616461 97616562
refGene_gn=PEX1;refGene_tx=XM_023250341.1;refGene_gn=PEX1;refGene_tx=XM_023250342.1;refGene_gn=PEX1;refGene_tx=XM_023250343.1
A2	97616811 97616935
refGene_gn=PEX1;refGene_tx=XM_023250341.1;refGene_gn=PEX1;refGene_tx=XM_023250342.1;refGene_gn=PEX1;refGene_tx=XM_023250343.1
A2	97620009 97620129
refGene_gn=PEX1;refGene_tx=XM_023250341.1;refGene_gn=PEX1;refGene_tx=XM_023250342.1;refGene_gn=PEX1;refGene_tx=XM_023250343.1
A2	97622004 97622690
refGene_gn=PEX1;refGene_tx=XM_023250341.1;refGene_gn=PEX1;refGene_tx=XM_023250342.1;refGene_gn=PEX1;refGene_tx=XM_023250343.1
A2	97622781 97622896
refGene_gn=PEX1;refGene_tx=XM_023250341.1;refGene_gn=PEX1;refGene_tx=XM_023250342.1;refGene_gn=PEX1;refGene_tx=XM_023250343.1
A2	97623314 97623398
refGene_gn=PEX1;refGene_tx=XM_023250341.1;refGene_gn=PEX1;refGene_tx=XM_023250342.1;refGene_gn=PEX1;refGene_tx=XM_023250343.1
A2	97626199 97626343

refGene_gn=PEX1;refGene_tx=XM_023250341.1;refGene_gn=PEX1;refGene_tx=XM_023250342.1;refGene_gn=PEX1;refGene_tx=XM_023250343.1
A2 97631172 97631301
refGene_gn=PEX1;refGene_tx=XM_023250341.1;refGene_gn=PEX1;refGene_tx=XM_023250342.1;refGene_gn=PEX1;refGene_tx=XM_023250343.1
A2 97631659 97631770 refGene_gn=RBM48;refGene_tx=XM_003982745.5
A2 97632422 97632613 refGene_gn=RBM48;refGene_tx=XM_003982745.5
A2 97634319 97634465 refGene_gn=RBM48;refGene_tx=XM_003982745.5
A2 97635938 97636504
refGene_gn=LOC101086547;refGene_tx=XM_023250345.1;refGene_gn=RBM48;refGene_tx=XM_003982745.5
A2 97638371 97638458 refGene_gn=RBM48;refGene_tx=XM_003982745.5
A2 97639056 97639113
refGene_gn=LOC101086547;refGene_tx=XM_023250344.1
A2 97641036 97641141
refGene_gn=LOC101086547;refGene_tx=XM_023250344.1;refGene_gn=LOC101086547;refGene_tx=XM_023250345.1
A2 97644605 97644757
refGene_gn=LOC101086547;refGene_tx=XM_023250344.1;refGene_gn=LOC101086547;refGene_tx=XM_023250345.1
A2 97651622 97651727
refGene_gn=LOC101086547;refGene_tx=XM_023250344.1;refGene_gn=LOC101086547;refGene_tx=XM_023250345.1
A2 97652027 97652183
refGene_gn=LOC101086547;refGene_tx=XM_023250344.1;refGene_gn=LOC101086547;refGene_tx=XM_023250345.1
A2 97661955 97662103
refGene_gn=LOC101086547;refGene_tx=XM_023250344.1;refGene_gn=LOC101086547;refGene_tx=XM_023250345.1
A2 97665678 97665765
refGene_gn=FAM133B;refGene_tx=XM_011280385.3;refGene_gn=FAM133B;refGene_tx=XM_023250346.1
A2 97668867 97668909
refGene_gn=FAM133B;refGene_tx=XM_011280385.3;refGene_gn=FAM133B;refGene_tx=XM_023250346.1
A2 97670485 97670578
refGene_gn=FAM133B;refGene_tx=XM_011280385.3;refGene_gn=FAM133B;refGene_tx=XM_023250346.1
A2 97677827 97677878
refGene_gn=FAM133B;refGene_tx=XM_011280385.3;refGene_gn=FAM133B;refGene_tx=XM_023250346.1
A2 97679182 97679275
refGene_gn=FAM133B;refGene_tx=XM_011280385.3;refGene_gn=FAM133B;refGene_tx=XM_023250346.1
A2 97679841 97679904
refGene_gn=FAM133B;refGene_tx=XM_011280385.3;refGene_gn=FAM133B;refGene_tx=XM_023250346.1
A2 97680373 97680406
refGene_gn=FAM133B;refGene_tx=XM_011280385.3;refGene_gn=FAM133B;refGene_tx=XM_023250346.1

A2 97680518 97680593
refGene_gn=FAM133B;refGene_tx=XM_011280385.3;refGene_gn=FAM133B;refGene_tx=XM_023250346.1
A2 97681491 97681570
refGene_gn=FAM133B;refGene_tx=XM_011280385.3;refGene_gn=FAM133B;refGene_tx=XM_023250346.1
A2 97683390 97683488
refGene_gn=FAM133B;refGene_tx=XM_011280385.3;refGene_gn=FAM133B;refGene_tx=XM_023250346.1
A2 97693687 97693711 refGene_gn=FAM133B;refGene_tx=XM_011280385.3
A2 97719780 97719927 refGene_gn=CDK6;refGene_tx=XM_003982756.4
A2 97725265 97725401 refGene_gn=CDK6;refGene_tx=XM_003982756.4
A2 97730849 97730900 refGene_gn=CDK6;refGene_tx=XM_003982756.4
A2 97785549 97785659 refGene_gn=CDK6;refGene_tx=XM_003982756.4
A2 97847502 97847670 refGene_gn=CDK6;refGene_tx=XM_003982756.4
A2 97898315 97898451 refGene_gn=CDK6;refGene_tx=XM_003982756.4
A2 97964681 97964914 refGene_gn=CDK6;refGene_tx=XM_003982756.4
A2 98144969 98144977 refGene_gn=SAMD9;refGene_tx=XM_023242079.1
A2 98162815 98162977 refGene_gn=SAMD9;refGene_tx=XM_023242079.1
A2 98163300 98163441 refGene_gn=SAMD9;refGene_tx=XM_023242079.1
A2 98163443 98163584 refGene_gn=SAMD9;refGene_tx=XM_023242079.1
A2 98163586 98163760 refGene_gn=SAMD9;refGene_tx=XM_023242079.1
A2 98163762 98163911 refGene_gn=SAMD9;refGene_tx=XM_023242079.1
A2 98163912 98164205 refGene_gn=SAMD9;refGene_tx=XM_023242079.1
A2 98164206 98164667 refGene_gn=SAMD9;refGene_tx=XM_023242079.1
A2 98164669 98165134 refGene_gn=SAMD9;refGene_tx=XM_023242079.1
A2 98165136 98165362 refGene_gn=SAMD9;refGene_tx=XM_023242079.1
A2 98165364 98165543 refGene_gn=SAMD9;refGene_tx=XM_023242079.1
A2 98165545 98165631 refGene_gn=SAMD9;refGene_tx=XM_023242079.1
A2 98165632 98165777 refGene_gn=SAMD9;refGene_tx=XM_023242079.1
A2 98165779 98165894 refGene_gn=SAMD9;refGene_tx=XM_023242079.1
A2 98165895 98166030 refGene_gn=SAMD9;refGene_tx=XM_023242079.1
A2 98188605 98193378 refGene_gn=SAMD9L;refGene_tx=XM_003982747.4
A2 98242713 98242827
refGene_gn=HEPACAM2;refGene_tx=XM_023250347.1;refGene_gn=HEPACAM2;refGene_tx=XM_023250348.1
A2 98246568 98246642
refGene_gn=HEPACAM2;refGene_tx=XM_023250347.1;refGene_gn=HEPACAM2;refGene_tx=XM_023250348.1
A2 98247974 98248012
refGene_gn=HEPACAM2;refGene_tx=XM_023250347.1;refGene_gn=HEPACAM2;refGene_tx=XM_023250348.1
A2 98248103 98248128
refGene_gn=HEPACAM2;refGene_tx=XM_023250347.1;refGene_gn=HEPACAM2;refGene_tx=XM_023250348.1
A2 98248217 98248334
refGene_gn=HEPACAM2;refGene_tx=XM_023250347.1;refGene_gn=HEPACAM2;refGene_tx=XM_023250348.1
A2 98255958 98255967
refGene_gn=HEPACAM2;refGene_tx=XM_023250347.1;refGene_gn=HEPACAM2;refG

ene_tx=XM_023250348.1
A2 98258855 98259152
refGene_gn=HEPACAM2;refGene_tx=XM_023250347.1;refGene_gn=HEPACAM2;refGene_tx=XM_023250348.1
A2 98269636 98269921
refGene_gn=HEPACAM2;refGene_tx=XM_023250347.1;refGene_gn=HEPACAM2;refGene_tx=XM_023250348.1
A2 98273446 98273800
refGene_gn=HEPACAM2;refGene_tx=XM_023250347.1;refGene_gn=HEPACAM2;refGene_tx=XM_023250348.1
A2 98276629 98276635 refGene_gn=HEPACAM2;refGene_tx=XM_023250348.1
A2 98280741 98280807 refGene_gn=HEPACAM2;refGene_tx=XM_023250348.1
A2 98281204 98281283
refGene_gn=HEPACAM2;refGene_tx=XM_023250347.1;refGene_gn=HEPACAM2;refGene_tx=XM_023250348.1
A2 98287708 98287741
refGene_gn=VPS50;refGene_tx=XM_011280388.3;refGene_gn=VPS50;refGene_tx=XM_019825464.2
A2 98294951 98295020 refGene_gn=VPS50;refGene_tx=XM_011280388.3
A2 98309823 98309946
refGene_gn=VPS50;refGene_tx=XM_011280388.3;refGene_gn=VPS50;refGene_tx=XM_019825464.2
A2 98311256 98311328
refGene_gn=VPS50;refGene_tx=XM_011280388.3;refGene_gn=VPS50;refGene_tx=XM_019825464.2
A2 98313564 98313618
refGene_gn=VPS50;refGene_tx=XM_011280388.3;refGene_gn=VPS50;refGene_tx=XM_019825464.2
A2 98314454 98314525
refGene_gn=VPS50;refGene_tx=XM_011280388.3;refGene_gn=VPS50;refGene_tx=XM_019825464.2
A2 98315214 98315332
refGene_gn=VPS50;refGene_tx=XM_011280388.3;refGene_gn=VPS50;refGene_tx=XM_019825464.2
A2 98315646 98315682
refGene_gn=VPS50;refGene_tx=XM_011280388.3;refGene_gn=VPS50;refGene_tx=XM_019825464.2
A2 98316859 98316942
refGene_gn=VPS50;refGene_tx=XM_011280388.3;refGene_gn=VPS50;refGene_tx=XM_019825464.2
A2 98331403 98331446
refGene_gn=VPS50;refGene_tx=XM_011280388.3;refGene_gn=VPS50;refGene_tx=XM_019825464.2
A2 98333463 98333562
refGene_gn=VPS50;refGene_tx=XM_011280388.3;refGene_gn=VPS50;refGene_tx=XM_019825464.2
A2 98335186 98335327
refGene_gn=VPS50;refGene_tx=XM_011280388.3;refGene_gn=VPS50;refGene_tx=XM_019825464.2
A2 98350709 98350842

refGene_gn=VPS50;refGene_tx=XM_011280388.3;refGene_gn=VPS50;refGene_tx=XM_019825464.2
A2 98354429 98354518 refGene_gn=VPS50;refGene_tx=XM_011280388.3;refGene_gn=VPS50;refGene_tx=XM_019825464.2
A2 98361452 98361547 refGene_gn=VPS50;refGene_tx=XM_011280388.3;refGene_gn=VPS50;refGene_tx=XM_019825464.2
A2 98361856 98361955 refGene_gn=VPS50;refGene_tx=XM_011280388.3;refGene_gn=VPS50;refGene_tx=XM_019825464.2
A2 98373486 98373577 refGene_gn=VPS50;refGene_tx=XM_011280388.3;refGene_gn=VPS50;refGene_tx=XM_019825464.2
A2 98376658 98376835 refGene_gn=VPS50;refGene_tx=XM_011280388.3;refGene_gn=VPS50;refGene_tx=XM_019825464.2
A2 98380179 98380298 refGene_gn=VPS50;refGene_tx=XM_011280388.3;refGene_gn=VPS50;refGene_tx=XM_019825464.2
A2 98382455 98382562 refGene_gn=VPS50;refGene_tx=XM_011280388.3;refGene_gn=VPS50;refGene_tx=XM_019825464.2
A2 98392326 98392448 refGene_gn=VPS50;refGene_tx=XM_011280388.3;refGene_gn=VPS50;refGene_tx=XM_019825464.2
A2 98400149 98400230 refGene_gn=VPS50;refGene_tx=XM_011280388.3;refGene_gn=VPS50;refGene_tx=XM_019825464.2
A2 98408707 98408856 refGene_gn=VPS50;refGene_tx=XM_011280388.3;refGene_gn=VPS50;refGene_tx=XM_019825464.2
A2 98409867 98409964 refGene_gn=VPS50;refGene_tx=XM_011280388.3;refGene_gn=VPS50;refGene_tx=XM_019825464.2
A2 98410995 98411154 refGene_gn=VPS50;refGene_tx=XM_011280388.3;refGene_gn=VPS50;refGene_tx=XM_019825464.2
A2 98414582 98414704 refGene_gn=VPS50;refGene_tx=XM_011280388.3;refGene_gn=VPS50;refGene_tx=XM_019825464.2
A2 98416472 98416662 refGene_gn=VPS50;refGene_tx=XM_011280388.3;refGene_gn=VPS50;refGene_tx=XM_019825464.2
A2 98419634 98419754 refGene_gn=VPS50;refGene_tx=XM_011280388.3;refGene_gn=VPS50;refGene_tx=XM_019825464.2
A2 98474896 98475154 refGene_gn=CALCR;refGene_tx=XM_003982757.4
A2 98482694 98482736 refGene_gn=CALCR;refGene_tx=XM_003982757.4
A2 98483429 98483648 refGene_gn=CALCR;refGene_tx=XM_003982757.4

A2 98485991 98486058 refGene_gn=CALCR;refGene_tx=XM_003982757.4
A2 98486165 98486226 refGene_gn=CALCR;refGene_tx=XM_003982757.4
A2 98491827 98491981 refGene_gn=CALCR;refGene_tx=XM_003982757.4
A2 98508303 98508430 refGene_gn=CALCR;refGene_tx=XM_003982757.4
A2 98516024 98516116 refGene_gn=CALCR;refGene_tx=XM_003982757.4
A2 98518974 98519087 refGene_gn=CALCR;refGene_tx=XM_003982757.4
A2 98525446 98525557 refGene_gn=CALCR;refGene_tx=XM_003982757.4
A2 98527687 98527844 refGene_gn=CALCR;refGene_tx=XM_003982757.4
A2 98534439 98534490 refGene_gn=CALCR;refGene_tx=XM_003982757.4
A2 98945642 98945725 refGene_gn=TFPI2;refGene_tx=XM_003982759.5
A2 98946325 98946496 refGene_gn=TFPI2;refGene_tx=XM_003982759.5
A2 98947718 98947898 refGene_gn=TFPI2;refGene_tx=XM_003982759.5
A2 98948535 98948718 refGene_gn=TFPI2;refGene_tx=XM_003982759.5
A2 98949021 98949109 refGene_gn=TFPI2;refGene_tx=XM_003982759.5
A2 98962863 98962959
refGene_gn=GNGT1;refGene_tx=XM_019825466.1;refGene_gn=GNGT1;refGene_tx
=XM_019825467.1
A2 98967638 98967767
refGene_gn=GNGT1;refGene_tx=XM_019825466.1;refGene_gn=GNGT1;refGene_tx
=XM_019825467.1
A2 98975211 98975307 refGene_gn=GNG11;refGene_tx=XM_003982762.5
A2 98978804 98978930 refGene_gn=GNG11;refGene_tx=XM_003982762.5
A2 99040401 99040460 refGene_gn=BET1;refGene_tx=XM_006929186.4
A2 99042047 99042203
refGene_gn=BET1;refGene_tx=XM_003982763.4;refGene_gn=BET1;refGene_tx=X
M_006929186.4
A2 99043747 99043804
refGene_gn=BET1;refGene_tx=XM_003982763.4;refGene_gn=BET1;refGene_tx=X
M_006929186.4
A2 99046374 99046499
refGene_gn=BET1;refGene_tx=XM_003982763.4;refGene_gn=BET1;refGene_tx=X
M_006929186.4
A2 99050587 99050606
refGene_gn=BET1;refGene_tx=XM_003982763.4;refGene_gn=BET1;refGene_tx=X
M_006929186.4
A2 99052655 99053306
refGene_gn=LOC111559500;refGene_tx=XM_023250350.1
A2 99442596 99442666 refGene_gn=COL1A2;refGene_tx=XM_003982764.5
A2 99445185 99445196 refGene_gn=COL1A2;refGene_tx=XM_003982764.5
A2 99445859 99445874 refGene_gn=COL1A2;refGene_tx=XM_003982764.5
A2 99446513 99446549 refGene_gn=COL1A2;refGene_tx=XM_003982764.5
A2 99447736 99447829 refGene_gn=COL1A2;refGene_tx=XM_003982764.5
A2 99449107 99449164 refGene_gn=COL1A2;refGene_tx=XM_003982764.5
A2 99451950 99451995 refGene_gn=COL1A2;refGene_tx=XM_003982764.5
A2 99452082 99452136 refGene_gn=COL1A2;refGene_tx=XM_003982764.5
A2 99452225 99452279 refGene_gn=COL1A2;refGene_tx=XM_003982764.5
A2 99452596 99452650 refGene_gn=COL1A2;refGene_tx=XM_003982764.5
A2 99453070 99453124 refGene_gn=COL1A2;refGene_tx=XM_003982764.5
A2 99453618 99453672 refGene_gn=COL1A2;refGene_tx=XM_003982764.5
A2 99455403 99455448 refGene_gn=COL1A2;refGene_tx=XM_003982764.5

A2 99455741 99455795 refGene_gn=COL1A2;refGene_tx=XM_003982764.5
A2 99455897 99455942 refGene_gn=COL1A2;refGene_tx=XM_003982764.5
A2 99456303 99456357 refGene_gn=COL1A2;refGene_tx=XM_003982764.5
A2 99456862 99456961 refGene_gn=COL1A2;refGene_tx=XM_003982764.5
A2 99457083 99457128 refGene_gn=COL1A2;refGene_tx=XM_003982764.5
A2 99457235 99457334 refGene_gn=COL1A2;refGene_tx=XM_003982764.5
A2 99457772 99457826 refGene_gn=COL1A2;refGene_tx=XM_003982764.5
A2 99457989 99458097 refGene_gn=COL1A2;refGene_tx=XM_003982764.5
A2 99458441 99458495 refGene_gn=COL1A2;refGene_tx=XM_003982764.5
A2 99458584 99458683 refGene_gn=COL1A2;refGene_tx=XM_003982764.5
A2 99459586 99459640 refGene_gn=COL1A2;refGene_tx=XM_003982764.5
A2 99460105 99460204 refGene_gn=COL1A2;refGene_tx=XM_003982764.5
A2 99460595 99460649 refGene_gn=COL1A2;refGene_tx=XM_003982764.5
A2 99461113 99461167 refGene_gn=COL1A2;refGene_tx=XM_003982764.5
A2 99461319 99461373 refGene_gn=COL1A2;refGene_tx=XM_003982764.5
A2 99461647 99461701 refGene_gn=COL1A2;refGene_tx=XM_003982764.5
A2 99462678 99462723 refGene_gn=COL1A2;refGene_tx=XM_003982764.5
A2 99463917 99464016 refGene_gn=COL1A2;refGene_tx=XM_003982764.5
A2 99465410 99465518 refGene_gn=COL1A2;refGene_tx=XM_003982764.5
A2 99466169 99466223 refGene_gn=COL1A2;refGene_tx=XM_003982764.5
A2 99467186 99467240 refGene_gn=COL1A2;refGene_tx=XM_003982764.5
A2 99467936 99467990 refGene_gn=COL1A2;refGene_tx=XM_003982764.5
A2 99468087 99468141 refGene_gn=COL1A2;refGene_tx=XM_003982764.5
A2 99468237 99468345 refGene_gn=COL1A2;refGene_tx=XM_003982764.5
A2 99468704 99468758 refGene_gn=COL1A2;refGene_tx=XM_003982764.5
A2 99469878 99469932 refGene_gn=COL1A2;refGene_tx=XM_003982764.5
A2 99470907 99471069 refGene_gn=COL1A2;refGene_tx=XM_003982764.5
A2 99471776 99471884 refGene_gn=COL1A2;refGene_tx=XM_003982764.5
A2 99472557 99472665 refGene_gn=COL1A2;refGene_tx=XM_003982764.5
A2 99473100 99473154 refGene_gn=COL1A2;refGene_tx=XM_003982764.5
A2 99473241 99473349 refGene_gn=COL1A2;refGene_tx=XM_003982764.5
A2 99473493 99473547 refGene_gn=COL1A2;refGene_tx=XM_003982764.5
A2 99473928 99474036 refGene_gn=COL1A2;refGene_tx=XM_003982764.5
A2 99474494 99474548 refGene_gn=COL1A2;refGene_tx=XM_003982764.5
A2 99474670 99474778 refGene_gn=COL1A2;refGene_tx=XM_003982764.5
A2 99475113 99475372 refGene_gn=COL1A2;refGene_tx=XM_003982764.5
A2 99475753 99475938 refGene_gn=COL1A2;refGene_tx=XM_003982764.5
A2 99476632 99476875 refGene_gn=COL1A2;refGene_tx=XM_003982764.5
A2 99477842 99477989 refGene_gn=COL1A2;refGene_tx=XM_003982764.5
A2 99536617 99536750
refGene_gn=CASD1;refGene_tx=XM_023250351.1;refGene_gn=CASD1;refGene_tx=XM_023250352.1;refGene_gn=CASD1;refGene_tx=XM_023250354.1;refGene_gn=CASD1;refGene_tx=XM_023250355.1;refGene_gn=CASD1;refGene_tx=XM_023250356.1;refGene_gn=CASD1;refGene_tx=XM_023250357.1;refGene_gn=CASD1;refGene_tx=XM_023250358.1
A2 99545078 99545175
refGene_gn=CASD1;refGene_tx=XM_023250351.1;refGene_gn=CASD1;refGene_tx=XM_023250352.1;refGene_gn=CASD1;refGene_tx=XM_023250354.1;refGene_gn=CASD1;refGene_tx=XM_023250355.1;refGene_gn=CASD1;refGene_tx=XM_023250356.1;refGene_gn=CASD1;refGene_tx=XM_023250357.1;refGene_gn=CASD1;refGe

ne_tx=XM_023250358.1
A2 99545972 99546093
refGene_gn=CASD1;refGene_tx=XM_023250351.1;refGene_gn=CASD1;refGene_tx=XM_023250352.1;refGene_gn=CASD1;refGene_tx=XM_023250354.1;refGene_gn=CASD1;refGene_tx=XM_023250355.1;refGene_gn=CASD1;refGene_tx=XM_023250356.1;refGene_gn=CASD1;refGene_tx=XM_023250357.1;refGene_gn=CASD1;refGene_tx=XM_023250358.1
A2 99553983 99554028
refGene_gn=CASD1;refGene_tx=XM_023250351.1;refGene_gn=CASD1;refGene_tx=XM_023250352.1;refGene_gn=CASD1;refGene_tx=XM_023250354.1;refGene_gn=CASD1;refGene_tx=XM_023250355.1;refGene_gn=CASD1;refGene_tx=XM_023250356.1;refGene_gn=CASD1;refGene_tx=XM_023250357.1;refGene_gn=CASD1;refGene_tx=XM_023250358.1
A2 99555505 99555568
refGene_gn=CASD1;refGene_tx=XM_023250351.1;refGene_gn=CASD1;refGene_tx=XM_023250352.1;refGene_gn=CASD1;refGene_tx=XM_023250354.1;refGene_gn=CASD1;refGene_tx=XM_023250355.1;refGene_gn=CASD1;refGene_tx=XM_023250356.1;refGene_gn=CASD1;refGene_tx=XM_023250357.1;refGene_gn=CASD1;refGene_tx=XM_023250358.1
A2 99559877 99559922
refGene_gn=CASD1;refGene_tx=XM_023250351.1;refGene_gn=CASD1;refGene_tx=XM_023250352.1;refGene_gn=CASD1;refGene_tx=XM_023250354.1;refGene_gn=CASD1;refGene_tx=XM_023250355.1;refGene_gn=CASD1;refGene_tx=XM_023250356.1;refGene_gn=CASD1;refGene_tx=XM_023250357.1;refGene_gn=CASD1;refGene_tx=XM_023250358.1
A2 99560353 99560477
refGene_gn=CASD1;refGene_tx=XM_023250351.1;refGene_gn=CASD1;refGene_tx=XM_023250352.1;refGene_gn=CASD1;refGene_tx=XM_023250354.1;refGene_gn=CASD1;refGene_tx=XM_023250355.1;refGene_gn=CASD1;refGene_tx=XM_023250356.1;refGene_gn=CASD1;refGene_tx=XM_023250357.1;refGene_gn=CASD1;refGene_tx=XM_023250358.1
A2 99561847 99562062
refGene_gn=CASD1;refGene_tx=XM_023250351.1;refGene_gn=CASD1;refGene_tx=XM_023250352.1;refGene_gn=CASD1;refGene_tx=XM_023250354.1;refGene_gn=CASD1;refGene_tx=XM_023250355.1;refGene_gn=CASD1;refGene_tx=XM_023250356.1;refGene_gn=CASD1;refGene_tx=XM_023250357.1;refGene_gn=CASD1;refGene_tx=XM_023250358.1
A2 99563656 99564079
refGene_gn=CASD1;refGene_tx=XM_023250351.1;refGene_gn=CASD1;refGene_tx=XM_023250352.1;refGene_gn=CASD1;refGene_tx=XM_023250354.1;refGene_gn=CASD1;refGene_tx=XM_023250355.1;refGene_gn=CASD1;refGene_tx=XM_023250356.1;refGene_gn=CASD1;refGene_tx=XM_023250357.1;refGene_gn=CASD1;refGene_tx=XM_023250358.1
A2 99565116 99565206
refGene_gn=CASD1;refGene_tx=XM_023250351.1;refGene_gn=CASD1;refGene_tx=XM_023250352.1;refGene_gn=CASD1;refGene_tx=XM_023250354.1;refGene_gn=CASD1;refGene_tx=XM_023250355.1;refGene_gn=CASD1;refGene_tx=XM_023250356.1;refGene_gn=CASD1;refGene_tx=XM_023250357.1;refGene_gn=CASD1;refGene_tx=XM_023250358.1
A2 99572941 99573061

refGene_gn=CASD1;refGene_tx=XM_023250351.1;refGene_gn=CASD1;refGene_tx=XM_023250352.1;refGene_gn=CASD1;refGene_tx=XM_023250354.1;refGene_gn=CASD1;refGene_tx=XM_023250355.1;refGene_gn=CASD1;refGene_tx=XM_023250356.1;refGene_gn=CASD1;refGene_tx=XM_023250357.1
A2 99574029 99574186
refGene_gn=CASD1;refGene_tx=XM_023250351.1;refGene_gn=CASD1;refGene_tx=XM_023250352.1;refGene_gn=CASD1;refGene_tx=XM_023250354.1;refGene_gn=CASD1;refGene_tx=XM_023250355.1;refGene_gn=CASD1;refGene_tx=XM_023250356.1;refGene_gn=CASD1;refGene_tx=XM_023250357.1
A2 99575604 99575684
refGene_gn=CASD1;refGene_tx=XM_023250351.1;refGene_gn=CASD1;refGene_tx=XM_023250352.1;refGene_gn=CASD1;refGene_tx=XM_023250354.1;refGene_gn=CASD1;refGene_tx=XM_023250355.1;refGene_gn=CASD1;refGene_tx=XM_023250356.1;refGene_gn=CASD1;refGene_tx=XM_023250357.1
A2 99578018 99578120
refGene_gn=CASD1;refGene_tx=XM_023250351.1;refGene_gn=CASD1;refGene_tx=XM_023250352.1;refGene_gn=CASD1;refGene_tx=XM_023250354.1;refGene_gn=CASD1;refGene_tx=XM_023250355.1;refGene_gn=CASD1;refGene_tx=XM_023250356.1;refGene_gn=CASD1;refGene_tx=XM_023250357.1
A2 99580593 99580734
refGene_gn=CASD1;refGene_tx=XM_023250351.1;refGene_gn=CASD1;refGene_tx=XM_023250352.1;refGene_gn=CASD1;refGene_tx=XM_023250356.1;refGene_gn=CASD1;refGene_tx=XM_023250357.1;refGene_gn=CASD1;refGene_tx=XM_023250358.1
A2 99581287 99581365
refGene_gn=CASD1;refGene_tx=XM_023250351.1;refGene_gn=CASD1;refGene_tx=XM_023250352.1;refGene_gn=CASD1;refGene_tx=XM_023250354.1;refGene_gn=CASD1;refGene_tx=XM_023250355.1;refGene_gn=CASD1;refGene_tx=XM_023250356.1;refGene_gn=CASD1;refGene_tx=XM_023250357.1;refGene_gn=CASD1;refGene_tx=XM_023250358.1
A2 99585092 99585185
refGene_gn=CASD1;refGene_tx=XM_023250351.1;refGene_gn=CASD1;refGene_tx=XM_023250354.1;refGene_gn=CASD1;refGene_tx=XM_023250356.1;refGene_gn=CASD1;refGene_tx=XM_023250358.1
A2 99587504 99587771
refGene_gn=CASD1;refGene_tx=XM_023250351.1;refGene_gn=CASD1;refGene_tx=XM_023250352.1;refGene_gn=CASD1;refGene_tx=XM_023250354.1;refGene_gn=CASD1;refGene_tx=XM_023250355.1;refGene_gn=CASD1;refGene_tx=XM_023250358.1
A2 99591484 99591511
refGene_gn=CASD1;refGene_tx=XM_023250356.1;refGene_gn=CASD1;refGene_tx=XM_023250357.1
A2 99615729 99615780
refGene_gn=SGCE;refGene_tx=XM_006929187.3;refGene_gn=SGCE;refGene_tx=XM_019825476.2;refGene_gn=SGCE;refGene_tx=XM_019825477.2;refGene_gn=SGCE;refGene_tx=XM_019825480.2;refGene_gn=SGCE;refGene_tx=XM_019825481.2;refGene_gn=SGCE;refGene_tx=XM_019825482.2
A2 99617726 99617761
refGene_gn=SGCE;refGene_tx=XM_019825476.2;refGene_gn=SGCE;refGene_tx=XM_019825477.2;refGene_gn=SGCE;refGene_tx=XM_019825480.2;refGene_gn=SGC

E;refGene_tx=XM_019825481.2;refGene_gn=SGCE;refGene_tx=XM_019825482.2
A2 99618632 99618687
refGene_gn=SGCE;refGene_tx=XM_006929187.3;refGene_gn=SGCE;refGene_tx=XM_011280390.3;refGene_gn=SGCE;refGene_tx=XM_019825476.2;refGene_gn=SGCE;refGene_tx=XM_019825477.2;refGene_gn=SGCE;refGene_tx=XM_019825478.2;refGene_gn=SGCE;refGene_tx=XM_019825479.2;refGene_gn=SGCE;refGene_tx=XM_019825480.2;refGene_gn=SGCE;refGene_tx=XM_019825481.2;refGene_gn=SGCE;refGene_tx=XM_019825482.2
A2 99628496 99628685
refGene_gn=SGCE;refGene_tx=XM_006929187.3;refGene_gn=SGCE;refGene_tx=XM_011280390.3;refGene_gn=SGCE;refGene_tx=XM_019825476.2;refGene_gn=SGCE;refGene_tx=XM_019825477.2;refGene_gn=SGCE;refGene_tx=XM_019825478.2;refGene_gn=SGCE;refGene_tx=XM_019825479.2;refGene_gn=SGCE;refGene_tx=XM_019825480.2;refGene_gn=SGCE;refGene_tx=XM_019825481.2;refGene_gn=SGCE;refGene_tx=XM_019825482.2
A2 99629118 99629122 refGene_gn=SGCE;refGene_tx=XM_019825483.2
A2 99629421 99629448
refGene_gn=SGCE;refGene_tx=XM_006929187.3;refGene_gn=SGCE;refGene_tx=XM_011280390.3;refGene_gn=SGCE;refGene_tx=XM_019825476.2;refGene_gn=SGCE;refGene_tx=XM_019825478.2;refGene_gn=SGCE;refGene_tx=XM_019825480.2;refGene_gn=SGCE;refGene_tx=XM_019825482.2;refGene_gn=SGCE;refGene_tx=XM_019825483.2
A2 99630381 99630593
refGene_gn=SGCE;refGene_tx=XM_006929187.3;refGene_gn=SGCE;refGene_tx=XM_011280390.3;refGene_gn=SGCE;refGene_tx=XM_019825476.2;refGene_gn=SGCE;refGene_tx=XM_019825477.2;refGene_gn=SGCE;refGene_tx=XM_019825478.2;refGene_gn=SGCE;refGene_tx=XM_019825479.2;refGene_gn=SGCE;refGene_tx=XM_019825480.2;refGene_gn=SGCE;refGene_tx=XM_019825481.2;refGene_gn=SGCE;refGene_tx=XM_019825482.2;refGene_gn=SGCE;refGene_tx=XM_019825483.2
A2 99633381 99633544
refGene_gn=SGCE;refGene_tx=XM_006929187.3;refGene_gn=SGCE;refGene_tx=XM_011280390.3;refGene_gn=SGCE;refGene_tx=XM_019825476.2;refGene_gn=SGCE;refGene_tx=XM_019825477.2;refGene_gn=SGCE;refGene_tx=XM_019825478.2;refGene_gn=SGCE;refGene_tx=XM_019825479.2;refGene_gn=SGCE;refGene_tx=XM_019825480.2;refGene_gn=SGCE;refGene_tx=XM_019825481.2;refGene_gn=SGCE;refGene_tx=XM_019825482.2;refGene_gn=SGCE;refGene_tx=XM_019825483.2
A2 99643961 99644160
refGene_gn=SGCE;refGene_tx=XM_006929187.3;refGene_gn=SGCE;refGene_tx=XM_011280390.3;refGene_gn=SGCE;refGene_tx=XM_019825476.2;refGene_gn=SGCE;refGene_tx=XM_019825477.2;refGene_gn=SGCE;refGene_tx=XM_019825478.2;refGene_gn=SGCE;refGene_tx=XM_019825479.2;refGene_gn=SGCE;refGene_tx=XM_019825480.2;refGene_gn=SGCE;refGene_tx=XM_019825481.2;refGene_gn=SGCE;refGene_tx=XM_019825482.2;refGene_gn=SGCE;refGene_tx=XM_019825483.2
A2 99649396 99649469
refGene_gn=SGCE;refGene_tx=XM_006929187.3;refGene_gn=SGCE;refGene_tx=XM_011280390.3;refGene_gn=SGCE;refGene_tx=XM_019825476.2;refGene_gn=SGCE;refGene_tx=XM_019825477.2;refGene_gn=SGCE;refGene_tx=XM_019825478.2;refGene_gn=SGCE;refGene_tx=XM_019825479.2;refGene_gn=SGCE;refGene_tx=XM_019825480.2;refGene_gn=SGCE;refGene_tx=XM_019825481.2;refGene_gn=SGCE;refGene_tx=XM_019825482.2;refGene_gn=SGCE;refGene_tx=XM_019825483.2

A2	99655425	99655583
refGene_gn=SGCE;refGene_tx=XM_006929187.3;refGene_gn=SGCE;refGene_tx=XM_011280390.3;refGene_gn=SGCE;refGene_tx=XM_019825476.2;refGene_gn=SGCE;refGene_tx=XM_019825477.2;refGene_gn=SGCE;refGene_tx=XM_019825478.2;refGene_gn=SGCE;refGene_tx=XM_019825479.2;refGene_gn=SGCE;refGene_tx=XM_019825480.2;refGene_gn=SGCE;refGene_tx=XM_019825481.2;refGene_gn=SGCE;refGene_tx=XM_019825482.2;refGene_gn=SGCE;refGene_tx=XM_019825483.2
A2	99657559	99657682
refGene_gn=SGCE;refGene_tx=XM_006929187.3;refGene_gn=SGCE;refGene_tx=XM_019825476.2;refGene_gn=SGCE;refGene_tx=XM_019825477.2;refGene_gn=SGCE;refGene_tx=XM_019825478.2;refGene_gn=SGCE;refGene_tx=XM_019825479.2;refGene_gn=SGCE;refGene_tx=XM_019825480.2;refGene_gn=SGCE;refGene_tx=XM_019825481.2;refGene_gn=SGCE;refGene_tx=XM_019825483.2
A2	99668089	99668197
refGene_gn=SGCE;refGene_tx=XM_006929187.3;refGene_gn=SGCE;refGene_tx=XM_019825476.2;refGene_gn=SGCE;refGene_tx=XM_019825477.2;refGene_gn=SGCE;refGene_tx=XM_019825478.2;refGene_gn=SGCE;refGene_tx=XM_019825479.2;refGene_gn=SGCE;refGene_tx=XM_019825483.2
A2	99685000	99685109
refGene_gn=SGCE;refGene_tx=XM_006929187.3;refGene_gn=SGCE;refGene_tx=XM_011280390.3;refGene_gn=SGCE;refGene_tx=XM_019825476.2;refGene_gn=SGCE;refGene_tx=XM_019825477.2;refGene_gn=SGCE;refGene_tx=XM_019825478.2;refGene_gn=SGCE;refGene_tx=XM_019825479.2;refGene_gn=SGCE;refGene_tx=XM_019825480.2;refGene_gn=SGCE;refGene_tx=XM_019825481.2;refGene_gn=SGCE;refGene_tx=XM_019825482.2;refGene_gn=SGCE;refGene_tx=XM_019825483.2
A2	99685611	99685616
refGene_gn=PEG10;refGene_tx=NM_001291089.1;refGene_gn=PEG10;refGene_tx=NM_001291090.1
A2	99692107	99694486
refGene_gn=PEG10;refGene_tx=NM_001291089.1;refGene_gn=PEG10;refGene_tx=NM_001291090.1
A2	99837601	99838993
refGene_gn=PPP1R9A;refGene_tx=XM_011280391.3;refGene_gn=PPP1R9A;refGene_tx=XM_011280392.3;refGene_gn=PPP1R9A;refGene_tx=XM_011280393.3;refGene_gn=PPP1R9A;refGene_tx=XM_019825492.2;refGene_gn=PPP1R9A;refGene_tx=XM_019825494.2;refGene_gn=PPP1R9A;refGene_tx=XM_019825496.2;refGene_gn=PPP1R9A;refGene_tx=XM_019825497.2;refGene_gn=PPP1R9A;refGene_tx=XM_019825499.2;refGene_gn=PPP1R9A;refGene_tx=XM_019825500.2;refGene_gn=PPP1R9A;refGene_tx=XM_019825501.2;refGene_gn=PPP1R9A;refGene_tx=XM_023250361.1;refGene_gn=PPP1R9A;refGene_tx=XM_023250362.1;refGene_gn=PPP1R9A;refGene_tx=XM_023250363.1;refGene_gn=PPP1R9A;refGene_tx=XM_023250364.1;refGene_gn=PPP1R9A;refGene_tx=XM_023250365.1
A2	100007690	100007823
refGene_gn=PPP1R9A;refGene_tx=XM_011280391.3;refGene_gn=PPP1R9A;refGene_tx=XM_011280392.3;refGene_gn=PPP1R9A;refGene_tx=XM_011280393.3;refGene_gn=PPP1R9A;refGene_tx=XM_019825492.2;refGene_gn=PPP1R9A;refGene_tx=XM_019825494.2;refGene_gn=PPP1R9A;refGene_tx=XM_019825496.2;refGene_gn=PPP1R9A;refGene_tx=XM_019825497.2;refGene_gn=PPP1R9A;refGene_tx=XM_019825499.2;refGene_gn=PPP1R9A;refGene_tx=XM_019825500.2;refGene_gn=PPP1R9A;refGene_tx=XM_019825501.2;refGene_gn=PPP1R9A;refGene_tx=XM_0232503

61.1;refGene_gn=PPP1R9A;refGene_tx=XM_023250362.1;refGene_gn=PPP1R9A;refGene_tx=XM_023250363.1;refGene_gn=PPP1R9A;refGene_tx=XM_023250364.1;refGene_gn=PPP1R9A;refGene_tx=XM_023250365.1
A2 100017154 100017275
refGene_gn=PPP1R9A;refGene_tx=XM_011280391.3;refGene_gn=PPP1R9A;refGene_tx=XM_011280392.3;refGene_gn=PPP1R9A;refGene_tx=XM_011280393.3;refGene_gn=PPP1R9A;refGene_tx=XM_019825492.2;refGene_gn=PPP1R9A;refGene_tx=XM_019825494.2;refGene_gn=PPP1R9A;refGene_tx=XM_019825496.2;refGene_gn=PPP1R9A;refGene_tx=XM_019825497.2;refGene_gn=PPP1R9A;refGene_tx=XM_019825499.2;refGene_gn=PPP1R9A;refGene_tx=XM_019825500.2;refGene_gn=PPP1R9A;refGene_tx=XM_019825501.2;refGene_gn=PPP1R9A;refGene_tx=XM_023250361.1;refGene_gn=PPP1R9A;refGene_tx=XM_023250362.1;refGene_gn=PPP1R9A;refGene_tx=XM_023250363.1;refGene_gn=PPP1R9A;refGene_tx=XM_023250364.1;refGene_gn=PPP1R9A;refGene_tx=XM_023250365.1
A2 100040024 100040129
refGene_gn=PPP1R9A;refGene_tx=XM_011280391.3;refGene_gn=PPP1R9A;refGene_tx=XM_011280392.3;refGene_gn=PPP1R9A;refGene_tx=XM_011280393.3;refGene_gn=PPP1R9A;refGene_tx=XM_019825492.2;refGene_gn=PPP1R9A;refGene_tx=XM_019825494.2;refGene_gn=PPP1R9A;refGene_tx=XM_019825496.2;refGene_gn=PPP1R9A;refGene_tx=XM_019825497.2;refGene_gn=PPP1R9A;refGene_tx=XM_019825499.2;refGene_gn=PPP1R9A;refGene_tx=XM_019825500.2;refGene_gn=PPP1R9A;refGene_tx=XM_019825501.2;refGene_gn=PPP1R9A;refGene_tx=XM_023250361.1;refGene_gn=PPP1R9A;refGene_tx=XM_023250362.1;refGene_gn=PPP1R9A;refGene_tx=XM_023250363.1;refGene_gn=PPP1R9A;refGene_tx=XM_023250364.1;refGene_gn=PPP1R9A;refGene_tx=XM_023250365.1
A2 100065424 100065560
refGene_gn=PPP1R9A;refGene_tx=XM_011280391.3;refGene_gn=PPP1R9A;refGene_tx=XM_011280392.3;refGene_gn=PPP1R9A;refGene_tx=XM_011280393.3;refGene_gn=PPP1R9A;refGene_tx=XM_019825492.2;refGene_gn=PPP1R9A;refGene_tx=XM_019825494.2;refGene_gn=PPP1R9A;refGene_tx=XM_019825496.2;refGene_gn=PPP1R9A;refGene_tx=XM_019825497.2;refGene_gn=PPP1R9A;refGene_tx=XM_019825499.2;refGene_gn=PPP1R9A;refGene_tx=XM_019825500.2;refGene_gn=PPP1R9A;refGene_tx=XM_019825501.2;refGene_gn=PPP1R9A;refGene_tx=XM_023250361.1;refGene_gn=PPP1R9A;refGene_tx=XM_023250362.1;refGene_gn=PPP1R9A;refGene_tx=XM_023250363.1;refGene_gn=PPP1R9A;refGene_tx=XM_023250364.1;refGene_gn=PPP1R9A;refGene_tx=XM_023250365.1
A2 100069427 100069490
refGene_gn=PPP1R9A;refGene_tx=XM_019825492.2;refGene_gn=PPP1R9A;refGene_tx=XM_019825494.2;refGene_gn=PPP1R9A;refGene_tx=XM_019825497.2;refGene_gn=PPP1R9A;refGene_tx=XM_019825499.2;refGene_gn=PPP1R9A;refGene_tx=XM_023250361.1;refGene_gn=PPP1R9A;refGene_tx=XM_023250362.1;refGene_gn=PPP1R9A;refGene_tx=XM_023250363.1;refGene_gn=PPP1R9A;refGene_tx=XM_023250364.1;refGene_gn=PPP1R9A;refGene_tx=XM_023250365.1
A2 100070513 100070579
refGene_gn=PPP1R9A;refGene_tx=XM_019825492.2;refGene_gn=PPP1R9A;refGene_tx=XM_019825494.2;refGene_gn=PPP1R9A;refGene_tx=XM_019825496.2;refGene_gn=PPP1R9A;refGene_tx=XM_019825497.2;refGene_gn=PPP1R9A;refGene_tx=XM_019825499.2;refGene_gn=PPP1R9A;refGene_tx=XM_019825500.2;refGene_gn=PPP1R9A;refGene_tx=XM_023250361.1;refGene_gn=PPP1R9A;refGene_tx=XM_023250362.1;refGene_gn=PPP1R9A;refGene_tx=XM_023250364.1;refGene_gn=PPP1

R9A;refGene_tx=XM_023250365.1
A2 100091296 100091452
refGene_gn=PPP1R9A;refGene_tx=XM_011280391.3;refGene_gn=PPP1R9A;refGene_tx=XM_011280392.3;refGene_gn=PPP1R9A;refGene_tx=XM_011280393.3;refGene_gn=PPP1R9A;refGene_tx=XM_019825492.2;refGene_gn=PPP1R9A;refGene_tx=XM_019825494.2;refGene_gn=PPP1R9A;refGene_tx=XM_019825496.2;refGene_gn=PPP1R9A;refGene_tx=XM_019825497.2;refGene_gn=PPP1R9A;refGene_tx=XM_019825499.2;refGene_gn=PPP1R9A;refGene_tx=XM_019825500.2;refGene_gn=PPP1R9A;refGene_tx=XM_019825501.2;refGene_gn=PPP1R9A;refGene_tx=XM_023250361.1;refGene_gn=PPP1R9A;refGene_tx=XM_023250362.1;refGene_gn=PPP1R9A;refGene_tx=XM_023250363.1;refGene_gn=PPP1R9A;refGene_tx=XM_023250364.1;refGene_gn=PPP1R9A;refGene_tx=XM_023250365.1
A2 100106972 100107026
refGene_gn=PPP1R9A;refGene_tx=XM_011280391.3;refGene_gn=PPP1R9A;refGene_tx=XM_011280392.3;refGene_gn=PPP1R9A;refGene_tx=XM_011280393.3;refGene_gn=PPP1R9A;refGene_tx=XM_019825492.2;refGene_gn=PPP1R9A;refGene_tx=XM_019825494.2;refGene_gn=PPP1R9A;refGene_tx=XM_019825496.2;refGene_gn=PPP1R9A;refGene_tx=XM_019825497.2;refGene_gn=PPP1R9A;refGene_tx=XM_019825499.2;refGene_gn=PPP1R9A;refGene_tx=XM_019825500.2;refGene_gn=PPP1R9A;refGene_tx=XM_019825501.2;refGene_gn=PPP1R9A;refGene_tx=XM_023250361.1;refGene_gn=PPP1R9A;refGene_tx=XM_023250362.1;refGene_gn=PPP1R9A;refGene_tx=XM_023250363.1;refGene_gn=PPP1R9A;refGene_tx=XM_023250364.1;refGene_gn=PPP1R9A;refGene_tx=XM_023250365.1
A2 100109619 100109849
refGene_gn=PPP1R9A;refGene_tx=XM_011280391.3;refGene_gn=PPP1R9A;refGene_tx=XM_011280392.3;refGene_gn=PPP1R9A;refGene_tx=XM_011280393.3;refGene_gn=PPP1R9A;refGene_tx=XM_019825492.2;refGene_gn=PPP1R9A;refGene_tx=XM_019825494.2;refGene_gn=PPP1R9A;refGene_tx=XM_019825496.2;refGene_gn=PPP1R9A;refGene_tx=XM_019825497.2;refGene_gn=PPP1R9A;refGene_tx=XM_019825499.2;refGene_gn=PPP1R9A;refGene_tx=XM_019825500.2;refGene_gn=PPP1R9A;refGene_tx=XM_019825501.2;refGene_gn=PPP1R9A;refGene_tx=XM_023250361.1;refGene_gn=PPP1R9A;refGene_tx=XM_023250362.1;refGene_gn=PPP1R9A;refGene_tx=XM_023250363.1;refGene_gn=PPP1R9A;refGene_tx=XM_023250364.1;refGene_gn=PPP1R9A;refGene_tx=XM_023250365.1
A2 100111353 100111450
refGene_gn=PPP1R9A;refGene_tx=XM_011280391.3;refGene_gn=PPP1R9A;refGene_tx=XM_011280392.3;refGene_gn=PPP1R9A;refGene_tx=XM_011280393.3;refGene_gn=PPP1R9A;refGene_tx=XM_019825492.2;refGene_gn=PPP1R9A;refGene_tx=XM_019825494.2;refGene_gn=PPP1R9A;refGene_tx=XM_019825496.2;refGene_gn=PPP1R9A;refGene_tx=XM_019825497.2;refGene_gn=PPP1R9A;refGene_tx=XM_019825499.2;refGene_gn=PPP1R9A;refGene_tx=XM_019825500.2;refGene_gn=PPP1R9A;refGene_tx=XM_019825501.2;refGene_gn=PPP1R9A;refGene_tx=XM_023250361.1;refGene_gn=PPP1R9A;refGene_tx=XM_023250362.1;refGene_gn=PPP1R9A;refGene_tx=XM_023250363.1;refGene_gn=PPP1R9A;refGene_tx=XM_023250364.1;refGene_gn=PPP1R9A;refGene_tx=XM_023250365.1
A2 100111553 100111725
refGene_gn=PPP1R9A;refGene_tx=XM_011280391.3;refGene_gn=PPP1R9A;refGene_tx=XM_011280392.3;refGene_gn=PPP1R9A;refGene_tx=XM_011280393.3;refGene_gn=PPP1R9A;refGene_tx=XM_019825492.2;refGene_gn=PPP1R9A;refGene_tx=XM_019825494.2;refGene_gn=PPP1R9A;refGene_tx=XM_019825496.2;refGene_gn

=PPP1R9A;refGene_tx=XM_019825497.2;refGene_gn=PPP1R9A;refGene_tx=XM_01
9825499.2;refGene_gn=PPP1R9A;refGene_tx=XM_019825500.2;refGene_gn=PPP1
R9A;refGene_tx=XM_019825501.2;refGene_gn=PPP1R9A;refGene_tx=XM_0232503
61.1;refGene_gn=PPP1R9A;refGene_tx=XM_023250362.1;refGene_gn=PPP1R9A;r
efGene_tx=XM_023250363.1;refGene_gn=PPP1R9A;refGene_tx=XM_023250364.1;
refGene_gn=PPP1R9A;refGene_tx=XM_023250365.1
A2 100129824 100129982
refGene_gn=PPP1R9A;refGene_tx=XM_011280391.3;refGene_gn=PPP1R9A;refGen
e_tx=XM_011280392.3;refGene_gn=PPP1R9A;refGene_tx=XM_011280393.3;refGe
ne_gn=PPP1R9A;refGene_tx=XM_019825492.2;refGene_gn=PPP1R9A;refGene_tx=
XM_019825494.2;refGene_gn=PPP1R9A;refGene_tx=XM_019825496.2;refGene_gn
=PPP1R9A;refGene_tx=XM_019825497.2;refGene_gn=PPP1R9A;refGene_tx=XM_01
9825499.2;refGene_gn=PPP1R9A;refGene_tx=XM_019825500.2;refGene_gn=PPP1
R9A;refGene_tx=XM_019825501.2;refGene_gn=PPP1R9A;refGene_tx=XM_0232503
61.1;refGene_gn=PPP1R9A;refGene_tx=XM_023250362.1;refGene_gn=PPP1R9A;r
efGene_tx=XM_023250363.1;refGene_gn=PPP1R9A;refGene_tx=XM_023250364.1;
refGene_gn=PPP1R9A;refGene_tx=XM_023250365.1
A2 100130483 100130781
refGene_gn=PPP1R9A;refGene_tx=XM_011280391.3;refGene_gn=PPP1R9A;refGen
e_tx=XM_019825492.2;refGene_gn=PPP1R9A;refGene_tx=XM_019825494.2;refGe
ne_gn=PPP1R9A;refGene_tx=XM_019825496.2;refGene_gn=PPP1R9A;refGene_tx=
XM_019825499.2;refGene_gn=PPP1R9A;refGene_tx=XM_019825500.2;refGene_gn
=PPP1R9A;refGene_tx=XM_019825501.2;refGene_gn=PPP1R9A;refGene_tx=XM_02
3250361.1;refGene_gn=PPP1R9A;refGene_tx=XM_023250362.1;refGene_gn=PPP1
R9A;refGene_tx=XM_023250363.1;refGene_gn=PPP1R9A;refGene_tx=XM_0232503
64.1;refGene_gn=PPP1R9A;refGene_tx=XM_023250365.1
A2 100143146 100143234
refGene_gn=PPP1R9A;refGene_tx=XM_011280391.3;refGene_gn=PPP1R9A;refGen
e_tx=XM_019825492.2;refGene_gn=PPP1R9A;refGene_tx=XM_019825494.2;refGe
ne_gn=PPP1R9A;refGene_tx=XM_019825496.2;refGene_gn=PPP1R9A;refGene_tx=
XM_019825500.2;refGene_gn=PPP1R9A;refGene_tx=XM_019825501.2;refGene_gn
=PPP1R9A;refGene_tx=XM_023250361.1;refGene_gn=PPP1R9A;refGene_tx=XM_02
3250362.1;refGene_gn=PPP1R9A;refGene_tx=XM_023250363.1;refGene_gn=PPP1
R9A;refGene_tx=XM_023250364.1;refGene_gn=PPP1R9A;refGene_tx=XM_0232503
65.1
A2 100143333 100143411
refGene_gn=PPP1R9A;refGene_tx=XM_011280391.3;refGene_gn=PPP1R9A;refGen
e_tx=XM_019825492.2;refGene_gn=PPP1R9A;refGene_tx=XM_019825494.2;refGe
ne_gn=PPP1R9A;refGene_tx=XM_019825496.2;refGene_gn=PPP1R9A;refGene_tx=
XM_019825500.2;refGene_gn=PPP1R9A;refGene_tx=XM_019825501.2;refGene_gn
=PPP1R9A;refGene_tx=XM_023250361.1;refGene_gn=PPP1R9A;refGene_tx=XM_02
3250362.1;refGene_gn=PPP1R9A;refGene_tx=XM_023250363.1;refGene_gn=PPP1
R9A;refGene_tx=XM_023250364.1;refGene_gn=PPP1R9A;refGene_tx=XM_0232503
65.1
A2 100145532 100145701
refGene_gn=PPP1R9A;refGene_tx=XM_023250365.1
A2 100155478 100155677
refGene_gn=PPP1R9A;refGene_tx=XM_011280391.3;refGene_gn=PPP1R9A;refGen
e_tx=XM_019825492.2;refGene_gn=PPP1R9A;refGene_tx=XM_019825494.2;refGe
ne_gn=PPP1R9A;refGene_tx=XM_019825496.2;refGene_gn=PPP1R9A;refGene_tx=

XM_019825499.2;refGene_gn=PPP1R9A;refGene_tx=XM_019825500.2;refGene_gn
=PPP1R9A;refGene_tx=XM_019825501.2;refGene_gn=PPP1R9A;refGene_tx=XM_02
3250361.1;refGene_gn=PPP1R9A;refGene_tx=XM_023250362.1;refGene_gn=PPP1
R9A;refGene_tx=XM_023250363.1;refGene_gn=PPP1R9A;refGene_tx=XM_0232503
64.1
A2 100157705 100157825
refGene_gn=PPP1R9A;refGene_tx=XM_011280391.3;refGene_gn=PPP1R9A;refGen
e_tx=XM_011280392.3;refGene_gn=PPP1R9A;refGene_tx=XM_011280393.3;refGe
ne_gn=PPP1R9A;refGene_tx=XM_019825492.2;refGene_gn=PPP1R9A;refGene_tx=
XM_019825494.2;refGene_gn=PPP1R9A;refGene_tx=XM_019825496.2;refGene_gn
=PPP1R9A;refGene_tx=XM_019825497.2;refGene_gn=PPP1R9A;refGene_tx=XM_01
9825500.2;refGene_gn=PPP1R9A;refGene_tx=XM_019825501.2;refGene_gn=PPP1
R9A;refGene_tx=XM_023250361.1;refGene_gn=PPP1R9A;refGene_tx=XM_0232503
62.1;refGene_gn=PPP1R9A;refGene_tx=XM_023250363.1;refGene_gn=PPP1R9A;r
efGene_tx=XM_023250364.1
A2 100158671 100158695
refGene_gn=PPP1R9A;refGene_tx=XM_011280391.3;refGene_gn=PPP1R9A;refGen
e_tx=XM_011280392.3;refGene_gn=PPP1R9A;refGene_tx=XM_019825492.2;refGe
ne_gn=PPP1R9A;refGene_tx=XM_019825496.2;refGene_gn=PPP1R9A;refGene_tx=
XM_019825497.2;refGene_gn=PPP1R9A;refGene_tx=XM_023250361.1;refGene_gn
=PPP1R9A;refGene_tx=XM_023250362.1;refGene_gn=PPP1R9A;refGene_tx=XM_02
3250363.1;refGene_gn=PPP1R9A;refGene_tx=XM_023250364.1
A2 100159802 100159985
refGene_gn=PPP1R9A;refGene_tx=XM_011280391.3;refGene_gn=PPP1R9A;refGen
e_tx=XM_011280392.3;refGene_gn=PPP1R9A;refGene_tx=XM_011280393.3;refGe
ne_gn=PPP1R9A;refGene_tx=XM_019825492.2;refGene_gn=PPP1R9A;refGene_tx=
XM_019825494.2;refGene_gn=PPP1R9A;refGene_tx=XM_019825496.2;refGene_gn
=PPP1R9A;refGene_tx=XM_019825497.2;refGene_gn=PPP1R9A;refGene_tx=XM_01
9825500.2;refGene_gn=PPP1R9A;refGene_tx=XM_019825501.2;refGene_gn=PPP1
R9A;refGene_tx=XM_023250361.1;refGene_gn=PPP1R9A;refGene_tx=XM_0232503
62.1;refGene_gn=PPP1R9A;refGene_tx=XM_023250363.1
A2 100161343 100161553
refGene_gn=PPP1R9A;refGene_tx=XM_011280391.3;refGene_gn=PPP1R9A;refGen
e_tx=XM_011280392.3;refGene_gn=PPP1R9A;refGene_tx=XM_011280393.3;refGe
ne_gn=PPP1R9A;refGene_tx=XM_019825492.2;refGene_gn=PPP1R9A;refGene_tx=
XM_019825494.2;refGene_gn=PPP1R9A;refGene_tx=XM_019825496.2;refGene_gn
=PPP1R9A;refGene_tx=XM_019825497.2;refGene_gn=PPP1R9A;refGene_tx=XM_01
9825500.2;refGene_gn=PPP1R9A;refGene_tx=XM_019825501.2;refGene_gn=PPP1
R9A;refGene_tx=XM_023250361.1;refGene_gn=PPP1R9A;refGene_tx=XM_0232503
62.1;refGene_gn=PPP1R9A;refGene_tx=XM_023250363.1;refGene_gn=PPP1R9A;r
efGene_tx=XM_023250364.1
A2 100169930 100170089
refGene_gn=PON1;refGene_tx=XM_003982775.3
A2 100172572 100172701
refGene_gn=PON1;refGene_tx=XM_003982775.3
A2 100183483 100183565
refGene_gn=PON1;refGene_tx=XM_003982775.3
A2 100184747 100184948
refGene_gn=PON1;refGene_tx=XM_003982775.3
A2 100188597 100188724

refGene_gn=PON1;refGene_tx=XM_003982775.3
A2 100194155 100194324
refGene_gn=PON1;refGene_tx=XM_003982775.3
A2 100195557 100195613
refGene_gn=PON1;refGene_tx=XM_003982775.3
A2 100196998 100197069
refGene_gn=PON1;refGene_tx=XM_003982775.3
A2 100204037 100204111
refGene_gn=PON1;refGene_tx=XM_003982775.3
A2 100224509 100224668
refGene_gn=PON3;refGene_tx=XM_003982776.5
A2 100227558 100227687
refGene_gn=PON3;refGene_tx=XM_003982776.5
A2 100227939 100228021
refGene_gn=PON3;refGene_tx=XM_003982776.5
A2 100229283 100229484
refGene_gn=PON3;refGene_tx=XM_003982776.5
A2 100233443 100233570
refGene_gn=PON3;refGene_tx=XM_003982776.5
A2 100235794 100235960
refGene_gn=PON3;refGene_tx=XM_003982776.5
A2 100241500 100241556
refGene_gn=PON3;refGene_tx=XM_003982776.5
A2 100249008 100249079
refGene_gn=PON3;refGene_tx=XM_003982776.5
A2 100250598 100250672
refGene_gn=PON3;refGene_tx=XM_003982776.5
A2 100259442 100259601
refGene_gn=PON2;refGene_tx=XM_023250366.1
A2 100260238 100260367
refGene_gn=PON2;refGene_tx=XM_023250366.1
A2 100261108 100261190
refGene_gn=PON2;refGene_tx=XM_023250366.1
A2 100263766 100263967
refGene_gn=PON2;refGene_tx=XM_023250366.1
A2 100265444 100265571
refGene_gn=PON2;refGene_tx=XM_023250366.1
A2 100266107 100266273
refGene_gn=PON2;refGene_tx=XM_023250366.1
A2 100271376 100271432
refGene_gn=PON2;refGene_tx=XM_023250366.1
A2 100277628 100277699
refGene_gn=PON2;refGene_tx=XM_023250366.1
A2 100286754 100286828
refGene_gn=PON2;refGene_tx=XM_023250366.1
A2 100339931 100340118
refGene_gn=ASB4;refGene_tx=XM_003982777.5
A2 100349876 100350176
refGene_gn=ASB4;refGene_tx=XM_003982777.5
A2 100383104 100383595

refGene_gn=ASB4;refGene_tx=XM_003982777.5
A2	100393078	100393192
refGene_gn=ASB4;refGene_tx=XM_003982777.5
A2	100394271	100394460
refGene_gn=ASB4;refGene_tx=XM_003982777.5
A2	100434421	100434565
refGene_gn=PDK4;refGene_tx=XM_003982778.5
A2	100435701	100435815
refGene_gn=PDK4;refGene_tx=XM_003982778.5
A2	100436117	100436228
refGene_gn=PDK4;refGene_tx=XM_003982778.5
A2	100436428	100436527
refGene_gn=PDK4;refGene_tx=XM_003982778.5
A2	100438532	100438609
refGene_gn=PDK4;refGene_tx=XM_003982778.5
A2	100440334	100440412
refGene_gn=PDK4;refGene_tx=XM_003982778.5
A2	100440895	100440982
refGene_gn=PDK4;refGene_tx=XM_003982778.5
A2	100441144	100441329
refGene_gn=PDK4;refGene_tx=XM_003982778.5
A2	100442096	100442168
refGene_gn=PDK4;refGene_tx=XM_003982778.5
A2	100443571	100443713
refGene_gn=PDK4;refGene_tx=XM_003982778.5
A2	100444650	100444780
refGene_gn=PDK4;refGene_tx=XM_003982778.5
A2	100614905	100615013
refGene_gn=DYNC1I1;refGene_tx=XM_003982780.4;refGene_gn=DYNC1I1;refGene_tx=XM_006929189.3;refGene_gn=DYNC1I1;refGene_tx=XM_019825503.1;refGene_gn=DYNC1I1;refGene_tx=XM_019825504.1;refGene_gn=DYNC1I1;refGene_tx=XM_023250368.1
A2	100622263	100622378
refGene_gn=DYNC1I1;refGene_tx=XM_003982780.4;refGene_gn=DYNC1I1;refGene_tx=XM_006929189.3;refGene_gn=DYNC1I1;refGene_tx=XM_019825503.1;refGene_gn=DYNC1I1;refGene_tx=XM_019825504.1;refGene_gn=DYNC1I1;refGene_tx=XM_023250368.1
A2	100625075	100625217
refGene_gn=DYNC1I1;refGene_tx=XM_003982780.4;refGene_gn=DYNC1I1;refGene_tx=XM_006929189.3;refGene_gn=DYNC1I1;refGene_tx=XM_019825503.1;refGene_gn=DYNC1I1;refGene_tx=XM_019825504.1;refGene_gn=DYNC1I1;refGene_tx=XM_023250368.1
A2	100639605	100639665
refGene_gn=DYNC1I1;refGene_tx=XM_006929189.3;refGene_gn=DYNC1I1;refGene_tx=XM_019825503.1;refGene_gn=DYNC1I1;refGene_tx=XM_023250368.1
A2	100657938	100658054
refGene_gn=DYNC1I1;refGene_tx=XM_003982780.4;refGene_gn=DYNC1I1;refGene_tx=XM_006929189.3;refGene_gn=DYNC1I1;refGene_tx=XM_019825503.1;refGene_gn=DYNC1I1;refGene_tx=XM_019825504.1;refGene_gn=DYNC1I1;refGene_tx=XM_023250368.1

A2 100770543 100770633
refGene_gn=DYNC1I1;refGene_tx=XM_003982780.4;refGene_gn=DYNC1I1;refGene_tx=XM_006929189.3;refGene_gn=DYNC1I1;refGene_tx=XM_019825503.1;refGene_gn=DYNC1I1;refGene_tx=XM_019825504.1;refGene_gn=DYNC1I1;refGene_tx=XM_023250368.1
A2 100777977 100778140
refGene_gn=DYNC1I1;refGene_tx=XM_003982780.4;refGene_gn=DYNC1I1;refGene_tx=XM_006929189.3;refGene_gn=DYNC1I1;refGene_tx=XM_019825503.1;refGene_gn=DYNC1I1;refGene_tx=XM_019825504.1;refGene_gn=DYNC1I1;refGene_tx=XM_023250368.1
A2 100780172 100780272
refGene_gn=DYNC1I1;refGene_tx=XM_003982780.4;refGene_gn=DYNC1I1;refGene_tx=XM_006929189.3;refGene_gn=DYNC1I1;refGene_tx=XM_019825503.1;refGene_gn=DYNC1I1;refGene_tx=XM_019825504.1;refGene_gn=DYNC1I1;refGene_tx=XM_023250368.1
A2 100788960 100789086
refGene_gn=DYNC1I1;refGene_tx=XM_003982780.4;refGene_gn=DYNC1I1;refGene_tx=XM_006929189.3;refGene_gn=DYNC1I1;refGene_tx=XM_019825503.1;refGene_gn=DYNC1I1;refGene_tx=XM_019825504.1;refGene_gn=DYNC1I1;refGene_tx=XM_023250368.1
A2 100825236 100825383
refGene_gn=DYNC1I1;refGene_tx=XM_003982780.4;refGene_gn=DYNC1I1;refGene_tx=XM_006929189.3;refGene_gn=DYNC1I1;refGene_tx=XM_019825503.1;refGene_gn=DYNC1I1;refGene_tx=XM_019825504.1;refGene_gn=DYNC1I1;refGene_tx=XM_023250368.1
A2 100829558 100829672
refGene_gn=DYNC1I1;refGene_tx=XM_003982780.4;refGene_gn=DYNC1I1;refGene_tx=XM_006929189.3;refGene_gn=DYNC1I1;refGene_tx=XM_019825503.1;refGene_gn=DYNC1I1;refGene_tx=XM_019825504.1;refGene_gn=DYNC1I1;refGene_tx=XM_023250368.1
A2 100832146 100832280
refGene_gn=DYNC1I1;refGene_tx=XM_003982780.4;refGene_gn=DYNC1I1;refGene_tx=XM_006929189.3;refGene_gn=DYNC1I1;refGene_tx=XM_019825503.1;refGene_gn=DYNC1I1;refGene_tx=XM_019825504.1;refGene_gn=DYNC1I1;refGene_tx=XM_023250368.1
A2 100835770 100835915
refGene_gn=DYNC1I1;refGene_tx=XM_003982780.4;refGene_gn=DYNC1I1;refGene_tx=XM_006929189.3;refGene_gn=DYNC1I1;refGene_tx=XM_019825503.1;refGene_gn=DYNC1I1;refGene_tx=XM_019825504.1;refGene_gn=DYNC1I1;refGene_tx=XM_023250368.1
A2 100874602 100874743
refGene_gn=DYNC1I1;refGene_tx=XM_003982780.4;refGene_gn=DYNC1I1;refGene_tx=XM_006929189.3;refGene_gn=DYNC1I1;refGene_tx=XM_019825503.1;refGene_gn=DYNC1I1;refGene_tx=XM_019825504.1;refGene_gn=DYNC1I1;refGene_tx=XM_023250368.1
A2 100878903 100879029
refGene_gn=DYNC1I1;refGene_tx=XM_003982780.4;refGene_gn=DYNC1I1;refGene_tx=XM_006929189.3;refGene_gn=DYNC1I1;refGene_tx=XM_019825503.1;refGene_gn=DYNC1I1;refGene_tx=XM_019825504.1;refGene_gn=DYNC1I1;refGene_tx=XM_023250368.1

A2	100896084	100896195
refGene_gn=DYNC1I1;refGene_tx=XM_003982780.4;refGene_gn=DYNC1I1;refGen
e_tx=XM_006929189.3;refGene_gn=DYNC1I1;refGene_tx=XM_019825503.1;refGe
ne_gn=DYNC1I1;refGene_tx=XM_019825504.1;refGene_gn=DYNC1I1;refGene_tx=
XM_023250368.1
A2	100914019	100914206
refGene_gn=SLC25A13;refGene_tx=XM_003982783.4;refGene_gn=SLC25A13;refG
ene_tx=XM_006929191.2;refGene_gn=SLC25A13;refGene_tx=XM_019825505.2;re
fGene_gn=SLC25A13;refGene_tx=XM_019825506.2;refGene_gn=SLC25A13;refGen
e_tx=XM_019825507.2
A2	100914721	100914812
refGene_gn=SLC25A13;refGene_tx=XM_003982783.4;refGene_gn=SLC25A13;refG
ene_tx=XM_006929191.2;refGene_gn=SLC25A13;refGene_tx=XM_019825505.2;re
fGene_gn=SLC25A13;refGene_tx=XM_019825506.2;refGene_gn=SLC25A13;refGen
e_tx=XM_019825507.2
A2	100914905	100915064
refGene_gn=SLC25A13;refGene_tx=XM_003982783.4;refGene_gn=SLC25A13;refG
ene_tx=XM_006929191.2;refGene_gn=SLC25A13;refGene_tx=XM_019825505.2;re
fGene_gn=SLC25A13;refGene_tx=XM_019825506.2;refGene_gn=SLC25A13;refGen
e_tx=XM_019825507.2
A2	100925790	100925929
refGene_gn=SLC25A13;refGene_tx=XM_003982783.4;refGene_gn=SLC25A13;refG
ene_tx=XM_006929191.2;refGene_gn=SLC25A13;refGene_tx=XM_019825505.2;re
fGene_gn=SLC25A13;refGene_tx=XM_019825506.2;refGene_gn=SLC25A13;refGen
e_tx=XM_019825507.2
A2	100944262	100944403
refGene_gn=SLC25A13;refGene_tx=XM_003982783.4;refGene_gn=SLC25A13;refG
ene_tx=XM_006929191.2;refGene_gn=SLC25A13;refGene_tx=XM_019825505.2;re
fGene_gn=SLC25A13;refGene_tx=XM_019825506.2;refGene_gn=SLC25A13;refGen
e_tx=XM_019825507.2
A2	100965835	100965916
refGene_gn=SLC25A13;refGene_tx=XM_003982783.4;refGene_gn=SLC25A13;refG
ene_tx=XM_006929191.2;refGene_gn=SLC25A13;refGene_tx=XM_019825505.2;re
fGene_gn=SLC25A13;refGene_tx=XM_019825506.2;refGene_gn=SLC25A13;refGen
e_tx=XM_019825507.2
A2	100967707	100967760
refGene_gn=SLC25A13;refGene_tx=XM_003982783.4;refGene_gn=SLC25A13;refG
ene_tx=XM_006929191.2;refGene_gn=SLC25A13;refGene_tx=XM_019825505.2;re
fGene_gn=SLC25A13;refGene_tx=XM_019825506.2;refGene_gn=SLC25A13;refGen
e_tx=XM_019825507.2
A2	100981346	100981505
refGene_gn=SLC25A13;refGene_tx=XM_003982783.4;refGene_gn=SLC25A13;refG
ene_tx=XM_006929191.2;refGene_gn=SLC25A13;refGene_tx=XM_019825505.2;re
fGene_gn=SLC25A13;refGene_tx=XM_019825506.2;refGene_gn=SLC25A13;refGen
e_tx=XM_019825507.2
A2	100982257	100982342
refGene_gn=SLC25A13;refGene_tx=XM_003982783.4;refGene_gn=SLC25A13;refG
ene_tx=XM_006929191.2;refGene_gn=SLC25A13;refGene_tx=XM_019825505.2;re
fGene_gn=SLC25A13;refGene_tx=XM_019825506.2;refGene_gn=SLC25A13;refGen
e_tx=XM_019825507.2

A2	100986127	100986212
refGene_gn=SLC25A13;refGene_tx=XM_003982783.4;refGene_gn=SLC25A13;refGene_tx=XM_006929191.2;refGene_gn=SLC25A13;refGene_tx=XM_019825505.2;refGene_gn=SLC25A13;refGene_tx=XM_019825506.2;refGene_gn=SLC25A13;refGene_tx=XM_019825507.2
A2	100986438	100986532
refGene_gn=SLC25A13;refGene_tx=XM_003982783.4;refGene_gn=SLC25A13;refGene_tx=XM_006929191.2;refGene_gn=SLC25A13;refGene_tx=XM_019825505.2;refGene_gn=SLC25A13;refGene_tx=XM_019825506.2;refGene_gn=SLC25A13;refGene_tx=XM_019825507.2
A2	100987279	100987418
refGene_gn=SLC25A13;refGene_tx=XM_003982783.4;refGene_gn=SLC25A13;refGene_tx=XM_006929191.2;refGene_gn=SLC25A13;refGene_tx=XM_019825505.2;refGene_gn=SLC25A13;refGene_tx=XM_019825506.2;refGene_gn=SLC25A13;refGene_tx=XM_019825507.2
A2	100989502	100989649
refGene_gn=SLC25A13;refGene_tx=XM_003982783.4;refGene_gn=SLC25A13;refGene_tx=XM_006929191.2;refGene_gn=SLC25A13;refGene_tx=XM_019825505.2;refGene_gn=SLC25A13;refGene_tx=XM_019825506.2;refGene_gn=SLC25A13;refGene_tx=XM_019825507.2
A2	101005524	101005664
refGene_gn=SLC25A13;refGene_tx=XM_003982783.4;refGene_gn=SLC25A13;refGene_tx=XM_006929191.2;refGene_gn=SLC25A13;refGene_tx=XM_019825505.2;refGene_gn=SLC25A13;refGene_tx=XM_019825506.2;refGene_gn=SLC25A13;refGene_tx=XM_019825507.2
A2	101027829	101027945
refGene_gn=SLC25A13;refGene_tx=XM_003982783.4;refGene_gn=SLC25A13;refGene_tx=XM_019825505.2;refGene_gn=SLC25A13;refGene_tx=XM_019825506.2;refGene_gn=SLC25A13;refGene_tx=XM_019825507.2
A2	101058096	101058239
refGene_gn=SLC25A13;refGene_tx=XM_003982783.4;refGene_gn=SLC25A13;refGene_tx=XM_006929191.2;refGene_gn=SLC25A13;refGene_tx=XM_019825505.2;refGene_gn=SLC25A13;refGene_tx=XM_019825506.2
A2	101067704	101067758
refGene_gn=SLC25A13;refGene_tx=XM_003982783.4
A2	101103152	101103167
refGene_gn=SLC25A13;refGene_tx=XM_003982783.4
A2	101428684	101428727
refGene_gn=SEM1;refGene_tx=XM_023250369.1
A2	101434854	101434948
refGene_gn=SEM1;refGene_tx=XM_023250369.1
A2	101452728	101452804
refGene_gn=SEM1;refGene_tx=XM_023250369.1
A2	101759040	101759464
refGene_gn=DLX6;refGene_tx=XM_011280395.3
A2	101760759	101760953
refGene_gn=DLX6;refGene_tx=XM_011280395.3
A2	101762805	101763057
refGene_gn=DLX6;refGene_tx=XM_011280395.3
A2	101774078	101774408

refGene_gn=DLX5;refGene_tx=XM_003982789.4
A2 101775509 101775694
refGene_gn=DLX5;refGene_tx=XM_003982789.4
A2 101777636 101777991
refGene_gn=DLX5;refGene_tx=XM_003982789.4
A2 101836944 101837118
refGene_gn=SDHAF3;refGene_tx=XM_003982790.5
A2 101906260 101906464
refGene_gn=SDHAF3;refGene_tx=XM_003982790.5
A2 102359255 102359348
refGene_gn=TAC1;refGene_tx=XM_011280396.3;refGene_gn=TAC1;refGene_tx=XM_011280397.3;refGene_gn=TAC1;refGene_tx=XM_011280398.3;refGene_gn=TAC1;refGene_tx=XM_019825509.2
A2 102359575 102359707
refGene_gn=TAC1;refGene_tx=XM_003982791.5;refGene_gn=TAC1;refGene_tx=XM_011280396.3;refGene_gn=TAC1;refGene_tx=XM_011280397.3;refGene_gn=TAC1;refGene_tx=XM_011280398.3;refGene_gn=TAC1;refGene_tx=XM_019825509.2
A2 102360767 102360864
refGene_gn=TAC1;refGene_tx=XM_003982791.5;refGene_gn=TAC1;refGene_tx=XM_011280396.3;refGene_gn=TAC1;refGene_tx=XM_011280397.3;refGene_gn=TAC1;refGene_tx=XM_011280398.3;refGene_gn=TAC1;refGene_tx=XM_019825509.2
A2 102361324 102361369
refGene_gn=TAC1;refGene_tx=XM_003982791.5;refGene_gn=TAC1;refGene_tx=XM_011280397.3;refGene_gn=TAC1;refGene_tx=XM_019825509.2
A2 102361846 102361870
refGene_gn=TAC1;refGene_tx=XM_003982791.5;refGene_gn=TAC1;refGene_tx=XM_011280396.3;refGene_gn=TAC1;refGene_tx=XM_011280397.3;refGene_gn=TAC1;refGene_tx=XM_011280398.3;refGene_gn=TAC1;refGene_tx=XM_019825509.2
A2 102363410 102363464
refGene_gn=TAC1;refGene_tx=XM_003982791.5;refGene_gn=TAC1;refGene_tx=XM_011280396.3;refGene_gn=TAC1;refGene_tx=XM_019825509.2
A2 102366519 102366569
refGene_gn=TAC1;refGene_tx=XM_003982791.5;refGene_gn=TAC1;refGene_tx=XM_011280396.3;refGene_gn=TAC1;refGene_tx=XM_011280397.3;refGene_gn=TAC1;refGene_tx=XM_011280398.3;refGene_gn=TAC1;refGene_tx=XM_019825509.2
A2 102450011 102450221
refGene_gn=ASNS;refGene_tx=XM_006929200.4;refGene_gn=ASNS;refGene_tx=XM_006929201.4;refGene_gn=ASNS;refGene_tx=XM_023250370.1
A2 102450939 102451095
refGene_gn=ASNS;refGene_tx=XM_006929200.4;refGene_gn=ASNS;refGene_tx=XM_006929201.4;refGene_gn=ASNS;refGene_tx=XM_023250370.1
A2 102451182 102451264
refGene_gn=ASNS;refGene_tx=XM_006929200.4;refGene_gn=ASNS;refGene_tx=XM_006929201.4;refGene_gn=ASNS;refGene_tx=XM_023250370.1
A2 102452540 102452641
refGene_gn=ASNS;refGene_tx=XM_006929200.4;refGene_gn=ASNS;refGene_tx=XM_006929201.4;refGene_gn=ASNS;refGene_tx=XM_023250370.1
A2 102453096 102453203
refGene_gn=ASNS;refGene_tx=XM_006929200.4;refGene_gn=ASNS;refGene_tx=XM_006929201.4;refGene_gn=ASNS;refGene_tx=XM_023250370.1

A2 102455708 102455835
refGene_gn=ASNS;refGene_tx=XM_006929200.4;refGene_gn=ASNS;refGene_tx=XM_006929201.4;refGene_gn=ASNS;refGene_tx=XM_023250370.1
A2 102456891 102457019
refGene_gn=ASNS;refGene_tx=XM_006929200.4;refGene_gn=ASNS;refGene_tx=XM_006929201.4;refGene_gn=ASNS;refGene_tx=XM_023250370.1
A2 102457669 102457771
refGene_gn=ASNS;refGene_tx=XM_006929200.4;refGene_gn=ASNS;refGene_tx=XM_006929201.4;refGene_gn=ASNS;refGene_tx=XM_023250370.1
A2 102458031 102458217
refGene_gn=ASNS;refGene_tx=XM_006929200.4;refGene_gn=ASNS;refGene_tx=XM_006929201.4;refGene_gn=ASNS;refGene_tx=XM_023250370.1
A2 102459126 102459364
refGene_gn=ASNS;refGene_tx=XM_006929200.4;refGene_gn=ASNS;refGene_tx=XM_006929201.4;refGene_gn=ASNS;refGene_tx=XM_023250370.1
A2 102463214 102463463
refGene_gn=ASNS;refGene_tx=XM_006929200.4;refGene_gn=ASNS;refGene_tx=XM_006929201.4;refGene_gn=ASNS;refGene_tx=XM_023250370.1
A2 102613118 102613125
refGene_gn=C1GALT1;refGene_tx=XM_023250371.1
A2 102639471 102639708
refGene_gn=C1GALT1;refGene_tx=XM_003982796.5;refGene_gn=C1GALT1;refGene_tx=XM_023250371.1;refGene_gn=C1GALT1;refGene_tx=XM_023250372.1
A2 102643126 102643794
refGene_gn=C1GALT1;refGene_tx=XM_003982796.5;refGene_gn=C1GALT1;refGene_tx=XM_023250371.1;refGene_gn=C1GALT1;refGene_tx=XM_023250372.1
A2 102650416 102650608
refGene_gn=C1GALT1;refGene_tx=XM_003982796.5;refGene_gn=C1GALT1;refGene_tx=XM_023250371.1;refGene_gn=C1GALT1;refGene_tx=XM_023250372.1
A2 102747098 102747271
refGene_gn=COL28A1;refGene_tx=XM_006929203.4
A2 102749723 102749916
refGene_gn=COL28A1;refGene_tx=XM_006929203.4
A2 102761243 102761401
refGene_gn=COL28A1;refGene_tx=XM_006929203.4
A2 102763623 102764178
refGene_gn=COL28A1;refGene_tx=XM_006929203.4
A2 102765804 102765841
refGene_gn=COL28A1;refGene_tx=XM_006929203.4
A2 102771931 102771967
refGene_gn=COL28A1;refGene_tx=XM_006929203.4
A2 102772056 102772137
refGene_gn=COL28A1;refGene_tx=XM_006929203.4
A2 102772684 102772753
refGene_gn=COL28A1;refGene_tx=XM_006929203.4
A2 102808992 102809061
refGene_gn=COL28A1;refGene_tx=XM_006929203.4
A2 102813581 102813650
refGene_gn=COL28A1;refGene_tx=XM_006929203.4
A2 102828161 102828230

refGene_gn=COL28A1;refGene_tx=XM_006929203.4
A2 102828401 102828470
refGene_gn=COL28A1;refGene_tx=XM_006929203.4
A2 102832048 102832117
refGene_gn=COL28A1;refGene_tx=XM_006929203.4
A2 102832766 102832835
refGene_gn=COL28A1;refGene_tx=XM_006929203.4
A2 102836324 102836396
refGene_gn=COL28A1;refGene_tx=XM_006929203.4
A2 102839092 102839161
refGene_gn=COL28A1;refGene_tx=XM_006929203.4
A2 102839912 102839984
refGene_gn=COL28A1;refGene_tx=XM_006929203.4
A2 102848677 102848746
refGene_gn=COL28A1;refGene_tx=XM_006929203.4
A2 102850244 102850313
refGene_gn=COL28A1;refGene_tx=XM_006929203.4
A2 102854387 102854456
refGene_gn=COL28A1;refGene_tx=XM_006929203.4
A2 102857471 102857540
refGene_gn=COL28A1;refGene_tx=XM_006929203.4
A2 102861187 102861256
refGene_gn=COL28A1;refGene_tx=XM_006929203.4
A2 102868297 102868366
refGene_gn=COL28A1;refGene_tx=XM_006929203.4
A2 102869435 102869504
refGene_gn=COL28A1;refGene_tx=XM_006929203.4
A2 102885022 102885076
refGene_gn=COL28A1;refGene_tx=XM_006929203.4
A2 102886171 102886216
refGene_gn=COL28A1;refGene_tx=XM_006929203.4
A2 102888769 102888814
refGene_gn=COL28A1;refGene_tx=XM_006929203.4
A2 102895611 102895638
refGene_gn=COL28A1;refGene_tx=XM_006929203.4
A2 102899963 102900005
refGene_gn=COL28A1;refGene_tx=XM_006929203.4
A2 102904308 102904362
refGene_gn=COL28A1;refGene_tx=XM_006929203.4
A2 102908249 102908306
refGene_gn=COL28A1;refGene_tx=XM_006929203.4
A2 102910001 102910022
refGene_gn=COL28A1;refGene_tx=XM_006929203.4
A2 102916643 102917200
refGene_gn=COL28A1;refGene_tx=XM_006929203.4
A2 102918034 102918161
refGene_gn=COL28A1;refGene_tx=XM_006929203.4
A2 102946405 102947699
refGene_gn=MIOS;refGene_tx=XM_023250375.1;refGene_gn=MIOS;refGene_tx=XM_023250376.1;refGene_gn=MIOS;refGene_tx=XM_023250377.1;refGene_gn=MIO

S;refGene_tx=XM_023250378.1;refGene_gn=MIOS;refGene_tx=XM_023250379.1
A2 102948021 102948120
refGene_gn=MIOS;refGene_tx=XM_023250375.1;refGene_gn=MIOS;refGene_tx=XM_023250376.1;refGene_gn=MIOS;refGene_tx=XM_023250377.1;refGene_gn=MIOS;refGene_tx=XM_023250378.1;refGene_gn=MIOS;refGene_tx=XM_023250379.1
A2 102966024 102966279
refGene_gn=MIOS;refGene_tx=XM_023250375.1;refGene_gn=MIOS;refGene_tx=XM_023250376.1;refGene_gn=MIOS;refGene_tx=XM_023250377.1;refGene_gn=MIOS;refGene_tx=XM_023250378.1;refGene_gn=MIOS;refGene_tx=XM_023250379.1
A2 102968772 102968942
refGene_gn=MIOS;refGene_tx=XM_023250375.1;refGene_gn=MIOS;refGene_tx=XM_023250376.1;refGene_gn=MIOS;refGene_tx=XM_023250377.1;refGene_gn=MIOS;refGene_tx=XM_023250378.1;refGene_gn=MIOS;refGene_tx=XM_023250379.1
A2 102971892 102971958
refGene_gn=MIOS;refGene_tx=XM_023250375.1;refGene_gn=MIOS;refGene_tx=XM_023250376.1;refGene_gn=MIOS;refGene_tx=XM_023250377.1;refGene_gn=MIOS;refGene_tx=XM_023250378.1;refGene_gn=MIOS;refGene_tx=XM_023250379.1
A2 102972793 102972952
refGene_gn=MIOS;refGene_tx=XM_023250375.1;refGene_gn=MIOS;refGene_tx=XM_023250376.1;refGene_gn=MIOS;refGene_tx=XM_023250377.1;refGene_gn=MIOS;refGene_tx=XM_023250378.1;refGene_gn=MIOS;refGene_tx=XM_023250379.1
A2 102976976 102977129
refGene_gn=MIOS;refGene_tx=XM_023250375.1;refGene_gn=MIOS;refGene_tx=XM_023250376.1;refGene_gn=MIOS;refGene_tx=XM_023250377.1;refGene_gn=MIOS;refGene_tx=XM_023250378.1;refGene_gn=MIOS;refGene_tx=XM_023250379.1
A2 102978107 102978312
refGene_gn=MIOS;refGene_tx=XM_023250375.1;refGene_gn=MIOS;refGene_tx=XM_023250376.1;refGene_gn=MIOS;refGene_tx=XM_023250377.1;refGene_gn=MIOS;refGene_tx=XM_023250378.1;refGene_gn=MIOS;refGene_tx=XM_023250379.1
A2 102985020 102985150
refGene_gn=MIOS;refGene_tx=XM_023250375.1;refGene_gn=MIOS;refGene_tx=XM_023250376.1;refGene_gn=MIOS;refGene_tx=XM_023250377.1;refGene_gn=MIOS;refGene_tx=XM_023250378.1;refGene_gn=MIOS;refGene_tx=XM_023250379.1
A2 102985888 102985985
refGene_gn=MIOS;refGene_tx=XM_023250375.1;refGene_gn=MIOS;refGene_tx=XM_023250376.1;refGene_gn=MIOS;refGene_tx=XM_023250377.1;refGene_gn=MIOS;refGene_tx=XM_023250378.1;refGene_gn=MIOS;refGene_tx=XM_023250379.1
A2 103007750 103007833
refGene_gn=RPA3;refGene_tx=XM_004001404.5
A2 103008620 103008729
refGene_gn=RPA3;refGene_tx=XM_004001404.5
A2 103009877 103009952
refGene_gn=RPA3;refGene_tx=XM_004001404.5
A2 103011269 103011368
refGene_gn=RPA3;refGene_tx=XM_004001404.5
A2 103034135 103034217
refGene_gn=UMAD1;refGene_tx=XM_011280402.2
A2 103156043 103156117
refGene_gn=UMAD1;refGene_tx=XM_011280402.2
A2 103230409 103230667

refGene_gn=UMAD1;refGene_tx=XM_011280402.2
A2	103319513	103319568
refGene_gn=GLCCI1;refGene_tx=XM_023242103.1
A2	103322113	103323196
refGene_gn=GLCCI1;refGene_tx=XM_023242103.1
A2	103355580	103355732
refGene_gn=GLCCI1;refGene_tx=XM_023242103.1
A2	103377574	103377661
refGene_gn=GLCCI1;refGene_tx=XM_023242103.1
A2	103399958	103400075
refGene_gn=GLCCI1;refGene_tx=XM_023242103.1
A2	103404979	103405132
refGene_gn=GLCCI1;refGene_tx=XM_023242103.1
A2	103410295	103410506
refGene_gn=GLCCI1;refGene_tx=XM_023242103.1
A2	103424611	103424732
refGene_gn=GLCCI1;refGene_tx=XM_023242103.1
A2	103425881	103426224
refGene_gn=GLCCI1;refGene_tx=XM_023242103.1
A2	103444687	103444812
refGene_gn=ICA1;refGene_tx=XM_023250380.1;refGene_gn=ICA1;refGene_tx=XM_023250381.1;refGene_gn=ICA1;refGene_tx=XM_023250382.1;refGene_gn=ICA1;refGene_tx=XM_023250384.1;refGene_gn=ICA1;refGene_tx=XM_023250385.1;refGene_gn=ICA1;refGene_tx=XM_023250386.1;refGene_gn=ICA1;refGene_tx=XM_023250387.1;refGene_gn=ICA1;refGene_tx=XM_023250388.1;refGene_gn=ICA1;refGene_tx=XM_023250389.1;refGene_gn=ICA1;refGene_tx=XM_023250390.1;refGene_gn=ICA1;refGene_tx=XM_023250391.1;refGene_gn=ICA1;refGene_tx=XM_023250393.1;refGene_gn=ICA1;refGene_tx=XM_023250396.1
A2	103447041	103447131
refGene_gn=ICA1;refGene_tx=XM_023250380.1;refGene_gn=ICA1;refGene_tx=XM_023250381.1;refGene_gn=ICA1;refGene_tx=XM_023250384.1;refGene_gn=ICA1;refGene_tx=XM_023250385.1;refGene_gn=ICA1;refGene_tx=XM_023250386.1;refGene_gn=ICA1;refGene_tx=XM_023250387.1;refGene_gn=ICA1;refGene_tx=XM_023250388.1;refGene_gn=ICA1;refGene_tx=XM_023250389.1;refGene_gn=ICA1;refGene_tx=XM_023250390.1;refGene_gn=ICA1;refGene_tx=XM_023250391.1;refGene_gn=ICA1;refGene_tx=XM_023250393.1
A2	103454215	103454226
refGene_gn=ICA1;refGene_tx=XM_023250392.1
A2	103454362	103454412
refGene_gn=ICA1;refGene_tx=XM_023250383.1;refGene_gn=ICA1;refGene_tx=XM_023250395.1
A2	103456951	103457221
refGene_gn=ICA1;refGene_tx=XM_023250380.1;refGene_gn=ICA1;refGene_tx=XM_023250381.1;refGene_gn=ICA1;refGene_tx=XM_023250382.1;refGene_gn=ICA1;refGene_tx=XM_023250383.1;refGene_gn=ICA1;refGene_tx=XM_023250384.1;refGene_gn=ICA1;refGene_tx=XM_023250385.1;refGene_gn=ICA1;refGene_tx=XM_023250386.1;refGene_gn=ICA1;refGene_tx=XM_023250387.1;refGene_gn=ICA1;refGene_tx=XM_023250388.1;refGene_gn=ICA1;refGene_tx=XM_023250389.1;refGene_gn=ICA1;refGene_tx=XM_023250390.1;refGene_gn=ICA1;refGene_tx=XM_023250391.1;refGene_gn=ICA1;refGene_tx=XM_023250392.1;refGene_gn=ICA

1;refGene_tx=XM_023250396.1
A2	103461476	103461529
refGene_gn=ICA1;refGene_tx=XM_023250394.1
A2	103467094	103467127
refGene_gn=ICA1;refGene_tx=XM_023250380.1;refGene_gn=ICA1;refGene_tx=XM_023250381.1;refGene_gn=ICA1;refGene_tx=XM_023250382.1;refGene_gn=ICA1;refGene_tx=XM_023250383.1;refGene_gn=ICA1;refGene_tx=XM_023250384.1;refGene_gn=ICA1;refGene_tx=XM_023250385.1;refGene_gn=ICA1;refGene_tx=XM_023250386.1;refGene_gn=ICA1;refGene_tx=XM_023250387.1;refGene_gn=ICA1;refGene_tx=XM_023250388.1;refGene_gn=ICA1;refGene_tx=XM_023250389.1;refGene_gn=ICA1;refGene_tx=XM_023250390.1;refGene_gn=ICA1;refGene_tx=XM_023250391.1;refGene_gn=ICA1;refGene_tx=XM_023250392.1;refGene_gn=ICA1;refGene_tx=XM_023250393.1;refGene_gn=ICA1;refGene_tx=XM_023250394.1;refGene_gn=ICA1;refGene_tx=XM_023250395.1;refGene_gn=ICA1;refGene_tx=XM_023250396.1
A2	103467237	103467300
refGene_gn=ICA1;refGene_tx=XM_023250380.1;refGene_gn=ICA1;refGene_tx=XM_023250381.1;refGene_gn=ICA1;refGene_tx=XM_023250382.1;refGene_gn=ICA1;refGene_tx=XM_023250383.1;refGene_gn=ICA1;refGene_tx=XM_023250384.1;refGene_gn=ICA1;refGene_tx=XM_023250385.1;refGene_gn=ICA1;refGene_tx=XM_023250386.1;refGene_gn=ICA1;refGene_tx=XM_023250387.1;refGene_gn=ICA1;refGene_tx=XM_023250388.1;refGene_gn=ICA1;refGene_tx=XM_023250389.1;refGene_gn=ICA1;refGene_tx=XM_023250390.1;refGene_gn=ICA1;refGene_tx=XM_023250391.1;refGene_gn=ICA1;refGene_tx=XM_023250392.1;refGene_gn=ICA1;refGene_tx=XM_023250393.1;refGene_gn=ICA1;refGene_tx=XM_023250394.1;refGene_gn=ICA1;refGene_tx=XM_023250395.1;refGene_gn=ICA1;refGene_tx=XM_023250396.1
A2	103470563	103470616
refGene_gn=ICA1;refGene_tx=XM_023250380.1;refGene_gn=ICA1;refGene_tx=XM_023250381.1;refGene_gn=ICA1;refGene_tx=XM_023250382.1;refGene_gn=ICA1;refGene_tx=XM_023250383.1;refGene_gn=ICA1;refGene_tx=XM_023250384.1;refGene_gn=ICA1;refGene_tx=XM_023250385.1;refGene_gn=ICA1;refGene_tx=XM_023250386.1;refGene_gn=ICA1;refGene_tx=XM_023250387.1;refGene_gn=ICA1;refGene_tx=XM_023250388.1;refGene_gn=ICA1;refGene_tx=XM_023250389.1;refGene_gn=ICA1;refGene_tx=XM_023250390.1;refGene_gn=ICA1;refGene_tx=XM_023250391.1;refGene_gn=ICA1;refGene_tx=XM_023250392.1;refGene_gn=ICA1;refGene_tx=XM_023250393.1;refGene_gn=ICA1;refGene_tx=XM_023250394.1;refGene_gn=ICA1;refGene_tx=XM_023250395.1;refGene_gn=ICA1;refGene_tx=XM_023250396.1
A2	103472624	103472722
refGene_gn=ICA1;refGene_tx=XM_023250380.1;refGene_gn=ICA1;refGene_tx=XM_023250381.1;refGene_gn=ICA1;refGene_tx=XM_023250382.1;refGene_gn=ICA1;refGene_tx=XM_023250383.1;refGene_gn=ICA1;refGene_tx=XM_023250384.1;refGene_gn=ICA1;refGene_tx=XM_023250385.1;refGene_gn=ICA1;refGene_tx=XM_023250386.1;refGene_gn=ICA1;refGene_tx=XM_023250387.1;refGene_gn=ICA1;refGene_tx=XM_023250388.1;refGene_gn=ICA1;refGene_tx=XM_023250389.1;refGene_gn=ICA1;refGene_tx=XM_023250390.1;refGene_gn=ICA1;refGene_tx=XM_023250391.1;refGene_gn=ICA1;refGene_tx=XM_023250392.1;refGene_gn=ICA1;refGene_tx=XM_023250393.1;refGene_gn=ICA1;refGene_tx=XM_023250394.1;refGene_gn=ICA1;refGene_tx=XM_023250395.1;refGene_gn=ICA1;refGene_tx=X

M_023250396.1
A2	103485092	103485191
refGene_gn=ICA1;refGene_tx=XM_023250380.1;refGene_gn=ICA1;refGene_tx=XM_023250381.1;refGene_gn=ICA1;refGene_tx=XM_023250382.1;refGene_gn=ICA1;refGene_tx=XM_023250383.1;refGene_gn=ICA1;refGene_tx=XM_023250384.1;refGene_gn=ICA1;refGene_tx=XM_023250385.1;refGene_gn=ICA1;refGene_tx=XM_023250386.1;refGene_gn=ICA1;refGene_tx=XM_023250387.1;refGene_gn=ICA1;refGene_tx=XM_023250388.1;refGene_gn=ICA1;refGene_tx=XM_023250389.1;refGene_gn=ICA1;refGene_tx=XM_023250390.1;refGene_gn=ICA1;refGene_tx=XM_023250391.1;refGene_gn=ICA1;refGene_tx=XM_023250392.1;refGene_gn=ICA1;refGene_tx=XM_023250393.1;refGene_gn=ICA1;refGene_tx=XM_023250394.1;refGene_gn=ICA1;refGene_tx=XM_023250395.1;refGene_gn=ICA1;refGene_tx=XM_023250396.1
A2	103486196	103486322
refGene_gn=ICA1;refGene_tx=XM_023250380.1;refGene_gn=ICA1;refGene_tx=XM_023250381.1;refGene_gn=ICA1;refGene_tx=XM_023250382.1;refGene_gn=ICA1;refGene_tx=XM_023250383.1;refGene_gn=ICA1;refGene_tx=XM_023250384.1;refGene_gn=ICA1;refGene_tx=XM_023250385.1;refGene_gn=ICA1;refGene_tx=XM_023250386.1;refGene_gn=ICA1;refGene_tx=XM_023250387.1;refGene_gn=ICA1;refGene_tx=XM_023250388.1;refGene_gn=ICA1;refGene_tx=XM_023250389.1;refGene_gn=ICA1;refGene_tx=XM_023250390.1;refGene_gn=ICA1;refGene_tx=XM_023250391.1;refGene_gn=ICA1;refGene_tx=XM_023250392.1;refGene_gn=ICA1;refGene_tx=XM_023250393.1;refGene_gn=ICA1;refGene_tx=XM_023250394.1;refGene_gn=ICA1;refGene_tx=XM_023250395.1;refGene_gn=ICA1;refGene_tx=XM_023250396.1
A2	103555759	103555958
refGene_gn=ICA1;refGene_tx=XM_023250380.1;refGene_gn=ICA1;refGene_tx=XM_023250381.1;refGene_gn=ICA1;refGene_tx=XM_023250382.1;refGene_gn=ICA1;refGene_tx=XM_023250383.1;refGene_gn=ICA1;refGene_tx=XM_023250384.1;refGene_gn=ICA1;refGene_tx=XM_023250385.1;refGene_gn=ICA1;refGene_tx=XM_023250386.1;refGene_gn=ICA1;refGene_tx=XM_023250387.1;refGene_gn=ICA1;refGene_tx=XM_023250388.1;refGene_gn=ICA1;refGene_tx=XM_023250389.1;refGene_gn=ICA1;refGene_tx=XM_023250390.1;refGene_gn=ICA1;refGene_tx=XM_023250391.1;refGene_gn=ICA1;refGene_tx=XM_023250392.1;refGene_gn=ICA1;refGene_tx=XM_023250393.1;refGene_gn=ICA1;refGene_tx=XM_023250394.1;refGene_gn=ICA1;refGene_tx=XM_023250395.1;refGene_gn=ICA1;refGene_tx=XM_023250396.1
A2	103559881	103560005
refGene_gn=ICA1;refGene_tx=XM_023250380.1;refGene_gn=ICA1;refGene_tx=XM_023250381.1;refGene_gn=ICA1;refGene_tx=XM_023250382.1;refGene_gn=ICA1;refGene_tx=XM_023250383.1;refGene_gn=ICA1;refGene_tx=XM_023250384.1;refGene_gn=ICA1;refGene_tx=XM_023250385.1;refGene_gn=ICA1;refGene_tx=XM_023250386.1;refGene_gn=ICA1;refGene_tx=XM_023250387.1;refGene_gn=ICA1;refGene_tx=XM_023250388.1;refGene_gn=ICA1;refGene_tx=XM_023250389.1;refGene_gn=ICA1;refGene_tx=XM_023250390.1;refGene_gn=ICA1;refGene_tx=XM_023250391.1;refGene_gn=ICA1;refGene_tx=XM_023250392.1;refGene_gn=ICA1;refGene_tx=XM_023250393.1;refGene_gn=ICA1;refGene_tx=XM_023250394.1;refGene_gn=ICA1;refGene_tx=XM_023250395.1;refGene_gn=ICA1;refGene_tx=XM_023250396.1
A2	103565276	103565349

refGene_gn=ICA1;refGene_tx=XM_023250380.1;refGene_gn=ICA1;refGene_tx=XM_023250381.1;refGene_gn=ICA1;refGene_tx=XM_023250382.1;refGene_gn=ICA1;refGene_tx=XM_023250383.1;refGene_gn=ICA1;refGene_tx=XM_023250384.1;refGene_gn=ICA1;refGene_tx=XM_023250385.1;refGene_gn=ICA1;refGene_tx=XM_023250386.1;refGene_gn=ICA1;refGene_tx=XM_023250387.1;refGene_gn=ICA1;refGene_tx=XM_023250388.1;refGene_gn=ICA1;refGene_tx=XM_023250389.1;refGene_gn=ICA1;refGene_tx=XM_023250390.1;refGene_gn=ICA1;refGene_tx=XM_023250391.1;refGene_gn=ICA1;refGene_tx=XM_023250392.1;refGene_gn=ICA1;refGene_tx=XM_023250393.1;refGene_gn=ICA1;refGene_tx=XM_023250394.1;refGene_gn=ICA1;refGene_tx=XM_023250395.1;refGene_gn=ICA1;refGene_tx=XM_023250396.1
A2 103569582 103569748
refGene_gn=ICA1;refGene_tx=XM_023250380.1;refGene_gn=ICA1;refGene_tx=XM_023250381.1;refGene_gn=ICA1;refGene_tx=XM_023250382.1;refGene_gn=ICA1;refGene_tx=XM_023250383.1;refGene_gn=ICA1;refGene_tx=XM_023250384.1;refGene_gn=ICA1;refGene_tx=XM_023250385.1;refGene_gn=ICA1;refGene_tx=XM_023250386.1;refGene_gn=ICA1;refGene_tx=XM_023250387.1;refGene_gn=ICA1;refGene_tx=XM_023250388.1;refGene_gn=ICA1;refGene_tx=XM_023250389.1;refGene_gn=ICA1;refGene_tx=XM_023250390.1;refGene_gn=ICA1;refGene_tx=XM_023250391.1;refGene_gn=ICA1;refGene_tx=XM_023250392.1;refGene_gn=ICA1;refGene_tx=XM_023250393.1;refGene_gn=ICA1;refGene_tx=XM_023250394.1;refGene_gn=ICA1;refGene_tx=XM_023250395.1;refGene_gn=ICA1;refGene_tx=XM_023250396.1
A2 103573431 103573469
refGene_gn=ICA1;refGene_tx=XM_023250380.1;refGene_gn=ICA1;refGene_tx=XM_023250381.1;refGene_gn=ICA1;refGene_tx=XM_023250382.1;refGene_gn=ICA1;refGene_tx=XM_023250383.1;refGene_gn=ICA1;refGene_tx=XM_023250384.1;refGene_gn=ICA1;refGene_tx=XM_023250385.1;refGene_gn=ICA1;refGene_tx=XM_023250386.1;refGene_gn=ICA1;refGene_tx=XM_023250387.1;refGene_gn=ICA1;refGene_tx=XM_023250388.1;refGene_gn=ICA1;refGene_tx=XM_023250389.1;refGene_gn=ICA1;refGene_tx=XM_023250390.1;refGene_gn=ICA1;refGene_tx=XM_023250391.1;refGene_gn=ICA1;refGene_tx=XM_023250392.1;refGene_gn=ICA1;refGene_tx=XM_023250393.1;refGene_gn=ICA1;refGene_tx=XM_023250394.1;refGene_gn=ICA1;refGene_tx=XM_023250395.1;refGene_gn=ICA1;refGene_tx=XM_023250396.1
A2 103594989 103595148
refGene_gn=ICA1;refGene_tx=XM_023250380.1;refGene_gn=ICA1;refGene_tx=XM_023250381.1;refGene_gn=ICA1;refGene_tx=XM_023250382.1;refGene_gn=ICA1;refGene_tx=XM_023250383.1;refGene_gn=ICA1;refGene_tx=XM_023250392.1;refGene_gn=ICA1;refGene_tx=XM_023250393.1;refGene_gn=ICA1;refGene_tx=XM_023250394.1;refGene_gn=ICA1;refGene_tx=XM_023250395.1;refGene_gn=ICA1;refGene_tx=XM_023250396.1
A2 103744033 103744087
refGene_gn=NXPH1;refGene_tx=XM_019825521.2
A2 104036543 104036675
refGene_gn=NXPH1;refGene_tx=XM_019825521.2
A2 104036676 104037306
refGene_gn=NXPH1;refGene_tx=XM_019825521.2
A2 105784792 105784849
refGene_gn=NDUFA4;refGene_tx=XM_003982803.4

A2 105788554 105788615
refGene_gn=NDUFA4;refGene_tx=XM_003982803.4
A2 105789510 105789599
refGene_gn=NDUFA4;refGene_tx=XM_003982803.4
A2 105790696 105790738
refGene_gn=NDUFA4;refGene_tx=XM_003982803.4
A2 105819427 105819428
refGene_gn=PHF14;refGene_tx=XM_011280403.2;refGene_gn=PHF14;refGene_tx
=XM_011280405.3;refGene_gn=PHF14;refGene_tx=XM_011280406.2;refGene_gn=
PHF14;refGene_tx=XM_011280407.3;refGene_gn=PHF14;refGene_tx=XM_0198255
23.2;refGene_gn=PHF14;refGene_tx=XM_019825526.1;refGene_gn=PHF14;refGe
ne_tx=XM_019825527.2;refGene_gn=PHF14;refGene_tx=XM_023250397.1
A2 105819918 105820029
refGene_gn=PHF14;refGene_tx=XM_011280403.2;refGene_gn=PHF14;refGene_tx
=XM_011280405.3;refGene_gn=PHF14;refGene_tx=XM_011280406.2;refGene_gn=
PHF14;refGene_tx=XM_011280407.3;refGene_gn=PHF14;refGene_tx=XM_0198255
23.2;refGene_gn=PHF14;refGene_tx=XM_019825526.1;refGene_gn=PHF14;refGe
ne_tx=XM_019825527.2;refGene_gn=PHF14;refGene_tx=XM_023250397.1
A2 105827666 105828463
refGene_gn=PHF14;refGene_tx=XM_011280403.2;refGene_gn=PHF14;refGene_tx
=XM_011280405.3;refGene_gn=PHF14;refGene_tx=XM_011280406.2;refGene_gn=
PHF14;refGene_tx=XM_011280407.3;refGene_gn=PHF14;refGene_tx=XM_0198255
23.2;refGene_gn=PHF14;refGene_tx=XM_019825526.1;refGene_gn=PHF14;refGe
ne_tx=XM_019825527.2;refGene_gn=PHF14;refGene_tx=XM_023250397.1
A2 105835336 105835481
refGene_gn=PHF14;refGene_tx=XM_011280403.2;refGene_gn=PHF14;refGene_tx
=XM_011280405.3;refGene_gn=PHF14;refGene_tx=XM_011280406.2;refGene_gn=
PHF14;refGene_tx=XM_011280407.3;refGene_gn=PHF14;refGene_tx=XM_0198255
23.2;refGene_gn=PHF14;refGene_tx=XM_019825526.1;refGene_gn=PHF14;refGe
ne_tx=XM_019825527.2;refGene_gn=PHF14;refGene_tx=XM_023250397.1
A2 105856186 105856346
refGene_gn=PHF14;refGene_tx=XM_011280403.2;refGene_gn=PHF14;refGene_tx
=XM_011280405.3;refGene_gn=PHF14;refGene_tx=XM_011280406.2;refGene_gn=
PHF14;refGene_tx=XM_011280407.3;refGene_gn=PHF14;refGene_tx=XM_0198255
23.2;refGene_gn=PHF14;refGene_tx=XM_019825526.1;refGene_gn=PHF14;refGe
ne_tx=XM_019825527.2;refGene_gn=PHF14;refGene_tx=XM_023250397.1
A2 105861956 105862068
refGene_gn=PHF14;refGene_tx=XM_011280403.2;refGene_gn=PHF14;refGene_tx
=XM_011280405.3;refGene_gn=PHF14;refGene_tx=XM_011280406.2;refGene_gn=
PHF14;refGene_tx=XM_011280407.3;refGene_gn=PHF14;refGene_tx=XM_0198255
23.2;refGene_gn=PHF14;refGene_tx=XM_019825526.1;refGene_gn=PHF14;refGe
ne_tx=XM_019825527.2;refGene_gn=PHF14;refGene_tx=XM_023250397.1
A2 105865216 105865354
refGene_gn=PHF14;refGene_tx=XM_011280403.2;refGene_gn=PHF14;refGene_tx
=XM_011280405.3;refGene_gn=PHF14;refGene_tx=XM_011280406.2;refGene_gn=
PHF14;refGene_tx=XM_011280407.3;refGene_gn=PHF14;refGene_tx=XM_0198255
23.2;refGene_gn=PHF14;refGene_tx=XM_019825526.1;refGene_gn=PHF14;refGe
ne_tx=XM_019825527.2;refGene_gn=PHF14;refGene_tx=XM_023250397.1
A2 105869125 105869272
refGene_gn=PHF14;refGene_tx=XM_011280403.2;refGene_gn=PHF14;refGene_tx

=XM_011280405.3;refGene_gn=PHF14;refGene_tx=XM_011280407.3;refGene_gn=PHF14;refGene_tx=XM_019825523.2;refGene_gn=PHF14;refGene_tx=XM_019825526.1;refGene_gn=PHF14;refGene_tx=XM_019825527.2;refGene_gn=PHF14;refGene_tx=XM_023250397.1
A2 105869898 105870169
refGene_gn=PHF14;refGene_tx=XM_011280403.2;refGene_gn=PHF14;refGene_tx=XM_011280405.3;refGene_gn=PHF14;refGene_tx=XM_011280406.2;refGene_gn=PHF14;refGene_tx=XM_011280407.3;refGene_gn=PHF14;refGene_tx=XM_019825523.2;refGene_gn=PHF14;refGene_tx=XM_019825526.1;refGene_gn=PHF14;refGene_tx=XM_019825527.2;refGene_gn=PHF14;refGene_tx=XM_023250397.1
A2 105870723 105870830
refGene_gn=PHF14;refGene_tx=XM_011280403.2;refGene_gn=PHF14;refGene_tx=XM_011280405.3;refGene_gn=PHF14;refGene_tx=XM_011280406.2;refGene_gn=PHF14;refGene_tx=XM_011280407.3;refGene_gn=PHF14;refGene_tx=XM_019825523.2;refGene_gn=PHF14;refGene_tx=XM_019825526.1;refGene_gn=PHF14;refGene_tx=XM_019825527.2;refGene_gn=PHF14;refGene_tx=XM_023250397.1
A2 105871955 105872051
refGene_gn=PHF14;refGene_tx=XM_011280403.2;refGene_gn=PHF14;refGene_tx=XM_011280405.3;refGene_gn=PHF14;refGene_tx=XM_011280406.2;refGene_gn=PHF14;refGene_tx=XM_011280407.3;refGene_gn=PHF14;refGene_tx=XM_019825523.2;refGene_gn=PHF14;refGene_tx=XM_019825526.1;refGene_gn=PHF14;refGene_tx=XM_019825527.2;refGene_gn=PHF14;refGene_tx=XM_023250397.1
A2 105873885 105873989
refGene_gn=PHF14;refGene_tx=XM_011280403.2;refGene_gn=PHF14;refGene_tx=XM_011280405.3;refGene_gn=PHF14;refGene_tx=XM_011280406.2;refGene_gn=PHF14;refGene_tx=XM_011280407.3;refGene_gn=PHF14;refGene_tx=XM_019825523.2;refGene_gn=PHF14;refGene_tx=XM_019825526.1;refGene_gn=PHF14;refGene_tx=XM_019825527.2;refGene_gn=PHF14;refGene_tx=XM_023250397.1
A2 105875788 105875920
refGene_gn=PHF14;refGene_tx=XM_011280403.2;refGene_gn=PHF14;refGene_tx=XM_011280405.3;refGene_gn=PHF14;refGene_tx=XM_011280406.2;refGene_gn=PHF14;refGene_tx=XM_011280407.3;refGene_gn=PHF14;refGene_tx=XM_019825523.2;refGene_gn=PHF14;refGene_tx=XM_019825526.1;refGene_gn=PHF14;refGene_tx=XM_019825527.2;refGene_gn=PHF14;refGene_tx=XM_023250397.1
A2 105896276 105896445
refGene_gn=PHF14;refGene_tx=XM_011280403.2;refGene_gn=PHF14;refGene_tx=XM_011280405.3;refGene_gn=PHF14;refGene_tx=XM_011280406.2;refGene_gn=PHF14;refGene_tx=XM_011280407.3;refGene_gn=PHF14;refGene_tx=XM_019825523.2;refGene_gn=PHF14;refGene_tx=XM_019825526.1;refGene_gn=PHF14;refGene_tx=XM_019825527.2;refGene_gn=PHF14;refGene_tx=XM_023250397.1
A2 105921592 105921643
refGene_gn=PHF14;refGene_tx=XM_011280403.2;refGene_gn=PHF14;refGene_tx=XM_011280405.3;refGene_gn=PHF14;refGene_tx=XM_011280406.2;refGene_gn=PHF14;refGene_tx=XM_011280407.3;refGene_gn=PHF14;refGene_tx=XM_019825523.2;refGene_gn=PHF14;refGene_tx=XM_019825526.1;refGene_gn=PHF14;refGene_tx=XM_019825527.2;refGene_gn=PHF14;refGene_tx=XM_023250397.1
A2 105921774 105921900
refGene_gn=PHF14;refGene_tx=XM_011280403.2;refGene_gn=PHF14;refGene_tx=XM_011280405.3;refGene_gn=PHF14;refGene_tx=XM_011280406.2;refGene_gn=PHF14;refGene_tx=XM_011280407.3;refGene_gn=PHF14;refGene_tx=XM_0198255

23.2;refGene_gn=PHF14;refGene_tx=XM_019825526.1;refGene_gn=PHF14;refGene_tx=XM_019825527.2;refGene_gn=PHF14;refGene_tx=XM_023250397.1
A2 105924919 105924959
refGene_gn=PHF14;refGene_tx=XM_019825526.1
A2 105965326 105965339
refGene_gn=PHF14;refGene_tx=XM_019825527.2
A2 105974231 105974394
refGene_gn=PHF14;refGene_tx=XM_011280403.2;refGene_gn=PHF14;refGene_tx=XM_011280405.3;refGene_gn=PHF14;refGene_tx=XM_011280406.2;refGene_gn=PHF14;refGene_tx=XM_019825523.2;refGene_gn=PHF14;refGene_tx=XM_023250397.1
A2 105999694 105999724
refGene_gn=PHF14;refGene_tx=XM_019825523.2
A2 106000602 106000614
refGene_gn=PHF14;refGene_tx=XM_011280405.3
A2 106036397 106036472
refGene_gn=PHF14;refGene_tx=XM_011280403.2;refGene_gn=PHF14;refGene_tx=XM_011280406.2
A2 106266937 106267022
refGene_gn=THSD7A;refGene_tx=XM_019825528.2;refGene_gn=THSD7A;refGene_tx=XM_019825529.2;refGene_gn=THSD7A;refGene_tx=XM_019825530.2
A2 106267702 106267790
refGene_gn=THSD7A;refGene_tx=XM_019825528.2;refGene_gn=THSD7A;refGene_tx=XM_019825529.2
A2 106270202 106270413
refGene_gn=THSD7A;refGene_tx=XM_019825528.2;refGene_gn=THSD7A;refGene_tx=XM_019825529.2;refGene_gn=THSD7A;refGene_tx=XM_019825530.2
A2 106270765 106270848
refGene_gn=THSD7A;refGene_tx=XM_019825528.2;refGene_gn=THSD7A;refGene_tx=XM_019825529.2;refGene_gn=THSD7A;refGene_tx=XM_019825530.2
A2 106274103 106274199
refGene_gn=THSD7A;refGene_tx=XM_019825528.2;refGene_gn=THSD7A;refGene_tx=XM_019825529.2;refGene_gn=THSD7A;refGene_tx=XM_019825530.2
A2 106284477 106284651
refGene_gn=THSD7A;refGene_tx=XM_019825528.2;refGene_gn=THSD7A;refGene_tx=XM_019825529.2;refGene_gn=THSD7A;refGene_tx=XM_019825530.2
A2 106289079 106289254
refGene_gn=THSD7A;refGene_tx=XM_019825528.2;refGene_gn=THSD7A;refGene_tx=XM_019825529.2;refGene_gn=THSD7A;refGene_tx=XM_019825530.2
A2 106290068 106290214
refGene_gn=THSD7A;refGene_tx=XM_019825528.2;refGene_gn=THSD7A;refGene_tx=XM_019825529.2;refGene_gn=THSD7A;refGene_tx=XM_019825530.2
A2 106290444 106290562
refGene_gn=THSD7A;refGene_tx=XM_019825528.2;refGene_gn=THSD7A;refGene_tx=XM_019825529.2;refGene_gn=THSD7A;refGene_tx=XM_019825530.2
A2 106294840 106294956
refGene_gn=THSD7A;refGene_tx=XM_019825528.2;refGene_gn=THSD7A;refGene_tx=XM_019825529.2;refGene_gn=THSD7A;refGene_tx=XM_019825530.2
A2 106296267 106296412
refGene_gn=THSD7A;refGene_tx=XM_019825528.2;refGene_gn=THSD7A;refGene_

tx=XM_019825529.2;refGene_gn=THSD7A;refGene_tx=XM_019825530.2
A2 106298064 106298218
refGene_gn=THSD7A;refGene_tx=XM_019825528.2;refGene_gn=THSD7A;refGene_tx=XM_019825529.2;refGene_gn=THSD7A;refGene_tx=XM_019825530.2
A2 106303152 106303286
refGene_gn=THSD7A;refGene_tx=XM_019825528.2;refGene_gn=THSD7A;refGene_tx=XM_019825529.2;refGene_gn=THSD7A;refGene_tx=XM_019825530.2
A2 106304976 106304982
refGene_gn=THSD7A;refGene_tx=XM_019825528.2;refGene_gn=THSD7A;refGene_tx=XM_019825529.2;refGene_gn=THSD7A;refGene_tx=XM_019825530.2
A2 106307623 106307802
refGene_gn=THSD7A;refGene_tx=XM_019825528.2;refGene_gn=THSD7A;refGene_tx=XM_019825529.2;refGene_gn=THSD7A;refGene_tx=XM_019825530.2
A2 106328589 106328853
refGene_gn=THSD7A;refGene_tx=XM_019825528.2;refGene_gn=THSD7A;refGene_tx=XM_019825529.2;refGene_gn=THSD7A;refGene_tx=XM_019825530.2
A2 106330318 106330513
refGene_gn=THSD7A;refGene_tx=XM_019825528.2;refGene_gn=THSD7A;refGene_tx=XM_019825529.2;refGene_gn=THSD7A;refGene_tx=XM_019825530.2
A2 106343064 106343168
refGene_gn=THSD7A;refGene_tx=XM_019825528.2;refGene_gn=THSD7A;refGene_tx=XM_019825529.2;refGene_gn=THSD7A;refGene_tx=XM_019825530.2
A2 106344723 106344856
refGene_gn=THSD7A;refGene_tx=XM_019825528.2;refGene_gn=THSD7A;refGene_tx=XM_019825529.2;refGene_gn=THSD7A;refGene_tx=XM_019825530.2
A2 106353931 106354047
refGene_gn=THSD7A;refGene_tx=XM_019825528.2;refGene_gn=THSD7A;refGene_tx=XM_019825529.2;refGene_gn=THSD7A;refGene_tx=XM_019825530.2
A2 106358702 106358937
refGene_gn=THSD7A;refGene_tx=XM_019825528.2;refGene_gn=THSD7A;refGene_tx=XM_019825529.2;refGene_gn=THSD7A;refGene_tx=XM_019825530.2
A2 106367818 106368013
refGene_gn=THSD7A;refGene_tx=XM_019825528.2;refGene_gn=THSD7A;refGene_tx=XM_019825529.2;refGene_gn=THSD7A;refGene_tx=XM_019825530.2
A2 106434567 106434780
refGene_gn=THSD7A;refGene_tx=XM_019825528.2;refGene_gn=THSD7A;refGene_tx=XM_019825529.2;refGene_gn=THSD7A;refGene_tx=XM_019825530.2
A2 106436037 106436193
refGene_gn=THSD7A;refGene_tx=XM_019825528.2;refGene_gn=THSD7A;refGene_tx=XM_019825529.2;refGene_gn=THSD7A;refGene_tx=XM_019825530.2
A2 106475199 106475268
refGene_gn=THSD7A;refGene_tx=XM_019825528.2;refGene_gn=THSD7A;refGene_tx=XM_019825530.2
A2 106479352 106479534
refGene_gn=THSD7A;refGene_tx=XM_019825528.2;refGene_gn=THSD7A;refGene_tx=XM_019825529.2;refGene_gn=THSD7A;refGene_tx=XM_019825530.2
A2 106481004 106481253
refGene_gn=THSD7A;refGene_tx=XM_019825528.2;refGene_gn=THSD7A;refGene_tx=XM_019825529.2;refGene_gn=THSD7A;refGene_tx=XM_019825530.2
A2 106521127 106521959

refGene_gn=THSD7A;refGene_tx=XM_019825528.2;refGene_gn=THSD7A;refGene_tx=XM_019825529.2;refGene_gn=THSD7A;refGene_tx=XM_019825530.2
A2 106691684 106692009
refGene_gn=THSD7A;refGene_tx=XM_019825528.2;refGene_gn=THSD7A;refGene_tx=XM_019825529.2;refGene_gn=THSD7A;refGene_tx=XM_019825530.2
A2 107037442 107037662
refGene_gn=TMEM106B;refGene_tx=XM_003982805.5;refGene_gn=TMEM106B;refGene_tx=XM_006929217.4;refGene_gn=TMEM106B;refGene_tx=XM_011280408.3
A2 107040915 107040979
refGene_gn=TMEM106B;refGene_tx=XM_003982805.5;refGene_gn=TMEM106B;refGene_tx=XM_006929217.4;refGene_gn=TMEM106B;refGene_tx=XM_011280408.3
A2 107047512 107047672
refGene_gn=TMEM106B;refGene_tx=XM_003982805.5;refGene_gn=TMEM106B;refGene_tx=XM_006929217.4;refGene_gn=TMEM106B;refGene_tx=XM_011280408.3
A2 107050752 107050893
refGene_gn=TMEM106B;refGene_tx=XM_003982805.5;refGene_gn=TMEM106B;refGene_tx=XM_006929217.4;refGene_gn=TMEM106B;refGene_tx=XM_011280408.3
A2 107051143 107051193
refGene_gn=TMEM106B;refGene_tx=XM_003982805.5;refGene_gn=TMEM106B;refGene_tx=XM_006929217.4;refGene_gn=TMEM106B;refGene_tx=XM_011280408.3
A2 107057449 107057503
refGene_gn=TMEM106B;refGene_tx=XM_003982805.5;refGene_gn=TMEM106B;refGene_tx=XM_006929217.4;refGene_gn=TMEM106B;refGene_tx=XM_011280408.3
A2 107058273 107058412
refGene_gn=TMEM106B;refGene_tx=XM_003982805.5;refGene_gn=TMEM106B;refGene_tx=XM_006929217.4;refGene_gn=TMEM106B;refGene_tx=XM_011280408.3
A2 107107273 107107288
refGene_gn=VWDE;refGene_tx=XM_023250399.1;refGene_gn=VWDE;refGene_tx=XM_023250400.1;refGene_gn=VWDE;refGene_tx=XM_023250401.1;refGene_gn=VWDE;refGene_tx=XM_023250402.1;refGene_gn=VWDE;refGene_tx=XM_023250403.1
A2 107111167 107111271
refGene_gn=VWDE;refGene_tx=XM_023250399.1;refGene_gn=VWDE;refGene_tx=XM_023250400.1;refGene_gn=VWDE;refGene_tx=XM_023250401.1;refGene_gn=VWDE;refGene_tx=XM_023250402.1;refGene_gn=VWDE;refGene_tx=XM_023250403.1
A2 107113786 107113882
refGene_gn=VWDE;refGene_tx=XM_023250399.1;refGene_gn=VWDE;refGene_tx=XM_023250400.1;refGene_gn=VWDE;refGene_tx=XM_023250401.1;refGene_gn=VWDE;refGene_tx=XM_023250402.1;refGene_gn=VWDE;refGene_tx=XM_023250403.1
A2 107115089 107115185
refGene_gn=VWDE;refGene_tx=XM_023250399.1;refGene_gn=VWDE;refGene_tx=XM_023250400.1;refGene_gn=VWDE;refGene_tx=XM_023250401.1;refGene_gn=VWDE;refGene_tx=XM_023250403.1
A2 107115289 107115385
refGene_gn=VWDE;refGene_tx=XM_023250399.1;refGene_gn=VWDE;refGene_tx=XM_023250400.1;refGene_gn=VWDE;refGene_tx=XM_023250403.1
A2 107118769 107118865
refGene_gn=VWDE;refGene_tx=XM_023250399.1;refGene_gn=VWDE;refGene_tx=XM_023250400.1;refGene_gn=VWDE;refGene_tx=XM_023250401.1;refGene_gn=VWDE;refGene_tx=XM_023250402.1;refGene_gn=VWDE;refGene_tx=XM_023250403.1
A2 107120104 107120200

refGene_gn=VWDE;refGene_tx=XM_023250399.1;refGene_gn=VWDE;refGene_tx=XM_023250400.1;refGene_gn=VWDE;refGene_tx=XM_023250401.1;refGene_gn=VWDE;refGene_tx=XM_023250402.1;refGene_gn=VWDE;refGene_tx=XM_023250403.1
A2 107120993 107121089
refGene_gn=VWDE;refGene_tx=XM_023250399.1;refGene_gn=VWDE;refGene_tx=XM_023250400.1;refGene_gn=VWDE;refGene_tx=XM_023250401.1;refGene_gn=VWDE;refGene_tx=XM_023250402.1;refGene_gn=VWDE;refGene_tx=XM_023250403.1
A2 107122086 107122182
refGene_gn=VWDE;refGene_tx=XM_023250399.1;refGene_gn=VWDE;refGene_tx=XM_023250400.1;refGene_gn=VWDE;refGene_tx=XM_023250401.1;refGene_gn=VWDE;refGene_tx=XM_023250402.1;refGene_gn=VWDE;refGene_tx=XM_023250403.1
A2 107122263 107122359
refGene_gn=VWDE;refGene_tx=XM_023250399.1;refGene_gn=VWDE;refGene_tx=XM_023250400.1;refGene_gn=VWDE;refGene_tx=XM_023250401.1;refGene_gn=VWDE;refGene_tx=XM_023250402.1;refGene_gn=VWDE;refGene_tx=XM_023250403.1
A2 107123723 107124293
refGene_gn=VWDE;refGene_tx=XM_023250399.1;refGene_gn=VWDE;refGene_tx=XM_023250400.1;refGene_gn=VWDE;refGene_tx=XM_023250401.1;refGene_gn=VWDE;refGene_tx=XM_023250402.1;refGene_gn=VWDE;refGene_tx=XM_023250403.1
A2 107126575 107126716
refGene_gn=VWDE;refGene_tx=XM_023250399.1;refGene_gn=VWDE;refGene_tx=XM_023250400.1;refGene_gn=VWDE;refGene_tx=XM_023250401.1;refGene_gn=VWDE;refGene_tx=XM_023250402.1;refGene_gn=VWDE;refGene_tx=XM_023250403.1
A2 107130571 107130709
refGene_gn=VWDE;refGene_tx=XM_023250399.1;refGene_gn=VWDE;refGene_tx=XM_023250400.1;refGene_gn=VWDE;refGene_tx=XM_023250401.1;refGene_gn=VWDE;refGene_tx=XM_023250402.1;refGene_gn=VWDE;refGene_tx=XM_023250403.1
A2 107132510 107132730
refGene_gn=VWDE;refGene_tx=XM_023250399.1;refGene_gn=VWDE;refGene_tx=XM_023250400.1;refGene_gn=VWDE;refGene_tx=XM_023250401.1;refGene_gn=VWDE;refGene_tx=XM_023250402.1;refGene_gn=VWDE;refGene_tx=XM_023250403.1
A2 107133794 107134045
refGene_gn=VWDE;refGene_tx=XM_023250399.1;refGene_gn=VWDE;refGene_tx=XM_023250400.1;refGene_gn=VWDE;refGene_tx=XM_023250401.1;refGene_gn=VWDE;refGene_tx=XM_023250402.1;refGene_gn=VWDE;refGene_tx=XM_023250403.1
A2 107140408 107140523
refGene_gn=VWDE;refGene_tx=XM_023250399.1;refGene_gn=VWDE;refGene_tx=XM_023250400.1;refGene_gn=VWDE;refGene_tx=XM_023250401.1;refGene_gn=VWDE;refGene_tx=XM_023250402.1;refGene_gn=VWDE;refGene_tx=XM_023250403.1
A2 107142352 107142457
refGene_gn=VWDE;refGene_tx=XM_023250399.1;refGene_gn=VWDE;refGene_tx=XM_023250400.1;refGene_gn=VWDE;refGene_tx=XM_023250401.1;refGene_gn=VWDE;refGene_tx=XM_023250402.1;refGene_gn=VWDE;refGene_tx=XM_023250403.1
A2 107142581 107142737
refGene_gn=VWDE;refGene_tx=XM_023250399.1;refGene_gn=VWDE;refGene_tx=XM_023250400.1;refGene_gn=VWDE;refGene_tx=XM_023250401.1;refGene_gn=VWDE;refGene_tx=XM_023250402.1;refGene_gn=VWDE;refGene_tx=XM_023250403.1
A2 107146402 107146539
refGene_gn=VWDE;refGene_tx=XM_023250399.1;refGene_gn=VWDE;refGene_tx=XM_023250400.1;refGene_gn=VWDE;refGene_tx=XM_023250401.1;refGene_gn=VWD

E;refGene_tx=XM_023250402.1;refGene_gn=VWDE;refGene_tx=XM_023250403.1
A2 107148773 107149735
refGene_gn=VWDE;refGene_tx=XM_023250399.1;refGene_gn=VWDE;refGene_tx=XM_023250400.1;refGene_gn=VWDE;refGene_tx=XM_023250401.1;refGene_gn=VWDE;refGene_tx=XM_023250402.1;refGene_gn=VWDE;refGene_tx=XM_023250403.1
A2 107150156 107150365
refGene_gn=VWDE;refGene_tx=XM_023250399.1;refGene_gn=VWDE;refGene_tx=XM_023250400.1;refGene_gn=VWDE;refGene_tx=XM_023250401.1;refGene_gn=VWDE;refGene_tx=XM_023250402.1;refGene_gn=VWDE;refGene_tx=XM_023250403.1
A2 107150871 107151142
refGene_gn=VWDE;refGene_tx=XM_023250399.1;refGene_gn=VWDE;refGene_tx=XM_023250400.1;refGene_gn=VWDE;refGene_tx=XM_023250401.1;refGene_gn=VWDE;refGene_tx=XM_023250402.1;refGene_gn=VWDE;refGene_tx=XM_023250403.1
A2 107152566 107152640
refGene_gn=VWDE;refGene_tx=XM_023250399.1;refGene_gn=VWDE;refGene_tx=XM_023250400.1;refGene_gn=VWDE;refGene_tx=XM_023250401.1;refGene_gn=VWDE;refGene_tx=XM_023250402.1;refGene_gn=VWDE;refGene_tx=XM_023250403.1
A2 107152931 107153149
refGene_gn=VWDE;refGene_tx=XM_023250399.1;refGene_gn=VWDE;refGene_tx=XM_023250400.1;refGene_gn=VWDE;refGene_tx=XM_023250401.1;refGene_gn=VWDE;refGene_tx=XM_023250402.1;refGene_gn=VWDE;refGene_tx=XM_023250403.1
A2 107155987 107156132
refGene_gn=VWDE;refGene_tx=XM_023250399.1;refGene_gn=VWDE;refGene_tx=XM_023250400.1;refGene_gn=VWDE;refGene_tx=XM_023250401.1;refGene_gn=VWDE;refGene_tx=XM_023250402.1;refGene_gn=VWDE;refGene_tx=XM_023250403.1
A2 107157862 107157952
refGene_gn=VWDE;refGene_tx=XM_023250399.1;refGene_gn=VWDE;refGene_tx=XM_023250400.1;refGene_gn=VWDE;refGene_tx=XM_023250401.1;refGene_gn=VWDE;refGene_tx=XM_023250402.1;refGene_gn=VWDE;refGene_tx=XM_023250403.1
A2 107158509 107158757
refGene_gn=VWDE;refGene_tx=XM_023250399.1;refGene_gn=VWDE;refGene_tx=XM_023250400.1;refGene_gn=VWDE;refGene_tx=XM_023250401.1;refGene_gn=VWDE;refGene_tx=XM_023250402.1;refGene_gn=VWDE;refGene_tx=XM_023250403.1
A2 107162743 107162809
refGene_gn=VWDE;refGene_tx=XM_023250399.1;refGene_gn=VWDE;refGene_tx=XM_023250401.1;refGene_gn=VWDE;refGene_tx=XM_023250402.1;refGene_gn=VWDE;refGene_tx=XM_023250403.1
A2 107172460 107172692
refGene_gn=VWDE;refGene_tx=XM_023250399.1;refGene_gn=VWDE;refGene_tx=XM_023250400.1;refGene_gn=VWDE;refGene_tx=XM_023250401.1;refGene_gn=VWDE;refGene_tx=XM_023250402.1;refGene_gn=VWDE;refGene_tx=XM_023250403.1
A2 107175540 107175725
refGene_gn=VWDE;refGene_tx=XM_023250399.1;refGene_gn=VWDE;refGene_tx=XM_023250400.1;refGene_gn=VWDE;refGene_tx=XM_023250401.1;refGene_gn=VWDE;refGene_tx=XM_023250402.1;refGene_gn=VWDE;refGene_tx=XM_023250403.1
A2 107184924 107184991
refGene_gn=VWDE;refGene_tx=XM_023250399.1;refGene_gn=VWDE;refGene_tx=XM_023250400.1;refGene_gn=VWDE;refGene_tx=XM_023250401.1;refGene_gn=VWDE;refGene_tx=XM_023250402.1
A2 107255290 107256205

refGene_gn=LOC101100954;refGene_tx=XM_003982806.3
A2 107308253 107308452
refGene_gn=SCIN;refGene_tx=XM_003982807.5
A2 107314104 107314259
refGene_gn=SCIN;refGene_tx=XM_003982807.5
A2 107318489 107318651
refGene_gn=SCIN;refGene_tx=XM_003982807.5
A2 107334918 107335068
refGene_gn=SCIN;refGene_tx=XM_003982807.5
A2 107358036 107358129
refGene_gn=SCIN;refGene_tx=XM_003982807.5
A2 107360815 107360948
refGene_gn=SCIN;refGene_tx=XM_003982807.5
A2 107361900 107361989
refGene_gn=SCIN;refGene_tx=XM_003982807.5
A2 107362836 107363052
refGene_gn=SCIN;refGene_tx=XM_003982807.5
A2 107366383 107366505
refGene_gn=SCIN;refGene_tx=XM_003982807.5
A2 107371923 107372014
refGene_gn=SCIN;refGene_tx=XM_003982807.5
A2 107374675 107374846
refGene_gn=SCIN;refGene_tx=XM_003982807.5
A2 107380945 107381123
refGene_gn=SCIN;refGene_tx=XM_003982807.5
A2 107381380 107381502
refGene_gn=SCIN;refGene_tx=XM_003982807.5
A2 107385555 107385633
refGene_gn=SCIN;refGene_tx=XM_003982807.5
A2 107388508 107388569
refGene_gn=SCIN;refGene_tx=XM_003982807.5
A2 107389200 107389328
refGene_gn=SCIN;refGene_tx=XM_003982807.5
A2 107437715 107438318
refGene_gn=ARL4A;refGene_tx=XM_003982808.5;refGene_gn=ARL4A;refGene_tx=XM_006929218.4;refGene_gn=ARL4A;refGene_tx=XM_006929219.3;refGene_gn=ARL4A;refGene_tx=XM_006929220.3;refGene_gn=ARL4A;refGene_tx=XM_006929221.3;refGene_gn=ARL4A;refGene_tx=XM_006929222.3
A2 108546644 108546866
refGene_gn=ETV1;refGene_tx=XM_006929226.4;refGene_gn=ETV1;refGene_tx=XM_011280409.3;refGene_gn=ETV1;refGene_tx=XM_011280410.3;refGene_gn=ETV1;refGene_tx=XM_011280411.3;refGene_gn=ETV1;refGene_tx=XM_011280412.3;refGene_gn=ETV1;refGene_tx=XM_019825531.2;refGene_gn=ETV1;refGene_tx=XM_019825532.2;refGene_gn=ETV1;refGene_tx=XM_019825533.2;refGene_gn=ETV1;refGene_tx=XM_019825534.2;refGene_gn=ETV1;refGene_tx=XM_019825535.2;refGene_gn=ETV1;refGene_tx=XM_019825536.2
A2 108551864 108551966
refGene_gn=ETV1;refGene_tx=XM_006929226.4;refGene_gn=ETV1;refGene_tx=XM_011280409.3;refGene_gn=ETV1;refGene_tx=XM_011280410.3;refGene_gn=ETV1;refGene_tx=XM_011280411.3;refGene_gn=ETV1;refGene_tx=XM_011280412.3;

refGene_gn=ETV1;refGene_tx=XM_019825531.2;refGene_gn=ETV1;refGene_tx=XM_019825532.2;refGene_gn=ETV1;refGene_tx=XM_019825533.2;refGene_gn=ETV1;refGene_tx=XM_019825534.2;refGene_gn=ETV1;refGene_tx=XM_019825535.2;refGene_gn=ETV1;refGene_tx=XM_019825536.2
A2 108556125 108556295
refGene_gn=ETV1;refGene_tx=XM_006929226.4;refGene_gn=ETV1;refGene_tx=XM_011280409.3;refGene_gn=ETV1;refGene_tx=XM_011280410.3;refGene_gn=ETV1;refGene_tx=XM_011280411.3;refGene_gn=ETV1;refGene_tx=XM_011280412.3;refGene_gn=ETV1;refGene_tx=XM_019825531.2;refGene_gn=ETV1;refGene_tx=XM_019825532.2;refGene_gn=ETV1;refGene_tx=XM_019825533.2;refGene_gn=ETV1;refGene_tx=XM_019825534.2;refGene_gn=ETV1;refGene_tx=XM_019825535.2;refGene_gn=ETV1;refGene_tx=XM_019825536.2
A2 108559407 108559476
refGene_gn=ETV1;refGene_tx=XM_006929226.4;refGene_gn=ETV1;refGene_tx=XM_011280409.3;refGene_gn=ETV1;refGene_tx=XM_011280410.3;refGene_gn=ETV1;refGene_tx=XM_011280411.3;refGene_gn=ETV1;refGene_tx=XM_011280412.3;refGene_gn=ETV1;refGene_tx=XM_019825531.2;refGene_gn=ETV1;refGene_tx=XM_019825532.2;refGene_gn=ETV1;refGene_tx=XM_019825533.2;refGene_gn=ETV1;refGene_tx=XM_019825534.2;refGene_gn=ETV1;refGene_tx=XM_019825535.2;refGene_gn=ETV1;refGene_tx=XM_019825536.2
A2 108561062 108561131
refGene_gn=ETV1;refGene_tx=XM_006929226.4;refGene_gn=ETV1;refGene_tx=XM_011280409.3;refGene_gn=ETV1;refGene_tx=XM_011280410.3;refGene_gn=ETV1;refGene_tx=XM_011280411.3;refGene_gn=ETV1;refGene_tx=XM_011280412.3;refGene_gn=ETV1;refGene_tx=XM_019825531.2;refGene_gn=ETV1;refGene_tx=XM_019825532.2;refGene_gn=ETV1;refGene_tx=XM_019825533.2;refGene_gn=ETV1;refGene_tx=XM_019825534.2;refGene_gn=ETV1;refGene_tx=XM_019825535.2;refGene_gn=ETV1;refGene_tx=XM_019825536.2
A2 108579323 108579571
refGene_gn=ETV1;refGene_tx=XM_006929226.4;refGene_gn=ETV1;refGene_tx=XM_011280409.3;refGene_gn=ETV1;refGene_tx=XM_011280410.3;refGene_gn=ETV1;refGene_tx=XM_011280411.3;refGene_gn=ETV1;refGene_tx=XM_011280412.3;refGene_gn=ETV1;refGene_tx=XM_019825531.2;refGene_gn=ETV1;refGene_tx=XM_019825532.2;refGene_gn=ETV1;refGene_tx=XM_019825533.2;refGene_gn=ETV1;refGene_tx=XM_019825534.2;refGene_gn=ETV1;refGene_tx=XM_019825535.2;refGene_gn=ETV1;refGene_tx=XM_019825536.2
A2 108583552 108583741
refGene_gn=ETV1;refGene_tx=XM_006929226.4;refGene_gn=ETV1;refGene_tx=XM_011280409.3;refGene_gn=ETV1;refGene_tx=XM_011280410.3;refGene_gn=ETV1;refGene_tx=XM_011280411.3;refGene_gn=ETV1;refGene_tx=XM_011280412.3;refGene_gn=ETV1;refGene_tx=XM_019825531.2;refGene_gn=ETV1;refGene_tx=XM_019825532.2;refGene_gn=ETV1;refGene_tx=XM_019825533.2;refGene_gn=ETV1;refGene_tx=XM_019825534.2;refGene_gn=ETV1;refGene_tx=XM_019825535.2;refGene_gn=ETV1;refGene_tx=XM_019825536.2
A2 108586487 108586617
refGene_gn=ETV1;refGene_tx=XM_006929226.4;refGene_gn=ETV1;refGene_tx=XM_011280409.3;refGene_gn=ETV1;refGene_tx=XM_011280410.3;refGene_gn=ETV1;refGene_tx=XM_011280411.3;refGene_gn=ETV1;refGene_tx=XM_011280412.3;refGene_gn=ETV1;refGene_tx=XM_019825531.2;refGene_gn=ETV1;refGene_tx=XM_019825532.2;refGene_gn=ETV1;refGene_tx=XM_019825533.2;refGene_gn=ETV

1;refGene_tx=XM_019825534.2;refGene_gn=ETV1;refGene_tx=XM_019825535.2;
refGene_gn=ETV1;refGene_tx=XM_019825536.2
A2 108623392 108623446
refGene_gn=ETV1;refGene_tx=XM_006929226.4;refGene_gn=ETV1;refGene_tx=X
M_011280409.3;refGene_gn=ETV1;refGene_tx=XM_011280410.3;refGene_gn=ETV
1;refGene_tx=XM_011280411.3;refGene_gn=ETV1;refGene_tx=XM_011280412.3;
refGene_gn=ETV1;refGene_tx=XM_019825531.2;refGene_gn=ETV1;refGene_tx=X
M_019825532.2;refGene_gn=ETV1;refGene_tx=XM_019825533.2;refGene_gn=ETV
1;refGene_tx=XM_019825536.2
A2 108631402 108631415
refGene_gn=ETV1;refGene_tx=XM_019825536.2
A2 108632176 108632237
refGene_gn=ETV1;refGene_tx=XM_011280412.3
A2 108632705 108632753
refGene_gn=ETV1;refGene_tx=XM_011280409.3;refGene_gn=ETV1;refGene_tx=X
M_011280410.3;refGene_gn=ETV1;refGene_tx=XM_011280411.3;refGene_gn=ETV
1;refGene_tx=XM_019825531.2;refGene_gn=ETV1;refGene_tx=XM_019825532.2;
refGene_gn=ETV1;refGene_tx=XM_019825534.2;refGene_gn=ETV1;refGene_tx=X
M_019825535.2
A2 108634146 108634234
refGene_gn=ETV1;refGene_tx=XM_006929226.4;refGene_gn=ETV1;refGene_tx=X
M_011280409.3;refGene_gn=ETV1;refGene_tx=XM_011280410.3;refGene_gn=ETV
1;refGene_tx=XM_011280411.3;refGene_gn=ETV1;refGene_tx=XM_019825531.2;
refGene_gn=ETV1;refGene_tx=XM_019825532.2;refGene_gn=ETV1;refGene_tx=X
M_019825533.2;refGene_gn=ETV1;refGene_tx=XM_019825534.2;refGene_gn=ETV
1;refGene_tx=XM_019825535.2
A2 108634696 108634780
refGene_gn=ETV1;refGene_tx=XM_019825531.2;refGene_gn=ETV1;refGene_tx=X
M_019825533.2;refGene_gn=ETV1;refGene_tx=XM_019825534.2
A2 108635039 108635084
refGene_gn=ETV1;refGene_tx=XM_006929226.4;refGene_gn=ETV1;refGene_tx=X
M_011280409.3;refGene_gn=ETV1;refGene_tx=XM_011280410.3;refGene_gn=ETV
1;refGene_tx=XM_011280411.3;refGene_gn=ETV1;refGene_tx=XM_019825532.2;
refGene_gn=ETV1;refGene_tx=XM_019825535.2
A2 108792619 108792727
refGene_gn=DGKB;refGene_tx=XM_023250405.1;refGene_gn=DGKB;refGene_tx=X
M_023250406.1;refGene_gn=DGKB;refGene_tx=XM_023250407.1;refGene_gn=DGK
B;refGene_tx=XM_023250408.1;refGene_gn=DGKB;refGene_tx=XM_023250409.1;
refGene_gn=DGKB;refGene_tx=XM_023250410.1;refGene_gn=DGKB;refGene_tx=X
M_023250411.1;refGene_gn=DGKB;refGene_tx=XM_023250412.1;refGene_gn=DGK
B;refGene_tx=XM_023250413.1
A2 108827445 108827506
refGene_gn=DGKB;refGene_tx=XM_023250405.1;refGene_gn=DGKB;refGene_tx=X
M_023250406.1;refGene_gn=DGKB;refGene_tx=XM_023250407.1;refGene_gn=DGK
B;refGene_tx=XM_023250408.1;refGene_gn=DGKB;refGene_tx=XM_023250409.1;
refGene_gn=DGKB;refGene_tx=XM_023250410.1;refGene_gn=DGKB;refGene_tx=X
M_023250411.1;refGene_gn=DGKB;refGene_tx=XM_023250412.1;refGene_gn=DGK
B;refGene_tx=XM_023250413.1
A2 108828454 108828575
refGene_gn=DGKB;refGene_tx=XM_023250405.1;refGene_gn=DGKB;refGene_tx=X

M_023250406.1;refGene_gn=DGKB;refGene_tx=XM_023250407.1;refGene_gn=DGK
B;refGene_tx=XM_023250408.1;refGene_gn=DGKB;refGene_tx=XM_023250409.1;
refGene_gn=DGKB;refGene_tx=XM_023250410.1;refGene_gn=DGKB;refGene_tx=X
M_023250411.1;refGene_gn=DGKB;refGene_tx=XM_023250412.1;refGene_gn=DGK
B;refGene_tx=XM_023250413.1
A2 108969634 108969830
refGene_gn=DGKB;refGene_tx=XM_023250405.1;refGene_gn=DGKB;refGene_tx=X
M_023250406.1;refGene_gn=DGKB;refGene_tx=XM_023250407.1;refGene_gn=DGK
B;refGene_tx=XM_023250408.1;refGene_gn=DGKB;refGene_tx=XM_023250409.1;
refGene_gn=DGKB;refGene_tx=XM_023250410.1;refGene_gn=DGKB;refGene_tx=X
M_023250411.1;refGene_gn=DGKB;refGene_tx=XM_023250412.1;refGene_gn=DGK
B;refGene_tx=XM_023250413.1
A2 108980185 108980276
refGene_gn=DGKB;refGene_tx=XM_023250405.1;refGene_gn=DGKB;refGene_tx=X
M_023250406.1;refGene_gn=DGKB;refGene_tx=XM_023250407.1;refGene_gn=DGK
B;refGene_tx=XM_023250408.1;refGene_gn=DGKB;refGene_tx=XM_023250409.1;
refGene_gn=DGKB;refGene_tx=XM_023250410.1;refGene_gn=DGKB;refGene_tx=X
M_023250411.1;refGene_gn=DGKB;refGene_tx=XM_023250412.1;refGene_gn=DGK
B;refGene_tx=XM_023250413.1
A2 109106103 109106168
refGene_gn=DGKB;refGene_tx=XM_023250405.1;refGene_gn=DGKB;refGene_tx=X
M_023250406.1;refGene_gn=DGKB;refGene_tx=XM_023250407.1;refGene_gn=DGK
B;refGene_tx=XM_023250408.1;refGene_gn=DGKB;refGene_tx=XM_023250409.1;
refGene_gn=DGKB;refGene_tx=XM_023250410.1;refGene_gn=DGKB;refGene_tx=X
M_023250411.1;refGene_gn=DGKB;refGene_tx=XM_023250412.1;refGene_gn=DGK
B;refGene_tx=XM_023250413.1
A2 109204392 109204553
refGene_gn=DGKB;refGene_tx=XM_023250405.1;refGene_gn=DGKB;refGene_tx=X
M_023250406.1;refGene_gn=DGKB;refGene_tx=XM_023250407.1;refGene_gn=DGK
B;refGene_tx=XM_023250408.1;refGene_gn=DGKB;refGene_tx=XM_023250409.1;
refGene_gn=DGKB;refGene_tx=XM_023250410.1;refGene_gn=DGKB;refGene_tx=X
M_023250411.1;refGene_gn=DGKB;refGene_tx=XM_023250412.1;refGene_gn=DGK
B;refGene_tx=XM_023250413.1
A2 109217494 109217584
refGene_gn=DGKB;refGene_tx=XM_023250405.1;refGene_gn=DGKB;refGene_tx=X
M_023250406.1;refGene_gn=DGKB;refGene_tx=XM_023250407.1;refGene_gn=DGK
B;refGene_tx=XM_023250408.1;refGene_gn=DGKB;refGene_tx=XM_023250409.1;
refGene_gn=DGKB;refGene_tx=XM_023250410.1;refGene_gn=DGKB;refGene_tx=X
M_023250411.1;refGene_gn=DGKB;refGene_tx=XM_023250412.1;refGene_gn=DGK
B;refGene_tx=XM_023250413.1
A2 109219718 109219804
refGene_gn=DGKB;refGene_tx=XM_023250405.1;refGene_gn=DGKB;refGene_tx=X
M_023250406.1;refGene_gn=DGKB;refGene_tx=XM_023250407.1;refGene_gn=DGK
B;refGene_tx=XM_023250408.1;refGene_gn=DGKB;refGene_tx=XM_023250409.1;
refGene_gn=DGKB;refGene_tx=XM_023250410.1;refGene_gn=DGKB;refGene_tx=X
M_023250411.1;refGene_gn=DGKB;refGene_tx=XM_023250412.1;refGene_gn=DGK
B;refGene_tx=XM_023250413.1
A2 109240845 109240920
refGene_gn=DGKB;refGene_tx=XM_023250405.1;refGene_gn=DGKB;refGene_tx=X
M_023250406.1;refGene_gn=DGKB;refGene_tx=XM_023250407.1;refGene_gn=DGK

B;refGene_tx=XM_023250408.1;refGene_gn=DGKB;refGene_tx=XM_023250409.1;
refGene_gn=DGKB;refGene_tx=XM_023250410.1;refGene_gn=DGKB;refGene_tx=X
M_023250411.1;refGene_gn=DGKB;refGene_tx=XM_023250412.1;refGene_gn=DGK
B;refGene_tx=XM_023250413.1
A2 109246562 109246636
refGene_gn=DGKB;refGene_tx=XM_023250405.1;refGene_gn=DGKB;refGene_tx=X
M_023250406.1;refGene_gn=DGKB;refGene_tx=XM_023250407.1;refGene_gn=DGK
B;refGene_tx=XM_023250408.1;refGene_gn=DGKB;refGene_tx=XM_023250409.1;
refGene_gn=DGKB;refGene_tx=XM_023250410.1;refGene_gn=DGKB;refGene_tx=X
M_023250411.1;refGene_gn=DGKB;refGene_tx=XM_023250412.1;refGene_gn=DGK
B;refGene_tx=XM_023250413.1
A2 109256356 109256473
refGene_gn=DGKB;refGene_tx=XM_023250405.1;refGene_gn=DGKB;refGene_tx=X
M_023250406.1;refGene_gn=DGKB;refGene_tx=XM_023250407.1;refGene_gn=DGK
B;refGene_tx=XM_023250408.1;refGene_gn=DGKB;refGene_tx=XM_023250409.1;
refGene_gn=DGKB;refGene_tx=XM_023250410.1;refGene_gn=DGKB;refGene_tx=X
M_023250411.1;refGene_gn=DGKB;refGene_tx=XM_023250412.1;refGene_gn=DGK
B;refGene_tx=XM_023250413.1
A2 109265559 109265595
refGene_gn=DGKB;refGene_tx=XM_023250405.1;refGene_gn=DGKB;refGene_tx=X
M_023250406.1;refGene_gn=DGKB;refGene_tx=XM_023250407.1;refGene_gn=DGK
B;refGene_tx=XM_023250408.1;refGene_gn=DGKB;refGene_tx=XM_023250409.1;
refGene_gn=DGKB;refGene_tx=XM_023250412.1;refGene_gn=DGKB;refGene_tx=X
M_023250413.1
A2 109290796 109290895
refGene_gn=DGKB;refGene_tx=XM_023250405.1;refGene_gn=DGKB;refGene_tx=X
M_023250406.1;refGene_gn=DGKB;refGene_tx=XM_023250407.1;refGene_gn=DGK
B;refGene_tx=XM_023250408.1;refGene_gn=DGKB;refGene_tx=XM_023250409.1;
refGene_gn=DGKB;refGene_tx=XM_023250410.1;refGene_gn=DGKB;refGene_tx=X
M_023250411.1;refGene_gn=DGKB;refGene_tx=XM_023250412.1;refGene_gn=DGK
B;refGene_tx=XM_023250413.1
A2 109302584 109302701
refGene_gn=DGKB;refGene_tx=XM_023250405.1;refGene_gn=DGKB;refGene_tx=X
M_023250406.1;refGene_gn=DGKB;refGene_tx=XM_023250407.1;refGene_gn=DGK
B;refGene_tx=XM_023250408.1;refGene_gn=DGKB;refGene_tx=XM_023250409.1;
refGene_gn=DGKB;refGene_tx=XM_023250410.1;refGene_gn=DGKB;refGene_tx=X
M_023250411.1;refGene_gn=DGKB;refGene_tx=XM_023250412.1;refGene_gn=DGK
B;refGene_tx=XM_023250413.1
A2 109302784 109302873
refGene_gn=DGKB;refGene_tx=XM_023250405.1;refGene_gn=DGKB;refGene_tx=X
M_023250406.1;refGene_gn=DGKB;refGene_tx=XM_023250407.1;refGene_gn=DGK
B;refGene_tx=XM_023250408.1;refGene_gn=DGKB;refGene_tx=XM_023250409.1;
refGene_gn=DGKB;refGene_tx=XM_023250410.1;refGene_gn=DGKB;refGene_tx=X
M_023250411.1;refGene_gn=DGKB;refGene_tx=XM_023250412.1;refGene_gn=DGK
B;refGene_tx=XM_023250413.1
A2 109305473 109305591
refGene_gn=DGKB;refGene_tx=XM_023250405.1;refGene_gn=DGKB;refGene_tx=X
M_023250406.1;refGene_gn=DGKB;refGene_tx=XM_023250407.1;refGene_gn=DGK
B;refGene_tx=XM_023250408.1;refGene_gn=DGKB;refGene_tx=XM_023250409.1;
refGene_gn=DGKB;refGene_tx=XM_023250410.1;refGene_gn=DGKB;refGene_tx=X

M_023250411.1;refGene_gn=DGKB;refGene_tx=XM_023250412.1;refGene_gn=DGK
B;refGene_tx=XM_023250413.1
A2 109314406 109314526
refGene_gn=DGKB;refGene_tx=XM_023250405.1;refGene_gn=DGKB;refGene_tx=X
M_023250406.1;refGene_gn=DGKB;refGene_tx=XM_023250407.1;refGene_gn=DGK
B;refGene_tx=XM_023250408.1;refGene_gn=DGKB;refGene_tx=XM_023250409.1;
refGene_gn=DGKB;refGene_tx=XM_023250410.1;refGene_gn=DGKB;refGene_tx=X
M_023250411.1;refGene_gn=DGKB;refGene_tx=XM_023250412.1;refGene_gn=DGK
B;refGene_tx=XM_023250413.1
A2 109317841 109317916
refGene_gn=DGKB;refGene_tx=XM_023250405.1;refGene_gn=DGKB;refGene_tx=X
M_023250406.1;refGene_gn=DGKB;refGene_tx=XM_023250407.1;refGene_gn=DGK
B;refGene_tx=XM_023250408.1;refGene_gn=DGKB;refGene_tx=XM_023250409.1;
refGene_gn=DGKB;refGene_tx=XM_023250410.1;refGene_gn=DGKB;refGene_tx=X
M_023250411.1;refGene_gn=DGKB;refGene_tx=XM_023250412.1;refGene_gn=DGK
B;refGene_tx=XM_023250413.1
A2 109321216 109321266
refGene_gn=DGKB;refGene_tx=XM_023250405.1;refGene_gn=DGKB;refGene_tx=X
M_023250406.1;refGene_gn=DGKB;refGene_tx=XM_023250407.1;refGene_gn=DGK
B;refGene_tx=XM_023250408.1;refGene_gn=DGKB;refGene_tx=XM_023250409.1;
refGene_gn=DGKB;refGene_tx=XM_023250410.1;refGene_gn=DGKB;refGene_tx=X
M_023250411.1;refGene_gn=DGKB;refGene_tx=XM_023250412.1;refGene_gn=DGK
B;refGene_tx=XM_023250413.1
A2 109332196 109332337
refGene_gn=DGKB;refGene_tx=XM_023250405.1;refGene_gn=DGKB;refGene_tx=X
M_023250406.1;refGene_gn=DGKB;refGene_tx=XM_023250407.1;refGene_gn=DGK
B;refGene_tx=XM_023250408.1;refGene_gn=DGKB;refGene_tx=XM_023250409.1;
refGene_gn=DGKB;refGene_tx=XM_023250410.1;refGene_gn=DGKB;refGene_tx=X
M_023250411.1;refGene_gn=DGKB;refGene_tx=XM_023250412.1;refGene_gn=DGK
B;refGene_tx=XM_023250413.1
A2 109348766 109348920
refGene_gn=DGKB;refGene_tx=XM_023250405.1;refGene_gn=DGKB;refGene_tx=X
M_023250406.1;refGene_gn=DGKB;refGene_tx=XM_023250407.1;refGene_gn=DGK
B;refGene_tx=XM_023250408.1;refGene_gn=DGKB;refGene_tx=XM_023250409.1;
refGene_gn=DGKB;refGene_tx=XM_023250410.1;refGene_gn=DGKB;refGene_tx=X
M_023250411.1;refGene_gn=DGKB;refGene_tx=XM_023250412.1;refGene_gn=DGK
B;refGene_tx=XM_023250413.1
A2 109357191 109357212
refGene_gn=DGKB;refGene_tx=XM_023250405.1;refGene_gn=DGKB;refGene_tx=X
M_023250406.1;refGene_gn=DGKB;refGene_tx=XM_023250407.1;refGene_gn=DGK
B;refGene_tx=XM_023250410.1;refGene_gn=DGKB;refGene_tx=XM_023250412.1
A2 109361044 109361121
refGene_gn=DGKB;refGene_tx=XM_023250405.1;refGene_gn=DGKB;refGene_tx=X
M_023250406.1;refGene_gn=DGKB;refGene_tx=XM_023250407.1;refGene_gn=DGK
B;refGene_tx=XM_023250408.1;refGene_gn=DGKB;refGene_tx=XM_023250409.1;
refGene_gn=DGKB;refGene_tx=XM_023250410.1;refGene_gn=DGKB;refGene_tx=X
M_023250411.1;refGene_gn=DGKB;refGene_tx=XM_023250412.1
A2 109450832 109450902
refGene_gn=DGKB;refGene_tx=XM_023250405.1;refGene_gn=DGKB;refGene_tx=X
M_023250406.1;refGene_gn=DGKB;refGene_tx=XM_023250407.1;refGene_gn=DGK

B;refGene_tx=XM_023250408.1;refGene_gn=DGKB;refGene_tx=XM_023250409.1;
refGene_gn=DGKB;refGene_tx=XM_023250410.1;refGene_gn=DGKB;refGene_tx=X
M_023250411.1;refGene_gn=DGKB;refGene_tx=XM_023250413.1
A2 109777890 109777965
refGene_gn=AGMO;refGene_tx=XM_006929227.4;refGene_gn=AGMO;refGene_tx=X
M_011280414.3;refGene_gn=AGMO;refGene_tx=XM_011280416.3;refGene_gn=AGM
O;refGene_tx=XM_019825539.2;refGene_gn=AGMO;refGene_tx=XM_023250414.1
A2 109873415 109873421
refGene_gn=AGMO;refGene_tx=XM_019825538.2
A2 109925360 109925466
refGene_gn=AGMO;refGene_tx=XM_006929227.4;refGene_gn=AGMO;refGene_tx=X
M_011280414.3;refGene_gn=AGMO;refGene_tx=XM_011280416.3;refGene_gn=AGM
O;refGene_tx=XM_019825538.2;refGene_gn=AGMO;refGene_tx=XM_019825539.2;
refGene_gn=AGMO;refGene_tx=XM_023250414.1
A2 109935635 109935718
refGene_gn=AGMO;refGene_tx=XM_006929227.4;refGene_gn=AGMO;refGene_tx=X
M_011280414.3;refGene_gn=AGMO;refGene_tx=XM_011280416.3;refGene_gn=AGM
O;refGene_tx=XM_019825538.2;refGene_gn=AGMO;refGene_tx=XM_019825539.2;
refGene_gn=AGMO;refGene_tx=XM_023250414.1
A2 109945107 109945224
refGene_gn=AGMO;refGene_tx=XM_006929227.4;refGene_gn=AGMO;refGene_tx=X
M_011280414.3;refGene_gn=AGMO;refGene_tx=XM_011280416.3;refGene_gn=AGM
O;refGene_tx=XM_019825538.2;refGene_gn=AGMO;refGene_tx=XM_019825539.2;
refGene_gn=AGMO;refGene_tx=XM_023250414.1
A2 109947382 109947517
refGene_gn=AGMO;refGene_tx=XM_006929227.4;refGene_gn=AGMO;refGene_tx=X
M_011280414.3;refGene_gn=AGMO;refGene_tx=XM_011280416.3;refGene_gn=AGM
O;refGene_tx=XM_019825538.2;refGene_gn=AGMO;refGene_tx=XM_019825539.2;
refGene_gn=AGMO;refGene_tx=XM_023250414.1
A2 109950752 109950832
refGene_gn=AGMO;refGene_tx=XM_006929227.4;refGene_gn=AGMO;refGene_tx=X
M_011280414.3;refGene_gn=AGMO;refGene_tx=XM_011280416.3;refGene_gn=AGM
O;refGene_tx=XM_019825538.2;refGene_gn=AGMO;refGene_tx=XM_019825539.2;
refGene_gn=AGMO;refGene_tx=XM_023250414.1
A2 109950923 109950989
refGene_gn=AGMO;refGene_tx=XM_006929227.4;refGene_gn=AGMO;refGene_tx=X
M_011280414.3;refGene_gn=AGMO;refGene_tx=XM_011280416.3;refGene_gn=AGM
O;refGene_tx=XM_019825538.2;refGene_gn=AGMO;refGene_tx=XM_019825539.2;
refGene_gn=AGMO;refGene_tx=XM_023250414.1
A2 109953848 109953915
refGene_gn=AGMO;refGene_tx=XM_006929227.4;refGene_gn=AGMO;refGene_tx=X
M_011280414.3;refGene_gn=AGMO;refGene_tx=XM_011280416.3;refGene_gn=AGM
O;refGene_tx=XM_019825538.2;refGene_gn=AGMO;refGene_tx=XM_019825539.2;
refGene_gn=AGMO;refGene_tx=XM_023250414.1
A2 109973681 109973777
refGene_gn=AGMO;refGene_tx=XM_006929227.4;refGene_gn=AGMO;refGene_tx=X
M_011280414.3;refGene_gn=AGMO;refGene_tx=XM_011280416.3;refGene_gn=AGM
O;refGene_tx=XM_019825538.2;refGene_gn=AGMO;refGene_tx=XM_019825539.2;
refGene_gn=AGMO;refGene_tx=XM_023250414.1
A2 109988869 109988973

refGene_gn=AGMO;refGene_tx=XM_006929227.4;refGene_gn=AGMO;refGene_tx=XM_011280414.3;refGene_gn=AGMO;refGene_tx=XM_011280416.3;refGene_gn=AGMO;refGene_tx=XM_019825538.2;refGene_gn=AGMO;refGene_tx=XM_019825539.2;refGene_gn=AGMO;refGene_tx=XM_023250414.1
A2 110090098 110090250
refGene_gn=AGMO;refGene_tx=XM_006929227.4;refGene_gn=AGMO;refGene_tx=XM_011280414.3;refGene_gn=AGMO;refGene_tx=XM_011280416.3;refGene_gn=AGMO;refGene_tx=XM_019825538.2;refGene_gn=AGMO;refGene_tx=XM_019825539.2;refGene_gn=AGMO;refGene_tx=XM_023250414.1
A2 110101497 110101628
refGene_gn=AGMO;refGene_tx=XM_006929227.4;refGene_gn=AGMO;refGene_tx=XM_011280414.3;refGene_gn=AGMO;refGene_tx=XM_011280416.3;refGene_gn=AGMO;refGene_tx=XM_019825538.2;refGene_gn=AGMO;refGene_tx=XM_019825539.2;refGene_gn=AGMO;refGene_tx=XM_023250414.1
A2 110103283 110103490
refGene_gn=AGMO;refGene_tx=XM_006929227.4;refGene_gn=AGMO;refGene_tx=XM_011280414.3;refGene_gn=AGMO;refGene_tx=XM_011280416.3;refGene_gn=AGMO;refGene_tx=XM_019825538.2;refGene_gn=AGMO;refGene_tx=XM_023250414.1
A2 110159123 110159192
refGene_gn=AGMO;refGene_tx=XM_006929227.4;refGene_gn=AGMO;refGene_tx=XM_019825538.2;refGene_gn=AGMO;refGene_tx=XM_019825539.2
A2 110168817 110169042
refGene_gn=MEOX2;refGene_tx=XM_003982817.5
A2 110184865 110185038
refGene_gn=MEOX2;refGene_tx=XM_003982817.5
A2 110232080 110232594
refGene_gn=MEOX2;refGene_tx=XM_003982817.5
A2 110633048 110633153
refGene_gn=ISPD;refGene_tx=XM_023250417.1
A2 110692437 110692476
refGene_gn=ISPD;refGene_tx=XM_023250415.1;refGene_gn=ISPD;refGene_tx=XM_023250416.1
A2 110754319 110754451
refGene_gn=ISPD;refGene_tx=XM_023250415.1;refGene_gn=ISPD;refGene_tx=XM_023250416.1;refGene_gn=ISPD;refGene_tx=XM_023250417.1
A2 110778325 110778418
refGene_gn=ISPD;refGene_tx=XM_023250415.1;refGene_gn=ISPD;refGene_tx=XM_023250416.1;refGene_gn=ISPD;refGene_tx=XM_023250417.1
A2 110778899 110778992
refGene_gn=ISPD;refGene_tx=XM_023250415.1;refGene_gn=ISPD;refGene_tx=XM_023250416.1;refGene_gn=ISPD;refGene_tx=XM_023250417.1
A2 110810189 110810284
refGene_gn=ISPD;refGene_tx=XM_023250415.1;refGene_gn=ISPD;refGene_tx=XM_023250416.1;refGene_gn=ISPD;refGene_tx=XM_023250417.1
A2 110833300 110833346
refGene_gn=ISPD;refGene_tx=XM_023250415.1;refGene_gn=ISPD;refGene_tx=XM_023250416.1;refGene_gn=ISPD;refGene_tx=XM_023250417.1
A2 110840487 110840592
refGene_gn=ISPD;refGene_tx=XM_023250415.1;refGene_gn=ISPD;refGene_tx=XM_023250416.1;refGene_gn=ISPD;refGene_tx=XM_023250417.1

A2 110914384 110914534
refGene_gn=ISPD;refGene_tx=XM_023250415.1;refGene_gn=ISPD;refGene_tx=XM_023250416.1;refGene_gn=ISPD;refGene_tx=XM_023250417.1
A2 110934582 110934859
refGene_gn=ISPD;refGene_tx=XM_023250415.1;refGene_gn=ISPD;refGene_tx=XM_023250416.1;refGene_gn=ISPD;refGene_tx=XM_023250417.1
A2 110953676 110953924
refGene_gn=ISPD;refGene_tx=XM_023250415.1;refGene_gn=ISPD;refGene_tx=XM_023250416.1;refGene_gn=ISPD;refGene_tx=XM_023250417.1
A2 110991609 110992025
refGene_gn=SOSTDC1;refGene_tx=XM_003982819.5;refGene_gn=SOSTDC1;refGene_tx=XM_006929228.4;refGene_gn=SOSTDC1;refGene_tx=XM_006929229.4;refGene_gn=SOSTDC1;refGene_tx=XM_019825544.2
A2 110994532 110994737
refGene_gn=SOSTDC1;refGene_tx=XM_003982819.5;refGene_gn=SOSTDC1;refGene_tx=XM_006929228.4;refGene_gn=SOSTDC1;refGene_tx=XM_006929229.4;refGene_gn=SOSTDC1;refGene_tx=XM_019825544.2
A2 111057340 111057344
refGene_gn=SOSTDC1;refGene_tx=XM_006929228.4;refGene_gn=SOSTDC1;refGene_tx=XM_006929229.4;refGene_gn=SOSTDC1;refGene_tx=XM_019825544.2
A2 111058100 111058193
refGene_gn=LRRC72;refGene_tx=XM_006929230.4;refGene_gn=LRRC72;refGene_tx=XM_023250419.1
A2 111063950 111064057
refGene_gn=LRRC72;refGene_tx=XM_006929230.4;refGene_gn=LRRC72;refGene_tx=XM_023250418.1;refGene_gn=LRRC72;refGene_tx=XM_023250419.1
A2 111068733 111068803
refGene_gn=LRRC72;refGene_tx=XM_006929230.4;refGene_gn=LRRC72;refGene_tx=XM_023250418.1;refGene_gn=LRRC72;refGene_tx=XM_023250419.1
A2 111079718 111079800
refGene_gn=LRRC72;refGene_tx=XM_006929230.4;refGene_gn=LRRC72;refGene_tx=XM_023250418.1;refGene_gn=LRRC72;refGene_tx=XM_023250419.1
A2 111082190 111082301
refGene_gn=LRRC72;refGene_tx=XM_006929230.4;refGene_gn=LRRC72;refGene_tx=XM_023250418.1;refGene_gn=LRRC72;refGene_tx=XM_023250419.1
A2 111090902 111090992
refGene_gn=LRRC72;refGene_tx=XM_006929230.4;refGene_gn=LRRC72;refGene_tx=XM_023250418.1
A2 111091659 111091812
refGene_gn=LRRC72;refGene_tx=XM_006929230.4;refGene_gn=LRRC72;refGene_tx=XM_023250418.1
A2 111098368 111098396
refGene_gn=LRRC72;refGene_tx=XM_006929230.4;refGene_gn=LRRC72;refGene_tx=XM_023250418.1;refGene_gn=LRRC72;refGene_tx=XM_023250419.1
A2 111099899 111100065
refGene_gn=LRRC72;refGene_tx=XM_006929230.4;refGene_gn=LRRC72;refGene_tx=XM_023250418.1;refGene_gn=LRRC72;refGene_tx=XM_023250419.1
A2 111107277 111107462
refGene_gn=ANKMY2;refGene_tx=XM_011280417.3;refGene_gn=ANKMY2;refGene_tx=XM_011280418.3

A2	111108900	111109030
refGene_gn=ANKMY2;refGene_tx=XM_011280417.3;refGene_gn=ANKMY2;refGene_tx=XM_011280418.3
A2	111110994	111111123
refGene_gn=ANKMY2;refGene_tx=XM_011280417.3
A2	111114990	111115126
refGene_gn=ANKMY2;refGene_tx=XM_011280417.3;refGene_gn=ANKMY2;refGene_tx=XM_011280418.3
A2	111115929	111116144
refGene_gn=ANKMY2;refGene_tx=XM_011280417.3;refGene_gn=ANKMY2;refGene_tx=XM_011280418.3
A2	111120995	111121156
refGene_gn=ANKMY2;refGene_tx=XM_011280417.3;refGene_gn=ANKMY2;refGene_tx=XM_011280418.3
A2	111130175	111130274
refGene_gn=ANKMY2;refGene_tx=XM_011280417.3;refGene_gn=ANKMY2;refGene_tx=XM_011280418.3
A2	111133515	111133654
refGene_gn=ANKMY2;refGene_tx=XM_011280417.3;refGene_gn=ANKMY2;refGene_tx=XM_011280418.3
A2	111140405	111140470
refGene_gn=ANKMY2;refGene_tx=XM_011280417.3;refGene_gn=ANKMY2;refGene_tx=XM_011280418.3
A2	111145958	111146025
refGene_gn=ANKMY2;refGene_tx=XM_011280417.3;refGene_gn=ANKMY2;refGene_tx=XM_011280418.3
A2	111169446	111169504
refGene_gn=BZW2;refGene_tx=XM_003982820.4;refGene_gn=BZW2;refGene_tx=XM_006929231.3
A2	111177793	111177970
refGene_gn=BZW2;refGene_tx=XM_003982820.4;refGene_gn=BZW2;refGene_tx=XM_006929231.3
A2	111182533	111182637
refGene_gn=BZW2;refGene_tx=XM_003982820.4;refGene_gn=BZW2;refGene_tx=XM_006929231.3
A2	111185038	111185104
refGene_gn=BZW2;refGene_tx=XM_003982820.4;refGene_gn=BZW2;refGene_tx=XM_006929231.3
A2	111188668	111188804
refGene_gn=BZW2;refGene_tx=XM_003982820.4;refGene_gn=BZW2;refGene_tx=XM_006929231.3
A2	111191837	111191947
refGene_gn=BZW2;refGene_tx=XM_003982820.4;refGene_gn=BZW2;refGene_tx=XM_006929231.3
A2	111196852	111197023
refGene_gn=BZW2;refGene_tx=XM_003982820.4;refGene_gn=BZW2;refGene_tx=XM_006929231.3
A2	111198890	111199037
refGene_gn=BZW2;refGene_tx=XM_003982820.4;refGene_gn=BZW2;refGene_tx=XM_006929231.3

A2 111200356 111200495
refGene_gn=BZW2;refGene_tx=XM_003982820.4;refGene_gn=BZW2;refGene_tx=X
M_006929231.3
A2 111207996 111208119
refGene_gn=BZW2;refGene_tx=XM_003982820.4;refGene_gn=BZW2;refGene_tx=X
M_006929231.3
A2 111209573 111209602
refGene_gn=BZW2;refGene_tx=XM_003982820.4;refGene_gn=BZW2;refGene_tx=X
M_006929231.3
A2 111253428 111253491
refGene_gn=TSPAN13;refGene_tx=XM_003982822.4
A2 111276541 111276709
refGene_gn=TSPAN13;refGene_tx=XM_003982822.4
A2 111277411 111277492
refGene_gn=TSPAN13;refGene_tx=XM_003982822.4
A2 111278125 111278239
refGene_gn=TSPAN13;refGene_tx=XM_003982822.4
A2 111279469 111279583
refGene_gn=TSPAN13;refGene_tx=XM_003982822.4
A2 111284951 111285026
refGene_gn=TSPAN13;refGene_tx=XM_003982822.4
A2 111292364 111292414
refGene_gn=AGR2;refGene_tx=XM_006929232.3;refGene_gn=AGR2;refGene_tx=X
M_019825546.2;refGene_gn=AGR2;refGene_tx=XM_019825548.2;refGene_gn=AGR
2;refGene_tx=XM_019825549.2
A2 111293601 111293685
refGene_gn=AGR2;refGene_tx=XM_006929232.3;refGene_gn=AGR2;refGene_tx=X
M_019825546.2;refGene_gn=AGR2;refGene_tx=XM_019825548.2;refGene_gn=AGR
2;refGene_tx=XM_019825549.2
A2 111296058 111296122
refGene_gn=AGR2;refGene_tx=XM_006929232.3;refGene_gn=AGR2;refGene_tx=X
M_019825546.2;refGene_gn=AGR2;refGene_tx=XM_019825548.2;refGene_gn=AGR
2;refGene_tx=XM_019825549.2
A2 111298606 111298680
refGene_gn=AGR2;refGene_tx=XM_006929232.3;refGene_gn=AGR2;refGene_tx=X
M_019825546.2;refGene_gn=AGR2;refGene_tx=XM_019825548.2;refGene_gn=AGR
2;refGene_tx=XM_019825549.2
A2 111299160 111299213
refGene_gn=AGR2;refGene_tx=XM_006929232.3;refGene_gn=AGR2;refGene_tx=X
M_019825546.2;refGene_gn=AGR2;refGene_tx=XM_019825548.2;refGene_gn=AGR
2;refGene_tx=XM_019825549.2
A2 111299331 111299395
refGene_gn=AGR2;refGene_tx=XM_006929232.3;refGene_gn=AGR2;refGene_tx=X
M_019825546.2;refGene_gn=AGR2;refGene_tx=XM_019825548.2;refGene_gn=AGR
2;refGene_tx=XM_019825549.2
A2 111299608 111299747
refGene_gn=AGR2;refGene_tx=XM_006929232.3;refGene_gn=AGR2;refGene_tx=X
M_019825546.2;refGene_gn=AGR2;refGene_tx=XM_019825548.2;refGene_gn=AGR
2;refGene_tx=XM_019825549.2
A2 111733659 111733724

refGene_gn=AHR;refGene_tx=XM_011280419.3;refGene_gn=AHR;refGene_tx=XM_019825550.2
A2 111745053 111745241
refGene_gn=AHR;refGene_tx=XM_011280419.3;refGene_gn=AHR;refGene_tx=XM_019825550.2
A2 111763413 111763517
refGene_gn=AHR;refGene_tx=XM_011280419.3;refGene_gn=AHR;refGene_tx=XM_019825550.2
A2 111766465 111766528
refGene_gn=AHR;refGene_tx=XM_019825551.2
A2 111767987 111768077
refGene_gn=AHR;refGene_tx=XM_011280419.3;refGene_gn=AHR;refGene_tx=XM_019825550.2;refGene_gn=AHR;refGene_tx=XM_019825551.2
A2 111769688 111769812
refGene_gn=AHR;refGene_tx=XM_011280419.3;refGene_gn=AHR;refGene_tx=XM_019825550.2;refGene_gn=AHR;refGene_tx=XM_019825551.2
A2 111770463 111770594
refGene_gn=AHR;refGene_tx=XM_011280419.3;refGene_gn=AHR;refGene_tx=XM_019825550.2;refGene_gn=AHR;refGene_tx=XM_019825551.2
A2 111772247 111772450
refGene_gn=AHR;refGene_tx=XM_011280419.3;refGene_gn=AHR;refGene_tx=XM_019825550.2;refGene_gn=AHR;refGene_tx=XM_019825551.2
A2 111773200 111773310
refGene_gn=AHR;refGene_tx=XM_011280419.3;refGene_gn=AHR;refGene_tx=XM_019825550.2;refGene_gn=AHR;refGene_tx=XM_019825551.2
A2 111774428 111774570
refGene_gn=AHR;refGene_tx=XM_011280419.3;refGene_gn=AHR;refGene_tx=XM_019825550.2;refGene_gn=AHR;refGene_tx=XM_019825551.2
A2 111781642 111782888
refGene_gn=AHR;refGene_tx=XM_011280419.3;refGene_gn=AHR;refGene_tx=XM_019825550.2;refGene_gn=AHR;refGene_tx=XM_019825551.2
A2 111785604 111785751
refGene_gn=AHR;refGene_tx=XM_011280419.3;refGene_gn=AHR;refGene_tx=XM_019825550.2;refGene_gn=AHR;refGene_tx=XM_019825551.2
A2 112189648 112189896
refGene_gn=SNX13;refGene_tx=XM_023250420.1
A2 112192337 112192450
refGene_gn=SNX13;refGene_tx=XM_023250420.1
A2 112194101 112194170
refGene_gn=SNX13;refGene_tx=XM_023250420.1
A2 112194452 112194598
refGene_gn=SNX13;refGene_tx=XM_023250420.1
A2 112197546 112197618
refGene_gn=SNX13;refGene_tx=XM_023250420.1
A2 112199114 112199276
refGene_gn=SNX13;refGene_tx=XM_023250420.1
A2 112204141 112204230
refGene_gn=SNX13;refGene_tx=XM_023250420.1
A2 112204232 112204251
refGene_gn=SNX13;refGene_tx=XM_023250420.1

A2 112206423 112206531
refGene_gn=SNX13;refGene_tx=XM_023250420.1
A2 112217461 112217601
refGene_gn=SNX13;refGene_tx=XM_023250420.1
A2 112221907 112221977
refGene_gn=SNX13;refGene_tx=XM_023250420.1
A2 112226909 112226947
refGene_gn=SNX13;refGene_tx=XM_023250420.1
A2 112229557 112229690
refGene_gn=SNX13;refGene_tx=XM_023250420.1
A2 112230292 112230397
refGene_gn=SNX13;refGene_tx=XM_023250420.1
A2 112235533 112235727
refGene_gn=SNX13;refGene_tx=XM_023250420.1
A2 112237506 112237606
refGene_gn=SNX13;refGene_tx=XM_023250420.1
A2 112241945 112242034
refGene_gn=SNX13;refGene_tx=XM_023250420.1
A2 112242402 112242541
refGene_gn=SNX13;refGene_tx=XM_023250420.1
A2 112246065 112246149
refGene_gn=SNX13;refGene_tx=XM_023250420.1
A2 112250837 112250926
refGene_gn=SNX13;refGene_tx=XM_023250420.1
A2 112252170 112252272
refGene_gn=SNX13;refGene_tx=XM_023250420.1
A2 112252364 112252486
refGene_gn=SNX13;refGene_tx=XM_023250420.1
A2 112261786 112261858
refGene_gn=SNX13;refGene_tx=XM_023250420.1
A2 112267663 112267785
refGene_gn=SNX13;refGene_tx=XM_023250420.1
A2 112268729 112268819
refGene_gn=SNX13;refGene_tx=XM_023250420.1
A2 112270524 112270627
refGene_gn=SNX13;refGene_tx=XM_023250420.1
A2 112274769 112274882
refGene_gn=SNX13;refGene_tx=XM_023250420.1
A2 112317652 112318450
refGene_gn=SNX13;refGene_tx=XM_023250420.1
A2 112374456 112375413
refGene_gn=PRPS1L1;refGene_tx=XM_019825553.2
A2 112494178 112494200
refGene_gn=HDAC9;refGene_tx=XM_019825556.1;refGene_gn=HDAC9;refGene_tx
=XM_019825557.2;refGene_gn=HDAC9;refGene_tx=XM_019825558.2;refGene_gn=
HDAC9;refGene_tx=XM_019825560.2;refGene_gn=HDAC9;refGene_tx=XM_0198255
61.2;refGene_gn=HDAC9;refGene_tx=XM_019825562.2;refGene_gn=HDAC9;refGe
ne_tx=XM_019825563.2;refGene_gn=HDAC9;refGene_tx=XM_019825564.2;refGen
e_gn=HDAC9;refGene_tx=XM_019825565.2;refGene_gn=HDAC9;refGene_tx=XM_01
9825573.2;refGene_gn=HDAC9;refGene_tx=XM_019825574.2;refGene_gn=HDAC9;

refGene_tx=XM_019825576.2;refGene_gn=HDAC9;refGene_tx=XM_019825577.2;refGene_gn=HDAC9;refGene_tx=XM_019825580.2;refGene_gn=HDAC9;refGene_tx=XM_019825581.2;refGene_gn=HDAC9;refGene_tx=XM_019825582.2;refGene_gn=HDAC9;refGene_tx=XM_023250421.1;refGene_gn=HDAC9;refGene_tx=XM_023250422.1;refGene_gn=HDAC9;refGene_tx=XM_023250423.1
A2 112819591 112819654
refGene_gn=HDAC9;refGene_tx=XM_019825556.1;refGene_gn=HDAC9;refGene_tx=XM_019825557.2;refGene_gn=HDAC9;refGene_tx=XM_019825558.2;refGene_gn=HDAC9;refGene_tx=XM_019825560.2;refGene_gn=HDAC9;refGene_tx=XM_019825561.2;refGene_gn=HDAC9;refGene_tx=XM_019825562.2;refGene_gn=HDAC9;refGene_tx=XM_019825563.2;refGene_gn=HDAC9;refGene_tx=XM_019825564.2;refGene_gn=HDAC9;refGene_tx=XM_019825565.2;refGene_gn=HDAC9;refGene_tx=XM_019825566.1;refGene_gn=HDAC9;refGene_tx=XM_019825567.1;refGene_gn=HDAC9;refGene_tx=XM_019825568.1;refGene_gn=HDAC9;refGene_tx=XM_019825573.2;refGene_gn=HDAC9;refGene_tx=XM_019825574.2;refGene_gn=HDAC9;refGene_tx=XM_019825576.2;refGene_gn=HDAC9;refGene_tx=XM_019825577.2;refGene_gn=HDAC9;refGene_tx=XM_019825580.2;refGene_gn=HDAC9;refGene_tx=XM_019825581.2;refGene_gn=HDAC9;refGene_tx=XM_019825582.2;refGene_gn=HDAC9;refGene_tx=XM_023250421.1;refGene_gn=HDAC9;refGene_tx=XM_023250422.1;refGene_gn=HDAC9;refGene_tx=XM_023250423.1
A2 112853077 112853096
refGene_gn=HDAC9;refGene_tx=XM_019825569.1;refGene_gn=HDAC9;refGene_tx=XM_019825570.2
A2 112905662 112905904
refGene_gn=HDAC9;refGene_tx=XM_019825556.1;refGene_gn=HDAC9;refGene_tx=XM_019825557.2;refGene_gn=HDAC9;refGene_tx=XM_019825558.2;refGene_gn=HDAC9;refGene_tx=XM_019825560.2;refGene_gn=HDAC9;refGene_tx=XM_019825561.2;refGene_gn=HDAC9;refGene_tx=XM_019825562.2;refGene_gn=HDAC9;refGene_tx=XM_019825563.2;refGene_gn=HDAC9;refGene_tx=XM_019825564.2;refGene_gn=HDAC9;refGene_tx=XM_019825565.2;refGene_gn=HDAC9;refGene_tx=XM_019825566.1;refGene_gn=HDAC9;refGene_tx=XM_019825567.1;refGene_gn=HDAC9;refGene_tx=XM_019825568.1;refGene_gn=HDAC9;refGene_tx=XM_019825569.1;refGene_gn=HDAC9;refGene_tx=XM_019825570.2;refGene_gn=HDAC9;refGene_tx=XM_019825573.2;refGene_gn=HDAC9;refGene_tx=XM_019825574.2;refGene_gn=HDAC9;refGene_tx=XM_019825576.2;refGene_gn=HDAC9;refGene_tx=XM_019825577.2;refGene_gn=HDAC9;refGene_tx=XM_019825580.2;refGene_gn=HDAC9;refGene_tx=XM_019825581.2;refGene_gn=HDAC9;refGene_tx=XM_019825582.2;refGene_gn=HDAC9;refGene_tx=XM_023250421.1;refGene_gn=HDAC9;refGene_tx=XM_023250422.1;refGene_gn=HDAC9;refGene_tx=XM_023250423.1
A2 112910923 112911089
refGene_gn=HDAC9;refGene_tx=XM_019825556.1;refGene_gn=HDAC9;refGene_tx=XM_019825557.2;refGene_gn=HDAC9;refGene_tx=XM_019825558.2;refGene_gn=HDAC9;refGene_tx=XM_019825560.2;refGene_gn=HDAC9;refGene_tx=XM_019825561.2;refGene_gn=HDAC9;refGene_tx=XM_019825562.2;refGene_gn=HDAC9;refGene_tx=XM_019825563.2;refGene_gn=HDAC9;refGene_tx=XM_019825564.2;refGene_gn=HDAC9;refGene_tx=XM_019825565.2;refGene_gn=HDAC9;refGene_tx=XM_019825566.1;refGene_gn=HDAC9;refGene_tx=XM_019825567.1;refGene_gn=HDAC9;refGene_tx=XM_019825568.1;refGene_gn=HDAC9;refGene_tx=XM_019825569.1;refGene_gn=HDAC9;refGene_tx=XM_019825570.2;refGene_gn=HDAC9;refGene_tx=XM_019825573.2;refGene_gn=HDAC9;refGene_tx=XM_019825574.2;refGene_gn=H

DAC9;refGene_tx=XM_019825576.2;refGene_gn=HDAC9;refGene_tx=XM_01982557
7.2;refGene_gn=HDAC9;refGene_tx=XM_019825580.2;refGene_gn=HDAC9;refGen
e_tx=XM_019825581.2;refGene_gn=HDAC9;refGene_tx=XM_019825582.2;refGene
_gn=HDAC9;refGene_tx=XM_023250421.1;refGene_gn=HDAC9;refGene_tx=XM_023
250422.1;refGene_gn=HDAC9;refGene_tx=XM_023250423.1
A2 112912060 112912187
refGene_gn=HDAC9;refGene_tx=XM_019825556.1;refGene_gn=HDAC9;refGene_tx
=XM_019825557.2;refGene_gn=HDAC9;refGene_tx=XM_019825558.2;refGene_gn=
HDAC9;refGene_tx=XM_019825560.2;refGene_gn=HDAC9;refGene_tx=XM_0198255
61.2;refGene_gn=HDAC9;refGene_tx=XM_019825562.2;refGene_gn=HDAC9;refGe
ne_tx=XM_019825563.2;refGene_gn=HDAC9;refGene_tx=XM_019825564.2;refGen
e_gn=HDAC9;refGene_tx=XM_019825565.2;refGene_gn=HDAC9;refGene_tx=XM_01
9825566.1;refGene_gn=HDAC9;refGene_tx=XM_019825567.1;refGene_gn=HDAC9;
refGene_tx=XM_019825568.1;refGene_gn=HDAC9;refGene_tx=XM_019825569.1;r
efGene_gn=HDAC9;refGene_tx=XM_019825570.2;refGene_gn=HDAC9;refGene_tx=
XM_019825573.2;refGene_gn=HDAC9;refGene_tx=XM_019825574.2;refGene_gn=H
DAC9;refGene_tx=XM_019825576.2;refGene_gn=HDAC9;refGene_tx=XM_01982557
7.2;refGene_gn=HDAC9;refGene_tx=XM_019825580.2;refGene_gn=HDAC9;refGen
e_tx=XM_019825581.2;refGene_gn=HDAC9;refGene_tx=XM_019825582.2;refGene
_gn=HDAC9;refGene_tx=XM_023250421.1;refGene_gn=HDAC9;refGene_tx=XM_023
250422.1;refGene_gn=HDAC9;refGene_tx=XM_023250423.1
A2 112914446 112914568
refGene_gn=HDAC9;refGene_tx=XM_019825556.1;refGene_gn=HDAC9;refGene_tx
=XM_019825557.2;refGene_gn=HDAC9;refGene_tx=XM_019825558.2;refGene_gn=
HDAC9;refGene_tx=XM_019825560.2;refGene_gn=HDAC9;refGene_tx=XM_0198255
61.2;refGene_gn=HDAC9;refGene_tx=XM_019825562.2;refGene_gn=HDAC9;refGe
ne_tx=XM_019825563.2;refGene_gn=HDAC9;refGene_tx=XM_019825564.2;refGen
e_gn=HDAC9;refGene_tx=XM_019825565.2;refGene_gn=HDAC9;refGene_tx=XM_01
9825566.1;refGene_gn=HDAC9;refGene_tx=XM_019825567.1;refGene_gn=HDAC9;
refGene_tx=XM_019825568.1;refGene_gn=HDAC9;refGene_tx=XM_019825569.1;r
efGene_gn=HDAC9;refGene_tx=XM_019825570.2;refGene_gn=HDAC9;refGene_tx=
XM_019825573.2;refGene_gn=HDAC9;refGene_tx=XM_019825574.2;refGene_gn=H
DAC9;refGene_tx=XM_019825576.2;refGene_gn=HDAC9;refGene_tx=XM_01982557
7.2;refGene_gn=HDAC9;refGene_tx=XM_019825580.2;refGene_gn=HDAC9;refGen
e_tx=XM_019825581.2;refGene_gn=HDAC9;refGene_tx=XM_019825582.2;refGene
_gn=HDAC9;refGene_tx=XM_023250421.1;refGene_gn=HDAC9;refGene_tx=XM_023
250422.1;refGene_gn=HDAC9;refGene_tx=XM_023250423.1
A2 112950901 112951033
refGene_gn=HDAC9;refGene_tx=XM_019825556.1;refGene_gn=HDAC9;refGene_tx
=XM_019825557.2;refGene_gn=HDAC9;refGene_tx=XM_019825558.2;refGene_gn=
HDAC9;refGene_tx=XM_019825560.2;refGene_gn=HDAC9;refGene_tx=XM_0198255
61.2;refGene_gn=HDAC9;refGene_tx=XM_019825562.2;refGene_gn=HDAC9;refGe
ne_tx=XM_019825563.2;refGene_gn=HDAC9;refGene_tx=XM_019825564.2;refGen
e_gn=HDAC9;refGene_tx=XM_019825565.2;refGene_gn=HDAC9;refGene_tx=XM_01
9825566.1;refGene_gn=HDAC9;refGene_tx=XM_019825567.1;refGene_gn=HDAC9;
refGene_tx=XM_019825568.1;refGene_gn=HDAC9;refGene_tx=XM_019825569.1;r
efGene_gn=HDAC9;refGene_tx=XM_019825570.2;refGene_gn=HDAC9;refGene_tx=
XM_019825576.2;refGene_gn=HDAC9;refGene_tx=XM_019825577.2;refGene_gn=H
DAC9;refGene_tx=XM_023250421.1
A2 112954417 112954533

refGene_gn=HDAC9;refGene_tx=XM_019825556.1;refGene_gn=HDAC9;refGene_tx
=XM_019825557.2;refGene_gn=HDAC9;refGene_tx=XM_019825558.2;refGene_gn=
HDAC9;refGene_tx=XM_019825560.2;refGene_gn=HDAC9;refGene_tx=XM_0198255
61.2;refGene_gn=HDAC9;refGene_tx=XM_019825562.2;refGene_gn=HDAC9;refGe
ne_tx=XM_019825563.2;refGene_gn=HDAC9;refGene_tx=XM_019825564.2;refGen
e_gn=HDAC9;refGene_tx=XM_019825565.2;refGene_gn=HDAC9;refGene_tx=XM_01
9825566.1;refGene_gn=HDAC9;refGene_tx=XM_019825567.1;refGene_gn=HDAC9;
refGene_tx=XM_019825568.1;refGene_gn=HDAC9;refGene_tx=XM_019825569.1;r
efGene_gn=HDAC9;refGene_tx=XM_019825570.2;refGene_gn=HDAC9;refGene_tx=
XM_019825573.2;refGene_gn=HDAC9;refGene_tx=XM_019825574.2;refGene_gn=H
DAC9;refGene_tx=XM_019825576.2;refGene_gn=HDAC9;refGene_tx=XM_01982557
7.2;refGene_gn=HDAC9;refGene_tx=XM_019825580.2;refGene_gn=HDAC9;refGen
e_tx=XM_019825581.2;refGene_gn=HDAC9;refGene_tx=XM_019825582.2;refGene
_gn=HDAC9;refGene_tx=XM_023250421.1;refGene_gn=HDAC9;refGene_tx=XM_023
250422.1;refGene_gn=HDAC9;refGene_tx=XM_023250423.1
A2 112965130 112965253
refGene_gn=HDAC9;refGene_tx=XM_019825556.1;refGene_gn=HDAC9;refGene_tx
=XM_019825557.2;refGene_gn=HDAC9;refGene_tx=XM_019825558.2;refGene_gn=
HDAC9;refGene_tx=XM_019825560.2;refGene_gn=HDAC9;refGene_tx=XM_0198255
61.2;refGene_gn=HDAC9;refGene_tx=XM_019825562.2;refGene_gn=HDAC9;refGe
ne_tx=XM_019825563.2;refGene_gn=HDAC9;refGene_tx=XM_019825564.2;refGen
e_gn=HDAC9;refGene_tx=XM_019825565.2;refGene_gn=HDAC9;refGene_tx=XM_01
9825566.1;refGene_gn=HDAC9;refGene_tx=XM_019825567.1;refGene_gn=HDAC9;
refGene_tx=XM_019825568.1;refGene_gn=HDAC9;refGene_tx=XM_019825569.1;r
efGene_gn=HDAC9;refGene_tx=XM_019825570.2;refGene_gn=HDAC9;refGene_tx=
XM_019825573.2;refGene_gn=HDAC9;refGene_tx=XM_019825574.2;refGene_gn=H
DAC9;refGene_tx=XM_019825576.2;refGene_gn=HDAC9;refGene_tx=XM_01982557
7.2;refGene_gn=HDAC9;refGene_tx=XM_019825580.2;refGene_gn=HDAC9;refGen
e_tx=XM_019825581.2;refGene_gn=HDAC9;refGene_tx=XM_019825582.2;refGene
_gn=HDAC9;refGene_tx=XM_023250422.1;refGene_gn=HDAC9;refGene_tx=XM_023
250423.1
A2 112968487 112968698
refGene_gn=HDAC9;refGene_tx=XM_019825556.1;refGene_gn=HDAC9;refGene_tx
=XM_019825557.2;refGene_gn=HDAC9;refGene_tx=XM_019825558.2;refGene_gn=
HDAC9;refGene_tx=XM_019825560.2;refGene_gn=HDAC9;refGene_tx=XM_0198255
61.2;refGene_gn=HDAC9;refGene_tx=XM_019825562.2;refGene_gn=HDAC9;refGe
ne_tx=XM_019825563.2;refGene_gn=HDAC9;refGene_tx=XM_019825564.2;refGen
e_gn=HDAC9;refGene_tx=XM_019825565.2;refGene_gn=HDAC9;refGene_tx=XM_01
9825566.1;refGene_gn=HDAC9;refGene_tx=XM_019825567.1;refGene_gn=HDAC9;
refGene_tx=XM_019825568.1;refGene_gn=HDAC9;refGene_tx=XM_019825569.1;r
efGene_gn=HDAC9;refGene_tx=XM_019825570.2;refGene_gn=HDAC9;refGene_tx=
XM_019825573.2;refGene_gn=HDAC9;refGene_tx=XM_019825574.2;refGene_gn=H
DAC9;refGene_tx=XM_019825576.2;refGene_gn=HDAC9;refGene_tx=XM_01982557
7.2;refGene_gn=HDAC9;refGene_tx=XM_019825580.2;refGene_gn=HDAC9;refGen
e_tx=XM_019825581.2;refGene_gn=HDAC9;refGene_tx=XM_019825582.2;refGene
_gn=HDAC9;refGene_tx=XM_023250421.1;refGene_gn=HDAC9;refGene_tx=XM_023
250422.1;refGene_gn=HDAC9;refGene_tx=XM_023250423.1
A2 112969116 112969334
refGene_gn=HDAC9;refGene_tx=XM_019825556.1;refGene_gn=HDAC9;refGene_tx
=XM_019825557.2;refGene_gn=HDAC9;refGene_tx=XM_019825558.2;refGene_gn=

HDAC9;refGene_tx=XM_019825560.2;refGene_gn=HDAC9;refGene_tx=XM_019825561.2;refGene_gn=HDAC9;refGene_tx=XM_019825562.2;refGene_gn=HDAC9;refGene_tx=XM_019825563.2;refGene_gn=HDAC9;refGene_tx=XM_019825564.2;refGene_gn=HDAC9;refGene_tx=XM_019825565.2;refGene_gn=HDAC9;refGene_tx=XM_019825566.1;refGene_gn=HDAC9;refGene_tx=XM_019825567.1;refGene_gn=HDAC9;refGene_tx=XM_019825568.1;refGene_gn=HDAC9;refGene_tx=XM_019825569.1;refGene_gn=HDAC9;refGene_tx=XM_019825570.2;refGene_gn=HDAC9;refGene_tx=XM_019825573.2;refGene_gn=HDAC9;refGene_tx=XM_019825574.2;refGene_gn=HDAC9;refGene_tx=XM_019825576.2;refGene_gn=HDAC9;refGene_tx=XM_019825577.2;refGene_gn=HDAC9;refGene_tx=XM_019825580.2;refGene_gn=HDAC9;refGene_tx=XM_019825581.2;refGene_gn=HDAC9;refGene_tx=XM_019825582.2;refGene_gn=HDAC9;refGene_tx=XM_023250421.1;refGene_gn=HDAC9;refGene_tx=XM_023250422.1;refGene_gn=HDAC9;refGene_tx=XM_023250423.1
A2 112987229 112987544
refGene_gn=HDAC9;refGene_tx=XM_019825556.1;refGene_gn=HDAC9;refGene_tx=XM_019825557.2;refGene_gn=HDAC9;refGene_tx=XM_019825558.2;refGene_gn=HDAC9;refGene_tx=XM_019825560.2;refGene_gn=HDAC9;refGene_tx=XM_019825561.2;refGene_gn=HDAC9;refGene_tx=XM_019825562.2;refGene_gn=HDAC9;refGene_tx=XM_019825563.2;refGene_gn=HDAC9;refGene_tx=XM_019825564.2;refGene_gn=HDAC9;refGene_tx=XM_019825565.2;refGene_gn=HDAC9;refGene_tx=XM_019825566.1;refGene_gn=HDAC9;refGene_tx=XM_019825567.1;refGene_gn=HDAC9;refGene_tx=XM_019825568.1;refGene_gn=HDAC9;refGene_tx=XM_019825569.1;refGene_gn=HDAC9;refGene_tx=XM_019825570.2;refGene_gn=HDAC9;refGene_tx=XM_019825573.2;refGene_gn=HDAC9;refGene_tx=XM_019825574.2;refGene_gn=HDAC9;refGene_tx=XM_019825576.2;refGene_gn=HDAC9;refGene_tx=XM_019825577.2;refGene_gn=HDAC9;refGene_tx=XM_019825580.2;refGene_gn=HDAC9;refGene_tx=XM_019825581.2;refGene_gn=HDAC9;refGene_tx=XM_019825582.2;refGene_gn=HDAC9;refGene_tx=XM_023250421.1;refGene_gn=HDAC9;refGene_tx=XM_023250422.1;refGene_gn=HDAC9;refGene_tx=XM_023250423.1
A2 113060510 113060688
refGene_gn=HDAC9;refGene_tx=XM_019825556.1;refGene_gn=HDAC9;refGene_tx=XM_019825557.2;refGene_gn=HDAC9;refGene_tx=XM_019825558.2;refGene_gn=HDAC9;refGene_tx=XM_019825560.2;refGene_gn=HDAC9;refGene_tx=XM_019825561.2;refGene_gn=HDAC9;refGene_tx=XM_019825562.2;refGene_gn=HDAC9;refGene_tx=XM_019825563.2;refGene_gn=HDAC9;refGene_tx=XM_019825564.2;refGene_gn=HDAC9;refGene_tx=XM_019825565.2;refGene_gn=HDAC9;refGene_tx=XM_019825566.1;refGene_gn=HDAC9;refGene_tx=XM_019825567.1;refGene_gn=HDAC9;refGene_tx=XM_019825568.1;refGene_gn=HDAC9;refGene_tx=XM_019825569.1;refGene_gn=HDAC9;refGene_tx=XM_019825570.2;refGene_gn=HDAC9;refGene_tx=XM_019825573.2;refGene_gn=HDAC9;refGene_tx=XM_019825574.2;refGene_gn=HDAC9;refGene_tx=XM_023250421.1;refGene_gn=HDAC9;refGene_tx=XM_023250422.1
A2 113076075 113076209
refGene_gn=HDAC9;refGene_tx=XM_019825556.1;refGene_gn=HDAC9;refGene_tx=XM_019825557.2;refGene_gn=HDAC9;refGene_tx=XM_019825558.2;refGene_gn=HDAC9;refGene_tx=XM_019825560.2;refGene_gn=HDAC9;refGene_tx=XM_019825561.2;refGene_gn=HDAC9;refGene_tx=XM_019825562.2;refGene_gn=HDAC9;refGene_tx=XM_019825563.2;refGene_gn=HDAC9;refGene_tx=XM_019825564.2;refGene_gn=HDAC9;refGene_tx=XM_019825565.2;refGene_gn=HDAC9;refGene_tx=XM_019825566.1;refGene_gn=HDAC9;refGene_tx=XM_019825567.1;refGene_gn=HDAC9;

refGene_tx=XM_019825568.1;refGene_gn=HDAC9;refGene_tx=XM_019825569.1;r
efGene_gn=HDAC9;refGene_tx=XM_019825570.2;refGene_gn=HDAC9;refGene_tx=
XM_019825573.2;refGene_gn=HDAC9;refGene_tx=XM_019825574.2;refGene_gn=H
DAC9;refGene_tx=XM_023250421.1;refGene_gn=HDAC9;refGene_tx=XM_02325042
2.1
A2 113090360 113090481
refGene_gn=HDAC9;refGene_tx=XM_019825556.1;refGene_gn=HDAC9;refGene_tx
=XM_019825557.2;refGene_gn=HDAC9;refGene_tx=XM_019825558.2;refGene_gn=
HDAC9;refGene_tx=XM_019825560.2;refGene_gn=HDAC9;refGene_tx=XM_0198255
61.2;refGene_gn=HDAC9;refGene_tx=XM_019825562.2;refGene_gn=HDAC9;refGe
ne_tx=XM_019825563.2;refGene_gn=HDAC9;refGene_tx=XM_019825564.2;refGen
e_gn=HDAC9;refGene_tx=XM_019825565.2;refGene_gn=HDAC9;refGene_tx=XM_01
9825566.1;refGene_gn=HDAC9;refGene_tx=XM_019825567.1;refGene_gn=HDAC9;
refGene_tx=XM_019825568.1;refGene_gn=HDAC9;refGene_tx=XM_019825569.1;r
efGene_gn=HDAC9;refGene_tx=XM_019825570.2;refGene_gn=HDAC9;refGene_tx=
XM_019825573.2;refGene_gn=HDAC9;refGene_tx=XM_019825574.2;refGene_gn=H
DAC9;refGene_tx=XM_023250421.1;refGene_gn=HDAC9;refGene_tx=XM_02325042
2.1
A2 113095890 113095940
refGene_gn=HDAC9;refGene_tx=XM_019825556.1;refGene_gn=HDAC9;refGene_tx
=XM_019825557.2;refGene_gn=HDAC9;refGene_tx=XM_019825558.2;refGene_gn=
HDAC9;refGene_tx=XM_019825560.2;refGene_gn=HDAC9;refGene_tx=XM_0198255
61.2;refGene_gn=HDAC9;refGene_tx=XM_019825562.2;refGene_gn=HDAC9;refGe
ne_tx=XM_019825563.2;refGene_gn=HDAC9;refGene_tx=XM_019825564.2;refGen
e_gn=HDAC9;refGene_tx=XM_019825565.2;refGene_gn=HDAC9;refGene_tx=XM_01
9825566.1;refGene_gn=HDAC9;refGene_tx=XM_019825567.1;refGene_gn=HDAC9;
refGene_tx=XM_019825568.1;refGene_gn=HDAC9;refGene_tx=XM_019825569.1;r
efGene_gn=HDAC9;refGene_tx=XM_019825570.2;refGene_gn=HDAC9;refGene_tx=
XM_019825573.2;refGene_gn=HDAC9;refGene_tx=XM_019825574.2;refGene_gn=H
DAC9;refGene_tx=XM_023250421.1;refGene_gn=HDAC9;refGene_tx=XM_02325042
2.1
A2 113132303 113132411
refGene_gn=HDAC9;refGene_tx=XM_019825556.1;refGene_gn=HDAC9;refGene_tx
=XM_019825557.2;refGene_gn=HDAC9;refGene_tx=XM_019825558.2;refGene_gn=
HDAC9;refGene_tx=XM_019825560.2;refGene_gn=HDAC9;refGene_tx=XM_0198255
61.2;refGene_gn=HDAC9;refGene_tx=XM_019825562.2;refGene_gn=HDAC9;refGe
ne_tx=XM_019825563.2;refGene_gn=HDAC9;refGene_tx=XM_019825564.2;refGen
e_gn=HDAC9;refGene_tx=XM_019825565.2;refGene_gn=HDAC9;refGene_tx=XM_01
9825566.1;refGene_gn=HDAC9;refGene_tx=XM_019825567.1;refGene_gn=HDAC9;
refGene_tx=XM_019825568.1;refGene_gn=HDAC9;refGene_tx=XM_019825569.1;r
efGene_gn=HDAC9;refGene_tx=XM_019825570.2;refGene_gn=HDAC9;refGene_tx=
XM_019825573.2;refGene_gn=HDAC9;refGene_tx=XM_019825574.2;refGene_gn=H
DAC9;refGene_tx=XM_023250421.1;refGene_gn=HDAC9;refGene_tx=XM_02325042
2.1
A2 113165399 113165455
refGene_gn=HDAC9;refGene_tx=XM_019825556.1;refGene_gn=HDAC9;refGene_tx
=XM_019825557.2;refGene_gn=HDAC9;refGene_tx=XM_019825558.2;refGene_gn=
HDAC9;refGene_tx=XM_019825560.2;refGene_gn=HDAC9;refGene_tx=XM_0198255
61.2;refGene_gn=HDAC9;refGene_tx=XM_019825562.2;refGene_gn=HDAC9;refGe
ne_tx=XM_019825563.2;refGene_gn=HDAC9;refGene_tx=XM_019825564.2;refGen

e_gn=HDAC9;refGene_tx=XM_019825565.2;refGene_gn=HDAC9;refGene_tx=XM_01
9825566.1;refGene_gn=HDAC9;refGene_tx=XM_019825567.1;refGene_gn=HDAC9;
refGene_tx=XM_019825568.1;refGene_gn=HDAC9;refGene_tx=XM_019825569.1;r
efGene_gn=HDAC9;refGene_tx=XM_019825570.2;refGene_gn=HDAC9;refGene_tx=
XM_019825573.2;refGene_gn=HDAC9;refGene_tx=XM_019825574.2;refGene_gn=H
DAC9;refGene_tx=XM_023250421.1;refGene_gn=HDAC9;refGene_tx=XM_02325042
2.1
A2 113165699 113165787
refGene_gn=HDAC9;refGene_tx=XM_019825556.1;refGene_gn=HDAC9;refGene_tx
=XM_019825557.2;refGene_gn=HDAC9;refGene_tx=XM_019825558.2;refGene_gn=
HDAC9;refGene_tx=XM_019825560.2;refGene_gn=HDAC9;refGene_tx=XM_0198255
61.2;refGene_gn=HDAC9;refGene_tx=XM_019825562.2;refGene_gn=HDAC9;refGe
ne_tx=XM_019825563.2;refGene_gn=HDAC9;refGene_tx=XM_019825564.2;refGen
e_gn=HDAC9;refGene_tx=XM_019825565.2;refGene_gn=HDAC9;refGene_tx=XM_01
9825566.1;refGene_gn=HDAC9;refGene_tx=XM_019825567.1;refGene_gn=HDAC9;
refGene_tx=XM_019825568.1;refGene_gn=HDAC9;refGene_tx=XM_019825569.1;r
efGene_gn=HDAC9;refGene_tx=XM_019825570.2;refGene_gn=HDAC9;refGene_tx=
XM_019825573.2;refGene_gn=HDAC9;refGene_tx=XM_019825574.2;refGene_gn=H
DAC9;refGene_tx=XM_023250421.1;refGene_gn=HDAC9;refGene_tx=XM_02325042
2.1
A2 113171469 113171589
refGene_gn=HDAC9;refGene_tx=XM_019825556.1;refGene_gn=HDAC9;refGene_tx
=XM_019825557.2;refGene_gn=HDAC9;refGene_tx=XM_019825558.2;refGene_gn=
HDAC9;refGene_tx=XM_019825560.2;refGene_gn=HDAC9;refGene_tx=XM_0198255
61.2;refGene_gn=HDAC9;refGene_tx=XM_019825562.2;refGene_gn=HDAC9;refGe
ne_tx=XM_019825563.2;refGene_gn=HDAC9;refGene_tx=XM_019825564.2;refGen
e_gn=HDAC9;refGene_tx=XM_019825565.2;refGene_gn=HDAC9;refGene_tx=XM_01
9825566.1;refGene_gn=HDAC9;refGene_tx=XM_019825567.1;refGene_gn=HDAC9;
refGene_tx=XM_019825568.1;refGene_gn=HDAC9;refGene_tx=XM_019825569.1;r
efGene_gn=HDAC9;refGene_tx=XM_019825570.2;refGene_gn=HDAC9;refGene_tx=
XM_019825573.2;refGene_gn=HDAC9;refGene_tx=XM_019825574.2;refGene_gn=H
DAC9;refGene_tx=XM_023250421.1;refGene_gn=HDAC9;refGene_tx=XM_02325042
2.1
A2 113171904 113172002
refGene_gn=HDAC9;refGene_tx=XM_019825556.1;refGene_gn=HDAC9;refGene_tx
=XM_019825557.2;refGene_gn=HDAC9;refGene_tx=XM_019825558.2;refGene_gn=
HDAC9;refGene_tx=XM_019825560.2;refGene_gn=HDAC9;refGene_tx=XM_0198255
61.2;refGene_gn=HDAC9;refGene_tx=XM_019825562.2;refGene_gn=HDAC9;refGe
ne_tx=XM_019825563.2;refGene_gn=HDAC9;refGene_tx=XM_019825564.2;refGen
e_gn=HDAC9;refGene_tx=XM_019825565.2;refGene_gn=HDAC9;refGene_tx=XM_01
9825566.1;refGene_gn=HDAC9;refGene_tx=XM_019825567.1;refGene_gn=HDAC9;
refGene_tx=XM_019825568.1;refGene_gn=HDAC9;refGene_tx=XM_019825569.1;r
efGene_gn=HDAC9;refGene_tx=XM_019825570.2;refGene_gn=HDAC9;refGene_tx=
XM_019825573.2;refGene_gn=HDAC9;refGene_tx=XM_019825574.2;refGene_gn=H
DAC9;refGene_tx=XM_023250421.1;refGene_gn=HDAC9;refGene_tx=XM_02325042
2.1
A2 113223923 113224042
refGene_gn=HDAC9;refGene_tx=XM_019825556.1;refGene_gn=HDAC9;refGene_tx
=XM_019825557.2;refGene_gn=HDAC9;refGene_tx=XM_019825558.2;refGene_gn=
HDAC9;refGene_tx=XM_019825560.2;refGene_gn=HDAC9;refGene_tx=XM_0198255

61.2;refGene_gn=HDAC9;refGene_tx=XM_019825562.2;refGene_gn=HDAC9;refGene_tx=XM_019825563.2;refGene_gn=HDAC9;refGene_tx=XM_019825564.2;refGene_gn=HDAC9;refGene_tx=XM_019825565.2;refGene_gn=HDAC9;refGene_tx=XM_019825566.1;refGene_gn=HDAC9;refGene_tx=XM_019825567.1;refGene_gn=HDAC9;refGene_tx=XM_019825568.1;refGene_gn=HDAC9;refGene_tx=XM_019825569.1;refGene_gn=HDAC9;refGene_tx=XM_019825570.2;refGene_gn=HDAC9;refGene_tx=XM_019825573.2;refGene_gn=HDAC9;refGene_tx=XM_019825574.2;refGene_gn=HDAC9;refGene_tx=XM_023250421.1;refGene_gn=HDAC9;refGene_tx=XM_023250422.1
A2 113285090 113285224
refGene_gn=HDAC9;refGene_tx=XM_019825556.1;refGene_gn=HDAC9;refGene_tx=XM_019825557.2;refGene_gn=HDAC9;refGene_tx=XM_019825558.2;refGene_gn=HDAC9;refGene_tx=XM_019825560.2;refGene_gn=HDAC9;refGene_tx=XM_019825561.2;refGene_gn=HDAC9;refGene_tx=XM_019825562.2;refGene_gn=HDAC9;refGene_tx=XM_019825563.2;refGene_gn=HDAC9;refGene_tx=XM_019825564.2;refGene_gn=HDAC9;refGene_tx=XM_019825565.2;refGene_gn=HDAC9;refGene_tx=XM_019825566.1;refGene_gn=HDAC9;refGene_tx=XM_019825567.1;refGene_gn=HDAC9;refGene_tx=XM_019825568.1;refGene_gn=HDAC9;refGene_tx=XM_019825569.1;refGene_gn=HDAC9;refGene_tx=XM_019825570.2;refGene_gn=HDAC9;refGene_tx=XM_019825573.2;refGene_gn=HDAC9;refGene_tx=XM_019825574.2;refGene_gn=HDAC9;refGene_tx=XM_023250421.1;refGene_gn=HDAC9;refGene_tx=XM_023250422.1
A2 113308982 113309067
refGene_gn=HDAC9;refGene_tx=XM_019825556.1;refGene_gn=HDAC9;refGene_tx=XM_019825557.2;refGene_gn=HDAC9;refGene_tx=XM_019825558.2;refGene_gn=HDAC9;refGene_tx=XM_019825560.2;refGene_gn=HDAC9;refGene_tx=XM_019825561.2;refGene_gn=HDAC9;refGene_tx=XM_019825562.2;refGene_gn=HDAC9;refGene_tx=XM_019825563.2;refGene_gn=HDAC9;refGene_tx=XM_019825564.2;refGene_gn=HDAC9;refGene_tx=XM_019825565.2;refGene_gn=HDAC9;refGene_tx=XM_019825566.1;refGene_gn=HDAC9;refGene_tx=XM_019825567.1;refGene_gn=HDAC9;refGene_tx=XM_019825568.1;refGene_gn=HDAC9;refGene_tx=XM_019825569.1;refGene_gn=HDAC9;refGene_tx=XM_019825570.2;refGene_gn=HDAC9;refGene_tx=XM_019825573.2;refGene_gn=HDAC9;refGene_tx=XM_019825574.2;refGene_gn=HDAC9;refGene_tx=XM_023250421.1;refGene_gn=HDAC9;refGene_tx=XM_023250422.1
A2 113340972 113341150
refGene_gn=HDAC9;refGene_tx=XM_019825556.1;refGene_gn=HDAC9;refGene_tx=XM_019825557.2;refGene_gn=HDAC9;refGene_tx=XM_019825558.2;refGene_gn=HDAC9;refGene_tx=XM_019825560.2;refGene_gn=HDAC9;refGene_tx=XM_019825561.2;refGene_gn=HDAC9;refGene_tx=XM_019825562.2;refGene_gn=HDAC9;refGene_tx=XM_019825563.2;refGene_gn=HDAC9;refGene_tx=XM_019825564.2;refGene_gn=HDAC9;refGene_tx=XM_019825565.2;refGene_gn=HDAC9;refGene_tx=XM_019825566.1;refGene_gn=HDAC9;refGene_tx=XM_019825567.1;refGene_gn=HDAC9;refGene_tx=XM_019825568.1;refGene_gn=HDAC9;refGene_tx=XM_019825569.1;refGene_gn=HDAC9;refGene_tx=XM_019825570.2;refGene_gn=HDAC9;refGene_tx=XM_019825573.2;refGene_gn=HDAC9;refGene_tx=XM_019825574.2;refGene_gn=HDAC9;refGene_tx=XM_023250421.1;refGene_gn=HDAC9;refGene_tx=XM_023250422.1
A2 113361776 113361816
refGene_gn=HDAC9;refGene_tx=XM_019825556.1;refGene_gn=HDAC9;refGene_tx

=XM_019825557.2;refGene_gn=HDAC9;refGene_tx=XM_019825558.2;refGene_gn=HDAC9;refGene_tx=XM_019825560.2;refGene_gn=HDAC9;refGene_tx=XM_019825561.2;refGene_gn=HDAC9;refGene_tx=XM_019825562.2;refGene_gn=HDAC9;refGene_tx=XM_019825563.2;refGene_gn=HDAC9;refGene_tx=XM_019825564.2;refGene_gn=HDAC9;refGene_tx=XM_019825565.2;refGene_gn=HDAC9;refGene_tx=XM_019825566.1;refGene_gn=HDAC9;refGene_tx=XM_019825567.1;refGene_gn=HDAC9;refGene_tx=XM_019825568.1;refGene_gn=HDAC9;refGene_tx=XM_019825569.1;refGene_gn=HDAC9;refGene_tx=XM_019825570.2;refGene_gn=HDAC9;refGene_tx=XM_019825573.2;refGene_gn=HDAC9;refGene_tx=XM_019825574.2;refGene_gn=HDAC9;refGene_tx=XM_023250421.1;refGene_gn=HDAC9;refGene_tx=XM_023250422.1
A2 113454010 113454613
refGene_gn=TWIST1;refGene_tx=XM_003982857.5
A2 113486300 113486795
refGene_gn=FERD3L;refGene_tx=XM_019826099.1
A2 113917574 113917971
refGene_gn=TWISTNB;refGene_tx=XM_003982831.4
A2 113918856 113919065
refGene_gn=TWISTNB;refGene_tx=XM_003982831.4
A2 113922231 113922373
refGene_gn=TWISTNB;refGene_tx=XM_003982831.4
A2 113947135 113947389
refGene_gn=TWISTNB;refGene_tx=XM_003982831.4
A2 113961595 113961655
refGene_gn=TMEM196;refGene_tx=XM_003982832.4;refGene_gn=TMEM196;refGene_tx=XM_019825583.2;refGene_gn=TMEM196;refGene_tx=XM_019825584.2;refGene_gn=TMEM196;refGene_tx=XM_019825585.2
A2 113964804 113964878
refGene_gn=TMEM196;refGene_tx=XM_019825583.2;refGene_gn=TMEM196;refGene_tx=XM_019825584.2
A2 113966059 113966314
refGene_gn=TMEM196;refGene_tx=XM_003982832.4;refGene_gn=TMEM196;refGene_tx=XM_019825583.2;refGene_gn=TMEM196;refGene_tx=XM_019825584.2;refGene_gn=TMEM196;refGene_tx=XM_019825585.2
A2 113970377 113970434
refGene_gn=TMEM196;refGene_tx=XM_003982832.4;refGene_gn=TMEM196;refGene_tx=XM_019825583.2;refGene_gn=TMEM196;refGene_tx=XM_019825584.2;refGene_gn=TMEM196;refGene_tx=XM_019825585.2
A2 114010573 114010738
refGene_gn=TMEM196;refGene_tx=XM_003982832.4;refGene_gn=TMEM196;refGene_tx=XM_019825583.2;refGene_gn=TMEM196;refGene_tx=XM_019825584.2;refGene_gn=TMEM196;refGene_tx=XM_019825585.2
A2 114277467 114277680
refGene_gn=MACC1;refGene_tx=XM_003982833.4
A2 114291487 114291676
refGene_gn=MACC1;refGene_tx=XM_003982833.4
A2 114295141 114297183
refGene_gn=MACC1;refGene_tx=XM_003982833.4
A2 114298659 114298774
refGene_gn=MACC1;refGene_tx=XM_003982833.4

A2 114466749 114466876
refGene_gn=ITGB8;refGene_tx=XM_003982834.5
A2 114492794 114492880
refGene_gn=ITGB8;refGene_tx=XM_003982834.5
A2 114498426 114498598
refGene_gn=ITGB8;refGene_tx=XM_003982834.5
A2 114513344 114513591
refGene_gn=ITGB8;refGene_tx=XM_003982834.5
A2 114514902 114515068
refGene_gn=ITGB8;refGene_tx=XM_003982834.5
A2 114515663 114515822
refGene_gn=ITGB8;refGene_tx=XM_003982834.5
A2 114526051 114526147
refGene_gn=ITGB8;refGene_tx=XM_003982834.5
A2 114529404 114529494
refGene_gn=ITGB8;refGene_tx=XM_003982834.5
A2 114533547 114533682
refGene_gn=ITGB8;refGene_tx=XM_003982834.5
A2 114536224 114536630
refGene_gn=ITGB8;refGene_tx=XM_003982834.5
A2 114539128 114539354
refGene_gn=ITGB8;refGene_tx=XM_003982834.5
A2 114542029 114542136
refGene_gn=ITGB8;refGene_tx=XM_003982834.5
A2 114545088 114545252
refGene_gn=ITGB8;refGene_tx=XM_003982834.5
A2 114545348 114545471
refGene_gn=ITGB8;refGene_tx=XM_003982834.5
A2 114759236 114759289
refGene_gn=ABCB5;refGene_tx=XM_023250426.1
A2 114761514 114761569
refGene_gn=ABCB5;refGene_tx=XM_023250426.1
A2 114763169 114763320
refGene_gn=ABCB5;refGene_tx=XM_023250426.1
A2 114766987 114767042
refGene_gn=ABCB5;refGene_tx=XM_023250426.1
A2 114775970 114776162
refGene_gn=ABCB5;refGene_tx=XM_023250426.1
A2 114776247 114776419
refGene_gn=ABCB5;refGene_tx=XM_023250426.1
A2 114778816 114778941
refGene_gn=ABCB5;refGene_tx=XM_023250426.1
A2 114779021 114779199
refGene_gn=ABCB5;refGene_tx=XM_023250426.1
A2 114780507 114780621
refGene_gn=ABCB5;refGene_tx=XM_023250426.1
A2 114780931 114781042
refGene_gn=ABCB5;refGene_tx=XM_023250426.1
A2 114784280 114784406
refGene_gn=ABCB5;refGene_tx=XM_023250426.1

A2 114785697 114785901
refGene_gn=ABCB5;refGene_tx=XM_023250426.1
A2 114792459 114792630
refGene_gn=ABCB5;refGene_tx=XM_023250426.1
A2 114810671 114810833
refGene_gn=ABCB5;refGene_tx=XM_023250426.1
A2 114814369 114814510
refGene_gn=ABCB5;refGene_tx=XM_023250426.1
A2 114827056 114827200
refGene_gn=ABCB5;refGene_tx=XM_023250426.1
A2 114828114 114828219
refGene_gn=ABCB5;refGene_tx=XM_023250426.1
A2 114828395 114828473
refGene_gn=ABCB5;refGene_tx=XM_023250426.1
A2 114832680 114832764
refGene_gn=ABCB5;refGene_tx=XM_023250426.1
A2 114841020 114841224
refGene_gn=ABCB5;refGene_tx=XM_023250426.1
A2 114844135 114844236
refGene_gn=ABCB5;refGene_tx=XM_023250426.1
A2 114845111 114845252
refGene_gn=ABCB5;refGene_tx=XM_023250426.1
A2 114857803 114857960
refGene_gn=ABCB5;refGene_tx=XM_023250426.1
A2 114860627 114860825
refGene_gn=ABCB5;refGene_tx=XM_023250426.1
A2 114865496 114865703
refGene_gn=ABCB5;refGene_tx=XM_023250426.1
A2 114871760 114871907
refGene_gn=ABCB5;refGene_tx=XM_023250426.1
A2 114873441 114873639
refGene_gn=ABCB5;refGene_tx=XM_023250426.1
A2 114898495 114899998
refGene_gn=SP8;refGene_tx=XM_023250427.1;refGene_gn=SP8;refGene_tx=XM_
023250428.1
A2 114901012 114901033
refGene_gn=SP8;refGene_tx=XM_023250427.1
A2 115485078 115485085
refGene_gn=SP4;refGene_tx=XM_023250429.1;refGene_gn=SP4;refGene_tx=XM_
023250431.1
A2 115485495 115485620
refGene_gn=SP4;refGene_tx=XM_023250429.1;refGene_gn=SP4;refGene_tx=XM_
023250431.1
A2 115486146 115487701
refGene_gn=SP4;refGene_tx=XM_023250429.1;refGene_gn=SP4;refGene_tx=XM_
023250431.1
A2 115532676 115532905
refGene_gn=SP4;refGene_tx=XM_023250429.1;refGene_gn=SP4;refGene_tx=XM_
023250431.1
A2 115540738 115540938

refGene_gn=SP4;refGene_tx=XM_023250429.1;refGene_gn=SP4;refGene_tx=XM_023250431.1
A2	115548566	115548649
refGene_gn=SP4;refGene_tx=XM_023250431.1
A2	115564237	115564485
refGene_gn=SP4;refGene_tx=XM_023250429.1
A2	115591829	115592195
refGene_gn=DNAH11;refGene_tx=XM_023250435.1
A2	115593711	115593855
refGene_gn=DNAH11;refGene_tx=XM_023250435.1
A2	115610003	115610200
refGene_gn=DNAH11;refGene_tx=XM_023250435.1
A2	115611001	115611200
refGene_gn=DNAH11;refGene_tx=XM_023250435.1
A2	115611936	115612036
refGene_gn=DNAH11;refGene_tx=XM_023250435.1
A2	115615595	115615807
refGene_gn=DNAH11;refGene_tx=XM_023250435.1
A2	115623531	115623762
refGene_gn=DNAH11;refGene_tx=XM_023250435.1
A2	115625391	115625513
refGene_gn=DNAH11;refGene_tx=XM_023250435.1
A2	115625514	115625557
refGene_gn=DNAH11;refGene_tx=XM_023250435.1
A2	115648147	115648264
refGene_gn=DNAH11;refGene_tx=XM_023250435.1
A2	115654663	115654801
refGene_gn=DNAH11;refGene_tx=XM_023250435.1
A2	115655112	115655237
refGene_gn=DNAH11;refGene_tx=XM_023250435.1
A2	115655786	115655982
refGene_gn=DNAH11;refGene_tx=XM_023250435.1
A2	115662100	115662205
refGene_gn=DNAH11;refGene_tx=XM_023250435.1
A2	115662578	115662971
refGene_gn=DNAH11;refGene_tx=XM_023250435.1
A2	115670642	115670975
refGene_gn=DNAH11;refGene_tx=XM_023250435.1
A2	115671198	115671453
refGene_gn=DNAH11;refGene_tx=XM_023250435.1
A2	115671533	115671703
refGene_gn=DNAH11;refGene_tx=XM_023250435.1
A2	115671958	115672181
refGene_gn=DNAH11;refGene_tx=XM_023250435.1
A2	115676233	115676350
refGene_gn=DNAH11;refGene_tx=XM_023250435.1
A2	115676479	115676566
refGene_gn=DNAH11;refGene_tx=XM_023250435.1
A2	115680407	115680566
refGene_gn=DNAH11;refGene_tx=XM_023250435.1

A2 115681931 115682015
refGene_gn=DNAH11;refGene_tx=XM_023250435.1
A2 115683807 115683966
refGene_gn=DNAH11;refGene_tx=XM_023250435.1
A2 115686858 115686981
refGene_gn=DNAH11;refGene_tx=XM_023250435.1
A2 115687778 115687901
refGene_gn=DNAH11;refGene_tx=XM_023250435.1
A2 115689177 115689402
refGene_gn=DNAH11;refGene_tx=XM_023250435.1
A2 115690898 115690990
refGene_gn=DNAH11;refGene_tx=XM_023250435.1
A2 115692531 115692658
refGene_gn=DNAH11;refGene_tx=XM_023250435.1
A2 115710525 115710672
refGene_gn=DNAH11;refGene_tx=XM_023250435.1
A2 115713943 115714177
refGene_gn=DNAH11;refGene_tx=XM_023250435.1
A2 115738905 115739037
refGene_gn=DNAH11;refGene_tx=XM_023250435.1
A2 115740865 115741026
refGene_gn=DNAH11;refGene_tx=XM_023250435.1
A2 115744628 115744785
refGene_gn=DNAH11;refGene_tx=XM_023250435.1
A2 115744906 115745052
refGene_gn=DNAH11;refGene_tx=XM_023250435.1
A2 115748221 115748338
refGene_gn=DNAH11;refGene_tx=XM_023250435.1
A2 115753476 115753615
refGene_gn=DNAH11;refGene_tx=XM_023250435.1
A2 115756728 115756821
refGene_gn=DNAH11;refGene_tx=XM_023250435.1
A2 115758552 115758747
refGene_gn=DNAH11;refGene_tx=XM_023250435.1
A2 115760117 115760195
refGene_gn=DNAH11;refGene_tx=XM_023250435.1
A2 115762124 115762261
refGene_gn=DNAH11;refGene_tx=XM_023250435.1
A2 115764491 115764642
refGene_gn=DNAH11;refGene_tx=XM_023250435.1
A2 115766052 115766201
refGene_gn=DNAH11;refGene_tx=XM_023250435.1
A2 115774771 115774922
refGene_gn=DNAH11;refGene_tx=XM_023250435.1
A2 115777773 115777905
refGene_gn=DNAH11;refGene_tx=XM_023250435.1
A2 115782776 115782954
refGene_gn=DNAH11;refGene_tx=XM_023250435.1
A2 115787975 115788034
refGene_gn=DNAH11;refGene_tx=XM_023250435.1

A2 115788035 115788177
refGene_gn=DNAH11;refGene_tx=XM_023250435.1
A2 115790824 115790990
refGene_gn=DNAH11;refGene_tx=XM_023250435.1
A2 115791650 115791753
refGene_gn=DNAH11;refGene_tx=XM_023250435.1
A2 115793451 115793691
refGene_gn=DNAH11;refGene_tx=XM_023250435.1
A2 115795091 115795253
refGene_gn=DNAH11;refGene_tx=XM_023250435.1
A2 115795524 115795718
refGene_gn=DNAH11;refGene_tx=XM_023250435.1
A2 115800641 115800804
refGene_gn=DNAH11;refGene_tx=XM_023250435.1
A2 115802571 115802695
refGene_gn=DNAH11;refGene_tx=XM_023250435.1
A2 115803144 115803287
refGene_gn=DNAH11;refGene_tx=XM_023250435.1
A2 115822192 115822354
refGene_gn=DNAH11;refGene_tx=XM_023250435.1
A2 115830636 115830870
refGene_gn=DNAH11;refGene_tx=XM_023250435.1
A2 115834597 115834744
refGene_gn=DNAH11;refGene_tx=XM_023250435.1
A2 115840907 115841021
refGene_gn=DNAH11;refGene_tx=XM_023250435.1
A2 115843280 115843424
refGene_gn=DNAH11;refGene_tx=XM_023250435.1
A2 115844037 115844220
refGene_gn=DNAH11;refGene_tx=XM_023250435.1
A2 115846182 115846284
refGene_gn=DNAH11;refGene_tx=XM_023250435.1
A2 115851382 115851517
refGene_gn=DNAH11;refGene_tx=XM_023250435.1
A2 115856570 115856741
refGene_gn=DNAH11;refGene_tx=XM_023250435.1
A2 115866122 115866358
refGene_gn=DNAH11;refGene_tx=XM_023250435.1
A2 115867562 115867685
refGene_gn=DNAH11;refGene_tx=XM_023250435.1
A2 115888058 115888263
refGene_gn=DNAH11;refGene_tx=XM_023250435.1
A2 115899658 115899823
refGene_gn=DNAH11;refGene_tx=XM_023250435.1
A2 115901777 115901918
refGene_gn=DNAH11;refGene_tx=XM_023250435.1
A2 115905298 115905469
refGene_gn=DNAH11;refGene_tx=XM_023250435.1
A2 115907997 115908120
refGene_gn=DNAH11;refGene_tx=XM_023250435.1

A2	115909534	115909728
refGene_gn=DNAH11;refGene_tx=XM_023250435.1
A2	115910889	115911038
refGene_gn=DNAH11;refGene_tx=XM_023250435.1
A2	115911889	115912017
refGene_gn=DNAH11;refGene_tx=XM_023250435.1
A2	115915887	115916115
refGene_gn=DNAH11;refGene_tx=XM_023250435.1
A2	115921915	115922107
refGene_gn=DNAH11;refGene_tx=XM_023250435.1
A2	115924985	115925105
refGene_gn=DNAH11;refGene_tx=XM_023250435.1
A2	115934777	115935020
refGene_gn=DNAH11;refGene_tx=XM_023250435.1
A2	115937859	115938042
refGene_gn=DNAH11;refGene_tx=XM_023250435.1
A2	115938121	115938237
refGene_gn=DNAH11;refGene_tx=XM_023250435.1
A2	115945757	115945870
refGene_gn=DNAH11;refGene_tx=XM_023250435.1
A2	115946326	115946467
refGene_gn=DNAH11;refGene_tx=XM_023250435.1
A2	115947137	115947385
refGene_gn=DNAH11;refGene_tx=XM_023250435.1
A2	115948184	115948215
refGene_gn=CDCA7L;refGene_tx=XM_019825596.2;refGene_gn=CDCA7L;refGene_tx=XM_019825597.2;refGene_gn=CDCA7L;refGene_tx=XM_023250432.1;refGene_gn=CDCA7L;refGene_tx=XM_023250433.1;refGene_gn=CDCA7L;refGene_tx=XM_023250434.1
A2	115948718	115948855
refGene_gn=CDCA7L;refGene_tx=XM_019825596.2;refGene_gn=CDCA7L;refGene_tx=XM_019825597.2;refGene_gn=CDCA7L;refGene_tx=XM_023250432.1;refGene_gn=CDCA7L;refGene_tx=XM_023250433.1;refGene_gn=CDCA7L;refGene_tx=XM_023250434.1
A2	115949728	115949878
refGene_gn=CDCA7L;refGene_tx=XM_019825596.2;refGene_gn=CDCA7L;refGene_tx=XM_019825597.2;refGene_gn=CDCA7L;refGene_tx=XM_023250432.1;refGene_gn=CDCA7L;refGene_tx=XM_023250433.1;refGene_gn=CDCA7L;refGene_tx=XM_023250434.1
A2	115951397	115951523
refGene_gn=CDCA7L;refGene_tx=XM_019825596.2;refGene_gn=CDCA7L;refGene_tx=XM_019825597.2;refGene_gn=CDCA7L;refGene_tx=XM_023250432.1;refGene_gn=CDCA7L;refGene_tx=XM_023250433.1;refGene_gn=CDCA7L;refGene_tx=XM_023250434.1
A2	115952184	115952355
refGene_gn=CDCA7L;refGene_tx=XM_019825596.2;refGene_gn=CDCA7L;refGene_tx=XM_019825597.2;refGene_gn=CDCA7L;refGene_tx=XM_023250432.1;refGene_gn=CDCA7L;refGene_tx=XM_023250433.1;refGene_gn=CDCA7L;refGene_tx=XM_023250434.1
A2	115952470	115952542

refGene_gn=CDCA7L;refGene_tx=XM_019825596.2;refGene_gn=CDCA7L;refGene_tx=XM_019825597.2;refGene_gn=CDCA7L;refGene_tx=XM_023250432.1;refGene_gn=CDCA7L;refGene_tx=XM_023250433.1;refGene_gn=CDCA7L;refGene_tx=XM_023250434.1
A2 115954569 115954938
refGene_gn=CDCA7L;refGene_tx=XM_019825596.2;refGene_gn=CDCA7L;refGene_tx=XM_019825597.2;refGene_gn=CDCA7L;refGene_tx=XM_023250432.1;refGene_gn=CDCA7L;refGene_tx=XM_023250433.1;refGene_gn=CDCA7L;refGene_tx=XM_023250434.1
A2 115958734 115958872
refGene_gn=CDCA7L;refGene_tx=XM_019825596.2;refGene_gn=CDCA7L;refGene_tx=XM_019825597.2;refGene_gn=CDCA7L;refGene_tx=XM_023250432.1;refGene_gn=CDCA7L;refGene_tx=XM_023250433.1;refGene_gn=CDCA7L;refGene_tx=XM_023250434.1
A2 115964099 115964240
refGene_gn=CDCA7L;refGene_tx=XM_019825596.2;refGene_gn=CDCA7L;refGene_tx=XM_019825597.2;refGene_gn=CDCA7L;refGene_tx=XM_023250432.1;refGene_gn=CDCA7L;refGene_tx=XM_023250433.1
A2 115979954 115980002
refGene_gn=CDCA7L;refGene_tx=XM_019825596.2;refGene_gn=CDCA7L;refGene_tx=XM_019825597.2;refGene_gn=CDCA7L;refGene_tx=XM_023250432.1
A2 115985496 115985520
refGene_gn=CDCA7L;refGene_tx=XM_023250433.1;refGene_gn=CDCA7L;refGene_tx=XM_023250434.1
A2 116142267 116142386
refGene_gn=RAPGEF5;refGene_tx=XM_023250436.1;refGene_gn=RAPGEF5;refGene_tx=XM_023250437.1
A2 116146297 116146352
refGene_gn=RAPGEF5;refGene_tx=XM_023250436.1;refGene_gn=RAPGEF5;refGene_tx=XM_023250437.1
A2 116151489 116151554
refGene_gn=RAPGEF5;refGene_tx=XM_023250436.1;refGene_gn=RAPGEF5;refGene_tx=XM_023250437.1
A2 116157201 116157289
refGene_gn=RAPGEF5;refGene_tx=XM_023250436.1;refGene_gn=RAPGEF5;refGene_tx=XM_023250437.1
A2 116158079 116158130
refGene_gn=RAPGEF5;refGene_tx=XM_023250436.1;refGene_gn=RAPGEF5;refGene_tx=XM_023250437.1
A2 116160509 116160600
refGene_gn=RAPGEF5;refGene_tx=XM_023250436.1;refGene_gn=RAPGEF5;refGene_tx=XM_023250437.1
A2 116165826 116166005
refGene_gn=RAPGEF5;refGene_tx=XM_023250436.1;refGene_gn=RAPGEF5;refGene_tx=XM_023250437.1
A2 116167987 116168110
refGene_gn=RAPGEF5;refGene_tx=XM_023250436.1;refGene_gn=RAPGEF5;refGene_tx=XM_023250437.1
A2 116171291 116171389
refGene_gn=RAPGEF5;refGene_tx=XM_023250436.1;refGene_gn=RAPGEF5;refGen

e_tx=XM_023250437.1
A2	116175107	116175257
refGene_gn=RAPGEF5;refGene_tx=XM_023250436.1;refGene_gn=RAPGEF5;refGene_tx=XM_023250437.1
A2	116177585	116177664
refGene_gn=RAPGEF5;refGene_tx=XM_023250436.1;refGene_gn=RAPGEF5;refGene_tx=XM_023250437.1
A2	116178635	116178666
refGene_gn=RAPGEF5;refGene_tx=XM_023250436.1;refGene_gn=RAPGEF5;refGene_tx=XM_023250437.1
A2	116180836	116180934
refGene_gn=RAPGEF5;refGene_tx=XM_023250436.1;refGene_gn=RAPGEF5;refGene_tx=XM_023250437.1
A2	116182114	116182259
refGene_gn=RAPGEF5;refGene_tx=XM_023250436.1;refGene_gn=RAPGEF5;refGene_tx=XM_023250437.1
A2	116187777	116187856
refGene_gn=RAPGEF5;refGene_tx=XM_023250436.1;refGene_gn=RAPGEF5;refGene_tx=XM_023250437.1
A2	116211624	116211713
refGene_gn=RAPGEF5;refGene_tx=XM_023250436.1;refGene_gn=RAPGEF5;refGene_tx=XM_023250437.1
A2	116212158	116212292
refGene_gn=RAPGEF5;refGene_tx=XM_023250436.1;refGene_gn=RAPGEF5;refGene_tx=XM_023250437.1
A2	116233117	116233192
refGene_gn=RAPGEF5;refGene_tx=XM_023250436.1
A2	116240160	116240286
refGene_gn=RAPGEF5;refGene_tx=XM_023250436.1;refGene_gn=RAPGEF5;refGene_tx=XM_023250437.1
A2	116255138	116255212
refGene_gn=RAPGEF5;refGene_tx=XM_023250436.1;refGene_gn=RAPGEF5;refGene_tx=XM_023250437.1
A2	116279893	116279942
refGene_gn=RAPGEF5;refGene_tx=XM_023250436.1;refGene_gn=RAPGEF5;refGene_tx=XM_023250437.1
A2	116302865	116302932
refGene_gn=RAPGEF5;refGene_tx=XM_023250436.1;refGene_gn=RAPGEF5;refGene_tx=XM_023250437.1
A2	116319079	116319248
refGene_gn=RAPGEF5;refGene_tx=XM_023250436.1;refGene_gn=RAPGEF5;refGene_tx=XM_023250437.1
A2	116320923	116320975
refGene_gn=RAPGEF5;refGene_tx=XM_023250436.1;refGene_gn=RAPGEF5;refGene_tx=XM_023250437.1
A2	116632391	116632402
refGene_gn=IL6;refGene_tx=XM_019814895.1
A2	116633437	116633496
refGene_gn=IL6;refGene_tx=NM_001009211.1;refGene_gn=IL6;refGene_tx=XM_019814895.1

A2 116633660 116633839
refGene_gn=IL6;refGene_tx=NM_001009211.1;refGene_gn=IL6;refGene_tx=XM_019814895.1
A2 116634939 116635053
refGene_gn=IL6;refGene_tx=NM_001009211.1;refGene_gn=IL6;refGene_tx=XM_019814895.1
A2 116635769 116635916
refGene_gn=IL6;refGene_tx=NM_001009211.1;refGene_gn=IL6;refGene_tx=XM_019814895.1
A2 116637127 116637295
refGene_gn=IL6;refGene_tx=NM_001009211.1;refGene_gn=IL6;refGene_tx=XM_019814895.1
A2 116688204 116688220
refGene_gn=TOMM7;refGene_tx=XM_011280427.3
A2 116692552 116692601
refGene_gn=TOMM7;refGene_tx=XM_011280427.3
A2 116696454 116696557
refGene_gn=TOMM7;refGene_tx=XM_011280427.3
A2 116796664 116797239
refGene_gn=FAM126A;refGene_tx=XM_003982838.5
A2 116798060 116798329
refGene_gn=FAM126A;refGene_tx=XM_006929237.3
A2 116811509 116811669
refGene_gn=FAM126A;refGene_tx=XM_003982838.5;refGene_gn=FAM126A;refGene_tx=XM_006929237.3
A2 116812408 116812496
refGene_gn=FAM126A;refGene_tx=XM_003982838.5;refGene_gn=FAM126A;refGene_tx=XM_006929237.3
A2 116815555 116815672
refGene_gn=FAM126A;refGene_tx=XM_003982838.5;refGene_gn=FAM126A;refGene_tx=XM_006929237.3
A2 116829035 116829131
refGene_gn=FAM126A;refGene_tx=XM_003982838.5;refGene_gn=FAM126A;refGene_tx=XM_006929237.3
A2 116829554 116829670
refGene_gn=FAM126A;refGene_tx=XM_003982838.5;refGene_gn=FAM126A;refGene_tx=XM_006929237.3
A2 116830226 116830307
refGene_gn=FAM126A;refGene_tx=XM_003982838.5;refGene_gn=FAM126A;refGene_tx=XM_006929237.3
A2 116831423 116831603
refGene_gn=FAM126A;refGene_tx=XM_003982838.5;refGene_gn=FAM126A;refGene_tx=XM_006929237.3
A2 116838876 116838978
refGene_gn=FAM126A;refGene_tx=XM_003982838.5;refGene_gn=FAM126A;refGene_tx=XM_006929237.3
A2 116845679 116845730
refGene_gn=FAM126A;refGene_tx=XM_003982838.5;refGene_gn=FAM126A;refGene_tx=XM_006929237.3
A2 116965419 116965539

refGene_gn=KLHL7;refGene_tx=XM_003982839.5;refGene_gn=KLHL7;refGene_tx=XM_023250441.1
A2 116966647 116966665
refGene_gn=KLHL7;refGene_tx=XM_019825599.1
A2 116986121 116986224
refGene_gn=KLHL7;refGene_tx=XM_003982839.5;refGene_gn=KLHL7;refGene_tx=XM_003982840.4;refGene_gn=KLHL7;refGene_tx=XM_011280428.3;refGene_gn=KLHL7;refGene_tx=XM_011280429.3;refGene_gn=KLHL7;refGene_tx=XM_019825599.1;refGene_gn=KLHL7;refGene_tx=XM_023250440.1;refGene_gn=KLHL7;refGene_tx=XM_023250441.1
A2 116987040 116987134
refGene_gn=KLHL7;refGene_tx=XM_003982839.5;refGene_gn=KLHL7;refGene_tx=XM_003982840.4;refGene_gn=KLHL7;refGene_tx=XM_011280428.3;refGene_gn=KLHL7;refGene_tx=XM_011280429.3;refGene_gn=KLHL7;refGene_tx=XM_019825599.1;refGene_gn=KLHL7;refGene_tx=XM_023250440.1;refGene_gn=KLHL7;refGene_tx=XM_023250441.1
A2 116987413 116987538
refGene_gn=KLHL7;refGene_tx=XM_003982839.5;refGene_gn=KLHL7;refGene_tx=XM_003982840.4;refGene_gn=KLHL7;refGene_tx=XM_011280428.3;refGene_gn=KLHL7;refGene_tx=XM_011280429.3;refGene_gn=KLHL7;refGene_tx=XM_019825599.1;refGene_gn=KLHL7;refGene_tx=XM_023250440.1;refGene_gn=KLHL7;refGene_tx=XM_023250441.1
A2 116988569 116988631
refGene_gn=KLHL7;refGene_tx=XM_023250441.1
A2 116995250 116995426
refGene_gn=KLHL7;refGene_tx=XM_003982839.5;refGene_gn=KLHL7;refGene_tx=XM_003982840.4;refGene_gn=KLHL7;refGene_tx=XM_011280428.3;refGene_gn=KLHL7;refGene_tx=XM_011280429.3;refGene_gn=KLHL7;refGene_tx=XM_019825599.1;refGene_gn=KLHL7;refGene_tx=XM_023250440.1
A2 116997878 116998053
refGene_gn=KLHL7;refGene_tx=XM_003982839.5;refGene_gn=KLHL7;refGene_tx=XM_003982840.4;refGene_gn=KLHL7;refGene_tx=XM_011280428.3;refGene_gn=KLHL7;refGene_tx=XM_011280429.3;refGene_gn=KLHL7;refGene_tx=XM_019825599.1;refGene_gn=KLHL7;refGene_tx=XM_023250440.1
A2 117009827 117009970
refGene_gn=KLHL7;refGene_tx=XM_003982839.5;refGene_gn=KLHL7;refGene_tx=XM_003982840.4;refGene_gn=KLHL7;refGene_tx=XM_011280428.3;refGene_gn=KLHL7;refGene_tx=XM_011280429.3;refGene_gn=KLHL7;refGene_tx=XM_019825599.1;refGene_gn=KLHL7;refGene_tx=XM_023250440.1
A2 117024997 117025238
refGene_gn=KLHL7;refGene_tx=XM_003982839.5;refGene_gn=KLHL7;refGene_tx=XM_003982840.4;refGene_gn=KLHL7;refGene_tx=XM_011280428.3;refGene_gn=KLHL7;refGene_tx=XM_011280429.3;refGene_gn=KLHL7;refGene_tx=XM_019825599.1;refGene_gn=KLHL7;refGene_tx=XM_023250440.1
A2 117027652 117027854
refGene_gn=KLHL7;refGene_tx=XM_003982839.5;refGene_gn=KLHL7;refGene_tx=XM_003982840.4;refGene_gn=KLHL7;refGene_tx=XM_011280428.3;refGene_gn=KLHL7;refGene_tx=XM_011280429.3;refGene_gn=KLHL7;refGene_tx=XM_019825599.1;refGene_gn=KLHL7;refGene_tx=XM_023250440.1
A2 117032773 117032871

refGene_gn=KLHL7;refGene_tx=XM_003982839.5;refGene_gn=KLHL7;refGene_tx=XM_003982840.4;refGene_gn=KLHL7;refGene_tx=XM_011280428.3;refGene_gn=KLHL7;refGene_tx=XM_011280429.3;refGene_gn=KLHL7;refGene_tx=XM_019825599.1;refGene_gn=KLHL7;refGene_tx=XM_023250440.1
A2 117033773 117034057
refGene_gn=KLHL7;refGene_tx=XM_003982839.5;refGene_gn=KLHL7;refGene_tx=XM_003982840.4;refGene_gn=KLHL7;refGene_tx=XM_011280428.3;refGene_gn=KLHL7;refGene_tx=XM_011280429.3;refGene_gn=KLHL7;refGene_tx=XM_019825599.1;refGene_gn=KLHL7;refGene_tx=XM_023250440.1
A2 117041898 117042010
refGene_gn=NUPL2;refGene_tx=XM_023250442.1
A2 117044212 117044441
refGene_gn=NUPL2;refGene_tx=XM_023250442.1
A2 117047829 117047924
refGene_gn=NUPL2;refGene_tx=XM_023250442.1
A2 117055690 117055767
refGene_gn=NUPL2;refGene_tx=XM_023250442.1
A2 117056501 117056588
refGene_gn=NUPL2;refGene_tx=XM_023250442.1
A2 117059113 117059198
refGene_gn=NUPL2;refGene_tx=XM_023250442.1
A2 117060417 117060986
refGene_gn=NUPL2;refGene_tx=XM_023250442.1
A2 117103884 117103957
refGene_gn=GPNMB;refGene_tx=XM_003982845.4
A2 117112151 117112304
refGene_gn=GPNMB;refGene_tx=XM_003982845.4
A2 117113174 117113318
refGene_gn=GPNMB;refGene_tx=XM_003982845.4
A2 117115683 117115848
refGene_gn=GPNMB;refGene_tx=XM_003982845.4
A2 117118049 117118208
refGene_gn=GPNMB;refGene_tx=XM_003982845.4
A2 117118505 117118817
refGene_gn=GPNMB;refGene_tx=XM_003982845.4
A2 117122508 117122607
refGene_gn=GPNMB;refGene_tx=XM_003982845.4
A2 117123395 117123498
refGene_gn=GPNMB;refGene_tx=XM_003982845.4
A2 117124804 117125019
refGene_gn=GPNMB;refGene_tx=XM_003982845.4
A2 117128357 117128451
refGene_gn=GPNMB;refGene_tx=XM_003982845.4
A2 117128896 117129050
refGene_gn=GPNMB;refGene_tx=XM_003982845.4
A2 117144398 117144654
refGene_gn=MALSU1;refGene_tx=XM_003982861.5
A2 117146142 117146321
refGene_gn=MALSU1;refGene_tx=XM_003982861.5
A2 117153012 117153094

refGene_gn=MALSU1;refGene_tx=XM_003982861.5
A2 117154509 117154691
refGene_gn=MALSU1;refGene_tx=XM_003982861.5
A2 117157518 117157617
refGene_gn=IGF2BP3;refGene_tx=XM_003982846.4;refGene_gn=IGF2BP3;refGene_tx=XM_019825602.2
A2 117157895 117158009
refGene_gn=IGF2BP3;refGene_tx=XM_003982846.4;refGene_gn=IGF2BP3;refGene_tx=XM_019825601.2;refGene_gn=IGF2BP3;refGene_tx=XM_019825602.2
A2 117158623 117158808
refGene_gn=IGF2BP3;refGene_tx=XM_003982846.4;refGene_gn=IGF2BP3;refGene_tx=XM_019825601.2;refGene_gn=IGF2BP3;refGene_tx=XM_019825602.2
A2 117161947 117162022
refGene_gn=IGF2BP3;refGene_tx=XM_003982846.4;refGene_gn=IGF2BP3;refGene_tx=XM_019825601.2;refGene_gn=IGF2BP3;refGene_tx=XM_019825602.2
A2 117165159 117165276
refGene_gn=IGF2BP3;refGene_tx=XM_003982846.4;refGene_gn=IGF2BP3;refGene_tx=XM_019825601.2;refGene_gn=IGF2BP3;refGene_tx=XM_019825602.2
A2 117182380 117182506
refGene_gn=IGF2BP3;refGene_tx=XM_003982846.4;refGene_gn=IGF2BP3;refGene_tx=XM_019825601.2;refGene_gn=IGF2BP3;refGene_tx=XM_019825602.2
A2 117184084 117184220
refGene_gn=IGF2BP3;refGene_tx=XM_003982846.4;refGene_gn=IGF2BP3;refGene_tx=XM_019825601.2;refGene_gn=IGF2BP3;refGene_tx=XM_019825602.2
A2 117186349 117186472
refGene_gn=IGF2BP3;refGene_tx=XM_003982846.4;refGene_gn=IGF2BP3;refGene_tx=XM_019825601.2;refGene_gn=IGF2BP3;refGene_tx=XM_019825602.2
A2 117187732 117187867
refGene_gn=IGF2BP3;refGene_tx=XM_003982846.4;refGene_gn=IGF2BP3;refGene_tx=XM_019825601.2;refGene_gn=IGF2BP3;refGene_tx=XM_019825602.2
A2 117191918 117192200
refGene_gn=IGF2BP3;refGene_tx=XM_003982846.4;refGene_gn=IGF2BP3;refGene_tx=XM_019825601.2;refGene_gn=IGF2BP3;refGene_tx=XM_019825602.2
A2 117201477 117201541
refGene_gn=IGF2BP3;refGene_tx=XM_003982846.4;refGene_gn=IGF2BP3;refGene_tx=XM_019825601.2
A2 117201632 117201684
refGene_gn=IGF2BP3;refGene_tx=XM_003982846.4;refGene_gn=IGF2BP3;refGene_tx=XM_019825601.2
A2 117259001 117259050
refGene_gn=IGF2BP3;refGene_tx=XM_003982846.4;refGene_gn=IGF2BP3;refGene_tx=XM_019825601.2
A2 117268265 117268514
refGene_gn=LOC101091006;refGene_tx=XM_011280432.3
A2 117305722 117305783
refGene_gn=IGF2BP3;refGene_tx=XM_003982846.4;refGene_gn=IGF2BP3;refGene_tx=XM_019825601.2;refGene_gn=IGF2BP3;refGene_tx=XM_019825602.2
A2 117307144 117307319
refGene_gn=IGF2BP3;refGene_tx=XM_003982846.4;refGene_gn=IGF2BP3;refGene_tx=XM_019825601.2;refGene_gn=IGF2BP3;refGene_tx=XM_019825602.2

A2 117345251 117345262
refGene_gn=TRA2A;refGene_tx=XM_003982850.5
A2 117345418 117345486
refGene_gn=TRA2A;refGene_tx=XM_003982850.5
A2 117346212 117346338
refGene_gn=TRA2A;refGene_tx=XM_003982850.5
A2 117347410 117347526
refGene_gn=TRA2A;refGene_tx=XM_003982850.5
A2 117349982 117350171
refGene_gn=TRA2A;refGene_tx=XM_003982850.5
A2 117351714 117351880
refGene_gn=TRA2A;refGene_tx=XM_003982850.5
A2 117355507 117355641
refGene_gn=TRA2A;refGene_tx=XM_003982850.5
A2 117365960 117365996
refGene_gn=TRA2A;refGene_tx=XM_003982850.5
A2 117439478 117439716
refGene_gn=CCDC126;refGene_tx=XM_023250443.1
A2 117462462 117462647
refGene_gn=CCDC126;refGene_tx=XM_023250443.1
A2 117528381 117528440
refGene_gn=STK31;refGene_tx=XM_006929245.4
A2 117528518 117528568
refGene_gn=STK31;refGene_tx=XM_011280434.3
A2 117530102 117530149
refGene_gn=STK31;refGene_tx=XM_006929245.4;refGene_gn=STK31;refGene_tx
=XM_011280434.3;refGene_gn=STK31;refGene_tx=XM_011280435.3
A2 117530237 117530290
refGene_gn=STK31;refGene_tx=XM_006929245.4;refGene_gn=STK31;refGene_tx
=XM_011280434.3;refGene_gn=STK31;refGene_tx=XM_011280435.3
A2 117535019 117535118
refGene_gn=STK31;refGene_tx=XM_006929245.4;refGene_gn=STK31;refGene_tx
=XM_011280434.3;refGene_gn=STK31;refGene_tx=XM_011280435.3;refGene_gn=
STK31;refGene_tx=XM_019825604.2
A2 117540630 117540705
refGene_gn=STK31;refGene_tx=XM_006929245.4;refGene_gn=STK31;refGene_tx
=XM_011280434.3;refGene_gn=STK31;refGene_tx=XM_011280435.3;refGene_gn=
STK31;refGene_tx=XM_019825604.2
A2 117542656 117542815
refGene_gn=STK31;refGene_tx=XM_006929245.4;refGene_gn=STK31;refGene_tx
=XM_011280434.3;refGene_gn=STK31;refGene_tx=XM_011280435.3;refGene_gn=
STK31;refGene_tx=XM_019825604.2
A2 117547420 117547779
refGene_gn=STK31;refGene_tx=XM_006929245.4;refGene_gn=STK31;refGene_tx
=XM_011280434.3;refGene_gn=STK31;refGene_tx=XM_011280435.3;refGene_gn=
STK31;refGene_tx=XM_019825604.2
A2 117548915 117549090
refGene_gn=STK31;refGene_tx=XM_006929245.4;refGene_gn=STK31;refGene_tx
=XM_011280434.3;refGene_gn=STK31;refGene_tx=XM_011280435.3;refGene_gn=
STK31;refGene_tx=XM_019825604.2

A2	117555612	117555728
refGene_gn=STK31;refGene_tx=XM_006929245.4;refGene_gn=STK31;refGene_tx=XM_011280434.3;refGene_gn=STK31;refGene_tx=XM_011280435.3;refGene_gn=STK31;refGene_tx=XM_019825604.2
A2	117558100	117558260
refGene_gn=STK31;refGene_tx=XM_006929245.4;refGene_gn=STK31;refGene_tx=XM_011280434.3;refGene_gn=STK31;refGene_tx=XM_011280435.3;refGene_gn=STK31;refGene_tx=XM_019825604.2
A2	117561215	117561338
refGene_gn=STK31;refGene_tx=XM_006929245.4;refGene_gn=STK31;refGene_tx=XM_011280434.3;refGene_gn=STK31;refGene_tx=XM_011280435.3;refGene_gn=STK31;refGene_tx=XM_019825604.2
A2	117565055	117565235
refGene_gn=STK31;refGene_tx=XM_006929245.4;refGene_gn=STK31;refGene_tx=XM_011280434.3;refGene_gn=STK31;refGene_tx=XM_011280435.3;refGene_gn=STK31;refGene_tx=XM_019825604.2
A2	117567280	117567397
refGene_gn=STK31;refGene_tx=XM_006929245.4;refGene_gn=STK31;refGene_tx=XM_011280434.3;refGene_gn=STK31;refGene_tx=XM_011280435.3;refGene_gn=STK31;refGene_tx=XM_019825604.2
A2	117568697	117568817
refGene_gn=STK31;refGene_tx=XM_006929245.4;refGene_gn=STK31;refGene_tx=XM_011280434.3;refGene_gn=STK31;refGene_tx=XM_011280435.3;refGene_gn=STK31;refGene_tx=XM_019825604.2
A2	117571405	117571537
refGene_gn=STK31;refGene_tx=XM_006929245.4;refGene_gn=STK31;refGene_tx=XM_011280434.3;refGene_gn=STK31;refGene_tx=XM_011280435.3;refGene_gn=STK31;refGene_tx=XM_019825604.2
A2	117572784	117572886
refGene_gn=STK31;refGene_tx=XM_006929245.4;refGene_gn=STK31;refGene_tx=XM_011280434.3;refGene_gn=STK31;refGene_tx=XM_011280435.3;refGene_gn=STK31;refGene_tx=XM_019825604.2
A2	117574311	117574392
refGene_gn=STK31;refGene_tx=XM_006929245.4;refGene_gn=STK31;refGene_tx=XM_011280434.3;refGene_gn=STK31;refGene_tx=XM_011280435.3;refGene_gn=STK31;refGene_tx=XM_019825604.2
A2	117576236	117576362
refGene_gn=STK31;refGene_tx=XM_006929245.4;refGene_gn=STK31;refGene_tx=XM_011280434.3;refGene_gn=STK31;refGene_tx=XM_011280435.3;refGene_gn=STK31;refGene_tx=XM_019825604.2
A2	117576974	117577100
refGene_gn=STK31;refGene_tx=XM_006929245.4;refGene_gn=STK31;refGene_tx=XM_011280434.3;refGene_gn=STK31;refGene_tx=XM_011280435.3;refGene_gn=STK31;refGene_tx=XM_019825604.2
A2	117577488	117577575
refGene_gn=STK31;refGene_tx=XM_006929245.4;refGene_gn=STK31;refGene_tx=XM_011280434.3;refGene_gn=STK31;refGene_tx=XM_011280435.3;refGene_gn=STK31;refGene_tx=XM_019825604.2
A2	117578230	117578380
refGene_gn=STK31;refGene_tx=XM_006929245.4;refGene_gn=STK31;refGene_tx

=XM_011280434.3;refGene_gn=STK31;refGene_tx=XM_011280435.3;refGene_gn=STK31;refGene_tx=XM_019825604.2
A2 117583311 117583434
refGene_gn=STK31;refGene_tx=XM_006929245.4;refGene_gn=STK31;refGene_tx=XM_011280434.3;refGene_gn=STK31;refGene_tx=XM_011280435.3;refGene_gn=STK31;refGene_tx=XM_019825604.2
A2 117593779 117593848
refGene_gn=STK31;refGene_tx=XM_006929245.4;refGene_gn=STK31;refGene_tx=XM_011280434.3;refGene_gn=STK31;refGene_tx=XM_011280435.3;refGene_gn=STK31;refGene_tx=XM_019825604.2
A2 117604151 117604379
refGene_gn=STK31;refGene_tx=XM_006929245.4;refGene_gn=STK31;refGene_tx=XM_011280434.3;refGene_gn=STK31;refGene_tx=XM_011280435.3;refGene_gn=STK31;refGene_tx=XM_019825604.2
A2 118012377 118012565
refGene_gn=NPY;refGene_tx=XM_019825605.2
A2 118016635 118016716
refGene_gn=NPY;refGene_tx=XM_019825605.2
A2 118018567 118018592
refGene_gn=NPY;refGene_tx=XM_019825605.2
A2 118317926 118317948
refGene_gn=MPP6;refGene_tx=XM_006929252.3;refGene_gn=MPP6;refGene_tx=XM_019825607.2;refGene_gn=MPP6;refGene_tx=XM_023250447.1
A2 118343068 118343190
refGene_gn=MPP6;refGene_tx=XM_006929252.3;refGene_gn=MPP6;refGene_tx=XM_006929253.4;refGene_gn=MPP6;refGene_tx=XM_019825607.2;refGene_gn=MPP6;refGene_tx=XM_023250444.1;refGene_gn=MPP6;refGene_tx=XM_023250445.1;refGene_gn=MPP6;refGene_tx=XM_023250446.1;refGene_gn=MPP6;refGene_tx=XM_023250447.1
A2 118362320 118362473
refGene_gn=MPP6;refGene_tx=XM_006929252.3;refGene_gn=MPP6;refGene_tx=XM_006929253.4;refGene_gn=MPP6;refGene_tx=XM_019825607.2;refGene_gn=MPP6;refGene_tx=XM_023250444.1;refGene_gn=MPP6;refGene_tx=XM_023250445.1;refGene_gn=MPP6;refGene_tx=XM_023250446.1;refGene_gn=MPP6;refGene_tx=XM_023250447.1
A2 118366060 118366213
refGene_gn=MPP6;refGene_tx=XM_006929252.3;refGene_gn=MPP6;refGene_tx=XM_006929253.4;refGene_gn=MPP6;refGene_tx=XM_019825607.2;refGene_gn=MPP6;refGene_tx=XM_023250444.1;refGene_gn=MPP6;refGene_tx=XM_023250445.1;refGene_gn=MPP6;refGene_tx=XM_023250446.1;refGene_gn=MPP6;refGene_tx=XM_023250447.1
A2 118367001 118367229
refGene_gn=MPP6;refGene_tx=XM_006929252.3;refGene_gn=MPP6;refGene_tx=XM_006929253.4;refGene_gn=MPP6;refGene_tx=XM_019825607.2;refGene_gn=MPP6;refGene_tx=XM_023250444.1;refGene_gn=MPP6;refGene_tx=XM_023250445.1;refGene_gn=MPP6;refGene_tx=XM_023250446.1;refGene_gn=MPP6;refGene_tx=XM_023250447.1
A2 118377691 118377733
refGene_gn=MPP6;refGene_tx=XM_006929252.3;refGene_gn=MPP6;refGene_tx=XM_006929253.4;refGene_gn=MPP6;refGene_tx=XM_019825607.2;refGene_gn=MPP

6;refGene_tx=XM_023250444.1;refGene_gn=MPP6;refGene_tx=XM_023250445.1;
refGene_gn=MPP6;refGene_tx=XM_023250446.1
A2 118379233 118379365
refGene_gn=MPP6;refGene_tx=XM_006929252.3;refGene_gn=MPP6;refGene_tx=X
M_006929253.4;refGene_gn=MPP6;refGene_tx=XM_019825607.2;refGene_gn=MPP
6;refGene_tx=XM_023250444.1;refGene_gn=MPP6;refGene_tx=XM_023250445.1;
refGene_gn=MPP6;refGene_tx=XM_023250446.1;refGene_gn=MPP6;refGene_tx=X
M_023250447.1
A2 118381422 118381522
refGene_gn=MPP6;refGene_tx=XM_006929252.3;refGene_gn=MPP6;refGene_tx=X
M_006929253.4;refGene_gn=MPP6;refGene_tx=XM_019825607.2;refGene_gn=MPP
6;refGene_tx=XM_023250444.1;refGene_gn=MPP6;refGene_tx=XM_023250445.1;
refGene_gn=MPP6;refGene_tx=XM_023250446.1;refGene_gn=MPP6;refGene_tx=X
M_023250447.1
A2 118381844 118381913
refGene_gn=MPP6;refGene_tx=XM_006929252.3;refGene_gn=MPP6;refGene_tx=X
M_006929253.4;refGene_gn=MPP6;refGene_tx=XM_019825607.2;refGene_gn=MPP
6;refGene_tx=XM_023250444.1;refGene_gn=MPP6;refGene_tx=XM_023250445.1;
refGene_gn=MPP6;refGene_tx=XM_023250446.1;refGene_gn=MPP6;refGene_tx=X
M_023250447.1
A2 118383650 118383812
refGene_gn=MPP6;refGene_tx=XM_006929252.3;refGene_gn=MPP6;refGene_tx=X
M_006929253.4;refGene_gn=MPP6;refGene_tx=XM_019825607.2;refGene_gn=MPP
6;refGene_tx=XM_023250444.1;refGene_gn=MPP6;refGene_tx=XM_023250445.1;
refGene_gn=MPP6;refGene_tx=XM_023250446.1;refGene_gn=MPP6;refGene_tx=X
M_023250447.1
A2 118395005 118395208
refGene_gn=MPP6;refGene_tx=XM_006929252.3;refGene_gn=MPP6;refGene_tx=X
M_006929253.4;refGene_gn=MPP6;refGene_tx=XM_019825607.2;refGene_gn=MPP
6;refGene_tx=XM_023250444.1;refGene_gn=MPP6;refGene_tx=XM_023250445.1;
refGene_gn=MPP6;refGene_tx=XM_023250446.1;refGene_gn=MPP6;refGene_tx=X
M_023250447.1
A2 118396197 118396326
refGene_gn=MPP6;refGene_tx=XM_006929252.3;refGene_gn=MPP6;refGene_tx=X
M_006929253.4;refGene_gn=MPP6;refGene_tx=XM_019825607.2;refGene_gn=MPP
6;refGene_tx=XM_023250444.1;refGene_gn=MPP6;refGene_tx=XM_023250445.1;
refGene_gn=MPP6;refGene_tx=XM_023250446.1;refGene_gn=MPP6;refGene_tx=X
M_023250447.1
A2 118403236 118403413
refGene_gn=MPP6;refGene_tx=XM_006929252.3;refGene_gn=MPP6;refGene_tx=X
M_006929253.4;refGene_gn=MPP6;refGene_tx=XM_019825607.2;refGene_gn=MPP
6;refGene_tx=XM_023250444.1;refGene_gn=MPP6;refGene_tx=XM_023250445.1;
refGene_gn=MPP6;refGene_tx=XM_023250446.1;refGene_gn=MPP6;refGene_tx=X
M_023250447.1
A2 118412759 118412990
refGene_gn=GSDME;refGene_tx=XM_003982887.5;refGene_gn=GSDME;refGene_tx
=XM_006929254.3
A2 118415698 118415772
refGene_gn=GSDME;refGene_tx=XM_003982887.5;refGene_gn=GSDME;refGene_tx
=XM_006929254.3

A2 118426043 118426239
refGene_gn=GSDME;refGene_tx=XM_003982887.5;refGene_gn=GSDME;refGene_tx
=XM_006929254.3
A2 118427976 118428104
refGene_gn=GSDME;refGene_tx=XM_003982887.5;refGene_gn=GSDME;refGene_tx
=XM_006929254.3
A2 118430155 118430317
refGene_gn=GSDME;refGene_tx=XM_003982887.5;refGene_gn=GSDME;refGene_tx
=XM_006929254.3
A2 118435198 118435319
refGene_gn=GSDME;refGene_tx=XM_003982887.5;refGene_gn=GSDME;refGene_tx
=XM_006929254.3
A2 118436947 118437119
refGene_gn=GSDME;refGene_tx=XM_003982887.5;refGene_gn=GSDME;refGene_tx
=XM_006929254.3
A2 118464919 118465112
refGene_gn=GSDME;refGene_tx=XM_003982887.5;refGene_gn=GSDME;refGene_tx
=XM_006929254.3
A2 118467325 118467536
refGene_gn=GSDME;refGene_tx=XM_003982887.5;refGene_gn=GSDME;refGene_tx
=XM_006929254.3
A2 118509696 118509793
refGene_gn=OSBPL3;refGene_tx=XM_003982866.5;refGene_gn=OSBPL3;refGene_
tx=XM_003982867.5;refGene_gn=OSBPL3;refGene_tx=XM_006929256.4;refGene_
gn=OSBPL3;refGene_tx=XM_019825608.2;refGene_gn=OSBPL3;refGene_tx=XM_01
9825609.2;refGene_gn=OSBPL3;refGene_tx=XM_019825610.2
A2 118513936 118514059
refGene_gn=OSBPL3;refGene_tx=XM_003982866.5;refGene_gn=OSBPL3;refGene_
tx=XM_003982867.5;refGene_gn=OSBPL3;refGene_tx=XM_006929256.4;refGene_
gn=OSBPL3;refGene_tx=XM_019825608.2;refGene_gn=OSBPL3;refGene_tx=XM_01
9825609.2;refGene_gn=OSBPL3;refGene_tx=XM_019825610.2
A2 118515659 118515786
refGene_gn=OSBPL3;refGene_tx=XM_003982866.5;refGene_gn=OSBPL3;refGene_
tx=XM_003982867.5;refGene_gn=OSBPL3;refGene_tx=XM_006929256.4;refGene_
gn=OSBPL3;refGene_tx=XM_019825608.2;refGene_gn=OSBPL3;refGene_tx=XM_01
9825609.2;refGene_gn=OSBPL3;refGene_tx=XM_019825610.2
A2 118517472 118517617
refGene_gn=OSBPL3;refGene_tx=XM_003982866.5;refGene_gn=OSBPL3;refGene_
tx=XM_003982867.5;refGene_gn=OSBPL3;refGene_tx=XM_006929256.4;refGene_
gn=OSBPL3;refGene_tx=XM_019825608.2;refGene_gn=OSBPL3;refGene_tx=XM_01
9825609.2;refGene_gn=OSBPL3;refGene_tx=XM_019825610.2
A2 118520405 118520550
refGene_gn=OSBPL3;refGene_tx=XM_003982866.5;refGene_gn=OSBPL3;refGene_
tx=XM_003982867.5;refGene_gn=OSBPL3;refGene_tx=XM_006929256.4;refGene_
gn=OSBPL3;refGene_tx=XM_019825608.2;refGene_gn=OSBPL3;refGene_tx=XM_01
9825609.2;refGene_gn=OSBPL3;refGene_tx=XM_019825610.2
A2 118522210 118522289
refGene_gn=OSBPL3;refGene_tx=XM_003982866.5;refGene_gn=OSBPL3;refGene_
tx=XM_003982867.5;refGene_gn=OSBPL3;refGene_tx=XM_006929256.4;refGene_
gn=OSBPL3;refGene_tx=XM_019825608.2;refGene_gn=OSBPL3;refGene_tx=XM_01

9825609.2;refGene_gn=OSBPL3;refGene_tx=XM_019825610.2
A2 118526476 118526540
refGene_gn=OSBPL3;refGene_tx=XM_003982866.5;refGene_gn=OSBPL3;refGene_tx=XM_003982867.5;refGene_gn=OSBPL3;refGene_tx=XM_006929256.4;refGene_gn=OSBPL3;refGene_tx=XM_019825608.2;refGene_gn=OSBPL3;refGene_tx=XM_019825609.2;refGene_gn=OSBPL3;refGene_tx=XM_019825610.2
A2 118537326 118537464
refGene_gn=OSBPL3;refGene_tx=XM_003982866.5;refGene_gn=OSBPL3;refGene_tx=XM_003982867.5;refGene_gn=OSBPL3;refGene_tx=XM_006929256.4;refGene_gn=OSBPL3;refGene_tx=XM_019825608.2;refGene_gn=OSBPL3;refGene_tx=XM_019825609.2;refGene_gn=OSBPL3;refGene_tx=XM_019825610.2
A2 118540150 118540401
refGene_gn=OSBPL3;refGene_tx=XM_003982866.5;refGene_gn=OSBPL3;refGene_tx=XM_003982867.5;refGene_gn=OSBPL3;refGene_tx=XM_006929256.4;refGene_gn=OSBPL3;refGene_tx=XM_019825608.2;refGene_gn=OSBPL3;refGene_tx=XM_019825609.2;refGene_gn=OSBPL3;refGene_tx=XM_019825610.2
A2 118544279 118544373
refGene_gn=OSBPL3;refGene_tx=XM_003982866.5;refGene_gn=OSBPL3;refGene_tx=XM_003982867.5;refGene_gn=OSBPL3;refGene_tx=XM_006929256.4;refGene_gn=OSBPL3;refGene_tx=XM_019825608.2;refGene_gn=OSBPL3;refGene_tx=XM_019825609.2;refGene_gn=OSBPL3;refGene_tx=XM_019825610.2
A2 118545292 118545427
refGene_gn=OSBPL3;refGene_tx=XM_003982866.5;refGene_gn=OSBPL3;refGene_tx=XM_003982867.5;refGene_gn=OSBPL3;refGene_tx=XM_006929256.4;refGene_gn=OSBPL3;refGene_tx=XM_019825608.2;refGene_gn=OSBPL3;refGene_tx=XM_019825609.2;refGene_gn=OSBPL3;refGene_tx=XM_019825610.2
A2 118552795 118552903
refGene_gn=OSBPL3;refGene_tx=XM_006929256.4;refGene_gn=OSBPL3;refGene_tx=XM_019825608.2;refGene_gn=OSBPL3;refGene_tx=XM_019825609.2;refGene_gn=OSBPL3;refGene_tx=XM_019825610.2
A2 118556180 118556311
refGene_gn=OSBPL3;refGene_tx=XM_003982866.5;refGene_gn=OSBPL3;refGene_tx=XM_003982867.5;refGene_gn=OSBPL3;refGene_tx=XM_006929256.4;refGene_gn=OSBPL3;refGene_tx=XM_019825608.2;refGene_gn=OSBPL3;refGene_tx=XM_019825609.2;refGene_gn=OSBPL3;refGene_tx=XM_019825610.2
A2 118566099 118566256
refGene_gn=OSBPL3;refGene_tx=XM_003982866.5;refGene_gn=OSBPL3;refGene_tx=XM_003982867.5;refGene_gn=OSBPL3;refGene_tx=XM_006929256.4;refGene_gn=OSBPL3;refGene_tx=XM_019825608.2;refGene_gn=OSBPL3;refGene_tx=XM_019825609.2;refGene_gn=OSBPL3;refGene_tx=XM_019825610.2
A2 118567299 118567392
refGene_gn=OSBPL3;refGene_tx=XM_003982867.5;refGene_gn=OSBPL3;refGene_tx=XM_006929256.4;refGene_gn=OSBPL3;refGene_tx=XM_019825608.2;refGene_gn=OSBPL3;refGene_tx=XM_019825609.2
A2 118567580 118567684
refGene_gn=OSBPL3;refGene_tx=XM_003982866.5;refGene_gn=OSBPL3;refGene_tx=XM_003982867.5;refGene_gn=OSBPL3;refGene_tx=XM_006929256.4;refGene_gn=OSBPL3;refGene_tx=XM_019825608.2;refGene_gn=OSBPL3;refGene_tx=XM_019825609.2;refGene_gn=OSBPL3;refGene_tx=XM_019825610.2
A2 118569747 118569871

refGene_gn=OSBPL3;refGene_tx=XM_003982866.5;refGene_gn=OSBPL3;refGene_tx=XM_003982867.5;refGene_gn=OSBPL3;refGene_tx=XM_006929256.4;refGene_gn=OSBPL3;refGene_tx=XM_019825608.2;refGene_gn=OSBPL3;refGene_tx=XM_019825609.2;refGene_gn=OSBPL3;refGene_tx=XM_019825610.2
A2 118570426 118570594
refGene_gn=OSBPL3;refGene_tx=XM_003982866.5;refGene_gn=OSBPL3;refGene_tx=XM_003982867.5;refGene_gn=OSBPL3;refGene_tx=XM_006929256.4;refGene_gn=OSBPL3;refGene_tx=XM_019825608.2;refGene_gn=OSBPL3;refGene_tx=XM_019825609.2;refGene_gn=OSBPL3;refGene_tx=XM_019825610.2
A2 118574149 118574263
refGene_gn=OSBPL3;refGene_tx=XM_003982866.5;refGene_gn=OSBPL3;refGene_tx=XM_003982867.5;refGene_gn=OSBPL3;refGene_tx=XM_006929256.4;refGene_gn=OSBPL3;refGene_tx=XM_019825608.2;refGene_gn=OSBPL3;refGene_tx=XM_019825609.2;refGene_gn=OSBPL3;refGene_tx=XM_019825610.2
A2 118575238 118575292
refGene_gn=OSBPL3;refGene_tx=XM_003982866.5;refGene_gn=OSBPL3;refGene_tx=XM_003982867.5;refGene_gn=OSBPL3;refGene_tx=XM_006929256.4;refGene_gn=OSBPL3;refGene_tx=XM_019825608.2;refGene_gn=OSBPL3;refGene_tx=XM_019825609.2;refGene_gn=OSBPL3;refGene_tx=XM_019825610.2
A2 118575472 118575589
refGene_gn=OSBPL3;refGene_tx=XM_003982866.5;refGene_gn=OSBPL3;refGene_tx=XM_003982867.5;refGene_gn=OSBPL3;refGene_tx=XM_006929256.4;refGene_gn=OSBPL3;refGene_tx=XM_019825608.2;refGene_gn=OSBPL3;refGene_tx=XM_019825609.2;refGene_gn=OSBPL3;refGene_tx=XM_019825610.2
A2 118597758 118597851
refGene_gn=OSBPL3;refGene_tx=XM_003982866.5;refGene_gn=OSBPL3;refGene_tx=XM_003982867.5;refGene_gn=OSBPL3;refGene_tx=XM_006929256.4;refGene_gn=OSBPL3;refGene_tx=XM_019825608.2;refGene_gn=OSBPL3;refGene_tx=XM_019825609.2;refGene_gn=OSBPL3;refGene_tx=XM_019825610.2
A2 118772951 118773100
refGene_gn=LOC101098213;refGene_tx=XM_006929257.2;refGene_gn=LOC101098213;refGene_tx=XM_019825614.1
A2 118773211 118773380
refGene_gn=LOC101098213;refGene_tx=XM_006929257.2;refGene_gn=LOC101098213;refGene_tx=XM_019825614.1
A2 118782139 118782961
refGene_gn=CA2H7orf31;refGene_tx=XM_003982869.4;refGene_gn=CA2H7orf31;refGene_tx=XM_019825612.2
A2 118786389 118786481
refGene_gn=CA2H7orf31;refGene_tx=XM_003982869.4;refGene_gn=CA2H7orf31;refGene_tx=XM_019825612.2
A2 118786773 118786948
refGene_gn=CA2H7orf31;refGene_tx=XM_003982869.4;refGene_gn=CA2H7orf31;refGene_tx=XM_019825612.2
A2 118793857 118793946
refGene_gn=CA2H7orf31;refGene_tx=XM_003982869.4;refGene_gn=CA2H7orf31;refGene_tx=XM_019825612.2
A2 118797763 118797945
refGene_gn=CA2H7orf31;refGene_tx=XM_003982869.4;refGene_gn=CA2H7orf31;refGene_tx=XM_019825612.2

A2 118801016 118801046
refGene_gn=CA2H7orf31;refGene_tx=XM_003982869.4;refGene_gn=CA2H7orf31;
refGene_tx=XM_019825612.2
A2 118803100 118803198
refGene_gn=CA2H7orf31;refGene_tx=XM_003982869.4;refGene_gn=CA2H7orf31;
refGene_tx=XM_019825612.2
A2 118813763 118813940
refGene_gn=CA2H7orf31;refGene_tx=XM_003982869.4
A2 118824387 118824521
refGene_gn=CA2H7orf31;refGene_tx=XM_003982869.4
A2 118854167 118854264
refGene_gn=NPVF;refGene_tx=XM_023242159.1
A2 118855288 118855692
refGene_gn=NPVF;refGene_tx=XM_023242159.1
A2 118857361 118857499
refGene_gn=NPVF;refGene_tx=XM_023242159.1
A2 119679230 119679800
refGene_gn=NFE2L3;refGene_tx=XM_019825057.2;refGene_gn=NFE2L3;refGene_
tx=XM_023250448.1
A2 119704243 119704423
refGene_gn=NFE2L3;refGene_tx=XM_019825057.2;refGene_gn=NFE2L3;refGene_
tx=XM_023250448.1
A2 119708409 119708493
refGene_gn=NFE2L3;refGene_tx=XM_019825057.2
A2 119709109 119710348
refGene_gn=NFE2L3;refGene_tx=XM_019825057.2;refGene_gn=NFE2L3;refGene_
tx=XM_023250448.1
A2 119717261 119717323
refGene_gn=HNRNPA2B1;refGene_tx=XM_011280438.3
A2 119717924 119718047
refGene_gn=HNRNPA2B1;refGene_tx=XM_011280438.3
A2 119718301 119718421
refGene_gn=HNRNPA2B1;refGene_tx=XM_011280438.3
A2 119719696 119719759
refGene_gn=HNRNPA2B1;refGene_tx=XM_011280438.3
A2 119720434 119720515
refGene_gn=HNRNPA2B1;refGene_tx=XM_011280438.3
A2 119720610 119720712
refGene_gn=HNRNPA2B1;refGene_tx=XM_011280438.3
A2 119720840 119721051
refGene_gn=HNRNPA2B1;refGene_tx=XM_011280438.3
A2 119721320 119721467
refGene_gn=HNRNPA2B1;refGene_tx=XM_011280438.3
A2 119721675 119721786
refGene_gn=HNRNPA2B1;refGene_tx=XM_011280438.3
A2 119721883 119721919
refGene_gn=HNRNPA2B1;refGene_tx=XM_011280438.3
A2 119724821 119724827
refGene_gn=HNRNPA2B1;refGene_tx=XM_011280438.3
A2 119727351 119727375

refGene_gn=CBX3;refGene_tx=XM_011280439.3;refGene_gn=CBX3;refGene_tx=XM_023250449.1
A2 119730206 119730349
refGene_gn=CBX3;refGene_tx=XM_011280439.3;refGene_gn=CBX3;refGene_tx=XM_011280440.3;refGene_gn=CBX3;refGene_tx=XM_023250449.1;refGene_gn=CBX3;refGene_tx=XM_023250451.1
A2 119733173 119733336
refGene_gn=CBX3;refGene_tx=XM_011280439.3;refGene_gn=CBX3;refGene_tx=XM_011280440.3;refGene_gn=CBX3;refGene_tx=XM_023250449.1;refGene_gn=CBX3;refGene_tx=XM_023250451.1
A2 119735519 119735614
refGene_gn=CBX3;refGene_tx=XM_011280439.3;refGene_gn=CBX3;refGene_tx=XM_011280440.3;refGene_gn=CBX3;refGene_tx=XM_023250449.1;refGene_gn=CBX3;refGene_tx=XM_023250451.1
A2 119735959 119736086
refGene_gn=CBX3;refGene_tx=XM_011280439.3;refGene_gn=CBX3;refGene_tx=XM_011280440.3;refGene_gn=CBX3;refGene_tx=XM_023250449.1;refGene_gn=CBX3;refGene_tx=XM_023250451.1
A2 119828070 119828094
refGene_gn=SNX10;refGene_tx=XM_006929264.3;refGene_gn=SNX10;refGene_tx=XM_019825617.2;refGene_gn=SNX10;refGene_tx=XM_019825618.2;refGene_gn=SNX10;refGene_tx=XM_019825619.2
A2 119846158 119846245
refGene_gn=SNX10;refGene_tx=XM_006929264.3;refGene_gn=SNX10;refGene_tx=XM_019825617.2;refGene_gn=SNX10;refGene_tx=XM_019825618.2;refGene_gn=SNX10;refGene_tx=XM_019825619.2
A2 119849849 119849950
refGene_gn=SNX10;refGene_tx=XM_006929264.3;refGene_gn=SNX10;refGene_tx=XM_019825617.2;refGene_gn=SNX10;refGene_tx=XM_019825618.2;refGene_gn=SNX10;refGene_tx=XM_019825619.2
A2 119850339 119850438
refGene_gn=SNX10;refGene_tx=XM_006929264.3;refGene_gn=SNX10;refGene_tx=XM_019825617.2;refGene_gn=SNX10;refGene_tx=XM_019825618.2;refGene_gn=SNX10;refGene_tx=XM_019825619.2
A2 119857642 119857855
refGene_gn=SNX10;refGene_tx=XM_006929264.3;refGene_gn=SNX10;refGene_tx=XM_019825617.2;refGene_gn=SNX10;refGene_tx=XM_019825618.2;refGene_gn=SNX10;refGene_tx=XM_019825619.2
A2 119858327 119858409
refGene_gn=SNX10;refGene_tx=XM_006929264.3;refGene_gn=SNX10;refGene_tx=XM_019825617.2;refGene_gn=SNX10;refGene_tx=XM_019825618.2;refGene_gn=SNX10;refGene_tx=XM_019825619.2
A2 120144387 120144480
refGene_gn=SKAP2;refGene_tx=XM_003982875.5;refGene_gn=SKAP2;refGene_tx=XM_006929265.4
A2 120157895 120158008
refGene_gn=SKAP2;refGene_tx=XM_003982875.5;refGene_gn=SKAP2;refGene_tx=XM_006929265.4
A2 120164058 120164136
refGene_gn=SKAP2;refGene_tx=XM_003982875.5;refGene_gn=SKAP2;refGene_tx

=XM_006929265.4
A2	120199565	120199688
refGene_gn=SKAP2;refGene_tx=XM_003982875.5;refGene_gn=SKAP2;refGene_tx
=XM_006929265.4
A2	120200043	120200107
refGene_gn=SKAP2;refGene_tx=XM_003982875.5;refGene_gn=SKAP2;refGene_tx
=XM_006929265.4
A2	120200987	120201112
refGene_gn=SKAP2;refGene_tx=XM_003982875.5;refGene_gn=SKAP2;refGene_tx
=XM_006929265.4
A2	120211562	120211646
refGene_gn=SKAP2;refGene_tx=XM_003982875.5;refGene_gn=SKAP2;refGene_tx
=XM_006929265.4
A2	120212918	120212996
refGene_gn=SKAP2;refGene_tx=XM_003982875.5;refGene_gn=SKAP2;refGene_tx
=XM_006929265.4
A2	120304628	120304736
refGene_gn=SKAP2;refGene_tx=XM_003982875.5;refGene_gn=SKAP2;refGene_tx
=XM_006929265.4
A2	120313952	120313978
refGene_gn=SKAP2;refGene_tx=XM_003982875.5;refGene_gn=SKAP2;refGene_tx
=XM_006929265.4
A2	120314574	120314680
refGene_gn=SKAP2;refGene_tx=XM_003982875.5;refGene_gn=SKAP2;refGene_tx
=XM_006929265.4
A2	120323801	120323868
refGene_gn=SKAP2;refGene_tx=XM_003982875.5;refGene_gn=SKAP2;refGene_tx
=XM_006929265.4
A2	120457481	120457546
refGene_gn=LOC102901262;refGene_tx=XM_023242167.1
A2	120494120	120494162
refGene_gn=LOC102901262;refGene_tx=XM_023242167.1
A2	120496063	120496626
refGene_gn=LOC102901262;refGene_tx=XM_023242167.1
A2	120498185	120498211
refGene_gn=LOC102901262;refGene_tx=XM_023242167.1
A2	120502667	120503023
refGene_gn=HOXA1;refGene_tx=XM_019825620.1
A2	120503482	120504134
refGene_gn=HOXA1;refGene_tx=XM_019825620.1;refGene_gn=HOXA1;refGene_tx
=XM_019825621.1
A2	120508903	120509643
refGene_gn=HOXA2;refGene_tx=XM_003982877.4
A2	120510293	120510684
refGene_gn=HOXA2;refGene_tx=XM_003982877.4
A2	120516238	120517038
refGene_gn=HOXA3;refGene_tx=XM_006929270.3;refGene_gn=HOXA3;refGene_tx
=XM_006929271.3;refGene_gn=HOXA3;refGene_tx=XM_023250452.1
A2	120518422	120518948
refGene_gn=HOXA3;refGene_tx=XM_006929270.3;refGene_gn=HOXA3;refGene_tx

=XM_006929271.3;refGene_gn=HOXA3;refGene_tx=XM_023250452.1
A2	120537061	120537408
refGene_gn=HOXA4;refGene_tx=XM_003982889.5
A2	120537960	120538594
refGene_gn=HOXA4;refGene_tx=XM_003982889.5
A2	120549790	120550041
refGene_gn=HOXA5;refGene_tx=XM_023250453.1
A2	120551026	120551921
refGene_gn=HOXA5;refGene_tx=XM_023250453.1
A2	120553616	120553876
refGene_gn=HOXA6;refGene_tx=XM_003982880.5
A2	120555301	120555743
refGene_gn=HOXA6;refGene_tx=XM_003982880.5
A2	120562962	120563288
refGene_gn=HOXA7;refGene_tx=XM_023250454.1
A2	120564261	120564640
refGene_gn=HOXA7;refGene_tx=XM_023250454.1
A2	120571785	120572024
refGene_gn=HOXA9;refGene_tx=XM_003982882.5
A2	120573129	120573709
refGene_gn=HOXA9;refGene_tx=XM_003982882.5
A2	120580333	120580608
refGene_gn=HOXA10;refGene_tx=XM_023250455.1
A2	120581813	120582777
refGene_gn=HOXA10;refGene_tx=XM_023250455.1
A2	120591452	120591685
refGene_gn=HOXA11;refGene_tx=XM_023250456.1
A2	120593158	120593909
refGene_gn=HOXA11;refGene_tx=XM_023250456.1
A2	120607231	120607476
refGene_gn=HOXA13;refGene_tx=XM_019825059.2
A2	120608194	120609119
refGene_gn=HOXA13;refGene_tx=XM_019825059.2
A2	120652723	120653150
refGene_gn=EVX1;refGene_tx=XM_023250457.1
A2	120654754	120655002
refGene_gn=EVX1;refGene_tx=XM_023250457.1
A2	120655630	120656170
refGene_gn=EVX1;refGene_tx=XM_023250457.1
A2	120919397	120919556
refGene_gn=HIBADH;refGene_tx=XM_003982884.4
A2	120923964	120924121
refGene_gn=HIBADH;refGene_tx=XM_003982884.4
A2	120931548	120931625
refGene_gn=HIBADH;refGene_tx=XM_003982884.4
A2	120933687	120933821
refGene_gn=HIBADH;refGene_tx=XM_003982884.4
A2	121003496	121003618
refGene_gn=HIBADH;refGene_tx=XM_003982884.4
A2	121006340	121006450

refGene_gn=HIBADH;refGene_tx=XM_003982884.4
A2	121015677	121015838
refGene_gn=HIBADH;refGene_tx=XM_003982884.4
A2	121028420	121028511
refGene_gn=HIBADH;refGene_tx=XM_003982884.4
A2	121105033	121105195
refGene_gn=TAX1BP1;refGene_tx=XM_006929273.4;refGene_gn=TAX1BP1;refGene_tx=XM_006929274.4
A2	121119934	121120037
refGene_gn=TAX1BP1;refGene_tx=XM_006929273.4;refGene_gn=TAX1BP1;refGene_tx=XM_006929274.4
A2	121124903	121125091
refGene_gn=TAX1BP1;refGene_tx=XM_006929273.4;refGene_gn=TAX1BP1;refGene_tx=XM_006929274.4
A2	121128435	121128594
refGene_gn=TAX1BP1;refGene_tx=XM_006929273.4;refGene_gn=TAX1BP1;refGene_tx=XM_006929274.4
A2	121138619	121138768
refGene_gn=TAX1BP1;refGene_tx=XM_006929273.4;refGene_gn=TAX1BP1;refGene_tx=XM_006929274.4
A2	121138860	121138951
refGene_gn=TAX1BP1;refGene_tx=XM_006929273.4;refGene_gn=TAX1BP1;refGene_tx=XM_006929274.4
A2	121142980	121143166
refGene_gn=TAX1BP1;refGene_tx=XM_006929273.4;refGene_gn=TAX1BP1;refGene_tx=XM_006929274.4
A2	121147421	121147646
refGene_gn=TAX1BP1;refGene_tx=XM_006929273.4;refGene_gn=TAX1BP1;refGene_tx=XM_006929274.4
A2	121148179	121148326
refGene_gn=TAX1BP1;refGene_tx=XM_006929273.4;refGene_gn=TAX1BP1;refGene_tx=XM_006929274.4
A2	121149701	121149816
refGene_gn=TAX1BP1;refGene_tx=XM_006929273.4;refGene_gn=TAX1BP1;refGene_tx=XM_006929274.4
A2	121151056	121151160
refGene_gn=TAX1BP1;refGene_tx=XM_006929273.4;refGene_gn=TAX1BP1;refGene_tx=XM_006929274.4
A2	121153981	121154107
refGene_gn=TAX1BP1;refGene_tx=XM_006929273.4;refGene_gn=TAX1BP1;refGene_tx=XM_006929274.4
A2	121158029	121158101
refGene_gn=TAX1BP1;refGene_tx=XM_006929273.4;refGene_gn=TAX1BP1;refGene_tx=XM_006929274.4
A2	121176909	121177093
refGene_gn=TAX1BP1;refGene_tx=XM_006929273.4;refGene_gn=TAX1BP1;refGene_tx=XM_006929274.4
A2	121177697	121177846
refGene_gn=TAX1BP1;refGene_tx=XM_006929273.4;refGene_gn=TAX1BP1;refGene_tx=XM_006929274.4

A2	121187627	121187710
refGene_gn=TAX1BP1;refGene_tx=XM_006929273.4;refGene_gn=TAX1BP1;refGene_tx=XM_006929274.4
A2	121188332	121188534
refGene_gn=TAX1BP1;refGene_tx=XM_006929273.4;refGene_gn=TAX1BP1;refGene_tx=XM_006929274.4
A2	121192197	121192374
refGene_gn=JAZF1;refGene_tx=XM_011280443.3;refGene_gn=JAZF1;refGene_tx=XM_019825625.2
A2	121200823	121200993
refGene_gn=JAZF1;refGene_tx=XM_011280443.3;refGene_gn=JAZF1;refGene_tx=XM_019825625.2
A2	121242428	121242625
refGene_gn=JAZF1;refGene_tx=XM_011280443.3;refGene_gn=JAZF1;refGene_tx=XM_019825625.2
A2	121345069	121345142
refGene_gn=JAZF1;refGene_tx=XM_011280443.3;refGene_gn=JAZF1;refGene_tx=XM_019825625.2
A2	121356470	121356543
refGene_gn=JAZF1;refGene_tx=XM_019825625.2
A2	121545106	121545221
refGene_gn=JAZF1;refGene_tx=XM_011280443.3
A2	121770455	121770458
refGene_gn=CREB5;refGene_tx=XM_019825626.2;refGene_gn=CREB5;refGene_tx=XM_019825627.2
A2	121833722	121833740
refGene_gn=CREB5;refGene_tx=XM_019825628.2;refGene_gn=CREB5;refGene_tx=XM_019825629.2;refGene_gn=CREB5;refGene_tx=XM_023250458.1
A2	121840405	121840477
refGene_gn=CREB5;refGene_tx=XM_006929280.4;refGene_gn=CREB5;refGene_tx=XM_019825626.2;refGene_gn=CREB5;refGene_tx=XM_019825627.2;refGene_gn=CREB5;refGene_tx=XM_019825628.2;refGene_gn=CREB5;refGene_tx=XM_019825629.2;refGene_gn=CREB5;refGene_tx=XM_023250458.1
A2	121844806	121844900
refGene_gn=CREB5;refGene_tx=XM_006929280.4;refGene_gn=CREB5;refGene_tx=XM_019825626.2;refGene_gn=CREB5;refGene_tx=XM_019825627.2;refGene_gn=CREB5;refGene_tx=XM_019825628.2;refGene_gn=CREB5;refGene_tx=XM_019825629.2;refGene_gn=CREB5;refGene_tx=XM_023250458.1
A2	121857297	121857419
refGene_gn=CREB5;refGene_tx=XM_006929280.4;refGene_gn=CREB5;refGene_tx=XM_019825626.2;refGene_gn=CREB5;refGene_tx=XM_019825627.2;refGene_gn=CREB5;refGene_tx=XM_019825628.2;refGene_gn=CREB5;refGene_tx=XM_019825629.2;refGene_gn=CREB5;refGene_tx=XM_023250458.1
A2	121922082	121922255
refGene_gn=CREB5;refGene_tx=XM_006929280.4;refGene_gn=CREB5;refGene_tx=XM_019825626.2;refGene_gn=CREB5;refGene_tx=XM_019825627.2;refGene_gn=CREB5;refGene_tx=XM_019825628.2;refGene_gn=CREB5;refGene_tx=XM_019825629.2;refGene_gn=CREB5;refGene_tx=XM_019825630.2;refGene_gn=CREB5;refGene_tx=XM_023250458.1
A2	122041853	122041900

refGene_gn=CREB5;refGene_tx=XM_003982897.5
A2 122071531 122071658
refGene_gn=CREB5;refGene_tx=XM_003982897.5;refGene_gn=CREB5;refGene_tx=XM_006929280.4;refGene_gn=CREB5;refGene_tx=XM_019825626.2;refGene_gn=CREB5;refGene_tx=XM_019825627.2;refGene_gn=CREB5;refGene_tx=XM_019825628.2;refGene_gn=CREB5;refGene_tx=XM_019825629.2;refGene_gn=CREB5;refGene_tx=XM_019825630.2;refGene_gn=CREB5;refGene_tx=XM_023250458.1
A2 122077308 122077419
refGene_gn=CREB5;refGene_tx=XM_003982897.5;refGene_gn=CREB5;refGene_tx=XM_006929280.4;refGene_gn=CREB5;refGene_tx=XM_019825626.2;refGene_gn=CREB5;refGene_tx=XM_019825627.2;refGene_gn=CREB5;refGene_tx=XM_019825629.2;refGene_gn=CREB5;refGene_tx=XM_019825630.2;refGene_gn=CREB5;refGene_tx=XM_023250458.1
A2 122154157 122154475
refGene_gn=CREB5;refGene_tx=XM_003982897.5;refGene_gn=CREB5;refGene_tx=XM_006929280.4;refGene_gn=CREB5;refGene_tx=XM_019825626.2;refGene_gn=CREB5;refGene_tx=XM_019825627.2;refGene_gn=CREB5;refGene_tx=XM_019825628.2;refGene_gn=CREB5;refGene_tx=XM_019825629.2;refGene_gn=CREB5;refGene_tx=XM_019825630.2;refGene_gn=CREB5;refGene_tx=XM_023250458.1
A2 122158544 122158772
refGene_gn=CREB5;refGene_tx=XM_003982897.5;refGene_gn=CREB5;refGene_tx=XM_006929280.4;refGene_gn=CREB5;refGene_tx=XM_019825626.2;refGene_gn=CREB5;refGene_tx=XM_019825627.2;refGene_gn=CREB5;refGene_tx=XM_019825628.2;refGene_gn=CREB5;refGene_tx=XM_019825629.2;refGene_gn=CREB5;refGene_tx=XM_019825630.2;refGene_gn=CREB5;refGene_tx=XM_023250458.1
A2 122167063 122167172
refGene_gn=CREB5;refGene_tx=XM_003982897.5;refGene_gn=CREB5;refGene_tx=XM_006929280.4;refGene_gn=CREB5;refGene_tx=XM_019825626.2;refGene_gn=CREB5;refGene_tx=XM_019825627.2;refGene_gn=CREB5;refGene_tx=XM_019825628.2;refGene_gn=CREB5;refGene_tx=XM_019825629.2;refGene_gn=CREB5;refGene_tx=XM_019825630.2;refGene_gn=CREB5;refGene_tx=XM_023250458.1
A2 122167763 122167768
refGene_gn=CREB5;refGene_tx=XM_019825629.2
A2 122168103 122168267
refGene_gn=CREB5;refGene_tx=XM_003982897.5;refGene_gn=CREB5;refGene_tx=XM_006929280.4;refGene_gn=CREB5;refGene_tx=XM_019825626.2;refGene_gn=CREB5;refGene_tx=XM_019825627.2;refGene_gn=CREB5;refGene_tx=XM_019825628.2;refGene_gn=CREB5;refGene_tx=XM_019825630.2;refGene_gn=CREB5;refGene_tx=XM_023250458.1
A2 122290380 122292861
refGene_gn=TRIL;refGene_tx=XM_019825631.2;refGene_gn=TRIL;refGene_tx=XM_023250459.1;refGene_gn=TRIL;refGene_tx=XM_023250460.1
A2 122335322 122335427
refGene_gn=CPVL;refGene_tx=XM_019825633.2;refGene_gn=CPVL;refGene_tx=XM_023250461.1
A2 122369280 122369463
refGene_gn=CPVL;refGene_tx=XM_019825633.2;refGene_gn=CPVL;refGene_tx=XM_023250461.1
A2 122396709 122396883
refGene_gn=CPVL;refGene_tx=XM_019825633.2;refGene_gn=CPVL;refGene_tx=X

M_023250461.1
A2	122398606	122398705
refGene_gn=CPVL;refGene_tx=XM_019825633.2;refGene_gn=CPVL;refGene_tx=X
M_023250461.1
A2	122403947	122404079
refGene_gn=CPVL;refGene_tx=XM_019825633.2
A2	122404613	122404736
refGene_gn=CPVL;refGene_tx=XM_019825633.2
A2	122419434	122419501
refGene_gn=CPVL;refGene_tx=XM_019825633.2;refGene_gn=CPVL;refGene_tx=X
M_023250461.1
A2	122425630	122425710
refGene_gn=CPVL;refGene_tx=XM_019825633.2;refGene_gn=CPVL;refGene_tx=X
M_023250461.1
A2	122427590	122427649
refGene_gn=CPVL;refGene_tx=XM_019825633.2;refGene_gn=CPVL;refGene_tx=X
M_023250461.1
A2	122428610	122428725
refGene_gn=CPVL;refGene_tx=XM_019825633.2;refGene_gn=CPVL;refGene_tx=X
M_023250461.1
A2	122443118	122443237
refGene_gn=CPVL;refGene_tx=XM_019825633.2;refGene_gn=CPVL;refGene_tx=X
M_023250461.1
A2	122451230	122451393
refGene_gn=CPVL;refGene_tx=XM_019825633.2;refGene_gn=CPVL;refGene_tx=X
M_023250461.1
A2	122510202	122510251
refGene_gn=CHN2;refGene_tx=XM_023250462.1
A2	122666216	122666255
refGene_gn=CHN2;refGene_tx=XM_023250462.1
A2	122675838	122675894
refGene_gn=CHN2;refGene_tx=XM_023250462.1
A2	122703225	122703257
refGene_gn=CHN2;refGene_tx=XM_023250462.1
A2	122708128	122708242
refGene_gn=CHN2;refGene_tx=XM_023250462.1
A2	122710037	122710323
refGene_gn=CHN2;refGene_tx=XM_023250462.1
A2	122775399	122775402
refGene_gn=CHN2;refGene_tx=XM_023250463.1
A2	122782499	122782577
refGene_gn=CHN2;refGene_tx=XM_023250462.1;refGene_gn=CHN2;refGene_tx=X
M_023250463.1
A2	122800305	122800390
refGene_gn=CHN2;refGene_tx=XM_023250462.1;refGene_gn=CHN2;refGene_tx=X
M_023250463.1
A2	122804676	122804850
refGene_gn=CHN2;refGene_tx=XM_023250462.1;refGene_gn=CHN2;refGene_tx=X
M_023250463.1
A2	122808275	122808353

refGene_gn=CHN2;refGene_tx=XM_023250462.1;refGene_gn=CHN2;refGene_tx=XM_023250463.1
A2 122811000 122811138
refGene_gn=CHN2;refGene_tx=XM_023250462.1;refGene_gn=CHN2;refGene_tx=XM_023250463.1
A2 122813038 122813144
refGene_gn=CHN2;refGene_tx=XM_023250462.1;refGene_gn=CHN2;refGene_tx=XM_023250463.1
A2 122815910 122816082
refGene_gn=CHN2;refGene_tx=XM_023250462.1;refGene_gn=CHN2;refGene_tx=XM_023250463.1
A2 122842770 122843160
refGene_gn=PRR15;refGene_tx=XM_006929282.4;refGene_gn=PRR15;refGene_tx=XM_006929283.3;refGene_gn=PRR15;refGene_tx=XM_006929285.3;refGene_gn=PRR15;refGene_tx=XM_011280445.3
A2 122995692 122995863
refGene_gn=WIPF3;refGene_tx=XM_023242173.1
A2 123005393 123005457
refGene_gn=WIPF3;refGene_tx=XM_023242173.1
A2 123006635 123006797
refGene_gn=WIPF3;refGene_tx=XM_023242173.1
A2 123018097 123018283
refGene_gn=WIPF3;refGene_tx=XM_023242173.1
A2 123020339 123020471
refGene_gn=WIPF3;refGene_tx=XM_023242173.1
A2 123025523 123026276
refGene_gn=WIPF3;refGene_tx=XM_023242173.1
A2 123029209 123029359
refGene_gn=WIPF3;refGene_tx=XM_023242173.1
A2 123030369 123030471
refGene_gn=WIPF3;refGene_tx=XM_023242173.1
A2 123041315 123041392
refGene_gn=WIPF3;refGene_tx=XM_023242173.1
A2 123043433 123043457
refGene_gn=WIPF3;refGene_tx=XM_023242173.1
A2 123053189 123053348
refGene_gn=SCRN1;refGene_tx=XM_019825641.2;refGene_gn=SCRN1;refGene_tx=XM_023250464.1
A2 123055691 123055872
refGene_gn=SCRN1;refGene_tx=XM_019825641.2;refGene_gn=SCRN1;refGene_tx=XM_023250464.1
A2 123064760 123064926
refGene_gn=SCRN1;refGene_tx=XM_019825641.2;refGene_gn=SCRN1;refGene_tx=XM_023250464.1
A2 123067663 123067858
refGene_gn=SCRN1;refGene_tx=XM_019825641.2;refGene_gn=SCRN1;refGene_tx=XM_023250464.1
A2 123071031 123071234
refGene_gn=SCRN1;refGene_tx=XM_019825641.2;refGene_gn=SCRN1;refGene_tx=XM_023250464.1

A2 123075398 123075580
refGene_gn=SCRN1;refGene_tx=XM_019825641.2;refGene_gn=SCRN1;refGene_tx=XM_023250464.1
A2 123089583 123089743
refGene_gn=SCRN1;refGene_tx=XM_019825641.2;refGene_gn=SCRN1;refGene_tx=XM_023250464.1
A2 123090205 123090216
refGene_gn=SCRN1;refGene_tx=XM_019825641.2
A2 123120686 123120725
refGene_gn=FKBP14;refGene_tx=XM_019825642.2
A2 123124232 123124391
refGene_gn=FKBP14;refGene_tx=XM_003982902.4
A2 123127260 123127388
refGene_gn=FKBP14;refGene_tx=XM_003982902.4;refGene_gn=FKBP14;refGene_tx=XM_019825642.2
A2 123130724 123130876
refGene_gn=FKBP14;refGene_tx=XM_003982902.4;refGene_gn=FKBP14;refGene_tx=XM_019825642.2
A2 123133971 123134168
refGene_gn=FKBP14;refGene_tx=XM_003982902.4;refGene_gn=FKBP14;refGene_tx=XM_019825642.2
A2 123136482 123136522
refGene_gn=PLEKHA8;refGene_tx=XM_003982903.5;refGene_gn=PLEKHA8;refGene_tx=XM_019825643.2;refGene_gn=PLEKHA8;refGene_tx=XM_019825644.2;refGene_gn=PLEKHA8;refGene_tx=XM_019825645.2
A2 123158233 123158350
refGene_gn=PLEKHA8;refGene_tx=XM_003982903.5;refGene_gn=PLEKHA8;refGene_tx=XM_019825643.2;refGene_gn=PLEKHA8;refGene_tx=XM_019825644.2;refGene_gn=PLEKHA8;refGene_tx=XM_019825645.2
A2 123159588 123159744
refGene_gn=PLEKHA8;refGene_tx=XM_003982903.5;refGene_gn=PLEKHA8;refGene_tx=XM_019825643.2;refGene_gn=PLEKHA8;refGene_tx=XM_019825644.2;refGene_gn=PLEKHA8;refGene_tx=XM_019825645.2
A2 123161123 123161248
refGene_gn=PLEKHA8;refGene_tx=XM_003982903.5;refGene_gn=PLEKHA8;refGene_tx=XM_019825643.2;refGene_gn=PLEKHA8;refGene_tx=XM_019825644.2;refGene_gn=PLEKHA8;refGene_tx=XM_019825645.2
A2 123164884 123165043
refGene_gn=PLEKHA8;refGene_tx=XM_003982903.5;refGene_gn=PLEKHA8;refGene_tx=XM_019825643.2;refGene_gn=PLEKHA8;refGene_tx=XM_019825644.2;refGene_gn=PLEKHA8;refGene_tx=XM_019825645.2
A2 123166028 123166069
refGene_gn=PLEKHA8;refGene_tx=XM_003982903.5;refGene_gn=PLEKHA8;refGene_tx=XM_019825643.2;refGene_gn=PLEKHA8;refGene_tx=XM_019825644.2;refGene_gn=PLEKHA8;refGene_tx=XM_019825645.2
A2 123167806 123167961
refGene_gn=PLEKHA8;refGene_tx=XM_003982903.5;refGene_gn=PLEKHA8;refGene_tx=XM_019825643.2;refGene_gn=PLEKHA8;refGene_tx=XM_019825644.2;refGene_gn=PLEKHA8;refGene_tx=XM_019825645.2
A2 123169842 123170002

refGene_gn=PLEKHA8;refGene_tx=XM_003982903.5;refGene_gn=PLEKHA8;refGene_tx=XM_019825643.2;refGene_gn=PLEKHA8;refGene_tx=XM_019825644.2;refGene_gn=PLEKHA8;refGene_tx=XM_019825645.2
A2 123170393 123170479
refGene_gn=PLEKHA8;refGene_tx=XM_003982903.5;refGene_gn=PLEKHA8;refGene_tx=XM_019825643.2;refGene_gn=PLEKHA8;refGene_tx=XM_019825644.2;refGene_gn=PLEKHA8;refGene_tx=XM_019825645.2
A2 123174904 123174963
refGene_gn=PLEKHA8;refGene_tx=XM_003982903.5;refGene_gn=PLEKHA8;refGene_tx=XM_019825643.2;refGene_gn=PLEKHA8;refGene_tx=XM_019825644.2;refGene_gn=PLEKHA8;refGene_tx=XM_019825645.2
A2 123175795 123175926
refGene_gn=PLEKHA8;refGene_tx=XM_003982903.5;refGene_gn=PLEKHA8;refGene_tx=XM_019825643.2;refGene_gn=PLEKHA8;refGene_tx=XM_019825644.2;refGene_gn=PLEKHA8;refGene_tx=XM_019825645.2
A2 123176557 123176628
refGene_gn=PLEKHA8;refGene_tx=XM_003982903.5;refGene_gn=PLEKHA8;refGene_tx=XM_019825643.2;refGene_gn=PLEKHA8;refGene_tx=XM_019825644.2;refGene_gn=PLEKHA8;refGene_tx=XM_019825645.2
A2 123177730 123177936
refGene_gn=PLEKHA8;refGene_tx=XM_019825643.2;refGene_gn=PLEKHA8;refGene_tx=XM_019825645.2
A2 123186615 123186677
refGene_gn=PLEKHA8;refGene_tx=XM_003982903.5;refGene_gn=PLEKHA8;refGene_tx=XM_019825644.2
A2 123192075 123192273
refGene_gn=PLEKHA8;refGene_tx=XM_003982903.5;refGene_gn=PLEKHA8;refGene_tx=XM_019825644.2
A2 123236714 123236876
refGene_gn=MTURN;refGene_tx=XM_023250465.1
A2 123246917 123247040
refGene_gn=MTURN;refGene_tx=XM_023250465.1
A2 123261193 123261304
refGene_gn=MTURN;refGene_tx=XM_023250465.1
A2 123376933 123377405
refGene_gn=ZNRF2;refGene_tx=XM_023250466.1;refGene_gn=ZNRF2;refGene_tx=XM_023250467.1
A2 123431650 123431746
refGene_gn=ZNRF2;refGene_tx=XM_023250466.1;refGene_gn=ZNRF2;refGene_tx=XM_023250467.1
A2 123458548 123458700
refGene_gn=ZNRF2;refGene_tx=XM_023250467.1
A2 123464365 123464471
refGene_gn=ZNRF2;refGene_tx=XM_023250466.1
A2 123471397 123471455
refGene_gn=ZNRF2;refGene_tx=XM_023250466.1
A2 123521126 123521199
refGene_gn=NOD1;refGene_tx=XM_003982904.5;refGene_gn=NOD1;refGene_tx=XM_011280447.3;refGene_gn=NOD1;refGene_tx=XM_011280448.3;refGene_gn=NOD1;refGene_tx=XM_011280449.3;refGene_gn=NOD1;refGene_tx=XM_019825648.2;

refGene_gn=NOD1;refGene_tx=XM_019825649.2;refGene_gn=NOD1;refGene_tx=XM_023250468.1
A2 123531925 123532009
refGene_gn=NOD1;refGene_tx=XM_003982904.5;refGene_gn=NOD1;refGene_tx=XM_011280447.3;refGene_gn=NOD1;refGene_tx=XM_011280448.3;refGene_gn=NOD1;refGene_tx=XM_011280449.3;refGene_gn=NOD1;refGene_tx=XM_019825648.2;refGene_gn=NOD1;refGene_tx=XM_019825649.2;refGene_gn=NOD1;refGene_tx=XM_023250468.1
A2 123539240 123539324
refGene_gn=NOD1;refGene_tx=XM_003982904.5;refGene_gn=NOD1;refGene_tx=XM_011280447.3;refGene_gn=NOD1;refGene_tx=XM_011280448.3;refGene_gn=NOD1;refGene_tx=XM_011280449.3;refGene_gn=NOD1;refGene_tx=XM_019825648.2;refGene_gn=NOD1;refGene_tx=XM_019825649.2;refGene_gn=NOD1;refGene_tx=XM_023250468.1
A2 123541665 123541749
refGene_gn=NOD1;refGene_tx=XM_003982904.5;refGene_gn=NOD1;refGene_tx=XM_011280447.3;refGene_gn=NOD1;refGene_tx=XM_011280448.3;refGene_gn=NOD1;refGene_tx=XM_011280449.3;refGene_gn=NOD1;refGene_tx=XM_019825648.2;refGene_gn=NOD1;refGene_tx=XM_019825649.2;refGene_gn=NOD1;refGene_tx=XM_023250468.1
A2 123542639 123542723
refGene_gn=NOD1;refGene_tx=XM_003982904.5;refGene_gn=NOD1;refGene_tx=XM_011280447.3;refGene_gn=NOD1;refGene_tx=XM_011280448.3;refGene_gn=NOD1;refGene_tx=XM_011280449.3;refGene_gn=NOD1;refGene_tx=XM_019825648.2;refGene_gn=NOD1;refGene_tx=XM_019825649.2;refGene_gn=NOD1;refGene_tx=XM_023250468.1
A2 123545902 123545939
refGene_gn=NOD1;refGene_tx=XM_019825650.2
A2 123546558 123546642
refGene_gn=NOD1;refGene_tx=XM_003982904.5;refGene_gn=NOD1;refGene_tx=XM_011280447.3;refGene_gn=NOD1;refGene_tx=XM_011280448.3;refGene_gn=NOD1;refGene_tx=XM_011280449.3;refGene_gn=NOD1;refGene_tx=XM_019825648.2;refGene_gn=NOD1;refGene_tx=XM_019825649.2;refGene_gn=NOD1;refGene_tx=XM_019825650.2;refGene_gn=NOD1;refGene_tx=XM_023250468.1
A2 123547452 123547536
refGene_gn=NOD1;refGene_tx=XM_003982904.5;refGene_gn=NOD1;refGene_tx=XM_011280447.3;refGene_gn=NOD1;refGene_tx=XM_011280448.3;refGene_gn=NOD1;refGene_tx=XM_011280449.3;refGene_gn=NOD1;refGene_tx=XM_019825648.2;refGene_gn=NOD1;refGene_tx=XM_019825649.2;refGene_gn=NOD1;refGene_tx=XM_019825650.2;refGene_gn=NOD1;refGene_tx=XM_023250468.1
A2 123547730 123547836
refGene_gn=NOD1;refGene_tx=XM_019825651.2
A2 123548704 123548788
refGene_gn=NOD1;refGene_tx=XM_003982904.5;refGene_gn=NOD1;refGene_tx=XM_011280447.3;refGene_gn=NOD1;refGene_tx=XM_011280448.3;refGene_gn=NOD1;refGene_tx=XM_011280449.3;refGene_gn=NOD1;refGene_tx=XM_019825648.2;refGene_gn=NOD1;refGene_tx=XM_019825649.2;refGene_gn=NOD1;refGene_tx=XM_019825650.2;refGene_gn=NOD1;refGene_tx=XM_019825651.2;refGene_gn=NOD1;refGene_tx=XM_023250468.1
A2 123551479 123553304

refGene_gn=NOD1;refGene_tx=XM_003982904.5;refGene_gn=NOD1;refGene_tx=XM_011280447.3;refGene_gn=NOD1;refGene_tx=XM_011280448.3;refGene_gn=NOD1;refGene_tx=XM_011280449.3;refGene_gn=NOD1;refGene_tx=XM_019825648.2;refGene_gn=NOD1;refGene_tx=XM_019825649.2;refGene_gn=NOD1;refGene_tx=XM_019825650.2;refGene_gn=NOD1;refGene_tx=XM_019825651.2;refGene_gn=NOD1;refGene_tx=XM_023250468.1
A2 123557172 123557347
refGene_gn=NOD1;refGene_tx=XM_003982904.5;refGene_gn=NOD1;refGene_tx=XM_011280447.3;refGene_gn=NOD1;refGene_tx=XM_011280448.3;refGene_gn=NOD1;refGene_tx=XM_011280449.3;refGene_gn=NOD1;refGene_tx=XM_019825648.2;refGene_gn=NOD1;refGene_tx=XM_019825649.2;refGene_gn=NOD1;refGene_tx=XM_019825650.2;refGene_gn=NOD1;refGene_tx=XM_019825651.2;refGene_gn=NOD1;refGene_tx=XM_023250468.1
A2 123558255 123558456
refGene_gn=NOD1;refGene_tx=XM_003982904.5;refGene_gn=NOD1;refGene_tx=XM_011280447.3;refGene_gn=NOD1;refGene_tx=XM_011280448.3;refGene_gn=NOD1;refGene_tx=XM_011280449.3;refGene_gn=NOD1;refGene_tx=XM_019825648.2;refGene_gn=NOD1;refGene_tx=XM_019825649.2;refGene_gn=NOD1;refGene_tx=XM_019825650.2;refGene_gn=NOD1;refGene_tx=XM_019825651.2;refGene_gn=NOD1;refGene_tx=XM_023250468.1
A2 123614474 123614618
refGene_gn=GGCT;refGene_tx=XM_003982905.5
A2 123616975 123617111
refGene_gn=GGCT;refGene_tx=XM_003982905.5
A2 123618510 123618656
refGene_gn=GGCT;refGene_tx=XM_003982905.5
A2 123622887 123623028
refGene_gn=GGCT;refGene_tx=XM_003982905.5
A2 123686317 123686539
refGene_gn=GARS;refGene_tx=XM_003982939.5
A2 123698185 123698287
refGene_gn=GARS;refGene_tx=XM_003982939.5
A2 123699010 123699113
refGene_gn=GARS;refGene_tx=XM_003982939.5
A2 123700406 123700548
refGene_gn=GARS;refGene_tx=XM_003982939.5
A2 123701462 123701551
refGene_gn=GARS;refGene_tx=XM_003982939.5
A2 123702452 123702529
refGene_gn=GARS;refGene_tx=XM_003982939.5
A2 123705856 123706002
refGene_gn=GARS;refGene_tx=XM_003982939.5
A2 123708359 123708509
refGene_gn=GARS;refGene_tx=XM_003982939.5
A2 123712787 123712950
refGene_gn=GARS;refGene_tx=XM_003982939.5
A2 123714268 123714433
refGene_gn=GARS;refGene_tx=XM_003982939.5
A2 123718534 123718642
refGene_gn=GARS;refGene_tx=XM_003982939.5

A2 123719454 123719600
refGene_gn=GARS;refGene_tx=XM_003982939.5
A2 123723932 123724018
refGene_gn=GARS;refGene_tx=XM_003982939.5
A2 123725813 123725923
refGene_gn=GARS;refGene_tx=XM_003982939.5
A2 123728945 123729039
refGene_gn=GARS;refGene_tx=XM_003982939.5
A2 123729720 123729911
refGene_gn=GARS;refGene_tx=XM_003982939.5
A2 123731500 123731626
refGene_gn=GARS;refGene_tx=XM_003982939.5
A2 123748175 123748316
refGene_gn=CRHR2;refGene_tx=XM_023250469.1;refGene_gn=CRHR2;refGene_tx
=XM_023250470.1;refGene_gn=CRHR2;refGene_tx=XM_023250471.1
A2 123749657 123749699
refGene_gn=CRHR2;refGene_tx=XM_023250469.1;refGene_gn=CRHR2;refGene_tx
=XM_023250470.1;refGene_gn=CRHR2;refGene_tx=XM_023250471.1
A2 123750105 123750241
refGene_gn=CRHR2;refGene_tx=XM_023250469.1;refGene_gn=CRHR2;refGene_tx
=XM_023250470.1;refGene_gn=CRHR2;refGene_tx=XM_023250471.1
A2 123750421 123750507
refGene_gn=CRHR2;refGene_tx=XM_023250469.1;refGene_gn=CRHR2;refGene_tx
=XM_023250470.1;refGene_gn=CRHR2;refGene_tx=XM_023250471.1
A2 123756669 123756742
refGene_gn=CRHR2;refGene_tx=XM_023250469.1;refGene_gn=CRHR2;refGene_tx
=XM_023250470.1;refGene_gn=CRHR2;refGene_tx=XM_023250471.1
A2 123758242 123758303
refGene_gn=CRHR2;refGene_tx=XM_023250469.1;refGene_gn=CRHR2;refGene_tx
=XM_023250470.1;refGene_gn=CRHR2;refGene_tx=XM_023250471.1
A2 123758799 123758953
refGene_gn=CRHR2;refGene_tx=XM_023250469.1;refGene_gn=CRHR2;refGene_tx
=XM_023250470.1;refGene_gn=CRHR2;refGene_tx=XM_023250471.1
A2 123761076 123761194
refGene_gn=CRHR2;refGene_tx=XM_023250469.1;refGene_gn=CRHR2;refGene_tx
=XM_023250470.1;refGene_gn=CRHR2;refGene_tx=XM_023250471.1
A2 123761554 123761664
refGene_gn=CRHR2;refGene_tx=XM_023250469.1;refGene_gn=CRHR2;refGene_tx
=XM_023250470.1;refGene_gn=CRHR2;refGene_tx=XM_023250471.1
A2 123763309 123763395
refGene_gn=CRHR2;refGene_tx=XM_023250469.1;refGene_gn=CRHR2;refGene_tx
=XM_023250470.1;refGene_gn=CRHR2;refGene_tx=XM_023250471.1
A2 123776900 123777026
refGene_gn=CRHR2;refGene_tx=XM_023250469.1;refGene_gn=CRHR2;refGene_tx
=XM_023250470.1;refGene_gn=CRHR2;refGene_tx=XM_023250471.1
A2 123777162 123777265
refGene_gn=CRHR2;refGene_tx=XM_023250470.1
A2 123783975 123784069
refGene_gn=CRHR2;refGene_tx=XM_023250469.1;refGene_gn=CRHR2;refGene_tx
=XM_023250471.1

A2	123793233	123793320
refGene_gn=CRHR2;refGene_tx=XM_023250469.1
A2	123853010	123853073
refGene_gn=MINDY4;refGene_tx=XM_011280450.3;refGene_gn=MINDY4;refGene_tx=XM_019825654.2
A2	123860986	123861106
refGene_gn=MINDY4;refGene_tx=XM_011280450.3;refGene_gn=MINDY4;refGene_tx=XM_019825653.2;refGene_gn=MINDY4;refGene_tx=XM_019825654.2
A2	123865034	123865270
refGene_gn=MINDY4;refGene_tx=XM_011280450.3;refGene_gn=MINDY4;refGene_tx=XM_019825653.2;refGene_gn=MINDY4;refGene_tx=XM_019825654.2
A2	123868243	123868490
refGene_gn=MINDY4;refGene_tx=XM_011280450.3;refGene_gn=MINDY4;refGene_tx=XM_019825653.2;refGene_gn=MINDY4;refGene_tx=XM_019825654.2
A2	123873974	123874405
refGene_gn=MINDY4;refGene_tx=XM_011280450.3;refGene_gn=MINDY4;refGene_tx=XM_019825653.2;refGene_gn=MINDY4;refGene_tx=XM_019825654.2
A2	123903660	123903719
refGene_gn=MINDY4;refGene_tx=XM_011280450.3;refGene_gn=MINDY4;refGene_tx=XM_019825653.2;refGene_gn=MINDY4;refGene_tx=XM_019825654.2
A2	123908917	123909024
refGene_gn=MINDY4;refGene_tx=XM_011280450.3;refGene_gn=MINDY4;refGene_tx=XM_019825653.2;refGene_gn=MINDY4;refGene_tx=XM_019825654.2
A2	123911496	123911613
refGene_gn=MINDY4;refGene_tx=XM_011280450.3;refGene_gn=MINDY4;refGene_tx=XM_019825653.2;refGene_gn=MINDY4;refGene_tx=XM_019825654.2
A2	123912838	123912927
refGene_gn=MINDY4;refGene_tx=XM_011280450.3;refGene_gn=MINDY4;refGene_tx=XM_019825653.2;refGene_gn=MINDY4;refGene_tx=XM_019825654.2
A2	123921825	123921927
refGene_gn=MINDY4;refGene_tx=XM_011280450.3;refGene_gn=MINDY4;refGene_tx=XM_019825653.2;refGene_gn=MINDY4;refGene_tx=XM_019825654.2
A2	123923759	123923823
refGene_gn=MINDY4;refGene_tx=XM_011280450.3;refGene_gn=MINDY4;refGene_tx=XM_019825653.2;refGene_gn=MINDY4;refGene_tx=XM_019825654.2
A2	123924892	123924958
refGene_gn=MINDY4;refGene_tx=XM_011280450.3;refGene_gn=MINDY4;refGene_tx=XM_019825653.2;refGene_gn=MINDY4;refGene_tx=XM_019825654.2
A2	123930387	123930455
refGene_gn=MINDY4;refGene_tx=XM_011280450.3;refGene_gn=MINDY4;refGene_tx=XM_019825653.2;refGene_gn=MINDY4;refGene_tx=XM_019825654.2
A2	123943029	123943093
refGene_gn=MINDY4;refGene_tx=XM_011280450.3;refGene_gn=MINDY4;refGene_tx=XM_019825653.2;refGene_gn=MINDY4;refGene_tx=XM_019825654.2
A2	123946183	123946397
refGene_gn=MINDY4;refGene_tx=XM_011280450.3;refGene_gn=MINDY4;refGene_tx=XM_019825653.2;refGene_gn=MINDY4;refGene_tx=XM_019825654.2
A2	123952719	123952900
refGene_gn=MINDY4;refGene_tx=XM_011280450.3;refGene_gn=MINDY4;refGene_tx=XM_019825653.2;refGene_gn=MINDY4;refGene_tx=XM_019825654.2

A2	123953433	123953506
refGene_gn=MINDY4;refGene_tx=XM_011280450.3;refGene_gn=MINDY4;refGene_tx=XM_019825653.2
A2	123962917	123962966
refGene_gn=MINDY4;refGene_tx=XM_011280450.3;refGene_gn=MINDY4;refGene_tx=XM_019825653.2
A2	123982439	123982829
refGene_gn=AQP1;refGene_tx=XM_003982907.4
A2	123992092	123992257
refGene_gn=AQP1;refGene_tx=XM_003982907.4
A2	123992563	123992644
refGene_gn=AQP1;refGene_tx=XM_003982907.4
A2	123993384	123993564
refGene_gn=AQP1;refGene_tx=XM_003982907.4
A2	124018061	124018118
refGene_gn=GHRHR;refGene_tx=XM_003982909.2
A2	124020219	124020322
refGene_gn=GHRHR;refGene_tx=XM_003982909.2
A2	124020439	124020547
refGene_gn=GHRHR;refGene_tx=XM_003982909.2;refGene_gn=GHRHR;refGene_tx=XM_023250472.1
A2	124021095	124021193
refGene_gn=GHRHR;refGene_tx=XM_003982909.2;refGene_gn=GHRHR;refGene_tx=XM_023250472.1
A2	124022247	124022345
refGene_gn=GHRHR;refGene_tx=XM_003982909.2;refGene_gn=GHRHR;refGene_tx=XM_023250472.1
A2	124023046	124023179
refGene_gn=GHRHR;refGene_tx=XM_003982909.2;refGene_gn=GHRHR;refGene_tx=XM_023250472.1
A2	124024411	124024565
refGene_gn=GHRHR;refGene_tx=XM_003982909.2;refGene_gn=GHRHR;refGene_tx=XM_023250472.1
A2	124024845	124024906
refGene_gn=GHRHR;refGene_tx=XM_003982909.2;refGene_gn=GHRHR;refGene_tx=XM_023250472.1
A2	124025421	124025491
refGene_gn=GHRHR;refGene_tx=XM_003982909.2;refGene_gn=GHRHR;refGene_tx=XM_023250472.1
A2	124026191	124026283
refGene_gn=GHRHR;refGene_tx=XM_003982909.2;refGene_gn=GHRHR;refGene_tx=XM_023250472.1
A2	124026848	124026978
refGene_gn=GHRHR;refGene_tx=XM_003982909.2;refGene_gn=GHRHR;refGene_tx=XM_023250472.1
A2	124027653	124027695
refGene_gn=GHRHR;refGene_tx=XM_003982909.2;refGene_gn=GHRHR;refGene_tx=XM_023250472.1
A2	124029475	124029601
refGene_gn=GHRHR;refGene_tx=XM_003982909.2;refGene_gn=GHRHR;refGene_tx

=XM_023250472.1
A2	124102878	124102929
refGene_gn=ADCYAP1R1;refGene_tx=XM_003982910.5;refGene_gn=ADCYAP1R1;refGene_tx=XM_003982912.5;refGene_gn=ADCYAP1R1;refGene_tx=XM_006929290.4;refGene_gn=ADCYAP1R1;refGene_tx=XM_019825655.2;refGene_gn=ADCYAP1R1;refGene_tx=XM_019825656.2;refGene_gn=ADCYAP1R1;refGene_tx=XM_019825657.2
A2	124104446	124104552
refGene_gn=ADCYAP1R1;refGene_tx=XM_003982910.5;refGene_gn=ADCYAP1R1;refGene_tx=XM_003982912.5;refGene_gn=ADCYAP1R1;refGene_tx=XM_006929290.4;refGene_gn=ADCYAP1R1;refGene_tx=XM_019825655.2;refGene_gn=ADCYAP1R1;refGene_tx=XM_019825656.2;refGene_gn=ADCYAP1R1;refGene_tx=XM_019825657.2
A2	124118475	124118583
refGene_gn=ADCYAP1R1;refGene_tx=XM_003982910.5;refGene_gn=ADCYAP1R1;refGene_tx=XM_003982912.5;refGene_gn=ADCYAP1R1;refGene_tx=XM_006929290.4;refGene_gn=ADCYAP1R1;refGene_tx=XM_019825655.2;refGene_gn=ADCYAP1R1;refGene_tx=XM_019825656.2;refGene_gn=ADCYAP1R1;refGene_tx=XM_019825657.2
A2	124121145	124121166
refGene_gn=ADCYAP1R1;refGene_tx=XM_003982910.5;refGene_gn=ADCYAP1R1;refGene_tx=XM_003982912.5;refGene_gn=ADCYAP1R1;refGene_tx=XM_006929290.4;refGene_gn=ADCYAP1R1;refGene_tx=XM_019825655.2;refGene_gn=ADCYAP1R1;refGene_tx=XM_019825656.2;refGene_gn=ADCYAP1R1;refGene_tx=XM_019825657.2
A2	124121345	124121358
refGene_gn=ADCYAP1R1;refGene_tx=XM_023250473.1
A2	124122207	124122249
refGene_gn=ADCYAP1R1;refGene_tx=XM_003982910.5;refGene_gn=ADCYAP1R1;refGene_tx=XM_003982912.5;refGene_gn=ADCYAP1R1;refGene_tx=XM_006929290.4;refGene_gn=ADCYAP1R1;refGene_tx=XM_019825655.2;refGene_gn=ADCYAP1R1;refGene_tx=XM_019825656.2;refGene_gn=ADCYAP1R1;refGene_tx=XM_019825657.2;refGene_gn=ADCYAP1R1;refGene_tx=XM_023250473.1
A2	124124640	124124750
refGene_gn=ADCYAP1R1;refGene_tx=XM_003982910.5;refGene_gn=ADCYAP1R1;refGene_tx=XM_003982912.5;refGene_gn=ADCYAP1R1;refGene_tx=XM_006929290.4;refGene_gn=ADCYAP1R1;refGene_tx=XM_019825655.2;refGene_gn=ADCYAP1R1;refGene_tx=XM_019825656.2;refGene_gn=ADCYAP1R1;refGene_tx=XM_019825657.2;refGene_gn=ADCYAP1R1;refGene_tx=XM_023250473.1
A2	124125357	124125455
refGene_gn=ADCYAP1R1;refGene_tx=XM_003982910.5;refGene_gn=ADCYAP1R1;refGene_tx=XM_003982912.5;refGene_gn=ADCYAP1R1;refGene_tx=XM_006929290.4;refGene_gn=ADCYAP1R1;refGene_tx=XM_019825655.2;refGene_gn=ADCYAP1R1;refGene_tx=XM_019825656.2;refGene_gn=ADCYAP1R1;refGene_tx=XM_019825657.2;refGene_gn=ADCYAP1R1;refGene_tx=XM_023250473.1
A2	124125907	124126040
refGene_gn=ADCYAP1R1;refGene_tx=XM_003982910.5;refGene_gn=ADCYAP1R1;refGene_tx=XM_003982912.5;refGene_gn=ADCYAP1R1;refGene_tx=XM_006929290.4;refGene_gn=ADCYAP1R1;refGene_tx=XM_019825655.2;refGene_gn=ADCYAP1R1;refGene_tx=XM_019825656.2;refGene_gn=ADCYAP1R1;refGene_tx=XM_019825657.

2;refGene_gn=ADCYAP1R1;refGene_tx=XM_023250473.1
A2 124126974 124127128
refGene_gn=ADCYAP1R1;refGene_tx=XM_003982910.5;refGene_gn=ADCYAP1R1;refGene_tx=XM_003982912.5;refGene_gn=ADCYAP1R1;refGene_tx=XM_006929290.4;refGene_gn=ADCYAP1R1;refGene_tx=XM_019825655.2;refGene_gn=ADCYAP1R1;refGene_tx=XM_019825656.2;refGene_gn=ADCYAP1R1;refGene_tx=XM_019825657.2;refGene_gn=ADCYAP1R1;refGene_tx=XM_023250473.1
A2 124127564 124127625
refGene_gn=ADCYAP1R1;refGene_tx=XM_003982910.5;refGene_gn=ADCYAP1R1;refGene_tx=XM_003982912.5;refGene_gn=ADCYAP1R1;refGene_tx=XM_006929290.4;refGene_gn=ADCYAP1R1;refGene_tx=XM_019825655.2;refGene_gn=ADCYAP1R1;refGene_tx=XM_019825656.2;refGene_gn=ADCYAP1R1;refGene_tx=XM_019825657.2;refGene_gn=ADCYAP1R1;refGene_tx=XM_023250473.1
A2 124128158 124128228
refGene_gn=ADCYAP1R1;refGene_tx=XM_003982910.5;refGene_gn=ADCYAP1R1;refGene_tx=XM_003982912.5;refGene_gn=ADCYAP1R1;refGene_tx=XM_006929290.4;refGene_gn=ADCYAP1R1;refGene_tx=XM_019825655.2;refGene_gn=ADCYAP1R1;refGene_tx=XM_019825656.2;refGene_gn=ADCYAP1R1;refGene_tx=XM_019825657.2;refGene_gn=ADCYAP1R1;refGene_tx=XM_023250473.1
A2 124132727 124132819
refGene_gn=ADCYAP1R1;refGene_tx=XM_003982910.5;refGene_gn=ADCYAP1R1;refGene_tx=XM_003982912.5;refGene_gn=ADCYAP1R1;refGene_tx=XM_006929290.4;refGene_gn=ADCYAP1R1;refGene_tx=XM_019825655.2;refGene_gn=ADCYAP1R1;refGene_tx=XM_019825656.2;refGene_gn=ADCYAP1R1;refGene_tx=XM_019825657.2;refGene_gn=ADCYAP1R1;refGene_tx=XM_023250473.1
A2 124135719 124135803
refGene_gn=ADCYAP1R1;refGene_tx=XM_019825655.2;refGene_gn=ADCYAP1R1;refGene_tx=XM_019825656.2;refGene_gn=ADCYAP1R1;refGene_tx=XM_019825657.2;refGene_gn=ADCYAP1R1;refGene_tx=XM_023250473.1
A2 124139828 124139912
refGene_gn=ADCYAP1R1;refGene_tx=XM_003982912.5;refGene_gn=ADCYAP1R1;refGene_tx=XM_006929290.4;refGene_gn=ADCYAP1R1;refGene_tx=XM_019825655.2;refGene_gn=ADCYAP1R1;refGene_tx=XM_019825656.2;refGene_gn=ADCYAP1R1;refGene_tx=XM_023250473.1
A2 124143038 124143168
refGene_gn=ADCYAP1R1;refGene_tx=XM_003982910.5;refGene_gn=ADCYAP1R1;refGene_tx=XM_003982912.5;refGene_gn=ADCYAP1R1;refGene_tx=XM_006929290.4;refGene_gn=ADCYAP1R1;refGene_tx=XM_019825655.2;refGene_gn=ADCYAP1R1;refGene_tx=XM_019825656.2;refGene_gn=ADCYAP1R1;refGene_tx=XM_019825657.2;refGene_gn=ADCYAP1R1;refGene_tx=XM_023250473.1
A2 124144540 124144582
refGene_gn=ADCYAP1R1;refGene_tx=XM_003982910.5;refGene_gn=ADCYAP1R1;refGene_tx=XM_003982912.5;refGene_gn=ADCYAP1R1;refGene_tx=XM_006929290.4;refGene_gn=ADCYAP1R1;refGene_tx=XM_019825655.2;refGene_gn=ADCYAP1R1;refGene_tx=XM_019825656.2;refGene_gn=ADCYAP1R1;refGene_tx=XM_019825657.2;refGene_gn=ADCYAP1R1;refGene_tx=XM_023250473.1
A2 124145999 124146188
refGene_gn=ADCYAP1R1;refGene_tx=XM_003982910.5;refGene_gn=ADCYAP1R1;refGene_tx=XM_003982912.5;refGene_gn=ADCYAP1R1;refGene_tx=XM_006929290.4;refGene_gn=ADCYAP1R1;refGene_tx=XM_019825655.2;refGene_gn=ADCYAP1R1;r

efGene_tx=XM_019825656.2;refGene_gn=ADCYAP1R1;refGene_tx=XM_019825657.2;refGene_gn=ADCYAP1R1;refGene_tx=XM_023250473.1
A2 124312890 124313859
refGene_gn=NEUROD6;refGene_tx=XM_023250474.1
A2 124313860 124313905
refGene_gn=NEUROD6;refGene_tx=XM_023250474.1
A2 124480781 124480819
refGene_gn=CCDC129;refGene_tx=XM_023242186.1
A2 124492016 124492198
refGene_gn=CCDC129;refGene_tx=XM_023242186.1
A2 124493589 124493638
refGene_gn=CCDC129;refGene_tx=XM_023242186.1
A2 124493967 124494011
refGene_gn=CCDC129;refGene_tx=XM_023242186.1
A2 124503172 124503224
refGene_gn=CCDC129;refGene_tx=XM_023242186.1
A2 124506127 124506214
refGene_gn=CCDC129;refGene_tx=XM_023242186.1
A2 124508225 124508428
refGene_gn=CCDC129;refGene_tx=XM_023242186.1
A2 124511529 124512104
refGene_gn=CCDC129;refGene_tx=XM_023242186.1
A2 124518047 124518105
refGene_gn=CCDC129;refGene_tx=XM_023242186.1
A2 124557480 124557563
refGene_gn=CCDC129;refGene_tx=XM_023242186.1
A2 124558620 124559910
refGene_gn=CCDC129;refGene_tx=XM_023242186.1
A2 124564516 124564644
refGene_gn=CCDC129;refGene_tx=XM_023242186.1
A2 124565876 124565988
refGene_gn=CCDC129;refGene_tx=XM_023242186.1
A2 124566460 124566766
refGene_gn=CCDC129;refGene_tx=XM_023242186.1
A2 124609841 124609983
refGene_gn=PPP1R17;refGene_tx=XM_019825659.2
A2 124613879 124613937
refGene_gn=PPP1R17;refGene_tx=XM_019825659.2
A2 124615116 124615223
refGene_gn=PPP1R17;refGene_tx=XM_006929292.3;refGene_gn=PPP1R17;refGene_tx=XM_019825659.2
A2 124618607 124618760
refGene_gn=PPP1R17;refGene_tx=XM_006929292.3;refGene_gn=PPP1R17;refGene_tx=XM_019825659.2
A2 124620135 124620288
refGene_gn=PPP1R17;refGene_tx=XM_006929292.3;refGene_gn=PPP1R17;refGene_tx=XM_019825659.2
A2 124626494 124626574
refGene_gn=PPP1R17;refGene_tx=XM_006929292.3;refGene_gn=PPP1R17;refGene_tx=XM_019825659.2

A2	124662469	124662639
refGene_gn=PDE1C;refGene_tx=XM_019825660.1;refGene_gn=PDE1C;refGene_tx=XM_019825661.1;refGene_gn=PDE1C;refGene_tx=XM_023250475.1;refGene_gn=PDE1C;refGene_tx=XM_023250476.1
A2	124673400	124673414
refGene_gn=PDE1C;refGene_tx=XM_019825662.2
A2	124677860	124677868
refGene_gn=PDE1C;refGene_tx=XM_019825663.2
A2	124680602	124680671
refGene_gn=PDE1C;refGene_tx=XM_019825660.1;refGene_gn=PDE1C;refGene_tx=XM_019825662.2;refGene_gn=PDE1C;refGene_tx=XM_019825663.2;refGene_gn=PDE1C;refGene_tx=XM_023250475.1;refGene_gn=PDE1C;refGene_tx=XM_023250476.1
A2	124694233	124694247
refGene_gn=PDE1C;refGene_tx=XM_019825664.2
A2	124711810	124711888
refGene_gn=PDE1C;refGene_tx=XM_019825660.1;refGene_gn=PDE1C;refGene_tx=XM_019825661.1;refGene_gn=PDE1C;refGene_tx=XM_019825662.2;refGene_gn=PDE1C;refGene_tx=XM_019825663.2;refGene_gn=PDE1C;refGene_tx=XM_019825664.2;refGene_gn=PDE1C;refGene_tx=XM_023250475.1;refGene_gn=PDE1C;refGene_tx=XM_023250476.1
A2	124718482	124718713
refGene_gn=PDE1C;refGene_tx=XM_019825660.1;refGene_gn=PDE1C;refGene_tx=XM_019825661.1;refGene_gn=PDE1C;refGene_tx=XM_019825662.2;refGene_gn=PDE1C;refGene_tx=XM_019825663.2;refGene_gn=PDE1C;refGene_tx=XM_019825664.2;refGene_gn=PDE1C;refGene_tx=XM_023250475.1;refGene_gn=PDE1C;refGene_tx=XM_023250476.1
A2	124728559	124728735
refGene_gn=PDE1C;refGene_tx=XM_019825660.1;refGene_gn=PDE1C;refGene_tx=XM_019825661.1;refGene_gn=PDE1C;refGene_tx=XM_019825662.2;refGene_gn=PDE1C;refGene_tx=XM_019825663.2;refGene_gn=PDE1C;refGene_tx=XM_019825664.2;refGene_gn=PDE1C;refGene_tx=XM_023250475.1;refGene_gn=PDE1C;refGene_tx=XM_023250476.1
A2	124730371	124730492
refGene_gn=PDE1C;refGene_tx=XM_019825660.1;refGene_gn=PDE1C;refGene_tx=XM_019825661.1;refGene_gn=PDE1C;refGene_tx=XM_019825662.2;refGene_gn=PDE1C;refGene_tx=XM_019825663.2;refGene_gn=PDE1C;refGene_tx=XM_019825664.2;refGene_gn=PDE1C;refGene_tx=XM_023250475.1;refGene_gn=PDE1C;refGene_tx=XM_023250476.1
A2	124733848	124733930
refGene_gn=PDE1C;refGene_tx=XM_019825660.1;refGene_gn=PDE1C;refGene_tx=XM_019825661.1;refGene_gn=PDE1C;refGene_tx=XM_019825662.2;refGene_gn=PDE1C;refGene_tx=XM_019825663.2;refGene_gn=PDE1C;refGene_tx=XM_019825664.2;refGene_gn=PDE1C;refGene_tx=XM_023250475.1;refGene_gn=PDE1C;refGene_tx=XM_023250476.1
A2	124742153	124742274
refGene_gn=PDE1C;refGene_tx=XM_019825660.1;refGene_gn=PDE1C;refGene_tx=XM_019825661.1;refGene_gn=PDE1C;refGene_tx=XM_019825662.2;refGene_gn=PDE1C;refGene_tx=XM_019825663.2;refGene_gn=PDE1C;refGene_tx=XM_019825664.2;refGene_gn=PDE1C;refGene_tx=XM_023250475.1;refGene_gn=PDE1C;refGe

ne_tx=XM_023250476.1
A2 124742822 124742924
refGene_gn=PDE1C;refGene_tx=XM_019825660.1;refGene_gn=PDE1C;refGene_tx=XM_019825661.1;refGene_gn=PDE1C;refGene_tx=XM_019825662.2;refGene_gn=PDE1C;refGene_tx=XM_019825663.2;refGene_gn=PDE1C;refGene_tx=XM_019825664.2;refGene_gn=PDE1C;refGene_tx=XM_023250475.1;refGene_gn=PDE1C;refGene_tx=XM_023250476.1
A2 124744966 124745095
refGene_gn=PDE1C;refGene_tx=XM_019825660.1;refGene_gn=PDE1C;refGene_tx=XM_019825661.1;refGene_gn=PDE1C;refGene_tx=XM_019825662.2;refGene_gn=PDE1C;refGene_tx=XM_019825663.2;refGene_gn=PDE1C;refGene_tx=XM_019825664.2;refGene_gn=PDE1C;refGene_tx=XM_023250475.1;refGene_gn=PDE1C;refGene_tx=XM_023250476.1
A2 124747573 124747674
refGene_gn=PDE1C;refGene_tx=XM_019825660.1;refGene_gn=PDE1C;refGene_tx=XM_019825661.1;refGene_gn=PDE1C;refGene_tx=XM_019825662.2;refGene_gn=PDE1C;refGene_tx=XM_019825663.2;refGene_gn=PDE1C;refGene_tx=XM_019825664.2;refGene_gn=PDE1C;refGene_tx=XM_023250475.1;refGene_gn=PDE1C;refGene_tx=XM_023250476.1
A2 124759410 124759551
refGene_gn=PDE1C;refGene_tx=XM_019825660.1;refGene_gn=PDE1C;refGene_tx=XM_019825661.1;refGene_gn=PDE1C;refGene_tx=XM_019825662.2;refGene_gn=PDE1C;refGene_tx=XM_019825663.2;refGene_gn=PDE1C;refGene_tx=XM_019825664.2;refGene_gn=PDE1C;refGene_tx=XM_023250475.1;refGene_gn=PDE1C;refGene_tx=XM_023250476.1
A2 124761946 124762063
refGene_gn=PDE1C;refGene_tx=XM_019825660.1;refGene_gn=PDE1C;refGene_tx=XM_019825661.1;refGene_gn=PDE1C;refGene_tx=XM_019825662.2;refGene_gn=PDE1C;refGene_tx=XM_019825663.2;refGene_gn=PDE1C;refGene_tx=XM_019825664.2;refGene_gn=PDE1C;refGene_tx=XM_023250475.1;refGene_gn=PDE1C;refGene_tx=XM_023250476.1
A2 124767687 124767754
refGene_gn=PDE1C;refGene_tx=XM_019825660.1;refGene_gn=PDE1C;refGene_tx=XM_019825661.1;refGene_gn=PDE1C;refGene_tx=XM_019825662.2;refGene_gn=PDE1C;refGene_tx=XM_019825663.2;refGene_gn=PDE1C;refGene_tx=XM_019825664.2;refGene_gn=PDE1C;refGene_tx=XM_023250475.1;refGene_gn=PDE1C;refGene_tx=XM_023250476.1
A2 124768742 124768925
refGene_gn=PDE1C;refGene_tx=XM_019825660.1;refGene_gn=PDE1C;refGene_tx=XM_019825661.1;refGene_gn=PDE1C;refGene_tx=XM_019825662.2;refGene_gn=PDE1C;refGene_tx=XM_019825663.2;refGene_gn=PDE1C;refGene_tx=XM_019825664.2;refGene_gn=PDE1C;refGene_tx=XM_023250475.1;refGene_gn=PDE1C;refGene_tx=XM_023250476.1
A2 124771033 124771147
refGene_gn=PDE1C;refGene_tx=XM_019825660.1;refGene_gn=PDE1C;refGene_tx=XM_019825661.1;refGene_gn=PDE1C;refGene_tx=XM_019825662.2;refGene_gn=PDE1C;refGene_tx=XM_019825663.2;refGene_gn=PDE1C;refGene_tx=XM_019825664.2;refGene_gn=PDE1C;refGene_tx=XM_023250475.1;refGene_gn=PDE1C;refGene_tx=XM_023250476.1
A2 124948118 124948145

refGene_gn=PDE1C;refGene_tx=XM_023250475.1;refGene_gn=PDE1C;refGene_tx
=XM_023250476.1
A2 124967509 124967610
refGene_gn=PDE1C;refGene_tx=XM_023250475.1;refGene_gn=PDE1C;refGene_tx
=XM_023250476.1
A2 125053345 125053517
refGene_gn=PDE1C;refGene_tx=XM_019825660.1;refGene_gn=PDE1C;refGene_tx
=XM_019825661.1;refGene_gn=PDE1C;refGene_tx=XM_019825662.2;refGene_gn=
PDE1C;refGene_tx=XM_019825663.2;refGene_gn=PDE1C;refGene_tx=XM_0198256
64.2
A2 125082243 125082294
refGene_gn=PDE1C;refGene_tx=XM_019825660.1;refGene_gn=PDE1C;refGene_tx
=XM_019825661.1;refGene_gn=PDE1C;refGene_tx=XM_019825662.2;refGene_gn=
PDE1C;refGene_tx=XM_019825663.2;refGene_gn=PDE1C;refGene_tx=XM_0198256
64.2
A2 125173690 125173775
refGene_gn=PDE1C;refGene_tx=XM_019825660.1;refGene_gn=PDE1C;refGene_tx
=XM_019825661.1;refGene_gn=PDE1C;refGene_tx=XM_019825662.2;refGene_gn=
PDE1C;refGene_tx=XM_019825663.2;refGene_gn=PDE1C;refGene_tx=XM_0198256
64.2
A2 125313553 125313586
refGene_gn=LSM5;refGene_tx=XM_003982919.4
A2 125313987 125314060
refGene_gn=LSM5;refGene_tx=XM_003982919.4
A2 125315006 125315034
refGene_gn=LSM5;refGene_tx=XM_003982919.4
A2 125315323 125315419
refGene_gn=LSM5;refGene_tx=XM_003982919.4
A2 125316546 125316592
refGene_gn=LSM5;refGene_tx=XM_003982919.4
A2 125323087 125323385
refGene_gn=AVL9;refGene_tx=XM_003982920.4;refGene_gn=AVL9;refGene_tx=X
M_023250477.1
A2 125344829 125344950
refGene_gn=AVL9;refGene_tx=XM_003982920.4;refGene_gn=AVL9;refGene_tx=X
M_023250477.1
A2 125346373 125346459
refGene_gn=AVL9;refGene_tx=XM_003982920.4;refGene_gn=AVL9;refGene_tx=X
M_023250477.1
A2 125348919 125348991
refGene_gn=AVL9;refGene_tx=XM_003982920.4;refGene_gn=AVL9;refGene_tx=X
M_023250477.1
A2 125359048 125359138
refGene_gn=AVL9;refGene_tx=XM_003982920.4;refGene_gn=AVL9;refGene_tx=X
M_023250477.1
A2 125359879 125359946
refGene_gn=AVL9;refGene_tx=XM_003982920.4;refGene_gn=AVL9;refGene_tx=X
M_023250477.1
A2 125361561 125361602
refGene_gn=AVL9;refGene_tx=XM_003982920.4;refGene_gn=AVL9;refGene_tx=X

M_023250477.1
A2	125362581	125362620
refGene_gn=AVL9;refGene_tx=XM_003982920.4;refGene_gn=AVL9;refGene_tx=X
M_023250477.1
A2	125367063	125367133
refGene_gn=AVL9;refGene_tx=XM_003982920.4;refGene_gn=AVL9;refGene_tx=X
M_023250477.1
A2	125367666	125368205
refGene_gn=AVL9;refGene_tx=XM_003982920.4;refGene_gn=AVL9;refGene_tx=X
M_023250477.1
A2	125376302	125376437
refGene_gn=AVL9;refGene_tx=XM_003982920.4;refGene_gn=AVL9;refGene_tx=X
M_023250477.1
A2	125378236	125378456
refGene_gn=AVL9;refGene_tx=XM_003982920.4;refGene_gn=AVL9;refGene_tx=X
M_023250477.1
A2	125380930	125381048
refGene_gn=AVL9;refGene_tx=XM_003982920.4;refGene_gn=AVL9;refGene_tx=X
M_023250477.1
A2	125383708	125383762
refGene_gn=AVL9;refGene_tx=XM_003982920.4;refGene_gn=AVL9;refGene_tx=X
M_023250477.1
A2	125384241	125384330
refGene_gn=AVL9;refGene_tx=XM_003982920.4;refGene_gn=AVL9;refGene_tx=X
M_023250477.1
A2	125387287	125387403
refGene_gn=AVL9;refGene_tx=XM_003982920.4;refGene_gn=AVL9;refGene_tx=X
M_023250477.1
A2	125414867	125416403
refGene_gn=KBTBD2;refGene_tx=XM_023250478.1
A2	125418795	125418961
refGene_gn=KBTBD2;refGene_tx=XM_023250478.1
A2	125425735	125425905
refGene_gn=KBTBD2;refGene_tx=XM_023250478.1
A2	125516064	125516206
refGene_gn=LOC109497530;refGene_tx=XM_019825666.2
A2	125524453	125525379
refGene_gn=LOC109497530;refGene_tx=XM_019825666.2
A2	125527313	125527379
refGene_gn=LOC101095968;refGene_tx=XM_023250487.1;refGene_gn=LOC101095
968;refGene_tx=XM_023250488.1;refGene_gn=LOC101095968;refGene_tx=XM_02
3250489.1
A2	125534807	125535006
refGene_gn=LOC101095968;refGene_tx=XM_011280504.3;refGene_gn=LOC101095
968;refGene_tx=XM_023250479.1;refGene_gn=LOC101095968;refGene_tx=XM_02
3250480.1;refGene_gn=LOC101095968;refGene_tx=XM_023250481.1;refGene_gn
=LOC101095968;refGene_tx=XM_023250482.1;refGene_gn=LOC101095968;refGen
e_tx=XM_023250483.1;refGene_gn=LOC101095968;refGene_tx=XM_023250484.1;
refGene_gn=LOC101095968;refGene_tx=XM_023250485.1;refGene_gn=LOC101095
968;refGene_tx=XM_023250486.1

A2	125535725	125535787
refGene_gn=LOC101095968;refGene_tx=XM_011280504.3;refGene_gn=LOC101095968;refGene_tx=XM_023250479.1;refGene_gn=LOC101095968;refGene_tx=XM_023250480.1;refGene_gn=LOC101095968;refGene_tx=XM_023250481.1;refGene_gn=LOC101095968;refGene_tx=XM_023250482.1;refGene_gn=LOC101095968;refGene_tx=XM_023250483.1;refGene_gn=LOC101095968;refGene_tx=XM_023250484.1;refGene_gn=LOC101095968;refGene_tx=XM_023250485.1;refGene_gn=LOC101095968;refGene_tx=XM_023250486.1
A2	125542308	125542400
refGene_gn=LOC101095968;refGene_tx=XM_011280504.3;refGene_gn=LOC101095968;refGene_tx=XM_023250479.1;refGene_gn=LOC101095968;refGene_tx=XM_023250480.1;refGene_gn=LOC101095968;refGene_tx=XM_023250481.1;refGene_gn=LOC101095968;refGene_tx=XM_023250482.1;refGene_gn=LOC101095968;refGene_tx=XM_023250483.1;refGene_gn=LOC101095968;refGene_tx=XM_023250484.1;refGene_gn=LOC101095968;refGene_tx=XM_023250485.1;refGene_gn=LOC101095968;refGene_tx=XM_023250486.1;refGene_gn=LOC101095968;refGene_tx=XM_023250487.1;refGene_gn=LOC101095968;refGene_tx=XM_023250488.1;refGene_gn=LOC101095968;refGene_tx=XM_023250489.1
A2	125548063	125548193
refGene_gn=LOC101095968;refGene_tx=XM_011280504.3;refGene_gn=LOC101095968;refGene_tx=XM_023250479.1;refGene_gn=LOC101095968;refGene_tx=XM_023250480.1;refGene_gn=LOC101095968;refGene_tx=XM_023250481.1;refGene_gn=LOC101095968;refGene_tx=XM_023250482.1;refGene_gn=LOC101095968;refGene_tx=XM_023250483.1;refGene_gn=LOC101095968;refGene_tx=XM_023250484.1;refGene_gn=LOC101095968;refGene_tx=XM_023250485.1;refGene_gn=LOC101095968;refGene_tx=XM_023250486.1;refGene_gn=LOC101095968;refGene_tx=XM_023250487.1;refGene_gn=LOC101095968;refGene_tx=XM_023250488.1;refGene_gn=LOC101095968;refGene_tx=XM_023250489.1
A2	125550007	125550038
refGene_gn=LOC101095968;refGene_tx=XM_011280504.3;refGene_gn=LOC101095968;refGene_tx=XM_023250479.1;refGene_gn=LOC101095968;refGene_tx=XM_023250480.1;refGene_gn=LOC101095968;refGene_tx=XM_023250481.1;refGene_gn=LOC101095968;refGene_tx=XM_023250482.1;refGene_gn=LOC101095968;refGene_tx=XM_023250483.1;refGene_gn=LOC101095968;refGene_tx=XM_023250484.1;refGene_gn=LOC101095968;refGene_tx=XM_023250485.1;refGene_gn=LOC101095968;refGene_tx=XM_023250486.1;refGene_gn=LOC101095968;refGene_tx=XM_023250487.1;refGene_gn=LOC101095968;refGene_tx=XM_023250488.1;refGene_gn=LOC101095968;refGene_tx=XM_023250489.1
A2	125553887	125554102
refGene_gn=LOC101095968;refGene_tx=XM_011280504.3;refGene_gn=LOC101095968;refGene_tx=XM_023250479.1;refGene_gn=LOC101095968;refGene_tx=XM_023250480.1;refGene_gn=LOC101095968;refGene_tx=XM_023250481.1;refGene_gn=LOC101095968;refGene_tx=XM_023250482.1;refGene_gn=LOC101095968;refGene_tx=XM_023250483.1;refGene_gn=LOC101095968;refGene_tx=XM_023250484.1;refGene_gn=LOC101095968;refGene_tx=XM_023250485.1;refGene_gn=LOC101095968;refGene_tx=XM_023250487.1;refGene_gn=LOC101095968;refGene_tx=XM_023250488.1;refGene_gn=LOC101095968;refGene_tx=XM_023250489.1
A2	125554756	125554908
refGene_gn=LOC101095968;refGene_tx=XM_023250486.1
A2	125564643	125564870

refGene_gn=FKBP9;refGene_tx=XM_019825670.2
A2 125577706 125577852
refGene_gn=FKBP9;refGene_tx=XM_019825670.2
A2 125578238 125578428
refGene_gn=FKBP9;refGene_tx=XM_019825670.2
A2 125579081 125579227
refGene_gn=FKBP9;refGene_tx=XM_019825670.2
A2 125582712 125582902
refGene_gn=FKBP9;refGene_tx=XM_019825670.2
A2 125589759 125589905
refGene_gn=FKBP9;refGene_tx=XM_019825670.2
A2 125596054 125596241
refGene_gn=FKBP9;refGene_tx=XM_019825670.2
A2 125599942 125600088
refGene_gn=FKBP9;refGene_tx=XM_019825670.2
A2 125602626 125602790
refGene_gn=FKBP9;refGene_tx=XM_019825670.2
A2 125605063 125605240
refGene_gn=FKBP9;refGene_tx=XM_019825670.2
A2 125614505 125614607
refGene_gn=NT5C3A;refGene_tx=XM_003982923.5;refGene_gn=NT5C3A;refGene_tx=XM_003982924.5;refGene_gn=NT5C3A;refGene_tx=XM_003982925.4;refGene_gn=NT5C3A;refGene_tx=XM_006929293.3;refGene_gn=NT5C3A;refGene_tx=XM_006929294.4
A2 125615104 125615305
refGene_gn=NT5C3A;refGene_tx=XM_003982923.5;refGene_gn=NT5C3A;refGene_tx=XM_003982924.5;refGene_gn=NT5C3A;refGene_tx=XM_003982925.4;refGene_gn=NT5C3A;refGene_tx=XM_006929293.3;refGene_gn=NT5C3A;refGene_tx=XM_006929294.4
A2 125616203 125616366
refGene_gn=NT5C3A;refGene_tx=XM_003982923.5;refGene_gn=NT5C3A;refGene_tx=XM_003982924.5;refGene_gn=NT5C3A;refGene_tx=XM_003982925.4;refGene_gn=NT5C3A;refGene_tx=XM_006929293.3;refGene_gn=NT5C3A;refGene_tx=XM_006929294.4
A2 125617978 125618068
refGene_gn=NT5C3A;refGene_tx=XM_003982923.5;refGene_gn=NT5C3A;refGene_tx=XM_003982924.5;refGene_gn=NT5C3A;refGene_tx=XM_003982925.4;refGene_gn=NT5C3A;refGene_tx=XM_006929293.3;refGene_gn=NT5C3A;refGene_tx=XM_006929294.4
A2 125619453 125619539
refGene_gn=NT5C3A;refGene_tx=XM_003982923.5;refGene_gn=NT5C3A;refGene_tx=XM_003982924.5;refGene_gn=NT5C3A;refGene_tx=XM_003982925.4;refGene_gn=NT5C3A;refGene_tx=XM_006929293.3;refGene_gn=NT5C3A;refGene_tx=XM_006929294.4
A2 125620511 125620558
refGene_gn=NT5C3A;refGene_tx=XM_003982923.5;refGene_gn=NT5C3A;refGene_tx=XM_003982924.5;refGene_gn=NT5C3A;refGene_tx=XM_003982925.4;refGene_gn=NT5C3A;refGene_tx=XM_006929293.3;refGene_gn=NT5C3A;refGene_tx=XM_006929294.4
A2 125621381 125621451

refGene_gn=NT5C3A;refGene_tx=XM_003982923.5;refGene_gn=NT5C3A;refGene_tx=XM_003982924.5;refGene_gn=NT5C3A;refGene_tx=XM_003982925.4;refGene_gn=NT5C3A;refGene_tx=XM_006929293.3;refGene_gn=NT5C3A;refGene_tx=XM_006929294.4
A2 125623844 125623943
refGene_gn=NT5C3A;refGene_tx=XM_003982923.5;refGene_gn=NT5C3A;refGene_tx=XM_003982924.5;refGene_gn=NT5C3A;refGene_tx=XM_003982925.4;refGene_gn=NT5C3A;refGene_tx=XM_006929293.3;refGene_gn=NT5C3A;refGene_tx=XM_006929294.4
A2 125629949 125629985
refGene_gn=NT5C3A;refGene_tx=XM_003982924.5;refGene_gn=NT5C3A;refGene_tx=XM_003982925.4;refGene_gn=NT5C3A;refGene_tx=XM_006929293.3;refGene_gn=NT5C3A;refGene_tx=XM_006929294.4
A2 125667978 125668116
refGene_gn=NT5C3A;refGene_tx=XM_003982923.5
A2 125689145 125689904
refGene_gn=LOC111557411;refGene_tx=XM_023242187.1
A2 125717737 125717939
refGene_gn=LOC101096219;refGene_tx=XM_023250490.1
A2 125718647 125718709
refGene_gn=LOC101096219;refGene_tx=XM_023250490.1
A2 125720679 125720771
refGene_gn=LOC101096219;refGene_tx=XM_023250490.1
A2 125723097 125723227
refGene_gn=LOC101096219;refGene_tx=XM_023250490.1
A2 125724279 125724310
refGene_gn=LOC101096219;refGene_tx=XM_023250490.1
A2 125736490 125736642
refGene_gn=LOC101096219;refGene_tx=XM_023250490.1
A2 125759873 125759985
refGene_gn=BBS9;refGene_tx=XM_006929295.4;refGene_gn=BBS9;refGene_tx=XM_006929296.4;refGene_gn=BBS9;refGene_tx=XM_019825674.2;refGene_gn=BBS9;refGene_tx=XM_019825680.2;refGene_gn=BBS9;refGene_tx=XM_023250491.1
A2 125768791 125768942
refGene_gn=BBS9;refGene_tx=XM_006929295.4;refGene_gn=BBS9;refGene_tx=XM_006929296.4;refGene_gn=BBS9;refGene_tx=XM_019825674.2;refGene_gn=BBS9;refGene_tx=XM_019825680.2;refGene_gn=BBS9;refGene_tx=XM_023250491.1
A2 125771559 125771624
refGene_gn=BBS9;refGene_tx=XM_006929295.4;refGene_gn=BBS9;refGene_tx=XM_006929296.4;refGene_gn=BBS9;refGene_tx=XM_019825674.2;refGene_gn=BBS9;refGene_tx=XM_019825676.2;refGene_gn=BBS9;refGene_tx=XM_019825680.2;refGene_gn=BBS9;refGene_tx=XM_023250491.1
A2 125778683 125778797
refGene_gn=BBS9;refGene_tx=XM_006929295.4;refGene_gn=BBS9;refGene_tx=XM_006929296.4;refGene_gn=BBS9;refGene_tx=XM_019825674.2;refGene_gn=BBS9;refGene_tx=XM_019825676.2;refGene_gn=BBS9;refGene_tx=XM_019825677.2;refGene_gn=BBS9;refGene_tx=XM_019825680.2;refGene_gn=BBS9;refGene_tx=XM_023250491.1
A2 125853329 125853504
refGene_gn=BBS9;refGene_tx=XM_006929295.4;refGene_gn=BBS9;refGene_tx=X

M_006929296.4;refGene_gn=BBS9;refGene_tx=XM_019825674.2;refGene_gn=BBS9;refGene_tx=XM_019825676.2;refGene_gn=BBS9;refGene_tx=XM_019825677.2;refGene_gn=BBS9;refGene_tx=XM_019825680.2;refGene_gn=BBS9;refGene_tx=XM_023250491.1;refGene_gn=BBS9;refGene_tx=XM_023250492.1
A2 125863199 125863284
refGene_gn=BBS9;refGene_tx=XM_006929295.4;refGene_gn=BBS9;refGene_tx=XM_006929296.4;refGene_gn=BBS9;refGene_tx=XM_019825674.2;refGene_gn=BBS9;refGene_tx=XM_019825676.2;refGene_gn=BBS9;refGene_tx=XM_019825677.2;refGene_gn=BBS9;refGene_tx=XM_019825680.2;refGene_gn=BBS9;refGene_tx=XM_023250491.1;refGene_gn=BBS9;refGene_tx=XM_023250492.1
A2 125873581 125873765
refGene_gn=BBS9;refGene_tx=XM_006929295.4;refGene_gn=BBS9;refGene_tx=XM_006929296.4;refGene_gn=BBS9;refGene_tx=XM_019825674.2;refGene_gn=BBS9;refGene_tx=XM_019825676.2;refGene_gn=BBS9;refGene_tx=XM_019825677.2;refGene_gn=BBS9;refGene_tx=XM_019825680.2;refGene_gn=BBS9;refGene_tx=XM_023250491.1;refGene_gn=BBS9;refGene_tx=XM_023250492.1
A2 125874421 125874551
refGene_gn=BBS9;refGene_tx=XM_006929295.4;refGene_gn=BBS9;refGene_tx=XM_006929296.4;refGene_gn=BBS9;refGene_tx=XM_019825674.2;refGene_gn=BBS9;refGene_tx=XM_019825676.2;refGene_gn=BBS9;refGene_tx=XM_019825677.2;refGene_gn=BBS9;refGene_tx=XM_019825680.2;refGene_gn=BBS9;refGene_tx=XM_023250491.1;refGene_gn=BBS9;refGene_tx=XM_023250492.1
A2 125924947 125925129
refGene_gn=BBS9;refGene_tx=XM_006929295.4;refGene_gn=BBS9;refGene_tx=XM_006929296.4;refGene_gn=BBS9;refGene_tx=XM_019825674.2;refGene_gn=BBS9;refGene_tx=XM_019825676.2;refGene_gn=BBS9;refGene_tx=XM_019825677.2;refGene_gn=BBS9;refGene_tx=XM_019825680.2;refGene_gn=BBS9;refGene_tx=XM_023250491.1;refGene_gn=BBS9;refGene_tx=XM_023250492.1
A2 125930298 125930375
refGene_gn=BBS9;refGene_tx=XM_006929295.4;refGene_gn=BBS9;refGene_tx=XM_006929296.4;refGene_gn=BBS9;refGene_tx=XM_019825674.2;refGene_gn=BBS9;refGene_tx=XM_019825676.2;refGene_gn=BBS9;refGene_tx=XM_019825677.2;refGene_gn=BBS9;refGene_tx=XM_019825680.2;refGene_gn=BBS9;refGene_tx=XM_023250491.1;refGene_gn=BBS9;refGene_tx=XM_023250492.1
A2 125934113 125934167
refGene_gn=BBS9;refGene_tx=XM_006929295.4;refGene_gn=BBS9;refGene_tx=XM_006929296.4;refGene_gn=BBS9;refGene_tx=XM_019825674.2;refGene_gn=BBS9;refGene_tx=XM_019825676.2;refGene_gn=BBS9;refGene_tx=XM_019825677.2;refGene_gn=BBS9;refGene_tx=XM_019825680.2;refGene_gn=BBS9;refGene_tx=XM_023250491.1;refGene_gn=BBS9;refGene_tx=XM_023250492.1
A2 125940319 125940422
refGene_gn=BBS9;refGene_tx=XM_006929295.4;refGene_gn=BBS9;refGene_tx=XM_006929296.4;refGene_gn=BBS9;refGene_tx=XM_019825674.2;refGene_gn=BBS9;refGene_tx=XM_019825676.2;refGene_gn=BBS9;refGene_tx=XM_019825677.2;refGene_gn=BBS9;refGene_tx=XM_019825680.2;refGene_gn=BBS9;refGene_tx=XM_023250491.1;refGene_gn=BBS9;refGene_tx=XM_023250492.1
A2 125941985 125942090
refGene_gn=BBS9;refGene_tx=XM_006929295.4;refGene_gn=BBS9;refGene_tx=XM_006929296.4;refGene_gn=BBS9;refGene_tx=XM_019825674.2;refGene_gn=BBS9;refGene_tx=XM_019825676.2;refGene_gn=BBS9;refGene_tx=XM_019825677.2;

refGene_gn=BBS9;refGene_tx=XM_019825680.2;refGene_gn=BBS9;refGene_tx=XM_023250491.1;refGene_gn=BBS9;refGene_tx=XM_023250492.1
A2	125944435	125944450
refGene_gn=BBS9;refGene_tx=XM_019825674.2;refGene_gn=BBS9;refGene_tx=XM_019825676.2;refGene_gn=BBS9;refGene_tx=XM_019825677.2;refGene_gn=BBS9;refGene_tx=XM_019825680.2;refGene_gn=BBS9;refGene_tx=XM_023250491.1;refGene_gn=BBS9;refGene_tx=XM_023250492.1
A2	125951816	125951957
refGene_gn=BBS9;refGene_tx=XM_006929295.4;refGene_gn=BBS9;refGene_tx=XM_006929296.4;refGene_gn=BBS9;refGene_tx=XM_019825674.2;refGene_gn=BBS9;refGene_tx=XM_019825676.2;refGene_gn=BBS9;refGene_tx=XM_019825677.2;refGene_gn=BBS9;refGene_tx=XM_019825680.2;refGene_gn=BBS9;refGene_tx=XM_023250491.1;refGene_gn=BBS9;refGene_tx=XM_023250492.1
A2	125969266	125969362
refGene_gn=BBS9;refGene_tx=XM_006929295.4;refGene_gn=BBS9;refGene_tx=XM_006929296.4;refGene_gn=BBS9;refGene_tx=XM_019825674.2;refGene_gn=BBS9;refGene_tx=XM_019825676.2;refGene_gn=BBS9;refGene_tx=XM_019825677.2;refGene_gn=BBS9;refGene_tx=XM_019825680.2;refGene_gn=BBS9;refGene_tx=XM_023250491.1;refGene_gn=BBS9;refGene_tx=XM_023250492.1
A2	125983539	125983712
refGene_gn=BBS9;refGene_tx=XM_006929295.4;refGene_gn=BBS9;refGene_tx=XM_006929296.4;refGene_gn=BBS9;refGene_tx=XM_019825674.2;refGene_gn=BBS9;refGene_tx=XM_019825676.2;refGene_gn=BBS9;refGene_tx=XM_019825677.2;refGene_gn=BBS9;refGene_tx=XM_019825680.2;refGene_gn=BBS9;refGene_tx=XM_023250491.1;refGene_gn=BBS9;refGene_tx=XM_023250492.1
A2	125987434	125987587
refGene_gn=BBS9;refGene_tx=XM_006929295.4;refGene_gn=BBS9;refGene_tx=XM_006929296.4;refGene_gn=BBS9;refGene_tx=XM_019825674.2;refGene_gn=BBS9;refGene_tx=XM_019825676.2;refGene_gn=BBS9;refGene_tx=XM_019825677.2;refGene_gn=BBS9;refGene_tx=XM_019825680.2;refGene_gn=BBS9;refGene_tx=XM_023250491.1;refGene_gn=BBS9;refGene_tx=XM_023250492.1
A2	126045590	126045650
refGene_gn=BBS9;refGene_tx=XM_019825680.2
A2	126105288	126105471
refGene_gn=BBS9;refGene_tx=XM_006929295.4;refGene_gn=BBS9;refGene_tx=XM_006929296.4;refGene_gn=BBS9;refGene_tx=XM_019825674.2;refGene_gn=BBS9;refGene_tx=XM_019825676.2;refGene_gn=BBS9;refGene_tx=XM_019825677.2;refGene_gn=BBS9;refGene_tx=XM_023250491.1;refGene_gn=BBS9;refGene_tx=XM_023250492.1
A2	126129493	126129716
refGene_gn=BBS9;refGene_tx=XM_006929295.4;refGene_gn=BBS9;refGene_tx=XM_006929296.4;refGene_gn=BBS9;refGene_tx=XM_019825674.2;refGene_gn=BBS9;refGene_tx=XM_019825676.2;refGene_gn=BBS9;refGene_tx=XM_019825677.2;refGene_gn=BBS9;refGene_tx=XM_023250491.1;refGene_gn=BBS9;refGene_tx=XM_023250492.1
A2	126192825	126192939
refGene_gn=BBS9;refGene_tx=XM_006929295.4;refGene_gn=BBS9;refGene_tx=XM_006929296.4;refGene_gn=BBS9;refGene_tx=XM_019825674.2;refGene_gn=BBS9;refGene_tx=XM_019825676.2;refGene_gn=BBS9;refGene_tx=XM_019825677.2;refGene_gn=BBS9;refGene_tx=XM_023250491.1;refGene_gn=BBS9;refGene_tx=X

M_023250492.1
A2	126193148	126193180
refGene_gn=BBS9;refGene_tx=XM_006929295.4;refGene_gn=BBS9;refGene_tx=XM_006929296.4;refGene_gn=BBS9;refGene_tx=XM_019825674.2;refGene_gn=BBS9;refGene_tx=XM_019825676.2;refGene_gn=BBS9;refGene_tx=XM_019825677.2;refGene_gn=BBS9;refGene_tx=XM_023250491.1;refGene_gn=BBS9;refGene_tx=XM_023250492.1
A2	126491831	126492596
refGene_gn=LOC111559526;refGene_tx=XM_023250493.1
A2	126492834	126492967
refGene_gn=BMPER;refGene_tx=XM_003982930.5;refGene_gn=BMPER;refGene_tx=XM_019825681.2;refGene_gn=BMPER;refGene_tx=XM_019825682.2;refGene_gn=LOC111559526;refGene_tx=XM_023250493.1
A2	126493999	126494085
refGene_gn=BMPER;refGene_tx=XM_003982930.5;refGene_gn=BMPER;refGene_tx=XM_006929299.4;refGene_gn=BMPER;refGene_tx=XM_019825681.2;refGene_gn=BMPER;refGene_tx=XM_019825682.2
A2	126523239	126523339
refGene_gn=BMPER;refGene_tx=XM_003982930.5;refGene_gn=BMPER;refGene_tx=XM_006929299.4;refGene_gn=BMPER;refGene_tx=XM_019825681.2;refGene_gn=BMPER;refGene_tx=XM_019825682.2
A2	126548912	126548995
refGene_gn=BMPER;refGene_tx=XM_003982930.5;refGene_gn=BMPER;refGene_tx=XM_006929299.4;refGene_gn=BMPER;refGene_tx=XM_019825681.2;refGene_gn=BMPER;refGene_tx=XM_019825682.2
A2	126552899	126552990
refGene_gn=BMPER;refGene_tx=XM_003982930.5;refGene_gn=BMPER;refGene_tx=XM_006929299.4;refGene_gn=BMPER;refGene_tx=XM_019825681.2;refGene_gn=BMPER;refGene_tx=XM_019825682.2
A2	126557162	126557245
refGene_gn=BMPER;refGene_tx=XM_003982930.5;refGene_gn=BMPER;refGene_tx=XM_006929299.4;refGene_gn=BMPER;refGene_tx=XM_019825681.2;refGene_gn=BMPER;refGene_tx=XM_019825682.2
A2	126625764	126625864
refGene_gn=BMPER;refGene_tx=XM_003982930.5;refGene_gn=BMPER;refGene_tx=XM_006929299.4;refGene_gn=BMPER;refGene_tx=XM_019825681.2;refGene_gn=BMPER;refGene_tx=XM_019825682.2
A2	126630809	126630919
refGene_gn=BMPER;refGene_tx=XM_003982930.5;refGene_gn=BMPER;refGene_tx=XM_006929299.4;refGene_gn=BMPER;refGene_tx=XM_019825681.2;refGene_gn=BMPER;refGene_tx=XM_019825682.2
A2	126634513	126634654
refGene_gn=BMPER;refGene_tx=XM_003982930.5;refGene_gn=BMPER;refGene_tx=XM_006929299.4;refGene_gn=BMPER;refGene_tx=XM_019825681.2;refGene_gn=BMPER;refGene_tx=XM_019825682.2
A2	126636839	126636944
refGene_gn=BMPER;refGene_tx=XM_003982930.5;refGene_gn=BMPER;refGene_tx=XM_006929299.4;refGene_gn=BMPER;refGene_tx=XM_019825681.2;refGene_gn=BMPER;refGene_tx=XM_019825682.2
A2	126640195	126640241

refGene_gn=BMPER;refGene_tx=XM_003982930.5;refGene_gn=BMPER;refGene_tx=XM_006929299.4;refGene_gn=BMPER;refGene_tx=XM_019825681.2;refGene_gn=BMPER;refGene_tx=XM_019825682.2
A2 126663053 126663383
refGene_gn=BMPER;refGene_tx=XM_003982930.5;refGene_gn=BMPER;refGene_tx=XM_006929299.4;refGene_gn=BMPER;refGene_tx=XM_019825681.2
A2 126669973 126670310
refGene_gn=BMPER;refGene_tx=XM_003982930.5;refGene_gn=BMPER;refGene_tx=XM_006929299.4;refGene_gn=BMPER;refGene_tx=XM_019825681.2;refGene_gn=BMPER;refGene_tx=XM_019825682.2
A2 126686662 126686678
refGene_gn=BMPER;refGene_tx=XM_019825681.2
A2 126729735 126729866
refGene_gn=BMPER;refGene_tx=XM_003982930.5;refGene_gn=BMPER;refGene_tx=XM_006929299.4;refGene_gn=BMPER;refGene_tx=XM_019825682.2
A2 126741415 126741597
refGene_gn=BMPER;refGene_tx=XM_003982930.5;refGene_gn=BMPER;refGene_tx=XM_006929299.4;refGene_gn=BMPER;refGene_tx=XM_019825682.2
A2 127080743 127080899
refGene_gn=NPSR1;refGene_tx=XM_011280501.2
A2 127109795 127109928
refGene_gn=NPSR1;refGene_tx=XM_011280501.2
A2 127176552 127176656
refGene_gn=NPSR1;refGene_tx=XM_011280501.2
A2 127186614 127186708
refGene_gn=NPSR1;refGene_tx=XM_011280501.2
A2 127192432 127192634
refGene_gn=NPSR1;refGene_tx=XM_011280501.2
A2 127204780 127204857
refGene_gn=NPSR1;refGene_tx=XM_011280501.2
A2 127221339 127221426
refGene_gn=NPSR1;refGene_tx=XM_011280501.2
A2 127231897 127232078
refGene_gn=NPSR1;refGene_tx=XM_011280501.2
A2 127232965 127233044
refGene_gn=NPSR1;refGene_tx=XM_011280501.2
A2 127292582 127292739
refGene_gn=DPY19L1;refGene_tx=XM_023250495.1;refGene_gn=DPY19L1;refGene_tx=XM_023250496.1;refGene_gn=DPY19L1;refGene_tx=XM_023250497.1
A2 127298979 127299105
refGene_gn=DPY19L1;refGene_tx=XM_023250495.1;refGene_gn=DPY19L1;refGene_tx=XM_023250496.1;refGene_gn=DPY19L1;refGene_tx=XM_023250497.1
A2 127300263 127300363
refGene_gn=DPY19L1;refGene_tx=XM_023250495.1;refGene_gn=DPY19L1;refGene_tx=XM_023250496.1;refGene_gn=DPY19L1;refGene_tx=XM_023250497.1
A2 127301091 127301266
refGene_gn=DPY19L1;refGene_tx=XM_023250495.1;refGene_gn=DPY19L1;refGene_tx=XM_023250496.1;refGene_gn=DPY19L1;refGene_tx=XM_023250497.1
A2 127302655 127302775
refGene_gn=DPY19L1;refGene_tx=XM_023250495.1;refGene_gn=DPY19L1;refGen

e_tx=XM_023250496.1;refGene_gn=DPY19L1;refGene_tx=XM_023250497.1
A2 127303912 127303937
refGene_gn=DPY19L1;refGene_tx=XM_023250495.1;refGene_gn=DPY19L1;refGene_tx=XM_023250496.1;refGene_gn=DPY19L1;refGene_tx=XM_023250497.1
A2 127306098 127306148
refGene_gn=DPY19L1;refGene_tx=XM_023250495.1;refGene_gn=DPY19L1;refGene_tx=XM_023250496.1;refGene_gn=DPY19L1;refGene_tx=XM_023250497.1
A2 127308112 127308184
refGene_gn=DPY19L1;refGene_tx=XM_023250495.1;refGene_gn=DPY19L1;refGene_tx=XM_023250496.1;refGene_gn=DPY19L1;refGene_tx=XM_023250497.1
A2 127312492 127312594
refGene_gn=DPY19L1;refGene_tx=XM_023250495.1;refGene_gn=DPY19L1;refGene_tx=XM_023250496.1;refGene_gn=DPY19L1;refGene_tx=XM_023250497.1
A2 127319007 127319088
refGene_gn=DPY19L1;refGene_tx=XM_023250495.1;refGene_gn=DPY19L1;refGene_tx=XM_023250496.1;refGene_gn=DPY19L1;refGene_tx=XM_023250497.1
A2 127319624 127319684
refGene_gn=DPY19L1;refGene_tx=XM_023250495.1;refGene_gn=DPY19L1;refGene_tx=XM_023250496.1;refGene_gn=DPY19L1;refGene_tx=XM_023250497.1
A2 127321796 127321883
refGene_gn=DPY19L1;refGene_tx=XM_023250495.1;refGene_gn=DPY19L1;refGene_tx=XM_023250496.1;refGene_gn=DPY19L1;refGene_tx=XM_023250497.1
A2 127330951 127331029
refGene_gn=DPY19L1;refGene_tx=XM_023250495.1;refGene_gn=DPY19L1;refGene_tx=XM_023250496.1;refGene_gn=DPY19L1;refGene_tx=XM_023250497.1
A2 127333193 127333293
refGene_gn=DPY19L1;refGene_tx=XM_023250495.1;refGene_gn=DPY19L1;refGene_tx=XM_023250496.1;refGene_gn=DPY19L1;refGene_tx=XM_023250497.1
A2 127335147 127335239
refGene_gn=DPY19L1;refGene_tx=XM_023250495.1;refGene_gn=DPY19L1;refGene_tx=XM_023250496.1;refGene_gn=DPY19L1;refGene_tx=XM_023250497.1
A2 127346558 127346616
refGene_gn=DPY19L1;refGene_tx=XM_023250495.1;refGene_gn=DPY19L1;refGene_tx=XM_023250496.1;refGene_gn=DPY19L1;refGene_tx=XM_023250497.1
A2 127359985 127360079
refGene_gn=DPY19L1;refGene_tx=XM_023250495.1;refGene_gn=DPY19L1;refGene_tx=XM_023250496.1;refGene_gn=DPY19L1;refGene_tx=XM_023250497.1
A2 127360897 127361018
refGene_gn=DPY19L1;refGene_tx=XM_023250495.1;refGene_gn=DPY19L1;refGene_tx=XM_023250496.1;refGene_gn=DPY19L1;refGene_tx=XM_023250497.1
A2 127364134 127364272
refGene_gn=DPY19L1;refGene_tx=XM_023250495.1;refGene_gn=DPY19L1;refGene_tx=XM_023250496.1;refGene_gn=DPY19L1;refGene_tx=XM_023250497.1
A2 127367789 127367877
refGene_gn=DPY19L1;refGene_tx=XM_023250495.1;refGene_gn=DPY19L1;refGene_tx=XM_023250496.1;refGene_gn=DPY19L1;refGene_tx=XM_023250497.1
A2 127368520 127368545
refGene_gn=DPY19L1;refGene_tx=XM_023250495.1;refGene_gn=DPY19L1;refGene_tx=XM_023250496.1;refGene_gn=DPY19L1;refGene_tx=XM_023250497.1
A2 127386691 127386989

refGene_gn=DPY19L1;refGene_tx=XM_023250495.1;refGene_gn=DPY19L1;refGene_tx=XM_023250496.1;refGene_gn=DPY19L1;refGene_tx=XM_023250497.1
A2 127422191 127422342
refGene_gn=DPY19L2;refGene_tx=XM_003982944.5
A2 127428119 127428245
refGene_gn=DPY19L2;refGene_tx=XM_003982944.5
A2 127429585 127429685
refGene_gn=DPY19L2;refGene_tx=XM_003982944.5
A2 127432341 127432516
refGene_gn=DPY19L2;refGene_tx=XM_003982944.5
A2 127433434 127433554
refGene_gn=DPY19L2;refGene_tx=XM_003982944.5
A2 127441196 127441221
refGene_gn=DPY19L2;refGene_tx=XM_003982944.5
A2 127452436 127452483
refGene_gn=DPY19L2;refGene_tx=XM_003982944.5
A2 127455893 127455965
refGene_gn=DPY19L2;refGene_tx=XM_003982944.5
A2 127457049 127457151
refGene_gn=DPY19L2;refGene_tx=XM_003982944.5
A2 127459344 127459425
refGene_gn=DPY19L2;refGene_tx=XM_003982944.5
A2 127464581 127464641
refGene_gn=DPY19L2;refGene_tx=XM_003982944.5
A2 127471316 127471403
refGene_gn=DPY19L2;refGene_tx=XM_003982944.5
A2 127472236 127472314
refGene_gn=DPY19L2;refGene_tx=XM_003982944.5
A2 127486223 127486323
refGene_gn=DPY19L2;refGene_tx=XM_003982944.5
A2 127487509 127487601
refGene_gn=DPY19L2;refGene_tx=XM_003982944.5
A2 127489264 127489322
refGene_gn=DPY19L2;refGene_tx=XM_003982944.5
A2 127494073 127494167
refGene_gn=DPY19L2;refGene_tx=XM_003982944.5
A2 127496481 127496602
refGene_gn=DPY19L2;refGene_tx=XM_003982944.5
A2 127508608 127508746
refGene_gn=DPY19L2;refGene_tx=XM_003982944.5
A2 127510777 127510865
refGene_gn=DPY19L2;refGene_tx=XM_003982944.5
A2 127512656 127512681
refGene_gn=DPY19L2;refGene_tx=XM_003982944.5
A2 127517104 127517480
refGene_gn=DPY19L2;refGene_tx=XM_003982944.5
A2 127533573 127533914
refGene_gn=TBX20;refGene_tx=XM_003982933.2
A2 127535610 127535723
refGene_gn=TBX20;refGene_tx=XM_003982933.2

A2	127567416	127567493
refGene_gn=TBX20;refGene_tx=XM_003982933.2
A2	127577586	127577745
refGene_gn=TBX20;refGene_tx=XM_003982933.2
A2	127581658	127581767
refGene_gn=TBX20;refGene_tx=XM_003982933.2
A2	127585816	127585981
refGene_gn=TBX20;refGene_tx=XM_003982933.2
A2	127587284	127587540
refGene_gn=TBX20;refGene_tx=XM_003982933.2
A2	127590901	127591028
refGene_gn=TBX20;refGene_tx=XM_003982933.2
A2	127903434	127903596
refGene_gn=HERPUD2;refGene_tx=XM_011280500.2;refGene_gn=HERPUD2;refGene_tx=XM_019825684.2;refGene_gn=HERPUD2;refGene_tx=XM_019825685.2;refGene_gn=HERPUD2;refGene_tx=XM_023250498.1
A2	127904083	127904201
refGene_gn=HERPUD2;refGene_tx=XM_011280500.2;refGene_gn=HERPUD2;refGene_tx=XM_019825684.2;refGene_gn=HERPUD2;refGene_tx=XM_019825685.2;refGene_gn=HERPUD2;refGene_tx=XM_023250498.1
A2	127904915	127905239
refGene_gn=HERPUD2;refGene_tx=XM_011280500.2;refGene_gn=HERPUD2;refGene_tx=XM_019825684.2;refGene_gn=HERPUD2;refGene_tx=XM_019825685.2;refGene_gn=HERPUD2;refGene_tx=XM_023250498.1
A2	127909366	127909489
refGene_gn=HERPUD2;refGene_tx=XM_011280500.2;refGene_gn=HERPUD2;refGene_tx=XM_019825684.2;refGene_gn=HERPUD2;refGene_tx=XM_019825685.2;refGene_gn=HERPUD2;refGene_tx=XM_023250498.1
A2	127921422	127921577
refGene_gn=HERPUD2;refGene_tx=XM_011280500.2;refGene_gn=HERPUD2;refGene_tx=XM_019825684.2;refGene_gn=HERPUD2;refGene_tx=XM_019825685.2;refGene_gn=HERPUD2;refGene_tx=XM_023250498.1
A2	127923602	127923716
refGene_gn=HERPUD2;refGene_tx=XM_011280500.2;refGene_gn=HERPUD2;refGene_tx=XM_019825684.2;refGene_gn=HERPUD2;refGene_tx=XM_019825685.2;refGene_gn=HERPUD2;refGene_tx=XM_023250498.1
A2	127927245	127927323
refGene_gn=HERPUD2;refGene_tx=XM_011280500.2;refGene_gn=HERPUD2;refGene_tx=XM_019825684.2
A2	127937051	127937198
refGene_gn=HERPUD2;refGene_tx=XM_011280500.2;refGene_gn=HERPUD2;refGene_tx=XM_019825684.2;refGene_gn=HERPUD2;refGene_tx=XM_019825685.2
A2	128026980	128027041
refGene_gn=SEPT7;refGene_tx=XM_003982945.5;refGene_gn=SEPT7;refGene_tx=XM_003982946.5;refGene_gn=SEPT7;refGene_tx=XM_006929309.4;refGene_gn=SEPT7;refGene_tx=XM_019825686.2
A2	128027141	128027166
refGene_gn=SEPT7;refGene_tx=XM_019825688.2;refGene_gn=SEPT7;refGene_tx=XM_019825690.2
A2	128028488	128028490

refGene_gn=SEPT7;refGene_tx=XM_003982946.5;refGene_gn=SEPT7;refGene_tx
=XM_006929309.4;refGene_gn=SEPT7;refGene_tx=XM_019825688.2;refGene_gn=
SEPT7;refGene_tx=XM_019825690.2
A2 128055968 128055984
refGene_gn=SEPT7;refGene_tx=XM_019825687.2
A2 128057550 128057552
refGene_gn=SEPT7;refGene_tx=XM_019825687.2
A2 128059671 128059676
refGene_gn=SEPT7;refGene_tx=XM_003982945.5;refGene_gn=SEPT7;refGene_tx
=XM_019825686.2
A2 128060200 128060224
refGene_gn=SEPT7;refGene_tx=XM_019825689.2
A2 128060978 128061081
refGene_gn=SEPT7;refGene_tx=XM_003982945.5;refGene_gn=SEPT7;refGene_tx
=XM_003982946.5;refGene_gn=SEPT7;refGene_tx=XM_006929309.4;refGene_gn=
SEPT7;refGene_tx=XM_019825686.2;refGene_gn=SEPT7;refGene_tx=XM_0198256
87.2;refGene_gn=SEPT7;refGene_tx=XM_019825688.2;refGene_gn=SEPT7;refGe
ne_tx=XM_019825689.2;refGene_gn=SEPT7;refGene_tx=XM_019825690.2
A2 128094293 128094400
refGene_gn=SEPT7;refGene_tx=XM_003982945.5;refGene_gn=SEPT7;refGene_tx
=XM_003982946.5;refGene_gn=SEPT7;refGene_tx=XM_006929309.4;refGene_gn=
SEPT7;refGene_tx=XM_019825686.2;refGene_gn=SEPT7;refGene_tx=XM_0198256
87.2;refGene_gn=SEPT7;refGene_tx=XM_019825688.2;refGene_gn=SEPT7;refGe
ne_tx=XM_019825689.2;refGene_gn=SEPT7;refGene_tx=XM_019825690.2
A2 128106276 128106377
refGene_gn=SEPT7;refGene_tx=XM_003982945.5;refGene_gn=SEPT7;refGene_tx
=XM_003982946.5;refGene_gn=SEPT7;refGene_tx=XM_006929309.4;refGene_gn=
SEPT7;refGene_tx=XM_019825686.2;refGene_gn=SEPT7;refGene_tx=XM_0198256
87.2;refGene_gn=SEPT7;refGene_tx=XM_019825688.2;refGene_gn=SEPT7;refGe
ne_tx=XM_019825689.2;refGene_gn=SEPT7;refGene_tx=XM_019825690.2
A2 128107201 128107336
refGene_gn=SEPT7;refGene_tx=XM_003982945.5;refGene_gn=SEPT7;refGene_tx
=XM_003982946.5;refGene_gn=SEPT7;refGene_tx=XM_006929309.4;refGene_gn=
SEPT7;refGene_tx=XM_019825686.2;refGene_gn=SEPT7;refGene_tx=XM_0198256
87.2;refGene_gn=SEPT7;refGene_tx=XM_019825688.2;refGene_gn=SEPT7;refGe
ne_tx=XM_019825689.2;refGene_gn=SEPT7;refGene_tx=XM_019825690.2
A2 128112871 128112989
refGene_gn=SEPT7;refGene_tx=XM_003982945.5;refGene_gn=SEPT7;refGene_tx
=XM_003982946.5;refGene_gn=SEPT7;refGene_tx=XM_006929309.4;refGene_gn=
SEPT7;refGene_tx=XM_019825686.2;refGene_gn=SEPT7;refGene_tx=XM_0198256
87.2;refGene_gn=SEPT7;refGene_tx=XM_019825688.2;refGene_gn=SEPT7;refGe
ne_tx=XM_019825689.2;refGene_gn=SEPT7;refGene_tx=XM_019825690.2
A2 128117510 128117603
refGene_gn=SEPT7;refGene_tx=XM_003982945.5;refGene_gn=SEPT7;refGene_tx
=XM_003982946.5;refGene_gn=SEPT7;refGene_tx=XM_006929309.4;refGene_gn=
SEPT7;refGene_tx=XM_019825686.2;refGene_gn=SEPT7;refGene_tx=XM_0198256
87.2;refGene_gn=SEPT7;refGene_tx=XM_019825688.2;refGene_gn=SEPT7;refGe
ne_tx=XM_019825689.2;refGene_gn=SEPT7;refGene_tx=XM_019825690.2
A2 128118930 128119027
refGene_gn=SEPT7;refGene_tx=XM_003982945.5;refGene_gn=SEPT7;refGene_tx

=XM_003982946.5;refGene_gn=SEPT7;refGene_tx=XM_006929309.4;refGene_gn=SEPT7;refGene_tx=XM_019825686.2;refGene_gn=SEPT7;refGene_tx=XM_019825687.2;refGene_gn=SEPT7;refGene_tx=XM_019825688.2;refGene_gn=SEPT7;refGene_tx=XM_019825689.2;refGene_gn=SEPT7;refGene_tx=XM_019825690.2
A2 128120403 128120455
refGene_gn=SEPT7;refGene_tx=XM_003982945.5;refGene_gn=SEPT7;refGene_tx=XM_003982946.5;refGene_gn=SEPT7;refGene_tx=XM_006929309.4;refGene_gn=SEPT7;refGene_tx=XM_019825686.2;refGene_gn=SEPT7;refGene_tx=XM_019825687.2;refGene_gn=SEPT7;refGene_tx=XM_019825688.2;refGene_gn=SEPT7;refGene_tx=XM_019825689.2;refGene_gn=SEPT7;refGene_tx=XM_019825690.2
A2 128126116 128126242
refGene_gn=SEPT7;refGene_tx=XM_003982945.5;refGene_gn=SEPT7;refGene_tx=XM_003982946.5;refGene_gn=SEPT7;refGene_tx=XM_006929309.4;refGene_gn=SEPT7;refGene_tx=XM_019825686.2;refGene_gn=SEPT7;refGene_tx=XM_019825687.2;refGene_gn=SEPT7;refGene_tx=XM_019825688.2;refGene_gn=SEPT7;refGene_tx=XM_019825689.2;refGene_gn=SEPT7;refGene_tx=XM_019825690.2
A2 128136564 128136700
refGene_gn=SEPT7;refGene_tx=XM_003982945.5;refGene_gn=SEPT7;refGene_tx=XM_003982946.5;refGene_gn=SEPT7;refGene_tx=XM_006929309.4;refGene_gn=SEPT7;refGene_tx=XM_019825686.2;refGene_gn=SEPT7;refGene_tx=XM_019825687.2;refGene_gn=SEPT7;refGene_tx=XM_019825688.2;refGene_gn=SEPT7;refGene_tx=XM_019825689.2;refGene_gn=SEPT7;refGene_tx=XM_019825690.2
A2 128141043 128141183
refGene_gn=SEPT7;refGene_tx=XM_003982945.5;refGene_gn=SEPT7;refGene_tx=XM_003982946.5;refGene_gn=SEPT7;refGene_tx=XM_006929309.4;refGene_gn=SEPT7;refGene_tx=XM_019825686.2;refGene_gn=SEPT7;refGene_tx=XM_019825687.2;refGene_gn=SEPT7;refGene_tx=XM_019825688.2;refGene_gn=SEPT7;refGene_tx=XM_019825689.2;refGene_gn=SEPT7;refGene_tx=XM_019825690.2
A2 128142209 128142249
refGene_gn=SEPT7;refGene_tx=XM_003982945.5;refGene_gn=SEPT7;refGene_tx=XM_003982946.5;refGene_gn=SEPT7;refGene_tx=XM_006929309.4;refGene_gn=SEPT7;refGene_tx=XM_019825686.2;refGene_gn=SEPT7;refGene_tx=XM_019825687.2;refGene_gn=SEPT7;refGene_tx=XM_019825688.2;refGene_gn=SEPT7;refGene_tx=XM_019825689.2;refGene_gn=SEPT7;refGene_tx=XM_019825690.2
A2 128257161 128258039
refGene_gn=EEPD1;refGene_tx=XM_003982948.3;refGene_gn=EEPD1;refGene_tx=XM_019825693.2;refGene_gn=EEPD1;refGene_tx=XM_019825694.2;refGene_gn=EEPD1;refGene_tx=XM_019825695.2
A2 128325533 128325585
refGene_gn=EEPD1;refGene_tx=XM_003982948.3;refGene_gn=EEPD1;refGene_tx=XM_019825693.2;refGene_gn=EEPD1;refGene_tx=XM_019825694.2;refGene_gn=EEPD1;refGene_tx=XM_019825695.2
A2 128354260 128354371
refGene_gn=EEPD1;refGene_tx=XM_003982948.3;refGene_gn=EEPD1;refGene_tx=XM_019825693.2;refGene_gn=EEPD1;refGene_tx=XM_019825694.2;refGene_gn=EEPD1;refGene_tx=XM_019825695.2
A2 128358047 128358182
refGene_gn=EEPD1;refGene_tx=XM_003982948.3;refGene_gn=EEPD1;refGene_tx=XM_019825693.2;refGene_gn=EEPD1;refGene_tx=XM_019825694.2;refGene_gn=EEPD1;refGene_tx=XM_019825695.2

A2 128360344 128360483
refGene_gn=EEPD1;refGene_tx=XM_003982948.3;refGene_gn=EEPD1;refGene_tx=XM_019825693.2;refGene_gn=EEPD1;refGene_tx=XM_019825694.2;refGene_gn=EEPD1;refGene_tx=XM_019825695.2
A2 128364993 128365188
refGene_gn=EEPD1;refGene_tx=XM_003982948.3;refGene_gn=EEPD1;refGene_tx=XM_019825693.2;refGene_gn=EEPD1;refGene_tx=XM_019825694.2;refGene_gn=EEPD1;refGene_tx=XM_019825695.2
A2 128367417 128367617
refGene_gn=EEPD1;refGene_tx=XM_003982948.3;refGene_gn=EEPD1;refGene_tx=XM_019825693.2;refGene_gn=EEPD1;refGene_tx=XM_019825694.2;refGene_gn=EEPD1;refGene_tx=XM_019825695.2
A2 128406783 128406945
refGene_gn=KIAA0895;refGene_tx=XM_003982951.5;refGene_gn=KIAA0895;refGene_tx=XM_006929310.4;refGene_gn=KIAA0895;refGene_tx=XM_011280498.3;refGene_gn=KIAA0895;refGene_tx=XM_019825701.2
A2 128409710 128409808
refGene_gn=KIAA0895;refGene_tx=XM_003982951.5;refGene_gn=KIAA0895;refGene_tx=XM_006929310.4;refGene_gn=KIAA0895;refGene_tx=XM_011280498.3;refGene_gn=KIAA0895;refGene_tx=XM_019825701.2;refGene_gn=KIAA0895;refGene_tx=XM_023250500.1
A2 128423570 128423876
refGene_gn=KIAA0895;refGene_tx=XM_003982951.5;refGene_gn=KIAA0895;refGene_tx=XM_006929310.4;refGene_gn=KIAA0895;refGene_tx=XM_011280498.3;refGene_gn=KIAA0895;refGene_tx=XM_019825701.2;refGene_gn=KIAA0895;refGene_tx=XM_023250500.1
A2 128424383 128424516
refGene_gn=KIAA0895;refGene_tx=XM_003982951.5;refGene_gn=KIAA0895;refGene_tx=XM_006929310.4;refGene_gn=KIAA0895;refGene_tx=XM_011280498.3;refGene_gn=KIAA0895;refGene_tx=XM_019825701.2;refGene_gn=KIAA0895;refGene_tx=XM_023250500.1
A2 128425127 128425271
refGene_gn=KIAA0895;refGene_tx=XM_003982951.5;refGene_gn=KIAA0895;refGene_tx=XM_006929310.4;refGene_gn=KIAA0895;refGene_tx=XM_011280498.3;refGene_gn=KIAA0895;refGene_tx=XM_019825701.2;refGene_gn=KIAA0895;refGene_tx=XM_023250500.1
A2 128452011 128452706
refGene_gn=KIAA0895;refGene_tx=XM_003982951.5;refGene_gn=KIAA0895;refGene_tx=XM_006929310.4;refGene_gn=KIAA0895;refGene_tx=XM_011280498.3;refGene_gn=KIAA0895;refGene_tx=XM_023250500.1
A2 128461889 128461914
refGene_gn=KIAA0895;refGene_tx=XM_003982951.5;refGene_gn=KIAA0895;refGene_tx=XM_006929310.4
A2 128471517 128471587
refGene_gn=KIAA0895;refGene_tx=XM_011280498.3;refGene_gn=KIAA0895;refGene_tx=XM_023250500.1
A2 128478192 128478306
refGene_gn=ANLN;refGene_tx=XM_003982949.4;refGene_gn=ANLN;refGene_tx=XM_019825696.1;refGene_gn=ANLN;refGene_tx=XM_019825698.1;refGene_gn=ANLN;refGene_tx=XM_019825699.1;refGene_gn=KIAA0895;refGene_tx=XM_01128049

8.3;refGene_gn=KIAA0895;refGene_tx=XM_023250500.1
A2 128484754 128484908
refGene_gn=ANLN;refGene_tx=XM_003982949.4;refGene_gn=ANLN;refGene_tx=XM_019825696.1;refGene_gn=ANLN;refGene_tx=XM_019825698.1;refGene_gn=ANLN;refGene_tx=XM_019825699.1
A2 128486397 128486712
refGene_gn=ANLN;refGene_tx=XM_003982949.4;refGene_gn=ANLN;refGene_tx=XM_019825696.1;refGene_gn=ANLN;refGene_tx=XM_019825698.1;refGene_gn=ANLN;refGene_tx=XM_019825699.1
A2 128490882 128491268
refGene_gn=ANLN;refGene_tx=XM_003982949.4;refGene_gn=ANLN;refGene_tx=XM_019825696.1;refGene_gn=ANLN;refGene_tx=XM_019825698.1;refGene_gn=ANLN;refGene_tx=XM_019825699.1
A2 128492635 128492864
refGene_gn=ANLN;refGene_tx=XM_003982949.4;refGene_gn=ANLN;refGene_tx=XM_019825696.1;refGene_gn=ANLN;refGene_tx=XM_019825698.1;refGene_gn=ANLN;refGene_tx=XM_019825699.1
A2 128495827 128496018
refGene_gn=ANLN;refGene_tx=XM_003982949.4;refGene_gn=ANLN;refGene_tx=XM_019825696.1;refGene_gn=ANLN;refGene_tx=XM_019825698.1;refGene_gn=ANLN;refGene_tx=XM_019825699.1
A2 128496295 128496403
refGene_gn=ANLN;refGene_tx=XM_003982949.4;refGene_gn=ANLN;refGene_tx=XM_019825696.1;refGene_gn=ANLN;refGene_tx=XM_019825698.1
A2 128499379 128499506
refGene_gn=ANLN;refGene_tx=XM_003982949.4;refGene_gn=ANLN;refGene_tx=XM_019825696.1;refGene_gn=ANLN;refGene_tx=XM_019825698.1;refGene_gn=ANLN;refGene_tx=XM_019825699.1
A2 128502572 128502683
refGene_gn=ANLN;refGene_tx=XM_003982949.4;refGene_gn=ANLN;refGene_tx=XM_019825696.1;refGene_gn=ANLN;refGene_tx=XM_019825698.1;refGene_gn=ANLN;refGene_tx=XM_019825699.1
A2 128503735 128503974
refGene_gn=ANLN;refGene_tx=XM_003982949.4;refGene_gn=ANLN;refGene_tx=XM_019825696.1;refGene_gn=ANLN;refGene_tx=XM_019825698.1;refGene_gn=ANLN;refGene_tx=XM_019825699.1
A2 128504678 128504824
refGene_gn=ANLN;refGene_tx=XM_003982949.4;refGene_gn=ANLN;refGene_tx=XM_019825696.1;refGene_gn=ANLN;refGene_tx=XM_019825698.1;refGene_gn=ANLN;refGene_tx=XM_019825699.1
A2 128505032 128505180
refGene_gn=ANLN;refGene_tx=XM_003982949.4;refGene_gn=ANLN;refGene_tx=XM_019825696.1;refGene_gn=ANLN;refGene_tx=XM_019825698.1;refGene_gn=ANLN;refGene_tx=XM_019825699.1
A2 128505805 128505941
refGene_gn=ANLN;refGene_tx=XM_003982949.4;refGene_gn=ANLN;refGene_tx=XM_019825696.1;refGene_gn=ANLN;refGene_tx=XM_019825698.1;refGene_gn=ANLN;refGene_tx=XM_019825699.1
A2 128506589 128506772
refGene_gn=ANLN;refGene_tx=XM_003982949.4;refGene_gn=ANLN;refGene_tx=X

M_019825696.1;refGene_gn=ANLN;refGene_tx=XM_019825698.1;refGene_gn=ANLN;refGene_tx=XM_019825699.1
A2 128508051 128508178
refGene_gn=ANLN;refGene_tx=XM_003982949.4;refGene_gn=ANLN;refGene_tx=XM_019825696.1;refGene_gn=ANLN;refGene_tx=XM_019825698.1;refGene_gn=ANLN;refGene_tx=XM_019825699.1
A2 128509069 128509118
refGene_gn=ANLN;refGene_tx=XM_003982949.4;refGene_gn=ANLN;refGene_tx=XM_019825696.1;refGene_gn=ANLN;refGene_tx=XM_019825698.1;refGene_gn=ANLN;refGene_tx=XM_019825699.1
A2 128509210 128509267
refGene_gn=ANLN;refGene_tx=XM_003982949.4;refGene_gn=ANLN;refGene_tx=XM_019825696.1;refGene_gn=ANLN;refGene_tx=XM_019825698.1;refGene_gn=ANLN;refGene_tx=XM_019825699.1
A2 128509710 128509749
refGene_gn=ANLN;refGene_tx=XM_003982949.4;refGene_gn=ANLN;refGene_tx=XM_019825696.1;refGene_gn=ANLN;refGene_tx=XM_019825698.1;refGene_gn=ANLN;refGene_tx=XM_019825699.1
A2 128510021 128510043
refGene_gn=ANLN;refGene_tx=XM_003982949.4;refGene_gn=ANLN;refGene_tx=XM_019825696.1;refGene_gn=ANLN;refGene_tx=XM_019825698.1;refGene_gn=ANLN;refGene_tx=XM_019825699.1
A2 128510950 128511063
refGene_gn=ANLN;refGene_tx=XM_003982949.4;refGene_gn=ANLN;refGene_tx=XM_019825696.1;refGene_gn=ANLN;refGene_tx=XM_019825698.1;refGene_gn=ANLN;refGene_tx=XM_019825699.1
A2 128511987 128512041
refGene_gn=ANLN;refGene_tx=XM_019825696.1;refGene_gn=ANLN;refGene_tx=XM_019825698.1;refGene_gn=ANLN;refGene_tx=XM_019825699.1
A2 128516519 128516606
refGene_gn=ANLN;refGene_tx=XM_003982949.4;refGene_gn=ANLN;refGene_tx=XM_019825696.1;refGene_gn=ANLN;refGene_tx=XM_019825698.1;refGene_gn=ANLN;refGene_tx=XM_019825699.1
A2 128521661 128521769
refGene_gn=ANLN;refGene_tx=XM_003982949.4;refGene_gn=ANLN;refGene_tx=XM_019825696.1;refGene_gn=ANLN;refGene_tx=XM_019825698.1;refGene_gn=ANLN;refGene_tx=XM_019825699.1
A2 128522609 128522782
refGene_gn=ANLN;refGene_tx=XM_003982949.4;refGene_gn=ANLN;refGene_tx=XM_019825696.1;refGene_gn=ANLN;refGene_tx=XM_019825698.1;refGene_gn=ANLN;refGene_tx=XM_019825699.1
A2 128524238 128524362
refGene_gn=ANLN;refGene_tx=XM_003982949.4;refGene_gn=ANLN;refGene_tx=XM_019825696.1;refGene_gn=ANLN;refGene_tx=XM_019825698.1;refGene_gn=ANLN;refGene_tx=XM_019825699.1
A2 128578122 128578251
refGene_gn=AOAH;refGene_tx=XM_003982953.5;refGene_gn=AOAH;refGene_tx=XM_023250501.1
A2 128587574 128587651
refGene_gn=AOAH;refGene_tx=XM_003982953.5;refGene_gn=AOAH;refGene_tx=X

M_023250501.1
A2	128592832	128592929
refGene_gn=AOAH;refGene_tx=XM_003982953.5;refGene_gn=AOAH;refGene_tx=X
M_023250501.1
A2	128594198	128594258
refGene_gn=AOAH;refGene_tx=XM_003982953.5;refGene_gn=AOAH;refGene_tx=X
M_023250501.1
A2	128594346	128594405
refGene_gn=AOAH;refGene_tx=XM_003982953.5;refGene_gn=AOAH;refGene_tx=X
M_023250501.1
A2	128601297	128601470
refGene_gn=AOAH;refGene_tx=XM_003982953.5;refGene_gn=AOAH;refGene_tx=X
M_023250501.1
A2	128609183	128609258
refGene_gn=AOAH;refGene_tx=XM_003982953.5;refGene_gn=AOAH;refGene_tx=X
M_023250501.1
A2	128609997	128610034
refGene_gn=AOAH;refGene_tx=XM_003982953.5;refGene_gn=AOAH;refGene_tx=X
M_023250501.1
A2	128626333	128626416
refGene_gn=AOAH;refGene_tx=XM_003982953.5;refGene_gn=AOAH;refGene_tx=X
M_023250501.1
A2	128646188	128646280
refGene_gn=AOAH;refGene_tx=XM_003982953.5;refGene_gn=AOAH;refGene_tx=X
M_023250501.1
A2	128668704	128668799
refGene_gn=AOAH;refGene_tx=XM_003982953.5;refGene_gn=AOAH;refGene_tx=X
M_023250501.1
A2	128670693	128670742
refGene_gn=AOAH;refGene_tx=XM_003982953.5;refGene_gn=AOAH;refGene_tx=X
M_023250501.1
A2	128671963	128672012
refGene_gn=AOAH;refGene_tx=XM_003982953.5;refGene_gn=AOAH;refGene_tx=X
M_023250501.1
A2	128672935	128673006
refGene_gn=AOAH;refGene_tx=XM_003982953.5;refGene_gn=AOAH;refGene_tx=X
M_023250501.1
A2	128674063	128674124
refGene_gn=AOAH;refGene_tx=XM_003982953.5;refGene_gn=AOAH;refGene_tx=X
M_023250501.1
A2	128680624	128680695
refGene_gn=AOAH;refGene_tx=XM_003982953.5;refGene_gn=AOAH;refGene_tx=X
M_023250501.1
A2	128685849	128685909
refGene_gn=AOAH;refGene_tx=XM_003982953.5;refGene_gn=AOAH;refGene_tx=X
M_023250501.1
A2	128702860	128702960
refGene_gn=AOAH;refGene_tx=XM_003982953.5;refGene_gn=AOAH;refGene_tx=X
M_023250501.1
A2	128710219	128710286

refGene_gn=AOAH;refGene_tx=XM_003982953.5;refGene_gn=AOAH;refGene_tx=XM_023250501.1
A2 128719526 128719622
refGene_gn=AOAH;refGene_tx=XM_003982953.5;refGene_gn=AOAH;refGene_tx=XM_023250501.1
A2 128744038 128744165
refGene_gn=AOAH;refGene_tx=XM_003982953.5;refGene_gn=AOAH;refGene_tx=XM_023250501.1
A2 128859434 128859635
refGene_gn=ELMO1;refGene_tx=XM_019825703.2;refGene_gn=ELMO1;refGene_tx=XM_019825705.2;refGene_gn=ELMO1;refGene_tx=XM_019825706.2;refGene_gn=ELMO1;refGene_tx=XM_019825707.2;refGene_gn=ELMO1;refGene_tx=XM_019825709.2;refGene_gn=ELMO1;refGene_tx=XM_019825710.2;refGene_gn=ELMO1;refGene_tx=XM_023250502.1;refGene_gn=ELMO1;refGene_tx=XM_023250503.1;refGene_gn=ELMO1;refGene_tx=XM_023250504.1;refGene_gn=ELMO1;refGene_tx=XM_023250505.1;refGene_gn=ELMO1;refGene_tx=XM_023250506.1;refGene_gn=ELMO1;refGene_tx=XM_023250507.1;refGene_gn=ELMO1;refGene_tx=XM_023250508.1;refGene_gn=ELMO1;refGene_tx=XM_023250509.1
A2 128865221 128865299
refGene_gn=ELMO1;refGene_tx=XM_019825703.2;refGene_gn=ELMO1;refGene_tx=XM_019825705.2;refGene_gn=ELMO1;refGene_tx=XM_019825706.2;refGene_gn=ELMO1;refGene_tx=XM_019825707.2;refGene_gn=ELMO1;refGene_tx=XM_019825709.2;refGene_gn=ELMO1;refGene_tx=XM_019825710.2;refGene_gn=ELMO1;refGene_tx=XM_023250502.1;refGene_gn=ELMO1;refGene_tx=XM_023250503.1;refGene_gn=ELMO1;refGene_tx=XM_023250504.1;refGene_gn=ELMO1;refGene_tx=XM_023250505.1;refGene_gn=ELMO1;refGene_tx=XM_023250506.1;refGene_gn=ELMO1;refGene_tx=XM_023250507.1;refGene_gn=ELMO1;refGene_tx=XM_023250508.1;refGene_gn=ELMO1;refGene_tx=XM_023250509.1
A2 128875669 128875752
refGene_gn=ELMO1;refGene_tx=XM_019825703.2;refGene_gn=ELMO1;refGene_tx=XM_019825705.2;refGene_gn=ELMO1;refGene_tx=XM_019825706.2;refGene_gn=ELMO1;refGene_tx=XM_019825707.2;refGene_gn=ELMO1;refGene_tx=XM_019825709.2;refGene_gn=ELMO1;refGene_tx=XM_019825710.2;refGene_gn=ELMO1;refGene_tx=XM_023250502.1;refGene_gn=ELMO1;refGene_tx=XM_023250503.1;refGene_gn=ELMO1;refGene_tx=XM_023250504.1;refGene_gn=ELMO1;refGene_tx=XM_023250505.1;refGene_gn=ELMO1;refGene_tx=XM_023250506.1;refGene_gn=ELMO1;refGene_tx=XM_023250507.1;refGene_gn=ELMO1;refGene_tx=XM_023250508.1;refGene_gn=ELMO1;refGene_tx=XM_023250509.1
A2 128883478 128883586
refGene_gn=ELMO1;refGene_tx=XM_019825703.2;refGene_gn=ELMO1;refGene_tx=XM_019825705.2;refGene_gn=ELMO1;refGene_tx=XM_019825706.2;refGene_gn=ELMO1;refGene_tx=XM_019825707.2;refGene_gn=ELMO1;refGene_tx=XM_019825709.2;refGene_gn=ELMO1;refGene_tx=XM_019825710.2;refGene_gn=ELMO1;refGene_tx=XM_023250502.1;refGene_gn=ELMO1;refGene_tx=XM_023250503.1;refGene_gn=ELMO1;refGene_tx=XM_023250504.1;refGene_gn=ELMO1;refGene_tx=XM_023250505.1;refGene_gn=ELMO1;refGene_tx=XM_023250506.1;refGene_gn=ELMO1;refGene_tx=XM_023250507.1;refGene_gn=ELMO1;refGene_tx=XM_023250508.1;refGene_gn=ELMO1;refGene_tx=XM_023250509.1
A2 128892723 128892836
refGene_gn=ELMO1;refGene_tx=XM_019825703.2;refGene_gn=ELMO1;refGene_tx

=XM_019825705.2;refGene_gn=ELMO1;refGene_tx=XM_019825706.2;refGene_gn=ELMO1;refGene_tx=XM_019825707.2;refGene_gn=ELMO1;refGene_tx=XM_019825709.2;refGene_gn=ELMO1;refGene_tx=XM_019825710.2;refGene_gn=ELMO1;refGene_tx=XM_023250502.1;refGene_gn=ELMO1;refGene_tx=XM_023250503.1;refGene_gn=ELMO1;refGene_tx=XM_023250504.1;refGene_gn=ELMO1;refGene_tx=XM_023250505.1;refGene_gn=ELMO1;refGene_tx=XM_023250506.1;refGene_gn=ELMO1;refGene_tx=XM_023250507.1;refGene_gn=ELMO1;refGene_tx=XM_023250508.1;refGene_gn=ELMO1;refGene_tx=XM_023250509.1
A2 128903508 128903672
refGene_gn=ELMO1;refGene_tx=XM_019825703.2;refGene_gn=ELMO1;refGene_tx=XM_019825705.2;refGene_gn=ELMO1;refGene_tx=XM_019825706.2;refGene_gn=ELMO1;refGene_tx=XM_019825707.2;refGene_gn=ELMO1;refGene_tx=XM_019825709.2;refGene_gn=ELMO1;refGene_tx=XM_019825710.2;refGene_gn=ELMO1;refGene_tx=XM_023250502.1;refGene_gn=ELMO1;refGene_tx=XM_023250503.1;refGene_gn=ELMO1;refGene_tx=XM_023250504.1;refGene_gn=ELMO1;refGene_tx=XM_023250505.1;refGene_gn=ELMO1;refGene_tx=XM_023250506.1;refGene_gn=ELMO1;refGene_tx=XM_023250507.1;refGene_gn=ELMO1;refGene_tx=XM_023250508.1;refGene_gn=ELMO1;refGene_tx=XM_023250509.1
A2 129022280 129022417
refGene_gn=ELMO1;refGene_tx=XM_019825703.2;refGene_gn=ELMO1;refGene_tx=XM_019825705.2;refGene_gn=ELMO1;refGene_tx=XM_019825706.2;refGene_gn=ELMO1;refGene_tx=XM_019825707.2;refGene_gn=ELMO1;refGene_tx=XM_019825709.2;refGene_gn=ELMO1;refGene_tx=XM_023250502.1;refGene_gn=ELMO1;refGene_tx=XM_023250503.1;refGene_gn=ELMO1;refGene_tx=XM_023250504.1;refGene_gn=ELMO1;refGene_tx=XM_023250505.1;refGene_gn=ELMO1;refGene_tx=XM_023250506.1;refGene_gn=ELMO1;refGene_tx=XM_023250507.1;refGene_gn=ELMO1;refGene_tx=XM_023250508.1;refGene_gn=ELMO1;refGene_tx=XM_023250509.1
A2 129099309 129099418
refGene_gn=ELMO1;refGene_tx=XM_019825703.2;refGene_gn=ELMO1;refGene_tx=XM_019825705.2;refGene_gn=ELMO1;refGene_tx=XM_019825706.2;refGene_gn=ELMO1;refGene_tx=XM_019825707.2;refGene_gn=ELMO1;refGene_tx=XM_019825709.2;refGene_gn=ELMO1;refGene_tx=XM_023250502.1;refGene_gn=ELMO1;refGene_tx=XM_023250503.1;refGene_gn=ELMO1;refGene_tx=XM_023250504.1;refGene_gn=ELMO1;refGene_tx=XM_023250505.1;refGene_gn=ELMO1;refGene_tx=XM_023250506.1;refGene_gn=ELMO1;refGene_tx=XM_023250507.1;refGene_gn=ELMO1;refGene_tx=XM_023250508.1;refGene_gn=ELMO1;refGene_tx=XM_023250509.1
A2 129133506 129133611
refGene_gn=ELMO1;refGene_tx=XM_019825703.2;refGene_gn=ELMO1;refGene_tx=XM_019825705.2;refGene_gn=ELMO1;refGene_tx=XM_019825706.2;refGene_gn=ELMO1;refGene_tx=XM_019825707.2;refGene_gn=ELMO1;refGene_tx=XM_019825709.2;refGene_gn=ELMO1;refGene_tx=XM_023250502.1;refGene_gn=ELMO1;refGene_tx=XM_023250503.1;refGene_gn=ELMO1;refGene_tx=XM_023250504.1;refGene_gn=ELMO1;refGene_tx=XM_023250505.1;refGene_gn=ELMO1;refGene_tx=XM_023250506.1;refGene_gn=ELMO1;refGene_tx=XM_023250507.1;refGene_gn=ELMO1;refGene_tx=XM_023250508.1;refGene_gn=ELMO1;refGene_tx=XM_023250509.1
A2 129182820 129182952
refGene_gn=ELMO1;refGene_tx=XM_019825703.2;refGene_gn=ELMO1;refGene_tx=XM_019825705.2;refGene_gn=ELMO1;refGene_tx=XM_019825706.2;refGene_gn=ELMO1;refGene_tx=XM_019825707.2;refGene_gn=ELMO1;refGene_tx=XM_019825709.2;refGene_gn=ELMO1;refGene_tx=XM_023250502.1;refGene_gn=ELMO1;refGe

ne_tx=XM_023250503.1;refGene_gn=ELMO1;refGene_tx=XM_023250504.1;refGen
e_gn=ELMO1;refGene_tx=XM_023250505.1;refGene_gn=ELMO1;refGene_tx=XM_02
3250506.1;refGene_gn=ELMO1;refGene_tx=XM_023250507.1;refGene_gn=ELMO1;
refGene_tx=XM_023250508.1;refGene_gn=ELMO1;refGene_tx=XM_023250509.1
A2 129184379 129184502
refGene_gn=ELMO1;refGene_tx=XM_019825703.2;refGene_gn=ELMO1;refGene_tx
=XM_019825705.2;refGene_gn=ELMO1;refGene_tx=XM_019825706.2;refGene_gn=
ELMO1;refGene_tx=XM_019825707.2;refGene_gn=ELMO1;refGene_tx=XM_0198257
09.2;refGene_gn=ELMO1;refGene_tx=XM_023250502.1;refGene_gn=ELMO1;refGe
ne_tx=XM_023250503.1;refGene_gn=ELMO1;refGene_tx=XM_023250504.1;refGen
e_gn=ELMO1;refGene_tx=XM_023250505.1;refGene_gn=ELMO1;refGene_tx=XM_02
3250506.1;refGene_gn=ELMO1;refGene_tx=XM_023250507.1;refGene_gn=ELMO1;
refGene_tx=XM_023250508.1;refGene_gn=ELMO1;refGene_tx=XM_023250509.1
A2 129187049 129187100
refGene_gn=ELMO1;refGene_tx=XM_019825703.2;refGene_gn=ELMO1;refGene_tx
=XM_019825705.2;refGene_gn=ELMO1;refGene_tx=XM_019825706.2;refGene_gn=
ELMO1;refGene_tx=XM_019825707.2;refGene_gn=ELMO1;refGene_tx=XM_0198257
09.2;refGene_gn=ELMO1;refGene_tx=XM_023250502.1;refGene_gn=ELMO1;refGe
ne_tx=XM_023250503.1;refGene_gn=ELMO1;refGene_tx=XM_023250504.1;refGen
e_gn=ELMO1;refGene_tx=XM_023250505.1;refGene_gn=ELMO1;refGene_tx=XM_02
3250506.1;refGene_gn=ELMO1;refGene_tx=XM_023250507.1;refGene_gn=ELMO1;
refGene_tx=XM_023250508.1;refGene_gn=ELMO1;refGene_tx=XM_023250509.1
A2 129188107 129188152
refGene_gn=ELMO1;refGene_tx=XM_019825703.2;refGene_gn=ELMO1;refGene_tx
=XM_019825705.2;refGene_gn=ELMO1;refGene_tx=XM_019825706.2;refGene_gn=
ELMO1;refGene_tx=XM_019825707.2;refGene_gn=ELMO1;refGene_tx=XM_0198257
09.2;refGene_gn=ELMO1;refGene_tx=XM_023250502.1;refGene_gn=ELMO1;refGe
ne_tx=XM_023250503.1;refGene_gn=ELMO1;refGene_tx=XM_023250504.1;refGen
e_gn=ELMO1;refGene_tx=XM_023250505.1;refGene_gn=ELMO1;refGene_tx=XM_02
3250506.1;refGene_gn=ELMO1;refGene_tx=XM_023250507.1;refGene_gn=ELMO1;
refGene_tx=XM_023250508.1
A2 129192700 129192779
refGene_gn=ELMO1;refGene_tx=XM_019825703.2;refGene_gn=ELMO1;refGene_tx
=XM_019825705.2;refGene_gn=ELMO1;refGene_tx=XM_019825706.2;refGene_gn=
ELMO1;refGene_tx=XM_019825707.2;refGene_gn=ELMO1;refGene_tx=XM_0198257
09.2;refGene_gn=ELMO1;refGene_tx=XM_023250502.1;refGene_gn=ELMO1;refGe
ne_tx=XM_023250503.1;refGene_gn=ELMO1;refGene_tx=XM_023250504.1;refGen
e_gn=ELMO1;refGene_tx=XM_023250505.1;refGene_gn=ELMO1;refGene_tx=XM_02
3250506.1;refGene_gn=ELMO1;refGene_tx=XM_023250507.1;refGene_gn=ELMO1;
refGene_tx=XM_023250508.1;refGene_gn=ELMO1;refGene_tx=XM_023250509.1
A2 129194922 129195074
refGene_gn=ELMO1;refGene_tx=XM_019825703.2;refGene_gn=ELMO1;refGene_tx
=XM_019825705.2;refGene_gn=ELMO1;refGene_tx=XM_019825706.2;refGene_gn=
ELMO1;refGene_tx=XM_019825707.2;refGene_gn=ELMO1;refGene_tx=XM_0198257
09.2;refGene_gn=ELMO1;refGene_tx=XM_023250502.1;refGene_gn=ELMO1;refGe
ne_tx=XM_023250503.1;refGene_gn=ELMO1;refGene_tx=XM_023250504.1;refGen
e_gn=ELMO1;refGene_tx=XM_023250505.1;refGene_gn=ELMO1;refGene_tx=XM_02
3250506.1;refGene_gn=ELMO1;refGene_tx=XM_023250507.1;refGene_gn=ELMO1;
refGene_tx=XM_023250508.1;refGene_gn=ELMO1;refGene_tx=XM_023250509.1
A2 129202228 129202328

refGene_gn=ELMO1;refGene_tx=XM_019825703.2;refGene_gn=ELMO1;refGene_tx
=XM_019825705.2;refGene_gn=ELMO1;refGene_tx=XM_019825706.2;refGene_gn=
ELMO1;refGene_tx=XM_019825707.2;refGene_gn=ELMO1;refGene_tx=XM_0198257
09.2;refGene_gn=ELMO1;refGene_tx=XM_023250502.1;refGene_gn=ELMO1;refGe
ne_tx=XM_023250503.1;refGene_gn=ELMO1;refGene_tx=XM_023250504.1;refGen
e_gn=ELMO1;refGene_tx=XM_023250505.1;refGene_gn=ELMO1;refGene_tx=XM_02
3250506.1;refGene_gn=ELMO1;refGene_tx=XM_023250507.1;refGene_gn=ELMO1;
refGene_tx=XM_023250508.1;refGene_gn=ELMO1;refGene_tx=XM_023250509.1
A2 129213982 129214018
refGene_gn=ELMO1;refGene_tx=XM_019825703.2;refGene_gn=ELMO1;refGene_tx
=XM_019825705.2;refGene_gn=ELMO1;refGene_tx=XM_019825706.2;refGene_gn=
ELMO1;refGene_tx=XM_019825707.2;refGene_gn=ELMO1;refGene_tx=XM_0198257
09.2;refGene_gn=ELMO1;refGene_tx=XM_023250502.1;refGene_gn=ELMO1;refGe
ne_tx=XM_023250503.1;refGene_gn=ELMO1;refGene_tx=XM_023250504.1;refGen
e_gn=ELMO1;refGene_tx=XM_023250505.1;refGene_gn=ELMO1;refGene_tx=XM_02
3250506.1;refGene_gn=ELMO1;refGene_tx=XM_023250507.1;refGene_gn=ELMO1;
refGene_tx=XM_023250508.1;refGene_gn=ELMO1;refGene_tx=XM_023250509.1
A2 129218088 129218108
refGene_gn=ELMO1;refGene_tx=XM_019825709.2
A2 129225465 129225635
refGene_gn=ELMO1;refGene_tx=XM_019825703.2;refGene_gn=ELMO1;refGene_tx
=XM_019825705.2;refGene_gn=ELMO1;refGene_tx=XM_019825706.2;refGene_gn=
ELMO1;refGene_tx=XM_019825707.2;refGene_gn=ELMO1;refGene_tx=XM_0232505
02.1;refGene_gn=ELMO1;refGene_tx=XM_023250503.1;refGene_gn=ELMO1;refGe
ne_tx=XM_023250504.1;refGene_gn=ELMO1;refGene_tx=XM_023250505.1;refGen
e_gn=ELMO1;refGene_tx=XM_023250506.1;refGene_gn=ELMO1;refGene_tx=XM_02
3250507.1;refGene_gn=ELMO1;refGene_tx=XM_023250508.1;refGene_gn=ELMO1;
refGene_tx=XM_023250509.1
A2 129229800 129229851
refGene_gn=ELMO1;refGene_tx=XM_019825703.2;refGene_gn=ELMO1;refGene_tx
=XM_019825705.2;refGene_gn=ELMO1;refGene_tx=XM_019825706.2;refGene_gn=
ELMO1;refGene_tx=XM_019825707.2;refGene_gn=ELMO1;refGene_tx=XM_0232505
02.1;refGene_gn=ELMO1;refGene_tx=XM_023250503.1;refGene_gn=ELMO1;refGe
ne_tx=XM_023250504.1;refGene_gn=ELMO1;refGene_tx=XM_023250505.1;refGen
e_gn=ELMO1;refGene_tx=XM_023250506.1;refGene_gn=ELMO1;refGene_tx=XM_02
3250507.1;refGene_gn=ELMO1;refGene_tx=XM_023250508.1;refGene_gn=ELMO1;
refGene_tx=XM_023250509.1
A2 129264994 129265067
refGene_gn=ELMO1;refGene_tx=XM_019825703.2;refGene_gn=ELMO1;refGene_tx
=XM_019825705.2;refGene_gn=ELMO1;refGene_tx=XM_019825706.2;refGene_gn=
ELMO1;refGene_tx=XM_019825707.2;refGene_gn=ELMO1;refGene_tx=XM_0232505
02.1;refGene_gn=ELMO1;refGene_tx=XM_023250503.1;refGene_gn=ELMO1;refGe
ne_tx=XM_023250504.1;refGene_gn=ELMO1;refGene_tx=XM_023250505.1;refGen
e_gn=ELMO1;refGene_tx=XM_023250506.1;refGene_gn=ELMO1;refGene_tx=XM_02
3250507.1;refGene_gn=ELMO1;refGene_tx=XM_023250508.1;refGene_gn=ELMO1;
refGene_tx=XM_023250509.1
A2 129266100 129266141
refGene_gn=ELMO1;refGene_tx=XM_019825703.2;refGene_gn=ELMO1;refGene_tx
=XM_019825705.2;refGene_gn=ELMO1;refGene_tx=XM_019825706.2;refGene_gn=
ELMO1;refGene_tx=XM_019825707.2;refGene_gn=ELMO1;refGene_tx=XM_0232505

02.1;refGene_gn=ELMO1;refGene_tx=XM_023250503.1;refGene_gn=ELMO1;refGene_tx=XM_023250504.1;refGene_gn=ELMO1;refGene_tx=XM_023250505.1;refGene_gn=ELMO1;refGene_tx=XM_023250506.1;refGene_gn=ELMO1;refGene_tx=XM_023250507.1;refGene_gn=ELMO1;refGene_tx=XM_023250508.1;refGene_gn=ELMO1;refGene_tx=XM_023250509.1
A2 129289582 129289660
refGene_gn=ELMO1;refGene_tx=XM_019825703.2;refGene_gn=ELMO1;refGene_tx=XM_019825705.2;refGene_gn=ELMO1;refGene_tx=XM_019825706.2;refGene_gn=ELMO1;refGene_tx=XM_019825707.2;refGene_gn=ELMO1;refGene_tx=XM_023250502.1;refGene_gn=ELMO1;refGene_tx=XM_023250503.1;refGene_gn=ELMO1;refGene_tx=XM_023250504.1;refGene_gn=ELMO1;refGene_tx=XM_023250505.1;refGene_gn=ELMO1;refGene_tx=XM_023250506.1;refGene_gn=ELMO1;refGene_tx=XM_023250507.1;refGene_gn=ELMO1;refGene_tx=XM_023250508.1;refGene_gn=ELMO1;refGene_tx=XM_023250509.1
A2 129510234 129510801
refGene_gn=LOC101081129;refGene_tx=XM_019825712.2
A2 129556490 129557396
refGene_gn=GPR141;refGene_tx=XM_003982959.4
A2 129583379 129584288
refGene_gn=THAP5;refGene_tx=XM_003982960.5
A2 129584775 129584968
refGene_gn=THAP5;refGene_tx=XM_003982960.5
A2 129588455 129588535
refGene_gn=THAP5;refGene_tx=XM_003982960.5
A2 129590946 129591163
refGene_gn=DNAJB9;refGene_tx=XM_003982962.5
A2 129592141 129592593
refGene_gn=DNAJB9;refGene_tx=XM_003982962.5
A2 131300992 131301112
refGene_gn=IMMP2L;refGene_tx=XM_003982963.5;refGene_gn=IMMP2L;refGene_tx=XM_019825713.2;refGene_gn=IMMP2L;refGene_tx=XM_019825714.2;refGene_gn=IMMP2L;refGene_tx=XM_019825715.2;refGene_gn=IMMP2L;refGene_tx=XM_019825716.2;refGene_gn=IMMP2L;refGene_tx=XM_023250510.1
A2 131514794 131514897
refGene_gn=IMMP2L;refGene_tx=XM_003982963.5;refGene_gn=IMMP2L;refGene_tx=XM_019825713.2;refGene_gn=IMMP2L;refGene_tx=XM_019825714.2;refGene_gn=IMMP2L;refGene_tx=XM_019825715.2;refGene_gn=IMMP2L;refGene_tx=XM_019825716.2;refGene_gn=IMMP2L;refGene_tx=XM_023250510.1
A2 131582303 131582369
refGene_gn=IMMP2L;refGene_tx=XM_003982963.5;refGene_gn=IMMP2L;refGene_tx=XM_019825713.2;refGene_gn=IMMP2L;refGene_tx=XM_019825714.2;refGene_gn=IMMP2L;refGene_tx=XM_019825715.2;refGene_gn=IMMP2L;refGene_tx=XM_019825716.2;refGene_gn=IMMP2L;refGene_tx=XM_023250510.1
A2 131752036 131754163
refGene_gn=LRRN3;refGene_tx=XM_006929314.4
A2 132112523 132112627
refGene_gn=IMMP2L;refGene_tx=XM_003982963.5;refGene_gn=IMMP2L;refGene_tx=XM_019825713.2;refGene_gn=IMMP2L;refGene_tx=XM_019825714.2;refGene_gn=IMMP2L;refGene_tx=XM_019825715.2;refGene_gn=IMMP2L;refGene_tx=XM_019825716.2;refGene_gn=IMMP2L;refGene_tx=XM_023250510.1

A2	132145277	132145412
refGene_gn=IMMP2L;refGene_tx=XM_003982963.5;refGene_gn=IMMP2L;refGene_tx=XM_019825713.2;refGene_gn=IMMP2L;refGene_tx=XM_019825714.2;refGene_gn=IMMP2L;refGene_tx=XM_019825715.2;refGene_gn=IMMP2L;refGene_tx=XM_019825716.2;refGene_gn=IMMP2L;refGene_tx=XM_023250510.1
A2	132300816	132301248
refGene_gn=DOCK4;refGene_tx=XM_011280513.3;refGene_gn=DOCK4;refGene_tx=XM_019825718.2;refGene_gn=DOCK4;refGene_tx=XM_019825719.2;refGene_gn=DOCK4;refGene_tx=XM_019825720.2
A2	132304401	132304463
refGene_gn=DOCK4;refGene_tx=XM_011280513.3;refGene_gn=DOCK4;refGene_tx=XM_019825718.2;refGene_gn=DOCK4;refGene_tx=XM_019825719.2;refGene_gn=DOCK4;refGene_tx=XM_019825720.2
A2	132307227	132307341
refGene_gn=DOCK4;refGene_tx=XM_011280513.3;refGene_gn=DOCK4;refGene_tx=XM_019825718.2;refGene_gn=DOCK4;refGene_tx=XM_019825719.2
A2	132309109	132309182
refGene_gn=DOCK4;refGene_tx=XM_011280513.3;refGene_gn=DOCK4;refGene_tx=XM_019825718.2;refGene_gn=DOCK4;refGene_tx=XM_019825719.2;refGene_gn=DOCK4;refGene_tx=XM_019825720.2
A2	132310885	132310995
refGene_gn=DOCK4;refGene_tx=XM_011280513.3;refGene_gn=DOCK4;refGene_tx=XM_019825718.2;refGene_gn=DOCK4;refGene_tx=XM_019825719.2;refGene_gn=DOCK4;refGene_tx=XM_019825720.2
A2	132311120	132311202
refGene_gn=DOCK4;refGene_tx=XM_011280513.3;refGene_gn=DOCK4;refGene_tx=XM_019825718.2;refGene_gn=DOCK4;refGene_tx=XM_019825719.2;refGene_gn=DOCK4;refGene_tx=XM_019825720.2
A2	132312242	132312363
refGene_gn=DOCK4;refGene_tx=XM_011280513.3;refGene_gn=DOCK4;refGene_tx=XM_019825718.2;refGene_gn=DOCK4;refGene_tx=XM_019825719.2;refGene_gn=DOCK4;refGene_tx=XM_019825720.2
A2	132312688	132312810
refGene_gn=DOCK4;refGene_tx=XM_011280513.3;refGene_gn=DOCK4;refGene_tx=XM_019825718.2;refGene_gn=DOCK4;refGene_tx=XM_019825719.2;refGene_gn=DOCK4;refGene_tx=XM_019825720.2
A2	132313140	132313260
refGene_gn=DOCK4;refGene_tx=XM_011280513.3;refGene_gn=DOCK4;refGene_tx=XM_019825718.2;refGene_gn=DOCK4;refGene_tx=XM_019825719.2;refGene_gn=DOCK4;refGene_tx=XM_019825720.2
A2	132316540	132316624
refGene_gn=DOCK4;refGene_tx=XM_011280513.3;refGene_gn=DOCK4;refGene_tx=XM_019825718.2;refGene_gn=DOCK4;refGene_tx=XM_019825719.2;refGene_gn=DOCK4;refGene_tx=XM_019825720.2
A2	132317522	132317699
refGene_gn=DOCK4;refGene_tx=XM_011280513.3;refGene_gn=DOCK4;refGene_tx=XM_019825718.2;refGene_gn=DOCK4;refGene_tx=XM_019825719.2;refGene_gn=DOCK4;refGene_tx=XM_019825720.2
A2	132325178	132325265
refGene_gn=DOCK4;refGene_tx=XM_011280513.3;refGene_gn=DOCK4;refGene_tx

=XM_019825718.2;refGene_gn=DOCK4;refGene_tx=XM_019825719.2;refGene_gn=DOCK4;refGene_tx=XM_019825720.2
A2 132327907 132328074
refGene_gn=DOCK4;refGene_tx=XM_011280513.3;refGene_gn=DOCK4;refGene_tx=XM_019825718.2;refGene_gn=DOCK4;refGene_tx=XM_019825719.2;refGene_gn=DOCK4;refGene_tx=XM_019825720.2
A2 132329465 132329607
refGene_gn=DOCK4;refGene_tx=XM_011280513.3;refGene_gn=DOCK4;refGene_tx=XM_019825718.2;refGene_gn=DOCK4;refGene_tx=XM_019825719.2;refGene_gn=DOCK4;refGene_tx=XM_019825720.2
A2 132334957 132335062
refGene_gn=DOCK4;refGene_tx=XM_011280513.3;refGene_gn=DOCK4;refGene_tx=XM_019825718.2;refGene_gn=DOCK4;refGene_tx=XM_019825719.2;refGene_gn=DOCK4;refGene_tx=XM_019825720.2
A2 132337096 132337183
refGene_gn=DOCK4;refGene_tx=XM_011280513.3;refGene_gn=DOCK4;refGene_tx=XM_019825718.2;refGene_gn=DOCK4;refGene_tx=XM_019825719.2;refGene_gn=DOCK4;refGene_tx=XM_019825720.2
A2 132338900 132339049
refGene_gn=DOCK4;refGene_tx=XM_011280513.3;refGene_gn=DOCK4;refGene_tx=XM_019825718.2;refGene_gn=DOCK4;refGene_tx=XM_019825719.2;refGene_gn=DOCK4;refGene_tx=XM_019825720.2
A2 132346866 132346960
refGene_gn=DOCK4;refGene_tx=XM_011280513.3;refGene_gn=DOCK4;refGene_tx=XM_019825718.2;refGene_gn=DOCK4;refGene_tx=XM_019825719.2;refGene_gn=DOCK4;refGene_tx=XM_019825720.2
A2 132350875 132350936
refGene_gn=DOCK4;refGene_tx=XM_011280513.3;refGene_gn=DOCK4;refGene_tx=XM_019825718.2;refGene_gn=DOCK4;refGene_tx=XM_019825719.2;refGene_gn=DOCK4;refGene_tx=XM_019825720.2
A2 132351786 132351882
refGene_gn=DOCK4;refGene_tx=XM_011280513.3;refGene_gn=DOCK4;refGene_tx=XM_019825718.2;refGene_gn=DOCK4;refGene_tx=XM_019825719.2;refGene_gn=DOCK4;refGene_tx=XM_019825720.2
A2 132352026 132352053
refGene_gn=DOCK4;refGene_tx=XM_011280513.3;refGene_gn=DOCK4;refGene_tx=XM_019825719.2;refGene_gn=DOCK4;refGene_tx=XM_019825720.2
A2 132357196 132357282
refGene_gn=DOCK4;refGene_tx=XM_011280513.3;refGene_gn=DOCK4;refGene_tx=XM_019825718.2;refGene_gn=DOCK4;refGene_tx=XM_019825719.2;refGene_gn=DOCK4;refGene_tx=XM_019825720.2
A2 132358996 132359145
refGene_gn=DOCK4;refGene_tx=XM_011280513.3;refGene_gn=DOCK4;refGene_tx=XM_019825718.2;refGene_gn=DOCK4;refGene_tx=XM_019825719.2;refGene_gn=DOCK4;refGene_tx=XM_019825720.2
A2 132366849 132366908
refGene_gn=DOCK4;refGene_tx=XM_011280513.3;refGene_gn=DOCK4;refGene_tx=XM_019825718.2;refGene_gn=DOCK4;refGene_tx=XM_019825719.2;refGene_gn=DOCK4;refGene_tx=XM_019825720.2
A2 132368460 132368561

refGene_gn=DOCK4;refGene_tx=XM_011280513.3;refGene_gn=DOCK4;refGene_tx=XM_019825718.2;refGene_gn=DOCK4;refGene_tx=XM_019825719.2;refGene_gn=DOCK4;refGene_tx=XM_019825720.2
A2 132370595 132370671 refGene_gn=DOCK4;refGene_tx=XM_011280513.3;refGene_gn=DOCK4;refGene_tx=XM_019825718.2;refGene_gn=DOCK4;refGene_tx=XM_019825719.2;refGene_gn=DOCK4;refGene_tx=XM_019825720.2
A2 132379804 132379899 refGene_gn=DOCK4;refGene_tx=XM_011280513.3;refGene_gn=DOCK4;refGene_tx=XM_019825718.2;refGene_gn=DOCK4;refGene_tx=XM_019825719.2;refGene_gn=DOCK4;refGene_tx=XM_019825720.2
A2 132390055 132390154 refGene_gn=DOCK4;refGene_tx=XM_011280513.3;refGene_gn=DOCK4;refGene_tx=XM_019825718.2;refGene_gn=DOCK4;refGene_tx=XM_019825719.2;refGene_gn=DOCK4;refGene_tx=XM_019825720.2
A2 132413409 132413544 refGene_gn=DOCK4;refGene_tx=XM_011280513.3;refGene_gn=DOCK4;refGene_tx=XM_019825718.2;refGene_gn=DOCK4;refGene_tx=XM_019825719.2;refGene_gn=DOCK4;refGene_tx=XM_019825720.2
A2 132415709 132415837 refGene_gn=DOCK4;refGene_tx=XM_011280513.3;refGene_gn=DOCK4;refGene_tx=XM_019825718.2;refGene_gn=DOCK4;refGene_tx=XM_019825719.2;refGene_gn=DOCK4;refGene_tx=XM_019825720.2
A2 132427621 132427814 refGene_gn=DOCK4;refGene_tx=XM_011280513.3;refGene_gn=DOCK4;refGene_tx=XM_019825718.2;refGene_gn=DOCK4;refGene_tx=XM_019825719.2;refGene_gn=DOCK4;refGene_tx=XM_019825720.2
A2 132432473 132432644 refGene_gn=DOCK4;refGene_tx=XM_011280513.3;refGene_gn=DOCK4;refGene_tx=XM_019825718.2;refGene_gn=DOCK4;refGene_tx=XM_019825719.2;refGene_gn=DOCK4;refGene_tx=XM_019825720.2
A2 132433977 132434059 refGene_gn=DOCK4;refGene_tx=XM_011280513.3;refGene_gn=DOCK4;refGene_tx=XM_019825718.2;refGene_gn=DOCK4;refGene_tx=XM_019825719.2;refGene_gn=DOCK4;refGene_tx=XM_019825720.2
A2 132436572 132436673 refGene_gn=DOCK4;refGene_tx=XM_011280513.3;refGene_gn=DOCK4;refGene_tx=XM_019825718.2;refGene_gn=DOCK4;refGene_tx=XM_019825719.2;refGene_gn=DOCK4;refGene_tx=XM_019825720.2
A2 132436849 132436933 refGene_gn=DOCK4;refGene_tx=XM_011280513.3;refGene_gn=DOCK4;refGene_tx=XM_019825718.2;refGene_gn=DOCK4;refGene_tx=XM_019825719.2;refGene_gn=DOCK4;refGene_tx=XM_019825720.2
A2 132437041 132437139 refGene_gn=DOCK4;refGene_tx=XM_011280513.3;refGene_gn=DOCK4;refGene_tx=XM_019825718.2;refGene_gn=DOCK4;refGene_tx=XM_019825719.2;refGene_gn=DOCK4;refGene_tx=XM_019825720.2
A2 132442085 132442242 refGene_gn=DOCK4;refGene_tx=XM_011280513.3;refGene_gn=DOCK4;refGene_tx=XM_019825718.2;refGene_gn=DOCK4;refGene_tx=XM_019825719.2;refGene_gn=

DOCK4;refGene_tx=XM_019825720.2
A2	132456984	132457091
refGene_gn=DOCK4;refGene_tx=XM_011280513.3;refGene_gn=DOCK4;refGene_tx=XM_019825718.2;refGene_gn=DOCK4;refGene_tx=XM_019825719.2;refGene_gn=DOCK4;refGene_tx=XM_019825720.2
A2	132462614	132462777
refGene_gn=DOCK4;refGene_tx=XM_011280513.3;refGene_gn=DOCK4;refGene_tx=XM_019825718.2;refGene_gn=DOCK4;refGene_tx=XM_019825719.2;refGene_gn=DOCK4;refGene_tx=XM_019825720.2
A2	132463578	132463703
refGene_gn=DOCK4;refGene_tx=XM_011280513.3;refGene_gn=DOCK4;refGene_tx=XM_019825718.2;refGene_gn=DOCK4;refGene_tx=XM_019825719.2;refGene_gn=DOCK4;refGene_tx=XM_019825720.2
A2	132473237	132473363
refGene_gn=DOCK4;refGene_tx=XM_011280513.3;refGene_gn=DOCK4;refGene_tx=XM_019825718.2;refGene_gn=DOCK4;refGene_tx=XM_019825719.2;refGene_gn=DOCK4;refGene_tx=XM_019825720.2
A2	132491048	132491137
refGene_gn=DOCK4;refGene_tx=XM_011280513.3;refGene_gn=DOCK4;refGene_tx=XM_019825718.2;refGene_gn=DOCK4;refGene_tx=XM_019825719.2;refGene_gn=DOCK4;refGene_tx=XM_019825720.2
A2	132494935	132495068
refGene_gn=DOCK4;refGene_tx=XM_011280513.3;refGene_gn=DOCK4;refGene_tx=XM_019825718.2;refGene_gn=DOCK4;refGene_tx=XM_019825719.2;refGene_gn=DOCK4;refGene_tx=XM_019825720.2
A2	132499122	132499183
refGene_gn=DOCK4;refGene_tx=XM_011280513.3;refGene_gn=DOCK4;refGene_tx=XM_019825718.2;refGene_gn=DOCK4;refGene_tx=XM_019825719.2;refGene_gn=DOCK4;refGene_tx=XM_019825720.2
A2	132500254	132500336
refGene_gn=DOCK4;refGene_tx=XM_011280513.3;refGene_gn=DOCK4;refGene_tx=XM_019825718.2;refGene_gn=DOCK4;refGene_tx=XM_019825719.2;refGene_gn=DOCK4;refGene_tx=XM_019825720.2
A2	132519617	132519769
refGene_gn=DOCK4;refGene_tx=XM_011280513.3;refGene_gn=DOCK4;refGene_tx=XM_019825718.2;refGene_gn=DOCK4;refGene_tx=XM_019825719.2;refGene_gn=DOCK4;refGene_tx=XM_019825720.2
A2	132527029	132527114
refGene_gn=DOCK4;refGene_tx=XM_011280513.3;refGene_gn=DOCK4;refGene_tx=XM_019825718.2;refGene_gn=DOCK4;refGene_tx=XM_019825719.2;refGene_gn=DOCK4;refGene_tx=XM_019825720.2
A2	132535384	132535533
refGene_gn=DOCK4;refGene_tx=XM_011280513.3;refGene_gn=DOCK4;refGene_tx=XM_019825718.2;refGene_gn=DOCK4;refGene_tx=XM_019825719.2;refGene_gn=DOCK4;refGene_tx=XM_019825720.2
A2	132540265	132540362
refGene_gn=DOCK4;refGene_tx=XM_011280513.3;refGene_gn=DOCK4;refGene_tx=XM_019825718.2;refGene_gn=DOCK4;refGene_tx=XM_019825719.2;refGene_gn=DOCK4;refGene_tx=XM_019825720.2
A2	132545173	132545229

refGene_gn=DOCK4;refGene_tx=XM_011280513.3;refGene_gn=DOCK4;refGene_tx=XM_019825718.2;refGene_gn=DOCK4;refGene_tx=XM_019825719.2;refGene_gn=DOCK4;refGene_tx=XM_019825720.2
A2 132547265 132547306
refGene_gn=DOCK4;refGene_tx=XM_011280513.3;refGene_gn=DOCK4;refGene_tx=XM_019825718.2;refGene_gn=DOCK4;refGene_tx=XM_019825719.2;refGene_gn=DOCK4;refGene_tx=XM_019825720.2
A2 132549747 132549831
refGene_gn=DOCK4;refGene_tx=XM_011280513.3;refGene_gn=DOCK4;refGene_tx=XM_019825718.2;refGene_gn=DOCK4;refGene_tx=XM_019825719.2;refGene_gn=DOCK4;refGene_tx=XM_019825720.2
A2 132734100 132734137
refGene_gn=DOCK4;refGene_tx=XM_011280513.3;refGene_gn=DOCK4;refGene_tx=XM_019825718.2;refGene_gn=DOCK4;refGene_tx=XM_019825719.2;refGene_gn=DOCK4;refGene_tx=XM_019825720.2
A2 132734728 132734819
refGene_gn=ZNF277;refGene_tx=XM_006929317.4
A2 132800957 132801159
refGene_gn=ZNF277;refGene_tx=XM_006929317.4;refGene_gn=ZNF277;refGene_tx=XM_006929318.3;refGene_gn=ZNF277;refGene_tx=XM_006929319.3
A2 132809004 132809093
refGene_gn=ZNF277;refGene_tx=XM_006929317.4;refGene_gn=ZNF277;refGene_tx=XM_006929318.3;refGene_gn=ZNF277;refGene_tx=XM_006929319.3
A2 132809483 132809566
refGene_gn=ZNF277;refGene_tx=XM_006929317.4;refGene_gn=ZNF277;refGene_tx=XM_006929318.3;refGene_gn=ZNF277;refGene_tx=XM_006929319.3
A2 132828142 132828234
refGene_gn=ZNF277;refGene_tx=XM_006929317.4;refGene_gn=ZNF277;refGene_tx=XM_006929318.3;refGene_gn=ZNF277;refGene_tx=XM_006929319.3
A2 132831574 132831685
refGene_gn=ZNF277;refGene_tx=XM_006929317.4;refGene_gn=ZNF277;refGene_tx=XM_006929318.3;refGene_gn=ZNF277;refGene_tx=XM_006929319.3
A2 132833354 132833487
refGene_gn=ZNF277;refGene_tx=XM_006929317.4;refGene_gn=ZNF277;refGene_tx=XM_006929318.3;refGene_gn=ZNF277;refGene_tx=XM_006929319.3
A2 132840404 132840472
refGene_gn=ZNF277;refGene_tx=XM_006929317.4;refGene_gn=ZNF277;refGene_tx=XM_006929318.3;refGene_gn=ZNF277;refGene_tx=XM_006929319.3
A2 132841612 132841709
refGene_gn=ZNF277;refGene_tx=XM_006929317.4;refGene_gn=ZNF277;refGene_tx=XM_006929318.3;refGene_gn=ZNF277;refGene_tx=XM_006929319.3
A2 132843438 132843481
refGene_gn=ZNF277;refGene_tx=XM_006929317.4;refGene_gn=ZNF277;refGene_tx=XM_006929318.3;refGene_gn=ZNF277;refGene_tx=XM_006929319.3
A2 132844443 132844618
refGene_gn=ZNF277;refGene_tx=XM_006929317.4;refGene_gn=ZNF277;refGene_tx=XM_006929318.3;refGene_gn=ZNF277;refGene_tx=XM_006929319.3
A2 132846057 132846244
refGene_gn=ZNF277;refGene_tx=XM_006929317.4;refGene_gn=ZNF277;refGene_tx=XM_006929318.3;refGene_gn=ZNF277;refGene_tx=XM_006929319.3

A2 132943512 132943603
refGene_gn=IFRD1;refGene_tx=XM_011280514.3
A2 132947382 132947487
refGene_gn=IFRD1;refGene_tx=XM_011280514.3;refGene_gn=IFRD1;refGene_tx=XM_019825722.2
A2 132947640 132947725
refGene_gn=IFRD1;refGene_tx=XM_011280514.3;refGene_gn=IFRD1;refGene_tx=XM_019825722.2
A2 132948480 132948605
refGene_gn=IFRD1;refGene_tx=XM_011280514.3;refGene_gn=IFRD1;refGene_tx=XM_019825722.2
A2 132950198 132950356
refGene_gn=IFRD1;refGene_tx=XM_011280514.3;refGene_gn=IFRD1;refGene_tx=XM_019825722.2
A2 132953047 132953098
refGene_gn=IFRD1;refGene_tx=XM_011280514.3;refGene_gn=IFRD1;refGene_tx=XM_019825722.2
A2 132953179 132953358
refGene_gn=IFRD1;refGene_tx=XM_011280514.3;refGene_gn=IFRD1;refGene_tx=XM_019825722.2
A2 132953444 132953553
refGene_gn=IFRD1;refGene_tx=XM_011280514.3;refGene_gn=IFRD1;refGene_tx=XM_019825722.2
A2 132957862 132957997
refGene_gn=IFRD1;refGene_tx=XM_011280514.3;refGene_gn=IFRD1;refGene_tx=XM_019825722.2
A2 132961780 132961909
refGene_gn=IFRD1;refGene_tx=XM_011280514.3;refGene_gn=IFRD1;refGene_tx=XM_019825722.2
A2 132962303 132962399
refGene_gn=IFRD1;refGene_tx=XM_011280514.3;refGene_gn=IFRD1;refGene_tx=XM_019825722.2
A2 132964146 132964236
refGene_gn=IFRD1;refGene_tx=XM_011280514.3;refGene_gn=IFRD1;refGene_tx=XM_019825722.2
A2 132973070 132973194
refGene_gn=LSMEM1;refGene_tx=XM_003982967.3
A2 132975167 132975296
refGene_gn=LSMEM1;refGene_tx=XM_003982967.3
A2 132979217 132979348
refGene_gn=LSMEM1;refGene_tx=XM_003982967.3
A2 133231300 133231848
refGene_gn=TMEM168;refGene_tx=XM_011280515.3;refGene_gn=TMEM168;refGene_tx=XM_019825724.2;refGene_gn=TMEM168;refGene_tx=XM_019825725.2
A2 133237390 133237665
refGene_gn=TMEM168;refGene_tx=XM_011280515.3;refGene_gn=TMEM168;refGene_tx=XM_019825724.2;refGene_gn=TMEM168;refGene_tx=XM_019825725.2
A2 133240476 133240619
refGene_gn=TMEM168;refGene_tx=XM_011280515.3;refGene_gn=TMEM168;refGene_tx=XM_019825724.2;refGene_gn=TMEM168;refGene_tx=XM_019825725.2

A2	133254529	133255657
refGene_gn=TMEM168;refGene_tx=XM_011280515.3;refGene_gn=TMEM168;refGene_tx=XM_019825724.2;refGene_gn=TMEM168;refGene_tx=XM_019825725.2
A2	133289363	133289834
refGene_gn=BMT2;refGene_tx=XM_003982969.5
A2	133289835	133289994
refGene_gn=BMT2;refGene_tx=XM_003982969.5
A2	133301684	133301799
refGene_gn=BMT2;refGene_tx=XM_003982969.5
A2	133342815	133342986
refGene_gn=BMT2;refGene_tx=XM_003982969.5
A2	133357420	133357562
refGene_gn=BMT2;refGene_tx=XM_003982969.5
A2	133373578	133373738
refGene_gn=BMT2;refGene_tx=XM_003982969.5
A2	133506946	133508059
refGene_gn=GPR85;refGene_tx=XM_011280516.3;refGene_gn=GPR85;refGene_tx=XM_011280517.3;refGene_gn=GPR85;refGene_tx=XM_011280518.3;refGene_gn=GPR85;refGene_tx=XM_019825726.2;refGene_gn=GPR85;refGene_tx=XM_023250513.1
A2	133533914	133534094
refGene_gn=LOC102899533;refGene_tx=XM_023250514.1
A2	134157858	134160255
refGene_gn=PPP1R3A;refGene_tx=XM_003982971.4
A2	134162173	134162298
refGene_gn=PPP1R3A;refGene_tx=XM_003982971.4
A2	134162411	134162470
refGene_gn=PPP1R3A;refGene_tx=XM_003982971.4
A2	134194865	134195653
refGene_gn=PPP1R3A;refGene_tx=XM_003982971.4
A2	134592693	134592861
refGene_gn=FOXP2;refGene_tx=NM_001113177.1;refGene_gn=FOXP2;refGene_tx=XM_023240626.1
A2	134699401	134699491
refGene_gn=FOXP2;refGene_tx=NM_001113177.1;refGene_gn=FOXP2;refGene_tx=XM_023240626.1
A2	134731894	134731969
refGene_gn=FOXP2;refGene_tx=XM_023240626.1
A2	134787925	134788063
refGene_gn=FOXP2;refGene_tx=NM_001113177.1;refGene_gn=FOXP2;refGene_tx=XM_023240626.1
A2	134789191	134789392
refGene_gn=FOXP2;refGene_tx=NM_001113177.1;refGene_gn=FOXP2;refGene_tx=XM_023240626.1
A2	134790958	134791130
refGene_gn=FOXP2;refGene_tx=NM_001113177.1;refGene_gn=FOXP2;refGene_tx=XM_023240626.1
A2	134802697	134802911
refGene_gn=FOXP2;refGene_tx=NM_001113177.1;refGene_gn=FOXP2;refGene_tx=XM_023240626.1

A2 134804699 134804804
refGene_gn=FOXP2;refGene_tx=NM_001113177.1;refGene_gn=FOXP2;refGene_tx=XM_023240626.1
A2 134811981 134812069
refGene_gn=FOXP2;refGene_tx=NM_001113177.1;refGene_gn=FOXP2;refGene_tx=XM_023240626.1
A2 134813680 134813764
refGene_gn=FOXP2;refGene_tx=NM_001113177.1;refGene_gn=FOXP2;refGene_tx=XM_023240626.1
A2 134817992 134818194
refGene_gn=FOXP2;refGene_tx=NM_001113177.1;refGene_gn=FOXP2;refGene_tx=XM_023240626.1
A2 134819290 134819367
refGene_gn=FOXP2;refGene_tx=NM_001113177.1;refGene_gn=FOXP2;refGene_tx=XM_023240626.1
A2 134819506 134819608
refGene_gn=FOXP2;refGene_tx=NM_001113177.1;refGene_gn=FOXP2;refGene_tx=XM_023240626.1
A2 134822010 134822132
refGene_gn=FOXP2;refGene_tx=NM_001113177.1;refGene_gn=FOXP2;refGene_tx=XM_023240626.1
A2 134823395 134823465
refGene_gn=FOXP2;refGene_tx=NM_001113177.1;refGene_gn=FOXP2;refGene_tx=XM_023240626.1
A2 134824240 134824404
refGene_gn=FOXP2;refGene_tx=NM_001113177.1;refGene_gn=FOXP2;refGene_tx=XM_023240626.1
A2 134852435 134852580
refGene_gn=FOXP2;refGene_tx=NM_001113177.1;refGene_gn=FOXP2;refGene_tx=XM_023240626.1
A2 135054509 135054718
refGene_gn=LOC111557449;refGene_tx=XM_023242189.1
A2 135055032 135055519
refGene_gn=LOC111557449;refGene_tx=XM_023242189.1
A2 135055634 135055975
refGene_gn=LOC111557449;refGene_tx=XM_023242189.1;refGene_gn=MDFIC;refGene_tx=XM_023250515.1
A2 135074118 135074241
refGene_gn=MDFIC;refGene_tx=XM_023250515.1
A2 135108009 135108288
refGene_gn=MDFIC;refGene_tx=XM_023250515.1
A2 135139762 135140010
refGene_gn=MDFIC;refGene_tx=XM_023250515.1
A2 136016164 136016545
refGene_gn=TFEC;refGene_tx=XM_003982972.4;refGene_gn=TFEC;refGene_tx=XM_019825731.1;refGene_gn=TFEC;refGene_tx=XM_019825733.1;refGene_gn=TFEC;refGene_tx=XM_019825735.1
A2 136017428 136017576
refGene_gn=TFEC;refGene_tx=XM_003982972.4;refGene_gn=TFEC;refGene_tx=XM_019825731.1;refGene_gn=TFEC;refGene_tx=XM_019825733.1;refGene_gn=TFE

C;refGene_tx=XM_019825735.1
A2 136023988 136024064
refGene_gn=TFEC;refGene_tx=XM_003982972.4;refGene_gn=TFEC;refGene_tx=XM_019825731.1;refGene_gn=TFEC;refGene_tx=XM_019825733.1;refGene_gn=TFEC;refGene_tx=XM_019825735.1
A2 136027645 136027702
refGene_gn=TFEC;refGene_tx=XM_003982972.4;refGene_gn=TFEC;refGene_tx=XM_019825731.1;refGene_gn=TFEC;refGene_tx=XM_019825733.1;refGene_gn=TFEC;refGene_tx=XM_019825735.1
A2 136029869 136029984
refGene_gn=TFEC;refGene_tx=XM_003982972.4;refGene_gn=TFEC;refGene_tx=XM_019825731.1;refGene_gn=TFEC;refGene_tx=XM_019825733.1;refGene_gn=TFEC;refGene_tx=XM_019825735.1
A2 136048508 136048595
refGene_gn=TFEC;refGene_tx=XM_003982972.4;refGene_gn=TFEC;refGene_tx=XM_019825731.1;refGene_gn=TFEC;refGene_tx=XM_019825735.1
A2 136060405 136060654
refGene_gn=TFEC;refGene_tx=XM_003982972.4;refGene_gn=TFEC;refGene_tx=XM_019825731.1;refGene_gn=TFEC;refGene_tx=XM_019825733.1;refGene_gn=TFEC;refGene_tx=XM_019825735.1
A2 136146316 136146514
refGene_gn=TFEC;refGene_tx=XM_019825731.1;refGene_gn=TFEC;refGene_tx=XM_019825733.1
A2 136235439 136235466
refGene_gn=TES;refGene_tx=NM_001168698.1
A2 136260093 136260179
refGene_gn=TES;refGene_tx=NM_001168698.1;refGene_gn=TES;refGene_tx=XM_019814600.2
A2 136271847 136272100
refGene_gn=TES;refGene_tx=NM_001168698.1;refGene_gn=TES;refGene_tx=XM_019814600.2
A2 136272986 136273322
refGene_gn=TES;refGene_tx=NM_001168698.1;refGene_gn=TES;refGene_tx=XM_019814600.2
A2 136274582 136274798
refGene_gn=TES;refGene_tx=NM_001168698.1;refGene_gn=TES;refGene_tx=XM_019814600.2
A2 136275200 136275359
refGene_gn=TES;refGene_tx=NM_001168698.1;refGene_gn=TES;refGene_tx=XM_019814600.2
A2 136280832 136281021
refGene_gn=TES;refGene_tx=NM_001168698.1;refGene_gn=TES;refGene_tx=XM_019814600.2
A2 136488633 136488783
refGene_gn=CAV2;refGene_tx=NM_001168694.1;refGene_gn=CAV2;refGene_tx=XM_023240768.1
A2 136489101 136489289
refGene_gn=CAV2;refGene_tx=NM_001168694.1
A2 136493999 136494188
refGene_gn=CAV2;refGene_tx=NM_001168694.1;refGene_gn=CAV2;refGene_tx=X

M_023240768.1
A2	136519731	136519761
refGene_gn=CAV1;refGene_tx=NM_001024163.1
A2	136521194	136521359
refGene_gn=CAV1;refGene_tx=NM_001024163.1;refGene_gn=CAV1;refGene_tx=X
M_023240783.1
A2	136552271	136552613
refGene_gn=CAV1;refGene_tx=NM_001024163.1;refGene_gn=CAV1;refGene_tx=X
M_023240783.1
A2	136687562	136688111
refGene_gn=MET;refGene_tx=NM_001168696.1
A2	136688112	136688766
refGene_gn=MET;refGene_tx=NM_001168696.1
A2	136715335	136715527
refGene_gn=MET;refGene_tx=NM_001168696.1
A2	136726786	136726921
refGene_gn=MET;refGene_tx=NM_001168696.1
A2	136727942	136728116
refGene_gn=MET;refGene_tx=NM_001168696.1
A2	136733038	136733199
refGene_gn=MET;refGene_tx=NM_001168696.1
A2	136734980	136735083
refGene_gn=MET;refGene_tx=NM_001168696.1
A2	136735193	136735330
refGene_gn=MET;refGene_tx=NM_001168696.1
A2	136736022	136736184
refGene_gn=MET;refGene_tx=NM_001168696.1
A2	136737211	136737311
refGene_gn=MET;refGene_tx=NM_001168696.1
A2	136746834	136747053
refGene_gn=MET;refGene_tx=NM_001168696.1
A2	136752728	136752875
refGene_gn=MET;refGene_tx=NM_001168696.1
A2	136754687	136754844
refGene_gn=MET;refGene_tx=NM_001168696.1
A2	136755046	136755187
refGene_gn=MET;refGene_tx=NM_001168696.1
A2	136758369	136758600
refGene_gn=MET;refGene_tx=NM_001168696.1
A2	136760510	136760591
refGene_gn=MET;refGene_tx=NM_001168696.1
A2	136762187	136762369
refGene_gn=MET;refGene_tx=NM_001168696.1
A2	136767251	136767361
refGene_gn=MET;refGene_tx=NM_001168696.1
A2	136768534	136768700
refGene_gn=MET;refGene_tx=NM_001168696.1
A2	136778180	136778317
refGene_gn=MET;refGene_tx=NM_001168696.1
A2	136778425	136778636

refGene_gn=MET;refGene_tx=NM_001168696.1
A2 136837783 136837822
refGene_gn=CAPZA2;refGene_tx=NM_001168692.1
A2 136867496 136867560
refGene_gn=CAPZA2;refGene_tx=NM_001168692.1
A2 136872165 136872217
refGene_gn=CAPZA2;refGene_tx=NM_001168692.1
A2 136878958 136879022
refGene_gn=CAPZA2;refGene_tx=NM_001168692.1
A2 136883135 136883342
refGene_gn=CAPZA2;refGene_tx=NM_001168692.1
A2 136885421 136885501
refGene_gn=CAPZA2;refGene_tx=NM_001168692.1
A2 136889334 136889413
refGene_gn=CAPZA2;refGene_tx=NM_001168692.1
A2 136891476 136891548
refGene_gn=CAPZA2;refGene_tx=NM_001168692.1
A2 136896048 136896111
refGene_gn=CAPZA2;refGene_tx=NM_001168692.1
A2 136896965 136897106
refGene_gn=CAPZA2;refGene_tx=NM_001168692.1
A2 136928940 136929091
refGene_gn=ST7;refGene_tx=NM_001168697.1;refGene_gn=ST7;refGene_tx=XM_006929305.4;refGene_gn=ST7;refGene_tx=XM_006929306.4;refGene_gn=ST7;refGene_tx=XM_006929307.4
A2 136983799 136983821
refGene_gn=ST7;refGene_tx=XM_006929308.4
A2 137057218 137057301
refGene_gn=ST7;refGene_tx=NM_001168697.1;refGene_gn=ST7;refGene_tx=XM_006929305.4;refGene_gn=ST7;refGene_tx=XM_006929306.4;refGene_gn=ST7;refGene_tx=XM_006929307.4;refGene_gn=ST7;refGene_tx=XM_006929308.4
A2 137068859 137069019
refGene_gn=ST7;refGene_tx=NM_001168697.1;refGene_gn=ST7;refGene_tx=XM_006929305.4;refGene_gn=ST7;refGene_tx=XM_006929306.4;refGene_gn=ST7;refGene_tx=XM_006929307.4;refGene_gn=ST7;refGene_tx=XM_006929308.4
A2 137078885 137078940
refGene_gn=ST7;refGene_tx=NM_001168697.1;refGene_gn=ST7;refGene_tx=XM_006929305.4;refGene_gn=ST7;refGene_tx=XM_006929306.4;refGene_gn=ST7;refGene_tx=XM_006929307.4;refGene_gn=ST7;refGene_tx=XM_006929308.4
A2 137079575 137079691
refGene_gn=ST7;refGene_tx=NM_001168697.1;refGene_gn=ST7;refGene_tx=XM_006929305.4;refGene_gn=ST7;refGene_tx=XM_006929306.4;refGene_gn=ST7;refGene_tx=XM_006929307.4;refGene_gn=ST7;refGene_tx=XM_006929308.4
A2 137081478 137081554
refGene_gn=ST7;refGene_tx=NM_001168697.1;refGene_gn=ST7;refGene_tx=XM_006929305.4;refGene_gn=ST7;refGene_tx=XM_006929306.4;refGene_gn=ST7;refGene_tx=XM_006929307.4;refGene_gn=ST7;refGene_tx=XM_006929308.4
A2 137083714 137083783
refGene_gn=ST7;refGene_tx=XM_006929305.4;refGene_gn=ST7;refGene_tx=XM_006929306.4;refGene_gn=ST7;refGene_tx=XM_006929308.4

A2 137085694 137085849
refGene_gn=ST7;refGene_tx=NM_001168697.1;refGene_gn=ST7;refGene_tx=XM_006929305.4;refGene_gn=ST7;refGene_tx=XM_006929306.4;refGene_gn=ST7;refGene_tx=XM_006929307.4;refGene_gn=ST7;refGene_tx=XM_006929308.4
A2 137088253 137088351
refGene_gn=ST7;refGene_tx=NM_001168697.1;refGene_gn=ST7;refGene_tx=XM_006929305.4;refGene_gn=ST7;refGene_tx=XM_006929306.4;refGene_gn=ST7;refGene_tx=XM_006929307.4;refGene_gn=ST7;refGene_tx=XM_006929308.4
A2 137118038 137118153
refGene_gn=ST7;refGene_tx=NM_001168697.1;refGene_gn=ST7;refGene_tx=XM_006929305.4;refGene_gn=ST7;refGene_tx=XM_006929306.4;refGene_gn=ST7;refGene_tx=XM_006929307.4;refGene_gn=ST7;refGene_tx=XM_006929308.4
A2 137138654 137138727
refGene_gn=ST7;refGene_tx=NM_001168697.1;refGene_gn=ST7;refGene_tx=XM_006929305.4;refGene_gn=ST7;refGene_tx=XM_006929306.4;refGene_gn=ST7;refGene_tx=XM_006929307.4;refGene_gn=ST7;refGene_tx=XM_006929308.4
A2 137140180 137140283
refGene_gn=ST7;refGene_tx=NM_001168697.1;refGene_gn=ST7;refGene_tx=XM_006929305.4;refGene_gn=ST7;refGene_tx=XM_006929306.4;refGene_gn=ST7;refGene_tx=XM_006929307.4;refGene_gn=ST7;refGene_tx=XM_006929308.4
A2 137157777 137157928
refGene_gn=ST7;refGene_tx=NM_001168697.1;refGene_gn=ST7;refGene_tx=XM_006929305.4;refGene_gn=ST7;refGene_tx=XM_006929306.4;refGene_gn=ST7;refGene_tx=XM_006929307.4;refGene_gn=ST7;refGene_tx=XM_006929308.4
A2 137166991 137167084
refGene_gn=ST7;refGene_tx=NM_001168697.1;refGene_gn=ST7;refGene_tx=XM_006929305.4;refGene_gn=ST7;refGene_tx=XM_006929306.4;refGene_gn=ST7;refGene_tx=XM_006929307.4;refGene_gn=ST7;refGene_tx=XM_006929308.4
A2 137169929 137170069
refGene_gn=ST7;refGene_tx=NM_001168697.1;refGene_gn=ST7;refGene_tx=XM_006929305.4;refGene_gn=ST7;refGene_tx=XM_006929306.4;refGene_gn=ST7;refGene_tx=XM_006929307.4;refGene_gn=ST7;refGene_tx=XM_006929308.4
A2 137171156 137171276
refGene_gn=ST7;refGene_tx=XM_006929305.4;refGene_gn=ST7;refGene_tx=XM_006929307.4;refGene_gn=ST7;refGene_tx=XM_006929308.4
A2 137177791 137177887
refGene_gn=ST7;refGene_tx=NM_001168697.1;refGene_gn=ST7;refGene_tx=XM_006929306.4
A2 137219007 137219237
refGene_gn=WNT2;refGene_tx=NM_001168699.1
A2 137236934 137237199
refGene_gn=WNT2;refGene_tx=NM_001168699.1
A2 137254442 137254720
refGene_gn=WNT2;refGene_tx=NM_001168699.1
A2 137259890 137260117
refGene_gn=WNT2;refGene_tx=NM_001168699.1
A2 137262484 137262567
refGene_gn=WNT2;refGene_tx=NM_001168699.1
A2 137291104 137291257
refGene_gn=ASZ1;refGene_tx=NM_001025078.2

A2 137294489 137294603
refGene_gn=ASZ1;refGene_tx=NM_001025078.2
A2 137295445 137295551
refGene_gn=ASZ1;refGene_tx=NM_001025078.2
A2 137299688 137299798
refGene_gn=ASZ1;refGene_tx=NM_001025078.2
A2 137300776 137300833
refGene_gn=ASZ1;refGene_tx=NM_001025078.2
A2 137302084 137302160
refGene_gn=ASZ1;refGene_tx=NM_001025078.2
A2 137302908 137303033
refGene_gn=ASZ1;refGene_tx=NM_001025078.2
A2 137304175 137304310
refGene_gn=ASZ1;refGene_tx=NM_001025078.2
A2 137304838 137304950
refGene_gn=ASZ1;refGene_tx=NM_001025078.2
A2 137335439 137335551
refGene_gn=ASZ1;refGene_tx=NM_001025078.2
A2 137338480 137338603
refGene_gn=ASZ1;refGene_tx=NM_001025078.2
A2 137342574 137342674
refGene_gn=ASZ1;refGene_tx=NM_001025078.2
A2 137343074 137343179
refGene_gn=ASZ1;refGene_tx=NM_001025078.2
A2 137407848 137407901
refGene_gn=CFTR;refGene_tx=NM_001048009.1
A2 137436744 137436855
refGene_gn=CFTR;refGene_tx=NM_001048009.1
A2 137441275 137441384
refGene_gn=CFTR;refGene_tx=NM_001048009.1
A2 137458969 137459185
refGene_gn=CFTR;refGene_tx=NM_001048009.1
A2 137462372 137462462
refGene_gn=CFTR;refGene_tx=NM_001048009.1
A2 137463313 137463477
refGene_gn=CFTR;refGene_tx=NM_001048009.1
A2 137464320 137464446
refGene_gn=CFTR;refGene_tx=NM_001048009.1
A2 137469123 137469370
refGene_gn=CFTR;refGene_tx=NM_001048009.1
A2 137471701 137471794
refGene_gn=CFTR;refGene_tx=NM_001048009.1
A2 137478791 137478974
refGene_gn=CFTR;refGene_tx=NM_001048009.1
A2 137482573 137482765
refGene_gn=CFTR;refGene_tx=NM_001048009.1
A2 137505216 137505311
refGene_gn=CFTR;refGene_tx=NM_001048009.1
A2 137508295 137508382
refGene_gn=CFTR;refGene_tx=NM_001048009.1

A2 137509872 137510593
refGene_gn=CFTR;refGene_tx=NM_001048009.1
A2 137513308 137513437
refGene_gn=CFTR;refGene_tx=NM_001048009.1
A2 137526153 137526191
refGene_gn=CFTR;refGene_tx=NM_001048009.1
A2 137526658 137526909
refGene_gn=CFTR;refGene_tx=NM_001048009.1
A2 137531002 137531082
refGene_gn=CFTR;refGene_tx=NM_001048009.1
A2 137536265 137536416
refGene_gn=CFTR;refGene_tx=NM_001048009.1
A2 137537805 137538033
refGene_gn=CFTR;refGene_tx=NM_001048009.1
A2 137540652 137540753
refGene_gn=CFTR;refGene_tx=NM_001048009.1
A2 137554709 137554964
refGene_gn=CFTR;refGene_tx=NM_001048009.1
A2 137570741 137570897
refGene_gn=CFTR;refGene_tx=NM_001048009.1
A2 137582680 137582770
refGene_gn=CFTR;refGene_tx=NM_001048009.1
A2 137592373 137592546
refGene_gn=CFTR;refGene_tx=NM_001048009.1
A2 137593151 137593257
refGene_gn=CFTR;refGene_tx=NM_001048009.1
A2 137594721 137594922
refGene_gn=CFTR;refGene_tx=NM_001048009.1
A2 137672187 137672433
refGene_gn=CTTNBP2;refGene_tx=NM_001168695.1;refGene_gn=CTTNBP2;refGene_tx=XM_023240808.1;refGene_gn=CTTNBP2;refGene_tx=XM_023240812.1;refGene_gn=CTTNBP2;refGene_tx=XM_023240816.1;refGene_gn=CTTNBP2;refGene_tx=XM_023240817.1
A2 137676387 137676483
refGene_gn=CTTNBP2;refGene_tx=NM_001168695.1;refGene_gn=CTTNBP2;refGene_tx=XM_023240808.1;refGene_gn=CTTNBP2;refGene_tx=XM_023240812.1;refGene_gn=CTTNBP2;refGene_tx=XM_023240816.1;refGene_gn=CTTNBP2;refGene_tx=XM_023240817.1
A2 137677755 137677888
refGene_gn=CTTNBP2;refGene_tx=NM_001168695.1;refGene_gn=CTTNBP2;refGene_tx=XM_023240808.1;refGene_gn=CTTNBP2;refGene_tx=XM_023240812.1;refGene_gn=CTTNBP2;refGene_tx=XM_023240816.1;refGene_gn=CTTNBP2;refGene_tx=XM_023240817.1
A2 137678617 137678636
refGene_gn=CTTNBP2;refGene_tx=XM_023240818.1;refGene_gn=CTTNBP2;refGene_tx=XM_023240820.1;refGene_gn=CTTNBP2;refGene_tx=XM_023240824.1
A2 137679440 137679504
refGene_gn=CTTNBP2;refGene_tx=NM_001168695.1;refGene_gn=CTTNBP2;refGene_tx=XM_023240808.1;refGene_gn=CTTNBP2;refGene_tx=XM_023240812.1;refGene_gn=CTTNBP2;refGene_tx=XM_023240816.1;refGene_gn=CTTNBP2;refGene_tx=

XM_023240817.1;refGene_gn=CTTNBP2;refGene_tx=XM_023240818.1;refGene_gn=CTTNBP2;refGene_tx=XM_023240820.1;refGene_gn=CTTNBP2;refGene_tx=XM_023240824.1
A2 137681878 137682064
refGene_gn=CTTNBP2;refGene_tx=NM_001168695.1;refGene_gn=CTTNBP2;refGene_tx=XM_023240808.1;refGene_gn=CTTNBP2;refGene_tx=XM_023240812.1;refGene_gn=CTTNBP2;refGene_tx=XM_023240816.1;refGene_gn=CTTNBP2;refGene_tx=XM_023240817.1;refGene_gn=CTTNBP2;refGene_tx=XM_023240818.1;refGene_gn=CTTNBP2;refGene_tx=XM_023240820.1;refGene_gn=CTTNBP2;refGene_tx=XM_023240824.1
A2 137682646 137682852
refGene_gn=CTTNBP2;refGene_tx=NM_001168695.1;refGene_gn=CTTNBP2;refGene_tx=XM_023240808.1;refGene_gn=CTTNBP2;refGene_tx=XM_023240812.1;refGene_gn=CTTNBP2;refGene_tx=XM_023240816.1;refGene_gn=CTTNBP2;refGene_tx=XM_023240817.1;refGene_gn=CTTNBP2;refGene_tx=XM_023240818.1;refGene_gn=CTTNBP2;refGene_tx=XM_023240820.1;refGene_gn=CTTNBP2;refGene_tx=XM_023240824.1
A2 137685587 137685766
refGene_gn=CTTNBP2;refGene_tx=NM_001168695.1;refGene_gn=CTTNBP2;refGene_tx=XM_023240808.1;refGene_gn=CTTNBP2;refGene_tx=XM_023240812.1;refGene_gn=CTTNBP2;refGene_tx=XM_023240816.1;refGene_gn=CTTNBP2;refGene_tx=XM_023240817.1;refGene_gn=CTTNBP2;refGene_tx=XM_023240818.1;refGene_gn=CTTNBP2;refGene_tx=XM_023240820.1;refGene_gn=CTTNBP2;refGene_tx=XM_023240824.1
A2 137689015 137689203
refGene_gn=CTTNBP2;refGene_tx=NM_001168695.1;refGene_gn=CTTNBP2;refGene_tx=XM_023240808.1;refGene_gn=CTTNBP2;refGene_tx=XM_023240812.1;refGene_gn=CTTNBP2;refGene_tx=XM_023240816.1;refGene_gn=CTTNBP2;refGene_tx=XM_023240817.1;refGene_gn=CTTNBP2;refGene_tx=XM_023240818.1;refGene_gn=CTTNBP2;refGene_tx=XM_023240820.1;refGene_gn=CTTNBP2;refGene_tx=XM_023240824.1
A2 137689378 137689531
refGene_gn=CTTNBP2;refGene_tx=NM_001168695.1;refGene_gn=CTTNBP2;refGene_tx=XM_023240808.1;refGene_gn=CTTNBP2;refGene_tx=XM_023240812.1;refGene_gn=CTTNBP2;refGene_tx=XM_023240816.1;refGene_gn=CTTNBP2;refGene_tx=XM_023240817.1;refGene_gn=CTTNBP2;refGene_tx=XM_023240818.1;refGene_gn=CTTNBP2;refGene_tx=XM_023240820.1;refGene_gn=CTTNBP2;refGene_tx=XM_023240824.1
A2 137699463 137699563
refGene_gn=CTTNBP2;refGene_tx=NM_001168695.1;refGene_gn=CTTNBP2;refGene_tx=XM_023240808.1;refGene_gn=CTTNBP2;refGene_tx=XM_023240812.1;refGene_gn=CTTNBP2;refGene_tx=XM_023240816.1;refGene_gn=CTTNBP2;refGene_tx=XM_023240817.1;refGene_gn=CTTNBP2;refGene_tx=XM_023240818.1;refGene_gn=CTTNBP2;refGene_tx=XM_023240820.1;refGene_gn=CTTNBP2;refGene_tx=XM_023240824.1
A2 137699637 137699724
refGene_gn=CTTNBP2;refGene_tx=NM_001168695.1;refGene_gn=CTTNBP2;refGene_tx=XM_023240808.1;refGene_gn=CTTNBP2;refGene_tx=XM_023240812.1;refGene_gn=CTTNBP2;refGene_tx=XM_023240816.1;refGene_gn=CTTNBP2;refGene_tx=XM_023240817.1;refGene_gn=CTTNBP2;refGene_tx=XM_023240818.1;refGene_gn

=CTTNBP2;refGene_tx=XM_023240820.1;refGene_gn=CTTNBP2;refGene_tx=XM_02
3240824.1
A2 137708630 137708710
refGene_gn=CTTNBP2;refGene_tx=NM_001168695.1;refGene_gn=CTTNBP2;refGen
e_tx=XM_023240808.1;refGene_gn=CTTNBP2;refGene_tx=XM_023240812.1;refGe
ne_gn=CTTNBP2;refGene_tx=XM_023240816.1;refGene_gn=CTTNBP2;refGene_tx=
XM_023240817.1;refGene_gn=CTTNBP2;refGene_tx=XM_023240818.1;refGene_gn
=CTTNBP2;refGene_tx=XM_023240820.1;refGene_gn=CTTNBP2;refGene_tx=XM_02
3240824.1
A2 137709962 137710058
refGene_gn=CTTNBP2;refGene_tx=NM_001168695.1;refGene_gn=CTTNBP2;refGen
e_tx=XM_023240808.1;refGene_gn=CTTNBP2;refGene_tx=XM_023240812.1;refGe
ne_gn=CTTNBP2;refGene_tx=XM_023240816.1;refGene_gn=CTTNBP2;refGene_tx=
XM_023240817.1;refGene_gn=CTTNBP2;refGene_tx=XM_023240818.1;refGene_gn
=CTTNBP2;refGene_tx=XM_023240820.1;refGene_gn=CTTNBP2;refGene_tx=XM_02
3240824.1
A2 137712203 137712479
refGene_gn=CTTNBP2;refGene_tx=NM_001168695.1;refGene_gn=CTTNBP2;refGen
e_tx=XM_023240808.1;refGene_gn=CTTNBP2;refGene_tx=XM_023240812.1;refGe
ne_gn=CTTNBP2;refGene_tx=XM_023240816.1;refGene_gn=CTTNBP2;refGene_tx=
XM_023240817.1;refGene_gn=CTTNBP2;refGene_tx=XM_023240818.1;refGene_gn
=CTTNBP2;refGene_tx=XM_023240820.1;refGene_gn=CTTNBP2;refGene_tx=XM_02
3240824.1
A2 137716897 137717015
refGene_gn=CTTNBP2;refGene_tx=NM_001168695.1;refGene_gn=CTTNBP2;refGen
e_tx=XM_023240808.1;refGene_gn=CTTNBP2;refGene_tx=XM_023240812.1;refGe
ne_gn=CTTNBP2;refGene_tx=XM_023240816.1;refGene_gn=CTTNBP2;refGene_tx=
XM_023240817.1;refGene_gn=CTTNBP2;refGene_tx=XM_023240818.1;refGene_gn
=CTTNBP2;refGene_tx=XM_023240820.1;refGene_gn=CTTNBP2;refGene_tx=XM_02
3240824.1
A2 137727036 137727291
refGene_gn=CTTNBP2;refGene_tx=NM_001168695.1;refGene_gn=CTTNBP2;refGen
e_tx=XM_023240808.1;refGene_gn=CTTNBP2;refGene_tx=XM_023240812.1;refGe
ne_gn=CTTNBP2;refGene_tx=XM_023240816.1;refGene_gn=CTTNBP2;refGene_tx=
XM_023240817.1;refGene_gn=CTTNBP2;refGene_tx=XM_023240818.1;refGene_gn
=CTTNBP2;refGene_tx=XM_023240820.1;refGene_gn=CTTNBP2;refGene_tx=XM_02
3240824.1
A2 137731607 137731758
refGene_gn=CTTNBP2;refGene_tx=NM_001168695.1;refGene_gn=CTTNBP2;refGen
e_tx=XM_023240808.1;refGene_gn=CTTNBP2;refGene_tx=XM_023240812.1;refGe
ne_gn=CTTNBP2;refGene_tx=XM_023240816.1;refGene_gn=CTTNBP2;refGene_tx=
XM_023240817.1;refGene_gn=CTTNBP2;refGene_tx=XM_023240818.1;refGene_gn
=CTTNBP2;refGene_tx=XM_023240820.1;refGene_gn=CTTNBP2;refGene_tx=XM_02
3240824.1
A2 137734028 137734128
refGene_gn=CTTNBP2;refGene_tx=NM_001168695.1;refGene_gn=CTTNBP2;refGen
e_tx=XM_023240808.1;refGene_gn=CTTNBP2;refGene_tx=XM_023240812.1;refGe
ne_gn=CTTNBP2;refGene_tx=XM_023240816.1;refGene_gn=CTTNBP2;refGene_tx=
XM_023240817.1;refGene_gn=CTTNBP2;refGene_tx=XM_023240818.1;refGene_gn
=CTTNBP2;refGene_tx=XM_023240820.1;refGene_gn=CTTNBP2;refGene_tx=XM_02

3240824.1
A2	137735485	137735689
refGene_gn=CTTNBP2;refGene_tx=NM_001168695.1;refGene_gn=CTTNBP2;refGene_tx=XM_023240808.1;refGene_gn=CTTNBP2;refGene_tx=XM_023240812.1;refGene_gn=CTTNBP2;refGene_tx=XM_023240816.1;refGene_gn=CTTNBP2;refGene_tx=XM_023240817.1;refGene_gn=CTTNBP2;refGene_tx=XM_023240818.1;refGene_gn=CTTNBP2;refGene_tx=XM_023240820.1;refGene_gn=CTTNBP2;refGene_tx=XM_023240824.1
A2	137737251	137737291
refGene_gn=CTTNBP2;refGene_tx=XM_023240812.1
A2	137741678	137743323
refGene_gn=CTTNBP2;refGene_tx=NM_001168695.1;refGene_gn=CTTNBP2;refGene_tx=XM_023240808.1;refGene_gn=CTTNBP2;refGene_tx=XM_023240812.1;refGene_gn=CTTNBP2;refGene_tx=XM_023240816.1;refGene_gn=CTTNBP2;refGene_tx=XM_023240817.1;refGene_gn=CTTNBP2;refGene_tx=XM_023240818.1;refGene_gn=CTTNBP2;refGene_tx=XM_023240820.1;refGene_gn=CTTNBP2;refGene_tx=XM_023240824.1
A2	137758340	137758565
refGene_gn=CTTNBP2;refGene_tx=NM_001168695.1;refGene_gn=CTTNBP2;refGene_tx=XM_023240808.1;refGene_gn=CTTNBP2;refGene_tx=XM_023240812.1;refGene_gn=CTTNBP2;refGene_tx=XM_023240816.1;refGene_gn=CTTNBP2;refGene_tx=XM_023240817.1;refGene_gn=CTTNBP2;refGene_tx=XM_023240818.1;refGene_gn=CTTNBP2;refGene_tx=XM_023240820.1;refGene_gn=CTTNBP2;refGene_tx=XM_023240824.1
A2	137810445	137810553
refGene_gn=CTTNBP2;refGene_tx=NM_001168695.1;refGene_gn=CTTNBP2;refGene_tx=XM_023240808.1;refGene_gn=CTTNBP2;refGene_tx=XM_023240812.1;refGene_gn=CTTNBP2;refGene_tx=XM_023240816.1;refGene_gn=CTTNBP2;refGene_tx=XM_023240818.1;refGene_gn=CTTNBP2;refGene_tx=XM_023240820.1;refGene_gn=CTTNBP2;refGene_tx=XM_023240824.1
A2	137811425	137811443
refGene_gn=CTTNBP2;refGene_tx=XM_023240808.1
A2	137822249	137822330
refGene_gn=CTTNBP2;refGene_tx=NM_001168695.1;refGene_gn=CTTNBP2;refGene_tx=XM_023240812.1;refGene_gn=CTTNBP2;refGene_tx=XM_023240818.1;refGene_gn=CTTNBP2;refGene_tx=XM_023240820.1;refGene_gn=CTTNBP2;refGene_tx=XM_023240824.1
A2	138086860	138086891
refGene_gn=LSM8;refGene_tx=XM_003982973.3
A2	138088037	138088078
refGene_gn=LSM8;refGene_tx=XM_003982973.3
A2	138091306	138091434
refGene_gn=LSM8;refGene_tx=XM_003982973.3
A2	138094153	138094244
refGene_gn=LSM8;refGene_tx=XM_003982973.3
A2	138116258	138116266
refGene_gn=ANKRD7;refGene_tx=XM_003982977.5
A2	138117698	138117808
refGene_gn=ANKRD7;refGene_tx=XM_003982977.5
A2	138126697	138126835

```
refGene_gn=ANKRD7;refGene_tx=XM_003982977.5
A2      138127500       138127607
refGene_gn=ANKRD7;refGene_tx=XM_003982977.5
A2      138129296       138129470
refGene_gn=ANKRD7;refGene_tx=XM_003982977.5
A2      138129623       138129738
refGene_gn=ANKRD7;refGene_tx=XM_003982977.5
A2      138132298       138132528
refGene_gn=ANKRD7;refGene_tx=XM_003982977.5
A2      139290559       139290951
refGene_gn=LOC101086129;refGene_tx=XM_023250516.1
A2      139290953       139291134
refGene_gn=LOC101086129;refGene_tx=XM_023250516.1
A2      139669394       139670509
refGene_gn=KCND2;refGene_tx=XM_011280521.2
A2      140142730       140142893
refGene_gn=KCND2;refGene_tx=XM_011280521.2
A2      140152074       140152170
refGene_gn=KCND2;refGene_tx=XM_011280521.2
A2      140153032       140153125
refGene_gn=KCND2;refGene_tx=XM_011280521.2
A2      140156629       140156877
refGene_gn=KCND2;refGene_tx=XM_011280521.2
A2      140158655       140158833
refGene_gn=KCND2;refGene_tx=XM_011280521.2
A2      140190420       140190726
refGene_gn=TSPAN12;refGene_tx=XM_003982978.3;refGene_gn=TSPAN12;refGen
e_tx=XM_023250517.1
A2      140211863       140212007
refGene_gn=TSPAN12;refGene_tx=XM_003982978.3;refGene_gn=TSPAN12;refGen
e_tx=XM_023250517.1
A2      140216831       140216939
refGene_gn=TSPAN12;refGene_tx=XM_003982978.3;refGene_gn=TSPAN12;refGen
e_tx=XM_023250517.1
A2      140220646       140220721
refGene_gn=TSPAN12;refGene_tx=XM_003982978.3;refGene_gn=TSPAN12;refGen
e_tx=XM_023250517.1
A2      140241506       140241642
refGene_gn=TSPAN12;refGene_tx=XM_003982978.3;refGene_gn=TSPAN12;refGen
e_tx=XM_023250517.1
A2      140242855       140242938
refGene_gn=TSPAN12;refGene_tx=XM_003982978.3;refGene_gn=TSPAN12;refGen
e_tx=XM_023250517.1
A2      140255145       140255211
refGene_gn=TSPAN12;refGene_tx=XM_003982978.3;refGene_gn=TSPAN12;refGen
e_tx=XM_023250517.1
A2      140323641       140323669
refGene_gn=ING3;refGene_tx=XM_003982979.5;refGene_gn=ING3;refGene_tx=X
M_006929324.4
A2      140323915       140323987
```

refGene_gn=ING3;refGene_tx=XM_003982979.5;refGene_gn=ING3;refGene_tx=XM_006929324.4
A2 140326669 140326770
refGene_gn=ING3;refGene_tx=XM_003982979.5;refGene_gn=ING3;refGene_tx=XM_006929324.4
A2 140329053 140329119
refGene_gn=ING3;refGene_tx=XM_003982979.5;refGene_gn=ING3;refGene_tx=XM_006929324.4
A2 140339442 140339539
refGene_gn=ING3;refGene_tx=XM_003982979.5;refGene_gn=ING3;refGene_tx=XM_006929324.4
A2 140341202 140341274
refGene_gn=ING3;refGene_tx=XM_003982979.5;refGene_gn=ING3;refGene_tx=XM_006929324.4
A2 140342164 140342284
refGene_gn=ING3;refGene_tx=XM_003982979.5;refGene_gn=ING3;refGene_tx=XM_006929324.4
A2 140342557 140342715
refGene_gn=ING3;refGene_tx=XM_003982979.5;refGene_gn=ING3;refGene_tx=XM_006929324.4
A2 140343293 140343487
refGene_gn=ING3;refGene_tx=XM_003982979.5;refGene_gn=ING3;refGene_tx=XM_006929324.4
A2 140345260 140345453
refGene_gn=ING3;refGene_tx=XM_003982979.5;refGene_gn=ING3;refGene_tx=XM_006929324.4
A2 140347979 140348018
refGene_gn=ING3;refGene_tx=XM_003982979.5;refGene_gn=ING3;refGene_tx=XM_006929324.4
A2 140349558 140349675
refGene_gn=ING3;refGene_tx=XM_003982979.5;refGene_gn=ING3;refGene_tx=XM_006929324.4
A2 140359719 140359968
refGene_gn=CPED1;refGene_tx=XM_003982981.5;refGene_gn=CPED1;refGene_tx=XM_006929325.4;refGene_gn=CPED1;refGene_tx=XM_006929326.4;refGene_gn=CPED1;refGene_tx=XM_019825739.2
A2 140386689 140386873
refGene_gn=CPED1;refGene_tx=XM_003982981.5;refGene_gn=CPED1;refGene_tx=XM_006929325.4;refGene_gn=CPED1;refGene_tx=XM_006929326.4;refGene_gn=CPED1;refGene_tx=XM_019825739.2
A2 140417373 140417480
refGene_gn=CPED1;refGene_tx=XM_003982981.5;refGene_gn=CPED1;refGene_tx=XM_006929325.4;refGene_gn=CPED1;refGene_tx=XM_006929326.4;refGene_gn=CPED1;refGene_tx=XM_019825740.2
A2 140423579 140423655
refGene_gn=CPED1;refGene_tx=XM_003982981.5;refGene_gn=CPED1;refGene_tx=XM_006929325.4;refGene_gn=CPED1;refGene_tx=XM_006929326.4;refGene_gn=CPED1;refGene_tx=XM_019825740.2
A2 140455833 140455957
refGene_gn=CPED1;refGene_tx=XM_003982981.5;refGene_gn=CPED1;refGene_tx

=XM_006929325.4;refGene_gn=CPED1;refGene_tx=XM_006929326.4;refGene_gn=CPED1;refGene_tx=XM_019825739.2;refGene_gn=CPED1;refGene_tx=XM_019825740.2
A2 140457515 140457684
refGene_gn=CPED1;refGene_tx=XM_003982981.5;refGene_gn=CPED1;refGene_tx=XM_006929325.4;refGene_gn=CPED1;refGene_tx=XM_006929326.4;refGene_gn=CPED1;refGene_tx=XM_019825739.2;refGene_gn=CPED1;refGene_tx=XM_019825740.2
A2 140486364 140486507
refGene_gn=CPED1;refGene_tx=XM_003982981.5;refGene_gn=CPED1;refGene_tx=XM_006929325.4;refGene_gn=CPED1;refGene_tx=XM_006929326.4;refGene_gn=CPED1;refGene_tx=XM_019825739.2;refGene_gn=CPED1;refGene_tx=XM_019825740.2
A2 140487289 140487362
refGene_gn=CPED1;refGene_tx=XM_003982981.5;refGene_gn=CPED1;refGene_tx=XM_006929325.4;refGene_gn=CPED1;refGene_tx=XM_006929326.4;refGene_gn=CPED1;refGene_tx=XM_019825739.2;refGene_gn=CPED1;refGene_tx=XM_019825740.2
A2 140488697 140488865
refGene_gn=CPED1;refGene_tx=XM_003982981.5;refGene_gn=CPED1;refGene_tx=XM_006929325.4;refGene_gn=CPED1;refGene_tx=XM_006929326.4;refGene_gn=CPED1;refGene_tx=XM_019825739.2;refGene_gn=CPED1;refGene_tx=XM_019825740.2
A2 140490370 140490475
refGene_gn=CPED1;refGene_tx=XM_003982981.5;refGene_gn=CPED1;refGene_tx=XM_006929325.4;refGene_gn=CPED1;refGene_tx=XM_019825739.2;refGene_gn=CPED1;refGene_tx=XM_019825740.2
A2 140492121 140492291
refGene_gn=CPED1;refGene_tx=XM_003982981.5;refGene_gn=CPED1;refGene_tx=XM_006929325.4;refGene_gn=CPED1;refGene_tx=XM_006929326.4;refGene_gn=CPED1;refGene_tx=XM_019825739.2;refGene_gn=CPED1;refGene_tx=XM_019825740.2
A2 140496464 140496535
refGene_gn=CPED1;refGene_tx=XM_003982981.5;refGene_gn=CPED1;refGene_tx=XM_006929325.4;refGene_gn=CPED1;refGene_tx=XM_006929326.4;refGene_gn=CPED1;refGene_tx=XM_019825739.2;refGene_gn=CPED1;refGene_tx=XM_019825740.2
A2 140498259 140498310
refGene_gn=CPED1;refGene_tx=XM_003982981.5;refGene_gn=CPED1;refGene_tx=XM_006929325.4;refGene_gn=CPED1;refGene_tx=XM_006929326.4;refGene_gn=CPED1;refGene_tx=XM_019825739.2;refGene_gn=CPED1;refGene_tx=XM_019825740.2
A2 140502964 140503148
refGene_gn=CPED1;refGene_tx=XM_003982981.5;refGene_gn=CPED1;refGene_tx=XM_006929325.4;refGene_gn=CPED1;refGene_tx=XM_006929326.4;refGene_gn=CPED1;refGene_tx=XM_019825739.2;refGene_gn=CPED1;refGene_tx=XM_019825740.2
A2 140504065 140504234
refGene_gn=CPED1;refGene_tx=XM_003982981.5;refGene_gn=CPED1;refGene_tx=XM_006929325.4;refGene_gn=CPED1;refGene_tx=XM_006929326.4;refGene_gn=

CPED1;refGene_tx=XM_019825739.2;refGene_gn=CPED1;refGene_tx=XM_0198257
40.2
A2 140513047 140513095
refGene_gn=CPED1;refGene_tx=XM_006929325.4;refGene_gn=CPED1;refGene_tx
=XM_006929326.4;refGene_gn=CPED1;refGene_tx=XM_019825739.2;refGene_gn=
CPED1;refGene_tx=XM_019825740.2
A2 140585953 140586071
refGene_gn=CPED1;refGene_tx=XM_003982981.5;refGene_gn=CPED1;refGene_tx
=XM_006929325.4;refGene_gn=CPED1;refGene_tx=XM_006929326.4;refGene_gn=
CPED1;refGene_tx=XM_019825739.2;refGene_gn=CPED1;refGene_tx=XM_0198257
40.2
A2 140592579 140592716
refGene_gn=CPED1;refGene_tx=XM_003982981.5;refGene_gn=CPED1;refGene_tx
=XM_006929325.4;refGene_gn=CPED1;refGene_tx=XM_006929326.4;refGene_gn=
CPED1;refGene_tx=XM_019825739.2;refGene_gn=CPED1;refGene_tx=XM_0198257
40.2
A2 140607272 140607493
refGene_gn=CPED1;refGene_tx=XM_003982981.5;refGene_gn=CPED1;refGene_tx
=XM_006929325.4;refGene_gn=CPED1;refGene_tx=XM_006929326.4;refGene_gn=
CPED1;refGene_tx=XM_019825739.2;refGene_gn=CPED1;refGene_tx=XM_0198257
40.2
A2 140607749 140607851
refGene_gn=CPED1;refGene_tx=XM_003982981.5;refGene_gn=CPED1;refGene_tx
=XM_006929325.4;refGene_gn=CPED1;refGene_tx=XM_006929326.4;refGene_gn=
CPED1;refGene_tx=XM_019825739.2;refGene_gn=CPED1;refGene_tx=XM_0198257
40.2
A2 140608255 140608343
refGene_gn=CPED1;refGene_tx=XM_003982981.5;refGene_gn=CPED1;refGene_tx
=XM_006929325.4;refGene_gn=CPED1;refGene_tx=XM_006929326.4;refGene_gn=
CPED1;refGene_tx=XM_019825739.2;refGene_gn=CPED1;refGene_tx=XM_0198257
40.2
A2 140612385 140612532
refGene_gn=CPED1;refGene_tx=XM_003982981.5;refGene_gn=CPED1;refGene_tx
=XM_006929325.4;refGene_gn=CPED1;refGene_tx=XM_006929326.4;refGene_gn=
CPED1;refGene_tx=XM_019825739.2;refGene_gn=CPED1;refGene_tx=XM_0198257
40.2
A2 140628812 140629025
refGene_gn=CPED1;refGene_tx=XM_003982981.5;refGene_gn=CPED1;refGene_tx
=XM_006929325.4;refGene_gn=CPED1;refGene_tx=XM_006929326.4;refGene_gn=
CPED1;refGene_tx=XM_019825739.2;refGene_gn=CPED1;refGene_tx=XM_0198257
40.2
A2 140652557 140652991
refGene_gn=WNT16;refGene_tx=XM_003982982.4
A2 140653174 140653416
refGene_gn=WNT16;refGene_tx=XM_003982982.4
A2 140655389 140655676
refGene_gn=WNT16;refGene_tx=XM_003982982.4
A2 140662418 140662883
refGene_gn=WNT16;refGene_tx=XM_003982982.4
A2 140675391 140675481

refGene_gn=FAM3C;refGene_tx=XM_003982983.5;refGene_gn=FAM3C;refGene_tx=XM_023250519.1
A2 140676074 140676201
refGene_gn=FAM3C;refGene_tx=XM_003982983.5;refGene_gn=FAM3C;refGene_tx=XM_023250519.1
A2 140687083 140687168
refGene_gn=FAM3C;refGene_tx=XM_003982983.5;refGene_gn=FAM3C;refGene_tx=XM_023250519.1
A2 140689137 140689188
refGene_gn=FAM3C;refGene_tx=XM_003982983.5;refGene_gn=FAM3C;refGene_tx=XM_023250519.1
A2 140690322 140690381
refGene_gn=FAM3C;refGene_tx=XM_003982983.5;refGene_gn=FAM3C;refGene_tx=XM_023250519.1
A2 140703149 140703273
refGene_gn=FAM3C;refGene_tx=XM_003982983.5;refGene_gn=FAM3C;refGene_tx=XM_023250519.1
A2 140703918 140703948
refGene_gn=FAM3C;refGene_tx=XM_003982983.5;refGene_gn=FAM3C;refGene_tx=XM_023250519.1
A2 140712669 140712774
refGene_gn=FAM3C;refGene_tx=XM_003982983.5;refGene_gn=FAM3C;refGene_tx=XM_023250519.1
A2 140716768 140716781
refGene_gn=FAM3C;refGene_tx=XM_003982983.5;refGene_gn=FAM3C;refGene_tx=XM_023250519.1
A2 141082012 141082070
refGene_gn=PTPRZ1;refGene_tx=XM_023250521.1;refGene_gn=PTPRZ1;refGene_tx=XM_023250522.1;refGene_gn=PTPRZ1;refGene_tx=XM_023250523.1;refGene_gn=PTPRZ1;refGene_tx=XM_023250524.1
A2 141137287 141137353
refGene_gn=PTPRZ1;refGene_tx=XM_023250521.1;refGene_gn=PTPRZ1;refGene_tx=XM_023250522.1;refGene_gn=PTPRZ1;refGene_tx=XM_023250524.1
A2 141175519 141175699
refGene_gn=PTPRZ1;refGene_tx=XM_023250521.1;refGene_gn=PTPRZ1;refGene_tx=XM_023250522.1;refGene_gn=PTPRZ1;refGene_tx=XM_023250523.1;refGene_gn=PTPRZ1;refGene_tx=XM_023250524.1
A2 141177968 141178120
refGene_gn=PTPRZ1;refGene_tx=XM_023250521.1;refGene_gn=PTPRZ1;refGene_tx=XM_023250522.1;refGene_gn=PTPRZ1;refGene_tx=XM_023250523.1;refGene_gn=PTPRZ1;refGene_tx=XM_023250524.1
A2 141181241 141181337
refGene_gn=PTPRZ1;refGene_tx=XM_023250521.1;refGene_gn=PTPRZ1;refGene_tx=XM_023250522.1;refGene_gn=PTPRZ1;refGene_tx=XM_023250523.1;refGene_gn=PTPRZ1;refGene_tx=XM_023250524.1
A2 141181839 141181906
refGene_gn=PTPRZ1;refGene_tx=XM_023250521.1;refGene_gn=PTPRZ1;refGene_tx=XM_023250522.1;refGene_gn=PTPRZ1;refGene_tx=XM_023250523.1;refGene_gn=PTPRZ1;refGene_tx=XM_023250524.1
A2 141195931 141196089

refGene_gn=PTPRZ1;refGene_tx=XM_023250521.1;refGene_gn=PTPRZ1;refGene_tx=XM_023250522.1;refGene_gn=PTPRZ1;refGene_tx=XM_023250523.1;refGene_gn=PTPRZ1;refGene_tx=XM_023250524.1
A2 141196241 141196392
refGene_gn=PTPRZ1;refGene_tx=XM_023250521.1;refGene_gn=PTPRZ1;refGene_tx=XM_023250522.1;refGene_gn=PTPRZ1;refGene_tx=XM_023250523.1;refGene_gn=PTPRZ1;refGene_tx=XM_023250524.1
A2 141206666 141206851
refGene_gn=PTPRZ1;refGene_tx=XM_023250521.1;refGene_gn=PTPRZ1;refGene_tx=XM_023250522.1;refGene_gn=PTPRZ1;refGene_tx=XM_023250523.1;refGene_gn=PTPRZ1;refGene_tx=XM_023250524.1
A2 141208483 141208610
refGene_gn=PTPRZ1;refGene_tx=XM_023250521.1;refGene_gn=PTPRZ1;refGene_tx=XM_023250522.1;refGene_gn=PTPRZ1;refGene_tx=XM_023250523.1;refGene_gn=PTPRZ1;refGene_tx=XM_023250524.1
A2 141214986 141215033
refGene_gn=PTPRZ1;refGene_tx=XM_023250521.1;refGene_gn=PTPRZ1;refGene_tx=XM_023250522.1;refGene_gn=PTPRZ1;refGene_tx=XM_023250523.1;refGene_gn=PTPRZ1;refGene_tx=XM_023250524.1
A2 141218481 141222067
refGene_gn=PTPRZ1;refGene_tx=XM_023250521.1;refGene_gn=PTPRZ1;refGene_tx=XM_023250522.1;refGene_gn=PTPRZ1;refGene_tx=XM_023250523.1;refGene_gn=PTPRZ1;refGene_tx=XM_023250524.1
A2 141227259 141227404
refGene_gn=PTPRZ1;refGene_tx=XM_023250521.1;refGene_gn=PTPRZ1;refGene_tx=XM_023250522.1;refGene_gn=PTPRZ1;refGene_tx=XM_023250523.1;refGene_gn=PTPRZ1;refGene_tx=XM_023250524.1
A2 141234992 141235084
refGene_gn=PTPRZ1;refGene_tx=XM_023250521.1;refGene_gn=PTPRZ1;refGene_tx=XM_023250522.1;refGene_gn=PTPRZ1;refGene_tx=XM_023250523.1;refGene_gn=PTPRZ1;refGene_tx=XM_023250524.1
A2 141237795 141237881
refGene_gn=PTPRZ1;refGene_tx=XM_023250521.1;refGene_gn=PTPRZ1;refGene_tx=XM_023250522.1;refGene_gn=PTPRZ1;refGene_tx=XM_023250523.1;refGene_gn=PTPRZ1;refGene_tx=XM_023250524.1
A2 141240761 141240782
refGene_gn=PTPRZ1;refGene_tx=XM_023250521.1;refGene_gn=PTPRZ1;refGene_tx=XM_023250523.1;refGene_gn=PTPRZ1;refGene_tx=XM_023250524.1
A2 141240947 141241044
refGene_gn=PTPRZ1;refGene_tx=XM_023250521.1;refGene_gn=PTPRZ1;refGene_tx=XM_023250522.1;refGene_gn=PTPRZ1;refGene_tx=XM_023250523.1;refGene_gn=PTPRZ1;refGene_tx=XM_023250524.1
A2 141245696 141245779
refGene_gn=PTPRZ1;refGene_tx=XM_023250521.1;refGene_gn=PTPRZ1;refGene_tx=XM_023250522.1;refGene_gn=PTPRZ1;refGene_tx=XM_023250523.1;refGene_gn=PTPRZ1;refGene_tx=XM_023250524.1
A2 141247304 141247439
refGene_gn=PTPRZ1;refGene_tx=XM_023250521.1;refGene_gn=PTPRZ1;refGene_tx=XM_023250522.1;refGene_gn=PTPRZ1;refGene_tx=XM_023250523.1;refGene_gn=PTPRZ1;refGene_tx=XM_023250524.1

A2 141247984 141248119
refGene_gn=PTPRZ1;refGene_tx=XM_023250521.1;refGene_gn=PTPRZ1;refGene_tx=XM_023250522.1;refGene_gn=PTPRZ1;refGene_tx=XM_023250523.1;refGene_gn=PTPRZ1;refGene_tx=XM_023250524.1
A2 141249104 141249268
refGene_gn=PTPRZ1;refGene_tx=XM_023250521.1;refGene_gn=PTPRZ1;refGene_tx=XM_023250522.1;refGene_gn=PTPRZ1;refGene_tx=XM_023250523.1;refGene_gn=PTPRZ1;refGene_tx=XM_023250524.1
A2 141251141 141251277
refGene_gn=PTPRZ1;refGene_tx=XM_023250521.1;refGene_gn=PTPRZ1;refGene_tx=XM_023250522.1;refGene_gn=PTPRZ1;refGene_tx=XM_023250523.1;refGene_gn=PTPRZ1;refGene_tx=XM_023250524.1
A2 141252740 141252887
refGene_gn=PTPRZ1;refGene_tx=XM_023250521.1;refGene_gn=PTPRZ1;refGene_tx=XM_023250522.1;refGene_gn=PTPRZ1;refGene_tx=XM_023250523.1;refGene_gn=PTPRZ1;refGene_tx=XM_023250524.1
A2 141260209 141260303
refGene_gn=PTPRZ1;refGene_tx=XM_023250521.1;refGene_gn=PTPRZ1;refGene_tx=XM_023250522.1;refGene_gn=PTPRZ1;refGene_tx=XM_023250523.1;refGene_gn=PTPRZ1;refGene_tx=XM_023250524.1
A2 141260584 141260658
refGene_gn=PTPRZ1;refGene_tx=XM_023250521.1;refGene_gn=PTPRZ1;refGene_tx=XM_023250522.1;refGene_gn=PTPRZ1;refGene_tx=XM_023250523.1;refGene_gn=PTPRZ1;refGene_tx=XM_023250524.1
A2 141262494 141262624
refGene_gn=PTPRZ1;refGene_tx=XM_023250521.1;refGene_gn=PTPRZ1;refGene_tx=XM_023250522.1;refGene_gn=PTPRZ1;refGene_tx=XM_023250523.1;refGene_gn=PTPRZ1;refGene_tx=XM_023250524.1
A2 141263409 141263560
refGene_gn=PTPRZ1;refGene_tx=XM_023250521.1;refGene_gn=PTPRZ1;refGene_tx=XM_023250522.1;refGene_gn=PTPRZ1;refGene_tx=XM_023250523.1;refGene_gn=PTPRZ1;refGene_tx=XM_023250524.1
A2 141264842 141264985
refGene_gn=PTPRZ1;refGene_tx=XM_023250524.1
A2 141265738 141265874
refGene_gn=PTPRZ1;refGene_tx=XM_023250524.1
A2 141267289 141267430
refGene_gn=PTPRZ1;refGene_tx=XM_023250524.1
A2 141281024 141281143
refGene_gn=AASS;refGene_tx=XM_003982984.3;refGene_gn=AASS;refGene_tx=XM_006929327.2;refGene_gn=AASS;refGene_tx=XM_019825742.1;refGene_gn=AASS;refGene_tx=XM_019825743.1
A2 141282276 141282453
refGene_gn=AASS;refGene_tx=XM_003982984.3;refGene_gn=AASS;refGene_tx=XM_006929327.2;refGene_gn=AASS;refGene_tx=XM_019825742.1;refGene_gn=AASS;refGene_tx=XM_019825743.1
A2 141282983 141283072
refGene_gn=AASS;refGene_tx=XM_003982984.3;refGene_gn=AASS;refGene_tx=XM_006929327.2;refGene_gn=AASS;refGene_tx=XM_019825742.1;refGene_gn=AASS;refGene_tx=XM_019825743.1

A2 141283750 141283866
refGene_gn=AASS;refGene_tx=XM_003982984.3;refGene_gn=AASS;refGene_tx=XM_006929327.2;refGene_gn=AASS;refGene_tx=XM_019825742.1;refGene_gn=AASS;refGene_tx=XM_019825743.1
A2 141285956 141286052
refGene_gn=AASS;refGene_tx=XM_003982984.3;refGene_gn=AASS;refGene_tx=XM_006929327.2;refGene_gn=AASS;refGene_tx=XM_019825742.1;refGene_gn=AASS;refGene_tx=XM_019825743.1
A2 141288912 141289080
refGene_gn=AASS;refGene_tx=XM_003982984.3;refGene_gn=AASS;refGene_tx=XM_006929327.2;refGene_gn=AASS;refGene_tx=XM_019825742.1;refGene_gn=AASS;refGene_tx=XM_019825743.1
A2 141295031 141295172
refGene_gn=AASS;refGene_tx=XM_003982984.3;refGene_gn=AASS;refGene_tx=XM_006929327.2;refGene_gn=AASS;refGene_tx=XM_019825742.1;refGene_gn=AASS;refGene_tx=XM_019825743.1
A2 141296048 141296157
refGene_gn=AASS;refGene_tx=XM_003982984.3;refGene_gn=AASS;refGene_tx=XM_006929327.2;refGene_gn=AASS;refGene_tx=XM_019825742.1;refGene_gn=AASS;refGene_tx=XM_019825743.1
A2 141296249 141296360
refGene_gn=AASS;refGene_tx=XM_003982984.3;refGene_gn=AASS;refGene_tx=XM_006929327.2;refGene_gn=AASS;refGene_tx=XM_019825742.1;refGene_gn=AASS;refGene_tx=XM_019825743.1
A2 141300183 141300310
refGene_gn=AASS;refGene_tx=XM_003982984.3;refGene_gn=AASS;refGene_tx=XM_006929327.2;refGene_gn=AASS;refGene_tx=XM_019825743.1
A2 141300497 141300619
refGene_gn=AASS;refGene_tx=XM_003982984.3;refGene_gn=AASS;refGene_tx=XM_006929327.2;refGene_gn=AASS;refGene_tx=XM_019825743.1
A2 141303262 141303330
refGene_gn=AASS;refGene_tx=XM_003982984.3;refGene_gn=AASS;refGene_tx=XM_006929327.2;refGene_gn=AASS;refGene_tx=XM_019825742.1;refGene_gn=AASS;refGene_tx=XM_019825743.1
A2 141303493 141303553
refGene_gn=AASS;refGene_tx=XM_003982984.3;refGene_gn=AASS;refGene_tx=XM_006929327.2;refGene_gn=AASS;refGene_tx=XM_019825742.1;refGene_gn=AASS;refGene_tx=XM_019825743.1
A2 141318696 141318808
refGene_gn=AASS;refGene_tx=XM_003982984.3;refGene_gn=AASS;refGene_tx=XM_006929327.2;refGene_gn=AASS;refGene_tx=XM_019825742.1;refGene_gn=AASS;refGene_tx=XM_019825743.1
A2 141319198 141319321
refGene_gn=AASS;refGene_tx=XM_003982984.3;refGene_gn=AASS;refGene_tx=XM_006929327.2;refGene_gn=AASS;refGene_tx=XM_019825742.1;refGene_gn=AASS;refGene_tx=XM_019825743.1
A2 141320726 141320875
refGene_gn=AASS;refGene_tx=XM_003982984.3;refGene_gn=AASS;refGene_tx=XM_006929327.2;refGene_gn=AASS;refGene_tx=XM_019825742.1;refGene_gn=AASS;refGene_tx=XM_019825743.1

A2 141322265 141322393
refGene_gn=AASS;refGene_tx=XM_003982984.3;refGene_gn=AASS;refGene_tx=XM_006929327.2;refGene_gn=AASS;refGene_tx=XM_019825742.1;refGene_gn=AASS;refGene_tx=XM_019825743.1
A2 141322520 141322599
refGene_gn=AASS;refGene_tx=XM_003982984.3;refGene_gn=AASS;refGene_tx=XM_006929327.2;refGene_gn=AASS;refGene_tx=XM_019825742.1;refGene_gn=AASS;refGene_tx=XM_019825743.1
A2 141323532 141323679
refGene_gn=AASS;refGene_tx=XM_003982984.3;refGene_gn=AASS;refGene_tx=XM_006929327.2;refGene_gn=AASS;refGene_tx=XM_019825742.1;refGene_gn=AASS;refGene_tx=XM_019825743.1
A2 141323787 141323855
refGene_gn=AASS;refGene_tx=XM_003982984.3;refGene_gn=AASS;refGene_tx=XM_006929327.2;refGene_gn=AASS;refGene_tx=XM_019825742.1
A2 141326880 141326965
refGene_gn=AASS;refGene_tx=XM_003982984.3;refGene_gn=AASS;refGene_tx=XM_006929327.2;refGene_gn=AASS;refGene_tx=XM_019825742.1
A2 141329671 141329848
refGene_gn=AASS;refGene_tx=XM_003982984.3;refGene_gn=AASS;refGene_tx=XM_006929327.2;refGene_gn=AASS;refGene_tx=XM_019825742.1
A2 141334008 141334218
refGene_gn=AASS;refGene_tx=XM_003982984.3;refGene_gn=AASS;refGene_tx=XM_006929327.2;refGene_gn=AASS;refGene_tx=XM_019825742.1
A2 141456027 141456386
refGene_gn=FEZF1;refGene_tx=XM_003982993.5;refGene_gn=FEZF1;refGene_tx=XM_011280523.3
A2 141456813 141456946
refGene_gn=FEZF1;refGene_tx=XM_003982993.5;refGene_gn=FEZF1;refGene_tx=XM_011280523.3
A2 141457149 141457284
refGene_gn=FEZF1;refGene_tx=XM_003982993.5;refGene_gn=FEZF1;refGene_tx=XM_011280523.3
A2 141457580 141458454
refGene_gn=FEZF1;refGene_tx=XM_003982993.5;refGene_gn=FEZF1;refGene_tx=XM_011280523.3
A2 141460989 141461006
refGene_gn=FEZF1;refGene_tx=XM_003982993.5
A2 141474535 141474706
refGene_gn=CADPS2;refGene_tx=XM_023250526.1
A2 141480609 141480714
refGene_gn=CADPS2;refGene_tx=XM_023250526.1
A2 141500020 141500128
refGene_gn=CADPS2;refGene_tx=XM_023250526.1
A2 141511430 141511463
refGene_gn=CADPS2;refGene_tx=XM_023250526.1
A2 141511561 141511645
refGene_gn=CADPS2;refGene_tx=XM_023250526.1
A2 141524405 141524480
refGene_gn=CADPS2;refGene_tx=XM_023250526.1

A2 141531975 141532123
refGene_gn=CADPS2;refGene_tx=XM_023250526.1
A2 141533508 141533664
refGene_gn=CADPS2;refGene_tx=XM_023250526.1
A2 141538394 141538514
refGene_gn=CADPS2;refGene_tx=XM_023250526.1
A2 141538639 141538781
refGene_gn=CADPS2;refGene_tx=XM_023250526.1
A2 141553454 141553611
refGene_gn=CADPS2;refGene_tx=XM_023250526.1
A2 141566662 141566671
refGene_gn=CADPS2;refGene_tx=XM_023250526.1
A2 141568483 141568587
refGene_gn=CADPS2;refGene_tx=XM_023250526.1
A2 141589817 141589838
refGene_gn=CADPS2;refGene_tx=XM_023250526.1
A2 141591874 141591998
refGene_gn=CADPS2;refGene_tx=XM_023250526.1
A2 141594967 141595031
refGene_gn=CADPS2;refGene_tx=XM_023250526.1
A2 141602130 141602232
refGene_gn=CADPS2;refGene_tx=XM_023250526.1
A2 141623136 141623324
refGene_gn=CADPS2;refGene_tx=XM_023250526.1
A2 141625596 141625733
refGene_gn=CADPS2;refGene_tx=XM_023250526.1
A2 141632913 141633114
refGene_gn=CADPS2;refGene_tx=XM_023250526.1
A2 141634115 141634224
refGene_gn=CADPS2;refGene_tx=XM_023250526.1
A2 141664335 141664402
refGene_gn=CADPS2;refGene_tx=XM_023250526.1
A2 141714332 141714472
refGene_gn=CADPS2;refGene_tx=XM_023250526.1
A2 141745265 141745307
refGene_gn=CADPS2;refGene_tx=XM_023250526.1
A2 141745308 141745378
refGene_gn=CADPS2;refGene_tx=XM_023250526.1
A2 141790637 141790759
refGene_gn=CADPS2;refGene_tx=XM_023250526.1
A2 141796157 141796394
refGene_gn=CADPS2;refGene_tx=XM_023250526.1
A2 141802886 141802967
refGene_gn=CADPS2;refGene_tx=XM_023250526.1
A2 141829666 141829999
refGene_gn=CADPS2;refGene_tx=XM_023250526.1
A2 141851932 141853069
refGene_gn=RNF133;refGene_tx=XM_023250527.1
A2 141855905 141856835
refGene_gn=RNF148;refGene_tx=XM_003982985.5

A2 141896799 141896913
refGene_gn=CADPS2;refGene_tx=XM_023250526.1
A2 142022358 142022697
refGene_gn=CADPS2;refGene_tx=XM_023250526.1
A2 142107712 142107919
refGene_gn=TAS2R16;refGene_tx=XM_011280553.2
A2 142107921 142108206
refGene_gn=TAS2R16;refGene_tx=XM_011280553.2
A2 142108207 142108237
refGene_gn=TAS2R16;refGene_tx=XM_011280553.2
A2 142108238 142108443
refGene_gn=TAS2R16;refGene_tx=XM_011280553.2
A2 142108444 142108559
refGene_gn=TAS2R16;refGene_tx=XM_011280553.2
A2 142109785 142110035
refGene_gn=TAS2R16;refGene_tx=XM_011280553.2
A2 142111024 142111157
refGene_gn=TAS2R16;refGene_tx=XM_011280553.2
A2 142181276 142181353
refGene_gn=TAS2R16;refGene_tx=XM_011280553.2
A2 142185350 142185438
refGene_gn=TAS2R16;refGene_tx=XM_011280553.2
A2 142202761 142202899
refGene_gn=SLC13A1;refGene_tx=XM_019825748.1;refGene_gn=SLC13A1;refGen
e_tx=XM_019825749.1;refGene_gn=SLC13A1;refGene_tx=XM_019825750.2
A2 142204930 142205068
refGene_gn=SLC13A1;refGene_tx=XM_019825748.1;refGene_gn=SLC13A1;refGen
e_tx=XM_019825749.1;refGene_gn=SLC13A1;refGene_tx=XM_019825750.2
A2 142206491 142206653
refGene_gn=SLC13A1;refGene_tx=XM_019825748.1;refGene_gn=SLC13A1;refGen
e_tx=XM_019825749.1;refGene_gn=SLC13A1;refGene_tx=XM_019825750.2
A2 142209766 142209876
refGene_gn=SLC13A1;refGene_tx=XM_019825748.1;refGene_gn=SLC13A1;refGen
e_tx=XM_019825749.1;refGene_gn=SLC13A1;refGene_tx=XM_019825750.2
A2 142213106 142213213
refGene_gn=SLC13A1;refGene_tx=XM_019825748.1;refGene_gn=SLC13A1;refGen
e_tx=XM_019825749.1;refGene_gn=SLC13A1;refGene_tx=XM_019825750.2
A2 142216541 142216643
refGene_gn=SLC13A1;refGene_tx=XM_019825748.1;refGene_gn=SLC13A1;refGen
e_tx=XM_019825749.1;refGene_gn=SLC13A1;refGene_tx=XM_019825750.2
A2 142217060 142217159
refGene_gn=SLC13A1;refGene_tx=XM_019825748.1;refGene_gn=SLC13A1;refGen
e_tx=XM_019825749.1;refGene_gn=SLC13A1;refGene_tx=XM_019825750.2
A2 142221760 142221880
refGene_gn=SLC13A1;refGene_tx=XM_019825748.1;refGene_gn=SLC13A1;refGen
e_tx=XM_019825749.1;refGene_gn=SLC13A1;refGene_tx=XM_019825750.2
A2 142235174 142235326
refGene_gn=SLC13A1;refGene_tx=XM_019825748.1;refGene_gn=SLC13A1;refGen
e_tx=XM_019825749.1;refGene_gn=SLC13A1;refGene_tx=XM_019825750.2
A2 142265136 142265185

refGene_gn=SLC13A1;refGene_tx=XM_019825748.1;refGene_gn=SLC13A1;refGene_tx=XM_019825749.1;refGene_gn=SLC13A1;refGene_tx=XM_019825750.2
A2 142265266 142265324
refGene_gn=SLC13A1;refGene_tx=XM_019825748.1;refGene_gn=SLC13A1;refGene_tx=XM_019825749.1;refGene_gn=SLC13A1;refGene_tx=XM_019825750.2
A2 142265947 142266135
refGene_gn=SLC13A1;refGene_tx=XM_019825748.1;refGene_gn=SLC13A1;refGene_tx=XM_019825749.1;refGene_gn=SLC13A1;refGene_tx=XM_019825750.2
A2 142268710 142268847
refGene_gn=SLC13A1;refGene_tx=XM_019825748.1;refGene_gn=SLC13A1;refGene_tx=XM_019825749.1;refGene_gn=SLC13A1;refGene_tx=XM_019825750.2
A2 142278431 142278560
refGene_gn=SLC13A1;refGene_tx=XM_019825748.1;refGene_gn=SLC13A1;refGene_tx=XM_019825749.1;refGene_gn=SLC13A1;refGene_tx=XM_019825750.2
A2 142300105 142300204
refGene_gn=SLC13A1;refGene_tx=XM_019825748.1;refGene_gn=SLC13A1;refGene_tx=XM_019825749.1;refGene_gn=SLC13A1;refGene_tx=XM_019825750.2
A2 142416477 142416486
refGene_gn=IQUB;refGene_tx=XM_019825746.2
A2 142482980 142483163
refGene_gn=IQUB;refGene_tx=XM_003982987.4
A2 142486892 142487078
refGene_gn=IQUB;refGene_tx=XM_003982987.4;refGene_gn=IQUB;refGene_tx=XM_019825746.2
A2 142487986 142488235
refGene_gn=IQUB;refGene_tx=XM_003982987.4;refGene_gn=IQUB;refGene_tx=XM_019825746.2
A2 142492279 142492456
refGene_gn=IQUB;refGene_tx=XM_003982987.4;refGene_gn=IQUB;refGene_tx=XM_019825746.2
A2 142495097 142495268
refGene_gn=IQUB;refGene_tx=XM_003982987.4;refGene_gn=IQUB;refGene_tx=XM_019825746.2
A2 142503832 142504008
refGene_gn=IQUB;refGene_tx=XM_003982987.4;refGene_gn=IQUB;refGene_tx=XM_019825746.2
A2 142515297 142515508
refGene_gn=IQUB;refGene_tx=XM_003982987.4;refGene_gn=IQUB;refGene_tx=XM_019825746.2
A2 142528128 142528284
refGene_gn=IQUB;refGene_tx=XM_003982987.4;refGene_gn=IQUB;refGene_tx=XM_019825746.2
A2 142528840 142529013
refGene_gn=IQUB;refGene_tx=XM_003982987.4;refGene_gn=IQUB;refGene_tx=XM_019825746.2
A2 142529100 142529262
refGene_gn=IQUB;refGene_tx=XM_003982987.4;refGene_gn=IQUB;refGene_tx=XM_019825746.2
A2 142532197 142532332
refGene_gn=IQUB;refGene_tx=XM_003982987.4;refGene_gn=IQUB;refGene_tx=X

M_019825746.2
A2	142535233	142535657
refGene_gn=IQUB;refGene_tx=XM_003982987.4;refGene_gn=IQUB;refGene_tx=XM_019825746.2
A2	142547873	142547975
refGene_gn=NDUFA5;refGene_tx=XM_003982988.5;refGene_gn=NDUFA5;refGene_tx=XM_006929330.4
A2	142550665	142550731
refGene_gn=NDUFA5;refGene_tx=XM_003982988.5;refGene_gn=NDUFA5;refGene_tx=XM_006929330.4
A2	142555338	142555455
refGene_gn=NDUFA5;refGene_tx=XM_003982988.5
A2	142559730	142559775
refGene_gn=NDUFA5;refGene_tx=XM_003982988.5;refGene_gn=NDUFA5;refGene_tx=XM_006929330.4
A2	142560100	142560121
refGene_gn=NDUFA5;refGene_tx=XM_003982988.5;refGene_gn=NDUFA5;refGene_tx=XM_006929330.4
A2	142600476	142600583
refGene_gn=ASB15;refGene_tx=XM_003982989.4;refGene_gn=ASB15;refGene_tx=XM_006929333.3;refGene_gn=ASB15;refGene_tx=XM_019825752.2;refGene_gn=ASB15;refGene_tx=XM_019825753.2
A2	142602399	142602452
refGene_gn=ASB15;refGene_tx=XM_003982989.4;refGene_gn=ASB15;refGene_tx=XM_006929333.3;refGene_gn=ASB15;refGene_tx=XM_019825752.2;refGene_gn=ASB15;refGene_tx=XM_019825753.2
A2	142602543	142602675
refGene_gn=ASB15;refGene_tx=XM_003982989.4;refGene_gn=ASB15;refGene_tx=XM_006929333.3;refGene_gn=ASB15;refGene_tx=XM_019825752.2;refGene_gn=ASB15;refGene_tx=XM_019825753.2
A2	142604229	142604388
refGene_gn=ASB15;refGene_tx=XM_003982989.4;refGene_gn=ASB15;refGene_tx=XM_006929333.3;refGene_gn=ASB15;refGene_tx=XM_019825752.2;refGene_gn=ASB15;refGene_tx=XM_019825753.2
A2	142609268	142609514
refGene_gn=ASB15;refGene_tx=XM_003982989.4;refGene_gn=ASB15;refGene_tx=XM_006929333.3;refGene_gn=ASB15;refGene_tx=XM_019825752.2;refGene_gn=ASB15;refGene_tx=XM_019825753.2
A2	142610450	142610622
refGene_gn=ASB15;refGene_tx=XM_003982989.4;refGene_gn=ASB15;refGene_tx=XM_006929333.3;refGene_gn=ASB15;refGene_tx=XM_019825752.2;refGene_gn=ASB15;refGene_tx=XM_019825753.2
A2	142612244	142612815
refGene_gn=ASB15;refGene_tx=XM_003982989.4;refGene_gn=ASB15;refGene_tx=XM_006929333.3;refGene_gn=ASB15;refGene_tx=XM_019825752.2;refGene_gn=ASB15;refGene_tx=XM_019825753.2
A2	142613337	142613491
refGene_gn=ASB15;refGene_tx=XM_003982989.4;refGene_gn=ASB15;refGene_tx=XM_019825752.2;refGene_gn=ASB15;refGene_tx=XM_019825753.2
A2	142618740	142618913

refGene_gn=ASB15;refGene_tx=XM_003982989.4;refGene_gn=ASB15;refGene_tx=XM_006929333.3;refGene_gn=ASB15;refGene_tx=XM_019825752.2;refGene_gn=ASB15;refGene_tx=XM_019825753.2
A2 142633839 142634112
refGene_gn=LMOD2;refGene_tx=XM_011280526.3
A2 142639470 142640871
refGene_gn=LMOD2;refGene_tx=XM_011280526.3
A2 142641633 142641660
refGene_gn=LMOD2;refGene_tx=XM_011280526.3
A2 142662044 142662106
refGene_gn=WASL;refGene_tx=XM_011280527.2
A2 142664547 142664656
refGene_gn=WASL;refGene_tx=XM_011280527.2
A2 142668620 142669141
refGene_gn=WASL;refGene_tx=XM_011280527.2
A2 142674761 142674915
refGene_gn=WASL;refGene_tx=XM_011280527.2
A2 142675921 142675964
refGene_gn=WASL;refGene_tx=XM_011280527.2
A2 142677225 142677394
refGene_gn=WASL;refGene_tx=XM_011280527.2
A2 142684213 142684237
refGene_gn=WASL;refGene_tx=XM_011280527.2
A2 142687568 142687665
refGene_gn=WASL;refGene_tx=XM_011280527.2
A2 142688871 142688958
refGene_gn=WASL;refGene_tx=XM_011280527.2
A2 142690770 142690905
refGene_gn=WASL;refGene_tx=XM_011280527.2
A2 142721690 142721807
refGene_gn=WASL;refGene_tx=XM_011280527.2
A2 142782460 142783381
refGene_gn=HYAL4;refGene_tx=XM_006929334.3
A2 142787448 142787538
refGene_gn=HYAL4;refGene_tx=XM_006929334.3
A2 142789475 142789922
refGene_gn=HYAL4;refGene_tx=XM_006929334.3
A2 142807878 142808832
refGene_gn=LOC101093011;refGene_tx=XM_023250528.1
A2 142817654 142817744
refGene_gn=LOC101093011;refGene_tx=XM_023250528.1
A2 142819698 142820109
refGene_gn=LOC101093011;refGene_tx=XM_023250528.1
A2 142845013 142845166
refGene_gn=SPAM1;refGene_tx=XM_011280528.3
A2 142845167 142845971
refGene_gn=SPAM1;refGene_tx=XM_011280528.3
A2 142846446 142846536
refGene_gn=SPAM1;refGene_tx=XM_011280528.3
A2 142859139 142859691

refGene_gn=SPAM1;refGene_tx=XM_011280528.3
A2	142919680	142920805
refGene_gn=TMEM229A;refGene_tx=XM_023250529.1
A2	143475238	143476057
refGene_gn=GPR37;refGene_tx=XM_006929340.4
A2	143492755	143493778
refGene_gn=GPR37;refGene_tx=XM_006929340.4
A2	143543358	143543471
refGene_gn=POT1;refGene_tx=XM_006929341.4;refGene_gn=POT1;refGene_tx=XM_006929342.4;refGene_gn=POT1;refGene_tx=XM_006929343.4
A2	143545714	143545820
refGene_gn=POT1;refGene_tx=XM_006929341.4;refGene_gn=POT1;refGene_tx=XM_006929342.4;refGene_gn=POT1;refGene_tx=XM_006929343.4
A2	143547431	143547523
refGene_gn=POT1;refGene_tx=XM_006929341.4;refGene_gn=POT1;refGene_tx=XM_006929342.4;refGene_gn=POT1;refGene_tx=XM_006929343.4
A2	143549378	143549467
refGene_gn=POT1;refGene_tx=XM_006929341.4;refGene_gn=POT1;refGene_tx=XM_006929342.4;refGene_gn=POT1;refGene_tx=XM_006929343.4
A2	143553811	143553947
refGene_gn=POT1;refGene_tx=XM_006929341.4;refGene_gn=POT1;refGene_tx=XM_006929342.4;refGene_gn=POT1;refGene_tx=XM_006929343.4
A2	143557214	143557420
refGene_gn=POT1;refGene_tx=XM_006929341.4;refGene_gn=POT1;refGene_tx=XM_006929342.4;refGene_gn=POT1;refGene_tx=XM_006929343.4
A2	143558779	143558936
refGene_gn=POT1;refGene_tx=XM_006929341.4;refGene_gn=POT1;refGene_tx=XM_006929342.4;refGene_gn=POT1;refGene_tx=XM_006929343.4
A2	143561037	143561094
refGene_gn=POT1;refGene_tx=XM_006929341.4;refGene_gn=POT1;refGene_tx=XM_006929342.4;refGene_gn=POT1;refGene_tx=XM_006929343.4
A2	143565697	143565786
refGene_gn=POT1;refGene_tx=XM_006929341.4;refGene_gn=POT1;refGene_tx=XM_006929342.4;refGene_gn=POT1;refGene_tx=XM_006929343.4
A2	143566909	143567079
refGene_gn=POT1;refGene_tx=XM_006929341.4;refGene_gn=POT1;refGene_tx=XM_006929342.4;refGene_gn=POT1;refGene_tx=XM_006929343.4
A2	143572232	143572388
refGene_gn=POT1;refGene_tx=XM_006929341.4;refGene_gn=POT1;refGene_tx=XM_006929342.4;refGene_gn=POT1;refGene_tx=XM_006929343.4
A2	143577039	143577330
refGene_gn=POT1;refGene_tx=XM_006929341.4;refGene_gn=POT1;refGene_tx=XM_006929342.4;refGene_gn=POT1;refGene_tx=XM_006929343.4
A2	143580736	143580867
refGene_gn=POT1;refGene_tx=XM_006929341.4;refGene_gn=POT1;refGene_tx=XM_006929342.4;refGene_gn=POT1;refGene_tx=XM_006929343.4
A2	143595051	143595166
refGene_gn=POT1;refGene_tx=XM_006929341.4;refGene_gn=POT1;refGene_tx=XM_006929342.4;refGene_gn=POT1;refGene_tx=XM_006929343.4
A2	143597774	143597822

refGene_gn=POT1;refGene_tx=XM_006929341.4;refGene_gn=POT1;refGene_tx=XM_006929342.4;refGene_gn=POT1;refGene_tx=XM_006929343.4
A2	143598585	143598699
refGene_gn=POT1;refGene_tx=XM_006929341.4;refGene_gn=POT1;refGene_tx=XM_006929342.4;refGene_gn=POT1;refGene_tx=XM_006929343.4
A2	143609434	143609526
refGene_gn=POT1;refGene_tx=XM_006929341.4;refGene_gn=POT1;refGene_tx=XM_006929342.4;refGene_gn=POT1;refGene_tx=XM_006929343.4
A2	143613549	143613622
refGene_gn=POT1;refGene_tx=XM_006929341.4;refGene_gn=POT1;refGene_tx=XM_006929342.4;refGene_gn=POT1;refGene_tx=XM_006929343.4
A2	143621728	143621851
refGene_gn=POT1;refGene_tx=XM_006929341.4;refGene_gn=POT1;refGene_tx=XM_006929342.4;refGene_gn=POT1;refGene_tx=XM_006929343.4
A2	144934978	144935028
refGene_gn=GRM8;refGene_tx=XM_011280531.3;refGene_gn=GRM8;refGene_tx=XM_019825760.2;refGene_gn=GRM8;refGene_tx=XM_019825761.2
A2	144941228	144941525
refGene_gn=GRM8;refGene_tx=XM_011280531.3;refGene_gn=GRM8;refGene_tx=XM_019825760.2;refGene_gn=GRM8;refGene_tx=XM_019825761.2;refGene_gn=GRM8;refGene_tx=XM_019825764.2
A2	144952945	144952975
refGene_gn=GRM8;refGene_tx=XM_023250530.1
A2	145014636	145015572
refGene_gn=GRM8;refGene_tx=XM_011280531.3;refGene_gn=GRM8;refGene_tx=XM_019825760.2;refGene_gn=GRM8;refGene_tx=XM_019825761.2;refGene_gn=GRM8;refGene_tx=XM_019825764.2;refGene_gn=GRM8;refGene_tx=XM_023250530.1
A2	145071319	145071456
refGene_gn=GRM8;refGene_tx=XM_011280531.3;refGene_gn=GRM8;refGene_tx=XM_019825760.2;refGene_gn=GRM8;refGene_tx=XM_019825761.2;refGene_gn=GRM8;refGene_tx=XM_019825764.2;refGene_gn=GRM8;refGene_tx=XM_023250530.1
A2	145211771	145211972
refGene_gn=GRM8;refGene_tx=XM_011280531.3;refGene_gn=GRM8;refGene_tx=XM_019825760.2;refGene_gn=GRM8;refGene_tx=XM_019825761.2;refGene_gn=GRM8;refGene_tx=XM_019825764.2;refGene_gn=GRM8;refGene_tx=XM_023250530.1
A2	145339342	145339480
refGene_gn=GRM8;refGene_tx=XM_011280531.3;refGene_gn=GRM8;refGene_tx=XM_019825760.2;refGene_gn=GRM8;refGene_tx=XM_019825761.2;refGene_gn=GRM8;refGene_tx=XM_019825764.2;refGene_gn=GRM8;refGene_tx=XM_023250530.1
A2	145340576	145340731
refGene_gn=GRM8;refGene_tx=XM_011280531.3;refGene_gn=GRM8;refGene_tx=XM_019825760.2;refGene_gn=GRM8;refGene_tx=XM_019825761.2;refGene_gn=GRM8;refGene_tx=XM_019825764.2;refGene_gn=GRM8;refGene_tx=XM_023250530.1
A2	145341188	145341324
refGene_gn=GRM8;refGene_tx=XM_011280531.3;refGene_gn=GRM8;refGene_tx=XM_019825760.2;refGene_gn=GRM8;refGene_tx=XM_019825761.2;refGene_gn=GRM8;refGene_tx=XM_019825764.2;refGene_gn=GRM8;refGene_tx=XM_023250530.1
A2	145537927	145538144
refGene_gn=GRM8;refGene_tx=XM_011280531.3;refGene_gn=GRM8;refGene_tx=XM_019825760.2;refGene_gn=GRM8;refGene_tx=XM_019825761.2;refGene_gn=GRM

8;refGene_tx=XM_019825764.2;refGene_gn=GRM8;refGene_tx=XM_023250530.1
A2	145687911	145688421
refGene_gn=GRM8;refGene_tx=XM_011280531.3;refGene_gn=GRM8;refGene_tx=XM_019825760.2;refGene_gn=GRM8;refGene_tx=XM_019825761.2;refGene_gn=GRM8;refGene_tx=XM_019825764.2;refGene_gn=GRM8;refGene_tx=XM_023250530.1
A2	145793995	145793996
refGene_gn=ZNF800;refGene_tx=XM_023250532.1
A2	145795220	145796917
refGene_gn=ZNF800;refGene_tx=XM_023250531.1;refGene_gn=ZNF800;refGene_tx=XM_023250532.1
A2	145799332	145799476
refGene_gn=ZNF800;refGene_tx=XM_023250531.1;refGene_gn=ZNF800;refGene_tx=XM_023250532.1
A2	145808409	145808505
refGene_gn=ZNF800;refGene_tx=XM_023250531.1;refGene_gn=ZNF800;refGene_tx=XM_023250532.1
A2	145813574	145813635
refGene_gn=ZNF800;refGene_tx=XM_023250531.1;refGene_gn=ZNF800;refGene_tx=XM_023250532.1
A2	145940647	145941943
refGene_gn=GCC1;refGene_tx=XM_003983006.5
A2	145942930	145943962
refGene_gn=GCC1;refGene_tx=XM_003983006.5
A2	145947595	145947662
refGene_gn=ARF5;refGene_tx=XM_019825068.2
A2	145948200	145948281
refGene_gn=ARF5;refGene_tx=XM_019825068.2
A2	145948632	145948742
refGene_gn=ARF5;refGene_tx=XM_019825068.2
A2	145949513	145949585
refGene_gn=ARF5;refGene_tx=XM_019825068.2
A2	145950127	145950253
refGene_gn=ARF5;refGene_tx=XM_019825068.2
A2	145950355	145950442
refGene_gn=ARF5;refGene_tx=XM_019825068.2
A2	145952482	145952623
refGene_gn=FSCN3;refGene_tx=XM_003983007.4
A2	145953675	145954372
refGene_gn=FSCN3;refGene_tx=XM_003983007.4
A2	145954689	145954808
refGene_gn=FSCN3;refGene_tx=XM_003983007.4
A2	145956363	145956523
refGene_gn=FSCN3;refGene_tx=XM_003983007.4
A2	145956994	145957165
refGene_gn=FSCN3;refGene_tx=XM_003983007.4
A2	145957501	145957707
refGene_gn=FSCN3;refGene_tx=XM_003983007.4
A2	145963469	145963691
refGene_gn=PAX4;refGene_tx=XM_004001293.3
A2	145963692	145963706

refGene_gn=PAX4;refGene_tx=XM_004001293.3
A2	145964035	145964174
refGene_gn=PAX4;refGene_tx=XM_004001293.3
A2	145964425	145964481
refGene_gn=PAX4;refGene_tx=XM_004001293.3
A2	145964945	145965015
refGene_gn=PAX4;refGene_tx=XM_004001293.3
A2	145965375	145965458
refGene_gn=PAX4;refGene_tx=XM_004001293.3
A2	145965681	145965807
refGene_gn=PAX4;refGene_tx=XM_004001293.3
A2	145966413	145966489
refGene_gn=PAX4;refGene_tx=XM_004001293.3
A2	145966770	145966986
refGene_gn=PAX4;refGene_tx=XM_004001293.3
A2	145967276	145967456
refGene_gn=PAX4;refGene_tx=XM_004001293.3
A2	145967806	145967995
refGene_gn=PAX4;refGene_tx=XM_004001293.3
A2	146002135	146002213
refGene_gn=SND1;refGene_tx=XM_011280535.3
A2	146030962	146031112
refGene_gn=SND1;refGene_tx=XM_011280535.3
A2	146037293	146037414
refGene_gn=SND1;refGene_tx=XM_011280535.3
A2	146042569	146042648
refGene_gn=SND1;refGene_tx=XM_011280535.3;refGene_gn=SND1;refGene_tx=XM_023250533.1
A2	146045023	146045184
refGene_gn=SND1;refGene_tx=XM_011280535.3;refGene_gn=SND1;refGene_tx=XM_023250533.1
A2	146045906	146045998
refGene_gn=SND1;refGene_tx=XM_011280535.3;refGene_gn=SND1;refGene_tx=XM_023250533.1
A2	146046652	146046811
refGene_gn=SND1;refGene_tx=XM_011280535.3;refGene_gn=SND1;refGene_tx=XM_023250533.1
A2	146048169	146048276
refGene_gn=SND1;refGene_tx=XM_011280535.3;refGene_gn=SND1;refGene_tx=XM_023250533.1
A2	146050560	146050651
refGene_gn=SND1;refGene_tx=XM_011280535.3;refGene_gn=SND1;refGene_tx=XM_023250533.1
A2	146063421	146063535
refGene_gn=SND1;refGene_tx=XM_011280535.3;refGene_gn=SND1;refGene_tx=XM_023250533.1
A2	146155114	146155204
refGene_gn=SND1;refGene_tx=XM_011280535.3;refGene_gn=SND1;refGene_tx=XM_023250533.1
A2	146193825	146193926

refGene_gn=SND1;refGene_tx=XM_011280535.3;refGene_gn=SND1;refGene_tx=XM_023250533.1
A2 146235683 146235794
refGene_gn=SND1;refGene_tx=XM_011280535.3;refGene_gn=SND1;refGene_tx=XM_023250533.1
A2 146254282 146254355
refGene_gn=SND1;refGene_tx=XM_011280535.3;refGene_gn=SND1;refGene_tx=XM_023250533.1
A2 146276487 146276629
refGene_gn=SND1;refGene_tx=XM_011280535.3;refGene_gn=SND1;refGene_tx=XM_023250533.1
A2 146330396 146330506
refGene_gn=SND1;refGene_tx=XM_011280535.3;refGene_gn=SND1;refGene_tx=XM_023250533.1
A2 146366425 146368378
refGene_gn=LRRC4;refGene_tx=XM_003983010.5
A2 146411394 146411583
refGene_gn=SND1;refGene_tx=XM_011280535.3;refGene_gn=SND1;refGene_tx=XM_023250533.1
A2 146417896 146418038
refGene_gn=SND1;refGene_tx=XM_011280535.3;refGene_gn=SND1;refGene_tx=XM_023250533.1
A2 146421258 146421382
refGene_gn=SND1;refGene_tx=XM_011280535.3;refGene_gn=SND1;refGene_tx=XM_023250533.1
A2 146422280 146422350
refGene_gn=SND1;refGene_tx=XM_011280535.3;refGene_gn=SND1;refGene_tx=XM_023250533.1
A2 146423433 146423547
refGene_gn=SND1;refGene_tx=XM_011280535.3;refGene_gn=SND1;refGene_tx=XM_023250533.1
A2 146425124 146425328
refGene_gn=SND1;refGene_tx=XM_011280535.3;refGene_gn=SND1;refGene_tx=XM_023250533.1
A2 146426929 146426974
refGene_gn=SND1;refGene_tx=XM_011280535.3;refGene_gn=SND1;refGene_tx=XM_023250533.1
A2 146427067 146427133
refGene_gn=SND1;refGene_tx=XM_011280535.3;refGene_gn=SND1;refGene_tx=XM_023250533.1
A2 146571415 146571559
refGene_gn=LEP;refGene_tx=NM_001009850.1;refGene_gn=LEP;refGene_tx=XM_006929350.4;refGene_gn=LEP;refGene_tx=XM_006929351.4;refGene_gn=LEP;refGene_tx=XM_006929352.4;refGene_gn=LEP;refGene_tx=XM_011280510.3;refGene_gn=LEP;refGene_tx=XM_011280511.3
A2 146572636 146572996
refGene_gn=LEP;refGene_tx=NM_001009850.1;refGene_gn=LEP;refGene_tx=XM_006929350.4;refGene_gn=LEP;refGene_tx=XM_006929351.4;refGene_gn=LEP;refGene_tx=XM_006929352.4;refGene_gn=LEP;refGene_tx=XM_011280510.3;refGene_gn=LEP;refGene_tx=XM_011280511.3

A2 146620885 146621020
refGene_gn=RBM28;refGene_tx=XM_011280536.3;refGene_gn=RBM28;refGene_tx=XM_019825765.2
A2 146623628 146623728
refGene_gn=RBM28;refGene_tx=XM_011280536.3;refGene_gn=RBM28;refGene_tx=XM_019825765.2
A2 146624349 146624606
refGene_gn=RBM28;refGene_tx=XM_011280536.3;refGene_gn=RBM28;refGene_tx=XM_019825765.2
A2 146627144 146627219
refGene_gn=RBM28;refGene_tx=XM_011280536.3;refGene_gn=RBM28;refGene_tx=XM_019825765.2
A2 146627531 146627681
refGene_gn=RBM28;refGene_tx=XM_011280536.3;refGene_gn=RBM28;refGene_tx=XM_019825765.2
A2 146629816 146629975
refGene_gn=RBM28;refGene_tx=XM_011280536.3;refGene_gn=RBM28;refGene_tx=XM_019825765.2
A2 146632010 146632075
refGene_gn=RBM28;refGene_tx=XM_011280536.3;refGene_gn=RBM28;refGene_tx=XM_019825765.2
A2 146633102 146633238
refGene_gn=RBM28;refGene_tx=XM_011280536.3;refGene_gn=RBM28;refGene_tx=XM_019825765.2
A2 146634150 146634224
refGene_gn=RBM28;refGene_tx=XM_011280536.3;refGene_gn=RBM28;refGene_tx=XM_019825765.2
A2 146636804 146636914
refGene_gn=RBM28;refGene_tx=XM_011280536.3;refGene_gn=RBM28;refGene_tx=XM_019825765.2
A2 146639741 146639814
refGene_gn=RBM28;refGene_tx=XM_011280536.3;refGene_gn=RBM28;refGene_tx=XM_019825765.2
A2 146641987 146642151
refGene_gn=RBM28;refGene_tx=XM_011280536.3;refGene_gn=RBM28;refGene_tx=XM_019825765.2
A2 146642292 146642443
refGene_gn=RBM28;refGene_tx=XM_011280536.3;refGene_gn=RBM28;refGene_tx=XM_019825765.2
A2 146643272 146643344
refGene_gn=RBM28;refGene_tx=XM_011280536.3;refGene_gn=RBM28;refGene_tx=XM_019825765.2
A2 146644444 146644537
refGene_gn=RBM28;refGene_tx=XM_011280536.3;refGene_gn=RBM28;refGene_tx=XM_019825765.2
A2 146645049 146645125
refGene_gn=RBM28;refGene_tx=XM_011280536.3;refGene_gn=RBM28;refGene_tx=XM_019825765.2
A2 146645547 146645642
refGene_gn=RBM28;refGene_tx=XM_011280536.3;refGene_gn=RBM28;refGene_tx

=XM_019825765.2
A2 146645954 146646113
refGene_gn=RBM28;refGene_tx=XM_011280536.3;refGene_gn=RBM28;refGene_tx
=XM_019825765.2
A2 146650014 146650132
refGene_gn=RBM28;refGene_tx=XM_011280536.3;refGene_gn=RBM28;refGene_tx
=XM_019825765.2
A2 146656642 146658468
refGene_gn=PRRT4;refGene_tx=XM_023250534.1;refGene_gn=PRRT4;refGene_tx
=XM_023250535.1;refGene_gn=PRRT4;refGene_tx=XM_023250536.1
A2 146663336 146663456
refGene_gn=PRRT4;refGene_tx=XM_023250534.1;refGene_gn=PRRT4;refGene_tx
=XM_023250535.1;refGene_gn=PRRT4;refGene_tx=XM_023250536.1
A2 146663805 146663910
refGene_gn=PRRT4;refGene_tx=XM_023250534.1;refGene_gn=PRRT4;refGene_tx
=XM_023250535.1;refGene_gn=PRRT4;refGene_tx=XM_023250536.1
A2 146663995 146664641
refGene_gn=PRRT4;refGene_tx=XM_023250534.1;refGene_gn=PRRT4;refGene_tx
=XM_023250535.1;refGene_gn=PRRT4;refGene_tx=XM_023250536.1
A2 146683857 146683958
refGene_gn=IMPDH1;refGene_tx=XM_023250537.1;refGene_gn=IMPDH1;refGene_
tx=XM_023250538.1;refGene_gn=IMPDH1;refGene_tx=XM_023250539.1;refGene_
gn=IMPDH1;refGene_tx=XM_023250540.1
A2 146684571 146684588
refGene_gn=IMPDH1;refGene_tx=XM_023250537.1;refGene_gn=IMPDH1;refGene_
tx=XM_023250538.1
A2 146685154 146685238
refGene_gn=IMPDH1;refGene_tx=XM_023250537.1;refGene_gn=IMPDH1;refGene_
tx=XM_023250538.1;refGene_gn=IMPDH1;refGene_tx=XM_023250539.1;refGene_
gn=IMPDH1;refGene_tx=XM_023250540.1
A2 146685368 146685512
refGene_gn=IMPDH1;refGene_tx=XM_023250537.1;refGene_gn=IMPDH1;refGene_
tx=XM_023250538.1;refGene_gn=IMPDH1;refGene_tx=XM_023250539.1;refGene_
gn=IMPDH1;refGene_tx=XM_023250540.1
A2 146685828 146685973
refGene_gn=IMPDH1;refGene_tx=XM_023250537.1;refGene_gn=IMPDH1;refGene_
tx=XM_023250538.1;refGene_gn=IMPDH1;refGene_tx=XM_023250539.1;refGene_
gn=IMPDH1;refGene_tx=XM_023250540.1
A2 146686079 146686223
refGene_gn=IMPDH1;refGene_tx=XM_023250537.1;refGene_gn=IMPDH1;refGene_
tx=XM_023250538.1;refGene_gn=IMPDH1;refGene_tx=XM_023250539.1;refGene_
gn=IMPDH1;refGene_tx=XM_023250540.1
A2 146687478 146687574
refGene_gn=IMPDH1;refGene_tx=XM_023250537.1;refGene_gn=IMPDH1;refGene_
tx=XM_023250538.1;refGene_gn=IMPDH1;refGene_tx=XM_023250539.1;refGene_
gn=IMPDH1;refGene_tx=XM_023250540.1
A2 146687822 146687913
refGene_gn=IMPDH1;refGene_tx=XM_023250537.1;refGene_gn=IMPDH1;refGene_
tx=XM_023250538.1;refGene_gn=IMPDH1;refGene_tx=XM_023250539.1;refGene_
gn=IMPDH1;refGene_tx=XM_023250540.1

A2	146688560	146688760
refGene_gn=IMPDH1;refGene_tx=XM_023250537.1;refGene_gn=IMPDH1;refGene_tx=XM_023250538.1;refGene_gn=IMPDH1;refGene_tx=XM_023250539.1;refGene_gn=IMPDH1;refGene_tx=XM_023250540.1
A2	146689558	146689646
refGene_gn=IMPDH1;refGene_tx=XM_023250537.1;refGene_gn=IMPDH1;refGene_tx=XM_023250538.1;refGene_gn=IMPDH1;refGene_tx=XM_023250539.1;refGene_gn=IMPDH1;refGene_tx=XM_023250540.1
A2	146689793	146690000
refGene_gn=IMPDH1;refGene_tx=XM_023250537.1;refGene_gn=IMPDH1;refGene_tx=XM_023250538.1;refGene_gn=IMPDH1;refGene_tx=XM_023250539.1;refGene_gn=IMPDH1;refGene_tx=XM_023250540.1
A2	146690275	146690350
refGene_gn=IMPDH1;refGene_tx=XM_023250537.1;refGene_gn=IMPDH1;refGene_tx=XM_023250538.1;refGene_gn=IMPDH1;refGene_tx=XM_023250539.1;refGene_gn=IMPDH1;refGene_tx=XM_023250540.1
A2	146690478	146690580
refGene_gn=IMPDH1;refGene_tx=XM_023250537.1;refGene_gn=IMPDH1;refGene_tx=XM_023250538.1;refGene_gn=IMPDH1;refGene_tx=XM_023250539.1;refGene_gn=IMPDH1;refGene_tx=XM_023250540.1
A2	146692570	146692619
refGene_gn=IMPDH1;refGene_tx=XM_023250537.1;refGene_gn=IMPDH1;refGene_tx=XM_023250538.1;refGene_gn=IMPDH1;refGene_tx=XM_023250539.1;refGene_gn=IMPDH1;refGene_tx=XM_023250540.1
A2	146694721	146694823
refGene_gn=IMPDH1;refGene_tx=XM_023250537.1;refGene_gn=IMPDH1;refGene_tx=XM_023250538.1;refGene_gn=IMPDH1;refGene_tx=XM_023250539.1;refGene_gn=IMPDH1;refGene_tx=XM_023250540.1
A2	146698965	146699135
refGene_gn=IMPDH1;refGene_tx=XM_023250537.1;refGene_gn=IMPDH1;refGene_tx=XM_023250538.1;refGene_gn=IMPDH1;refGene_tx=XM_023250539.1;refGene_gn=IMPDH1;refGene_tx=XM_023250540.1
A2	146736026	146736223
refGene_gn=HILPDA;refGene_tx=XM_003983035.5
A2	146737323	146737547
refGene_gn=HILPDA;refGene_tx=XM_003983035.5
A2	146737946	146737949
refGene_gn=FAM71F2;refGene_tx=XM_019825771.2
A2	146749811	146749964
refGene_gn=FAM71F2;refGene_tx=XM_003983013.5;refGene_gn=FAM71F2;refGene_tx=XM_006929354.4
A2	146752601	146752786
refGene_gn=FAM71F2;refGene_tx=XM_003983013.5;refGene_gn=FAM71F2;refGene_tx=XM_006929354.4;refGene_gn=FAM71F2;refGene_tx=XM_006929356.3;refGene_gn=FAM71F2;refGene_tx=XM_019825771.2
A2	146753616	146753866
refGene_gn=FAM71F2;refGene_tx=XM_003983013.5;refGene_gn=FAM71F2;refGene_tx=XM_006929354.4;refGene_gn=FAM71F2;refGene_tx=XM_006929356.3;refGene_gn=FAM71F2;refGene_tx=XM_019825771.2
A2	146760284	146760403

refGene_gn=FAM71F2;refGene_tx=XM_003983013.5;refGene_gn=FAM71F2;refGene_tx=XM_006929354.4;refGene_gn=FAM71F2;refGene_tx=XM_006929356.3;refGene_gn=FAM71F2;refGene_tx=XM_019825771.2
A2 146765168 146765364
refGene_gn=FAM71F2;refGene_tx=XM_003983013.5;refGene_gn=FAM71F2;refGene_tx=XM_006929354.4;refGene_gn=FAM71F2;refGene_tx=XM_006929356.3;refGene_gn=FAM71F2;refGene_tx=XM_019825771.2
A2 146780799 146780901
refGene_gn=FAM71F1;refGene_tx=XM_019825770.2
A2 146780925 146781174
refGene_gn=FAM71F1;refGene_tx=XM_006929357.3;refGene_gn=FAM71F1;refGene_tx=XM_006929360.3;refGene_gn=FAM71F1;refGene_tx=XM_019825768.2
A2 146782245 146782430
refGene_gn=FAM71F1;refGene_tx=XM_006929357.3;refGene_gn=FAM71F1;refGene_tx=XM_006929360.3;refGene_gn=FAM71F1;refGene_tx=XM_019825768.2;refGene_gn=FAM71F1;refGene_tx=XM_019825770.2
A2 146783770 146784023
refGene_gn=FAM71F1;refGene_tx=XM_006929357.3;refGene_gn=FAM71F1;refGene_tx=XM_006929360.3;refGene_gn=FAM71F1;refGene_tx=XM_019825768.2;refGene_gn=FAM71F1;refGene_tx=XM_019825770.2
A2 146788725 146788847
refGene_gn=FAM71F1;refGene_tx=XM_006929357.3;refGene_gn=FAM71F1;refGene_tx=XM_006929360.3;refGene_gn=FAM71F1;refGene_tx=XM_019825768.2;refGene_gn=FAM71F1;refGene_tx=XM_019825770.2
A2 146791499 146791543
refGene_gn=FAM71F1;refGene_tx=XM_006929357.3;refGene_gn=FAM71F1;refGene_tx=XM_019825768.2;refGene_gn=FAM71F1;refGene_tx=XM_019825770.2
A2 146794890 146795058
refGene_gn=FAM71F1;refGene_tx=XM_006929357.3;refGene_gn=FAM71F1;refGene_tx=XM_006929360.3;refGene_gn=FAM71F1;refGene_tx=XM_019825768.2;refGene_gn=FAM71F1;refGene_tx=XM_019825770.2
A2 146795815 146795829
refGene_gn=FAM71F1;refGene_tx=XM_006929357.3;refGene_gn=FAM71F1;refGene_tx=XM_019825768.2;refGene_gn=FAM71F1;refGene_tx=XM_019825770.2
A2 146814281 146814502
refGene_gn=CALU;refGene_tx=XM_003983016.5;refGene_gn=CALU;refGene_tx=XM_003983017.5
A2 146819889 146820083
refGene_gn=CALU;refGene_tx=XM_003983016.5
A2 146820156 146820350
refGene_gn=CALU;refGene_tx=XM_003983017.5
A2 146823775 146823942
refGene_gn=CALU;refGene_tx=XM_003983016.5;refGene_gn=CALU;refGene_tx=XM_003983017.5
A2 146825303 146825364
refGene_gn=CALU;refGene_tx=XM_003983016.5;refGene_gn=CALU;refGene_tx=XM_003983017.5
A2 146830461 146830661
refGene_gn=CALU;refGene_tx=XM_003983016.5;refGene_gn=CALU;refGene_tx=XM_003983017.5

A2 146832157 146832262
refGene_gn=CALU;refGene_tx=XM_003983016.5;refGene_gn=CALU;refGene_tx=X
M_003983017.5
A2 146835615 146835735
refGene_gn=OPN1SW;refGene_tx=XM_003983019.5
A2 146836669 146836909
refGene_gn=OPN1SW;refGene_tx=XM_003983019.5
A2 146837445 146837611
refGene_gn=OPN1SW;refGene_tx=XM_003983019.5
A2 146837957 146838126
refGene_gn=OPN1SW;refGene_tx=XM_003983019.5
A2 146838444 146838799
refGene_gn=OPN1SW;refGene_tx=XM_003983019.5
A2 146851698 146851774
refGene_gn=CCDC136;refGene_tx=XM_023250541.1;refGene_gn=CCDC136;refGen
e_tx=XM_023250542.1;refGene_gn=CCDC136;refGene_tx=XM_023250543.1;refGe
ne_gn=CCDC136;refGene_tx=XM_023250544.1;refGene_gn=CCDC136;refGene_tx=
XM_023250545.1;refGene_gn=CCDC136;refGene_tx=XM_023250546.1
A2 146852564 146852672
refGene_gn=CCDC136;refGene_tx=XM_023250541.1;refGene_gn=CCDC136;refGen
e_tx=XM_023250542.1;refGene_gn=CCDC136;refGene_tx=XM_023250543.1;refGe
ne_gn=CCDC136;refGene_tx=XM_023250544.1;refGene_gn=CCDC136;refGene_tx=
XM_023250545.1;refGene_gn=CCDC136;refGene_tx=XM_023250546.1
A2 146854512 146854767
refGene_gn=CCDC136;refGene_tx=XM_023250541.1;refGene_gn=CCDC136;refGen
e_tx=XM_023250542.1;refGene_gn=CCDC136;refGene_tx=XM_023250543.1;refGe
ne_gn=CCDC136;refGene_tx=XM_023250544.1;refGene_gn=CCDC136;refGene_tx=
XM_023250545.1;refGene_gn=CCDC136;refGene_tx=XM_023250546.1
A2 146854844 146854919
refGene_gn=CCDC136;refGene_tx=XM_023250541.1;refGene_gn=CCDC136;refGen
e_tx=XM_023250542.1;refGene_gn=CCDC136;refGene_tx=XM_023250543.1;refGe
ne_gn=CCDC136;refGene_tx=XM_023250544.1;refGene_gn=CCDC136;refGene_tx=
XM_023250545.1;refGene_gn=CCDC136;refGene_tx=XM_023250546.1
A2 146858192 146858208
refGene_gn=CCDC136;refGene_tx=XM_019825773.2
A2 146859146 146859470
refGene_gn=CCDC136;refGene_tx=XM_019825773.2;refGene_gn=CCDC136;refGen
e_tx=XM_023250541.1;refGene_gn=CCDC136;refGene_tx=XM_023250542.1;refGe
ne_gn=CCDC136;refGene_tx=XM_023250543.1;refGene_gn=CCDC136;refGene_tx=
XM_023250544.1;refGene_gn=CCDC136;refGene_tx=XM_023250545.1;refGene_gn
=CCDC136;refGene_tx=XM_023250546.1
A2 146862553 146862665
refGene_gn=CCDC136;refGene_tx=XM_019825773.2;refGene_gn=CCDC136;refGen
e_tx=XM_023250541.1;refGene_gn=CCDC136;refGene_tx=XM_023250542.1;refGe
ne_gn=CCDC136;refGene_tx=XM_023250543.1;refGene_gn=CCDC136;refGene_tx=
XM_023250544.1;refGene_gn=CCDC136;refGene_tx=XM_023250545.1;refGene_gn
=CCDC136;refGene_tx=XM_023250546.1
A2 146863229 146863398
refGene_gn=CCDC136;refGene_tx=XM_019825773.2;refGene_gn=CCDC136;refGen
e_tx=XM_023250541.1;refGene_gn=CCDC136;refGene_tx=XM_023250542.1;refGe

ne_gn=CCDC136;refGene_tx=XM_023250543.1;refGene_gn=CCDC136;refGene_tx=XM_023250544.1;refGene_gn=CCDC136;refGene_tx=XM_023250545.1;refGene_gn=CCDC136;refGene_tx=XM_023250546.1
A2 146863639 146863780
refGene_gn=CCDC136;refGene_tx=XM_019825773.2;refGene_gn=CCDC136;refGene_tx=XM_023250541.1;refGene_gn=CCDC136;refGene_tx=XM_023250542.1;refGene_gn=CCDC136;refGene_tx=XM_023250543.1;refGene_gn=CCDC136;refGene_tx=XM_023250544.1;refGene_gn=CCDC136;refGene_tx=XM_023250545.1;refGene_gn=CCDC136;refGene_tx=XM_023250546.1
A2 146864122 146864281
refGene_gn=CCDC136;refGene_tx=XM_019825773.2;refGene_gn=CCDC136;refGene_tx=XM_023250541.1;refGene_gn=CCDC136;refGene_tx=XM_023250542.1;refGene_gn=CCDC136;refGene_tx=XM_023250543.1;refGene_gn=CCDC136;refGene_tx=XM_023250544.1;refGene_gn=CCDC136;refGene_tx=XM_023250545.1;refGene_gn=CCDC136;refGene_tx=XM_023250546.1
A2 146864544 146864715
refGene_gn=CCDC136;refGene_tx=XM_019825773.2;refGene_gn=CCDC136;refGene_tx=XM_023250541.1;refGene_gn=CCDC136;refGene_tx=XM_023250542.1;refGene_gn=CCDC136;refGene_tx=XM_023250543.1;refGene_gn=CCDC136;refGene_tx=XM_023250544.1;refGene_gn=CCDC136;refGene_tx=XM_023250545.1;refGene_gn=CCDC136;refGene_tx=XM_023250546.1
A2 146865330 146865516
refGene_gn=CCDC136;refGene_tx=XM_019825773.2;refGene_gn=CCDC136;refGene_tx=XM_023250541.1;refGene_gn=CCDC136;refGene_tx=XM_023250542.1;refGene_gn=CCDC136;refGene_tx=XM_023250543.1;refGene_gn=CCDC136;refGene_tx=XM_023250544.1;refGene_gn=CCDC136;refGene_tx=XM_023250545.1
A2 146867153 146867348
refGene_gn=CCDC136;refGene_tx=XM_019825773.2;refGene_gn=CCDC136;refGene_tx=XM_023250541.1;refGene_gn=CCDC136;refGene_tx=XM_023250542.1;refGene_gn=CCDC136;refGene_tx=XM_023250543.1;refGene_gn=CCDC136;refGene_tx=XM_023250544.1
A2 146867837 146868065
refGene_gn=CCDC136;refGene_tx=XM_019825773.2;refGene_gn=CCDC136;refGene_tx=XM_023250541.1;refGene_gn=CCDC136;refGene_tx=XM_023250542.1;refGene_gn=CCDC136;refGene_tx=XM_023250543.1;refGene_gn=CCDC136;refGene_tx=XM_023250544.1
A2 146869511 146870021
refGene_gn=CCDC136;refGene_tx=XM_019825773.2;refGene_gn=CCDC136;refGene_tx=XM_023250541.1;refGene_gn=CCDC136;refGene_tx=XM_023250542.1;refGene_gn=CCDC136;refGene_tx=XM_023250543.1
A2 146870358 146870580
refGene_gn=CCDC136;refGene_tx=XM_019825773.2;refGene_gn=CCDC136;refGene_tx=XM_023250541.1;refGene_gn=CCDC136;refGene_tx=XM_023250542.1;refGene_gn=CCDC136;refGene_tx=XM_023250543.1
A2 146871649 146871928
refGene_gn=CCDC136;refGene_tx=XM_019825773.2;refGene_gn=CCDC136;refGene_tx=XM_023250541.1;refGene_gn=CCDC136;refGene_tx=XM_023250542.1;refGene_gn=CCDC136;refGene_tx=XM_023250543.1
A2 146872633 146872933
refGene_gn=CCDC136;refGene_tx=XM_019825773.2;refGene_gn=CCDC136;refGen

e_tx=XM_023250541.1;refGene_gn=CCDC136;refGene_tx=XM_023250542.1;refGene_gn=CCDC136;refGene_tx=XM_023250543.1
A2	146874768	146874870
refGene_gn=CCDC136;refGene_tx=XM_023250543.1;refGene_gn=CCDC136;refGene_tx=XM_023250544.1;refGene_gn=CCDC136;refGene_tx=XM_023250545.1;refGene_gn=CCDC136;refGene_tx=XM_023250546.1
A2	146879198	146879309
refGene_gn=CCDC136;refGene_tx=XM_023250542.1
A2	146879643	146879796
refGene_gn=CCDC136;refGene_tx=XM_019825773.2;refGene_gn=CCDC136;refGene_tx=XM_023250541.1
A2	146885673	146886028
refGene_gn=FLNC;refGene_tx=XM_003983022.5
A2	146890363	146890612
refGene_gn=FLNC;refGene_tx=XM_003983022.5
A2	146892168	146892266
refGene_gn=FLNC;refGene_tx=XM_003983022.5
A2	146892414	146892565
refGene_gn=FLNC;refGene_tx=XM_003983022.5
A2	146892660	146892779
refGene_gn=FLNC;refGene_tx=XM_003983022.5
A2	146892987	146893065
refGene_gn=FLNC;refGene_tx=XM_003983022.5
A2	146893260	146893423
refGene_gn=FLNC;refGene_tx=XM_003983022.5
A2	146893593	146893794
refGene_gn=FLNC;refGene_tx=XM_003983022.5
A2	146895004	146895142
refGene_gn=FLNC;refGene_tx=XM_003983022.5
A2	146895539	146895666
refGene_gn=FLNC;refGene_tx=XM_003983022.5
A2	146895809	146895946
refGene_gn=FLNC;refGene_tx=XM_003983022.5
A2	146896155	146896349
refGene_gn=FLNC;refGene_tx=XM_003983022.5
A2	146896442	146896512
refGene_gn=FLNC;refGene_tx=XM_003983022.5
A2	146896514	146896555
refGene_gn=FLNC;refGene_tx=XM_003983022.5
A2	146897157	146897301
refGene_gn=FLNC;refGene_tx=XM_003983022.5
A2	146897520	146897644
refGene_gn=FLNC;refGene_tx=XM_003983022.5
A2	146897779	146897940
refGene_gn=FLNC;refGene_tx=XM_003983022.5
A2	146898146	146898237
refGene_gn=FLNC;refGene_tx=XM_003983022.5
A2	146898326	146898496
refGene_gn=FLNC;refGene_tx=XM_003983022.5
A2	146898716	146898834

refGene_gn=FLNC;refGene_tx=XM_003983022.5
A2	146898923	146899186
refGene_gn=FLNC;refGene_tx=XM_003983022.5
A2	146899577	146900175
refGene_gn=FLNC;refGene_tx=XM_003983022.5
A2	146900837	146901011
refGene_gn=FLNC;refGene_tx=XM_003983022.5
A2	146901118	146901281
refGene_gn=FLNC;refGene_tx=XM_003983022.5
A2	146901561	146901722
refGene_gn=FLNC;refGene_tx=XM_003983022.5
A2	146902391	146902559
refGene_gn=FLNC;refGene_tx=XM_003983022.5
A2	146902638	146902762
refGene_gn=FLNC;refGene_tx=XM_003983022.5
A2	146903198	146903355
refGene_gn=FLNC;refGene_tx=XM_003983022.5
A2	146903430	146903620
refGene_gn=FLNC;refGene_tx=XM_003983022.5
A2	146903805	146903829
refGene_gn=FLNC;refGene_tx=XM_003983022.5
A2	146903951	146904199
refGene_gn=FLNC;refGene_tx=XM_003983022.5
A2	146904645	146904726
refGene_gn=FLNC;refGene_tx=XM_003983022.5
A2	146905027	146905127
refGene_gn=FLNC;refGene_tx=XM_003983022.5
A2	146905376	146905517
refGene_gn=FLNC;refGene_tx=XM_003983022.5
A2	146905786	146905915
refGene_gn=FLNC;refGene_tx=XM_003983022.5
A2	146906007	146906181
refGene_gn=FLNC;refGene_tx=XM_003983022.5
A2	146906795	146906957
refGene_gn=FLNC;refGene_tx=XM_003983022.5
A2	146907070	146907274
refGene_gn=FLNC;refGene_tx=XM_003983022.5
A2	146907692	146907845
refGene_gn=FLNC;refGene_tx=XM_003983022.5
A2	146907930	146908053
refGene_gn=FLNC;refGene_tx=XM_003983022.5
A2	146908166	146908409
refGene_gn=FLNC;refGene_tx=XM_003983022.5
A2	146908601	146908871
refGene_gn=FLNC;refGene_tx=XM_003983022.5
A2	146908973	146909111
refGene_gn=FLNC;refGene_tx=XM_003983022.5
A2	146909338	146909454
refGene_gn=FLNC;refGene_tx=XM_003983022.5
A2	146910026	146910159

refGene_gn=FLNC;refGene_tx=XM_003983022.5
A2 146910253 146910430
refGene_gn=FLNC;refGene_tx=XM_003983022.5
A2 146910580 146910799
refGene_gn=FLNC;refGene_tx=XM_003983022.5
A2 146911412 146911622
refGene_gn=FLNC;refGene_tx=XM_003983022.5
A2 146911746 146911934
refGene_gn=FLNC;refGene_tx=XM_003983022.5
A2 146917034 146917192
refGene_gn=ATP6V1F;refGene_tx=XM_003983025.5
A2 146919660 146919862
refGene_gn=ATP6V1F;refGene_tx=XM_003983025.5
A2 146920547 146921115
refGene_gn=LOC102900188;refGene_tx=XM_006929363.2
A2 146922679 146922789
refGene_gn=LOC102900188;refGene_tx=XM_006929363.2
A2 146923035 146923077
refGene_gn=LOC111559540;refGene_tx=XM_023250548.1
A2 146923542 146923632
refGene_gn=LOC111559540;refGene_tx=XM_023250548.1
A2 146923733 146924879
refGene_gn=LOC111559540;refGene_tx=XM_023250548.1
A2 146925188 146925250
refGene_gn=LOC111559540;refGene_tx=XM_023250548.1
A2 146926830 146926934
refGene_gn=LOC111559540;refGene_tx=XM_023250548.1
A2 146927240 146927308
refGene_gn=LOC111559540;refGene_tx=XM_023250548.1
A2 146928373 146928535
refGene_gn=LOC111559540;refGene_tx=XM_023250548.1
A2 146928611 146928641
refGene_gn=LOC111559540;refGene_tx=XM_023250548.1
A2 146930771 146931022
refGene_gn=KCP;refGene_tx=XM_019825774.2
A2 146931175 146931482
refGene_gn=KCP;refGene_tx=XM_019825774.2
A2 146931914 146932079
refGene_gn=KCP;refGene_tx=XM_019825774.2
A2 146932858 146932960
refGene_gn=KCP;refGene_tx=XM_019825774.2
A2 146933046 146933158
refGene_gn=KCP;refGene_tx=XM_019825774.2
A2 146933239 146933412
refGene_gn=KCP;refGene_tx=XM_019825774.2
A2 146933851 146933994
refGene_gn=KCP;refGene_tx=XM_019825774.2
A2 146934082 146934185
refGene_gn=KCP;refGene_tx=XM_019825774.2
A2 146934528 146934617

refGene_gn=KCP;refGene_tx=XM_019825774.2
A2 146934757 146934802
refGene_gn=KCP;refGene_tx=XM_019825774.2
A2 146935114 146935214
refGene_gn=KCP;refGene_tx=XM_019825774.2
A2 146935412 146935492
refGene_gn=KCP;refGene_tx=XM_019825774.2
A2 146936430 146936551
refGene_gn=KCP;refGene_tx=XM_019825774.2
A2 146937125 146937208
refGene_gn=KCP;refGene_tx=XM_019825774.2
A2 146937395 146937569
refGene_gn=KCP;refGene_tx=XM_019825774.2
A2 146938537 146938631
refGene_gn=KCP;refGene_tx=XM_019825774.2
A2 146938738 146938821
refGene_gn=KCP;refGene_tx=XM_019825774.2
A2 146938953 146939044
refGene_gn=KCP;refGene_tx=XM_019825774.2
A2 146939304 146939390
refGene_gn=KCP;refGene_tx=XM_019825774.2
A2 146940009 146940186
refGene_gn=KCP;refGene_tx=XM_019825774.2
A2 146941923 146942094
refGene_gn=KCP;refGene_tx=XM_019825774.2
A2 146942392 146942581
refGene_gn=KCP;refGene_tx=XM_019825774.2
A2 146942681 146942775
refGene_gn=KCP;refGene_tx=XM_019825774.2
A2 146942927 146943010
refGene_gn=KCP;refGene_tx=XM_019825774.2
A2 146943145 146943319
refGene_gn=KCP;refGene_tx=XM_019825774.2
A2 146943985 146944079
refGene_gn=KCP;refGene_tx=XM_019825774.2
A2 146944156 146944263
refGene_gn=KCP;refGene_tx=XM_019825774.2
A2 146944352 146944505
refGene_gn=KCP;refGene_tx=XM_019825774.2
A2 146944739 146944821
refGene_gn=KCP;refGene_tx=XM_019825774.2
A2 146944895 146944981
refGene_gn=KCP;refGene_tx=XM_019825774.2
A2 146945314 146945408
refGene_gn=KCP;refGene_tx=XM_019825774.2
A2 146945485 146945565
refGene_gn=KCP;refGene_tx=XM_019825774.2
A2 146945671 146945765
refGene_gn=KCP;refGene_tx=XM_019825774.2
A2 146948480 146948563

refGene_gn=KCP;refGene_tx=XM_019825774.2
A2 146949424 146949518
refGene_gn=KCP;refGene_tx=XM_019825774.2
A2 146949746 146949829
refGene_gn=KCP;refGene_tx=XM_019825774.2
A2 146949971 146950068
refGene_gn=KCP;refGene_tx=XM_019825774.2
A2 146950265 146950333
refGene_gn=KCP;refGene_tx=XM_019825774.2
A2 146951920 146952005
refGene_gn=KCP;refGene_tx=XM_019825774.2
A2 146952702 146952779
refGene_gn=KCP;refGene_tx=XM_019825774.2
A2 146952868 146953058
refGene_gn=KCP;refGene_tx=XM_019825774.2
A2 146953890 146954057
refGene_gn=KCP;refGene_tx=XM_019825774.2
A2 146956577 146956656
refGene_gn=KCP;refGene_tx=XM_019825774.2
A2 146978822 146979017
refGene_gn=IRF5;refGene_tx=XM_006929365.4;refGene_gn=IRF5;refGene_tx=XM_023250551.1
A2 146981092 146981282
refGene_gn=IRF5;refGene_tx=XM_006929365.4;refGene_gn=IRF5;refGene_tx=XM_023250551.1
A2 146981696 146981761
refGene_gn=IRF5;refGene_tx=XM_006929365.4;refGene_gn=IRF5;refGene_tx=XM_023250551.1
A2 146982263 146982297
refGene_gn=IRF5;refGene_tx=XM_006929365.4;refGene_gn=IRF5;refGene_tx=XM_023250551.1
A2 146982464 146982734
refGene_gn=IRF5;refGene_tx=XM_006929365.4;refGene_gn=IRF5;refGene_tx=XM_023250551.1
A2 146983000 146983393
refGene_gn=IRF5;refGene_tx=XM_006929365.4;refGene_gn=IRF5;refGene_tx=XM_023250551.1
A2 146983478 146983597
refGene_gn=IRF5;refGene_tx=XM_006929365.4;refGene_gn=IRF5;refGene_tx=XM_023250551.1
A2 146983869 146984118
refGene_gn=IRF5;refGene_tx=XM_006929365.4;refGene_gn=IRF5;refGene_tx=XM_023250551.1
A2 146988444 146988505
refGene_gn=TNPO3;refGene_tx=XM_023250549.1;refGene_gn=TNPO3;refGene_tx=XM_023250550.1
A2 146994768 146994881
refGene_gn=TNPO3;refGene_tx=XM_023250549.1;refGene_gn=TNPO3;refGene_tx=XM_023250550.1
A2 146998543 146998711

refGene_gn=TNPO3;refGene_tx=XM_023250549.1;refGene_gn=TNPO3;refGene_tx=XM_023250550.1
A2 146999549 146999706
refGene_gn=TNPO3;refGene_tx=XM_023250549.1;refGene_gn=TNPO3;refGene_tx=XM_023250550.1
A2 147001265 147001360
refGene_gn=TNPO3;refGene_tx=XM_023250549.1;refGene_gn=TNPO3;refGene_tx=XM_023250550.1
A2 147002453 147002570
refGene_gn=TNPO3;refGene_tx=XM_023250549.1;refGene_gn=TNPO3;refGene_tx=XM_023250550.1
A2 147005453 147005594
refGene_gn=TNPO3;refGene_tx=XM_023250549.1;refGene_gn=TNPO3;refGene_tx=XM_023250550.1
A2 147006412 147006473
refGene_gn=TNPO3;refGene_tx=XM_023250549.1;refGene_gn=TNPO3;refGene_tx=XM_023250550.1
A2 147008165 147008242
refGene_gn=TNPO3;refGene_tx=XM_023250549.1;refGene_gn=TNPO3;refGene_tx=XM_023250550.1
A2 147008957 147009049
refGene_gn=TNPO3;refGene_tx=XM_023250549.1;refGene_gn=TNPO3;refGene_tx=XM_023250550.1
A2 147011505 147011697
refGene_gn=TNPO3;refGene_tx=XM_023250549.1;refGene_gn=TNPO3;refGene_tx=XM_023250550.1
A2 147015136 147015276
refGene_gn=TNPO3;refGene_tx=XM_023250549.1;refGene_gn=TNPO3;refGene_tx=XM_023250550.1
A2 147017212 147017304
refGene_gn=TNPO3;refGene_tx=XM_023250549.1;refGene_gn=TNPO3;refGene_tx=XM_023250550.1
A2 147019043 147019151
refGene_gn=TNPO3;refGene_tx=XM_023250549.1;refGene_gn=TNPO3;refGene_tx=XM_023250550.1
A2 147021367 147021514
refGene_gn=TNPO3;refGene_tx=XM_023250549.1;refGene_gn=TNPO3;refGene_tx=XM_023250550.1
A2 147023163 147023302
refGene_gn=TNPO3;refGene_tx=XM_023250549.1;refGene_gn=TNPO3;refGene_tx=XM_023250550.1
A2 147023767 147023943
refGene_gn=TNPO3;refGene_tx=XM_023250549.1;refGene_gn=TNPO3;refGene_tx=XM_023250550.1
A2 147026372 147026516
refGene_gn=TNPO3;refGene_tx=XM_023250549.1;refGene_gn=TNPO3;refGene_tx=XM_023250550.1
A2 147031651 147031808
refGene_gn=TNPO3;refGene_tx=XM_023250549.1;refGene_gn=TNPO3;refGene_tx=XM_023250550.1

A2 147033277 147033351
refGene_gn=TNPO3;refGene_tx=XM_023250549.1;refGene_gn=TNPO3;refGene_tx=XM_023250550.1
A2 147034317 147034518
refGene_gn=TNPO3;refGene_tx=XM_023250549.1;refGene_gn=TNPO3;refGene_tx=XM_023250550.1
A2 147063774 147063894
refGene_gn=TNPO3;refGene_tx=XM_023250549.1
A2 147137889 147137991
refGene_gn=TSPAN33;refGene_tx=XM_003983031.4;refGene_gn=TSPAN33;refGene_tx=XM_023250552.1
A2 147152004 147152062
refGene_gn=TSPAN33;refGene_tx=XM_003983031.4;refGene_gn=TSPAN33;refGene_tx=XM_023250552.1
A2 147152713 147152841
refGene_gn=TSPAN33;refGene_tx=XM_003983031.4;refGene_gn=TSPAN33;refGene_tx=XM_023250552.1
A2 147153175 147153250
refGene_gn=TSPAN33;refGene_tx=XM_003983031.4;refGene_gn=TSPAN33;refGene_tx=XM_023250552.1
A2 147154300 147154396
refGene_gn=TSPAN33;refGene_tx=XM_003983031.4;refGene_gn=TSPAN33;refGene_tx=XM_023250552.1
A2 147156202 147156331
refGene_gn=TSPAN33;refGene_tx=XM_003983031.4;refGene_gn=TSPAN33;refGene_tx=XM_023250552.1
A2 147156733 147156895
refGene_gn=TSPAN33;refGene_tx=XM_003983031.4;refGene_gn=TSPAN33;refGene_tx=XM_023250552.1
A2 147157105 147157207
refGene_gn=TSPAN33;refGene_tx=XM_003983031.4;refGene_gn=TSPAN33;refGene_tx=XM_023250552.1
A2 147173411 147173745
refGene_gn=SMO;refGene_tx=XM_011280558.2
A2 147186176 147186382
refGene_gn=SMO;refGene_tx=XM_011280558.2
A2 147187128 147187338
refGene_gn=SMO;refGene_tx=XM_011280558.2
A2 147187586 147187759
refGene_gn=SMO;refGene_tx=XM_011280558.2
A2 147188091 147188311
refGene_gn=SMO;refGene_tx=XM_011280558.2
A2 147188417 147188541
refGene_gn=SMO;refGene_tx=XM_011280558.2
A2 147189708 147189801
refGene_gn=SMO;refGene_tx=XM_011280558.2
A2 147190253 147190362
refGene_gn=SMO;refGene_tx=XM_011280558.2
A2 147191183 147191369
refGene_gn=SMO;refGene_tx=XM_011280558.2

A2	147191828	147191977
refGene_gn=SMO;refGene_tx=XM_011280558.2
A2	147192438	147192573
refGene_gn=SMO;refGene_tx=XM_011280558.2
A2	147192811	147193254
refGene_gn=SMO;refGene_tx=XM_011280558.2
A2	147203695	147204058
refGene_gn=AHCYL2;refGene_tx=XM_023250553.1;refGene_gn=AHCYL2;refGene_
tx=XM_023250554.1
A2	147317919	147317976
refGene_gn=AHCYL2;refGene_tx=XM_006929367.3;refGene_gn=AHCYL2;refGene_
tx=XM_019825777.2
A2	147323822	147323879
refGene_gn=AHCYL2;refGene_tx=XM_019825775.2;refGene_gn=AHCYL2;refGene_
tx=XM_019825776.2
A2	147326949	147327061
refGene_gn=AHCYL2;refGene_tx=XM_006929367.3;refGene_gn=AHCYL2;refGene_
tx=XM_019825775.2;refGene_gn=AHCYL2;refGene_tx=XM_019825776.2;refGene_
gn=AHCYL2;refGene_tx=XM_019825777.2;refGene_gn=AHCYL2;refGene_tx=XM_02
3250553.1;refGene_gn=AHCYL2;refGene_tx=XM_023250554.1
A2	147335452	147335596
refGene_gn=AHCYL2;refGene_tx=XM_006929367.3;refGene_gn=AHCYL2;refGene_
tx=XM_019825775.2;refGene_gn=AHCYL2;refGene_tx=XM_019825776.2;refGene_
gn=AHCYL2;refGene_tx=XM_019825777.2;refGene_gn=AHCYL2;refGene_tx=XM_02
3250553.1;refGene_gn=AHCYL2;refGene_tx=XM_023250554.1
A2	147336255	147336356
refGene_gn=AHCYL2;refGene_tx=XM_006929367.3;refGene_gn=AHCYL2;refGene_
tx=XM_019825775.2;refGene_gn=AHCYL2;refGene_tx=XM_019825776.2;refGene_
gn=AHCYL2;refGene_tx=XM_019825777.2;refGene_gn=AHCYL2;refGene_tx=XM_01
9825780.2;refGene_gn=AHCYL2;refGene_tx=XM_019825781.2;refGene_gn=AHCYL
2;refGene_tx=XM_023250553.1;refGene_gn=AHCYL2;refGene_tx=XM_023250554.
1
A2	147343669	147343772
refGene_gn=AHCYL2;refGene_tx=XM_006929367.3;refGene_gn=AHCYL2;refGene_
tx=XM_019825775.2;refGene_gn=AHCYL2;refGene_tx=XM_019825776.2;refGene_
gn=AHCYL2;refGene_tx=XM_019825777.2;refGene_gn=AHCYL2;refGene_tx=XM_01
9825780.2;refGene_gn=AHCYL2;refGene_tx=XM_019825781.2;refGene_gn=AHCYL
2;refGene_tx=XM_023250553.1;refGene_gn=AHCYL2;refGene_tx=XM_023250554.
1
A2	147345224	147345319
refGene_gn=AHCYL2;refGene_tx=XM_006929367.3;refGene_gn=AHCYL2;refGene_
tx=XM_019825775.2;refGene_gn=AHCYL2;refGene_tx=XM_019825776.2;refGene_
gn=AHCYL2;refGene_tx=XM_019825777.2;refGene_gn=AHCYL2;refGene_tx=XM_01
9825780.2;refGene_gn=AHCYL2;refGene_tx=XM_019825781.2;refGene_gn=AHCYL
2;refGene_tx=XM_023250553.1;refGene_gn=AHCYL2;refGene_tx=XM_023250554.
1
A2	147348107	147348214
refGene_gn=AHCYL2;refGene_tx=XM_006929367.3;refGene_gn=AHCYL2;refGene_
tx=XM_019825775.2;refGene_gn=AHCYL2;refGene_tx=XM_019825776.2;refGene_
gn=AHCYL2;refGene_tx=XM_019825777.2;refGene_gn=AHCYL2;refGene_tx=XM_01

9825780.2;refGene_gn=AHCYL2;refGene_tx=XM_019825781.2;refGene_gn=AHCYL2;refGene_tx=XM_023250553.1;refGene_gn=AHCYL2;refGene_tx=XM_023250554.1
A2 147349748 147349865
refGene_gn=AHCYL2;refGene_tx=XM_006929367.3;refGene_gn=AHCYL2;refGene_tx=XM_019825775.2;refGene_gn=AHCYL2;refGene_tx=XM_019825776.2;refGene_gn=AHCYL2;refGene_tx=XM_019825777.2;refGene_gn=AHCYL2;refGene_tx=XM_019825780.2;refGene_gn=AHCYL2;refGene_tx=XM_019825781.2;refGene_gn=AHCYL2;refGene_tx=XM_023250553.1;refGene_gn=AHCYL2;refGene_tx=XM_023250554.1
A2 147353012 147353076
refGene_gn=AHCYL2;refGene_tx=XM_006929367.3;refGene_gn=AHCYL2;refGene_tx=XM_019825775.2;refGene_gn=AHCYL2;refGene_tx=XM_019825776.2;refGene_gn=AHCYL2;refGene_tx=XM_019825777.2;refGene_gn=AHCYL2;refGene_tx=XM_019825780.2;refGene_gn=AHCYL2;refGene_tx=XM_019825781.2;refGene_gn=AHCYL2;refGene_tx=XM_023250553.1;refGene_gn=AHCYL2;refGene_tx=XM_023250554.1
A2 147353561 147353650
refGene_gn=AHCYL2;refGene_tx=XM_006929367.3;refGene_gn=AHCYL2;refGene_tx=XM_019825775.2;refGene_gn=AHCYL2;refGene_tx=XM_019825776.2;refGene_gn=AHCYL2;refGene_tx=XM_019825777.2;refGene_gn=AHCYL2;refGene_tx=XM_019825780.2;refGene_gn=AHCYL2;refGene_tx=XM_019825781.2;refGene_gn=AHCYL2;refGene_tx=XM_023250553.1;refGene_gn=AHCYL2;refGene_tx=XM_023250554.1
A2 147356543 147356614
refGene_gn=AHCYL2;refGene_tx=XM_006929367.3;refGene_gn=AHCYL2;refGene_tx=XM_019825775.2;refGene_gn=AHCYL2;refGene_tx=XM_019825776.2;refGene_gn=AHCYL2;refGene_tx=XM_019825777.2;refGene_gn=AHCYL2;refGene_tx=XM_019825780.2;refGene_gn=AHCYL2;refGene_tx=XM_019825781.2;refGene_gn=AHCYL2;refGene_tx=XM_023250553.1;refGene_gn=AHCYL2;refGene_tx=XM_023250554.1
A2 147359928 147360023
refGene_gn=AHCYL2;refGene_tx=XM_006929367.3;refGene_gn=AHCYL2;refGene_tx=XM_019825775.2;refGene_gn=AHCYL2;refGene_tx=XM_019825776.2;refGene_gn=AHCYL2;refGene_tx=XM_019825777.2;refGene_gn=AHCYL2;refGene_tx=XM_019825780.2;refGene_gn=AHCYL2;refGene_tx=XM_019825781.2;refGene_gn=AHCYL2;refGene_tx=XM_023250553.1;refGene_gn=AHCYL2;refGene_tx=XM_023250554.1
A2 147366620 147366719
refGene_gn=AHCYL2;refGene_tx=XM_006929367.3;refGene_gn=AHCYL2;refGene_tx=XM_019825775.2;refGene_gn=AHCYL2;refGene_tx=XM_019825776.2;refGene_gn=AHCYL2;refGene_tx=XM_019825777.2;refGene_gn=AHCYL2;refGene_tx=XM_019825780.2;refGene_gn=AHCYL2;refGene_tx=XM_019825781.2;refGene_gn=AHCYL2;refGene_tx=XM_023250553.1;refGene_gn=AHCYL2;refGene_tx=XM_023250554.1
A2 147368391 147368460
refGene_gn=AHCYL2;refGene_tx=XM_006929367.3;refGene_gn=AHCYL2;refGene_tx=XM_019825775.2;refGene_gn=AHCYL2;refGene_tx=XM_019825776.2;refGene_gn=AHCYL2;refGene_tx=XM_019825777.2;refGene_gn=AHCYL2;refGene_tx=XM_019825780.2;refGene_gn=AHCYL2;refGene_tx=XM_019825781.2;refGene_gn=AHCYL

2;refGene_tx=XM_023250553.1;refGene_gn=AHCYL2;refGene_tx=XM_023250554.
1
A2 147368562 147368641
refGene_gn=AHCYL2;refGene_tx=XM_006929367.3;refGene_gn=AHCYL2;refGene_
tx=XM_019825775.2;refGene_gn=AHCYL2;refGene_tx=XM_019825776.2;refGene_
gn=AHCYL2;refGene_tx=XM_019825777.2;refGene_gn=AHCYL2;refGene_tx=XM_01
9825780.2;refGene_gn=AHCYL2;refGene_tx=XM_019825781.2;refGene_gn=AHCYL
2;refGene_tx=XM_023250553.1;refGene_gn=AHCYL2;refGene_tx=XM_023250554.
1
A2 147369898 147370019
refGene_gn=AHCYL2;refGene_tx=XM_006929367.3;refGene_gn=AHCYL2;refGene_
tx=XM_019825775.2;refGene_gn=AHCYL2;refGene_tx=XM_019825776.2;refGene_
gn=AHCYL2;refGene_tx=XM_019825777.2;refGene_gn=AHCYL2;refGene_tx=XM_01
9825780.2;refGene_gn=AHCYL2;refGene_tx=XM_019825781.2;refGene_gn=AHCYL
2;refGene_tx=XM_023250553.1;refGene_gn=AHCYL2;refGene_tx=XM_023250554.
1
A2 147370425 147370432
refGene_gn=AHCYL2;refGene_tx=XM_006929367.3;refGene_gn=AHCYL2;refGene_
tx=XM_019825775.2;refGene_gn=AHCYL2;refGene_tx=XM_019825776.2;refGene_
gn=AHCYL2;refGene_tx=XM_019825777.2;refGene_gn=AHCYL2;refGene_tx=XM_01
9825780.2;refGene_gn=AHCYL2;refGene_tx=XM_019825781.2;refGene_gn=AHCYL
2;refGene_tx=XM_023250553.1;refGene_gn=AHCYL2;refGene_tx=XM_023250554.
1
A2 147378094 147378223
refGene_gn=STRIP2;refGene_tx=XM_003983039.5;refGene_gn=STRIP2;refGene_
tx=XM_019825782.2;refGene_gn=STRIP2;refGene_tx=XM_019825783.2
A2 147384120 147384190
refGene_gn=STRIP2;refGene_tx=XM_003983039.5;refGene_gn=STRIP2;refGene_
tx=XM_019825782.2;refGene_gn=STRIP2;refGene_tx=XM_019825783.2
A2 147387167 147387242
refGene_gn=STRIP2;refGene_tx=XM_003983039.5;refGene_gn=STRIP2;refGene_
tx=XM_019825782.2;refGene_gn=STRIP2;refGene_tx=XM_019825783.2
A2 147387683 147387822
refGene_gn=STRIP2;refGene_tx=XM_019825782.2;refGene_gn=STRIP2;refGene_
tx=XM_019825783.2
A2 147388227 147388362
refGene_gn=STRIP2;refGene_tx=XM_003983039.5;refGene_gn=STRIP2;refGene_
tx=XM_019825782.2;refGene_gn=STRIP2;refGene_tx=XM_019825783.2
A2 147389715 147389836
refGene_gn=STRIP2;refGene_tx=XM_003983039.5;refGene_gn=STRIP2;refGene_
tx=XM_019825782.2;refGene_gn=STRIP2;refGene_tx=XM_019825783.2
A2 147390601 147390670
refGene_gn=STRIP2;refGene_tx=XM_003983039.5;refGene_gn=STRIP2;refGene_
tx=XM_019825782.2;refGene_gn=STRIP2;refGene_tx=XM_019825783.2
A2 147390869 147390976
refGene_gn=STRIP2;refGene_tx=XM_003983039.5;refGene_gn=STRIP2;refGene_
tx=XM_019825782.2;refGene_gn=STRIP2;refGene_tx=XM_019825783.2
A2 147392401 147392529
refGene_gn=STRIP2;refGene_tx=XM_003983039.5;refGene_gn=STRIP2;refGene_
tx=XM_019825782.2;refGene_gn=STRIP2;refGene_tx=XM_019825783.2

A2 147393421 147393625
refGene_gn=STRIP2;refGene_tx=XM_003983039.5;refGene_gn=STRIP2;refGene_tx=XM_019825782.2;refGene_gn=STRIP2;refGene_tx=XM_019825783.2
A2 147394926 147395162
refGene_gn=STRIP2;refGene_tx=XM_003983039.5;refGene_gn=STRIP2;refGene_tx=XM_019825782.2;refGene_gn=STRIP2;refGene_tx=XM_019825783.2
A2 147395422 147395488
refGene_gn=STRIP2;refGene_tx=XM_003983039.5;refGene_gn=STRIP2;refGene_tx=XM_019825782.2;refGene_gn=STRIP2;refGene_tx=XM_019825783.2
A2 147396188 147396252
refGene_gn=STRIP2;refGene_tx=XM_003983039.5;refGene_gn=STRIP2;refGene_tx=XM_019825782.2;refGene_gn=STRIP2;refGene_tx=XM_019825783.2
A2 147397205 147397277
refGene_gn=STRIP2;refGene_tx=XM_003983039.5;refGene_gn=STRIP2;refGene_tx=XM_019825782.2;refGene_gn=STRIP2;refGene_tx=XM_019825783.2
A2 147400108 147400183
refGene_gn=STRIP2;refGene_tx=XM_003983039.5;refGene_gn=STRIP2;refGene_tx=XM_019825782.2;refGene_gn=STRIP2;refGene_tx=XM_019825783.2
A2 147400876 147400974
refGene_gn=STRIP2;refGene_tx=XM_003983039.5;refGene_gn=STRIP2;refGene_tx=XM_019825782.2;refGene_gn=STRIP2;refGene_tx=XM_019825783.2
A2 147401445 147401572
refGene_gn=STRIP2;refGene_tx=XM_003983039.5;refGene_gn=STRIP2;refGene_tx=XM_019825782.2;refGene_gn=STRIP2;refGene_tx=XM_019825783.2
A2 147404145 147404246
refGene_gn=STRIP2;refGene_tx=XM_003983039.5;refGene_gn=STRIP2;refGene_tx=XM_019825782.2;refGene_gn=STRIP2;refGene_tx=XM_019825783.2
A2 147406956 147407023
refGene_gn=STRIP2;refGene_tx=XM_003983039.5;refGene_gn=STRIP2;refGene_tx=XM_019825782.2;refGene_gn=STRIP2;refGene_tx=XM_019825783.2
A2 147407093 147407132
refGene_gn=STRIP2;refGene_tx=XM_019825783.2
A2 147422135 147422240
refGene_gn=STRIP2;refGene_tx=XM_003983039.5;refGene_gn=STRIP2;refGene_tx=XM_019825782.2
A2 147424211 147424416
refGene_gn=STRIP2;refGene_tx=XM_003983039.5;refGene_gn=STRIP2;refGene_tx=XM_019825782.2
A2 147427863 147428114
refGene_gn=STRIP2;refGene_tx=XM_003983039.5;refGene_gn=STRIP2;refGene_tx=XM_019825782.2
A2 147438219 147438304
refGene_gn=SMKR1;refGene_tx=XM_019825784.2;refGene_gn=SMKR1;refGene_tx=XM_023250555.1;refGene_gn=SMKR1;refGene_tx=XM_023250556.1;refGene_gn=SMKR1;refGene_tx=XM_023250557.1;refGene_gn=SMKR1;refGene_tx=XM_023250558.1;refGene_gn=SMKR1;refGene_tx=XM_023250559.1
A2 147443544 147443589
refGene_gn=SMKR1;refGene_tx=XM_019825785.2
A2 147445201 147445396
refGene_gn=SMKR1;refGene_tx=XM_019825784.2;refGene_gn=SMKR1;refGene_tx

=XM_019825785.2;refGene_gn=SMKR1;refGene_tx=XM_023250555.1;refGene_gn=SMKR1;refGene_tx=XM_023250556.1;refGene_gn=SMKR1;refGene_tx=XM_023250557.1;refGene_gn=SMKR1;refGene_tx=XM_023250558.1;refGene_gn=SMKR1;refGene_tx=XM_023250559.1
A2 147557111 147557334
refGene_gn=NRF1;refGene_tx=XM_023250560.1;refGene_gn=NRF1;refGene_tx=XM_023250561.1;refGene_gn=NRF1;refGene_tx=XM_023250562.1;refGene_gn=NRF1;refGene_tx=XM_023250563.1;refGene_gn=NRF1;refGene_tx=XM_023250564.1
A2 147566039 147566154
refGene_gn=NRF1;refGene_tx=XM_023250560.1;refGene_gn=NRF1;refGene_tx=XM_023250561.1;refGene_gn=NRF1;refGene_tx=XM_023250562.1;refGene_gn=NRF1;refGene_tx=XM_023250563.1;refGene_gn=NRF1;refGene_tx=XM_023250564.1
A2 147568227 147568354
refGene_gn=NRF1;refGene_tx=XM_023250560.1;refGene_gn=NRF1;refGene_tx=XM_023250561.1;refGene_gn=NRF1;refGene_tx=XM_023250562.1;refGene_gn=NRF1;refGene_tx=XM_023250563.1;refGene_gn=NRF1;refGene_tx=XM_023250564.1
A2 147587816 147587957
refGene_gn=NRF1;refGene_tx=XM_023250560.1;refGene_gn=NRF1;refGene_tx=XM_023250561.1;refGene_gn=NRF1;refGene_tx=XM_023250562.1;refGene_gn=NRF1;refGene_tx=XM_023250563.1;refGene_gn=NRF1;refGene_tx=XM_023250564.1
A2 147603938 147604097
refGene_gn=NRF1;refGene_tx=XM_023250560.1;refGene_gn=NRF1;refGene_tx=XM_023250561.1;refGene_gn=NRF1;refGene_tx=XM_023250562.1;refGene_gn=NRF1;refGene_tx=XM_023250563.1;refGene_gn=NRF1;refGene_tx=XM_023250564.1
A2 147604871 147605069
refGene_gn=NRF1;refGene_tx=XM_023250560.1;refGene_gn=NRF1;refGene_tx=XM_023250561.1;refGene_gn=NRF1;refGene_tx=XM_023250562.1;refGene_gn=NRF1;refGene_tx=XM_023250563.1;refGene_gn=NRF1;refGene_tx=XM_023250564.1
A2 147605679 147605781
refGene_gn=NRF1;refGene_tx=XM_023250560.1;refGene_gn=NRF1;refGene_tx=XM_023250561.1;refGene_gn=NRF1;refGene_tx=XM_023250562.1;refGene_gn=NRF1;refGene_tx=XM_023250563.1;refGene_gn=NRF1;refGene_tx=XM_023250564.1
A2 147609579 147609738
refGene_gn=NRF1;refGene_tx=XM_023250560.1;refGene_gn=NRF1;refGene_tx=XM_023250561.1;refGene_gn=NRF1;refGene_tx=XM_023250562.1;refGene_gn=NRF1;refGene_tx=XM_023250564.1
A2 147612139 147612297
refGene_gn=NRF1;refGene_tx=XM_023250560.1;refGene_gn=NRF1;refGene_tx=XM_023250561.1;refGene_gn=NRF1;refGene_tx=XM_023250562.1;refGene_gn=NRF1;refGene_tx=XM_023250563.1;refGene_gn=NRF1;refGene_tx=XM_023250564.1
A2 147612379 147612407
refGene_gn=NRF1;refGene_tx=XM_023250564.1
A2 147619842 147619967
refGene_gn=NRF1;refGene_tx=XM_023250560.1;refGene_gn=NRF1;refGene_tx=XM_023250561.1;refGene_gn=NRF1;refGene_tx=XM_023250562.1;refGene_gn=NRF1;refGene_tx=XM_023250563.1
A2 147627081 147627167
refGene_gn=NRF1;refGene_tx=XM_023250562.1
A2 147652075 147652239
refGene_gn=NRF1;refGene_tx=XM_023250560.1;refGene_gn=NRF1;refGene_tx=X

M_023250561.1;refGene_gn=NRF1;refGene_tx=XM_023250563.1
A2	147726121	147726246
refGene_gn=UBE2H;refGene_tx=XM_003983041.5;refGene_gn=UBE2H;refGene_tx=XM_019825792.2;refGene_gn=UBE2H;refGene_tx=XM_019825793.2;refGene_gn=UBE2H;refGene_tx=XM_019825794.2;refGene_gn=UBE2H;refGene_tx=XM_023250565.1
A2	147730551	147730680
refGene_gn=UBE2H;refGene_tx=XM_003983041.5;refGene_gn=UBE2H;refGene_tx=XM_019825792.2;refGene_gn=UBE2H;refGene_tx=XM_019825793.2;refGene_gn=UBE2H;refGene_tx=XM_019825794.2;refGene_gn=UBE2H;refGene_tx=XM_023250565.1
A2	147745253	147745306
refGene_gn=UBE2H;refGene_tx=XM_003983041.5;refGene_gn=UBE2H;refGene_tx=XM_019825792.2;refGene_gn=UBE2H;refGene_tx=XM_023250565.1
A2	147746755	147746795
refGene_gn=UBE2H;refGene_tx=XM_003983041.5;refGene_gn=UBE2H;refGene_tx=XM_019825792.2;refGene_gn=UBE2H;refGene_tx=XM_023250565.1
A2	147766437	147766512
refGene_gn=UBE2H;refGene_tx=XM_003983041.5;refGene_gn=UBE2H;refGene_tx=XM_019825792.2;refGene_gn=UBE2H;refGene_tx=XM_019825793.2;refGene_gn=UBE2H;refGene_tx=XM_019825794.2;refGene_gn=UBE2H;refGene_tx=XM_023250565.1
A2	147767614	147767691
refGene_gn=UBE2H;refGene_tx=XM_003983041.5;refGene_gn=UBE2H;refGene_tx=XM_019825792.2;refGene_gn=UBE2H;refGene_tx=XM_019825793.2;refGene_gn=UBE2H;refGene_tx=XM_019825794.2;refGene_gn=UBE2H;refGene_tx=XM_023250565.1
A2	147772150	147772192
refGene_gn=UBE2H;refGene_tx=XM_019825792.2;refGene_gn=UBE2H;refGene_tx=XM_019825793.2
A2	147793586	147793640
refGene_gn=UBE2H;refGene_tx=XM_023250565.1
A2	147793721	147793759
refGene_gn=UBE2H;refGene_tx=XM_023250565.1
A2	147825107	147825160
refGene_gn=UBE2H;refGene_tx=XM_003983041.5;refGene_gn=UBE2H;refGene_tx=XM_019825792.2;refGene_gn=UBE2H;refGene_tx=XM_019825793.2;refGene_gn=UBE2H;refGene_tx=XM_019825794.2
A2	147890649	147890718
refGene_gn=ZC3HC1;refGene_tx=XM_003983043.4
A2	147892851	147893058
refGene_gn=ZC3HC1;refGene_tx=XM_003983043.4
A2	147893718	147893931
refGene_gn=ZC3HC1;refGene_tx=XM_003983043.4
A2	147894750	147894994
refGene_gn=ZC3HC1;refGene_tx=XM_003983043.4
A2	147896031	147896186
refGene_gn=ZC3HC1;refGene_tx=XM_003983043.4
A2	147897575	147897703
refGene_gn=ZC3HC1;refGene_tx=XM_003983043.4

A2 147907123 147907207
refGene_gn=ZC3HC1;refGene_tx=XM_003983043.4
A2 147907746 147907897
refGene_gn=ZC3HC1;refGene_tx=XM_003983043.4
A2 147912269 147912381
refGene_gn=ZC3HC1;refGene_tx=XM_003983043.4
A2 147913927 147914073
refGene_gn=ZC3HC1;refGene_tx=XM_003983043.4
A2 147930094 147930266
refGene_gn=KLHDC10;refGene_tx=XM_011280559.3;refGene_gn=KLHDC10;refGene_tx=XM_023250566.1;refGene_gn=KLHDC10;refGene_tx=XM_023250567.1;refGene_gn=KLHDC10;refGene_tx=XM_023250568.1
A2 147959693 147959780
refGene_gn=KLHDC10;refGene_tx=XM_011280559.3;refGene_gn=KLHDC10;refGene_tx=XM_023250566.1
A2 147975126 147975348
refGene_gn=KLHDC10;refGene_tx=XM_011280559.3;refGene_gn=KLHDC10;refGene_tx=XM_023250566.1;refGene_gn=KLHDC10;refGene_tx=XM_023250567.1;refGene_gn=KLHDC10;refGene_tx=XM_023250568.1
A2 147977440 147977595
refGene_gn=KLHDC10;refGene_tx=XM_011280559.3;refGene_gn=KLHDC10;refGene_tx=XM_023250566.1;refGene_gn=KLHDC10;refGene_tx=XM_023250567.1;refGene_gn=KLHDC10;refGene_tx=XM_023250568.1
A2 147978769 147978918
refGene_gn=KLHDC10;refGene_tx=XM_011280559.3;refGene_gn=KLHDC10;refGene_tx=XM_023250566.1;refGene_gn=KLHDC10;refGene_tx=XM_023250567.1;refGene_gn=KLHDC10;refGene_tx=XM_023250568.1
A2 147980834 147980919
refGene_gn=KLHDC10;refGene_tx=XM_011280559.3;refGene_gn=KLHDC10;refGene_tx=XM_023250566.1;refGene_gn=KLHDC10;refGene_tx=XM_023250567.1;refGene_gn=KLHDC10;refGene_tx=XM_023250568.1
A2 147981979 147982046
refGene_gn=KLHDC10;refGene_tx=XM_011280559.3;refGene_gn=KLHDC10;refGene_tx=XM_023250566.1;refGene_gn=KLHDC10;refGene_tx=XM_023250567.1;refGene_gn=KLHDC10;refGene_tx=XM_023250568.1
A2 147982576 147982624
refGene_gn=KLHDC10;refGene_tx=XM_011280559.3;refGene_gn=KLHDC10;refGene_tx=XM_023250566.1;refGene_gn=KLHDC10;refGene_tx=XM_023250567.1;refGene_gn=KLHDC10;refGene_tx=XM_023250568.1
A2 147984660 147984800
refGene_gn=KLHDC10;refGene_tx=XM_011280559.3;refGene_gn=KLHDC10;refGene_tx=XM_023250566.1;refGene_gn=KLHDC10;refGene_tx=XM_023250567.1;refGene_gn=KLHDC10;refGene_tx=XM_023250568.1
A2 147985437 147985647
refGene_gn=KLHDC10;refGene_tx=XM_011280559.3;refGene_gn=KLHDC10;refGene_tx=XM_023250568.1
A2 148001041 148001257
refGene_gn=KLHDC10;refGene_tx=XM_023250566.1;refGene_gn=KLHDC10;refGene_tx=XM_023250567.1
A2 148014701 148014756

refGene_gn=TMEM209;refGene_tx=XM_011280543.3;refGene_gn=TMEM209;refGen
e_tx=XM_011280544.3;refGene_gn=TMEM209;refGene_tx=XM_023250569.1
A2 148015370 148015380
refGene_gn=TMEM209;refGene_tx=XM_019825796.2
A2 148019293 148019367
refGene_gn=TMEM209;refGene_tx=XM_011280543.3;refGene_gn=TMEM209;refGen
e_tx=XM_011280544.3;refGene_gn=TMEM209;refGene_tx=XM_019825796.2;refGe
ne_gn=TMEM209;refGene_tx=XM_023250569.1
A2 148022023 148022121
refGene_gn=TMEM209;refGene_tx=XM_011280543.3;refGene_gn=TMEM209;refGen
e_tx=XM_011280544.3;refGene_gn=TMEM209;refGene_tx=XM_019825796.2;refGe
ne_gn=TMEM209;refGene_tx=XM_023250569.1
A2 148022214 148022329
refGene_gn=TMEM209;refGene_tx=XM_011280543.3;refGene_gn=TMEM209;refGen
e_tx=XM_011280544.3;refGene_gn=TMEM209;refGene_tx=XM_019825796.2;refGe
ne_gn=TMEM209;refGene_tx=XM_023250569.1
A2 148023804 148023902
refGene_gn=TMEM209;refGene_tx=XM_011280543.3;refGene_gn=TMEM209;refGen
e_tx=XM_011280544.3;refGene_gn=TMEM209;refGene_tx=XM_019825796.2;refGe
ne_gn=TMEM209;refGene_tx=XM_023250569.1
A2 148026465 148026591
refGene_gn=TMEM209;refGene_tx=XM_011280543.3;refGene_gn=TMEM209;refGen
e_tx=XM_011280544.3;refGene_gn=TMEM209;refGene_tx=XM_019825796.2;refGe
ne_gn=TMEM209;refGene_tx=XM_023250569.1
A2 148032078 148032175
refGene_gn=TMEM209;refGene_tx=XM_011280543.3;refGene_gn=TMEM209;refGen
e_tx=XM_011280544.3;refGene_gn=TMEM209;refGene_tx=XM_019825796.2;refGe
ne_gn=TMEM209;refGene_tx=XM_023250569.1
A2 148034160 148034232
refGene_gn=TMEM209;refGene_tx=XM_011280543.3;refGene_gn=TMEM209;refGen
e_tx=XM_011280544.3;refGene_gn=TMEM209;refGene_tx=XM_019825796.2;refGe
ne_gn=TMEM209;refGene_tx=XM_023250569.1
A2 148034850 148035026
refGene_gn=TMEM209;refGene_tx=XM_011280543.3;refGene_gn=TMEM209;refGen
e_tx=XM_011280544.3;refGene_gn=TMEM209;refGene_tx=XM_019825796.2;refGe
ne_gn=TMEM209;refGene_tx=XM_023250569.1
A2 148039887 148040089
refGene_gn=TMEM209;refGene_tx=XM_011280543.3;refGene_gn=TMEM209;refGen
e_tx=XM_011280544.3;refGene_gn=TMEM209;refGene_tx=XM_019825796.2;refGe
ne_gn=TMEM209;refGene_tx=XM_023250569.1
A2 148044485 148044727
refGene_gn=TMEM209;refGene_tx=XM_011280543.3;refGene_gn=TMEM209;refGen
e_tx=XM_011280544.3;refGene_gn=TMEM209;refGene_tx=XM_019825796.2;refGe
ne_gn=TMEM209;refGene_tx=XM_023250569.1
A2 148045224 148045356
refGene_gn=TMEM209;refGene_tx=XM_011280543.3;refGene_gn=TMEM209;refGen
e_tx=XM_011280544.3;refGene_gn=TMEM209;refGene_tx=XM_019825796.2;refGe
ne_gn=TMEM209;refGene_tx=XM_023250569.1
A2 148046551 148046610
refGene_gn=TMEM209;refGene_tx=XM_011280543.3;refGene_gn=TMEM209;refGen

e_tx=XM_011280544.3;refGene_gn=TMEM209;refGene_tx=XM_019825796.2;refGene_gn=TMEM209;refGene_tx=XM_023250569.1
A2 148046766 148046903
refGene_gn=TMEM209;refGene_tx=XM_011280543.3;refGene_gn=TMEM209;refGene_tx=XM_011280544.3;refGene_gn=TMEM209;refGene_tx=XM_019825796.2;refGene_gn=TMEM209;refGene_tx=XM_023250569.1
A2 148048055 148048058
refGene_gn=TMEM209;refGene_tx=XM_011280543.3;refGene_gn=TMEM209;refGene_tx=XM_019825796.2
A2 148048748 148048769
refGene_gn=SSMEM1;refGene_tx=XM_019825797.2
A2 148049160 148049178
refGene_gn=SSMEM1;refGene_tx=XM_006929372.2
A2 148050934 148051117
refGene_gn=SSMEM1;refGene_tx=XM_003983069.4
A2 148057343 148057398
refGene_gn=SSMEM1;refGene_tx=XM_003983069.4;refGene_gn=SSMEM1;refGene_tx=XM_006929372.2;refGene_gn=SSMEM1;refGene_tx=XM_019825797.2
A2 148059871 148060368
refGene_gn=SSMEM1;refGene_tx=XM_003983069.4;refGene_gn=SSMEM1;refGene_tx=XM_006929372.2;refGene_gn=SSMEM1;refGene_tx=XM_019825797.2
A2 148084313 148084378
refGene_gn=CPA2;refGene_tx=XM_003983050.5
A2 148086481 148086563
refGene_gn=CPA2;refGene_tx=XM_003983050.5
A2 148087234 148087369
refGene_gn=CPA2;refGene_tx=XM_003983050.5
A2 148088167 148088263
refGene_gn=CPA2;refGene_tx=XM_003983050.5
A2 148089960 148090062
refGene_gn=CPA2;refGene_tx=XM_003983050.5
A2 148092594 148092693
refGene_gn=CPA2;refGene_tx=XM_003983050.5
A2 148094043 148094154
refGene_gn=CPA2;refGene_tx=XM_003983050.5
A2 148095284 148095375
refGene_gn=CPA2;refGene_tx=XM_003983050.5
A2 148096942 148097142
refGene_gn=CPA2;refGene_tx=XM_003983050.5
A2 148099220 148099305
refGene_gn=CPA2;refGene_tx=XM_003983050.5
A2 148105761 148105949
refGene_gn=CPA2;refGene_tx=XM_003983050.5
A2 148108579 148108647
refGene_gn=CPA4;refGene_tx=XM_003983051.4;refGene_gn=CPA4;refGene_tx=XM_019825803.2
A2 148113203 148113285
refGene_gn=CPA4;refGene_tx=XM_003983051.4;refGene_gn=CPA4;refGene_tx=XM_019825803.2
A2 148113689 148113824

refGene_gn=CPA4;refGene_tx=XM_003983051.4;refGene_gn=CPA4;refGene_tx=XM_019825803.2
A2 148114922 148115021
refGene_gn=CPA4;refGene_tx=XM_003983051.4;refGene_gn=CPA4;refGene_tx=XM_019825803.2
A2 148117824 148117926
refGene_gn=CPA4;refGene_tx=XM_003983051.4;refGene_gn=CPA4;refGene_tx=XM_019825803.2
A2 148119244 148119349
refGene_gn=CPA4;refGene_tx=XM_003983051.4;refGene_gn=CPA4;refGene_tx=XM_019825803.2
A2 148120059 148120170
refGene_gn=CPA4;refGene_tx=XM_003983051.4;refGene_gn=CPA4;refGene_tx=XM_019825803.2
A2 148121836 148121927
refGene_gn=CPA4;refGene_tx=XM_003983051.4;refGene_gn=CPA4;refGene_tx=XM_019825803.2
A2 148123145 148123345
refGene_gn=CPA4;refGene_tx=XM_003983051.4;refGene_gn=CPA4;refGene_tx=XM_019825803.2
A2 148124321 148124406
refGene_gn=CPA4;refGene_tx=XM_003983051.4;refGene_gn=CPA4;refGene_tx=XM_019825803.2
A2 148127252 148127326
refGene_gn=CPA4;refGene_tx=XM_019825803.2
A2 148129417 148129605
refGene_gn=CPA4;refGene_tx=XM_003983051.4
A2 148147952 148148068
refGene_gn=CPA5;refGene_tx=XM_006929376.4;refGene_gn=CPA5;refGene_tx=XM_011280548.3
A2 148149923 148150005
refGene_gn=CPA5;refGene_tx=XM_006929376.4;refGene_gn=CPA5;refGene_tx=XM_011280548.3
A2 148151041 148151176
refGene_gn=CPA5;refGene_tx=XM_006929376.4;refGene_gn=CPA5;refGene_tx=XM_011280548.3
A2 148159659 148159758
refGene_gn=CPA5;refGene_tx=XM_006929376.4;refGene_gn=CPA5;refGene_tx=XM_011280548.3
A2 148160908 148161010
refGene_gn=CPA5;refGene_tx=XM_006929376.4;refGene_gn=CPA5;refGene_tx=XM_011280548.3
A2 148162190 148162292
refGene_gn=CPA5;refGene_tx=XM_006929376.4;refGene_gn=CPA5;refGene_tx=XM_011280548.3
A2 148162636 148162747
refGene_gn=CPA5;refGene_tx=XM_006929376.4;refGene_gn=CPA5;refGene_tx=XM_011280548.3
A2 148163183 148163274
refGene_gn=CPA5;refGene_tx=XM_006929376.4;refGene_gn=CPA5;refGene_tx=X

M_011280548.3
A2	148166524	148166724
refGene_gn=CPA5;refGene_tx=XM_006929376.4;refGene_gn=CPA5;refGene_tx=X
M_011280548.3
A2	148167017	148167102
refGene_gn=CPA5;refGene_tx=XM_006929376.4;refGene_gn=CPA5;refGene_tx=X
M_011280548.3
A2	148167615	148167803
refGene_gn=CPA5;refGene_tx=XM_006929376.4;refGene_gn=CPA5;refGene_tx=X
M_011280548.3
A2	148178413	148178478
refGene_gn=CPA1;refGene_tx=XM_003983055.5;refGene_gn=CPA1;refGene_tx=X
M_023250570.1
A2	148178975	148179057
refGene_gn=CPA1;refGene_tx=XM_003983055.5;refGene_gn=CPA1;refGene_tx=X
M_023250570.1
A2	148179509	148179743
refGene_gn=CPA1;refGene_tx=XM_003983055.5;refGene_gn=CPA1;refGene_tx=X
M_023250570.1
A2	148179976	148180078
refGene_gn=CPA1;refGene_tx=XM_003983055.5;refGene_gn=CPA1;refGene_tx=X
M_023250570.1
A2	148180619	148180721
refGene_gn=CPA1;refGene_tx=XM_003983055.5;refGene_gn=CPA1;refGene_tx=X
M_023250570.1
A2	148180896	148181007
refGene_gn=CPA1;refGene_tx=XM_003983055.5;refGene_gn=CPA1;refGene_tx=X
M_023250570.1
A2	148181882	148181985
refGene_gn=CPA1;refGene_tx=XM_003983055.5;refGene_gn=CPA1;refGene_tx=X
M_023250570.1
A2	148182483	148182684
refGene_gn=CPA1;refGene_tx=XM_003983055.5;refGene_gn=CPA1;refGene_tx=X
M_023250570.1
A2	148183069	148183154
refGene_gn=CPA1;refGene_tx=XM_003983055.5;refGene_gn=CPA1;refGene_tx=X
M_023250570.1
A2	148184846	148185034
refGene_gn=CPA1;refGene_tx=XM_003983055.5;refGene_gn=CPA1;refGene_tx=X
M_023250570.1
A2	148195469	148195618
refGene_gn=CEP41;refGene_tx=XM_003983057.5;refGene_gn=CEP41;refGene_tx
=XM_006929373.4;refGene_gn=CEP41;refGene_tx=XM_023250571.1
A2	148196318	148196534
refGene_gn=CEP41;refGene_tx=XM_003983057.5;refGene_gn=CEP41;refGene_tx
=XM_006929373.4;refGene_gn=CEP41;refGene_tx=XM_023250571.1
A2	148197008	148197123
refGene_gn=CEP41;refGene_tx=XM_003983057.5;refGene_gn=CEP41;refGene_tx
=XM_006929373.4;refGene_gn=CEP41;refGene_tx=XM_023250571.1
A2	148198202	148198270

refGene_gn=CEP41;refGene_tx=XM_003983057.5;refGene_gn=CEP41;refGene_tx=XM_006929373.4;refGene_gn=CEP41;refGene_tx=XM_023250571.1
A2 148198920 148199072
refGene_gn=CEP41;refGene_tx=XM_003983057.5;refGene_gn=CEP41;refGene_tx=XM_006929373.4;refGene_gn=CEP41;refGene_tx=XM_023250571.1
A2 148200764 148200909
refGene_gn=CEP41;refGene_tx=XM_003983057.5;refGene_gn=CEP41;refGene_tx=XM_006929373.4;refGene_gn=CEP41;refGene_tx=XM_023250571.1
A2 148207686 148207756
refGene_gn=CEP41;refGene_tx=XM_003983057.5;refGene_gn=CEP41;refGene_tx=XM_006929373.4;refGene_gn=CEP41;refGene_tx=XM_023250571.1
A2 148208797 148208859
refGene_gn=CEP41;refGene_tx=XM_003983057.5;refGene_gn=CEP41;refGene_tx=XM_006929373.4;refGene_gn=CEP41;refGene_tx=XM_023250571.1
A2 148213072 148213120
refGene_gn=CEP41;refGene_tx=XM_003983057.5;refGene_gn=CEP41;refGene_tx=XM_006929373.4;refGene_gn=CEP41;refGene_tx=XM_023250571.1
A2 148227137 148227201
refGene_gn=CEP41;refGene_tx=XM_003983057.5;refGene_gn=CEP41;refGene_tx=XM_006929373.4;refGene_gn=CEP41;refGene_tx=XM_023250571.1
A2 148240212 148240254
refGene_gn=CEP41;refGene_tx=XM_023250571.1
A2 148240816 148240867
refGene_gn=CEP41;refGene_tx=XM_006929373.4
A2 148241266 148241299
refGene_gn=CEP41;refGene_tx=XM_003983057.5
A2 148297876 148297902
refGene_gn=MEST;refGene_tx=XM_023250573.1
A2 148301055 148301210
refGene_gn=MEST;refGene_tx=XM_023250573.1
A2 148302885 148302956
refGene_gn=MEST;refGene_tx=XM_023250573.1
A2 148302957 148302966
refGene_gn=MEST;refGene_tx=XM_023250573.1
A2 148303350 148303428
refGene_gn=MEST;refGene_tx=XM_023250573.1
A2 148303551 148303688
refGene_gn=MEST;refGene_tx=XM_023250573.1
A2 148303865 148303924
refGene_gn=MEST;refGene_tx=XM_023250573.1
A2 148305189 148305230
refGene_gn=MEST;refGene_tx=XM_023250573.1
A2 148306002 148306073
refGene_gn=MEST;refGene_tx=XM_023250573.1
A2 148306314 148306416
refGene_gn=MEST;refGene_tx=XM_023250573.1
A2 148306962 148307039
refGene_gn=MEST;refGene_tx=XM_023250573.1
A2 148309270 148309334
refGene_gn=MEST;refGene_tx=XM_023250573.1

A2 148310264 148310382
refGene_gn=MEST;refGene_tx=XM_023250573.1
A2 148312005 148312136
refGene_gn=COPG2;refGene_tx=XM_019825808.1;refGene_gn=COPG2;refGene_tx
=XM_023250572.1
A2 148312613 148312712
refGene_gn=COPG2;refGene_tx=XM_019825808.1;refGene_gn=COPG2;refGene_tx
=XM_023250572.1
A2 148312967 148313106
refGene_gn=COPG2;refGene_tx=XM_019825808.1;refGene_gn=COPG2;refGene_tx
=XM_023250572.1
A2 148313835 148313933
refGene_gn=COPG2;refGene_tx=XM_019825808.1;refGene_gn=COPG2;refGene_tx
=XM_023250572.1
A2 148358306 148358478
refGene_gn=COPG2;refGene_tx=XM_019825808.1;refGene_gn=COPG2;refGene_tx
=XM_023250572.1
A2 148359149 148359289
refGene_gn=COPG2;refGene_tx=XM_019825808.1;refGene_gn=COPG2;refGene_tx
=XM_023250572.1
A2 148359721 148359784
refGene_gn=COPG2;refGene_tx=XM_019825808.1;refGene_gn=COPG2;refGene_tx
=XM_023250572.1
A2 148361260 148361386
refGene_gn=COPG2;refGene_tx=XM_019825808.1;refGene_gn=COPG2;refGene_tx
=XM_023250572.1
A2 148361891 148361995
refGene_gn=COPG2;refGene_tx=XM_019825808.1;refGene_gn=COPG2;refGene_tx
=XM_023250572.1
A2 148362823 148362899
refGene_gn=COPG2;refGene_tx=XM_019825808.1;refGene_gn=COPG2;refGene_tx
=XM_023250572.1
A2 148365487 148365731
refGene_gn=COPG2;refGene_tx=XM_019825808.1;refGene_gn=COPG2;refGene_tx
=XM_023250572.1
A2 148366050 148366146
refGene_gn=COPG2;refGene_tx=XM_019825808.1;refGene_gn=COPG2;refGene_tx
=XM_023250572.1
A2 148370294 148370483
refGene_gn=COPG2;refGene_tx=XM_019825808.1;refGene_gn=COPG2;refGene_tx
=XM_023250572.1
A2 148373240 148373308
refGene_gn=COPG2;refGene_tx=XM_019825808.1;refGene_gn=COPG2;refGene_tx
=XM_023250572.1
A2 148373861 148373995
refGene_gn=COPG2;refGene_tx=XM_019825808.1;refGene_gn=COPG2;refGene_tx
=XM_023250572.1
A2 148396961 148397119
refGene_gn=COPG2;refGene_tx=XM_019825808.1;refGene_gn=COPG2;refGene_tx
=XM_023250572.1

A2	148398159	148398246
refGene_gn=COPG2;refGene_tx=XM_019825808.1;refGene_gn=COPG2;refGene_tx=XM_023250572.1
A2	148399412	148399505
refGene_gn=COPG2;refGene_tx=XM_019825808.1;refGene_gn=COPG2;refGene_tx=XM_023250572.1
A2	148400832	148400908
refGene_gn=COPG2;refGene_tx=XM_019825808.1;refGene_gn=COPG2;refGene_tx=XM_023250572.1
A2	148409301	148409381
refGene_gn=COPG2;refGene_tx=XM_019825808.1;refGene_gn=COPG2;refGene_tx=XM_023250572.1
A2	148423440	148423512
refGene_gn=COPG2;refGene_tx=XM_023250572.1
A2	148427240	148427321
refGene_gn=COPG2;refGene_tx=XM_023250572.1
A2	148428089	148428142
refGene_gn=COPG2;refGene_tx=XM_023250572.1
A2	148429295	148429332
refGene_gn=COPG2;refGene_tx=XM_023250572.1
A2	148429766	148429936
refGene_gn=TSGA13;refGene_tx=XM_006929378.3;refGene_gn=TSGA13;refGene_tx=XM_006929379.4;refGene_gn=TSGA13;refGene_tx=XM_019825810.2
A2	148431553	148431681
refGene_gn=TSGA13;refGene_tx=XM_006929378.3;refGene_gn=TSGA13;refGene_tx=XM_006929379.4;refGene_gn=TSGA13;refGene_tx=XM_019825810.2
A2	148432214	148432354
refGene_gn=TSGA13;refGene_tx=XM_006929378.3;refGene_gn=TSGA13;refGene_tx=XM_006929379.4;refGene_gn=TSGA13;refGene_tx=XM_019825810.2
A2	148437273	148437480
refGene_gn=TSGA13;refGene_tx=XM_006929378.3;refGene_gn=TSGA13;refGene_tx=XM_006929379.4;refGene_gn=TSGA13;refGene_tx=XM_019825810.2
A2	148438391	148438463
refGene_gn=TSGA13;refGene_tx=XM_006929378.3;refGene_gn=TSGA13;refGene_tx=XM_006929379.4;refGene_gn=TSGA13;refGene_tx=XM_019825810.2
A2	148440367	148440446
refGene_gn=TSGA13;refGene_tx=XM_006929378.3;refGene_gn=TSGA13;refGene_tx=XM_006929379.4;refGene_gn=TSGA13;refGene_tx=XM_019825810.2
A2	148442482	148442505
refGene_gn=TSGA13;refGene_tx=XM_006929378.3;refGene_gn=TSGA13;refGene_tx=XM_006929379.4;refGene_gn=TSGA13;refGene_tx=XM_019825810.2
A2	148486373	148487357
refGene_gn=KLF14;refGene_tx=XM_023250574.1
A2	148743704	148743860
refGene_gn=LOC101093924;refGene_tx=XM_019825071.1
A2	148744805	148744901
refGene_gn=LOC101093924;refGene_tx=XM_019825071.1
A2	148755200	148756133
refGene_gn=LOC101093924;refGene_tx=XM_019825071.1
A2	148849176	148849193

refGene_gn=MKLN1;refGene_tx=XM_003983064.5
A2 148983386 148983394
refGene_gn=MKLN1;refGene_tx=XM_006929380.3
A2 149015582 149015680
refGene_gn=MKLN1;refGene_tx=XM_003983063.5
A2 149063258 149063328
refGene_gn=MKLN1;refGene_tx=XM_003983063.5;refGene_gn=MKLN1;refGene_tx=XM_003983064.5;refGene_gn=MKLN1;refGene_tx=XM_006929380.3
A2 149087773 149087916
refGene_gn=MKLN1;refGene_tx=XM_003983063.5;refGene_gn=MKLN1;refGene_tx=XM_003983064.5;refGene_gn=MKLN1;refGene_tx=XM_006929380.3
A2 149089137 149089226
refGene_gn=MKLN1;refGene_tx=XM_003983063.5;refGene_gn=MKLN1;refGene_tx=XM_003983064.5;refGene_gn=MKLN1;refGene_tx=XM_006929380.3
A2 149096744 149096854
refGene_gn=MKLN1;refGene_tx=XM_003983063.5;refGene_gn=MKLN1;refGene_tx=XM_003983064.5;refGene_gn=MKLN1;refGene_tx=XM_006929380.3
A2 149099456 149099649
refGene_gn=MKLN1;refGene_tx=XM_003983063.5;refGene_gn=MKLN1;refGene_tx=XM_003983064.5;refGene_gn=MKLN1;refGene_tx=XM_006929380.3
A2 149117479 149117557
refGene_gn=MKLN1;refGene_tx=XM_003983063.5;refGene_gn=MKLN1;refGene_tx=XM_003983064.5;refGene_gn=MKLN1;refGene_tx=XM_006929380.3
A2 149119772 149119838
refGene_gn=MKLN1;refGene_tx=XM_003983063.5;refGene_gn=MKLN1;refGene_tx=XM_003983064.5;refGene_gn=MKLN1;refGene_tx=XM_006929380.3
A2 149130004 149130117
refGene_gn=MKLN1;refGene_tx=XM_003983063.5;refGene_gn=MKLN1;refGene_tx=XM_003983064.5;refGene_gn=MKLN1;refGene_tx=XM_006929380.3
A2 149140083 149140296
refGene_gn=MKLN1;refGene_tx=XM_003983063.5;refGene_gn=MKLN1;refGene_tx=XM_003983064.5;refGene_gn=MKLN1;refGene_tx=XM_006929380.3
A2 149147379 149147601
refGene_gn=MKLN1;refGene_tx=XM_003983063.5;refGene_gn=MKLN1;refGene_tx=XM_003983064.5;refGene_gn=MKLN1;refGene_tx=XM_006929380.3
A2 149150299 149150429
refGene_gn=MKLN1;refGene_tx=XM_003983063.5;refGene_gn=MKLN1;refGene_tx=XM_003983064.5;refGene_gn=MKLN1;refGene_tx=XM_006929380.3
A2 149169127 149169275
refGene_gn=MKLN1;refGene_tx=XM_003983063.5;refGene_gn=MKLN1;refGene_tx=XM_003983064.5;refGene_gn=MKLN1;refGene_tx=XM_006929380.3
A2 149173409 149173524
refGene_gn=MKLN1;refGene_tx=XM_003983063.5;refGene_gn=MKLN1;refGene_tx=XM_003983064.5;refGene_gn=MKLN1;refGene_tx=XM_006929380.3
A2 149175040 149175180
refGene_gn=MKLN1;refGene_tx=XM_003983063.5;refGene_gn=MKLN1;refGene_tx=XM_003983064.5;refGene_gn=MKLN1;refGene_tx=XM_006929380.3
A2 149182072 149182175
refGene_gn=MKLN1;refGene_tx=XM_003983063.5;refGene_gn=MKLN1;refGene_tx=XM_003983064.5;refGene_gn=MKLN1;refGene_tx=XM_006929380.3

A2 149193311 149193366
refGene_gn=MKLN1;refGene_tx=XM_003983063.5;refGene_gn=MKLN1;refGene_tx=XM_003983064.5;refGene_gn=MKLN1;refGene_tx=XM_006929380.3
A2 149204706 149204828
refGene_gn=MKLN1;refGene_tx=XM_003983063.5;refGene_gn=MKLN1;refGene_tx=XM_003983064.5;refGene_gn=MKLN1;refGene_tx=XM_006929380.3
A2 149221057 149221255
refGene_gn=PODXL;refGene_tx=XM_003983072.4
A2 149222275 149222443
refGene_gn=PODXL;refGene_tx=XM_003983072.4
A2 149222692 149222754
refGene_gn=PODXL;refGene_tx=XM_003983072.4
A2 149222992 149223140
refGene_gn=PODXL;refGene_tx=XM_003983072.4
A2 149225310 149225382
refGene_gn=PODXL;refGene_tx=XM_003983072.4
A2 149225670 149226038
refGene_gn=PODXL;refGene_tx=XM_003983072.4
A2 149228549 149229203
refGene_gn=PODXL;refGene_tx=XM_003983072.4
A2 149270609 149270676
refGene_gn=PODXL;refGene_tx=XM_003983072.4
A2 149752565 149752661
refGene_gn=PLXNA4;refGene_tx=XM_023250576.1
A2 149754694 149754845
refGene_gn=PLXNA4;refGene_tx=XM_023250576.1
A2 149761456 149761669
refGene_gn=PLXNA4;refGene_tx=XM_023250576.1
A2 149765511 149765681
refGene_gn=PLXNA4;refGene_tx=XM_023250576.1
A2 149766795 149766986
refGene_gn=PLXNA4;refGene_tx=XM_023250576.1
A2 149768122 149768222
refGene_gn=PLXNA4;refGene_tx=XM_023250576.1
A2 149768783 149768887
refGene_gn=PLXNA4;refGene_tx=XM_023250576.1
A2 149780734 149780894
refGene_gn=PLXNA4;refGene_tx=XM_023250576.1
A2 149784654 149784801
refGene_gn=PLXNA4;refGene_tx=XM_023250576.1
A2 149785556 149785623
refGene_gn=PLXNA4;refGene_tx=XM_023250576.1
A2 149788562 149788831
refGene_gn=PLXNA4;refGene_tx=XM_023250576.1
A2 149796260 149796403
refGene_gn=PLXNA4;refGene_tx=XM_023250576.1
A2 149799226 149799461
refGene_gn=PLXNA4;refGene_tx=XM_023250576.1
A2 149800121 149800268
refGene_gn=PLXNA4;refGene_tx=XM_023250576.1

A2 149800940 149801180
refGene_gn=PLXNA4;refGene_tx=XM_023250576.1
A2 149801632 149801726
refGene_gn=PLXNA4;refGene_tx=XM_023250576.1
A2 149804519 149804684
refGene_gn=PLXNA4;refGene_tx=XM_023250576.1
A2 149805859 149805996
refGene_gn=PLXNA4;refGene_tx=XM_023250576.1
A2 149811116 149811234
refGene_gn=PLXNA4;refGene_tx=XM_023250576.1
A2 149815388 149815540
refGene_gn=PLXNA4;refGene_tx=XM_023250576.1
A2 149819390 149819581
refGene_gn=PLXNA4;refGene_tx=XM_023250576.1
A2 149819997 149820094
refGene_gn=PLXNA4;refGene_tx=XM_023250576.1
A2 149827309 149827510
refGene_gn=PLXNA4;refGene_tx=XM_023250576.1
A2 149838194 149838309
refGene_gn=PLXNA4;refGene_tx=XM_023250576.1
A2 149840786 149840886
refGene_gn=PLXNA4;refGene_tx=XM_023250576.1
A2 149842121 149842275
refGene_gn=PLXNA4;refGene_tx=XM_023250576.1
A2 149842979 149843103
refGene_gn=PLXNA4;refGene_tx=XM_023250576.1
A2 149857591 149857692
refGene_gn=PLXNA4;refGene_tx=XM_023250576.1
A2 149907978 149908110
refGene_gn=PLXNA4;refGene_tx=XM_023250576.1
A2 150059740 150059839
refGene_gn=PLXNA4;refGene_tx=XM_023250577.1
A2 150096841 150097024
refGene_gn=PLXNA4;refGene_tx=XM_023250576.1;refGene_gn=PLXNA4;refGene_tx=XM_023250577.1
A2 150113837 150115022
refGene_gn=PLXNA4;refGene_tx=XM_023250576.1;refGene_gn=PLXNA4;refGene_tx=XM_023250577.1
A2 150365832 150365869
refGene_gn=CHCHD3;refGene_tx=XM_006929382.3;refGene_gn=CHCHD3;refGene_tx=XM_019825818.2;refGene_gn=CHCHD3;refGene_tx=XM_023250578.1;refGene_gn=CHCHD3;refGene_tx=XM_023250579.1
A2 150377386 150377522
refGene_gn=CHCHD3;refGene_tx=XM_006929382.3;refGene_gn=CHCHD3;refGene_tx=XM_019825818.2;refGene_gn=CHCHD3;refGene_tx=XM_023250578.1;refGene_gn=CHCHD3;refGene_tx=XM_023250579.1
A2 150402531 150402617
refGene_gn=CHCHD3;refGene_tx=XM_023250580.1
A2 150415584 150415664
refGene_gn=CHCHD3;refGene_tx=XM_023250578.1;refGene_gn=CHCHD3;refGene_

tx=XM_023250579.1;refGene_gn=CHCHD3;refGene_tx=XM_023250580.1
A2	150420574	150420645
refGene_gn=CHCHD3;refGene_tx=XM_006929382.3;refGene_gn=CHCHD3;refGene_tx=XM_019825818.2;refGene_gn=CHCHD3;refGene_tx=XM_023250578.1;refGene_gn=CHCHD3;refGene_tx=XM_023250579.1;refGene_gn=CHCHD3;refGene_tx=XM_023250580.1
A2	150463923	150464007
refGene_gn=CHCHD3;refGene_tx=XM_006929382.3;refGene_gn=CHCHD3;refGene_tx=XM_019825818.2;refGene_gn=CHCHD3;refGene_tx=XM_023250578.1;refGene_gn=CHCHD3;refGene_tx=XM_023250579.1;refGene_gn=CHCHD3;refGene_tx=XM_023250580.1
A2	150465056	150465071
refGene_gn=CHCHD3;refGene_tx=XM_019825818.2;refGene_gn=CHCHD3;refGene_tx=XM_023250578.1;refGene_gn=CHCHD3;refGene_tx=XM_023250580.1
A2	150542216	150542334
refGene_gn=CHCHD3;refGene_tx=XM_006929382.3;refGene_gn=CHCHD3;refGene_tx=XM_019825818.2;refGene_gn=CHCHD3;refGene_tx=XM_023250578.1;refGene_gn=CHCHD3;refGene_tx=XM_023250579.1;refGene_gn=CHCHD3;refGene_tx=XM_023250580.1
A2	150585556	150585638
refGene_gn=CHCHD3;refGene_tx=XM_006929382.3;refGene_gn=CHCHD3;refGene_tx=XM_019825818.2;refGene_gn=CHCHD3;refGene_tx=XM_023250578.1;refGene_gn=CHCHD3;refGene_tx=XM_023250579.1;refGene_gn=CHCHD3;refGene_tx=XM_023250580.1
A2	150634741	150634829
refGene_gn=CHCHD3;refGene_tx=XM_006929382.3;refGene_gn=CHCHD3;refGene_tx=XM_019825818.2;refGene_gn=CHCHD3;refGene_tx=XM_023250578.1;refGene_gn=CHCHD3;refGene_tx=XM_023250579.1;refGene_gn=CHCHD3;refGene_tx=XM_023250580.1
A2	150646017	150646098
refGene_gn=CHCHD3;refGene_tx=XM_006929382.3;refGene_gn=CHCHD3;refGene_tx=XM_019825818.2;refGene_gn=CHCHD3;refGene_tx=XM_023250578.1;refGene_gn=CHCHD3;refGene_tx=XM_023250579.1;refGene_gn=CHCHD3;refGene_tx=XM_023250580.1
A2	150797509	150797595
refGene_gn=EXOC4;refGene_tx=XM_003983067.4
A2	150821238	150821428
refGene_gn=EXOC4;refGene_tx=XM_003983067.4
A2	150831658	150831853
refGene_gn=EXOC4;refGene_tx=XM_003983067.4
A2	150850465	150850650
refGene_gn=EXOC4;refGene_tx=XM_003983067.4
A2	150861800	150861910
refGene_gn=EXOC4;refGene_tx=XM_003983067.4
A2	150878896	150879140
refGene_gn=EXOC4;refGene_tx=XM_003983067.4
A2	150898497	150898672
refGene_gn=EXOC4;refGene_tx=XM_003983067.4
A2	150996143	150996289
refGene_gn=EXOC4;refGene_tx=XM_003983067.4

A2 151000969 151001058
refGene_gn=EXOC4;refGene_tx=XM_003983067.4
A2 151136506 151136603
refGene_gn=EXOC4;refGene_tx=XM_003983067.4
A2 151331624 151331844
refGene_gn=EXOC4;refGene_tx=XM_003983067.4
A2 151388801 151388938
refGene_gn=EXOC4;refGene_tx=XM_003983067.4
A2 151402603 151402759
refGene_gn=EXOC4;refGene_tx=XM_003983067.4
A2 151429752 151429931
refGene_gn=EXOC4;refGene_tx=XM_003983067.4
A2 151476306 151476448
refGene_gn=EXOC4;refGene_tx=XM_003983067.4
A2 151486141 151486320
refGene_gn=EXOC4;refGene_tx=XM_003983067.4
A2 151490002 151490162
refGene_gn=EXOC4;refGene_tx=XM_003983067.4
A2 151547139 151547377
refGene_gn=EXOC4;refGene_tx=XM_003983067.4
A2 151586869 151587166
refGene_gn=LRGUK;refGene_tx=XM_023250581.1;refGene_gn=LRGUK;refGene_tx=XM_023250582.1;refGene_gn=LRGUK;refGene_tx=XM_023250584.1
A2 151594414 151594522
refGene_gn=LRGUK;refGene_tx=XM_023250581.1;refGene_gn=LRGUK;refGene_tx=XM_023250582.1;refGene_gn=LRGUK;refGene_tx=XM_023250584.1
A2 151597046 151597128
refGene_gn=LRGUK;refGene_tx=XM_023250581.1;refGene_gn=LRGUK;refGene_tx=XM_023250582.1;refGene_gn=LRGUK;refGene_tx=XM_023250584.1
A2 151600756 151600857
refGene_gn=LRGUK;refGene_tx=XM_023250581.1;refGene_gn=LRGUK;refGene_tx=XM_023250582.1;refGene_gn=LRGUK;refGene_tx=XM_023250584.1
A2 151602090 151602171
refGene_gn=LRGUK;refGene_tx=XM_023250583.1
A2 151605740 151605822
refGene_gn=LRGUK;refGene_tx=XM_023250581.1;refGene_gn=LRGUK;refGene_tx=XM_023250582.1;refGene_gn=LRGUK;refGene_tx=XM_023250583.1;refGene_gn=LRGUK;refGene_tx=XM_023250584.1
A2 151616283 151616408
refGene_gn=LRGUK;refGene_tx=XM_023250581.1;refGene_gn=LRGUK;refGene_tx=XM_023250582.1;refGene_gn=LRGUK;refGene_tx=XM_023250583.1;refGene_gn=LRGUK;refGene_tx=XM_023250584.1
A2 151620628 151620772
refGene_gn=LRGUK;refGene_tx=XM_023250581.1;refGene_gn=LRGUK;refGene_tx=XM_023250582.1;refGene_gn=LRGUK;refGene_tx=XM_023250583.1;refGene_gn=LRGUK;refGene_tx=XM_023250584.1
A2 151627062 151627143
refGene_gn=LRGUK;refGene_tx=XM_023250581.1;refGene_gn=LRGUK;refGene_tx=XM_023250582.1;refGene_gn=LRGUK;refGene_tx=XM_023250583.1;refGene_gn=LRGUK;refGene_tx=XM_023250584.1

A2 151628779 151628866
refGene_gn=LRGUK;refGene_tx=XM_023250581.1;refGene_gn=LRGUK;refGene_tx=XM_023250582.1;refGene_gn=LRGUK;refGene_tx=XM_023250583.1;refGene_gn=LRGUK;refGene_tx=XM_023250584.1
A2 151630411 151630518
refGene_gn=LRGUK;refGene_tx=XM_023250581.1;refGene_gn=LRGUK;refGene_tx=XM_023250582.1;refGene_gn=LRGUK;refGene_tx=XM_023250583.1;refGene_gn=LRGUK;refGene_tx=XM_023250584.1
A2 151635820 151635940
refGene_gn=LRGUK;refGene_tx=XM_023250581.1;refGene_gn=LRGUK;refGene_tx=XM_023250582.1;refGene_gn=LRGUK;refGene_tx=XM_023250583.1;refGene_gn=LRGUK;refGene_tx=XM_023250584.1
A2 151641603 151641700
refGene_gn=LRGUK;refGene_tx=XM_023250581.1;refGene_gn=LRGUK;refGene_tx=XM_023250582.1;refGene_gn=LRGUK;refGene_tx=XM_023250583.1;refGene_gn=LRGUK;refGene_tx=XM_023250584.1
A2 151645355 151645469
refGene_gn=LRGUK;refGene_tx=XM_023250581.1;refGene_gn=LRGUK;refGene_tx=XM_023250582.1;refGene_gn=LRGUK;refGene_tx=XM_023250583.1;refGene_gn=LRGUK;refGene_tx=XM_023250584.1
A2 151648349 151648551
refGene_gn=LRGUK;refGene_tx=XM_023250581.1;refGene_gn=LRGUK;refGene_tx=XM_023250582.1;refGene_gn=LRGUK;refGene_tx=XM_023250583.1;refGene_gn=LRGUK;refGene_tx=XM_023250584.1
A2 151649412 151649508
refGene_gn=LRGUK;refGene_tx=XM_023250581.1;refGene_gn=LRGUK;refGene_tx=XM_023250582.1;refGene_gn=LRGUK;refGene_tx=XM_023250583.1;refGene_gn=LRGUK;refGene_tx=XM_023250584.1
A2 151655582 151657762
refGene_gn=LRGUK;refGene_tx=XM_023250581.1;refGene_gn=LRGUK;refGene_tx=XM_023250582.1;refGene_gn=LRGUK;refGene_tx=XM_023250583.1
A2 151670285 151670425
refGene_gn=LRGUK;refGene_tx=XM_023250584.1
A2 151682305 151682394
refGene_gn=LRGUK;refGene_tx=XM_023250584.1
A2 151684497 151684620
refGene_gn=LRGUK;refGene_tx=XM_023250584.1
A2 151691357 151691509
refGene_gn=LRGUK;refGene_tx=XM_023250584.1
A2 151695311 151695457
refGene_gn=LRGUK;refGene_tx=XM_023250584.1
A2 151744986 151745233
refGene_gn=SLC35B4;refGene_tx=XM_011280550.3
A2 151745312 151745383
refGene_gn=SLC35B4;refGene_tx=XM_019825822.2;refGene_gn=SLC35B4;refGene_tx=XM_019825823.2
A2 151746024 151746157
refGene_gn=SLC35B4;refGene_tx=XM_011280550.3;refGene_gn=SLC35B4;refGene_tx=XM_019825822.2;refGene_gn=SLC35B4;refGene_tx=XM_019825823.2
A2 151749755 151749831

refGene_gn=SLC35B4;refGene_tx=XM_011280550.3;refGene_gn=SLC35B4;refGene_tx=XM_019825822.2;refGene_gn=SLC35B4;refGene_tx=XM_019825823.2
A2 151751131 151751241
refGene_gn=SLC35B4;refGene_tx=XM_011280550.3;refGene_gn=SLC35B4;refGene_tx=XM_019825822.2;refGene_gn=SLC35B4;refGene_tx=XM_019825823.2
A2 151752900 151752961
refGene_gn=SLC35B4;refGene_tx=XM_011280550.3;refGene_gn=SLC35B4;refGene_tx=XM_019825822.2;refGene_gn=SLC35B4;refGene_tx=XM_019825823.2
A2 151753179 151753261
refGene_gn=SLC35B4;refGene_tx=XM_011280550.3;refGene_gn=SLC35B4;refGene_tx=XM_019825822.2;refGene_gn=SLC35B4;refGene_tx=XM_019825823.2
A2 151755840 151755890
refGene_gn=SLC35B4;refGene_tx=XM_011280550.3;refGene_gn=SLC35B4;refGene_tx=XM_019825822.2;refGene_gn=SLC35B4;refGene_tx=XM_019825823.2
A2 151757187 151757290
refGene_gn=SLC35B4;refGene_tx=XM_011280550.3;refGene_gn=SLC35B4;refGene_tx=XM_019825822.2;refGene_gn=SLC35B4;refGene_tx=XM_019825823.2
A2 151775010 151775124
refGene_gn=SLC35B4;refGene_tx=XM_011280550.3;refGene_gn=SLC35B4;refGene_tx=XM_019825822.2;refGene_gn=SLC35B4;refGene_tx=XM_019825823.2
A2 151781962 151782039
refGene_gn=SLC35B4;refGene_tx=XM_011280550.3;refGene_gn=SLC35B4;refGene_tx=XM_019825822.2;refGene_gn=SLC35B4;refGene_tx=XM_019825823.2
A2 151900132 151900175
refGene_gn=AKR1B1;refGene_tx=XM_003983075.4
A2 151902302 151902385
refGene_gn=AKR1B1;refGene_tx=XM_003983075.4
A2 151904282 151904366
refGene_gn=AKR1B1;refGene_tx=XM_003983075.4
A2 151904882 151904964
refGene_gn=AKR1B1;refGene_tx=XM_003983075.4
A2 151905276 151905383
refGene_gn=AKR1B1;refGene_tx=XM_003983075.4
A2 151905846 151905969
refGene_gn=AKR1B1;refGene_tx=XM_003983075.4
A2 151906271 151906349
refGene_gn=AKR1B1;refGene_tx=XM_003983075.4
A2 151907291 151907408
refGene_gn=AKR1B1;refGene_tx=XM_003983075.4
A2 151908104 151908272
refGene_gn=AKR1B1;refGene_tx=XM_003983075.4
A2 151916215 151916281
refGene_gn=AKR1B1;refGene_tx=XM_003983075.4
A2 152000400 152000519
refGene_gn=LOC102902631;refGene_tx=XM_023242242.1
A2 152008220 152008299
refGene_gn=LOC102902631;refGene_tx=XM_023242242.1
A2 152030213 152030731
refGene_gn=LOC102902631;refGene_tx=XM_023242242.1
A2 152035580 152035667

refGene_gn=LOC102902631;refGene_tx=XM_023242242.1
A2	152036377	152036402
refGene_gn=LOC102902631;refGene_tx=XM_023242242.1
A2	152044942	152045543
refGene_gn=BPGM;refGene_tx=XM_003983076.5;refGene_gn=BPGM;refGene_tx=XM_006929384.4;refGene_gn=BPGM;refGene_tx=XM_006929385.4;refGene_gn=BPGM;refGene_tx=XM_006929386.4
A2	152057383	152057565
refGene_gn=BPGM;refGene_tx=XM_003983076.5;refGene_gn=BPGM;refGene_tx=XM_006929384.4;refGene_gn=BPGM;refGene_tx=XM_006929385.4;refGene_gn=BPGM;refGene_tx=XM_006929386.4
A2	152246778	152246849
refGene_gn=CALD1;refGene_tx=XM_019825825.2;refGene_gn=CALD1;refGene_tx=XM_019825826.2;refGene_gn=CALD1;refGene_tx=XM_019825827.2;refGene_gn=CALD1;refGene_tx=XM_019825828.2;refGene_gn=CALD1;refGene_tx=XM_019825829.2;refGene_gn=CALD1;refGene_tx=XM_019825831.2
A2	152273953	152274006
refGene_gn=CALD1;refGene_tx=XM_019825832.2
A2	152308950	152309097
refGene_gn=CALD1;refGene_tx=XM_019825825.2;refGene_gn=CALD1;refGene_tx=XM_019825826.2;refGene_gn=CALD1;refGene_tx=XM_019825827.2;refGene_gn=CALD1;refGene_tx=XM_019825828.2;refGene_gn=CALD1;refGene_tx=XM_019825829.2;refGene_gn=CALD1;refGene_tx=XM_019825831.2;refGene_gn=CALD1;refGene_tx=XM_019825832.2
A2	152313080	152314029
refGene_gn=CALD1;refGene_tx=XM_019825825.2;refGene_gn=CALD1;refGene_tx=XM_019825826.2;refGene_gn=CALD1;refGene_tx=XM_019825827.2;refGene_gn=CALD1;refGene_tx=XM_019825828.2;refGene_gn=CALD1;refGene_tx=XM_019825829.2;refGene_gn=CALD1;refGene_tx=XM_019825831.2;refGene_gn=CALD1;refGene_tx=XM_019825832.2
A2	152315314	152315389
refGene_gn=CALD1;refGene_tx=XM_019825825.2;refGene_gn=CALD1;refGene_tx=XM_019825826.2;refGene_gn=CALD1;refGene_tx=XM_019825827.2;refGene_gn=CALD1;refGene_tx=XM_019825828.2
A2	152320630	152320773
refGene_gn=CALD1;refGene_tx=XM_019825825.2;refGene_gn=CALD1;refGene_tx=XM_019825826.2;refGene_gn=CALD1;refGene_tx=XM_019825827.2;refGene_gn=CALD1;refGene_tx=XM_019825828.2;refGene_gn=CALD1;refGene_tx=XM_019825829.2;refGene_gn=CALD1;refGene_tx=XM_019825831.2;refGene_gn=CALD1;refGene_tx=XM_019825832.2
A2	152327899	152328164
refGene_gn=CALD1;refGene_tx=XM_019825825.2;refGene_gn=CALD1;refGene_tx=XM_019825826.2;refGene_gn=CALD1;refGene_tx=XM_019825827.2;refGene_gn=CALD1;refGene_tx=XM_019825828.2;refGene_gn=CALD1;refGene_tx=XM_019825829.2;refGene_gn=CALD1;refGene_tx=XM_019825831.2;refGene_gn=CALD1;refGene_tx=XM_019825832.2
A2	152330683	152330824
refGene_gn=CALD1;refGene_tx=XM_019825825.2;refGene_gn=CALD1;refGene_tx=XM_019825826.2;refGene_gn=CALD1;refGene_tx=XM_019825827.2;refGene_gn=CALD1;refGene_tx=XM_019825828.2;refGene_gn=CALD1;refGene_tx=XM_0198258

29.2;refGene_gn=CALD1;refGene_tx=XM_019825831.2;refGene_gn=CALD1;refGene_tx=XM_019825832.2
A2 152336128 152336172
refGene_gn=CALD1;refGene_tx=XM_019825825.2;refGene_gn=CALD1;refGene_tx=XM_019825826.2;refGene_gn=CALD1;refGene_tx=XM_019825827.2;refGene_gn=CALD1;refGene_tx=XM_019825828.2;refGene_gn=CALD1;refGene_tx=XM_019825829.2;refGene_gn=CALD1;refGene_tx=XM_019825831.2;refGene_gn=CALD1;refGene_tx=XM_019825832.2
A2 152336276 152336358
refGene_gn=CALD1;refGene_tx=XM_019825825.2;refGene_gn=CALD1;refGene_tx=XM_019825826.2;refGene_gn=CALD1;refGene_tx=XM_019825827.2;refGene_gn=CALD1;refGene_tx=XM_019825828.2;refGene_gn=CALD1;refGene_tx=XM_019825829.2;refGene_gn=CALD1;refGene_tx=XM_019825831.2;refGene_gn=CALD1;refGene_tx=XM_019825832.2
A2 152338502 152338640
refGene_gn=CALD1;refGene_tx=XM_019825825.2;refGene_gn=CALD1;refGene_tx=XM_019825826.2;refGene_gn=CALD1;refGene_tx=XM_019825827.2;refGene_gn=CALD1;refGene_tx=XM_019825828.2;refGene_gn=CALD1;refGene_tx=XM_019825829.2;refGene_gn=CALD1;refGene_tx=XM_019825831.2;refGene_gn=CALD1;refGene_tx=XM_019825832.2
A2 152339145 152339241
refGene_gn=CALD1;refGene_tx=XM_019825825.2;refGene_gn=CALD1;refGene_tx=XM_019825826.2;refGene_gn=CALD1;refGene_tx=XM_019825827.2;refGene_gn=CALD1;refGene_tx=XM_019825828.2;refGene_gn=CALD1;refGene_tx=XM_019825829.2;refGene_gn=CALD1;refGene_tx=XM_019825831.2;refGene_gn=CALD1;refGene_tx=XM_019825832.2
A2 152344574 152344655
refGene_gn=CALD1;refGene_tx=XM_019825825.2;refGene_gn=CALD1;refGene_tx=XM_019825826.2;refGene_gn=CALD1;refGene_tx=XM_019825827.2;refGene_gn=CALD1;refGene_tx=XM_019825828.2;refGene_gn=CALD1;refGene_tx=XM_019825829.2;refGene_gn=CALD1;refGene_tx=XM_019825831.2;refGene_gn=CALD1;refGene_tx=XM_019825832.2
A2 152347588 152347597
refGene_gn=CALD1;refGene_tx=XM_019825825.2;refGene_gn=CALD1;refGene_tx=XM_019825826.2;refGene_gn=CALD1;refGene_tx=XM_019825827.2;refGene_gn=CALD1;refGene_tx=XM_019825828.2;refGene_gn=CALD1;refGene_tx=XM_019825829.2;refGene_gn=CALD1;refGene_tx=XM_019825831.2;refGene_gn=CALD1;refGene_tx=XM_019825832.2
A2 152361035 152361134
refGene_gn=AGBL3;refGene_tx=XM_019825833.2;refGene_gn=AGBL3;refGene_tx=XM_019825834.2;refGene_gn=AGBL3;refGene_tx=XM_019825836.2;refGene_gn=AGBL3;refGene_tx=XM_019825837.2;refGene_gn=AGBL3;refGene_tx=XM_019825839.2;refGene_gn=AGBL3;refGene_tx=XM_019825840.2;refGene_gn=AGBL3;refGene_tx=XM_019825843.2;refGene_gn=AGBL3;refGene_tx=XM_019825844.2;refGene_gn=AGBL3;refGene_tx=XM_019825846.2
A2 152362048 152362109
refGene_gn=AGBL3;refGene_tx=XM_019825833.2;refGene_gn=AGBL3;refGene_tx=XM_019825834.2;refGene_gn=AGBL3;refGene_tx=XM_019825836.2;refGene_gn=AGBL3;refGene_tx=XM_019825837.2;refGene_gn=AGBL3;refGene_tx=XM_019825839.2;refGene_gn=AGBL3;refGene_tx=XM_019825840.2;refGene_gn=AGBL3;refGe

ne_tx=XM_019825841.2;refGene_gn=AGBL3;refGene_tx=XM_019825842.2;refGene_gn=AGBL3;refGene_tx=XM_019825843.2;refGene_gn=AGBL3;refGene_tx=XM_019825844.2;refGene_gn=AGBL3;refGene_tx=XM_019825846.2
A2 152363190 152363220
refGene_gn=AGBL3;refGene_tx=XM_019825833.2;refGene_gn=AGBL3;refGene_tx=XM_019825834.2;refGene_gn=AGBL3;refGene_tx=XM_019825836.2;refGene_gn=AGBL3;refGene_tx=XM_019825839.2;refGene_gn=AGBL3;refGene_tx=XM_019825840.2;refGene_gn=AGBL3;refGene_tx=XM_019825841.2;refGene_gn=AGBL3;refGene_tx=XM_019825843.2;refGene_gn=AGBL3;refGene_tx=XM_019825844.2
A2 152364856 152365042
refGene_gn=AGBL3;refGene_tx=XM_019825833.2;refGene_gn=AGBL3;refGene_tx=XM_019825834.2;refGene_gn=AGBL3;refGene_tx=XM_019825836.2;refGene_gn=AGBL3;refGene_tx=XM_019825837.2;refGene_gn=AGBL3;refGene_tx=XM_019825839.2;refGene_gn=AGBL3;refGene_tx=XM_019825840.2;refGene_gn=AGBL3;refGene_tx=XM_019825841.2;refGene_gn=AGBL3;refGene_tx=XM_019825842.2;refGene_gn=AGBL3;refGene_tx=XM_019825843.2;refGene_gn=AGBL3;refGene_tx=XM_019825844.2;refGene_gn=AGBL3;refGene_tx=XM_019825846.2
A2 152379872 152379980
refGene_gn=AGBL3;refGene_tx=XM_019825833.2;refGene_gn=AGBL3;refGene_tx=XM_019825834.2;refGene_gn=AGBL3;refGene_tx=XM_019825836.2;refGene_gn=AGBL3;refGene_tx=XM_019825837.2;refGene_gn=AGBL3;refGene_tx=XM_019825839.2;refGene_gn=AGBL3;refGene_tx=XM_019825840.2;refGene_gn=AGBL3;refGene_tx=XM_019825841.2;refGene_gn=AGBL3;refGene_tx=XM_019825842.2;refGene_gn=AGBL3;refGene_tx=XM_019825843.2;refGene_gn=AGBL3;refGene_tx=XM_019825844.2;refGene_gn=AGBL3;refGene_tx=XM_019825846.2
A2 152383058 152383197
refGene_gn=AGBL3;refGene_tx=XM_019825833.2;refGene_gn=AGBL3;refGene_tx=XM_019825834.2;refGene_gn=AGBL3;refGene_tx=XM_019825836.2;refGene_gn=AGBL3;refGene_tx=XM_019825837.2;refGene_gn=AGBL3;refGene_tx=XM_019825839.2;refGene_gn=AGBL3;refGene_tx=XM_019825840.2;refGene_gn=AGBL3;refGene_tx=XM_019825841.2;refGene_gn=AGBL3;refGene_tx=XM_019825842.2;refGene_gn=AGBL3;refGene_tx=XM_019825843.2;refGene_gn=AGBL3;refGene_tx=XM_019825844.2;refGene_gn=AGBL3;refGene_tx=XM_019825846.2
A2 152384396 152385176
refGene_gn=AGBL3;refGene_tx=XM_019825833.2;refGene_gn=AGBL3;refGene_tx=XM_019825834.2;refGene_gn=AGBL3;refGene_tx=XM_019825836.2;refGene_gn=AGBL3;refGene_tx=XM_019825837.2;refGene_gn=AGBL3;refGene_tx=XM_019825839.2;refGene_gn=AGBL3;refGene_tx=XM_019825840.2;refGene_gn=AGBL3;refGene_tx=XM_019825841.2;refGene_gn=AGBL3;refGene_tx=XM_019825842.2;refGene_gn=AGBL3;refGene_tx=XM_019825843.2;refGene_gn=AGBL3;refGene_tx=XM_019825844.2;refGene_gn=AGBL3;refGene_tx=XM_019825845.1;refGene_gn=AGBL3;refGene_tx=XM_019825846.2
A2 152389242 152389405
refGene_gn=AGBL3;refGene_tx=XM_019825833.2;refGene_gn=AGBL3;refGene_tx=XM_019825834.2;refGene_gn=AGBL3;refGene_tx=XM_019825836.2;refGene_gn=AGBL3;refGene_tx=XM_019825837.2;refGene_gn=AGBL3;refGene_tx=XM_019825839.2;refGene_gn=AGBL3;refGene_tx=XM_019825840.2;refGene_gn=AGBL3;refGene_tx=XM_019825841.2;refGene_gn=AGBL3;refGene_tx=XM_019825842.2;refGene_gn=AGBL3;refGene_tx=XM_019825843.2;refGene_gn=AGBL3;refGene_tx=XM_019825844.2;refGene_gn=AGBL3;refGene_tx=XM_019825845.1;refGene_gn=AGBL3;

refGene_tx=XM_019825846.2
A2 152390989 152391116
refGene_gn=AGBL3;refGene_tx=XM_019825833.2;refGene_gn=AGBL3;refGene_tx=XM_019825834.2;refGene_gn=AGBL3;refGene_tx=XM_019825836.2;refGene_gn=AGBL3;refGene_tx=XM_019825837.2;refGene_gn=AGBL3;refGene_tx=XM_019825839.2;refGene_gn=AGBL3;refGene_tx=XM_019825840.2;refGene_gn=AGBL3;refGene_tx=XM_019825841.2;refGene_gn=AGBL3;refGene_tx=XM_019825842.2;refGene_gn=AGBL3;refGene_tx=XM_019825843.2;refGene_gn=AGBL3;refGene_tx=XM_019825844.2;refGene_gn=AGBL3;refGene_tx=XM_019825845.1;refGene_gn=AGBL3;refGene_tx=XM_019825846.2
A2 152392302 152392403
refGene_gn=AGBL3;refGene_tx=XM_019825833.2;refGene_gn=AGBL3;refGene_tx=XM_019825834.2;refGene_gn=AGBL3;refGene_tx=XM_019825836.2;refGene_gn=AGBL3;refGene_tx=XM_019825837.2;refGene_gn=AGBL3;refGene_tx=XM_019825839.2;refGene_gn=AGBL3;refGene_tx=XM_019825840.2;refGene_gn=AGBL3;refGene_tx=XM_019825841.2;refGene_gn=AGBL3;refGene_tx=XM_019825842.2;refGene_gn=AGBL3;refGene_tx=XM_019825843.2;refGene_gn=AGBL3;refGene_tx=XM_019825844.2;refGene_gn=AGBL3;refGene_tx=XM_019825845.1;refGene_gn=AGBL3;refGene_tx=XM_019825846.2
A2 152392623 152392736
refGene_gn=AGBL3;refGene_tx=XM_019825833.2;refGene_gn=AGBL3;refGene_tx=XM_019825834.2;refGene_gn=AGBL3;refGene_tx=XM_019825836.2;refGene_gn=AGBL3;refGene_tx=XM_019825837.2;refGene_gn=AGBL3;refGene_tx=XM_019825839.2;refGene_gn=AGBL3;refGene_tx=XM_019825840.2;refGene_gn=AGBL3;refGene_tx=XM_019825841.2;refGene_gn=AGBL3;refGene_tx=XM_019825842.2;refGene_gn=AGBL3;refGene_tx=XM_019825843.2;refGene_gn=AGBL3;refGene_tx=XM_019825844.2;refGene_gn=AGBL3;refGene_tx=XM_019825845.1;refGene_gn=AGBL3;refGene_tx=XM_019825846.2
A2 152393242 152393309
refGene_gn=AGBL3;refGene_tx=XM_019825833.2;refGene_gn=AGBL3;refGene_tx=XM_019825834.2;refGene_gn=AGBL3;refGene_tx=XM_019825836.2;refGene_gn=AGBL3;refGene_tx=XM_019825837.2;refGene_gn=AGBL3;refGene_tx=XM_019825839.2;refGene_gn=AGBL3;refGene_tx=XM_019825840.2;refGene_gn=AGBL3;refGene_tx=XM_019825841.2;refGene_gn=AGBL3;refGene_tx=XM_019825842.2;refGene_gn=AGBL3;refGene_tx=XM_019825843.2;refGene_gn=AGBL3;refGene_tx=XM_019825844.2;refGene_gn=AGBL3;refGene_tx=XM_019825845.1;refGene_gn=AGBL3;refGene_tx=XM_019825846.2
A2 152398603 152398690
refGene_gn=AGBL3;refGene_tx=XM_019825833.2;refGene_gn=AGBL3;refGene_tx=XM_019825834.2;refGene_gn=AGBL3;refGene_tx=XM_019825836.2;refGene_gn=AGBL3;refGene_tx=XM_019825837.2;refGene_gn=AGBL3;refGene_tx=XM_019825841.2;refGene_gn=AGBL3;refGene_tx=XM_019825842.2;refGene_gn=AGBL3;refGene_tx=XM_019825843.2;refGene_gn=AGBL3;refGene_tx=XM_019825844.2;refGene_gn=AGBL3;refGene_tx=XM_019825845.1;refGene_gn=AGBL3;refGene_tx=XM_019825846.2
A2 152402025 152402083
refGene_gn=AGBL3;refGene_tx=XM_019825833.2;refGene_gn=AGBL3;refGene_tx=XM_019825834.2;refGene_gn=AGBL3;refGene_tx=XM_019825836.2;refGene_gn=AGBL3;refGene_tx=XM_019825837.2;refGene_gn=AGBL3;refGene_tx=XM_019825839.2;refGene_gn=AGBL3;refGene_tx=XM_019825840.2;refGene_gn=AGBL3;refGe

ne_tx=XM_019825841.2;refGene_gn=AGBL3;refGene_tx=XM_019825842.2;refGene_gn=AGBL3;refGene_tx=XM_019825843.2;refGene_gn=AGBL3;refGene_tx=XM_019825844.2;refGene_gn=AGBL3;refGene_tx=XM_019825845.1;refGene_gn=AGBL3;refGene_tx=XM_019825846.2
A2 152403572 152403644
refGene_gn=AGBL3;refGene_tx=XM_019825833.2;refGene_gn=AGBL3;refGene_tx=XM_019825834.2;refGene_gn=AGBL3;refGene_tx=XM_019825836.2;refGene_gn=AGBL3;refGene_tx=XM_019825837.2;refGene_gn=AGBL3;refGene_tx=XM_019825839.2;refGene_gn=AGBL3;refGene_tx=XM_019825840.2;refGene_gn=AGBL3;refGene_tx=XM_019825841.2;refGene_gn=AGBL3;refGene_tx=XM_019825842.2;refGene_gn=AGBL3;refGene_tx=XM_019825843.2;refGene_gn=AGBL3;refGene_tx=XM_019825844.2;refGene_gn=AGBL3;refGene_tx=XM_019825845.1;refGene_gn=AGBL3;refGene_tx=XM_019825846.2
A2 152417691 152417846
refGene_gn=AGBL3;refGene_tx=XM_019825844.2
A2 152420996 152421045
refGene_gn=AGBL3;refGene_tx=XM_019825833.2;refGene_gn=AGBL3;refGene_tx=XM_019825834.2;refGene_gn=AGBL3;refGene_tx=XM_019825836.2;refGene_gn=AGBL3;refGene_tx=XM_019825837.2;refGene_gn=AGBL3;refGene_tx=XM_019825839.2;refGene_gn=AGBL3;refGene_tx=XM_019825840.2;refGene_gn=AGBL3;refGene_tx=XM_019825841.2;refGene_gn=AGBL3;refGene_tx=XM_019825842.2;refGene_gn=AGBL3;refGene_tx=XM_019825845.1
A2 152422219 152422416
refGene_gn=AGBL3;refGene_tx=XM_019825833.2;refGene_gn=AGBL3;refGene_tx=XM_019825834.2;refGene_gn=AGBL3;refGene_tx=XM_019825836.2;refGene_gn=AGBL3;refGene_tx=XM_019825837.2;refGene_gn=AGBL3;refGene_tx=XM_019825839.2;refGene_gn=AGBL3;refGene_tx=XM_019825840.2;refGene_gn=AGBL3;refGene_tx=XM_019825841.2;refGene_gn=AGBL3;refGene_tx=XM_019825842.2;refGene_gn=AGBL3;refGene_tx=XM_019825845.1
A2 152426982 152427214
refGene_gn=AGBL3;refGene_tx=XM_019825833.2;refGene_gn=AGBL3;refGene_tx=XM_019825834.2;refGene_gn=AGBL3;refGene_tx=XM_019825836.2;refGene_gn=AGBL3;refGene_tx=XM_019825837.2;refGene_gn=AGBL3;refGene_tx=XM_019825839.2;refGene_gn=AGBL3;refGene_tx=XM_019825840.2;refGene_gn=AGBL3;refGene_tx=XM_019825841.2;refGene_gn=AGBL3;refGene_tx=XM_019825842.2;refGene_gn=AGBL3;refGene_tx=XM_019825843.2;refGene_gn=AGBL3;refGene_tx=XM_019825845.1;refGene_gn=AGBL3;refGene_tx=XM_019825846.2
A2 152430916 152431316
refGene_gn=AGBL3;refGene_tx=XM_019825833.2;refGene_gn=AGBL3;refGene_tx=XM_019825834.2;refGene_gn=AGBL3;refGene_tx=XM_019825836.2;refGene_gn=AGBL3;refGene_tx=XM_019825837.2;refGene_gn=AGBL3;refGene_tx=XM_019825839.2;refGene_gn=AGBL3;refGene_tx=XM_019825840.2;refGene_gn=AGBL3;refGene_tx=XM_019825841.2;refGene_gn=AGBL3;refGene_tx=XM_019825842.2;refGene_gn=AGBL3;refGene_tx=XM_019825843.2;refGene_gn=AGBL3;refGene_tx=XM_019825845.1;refGene_gn=AGBL3;refGene_tx=XM_019825846.2
A2 152461101 152461114
refGene_gn=TMEM140;refGene_tx=XM_006929395.3
A2 152463028 152463621
refGene_gn=TMEM140;refGene_tx=XM_006929395.3;refGene_gn=TMEM140;refGene_tx=XM_006929396.3;refGene_gn=TMEM140;refGene_tx=XM_006929397.3

A2 152465137 152465395
refGene_gn=CA2H7orf49;refGene_tx=XM_003983080.5;refGene_gn=CA2H7orf49;
refGene_tx=XM_011280583.3;refGene_gn=CA2H7orf49;refGene_tx=XM_01128058
4.3;refGene_gn=CA2H7orf49;refGene_tx=XM_011280585.3;refGene_gn=CA2H7or
f49;refGene_tx=XM_011280586.3;refGene_gn=CA2H7orf49;refGene_tx=XM_0198
25848.2;refGene_gn=CA2H7orf49;refGene_tx=XM_019825850.2;refGene_gn=CA2
H7orf49;refGene_tx=XM_019825851.2;refGene_gn=CA2H7orf49;refGene_tx=XM_
019825852.2;refGene_gn=CA2H7orf49;refGene_tx=XM_019825853.1;refGene_gn
=CA2H7orf49;refGene_tx=XM_023250586.1
A2 152466296 152466372
refGene_gn=CA2H7orf49;refGene_tx=XM_003983080.5;refGene_gn=CA2H7orf49;
refGene_tx=XM_011280583.3;refGene_gn=CA2H7orf49;refGene_tx=XM_01128058
4.3;refGene_gn=CA2H7orf49;refGene_tx=XM_011280585.3;refGene_gn=CA2H7or
f49;refGene_tx=XM_011280586.3;refGene_gn=CA2H7orf49;refGene_tx=XM_0198
25848.2;refGene_gn=CA2H7orf49;refGene_tx=XM_019825850.2;refGene_gn=CA2
H7orf49;refGene_tx=XM_019825851.2;refGene_gn=CA2H7orf49;refGene_tx=XM_
019825852.2;refGene_gn=CA2H7orf49;refGene_tx=XM_019825853.1;refGene_gn
=CA2H7orf49;refGene_tx=XM_023250586.1
A2 152466626 152466766
refGene_gn=CA2H7orf49;refGene_tx=XM_003983080.5;refGene_gn=CA2H7orf49;
refGene_tx=XM_011280583.3;refGene_gn=CA2H7orf49;refGene_tx=XM_01128058
4.3;refGene_gn=CA2H7orf49;refGene_tx=XM_011280585.3;refGene_gn=CA2H7or
f49;refGene_tx=XM_011280586.3;refGene_gn=CA2H7orf49;refGene_tx=XM_0198
25848.2;refGene_gn=CA2H7orf49;refGene_tx=XM_019825850.2;refGene_gn=CA2
H7orf49;refGene_tx=XM_019825851.2;refGene_gn=CA2H7orf49;refGene_tx=XM_
019825852.2;refGene_gn=CA2H7orf49;refGene_tx=XM_023250586.1
A2 152472428 152472593
refGene_gn=WDR91;refGene_tx=XM_011280581.3
A2 152473386 152473464
refGene_gn=WDR91;refGene_tx=XM_011280582.3
A2 152475014 152475212
refGene_gn=WDR91;refGene_tx=XM_011280581.3;refGene_gn=WDR91;refGene_tx
=XM_011280582.3
A2 152476479 152476592
refGene_gn=WDR91;refGene_tx=XM_011280581.3;refGene_gn=WDR91;refGene_tx
=XM_011280582.3
A2 152477617 152477726
refGene_gn=WDR91;refGene_tx=XM_011280581.3;refGene_gn=WDR91;refGene_tx
=XM_011280582.3
A2 152479745 152479914
refGene_gn=WDR91;refGene_tx=XM_011280581.3;refGene_gn=WDR91;refGene_tx
=XM_011280582.3
A2 152480055 152480150
refGene_gn=WDR91;refGene_tx=XM_011280581.3;refGene_gn=WDR91;refGene_tx
=XM_011280582.3
A2 152481729 152481880
refGene_gn=WDR91;refGene_tx=XM_011280581.3;refGene_gn=WDR91;refGene_tx
=XM_011280582.3
A2 152482611 152482805
refGene_gn=WDR91;refGene_tx=XM_011280581.3;refGene_gn=WDR91;refGene_tx

=XM_011280582.3
A2	152484311	152484467
refGene_gn=WDR91;refGene_tx=XM_011280581.3;refGene_gn=WDR91;refGene_tx
=XM_011280582.3
A2	152493512	152493678
refGene_gn=WDR91;refGene_tx=XM_011280581.3;refGene_gn=WDR91;refGene_tx
=XM_011280582.3
A2	152494764	152494895
refGene_gn=WDR91;refGene_tx=XM_011280581.3;refGene_gn=WDR91;refGene_tx
=XM_011280582.3
A2	152495879	152495962
refGene_gn=WDR91;refGene_tx=XM_011280581.3;refGene_gn=WDR91;refGene_tx
=XM_011280582.3
A2	152497533	152497741
refGene_gn=WDR91;refGene_tx=XM_011280581.3;refGene_gn=WDR91;refGene_tx
=XM_011280582.3
A2	152498292	152498472
refGene_gn=WDR91;refGene_tx=XM_011280581.3;refGene_gn=WDR91;refGene_tx
=XM_011280582.3
A2	152500003	152500126
refGene_gn=WDR91;refGene_tx=XM_011280581.3;refGene_gn=WDR91;refGene_tx
=XM_011280582.3
A2	152519569	152519761
refGene_gn=STRA8;refGene_tx=XM_023250590.1
A2	152521845	152521921
refGene_gn=STRA8;refGene_tx=XM_023250590.1
A2	152522507	152522592
refGene_gn=STRA8;refGene_tx=XM_023250590.1
A2	152524686	152524908
refGene_gn=STRA8;refGene_tx=XM_023250590.1
A2	152525787	152526073
refGene_gn=STRA8;refGene_tx=XM_023250590.1
A2	152530307	152530381
refGene_gn=STRA8;refGene_tx=XM_023250590.1
A2	152530990	152531028
refGene_gn=LOC102900083;refGene_tx=XM_019825862.2;refGene_gn=LOC102900
083;refGene_tx=XM_023250589.1
A2	152532003	152532115
refGene_gn=STRA8;refGene_tx=XM_023250590.1
A2	152535176	152535251
refGene_gn=STRA8;refGene_tx=XM_023250590.1
A2	152537177	152537269
refGene_gn=LOC102900083;refGene_tx=XM_011280601.3;refGene_gn=LOC102900
083;refGene_tx=XM_019825860.2
A2	152537973	152538144
refGene_gn=LOC102900083;refGene_tx=XM_011280601.3;refGene_gn=LOC102900
083;refGene_tx=XM_019825860.2;refGene_gn=LOC102900083;refGene_tx=XM_01
9825862.2
A2	152539499	152539595
refGene_gn=LOC102900083;refGene_tx=XM_011280601.3;refGene_gn=LOC102900

083;refGene_tx=XM_019825860.2;refGene_gn=LOC102900083;refGene_tx=XM_019825862.2;refGene_gn=LOC102900083;refGene_tx=XM_023250589.1
A2 152539690 152539799
refGene_gn=LOC102900083;refGene_tx=XM_011280601.3;refGene_gn=LOC102900083;refGene_tx=XM_019825860.2;refGene_gn=LOC102900083;refGene_tx=XM_019825862.2;refGene_gn=LOC102900083;refGene_tx=XM_023250589.1
A2 152540204 152540369
refGene_gn=LOC102900083;refGene_tx=XM_011280601.3;refGene_gn=LOC102900083;refGene_tx=XM_019825860.2;refGene_gn=LOC102900083;refGene_tx=XM_019825862.2;refGene_gn=LOC102900083;refGene_tx=XM_023250589.1
A2 152544777 152545031
refGene_gn=LOC102900083;refGene_tx=XM_011280601.3;refGene_gn=LOC102900083;refGene_tx=XM_019825860.2;refGene_gn=LOC102900083;refGene_tx=XM_019825862.2;refGene_gn=LOC102900083;refGene_tx=XM_023250589.1
A2 152546326 152546483
refGene_gn=LOC102900083;refGene_tx=XM_011280601.3;refGene_gn=LOC102900083;refGene_tx=XM_019825860.2;refGene_gn=LOC102900083;refGene_tx=XM_019825862.2;refGene_gn=LOC102900083;refGene_tx=XM_023250589.1
A2 152547576 152547697
refGene_gn=LOC102900083;refGene_tx=XM_011280601.3;refGene_gn=LOC102900083;refGene_tx=XM_019825860.2;refGene_gn=LOC102900083;refGene_tx=XM_019825862.2;refGene_gn=LOC102900083;refGene_tx=XM_023250589.1
A2 152548832 152549014
refGene_gn=LOC102900083;refGene_tx=XM_011280601.3;refGene_gn=LOC102900083;refGene_tx=XM_019825860.2;refGene_gn=LOC102900083;refGene_tx=XM_019825862.2;refGene_gn=LOC102900083;refGene_tx=XM_023250589.1
A2 152555632 152555792
refGene_gn=LOC102900083;refGene_tx=XM_011280601.3;refGene_gn=LOC102900083;refGene_tx=XM_019825860.2;refGene_gn=LOC102900083;refGene_tx=XM_019825862.2;refGene_gn=LOC102900083;refGene_tx=XM_023250589.1
A2 152573090 152573248
refGene_gn=LOC102900083;refGene_tx=XM_011280601.3;refGene_gn=LOC102900083;refGene_tx=XM_019825860.2;refGene_gn=LOC102900083;refGene_tx=XM_019825862.2;refGene_gn=LOC102900083;refGene_tx=XM_023250589.1
A2 152583430 152583577
refGene_gn=LOC102900083;refGene_tx=XM_011280601.3;refGene_gn=LOC102900083;refGene_tx=XM_019825860.2;refGene_gn=LOC102900083;refGene_tx=XM_019825862.2;refGene_gn=LOC102900083;refGene_tx=XM_023250589.1
A2 152585919 152585961
refGene_gn=LOC102900083;refGene_tx=XM_011280601.3;refGene_gn=LOC102900083;refGene_tx=XM_019825860.2;refGene_gn=LOC102900083;refGene_tx=XM_019825862.2;refGene_gn=LOC102900083;refGene_tx=XM_023250589.1
A2 152615436 152615738
refGene_gn=CNOT4;refGene_tx=XM_006929399.3;refGene_gn=CNOT4;refGene_tx=XM_006929400.3;refGene_gn=CNOT4;refGene_tx=XM_011280578.3;refGene_gn=CNOT4;refGene_tx=XM_011280579.3
A2 152616436 152616649
refGene_gn=CNOT4;refGene_tx=XM_006929399.3;refGene_gn=CNOT4;refGene_tx=XM_011280578.3
A2 152641588 152641689

refGene_gn=CNOT4;refGene_tx=XM_006929401.3;refGene_gn=CNOT4;refGene_tx
=XM_011280580.2
A2	152646689	152647187
refGene_gn=CNOT4;refGene_tx=XM_006929399.3;refGene_gn=CNOT4;refGene_tx
=XM_006929400.3;refGene_gn=CNOT4;refGene_tx=XM_006929401.3;refGene_gn=
CNOT4;refGene_tx=XM_011280578.3;refGene_gn=CNOT4;refGene_tx=XM_0112805
79.3;refGene_gn=CNOT4;refGene_tx=XM_011280580.2
A2	152648379	152648629
refGene_gn=CNOT4;refGene_tx=XM_006929399.3;refGene_gn=CNOT4;refGene_tx
=XM_006929400.3;refGene_gn=CNOT4;refGene_tx=XM_006929401.3;refGene_gn=
CNOT4;refGene_tx=XM_011280578.3;refGene_gn=CNOT4;refGene_tx=XM_0112805
79.3;refGene_gn=CNOT4;refGene_tx=XM_011280580.2
A2	152651513	152651571
refGene_gn=CNOT4;refGene_tx=XM_006929399.3;refGene_gn=CNOT4;refGene_tx
=XM_006929400.3;refGene_gn=CNOT4;refGene_tx=XM_006929401.3;refGene_gn=
CNOT4;refGene_tx=XM_011280578.3;refGene_gn=CNOT4;refGene_tx=XM_0112805
79.3;refGene_gn=CNOT4;refGene_tx=XM_011280580.2
A2	152670827	152670961
refGene_gn=CNOT4;refGene_tx=XM_006929399.3;refGene_gn=CNOT4;refGene_tx
=XM_006929400.3;refGene_gn=CNOT4;refGene_tx=XM_006929401.3;refGene_gn=
CNOT4;refGene_tx=XM_011280578.3;refGene_gn=CNOT4;refGene_tx=XM_0112805
79.3;refGene_gn=CNOT4;refGene_tx=XM_011280580.2
A2	152674181	152674307
refGene_gn=CNOT4;refGene_tx=XM_006929399.3;refGene_gn=CNOT4;refGene_tx
=XM_006929400.3;refGene_gn=CNOT4;refGene_tx=XM_006929401.3;refGene_gn=
CNOT4;refGene_tx=XM_011280578.3;refGene_gn=CNOT4;refGene_tx=XM_0112805
79.3;refGene_gn=CNOT4;refGene_tx=XM_011280580.2
A2	152675076	152675178
refGene_gn=CNOT4;refGene_tx=XM_006929399.3;refGene_gn=CNOT4;refGene_tx
=XM_006929400.3;refGene_gn=CNOT4;refGene_tx=XM_006929401.3;refGene_gn=
CNOT4;refGene_tx=XM_011280578.3;refGene_gn=CNOT4;refGene_tx=XM_0112805
79.3;refGene_gn=CNOT4;refGene_tx=XM_011280580.2
A2	152675919	152676006
refGene_gn=CNOT4;refGene_tx=XM_006929399.3;refGene_gn=CNOT4;refGene_tx
=XM_006929400.3;refGene_gn=CNOT4;refGene_tx=XM_006929401.3;refGene_gn=
CNOT4;refGene_tx=XM_011280578.3;refGene_gn=CNOT4;refGene_tx=XM_0112805
79.3;refGene_gn=CNOT4;refGene_tx=XM_011280580.2
A2	152683516	152683714
refGene_gn=CNOT4;refGene_tx=XM_006929399.3;refGene_gn=CNOT4;refGene_tx
=XM_006929400.3;refGene_gn=CNOT4;refGene_tx=XM_006929401.3;refGene_gn=
CNOT4;refGene_tx=XM_011280578.3;refGene_gn=CNOT4;refGene_tx=XM_0112805
79.3;refGene_gn=CNOT4;refGene_tx=XM_011280580.2
A2	152697375	152697549
refGene_gn=CNOT4;refGene_tx=XM_006929399.3;refGene_gn=CNOT4;refGene_tx
=XM_006929400.3;refGene_gn=CNOT4;refGene_tx=XM_006929401.3;refGene_gn=
CNOT4;refGene_tx=XM_011280578.3;refGene_gn=CNOT4;refGene_tx=XM_0112805
79.3;refGene_gn=CNOT4;refGene_tx=XM_011280580.2
A2	152795956	152795984
refGene_gn=NUP205;refGene_tx=XM_003983084.5
A2	152804235	152804378

refGene_gn=NUP205;refGene_tx=XM_003983084.5
A2 152805599 152805771
refGene_gn=NUP205;refGene_tx=XM_003983084.5
A2 152808412 152808557
refGene_gn=NUP205;refGene_tx=XM_003983084.5
A2 152808939 152809099
refGene_gn=NUP205;refGene_tx=XM_003983084.5
A2 152809531 152809760
refGene_gn=NUP205;refGene_tx=XM_003983084.5
A2 152812350 152812515
refGene_gn=NUP205;refGene_tx=XM_003983084.5
A2 152817333 152817509
refGene_gn=NUP205;refGene_tx=XM_003983084.5
A2 152821688 152821805
refGene_gn=NUP205;refGene_tx=XM_003983084.5
A2 152821985 152822123
refGene_gn=NUP205;refGene_tx=XM_003983084.5
A2 152823646 152823797
refGene_gn=NUP205;refGene_tx=XM_003983084.5
A2 152824964 152825170
refGene_gn=NUP205;refGene_tx=XM_003983084.5
A2 152827178 152827361
refGene_gn=NUP205;refGene_tx=XM_003983084.5
A2 152828756 152828807
refGene_gn=NUP205;refGene_tx=XM_003983084.5
A2 152829416 152829626
refGene_gn=NUP205;refGene_tx=XM_003983084.5
A2 152832653 152832753
refGene_gn=NUP205;refGene_tx=XM_003983084.5
A2 152833170 152833308
refGene_gn=NUP205;refGene_tx=XM_003983084.5
A2 152834438 152834628
refGene_gn=NUP205;refGene_tx=XM_003983084.5
A2 152835371 152835492
refGene_gn=NUP205;refGene_tx=XM_003983084.5
A2 152838040 152838122
refGene_gn=NUP205;refGene_tx=XM_003983084.5
A2 152838666 152838831
refGene_gn=NUP205;refGene_tx=XM_003983084.5
A2 152839141 152839266
refGene_gn=NUP205;refGene_tx=XM_003983084.5
A2 152843783 152843898
refGene_gn=NUP205;refGene_tx=XM_003983084.5
A2 152845795 152845945
refGene_gn=NUP205;refGene_tx=XM_003983084.5
A2 152846509 152846581
refGene_gn=NUP205;refGene_tx=XM_003983084.5
A2 152847165 152847323
refGene_gn=NUP205;refGene_tx=XM_003983084.5
A2 152847826 152847907

refGene_gn=NUP205;refGene_tx=XM_003983084.5
A2 152848693 152848885
refGene_gn=NUP205;refGene_tx=XM_003983084.5
A2 152849409 152849677
refGene_gn=NUP205;refGene_tx=XM_003983084.5
A2 152849767 152849866
refGene_gn=NUP205;refGene_tx=XM_003983084.5
A2 152857911 152858060
refGene_gn=NUP205;refGene_tx=XM_003983084.5
A2 152859513 152859705
refGene_gn=NUP205;refGene_tx=XM_003983084.5
A2 152860589 152860711
refGene_gn=NUP205;refGene_tx=XM_003983084.5
A2 152862342 152862481
refGene_gn=NUP205;refGene_tx=XM_003983084.5
A2 152864589 152864716
refGene_gn=NUP205;refGene_tx=XM_003983084.5
A2 152870234 152870311
refGene_gn=NUP205;refGene_tx=XM_003983084.5
A2 152872422 152872551
refGene_gn=NUP205;refGene_tx=XM_003983084.5
A2 152872770 152872897
refGene_gn=NUP205;refGene_tx=XM_003983084.5
A2 152876402 152876569
refGene_gn=NUP205;refGene_tx=XM_003983084.5
A2 152878085 152878209
refGene_gn=NUP205;refGene_tx=XM_003983084.5
A2 152878681 152878810
refGene_gn=NUP205;refGene_tx=XM_003983084.5
A2 152879362 152879436
refGene_gn=NUP205;refGene_tx=XM_003983084.5
A2 152881979 152882132
refGene_gn=NUP205;refGene_tx=XM_003983084.5
A2 152889688 152889916
refGene_gn=STMP1;refGene_tx=XM_023242248.1
A2 152902731 152902785
refGene_gn=STMP1;refGene_tx=XM_023242248.1
A2 152903639 152903714
refGene_gn=STMP1;refGene_tx=XM_023242248.1
A2 152911010 152911148
refGene_gn=SLC13A4;refGene_tx=XM_019825865.2
A2 152912852 152912990
refGene_gn=SLC13A4;refGene_tx=XM_019825865.2
A2 152914194 152914356
refGene_gn=SLC13A4;refGene_tx=XM_019825865.2
A2 152917569 152917634
refGene_gn=SLC13A4;refGene_tx=XM_019825865.2
A2 152917636 152917693
refGene_gn=SLC13A4;refGene_tx=XM_019825865.2
A2 152917907 152918005

refGene_gn=SLC13A4;refGene_tx=XM_019825865.2
A2 152918969 152919071
refGene_gn=SLC13A4;refGene_tx=XM_019825865.2
A2 152920655 152920757
refGene_gn=SLC13A4;refGene_tx=XM_019825865.2
A2 152921976 152922096
refGene_gn=SLC13A4;refGene_tx=XM_019825865.2
A2 152926499 152926684
refGene_gn=SLC13A4;refGene_tx=XM_019825865.2
A2 152928607 152928685
refGene_gn=SLC13A4;refGene_tx=XM_019825865.2
A2 152929458 152929498
refGene_gn=SLC13A4;refGene_tx=XM_019825865.2
A2 152932204 152932259
refGene_gn=SLC13A4;refGene_tx=XM_019825865.2
A2 152932666 152932839
refGene_gn=SLC13A4;refGene_tx=XM_019825865.2
A2 152934810 152934947
refGene_gn=SLC13A4;refGene_tx=XM_019825865.2
A2 152948157 152948286
refGene_gn=SLC13A4;refGene_tx=XM_019825865.2
A2 152953547 152953646
refGene_gn=SLC13A4;refGene_tx=XM_019825865.2
A2 152958823 152959168
refGene_gn=FAM180A;refGene_tx=XM_003983091.4
A2 152960639 152960740
refGene_gn=FAM180A;refGene_tx=XM_003983091.4
A2 152971218 152971294
refGene_gn=FAM180A;refGene_tx=XM_003983091.4
A2 153133594 153133681
refGene_gn=MTPN;refGene_tx=XM_003983086.5
A2 153153446 153153530
refGene_gn=MTPN;refGene_tx=XM_003983086.5
A2 153154380 153154494
refGene_gn=MTPN;refGene_tx=XM_003983086.5
A2 153189068 153189140
refGene_gn=MTPN;refGene_tx=XM_003983086.5
A2 153960345 153961746
refGene_gn=CHRM2;refGene_tx=XM_019825870.1
A2 154096005 154096061
refGene_gn=PTN;refGene_tx=XM_006929409.3;refGene_gn=PTN;refGene_tx=XM_
023250600.1
A2 154116418 154116580
refGene_gn=PTN;refGene_tx=XM_006929409.3;refGene_gn=PTN;refGene_tx=XM_
023250600.1
A2 154119103 154119277
refGene_gn=PTN;refGene_tx=XM_006929409.3;refGene_gn=PTN;refGene_tx=XM_
023250600.1
A2 154120236 154120351
refGene_gn=PTN;refGene_tx=XM_006929409.3;refGene_gn=PTN;refGene_tx=XM_

023250600.1
A2 154231568 154231685
refGene_gn=DGKI;refGene_tx=XM_019825868.2;refGene_gn=DGKI;refGene_tx=XM_019825869.2;refGene_gn=DGKI;refGene_tx=XM_023250594.1;refGene_gn=DGKI;refGene_tx=XM_023250595.1;refGene_gn=DGKI;refGene_tx=XM_023250596.1;refGene_gn=DGKI;refGene_tx=XM_023250598.1;refGene_gn=DGKI;refGene_tx=XM_023250599.1
A2 154235562 154235662
refGene_gn=DGKI;refGene_tx=XM_019825868.2;refGene_gn=DGKI;refGene_tx=XM_019825869.2;refGene_gn=DGKI;refGene_tx=XM_023250594.1;refGene_gn=DGKI;refGene_tx=XM_023250595.1;refGene_gn=DGKI;refGene_tx=XM_023250596.1;refGene_gn=DGKI;refGene_tx=XM_023250598.1;refGene_gn=DGKI;refGene_tx=XM_023250599.1
A2 154237617 154237654
refGene_gn=DGKI;refGene_tx=XM_019825868.2;refGene_gn=DGKI;refGene_tx=XM_019825869.2;refGene_gn=DGKI;refGene_tx=XM_023250594.1;refGene_gn=DGKI;refGene_tx=XM_023250595.1;refGene_gn=DGKI;refGene_tx=XM_023250596.1;refGene_gn=DGKI;refGene_tx=XM_023250598.1;refGene_gn=DGKI;refGene_tx=XM_023250599.1
A2 154248062 154248183
refGene_gn=DGKI;refGene_tx=XM_019825868.2;refGene_gn=DGKI;refGene_tx=XM_019825869.2;refGene_gn=DGKI;refGene_tx=XM_023250594.1;refGene_gn=DGKI;refGene_tx=XM_023250595.1;refGene_gn=DGKI;refGene_tx=XM_023250596.1;refGene_gn=DGKI;refGene_tx=XM_023250598.1;refGene_gn=DGKI;refGene_tx=XM_023250599.1
A2 154251630 154251668
refGene_gn=DGKI;refGene_tx=XM_019825868.2;refGene_gn=DGKI;refGene_tx=XM_019825869.2;refGene_gn=DGKI;refGene_tx=XM_023250594.1;refGene_gn=DGKI;refGene_tx=XM_023250595.1;refGene_gn=DGKI;refGene_tx=XM_023250596.1;refGene_gn=DGKI;refGene_tx=XM_023250598.1;refGene_gn=DGKI;refGene_tx=XM_023250599.1
A2 154280246 154280272
refGene_gn=DGKI;refGene_tx=XM_019825868.2;refGene_gn=DGKI;refGene_tx=XM_019825869.2;refGene_gn=DGKI;refGene_tx=XM_023250594.1;refGene_gn=DGKI;refGene_tx=XM_023250595.1;refGene_gn=DGKI;refGene_tx=XM_023250596.1;refGene_gn=DGKI;refGene_tx=XM_023250598.1;refGene_gn=DGKI;refGene_tx=XM_023250599.1
A2 154306598 154306721
refGene_gn=DGKI;refGene_tx=XM_019825868.2;refGene_gn=DGKI;refGene_tx=XM_019825869.2;refGene_gn=DGKI;refGene_tx=XM_023250594.1;refGene_gn=DGKI;refGene_tx=XM_023250595.1;refGene_gn=DGKI;refGene_tx=XM_023250596.1;refGene_gn=DGKI;refGene_tx=XM_023250598.1;refGene_gn=DGKI;refGene_tx=XM_023250599.1
A2 154308762 154308890
refGene_gn=DGKI;refGene_tx=XM_019825868.2;refGene_gn=DGKI;refGene_tx=XM_019825869.2;refGene_gn=DGKI;refGene_tx=XM_023250594.1;refGene_gn=DGKI;refGene_tx=XM_023250595.1;refGene_gn=DGKI;refGene_tx=XM_023250596.1;refGene_gn=DGKI;refGene_tx=XM_023250598.1;refGene_gn=DGKI;refGene_tx=XM_023250599.1
A2 154309806 154309847

refGene_gn=DGKI;refGene_tx=XM_019825868.2;refGene_gn=DGKI;refGene_tx=XM_019825869.2;refGene_gn=DGKI;refGene_tx=XM_023250594.1;refGene_gn=DGKI;refGene_tx=XM_023250595.1;refGene_gn=DGKI;refGene_tx=XM_023250596.1;refGene_gn=DGKI;refGene_tx=XM_023250598.1;refGene_gn=DGKI;refGene_tx=XM_023250599.1
A2 154311922 154311992
refGene_gn=DGKI;refGene_tx=XM_019825868.2;refGene_gn=DGKI;refGene_tx=XM_019825869.2;refGene_gn=DGKI;refGene_tx=XM_023250594.1;refGene_gn=DGKI;refGene_tx=XM_023250595.1;refGene_gn=DGKI;refGene_tx=XM_023250596.1;refGene_gn=DGKI;refGene_tx=XM_023250598.1;refGene_gn=DGKI;refGene_tx=XM_023250599.1
A2 154327058 154327127
refGene_gn=DGKI;refGene_tx=XM_019825868.2;refGene_gn=DGKI;refGene_tx=XM_023250594.1;refGene_gn=DGKI;refGene_tx=XM_023250595.1;refGene_gn=DGKI;refGene_tx=XM_023250596.1;refGene_gn=DGKI;refGene_tx=XM_023250599.1
A2 154327570 154327615
refGene_gn=DGKI;refGene_tx=XM_019825868.2;refGene_gn=DGKI;refGene_tx=XM_019825869.2;refGene_gn=DGKI;refGene_tx=XM_023250594.1;refGene_gn=DGKI;refGene_tx=XM_023250595.1;refGene_gn=DGKI;refGene_tx=XM_023250596.1;refGene_gn=DGKI;refGene_tx=XM_023250598.1;refGene_gn=DGKI;refGene_tx=XM_023250599.1
A2 154329857 154329937
refGene_gn=DGKI;refGene_tx=XM_019825868.2;refGene_gn=DGKI;refGene_tx=XM_019825869.2;refGene_gn=DGKI;refGene_tx=XM_023250594.1;refGene_gn=DGKI;refGene_tx=XM_023250595.1;refGene_gn=DGKI;refGene_tx=XM_023250596.1;refGene_gn=DGKI;refGene_tx=XM_023250598.1;refGene_gn=DGKI;refGene_tx=XM_023250599.1
A2 154351083 154351184
refGene_gn=DGKI;refGene_tx=XM_019825868.2;refGene_gn=DGKI;refGene_tx=XM_019825869.2;refGene_gn=DGKI;refGene_tx=XM_023250594.1;refGene_gn=DGKI;refGene_tx=XM_023250595.1;refGene_gn=DGKI;refGene_tx=XM_023250596.1;refGene_gn=DGKI;refGene_tx=XM_023250598.1;refGene_gn=DGKI;refGene_tx=XM_023250599.1
A2 154378686 154378749
refGene_gn=DGKI;refGene_tx=XM_023250594.1;refGene_gn=DGKI;refGene_tx=XM_023250595.1;refGene_gn=DGKI;refGene_tx=XM_023250596.1;refGene_gn=DGKI;refGene_tx=XM_023250598.1;refGene_gn=DGKI;refGene_tx=XM_023250599.1
A2 154384243 154384443
refGene_gn=DGKI;refGene_tx=XM_019825868.2;refGene_gn=DGKI;refGene_tx=XM_019825869.2;refGene_gn=DGKI;refGene_tx=XM_023250594.1;refGene_gn=DGKI;refGene_tx=XM_023250595.1;refGene_gn=DGKI;refGene_tx=XM_023250596.1;refGene_gn=DGKI;refGene_tx=XM_023250598.1;refGene_gn=DGKI;refGene_tx=XM_023250599.1
A2 154395471 154395583
refGene_gn=DGKI;refGene_tx=XM_019825868.2;refGene_gn=DGKI;refGene_tx=XM_019825869.2;refGene_gn=DGKI;refGene_tx=XM_023250594.1;refGene_gn=DGKI;refGene_tx=XM_023250595.1;refGene_gn=DGKI;refGene_tx=XM_023250596.1;refGene_gn=DGKI;refGene_tx=XM_023250598.1;refGene_gn=DGKI;refGene_tx=XM_023250599.1
A2 154397229 154397303

refGene_gn=DGKI;refGene_tx=XM_019825868.2;refGene_gn=DGKI;refGene_tx=XM_019825869.2;refGene_gn=DGKI;refGene_tx=XM_023250594.1;refGene_gn=DGKI;refGene_tx=XM_023250595.1;refGene_gn=DGKI;refGene_tx=XM_023250596.1;refGene_gn=DGKI;refGene_tx=XM_023250598.1;refGene_gn=DGKI;refGene_tx=XM_023250599.1
A2 154402064 154402127
refGene_gn=DGKI;refGene_tx=XM_019825868.2;refGene_gn=DGKI;refGene_tx=XM_019825869.2;refGene_gn=DGKI;refGene_tx=XM_023250594.1;refGene_gn=DGKI;refGene_tx=XM_023250595.1;refGene_gn=DGKI;refGene_tx=XM_023250596.1;refGene_gn=DGKI;refGene_tx=XM_023250598.1;refGene_gn=DGKI;refGene_tx=XM_023250599.1
A2 154403059 154403115
refGene_gn=DGKI;refGene_tx=XM_019825868.2;refGene_gn=DGKI;refGene_tx=XM_019825869.2;refGene_gn=DGKI;refGene_tx=XM_023250594.1;refGene_gn=DGKI;refGene_tx=XM_023250595.1;refGene_gn=DGKI;refGene_tx=XM_023250596.1;refGene_gn=DGKI;refGene_tx=XM_023250598.1;refGene_gn=DGKI;refGene_tx=XM_023250599.1
A2 154412266 154412345
refGene_gn=DGKI;refGene_tx=XM_019825868.2;refGene_gn=DGKI;refGene_tx=XM_019825869.2;refGene_gn=DGKI;refGene_tx=XM_023250594.1;refGene_gn=DGKI;refGene_tx=XM_023250595.1;refGene_gn=DGKI;refGene_tx=XM_023250596.1;refGene_gn=DGKI;refGene_tx=XM_023250598.1;refGene_gn=DGKI;refGene_tx=XM_023250599.1
A2 154416590 154416728
refGene_gn=DGKI;refGene_tx=XM_019825868.2;refGene_gn=DGKI;refGene_tx=XM_019825869.2;refGene_gn=DGKI;refGene_tx=XM_023250594.1;refGene_gn=DGKI;refGene_tx=XM_023250595.1;refGene_gn=DGKI;refGene_tx=XM_023250596.1;refGene_gn=DGKI;refGene_tx=XM_023250598.1;refGene_gn=DGKI;refGene_tx=XM_023250599.1
A2 154418125 154418239
refGene_gn=DGKI;refGene_tx=XM_019825868.2;refGene_gn=DGKI;refGene_tx=XM_019825869.2;refGene_gn=DGKI;refGene_tx=XM_023250594.1;refGene_gn=DGKI;refGene_tx=XM_023250595.1;refGene_gn=DGKI;refGene_tx=XM_023250596.1;refGene_gn=DGKI;refGene_tx=XM_023250598.1;refGene_gn=DGKI;refGene_tx=XM_023250599.1
A2 154428376 154428437
refGene_gn=DGKI;refGene_tx=XM_019825868.2;refGene_gn=DGKI;refGene_tx=XM_019825869.2;refGene_gn=DGKI;refGene_tx=XM_023250594.1;refGene_gn=DGKI;refGene_tx=XM_023250595.1;refGene_gn=DGKI;refGene_tx=XM_023250596.1;refGene_gn=DGKI;refGene_tx=XM_023250598.1;refGene_gn=DGKI;refGene_tx=XM_023250599.1
A2 154430603 154430686
refGene_gn=DGKI;refGene_tx=XM_019825868.2;refGene_gn=DGKI;refGene_tx=XM_019825869.2;refGene_gn=DGKI;refGene_tx=XM_023250594.1;refGene_gn=DGKI;refGene_tx=XM_023250595.1;refGene_gn=DGKI;refGene_tx=XM_023250596.1;refGene_gn=DGKI;refGene_tx=XM_023250598.1;refGene_gn=DGKI;refGene_tx=XM_023250599.1
A2 154440811 154440910
refGene_gn=DGKI;refGene_tx=XM_019825868.2;refGene_gn=DGKI;refGene_tx=XM_019825869.2;refGene_gn=DGKI;refGene_tx=XM_023250594.1;refGene_gn=DGK

I;refGene_tx=XM_023250595.1;refGene_gn=DGKI;refGene_tx=XM_023250596.1;
refGene_gn=DGKI;refGene_tx=XM_023250598.1;refGene_gn=DGKI;refGene_tx=X
M_023250599.1
A2 154441442 154441517
refGene_gn=DGKI;refGene_tx=XM_019825868.2;refGene_gn=DGKI;refGene_tx=X
M_019825869.2;refGene_gn=DGKI;refGene_tx=XM_023250594.1;refGene_gn=DGK
I;refGene_tx=XM_023250595.1;refGene_gn=DGKI;refGene_tx=XM_023250596.1;
refGene_gn=DGKI;refGene_tx=XM_023250598.1;refGene_gn=DGKI;refGene_tx=X
M_023250599.1
A2 154450457 154450574
refGene_gn=DGKI;refGene_tx=XM_019825868.2;refGene_gn=DGKI;refGene_tx=X
M_019825869.2;refGene_gn=DGKI;refGene_tx=XM_023250594.1;refGene_gn=DGK
I;refGene_tx=XM_023250595.1;refGene_gn=DGKI;refGene_tx=XM_023250596.1;
refGene_gn=DGKI;refGene_tx=XM_023250598.1;refGene_gn=DGKI;refGene_tx=X
M_023250599.1
A2 154453765 154453837
refGene_gn=DGKI;refGene_tx=XM_019825868.2;refGene_gn=DGKI;refGene_tx=X
M_019825869.2;refGene_gn=DGKI;refGene_tx=XM_023250594.1;refGene_gn=DGK
I;refGene_tx=XM_023250595.1;refGene_gn=DGKI;refGene_tx=XM_023250596.1;
refGene_gn=DGKI;refGene_tx=XM_023250598.1;refGene_gn=DGKI;refGene_tx=X
M_023250599.1
A2 154471865 154471931
refGene_gn=DGKI;refGene_tx=XM_019825868.2;refGene_gn=DGKI;refGene_tx=X
M_019825869.2;refGene_gn=DGKI;refGene_tx=XM_023250594.1;refGene_gn=DGK
I;refGene_tx=XM_023250595.1;refGene_gn=DGKI;refGene_tx=XM_023250596.1;
refGene_gn=DGKI;refGene_tx=XM_023250598.1;refGene_gn=DGKI;refGene_tx=X
M_023250599.1
A2 154479760 154479817
refGene_gn=DGKI;refGene_tx=XM_019825868.2;refGene_gn=DGKI;refGene_tx=X
M_019825869.2;refGene_gn=DGKI;refGene_tx=XM_023250594.1;refGene_gn=DGK
I;refGene_tx=XM_023250595.1;refGene_gn=DGKI;refGene_tx=XM_023250596.1;
refGene_gn=DGKI;refGene_tx=XM_023250598.1
A2 154481439 154481514
refGene_gn=DGKI;refGene_tx=XM_019825868.2;refGene_gn=DGKI;refGene_tx=X
M_019825869.2;refGene_gn=DGKI;refGene_tx=XM_023250594.1;refGene_gn=DGK
I;refGene_tx=XM_023250595.1;refGene_gn=DGKI;refGene_tx=XM_023250596.1;
refGene_gn=DGKI;refGene_tx=XM_023250598.1
A2 154503969 154504065
refGene_gn=DGKI;refGene_tx=XM_019825868.2;refGene_gn=DGKI;refGene_tx=X
M_019825869.2;refGene_gn=DGKI;refGene_tx=XM_023250594.1;refGene_gn=DGK
I;refGene_tx=XM_023250595.1;refGene_gn=DGKI;refGene_tx=XM_023250596.1;
refGene_gn=DGKI;refGene_tx=XM_023250598.1;refGene_gn=DGKI;refGene_tx=X
M_023250599.1
A2 154517205 154517314
refGene_gn=DGKI;refGene_tx=XM_019825868.2;refGene_gn=DGKI;refGene_tx=X
M_019825869.2;refGene_gn=DGKI;refGene_tx=XM_023250594.1;refGene_gn=DGK
I;refGene_tx=XM_023250595.1;refGene_gn=DGKI;refGene_tx=XM_023250596.1;
refGene_gn=DGKI;refGene_tx=XM_023250598.1;refGene_gn=DGKI;refGene_tx=X
M_023250599.1
A2 154668028 154668441

refGene_gn=DGKI;refGene_tx=XM_019825868.2;refGene_gn=DGKI;refGene_tx=XM_019825869.2;refGene_gn=DGKI;refGene_tx=XM_023250594.1;refGene_gn=DGKI;refGene_tx=XM_023250595.1;refGene_gn=DGKI;refGene_tx=XM_023250596.1;refGene_gn=DGKI;refGene_tx=XM_023250598.1;refGene_gn=DGKI;refGene_tx=XM_023250599.1
A2 154702712 154702788
refGene_gn=CREB3L2;refGene_tx=XM_003983094.5;refGene_gn=CREB3L2;refGene_tx=XM_019825875.2
A2 154704394 154704615
refGene_gn=CREB3L2;refGene_tx=XM_003983094.5;refGene_gn=CREB3L2;refGene_tx=XM_019825874.2;refGene_gn=CREB3L2;refGene_tx=XM_019825875.2
A2 154706018 154706145
refGene_gn=CREB3L2;refGene_tx=XM_003983094.5;refGene_gn=CREB3L2;refGene_tx=XM_019825874.2;refGene_gn=CREB3L2;refGene_tx=XM_019825875.2
A2 154706420 154706520
refGene_gn=CREB3L2;refGene_tx=XM_003983094.5;refGene_gn=CREB3L2;refGene_tx=XM_019825874.2;refGene_gn=CREB3L2;refGene_tx=XM_019825875.2
A2 154718715 154718784
refGene_gn=CREB3L2;refGene_tx=XM_003983094.5;refGene_gn=CREB3L2;refGene_tx=XM_019825874.2;refGene_gn=CREB3L2;refGene_tx=XM_019825875.2
A2 154720118 154720177
refGene_gn=CREB3L2;refGene_tx=XM_003983094.5;refGene_gn=CREB3L2;refGene_tx=XM_019825874.2;refGene_gn=CREB3L2;refGene_tx=XM_019825875.2
A2 154721390 154721537
refGene_gn=CREB3L2;refGene_tx=XM_003983094.5;refGene_gn=CREB3L2;refGene_tx=XM_019825874.2;refGene_gn=CREB3L2;refGene_tx=XM_019825875.2
A2 154723785 154723970
refGene_gn=CREB3L2;refGene_tx=XM_003983094.5;refGene_gn=CREB3L2;refGene_tx=XM_019825874.2;refGene_gn=CREB3L2;refGene_tx=XM_019825875.2
A2 154728654 154728742
refGene_gn=CREB3L2;refGene_tx=XM_003983094.5;refGene_gn=CREB3L2;refGene_tx=XM_019825874.2
A2 154730205 154730381
refGene_gn=CREB3L2;refGene_tx=XM_003983094.5;refGene_gn=CREB3L2;refGene_tx=XM_019825874.2
A2 154745530 154745747
refGene_gn=CREB3L2;refGene_tx=XM_003983094.5;refGene_gn=CREB3L2;refGene_tx=XM_019825874.2;refGene_gn=CREB3L2;refGene_tx=XM_019825875.2
A2 154818670 154818772
refGene_gn=CREB3L2;refGene_tx=XM_003983094.5;refGene_gn=CREB3L2;refGene_tx=XM_019825874.2;refGene_gn=CREB3L2;refGene_tx=XM_019825875.2
A2 154821528 154821591
refGene_gn=AKR1D1;refGene_tx=XM_006929410.4
A2 154867828 154867921
refGene_gn=AKR1D1;refGene_tx=XM_019825876.2
A2 154868897 154869076
refGene_gn=LOC101100956;refGene_tx=XM_023250602.1
A2 154869980 154870208
refGene_gn=LOC101100956;refGene_tx=XM_023250602.1;refGene_gn=LOC101100956;refGene_tx=XM_023250603.1

A2	154878747	154878915
refGene_gn=AKR1D1;refGene_tx=XM_006929410.4;refGene_gn=AKR1D1;refGene_tx=XM_019825876.2
A2	154882141	154882258
refGene_gn=AKR1D1;refGene_tx=XM_006929410.4;refGene_gn=AKR1D1;refGene_tx=XM_019825876.2
A2	154887914	154887992
refGene_gn=AKR1D1;refGene_tx=XM_006929410.4;refGene_gn=AKR1D1;refGene_tx=XM_019825876.2
A2	154892496	154892619
refGene_gn=AKR1D1;refGene_tx=XM_006929410.4;refGene_gn=AKR1D1;refGene_tx=XM_019825876.2
A2	154897561	154897671
refGene_gn=AKR1D1;refGene_tx=XM_006929410.4;refGene_gn=AKR1D1;refGene_tx=XM_019825876.2
A2	154898865	154899031
refGene_gn=AKR1D1;refGene_tx=XM_006929410.4;refGene_gn=AKR1D1;refGene_tx=XM_019825876.2
A2	154908171	154908254
refGene_gn=AKR1D1;refGene_tx=XM_006929410.4;refGene_gn=AKR1D1;refGene_tx=XM_019825876.2
A2	154910264	154910307
refGene_gn=AKR1D1;refGene_tx=XM_006929410.4;refGene_gn=AKR1D1;refGene_tx=XM_019825876.2
A2	155119573	155119946
refGene_gn=LOC111557457;refGene_tx=XM_023242259.1
A2	155119999	155120163
refGene_gn=LOC111557457;refGene_tx=XM_023242259.1
A2	155120318	155120682
refGene_gn=LOC101089090;refGene_tx=XM_019825074.2;refGene_gn=LOC101089090;refGene_tx=XM_023250604.1
A2	155180537	155180656
refGene_gn=LOC101089090;refGene_tx=XM_019825074.2;refGene_gn=LOC101089090;refGene_tx=XM_023250604.1
A2	155188839	155188987
refGene_gn=LOC101089090;refGene_tx=XM_019825074.2;refGene_gn=LOC101089090;refGene_tx=XM_023250604.1
A2	155191965	155192098
refGene_gn=LOC101089090;refGene_tx=XM_019825074.2;refGene_gn=LOC101089090;refGene_tx=XM_023250604.1
A2	155198176	155198293
refGene_gn=LOC101089090;refGene_tx=XM_019825074.2;refGene_gn=LOC101089090;refGene_tx=XM_023250604.1
A2	155201049	155201164
refGene_gn=LOC101089090;refGene_tx=XM_019825074.2;refGene_gn=LOC101089090;refGene_tx=XM_023250604.1
A2	155203313	155203460
refGene_gn=LOC101089090;refGene_tx=XM_019825074.2;refGene_gn=LOC101089090;refGene_tx=XM_023250604.1
A2	155212034	155212152

refGene_gn=LOC101089090;refGene_tx=XM_019825074.2;refGene_gn=LOC101089090;refGene_tx=XM_023250604.1
A2 155214799 155215068
refGene_gn=LOC101089090;refGene_tx=XM_019825074.2;refGene_gn=LOC101089090;refGene_tx=XM_023250604.1
A2 155219812 155219986
refGene_gn=LOC101089090;refGene_tx=XM_019825074.2;refGene_gn=LOC101089090;refGene_tx=XM_023250604.1
A2 155229474 155229648
refGene_gn=LOC101089090;refGene_tx=XM_019825074.2;refGene_gn=LOC101089090;refGene_tx=XM_023250604.1
A2 155232089 155232225
refGene_gn=LOC101089090;refGene_tx=XM_019825074.2;refGene_gn=LOC101089090;refGene_tx=XM_023250604.1
A2 155233980 155234053
refGene_gn=LOC101089090;refGene_tx=XM_019825074.2;refGene_gn=LOC101089090;refGene_tx=XM_023250604.1
A2 155235136 155235305
refGene_gn=LOC101089090;refGene_tx=XM_019825074.2;refGene_gn=LOC101089090;refGene_tx=XM_023250604.1
A2 155237586 155237915
refGene_gn=LOC101089090;refGene_tx=XM_019825074.2;refGene_gn=LOC101089090;refGene_tx=XM_023250604.1
A2 155239480 155239613
refGene_gn=LOC101089090;refGene_tx=XM_019825074.2;refGene_gn=LOC101089090;refGene_tx=XM_023250604.1
A2 155240851 155240926
refGene_gn=LOC101089090;refGene_tx=XM_019825074.2;refGene_gn=LOC101089090;refGene_tx=XM_023250604.1
A2 155241575 155241725
refGene_gn=LOC101089090;refGene_tx=XM_019825074.2;refGene_gn=LOC101089090;refGene_tx=XM_023250604.1
A2 155242211 155242421
refGene_gn=LOC101089090;refGene_tx=XM_019825074.2;refGene_gn=LOC101089090;refGene_tx=XM_023250604.1
A2 155258439 155258451
refGene_gn=SVOPL;refGene_tx=XM_023250606.1
A2 155258912 155259026
refGene_gn=SVOPL;refGene_tx=XM_023250606.1
A2 155272501 155272591
refGene_gn=SVOPL;refGene_tx=XM_023250606.1
A2 155280662 155280744
refGene_gn=SVOPL;refGene_tx=XM_023250606.1
A2 155281472 155281584
refGene_gn=SVOPL;refGene_tx=XM_023250606.1
A2 155282211 155282411
refGene_gn=SVOPL;refGene_tx=XM_023250606.1
A2 155292417 155292491
refGene_gn=SVOPL;refGene_tx=XM_023250606.1
A2 155302996 155303125

refGene_gn=SVOPL;refGene_tx=XM_023250606.1
A2 155305635 155305761
refGene_gn=SVOPL;refGene_tx=XM_023250606.1
A2 155309139 155309203
refGene_gn=SVOPL;refGene_tx=XM_023250606.1
A2 155312036 155312161
refGene_gn=SVOPL;refGene_tx=XM_023250606.1
A2 155313159 155313231
refGene_gn=SVOPL;refGene_tx=XM_023250606.1
A2 155321901 155322000
refGene_gn=SVOPL;refGene_tx=XM_023250606.1
A2 155326030 155326122
refGene_gn=SVOPL;refGene_tx=XM_023250606.1
A2 155326307 155326389
refGene_gn=SVOPL;refGene_tx=XM_023250606.1
A2 155352477 155352571
refGene_gn=ATP6V0A4;refGene_tx=XM_011280577.3;refGene_gn=ATP6V0A4;refGene_tx=XM_019825880.2;refGene_gn=ATP6V0A4;refGene_tx=XM_019825881.2
A2 155354039 155354211
refGene_gn=ATP6V0A4;refGene_tx=XM_011280577.3;refGene_gn=ATP6V0A4;refGene_tx=XM_019825880.2;refGene_gn=ATP6V0A4;refGene_tx=XM_019825881.2
A2 155357366 155357484
refGene_gn=ATP6V0A4;refGene_tx=XM_011280577.3;refGene_gn=ATP6V0A4;refGene_tx=XM_019825880.2;refGene_gn=ATP6V0A4;refGene_tx=XM_019825881.2
A2 155361163 155361292
refGene_gn=ATP6V0A4;refGene_tx=XM_011280577.3;refGene_gn=ATP6V0A4;refGene_tx=XM_019825880.2;refGene_gn=ATP6V0A4;refGene_tx=XM_019825881.2
A2 155388277 155388379
refGene_gn=ATP6V0A4;refGene_tx=XM_011280577.3;refGene_gn=ATP6V0A4;refGene_tx=XM_019825880.2;refGene_gn=ATP6V0A4;refGene_tx=XM_019825881.2
A2 155391635 155391852
refGene_gn=ATP6V0A4;refGene_tx=XM_011280577.3;refGene_gn=ATP6V0A4;refGene_tx=XM_019825880.2;refGene_gn=ATP6V0A4;refGene_tx=XM_019825881.2
A2 155393465 155393584
refGene_gn=ATP6V0A4;refGene_tx=XM_011280577.3;refGene_gn=ATP6V0A4;refGene_tx=XM_019825880.2;refGene_gn=ATP6V0A4;refGene_tx=XM_019825881.2
A2 155397649 155397743
refGene_gn=ATP6V0A4;refGene_tx=XM_011280577.3;refGene_gn=ATP6V0A4;refGene_tx=XM_019825880.2;refGene_gn=ATP6V0A4;refGene_tx=XM_019825881.2
A2 155400850 155401008
refGene_gn=ATP6V0A4;refGene_tx=XM_011280577.3;refGene_gn=ATP6V0A4;refGene_tx=XM_019825880.2;refGene_gn=ATP6V0A4;refGene_tx=XM_019825881.2
A2 155402352 155402492
refGene_gn=ATP6V0A4;refGene_tx=XM_011280577.3;refGene_gn=ATP6V0A4;refGene_tx=XM_019825880.2;refGene_gn=ATP6V0A4;refGene_tx=XM_019825881.2
A2 155403338 155403489
refGene_gn=ATP6V0A4;refGene_tx=XM_011280577.3;refGene_gn=ATP6V0A4;refGene_tx=XM_019825880.2;refGene_gn=ATP6V0A4;refGene_tx=XM_019825881.2
A2 155406991 155407204
refGene_gn=ATP6V0A4;refGene_tx=XM_011280577.3;refGene_gn=ATP6V0A4;refG

ene_tx=XM_019825880.2;refGene_gn=ATP6V0A4;refGene_tx=XM_019825881.2
A2 155409741 155409835
refGene_gn=ATP6V0A4;refGene_tx=XM_011280577.3;refGene_gn=ATP6V0A4;refGene_tx=XM_019825880.2;refGene_gn=ATP6V0A4;refGene_tx=XM_019825881.2
A2 155410973 155411056
refGene_gn=ATP6V0A4;refGene_tx=XM_011280577.3;refGene_gn=ATP6V0A4;refGene_tx=XM_019825880.2;refGene_gn=ATP6V0A4;refGene_tx=XM_019825881.2
A2 155416234 155416361
refGene_gn=ATP6V0A4;refGene_tx=XM_011280577.3;refGene_gn=ATP6V0A4;refGene_tx=XM_019825880.2;refGene_gn=ATP6V0A4;refGene_tx=XM_019825881.2
A2 155417694 155417789
refGene_gn=ATP6V0A4;refGene_tx=XM_011280577.3;refGene_gn=ATP6V0A4;refGene_tx=XM_019825880.2;refGene_gn=ATP6V0A4;refGene_tx=XM_019825881.2
A2 155418238 155418364
refGene_gn=ATP6V0A4;refGene_tx=XM_011280577.3;refGene_gn=ATP6V0A4;refGene_tx=XM_019825880.2;refGene_gn=ATP6V0A4;refGene_tx=XM_019825881.2
A2 155425403 155425498
refGene_gn=ATP6V0A4;refGene_tx=XM_011280577.3;refGene_gn=ATP6V0A4;refGene_tx=XM_019825880.2;refGene_gn=ATP6V0A4;refGene_tx=XM_019825881.2
A2 155425795 155425874
refGene_gn=ATP6V0A4;refGene_tx=XM_011280577.3;refGene_gn=ATP6V0A4;refGene_tx=XM_019825880.2;refGene_gn=ATP6V0A4;refGene_tx=XM_019825881.2
A2 155426957 155427074
refGene_gn=ATP6V0A4;refGene_tx=XM_011280577.3;refGene_gn=ATP6V0A4;refGene_tx=XM_019825880.2;refGene_gn=ATP6V0A4;refGene_tx=XM_019825881.2
A2 155452357 155452439
refGene_gn=TMEM213;refGene_tx=XM_003983098.4;refGene_gn=TMEM213;refGene_tx=XM_019825883.2;refGene_gn=TMEM213;refGene_tx=XM_019825884.2
A2 155455991 155456063
refGene_gn=TMEM213;refGene_tx=XM_003983098.4;refGene_gn=TMEM213;refGene_tx=XM_019825883.2;refGene_gn=TMEM213;refGene_tx=XM_019825884.2
A2 155456908 155457078
refGene_gn=TMEM213;refGene_tx=XM_003983098.4;refGene_gn=TMEM213;refGene_tx=XM_019825883.2;refGene_gn=TMEM213;refGene_tx=XM_019825884.2
A2 155481846 155482101
refGene_gn=KIAA1549;refGene_tx=XM_023242271.1
A2 155484533 155484679
refGene_gn=KIAA1549;refGene_tx=XM_023242271.1
A2 155495732 155495890
refGene_gn=KIAA1549;refGene_tx=XM_023242271.1
A2 155502149 155502196
refGene_gn=KIAA1549;refGene_tx=XM_023242271.1
A2 155513008 155513326
refGene_gn=KIAA1549;refGene_tx=XM_023242271.1
A2 155518934 155519088
refGene_gn=KIAA1549;refGene_tx=XM_023242271.1
A2 155520046 155520270
refGene_gn=KIAA1549;refGene_tx=XM_023242271.1
A2 155521790 155521996
refGene_gn=KIAA1549;refGene_tx=XM_023242271.1

A2	155530441	155530557
refGene_gn=KIAA1549;refGene_tx=XM_023242271.1
A2	155532266	155532463
refGene_gn=KIAA1549;refGene_tx=XM_023242271.1
A2	155539657	155539842
refGene_gn=KIAA1549;refGene_tx=XM_023242271.1
A2	155542806	155542984
refGene_gn=KIAA1549;refGene_tx=XM_023242271.1
A2	155552989	155553138
refGene_gn=KIAA1549;refGene_tx=XM_023242271.1
A2	155554605	155554665
refGene_gn=KIAA1549;refGene_tx=XM_023242271.1
A2	155555894	155556078
refGene_gn=KIAA1549;refGene_tx=XM_023242271.1
A2	155558473	155558604
refGene_gn=KIAA1549;refGene_tx=XM_023242271.1
A2	155560315	155560493
refGene_gn=KIAA1549;refGene_tx=XM_023242271.1
A2	155561588	155561677
refGene_gn=KIAA1549;refGene_tx=XM_023242271.1
A2	155566364	155567127
refGene_gn=KIAA1549;refGene_tx=XM_023242271.1
A2	155567129	155569036
refGene_gn=KIAA1549;refGene_tx=XM_023242271.1
A2	155624035	155624153
refGene_gn=KIAA1549;refGene_tx=XM_023242271.1
A2	155624154	155624267
refGene_gn=KIAA1549;refGene_tx=XM_023242271.1
A2	155624325	155624893
refGene_gn=KIAA1549;refGene_tx=XM_023242271.1
A2	155626870	155626915
refGene_gn=KIAA1549;refGene_tx=XM_023242271.1
A2	155647012	155647146
refGene_gn=ZC3HAV1L;refGene_tx=XM_023250607.1
A2	155649104	155649363
refGene_gn=ZC3HAV1L;refGene_tx=XM_023250607.1
A2	155654718	155654854
refGene_gn=ZC3HAV1L;refGene_tx=XM_023250607.1
A2	155655753	155656121
refGene_gn=ZC3HAV1L;refGene_tx=XM_023250607.1
A2	155666590	155666856
refGene_gn=ZC3HAV1;refGene_tx=XM_006929411.4
A2	155678304	155678438
refGene_gn=ZC3HAV1;refGene_tx=XM_006929411.4
A2	155679704	155679835
refGene_gn=ZC3HAV1;refGene_tx=XM_006929411.4
A2	155680597	155680679
refGene_gn=ZC3HAV1;refGene_tx=XM_006929411.4
A2	155683427	155683530
refGene_gn=ZC3HAV1;refGene_tx=XM_006929411.4

A2	155688199	155688320
refGene_gn=ZC3HAV1;refGene_tx=XM_006929411.4
A2	155691115	155691290
refGene_gn=ZC3HAV1;refGene_tx=XM_006929411.4
A2	155694121	155694245
refGene_gn=ZC3HAV1;refGene_tx=XM_006929411.4
A2	155697119	155697218
refGene_gn=ZC3HAV1;refGene_tx=XM_006929411.4
A2	155697661	155698798
refGene_gn=ZC3HAV1;refGene_tx=XM_006929411.4
A2	155702331	155702584
refGene_gn=ZC3HAV1;refGene_tx=XM_006929411.4
A2	155707662	155707798
refGene_gn=ZC3HAV1;refGene_tx=XM_006929411.4
A2	155725092	155725400
refGene_gn=ZC3HAV1;refGene_tx=XM_006929411.4
A2	155743798	155743801
refGene_gn=TTC26;refGene_tx=XM_011280575.3
A2	155744307	155744445
refGene_gn=TTC26;refGene_tx=XM_011280575.3
A2	155747769	155747862
refGene_gn=TTC26;refGene_tx=XM_011280575.3
A2	155750944	155751059
refGene_gn=TTC26;refGene_tx=XM_011280575.3;refGene_gn=TTC26;refGene_tx=XM_011280576.3;refGene_gn=TTC26;refGene_tx=XM_019825885.2
A2	155753272	155753322
refGene_gn=TTC26;refGene_tx=XM_011280575.3;refGene_gn=TTC26;refGene_tx=XM_011280576.3;refGene_gn=TTC26;refGene_tx=XM_019825885.2
A2	155759544	155759690
refGene_gn=TTC26;refGene_tx=XM_011280575.3;refGene_gn=TTC26;refGene_tx=XM_011280576.3;refGene_gn=TTC26;refGene_tx=XM_019825885.2;refGene_gn=TTC26;refGene_tx=XM_019825886.2
A2	155760596	155760768
refGene_gn=TTC26;refGene_tx=XM_011280575.3;refGene_gn=TTC26;refGene_tx=XM_011280576.3;refGene_gn=TTC26;refGene_tx=XM_019825885.2;refGene_gn=TTC26;refGene_tx=XM_019825886.2
A2	155768530	155768605
refGene_gn=TTC26;refGene_tx=XM_011280575.3;refGene_gn=TTC26;refGene_tx=XM_011280576.3;refGene_gn=TTC26;refGene_tx=XM_019825885.2;refGene_gn=TTC26;refGene_tx=XM_019825886.2
A2	155773221	155773324
refGene_gn=TTC26;refGene_tx=XM_011280575.3;refGene_gn=TTC26;refGene_tx=XM_011280576.3;refGene_gn=TTC26;refGene_tx=XM_019825885.2;refGene_gn=TTC26;refGene_tx=XM_019825886.2
A2	155774557	155774613
refGene_gn=TTC26;refGene_tx=XM_011280575.3;refGene_gn=TTC26;refGene_tx=XM_011280576.3;refGene_gn=TTC26;refGene_tx=XM_019825885.2;refGene_gn=TTC26;refGene_tx=XM_019825886.2
A2	155776172	155776226
refGene_gn=TTC26;refGene_tx=XM_011280575.3;refGene_gn=TTC26;refGene_tx

=XM_011280576.3;refGene_gn=TTC26;refGene_tx=XM_019825885.2;refGene_gn=TTC26;refGene_tx=XM_019825886.2
A2 155777126 155777190
refGene_gn=TTC26;refGene_tx=XM_011280575.3;refGene_gn=TTC26;refGene_tx=XM_011280576.3;refGene_gn=TTC26;refGene_tx=XM_019825885.2;refGene_gn=TTC26;refGene_tx=XM_019825886.2
A2 155779150 155779239
refGene_gn=TTC26;refGene_tx=XM_011280575.3;refGene_gn=TTC26;refGene_tx=XM_011280576.3;refGene_gn=TTC26;refGene_tx=XM_019825885.2;refGene_gn=TTC26;refGene_tx=XM_019825886.2
A2 155779453 155779537
refGene_gn=TTC26;refGene_tx=XM_011280575.3;refGene_gn=TTC26;refGene_tx=XM_011280576.3;refGene_gn=TTC26;refGene_tx=XM_019825885.2;refGene_gn=TTC26;refGene_tx=XM_019825886.2
A2 155781706 155781776
refGene_gn=TTC26;refGene_tx=XM_011280575.3;refGene_gn=TTC26;refGene_tx=XM_011280576.3;refGene_gn=TTC26;refGene_tx=XM_019825885.2;refGene_gn=TTC26;refGene_tx=XM_019825886.2
A2 155784161 155784268
refGene_gn=TTC26;refGene_tx=XM_011280575.3;refGene_gn=TTC26;refGene_tx=XM_011280576.3;refGene_gn=TTC26;refGene_tx=XM_019825885.2;refGene_gn=TTC26;refGene_tx=XM_019825886.2
A2 155789384 155789518
refGene_gn=TTC26;refGene_tx=XM_011280575.3;refGene_gn=TTC26;refGene_tx=XM_011280576.3;refGene_gn=TTC26;refGene_tx=XM_019825885.2;refGene_gn=TTC26;refGene_tx=XM_019825886.2
A2 155792159 155792271
refGene_gn=TTC26;refGene_tx=XM_011280575.3;refGene_gn=TTC26;refGene_tx=XM_011280576.3;refGene_gn=TTC26;refGene_tx=XM_019825885.2;refGene_gn=TTC26;refGene_tx=XM_019825886.2
A2 155813496 155813967
refGene_gn=UBN2;refGene_tx=XM_011280574.3;refGene_gn=UBN2;refGene_tx=XM_023250608.1;refGene_gn=UBN2;refGene_tx=XM_023250609.1;refGene_gn=UBN2;refGene_tx=XM_023250610.1;refGene_gn=UBN2;refGene_tx=XM_023250611.1
A2 155819635 155819728
refGene_gn=UBN2;refGene_tx=XM_011280574.3;refGene_gn=UBN2;refGene_tx=XM_023250608.1;refGene_gn=UBN2;refGene_tx=XM_023250609.1;refGene_gn=UBN2;refGene_tx=XM_023250610.1;refGene_gn=UBN2;refGene_tx=XM_023250611.1
A2 155841164 155841266
refGene_gn=UBN2;refGene_tx=XM_011280574.3;refGene_gn=UBN2;refGene_tx=XM_023250608.1;refGene_gn=UBN2;refGene_tx=XM_023250610.1;refGene_gn=UBN2;refGene_tx=XM_023250611.1
A2 155844932 155845070
refGene_gn=UBN2;refGene_tx=XM_011280574.3;refGene_gn=UBN2;refGene_tx=XM_023250608.1;refGene_gn=UBN2;refGene_tx=XM_023250609.1;refGene_gn=UBN2;refGene_tx=XM_023250610.1;refGene_gn=UBN2;refGene_tx=XM_023250611.1
A2 155845677 155845781
refGene_gn=UBN2;refGene_tx=XM_011280574.3;refGene_gn=UBN2;refGene_tx=XM_023250608.1;refGene_gn=UBN2;refGene_tx=XM_023250609.1;refGene_gn=UBN2;refGene_tx=XM_023250610.1;refGene_gn=UBN2;refGene_tx=XM_023250611.1

A2	155847772	155848262
refGene_gn=UBN2;refGene_tx=XM_011280574.3;refGene_gn=UBN2;refGene_tx=XM_023250608.1;refGene_gn=UBN2;refGene_tx=XM_023250609.1;refGene_gn=UBN2;refGene_tx=XM_023250610.1;refGene_gn=UBN2;refGene_tx=XM_023250611.1
A2	155853293	155853364
refGene_gn=UBN2;refGene_tx=XM_011280574.3;refGene_gn=UBN2;refGene_tx=XM_023250608.1;refGene_gn=UBN2;refGene_tx=XM_023250609.1;refGene_gn=UBN2;refGene_tx=XM_023250610.1;refGene_gn=UBN2;refGene_tx=XM_023250611.1
A2	155858505	155858635
refGene_gn=UBN2;refGene_tx=XM_011280574.3;refGene_gn=UBN2;refGene_tx=XM_023250608.1;refGene_gn=UBN2;refGene_tx=XM_023250609.1;refGene_gn=UBN2;refGene_tx=XM_023250610.1;refGene_gn=UBN2;refGene_tx=XM_023250611.1
A2	155861671	155861790
refGene_gn=UBN2;refGene_tx=XM_011280574.3;refGene_gn=UBN2;refGene_tx=XM_023250608.1;refGene_gn=UBN2;refGene_tx=XM_023250609.1;refGene_gn=UBN2;refGene_tx=XM_023250610.1;refGene_gn=UBN2;refGene_tx=XM_023250611.1
A2	155862296	155862410
refGene_gn=UBN2;refGene_tx=XM_011280574.3;refGene_gn=UBN2;refGene_tx=XM_023250608.1;refGene_gn=UBN2;refGene_tx=XM_023250609.1;refGene_gn=UBN2;refGene_tx=XM_023250610.1;refGene_gn=UBN2;refGene_tx=XM_023250611.1
A2	155862991	155863135
refGene_gn=UBN2;refGene_tx=XM_011280574.3;refGene_gn=UBN2;refGene_tx=XM_023250608.1;refGene_gn=UBN2;refGene_tx=XM_023250609.1;refGene_gn=UBN2;refGene_tx=XM_023250611.1
A2	155863867	155863918
refGene_gn=UBN2;refGene_tx=XM_011280574.3;refGene_gn=UBN2;refGene_tx=XM_023250608.1;refGene_gn=UBN2;refGene_tx=XM_023250609.1;refGene_gn=UBN2;refGene_tx=XM_023250610.1;refGene_gn=UBN2;refGene_tx=XM_023250611.1
A2	155866260	155866303
refGene_gn=UBN2;refGene_tx=XM_011280574.3;refGene_gn=UBN2;refGene_tx=XM_023250608.1;refGene_gn=UBN2;refGene_tx=XM_023250609.1;refGene_gn=UBN2;refGene_tx=XM_023250610.1;refGene_gn=UBN2;refGene_tx=XM_023250611.1
A2	155870510	155872061
refGene_gn=UBN2;refGene_tx=XM_011280574.3;refGene_gn=UBN2;refGene_tx=XM_023250608.1;refGene_gn=UBN2;refGene_tx=XM_023250609.1;refGene_gn=UBN2;refGene_tx=XM_023250610.1;refGene_gn=UBN2;refGene_tx=XM_023250611.1
A2	155878150	155878382
refGene_gn=UBN2;refGene_tx=XM_011280574.3;refGene_gn=UBN2;refGene_tx=XM_023250608.1;refGene_gn=UBN2;refGene_tx=XM_023250609.1;refGene_gn=UBN2;refGene_tx=XM_023250610.1
A2	155878819	155878909
refGene_gn=UBN2;refGene_tx=XM_011280574.3;refGene_gn=UBN2;refGene_tx=XM_023250608.1;refGene_gn=UBN2;refGene_tx=XM_023250609.1;refGene_gn=UBN2;refGene_tx=XM_023250610.1
A2	155881957	155882007
refGene_gn=UBN2;refGene_tx=XM_011280574.3;refGene_gn=UBN2;refGene_tx=XM_023250608.1;refGene_gn=UBN2;refGene_tx=XM_023250609.1;refGene_gn=UBN2;refGene_tx=XM_023250610.1
A2	155901181	155901220
refGene_gn=UBN2;refGene_tx=XM_023250611.1

A2	155909946	155910090
refGene_gn=FMC1;refGene_tx=XM_006929412.4
A2	155913682	155913886
refGene_gn=FMC1;refGene_tx=XM_006929412.4
A2	155921903	155921964
refGene_gn=LOC101080269;refGene_tx=XM_003983100.4;refGene_gn=LOC101080269;refGene_tx=XM_023250613.1
A2	155924959	155924997
refGene_gn=LOC101080269;refGene_tx=XM_006929413.2;refGene_gn=LOC101080269;refGene_tx=XM_023250615.1
A2	155928030	155928037
refGene_gn=LOC101080269;refGene_tx=XM_023250614.1
A2	155932106	155932250
refGene_gn=LOC101080269;refGene_tx=XM_019825888.2;refGene_gn=LOC101080269;refGene_tx=XM_019825889.2;refGene_gn=LOC101080269;refGene_tx=XM_023250612.1;refGene_gn=LOC101080269;refGene_tx=XM_023250613.1;refGene_gn=LOC101080269;refGene_tx=XM_023250614.1;refGene_gn=LOC101080269;refGene_tx=XM_023250616.1
A2	155935017	155935088
refGene_gn=LOC101080269;refGene_tx=XM_003983102.4;refGene_gn=LOC101080269;refGene_tx=XM_006929413.2;refGene_gn=LOC101080269;refGene_tx=XM_019825887.2;refGene_gn=LOC101080269;refGene_tx=XM_023250612.1;refGene_gn=LOC101080269;refGene_tx=XM_023250613.1;refGene_gn=LOC101080269;refGene_tx=XM_023250614.1;refGene_gn=LOC101080269;refGene_tx=XM_023250615.1
A2	155936095	155936169
refGene_gn=LOC101080269;refGene_tx=XM_023250612.1;refGene_gn=LOC101080269;refGene_tx=XM_023250613.1;refGene_gn=LOC101080269;refGene_tx=XM_023250614.1;refGene_gn=LOC101080269;refGene_tx=XM_023250615.1
A2	155936661	155936756
refGene_gn=LOC101080269;refGene_tx=XM_003983100.4;refGene_gn=LOC101080269;refGene_tx=XM_003983102.4;refGene_gn=LOC101080269;refGene_tx=XM_006929413.2;refGene_gn=LOC101080269;refGene_tx=XM_019825887.2;refGene_gn=LOC101080269;refGene_tx=XM_019825888.2;refGene_gn=LOC101080269;refGene_tx=XM_019825889.2;refGene_gn=LOC101080269;refGene_tx=XM_023250612.1;refGene_gn=LOC101080269;refGene_tx=XM_023250613.1;refGene_gn=LOC101080269;refGene_tx=XM_023250614.1;refGene_gn=LOC101080269;refGene_tx=XM_023250615.1;refGene_gn=LOC101080269;refGene_tx=XM_023250616.1
A2	155954180	155954279
refGene_gn=LOC101080269;refGene_tx=XM_003983100.4;refGene_gn=LOC101080269;refGene_tx=XM_003983102.4;refGene_gn=LOC101080269;refGene_tx=XM_006929413.2;refGene_gn=LOC101080269;refGene_tx=XM_011280572.3;refGene_gn=LOC101080269;refGene_tx=XM_019825887.2;refGene_gn=LOC101080269;refGene_tx=XM_019825888.2;refGene_gn=LOC101080269;refGene_tx=XM_019825889.2;refGene_gn=LOC101080269;refGene_tx=XM_023250612.1;refGene_gn=LOC101080269;refGene_tx=XM_023250613.1;refGene_gn=LOC101080269;refGene_tx=XM_023250614.1;refGene_gn=LOC101080269;refGene_tx=XM_023250615.1;refGene_gn=LOC101080269;refGene_tx=XM_023250616.1
A2	155955251	155955362
refGene_gn=LOC101080269;refGene_tx=XM_003983100.4;refGene_gn=LOC101080269;refGene_tx=XM_003983102.4;refGene_gn=LOC101080269;refGene_tx=XM_00

6929413.2;refGene_gn=LOC101080269;refGene_tx=XM_011280572.3;refGene_gn
=LOC101080269;refGene_tx=XM_019825887.2;refGene_gn=LOC101080269;refGen
e_tx=XM_019825888.2;refGene_gn=LOC101080269;refGene_tx=XM_019825889.2;
refGene_gn=LOC101080269;refGene_tx=XM_023250612.1;refGene_gn=LOC101080
269;refGene_tx=XM_023250613.1;refGene_gn=LOC101080269;refGene_tx=XM_02
3250614.1;refGene_gn=LOC101080269;refGene_tx=XM_023250615.1;refGene_gn
=LOC101080269;refGene_tx=XM_023250616.1
A2 155959064 155959208
refGene_gn=LOC101080269;refGene_tx=XM_003983100.4;refGene_gn=LOC101080
269;refGene_tx=XM_003983102.4;refGene_gn=LOC101080269;refGene_tx=XM_00
6929413.2;refGene_gn=LOC101080269;refGene_tx=XM_011280572.3;refGene_gn
=LOC101080269;refGene_tx=XM_019825887.2;refGene_gn=LOC101080269;refGen
e_tx=XM_019825888.2;refGene_gn=LOC101080269;refGene_tx=XM_019825889.2;
refGene_gn=LOC101080269;refGene_tx=XM_023250612.1;refGene_gn=LOC101080
269;refGene_tx=XM_023250613.1;refGene_gn=LOC101080269;refGene_tx=XM_02
3250614.1;refGene_gn=LOC101080269;refGene_tx=XM_023250615.1;refGene_gn
=LOC101080269;refGene_tx=XM_023250616.1
A2 155964705 155964882
refGene_gn=LOC101080269;refGene_tx=XM_003983100.4;refGene_gn=LOC101080
269;refGene_tx=XM_003983102.4;refGene_gn=LOC101080269;refGene_tx=XM_00
6929413.2;refGene_gn=LOC101080269;refGene_tx=XM_011280572.3;refGene_gn
=LOC101080269;refGene_tx=XM_019825887.2;refGene_gn=LOC101080269;refGen
e_tx=XM_019825888.2;refGene_gn=LOC101080269;refGene_tx=XM_019825889.2;
refGene_gn=LOC101080269;refGene_tx=XM_023250612.1;refGene_gn=LOC101080
269;refGene_tx=XM_023250613.1;refGene_gn=LOC101080269;refGene_tx=XM_02
3250614.1;refGene_gn=LOC101080269;refGene_tx=XM_023250615.1;refGene_gn
=LOC101080269;refGene_tx=XM_023250616.1
A2 155967130 155967222
refGene_gn=LOC101080269;refGene_tx=XM_003983100.4;refGene_gn=LOC101080
269;refGene_tx=XM_003983102.4;refGene_gn=LOC101080269;refGene_tx=XM_00
6929413.2;refGene_gn=LOC101080269;refGene_tx=XM_011280572.3;refGene_gn
=LOC101080269;refGene_tx=XM_019825887.2;refGene_gn=LOC101080269;refGen
e_tx=XM_019825888.2;refGene_gn=LOC101080269;refGene_tx=XM_019825889.2;
refGene_gn=LOC101080269;refGene_tx=XM_023250612.1;refGene_gn=LOC101080
269;refGene_tx=XM_023250613.1;refGene_gn=LOC101080269;refGene_tx=XM_02
3250614.1;refGene_gn=LOC101080269;refGene_tx=XM_023250615.1;refGene_gn
=LOC101080269;refGene_tx=XM_023250616.1
A2 155969902 155969932
refGene_gn=LOC101080269;refGene_tx=XM_003983100.4;refGene_gn=LOC101080
269;refGene_tx=XM_003983102.4;refGene_gn=LOC101080269;refGene_tx=XM_00
6929413.2;refGene_gn=LOC101080269;refGene_tx=XM_011280572.3;refGene_gn
=LOC101080269;refGene_tx=XM_019825887.2;refGene_gn=LOC101080269;refGen
e_tx=XM_019825888.2;refGene_gn=LOC101080269;refGene_tx=XM_019825889.2;
refGene_gn=LOC101080269;refGene_tx=XM_023250612.1;refGene_gn=LOC101080
269;refGene_tx=XM_023250613.1;refGene_gn=LOC101080269;refGene_tx=XM_02
3250614.1;refGene_gn=LOC101080269;refGene_tx=XM_023250615.1;refGene_gn
=LOC101080269;refGene_tx=XM_023250616.1
A2 155973207 155973399
refGene_gn=LOC101080269;refGene_tx=XM_003983100.4;refGene_gn=LOC101080
269;refGene_tx=XM_003983102.4;refGene_gn=LOC101080269;refGene_tx=XM_00

6929413.2;refGene_gn=LOC101080269;refGene_tx=XM_011280572.3;refGene_gn
=LOC101080269;refGene_tx=XM_019825887.2;refGene_gn=LOC101080269;refGen
e_tx=XM_019825888.2;refGene_gn=LOC101080269;refGene_tx=XM_019825889.2;
refGene_gn=LOC101080269;refGene_tx=XM_023250612.1;refGene_gn=LOC101080
269;refGene_tx=XM_023250613.1;refGene_gn=LOC101080269;refGene_tx=XM_02
3250614.1;refGene_gn=LOC101080269;refGene_tx=XM_023250615.1;refGene_gn
=LOC101080269;refGene_tx=XM_023250616.1
A2 155977064 155977242
refGene_gn=LOC101080269;refGene_tx=XM_003983100.4;refGene_gn=LOC101080
269;refGene_tx=XM_003983102.4;refGene_gn=LOC101080269;refGene_tx=XM_00
6929413.2;refGene_gn=LOC101080269;refGene_tx=XM_011280572.3;refGene_gn
=LOC101080269;refGene_tx=XM_019825887.2;refGene_gn=LOC101080269;refGen
e_tx=XM_019825888.2;refGene_gn=LOC101080269;refGene_tx=XM_019825889.2;
refGene_gn=LOC101080269;refGene_tx=XM_023250612.1;refGene_gn=LOC101080
269;refGene_tx=XM_023250613.1;refGene_gn=LOC101080269;refGene_tx=XM_02
3250614.1;refGene_gn=LOC101080269;refGene_tx=XM_023250615.1;refGene_gn
=LOC101080269;refGene_tx=XM_023250616.1
A2 155999269 155999390
refGene_gn=KLRG2;refGene_tx=XM_023250617.1
A2 155999892 155999996
refGene_gn=KLRG2;refGene_tx=XM_023250617.1
A2 156003931 156004077
refGene_gn=KLRG2;refGene_tx=XM_023250617.1
A2 156004522 156004624
refGene_gn=KLRG2;refGene_tx=XM_023250617.1
A2 156010248 156010996
refGene_gn=KLRG2;refGene_tx=XM_023250617.1
A2 156031696 156031886
refGene_gn=CLEC2L;refGene_tx=XM_006929449.4
A2 156040910 156040985
refGene_gn=CLEC2L;refGene_tx=XM_006929449.4
A2 156043150 156043317
refGene_gn=CLEC2L;refGene_tx=XM_006929449.4
A2 156044259 156044360
refGene_gn=CLEC2L;refGene_tx=XM_006929449.4
A2 156046290 156046402
refGene_gn=CLEC2L;refGene_tx=XM_006929449.4
A2 156064728 156065385
refGene_gn=HIPK2;refGene_tx=XM_023250618.1;refGene_gn=HIPK2;refGene_tx
=XM_023250619.1;refGene_gn=HIPK2;refGene_tx=XM_023250620.1;refGene_gn=
HIPK2;refGene_tx=XM_023250621.1
A2 156066770 156066928
refGene_gn=HIPK2;refGene_tx=XM_023250618.1;refGene_gn=HIPK2;refGene_tx
=XM_023250619.1;refGene_gn=HIPK2;refGene_tx=XM_023250620.1;refGene_gn=
HIPK2;refGene_tx=XM_023250621.1
A2 156073099 156073344
refGene_gn=HIPK2;refGene_tx=XM_023250618.1;refGene_gn=HIPK2;refGene_tx
=XM_023250619.1;refGene_gn=HIPK2;refGene_tx=XM_023250620.1;refGene_gn=
HIPK2;refGene_tx=XM_023250621.1
A2 156085748 156086030

refGene_gn=HIPK2;refGene_tx=XM_023250618.1;refGene_gn=HIPK2;refGene_tx=XM_023250619.1;refGene_gn=HIPK2;refGene_tx=XM_023250620.1;refGene_gn=HIPK2;refGene_tx=XM_023250621.1
A2	156088855	156089035
refGene_gn=HIPK2;refGene_tx=XM_023250618.1;refGene_gn=HIPK2;refGene_tx=XM_023250619.1;refGene_gn=HIPK2;refGene_tx=XM_023250620.1;refGene_gn=HIPK2;refGene_tx=XM_023250621.1
A2	156091758	156091901
refGene_gn=HIPK2;refGene_tx=XM_023250618.1;refGene_gn=HIPK2;refGene_tx=XM_023250619.1;refGene_gn=HIPK2;refGene_tx=XM_023250620.1;refGene_gn=HIPK2;refGene_tx=XM_023250621.1
A2	156102079	156102201
refGene_gn=HIPK2;refGene_tx=XM_023250618.1;refGene_gn=HIPK2;refGene_tx=XM_023250619.1;refGene_gn=HIPK2;refGene_tx=XM_023250620.1;refGene_gn=HIPK2;refGene_tx=XM_023250621.1
A2	156103156	156103364
refGene_gn=HIPK2;refGene_tx=XM_023250618.1;refGene_gn=HIPK2;refGene_tx=XM_023250619.1;refGene_gn=HIPK2;refGene_tx=XM_023250620.1;refGene_gn=HIPK2;refGene_tx=XM_023250621.1
A2	156106631	156106794
refGene_gn=HIPK2;refGene_tx=XM_023250618.1;refGene_gn=HIPK2;refGene_tx=XM_023250619.1;refGene_gn=HIPK2;refGene_tx=XM_023250620.1;refGene_gn=HIPK2;refGene_tx=XM_023250621.1
A2	156111970	156112155
refGene_gn=HIPK2;refGene_tx=XM_023250618.1;refGene_gn=HIPK2;refGene_tx=XM_023250619.1;refGene_gn=HIPK2;refGene_tx=XM_023250620.1;refGene_gn=HIPK2;refGene_tx=XM_023250621.1
A2	156114188	156114275
refGene_gn=HIPK2;refGene_tx=XM_023250618.1;refGene_gn=HIPK2;refGene_tx=XM_023250619.1;refGene_gn=HIPK2;refGene_tx=XM_023250620.1;refGene_gn=HIPK2;refGene_tx=XM_023250621.1
A2	156116103	156116223
refGene_gn=HIPK2;refGene_tx=XM_023250618.1;refGene_gn=HIPK2;refGene_tx=XM_023250619.1;refGene_gn=HIPK2;refGene_tx=XM_023250620.1;refGene_gn=HIPK2;refGene_tx=XM_023250621.1
A2	156116459	156116583
refGene_gn=HIPK2;refGene_tx=XM_023250618.1;refGene_gn=HIPK2;refGene_tx=XM_023250619.1;refGene_gn=HIPK2;refGene_tx=XM_023250620.1;refGene_gn=HIPK2;refGene_tx=XM_023250621.1
A2	156182502	156183586
refGene_gn=HIPK2;refGene_tx=XM_023250618.1;refGene_gn=HIPK2;refGene_tx=XM_023250619.1;refGene_gn=HIPK2;refGene_tx=XM_023250620.1;refGene_gn=HIPK2;refGene_tx=XM_023250621.1
A2	156243316	156243335
refGene_gn=HIPK2;refGene_tx=XM_023250618.1;refGene_gn=HIPK2;refGene_tx=XM_023250619.1;refGene_gn=HIPK2;refGene_tx=XM_023250620.1;refGene_gn=HIPK2;refGene_tx=XM_023250621.1
A2	156286176	156286265
refGene_gn=TBXAS1;refGene_tx=XM_011280571.3
A2	156321898	156321992

refGene_gn=TBXAS1;refGene_tx=XM_011280571.3
A2 156324637 156324690
refGene_gn=TBXAS1;refGene_tx=XM_011280571.3
A2 156355479 156355576
refGene_gn=TBXAS1;refGene_tx=XM_011280571.3
A2 156367549 156367666
refGene_gn=TBXAS1;refGene_tx=XM_011280571.3
A2 156383902 156383991
refGene_gn=TBXAS1;refGene_tx=XM_011280571.3
A2 156385741 156385890
refGene_gn=TBXAS1;refGene_tx=XM_011280571.3
A2 156387644 156387775
refGene_gn=TBXAS1;refGene_tx=XM_011280571.3
A2 156391476 156391788
refGene_gn=TBXAS1;refGene_tx=XM_011280571.3
A2 156432536 156432628
refGene_gn=TBXAS1;refGene_tx=XM_011280571.3
A2 156436528 156436666
refGene_gn=TBXAS1;refGene_tx=XM_011280571.3
A2 156438200 156438363
refGene_gn=TBXAS1;refGene_tx=XM_011280571.3
A2 156440597 156440672
refGene_gn=TBXAS1;refGene_tx=XM_011280571.3
A2 156444479 156444805
refGene_gn=PARP12;refGene_tx=XM_023250622.1
A2 156446200 156446352
refGene_gn=PARP12;refGene_tx=XM_023250622.1
A2 156447166 156447297
refGene_gn=PARP12;refGene_tx=XM_023250622.1
A2 156449087 156449163
refGene_gn=PARP12;refGene_tx=XM_023250622.1
A2 156455329 156455426
refGene_gn=PARP12;refGene_tx=XM_023250622.1
A2 156458722 156458864
refGene_gn=PARP12;refGene_tx=XM_023250622.1
A2 156462480 156462676
refGene_gn=PARP12;refGene_tx=XM_023250622.1
A2 156467150 156467274
refGene_gn=PARP12;refGene_tx=XM_023250622.1
A2 156474008 156474110
refGene_gn=PARP12;refGene_tx=XM_023250622.1
A2 156475609 156475910
refGene_gn=PARP12;refGene_tx=XM_023250622.1
A2 156478618 156478754
refGene_gn=PARP12;refGene_tx=XM_023250622.1
A2 156482275 156482601
refGene_gn=PARP12;refGene_tx=XM_023250622.1
A2 156497530 156497625
refGene_gn=KDM7A;refGene_tx=XM_023250623.1
A2 156498289 156498563

refGene_gn=KDM7A;refGene_tx=XM_023250623.1
A2 156501078 156501164
refGene_gn=KDM7A;refGene_tx=XM_023250623.1
A2 156502543 156502752
refGene_gn=KDM7A;refGene_tx=XM_023250623.1
A2 156503082 156503231
refGene_gn=KDM7A;refGene_tx=XM_023250623.1
A2 156503720 156503818
refGene_gn=KDM7A;refGene_tx=XM_023250623.1
A2 156505036 156505191
refGene_gn=KDM7A;refGene_tx=XM_023250623.1
A2 156505404 156505529
refGene_gn=KDM7A;refGene_tx=XM_023250623.1
A2 156507030 156507240
refGene_gn=KDM7A;refGene_tx=XM_023250623.1
A2 156514213 156514303
refGene_gn=KDM7A;refGene_tx=XM_023250623.1
A2 156516608 156516700
refGene_gn=KDM7A;refGene_tx=XM_023250623.1
A2 156520074 156520181
refGene_gn=KDM7A;refGene_tx=XM_023250623.1
A2 156521537 156521625
refGene_gn=KDM7A;refGene_tx=XM_023250623.1
A2 156525315 156525478
refGene_gn=KDM7A;refGene_tx=XM_023250623.1
A2 156526695 156526882
refGene_gn=KDM7A;refGene_tx=XM_023250623.1
A2 156527187 156527329
refGene_gn=KDM7A;refGene_tx=XM_023250623.1
A2 156529145 156529306
refGene_gn=KDM7A;refGene_tx=XM_023250623.1
A2 156532588 156532706
refGene_gn=KDM7A;refGene_tx=XM_023250623.1
A2 156537598 156537684
refGene_gn=KDM7A;refGene_tx=XM_023250623.1
A2 156573593 156573787
refGene_gn=KDM7A;refGene_tx=XM_023250623.1
A2 156699167 156699260
refGene_gn=SLC37A3;refGene_tx=XM_023250629.1;refGene_gn=SLC37A3;refGene_tx=XM_023250630.1
A2 156699821 156699970
refGene_gn=SLC37A3;refGene_tx=XM_023250624.1;refGene_gn=SLC37A3;refGene_tx=XM_023250625.1;refGene_gn=SLC37A3;refGene_tx=XM_023250626.1;refGene_gn=SLC37A3;refGene_tx=XM_023250627.1;refGene_gn=SLC37A3;refGene_tx=XM_023250628.1
A2 156700653 156700719
refGene_gn=SLC37A3;refGene_tx=XM_023250624.1;refGene_gn=SLC37A3;refGene_tx=XM_023250625.1;refGene_gn=SLC37A3;refGene_tx=XM_023250626.1;refGene_gn=SLC37A3;refGene_tx=XM_023250627.1;refGene_gn=SLC37A3;refGene_tx=XM_023250628.1;refGene_gn=SLC37A3;refGene_tx=XM_023250629.1

A2 156701208 156701214
refGene_gn=SLC37A3;refGene_tx=XM_023250631.1
A2 156705719 156705871
refGene_gn=SLC37A3;refGene_tx=XM_023250624.1;refGene_gn=SLC37A3;refGene_tx=XM_023250625.1;refGene_gn=SLC37A3;refGene_tx=XM_023250626.1;refGene_gn=SLC37A3;refGene_tx=XM_023250627.1;refGene_gn=SLC37A3;refGene_tx=XM_023250628.1;refGene_gn=SLC37A3;refGene_tx=XM_023250629.1;refGene_gn=SLC37A3;refGene_tx=XM_023250630.1;refGene_gn=SLC37A3;refGene_tx=XM_023250631.1
A2 156707615 156707663
refGene_gn=SLC37A3;refGene_tx=XM_023250624.1;refGene_gn=SLC37A3;refGene_tx=XM_023250625.1;refGene_gn=SLC37A3;refGene_tx=XM_023250626.1;refGene_gn=SLC37A3;refGene_tx=XM_023250627.1;refGene_gn=SLC37A3;refGene_tx=XM_023250628.1;refGene_gn=SLC37A3;refGene_tx=XM_023250629.1;refGene_gn=SLC37A3;refGene_tx=XM_023250630.1;refGene_gn=SLC37A3;refGene_tx=XM_023250631.1
A2 156708266 156708368
refGene_gn=SLC37A3;refGene_tx=XM_023250624.1;refGene_gn=SLC37A3;refGene_tx=XM_023250625.1;refGene_gn=SLC37A3;refGene_tx=XM_023250626.1;refGene_gn=SLC37A3;refGene_tx=XM_023250627.1;refGene_gn=SLC37A3;refGene_tx=XM_023250628.1;refGene_gn=SLC37A3;refGene_tx=XM_023250629.1;refGene_gn=SLC37A3;refGene_tx=XM_023250630.1;refGene_gn=SLC37A3;refGene_tx=XM_023250631.1
A2 156710212 156710354
refGene_gn=SLC37A3;refGene_tx=XM_023250624.1;refGene_gn=SLC37A3;refGene_tx=XM_023250625.1;refGene_gn=SLC37A3;refGene_tx=XM_023250626.1;refGene_gn=SLC37A3;refGene_tx=XM_023250627.1;refGene_gn=SLC37A3;refGene_tx=XM_023250628.1;refGene_gn=SLC37A3;refGene_tx=XM_023250629.1;refGene_gn=SLC37A3;refGene_tx=XM_023250630.1;refGene_gn=SLC37A3;refGene_tx=XM_023250631.1
A2 156712821 156713000
refGene_gn=SLC37A3;refGene_tx=XM_023250624.1;refGene_gn=SLC37A3;refGene_tx=XM_023250625.1;refGene_gn=SLC37A3;refGene_tx=XM_023250626.1;refGene_gn=SLC37A3;refGene_tx=XM_023250627.1;refGene_gn=SLC37A3;refGene_tx=XM_023250628.1;refGene_gn=SLC37A3;refGene_tx=XM_023250629.1;refGene_gn=SLC37A3;refGene_tx=XM_023250630.1;refGene_gn=SLC37A3;refGene_tx=XM_023250631.1
A2 156713742 156713827
refGene_gn=SLC37A3;refGene_tx=XM_023250624.1;refGene_gn=SLC37A3;refGene_tx=XM_023250625.1;refGene_gn=SLC37A3;refGene_tx=XM_023250626.1;refGene_gn=SLC37A3;refGene_tx=XM_023250627.1;refGene_gn=SLC37A3;refGene_tx=XM_023250628.1;refGene_gn=SLC37A3;refGene_tx=XM_023250629.1;refGene_gn=SLC37A3;refGene_tx=XM_023250630.1;refGene_gn=SLC37A3;refGene_tx=XM_023250631.1
A2 156716198 156716295
refGene_gn=SLC37A3;refGene_tx=XM_023250624.1;refGene_gn=SLC37A3;refGene_tx=XM_023250625.1;refGene_gn=SLC37A3;refGene_tx=XM_023250626.1;refGene_gn=SLC37A3;refGene_tx=XM_023250627.1;refGene_gn=SLC37A3;refGene_tx=XM_023250628.1;refGene_gn=SLC37A3;refGene_tx=XM_023250629.1;refGene_gn=SLC37A3;refGene_tx=XM_023250630.1;refGene_gn=SLC37A3;refGene_tx=XM_02

3250631.1
A2	156720108	156720254
refGene_gn=SLC37A3;refGene_tx=XM_023250624.1;refGene_gn=SLC37A3;refGene_tx=XM_023250625.1;refGene_gn=SLC37A3;refGene_tx=XM_023250626.1;refGene_gn=SLC37A3;refGene_tx=XM_023250627.1;refGene_gn=SLC37A3;refGene_tx=XM_023250628.1;refGene_gn=SLC37A3;refGene_tx=XM_023250629.1;refGene_gn=SLC37A3;refGene_tx=XM_023250630.1;refGene_gn=SLC37A3;refGene_tx=XM_023250631.1
A2	156722329	156722413
refGene_gn=SLC37A3;refGene_tx=XM_023250624.1;refGene_gn=SLC37A3;refGene_tx=XM_023250625.1;refGene_gn=SLC37A3;refGene_tx=XM_023250626.1;refGene_gn=SLC37A3;refGene_tx=XM_023250627.1;refGene_gn=SLC37A3;refGene_tx=XM_023250628.1;refGene_gn=SLC37A3;refGene_tx=XM_023250629.1;refGene_gn=SLC37A3;refGene_tx=XM_023250630.1;refGene_gn=SLC37A3;refGene_tx=XM_023250631.1
A2	156725351	156725444
refGene_gn=SLC37A3;refGene_tx=XM_023250624.1;refGene_gn=SLC37A3;refGene_tx=XM_023250625.1;refGene_gn=SLC37A3;refGene_tx=XM_023250626.1;refGene_gn=SLC37A3;refGene_tx=XM_023250627.1;refGene_gn=SLC37A3;refGene_tx=XM_023250628.1;refGene_gn=SLC37A3;refGene_tx=XM_023250629.1;refGene_gn=SLC37A3;refGene_tx=XM_023250630.1;refGene_gn=SLC37A3;refGene_tx=XM_023250631.1
A2	156731803	156731912
refGene_gn=SLC37A3;refGene_tx=XM_023250624.1;refGene_gn=SLC37A3;refGene_tx=XM_023250625.1;refGene_gn=SLC37A3;refGene_tx=XM_023250626.1;refGene_gn=SLC37A3;refGene_tx=XM_023250627.1;refGene_gn=SLC37A3;refGene_tx=XM_023250628.1;refGene_gn=SLC37A3;refGene_tx=XM_023250629.1;refGene_gn=SLC37A3;refGene_tx=XM_023250630.1;refGene_gn=SLC37A3;refGene_tx=XM_023250631.1
A2	156733313	156733402
refGene_gn=SLC37A3;refGene_tx=XM_023250624.1;refGene_gn=SLC37A3;refGene_tx=XM_023250625.1;refGene_gn=SLC37A3;refGene_tx=XM_023250626.1;refGene_gn=SLC37A3;refGene_tx=XM_023250627.1;refGene_gn=SLC37A3;refGene_tx=XM_023250628.1;refGene_gn=SLC37A3;refGene_tx=XM_023250629.1;refGene_gn=SLC37A3;refGene_tx=XM_023250630.1;refGene_gn=SLC37A3;refGene_tx=XM_023250631.1
A2	156748343	156748544
refGene_gn=RAB19;refGene_tx=XM_011280594.3;refGene_gn=RAB19;refGene_tx=XM_019825898.2
A2	156751119	156751303
refGene_gn=RAB19;refGene_tx=XM_011280594.3;refGene_gn=RAB19;refGene_tx=XM_019825898.2
A2	156755678	156755947
refGene_gn=RAB19;refGene_tx=XM_011280594.3;refGene_gn=RAB19;refGene_tx=XM_019825898.2
A2	156771448	156771661
refGene_gn=MKRN1;refGene_tx=XM_003983147.4;refGene_gn=MKRN1;refGene_tx=XM_011280570.3
A2	156772024	156772163
refGene_gn=MKRN1;refGene_tx=XM_003983147.4;refGene_gn=MKRN1;refGene_tx

=XM_011280570.3
A2	156772722	156772833
refGene_gn=MKRN1;refGene_tx=XM_003983147.4;refGene_gn=MKRN1;refGene_tx=XM_011280570.3
A2	156773643	156773858
refGene_gn=MKRN1;refGene_tx=XM_003983147.4;refGene_gn=MKRN1;refGene_tx=XM_011280570.3
A2	156776076	156776303
refGene_gn=MKRN1;refGene_tx=XM_003983147.4;refGene_gn=MKRN1;refGene_tx=XM_011280570.3
A2	156776988	156777218
refGene_gn=MKRN1;refGene_tx=XM_003983147.4;refGene_gn=MKRN1;refGene_tx=XM_011280570.3
A2	156782503	156782632
refGene_gn=MKRN1;refGene_tx=XM_003983147.4;refGene_gn=MKRN1;refGene_tx=XM_011280570.3
A2	156790325	156790510
refGene_gn=MKRN1;refGene_tx=XM_003983147.4
A2	156822828	156822860
refGene_gn=DENND2A;refGene_tx=XM_019825902.2;refGene_gn=DENND2A;refGene_tx=XM_023250632.1;refGene_gn=DENND2A;refGene_tx=XM_023250633.1
A2	156823980	156824067
refGene_gn=DENND2A;refGene_tx=XM_019825902.2;refGene_gn=DENND2A;refGene_tx=XM_023250632.1;refGene_gn=DENND2A;refGene_tx=XM_023250633.1
A2	156825669	156825915
refGene_gn=DENND2A;refGene_tx=XM_019825902.2;refGene_gn=DENND2A;refGene_tx=XM_023250632.1;refGene_gn=DENND2A;refGene_tx=XM_023250633.1
A2	156826229	156826347
refGene_gn=DENND2A;refGene_tx=XM_019825902.2;refGene_gn=DENND2A;refGene_tx=XM_023250632.1;refGene_gn=DENND2A;refGene_tx=XM_023250633.1
A2	156830320	156830362
refGene_gn=DENND2A;refGene_tx=XM_019825902.2;refGene_gn=DENND2A;refGene_tx=XM_023250632.1;refGene_gn=DENND2A;refGene_tx=XM_023250633.1
A2	156831984	156832162
refGene_gn=DENND2A;refGene_tx=XM_019825902.2;refGene_gn=DENND2A;refGene_tx=XM_023250632.1;refGene_gn=DENND2A;refGene_tx=XM_023250633.1
A2	156845103	156845252
refGene_gn=DENND2A;refGene_tx=XM_019825902.2;refGene_gn=DENND2A;refGene_tx=XM_023250632.1;refGene_gn=DENND2A;refGene_tx=XM_023250633.1
A2	156846965	156847106
refGene_gn=DENND2A;refGene_tx=XM_019825902.2;refGene_gn=DENND2A;refGene_tx=XM_023250632.1;refGene_gn=DENND2A;refGene_tx=XM_023250633.1
A2	156853664	156853742
refGene_gn=DENND2A;refGene_tx=XM_019825902.2;refGene_gn=DENND2A;refGene_tx=XM_023250632.1;refGene_gn=DENND2A;refGene_tx=XM_023250633.1
A2	156855039	156855109
refGene_gn=DENND2A;refGene_tx=XM_019825902.2;refGene_gn=DENND2A;refGene_tx=XM_023250632.1;refGene_gn=DENND2A;refGene_tx=XM_023250633.1
A2	156856070	156856180
refGene_gn=DENND2A;refGene_tx=XM_019825902.2;refGene_gn=DENND2A;refGen

e_tx=XM_023250632.1;refGene_gn=DENND2A;refGene_tx=XM_023250633.1
A2 156860770 156860958
refGene_gn=DENND2A;refGene_tx=XM_019825902.2;refGene_gn=DENND2A;refGene_tx=XM_023250632.1;refGene_gn=DENND2A;refGene_tx=XM_023250633.1
A2 156862065 156862116
refGene_gn=DENND2A;refGene_tx=XM_019825902.2;refGene_gn=DENND2A;refGene_tx=XM_023250632.1;refGene_gn=DENND2A;refGene_tx=XM_023250633.1
A2 156862648 156862742
refGene_gn=DENND2A;refGene_tx=XM_019825902.2;refGene_gn=DENND2A;refGene_tx=XM_023250632.1;refGene_gn=DENND2A;refGene_tx=XM_023250633.1
A2 156867419 156867620
refGene_gn=DENND2A;refGene_tx=XM_019825902.2;refGene_gn=DENND2A;refGene_tx=XM_023250632.1;refGene_gn=DENND2A;refGene_tx=XM_023250633.1
A2 156876397 156876519
refGene_gn=DENND2A;refGene_tx=XM_019825902.2;refGene_gn=DENND2A;refGene_tx=XM_023250632.1;refGene_gn=DENND2A;refGene_tx=XM_023250633.1
A2 156878983 156879111
refGene_gn=DENND2A;refGene_tx=XM_019825902.2;refGene_gn=DENND2A;refGene_tx=XM_023250632.1;refGene_gn=DENND2A;refGene_tx=XM_023250633.1
A2 156893195 156894350
refGene_gn=DENND2A;refGene_tx=XM_019825902.2;refGene_gn=DENND2A;refGene_tx=XM_023250632.1;refGene_gn=DENND2A;refGene_tx=XM_023250633.1
A2 156896902 156897004
refGene_gn=DENND2A;refGene_tx=XM_019825902.2;refGene_gn=DENND2A;refGene_tx=XM_023250632.1
A2 156910709 156910783
refGene_gn=DENND2A;refGene_tx=XM_019825902.2
A2 156937145 156938078
refGene_gn=ADCK2;refGene_tx=XM_003983109.5;refGene_gn=ADCK2;refGene_tx=XM_006929422.4
A2 156939983 156940130
refGene_gn=ADCK2;refGene_tx=XM_003983109.5;refGene_gn=ADCK2;refGene_tx=XM_006929422.4
A2 156942571 156942700
refGene_gn=ADCK2;refGene_tx=XM_003983109.5;refGene_gn=ADCK2;refGene_tx=XM_006929422.4
A2 156944732 156944828
refGene_gn=ADCK2;refGene_tx=XM_003983109.5;refGene_gn=ADCK2;refGene_tx=XM_006929422.4
A2 156947107 156947359
refGene_gn=ADCK2;refGene_tx=XM_003983109.5;refGene_gn=ADCK2;refGene_tx=XM_006929422.4
A2 156948999 156949128
refGene_gn=ADCK2;refGene_tx=XM_003983109.5;refGene_gn=ADCK2;refGene_tx=XM_006929422.4
A2 156949409 156949454
refGene_gn=ADCK2;refGene_tx=XM_006929422.4
A2 156950658 156950712
refGene_gn=ADCK2;refGene_tx=XM_003983109.5
A2 156954837 156954978

refGene_gn=ADCK2;refGene_tx=XM_003983109.5
A2	156959992	156960090
refGene_gn=NDUFB2;refGene_tx=XM_003983110.5;refGene_gn=NDUFB2;refGene_tx=XM_006929423.4
A2	156964882	156965027
refGene_gn=NDUFB2;refGene_tx=XM_003983110.5;refGene_gn=NDUFB2;refGene_tx=XM_006929423.4
A2	156966183	156966258
refGene_gn=NDUFB2;refGene_tx=XM_003983110.5;refGene_gn=NDUFB2;refGene_tx=XM_006929423.4
A2	156980180	156980203
refGene_gn=BRAF;refGene_tx=XM_006929424.4;refGene_gn=BRAF;refGene_tx=XM_019825906.2;refGene_gn=BRAF;refGene_tx=XM_019825907.2;refGene_gn=BRAF;refGene_tx=XM_019825910.2
A2	156986496	156986650
refGene_gn=BRAF;refGene_tx=XM_006929424.4;refGene_gn=BRAF;refGene_tx=XM_019825906.2;refGene_gn=BRAF;refGene_tx=XM_019825907.2;refGene_gn=BRAF;refGene_tx=XM_019825910.2
A2	156987551	156987602
refGene_gn=BRAF;refGene_tx=XM_019825906.2;refGene_gn=BRAF;refGene_tx=XM_019825910.2
A2	156990558	156990693
refGene_gn=BRAF;refGene_tx=XM_006929424.4;refGene_gn=BRAF;refGene_tx=XM_019825906.2;refGene_gn=BRAF;refGene_tx=XM_019825907.2;refGene_gn=BRAF;refGene_tx=XM_019825910.2
A2	156997467	156997599
refGene_gn=BRAF;refGene_tx=XM_006929424.4;refGene_gn=BRAF;refGene_tx=XM_019825906.2;refGene_gn=BRAF;refGene_tx=XM_019825907.2;refGene_gn=BRAF;refGene_tx=XM_019825910.2
A2	157001175	157001294
refGene_gn=BRAF;refGene_tx=XM_006929424.4;refGene_gn=BRAF;refGene_tx=XM_019825906.2;refGene_gn=BRAF;refGene_tx=XM_019825907.2;refGene_gn=BRAF;refGene_tx=XM_019825910.2
A2	157002117	157002164
refGene_gn=BRAF;refGene_tx=XM_006929424.4;refGene_gn=BRAF;refGene_tx=XM_019825906.2;refGene_gn=BRAF;refGene_tx=XM_019825907.2;refGene_gn=BRAF;refGene_tx=XM_019825910.2
A2	157019912	157020089
refGene_gn=BRAF;refGene_tx=XM_006929424.4;refGene_gn=BRAF;refGene_tx=XM_019825906.2;refGene_gn=BRAF;refGene_tx=XM_019825907.2;refGene_gn=BRAF;refGene_tx=XM_019825910.2
A2	157020931	157021016
refGene_gn=BRAF;refGene_tx=XM_006929424.4;refGene_gn=BRAF;refGene_tx=XM_019825906.2;refGene_gn=BRAF;refGene_tx=XM_019825907.2;refGene_gn=BRAF;refGene_tx=XM_019825910.2
A2	157023033	157023151
refGene_gn=BRAF;refGene_tx=XM_006929424.4;refGene_gn=BRAF;refGene_tx=XM_019825906.2;refGene_gn=BRAF;refGene_tx=XM_019825907.2;refGene_gn=BRAF;refGene_tx=XM_019825910.2
A2	157023398	157023535

refGene_gn=BRAF;refGene_tx=XM_006929424.4;refGene_gn=BRAF;refGene_tx=XM_019825906.2;refGene_gn=BRAF;refGene_tx=XM_019825907.2;refGene_gn=BRAF;refGene_tx=XM_019825910.2
A2 157025670 157025790
refGene_gn=BRAF;refGene_tx=XM_019825906.2;refGene_gn=BRAF;refGene_tx=XM_019825907.2
A2 157027225 157027262
refGene_gn=BRAF;refGene_tx=XM_006929424.4;refGene_gn=BRAF;refGene_tx=XM_019825906.2;refGene_gn=BRAF;refGene_tx=XM_019825907.2;refGene_gn=BRAF;refGene_tx=XM_019825910.2
A2 157035265 157035425
refGene_gn=BRAF;refGene_tx=XM_006929424.4;refGene_gn=BRAF;refGene_tx=XM_019825906.2;refGene_gn=BRAF;refGene_tx=XM_019825907.2;refGene_gn=BRAF;refGene_tx=XM_019825910.2
A2 157041431 157041551
refGene_gn=BRAF;refGene_tx=XM_006929424.4;refGene_gn=BRAF;refGene_tx=XM_019825906.2;refGene_gn=BRAF;refGene_tx=XM_019825907.2;refGene_gn=BRAF;refGene_tx=XM_019825910.2
A2 157042509 157042658
refGene_gn=BRAF;refGene_tx=XM_006929424.4;refGene_gn=BRAF;refGene_tx=XM_019825906.2;refGene_gn=BRAF;refGene_tx=XM_019825907.2;refGene_gn=BRAF;refGene_tx=XM_019825910.2
A2 157047618 157047721
refGene_gn=BRAF;refGene_tx=XM_006929424.4;refGene_gn=BRAF;refGene_tx=XM_019825906.2;refGene_gn=BRAF;refGene_tx=XM_019825907.2;refGene_gn=BRAF;refGene_tx=XM_019825910.2
A2 157048570 157048674
refGene_gn=BRAF;refGene_tx=XM_006929424.4;refGene_gn=BRAF;refGene_tx=XM_019825906.2;refGene_gn=BRAF;refGene_tx=XM_019825907.2;refGene_gn=BRAF;refGene_tx=XM_019825910.2
A2 157054864 157055128
refGene_gn=BRAF;refGene_tx=XM_006929424.4;refGene_gn=BRAF;refGene_tx=XM_019825906.2;refGene_gn=BRAF;refGene_tx=XM_019825907.2;refGene_gn=BRAF;refGene_tx=XM_019825910.2
A2 157069781 157069883
refGene_gn=BRAF;refGene_tx=XM_006929424.4;refGene_gn=BRAF;refGene_tx=XM_019825906.2;refGene_gn=BRAF;refGene_tx=XM_019825907.2;refGene_gn=BRAF;refGene_tx=XM_019825910.2
A2 157139253 157139382
refGene_gn=BRAF;refGene_tx=XM_006929424.4;refGene_gn=BRAF;refGene_tx=XM_019825906.2;refGene_gn=BRAF;refGene_tx=XM_019825907.2;refGene_gn=BRAF;refGene_tx=XM_019825910.2
A2 157192443 157192549
refGene_gn=MRPS33;refGene_tx=XM_003983111.4
A2 157196344 157196559
refGene_gn=MRPS33;refGene_tx=XM_003983111.4
A2 157246711 157247093
refGene_gn=TMEM178B;refGene_tx=XM_003983112.5
A2 157366527 157366641
refGene_gn=TMEM178B;refGene_tx=XM_003983112.5

A2 157570784 157570922
refGene_gn=TMEM178B;refGene_tx=XM_003983112.5
A2 157603132 157603383
refGene_gn=TMEM178B;refGene_tx=XM_003983112.5
A2 157680986 157681087
refGene_gn=AGK;refGene_tx=XM_023250638.1
A2 157705125 157705165
refGene_gn=AGK;refGene_tx=XM_023250638.1
A2 157708663 157708713
refGene_gn=AGK;refGene_tx=XM_023250638.1
A2 157708714 157708741
refGene_gn=AGK;refGene_tx=XM_023250638.1
A2 157713670 157713746
refGene_gn=AGK;refGene_tx=XM_023250638.1
A2 157718514 157718607
refGene_gn=AGK;refGene_tx=XM_023250638.1
A2 157720303 157720336
refGene_gn=AGK;refGene_tx=XM_023250638.1
A2 157721421 157721516
refGene_gn=AGK;refGene_tx=XM_023250638.1
A2 157729112 157729182
refGene_gn=AGK;refGene_tx=XM_023250638.1
A2 157736286 157736366
refGene_gn=AGK;refGene_tx=XM_023250638.1
A2 157742733 157742791
refGene_gn=AGK;refGene_tx=XM_023250638.1
A2 157747196 157747344
refGene_gn=AGK;refGene_tx=XM_023250638.1
A2 157747784 157747882
refGene_gn=AGK;refGene_tx=XM_023250638.1
A2 157753926 157753997
refGene_gn=AGK;refGene_tx=XM_023250638.1
A2 157755906 157755991
refGene_gn=AGK;refGene_tx=XM_023250638.1
A2 157757040 157757178
refGene_gn=AGK;refGene_tx=XM_023250638.1
A2 157765506 157765702
refGene_gn=KIAA1147;refGene_tx=XM_023250635.1;refGene_gn=KIAA1147;refG
ene_tx=XM_023250636.1;refGene_gn=KIAA1147;refGene_tx=XM_023250637.1
A2 157767389 157767458
refGene_gn=KIAA1147;refGene_tx=XM_023250635.1;refGene_gn=KIAA1147;refG
ene_tx=XM_023250636.1;refGene_gn=KIAA1147;refGene_tx=XM_023250637.1
A2 157768096 157768247
refGene_gn=KIAA1147;refGene_tx=XM_023250635.1;refGene_gn=KIAA1147;refG
ene_tx=XM_023250636.1;refGene_gn=KIAA1147;refGene_tx=XM_023250637.1
A2 157768476 157768608
refGene_gn=KIAA1147;refGene_tx=XM_023250635.1;refGene_gn=KIAA1147;refG
ene_tx=XM_023250636.1;refGene_gn=KIAA1147;refGene_tx=XM_023250637.1
A2 157769571 157769710
refGene_gn=KIAA1147;refGene_tx=XM_023250635.1;refGene_gn=KIAA1147;refG

ene_tx=XM_023250636.1;refGene_gn=KIAA1147;refGene_tx=XM_023250637.1
A2 157776980 157777134
refGene_gn=KIAA1147;refGene_tx=XM_023250635.1;refGene_gn=KIAA1147;refGene_tx=XM_023250636.1;refGene_gn=KIAA1147;refGene_tx=XM_023250637.1
A2 157787900 157788059
refGene_gn=KIAA1147;refGene_tx=XM_023250635.1;refGene_gn=KIAA1147;refGene_tx=XM_023250636.1;refGene_gn=KIAA1147;refGene_tx=XM_023250637.1
A2 157789274 157789374
refGene_gn=KIAA1147;refGene_tx=XM_023250635.1;refGene_gn=KIAA1147;refGene_tx=XM_023250636.1;refGene_gn=KIAA1147;refGene_tx=XM_023250637.1
A2 157805800 157806068
refGene_gn=KIAA1147;refGene_tx=XM_023250635.1;refGene_gn=KIAA1147;refGene_tx=XM_023250636.1;refGene_gn=KIAA1147;refGene_tx=XM_023250637.1
A2 157820262 157820607
refGene_gn=WEE2;refGene_tx=XM_023250639.1;refGene_gn=WEE2;refGene_tx=XM_023250640.1
A2 157828053 157828250
refGene_gn=WEE2;refGene_tx=XM_023250639.1;refGene_gn=WEE2;refGene_tx=XM_023250640.1
A2 157829997 157830049
refGene_gn=WEE2;refGene_tx=XM_023250639.1;refGene_gn=WEE2;refGene_tx=XM_023250640.1
A2 157832836 157833009
refGene_gn=WEE2;refGene_tx=XM_023250639.1;refGene_gn=WEE2;refGene_tx=XM_023250640.1
A2 157834628 157834750
refGene_gn=WEE2;refGene_tx=XM_023250639.1;refGene_gn=WEE2;refGene_tx=XM_023250640.1
A2 157839914 157840061
refGene_gn=WEE2;refGene_tx=XM_023250639.1;refGene_gn=WEE2;refGene_tx=XM_023250640.1
A2 157840685 157840793
refGene_gn=WEE2;refGene_tx=XM_023250639.1;refGene_gn=WEE2;refGene_tx=XM_023250640.1
A2 157840936 157841022
refGene_gn=WEE2;refGene_tx=XM_023250639.1;refGene_gn=WEE2;refGene_tx=XM_023250640.1
A2 157841755 157841926
refGene_gn=WEE2;refGene_tx=XM_023250639.1;refGene_gn=WEE2;refGene_tx=XM_023250640.1
A2 157843577 157843720
refGene_gn=WEE2;refGene_tx=XM_023250639.1
A2 157847880 157848037
refGene_gn=WEE2;refGene_tx=XM_023250639.1;refGene_gn=WEE2;refGene_tx=XM_023250640.1
A2 157855085 157855109
refGene_gn=SSBP1;refGene_tx=XM_006929427.3;refGene_gn=SSBP1;refGene_tx=XM_006929429.3;refGene_gn=SSBP1;refGene_tx=XM_006929432.3;refGene_gn=SSBP1;refGene_tx=XM_019825921.2;refGene_gn=SSBP1;refGene_tx=XM_019825923.2;refGene_gn=SSBP1;refGene_tx=XM_019825924.1;refGene_gn=SSBP1;refGe

ne_tx=XM_019825925.1;refGene_gn=SSBP1;refGene_tx=XM_019825926.2;refGene_gn=SSBP1;refGene_tx=XM_023250641.1
A2	157857545	157857606
refGene_gn=SSBP1;refGene_tx=XM_006929427.3;refGene_gn=SSBP1;refGene_tx=XM_006929429.3;refGene_gn=SSBP1;refGene_tx=XM_006929432.3;refGene_gn=SSBP1;refGene_tx=XM_019825921.2;refGene_gn=SSBP1;refGene_tx=XM_019825923.2;refGene_gn=SSBP1;refGene_tx=XM_019825924.1;refGene_gn=SSBP1;refGene_tx=XM_019825925.1;refGene_gn=SSBP1;refGene_tx=XM_019825926.2;refGene_gn=SSBP1;refGene_tx=XM_023250641.1
A2	157858194	157858335
refGene_gn=SSBP1;refGene_tx=XM_006929427.3;refGene_gn=SSBP1;refGene_tx=XM_006929429.3;refGene_gn=SSBP1;refGene_tx=XM_006929432.3;refGene_gn=SSBP1;refGene_tx=XM_019825921.2;refGene_gn=SSBP1;refGene_tx=XM_019825923.2;refGene_gn=SSBP1;refGene_tx=XM_019825924.1;refGene_gn=SSBP1;refGene_tx=XM_019825925.1;refGene_gn=SSBP1;refGene_tx=XM_019825926.2;refGene_gn=SSBP1;refGene_tx=XM_023250641.1
A2	157858540	157858634
refGene_gn=SSBP1;refGene_tx=XM_006929427.3;refGene_gn=SSBP1;refGene_tx=XM_006929429.3;refGene_gn=SSBP1;refGene_tx=XM_006929432.3;refGene_gn=SSBP1;refGene_tx=XM_019825921.2;refGene_gn=SSBP1;refGene_tx=XM_019825923.2;refGene_gn=SSBP1;refGene_tx=XM_019825924.1;refGene_gn=SSBP1;refGene_tx=XM_019825925.1;refGene_gn=SSBP1;refGene_tx=XM_019825926.2;refGene_gn=SSBP1;refGene_tx=XM_023250641.1
A2	157860321	157860410
refGene_gn=SSBP1;refGene_tx=XM_006929427.3;refGene_gn=SSBP1;refGene_tx=XM_006929429.3;refGene_gn=SSBP1;refGene_tx=XM_006929432.3;refGene_gn=SSBP1;refGene_tx=XM_019825921.2;refGene_gn=SSBP1;refGene_tx=XM_019825923.2;refGene_gn=SSBP1;refGene_tx=XM_019825924.1;refGene_gn=SSBP1;refGene_tx=XM_019825925.1;refGene_gn=SSBP1;refGene_tx=XM_019825926.2;refGene_gn=SSBP1;refGene_tx=XM_023250641.1
A2	157864790	157864834
refGene_gn=SSBP1;refGene_tx=XM_006929427.3;refGene_gn=SSBP1;refGene_tx=XM_006929429.3;refGene_gn=SSBP1;refGene_tx=XM_019825921.2;refGene_gn=SSBP1;refGene_tx=XM_019825923.2;refGene_gn=SSBP1;refGene_tx=XM_019825924.1;refGene_gn=SSBP1;refGene_tx=XM_019825925.1;refGene_gn=SSBP1;refGene_tx=XM_019825926.2;refGene_gn=SSBP1;refGene_tx=XM_023250641.1
A2	157870855	157870869
refGene_gn=SSBP1;refGene_tx=XM_006929432.3
A2	157878201	157879152
refGene_gn=TAS2R3;refGene_tx=XM_011280593.1
A2	157888070	157888991
refGene_gn=TAS2R4;refGene_tx=XM_011280592.1
A2	157909813	157910378
refGene_gn=TAS2R5;refGene_tx=XM_011280591.3
A2	157910380	157910508
refGene_gn=TAS2R5;refGene_tx=XM_011280591.3
A2	157910509	157910561
refGene_gn=TAS2R5;refGene_tx=XM_011280591.3
A2	157910563	157910829
refGene_gn=TAS2R5;refGene_tx=XM_011280591.3

A2 157927024 157927165
refGene_gn=PRSS37;refGene_tx=XM_003983114.4
A2 157927709 157927846
refGene_gn=PRSS37;refGene_tx=XM_003983114.4
A2 157928425 157928679
refGene_gn=PRSS37;refGene_tx=XM_003983114.4
A2 157929940 157930082
refGene_gn=PRSS37;refGene_tx=XM_003983114.4
A2 157932126 157932160
refGene_gn=PRSS37;refGene_tx=XM_003983114.4
A2 157935510 157935547
refGene_gn=LOC101094087;refGene_tx=XM_006929433.3
A2 157945972 157946114
refGene_gn=LOC101094087;refGene_tx=XM_006929433.3
A2 157951800 157952048
refGene_gn=LOC101094087;refGene_tx=XM_006929433.3
A2 157952867 157953004
refGene_gn=LOC101094087;refGene_tx=XM_006929433.3
A2 157953386 157953536
refGene_gn=LOC101094087;refGene_tx=XM_006929433.3
A2 157979973 157980918
refGene_gn=LOC101083668;refGene_tx=XM_003983115.3
A2 157997026 157997633
refGene_gn=LOC101083922;refGene_tx=XM_003983116.4
A2 157997635 157997970
refGene_gn=LOC101083922;refGene_tx=XM_003983116.4
A2 158009754 158010699
refGene_gn=LOC101084183;refGene_tx=XM_003983117.3
A2 158024157 158024272
refGene_gn=CLEC5A;refGene_tx=XM_003983118.4;refGene_gn=CLEC5A;refGene_tx=XM_019825931.2
A2 158025452 158025556
refGene_gn=CLEC5A;refGene_tx=XM_003983118.4;refGene_gn=CLEC5A;refGene_tx=XM_019825931.2
A2 158028675 158028812
refGene_gn=CLEC5A;refGene_tx=XM_003983118.4;refGene_gn=CLEC5A;refGene_tx=XM_019825931.2
A2 158029126 158029186
refGene_gn=CLEC5A;refGene_tx=XM_003983118.4;refGene_gn=CLEC5A;refGene_tx=XM_019825931.2
A2 158031082 158031142
refGene_gn=CLEC5A;refGene_tx=XM_003983118.4
A2 158031973 158032052
refGene_gn=CLEC5A;refGene_tx=XM_003983118.4;refGene_gn=CLEC5A;refGene_tx=XM_019825931.2
A2 158053038 158054043
refGene_gn=TAS2R38;refGene_tx=XM_003983119.1
A2 158082936 158083063
refGene_gn=MGAM;refGene_tx=XM_011280590.3;refGene_gn=MGAM;refGene_tx=XM_019825927.1

A2 158084659 158084754
refGene_gn=MGAM;refGene_tx=XM_011280590.3;refGene_gn=MGAM;refGene_tx=XM_019825927.1
A2 158092177 158092298
refGene_gn=MGAM;refGene_tx=XM_011280590.3;refGene_gn=MGAM;refGene_tx=XM_019825927.1
A2 158093514 158093624
refGene_gn=MGAM;refGene_tx=XM_011280590.3;refGene_gn=MGAM;refGene_tx=XM_019825927.1
A2 158094179 158094331
refGene_gn=MGAM;refGene_tx=XM_011280590.3;refGene_gn=MGAM;refGene_tx=XM_019825927.1
A2 158094922 158095094
refGene_gn=MGAM;refGene_tx=XM_011280590.3;refGene_gn=MGAM;refGene_tx=XM_019825927.1
A2 158097295 158097395
refGene_gn=MGAM;refGene_tx=XM_011280590.3;refGene_gn=MGAM;refGene_tx=XM_019825927.1
A2 158099520 158099633
refGene_gn=MGAM;refGene_tx=XM_011280590.3;refGene_gn=MGAM;refGene_tx=XM_019825927.1
A2 158100162 158100288
refGene_gn=MGAM;refGene_tx=XM_011280590.3;refGene_gn=MGAM;refGene_tx=XM_019825927.1
A2 158101582 158101714
refGene_gn=MGAM;refGene_tx=XM_011280590.3;refGene_gn=MGAM;refGene_tx=XM_019825927.1
A2 158101877 158101994
refGene_gn=MGAM;refGene_tx=XM_011280590.3;refGene_gn=MGAM;refGene_tx=XM_019825927.1
A2 158103676 158103790
refGene_gn=MGAM;refGene_tx=XM_011280590.3;refGene_gn=MGAM;refGene_tx=XM_019825927.1
A2 158104877 158104962
refGene_gn=MGAM;refGene_tx=XM_011280590.3;refGene_gn=MGAM;refGene_tx=XM_019825927.1
A2 158106679 158106797
refGene_gn=MGAM;refGene_tx=XM_011280590.3;refGene_gn=MGAM;refGene_tx=XM_019825927.1
A2 158107918 158108090
refGene_gn=MGAM;refGene_tx=XM_011280590.3;refGene_gn=MGAM;refGene_tx=XM_019825927.1
A2 158109390 158109507
refGene_gn=MGAM;refGene_tx=XM_011280590.3;refGene_gn=MGAM;refGene_tx=XM_019825927.1
A2 158110059 158110214
refGene_gn=MGAM;refGene_tx=XM_011280590.3;refGene_gn=MGAM;refGene_tx=XM_019825927.1
A2 158111526 158111611
refGene_gn=MGAM;refGene_tx=XM_011280590.3;refGene_gn=MGAM;refGene_tx=X

M_019825927.1
A2 158112352 158112409
refGene_gn=MGAM;refGene_tx=XM_011280590.3;refGene_gn=MGAM;refGene_tx=X
M_019825927.1
A2 158112912 158113037
refGene_gn=MGAM;refGene_tx=XM_011280590.3;refGene_gn=MGAM;refGene_tx=X
M_019825927.1
A2 158116479 158116568
refGene_gn=MGAM;refGene_tx=XM_011280590.3;refGene_gn=MGAM;refGene_tx=X
M_019825927.1
A2 158117233 158117283
refGene_gn=MGAM;refGene_tx=XM_011280590.3;refGene_gn=MGAM;refGene_tx=X
M_019825927.1
A2 158118527 158118695
refGene_gn=MGAM;refGene_tx=XM_011280590.3;refGene_gn=MGAM;refGene_tx=X
M_019825927.1
A2 158121661 158121814
refGene_gn=MGAM;refGene_tx=XM_011280590.3;refGene_gn=MGAM;refGene_tx=X
M_019825927.1
A2 158122152 158122353
refGene_gn=MGAM;refGene_tx=XM_011280590.3;refGene_gn=MGAM;refGene_tx=X
M_019825927.1
A2 158124333 158124488
refGene_gn=MGAM;refGene_tx=XM_011280590.3;refGene_gn=MGAM;refGene_tx=X
M_019825927.1
A2 158125467 158125636
refGene_gn=MGAM;refGene_tx=XM_011280590.3;refGene_gn=MGAM;refGene_tx=X
M_019825927.1
A2 158125891 158125988
refGene_gn=MGAM;refGene_tx=XM_011280590.3;refGene_gn=MGAM;refGene_tx=X
M_019825927.1
A2 158127591 158127704
refGene_gn=MGAM;refGene_tx=XM_011280590.3;refGene_gn=MGAM;refGene_tx=X
M_019825927.1
A2 158128667 158128793
refGene_gn=MGAM;refGene_tx=XM_011280590.3;refGene_gn=MGAM;refGene_tx=X
M_019825927.1
A2 158129734 158129863
refGene_gn=MGAM;refGene_tx=XM_011280590.3;refGene_gn=MGAM;refGene_tx=X
M_019825927.1
A2 158130123 158130234
refGene_gn=MGAM;refGene_tx=XM_011280590.3;refGene_gn=MGAM;refGene_tx=X
M_019825927.1
A2 158130548 158130611
refGene_gn=MGAM;refGene_tx=XM_011280590.3;refGene_gn=MGAM;refGene_tx=X
M_019825927.1
A2 158136095 158136230
refGene_gn=MGAM;refGene_tx=XM_011280590.3;refGene_gn=MGAM;refGene_tx=X
M_019825927.1
A2 158136659 158136747

refGene_gn=MGAM;refGene_tx=XM_011280590.3;refGene_gn=MGAM;refGene_tx=XM_019825927.1
A2 158137810 158137949
refGene_gn=MGAM;refGene_tx=XM_011280590.3;refGene_gn=MGAM;refGene_tx=XM_019825927.1
A2 158138787 158138921
refGene_gn=MGAM;refGene_tx=XM_011280590.3;refGene_gn=MGAM;refGene_tx=XM_019825927.1
A2 158139053 158139088
refGene_gn=MGAM;refGene_tx=XM_011280590.3;refGene_gn=MGAM;refGene_tx=XM_019825927.1
A2 158139181 158139298
refGene_gn=MGAM;refGene_tx=XM_011280590.3;refGene_gn=MGAM;refGene_tx=XM_019825927.1
A2 158142797 158142946
refGene_gn=MGAM;refGene_tx=XM_011280590.3;refGene_gn=MGAM;refGene_tx=XM_019825927.1
A2 158143781 158143866
refGene_gn=MGAM;refGene_tx=XM_011280590.3;refGene_gn=MGAM;refGene_tx=XM_019825927.1
A2 158144495 158144552
refGene_gn=MGAM;refGene_tx=XM_011280590.3;refGene_gn=MGAM;refGene_tx=XM_019825927.1
A2 158152192 158152317
refGene_gn=MGAM;refGene_tx=XM_011280590.3;refGene_gn=MGAM;refGene_tx=XM_019825927.1
A2 158153430 158153519
refGene_gn=MGAM;refGene_tx=XM_011280590.3;refGene_gn=MGAM;refGene_tx=XM_019825927.1
A2 158155014 158155064
refGene_gn=MGAM;refGene_tx=XM_011280590.3;refGene_gn=MGAM;refGene_tx=XM_019825927.1
A2 158155828 158155999
refGene_gn=MGAM;refGene_tx=XM_011280590.3;refGene_gn=MGAM;refGene_tx=XM_019825927.1
A2 158157968 158158052
refGene_gn=MGAM;refGene_tx=XM_011280590.3;refGene_gn=MGAM;refGene_tx=XM_019825927.1
A2 158172360 158172466
refGene_gn=LOC101094837;refGene_tx=XM_023242290.1
A2 158176661 158176741
refGene_gn=LOC101094837;refGene_tx=XM_023242290.1
A2 158184606 158184727
refGene_gn=LOC101094837;refGene_tx=XM_023242290.1
A2 158184832 158184942
refGene_gn=LOC101094837;refGene_tx=XM_023242290.1
A2 158185495 158185650
refGene_gn=LOC101094837;refGene_tx=XM_023242290.1
A2 158186501 158186673
refGene_gn=LOC101094837;refGene_tx=XM_023242290.1

A2 158188902 158189002
refGene_gn=LOC101094837;refGene_tx=XM_023242290.1
A2 158189761 158189874
refGene_gn=LOC101094837;refGene_tx=XM_023242290.1
A2 158191864 158191990
refGene_gn=LOC101094837;refGene_tx=XM_023242290.1
A2 158194077 158194209
refGene_gn=LOC101094837;refGene_tx=XM_023242290.1
A2 158194298 158194397
refGene_gn=LOC101094837;refGene_tx=XM_023242290.1
A2 158196782 158196896
refGene_gn=LOC101094837;refGene_tx=XM_023242290.1
A2 158198244 158198329
refGene_gn=LOC101094837;refGene_tx=XM_023242290.1
A2 158201194 158201312
refGene_gn=LOC101094837;refGene_tx=XM_023242290.1
A2 158204580 158204752
refGene_gn=LOC101094837;refGene_tx=XM_023242290.1
A2 158205514 158205631
refGene_gn=LOC101094837;refGene_tx=XM_023242290.1
A2 158206345 158206500
refGene_gn=LOC101094837;refGene_tx=XM_023242290.1
A2 158206627 158206712
refGene_gn=LOC101094837;refGene_tx=XM_023242290.1
A2 158207834 158207891
refGene_gn=LOC101094837;refGene_tx=XM_023242290.1
A2 158208703 158208828
refGene_gn=LOC101094837;refGene_tx=XM_023242290.1
A2 158209691 158209780
refGene_gn=LOC101094837;refGene_tx=XM_023242290.1
A2 158210578 158210616
refGene_gn=LOC101094837;refGene_tx=XM_023242290.1
A2 158212777 158212945
refGene_gn=LOC101094837;refGene_tx=XM_023242290.1
A2 158214130 158214286
refGene_gn=LOC101094837;refGene_tx=XM_023242290.1
A2 158215426 158215648
refGene_gn=LOC101094837;refGene_tx=XM_023242290.1
A2 158217710 158217865
refGene_gn=LOC101094837;refGene_tx=XM_023242290.1
A2 158219368 158219537
refGene_gn=LOC101094837;refGene_tx=XM_023242290.1
A2 158220070 158220167
refGene_gn=LOC101094837;refGene_tx=XM_023242290.1
A2 158220608 158220721
refGene_gn=LOC101094837;refGene_tx=XM_023242290.1
A2 158221122 158221248
refGene_gn=LOC101094837;refGene_tx=XM_023242290.1
A2 158222791 158222920
refGene_gn=LOC101094837;refGene_tx=XM_023242290.1

A2	158224347	158224455
refGene_gn=LOC101094837;refGene_tx=XM_023242290.1
A2	158226193	158226256
refGene_gn=LOC101094837;refGene_tx=XM_023242290.1
A2	158227305	158227440
refGene_gn=LOC101094837;refGene_tx=XM_023242290.1
A2	158228978	158229063
refGene_gn=LOC101094837;refGene_tx=XM_023242290.1
A2	158230128	158230267
refGene_gn=LOC101094837;refGene_tx=XM_023242290.1
A2	158232221	158232355
refGene_gn=LOC101094837;refGene_tx=XM_023242290.1
A2	158232634	158232669
refGene_gn=LOC101094837;refGene_tx=XM_023242290.1
A2	158232769	158232886
refGene_gn=LOC101094837;refGene_tx=XM_023242290.1
A2	158233446	158233595
refGene_gn=LOC101094837;refGene_tx=XM_023242290.1
A2	158233714	158233799
refGene_gn=LOC101094837;refGene_tx=XM_023242290.1
A2	158234539	158234596
refGene_gn=LOC101094837;refGene_tx=XM_023242290.1
A2	158235019	158235144
refGene_gn=LOC101094837;refGene_tx=XM_023242290.1
A2	158237231	158237320
refGene_gn=LOC101094837;refGene_tx=XM_023242290.1
A2	158246407	158246457
refGene_gn=LOC101094837;refGene_tx=XM_023242290.1
A2	158249509	158249683
refGene_gn=LOC101094837;refGene_tx=XM_023242290.1
A2	158251016	158251278
refGene_gn=LOC101094837;refGene_tx=XM_023242290.1
A2	158251488	158251783
refGene_gn=LOC101094837;refGene_tx=XM_023242290.1
A2	158251898	158252089
refGene_gn=LOC101094837;refGene_tx=XM_023242290.1
A2	158252199	158252554
refGene_gn=LOC101094837;refGene_tx=XM_023242290.1
A2	158253137	158254216
refGene_gn=LOC101094837;refGene_tx=XM_023242290.1
A2	158254732	158255040
refGene_gn=LOC101094837;refGene_tx=XM_023242290.1
A2	158266273	158266516
refGene_gn=LOC101084948;refGene_tx=XM_003983120.4
A2	158266828	158266943
refGene_gn=LOC101084948;refGene_tx=XM_003983120.4
A2	158267688	158267819
refGene_gn=LOC101084948;refGene_tx=XM_003983120.4
A2	158269350	158269410
refGene_gn=LOC101084948;refGene_tx=XM_003983120.4

A2	158269570	158269667
refGene_gn=LOC101084948;refGene_tx=XM_003983120.4
A2	158269896	158269988
refGene_gn=LOC101084948;refGene_tx=XM_003983120.4
A2	158270323	158270490
refGene_gn=LOC101084948;refGene_tx=XM_003983120.4
A2	158270675	158270778
refGene_gn=LOC101084948;refGene_tx=XM_003983120.4
A2	158270967	158271135
refGene_gn=LOC101084948;refGene_tx=XM_003983120.4
A2	158271911	158271995
refGene_gn=LOC101084948;refGene_tx=XM_003983120.4
A2	158272208	158272370
refGene_gn=LOC101084948;refGene_tx=XM_003983120.4
A2	158272886	158273033
refGene_gn=LOC101084948;refGene_tx=XM_003983120.4
A2	158274142	158274430
refGene_gn=LOC101084948;refGene_tx=XM_003983120.4
A2	158289386	158289536
refGene_gn=LOC101085196;refGene_tx=XM_003983121.4
A2	158289640	158289780
refGene_gn=LOC101085196;refGene_tx=XM_003983121.4
A2	158292204	158292461
refGene_gn=LOC101085196;refGene_tx=XM_003983121.4
A2	158292685	158292824
refGene_gn=LOC101085196;refGene_tx=XM_003983121.4
A2	158294501	158294541
refGene_gn=LOC101085196;refGene_tx=XM_003983121.4
A2	158300531	158300681
refGene_gn=LOC102902015;refGene_tx=XM_006929434.3
A2	158301072	158301209
refGene_gn=LOC102902015;refGene_tx=XM_006929434.3
A2	158303524	158303775
refGene_gn=LOC102902015;refGene_tx=XM_006929434.3
A2	158309211	158309368
refGene_gn=LOC102902015;refGene_tx=XM_006929434.3
A2	158310144	158310184
refGene_gn=LOC102902015;refGene_tx=XM_006929434.3
A2	158317835	158317985
refGene_gn=LOC101085453;refGene_tx=XM_003983122.3
A2	158318532	158318669
refGene_gn=LOC101085453;refGene_tx=XM_003983122.3
A2	158319175	158319429
refGene_gn=LOC101085453;refGene_tx=XM_003983122.3
A2	158320205	158320365
refGene_gn=LOC101085453;refGene_tx=XM_003983122.3
A2	158322372	158322412
refGene_gn=LOC101085453;refGene_tx=XM_003983122.3
A2	158332885	158335049	refGene_gn=LOC101095329
A2	158344426	158346514	refGene_gn=LOC105260394

A2 158363648 158363697
refGene_gn=LOC111559549;refGene_tx=XM_023250642.1
A2 158363800 158364495
refGene_gn=LOC111559549;refGene_tx=XM_023250642.1
A2 158365402 158366447 refGene_gn=LOC109497660
A2 158374929 158375885 refGene_gn=LOC111557462
A2 158387102 158389298 refGene_gn=LOC109497661
A2 158478139 158538534
refGene_gn=LOC101085707;refGene_tx=XM_003983123.3;refGene_gn=LOC101097308;refGene_gn=LOC111559550
A2 158540508 158547370 refGene_gn=LOC101097553
A2 158553219 158553370
refGene_gn=LOC101097801;refGene_tx=XM_019825942.2;refGene_gn=LOC101097801;refGene_tx=XM_023250645.1;refGene_gn=LOC101097801;refGene_tx=XM_023250647.1
A2 158556728 158556840
refGene_gn=LOC101097801;refGene_tx=XM_019825944.2;refGene_gn=LOC101097801;refGene_tx=XM_023250646.1
A2 158557840 158558150
refGene_gn=LOC101097801;refGene_tx=XM_019825942.2;refGene_gn=LOC101097801;refGene_tx=XM_019825944.2;refGene_gn=LOC101097801;refGene_tx=XM_023250645.1;refGene_gn=LOC101097801;refGene_tx=XM_023250646.1;refGene_gn=LOC101097801;refGene_tx=XM_023250647.1
A2 158558521 158558564
refGene_gn=LOC101097801;refGene_tx=XM_023250647.1
A2 158559385 158559560
refGene_gn=LOC101097801;refGene_tx=XM_019825942.2;refGene_gn=LOC101097801;refGene_tx=XM_019825944.2;refGene_gn=LOC101097801;refGene_tx=XM_023250645.1;refGene_gn=LOC101097801;refGene_tx=XM_023250646.1
A2 158601254 158601354
refGene_gn=EPHB6;refGene_tx=XM_003983124.5;refGene_gn=EPHB6;refGene_tx=XM_023250648.1;refGene_gn=EPHB6;refGene_tx=XM_023250649.1;refGene_gn=EPHB6;refGene_tx=XM_023250650.1;refGene_gn=EPHB6;refGene_tx=XM_023250651.1
A2 158601628 158601693
refGene_gn=EPHB6;refGene_tx=XM_003983124.5;refGene_gn=EPHB6;refGene_tx=XM_023250648.1;refGene_gn=EPHB6;refGene_tx=XM_023250649.1;refGene_gn=EPHB6;refGene_tx=XM_023250650.1;refGene_gn=EPHB6;refGene_tx=XM_023250651.1
A2 158601880 158602634
refGene_gn=EPHB6;refGene_tx=XM_003983124.5;refGene_gn=EPHB6;refGene_tx=XM_023250648.1;refGene_gn=EPHB6;refGene_tx=XM_023250649.1;refGene_gn=EPHB6;refGene_tx=XM_023250650.1;refGene_gn=EPHB6;refGene_tx=XM_023250651.1
A2 158603366 158603522
refGene_gn=EPHB6;refGene_tx=XM_003983124.5;refGene_gn=EPHB6;refGene_tx=XM_023250648.1;refGene_gn=EPHB6;refGene_tx=XM_023250649.1;refGene_gn=EPHB6;refGene_tx=XM_023250650.1;refGene_gn=EPHB6;refGene_tx=XM_023250651.1
A2 158603893 158604250

refGene_gn=EPHB6;refGene_tx=XM_003983124.5;refGene_gn=EPHB6;refGene_tx=XM_023250648.1;refGene_gn=EPHB6;refGene_tx=XM_023250649.1;refGene_gn=EPHB6;refGene_tx=XM_023250650.1;refGene_gn=EPHB6;refGene_tx=XM_023250651.1
A2 158604362 158604487
refGene_gn=EPHB6;refGene_tx=XM_003983124.5;refGene_gn=EPHB6;refGene_tx=XM_023250648.1;refGene_gn=EPHB6;refGene_tx=XM_023250649.1;refGene_gn=EPHB6;refGene_tx=XM_023250650.1;refGene_gn=EPHB6;refGene_tx=XM_023250651.1
A2 158604614 158604777
refGene_gn=EPHB6;refGene_tx=XM_003983124.5;refGene_gn=EPHB6;refGene_tx=XM_023250648.1;refGene_gn=EPHB6;refGene_tx=XM_023250649.1;refGene_gn=EPHB6;refGene_tx=XM_023250650.1;refGene_gn=EPHB6;refGene_tx=XM_023250651.1
A2 158605046 158605161
refGene_gn=EPHB6;refGene_tx=XM_003983124.5;refGene_gn=EPHB6;refGene_tx=XM_023250648.1;refGene_gn=EPHB6;refGene_tx=XM_023250649.1;refGene_gn=EPHB6;refGene_tx=XM_023250650.1;refGene_gn=EPHB6;refGene_tx=XM_023250651.1
A2 158605408 158605461
refGene_gn=EPHB6;refGene_tx=XM_003983124.5;refGene_gn=EPHB6;refGene_tx=XM_023250648.1;refGene_gn=EPHB6;refGene_tx=XM_023250649.1;refGene_gn=EPHB6;refGene_tx=XM_023250650.1;refGene_gn=EPHB6;refGene_tx=XM_023250651.1
A2 158605692 158605812
refGene_gn=EPHB6;refGene_tx=XM_003983124.5;refGene_gn=EPHB6;refGene_tx=XM_023250648.1;refGene_gn=EPHB6;refGene_tx=XM_023250649.1;refGene_gn=EPHB6;refGene_tx=XM_023250650.1;refGene_gn=EPHB6;refGene_tx=XM_023250651.1
A2 158605927 158606175
refGene_gn=EPHB6;refGene_tx=XM_003983124.5;refGene_gn=EPHB6;refGene_tx=XM_023250648.1;refGene_gn=EPHB6;refGene_tx=XM_023250649.1;refGene_gn=EPHB6;refGene_tx=XM_023250650.1;refGene_gn=EPHB6;refGene_tx=XM_023250651.1
A2 158606440 158606614
refGene_gn=EPHB6;refGene_tx=XM_003983124.5;refGene_gn=EPHB6;refGene_tx=XM_023250648.1;refGene_gn=EPHB6;refGene_tx=XM_023250649.1;refGene_gn=EPHB6;refGene_tx=XM_023250650.1;refGene_gn=EPHB6;refGene_tx=XM_023250651.1
A2 158606843 158606993
refGene_gn=EPHB6;refGene_tx=XM_003983124.5;refGene_gn=EPHB6;refGene_tx=XM_023250648.1;refGene_gn=EPHB6;refGene_tx=XM_023250649.1;refGene_gn=EPHB6;refGene_tx=XM_023250650.1;refGene_gn=EPHB6;refGene_tx=XM_023250651.1
A2 158607252 158607446
refGene_gn=EPHB6;refGene_tx=XM_003983124.5;refGene_gn=EPHB6;refGene_tx=XM_023250648.1;refGene_gn=EPHB6;refGene_tx=XM_023250649.1;refGene_gn=EPHB6;refGene_tx=XM_023250650.1;refGene_gn=EPHB6;refGene_tx=XM_023250651.1
A2 158607566 158607722

refGene_gn=EPHB6;refGene_tx=XM_003983124.5;refGene_gn=EPHB6;refGene_tx=XM_023250648.1;refGene_gn=EPHB6;refGene_tx=XM_023250649.1;refGene_gn=EPHB6;refGene_tx=XM_023250650.1;refGene_gn=EPHB6;refGene_tx=XM_023250651.1
A2	158607830	158607939
refGene_gn=EPHB6;refGene_tx=XM_003983124.5;refGene_gn=EPHB6;refGene_tx=XM_023250648.1;refGene_gn=EPHB6;refGene_tx=XM_023250649.1;refGene_gn=EPHB6;refGene_tx=XM_023250650.1;refGene_gn=EPHB6;refGene_tx=XM_023250651.1
A2	158608790	158609070
refGene_gn=TRPV6;refGene_tx=XM_019825946.2
A2	158609363	158609470
refGene_gn=TRPV6;refGene_tx=XM_019825946.2
A2	158610448	158610717
refGene_gn=TRPV6;refGene_tx=XM_019825946.2
A2	158611089	158611156
refGene_gn=TRPV6;refGene_tx=XM_019825946.2
A2	158611542	158611708
refGene_gn=TRPV6;refGene_tx=XM_019825946.2
A2	158611920	158611997
refGene_gn=TRPV6;refGene_tx=XM_019825946.2
A2	158612096	158612183
refGene_gn=TRPV6;refGene_tx=XM_019825946.2
A2	158612497	158612710
refGene_gn=TRPV6;refGene_tx=XM_019825946.2
A2	158612792	158612939
refGene_gn=TRPV6;refGene_tx=XM_019825946.2
A2	158613309	158613485
refGene_gn=TRPV6;refGene_tx=XM_019825946.2
A2	158613587	158613686
refGene_gn=TRPV6;refGene_tx=XM_019825946.2
A2	158613982	158614120
refGene_gn=TRPV6;refGene_tx=XM_019825946.2
A2	158614435	158614558
refGene_gn=TRPV6;refGene_tx=XM_019825946.2
A2	158614707	158614805
refGene_gn=TRPV6;refGene_tx=XM_019825946.2
A2	158622350	158622478
refGene_gn=TRPV6;refGene_tx=XM_019825946.2
A2	158637657	158637964
refGene_gn=TRPV5;refGene_tx=XM_023250652.1
A2	158638651	158638758
refGene_gn=TRPV5;refGene_tx=XM_023250652.1
A2	158642250	158642519
refGene_gn=TRPV5;refGene_tx=XM_023250652.1
A2	158644604	158644671
refGene_gn=TRPV5;refGene_tx=XM_023250652.1
A2	158644830	158644996
refGene_gn=TRPV5;refGene_tx=XM_023250652.1
A2	158645241	158645318

refGene_gn=TRPV5;refGene_tx=XM_023250652.1
A2 158645416 158645503
refGene_gn=TRPV5;refGene_tx=XM_023250652.1
A2 158656468 158656681
refGene_gn=TRPV5;refGene_tx=XM_023250652.1
A2 158658524 158658671
refGene_gn=TRPV5;refGene_tx=XM_023250652.1
A2 158659041 158659217
refGene_gn=TRPV5;refGene_tx=XM_023250652.1
A2 158659319 158659418
refGene_gn=TRPV5;refGene_tx=XM_023250652.1
A2 158659697 158659835
refGene_gn=TRPV5;refGene_tx=XM_023250652.1
A2 158662366 158662489
refGene_gn=TRPV5;refGene_tx=XM_023250652.1
A2 158662643 158662741
refGene_gn=TRPV5;refGene_tx=XM_023250652.1
A2 158665004 158665132
refGene_gn=TRPV5;refGene_tx=XM_023250652.1
A2 158670255 158670291
refGene_gn=LLCFC1;refGene_tx=XM_011280613.3
A2 158670406 158670527
refGene_gn=LLCFC1;refGene_tx=XM_011280613.3
A2 158671114 158671293
refGene_gn=LLCFC1;refGene_tx=XM_011280613.3
A2 158672004 158672166
refGene_gn=KEL;refGene_tx=XM_023250653.1;refGene_gn=KEL;refGene_tx=XM_023250654.1;refGene_gn=KEL;refGene_tx=XM_023250655.1;refGene_gn=KEL;refGene_tx=XM_023250656.1;refGene_gn=KEL;refGene_tx=XM_023250657.1;refGene_gn=KEL;refGene_tx=XM_023250658.1;refGene_gn=KEL;refGene_tx=XM_023250659.1;refGene_gn=KEL;refGene_tx=XM_023250660.1;refGene_gn=KEL;refGene_tx=XM_023250661.1;refGene_gn=KEL;refGene_tx=XM_023250663.1;refGene_gn=KEL;refGene_tx=XM_023250664.1;refGene_gn=KEL;refGene_tx=XM_023250665.1
A2 158673249 158673345
refGene_gn=KEL;refGene_tx=XM_023250653.1;refGene_gn=KEL;refGene_tx=XM_023250654.1;refGene_gn=KEL;refGene_tx=XM_023250655.1;refGene_gn=KEL;refGene_tx=XM_023250656.1;refGene_gn=KEL;refGene_tx=XM_023250657.1;refGene_gn=KEL;refGene_tx=XM_023250658.1;refGene_gn=KEL;refGene_tx=XM_023250659.1;refGene_gn=KEL;refGene_tx=XM_023250660.1;refGene_gn=KEL;refGene_tx=XM_023250661.1;refGene_gn=KEL;refGene_tx=XM_023250663.1;refGene_gn=KEL;refGene_tx=XM_023250664.1;refGene_gn=KEL;refGene_tx=XM_023250665.1
A2 158673694 158673864
refGene_gn=KEL;refGene_tx=XM_023250653.1;refGene_gn=KEL;refGene_tx=XM_023250654.1;refGene_gn=KEL;refGene_tx=XM_023250655.1;refGene_gn=KEL;refGene_tx=XM_023250656.1;refGene_gn=KEL;refGene_tx=XM_023250657.1;refGene_gn=KEL;refGene_tx=XM_023250658.1;refGene_gn=KEL;refGene_tx=XM_023250659.1;refGene_gn=KEL;refGene_tx=XM_023250660.1;refGene_gn=KEL;refGene_tx=XM_023250661.1;refGene_gn=KEL;refGene_tx=XM_023250662.1;refGene_gn

=KEL;refGene_tx=XM_023250663.1;refGene_gn=KEL;refGene_tx=XM_023250664.
1;refGene_gn=KEL;refGene_tx=XM_023250665.1
A2 158674072 158674144
refGene_gn=KEL;refGene_tx=XM_023250653.1;refGene_gn=KEL;refGene_tx=XM_
023250654.1;refGene_gn=KEL;refGene_tx=XM_023250655.1;refGene_gn=KEL;re
fGene_tx=XM_023250656.1;refGene_gn=KEL;refGene_tx=XM_023250657.1;refGe
ne_gn=KEL;refGene_tx=XM_023250658.1;refGene_gn=KEL;refGene_tx=XM_02325
0659.1;refGene_gn=KEL;refGene_tx=XM_023250660.1;refGene_gn=KEL;refGene
_tx=XM_023250661.1;refGene_gn=KEL;refGene_tx=XM_023250662.1;refGene_gn
=KEL;refGene_tx=XM_023250663.1;refGene_gn=KEL;refGene_tx=XM_023250664.
1;refGene_gn=KEL;refGene_tx=XM_023250665.1
A2 158674286 158674397
refGene_gn=KEL;refGene_tx=XM_023250653.1;refGene_gn=KEL;refGene_tx=XM_
023250654.1;refGene_gn=KEL;refGene_tx=XM_023250655.1;refGene_gn=KEL;re
fGene_tx=XM_023250656.1;refGene_gn=KEL;refGene_tx=XM_023250657.1;refGe
ne_gn=KEL;refGene_tx=XM_023250658.1;refGene_gn=KEL;refGene_tx=XM_02325
0659.1;refGene_gn=KEL;refGene_tx=XM_023250660.1;refGene_gn=KEL;refGene
_tx=XM_023250661.1;refGene_gn=KEL;refGene_tx=XM_023250662.1;refGene_gn
=KEL;refGene_tx=XM_023250663.1;refGene_gn=KEL;refGene_tx=XM_023250664.
1;refGene_gn=KEL;refGene_tx=XM_023250665.1
A2 158674583 158674684
refGene_gn=KEL;refGene_tx=XM_023250653.1;refGene_gn=KEL;refGene_tx=XM_
023250654.1;refGene_gn=KEL;refGene_tx=XM_023250655.1;refGene_gn=KEL;re
fGene_tx=XM_023250656.1;refGene_gn=KEL;refGene_tx=XM_023250657.1;refGe
ne_gn=KEL;refGene_tx=XM_023250658.1;refGene_gn=KEL;refGene_tx=XM_02325
0659.1;refGene_gn=KEL;refGene_tx=XM_023250660.1;refGene_gn=KEL;refGene
_tx=XM_023250661.1;refGene_gn=KEL;refGene_tx=XM_023250662.1;refGene_gn
=KEL;refGene_tx=XM_023250663.1;refGene_gn=KEL;refGene_tx=XM_023250664.
1;refGene_gn=KEL;refGene_tx=XM_023250665.1
A2 158675124 158675202
refGene_gn=KEL;refGene_tx=XM_023250653.1;refGene_gn=KEL;refGene_tx=XM_
023250654.1;refGene_gn=KEL;refGene_tx=XM_023250655.1;refGene_gn=KEL;re
fGene_tx=XM_023250656.1;refGene_gn=KEL;refGene_tx=XM_023250657.1;refGe
ne_gn=KEL;refGene_tx=XM_023250658.1;refGene_gn=KEL;refGene_tx=XM_02325
0659.1;refGene_gn=KEL;refGene_tx=XM_023250660.1;refGene_gn=KEL;refGene
_tx=XM_023250661.1;refGene_gn=KEL;refGene_tx=XM_023250662.1;refGene_gn
=KEL;refGene_tx=XM_023250663.1;refGene_gn=KEL;refGene_tx=XM_023250664.
1;refGene_gn=KEL;refGene_tx=XM_023250665.1
A2 158675438 158675537
refGene_gn=KEL;refGene_tx=XM_023250653.1;refGene_gn=KEL;refGene_tx=XM_
023250654.1;refGene_gn=KEL;refGene_tx=XM_023250655.1;refGene_gn=KEL;re
fGene_tx=XM_023250656.1;refGene_gn=KEL;refGene_tx=XM_023250657.1;refGe
ne_gn=KEL;refGene_tx=XM_023250658.1;refGene_gn=KEL;refGene_tx=XM_02325
0659.1;refGene_gn=KEL;refGene_tx=XM_023250660.1;refGene_gn=KEL;refGene
_tx=XM_023250661.1;refGene_gn=KEL;refGene_tx=XM_023250662.1;refGene_gn
=KEL;refGene_tx=XM_023250663.1;refGene_gn=KEL;refGene_tx=XM_023250664.
1;refGene_gn=KEL;refGene_tx=XM_023250665.1
A2 158676703 158676814
refGene_gn=KEL;refGene_tx=XM_023250653.1;refGene_gn=KEL;refGene_tx=XM_
023250654.1;refGene_gn=KEL;refGene_tx=XM_023250655.1;refGene_gn=KEL;re

fGene_tx=XM_023250656.1;refGene_gn=KEL;refGene_tx=XM_023250657.1;refGene_gn=KEL;refGene_tx=XM_023250658.1;refGene_gn=KEL;refGene_tx=XM_023250659.1;refGene_gn=KEL;refGene_tx=XM_023250660.1;refGene_gn=KEL;refGene_tx=XM_023250661.1;refGene_gn=KEL;refGene_tx=XM_023250662.1;refGene_gn=KEL;refGene_tx=XM_023250663.1;refGene_gn=KEL;refGene_tx=XM_023250664.1;refGene_gn=KEL;refGene_tx=XM_023250665.1
A2 158684139 158684269
refGene_gn=KEL;refGene_tx=XM_023250653.1;refGene_gn=KEL;refGene_tx=XM_023250654.1;refGene_gn=KEL;refGene_tx=XM_023250655.1;refGene_gn=KEL;refGene_tx=XM_023250656.1;refGene_gn=KEL;refGene_tx=XM_023250657.1;refGene_gn=KEL;refGene_tx=XM_023250658.1;refGene_gn=KEL;refGene_tx=XM_023250659.1;refGene_gn=KEL;refGene_tx=XM_023250660.1;refGene_gn=KEL;refGene_tx=XM_023250661.1;refGene_gn=KEL;refGene_tx=XM_023250662.1;refGene_gn=KEL;refGene_tx=XM_023250663.1;refGene_gn=KEL;refGene_tx=XM_023250664.1;refGene_gn=KEL;refGene_tx=XM_023250665.1
A2 158685369 158685518
refGene_gn=KEL;refGene_tx=XM_023250653.1;refGene_gn=KEL;refGene_tx=XM_023250654.1;refGene_gn=KEL;refGene_tx=XM_023250655.1;refGene_gn=KEL;refGene_tx=XM_023250656.1;refGene_gn=KEL;refGene_tx=XM_023250657.1;refGene_gn=KEL;refGene_tx=XM_023250658.1;refGene_gn=KEL;refGene_tx=XM_023250659.1;refGene_gn=KEL;refGene_tx=XM_023250660.1;refGene_gn=KEL;refGene_tx=XM_023250661.1;refGene_gn=KEL;refGene_tx=XM_023250662.1;refGene_gn=KEL;refGene_tx=XM_023250663.1;refGene_gn=KEL;refGene_tx=XM_023250664.1;refGene_gn=KEL;refGene_tx=XM_023250665.1
A2 158685760 158685949
refGene_gn=KEL;refGene_tx=XM_023250653.1;refGene_gn=KEL;refGene_tx=XM_023250654.1;refGene_gn=KEL;refGene_tx=XM_023250655.1;refGene_gn=KEL;refGene_tx=XM_023250656.1;refGene_gn=KEL;refGene_tx=XM_023250657.1;refGene_gn=KEL;refGene_tx=XM_023250658.1;refGene_gn=KEL;refGene_tx=XM_023250659.1;refGene_gn=KEL;refGene_tx=XM_023250660.1;refGene_gn=KEL;refGene_tx=XM_023250662.1;refGene_gn=KEL;refGene_tx=XM_023250663.1;refGene_gn=KEL;refGene_tx=XM_023250664.1;refGene_gn=KEL;refGene_tx=XM_023250665.1
A2 158686035 158686188
refGene_gn=KEL;refGene_tx=XM_023250653.1;refGene_gn=KEL;refGene_tx=XM_023250654.1;refGene_gn=KEL;refGene_tx=XM_023250655.1;refGene_gn=KEL;refGene_tx=XM_023250656.1;refGene_gn=KEL;refGene_tx=XM_023250657.1;refGene_gn=KEL;refGene_tx=XM_023250658.1;refGene_gn=KEL;refGene_tx=XM_023250659.1;refGene_gn=KEL;refGene_tx=XM_023250660.1;refGene_gn=KEL;refGene_tx=XM_023250662.1;refGene_gn=KEL;refGene_tx=XM_023250663.1;refGene_gn=KEL;refGene_tx=XM_023250664.1;refGene_gn=KEL;refGene_tx=XM_023250665.1
A2 158689135 158689282
refGene_gn=KEL;refGene_tx=XM_023250653.1;refGene_gn=KEL;refGene_tx=XM_023250654.1;refGene_gn=KEL;refGene_tx=XM_023250655.1;refGene_gn=KEL;refGene_tx=XM_023250657.1;refGene_gn=KEL;refGene_tx=XM_023250658.1;refGene_gn=KEL;refGene_tx=XM_023250659.1;refGene_gn=KEL;refGene_tx=XM_023250661.1;refGene_gn=KEL;refGene_tx=XM_023250662.1;refGene_gn=KEL;refGene_tx=XM_023250663.1;refGene_gn=KEL;refGene_tx=XM_023250664.1
A2 158689632 158689757

refGene_gn=KEL;refGene_tx=XM_023250653.1;refGene_gn=KEL;refGene_tx=XM_023250654.1;refGene_gn=KEL;refGene_tx=XM_023250655.1;refGene_gn=KEL;refGene_tx=XM_023250656.1;refGene_gn=KEL;refGene_tx=XM_023250657.1;refGene_gn=KEL;refGene_tx=XM_023250658.1;refGene_gn=KEL;refGene_tx=XM_023250659.1;refGene_gn=KEL;refGene_tx=XM_023250660.1;refGene_gn=KEL;refGene_tx=XM_023250661.1;refGene_gn=KEL;refGene_tx=XM_023250662.1;refGene_gn=KEL;refGene_tx=XM_023250663.1;refGene_gn=KEL;refGene_tx=XM_023250664.1;refGene_gn=KEL;refGene_tx=XM_023250665.1
A2	158691217	158691394
refGene_gn=KEL;refGene_tx=XM_023250653.1;refGene_gn=KEL;refGene_tx=XM_023250654.1;refGene_gn=KEL;refGene_tx=XM_023250655.1;refGene_gn=KEL;refGene_tx=XM_023250656.1;refGene_gn=KEL;refGene_tx=XM_023250657.1;refGene_gn=KEL;refGene_tx=XM_023250658.1;refGene_gn=KEL;refGene_tx=XM_023250659.1;refGene_gn=KEL;refGene_tx=XM_023250660.1;refGene_gn=KEL;refGene_tx=XM_023250661.1;refGene_gn=KEL;refGene_tx=XM_023250662.1;refGene_gn=KEL;refGene_tx=XM_023250663.1;refGene_gn=KEL;refGene_tx=XM_023250664.1;refGene_gn=KEL;refGene_tx=XM_023250665.1
A2	158691678	158691841
refGene_gn=KEL;refGene_tx=XM_023250653.1;refGene_gn=KEL;refGene_tx=XM_023250654.1;refGene_gn=KEL;refGene_tx=XM_023250655.1;refGene_gn=KEL;refGene_tx=XM_023250656.1;refGene_gn=KEL;refGene_tx=XM_023250657.1;refGene_gn=KEL;refGene_tx=XM_023250658.1;refGene_gn=KEL;refGene_tx=XM_023250659.1;refGene_gn=KEL;refGene_tx=XM_023250660.1;refGene_gn=KEL;refGene_tx=XM_023250661.1;refGene_gn=KEL;refGene_tx=XM_023250662.1
A2	158692129	158692264
refGene_gn=KEL;refGene_tx=XM_023250653.1;refGene_gn=KEL;refGene_tx=XM_023250654.1;refGene_gn=KEL;refGene_tx=XM_023250655.1;refGene_gn=KEL;refGene_tx=XM_023250656.1;refGene_gn=KEL;refGene_tx=XM_023250657.1;refGene_gn=KEL;refGene_tx=XM_023250659.1;refGene_gn=KEL;refGene_tx=XM_023250660.1;refGene_gn=KEL;refGene_tx=XM_023250661.1;refGene_gn=KEL;refGene_tx=XM_023250662.1
A2	158741297	158742242
refGene_gn=LOC101086714;refGene_tx=XM_003983127.3
A2	158756725	158757670
refGene_gn=LOC101098529;refGene_tx=XM_003983163.5
A2	158772767	158773712
refGene_gn=LOC101087220;refGene_tx=XM_003983129.3
A2	158783708	158784400
refGene_gn=LOC111557463;refGene_tx=XM_023242295.1
A2	158784402	158784679
refGene_gn=LOC111557463;refGene_tx=XM_023242295.1
A2	158791578	158791637
refGene_gn=LOC111557463;refGene_tx=XM_023242295.1
A2	158792258	158792304
refGene_gn=LOC111557463;refGene_tx=XM_023242295.1
A2	158836207	158836302
refGene_gn=PIP;refGene_tx=XM_011280701.2
A2	158838329	158838438
refGene_gn=PIP;refGene_tx=XM_011280701.2
A2	158843964	158844079

refGene_gn=PIP;refGene_tx=XM_011280701.2
A2 158844628 158844753
refGene_gn=PIP;refGene_tx=XM_011280701.2
A2 158874284 158874360
refGene_gn=TAS2R39;refGene_tx=XM_023250666.1
A2 158874362 158875024
refGene_gn=TAS2R39;refGene_tx=XM_023250666.1
A2 158898024 158898732
refGene_gn=TAS2R40;refGene_tx=XM_019825957.2
A2 158898734 158899016
refGene_gn=TAS2R40;refGene_tx=XM_019825957.2
A2 158920701 158920773
refGene_gn=GSTK1;refGene_tx=XM_003983130.5
A2 158921258 158921340
refGene_gn=GSTK1;refGene_tx=XM_003983130.5
A2 158921516 158921645
refGene_gn=GSTK1;refGene_tx=XM_003983130.5
A2 158921950 158922051
refGene_gn=GSTK1;refGene_tx=XM_003983130.5
A2 158922224 158922260
refGene_gn=GSTK1;refGene_tx=XM_003983130.5
A2 158923890 158924007
refGene_gn=GSTK1;refGene_tx=XM_003983130.5
A2 158924359 158924453
refGene_gn=GSTK1;refGene_tx=XM_003983130.5
A2 158924713 158924763
refGene_gn=GSTK1;refGene_tx=XM_003983130.5
A2 158936880 158936990
refGene_gn=TMEM139;refGene_tx=XM_019825961.2
A2 158937997 158938230
refGene_gn=TMEM139;refGene_tx=XM_019825959.1;refGene_gn=TMEM139;refGen
e_tx=XM_019825960.2
A2 158938450 158938841
refGene_gn=TMEM139;refGene_tx=XM_019825959.1;refGene_gn=TMEM139;refGen
e_tx=XM_019825960.2;refGene_gn=TMEM139;refGene_tx=XM_019825961.2
A2 158940429 158940503
refGene_gn=CASP2;refGene_tx=XM_003983133.4;refGene_gn=CASP2;refGene_tx
=XM_006929443.3;refGene_gn=CASP2;refGene_tx=XM_019825963.2
A2 158943872 158944023
refGene_gn=CASP2;refGene_tx=XM_003983133.4;refGene_gn=CASP2;refGene_tx
=XM_006929443.3;refGene_gn=CASP2;refGene_tx=XM_019825963.2
A2 158944310 158944478
refGene_gn=CASP2;refGene_tx=XM_003983133.4;refGene_gn=CASP2;refGene_tx
=XM_006929443.3;refGene_gn=CASP2;refGene_tx=XM_019825963.2
A2 158944628 158944710
refGene_gn=CASP2;refGene_tx=XM_003983133.4;refGene_gn=CASP2;refGene_tx
=XM_006929443.3;refGene_gn=CASP2;refGene_tx=XM_019825963.2
A2 158945265 158945360
refGene_gn=CASP2;refGene_tx=XM_003983133.4;refGene_gn=CASP2;refGene_tx
=XM_006929443.3;refGene_gn=CASP2;refGene_tx=XM_019825963.2

A2 158945642 158945819
refGene_gn=CASP2;refGene_tx=XM_003983133.4;refGene_gn=CASP2;refGene_tx=XM_006929443.3
A2 158952609 158952738
refGene_gn=CASP2;refGene_tx=XM_003983133.4;refGene_gn=CASP2;refGene_tx=XM_006929443.3;refGene_gn=CASP2;refGene_tx=XM_019825963.2
A2 158952893 158952984
refGene_gn=CASP2;refGene_tx=XM_003983133.4;refGene_gn=CASP2;refGene_tx=XM_006929443.3;refGene_gn=CASP2;refGene_tx=XM_019825963.2
A2 158953069 158953134
refGene_gn=CASP2;refGene_tx=XM_006929443.3
A2 158956203 158956350
refGene_gn=CASP2;refGene_tx=XM_003983133.4;refGene_gn=CASP2;refGene_tx=XM_019825963.2
A2 158956526 158956636
refGene_gn=CASP2;refGene_tx=XM_003983133.4;refGene_gn=CASP2;refGene_tx=XM_019825963.2
A2 158956769 158956901
refGene_gn=CASP2;refGene_tx=XM_003983133.4;refGene_gn=CASP2;refGene_tx=XM_019825963.2
A2 158963697 158963877
refGene_gn=CLCN1;refGene_tx=NM_001305027.1
A2 158966469 158966590
refGene_gn=CLCN1;refGene_tx=NM_001305027.1
A2 158966958 158967090
refGene_gn=CLCN1;refGene_tx=NM_001305027.1
A2 158967660 158967789
refGene_gn=CLCN1;refGene_tx=NM_001305027.1
A2 158968042 158968176
refGene_gn=CLCN1;refGene_tx=NM_001305027.1
A2 158969331 158969409
refGene_gn=CLCN1;refGene_tx=NM_001305027.1
A2 158970495 158970574
refGene_gn=CLCN1;refGene_tx=NM_001305027.1
A2 158975889 158976015
refGene_gn=CLCN1;refGene_tx=NM_001305027.1
A2 158976302 158976387
refGene_gn=CLCN1;refGene_tx=NM_001305027.1
A2 158976593 158976695
refGene_gn=CLCN1;refGene_tx=NM_001305027.1
A2 158977183 158977268
refGene_gn=CLCN1;refGene_tx=NM_001305027.1
A2 158977721 158977871
refGene_gn=CLCN1;refGene_tx=NM_001305027.1
A2 158983903 158983973
refGene_gn=CLCN1;refGene_tx=NM_001305027.1
A2 158984162 158984273
refGene_gn=CLCN1;refGene_tx=NM_001305027.1
A2 158985932 158986146
refGene_gn=CLCN1;refGene_tx=NM_001305027.1

A2 158986363 158986497
refGene_gn=CLCN1;refGene_tx=NM_001305027.1
A2 158988945 158989190
refGene_gn=CLCN1;refGene_tx=NM_001305027.1
A2 158989478 158989590
refGene_gn=CLCN1;refGene_tx=NM_001305027.1
A2 158989915 158989995
refGene_gn=CLCN1;refGene_tx=NM_001305027.1
A2 158990236 158990275
refGene_gn=CLCN1;refGene_tx=NM_001305027.1
A2 158993933 158994038
refGene_gn=CLCN1;refGene_tx=NM_001305027.1
A2 158994127 158994214
refGene_gn=CLCN1;refGene_tx=NM_001305027.1
A2 158994702 158995074
refGene_gn=CLCN1;refGene_tx=NM_001305027.1
A2 158999386 158999859
refGene_gn=FAM131B;refGene_tx=XM_023250667.1;refGene_gn=FAM131B;refGene_tx=XM_023250668.1
A2 159000057 159000201
refGene_gn=FAM131B;refGene_tx=XM_023250667.1;refGene_gn=FAM131B;refGene_tx=XM_023250668.1
A2 159002323 159002521
refGene_gn=FAM131B;refGene_tx=XM_023250667.1;refGene_gn=FAM131B;refGene_tx=XM_023250668.1
A2 159002805 159002899
refGene_gn=FAM131B;refGene_tx=XM_023250667.1;refGene_gn=FAM131B;refGene_tx=XM_023250668.1
A2 159003154 159003190
refGene_gn=FAM131B;refGene_tx=XM_023250667.1;refGene_gn=FAM131B;refGene_tx=XM_023250668.1
A2 159003553 159003663
refGene_gn=FAM131B;refGene_tx=XM_023250667.1;refGene_gn=FAM131B;refGene_tx=XM_023250668.1
A2 159006030 159006058
refGene_gn=FAM131B;refGene_tx=XM_023250667.1
A2 159006418 159006446
refGene_gn=FAM131B;refGene_tx=XM_023250668.1
A2 159030566 159030774
refGene_gn=ZYX;refGene_tx=XM_023250674.1
A2 159031269 159031469
refGene_gn=ZYX;refGene_tx=XM_023250674.1
A2 159031621 159031714
refGene_gn=ZYX;refGene_tx=XM_023250674.1
A2 159031810 159032347
refGene_gn=ZYX;refGene_tx=XM_023250674.1
A2 159036285 159036406
refGene_gn=ZYX;refGene_tx=XM_023250674.1
A2 159036535 159036705
refGene_gn=ZYX;refGene_tx=XM_023250674.1

A2 159036842 159037021
refGene_gn=ZYX;refGene_tx=XM_023250674.1
A2 159037890 159038011
refGene_gn=ZYX;refGene_tx=XM_023250674.1
A2 159038618 159038723
refGene_gn=ZYX;refGene_tx=XM_023250674.1
A2 159039569 159039648
refGene_gn=EPHA1;refGene_tx=XM_023250669.1
A2 159039737 159039893
refGene_gn=EPHA1;refGene_tx=XM_023250669.1
A2 159040101 159040162
refGene_gn=EPHA1;refGene_tx=XM_023250670.1
A2 159041286 159041480
refGene_gn=EPHA1;refGene_tx=XM_023250669.1;refGene_gn=EPHA1;refGene_tx
=XM_023250670.1
A2 159041721 159041871
refGene_gn=EPHA1;refGene_tx=XM_023250669.1;refGene_gn=EPHA1;refGene_tx
=XM_023250670.1
A2 159042060 159042267
refGene_gn=EPHA1;refGene_tx=XM_023250669.1;refGene_gn=EPHA1;refGene_tx
=XM_023250670.1
A2 159042388 159042450
refGene_gn=EPHA1;refGene_tx=XM_023250669.1;refGene_gn=EPHA1;refGene_tx
=XM_023250670.1
A2 159042611 159042797
refGene_gn=EPHA1;refGene_tx=XM_023250669.1;refGene_gn=EPHA1;refGene_tx
=XM_023250670.1
A2 159043625 159043751
refGene_gn=EPHA1;refGene_tx=XM_023250669.1;refGene_gn=EPHA1;refGene_tx
=XM_023250670.1
A2 159044505 159044564
refGene_gn=EPHA1;refGene_tx=XM_023250669.1;refGene_gn=EPHA1;refGene_tx
=XM_023250670.1
A2 159044786 159044883
refGene_gn=EPHA1;refGene_tx=XM_023250669.1;refGene_gn=EPHA1;refGene_tx
=XM_023250670.1
A2 159045793 159045944
refGene_gn=EPHA1;refGene_tx=XM_023250669.1;refGene_gn=EPHA1;refGene_tx
=XM_023250670.1
A2 159046201 159046329
refGene_gn=EPHA1;refGene_tx=XM_023250669.1;refGene_gn=EPHA1;refGene_tx
=XM_023250670.1
A2 159046482 159046827
refGene_gn=EPHA1;refGene_tx=XM_023250669.1;refGene_gn=EPHA1;refGene_tx
=XM_023250670.1
A2 159047112 159047268
refGene_gn=EPHA1;refGene_tx=XM_023250669.1;refGene_gn=EPHA1;refGene_tx
=XM_023250670.1
A2 159047510 159047913
refGene_gn=EPHA1;refGene_tx=XM_023250669.1;refGene_gn=EPHA1;refGene_tx

=XM_023250670.1
A2	159049855	159050137
refGene_gn=EPHA1;refGene_tx=XM_023250669.1;refGene_gn=EPHA1;refGene_tx
=XM_023250670.1
A2	159054540	159054608
refGene_gn=EPHA1;refGene_tx=XM_023250669.1;refGene_gn=EPHA1;refGene_tx
=XM_023250670.1
A2	159055687	159055769
refGene_gn=EPHA1;refGene_tx=XM_023250669.1;refGene_gn=EPHA1;refGene_tx
=XM_023250670.1
A2	159082395	159082762
refGene_gn=LOC111559554;refGene_tx=XM_023250675.1
A2	159082764	159083513
refGene_gn=LOC111559554;refGene_tx=XM_023250675.1
A2	159088519	159088727
refGene_gn=TAS2R60;refGene_tx=XM_011280699.3
A2	159088729	159088942
refGene_gn=TAS2R60;refGene_tx=XM_011280699.3
A2	159088944	159088974
refGene_gn=TAS2R60;refGene_tx=XM_011280699.3
A2	159088976	159089135
refGene_gn=TAS2R60;refGene_tx=XM_011280699.3
A2	159089137	159089182
refGene_gn=TAS2R60;refGene_tx=XM_011280699.3
A2	159089184	159089450
refGene_gn=TAS2R60;refGene_tx=XM_011280699.3
A2	159113347	159113386
refGene_gn=TAS2R41;refGene_tx=XM_011280698.3
A2	159113528	159113691
refGene_gn=TAS2R41;refGene_tx=XM_011280698.3
A2	159113693	159113822
refGene_gn=TAS2R41;refGene_tx=XM_011280698.3
A2	159113824	159114234
refGene_gn=TAS2R41;refGene_tx=XM_011280698.3
A2	159114236	159114530
refGene_gn=TAS2R41;refGene_tx=XM_011280698.3
A2	159121860	159122349
refGene_gn=LOC101082225;refGene_tx=XM_019825968.2
A2	159128751	159129693
refGene_gn=LOC101082481;refGene_tx=XM_003983180.3
A2	159154054	159155020
refGene_gn=LOC101082738;refGene_tx=XM_003983181.3
A2	159216571	159216826
refGene_gn=TCAF2;refGene_tx=XM_023250671.1;refGene_gn=TCAF2;refGene_tx
=XM_023250672.1;refGene_gn=TCAF2;refGene_tx=XM_023250673.1
A2	159219136	159219474
refGene_gn=TCAF2;refGene_tx=XM_023250671.1;refGene_gn=TCAF2;refGene_tx
=XM_023250672.1;refGene_gn=TCAF2;refGene_tx=XM_023250673.1
A2	159220317	159220557
refGene_gn=TCAF2;refGene_tx=XM_023250671.1;refGene_gn=TCAF2;refGene_tx

=XM_023250672.1;refGene_gn=TCAF2;refGene_tx=XM_023250673.1
A2 159220862 159221076
refGene_gn=TCAF2;refGene_tx=XM_023250671.1;refGene_gn=TCAF2;refGene_tx=XM_023250672.1;refGene_gn=TCAF2;refGene_tx=XM_023250673.1
A2 159222103 159222201
refGene_gn=TCAF2;refGene_tx=XM_023250671.1;refGene_gn=TCAF2;refGene_tx=XM_023250672.1;refGene_gn=TCAF2;refGene_tx=XM_023250673.1
A2 159222495 159223487
refGene_gn=TCAF2;refGene_tx=XM_023250671.1;refGene_gn=TCAF2;refGene_tx=XM_023250672.1;refGene_gn=TCAF2;refGene_tx=XM_023250673.1
A2 159236713 159237336
refGene_gn=TCAF2;refGene_tx=XM_023250671.1;refGene_gn=TCAF2;refGene_tx=XM_023250672.1;refGene_gn=TCAF2;refGene_tx=XM_023250673.1
A2 159253820 159253830
refGene_gn=TCAF1;refGene_tx=XM_006929456.2;refGene_gn=TCAF1;refGene_tx=XM_011280675.2;refGene_gn=TCAF1;refGene_tx=XM_023250676.1
A2 159256396 159256651
refGene_gn=TCAF1;refGene_tx=XM_006929456.2;refGene_gn=TCAF1;refGene_tx=XM_011280675.2;refGene_gn=TCAF1;refGene_tx=XM_023250676.1;refGene_gn=TCAF1;refGene_tx=XM_023250677.1
A2 159258038 159258376
refGene_gn=TCAF1;refGene_tx=XM_006929456.2;refGene_gn=TCAF1;refGene_tx=XM_011280675.2;refGene_gn=TCAF1;refGene_tx=XM_023250676.1;refGene_gn=TCAF1;refGene_tx=XM_023250677.1
A2 159259649 159259889
refGene_gn=TCAF1;refGene_tx=XM_006929456.2;refGene_gn=TCAF1;refGene_tx=XM_011280675.2;refGene_gn=TCAF1;refGene_tx=XM_023250676.1;refGene_gn=TCAF1;refGene_tx=XM_023250677.1
A2 159260147 159260361
refGene_gn=TCAF1;refGene_tx=XM_006929456.2;refGene_gn=TCAF1;refGene_tx=XM_011280675.2;refGene_gn=TCAF1;refGene_tx=XM_023250676.1;refGene_gn=TCAF1;refGene_tx=XM_023250677.1
A2 159261641 159261739
refGene_gn=TCAF1;refGene_tx=XM_006929456.2;refGene_gn=TCAF1;refGene_tx=XM_011280675.2;refGene_gn=TCAF1;refGene_tx=XM_023250676.1;refGene_gn=TCAF1;refGene_tx=XM_023250677.1
A2 159262002 159262997
refGene_gn=TCAF1;refGene_tx=XM_006929456.2;refGene_gn=TCAF1;refGene_tx=XM_011280675.2;refGene_gn=TCAF1;refGene_tx=XM_023250676.1;refGene_gn=TCAF1;refGene_tx=XM_023250677.1
A2 159284872 159285492
refGene_gn=TCAF1;refGene_tx=XM_006929456.2;refGene_gn=TCAF1;refGene_tx=XM_011280675.2;refGene_gn=TCAF1;refGene_tx=XM_023250676.1;refGene_gn=TCAF1;refGene_tx=XM_023250677.1
A2 159336214 159337168
refGene_gn=LOC101083254;refGene_tx=XM_023250678.1
A2 159362943 159363897
refGene_gn=LOC101100878;refGene_tx=XM_003983171.3
A2 159386028 159386982
refGene_gn=LOC101101123;refGene_tx=XM_023241745.1

A2 159407869 159408823
refGene_gn=LOC101101377;refGene_tx=XM_003983173.1
A2 159427853 159428789
refGene_gn=LOC101083503;refGene_tx=XM_003983183.1
A2 159442910 159443840
refGene_gn=LOC101101632;refGene_tx=XM_003983174.1
A2 159463687 159464623
refGene_gn=LOC101083761;refGene_tx=XM_006929462.1
A2 159509513 159510446
refGene_gn=LOC101080445;refGene_tx=XM_003983175.2
A2 159521671 159522604
refGene_gn=LOC101080702;refGene_tx=XM_003983176.2
A2 159532913 159533846
refGene_gn=LOC101084019;refGene_tx=XM_003983185.1
A2 159548624 159549557
refGene_gn=LOC101084275;refGene_tx=XM_003983186.1
A2 159563123 159564053
refGene_gn=LOC101080959;refGene_tx=XM_003983177.1
A2 159594337 159594357
refGene_gn=LOC101084529;refGene_tx=XM_023241799.1
A2 159594358 159595140
refGene_gn=LOC101084529;refGene_tx=XM_023241799.1
A2 159595142 159595243
refGene_gn=LOC101084529;refGene_tx=XM_023241799.1
A2 159621660 159622585
refGene_gn=LOC101084786;refGene_tx=XM_023242306.1
A2 159631020 159631145
refGene_gn=LOC101084786;refGene_tx=XM_023242306.1
A2 159637767 159638709
refGene_gn=LOC101085032;refGene_tx=XM_011279609.1
A2 159656001 159656934
refGene_gn=LOC101081211;refGene_tx=XM_003983178.2
A2 159683542 159683957
refGene_gn=ARHGEF5;refGene_tx=XM_023250679.1;refGene_gn=ARHGEF5;refGen
e_tx=XM_023250680.1
A2 159684563 159686471
refGene_gn=ARHGEF5;refGene_tx=XM_023250679.1;refGene_gn=ARHGEF5;refGen
e_tx=XM_023250680.1
A2 159686948 159687006
refGene_gn=ARHGEF5;refGene_tx=XM_023250679.1;refGene_gn=ARHGEF5;refGen
e_tx=XM_023250680.1
A2 159687436 159687501
refGene_gn=ARHGEF5;refGene_tx=XM_023250679.1;refGene_gn=ARHGEF5;refGen
e_tx=XM_023250680.1
A2 159687706 159687960
refGene_gn=ARHGEF5;refGene_tx=XM_023250679.1;refGene_gn=ARHGEF5;refGen
e_tx=XM_023250680.1
A2 159690773 159690934
refGene_gn=ARHGEF5;refGene_tx=XM_023250679.1;refGene_gn=ARHGEF5;refGen
e_tx=XM_023250680.1

A2 159691437 159691590
refGene_gn=ARHGEF5;refGene_tx=XM_023250679.1;refGene_gn=ARHGEF5;refGene_tx=XM_023250680.1
A2 159691870 159692000
refGene_gn=ARHGEF5;refGene_tx=XM_023250679.1;refGene_gn=ARHGEF5;refGene_tx=XM_023250680.1
A2 159692307 159692382
refGene_gn=ARHGEF5;refGene_tx=XM_023250679.1;refGene_gn=ARHGEF5;refGene_tx=XM_023250680.1
A2 159692826 159692916
refGene_gn=ARHGEF5;refGene_tx=XM_023250679.1;refGene_gn=ARHGEF5;refGene_tx=XM_023250680.1
A2 159694308 159694466
refGene_gn=ARHGEF5;refGene_tx=XM_023250679.1;refGene_gn=ARHGEF5;refGene_tx=XM_023250680.1
A2 159695067 159695229
refGene_gn=ARHGEF5;refGene_tx=XM_023250679.1;refGene_gn=ARHGEF5;refGene_tx=XM_023250680.1
A2 159696784 159696855
refGene_gn=ARHGEF5;refGene_tx=XM_023250679.1;refGene_gn=ARHGEF5;refGene_tx=XM_023250680.1
A2 159698626 159698731
refGene_gn=ARHGEF5;refGene_tx=XM_023250679.1;refGene_gn=ARHGEF5;refGene_tx=XM_023250680.1
A2 159699442 159699600
refGene_gn=ARHGEF5;refGene_tx=XM_023250679.1;refGene_gn=ARHGEF5;refGene_tx=XM_023250680.1
A2 159710302 159710604
refGene_gn=NOBOX;refGene_tx=XM_023242315.1
A2 159711293 159711604
refGene_gn=NOBOX;refGene_tx=XM_023242315.1
A2 159711928 159712151
refGene_gn=NOBOX;refGene_tx=XM_023242315.1
A2 159712363 159712449
refGene_gn=NOBOX;refGene_tx=XM_023242315.1
A2 159712728 159712892
refGene_gn=NOBOX;refGene_tx=XM_023242315.1
A2 159713134 159713292
refGene_gn=NOBOX;refGene_tx=XM_023242315.1
A2 159713892 159714462
refGene_gn=NOBOX;refGene_tx=XM_023242315.1
A2 159715800 159716311
refGene_gn=NOBOX;refGene_tx=XM_023242315.1
A2 159719239 159719263
refGene_gn=NOBOX;refGene_tx=XM_023242315.1
A2 159719389 159719634
refGene_gn=NOBOX;refGene_tx=XM_023242315.1
A2 159723360 159723616
refGene_gn=NOBOX;refGene_tx=XM_023242315.1
A2 159761355 159761474

refGene_gn=TPK1;refGene_tx=XM_023250681.1;refGene_gn=TPK1;refGene_tx=XM_023250682.1;refGene_gn=TPK1;refGene_tx=XM_023250683.1;refGene_gn=TPK1;refGene_tx=XM_023250684.1;refGene_gn=TPK1;refGene_tx=XM_023250685.1;refGene_gn=TPK1;refGene_tx=XM_023250686.1;refGene_gn=TPK1;refGene_tx=XM_023250687.1
A2 159853886 159853998
refGene_gn=TPK1;refGene_tx=XM_023250681.1;refGene_gn=TPK1;refGene_tx=XM_023250682.1;refGene_gn=TPK1;refGene_tx=XM_023250683.1;refGene_gn=TPK1;refGene_tx=XM_023250684.1;refGene_gn=TPK1;refGene_tx=XM_023250685.1;refGene_gn=TPK1;refGene_tx=XM_023250686.1;refGene_gn=TPK1;refGene_tx=XM_023250687.1
A2 159897023 159897170
refGene_gn=TPK1;refGene_tx=XM_023250681.1;refGene_gn=TPK1;refGene_tx=XM_023250682.1;refGene_gn=TPK1;refGene_tx=XM_023250683.1;refGene_gn=TPK1;refGene_tx=XM_023250686.1;refGene_gn=TPK1;refGene_tx=XM_023250687.1
A2 159924011 159924107
refGene_gn=TPK1;refGene_tx=XM_023250681.1;refGene_gn=TPK1;refGene_tx=XM_023250682.1;refGene_gn=TPK1;refGene_tx=XM_023250683.1;refGene_gn=TPK1;refGene_tx=XM_023250684.1;refGene_gn=TPK1;refGene_tx=XM_023250685.1;refGene_gn=TPK1;refGene_tx=XM_023250686.1;refGene_gn=TPK1;refGene_tx=XM_023250687.1
A2 159944901 159944974
refGene_gn=TPK1;refGene_tx=XM_023250681.1;refGene_gn=TPK1;refGene_tx=XM_023250682.1;refGene_gn=TPK1;refGene_tx=XM_023250683.1;refGene_gn=TPK1;refGene_tx=XM_023250684.1;refGene_gn=TPK1;refGene_tx=XM_023250685.1;refGene_gn=TPK1;refGene_tx=XM_023250686.1;refGene_gn=TPK1;refGene_tx=XM_023250687.1
A2 159947819 159947830
refGene_gn=TPK1;refGene_tx=XM_023250687.1
A2 159967864 159967934
refGene_gn=TPK1;refGene_tx=XM_023250682.1;refGene_gn=TPK1;refGene_tx=XM_023250683.1;refGene_gn=TPK1;refGene_tx=XM_023250685.1
A2 160032044 160032116
refGene_gn=TPK1;refGene_tx=XM_023250681.1;refGene_gn=TPK1;refGene_tx=XM_023250682.1;refGene_gn=TPK1;refGene_tx=XM_023250683.1;refGene_gn=TPK1;refGene_tx=XM_023250684.1;refGene_gn=TPK1;refGene_tx=XM_023250685.1;refGene_gn=TPK1;refGene_tx=XM_023250686.1
A2 160111155 160111214
refGene_gn=TPK1;refGene_tx=XM_023250681.1;refGene_gn=TPK1;refGene_tx=XM_023250682.1;refGene_gn=TPK1;refGene_tx=XM_023250683.1;refGene_gn=TPK1;refGene_tx=XM_023250684.1;refGene_gn=TPK1;refGene_tx=XM_023250685.1
A2 160112012 160112072
refGene_gn=TPK1;refGene_tx=XM_023250681.1;refGene_gn=TPK1;refGene_tx=XM_023250684.1
A2 161247427 161247524
refGene_gn=CNTNAP2;refGene_tx=XM_023250688.1
A2 161835355 161835466
refGene_gn=CNTNAP2;refGene_tx=XM_023250688.1
A2 161893766 161893960
refGene_gn=CNTNAP2;refGene_tx=XM_023250688.1

A2 162078945 162079093
refGene_gn=CNTNAP2;refGene_tx=XM_023250688.1
A2 162143109 162143313
refGene_gn=CNTNAP2;refGene_tx=XM_023250688.1
A2 162147250 162147435
refGene_gn=CNTNAP2;refGene_tx=XM_023250688.1
A2 162153683 162153827
refGene_gn=CNTNAP2;refGene_tx=XM_023250688.1
A2 162157436 162157701
refGene_gn=CNTNAP2;refGene_tx=XM_023250688.1
A2 162302305 162302455
refGene_gn=CNTNAP2;refGene_tx=XM_023250688.1
A2 162383760 162383932
refGene_gn=CNTNAP2;refGene_tx=XM_023250688.1
A2 162451139 162451246
refGene_gn=CNTNAP2;refGene_tx=XM_023250688.1
A2 162507993 162508113
refGene_gn=CNTNAP2;refGene_tx=XM_023250688.1
A2 162557352 162557553
refGene_gn=CNTNAP2;refGene_tx=XM_023250688.1
A2 162785444 162785601
refGene_gn=CNTNAP2;refGene_tx=XM_023250688.1
A2 162857375 162857503
refGene_gn=CNTNAP2;refGene_tx=XM_023250688.1
A2 162965289 162965460
refGene_gn=CNTNAP2;refGene_tx=XM_023250688.1
A2 162989460 162989679
refGene_gn=CNTNAP2;refGene_tx=XM_023250688.1
A2 163011875 163012112
refGene_gn=CNTNAP2;refGene_tx=XM_023250688.1
A2 163064630 163064867
refGene_gn=CNTNAP2;refGene_tx=XM_023250688.1
A2 163073401 163073535
refGene_gn=CNTNAP2;refGene_tx=XM_023250688.1
A2 163099116 163099210
refGene_gn=CNTNAP2;refGene_tx=XM_023250688.1
A2 163186997 163187237
refGene_gn=CNTNAP2;refGene_tx=XM_023250688.1
A2 163211415 163211496
refGene_gn=CNTNAP2;refGene_tx=XM_023250688.1
A2 163215672 163215872
refGene_gn=CNTNAP2;refGene_tx=XM_023250688.1
A2 163338789 163339600
refGene_gn=LOC109497681;refGene_tx=XM_019825981.2;refGene_gn=LOC109497
681;refGene_tx=XM_023250689.1
A2 163340846 163340915
refGene_gn=LOC109497681;refGene_tx=XM_019825981.2;refGene_gn=LOC109497
681;refGene_tx=XM_023250689.1
A2 163342538 163342666
refGene_gn=LOC109497681;refGene_tx=XM_019825981.2;refGene_gn=LOC109497

681;refGene_tx=XM_023250689.1
A2 163401620 163401760
refGene_gn=CUL1;refGene_tx=XM_023250690.1;refGene_gn=CUL1;refGene_tx=XM_023250691.1;refGene_gn=CUL1;refGene_tx=XM_023250692.1
A2 163427459 163427634
refGene_gn=CUL1;refGene_tx=XM_023250690.1;refGene_gn=CUL1;refGene_tx=XM_023250691.1;refGene_gn=CUL1;refGene_tx=XM_023250692.1
A2 163429692 163429860
refGene_gn=CUL1;refGene_tx=XM_023250690.1;refGene_gn=CUL1;refGene_tx=XM_023250691.1;refGene_gn=CUL1;refGene_tx=XM_023250692.1
A2 163433721 163433772
refGene_gn=CUL1;refGene_tx=XM_023250690.1;refGene_gn=CUL1;refGene_tx=XM_023250691.1;refGene_gn=CUL1;refGene_tx=XM_023250692.1
A2 163433982 163434073
refGene_gn=CUL1;refGene_tx=XM_023250690.1;refGene_gn=CUL1;refGene_tx=XM_023250691.1;refGene_gn=CUL1;refGene_tx=XM_023250692.1
A2 163435263 163435427
refGene_gn=CUL1;refGene_tx=XM_023250690.1;refGene_gn=CUL1;refGene_tx=XM_023250691.1;refGene_gn=CUL1;refGene_tx=XM_023250692.1
A2 163441023 163441186
refGene_gn=CUL1;refGene_tx=XM_023250690.1;refGene_gn=CUL1;refGene_tx=XM_023250691.1;refGene_gn=CUL1;refGene_tx=XM_023250692.1
A2 163442054 163442185
refGene_gn=CUL1;refGene_tx=XM_023250690.1;refGene_gn=CUL1;refGene_tx=XM_023250691.1;refGene_gn=CUL1;refGene_tx=XM_023250692.1
A2 163456492 163456600
refGene_gn=CUL1;refGene_tx=XM_023250690.1;refGene_gn=CUL1;refGene_tx=XM_023250691.1;refGene_gn=CUL1;refGene_tx=XM_023250692.1
A2 163456684 163456791
refGene_gn=CUL1;refGene_tx=XM_023250690.1;refGene_gn=CUL1;refGene_tx=XM_023250691.1;refGene_gn=CUL1;refGene_tx=XM_023250692.1
A2 163459185 163459234
refGene_gn=CUL1;refGene_tx=XM_023250690.1;refGene_gn=CUL1;refGene_tx=XM_023250691.1;refGene_gn=CUL1;refGene_tx=XM_023250692.1
A2 163459603 163459735
refGene_gn=CUL1;refGene_tx=XM_023250690.1;refGene_gn=CUL1;refGene_tx=XM_023250691.1;refGene_gn=CUL1;refGene_tx=XM_023250692.1
A2 163460881 163460999
refGene_gn=CUL1;refGene_tx=XM_023250690.1;refGene_gn=CUL1;refGene_tx=XM_023250691.1;refGene_gn=CUL1;refGene_tx=XM_023250692.1
A2 163462129 163462206
refGene_gn=CUL1;refGene_tx=XM_023250690.1;refGene_gn=CUL1;refGene_tx=XM_023250691.1;refGene_gn=CUL1;refGene_tx=XM_023250692.1
A2 163462629 163462761
refGene_gn=CUL1;refGene_tx=XM_023250690.1;refGene_gn=CUL1;refGene_tx=XM_023250691.1;refGene_gn=CUL1;refGene_tx=XM_023250692.1
A2 163463627 163463732
refGene_gn=CUL1;refGene_tx=XM_023250690.1;refGene_gn=CUL1;refGene_tx=XM_023250691.1;refGene_gn=CUL1;refGene_tx=XM_023250692.1
A2 163467380 163467428

refGene_gn=CUL1;refGene_tx=XM_023250690.1;refGene_gn=CUL1;refGene_tx=XM_023250691.1;refGene_gn=CUL1;refGene_tx=XM_023250692.1
A2 163467507 163467590
refGene_gn=CUL1;refGene_tx=XM_023250690.1;refGene_gn=CUL1;refGene_tx=XM_023250691.1;refGene_gn=CUL1;refGene_tx=XM_023250692.1
A2 163468083 163468189
refGene_gn=CUL1;refGene_tx=XM_023250690.1;refGene_gn=CUL1;refGene_tx=XM_023250691.1;refGene_gn=CUL1;refGene_tx=XM_023250692.1
A2 163468767 163468881
refGene_gn=CUL1;refGene_tx=XM_023250690.1;refGene_gn=CUL1;refGene_tx=XM_023250691.1;refGene_gn=CUL1;refGene_tx=XM_023250692.1
A2 163469995 163470076
refGene_gn=CUL1;refGene_tx=XM_023250690.1;refGene_gn=CUL1;refGene_tx=XM_023250691.1;refGene_gn=CUL1;refGene_tx=XM_023250692.1
A2 163471283 163471389
refGene_gn=LOC109497680;refGene_tx=XM_019825978.2;refGene_gn=LOC109497680;refGene_tx=XM_019825979.2;refGene_gn=LOC109497680;refGene_tx=XM_019825980.2
A2 163472270 163472423
refGene_gn=LOC109497680;refGene_tx=XM_019825978.2;refGene_gn=LOC109497680;refGene_tx=XM_019825979.2;refGene_gn=LOC109497680;refGene_tx=XM_019825980.2
A2 163475380 163475441
refGene_gn=EZH2;refGene_tx=XM_023250693.1
A2 163475928 163476023
refGene_gn=LOC109497680;refGene_tx=XM_019825978.2;refGene_gn=LOC109497680;refGene_tx=XM_019825980.2
A2 163476664 163476697
refGene_gn=LOC109497680;refGene_tx=XM_019825980.2
A2 163477870 163477955
refGene_gn=EZH2;refGene_tx=XM_023250693.1
A2 163478113 163478194
refGene_gn=EZH2;refGene_tx=XM_023250693.1
A2 163478366 163478776
refGene_gn=LOC109497680;refGene_tx=XM_019825978.2;refGene_gn=LOC109497680;refGene_tx=XM_019825979.2
A2 163478842 163478924
refGene_gn=EZH2;refGene_tx=XM_023250693.1
A2 163479566 163479662
refGene_gn=EZH2;refGene_tx=XM_023250693.1
A2 163480846 163481025
refGene_gn=EZH2;refGene_tx=XM_023250693.1
A2 163481984 163482110
refGene_gn=EZH2;refGene_tx=XM_023250693.1
A2 163482551 163482592
refGene_gn=EZH2;refGene_tx=XM_023250693.1
A2 163484229 163484324
refGene_gn=EZH2;refGene_tx=XM_023250693.1
A2 163484704 163484874
refGene_gn=EZH2;refGene_tx=XM_023250693.1

A2 163485682 163485710
refGene_gn=EZH2;refGene_tx=XM_023250693.1
A2 163485711 163485921
refGene_gn=EZH2;refGene_tx=XM_023250693.1
A2 163487279 163487371
refGene_gn=EZH2;refGene_tx=XM_023250693.1
A2 163495122 163495286
refGene_gn=EZH2;refGene_tx=XM_023250693.1
A2 163496209 163496312
refGene_gn=EZH2;refGene_tx=XM_023250693.1
A2 163497168 163497309
refGene_gn=EZH2;refGene_tx=XM_023250693.1
A2 163498128 163498249
refGene_gn=EZH2;refGene_tx=XM_023250693.1
A2 163501401 163501518
refGene_gn=EZH2;refGene_tx=XM_023250693.1
A2 163504900 163504948
refGene_gn=EZH2;refGene_tx=XM_023250693.1
A2 163516715 163516817
refGene_gn=EZH2;refGene_tx=XM_023250693.1
A2 163517375 163517492
refGene_gn=EZH2;refGene_tx=XM_023250693.1
A2 163645823 163646239
refGene_gn=PDIA4;refGene_tx=XM_023250695.1
A2 163646994 163647228
refGene_gn=PDIA4;refGene_tx=XM_023250695.1
A2 163647694 163647851
refGene_gn=PDIA4;refGene_tx=XM_023250695.1
A2 163651157 163651309
refGene_gn=PDIA4;refGene_tx=XM_023250695.1
A2 163653330 163653492
refGene_gn=PDIA4;refGene_tx=XM_023250695.1
A2 163653622 163653828
refGene_gn=PDIA4;refGene_tx=XM_023250695.1
A2 163656065 163656204
refGene_gn=PDIA4;refGene_tx=XM_023250695.1
A2 163658722 163658928
refGene_gn=PDIA4;refGene_tx=XM_023250695.1
A2 163660930 163661111
refGene_gn=PDIA4;refGene_tx=XM_023250695.1
A2 163667796 163667884
refGene_gn=PDIA4;refGene_tx=XM_023250695.1
A2 163688274 163690340
refGene_gn=ZNF786;refGene_tx=XM_011280661.3;refGene_gn=ZNF786;refGene_
tx=XM_023250700.1
A2 163691768 163691921
refGene_gn=ZNF786;refGene_tx=XM_011280661.3;refGene_gn=ZNF786;refGene_
tx=XM_023250700.1
A2 163695470 163695597
refGene_gn=ZNF786;refGene_tx=XM_011280661.3;refGene_gn=ZNF786;refGene_

tx=XM_023250700.1
A2	163703468	163703486
refGene_gn=ZNF786;refGene_tx=XM_023250700.1
A2	163715565	163717520
refGene_gn=ZNF425;refGene_tx=XM_003983195.4;refGene_gn=ZNF425;refGene_
tx=XM_023250702.1
A2	163720183	163720339
refGene_gn=ZNF425;refGene_tx=XM_003983195.4;refGene_gn=ZNF425;refGene_
tx=XM_023250702.1
A2	163723280	163723407
refGene_gn=ZNF425;refGene_tx=XM_003983195.4
A2	163724774	163724792
refGene_gn=ZNF425;refGene_tx=XM_003983195.4
A2	163747869	163747893
refGene_gn=ZNF398;refGene_tx=XM_023250704.1
A2	163752380	163753085
refGene_gn=ZNF398;refGene_tx=XM_006929467.4;refGene_gn=ZNF398;refGene_
tx=XM_023250704.1
A2	163761109	163761236
refGene_gn=ZNF398;refGene_tx=XM_006929467.4;refGene_gn=ZNF398;refGene_
tx=XM_023250703.1;refGene_gn=ZNF398;refGene_tx=XM_023250704.1
A2	163761784	163761898
refGene_gn=ZNF398;refGene_tx=XM_006929467.4;refGene_gn=ZNF398;refGene_
tx=XM_023250703.1;refGene_gn=ZNF398;refGene_tx=XM_023250704.1
A2	163771908	163772022
refGene_gn=ZNF398;refGene_tx=XM_006929467.4;refGene_gn=ZNF398;refGene_
tx=XM_023250703.1;refGene_gn=ZNF398;refGene_tx=XM_023250704.1
A2	163773095	163774249
refGene_gn=ZNF398;refGene_tx=XM_006929467.4;refGene_gn=ZNF398;refGene_
tx=XM_023250703.1;refGene_gn=ZNF398;refGene_tx=XM_023250704.1
A2	163784486	163784651
refGene_gn=ZNF282;refGene_tx=XM_023250699.1
A2	163786940	163787360
refGene_gn=ZNF282;refGene_tx=XM_023250699.1
A2	163796236	163796363
refGene_gn=ZNF282;refGene_tx=XM_023250699.1
A2	163796902	163797022
refGene_gn=ZNF282;refGene_tx=XM_023250699.1
A2	163799199	163799319
refGene_gn=ZNF282;refGene_tx=XM_023250699.1
A2	163800807	163800921
refGene_gn=ZNF282;refGene_tx=XM_023250699.1
A2	163802373	163802487
refGene_gn=ZNF282;refGene_tx=XM_023250699.1
A2	163808616	163809383
refGene_gn=ZNF282;refGene_tx=XM_023250699.1
A2	163817794	163817818
refGene_gn=ZNF212;refGene_tx=XM_003983196.5
A2	163817858	163818227
refGene_gn=ZNF212;refGene_tx=XM_019825985.2;refGene_gn=ZNF212;refGene_

tx=XM_019825986.2
A2	163828420	163828810
refGene_gn=ZNF212;refGene_tx=XM_003983196.5;refGene_gn=ZNF212;refGene_tx=XM_019825985.2;refGene_gn=ZNF212;refGene_tx=XM_019825986.2
A2	163828942	163829069
refGene_gn=ZNF212;refGene_tx=XM_003983196.5;refGene_gn=ZNF212;refGene_tx=XM_019825985.2;refGene_gn=ZNF212;refGene_tx=XM_019825986.2
A2	163830654	163830744
refGene_gn=ZNF212;refGene_tx=XM_003983196.5;refGene_gn=ZNF212;refGene_tx=XM_019825985.2;refGene_gn=ZNF212;refGene_tx=XM_019825986.2
A2	163831507	163832364
refGene_gn=ZNF212;refGene_tx=XM_003983196.5;refGene_gn=ZNF212;refGene_tx=XM_019825985.2;refGene_gn=ZNF212;refGene_tx=XM_019825986.2
A2	163837629	163837635
refGene_gn=ZNF783;refGene_tx=XM_006929466.4
A2	163840468	163840492
refGene_gn=ZNF783;refGene_tx=XM_023250705.1
A2	163843795	163844197
refGene_gn=ZNF783;refGene_tx=XM_006929466.4;refGene_gn=ZNF783;refGene_tx=XM_023250705.1
A2	163844285	163844412
refGene_gn=ZNF783;refGene_tx=XM_006929466.4;refGene_gn=ZNF783;refGene_tx=XM_023250705.1
A2	163844576	163844705
refGene_gn=ZNF783;refGene_tx=XM_006929466.4;refGene_gn=ZNF783;refGene_tx=XM_023250705.1
A2	163851924	163852053
refGene_gn=ZNF783;refGene_tx=XM_006929466.4;refGene_gn=ZNF783;refGene_tx=XM_023250705.1
A2	163853079	163853172
refGene_gn=ZNF783;refGene_tx=XM_006929466.4;refGene_gn=ZNF783;refGene_tx=XM_023250705.1
A2	163854974	163855813
refGene_gn=ZNF783;refGene_tx=XM_006929466.4;refGene_gn=ZNF783;refGene_tx=XM_023250705.1
A2	163905037	163906194
refGene_gn=ZNF777;refGene_tx=XM_023250696.1;refGene_gn=ZNF777;refGene_tx=XM_023250697.1;refGene_gn=ZNF777;refGene_tx=XM_023250698.1
A2	163909613	163909865
refGene_gn=ZNF777;refGene_tx=XM_023250696.1;refGene_gn=ZNF777;refGene_tx=XM_023250697.1;refGene_gn=ZNF777;refGene_tx=XM_023250698.1
A2	163923868	163923982
refGene_gn=ZNF777;refGene_tx=XM_023250696.1;refGene_gn=ZNF777;refGene_tx=XM_023250697.1;refGene_gn=ZNF777;refGene_tx=XM_023250698.1
A2	163926040	163926167
refGene_gn=ZNF777;refGene_tx=XM_023250696.1;refGene_gn=ZNF777;refGene_tx=XM_023250697.1;refGene_gn=ZNF777;refGene_tx=XM_023250698.1
A2	163927414	163928266
refGene_gn=ZNF777;refGene_tx=XM_023250696.1;refGene_gn=ZNF777;refGene_tx=XM_023250697.1;refGene_gn=ZNF777;refGene_tx=XM_023250698.1

A2 163932271 163932388
refGene_gn=ZNF777;refGene_tx=XM_023250696.1
A2 163947203 163948261
refGene_gn=ZNF746;refGene_tx=XM_023250706.1;refGene_gn=ZNF746;refGene_tx=XM_023250707.1;refGene_gn=ZNF746;refGene_tx=XM_023250708.1;refGene_gn=ZNF746;refGene_tx=XM_023250709.1
A2 163949200 163949326
refGene_gn=ZNF746;refGene_tx=XM_023250706.1;refGene_gn=ZNF746;refGene_tx=XM_023250707.1;refGene_gn=ZNF746;refGene_tx=XM_023250708.1;refGene_gn=ZNF746;refGene_tx=XM_023250709.1
A2 163949814 163950006
refGene_gn=ZNF746;refGene_tx=XM_023250706.1;refGene_gn=ZNF746;refGene_tx=XM_023250707.1;refGene_gn=ZNF746;refGene_tx=XM_023250708.1;refGene_gn=ZNF746;refGene_tx=XM_023250709.1
A2 163965662 163965776
refGene_gn=ZNF746;refGene_tx=XM_023250706.1;refGene_gn=ZNF746;refGene_tx=XM_023250707.1;refGene_gn=ZNF746;refGene_tx=XM_023250708.1;refGene_gn=ZNF746;refGene_tx=XM_023250709.1
A2 163966777 163966904
refGene_gn=ZNF746;refGene_tx=XM_023250706.1;refGene_gn=ZNF746;refGene_tx=XM_023250707.1;refGene_gn=ZNF746;refGene_tx=XM_023250708.1;refGene_gn=ZNF746;refGene_tx=XM_023250709.1
A2 163966991 163967291
refGene_gn=ZNF746;refGene_tx=XM_023250706.1;refGene_gn=ZNF746;refGene_tx=XM_023250707.1;refGene_gn=ZNF746;refGene_tx=XM_023250708.1;refGene_gn=ZNF746;refGene_tx=XM_023250709.1
A2 163970598 163970622
refGene_gn=ZNF746;refGene_tx=XM_023250706.1;refGene_gn=ZNF746;refGene_tx=XM_023250707.1;refGene_gn=ZNF746;refGene_tx=XM_023250708.1;refGene_gn=ZNF746;refGene_tx=XM_023250709.1
A2 164031207 164031219
refGene_gn=KRBA1;refGene_tx=XM_011280654.3;refGene_gn=KRBA1;refGene_tx=XM_011280655.3;refGene_gn=KRBA1;refGene_tx=XM_011280656.3;refGene_gn=KRBA1;refGene_tx=XM_011280657.3;refGene_gn=KRBA1;refGene_tx=XM_011280658.3;refGene_gn=KRBA1;refGene_tx=XM_019825987.2;refGene_gn=KRBA1;refGene_tx=XM_019825988.2
A2 164037944 164038071
refGene_gn=KRBA1;refGene_tx=XM_011280654.3;refGene_gn=KRBA1;refGene_tx=XM_011280655.3;refGene_gn=KRBA1;refGene_tx=XM_011280656.3;refGene_gn=KRBA1;refGene_tx=XM_011280657.3;refGene_gn=KRBA1;refGene_tx=XM_011280658.3;refGene_gn=KRBA1;refGene_tx=XM_019825987.2;refGene_gn=KRBA1;refGene_tx=XM_019825988.2
A2 164038414 164038534
refGene_gn=KRBA1;refGene_tx=XM_011280654.3;refGene_gn=KRBA1;refGene_tx=XM_011280655.3;refGene_gn=KRBA1;refGene_tx=XM_011280656.3;refGene_gn=KRBA1;refGene_tx=XM_011280657.3;refGene_gn=KRBA1;refGene_tx=XM_011280658.3;refGene_gn=KRBA1;refGene_tx=XM_019825987.2;refGene_gn=KRBA1;refGene_tx=XM_019825988.2
A2 164039228 164039369
refGene_gn=KRBA1;refGene_tx=XM_011280654.3;refGene_gn=KRBA1;refGene_tx

=XM_011280655.3;refGene_gn=KRBA1;refGene_tx=XM_011280656.3;refGene_gn=KRBA1;refGene_tx=XM_011280657.3;refGene_gn=KRBA1;refGene_tx=XM_011280658.3;refGene_gn=KRBA1;refGene_tx=XM_019825987.2;refGene_gn=KRBA1;refGene_tx=XM_019825988.2
A2 164039994 164040174
refGene_gn=KRBA1;refGene_tx=XM_011280654.3;refGene_gn=KRBA1;refGene_tx=XM_011280655.3;refGene_gn=KRBA1;refGene_tx=XM_011280656.3;refGene_gn=KRBA1;refGene_tx=XM_011280657.3;refGene_gn=KRBA1;refGene_tx=XM_011280658.3;refGene_gn=KRBA1;refGene_tx=XM_019825987.2;refGene_gn=KRBA1;refGene_tx=XM_019825988.2
A2 164041021 164041153
refGene_gn=KRBA1;refGene_tx=XM_011280654.3;refGene_gn=KRBA1;refGene_tx=XM_011280655.3;refGene_gn=KRBA1;refGene_tx=XM_011280656.3;refGene_gn=KRBA1;refGene_tx=XM_011280657.3;refGene_gn=KRBA1;refGene_tx=XM_011280658.3;refGene_gn=KRBA1;refGene_tx=XM_019825987.2;refGene_gn=KRBA1;refGene_tx=XM_019825988.2
A2 164041359 164041542
refGene_gn=KRBA1;refGene_tx=XM_011280654.3;refGene_gn=KRBA1;refGene_tx=XM_011280655.3;refGene_gn=KRBA1;refGene_tx=XM_011280656.3;refGene_gn=KRBA1;refGene_tx=XM_011280657.3;refGene_gn=KRBA1;refGene_tx=XM_019825987.2;refGene_gn=KRBA1;refGene_tx=XM_019825988.2
A2 164042420 164042561
refGene_gn=KRBA1;refGene_tx=XM_011280654.3;refGene_gn=KRBA1;refGene_tx=XM_011280655.3;refGene_gn=KRBA1;refGene_tx=XM_011280656.3;refGene_gn=KRBA1;refGene_tx=XM_011280657.3;refGene_gn=KRBA1;refGene_tx=XM_011280658.3;refGene_gn=KRBA1;refGene_tx=XM_019825987.2;refGene_gn=KRBA1;refGene_tx=XM_019825988.2
A2 164043231 164043450
refGene_gn=KRBA1;refGene_tx=XM_011280654.3;refGene_gn=KRBA1;refGene_tx=XM_011280655.3;refGene_gn=KRBA1;refGene_tx=XM_011280656.3;refGene_gn=KRBA1;refGene_tx=XM_011280657.3;refGene_gn=KRBA1;refGene_tx=XM_011280658.3;refGene_gn=KRBA1;refGene_tx=XM_019825987.2;refGene_gn=KRBA1;refGene_tx=XM_019825988.2
A2 164043953 164044127
refGene_gn=KRBA1;refGene_tx=XM_011280654.3;refGene_gn=KRBA1;refGene_tx=XM_011280655.3;refGene_gn=KRBA1;refGene_tx=XM_011280656.3;refGene_gn=KRBA1;refGene_tx=XM_011280657.3;refGene_gn=KRBA1;refGene_tx=XM_011280658.3;refGene_gn=KRBA1;refGene_tx=XM_019825987.2;refGene_gn=KRBA1;refGene_tx=XM_019825988.2
A2 164044531 164044705
refGene_gn=KRBA1;refGene_tx=XM_011280654.3;refGene_gn=KRBA1;refGene_tx=XM_011280655.3;refGene_gn=KRBA1;refGene_tx=XM_011280656.3;refGene_gn=KRBA1;refGene_tx=XM_011280657.3;refGene_gn=KRBA1;refGene_tx=XM_011280658.3;refGene_gn=KRBA1;refGene_tx=XM_019825987.2;refGene_gn=KRBA1;refGene_tx=XM_019825988.2
A2 164046841 164046994
refGene_gn=KRBA1;refGene_tx=XM_011280654.3;refGene_gn=KRBA1;refGene_tx=XM_011280655.3;refGene_gn=KRBA1;refGene_tx=XM_011280656.3;refGene_gn=KRBA1;refGene_tx=XM_011280657.3;refGene_gn=KRBA1;refGene_tx=XM_011280658.3;refGene_gn=KRBA1;refGene_tx=XM_019825987.2;refGene_gn=KRBA1;refGe

ne_tx=XM_019825988.2
A2 164047496 164047704
refGene_gn=KRBA1;refGene_tx=XM_011280654.3;refGene_gn=KRBA1;refGene_tx=XM_011280655.3;refGene_gn=KRBA1;refGene_tx=XM_011280656.3;refGene_gn=KRBA1;refGene_tx=XM_011280657.3;refGene_gn=KRBA1;refGene_tx=XM_011280658.3;refGene_gn=KRBA1;refGene_tx=XM_019825987.2;refGene_gn=KRBA1;refGene_tx=XM_019825988.2
A2 164048490 164048703
refGene_gn=KRBA1;refGene_tx=XM_011280654.3;refGene_gn=KRBA1;refGene_tx=XM_011280655.3;refGene_gn=KRBA1;refGene_tx=XM_011280656.3;refGene_gn=KRBA1;refGene_tx=XM_011280657.3;refGene_gn=KRBA1;refGene_tx=XM_011280658.3;refGene_gn=KRBA1;refGene_tx=XM_019825987.2;refGene_gn=KRBA1;refGene_tx=XM_019825988.2
A2 164048794 164048926
refGene_gn=KRBA1;refGene_tx=XM_011280654.3;refGene_gn=KRBA1;refGene_tx=XM_011280655.3;refGene_gn=KRBA1;refGene_tx=XM_011280656.3;refGene_gn=KRBA1;refGene_tx=XM_011280657.3;refGene_gn=KRBA1;refGene_tx=XM_011280658.3;refGene_gn=KRBA1;refGene_tx=XM_019825987.2;refGene_gn=KRBA1;refGene_tx=XM_019825988.2
A2 164049933 164050041
refGene_gn=KRBA1;refGene_tx=XM_011280654.3;refGene_gn=KRBA1;refGene_tx=XM_011280655.3;refGene_gn=KRBA1;refGene_tx=XM_011280656.3;refGene_gn=KRBA1;refGene_tx=XM_011280657.3;refGene_gn=KRBA1;refGene_tx=XM_011280658.3;refGene_gn=KRBA1;refGene_tx=XM_019825987.2;refGene_gn=KRBA1;refGene_tx=XM_019825988.2
A2 164050582 164050744
refGene_gn=KRBA1;refGene_tx=XM_011280654.3;refGene_gn=KRBA1;refGene_tx=XM_011280655.3;refGene_gn=KRBA1;refGene_tx=XM_011280656.3;refGene_gn=KRBA1;refGene_tx=XM_011280657.3;refGene_gn=KRBA1;refGene_tx=XM_011280658.3;refGene_gn=KRBA1;refGene_tx=XM_019825987.2;refGene_gn=KRBA1;refGene_tx=XM_019825988.2
A2 164052459 164053385
refGene_gn=KRBA1;refGene_tx=XM_011280654.3;refGene_gn=KRBA1;refGene_tx=XM_011280655.3;refGene_gn=KRBA1;refGene_tx=XM_011280656.3;refGene_gn=KRBA1;refGene_tx=XM_011280657.3;refGene_gn=KRBA1;refGene_tx=XM_011280658.3;refGene_gn=KRBA1;refGene_tx=XM_019825987.2;refGene_gn=KRBA1;refGene_tx=XM_019825988.2
A2 164077156 164078658
refGene_gn=ZNF467;refGene_tx=XM_023250710.1;refGene_gn=ZNF467;refGene_tx=XM_023250711.1;refGene_gn=ZNF467;refGene_tx=XM_023250712.1;refGene_gn=ZNF467;refGene_tx=XM_023250713.1;refGene_gn=ZNF467;refGene_tx=XM_023250714.1
A2 164081496 164081610
refGene_gn=ZNF467;refGene_tx=XM_023250710.1;refGene_gn=ZNF467;refGene_tx=XM_023250711.1;refGene_gn=ZNF467;refGene_tx=XM_023250712.1;refGene_gn=ZNF467;refGene_tx=XM_023250713.1;refGene_gn=ZNF467;refGene_tx=XM_023250714.1
A2 164082418 164082535
refGene_gn=ZNF467;refGene_tx=XM_023250710.1;refGene_gn=ZNF467;refGene_tx=XM_023250711.1;refGene_gn=ZNF467;refGene_tx=XM_023250712.1;refGene_

gn=ZNF467;refGene_tx=XM_023250713.1;refGene_gn=ZNF467;refGene_tx=XM_023250714.1
A2 164082997 164083074
refGene_gn=ZNF467;refGene_tx=XM_023250710.1;refGene_gn=ZNF467;refGene_tx=XM_023250711.1;refGene_gn=ZNF467;refGene_tx=XM_023250712.1
A2 164083386 164083466
refGene_gn=ZNF467;refGene_tx=XM_023250710.1
A2 164086562 164086679
refGene_gn=SSPO;refGene_tx=XM_023242329.1
A2 164088335 164088640
refGene_gn=SSPO;refGene_tx=XM_023242329.1
A2 164089057 164089140
refGene_gn=SSPO;refGene_tx=XM_023242329.1
A2 164089254 164089434
refGene_gn=SSPO;refGene_tx=XM_023242329.1
A2 164089688 164089933
refGene_gn=SSPO;refGene_tx=XM_023242329.1
A2 164090032 164090132
refGene_gn=SSPO;refGene_tx=XM_023242329.1
A2 164091083 164091242
refGene_gn=SSPO;refGene_tx=XM_023242329.1
A2 164091328 164091408
refGene_gn=SSPO;refGene_tx=XM_023242329.1
A2 164091887 164092040
refGene_gn=SSPO;refGene_tx=XM_023242329.1
A2 164092269 164092441
refGene_gn=SSPO;refGene_tx=XM_023242329.1
A2 164092643 164092884
refGene_gn=SSPO;refGene_tx=XM_023242329.1
A2 164093186 164093331
refGene_gn=SSPO;refGene_tx=XM_023242329.1
A2 164094242 164094360
refGene_gn=SSPO;refGene_tx=XM_023242329.1
A2 164094445 164094621
refGene_gn=SSPO;refGene_tx=XM_023242329.1
A2 164094703 164094931
refGene_gn=SSPO;refGene_tx=XM_023242329.1
A2 164095134 164095300
refGene_gn=SSPO;refGene_tx=XM_023242329.1
A2 164095529 164095764
refGene_gn=SSPO;refGene_tx=XM_023242329.1
A2 164096515 164096705
refGene_gn=SSPO;refGene_tx=XM_023242329.1
A2 164096846 164096941
refGene_gn=SSPO;refGene_tx=XM_023242329.1
A2 164097130 164097387
refGene_gn=SSPO;refGene_tx=XM_023242329.1
A2 164097704 164097839
refGene_gn=SSPO;refGene_tx=XM_023242329.1
A2 164098468 164098665

refGene_gn=SSPO;refGene_tx=XM_023242329.1
A2	164098795	164098923
refGene_gn=SSPO;refGene_tx=XM_023242329.1
A2	164099010	164099120
refGene_gn=SSPO;refGene_tx=XM_023242329.1
A2	164099448	164099598
refGene_gn=SSPO;refGene_tx=XM_023242329.1
A2	164099734	164099828
refGene_gn=SSPO;refGene_tx=XM_023242329.1
A2	164099922	164100092
refGene_gn=SSPO;refGene_tx=XM_023242329.1
A2	164100304	164100538
refGene_gn=SSPO;refGene_tx=XM_023242329.1
A2	164100747	164100969
refGene_gn=SSPO;refGene_tx=XM_023242329.1
A2	164101397	164101634
refGene_gn=SSPO;refGene_tx=XM_023242329.1
A2	164102225	164102354
refGene_gn=SSPO;refGene_tx=XM_023242329.1
A2	164102452	164102544
refGene_gn=SSPO;refGene_tx=XM_023242329.1
A2	164102641	164102873
refGene_gn=SSPO;refGene_tx=XM_023242329.1
A2	164103010	164103163
refGene_gn=SSPO;refGene_tx=XM_023242329.1
A2	164103257	164103398
refGene_gn=SSPO;refGene_tx=XM_023242329.1
A2	164103500	164103683
refGene_gn=SSPO;refGene_tx=XM_023242329.1
A2	164103789	164103940
refGene_gn=SSPO;refGene_tx=XM_023242329.1
A2	164104563	164104717
refGene_gn=SSPO;refGene_tx=XM_023242329.1
A2	164104858	164104916
refGene_gn=SSPO;refGene_tx=XM_023242329.1
A2	164105717	164105822
refGene_gn=SSPO;refGene_tx=XM_023242329.1
A2	164106120	164106324
refGene_gn=SSPO;refGene_tx=XM_023242329.1
A2	164106636	164106746
refGene_gn=SSPO;refGene_tx=XM_023242329.1
A2	164107634	164107788
refGene_gn=SSPO;refGene_tx=XM_023242329.1
A2	164107868	164108022
refGene_gn=SSPO;refGene_tx=XM_023242329.1
A2	164108511	164108584
refGene_gn=SSPO;refGene_tx=XM_023242329.1
A2	164109380	164109438
refGene_gn=SSPO;refGene_tx=XM_023242329.1
A2	164110805	164110982

refGene_gn=SSPO;refGene_tx=XM_023242329.1
A2	164111281	164111425
refGene_gn=SSPO;refGene_tx=XM_023242329.1
A2	164112497	164112668
refGene_gn=SSPO;refGene_tx=XM_023242329.1
A2	164112752	164112917
refGene_gn=SSPO;refGene_tx=XM_023242329.1
A2	164113634	164113814
refGene_gn=SSPO;refGene_tx=XM_023242329.1
A2	164113932	164114055
refGene_gn=SSPO;refGene_tx=XM_023242329.1
A2	164114190	164114320
refGene_gn=SSPO;refGene_tx=XM_023242329.1
A2	164114424	164114550
refGene_gn=SSPO;refGene_tx=XM_023242329.1
A2	164114726	164114818
refGene_gn=SSPO;refGene_tx=XM_023242329.1
A2	164115691	164115865
refGene_gn=SSPO;refGene_tx=XM_023242329.1
A2	164116135	164116300
refGene_gn=SSPO;refGene_tx=XM_023242329.1
A2	164116936	164117189
refGene_gn=SSPO;refGene_tx=XM_023242329.1
A2	164118139	164118186
refGene_gn=SSPO;refGene_tx=XM_023242329.1
A2	164118359	164118530
refGene_gn=SSPO;refGene_tx=XM_023242329.1
A2	164118835	164118967
refGene_gn=SSPO;refGene_tx=XM_023242329.1
A2	164119194	164119340
refGene_gn=SSPO;refGene_tx=XM_023242329.1
A2	164120684	164120779
refGene_gn=SSPO;refGene_tx=XM_023242329.1
A2	164121765	164121818
refGene_gn=SSPO;refGene_tx=XM_023242329.1
A2	164121978	164122185
refGene_gn=SSPO;refGene_tx=XM_023242329.1
A2	164122306	164122477
refGene_gn=SSPO;refGene_tx=XM_023242329.1
A2	164122582	164122662
refGene_gn=SSPO;refGene_tx=XM_023242329.1
A2	164123856	164124026
refGene_gn=SSPO;refGene_tx=XM_023242329.1
A2	164124566	164124613
refGene_gn=SSPO;refGene_tx=XM_023242329.1
A2	164124729	164124921
refGene_gn=SSPO;refGene_tx=XM_023242329.1
A2	164125011	164125182
refGene_gn=SSPO;refGene_tx=XM_023242329.1
A2	164125443	164125644

refGene_gn=SSPO;refGene_tx=XM_023242329.1
A2	164125951	164126048
refGene_gn=SSPO;refGene_tx=XM_023242329.1
A2	164126156	164126356
refGene_gn=SSPO;refGene_tx=XM_023242329.1
A2	164126701	164126868
refGene_gn=SSPO;refGene_tx=XM_023242329.1
A2	164127984	164128145
refGene_gn=SSPO;refGene_tx=XM_023242329.1
A2	164128335	164128559
refGene_gn=SSPO;refGene_tx=XM_023242329.1
A2	164129164	164129206
refGene_gn=SSPO;refGene_tx=XM_023242329.1
A2	164129374	164129530
refGene_gn=SSPO;refGene_tx=XM_023242329.1
A2	164130024	164130234
refGene_gn=SSPO;refGene_tx=XM_023242329.1
A2	164130396	164130564
refGene_gn=SSPO;refGene_tx=XM_023242329.1
A2	164131292	164131424
refGene_gn=SSPO;refGene_tx=XM_023242329.1
A2	164131525	164131772
refGene_gn=SSPO;refGene_tx=XM_023242329.1
A2	164132053	164132206
refGene_gn=SSPO;refGene_tx=XM_023242329.1
A2	164132285	164132332
refGene_gn=SSPO;refGene_tx=XM_023242329.1
A2	164132490	164132835
refGene_gn=SSPO;refGene_tx=XM_023242329.1
A2	164133178	164133352
refGene_gn=SSPO;refGene_tx=XM_023242329.1
A2	164133886	164134017
refGene_gn=SSPO;refGene_tx=XM_023242329.1
A2	164134279	164134404
refGene_gn=SSPO;refGene_tx=XM_023242329.1
A2	164134624	164134671
refGene_gn=SSPO;refGene_tx=XM_023242329.1
A2	164134925	164135204
refGene_gn=SSPO;refGene_tx=XM_023242329.1
A2	164135569	164135728
refGene_gn=SSPO;refGene_tx=XM_023242329.1
A2	164135897	164136014
refGene_gn=SSPO;refGene_tx=XM_023242329.1
A2	164136475	164136614
refGene_gn=SSPO;refGene_tx=XM_023242329.1
A2	164136751	164136954
refGene_gn=SSPO;refGene_tx=XM_023242329.1
A2	164137101	164137229
refGene_gn=SSPO;refGene_tx=XM_023242329.1
A2	164137327	164137470

refGene_gn=SSPO;refGene_tx=XM_023242329.1
A2 164137604 164137651
refGene_gn=SSPO;refGene_tx=XM_023242329.1
A2 164138417 164138635
refGene_gn=SSPO;refGene_tx=XM_023242329.1
A2 164138937 164139033
refGene_gn=SSPO;refGene_tx=XM_023242329.1
A2 164139164 164139225
refGene_gn=SSPO;refGene_tx=XM_023242329.1
A2 164139512 164139643
refGene_gn=SSPO;refGene_tx=XM_023242329.1
A2 164141188 164141339
refGene_gn=SSPO;refGene_tx=XM_023242329.1
A2 164141937 164141960
refGene_gn=SSPO;refGene_tx=XM_023242329.1
A2 164148220 164148244
refGene_gn=ZNF862;refGene_tx=XM_003983202.5
A2 164157755 164157867
refGene_gn=ZNF862;refGene_tx=XM_003983202.5;refGene_gn=ZNF862;refGene_tx=XM_006929477.4;refGene_gn=ZNF862;refGene_tx=XM_006929478.4
A2 164159595 164159700
refGene_gn=ZNF862;refGene_tx=XM_003983202.5;refGene_gn=ZNF862;refGene_tx=XM_006929477.4;refGene_gn=ZNF862;refGene_tx=XM_006929478.4
A2 164162001 164162699
refGene_gn=ZNF862;refGene_tx=XM_003983202.5;refGene_gn=ZNF862;refGene_tx=XM_006929477.4;refGene_gn=ZNF862;refGene_tx=XM_006929478.4
A2 164164196 164164374
refGene_gn=ZNF862;refGene_tx=XM_003983202.5;refGene_gn=ZNF862;refGene_tx=XM_006929477.4;refGene_gn=ZNF862;refGene_tx=XM_006929478.4
A2 164177131 164177236
refGene_gn=ZNF862;refGene_tx=XM_003983202.5;refGene_gn=ZNF862;refGene_tx=XM_006929477.4;refGene_gn=ZNF862;refGene_tx=XM_006929478.4
A2 164178548 164180663
refGene_gn=ZNF862;refGene_tx=XM_003983202.5;refGene_gn=ZNF862;refGene_tx=XM_006929477.4;refGene_gn=ZNF862;refGene_tx=XM_006929478.4
A2 164182200 164182376
refGene_gn=ZNF862;refGene_tx=XM_003983202.5;refGene_gn=ZNF862;refGene_tx=XM_006929477.4;refGene_gn=ZNF862;refGene_tx=XM_006929478.4
A2 164185497 164185802
refGene_gn=ATP6V0E2;refGene_tx=XM_023250715.1
A2 164187173 164187221
refGene_gn=ATP6V0E2;refGene_tx=XM_023250715.1
A2 164189755 164189849
refGene_gn=ATP6V0E2;refGene_tx=XM_023250715.1
A2 164202587 164205818
refGene_gn=ZBED6CL;refGene_tx=XM_006929480.4;refGene_gn=ZBED6CL;refGene_tx=XM_023250716.1
A2 164209781 164210561
refGene_gn=LRRC61;refGene_tx=XM_003983203.5
A2 164211085 164211439

refGene_gn=RARRES2;refGene_tx=XM_019825990.2;refGene_gn=RARRES2;refGene_tx=XM_019825991.2;refGene_gn=RARRES2;refGene_tx=XM_019825992.2;refGene_gn=RARRES2;refGene_tx=XM_019825993.2
A2 164211629 164211746
refGene_gn=RARRES2;refGene_tx=XM_006929482.4
A2 164211970 164212066
refGene_gn=RARRES2;refGene_tx=XM_006929482.4;refGene_gn=RARRES2;refGene_tx=XM_019825990.2;refGene_gn=RARRES2;refGene_tx=XM_019825991.2;refGene_gn=RARRES2;refGene_tx=XM_019825992.2;refGene_gn=RARRES2;refGene_tx=XM_019825993.2
A2 164213018 164213123
refGene_gn=RARRES2;refGene_tx=XM_006929482.4;refGene_gn=RARRES2;refGene_tx=XM_019825990.2;refGene_gn=RARRES2;refGene_tx=XM_019825991.2;refGene_gn=RARRES2;refGene_tx=XM_019825992.2;refGene_gn=RARRES2;refGene_tx=XM_019825993.2
A2 164213326 164213500
refGene_gn=RARRES2;refGene_tx=XM_006929482.4;refGene_gn=RARRES2;refGene_tx=XM_019825990.2;refGene_gn=RARRES2;refGene_tx=XM_019825991.2;refGene_gn=RARRES2;refGene_tx=XM_019825992.2;refGene_gn=RARRES2;refGene_tx=XM_019825993.2
A2 164225644 164225683
refGene_gn=REPIN1;refGene_tx=XM_003983205.5;refGene_gn=REPIN1;refGene_tx=XM_023250718.1;refGene_gn=REPIN1;refGene_tx=XM_023250719.1
A2 164226414 164226571
refGene_gn=REPIN1;refGene_tx=XM_003983205.5;refGene_gn=REPIN1;refGene_tx=XM_006929486.4;refGene_gn=REPIN1;refGene_tx=XM_019825995.2;refGene_gn=REPIN1;refGene_tx=XM_019825996.2;refGene_gn=REPIN1;refGene_tx=XM_023250718.1;refGene_gn=REPIN1;refGene_tx=XM_023250719.1
A2 164227982 164229697
refGene_gn=REPIN1;refGene_tx=XM_003983205.5;refGene_gn=REPIN1;refGene_tx=XM_006929486.4;refGene_gn=REPIN1;refGene_tx=XM_006929487.4;refGene_gn=REPIN1;refGene_tx=XM_006929488.4;refGene_gn=REPIN1;refGene_tx=XM_019825995.2;refGene_gn=REPIN1;refGene_tx=XM_019825996.2;refGene_gn=REPIN1;refGene_tx=XM_019825997.2;refGene_gn=REPIN1;refGene_tx=XM_023250718.1;refGene_gn=REPIN1;refGene_tx=XM_023250719.1
A2 164245455 164245486
refGene_gn=ZNF775;refGene_tx=XM_023250720.1
A2 164252563 164252740
refGene_gn=ZNF775;refGene_tx=XM_023250720.1
A2 164252742 164253059
refGene_gn=ZNF775;refGene_tx=XM_023250720.1
A2 164253060 164254131
refGene_gn=ZNF775;refGene_tx=XM_023250720.1
A2 164264133 164264745
refGene_gn=LOC111559559;refGene_tx=XM_023250722.1
A2 164288122 164288157
refGene_gn=GIMAP8;refGene_tx=XM_023250723.1
A2 164293470 164294131
refGene_gn=GIMAP8;refGene_tx=XM_023250723.1
A2 164298118 164298164

refGene_gn=GIMAP8;refGene_tx=XM_023250723.1
A2 164300827 164301454
refGene_gn=GIMAP8;refGene_tx=XM_023250723.1
A2 164303751 164304473
refGene_gn=GIMAP8;refGene_tx=XM_023250723.1
A2 164779292 164780074
refGene_gn=GIMAP6;refGene_tx=XM_006929496.4
A2 164782678 164782763
refGene_gn=GIMAP6;refGene_tx=XM_006929496.4
A2 164785417 164785591
refGene_gn=GIMAP6;refGene_tx=XM_006929496.4
A2 164812500 164812528
refGene_gn=GIMAP2;refGene_tx=XM_003983236.4
A2 164813361 164814341
refGene_gn=GIMAP2;refGene_tx=XM_003983236.4
A2 164825452 164825976
refGene_gn=LOC101101043;refGene_tx=XM_023242338.1
A2 164825978 164826209
refGene_gn=LOC101101043;refGene_tx=XM_023242338.1
A2 164828097 164828149
refGene_gn=LOC101101043;refGene_tx=XM_023242338.1
A2 164939885 164939928
refGene_gn=LOC101081130;refGene_tx=XM_006929498.4;refGene_gn=LOC101081130;refGene_tx=XM_019825999.2
A2 164941838 164942725
refGene_gn=LOC101081130;refGene_tx=XM_006929498.4;refGene_gn=LOC101081130;refGene_tx=XM_019825999.2
A2 164969227 164969320
refGene_gn=TMEM176B;refGene_tx=XM_003983215.5;refGene_gn=TMEM176B;refGene_tx=XM_011280643.3;refGene_gn=TMEM176B;refGene_tx=XM_011280644.3;refGene_gn=TMEM176B;refGene_tx=XM_019826003.2;refGene_gn=TMEM176B;refGene_tx=XM_019826004.2;refGene_gn=TMEM176B;refGene_tx=XM_023250725.1;refGene_gn=TMEM176B;refGene_tx=XM_023250726.1
A2 164969659 164969779
refGene_gn=TMEM176B;refGene_tx=XM_003983215.5;refGene_gn=TMEM176B;refGene_tx=XM_011280643.3;refGene_gn=TMEM176B;refGene_tx=XM_011280644.3;refGene_gn=TMEM176B;refGene_tx=XM_019826003.2;refGene_gn=TMEM176B;refGene_tx=XM_019826004.2;refGene_gn=TMEM176B;refGene_tx=XM_023250725.1;refGene_gn=TMEM176B;refGene_tx=XM_023250726.1
A2 164970682 164970913
refGene_gn=TMEM176B;refGene_tx=XM_003983215.5;refGene_gn=TMEM176B;refGene_tx=XM_011280643.3;refGene_gn=TMEM176B;refGene_tx=XM_011280644.3;refGene_gn=TMEM176B;refGene_tx=XM_019826003.2;refGene_gn=TMEM176B;refGene_tx=XM_019826004.2;refGene_gn=TMEM176B;refGene_tx=XM_023250725.1;refGene_gn=TMEM176B;refGene_tx=XM_023250726.1
A2 164971135 164971192
refGene_gn=TMEM176B;refGene_tx=XM_003983215.5;refGene_gn=TMEM176B;refGene_tx=XM_011280643.3;refGene_gn=TMEM176B;refGene_tx=XM_011280644.3;refGene_gn=TMEM176B;refGene_tx=XM_019826003.2;refGene_gn=TMEM176B;refGene_tx=XM_019826004.2;refGene_gn=TMEM176B;refGene_tx=XM_023250725.1;refG

ene_gn=TMEM176B;refGene_tx=XM_023250726.1
A2	164971550	164971661
refGene_gn=TMEM176B;refGene_tx=XM_003983215.5;refGene_gn=TMEM176B;refGene_tx=XM_011280643.3;refGene_gn=TMEM176B;refGene_tx=XM_011280644.3;refGene_gn=TMEM176B;refGene_tx=XM_019826003.2;refGene_gn=TMEM176B;refGene_tx=XM_019826004.2;refGene_gn=TMEM176B;refGene_tx=XM_023250725.1;refGene_gn=TMEM176B;refGene_tx=XM_023250726.1
A2	164973386	164973590
refGene_gn=TMEM176B;refGene_tx=XM_003983215.5;refGene_gn=TMEM176B;refGene_tx=XM_011280643.3;refGene_gn=TMEM176B;refGene_tx=XM_011280644.3;refGene_gn=TMEM176B;refGene_tx=XM_019826003.2;refGene_gn=TMEM176B;refGene_tx=XM_019826004.2;refGene_gn=TMEM176B;refGene_tx=XM_023250725.1;refGene_gn=TMEM176B;refGene_tx=XM_023250726.1
A2	164977962	164978160
refGene_gn=TMEM176A;refGene_tx=XM_006929500.4;refGene_gn=TMEM176A;refGene_tx=XM_019826006.2;refGene_gn=TMEM176A;refGene_tx=XM_019826007.2;refGene_gn=TMEM176A;refGene_tx=XM_019826008.2
A2	164978621	164978732
refGene_gn=TMEM176A;refGene_tx=XM_006929500.4;refGene_gn=TMEM176A;refGene_tx=XM_019826006.2;refGene_gn=TMEM176A;refGene_tx=XM_019826007.2
A2	164979829	164979886
refGene_gn=TMEM176A;refGene_tx=XM_006929500.4;refGene_gn=TMEM176A;refGene_tx=XM_019826006.2;refGene_gn=TMEM176A;refGene_tx=XM_019826007.2;refGene_gn=TMEM176A;refGene_tx=XM_019826008.2
A2	164980057	164980279
refGene_gn=TMEM176A;refGene_tx=XM_006929500.4;refGene_gn=TMEM176A;refGene_tx=XM_019826006.2;refGene_gn=TMEM176A;refGene_tx=XM_019826007.2;refGene_gn=TMEM176A;refGene_tx=XM_019826008.2
A2	164980817	164980925
refGene_gn=TMEM176A;refGene_tx=XM_006929500.4;refGene_gn=TMEM176A;refGene_tx=XM_019826006.2;refGene_gn=TMEM176A;refGene_tx=XM_019826007.2;refGene_gn=TMEM176A;refGene_tx=XM_019826008.2
A2	164981289	164981331
refGene_gn=TMEM176A;refGene_tx=XM_006929500.4;refGene_gn=TMEM176A;refGene_tx=XM_019826007.2;refGene_gn=TMEM176A;refGene_tx=XM_019826008.2
A2	164984457	164984499
refGene_gn=TMEM176A;refGene_tx=XM_019826006.2
A2	165017987	165019572
refGene_gn=AOC1;refGene_tx=XM_019826009.2
A2	165020144	165020430
refGene_gn=AOC1;refGene_tx=XM_019826009.2
A2	165020763	165020896
refGene_gn=AOC1;refGene_tx=XM_019826009.2
A2	165021316	165021586
refGene_gn=AOC1;refGene_tx=XM_019826009.2
A2	165067964	165068114
refGene_gn=KCNH2;refGene_tx=XM_023250727.1;refGene_gn=KCNH2;refGene_tx=XM_023250728.1;refGene_gn=KCNH2;refGene_tx=XM_023250729.1;refGene_gn=KCNH2;refGene_tx=XM_023250730.1
A2	165069396	165069574

refGene_gn=KCNH2;refGene_tx=XM_023250727.1;refGene_gn=KCNH2;refGene_tx=XM_023250728.1;refGene_gn=KCNH2;refGene_tx=XM_023250729.1;refGene_gn=KCNH2;refGene_tx=XM_023250730.1
A2 165069898 165070085
refGene_gn=KCNH2;refGene_tx=XM_023250727.1;refGene_gn=KCNH2;refGene_tx=XM_023250728.1;refGene_gn=KCNH2;refGene_tx=XM_023250729.1;refGene_gn=KCNH2;refGene_tx=XM_023250730.1
A2 165070159 165070432
refGene_gn=KCNH2;refGene_tx=XM_023250727.1;refGene_gn=KCNH2;refGene_tx=XM_023250728.1;refGene_gn=KCNH2;refGene_tx=XM_023250729.1;refGene_gn=KCNH2;refGene_tx=XM_023250730.1
A2 165071018 165071118
refGene_gn=KCNH2;refGene_tx=XM_023250727.1;refGene_gn=KCNH2;refGene_tx=XM_023250728.1;refGene_gn=KCNH2;refGene_tx=XM_023250729.1;refGene_gn=KCNH2;refGene_tx=XM_023250730.1
A2 165071395 165071589
refGene_gn=KCNH2;refGene_tx=XM_023250727.1;refGene_gn=KCNH2;refGene_tx=XM_023250728.1;refGene_gn=KCNH2;refGene_tx=XM_023250729.1;refGene_gn=KCNH2;refGene_tx=XM_023250730.1
A2 165072718 165072971
refGene_gn=KCNH2;refGene_tx=XM_023250727.1;refGene_gn=KCNH2;refGene_tx=XM_023250728.1;refGene_gn=KCNH2;refGene_tx=XM_023250730.1
A2 165073413 165073613
refGene_gn=KCNH2;refGene_tx=XM_023250727.1;refGene_gn=KCNH2;refGene_tx=XM_023250728.1;refGene_gn=KCNH2;refGene_tx=XM_023250730.1
A2 165073911 165074299
refGene_gn=KCNH2;refGene_tx=XM_023250727.1;refGene_gn=KCNH2;refGene_tx=XM_023250728.1;refGene_gn=KCNH2;refGene_tx=XM_023250729.1;refGene_gn=KCNH2;refGene_tx=XM_023250730.1
A2 165074762 165075191
refGene_gn=KCNH2;refGene_tx=XM_023250727.1;refGene_gn=KCNH2;refGene_tx=XM_023250728.1;refGene_gn=KCNH2;refGene_tx=XM_023250729.1;refGene_gn=KCNH2;refGene_tx=XM_023250730.1
A2 165077608 165077716
refGene_gn=KCNH2;refGene_tx=XM_023250730.1
A2 165079561 165079773
refGene_gn=KCNH2;refGene_tx=XM_023250727.1;refGene_gn=KCNH2;refGene_tx=XM_023250728.1;refGene_gn=KCNH2;refGene_tx=XM_023250729.1
A2 165080335 165080803
refGene_gn=KCNH2;refGene_tx=XM_023250727.1;refGene_gn=KCNH2;refGene_tx=XM_023250728.1;refGene_gn=KCNH2;refGene_tx=XM_023250729.1
A2 165081889 165082054
refGene_gn=KCNH2;refGene_tx=XM_023250727.1;refGene_gn=KCNH2;refGene_tx=XM_023250728.1;refGene_gn=KCNH2;refGene_tx=XM_023250729.1
A2 165096452 165096683
refGene_gn=KCNH2;refGene_tx=XM_023250727.1;refGene_gn=KCNH2;refGene_tx=XM_023250728.1;refGene_gn=KCNH2;refGene_tx=XM_023250729.1
A2 165099448 165099524
refGene_gn=KCNH2;refGene_tx=XM_023250727.1;refGene_gn=KCNH2;refGene_tx=XM_023250728.1;refGene_gn=KCNH2;refGene_tx=XM_023250729.1

A2	165112538	165112702
refGene_gn=NOS3;refGene_tx=XM_003983218.5;refGene_gn=NOS3;refGene_tx=XM_006929504.4
A2	165114004	165114110
refGene_gn=NOS3;refGene_tx=XM_003983218.5;refGene_gn=NOS3;refGene_tx=XM_006929504.4
A2	165114455	165114604
refGene_gn=NOS3;refGene_tx=XM_003983218.5;refGene_gn=NOS3;refGene_tx=XM_006929504.4
A2	165114813	165114976
refGene_gn=NOS3;refGene_tx=XM_003983218.5;refGene_gn=NOS3;refGene_tx=XM_006929504.4;refGene_gn=NOS3;refGene_tx=XM_019826010.2;refGene_gn=NOS3;refGene_tx=XM_019826011.2
A2	165116187	165116333
refGene_gn=NOS3;refGene_tx=XM_003983218.5;refGene_gn=NOS3;refGene_tx=XM_006929504.4;refGene_gn=NOS3;refGene_tx=XM_019826010.2;refGene_gn=NOS3;refGene_tx=XM_019826011.2
A2	165116427	165116569
refGene_gn=NOS3;refGene_tx=XM_003983218.5;refGene_gn=NOS3;refGene_tx=XM_006929504.4;refGene_gn=NOS3;refGene_tx=XM_019826010.2;refGene_gn=NOS3;refGene_tx=XM_019826011.2
A2	165116766	165116906
refGene_gn=NOS3;refGene_tx=XM_003983218.5;refGene_gn=NOS3;refGene_tx=XM_006929504.4;refGene_gn=NOS3;refGene_tx=XM_019826010.2;refGene_gn=NOS3;refGene_tx=XM_019826011.2
A2	165117014	165117189
refGene_gn=NOS3;refGene_tx=XM_003983218.5;refGene_gn=NOS3;refGene_tx=XM_006929504.4;refGene_gn=NOS3;refGene_tx=XM_019826010.2;refGene_gn=NOS3;refGene_tx=XM_019826011.2
A2	165117678	165117780
refGene_gn=NOS3;refGene_tx=XM_003983218.5;refGene_gn=NOS3;refGene_tx=XM_006929504.4;refGene_gn=NOS3;refGene_tx=XM_019826010.2;refGene_gn=NOS3;refGene_tx=XM_019826011.2
A2	165118385	165118580
refGene_gn=NOS3;refGene_tx=XM_003983218.5;refGene_gn=NOS3;refGene_tx=XM_006929504.4;refGene_gn=NOS3;refGene_tx=XM_019826010.2;refGene_gn=NOS3;refGene_tx=XM_019826011.2
A2	165118691	165118765
refGene_gn=NOS3;refGene_tx=XM_003983218.5;refGene_gn=NOS3;refGene_tx=XM_006929504.4;refGene_gn=NOS3;refGene_tx=XM_019826010.2;refGene_gn=NOS3;refGene_tx=XM_019826011.2
A2	165118948	165119093
refGene_gn=NOS3;refGene_tx=XM_003983218.5;refGene_gn=NOS3;refGene_tx=XM_006929504.4;refGene_gn=NOS3;refGene_tx=XM_019826010.2;refGene_gn=NOS3;refGene_tx=XM_019826011.2
A2	165119355	165119460
refGene_gn=NOS3;refGene_tx=XM_003983218.5;refGene_gn=NOS3;refGene_tx=XM_006929504.4;refGene_gn=NOS3;refGene_tx=XM_019826010.2;refGene_gn=NOS3;refGene_tx=XM_019826011.2
A2	165122618	165122686

refGene_gn=NOS3;refGene_tx=XM_003983218.5;refGene_gn=NOS3;refGene_tx=XM_006929504.4;refGene_gn=NOS3;refGene_tx=XM_019826010.2;refGene_gn=NOS3;refGene_tx=XM_019826011.2
A2 165123084 165123201
refGene_gn=NOS3;refGene_tx=XM_003983218.5;refGene_gn=NOS3;refGene_tx=XM_006929504.4;refGene_gn=NOS3;refGene_tx=XM_019826010.2;refGene_gn=NOS3;refGene_tx=XM_019826011.2
A2 165123310 165123485
refGene_gn=NOS3;refGene_tx=XM_003983218.5;refGene_gn=NOS3;refGene_tx=XM_006929504.4;refGene_gn=NOS3;refGene_tx=XM_019826010.2;refGene_gn=NOS3;refGene_tx=XM_019826011.2
A2 165124829 165124962
refGene_gn=NOS3;refGene_tx=XM_003983218.5;refGene_gn=NOS3;refGene_tx=XM_006929504.4;refGene_gn=NOS3;refGene_tx=XM_019826010.2;refGene_gn=NOS3;refGene_tx=XM_019826011.2
A2 165125075 165125154
refGene_gn=NOS3;refGene_tx=XM_003983218.5;refGene_gn=NOS3;refGene_tx=XM_006929504.4;refGene_gn=NOS3;refGene_tx=XM_019826010.2;refGene_gn=NOS3;refGene_tx=XM_019826011.2
A2 165125589 165125777
refGene_gn=NOS3;refGene_tx=XM_003983218.5;refGene_gn=NOS3;refGene_tx=XM_006929504.4;refGene_gn=NOS3;refGene_tx=XM_019826010.2;refGene_gn=NOS3;refGene_tx=XM_019826011.2
A2 165126357 165126530
refGene_gn=NOS3;refGene_tx=XM_003983218.5;refGene_gn=NOS3;refGene_tx=XM_006929504.4;refGene_gn=NOS3;refGene_tx=XM_019826010.2;refGene_gn=NOS3;refGene_tx=XM_019826011.2
A2 165126796 165127007
refGene_gn=NOS3;refGene_tx=XM_003983218.5;refGene_gn=NOS3;refGene_tx=XM_006929504.4;refGene_gn=NOS3;refGene_tx=XM_019826010.2;refGene_gn=NOS3;refGene_tx=XM_019826011.2
A2 165127093 165127181
refGene_gn=NOS3;refGene_tx=XM_003983218.5;refGene_gn=NOS3;refGene_tx=XM_006929504.4;refGene_gn=NOS3;refGene_tx=XM_019826010.2;refGene_gn=NOS3;refGene_tx=XM_019826011.2
A2 165128401 165128523
refGene_gn=NOS3;refGene_tx=XM_003983218.5;refGene_gn=NOS3;refGene_tx=XM_006929504.4;refGene_gn=NOS3;refGene_tx=XM_019826010.2;refGene_gn=NOS3;refGene_tx=XM_019826011.2
A2 165129283 165129432
refGene_gn=NOS3;refGene_tx=XM_003983218.5;refGene_gn=NOS3;refGene_tx=XM_006929504.4;refGene_gn=NOS3;refGene_tx=XM_019826010.2;refGene_gn=NOS3;refGene_tx=XM_019826011.2
A2 165129794 165129989
refGene_gn=NOS3;refGene_tx=XM_003983218.5;refGene_gn=NOS3;refGene_tx=XM_006929504.4;refGene_gn=NOS3;refGene_tx=XM_019826010.2;refGene_gn=NOS3;refGene_tx=XM_019826011.2
A2 165130099 165130261
refGene_gn=NOS3;refGene_tx=XM_003983218.5;refGene_gn=NOS3;refGene_tx=XM_006929504.4;refGene_gn=NOS3;refGene_tx=XM_019826010.2;refGene_gn=NOS

3;refGene_tx=XM_019826011.2
A2 165131634 165131762
refGene_gn=ATG9B;refGene_tx=XM_003983219.4;refGene_gn=ATG9B;refGene_tx=XM_019826012.2
A2 165131848 165131972
refGene_gn=ATG9B;refGene_tx=XM_003983219.4;refGene_gn=ATG9B;refGene_tx=XM_019826012.2
A2 165132173 165132270
refGene_gn=ATG9B;refGene_tx=XM_003983219.4;refGene_gn=ATG9B;refGene_tx=XM_019826012.2
A2 165132451 165132585
refGene_gn=ATG9B;refGene_tx=XM_003983219.4;refGene_gn=ATG9B;refGene_tx=XM_019826012.2
A2 165132754 165132991
refGene_gn=ATG9B;refGene_tx=XM_003983219.4;refGene_gn=ATG9B;refGene_tx=XM_019826012.2
A2 165133579 165133759
refGene_gn=ATG9B;refGene_tx=XM_003983219.4;refGene_gn=ATG9B;refGene_tx=XM_019826012.2
A2 165133975 165134129
refGene_gn=ATG9B;refGene_tx=XM_003983219.4;refGene_gn=ATG9B;refGene_tx=XM_019826012.2
A2 165134315 165135070
refGene_gn=ATG9B;refGene_tx=XM_003983219.4;refGene_gn=ATG9B;refGene_tx=XM_019826012.2
A2 165136635 165136777
refGene_gn=ATG9B;refGene_tx=XM_003983219.4;refGene_gn=ATG9B;refGene_tx=XM_019826012.2
A2 165137310 165137472
refGene_gn=ATG9B;refGene_tx=XM_003983219.4;refGene_gn=ATG9B;refGene_tx=XM_019826012.2
A2 165137669 165137734
refGene_gn=ATG9B;refGene_tx=XM_003983219.4;refGene_gn=ATG9B;refGene_tx=XM_019826012.2
A2 165137900 165137944
refGene_gn=ATG9B;refGene_tx=XM_003983219.4;refGene_gn=ATG9B;refGene_tx=XM_019826012.2
A2 165138063 165138604
refGene_gn=ATG9B;refGene_tx=XM_003983219.4;refGene_gn=ATG9B;refGene_tx=XM_019826012.2
A2 165140994 165141138
refGene_gn=ABCB8;refGene_tx=XM_003983242.5;refGene_gn=ABCB8;refGene_tx=XM_006929505.4;refGene_gn=ABCB8;refGene_tx=XM_006929506.4
A2 165145042 165145355
refGene_gn=ABCB8;refGene_tx=XM_003983242.5;refGene_gn=ABCB8;refGene_tx=XM_006929505.4
A2 165145719 165145875
refGene_gn=ABCB8;refGene_tx=XM_003983242.5;refGene_gn=ABCB8;refGene_tx=XM_006929505.4;refGene_gn=ABCB8;refGene_tx=XM_006929506.4
A2 165145952 165146047

refGene_gn=ABCB8;refGene_tx=XM_003983242.5;refGene_gn=ABCB8;refGene_tx=XM_006929505.4;refGene_gn=ABCB8;refGene_tx=XM_006929506.4
A2 165146180 165146286
refGene_gn=ABCB8;refGene_tx=XM_003983242.5;refGene_gn=ABCB8;refGene_tx=XM_006929505.4;refGene_gn=ABCB8;refGene_tx=XM_006929506.4
A2 165147082 165147244
refGene_gn=ABCB8;refGene_tx=XM_003983242.5;refGene_gn=ABCB8;refGene_tx=XM_006929505.4;refGene_gn=ABCB8;refGene_tx=XM_006929506.4
A2 165147343 165147429
refGene_gn=ABCB8;refGene_tx=XM_003983242.5;refGene_gn=ABCB8;refGene_tx=XM_006929505.4;refGene_gn=ABCB8;refGene_tx=XM_006929506.4
A2 165147544 165147642
refGene_gn=ABCB8;refGene_tx=XM_003983242.5;refGene_gn=ABCB8;refGene_tx=XM_006929505.4;refGene_gn=ABCB8;refGene_tx=XM_006929506.4
A2 165147925 165148031
refGene_gn=ABCB8;refGene_tx=XM_003983242.5;refGene_gn=ABCB8;refGene_tx=XM_006929505.4;refGene_gn=ABCB8;refGene_tx=XM_006929506.4
A2 165151014 165151048
refGene_gn=ABCB8;refGene_tx=XM_003983242.5;refGene_gn=ABCB8;refGene_tx=XM_006929505.4;refGene_gn=ABCB8;refGene_tx=XM_006929506.4
A2 165151248 165151385
refGene_gn=ABCB8;refGene_tx=XM_003983242.5;refGene_gn=ABCB8;refGene_tx=XM_006929505.4;refGene_gn=ABCB8;refGene_tx=XM_006929506.4
A2 165151547 165151642
refGene_gn=ABCB8;refGene_tx=XM_003983242.5;refGene_gn=ABCB8;refGene_tx=XM_006929505.4;refGene_gn=ABCB8;refGene_tx=XM_006929506.4
A2 165151819 165151953
refGene_gn=ABCB8;refGene_tx=XM_003983242.5;refGene_gn=ABCB8;refGene_tx=XM_006929505.4;refGene_gn=ABCB8;refGene_tx=XM_006929506.4
A2 165152732 165152880
refGene_gn=ABCB8;refGene_tx=XM_003983242.5;refGene_gn=ABCB8;refGene_tx=XM_006929505.4;refGene_gn=ABCB8;refGene_tx=XM_006929506.4
A2 165154239 165154490
refGene_gn=ABCB8;refGene_tx=XM_003983242.5;refGene_gn=ABCB8;refGene_tx=XM_006929505.4;refGene_gn=ABCB8;refGene_tx=XM_006929506.4
A2 165155121 165155247
refGene_gn=ABCB8;refGene_tx=XM_003983242.5;refGene_gn=ABCB8;refGene_tx=XM_006929505.4;refGene_gn=ABCB8;refGene_tx=XM_006929506.4
A2 165159319 165159853
refGene_gn=ASIC3;refGene_tx=XM_011280641.2;refGene_gn=ASIC3;refGene_tx=XM_019826014.1;refGene_gn=ASIC3;refGene_tx=XM_019826015.2;refGene_gn=ASIC3;refGene_tx=XM_019826016.1;refGene_gn=ASIC3;refGene_tx=XM_019826019.1;refGene_gn=ASIC3;refGene_tx=XM_019826020.1
A2 165160593 165160744
refGene_gn=ASIC3;refGene_tx=XM_011280641.2;refGene_gn=ASIC3;refGene_tx=XM_019826014.1;refGene_gn=ASIC3;refGene_tx=XM_019826015.2;refGene_gn=ASIC3;refGene_tx=XM_019826016.1;refGene_gn=ASIC3;refGene_tx=XM_019826019.1;refGene_gn=ASIC3;refGene_tx=XM_019826020.1
A2 165160968 165161096
refGene_gn=ASIC3;refGene_tx=XM_011280641.2;refGene_gn=ASIC3;refGene_tx

=XM_019826014.1;refGene_gn=ASIC3;refGene_tx=XM_019826015.2;refGene_gn=
ASIC3;refGene_tx=XM_019826016.1;refGene_gn=ASIC3;refGene_tx=XM_0198260
19.1;refGene_gn=ASIC3;refGene_tx=XM_019826020.1
A2 165161252 165161454
refGene_gn=ASIC3;refGene_tx=XM_011280641.2;refGene_gn=ASIC3;refGene_tx
=XM_019826014.1;refGene_gn=ASIC3;refGene_tx=XM_019826015.2;refGene_gn=
ASIC3;refGene_tx=XM_019826016.1;refGene_gn=ASIC3;refGene_tx=XM_0198260
19.1;refGene_gn=ASIC3;refGene_tx=XM_019826020.1
A2 165161547 165161604
refGene_gn=ASIC3;refGene_tx=XM_011280641.2;refGene_gn=ASIC3;refGene_tx
=XM_019826014.1;refGene_gn=ASIC3;refGene_tx=XM_019826015.2;refGene_gn=
ASIC3;refGene_tx=XM_019826016.1;refGene_gn=ASIC3;refGene_tx=XM_0198260
19.1;refGene_gn=ASIC3;refGene_tx=XM_019826020.1
A2 165161705 165161853
refGene_gn=ASIC3;refGene_tx=XM_011280641.2;refGene_gn=ASIC3;refGene_tx
=XM_019826014.1;refGene_gn=ASIC3;refGene_tx=XM_019826015.2;refGene_gn=
ASIC3;refGene_tx=XM_019826016.1;refGene_gn=ASIC3;refGene_tx=XM_0198260
19.1;refGene_gn=ASIC3;refGene_tx=XM_019826020.1
A2 165162155 165162247
refGene_gn=ASIC3;refGene_tx=XM_011280641.2;refGene_gn=ASIC3;refGene_tx
=XM_019826014.1;refGene_gn=ASIC3;refGene_tx=XM_019826015.2;refGene_gn=
ASIC3;refGene_tx=XM_019826016.1;refGene_gn=ASIC3;refGene_tx=XM_0198260
19.1;refGene_gn=ASIC3;refGene_tx=XM_019826020.1
A2 165162385 165162465
refGene_gn=ASIC3;refGene_tx=XM_011280641.2;refGene_gn=ASIC3;refGene_tx
=XM_019826014.1;refGene_gn=ASIC3;refGene_tx=XM_019826015.2;refGene_gn=
ASIC3;refGene_tx=XM_019826016.1;refGene_gn=ASIC3;refGene_tx=XM_0198260
20.1
A2 165162562 165162634
refGene_gn=ASIC3;refGene_tx=XM_011280641.2;refGene_gn=ASIC3;refGene_tx
=XM_019826014.1;refGene_gn=ASIC3;refGene_tx=XM_019826015.2;refGene_gn=
ASIC3;refGene_tx=XM_019826016.1;refGene_gn=ASIC3;refGene_tx=XM_0198260
19.1;refGene_gn=ASIC3;refGene_tx=XM_019826020.1
A2 165162805 165162864
refGene_gn=ASIC3;refGene_tx=XM_011280641.2;refGene_gn=ASIC3;refGene_tx
=XM_019826016.1;refGene_gn=ASIC3;refGene_tx=XM_019826020.1
A2 165162973 165163170
refGene_gn=ASIC3;refGene_tx=XM_011280641.2;refGene_gn=ASIC3;refGene_tx
=XM_019826014.1;refGene_gn=ASIC3;refGene_tx=XM_019826015.2;refGene_gn=
ASIC3;refGene_tx=XM_019826016.1;refGene_gn=ASIC3;refGene_tx=XM_0198260
20.1
A2 165164401 165164488
refGene_gn=CDK5;refGene_tx=XM_003983221.4
A2 165164614 165164695
refGene_gn=CDK5;refGene_tx=XM_003983221.4
A2 165164797 165164858
refGene_gn=CDK5;refGene_tx=XM_003983221.4
A2 165165519 165165589
refGene_gn=CDK5;refGene_tx=XM_003983221.4
A2 165165758 165165855

refGene_gn=CDK5;refGene_tx=XM_003983221.4
A2	165166019	165166094
refGene_gn=CDK5;refGene_tx=XM_003983221.4
A2	165166242	165166338
refGene_gn=CDK5;refGene_tx=XM_003983221.4
A2	165167074	165167131
refGene_gn=CDK5;refGene_tx=XM_003983221.4
A2	165167230	165167291
refGene_gn=CDK5;refGene_tx=XM_003983221.4
A2	165167459	165167527
refGene_gn=CDK5;refGene_tx=XM_003983221.4
A2	165167619	165167708
refGene_gn=CDK5;refGene_tx=XM_003983221.4
A2	165168364	165168401
refGene_gn=CDK5;refGene_tx=XM_003983221.4
A2	165172732	165172783
refGene_gn=SLC4A2;refGene_tx=XM_023250732.1
A2	165173434	165173458
refGene_gn=SLC4A2;refGene_tx=XM_019826021.2
A2	165174473	165174639
refGene_gn=SLC4A2;refGene_tx=XM_019826021.2;refGene_gn=SLC4A2;refGene_
tx=XM_023250732.1
A2	165174776	165175018
refGene_gn=SLC4A2;refGene_tx=XM_019826021.2;refGene_gn=SLC4A2;refGene_
tx=XM_023250732.1
A2	165175098	165175217
refGene_gn=SLC4A2;refGene_tx=XM_019826021.2;refGene_gn=SLC4A2;refGene_
tx=XM_023250732.1
A2	165176632	165176865
refGene_gn=SLC4A2;refGene_tx=XM_019826021.2;refGene_gn=SLC4A2;refGene_
tx=XM_023250732.1
A2	165176956	165177099
refGene_gn=SLC4A2;refGene_tx=XM_019826021.2;refGene_gn=SLC4A2;refGene_
tx=XM_023250732.1
A2	165178348	165178529
refGene_gn=SLC4A2;refGene_tx=XM_019826021.2;refGene_gn=SLC4A2;refGene_
tx=XM_023250732.1
A2	165180447	165180583
refGene_gn=SLC4A2;refGene_tx=XM_019826021.2;refGene_gn=SLC4A2;refGene_
tx=XM_023250732.1
A2	165180674	165180840
refGene_gn=SLC4A2;refGene_tx=XM_019826021.2;refGene_gn=SLC4A2;refGene_
tx=XM_023250732.1
A2	165180993	165181108
refGene_gn=SLC4A2;refGene_tx=XM_019826021.2;refGene_gn=SLC4A2;refGene_
tx=XM_023250732.1
A2	165181238	165181423
refGene_gn=SLC4A2;refGene_tx=XM_019826021.2;refGene_gn=SLC4A2;refGene_
tx=XM_023250732.1
A2	165181535	165181761

refGene_gn=SLC4A2;refGene_tx=XM_019826021.2;refGene_gn=SLC4A2;refGene_tx=XM_023250732.1
A2 165181904 165182120
refGene_gn=SLC4A2;refGene_tx=XM_019826021.2;refGene_gn=SLC4A2;refGene_tx=XM_023250732.1
A2 165182200 165182349
refGene_gn=SLC4A2;refGene_tx=XM_019826021.2;refGene_gn=SLC4A2;refGene_tx=XM_023250732.1
A2 165182473 165182668
refGene_gn=SLC4A2;refGene_tx=XM_019826021.2;refGene_gn=SLC4A2;refGene_tx=XM_023250732.1
A2 165183525 165183780
refGene_gn=SLC4A2;refGene_tx=XM_019826021.2;refGene_gn=SLC4A2;refGene_tx=XM_023250732.1
A2 165183887 165183977
refGene_gn=SLC4A2;refGene_tx=XM_019826021.2;refGene_gn=SLC4A2;refGene_tx=XM_023250732.1
A2 165184176 165184343
refGene_gn=SLC4A2;refGene_tx=XM_019826021.2;refGene_gn=SLC4A2;refGene_tx=XM_023250732.1
A2 165184729 165184983
refGene_gn=SLC4A2;refGene_tx=XM_019826021.2;refGene_gn=SLC4A2;refGene_tx=XM_023250732.1
A2 165185078 165185248
refGene_gn=SLC4A2;refGene_tx=XM_019826021.2;refGene_gn=SLC4A2;refGene_tx=XM_023250732.1
A2 165185518 165185692
refGene_gn=SLC4A2;refGene_tx=XM_019826021.2;refGene_gn=SLC4A2;refGene_tx=XM_023250732.1
A2 165185798 165185879
refGene_gn=SLC4A2;refGene_tx=XM_019826021.2;refGene_gn=SLC4A2;refGene_tx=XM_023250732.1
A2 165186240 165186348
refGene_gn=FASTK;refGene_tx=XM_011280640.2
A2 165186454 165186538
refGene_gn=FASTK;refGene_tx=XM_011280640.2
A2 165186539 165186570
refGene_gn=FASTK;refGene_tx=XM_011280640.2
A2 165186643 165186779
refGene_gn=FASTK;refGene_tx=XM_011280640.2
A2 165186863 165186955
refGene_gn=FASTK;refGene_tx=XM_011280640.2
A2 165187188 165187348
refGene_gn=FASTK;refGene_tx=XM_011280640.2
A2 165187452 165187666
refGene_gn=FASTK;refGene_tx=XM_011280640.2
A2 165188153 165188293
refGene_gn=FASTK;refGene_tx=XM_011280640.2
A2 165188419 165188599
refGene_gn=FASTK;refGene_tx=XM_011280640.2

A2	165189092	165189515
refGene_gn=FASTK;refGene_tx=XM_011280640.2
A2	165190298	165190380
refGene_gn=FASTK;refGene_tx=XM_011280640.2
A2	165191264	165191616
refGene_gn=TMUB1;refGene_tx=XM_003983225.4
A2	165191898	165192317
refGene_gn=TMUB1;refGene_tx=XM_003983225.4
A2	165193199	165193313
refGene_gn=TMUB1;refGene_tx=XM_003983225.4
A2	165196450	165196781
refGene_gn=AGAP3;refGene_tx=XM_023250734.1;refGene_gn=AGAP3;refGene_tx=XM_023250735.1;refGene_gn=AGAP3;refGene_tx=XM_023250736.1;refGene_gn=AGAP3;refGene_tx=XM_023250739.1;refGene_gn=AGAP3;refGene_tx=XM_023250740.1
A2	165220506	165221425
refGene_gn=AGAP3;refGene_tx=XM_023250738.1
A2	165222681	165222740
refGene_gn=AGAP3;refGene_tx=XM_023250734.1;refGene_gn=AGAP3;refGene_tx=XM_023250735.1;refGene_gn=AGAP3;refGene_tx=XM_023250736.1;refGene_gn=AGAP3;refGene_tx=XM_023250738.1;refGene_gn=AGAP3;refGene_tx=XM_023250739.1;refGene_gn=AGAP3;refGene_tx=XM_023250740.1
A2	165223007	165223095
refGene_gn=AGAP3;refGene_tx=XM_023250734.1;refGene_gn=AGAP3;refGene_tx=XM_023250735.1;refGene_gn=AGAP3;refGene_tx=XM_023250736.1;refGene_gn=AGAP3;refGene_tx=XM_023250738.1;refGene_gn=AGAP3;refGene_tx=XM_023250739.1;refGene_gn=AGAP3;refGene_tx=XM_023250740.1
A2	165223298	165223384
refGene_gn=AGAP3;refGene_tx=XM_023250734.1;refGene_gn=AGAP3;refGene_tx=XM_023250735.1;refGene_gn=AGAP3;refGene_tx=XM_023250736.1;refGene_gn=AGAP3;refGene_tx=XM_023250738.1;refGene_gn=AGAP3;refGene_tx=XM_023250739.1;refGene_gn=AGAP3;refGene_tx=XM_023250740.1
A2	165223528	165223670
refGene_gn=AGAP3;refGene_tx=XM_023250734.1;refGene_gn=AGAP3;refGene_tx=XM_023250735.1;refGene_gn=AGAP3;refGene_tx=XM_023250736.1;refGene_gn=AGAP3;refGene_tx=XM_023250737.1;refGene_gn=AGAP3;refGene_tx=XM_023250738.1;refGene_gn=AGAP3;refGene_tx=XM_023250739.1;refGene_gn=AGAP3;refGene_tx=XM_023250740.1
A2	165224059	165224194
refGene_gn=AGAP3;refGene_tx=XM_023250734.1;refGene_gn=AGAP3;refGene_tx=XM_023250735.1;refGene_gn=AGAP3;refGene_tx=XM_023250736.1;refGene_gn=AGAP3;refGene_tx=XM_023250737.1;refGene_gn=AGAP3;refGene_tx=XM_023250738.1;refGene_gn=AGAP3;refGene_tx=XM_023250739.1;refGene_gn=AGAP3;refGene_tx=XM_023250740.1
A2	165224354	165224482
refGene_gn=AGAP3;refGene_tx=XM_023250734.1;refGene_gn=AGAP3;refGene_tx=XM_023250735.1;refGene_gn=AGAP3;refGene_tx=XM_023250736.1;refGene_gn=AGAP3;refGene_tx=XM_023250737.1;refGene_gn=AGAP3;refGene_tx=XM_023250738.1;refGene_gn=AGAP3;refGene_tx=XM_023250739.1;refGene_gn=AGAP3;refGene_tx=XM_023250740.1

A2	165226080	165226239
refGene_gn=AGAP3;refGene_tx=XM_023250734.1;refGene_gn=AGAP3;refGene_tx=XM_023250735.1;refGene_gn=AGAP3;refGene_tx=XM_023250736.1;refGene_gn=AGAP3;refGene_tx=XM_023250737.1;refGene_gn=AGAP3;refGene_tx=XM_023250738.1;refGene_gn=AGAP3;refGene_tx=XM_023250739.1
A2	165228697	165228760
refGene_gn=AGAP3;refGene_tx=XM_023250738.1;refGene_gn=AGAP3;refGene_tx=XM_023250739.1;refGene_gn=AGAP3;refGene_tx=XM_023250740.1
A2	165229801	165229894
refGene_gn=AGAP3;refGene_tx=XM_023250734.1;refGene_gn=AGAP3;refGene_tx=XM_023250735.1;refGene_gn=AGAP3;refGene_tx=XM_023250736.1;refGene_gn=AGAP3;refGene_tx=XM_023250737.1
A2	165234463	165234568
refGene_gn=AGAP3;refGene_tx=XM_023250734.1;refGene_gn=AGAP3;refGene_tx=XM_023250735.1;refGene_gn=AGAP3;refGene_tx=XM_023250736.1;refGene_gn=AGAP3;refGene_tx=XM_023250737.1
A2	165240875	165241044
refGene_gn=AGAP3;refGene_tx=XM_023250734.1;refGene_gn=AGAP3;refGene_tx=XM_023250735.1;refGene_gn=AGAP3;refGene_tx=XM_023250736.1;refGene_gn=AGAP3;refGene_tx=XM_023250737.1
A2	165244793	165244907
refGene_gn=AGAP3;refGene_tx=XM_023250734.1;refGene_gn=AGAP3;refGene_tx=XM_023250736.1;refGene_gn=AGAP3;refGene_tx=XM_023250737.1
A2	165246646	165246784
refGene_gn=AGAP3;refGene_tx=XM_023250734.1;refGene_gn=AGAP3;refGene_tx=XM_023250736.1;refGene_gn=AGAP3;refGene_tx=XM_023250737.1
A2	165248483	165248638
refGene_gn=AGAP3;refGene_tx=XM_023250734.1;refGene_gn=AGAP3;refGene_tx=XM_023250735.1;refGene_gn=AGAP3;refGene_tx=XM_023250736.1;refGene_gn=AGAP3;refGene_tx=XM_023250737.1
A2	165248729	165248820
refGene_gn=AGAP3;refGene_tx=XM_023250734.1;refGene_gn=AGAP3;refGene_tx=XM_023250735.1;refGene_gn=AGAP3;refGene_tx=XM_023250736.1;refGene_gn=AGAP3;refGene_tx=XM_023250737.1
A2	165248966	165249189
refGene_gn=AGAP3;refGene_tx=XM_023250734.1;refGene_gn=AGAP3;refGene_tx=XM_023250735.1;refGene_gn=AGAP3;refGene_tx=XM_023250736.1;refGene_gn=AGAP3;refGene_tx=XM_023250737.1
A2	165249722	165249798
refGene_gn=AGAP3;refGene_tx=XM_023250736.1
A2	165249854	165250110
refGene_gn=AGAP3;refGene_tx=XM_023250734.1;refGene_gn=AGAP3;refGene_tx=XM_023250735.1;refGene_gn=AGAP3;refGene_tx=XM_023250737.1
A2	165250252	165250459
refGene_gn=AGAP3;refGene_tx=XM_023250734.1;refGene_gn=AGAP3;refGene_tx=XM_023250735.1;refGene_gn=AGAP3;refGene_tx=XM_023250737.1
A2	165254278	165254832
refGene_gn=GBX1;refGene_tx=XM_019826026.2
A2	165271413	165271948
refGene_gn=GBX1;refGene_tx=XM_019826026.2

A2	165278578	165278764
refGene_gn=ASB10;refGene_tx=XM_003983227.5
A2	165279044	165279158
refGene_gn=ASB10;refGene_tx=XM_003983227.5
A2	165282582	165282819
refGene_gn=ASB10;refGene_tx=XM_023250741.1
A2	165282899	165283419
refGene_gn=ASB10;refGene_tx=XM_003983227.5;refGene_gn=ASB10;refGene_tx=XM_023250741.1
A2	165286143	165286411
refGene_gn=ASB10;refGene_tx=XM_003983227.5;refGene_gn=ASB10;refGene_tx=XM_023250741.1
A2	165286559	165286869
refGene_gn=ASB10;refGene_tx=XM_023250741.1
A2	165287207	165287478
refGene_gn=ASB10;refGene_tx=XM_003983227.5
A2	165289905	165290007
refGene_gn=IQCA1L;refGene_tx=XM_011280634.3;refGene_gn=IQCA1L;refGene_tx=XM_019826028.2
A2	165290563	165290802
refGene_gn=IQCA1L;refGene_tx=XM_011280634.3;refGene_gn=IQCA1L;refGene_tx=XM_019826028.2
A2	165291176	165291354
refGene_gn=IQCA1L;refGene_tx=XM_011280634.3;refGene_gn=IQCA1L;refGene_tx=XM_019826028.2
A2	165291643	165291730
refGene_gn=IQCA1L;refGene_tx=XM_011280634.3;refGene_gn=IQCA1L;refGene_tx=XM_019826028.2
A2	165292113	165292313
refGene_gn=IQCA1L;refGene_tx=XM_011280634.3;refGene_gn=IQCA1L;refGene_tx=XM_019826028.2
A2	165292906	165293014
refGene_gn=IQCA1L;refGene_tx=XM_011280634.3;refGene_gn=IQCA1L;refGene_tx=XM_019826028.2
A2	165293169	165293252
refGene_gn=IQCA1L;refGene_tx=XM_011280634.3;refGene_gn=IQCA1L;refGene_tx=XM_019826028.2
A2	165294033	165294089
refGene_gn=IQCA1L;refGene_tx=XM_011280634.3;refGene_gn=IQCA1L;refGene_tx=XM_019826028.2
A2	165294177	165294261
refGene_gn=IQCA1L;refGene_tx=XM_011280634.3;refGene_gn=IQCA1L;refGene_tx=XM_019826028.2
A2	165295035	165295142
refGene_gn=IQCA1L;refGene_tx=XM_011280634.3;refGene_gn=IQCA1L;refGene_tx=XM_019826028.2
A2	165295549	165295625
refGene_gn=IQCA1L;refGene_tx=XM_011280634.3;refGene_gn=IQCA1L;refGene_tx=XM_019826028.2
A2	165295764	165295890

refGene_gn=IQCA1L;refGene_tx=XM_011280634.3;refGene_gn=IQCA1L;refGene_tx=XM_019826028.2
A2 165296353 165296438
refGene_gn=IQCA1L;refGene_tx=XM_011280634.3;refGene_gn=IQCA1L;refGene_tx=XM_019826028.2
A2 165297029 165297235
refGene_gn=IQCA1L;refGene_tx=XM_011280634.3;refGene_gn=IQCA1L;refGene_tx=XM_019826028.2
A2 165298773 165298845
refGene_gn=IQCA1L;refGene_tx=XM_011280634.3;refGene_gn=IQCA1L;refGene_tx=XM_019826028.2
A2 165300051 165300237
refGene_gn=IQCA1L;refGene_tx=XM_011280634.3;refGene_gn=IQCA1L;refGene_tx=XM_019826028.2
A2 165300548 165300668
refGene_gn=IQCA1L;refGene_tx=XM_011280634.3;refGene_gn=IQCA1L;refGene_tx=XM_019826028.2
A2 165301378 165301706
refGene_gn=IQCA1L;refGene_tx=XM_011280634.3
A2 165302372 165302383
refGene_gn=IQCA1L;refGene_tx=XM_011280634.3
A2 165304069 165304306
refGene_gn=LOC102899259;refGene_tx=XM_019826036.2
A2 165306249 165306381
refGene_gn=LOC102899259;refGene_tx=XM_019826036.2
A2 165309556 165309694
refGene_gn=ABCF2;refGene_tx=XM_006929511.4;refGene_gn=ABCF2;refGene_tx=XM_006929512.4
A2 165310401 165310605
refGene_gn=ABCF2;refGene_tx=XM_006929511.4;refGene_gn=ABCF2;refGene_tx=XM_006929512.4
A2 165311291 165311420
refGene_gn=ABCF2;refGene_tx=XM_006929511.4;refGene_gn=ABCF2;refGene_tx=XM_006929512.4
A2 165311620 165311683
refGene_gn=ABCF2;refGene_tx=XM_006929511.4;refGene_gn=ABCF2;refGene_tx=XM_006929512.4
A2 165312355 165312466
refGene_gn=ABCF2;refGene_tx=XM_006929511.4;refGene_gn=ABCF2;refGene_tx=XM_006929512.4
A2 165312830 165312920
refGene_gn=ABCF2;refGene_tx=XM_006929511.4;refGene_gn=ABCF2;refGene_tx=XM_006929512.4
A2 165313005 165313125
refGene_gn=ABCF2;refGene_tx=XM_006929511.4;refGene_gn=ABCF2;refGene_tx=XM_006929512.4
A2 165313296 165313392
refGene_gn=ABCF2;refGene_tx=XM_006929511.4;refGene_gn=ABCF2;refGene_tx=XM_006929512.4
A2 165314581 165314684

refGene_gn=ABCF2;refGene_tx=XM_006929511.4;refGene_gn=ABCF2;refGene_tx=XM_006929512.4
A2	165315514	165315610
refGene_gn=ABCF2;refGene_tx=XM_006929511.4;refGene_gn=ABCF2;refGene_tx=XM_006929512.4
A2	165316770	165316942
refGene_gn=ABCF2;refGene_tx=XM_006929511.4;refGene_gn=ABCF2;refGene_tx=XM_006929512.4
A2	165317033	165317216
refGene_gn=ABCF2;refGene_tx=XM_006929511.4;refGene_gn=ABCF2;refGene_tx=XM_006929512.4
A2	165317710	165317923
refGene_gn=ABCF2;refGene_tx=XM_006929511.4;refGene_gn=ABCF2;refGene_tx=XM_006929512.4
A2	165323670	165323827
refGene_gn=ABCF2;refGene_tx=XM_006929511.4;refGene_gn=ABCF2;refGene_tx=XM_006929512.4
A2	165333164	165333451
refGene_gn=CHPF2;refGene_tx=XM_003983230.5
A2	165333884	165334449
refGene_gn=CHPF2;refGene_tx=XM_003983230.5
A2	165335039	165335222
refGene_gn=CHPF2;refGene_tx=XM_003983230.5
A2	165335931	165337239
refGene_gn=CHPF2;refGene_tx=XM_003983230.5
A2	165337660	165337714
refGene_gn=SMARCD3;refGene_tx=XM_003983231.5;refGene_gn=SMARCD3;refGene_tx=XM_019826029.2;refGene_gn=SMARCD3;refGene_tx=XM_019826030.2;refGene_gn=SMARCD3;refGene_tx=XM_019826032.2;refGene_gn=SMARCD3;refGene_tx=XM_019826033.2;refGene_gn=SMARCD3;refGene_tx=XM_019826034.2;refGene_gn=SMARCD3;refGene_tx=XM_019826035.2
A2	165337911	165338013
refGene_gn=SMARCD3;refGene_tx=XM_003983231.5;refGene_gn=SMARCD3;refGene_tx=XM_019826029.2;refGene_gn=SMARCD3;refGene_tx=XM_019826030.2;refGene_gn=SMARCD3;refGene_tx=XM_019826032.2;refGene_gn=SMARCD3;refGene_tx=XM_019826033.2;refGene_gn=SMARCD3;refGene_tx=XM_019826034.2;refGene_gn=SMARCD3;refGene_tx=XM_019826035.2
A2	165338132	165338255
refGene_gn=SMARCD3;refGene_tx=XM_003983231.5;refGene_gn=SMARCD3;refGene_tx=XM_019826029.2;refGene_gn=SMARCD3;refGene_tx=XM_019826030.2;refGene_gn=SMARCD3;refGene_tx=XM_019826032.2;refGene_gn=SMARCD3;refGene_tx=XM_019826033.2;refGene_gn=SMARCD3;refGene_tx=XM_019826034.2;refGene_gn=SMARCD3;refGene_tx=XM_019826035.2
A2	165338533	165338669
refGene_gn=SMARCD3;refGene_tx=XM_003983231.5;refGene_gn=SMARCD3;refGene_tx=XM_019826029.2;refGene_gn=SMARCD3;refGene_tx=XM_019826030.2;refGene_gn=SMARCD3;refGene_tx=XM_019826032.2;refGene_gn=SMARCD3;refGene_tx=XM_019826033.2;refGene_gn=SMARCD3;refGene_tx=XM_019826034.2;refGene_gn=SMARCD3;refGene_tx=XM_019826035.2
A2	165338815	165338913

refGene_gn=SMARCD3;refGene_tx=XM_003983231.5;refGene_gn=SMARCD3;refGene_tx=XM_019826029.2;refGene_gn=SMARCD3;refGene_tx=XM_019826030.2;refGene_gn=SMARCD3;refGene_tx=XM_019826032.2;refGene_gn=SMARCD3;refGene_tx=XM_019826033.2;refGene_gn=SMARCD3;refGene_tx=XM_019826034.2;refGene_gn=SMARCD3;refGene_tx=XM_019826035.2
A2 165339627 165339789
refGene_gn=SMARCD3;refGene_tx=XM_003983231.5;refGene_gn=SMARCD3;refGene_tx=XM_019826029.2;refGene_gn=SMARCD3;refGene_tx=XM_019826030.2;refGene_gn=SMARCD3;refGene_tx=XM_019826032.2;refGene_gn=SMARCD3;refGene_tx=XM_019826033.2;refGene_gn=SMARCD3;refGene_tx=XM_019826034.2;refGene_gn=SMARCD3;refGene_tx=XM_019826035.2
A2 165340016 165340118
refGene_gn=SMARCD3;refGene_tx=XM_003983231.5;refGene_gn=SMARCD3;refGene_tx=XM_019826029.2;refGene_gn=SMARCD3;refGene_tx=XM_019826030.2;refGene_gn=SMARCD3;refGene_tx=XM_019826032.2;refGene_gn=SMARCD3;refGene_tx=XM_019826033.2;refGene_gn=SMARCD3;refGene_tx=XM_019826034.2;refGene_gn=SMARCD3;refGene_tx=XM_019826035.2
A2 165340292 165340462
refGene_gn=SMARCD3;refGene_tx=XM_003983231.5;refGene_gn=SMARCD3;refGene_tx=XM_019826029.2;refGene_gn=SMARCD3;refGene_tx=XM_019826030.2;refGene_gn=SMARCD3;refGene_tx=XM_019826032.2;refGene_gn=SMARCD3;refGene_tx=XM_019826033.2;refGene_gn=SMARCD3;refGene_tx=XM_019826034.2;refGene_gn=SMARCD3;refGene_tx=XM_019826035.2
A2 165340603 165340762
refGene_gn=SMARCD3;refGene_tx=XM_003983231.5;refGene_gn=SMARCD3;refGene_tx=XM_019826029.2;refGene_gn=SMARCD3;refGene_tx=XM_019826030.2;refGene_gn=SMARCD3;refGene_tx=XM_019826032.2;refGene_gn=SMARCD3;refGene_tx=XM_019826033.2;refGene_gn=SMARCD3;refGene_tx=XM_019826034.2;refGene_gn=SMARCD3;refGene_tx=XM_019826035.2
A2 165340882 165341005
refGene_gn=SMARCD3;refGene_tx=XM_003983231.5;refGene_gn=SMARCD3;refGene_tx=XM_019826029.2;refGene_gn=SMARCD3;refGene_tx=XM_019826030.2;refGene_gn=SMARCD3;refGene_tx=XM_019826032.2;refGene_gn=SMARCD3;refGene_tx=XM_019826033.2;refGene_gn=SMARCD3;refGene_tx=XM_019826034.2;refGene_gn=SMARCD3;refGene_tx=XM_019826035.2
A2 165341844 165341887
refGene_gn=SMARCD3;refGene_tx=XM_003983231.5;refGene_gn=SMARCD3;refGene_tx=XM_019826029.2;refGene_gn=SMARCD3;refGene_tx=XM_019826030.2;refGene_gn=SMARCD3;refGene_tx=XM_019826032.2;refGene_gn=SMARCD3;refGene_tx=XM_019826033.2;refGene_gn=SMARCD3;refGene_tx=XM_019826034.2;refGene_gn=SMARCD3;refGene_tx=XM_019826035.2
A2 165343615 165343827
refGene_gn=SMARCD3;refGene_tx=XM_003983231.5;refGene_gn=SMARCD3;refGene_tx=XM_019826029.2;refGene_gn=SMARCD3;refGene_tx=XM_019826030.2;refGene_gn=SMARCD3;refGene_tx=XM_019826032.2;refGene_gn=SMARCD3;refGene_tx=XM_019826033.2;refGene_gn=SMARCD3;refGene_tx=XM_019826034.2
A2 165346769 165346847
refGene_gn=SMARCD3;refGene_tx=XM_003983231.5;refGene_gn=SMARCD3;refGene_tx=XM_019826029.2;refGene_gn=SMARCD3;refGene_tx=XM_019826033.2
A2 165365637 165365676

refGene_gn=SMARCD3;refGene_tx=XM_019826030.2;refGene_gn=SMARCD3;refGene_tx=XM_019826032.2
A2 165413280 165413351
refGene_gn=NUB1;refGene_tx=XM_023250743.1;refGene_gn=NUB1;refGene_tx=XM_023250746.1
A2 165415796 165415914
refGene_gn=NUB1;refGene_tx=XM_023250743.1;refGene_gn=NUB1;refGene_tx=XM_023250744.1;refGene_gn=NUB1;refGene_tx=XM_023250745.1;refGene_gn=NUB1;refGene_tx=XM_023250746.1
A2 165418070 165418238
refGene_gn=NUB1;refGene_tx=XM_023250743.1;refGene_gn=NUB1;refGene_tx=XM_023250744.1;refGene_gn=NUB1;refGene_tx=XM_023250745.1;refGene_gn=NUB1;refGene_tx=XM_023250746.1
A2 165420205 165420264
refGene_gn=NUB1;refGene_tx=XM_023250743.1;refGene_gn=NUB1;refGene_tx=XM_023250744.1;refGene_gn=NUB1;refGene_tx=XM_023250745.1;refGene_gn=NUB1;refGene_tx=XM_023250746.1
A2 165421250 165421321
refGene_gn=NUB1;refGene_tx=XM_023250743.1;refGene_gn=NUB1;refGene_tx=XM_023250744.1;refGene_gn=NUB1;refGene_tx=XM_023250745.1;refGene_gn=NUB1;refGene_tx=XM_023250746.1
A2 165423977 165424160
refGene_gn=NUB1;refGene_tx=XM_023250743.1;refGene_gn=NUB1;refGene_tx=XM_023250744.1;refGene_gn=NUB1;refGene_tx=XM_023250745.1;refGene_gn=NUB1;refGene_tx=XM_023250746.1
A2 165424356 165424448
refGene_gn=NUB1;refGene_tx=XM_023250743.1;refGene_gn=NUB1;refGene_tx=XM_023250744.1;refGene_gn=NUB1;refGene_tx=XM_023250745.1;refGene_gn=NUB1;refGene_tx=XM_023250746.1
A2 165426301 165426408
refGene_gn=NUB1;refGene_tx=XM_023250743.1;refGene_gn=NUB1;refGene_tx=XM_023250744.1;refGene_gn=NUB1;refGene_tx=XM_023250745.1;refGene_gn=NUB1;refGene_tx=XM_023250746.1
A2 165431069 165431096
refGene_gn=WDR86;refGene_tx=XM_023250750.1
A2 165431922 165432109
refGene_gn=NUB1;refGene_tx=XM_023250743.1;refGene_gn=NUB1;refGene_tx=XM_023250744.1;refGene_gn=NUB1;refGene_tx=XM_023250745.1;refGene_gn=NUB1;refGene_tx=XM_023250746.1
A2 165432710 165432818
refGene_gn=NUB1;refGene_tx=XM_023250743.1;refGene_gn=NUB1;refGene_tx=XM_023250744.1;refGene_gn=NUB1;refGene_tx=XM_023250745.1;refGene_gn=NUB1;refGene_tx=XM_023250746.1
A2 165433625 165433778
refGene_gn=NUB1;refGene_tx=XM_023250743.1;refGene_gn=NUB1;refGene_tx=XM_023250744.1;refGene_gn=NUB1;refGene_tx=XM_023250745.1;refGene_gn=NUB1;refGene_tx=XM_023250746.1
A2 165434132 165434192
refGene_gn=WDR86;refGene_tx=XM_023250748.1
A2 165438011 165438053

refGene_gn=WDR86;refGene_tx=XM_023250749.1
A2 165438059 165438206
refGene_gn=NUB1;refGene_tx=XM_023250743.1;refGene_gn=NUB1;refGene_tx=XM_023250744.1;refGene_gn=NUB1;refGene_tx=XM_023250745.1;refGene_gn=NUB1;refGene_tx=XM_023250746.1
A2 165439717 165439813
refGene_gn=NUB1;refGene_tx=XM_023250743.1;refGene_gn=NUB1;refGene_tx=XM_023250744.1;refGene_gn=NUB1;refGene_tx=XM_023250745.1;refGene_gn=NUB1;refGene_tx=XM_023250746.1
A2 165440490 165440668
refGene_gn=NUB1;refGene_tx=XM_023250743.1;refGene_gn=NUB1;refGene_tx=XM_023250744.1;refGene_gn=NUB1;refGene_tx=XM_023250745.1;refGene_gn=NUB1;refGene_tx=XM_023250746.1
A2 165440864 165441068
refGene_gn=NUB1;refGene_tx=XM_023250743.1;refGene_gn=NUB1;refGene_tx=XM_023250744.1;refGene_gn=NUB1;refGene_tx=XM_023250745.1;refGene_gn=NUB1;refGene_tx=XM_023250746.1
A2 165444484 165444658
refGene_gn=WDR86;refGene_tx=XM_023250747.1
A2 165444886 165444990
refGene_gn=WDR86;refGene_tx=XM_023250747.1;refGene_gn=WDR86;refGene_tx=XM_023250748.1;refGene_gn=WDR86;refGene_tx=XM_023250749.1;refGene_gn=WDR86;refGene_tx=XM_023250750.1
A2 165448321 165448457
refGene_gn=WDR86;refGene_tx=XM_023250747.1;refGene_gn=WDR86;refGene_tx=XM_023250748.1;refGene_gn=WDR86;refGene_tx=XM_023250749.1;refGene_gn=WDR86;refGene_tx=XM_023250750.1
A2 165451732 165452147
refGene_gn=WDR86;refGene_tx=XM_023250747.1;refGene_gn=WDR86;refGene_tx=XM_023250748.1;refGene_gn=WDR86;refGene_tx=XM_023250749.1;refGene_gn=WDR86;refGene_tx=XM_023250750.1
A2 165456874 165457016
refGene_gn=WDR86;refGene_tx=XM_023250747.1;refGene_gn=WDR86;refGene_tx=XM_023250748.1;refGene_gn=WDR86;refGene_tx=XM_023250749.1;refGene_gn=WDR86;refGene_tx=XM_023250750.1
A2 165466100 165466323
refGene_gn=WDR86;refGene_tx=XM_023250747.1;refGene_gn=WDR86;refGene_tx=XM_023250748.1;refGene_gn=WDR86;refGene_tx=XM_023250749.1;refGene_gn=WDR86;refGene_tx=XM_023250750.1
A2 165480166 165480302
refGene_gn=CRYGN;refGene_tx=XM_003983232.5
A2 165482833 165482979
refGene_gn=CRYGN;refGene_tx=XM_003983232.5
A2 165485299 165485548
refGene_gn=CRYGN;refGene_tx=XM_003983232.5
A2 165487188 165487209
refGene_gn=CRYGN;refGene_tx=XM_003983232.5
A2 165501761 165501854
refGene_gn=RHEB;refGene_tx=XM_023250751.1
A2 165507535 165507617

refGene_gn=RHEB;refGene_tx=XM_023250751.1
A2 165508242 165508290
refGene_gn=RHEB;refGene_tx=XM_023250751.1
A2 165508405 165508462
refGene_gn=RHEB;refGene_tx=XM_023250751.1
A2 165509352 165509435
refGene_gn=RHEB;refGene_tx=XM_023250751.1
A2 165516273 165516341
refGene_gn=RHEB;refGene_tx=XM_023250751.1
A2 165522869 165522941
refGene_gn=RHEB;refGene_tx=XM_023250751.1
A2 165545226 165545278
refGene_gn=RHEB;refGene_tx=XM_023250751.1
A2 165581662 165581694
refGene_gn=PRKAG2;refGene_tx=XM_023250752.1;refGene_gn=PRKAG2;refGene_tx=XM_023250753.1;refGene_gn=PRKAG2;refGene_tx=XM_023250754.1;refGene_gn=PRKAG2;refGene_tx=XM_023250755.1;refGene_gn=PRKAG2;refGene_tx=XM_023250756.1;refGene_gn=PRKAG2;refGene_tx=XM_023250757.1;refGene_gn=PRKAG2;refGene_tx=XM_023250758.1;refGene_gn=PRKAG2;refGene_tx=XM_023250759.1
A2 165588593 165588687
refGene_gn=PRKAG2;refGene_tx=XM_023250752.1;refGene_gn=PRKAG2;refGene_tx=XM_023250753.1;refGene_gn=PRKAG2;refGene_tx=XM_023250754.1;refGene_gn=PRKAG2;refGene_tx=XM_023250755.1;refGene_gn=PRKAG2;refGene_tx=XM_023250756.1;refGene_gn=PRKAG2;refGene_tx=XM_023250757.1;refGene_gn=PRKAG2;refGene_tx=XM_023250758.1;refGene_gn=PRKAG2;refGene_tx=XM_023250759.1
A2 165591258 165591405
refGene_gn=PRKAG2;refGene_tx=XM_023250752.1;refGene_gn=PRKAG2;refGene_tx=XM_023250753.1;refGene_gn=PRKAG2;refGene_tx=XM_023250754.1;refGene_gn=PRKAG2;refGene_tx=XM_023250755.1;refGene_gn=PRKAG2;refGene_tx=XM_023250756.1;refGene_gn=PRKAG2;refGene_tx=XM_023250757.1;refGene_gn=PRKAG2;refGene_tx=XM_023250758.1;refGene_gn=PRKAG2;refGene_tx=XM_023250759.1
A2 165593006 165593044
refGene_gn=PRKAG2;refGene_tx=XM_023250752.1;refGene_gn=PRKAG2;refGene_tx=XM_023250753.1;refGene_gn=PRKAG2;refGene_tx=XM_023250754.1;refGene_gn=PRKAG2;refGene_tx=XM_023250755.1;refGene_gn=PRKAG2;refGene_tx=XM_023250756.1;refGene_gn=PRKAG2;refGene_tx=XM_023250757.1;refGene_gn=PRKAG2;refGene_tx=XM_023250758.1;refGene_gn=PRKAG2;refGene_tx=XM_023250759.1
A2 165593385 165593551
refGene_gn=PRKAG2;refGene_tx=XM_023250752.1;refGene_gn=PRKAG2;refGene_tx=XM_023250753.1;refGene_gn=PRKAG2;refGene_tx=XM_023250754.1;refGene_gn=PRKAG2;refGene_tx=XM_023250755.1;refGene_gn=PRKAG2;refGene_tx=XM_023250756.1;refGene_gn=PRKAG2;refGene_tx=XM_023250757.1;refGene_gn=PRKAG2;refGene_tx=XM_023250758.1;refGene_gn=PRKAG2;refGene_tx=XM_023250759.1
A2 165596290 165596417
refGene_gn=PRKAG2;refGene_tx=XM_023250752.1;refGene_gn=PRKAG2;refGene_

tx=XM_023250753.1;refGene_gn=PRKAG2;refGene_tx=XM_023250754.1;refGene_gn=PRKAG2;refGene_tx=XM_023250755.1;refGene_gn=PRKAG2;refGene_tx=XM_023250756.1;refGene_gn=PRKAG2;refGene_tx=XM_023250757.1;refGene_gn=PRKAG2;refGene_tx=XM_023250758.1;refGene_gn=PRKAG2;refGene_tx=XM_023250759.1
A2 165597257 165597312
refGene_gn=PRKAG2;refGene_tx=XM_023250752.1;refGene_gn=PRKAG2;refGene_tx=XM_023250753.1;refGene_gn=PRKAG2;refGene_tx=XM_023250754.1;refGene_gn=PRKAG2;refGene_tx=XM_023250755.1;refGene_gn=PRKAG2;refGene_tx=XM_023250756.1;refGene_gn=PRKAG2;refGene_tx=XM_023250757.1;refGene_gn=PRKAG2;refGene_tx=XM_023250758.1;refGene_gn=PRKAG2;refGene_tx=XM_023250759.1
A2 165598957 165599003
refGene_gn=PRKAG2;refGene_tx=XM_023250752.1;refGene_gn=PRKAG2;refGene_tx=XM_023250753.1;refGene_gn=PRKAG2;refGene_tx=XM_023250754.1;refGene_gn=PRKAG2;refGene_tx=XM_023250755.1;refGene_gn=PRKAG2;refGene_tx=XM_023250756.1;refGene_gn=PRKAG2;refGene_tx=XM_023250757.1;refGene_gn=PRKAG2;refGene_tx=XM_023250758.1;refGene_gn=PRKAG2;refGene_tx=XM_023250759.1
A2 165606120 165606179
refGene_gn=PRKAG2;refGene_tx=XM_023250752.1;refGene_gn=PRKAG2;refGene_tx=XM_023250753.1;refGene_gn=PRKAG2;refGene_tx=XM_023250754.1;refGene_gn=PRKAG2;refGene_tx=XM_023250755.1;refGene_gn=PRKAG2;refGene_tx=XM_023250756.1;refGene_gn=PRKAG2;refGene_tx=XM_023250757.1;refGene_gn=PRKAG2;refGene_tx=XM_023250758.1;refGene_gn=PRKAG2;refGene_tx=XM_023250759.1
A2 165607269 165607351
refGene_gn=PRKAG2;refGene_tx=XM_023250752.1;refGene_gn=PRKAG2;refGene_tx=XM_023250753.1;refGene_gn=PRKAG2;refGene_tx=XM_023250754.1;refGene_gn=PRKAG2;refGene_tx=XM_023250755.1;refGene_gn=PRKAG2;refGene_tx=XM_023250756.1;refGene_gn=PRKAG2;refGene_tx=XM_023250757.1;refGene_gn=PRKAG2;refGene_tx=XM_023250758.1;refGene_gn=PRKAG2;refGene_tx=XM_023250759.1
A2 165620340 165620447
refGene_gn=PRKAG2;refGene_tx=XM_023250752.1;refGene_gn=PRKAG2;refGene_tx=XM_023250753.1;refGene_gn=PRKAG2;refGene_tx=XM_023250754.1;refGene_gn=PRKAG2;refGene_tx=XM_023250755.1;refGene_gn=PRKAG2;refGene_tx=XM_023250756.1;refGene_gn=PRKAG2;refGene_tx=XM_023250757.1;refGene_gn=PRKAG2;refGene_tx=XM_023250758.1;refGene_gn=PRKAG2;refGene_tx=XM_023250759.1
A2 165640319 165640389
refGene_gn=PRKAG2;refGene_tx=XM_023250752.1;refGene_gn=PRKAG2;refGene_tx=XM_023250753.1;refGene_gn=PRKAG2;refGene_tx=XM_023250754.1;refGene_gn=PRKAG2;refGene_tx=XM_023250755.1;refGene_gn=PRKAG2;refGene_tx=XM_023250756.1;refGene_gn=PRKAG2;refGene_tx=XM_023250757.1;refGene_gn=PRKAG2;refGene_tx=XM_023250758.1;refGene_gn=PRKAG2;refGene_tx=XM_023250759.1
A2 165678343 165678561
refGene_gn=PRKAG2;refGene_tx=XM_023250752.1;refGene_gn=PRKAG2;refGene_tx=XM_023250753.1;refGene_gn=PRKAG2;refGene_tx=XM_023250754.1;refGene_

gn=PRKAG2;refGene_tx=XM_023250755.1;refGene_gn=PRKAG2;refGene_tx=XM_023250756.1;refGene_gn=PRKAG2;refGene_tx=XM_023250757.1
A2 165720026 165720107
refGene_gn=PRKAG2;refGene_tx=XM_023250752.1;refGene_gn=PRKAG2;refGene_tx=XM_023250753.1;refGene_gn=PRKAG2;refGene_tx=XM_023250756.1
A2 165728057 165728151
refGene_gn=PRKAG2;refGene_tx=XM_023250757.1
A2 165759808 165760088
refGene_gn=PRKAG2;refGene_tx=XM_023250752.1;refGene_gn=PRKAG2;refGene_tx=XM_023250753.1;refGene_gn=PRKAG2;refGene_tx=XM_023250754.1;refGene_gn=PRKAG2;refGene_tx=XM_023250755.1;refGene_gn=PRKAG2;refGene_tx=XM_023250756.1
A2 165766708 165766780
refGene_gn=PRKAG2;refGene_tx=XM_023250752.1;refGene_gn=PRKAG2;refGene_tx=XM_023250753.1;refGene_gn=PRKAG2;refGene_tx=XM_023250754.1;refGene_gn=PRKAG2;refGene_tx=XM_023250755.1;refGene_gn=PRKAG2;refGene_tx=XM_023250756.1
A2 165842421 165842535
refGene_gn=PRKAG2;refGene_tx=XM_023250752.1;refGene_gn=PRKAG2;refGene_tx=XM_023250753.1;refGene_gn=PRKAG2;refGene_tx=XM_023250754.1;refGene_gn=PRKAG2;refGene_tx=XM_023250755.1
A2 165899536 165899547
refGene_gn=GALNTL5;refGene_tx=XM_019826048.2
A2 165904495 165904757
refGene_gn=GALNTL5;refGene_tx=XM_011280625.3;refGene_gn=GALNTL5;refGene_tx=XM_011280626.3;refGene_gn=GALNTL5;refGene_tx=XM_019826046.2;refGene_gn=GALNTL5;refGene_tx=XM_019826049.1;refGene_gn=GALNTL5;refGene_tx=XM_023250760.1
A2 165910298 165910534
refGene_gn=GALNTL5;refGene_tx=XM_011280628.2
A2 165913446 165913567
refGene_gn=GALNTL5;refGene_tx=XM_011280625.3;refGene_gn=GALNTL5;refGene_tx=XM_011280626.3;refGene_gn=GALNTL5;refGene_tx=XM_019826046.2;refGene_gn=GALNTL5;refGene_tx=XM_023250760.1
A2 165919840 165920007
refGene_gn=GALNTL5;refGene_tx=XM_011280625.3;refGene_gn=GALNTL5;refGene_tx=XM_011280626.3;refGene_gn=GALNTL5;refGene_tx=XM_011280628.2;refGene_gn=GALNTL5;refGene_tx=XM_019826046.2;refGene_gn=GALNTL5;refGene_tx=XM_019826048.2;refGene_gn=GALNTL5;refGene_tx=XM_019826049.1;refGene_gn=GALNTL5;refGene_tx=XM_023250760.1
A2 165921646 165921769
refGene_gn=GALNTL5;refGene_tx=XM_011280625.3;refGene_gn=GALNTL5;refGene_tx=XM_011280626.3;refGene_gn=GALNTL5;refGene_tx=XM_011280628.2;refGene_gn=GALNTL5;refGene_tx=XM_019826048.2;refGene_gn=GALNTL5;refGene_tx=XM_019826049.1;refGene_gn=GALNTL5;refGene_tx=XM_023250760.1
A2 165936661 165936911
refGene_gn=GALNTL5;refGene_tx=XM_011280625.3;refGene_gn=GALNTL5;refGene_tx=XM_011280626.3;refGene_gn=GALNTL5;refGene_tx=XM_011280628.2;refGene_gn=GALNTL5;refGene_tx=XM_019826046.2;refGene_gn=GALNTL5;refGene_tx=XM_019826048.2;refGene_gn=GALNTL5;refGene_tx=XM_019826049.1;refGene_gn

=GALNTL5;refGene_tx=XM_023250760.1
A2	165942996	165943114
refGene_gn=GALNTL5;refGene_tx=XM_011280625.3;refGene_gn=GALNTL5;refGene_tx=XM_011280626.3;refGene_gn=GALNTL5;refGene_tx=XM_011280628.2;refGene_gn=GALNTL5;refGene_tx=XM_019826046.2;refGene_gn=GALNTL5;refGene_tx=XM_019826048.2;refGene_gn=GALNTL5;refGene_tx=XM_019826049.1;refGene_gn=GALNTL5;refGene_tx=XM_023250760.1
A2	165954782	165954932
refGene_gn=GALNTL5;refGene_tx=XM_011280625.3;refGene_gn=GALNTL5;refGene_tx=XM_011280626.3;refGene_gn=GALNTL5;refGene_tx=XM_011280628.2;refGene_gn=GALNTL5;refGene_tx=XM_019826046.2;refGene_gn=GALNTL5;refGene_tx=XM_019826048.2;refGene_gn=GALNTL5;refGene_tx=XM_019826049.1;refGene_gn=GALNTL5;refGene_tx=XM_023250760.1
A2	165957458	165957614
refGene_gn=GALNTL5;refGene_tx=XM_011280625.3;refGene_gn=GALNTL5;refGene_tx=XM_011280626.3;refGene_gn=GALNTL5;refGene_tx=XM_011280628.2;refGene_gn=GALNTL5;refGene_tx=XM_019826046.2;refGene_gn=GALNTL5;refGene_tx=XM_019826048.2;refGene_gn=GALNTL5;refGene_tx=XM_019826049.1;refGene_gn=GALNTL5;refGene_tx=XM_023250760.1
A2	165983495	165983790
refGene_gn=GALNT11;refGene_tx=XM_006929523.4;refGene_gn=GALNT11;refGene_tx=XM_006929524.4;refGene_gn=GALNT11;refGene_tx=XM_006929525.4;refGene_gn=GALNT11;refGene_tx=XM_006929526.4;refGene_gn=GALNT11;refGene_tx=XM_019826051.2;refGene_gn=GALNT11;refGene_tx=XM_019826052.2;refGene_gn=GALNT11;refGene_tx=XM_019826053.2
A2	165984827	165984951
refGene_gn=GALNT11;refGene_tx=XM_006929523.4;refGene_gn=GALNT11;refGene_tx=XM_006929524.4;refGene_gn=GALNT11;refGene_tx=XM_006929525.4;refGene_gn=GALNT11;refGene_tx=XM_006929526.4;refGene_gn=GALNT11;refGene_tx=XM_019826051.2;refGene_gn=GALNT11;refGene_tx=XM_019826052.2;refGene_gn=GALNT11;refGene_tx=XM_019826053.2
A2	165986374	165986541
refGene_gn=GALNT11;refGene_tx=XM_006929523.4;refGene_gn=GALNT11;refGene_tx=XM_006929524.4;refGene_gn=GALNT11;refGene_tx=XM_006929525.4;refGene_gn=GALNT11;refGene_tx=XM_006929526.4;refGene_gn=GALNT11;refGene_tx=XM_019826051.2;refGene_gn=GALNT11;refGene_tx=XM_019826052.2;refGene_gn=GALNT11;refGene_tx=XM_019826053.2
A2	165989152	165989278
refGene_gn=GALNT11;refGene_tx=XM_006929523.4;refGene_gn=GALNT11;refGene_tx=XM_006929524.4;refGene_gn=GALNT11;refGene_tx=XM_006929525.4;refGene_gn=GALNT11;refGene_tx=XM_006929526.4;refGene_gn=GALNT11;refGene_tx=XM_019826051.2;refGene_gn=GALNT11;refGene_tx=XM_019826052.2;refGene_gn=GALNT11;refGene_tx=XM_019826053.2
A2	165991191	165991441
refGene_gn=GALNT11;refGene_tx=XM_006929523.4;refGene_gn=GALNT11;refGene_tx=XM_006929524.4;refGene_gn=GALNT11;refGene_tx=XM_006929525.4;refGene_gn=GALNT11;refGene_tx=XM_006929526.4;refGene_gn=GALNT11;refGene_tx=XM_019826051.2;refGene_gn=GALNT11;refGene_tx=XM_019826052.2;refGene_gn=GALNT11;refGene_tx=XM_019826053.2
A2	165996804	165996922

refGene_gn=GALNT11;refGene_tx=XM_006929523.4;refGene_gn=GALNT11;refGene_tx=XM_006929524.4;refGene_gn=GALNT11;refGene_tx=XM_006929525.4;refGene_gn=GALNT11;refGene_tx=XM_006929526.4;refGene_gn=GALNT11;refGene_tx=XM_019826051.2;refGene_gn=GALNT11;refGene_tx=XM_019826052.2;refGene_gn=GALNT11;refGene_tx=XM_019826053.2
A2 165998221 165998374
refGene_gn=GALNT11;refGene_tx=XM_006929523.4;refGene_gn=GALNT11;refGene_tx=XM_006929524.4;refGene_gn=GALNT11;refGene_tx=XM_006929525.4;refGene_gn=GALNT11;refGene_tx=XM_006929526.4;refGene_gn=GALNT11;refGene_tx=XM_019826051.2;refGene_gn=GALNT11;refGene_tx=XM_019826052.2;refGene_gn=GALNT11;refGene_tx=XM_019826053.2
A2 166000721 166000940
refGene_gn=GALNT11;refGene_tx=XM_006929523.4;refGene_gn=GALNT11;refGene_tx=XM_006929524.4;refGene_gn=GALNT11;refGene_tx=XM_006929525.4;refGene_gn=GALNT11;refGene_tx=XM_006929526.4;refGene_gn=GALNT11;refGene_tx=XM_019826051.2;refGene_gn=GALNT11;refGene_tx=XM_019826052.2;refGene_gn=GALNT11;refGene_tx=XM_019826053.2
A2 166002413 166002518
refGene_gn=GALNT11;refGene_tx=XM_006929523.4;refGene_gn=GALNT11;refGene_tx=XM_006929524.4;refGene_gn=GALNT11;refGene_tx=XM_006929525.4;refGene_gn=GALNT11;refGene_tx=XM_006929526.4;refGene_gn=GALNT11;refGene_tx=XM_019826051.2;refGene_gn=GALNT11;refGene_tx=XM_019826052.2;refGene_gn=GALNT11;refGene_tx=XM_019826053.2
A2 166004324 166004462
refGene_gn=GALNT11;refGene_tx=XM_006929523.4;refGene_gn=GALNT11;refGene_tx=XM_006929524.4;refGene_gn=GALNT11;refGene_tx=XM_006929525.4;refGene_gn=GALNT11;refGene_tx=XM_006929526.4;refGene_gn=GALNT11;refGene_tx=XM_019826051.2;refGene_gn=GALNT11;refGene_tx=XM_019826052.2;refGene_gn=GALNT11;refGene_tx=XM_019826053.2
A2 166004675 166004807
refGene_gn=GALNT11;refGene_tx=XM_006929523.4;refGene_gn=GALNT11;refGene_tx=XM_006929524.4;refGene_gn=GALNT11;refGene_tx=XM_006929525.4;refGene_gn=GALNT11;refGene_tx=XM_006929526.4;refGene_gn=GALNT11;refGene_tx=XM_019826051.2;refGene_gn=GALNT11;refGene_tx=XM_019826052.2;refGene_gn=GALNT11;refGene_tx=XM_019826053.2
A2 166022994 166023087
refGene_gn=KMT2C;refGene_tx=XM_023250761.1;refGene_gn=KMT2C;refGene_tx=XM_023250762.1;refGene_gn=KMT2C;refGene_tx=XM_023250763.1;refGene_gn=KMT2C;refGene_tx=XM_023250764.1;refGene_gn=KMT2C;refGene_tx=XM_023250765.1;refGene_gn=KMT2C;refGene_tx=XM_023250766.1;refGene_gn=KMT2C;refGene_tx=XM_023250767.1;refGene_gn=KMT2C;refGene_tx=XM_023250768.1;refGene_gn=KMT2C;refGene_tx=XM_023250769.1;refGene_gn=KMT2C;refGene_tx=XM_023250770.1;refGene_gn=KMT2C;refGene_tx=XM_023250771.1;refGene_gn=KMT2C;refGene_tx=XM_023250772.1;refGene_gn=KMT2C;refGene_tx=XM_023250773.1;refGene_gn=KMT2C;refGene_tx=XM_023250774.1;refGene_gn=KMT2C;refGene_tx=XM_023250775.1
A2 166024752 166024861
refGene_gn=KMT2C;refGene_tx=XM_023250761.1;refGene_gn=KMT2C;refGene_tx=XM_023250762.1;refGene_gn=KMT2C;refGene_tx=XM_023250763.1;refGene_gn=KMT2C;refGene_tx=XM_023250764.1;refGene_gn=KMT2C;refGene_tx=XM_0232507

65.1;refGene_gn=KMT2C;refGene_tx=XM_023250766.1;refGene_gn=KMT2C;refGene_tx=XM_023250767.1;refGene_gn=KMT2C;refGene_tx=XM_023250768.1;refGene_gn=KMT2C;refGene_tx=XM_023250769.1;refGene_gn=KMT2C;refGene_tx=XM_023250770.1;refGene_gn=KMT2C;refGene_tx=XM_023250771.1;refGene_gn=KMT2C;refGene_tx=XM_023250772.1;refGene_gn=KMT2C;refGene_tx=XM_023250773.1;refGene_gn=KMT2C;refGene_tx=XM_023250774.1;refGene_gn=KMT2C;refGene_tx=XM_023250775.1
A2 166025112 166025186
refGene_gn=KMT2C;refGene_tx=XM_023250761.1;refGene_gn=KMT2C;refGene_tx=XM_023250762.1;refGene_gn=KMT2C;refGene_tx=XM_023250763.1;refGene_gn=KMT2C;refGene_tx=XM_023250764.1;refGene_gn=KMT2C;refGene_tx=XM_023250765.1;refGene_gn=KMT2C;refGene_tx=XM_023250766.1;refGene_gn=KMT2C;refGene_tx=XM_023250767.1;refGene_gn=KMT2C;refGene_tx=XM_023250768.1;refGene_gn=KMT2C;refGene_tx=XM_023250769.1;refGene_gn=KMT2C;refGene_tx=XM_023250770.1;refGene_gn=KMT2C;refGene_tx=XM_023250771.1;refGene_gn=KMT2C;refGene_tx=XM_023250772.1;refGene_gn=KMT2C;refGene_tx=XM_023250773.1;refGene_gn=KMT2C;refGene_tx=XM_023250774.1;refGene_gn=KMT2C;refGene_tx=XM_023250775.1
A2 166025545 166025662
refGene_gn=KMT2C;refGene_tx=XM_023250761.1;refGene_gn=KMT2C;refGene_tx=XM_023250762.1;refGene_gn=KMT2C;refGene_tx=XM_023250763.1;refGene_gn=KMT2C;refGene_tx=XM_023250764.1;refGene_gn=KMT2C;refGene_tx=XM_023250765.1;refGene_gn=KMT2C;refGene_tx=XM_023250766.1;refGene_gn=KMT2C;refGene_tx=XM_023250767.1;refGene_gn=KMT2C;refGene_tx=XM_023250768.1;refGene_gn=KMT2C;refGene_tx=XM_023250769.1;refGene_gn=KMT2C;refGene_tx=XM_023250770.1;refGene_gn=KMT2C;refGene_tx=XM_023250771.1;refGene_gn=KMT2C;refGene_tx=XM_023250772.1;refGene_gn=KMT2C;refGene_tx=XM_023250773.1;refGene_gn=KMT2C;refGene_tx=XM_023250774.1;refGene_gn=KMT2C;refGene_tx=XM_023250775.1
A2 166029882 166030051
refGene_gn=KMT2C;refGene_tx=XM_023250761.1;refGene_gn=KMT2C;refGene_tx=XM_023250762.1;refGene_gn=KMT2C;refGene_tx=XM_023250763.1;refGene_gn=KMT2C;refGene_tx=XM_023250764.1;refGene_gn=KMT2C;refGene_tx=XM_023250765.1;refGene_gn=KMT2C;refGene_tx=XM_023250766.1;refGene_gn=KMT2C;refGene_tx=XM_023250767.1;refGene_gn=KMT2C;refGene_tx=XM_023250768.1;refGene_gn=KMT2C;refGene_tx=XM_023250769.1;refGene_gn=KMT2C;refGene_tx=XM_023250770.1;refGene_gn=KMT2C;refGene_tx=XM_023250771.1;refGene_gn=KMT2C;refGene_tx=XM_023250772.1;refGene_gn=KMT2C;refGene_tx=XM_023250773.1;refGene_gn=KMT2C;refGene_tx=XM_023250774.1;refGene_gn=KMT2C;refGene_tx=XM_023250775.1
A2 166030318 166030461
refGene_gn=KMT2C;refGene_tx=XM_023250761.1;refGene_gn=KMT2C;refGene_tx=XM_023250762.1;refGene_gn=KMT2C;refGene_tx=XM_023250763.1;refGene_gn=KMT2C;refGene_tx=XM_023250764.1;refGene_gn=KMT2C;refGene_tx=XM_023250765.1;refGene_gn=KMT2C;refGene_tx=XM_023250766.1;refGene_gn=KMT2C;refGene_tx=XM_023250767.1;refGene_gn=KMT2C;refGene_tx=XM_023250768.1;refGene_gn=KMT2C;refGene_tx=XM_023250769.1;refGene_gn=KMT2C;refGene_tx=XM_023250770.1;refGene_gn=KMT2C;refGene_tx=XM_023250771.1;refGene_gn=KMT2C;refGene_tx=XM_023250772.1;refGene_gn=KMT2C;refGene_tx=XM_023250773.1;refGene_gn=KMT2C;refGene_tx=XM_023250774.1;refGene_gn=KMT2C;refGene_tx=

XM_023250775.1
A2 166031736 166031873
refGene_gn=KMT2C;refGene_tx=XM_023250761.1;refGene_gn=KMT2C;refGene_tx=XM_023250762.1;refGene_gn=KMT2C;refGene_tx=XM_023250763.1;refGene_gn=KMT2C;refGene_tx=XM_023250764.1;refGene_gn=KMT2C;refGene_tx=XM_023250765.1;refGene_gn=KMT2C;refGene_tx=XM_023250766.1;refGene_gn=KMT2C;refGene_tx=XM_023250767.1;refGene_gn=KMT2C;refGene_tx=XM_023250768.1;refGene_gn=KMT2C;refGene_tx=XM_023250769.1;refGene_gn=KMT2C;refGene_tx=XM_023250770.1;refGene_gn=KMT2C;refGene_tx=XM_023250771.1;refGene_gn=KMT2C;refGene_tx=XM_023250772.1;refGene_gn=KMT2C;refGene_tx=XM_023250773.1;refGene_gn=KMT2C;refGene_tx=XM_023250774.1;refGene_gn=KMT2C;refGene_tx=XM_023250775.1
A2 166032794 166033914
refGene_gn=KMT2C;refGene_tx=XM_023250761.1;refGene_gn=KMT2C;refGene_tx=XM_023250762.1;refGene_gn=KMT2C;refGene_tx=XM_023250763.1;refGene_gn=KMT2C;refGene_tx=XM_023250764.1;refGene_gn=KMT2C;refGene_tx=XM_023250765.1;refGene_gn=KMT2C;refGene_tx=XM_023250766.1;refGene_gn=KMT2C;refGene_tx=XM_023250767.1;refGene_gn=KMT2C;refGene_tx=XM_023250768.1;refGene_gn=KMT2C;refGene_tx=XM_023250769.1;refGene_gn=KMT2C;refGene_tx=XM_023250770.1;refGene_gn=KMT2C;refGene_tx=XM_023250771.1;refGene_gn=KMT2C;refGene_tx=XM_023250772.1;refGene_gn=KMT2C;refGene_tx=XM_023250773.1;refGene_gn=KMT2C;refGene_tx=XM_023250774.1;refGene_gn=KMT2C;refGene_tx=XM_023250775.1
A2 166035299 166035407
refGene_gn=KMT2C;refGene_tx=XM_023250761.1;refGene_gn=KMT2C;refGene_tx=XM_023250762.1;refGene_gn=KMT2C;refGene_tx=XM_023250763.1;refGene_gn=KMT2C;refGene_tx=XM_023250764.1;refGene_gn=KMT2C;refGene_tx=XM_023250765.1;refGene_gn=KMT2C;refGene_tx=XM_023250766.1;refGene_gn=KMT2C;refGene_tx=XM_023250767.1;refGene_gn=KMT2C;refGene_tx=XM_023250768.1;refGene_gn=KMT2C;refGene_tx=XM_023250769.1;refGene_gn=KMT2C;refGene_tx=XM_023250770.1;refGene_gn=KMT2C;refGene_tx=XM_023250771.1;refGene_gn=KMT2C;refGene_tx=XM_023250772.1;refGene_gn=KMT2C;refGene_tx=XM_023250773.1;refGene_gn=KMT2C;refGene_tx=XM_023250774.1;refGene_gn=KMT2C;refGene_tx=XM_023250775.1
A2 166035848 166035982
refGene_gn=KMT2C;refGene_tx=XM_023250761.1;refGene_gn=KMT2C;refGene_tx=XM_023250762.1;refGene_gn=KMT2C;refGene_tx=XM_023250763.1;refGene_gn=KMT2C;refGene_tx=XM_023250764.1;refGene_gn=KMT2C;refGene_tx=XM_023250765.1;refGene_gn=KMT2C;refGene_tx=XM_023250766.1;refGene_gn=KMT2C;refGene_tx=XM_023250767.1;refGene_gn=KMT2C;refGene_tx=XM_023250768.1;refGene_gn=KMT2C;refGene_tx=XM_023250769.1;refGene_gn=KMT2C;refGene_tx=XM_023250770.1;refGene_gn=KMT2C;refGene_tx=XM_023250771.1;refGene_gn=KMT2C;refGene_tx=XM_023250772.1;refGene_gn=KMT2C;refGene_tx=XM_023250773.1;refGene_gn=KMT2C;refGene_tx=XM_023250774.1;refGene_gn=KMT2C;refGene_tx=XM_023250775.1
A2 166036994 166037244
refGene_gn=KMT2C;refGene_tx=XM_023250761.1;refGene_gn=KMT2C;refGene_tx=XM_023250762.1;refGene_gn=KMT2C;refGene_tx=XM_023250763.1;refGene_gn=KMT2C;refGene_tx=XM_023250764.1;refGene_gn=KMT2C;refGene_tx=XM_023250765.1;refGene_gn=KMT2C;refGene_tx=XM_023250766.1;refGene_gn=KMT2C;refGe

ne_tx=XM_023250767.1;refGene_gn=KMT2C;refGene_tx=XM_023250768.1;refGen
e_gn=KMT2C;refGene_tx=XM_023250769.1;refGene_gn=KMT2C;refGene_tx=XM_02
3250770.1;refGene_gn=KMT2C;refGene_tx=XM_023250771.1;refGene_gn=KMT2C;
refGene_tx=XM_023250772.1;refGene_gn=KMT2C;refGene_tx=XM_023250773.1;r
efGene_gn=KMT2C;refGene_tx=XM_023250774.1;refGene_gn=KMT2C;refGene_tx=
XM_023250775.1
A2 166038058 166038195
refGene_gn=KMT2C;refGene_tx=XM_023250761.1;refGene_gn=KMT2C;refGene_tx
=XM_023250762.1;refGene_gn=KMT2C;refGene_tx=XM_023250763.1;refGene_gn=
KMT2C;refGene_tx=XM_023250764.1;refGene_gn=KMT2C;refGene_tx=XM_0232507
65.1;refGene_gn=KMT2C;refGene_tx=XM_023250766.1;refGene_gn=KMT2C;refGe
ne_tx=XM_023250767.1;refGene_gn=KMT2C;refGene_tx=XM_023250768.1;refGen
e_gn=KMT2C;refGene_tx=XM_023250769.1;refGene_gn=KMT2C;refGene_tx=XM_02
3250770.1;refGene_gn=KMT2C;refGene_tx=XM_023250771.1;refGene_gn=KMT2C;
refGene_tx=XM_023250772.1;refGene_gn=KMT2C;refGene_tx=XM_023250773.1;r
efGene_gn=KMT2C;refGene_tx=XM_023250774.1;refGene_gn=KMT2C;refGene_tx=
XM_023250775.1
A2 166038375 166038554
refGene_gn=KMT2C;refGene_tx=XM_023250761.1;refGene_gn=KMT2C;refGene_tx
=XM_023250762.1;refGene_gn=KMT2C;refGene_tx=XM_023250763.1;refGene_gn=
KMT2C;refGene_tx=XM_023250764.1;refGene_gn=KMT2C;refGene_tx=XM_0232507
65.1;refGene_gn=KMT2C;refGene_tx=XM_023250766.1;refGene_gn=KMT2C;refGe
ne_tx=XM_023250767.1;refGene_gn=KMT2C;refGene_tx=XM_023250768.1;refGen
e_gn=KMT2C;refGene_tx=XM_023250769.1;refGene_gn=KMT2C;refGene_tx=XM_02
3250770.1;refGene_gn=KMT2C;refGene_tx=XM_023250771.1;refGene_gn=KMT2C;
refGene_tx=XM_023250772.1;refGene_gn=KMT2C;refGene_tx=XM_023250773.1;r
efGene_gn=KMT2C;refGene_tx=XM_023250774.1;refGene_gn=KMT2C;refGene_tx=
XM_023250775.1
A2 166040105 166040253
refGene_gn=KMT2C;refGene_tx=XM_023250761.1;refGene_gn=KMT2C;refGene_tx
=XM_023250762.1;refGene_gn=KMT2C;refGene_tx=XM_023250763.1;refGene_gn=
KMT2C;refGene_tx=XM_023250764.1;refGene_gn=KMT2C;refGene_tx=XM_0232507
65.1;refGene_gn=KMT2C;refGene_tx=XM_023250766.1;refGene_gn=KMT2C;refGe
ne_tx=XM_023250767.1;refGene_gn=KMT2C;refGene_tx=XM_023250768.1;refGen
e_gn=KMT2C;refGene_tx=XM_023250769.1;refGene_gn=KMT2C;refGene_tx=XM_02
3250770.1;refGene_gn=KMT2C;refGene_tx=XM_023250771.1;refGene_gn=KMT2C;
refGene_tx=XM_023250772.1;refGene_gn=KMT2C;refGene_tx=XM_023250773.1;r
efGene_gn=KMT2C;refGene_tx=XM_023250774.1;refGene_gn=KMT2C;refGene_tx=
XM_023250775.1
A2 166040400 166040542
refGene_gn=KMT2C;refGene_tx=XM_023250761.1;refGene_gn=KMT2C;refGene_tx
=XM_023250762.1;refGene_gn=KMT2C;refGene_tx=XM_023250763.1;refGene_gn=
KMT2C;refGene_tx=XM_023250764.1;refGene_gn=KMT2C;refGene_tx=XM_0232507
65.1;refGene_gn=KMT2C;refGene_tx=XM_023250766.1;refGene_gn=KMT2C;refGe
ne_tx=XM_023250767.1;refGene_gn=KMT2C;refGene_tx=XM_023250768.1;refGen
e_gn=KMT2C;refGene_tx=XM_023250769.1;refGene_gn=KMT2C;refGene_tx=XM_02
3250770.1;refGene_gn=KMT2C;refGene_tx=XM_023250771.1;refGene_gn=KMT2C;
refGene_tx=XM_023250772.1;refGene_gn=KMT2C;refGene_tx=XM_023250773.1;r
efGene_gn=KMT2C;refGene_tx=XM_023250774.1;refGene_gn=KMT2C;refGene_tx=
XM_023250775.1

A2 166041970 166042135
refGene_gn=KMT2C;refGene_tx=XM_023250761.1;refGene_gn=KMT2C;refGene_tx=XM_023250762.1;refGene_gn=KMT2C;refGene_tx=XM_023250763.1;refGene_gn=KMT2C;refGene_tx=XM_023250764.1;refGene_gn=KMT2C;refGene_tx=XM_023250765.1;refGene_gn=KMT2C;refGene_tx=XM_023250766.1;refGene_gn=KMT2C;refGene_tx=XM_023250767.1;refGene_gn=KMT2C;refGene_tx=XM_023250768.1;refGene_gn=KMT2C;refGene_tx=XM_023250769.1;refGene_gn=KMT2C;refGene_tx=XM_023250770.1
A2 166042824 166043037
refGene_gn=KMT2C;refGene_tx=XM_023250761.1;refGene_gn=KMT2C;refGene_tx=XM_023250762.1;refGene_gn=KMT2C;refGene_tx=XM_023250763.1;refGene_gn=KMT2C;refGene_tx=XM_023250764.1;refGene_gn=KMT2C;refGene_tx=XM_023250765.1;refGene_gn=KMT2C;refGene_tx=XM_023250766.1;refGene_gn=KMT2C;refGene_tx=XM_023250767.1;refGene_gn=KMT2C;refGene_tx=XM_023250768.1;refGene_gn=KMT2C;refGene_tx=XM_023250769.1;refGene_gn=KMT2C;refGene_tx=XM_023250770.1;refGene_gn=KMT2C;refGene_tx=XM_023250771.1;refGene_gn=KMT2C;refGene_tx=XM_023250772.1;refGene_gn=KMT2C;refGene_tx=XM_023250773.1;refGene_gn=KMT2C;refGene_tx=XM_023250774.1;refGene_gn=KMT2C;refGene_tx=XM_023250775.1
A2 166047451 166049122
refGene_gn=KMT2C;refGene_tx=XM_023250761.1;refGene_gn=KMT2C;refGene_tx=XM_023250762.1;refGene_gn=KMT2C;refGene_tx=XM_023250763.1;refGene_gn=KMT2C;refGene_tx=XM_023250764.1;refGene_gn=KMT2C;refGene_tx=XM_023250765.1;refGene_gn=KMT2C;refGene_tx=XM_023250766.1;refGene_gn=KMT2C;refGene_tx=XM_023250767.1;refGene_gn=KMT2C;refGene_tx=XM_023250768.1;refGene_gn=KMT2C;refGene_tx=XM_023250769.1;refGene_gn=KMT2C;refGene_tx=XM_023250770.1;refGene_gn=KMT2C;refGene_tx=XM_023250771.1;refGene_gn=KMT2C;refGene_tx=XM_023250772.1;refGene_gn=KMT2C;refGene_tx=XM_023250773.1;refGene_gn=KMT2C;refGene_tx=XM_023250774.1;refGene_gn=KMT2C;refGene_tx=XM_023250775.1
A2 166051603 166051836
refGene_gn=KMT2C;refGene_tx=XM_023250761.1;refGene_gn=KMT2C;refGene_tx=XM_023250762.1;refGene_gn=KMT2C;refGene_tx=XM_023250763.1;refGene_gn=KMT2C;refGene_tx=XM_023250764.1;refGene_gn=KMT2C;refGene_tx=XM_023250765.1;refGene_gn=KMT2C;refGene_tx=XM_023250766.1;refGene_gn=KMT2C;refGene_tx=XM_023250767.1;refGene_gn=KMT2C;refGene_tx=XM_023250768.1;refGene_gn=KMT2C;refGene_tx=XM_023250769.1;refGene_gn=KMT2C;refGene_tx=XM_023250770.1;refGene_gn=KMT2C;refGene_tx=XM_023250771.1;refGene_gn=KMT2C;refGene_tx=XM_023250772.1;refGene_gn=KMT2C;refGene_tx=XM_023250773.1;refGene_gn=KMT2C;refGene_tx=XM_023250774.1;refGene_gn=KMT2C;refGene_tx=XM_023250775.1
A2 166053725 166053789
refGene_gn=KMT2C;refGene_tx=XM_023250761.1;refGene_gn=KMT2C;refGene_tx=XM_023250762.1;refGene_gn=KMT2C;refGene_tx=XM_023250763.1;refGene_gn=KMT2C;refGene_tx=XM_023250764.1;refGene_gn=KMT2C;refGene_tx=XM_023250765.1;refGene_gn=KMT2C;refGene_tx=XM_023250766.1;refGene_gn=KMT2C;refGene_tx=XM_023250767.1;refGene_gn=KMT2C;refGene_tx=XM_023250768.1;refGene_gn=KMT2C;refGene_tx=XM_023250769.1;refGene_gn=KMT2C;refGene_tx=XM_023250770.1;refGene_gn=KMT2C;refGene_tx=XM_023250771.1;refGene_gn=KMT2C;refGene_tx=XM_023250772.1;refGene_gn=KMT2C;refGene_tx=XM_023250773.1;r

efGene_gn=KMT2C;refGene_tx=XM_023250774.1;refGene_gn=KMT2C;refGene_tx=
XM_023250775.1
A2 166055363 166055442
refGene_gn=KMT2C;refGene_tx=XM_023250761.1;refGene_gn=KMT2C;refGene_tx
=XM_023250762.1;refGene_gn=KMT2C;refGene_tx=XM_023250763.1;refGene_gn=
KMT2C;refGene_tx=XM_023250764.1;refGene_gn=KMT2C;refGene_tx=XM_0232507
65.1;refGene_gn=KMT2C;refGene_tx=XM_023250766.1;refGene_gn=KMT2C;refGe
ne_tx=XM_023250767.1;refGene_gn=KMT2C;refGene_tx=XM_023250768.1;refGen
e_gn=KMT2C;refGene_tx=XM_023250769.1;refGene_gn=KMT2C;refGene_tx=XM_02
3250770.1;refGene_gn=KMT2C;refGene_tx=XM_023250771.1;refGene_gn=KMT2C;
refGene_tx=XM_023250772.1;refGene_gn=KMT2C;refGene_tx=XM_023250773.1;r
efGene_gn=KMT2C;refGene_tx=XM_023250774.1;refGene_gn=KMT2C;refGene_tx=
XM_023250775.1
A2 166058658 166058770
refGene_gn=KMT2C;refGene_tx=XM_023250761.1;refGene_gn=KMT2C;refGene_tx
=XM_023250762.1;refGene_gn=KMT2C;refGene_tx=XM_023250763.1;refGene_gn=
KMT2C;refGene_tx=XM_023250764.1;refGene_gn=KMT2C;refGene_tx=XM_0232507
65.1;refGene_gn=KMT2C;refGene_tx=XM_023250766.1;refGene_gn=KMT2C;refGe
ne_tx=XM_023250767.1;refGene_gn=KMT2C;refGene_tx=XM_023250768.1;refGen
e_gn=KMT2C;refGene_tx=XM_023250769.1;refGene_gn=KMT2C;refGene_tx=XM_02
3250770.1;refGene_gn=KMT2C;refGene_tx=XM_023250771.1;refGene_gn=KMT2C;
refGene_tx=XM_023250772.1;refGene_gn=KMT2C;refGene_tx=XM_023250773.1;r
efGene_gn=KMT2C;refGene_tx=XM_023250774.1;refGene_gn=KMT2C;refGene_tx=
XM_023250775.1
A2 166059846 166061651
refGene_gn=KMT2C;refGene_tx=XM_023250761.1;refGene_gn=KMT2C;refGene_tx
=XM_023250762.1;refGene_gn=KMT2C;refGene_tx=XM_023250763.1;refGene_gn=
KMT2C;refGene_tx=XM_023250764.1;refGene_gn=KMT2C;refGene_tx=XM_0232507
65.1;refGene_gn=KMT2C;refGene_tx=XM_023250766.1;refGene_gn=KMT2C;refGe
ne_tx=XM_023250767.1;refGene_gn=KMT2C;refGene_tx=XM_023250768.1;refGen
e_gn=KMT2C;refGene_tx=XM_023250769.1;refGene_gn=KMT2C;refGene_tx=XM_02
3250770.1;refGene_gn=KMT2C;refGene_tx=XM_023250771.1;refGene_gn=KMT2C;
refGene_tx=XM_023250772.1;refGene_gn=KMT2C;refGene_tx=XM_023250773.1;r
efGene_gn=KMT2C;refGene_tx=XM_023250774.1;refGene_gn=KMT2C;refGene_tx=
XM_023250775.1
A2 166063217 166063510
refGene_gn=KMT2C;refGene_tx=XM_023250761.1;refGene_gn=KMT2C;refGene_tx
=XM_023250762.1;refGene_gn=KMT2C;refGene_tx=XM_023250763.1;refGene_gn=
KMT2C;refGene_tx=XM_023250764.1;refGene_gn=KMT2C;refGene_tx=XM_0232507
65.1;refGene_gn=KMT2C;refGene_tx=XM_023250766.1;refGene_gn=KMT2C;refGe
ne_tx=XM_023250767.1;refGene_gn=KMT2C;refGene_tx=XM_023250768.1;refGen
e_gn=KMT2C;refGene_tx=XM_023250769.1;refGene_gn=KMT2C;refGene_tx=XM_02
3250770.1;refGene_gn=KMT2C;refGene_tx=XM_023250771.1;refGene_gn=KMT2C;
refGene_tx=XM_023250772.1;refGene_gn=KMT2C;refGene_tx=XM_023250773.1;r
efGene_gn=KMT2C;refGene_tx=XM_023250774.1;refGene_gn=KMT2C;refGene_tx=
XM_023250775.1
A2 166063974 166065834
refGene_gn=KMT2C;refGene_tx=XM_023250761.1;refGene_gn=KMT2C;refGene_tx
=XM_023250762.1;refGene_gn=KMT2C;refGene_tx=XM_023250763.1;refGene_gn=
KMT2C;refGene_tx=XM_023250764.1;refGene_gn=KMT2C;refGene_tx=XM_0232507

65.1;refGene_gn=KMT2C;refGene_tx=XM_023250766.1;refGene_gn=KMT2C;refGene_tx=XM_023250767.1;refGene_gn=KMT2C;refGene_tx=XM_023250768.1;refGene_gn=KMT2C;refGene_tx=XM_023250769.1;refGene_gn=KMT2C;refGene_tx=XM_023250770.1;refGene_gn=KMT2C;refGene_tx=XM_023250771.1;refGene_gn=KMT2C;refGene_tx=XM_023250772.1;refGene_gn=KMT2C;refGene_tx=XM_023250773.1;refGene_gn=KMT2C;refGene_tx=XM_023250774.1;refGene_gn=KMT2C;refGene_tx=XM_023250775.1
A2	166066452	166066635
refGene_gn=KMT2C;refGene_tx=XM_023250761.1;refGene_gn=KMT2C;refGene_tx=XM_023250762.1;refGene_gn=KMT2C;refGene_tx=XM_023250763.1;refGene_gn=KMT2C;refGene_tx=XM_023250764.1;refGene_gn=KMT2C;refGene_tx=XM_023250765.1;refGene_gn=KMT2C;refGene_tx=XM_023250766.1;refGene_gn=KMT2C;refGene_tx=XM_023250767.1;refGene_gn=KMT2C;refGene_tx=XM_023250768.1;refGene_gn=KMT2C;refGene_tx=XM_023250769.1;refGene_gn=KMT2C;refGene_tx=XM_023250770.1;refGene_gn=KMT2C;refGene_tx=XM_023250771.1;refGene_gn=KMT2C;refGene_tx=XM_023250772.1;refGene_gn=KMT2C;refGene_tx=XM_023250773.1;refGene_gn=KMT2C;refGene_tx=XM_023250774.1;refGene_gn=KMT2C;refGene_tx=XM_023250775.1
A2	166067754	166067828
refGene_gn=KMT2C;refGene_tx=XM_023250761.1;refGene_gn=KMT2C;refGene_tx=XM_023250762.1;refGene_gn=KMT2C;refGene_tx=XM_023250763.1;refGene_gn=KMT2C;refGene_tx=XM_023250764.1;refGene_gn=KMT2C;refGene_tx=XM_023250765.1;refGene_gn=KMT2C;refGene_tx=XM_023250766.1;refGene_gn=KMT2C;refGene_tx=XM_023250767.1;refGene_gn=KMT2C;refGene_tx=XM_023250768.1;refGene_gn=KMT2C;refGene_tx=XM_023250769.1;refGene_gn=KMT2C;refGene_tx=XM_023250770.1;refGene_gn=KMT2C;refGene_tx=XM_023250771.1;refGene_gn=KMT2C;refGene_tx=XM_023250772.1;refGene_gn=KMT2C;refGene_tx=XM_023250773.1;refGene_gn=KMT2C;refGene_tx=XM_023250774.1;refGene_gn=KMT2C;refGene_tx=XM_023250775.1
A2	166068850	166069065
refGene_gn=KMT2C;refGene_tx=XM_023250761.1;refGene_gn=KMT2C;refGene_tx=XM_023250762.1;refGene_gn=KMT2C;refGene_tx=XM_023250763.1;refGene_gn=KMT2C;refGene_tx=XM_023250764.1;refGene_gn=KMT2C;refGene_tx=XM_023250765.1;refGene_gn=KMT2C;refGene_tx=XM_023250766.1;refGene_gn=KMT2C;refGene_tx=XM_023250767.1;refGene_gn=KMT2C;refGene_tx=XM_023250768.1;refGene_gn=KMT2C;refGene_tx=XM_023250769.1;refGene_gn=KMT2C;refGene_tx=XM_023250770.1;refGene_gn=KMT2C;refGene_tx=XM_023250771.1;refGene_gn=KMT2C;refGene_tx=XM_023250772.1;refGene_gn=KMT2C;refGene_tx=XM_023250773.1;refGene_gn=KMT2C;refGene_tx=XM_023250774.1;refGene_gn=KMT2C;refGene_tx=XM_023250775.1
A2	166069303	166069436
refGene_gn=KMT2C;refGene_tx=XM_023250761.1;refGene_gn=KMT2C;refGene_tx=XM_023250762.1;refGene_gn=KMT2C;refGene_tx=XM_023250763.1;refGene_gn=KMT2C;refGene_tx=XM_023250764.1;refGene_gn=KMT2C;refGene_tx=XM_023250765.1;refGene_gn=KMT2C;refGene_tx=XM_023250766.1;refGene_gn=KMT2C;refGene_tx=XM_023250767.1;refGene_gn=KMT2C;refGene_tx=XM_023250768.1;refGene_gn=KMT2C;refGene_tx=XM_023250769.1;refGene_gn=KMT2C;refGene_tx=XM_023250770.1;refGene_gn=KMT2C;refGene_tx=XM_023250771.1;refGene_gn=KMT2C;refGene_tx=XM_023250772.1;refGene_gn=KMT2C;refGene_tx=XM_023250773.1;refGene_gn=KMT2C;refGene_tx=XM_023250774.1;refGene_gn=KMT2C;refGene_tx=

XM_023250775.1
A2 166074506 166074656
refGene_gn=KMT2C;refGene_tx=XM_023250761.1;refGene_gn=KMT2C;refGene_tx=XM_023250762.1;refGene_gn=KMT2C;refGene_tx=XM_023250763.1;refGene_gn=KMT2C;refGene_tx=XM_023250764.1;refGene_gn=KMT2C;refGene_tx=XM_023250765.1;refGene_gn=KMT2C;refGene_tx=XM_023250766.1;refGene_gn=KMT2C;refGene_tx=XM_023250767.1;refGene_gn=KMT2C;refGene_tx=XM_023250768.1;refGene_gn=KMT2C;refGene_tx=XM_023250769.1;refGene_gn=KMT2C;refGene_tx=XM_023250770.1;refGene_gn=KMT2C;refGene_tx=XM_023250771.1;refGene_gn=KMT2C;refGene_tx=XM_023250772.1;refGene_gn=KMT2C;refGene_tx=XM_023250773.1;refGene_gn=KMT2C;refGene_tx=XM_023250774.1;refGene_gn=KMT2C;refGene_tx=XM_023250775.1
A2 166074755 166074788
refGene_gn=KMT2C;refGene_tx=XM_023250761.1;refGene_gn=KMT2C;refGene_tx=XM_023250762.1;refGene_gn=KMT2C;refGene_tx=XM_023250763.1;refGene_gn=KMT2C;refGene_tx=XM_023250764.1;refGene_gn=KMT2C;refGene_tx=XM_023250765.1;refGene_gn=KMT2C;refGene_tx=XM_023250766.1;refGene_gn=KMT2C;refGene_tx=XM_023250767.1;refGene_gn=KMT2C;refGene_tx=XM_023250768.1;refGene_gn=KMT2C;refGene_tx=XM_023250769.1;refGene_gn=KMT2C;refGene_tx=XM_023250770.1;refGene_gn=KMT2C;refGene_tx=XM_023250771.1;refGene_gn=KMT2C;refGene_tx=XM_023250772.1;refGene_gn=KMT2C;refGene_tx=XM_023250773.1;refGene_gn=KMT2C;refGene_tx=XM_023250774.1;refGene_gn=KMT2C;refGene_tx=XM_023250775.1
A2 166074970 166075108
refGene_gn=KMT2C;refGene_tx=XM_023250761.1;refGene_gn=KMT2C;refGene_tx=XM_023250762.1;refGene_gn=KMT2C;refGene_tx=XM_023250763.1;refGene_gn=KMT2C;refGene_tx=XM_023250764.1;refGene_gn=KMT2C;refGene_tx=XM_023250765.1;refGene_gn=KMT2C;refGene_tx=XM_023250766.1;refGene_gn=KMT2C;refGene_tx=XM_023250767.1;refGene_gn=KMT2C;refGene_tx=XM_023250768.1;refGene_gn=KMT2C;refGene_tx=XM_023250769.1;refGene_gn=KMT2C;refGene_tx=XM_023250770.1;refGene_gn=KMT2C;refGene_tx=XM_023250771.1;refGene_gn=KMT2C;refGene_tx=XM_023250772.1;refGene_gn=KMT2C;refGene_tx=XM_023250773.1;refGene_gn=KMT2C;refGene_tx=XM_023250774.1;refGene_gn=KMT2C;refGene_tx=XM_023250775.1
A2 166076113 166076161
refGene_gn=KMT2C;refGene_tx=XM_023250761.1;refGene_gn=KMT2C;refGene_tx=XM_023250762.1;refGene_gn=KMT2C;refGene_tx=XM_023250763.1;refGene_gn=KMT2C;refGene_tx=XM_023250764.1;refGene_gn=KMT2C;refGene_tx=XM_023250765.1;refGene_gn=KMT2C;refGene_tx=XM_023250766.1;refGene_gn=KMT2C;refGene_tx=XM_023250768.1;refGene_gn=KMT2C;refGene_tx=XM_023250769.1;refGene_gn=KMT2C;refGene_tx=XM_023250770.1;refGene_gn=KMT2C;refGene_tx=XM_023250771.1;refGene_gn=KMT2C;refGene_tx=XM_023250772.1;refGene_gn=KMT2C;refGene_tx=XM_023250773.1;refGene_gn=KMT2C;refGene_tx=XM_023250774.1;refGene_gn=KMT2C;refGene_tx=XM_023250775.1
A2 166076490 166076595
refGene_gn=KMT2C;refGene_tx=XM_023250761.1;refGene_gn=KMT2C;refGene_tx=XM_023250762.1;refGene_gn=KMT2C;refGene_tx=XM_023250763.1;refGene_gn=KMT2C;refGene_tx=XM_023250764.1;refGene_gn=KMT2C;refGene_tx=XM_023250765.1;refGene_gn=KMT2C;refGene_tx=XM_023250766.1;refGene_gn=KMT2C;refGene_tx=XM_023250767.1;refGene_gn=KMT2C;refGene_tx=XM_023250768.1;refGen

e_gn=KMT2C;refGene_tx=XM_023250769.1;refGene_gn=KMT2C;refGene_tx=XM_023250770.1;refGene_gn=KMT2C;refGene_tx=XM_023250771.1;refGene_gn=KMT2C;refGene_tx=XM_023250772.1;refGene_gn=KMT2C;refGene_tx=XM_023250773.1;refGene_gn=KMT2C;refGene_tx=XM_023250774.1;refGene_gn=KMT2C;refGene_tx=XM_023250775.1
A2 166079822 166080003
refGene_gn=KMT2C;refGene_tx=XM_023250761.1;refGene_gn=KMT2C;refGene_tx=XM_023250762.1;refGene_gn=KMT2C;refGene_tx=XM_023250763.1;refGene_gn=KMT2C;refGene_tx=XM_023250764.1;refGene_gn=KMT2C;refGene_tx=XM_023250765.1;refGene_gn=KMT2C;refGene_tx=XM_023250766.1;refGene_gn=KMT2C;refGene_tx=XM_023250767.1;refGene_gn=KMT2C;refGene_tx=XM_023250768.1;refGene_gn=KMT2C;refGene_tx=XM_023250769.1;refGene_gn=KMT2C;refGene_tx=XM_023250770.1;refGene_gn=KMT2C;refGene_tx=XM_023250771.1;refGene_gn=KMT2C;refGene_tx=XM_023250772.1;refGene_gn=KMT2C;refGene_tx=XM_023250773.1;refGene_gn=KMT2C;refGene_tx=XM_023250774.1;refGene_gn=KMT2C;refGene_tx=XM_023250775.1
A2 166084284 166084415
refGene_gn=KMT2C;refGene_tx=XM_023250761.1;refGene_gn=KMT2C;refGene_tx=XM_023250762.1;refGene_gn=KMT2C;refGene_tx=XM_023250763.1;refGene_gn=KMT2C;refGene_tx=XM_023250764.1;refGene_gn=KMT2C;refGene_tx=XM_023250765.1;refGene_gn=KMT2C;refGene_tx=XM_023250766.1;refGene_gn=KMT2C;refGene_tx=XM_023250767.1;refGene_gn=KMT2C;refGene_tx=XM_023250768.1;refGene_gn=KMT2C;refGene_tx=XM_023250769.1;refGene_gn=KMT2C;refGene_tx=XM_023250770.1;refGene_gn=KMT2C;refGene_tx=XM_023250771.1;refGene_gn=KMT2C;refGene_tx=XM_023250772.1;refGene_gn=KMT2C;refGene_tx=XM_023250773.1;refGene_gn=KMT2C;refGene_tx=XM_023250774.1;refGene_gn=KMT2C;refGene_tx=XM_023250775.1
A2 166087106 166087226
refGene_gn=KMT2C;refGene_tx=XM_023250761.1;refGene_gn=KMT2C;refGene_tx=XM_023250762.1;refGene_gn=KMT2C;refGene_tx=XM_023250763.1;refGene_gn=KMT2C;refGene_tx=XM_023250764.1;refGene_gn=KMT2C;refGene_tx=XM_023250765.1;refGene_gn=KMT2C;refGene_tx=XM_023250766.1;refGene_gn=KMT2C;refGene_tx=XM_023250767.1;refGene_gn=KMT2C;refGene_tx=XM_023250768.1;refGene_gn=KMT2C;refGene_tx=XM_023250769.1;refGene_gn=KMT2C;refGene_tx=XM_023250770.1;refGene_gn=KMT2C;refGene_tx=XM_023250771.1;refGene_gn=KMT2C;refGene_tx=XM_023250772.1;refGene_gn=KMT2C;refGene_tx=XM_023250773.1;refGene_gn=KMT2C;refGene_tx=XM_023250774.1;refGene_gn=KMT2C;refGene_tx=XM_023250775.1
A2 166089551 166089680
refGene_gn=KMT2C;refGene_tx=XM_023250761.1;refGene_gn=KMT2C;refGene_tx=XM_023250762.1;refGene_gn=KMT2C;refGene_tx=XM_023250763.1;refGene_gn=KMT2C;refGene_tx=XM_023250764.1;refGene_gn=KMT2C;refGene_tx=XM_023250765.1;refGene_gn=KMT2C;refGene_tx=XM_023250766.1;refGene_gn=KMT2C;refGene_tx=XM_023250767.1;refGene_gn=KMT2C;refGene_tx=XM_023250768.1;refGene_gn=KMT2C;refGene_tx=XM_023250769.1;refGene_gn=KMT2C;refGene_tx=XM_023250770.1;refGene_gn=KMT2C;refGene_tx=XM_023250771.1;refGene_gn=KMT2C;refGene_tx=XM_023250772.1;refGene_gn=KMT2C;refGene_tx=XM_023250773.1;refGene_gn=KMT2C;refGene_tx=XM_023250774.1;refGene_gn=KMT2C;refGene_tx=XM_023250775.1
A2 166094544 166094757

refGene_gn=KMT2C;refGene_tx=XM_023250761.1;refGene_gn=KMT2C;refGene_tx=XM_023250762.1;refGene_gn=KMT2C;refGene_tx=XM_023250763.1;refGene_gn=KMT2C;refGene_tx=XM_023250764.1;refGene_gn=KMT2C;refGene_tx=XM_023250765.1;refGene_gn=KMT2C;refGene_tx=XM_023250766.1;refGene_gn=KMT2C;refGene_tx=XM_023250767.1;refGene_gn=KMT2C;refGene_tx=XM_023250768.1;refGene_gn=KMT2C;refGene_tx=XM_023250769.1;refGene_gn=KMT2C;refGene_tx=XM_023250770.1;refGene_gn=KMT2C;refGene_tx=XM_023250771.1;refGene_gn=KMT2C;refGene_tx=XM_023250772.1;refGene_gn=KMT2C;refGene_tx=XM_023250773.1;refGene_gn=KMT2C;refGene_tx=XM_023250774.1;refGene_gn=KMT2C;refGene_tx=XM_023250775.1
A2	166095709	166095775
refGene_gn=KMT2C;refGene_tx=XM_023250761.1;refGene_gn=KMT2C;refGene_tx=XM_023250762.1;refGene_gn=KMT2C;refGene_tx=XM_023250763.1;refGene_gn=KMT2C;refGene_tx=XM_023250764.1;refGene_gn=KMT2C;refGene_tx=XM_023250765.1;refGene_gn=KMT2C;refGene_tx=XM_023250766.1;refGene_gn=KMT2C;refGene_tx=XM_023250767.1;refGene_gn=KMT2C;refGene_tx=XM_023250768.1;refGene_gn=KMT2C;refGene_tx=XM_023250769.1;refGene_gn=KMT2C;refGene_tx=XM_023250770.1;refGene_gn=KMT2C;refGene_tx=XM_023250771.1;refGene_gn=KMT2C;refGene_tx=XM_023250772.1;refGene_gn=KMT2C;refGene_tx=XM_023250773.1;refGene_gn=KMT2C;refGene_tx=XM_023250774.1;refGene_gn=KMT2C;refGene_tx=XM_023250775.1
A2	166096250	166096360
refGene_gn=KMT2C;refGene_tx=XM_023250761.1;refGene_gn=KMT2C;refGene_tx=XM_023250762.1;refGene_gn=KMT2C;refGene_tx=XM_023250763.1;refGene_gn=KMT2C;refGene_tx=XM_023250764.1;refGene_gn=KMT2C;refGene_tx=XM_023250765.1;refGene_gn=KMT2C;refGene_tx=XM_023250766.1;refGene_gn=KMT2C;refGene_tx=XM_023250767.1;refGene_gn=KMT2C;refGene_tx=XM_023250768.1;refGene_gn=KMT2C;refGene_tx=XM_023250769.1;refGene_gn=KMT2C;refGene_tx=XM_023250770.1;refGene_gn=KMT2C;refGene_tx=XM_023250771.1;refGene_gn=KMT2C;refGene_tx=XM_023250772.1;refGene_gn=KMT2C;refGene_tx=XM_023250773.1;refGene_gn=KMT2C;refGene_tx=XM_023250774.1;refGene_gn=KMT2C;refGene_tx=XM_023250775.1
A2	166097624	166097789
refGene_gn=KMT2C;refGene_tx=XM_023250761.1;refGene_gn=KMT2C;refGene_tx=XM_023250762.1;refGene_gn=KMT2C;refGene_tx=XM_023250763.1;refGene_gn=KMT2C;refGene_tx=XM_023250764.1;refGene_gn=KMT2C;refGene_tx=XM_023250765.1;refGene_gn=KMT2C;refGene_tx=XM_023250766.1;refGene_gn=KMT2C;refGene_tx=XM_023250767.1;refGene_gn=KMT2C;refGene_tx=XM_023250768.1;refGene_gn=KMT2C;refGene_tx=XM_023250769.1;refGene_gn=KMT2C;refGene_tx=XM_023250770.1;refGene_gn=KMT2C;refGene_tx=XM_023250771.1;refGene_gn=KMT2C;refGene_tx=XM_023250772.1;refGene_gn=KMT2C;refGene_tx=XM_023250773.1;refGene_gn=KMT2C;refGene_tx=XM_023250774.1;refGene_gn=KMT2C;refGene_tx=XM_023250775.1
A2	166098039	166098221
refGene_gn=KMT2C;refGene_tx=XM_023250761.1;refGene_gn=KMT2C;refGene_tx=XM_023250762.1;refGene_gn=KMT2C;refGene_tx=XM_023250763.1;refGene_gn=KMT2C;refGene_tx=XM_023250764.1;refGene_gn=KMT2C;refGene_tx=XM_023250765.1;refGene_gn=KMT2C;refGene_tx=XM_023250766.1;refGene_gn=KMT2C;refGene_tx=XM_023250767.1;refGene_gn=KMT2C;refGene_tx=XM_023250768.1;refGene_gn=KMT2C;refGene_tx=XM_023250769.1;refGene_gn=KMT2C;refGene_tx=XM_02

3250770.1;refGene_gn=KMT2C;refGene_tx=XM_023250771.1;refGene_gn=KMT2C;
refGene_tx=XM_023250772.1;refGene_gn=KMT2C;refGene_tx=XM_023250773.1;r
efGene_gn=KMT2C;refGene_tx=XM_023250774.1;refGene_gn=KMT2C;refGene_tx=
XM_023250775.1
A2 166105381 166105486
refGene_gn=KMT2C;refGene_tx=XM_023250761.1;refGene_gn=KMT2C;refGene_tx
=XM_023250762.1;refGene_gn=KMT2C;refGene_tx=XM_023250763.1;refGene_gn=
KMT2C;refGene_tx=XM_023250764.1;refGene_gn=KMT2C;refGene_tx=XM_0232507
65.1;refGene_gn=KMT2C;refGene_tx=XM_023250766.1;refGene_gn=KMT2C;refGe
ne_tx=XM_023250767.1;refGene_gn=KMT2C;refGene_tx=XM_023250768.1;refGen
e_gn=KMT2C;refGene_tx=XM_023250769.1;refGene_gn=KMT2C;refGene_tx=XM_02
3250770.1;refGene_gn=KMT2C;refGene_tx=XM_023250771.1;refGene_gn=KMT2C;
refGene_tx=XM_023250772.1;refGene_gn=KMT2C;refGene_tx=XM_023250773.1;r
efGene_gn=KMT2C;refGene_tx=XM_023250774.1;refGene_gn=KMT2C;refGene_tx=
XM_023250775.1
A2 166105651 166105753
refGene_gn=KMT2C;refGene_tx=XM_023250761.1;refGene_gn=KMT2C;refGene_tx
=XM_023250762.1;refGene_gn=KMT2C;refGene_tx=XM_023250763.1;refGene_gn=
KMT2C;refGene_tx=XM_023250764.1;refGene_gn=KMT2C;refGene_tx=XM_0232507
65.1;refGene_gn=KMT2C;refGene_tx=XM_023250766.1;refGene_gn=KMT2C;refGe
ne_tx=XM_023250767.1;refGene_gn=KMT2C;refGene_tx=XM_023250768.1;refGen
e_gn=KMT2C;refGene_tx=XM_023250769.1;refGene_gn=KMT2C;refGene_tx=XM_02
3250770.1;refGene_gn=KMT2C;refGene_tx=XM_023250771.1;refGene_gn=KMT2C;
refGene_tx=XM_023250772.1;refGene_gn=KMT2C;refGene_tx=XM_023250773.1;r
efGene_gn=KMT2C;refGene_tx=XM_023250774.1;refGene_gn=KMT2C;refGene_tx=
XM_023250775.1
A2 166114316 166114433
refGene_gn=KMT2C;refGene_tx=XM_023250761.1;refGene_gn=KMT2C;refGene_tx
=XM_023250762.1;refGene_gn=KMT2C;refGene_tx=XM_023250763.1;refGene_gn=
KMT2C;refGene_tx=XM_023250764.1;refGene_gn=KMT2C;refGene_tx=XM_0232507
65.1;refGene_gn=KMT2C;refGene_tx=XM_023250766.1;refGene_gn=KMT2C;refGe
ne_tx=XM_023250767.1;refGene_gn=KMT2C;refGene_tx=XM_023250768.1;refGen
e_gn=KMT2C;refGene_tx=XM_023250769.1;refGene_gn=KMT2C;refGene_tx=XM_02
3250770.1;refGene_gn=KMT2C;refGene_tx=XM_023250771.1;refGene_gn=KMT2C;
refGene_tx=XM_023250772.1;refGene_gn=KMT2C;refGene_tx=XM_023250773.1;r
efGene_gn=KMT2C;refGene_tx=XM_023250774.1;refGene_gn=KMT2C;refGene_tx=
XM_023250775.1
A2 166116671 166116791
refGene_gn=KMT2C;refGene_tx=XM_023250761.1;refGene_gn=KMT2C;refGene_tx
=XM_023250762.1;refGene_gn=KMT2C;refGene_tx=XM_023250763.1;refGene_gn=
KMT2C;refGene_tx=XM_023250764.1;refGene_gn=KMT2C;refGene_tx=XM_0232507
65.1;refGene_gn=KMT2C;refGene_tx=XM_023250766.1;refGene_gn=KMT2C;refGe
ne_tx=XM_023250767.1;refGene_gn=KMT2C;refGene_tx=XM_023250768.1;refGen
e_gn=KMT2C;refGene_tx=XM_023250769.1;refGene_gn=KMT2C;refGene_tx=XM_02
3250771.1;refGene_gn=KMT2C;refGene_tx=XM_023250772.1;refGene_gn=KMT2C;
refGene_tx=XM_023250773.1;refGene_gn=KMT2C;refGene_tx=XM_023250774.1;r
efGene_gn=KMT2C;refGene_tx=XM_023250775.1
A2 166127005 166127718
refGene_gn=KMT2C;refGene_tx=XM_023250761.1;refGene_gn=KMT2C;refGene_tx
=XM_023250762.1;refGene_gn=KMT2C;refGene_tx=XM_023250763.1;refGene_gn=

KMT2C;refGene_tx=XM_023250764.1;refGene_gn=KMT2C;refGene_tx=XM_0232507
65.1;refGene_gn=KMT2C;refGene_tx=XM_023250766.1;refGene_gn=KMT2C;refGe
ne_tx=XM_023250767.1;refGene_gn=KMT2C;refGene_tx=XM_023250768.1;refGen
e_gn=KMT2C;refGene_tx=XM_023250769.1;refGene_gn=KMT2C;refGene_tx=XM_02
3250770.1;refGene_gn=KMT2C;refGene_tx=XM_023250771.1;refGene_gn=KMT2C;
refGene_tx=XM_023250772.1;refGene_gn=KMT2C;refGene_tx=XM_023250773.1;r
efGene_gn=KMT2C;refGene_tx=XM_023250774.1;refGene_gn=KMT2C;refGene_tx=
XM_023250775.1
A2 166130210 166130288
refGene_gn=KMT2C;refGene_tx=XM_023250761.1;refGene_gn=KMT2C;refGene_tx
=XM_023250762.1;refGene_gn=KMT2C;refGene_tx=XM_023250763.1;refGene_gn=
KMT2C;refGene_tx=XM_023250764.1;refGene_gn=KMT2C;refGene_tx=XM_0232507
65.1;refGene_gn=KMT2C;refGene_tx=XM_023250766.1;refGene_gn=KMT2C;refGe
ne_tx=XM_023250767.1;refGene_gn=KMT2C;refGene_tx=XM_023250768.1;refGen
e_gn=KMT2C;refGene_tx=XM_023250769.1;refGene_gn=KMT2C;refGene_tx=XM_02
3250770.1;refGene_gn=KMT2C;refGene_tx=XM_023250771.1;refGene_gn=KMT2C;
refGene_tx=XM_023250772.1;refGene_gn=KMT2C;refGene_tx=XM_023250773.1;r
efGene_gn=KMT2C;refGene_tx=XM_023250774.1;refGene_gn=KMT2C;refGene_tx=
XM_023250775.1
A2 166131063 166131177
refGene_gn=KMT2C;refGene_tx=XM_023250761.1;refGene_gn=KMT2C;refGene_tx
=XM_023250762.1;refGene_gn=KMT2C;refGene_tx=XM_023250763.1;refGene_gn=
KMT2C;refGene_tx=XM_023250764.1;refGene_gn=KMT2C;refGene_tx=XM_0232507
65.1;refGene_gn=KMT2C;refGene_tx=XM_023250766.1;refGene_gn=KMT2C;refGe
ne_tx=XM_023250767.1;refGene_gn=KMT2C;refGene_tx=XM_023250768.1;refGen
e_gn=KMT2C;refGene_tx=XM_023250769.1;refGene_gn=KMT2C;refGene_tx=XM_02
3250770.1;refGene_gn=KMT2C;refGene_tx=XM_023250771.1;refGene_gn=KMT2C;
refGene_tx=XM_023250772.1;refGene_gn=KMT2C;refGene_tx=XM_023250773.1;r
efGene_gn=KMT2C;refGene_tx=XM_023250774.1;refGene_gn=KMT2C;refGene_tx=
XM_023250775.1
A2 166132262 166132414
refGene_gn=KMT2C;refGene_tx=XM_023250761.1;refGene_gn=KMT2C;refGene_tx
=XM_023250762.1;refGene_gn=KMT2C;refGene_tx=XM_023250763.1;refGene_gn=
KMT2C;refGene_tx=XM_023250764.1;refGene_gn=KMT2C;refGene_tx=XM_0232507
65.1;refGene_gn=KMT2C;refGene_tx=XM_023250766.1;refGene_gn=KMT2C;refGe
ne_tx=XM_023250767.1;refGene_gn=KMT2C;refGene_tx=XM_023250768.1;refGen
e_gn=KMT2C;refGene_tx=XM_023250769.1;refGene_gn=KMT2C;refGene_tx=XM_02
3250770.1;refGene_gn=KMT2C;refGene_tx=XM_023250771.1;refGene_gn=KMT2C;
refGene_tx=XM_023250772.1;refGene_gn=KMT2C;refGene_tx=XM_023250773.1;r
efGene_gn=KMT2C;refGene_tx=XM_023250774.1;refGene_gn=KMT2C;refGene_tx=
XM_023250775.1
A2 166132871 166133041
refGene_gn=KMT2C;refGene_tx=XM_023250761.1;refGene_gn=KMT2C;refGene_tx
=XM_023250762.1;refGene_gn=KMT2C;refGene_tx=XM_023250763.1;refGene_gn=
KMT2C;refGene_tx=XM_023250764.1;refGene_gn=KMT2C;refGene_tx=XM_0232507
65.1;refGene_gn=KMT2C;refGene_tx=XM_023250766.1;refGene_gn=KMT2C;refGe
ne_tx=XM_023250767.1;refGene_gn=KMT2C;refGene_tx=XM_023250768.1;refGen
e_gn=KMT2C;refGene_tx=XM_023250769.1;refGene_gn=KMT2C;refGene_tx=XM_02
3250770.1;refGene_gn=KMT2C;refGene_tx=XM_023250771.1;refGene_gn=KMT2C;
refGene_tx=XM_023250772.1;refGene_gn=KMT2C;refGene_tx=XM_023250773.1;r

efGene_gn=KMT2C;refGene_tx=XM_023250774.1;refGene_gn=KMT2C;refGene_tx=XM_023250775.1
A2 166141924 166142039
refGene_gn=KMT2C;refGene_tx=XM_023250761.1;refGene_gn=KMT2C;refGene_tx=XM_023250762.1;refGene_gn=KMT2C;refGene_tx=XM_023250763.1;refGene_gn=KMT2C;refGene_tx=XM_023250764.1;refGene_gn=KMT2C;refGene_tx=XM_023250765.1;refGene_gn=KMT2C;refGene_tx=XM_023250766.1;refGene_gn=KMT2C;refGene_tx=XM_023250767.1;refGene_gn=KMT2C;refGene_tx=XM_023250768.1;refGene_gn=KMT2C;refGene_tx=XM_023250769.1;refGene_gn=KMT2C;refGene_tx=XM_023250770.1;refGene_gn=KMT2C;refGene_tx=XM_023250771.1;refGene_gn=KMT2C;refGene_tx=XM_023250772.1;refGene_gn=KMT2C;refGene_tx=XM_023250773.1;refGene_gn=KMT2C;refGene_tx=XM_023250774.1;refGene_gn=KMT2C;refGene_tx=XM_023250775.1
A2 166143968 166144140
refGene_gn=KMT2C;refGene_tx=XM_023250761.1;refGene_gn=KMT2C;refGene_tx=XM_023250762.1;refGene_gn=KMT2C;refGene_tx=XM_023250763.1;refGene_gn=KMT2C;refGene_tx=XM_023250764.1;refGene_gn=KMT2C;refGene_tx=XM_023250765.1;refGene_gn=KMT2C;refGene_tx=XM_023250766.1;refGene_gn=KMT2C;refGene_tx=XM_023250767.1;refGene_gn=KMT2C;refGene_tx=XM_023250768.1;refGene_gn=KMT2C;refGene_tx=XM_023250769.1;refGene_gn=KMT2C;refGene_tx=XM_023250770.1;refGene_gn=KMT2C;refGene_tx=XM_023250771.1;refGene_gn=KMT2C;refGene_tx=XM_023250772.1;refGene_gn=KMT2C;refGene_tx=XM_023250773.1;refGene_gn=KMT2C;refGene_tx=XM_023250774.1;refGene_gn=KMT2C;refGene_tx=XM_023250775.1
A2 166151356 166151519
refGene_gn=KMT2C;refGene_tx=XM_023250761.1;refGene_gn=KMT2C;refGene_tx=XM_023250762.1;refGene_gn=KMT2C;refGene_tx=XM_023250763.1;refGene_gn=KMT2C;refGene_tx=XM_023250764.1;refGene_gn=KMT2C;refGene_tx=XM_023250765.1;refGene_gn=KMT2C;refGene_tx=XM_023250766.1;refGene_gn=KMT2C;refGene_tx=XM_023250767.1;refGene_gn=KMT2C;refGene_tx=XM_023250768.1;refGene_gn=KMT2C;refGene_tx=XM_023250769.1;refGene_gn=KMT2C;refGene_tx=XM_023250770.1;refGene_gn=KMT2C;refGene_tx=XM_023250771.1;refGene_gn=KMT2C;refGene_tx=XM_023250772.1;refGene_gn=KMT2C;refGene_tx=XM_023250773.1;refGene_gn=KMT2C;refGene_tx=XM_023250774.1;refGene_gn=KMT2C;refGene_tx=XM_023250775.1
A2 166184983 166185093
refGene_gn=KMT2C;refGene_tx=XM_023250761.1;refGene_gn=KMT2C;refGene_tx=XM_023250762.1;refGene_gn=KMT2C;refGene_tx=XM_023250763.1;refGene_gn=KMT2C;refGene_tx=XM_023250764.1;refGene_gn=KMT2C;refGene_tx=XM_023250765.1;refGene_gn=KMT2C;refGene_tx=XM_023250766.1;refGene_gn=KMT2C;refGene_tx=XM_023250767.1;refGene_gn=KMT2C;refGene_tx=XM_023250768.1;refGene_gn=KMT2C;refGene_tx=XM_023250769.1;refGene_gn=KMT2C;refGene_tx=XM_023250770.1;refGene_gn=KMT2C;refGene_tx=XM_023250771.1;refGene_gn=KMT2C;refGene_tx=XM_023250772.1;refGene_gn=KMT2C;refGene_tx=XM_023250773.1;refGene_gn=KMT2C;refGene_tx=XM_023250774.1;refGene_gn=KMT2C;refGene_tx=XM_023250775.1
A2 166186123 166186272
refGene_gn=KMT2C;refGene_tx=XM_023250761.1;refGene_gn=KMT2C;refGene_tx=XM_023250762.1;refGene_gn=KMT2C;refGene_tx=XM_023250763.1;refGene_gn=KMT2C;refGene_tx=XM_023250764.1;refGene_gn=KMT2C;refGene_tx=XM_0232507

65.1;refGene_gn=KMT2C;refGene_tx=XM_023250766.1;refGene_gn=KMT2C;refGene_tx=XM_023250767.1;refGene_gn=KMT2C;refGene_tx=XM_023250768.1;refGene_gn=KMT2C;refGene_tx=XM_023250769.1;refGene_gn=KMT2C;refGene_tx=XM_023250770.1;refGene_gn=KMT2C;refGene_tx=XM_023250771.1;refGene_gn=KMT2C;refGene_tx=XM_023250772.1;refGene_gn=KMT2C;refGene_tx=XM_023250773.1;refGene_gn=KMT2C;refGene_tx=XM_023250774.1;refGene_gn=KMT2C;refGene_tx=XM_023250775.1
A2 166189539 166189740
refGene_gn=KMT2C;refGene_tx=XM_023250761.1;refGene_gn=KMT2C;refGene_tx=XM_023250762.1;refGene_gn=KMT2C;refGene_tx=XM_023250763.1;refGene_gn=KMT2C;refGene_tx=XM_023250764.1;refGene_gn=KMT2C;refGene_tx=XM_023250765.1;refGene_gn=KMT2C;refGene_tx=XM_023250766.1;refGene_gn=KMT2C;refGene_tx=XM_023250767.1;refGene_gn=KMT2C;refGene_tx=XM_023250768.1;refGene_gn=KMT2C;refGene_tx=XM_023250769.1;refGene_gn=KMT2C;refGene_tx=XM_023250770.1;refGene_gn=KMT2C;refGene_tx=XM_023250771.1;refGene_gn=KMT2C;refGene_tx=XM_023250772.1;refGene_gn=KMT2C;refGene_tx=XM_023250773.1;refGene_gn=KMT2C;refGene_tx=XM_023250774.1;refGene_gn=KMT2C;refGene_tx=XM_023250775.1
A2 166196304 166196446
refGene_gn=KMT2C;refGene_tx=XM_023250761.1;refGene_gn=KMT2C;refGene_tx=XM_023250762.1;refGene_gn=KMT2C;refGene_tx=XM_023250763.1;refGene_gn=KMT2C;refGene_tx=XM_023250764.1;refGene_gn=KMT2C;refGene_tx=XM_023250765.1;refGene_gn=KMT2C;refGene_tx=XM_023250766.1;refGene_gn=KMT2C;refGene_tx=XM_023250767.1;refGene_gn=KMT2C;refGene_tx=XM_023250768.1;refGene_gn=KMT2C;refGene_tx=XM_023250769.1;refGene_gn=KMT2C;refGene_tx=XM_023250770.1;refGene_gn=KMT2C;refGene_tx=XM_023250771.1;refGene_gn=KMT2C;refGene_tx=XM_023250772.1;refGene_gn=KMT2C;refGene_tx=XM_023250773.1;refGene_gn=KMT2C;refGene_tx=XM_023250774.1;refGene_gn=KMT2C;refGene_tx=XM_023250775.1
A2 166229823 166229912
refGene_gn=KMT2C;refGene_tx=XM_023250761.1;refGene_gn=KMT2C;refGene_tx=XM_023250762.1;refGene_gn=KMT2C;refGene_tx=XM_023250763.1;refGene_gn=KMT2C;refGene_tx=XM_023250764.1;refGene_gn=KMT2C;refGene_tx=XM_023250765.1;refGene_gn=KMT2C;refGene_tx=XM_023250766.1;refGene_gn=KMT2C;refGene_tx=XM_023250767.1;refGene_gn=KMT2C;refGene_tx=XM_023250768.1;refGene_gn=KMT2C;refGene_tx=XM_023250769.1;refGene_gn=KMT2C;refGene_tx=XM_023250770.1;refGene_gn=KMT2C;refGene_tx=XM_023250771.1;refGene_gn=KMT2C;refGene_tx=XM_023250772.1;refGene_gn=KMT2C;refGene_tx=XM_023250773.1;refGene_gn=KMT2C;refGene_tx=XM_023250774.1;refGene_gn=KMT2C;refGene_tx=XM_023250775.1
A2 166307034 166307147
refGene_gn=KMT2C;refGene_tx=XM_023250768.1
A2 166308885 166309046
refGene_gn=KMT2C;refGene_tx=XM_023250761.1;refGene_gn=KMT2C;refGene_tx=XM_023250762.1;refGene_gn=KMT2C;refGene_tx=XM_023250763.1;refGene_gn=KMT2C;refGene_tx=XM_023250764.1;refGene_gn=KMT2C;refGene_tx=XM_023250765.1;refGene_gn=KMT2C;refGene_tx=XM_023250766.1;refGene_gn=KMT2C;refGene_tx=XM_023250767.1;refGene_gn=KMT2C;refGene_tx=XM_023250769.1;refGene_gn=KMT2C;refGene_tx=XM_023250770.1;refGene_gn=KMT2C;refGene_tx=XM_023250771.1;refGene_gn=KMT2C;refGene_tx=XM_023250772.1;refGene_gn=KMT2C;

refGene_tx=XM_023250773.1;refGene_gn=KMT2C;refGene_tx=XM_023250774.1;refGene_gn=KMT2C;refGene_tx=XM_023250775.1
A2	166317677	166319255
refGene_gn=CCT8L2;refGene_tx=XM_006944157.4
A2	166465470	166466180
refGene_gn=XRCC2;refGene_tx=XM_011280622.3
A2	166475967	166476049
refGene_gn=XRCC2;refGene_tx=XM_011280622.3
A2	166489826	166489865
refGene_gn=XRCC2;refGene_tx=XM_011280622.3
A2	166546306	166546376
refGene_gn=ACTR3B;refGene_tx=XM_023250779.1
A2	166547274	166547318
refGene_gn=ACTR3B;refGene_tx=XM_023250777.1
A2	166548785	166548822
refGene_gn=ACTR3B;refGene_tx=XM_023250778.1;refGene_gn=ACTR3B;refGene_tx=XM_023250780.1
A2	166566944	166567000
refGene_gn=ACTR3B;refGene_tx=XM_023250777.1;refGene_gn=ACTR3B;refGene_tx=XM_023250778.1
A2	166567418	166567605
refGene_gn=ACTR3B;refGene_tx=XM_023250776.1
A2	166581063	166581188
refGene_gn=ACTR3B;refGene_tx=XM_023250776.1;refGene_gn=ACTR3B;refGene_tx=XM_023250777.1;refGene_gn=ACTR3B;refGene_tx=XM_023250778.1;refGene_gn=ACTR3B;refGene_tx=XM_023250779.1;refGene_gn=ACTR3B;refGene_tx=XM_023250780.1
A2	166582069	166582180
refGene_gn=ACTR3B;refGene_tx=XM_023250776.1;refGene_gn=ACTR3B;refGene_tx=XM_023250777.1;refGene_gn=ACTR3B;refGene_tx=XM_023250778.1;refGene_gn=ACTR3B;refGene_tx=XM_023250779.1;refGene_gn=ACTR3B;refGene_tx=XM_023250780.1;refGene_gn=ACTR3B;refGene_tx=XM_023250782.1;refGene_gn=ACTR3B;refGene_tx=XM_023250784.1
A2	166593263	166593359
refGene_gn=ACTR3B;refGene_tx=XM_023250776.1;refGene_gn=ACTR3B;refGene_tx=XM_023250777.1;refGene_gn=ACTR3B;refGene_tx=XM_023250778.1;refGene_gn=ACTR3B;refGene_tx=XM_023250779.1;refGene_gn=ACTR3B;refGene_tx=XM_023250780.1;refGene_gn=ACTR3B;refGene_tx=XM_023250782.1;refGene_gn=ACTR3B;refGene_tx=XM_023250783.1;refGene_gn=ACTR3B;refGene_tx=XM_023250784.1
A2	166595093	166595201
refGene_gn=ACTR3B;refGene_tx=XM_023250776.1;refGene_gn=ACTR3B;refGene_tx=XM_023250777.1;refGene_gn=ACTR3B;refGene_tx=XM_023250778.1;refGene_gn=ACTR3B;refGene_tx=XM_023250779.1;refGene_gn=ACTR3B;refGene_tx=XM_023250780.1;refGene_gn=ACTR3B;refGene_tx=XM_023250782.1;refGene_gn=ACTR3B;refGene_tx=XM_023250783.1;refGene_gn=ACTR3B;refGene_tx=XM_023250784.1
A2	166597869	166598013
refGene_gn=ACTR3B;refGene_tx=XM_023250776.1;refGene_gn=ACTR3B;refGene_tx=XM_023250777.1;refGene_gn=ACTR3B;refGene_tx=XM_023250778.1;refGene_

gn=ACTR3B;refGene_tx=XM_023250779.1;refGene_gn=ACTR3B;refGene_tx=XM_023250780.1;refGene_gn=ACTR3B;refGene_tx=XM_023250782.1;refGene_gn=ACTR3B;refGene_tx=XM_023250783.1;refGene_gn=ACTR3B;refGene_tx=XM_023250784.1
A2	166600929	166601103
refGene_gn=ACTR3B;refGene_tx=XM_023250776.1;refGene_gn=ACTR3B;refGene_tx=XM_023250777.1;refGene_gn=ACTR3B;refGene_tx=XM_023250778.1;refGene_gn=ACTR3B;refGene_tx=XM_023250779.1;refGene_gn=ACTR3B;refGene_tx=XM_023250780.1;refGene_gn=ACTR3B;refGene_tx=XM_023250782.1;refGene_gn=ACTR3B;refGene_tx=XM_023250783.1;refGene_gn=ACTR3B;refGene_tx=XM_023250784.1
A2	166602660	166602753
refGene_gn=ACTR3B;refGene_tx=XM_023250776.1;refGene_gn=ACTR3B;refGene_tx=XM_023250777.1;refGene_gn=ACTR3B;refGene_tx=XM_023250778.1;refGene_gn=ACTR3B;refGene_tx=XM_023250779.1;refGene_gn=ACTR3B;refGene_tx=XM_023250780.1;refGene_gn=ACTR3B;refGene_tx=XM_023250782.1;refGene_gn=ACTR3B;refGene_tx=XM_023250783.1;refGene_gn=ACTR3B;refGene_tx=XM_023250784.1
A2	166634637	166634763
refGene_gn=ACTR3B;refGene_tx=XM_023250776.1;refGene_gn=ACTR3B;refGene_tx=XM_023250777.1;refGene_gn=ACTR3B;refGene_tx=XM_023250778.1;refGene_gn=ACTR3B;refGene_tx=XM_023250779.1;refGene_gn=ACTR3B;refGene_tx=XM_023250780.1;refGene_gn=ACTR3B;refGene_tx=XM_023250782.1;refGene_gn=ACTR3B;refGene_tx=XM_023250783.1;refGene_gn=ACTR3B;refGene_tx=XM_023250784.1
A2	166635663	166635747
refGene_gn=ACTR3B;refGene_tx=XM_023250776.1;refGene_gn=ACTR3B;refGene_tx=XM_023250777.1;refGene_gn=ACTR3B;refGene_tx=XM_023250778.1;refGene_gn=ACTR3B;refGene_tx=XM_023250779.1;refGene_gn=ACTR3B;refGene_tx=XM_023250780.1;refGene_gn=ACTR3B;refGene_tx=XM_023250782.1;refGene_gn=ACTR3B;refGene_tx=XM_023250783.1;refGene_gn=ACTR3B;refGene_tx=XM_023250784.1
A2	166636225	166636321
refGene_gn=ACTR3B;refGene_tx=XM_023250776.1;refGene_gn=ACTR3B;refGene_tx=XM_023250777.1;refGene_gn=ACTR3B;refGene_tx=XM_023250778.1;refGene_gn=ACTR3B;refGene_tx=XM_023250779.1;refGene_gn=ACTR3B;refGene_tx=XM_023250780.1;refGene_gn=ACTR3B;refGene_tx=XM_023250782.1;refGene_gn=ACTR3B;refGene_tx=XM_023250783.1;refGene_gn=ACTR3B;refGene_tx=XM_023250784.1
A2	167237299	167237359
refGene_gn=DPP6;refGene_tx=XM_023250893.1
A2	167327314	167327365
refGene_gn=DPP6;refGene_tx=XM_023250891.1
A2	167475482	167475719
refGene_gn=DPP6;refGene_tx=XM_023250889.1;refGene_gn=DPP6;refGene_tx=XM_023250895.1
A2	167590784	167590835
refGene_gn=DPP6;refGene_tx=XM_023250892.1
A2	167696030	167696087
refGene_gn=DPP6;refGene_tx=XM_023250890.1

A2 167820700 167820815
refGene_gn=DPP6;refGene_tx=XM_023250889.1;refGene_gn=DPP6;refGene_tx=XM_023250890.1;refGene_gn=DPP6;refGene_tx=XM_023250891.1;refGene_gn=DPP6;refGene_tx=XM_023250892.1;refGene_gn=DPP6;refGene_tx=XM_023250893.1;refGene_gn=DPP6;refGene_tx=XM_023250895.1
A2 167843835 167843934
refGene_gn=DPP6;refGene_tx=XM_023250889.1;refGene_gn=DPP6;refGene_tx=XM_023250890.1;refGene_gn=DPP6;refGene_tx=XM_023250891.1;refGene_gn=DPP6;refGene_tx=XM_023250892.1;refGene_gn=DPP6;refGene_tx=XM_023250893.1;refGene_gn=DPP6;refGene_tx=XM_023250895.1
A2 167900792 167900887
refGene_gn=DPP6;refGene_tx=XM_023250889.1;refGene_gn=DPP6;refGene_tx=XM_023250890.1;refGene_gn=DPP6;refGene_tx=XM_023250891.1;refGene_gn=DPP6;refGene_tx=XM_023250892.1;refGene_gn=DPP6;refGene_tx=XM_023250893.1;refGene_gn=DPP6;refGene_tx=XM_023250895.1
A2 167913797 167913872
refGene_gn=DPP6;refGene_tx=XM_023250889.1;refGene_gn=DPP6;refGene_tx=XM_023250890.1;refGene_gn=DPP6;refGene_tx=XM_023250891.1;refGene_gn=DPP6;refGene_tx=XM_023250892.1;refGene_gn=DPP6;refGene_tx=XM_023250893.1;refGene_gn=DPP6;refGene_tx=XM_023250895.1
A2 167980912 167980965
refGene_gn=DPP6;refGene_tx=XM_023250889.1;refGene_gn=DPP6;refGene_tx=XM_023250890.1;refGene_gn=DPP6;refGene_tx=XM_023250891.1;refGene_gn=DPP6;refGene_tx=XM_023250892.1;refGene_gn=DPP6;refGene_tx=XM_023250893.1;refGene_gn=DPP6;refGene_tx=XM_023250895.1
A2 168020842 168020924
refGene_gn=DPP6;refGene_tx=XM_023250889.1;refGene_gn=DPP6;refGene_tx=XM_023250890.1;refGene_gn=DPP6;refGene_tx=XM_023250891.1;refGene_gn=DPP6;refGene_tx=XM_023250892.1;refGene_gn=DPP6;refGene_tx=XM_023250893.1;refGene_gn=DPP6;refGene_tx=XM_023250895.1
A2 168062970 168063091
refGene_gn=DPP6;refGene_tx=XM_023250889.1;refGene_gn=DPP6;refGene_tx=XM_023250890.1;refGene_gn=DPP6;refGene_tx=XM_023250891.1;refGene_gn=DPP6;refGene_tx=XM_023250892.1;refGene_gn=DPP6;refGene_tx=XM_023250893.1;refGene_gn=DPP6;refGene_tx=XM_023250895.1
A2 168096999 168097154
refGene_gn=DPP6;refGene_tx=XM_023250889.1;refGene_gn=DPP6;refGene_tx=XM_023250890.1;refGene_gn=DPP6;refGene_tx=XM_023250891.1;refGene_gn=DPP6;refGene_tx=XM_023250892.1;refGene_gn=DPP6;refGene_tx=XM_023250893.1;refGene_gn=DPP6;refGene_tx=XM_023250895.1
A2 168099799 168099897
refGene_gn=DPP6;refGene_tx=XM_023250889.1;refGene_gn=DPP6;refGene_tx=XM_023250890.1;refGene_gn=DPP6;refGene_tx=XM_023250891.1;refGene_gn=DPP6;refGene_tx=XM_023250892.1;refGene_gn=DPP6;refGene_tx=XM_023250893.1;refGene_gn=DPP6;refGene_tx=XM_023250895.1
A2 168119534 168119658
refGene_gn=DPP6;refGene_tx=XM_023250889.1;refGene_gn=DPP6;refGene_tx=XM_023250890.1;refGene_gn=DPP6;refGene_tx=XM_023250891.1;refGene_gn=DPP6;refGene_tx=XM_023250892.1;refGene_gn=DPP6;refGene_tx=XM_023250893.1;refGene_gn=DPP6;refGene_tx=XM_023250895.1

A2 168121225 168121264
refGene_gn=DPP6;refGene_tx=XM_023250889.1;refGene_gn=DPP6;refGene_tx=XM_023250890.1;refGene_gn=DPP6;refGene_tx=XM_023250891.1;refGene_gn=DPP6;refGene_tx=XM_023250892.1;refGene_gn=DPP6;refGene_tx=XM_023250893.1;refGene_gn=DPP6;refGene_tx=XM_023250895.1
A2 168125579 168125687
refGene_gn=DPP6;refGene_tx=XM_023250889.1;refGene_gn=DPP6;refGene_tx=XM_023250890.1;refGene_gn=DPP6;refGene_tx=XM_023250891.1;refGene_gn=DPP6;refGene_tx=XM_023250892.1;refGene_gn=DPP6;refGene_tx=XM_023250893.1;refGene_gn=DPP6;refGene_tx=XM_023250895.1
A2 168127943 168128035
refGene_gn=DPP6;refGene_tx=XM_023250889.1;refGene_gn=DPP6;refGene_tx=XM_023250890.1;refGene_gn=DPP6;refGene_tx=XM_023250891.1;refGene_gn=DPP6;refGene_tx=XM_023250892.1;refGene_gn=DPP6;refGene_tx=XM_023250893.1;refGene_gn=DPP6;refGene_tx=XM_023250895.1
A2 168128810 168128858
refGene_gn=DPP6;refGene_tx=XM_023250889.1;refGene_gn=DPP6;refGene_tx=XM_023250890.1;refGene_gn=DPP6;refGene_tx=XM_023250891.1;refGene_gn=DPP6;refGene_tx=XM_023250892.1;refGene_gn=DPP6;refGene_tx=XM_023250893.1;refGene_gn=DPP6;refGene_tx=XM_023250895.1
A2 168130605 168130724
refGene_gn=DPP6;refGene_tx=XM_023250889.1;refGene_gn=DPP6;refGene_tx=XM_023250890.1;refGene_gn=DPP6;refGene_tx=XM_023250891.1;refGene_gn=DPP6;refGene_tx=XM_023250892.1;refGene_gn=DPP6;refGene_tx=XM_023250893.1;refGene_gn=DPP6;refGene_tx=XM_023250895.1
A2 168160436 168160484
refGene_gn=DPP6;refGene_tx=XM_023250889.1;refGene_gn=DPP6;refGene_tx=XM_023250890.1;refGene_gn=DPP6;refGene_tx=XM_023250891.1;refGene_gn=DPP6;refGene_tx=XM_023250892.1;refGene_gn=DPP6;refGene_tx=XM_023250893.1;refGene_gn=DPP6;refGene_tx=XM_023250895.1
A2 168172990 168173089
refGene_gn=DPP6;refGene_tx=XM_023250889.1;refGene_gn=DPP6;refGene_tx=XM_023250890.1;refGene_gn=DPP6;refGene_tx=XM_023250891.1;refGene_gn=DPP6;refGene_tx=XM_023250892.1;refGene_gn=DPP6;refGene_tx=XM_023250893.1;refGene_gn=DPP6;refGene_tx=XM_023250895.1
A2 168177766 168177836
refGene_gn=DPP6;refGene_tx=XM_023250889.1;refGene_gn=DPP6;refGene_tx=XM_023250890.1;refGene_gn=DPP6;refGene_tx=XM_023250891.1;refGene_gn=DPP6;refGene_tx=XM_023250892.1;refGene_gn=DPP6;refGene_tx=XM_023250893.1;refGene_gn=DPP6;refGene_tx=XM_023250895.1
A2 168179068 168179263
refGene_gn=DPP6;refGene_tx=XM_023250889.1;refGene_gn=DPP6;refGene_tx=XM_023250890.1;refGene_gn=DPP6;refGene_tx=XM_023250891.1;refGene_gn=DPP6;refGene_tx=XM_023250892.1;refGene_gn=DPP6;refGene_tx=XM_023250893.1;refGene_gn=DPP6;refGene_tx=XM_023250895.1
A2 168185581 168185636
refGene_gn=DPP6;refGene_tx=XM_023250889.1;refGene_gn=DPP6;refGene_tx=XM_023250890.1;refGene_gn=DPP6;refGene_tx=XM_023250891.1;refGene_gn=DPP6;refGene_tx=XM_023250892.1;refGene_gn=DPP6;refGene_tx=XM_023250893.1;refGene_gn=DPP6;refGene_tx=XM_023250895.1

A2	168188215	168188327
refGene_gn=DPP6;refGene_tx=XM_023250889.1;refGene_gn=DPP6;refGene_tx=XM_023250890.1;refGene_gn=DPP6;refGene_tx=XM_023250891.1;refGene_gn=DPP6;refGene_tx=XM_023250892.1;refGene_gn=DPP6;refGene_tx=XM_023250893.1;refGene_gn=DPP6;refGene_tx=XM_023250895.1
A2	168189698	168189757
refGene_gn=DPP6;refGene_tx=XM_023250889.1;refGene_gn=DPP6;refGene_tx=XM_023250890.1;refGene_gn=DPP6;refGene_tx=XM_023250891.1;refGene_gn=DPP6;refGene_tx=XM_023250892.1;refGene_gn=DPP6;refGene_tx=XM_023250893.1;refGene_gn=DPP6;refGene_tx=XM_023250895.1
A2	168190559	168190632
refGene_gn=DPP6;refGene_tx=XM_023250889.1;refGene_gn=DPP6;refGene_tx=XM_023250890.1;refGene_gn=DPP6;refGene_tx=XM_023250891.1;refGene_gn=DPP6;refGene_tx=XM_023250892.1;refGene_gn=DPP6;refGene_tx=XM_023250893.1;refGene_gn=DPP6;refGene_tx=XM_023250895.1
A2	168190718	168190795
refGene_gn=DPP6;refGene_tx=XM_023250889.1;refGene_gn=DPP6;refGene_tx=XM_023250890.1;refGene_gn=DPP6;refGene_tx=XM_023250891.1;refGene_gn=DPP6;refGene_tx=XM_023250892.1;refGene_gn=DPP6;refGene_tx=XM_023250893.1;refGene_gn=DPP6;refGene_tx=XM_023250895.1
A2	168192835	168192982
refGene_gn=DPP6;refGene_tx=XM_023250895.1
A2	168223183	168223199
refGene_gn=PAXIP1;refGene_tx=XM_023242347.1
A2	168223882	168223943
refGene_gn=PAXIP1;refGene_tx=XM_023242347.1
A2	168224020	168224096
refGene_gn=PAXIP1;refGene_tx=XM_023242347.1
A2	168224183	168224318
refGene_gn=PAXIP1;refGene_tx=XM_023242347.1
A2	168225345	168225446
refGene_gn=PAXIP1;refGene_tx=XM_023242347.1
A2	168231568	168231737
refGene_gn=PAXIP1;refGene_tx=XM_023242347.1
A2	168232719	168232822
refGene_gn=PAXIP1;refGene_tx=XM_023242347.1
A2	168234330	168234401
refGene_gn=PAXIP1;refGene_tx=XM_023242347.1
A2	168236130	168236174
refGene_gn=PAXIP1;refGene_tx=XM_023242347.1
A2	168237180	168237365
refGene_gn=PAXIP1;refGene_tx=XM_023242347.1
A2	168237805	168237927
refGene_gn=PAXIP1;refGene_tx=XM_023242347.1
A2	168238545	168238683
refGene_gn=PAXIP1;refGene_tx=XM_023242347.1
A2	168239517	168239613
refGene_gn=PAXIP1;refGene_tx=XM_023242347.1
A2	168243084	168243179
refGene_gn=PAXIP1;refGene_tx=XM_023242347.1

A2	168243584	168243795
refGene_gn=PAXIP1;refGene_tx=XM_023242347.1
A2	168243869	168244148
refGene_gn=PAXIP1;refGene_tx=XM_023242347.1
A2	168249771	168250410
refGene_gn=PAXIP1;refGene_tx=XM_023242347.1
A2	168256213	168256327
refGene_gn=PAXIP1;refGene_tx=XM_023242347.1
A2	168262951	168263015
refGene_gn=PAXIP1;refGene_tx=XM_023242347.1
A2	168265348	168265392
refGene_gn=PAXIP1;refGene_tx=XM_023242347.1
A2	168269968	168270103
refGene_gn=PAXIP1;refGene_tx=XM_023242347.1
A2	168274049	168274130
refGene_gn=PAXIP1;refGene_tx=XM_023242347.1
A2	168307448	168308186
refGene_gn=HTR5A;refGene_tx=XM_003983252.5
A2	168319024	168319357
refGene_gn=HTR5A;refGene_tx=XM_003983252.5
A2	168419735	168420138
refGene_gn=INSIG1;refGene_tx=XM_023250896.1
A2	168422495	168422620
refGene_gn=INSIG1;refGene_tx=XM_023250896.1
A2	168424053	168424220
refGene_gn=INSIG1;refGene_tx=XM_023250896.1
A2	168424563	168424663
refGene_gn=INSIG1;refGene_tx=XM_023250896.1
A2	168430025	168430055
refGene_gn=INSIG1;refGene_tx=XM_023250896.1
A2	168516695	168517392
refGene_gn=EN2;refGene_tx=XM_006929557.4
A2	168520694	168521011
refGene_gn=EN2;refGene_tx=XM_006929557.4
A2	168554696	168554797
refGene_gn=CNPY1;refGene_tx=XM_023250786.1
A2	168557671	168557851
refGene_gn=CNPY1;refGene_tx=XM_006929537.3;refGene_gn=CNPY1;refGene_tx
=XM_006929538.3;refGene_gn=CNPY1;refGene_tx=XM_023250787.1
A2	168559306	168559338
refGene_gn=CNPY1;refGene_tx=XM_023250788.1
A2	168561511	168561608
refGene_gn=CNPY1;refGene_tx=XM_006929537.3;refGene_gn=CNPY1;refGene_tx
=XM_006929538.3;refGene_gn=CNPY1;refGene_tx=XM_023250786.1;refGene_gn=
CNPY1;refGene_tx=XM_023250788.1
A2	168563364	168563568
refGene_gn=CNPY1;refGene_tx=XM_006929537.3;refGene_gn=CNPY1;refGene_tx
=XM_006929538.3;refGene_gn=CNPY1;refGene_tx=XM_023250786.1;refGene_gn=
CNPY1;refGene_tx=XM_023250787.1;refGene_gn=CNPY1;refGene_tx=XM_0232507
88.1

A2 168593724 168593823
refGene_gn=CNPY1;refGene_tx=XM_006929537.3;refGene_gn=CNPY1;refGene_tx
=XM_006929538.3;refGene_gn=CNPY1;refGene_tx=XM_023250786.1;refGene_gn=
CNPY1;refGene_tx=XM_023250787.1;refGene_gn=CNPY1;refGene_tx=XM_0232507
88.1
A2 168647654 168647668
refGene_gn=LOC111556233;refGene_tx=XM_023250800.1;refGene_gn=LOC111556
233;refGene_tx=XM_023250801.1
A2 168648083 168648812
refGene_gn=LOC111556233;refGene_tx=XM_023250800.1;refGene_gn=LOC111556
233;refGene_tx=XM_023250801.1;refGene_gn=RBM33;refGene_tx=XM_023250789
.
1;refGene_gn=RBM33;refGene_tx=XM_023250790.1;refGene_gn=RBM33;refGene_
tx=XM_023250791.1;refGene_gn=RBM33;refGene_tx=XM_023250792.1;refGene_g
n=RBM33;refGene_tx=XM_023250795.1;refGene_gn=RBM33;refGene_tx=XM_02325
0797.1
A2 168675045 168675124
refGene_gn=RBM33;refGene_tx=XM_023250789.1;refGene_gn=RBM33;refGene_tx
=XM_023250790.1;refGene_gn=RBM33;refGene_tx=XM_023250791.1;refGene_gn=
RBM33;refGene_tx=XM_023250792.1;refGene_gn=RBM33;refGene_tx=XM_0232507
94.1;refGene_gn=RBM33;refGene_tx=XM_023250795.1;refGene_gn=RBM33;refGe
ne_tx=XM_023250797.1
A2 168675242 168675243
refGene_gn=RBM33;refGene_tx=XM_023250796.1
A2 168682609 168682658
refGene_gn=RBM33;refGene_tx=XM_023250789.1;refGene_gn=RBM33;refGene_tx
=XM_023250790.1;refGene_gn=RBM33;refGene_tx=XM_023250791.1;refGene_gn=
RBM33;refGene_tx=XM_023250792.1;refGene_gn=RBM33;refGene_tx=XM_0232507
94.1;refGene_gn=RBM33;refGene_tx=XM_023250795.1;refGene_gn=RBM33;refGe
ne_tx=XM_023250796.1;refGene_gn=RBM33;refGene_tx=XM_023250797.1
A2 168689879 168689956
refGene_gn=RBM33;refGene_tx=XM_023250789.1;refGene_gn=RBM33;refGene_tx
=XM_023250790.1;refGene_gn=RBM33;refGene_tx=XM_023250791.1;refGene_gn=
RBM33;refGene_tx=XM_023250792.1;refGene_gn=RBM33;refGene_tx=XM_0232507
95.1;refGene_gn=RBM33;refGene_tx=XM_023250797.1
A2 168692296 168692609
refGene_gn=RBM33;refGene_tx=XM_023250789.1;refGene_gn=RBM33;refGene_tx
=XM_023250790.1;refGene_gn=RBM33;refGene_tx=XM_023250791.1;refGene_gn=
RBM33;refGene_tx=XM_023250792.1;refGene_gn=RBM33;refGene_tx=XM_0232507
94.1;refGene_gn=RBM33;refGene_tx=XM_023250795.1;refGene_gn=RBM33;refGe
ne_tx=XM_023250796.1;refGene_gn=RBM33;refGene_tx=XM_023250797.1
A2 168696602 168696755
refGene_gn=RBM33;refGene_tx=XM_023250798.1
A2 168709713 168709885
refGene_gn=RBM33;refGene_tx=XM_023250789.1;refGene_gn=RBM33;refGene_tx
=XM_023250790.1;refGene_gn=RBM33;refGene_tx=XM_023250791.1;refGene_gn=
RBM33;refGene_tx=XM_023250792.1;refGene_gn=RBM33;refGene_tx=XM_0232507
94.1;refGene_gn=RBM33;refGene_tx=XM_023250795.1;refGene_gn=RBM33;refGe
ne_tx=XM_023250796.1;refGene_gn=RBM33;refGene_tx=XM_023250797.1;refGen
e_gn=RBM33;refGene_tx=XM_023250798.1;refGene_gn=RBM33;refGene_tx=XM_02

3250799.1
A2	168714693	168714902
refGene_gn=RBM33;refGene_tx=XM_023250789.1;refGene_gn=RBM33;refGene_tx=XM_023250790.1;refGene_gn=RBM33;refGene_tx=XM_023250791.1;refGene_gn=RBM33;refGene_tx=XM_023250792.1;refGene_gn=RBM33;refGene_tx=XM_023250794.1;refGene_gn=RBM33;refGene_tx=XM_023250795.1;refGene_gn=RBM33;refGene_tx=XM_023250796.1;refGene_gn=RBM33;refGene_tx=XM_023250797.1;refGene_gn=RBM33;refGene_tx=XM_023250798.1;refGene_gn=RBM33;refGene_tx=XM_023250799.1
A2	168718350	168718636
refGene_gn=RBM33;refGene_tx=XM_023250789.1;refGene_gn=RBM33;refGene_tx=XM_023250790.1;refGene_gn=RBM33;refGene_tx=XM_023250791.1;refGene_gn=RBM33;refGene_tx=XM_023250792.1;refGene_gn=RBM33;refGene_tx=XM_023250794.1;refGene_gn=RBM33;refGene_tx=XM_023250795.1;refGene_gn=RBM33;refGene_tx=XM_023250796.1;refGene_gn=RBM33;refGene_tx=XM_023250797.1;refGene_gn=RBM33;refGene_tx=XM_023250798.1;refGene_gn=RBM33;refGene_tx=XM_023250799.1
A2	168724713	168724772
refGene_gn=RBM33;refGene_tx=XM_023250789.1;refGene_gn=RBM33;refGene_tx=XM_023250790.1;refGene_gn=RBM33;refGene_tx=XM_023250791.1;refGene_gn=RBM33;refGene_tx=XM_023250792.1;refGene_gn=RBM33;refGene_tx=XM_023250794.1;refGene_gn=RBM33;refGene_tx=XM_023250795.1;refGene_gn=RBM33;refGene_tx=XM_023250796.1;refGene_gn=RBM33;refGene_tx=XM_023250797.1;refGene_gn=RBM33;refGene_tx=XM_023250798.1;refGene_gn=RBM33;refGene_tx=XM_023250799.1
A2	168743042	168743175
refGene_gn=RBM33;refGene_tx=XM_023250789.1;refGene_gn=RBM33;refGene_tx=XM_023250790.1;refGene_gn=RBM33;refGene_tx=XM_023250791.1;refGene_gn=RBM33;refGene_tx=XM_023250792.1;refGene_gn=RBM33;refGene_tx=XM_023250794.1;refGene_gn=RBM33;refGene_tx=XM_023250795.1;refGene_gn=RBM33;refGene_tx=XM_023250796.1;refGene_gn=RBM33;refGene_tx=XM_023250797.1;refGene_gn=RBM33;refGene_tx=XM_023250798.1;refGene_gn=RBM33;refGene_tx=XM_023250799.1
A2	168743544	168743888
refGene_gn=RBM33;refGene_tx=XM_023250789.1;refGene_gn=RBM33;refGene_tx=XM_023250790.1;refGene_gn=RBM33;refGene_tx=XM_023250791.1;refGene_gn=RBM33;refGene_tx=XM_023250792.1;refGene_gn=RBM33;refGene_tx=XM_023250794.1;refGene_gn=RBM33;refGene_tx=XM_023250795.1;refGene_gn=RBM33;refGene_tx=XM_023250796.1;refGene_gn=RBM33;refGene_tx=XM_023250797.1;refGene_gn=RBM33;refGene_tx=XM_023250798.1;refGene_gn=RBM33;refGene_tx=XM_023250799.1
A2	168745007	168745310
refGene_gn=RBM33;refGene_tx=XM_023250789.1;refGene_gn=RBM33;refGene_tx=XM_023250790.1;refGene_gn=RBM33;refGene_tx=XM_023250791.1;refGene_gn=RBM33;refGene_tx=XM_023250792.1;refGene_gn=RBM33;refGene_tx=XM_023250794.1;refGene_gn=RBM33;refGene_tx=XM_023250795.1;refGene_gn=RBM33;refGene_tx=XM_023250796.1;refGene_gn=RBM33;refGene_tx=XM_023250797.1;refGene_gn=RBM33;refGene_tx=XM_023250798.1;refGene_gn=RBM33;refGene_tx=XM_023250799.1
A2	168747313	168747598

refGene_gn=RBM33;refGene_tx=XM_023250789.1;refGene_gn=RBM33;refGene_tx=XM_023250790.1;refGene_gn=RBM33;refGene_tx=XM_023250791.1;refGene_gn=RBM33;refGene_tx=XM_023250792.1;refGene_gn=RBM33;refGene_tx=XM_023250794.1;refGene_gn=RBM33;refGene_tx=XM_023250795.1;refGene_gn=RBM33;refGene_tx=XM_023250796.1;refGene_gn=RBM33;refGene_tx=XM_023250797.1;refGene_gn=RBM33;refGene_tx=XM_023250798.1;refGene_gn=RBM33;refGene_tx=XM_023250799.1
A2 168750987 168751617
refGene_gn=RBM33;refGene_tx=XM_023250789.1;refGene_gn=RBM33;refGene_tx=XM_023250790.1;refGene_gn=RBM33;refGene_tx=XM_023250791.1;refGene_gn=RBM33;refGene_tx=XM_023250792.1;refGene_gn=RBM33;refGene_tx=XM_023250794.1;refGene_gn=RBM33;refGene_tx=XM_023250795.1;refGene_gn=RBM33;refGene_tx=XM_023250796.1;refGene_gn=RBM33;refGene_tx=XM_023250797.1;refGene_gn=RBM33;refGene_tx=XM_023250798.1;refGene_gn=RBM33;refGene_tx=XM_023250799.1
A2 168768917 168769127
refGene_gn=RBM33;refGene_tx=XM_023250789.1;refGene_gn=RBM33;refGene_tx=XM_023250790.1;refGene_gn=RBM33;refGene_tx=XM_023250791.1;refGene_gn=RBM33;refGene_tx=XM_023250792.1;refGene_gn=RBM33;refGene_tx=XM_023250794.1;refGene_gn=RBM33;refGene_tx=XM_023250796.1;refGene_gn=RBM33;refGene_tx=XM_023250798.1;refGene_gn=RBM33;refGene_tx=XM_023250799.1
A2 168773960 168774149
refGene_gn=RBM33;refGene_tx=XM_023250789.1;refGene_gn=RBM33;refGene_tx=XM_023250790.1;refGene_gn=RBM33;refGene_tx=XM_023250791.1;refGene_gn=RBM33;refGene_tx=XM_023250792.1;refGene_gn=RBM33;refGene_tx=XM_023250794.1;refGene_gn=RBM33;refGene_tx=XM_023250795.1;refGene_gn=RBM33;refGene_tx=XM_023250796.1;refGene_gn=RBM33;refGene_tx=XM_023250798.1;refGene_gn=RBM33;refGene_tx=XM_023250799.1
A2 168782235 168782324
refGene_gn=RBM33;refGene_tx=XM_023250789.1;refGene_gn=RBM33;refGene_tx=XM_023250790.1;refGene_gn=RBM33;refGene_tx=XM_023250791.1;refGene_gn=RBM33;refGene_tx=XM_023250792.1;refGene_gn=RBM33;refGene_tx=XM_023250794.1;refGene_gn=RBM33;refGene_tx=XM_023250795.1;refGene_gn=RBM33;refGene_tx=XM_023250796.1;refGene_gn=RBM33;refGene_tx=XM_023250797.1;refGene_gn=RBM33;refGene_tx=XM_023250798.1;refGene_gn=RBM33;refGene_tx=XM_023250799.1
A2 168782663 168782712
refGene_gn=RBM33;refGene_tx=XM_023250789.1;refGene_gn=RBM33;refGene_tx=XM_023250790.1;refGene_gn=RBM33;refGene_tx=XM_023250791.1;refGene_gn=RBM33;refGene_tx=XM_023250792.1;refGene_gn=RBM33;refGene_tx=XM_023250794.1;refGene_gn=RBM33;refGene_tx=XM_023250795.1;refGene_gn=RBM33;refGene_tx=XM_023250796.1;refGene_gn=RBM33;refGene_tx=XM_023250797.1;refGene_gn=RBM33;refGene_tx=XM_023250798.1;refGene_gn=RBM33;refGene_tx=XM_023250799.1
A2 168807637 168808461
refGene_gn=SHH;refGene_tx=XM_023242357.1
A2 168811191 168811453
refGene_gn=SHH;refGene_tx=XM_023242357.1
A2 168816434 168817046
refGene_gn=SHH;refGene_tx=XM_023242357.1

A2	168820334	168820531
refGene_gn=SHH;refGene_tx=XM_023242357.1
A2	168820850	168820920
refGene_gn=SHH;refGene_tx=XM_023242357.1
A2	168826098	168826262
refGene_gn=SHH;refGene_tx=XM_023242357.1
A2	168829869	168830008
refGene_gn=SHH;refGene_tx=XM_023242357.1
A2	169194233	169194733
refGene_gn=LOC102900227;refGene_tx=XM_023250803.1;refGene_gn=LOC102900227;refGene_tx=XM_023250804.1;refGene_gn=LOC102900227;refGene_tx=XM_023250805.1
A2	169195059	169195075
refGene_gn=LOC102900227;refGene_tx=XM_023250805.1
A2	169209056	169209186
refGene_gn=LOC102900227;refGene_tx=XM_023250803.1;refGene_gn=LOC102900227;refGene_tx=XM_023250804.1
A2	169401929	169401944
refGene_gn=RNF32;refGene_tx=XM_003983253.5;refGene_gn=RNF32;refGene_tx=XM_006929541.4;refGene_gn=RNF32;refGene_tx=XM_006929542.4;refGene_gn=RNF32;refGene_tx=XM_019826061.2;refGene_gn=RNF32;refGene_tx=XM_019826062.2;refGene_gn=RNF32;refGene_tx=XM_019826063.2;refGene_gn=RNF32;refGene_tx=XM_019826064.2;refGene_gn=RNF32;refGene_tx=XM_019826065.2;refGene_gn=RNF32;refGene_tx=XM_019826066.2;refGene_gn=RNF32;refGene_tx=XM_019826067.2;refGene_gn=RNF32;refGene_tx=XM_019826068.2;refGene_gn=RNF32;refGene_tx=XM_023250811.1;refGene_gn=RNF32;refGene_tx=XM_023250812.1;refGene_gn=RNF32;refGene_tx=XM_023250813.1
A2	169402462	169402727
refGene_gn=RNF32;refGene_tx=XM_003983253.5;refGene_gn=RNF32;refGene_tx=XM_006929541.4;refGene_gn=RNF32;refGene_tx=XM_006929542.4;refGene_gn=RNF32;refGene_tx=XM_019826061.2;refGene_gn=RNF32;refGene_tx=XM_019826062.2;refGene_gn=RNF32;refGene_tx=XM_019826063.2;refGene_gn=RNF32;refGene_tx=XM_019826064.2;refGene_gn=RNF32;refGene_tx=XM_019826065.2;refGene_gn=RNF32;refGene_tx=XM_019826066.2;refGene_gn=RNF32;refGene_tx=XM_019826067.2;refGene_gn=RNF32;refGene_tx=XM_019826068.2;refGene_gn=RNF32;refGene_tx=XM_023250811.1;refGene_gn=RNF32;refGene_tx=XM_023250812.1;refGene_gn=RNF32;refGene_tx=XM_023250813.1
A2	169406040	169406183
refGene_gn=RNF32;refGene_tx=XM_003983253.5;refGene_gn=RNF32;refGene_tx=XM_006929541.4;refGene_gn=RNF32;refGene_tx=XM_006929542.4;refGene_gn=RNF32;refGene_tx=XM_019826062.2;refGene_gn=RNF32;refGene_tx=XM_019826063.2;refGene_gn=RNF32;refGene_tx=XM_019826064.2;refGene_gn=RNF32;refGene_tx=XM_019826065.2;refGene_gn=RNF32;refGene_tx=XM_019826066.2;refGene_gn=RNF32;refGene_tx=XM_019826067.2;refGene_gn=RNF32;refGene_tx=XM_019826068.2;refGene_gn=RNF32;refGene_tx=XM_023250811.1;refGene_gn=RNF32;refGene_tx=XM_023250812.1;refGene_gn=RNF32;refGene_tx=XM_023250813.1
A2	169406797	169406871
refGene_gn=LMBR1;refGene_tx=XM_023250808.1
A2	169407491	169407619
refGene_gn=RNF32;refGene_tx=XM_019826061.2

A2 169409018 169409051
refGene_gn=RNF32;refGene_tx=XM_003983253.5;refGene_gn=RNF32;refGene_tx=XM_006929541.4;refGene_gn=RNF32;refGene_tx=XM_006929542.4;refGene_gn=RNF32;refGene_tx=XM_019826061.2;refGene_gn=RNF32;refGene_tx=XM_019826063.2;refGene_gn=RNF32;refGene_tx=XM_019826064.2;refGene_gn=RNF32;refGene_tx=XM_019826065.2;refGene_gn=RNF32;refGene_tx=XM_019826066.2;refGene_gn=RNF32;refGene_tx=XM_019826067.2;refGene_gn=RNF32;refGene_tx=XM_019826068.2;refGene_gn=RNF32;refGene_tx=XM_023250811.1;refGene_gn=RNF32;refGene_tx=XM_023250812.1;refGene_gn=RNF32;refGene_tx=XM_023250813.1
A2 169409586 169409711
refGene_gn=RNF32;refGene_tx=XM_003983253.5;refGene_gn=RNF32;refGene_tx=XM_006929541.4;refGene_gn=RNF32;refGene_tx=XM_006929542.4;refGene_gn=RNF32;refGene_tx=XM_019826061.2;refGene_gn=RNF32;refGene_tx=XM_019826062.2;refGene_gn=RNF32;refGene_tx=XM_019826063.2;refGene_gn=RNF32;refGene_tx=XM_019826064.2;refGene_gn=RNF32;refGene_tx=XM_019826065.2;refGene_gn=RNF32;refGene_tx=XM_019826066.2;refGene_gn=RNF32;refGene_tx=XM_019826067.2;refGene_gn=RNF32;refGene_tx=XM_019826068.2;refGene_gn=RNF32;refGene_tx=XM_023250811.1;refGene_gn=RNF32;refGene_tx=XM_023250812.1;refGene_gn=RNF32;refGene_tx=XM_023250813.1
A2 169409921 169410030
refGene_gn=RNF32;refGene_tx=XM_003983253.5;refGene_gn=RNF32;refGene_tx=XM_006929541.4;refGene_gn=RNF32;refGene_tx=XM_006929542.4;refGene_gn=RNF32;refGene_tx=XM_019826061.2;refGene_gn=RNF32;refGene_tx=XM_019826062.2;refGene_gn=RNF32;refGene_tx=XM_019826063.2;refGene_gn=RNF32;refGene_tx=XM_019826064.2;refGene_gn=RNF32;refGene_tx=XM_019826065.2;refGene_gn=RNF32;refGene_tx=XM_019826066.2;refGene_gn=RNF32;refGene_tx=XM_019826067.2;refGene_gn=RNF32;refGene_tx=XM_019826068.2;refGene_gn=RNF32;refGene_tx=XM_023250811.1;refGene_gn=RNF32;refGene_tx=XM_023250812.1;refGene_gn=RNF32;refGene_tx=XM_023250813.1
A2 169410761 169410884
refGene_gn=RNF32;refGene_tx=XM_019826066.2
A2 169411795 169411939
refGene_gn=RNF32;refGene_tx=XM_019826063.2
A2 169413971 169414034
refGene_gn=RNF32;refGene_tx=XM_019826063.2;refGene_gn=RNF32;refGene_tx=XM_019826068.2
A2 169423491 169423557
refGene_gn=RNF32;refGene_tx=XM_019826067.2
A2 169423988 169424033
refGene_gn=RNF32;refGene_tx=XM_023250813.1
A2 169435484 169435652
refGene_gn=RNF32;refGene_tx=XM_003983253.5;refGene_gn=RNF32;refGene_tx=XM_006929541.4;refGene_gn=RNF32;refGene_tx=XM_006929542.4;refGene_gn=RNF32;refGene_tx=XM_019826061.2;refGene_gn=RNF32;refGene_tx=XM_019826062.2;refGene_gn=RNF32;refGene_tx=XM_019826064.2;refGene_gn=RNF32;refGene_tx=XM_019826065.2;refGene_gn=RNF32;refGene_tx=XM_023250811.1
A2 169435878 169435881
refGene_gn=RNF32;refGene_tx=XM_019826065.2
A2 169435936 169435966
refGene_gn=RNF32;refGene_tx=XM_019826064.2

A2 169436701 169436953
refGene_gn=RNF32;refGene_tx=XM_003983253.5;refGene_gn=RNF32;refGene_tx
=XM_019826061.2;refGene_gn=RNF32;refGene_tx=XM_019826062.2;refGene_gn=
RNF32;refGene_tx=XM_023250811.1
A2 169437252 169437345
refGene_gn=RNF32;refGene_tx=XM_006929542.4
A2 169449565 169449718
refGene_gn=RNF32;refGene_tx=XM_006929541.4;refGene_gn=RNF32;refGene_tx
=XM_023250812.1
A2 169451926 169452012
refGene_gn=LMBR1;refGene_tx=XM_023250806.1;refGene_gn=LMBR1;refGene_tx
=XM_023250807.1;refGene_gn=LMBR1;refGene_tx=XM_023250809.1
A2 169455113 169455275
refGene_gn=LMBR1;refGene_tx=XM_023250806.1;refGene_gn=LMBR1;refGene_tx
=XM_023250809.1
A2 169467817 169467840
refGene_gn=LMBR1;refGene_tx=XM_023250810.1
A2 169475534 169475601
refGene_gn=LMBR1;refGene_tx=XM_023250806.1;refGene_gn=LMBR1;refGene_tx
=XM_023250807.1;refGene_gn=LMBR1;refGene_tx=XM_023250808.1;refGene_gn=
LMBR1;refGene_tx=XM_023250809.1;refGene_gn=LMBR1;refGene_tx=XM_0232508
10.1
A2 169476854 169476945
refGene_gn=LMBR1;refGene_tx=XM_023250806.1;refGene_gn=LMBR1;refGene_tx
=XM_023250807.1;refGene_gn=LMBR1;refGene_tx=XM_023250808.1;refGene_gn=
LMBR1;refGene_tx=XM_023250809.1;refGene_gn=LMBR1;refGene_tx=XM_0232508
10.1
A2 169477173 169477247
refGene_gn=LMBR1;refGene_tx=XM_023250806.1;refGene_gn=LMBR1;refGene_tx
=XM_023250807.1;refGene_gn=LMBR1;refGene_tx=XM_023250808.1;refGene_gn=
LMBR1;refGene_tx=XM_023250809.1;refGene_gn=LMBR1;refGene_tx=XM_0232508
10.1
A2 169479926 169480004
refGene_gn=LMBR1;refGene_tx=XM_023250806.1;refGene_gn=LMBR1;refGene_tx
=XM_023250807.1;refGene_gn=LMBR1;refGene_tx=XM_023250808.1;refGene_gn=
LMBR1;refGene_tx=XM_023250809.1;refGene_gn=LMBR1;refGene_tx=XM_0232508
10.1
A2 169480647 169480724
refGene_gn=LMBR1;refGene_tx=XM_023250806.1;refGene_gn=LMBR1;refGene_tx
=XM_023250807.1;refGene_gn=LMBR1;refGene_tx=XM_023250808.1;refGene_gn=
LMBR1;refGene_tx=XM_023250809.1;refGene_gn=LMBR1;refGene_tx=XM_0232508
10.1
A2 169486012 169486093
refGene_gn=LMBR1;refGene_tx=XM_023250806.1;refGene_gn=LMBR1;refGene_tx
=XM_023250807.1;refGene_gn=LMBR1;refGene_tx=XM_023250808.1;refGene_gn=
LMBR1;refGene_tx=XM_023250809.1;refGene_gn=LMBR1;refGene_tx=XM_0232508
10.1
A2 169504357 169504430
refGene_gn=LMBR1;refGene_tx=XM_023250806.1;refGene_gn=LMBR1;refGene_tx
=XM_023250807.1;refGene_gn=LMBR1;refGene_tx=XM_023250808.1;refGene_gn=

LMBR1;refGene_tx=XM_023250809.1;refGene_gn=LMBR1;refGene_tx=XM_0232508
10.1
A2 169508025 169508090
refGene_gn=LMBR1;refGene_tx=XM_023250806.1;refGene_gn=LMBR1;refGene_tx
=XM_023250807.1;refGene_gn=LMBR1;refGene_tx=XM_023250808.1;refGene_gn=
LMBR1;refGene_tx=XM_023250809.1;refGene_gn=LMBR1;refGene_tx=XM_0232508
10.1
A2 169509337 169509406
refGene_gn=LMBR1;refGene_tx=XM_023250806.1;refGene_gn=LMBR1;refGene_tx
=XM_023250807.1;refGene_gn=LMBR1;refGene_tx=XM_023250808.1;refGene_gn=
LMBR1;refGene_tx=XM_023250809.1;refGene_gn=LMBR1;refGene_tx=XM_0232508
10.1
A2 169509889 169510016
refGene_gn=LMBR1;refGene_tx=XM_023250806.1;refGene_gn=LMBR1;refGene_tx
=XM_023250807.1;refGene_gn=LMBR1;refGene_tx=XM_023250808.1;refGene_gn=
LMBR1;refGene_tx=XM_023250809.1;refGene_gn=LMBR1;refGene_tx=XM_0232508
10.1
A2 169539290 169539394
refGene_gn=LMBR1;refGene_tx=XM_023250806.1;refGene_gn=LMBR1;refGene_tx
=XM_023250807.1;refGene_gn=LMBR1;refGene_tx=XM_023250808.1;refGene_gn=
LMBR1;refGene_tx=XM_023250809.1;refGene_gn=LMBR1;refGene_tx=XM_0232508
10.1
A2 169562920 169563060
refGene_gn=LMBR1;refGene_tx=XM_023250806.1;refGene_gn=LMBR1;refGene_tx
=XM_023250807.1;refGene_gn=LMBR1;refGene_tx=XM_023250808.1;refGene_gn=
LMBR1;refGene_tx=XM_023250809.1;refGene_gn=LMBR1;refGene_tx=XM_0232508
10.1
A2 169563143 169563145
refGene_gn=LMBR1;refGene_tx=XM_023250809.1
A2 169569683 169569723
refGene_gn=LMBR1;refGene_tx=XM_023250806.1;refGene_gn=LMBR1;refGene_tx
=XM_023250807.1;refGene_gn=LMBR1;refGene_tx=XM_023250808.1;refGene_gn=
LMBR1;refGene_tx=XM_023250810.1
A2 169572106 169572179
refGene_gn=LMBR1;refGene_tx=XM_023250806.1;refGene_gn=LMBR1;refGene_tx
=XM_023250807.1;refGene_gn=LMBR1;refGene_tx=XM_023250808.1;refGene_gn=
LMBR1;refGene_tx=XM_023250810.1
A2 169603968 169604034
refGene_gn=LMBR1;refGene_tx=XM_023250806.1;refGene_gn=LMBR1;refGene_tx
=XM_023250807.1;refGene_gn=LMBR1;refGene_tx=XM_023250808.1;refGene_gn=
LMBR1;refGene_tx=XM_023250810.1
A2 169637102 169638092
refGene_gn=NOM1;refGene_tx=XM_019825089.2
A2 169639650 169639775
refGene_gn=NOM1;refGene_tx=XM_019825089.2
A2 169640996 169641192
refGene_gn=NOM1;refGene_tx=XM_019825089.2
A2 169643026 169643350
refGene_gn=NOM1;refGene_tx=XM_019825089.2
A2 169645288 169645399

refGene_gn=NOM1;refGene_tx=XM_019825089.2
A2	169646561	169646729
refGene_gn=NOM1;refGene_tx=XM_019825089.2
A2	169647416	169647538
refGene_gn=NOM1;refGene_tx=XM_019825089.2
A2	169648115	169648248
refGene_gn=NOM1;refGene_tx=XM_019825089.2
A2	169648750	169648882
refGene_gn=NOM1;refGene_tx=XM_019825089.2
A2	169650904	169651014
refGene_gn=NOM1;refGene_tx=XM_019825089.2
A2	169651289	169651464
refGene_gn=NOM1;refGene_tx=XM_019825089.2
A2	169661146	169661409
refGene_gn=LOC109497725;refGene_tx=XM_019826070.2
A2	169662486	169662571
refGene_gn=LOC109497725;refGene_tx=XM_019826070.2
A2	169662665	169663010
refGene_gn=LOC109497725;refGene_tx=XM_019826070.2
A2	169670382	169670742
refGene_gn=MNX1;refGene_tx=XM_023250815.1
A2	169671350	169671511
refGene_gn=MNX1;refGene_tx=XM_023250815.1
A2	169674382	169675106
refGene_gn=MNX1;refGene_tx=XM_023250815.1
A2	169753739	169753805
refGene_gn=UBE3C;refGene_tx=XM_003983254.5
A2	169780467	169780521
refGene_gn=UBE3C;refGene_tx=XM_003983254.5
A2	169783992	169784067
refGene_gn=UBE3C;refGene_tx=XM_003983254.5
A2	169784536	169784683
refGene_gn=UBE3C;refGene_tx=XM_003983254.5
A2	169788426	169788542
refGene_gn=UBE3C;refGene_tx=XM_003983254.5
A2	169788948	169789106
refGene_gn=UBE3C;refGene_tx=XM_003983254.5
A2	169793457	169793611
refGene_gn=UBE3C;refGene_tx=XM_003983254.5
A2	169794025	169794185
refGene_gn=UBE3C;refGene_tx=XM_003983254.5
A2	169794186	169794247
refGene_gn=UBE3C;refGene_tx=XM_003983254.5
A2	169795701	169795856
refGene_gn=UBE3C;refGene_tx=XM_003983254.5
A2	169799337	169799525
refGene_gn=UBE3C;refGene_tx=XM_003983254.5
A2	169811231	169811318
refGene_gn=UBE3C;refGene_tx=XM_003983254.5
A2	169816270	169816428

refGene_gn=UBE3C;refGene_tx=XM_003983254.5
A2 169816577 169816810
refGene_gn=UBE3C;refGene_tx=XM_003983254.5
A2 169825806 169825911
refGene_gn=UBE3C;refGene_tx=XM_003983254.5
A2 169828197 169828285
refGene_gn=UBE3C;refGene_tx=XM_003983254.5
A2 169829238 169829336
refGene_gn=UBE3C;refGene_tx=XM_003983254.5
A2 169830879 169831012
refGene_gn=UBE3C;refGene_tx=XM_003983254.5
A2 169835878 169836126
refGene_gn=UBE3C;refGene_tx=XM_003983254.5
A2 169850013 169850226
refGene_gn=UBE3C;refGene_tx=XM_003983254.5
A2 169857347 169857536
refGene_gn=UBE3C;refGene_tx=XM_003983254.5
A2 169857644 169857711
refGene_gn=UBE3C;refGene_tx=XM_003983254.5
A2 169865094 169865225
refGene_gn=UBE3C;refGene_tx=XM_003983254.5
A2 169872935 169873106
refGene_gn=UBE3C;refGene_tx=XM_003983254.5
A2 169947408 169947473
refGene_gn=DNAJB6;refGene_tx=XM_003983255.5;refGene_gn=DNAJB6;refGene_tx=XM_023250816.1;refGene_gn=DNAJB6;refGene_tx=XM_023250817.1;refGene_gn=DNAJB6;refGene_tx=XM_023250818.1;refGene_gn=DNAJB6;refGene_tx=XM_023250819.1
A2 169949749 169949859
refGene_gn=DNAJB6;refGene_tx=XM_003983255.5;refGene_gn=DNAJB6;refGene_tx=XM_023250816.1;refGene_gn=DNAJB6;refGene_tx=XM_023250817.1;refGene_gn=DNAJB6;refGene_tx=XM_023250818.1;refGene_gn=DNAJB6;refGene_tx=XM_023250819.1
A2 169951750 169951810
refGene_gn=DNAJB6;refGene_tx=XM_003983255.5;refGene_gn=DNAJB6;refGene_tx=XM_023250816.1;refGene_gn=DNAJB6;refGene_tx=XM_023250817.1;refGene_gn=DNAJB6;refGene_tx=XM_023250818.1;refGene_gn=DNAJB6;refGene_tx=XM_023250819.1
A2 169952653 169952764
refGene_gn=DNAJB6;refGene_tx=XM_003983255.5;refGene_gn=DNAJB6;refGene_tx=XM_023250816.1;refGene_gn=DNAJB6;refGene_tx=XM_023250817.1;refGene_gn=DNAJB6;refGene_tx=XM_023250818.1;refGene_gn=DNAJB6;refGene_tx=XM_023250819.1
A2 169965332 169965464
refGene_gn=DNAJB6;refGene_tx=XM_003983255.5;refGene_gn=DNAJB6;refGene_tx=XM_023250816.1;refGene_gn=DNAJB6;refGene_tx=XM_023250817.1;refGene_gn=DNAJB6;refGene_tx=XM_023250818.1;refGene_gn=DNAJB6;refGene_tx=XM_023250819.1
A2 169967530 169967672
refGene_gn=DNAJB6;refGene_tx=XM_003983255.5;refGene_gn=DNAJB6;refGene_

tx=XM_023250816.1;refGene_gn=DNAJB6;refGene_tx=XM_023250817.1;refGene_gn=DNAJB6;refGene_tx=XM_023250818.1;refGene_gn=DNAJB6;refGene_tx=XM_023250819.1
A2	169968196	169968302
refGene_gn=DNAJB6;refGene_tx=XM_003983255.5;refGene_gn=DNAJB6;refGene_tx=XM_023250816.1;refGene_gn=DNAJB6;refGene_tx=XM_023250817.1;refGene_gn=DNAJB6;refGene_tx=XM_023250818.1;refGene_gn=DNAJB6;refGene_tx=XM_023250819.1
A2	169983999	169984215
refGene_gn=DNAJB6;refGene_tx=XM_003983255.5;refGene_gn=DNAJB6;refGene_tx=XM_023250816.1;refGene_gn=DNAJB6;refGene_tx=XM_023250817.1;refGene_gn=DNAJB6;refGene_tx=XM_023250818.1
A2	169990148	169990231
refGene_gn=DNAJB6;refGene_tx=XM_003983255.5;refGene_gn=DNAJB6;refGene_tx=XM_023250816.1;refGene_gn=DNAJB6;refGene_tx=XM_023250817.1;refGene_gn=DNAJB6;refGene_tx=XM_023250818.1
A2	170056397	170056469
refGene_gn=PTPRN2;refGene_tx=XM_023250820.1;refGene_gn=PTPRN2;refGene_tx=XM_023250821.1;refGene_gn=PTPRN2;refGene_tx=XM_023250822.1;refGene_gn=PTPRN2;refGene_tx=XM_023250823.1
A2	170058077	170058151
refGene_gn=PTPRN2;refGene_tx=XM_023250820.1;refGene_gn=PTPRN2;refGene_tx=XM_023250821.1;refGene_gn=PTPRN2;refGene_tx=XM_023250822.1;refGene_gn=PTPRN2;refGene_tx=XM_023250823.1
A2	170071384	170071449
refGene_gn=PTPRN2;refGene_tx=XM_023250820.1;refGene_gn=PTPRN2;refGene_tx=XM_023250821.1;refGene_gn=PTPRN2;refGene_tx=XM_023250822.1;refGene_gn=PTPRN2;refGene_tx=XM_023250823.1
A2	170072859	170072913
refGene_gn=PTPRN2;refGene_tx=XM_023250820.1;refGene_gn=PTPRN2;refGene_tx=XM_023250821.1;refGene_gn=PTPRN2;refGene_tx=XM_023250822.1;refGene_gn=PTPRN2;refGene_tx=XM_023250823.1
A2	170079282	170079449
refGene_gn=PTPRN2;refGene_tx=XM_023250820.1;refGene_gn=PTPRN2;refGene_tx=XM_023250821.1;refGene_gn=PTPRN2;refGene_tx=XM_023250822.1;refGene_gn=PTPRN2;refGene_tx=XM_023250823.1
A2	170080446	170080566
refGene_gn=PTPRN2;refGene_tx=XM_023250820.1;refGene_gn=PTPRN2;refGene_tx=XM_023250821.1;refGene_gn=PTPRN2;refGene_tx=XM_023250822.1;refGene_gn=PTPRN2;refGene_tx=XM_023250823.1
A2	170091086	170091164
refGene_gn=PTPRN2;refGene_tx=XM_023250820.1;refGene_gn=PTPRN2;refGene_tx=XM_023250821.1;refGene_gn=PTPRN2;refGene_tx=XM_023250822.1;refGene_gn=PTPRN2;refGene_tx=XM_023250823.1
A2	170097342	170097416
refGene_gn=PTPRN2;refGene_tx=XM_023250820.1;refGene_gn=PTPRN2;refGene_tx=XM_023250821.1;refGene_gn=PTPRN2;refGene_tx=XM_023250822.1;refGene_gn=PTPRN2;refGene_tx=XM_023250823.1
A2	170112757	170112905
refGene_gn=PTPRN2;refGene_tx=XM_023250820.1;refGene_gn=PTPRN2;refGene_

tx=XM_023250821.1;refGene_gn=PTPRN2;refGene_tx=XM_023250822.1;refGene_gn=PTPRN2;refGene_tx=XM_023250823.1
A2 170137027 170137222
refGene_gn=PTPRN2;refGene_tx=XM_023250820.1;refGene_gn=PTPRN2;refGene_tx=XM_023250821.1;refGene_gn=PTPRN2;refGene_tx=XM_023250823.1
A2 170158861 170159074
refGene_gn=PTPRN2;refGene_tx=XM_023250820.1;refGene_gn=PTPRN2;refGene_tx=XM_023250821.1;refGene_gn=PTPRN2;refGene_tx=XM_023250822.1;refGene_gn=PTPRN2;refGene_tx=XM_023250823.1
A2 170320648 170320713
refGene_gn=PTPRN2;refGene_tx=XM_023250820.1;refGene_gn=PTPRN2;refGene_tx=XM_023250821.1;refGene_gn=PTPRN2;refGene_tx=XM_023250822.1;refGene_gn=PTPRN2;refGene_tx=XM_023250823.1
A2 170482461 170482541
refGene_gn=PTPRN2;refGene_tx=XM_023250820.1;refGene_gn=PTPRN2;refGene_tx=XM_023250821.1;refGene_gn=PTPRN2;refGene_tx=XM_023250822.1;refGene_gn=PTPRN2;refGene_tx=XM_023250823.1
A2 170498551 170498638
refGene_gn=PTPRN2;refGene_tx=XM_023250821.1;refGene_gn=PTPRN2;refGene_tx=XM_023250822.1;refGene_gn=PTPRN2;refGene_tx=XM_023250823.1
A2 170519723 170520103
refGene_gn=PTPRN2;refGene_tx=XM_023250820.1;refGene_gn=PTPRN2;refGene_tx=XM_023250821.1;refGene_gn=PTPRN2;refGene_tx=XM_023250822.1;refGene_gn=PTPRN2;refGene_tx=XM_023250823.1
A2 170522679 170522720
refGene_gn=PTPRN2;refGene_tx=XM_023250820.1;refGene_gn=PTPRN2;refGene_tx=XM_023250821.1;refGene_gn=PTPRN2;refGene_tx=XM_023250822.1;refGene_gn=PTPRN2;refGene_tx=XM_023250823.1
A2 170524280 170524463
refGene_gn=PTPRN2;refGene_tx=XM_023250820.1;refGene_gn=PTPRN2;refGene_tx=XM_023250821.1;refGene_gn=PTPRN2;refGene_tx=XM_023250822.1;refGene_gn=PTPRN2;refGene_tx=XM_023250823.1
A2 170541172 170541533
refGene_gn=PTPRN2;refGene_tx=XM_023250820.1;refGene_gn=PTPRN2;refGene_tx=XM_023250821.1;refGene_gn=PTPRN2;refGene_tx=XM_023250822.1;refGene_gn=PTPRN2;refGene_tx=XM_023250823.1
A2 170555493 170555656
refGene_gn=PTPRN2;refGene_tx=XM_023250820.1;refGene_gn=PTPRN2;refGene_tx=XM_023250821.1;refGene_gn=PTPRN2;refGene_tx=XM_023250822.1;refGene_gn=PTPRN2;refGene_tx=XM_023250823.1
A2 170565048 170565145
refGene_gn=PTPRN2;refGene_tx=XM_023250820.1;refGene_gn=PTPRN2;refGene_tx=XM_023250821.1;refGene_gn=PTPRN2;refGene_tx=XM_023250822.1;refGene_gn=PTPRN2;refGene_tx=XM_023250823.1
A2 170632033 170632147
refGene_gn=PTPRN2;refGene_tx=XM_023250820.1;refGene_gn=PTPRN2;refGene_tx=XM_023250821.1;refGene_gn=PTPRN2;refGene_tx=XM_023250822.1;refGene_gn=PTPRN2;refGene_tx=XM_023250823.1
A2 170696512 170696534
refGene_gn=PTPRN2;refGene_tx=XM_023250821.1

A2 170770690 170770741
refGene_gn=PTPRN2;refGene_tx=XM_023250820.1;refGene_gn=PTPRN2;refGene_tx=XM_023250822.1;refGene_gn=PTPRN2;refGene_tx=XM_023250823.1
A2 170860319 170860413
refGene_gn=PTPRN2;refGene_tx=XM_023250820.1;refGene_gn=PTPRN2;refGene_tx=XM_023250822.1;refGene_gn=PTPRN2;refGene_tx=XM_023250823.1
A2 170880600 170880652
refGene_gn=NCAPG2;refGene_tx=XM_019826078.2;refGene_gn=NCAPG2;refGene_tx=XM_023250824.1;refGene_gn=NCAPG2;refGene_tx=XM_023250825.1;refGene_gn=NCAPG2;refGene_tx=XM_023250826.1;refGene_gn=NCAPG2;refGene_tx=XM_023250827.1;refGene_gn=NCAPG2;refGene_tx=XM_023250828.1;refGene_gn=NCAPG2;refGene_tx=XM_023250829.1;refGene_gn=NCAPG2;refGene_tx=XM_023250830.1;refGene_gn=NCAPG2;refGene_tx=XM_023250831.1
A2 170886991 170887091
refGene_gn=NCAPG2;refGene_tx=XM_019826078.2;refGene_gn=NCAPG2;refGene_tx=XM_023250824.1;refGene_gn=NCAPG2;refGene_tx=XM_023250825.1;refGene_gn=NCAPG2;refGene_tx=XM_023250826.1;refGene_gn=NCAPG2;refGene_tx=XM_023250827.1;refGene_gn=NCAPG2;refGene_tx=XM_023250828.1;refGene_gn=NCAPG2;refGene_tx=XM_023250829.1;refGene_gn=NCAPG2;refGene_tx=XM_023250830.1;refGene_gn=NCAPG2;refGene_tx=XM_023250831.1;refGene_gn=NCAPG2;refGene_tx=XM_023250832.1
A2 170888258 170888359
refGene_gn=NCAPG2;refGene_tx=XM_019826078.2;refGene_gn=NCAPG2;refGene_tx=XM_023250824.1;refGene_gn=NCAPG2;refGene_tx=XM_023250825.1;refGene_gn=NCAPG2;refGene_tx=XM_023250826.1;refGene_gn=NCAPG2;refGene_tx=XM_023250827.1;refGene_gn=NCAPG2;refGene_tx=XM_023250828.1;refGene_gn=NCAPG2;refGene_tx=XM_023250829.1;refGene_gn=NCAPG2;refGene_tx=XM_023250830.1;refGene_gn=NCAPG2;refGene_tx=XM_023250831.1
A2 170889180 170889284
refGene_gn=NCAPG2;refGene_tx=XM_019826078.2;refGene_gn=NCAPG2;refGene_tx=XM_023250824.1;refGene_gn=NCAPG2;refGene_tx=XM_023250825.1;refGene_gn=NCAPG2;refGene_tx=XM_023250826.1;refGene_gn=NCAPG2;refGene_tx=XM_023250827.1;refGene_gn=NCAPG2;refGene_tx=XM_023250828.1;refGene_gn=NCAPG2;refGene_tx=XM_023250829.1;refGene_gn=NCAPG2;refGene_tx=XM_023250830.1;refGene_gn=NCAPG2;refGene_tx=XM_023250831.1;refGene_gn=NCAPG2;refGene_tx=XM_023250832.1
A2 170892674 170892815
refGene_gn=NCAPG2;refGene_tx=XM_019826078.2;refGene_gn=NCAPG2;refGene_tx=XM_023250824.1;refGene_gn=NCAPG2;refGene_tx=XM_023250825.1;refGene_gn=NCAPG2;refGene_tx=XM_023250826.1;refGene_gn=NCAPG2;refGene_tx=XM_023250827.1;refGene_gn=NCAPG2;refGene_tx=XM_023250828.1;refGene_gn=NCAPG2;refGene_tx=XM_023250829.1;refGene_gn=NCAPG2;refGene_tx=XM_023250831.1;refGene_gn=NCAPG2;refGene_tx=XM_023250832.1
A2 170895175 170895363
refGene_gn=NCAPG2;refGene_tx=XM_019826078.2;refGene_gn=NCAPG2;refGene_tx=XM_023250824.1;refGene_gn=NCAPG2;refGene_tx=XM_023250825.1;refGene_gn=NCAPG2;refGene_tx=XM_023250826.1;refGene_gn=NCAPG2;refGene_tx=XM_023250827.1;refGene_gn=NCAPG2;refGene_tx=XM_023250828.1;refGene_gn=NCAPG2;refGene_tx=XM_023250829.1;refGene_gn=NCAPG2;refGene_tx=XM_023250830.1;refGene_gn=NCAPG2;refGene_tx=XM_023250831.1;refGene_gn=NCAPG2;refGen

e_tx=XM_023250832.1
A2	170896719	170896819
refGene_gn=NCAPG2;refGene_tx=XM_019826078.2;refGene_gn=NCAPG2;refGene_tx=XM_023250824.1;refGene_gn=NCAPG2;refGene_tx=XM_023250825.1;refGene_gn=NCAPG2;refGene_tx=XM_023250826.1;refGene_gn=NCAPG2;refGene_tx=XM_023250827.1;refGene_gn=NCAPG2;refGene_tx=XM_023250828.1;refGene_gn=NCAPG2;refGene_tx=XM_023250829.1;refGene_gn=NCAPG2;refGene_tx=XM_023250830.1;refGene_gn=NCAPG2;refGene_tx=XM_023250831.1;refGene_gn=NCAPG2;refGene_tx=XM_023250832.1
A2	170897260	170897401
refGene_gn=NCAPG2;refGene_tx=XM_019826078.2;refGene_gn=NCAPG2;refGene_tx=XM_023250824.1;refGene_gn=NCAPG2;refGene_tx=XM_023250825.1;refGene_gn=NCAPG2;refGene_tx=XM_023250826.1;refGene_gn=NCAPG2;refGene_tx=XM_023250827.1;refGene_gn=NCAPG2;refGene_tx=XM_023250828.1;refGene_gn=NCAPG2;refGene_tx=XM_023250829.1;refGene_gn=NCAPG2;refGene_tx=XM_023250830.1;refGene_gn=NCAPG2;refGene_tx=XM_023250831.1;refGene_gn=NCAPG2;refGene_tx=XM_023250832.1
A2	170897487	170897604
refGene_gn=NCAPG2;refGene_tx=XM_019826078.2;refGene_gn=NCAPG2;refGene_tx=XM_023250824.1;refGene_gn=NCAPG2;refGene_tx=XM_023250825.1;refGene_gn=NCAPG2;refGene_tx=XM_023250826.1;refGene_gn=NCAPG2;refGene_tx=XM_023250827.1;refGene_gn=NCAPG2;refGene_tx=XM_023250828.1;refGene_gn=NCAPG2;refGene_tx=XM_023250829.1;refGene_gn=NCAPG2;refGene_tx=XM_023250830.1;refGene_gn=NCAPG2;refGene_tx=XM_023250831.1;refGene_gn=NCAPG2;refGene_tx=XM_023250832.1
A2	170898876	170899050
refGene_gn=NCAPG2;refGene_tx=XM_019826078.2;refGene_gn=NCAPG2;refGene_tx=XM_023250824.1;refGene_gn=NCAPG2;refGene_tx=XM_023250825.1;refGene_gn=NCAPG2;refGene_tx=XM_023250826.1;refGene_gn=NCAPG2;refGene_tx=XM_023250827.1;refGene_gn=NCAPG2;refGene_tx=XM_023250828.1;refGene_gn=NCAPG2;refGene_tx=XM_023250829.1;refGene_gn=NCAPG2;refGene_tx=XM_023250830.1;refGene_gn=NCAPG2;refGene_tx=XM_023250831.1;refGene_gn=NCAPG2;refGene_tx=XM_023250832.1
A2	170899168	170899319
refGene_gn=NCAPG2;refGene_tx=XM_019826078.2;refGene_gn=NCAPG2;refGene_tx=XM_023250824.1;refGene_gn=NCAPG2;refGene_tx=XM_023250825.1;refGene_gn=NCAPG2;refGene_tx=XM_023250826.1;refGene_gn=NCAPG2;refGene_tx=XM_023250827.1;refGene_gn=NCAPG2;refGene_tx=XM_023250828.1;refGene_gn=NCAPG2;refGene_tx=XM_023250829.1;refGene_gn=NCAPG2;refGene_tx=XM_023250830.1;refGene_gn=NCAPG2;refGene_tx=XM_023250831.1;refGene_gn=NCAPG2;refGene_tx=XM_023250832.1
A2	170900294	170900365
refGene_gn=NCAPG2;refGene_tx=XM_019826078.2;refGene_gn=NCAPG2;refGene_tx=XM_023250824.1;refGene_gn=NCAPG2;refGene_tx=XM_023250825.1;refGene_gn=NCAPG2;refGene_tx=XM_023250826.1;refGene_gn=NCAPG2;refGene_tx=XM_023250827.1;refGene_gn=NCAPG2;refGene_tx=XM_023250828.1;refGene_gn=NCAPG2;refGene_tx=XM_023250829.1;refGene_gn=NCAPG2;refGene_tx=XM_023250830.1;refGene_gn=NCAPG2;refGene_tx=XM_023250831.1;refGene_gn=NCAPG2;refGene_tx=XM_023250832.1
A2	170902629	170902803

refGene_gn=NCAPG2;refGene_tx=XM_019826078.2;refGene_gn=NCAPG2;refGene_
tx=XM_023250824.1;refGene_gn=NCAPG2;refGene_tx=XM_023250825.1;refGene_
gn=NCAPG2;refGene_tx=XM_023250826.1;refGene_gn=NCAPG2;refGene_tx=XM_02
3250827.1;refGene_gn=NCAPG2;refGene_tx=XM_023250828.1;refGene_gn=NCAPG
2;refGene_tx=XM_023250829.1;refGene_gn=NCAPG2;refGene_tx=XM_023250830.
1;refGene_gn=NCAPG2;refGene_tx=XM_023250831.1;refGene_gn=NCAPG2;refGen
e_tx=XM_023250832.1
A2 170907089 170907187
refGene_gn=NCAPG2;refGene_tx=XM_019826078.2;refGene_gn=NCAPG2;refGene_
tx=XM_023250824.1;refGene_gn=NCAPG2;refGene_tx=XM_023250825.1;refGene_
gn=NCAPG2;refGene_tx=XM_023250826.1;refGene_gn=NCAPG2;refGene_tx=XM_02
3250827.1;refGene_gn=NCAPG2;refGene_tx=XM_023250828.1;refGene_gn=NCAPG
2;refGene_tx=XM_023250829.1;refGene_gn=NCAPG2;refGene_tx=XM_023250830.
1;refGene_gn=NCAPG2;refGene_tx=XM_023250831.1;refGene_gn=NCAPG2;refGen
e_tx=XM_023250832.1
A2 170907415 170907638
refGene_gn=NCAPG2;refGene_tx=XM_019826078.2;refGene_gn=NCAPG2;refGene_
tx=XM_023250824.1;refGene_gn=NCAPG2;refGene_tx=XM_023250825.1;refGene_
gn=NCAPG2;refGene_tx=XM_023250826.1;refGene_gn=NCAPG2;refGene_tx=XM_02
3250827.1;refGene_gn=NCAPG2;refGene_tx=XM_023250828.1;refGene_gn=NCAPG
2;refGene_tx=XM_023250829.1;refGene_gn=NCAPG2;refGene_tx=XM_023250830.
1;refGene_gn=NCAPG2;refGene_tx=XM_023250831.1;refGene_gn=NCAPG2;refGen
e_tx=XM_023250832.1
A2 170913015 170913168
refGene_gn=NCAPG2;refGene_tx=XM_019826078.2;refGene_gn=NCAPG2;refGene_
tx=XM_023250824.1;refGene_gn=NCAPG2;refGene_tx=XM_023250825.1;refGene_
gn=NCAPG2;refGene_tx=XM_023250826.1;refGene_gn=NCAPG2;refGene_tx=XM_02
3250827.1;refGene_gn=NCAPG2;refGene_tx=XM_023250828.1;refGene_gn=NCAPG
2;refGene_tx=XM_023250829.1;refGene_gn=NCAPG2;refGene_tx=XM_023250830.
1;refGene_gn=NCAPG2;refGene_tx=XM_023250831.1;refGene_gn=NCAPG2;refGen
e_tx=XM_023250832.1
A2 170914834 170915014
refGene_gn=NCAPG2;refGene_tx=XM_019826078.2;refGene_gn=NCAPG2;refGene_
tx=XM_023250824.1;refGene_gn=NCAPG2;refGene_tx=XM_023250825.1;refGene_
gn=NCAPG2;refGene_tx=XM_023250826.1;refGene_gn=NCAPG2;refGene_tx=XM_02
3250827.1;refGene_gn=NCAPG2;refGene_tx=XM_023250828.1;refGene_gn=NCAPG
2;refGene_tx=XM_023250829.1;refGene_gn=NCAPG2;refGene_tx=XM_023250830.
1;refGene_gn=NCAPG2;refGene_tx=XM_023250831.1;refGene_gn=NCAPG2;refGen
e_tx=XM_023250832.1
A2 170921018 170921144
refGene_gn=NCAPG2;refGene_tx=XM_019826078.2;refGene_gn=NCAPG2;refGene_
tx=XM_023250824.1;refGene_gn=NCAPG2;refGene_tx=XM_023250825.1;refGene_
gn=NCAPG2;refGene_tx=XM_023250826.1;refGene_gn=NCAPG2;refGene_tx=XM_02
3250827.1;refGene_gn=NCAPG2;refGene_tx=XM_023250828.1;refGene_gn=NCAPG
2;refGene_tx=XM_023250829.1;refGene_gn=NCAPG2;refGene_tx=XM_023250830.
1;refGene_gn=NCAPG2;refGene_tx=XM_023250831.1;refGene_gn=NCAPG2;refGen
e_tx=XM_023250832.1
A2 170921731 170921827
refGene_gn=NCAPG2;refGene_tx=XM_019826078.2;refGene_gn=NCAPG2;refGene_
tx=XM_023250824.1;refGene_gn=NCAPG2;refGene_tx=XM_023250825.1;refGene_

gn=NCAPG2;refGene_tx=XM_023250826.1;refGene_gn=NCAPG2;refGene_tx=XM_023250827.1;refGene_gn=NCAPG2;refGene_tx=XM_023250828.1;refGene_gn=NCAPG2;refGene_tx=XM_023250829.1;refGene_gn=NCAPG2;refGene_tx=XM_023250830.1;refGene_gn=NCAPG2;refGene_tx=XM_023250831.1;refGene_gn=NCAPG2;refGene_tx=XM_023250832.1
A2 170925157 170925244
refGene_gn=NCAPG2;refGene_tx=XM_019826078.2;refGene_gn=NCAPG2;refGene_tx=XM_023250824.1;refGene_gn=NCAPG2;refGene_tx=XM_023250825.1;refGene_gn=NCAPG2;refGene_tx=XM_023250826.1;refGene_gn=NCAPG2;refGene_tx=XM_023250827.1;refGene_gn=NCAPG2;refGene_tx=XM_023250828.1;refGene_gn=NCAPG2;refGene_tx=XM_023250829.1;refGene_gn=NCAPG2;refGene_tx=XM_023250830.1;refGene_gn=NCAPG2;refGene_tx=XM_023250831.1;refGene_gn=NCAPG2;refGene_tx=XM_023250832.1
A2 170928331 170928401
refGene_gn=NCAPG2;refGene_tx=XM_019826078.2;refGene_gn=NCAPG2;refGene_tx=XM_023250824.1;refGene_gn=NCAPG2;refGene_tx=XM_023250825.1;refGene_gn=NCAPG2;refGene_tx=XM_023250826.1;refGene_gn=NCAPG2;refGene_tx=XM_023250827.1;refGene_gn=NCAPG2;refGene_tx=XM_023250828.1;refGene_gn=NCAPG2;refGene_tx=XM_023250829.1;refGene_gn=NCAPG2;refGene_tx=XM_023250830.1;refGene_gn=NCAPG2;refGene_tx=XM_023250831.1;refGene_gn=NCAPG2;refGene_tx=XM_023250832.1
A2 170930149 170930244
refGene_gn=NCAPG2;refGene_tx=XM_019826078.2;refGene_gn=NCAPG2;refGene_tx=XM_023250824.1;refGene_gn=NCAPG2;refGene_tx=XM_023250825.1;refGene_gn=NCAPG2;refGene_tx=XM_023250826.1;refGene_gn=NCAPG2;refGene_tx=XM_023250827.1;refGene_gn=NCAPG2;refGene_tx=XM_023250828.1;refGene_gn=NCAPG2;refGene_tx=XM_023250829.1;refGene_gn=NCAPG2;refGene_tx=XM_023250830.1;refGene_gn=NCAPG2;refGene_tx=XM_023250831.1;refGene_gn=NCAPG2;refGene_tx=XM_023250832.1
A2 170932877 170933012
refGene_gn=NCAPG2;refGene_tx=XM_019826078.2;refGene_gn=NCAPG2;refGene_tx=XM_023250824.1;refGene_gn=NCAPG2;refGene_tx=XM_023250825.1;refGene_gn=NCAPG2;refGene_tx=XM_023250826.1;refGene_gn=NCAPG2;refGene_tx=XM_023250827.1;refGene_gn=NCAPG2;refGene_tx=XM_023250828.1;refGene_gn=NCAPG2;refGene_tx=XM_023250829.1;refGene_gn=NCAPG2;refGene_tx=XM_023250830.1;refGene_gn=NCAPG2;refGene_tx=XM_023250831.1;refGene_gn=NCAPG2;refGene_tx=XM_023250832.1
A2 170935330 170935485
refGene_gn=NCAPG2;refGene_tx=XM_019826078.2;refGene_gn=NCAPG2;refGene_tx=XM_023250824.1;refGene_gn=NCAPG2;refGene_tx=XM_023250825.1;refGene_gn=NCAPG2;refGene_tx=XM_023250826.1;refGene_gn=NCAPG2;refGene_tx=XM_023250827.1;refGene_gn=NCAPG2;refGene_tx=XM_023250828.1;refGene_gn=NCAPG2;refGene_tx=XM_023250829.1;refGene_gn=NCAPG2;refGene_tx=XM_023250830.1;refGene_gn=NCAPG2;refGene_tx=XM_023250831.1;refGene_gn=NCAPG2;refGene_tx=XM_023250832.1
A2 170937239 170937354
refGene_gn=NCAPG2;refGene_tx=XM_019826078.2;refGene_gn=NCAPG2;refGene_tx=XM_023250824.1;refGene_gn=NCAPG2;refGene_tx=XM_023250825.1;refGene_gn=NCAPG2;refGene_tx=XM_023250826.1;refGene_gn=NCAPG2;refGene_tx=XM_023250827.1;refGene_gn=NCAPG2;refGene_tx=XM_023250828.1;refGene_gn=NCAPG

2;refGene_tx=XM_023250829.1;refGene_gn=NCAPG2;refGene_tx=XM_023250830.1;refGene_gn=NCAPG2;refGene_tx=XM_023250831.1;refGene_gn=NCAPG2;refGene_tx=XM_023250832.1
A2 170937952 170938141
refGene_gn=NCAPG2;refGene_tx=XM_019826078.2;refGene_gn=NCAPG2;refGene_tx=XM_023250824.1;refGene_gn=NCAPG2;refGene_tx=XM_023250825.1;refGene_gn=NCAPG2;refGene_tx=XM_023250826.1;refGene_gn=NCAPG2;refGene_tx=XM_023250827.1;refGene_gn=NCAPG2;refGene_tx=XM_023250828.1;refGene_gn=NCAPG2;refGene_tx=XM_023250829.1;refGene_gn=NCAPG2;refGene_tx=XM_023250830.1;refGene_gn=NCAPG2;refGene_tx=XM_023250831.1;refGene_gn=NCAPG2;refGene_tx=XM_023250832.1
A2 170942729 170942807
refGene_gn=NCAPG2;refGene_tx=XM_019826078.2;refGene_gn=NCAPG2;refGene_tx=XM_023250824.1;refGene_gn=NCAPG2;refGene_tx=XM_023250825.1;refGene_gn=NCAPG2;refGene_tx=XM_023250826.1;refGene_gn=NCAPG2;refGene_tx=XM_023250827.1;refGene_gn=NCAPG2;refGene_tx=XM_023250828.1;refGene_gn=NCAPG2;refGene_tx=XM_023250829.1;refGene_gn=NCAPG2;refGene_tx=XM_023250830.1;refGene_gn=NCAPG2;refGene_tx=XM_023250832.1
A2 170962437 170962483
refGene_gn=ESYT2;refGene_tx=XM_023250833.1;refGene_gn=ESYT2;refGene_tx=XM_023250834.1;refGene_gn=ESYT2;refGene_tx=XM_023250835.1;refGene_gn=ESYT2;refGene_tx=XM_023250836.1
A2 170962579 170962629
refGene_gn=ESYT2;refGene_tx=XM_023250833.1;refGene_gn=ESYT2;refGene_tx=XM_023250834.1;refGene_gn=ESYT2;refGene_tx=XM_023250835.1;refGene_gn=ESYT2;refGene_tx=XM_023250836.1
A2 170963435 170963541
refGene_gn=ESYT2;refGene_tx=XM_023250833.1;refGene_gn=ESYT2;refGene_tx=XM_023250834.1;refGene_gn=ESYT2;refGene_tx=XM_023250835.1;refGene_gn=ESYT2;refGene_tx=XM_023250836.1
A2 170965217 170965349
refGene_gn=ESYT2;refGene_tx=XM_023250833.1;refGene_gn=ESYT2;refGene_tx=XM_023250834.1;refGene_gn=ESYT2;refGene_tx=XM_023250835.1;refGene_gn=ESYT2;refGene_tx=XM_023250836.1
A2 170967163 170967262
refGene_gn=ESYT2;refGene_tx=XM_023250833.1;refGene_gn=ESYT2;refGene_tx=XM_023250834.1;refGene_gn=ESYT2;refGene_tx=XM_023250835.1;refGene_gn=ESYT2;refGene_tx=XM_023250836.1
A2 170968408 170968794
refGene_gn=ESYT2;refGene_tx=XM_023250833.1;refGene_gn=ESYT2;refGene_tx=XM_023250834.1;refGene_gn=ESYT2;refGene_tx=XM_023250835.1;refGene_gn=ESYT2;refGene_tx=XM_023250836.1
A2 170970540 170970690
refGene_gn=ESYT2;refGene_tx=XM_023250833.1;refGene_gn=ESYT2;refGene_tx=XM_023250834.1;refGene_gn=ESYT2;refGene_tx=XM_023250835.1;refGene_gn=ESYT2;refGene_tx=XM_023250836.1
A2 170972770 170972857
refGene_gn=ESYT2;refGene_tx=XM_023250833.1;refGene_gn=ESYT2;refGene_tx=XM_023250834.1;refGene_gn=ESYT2;refGene_tx=XM_023250835.1;refGene_gn=ESYT2;refGene_tx=XM_023250836.1

A2 170973761 170973836
refGene_gn=ESYT2;refGene_tx=XM_023250833.1;refGene_gn=ESYT2;refGene_tx=XM_023250834.1;refGene_gn=ESYT2;refGene_tx=XM_023250835.1;refGene_gn=ESYT2;refGene_tx=XM_023250836.1
A2 170977044 170977107
refGene_gn=ESYT2;refGene_tx=XM_023250833.1;refGene_gn=ESYT2;refGene_tx=XM_023250835.1
A2 170980853 170980955
refGene_gn=ESYT2;refGene_tx=XM_023250833.1;refGene_gn=ESYT2;refGene_tx=XM_023250834.1
A2 170982764 170982860
refGene_gn=ESYT2;refGene_tx=XM_023250833.1;refGene_gn=ESYT2;refGene_tx=XM_023250834.1;refGene_gn=ESYT2;refGene_tx=XM_023250835.1;refGene_gn=ESYT2;refGene_tx=XM_023250836.1
A2 170983527 170983617
refGene_gn=ESYT2;refGene_tx=XM_023250833.1;refGene_gn=ESYT2;refGene_tx=XM_023250834.1;refGene_gn=ESYT2;refGene_tx=XM_023250835.1;refGene_gn=ESYT2;refGene_tx=XM_023250836.1
A2 170984685 170984734
refGene_gn=ESYT2;refGene_tx=XM_023250833.1;refGene_gn=ESYT2;refGene_tx=XM_023250834.1;refGene_gn=ESYT2;refGene_tx=XM_023250835.1;refGene_gn=ESYT2;refGene_tx=XM_023250836.1
A2 170986154 170986237
refGene_gn=ESYT2;refGene_tx=XM_023250833.1;refGene_gn=ESYT2;refGene_tx=XM_023250834.1;refGene_gn=ESYT2;refGene_tx=XM_023250835.1;refGene_gn=ESYT2;refGene_tx=XM_023250836.1
A2 170987263 170987440
refGene_gn=ESYT2;refGene_tx=XM_023250833.1;refGene_gn=ESYT2;refGene_tx=XM_023250834.1;refGene_gn=ESYT2;refGene_tx=XM_023250835.1;refGene_gn=ESYT2;refGene_tx=XM_023250836.1
A2 170990949 170991070
refGene_gn=ESYT2;refGene_tx=XM_023250833.1;refGene_gn=ESYT2;refGene_tx=XM_023250834.1;refGene_gn=ESYT2;refGene_tx=XM_023250835.1;refGene_gn=ESYT2;refGene_tx=XM_023250836.1
A2 170996923 170996979
refGene_gn=ESYT2;refGene_tx=XM_023250833.1;refGene_gn=ESYT2;refGene_tx=XM_023250834.1;refGene_gn=ESYT2;refGene_tx=XM_023250835.1;refGene_gn=ESYT2;refGene_tx=XM_023250836.1
A2 171005641 171005731
refGene_gn=ESYT2;refGene_tx=XM_023250833.1;refGene_gn=ESYT2;refGene_tx=XM_023250834.1;refGene_gn=ESYT2;refGene_tx=XM_023250835.1;refGene_gn=ESYT2;refGene_tx=XM_023250836.1
A2 171005980 171006053
refGene_gn=ESYT2;refGene_tx=XM_023250833.1;refGene_gn=ESYT2;refGene_tx=XM_023250834.1;refGene_gn=ESYT2;refGene_tx=XM_023250835.1;refGene_gn=ESYT2;refGene_tx=XM_023250836.1
A2 171010188 171010265
refGene_gn=ESYT2;refGene_tx=XM_023250833.1;refGene_gn=ESYT2;refGene_tx=XM_023250834.1;refGene_gn=ESYT2;refGene_tx=XM_023250835.1;refGene_gn=ESYT2;refGene_tx=XM_023250836.1

A2	171013009	171013144
refGene_gn=ESYT2;refGene_tx=XM_023250833.1;refGene_gn=ESYT2;refGene_tx=XM_023250834.1;refGene_gn=ESYT2;refGene_tx=XM_023250835.1;refGene_gn=ESYT2;refGene_tx=XM_023250836.1
A2	171013991	171014033
refGene_gn=ESYT2;refGene_tx=XM_023250833.1;refGene_gn=ESYT2;refGene_tx=XM_023250834.1;refGene_gn=ESYT2;refGene_tx=XM_023250835.1;refGene_gn=ESYT2;refGene_tx=XM_023250836.1
A2	171041582	171041894
refGene_gn=ESYT2;refGene_tx=XM_023250833.1;refGene_gn=ESYT2;refGene_tx=XM_023250834.1;refGene_gn=ESYT2;refGene_tx=XM_023250835.1;refGene_gn=ESYT2;refGene_tx=XM_023250836.1
A2	171052038	171052053
refGene_gn=WDR60;refGene_tx=XM_003983271.5;refGene_gn=WDR60;refGene_tx=XM_006929551.4;refGene_gn=WDR60;refGene_tx=XM_006929552.4;refGene_gn=WDR60;refGene_tx=XM_006929553.4;refGene_gn=WDR60;refGene_tx=XM_019826089.2;refGene_gn=WDR60;refGene_tx=XM_019826090.2
A2	171059113	171059167
refGene_gn=WDR60;refGene_tx=XM_003983271.5;refGene_gn=WDR60;refGene_tx=XM_006929551.4;refGene_gn=WDR60;refGene_tx=XM_006929552.4;refGene_gn=WDR60;refGene_tx=XM_006929553.4;refGene_gn=WDR60;refGene_tx=XM_019826089.2;refGene_gn=WDR60;refGene_tx=XM_019826090.2
A2	171060328	171060743
refGene_gn=WDR60;refGene_tx=XM_003983271.5;refGene_gn=WDR60;refGene_tx=XM_006929551.4;refGene_gn=WDR60;refGene_tx=XM_006929552.4;refGene_gn=WDR60;refGene_tx=XM_006929553.4;refGene_gn=WDR60;refGene_tx=XM_019826089.2;refGene_gn=WDR60;refGene_tx=XM_019826090.2
A2	171063562	171063654
refGene_gn=WDR60;refGene_tx=XM_003983271.5;refGene_gn=WDR60;refGene_tx=XM_006929551.4;refGene_gn=WDR60;refGene_tx=XM_006929552.4;refGene_gn=WDR60;refGene_tx=XM_006929553.4;refGene_gn=WDR60;refGene_tx=XM_019826089.2;refGene_gn=WDR60;refGene_tx=XM_019826090.2
A2	171065568	171065877
refGene_gn=WDR60;refGene_tx=XM_003983271.5;refGene_gn=WDR60;refGene_tx=XM_006929551.4;refGene_gn=WDR60;refGene_tx=XM_006929552.4;refGene_gn=WDR60;refGene_tx=XM_006929553.4;refGene_gn=WDR60;refGene_tx=XM_019826089.2;refGene_gn=WDR60;refGene_tx=XM_019826090.2
A2	171070391	171070468
refGene_gn=WDR60;refGene_tx=XM_003983271.5;refGene_gn=WDR60;refGene_tx=XM_006929551.4;refGene_gn=WDR60;refGene_tx=XM_006929552.4;refGene_gn=WDR60;refGene_tx=XM_006929553.4;refGene_gn=WDR60;refGene_tx=XM_019826089.2;refGene_gn=WDR60;refGene_tx=XM_019826090.2
A2	171072842	171072897
refGene_gn=WDR60;refGene_tx=XM_003983271.5;refGene_gn=WDR60;refGene_tx=XM_006929551.4;refGene_gn=WDR60;refGene_tx=XM_006929552.4;refGene_gn=WDR60;refGene_tx=XM_006929553.4;refGene_gn=WDR60;refGene_tx=XM_019826089.2;refGene_gn=WDR60;refGene_tx=XM_019826090.2
A2	171075695	171075758
refGene_gn=WDR60;refGene_tx=XM_003983271.5;refGene_gn=WDR60;refGene_tx=XM_006929551.4;refGene_gn=WDR60;refGene_tx=XM_006929552.4;refGene_gn=

WDR60;refGene_tx=XM_019826089.2;refGene_gn=WDR60;refGene_tx=XM_0198260
90.2
A2	171079817	171079895
refGene_gn=WDR60;refGene_tx=XM_003983271.5;refGene_gn=WDR60;refGene_tx
=XM_006929551.4;refGene_gn=WDR60;refGene_tx=XM_006929552.4;refGene_gn=
WDR60;refGene_tx=XM_006929553.4;refGene_gn=WDR60;refGene_tx=XM_0198260
89.2;refGene_gn=WDR60;refGene_tx=XM_019826090.2
A2	171080459	171080676
refGene_gn=WDR60;refGene_tx=XM_003983271.5;refGene_gn=WDR60;refGene_tx
=XM_006929551.4;refGene_gn=WDR60;refGene_tx=XM_006929552.4;refGene_gn=
WDR60;refGene_tx=XM_006929553.4;refGene_gn=WDR60;refGene_tx=XM_0198260
89.2;refGene_gn=WDR60;refGene_tx=XM_019826090.2
A2	171081511	171081614
refGene_gn=WDR60;refGene_tx=XM_003983271.5;refGene_gn=WDR60;refGene_tx
=XM_006929551.4;refGene_gn=WDR60;refGene_tx=XM_006929552.4;refGene_gn=
WDR60;refGene_tx=XM_006929553.4;refGene_gn=WDR60;refGene_tx=XM_0198260
89.2;refGene_gn=WDR60;refGene_tx=XM_019826090.2
A2	171090201	171090331
refGene_gn=WDR60;refGene_tx=XM_003983271.5;refGene_gn=WDR60;refGene_tx
=XM_006929551.4;refGene_gn=WDR60;refGene_tx=XM_006929552.4;refGene_gn=
WDR60;refGene_tx=XM_006929553.4;refGene_gn=WDR60;refGene_tx=XM_0198260
89.2;refGene_gn=WDR60;refGene_tx=XM_019826090.2
A2	171092631	171092743
refGene_gn=WDR60;refGene_tx=XM_003983271.5;refGene_gn=WDR60;refGene_tx
=XM_006929551.4;refGene_gn=WDR60;refGene_tx=XM_006929552.4;refGene_gn=
WDR60;refGene_tx=XM_006929553.4;refGene_gn=WDR60;refGene_tx=XM_0198260
89.2;refGene_gn=WDR60;refGene_tx=XM_019826090.2
A2	171093496	171093582
refGene_gn=WDR60;refGene_tx=XM_003983271.5;refGene_gn=WDR60;refGene_tx
=XM_006929551.4;refGene_gn=WDR60;refGene_tx=XM_006929552.4;refGene_gn=
WDR60;refGene_tx=XM_006929553.4;refGene_gn=WDR60;refGene_tx=XM_0198260
89.2;refGene_gn=WDR60;refGene_tx=XM_019826090.2
A2	171094829	171094959
refGene_gn=WDR60;refGene_tx=XM_003983271.5;refGene_gn=WDR60;refGene_tx
=XM_006929551.4;refGene_gn=WDR60;refGene_tx=XM_006929552.4;refGene_gn=
WDR60;refGene_tx=XM_006929553.4;refGene_gn=WDR60;refGene_tx=XM_0198260
89.2;refGene_gn=WDR60;refGene_tx=XM_019826090.2
A2	171097104	171097277
refGene_gn=WDR60;refGene_tx=XM_003983271.5;refGene_gn=WDR60;refGene_tx
=XM_006929551.4;refGene_gn=WDR60;refGene_tx=XM_006929552.4;refGene_gn=
WDR60;refGene_tx=XM_006929553.4;refGene_gn=WDR60;refGene_tx=XM_0198260
89.2;refGene_gn=WDR60;refGene_tx=XM_019826090.2
A2	171098134	171098297
refGene_gn=WDR60;refGene_tx=XM_003983271.5;refGene_gn=WDR60;refGene_tx
=XM_006929551.4;refGene_gn=WDR60;refGene_tx=XM_006929552.4;refGene_gn=
WDR60;refGene_tx=XM_006929553.4;refGene_gn=WDR60;refGene_tx=XM_0198260
89.2;refGene_gn=WDR60;refGene_tx=XM_019826090.2
A2	171103322	171103436
refGene_gn=WDR60;refGene_tx=XM_003983271.5;refGene_gn=WDR60;refGene_tx
=XM_006929551.4;refGene_gn=WDR60;refGene_tx=XM_006929552.4;refGene_gn=

WDR60;refGene_tx=XM_006929553.4;refGene_gn=WDR60;refGene_tx=XM_019826089.2;refGene_gn=WDR60;refGene_tx=XM_019826090.2
A2 171103531 171103593
refGene_gn=WDR60;refGene_tx=XM_003983271.5;refGene_gn=WDR60;refGene_tx=XM_006929551.4;refGene_gn=WDR60;refGene_tx=XM_006929552.4;refGene_gn=WDR60;refGene_tx=XM_006929553.4;refGene_gn=WDR60;refGene_tx=XM_019826089.2;refGene_gn=WDR60;refGene_tx=XM_019826090.2
A2 171104034 171104086
refGene_gn=WDR60;refGene_tx=XM_003983271.5;refGene_gn=WDR60;refGene_tx=XM_006929551.4;refGene_gn=WDR60;refGene_tx=XM_006929552.4;refGene_gn=WDR60;refGene_tx=XM_006929553.4;refGene_gn=WDR60;refGene_tx=XM_019826089.2;refGene_gn=WDR60;refGene_tx=XM_019826090.2
A2 171107113 171107174
refGene_gn=WDR60;refGene_tx=XM_003983271.5;refGene_gn=WDR60;refGene_tx=XM_006929551.4;refGene_gn=WDR60;refGene_tx=XM_006929552.4;refGene_gn=WDR60;refGene_tx=XM_006929553.4;refGene_gn=WDR60;refGene_tx=XM_019826089.2;refGene_gn=WDR60;refGene_tx=XM_019826090.2
A2 171110324 171110424
refGene_gn=WDR60;refGene_tx=XM_003983271.5;refGene_gn=WDR60;refGene_tx=XM_006929551.4;refGene_gn=WDR60;refGene_tx=XM_006929552.4;refGene_gn=WDR60;refGene_tx=XM_006929553.4;refGene_gn=WDR60;refGene_tx=XM_019826089.2;refGene_gn=WDR60;refGene_tx=XM_019826090.2
A2 171110591 171110723
refGene_gn=WDR60;refGene_tx=XM_003983271.5;refGene_gn=WDR60;refGene_tx=XM_006929551.4;refGene_gn=WDR60;refGene_tx=XM_006929552.4;refGene_gn=WDR60;refGene_tx=XM_006929553.4;refGene_gn=WDR60;refGene_tx=XM_019826089.2;refGene_gn=WDR60;refGene_tx=XM_019826090.2
A2 171111093 171111306
refGene_gn=WDR60;refGene_tx=XM_019826090.2
A2 171112511 171112735
refGene_gn=WDR60;refGene_tx=XM_003983271.5;refGene_gn=WDR60;refGene_tx=XM_006929551.4;refGene_gn=WDR60;refGene_tx=XM_006929552.4;refGene_gn=WDR60;refGene_tx=XM_006929553.4;refGene_gn=WDR60;refGene_tx=XM_019826089.2
A2 171115309 171115484
refGene_gn=WDR60;refGene_tx=XM_003983271.5;refGene_gn=WDR60;refGene_tx=XM_006929551.4;refGene_gn=WDR60;refGene_tx=XM_006929552.4;refGene_gn=WDR60;refGene_tx=XM_006929553.4
A2 171127556 171127692
refGene_gn=WDR60;refGene_tx=XM_019826089.2
A2 171193227 171193401
refGene_gn=VIPR2;refGene_tx=XM_003983257.4;refGene_gn=VIPR2;refGene_tx=XM_011280616.2
A2 171194091 171194133
refGene_gn=VIPR2;refGene_tx=XM_003983257.4;refGene_gn=VIPR2;refGene_tx=XM_011280616.2
A2 171194206 171194336
refGene_gn=VIPR2;refGene_tx=XM_003983257.4;refGene_gn=VIPR2;refGene_tx=XM_011280616.2
A2 171197351 171197443

refGene_gn=VIPR2;refGene_tx=XM_003983257.4;refGene_gn=VIPR2;refGene_tx=XM_011280616.2
A2 171197959 171198029
refGene_gn=VIPR2;refGene_tx=XM_003983257.4;refGene_gn=VIPR2;refGene_tx=XM_011280616.2
A2 171199502 171199563
refGene_gn=VIPR2;refGene_tx=XM_003983257.4;refGene_gn=VIPR2;refGene_tx=XM_011280616.2
A2 171200239 171200390
refGene_gn=VIPR2;refGene_tx=XM_003983257.4;refGene_gn=VIPR2;refGene_tx=XM_011280616.2
A2 171205623 171205765
refGene_gn=VIPR2;refGene_tx=XM_003983257.4;refGene_gn=VIPR2;refGene_tx=XM_011280616.2
A2 171211251 171211349
refGene_gn=VIPR2;refGene_tx=XM_003983257.4;refGene_gn=VIPR2;refGene_tx=XM_011280616.2
A2 171232632 171232730
refGene_gn=VIPR2;refGene_tx=XM_003983257.4;refGene_gn=VIPR2;refGene_tx=XM_011280616.2
A2 171238332 171238440
refGene_gn=VIPR2;refGene_tx=XM_003983257.4
A2 171260244 171260344
refGene_gn=VIPR2;refGene_tx=XM_003983257.4;refGene_gn=VIPR2;refGene_tx=XM_011280616.2
A2 171262430 171262493
refGene_gn=VIPR2;refGene_tx=XM_003983257.4;refGene_gn=VIPR2;refGene_tx=XM_011280616.2
A3 160352 160770
refGene_gn=LOC111559734;refGene_tx=XM_023250927.1
A3 193494 193700
refGene_gn=LOC111559734;refGene_tx=XM_023250927.1
A3 336905 337285
refGene_gn=PCMTD2;refGene_tx=XM_019826177.1;refGene_gn=PCMTD2;refGene_tx=XM_023250928.1;refGene_gn=PCMTD2;refGene_tx=XM_023250929.1
A3 340463 340587
refGene_gn=PCMTD2;refGene_tx=XM_019826177.1;refGene_gn=PCMTD2;refGene_tx=XM_023250928.1;refGene_gn=PCMTD2;refGene_tx=XM_023250929.1
A3 346706 346878
refGene_gn=PCMTD2;refGene_tx=XM_019826177.1;refGene_gn=PCMTD2;refGene_tx=XM_023250928.1;refGene_gn=PCMTD2;refGene_tx=XM_023250929.1
A3 347885 347988
refGene_gn=PCMTD2;refGene_tx=XM_019826177.1;refGene_gn=PCMTD2;refGene_tx=XM_023250928.1;refGene_gn=PCMTD2;refGene_tx=XM_023250929.1
A3 351656 351987
refGene_gn=PCMTD2;refGene_tx=XM_019826177.1;refGene_gn=PCMTD2;refGene_tx=XM_023250928.1;refGene_gn=PCMTD2;refGene_tx=XM_023250929.1
A3 355828 355906 refGene_gn=PCMTD2;refGene_tx=XM_019826177.1
A3 369290 369431
refGene_gn=MYT1;refGene_tx=XM_011280801.3;refGene_gn=MYT1;refGene_tx=X

M_019826178.2;refGene_gn=MYT1;refGene_tx=XM_019826179.1;refGene_gn=MYT1;refGene_tx=XM_019826181.1;refGene_gn=MYT1;refGene_tx=XM_019826182.1;refGene_gn=MYT1;refGene_tx=XM_023250930.1
A3 369788 369932
refGene_gn=MYT1;refGene_tx=XM_011280801.3;refGene_gn=MYT1;refGene_tx=XM_019826178.2;refGene_gn=MYT1;refGene_tx=XM_019826179.1;refGene_gn=MYT1;refGene_tx=XM_019826181.1;refGene_gn=MYT1;refGene_tx=XM_019826182.1;refGene_gn=MYT1;refGene_tx=XM_023250930.1
A3 372034 372138
refGene_gn=MYT1;refGene_tx=XM_011280801.3;refGene_gn=MYT1;refGene_tx=XM_019826178.2;refGene_gn=MYT1;refGene_tx=XM_019826179.1;refGene_gn=MYT1;refGene_tx=XM_019826181.1;refGene_gn=MYT1;refGene_tx=XM_019826182.1;refGene_gn=MYT1;refGene_tx=XM_023250930.1
A3 372773 372865
refGene_gn=MYT1;refGene_tx=XM_011280801.3;refGene_gn=MYT1;refGene_tx=XM_019826178.2;refGene_gn=MYT1;refGene_tx=XM_019826179.1;refGene_gn=MYT1;refGene_tx=XM_019826181.1;refGene_gn=MYT1;refGene_tx=XM_019826182.1;refGene_gn=MYT1;refGene_tx=XM_023250930.1
A3 373918 374140
refGene_gn=MYT1;refGene_tx=XM_011280801.3;refGene_gn=MYT1;refGene_tx=XM_019826178.2;refGene_gn=MYT1;refGene_tx=XM_019826179.1;refGene_gn=MYT1;refGene_tx=XM_019826181.1;refGene_gn=MYT1;refGene_tx=XM_019826182.1;refGene_gn=MYT1;refGene_tx=XM_023250930.1
A3 378330 378414
refGene_gn=MYT1;refGene_tx=XM_011280801.3;refGene_gn=MYT1;refGene_tx=XM_019826178.2;refGene_gn=MYT1;refGene_tx=XM_019826179.1;refGene_gn=MYT1;refGene_tx=XM_019826181.1;refGene_gn=MYT1;refGene_tx=XM_019826182.1;refGene_gn=MYT1;refGene_tx=XM_023250930.1
A3 378777 378840
refGene_gn=MYT1;refGene_tx=XM_011280801.3;refGene_gn=MYT1;refGene_tx=XM_019826178.2;refGene_gn=MYT1;refGene_tx=XM_019826179.1;refGene_gn=MYT1;refGene_tx=XM_019826181.1;refGene_gn=MYT1;refGene_tx=XM_019826182.1;refGene_gn=MYT1;refGene_tx=XM_023250930.1
A3 383527 383596
refGene_gn=MYT1;refGene_tx=XM_011280801.3;refGene_gn=MYT1;refGene_tx=XM_019826178.2;refGene_gn=MYT1;refGene_tx=XM_019826179.1;refGene_gn=MYT1;refGene_tx=XM_019826181.1;refGene_gn=MYT1;refGene_tx=XM_019826182.1;refGene_gn=MYT1;refGene_tx=XM_023250930.1
A3 383707 383770
refGene_gn=MYT1;refGene_tx=XM_011280801.3;refGene_gn=MYT1;refGene_tx=XM_019826178.2;refGene_gn=MYT1;refGene_tx=XM_019826179.1;refGene_gn=MYT1;refGene_tx=XM_019826181.1;refGene_gn=MYT1;refGene_tx=XM_019826182.1;refGene_gn=MYT1;refGene_tx=XM_023250930.1
A3 384799 384954
refGene_gn=MYT1;refGene_tx=XM_011280801.3;refGene_gn=MYT1;refGene_tx=XM_019826178.2;refGene_gn=MYT1;refGene_tx=XM_019826179.1;refGene_gn=MYT1;refGene_tx=XM_019826181.1;refGene_gn=MYT1;refGene_tx=XM_019826182.1;refGene_gn=MYT1;refGene_tx=XM_023250930.1
A3 387849 388119
refGene_gn=MYT1;refGene_tx=XM_011280801.3;refGene_gn=MYT1;refGene_tx=X

M_019826178.2;refGene_gn=MYT1;refGene_tx=XM_019826179.1;refGene_gn=MYT1;refGene_tx=XM_019826181.1;refGene_gn=MYT1;refGene_tx=XM_019826182.1;refGene_gn=MYT1;refGene_tx=XM_023250930.1
A3 388646 388834
refGene_gn=MYT1;refGene_tx=XM_011280801.3;refGene_gn=MYT1;refGene_tx=XM_019826178.2;refGene_gn=MYT1;refGene_tx=XM_019826179.1;refGene_gn=MYT1;refGene_tx=XM_019826181.1;refGene_gn=MYT1;refGene_tx=XM_019826182.1;refGene_gn=MYT1;refGene_tx=XM_023250930.1
A3 390214 390429
refGene_gn=MYT1;refGene_tx=XM_011280801.3;refGene_gn=MYT1;refGene_tx=XM_019826178.2;refGene_gn=MYT1;refGene_tx=XM_019826179.1;refGene_gn=MYT1;refGene_tx=XM_019826181.1;refGene_gn=MYT1;refGene_tx=XM_019826182.1;refGene_gn=MYT1;refGene_tx=XM_023250930.1
A3 393277 393391
refGene_gn=MYT1;refGene_tx=XM_011280801.3;refGene_gn=MYT1;refGene_tx=XM_019826178.2;refGene_gn=MYT1;refGene_tx=XM_019826179.1;refGene_gn=MYT1;refGene_tx=XM_019826181.1;refGene_gn=MYT1;refGene_tx=XM_019826182.1;refGene_gn=MYT1;refGene_tx=XM_023250930.1
A3 394424 394515
refGene_gn=MYT1;refGene_tx=XM_011280801.3;refGene_gn=MYT1;refGene_tx=XM_019826178.2;refGene_gn=MYT1;refGene_tx=XM_019826179.1;refGene_gn=MYT1;refGene_tx=XM_019826181.1;refGene_gn=MYT1;refGene_tx=XM_019826182.1;refGene_gn=MYT1;refGene_tx=XM_023250930.1
A3 395069 395204
refGene_gn=MYT1;refGene_tx=XM_011280801.3;refGene_gn=MYT1;refGene_tx=XM_019826178.2;refGene_gn=MYT1;refGene_tx=XM_019826179.1;refGene_gn=MYT1;refGene_tx=XM_019826181.1;refGene_gn=MYT1;refGene_tx=XM_019826182.1;refGene_gn=MYT1;refGene_tx=XM_023250930.1
A3 397210 398140
refGene_gn=MYT1;refGene_tx=XM_011280801.3;refGene_gn=MYT1;refGene_tx=XM_019826178.2;refGene_gn=MYT1;refGene_tx=XM_019826179.1;refGene_gn=MYT1;refGene_tx=XM_019826181.1;refGene_gn=MYT1;refGene_tx=XM_019826182.1;refGene_gn=MYT1;refGene_tx=XM_023250930.1
A3 399455 399703
refGene_gn=MYT1;refGene_tx=XM_011280801.3;refGene_gn=MYT1;refGene_tx=XM_019826178.2;refGene_gn=MYT1;refGene_tx=XM_019826179.1;refGene_gn=MYT1;refGene_tx=XM_019826181.1;refGene_gn=MYT1;refGene_tx=XM_019826182.1;refGene_gn=MYT1;refGene_tx=XM_023250930.1
A3 400390 400453
refGene_gn=MYT1;refGene_tx=XM_011280801.3;refGene_gn=MYT1;refGene_tx=XM_019826178.2;refGene_gn=MYT1;refGene_tx=XM_019826179.1;refGene_gn=MYT1;refGene_tx=XM_019826181.1;refGene_gn=MYT1;refGene_tx=XM_019826182.1
A3 406634 406665
refGene_gn=MYT1;refGene_tx=XM_011280801.3;refGene_gn=MYT1;refGene_tx=XM_019826178.2;refGene_gn=MYT1;refGene_tx=XM_019826179.1;refGene_gn=MYT1;refGene_tx=XM_019826181.1;refGene_gn=MYT1;refGene_tx=XM_019826182.1
A3 407654 407709
refGene_gn=MYT1;refGene_tx=XM_011280801.3;refGene_gn=MYT1;refGene_tx=XM_019826178.2;refGene_gn=MYT1;refGene_tx=XM_019826179.1;refGene_gn=MYT1;refGene_tx=XM_019826181.1;refGene_gn=MYT1;refGene_tx=XM_019826182.1

A3 416184 416285
refGene_gn=MYT1;refGene_tx=XM_011280801.3;refGene_gn=MYT1;refGene_tx=XM_019826178.2
A3 438776 438801
refGene_gn=MYT1;refGene_tx=XM_011280801.3;refGene_gn=MYT1;refGene_tx=XM_019826178.2
A3 473822 474857 refGene_gn=NPBWR2;refGene_tx=XM_003983288.1
A3 480405 480929
refGene_gn=OPRL1;refGene_tx=XM_006929564.2;refGene_gn=OPRL1;refGene_tx=XM_019826183.1;refGene_gn=OPRL1;refGene_tx=XM_019826185.2;refGene_gn=OPRL1;refGene_tx=XM_019826187.1;refGene_gn=OPRL1;refGene_tx=XM_019826188.1;refGene_gn=OPRL1;refGene_tx=XM_019826190.1;refGene_gn=OPRL1;refGene_tx=XM_019826191.1
A3 481025 481381
refGene_gn=OPRL1;refGene_tx=XM_006929564.2;refGene_gn=OPRL1;refGene_tx=XM_019826183.1;refGene_gn=OPRL1;refGene_tx=XM_019826185.2;refGene_gn=OPRL1;refGene_tx=XM_019826187.1;refGene_gn=OPRL1;refGene_tx=XM_019826188.1;refGene_gn=OPRL1;refGene_tx=XM_019826190.1;refGene_gn=OPRL1;refGene_tx=XM_019826191.1
A3 486156 486453
refGene_gn=OPRL1;refGene_tx=XM_006929564.2;refGene_gn=OPRL1;refGene_tx=XM_019826183.1;refGene_gn=OPRL1;refGene_tx=XM_019826185.2;refGene_gn=OPRL1;refGene_tx=XM_019826187.1;refGene_gn=OPRL1;refGene_tx=XM_019826188.1;refGene_gn=OPRL1;refGene_tx=XM_019826190.1
A3 487726 487793
refGene_gn=OPRL1;refGene_tx=XM_019826183.1;refGene_gn=OPRL1;refGene_tx=XM_019826185.2;refGene_gn=OPRL1;refGene_tx=XM_019826187.1
A3 488059 488084 refGene_gn=OPRL1;refGene_tx=XM_019826185.2
A3 497001 497119 refGene_gn=OPRL1;refGene_tx=XM_019826183.1
A3 501265 501295
refGene_gn=RGS19;refGene_tx=XM_003983275.4;refGene_gn=RGS19;refGene_tx=XM_006929566.3;refGene_gn=RGS19;refGene_tx=XM_023250931.1
A3 501463 501585
refGene_gn=RGS19;refGene_tx=XM_003983275.4;refGene_gn=RGS19;refGene_tx=XM_006929566.3;refGene_gn=RGS19;refGene_tx=XM_023250931.1
A3 502923 502998
refGene_gn=RGS19;refGene_tx=XM_003983275.4;refGene_gn=RGS19;refGene_tx=XM_023250931.1
A3 503081 503316
refGene_gn=RGS19;refGene_tx=XM_003983275.4;refGene_gn=RGS19;refGene_tx=XM_006929566.3;refGene_gn=RGS19;refGene_tx=XM_023250931.1
A3 503399 503591
refGene_gn=RGS19;refGene_tx=XM_003983275.4;refGene_gn=RGS19;refGene_tx=XM_006929566.3;refGene_gn=RGS19;refGene_tx=XM_023250931.1
A3 505120 505129
refGene_gn=TCEA2;refGene_tx=XM_003983289.5;refGene_gn=TCEA2;refGene_tx=XM_006929567.4
A3 505323 505395
refGene_gn=TCEA2;refGene_tx=XM_003983289.5;refGene_gn=TCEA2;refGene_tx=XM_006929567.4

A3 506007 506154
refGene_gn=TCEA2;refGene_tx=XM_003983289.5;refGene_gn=TCEA2;refGene_tx=XM_006929567.4
A3 506223 506378
refGene_gn=TCEA2;refGene_tx=XM_003983289.5;refGene_gn=TCEA2;refGene_tx=XM_006929567.4
A3 506839 506896
refGene_gn=TCEA2;refGene_tx=XM_003983289.5;refGene_gn=TCEA2;refGene_tx=XM_006929567.4
A3 507175 507306
refGene_gn=TCEA2;refGene_tx=XM_003983289.5;refGene_gn=TCEA2;refGene_tx=XM_006929567.4
A3 508498 508589
refGene_gn=TCEA2;refGene_tx=XM_003983289.5;refGene_gn=TCEA2;refGene_tx=XM_006929567.4
A3 509672 509778
refGene_gn=TCEA2;refGene_tx=XM_003983289.5;refGene_gn=TCEA2;refGene_tx=XM_006929567.4
A3 510102 510165
refGene_gn=TCEA2;refGene_tx=XM_003983289.5;refGene_gn=TCEA2;refGene_tx=XM_006929567.4
A3 513162 513234
refGene_gn=TCEA2;refGene_tx=XM_003983289.5;refGene_gn=TCEA2;refGene_tx=XM_006929567.4
A3 524636 524994 refGene_gn=SOX18;refGene_tx=XM_023250932.1
A3 525115 525441 refGene_gn=SOX18;refGene_tx=XM_023250932.1
A3 525442 525943 refGene_gn=SOX18;refGene_tx=XM_023250932.1
A3 539310 539463 refGene_gn=PRPF6;refGene_tx=XM_023250933.1
A3 540086 540213 refGene_gn=PRPF6;refGene_tx=XM_023250933.1
A3 542234 542349 refGene_gn=PRPF6;refGene_tx=XM_023250933.1
A3 543605 543697 refGene_gn=PRPF6;refGene_tx=XM_023250933.1
A3 544426 544560 refGene_gn=PRPF6;refGene_tx=XM_023250933.1
A3 545032 545209 refGene_gn=PRPF6;refGene_tx=XM_023250933.1
A3 545766 545886 refGene_gn=PRPF6;refGene_tx=XM_023250933.1
A3 547659 547798 refGene_gn=PRPF6;refGene_tx=XM_023250933.1
A3 549465 549587 refGene_gn=PRPF6;refGene_tx=XM_023250933.1
A3 554666 554789 refGene_gn=PRPF6;refGene_tx=XM_023250933.1
A3 561601 561820 refGene_gn=PRPF6;refGene_tx=XM_023250933.1
A3 562774 562893 refGene_gn=PRPF6;refGene_tx=XM_023250933.1
A3 567391 567554 refGene_gn=PRPF6;refGene_tx=XM_023250933.1
A3 568396 568553 refGene_gn=PRPF6;refGene_tx=XM_023250933.1
A3 569009 569104 refGene_gn=PRPF6;refGene_tx=XM_023250933.1
A3 576856 577012 refGene_gn=PRPF6;refGene_tx=XM_023250933.1
A3 577243 577420 refGene_gn=PRPF6;refGene_tx=XM_023250933.1
A3 578357 578436 refGene_gn=PRPF6;refGene_tx=XM_023250933.1
A3 582406 582525 refGene_gn=PRPF6;refGene_tx=XM_023250933.1
A3 583627 583796 refGene_gn=PRPF6;refGene_tx=XM_023250933.1
A3 585962 586033 refGene_gn=PRPF6;refGene_tx=XM_023250933.1
A3 587617 587981
refGene_gn=SAMD10;refGene_tx=XM_003983277.5;refGene_gn=SAMD10;refGene_

tx=XM_019826194.2;refGene_gn=SAMD10;refGene_tx=XM_023250934.1
A3 588619 588696 refGene_gn=SAMD10;refGene_tx=XM_019826194.2
A3 589439 589446 refGene_gn=SAMD10;refGene_tx=XM_019826195.2
A3 589719 590388
refGene_gn=SAMD10;refGene_tx=XM_003983277.5;refGene_gn=SAMD10;refGene_tx=XM_011280798.3;refGene_gn=SAMD10;refGene_tx=XM_019826194.2;refGene_gn=SAMD10;refGene_tx=XM_019826195.2;refGene_gn=SAMD10;refGene_tx=XM_023250934.1
A3 590769 590910
refGene_gn=SAMD10;refGene_tx=XM_003983277.5;refGene_gn=SAMD10;refGene_tx=XM_011280798.3;refGene_gn=SAMD10;refGene_tx=XM_019826194.2;refGene_gn=SAMD10;refGene_tx=XM_019826195.2
A3 591110 591133
refGene_gn=SAMD10;refGene_tx=XM_003983277.5;refGene_gn=SAMD10;refGene_tx=XM_011280798.3;refGene_gn=SAMD10;refGene_tx=XM_019826194.2;refGene_gn=SAMD10;refGene_tx=XM_019826195.2
A3 596731 596855 refGene_gn=ZNF512B;refGene_tx=XM_023250935.1
A3 597144 597287 refGene_gn=ZNF512B;refGene_tx=XM_023250935.1
A3 597654 597783 refGene_gn=ZNF512B;refGene_tx=XM_023250935.1
A3 597902 598597 refGene_gn=ZNF512B;refGene_tx=XM_023250935.1
A3 598679 598906 refGene_gn=ZNF512B;refGene_tx=XM_023250935.1
A3 599022 599094 refGene_gn=ZNF512B;refGene_tx=XM_023250935.1
A3 599191 599338 refGene_gn=ZNF512B;refGene_tx=XM_023250935.1
A3 599501 599626 refGene_gn=ZNF512B;refGene_tx=XM_023250935.1
A3 599697 599790 refGene_gn=ZNF512B;refGene_tx=XM_023250935.1
A3 599996 600095 refGene_gn=ZNF512B;refGene_tx=XM_023250935.1
A3 600194 600365 refGene_gn=ZNF512B;refGene_tx=XM_023250935.1
A3 600594 600789 refGene_gn=ZNF512B;refGene_tx=XM_023250935.1
A3 601015 601117 refGene_gn=ZNF512B;refGene_tx=XM_023250935.1
A3 601419 601482 refGene_gn=ZNF512B;refGene_tx=XM_023250935.1
A3 602043 602142 refGene_gn=ZNF512B;refGene_tx=XM_023250935.1
A3 602504 602756 refGene_gn=ZNF512B;refGene_tx=XM_023250935.1
A3 606321 606434
refGene_gn=UCKL1;refGene_tx=XM_023250936.1;refGene_gn=UCKL1;refGene_tx=XM_023250938.1;refGene_gn=UCKL1;refGene_tx=XM_023250939.1
A3 607227 607256 refGene_gn=UCKL1;refGene_tx=XM_023250937.1
A3 607362 607464
refGene_gn=UCKL1;refGene_tx=XM_023250936.1;refGene_gn=UCKL1;refGene_tx=XM_023250937.1
A3 609316 609621
refGene_gn=UCKL1;refGene_tx=XM_006929569.4;refGene_gn=UCKL1;refGene_tx=XM_006929570.4
A3 612637 612825
refGene_gn=UCKL1;refGene_tx=XM_006929569.4;refGene_gn=UCKL1;refGene_tx=XM_006929570.4;refGene_gn=UCKL1;refGene_tx=XM_023250936.1;refGene_gn=UCKL1;refGene_tx=XM_023250937.1;refGene_gn=UCKL1;refGene_tx=XM_023250938.1;refGene_gn=UCKL1;refGene_tx=XM_023250939.1
A3 612996 613103
refGene_gn=UCKL1;refGene_tx=XM_006929569.4;refGene_gn=UCKL1;refGene_tx=XM_006929570.4;refGene_gn=UCKL1;refGene_tx=XM_023250936.1;refGene_gn=

UCKL1;refGene_tx=XM_023250937.1;refGene_gn=UCKL1;refGene_tx=XM_0232509
38.1;refGene_gn=UCKL1;refGene_tx=XM_023250939.1
A3 613218 613389
refGene_gn=UCKL1;refGene_tx=XM_006929569.4;refGene_gn=UCKL1;refGene_tx
=XM_006929570.4;refGene_gn=UCKL1;refGene_tx=XM_023250936.1;refGene_gn=
UCKL1;refGene_tx=XM_023250937.1;refGene_gn=UCKL1;refGene_tx=XM_0232509
38.1;refGene_gn=UCKL1;refGene_tx=XM_023250939.1
A3 613474 613546
refGene_gn=UCKL1;refGene_tx=XM_006929569.4;refGene_gn=UCKL1;refGene_tx
=XM_006929570.4;refGene_gn=UCKL1;refGene_tx=XM_023250936.1;refGene_gn=
UCKL1;refGene_tx=XM_023250937.1;refGene_gn=UCKL1;refGene_tx=XM_0232509
38.1;refGene_gn=UCKL1;refGene_tx=XM_023250939.1
A3 614688 614878
refGene_gn=UCKL1;refGene_tx=XM_006929569.4;refGene_gn=UCKL1;refGene_tx
=XM_006929570.4;refGene_gn=UCKL1;refGene_tx=XM_023250936.1;refGene_gn=
UCKL1;refGene_tx=XM_023250937.1;refGene_gn=UCKL1;refGene_tx=XM_0232509
38.1;refGene_gn=UCKL1;refGene_tx=XM_023250939.1
A3 614970 615032
refGene_gn=UCKL1;refGene_tx=XM_006929569.4;refGene_gn=UCKL1;refGene_tx
=XM_006929570.4;refGene_gn=UCKL1;refGene_tx=XM_023250936.1;refGene_gn=
UCKL1;refGene_tx=XM_023250937.1;refGene_gn=UCKL1;refGene_tx=XM_0232509
38.1;refGene_gn=UCKL1;refGene_tx=XM_023250939.1
A3 615769 615786
refGene_gn=UCKL1;refGene_tx=XM_006929570.4;refGene_gn=UCKL1;refGene_tx
=XM_023250939.1
A3 617032 617052
refGene_gn=UCKL1;refGene_tx=XM_006929569.4;refGene_gn=UCKL1;refGene_tx
=XM_023250936.1;refGene_gn=UCKL1;refGene_tx=XM_023250937.1;refGene_gn=
UCKL1;refGene_tx=XM_023250938.1
A3 617888 617987
refGene_gn=UCKL1;refGene_tx=XM_006929569.4;refGene_gn=UCKL1;refGene_tx
=XM_006929570.4;refGene_gn=UCKL1;refGene_tx=XM_023250936.1;refGene_gn=
UCKL1;refGene_tx=XM_023250937.1;refGene_gn=UCKL1;refGene_tx=XM_0232509
38.1;refGene_gn=UCKL1;refGene_tx=XM_023250939.1
A3 618059 618153
refGene_gn=UCKL1;refGene_tx=XM_006929569.4;refGene_gn=UCKL1;refGene_tx
=XM_006929570.4;refGene_gn=UCKL1;refGene_tx=XM_023250936.1;refGene_gn=
UCKL1;refGene_tx=XM_023250937.1;refGene_gn=UCKL1;refGene_tx=XM_0232509
38.1;refGene_gn=UCKL1;refGene_tx=XM_023250939.1
A3 618256 618319
refGene_gn=UCKL1;refGene_tx=XM_006929569.4;refGene_gn=UCKL1;refGene_tx
=XM_006929570.4;refGene_gn=UCKL1;refGene_tx=XM_023250936.1;refGene_gn=
UCKL1;refGene_tx=XM_023250937.1;refGene_gn=UCKL1;refGene_tx=XM_0232509
38.1;refGene_gn=UCKL1;refGene_tx=XM_023250939.1
A3 618423 618546
refGene_gn=UCKL1;refGene_tx=XM_006929569.4;refGene_gn=UCKL1;refGene_tx
=XM_006929570.4;refGene_gn=UCKL1;refGene_tx=XM_023250936.1;refGene_gn=
UCKL1;refGene_tx=XM_023250937.1;refGene_gn=UCKL1;refGene_tx=XM_0232509
38.1;refGene_gn=UCKL1;refGene_tx=XM_023250939.1
A3 618710 618818

refGene_gn=UCKL1;refGene_tx=XM_006929569.4;refGene_gn=UCKL1;refGene_tx=XM_006929570.4;refGene_gn=UCKL1;refGene_tx=XM_023250936.1;refGene_gn=UCKL1;refGene_tx=XM_023250937.1;refGene_gn=UCKL1;refGene_tx=XM_023250938.1;refGene_gn=UCKL1;refGene_tx=XM_023250939.1
A3 618886 619043
refGene_gn=UCKL1;refGene_tx=XM_006929569.4;refGene_gn=UCKL1;refGene_tx=XM_006929570.4;refGene_gn=UCKL1;refGene_tx=XM_023250936.1;refGene_gn=UCKL1;refGene_tx=XM_023250937.1;refGene_gn=UCKL1;refGene_tx=XM_023250938.1;refGene_gn=UCKL1;refGene_tx=XM_023250939.1
A3 619135 619239
refGene_gn=UCKL1;refGene_tx=XM_006929569.4;refGene_gn=UCKL1;refGene_tx=XM_006929570.4;refGene_gn=UCKL1;refGene_tx=XM_023250936.1;refGene_gn=UCKL1;refGene_tx=XM_023250937.1;refGene_gn=UCKL1;refGene_tx=XM_023250938.1;refGene_gn=UCKL1;refGene_tx=XM_023250939.1
A3 628546 628650 refGene_gn=DNAJC5;refGene_tx=XM_003983279.4
A3 629104 629276 refGene_gn=DNAJC5;refGene_tx=XM_003983279.4
A3 629919 630133 refGene_gn=DNAJC5;refGene_tx=XM_003983279.4
A3 630932 631039 refGene_gn=DNAJC5;refGene_tx=XM_003983279.4
A3 669512 669608
refGene_gn=TPD52L2;refGene_tx=XM_003983280.4;refGene_gn=TPD52L2;refGene_tx=XM_003983281.5;refGene_gn=TPD52L2;refGene_tx=XM_006929573.4;refGene_gn=TPD52L2;refGene_tx=XM_006929574.4;refGene_gn=TPD52L2;refGene_tx=XM_006929575.4;refGene_gn=TPD52L2;refGene_tx=XM_006929576.4;refGene_gn=TPD52L2;refGene_tx=XM_019826198.2;refGene_gn=TPD52L2;refGene_tx=XM_019826199.2
A3 670105 670154
refGene_gn=TPD52L2;refGene_tx=XM_003983280.4;refGene_gn=TPD52L2;refGene_tx=XM_003983281.5;refGene_gn=TPD52L2;refGene_tx=XM_006929573.4;refGene_gn=TPD52L2;refGene_tx=XM_006929574.4;refGene_gn=TPD52L2;refGene_tx=XM_006929575.4;refGene_gn=TPD52L2;refGene_tx=XM_006929576.4;refGene_gn=TPD52L2;refGene_tx=XM_019826198.2;refGene_gn=TPD52L2;refGene_tx=XM_019826199.2
A3 671192 671237
refGene_gn=TPD52L2;refGene_tx=XM_006929573.4;refGene_gn=TPD52L2;refGene_tx=XM_006929575.4;refGene_gn=TPD52L2;refGene_tx=XM_019826198.2;refGene_gn=TPD52L2;refGene_tx=XM_019826199.2
A3 672125 672152
refGene_gn=TPD52L2;refGene_tx=XM_006929573.4;refGene_gn=TPD52L2;refGene_tx=XM_006929574.4;refGene_gn=TPD52L2;refGene_tx=XM_006929575.4;refGene_gn=TPD52L2;refGene_tx=XM_006929576.4;refGene_gn=TPD52L2;refGene_tx=XM_019826198.2;refGene_gn=TPD52L2;refGene_tx=XM_019826199.2;refGene_gn=TPD52L2;refGene_tx=XM_019826201.1;refGene_gn=TPD52L2;refGene_tx=XM_019826202.1
A3 673652 673713
refGene_gn=TPD52L2;refGene_tx=XM_019826201.1;refGene_gn=TPD52L2;refGene_tx=XM_019826202.1
A3 675181 675287
refGene_gn=TPD52L2;refGene_tx=XM_003983280.4;refGene_gn=TPD52L2;refGene_tx=XM_003983281.5;refGene_gn=TPD52L2;refGene_tx=XM_006929573.4;refGene_gn=TPD52L2;refGene_tx=XM_006929574.4;refGene_gn=TPD52L2;refGene_tx=

XM_006929575.4;refGene_gn=TPD52L2;refGene_tx=XM_006929576.4;refGene_gn
=TPD52L2;refGene_tx=XM_019826198.2;refGene_gn=TPD52L2;refGene_tx=XM_01
9826199.2;refGene_gn=TPD52L2;refGene_tx=XM_019826201.1;refGene_gn=TPD5
2L2;refGene_tx=XM_019826202.1;refGene_gn=TPD52L2;refGene_tx=XM_0198262
03.1
A3 680027 680087
refGene_gn=TPD52L2;refGene_tx=XM_003983280.4;refGene_gn=TPD52L2;refGen
e_tx=XM_006929573.4;refGene_gn=TPD52L2;refGene_tx=XM_006929574.4;refGe
ne_gn=TPD52L2;refGene_tx=XM_019826198.2;refGene_gn=TPD52L2;refGene_tx=
XM_019826201.1;refGene_gn=TPD52L2;refGene_tx=XM_019826202.1
A3 681051 681200
refGene_gn=TPD52L2;refGene_tx=XM_003983280.4;refGene_gn=TPD52L2;refGen
e_tx=XM_003983281.5;refGene_gn=TPD52L2;refGene_tx=XM_006929573.4;refGe
ne_gn=TPD52L2;refGene_tx=XM_006929574.4;refGene_gn=TPD52L2;refGene_tx=
XM_006929575.4;refGene_gn=TPD52L2;refGene_tx=XM_006929576.4;refGene_gn
=TPD52L2;refGene_tx=XM_019826198.2;refGene_gn=TPD52L2;refGene_tx=XM_01
9826199.2;refGene_gn=TPD52L2;refGene_tx=XM_019826201.1;refGene_gn=TPD5
2L2;refGene_tx=XM_019826202.1;refGene_gn=TPD52L2;refGene_tx=XM_0198262
03.1
A3 683200 683346
refGene_gn=TPD52L2;refGene_tx=XM_003983280.4;refGene_gn=TPD52L2;refGen
e_tx=XM_003983281.5;refGene_gn=TPD52L2;refGene_tx=XM_006929573.4;refGe
ne_gn=TPD52L2;refGene_tx=XM_006929574.4;refGene_gn=TPD52L2;refGene_tx=
XM_006929575.4;refGene_gn=TPD52L2;refGene_tx=XM_006929576.4;refGene_gn
=TPD52L2;refGene_tx=XM_019826198.2;refGene_gn=TPD52L2;refGene_tx=XM_01
9826199.2;refGene_gn=TPD52L2;refGene_tx=XM_019826201.1;refGene_gn=TPD5
2L2;refGene_tx=XM_019826202.1;refGene_gn=TPD52L2;refGene_tx=XM_0198262
03.1
A3 687671 687690
refGene_gn=TPD52L2;refGene_tx=XM_003983280.4;refGene_gn=TPD52L2;refGen
e_tx=XM_003983281.5;refGene_gn=TPD52L2;refGene_tx=XM_006929573.4;refGe
ne_gn=TPD52L2;refGene_tx=XM_006929574.4;refGene_gn=TPD52L2;refGene_tx=
XM_006929575.4;refGene_gn=TPD52L2;refGene_tx=XM_006929576.4;refGene_gn
=TPD52L2;refGene_tx=XM_019826198.2;refGene_gn=TPD52L2;refGene_tx=XM_01
9826199.2;refGene_gn=TPD52L2;refGene_tx=XM_019826201.1;refGene_gn=TPD5
2L2;refGene_tx=XM_019826202.1;refGene_gn=TPD52L2;refGene_tx=XM_0198262
03.1
A3 689798 691214 refGene_gn=ABHD16B;refGene_tx=XM_003983292.5
A3 747531 748468
refGene_gn=ZBTB46;refGene_tx=XM_003983282.5;refGene_gn=ZBTB46;refGene_
tx=XM_006929577.4;refGene_gn=ZBTB46;refGene_tx=XM_023250941.1
A3 758989 759274
refGene_gn=ZBTB46;refGene_tx=XM_003983282.5;refGene_gn=ZBTB46;refGene_
tx=XM_006929577.4
A3 778106 778282
refGene_gn=ZBTB46;refGene_tx=XM_003983282.5;refGene_gn=ZBTB46;refGene_
tx=XM_006929577.4;refGene_gn=ZBTB46;refGene_tx=XM_023250941.1
A3 781001 781373
refGene_gn=ZBTB46;refGene_tx=XM_003983282.5;refGene_gn=ZBTB46;refGene_
tx=XM_006929577.4;refGene_gn=ZBTB46;refGene_tx=XM_023250941.1

A3 785561 785673
refGene_gn=SLC2A4RG;refGene_tx=XM_023250942.1;refGene_gn=SLC2A4RG;refGene_tx=XM_023250943.1
A3 785762 785854
refGene_gn=SLC2A4RG;refGene_tx=XM_023250942.1;refGene_gn=SLC2A4RG;refGene_tx=XM_023250943.1
A3 785938 786206
refGene_gn=SLC2A4RG;refGene_tx=XM_023250942.1;refGene_gn=SLC2A4RG;refGene_tx=XM_023250943.1
A3 786296 786397
refGene_gn=SLC2A4RG;refGene_tx=XM_023250942.1;refGene_gn=SLC2A4RG;refGene_tx=XM_023250943.1
A3 786469 786648
refGene_gn=SLC2A4RG;refGene_tx=XM_023250942.1;refGene_gn=SLC2A4RG;refGene_tx=XM_023250943.1
A3 787078 787188
refGene_gn=SLC2A4RG;refGene_tx=XM_023250942.1;refGene_gn=SLC2A4RG;refGene_tx=XM_023250943.1
A3 788107 788268
refGene_gn=SLC2A4RG;refGene_tx=XM_023250942.1;refGene_gn=SLC2A4RG;refGene_tx=XM_023250943.1
A3 788591 788702
refGene_gn=SLC2A4RG;refGene_tx=XM_023250942.1;refGene_gn=SLC2A4RG;refGene_tx=XM_023250943.1
A3 789829 790066
refGene_gn=LIME1;refGene_tx=XM_023250947.1;refGene_gn=LIME1;refGene_tx=XM_023250948.1;refGene_gn=LIME1;refGene_tx=XM_023250950.1
A3 790148 790489
refGene_gn=LIME1;refGene_tx=XM_023250947.1;refGene_gn=LIME1;refGene_tx=XM_023250948.1;refGene_gn=LIME1;refGene_tx=XM_023250950.1
A3 790613 790701
refGene_gn=LIME1;refGene_tx=XM_023250947.1;refGene_gn=LIME1;refGene_tx=XM_023250950.1
A3 790771 790829
refGene_gn=LIME1;refGene_tx=XM_023250947.1;refGene_gn=LIME1;refGene_tx=XM_023250950.1
A3 790979 791077 refGene_gn=LIME1;refGene_tx=XM_023250947.1
A3 791985 792121 refGene_gn=LIME1;refGene_tx=XM_023250948.1
A3 792964 793103
refGene_gn=ZGPAT;refGene_tx=XM_023250945.1;refGene_gn=ZGPAT;refGene_tx=XM_023250946.1
A3 793316 793716
refGene_gn=ZGPAT;refGene_tx=XM_023250945.1;refGene_gn=ZGPAT;refGene_tx=XM_023250946.1
A3 793802 793922
refGene_gn=ZGPAT;refGene_tx=XM_023250945.1;refGene_gn=ZGPAT;refGene_tx=XM_023250946.1
A3 794792 794948
refGene_gn=ZGPAT;refGene_tx=XM_023250945.1;refGene_gn=ZGPAT;refGene_tx=XM_023250946.1

A3	795160	795294
refGene_gn=ZGPAT;refGene_tx=XM_023250945.1;refGene_gn=ZGPAT;refGene_tx
=XM_023250946.1
A3	808265	808849
refGene_gn=ZGPAT;refGene_tx=XM_023250945.1;refGene_gn=ZGPAT;refGene_tx
=XM_023250946.1
A3	810109	810202
refGene_gn=ARFRP1;refGene_tx=XM_023250951.1;refGene_gn=ARFRP1;refGene_
tx=XM_023250952.1;refGene_gn=ARFRP1;refGene_tx=XM_023250953.1;refGene_
gn=ARFRP1;refGene_tx=XM_023250954.1;refGene_gn=ARFRP1;refGene_tx=XM_02
3250955.1;refGene_gn=ARFRP1;refGene_tx=XM_023250956.1;refGene_gn=ARFRP
1;refGene_tx=XM_023250957.1;refGene_gn=ARFRP1;refGene_tx=XM_023250958.
1;refGene_gn=ARFRP1;refGene_tx=XM_023250959.1;refGene_gn=ARFRP1;refGen
e_tx=XM_023250960.1
A3	810427	810515
refGene_gn=ARFRP1;refGene_tx=XM_023250951.1;refGene_gn=ARFRP1;refGene_
tx=XM_023250952.1;refGene_gn=ARFRP1;refGene_tx=XM_023250953.1;refGene_
gn=ARFRP1;refGene_tx=XM_023250954.1;refGene_gn=ARFRP1;refGene_tx=XM_02
3250955.1;refGene_gn=ARFRP1;refGene_tx=XM_023250956.1;refGene_gn=ARFRP
1;refGene_tx=XM_023250957.1;refGene_gn=ARFRP1;refGene_tx=XM_023250958.
1;refGene_gn=ARFRP1;refGene_tx=XM_023250959.1;refGene_gn=ARFRP1;refGen
e_tx=XM_023250960.1
A3	810742	810825
refGene_gn=ARFRP1;refGene_tx=XM_023250951.1;refGene_gn=ARFRP1;refGene_
tx=XM_023250952.1;refGene_gn=ARFRP1;refGene_tx=XM_023250953.1;refGene_
gn=ARFRP1;refGene_tx=XM_023250954.1;refGene_gn=ARFRP1;refGene_tx=XM_02
3250955.1;refGene_gn=ARFRP1;refGene_tx=XM_023250956.1;refGene_gn=ARFRP
1;refGene_tx=XM_023250957.1;refGene_gn=ARFRP1;refGene_tx=XM_023250958.
1;refGene_gn=ARFRP1;refGene_tx=XM_023250959.1;refGene_gn=ARFRP1;refGen
e_tx=XM_023250960.1
A3	813992	814161
refGene_gn=ARFRP1;refGene_tx=XM_023250951.1;refGene_gn=ARFRP1;refGene_
tx=XM_023250952.1;refGene_gn=ARFRP1;refGene_tx=XM_023250953.1;refGene_
gn=ARFRP1;refGene_tx=XM_023250954.1;refGene_gn=ARFRP1;refGene_tx=XM_02
3250955.1;refGene_gn=ARFRP1;refGene_tx=XM_023250956.1;refGene_gn=ARFRP
1;refGene_tx=XM_023250957.1;refGene_gn=ARFRP1;refGene_tx=XM_023250958.
1;refGene_gn=ARFRP1;refGene_tx=XM_023250959.1;refGene_gn=ARFRP1;refGen
e_tx=XM_023250960.1
A3	814375	814446
refGene_gn=ARFRP1;refGene_tx=XM_023250951.1;refGene_gn=ARFRP1;refGene_
tx=XM_023250952.1;refGene_gn=ARFRP1;refGene_tx=XM_023250953.1;refGene_
gn=ARFRP1;refGene_tx=XM_023250954.1;refGene_gn=ARFRP1;refGene_tx=XM_02
3250955.1;refGene_gn=ARFRP1;refGene_tx=XM_023250956.1;refGene_gn=ARFRP
1;refGene_tx=XM_023250957.1;refGene_gn=ARFRP1;refGene_tx=XM_023250958.
1;refGene_gn=ARFRP1;refGene_tx=XM_023250959.1;refGene_gn=ARFRP1;refGen
e_tx=XM_023250960.1
A3	814849	814906	refGene_gn=ARFRP1;refGene_tx=XM_023250960.1
A3	815000	815117
refGene_gn=ARFRP1;refGene_tx=XM_023250953.1;refGene_gn=ARFRP1;refGene_
tx=XM_023250954.1;refGene_gn=ARFRP1;refGene_tx=XM_023250956.1;refGene_

gn=ARFRP1;refGene_tx=XM_023250957.1;refGene_gn=ARFRP1;refGene_tx=XM_023250959.1
A3 815187 815412
refGene_gn=ARFRP1;refGene_tx=XM_023250951.1;refGene_gn=ARFRP1;refGene_tx=XM_023250952.1;refGene_gn=ARFRP1;refGene_tx=XM_023250953.1;refGene_gn=ARFRP1;refGene_tx=XM_023250954.1;refGene_gn=ARFRP1;refGene_tx=XM_023250955.1;refGene_gn=ARFRP1;refGene_tx=XM_023250956.1;refGene_gn=ARFRP1;refGene_tx=XM_023250957.1;refGene_gn=ARFRP1;refGene_tx=XM_023250958.1
A3 817299 817583 refGene_gn=TNFRSF6B;refGene_tx=XM_019826214.2
A3 817989 818193 refGene_gn=TNFRSF6B;refGene_tx=XM_019826214.2
A3 818294 818727 refGene_gn=TNFRSF6B;refGene_tx=XM_019826214.2
A3 819691 819772 refGene_gn=RTEL1;refGene_tx=XM_023250944.1
A3 819882 819983 refGene_gn=RTEL1;refGene_tx=XM_023250944.1
A3 820070 820205 refGene_gn=RTEL1;refGene_tx=XM_023250944.1
A3 820318 820480 refGene_gn=RTEL1;refGene_tx=XM_023250944.1
A3 820570 820807 refGene_gn=RTEL1;refGene_tx=XM_023250944.1
A3 820988 821105 refGene_gn=RTEL1;refGene_tx=XM_023250944.1
A3 821662 821803 refGene_gn=RTEL1;refGene_tx=XM_023250944.1
A3 821997 822196 refGene_gn=RTEL1;refGene_tx=XM_023250944.1
A3 823480 823576 refGene_gn=RTEL1;refGene_tx=XM_023250944.1
A3 824174 824332 refGene_gn=RTEL1;refGene_tx=XM_023250944.1
A3 824544 824692 refGene_gn=RTEL1;refGene_tx=XM_023250944.1
A3 824782 824906 refGene_gn=RTEL1;refGene_tx=XM_023250944.1
A3 825043 825159 refGene_gn=RTEL1;refGene_tx=XM_023250944.1
A3 825234 825381 refGene_gn=RTEL1;refGene_tx=XM_023250944.1
A3 825813 825891 refGene_gn=RTEL1;refGene_tx=XM_023250944.1
A3 826242 826320 refGene_gn=RTEL1;refGene_tx=XM_023250944.1
A3 826543 826629 refGene_gn=RTEL1;refGene_tx=XM_023250944.1
A3 826782 826823 refGene_gn=RTEL1;refGene_tx=XM_023250944.1
A3 826909 827023 refGene_gn=RTEL1;refGene_tx=XM_023250944.1
A3 827126 827259 refGene_gn=RTEL1;refGene_tx=XM_023250944.1
A3 829245 829327 refGene_gn=RTEL1;refGene_tx=XM_023250944.1
A3 829399 829474 refGene_gn=RTEL1;refGene_tx=XM_023250944.1
A3 832113 832169 refGene_gn=RTEL1;refGene_tx=XM_023250944.1
A3 832771 832869 refGene_gn=RTEL1;refGene_tx=XM_023250944.1
A3 833449 833528 refGene_gn=RTEL1;refGene_tx=XM_023250944.1
A3 833595 833634 refGene_gn=RTEL1;refGene_tx=XM_023250944.1
A3 841437 841552 refGene_gn=RTEL1;refGene_tx=XM_023250944.1
A3 841553 841592 refGene_gn=RTEL1;refGene_tx=XM_023250944.1
A3 842643 842709 refGene_gn=RTEL1;refGene_tx=XM_023250944.1
A3 843258 843343 refGene_gn=RTEL1;refGene_tx=XM_023250944.1
A3 844197 844273 refGene_gn=RTEL1;refGene_tx=XM_023250944.1
A3 848734 848795 refGene_gn=RTEL1;refGene_tx=XM_023250944.1
A3 848981 849063 refGene_gn=RTEL1;refGene_tx=XM_023250944.1
A3 849638 849732 refGene_gn=RTEL1;refGene_tx=XM_023250944.1
A3 850003 850196 refGene_gn=RTEL1;refGene_tx=XM_023250944.1
A3 851100 851202 refGene_gn=RTEL1;refGene_tx=XM_023250944.1
A3 854710 854729 refGene_gn=STMN3;refGene_tx=XM_023250961.1
A3 860760 860856 refGene_gn=STMN3;refGene_tx=XM_023250961.1

A3 861118 861294 refGene_gn=STMN3;refGene_tx=XM_023250961.1
A3 862564 862756 refGene_gn=STMN3;refGene_tx=XM_023250961.1
A3 863540 863600 refGene_gn=STMN3;refGene_tx=XM_023250961.1
A3 879242 879373
refGene_gn=GMEB2;refGene_tx=XM_019826227.2;refGene_gn=GMEB2;refGene_tx=XM_023250962.1;refGene_gn=GMEB2;refGene_tx=XM_023250963.1;refGene_gn=GMEB2;refGene_tx=XM_023250964.1
A3 882798 882896
refGene_gn=GMEB2;refGene_tx=XM_019826227.2;refGene_gn=GMEB2;refGene_tx=XM_023250962.1;refGene_gn=GMEB2;refGene_tx=XM_023250963.1;refGene_gn=GMEB2;refGene_tx=XM_023250964.1
A3 883391 883522
refGene_gn=GMEB2;refGene_tx=XM_019826227.2;refGene_gn=GMEB2;refGene_tx=XM_023250962.1;refGene_gn=GMEB2;refGene_tx=XM_023250963.1;refGene_gn=GMEB2;refGene_tx=XM_023250964.1
A3 888191 888295
refGene_gn=GMEB2;refGene_tx=XM_019826227.2;refGene_gn=GMEB2;refGene_tx=XM_023250962.1;refGene_gn=GMEB2;refGene_tx=XM_023250963.1;refGene_gn=GMEB2;refGene_tx=XM_023250964.1
A3 890619 890777
refGene_gn=GMEB2;refGene_tx=XM_019826227.2;refGene_gn=GMEB2;refGene_tx=XM_023250962.1;refGene_gn=GMEB2;refGene_tx=XM_023250963.1;refGene_gn=GMEB2;refGene_tx=XM_023250964.1
A3 892914 892986
refGene_gn=GMEB2;refGene_tx=XM_019826227.2;refGene_gn=GMEB2;refGene_tx=XM_023250962.1;refGene_gn=GMEB2;refGene_tx=XM_023250963.1;refGene_gn=GMEB2;refGene_tx=XM_023250964.1
A3 893234 893372
refGene_gn=GMEB2;refGene_tx=XM_019826227.2;refGene_gn=GMEB2;refGene_tx=XM_023250962.1;refGene_gn=GMEB2;refGene_tx=XM_023250963.1;refGene_gn=GMEB2;refGene_tx=XM_023250964.1
A3 893593 893716
refGene_gn=GMEB2;refGene_tx=XM_019826227.2;refGene_gn=GMEB2;refGene_tx=XM_023250962.1;refGene_gn=GMEB2;refGene_tx=XM_023250963.1;refGene_gn=GMEB2;refGene_tx=XM_023250964.1
A3 894350 895009
refGene_gn=GMEB2;refGene_tx=XM_019826227.2;refGene_gn=GMEB2;refGene_tx=XM_023250962.1;refGene_gn=GMEB2;refGene_tx=XM_023250963.1;refGene_gn=GMEB2;refGene_tx=XM_023250964.1
A3 911855 912874
refGene_gn=HELZ2;refGene_tx=XM_023250965.1;refGene_gn=HELZ2;refGene_tx=XM_023250966.1;refGene_gn=HELZ2;refGene_tx=XM_023250967.1
A3 913143 913327
refGene_gn=HELZ2;refGene_tx=XM_023250965.1;refGene_gn=HELZ2;refGene_tx=XM_023250966.1;refGene_gn=HELZ2;refGene_tx=XM_023250967.1
A3 913403 913511
refGene_gn=HELZ2;refGene_tx=XM_023250965.1;refGene_gn=HELZ2;refGene_tx=XM_023250966.1;refGene_gn=HELZ2;refGene_tx=XM_023250967.1
A3 913873 914394
refGene_gn=HELZ2;refGene_tx=XM_023250965.1;refGene_gn=HELZ2;refGene_tx

=XM_023250966.1;refGene_gn=HELZ2;refGene_tx=XM_023250967.1
A3	914488	915130
refGene_gn=HELZ2;refGene_tx=XM_023250965.1;refGene_gn=HELZ2;refGene_tx=XM_023250966.1;refGene_gn=HELZ2;refGene_tx=XM_023250967.1
A3	916193	916977
refGene_gn=HELZ2;refGene_tx=XM_023250965.1;refGene_gn=HELZ2;refGene_tx=XM_023250966.1;refGene_gn=HELZ2;refGene_tx=XM_023250967.1
A3	917269	917345
refGene_gn=HELZ2;refGene_tx=XM_023250965.1;refGene_gn=HELZ2;refGene_tx=XM_023250966.1;refGene_gn=HELZ2;refGene_tx=XM_023250967.1
A3	917428	921218
refGene_gn=HELZ2;refGene_tx=XM_023250965.1;refGene_gn=HELZ2;refGene_tx=XM_023250966.1;refGene_gn=HELZ2;refGene_tx=XM_023250967.1
A3	921376	921483
refGene_gn=HELZ2;refGene_tx=XM_023250965.1;refGene_gn=HELZ2;refGene_tx=XM_023250966.1;refGene_gn=HELZ2;refGene_tx=XM_023250967.1
A3	921571	921703
refGene_gn=HELZ2;refGene_tx=XM_023250965.1;refGene_gn=HELZ2;refGene_tx=XM_023250966.1;refGene_gn=HELZ2;refGene_tx=XM_023250967.1
A3	921788	921950
refGene_gn=HELZ2;refGene_tx=XM_023250965.1;refGene_gn=HELZ2;refGene_tx=XM_023250966.1;refGene_gn=HELZ2;refGene_tx=XM_023250967.1
A3	922031	922179
refGene_gn=HELZ2;refGene_tx=XM_023250965.1;refGene_gn=HELZ2;refGene_tx=XM_023250966.1;refGene_gn=HELZ2;refGene_tx=XM_023250967.1
A3	922291	922408
refGene_gn=HELZ2;refGene_tx=XM_023250965.1;refGene_gn=HELZ2;refGene_tx=XM_023250966.1;refGene_gn=HELZ2;refGene_tx=XM_023250967.1
A3	922513	922706
refGene_gn=HELZ2;refGene_tx=XM_023250965.1;refGene_gn=HELZ2;refGene_tx=XM_023250966.1;refGene_gn=HELZ2;refGene_tx=XM_023250967.1
A3	922869	923004
refGene_gn=HELZ2;refGene_tx=XM_023250965.1;refGene_gn=HELZ2;refGene_tx=XM_023250966.1;refGene_gn=HELZ2;refGene_tx=XM_023250967.1
A3	923206	923425
refGene_gn=HELZ2;refGene_tx=XM_023250965.1;refGene_gn=HELZ2;refGene_tx=XM_023250966.1;refGene_gn=HELZ2;refGene_tx=XM_023250967.1
A3	923561	923718
refGene_gn=HELZ2;refGene_tx=XM_023250965.1;refGene_gn=HELZ2;refGene_tx=XM_023250966.1;refGene_gn=HELZ2;refGene_tx=XM_023250967.1
A3	923833	924001
refGene_gn=HELZ2;refGene_tx=XM_023250965.1;refGene_gn=HELZ2;refGene_tx=XM_023250966.1;refGene_gn=HELZ2;refGene_tx=XM_023250967.1
A3	924367	924492
refGene_gn=HELZ2;refGene_tx=XM_023250965.1;refGene_gn=HELZ2;refGene_tx=XM_023250966.1;refGene_gn=HELZ2;refGene_tx=XM_023250967.1
A3	928593	929598
refGene_gn=FNDC11;refGene_tx=XM_006929587.4;refGene_gn=FNDC11;refGene_tx=XM_019826228.2;refGene_gn=FNDC11;refGene_tx=XM_019826229.2;refGene_gn=FNDC11;refGene_tx=XM_019826230.2;refGene_gn=FNDC11;refGene_tx=XM_01

9826232.2
A3	931023	931125	refGene_gn=FNDC11;refGene_tx=XM_006929587.4
A3	935409	935771	refGene_gn=SRMS;refGene_tx=XM_019828008.2
A3	939025	939147	refGene_gn=SRMS;refGene_tx=XM_019828008.2
A3	939615	939779	refGene_gn=SRMS;refGene_tx=XM_019828008.2
A3	940416	940600	refGene_gn=SRMS;refGene_tx=XM_019828008.2
A3	940707	940866	refGene_gn=SRMS;refGene_tx=XM_019828008.2
A3	941325	941507	refGene_gn=SRMS;refGene_tx=XM_019828008.2
A3	941597	941754	refGene_gn=SRMS;refGene_tx=XM_019828008.2
A3	941880	942074	refGene_gn=SRMS;refGene_tx=XM_019828008.2
A3	945410	945640
refGene_gn=PTK6;refGene_tx=XM_011280791.3;refGene_gn=PTK6;refGene_tx=XM_019826235.2
A3	946133	946445
refGene_gn=PTK6;refGene_tx=XM_019826233.2;refGene_gn=PTK6;refGene_tx=XM_019826234.2;refGene_gn=PTK6;refGene_tx=XM_019826236.2
A3	947103	947225
refGene_gn=PTK6;refGene_tx=XM_011280791.3;refGene_gn=PTK6;refGene_tx=XM_019826233.2;refGene_gn=PTK6;refGene_tx=XM_019826234.2;refGene_gn=PTK6;refGene_tx=XM_019826235.2;refGene_gn=PTK6;refGene_tx=XM_019826236.2
A3	947538	947699
refGene_gn=PTK6;refGene_tx=XM_011280791.3;refGene_gn=PTK6;refGene_tx=XM_019826233.2;refGene_gn=PTK6;refGene_tx=XM_019826234.2;refGene_gn=PTK6;refGene_tx=XM_019826235.2;refGene_gn=PTK6;refGene_tx=XM_019826236.2
A3	948171	948325
refGene_gn=PTK6;refGene_tx=XM_011280791.3;refGene_gn=PTK6;refGene_tx=XM_019826233.2;refGene_gn=PTK6;refGene_tx=XM_019826234.2;refGene_gn=PTK6;refGene_tx=XM_019826235.2;refGene_gn=PTK6;refGene_tx=XM_019826236.2
A3	949377	949539
refGene_gn=PTK6;refGene_tx=XM_011280791.3;refGene_gn=PTK6;refGene_tx=XM_019826233.2;refGene_gn=PTK6;refGene_tx=XM_019826234.2;refGene_gn=PTK6;refGene_tx=XM_019826235.2
A3	950661	950843
refGene_gn=PTK6;refGene_tx=XM_011280791.3;refGene_gn=PTK6;refGene_tx=XM_019826233.2;refGene_gn=PTK6;refGene_tx=XM_019826234.2;refGene_gn=PTK6;refGene_tx=XM_019826235.2;refGene_gn=PTK6;refGene_tx=XM_019826236.2
A3	952083	952237
refGene_gn=PTK6;refGene_tx=XM_011280791.3;refGene_gn=PTK6;refGene_tx=XM_019826233.2;refGene_gn=PTK6;refGene_tx=XM_019826234.2;refGene_gn=PTK6;refGene_tx=XM_019826235.2;refGene_gn=PTK6;refGene_tx=XM_019826236.2
A3	952471	952789
refGene_gn=PTK6;refGene_tx=XM_011280791.3;refGene_gn=PTK6;refGene_tx=XM_019826233.2;refGene_gn=PTK6;refGene_tx=XM_019826234.2;refGene_gn=PTK6;refGene_tx=XM_019826235.2;refGene_gn=PTK6;refGene_tx=XM_019826236.2
A3	954648	954763	refGene_gn=PPDPF;refGene_tx=XM_003983311.5
A3	954840	954937	refGene_gn=PPDPF;refGene_tx=XM_003983311.5
A3	955035	955113	refGene_gn=PPDPF;refGene_tx=XM_003983311.5
A3	955281	955336	refGene_gn=PPDPF;refGene_tx=XM_003983311.5
A3	975359	975503	refGene_gn=EEF1A2;refGene_tx=XM_023250968.1
A3	977164	977344	refGene_gn=EEF1A2;refGene_tx=XM_023250968.1

A3	977760	978057	refGene_gn=EEF1A2;refGene_tx=XM_023250968.1
A3	979531	979682	refGene_gn=EEF1A2;refGene_tx=XM_023250968.1
A3	980945	981202	refGene_gn=EEF1A2;refGene_tx=XM_023250968.1
A3	981850	982085	refGene_gn=EEF1A2;refGene_tx=XM_023250968.1
A3	982709	982837	refGene_gn=EEF1A2;refGene_tx=XM_023250968.1
A3	994433	994729
refGene_gn=KCNQ2;refGene_tx=XM_023250969.1;refGene_gn=KCNQ2;refGene_tx=XM_023250970.1;refGene_gn=KCNQ2;refGene_tx=XM_023250971.1;refGene_gn=KCNQ2;refGene_tx=XM_023250972.1;refGene_gn=KCNQ2;refGene_tx=XM_023250974.1;refGene_gn=KCNQ2;refGene_tx=XM_023250975.1;refGene_gn=KCNQ2;refGene_tx=XM_023250976.1;refGene_gn=KCNQ2;refGene_tx=XM_023250977.1;refGene_gn=KCNQ2;refGene_tx=XM_023250978.1;refGene_gn=KCNQ2;refGene_tx=XM_023250979.1;refGene_gn=KCNQ2;refGene_tx=XM_023250980.1;refGene_gn=KCNQ2;refGene_tx=XM_023250981.1;refGene_gn=KCNQ2;refGene_tx=XM_023250982.1;refGene_gn=KCNQ2;refGene_tx=XM_023250983.1;refGene_gn=KCNQ2;refGene_tx=XM_023250984.1;refGene_gn=KCNQ2;refGene_tx=XM_023250985.1;refGene_gn=KCNQ2;refGene_tx=XM_023250986.1
A3	1011106	1011197
refGene_gn=KCNQ2;refGene_tx=XM_023250969.1;refGene_gn=KCNQ2;refGene_tx=XM_023250970.1;refGene_gn=KCNQ2;refGene_tx=XM_023250971.1;refGene_gn=KCNQ2;refGene_tx=XM_023250972.1;refGene_gn=KCNQ2;refGene_tx=XM_023250974.1;refGene_gn=KCNQ2;refGene_tx=XM_023250975.1;refGene_gn=KCNQ2;refGene_tx=XM_023250976.1;refGene_gn=KCNQ2;refGene_tx=XM_023250977.1;refGene_gn=KCNQ2;refGene_tx=XM_023250978.1;refGene_gn=KCNQ2;refGene_tx=XM_023250979.1;refGene_gn=KCNQ2;refGene_tx=XM_023250980.1;refGene_gn=KCNQ2;refGene_tx=XM_023250981.1;refGene_gn=KCNQ2;refGene_tx=XM_023250982.1;refGene_gn=KCNQ2;refGene_tx=XM_023250983.1;refGene_gn=KCNQ2;refGene_tx=XM_023250984.1;refGene_gn=KCNQ2;refGene_tx=XM_023250985.1;refGene_gn=KCNQ2;refGene_tx=XM_023250986.1
A3	1011874	1012001
refGene_gn=KCNQ2;refGene_tx=XM_023250969.1;refGene_gn=KCNQ2;refGene_tx=XM_023250970.1;refGene_gn=KCNQ2;refGene_tx=XM_023250971.1;refGene_gn=KCNQ2;refGene_tx=XM_023250972.1;refGene_gn=KCNQ2;refGene_tx=XM_023250974.1;refGene_gn=KCNQ2;refGene_tx=XM_023250975.1;refGene_gn=KCNQ2;refGene_tx=XM_023250976.1;refGene_gn=KCNQ2;refGene_tx=XM_023250977.1;refGene_gn=KCNQ2;refGene_tx=XM_023250978.1;refGene_gn=KCNQ2;refGene_tx=XM_023250979.1;refGene_gn=KCNQ2;refGene_tx=XM_023250980.1;refGene_gn=KCNQ2;refGene_tx=XM_023250981.1;refGene_gn=KCNQ2;refGene_tx=XM_023250982.1;refGene_gn=KCNQ2;refGene_tx=XM_023250983.1;refGene_gn=KCNQ2;refGene_tx=XM_023250984.1;refGene_gn=KCNQ2;refGene_tx=XM_023250985.1;refGene_gn=KCNQ2;refGene_tx=XM_023250986.1
A3	1012416	1012592
refGene_gn=KCNQ2;refGene_tx=XM_023250969.1;refGene_gn=KCNQ2;refGene_tx=XM_023250970.1;refGene_gn=KCNQ2;refGene_tx=XM_023250971.1;refGene_gn=KCNQ2;refGene_tx=XM_023250972.1;refGene_gn=KCNQ2;refGene_tx=XM_023250974.1;refGene_gn=KCNQ2;refGene_tx=XM_023250975.1;refGene_gn=KCNQ2;refGene_tx=XM_023250976.1;refGene_gn=KCNQ2;refGene_tx=XM_023250977.1;refGene_gn=KCNQ2;refGene_tx=XM_023250978.1;refGene_gn=KCNQ2;refGene_tx=XM_023250979.1;refGene_gn=KCNQ2;refGene_tx=XM_023250980.1;refGene_gn=KCNQ2;refGene_tx=XM_023250981.1;refGene_gn=KCNQ2;refGene_tx=XM_023250982.1;r

efGene_gn=KCNQ2;refGene_tx=XM_023250983.1;refGene_gn=KCNQ2;refGene_tx=XM_023250984.1;refGene_gn=KCNQ2;refGene_tx=XM_023250985.1;refGene_gn=KCNQ2;refGene_tx=XM_023250986.1
A3 1013190 1013316
refGene_gn=KCNQ2;refGene_tx=XM_023250969.1;refGene_gn=KCNQ2;refGene_tx=XM_023250970.1;refGene_gn=KCNQ2;refGene_tx=XM_023250971.1;refGene_gn=KCNQ2;refGene_tx=XM_023250972.1;refGene_gn=KCNQ2;refGene_tx=XM_023250974.1;refGene_gn=KCNQ2;refGene_tx=XM_023250975.1;refGene_gn=KCNQ2;refGene_tx=XM_023250976.1;refGene_gn=KCNQ2;refGene_tx=XM_023250977.1;refGene_gn=KCNQ2;refGene_tx=XM_023250978.1;refGene_gn=KCNQ2;refGene_tx=XM_023250979.1;refGene_gn=KCNQ2;refGene_tx=XM_023250980.1;refGene_gn=KCNQ2;refGene_tx=XM_023250981.1;refGene_gn=KCNQ2;refGene_tx=XM_023250982.1;refGene_gn=KCNQ2;refGene_tx=XM_023250983.1;refGene_gn=KCNQ2;refGene_tx=XM_023250984.1;refGene_gn=KCNQ2;refGene_tx=XM_023250985.1;refGene_gn=KCNQ2;refGene_tx=XM_023250986.1
A3 1016169 1016280
refGene_gn=KCNQ2;refGene_tx=XM_023250969.1;refGene_gn=KCNQ2;refGene_tx=XM_023250970.1;refGene_gn=KCNQ2;refGene_tx=XM_023250971.1;refGene_gn=KCNQ2;refGene_tx=XM_023250972.1;refGene_gn=KCNQ2;refGene_tx=XM_023250974.1;refGene_gn=KCNQ2;refGene_tx=XM_023250975.1;refGene_gn=KCNQ2;refGene_tx=XM_023250976.1;refGene_gn=KCNQ2;refGene_tx=XM_023250977.1;refGene_gn=KCNQ2;refGene_tx=XM_023250978.1;refGene_gn=KCNQ2;refGene_tx=XM_023250979.1;refGene_gn=KCNQ2;refGene_tx=XM_023250980.1;refGene_gn=KCNQ2;refGene_tx=XM_023250981.1;refGene_gn=KCNQ2;refGene_tx=XM_023250982.1;refGene_gn=KCNQ2;refGene_tx=XM_023250983.1;refGene_gn=KCNQ2;refGene_tx=XM_023250984.1;refGene_gn=KCNQ2;refGene_tx=XM_023250985.1;refGene_gn=KCNQ2;refGene_tx=XM_023250986.1
A3 1016981 1017077
refGene_gn=KCNQ2;refGene_tx=XM_023250969.1;refGene_gn=KCNQ2;refGene_tx=XM_023250970.1;refGene_gn=KCNQ2;refGene_tx=XM_023250971.1;refGene_gn=KCNQ2;refGene_tx=XM_023250972.1;refGene_gn=KCNQ2;refGene_tx=XM_023250974.1;refGene_gn=KCNQ2;refGene_tx=XM_023250975.1;refGene_gn=KCNQ2;refGene_tx=XM_023250976.1;refGene_gn=KCNQ2;refGene_tx=XM_023250977.1;refGene_gn=KCNQ2;refGene_tx=XM_023250978.1;refGene_gn=KCNQ2;refGene_tx=XM_023250979.1;refGene_gn=KCNQ2;refGene_tx=XM_023250980.1;refGene_gn=KCNQ2;refGene_tx=XM_023250981.1;refGene_gn=KCNQ2;refGene_tx=XM_023250982.1;refGene_gn=KCNQ2;refGene_tx=XM_023250983.1;refGene_gn=KCNQ2;refGene_tx=XM_023250984.1;refGene_gn=KCNQ2;refGene_tx=XM_023250985.1;refGene_gn=KCNQ2;refGene_tx=XM_023250986.1
A3 1021591 1021686
refGene_gn=KCNQ2;refGene_tx=XM_023250969.1;refGene_gn=KCNQ2;refGene_tx=XM_023250970.1;refGene_gn=KCNQ2;refGene_tx=XM_023250971.1;refGene_gn=KCNQ2;refGene_tx=XM_023250972.1;refGene_gn=KCNQ2;refGene_tx=XM_023250974.1;refGene_gn=KCNQ2;refGene_tx=XM_023250975.1;refGene_gn=KCNQ2;refGene_tx=XM_023250976.1;refGene_gn=KCNQ2;refGene_tx=XM_023250977.1;refGene_gn=KCNQ2;refGene_tx=XM_023250978.1;refGene_gn=KCNQ2;refGene_tx=XM_023250979.1;refGene_gn=KCNQ2;refGene_tx=XM_023250980.1;refGene_gn=KCNQ2;refGene_tx=XM_023250981.1;refGene_gn=KCNQ2;refGene_tx=XM_023250982.1;refGene_gn=KCNQ2;refGene_tx=XM_023250983.1;refGene_gn=KCNQ2;refGene_tx=XM_023250984.1;refGene_gn=KCNQ2;refGene_tx=XM_023250985.1;refGene_gn=K

CNQ2;refGene_tx=XM_023250986.1
A3 1022831 1022861
refGene_gn=KCNQ2;refGene_tx=XM_023250969.1;refGene_gn=KCNQ2;refGene_tx=XM_023250970.1;refGene_gn=KCNQ2;refGene_tx=XM_023250972.1;refGene_gn=KCNQ2;refGene_tx=XM_023250974.1;refGene_gn=KCNQ2;refGene_tx=XM_023250976.1;refGene_gn=KCNQ2;refGene_tx=XM_023250977.1;refGene_gn=KCNQ2;refGene_tx=XM_023250978.1;refGene_gn=KCNQ2;refGene_tx=XM_023250980.1;refGene_gn=KCNQ2;refGene_tx=XM_023250981.1;refGene_gn=KCNQ2;refGene_tx=XM_023250984.1;refGene_gn=KCNQ2;refGene_tx=XM_023250985.1;refGene_gn=KCNQ2;refGene_tx=XM_023250986.1
A3 1025659 1025728
refGene_gn=KCNQ2;refGene_tx=XM_023250969.1;refGene_gn=KCNQ2;refGene_tx=XM_023250970.1;refGene_gn=KCNQ2;refGene_tx=XM_023250971.1;refGene_gn=KCNQ2;refGene_tx=XM_023250972.1;refGene_gn=KCNQ2;refGene_tx=XM_023250974.1;refGene_gn=KCNQ2;refGene_tx=XM_023250975.1;refGene_gn=KCNQ2;refGene_tx=XM_023250976.1;refGene_gn=KCNQ2;refGene_tx=XM_023250977.1;refGene_gn=KCNQ2;refGene_tx=XM_023250978.1;refGene_gn=KCNQ2;refGene_tx=XM_023250979.1;refGene_gn=KCNQ2;refGene_tx=XM_023250980.1;refGene_gn=KCNQ2;refGene_tx=XM_023250981.1;refGene_gn=KCNQ2;refGene_tx=XM_023250982.1;refGene_gn=KCNQ2;refGene_tx=XM_023250983.1;refGene_gn=KCNQ2;refGene_tx=XM_023250984.1;refGene_gn=KCNQ2;refGene_tx=XM_023250985.1;refGene_gn=KCNQ2;refGene_tx=XM_023250986.1
A3 1028934 1028964
refGene_gn=KCNQ2;refGene_tx=XM_023250969.1;refGene_gn=KCNQ2;refGene_tx=XM_023250970.1;refGene_gn=KCNQ2;refGene_tx=XM_023250971.1;refGene_gn=KCNQ2;refGene_tx=XM_023250972.1;refGene_gn=KCNQ2;refGene_tx=XM_023250974.1;refGene_gn=KCNQ2;refGene_tx=XM_023250975.1;refGene_gn=KCNQ2;refGene_tx=XM_023250976.1;refGene_gn=KCNQ2;refGene_tx=XM_023250977.1;refGene_gn=KCNQ2;refGene_tx=XM_023250978.1;refGene_gn=KCNQ2;refGene_tx=XM_023250979.1;refGene_gn=KCNQ2;refGene_tx=XM_023250980.1;refGene_gn=KCNQ2;refGene_tx=XM_023250981.1;refGene_gn=KCNQ2;refGene_tx=XM_023250982.1;refGene_gn=KCNQ2;refGene_tx=XM_023250983.1;refGene_gn=KCNQ2;refGene_tx=XM_023250984.1;refGene_gn=KCNQ2;refGene_tx=XM_023250985.1;refGene_gn=KCNQ2;refGene_tx=XM_023250986.1
A3 1032760 1032850 refGene_gn=KCNQ2;refGene_tx=XM_023250986.1
A3 1032950 1033004
refGene_gn=KCNQ2;refGene_tx=XM_023250969.1;refGene_gn=KCNQ2;refGene_tx=XM_023250970.1;refGene_gn=KCNQ2;refGene_tx=XM_023250971.1;refGene_gn=KCNQ2;refGene_tx=XM_023250978.1;refGene_gn=KCNQ2;refGene_tx=XM_023250986.1
A3 1036173 1036397
refGene_gn=KCNQ2;refGene_tx=XM_023250969.1;refGene_gn=KCNQ2;refGene_tx=XM_023250970.1;refGene_gn=KCNQ2;refGene_tx=XM_023250971.1;refGene_gn=KCNQ2;refGene_tx=XM_023250972.1;refGene_gn=KCNQ2;refGene_tx=XM_023250974.1;refGene_gn=KCNQ2;refGene_tx=XM_023250975.1;refGene_gn=KCNQ2;refGene_tx=XM_023250976.1;refGene_gn=KCNQ2;refGene_tx=XM_023250977.1;refGene_gn=KCNQ2;refGene_tx=XM_023250978.1;refGene_gn=KCNQ2;refGene_tx=XM_023250979.1;refGene_gn=KCNQ2;refGene_tx=XM_023250980.1;refGene_gn=KCNQ2;refGene_tx=XM_023250981.1;refGene_gn=KCNQ2;refGene_tx=XM_023250982.1;refGene_gn=KCNQ2;refGene_tx=XM_023250983.1;refGene_gn=KCNQ2;refGene_tx=

XM_023250984.1;refGene_gn=KCNQ2;refGene_tx=XM_023250985.1;refGene_gn=K
CNQ2;refGene_tx=XM_023250986.1
A3 1036939 1037048
refGene_gn=KCNQ2;refGene_tx=XM_023250969.1;refGene_gn=KCNQ2;refGene_tx
=XM_023250970.1;refGene_gn=KCNQ2;refGene_tx=XM_023250971.1;refGene_gn=
KCNQ2;refGene_tx=XM_023250972.1;refGene_gn=KCNQ2;refGene_tx=XM_0232509
74.1;refGene_gn=KCNQ2;refGene_tx=XM_023250975.1;refGene_gn=KCNQ2;refGe
ne_tx=XM_023250976.1;refGene_gn=KCNQ2;refGene_tx=XM_023250977.1;refGen
e_gn=KCNQ2;refGene_tx=XM_023250978.1;refGene_gn=KCNQ2;refGene_tx=XM_02
3250979.1;refGene_gn=KCNQ2;refGene_tx=XM_023250980.1;refGene_gn=KCNQ2;
refGene_tx=XM_023250981.1;refGene_gn=KCNQ2;refGene_tx=XM_023250982.1;r
efGene_gn=KCNQ2;refGene_tx=XM_023250983.1;refGene_gn=KCNQ2;refGene_tx=
XM_023250984.1;refGene_gn=KCNQ2;refGene_tx=XM_023250985.1;refGene_gn=K
CNQ2;refGene_tx=XM_023250986.1
A3 1037381 1037513
refGene_gn=KCNQ2;refGene_tx=XM_023250969.1;refGene_gn=KCNQ2;refGene_tx
=XM_023250970.1;refGene_gn=KCNQ2;refGene_tx=XM_023250971.1;refGene_gn=
KCNQ2;refGene_tx=XM_023250972.1;refGene_gn=KCNQ2;refGene_tx=XM_0232509
74.1;refGene_gn=KCNQ2;refGene_tx=XM_023250975.1;refGene_gn=KCNQ2;refGe
ne_tx=XM_023250976.1;refGene_gn=KCNQ2;refGene_tx=XM_023250977.1;refGen
e_gn=KCNQ2;refGene_tx=XM_023250978.1;refGene_gn=KCNQ2;refGene_tx=XM_02
3250979.1;refGene_gn=KCNQ2;refGene_tx=XM_023250980.1;refGene_gn=KCNQ2;
refGene_tx=XM_023250981.1;refGene_gn=KCNQ2;refGene_tx=XM_023250982.1;r
efGene_gn=KCNQ2;refGene_tx=XM_023250983.1;refGene_gn=KCNQ2;refGene_tx=
XM_023250984.1;refGene_gn=KCNQ2;refGene_tx=XM_023250985.1;refGene_gn=K
CNQ2;refGene_tx=XM_023250986.1
A3 1039220 1039328
refGene_gn=KCNQ2;refGene_tx=XM_023250969.1;refGene_gn=KCNQ2;refGene_tx
=XM_023250970.1;refGene_gn=KCNQ2;refGene_tx=XM_023250971.1;refGene_gn=
KCNQ2;refGene_tx=XM_023250972.1;refGene_gn=KCNQ2;refGene_tx=XM_0232509
74.1;refGene_gn=KCNQ2;refGene_tx=XM_023250975.1;refGene_gn=KCNQ2;refGe
ne_tx=XM_023250976.1;refGene_gn=KCNQ2;refGene_tx=XM_023250977.1;refGen
e_gn=KCNQ2;refGene_tx=XM_023250979.1;refGene_gn=KCNQ2;refGene_tx=XM_02
3250986.1
A3 1041871 1041995
refGene_gn=KCNQ2;refGene_tx=XM_023250969.1;refGene_gn=KCNQ2;refGene_tx
=XM_023250970.1;refGene_gn=KCNQ2;refGene_tx=XM_023250971.1;refGene_gn=
KCNQ2;refGene_tx=XM_023250972.1;refGene_gn=KCNQ2;refGene_tx=XM_0232509
74.1;refGene_gn=KCNQ2;refGene_tx=XM_023250975.1;refGene_gn=KCNQ2;refGe
ne_tx=XM_023250976.1;refGene_gn=KCNQ2;refGene_tx=XM_023250977.1;refGen
e_gn=KCNQ2;refGene_tx=XM_023250978.1;refGene_gn=KCNQ2;refGene_tx=XM_02
3250979.1;refGene_gn=KCNQ2;refGene_tx=XM_023250980.1;refGene_gn=KCNQ2;
refGene_tx=XM_023250981.1;refGene_gn=KCNQ2;refGene_tx=XM_023250982.1;r
efGene_gn=KCNQ2;refGene_tx=XM_023250983.1;refGene_gn=KCNQ2;refGene_tx=
XM_023250984.1;refGene_gn=KCNQ2;refGene_tx=XM_023250985.1;refGene_gn=K
CNQ2;refGene_tx=XM_023250986.1
A3 1042247 1042940
refGene_gn=KCNQ2;refGene_tx=XM_023250969.1;refGene_gn=KCNQ2;refGene_tx
=XM_023250970.1;refGene_gn=KCNQ2;refGene_tx=XM_023250971.1;refGene_gn=
KCNQ2;refGene_tx=XM_023250972.1;refGene_gn=KCNQ2;refGene_tx=XM_0232509

74.1;refGene_gn=KCNQ2;refGene_tx=XM_023250975.1;refGene_gn=KCNQ2;refGene_tx=XM_023250976.1;refGene_gn=KCNQ2;refGene_tx=XM_023250977.1;refGene_gn=KCNQ2;refGene_tx=XM_023250978.1;refGene_gn=KCNQ2;refGene_tx=XM_023250979.1;refGene_gn=KCNQ2;refGene_tx=XM_023250980.1;refGene_gn=KCNQ2;refGene_tx=XM_023250981.1;refGene_gn=KCNQ2;refGene_tx=XM_023250982.1;refGene_gn=KCNQ2;refGene_tx=XM_023250983.1;refGene_gn=KCNQ2;refGene_tx=XM_023250984.1;refGene_gn=KCNQ2;refGene_tx=XM_023250985.1;refGene_gn=KCNQ2;refGene_tx=XM_023250986.1
A3 1075908 1075990 refGene_gn=CHRNA4;refGene_tx=XM_023250987.1
A3 1077387 1077539 refGene_gn=CHRNA4;refGene_tx=XM_023250987.1
A3 1080579 1080624 refGene_gn=CHRNA4;refGene_tx=XM_023250987.1
A3 1080789 1080899 refGene_gn=CHRNA4;refGene_tx=XM_023250987.1
A3 1084779 1086166
refGene_gn=CHRNA4;refGene_tx=XM_019826238.2;refGene_gn=CHRNA4;refGene_tx=XM_023250987.1
A3 1089204 1089330
refGene_gn=CHRNA4;refGene_tx=XM_019826238.2;refGene_gn=CHRNA4;refGene_tx=XM_023250987.1
A3 1097698 1097724
refGene_gn=COL20A1;refGene_tx=XM_023250988.1;refGene_gn=COL20A1;refGene_tx=XM_023250989.1;refGene_gn=COL20A1;refGene_tx=XM_023250990.1;refGene_gn=COL20A1;refGene_tx=XM_023250991.1;refGene_gn=COL20A1;refGene_tx=XM_023250992.1;refGene_gn=COL20A1;refGene_tx=XM_023250993.1;refGene_gn=COL20A1;refGene_tx=XM_023250994.1;refGene_gn=COL20A1;refGene_tx=XM_023250995.1;refGene_gn=COL20A1;refGene_tx=XM_023250996.1;refGene_gn=COL20A1;refGene_tx=XM_023250997.1
A3 1098139 1098217
refGene_gn=COL20A1;refGene_tx=XM_023250988.1;refGene_gn=COL20A1;refGene_tx=XM_023250989.1;refGene_gn=COL20A1;refGene_tx=XM_023250990.1;refGene_gn=COL20A1;refGene_tx=XM_023250991.1;refGene_gn=COL20A1;refGene_tx=XM_023250992.1;refGene_gn=COL20A1;refGene_tx=XM_023250993.1;refGene_gn=COL20A1;refGene_tx=XM_023250995.1;refGene_gn=COL20A1;refGene_tx=XM_023250996.1;refGene_gn=COL20A1;refGene_tx=XM_023250997.1
A3 1098998 1099136
refGene_gn=COL20A1;refGene_tx=XM_023250988.1;refGene_gn=COL20A1;refGene_tx=XM_023250989.1;refGene_gn=COL20A1;refGene_tx=XM_023250990.1;refGene_gn=COL20A1;refGene_tx=XM_023250991.1;refGene_gn=COL20A1;refGene_tx=XM_023250992.1;refGene_gn=COL20A1;refGene_tx=XM_023250993.1;refGene_gn=COL20A1;refGene_tx=XM_023250994.1;refGene_gn=COL20A1;refGene_tx=XM_023250995.1;refGene_gn=COL20A1;refGene_tx=XM_023250996.1;refGene_gn=COL20A1;refGene_tx=XM_023250997.1;refGene_gn=COL20A1;refGene_tx=XM_023250998.1
A3 1099342 1099409
refGene_gn=COL20A1;refGene_tx=XM_023250988.1;refGene_gn=COL20A1;refGene_tx=XM_023250989.1;refGene_gn=COL20A1;refGene_tx=XM_023250990.1;refGene_gn=COL20A1;refGene_tx=XM_023250991.1;refGene_gn=COL20A1;refGene_tx=XM_023250992.1;refGene_gn=COL20A1;refGene_tx=XM_023250993.1;refGene_gn=COL20A1;refGene_tx=XM_023250994.1;refGene_gn=COL20A1;refGene_tx=XM_023250995.1;refGene_gn=COL20A1;refGene_tx=XM_023250996.1;refGene_gn=COL20A1;refGene_tx=XM_023250997.1

A3 1099521 1099557
refGene_gn=COL20A1;refGene_tx=XM_023250988.1;refGene_gn=COL20A1;refGen
e_tx=XM_023250989.1;refGene_gn=COL20A1;refGene_tx=XM_023250990.1;refGe
ne_gn=COL20A1;refGene_tx=XM_023250991.1;refGene_gn=COL20A1;refGene_tx=
XM_023250992.1;refGene_gn=COL20A1;refGene_tx=XM_023250993.1;refGene_gn
=COL20A1;refGene_tx=XM_023250994.1;refGene_gn=COL20A1;refGene_tx=XM_02
3250995.1;refGene_gn=COL20A1;refGene_tx=XM_023250996.1;refGene_gn=COL2
0A1;refGene_tx=XM_023250997.1;refGene_gn=COL20A1;refGene_tx=XM_0232509
98.1
A3 1099982 1100054
refGene_gn=COL20A1;refGene_tx=XM_023250988.1;refGene_gn=COL20A1;refGen
e_tx=XM_023250989.1;refGene_gn=COL20A1;refGene_tx=XM_023250990.1;refGe
ne_gn=COL20A1;refGene_tx=XM_023250991.1;refGene_gn=COL20A1;refGene_tx=
XM_023250992.1;refGene_gn=COL20A1;refGene_tx=XM_023250993.1;refGene_gn
=COL20A1;refGene_tx=XM_023250994.1;refGene_gn=COL20A1;refGene_tx=XM_02
3250995.1;refGene_gn=COL20A1;refGene_tx=XM_023250996.1;refGene_gn=COL2
0A1;refGene_tx=XM_023250997.1;refGene_gn=COL20A1;refGene_tx=XM_0232509
98.1
A3 1100705 1100759
refGene_gn=COL20A1;refGene_tx=XM_023250988.1;refGene_gn=COL20A1;refGen
e_tx=XM_023250989.1;refGene_gn=COL20A1;refGene_tx=XM_023250990.1;refGe
ne_gn=COL20A1;refGene_tx=XM_023250992.1;refGene_gn=COL20A1;refGene_tx=
XM_023250993.1;refGene_gn=COL20A1;refGene_tx=XM_023250994.1;refGene_gn
=COL20A1;refGene_tx=XM_023250995.1;refGene_gn=COL20A1;refGene_tx=XM_02
3250996.1;refGene_gn=COL20A1;refGene_tx=XM_023250997.1;refGene_gn=COL2
0A1;refGene_tx=XM_023250998.1
A3 1101034 1101088
refGene_gn=COL20A1;refGene_tx=XM_023250988.1;refGene_gn=COL20A1;refGen
e_tx=XM_023250989.1;refGene_gn=COL20A1;refGene_tx=XM_023250990.1;refGe
ne_gn=COL20A1;refGene_tx=XM_023250991.1;refGene_gn=COL20A1;refGene_tx=
XM_023250992.1;refGene_gn=COL20A1;refGene_tx=XM_023250993.1;refGene_gn
=COL20A1;refGene_tx=XM_023250994.1;refGene_gn=COL20A1;refGene_tx=XM_02
3250995.1;refGene_gn=COL20A1;refGene_tx=XM_023250996.1;refGene_gn=COL2
0A1;refGene_tx=XM_023250997.1;refGene_gn=COL20A1;refGene_tx=XM_0232509
98.1
A3 1101197 1101251
refGene_gn=COL20A1;refGene_tx=XM_023250988.1;refGene_gn=COL20A1;refGen
e_tx=XM_023250989.1;refGene_gn=COL20A1;refGene_tx=XM_023250990.1;refGe
ne_gn=COL20A1;refGene_tx=XM_023250991.1;refGene_gn=COL20A1;refGene_tx=
XM_023250992.1;refGene_gn=COL20A1;refGene_tx=XM_023250993.1;refGene_gn
=COL20A1;refGene_tx=XM_023250994.1;refGene_gn=COL20A1;refGene_tx=XM_02
3250995.1;refGene_gn=COL20A1;refGene_tx=XM_023250996.1;refGene_gn=COL2
0A1;refGene_tx=XM_023250997.1;refGene_gn=COL20A1;refGene_tx=XM_0232509
98.1
A3 1101330 1101426
refGene_gn=COL20A1;refGene_tx=XM_023250988.1;refGene_gn=COL20A1;refGen
e_tx=XM_023250989.1;refGene_gn=COL20A1;refGene_tx=XM_023250990.1;refGe
ne_gn=COL20A1;refGene_tx=XM_023250991.1;refGene_gn=COL20A1;refGene_tx=
XM_023250992.1;refGene_gn=COL20A1;refGene_tx=XM_023250993.1;refGene_gn
=COL20A1;refGene_tx=XM_023250994.1;refGene_gn=COL20A1;refGene_tx=XM_02

3250996.1;refGene_gn=COL20A1;refGene_tx=XM_023250998.1
A3 1102529 1102583
refGene_gn=COL20A1;refGene_tx=XM_023250988.1;refGene_gn=COL20A1;refGen
e_tx=XM_023250989.1;refGene_gn=COL20A1;refGene_tx=XM_023250990.1;refGe
ne_gn=COL20A1;refGene_tx=XM_023250991.1;refGene_gn=COL20A1;refGene_tx=
XM_023250992.1;refGene_gn=COL20A1;refGene_tx=XM_023250993.1;refGene_gn
=COL20A1;refGene_tx=XM_023250994.1;refGene_gn=COL20A1;refGene_tx=XM_02
3250995.1;refGene_gn=COL20A1;refGene_tx=XM_023250996.1;refGene_gn=COL2
0A1;refGene_tx=XM_023250997.1;refGene_gn=COL20A1;refGene_tx=XM_0232509
98.1
A3 1103255 1103342
refGene_gn=COL20A1;refGene_tx=XM_023250988.1;refGene_gn=COL20A1;refGen
e_tx=XM_023250989.1;refGene_gn=COL20A1;refGene_tx=XM_023250990.1;refGe
ne_gn=COL20A1;refGene_tx=XM_023250991.1;refGene_gn=COL20A1;refGene_tx=
XM_023250992.1;refGene_gn=COL20A1;refGene_tx=XM_023250993.1;refGene_gn
=COL20A1;refGene_tx=XM_023250994.1;refGene_gn=COL20A1;refGene_tx=XM_02
3250995.1;refGene_gn=COL20A1;refGene_tx=XM_023250996.1;refGene_gn=COL2
0A1;refGene_tx=XM_023250997.1;refGene_gn=COL20A1;refGene_tx=XM_0232509
98.1
A3 1103935 1104013
refGene_gn=COL20A1;refGene_tx=XM_023250988.1;refGene_gn=COL20A1;refGen
e_tx=XM_023250989.1;refGene_gn=COL20A1;refGene_tx=XM_023250990.1;refGe
ne_gn=COL20A1;refGene_tx=XM_023250991.1;refGene_gn=COL20A1;refGene_tx=
XM_023250992.1;refGene_gn=COL20A1;refGene_tx=XM_023250993.1;refGene_gn
=COL20A1;refGene_tx=XM_023250994.1;refGene_gn=COL20A1;refGene_tx=XM_02
3250995.1;refGene_gn=COL20A1;refGene_tx=XM_023250996.1;refGene_gn=COL2
0A1;refGene_tx=XM_023250997.1;refGene_gn=COL20A1;refGene_tx=XM_0232509
98.1
A3 1104103 1104262
refGene_gn=COL20A1;refGene_tx=XM_023250988.1;refGene_gn=COL20A1;refGen
e_tx=XM_023250989.1;refGene_gn=COL20A1;refGene_tx=XM_023250990.1;refGe
ne_gn=COL20A1;refGene_tx=XM_023250991.1;refGene_gn=COL20A1;refGene_tx=
XM_023250992.1;refGene_gn=COL20A1;refGene_tx=XM_023250993.1;refGene_gn
=COL20A1;refGene_tx=XM_023250994.1;refGene_gn=COL20A1;refGene_tx=XM_02
3250995.1;refGene_gn=COL20A1;refGene_tx=XM_023250996.1;refGene_gn=COL2
0A1;refGene_tx=XM_023250997.1;refGene_gn=COL20A1;refGene_tx=XM_0232509
98.1
A3 1104668 1104778
refGene_gn=COL20A1;refGene_tx=XM_023250988.1;refGene_gn=COL20A1;refGen
e_tx=XM_023250989.1;refGene_gn=COL20A1;refGene_tx=XM_023250990.1;refGe
ne_gn=COL20A1;refGene_tx=XM_023250991.1;refGene_gn=COL20A1;refGene_tx=
XM_023250992.1;refGene_gn=COL20A1;refGene_tx=XM_023250993.1;refGene_gn
=COL20A1;refGene_tx=XM_023250994.1;refGene_gn=COL20A1;refGene_tx=XM_02
3250995.1;refGene_gn=COL20A1;refGene_tx=XM_023250996.1;refGene_gn=COL2
0A1;refGene_tx=XM_023250997.1;refGene_gn=COL20A1;refGene_tx=XM_0232509
98.1
A3 1105009 1105152
refGene_gn=COL20A1;refGene_tx=XM_023250988.1;refGene_gn=COL20A1;refGen
e_tx=XM_023250989.1;refGene_gn=COL20A1;refGene_tx=XM_023250990.1;refGe
ne_gn=COL20A1;refGene_tx=XM_023250991.1;refGene_gn=COL20A1;refGene_tx=

XM_023250992.1;refGene_gn=COL20A1;refGene_tx=XM_023250993.1;refGene_gn=COL20A1;refGene_tx=XM_023250994.1;refGene_gn=COL20A1;refGene_tx=XM_023250995.1;refGene_gn=COL20A1;refGene_tx=XM_023250996.1;refGene_gn=COL20A1;refGene_tx=XM_023250997.1;refGene_gn=COL20A1;refGene_tx=XM_023250998.1
A3 1105571 1105643
refGene_gn=COL20A1;refGene_tx=XM_023250988.1;refGene_gn=COL20A1;refGene_tx=XM_023250989.1;refGene_gn=COL20A1;refGene_tx=XM_023250990.1;refGene_gn=COL20A1;refGene_tx=XM_023250991.1;refGene_gn=COL20A1;refGene_tx=XM_023250992.1;refGene_gn=COL20A1;refGene_tx=XM_023250994.1;refGene_gn=COL20A1;refGene_tx=XM_023250995.1;refGene_gn=COL20A1;refGene_tx=XM_023250996.1;refGene_gn=COL20A1;refGene_tx=XM_023250998.1
A3 1106989 1107128
refGene_gn=COL20A1;refGene_tx=XM_023250988.1;refGene_gn=COL20A1;refGene_tx=XM_023250989.1;refGene_gn=COL20A1;refGene_tx=XM_023250990.1;refGene_gn=COL20A1;refGene_tx=XM_023250991.1;refGene_gn=COL20A1;refGene_tx=XM_023250992.1;refGene_gn=COL20A1;refGene_tx=XM_023250993.1;refGene_gn=COL20A1;refGene_tx=XM_023250994.1;refGene_gn=COL20A1;refGene_tx=XM_023250995.1;refGene_gn=COL20A1;refGene_tx=XM_023250996.1;refGene_gn=COL20A1;refGene_tx=XM_023250997.1;refGene_gn=COL20A1;refGene_tx=XM_023250998.1
A3 1108036 1108075
refGene_gn=COL20A1;refGene_tx=XM_023250988.1;refGene_gn=COL20A1;refGene_tx=XM_023250989.1;refGene_gn=COL20A1;refGene_tx=XM_023250990.1;refGene_gn=COL20A1;refGene_tx=XM_023250991.1;refGene_gn=COL20A1;refGene_tx=XM_023250992.1;refGene_gn=COL20A1;refGene_tx=XM_023250993.1;refGene_gn=COL20A1;refGene_tx=XM_023250994.1;refGene_gn=COL20A1;refGene_tx=XM_023250995.1;refGene_gn=COL20A1;refGene_tx=XM_023250996.1;refGene_gn=COL20A1;refGene_tx=XM_023250997.1;refGene_gn=COL20A1;refGene_tx=XM_023250998.1
A3 1110896 1111026
refGene_gn=COL20A1;refGene_tx=XM_023250988.1;refGene_gn=COL20A1;refGene_tx=XM_023250989.1;refGene_gn=COL20A1;refGene_tx=XM_023250990.1;refGene_gn=COL20A1;refGene_tx=XM_023250991.1;refGene_gn=COL20A1;refGene_tx=XM_023250992.1;refGene_gn=COL20A1;refGene_tx=XM_023250993.1;refGene_gn=COL20A1;refGene_tx=XM_023250994.1;refGene_gn=COL20A1;refGene_tx=XM_023250995.1;refGene_gn=COL20A1;refGene_tx=XM_023250996.1;refGene_gn=COL20A1;refGene_tx=XM_023250997.1;refGene_gn=COL20A1;refGene_tx=XM_023250998.1
A3 1111205 1111348
refGene_gn=COL20A1;refGene_tx=XM_023250988.1;refGene_gn=COL20A1;refGene_tx=XM_023250989.1;refGene_gn=COL20A1;refGene_tx=XM_023250990.1;refGene_gn=COL20A1;refGene_tx=XM_023250991.1;refGene_gn=COL20A1;refGene_tx=XM_023250992.1;refGene_gn=COL20A1;refGene_tx=XM_023250993.1;refGene_gn=COL20A1;refGene_tx=XM_023250994.1;refGene_gn=COL20A1;refGene_tx=XM_023250995.1;refGene_gn=COL20A1;refGene_tx=XM_023250996.1;refGene_gn=COL20A1;refGene_tx=XM_023250997.1;refGene_gn=COL20A1;refGene_tx=XM_023250998.1
A3 1111747 1111880
refGene_gn=COL20A1;refGene_tx=XM_023250988.1;refGene_gn=COL20A1;refGen

e_tx=XM_023250989.1;refGene_gn=COL20A1;refGene_tx=XM_023250990.1;refGene_gn=COL20A1;refGene_tx=XM_023250991.1;refGene_gn=COL20A1;refGene_tx=XM_023250992.1;refGene_gn=COL20A1;refGene_tx=XM_023250993.1;refGene_gn=COL20A1;refGene_tx=XM_023250994.1;refGene_gn=COL20A1;refGene_tx=XM_023250995.1;refGene_gn=COL20A1;refGene_tx=XM_023250996.1;refGene_gn=COL20A1;refGene_tx=XM_023250997.1;refGene_gn=COL20A1;refGene_tx=XM_023250998.1
A3 1112044 1112187
refGene_gn=COL20A1;refGene_tx=XM_023250988.1;refGene_gn=COL20A1;refGene_tx=XM_023250989.1;refGene_gn=COL20A1;refGene_tx=XM_023250990.1;refGene_gn=COL20A1;refGene_tx=XM_023250991.1;refGene_gn=COL20A1;refGene_tx=XM_023250992.1;refGene_gn=COL20A1;refGene_tx=XM_023250993.1;refGene_gn=COL20A1;refGene_tx=XM_023250994.1;refGene_gn=COL20A1;refGene_tx=XM_023250995.1;refGene_gn=COL20A1;refGene_tx=XM_023250996.1;refGene_gn=COL20A1;refGene_tx=XM_023250997.1;refGene_gn=COL20A1;refGene_tx=XM_023250998.1
A3 1112631 1112761
refGene_gn=COL20A1;refGene_tx=XM_023250988.1;refGene_gn=COL20A1;refGene_tx=XM_023250989.1;refGene_gn=COL20A1;refGene_tx=XM_023250990.1;refGene_gn=COL20A1;refGene_tx=XM_023250991.1;refGene_gn=COL20A1;refGene_tx=XM_023250992.1;refGene_gn=COL20A1;refGene_tx=XM_023250993.1;refGene_gn=COL20A1;refGene_tx=XM_023250994.1;refGene_gn=COL20A1;refGene_tx=XM_023250995.1;refGene_gn=COL20A1;refGene_tx=XM_023250996.1;refGene_gn=COL20A1;refGene_tx=XM_023250997.1;refGene_gn=COL20A1;refGene_tx=XM_023250998.1
A3 1113251 1113391
refGene_gn=COL20A1;refGene_tx=XM_023250988.1;refGene_gn=COL20A1;refGene_tx=XM_023250989.1;refGene_gn=COL20A1;refGene_tx=XM_023250990.1;refGene_gn=COL20A1;refGene_tx=XM_023250991.1;refGene_gn=COL20A1;refGene_tx=XM_023250992.1;refGene_gn=COL20A1;refGene_tx=XM_023250993.1;refGene_gn=COL20A1;refGene_tx=XM_023250994.1;refGene_gn=COL20A1;refGene_tx=XM_023250995.1;refGene_gn=COL20A1;refGene_tx=XM_023250996.1;refGene_gn=COL20A1;refGene_tx=XM_023250997.1;refGene_gn=COL20A1;refGene_tx=XM_023250998.1
A3 1113565 1113689
refGene_gn=COL20A1;refGene_tx=XM_023250988.1;refGene_gn=COL20A1;refGene_tx=XM_023250989.1;refGene_gn=COL20A1;refGene_tx=XM_023250990.1;refGene_gn=COL20A1;refGene_tx=XM_023250991.1;refGene_gn=COL20A1;refGene_tx=XM_023250992.1;refGene_gn=COL20A1;refGene_tx=XM_023250993.1;refGene_gn=COL20A1;refGene_tx=XM_023250994.1;refGene_gn=COL20A1;refGene_tx=XM_023250995.1;refGene_gn=COL20A1;refGene_tx=XM_023250996.1;refGene_gn=COL20A1;refGene_tx=XM_023250997.1;refGene_gn=COL20A1;refGene_tx=XM_023250998.1
A3 1113797 1113943
refGene_gn=COL20A1;refGene_tx=XM_023250988.1;refGene_gn=COL20A1;refGene_tx=XM_023250989.1;refGene_gn=COL20A1;refGene_tx=XM_023250990.1;refGene_gn=COL20A1;refGene_tx=XM_023250991.1;refGene_gn=COL20A1;refGene_tx=XM_023250992.1;refGene_gn=COL20A1;refGene_tx=XM_023250993.1;refGene_gn=COL20A1;refGene_tx=XM_023250994.1;refGene_gn=COL20A1;refGene_tx=XM_023250995.1;refGene_gn=COL20A1;refGene_tx=XM_023250996.1;refGene_gn=COL2

0A1;refGene_tx=XM_023250997.1;refGene_gn=COL20A1;refGene_tx=XM_023250998.1
A3 1114697 1114827
refGene_gn=COL20A1;refGene_tx=XM_023250988.1;refGene_gn=COL20A1;refGene_tx=XM_023250989.1;refGene_gn=COL20A1;refGene_tx=XM_023250990.1;refGene_gn=COL20A1;refGene_tx=XM_023250991.1;refGene_gn=COL20A1;refGene_tx=XM_023250992.1;refGene_gn=COL20A1;refGene_tx=XM_023250993.1;refGene_gn=COL20A1;refGene_tx=XM_023250994.1;refGene_gn=COL20A1;refGene_tx=XM_023250995.1;refGene_gn=COL20A1;refGene_tx=XM_023250996.1;refGene_gn=COL20A1;refGene_tx=XM_023250997.1;refGene_gn=COL20A1;refGene_tx=XM_023250998.1
A3 1115470 1115643
refGene_gn=COL20A1;refGene_tx=XM_023250988.1;refGene_gn=COL20A1;refGene_tx=XM_023250989.1;refGene_gn=COL20A1;refGene_tx=XM_023250990.1;refGene_gn=COL20A1;refGene_tx=XM_023250991.1;refGene_gn=COL20A1;refGene_tx=XM_023250992.1;refGene_gn=COL20A1;refGene_tx=XM_023250993.1;refGene_gn=COL20A1;refGene_tx=XM_023250994.1;refGene_gn=COL20A1;refGene_tx=XM_023250995.1;refGene_gn=COL20A1;refGene_tx=XM_023250996.1;refGene_gn=COL20A1;refGene_tx=XM_023250997.1;refGene_gn=COL20A1;refGene_tx=XM_023250998.1
A3 1115916 1116075
refGene_gn=COL20A1;refGene_tx=XM_023250988.1;refGene_gn=COL20A1;refGene_tx=XM_023250989.1;refGene_gn=COL20A1;refGene_tx=XM_023250990.1;refGene_gn=COL20A1;refGene_tx=XM_023250991.1;refGene_gn=COL20A1;refGene_tx=XM_023250992.1;refGene_gn=COL20A1;refGene_tx=XM_023250993.1;refGene_gn=COL20A1;refGene_tx=XM_023250994.1;refGene_gn=COL20A1;refGene_tx=XM_023250995.1;refGene_gn=COL20A1;refGene_tx=XM_023250996.1;refGene_gn=COL20A1;refGene_tx=XM_023250997.1;refGene_gn=COL20A1;refGene_tx=XM_023250998.1
A3 1116823 1116988
refGene_gn=COL20A1;refGene_tx=XM_023250988.1;refGene_gn=COL20A1;refGene_tx=XM_023250989.1;refGene_gn=COL20A1;refGene_tx=XM_023250990.1;refGene_gn=COL20A1;refGene_tx=XM_023250991.1;refGene_gn=COL20A1;refGene_tx=XM_023250992.1;refGene_gn=COL20A1;refGene_tx=XM_023250993.1;refGene_gn=COL20A1;refGene_tx=XM_023250994.1;refGene_gn=COL20A1;refGene_tx=XM_023250995.1;refGene_gn=COL20A1;refGene_tx=XM_023250996.1;refGene_gn=COL20A1;refGene_tx=XM_023250997.1;refGene_gn=COL20A1;refGene_tx=XM_023250998.1
A3 1117384 1117504
refGene_gn=COL20A1;refGene_tx=XM_023250988.1;refGene_gn=COL20A1;refGene_tx=XM_023250989.1;refGene_gn=COL20A1;refGene_tx=XM_023250990.1;refGene_gn=COL20A1;refGene_tx=XM_023250991.1;refGene_gn=COL20A1;refGene_tx=XM_023250992.1;refGene_gn=COL20A1;refGene_tx=XM_023250993.1;refGene_gn=COL20A1;refGene_tx=XM_023250994.1;refGene_gn=COL20A1;refGene_tx=XM_023250995.1;refGene_gn=COL20A1;refGene_tx=XM_023250996.1;refGene_gn=COL20A1;refGene_tx=XM_023250997.1;refGene_gn=COL20A1;refGene_tx=XM_023250998.1
A3 1118052 1118211
refGene_gn=COL20A1;refGene_tx=XM_023250988.1;refGene_gn=COL20A1;refGene_tx=XM_023250989.1;refGene_gn=COL20A1;refGene_tx=XM_023250990.1;refGe

ne_gn=COL20A1;refGene_tx=XM_023250991.1;refGene_gn=COL20A1;refGene_tx=
XM_023250992.1;refGene_gn=COL20A1;refGene_tx=XM_023250993.1;refGene_gn
=COL20A1;refGene_tx=XM_023250994.1;refGene_gn=COL20A1;refGene_tx=XM_02
3250995.1;refGene_gn=COL20A1;refGene_tx=XM_023250996.1;refGene_gn=COL2
0A1;refGene_tx=XM_023250997.1;refGene_gn=COL20A1;refGene_tx=XM_0232509
98.1
A3	1118877	1119000
refGene_gn=COL20A1;refGene_tx=XM_023250988.1;refGene_gn=COL20A1;refGen
e_tx=XM_023250989.1;refGene_gn=COL20A1;refGene_tx=XM_023250990.1;refGe
ne_gn=COL20A1;refGene_tx=XM_023250991.1;refGene_gn=COL20A1;refGene_tx=
XM_023250992.1;refGene_gn=COL20A1;refGene_tx=XM_023250993.1;refGene_gn
=COL20A1;refGene_tx=XM_023250994.1;refGene_gn=COL20A1;refGene_tx=XM_02
3250995.1;refGene_gn=COL20A1;refGene_tx=XM_023250997.1;refGene_gn=COL2
0A1;refGene_tx=XM_023250998.1
A3	1119215	1119353
refGene_gn=COL20A1;refGene_tx=XM_023250988.1;refGene_gn=COL20A1;refGen
e_tx=XM_023250989.1;refGene_gn=COL20A1;refGene_tx=XM_023250990.1;refGe
ne_gn=COL20A1;refGene_tx=XM_023250991.1;refGene_gn=COL20A1;refGene_tx=
XM_023250992.1;refGene_gn=COL20A1;refGene_tx=XM_023250993.1;refGene_gn
=COL20A1;refGene_tx=XM_023250994.1;refGene_gn=COL20A1;refGene_tx=XM_02
3250995.1;refGene_gn=COL20A1;refGene_tx=XM_023250996.1;refGene_gn=COL2
0A1;refGene_tx=XM_023250997.1;refGene_gn=COL20A1;refGene_tx=XM_0232509
98.1
A3	1119509	1119786
refGene_gn=COL20A1;refGene_tx=XM_023250988.1;refGene_gn=COL20A1;refGen
e_tx=XM_023250989.1;refGene_gn=COL20A1;refGene_tx=XM_023250990.1;refGe
ne_gn=COL20A1;refGene_tx=XM_023250991.1;refGene_gn=COL20A1;refGene_tx=
XM_023250992.1;refGene_gn=COL20A1;refGene_tx=XM_023250993.1;refGene_gn
=COL20A1;refGene_tx=XM_023250994.1;refGene_gn=COL20A1;refGene_tx=XM_02
3250995.1;refGene_gn=COL20A1;refGene_tx=XM_023250996.1;refGene_gn=COL2
0A1;refGene_tx=XM_023250997.1;refGene_gn=COL20A1;refGene_tx=XM_0232509
98.1
A3	1125425	1125489	refGene_gn=COL20A1;refGene_tx=XM_023250992.1
A3	1133178	1133518
refGene_gn=ARFGAP1;refGene_tx=XM_023250999.1;refGene_gn=ARFGAP1;refGen
e_tx=XM_023251000.1;refGene_gn=ARFGAP1;refGene_tx=XM_023251001.1;refGe
ne_gn=ARFGAP1;refGene_tx=XM_023251002.1;refGene_gn=ARFGAP1;refGene_tx=
XM_023251003.1;refGene_gn=ARFGAP1;refGene_tx=XM_023251004.1;refGene_gn
=ARFGAP1;refGene_tx=XM_023251005.1;refGene_gn=ARFGAP1;refGene_tx=XM_02
3251006.1;refGene_gn=ARFGAP1;refGene_tx=XM_023251007.1
A3	1134220	1134297
refGene_gn=ARFGAP1;refGene_tx=XM_023250999.1;refGene_gn=ARFGAP1;refGen
e_tx=XM_023251000.1;refGene_gn=ARFGAP1;refGene_tx=XM_023251001.1;refGe
ne_gn=ARFGAP1;refGene_tx=XM_023251002.1;refGene_gn=ARFGAP1;refGene_tx=
XM_023251003.1;refGene_gn=ARFGAP1;refGene_tx=XM_023251004.1;refGene_gn
=ARFGAP1;refGene_tx=XM_023251005.1;refGene_gn=ARFGAP1;refGene_tx=XM_02
3251006.1;refGene_gn=ARFGAP1;refGene_tx=XM_023251007.1
A3	1134815	1134875
refGene_gn=ARFGAP1;refGene_tx=XM_023250999.1;refGene_gn=ARFGAP1;refGen
e_tx=XM_023251000.1;refGene_gn=ARFGAP1;refGene_tx=XM_023251001.1;refGe

ne_gn=ARFGAP1;refGene_tx=XM_023251002.1;refGene_gn=ARFGAP1;refGene_tx=XM_023251003.1
A3 1135302 1135359
refGene_gn=ARFGAP1;refGene_tx=XM_023250999.1;refGene_gn=ARFGAP1;refGene_tx=XM_023251000.1;refGene_gn=ARFGAP1;refGene_tx=XM_023251001.1;refGene_gn=ARFGAP1;refGene_tx=XM_023251002.1;refGene_gn=ARFGAP1;refGene_tx=XM_023251003.1;refGene_gn=ARFGAP1;refGene_tx=XM_023251004.1;refGene_gn=ARFGAP1;refGene_tx=XM_023251005.1;refGene_gn=ARFGAP1;refGene_tx=XM_023251006.1;refGene_gn=ARFGAP1;refGene_tx=XM_023251007.1
A3 1136194 1136224
refGene_gn=ARFGAP1;refGene_tx=XM_023250999.1;refGene_gn=ARFGAP1;refGene_tx=XM_023251000.1;refGene_gn=ARFGAP1;refGene_tx=XM_023251001.1;refGene_gn=ARFGAP1;refGene_tx=XM_023251002.1;refGene_gn=ARFGAP1;refGene_tx=XM_023251004.1;refGene_gn=ARFGAP1;refGene_tx=XM_023251005.1
A3 1137171 1137204
refGene_gn=ARFGAP1;refGene_tx=XM_023250999.1;refGene_gn=ARFGAP1;refGene_tx=XM_023251000.1;refGene_gn=ARFGAP1;refGene_tx=XM_023251001.1;refGene_gn=ARFGAP1;refGene_tx=XM_023251002.1;refGene_gn=ARFGAP1;refGene_tx=XM_023251003.1;refGene_gn=ARFGAP1;refGene_tx=XM_023251004.1;refGene_gn=ARFGAP1;refGene_tx=XM_023251005.1;refGene_gn=ARFGAP1;refGene_tx=XM_023251006.1;refGene_gn=ARFGAP1;refGene_tx=XM_023251007.1
A3 1138332 1138389
refGene_gn=ARFGAP1;refGene_tx=XM_023250999.1;refGene_gn=ARFGAP1;refGene_tx=XM_023251000.1;refGene_gn=ARFGAP1;refGene_tx=XM_023251001.1;refGene_gn=ARFGAP1;refGene_tx=XM_023251002.1;refGene_gn=ARFGAP1;refGene_tx=XM_023251003.1;refGene_gn=ARFGAP1;refGene_tx=XM_023251004.1;refGene_gn=ARFGAP1;refGene_tx=XM_023251005.1;refGene_gn=ARFGAP1;refGene_tx=XM_023251006.1;refGene_gn=ARFGAP1;refGene_tx=XM_023251007.1
A3 1139371 1139468
refGene_gn=ARFGAP1;refGene_tx=XM_023250999.1;refGene_gn=ARFGAP1;refGene_tx=XM_023251000.1;refGene_gn=ARFGAP1;refGene_tx=XM_023251001.1;refGene_gn=ARFGAP1;refGene_tx=XM_023251002.1;refGene_gn=ARFGAP1;refGene_tx=XM_023251003.1;refGene_gn=ARFGAP1;refGene_tx=XM_023251004.1;refGene_gn=ARFGAP1;refGene_tx=XM_023251005.1;refGene_gn=ARFGAP1;refGene_tx=XM_023251006.1;refGene_gn=ARFGAP1;refGene_tx=XM_023251007.1
A3 1139976 1140063
refGene_gn=ARFGAP1;refGene_tx=XM_023250999.1;refGene_gn=ARFGAP1;refGene_tx=XM_023251000.1;refGene_gn=ARFGAP1;refGene_tx=XM_023251001.1;refGene_gn=ARFGAP1;refGene_tx=XM_023251002.1;refGene_gn=ARFGAP1;refGene_tx=XM_023251003.1;refGene_gn=ARFGAP1;refGene_tx=XM_023251004.1;refGene_gn=ARFGAP1;refGene_tx=XM_023251005.1;refGene_gn=ARFGAP1;refGene_tx=XM_023251006.1;refGene_gn=ARFGAP1;refGene_tx=XM_023251007.1
A3 1140838 1140939
refGene_gn=ARFGAP1;refGene_tx=XM_023250999.1;refGene_gn=ARFGAP1;refGene_tx=XM_023251000.1;refGene_gn=ARFGAP1;refGene_tx=XM_023251001.1;refGene_gn=ARFGAP1;refGene_tx=XM_023251002.1;refGene_gn=ARFGAP1;refGene_tx=XM_023251003.1;refGene_gn=ARFGAP1;refGene_tx=XM_023251004.1;refGene_gn=ARFGAP1;refGene_tx=XM_023251005.1;refGene_gn=ARFGAP1;refGene_tx=XM_023251006.1;refGene_gn=ARFGAP1;refGene_tx=XM_023251007.1
A3 1141414 1141586

refGene_gn=ARFGAP1;refGene_tx=XM_023250999.1;refGene_gn=ARFGAP1;refGene_tx=XM_023251000.1;refGene_gn=ARFGAP1;refGene_tx=XM_023251001.1;refGene_gn=ARFGAP1;refGene_tx=XM_023251002.1;refGene_gn=ARFGAP1;refGene_tx=XM_023251003.1;refGene_gn=ARFGAP1;refGene_tx=XM_023251004.1;refGene_gn=ARFGAP1;refGene_tx=XM_023251005.1;refGene_gn=ARFGAP1;refGene_tx=XM_023251006.1;refGene_gn=ARFGAP1;refGene_tx=XM_023251007.1
A3 1141838 1141948
refGene_gn=ARFGAP1;refGene_tx=XM_023250999.1;refGene_gn=ARFGAP1;refGene_tx=XM_023251000.1;refGene_gn=ARFGAP1;refGene_tx=XM_023251001.1;refGene_gn=ARFGAP1;refGene_tx=XM_023251002.1;refGene_gn=ARFGAP1;refGene_tx=XM_023251003.1;refGene_gn=ARFGAP1;refGene_tx=XM_023251004.1;refGene_gn=ARFGAP1;refGene_tx=XM_023251005.1;refGene_gn=ARFGAP1;refGene_tx=XM_023251006.1;refGene_gn=ARFGAP1;refGene_tx=XM_023251007.1
A3 1142146 1142206
refGene_gn=ARFGAP1;refGene_tx=XM_023250999.1;refGene_gn=ARFGAP1;refGene_tx=XM_023251000.1;refGene_gn=ARFGAP1;refGene_tx=XM_023251001.1;refGene_gn=ARFGAP1;refGene_tx=XM_023251002.1;refGene_gn=ARFGAP1;refGene_tx=XM_023251003.1;refGene_gn=ARFGAP1;refGene_tx=XM_023251004.1;refGene_gn=ARFGAP1;refGene_tx=XM_023251005.1;refGene_gn=ARFGAP1;refGene_tx=XM_023251006.1;refGene_gn=ARFGAP1;refGene_tx=XM_023251007.1
A3 1172803 1172997
refGene_gn=BIRC7;refGene_tx=XM_019826258.2;refGene_gn=BIRC7;refGene_tx=XM_019826259.2
A3 1173163 1173640
refGene_gn=BIRC7;refGene_tx=XM_019826258.2;refGene_gn=BIRC7;refGene_tx=XM_019826259.2
A3 1174351 1174397
refGene_gn=BIRC7;refGene_tx=XM_019826258.2;refGene_gn=BIRC7;refGene_tx=XM_019826259.2
A3 1174504 1174586
refGene_gn=BIRC7;refGene_tx=XM_019826258.2;refGene_gn=BIRC7;refGene_tx=XM_019826259.2
A3 1174839 1174939
refGene_gn=BIRC7;refGene_tx=XM_019826258.2;refGene_gn=BIRC7;refGene_tx=XM_019826259.2
A3 1175694 1176064
refGene_gn=BIRC7;refGene_tx=XM_019826258.2;refGene_gn=BIRC7;refGene_tx=XM_019826259.2
A3 1188856 1188883 refGene_gn=YTHDF1;refGene_tx=XM_023251009.1
A3 1189054 1189079 refGene_gn=YTHDF1;refGene_tx=XM_023251009.1
A3 1190073 1190153 refGene_gn=YTHDF1;refGene_tx=XM_023251009.1
A3 1197822 1199343 refGene_gn=YTHDF1;refGene_tx=XM_023251009.1
A3 1201171 1201198 refGene_gn=YTHDF1;refGene_tx=XM_023251009.1
A3 1214929 1215806
refGene_gn=LOC111559738;refGene_tx=XM_023251010.1
A3 1216140 1216841
refGene_gn=LOC111559738;refGene_tx=XM_023251010.1
A3 1335991 1336019 refGene_gn=BHLHE23;refGene_tx=XM_023250923.1
A3 1337796 1337966 refGene_gn=BHLHE23;refGene_tx=XM_023250923.1
A3 1338303 1338476 refGene_gn=BHLHE23;refGene_tx=XM_023250923.1

A3 1340886 1341038 refGene_gn=BHLHE23;refGene_tx=XM_023250923.1
A3 1341952 1342128 refGene_gn=BHLHE23;refGene_tx=XM_023250923.1
A3 1342159 1343200 refGene_gn=BHLHE23;refGene_tx=XM_023250923.1
A3 1370877 1371041 refGene_gn=SLC17A9;refGene_tx=XM_023251011.1
A3 1371540 1371570 refGene_gn=SLC17A9;refGene_tx=XM_023251011.1
A3 1371727 1371783 refGene_gn=SLC17A9;refGene_tx=XM_023251011.1
A3 1372010 1372126 refGene_gn=SLC17A9;refGene_tx=XM_023251011.1
A3 1372456 1372491 refGene_gn=SLC17A9;refGene_tx=XM_023251011.1
A3 1373426 1373514 refGene_gn=SLC17A9;refGene_tx=XM_023251011.1
A3 1374135 1374232 refGene_gn=SLC17A9;refGene_tx=XM_023251011.1
A3 1374422 1374519 refGene_gn=SLC17A9;refGene_tx=XM_023251011.1
A3 1375089 1375220 refGene_gn=SLC17A9;refGene_tx=XM_023251011.1
A3 1376251 1376351 refGene_gn=SLC17A9;refGene_tx=XM_023251011.1
A3 1378259 1378399 refGene_gn=SLC17A9;refGene_tx=XM_023251011.1
A3 1378786 1378984 refGene_gn=SLC17A9;refGene_tx=XM_023251011.1
A3 1383897 1383959 refGene_gn=SLC17A9;refGene_tx=XM_023251011.1
A3 1390012 1390186
refGene_gn=GID8;refGene_tx=XM_003983298.5;refGene_gn=GID8;refGene_tx=XM_023251025.1
A3 1390959 1391157
refGene_gn=GID8;refGene_tx=XM_003983298.5;refGene_gn=GID8;refGene_tx=XM_023251025.1
A3 1391463 1391660
refGene_gn=GID8;refGene_tx=XM_003983298.5;refGene_gn=GID8;refGene_tx=XM_023251025.1
A3 1392689 1392807
refGene_gn=GID8;refGene_tx=XM_003983298.5;refGene_gn=GID8;refGene_tx=XM_023251025.1
A3 1415324 1416154
refGene_gn=DIDO1;refGene_tx=XM_023251012.1;refGene_gn=DIDO1;refGene_tx=XM_023251013.1;refGene_gn=DIDO1;refGene_tx=XM_023251014.1;refGene_gn=DIDO1;refGene_tx=XM_023251015.1;refGene_gn=DIDO1;refGene_tx=XM_023251016.1;refGene_gn=DIDO1;refGene_tx=XM_023251017.1
A3 1416571 1416893
refGene_gn=DIDO1;refGene_tx=XM_023251012.1;refGene_gn=DIDO1;refGene_tx=XM_023251013.1;refGene_gn=DIDO1;refGene_tx=XM_023251014.1;refGene_gn=DIDO1;refGene_tx=XM_023251015.1;refGene_gn=DIDO1;refGene_tx=XM_023251016.1;refGene_gn=DIDO1;refGene_tx=XM_023251017.1
A3 1419514 1419727
refGene_gn=DIDO1;refGene_tx=XM_023251012.1;refGene_gn=DIDO1;refGene_tx=XM_023251013.1;refGene_gn=DIDO1;refGene_tx=XM_023251014.1;refGene_gn=DIDO1;refGene_tx=XM_023251015.1;refGene_gn=DIDO1;refGene_tx=XM_023251016.1;refGene_gn=DIDO1;refGene_tx=XM_023251017.1
A3 1420551 1420762
refGene_gn=DIDO1;refGene_tx=XM_023251012.1;refGene_gn=DIDO1;refGene_tx=XM_023251013.1;refGene_gn=DIDO1;refGene_tx=XM_023251014.1;refGene_gn=DIDO1;refGene_tx=XM_023251015.1;refGene_gn=DIDO1;refGene_tx=XM_023251016.1;refGene_gn=DIDO1;refGene_tx=XM_023251017.1
A3 1429430 1429905
refGene_gn=DIDO1;refGene_tx=XM_023251012.1;refGene_gn=DIDO1;refGene_tx

=XM_023251013.1;refGene_gn=DIDO1;refGene_tx=XM_023251014.1;refGene_gn=DIDO1;refGene_tx=XM_023251015.1;refGene_gn=DIDO1;refGene_tx=XM_023251016.1;refGene_gn=DIDO1;refGene_tx=XM_023251017.1
A3 1430026 1430186
refGene_gn=DIDO1;refGene_tx=XM_023251012.1;refGene_gn=DIDO1;refGene_tx=XM_023251013.1;refGene_gn=DIDO1;refGene_tx=XM_023251014.1;refGene_gn=DIDO1;refGene_tx=XM_023251015.1;refGene_gn=DIDO1;refGene_tx=XM_023251016.1;refGene_gn=DIDO1;refGene_tx=XM_023251017.1
A3 1431429 1431546
refGene_gn=DIDO1;refGene_tx=XM_023251012.1;refGene_gn=DIDO1;refGene_tx=XM_023251013.1;refGene_gn=DIDO1;refGene_tx=XM_023251014.1;refGene_gn=DIDO1;refGene_tx=XM_023251015.1;refGene_gn=DIDO1;refGene_tx=XM_023251016.1;refGene_gn=DIDO1;refGene_tx=XM_023251017.1
A3 1431669 1431774
refGene_gn=DIDO1;refGene_tx=XM_023251012.1;refGene_gn=DIDO1;refGene_tx=XM_023251013.1;refGene_gn=DIDO1;refGene_tx=XM_023251014.1;refGene_gn=DIDO1;refGene_tx=XM_023251015.1;refGene_gn=DIDO1;refGene_tx=XM_023251016.1;refGene_gn=DIDO1;refGene_tx=XM_023251017.1
A3 1431951 1432087
refGene_gn=DIDO1;refGene_tx=XM_023251012.1;refGene_gn=DIDO1;refGene_tx=XM_023251013.1;refGene_gn=DIDO1;refGene_tx=XM_023251014.1;refGene_gn=DIDO1;refGene_tx=XM_023251015.1;refGene_gn=DIDO1;refGene_tx=XM_023251016.1;refGene_gn=DIDO1;refGene_tx=XM_023251017.1
A3 1432341 1432870
refGene_gn=DIDO1;refGene_tx=XM_023251012.1;refGene_gn=DIDO1;refGene_tx=XM_023251013.1;refGene_gn=DIDO1;refGene_tx=XM_023251014.1;refGene_gn=DIDO1;refGene_tx=XM_023251015.1;refGene_gn=DIDO1;refGene_tx=XM_023251016.1;refGene_gn=DIDO1;refGene_tx=XM_023251017.1
A3 1433521 1433675
refGene_gn=DIDO1;refGene_tx=XM_023251012.1;refGene_gn=DIDO1;refGene_tx=XM_023251013.1;refGene_gn=DIDO1;refGene_tx=XM_023251014.1;refGene_gn=DIDO1;refGene_tx=XM_023251015.1;refGene_gn=DIDO1;refGene_tx=XM_023251016.1;refGene_gn=DIDO1;refGene_tx=XM_023251017.1
A3 1434382 1434472
refGene_gn=DIDO1;refGene_tx=XM_023251012.1;refGene_gn=DIDO1;refGene_tx=XM_023251013.1;refGene_gn=DIDO1;refGene_tx=XM_023251014.1;refGene_gn=DIDO1;refGene_tx=XM_023251015.1;refGene_gn=DIDO1;refGene_tx=XM_023251016.1;refGene_gn=DIDO1;refGene_tx=XM_023251017.1
A3 1435286 1435482
refGene_gn=DIDO1;refGene_tx=XM_023251012.1;refGene_gn=DIDO1;refGene_tx=XM_023251013.1;refGene_gn=DIDO1;refGene_tx=XM_023251014.1;refGene_gn=DIDO1;refGene_tx=XM_023251015.1;refGene_gn=DIDO1;refGene_tx=XM_023251016.1;refGene_gn=DIDO1;refGene_tx=XM_023251017.1
A3 1442769 1445825
refGene_gn=DIDO1;refGene_tx=XM_023251012.1;refGene_gn=DIDO1;refGene_tx=XM_023251013.1;refGene_gn=DIDO1;refGene_tx=XM_023251014.1;refGene_gn=DIDO1;refGene_tx=XM_023251015.1;refGene_gn=DIDO1;refGene_tx=XM_023251016.1;refGene_gn=DIDO1;refGene_tx=XM_023251017.1
A3 1456929 1457597
refGene_gn=TCFL5;refGene_tx=XM_023251018.1;refGene_gn=TCFL5;refGene_tx

=XM_023251019.1;refGene_gn=TCFL5;refGene_tx=XM_023251020.1;refGene_gn=TCFL5;refGene_tx=XM_023251021.1;refGene_gn=TCFL5;refGene_tx=XM_023251022.1;refGene_gn=TCFL5;refGene_tx=XM_023251023.1;refGene_gn=TCFL5;refGene_tx=XM_023251024.1
A3 1458660 1458841
refGene_gn=TCFL5;refGene_tx=XM_023251018.1;refGene_gn=TCFL5;refGene_tx=XM_023251019.1;refGene_gn=TCFL5;refGene_tx=XM_023251020.1;refGene_gn=TCFL5;refGene_tx=XM_023251021.1;refGene_gn=TCFL5;refGene_tx=XM_023251022.1;refGene_gn=TCFL5;refGene_tx=XM_023251023.1;refGene_gn=TCFL5;refGene_tx=XM_023251024.1
A3 1459111 1459274
refGene_gn=TCFL5;refGene_tx=XM_023251018.1;refGene_gn=TCFL5;refGene_tx=XM_023251019.1;refGene_gn=TCFL5;refGene_tx=XM_023251020.1;refGene_gn=TCFL5;refGene_tx=XM_023251021.1;refGene_gn=TCFL5;refGene_tx=XM_023251022.1;refGene_gn=TCFL5;refGene_tx=XM_023251023.1;refGene_gn=TCFL5;refGene_tx=XM_023251024.1
A3 1460666 1460913
refGene_gn=TCFL5;refGene_tx=XM_023251018.1;refGene_gn=TCFL5;refGene_tx=XM_023251019.1;refGene_gn=TCFL5;refGene_tx=XM_023251020.1;refGene_gn=TCFL5;refGene_tx=XM_023251021.1;refGene_gn=TCFL5;refGene_tx=XM_023251022.1;refGene_gn=TCFL5;refGene_tx=XM_023251023.1;refGene_gn=TCFL5;refGene_tx=XM_023251024.1
A3 1463013 1463155
refGene_gn=TCFL5;refGene_tx=XM_023251018.1;refGene_gn=TCFL5;refGene_tx=XM_023251019.1;refGene_gn=TCFL5;refGene_tx=XM_023251020.1;refGene_gn=TCFL5;refGene_tx=XM_023251021.1;refGene_gn=TCFL5;refGene_tx=XM_023251022.1;refGene_gn=TCFL5;refGene_tx=XM_023251023.1;refGene_gn=TCFL5;refGene_tx=XM_023251024.1
A3 1469804 1469816 refGene_gn=TCFL5;refGene_tx=XM_023251024.1
A3 1472167 1472290
refGene_gn=TCFL5;refGene_tx=XM_023251018.1;refGene_gn=TCFL5;refGene_tx=XM_023251019.1;refGene_gn=TCFL5;refGene_tx=XM_023251020.1;refGene_gn=TCFL5;refGene_tx=XM_023251021.1;refGene_gn=TCFL5;refGene_tx=XM_023251022.1;refGene_gn=TCFL5;refGene_tx=XM_023251023.1
A3 1473511 1473702 refGene_gn=COL9A3;refGene_tx=XM_023250924.1
A3 1474406 1474484 refGene_gn=COL9A3;refGene_tx=XM_023250924.1
A3 1475422 1475605 refGene_gn=COL9A3;refGene_tx=XM_023250924.1
A3 1476046 1476101 refGene_gn=COL9A3;refGene_tx=XM_023250924.1
A3 1476233 1476380 refGene_gn=COL9A3;refGene_tx=XM_023250924.1
A3 1476523 1476556 refGene_gn=COL9A3;refGene_tx=XM_023250924.1
A3 1477641 1477686 refGene_gn=COL9A3;refGene_tx=XM_023250924.1
A3 1478136 1478172 refGene_gn=COL9A3;refGene_tx=XM_023250924.1
A3 1479132 1479204 refGene_gn=COL9A3;refGene_tx=XM_023250924.1
A3 1479314 1479368 refGene_gn=COL9A3;refGene_tx=XM_023250924.1
A3 1479826 1479880 refGene_gn=COL9A3;refGene_tx=XM_023250924.1
A3 1479962 1480016 refGene_gn=COL9A3;refGene_tx=XM_023250924.1
A3 1480116 1480143 refGene_gn=COL9A3;refGene_tx=XM_023250924.1
A3 1480144 1480162 refGene_gn=COL9A3;refGene_tx=XM_023250924.1
A3 1480325 1480379 refGene_gn=COL9A3;refGene_tx=XM_023250924.1
A3 1480464 1480518 refGene_gn=COL9A3;refGene_tx=XM_023250924.1

A3	1481250	1481304	refGene_gn=COL9A3;refGene_tx=XM_023250924.1
A3	1481931	1481985	refGene_gn=COL9A3;refGene_tx=XM_023250924.1
A3	1482347	1482401	refGene_gn=COL9A3;refGene_tx=XM_023250924.1
A3	1482636	1482690	refGene_gn=COL9A3;refGene_tx=XM_023250924.1
A3	1482976	1483030	refGene_gn=COL9A3;refGene_tx=XM_023250924.1
A3	1483652	1483706	refGene_gn=COL9A3;refGene_tx=XM_023250924.1
A3	1484093	1484150	refGene_gn=COL9A3;refGene_tx=XM_023250924.1
A3	1485288	1485330	refGene_gn=COL9A3;refGene_tx=XM_023250924.1
A3	1485661	1485715	refGene_gn=COL9A3;refGene_tx=XM_023250924.1
A3	1486320	1486344	refGene_gn=COL9A3;refGene_tx=XM_023250924.1
A3	1486652	1486688	refGene_gn=COL9A3;refGene_tx=XM_023250924.1
A3	1486797	1486881	refGene_gn=COL9A3;refGene_tx=XM_023250924.1
A3	1487415	1487422
refGene_gn=LOC111559739;refGene_tx=XM_023251026.1
A3	1488009	1488081
refGene_gn=LOC111559739;refGene_tx=XM_023251026.1
A3	1488681	1488716
refGene_gn=LOC111559739;refGene_tx=XM_023251026.1
A3	1489818	1489887
refGene_gn=LOC111559739;refGene_tx=XM_023251026.1
A3	1490442	1490514
refGene_gn=LOC111559739;refGene_tx=XM_023251026.1
A3	1495101	1496032	refGene_gn=OGFR;refGene_tx=XM_023251027.1
A3	1496758	1496907	refGene_gn=OGFR;refGene_tx=XM_023251027.1
A3	1497455	1497522	refGene_gn=OGFR;refGene_tx=XM_023251027.1
A3	1498532	1498611	refGene_gn=OGFR;refGene_tx=XM_023251027.1
A3	1499433	1499512	refGene_gn=OGFR;refGene_tx=XM_023251027.1
A3	1500137	1500206	refGene_gn=OGFR;refGene_tx=XM_023251027.1
A3	1502657	1502831	refGene_gn=OGFR;refGene_tx=XM_023251027.1
A3	1507364	1507546	refGene_gn=MRGBP;refGene_tx=XM_023251028.1
A3	1507978	1508053	refGene_gn=MRGBP;refGene_tx=XM_023251028.1
A3	1508366	1508448	refGene_gn=MRGBP;refGene_tx=XM_023251028.1
A3	1510424	1510546	refGene_gn=MRGBP;refGene_tx=XM_023251028.1
A3	1511157	1511305	refGene_gn=MRGBP;refGene_tx=XM_023251028.1
A3	1533366	1533616	refGene_gn=NTSR1;refGene_tx=XM_019828012.2
A3	1534490	1534581	refGene_gn=NTSR1;refGene_tx=XM_019828012.2
A3	1538465	1538667	refGene_gn=NTSR1;refGene_tx=XM_019828012.2
A3	1577841	1578555	refGene_gn=NTSR1;refGene_tx=XM_019828012.2
A3	1597189	1597354
refGene_gn=SLCO4A1;refGene_tx=XM_023251029.1;refGene_gn=SLCO4A1;refGen
e_tx=XM_023251030.1
A3	1599121	1599270
refGene_gn=SLCO4A1;refGene_tx=XM_023251029.1;refGene_gn=SLCO4A1;refGen
e_tx=XM_023251030.1
A3	1599670	1599735
refGene_gn=SLCO4A1;refGene_tx=XM_023251029.1;refGene_gn=SLCO4A1;refGen
e_tx=XM_023251030.1
A3	1600001	1600174
refGene_gn=SLCO4A1;refGene_tx=XM_023251029.1;refGene_gn=SLCO4A1;refGen
e_tx=XM_023251030.1

A3	1600264	1600430
refGene_gn=SLCO4A1;refGene_tx=XM_023251029.1;refGene_gn=SLCO4A1;refGene_tx=XM_023251030.1
A3	1601370	1601566
refGene_gn=SLCO4A1;refGene_tx=XM_023251029.1;refGene_gn=SLCO4A1;refGene_tx=XM_023251030.1
A3	1602798	1602953
refGene_gn=SLCO4A1;refGene_tx=XM_023251029.1;refGene_gn=SLCO4A1;refGene_tx=XM_023251030.1
A3	1606060	1606172
refGene_gn=SLCO4A1;refGene_tx=XM_023251029.1;refGene_gn=SLCO4A1;refGene_tx=XM_023251030.1
A3	1606371	1606493
refGene_gn=SLCO4A1;refGene_tx=XM_023251029.1;refGene_gn=SLCO4A1;refGene_tx=XM_023251030.1
A3	1607818	1607909
refGene_gn=SLCO4A1;refGene_tx=XM_023251029.1;refGene_gn=SLCO4A1;refGene_tx=XM_023251030.1
A3	1608956	1609819
refGene_gn=SLCO4A1;refGene_tx=XM_023251029.1;refGene_gn=SLCO4A1;refGene_tx=XM_023251030.1
A3	1620371	1620445	refGene_gn=SLCO4A1;refGene_tx=XM_023251029.1
A3	1745945	1746489	refGene_gn=GATA5;refGene_tx=XM_023251031.1
A3	1747693	1747869	refGene_gn=GATA5;refGene_tx=XM_023251031.1
A3	1752351	1752477	refGene_gn=GATA5;refGene_tx=XM_023251031.1
A3	1753268	1753356	refGene_gn=GATA5;refGene_tx=XM_023251031.1
A3	1753671	1753799	refGene_gn=GATA5;refGene_tx=XM_023251031.1
A3	1754151	1754307	refGene_gn=GATA5;refGene_tx=XM_023251031.1
A3	1785705	1785766	refGene_gn=RBBP8NL;refGene_tx=XM_019826267.2
A3	1786571	1786614	refGene_gn=RBBP8NL;refGene_tx=XM_019826267.2
A3	1787158	1787254	refGene_gn=RBBP8NL;refGene_tx=XM_019826267.2
A3	1787539	1787652	refGene_gn=RBBP8NL;refGene_tx=XM_019826267.2
A3	1788259	1788332	refGene_gn=RBBP8NL;refGene_tx=XM_019826267.2
A3	1788726	1788857	refGene_gn=RBBP8NL;refGene_tx=XM_019826267.2
A3	1789003	1789086	refGene_gn=RBBP8NL;refGene_tx=XM_019826267.2
A3	1789303	1789470	refGene_gn=RBBP8NL;refGene_tx=XM_019826267.2
A3	1789855	1790540	refGene_gn=RBBP8NL;refGene_tx=XM_019826267.2
A3	1790790	1790917	refGene_gn=RBBP8NL;refGene_tx=XM_019826267.2
A3	1791391	1791462	refGene_gn=RBBP8NL;refGene_tx=XM_019826267.2
A3	1791572	1791702	refGene_gn=RBBP8NL;refGene_tx=XM_019826267.2
A3	1793335	1793466	refGene_gn=RBBP8NL;refGene_tx=XM_019826267.2
A3	1796605	1797087	refGene_gn=CABLES2;refGene_tx=XM_023251032.1
A3	1804461	1804533	refGene_gn=CABLES2;refGene_tx=XM_023251032.1
A3	1804638	1804731	refGene_gn=CABLES2;refGene_tx=XM_023251032.1
A3	1805915	1805993	refGene_gn=CABLES2;refGene_tx=XM_023251032.1
A3	1806592	1806701	refGene_gn=CABLES2;refGene_tx=XM_023251032.1
A3	1807240	1807406	refGene_gn=CABLES2;refGene_tx=XM_023251032.1
A3	1807798	1807902	refGene_gn=CABLES2;refGene_tx=XM_023251032.1
A3	1808262	1808369	refGene_gn=CABLES2;refGene_tx=XM_023251032.1
A3	1809169	1809374	refGene_gn=CABLES2;refGene_tx=XM_023251032.1

| | | | |
|---|---|---|---|
| A3 | 1809575 | 1809716 | refGene_gn=CABLES2;refGene_tx=XM_023251032.1 |
| A3 | 1811918 | 1811928 | refGene_gn=RPS21;refGene_tx=XM_023251033.1 |
| A3 | 1812014 | 1812070 | refGene_gn=RPS21;refGene_tx=XM_023251033.1 |
| A3 | 1812367 | 1812439 | refGene_gn=RPS21;refGene_tx=XM_023251033.1 |
| A3 | 1812575 | 1812639 | refGene_gn=RPS21;refGene_tx=XM_023251033.1 |
| A3 | 1812731 | 1812781 | refGene_gn=RPS21;refGene_tx=XM_023251033.1 |
| A3 | 1830364 | 1830661 | refGene_gn=LAMA5;refGene_tx=XM_023251034.1 |
| A3 | 1834932 | 1835085 | refGene_gn=LAMA5;refGene_tx=XM_023251034.1 |
| A3 | 1844460 | 1844578 | refGene_gn=LAMA5;refGene_tx=XM_023251034.1 |
| A3 | 1845263 | 1845382 | refGene_gn=LAMA5;refGene_tx=XM_023251034.1 |
| A3 | 1845470 | 1845641 | refGene_gn=LAMA5;refGene_tx=XM_023251034.1 |
| A3 | 1845731 | 1845829 | refGene_gn=LAMA5;refGene_tx=XM_023251034.1 |
| A3 | 1851943 | 1852059 | refGene_gn=LAMA5;refGene_tx=XM_023251034.1 |
| A3 | 1852394 | 1852513 | refGene_gn=LAMA5;refGene_tx=XM_023251034.1 |
| A3 | 1852665 | 1852756 | refGene_gn=LAMA5;refGene_tx=XM_023251034.1 |
| A3 | 1852937 | 1853072 | refGene_gn=LAMA5;refGene_tx=XM_023251034.1 |
| A3 | 1853222 | 1853282 | refGene_gn=LAMA5;refGene_tx=XM_023251034.1 |
| A3 | 1857144 | 1857285 | refGene_gn=LAMA5;refGene_tx=XM_023251034.1 |
| A3 | 1857365 | 1857503 | refGene_gn=LAMA5;refGene_tx=XM_023251034.1 |
| A3 | 1857599 | 1857734 | refGene_gn=LAMA5;refGene_tx=XM_023251034.1 |
| A3 | 1857819 | 1857954 | refGene_gn=LAMA5;refGene_tx=XM_023251034.1 |
| A3 | 1858028 | 1858166 | refGene_gn=LAMA5;refGene_tx=XM_023251034.1 |
| A3 | 1858641 | 1858694 | refGene_gn=LAMA5;refGene_tx=XM_023251034.1 |
| A3 | 1858927 | 1859033 | refGene_gn=LAMA5;refGene_tx=XM_023251034.1 |
| A3 | 1859149 | 1859202 | refGene_gn=LAMA5;refGene_tx=XM_023251034.1 |
| A3 | 1859289 | 1859395 | refGene_gn=LAMA5;refGene_tx=XM_023251034.1 |
| A3 | 1859718 | 1859818 | refGene_gn=LAMA5;refGene_tx=XM_023251034.1 |
| A3 | 1859920 | 1860077 | refGene_gn=LAMA5;refGene_tx=XM_023251034.1 |
| A3 | 1860224 | 1860257 | refGene_gn=LAMA5;refGene_tx=XM_023251034.1 |
| A3 | 1860258 | 1860364 | refGene_gn=LAMA5;refGene_tx=XM_023251034.1 |
| A3 | 1860633 | 1860776 | refGene_gn=LAMA5;refGene_tx=XM_023251034.1 |
| A3 | 1860853 | 1860960 | refGene_gn=LAMA5;refGene_tx=XM_023251034.1 |
| A3 | 1861061 | 1861215 | refGene_gn=LAMA5;refGene_tx=XM_023251034.1 |
| A3 | 1861743 | 1861904 | refGene_gn=LAMA5;refGene_tx=XM_023251034.1 |
| A3 | 1861992 | 1862101 | refGene_gn=LAMA5;refGene_tx=XM_023251034.1 |
| A3 | 1863278 | 1863376 | refGene_gn=LAMA5;refGene_tx=XM_023251034.1 |
| A3 | 1863457 | 1863659 | refGene_gn=LAMA5;refGene_tx=XM_023251034.1 |
| A3 | 1863746 | 1863873 | refGene_gn=LAMA5;refGene_tx=XM_023251034.1 |
| A3 | 1864460 | 1864600 | refGene_gn=LAMA5;refGene_tx=XM_023251034.1 |
| A3 | 1865247 | 1865363 | refGene_gn=LAMA5;refGene_tx=XM_023251034.1 |
| A3 | 1865468 | 1865680 | refGene_gn=LAMA5;refGene_tx=XM_023251034.1 |
| A3 | 1866024 | 1866229 | refGene_gn=LAMA5;refGene_tx=XM_023251034.1 |
| A3 | 1866426 | 1866571 | refGene_gn=LAMA5;refGene_tx=XM_023251034.1 |
| A3 | 1866781 | 1866922 | refGene_gn=LAMA5;refGene_tx=XM_023251034.1 |
| A3 | 1867064 | 1867226 | refGene_gn=LAMA5;refGene_tx=XM_023251034.1 |
| A3 | 1867436 | 1867538 | refGene_gn=LAMA5;refGene_tx=XM_023251034.1 |
| A3 | 1867642 | 1867726 | refGene_gn=LAMA5;refGene_tx=XM_023251034.1 |
| A3 | 1868393 | 1868624 | refGene_gn=LAMA5;refGene_tx=XM_023251034.1 |
| A3 | 1869101 | 1869215 | refGene_gn=LAMA5;refGene_tx=XM_023251034.1 |
| A3 | 1869455 | 1869583 | refGene_gn=LAMA5;refGene_tx=XM_023251034.1 |

A3	1869785	1869866	refGene_gn=LAMA5;refGene_tx=XM_023251034.1
A3	1869990	1870205	refGene_gn=LAMA5;refGene_tx=XM_023251034.1
A3	1870387	1870488	refGene_gn=LAMA5;refGene_tx=XM_023251034.1
A3	1870603	1870784	refGene_gn=LAMA5;refGene_tx=XM_023251034.1
A3	1870870	1871020	refGene_gn=LAMA5;refGene_tx=XM_023251034.1
A3	1871559	1871711	refGene_gn=LAMA5;refGene_tx=XM_023251034.1
A3	1871779	1871890	refGene_gn=LAMA5;refGene_tx=XM_023251034.1
A3	1872073	1872185	refGene_gn=LAMA5;refGene_tx=XM_023251034.1
A3	1872816	1872987	refGene_gn=LAMA5;refGene_tx=XM_023251034.1
A3	1873166	1873363	refGene_gn=LAMA5;refGene_tx=XM_023251034.1
A3	1873476	1873593	refGene_gn=LAMA5;refGene_tx=XM_023251034.1
A3	1873731	1873886	refGene_gn=LAMA5;refGene_tx=XM_023251034.1
A3	1874182	1874324	refGene_gn=LAMA5;refGene_tx=XM_023251034.1
A3	1874428	1874531	refGene_gn=LAMA5;refGene_tx=XM_023251034.1
A3	1875106	1875217	refGene_gn=LAMA5;refGene_tx=XM_023251034.1
A3	1875940	1876120	refGene_gn=LAMA5;refGene_tx=XM_023251034.1
A3	1876210	1876359	refGene_gn=LAMA5;refGene_tx=XM_023251034.1
A3	1876439	1876610	refGene_gn=LAMA5;refGene_tx=XM_023251034.1
A3	1876747	1876884	refGene_gn=LAMA5;refGene_tx=XM_023251034.1
A3	1876958	1877112	refGene_gn=LAMA5;refGene_tx=XM_023251034.1
A3	1877248	1877382	refGene_gn=LAMA5;refGene_tx=XM_023251034.1
A3	1877457	1877620	refGene_gn=LAMA5;refGene_tx=XM_023251034.1
A3	1878414	1878549	refGene_gn=LAMA5;refGene_tx=XM_023251034.1
A3	1878749	1878898	refGene_gn=LAMA5;refGene_tx=XM_023251034.1
A3	1878967	1879100	refGene_gn=LAMA5;refGene_tx=XM_023251034.1
A3	1879186	1879330	refGene_gn=LAMA5;refGene_tx=XM_023251034.1
A3	1879433	1879564	refGene_gn=LAMA5;refGene_tx=XM_023251034.1
A3	1879666	1879837	refGene_gn=LAMA5;refGene_tx=XM_023251034.1
A3	1879926	1880065	refGene_gn=LAMA5;refGene_tx=XM_023251034.1
A3	1880176	1880322	refGene_gn=LAMA5;refGene_tx=XM_023251034.1
A3	1880390	1880583	refGene_gn=LAMA5;refGene_tx=XM_023251034.1
A3	1880657	1880822	refGene_gn=LAMA5;refGene_tx=XM_023251034.1
A3	1880947	1881104	refGene_gn=LAMA5;refGene_tx=XM_023251034.1
A3	1881214	1881348	refGene_gn=LAMA5;refGene_tx=XM_023251034.1
A3	1881444	1881538	refGene_gn=LAMA5;refGene_tx=XM_023251034.1
A3	1881697	1881820	refGene_gn=LAMA5;refGene_tx=XM_023251034.1
A3	1882001	1882141	refGene_gn=LAMA5;refGene_tx=XM_023251034.1
A3	1882817	1882921	refGene_gn=ADRM1;refGene_tx=XM_023251035.1
A3	1883105	1883208	refGene_gn=ADRM1;refGene_tx=XM_023251035.1
A3	1883365	1883523	refGene_gn=ADRM1;refGene_tx=XM_023251035.1
A3	1883728	1883961	refGene_gn=ADRM1;refGene_tx=XM_023251035.1
A3	1884220	1884302	refGene_gn=ADRM1;refGene_tx=XM_023251035.1
A3	1884746	1884833	refGene_gn=ADRM1;refGene_tx=XM_023251035.1
A3	1885151	1885275	refGene_gn=ADRM1;refGene_tx=XM_023251035.1
A3	1887229	1887346	refGene_gn=ADRM1;refGene_tx=XM_023251035.1
A3	1887943	1888156	refGene_gn=ADRM1;refGene_tx=XM_023251035.1
A3	1896334	1896437
refGene_gn=OSBPL2;refGene_tx=XM_011280784.2;refGene_gn=OSBPL2;refGene_
tx=XM_011280785.3;refGene_gn=OSBPL2;refGene_tx=XM_019826268.2;refGene_
gn=OSBPL2;refGene_tx=XM_019826269.2;refGene_gn=OSBPL2;refGene_tx=XM_01

9826271.1
A3	1898193	1898284
refGene_gn=OSBPL2;refGene_tx=XM_011280784.2;refGene_gn=OSBPL2;refGene_tx=XM_011280785.3;refGene_gn=OSBPL2;refGene_tx=XM_019826268.2;refGene_gn=OSBPL2;refGene_tx=XM_019826269.2;refGene_gn=OSBPL2;refGene_tx=XM_019826271.1
A3	1899193	1899317
refGene_gn=OSBPL2;refGene_tx=XM_011280784.2;refGene_gn=OSBPL2;refGene_tx=XM_011280785.3;refGene_gn=OSBPL2;refGene_tx=XM_019826268.2;refGene_gn=OSBPL2;refGene_tx=XM_019826269.2;refGene_gn=OSBPL2;refGene_tx=XM_019826271.1
A3	1899643	1899730	refGene_gn=OSBPL2;refGene_tx=XM_019826270.1
A3	1900326	1900455
refGene_gn=OSBPL2;refGene_tx=XM_011280784.2;refGene_gn=OSBPL2;refGene_tx=XM_011280785.3;refGene_gn=OSBPL2;refGene_tx=XM_019826268.2;refGene_gn=OSBPL2;refGene_tx=XM_019826269.2;refGene_gn=OSBPL2;refGene_tx=XM_019826270.1;refGene_gn=OSBPL2;refGene_tx=XM_019826271.1
A3	1901860	1901984
refGene_gn=OSBPL2;refGene_tx=XM_011280784.2;refGene_gn=OSBPL2;refGene_tx=XM_011280785.3;refGene_gn=OSBPL2;refGene_tx=XM_019826268.2;refGene_gn=OSBPL2;refGene_tx=XM_019826269.2;refGene_gn=OSBPL2;refGene_tx=XM_019826270.1;refGene_gn=OSBPL2;refGene_tx=XM_019826271.1
A3	1904402	1904492
refGene_gn=OSBPL2;refGene_tx=XM_011280784.2;refGene_gn=OSBPL2;refGene_tx=XM_011280785.3;refGene_gn=OSBPL2;refGene_tx=XM_019826268.2;refGene_gn=OSBPL2;refGene_tx=XM_019826269.2;refGene_gn=OSBPL2;refGene_tx=XM_019826270.1;refGene_gn=OSBPL2;refGene_tx=XM_019826271.1
A3	1905099	1905207
refGene_gn=OSBPL2;refGene_tx=XM_011280784.2;refGene_gn=OSBPL2;refGene_tx=XM_011280785.3;refGene_gn=OSBPL2;refGene_tx=XM_019826268.2;refGene_gn=OSBPL2;refGene_tx=XM_019826269.2;refGene_gn=OSBPL2;refGene_tx=XM_019826270.1;refGene_gn=OSBPL2;refGene_tx=XM_019826271.1
A3	1906816	1906999
refGene_gn=OSBPL2;refGene_tx=XM_011280784.2;refGene_gn=OSBPL2;refGene_tx=XM_011280785.3;refGene_gn=OSBPL2;refGene_tx=XM_019826268.2;refGene_gn=OSBPL2;refGene_tx=XM_019826269.2;refGene_gn=OSBPL2;refGene_tx=XM_019826270.1;refGene_gn=OSBPL2;refGene_tx=XM_019826271.1
A3	1909695	1909793
refGene_gn=OSBPL2;refGene_tx=XM_011280784.2;refGene_gn=OSBPL2;refGene_tx=XM_011280785.3;refGene_gn=OSBPL2;refGene_tx=XM_019826268.2;refGene_gn=OSBPL2;refGene_tx=XM_019826269.2;refGene_gn=OSBPL2;refGene_tx=XM_019826270.1;refGene_gn=OSBPL2;refGene_tx=XM_019826271.1
A3	1910247	1910382
refGene_gn=OSBPL2;refGene_tx=XM_011280784.2;refGene_gn=OSBPL2;refGene_tx=XM_011280785.3;refGene_gn=OSBPL2;refGene_tx=XM_019826268.2;refGene_gn=OSBPL2;refGene_tx=XM_019826269.2;refGene_gn=OSBPL2;refGene_tx=XM_019826270.1;refGene_gn=OSBPL2;refGene_tx=XM_019826271.1
A3	1913876	1913952
refGene_gn=OSBPL2;refGene_tx=XM_011280784.2;refGene_gn=OSBPL2;refGene_tx=XM_011280785.3;refGene_gn=OSBPL2;refGene_tx=XM_019826268.2;refGene_

gn=OSBPL2;refGene_tx=XM_019826269.2;refGene_gn=OSBPL2;refGene_tx=XM_01
9826270.1
A3 1915696 1915841
refGene_gn=OSBPL2;refGene_tx=XM_011280784.2;refGene_gn=OSBPL2;refGene_
tx=XM_011280785.3;refGene_gn=OSBPL2;refGene_tx=XM_019826268.2;refGene_
gn=OSBPL2;refGene_tx=XM_019826270.1
A3 1919130 1919167
refGene_gn=OSBPL2;refGene_tx=XM_011280784.2;refGene_gn=OSBPL2;refGene_
tx=XM_011280785.3;refGene_gn=OSBPL2;refGene_tx=XM_019826268.2;refGene_
gn=OSBPL2;refGene_tx=XM_019826270.1
A3 1948435 1948469 refGene_gn=HRH3;refGene_tx=XM_023251036.1
A3 1948471 1948684 refGene_gn=HRH3;refGene_tx=XM_023251036.1
A3 1949749 1949916 refGene_gn=HRH3;refGene_tx=XM_023251036.1
A3 1951059 1951980 refGene_gn=HRH3;refGene_tx=XM_023251036.1
A3 1964844 1965239
refGene_gn=MTG2;refGene_tx=XM_011280781.3;refGene_gn=MTG2;refGene_tx=X
M_011280782.3;refGene_gn=MTG2;refGene_tx=XM_011280783.3
A3 1965857 1965996
refGene_gn=MTG2;refGene_tx=XM_011280781.3;refGene_gn=MTG2;refGene_tx=X
M_011280782.3;refGene_gn=MTG2;refGene_tx=XM_011280783.3
A3 1966120 1966339
refGene_gn=MTG2;refGene_tx=XM_011280781.3;refGene_gn=MTG2;refGene_tx=X
M_011280782.3;refGene_gn=MTG2;refGene_tx=XM_011280783.3
A3 1967097 1967213
refGene_gn=MTG2;refGene_tx=XM_011280781.3;refGene_gn=MTG2;refGene_tx=X
M_011280782.3;refGene_gn=MTG2;refGene_tx=XM_011280783.3
A3 1968351 1968499
refGene_gn=MTG2;refGene_tx=XM_011280781.3;refGene_gn=MTG2;refGene_tx=X
M_011280782.3;refGene_gn=MTG2;refGene_tx=XM_011280783.3
A3 1970403 1970589
refGene_gn=MTG2;refGene_tx=XM_011280781.3;refGene_gn=MTG2;refGene_tx=X
M_011280782.3;refGene_gn=MTG2;refGene_tx=XM_011280783.3
A3 1982165 1982192
refGene_gn=SS18L1;refGene_tx=XM_011280780.3;refGene_gn=SS18L1;refGene_
tx=XM_019826273.2;refGene_gn=SS18L1;refGene_tx=XM_023251037.1;refGene_
gn=SS18L1;refGene_tx=XM_023251038.1;refGene_gn=SS18L1;refGene_tx=XM_02
3251039.1;refGene_gn=SS18L1;refGene_tx=XM_023251041.1
A3 1982740 1982868
refGene_gn=SS18L1;refGene_tx=XM_011280780.3;refGene_gn=SS18L1;refGene_
tx=XM_019826273.2;refGene_gn=SS18L1;refGene_tx=XM_023251037.1;refGene_
gn=SS18L1;refGene_tx=XM_023251038.1;refGene_gn=SS18L1;refGene_tx=XM_02
3251039.1;refGene_gn=SS18L1;refGene_tx=XM_023251041.1
A3 1986007 1986127
refGene_gn=SS18L1;refGene_tx=XM_011280780.3;refGene_gn=SS18L1;refGene_
tx=XM_019826273.2;refGene_gn=SS18L1;refGene_tx=XM_023251037.1;refGene_
gn=SS18L1;refGene_tx=XM_023251038.1;refGene_gn=SS18L1;refGene_tx=XM_02
3251039.1;refGene_gn=SS18L1;refGene_tx=XM_023251041.1
A3 1991360 1991527 refGene_gn=SS18L1;refGene_tx=XM_023251040.1
A3 1993774 1993864
refGene_gn=SS18L1;refGene_tx=XM_011280780.3;refGene_gn=SS18L1;refGene_

tx=XM_019826273.2;refGene_gn=SS18L1;refGene_tx=XM_023251037.1;refGene_gn=SS18L1;refGene_tx=XM_023251038.1;refGene_gn=SS18L1;refGene_tx=XM_023251039.1;refGene_gn=SS18L1;refGene_tx=XM_023251040.1;refGene_gn=SS18L1;refGene_tx=XM_023251041.1
A3 1994769 1994874
refGene_gn=SS18L1;refGene_tx=XM_011280780.3;refGene_gn=SS18L1;refGene_tx=XM_019826273.2;refGene_gn=SS18L1;refGene_tx=XM_023251037.1;refGene_gn=SS18L1;refGene_tx=XM_023251038.1;refGene_gn=SS18L1;refGene_tx=XM_023251039.1;refGene_gn=SS18L1;refGene_tx=XM_023251040.1;refGene_gn=SS18L1;refGene_tx=XM_023251041.1
A3 1995206 1995371
refGene_gn=SS18L1;refGene_tx=XM_011280780.3;refGene_gn=SS18L1;refGene_tx=XM_019826273.2;refGene_gn=SS18L1;refGene_tx=XM_023251037.1;refGene_gn=SS18L1;refGene_tx=XM_023251038.1;refGene_gn=SS18L1;refGene_tx=XM_023251039.1;refGene_gn=SS18L1;refGene_tx=XM_023251040.1;refGene_gn=SS18L1;refGene_tx=XM_023251041.1
A3 1996542 1996740
refGene_gn=SS18L1;refGene_tx=XM_011280780.3;refGene_gn=SS18L1;refGene_tx=XM_019826273.2;refGene_gn=SS18L1;refGene_tx=XM_023251037.1;refGene_gn=SS18L1;refGene_tx=XM_023251038.1;refGene_gn=SS18L1;refGene_tx=XM_023251039.1;refGene_gn=SS18L1;refGene_tx=XM_023251040.1;refGene_gn=SS18L1;refGene_tx=XM_023251041.1
A3 1997234 1997379
refGene_gn=SS18L1;refGene_tx=XM_011280780.3;refGene_gn=SS18L1;refGene_tx=XM_019826273.2;refGene_gn=SS18L1;refGene_tx=XM_023251037.1;refGene_gn=SS18L1;refGene_tx=XM_023251038.1;refGene_gn=SS18L1;refGene_tx=XM_023251039.1;refGene_gn=SS18L1;refGene_tx=XM_023251040.1;refGene_gn=SS18L1;refGene_tx=XM_023251041.1
A3 1998708 1998793
refGene_gn=SS18L1;refGene_tx=XM_011280780.3;refGene_gn=SS18L1;refGene_tx=XM_019826273.2;refGene_gn=SS18L1;refGene_tx=XM_023251037.1;refGene_gn=SS18L1;refGene_tx=XM_023251038.1;refGene_gn=SS18L1;refGene_tx=XM_023251039.1;refGene_gn=SS18L1;refGene_tx=XM_023251040.1;refGene_gn=SS18L1;refGene_tx=XM_023251041.1
A3 1998896 1998986
refGene_gn=SS18L1;refGene_tx=XM_023251037.1;refGene_gn=SS18L1;refGene_tx=XM_023251038.1;refGene_gn=SS18L1;refGene_tx=XM_023251040.1
A3 1999498 1999575
refGene_gn=SS18L1;refGene_tx=XM_019826273.2;refGene_gn=SS18L1;refGene_tx=XM_023251037.1;refGene_gn=SS18L1;refGene_tx=XM_023251038.1;refGene_gn=SS18L1;refGene_tx=XM_023251039.1;refGene_gn=SS18L1;refGene_tx=XM_023251040.1;refGene_gn=SS18L1;refGene_tx=XM_023251041.1
A3 1999890 1999893 refGene_gn=SS18L1;refGene_tx=XM_019826273.2
A3 2007306 2007375
refGene_gn=SS18L1;refGene_tx=XM_023251037.1;refGene_gn=SS18L1;refGene_tx=XM_023251039.1;refGene_gn=SS18L1;refGene_tx=XM_023251040.1
A3 2007893 2007989 refGene_gn=PSMA7;refGene_tx=XM_023251045.1
A3 2010145 2010272 refGene_gn=PSMA7;refGene_tx=XM_023251045.1
A3 2010824 2010949 refGene_gn=PSMA7;refGene_tx=XM_023251045.1
A3 2011645 2011768 refGene_gn=PSMA7;refGene_tx=XM_023251045.1

A3	2012225	2012345	refGene_gn=PSMA7;refGene_tx=XM_023251045.1
A3	2012901	2012964	refGene_gn=PSMA7;refGene_tx=XM_023251045.1
A3	2013363	2013456	refGene_gn=PSMA7;refGene_tx=XM_023251045.1
A3	2016985	2017157
refGene_gn=LSM14B;refGene_tx=XM_023251042.1;refGene_gn=LSM14B;refGene_tx=XM_023251043.1;refGene_gn=LSM14B;refGene_tx=XM_023251044.1
A3	2018926	2019077
refGene_gn=LSM14B;refGene_tx=XM_023251042.1;refGene_gn=LSM14B;refGene_tx=XM_023251043.1;refGene_gn=LSM14B;refGene_tx=XM_023251044.1
A3	2019393	2019555
refGene_gn=LSM14B;refGene_tx=XM_023251042.1;refGene_gn=LSM14B;refGene_tx=XM_023251043.1;refGene_gn=LSM14B;refGene_tx=XM_023251044.1
A3	2019778	2019856
refGene_gn=LSM14B;refGene_tx=XM_023251042.1;refGene_gn=LSM14B;refGene_tx=XM_023251044.1
A3	2020105	2020273
refGene_gn=LSM14B;refGene_tx=XM_023251042.1;refGene_gn=LSM14B;refGene_tx=XM_023251043.1;refGene_gn=LSM14B;refGene_tx=XM_023251044.1
A3	2022239	2022356
refGene_gn=LSM14B;refGene_tx=XM_023251042.1;refGene_gn=LSM14B;refGene_tx=XM_023251043.1
A3	2023117	2023253
refGene_gn=LSM14B;refGene_tx=XM_023251042.1;refGene_gn=LSM14B;refGene_tx=XM_023251043.1;refGene_gn=LSM14B;refGene_tx=XM_023251044.1
A3	2024349	2024513
refGene_gn=LSM14B;refGene_tx=XM_023251042.1;refGene_gn=LSM14B;refGene_tx=XM_023251043.1;refGene_gn=LSM14B;refGene_tx=XM_023251044.1
A3	2026234	2026361
refGene_gn=LSM14B;refGene_tx=XM_023251042.1;refGene_gn=LSM14B;refGene_tx=XM_023251043.1;refGene_gn=LSM14B;refGene_tx=XM_023251044.1
A3	2053309	2054660	refGene_gn=TAF4;refGene_tx=XM_023251046.1
A3	2089656	2089823	refGene_gn=TAF4;refGene_tx=XM_023251046.1
A3	2091021	2091141	refGene_gn=TAF4;refGene_tx=XM_023251046.1
A3	2094180	2094300	refGene_gn=TAF4;refGene_tx=XM_023251046.1
A3	2095064	2095187	refGene_gn=TAF4;refGene_tx=XM_023251046.1
A3	2096613	2096703	refGene_gn=TAF4;refGene_tx=XM_023251046.1
A3	2097273	2097519	refGene_gn=TAF4;refGene_tx=XM_023251046.1
A3	2099734	2099882	refGene_gn=TAF4;refGene_tx=XM_023251046.1
A3	2100293	2100408	refGene_gn=TAF4;refGene_tx=XM_023251046.1
A3	2102530	2102700	refGene_gn=TAF4;refGene_tx=XM_023251046.1
A3	2103023	2103154	refGene_gn=TAF4;refGene_tx=XM_023251046.1
A3	2103819	2103945	refGene_gn=TAF4;refGene_tx=XM_023251046.1
A3	2104380	2104437	refGene_gn=TAF4;refGene_tx=XM_023251046.1
A3	2104860	2104980	refGene_gn=TAF4;refGene_tx=XM_023251046.1
A3	2117555	2117723	refGene_gn=TAF4;refGene_tx=XM_023251046.1
A3	2133818	2134011
refGene_gn=CDH4;refGene_tx=XM_019826279.2;refGene_gn=CDH4;refGene_tx=XM_019826281.2;refGene_gn=CDH4;refGene_tx=XM_023251047.1;refGene_gn=CDH4;refGene_tx=XM_023251048.1
A3	2134784	2134966

refGene_gn=CDH4;refGene_tx=XM_019826279.2;refGene_gn=CDH4;refGene_tx=XM_019826281.2;refGene_gn=CDH4;refGene_tx=XM_023251047.1;refGene_gn=CDH4;refGene_tx=XM_023251048.1
A3 2136150 2136315
refGene_gn=CDH4;refGene_tx=XM_019826279.2;refGene_gn=CDH4;refGene_tx=XM_019826281.2;refGene_gn=CDH4;refGene_tx=XM_023251048.1
A3 2136916 2137056
refGene_gn=CDH4;refGene_tx=XM_019826279.2;refGene_gn=CDH4;refGene_tx=XM_019826281.2;refGene_gn=CDH4;refGene_tx=XM_023251047.1;refGene_gn=CDH4;refGene_tx=XM_023251048.1
A3 2137139 2137373
refGene_gn=CDH4;refGene_tx=XM_019826279.2;refGene_gn=CDH4;refGene_tx=XM_019826281.2;refGene_gn=CDH4;refGene_tx=XM_023251047.1;refGene_gn=CDH4;refGene_tx=XM_023251048.1
A3 2138467 2138701
refGene_gn=CDH4;refGene_tx=XM_019826279.2;refGene_gn=CDH4;refGene_tx=XM_019826281.2;refGene_gn=CDH4;refGene_tx=XM_023251047.1;refGene_gn=CDH4;refGene_tx=XM_023251048.1
A3 2141802 2141945
refGene_gn=CDH4;refGene_tx=XM_019826279.2;refGene_gn=CDH4;refGene_tx=XM_019826281.2;refGene_gn=CDH4;refGene_tx=XM_023251047.1;refGene_gn=CDH4;refGene_tx=XM_023251048.1
A3 2142412 2142666
refGene_gn=CDH4;refGene_tx=XM_019826279.2;refGene_gn=CDH4;refGene_tx=XM_019826281.2;refGene_gn=CDH4;refGene_tx=XM_023251047.1;refGene_gn=CDH4;refGene_tx=XM_023251048.1
A3 2148884 2149070
refGene_gn=CDH4;refGene_tx=XM_019826279.2;refGene_gn=CDH4;refGene_tx=XM_019826281.2;refGene_gn=CDH4;refGene_tx=XM_023251047.1;refGene_gn=CDH4;refGene_tx=XM_023251048.1
A3 2161447 2161585
refGene_gn=CDH4;refGene_tx=XM_019826279.2;refGene_gn=CDH4;refGene_tx=XM_019826281.2;refGene_gn=CDH4;refGene_tx=XM_023251047.1;refGene_gn=CDH4;refGene_tx=XM_023251048.1
A3 2180805 2180978
refGene_gn=CDH4;refGene_tx=XM_019826279.2;refGene_gn=CDH4;refGene_tx=XM_019826281.2;refGene_gn=CDH4;refGene_tx=XM_023251047.1;refGene_gn=CDH4;refGene_tx=XM_023251048.1
A3 2191233 2191378
refGene_gn=CDH4;refGene_tx=XM_019826279.2;refGene_gn=CDH4;refGene_tx=XM_019826281.2;refGene_gn=CDH4;refGene_tx=XM_023251047.1;refGene_gn=CDH4;refGene_tx=XM_023251048.1
A3 2198925 2199081
refGene_gn=CDH4;refGene_tx=XM_019826279.2;refGene_gn=CDH4;refGene_tx=XM_019826281.2;refGene_gn=CDH4;refGene_tx=XM_023251047.1;refGene_gn=CDH4;refGene_tx=XM_023251048.1
A3 2264070 2264244
refGene_gn=CDH4;refGene_tx=XM_019826279.2;refGene_gn=CDH4;refGene_tx=XM_019826281.2;refGene_gn=CDH4;refGene_tx=XM_023251047.1;refGene_gn=CDH4;refGene_tx=XM_023251048.1

A3	2285763	2285990
refGene_gn=CDH4;refGene_tx=XM_019826279.2;refGene_gn=CDH4;refGene_tx=XM_019826281.2;refGene_gn=CDH4;refGene_tx=XM_023251047.1;refGene_gn=CDH4;refGene_tx=XM_023251048.1
A3	2305881	2305936	refGene_gn=CDH4;refGene_tx=XM_019826281.2
A3	2670293	2670405
refGene_gn=CDH4;refGene_tx=XM_019826279.2;refGene_gn=CDH4;refGene_tx=XM_023251047.1;refGene_gn=CDH4;refGene_tx=XM_023251048.1
A3	2670827	2670869
refGene_gn=CDH4;refGene_tx=XM_019826279.2;refGene_gn=CDH4;refGene_tx=XM_023251047.1
A3	2672906	2672963	refGene_gn=CDH4;refGene_tx=XM_023251048.1
A3	3658978	3659125	refGene_gn=CDH26;refGene_tx=XM_006929597.3
A3	3661181	3661256	refGene_gn=CDH26;refGene_tx=XM_006929597.3
A3	3663589	3663643	refGene_gn=CDH26;refGene_tx=XM_006929597.3
A3	3665334	3665433	refGene_gn=CDH26;refGene_tx=XM_006929597.3
A3	3667016	3667094	refGene_gn=CDH26;refGene_tx=XM_006929597.3
A3	3668700	3668831	refGene_gn=CDH26;refGene_tx=XM_006929597.3
A3	3669372	3669594	refGene_gn=CDH26;refGene_tx=XM_006929597.3
A3	3670907	3671147	refGene_gn=CDH26;refGene_tx=XM_006929597.3
A3	3672976	3673119	refGene_gn=CDH26;refGene_tx=XM_006929597.3
A3	3676745	3677014	refGene_gn=CDH26;refGene_tx=XM_006929597.3
A3	3678253	3678439	refGene_gn=CDH26;refGene_tx=XM_006929597.3
A3	3680618	3680747	refGene_gn=CDH26;refGene_tx=XM_006929597.3
A3	3681181	3681348	refGene_gn=CDH26;refGene_tx=XM_006929597.3
A3	3683587	3683735	refGene_gn=CDH26;refGene_tx=XM_006929597.3
A3	3690067	3690229	refGene_gn=CDH26;refGene_tx=XM_006929597.3
A3	3691915	3692020	refGene_gn=CDH26;refGene_tx=XM_006929597.3
A3	3693539	3693596	refGene_gn=CDH26;refGene_tx=XM_006929597.3
A3	3700088	3700154	refGene_gn=CDH26;refGene_tx=XM_006929597.3
A3	3712107	3712550	refGene_gn=FAM217B;refGene_tx=XM_011280778.3
A3	3712552	3713222	refGene_gn=FAM217B;refGene_tx=XM_011280778.3
A3	3717603	3718497	refGene_gn=PPP1R3D;refGene_tx=XM_003983339.4
A3	3722132	3722403
refGene_gn=LOC111559732;refGene_tx=XM_023250925.1
A3	3722597	3723126
refGene_gn=LOC111559732;refGene_tx=XM_023250925.1
A3	3723264	3723370
refGene_gn=LOC111559732;refGene_tx=XM_023250925.1
A3	3734629	3734653	refGene_gn=SYCP2;refGene_tx=XM_023251049.1
A3	3736556	3736700	refGene_gn=SYCP2;refGene_tx=XM_023251049.1
A3	3737728	3737857	refGene_gn=SYCP2;refGene_tx=XM_023251049.1
A3	3738907	3739012	refGene_gn=SYCP2;refGene_tx=XM_023251049.1
A3	3739296	3739321	refGene_gn=SYCP2;refGene_tx=XM_023251049.1
A3	3742895	3742981	refGene_gn=SYCP2;refGene_tx=XM_023251049.1
A3	3743650	3743736	refGene_gn=SYCP2;refGene_tx=XM_023251049.1
A3	3744007	3744042	refGene_gn=SYCP2;refGene_tx=XM_023251049.1
A3	3744889	3745032	refGene_gn=SYCP2;refGene_tx=XM_023251049.1
A3	3745118	3745171	refGene_gn=SYCP2;refGene_tx=XM_023251049.1
A3	3746786	3746832	refGene_gn=SYCP2;refGene_tx=XM_023251049.1

A3	3747374	3747470	refGene_gn=SYCP2;refGene_tx=XM_023251049.1
A3	3751615	3751676	refGene_gn=SYCP2;refGene_tx=XM_023251049.1
A3	3764376	3764525	refGene_gn=SYCP2;refGene_tx=XM_023251049.1
A3	3765702	3765777	refGene_gn=SYCP2;refGene_tx=XM_023251049.1
A3	3766208	3766355	refGene_gn=SYCP2;refGene_tx=XM_023251049.1
A3	3769542	3769642	refGene_gn=SYCP2;refGene_tx=XM_023251049.1
A3	3770992	3771150	refGene_gn=SYCP2;refGene_tx=XM_023251049.1
A3	3772869	3772879	refGene_gn=SYCP2;refGene_tx=XM_023251049.1
A3	3772881	3772938	refGene_gn=SYCP2;refGene_tx=XM_023251049.1
A3	3773323	3773381	refGene_gn=SYCP2;refGene_tx=XM_023251049.1
A3	3773907	3774041	refGene_gn=SYCP2;refGene_tx=XM_023251049.1
A3	3774187	3774624	refGene_gn=SYCP2;refGene_tx=XM_023251049.1
A3	3782232	3782360	refGene_gn=SYCP2;refGene_tx=XM_023251049.1
A3	3783139	3783176	refGene_gn=SYCP2;refGene_tx=XM_023251049.1
A3	3785456	3785527	refGene_gn=SYCP2;refGene_tx=XM_023251049.1
A3	3785618	3785676	refGene_gn=SYCP2;refGene_tx=XM_023251049.1
A3	3786165	3786221	refGene_gn=SYCP2;refGene_tx=XM_023251049.1
A3	3786653	3786711	refGene_gn=SYCP2;refGene_tx=XM_023251049.1
A3	3787850	3788019	refGene_gn=SYCP2;refGene_tx=XM_023251049.1
A3	3789958	3789996	refGene_gn=SYCP2;refGene_tx=XM_023251049.1
A3	3790483	3790654	refGene_gn=SYCP2;refGene_tx=XM_023251049.1
A3	3792051	3792247	refGene_gn=SYCP2;refGene_tx=XM_023251049.1
A3	3793417	3793620	refGene_gn=SYCP2;refGene_tx=XM_023251049.1
A3	3797068	3797159	refGene_gn=SYCP2;refGene_tx=XM_023251049.1
A3	3797421	3797574	refGene_gn=SYCP2;refGene_tx=XM_023251049.1
A3	3798108	3798236	refGene_gn=SYCP2;refGene_tx=XM_023251049.1
A3	3799251	3799346	refGene_gn=SYCP2;refGene_tx=XM_023251049.1
A3	3799423	3799523	refGene_gn=SYCP2;refGene_tx=XM_023251049.1
A3	3799965	3800024	refGene_gn=SYCP2;refGene_tx=XM_023251049.1
A3	3800129	3800208	refGene_gn=SYCP2;refGene_tx=XM_023251049.1
A3	3800296	3800353	refGene_gn=SYCP2;refGene_tx=XM_023251049.1
A3	3801198	3801276	refGene_gn=SYCP2;refGene_tx=XM_023251049.1
A3	3813605	3813621	refGene_gn=PHACTR3;refGene_tx=XM_003983301.5
A3	3816301	3816378	refGene_gn=PHACTR3;refGene_tx=XM_003983301.5
A3	3818612	3818753	refGene_gn=PHACTR3;refGene_tx=XM_003983301.5
A3	3819750	3819812	refGene_gn=PHACTR3;refGene_tx=XM_003983301.5
A3	3823818	3823874	refGene_gn=PHACTR3;refGene_tx=XM_003983301.5
A3	3855950	3856104	refGene_gn=PHACTR3;refGene_tx=XM_003983301.5
A3	3889680	3889928	refGene_gn=PHACTR3;refGene_tx=XM_003983301.5
A3	3890650	3890825	refGene_gn=PHACTR3;refGene_tx=XM_003983301.5
A3	3895593	3895642	refGene_gn=PHACTR3;refGene_tx=XM_003983301.5
A3	3895643	3895807	refGene_gn=PHACTR3;refGene_tx=XM_003983301.5
A3	3904785	3904968	refGene_gn=PHACTR3;refGene_tx=XM_003983301.5
A3	3911555	3911633	refGene_gn=PHACTR3;refGene_tx=XM_003983301.5
A3	3915251	3915413	refGene_gn=PHACTR3;refGene_tx=XM_003983301.5
A3	4027648	4027766	refGene_gn=PHACTR3;refGene_tx=XM_003983301.5
A3	4222493	4222534	refGene_gn=EDN3;refGene_tx=XM_023251052.1;refGene_gn=EDN3;refGene_tx=XM_023251053.1;refGene_gn=EDN3;refGene_tx=XM_023251054.1;refGene_gn=EDN3;refGene_tx=XM_023251055.1

A3 4223634 4223822
refGene_gn=EDN3;refGene_tx=XM_023251050.1;refGene_gn=EDN3;refGene_tx=XM_023251051.1
A3 4224836 4224931
refGene_gn=EDN3;refGene_tx=XM_023251050.1;refGene_gn=EDN3;refGene_tx=XM_023251051.1;refGene_gn=EDN3;refGene_tx=XM_023251052.1;refGene_gn=EDN3;refGene_tx=XM_023251053.1;refGene_gn=EDN3;refGene_tx=XM_023251054.1;refGene_gn=EDN3;refGene_tx=XM_023251055.1
A3 4225868 4226042
refGene_gn=EDN3;refGene_tx=XM_023251050.1;refGene_gn=EDN3;refGene_tx=XM_023251051.1;refGene_gn=EDN3;refGene_tx=XM_023251052.1;refGene_gn=EDN3;refGene_tx=XM_023251053.1;refGene_gn=EDN3;refGene_tx=XM_023251054.1;refGene_gn=EDN3;refGene_tx=XM_023251055.1
A3 4241631 4241920
refGene_gn=EDN3;refGene_tx=XM_023251050.1;refGene_gn=EDN3;refGene_tx=XM_023251051.1;refGene_gn=EDN3;refGene_tx=XM_023251052.1;refGene_gn=EDN3;refGene_tx=XM_023251053.1;refGene_gn=EDN3;refGene_tx=XM_023251054.1;refGene_gn=EDN3;refGene_tx=XM_023251055.1
A3 4242547 4242599
refGene_gn=EDN3;refGene_tx=XM_023251050.1;refGene_gn=EDN3;refGene_tx=XM_023251051.1;refGene_gn=EDN3;refGene_tx=XM_023251052.1;refGene_gn=EDN3;refGene_tx=XM_023251053.1;refGene_gn=EDN3;refGene_tx=XM_023251054.1;refGene_gn=EDN3;refGene_tx=XM_023251055.1
A3 4286121 4286952
refGene_gn=ZNF831;refGene_tx=XM_019826290.2;refGene_gn=ZNF831;refGene_tx=XM_019826291.2;refGene_gn=ZNF831;refGene_tx=XM_023251056.1
A3 4287680 4287829
refGene_gn=ZNF831;refGene_tx=XM_019826290.2;refGene_gn=ZNF831;refGene_tx=XM_019826291.2;refGene_gn=ZNF831;refGene_tx=XM_023251056.1
A3 4329392 4329544
refGene_gn=ZNF831;refGene_tx=XM_019826290.2;refGene_gn=ZNF831;refGene_tx=XM_019826291.2;refGene_gn=ZNF831;refGene_tx=XM_023251056.1
A3 4345949 4346089
refGene_gn=ZNF831;refGene_tx=XM_019826290.2;refGene_gn=ZNF831;refGene_tx=XM_019826291.2;refGene_gn=ZNF831;refGene_tx=XM_023251056.1
A3 4347130 4350754
refGene_gn=ZNF831;refGene_tx=XM_019826290.2;refGene_gn=ZNF831;refGene_tx=XM_019826291.2;refGene_gn=ZNF831;refGene_tx=XM_023251056.1
A3 4487972 4488004 refGene_gn=PRELID3B;refGene_tx=XM_023251057.1
A3 4491837 4492006 refGene_gn=PRELID3B;refGene_tx=XM_023251057.1
A3 4493208 4493298 refGene_gn=PRELID3B;refGene_tx=XM_023251057.1
A3 4493732 4493803 refGene_gn=PRELID3B;refGene_tx=XM_023251057.1
A3 4493925 4494028 refGene_gn=PRELID3B;refGene_tx=XM_023251057.1
A3 4495665 4495785 refGene_gn=PRELID3B;refGene_tx=XM_023251057.1
A3 4498419 4498451
refGene_gn=ATP5F1E;refGene_tx=XM_023251058.1;refGene_gn=ATP5F1E;refGene_tx=XM_023251059.1;refGene_gn=ATP5F1E;refGene_tx=XM_023251060.1
A3 4500036 4500160
refGene_gn=ATP5F1E;refGene_tx=XM_023251058.1;refGene_gn=ATP5F1E;refGene_tx=XM_023251059.1;refGene_gn=ATP5F1E;refGene_tx=XM_023251060.1

A3 4505442 4506539
refGene_gn=TUBB1;refGene_tx=NM_001134989.1;refGene_gn=TUBB1;refGene_tx
=XM_023250905.1;refGene_gn=TUBB1;refGene_tx=XM_023250906.1
A3 4506657 4506768
refGene_gn=TUBB1;refGene_tx=NM_001134989.1;refGene_gn=TUBB1;refGene_tx
=XM_023250905.1;refGene_gn=TUBB1;refGene_tx=XM_023250906.1
A3 4507276 4507385 refGene_gn=TUBB1;refGene_tx=NM_001134989.1
A3 4511112 4511169 refGene_gn=TUBB1;refGene_tx=NM_001134989.1
A3 4524484 4524627 refGene_gn=CTSZ;refGene_tx=XM_011280775.3
A3 4525006 4525170 refGene_gn=CTSZ;refGene_tx=XM_011280775.3
A3 4529278 4529458 refGene_gn=CTSZ;refGene_tx=XM_011280775.3
A3 4531212 4531363 refGene_gn=CTSZ;refGene_tx=XM_011280775.3
A3 4531653 4531816 refGene_gn=CTSZ;refGene_tx=XM_011280775.3
A3 4532624 4532735 refGene_gn=CTSZ;refGene_tx=XM_011280775.3
A3 4533574 4533609
refGene_gn=NELFCD;refGene_tx=XM_023251061.1;refGene_gn=NELFCD;refGene_
tx=XM_023251062.1
A3 4533702 4533832
refGene_gn=NELFCD;refGene_tx=XM_023251061.1;refGene_gn=NELFCD;refGene_
tx=XM_023251062.1
A3 4534181 4534322
refGene_gn=NELFCD;refGene_tx=XM_023251061.1;refGene_gn=NELFCD;refGene_
tx=XM_023251062.1
A3 4534401 4534497
refGene_gn=NELFCD;refGene_tx=XM_023251061.1;refGene_gn=NELFCD;refGene_
tx=XM_023251062.1
A3 4535501 4535616
refGene_gn=NELFCD;refGene_tx=XM_023251061.1;refGene_gn=NELFCD;refGene_
tx=XM_023251062.1
A3 4536319 4536459
refGene_gn=NELFCD;refGene_tx=XM_023251061.1;refGene_gn=NELFCD;refGene_
tx=XM_023251062.1
A3 4536885 4537020
refGene_gn=NELFCD;refGene_tx=XM_023251061.1;refGene_gn=NELFCD;refGene_
tx=XM_023251062.1
A3 4537255 4537421
refGene_gn=NELFCD;refGene_tx=XM_023251061.1;refGene_gn=NELFCD;refGene_
tx=XM_023251062.1
A3 4538075 4538206
refGene_gn=NELFCD;refGene_tx=XM_023251061.1;refGene_gn=NELFCD;refGene_
tx=XM_023251062.1
A3 4538420 4538573
refGene_gn=NELFCD;refGene_tx=XM_023251061.1;refGene_gn=NELFCD;refGene_
tx=XM_023251062.1
A3 4539177 4539285
refGene_gn=NELFCD;refGene_tx=XM_023251061.1;refGene_gn=NELFCD;refGene_
tx=XM_023251062.1
A3 4540379 4540489
refGene_gn=NELFCD;refGene_tx=XM_023251061.1;refGene_gn=NELFCD;refGene_
tx=XM_023251062.1

A3	4541328	4541438
refGene_gn=NELFCD;refGene_tx=XM_023251061.1;refGene_gn=NELFCD;refGene_
tx=XM_023251062.1
A3	4541673	4541789
refGene_gn=NELFCD;refGene_tx=XM_023251061.1;refGene_gn=NELFCD;refGene_
tx=XM_023251062.1
A3	4545492	4545612
refGene_gn=NELFCD;refGene_tx=XM_023251061.1;refGene_gn=NELFCD;refGene_
tx=XM_023251062.1
A3	4611408	4611555
refGene_gn=LOC101098453;refGene_tx=XM_023251063.1
A3	4611781	4611849
refGene_gn=LOC101098453;refGene_tx=XM_023251063.1
A3	4612145	4612276
refGene_gn=LOC101098453;refGene_tx=XM_023251063.1
A3	4612384	4612505
refGene_gn=LOC101098453;refGene_tx=XM_023251063.1
A3	4612611	4612670
refGene_gn=LOC101098453;refGene_tx=XM_023251063.1
A3	4612785	4612859
refGene_gn=LOC101098453;refGene_tx=XM_023251063.1
A3	4612996	4613051
refGene_gn=LOC101098453;refGene_tx=XM_023251063.1
A3	4616184	4616282
refGene_gn=LOC101098453;refGene_tx=XM_023251063.1
A3	4618169	4618289
refGene_gn=LOC101098453;refGene_tx=XM_023251063.1
A3	4618379	4618434
refGene_gn=LOC101098453;refGene_tx=XM_023251063.1
A3	4622690	4622735
refGene_gn=LOC101098453;refGene_tx=XM_023251063.1
A3	4625891	4625964
refGene_gn=LOC101098453;refGene_tx=XM_023251063.1
A3	4663843	4664549
refGene_gn=LOC101098453;refGene_tx=XM_023251063.1
A3	4664550	4665708
refGene_gn=LOC101098453;refGene_tx=XM_023251063.1
A3	4678268	4679012
refGene_gn=LOC102900772;refGene_tx=XM_006929601.3
A3	4795230	4795389	refGene_gn=NPEPL1;refGene_tx=XM_023251064.1
A3	4795904	4796015	refGene_gn=NPEPL1;refGene_tx=XM_023251064.1
A3	4796550	4796727	refGene_gn=NPEPL1;refGene_tx=XM_023251064.1
A3	4797035	4797159	refGene_gn=NPEPL1;refGene_tx=XM_023251064.1
A3	4798109	4798210	refGene_gn=NPEPL1;refGene_tx=XM_023251064.1
A3	4801672	4801750	refGene_gn=NPEPL1;refGene_tx=XM_023251064.1
A3	4807671	4807814	refGene_gn=NPEPL1;refGene_tx=XM_023251064.1
A3	4810188	4810270	refGene_gn=NPEPL1;refGene_tx=XM_023251064.1
A3	4810726	4810816	refGene_gn=NPEPL1;refGene_tx=XM_023251064.1
A3	4814471	4814642	refGene_gn=NPEPL1;refGene_tx=XM_023251064.1
A3	4815088	4815274	refGene_gn=NPEPL1;refGene_tx=XM_023251064.1

A3 4815803 4815953 refGene_gn=NPEPL1;refGene_tx=XM_023251064.1
A3 4830294 4830399
refGene_gn=STX16;refGene_tx=XM_011280767.3;refGene_gn=STX16;refGene_tx=XM_011280769.3;refGene_gn=STX16;refGene_tx=XM_011280773.3;refGene_gn=STX16;refGene_tx=XM_019826299.2;refGene_gn=STX16;refGene_tx=XM_019826301.2;refGene_gn=STX16;refGene_tx=XM_023251065.1;refGene_gn=STX16;refGene_tx=XM_023251066.1;refGene_gn=STX16;refGene_tx=XM_023251067.1;refGene_gn=STX16;refGene_tx=XM_023251068.1
A3 4832469 4832533
refGene_gn=STX16;refGene_tx=XM_019826300.2;refGene_gn=STX16;refGene_tx=XM_019826302.2
A3 4832813 4832894
refGene_gn=STX16;refGene_tx=XM_011280767.3;refGene_gn=STX16;refGene_tx=XM_011280769.3;refGene_gn=STX16;refGene_tx=XM_011280773.3;refGene_gn=STX16;refGene_tx=XM_019826299.2;refGene_gn=STX16;refGene_tx=XM_019826300.2;refGene_gn=STX16;refGene_tx=XM_019826301.2;refGene_gn=STX16;refGene_tx=XM_019826302.2;refGene_gn=STX16;refGene_tx=XM_023251065.1;refGene_gn=STX16;refGene_tx=XM_023251066.1;refGene_gn=STX16;refGene_tx=XM_023251067.1;refGene_gn=STX16;refGene_tx=XM_023251068.1
A3 4833591 4833657 refGene_gn=STX16;refGene_tx=XM_019826303.2
A3 4834134 4834278
refGene_gn=STX16;refGene_tx=XM_011280767.3;refGene_gn=STX16;refGene_tx=XM_011280769.3;refGene_gn=STX16;refGene_tx=XM_011280773.3;refGene_gn=STX16;refGene_tx=XM_019826299.2;refGene_gn=STX16;refGene_tx=XM_019826300.2;refGene_gn=STX16;refGene_tx=XM_019826301.2;refGene_gn=STX16;refGene_tx=XM_019826302.2;refGene_gn=STX16;refGene_tx=XM_019826303.2;refGene_gn=STX16;refGene_tx=XM_023251065.1;refGene_gn=STX16;refGene_tx=XM_023251066.1;refGene_gn=STX16;refGene_tx=XM_023251067.1;refGene_gn=STX16;refGene_tx=XM_023251068.1
A3 4834834 4834926
refGene_gn=STX16;refGene_tx=XM_011280767.3;refGene_gn=STX16;refGene_tx=XM_011280769.3;refGene_gn=STX16;refGene_tx=XM_011280773.3;refGene_gn=STX16;refGene_tx=XM_019826299.2;refGene_gn=STX16;refGene_tx=XM_019826300.2;refGene_gn=STX16;refGene_tx=XM_019826301.2;refGene_gn=STX16;refGene_tx=XM_019826302.2;refGene_gn=STX16;refGene_tx=XM_019826303.2;refGene_gn=STX16;refGene_tx=XM_023251065.1;refGene_gn=STX16;refGene_tx=XM_023251066.1;refGene_gn=STX16;refGene_tx=XM_023251067.1;refGene_gn=STX16;refGene_tx=XM_023251068.1
A3 4836038 4836204
refGene_gn=STX16;refGene_tx=XM_011280767.3;refGene_gn=STX16;refGene_tx=XM_011280769.3;refGene_gn=STX16;refGene_tx=XM_011280773.3;refGene_gn=STX16;refGene_tx=XM_019826299.2;refGene_gn=STX16;refGene_tx=XM_019826300.2;refGene_gn=STX16;refGene_tx=XM_019826301.2;refGene_gn=STX16;refGene_tx=XM_019826302.2;refGene_gn=STX16;refGene_tx=XM_019826303.2;refGene_gn=STX16;refGene_tx=XM_023251065.1;refGene_gn=STX16;refGene_tx=XM_023251066.1;refGene_gn=STX16;refGene_tx=XM_023251067.1;refGene_gn=STX16;refGene_tx=XM_023251068.1
A3 4837415 4837556
refGene_gn=STX16;refGene_tx=XM_011280767.3;refGene_gn=STX16;refGene_tx=XM_011280769.3;refGene_gn=STX16;refGene_tx=XM_011280773.3;refGene_gn=

STX16;refGene_tx=XM_019826299.2;refGene_gn=STX16;refGene_tx=XM_0198263
00.2;refGene_gn=STX16;refGene_tx=XM_019826301.2;refGene_gn=STX16;refGe
ne_tx=XM_019826302.2;refGene_gn=STX16;refGene_tx=XM_019826303.2;refGen
e_gn=STX16;refGene_tx=XM_023251065.1;refGene_gn=STX16;refGene_tx=XM_02
3251066.1;refGene_gn=STX16;refGene_tx=XM_023251067.1;refGene_gn=STX16;
refGene_tx=XM_023251068.1
A3 4837966 4838074
refGene_gn=STX16;refGene_tx=XM_011280767.3;refGene_gn=STX16;refGene_tx
=XM_011280769.3;refGene_gn=STX16;refGene_tx=XM_011280773.3;refGene_gn=
STX16;refGene_tx=XM_019826299.2;refGene_gn=STX16;refGene_tx=XM_0198263
00.2;refGene_gn=STX16;refGene_tx=XM_019826301.2;refGene_gn=STX16;refGe
ne_tx=XM_019826302.2;refGene_gn=STX16;refGene_tx=XM_019826303.2;refGen
e_gn=STX16;refGene_tx=XM_023251065.1;refGene_gn=STX16;refGene_tx=XM_02
3251066.1;refGene_gn=STX16;refGene_tx=XM_023251067.1;refGene_gn=STX16;
refGene_tx=XM_023251068.1
A3 4838199 4838208
refGene_gn=STX16;refGene_tx=XM_011280773.3;refGene_gn=STX16;refGene_tx
=XM_023251067.1;refGene_gn=STX16;refGene_tx=XM_023251068.1
A3 4845400 4845412
refGene_gn=STX16;refGene_tx=XM_019826299.2;refGene_gn=STX16;refGene_tx
=XM_019826300.2;refGene_gn=STX16;refGene_tx=XM_019826301.2;refGene_gn=
STX16;refGene_tx=XM_019826302.2;refGene_gn=STX16;refGene_tx=XM_0198263
03.2;refGene_gn=STX16;refGene_tx=XM_023251065.1;refGene_gn=STX16;refGe
ne_tx=XM_023251066.1
A3 4845564 4845600
refGene_gn=STX16;refGene_tx=XM_019826299.2;refGene_gn=STX16;refGene_tx
=XM_019826300.2;refGene_gn=STX16;refGene_tx=XM_019826301.2;refGene_gn=
STX16;refGene_tx=XM_019826302.2;refGene_gn=STX16;refGene_tx=XM_0198263
03.2
A3 4853327 4853459
refGene_gn=STX16;refGene_tx=XM_011280767.3;refGene_gn=STX16;refGene_tx
=XM_011280769.3;refGene_gn=STX16;refGene_tx=XM_019826299.2;refGene_gn=
STX16;refGene_tx=XM_019826300.2;refGene_gn=STX16;refGene_tx=XM_0198263
01.2;refGene_gn=STX16;refGene_tx=XM_019826302.2;refGene_gn=STX16;refGe
ne_tx=XM_019826303.2;refGene_gn=STX16;refGene_tx=XM_023251065.1;refGen
e_gn=STX16;refGene_tx=XM_023251066.1
A3 5036148 5036306 refGene_gn=APCDD1L;refGene_tx=XM_023251069.1
A3 5038733 5039286 refGene_gn=APCDD1L;refGene_tx=XM_023251069.1
A3 5044501 5045248 refGene_gn=APCDD1L;refGene_tx=XM_023251069.1
A3 5063049 5063208 refGene_gn=VAPB;refGene_tx=XM_023251070.1
A3 5066203 5066380 refGene_gn=VAPB;refGene_tx=XM_023251070.1
A3 5068275 5068356 refGene_gn=VAPB;refGene_tx=XM_023251070.1
A3 5071707 5071811 refGene_gn=VAPB;refGene_tx=XM_023251070.1
A3 5085172 5085325 refGene_gn=VAPB;refGene_tx=XM_023251070.1
A3 5114260 5114318 refGene_gn=VAPB;refGene_tx=XM_023251070.1
A3 5139421 5139519
refGene_gn=RAB22A;refGene_tx=XM_011280763.3;refGene_gn=RAB22A;refGene_
tx=XM_011280764.3;refGene_gn=RAB22A;refGene_tx=XM_011280765.3
A3 5143923 5144033
refGene_gn=RAB22A;refGene_tx=XM_011280763.3;refGene_gn=RAB22A;refGene_

tx=XM_011280764.3;refGene_gn=RAB22A;refGene_tx=XM_011280765.3
A3	5144638	5144745
refGene_gn=RAB22A;refGene_tx=XM_011280763.3;refGene_gn=RAB22A;refGene_tx=XM_011280764.3;refGene_gn=RAB22A;refGene_tx=XM_011280765.3
A3	5144832	5144904
refGene_gn=RAB22A;refGene_tx=XM_011280763.3;refGene_gn=RAB22A;refGene_tx=XM_011280764.3
A3	5154994	5155076
refGene_gn=RAB22A;refGene_tx=XM_011280763.3;refGene_gn=RAB22A;refGene_tx=XM_011280764.3;refGene_gn=RAB22A;refGene_tx=XM_011280765.3
A3	5188722	5188802
refGene_gn=RAB22A;refGene_tx=XM_011280763.3;refGene_gn=RAB22A;refGene_tx=XM_011280764.3;refGene_gn=RAB22A;refGene_tx=XM_011280765.3
A3	5189787	5189823
refGene_gn=RAB22A;refGene_tx=XM_011280763.3;refGene_gn=RAB22A;refGene_tx=XM_011280764.3;refGene_gn=RAB22A;refGene_tx=XM_011280765.3
A3	5190264	5190271
refGene_gn=LOC101092674;refGene_tx=XM_023251071.1;refGene_gn=LOC101092674;refGene_tx=XM_023251073.1
A3	5190424	5190469
refGene_gn=LOC101092674;refGene_tx=XM_023251071.1;refGene_gn=LOC101092674;refGene_tx=XM_023251073.1
A3	5196809	5196889
refGene_gn=LOC101092674;refGene_tx=XM_019826311.2;refGene_gn=LOC101092674;refGene_tx=XM_023251072.1
A3	5218079	5218212
refGene_gn=LOC101092674;refGene_tx=XM_019826309.2;refGene_gn=LOC101092674;refGene_tx=XM_023251071.1;refGene_gn=LOC101092674;refGene_tx=XM_023251073.1
A3	5231000	5231056
refGene_gn=LOC101092674;refGene_tx=XM_019826312.2
A3	5231176	5231283
refGene_gn=LOC101092674;refGene_tx=XM_006929630.4;refGene_gn=LOC101092674;refGene_tx=XM_019826309.2;refGene_gn=LOC101092674;refGene_tx=XM_019826311.2;refGene_gn=LOC101092674;refGene_tx=XM_019826312.2;refGene_gn=LOC101092674;refGene_tx=XM_023251071.1;refGene_gn=LOC101092674;refGene_tx=XM_023251072.1;refGene_gn=LOC101092674;refGene_tx=XM_023251073.1
A3	5232581	5232724
refGene_gn=LOC101092674;refGene_tx=XM_006929630.4;refGene_gn=LOC101092674;refGene_tx=XM_019826309.2;refGene_gn=LOC101092674;refGene_tx=XM_019826311.2;refGene_gn=LOC101092674;refGene_tx=XM_019826312.2;refGene_gn=LOC101092674;refGene_tx=XM_023251071.1;refGene_gn=LOC101092674;refGene_tx=XM_023251072.1;refGene_gn=LOC101092674;refGene_tx=XM_023251073.1
A3	5241516	5241663
refGene_gn=LOC101092674;refGene_tx=XM_006929630.4;refGene_gn=LOC101092674;refGene_tx=XM_019826309.2;refGene_gn=LOC101092674;refGene_tx=XM_019826311.2;refGene_gn=LOC101092674;refGene_tx=XM_019826312.2;refGene_gn=LOC101092674;refGene_tx=XM_023251071.1;refGene_gn=LOC101092674;refGene_tx=XM_023251072.1;refGene_gn=LOC101092674;refGene_tx=XM_023251073.1
A3	5252444	5252552

refGene_gn=LOC101092674;refGene_tx=XM_006929630.4;refGene_gn=LOC101092674;refGene_tx=XM_019826309.2;refGene_gn=LOC101092674;refGene_tx=XM_019826311.2;refGene_gn=LOC101092674;refGene_tx=XM_019826312.2;refGene_gn=LOC101092674;refGene_tx=XM_023251071.1;refGene_gn=LOC101092674;refGene_tx=XM_023251072.1;refGene_gn=LOC101092674;refGene_tx=XM_023251073.1
A3 5253382 5253448
refGene_gn=LOC101092674;refGene_tx=XM_006929630.4;refGene_gn=LOC101092674;refGene_tx=XM_019826309.2;refGene_gn=LOC101092674;refGene_tx=XM_019826311.2;refGene_gn=LOC101092674;refGene_tx=XM_019826312.2;refGene_gn=LOC101092674;refGene_tx=XM_023251071.1;refGene_gn=LOC101092674;refGene_tx=XM_023251072.1;refGene_gn=LOC101092674;refGene_tx=XM_023251073.1
A3 5255325 5255484
refGene_gn=LOC101092674;refGene_tx=XM_006929630.4;refGene_gn=LOC101092674;refGene_tx=XM_019826309.2;refGene_gn=LOC101092674;refGene_tx=XM_019826311.2;refGene_gn=LOC101092674;refGene_tx=XM_019826312.2;refGene_gn=LOC101092674;refGene_tx=XM_023251071.1;refGene_gn=LOC101092674;refGene_tx=XM_023251072.1;refGene_gn=LOC101092674;refGene_tx=XM_023251073.1
A3 5256041 5256226
refGene_gn=LOC101092674;refGene_tx=XM_006929630.4;refGene_gn=LOC101092674;refGene_tx=XM_019826309.2;refGene_gn=LOC101092674;refGene_tx=XM_019826311.2;refGene_gn=LOC101092674;refGene_tx=XM_019826312.2;refGene_gn=LOC101092674;refGene_tx=XM_023251071.1;refGene_gn=LOC101092674;refGene_tx=XM_023251072.1;refGene_gn=LOC101092674;refGene_tx=XM_023251073.1
A3 5257358 5257819
refGene_gn=LOC101092674;refGene_tx=XM_006929630.4;refGene_gn=LOC101092674;refGene_tx=XM_019826309.2;refGene_gn=LOC101092674;refGene_tx=XM_019826311.2;refGene_gn=LOC101092674;refGene_tx=XM_019826312.2;refGene_gn=LOC101092674;refGene_tx=XM_023251071.1;refGene_gn=LOC101092674;refGene_tx=XM_023251072.1;refGene_gn=LOC101092674;refGene_tx=XM_023251073.1
A3 5259661 5259855
refGene_gn=LOC101092674;refGene_tx=XM_006929630.4;refGene_gn=LOC101092674;refGene_tx=XM_019826309.2;refGene_gn=LOC101092674;refGene_tx=XM_019826311.2;refGene_gn=LOC101092674;refGene_tx=XM_019826312.2;refGene_gn=LOC101092674;refGene_tx=XM_023251071.1;refGene_gn=LOC101092674;refGene_tx=XM_023251072.1;refGene_gn=LOC101092674;refGene_tx=XM_023251073.1
A3 5262570 5262666
refGene_gn=LOC101092674;refGene_tx=XM_006929630.4;refGene_gn=LOC101092674;refGene_tx=XM_019826309.2;refGene_gn=LOC101092674;refGene_tx=XM_019826311.2;refGene_gn=LOC101092674;refGene_tx=XM_019826312.2;refGene_gn=LOC101092674;refGene_tx=XM_023251071.1;refGene_gn=LOC101092674;refGene_tx=XM_023251072.1;refGene_gn=LOC101092674;refGene_tx=XM_023251073.1
A3 5263530 5263716
refGene_gn=LOC101092674;refGene_tx=XM_006929630.4;refGene_gn=LOC101092674;refGene_tx=XM_019826309.2;refGene_gn=LOC101092674;refGene_tx=XM_019826311.2;refGene_gn=LOC101092674;refGene_tx=XM_019826312.2;refGene_gn=LOC101092674;refGene_tx=XM_023251071.1;refGene_gn=LOC101092674;refGene_tx=XM_023251072.1;refGene_gn=LOC101092674;refGene_tx=XM_023251073.1
A3 5263990 5264152
refGene_gn=LOC101092674;refGene_tx=XM_006929630.4;refGene_gn=LOC101092674;refGene_tx=XM_019826309.2;refGene_gn=LOC101092674;refGene_tx=XM_01

9826311.2;refGene_gn=LOC101092674;refGene_tx=XM_019826312.2;refGene_gn=LOC101092674;refGene_tx=XM_023251071.1;refGene_gn=LOC101092674;refGene_tx=XM_023251072.1;refGene_gn=LOC101092674;refGene_tx=XM_023251073.1
A3 5265072 5265173
refGene_gn=LOC101092674;refGene_tx=XM_006929630.4;refGene_gn=LOC101092674;refGene_tx=XM_019826309.2;refGene_gn=LOC101092674;refGene_tx=XM_019826311.2;refGene_gn=LOC101092674;refGene_tx=XM_019826312.2;refGene_gn=LOC101092674;refGene_tx=XM_023251071.1;refGene_gn=LOC101092674;refGene_tx=XM_023251072.1;refGene_gn=LOC101092674;refGene_tx=XM_023251073.1
A3 5267357 5267478
refGene_gn=LOC101092674;refGene_tx=XM_006929630.4;refGene_gn=LOC101092674;refGene_tx=XM_019826309.2;refGene_gn=LOC101092674;refGene_tx=XM_019826311.2;refGene_gn=LOC101092674;refGene_tx=XM_019826312.2;refGene_gn=LOC101092674;refGene_tx=XM_023251071.1;refGene_gn=LOC101092674;refGene_tx=XM_023251072.1;refGene_gn=LOC101092674;refGene_tx=XM_023251073.1
A3 5268279 5268414
refGene_gn=LOC101092674;refGene_tx=XM_006929630.4;refGene_gn=LOC101092674;refGene_tx=XM_019826309.2;refGene_gn=LOC101092674;refGene_tx=XM_019826311.2;refGene_gn=LOC101092674;refGene_tx=XM_019826312.2;refGene_gn=LOC101092674;refGene_tx=XM_023251071.1;refGene_gn=LOC101092674;refGene_tx=XM_023251072.1;refGene_gn=LOC101092674;refGene_tx=XM_023251073.1
A3 5270449 5270591
refGene_gn=LOC101092674;refGene_tx=XM_006929630.4;refGene_gn=LOC101092674;refGene_tx=XM_019826309.2;refGene_gn=LOC101092674;refGene_tx=XM_019826311.2;refGene_gn=LOC101092674;refGene_tx=XM_019826312.2;refGene_gn=LOC101092674;refGene_tx=XM_023251071.1;refGene_gn=LOC101092674;refGene_tx=XM_023251072.1;refGene_gn=LOC101092674;refGene_tx=XM_023251073.1
A3 5271423 5271581
refGene_gn=LOC101092674;refGene_tx=XM_006929630.4;refGene_gn=LOC101092674;refGene_tx=XM_019826309.2;refGene_gn=LOC101092674;refGene_tx=XM_019826311.2;refGene_gn=LOC101092674;refGene_tx=XM_019826312.2;refGene_gn=LOC101092674;refGene_tx=XM_023251071.1;refGene_gn=LOC101092674;refGene_tx=XM_023251072.1;refGene_gn=LOC101092674;refGene_tx=XM_023251073.1
A3 5274252 5274607 refGene_gn=ANKRD60;refGene_tx=XM_023250926.1
A3 5278934 5279065 refGene_gn=ANKRD60;refGene_tx=XM_023250926.1
A3 5280503 5280669 refGene_gn=ANKRD60;refGene_tx=XM_023250926.1
A3 5283073 5283375 refGene_gn=ANKRD60;refGene_tx=XM_023250926.1
A3 5332544 5332706
refGene_gn=CA3H20orf85;refGene_tx=XM_003983347.4
A3 5334574 5334676
refGene_gn=CA3H20orf85;refGene_tx=XM_003983347.4
A3 5336847 5336929
refGene_gn=CA3H20orf85;refGene_tx=XM_003983347.4
A3 5339285 5339353
refGene_gn=CA3H20orf85;refGene_tx=XM_003983347.4
A3 5649323 5649432
refGene_gn=PMEPA1;refGene_tx=XM_023251074.1;refGene_gn=PMEPA1;refGene_tx=XM_023251075.1
A3 5695246 5695401
refGene_gn=PMEPA1;refGene_tx=XM_023251074.1;refGene_gn=PMEPA1;refGene_

tx=XM_023251075.1
A3 5700833 5700887
refGene_gn=PMEPA1;refGene_tx=XM_023251074.1;refGene_gn=PMEPA1;refGene_tx=XM_023251075.1
A3 5701348 5701897
refGene_gn=PMEPA1;refGene_tx=XM_023251074.1;refGene_gn=PMEPA1;refGene_tx=XM_023251075.1
A3 5717082 5717116
refGene_gn=ZBP1;refGene_tx=XM_006929633.3;refGene_gn=ZBP1;refGene_tx=XM_006929634.3
A3 5720451 5720673
refGene_gn=ZBP1;refGene_tx=XM_006929633.3;refGene_gn=ZBP1;refGene_tx=XM_006929634.3;refGene_gn=ZBP1;refGene_tx=XM_019826314.2;refGene_gn=ZBP1;refGene_tx=XM_019826315.2
A3 5721231 5721297
refGene_gn=ZBP1;refGene_tx=XM_006929633.3;refGene_gn=ZBP1;refGene_tx=XM_006929634.3;refGene_gn=ZBP1;refGene_tx=XM_019826314.2;refGene_gn=ZBP1;refGene_tx=XM_019826315.2
A3 5721674 5721848
refGene_gn=ZBP1;refGene_tx=XM_006929633.3;refGene_gn=ZBP1;refGene_tx=XM_006929634.3;refGene_gn=ZBP1;refGene_tx=XM_019826314.2;refGene_gn=ZBP1;refGene_tx=XM_019826315.2
A3 5723067 5723232
refGene_gn=ZBP1;refGene_tx=XM_006929633.3;refGene_gn=ZBP1;refGene_tx=XM_006929634.3;refGene_gn=ZBP1;refGene_tx=XM_019826314.2;refGene_gn=ZBP1;refGene_tx=XM_019826315.2
A3 5724827 5725034
refGene_gn=ZBP1;refGene_tx=XM_006929633.3;refGene_gn=ZBP1;refGene_tx=XM_006929634.3;refGene_gn=ZBP1;refGene_tx=XM_019826314.2;refGene_gn=ZBP1;refGene_tx=XM_019826315.2
A3 5726002 5726233
refGene_gn=ZBP1;refGene_tx=XM_006929633.3;refGene_gn=ZBP1;refGene_tx=XM_006929634.3;refGene_gn=ZBP1;refGene_tx=XM_019826314.2;refGene_gn=ZBP1;refGene_tx=XM_019826315.2
A3 5730696 5730938
refGene_gn=ZBP1;refGene_tx=XM_006929633.3;refGene_gn=ZBP1;refGene_tx=XM_006929634.3;refGene_gn=ZBP1;refGene_tx=XM_019826314.2;refGene_gn=ZBP1;refGene_tx=XM_019826315.2
A3 5766335 5766790 refGene_gn=PCK1;refGene_tx=XM_003983348.5
A3 5766935 5767031 refGene_gn=PCK1;refGene_tx=XM_003983348.5
A3 5767396 5767528 refGene_gn=PCK1;refGene_tx=XM_003983348.5
A3 5767622 5767847 refGene_gn=PCK1;refGene_tx=XM_003983348.5
A3 5768297 5768460 refGene_gn=PCK1;refGene_tx=XM_003983348.5
A3 5768801 5768989 refGene_gn=PCK1;refGene_tx=XM_003983348.5
A3 5769126 5769330 refGene_gn=PCK1;refGene_tx=XM_003983348.5
A3 5769515 5769597 refGene_gn=PCK1;refGene_tx=XM_003983348.5
A3 5769598 5769698 refGene_gn=PCK1;refGene_tx=XM_003983348.5
A3 5770179 5770403 refGene_gn=PCK1;refGene_tx=XM_003983348.5
A3 5802565 5803102 refGene_gn=CTCFL;refGene_tx=XM_023251076.1
A3 5803801 5804027 refGene_gn=CTCFL;refGene_tx=XM_023251076.1

A3 5808285 5808456 refGene_gn=CTCFL;refGene_tx=XM_023251076.1
A3 5808774 5808908 refGene_gn=CTCFL;refGene_tx=XM_023251076.1
A3 5811645 5811766 refGene_gn=CTCFL;refGene_tx=XM_023251076.1
A3 5813644 5813794 refGene_gn=CTCFL;refGene_tx=XM_023251076.1
A3 5815950 5816111 refGene_gn=CTCFL;refGene_tx=XM_023251076.1
A3 5818960 5819143 refGene_gn=CTCFL;refGene_tx=XM_023251076.1
A3 5825287 5825411 refGene_gn=CTCFL;refGene_tx=XM_023251076.1
A3 5830131 5830286 refGene_gn=CTCFL;refGene_tx=XM_023251076.1
A3 5898773 5899077 refGene_gn=RBM38;refGene_tx=XM_006929636.2
A3 5911471 5911526 refGene_gn=RBM38;refGene_tx=XM_006929636.2
A3 5912004 5912128 refGene_gn=RBM38;refGene_tx=XM_006929636.2
A3 5912935 5913172 refGene_gn=RBM38;refGene_tx=XM_006929636.2
A3 5927213 5927300
refGene_gn=RAE1;refGene_tx=XM_003983350.4;refGene_gn=RAE1;refGene_tx=XM_006929637.3;refGene_gn=RAE1;refGene_tx=XM_011280761.3;refGene_gn=RAE1;refGene_tx=XM_019826319.2
A3 5930612 5930807
refGene_gn=RAE1;refGene_tx=XM_003983350.4;refGene_gn=RAE1;refGene_tx=XM_006929637.3;refGene_gn=RAE1;refGene_tx=XM_011280761.3;refGene_gn=RAE1;refGene_tx=XM_019826319.2
A3 5931366 5931442
refGene_gn=RAE1;refGene_tx=XM_003983350.4;refGene_gn=RAE1;refGene_tx=XM_006929637.3;refGene_gn=RAE1;refGene_tx=XM_011280761.3;refGene_gn=RAE1;refGene_tx=XM_019826319.2
A3 5931523 5931630
refGene_gn=RAE1;refGene_tx=XM_003983350.4;refGene_gn=RAE1;refGene_tx=XM_006929637.3;refGene_gn=RAE1;refGene_tx=XM_011280761.3;refGene_gn=RAE1;refGene_tx=XM_019826319.2
A3 5935200 5935308
refGene_gn=RAE1;refGene_tx=XM_003983350.4;refGene_gn=RAE1;refGene_tx=XM_006929637.3;refGene_gn=RAE1;refGene_tx=XM_011280761.3;refGene_gn=RAE1;refGene_tx=XM_019826319.2
A3 5936289 5936361
refGene_gn=RAE1;refGene_tx=XM_003983350.4;refGene_gn=RAE1;refGene_tx=XM_006929637.3;refGene_gn=RAE1;refGene_tx=XM_011280761.3;refGene_gn=RAE1;refGene_tx=XM_019826319.2
A3 5936512 5936599
refGene_gn=RAE1;refGene_tx=XM_003983350.4;refGene_gn=RAE1;refGene_tx=XM_006929637.3;refGene_gn=RAE1;refGene_tx=XM_011280761.3;refGene_gn=RAE1;refGene_tx=XM_019826319.2
A3 5937345 5937432
refGene_gn=RAE1;refGene_tx=XM_003983350.4;refGene_gn=RAE1;refGene_tx=XM_006929637.3;refGene_gn=RAE1;refGene_tx=XM_011280761.3;refGene_gn=RAE1;refGene_tx=XM_019826319.2
A3 5942947 5943040
refGene_gn=RAE1;refGene_tx=XM_003983350.4;refGene_gn=RAE1;refGene_tx=XM_006929637.3;refGene_gn=RAE1;refGene_tx=XM_011280761.3;refGene_gn=RAE1;refGene_tx=XM_019826319.2
A3 5945090 5945195
refGene_gn=RAE1;refGene_tx=XM_003983350.4;refGene_gn=RAE1;refGene_tx=X

M_006929637.3;refGene_gn=RAE1;refGene_tx=XM_011280761.3;refGene_gn=RAE
1;refGene_tx=XM_019826319.2
A3 5945758 5945848
refGene_gn=RAE1;refGene_tx=XM_003983350.4;refGene_gn=RAE1;refGene_tx=X
M_006929637.3;refGene_gn=RAE1;refGene_tx=XM_011280761.3;refGene_gn=RAE
1;refGene_tx=XM_019826319.2
A3 5953733 5953853
refGene_gn=SPO11;refGene_tx=XM_003983351.3;refGene_gn=SPO11;refGene_tx
=XM_019826320.1
A3 5954354 5954466
refGene_gn=SPO11;refGene_tx=XM_003983351.3;refGene_gn=SPO11;refGene_tx
=XM_019826320.1
A3 5956648 5956725
refGene_gn=SPO11;refGene_tx=XM_003983351.3;refGene_gn=SPO11;refGene_tx
=XM_019826320.1
A3 5957661 5957699
refGene_gn=SPO11;refGene_tx=XM_003983351.3;refGene_gn=SPO11;refGene_tx
=XM_019826320.1
A3 5958013 5958113
refGene_gn=SPO11;refGene_tx=XM_003983351.3;refGene_gn=SPO11;refGene_tx
=XM_019826320.1
A3 5959557 5959667
refGene_gn=SPO11;refGene_tx=XM_003983351.3;refGene_gn=SPO11;refGene_tx
=XM_019826320.1
A3 5959997 5960034
refGene_gn=SPO11;refGene_tx=XM_003983351.3;refGene_gn=SPO11;refGene_tx
=XM_019826320.1
A3 5960742 5960829
refGene_gn=SPO11;refGene_tx=XM_003983351.3;refGene_gn=SPO11;refGene_tx
=XM_019826320.1
A3 5961382 5961491
refGene_gn=SPO11;refGene_tx=XM_003983351.3;refGene_gn=SPO11;refGene_tx
=XM_019826320.1
A3 5961713 5961780
refGene_gn=SPO11;refGene_tx=XM_003983351.3;refGene_gn=SPO11;refGene_tx
=XM_019826320.1
A3 5962194 5962283
refGene_gn=SPO11;refGene_tx=XM_003983351.3;refGene_gn=SPO11;refGene_tx
=XM_019826320.1
A3 5963561 5963675
refGene_gn=SPO11;refGene_tx=XM_003983351.3;refGene_gn=SPO11;refGene_tx
=XM_019826320.1
A3 5965837 5965968
refGene_gn=SPO11;refGene_tx=XM_003983351.3;refGene_gn=SPO11;refGene_tx
=XM_019826320.1
A3 6015795 6016213 refGene_gn=BMP7;refGene_tx=XM_011280760.3
A3 6048575 6048705 refGene_gn=BMP7;refGene_tx=XM_011280760.3
A3 6048706 6048766 refGene_gn=BMP7;refGene_tx=XM_011280760.3
A3 6077333 6077482 refGene_gn=BMP7;refGene_tx=XM_011280760.3
A3 6093072 6093231 refGene_gn=BMP7;refGene_tx=XM_011280760.3

A3	6093233	6093269	refGene_gn=BMP7;refGene_tx=XM_011280760.3
A3	6102152	6102229	refGene_gn=BMP7;refGene_tx=XM_011280760.3
A3	6103727	6103838	refGene_gn=BMP7;refGene_tx=XM_011280760.3
A3	6108367	6108517	refGene_gn=BMP7;refGene_tx=XM_011280760.3
A3	6518068	6518354
refGene_gn=TFAP2C;refGene_tx=XM_006929638.4;refGene_gn=TFAP2C;refGene_
tx=XM_019826322.2
A3	6519761	6519906
refGene_gn=TFAP2C;refGene_tx=XM_006929638.4;refGene_gn=TFAP2C;refGene_
tx=XM_019826322.2
A3	6521705	6521824
refGene_gn=TFAP2C;refGene_tx=XM_006929638.4;refGene_gn=TFAP2C;refGene_
tx=XM_019826322.2
A3	6522382	6522599
refGene_gn=TFAP2C;refGene_tx=XM_006929638.4;refGene_gn=TFAP2C;refGene_
tx=XM_019826322.2
A3	6523793	6523845
refGene_gn=TFAP2C;refGene_tx=XM_006929638.4;refGene_gn=TFAP2C;refGene_
tx=XM_019826322.2
A3	6523961	6524447
refGene_gn=TFAP2C;refGene_tx=XM_006929638.4;refGene_gn=TFAP2C;refGene_
tx=XM_019826322.2
A3	6524645	6524903	refGene_gn=TFAP2C;refGene_tx=XM_019826322.2
A3	6526060	6526108	refGene_gn=TFAP2C;refGene_tx=XM_006929638.4
A3	6603085	6603349
refGene_gn=LOC101094662;refGene_tx=XM_003983370.4
A3	6604515	6604764
refGene_gn=LOC101094662;refGene_tx=XM_003983370.4
A3	6608478	6609027
refGene_gn=RTFDC1;refGene_tx=XM_003983353.5;refGene_gn=RTFDC1;refGene_
tx=XM_019826324.2;refGene_gn=RTFDC1;refGene_tx=XM_019826325.2
A3	6610375	6610512
refGene_gn=RTFDC1;refGene_tx=XM_003983353.5;refGene_gn=RTFDC1;refGene_
tx=XM_019826324.2;refGene_gn=RTFDC1;refGene_tx=XM_019826325.2
A3	6610621	6610676
refGene_gn=RTFDC1;refGene_tx=XM_003983353.5;refGene_gn=RTFDC1;refGene_
tx=XM_019826324.2;refGene_gn=RTFDC1;refGene_tx=XM_019826325.2
A3	6613437	6613551
refGene_gn=RTFDC1;refGene_tx=XM_003983353.5;refGene_gn=RTFDC1;refGene_
tx=XM_019826324.2;refGene_gn=RTFDC1;refGene_tx=XM_019826325.2
A3	6624164	6625131
refGene_gn=GCNT7;refGene_tx=XM_003983354.5;refGene_gn=GCNT7;refGene_tx
=XM_019826328.2;refGene_gn=GCNT7;refGene_tx=XM_023251077.1;refGene_gn=
GCNT7;refGene_tx=XM_023251078.1
A3	6626090	6626115	refGene_gn=GCNT7;refGene_tx=XM_023251079.1
A3	6626243	6626339
refGene_gn=GCNT7;refGene_tx=XM_003983354.5;refGene_gn=GCNT7;refGene_tx
=XM_019826328.2;refGene_gn=GCNT7;refGene_tx=XM_023251077.1;refGene_gn=
GCNT7;refGene_tx=XM_023251078.1;refGene_gn=GCNT7;refGene_tx=XM_0232510
79.1

A3 6629444 6629668
refGene_gn=GCNT7;refGene_tx=XM_003983354.5;refGene_gn=GCNT7;refGene_tx
=XM_019826328.2;refGene_gn=GCNT7;refGene_tx=XM_023251079.1
A3 6631478 6631699
refGene_gn=GCNT7;refGene_tx=XM_023251077.1;refGene_gn=GCNT7;refGene_tx
=XM_023251078.1
A3 6639237 6639316
refGene_gn=RTFDC1;refGene_tx=XM_003983353.5;refGene_gn=RTFDC1;refGene_
tx=XM_019826324.2;refGene_gn=RTFDC1;refGene_tx=XM_019826325.2
A3 6642739 6642879
refGene_gn=RTFDC1;refGene_tx=XM_003983353.5;refGene_gn=RTFDC1;refGene_
tx=XM_019826324.2;refGene_gn=RTFDC1;refGene_tx=XM_019826325.2
A3 6644873 6644967
refGene_gn=RTFDC1;refGene_tx=XM_003983353.5;refGene_gn=RTFDC1;refGene_
tx=XM_019826324.2;refGene_gn=RTFDC1;refGene_tx=XM_019826325.2
A3 6645635 6645730
refGene_gn=RTFDC1;refGene_tx=XM_003983353.5;refGene_gn=RTFDC1;refGene_
tx=XM_019826324.2;refGene_gn=RTFDC1;refGene_tx=XM_019826325.2
A3 6649500 6649569
refGene_gn=RTFDC1;refGene_tx=XM_003983353.5;refGene_gn=RTFDC1;refGene_
tx=XM_019826324.2;refGene_gn=RTFDC1;refGene_tx=XM_019826325.2
A3 6657188 6657599 refGene_gn=CASS4;refGene_tx=XM_011280758.3
A3 6659708 6659786 refGene_gn=CASS4;refGene_tx=XM_011280758.3
A3 6661015 6662341 refGene_gn=CASS4;refGene_tx=XM_011280758.3
A3 6663224 6663305 refGene_gn=CASS4;refGene_tx=XM_011280758.3
A3 6664306 6664408 refGene_gn=CASS4;refGene_tx=XM_011280758.3
A3 6670101 6670524 refGene_gn=CASS4;refGene_tx=XM_011280758.3
A3 6692250 6692286 refGene_gn=CASS4;refGene_tx=XM_011280758.3
A3 6700280 6700540
refGene_gn=CSTF1;refGene_tx=XM_003983355.4;refGene_gn=CSTF1;refGene_tx
=XM_006929640.4
A3 6703689 6704080
refGene_gn=CSTF1;refGene_tx=XM_003983355.4;refGene_gn=CSTF1;refGene_tx
=XM_006929640.4
A3 6704893 6705091
refGene_gn=CSTF1;refGene_tx=XM_003983355.4;refGene_gn=CSTF1;refGene_tx
=XM_006929640.4
A3 6705243 6705521
refGene_gn=CSTF1;refGene_tx=XM_003983355.4;refGene_gn=CSTF1;refGene_tx
=XM_006929640.4
A3 6707707 6707876
refGene_gn=CSTF1;refGene_tx=XM_003983355.4;refGene_gn=CSTF1;refGene_tx
=XM_006929640.4
A3 6716353 6716395 refGene_gn=AURKA;refGene_tx=XM_006929641.4
A3 6719653 6719933 refGene_gn=AURKA;refGene_tx=XM_006929641.4
A3 6721473 6721528 refGene_gn=AURKA;refGene_tx=XM_006929641.4
A3 6722785 6722862 refGene_gn=AURKA;refGene_tx=XM_006929641.4
A3 6722864 6722976 refGene_gn=AURKA;refGene_tx=XM_006929641.4
A3 6724106 6724245 refGene_gn=AURKA;refGene_tx=XM_006929641.4
A3 6728695 6728844 refGene_gn=AURKA;refGene_tx=XM_006929641.4

A3	6731785	6731960	refGene_gn=AURKA;refGene_tx=XM_006929641.4
A3	6732092	6732278	refGene_gn=AURKA;refGene_tx=XM_006929641.4
A3	6736923	6737140
refGene_gn=FAM210B;refGene_tx=XM_023251081.1;refGene_gn=FAM210B;refGene_tx=XM_023251082.1
A3	6737596	6737853
refGene_gn=FAM210B;refGene_tx=XM_023251081.1;refGene_gn=FAM210B;refGene_tx=XM_023251082.1
A3	6744056	6744242
refGene_gn=FAM210B;refGene_tx=XM_023251081.1;refGene_gn=FAM210B;refGene_tx=XM_023251082.1
A3	6817423	6818395	refGene_gn=MC3R;refGene_tx=XM_023251083.1
A3	7004976	7005267
refGene_gn=CBLN4;refGene_tx=XM_003983358.5;refGene_gn=CBLN4;refGene_tx=XM_023251084.1
A3	7008358	7008475
refGene_gn=CBLN4;refGene_tx=XM_003983358.5;refGene_gn=CBLN4;refGene_tx=XM_023251084.1
A3	7010511	7010709
refGene_gn=CBLN4;refGene_tx=XM_003983358.5;refGene_gn=CBLN4;refGene_tx=XM_023251084.1
A3	7070018	7070104	refGene_gn=CBLN4;refGene_tx=XM_023251084.1
A3	8106346	8106411	refGene_gn=DOK5;refGene_tx=XM_006929650.3
A3	8113964	8114085	refGene_gn=DOK5;refGene_tx=XM_006929650.3
A3	8138241	8138377	refGene_gn=DOK5;refGene_tx=XM_006929650.3
A3	8152411	8152601	refGene_gn=DOK5;refGene_tx=XM_006929650.3
A3	8155003	8155123	refGene_gn=DOK5;refGene_tx=XM_006929650.3
A3	8155209	8155324	refGene_gn=DOK5;refGene_tx=XM_006929650.3
A3	8187526	8187634	refGene_gn=DOK5;refGene_tx=XM_006929650.3
A3	8252847	8252901	refGene_gn=DOK5;refGene_tx=XM_006929650.3
A3	8488682	8488814	refGene_gn=PFDN4;refGene_tx=XM_011280754.3
A3	8491781	8491922	refGene_gn=PFDN4;refGene_tx=XM_011280754.3
A3	8492761	8492869	refGene_gn=PFDN4;refGene_tx=XM_011280754.3
A3	8501201	8501225	refGene_gn=PFDN4;refGene_tx=XM_011280754.3
A3	8530821	8531079
refGene_gn=LOC101096141;refGene_tx=XM_023251085.1
A3	8531288	8531479
refGene_gn=LOC101096141;refGene_tx=XM_023251085.1
A3	8532707	8532801
refGene_gn=LOC101096141;refGene_tx=XM_023251085.1
A3	8534827	8534924
refGene_gn=LOC101096141;refGene_tx=XM_023251085.1
A3	8537745	8537837
refGene_gn=LOC101096141;refGene_tx=XM_023251085.1
A3	8538793	8538905
refGene_gn=LOC101096141;refGene_tx=XM_023251085.1
A3	8540309	8540455
refGene_gn=LOC101096141;refGene_tx=XM_023251085.1
A3	8544193	8544360
refGene_gn=LOC101096141;refGene_tx=XM_023251085.1

A3 8545175 8545254
refGene_gn=LOC101096141;refGene_tx=XM_023251085.1
A3 8545956 8546029
refGene_gn=LOC101096141;refGene_tx=XM_023251085.1
A3 8546031 8546153
refGene_gn=LOC101096141;refGene_tx=XM_023251085.1
A3 8546747 8546858
refGene_gn=LOC101096141;refGene_tx=XM_023251085.1
A3 8645753 8646269 refGene_gn=BCAS1;refGene_tx=XM_023251086.1
A3 8667251 8667410 refGene_gn=BCAS1;refGene_tx=XM_023251086.1
A3 8668403 8668448 refGene_gn=BCAS1;refGene_tx=XM_023251086.1
A3 8670854 8670989 refGene_gn=BCAS1;refGene_tx=XM_023251086.1
A3 8675831 8676008 refGene_gn=BCAS1;refGene_tx=XM_023251086.1
A3 8681418 8681460 refGene_gn=BCAS1;refGene_tx=XM_023251086.1
A3 8689169 8689343 refGene_gn=BCAS1;refGene_tx=XM_023251086.1
A3 8697032 8697098 refGene_gn=BCAS1;refGene_tx=XM_023251086.1
A3 8702593 8702857 refGene_gn=BCAS1;refGene_tx=XM_023251086.1
A3 8710208 8710283 refGene_gn=BCAS1;refGene_tx=XM_023251086.1
A3 8996110 8997503
refGene_gn=ZNF217;refGene_tx=XM_019828023.2;refGene_gn=ZNF217;refGene_tx=XM_023251087.1;refGene_gn=ZNF217;refGene_tx=XM_023251088.1;refGene_gn=ZNF217;refGene_tx=XM_023251089.1;refGene_gn=ZNF217;refGene_tx=XM_023251090.1;refGene_gn=ZNF217;refGene_tx=XM_023251091.1
A3 9000868 9000985
refGene_gn=ZNF217;refGene_tx=XM_019828023.2;refGene_gn=ZNF217;refGene_tx=XM_023251087.1;refGene_gn=ZNF217;refGene_tx=XM_023251088.1;refGene_gn=ZNF217;refGene_tx=XM_023251089.1;refGene_gn=ZNF217;refGene_tx=XM_023251090.1;refGene_gn=ZNF217;refGene_tx=XM_023251091.1
A3 9001576 9003004
refGene_gn=ZNF217;refGene_tx=XM_019828023.2;refGene_gn=ZNF217;refGene_tx=XM_023251087.1;refGene_gn=ZNF217;refGene_tx=XM_023251088.1;refGene_gn=ZNF217;refGene_tx=XM_023251089.1;refGene_gn=ZNF217;refGene_tx=XM_023251090.1;refGene_gn=ZNF217;refGene_tx=XM_023251091.1
A3 9004908 9005018
refGene_gn=ZNF217;refGene_tx=XM_019828023.2;refGene_gn=ZNF217;refGene_tx=XM_023251087.1;refGene_gn=ZNF217;refGene_tx=XM_023251088.1;refGene_gn=ZNF217;refGene_tx=XM_023251089.1;refGene_gn=ZNF217;refGene_tx=XM_023251090.1;refGene_gn=ZNF217;refGene_tx=XM_023251091.1
A3 9081039 9081041 refGene_gn=TSHZ2;refGene_tx=XM_019826348.2
A3 9237426 9237431 refGene_gn=TSHZ2;refGene_tx=XM_019826347.2
A3 9267599 9270664
refGene_gn=TSHZ2;refGene_tx=XM_003983378.5;refGene_gn=TSHZ2;refGene_tx=XM_006929653.4;refGene_gn=TSHZ2;refGene_tx=XM_011280753.3;refGene_gn=TSHZ2;refGene_tx=XM_019826344.2;refGene_gn=TSHZ2;refGene_tx=XM_019826347.2;refGene_gn=TSHZ2;refGene_tx=XM_019826348.2
A3 9543586 9543626
refGene_gn=TSHZ2;refGene_tx=XM_003983378.5;refGene_gn=TSHZ2;refGene_tx=XM_006929653.4;refGene_gn=TSHZ2;refGene_tx=XM_011280753.3;refGene_gn=TSHZ2;refGene_tx=XM_019826344.2;refGene_gn=TSHZ2;refGene_tx=XM_019826347.2;refGene_gn=TSHZ2;refGene_tx=XM_019826348.2

A3	10220289	10220335
refGene_gn=ZFP64;refGene_tx=XM_011280751.3;refGene_gn=ZFP64;refGene_tx=XM_011280752.3;refGene_gn=ZFP64;refGene_tx=XM_019826349.2;refGene_gn=ZFP64;refGene_tx=XM_019826350.2;refGene_gn=ZFP64;refGene_tx=XM_019826352.2
A3	10223819	10224059
refGene_gn=ZFP64;refGene_tx=XM_011280751.3;refGene_gn=ZFP64;refGene_tx=XM_011280752.3;refGene_gn=ZFP64;refGene_tx=XM_019826349.2;refGene_gn=ZFP64;refGene_tx=XM_019826350.2;refGene_gn=ZFP64;refGene_tx=XM_019826352.2
A3	10238270	10238432
refGene_gn=ZFP64;refGene_tx=XM_011280751.3;refGene_gn=ZFP64;refGene_tx=XM_011280752.3;refGene_gn=ZFP64;refGene_tx=XM_019826349.2;refGene_gn=ZFP64;refGene_tx=XM_019826350.2;refGene_gn=ZFP64;refGene_tx=XM_019826352.2
A3	10239502	10239565
refGene_gn=ZFP64;refGene_tx=XM_011280751.3;refGene_gn=ZFP64;refGene_tx=XM_011280752.3;refGene_gn=ZFP64;refGene_tx=XM_019826349.2;refGene_gn=ZFP64;refGene_tx=XM_019826350.2;refGene_gn=ZFP64;refGene_tx=XM_019826352.2
A3	10242949	10243201
refGene_gn=ZFP64;refGene_tx=XM_011280751.3;refGene_gn=ZFP64;refGene_tx=XM_011280752.3;refGene_gn=ZFP64;refGene_tx=XM_019826349.2
A3	10249111	10250394
refGene_gn=ZFP64;refGene_tx=XM_011280751.3;refGene_gn=ZFP64;refGene_tx=XM_011280752.3;refGene_gn=ZFP64;refGene_tx=XM_019826350.2
A3	10291622	10291728	refGene_gn=ZFP64;refGene_tx=XM_006929656.4
A3	10297920	10298067
refGene_gn=ZFP64;refGene_tx=XM_006929656.4;refGene_gn=ZFP64;refGene_tx=XM_019826349.2;refGene_gn=ZFP64;refGene_tx=XM_019826352.2
A3	10298507	10298570
refGene_gn=ZFP64;refGene_tx=XM_006929656.4;refGene_gn=ZFP64;refGene_tx=XM_019826349.2;refGene_gn=ZFP64;refGene_tx=XM_019826352.2
A3	10307295	10307547
refGene_gn=ZFP64;refGene_tx=XM_006929656.4;refGene_gn=ZFP64;refGene_tx=XM_019826349.2;refGene_gn=ZFP64;refGene_tx=XM_019826352.2
A3	10310263	10310973
refGene_gn=ZFP64;refGene_tx=XM_006929656.4;refGene_gn=ZFP64;refGene_tx=XM_019826349.2;refGene_gn=ZFP64;refGene_tx=XM_019826352.2
A3	10563861	10563991
refGene_gn=SALL4;refGene_tx=XM_003983380.5;refGene_gn=SALL4;refGene_tx=XM_019826343.2
A3	10575589	10577935
refGene_gn=SALL4;refGene_tx=XM_003983380.5;refGene_gn=SALL4;refGene_tx=XM_019826341.2;refGene_gn=SALL4;refGene_tx=XM_019826343.2
A3	10579052	10579333
refGene_gn=SALL4;refGene_tx=XM_003983380.5;refGene_gn=SALL4;refGene_tx=XM_019826341.2;refGene_gn=SALL4;refGene_tx=XM_019826343.2
A3	10582039	10582459
refGene_gn=SALL4;refGene_tx=XM_003983380.5;refGene_gn=SALL4;refGene_tx

=XM_019826341.2;refGene_gn=SALL4;refGene_tx=XM_019826343.2
A3 10595107 10595175 refGene_gn=ATP9A;refGene_tx=XM_023251092.1
A3 10605905 10605928 refGene_gn=ATP9A;refGene_tx=XM_019826353.1
A3 10613132 10613146 refGene_gn=ATP9A;refGene_tx=XM_006929657.2
A3 10626008 10626153
refGene_gn=ATP9A;refGene_tx=XM_006929657.2;refGene_gn=ATP9A;refGene_tx
=XM_019826353.1;refGene_gn=ATP9A;refGene_tx=XM_023251092.1
A3 10628038 10628152
refGene_gn=ATP9A;refGene_tx=XM_006929657.2;refGene_gn=ATP9A;refGene_tx
=XM_019826353.1;refGene_gn=ATP9A;refGene_tx=XM_023251092.1
A3 10633394 10633503
refGene_gn=ATP9A;refGene_tx=XM_006929657.2;refGene_gn=ATP9A;refGene_tx
=XM_019826353.1;refGene_gn=ATP9A;refGene_tx=XM_023251092.1
A3 10647212 10647271
refGene_gn=ATP9A;refGene_tx=XM_006929657.2;refGene_gn=ATP9A;refGene_tx
=XM_019826353.1;refGene_gn=ATP9A;refGene_tx=XM_023251092.1
A3 10648416 10648468
refGene_gn=ATP9A;refGene_tx=XM_006929657.2;refGene_gn=ATP9A;refGene_tx
=XM_019826353.1;refGene_gn=ATP9A;refGene_tx=XM_023251092.1
A3 10650729 10650824
refGene_gn=ATP9A;refGene_tx=XM_006929657.2;refGene_gn=ATP9A;refGene_tx
=XM_019826353.1;refGene_gn=ATP9A;refGene_tx=XM_023251092.1
A3 10655084 10655165
refGene_gn=ATP9A;refGene_tx=XM_006929657.2;refGene_gn=ATP9A;refGene_tx
=XM_019826353.1;refGene_gn=ATP9A;refGene_tx=XM_023251092.1
A3 10655954 10656030
refGene_gn=ATP9A;refGene_tx=XM_006929657.2;refGene_gn=ATP9A;refGene_tx
=XM_019826353.1;refGene_gn=ATP9A;refGene_tx=XM_023251092.1
A3 10661982 10662059
refGene_gn=ATP9A;refGene_tx=XM_006929657.2;refGene_gn=ATP9A;refGene_tx
=XM_019826353.1;refGene_gn=ATP9A;refGene_tx=XM_023251092.1
A3 10663964 10664125
refGene_gn=ATP9A;refGene_tx=XM_006929657.2;refGene_gn=ATP9A;refGene_tx
=XM_019826353.1;refGene_gn=ATP9A;refGene_tx=XM_023251092.1
A3 10665114 10665257
refGene_gn=ATP9A;refGene_tx=XM_006929657.2;refGene_gn=ATP9A;refGene_tx
=XM_019826353.1;refGene_gn=ATP9A;refGene_tx=XM_023251092.1
A3 10666210 10666323
refGene_gn=ATP9A;refGene_tx=XM_006929657.2;refGene_gn=ATP9A;refGene_tx
=XM_019826353.1;refGene_gn=ATP9A;refGene_tx=XM_023251092.1
A3 10674118 10674331
refGene_gn=ATP9A;refGene_tx=XM_006929657.2;refGene_gn=ATP9A;refGene_tx
=XM_019826353.1;refGene_gn=ATP9A;refGene_tx=XM_023251092.1
A3 10691368 10691530
refGene_gn=ATP9A;refGene_tx=XM_006929657.2;refGene_gn=ATP9A;refGene_tx
=XM_019826353.1;refGene_gn=ATP9A;refGene_tx=XM_023251092.1
A3 10702137 10702230
refGene_gn=ATP9A;refGene_tx=XM_006929657.2;refGene_gn=ATP9A;refGene_tx
=XM_019826353.1;refGene_gn=ATP9A;refGene_tx=XM_023251092.1
A3 10703288 10703372

refGene_gn=ATP9A;refGene_tx=XM_006929657.2;refGene_gn=ATP9A;refGene_tx=XM_019826353.1;refGene_gn=ATP9A;refGene_tx=XM_023251092.1
A3 10704733 10704904
refGene_gn=ATP9A;refGene_tx=XM_006929657.2;refGene_gn=ATP9A;refGene_tx=XM_019826353.1;refGene_gn=ATP9A;refGene_tx=XM_023251092.1
A3 10707140 10707239
refGene_gn=ATP9A;refGene_tx=XM_006929657.2;refGene_gn=ATP9A;refGene_tx=XM_019826353.1;refGene_gn=ATP9A;refGene_tx=XM_023251092.1
A3 10709597 10709687
refGene_gn=ATP9A;refGene_tx=XM_006929657.2;refGene_gn=ATP9A;refGene_tx=XM_019826353.1;refGene_gn=ATP9A;refGene_tx=XM_023251092.1
A3 10709802 10709947
refGene_gn=ATP9A;refGene_tx=XM_006929657.2;refGene_gn=ATP9A;refGene_tx=XM_019826353.1;refGene_gn=ATP9A;refGene_tx=XM_023251092.1
A3 10710766 10710831
refGene_gn=ATP9A;refGene_tx=XM_006929657.2;refGene_gn=ATP9A;refGene_tx=XM_019826353.1;refGene_gn=ATP9A;refGene_tx=XM_023251092.1
A3 10712147 10712303
refGene_gn=ATP9A;refGene_tx=XM_006929657.2;refGene_gn=ATP9A;refGene_tx=XM_019826353.1;refGene_gn=ATP9A;refGene_tx=XM_023251092.1
A3 10715847 10715912
refGene_gn=ATP9A;refGene_tx=XM_006929657.2;refGene_gn=ATP9A;refGene_tx=XM_019826353.1;refGene_gn=ATP9A;refGene_tx=XM_023251092.1
A3 10717745 10717854
refGene_gn=ATP9A;refGene_tx=XM_006929657.2;refGene_gn=ATP9A;refGene_tx=XM_019826353.1;refGene_gn=ATP9A;refGene_tx=XM_023251092.1
A3 10718492 10718550
refGene_gn=ATP9A;refGene_tx=XM_006929657.2;refGene_gn=ATP9A;refGene_tx=XM_019826353.1;refGene_gn=ATP9A;refGene_tx=XM_023251092.1
A3 10720727 10720931
refGene_gn=ATP9A;refGene_tx=XM_006929657.2;refGene_gn=ATP9A;refGene_tx=XM_019826353.1;refGene_gn=ATP9A;refGene_tx=XM_023251092.1
A3 10722188 10722325
refGene_gn=ATP9A;refGene_tx=XM_006929657.2;refGene_gn=ATP9A;refGene_tx=XM_019826353.1;refGene_gn=ATP9A;refGene_tx=XM_023251092.1
A3 10752685 10752755 refGene_gn=NFATC2;refGene_tx=XM_023251100.1
A3 10771143 10771273
refGene_gn=NFATC2;refGene_tx=XM_023251093.1;refGene_gn=NFATC2;refGene_tx=XM_023251094.1;refGene_gn=NFATC2;refGene_tx=XM_023251095.1;refGene_gn=NFATC2;refGene_tx=XM_023251096.1;refGene_gn=NFATC2;refGene_tx=XM_023251098.1;refGene_gn=NFATC2;refGene_tx=XM_023251099.1
A3 10791392 10792407
refGene_gn=NFATC2;refGene_tx=XM_023251093.1;refGene_gn=NFATC2;refGene_tx=XM_023251094.1;refGene_gn=NFATC2;refGene_tx=XM_023251095.1;refGene_gn=NFATC2;refGene_tx=XM_023251096.1;refGene_gn=NFATC2;refGene_tx=XM_023251098.1;refGene_gn=NFATC2;refGene_tx=XM_023251099.1;refGene_gn=NFATC2;refGene_tx=XM_023251100.1;refGene_gn=NFATC2;refGene_tx=XM_023251101.1;refGene_gn=NFATC2;refGene_tx=XM_023251102.1
A3 10796916 10797088
refGene_gn=NFATC2;refGene_tx=XM_023251093.1;refGene_gn=NFATC2;refGene_

tx=XM_023251094.1;refGene_gn=NFATC2;refGene_tx=XM_023251095.1;refGene_gn=NFATC2;refGene_tx=XM_023251096.1;refGene_gn=NFATC2;refGene_tx=XM_023251098.1;refGene_gn=NFATC2;refGene_tx=XM_023251099.1;refGene_gn=NFATC2;refGene_tx=XM_023251100.1;refGene_gn=NFATC2;refGene_tx=XM_023251101.1;refGene_gn=NFATC2;refGene_tx=XM_023251102.1
A3 10834218 10834421
refGene_gn=NFATC2;refGene_tx=XM_023251093.1;refGene_gn=NFATC2;refGene_tx=XM_023251094.1;refGene_gn=NFATC2;refGene_tx=XM_023251095.1;refGene_gn=NFATC2;refGene_tx=XM_023251096.1;refGene_gn=NFATC2;refGene_tx=XM_023251098.1;refGene_gn=NFATC2;refGene_tx=XM_023251099.1;refGene_gn=NFATC2;refGene_tx=XM_023251100.1;refGene_gn=NFATC2;refGene_tx=XM_023251101.1;refGene_gn=NFATC2;refGene_tx=XM_023251102.1
A3 10835435 10835608
refGene_gn=NFATC2;refGene_tx=XM_023251093.1;refGene_gn=NFATC2;refGene_tx=XM_023251094.1;refGene_gn=NFATC2;refGene_tx=XM_023251095.1;refGene_gn=NFATC2;refGene_tx=XM_023251096.1;refGene_gn=NFATC2;refGene_tx=XM_023251098.1;refGene_gn=NFATC2;refGene_tx=XM_023251099.1;refGene_gn=NFATC2;refGene_tx=XM_023251100.1;refGene_gn=NFATC2;refGene_tx=XM_023251101.1;refGene_gn=NFATC2;refGene_tx=XM_023251102.1
A3 10850400 10850541
refGene_gn=NFATC2;refGene_tx=XM_023251093.1;refGene_gn=NFATC2;refGene_tx=XM_023251094.1;refGene_gn=NFATC2;refGene_tx=XM_023251095.1;refGene_gn=NFATC2;refGene_tx=XM_023251096.1;refGene_gn=NFATC2;refGene_tx=XM_023251098.1;refGene_gn=NFATC2;refGene_tx=XM_023251099.1;refGene_gn=NFATC2;refGene_tx=XM_023251100.1;refGene_gn=NFATC2;refGene_tx=XM_023251101.1;refGene_gn=NFATC2;refGene_tx=XM_023251102.1
A3 10867854 10867910
refGene_gn=NFATC2;refGene_tx=XM_023251093.1;refGene_gn=NFATC2;refGene_tx=XM_023251094.1;refGene_gn=NFATC2;refGene_tx=XM_023251095.1;refGene_gn=NFATC2;refGene_tx=XM_023251096.1;refGene_gn=NFATC2;refGene_tx=XM_023251098.1;refGene_gn=NFATC2;refGene_tx=XM_023251099.1;refGene_gn=NFATC2;refGene_tx=XM_023251100.1;refGene_gn=NFATC2;refGene_tx=XM_023251101.1;refGene_gn=NFATC2;refGene_tx=XM_023251102.1
A3 10868302 10868429
refGene_gn=NFATC2;refGene_tx=XM_023251093.1;refGene_gn=NFATC2;refGene_tx=XM_023251094.1;refGene_gn=NFATC2;refGene_tx=XM_023251095.1;refGene_gn=NFATC2;refGene_tx=XM_023251096.1;refGene_gn=NFATC2;refGene_tx=XM_023251098.1;refGene_gn=NFATC2;refGene_tx=XM_023251099.1;refGene_gn=NFATC2;refGene_tx=XM_023251100.1;refGene_gn=NFATC2;refGene_tx=XM_023251101.1;refGene_gn=NFATC2;refGene_tx=XM_023251102.1
A3 10870678 10871368
refGene_gn=NFATC2;refGene_tx=XM_023251093.1;refGene_gn=NFATC2;refGene_tx=XM_023251094.1;refGene_gn=NFATC2;refGene_tx=XM_023251095.1;refGene_gn=NFATC2;refGene_tx=XM_023251096.1;refGene_gn=NFATC2;refGene_tx=XM_023251098.1;refGene_gn=NFATC2;refGene_tx=XM_023251099.1;refGene_gn=NFATC2;refGene_tx=XM_023251100.1;refGene_gn=NFATC2;refGene_tx=XM_023251101.1;refGene_gn=NFATC2;refGene_tx=XM_023251102.1
A3 10882321 10882359 refGene_gn=NFATC2;refGene_tx=XM_023251098.1
A3 10901250 10901294
refGene_gn=NFATC2;refGene_tx=XM_023251094.1;refGene_gn=NFATC2;refGene_

tx=XM_023251095.1;refGene_gn=NFATC2;refGene_tx=XM_023251096.1
A3 10908234 10908290
refGene_gn=NFATC2;refGene_tx=XM_023251093.1;refGene_gn=NFATC2;refGene_tx=XM_023251100.1;refGene_gn=NFATC2;refGene_tx=XM_023251101.1;refGene_gn=NFATC2;refGene_tx=XM_023251102.1
A3 10908700 10908729 refGene_gn=NFATC2;refGene_tx=XM_023251099.1
A3 11223512 11224298 refGene_gn=KCNG1;refGene_tx=XM_023251103.1
A3 11230380 11231151 refGene_gn=KCNG1;refGene_tx=XM_023251103.1
A3 11276821 11278231 refGene_gn=MOCS3;refGene_tx=XM_003983382.4
A3 11278521 11278682
refGene_gn=DPM1;refGene_tx=XM_003983383.4;refGene_gn=DPM1;refGene_tx=XM_019826360.2
A3 11282078 11282178
refGene_gn=DPM1;refGene_tx=XM_003983383.4;refGene_gn=DPM1;refGene_tx=XM_019826360.2
A3 11287843 11287877
refGene_gn=DPM1;refGene_tx=XM_003983383.4;refGene_gn=DPM1;refGene_tx=XM_019826360.2
A3 11289912 11289989
refGene_gn=DPM1;refGene_tx=XM_003983383.4;refGene_gn=DPM1;refGene_tx=XM_019826360.2
A3 11290073 11290099
refGene_gn=DPM1;refGene_tx=XM_003983383.4;refGene_gn=DPM1;refGene_tx=XM_019826360.2
A3 11292056 11292152
refGene_gn=DPM1;refGene_tx=XM_003983383.4;refGene_gn=DPM1;refGene_tx=XM_019826360.2
A3 11293117 11293127 refGene_gn=DPM1;refGene_tx=XM_019826360.2
A3 11295456 11295525 refGene_gn=DPM1;refGene_tx=XM_003983383.4
A3 11297018 11297133 refGene_gn=DPM1;refGene_tx=XM_003983383.4
A3 11297993 11298098 refGene_gn=DPM1;refGene_tx=XM_003983383.4
A3 11324214 11324322
refGene_gn=ADNP;refGene_tx=XM_003983384.5;refGene_gn=ADNP;refGene_tx=XM_006929663.4;refGene_gn=ADNP;refGene_tx=XM_019826358.2;refGene_gn=ADNP;refGene_tx=XM_023251104.1;refGene_gn=ADNP;refGene_tx=XM_023251105.1
A3 11326123 11326216
refGene_gn=ADNP;refGene_tx=XM_003983384.5;refGene_gn=ADNP;refGene_tx=XM_006929663.4;refGene_gn=ADNP;refGene_tx=XM_019826358.2;refGene_gn=ADNP;refGene_tx=XM_023251104.1;refGene_gn=ADNP;refGene_tx=XM_023251105.1
A3 11328002 11331110
refGene_gn=ADNP;refGene_tx=XM_003983384.5;refGene_gn=ADNP;refGene_tx=XM_006929663.4;refGene_gn=ADNP;refGene_tx=XM_019826358.2;refGene_gn=ADNP;refGene_tx=XM_023251104.1;refGene_gn=ADNP;refGene_tx=XM_023251105.1;refGene_gn=ADNP;refGene_tx=XM_023251106.1
A3 11344364 11344490
refGene_gn=BCAS4;refGene_tx=XM_023251107.1;refGene_gn=BCAS4;refGene_tx=XM_023251108.1;refGene_gn=BCAS4;refGene_tx=XM_023251109.1
A3 11368928 11369053 refGene_gn=BCAS4;refGene_tx=XM_023251108.1
A3 11371150 11371285
refGene_gn=BCAS4;refGene_tx=XM_023251107.1;refGene_gn=BCAS4;refGene_tx

=XM_023251108.1
A3 11379330 11379432
refGene_gn=BCAS4;refGene_tx=XM_023251107.1;refGene_gn=BCAS4;refGene_tx=XM_023251108.1;refGene_gn=BCAS4;refGene_tx=XM_023251109.1
A3 11389376 11389448
refGene_gn=BCAS4;refGene_tx=XM_023251107.1;refGene_gn=BCAS4;refGene_tx=XM_023251108.1;refGene_gn=BCAS4;refGene_tx=XM_023251109.1
A3 11405025 11405205
refGene_gn=BCAS4;refGene_tx=XM_023251107.1;refGene_gn=BCAS4;refGene_tx=XM_023251108.1;refGene_gn=BCAS4;refGene_tx=XM_023251109.1
A3 11433293 11434117 refGene_gn=PARD6B;refGene_tx=XM_023251110.1
A3 11442261 11442484 refGene_gn=PARD6B;refGene_tx=XM_023251110.1
A3 11446895 11446961 refGene_gn=PARD6B;refGene_tx=XM_023251110.1
A3 11554315 11554339
refGene_gn=PTPN1;refGene_tx=XM_003983386.5;refGene_gn=PTPN1;refGene_tx=XM_006929664.4
A3 11555594 11555766
refGene_gn=PTPN1;refGene_tx=XM_003983386.5;refGene_gn=PTPN1;refGene_tx=XM_006929664.4
A3 11558085 11558309
refGene_gn=PTPN1;refGene_tx=XM_003983386.5;refGene_gn=PTPN1;refGene_tx=XM_006929664.4
A3 11558683 11558845
refGene_gn=PTPN1;refGene_tx=XM_003983386.5;refGene_gn=PTPN1;refGene_tx=XM_006929664.4
A3 11559259 11559469
refGene_gn=PTPN1;refGene_tx=XM_003983386.5;refGene_gn=PTPN1;refGene_tx=XM_006929664.4
A3 11562655 11562793
refGene_gn=PTPN1;refGene_tx=XM_003983386.5;refGene_gn=PTPN1;refGene_tx=XM_006929664.4
A3 11568197 11568296
refGene_gn=PTPN1;refGene_tx=XM_003983386.5;refGene_gn=PTPN1;refGene_tx=XM_006929664.4
A3 11573054 11573155
refGene_gn=PTPN1;refGene_tx=XM_003983386.5;refGene_gn=PTPN1;refGene_tx=XM_006929664.4
A3 11576709 11576800 refGene_gn=PTPN1;refGene_tx=XM_003983386.5
A3 11621205 11621268 refGene_gn=PTPN1;refGene_tx=XM_003983386.5
A3 11899997 11901038 refGene_gn=CEBPB;refGene_tx=XM_023251111.1
A3 11928723 11928847
refGene_gn=TMEM189;refGene_tx=XM_011280810.2;refGene_gn=TMEM189;refGene_tx=XM_023251114.1
A3 11932202 11932254 refGene_gn=TMEM189;refGene_tx=XM_023251113.1
A3 11936570 11936690
refGene_gn=TMEM189;refGene_tx=XM_011280810.2;refGene_gn=TMEM189;refGene_tx=XM_023251113.1;refGene_gn=TMEM189;refGene_tx=XM_023251114.1
A3 11943286 11943378
refGene_gn=TMEM189;refGene_tx=XM_011280810.2;refGene_gn=TMEM189;refGene_tx=XM_023251113.1;refGene_gn=TMEM189;refGene_tx=XM_023251114.1

A3 11944532 11944677
refGene_gn=TMEM189;refGene_tx=XM_011280810.2;refGene_gn=TMEM189;refGene_tx=XM_023251113.1;refGene_gn=TMEM189;refGene_tx=XM_023251114.1
A3 11945558 11945758 refGene_gn=TMEM189;refGene_tx=XM_023251114.1
A3 11945984 11946197
refGene_gn=TMEM189;refGene_tx=XM_011280810.2;refGene_gn=TMEM189;refGene_tx=XM_023251113.1
A3 11948114 11948236
refGene_gn=TMEM189;refGene_tx=XM_011280810.2;refGene_gn=TMEM189;refGene_tx=XM_023251113.1
A3 11958748 11958770 refGene_gn=UBE2V1;refGene_tx=XM_003983388.5
A3 11959166 11959197 refGene_gn=UBE2V1;refGene_tx=XM_019826366.2
A3 11960100 11960127 refGene_gn=UBE2V1;refGene_tx=XM_019826367.2
A3 11977290 11977439
refGene_gn=UBE2V1;refGene_tx=XM_003983388.5;refGene_gn=UBE2V1;refGene_tx=XM_019826366.2
A3 11986694 11986820
refGene_gn=UBE2V1;refGene_tx=XM_003983388.5;refGene_gn=UBE2V1;refGene_tx=XM_019826366.2;refGene_gn=UBE2V1;refGene_tx=XM_019826367.2
A3 11988027 11988174
refGene_gn=UBE2V1;refGene_tx=XM_003983388.5;refGene_gn=UBE2V1;refGene_tx=XM_019826366.2;refGene_gn=UBE2V1;refGene_tx=XM_019826367.2
A3 12058870 12059055 refGene_gn=SNAI1;refGene_tx=XM_023251115.1
A3 12063019 12063547 refGene_gn=SNAI1;refGene_tx=XM_023251115.1
A3 12064134 12064216 refGene_gn=SNAI1;refGene_tx=XM_023251115.1
A3 12083463 12083529 refGene_gn=RNF114;refGene_tx=XM_003983392.5
A3 12084809 12084917 refGene_gn=RNF114;refGene_tx=XM_003983392.5
A3 12088421 12088536 refGene_gn=RNF114;refGene_tx=XM_003983392.5
A3 12088881 12088988 refGene_gn=RNF114;refGene_tx=XM_003983392.5
A3 12091347 12091498 refGene_gn=RNF114;refGene_tx=XM_003983392.5
A3 12097295 12097435 refGene_gn=RNF114;refGene_tx=XM_003983392.5
A3 12113843 12114179
refGene_gn=SPATA2;refGene_tx=XM_011280749.2;refGene_gn=SPATA2;refGene_tx=XM_019826369.2
A3 12115002 12116229
refGene_gn=SPATA2;refGene_tx=XM_011280749.2;refGene_gn=SPATA2;refGene_tx=XM_019826369.2
A3 12121105 12121678 refGene_gn=IL-29;refGene_tx=NM_001040680.1
A3 12132911 12133019
refGene_gn=SLC9A8;refGene_tx=XM_006929668.4;refGene_gn=SLC9A8;refGene_tx=XM_011280748.2;refGene_gn=SLC9A8;refGene_tx=XM_019826371.2;refGene_gn=SLC9A8;refGene_tx=XM_019826372.2;refGene_gn=SLC9A8;refGene_tx=XM_019826373.2
A3 12133935 12134082
refGene_gn=SLC9A8;refGene_tx=XM_006929668.4;refGene_gn=SLC9A8;refGene_tx=XM_011280748.2;refGene_gn=SLC9A8;refGene_tx=XM_019826371.2;refGene_gn=SLC9A8;refGene_tx=XM_019826372.2;refGene_gn=SLC9A8;refGene_tx=XM_019826373.2
A3 12135050 12135271
refGene_gn=SLC9A8;refGene_tx=XM_006929668.4;refGene_gn=SLC9A8;refGene_

tx=XM_011280748.2;refGene_gn=SLC9A8;refGene_tx=XM_019826371.2;refGene_gn=SLC9A8;refGene_tx=XM_019826372.2;refGene_gn=SLC9A8;refGene_tx=XM_019826373.2
A3 12138247 12138359
refGene_gn=SLC9A8;refGene_tx=XM_006929668.4;refGene_gn=SLC9A8;refGene_tx=XM_011280748.2;refGene_gn=SLC9A8;refGene_tx=XM_019826371.2;refGene_gn=SLC9A8;refGene_tx=XM_019826373.2
A3 12140912 12140995
refGene_gn=SLC9A8;refGene_tx=XM_006929668.4;refGene_gn=SLC9A8;refGene_tx=XM_011280748.2;refGene_gn=SLC9A8;refGene_tx=XM_019826371.2;refGene_gn=SLC9A8;refGene_tx=XM_019826373.2
A3 12143205 12143322
refGene_gn=SLC9A8;refGene_tx=XM_006929668.4;refGene_gn=SLC9A8;refGene_tx=XM_011280748.2;refGene_gn=SLC9A8;refGene_tx=XM_019826371.2;refGene_gn=SLC9A8;refGene_tx=XM_019826373.2
A3 12151657 12151763
refGene_gn=SLC9A8;refGene_tx=XM_006929668.4;refGene_gn=SLC9A8;refGene_tx=XM_011280748.2;refGene_gn=SLC9A8;refGene_tx=XM_019826371.2;refGene_gn=SLC9A8;refGene_tx=XM_019826372.2;refGene_gn=SLC9A8;refGene_tx=XM_019826373.2
A3 12153221 12153360
refGene_gn=SLC9A8;refGene_tx=XM_006929668.4;refGene_gn=SLC9A8;refGene_tx=XM_011280748.2;refGene_gn=SLC9A8;refGene_tx=XM_019826371.2;refGene_gn=SLC9A8;refGene_tx=XM_019826372.2;refGene_gn=SLC9A8;refGene_tx=XM_019826373.2
A3 12158991 12159135
refGene_gn=SLC9A8;refGene_tx=XM_006929668.4;refGene_gn=SLC9A8;refGene_tx=XM_011280748.2;refGene_gn=SLC9A8;refGene_tx=XM_019826371.2;refGene_gn=SLC9A8;refGene_tx=XM_019826372.2;refGene_gn=SLC9A8;refGene_tx=XM_019826373.2
A3 12164589 12164624
refGene_gn=SLC9A8;refGene_tx=XM_006929668.4;refGene_gn=SLC9A8;refGene_tx=XM_011280748.2;refGene_gn=SLC9A8;refGene_tx=XM_019826371.2;refGene_gn=SLC9A8;refGene_tx=XM_019826372.2;refGene_gn=SLC9A8;refGene_tx=XM_019826373.2
A3 12165749 12165851
refGene_gn=SLC9A8;refGene_tx=XM_006929668.4;refGene_gn=SLC9A8;refGene_tx=XM_011280748.2;refGene_gn=SLC9A8;refGene_tx=XM_019826371.2;refGene_gn=SLC9A8;refGene_tx=XM_019826372.2;refGene_gn=SLC9A8;refGene_tx=XM_019826373.2
A3 12170952 12171036
refGene_gn=SLC9A8;refGene_tx=XM_006929668.4;refGene_gn=SLC9A8;refGene_tx=XM_011280748.2;refGene_gn=SLC9A8;refGene_tx=XM_019826372.2;refGene_gn=SLC9A8;refGene_tx=XM_019826373.2
A3 12175815 12175874
refGene_gn=SLC9A8;refGene_tx=XM_006929668.4;refGene_gn=SLC9A8;refGene_tx=XM_011280748.2;refGene_gn=SLC9A8;refGene_tx=XM_019826371.2;refGene_gn=SLC9A8;refGene_tx=XM_019826372.2;refGene_gn=SLC9A8;refGene_tx=XM_019826373.2
A3 12189170 12189251

refGene_gn=SLC9A8;refGene_tx=XM_006929668.4;refGene_gn=SLC9A8;refGene_tx=XM_011280748.2;refGene_gn=SLC9A8;refGene_tx=XM_019826371.2;refGene_gn=SLC9A8;refGene_tx=XM_019826372.2
A3 12195704 12195883 refGene_gn=SLC9A8;refGene_tx=XM_006929668.4;refGene_gn=SLC9A8;refGene_tx=XM_011280748.2;refGene_gn=SLC9A8;refGene_tx=XM_019826371.2;refGene_gn=SLC9A8;refGene_tx=XM_019826372.2
A3 12197265 12197313 refGene_gn=SLC9A8;refGene_tx=XM_006929668.4;refGene_gn=SLC9A8;refGene_tx=XM_019826371.2;refGene_gn=SLC9A8;refGene_tx=XM_019826372.2
A3 12197766 12197792 refGene_gn=SLC9A8;refGene_tx=XM_006929668.4;refGene_gn=SLC9A8;refGene_tx=XM_011280748.2;refGene_gn=SLC9A8;refGene_tx=XM_019826371.2;refGene_gn=SLC9A8;refGene_tx=XM_019826372.2
A3 12264775 12264890 refGene_gn=B4GALT5;refGene_tx=XM_023251116.1
A3 12319999 12320134 refGene_gn=B4GALT5;refGene_tx=XM_023251116.1
A3 12330065 12330179 refGene_gn=B4GALT5;refGene_tx=XM_023251116.1
A3 12330799 12330924 refGene_gn=B4GALT5;refGene_tx=XM_023251116.1
A3 12331891 12332008 refGene_gn=B4GALT5;refGene_tx=XM_023251116.1
A3 12332599 12332787 refGene_gn=B4GALT5;refGene_tx=XM_023251116.1
A3 12333500 12333623 refGene_gn=B4GALT5;refGene_tx=XM_023251116.1
A3 12335814 12335916 refGene_gn=B4GALT5;refGene_tx=XM_023251116.1
A3 12336867 12337015 refGene_gn=B4GALT5;refGene_tx=XM_023251116.1
A3 12394314 12394388 refGene_gn=PTGIS;refGene_tx=XM_023251117.1;refGene_gn=PTGIS;refGene_tx=XM_023251118.1
A3 12402512 12402636 refGene_gn=PTGIS;refGene_tx=XM_023251117.1;refGene_gn=PTGIS;refGene_tx=XM_023251118.1
A3 12403851 12404030 refGene_gn=PTGIS;refGene_tx=XM_023251117.1;refGene_gn=PTGIS;refGene_tx=XM_023251118.1
A3 12405916 12406060 refGene_gn=PTGIS;refGene_tx=XM_023251117.1;refGene_gn=PTGIS;refGene_tx=XM_023251118.1
A3 12409406 12409558 refGene_gn=PTGIS;refGene_tx=XM_023251117.1;refGene_gn=PTGIS;refGene_tx=XM_023251118.1
A3 12417119 12417301 refGene_gn=PTGIS;refGene_tx=XM_023251117.1;refGene_gn=PTGIS;refGene_tx=XM_023251118.1
A3 12425456 12425625 refGene_gn=PTGIS;refGene_tx=XM_023251117.1;refGene_gn=PTGIS;refGene_tx=XM_023251118.1
A3 12426603 12426785 refGene_gn=PTGIS;refGene_tx=XM_023251117.1;refGene_gn=PTGIS;refGene_tx=XM_023251118.1
A3 12428670 12428822 refGene_gn=PTGIS;refGene_tx=XM_023251117.1;refGene_gn=PTGIS;refGene_tx=XM_023251118.1

A3 12429628 12429752 refGene_gn=PTGIS;refGene_tx=XM_023251117.1
A3 12431303 12431448 refGene_gn=PTGIS;refGene_tx=XM_023251118.1
A3 12444532 12445099 refGene_gn=KCNB1;refGene_tx=XM_003983394.5
A3 12534252 12536259 refGene_gn=KCNB1;refGene_tx=XM_003983394.5
A3 12605666 12605727 refGene_gn=ZNFX1;refGene_tx=XM_023251119.1
A3 12608752 12610561 refGene_gn=ZNFX1;refGene_tx=XM_023251119.1
A3 12615572 12615704 refGene_gn=ZNFX1;refGene_tx=XM_023251119.1
A3 12617128 12617277 refGene_gn=ZNFX1;refGene_tx=XM_023251119.1
A3 12617903 12618053 refGene_gn=ZNFX1;refGene_tx=XM_023251119.1
A3 12619127 12619242 refGene_gn=ZNFX1;refGene_tx=XM_023251119.1
A3 12620533 12620784 refGene_gn=ZNFX1;refGene_tx=XM_023251119.1
A3 12622618 12622758 refGene_gn=ZNFX1;refGene_tx=XM_023251119.1
A3 12624022 12624177 refGene_gn=ZNFX1;refGene_tx=XM_023251119.1
A3 12624545 12624691 refGene_gn=ZNFX1;refGene_tx=XM_023251119.1
A3 12625498 12625609 refGene_gn=ZNFX1;refGene_tx=XM_023251119.1
A3 12626516 12626612 refGene_gn=ZNFX1;refGene_tx=XM_023251119.1
A3 12627500 12629942 refGene_gn=ZNFX1;refGene_tx=XM_023251119.1
A3 12632544 12632563
refGene_gn=DDX27;refGene_tx=XM_011280746.3;refGene_gn=DDX27;refGene_tx
=XM_011280747.3
A3 12632689 12632764
refGene_gn=DDX27;refGene_tx=XM_011280746.3;refGene_gn=DDX27;refGene_tx
=XM_011280747.3
A3 12633770 12633858
refGene_gn=DDX27;refGene_tx=XM_011280746.3;refGene_gn=DDX27;refGene_tx
=XM_011280747.3
A3 12634392 12634516
refGene_gn=DDX27;refGene_tx=XM_011280746.3;refGene_gn=DDX27;refGene_tx
=XM_011280747.3
A3 12634610 12634705
refGene_gn=DDX27;refGene_tx=XM_011280746.3;refGene_gn=DDX27;refGene_tx
=XM_011280747.3
A3 12636468 12636571
refGene_gn=DDX27;refGene_tx=XM_011280746.3;refGene_gn=DDX27;refGene_tx
=XM_011280747.3
A3 12636789 12636896
refGene_gn=DDX27;refGene_tx=XM_011280746.3;refGene_gn=DDX27;refGene_tx
=XM_011280747.3
A3 12638074 12638252
refGene_gn=DDX27;refGene_tx=XM_011280746.3;refGene_gn=DDX27;refGene_tx
=XM_011280747.3
A3 12638351 12638433
refGene_gn=DDX27;refGene_tx=XM_011280746.3;refGene_gn=DDX27;refGene_tx
=XM_011280747.3
A3 12639281 12639435
refGene_gn=DDX27;refGene_tx=XM_011280746.3;refGene_gn=DDX27;refGene_tx
=XM_011280747.3
A3 12640309 12640451
refGene_gn=DDX27;refGene_tx=XM_011280746.3;refGene_gn=DDX27;refGene_tx
=XM_011280747.3

A3	12640622	12640722
refGene_gn=DDX27;refGene_tx=XM_011280746.3;refGene_gn=DDX27;refGene_tx=XM_011280747.3
A3	12643504	12643655
refGene_gn=DDX27;refGene_tx=XM_011280746.3;refGene_gn=DDX27;refGene_tx=XM_011280747.3
A3	12644274	12644448
refGene_gn=DDX27;refGene_tx=XM_011280746.3;refGene_gn=DDX27;refGene_tx=XM_011280747.3
A3	12645916	12646022
refGene_gn=DDX27;refGene_tx=XM_011280746.3;refGene_gn=DDX27;refGene_tx=XM_011280747.3
A3	12646683	12646770
refGene_gn=DDX27;refGene_tx=XM_011280746.3;refGene_gn=DDX27;refGene_tx=XM_011280747.3
A3	12646896	12646937
refGene_gn=DDX27;refGene_tx=XM_011280746.3;refGene_gn=DDX27;refGene_tx=XM_011280747.3
A3	12647481	12647647
refGene_gn=DDX27;refGene_tx=XM_011280746.3;refGene_gn=DDX27;refGene_tx=XM_011280747.3
A3	12647901	12647964
refGene_gn=DDX27;refGene_tx=XM_011280746.3;refGene_gn=DDX27;refGene_tx=XM_011280747.3
A3	12648649	12648796
refGene_gn=DDX27;refGene_tx=XM_011280746.3;refGene_gn=DDX27;refGene_tx=XM_011280747.3
A3	12650752	12650845
refGene_gn=DDX27;refGene_tx=XM_011280746.3;refGene_gn=DDX27;refGene_tx=XM_011280747.3
A3	12707646	12707851
refGene_gn=STAU1;refGene_tx=XM_006929670.2;refGene_gn=STAU1;refGene_tx=XM_023251120.1;refGene_gn=STAU1;refGene_tx=XM_023251121.1;refGene_gn=STAU1;refGene_tx=XM_023251122.1;refGene_gn=STAU1;refGene_tx=XM_023251123.1;refGene_gn=STAU1;refGene_tx=XM_023251124.1;refGene_gn=STAU1;refGene_tx=XM_023251125.1;refGene_gn=STAU1;refGene_tx=XM_023251126.1;refGene_gn=STAU1;refGene_tx=XM_023251127.1;refGene_gn=STAU1;refGene_tx=XM_023251128.1;refGene_gn=STAU1;refGene_tx=XM_023251132.1
A3	12709631	12709756
refGene_gn=STAU1;refGene_tx=XM_006929670.2;refGene_gn=STAU1;refGene_tx=XM_023251120.1;refGene_gn=STAU1;refGene_tx=XM_023251121.1;refGene_gn=STAU1;refGene_tx=XM_023251122.1;refGene_gn=STAU1;refGene_tx=XM_023251123.1;refGene_gn=STAU1;refGene_tx=XM_023251124.1;refGene_gn=STAU1;refGene_tx=XM_023251125.1;refGene_gn=STAU1;refGene_tx=XM_023251126.1;refGene_gn=STAU1;refGene_tx=XM_023251132.1
A3	12712729	12712934
refGene_gn=STAU1;refGene_tx=XM_006929670.2;refGene_gn=STAU1;refGene_tx=XM_023251120.1;refGene_gn=STAU1;refGene_tx=XM_023251121.1;refGene_gn=STAU1;refGene_tx=XM_023251122.1;refGene_gn=STAU1;refGene_tx=XM_023251123.1;refGene_gn=STAU1;refGene_tx=XM_023251124.1;refGene_gn=STAU1;refGe

ne_tx=XM_023251125.1;refGene_gn=STAU1;refGene_tx=XM_023251126.1;refGen
e_gn=STAU1;refGene_tx=XM_023251132.1
A3 12714791 12714930
refGene_gn=STAU1;refGene_tx=XM_006929670.2;refGene_gn=STAU1;refGene_tx
=XM_006929677.3;refGene_gn=STAU1;refGene_tx=XM_023251120.1;refGene_gn=
STAU1;refGene_tx=XM_023251121.1;refGene_gn=STAU1;refGene_tx=XM_0232511
22.1;refGene_gn=STAU1;refGene_tx=XM_023251123.1;refGene_gn=STAU1;refGe
ne_tx=XM_023251124.1;refGene_gn=STAU1;refGene_tx=XM_023251125.1;refGen
e_gn=STAU1;refGene_tx=XM_023251126.1;refGene_gn=STAU1;refGene_tx=XM_02
3251127.1;refGene_gn=STAU1;refGene_tx=XM_023251128.1;refGene_gn=STAU1;
refGene_tx=XM_023251130.1;refGene_gn=STAU1;refGene_tx=XM_023251131.1;r
efGene_gn=STAU1;refGene_tx=XM_023251132.1
A3 12716527 12716714
refGene_gn=STAU1;refGene_tx=XM_006929670.2;refGene_gn=STAU1;refGene_tx
=XM_006929677.3;refGene_gn=STAU1;refGene_tx=XM_023251120.1;refGene_gn=
STAU1;refGene_tx=XM_023251121.1;refGene_gn=STAU1;refGene_tx=XM_0232511
22.1;refGene_gn=STAU1;refGene_tx=XM_023251123.1;refGene_gn=STAU1;refGe
ne_tx=XM_023251124.1;refGene_gn=STAU1;refGene_tx=XM_023251125.1;refGen
e_gn=STAU1;refGene_tx=XM_023251126.1;refGene_gn=STAU1;refGene_tx=XM_02
3251127.1;refGene_gn=STAU1;refGene_tx=XM_023251128.1;refGene_gn=STAU1;
refGene_tx=XM_023251130.1;refGene_gn=STAU1;refGene_tx=XM_023251131.1;r
efGene_gn=STAU1;refGene_tx=XM_023251132.1
A3 12727523 12727622
refGene_gn=STAU1;refGene_tx=XM_006929670.2;refGene_gn=STAU1;refGene_tx
=XM_006929677.3;refGene_gn=STAU1;refGene_tx=XM_023251120.1;refGene_gn=
STAU1;refGene_tx=XM_023251121.1;refGene_gn=STAU1;refGene_tx=XM_0232511
22.1;refGene_gn=STAU1;refGene_tx=XM_023251123.1;refGene_gn=STAU1;refGe
ne_tx=XM_023251124.1;refGene_gn=STAU1;refGene_tx=XM_023251125.1;refGen
e_gn=STAU1;refGene_tx=XM_023251126.1;refGene_gn=STAU1;refGene_tx=XM_02
3251127.1;refGene_gn=STAU1;refGene_tx=XM_023251128.1;refGene_gn=STAU1;
refGene_tx=XM_023251130.1;refGene_gn=STAU1;refGene_tx=XM_023251131.1;r
efGene_gn=STAU1;refGene_tx=XM_023251132.1
A3 12735960 12736173
refGene_gn=STAU1;refGene_tx=XM_006929670.2;refGene_gn=STAU1;refGene_tx
=XM_006929677.3;refGene_gn=STAU1;refGene_tx=XM_023251120.1;refGene_gn=
STAU1;refGene_tx=XM_023251121.1;refGene_gn=STAU1;refGene_tx=XM_0232511
22.1;refGene_gn=STAU1;refGene_tx=XM_023251123.1;refGene_gn=STAU1;refGe
ne_tx=XM_023251124.1;refGene_gn=STAU1;refGene_tx=XM_023251125.1;refGen
e_gn=STAU1;refGene_tx=XM_023251126.1;refGene_gn=STAU1;refGene_tx=XM_02
3251127.1;refGene_gn=STAU1;refGene_tx=XM_023251128.1;refGene_gn=STAU1;
refGene_tx=XM_023251130.1;refGene_gn=STAU1;refGene_tx=XM_023251131.1;r
efGene_gn=STAU1;refGene_tx=XM_023251132.1
A3 12737683 12737827
refGene_gn=STAU1;refGene_tx=XM_006929670.2;refGene_gn=STAU1;refGene_tx
=XM_006929677.3;refGene_gn=STAU1;refGene_tx=XM_023251120.1;refGene_gn=
STAU1;refGene_tx=XM_023251121.1;refGene_gn=STAU1;refGene_tx=XM_0232511
22.1;refGene_gn=STAU1;refGene_tx=XM_023251123.1;refGene_gn=STAU1;refGe
ne_tx=XM_023251124.1;refGene_gn=STAU1;refGene_tx=XM_023251125.1;refGen
e_gn=STAU1;refGene_tx=XM_023251126.1;refGene_gn=STAU1;refGene_tx=XM_02
3251127.1;refGene_gn=STAU1;refGene_tx=XM_023251128.1;refGene_gn=STAU1;

refGene_tx=XM_023251130.1;refGene_gn=STAU1;refGene_tx=XM_023251131.1;refGene_gn=STAU1;refGene_tx=XM_023251132.1
A3 12739668 12739815
refGene_gn=STAU1;refGene_tx=XM_006929670.2;refGene_gn=STAU1;refGene_tx=XM_006929677.3;refGene_gn=STAU1;refGene_tx=XM_023251120.1;refGene_gn=STAU1;refGene_tx=XM_023251121.1;refGene_gn=STAU1;refGene_tx=XM_023251122.1;refGene_gn=STAU1;refGene_tx=XM_023251123.1;refGene_gn=STAU1;refGene_tx=XM_023251124.1;refGene_gn=STAU1;refGene_tx=XM_023251125.1;refGene_gn=STAU1;refGene_tx=XM_023251126.1;refGene_gn=STAU1;refGene_tx=XM_023251127.1;refGene_gn=STAU1;refGene_tx=XM_023251128.1;refGene_gn=STAU1;refGene_tx=XM_023251130.1;refGene_gn=STAU1;refGene_tx=XM_023251131.1;refGene_gn=STAU1;refGene_tx=XM_023251132.1
A3 12741042 12741118
refGene_gn=STAU1;refGene_tx=XM_006929670.2;refGene_gn=STAU1;refGene_tx=XM_006929677.3;refGene_gn=STAU1;refGene_tx=XM_023251120.1;refGene_gn=STAU1;refGene_tx=XM_023251121.1;refGene_gn=STAU1;refGene_tx=XM_023251122.1;refGene_gn=STAU1;refGene_tx=XM_023251123.1;refGene_gn=STAU1;refGene_tx=XM_023251124.1;refGene_gn=STAU1;refGene_tx=XM_023251125.1;refGene_gn=STAU1;refGene_tx=XM_023251126.1;refGene_gn=STAU1;refGene_tx=XM_023251127.1;refGene_gn=STAU1;refGene_tx=XM_023251128.1;refGene_gn=STAU1;refGene_tx=XM_023251130.1;refGene_gn=STAU1;refGene_tx=XM_023251131.1;refGene_gn=STAU1;refGene_tx=XM_023251132.1
A3 12741313 12741633
refGene_gn=STAU1;refGene_tx=XM_006929670.2;refGene_gn=STAU1;refGene_tx=XM_006929677.3;refGene_gn=STAU1;refGene_tx=XM_023251120.1;refGene_gn=STAU1;refGene_tx=XM_023251121.1;refGene_gn=STAU1;refGene_tx=XM_023251122.1;refGene_gn=STAU1;refGene_tx=XM_023251123.1;refGene_gn=STAU1;refGene_tx=XM_023251124.1;refGene_gn=STAU1;refGene_tx=XM_023251125.1;refGene_gn=STAU1;refGene_tx=XM_023251126.1;refGene_gn=STAU1;refGene_tx=XM_023251127.1;refGene_gn=STAU1;refGene_tx=XM_023251128.1;refGene_gn=STAU1;refGene_tx=XM_023251130.1;refGene_gn=STAU1;refGene_tx=XM_023251131.1;refGene_gn=STAU1;refGene_tx=XM_023251132.1
A3 12742225 12742348
refGene_gn=STAU1;refGene_tx=XM_006929670.2;refGene_gn=STAU1;refGene_tx=XM_006929677.3;refGene_gn=STAU1;refGene_tx=XM_023251120.1;refGene_gn=STAU1;refGene_tx=XM_023251121.1;refGene_gn=STAU1;refGene_tx=XM_023251122.1;refGene_gn=STAU1;refGene_tx=XM_023251123.1;refGene_gn=STAU1;refGene_tx=XM_023251124.1;refGene_gn=STAU1;refGene_tx=XM_023251125.1;refGene_gn=STAU1;refGene_tx=XM_023251126.1;refGene_gn=STAU1;refGene_tx=XM_023251127.1;refGene_gn=STAU1;refGene_tx=XM_023251128.1;refGene_gn=STAU1;refGene_tx=XM_023251130.1;refGene_gn=STAU1;refGene_tx=XM_023251131.1;refGene_gn=STAU1;refGene_tx=XM_023251132.1
A3 12743060 12743150
refGene_gn=STAU1;refGene_tx=XM_006929670.2;refGene_gn=STAU1;refGene_tx=XM_006929677.3;refGene_gn=STAU1;refGene_tx=XM_023251120.1;refGene_gn=STAU1;refGene_tx=XM_023251122.1;refGene_gn=STAU1;refGene_tx=XM_023251123.1;refGene_gn=STAU1;refGene_tx=XM_023251124.1;refGene_gn=STAU1;refGene_tx=XM_023251127.1;refGene_gn=STAU1;refGene_tx=XM_023251128.1;refGene_gn=STAU1;refGene_tx=XM_023251130.1;refGene_gn=STAU1;refGene_tx=XM_023251131.1

A3 12743931 12744015
refGene_gn=STAU1;refGene_tx=XM_006929670.2;refGene_gn=STAU1;refGene_tx=XM_006929677.3;refGene_gn=STAU1;refGene_tx=XM_023251120.1;refGene_gn=STAU1;refGene_tx=XM_023251121.1;refGene_gn=STAU1;refGene_tx=XM_023251122.1;refGene_gn=STAU1;refGene_tx=XM_023251123.1;refGene_gn=STAU1;refGene_tx=XM_023251127.1;refGene_gn=STAU1;refGene_tx=XM_023251128.1;refGene_gn=STAU1;refGene_tx=XM_023251130.1;refGene_gn=STAU1;refGene_tx=XM_023251131.1
A3 12755689 12755779
refGene_gn=CSE1L;refGene_tx=XM_006929678.4;refGene_gn=CSE1L;refGene_tx=XM_019826376.2
A3 12757156 12757388
refGene_gn=CSE1L;refGene_tx=XM_006929678.4;refGene_gn=CSE1L;refGene_tx=XM_019826376.2
A3 12758450 12758597
refGene_gn=CSE1L;refGene_tx=XM_006929678.4;refGene_gn=CSE1L;refGene_tx=XM_019826376.2
A3 12759958 12760040
refGene_gn=CSE1L;refGene_tx=XM_006929678.4;refGene_gn=CSE1L;refGene_tx=XM_019826376.2
A3 12760421 12760507
refGene_gn=CSE1L;refGene_tx=XM_006929678.4;refGene_gn=CSE1L;refGene_tx=XM_019826376.2
A3 12760608 12760706
refGene_gn=CSE1L;refGene_tx=XM_006929678.4;refGene_gn=CSE1L;refGene_tx=XM_019826376.2
A3 12761085 12761294
refGene_gn=CSE1L;refGene_tx=XM_006929678.4;refGene_gn=CSE1L;refGene_tx=XM_019826376.2
A3 12761656 12761807
refGene_gn=CSE1L;refGene_tx=XM_006929678.4;refGene_gn=CSE1L;refGene_tx=XM_019826376.2
A3 12763943 12764041
refGene_gn=CSE1L;refGene_tx=XM_006929678.4;refGene_gn=CSE1L;refGene_tx=XM_019826376.2
A3 12765695 12765799
refGene_gn=CSE1L;refGene_tx=XM_006929678.4;refGene_gn=CSE1L;refGene_tx=XM_019826376.2
A3 12766932 12767069
refGene_gn=CSE1L;refGene_tx=XM_006929678.4;refGene_gn=CSE1L;refGene_tx=XM_019826376.2
A3 12768233 12768295
refGene_gn=CSE1L;refGene_tx=XM_006929678.4;refGene_gn=CSE1L;refGene_tx=XM_019826376.2
A3 12770668 12770753
refGene_gn=CSE1L;refGene_tx=XM_006929678.4;refGene_gn=CSE1L;refGene_tx=XM_019826376.2
A3 12771471 12771674
refGene_gn=CSE1L;refGene_tx=XM_006929678.4;refGene_gn=CSE1L;refGene_tx=XM_019826376.2

A3 12772065 12772131
refGene_gn=CSE1L;refGene_tx=XM_006929678.4;refGene_gn=CSE1L;refGene_tx
=XM_019826376.2
A3 12774206 12774336
refGene_gn=CSE1L;refGene_tx=XM_006929678.4;refGene_gn=CSE1L;refGene_tx
=XM_019826376.2
A3 12774462 12774630
refGene_gn=CSE1L;refGene_tx=XM_006929678.4;refGene_gn=CSE1L;refGene_tx
=XM_019826376.2
A3 12776998 12777091
refGene_gn=CSE1L;refGene_tx=XM_006929678.4;refGene_gn=CSE1L;refGene_tx
=XM_019826376.2
A3 12778849 12778957
refGene_gn=CSE1L;refGene_tx=XM_006929678.4;refGene_gn=CSE1L;refGene_tx
=XM_019826376.2
A3 12780331 12780412 refGene_gn=CSE1L;refGene_tx=XM_019826376.2
A3 12781068 12781159
refGene_gn=CSE1L;refGene_tx=XM_006929678.4;refGene_gn=CSE1L;refGene_tx
=XM_019826376.2
A3 12781719 12781865
refGene_gn=CSE1L;refGene_tx=XM_006929678.4;refGene_gn=CSE1L;refGene_tx
=XM_019826376.2
A3 12781943 12782045
refGene_gn=CSE1L;refGene_tx=XM_006929678.4;refGene_gn=CSE1L;refGene_tx
=XM_019826376.2
A3 12784108 12784251
refGene_gn=CSE1L;refGene_tx=XM_006929678.4;refGene_gn=CSE1L;refGene_tx
=XM_019826376.2
A3 12789211 12789302
refGene_gn=CSE1L;refGene_tx=XM_006929678.4;refGene_gn=CSE1L;refGene_tx
=XM_019826376.2
A3 12816973 12817150 refGene_gn=ARFGEF2;refGene_tx=XM_023251133.1
A3 12817715 12817833 refGene_gn=ARFGEF2;refGene_tx=XM_023251133.1
A3 12821602 12821741 refGene_gn=ARFGEF2;refGene_tx=XM_023251133.1
A3 12822634 12822803 refGene_gn=ARFGEF2;refGene_tx=XM_023251133.1
A3 12827107 12827238 refGene_gn=ARFGEF2;refGene_tx=XM_023251133.1
A3 12829523 12829638 refGene_gn=ARFGEF2;refGene_tx=XM_023251133.1
A3 12831097 12831152 refGene_gn=ARFGEF2;refGene_tx=XM_023251133.1
A3 12831260 12831399 refGene_gn=ARFGEF2;refGene_tx=XM_023251133.1
A3 12833431 12833567 refGene_gn=ARFGEF2;refGene_tx=XM_023251133.1
A3 12836611 12836741 refGene_gn=ARFGEF2;refGene_tx=XM_023251133.1
A3 12836882 12837013 refGene_gn=ARFGEF2;refGene_tx=XM_023251133.1
A3 12838751 12838912 refGene_gn=ARFGEF2;refGene_tx=XM_023251133.1
A3 12840907 12841080 refGene_gn=ARFGEF2;refGene_tx=XM_023251133.1
A3 12846409 12846561 refGene_gn=ARFGEF2;refGene_tx=XM_023251133.1
A3 12851260 12851430 refGene_gn=ARFGEF2;refGene_tx=XM_023251133.1
A3 12851534 12851575 refGene_gn=ARFGEF2;refGene_tx=XM_023251133.1
A3 12856987 12857087 refGene_gn=ARFGEF2;refGene_tx=XM_023251133.1
A3 12858228 12858376 refGene_gn=ARFGEF2;refGene_tx=XM_023251133.1
A3 12862461 12862620 refGene_gn=ARFGEF2;refGene_tx=XM_023251133.1

A3	12863908	12864037	refGene_gn=ARFGEF2;refGene_tx=XM_023251133.1
A3	12865084	12865236	refGene_gn=ARFGEF2;refGene_tx=XM_023251133.1
A3	12865793	12865965	refGene_gn=ARFGEF2;refGene_tx=XM_023251133.1
A3	12866066	12866151	refGene_gn=ARFGEF2;refGene_tx=XM_023251133.1
A3	12868674	12868880	refGene_gn=ARFGEF2;refGene_tx=XM_023251133.1
A3	12869391	12869503	refGene_gn=ARFGEF2;refGene_tx=XM_023251133.1
A3	12871734	12871918	refGene_gn=ARFGEF2;refGene_tx=XM_023251133.1
A3	12872566	12872675	refGene_gn=ARFGEF2;refGene_tx=XM_023251133.1
A3	12874177	12874317	refGene_gn=ARFGEF2;refGene_tx=XM_023251133.1
A3	12875065	12875165	refGene_gn=ARFGEF2;refGene_tx=XM_023251133.1
A3	12875811	12876046	refGene_gn=ARFGEF2;refGene_tx=XM_023251133.1
A3	12877198	12877329	refGene_gn=ARFGEF2;refGene_tx=XM_023251133.1
A3	12879952	12880104	refGene_gn=ARFGEF2;refGene_tx=XM_023251133.1
A3	12881983	12882052	refGene_gn=ARFGEF2;refGene_tx=XM_023251133.1
A3	12891194	12891429	refGene_gn=ARFGEF2;refGene_tx=XM_023251133.1
A3	12891890	12892070	refGene_gn=ARFGEF2;refGene_tx=XM_023251133.1
A3	12893023	12893170	refGene_gn=ARFGEF2;refGene_tx=XM_023251133.1
A3	12903426	12903550	refGene_gn=ARFGEF2;refGene_tx=XM_023251133.1
A3	12904189	12904220	refGene_gn=ARFGEF2;refGene_tx=XM_023251133.1
A3	12918023	12918144	refGene_gn=ARFGEF2;refGene_tx=XM_023251133.1
A3	12980701	12980914	refGene_gn=PREX1;refGene_tx=XM_023251134.1
A3	13006562	13006577	refGene_gn=PREX1;refGene_tx=XM_023251136.1
A3	13060167	13060239
refGene_gn=PREX1;refGene_tx=XM_023251134.1;refGene_gn=PREX1;refGene_tx
=XM_023251136.1
A3	13065099	13065222
refGene_gn=PREX1;refGene_tx=XM_023251134.1;refGene_gn=PREX1;refGene_tx
=XM_023251136.1
A3	13071997	13072102
refGene_gn=PREX1;refGene_tx=XM_023251134.1;refGene_gn=PREX1;refGene_tx
=XM_023251136.1
A3	13077947	13078049
refGene_gn=PREX1;refGene_tx=XM_023251134.1;refGene_gn=PREX1;refGene_tx
=XM_023251136.1
A3	13086930	13087092
refGene_gn=PREX1;refGene_tx=XM_023251134.1;refGene_gn=PREX1;refGene_tx
=XM_023251136.1
A3	13093833	13093967
refGene_gn=PREX1;refGene_tx=XM_023251134.1;refGene_gn=PREX1;refGene_tx
=XM_023251136.1
A3	13100814	13100933
refGene_gn=PREX1;refGene_tx=XM_023251134.1;refGene_gn=PREX1;refGene_tx
=XM_023251136.1
A3	13101972	13102122
refGene_gn=PREX1;refGene_tx=XM_023251134.1;refGene_gn=PREX1;refGene_tx
=XM_023251136.1
A3	13104130	13104278
refGene_gn=PREX1;refGene_tx=XM_023251134.1;refGene_gn=PREX1;refGene_tx
=XM_023251136.1
A3	13111725	13111826

refGene_gn=PREX1;refGene_tx=XM_023251134.1;refGene_gn=PREX1;refGene_tx=XM_023251136.1
A3 13111953 13112057
refGene_gn=PREX1;refGene_tx=XM_023251134.1;refGene_gn=PREX1;refGene_tx=XM_023251136.1
A3 13112254 13112304
refGene_gn=PREX1;refGene_tx=XM_023251134.1;refGene_gn=PREX1;refGene_tx=XM_023251136.1
A3 13113871 13113947
refGene_gn=PREX1;refGene_tx=XM_023251134.1;refGene_gn=PREX1;refGene_tx=XM_023251136.1
A3 13122646 13122719
refGene_gn=PREX1;refGene_tx=XM_023251134.1;refGene_gn=PREX1;refGene_tx=XM_023251136.1
A3 13127509 13127652
refGene_gn=PREX1;refGene_tx=XM_023251134.1;refGene_gn=PREX1;refGene_tx=XM_023251136.1
A3 13129303 13129396
refGene_gn=PREX1;refGene_tx=XM_023251134.1;refGene_gn=PREX1;refGene_tx=XM_023251136.1
A3 13132174 13132323
refGene_gn=PREX1;refGene_tx=XM_023251134.1;refGene_gn=PREX1;refGene_tx=XM_023251136.1
A3 13134009 13134095
refGene_gn=PREX1;refGene_tx=XM_023251134.1;refGene_gn=PREX1;refGene_tx=XM_023251136.1
A3 13136817 13136954
refGene_gn=PREX1;refGene_tx=XM_023251134.1;refGene_gn=PREX1;refGene_tx=XM_023251136.1
A3 13137639 13137733
refGene_gn=PREX1;refGene_tx=XM_023251134.1;refGene_gn=PREX1;refGene_tx=XM_023251136.1
A3 13140419 13140607
refGene_gn=PREX1;refGene_tx=XM_023251134.1;refGene_gn=PREX1;refGene_tx=XM_023251136.1
A3 13140876 13141035
refGene_gn=PREX1;refGene_tx=XM_023251134.1;refGene_gn=PREX1;refGene_tx=XM_023251136.1
A3 13141691 13141902
refGene_gn=PREX1;refGene_tx=XM_023251134.1;refGene_gn=PREX1;refGene_tx=XM_023251136.1
A3 13142330 13142607
refGene_gn=PREX1;refGene_tx=XM_023251134.1;refGene_gn=PREX1;refGene_tx=XM_023251136.1
A3 13145028 13145235
refGene_gn=PREX1;refGene_tx=XM_023251134.1;refGene_gn=PREX1;refGene_tx=XM_023251136.1
A3 13146528 13146617
refGene_gn=PREX1;refGene_tx=XM_023251134.1;refGene_gn=PREX1;refGene_tx=XM_023251136.1

A3 13147897 13147980
refGene_gn=PREX1;refGene_tx=XM_023251134.1;refGene_gn=PREX1;refGene_tx
=XM_023251136.1
A3 13148099 13148190
refGene_gn=PREX1;refGene_tx=XM_023251134.1;refGene_gn=PREX1;refGene_tx
=XM_023251136.1
A3 13150651 13150780
refGene_gn=PREX1;refGene_tx=XM_023251134.1;refGene_gn=PREX1;refGene_tx
=XM_023251136.1
A3 13152448 13152490
refGene_gn=PREX1;refGene_tx=XM_023251134.1;refGene_gn=PREX1;refGene_tx
=XM_023251136.1
A3 13153545 13153766
refGene_gn=PREX1;refGene_tx=XM_023251134.1;refGene_gn=PREX1;refGene_tx
=XM_023251136.1
A3 13156204 13156304
refGene_gn=PREX1;refGene_tx=XM_023251134.1;refGene_gn=PREX1;refGene_tx
=XM_023251136.1
A3 13157848 13157992
refGene_gn=PREX1;refGene_tx=XM_023251134.1;refGene_gn=PREX1;refGene_tx
=XM_023251136.1
A3 13158090 13158205
refGene_gn=PREX1;refGene_tx=XM_023251134.1;refGene_gn=PREX1;refGene_tx
=XM_023251136.1
A3 13159892 13159959
refGene_gn=PREX1;refGene_tx=XM_023251134.1;refGene_gn=PREX1;refGene_tx
=XM_023251136.1
A3 13161255 13161428
refGene_gn=PREX1;refGene_tx=XM_023251134.1;refGene_gn=PREX1;refGene_tx
=XM_023251136.1
A3 13162882 13162985
refGene_gn=PREX1;refGene_tx=XM_023251134.1;refGene_gn=PREX1;refGene_tx
=XM_023251136.1
A3 13163208 13163276
refGene_gn=PREX1;refGene_tx=XM_023251134.1;refGene_gn=PREX1;refGene_tx
=XM_023251136.1
A3 13164150 13164193
refGene_gn=PREX1;refGene_tx=XM_023251134.1;refGene_gn=PREX1;refGene_tx
=XM_023251136.1
A3 13487442 13488193
refGene_gn=LOC109497960;refGene_tx=XM_019826388.2;refGene_gn=LOC109497
960;refGene_tx=XM_023251137.1
A3 13492518 13492616
refGene_gn=LOC109497960;refGene_tx=XM_019826388.2;refGene_gn=LOC109497
960;refGene_tx=XM_023251137.1
A3 13493709 13493772
refGene_gn=LOC109497960;refGene_tx=XM_019826388.2;refGene_gn=LOC109497
960;refGene_tx=XM_023251137.1
A3 13723959 13724134
refGene_gn=SULF2;refGene_tx=XM_023251149.1;refGene_gn=SULF2;refGene_tx

=XM_023251150.1
A3 13744292 13744532
refGene_gn=SULF2;refGene_tx=XM_023251149.1;refGene_gn=SULF2;refGene_tx
=XM_023251150.1
A3 13774589 13774741
refGene_gn=SULF2;refGene_tx=XM_023251149.1;refGene_gn=SULF2;refGene_tx
=XM_023251150.1
A3 13786188 13786358
refGene_gn=SULF2;refGene_tx=XM_023251149.1;refGene_gn=SULF2;refGene_tx
=XM_023251150.1
A3 13791097 13791248
refGene_gn=SULF2;refGene_tx=XM_023251149.1;refGene_gn=SULF2;refGene_tx
=XM_023251150.1
A3 13792447 13792623
refGene_gn=SULF2;refGene_tx=XM_023251149.1;refGene_gn=SULF2;refGene_tx
=XM_023251150.1
A3 13796218 13796347
refGene_gn=SULF2;refGene_tx=XM_023251149.1;refGene_gn=SULF2;refGene_tx
=XM_023251150.1
A3 13797315 13797372
refGene_gn=SULF2;refGene_tx=XM_023251149.1;refGene_gn=SULF2;refGene_tx
=XM_023251150.1
A3 13797866 13797996
refGene_gn=SULF2;refGene_tx=XM_023251149.1;refGene_gn=SULF2;refGene_tx
=XM_023251150.1
A3 13799380 13799573
refGene_gn=SULF2;refGene_tx=XM_023251149.1;refGene_gn=SULF2;refGene_tx
=XM_023251150.1
A3 13806840 13807069
refGene_gn=SULF2;refGene_tx=XM_023251149.1;refGene_gn=SULF2;refGene_tx
=XM_023251150.1
A3 13807191 13807288
refGene_gn=SULF2;refGene_tx=XM_023251149.1;refGene_gn=SULF2;refGene_tx
=XM_023251150.1
A3 13807639 13807734
refGene_gn=SULF2;refGene_tx=XM_023251149.1;refGene_gn=SULF2;refGene_tx
=XM_023251150.1
A3 13808701 13808761
refGene_gn=SULF2;refGene_tx=XM_023251149.1;refGene_gn=SULF2;refGene_tx
=XM_023251150.1
A3 13809103 13809273
refGene_gn=SULF2;refGene_tx=XM_023251149.1;refGene_gn=SULF2;refGene_tx
=XM_023251150.1
A3 13809449 13809592
refGene_gn=SULF2;refGene_tx=XM_023251149.1;refGene_gn=SULF2;refGene_tx
=XM_023251150.1
A3 13810716 13810840
refGene_gn=SULF2;refGene_tx=XM_023251149.1;refGene_gn=SULF2;refGene_tx
=XM_023251150.1
A3 13811511 13811545

refGene_gn=SULF2;refGene_tx=XM_023251149.1;refGene_gn=SULF2;refGene_tx=XM_023251150.1
A3 13811745 13811799
refGene_gn=SULF2;refGene_tx=XM_023251149.1;refGene_gn=SULF2;refGene_tx=XM_023251150.1
A3 13812677 13812708
refGene_gn=SULF2;refGene_tx=XM_023251149.1;refGene_gn=SULF2;refGene_tx=XM_023251150.1
A3 13817531 13817543
refGene_gn=NCOA3;refGene_tx=XM_019826389.2;refGene_gn=NCOA3;refGene_tx=XM_019826390.2;refGene_gn=NCOA3;refGene_tx=XM_023251138.1;refGene_gn=NCOA3;refGene_tx=XM_023251139.1;refGene_gn=NCOA3;refGene_tx=XM_023251140.1;refGene_gn=NCOA3;refGene_tx=XM_023251141.1;refGene_gn=NCOA3;refGene_tx=XM_023251142.1;refGene_gn=NCOA3;refGene_tx=XM_023251143.1;refGene_gn=NCOA3;refGene_tx=XM_023251144.1;refGene_gn=NCOA3;refGene_tx=XM_023251145.1;refGene_gn=NCOA3;refGene_tx=XM_023251146.1;refGene_gn=NCOA3;refGene_tx=XM_023251147.1
A3 13817848 13817990
refGene_gn=NCOA3;refGene_tx=XM_019826389.2;refGene_gn=NCOA3;refGene_tx=XM_019826390.2;refGene_gn=NCOA3;refGene_tx=XM_023251138.1;refGene_gn=NCOA3;refGene_tx=XM_023251139.1;refGene_gn=NCOA3;refGene_tx=XM_023251140.1;refGene_gn=NCOA3;refGene_tx=XM_023251141.1;refGene_gn=NCOA3;refGene_tx=XM_023251142.1;refGene_gn=NCOA3;refGene_tx=XM_023251143.1;refGene_gn=NCOA3;refGene_tx=XM_023251144.1;refGene_gn=NCOA3;refGene_tx=XM_023251145.1;refGene_gn=NCOA3;refGene_tx=XM_023251146.1;refGene_gn=NCOA3;refGene_tx=XM_023251147.1
A3 13818336 13818511
refGene_gn=NCOA3;refGene_tx=XM_019826389.2;refGene_gn=NCOA3;refGene_tx=XM_019826390.2;refGene_gn=NCOA3;refGene_tx=XM_023251138.1;refGene_gn=NCOA3;refGene_tx=XM_023251139.1;refGene_gn=NCOA3;refGene_tx=XM_023251140.1;refGene_gn=NCOA3;refGene_tx=XM_023251141.1;refGene_gn=NCOA3;refGene_tx=XM_023251142.1;refGene_gn=NCOA3;refGene_tx=XM_023251143.1;refGene_gn=NCOA3;refGene_tx=XM_023251144.1;refGene_gn=NCOA3;refGene_tx=XM_023251145.1;refGene_gn=NCOA3;refGene_tx=XM_023251146.1;refGene_gn=NCOA3;refGene_tx=XM_023251147.1
A3 13819593 13819837
refGene_gn=NCOA3;refGene_tx=XM_019826389.2;refGene_gn=NCOA3;refGene_tx=XM_019826390.2;refGene_gn=NCOA3;refGene_tx=XM_023251138.1;refGene_gn=NCOA3;refGene_tx=XM_023251139.1;refGene_gn=NCOA3;refGene_tx=XM_023251140.1;refGene_gn=NCOA3;refGene_tx=XM_023251141.1;refGene_gn=NCOA3;refGene_tx=XM_023251142.1;refGene_gn=NCOA3;refGene_tx=XM_023251143.1;refGene_gn=NCOA3;refGene_tx=XM_023251144.1;refGene_gn=NCOA3;refGene_tx=XM_023251145.1;refGene_gn=NCOA3;refGene_tx=XM_023251146.1;refGene_gn=NCOA3;refGene_tx=XM_023251147.1
A3 13820719 13820791 refGene_gn=NCOA3;refGene_tx=XM_023251148.1
A3 13821056 13821161
refGene_gn=NCOA3;refGene_tx=XM_019826389.2;refGene_gn=NCOA3;refGene_tx=XM_019826390.2;refGene_gn=NCOA3;refGene_tx=XM_023251138.1;refGene_gn=NCOA3;refGene_tx=XM_023251139.1;refGene_gn=NCOA3;refGene_tx=XM_023251140.1;refGene_gn=NCOA3;refGene_tx=XM_023251141.1;refGene_gn=NCOA3;refGe

ne_tx=XM_023251142.1;refGene_gn=NCOA3;refGene_tx=XM_023251143.1;refGen
e_gn=NCOA3;refGene_tx=XM_023251144.1;refGene_gn=NCOA3;refGene_tx=XM_02
3251145.1;refGene_gn=NCOA3;refGene_tx=XM_023251146.1;refGene_gn=NCOA3;
refGene_tx=XM_023251147.1;refGene_gn=NCOA3;refGene_tx=XM_023251148.1
A3 13823030 13823324
refGene_gn=NCOA3;refGene_tx=XM_019826389.2;refGene_gn=NCOA3;refGene_tx
=XM_019826390.2;refGene_gn=NCOA3;refGene_tx=XM_023251138.1;refGene_gn=
NCOA3;refGene_tx=XM_023251139.1;refGene_gn=NCOA3;refGene_tx=XM_0232511
40.1;refGene_gn=NCOA3;refGene_tx=XM_023251141.1;refGene_gn=NCOA3;refGe
ne_tx=XM_023251142.1;refGene_gn=NCOA3;refGene_tx=XM_023251143.1;refGen
e_gn=NCOA3;refGene_tx=XM_023251144.1;refGene_gn=NCOA3;refGene_tx=XM_02
3251145.1;refGene_gn=NCOA3;refGene_tx=XM_023251146.1;refGene_gn=NCOA3;
refGene_tx=XM_023251147.1;refGene_gn=NCOA3;refGene_tx=XM_023251148.1
A3 13829767 13829939
refGene_gn=NCOA3;refGene_tx=XM_019826389.2;refGene_gn=NCOA3;refGene_tx
=XM_019826390.2;refGene_gn=NCOA3;refGene_tx=XM_023251138.1;refGene_gn=
NCOA3;refGene_tx=XM_023251139.1;refGene_gn=NCOA3;refGene_tx=XM_0232511
40.1;refGene_gn=NCOA3;refGene_tx=XM_023251141.1;refGene_gn=NCOA3;refGe
ne_tx=XM_023251142.1;refGene_gn=NCOA3;refGene_tx=XM_023251143.1;refGen
e_gn=NCOA3;refGene_tx=XM_023251144.1;refGene_gn=NCOA3;refGene_tx=XM_02
3251145.1;refGene_gn=NCOA3;refGene_tx=XM_023251146.1;refGene_gn=NCOA3;
refGene_tx=XM_023251147.1;refGene_gn=NCOA3;refGene_tx=XM_023251148.1
A3 13830974 13831101
refGene_gn=NCOA3;refGene_tx=XM_019826389.2;refGene_gn=NCOA3;refGene_tx
=XM_019826390.2;refGene_gn=NCOA3;refGene_tx=XM_023251138.1;refGene_gn=
NCOA3;refGene_tx=XM_023251139.1;refGene_gn=NCOA3;refGene_tx=XM_0232511
40.1;refGene_gn=NCOA3;refGene_tx=XM_023251141.1;refGene_gn=NCOA3;refGe
ne_tx=XM_023251142.1;refGene_gn=NCOA3;refGene_tx=XM_023251143.1;refGen
e_gn=NCOA3;refGene_tx=XM_023251144.1;refGene_gn=NCOA3;refGene_tx=XM_02
3251145.1;refGene_gn=NCOA3;refGene_tx=XM_023251146.1;refGene_gn=NCOA3;
refGene_tx=XM_023251147.1;refGene_gn=NCOA3;refGene_tx=XM_023251148.1
A3 13831199 13831457
refGene_gn=NCOA3;refGene_tx=XM_019826389.2;refGene_gn=NCOA3;refGene_tx
=XM_019826390.2;refGene_gn=NCOA3;refGene_tx=XM_023251138.1;refGene_gn=
NCOA3;refGene_tx=XM_023251139.1;refGene_gn=NCOA3;refGene_tx=XM_0232511
40.1;refGene_gn=NCOA3;refGene_tx=XM_023251141.1;refGene_gn=NCOA3;refGe
ne_tx=XM_023251142.1;refGene_gn=NCOA3;refGene_tx=XM_023251143.1;refGen
e_gn=NCOA3;refGene_tx=XM_023251144.1;refGene_gn=NCOA3;refGene_tx=XM_02
3251145.1;refGene_gn=NCOA3;refGene_tx=XM_023251146.1;refGene_gn=NCOA3;
refGene_tx=XM_023251147.1;refGene_gn=NCOA3;refGene_tx=XM_023251148.1
A3 13832399 13832597
refGene_gn=NCOA3;refGene_tx=XM_019826389.2;refGene_gn=NCOA3;refGene_tx
=XM_019826390.2;refGene_gn=NCOA3;refGene_tx=XM_023251138.1;refGene_gn=
NCOA3;refGene_tx=XM_023251139.1;refGene_gn=NCOA3;refGene_tx=XM_0232511
40.1;refGene_gn=NCOA3;refGene_tx=XM_023251141.1;refGene_gn=NCOA3;refGe
ne_tx=XM_023251142.1;refGene_gn=NCOA3;refGene_tx=XM_023251143.1;refGen
e_gn=NCOA3;refGene_tx=XM_023251144.1;refGene_gn=NCOA3;refGene_tx=XM_02
3251145.1;refGene_gn=NCOA3;refGene_tx=XM_023251146.1;refGene_gn=NCOA3;
refGene_tx=XM_023251148.1
A3 13834154 13834290

refGene_gn=NCOA3;refGene_tx=XM_019826389.2;refGene_gn=NCOA3;refGene_tx
=XM_019826390.2;refGene_gn=NCOA3;refGene_tx=XM_023251138.1;refGene_gn=
NCOA3;refGene_tx=XM_023251139.1;refGene_gn=NCOA3;refGene_tx=XM_0232511
40.1;refGene_gn=NCOA3;refGene_tx=XM_023251141.1;refGene_gn=NCOA3;refGe
ne_tx=XM_023251142.1;refGene_gn=NCOA3;refGene_tx=XM_023251143.1;refGen
e_gn=NCOA3;refGene_tx=XM_023251144.1;refGene_gn=NCOA3;refGene_tx=XM_02
3251145.1;refGene_gn=NCOA3;refGene_tx=XM_023251146.1;refGene_gn=NCOA3;
refGene_tx=XM_023251147.1;refGene_gn=NCOA3;refGene_tx=XM_023251148.1
A3 13835509 13836378
refGene_gn=NCOA3;refGene_tx=XM_019826389.2;refGene_gn=NCOA3;refGene_tx
=XM_019826390.2;refGene_gn=NCOA3;refGene_tx=XM_023251138.1;refGene_gn=
NCOA3;refGene_tx=XM_023251139.1;refGene_gn=NCOA3;refGene_tx=XM_0232511
40.1;refGene_gn=NCOA3;refGene_tx=XM_023251141.1;refGene_gn=NCOA3;refGe
ne_tx=XM_023251142.1;refGene_gn=NCOA3;refGene_tx=XM_023251143.1;refGen
e_gn=NCOA3;refGene_tx=XM_023251144.1;refGene_gn=NCOA3;refGene_tx=XM_02
3251145.1;refGene_gn=NCOA3;refGene_tx=XM_023251146.1;refGene_gn=NCOA3;
refGene_tx=XM_023251147.1;refGene_gn=NCOA3;refGene_tx=XM_023251148.1
A3 13836551 13836943
refGene_gn=NCOA3;refGene_tx=XM_019826389.2;refGene_gn=NCOA3;refGene_tx
=XM_019826390.2;refGene_gn=NCOA3;refGene_tx=XM_023251138.1;refGene_gn=
NCOA3;refGene_tx=XM_023251139.1;refGene_gn=NCOA3;refGene_tx=XM_0232511
40.1;refGene_gn=NCOA3;refGene_tx=XM_023251141.1;refGene_gn=NCOA3;refGe
ne_tx=XM_023251142.1;refGene_gn=NCOA3;refGene_tx=XM_023251143.1;refGen
e_gn=NCOA3;refGene_tx=XM_023251144.1;refGene_gn=NCOA3;refGene_tx=XM_02
3251145.1;refGene_gn=NCOA3;refGene_tx=XM_023251146.1;refGene_gn=NCOA3;
refGene_tx=XM_023251147.1;refGene_gn=NCOA3;refGene_tx=XM_023251148.1
A3 13837909 13838057
refGene_gn=NCOA3;refGene_tx=XM_019826389.2;refGene_gn=NCOA3;refGene_tx
=XM_019826390.2;refGene_gn=NCOA3;refGene_tx=XM_023251138.1;refGene_gn=
NCOA3;refGene_tx=XM_023251139.1;refGene_gn=NCOA3;refGene_tx=XM_0232511
40.1;refGene_gn=NCOA3;refGene_tx=XM_023251141.1;refGene_gn=NCOA3;refGe
ne_tx=XM_023251142.1;refGene_gn=NCOA3;refGene_tx=XM_023251143.1;refGen
e_gn=NCOA3;refGene_tx=XM_023251144.1;refGene_gn=NCOA3;refGene_tx=XM_02
3251145.1;refGene_gn=NCOA3;refGene_tx=XM_023251146.1;refGene_gn=NCOA3;
refGene_tx=XM_023251147.1;refGene_gn=NCOA3;refGene_tx=XM_023251148.1
A3 13838317 13838458
refGene_gn=NCOA3;refGene_tx=XM_019826389.2;refGene_gn=NCOA3;refGene_tx
=XM_019826390.2;refGene_gn=NCOA3;refGene_tx=XM_023251138.1;refGene_gn=
NCOA3;refGene_tx=XM_023251139.1;refGene_gn=NCOA3;refGene_tx=XM_0232511
40.1;refGene_gn=NCOA3;refGene_tx=XM_023251141.1;refGene_gn=NCOA3;refGe
ne_tx=XM_023251142.1;refGene_gn=NCOA3;refGene_tx=XM_023251143.1;refGen
e_gn=NCOA3;refGene_tx=XM_023251144.1;refGene_gn=NCOA3;refGene_tx=XM_02
3251145.1;refGene_gn=NCOA3;refGene_tx=XM_023251146.1;refGene_gn=NCOA3;
refGene_tx=XM_023251147.1;refGene_gn=NCOA3;refGene_tx=XM_023251148.1
A3 13842010 13842112
refGene_gn=NCOA3;refGene_tx=XM_019826389.2;refGene_gn=NCOA3;refGene_tx
=XM_019826390.2;refGene_gn=NCOA3;refGene_tx=XM_023251138.1;refGene_gn=
NCOA3;refGene_tx=XM_023251139.1;refGene_gn=NCOA3;refGene_tx=XM_0232511
40.1;refGene_gn=NCOA3;refGene_tx=XM_023251141.1;refGene_gn=NCOA3;refGe
ne_tx=XM_023251142.1;refGene_gn=NCOA3;refGene_tx=XM_023251143.1;refGen

e_gn=NCOA3;refGene_tx=XM_023251144.1;refGene_gn=NCOA3;refGene_tx=XM_023251145.1;refGene_gn=NCOA3;refGene_tx=XM_023251146.1;refGene_gn=NCOA3;refGene_tx=XM_023251147.1;refGene_gn=NCOA3;refGene_tx=XM_023251148.1
A3 13842280 13842469
refGene_gn=NCOA3;refGene_tx=XM_019826389.2;refGene_gn=NCOA3;refGene_tx=XM_019826390.2;refGene_gn=NCOA3;refGene_tx=XM_023251138.1;refGene_gn=NCOA3;refGene_tx=XM_023251139.1;refGene_gn=NCOA3;refGene_tx=XM_023251140.1;refGene_gn=NCOA3;refGene_tx=XM_023251141.1;refGene_gn=NCOA3;refGene_tx=XM_023251142.1;refGene_gn=NCOA3;refGene_tx=XM_023251143.1;refGene_gn=NCOA3;refGene_tx=XM_023251144.1;refGene_gn=NCOA3;refGene_tx=XM_023251145.1;refGene_gn=NCOA3;refGene_tx=XM_023251146.1;refGene_gn=NCOA3;refGene_tx=XM_023251147.1;refGene_gn=NCOA3;refGene_tx=XM_023251148.1
A3 13842738 13842913
refGene_gn=NCOA3;refGene_tx=XM_019826389.2;refGene_gn=NCOA3;refGene_tx=XM_019826390.2;refGene_gn=NCOA3;refGene_tx=XM_023251138.1;refGene_gn=NCOA3;refGene_tx=XM_023251139.1;refGene_gn=NCOA3;refGene_tx=XM_023251140.1;refGene_gn=NCOA3;refGene_tx=XM_023251141.1;refGene_gn=NCOA3;refGene_tx=XM_023251142.1;refGene_gn=NCOA3;refGene_tx=XM_023251143.1;refGene_gn=NCOA3;refGene_tx=XM_023251144.1;refGene_gn=NCOA3;refGene_tx=XM_023251145.1;refGene_gn=NCOA3;refGene_tx=XM_023251146.1;refGene_gn=NCOA3;refGene_tx=XM_023251147.1;refGene_gn=NCOA3;refGene_tx=XM_023251148.1
A3 13843872 13843973
refGene_gn=NCOA3;refGene_tx=XM_019826389.2;refGene_gn=NCOA3;refGene_tx=XM_019826390.2;refGene_gn=NCOA3;refGene_tx=XM_023251138.1;refGene_gn=NCOA3;refGene_tx=XM_023251139.1;refGene_gn=NCOA3;refGene_tx=XM_023251140.1;refGene_gn=NCOA3;refGene_tx=XM_023251141.1;refGene_gn=NCOA3;refGene_tx=XM_023251142.1;refGene_gn=NCOA3;refGene_tx=XM_023251143.1;refGene_gn=NCOA3;refGene_tx=XM_023251144.1;refGene_gn=NCOA3;refGene_tx=XM_023251145.1;refGene_gn=NCOA3;refGene_tx=XM_023251146.1;refGene_gn=NCOA3;refGene_tx=XM_023251147.1;refGene_gn=NCOA3;refGene_tx=XM_023251148.1
A3 13844462 13844635
refGene_gn=NCOA3;refGene_tx=XM_019826389.2;refGene_gn=NCOA3;refGene_tx=XM_019826390.2;refGene_gn=NCOA3;refGene_tx=XM_023251138.1;refGene_gn=NCOA3;refGene_tx=XM_023251139.1;refGene_gn=NCOA3;refGene_tx=XM_023251140.1;refGene_gn=NCOA3;refGene_tx=XM_023251141.1;refGene_gn=NCOA3;refGene_tx=XM_023251142.1;refGene_gn=NCOA3;refGene_tx=XM_023251143.1;refGene_gn=NCOA3;refGene_tx=XM_023251144.1;refGene_gn=NCOA3;refGene_tx=XM_023251145.1;refGene_gn=NCOA3;refGene_tx=XM_023251146.1;refGene_gn=NCOA3;refGene_tx=XM_023251147.1;refGene_gn=NCOA3;refGene_tx=XM_023251148.1
A3 13845516 13845602
refGene_gn=NCOA3;refGene_tx=XM_019826389.2;refGene_gn=NCOA3;refGene_tx=XM_019826390.2;refGene_gn=NCOA3;refGene_tx=XM_023251138.1;refGene_gn=NCOA3;refGene_tx=XM_023251139.1;refGene_gn=NCOA3;refGene_tx=XM_023251140.1;refGene_gn=NCOA3;refGene_tx=XM_023251141.1;refGene_gn=NCOA3;refGene_tx=XM_023251142.1;refGene_gn=NCOA3;refGene_tx=XM_023251143.1;refGene_gn=NCOA3;refGene_tx=XM_023251144.1;refGene_gn=NCOA3;refGene_tx=XM_023251145.1;refGene_gn=NCOA3;refGene_tx=XM_023251146.1;refGene_gn=NCOA3;refGene_tx=XM_023251147.1;refGene_gn=NCOA3;refGene_tx=XM_023251148.1
A3 14100591 14100602 refGene_gn=ZMYND8;refGene_tx=XM_019826406.2
A3 14102709 14102723

refGene_gn=ZMYND8;refGene_tx=XM_003983402.5;refGene_gn=ZMYND8;refGene_tx=XM_003983403.5;refGene_gn=ZMYND8;refGene_tx=XM_011280734.3;refGene_gn=ZMYND8;refGene_tx=XM_011280738.3;refGene_gn=ZMYND8;refGene_tx=XM_019826405.2;refGene_gn=ZMYND8;refGene_tx=XM_019826413.2
A3 14106804 14106896 refGene_gn=ZMYND8;refGene_tx=XM_019826403.2
A3 14111538 14111609
refGene_gn=ZMYND8;refGene_tx=XM_003983402.5;refGene_gn=ZMYND8;refGene_tx=XM_003983403.5;refGene_gn=ZMYND8;refGene_tx=XM_011280734.3;refGene_gn=ZMYND8;refGene_tx=XM_011280738.3;refGene_gn=ZMYND8;refGene_tx=XM_019826396.2;refGene_gn=ZMYND8;refGene_tx=XM_019826398.2;refGene_gn=ZMYND8;refGene_tx=XM_019826399.2;refGene_gn=ZMYND8;refGene_tx=XM_019826400.2;refGene_gn=ZMYND8;refGene_tx=XM_019826401.2;refGene_gn=ZMYND8;refGene_tx=XM_019826402.2;refGene_gn=ZMYND8;refGene_tx=XM_019826403.2;refGene_gn=ZMYND8;refGene_tx=XM_019826404.2;refGene_gn=ZMYND8;refGene_tx=XM_019826405.2;refGene_gn=ZMYND8;refGene_tx=XM_019826406.2;refGene_gn=ZMYND8;refGene_tx=XM_019826409.2;refGene_gn=ZMYND8;refGene_tx=XM_019826410.2;refGene_gn=ZMYND8;refGene_tx=XM_019826411.2;refGene_gn=ZMYND8;refGene_tx=XM_019826412.2;refGene_gn=ZMYND8;refGene_tx=XM_019826413.2;refGene_gn=ZMYND8;refGene_tx=XM_019826419.2;refGene_gn=ZMYND8;refGene_tx=XM_023251152.1
A3 14132381 14132460 refGene_gn=ZMYND8;refGene_tx=XM_019826407.2
A3 14140041 14140190
refGene_gn=ZMYND8;refGene_tx=XM_003983402.5;refGene_gn=ZMYND8;refGene_tx=XM_003983403.5;refGene_gn=ZMYND8;refGene_tx=XM_011280734.3;refGene_gn=ZMYND8;refGene_tx=XM_011280738.3;refGene_gn=ZMYND8;refGene_tx=XM_019826396.2;refGene_gn=ZMYND8;refGene_tx=XM_019826398.2;refGene_gn=ZMYND8;refGene_tx=XM_019826399.2;refGene_gn=ZMYND8;refGene_tx=XM_019826400.2;refGene_gn=ZMYND8;refGene_tx=XM_019826401.2;refGene_gn=ZMYND8;refGene_tx=XM_019826402.2;refGene_gn=ZMYND8;refGene_tx=XM_019826403.2;refGene_gn=ZMYND8;refGene_tx=XM_019826404.2;refGene_gn=ZMYND8;refGene_tx=XM_019826405.2;refGene_gn=ZMYND8;refGene_tx=XM_019826406.2;refGene_gn=ZMYND8;refGene_tx=XM_019826407.2;refGene_gn=ZMYND8;refGene_tx=XM_019826409.2;refGene_gn=ZMYND8;refGene_tx=XM_019826410.2;refGene_gn=ZMYND8;refGene_tx=XM_019826411.2;refGene_gn=ZMYND8;refGene_tx=XM_019826412.2;refGene_gn=ZMYND8;refGene_tx=XM_019826413.2;refGene_gn=ZMYND8;refGene_tx=XM_019826419.2;refGene_gn=ZMYND8;refGene_tx=XM_023251152.1
A3 14147830 14148049
refGene_gn=ZMYND8;refGene_tx=XM_003983402.5;refGene_gn=ZMYND8;refGene_tx=XM_003983403.5;refGene_gn=ZMYND8;refGene_tx=XM_011280734.3;refGene_gn=ZMYND8;refGene_tx=XM_011280738.3;refGene_gn=ZMYND8;refGene_tx=XM_019826396.2;refGene_gn=ZMYND8;refGene_tx=XM_019826398.2;refGene_gn=ZMYND8;refGene_tx=XM_019826399.2;refGene_gn=ZMYND8;refGene_tx=XM_019826400.2;refGene_gn=ZMYND8;refGene_tx=XM_019826401.2;refGene_gn=ZMYND8;refGene_tx=XM_019826402.2;refGene_gn=ZMYND8;refGene_tx=XM_019826403.2;refGene_gn=ZMYND8;refGene_tx=XM_019826404.2;refGene_gn=ZMYND8;refGene_tx=XM_019826405.2;refGene_gn=ZMYND8;refGene_tx=XM_019826406.2;refGene_gn=ZMYND8;refGene_tx=XM_019826407.2;refGene_gn=ZMYND8;refGene_tx=XM_019826409.2;refGene_gn=ZMYND8;refGene_tx=XM_019826410.2;refGene_gn=ZMYND8;refGene_tx=XM_019826411.2;refGene_gn=ZMYND8;refGene_tx=XM_019826412.2;refGene_gn=ZMYND8;refGene_tx=XM_019826413.2;refGene_gn=ZMYND8;refGene_tx=X

M_019826416.2;refGene_gn=ZMYND8;refGene_tx=XM_019826417.2;refGene_gn=Z
MYND8;refGene_tx=XM_019826418.2;refGene_gn=ZMYND8;refGene_tx=XM_019826
419.2;refGene_gn=ZMYND8;refGene_tx=XM_023251152.1
A3 14150910 14151024
refGene_gn=ZMYND8;refGene_tx=XM_003983402.5;refGene_gn=ZMYND8;refGene_
tx=XM_003983403.5;refGene_gn=ZMYND8;refGene_tx=XM_011280734.3;refGene_
gn=ZMYND8;refGene_tx=XM_011280738.3;refGene_gn=ZMYND8;refGene_tx=XM_01
9826396.2;refGene_gn=ZMYND8;refGene_tx=XM_019826398.2;refGene_gn=ZMYND
8;refGene_tx=XM_019826399.2;refGene_gn=ZMYND8;refGene_tx=XM_019826400.
2;refGene_gn=ZMYND8;refGene_tx=XM_019826401.2;refGene_gn=ZMYND8;refGen
e_tx=XM_019826402.2;refGene_gn=ZMYND8;refGene_tx=XM_019826403.2;refGen
e_gn=ZMYND8;refGene_tx=XM_019826404.2;refGene_gn=ZMYND8;refGene_tx=XM_
019826405.2;refGene_gn=ZMYND8;refGene_tx=XM_019826406.2;refGene_gn=ZMY
ND8;refGene_tx=XM_019826407.2;refGene_gn=ZMYND8;refGene_tx=XM_01982640
9.2;refGene_gn=ZMYND8;refGene_tx=XM_019826410.2;refGene_gn=ZMYND8;refG
ene_tx=XM_019826411.2;refGene_gn=ZMYND8;refGene_tx=XM_019826412.2;refG
ene_gn=ZMYND8;refGene_tx=XM_019826413.2;refGene_gn=ZMYND8;refGene_tx=X
M_019826416.2;refGene_gn=ZMYND8;refGene_tx=XM_019826417.2;refGene_gn=Z
MYND8;refGene_tx=XM_019826418.2;refGene_gn=ZMYND8;refGene_tx=XM_019826
419.2;refGene_gn=ZMYND8;refGene_tx=XM_023251152.1
A3 14154455 14154548
refGene_gn=ZMYND8;refGene_tx=XM_003983402.5;refGene_gn=ZMYND8;refGene_
tx=XM_003983403.5;refGene_gn=ZMYND8;refGene_tx=XM_011280734.3;refGene_
gn=ZMYND8;refGene_tx=XM_011280738.3;refGene_gn=ZMYND8;refGene_tx=XM_01
9826396.2;refGene_gn=ZMYND8;refGene_tx=XM_019826398.2;refGene_gn=ZMYND
8;refGene_tx=XM_019826399.2;refGene_gn=ZMYND8;refGene_tx=XM_019826400.
2;refGene_gn=ZMYND8;refGene_tx=XM_019826401.2;refGene_gn=ZMYND8;refGen
e_tx=XM_019826402.2;refGene_gn=ZMYND8;refGene_tx=XM_019826403.2;refGen
e_gn=ZMYND8;refGene_tx=XM_019826404.2;refGene_gn=ZMYND8;refGene_tx=XM_
019826405.2;refGene_gn=ZMYND8;refGene_tx=XM_019826406.2;refGene_gn=ZMY
ND8;refGene_tx=XM_019826407.2;refGene_gn=ZMYND8;refGene_tx=XM_01982640
9.2;refGene_gn=ZMYND8;refGene_tx=XM_019826410.2;refGene_gn=ZMYND8;refG
ene_tx=XM_019826411.2;refGene_gn=ZMYND8;refGene_tx=XM_019826412.2;refG
ene_gn=ZMYND8;refGene_tx=XM_019826413.2;refGene_gn=ZMYND8;refGene_tx=X
M_019826416.2;refGene_gn=ZMYND8;refGene_tx=XM_019826417.2;refGene_gn=Z
MYND8;refGene_tx=XM_019826418.2;refGene_gn=ZMYND8;refGene_tx=XM_019826
419.2;refGene_gn=ZMYND8;refGene_tx=XM_023251152.1
A3 14154765 14154780 refGene_gn=ZMYND8;refGene_tx=XM_019826420.2
A3 14155720 14155808
refGene_gn=ZMYND8;refGene_tx=XM_003983402.5;refGene_gn=ZMYND8;refGene_
tx=XM_003983403.5;refGene_gn=ZMYND8;refGene_tx=XM_011280734.3;refGene_
gn=ZMYND8;refGene_tx=XM_011280738.3;refGene_gn=ZMYND8;refGene_tx=XM_01
9826396.2;refGene_gn=ZMYND8;refGene_tx=XM_019826398.2;refGene_gn=ZMYND
8;refGene_tx=XM_019826399.2;refGene_gn=ZMYND8;refGene_tx=XM_019826400.
2;refGene_gn=ZMYND8;refGene_tx=XM_019826401.2;refGene_gn=ZMYND8;refGen
e_tx=XM_019826402.2;refGene_gn=ZMYND8;refGene_tx=XM_019826403.2;refGen
e_gn=ZMYND8;refGene_tx=XM_019826404.2;refGene_gn=ZMYND8;refGene_tx=XM_
019826405.2;refGene_gn=ZMYND8;refGene_tx=XM_019826406.2;refGene_gn=ZMY
ND8;refGene_tx=XM_019826407.2;refGene_gn=ZMYND8;refGene_tx=XM_01982640
9.2;refGene_gn=ZMYND8;refGene_tx=XM_019826410.2;refGene_gn=ZMYND8;refG

ene_tx=XM_019826411.2;refGene_gn=ZMYND8;refGene_tx=XM_019826412.2;refGene_gn=ZMYND8;refGene_tx=XM_019826413.2;refGene_gn=ZMYND8;refGene_tx=XM_019826416.2;refGene_gn=ZMYND8;refGene_tx=XM_019826417.2;refGene_gn=ZMYND8;refGene_tx=XM_019826418.2;refGene_gn=ZMYND8;refGene_tx=XM_019826419.2;refGene_gn=ZMYND8;refGene_tx=XM_019826420.2;refGene_gn=ZMYND8;refGene_tx=XM_023251152.1
A3 14158267 14158323
refGene_gn=ZMYND8;refGene_tx=XM_003983402.5;refGene_gn=ZMYND8;refGene_tx=XM_003983403.5;refGene_gn=ZMYND8;refGene_tx=XM_011280734.3;refGene_gn=ZMYND8;refGene_tx=XM_011280738.3;refGene_gn=ZMYND8;refGene_tx=XM_019826396.2;refGene_gn=ZMYND8;refGene_tx=XM_019826398.2;refGene_gn=ZMYND8;refGene_tx=XM_019826399.2;refGene_gn=ZMYND8;refGene_tx=XM_019826400.2;refGene_gn=ZMYND8;refGene_tx=XM_019826401.2;refGene_gn=ZMYND8;refGene_tx=XM_019826402.2;refGene_gn=ZMYND8;refGene_tx=XM_019826403.2;refGene_gn=ZMYND8;refGene_tx=XM_019826404.2;refGene_gn=ZMYND8;refGene_tx=XM_019826405.2;refGene_gn=ZMYND8;refGene_tx=XM_019826406.2;refGene_gn=ZMYND8;refGene_tx=XM_019826407.2;refGene_gn=ZMYND8;refGene_tx=XM_019826409.2;refGene_gn=ZMYND8;refGene_tx=XM_019826410.2;refGene_gn=ZMYND8;refGene_tx=XM_019826411.2;refGene_gn=ZMYND8;refGene_tx=XM_019826412.2;refGene_gn=ZMYND8;refGene_tx=XM_019826413.2;refGene_gn=ZMYND8;refGene_tx=XM_019826416.2;refGene_gn=ZMYND8;refGene_tx=XM_019826417.2;refGene_gn=ZMYND8;refGene_tx=XM_019826418.2;refGene_gn=ZMYND8;refGene_tx=XM_019826419.2;refGene_gn=ZMYND8;refGene_tx=XM_019826420.2;refGene_gn=ZMYND8;refGene_tx=XM_023251152.1
A3 14162824 14162902
refGene_gn=ZMYND8;refGene_tx=XM_003983402.5;refGene_gn=ZMYND8;refGene_tx=XM_003983403.5;refGene_gn=ZMYND8;refGene_tx=XM_011280734.3;refGene_gn=ZMYND8;refGene_tx=XM_011280738.3;refGene_gn=ZMYND8;refGene_tx=XM_019826396.2;refGene_gn=ZMYND8;refGene_tx=XM_019826398.2;refGene_gn=ZMYND8;refGene_tx=XM_019826399.2;refGene_gn=ZMYND8;refGene_tx=XM_019826400.2;refGene_gn=ZMYND8;refGene_tx=XM_019826401.2;refGene_gn=ZMYND8;refGene_tx=XM_019826402.2;refGene_gn=ZMYND8;refGene_tx=XM_019826403.2;refGene_gn=ZMYND8;refGene_tx=XM_019826404.2;refGene_gn=ZMYND8;refGene_tx=XM_019826405.2;refGene_gn=ZMYND8;refGene_tx=XM_019826406.2;refGene_gn=ZMYND8;refGene_tx=XM_019826407.2;refGene_gn=ZMYND8;refGene_tx=XM_019826409.2;refGene_gn=ZMYND8;refGene_tx=XM_019826410.2;refGene_gn=ZMYND8;refGene_tx=XM_019826411.2;refGene_gn=ZMYND8;refGene_tx=XM_019826412.2;refGene_gn=ZMYND8;refGene_tx=XM_019826413.2;refGene_gn=ZMYND8;refGene_tx=XM_019826416.2;refGene_gn=ZMYND8;refGene_tx=XM_019826417.2;refGene_gn=ZMYND8;refGene_tx=XM_019826418.2;refGene_gn=ZMYND8;refGene_tx=XM_019826419.2;refGene_gn=ZMYND8;refGene_tx=XM_019826420.2;refGene_gn=ZMYND8;refGene_tx=XM_023251152.1
A3 14163924 14164040
refGene_gn=ZMYND8;refGene_tx=XM_003983402.5;refGene_gn=ZMYND8;refGene_tx=XM_003983403.5;refGene_gn=ZMYND8;refGene_tx=XM_011280734.3;refGene_gn=ZMYND8;refGene_tx=XM_011280738.3;refGene_gn=ZMYND8;refGene_tx=XM_019826396.2;refGene_gn=ZMYND8;refGene_tx=XM_019826398.2;refGene_gn=ZMYND8;refGene_tx=XM_019826399.2;refGene_gn=ZMYND8;refGene_tx=XM_019826400.2;refGene_gn=ZMYND8;refGene_tx=XM_019826401.2;refGene_gn=ZMYND8;refGene_tx=XM_019826402.2;refGene_gn=ZMYND8;refGene_tx=XM_019826403.2;refGen

e_gn=ZMYND8;refGene_tx=XM_019826404.2;refGene_gn=ZMYND8;refGene_tx=XM_019826405.2;refGene_gn=ZMYND8;refGene_tx=XM_019826406.2;refGene_gn=ZMYND8;refGene_tx=XM_019826407.2;refGene_gn=ZMYND8;refGene_tx=XM_019826409.2;refGene_gn=ZMYND8;refGene_tx=XM_019826410.2;refGene_gn=ZMYND8;refGene_tx=XM_019826411.2;refGene_gn=ZMYND8;refGene_tx=XM_019826412.2;refGene_gn=ZMYND8;refGene_tx=XM_019826413.2;refGene_gn=ZMYND8;refGene_tx=XM_019826416.2;refGene_gn=ZMYND8;refGene_tx=XM_019826417.2;refGene_gn=ZMYND8;refGene_tx=XM_019826418.2;refGene_gn=ZMYND8;refGene_tx=XM_019826419.2;refGene_gn=ZMYND8;refGene_tx=XM_019826420.2;refGene_gn=ZMYND8;refGene_tx=XM_023251152.1
A3 14170101 14170583
refGene_gn=ZMYND8;refGene_tx=XM_003983402.5;refGene_gn=ZMYND8;refGene_tx=XM_003983403.5;refGene_gn=ZMYND8;refGene_tx=XM_011280734.3;refGene_gn=ZMYND8;refGene_tx=XM_011280738.3;refGene_gn=ZMYND8;refGene_tx=XM_019826396.2;refGene_gn=ZMYND8;refGene_tx=XM_019826398.2;refGene_gn=ZMYND8;refGene_tx=XM_019826399.2;refGene_gn=ZMYND8;refGene_tx=XM_019826400.2;refGene_gn=ZMYND8;refGene_tx=XM_019826401.2;refGene_gn=ZMYND8;refGene_tx=XM_019826402.2;refGene_gn=ZMYND8;refGene_tx=XM_019826403.2;refGene_gn=ZMYND8;refGene_tx=XM_019826404.2;refGene_gn=ZMYND8;refGene_tx=XM_019826405.2;refGene_gn=ZMYND8;refGene_tx=XM_019826406.2;refGene_gn=ZMYND8;refGene_tx=XM_019826407.2;refGene_gn=ZMYND8;refGene_tx=XM_019826409.2;refGene_gn=ZMYND8;refGene_tx=XM_019826410.2;refGene_gn=ZMYND8;refGene_tx=XM_019826411.2;refGene_gn=ZMYND8;refGene_tx=XM_019826412.2;refGene_gn=ZMYND8;refGene_tx=XM_019826413.2;refGene_gn=ZMYND8;refGene_tx=XM_019826416.2;refGene_gn=ZMYND8;refGene_tx=XM_019826417.2;refGene_gn=ZMYND8;refGene_tx=XM_019826418.2;refGene_gn=ZMYND8;refGene_tx=XM_019826419.2;refGene_gn=ZMYND8;refGene_tx=XM_019826420.2;refGene_gn=ZMYND8;refGene_tx=XM_023251152.1
A3 14183849 14183990
refGene_gn=ZMYND8;refGene_tx=XM_003983402.5;refGene_gn=ZMYND8;refGene_tx=XM_003983403.5;refGene_gn=ZMYND8;refGene_tx=XM_011280734.3;refGene_gn=ZMYND8;refGene_tx=XM_011280738.3;refGene_gn=ZMYND8;refGene_tx=XM_019826396.2;refGene_gn=ZMYND8;refGene_tx=XM_019826398.2;refGene_gn=ZMYND8;refGene_tx=XM_019826399.2;refGene_gn=ZMYND8;refGene_tx=XM_019826400.2;refGene_gn=ZMYND8;refGene_tx=XM_019826401.2;refGene_gn=ZMYND8;refGene_tx=XM_019826402.2;refGene_gn=ZMYND8;refGene_tx=XM_019826403.2;refGene_gn=ZMYND8;refGene_tx=XM_019826404.2;refGene_gn=ZMYND8;refGene_tx=XM_019826405.2;refGene_gn=ZMYND8;refGene_tx=XM_019826406.2;refGene_gn=ZMYND8;refGene_tx=XM_019826407.2;refGene_gn=ZMYND8;refGene_tx=XM_019826409.2;refGene_gn=ZMYND8;refGene_tx=XM_019826410.2;refGene_gn=ZMYND8;refGene_tx=XM_019826411.2;refGene_gn=ZMYND8;refGene_tx=XM_019826412.2;refGene_gn=ZMYND8;refGene_tx=XM_019826413.2;refGene_gn=ZMYND8;refGene_tx=XM_019826416.2;refGene_gn=ZMYND8;refGene_tx=XM_019826417.2;refGene_gn=ZMYND8;refGene_tx=XM_019826418.2;refGene_gn=ZMYND8;refGene_tx=XM_019826419.2;refGene_gn=ZMYND8;refGene_tx=XM_019826420.2;refGene_gn=ZMYND8;refGene_tx=XM_023251152.1
A3 14194027 14194180
refGene_gn=ZMYND8;refGene_tx=XM_003983402.5;refGene_gn=ZMYND8;refGene_tx=XM_003983403.5;refGene_gn=ZMYND8;refGene_tx=XM_011280734.3;refGene_gn=ZMYND8;refGene_tx=XM_011280738.3;refGene_gn=ZMYND8;refGene_tx=XM_01

9826396.2;refGene_gn=ZMYND8;refGene_tx=XM_019826398.2;refGene_gn=ZMYND8;refGene_tx=XM_019826399.2;refGene_gn=ZMYND8;refGene_tx=XM_019826400.2;refGene_gn=ZMYND8;refGene_tx=XM_019826401.2;refGene_gn=ZMYND8;refGene_tx=XM_019826402.2;refGene_gn=ZMYND8;refGene_tx=XM_019826403.2;refGene_gn=ZMYND8;refGene_tx=XM_019826404.2;refGene_gn=ZMYND8;refGene_tx=XM_019826405.2;refGene_gn=ZMYND8;refGene_tx=XM_019826406.2;refGene_gn=ZMYND8;refGene_tx=XM_019826407.2;refGene_gn=ZMYND8;refGene_tx=XM_019826409.2;refGene_gn=ZMYND8;refGene_tx=XM_019826410.2;refGene_gn=ZMYND8;refGene_tx=XM_019826411.2;refGene_gn=ZMYND8;refGene_tx=XM_019826412.2;refGene_gn=ZMYND8;refGene_tx=XM_019826413.2;refGene_gn=ZMYND8;refGene_tx=XM_019826416.2;refGene_gn=ZMYND8;refGene_tx=XM_019826417.2;refGene_gn=ZMYND8;refGene_tx=XM_019826418.2;refGene_gn=ZMYND8;refGene_tx=XM_019826419.2;refGene_gn=ZMYND8;refGene_tx=XM_019826420.2;refGene_gn=ZMYND8;refGene_tx=XM_023251152.1

A3 14196767 14197277

refGene_gn=ZMYND8;refGene_tx=XM_003983402.5;refGene_gn=ZMYND8;refGene_tx=XM_003983403.5;refGene_gn=ZMYND8;refGene_tx=XM_011280734.3;refGene_gn=ZMYND8;refGene_tx=XM_011280738.3;refGene_gn=ZMYND8;refGene_tx=XM_019826396.2;refGene_gn=ZMYND8;refGene_tx=XM_019826398.2;refGene_gn=ZMYND8;refGene_tx=XM_019826399.2;refGene_gn=ZMYND8;refGene_tx=XM_019826400.2;refGene_gn=ZMYND8;refGene_tx=XM_019826401.2;refGene_gn=ZMYND8;refGene_tx=XM_019826402.2;refGene_gn=ZMYND8;refGene_tx=XM_019826403.2;refGene_gn=ZMYND8;refGene_tx=XM_019826404.2;refGene_gn=ZMYND8;refGene_tx=XM_019826405.2;refGene_gn=ZMYND8;refGene_tx=XM_019826406.2;refGene_gn=ZMYND8;refGene_tx=XM_019826407.2;refGene_gn=ZMYND8;refGene_tx=XM_019826409.2;refGene_gn=ZMYND8;refGene_tx=XM_019826410.2;refGene_gn=ZMYND8;refGene_tx=XM_019826411.2;refGene_gn=ZMYND8;refGene_tx=XM_019826412.2;refGene_gn=ZMYND8;refGene_tx=XM_019826413.2;refGene_gn=ZMYND8;refGene_tx=XM_019826416.2;refGene_gn=ZMYND8;refGene_tx=XM_019826417.2;refGene_gn=ZMYND8;refGene_tx=XM_019826418.2;refGene_gn=ZMYND8;refGene_tx=XM_019826420.2;refGene_gn=ZMYND8;refGene_tx=XM_023251152.1

A3 14202246 14202615

refGene_gn=ZMYND8;refGene_tx=XM_003983402.5;refGene_gn=ZMYND8;refGene_tx=XM_003983403.5;refGene_gn=ZMYND8;refGene_tx=XM_011280734.3;refGene_gn=ZMYND8;refGene_tx=XM_011280738.3;refGene_gn=ZMYND8;refGene_tx=XM_019826396.2;refGene_gn=ZMYND8;refGene_tx=XM_019826398.2;refGene_gn=ZMYND8;refGene_tx=XM_019826399.2;refGene_gn=ZMYND8;refGene_tx=XM_019826400.2;refGene_gn=ZMYND8;refGene_tx=XM_019826401.2;refGene_gn=ZMYND8;refGene_tx=XM_019826402.2;refGene_gn=ZMYND8;refGene_tx=XM_019826403.2;refGene_gn=ZMYND8;refGene_tx=XM_019826404.2;refGene_gn=ZMYND8;refGene_tx=XM_019826405.2;refGene_gn=ZMYND8;refGene_tx=XM_019826406.2;refGene_gn=ZMYND8;refGene_tx=XM_019826407.2;refGene_gn=ZMYND8;refGene_tx=XM_019826409.2;refGene_gn=ZMYND8;refGene_tx=XM_019826410.2;refGene_gn=ZMYND8;refGene_tx=XM_019826411.2;refGene_gn=ZMYND8;refGene_tx=XM_019826412.2;refGene_gn=ZMYND8;refGene_tx=XM_019826413.2;refGene_gn=ZMYND8;refGene_tx=XM_019826416.2;refGene_gn=ZMYND8;refGene_tx=XM_019826417.2;refGene_gn=ZMYND8;refGene_tx=XM_019826418.2;refGene_gn=ZMYND8;refGene_tx=XM_019826419.2;refGene_gn=ZMYND8;refGene_tx=XM_019826420.2;refGene_gn=ZMYND8;refGene_tx=XM_023251152.1

A3 14205330 14205521

refGene_gn=ZMYND8;refGene_tx=XM_003983402.5;refGene_gn=ZMYND8;refGene_tx=XM_003983403.5;refGene_gn=ZMYND8;refGene_tx=XM_011280734.3;refGene_gn=ZMYND8;refGene_tx=XM_011280738.3;refGene_gn=ZMYND8;refGene_tx=XM_019826396.2;refGene_gn=ZMYND8;refGene_tx=XM_019826398.2;refGene_gn=ZMYND8;refGene_tx=XM_019826399.2;refGene_gn=ZMYND8;refGene_tx=XM_019826400.2;refGene_gn=ZMYND8;refGene_tx=XM_019826401.2;refGene_gn=ZMYND8;refGene_tx=XM_019826402.2;refGene_gn=ZMYND8;refGene_tx=XM_019826403.2;refGene_gn=ZMYND8;refGene_tx=XM_019826404.2;refGene_gn=ZMYND8;refGene_tx=XM_019826405.2;refGene_gn=ZMYND8;refGene_tx=XM_019826406.2;refGene_gn=ZMYND8;refGene_tx=XM_019826407.2;refGene_gn=ZMYND8;refGene_tx=XM_019826409.2;refGene_gn=ZMYND8;refGene_tx=XM_019826410.2;refGene_gn=ZMYND8;refGene_tx=XM_019826411.2;refGene_gn=ZMYND8;refGene_tx=XM_019826412.2;refGene_gn=ZMYND8;refGene_tx=XM_019826413.2;refGene_gn=ZMYND8;refGene_tx=XM_019826416.2;refGene_gn=ZMYND8;refGene_tx=XM_019826417.2;refGene_gn=ZMYND8;refGene_tx=XM_019826418.2;refGene_gn=ZMYND8;refGene_tx=XM_019826419.2;refGene_gn=ZMYND8;refGene_tx=XM_019826420.2;refGene_gn=ZMYND8;refGene_tx=XM_023251152.1
A3 14210455 14210536
refGene_gn=ZMYND8;refGene_tx=XM_003983402.5;refGene_gn=ZMYND8;refGene_tx=XM_003983403.5;refGene_gn=ZMYND8;refGene_tx=XM_011280734.3;refGene_gn=ZMYND8;refGene_tx=XM_011280738.3;refGene_gn=ZMYND8;refGene_tx=XM_019826396.2;refGene_gn=ZMYND8;refGene_tx=XM_019826398.2;refGene_gn=ZMYND8;refGene_tx=XM_019826399.2;refGene_gn=ZMYND8;refGene_tx=XM_019826400.2;refGene_gn=ZMYND8;refGene_tx=XM_019826401.2;refGene_gn=ZMYND8;refGene_tx=XM_019826402.2;refGene_gn=ZMYND8;refGene_tx=XM_019826403.2;refGene_gn=ZMYND8;refGene_tx=XM_019826404.2;refGene_gn=ZMYND8;refGene_tx=XM_019826405.2;refGene_gn=ZMYND8;refGene_tx=XM_019826406.2;refGene_gn=ZMYND8;refGene_tx=XM_019826407.2;refGene_gn=ZMYND8;refGene_tx=XM_019826409.2;refGene_gn=ZMYND8;refGene_tx=XM_019826410.2;refGene_gn=ZMYND8;refGene_tx=XM_019826411.2;refGene_gn=ZMYND8;refGene_tx=XM_019826412.2;refGene_gn=ZMYND8;refGene_tx=XM_019826413.2;refGene_gn=ZMYND8;refGene_tx=XM_019826416.2;refGene_gn=ZMYND8;refGene_tx=XM_019826417.2;refGene_gn=ZMYND8;refGene_tx=XM_019826418.2;refGene_gn=ZMYND8;refGene_tx=XM_019826419.2;refGene_gn=ZMYND8;refGene_tx=XM_019826420.2;refGene_gn=ZMYND8;refGene_tx=XM_023251152.1
A3 14213781 14213860
refGene_gn=ZMYND8;refGene_tx=XM_003983402.5;refGene_gn=ZMYND8;refGene_tx=XM_003983403.5;refGene_gn=ZMYND8;refGene_tx=XM_011280734.3;refGene_gn=ZMYND8;refGene_tx=XM_011280738.3;refGene_gn=ZMYND8;refGene_tx=XM_019826396.2;refGene_gn=ZMYND8;refGene_tx=XM_019826398.2;refGene_gn=ZMYND8;refGene_tx=XM_019826399.2;refGene_gn=ZMYND8;refGene_tx=XM_019826400.2;refGene_gn=ZMYND8;refGene_tx=XM_019826401.2;refGene_gn=ZMYND8;refGene_tx=XM_019826402.2;refGene_gn=ZMYND8;refGene_tx=XM_019826403.2;refGene_gn=ZMYND8;refGene_tx=XM_019826404.2;refGene_gn=ZMYND8;refGene_tx=XM_019826405.2;refGene_gn=ZMYND8;refGene_tx=XM_019826406.2;refGene_gn=ZMYND8;refGene_tx=XM_019826407.2;refGene_gn=ZMYND8;refGene_tx=XM_019826409.2;refGene_gn=ZMYND8;refGene_tx=XM_019826410.2;refGene_gn=ZMYND8;refGene_tx=XM_019826411.2;refGene_gn=ZMYND8;refGene_tx=XM_019826412.2;refGene_gn=ZMYND8;refGene_tx=XM_019826413.2;refGene_gn=ZMYND8;refGene_tx=XM_019826416.2;refGene_gn=ZMYND8;refGene_tx=XM_019826417.2;refGene_gn=Z

MYND8;refGene_tx=XM_019826418.2;refGene_gn=ZMYND8;refGene_tx=XM_019826
419.2;refGene_gn=ZMYND8;refGene_tx=XM_019826420.2;refGene_gn=ZMYND8;re
fGene_tx=XM_023251152.1
A3 14216138 14216378
refGene_gn=ZMYND8;refGene_tx=XM_003983402.5;refGene_gn=ZMYND8;refGene_
tx=XM_003983403.5;refGene_gn=ZMYND8;refGene_tx=XM_011280734.3;refGene_
gn=ZMYND8;refGene_tx=XM_011280738.3;refGene_gn=ZMYND8;refGene_tx=XM_01
9826396.2;refGene_gn=ZMYND8;refGene_tx=XM_019826398.2;refGene_gn=ZMYND
8;refGene_tx=XM_019826399.2;refGene_gn=ZMYND8;refGene_tx=XM_019826400.
2;refGene_gn=ZMYND8;refGene_tx=XM_019826401.2;refGene_gn=ZMYND8;refGen
e_tx=XM_019826402.2;refGene_gn=ZMYND8;refGene_tx=XM_019826403.2;refGen
e_gn=ZMYND8;refGene_tx=XM_019826404.2;refGene_gn=ZMYND8;refGene_tx=XM_
019826405.2;refGene_gn=ZMYND8;refGene_tx=XM_019826406.2;refGene_gn=ZMY
ND8;refGene_tx=XM_019826407.2;refGene_gn=ZMYND8;refGene_tx=XM_01982640
9.2;refGene_gn=ZMYND8;refGene_tx=XM_019826410.2;refGene_gn=ZMYND8;refG
ene_tx=XM_019826411.2;refGene_gn=ZMYND8;refGene_tx=XM_019826412.2;refG
ene_gn=ZMYND8;refGene_tx=XM_019826413.2;refGene_gn=ZMYND8;refGene_tx=X
M_019826416.2;refGene_gn=ZMYND8;refGene_tx=XM_019826417.2;refGene_gn=Z
MYND8;refGene_tx=XM_019826418.2;refGene_gn=ZMYND8;refGene_tx=XM_019826
419.2;refGene_gn=ZMYND8;refGene_tx=XM_019826420.2;refGene_gn=ZMYND8;re
fGene_tx=XM_023251152.1
A3 14218313 14218459
refGene_gn=ZMYND8;refGene_tx=XM_003983402.5;refGene_gn=ZMYND8;refGene_
tx=XM_003983403.5;refGene_gn=ZMYND8;refGene_tx=XM_011280734.3;refGene_
gn=ZMYND8;refGene_tx=XM_011280738.3;refGene_gn=ZMYND8;refGene_tx=XM_01
9826396.2;refGene_gn=ZMYND8;refGene_tx=XM_019826398.2;refGene_gn=ZMYND
8;refGene_tx=XM_019826399.2;refGene_gn=ZMYND8;refGene_tx=XM_019826400.
2;refGene_gn=ZMYND8;refGene_tx=XM_019826401.2;refGene_gn=ZMYND8;refGen
e_tx=XM_019826402.2;refGene_gn=ZMYND8;refGene_tx=XM_019826403.2;refGen
e_gn=ZMYND8;refGene_tx=XM_019826404.2;refGene_gn=ZMYND8;refGene_tx=XM_
019826405.2;refGene_gn=ZMYND8;refGene_tx=XM_019826406.2;refGene_gn=ZMY
ND8;refGene_tx=XM_019826407.2;refGene_gn=ZMYND8;refGene_tx=XM_01982640
9.2;refGene_gn=ZMYND8;refGene_tx=XM_019826410.2;refGene_gn=ZMYND8;refG
ene_tx=XM_019826411.2;refGene_gn=ZMYND8;refGene_tx=XM_019826412.2;refG
ene_gn=ZMYND8;refGene_tx=XM_019826413.2;refGene_gn=ZMYND8;refGene_tx=X
M_019826416.2;refGene_gn=ZMYND8;refGene_tx=XM_019826417.2;refGene_gn=Z
MYND8;refGene_tx=XM_019826418.2;refGene_gn=ZMYND8;refGene_tx=XM_019826
419.2;refGene_gn=ZMYND8;refGene_tx=XM_019826420.2;refGene_gn=ZMYND8;re
fGene_tx=XM_023251152.1
A3 14219498 14219565
refGene_gn=ZMYND8;refGene_tx=XM_003983402.5;refGene_gn=ZMYND8;refGene_
tx=XM_003983403.5;refGene_gn=ZMYND8;refGene_tx=XM_011280734.3;refGene_
gn=ZMYND8;refGene_tx=XM_011280738.3;refGene_gn=ZMYND8;refGene_tx=XM_01
9826398.2;refGene_gn=ZMYND8;refGene_tx=XM_019826400.2;refGene_gn=ZMYND
8;refGene_tx=XM_019826403.2;refGene_gn=ZMYND8;refGene_tx=XM_019826409.
2
A3 14226590 14226815
refGene_gn=ZMYND8;refGene_tx=XM_003983402.5;refGene_gn=ZMYND8;refGene_
tx=XM_003983403.5;refGene_gn=ZMYND8;refGene_tx=XM_011280734.3;refGene_
gn=ZMYND8;refGene_tx=XM_011280738.3;refGene_gn=ZMYND8;refGene_tx=XM_01

9826396.2;refGene_gn=ZMYND8;refGene_tx=XM_019826398.2;refGene_gn=ZMYND8;refGene_tx=XM_019826399.2;refGene_gn=ZMYND8;refGene_tx=XM_019826400.2;refGene_gn=ZMYND8;refGene_tx=XM_019826401.2;refGene_gn=ZMYND8;refGene_tx=XM_019826402.2;refGene_gn=ZMYND8;refGene_tx=XM_019826403.2;refGene_gn=ZMYND8;refGene_tx=XM_019826404.2;refGene_gn=ZMYND8;refGene_tx=XM_019826405.2;refGene_gn=ZMYND8;refGene_tx=XM_019826406.2;refGene_gn=ZMYND8;refGene_tx=XM_019826407.2;refGene_gn=ZMYND8;refGene_tx=XM_019826409.2;refGene_gn=ZMYND8;refGene_tx=XM_019826410.2;refGene_gn=ZMYND8;refGene_tx=XM_019826411.2;refGene_gn=ZMYND8;refGene_tx=XM_019826412.2;refGene_gn=ZMYND8;refGene_tx=XM_019826413.2;refGene_gn=ZMYND8;refGene_tx=XM_019826416.2;refGene_gn=ZMYND8;refGene_tx=XM_019826417.2;refGene_gn=ZMYND8;refGene_tx=XM_019826418.2;refGene_gn=ZMYND8;refGene_tx=XM_019826419.2;refGene_gn=ZMYND8;refGene_tx=XM_019826420.2;refGene_gn=ZMYND8;refGene_tx=XM_023251152.1
A3 14240379 14240460
refGene_gn=EYA2;refGene_tx=XM_023251153.1;refGene_gn=EYA2;refGene_tx=XM_023251154.1;refGene_gn=EYA2;refGene_tx=XM_023251155.1
A3 14247293 14247394
refGene_gn=EYA2;refGene_tx=XM_023251153.1;refGene_gn=EYA2;refGene_tx=XM_023251154.1;refGene_gn=EYA2;refGene_tx=XM_023251155.1
A3 14252503 14252625
refGene_gn=EYA2;refGene_tx=XM_023251153.1;refGene_gn=EYA2;refGene_tx=XM_023251154.1;refGene_gn=EYA2;refGene_tx=XM_023251155.1
A3 14253560 14253675
refGene_gn=EYA2;refGene_tx=XM_023251153.1;refGene_gn=EYA2;refGene_tx=XM_023251154.1;refGene_gn=EYA2;refGene_tx=XM_023251155.1
A3 14257850 14258011
refGene_gn=EYA2;refGene_tx=XM_023251153.1;refGene_gn=EYA2;refGene_tx=XM_023251154.1;refGene_gn=EYA2;refGene_tx=XM_023251155.1
A3 14262510 14262569
refGene_gn=EYA2;refGene_tx=XM_023251153.1;refGene_gn=EYA2;refGene_tx=XM_023251154.1;refGene_gn=EYA2;refGene_tx=XM_023251155.1
A3 14285634 14285724
refGene_gn=EYA2;refGene_tx=XM_023251153.1;refGene_gn=EYA2;refGene_tx=XM_023251154.1;refGene_gn=EYA2;refGene_tx=XM_023251155.1
A3 14320501 14320585
refGene_gn=EYA2;refGene_tx=XM_023251153.1;refGene_gn=EYA2;refGene_tx=XM_023251154.1;refGene_gn=EYA2;refGene_tx=XM_023251155.1
A3 14327391 14327534
refGene_gn=EYA2;refGene_tx=XM_023251153.1;refGene_gn=EYA2;refGene_tx=XM_023251154.1;refGene_gn=EYA2;refGene_tx=XM_023251155.1
A3 14337966 14338144
refGene_gn=EYA2;refGene_tx=XM_023251153.1;refGene_gn=EYA2;refGene_tx=XM_023251154.1;refGene_gn=EYA2;refGene_tx=XM_023251155.1
A3 14340004 14340072
refGene_gn=EYA2;refGene_tx=XM_023251153.1;refGene_gn=EYA2;refGene_tx=XM_023251154.1;refGene_gn=EYA2;refGene_tx=XM_023251155.1
A3 14394357 14394474
refGene_gn=EYA2;refGene_tx=XM_023251153.1;refGene_gn=EYA2;refGene_tx=XM_023251154.1;refGene_gn=EYA2;refGene_tx=XM_023251155.1

A3 14403025 14403168
refGene_gn=EYA2;refGene_tx=XM_023251153.1;refGene_gn=EYA2;refGene_tx=XM_023251154.1;refGene_gn=EYA2;refGene_tx=XM_023251155.1
A3 14406440 14406486
refGene_gn=EYA2;refGene_tx=XM_023251153.1;refGene_gn=EYA2;refGene_tx=XM_023251154.1
A3 14421605 14421714
refGene_gn=EYA2;refGene_tx=XM_023251153.1;refGene_gn=EYA2;refGene_tx=XM_023251154.1
A3 14619602 14619637 refGene_gn=SLC2A10;refGene_tx=XM_023251158.1
A3 14631767 14631846
refGene_gn=SLC2A10;refGene_tx=XM_006929697.2;refGene_gn=SLC2A10;refGene_tx=XM_023251159.1
A3 14632419 14632597 refGene_gn=SLC2A10;refGene_tx=XM_023251157.1
A3 14634054 14634190
refGene_gn=SLC2A10;refGene_tx=XM_006929697.2;refGene_gn=SLC2A10;refGene_tx=XM_023251157.1
A3 14636128 14636251
refGene_gn=SLC2A10;refGene_tx=XM_006929697.2;refGene_gn=SLC2A10;refGene_tx=XM_023251157.1;refGene_gn=SLC2A10;refGene_tx=XM_023251158.1;refGene_gn=SLC2A10;refGene_tx=XM_023251159.1
A3 14636705 14637986
refGene_gn=SLC2A10;refGene_tx=XM_006929697.2;refGene_gn=SLC2A10;refGene_tx=XM_023251157.1;refGene_gn=SLC2A10;refGene_tx=XM_023251158.1;refGene_gn=SLC2A10;refGene_tx=XM_023251159.1
A3 14645861 14645865
refGene_gn=SLC2A10;refGene_tx=XM_006929697.2;refGene_gn=SLC2A10;refGene_tx=XM_023251157.1;refGene_gn=SLC2A10;refGene_tx=XM_023251158.1;refGene_gn=SLC2A10;refGene_tx=XM_023251159.1
A3 14651966 14652249 refGene_gn=TP53RK;refGene_tx=XM_023251164.1
A3 14653795 14654274 refGene_gn=TP53RK;refGene_tx=XM_023251164.1
A3 14667817 14667928
refGene_gn=SLC13A3;refGene_tx=XM_023251160.1;refGene_gn=SLC13A3;refGene_tx=XM_023251161.1;refGene_gn=SLC13A3;refGene_tx=XM_023251163.1
A3 14695602 14695868
refGene_gn=SLC13A3;refGene_tx=XM_023251160.1;refGene_gn=SLC13A3;refGene_tx=XM_023251161.1;refGene_gn=SLC13A3;refGene_tx=XM_023251163.1
A3 14698728 14698889
refGene_gn=SLC13A3;refGene_tx=XM_023251160.1;refGene_gn=SLC13A3;refGene_tx=XM_023251161.1;refGene_gn=SLC13A3;refGene_tx=XM_023251163.1
A3 14708851 14708918
refGene_gn=SLC13A3;refGene_tx=XM_023251160.1;refGene_gn=SLC13A3;refGene_tx=XM_023251161.1;refGene_gn=SLC13A3;refGene_tx=XM_023251163.1
A3 14712157 14712328
refGene_gn=SLC13A3;refGene_tx=XM_023251160.1;refGene_gn=SLC13A3;refGene_tx=XM_023251161.1
A3 14715210 14715336
refGene_gn=SLC13A3;refGene_tx=XM_023251160.1;refGene_gn=SLC13A3;refGene_tx=XM_023251161.1;refGene_gn=SLC13A3;refGene_tx=XM_023251163.1
A3 14717975 14718074

refGene_gn=SLC13A3;refGene_tx=XM_023251160.1;refGene_gn=SLC13A3;refGene_tx=XM_023251161.1;refGene_gn=SLC13A3;refGene_tx=XM_023251163.1
A3 14719046 14719151
refGene_gn=SLC13A3;refGene_tx=XM_023251160.1;refGene_gn=SLC13A3;refGene_tx=XM_023251161.1;refGene_gn=SLC13A3;refGene_tx=XM_023251163.1
A3 14723445 14723543
refGene_gn=SLC13A3;refGene_tx=XM_023251160.1;refGene_gn=SLC13A3;refGene_tx=XM_023251163.1
A3 14728595 14728708
refGene_gn=SLC13A3;refGene_tx=XM_023251160.1;refGene_gn=SLC13A3;refGene_tx=XM_023251161.1;refGene_gn=SLC13A3;refGene_tx=XM_023251163.1
A3 14735076 14735238
refGene_gn=SLC13A3;refGene_tx=XM_023251160.1;refGene_gn=SLC13A3;refGene_tx=XM_023251161.1;refGene_gn=SLC13A3;refGene_tx=XM_023251163.1
A3 14738173 14738311
refGene_gn=SLC13A3;refGene_tx=XM_023251160.1;refGene_gn=SLC13A3;refGene_tx=XM_023251161.1;refGene_gn=SLC13A3;refGene_tx=XM_023251163.1
A3 14741056 14741331
refGene_gn=SLC13A3;refGene_tx=XM_023251160.1;refGene_gn=SLC13A3;refGene_tx=XM_023251161.1;refGene_gn=SLC13A3;refGene_tx=XM_023251163.1
A3 14751806 14751819 refGene_gn=OCSTAMP;refGene_tx=XM_023251156.1
A3 14752011 14753096 refGene_gn=OCSTAMP;refGene_tx=XM_023251156.1
A3 14756023 14756427 refGene_gn=OCSTAMP;refGene_tx=XM_023251156.1
A3 14756428 14756639 refGene_gn=OCSTAMP;refGene_tx=XM_023251156.1
A3 14766428 14767193
refGene_gn=LOC101084866;refGene_tx=XM_003983507.3
A3 14769576 14769891
refGene_gn=LOC101084866;refGene_tx=XM_003983507.3
A3 14896606 14896684
refGene_gn=ELMO2;refGene_tx=XM_023251166.1;refGene_gn=ELMO2;refGene_tx=XM_023251167.1;refGene_gn=ELMO2;refGene_tx=XM_023251168.1;refGene_gn=ELMO2;refGene_tx=XM_023251169.1
A3 14896977 14897018
refGene_gn=ELMO2;refGene_tx=XM_023251166.1;refGene_gn=ELMO2;refGene_tx=XM_023251167.1;refGene_gn=ELMO2;refGene_tx=XM_023251168.1;refGene_gn=ELMO2;refGene_tx=XM_023251169.1
A3 14897417 14897490
refGene_gn=ELMO2;refGene_tx=XM_023251166.1;refGene_gn=ELMO2;refGene_tx=XM_023251167.1;refGene_gn=ELMO2;refGene_tx=XM_023251168.1;refGene_gn=ELMO2;refGene_tx=XM_023251169.1
A3 14897828 14897879
refGene_gn=ELMO2;refGene_tx=XM_023251166.1;refGene_gn=ELMO2;refGene_tx=XM_023251167.1;refGene_gn=ELMO2;refGene_tx=XM_023251168.1;refGene_gn=ELMO2;refGene_tx=XM_023251169.1
A3 14901359 14901541
refGene_gn=ELMO2;refGene_tx=XM_023251166.1;refGene_gn=ELMO2;refGene_tx=XM_023251167.1;refGene_gn=ELMO2;refGene_tx=XM_023251168.1;refGene_gn=ELMO2;refGene_tx=XM_023251169.1
A3 14903036 14903136
refGene_gn=ELMO2;refGene_tx=XM_023251166.1;refGene_gn=ELMO2;refGene_tx

=XM_023251167.1;refGene_gn=ELMO2;refGene_tx=XM_023251168.1;refGene_gn=ELMO2;refGene_tx=XM_023251169.1
A3 14904175 14904327
refGene_gn=ELMO2;refGene_tx=XM_023251166.1;refGene_gn=ELMO2;refGene_tx=XM_023251167.1;refGene_gn=ELMO2;refGene_tx=XM_023251168.1;refGene_gn=ELMO2;refGene_tx=XM_023251169.1
A3 14906821 14906900
refGene_gn=ELMO2;refGene_tx=XM_023251166.1;refGene_gn=ELMO2;refGene_tx=XM_023251167.1;refGene_gn=ELMO2;refGene_tx=XM_023251168.1;refGene_gn=ELMO2;refGene_tx=XM_023251169.1
A3 14908064 14908100
refGene_gn=ELMO2;refGene_tx=XM_023251166.1;refGene_gn=ELMO2;refGene_tx=XM_023251168.1;refGene_gn=ELMO2;refGene_tx=XM_023251169.1
A3 14910000 14910051
refGene_gn=ELMO2;refGene_tx=XM_023251166.1;refGene_gn=ELMO2;refGene_tx=XM_023251167.1;refGene_gn=ELMO2;refGene_tx=XM_023251168.1;refGene_gn=ELMO2;refGene_tx=XM_023251169.1
A3 14914343 14914466
refGene_gn=ELMO2;refGene_tx=XM_023251166.1;refGene_gn=ELMO2;refGene_tx=XM_023251167.1;refGene_gn=ELMO2;refGene_tx=XM_023251168.1;refGene_gn=ELMO2;refGene_tx=XM_023251169.1
A3 14914822 14914957
refGene_gn=ELMO2;refGene_tx=XM_023251166.1;refGene_gn=ELMO2;refGene_tx=XM_023251167.1;refGene_gn=ELMO2;refGene_tx=XM_023251168.1;refGene_gn=ELMO2;refGene_tx=XM_023251169.1
A3 14915542 14915647
refGene_gn=ELMO2;refGene_tx=XM_023251166.1;refGene_gn=ELMO2;refGene_tx=XM_023251167.1;refGene_gn=ELMO2;refGene_tx=XM_023251168.1;refGene_gn=ELMO2;refGene_tx=XM_023251169.1
A3 14915724 14915833
refGene_gn=ELMO2;refGene_tx=XM_023251166.1;refGene_gn=ELMO2;refGene_tx=XM_023251167.1;refGene_gn=ELMO2;refGene_tx=XM_023251168.1;refGene_gn=ELMO2;refGene_tx=XM_023251169.1
A3 14916270 14916407
refGene_gn=ELMO2;refGene_tx=XM_023251166.1;refGene_gn=ELMO2;refGene_tx=XM_023251167.1;refGene_gn=ELMO2;refGene_tx=XM_023251168.1;refGene_gn=ELMO2;refGene_tx=XM_023251169.1
A3 14917715 14917879
refGene_gn=ELMO2;refGene_tx=XM_023251166.1;refGene_gn=ELMO2;refGene_tx=XM_023251167.1;refGene_gn=ELMO2;refGene_tx=XM_023251168.1;refGene_gn=ELMO2;refGene_tx=XM_023251169.1
A3 14917965 14918078
refGene_gn=ELMO2;refGene_tx=XM_023251166.1;refGene_gn=ELMO2;refGene_tx=XM_023251167.1;refGene_gn=ELMO2;refGene_tx=XM_023251168.1;refGene_gn=ELMO2;refGene_tx=XM_023251169.1
A3 14918240 14918348
refGene_gn=ELMO2;refGene_tx=XM_023251166.1;refGene_gn=ELMO2;refGene_tx=XM_023251167.1;refGene_gn=ELMO2;refGene_tx=XM_023251168.1;refGene_gn=ELMO2;refGene_tx=XM_023251169.1
A3 14919105 14919188

refGene_gn=ELMO2;refGene_tx=XM_023251166.1;refGene_gn=ELMO2;refGene_tx=XM_023251167.1;refGene_gn=ELMO2;refGene_tx=XM_023251168.1;refGene_gn=ELMO2;refGene_tx=XM_023251169.1
A3 14920997 14921075
refGene_gn=ELMO2;refGene_tx=XM_023251166.1;refGene_gn=ELMO2;refGene_tx=XM_023251167.1;refGene_gn=ELMO2;refGene_tx=XM_023251168.1;refGene_gn=ELMO2;refGene_tx=XM_023251169.1
A3 14922648 14922849
refGene_gn=ELMO2;refGene_tx=XM_023251166.1;refGene_gn=ELMO2;refGene_tx=XM_023251167.1;refGene_gn=ELMO2;refGene_tx=XM_023251168.1;refGene_gn=ELMO2;refGene_tx=XM_023251169.1
A3 14931511 14931631 refGene_gn=SLC35C2;refGene_tx=XM_023251170.1
A3 14932211 14932362 refGene_gn=SLC35C2;refGene_tx=XM_023251170.1
A3 14933175 14933239 refGene_gn=SLC35C2;refGene_tx=XM_023251170.1
A3 14933946 14934019
refGene_gn=SLC35C2;refGene_tx=XM_006929705.4;refGene_gn=SLC35C2;refGene_tx=XM_019826426.2;refGene_gn=SLC35C2;refGene_tx=XM_019826427.2;refGene_gn=SLC35C2;refGene_tx=XM_023251170.1
A3 14934508 14934677
refGene_gn=SLC35C2;refGene_tx=XM_006929705.4;refGene_gn=SLC35C2;refGene_tx=XM_019826426.2;refGene_gn=SLC35C2;refGene_tx=XM_019826427.2;refGene_gn=SLC35C2;refGene_tx=XM_023251170.1
A3 14934761 14934845
refGene_gn=SLC35C2;refGene_tx=XM_006929705.4;refGene_gn=SLC35C2;refGene_tx=XM_019826426.2;refGene_gn=SLC35C2;refGene_tx=XM_019826427.2;refGene_gn=SLC35C2;refGene_tx=XM_023251170.1
A3 14936995 14937171
refGene_gn=SLC35C2;refGene_tx=XM_006929705.4;refGene_gn=SLC35C2;refGene_tx=XM_019826426.2;refGene_gn=SLC35C2;refGene_tx=XM_019826427.2;refGene_gn=SLC35C2;refGene_tx=XM_023251170.1
A3 14938303 14938433
refGene_gn=SLC35C2;refGene_tx=XM_006929705.4;refGene_gn=SLC35C2;refGene_tx=XM_019826426.2;refGene_gn=SLC35C2;refGene_tx=XM_019826427.2;refGene_gn=SLC35C2;refGene_tx=XM_023251170.1
A3 14938797 14938931
refGene_gn=SLC35C2;refGene_tx=XM_006929705.4;refGene_gn=SLC35C2;refGene_tx=XM_019826426.2;refGene_gn=SLC35C2;refGene_tx=XM_019826427.2;refGene_gn=SLC35C2;refGene_tx=XM_023251170.1
A3 15032006 15032261 refGene_gn=CDH22;refGene_tx=XM_023251172.1
A3 15040478 15040773 refGene_gn=CDH22;refGene_tx=XM_023251172.1
A3 15052940 15053060 refGene_gn=CDH22;refGene_tx=XM_023251172.1
A3 15061257 15061425 refGene_gn=CDH22;refGene_tx=XM_023251172.1
A3 15065037 15065231 refGene_gn=CDH22;refGene_tx=XM_023251172.1
A3 15067961 15068215 refGene_gn=CDH22;refGene_tx=XM_023251172.1
A3 15076941 15077078 refGene_gn=CDH22;refGene_tx=XM_023251172.1
A3 15087734 15087856 refGene_gn=CDH22;refGene_tx=XM_023251172.1
A3 15087973 15088091 refGene_gn=CDH22;refGene_tx=XM_023251172.1
A3 15094246 15094498 refGene_gn=CDH22;refGene_tx=XM_023251172.1
A3 15097269 15097841 refGene_gn=CDH22;refGene_tx=XM_023251172.1
A3 15139301 15139466

refGene_gn=CD40;refGene_tx=XM_003983509.5;refGene_gn=CD40;refGene_tx=XM_019826428.2
A3 15140097 15140126
refGene_gn=CD40;refGene_tx=XM_003983509.5;refGene_gn=CD40;refGene_tx=XM_019826428.2
A3 15140237 15140324
refGene_gn=CD40;refGene_tx=XM_003983509.5;refGene_gn=CD40;refGene_tx=XM_019826428.2
A3 15141974 15142036
refGene_gn=CD40;refGene_tx=XM_003983509.5;refGene_gn=CD40;refGene_tx=XM_019826428.2
A3 15144618 15144712
refGene_gn=CD40;refGene_tx=XM_003983509.5;refGene_gn=CD40;refGene_tx=XM_019826428.2
A3 15145065 15145212
refGene_gn=CD40;refGene_tx=XM_003983509.5;refGene_gn=CD40;refGene_tx=XM_019826428.2
A3 15145466 15145592
refGene_gn=CD40;refGene_tx=XM_003983509.5;refGene_gn=CD40;refGene_tx=XM_019826428.2
A3 15146152 15146231
refGene_gn=CD40;refGene_tx=XM_003983509.5;refGene_gn=CD40;refGene_tx=XM_019826428.2
A3 15148359 15148422 refGene_gn=CD40;refGene_tx=XM_019826428.2
A3 15150029 15150080 refGene_gn=CD40;refGene_tx=XM_003983509.5
A3 15180086 15180124
refGene_gn=NCOA5;refGene_tx=XM_023251173.1;refGene_gn=NCOA5;refGene_tx=XM_023251175.1
A3 15188830 15189157
refGene_gn=NCOA5;refGene_tx=XM_023251173.1;refGene_gn=NCOA5;refGene_tx=XM_023251174.1;refGene_gn=NCOA5;refGene_tx=XM_023251175.1
A3 15190517 15190654
refGene_gn=NCOA5;refGene_tx=XM_023251173.1;refGene_gn=NCOA5;refGene_tx=XM_023251174.1;refGene_gn=NCOA5;refGene_tx=XM_023251175.1
A3 15191860 15191987
refGene_gn=NCOA5;refGene_tx=XM_023251173.1;refGene_gn=NCOA5;refGene_tx=XM_023251174.1;refGene_gn=NCOA5;refGene_tx=XM_023251175.1
A3 15193823 15194023
refGene_gn=NCOA5;refGene_tx=XM_023251173.1;refGene_gn=NCOA5;refGene_tx=XM_023251174.1;refGene_gn=NCOA5;refGene_tx=XM_023251175.1
A3 15195372 15195716
refGene_gn=NCOA5;refGene_tx=XM_023251173.1;refGene_gn=NCOA5;refGene_tx=XM_023251174.1;refGene_gn=NCOA5;refGene_tx=XM_023251175.1
A3 15196155 15196745
refGene_gn=NCOA5;refGene_tx=XM_023251173.1;refGene_gn=NCOA5;refGene_tx=XM_023251174.1;refGene_gn=NCOA5;refGene_tx=XM_023251175.1
A3 15201395 15201487
refGene_gn=SLC12A5;refGene_tx=XM_023251177.1;refGene_gn=SLC12A5;refGene_tx=XM_023251178.1;refGene_gn=SLC12A5;refGene_tx=XM_023251179.1;refGene_gn=SLC12A5;refGene_tx=XM_023251180.1

A3 15201693 15201827
refGene_gn=SLC12A5;refGene_tx=XM_023251177.1;refGene_gn=SLC12A5;refGene_tx=XM_023251178.1;refGene_gn=SLC12A5;refGene_tx=XM_023251179.1;refGene_gn=SLC12A5;refGene_tx=XM_023251180.1
A3 15202067 15202082
refGene_gn=SLC12A5;refGene_tx=XM_023251177.1;refGene_gn=SLC12A5;refGene_tx=XM_023251178.1;refGene_gn=SLC12A5;refGene_tx=XM_023251179.1;refGene_gn=SLC12A5;refGene_tx=XM_023251180.1
A3 15202423 15202623
refGene_gn=SLC12A5;refGene_tx=XM_023251177.1;refGene_gn=SLC12A5;refGene_tx=XM_023251178.1;refGene_gn=SLC12A5;refGene_tx=XM_023251179.1;refGene_gn=SLC12A5;refGene_tx=XM_023251180.1
A3 15202721 15202844
refGene_gn=SLC12A5;refGene_tx=XM_023251177.1;refGene_gn=SLC12A5;refGene_tx=XM_023251178.1;refGene_gn=SLC12A5;refGene_tx=XM_023251179.1;refGene_gn=SLC12A5;refGene_tx=XM_023251180.1
A3 15203970 15204078
refGene_gn=SLC12A5;refGene_tx=XM_023251177.1;refGene_gn=SLC12A5;refGene_tx=XM_023251178.1;refGene_gn=SLC12A5;refGene_tx=XM_023251179.1;refGene_gn=SLC12A5;refGene_tx=XM_023251180.1
A3 15204994 15205126
refGene_gn=SLC12A5;refGene_tx=XM_023251177.1;refGene_gn=SLC12A5;refGene_tx=XM_023251178.1;refGene_gn=SLC12A5;refGene_tx=XM_023251179.1;refGene_gn=SLC12A5;refGene_tx=XM_023251180.1
A3 15205564 15205734
refGene_gn=SLC12A5;refGene_tx=XM_023251177.1;refGene_gn=SLC12A5;refGene_tx=XM_023251178.1;refGene_gn=SLC12A5;refGene_tx=XM_023251179.1;refGene_gn=SLC12A5;refGene_tx=XM_023251180.1
A3 15206606 15206802
refGene_gn=SLC12A5;refGene_tx=XM_023251177.1;refGene_gn=SLC12A5;refGene_tx=XM_023251178.1;refGene_gn=SLC12A5;refGene_tx=XM_023251179.1;refGene_gn=SLC12A5;refGene_tx=XM_023251180.1
A3 15208627 15208796
refGene_gn=SLC12A5;refGene_tx=XM_023251177.1;refGene_gn=SLC12A5;refGene_tx=XM_023251178.1;refGene_gn=SLC12A5;refGene_tx=XM_023251179.1;refGene_gn=SLC12A5;refGene_tx=XM_023251180.1
A3 15209941 15210046
refGene_gn=SLC12A5;refGene_tx=XM_023251177.1;refGene_gn=SLC12A5;refGene_tx=XM_023251178.1;refGene_gn=SLC12A5;refGene_tx=XM_023251179.1;refGene_gn=SLC12A5;refGene_tx=XM_023251180.1
A3 15210308 15210428
refGene_gn=SLC12A5;refGene_tx=XM_023251177.1;refGene_gn=SLC12A5;refGene_tx=XM_023251178.1;refGene_gn=SLC12A5;refGene_tx=XM_023251179.1;refGene_gn=SLC12A5;refGene_tx=XM_023251180.1
A3 15211360 15211459
refGene_gn=SLC12A5;refGene_tx=XM_023251177.1;refGene_gn=SLC12A5;refGene_tx=XM_023251178.1;refGene_gn=SLC12A5;refGene_tx=XM_023251179.1;refGene_gn=SLC12A5;refGene_tx=XM_023251180.1
A3 15211773 15211892
refGene_gn=SLC12A5;refGene_tx=XM_023251177.1;refGene_gn=SLC12A5;refGen

e_tx=XM_023251178.1;refGene_gn=SLC12A5;refGene_tx=XM_023251179.1;refGene_gn=SLC12A5;refGene_tx=XM_023251180.1
A3 15212499 15212674
refGene_gn=SLC12A5;refGene_tx=XM_023251177.1;refGene_gn=SLC12A5;refGene_tx=XM_023251178.1;refGene_gn=SLC12A5;refGene_tx=XM_023251179.1;refGene_gn=SLC12A5;refGene_tx=XM_023251180.1
A3 15214556 15214614
refGene_gn=SLC12A5;refGene_tx=XM_023251177.1;refGene_gn=SLC12A5;refGene_tx=XM_023251178.1;refGene_gn=SLC12A5;refGene_tx=XM_023251179.1;refGene_gn=SLC12A5;refGene_tx=XM_023251180.1
A3 15214755 15214854
refGene_gn=SLC12A5;refGene_tx=XM_023251177.1;refGene_gn=SLC12A5;refGene_tx=XM_023251178.1;refGene_gn=SLC12A5;refGene_tx=XM_023251179.1;refGene_gn=SLC12A5;refGene_tx=XM_023251180.1
A3 15215148 15215319
refGene_gn=SLC12A5;refGene_tx=XM_023251177.1;refGene_gn=SLC12A5;refGene_tx=XM_023251178.1;refGene_gn=SLC12A5;refGene_tx=XM_023251179.1;refGene_gn=SLC12A5;refGene_tx=XM_023251180.1
A3 15216931 15217143
refGene_gn=SLC12A5;refGene_tx=XM_023251177.1;refGene_gn=SLC12A5;refGene_tx=XM_023251178.1;refGene_gn=SLC12A5;refGene_tx=XM_023251179.1;refGene_gn=SLC12A5;refGene_tx=XM_023251180.1
A3 15217880 15218122
refGene_gn=SLC12A5;refGene_tx=XM_023251177.1;refGene_gn=SLC12A5;refGene_tx=XM_023251178.1;refGene_gn=SLC12A5;refGene_tx=XM_023251179.1;refGene_gn=SLC12A5;refGene_tx=XM_023251180.1
A3 15219230 15219361
refGene_gn=SLC12A5;refGene_tx=XM_023251177.1;refGene_gn=SLC12A5;refGene_tx=XM_023251178.1;refGene_gn=SLC12A5;refGene_tx=XM_023251179.1;refGene_gn=SLC12A5;refGene_tx=XM_023251180.1
A3 15219856 15219911
refGene_gn=SLC12A5;refGene_tx=XM_023251177.1;refGene_gn=SLC12A5;refGene_tx=XM_023251178.1;refGene_gn=SLC12A5;refGene_tx=XM_023251179.1;refGene_gn=SLC12A5;refGene_tx=XM_023251180.1
A3 15221331 15221478
refGene_gn=SLC12A5;refGene_tx=XM_023251177.1;refGene_gn=SLC12A5;refGene_tx=XM_023251178.1;refGene_gn=SLC12A5;refGene_tx=XM_023251179.1
A3 15221797 15221929
refGene_gn=SLC12A5;refGene_tx=XM_023251177.1;refGene_gn=SLC12A5;refGene_tx=XM_023251178.1;refGene_gn=SLC12A5;refGene_tx=XM_023251179.1
A3 15222234 15222329
refGene_gn=SLC12A5;refGene_tx=XM_023251177.1;refGene_gn=SLC12A5;refGene_tx=XM_023251178.1;refGene_gn=SLC12A5;refGene_tx=XM_023251179.1
A3 15227955 15228007
refGene_gn=SLC12A5;refGene_tx=XM_023251178.1;refGene_gn=SLC12A5;refGene_tx=XM_023251180.1
A3 15235447 15235568 refGene_gn=SLC12A5;refGene_tx=XM_023251177.1
A3 15240552 15240671 refGene_gn=MMP9;refGene_tx=XM_003983412.5
A3 15242044 15242148 refGene_gn=MMP9;refGene_tx=XM_003983412.5
A3 15242240 15242391 refGene_gn=MMP9;refGene_tx=XM_003983412.5

A3 15242708 15242848 refGene_gn=MMP9;refGene_tx=XM_003983412.5
A3 15242971 15243245 refGene_gn=MMP9;refGene_tx=XM_003983412.5
A3 15244068 15244224 refGene_gn=MMP9;refGene_tx=XM_003983412.5
A3 15244325 15244502 refGene_gn=MMP9;refGene_tx=XM_003983412.5
A3 15245148 15245322 refGene_gn=MMP9;refGene_tx=XM_003983412.5
A3 15245572 15245746 refGene_gn=MMP9;refGene_tx=XM_003983412.5
A3 15245834 15245963 refGene_gn=MMP9;refGene_tx=XM_003983412.5
A3 15246246 15246395 refGene_gn=MMP9;refGene_tx=XM_003983412.5
A3 15246754 15246987 refGene_gn=MMP9;refGene_tx=XM_003983412.5
A3 15247483 15247621 refGene_gn=MMP9;refGene_tx=XM_003983412.5
A3 15276794 15276995
refGene_gn=ZNF335;refGene_tx=XM_011280725.2;refGene_gn=ZNF335;refGene_tx=XM_011280727.3;refGene_gn=ZNF335;refGene_tx=XM_019826430.2;refGene_gn=ZNF335;refGene_tx=XM_023251181.1
A3 15278284 15278525
refGene_gn=ZNF335;refGene_tx=XM_011280725.2;refGene_gn=ZNF335;refGene_tx=XM_011280727.3;refGene_gn=ZNF335;refGene_tx=XM_019826430.2;refGene_gn=ZNF335;refGene_tx=XM_023251181.1
A3 15279364 15279442
refGene_gn=ZNF335;refGene_tx=XM_011280725.2;refGene_gn=ZNF335;refGene_tx=XM_011280727.3;refGene_gn=ZNF335;refGene_tx=XM_019826430.2;refGene_gn=ZNF335;refGene_tx=XM_023251181.1
A3 15279691 15279991
refGene_gn=ZNF335;refGene_tx=XM_011280725.2;refGene_gn=ZNF335;refGene_tx=XM_011280727.3;refGene_gn=ZNF335;refGene_tx=XM_019826430.2;refGene_gn=ZNF335;refGene_tx=XM_023251181.1
A3 15280085 15280232
refGene_gn=ZNF335;refGene_tx=XM_011280725.2;refGene_gn=ZNF335;refGene_tx=XM_011280727.3;refGene_gn=ZNF335;refGene_tx=XM_019826430.2;refGene_gn=ZNF335;refGene_tx=XM_023251181.1
A3 15281073 15281220
refGene_gn=ZNF335;refGene_tx=XM_011280725.2;refGene_gn=ZNF335;refGene_tx=XM_011280727.3;refGene_gn=ZNF335;refGene_tx=XM_019826430.2;refGene_gn=ZNF335;refGene_tx=XM_023251181.1
A3 15282510 15282763
refGene_gn=ZNF335;refGene_tx=XM_011280725.2;refGene_gn=ZNF335;refGene_tx=XM_011280727.3;refGene_gn=ZNF335;refGene_tx=XM_019826430.2;refGene_gn=ZNF335;refGene_tx=XM_023251181.1
A3 15282847 15283025
refGene_gn=ZNF335;refGene_tx=XM_011280725.2;refGene_gn=ZNF335;refGene_tx=XM_011280727.3;refGene_gn=ZNF335;refGene_tx=XM_019826430.2;refGene_gn=ZNF335;refGene_tx=XM_023251181.1
A3 15284005 15284118
refGene_gn=ZNF335;refGene_tx=XM_011280725.2;refGene_gn=ZNF335;refGene_tx=XM_011280727.3;refGene_gn=ZNF335;refGene_tx=XM_019826430.2;refGene_gn=ZNF335;refGene_tx=XM_023251181.1
A3 15284947 15284966
refGene_gn=ZNF335;refGene_tx=XM_011280725.2;refGene_gn=ZNF335;refGene_tx=XM_011280727.3;refGene_gn=ZNF335;refGene_tx=XM_019826430.2;refGene_gn=ZNF335;refGene_tx=XM_023251181.1

A3 15285095 15285212
refGene_gn=ZNF335;refGene_tx=XM_011280725.2;refGene_gn=ZNF335;refGene_tx=XM_011280727.3;refGene_gn=ZNF335;refGene_tx=XM_019826430.2;refGene_gn=ZNF335;refGene_tx=XM_023251181.1
A3 15285293 15285370
refGene_gn=ZNF335;refGene_tx=XM_011280725.2;refGene_gn=ZNF335;refGene_tx=XM_011280727.3;refGene_gn=ZNF335;refGene_tx=XM_019826430.2;refGene_gn=ZNF335;refGene_tx=XM_023251181.1
A3 15285452 15285613
refGene_gn=ZNF335;refGene_tx=XM_011280725.2;refGene_gn=ZNF335;refGene_tx=XM_011280727.3;refGene_gn=ZNF335;refGene_tx=XM_019826430.2;refGene_gn=ZNF335;refGene_tx=XM_023251181.1
A3 15286327 15286560
refGene_gn=ZNF335;refGene_tx=XM_011280725.2;refGene_gn=ZNF335;refGene_tx=XM_011280727.3;refGene_gn=ZNF335;refGene_tx=XM_019826430.2;refGene_gn=ZNF335;refGene_tx=XM_023251181.1
A3 15287474 15287568
refGene_gn=ZNF335;refGene_tx=XM_011280725.2;refGene_gn=ZNF335;refGene_tx=XM_011280727.3;refGene_gn=ZNF335;refGene_tx=XM_019826430.2;refGene_gn=ZNF335;refGene_tx=XM_023251181.1
A3 15287689 15287784
refGene_gn=ZNF335;refGene_tx=XM_011280725.2;refGene_gn=ZNF335;refGene_tx=XM_011280727.3;refGene_gn=ZNF335;refGene_tx=XM_019826430.2;refGene_gn=ZNF335;refGene_tx=XM_023251181.1
A3 15290325 15290585
refGene_gn=ZNF335;refGene_tx=XM_011280725.2;refGene_gn=ZNF335;refGene_tx=XM_011280727.3;refGene_gn=ZNF335;refGene_tx=XM_019826430.2;refGene_gn=ZNF335;refGene_tx=XM_023251181.1
A3 15291420 15291532
refGene_gn=ZNF335;refGene_tx=XM_011280725.2;refGene_gn=ZNF335;refGene_tx=XM_011280727.3;refGene_gn=ZNF335;refGene_tx=XM_019826430.2;refGene_gn=ZNF335;refGene_tx=XM_023251181.1
A3 15291629 15292004
refGene_gn=ZNF335;refGene_tx=XM_011280725.2;refGene_gn=ZNF335;refGene_tx=XM_011280727.3;refGene_gn=ZNF335;refGene_tx=XM_019826430.2;refGene_gn=ZNF335;refGene_tx=XM_023251181.1
A3 15292804 15292947
refGene_gn=ZNF335;refGene_tx=XM_011280725.2;refGene_gn=ZNF335;refGene_tx=XM_011280727.3;refGene_gn=ZNF335;refGene_tx=XM_019826430.2;refGene_gn=ZNF335;refGene_tx=XM_023251181.1
A3 15293026 15293181
refGene_gn=ZNF335;refGene_tx=XM_011280725.2;refGene_gn=ZNF335;refGene_tx=XM_011280727.3;refGene_gn=ZNF335;refGene_tx=XM_019826430.2;refGene_gn=ZNF335;refGene_tx=XM_023251181.1
A3 15293271 15293375
refGene_gn=ZNF335;refGene_tx=XM_011280725.2;refGene_gn=ZNF335;refGene_tx=XM_011280727.3;refGene_gn=ZNF335;refGene_tx=XM_019826430.2;refGene_gn=ZNF335;refGene_tx=XM_023251181.1
A3 15293465 15293543
refGene_gn=ZNF335;refGene_tx=XM_011280725.2;refGene_gn=ZNF335;refGene_

tx=XM_011280727.3;refGene_gn=ZNF335;refGene_tx=XM_019826430.2;refGene_gn=ZNF335;refGene_tx=XM_023251181.1
A3 15293741 15293825
refGene_gn=ZNF335;refGene_tx=XM_011280725.2;refGene_gn=ZNF335;refGene_tx=XM_011280727.3;refGene_gn=ZNF335;refGene_tx=XM_019826430.2;refGene_gn=ZNF335;refGene_tx=XM_023251181.1
A3 15293907 15293973
refGene_gn=ZNF335;refGene_tx=XM_011280725.2;refGene_gn=ZNF335;refGene_tx=XM_011280727.3;refGene_gn=ZNF335;refGene_tx=XM_019826430.2;refGene_gn=ZNF335;refGene_tx=XM_023251181.1
A3 15294056 15294138
refGene_gn=ZNF335;refGene_tx=XM_011280725.2;refGene_gn=ZNF335;refGene_tx=XM_011280727.3;refGene_gn=ZNF335;refGene_tx=XM_019826430.2;refGene_gn=ZNF335;refGene_tx=XM_023251181.1
A3 15294224 15294352
refGene_gn=ZNF335;refGene_tx=XM_011280725.2;refGene_gn=ZNF335;refGene_tx=XM_011280727.3;refGene_gn=ZNF335;refGene_tx=XM_019826430.2;refGene_gn=ZNF335;refGene_tx=XM_023251181.1
A3 15295686 15295915 refGene_gn=PCIF1;refGene_tx=XM_006929707.4
A3 15296000 15296176 refGene_gn=PCIF1;refGene_tx=XM_006929707.4
A3 15296278 15296372 refGene_gn=PCIF1;refGene_tx=XM_006929707.4
A3 15297073 15297258 refGene_gn=PCIF1;refGene_tx=XM_006929707.4
A3 15297344 15297431 refGene_gn=PCIF1;refGene_tx=XM_006929707.4
A3 15297576 15297749 refGene_gn=PCIF1;refGene_tx=XM_006929707.4
A3 15298423 15298586 refGene_gn=PCIF1;refGene_tx=XM_006929707.4
A3 15299658 15299758 refGene_gn=PCIF1;refGene_tx=XM_006929707.4
A3 15299971 15300055 refGene_gn=PCIF1;refGene_tx=XM_006929707.4
A3 15300141 15300289 refGene_gn=PCIF1;refGene_tx=XM_006929707.4
A3 15300869 15301024 refGene_gn=PCIF1;refGene_tx=XM_006929707.4
A3 15301161 15301292 refGene_gn=PCIF1;refGene_tx=XM_006929707.4
A3 15301484 15301507 refGene_gn=PCIF1;refGene_tx=XM_006929707.4
A3 15301509 15301621 refGene_gn=PCIF1;refGene_tx=XM_006929707.4
A3 15302792 15302917 refGene_gn=PCIF1;refGene_tx=XM_006929707.4
A3 15303007 15303150 refGene_gn=PCIF1;refGene_tx=XM_006929707.4
A3 15304110 15304133 refGene_gn=PCIF1;refGene_tx=XM_006929707.4
A3 15321549 15321649 refGene_gn=PLTP;refGene_tx=XM_006929709.4
A3 15321737 15321837 refGene_gn=PLTP;refGene_tx=XM_006929709.4
A3 15322508 15322637 refGene_gn=PLTP;refGene_tx=XM_006929709.4
A3 15322918 15323074 refGene_gn=PLTP;refGene_tx=XM_006929709.4
A3 15324481 15324545 refGene_gn=PLTP;refGene_tx=XM_006929709.4
A3 15324622 15324686 refGene_gn=PLTP;refGene_tx=XM_006929709.4
A3 15325380 15325472 refGene_gn=PLTP;refGene_tx=XM_006929709.4
A3 15326841 15327018 refGene_gn=PLTP;refGene_tx=XM_006929709.4
A3 15327107 15327167 refGene_gn=PLTP;refGene_tx=XM_006929709.4
A3 15328399 15328564 refGene_gn=PLTP;refGene_tx=XM_006929709.4
A3 15328661 15328729 refGene_gn=PLTP;refGene_tx=XM_006929709.4
A3 15329724 15329767 refGene_gn=PLTP;refGene_tx=XM_006929709.4
A3 15329871 15329935 refGene_gn=PLTP;refGene_tx=XM_006929709.4
A3 15330019 15330096 refGene_gn=PLTP;refGene_tx=XM_006929709.4
A3 15330508 15330640 refGene_gn=PLTP;refGene_tx=XM_006929709.4

A3 15331095 15331179
refGene_gn=CTSA;refGene_tx=XM_023251183.1;refGene_gn=CTSA;refGene_tx=XM_023251184.1
A3 15331432 15331570
refGene_gn=CTSA;refGene_tx=XM_023251183.1;refGene_gn=CTSA;refGene_tx=XM_023251184.1
A3 15331724 15331814
refGene_gn=CTSA;refGene_tx=XM_023251183.1;refGene_gn=CTSA;refGene_tx=XM_023251184.1
A3 15332464 15332540
refGene_gn=CTSA;refGene_tx=XM_023251183.1;refGene_gn=CTSA;refGene_tx=XM_023251184.1
A3 15333206 15333346
refGene_gn=CTSA;refGene_tx=XM_023251183.1;refGene_gn=CTSA;refGene_tx=XM_023251184.1
A3 15333445 15333524
refGene_gn=CTSA;refGene_tx=XM_023251183.1;refGene_gn=CTSA;refGene_tx=XM_023251184.1
A3 15333605 15333697
refGene_gn=CTSA;refGene_tx=XM_023251183.1;refGene_gn=CTSA;refGene_tx=XM_023251184.1
A3 15334290 15334375
refGene_gn=CTSA;refGene_tx=XM_023251183.1;refGene_gn=CTSA;refGene_tx=XM_023251184.1
A3 15334722 15334814
refGene_gn=CTSA;refGene_tx=XM_023251183.1;refGene_gn=CTSA;refGene_tx=XM_023251184.1
A3 15335127 15335283
refGene_gn=CTSA;refGene_tx=XM_023251183.1;refGene_gn=CTSA;refGene_tx=XM_023251184.1
A3 15335513 15335600
refGene_gn=CTSA;refGene_tx=XM_023251183.1;refGene_gn=CTSA;refGene_tx=XM_023251184.1
A3 15335691 15335742
refGene_gn=CTSA;refGene_tx=XM_023251183.1;refGene_gn=CTSA;refGene_tx=XM_023251184.1
A3 15336001 15336113
refGene_gn=CTSA;refGene_tx=XM_023251183.1;refGene_gn=CTSA;refGene_tx=XM_023251184.1
A3 15336263 15336454
refGene_gn=CTSA;refGene_tx=XM_023251183.1;refGene_gn=CTSA;refGene_tx=XM_023251184.1
A3 15336645 15336699
refGene_gn=CTSA;refGene_tx=XM_023251183.1;refGene_gn=CTSA;refGene_tx=XM_023251184.1
A3 15337079 15337821 refGene_gn=NEURL2;refGene_tx=XM_023251185.1
A3 15338582 15338698 refGene_gn=NEURL2;refGene_tx=XM_023251185.1
A3 15339917 15340038 refGene_gn=SPATA25;refGene_tx=XM_003983420.4
A3 15340411 15341043 refGene_gn=SPATA25;refGene_tx=XM_003983420.4
A3 15342190 15343561

refGene_gn=ZSWIM1;refGene_tx=XM_023251189.1;refGene_gn=ZSWIM1;refGene_tx=XM_023251190.1;refGene_gn=ZSWIM1;refGene_tx=XM_023251191.1;refGene_gn=ZSWIM1;refGene_tx=XM_023251192.1
A3 15346567 15348503
refGene_gn=ZSWIM3;refGene_tx=XM_003983511.5;refGene_gn=ZSWIM3;refGene_tx=XM_006929710.4
A3 15355173 15355328 refGene_gn=ZSWIM3;refGene_tx=XM_003983511.5
A3 15355831 15355950
refGene_gn=ACOT8;refGene_tx=XM_003983422.4;refGene_gn=ACOT8;refGene_tx=XM_019826434.2;refGene_gn=ACOT8;refGene_tx=XM_019826435.1;refGene_gn=ACOT8;refGene_tx=XM_019826436.1
A3 15356108 15356173 refGene_gn=ZSWIM3;refGene_tx=XM_006929710.4
A3 15357237 15357374
refGene_gn=ACOT8;refGene_tx=XM_003983422.4;refGene_gn=ACOT8;refGene_tx=XM_019826434.2;refGene_gn=ACOT8;refGene_tx=XM_019826435.1;refGene_gn=ACOT8;refGene_tx=XM_019826436.1
A3 15362726 15362952
refGene_gn=ACOT8;refGene_tx=XM_003983422.4;refGene_gn=ACOT8;refGene_tx=XM_019826434.2;refGene_gn=ACOT8;refGene_tx=XM_019826435.1;refGene_gn=ACOT8;refGene_tx=XM_019826436.1
A3 15364312 15364746 refGene_gn=ACOT8;refGene_tx=XM_019826433.1
A3 15364843 15365001
refGene_gn=ACOT8;refGene_tx=XM_003983422.4;refGene_gn=ACOT8;refGene_tx=XM_019826433.1;refGene_gn=ACOT8;refGene_tx=XM_019826434.2;refGene_gn=ACOT8;refGene_tx=XM_019826435.1;refGene_gn=ACOT8;refGene_tx=XM_019826436.1
A3 15365288 15365452 refGene_gn=ACOT8;refGene_tx=XM_019826435.1
A3 15365562 15365757
refGene_gn=ACOT8;refGene_tx=XM_003983422.4;refGene_gn=ACOT8;refGene_tx=XM_019826433.1;refGene_gn=ACOT8;refGene_tx=XM_019826434.2
A3 15365888 15365923 refGene_gn=ACOT8;refGene_tx=XM_019826434.2
A3 15367195 15367314
refGene_gn=ACOT8;refGene_tx=XM_003983422.4;refGene_gn=ACOT8;refGene_tx=XM_019826433.1;refGene_gn=ACOT8;refGene_tx=XM_019826436.1
A3 15367803 15368478
refGene_gn=SNX21;refGene_tx=XM_006929769.2;refGene_gn=SNX21;refGene_tx=XM_023251187.1;refGene_gn=SNX21;refGene_tx=XM_023251188.1
A3 15368652 15368663 refGene_gn=SNX21;refGene_tx=XM_023251187.1
A3 15371878 15372036
refGene_gn=SNX21;refGene_tx=XM_006929769.2;refGene_gn=SNX21;refGene_tx=XM_023251187.1;refGene_gn=SNX21;refGene_tx=XM_023251188.1
A3 15372464 15372732
refGene_gn=SNX21;refGene_tx=XM_006929769.2;refGene_gn=SNX21;refGene_tx=XM_023251187.1;refGene_gn=SNX21;refGene_tx=XM_023251188.1
A3 15373064 15373397
refGene_gn=SNX21;refGene_tx=XM_006929769.2;refGene_gn=SNX21;refGene_tx=XM_023251187.1;refGene_gn=SNX21;refGene_tx=XM_023251188.1
A3 15377596 15377599 refGene_gn=TNNC2;refGene_tx=XM_003983423.3
A3 15379033 15379085 refGene_gn=TNNC2;refGene_tx=XM_003983423.3
A3 15379220 15379364 refGene_gn=TNNC2;refGene_tx=XM_003983423.3

A3	15379456	15379571	refGene_gn=TNNC2;refGene_tx=XM_003983423.3
A3	15379739	15379876	refGene_gn=TNNC2;refGene_tx=XM_003983423.3
A3	15380041	15380073	refGene_gn=TNNC2;refGene_tx=XM_003983423.3
A3	15385008	15385067
refGene_gn=UBE2C;refGene_tx=XM_003983424.5;refGene_gn=UBE2C;refGene_tx=XM_006929712.4;refGene_gn=UBE2C;refGene_tx=XM_019826438.2
A3	15385531	15385591
refGene_gn=UBE2C;refGene_tx=XM_003983424.5;refGene_gn=UBE2C;refGene_tx=XM_006929712.4;refGene_gn=UBE2C;refGene_tx=XM_019826438.2
A3	15385706	15385911
refGene_gn=UBE2C;refGene_tx=XM_003983424.5;refGene_gn=UBE2C;refGene_tx=XM_006929712.4;refGene_gn=UBE2C;refGene_tx=XM_019826438.2
A3	15386919	15387006
refGene_gn=UBE2C;refGene_tx=XM_003983424.5;refGene_gn=UBE2C;refGene_tx=XM_006929712.4;refGene_gn=UBE2C;refGene_tx=XM_019826438.2
A3	15387774	15387802
refGene_gn=UBE2C;refGene_tx=XM_003983424.5;refGene_gn=UBE2C;refGene_tx=XM_006929712.4
A3	15388350	15388707
refGene_gn=UBE2C;refGene_tx=XM_003983424.5;refGene_gn=UBE2C;refGene_tx=XM_019826438.2
A3	15389622	15389761
refGene_gn=DNTTIP1;refGene_tx=XM_003983427.5;refGene_gn=DNTTIP1;refGene_tx=XM_011280716.3
A3	15389872	15389928
refGene_gn=DNTTIP1;refGene_tx=XM_003983427.5;refGene_gn=DNTTIP1;refGene_tx=XM_011280716.3
A3	15391256	15391355	refGene_gn=DNTTIP1;refGene_tx=XM_019826437.2
A3	15391661	15391733
refGene_gn=DNTTIP1;refGene_tx=XM_003983427.5;refGene_gn=DNTTIP1;refGene_tx=XM_011280716.3;refGene_gn=DNTTIP1;refGene_tx=XM_019826437.2
A3	15395241	15395302
refGene_gn=DNTTIP1;refGene_tx=XM_003983427.5;refGene_gn=DNTTIP1;refGene_tx=XM_011280716.3;refGene_gn=DNTTIP1;refGene_tx=XM_019826437.2
A3	15395404	15395463
refGene_gn=DNTTIP1;refGene_tx=XM_003983427.5;refGene_gn=DNTTIP1;refGene_tx=XM_011280716.3;refGene_gn=DNTTIP1;refGene_tx=XM_019826437.2
A3	15397693	15397739
refGene_gn=DNTTIP1;refGene_tx=XM_003983427.5;refGene_gn=DNTTIP1;refGene_tx=XM_011280716.3;refGene_gn=DNTTIP1;refGene_tx=XM_019826437.2
A3	15398605	15398664
refGene_gn=DNTTIP1;refGene_tx=XM_003983427.5;refGene_gn=DNTTIP1;refGene_tx=XM_011280716.3;refGene_gn=DNTTIP1;refGene_tx=XM_019826437.2
A3	15398914	15398971
refGene_gn=DNTTIP1;refGene_tx=XM_003983427.5;refGene_gn=DNTTIP1;refGene_tx=XM_011280716.3;refGene_gn=DNTTIP1;refGene_tx=XM_019826437.2
A3	15399232	15399301
refGene_gn=DNTTIP1;refGene_tx=XM_003983427.5;refGene_gn=DNTTIP1;refGene_tx=XM_011280716.3;refGene_gn=DNTTIP1;refGene_tx=XM_019826437.2
A3	15405746	15405845

refGene_gn=DNTTIP1;refGene_tx=XM_003983427.5;refGene_gn=DNTTIP1;refGene_tx=XM_011280716.3;refGene_gn=DNTTIP1;refGene_tx=XM_019826437.2
A3 15413149 15413246
refGene_gn=DNTTIP1;refGene_tx=XM_003983427.5;refGene_gn=DNTTIP1;refGene_tx=XM_011280716.3;refGene_gn=DNTTIP1;refGene_tx=XM_019826437.2
A3 15415623 15415694
refGene_gn=DNTTIP1;refGene_tx=XM_003983427.5;refGene_gn=DNTTIP1;refGene_tx=XM_011280716.3;refGene_gn=DNTTIP1;refGene_tx=XM_019826437.2
A3 15416252 15416357
refGene_gn=DNTTIP1;refGene_tx=XM_003983427.5;refGene_gn=DNTTIP1;refGene_tx=XM_019826437.2
A3 15418515 15418594
refGene_gn=WFDC3;refGene_tx=XM_006929713.3;refGene_gn=WFDC3;refGene_tx=XM_006929714.3
A3 15419055 15419187
refGene_gn=WFDC3;refGene_tx=XM_006929713.3;refGene_gn=WFDC3;refGene_tx=XM_006929714.3
A3 15421580 15421727
refGene_gn=WFDC3;refGene_tx=XM_006929713.3;refGene_gn=WFDC3;refGene_tx=XM_006929714.3
A3 15428725 15429089 refGene_gn=WFDC3;refGene_tx=XM_023251194.1
A3 15429360 15429492
refGene_gn=WFDC3;refGene_tx=XM_006929713.3;refGene_gn=WFDC3;refGene_tx=XM_023251194.1
A3 15431258 15431444
refGene_gn=WFDC3;refGene_tx=XM_006929713.3;refGene_gn=WFDC3;refGene_tx=XM_006929714.3;refGene_gn=WFDC3;refGene_tx=XM_023251194.1
A3 15432065 15432082
refGene_gn=WFDC3;refGene_tx=XM_006929713.3;refGene_gn=WFDC3;refGene_tx=XM_006929714.3;refGene_gn=WFDC3;refGene_tx=XM_023251194.1
A3 15462853 15462866 refGene_gn=SPINT4;refGene_tx=XM_023251197.1
A3 15463763 15463887 refGene_gn=SPINT4;refGene_tx=XM_023251197.1
A3 15463889 15463940 refGene_gn=SPINT4;refGene_tx=XM_023251197.1
A3 15464650 15464765 refGene_gn=SPINT4;refGene_tx=XM_023251197.1
A3 15479583 15479626 refGene_gn=WFDC13;refGene_tx=XM_019826443.2
A3 15481063 15481214 refGene_gn=WFDC13;refGene_tx=XM_019826443.2
A3 15483958 15484154 refGene_gn=WFDC13;refGene_tx=XM_019826443.2
A3 15489786 15489877 refGene_gn=WFDC10B;refGene_tx=XM_023251198.1
A3 15490786 15490920 refGene_gn=WFDC10B;refGene_tx=XM_023251198.1
A3 15498853 15499000
refGene_gn=WFDC11;refGene_tx=XM_023251195.1;refGene_gn=WFDC11;refGene_tx=XM_023251196.1
A3 15501021 15501101
refGene_gn=WFDC11;refGene_tx=XM_023251195.1;refGene_gn=WFDC11;refGene_tx=XM_023251196.1
A3 15504314 15504460
refGene_gn=WFDC11;refGene_tx=XM_023251195.1;refGene_gn=WFDC11;refGene_tx=XM_023251196.1
A3 15505420 15505563
refGene_gn=WFDC11;refGene_tx=XM_023251195.1;refGene_gn=WFDC11;refGene_

tx=XM_023251196.1
A3 15506066 15506096
refGene_gn=WFDC11;refGene_tx=XM_023251195.1;refGene_gn=WFDC11;refGene_
tx=XM_023251196.1
A3 15549141 15549337
refGene_gn=WFDC9;refGene_tx=XM_019826440.2;refGene_gn=WFDC9;refGene_tx
=XM_023251199.1
A3 15550282 15550430
refGene_gn=WFDC9;refGene_tx=XM_019826440.2;refGene_gn=WFDC9;refGene_tx
=XM_023251199.1
A3 15550941 15550972
refGene_gn=WFDC9;refGene_tx=XM_019826440.2;refGene_gn=WFDC9;refGene_tx
=XM_023251199.1
A3 15557740 15557846
refGene_gn=LOC109497976;refGene_tx=XM_019826441.2
A3 15558762 15558912
refGene_gn=LOC109497976;refGene_tx=XM_019826441.2
A3 15559449 15559535
refGene_gn=LOC109497976;refGene_tx=XM_019826441.2
A3 15566435 15566494 refGene_gn=WFDC8;refGene_tx=XM_023251200.1
A3 15572465 15572575 refGene_gn=WFDC8;refGene_tx=XM_023251200.1
A3 15575425 15575566 refGene_gn=WFDC8;refGene_tx=XM_023251200.1
A3 15576850 15577018 refGene_gn=WFDC8;refGene_tx=XM_023251200.1
A3 15578960 15579101 refGene_gn=WFDC8;refGene_tx=XM_023251200.1
A3 15580078 15580218 refGene_gn=WFDC8;refGene_tx=XM_023251200.1
A3 15587917 15588002
refGene_gn=LOC101087913;refGene_tx=XM_006929717.3
A3 15589726 15589858
refGene_gn=LOC101087913;refGene_tx=XM_006929717.3
A3 15593473 15593641
refGene_gn=LOC101087913;refGene_tx=XM_006929717.3
A3 15594124 15594132
refGene_gn=LOC101087913;refGene_tx=XM_006929717.3
A3 15596723 15596814
refGene_gn=LOC109497979;refGene_tx=XM_019826444.2
A3 15598000 15598132
refGene_gn=LOC109497979;refGene_tx=XM_019826444.2
A3 15599795 15599963
refGene_gn=LOC109497979;refGene_tx=XM_019826444.2
A3 15600613 15600615
refGene_gn=LOC109497979;refGene_tx=XM_019826444.2
A3 15622010 15622162
refGene_gn=WFDC2;refGene_tx=XM_003983428.5;refGene_gn=WFDC2;refGene_tx
=XM_011280714.2
A3 15624928 15625105
refGene_gn=WFDC2;refGene_tx=XM_003983428.5;refGene_gn=WFDC2;refGene_tx
=XM_011280714.2
A3 15625637 15625716
refGene_gn=WFDC2;refGene_tx=XM_003983428.5;refGene_gn=WFDC2;refGene_tx
=XM_011280714.2

A3 15655355 15655431
refGene_gn=LOC101088151;refGene_tx=XM_019828122.1
A3 15657783 15658079
refGene_gn=LOC101088151;refGene_tx=XM_019828122.1
A3 15662555 15662808
refGene_gn=PIGT;refGene_tx=XM_003983429.5;refGene_gn=PIGT;refGene_tx=XM_019826445.2;refGene_gn=PIGT;refGene_tx=XM_019826446.2
A3 15663520 15663604
refGene_gn=PIGT;refGene_tx=XM_003983429.5;refGene_gn=PIGT;refGene_tx=XM_019826445.2;refGene_gn=PIGT;refGene_tx=XM_019826446.2
A3 15663711 15663877
refGene_gn=PIGT;refGene_tx=XM_003983429.5;refGene_gn=PIGT;refGene_tx=XM_019826445.2;refGene_gn=PIGT;refGene_tx=XM_019826446.2
A3 15665597 15665798
refGene_gn=PIGT;refGene_tx=XM_003983429.5;refGene_gn=PIGT;refGene_tx=XM_019826445.2;refGene_gn=PIGT;refGene_tx=XM_019826446.2
A3 15666701 15666867
refGene_gn=PIGT;refGene_tx=XM_003983429.5;refGene_gn=PIGT;refGene_tx=XM_019826445.2;refGene_gn=PIGT;refGene_tx=XM_019826446.2
A3 15666957 15667055
refGene_gn=PIGT;refGene_tx=XM_003983429.5;refGene_gn=PIGT;refGene_tx=XM_019826445.2;refGene_gn=PIGT;refGene_tx=XM_019826446.2
A3 15667166 15667254
refGene_gn=PIGT;refGene_tx=XM_003983429.5;refGene_gn=PIGT;refGene_tx=XM_019826445.2;refGene_gn=PIGT;refGene_tx=XM_019826446.2
A3 15667766 15667853
refGene_gn=PIGT;refGene_tx=XM_003983429.5;refGene_gn=PIGT;refGene_tx=XM_019826445.2;refGene_gn=PIGT;refGene_tx=XM_019826446.2
A3 15668015 15668146
refGene_gn=PIGT;refGene_tx=XM_003983429.5;refGene_gn=PIGT;refGene_tx=XM_019826445.2;refGene_gn=PIGT;refGene_tx=XM_019826446.2
A3 15668448 15668576
refGene_gn=PIGT;refGene_tx=XM_003983429.5;refGene_gn=PIGT;refGene_tx=XM_019826445.2;refGene_gn=PIGT;refGene_tx=XM_019826446.2
A3 15670373 15670551
refGene_gn=PIGT;refGene_tx=XM_003983429.5;refGene_gn=PIGT;refGene_tx=XM_019826445.2;refGene_gn=PIGT;refGene_tx=XM_019826446.2
A3 15671098 15671285
refGene_gn=PIGT;refGene_tx=XM_003983429.5;refGene_gn=PIGT;refGene_tx=XM_019826445.2;refGene_gn=PIGT;refGene_tx=XM_019826446.2
A3 15677758 15677964
refGene_gn=DBNDD2;refGene_tx=XM_003983437.4;refGene_gn=DBNDD2;refGene_tx=XM_003983439.4;refGene_gn=DBNDD2;refGene_tx=XM_006929724.4;refGene_gn=DBNDD2;refGene_tx=XM_006929725.3;refGene_gn=DBNDD2;refGene_tx=XM_019826447.1
A3 15679346 15679484
refGene_gn=DBNDD2;refGene_tx=XM_003983437.4;refGene_gn=DBNDD2;refGene_tx=XM_003983439.4;refGene_gn=DBNDD2;refGene_tx=XM_006929724.4;refGene_gn=DBNDD2;refGene_tx=XM_006929725.3;refGene_gn=DBNDD2;refGene_tx=XM_019826447.1

A3 15679679 15679818
refGene_gn=DBNDD2;refGene_tx=XM_003983437.4;refGene_gn=DBNDD2;refGene_tx=XM_003983439.4;refGene_gn=DBNDD2;refGene_tx=XM_006929724.4;refGene_gn=DBNDD2;refGene_tx=XM_006929725.3;refGene_gn=DBNDD2;refGene_tx=XM_019826447.1
A3 15679819 15679826 refGene_gn=SYS1;refGene_tx=XM_006929719.3
A3 15702136 15702184 refGene_gn=TP53TG5;refGene_tx=XM_023251201.1
A3 15702464 15702539 refGene_gn=TP53TG5;refGene_tx=XM_023251201.1
A3 15702698 15702829 refGene_gn=TP53TG5;refGene_tx=XM_023251201.1
A3 15704290 15704930 refGene_gn=TP53TG5;refGene_tx=XM_023251201.1
A3 15711974 15712215 refGene_gn=SYS1;refGene_tx=XM_003983440.4
A3 15713422 15713490
refGene_gn=SYS1;refGene_tx=XM_003983440.4;refGene_gn=SYS1;refGene_tx=XM_006929719.3
A3 15714769 15714931
refGene_gn=SYS1;refGene_tx=XM_003983440.4;refGene_gn=SYS1;refGene_tx=XM_006929719.3
A3 15728579 15728648 refGene_gn=SDC4;refGene_tx=XM_003983441.4
A3 15740168 15740307 refGene_gn=SDC4;refGene_tx=XM_003983441.4
A3 15741354 15741401 refGene_gn=SDC4;refGene_tx=XM_003983441.4
A3 15742232 15742431 refGene_gn=SDC4;refGene_tx=XM_003983441.4
A3 15745354 15745506 refGene_gn=SDC4;refGene_tx=XM_003983441.4
A3 15754590 15754864
refGene_gn=RBPJL;refGene_tx=XM_023251206.1;refGene_gn=RBPJL;refGene_tx=XM_023251207.1
A3 15754953 15755057
refGene_gn=RBPJL;refGene_tx=XM_023251206.1;refGene_gn=RBPJL;refGene_tx=XM_023251207.1
A3 15755197 15755353
refGene_gn=RBPJL;refGene_tx=XM_023251206.1;refGene_gn=RBPJL;refGene_tx=XM_023251207.1
A3 15756245 15756398
refGene_gn=RBPJL;refGene_tx=XM_023251206.1;refGene_gn=RBPJL;refGene_tx=XM_023251207.1
A3 15756679 15756789
refGene_gn=RBPJL;refGene_tx=XM_023251206.1;refGene_gn=RBPJL;refGene_tx=XM_023251207.1
A3 15757188 15757326 refGene_gn=RBPJL;refGene_tx=XM_023251206.1
A3 15759352 15759527
refGene_gn=RBPJL;refGene_tx=XM_023251206.1;refGene_gn=RBPJL;refGene_tx=XM_023251207.1
A3 15759799 15759915
refGene_gn=RBPJL;refGene_tx=XM_023251206.1;refGene_gn=RBPJL;refGene_tx=XM_023251207.1
A3 15760084 15760155
refGene_gn=RBPJL;refGene_tx=XM_023251206.1;refGene_gn=RBPJL;refGene_tx=XM_023251207.1
A3 15761828 15761978
refGene_gn=RBPJL;refGene_tx=XM_023251206.1;refGene_gn=RBPJL;refGene_tx=XM_023251207.1

A3 15763168 15763277
refGene_gn=RBPJL;refGene_tx=XM_023251206.1;refGene_gn=RBPJL;refGene_tx=XM_023251207.1
A3 15764506 15764528
refGene_gn=RBPJL;refGene_tx=XM_023251206.1;refGene_gn=RBPJL;refGene_tx=XM_023251207.1
A3 15765581 15765660
refGene_gn=MATN4;refGene_tx=XM_023251203.1;refGene_gn=MATN4;refGene_tx=XM_023251204.1;refGene_gn=MATN4;refGene_tx=XM_023251205.1
A3 15766350 15766920
refGene_gn=MATN4;refGene_tx=XM_023251203.1;refGene_gn=MATN4;refGene_tx=XM_023251204.1;refGene_gn=MATN4;refGene_tx=XM_023251205.1
A3 15767617 15767740
refGene_gn=MATN4;refGene_tx=XM_023251203.1;refGene_gn=MATN4;refGene_tx=XM_023251204.1;refGene_gn=MATN4;refGene_tx=XM_023251205.1
A3 15769715 15769838
refGene_gn=MATN4;refGene_tx=XM_023251203.1;refGene_gn=MATN4;refGene_tx=XM_023251204.1;refGene_gn=MATN4;refGene_tx=XM_023251205.1
A3 15769928 15770051
refGene_gn=MATN4;refGene_tx=XM_023251203.1;refGene_gn=MATN4;refGene_tx=XM_023251204.1;refGene_gn=MATN4;refGene_tx=XM_023251205.1
A3 15770144 15770267
refGene_gn=MATN4;refGene_tx=XM_023251203.1;refGene_gn=MATN4;refGene_tx=XM_023251204.1;refGene_gn=MATN4;refGene_tx=XM_023251205.1
A3 15771719 15772133
refGene_gn=MATN4;refGene_tx=XM_023251203.1;refGene_gn=MATN4;refGene_tx=XM_023251204.1;refGene_gn=MATN4;refGene_tx=XM_023251205.1
A3 15772232 15772385
refGene_gn=MATN4;refGene_tx=XM_023251203.1;refGene_gn=MATN4;refGene_tx=XM_023251204.1;refGene_gn=MATN4;refGene_tx=XM_023251205.1
A3 15775343 15775363 refGene_gn=MATN4;refGene_tx=XM_023251205.1
A3 15775494 15775639
refGene_gn=MATN4;refGene_tx=XM_023251203.1;refGene_gn=MATN4;refGene_tx=XM_023251204.1
A3 15775721 15776445
refGene_gn=MATN4;refGene_tx=XM_023251203.1;refGene_gn=MATN4;refGene_tx=XM_023251204.1
A3 15808320 15808402 refGene_gn=SLPI;refGene_tx=XM_003983442.4
A3 15809308 15809467 refGene_gn=SLPI;refGene_tx=XM_003983442.4
A3 15809952 15810101 refGene_gn=SLPI;refGene_tx=XM_003983442.4
A3 15829536 15829565
refGene_gn=LOC101089181;refGene_tx=XM_011280711.3;refGene_gn=LOC101089181;refGene_tx=XM_023250912.1
A3 15830339 15830489
refGene_gn=LOC101089181;refGene_tx=XM_011280711.3;refGene_gn=LOC101089181;refGene_tx=XM_023250912.1
A3 15830760 15830922
refGene_gn=LOC101089181;refGene_tx=XM_011280711.3;refGene_gn=LOC101089181;refGene_tx=XM_023250912.1
A3 15831849 15831934

refGene_gn=LOC101089181;refGene_tx=XM_011280711.3
A3	15838735	15838760
refGene_gn=LOC101089181;refGene_tx=XM_023250912.1
A3	15870366	15870611	refGene_gn=PI3;refGene_tx=XM_006929726.2
A3	15871424	15871500	refGene_gn=PI3;refGene_tx=XM_006929726.2
A3	15881696	15881781	refGene_gn=WFDC5;refGene_tx=XM_006929727.3
A3	15885107	15885248	refGene_gn=WFDC5;refGene_tx=XM_006929727.3
A3	15885330	15885471	refGene_gn=WFDC5;refGene_tx=XM_006929727.3
A3	15885865	15885870	refGene_gn=WFDC5;refGene_tx=XM_006929727.3
A3	15895231	15896257	refGene_gn=KCNS1;refGene_tx=XM_023251208.1
A3	15898783	15899254	refGene_gn=KCNS1;refGene_tx=XM_023251208.1
A3	15914783	15914942	refGene_gn=STK4;refGene_tx=XM_003983444.5
A3	15953641	15953799	refGene_gn=STK4;refGene_tx=XM_003983444.5
A3	15973458	15973645	refGene_gn=STK4;refGene_tx=XM_003983444.5
A3	15974532	15974661	refGene_gn=STK4;refGene_tx=XM_003983444.5
A3	15977062	15977200	refGene_gn=STK4;refGene_tx=XM_003983444.5
A3	15978581	15978749	refGene_gn=STK4;refGene_tx=XM_003983444.5
A3	15983572	15983737	refGene_gn=STK4;refGene_tx=XM_003983444.5
A3	15990014	15990129	refGene_gn=STK4;refGene_tx=XM_003983444.5
A3	15992853	15992982	refGene_gn=STK4;refGene_tx=XM_003983444.5
A3	15998881	15998962	refGene_gn=STK4;refGene_tx=XM_003983444.5
A3	16004375	16004410	refGene_gn=STK4;refGene_tx=XM_003983444.5
A3	16012603	16012730	refGene_gn=TOMM34;refGene_tx=XM_003983445.5
A3	16016761	16016861	refGene_gn=TOMM34;refGene_tx=XM_003983445.5
A3	16018005	16018158	refGene_gn=TOMM34;refGene_tx=XM_003983445.5
A3	16020939	16021109	refGene_gn=TOMM34;refGene_tx=XM_003983445.5
A3	16024487	16024635	refGene_gn=TOMM34;refGene_tx=XM_003983445.5
A3	16028567	16028694	refGene_gn=TOMM34;refGene_tx=XM_003983445.5
A3	16028883	16028988	refGene_gn=TOMM34;refGene_tx=XM_003983445.5
A3	16032840	16032909	refGene_gn=PABPC1L;refGene_tx=XM_003983525.5
A3	16033360	16033491	refGene_gn=PABPC1L;refGene_tx=XM_003983525.5
A3	16034929	16035023	refGene_gn=PABPC1L;refGene_tx=XM_003983525.5
A3	16035887	16035991	refGene_gn=PABPC1L;refGene_tx=XM_003983525.5
A3	16036541	16036670	refGene_gn=PABPC1L;refGene_tx=XM_003983525.5
A3	16037303	16037388	refGene_gn=PABPC1L;refGene_tx=XM_003983525.5
A3	16038423	16038690	refGene_gn=PABPC1L;refGene_tx=XM_003983525.5
A3	16043344	16043440	refGene_gn=PABPC1L;refGene_tx=XM_003983525.5
A3	16043737	16043875	refGene_gn=PABPC1L;refGene_tx=XM_003983525.5
A3	16046939	16047034	refGene_gn=PABPC1L;refGene_tx=XM_003983525.5
A3	16047196	16047336	refGene_gn=PABPC1L;refGene_tx=XM_003983525.5
A3	16049459	16049575	refGene_gn=PABPC1L;refGene_tx=XM_003983525.5
A3	16051085	16051279	refGene_gn=PABPC1L;refGene_tx=XM_003983525.5
A3	16053319	16053512	refGene_gn=PABPC1L;refGene_tx=XM_003983525.5
A3	16056836	16056893
refGene_gn=YWHAB;refGene_tx=XM_019826454.2;refGene_gn=YWHAB;refGene_tx
=XM_023251209.1
A3	16057190	16057286
refGene_gn=YWHAB;refGene_tx=XM_019826454.2;refGene_gn=YWHAB;refGene_tx
=XM_023251209.1
A3	16058118	16058282

refGene_gn=YWHAB;refGene_tx=XM_019826454.2;refGene_gn=YWHAB;refGene_tx=XM_023251209.1
A3 16059161 16059285
refGene_gn=YWHAB;refGene_tx=XM_019826454.2;refGene_gn=YWHAB;refGene_tx=XM_023251209.1
A3 16061463 16061757
refGene_gn=YWHAB;refGene_tx=XM_019826454.2;refGene_gn=YWHAB;refGene_tx=XM_023251209.1
A3 16129349 16129446
refGene_gn=RIMS4;refGene_tx=XM_023251211.1;refGene_gn=RIMS4;refGene_tx=XM_023251212.1
A3 16170341 16170483
refGene_gn=RIMS4;refGene_tx=XM_023251211.1;refGene_gn=RIMS4;refGene_tx=XM_023251212.1
A3 16184572 16184685
refGene_gn=RIMS4;refGene_tx=XM_023251211.1;refGene_gn=RIMS4;refGene_tx=XM_023251212.1
A3 16184962 16185064
refGene_gn=RIMS4;refGene_tx=XM_023251211.1;refGene_gn=RIMS4;refGene_tx=XM_023251212.1
A3 16185707 16185847
refGene_gn=RIMS4;refGene_tx=XM_023251211.1;refGene_gn=RIMS4;refGene_tx=XM_023251212.1
A3 16188644 16188863
refGene_gn=RIMS4;refGene_tx=XM_023251211.1;refGene_gn=RIMS4;refGene_tx=XM_023251212.1
A3 16194168 16194878 refGene_gn=KCNK15;refGene_tx=XM_023251210.1
A3 16198759 16199042 refGene_gn=KCNK15;refGene_tx=XM_023251210.1
A3 16218182 16218403
refGene_gn=WISP2;refGene_tx=XM_003983447.4;refGene_gn=WISP2;refGene_tx=XM_006929730.3
A3 16220133 16220388
refGene_gn=WISP2;refGene_tx=XM_003983447.4;refGene_gn=WISP2;refGene_tx=XM_006929730.3
A3 16225953 16226170
refGene_gn=WISP2;refGene_tx=XM_003983447.4;refGene_gn=WISP2;refGene_tx=XM_006929730.3
A3 16230215 16230275
refGene_gn=WISP2;refGene_tx=XM_003983447.4;refGene_gn=WISP2;refGene_tx=XM_006929730.3
A3 16280477 16280510 refGene_gn=ADA;refGene_tx=XM_011280709.3
A3 16292196 16292258 refGene_gn=ADA;refGene_tx=XM_011280709.3
A3 16297929 16298052
refGene_gn=ADA;refGene_tx=XM_006929733.4;refGene_gn=ADA;refGene_tx=XM_011280709.3
A3 16301471 16301615
refGene_gn=ADA;refGene_tx=XM_006929733.4;refGene_gn=ADA;refGene_tx=XM_011280709.3
A3 16302386 16302502
refGene_gn=ADA;refGene_tx=XM_006929733.4;refGene_gn=ADA;refGene_tx=XM_

011280709.3
A3 16303443 16303571
refGene_gn=ADA;refGene_tx=XM_006929733.4;refGene_gn=ADA;refGene_tx=XM_011280709.3
A3 16304290 16304362
refGene_gn=ADA;refGene_tx=XM_006929733.4;refGene_gn=ADA;refGene_tx=XM_011280709.3
A3 16304442 16304544
refGene_gn=ADA;refGene_tx=XM_006929733.4;refGene_gn=ADA;refGene_tx=XM_011280709.3
A3 16304797 16304862
refGene_gn=ADA;refGene_tx=XM_006929733.4;refGene_gn=ADA;refGene_tx=XM_011280709.3
A3 16306574 16306704
refGene_gn=ADA;refGene_tx=XM_006929733.4;refGene_gn=ADA;refGene_tx=XM_011280709.3
A3 16307342 16307445
refGene_gn=ADA;refGene_tx=XM_006929733.4;refGene_gn=ADA;refGene_tx=XM_011280709.3
A3 16307922 16307936
refGene_gn=ADA;refGene_tx=XM_006929733.4;refGene_gn=ADA;refGene_tx=XM_011280709.3
A3 16309334 16309414
refGene_gn=PKIG;refGene_tx=XM_003983448.5;refGene_gn=PKIG;refGene_tx=XM_006929735.4;refGene_gn=PKIG;refGene_tx=XM_019826462.2
A3 16312711 16312859
refGene_gn=PKIG;refGene_tx=XM_003983448.5;refGene_gn=PKIG;refGene_tx=XM_006929735.4;refGene_gn=PKIG;refGene_tx=XM_019826462.2
A3 16410545 16410584 refGene_gn=SERINC3;refGene_tx=XM_011280708.3
A3 16416378 16416540 refGene_gn=SERINC3;refGene_tx=XM_011280708.3
A3 16417148 16417342 refGene_gn=SERINC3;refGene_tx=XM_011280708.3
A3 16418936 16419016 refGene_gn=SERINC3;refGene_tx=XM_011280708.3
A3 16419677 16419815 refGene_gn=SERINC3;refGene_tx=XM_011280708.3
A3 16422959 16423129 refGene_gn=SERINC3;refGene_tx=XM_011280708.3
A3 16423911 16424002 refGene_gn=SERINC3;refGene_tx=XM_011280708.3
A3 16424809 16424990 refGene_gn=SERINC3;refGene_tx=XM_011280708.3
A3 16429319 16429547 refGene_gn=SERINC3;refGene_tx=XM_011280708.3
A3 16430173 16430312 refGene_gn=SERINC3;refGene_tx=XM_011280708.3
A3 16437739 16438018
refGene_gn=TTPAL;refGene_tx=XM_003983449.4;refGene_gn=TTPAL;refGene_tx=XM_006929736.3;refGene_gn=TTPAL;refGene_tx=XM_019826456.2;refGene_gn=TTPAL;refGene_tx=XM_019826457.1;refGene_gn=TTPAL;refGene_tx=XM_019826458.1;refGene_gn=TTPAL;refGene_tx=XM_019826459.1;refGene_gn=TTPAL;refGene_tx=XM_019826460.1
A3 16440388 16440499
refGene_gn=TTPAL;refGene_tx=XM_003983449.4;refGene_gn=TTPAL;refGene_tx=XM_006929736.3;refGene_gn=TTPAL;refGene_tx=XM_019826456.2;refGene_gn=TTPAL;refGene_tx=XM_019826457.1;refGene_gn=TTPAL;refGene_tx=XM_019826458.1;refGene_gn=TTPAL;refGene_tx=XM_019826459.1;refGene_gn=TTPAL;refGene_tx=XM_019826460.1

A3	16442013	16442207
refGene_gn=TTPAL;refGene_tx=XM_003983449.4;refGene_gn=TTPAL;refGene_tx=XM_006929736.3;refGene_gn=TTPAL;refGene_tx=XM_019826456.2;refGene_gn=TTPAL;refGene_tx=XM_019826457.1;refGene_gn=TTPAL;refGene_tx=XM_019826458.1;refGene_gn=TTPAL;refGene_tx=XM_019826459.1;refGene_gn=TTPAL;refGene_tx=XM_019826460.1
A3	16444519	16444967
refGene_gn=TTPAL;refGene_tx=XM_003983449.4;refGene_gn=TTPAL;refGene_tx=XM_006929736.3;refGene_gn=TTPAL;refGene_tx=XM_019826456.2;refGene_gn=TTPAL;refGene_tx=XM_019826457.1;refGene_gn=TTPAL;refGene_tx=XM_019826458.1;refGene_gn=TTPAL;refGene_tx=XM_019826459.1;refGene_gn=TTPAL;refGene_tx=XM_019826460.1
A3	16483538	16483681
refGene_gn=HNF4A;refGene_tx=XM_006929740.4;refGene_gn=HNF4A;refGene_tx=XM_019826463.2;refGene_gn=HNF4A;refGene_tx=XM_019826464.1;refGene_gn=HNF4A;refGene_tx=XM_019826465.2;refGene_gn=HNF4A;refGene_tx=XM_019826466.1;refGene_gn=HNF4A;refGene_tx=XM_023251214.1;refGene_gn=HNF4A;refGene_tx=XM_023251215.1
A3	16484773	16484926
refGene_gn=HNF4A;refGene_tx=XM_006929740.4;refGene_gn=HNF4A;refGene_tx=XM_019826463.2;refGene_gn=HNF4A;refGene_tx=XM_019826464.1;refGene_gn=HNF4A;refGene_tx=XM_019826465.2;refGene_gn=HNF4A;refGene_tx=XM_019826466.1;refGene_gn=HNF4A;refGene_tx=XM_023251214.1;refGene_gn=HNF4A;refGene_tx=XM_023251215.1
A3	16487398	16487665
refGene_gn=HNF4A;refGene_tx=XM_006929740.4;refGene_gn=HNF4A;refGene_tx=XM_019826463.2;refGene_gn=HNF4A;refGene_tx=XM_019826464.1;refGene_gn=HNF4A;refGene_tx=XM_019826465.2;refGene_gn=HNF4A;refGene_tx=XM_019826466.1;refGene_gn=HNF4A;refGene_tx=XM_023251214.1;refGene_gn=HNF4A;refGene_tx=XM_023251215.1
A3	16490476	16490632
refGene_gn=HNF4A;refGene_tx=XM_006929740.4;refGene_gn=HNF4A;refGene_tx=XM_019826463.2;refGene_gn=HNF4A;refGene_tx=XM_019826464.1;refGene_gn=HNF4A;refGene_tx=XM_019826465.2;refGene_gn=HNF4A;refGene_tx=XM_019826466.1;refGene_gn=HNF4A;refGene_tx=XM_023251214.1;refGene_gn=HNF4A;refGene_tx=XM_023251215.1
A3	16491774	16491862
refGene_gn=HNF4A;refGene_tx=XM_006929740.4;refGene_gn=HNF4A;refGene_tx=XM_019826463.2;refGene_gn=HNF4A;refGene_tx=XM_019826464.1;refGene_gn=HNF4A;refGene_tx=XM_019826465.2;refGene_gn=HNF4A;refGene_tx=XM_019826466.1;refGene_gn=HNF4A;refGene_tx=XM_023251214.1;refGene_gn=HNF4A;refGene_tx=XM_023251215.1
A3	16495237	16495393
refGene_gn=HNF4A;refGene_tx=XM_006929740.4;refGene_gn=HNF4A;refGene_tx=XM_019826463.2;refGene_gn=HNF4A;refGene_tx=XM_019826464.1;refGene_gn=HNF4A;refGene_tx=XM_019826465.2;refGene_gn=HNF4A;refGene_tx=XM_019826466.1;refGene_gn=HNF4A;refGene_tx=XM_023251214.1;refGene_gn=HNF4A;refGene_tx=XM_023251215.1
A3	16496111	16496218
refGene_gn=HNF4A;refGene_tx=XM_006929740.4;refGene_gn=HNF4A;refGene_tx

=XM_019826463.2;refGene_gn=HNF4A;refGene_tx=XM_019826464.1;refGene_gn=HNF4A;refGene_tx=XM_019826465.2;refGene_gn=HNF4A;refGene_tx=XM_019826466.1;refGene_gn=HNF4A;refGene_tx=XM_023251214.1;refGene_gn=HNF4A;refGene_tx=XM_023251215.1
A3 16501346 16501441 refGene_gn=HNF4A;refGene_tx=XM_006929740.4;refGene_gn=HNF4A;refGene_tx=XM_019826463.2;refGene_gn=HNF4A;refGene_tx=XM_019826464.1;refGene_gn=HNF4A;refGene_tx=XM_019826465.2;refGene_gn=HNF4A;refGene_tx=XM_019826466.1;refGene_gn=HNF4A;refGene_tx=XM_023251214.1;refGene_gn=HNF4A;refGene_tx=XM_023251215.1
A3 16502574 16502749 refGene_gn=HNF4A;refGene_tx=XM_006929740.4;refGene_gn=HNF4A;refGene_tx=XM_019826463.2;refGene_gn=HNF4A;refGene_tx=XM_019826464.1;refGene_gn=HNF4A;refGene_tx=XM_019826465.2;refGene_gn=HNF4A;refGene_tx=XM_019826466.1;refGene_gn=HNF4A;refGene_tx=XM_023251214.1;refGene_gn=HNF4A;refGene_tx=XM_023251215.1
A3 16506556 16506671 refGene_gn=HNF4A;refGene_tx=XM_019826463.2;refGene_gn=HNF4A;refGene_tx=XM_019826464.1;refGene_gn=HNF4A;refGene_tx=XM_019826465.2;refGene_gn=HNF4A;refGene_tx=XM_019826466.1
A3 16516178 16516218 refGene_gn=HNF4A;refGene_tx=XM_006929740.4
A3 16547040 16547173 refGene_gn=R3HDML;refGene_tx=XM_003983453.3;refGene_gn=R3HDML;refGene_tx=XM_019826469.1
A3 16551517 16551633 refGene_gn=R3HDML;refGene_tx=XM_003983453.3;refGene_gn=R3HDML;refGene_tx=XM_019826469.1
A3 16553429 16553562 refGene_gn=R3HDML;refGene_tx=XM_003983453.3;refGene_gn=R3HDML;refGene_tx=XM_019826469.1
A3 16553685 16553804 refGene_gn=R3HDML;refGene_tx=XM_003983453.3;refGene_gn=R3HDML;refGene_tx=XM_019826469.1
A3 16556740 16556998 refGene_gn=R3HDML;refGene_tx=XM_003983453.3;refGene_gn=R3HDML;refGene_tx=XM_019826469.1
A3 16571808 16571981 refGene_gn=FITM2;refGene_tx=XM_003983454.5
A3 16573975 16574591 refGene_gn=FITM2;refGene_tx=XM_003983454.5
A3 16588550 16588894 refGene_gn=GDAP1L1;refGene_tx=XM_003983455.5;refGene_gn=GDAP1L1;refGene_tx=XM_006929741.4;refGene_gn=GDAP1L1;refGene_tx=XM_019826470.2;refGene_gn=GDAP1L1;refGene_tx=XM_019826471.2
A3 16599993 16600108 refGene_gn=GDAP1L1;refGene_tx=XM_003983455.5;refGene_gn=GDAP1L1;refGene_tx=XM_006929741.4;refGene_gn=GDAP1L1;refGene_tx=XM_019826470.2;refGene_gn=GDAP1L1;refGene_tx=XM_019826471.2
A3 16600893 16600991 refGene_gn=GDAP1L1;refGene_tx=XM_003983455.5;refGene_gn=GDAP1L1;refGene_tx=XM_006929741.4;refGene_gn=GDAP1L1;refGene_tx=XM_019826470.2;refGene_gn=GDAP1L1;refGene_tx=XM_019826471.2

A3 16605420 16605594
refGene_gn=GDAP1L1;refGene_tx=XM_003983455.5;refGene_gn=GDAP1L1;refGen
e_tx=XM_006929741.4;refGene_gn=GDAP1L1;refGene_tx=XM_019826470.2;refGe
ne_gn=GDAP1L1;refGene_tx=XM_019826471.2
A3 16606619 16606812
refGene_gn=GDAP1L1;refGene_tx=XM_003983455.5;refGene_gn=GDAP1L1;refGen
e_tx=XM_006929741.4;refGene_gn=GDAP1L1;refGene_tx=XM_019826470.2;refGe
ne_gn=GDAP1L1;refGene_tx=XM_019826471.2
A3 16614508 16614688 refGene_gn=GDAP1L1;refGene_tx=XM_003983455.5
A3 16642573 16642650 refGene_gn=OSER1;refGene_tx=XM_003983459.3
A3 16645570 16645684 refGene_gn=OSER1;refGene_tx=XM_003983459.3
A3 16651575 16652263 refGene_gn=OSER1;refGene_tx=XM_003983459.3
A3 16660214 16660593 refGene_gn=JPH2;refGene_tx=XM_023251216.1
A3 16681490 16682286 refGene_gn=JPH2;refGene_tx=XM_023251216.1
A3 16721936 16722055 refGene_gn=JPH2;refGene_tx=XM_023251216.1
A3 16723940 16724620 refGene_gn=JPH2;refGene_tx=XM_023251216.1
A3 16725411 16725492 refGene_gn=JPH2;refGene_tx=XM_023251216.1
A3 16737119 16737150 refGene_gn=TOX2;refGene_tx=XM_023251218.1
A3 16737175 16737212
refGene_gn=TOX2;refGene_tx=XM_023251217.1;refGene_gn=TOX2;refGene_tx=X
M_023251219.1;refGene_gn=TOX2;refGene_tx=XM_023251220.1;refGene_gn=TOX
2;refGene_tx=XM_023251221.1;refGene_gn=TOX2;refGene_tx=XM_023251222.1;
refGene_gn=TOX2;refGene_tx=XM_023251223.1;refGene_gn=TOX2;refGene_tx=X
M_023251224.1;refGene_gn=TOX2;refGene_tx=XM_023251225.1
A3 16739679 16739810
refGene_gn=TOX2;refGene_tx=XM_023251217.1;refGene_gn=TOX2;refGene_tx=X
M_023251218.1;refGene_gn=TOX2;refGene_tx=XM_023251219.1;refGene_gn=TOX
2;refGene_tx=XM_023251220.1;refGene_gn=TOX2;refGene_tx=XM_023251221.1;
refGene_gn=TOX2;refGene_tx=XM_023251222.1;refGene_gn=TOX2;refGene_tx=X
M_023251223.1;refGene_gn=TOX2;refGene_tx=XM_023251224.1;refGene_gn=TOX
2;refGene_tx=XM_023251225.1
A3 16740501 16740897
refGene_gn=TOX2;refGene_tx=XM_023251217.1;refGene_gn=TOX2;refGene_tx=X
M_023251218.1;refGene_gn=TOX2;refGene_tx=XM_023251219.1;refGene_gn=TOX
2;refGene_tx=XM_023251220.1;refGene_gn=TOX2;refGene_tx=XM_023251221.1;
refGene_gn=TOX2;refGene_tx=XM_023251222.1;refGene_gn=TOX2;refGene_tx=X
M_023251223.1;refGene_gn=TOX2;refGene_tx=XM_023251224.1;refGene_gn=TOX
2;refGene_tx=XM_023251225.1
A3 16741574 16741655
refGene_gn=TOX2;refGene_tx=XM_023251217.1;refGene_gn=TOX2;refGene_tx=X
M_023251218.1;refGene_gn=TOX2;refGene_tx=XM_023251219.1;refGene_gn=TOX
2;refGene_tx=XM_023251220.1;refGene_gn=TOX2;refGene_tx=XM_023251221.1;
refGene_gn=TOX2;refGene_tx=XM_023251222.1;refGene_gn=TOX2;refGene_tx=X
M_023251223.1;refGene_gn=TOX2;refGene_tx=XM_023251224.1;refGene_gn=TOX
2;refGene_tx=XM_023251225.1
A3 16742101 16742329
refGene_gn=TOX2;refGene_tx=XM_023251217.1;refGene_gn=TOX2;refGene_tx=X
M_023251218.1;refGene_gn=TOX2;refGene_tx=XM_023251219.1;refGene_gn=TOX
2;refGene_tx=XM_023251220.1;refGene_gn=TOX2;refGene_tx=XM_023251221.1;
refGene_gn=TOX2;refGene_tx=XM_023251222.1;refGene_gn=TOX2;refGene_tx=X

M_023251223.1;refGene_gn=TOX2;refGene_tx=XM_023251224.1;refGene_gn=TOX2;refGene_tx=XM_023251225.1
A3 16744728 16744968
refGene_gn=TOX2;refGene_tx=XM_023251217.1;refGene_gn=TOX2;refGene_tx=XM_023251218.1;refGene_gn=TOX2;refGene_tx=XM_023251219.1;refGene_gn=TOX2;refGene_tx=XM_023251220.1;refGene_gn=TOX2;refGene_tx=XM_023251221.1;refGene_gn=TOX2;refGene_tx=XM_023251222.1;refGene_gn=TOX2;refGene_tx=XM_023251223.1;refGene_gn=TOX2;refGene_tx=XM_023251224.1;refGene_gn=TOX2;refGene_tx=XM_023251225.1
A3 16785145 16785391
refGene_gn=TOX2;refGene_tx=XM_023251217.1;refGene_gn=TOX2;refGene_tx=XM_023251218.1;refGene_gn=TOX2;refGene_tx=XM_023251219.1;refGene_gn=TOX2;refGene_tx=XM_023251220.1;refGene_gn=TOX2;refGene_tx=XM_023251221.1;refGene_gn=TOX2;refGene_tx=XM_023251222.1;refGene_gn=TOX2;refGene_tx=XM_023251223.1;refGene_gn=TOX2;refGene_tx=XM_023251224.1;refGene_gn=TOX2;refGene_tx=XM_023251225.1
A3 16821400 16821466
refGene_gn=TOX2;refGene_tx=XM_023251217.1;refGene_gn=TOX2;refGene_tx=XM_023251218.1;refGene_gn=TOX2;refGene_tx=XM_023251219.1;refGene_gn=TOX2;refGene_tx=XM_023251220.1;refGene_gn=TOX2;refGene_tx=XM_023251222.1;refGene_gn=TOX2;refGene_tx=XM_023251223.1;refGene_gn=TOX2;refGene_tx=XM_023251224.1;refGene_gn=TOX2;refGene_tx=XM_023251225.1
A3 16821932 16821935 refGene_gn=TOX2;refGene_tx=XM_023251222.1
A3 16845735 16845861 refGene_gn=TOX2;refGene_tx=XM_023251219.1
A3 16863073 16863223
refGene_gn=TOX2;refGene_tx=XM_023251217.1;refGene_gn=TOX2;refGene_tx=XM_023251218.1
A3 16873030 16873129
refGene_gn=TOX2;refGene_tx=XM_023251220.1;refGene_gn=TOX2;refGene_tx=XM_023251221.1
A3 16999786 17000290 refGene_gn=GTSF1L;refGene_tx=XM_003983461.4
A3 17014736 17014865
refGene_gn=MYBL2;refGene_tx=XM_023251227.1;refGene_gn=MYBL2;refGene_tx=XM_023251228.1
A3 17015483 17015633
refGene_gn=MYBL2;refGene_tx=XM_023251227.1;refGene_gn=MYBL2;refGene_tx=XM_023251228.1
A3 17016996 17017101
refGene_gn=MYBL2;refGene_tx=XM_023251227.1;refGene_gn=MYBL2;refGene_tx=XM_023251228.1
A3 17018503 17018617
refGene_gn=MYBL2;refGene_tx=XM_023251227.1;refGene_gn=MYBL2;refGene_tx=XM_023251228.1
A3 17020013 17020113
refGene_gn=MYBL2;refGene_tx=XM_023251227.1;refGene_gn=MYBL2;refGene_tx=XM_023251228.1
A3 17024878 17025018
refGene_gn=MYBL2;refGene_tx=XM_023251227.1;refGene_gn=MYBL2;refGene_tx=XM_023251228.1
A3 17027225 17027645

refGene_gn=MYBL2;refGene_tx=XM_023251227.1;refGene_gn=MYBL2;refGene_tx=XM_023251228.1
A3 17030805 17031093
refGene_gn=MYBL2;refGene_tx=XM_023251227.1;refGene_gn=MYBL2;refGene_tx=XM_023251228.1
A3 17038050 17038213
refGene_gn=MYBL2;refGene_tx=XM_023251227.1;refGene_gn=MYBL2;refGene_tx=XM_023251228.1
A3 17041044 17041265
refGene_gn=MYBL2;refGene_tx=XM_023251227.1;refGene_gn=MYBL2;refGene_tx=XM_023251228.1
A3 17041468 17041561
refGene_gn=MYBL2;refGene_tx=XM_023251227.1;refGene_gn=MYBL2;refGene_tx=XM_023251228.1
A3 17042408 17042480
refGene_gn=MYBL2;refGene_tx=XM_023251227.1;refGene_gn=MYBL2;refGene_tx=XM_023251228.1
A3 17048142 17048236
refGene_gn=MYBL2;refGene_tx=XM_023251227.1;refGene_gn=MYBL2;refGene_tx=XM_023251228.1
A3 17051457 17051477
refGene_gn=MYBL2;refGene_tx=XM_023251227.1;refGene_gn=MYBL2;refGene_tx=XM_023251228.1
A3 17061047 17061095 refGene_gn=IFT52;refGene_tx=XM_003983463.5
A3 17062667 17062813 refGene_gn=IFT52;refGene_tx=XM_003983463.5
A3 17065282 17065391 refGene_gn=IFT52;refGene_tx=XM_003983463.5
A3 17066255 17066343 refGene_gn=IFT52;refGene_tx=XM_003983463.5
A3 17070680 17070835 refGene_gn=IFT52;refGene_tx=XM_003983463.5
A3 17074800 17074869 refGene_gn=IFT52;refGene_tx=XM_003983463.5
A3 17076636 17076723 refGene_gn=IFT52;refGene_tx=XM_003983463.5
A3 17079700 17079827 refGene_gn=IFT52;refGene_tx=XM_003983463.5
A3 17085597 17085669 refGene_gn=IFT52;refGene_tx=XM_003983463.5
A3 17086757 17086833 refGene_gn=IFT52;refGene_tx=XM_003983463.5
A3 17087117 17087247 refGene_gn=IFT52;refGene_tx=XM_003983463.5
A3 17090852 17090940 refGene_gn=IFT52;refGene_tx=XM_003983463.5
A3 17091796 17091921 refGene_gn=IFT52;refGene_tx=XM_003983463.5
A3 17093924 17093945 refGene_gn=IFT52;refGene_tx=XM_003983463.5
A3 17097255 17097420
refGene_gn=SGK2;refGene_tx=XM_019826482.2;refGene_gn=SGK2;refGene_tx=XM_019826483.2
A3 17100969 17101002 refGene_gn=SGK2;refGene_tx=XM_019826484.2
A3 17101086 17101176
refGene_gn=SGK2;refGene_tx=XM_019826482.2;refGene_gn=SGK2;refGene_tx=XM_019826483.2;refGene_gn=SGK2;refGene_tx=XM_019826484.2
A3 17103173 17103329
refGene_gn=SGK2;refGene_tx=XM_019826482.2;refGene_gn=SGK2;refGene_tx=XM_019826483.2;refGene_gn=SGK2;refGene_tx=XM_019826484.2
A3 17104225 17104321
refGene_gn=SGK2;refGene_tx=XM_019826482.2;refGene_gn=SGK2;refGene_tx=XM_019826483.2;refGene_gn=SGK2;refGene_tx=XM_019826484.2

A3 17107233 17107320
refGene_gn=SGK2;refGene_tx=XM_019826482.2;refGene_gn=SGK2;refGene_tx=XM_019826483.2;refGene_gn=SGK2;refGene_tx=XM_019826484.2
A3 17108313 17108350
refGene_gn=SGK2;refGene_tx=XM_019826482.2;refGene_gn=SGK2;refGene_tx=XM_019826483.2;refGene_gn=SGK2;refGene_tx=XM_019826484.2
A3 17108649 17108762
refGene_gn=SGK2;refGene_tx=XM_019826482.2;refGene_gn=SGK2;refGene_tx=XM_019826483.2;refGene_gn=SGK2;refGene_tx=XM_019826484.2
A3 17109821 17109953
refGene_gn=SGK2;refGene_tx=XM_019826482.2;refGene_gn=SGK2;refGene_tx=XM_019826483.2;refGene_gn=SGK2;refGene_tx=XM_019826484.2
A3 17110862 17110946
refGene_gn=SGK2;refGene_tx=XM_019826482.2;refGene_gn=SGK2;refGene_tx=XM_019826483.2;refGene_gn=SGK2;refGene_tx=XM_019826484.2
A3 17111148 17111206
refGene_gn=SGK2;refGene_tx=XM_019826482.2;refGene_gn=SGK2;refGene_tx=XM_019826483.2;refGene_gn=SGK2;refGene_tx=XM_019826484.2
A3 17111734 17111784
refGene_gn=SGK2;refGene_tx=XM_019826482.2;refGene_gn=SGK2;refGene_tx=XM_019826483.2;refGene_gn=SGK2;refGene_tx=XM_019826484.2
A3 17112295 17112331
refGene_gn=SGK2;refGene_tx=XM_019826482.2;refGene_gn=SGK2;refGene_tx=XM_019826483.2;refGene_gn=SGK2;refGene_tx=XM_019826484.2
A3 17127377 17127588 refGene_gn=L3MBTL1;refGene_tx=XM_023251230.1
A3 17129114 17129245 refGene_gn=L3MBTL1;refGene_tx=XM_023251230.1
A3 17132420 17132483 refGene_gn=L3MBTL1;refGene_tx=XM_023251230.1
A3 17132913 17133071 refGene_gn=L3MBTL1;refGene_tx=XM_023251230.1
A3 17137201 17137251 refGene_gn=L3MBTL1;refGene_tx=XM_023251230.1
A3 17137341 17137539 refGene_gn=L3MBTL1;refGene_tx=XM_023251230.1
A3 17137722 17137822 refGene_gn=L3MBTL1;refGene_tx=XM_023251230.1
A3 17138663 17138778 refGene_gn=L3MBTL1;refGene_tx=XM_023251230.1
A3 17139244 17139355 refGene_gn=L3MBTL1;refGene_tx=XM_023251230.1
A3 17139554 17139640 refGene_gn=L3MBTL1;refGene_tx=XM_023251230.1
A3 17140211 17140288 refGene_gn=L3MBTL1;refGene_tx=XM_023251230.1
A3 17140715 17140867 refGene_gn=L3MBTL1;refGene_tx=XM_023251230.1
A3 17142056 17142148 refGene_gn=L3MBTL1;refGene_tx=XM_023251230.1
A3 17142481 17142617 refGene_gn=L3MBTL1;refGene_tx=XM_023251230.1
A3 17143513 17143618 refGene_gn=L3MBTL1;refGene_tx=XM_023251230.1
A3 17144138 17144227 refGene_gn=L3MBTL1;refGene_tx=XM_023251230.1
A3 17151765 17151850 refGene_gn=L3MBTL1;refGene_tx=XM_023251230.1
A3 17152476 17152606 refGene_gn=L3MBTL1;refGene_tx=XM_023251230.1
A3 17152752 17152903 refGene_gn=L3MBTL1;refGene_tx=XM_023251230.1
A3 17153130 17153272 refGene_gn=L3MBTL1;refGene_tx=XM_023251230.1
A3 17153592 17153819 refGene_gn=L3MBTL1;refGene_tx=XM_023251230.1
A3 17153971 17154107 refGene_gn=L3MBTL1;refGene_tx=XM_023251230.1
A3 17171378 17171739 refGene_gn=SRSF6;refGene_tx=XM_023251231.1
A3 17171835 17171919 refGene_gn=SRSF6;refGene_tx=XM_023251231.1
A3 17172187 17172396 refGene_gn=SRSF6;refGene_tx=XM_023251231.1
A3 17172763 17172888 refGene_gn=SRSF6;refGene_tx=XM_023251231.1

A3 17174230 17174379 refGene_gn=SRSF6;refGene_tx=XM_023251231.1
A3 17174602 17174709 refGene_gn=SRSF6;refGene_tx=XM_023251231.1
A3 17384931 17385019
refGene_gn=PTPRT;refGene_tx=XM_023251232.1;refGene_gn=PTPRT;refGene_tx=XM_023251233.1;refGene_gn=PTPRT;refGene_tx=XM_023251234.1;refGene_gn=PTPRT;refGene_tx=XM_023251235.1;refGene_gn=PTPRT;refGene_tx=XM_023251236.1;refGene_gn=PTPRT;refGene_tx=XM_023251237.1;refGene_gn=PTPRT;refGene_tx=XM_023251238.1;refGene_gn=PTPRT;refGene_tx=XM_023251239.1;refGene_gn=PTPRT;refGene_tx=XM_023251240.1;refGene_gn=PTPRT;refGene_tx=XM_023251241.1;refGene_gn=PTPRT;refGene_tx=XM_023251242.1;refGene_gn=PTPRT;refGene_tx=XM_023251243.1
A3 17661224 17661350
refGene_gn=PTPRT;refGene_tx=XM_023251232.1;refGene_gn=PTPRT;refGene_tx=XM_023251233.1;refGene_gn=PTPRT;refGene_tx=XM_023251234.1;refGene_gn=PTPRT;refGene_tx=XM_023251235.1;refGene_gn=PTPRT;refGene_tx=XM_023251236.1;refGene_gn=PTPRT;refGene_tx=XM_023251237.1;refGene_gn=PTPRT;refGene_tx=XM_023251238.1;refGene_gn=PTPRT;refGene_tx=XM_023251239.1;refGene_gn=PTPRT;refGene_tx=XM_023251240.1;refGene_gn=PTPRT;refGene_tx=XM_023251241.1;refGene_gn=PTPRT;refGene_tx=XM_023251242.1;refGene_gn=PTPRT;refGene_tx=XM_023251243.1
A3 17743601 17743873
refGene_gn=PTPRT;refGene_tx=XM_023251232.1;refGene_gn=PTPRT;refGene_tx=XM_023251233.1;refGene_gn=PTPRT;refGene_tx=XM_023251234.1;refGene_gn=PTPRT;refGene_tx=XM_023251235.1;refGene_gn=PTPRT;refGene_tx=XM_023251236.1;refGene_gn=PTPRT;refGene_tx=XM_023251237.1;refGene_gn=PTPRT;refGene_tx=XM_023251238.1;refGene_gn=PTPRT;refGene_tx=XM_023251239.1;refGene_gn=PTPRT;refGene_tx=XM_023251240.1;refGene_gn=PTPRT;refGene_tx=XM_023251241.1;refGene_gn=PTPRT;refGene_tx=XM_023251242.1;refGene_gn=PTPRT;refGene_tx=XM_023251243.1
A3 17750364 17750446
refGene_gn=PTPRT;refGene_tx=XM_023251232.1;refGene_gn=PTPRT;refGene_tx=XM_023251233.1;refGene_gn=PTPRT;refGene_tx=XM_023251234.1;refGene_gn=PTPRT;refGene_tx=XM_023251235.1;refGene_gn=PTPRT;refGene_tx=XM_023251236.1;refGene_gn=PTPRT;refGene_tx=XM_023251237.1;refGene_gn=PTPRT;refGene_tx=XM_023251238.1;refGene_gn=PTPRT;refGene_tx=XM_023251239.1;refGene_gn=PTPRT;refGene_tx=XM_023251240.1;refGene_gn=PTPRT;refGene_tx=XM_023251241.1;refGene_gn=PTPRT;refGene_tx=XM_023251242.1;refGene_gn=PTPRT;refGene_tx=XM_023251243.1
A3 17759470 17759586
refGene_gn=PTPRT;refGene_tx=XM_023251232.1;refGene_gn=PTPRT;refGene_tx=XM_023251233.1;refGene_gn=PTPRT;refGene_tx=XM_023251234.1;refGene_gn=PTPRT;refGene_tx=XM_023251235.1;refGene_gn=PTPRT;refGene_tx=XM_023251236.1;refGene_gn=PTPRT;refGene_tx=XM_023251237.1;refGene_gn=PTPRT;refGene_tx=XM_023251238.1;refGene_gn=PTPRT;refGene_tx=XM_023251239.1;refGene_gn=PTPRT;refGene_tx=XM_023251240.1;refGene_gn=PTPRT;refGene_tx=XM_023251241.1;refGene_gn=PTPRT;refGene_tx=XM_023251242.1;refGene_gn=PTPRT;refGene_tx=XM_023251243.1
A3 17777917 17778092
refGene_gn=PTPRT;refGene_tx=XM_023251232.1;refGene_gn=PTPRT;refGene_tx=XM_023251233.1;refGene_gn=PTPRT;refGene_tx=XM_023251234.1;refGene_gn=

PTPRT;refGene_tx=XM_023251235.1;refGene_gn=PTPRT;refGene_tx=XM_023251236.1;refGene_gn=PTPRT;refGene_tx=XM_023251237.1;refGene_gn=PTPRT;refGene_tx=XM_023251238.1;refGene_gn=PTPRT;refGene_tx=XM_023251239.1;refGene_gn=PTPRT;refGene_tx=XM_023251240.1;refGene_gn=PTPRT;refGene_tx=XM_023251241.1;refGene_gn=PTPRT;refGene_tx=XM_023251242.1;refGene_gn=PTPRT;refGene_tx=XM_023251243.1
A3 17852094 17852388
refGene_gn=PTPRT;refGene_tx=XM_023251232.1;refGene_gn=PTPRT;refGene_tx=XM_023251233.1;refGene_gn=PTPRT;refGene_tx=XM_023251234.1;refGene_gn=PTPRT;refGene_tx=XM_023251235.1;refGene_gn=PTPRT;refGene_tx=XM_023251236.1;refGene_gn=PTPRT;refGene_tx=XM_023251237.1;refGene_gn=PTPRT;refGene_tx=XM_023251238.1;refGene_gn=PTPRT;refGene_tx=XM_023251239.1;refGene_gn=PTPRT;refGene_tx=XM_023251240.1;refGene_gn=PTPRT;refGene_tx=XM_023251241.1;refGene_gn=PTPRT;refGene_tx=XM_023251242.1;refGene_gn=PTPRT;refGene_tx=XM_023251243.1
A3 18039904 18040201
refGene_gn=PTPRT;refGene_tx=XM_023251232.1;refGene_gn=PTPRT;refGene_tx=XM_023251233.1;refGene_gn=PTPRT;refGene_tx=XM_023251234.1;refGene_gn=PTPRT;refGene_tx=XM_023251235.1;refGene_gn=PTPRT;refGene_tx=XM_023251236.1;refGene_gn=PTPRT;refGene_tx=XM_023251237.1;refGene_gn=PTPRT;refGene_tx=XM_023251238.1;refGene_gn=PTPRT;refGene_tx=XM_023251239.1;refGene_gn=PTPRT;refGene_tx=XM_023251240.1;refGene_gn=PTPRT;refGene_tx=XM_023251241.1;refGene_gn=PTPRT;refGene_tx=XM_023251242.1;refGene_gn=PTPRT;refGene_tx=XM_023251243.1
A3 18062928 18063038
refGene_gn=PTPRT;refGene_tx=XM_023251232.1;refGene_gn=PTPRT;refGene_tx=XM_023251233.1;refGene_gn=PTPRT;refGene_tx=XM_023251234.1;refGene_gn=PTPRT;refGene_tx=XM_023251235.1;refGene_gn=PTPRT;refGene_tx=XM_023251236.1;refGene_gn=PTPRT;refGene_tx=XM_023251237.1;refGene_gn=PTPRT;refGene_tx=XM_023251238.1;refGene_gn=PTPRT;refGene_tx=XM_023251239.1;refGene_gn=PTPRT;refGene_tx=XM_023251240.1;refGene_gn=PTPRT;refGene_tx=XM_023251241.1;refGene_gn=PTPRT;refGene_tx=XM_023251242.1;refGene_gn=PTPRT;refGene_tx=XM_023251243.1
A3 18152629 18152831
refGene_gn=PTPRT;refGene_tx=XM_023251232.1;refGene_gn=PTPRT;refGene_tx=XM_023251233.1;refGene_gn=PTPRT;refGene_tx=XM_023251234.1;refGene_gn=PTPRT;refGene_tx=XM_023251235.1;refGene_gn=PTPRT;refGene_tx=XM_023251236.1;refGene_gn=PTPRT;refGene_tx=XM_023251237.1;refGene_gn=PTPRT;refGene_tx=XM_023251238.1;refGene_gn=PTPRT;refGene_tx=XM_023251239.1;refGene_gn=PTPRT;refGene_tx=XM_023251240.1;refGene_gn=PTPRT;refGene_tx=XM_023251241.1;refGene_gn=PTPRT;refGene_tx=XM_023251242.1;refGene_gn=PTPRT;refGene_tx=XM_023251243.1
A3 18154158 18154261
refGene_gn=PTPRT;refGene_tx=XM_023251232.1;refGene_gn=PTPRT;refGene_tx=XM_023251233.1;refGene_gn=PTPRT;refGene_tx=XM_023251234.1;refGene_gn=PTPRT;refGene_tx=XM_023251235.1;refGene_gn=PTPRT;refGene_tx=XM_023251236.1;refGene_gn=PTPRT;refGene_tx=XM_023251237.1;refGene_gn=PTPRT;refGene_tx=XM_023251238.1;refGene_gn=PTPRT;refGene_tx=XM_023251239.1;refGene_gn=PTPRT;refGene_tx=XM_023251240.1;refGene_gn=PTPRT;refGene_tx=XM_023251241.1;refGene_gn=PTPRT;refGene_tx=XM_023251242.1;refGene_gn=PTPRT;

refGene_tx=XM_023251243.1
A3 18207780 18208054
refGene_gn=PTPRT;refGene_tx=XM_023251232.1;refGene_gn=PTPRT;refGene_tx
=XM_023251233.1;refGene_gn=PTPRT;refGene_tx=XM_023251234.1;refGene_gn=
PTPRT;refGene_tx=XM_023251235.1;refGene_gn=PTPRT;refGene_tx=XM_0232512
36.1;refGene_gn=PTPRT;refGene_tx=XM_023251237.1;refGene_gn=PTPRT;refGe
ne_tx=XM_023251238.1;refGene_gn=PTPRT;refGene_tx=XM_023251239.1;refGen
e_gn=PTPRT;refGene_tx=XM_023251240.1;refGene_gn=PTPRT;refGene_tx=XM_02
3251241.1;refGene_gn=PTPRT;refGene_tx=XM_023251242.1;refGene_gn=PTPRT;
refGene_tx=XM_023251243.1
A3 18239346 18239383
refGene_gn=PTPRT;refGene_tx=XM_023251232.1;refGene_gn=PTPRT;refGene_tx
=XM_023251233.1;refGene_gn=PTPRT;refGene_tx=XM_023251234.1;refGene_gn=
PTPRT;refGene_tx=XM_023251235.1;refGene_gn=PTPRT;refGene_tx=XM_0232512
36.1;refGene_gn=PTPRT;refGene_tx=XM_023251237.1;refGene_gn=PTPRT;refGe
ne_tx=XM_023251238.1;refGene_gn=PTPRT;refGene_tx=XM_023251239.1;refGen
e_gn=PTPRT;refGene_tx=XM_023251240.1;refGene_gn=PTPRT;refGene_tx=XM_02
3251241.1;refGene_gn=PTPRT;refGene_tx=XM_023251242.1;refGene_gn=PTPRT;
refGene_tx=XM_023251243.1
A3 18253784 18253841
refGene_gn=PTPRT;refGene_tx=XM_023251232.1;refGene_gn=PTPRT;refGene_tx
=XM_023251233.1;refGene_gn=PTPRT;refGene_tx=XM_023251234.1;refGene_gn=
PTPRT;refGene_tx=XM_023251235.1;refGene_gn=PTPRT;refGene_tx=XM_0232512
37.1;refGene_gn=PTPRT;refGene_tx=XM_023251238.1;refGene_gn=PTPRT;refGe
ne_tx=XM_023251240.1
A3 18274225 18274361
refGene_gn=PTPRT;refGene_tx=XM_023251232.1;refGene_gn=PTPRT;refGene_tx
=XM_023251233.1;refGene_gn=PTPRT;refGene_tx=XM_023251234.1;refGene_gn=
PTPRT;refGene_tx=XM_023251235.1;refGene_gn=PTPRT;refGene_tx=XM_0232512
36.1;refGene_gn=PTPRT;refGene_tx=XM_023251237.1;refGene_gn=PTPRT;refGe
ne_tx=XM_023251238.1;refGene_gn=PTPRT;refGene_tx=XM_023251239.1;refGen
e_gn=PTPRT;refGene_tx=XM_023251240.1;refGene_gn=PTPRT;refGene_tx=XM_02
3251241.1;refGene_gn=PTPRT;refGene_tx=XM_023251242.1;refGene_gn=PTPRT;
refGene_tx=XM_023251243.1
A3 18286363 18286393
refGene_gn=PTPRT;refGene_tx=XM_023251232.1;refGene_gn=PTPRT;refGene_tx
=XM_023251233.1;refGene_gn=PTPRT;refGene_tx=XM_023251234.1;refGene_gn=
PTPRT;refGene_tx=XM_023251236.1;refGene_gn=PTPRT;refGene_tx=XM_0232512
37.1;refGene_gn=PTPRT;refGene_tx=XM_023251238.1;refGene_gn=PTPRT;refGe
ne_tx=XM_023251241.1;refGene_gn=PTPRT;refGene_tx=XM_023251242.1
A3 18312491 18312649
refGene_gn=PTPRT;refGene_tx=XM_023251232.1;refGene_gn=PTPRT;refGene_tx
=XM_023251233.1;refGene_gn=PTPRT;refGene_tx=XM_023251234.1;refGene_gn=
PTPRT;refGene_tx=XM_023251235.1;refGene_gn=PTPRT;refGene_tx=XM_0232512
36.1;refGene_gn=PTPRT;refGene_tx=XM_023251237.1;refGene_gn=PTPRT;refGe
ne_tx=XM_023251238.1;refGene_gn=PTPRT;refGene_tx=XM_023251239.1;refGen
e_gn=PTPRT;refGene_tx=XM_023251240.1;refGene_gn=PTPRT;refGene_tx=XM_02
3251241.1;refGene_gn=PTPRT;refGene_tx=XM_023251242.1;refGene_gn=PTPRT;
refGene_tx=XM_023251243.1
A3 18349147 18349338

refGene_gn=PTPRT;refGene_tx=XM_023251232.1;refGene_gn=PTPRT;refGene_tx
=XM_023251233.1;refGene_gn=PTPRT;refGene_tx=XM_023251234.1;refGene_gn=
PTPRT;refGene_tx=XM_023251235.1;refGene_gn=PTPRT;refGene_tx=XM_0232512
36.1;refGene_gn=PTPRT;refGene_tx=XM_023251237.1;refGene_gn=PTPRT;refGe
ne_tx=XM_023251238.1;refGene_gn=PTPRT;refGene_tx=XM_023251239.1;refGen
e_gn=PTPRT;refGene_tx=XM_023251240.1;refGene_gn=PTPRT;refGene_tx=XM_02
3251241.1;refGene_gn=PTPRT;refGene_tx=XM_023251242.1;refGene_gn=PTPRT;
refGene_tx=XM_023251243.1
A3 18368039 18368127
refGene_gn=PTPRT;refGene_tx=XM_023251232.1;refGene_gn=PTPRT;refGene_tx
=XM_023251233.1;refGene_gn=PTPRT;refGene_tx=XM_023251234.1;refGene_gn=
PTPRT;refGene_tx=XM_023251235.1;refGene_gn=PTPRT;refGene_tx=XM_0232512
36.1;refGene_gn=PTPRT;refGene_tx=XM_023251237.1;refGene_gn=PTPRT;refGe
ne_tx=XM_023251238.1;refGene_gn=PTPRT;refGene_tx=XM_023251239.1;refGen
e_gn=PTPRT;refGene_tx=XM_023251240.1;refGene_gn=PTPRT;refGene_tx=XM_02
3251241.1;refGene_gn=PTPRT;refGene_tx=XM_023251242.1;refGene_gn=PTPRT;
refGene_tx=XM_023251243.1
A3 18382605 18382682
refGene_gn=PTPRT;refGene_tx=XM_023251232.1;refGene_gn=PTPRT;refGene_tx
=XM_023251233.1;refGene_gn=PTPRT;refGene_tx=XM_023251234.1;refGene_gn=
PTPRT;refGene_tx=XM_023251235.1;refGene_gn=PTPRT;refGene_tx=XM_0232512
36.1;refGene_gn=PTPRT;refGene_tx=XM_023251237.1;refGene_gn=PTPRT;refGe
ne_tx=XM_023251238.1;refGene_gn=PTPRT;refGene_tx=XM_023251239.1;refGen
e_gn=PTPRT;refGene_tx=XM_023251240.1;refGene_gn=PTPRT;refGene_tx=XM_02
3251241.1;refGene_gn=PTPRT;refGene_tx=XM_023251242.1;refGene_gn=PTPRT;
refGene_tx=XM_023251243.1
A3 18390706 18390743
refGene_gn=PTPRT;refGene_tx=XM_023251232.1;refGene_gn=PTPRT;refGene_tx
=XM_023251233.1;refGene_gn=PTPRT;refGene_tx=XM_023251234.1;refGene_gn=
PTPRT;refGene_tx=XM_023251235.1;refGene_gn=PTPRT;refGene_tx=XM_0232512
36.1;refGene_gn=PTPRT;refGene_tx=XM_023251237.1;refGene_gn=PTPRT;refGe
ne_tx=XM_023251238.1;refGene_gn=PTPRT;refGene_tx=XM_023251239.1;refGen
e_gn=PTPRT;refGene_tx=XM_023251240.1;refGene_gn=PTPRT;refGene_tx=XM_02
3251241.1;refGene_gn=PTPRT;refGene_tx=XM_023251242.1;refGene_gn=PTPRT;
refGene_tx=XM_023251243.1
A3 18392041 18392139
refGene_gn=PTPRT;refGene_tx=XM_023251232.1;refGene_gn=PTPRT;refGene_tx
=XM_023251233.1;refGene_gn=PTPRT;refGene_tx=XM_023251234.1;refGene_gn=
PTPRT;refGene_tx=XM_023251235.1;refGene_gn=PTPRT;refGene_tx=XM_0232512
36.1;refGene_gn=PTPRT;refGene_tx=XM_023251237.1;refGene_gn=PTPRT;refGe
ne_tx=XM_023251238.1;refGene_gn=PTPRT;refGene_tx=XM_023251239.1;refGen
e_gn=PTPRT;refGene_tx=XM_023251240.1;refGene_gn=PTPRT;refGene_tx=XM_02
3251241.1;refGene_gn=PTPRT;refGene_tx=XM_023251242.1;refGene_gn=PTPRT;
refGene_tx=XM_023251243.1
A3 18394641 18394701
refGene_gn=PTPRT;refGene_tx=XM_023251232.1;refGene_gn=PTPRT;refGene_tx
=XM_023251233.1;refGene_gn=PTPRT;refGene_tx=XM_023251234.1;refGene_gn=
PTPRT;refGene_tx=XM_023251235.1;refGene_gn=PTPRT;refGene_tx=XM_0232512
36.1;refGene_gn=PTPRT;refGene_tx=XM_023251239.1
A3 18395425 18395542

refGene_gn=PTPRT;refGene_tx=XM_023251232.1;refGene_gn=PTPRT;refGene_tx
=XM_023251233.1;refGene_gn=PTPRT;refGene_tx=XM_023251234.1;refGene_gn=
PTPRT;refGene_tx=XM_023251235.1;refGene_gn=PTPRT;refGene_tx=XM_0232512
36.1;refGene_gn=PTPRT;refGene_tx=XM_023251237.1;refGene_gn=PTPRT;refGe
ne_tx=XM_023251238.1;refGene_gn=PTPRT;refGene_tx=XM_023251239.1;refGen
e_gn=PTPRT;refGene_tx=XM_023251240.1;refGene_gn=PTPRT;refGene_tx=XM_02
3251241.1;refGene_gn=PTPRT;refGene_tx=XM_023251242.1;refGene_gn=PTPRT;
refGene_tx=XM_023251243.1
A3 18400098 18400253
refGene_gn=PTPRT;refGene_tx=XM_023251232.1;refGene_gn=PTPRT;refGene_tx
=XM_023251233.1;refGene_gn=PTPRT;refGene_tx=XM_023251234.1;refGene_gn=
PTPRT;refGene_tx=XM_023251235.1;refGene_gn=PTPRT;refGene_tx=XM_0232512
36.1;refGene_gn=PTPRT;refGene_tx=XM_023251237.1;refGene_gn=PTPRT;refGe
ne_tx=XM_023251238.1;refGene_gn=PTPRT;refGene_tx=XM_023251239.1;refGen
e_gn=PTPRT;refGene_tx=XM_023251240.1;refGene_gn=PTPRT;refGene_tx=XM_02
3251241.1;refGene_gn=PTPRT;refGene_tx=XM_023251242.1;refGene_gn=PTPRT;
refGene_tx=XM_023251243.1
A3 18403554 18403690
refGene_gn=PTPRT;refGene_tx=XM_023251232.1;refGene_gn=PTPRT;refGene_tx
=XM_023251233.1;refGene_gn=PTPRT;refGene_tx=XM_023251234.1;refGene_gn=
PTPRT;refGene_tx=XM_023251235.1;refGene_gn=PTPRT;refGene_tx=XM_0232512
36.1;refGene_gn=PTPRT;refGene_tx=XM_023251237.1;refGene_gn=PTPRT;refGe
ne_tx=XM_023251238.1;refGene_gn=PTPRT;refGene_tx=XM_023251239.1;refGen
e_gn=PTPRT;refGene_tx=XM_023251240.1;refGene_gn=PTPRT;refGene_tx=XM_02
3251241.1;refGene_gn=PTPRT;refGene_tx=XM_023251242.1;refGene_gn=PTPRT;
refGene_tx=XM_023251243.1
A3 18405770 18405920
refGene_gn=PTPRT;refGene_tx=XM_023251232.1;refGene_gn=PTPRT;refGene_tx
=XM_023251233.1;refGene_gn=PTPRT;refGene_tx=XM_023251234.1;refGene_gn=
PTPRT;refGene_tx=XM_023251235.1;refGene_gn=PTPRT;refGene_tx=XM_0232512
36.1;refGene_gn=PTPRT;refGene_tx=XM_023251237.1;refGene_gn=PTPRT;refGe
ne_tx=XM_023251238.1;refGene_gn=PTPRT;refGene_tx=XM_023251239.1;refGen
e_gn=PTPRT;refGene_tx=XM_023251240.1;refGene_gn=PTPRT;refGene_tx=XM_02
3251241.1;refGene_gn=PTPRT;refGene_tx=XM_023251242.1;refGene_gn=PTPRT;
refGene_tx=XM_023251243.1
A3 18407416 18407590
refGene_gn=PTPRT;refGene_tx=XM_023251232.1;refGene_gn=PTPRT;refGene_tx
=XM_023251233.1;refGene_gn=PTPRT;refGene_tx=XM_023251234.1;refGene_gn=
PTPRT;refGene_tx=XM_023251235.1;refGene_gn=PTPRT;refGene_tx=XM_0232512
36.1;refGene_gn=PTPRT;refGene_tx=XM_023251237.1;refGene_gn=PTPRT;refGe
ne_tx=XM_023251238.1;refGene_gn=PTPRT;refGene_tx=XM_023251239.1;refGen
e_gn=PTPRT;refGene_tx=XM_023251240.1;refGene_gn=PTPRT;refGene_tx=XM_02
3251241.1;refGene_gn=PTPRT;refGene_tx=XM_023251242.1;refGene_gn=PTPRT;
refGene_tx=XM_023251243.1
A3 18411174 18411306
refGene_gn=PTPRT;refGene_tx=XM_023251232.1;refGene_gn=PTPRT;refGene_tx
=XM_023251233.1;refGene_gn=PTPRT;refGene_tx=XM_023251234.1;refGene_gn=
PTPRT;refGene_tx=XM_023251235.1;refGene_gn=PTPRT;refGene_tx=XM_0232512
36.1;refGene_gn=PTPRT;refGene_tx=XM_023251237.1;refGene_gn=PTPRT;refGe
ne_tx=XM_023251238.1;refGene_gn=PTPRT;refGene_tx=XM_023251239.1;refGen

e_gn=PTPRT;refGene_tx=XM_023251240.1;refGene_gn=PTPRT;refGene_tx=XM_023251241.1;refGene_gn=PTPRT;refGene_tx=XM_023251242.1;refGene_gn=PTPRT;refGene_tx=XM_023251243.1
A3 18437328 18437454
refGene_gn=PTPRT;refGene_tx=XM_023251232.1;refGene_gn=PTPRT;refGene_tx=XM_023251233.1;refGene_gn=PTPRT;refGene_tx=XM_023251234.1;refGene_gn=PTPRT;refGene_tx=XM_023251235.1;refGene_gn=PTPRT;refGene_tx=XM_023251236.1;refGene_gn=PTPRT;refGene_tx=XM_023251237.1;refGene_gn=PTPRT;refGene_tx=XM_023251238.1;refGene_gn=PTPRT;refGene_tx=XM_023251239.1;refGene_gn=PTPRT;refGene_tx=XM_023251240.1;refGene_gn=PTPRT;refGene_tx=XM_023251241.1;refGene_gn=PTPRT;refGene_tx=XM_023251242.1;refGene_gn=PTPRT;refGene_tx=XM_023251243.1
A3 18438317 18438481
refGene_gn=PTPRT;refGene_tx=XM_023251232.1;refGene_gn=PTPRT;refGene_tx=XM_023251233.1;refGene_gn=PTPRT;refGene_tx=XM_023251234.1;refGene_gn=PTPRT;refGene_tx=XM_023251235.1;refGene_gn=PTPRT;refGene_tx=XM_023251236.1;refGene_gn=PTPRT;refGene_tx=XM_023251237.1;refGene_gn=PTPRT;refGene_tx=XM_023251238.1;refGene_gn=PTPRT;refGene_tx=XM_023251239.1;refGene_gn=PTPRT;refGene_tx=XM_023251240.1;refGene_gn=PTPRT;refGene_tx=XM_023251241.1;refGene_gn=PTPRT;refGene_tx=XM_023251242.1;refGene_gn=PTPRT;refGene_tx=XM_023251243.1
A3 18441659 18441795
refGene_gn=PTPRT;refGene_tx=XM_023251232.1;refGene_gn=PTPRT;refGene_tx=XM_023251233.1;refGene_gn=PTPRT;refGene_tx=XM_023251234.1;refGene_gn=PTPRT;refGene_tx=XM_023251235.1;refGene_gn=PTPRT;refGene_tx=XM_023251236.1;refGene_gn=PTPRT;refGene_tx=XM_023251237.1;refGene_gn=PTPRT;refGene_tx=XM_023251238.1;refGene_gn=PTPRT;refGene_tx=XM_023251239.1;refGene_gn=PTPRT;refGene_tx=XM_023251240.1;refGene_gn=PTPRT;refGene_tx=XM_023251241.1;refGene_gn=PTPRT;refGene_tx=XM_023251242.1;refGene_gn=PTPRT;refGene_tx=XM_023251243.1
A3 18442672 18442726
refGene_gn=PTPRT;refGene_tx=XM_023251232.1;refGene_gn=PTPRT;refGene_tx=XM_023251233.1;refGene_gn=PTPRT;refGene_tx=XM_023251234.1;refGene_gn=PTPRT;refGene_tx=XM_023251235.1;refGene_gn=PTPRT;refGene_tx=XM_023251236.1;refGene_gn=PTPRT;refGene_tx=XM_023251237.1;refGene_gn=PTPRT;refGene_tx=XM_023251238.1;refGene_gn=PTPRT;refGene_tx=XM_023251239.1;refGene_gn=PTPRT;refGene_tx=XM_023251240.1;refGene_gn=PTPRT;refGene_tx=XM_023251241.1;refGene_gn=PTPRT;refGene_tx=XM_023251242.1;refGene_gn=PTPRT;refGene_tx=XM_023251243.1
A3 18915396 18915429
refGene_gn=CHD6;refGene_tx=XM_019826498.2;refGene_gn=CHD6;refGene_tx=XM_019826499.2;refGene_gn=CHD6;refGene_tx=XM_023251244.1;refGene_gn=CHD6;refGene_tx=XM_023251245.1;refGene_gn=CHD6;refGene_tx=XM_023251247.1;refGene_gn=CHD6;refGene_tx=XM_023251248.1
A3 18927797 18928318
refGene_gn=CHD6;refGene_tx=XM_019826498.2;refGene_gn=CHD6;refGene_tx=XM_019826499.2;refGene_gn=CHD6;refGene_tx=XM_023251244.1;refGene_gn=CHD6;refGene_tx=XM_023251245.1;refGene_gn=CHD6;refGene_tx=XM_023251246.1;refGene_gn=CHD6;refGene_tx=XM_023251247.1;refGene_gn=CHD6;refGene_tx=XM_023251248.1

A3 18946003 18946151
refGene_gn=CHD6;refGene_tx=XM_019826498.2;refGene_gn=CHD6;refGene_tx=XM_019826499.2;refGene_gn=CHD6;refGene_tx=XM_023251244.1;refGene_gn=CHD6;refGene_tx=XM_023251245.1;refGene_gn=CHD6;refGene_tx=XM_023251246.1;refGene_gn=CHD6;refGene_tx=XM_023251247.1;refGene_gn=CHD6;refGene_tx=XM_023251248.1
A3 18947973 18948123
refGene_gn=CHD6;refGene_tx=XM_019826498.2;refGene_gn=CHD6;refGene_tx=XM_019826499.2;refGene_gn=CHD6;refGene_tx=XM_023251244.1;refGene_gn=CHD6;refGene_tx=XM_023251245.1;refGene_gn=CHD6;refGene_tx=XM_023251246.1;refGene_gn=CHD6;refGene_tx=XM_023251247.1;refGene_gn=CHD6;refGene_tx=XM_023251248.1
A3 18962560 18962623
refGene_gn=CHD6;refGene_tx=XM_019826498.2;refGene_gn=CHD6;refGene_tx=XM_019826499.2;refGene_gn=CHD6;refGene_tx=XM_023251244.1;refGene_gn=CHD6;refGene_tx=XM_023251245.1;refGene_gn=CHD6;refGene_tx=XM_023251246.1;refGene_gn=CHD6;refGene_tx=XM_023251247.1;refGene_gn=CHD6;refGene_tx=XM_023251248.1
A3 18965162 18965221
refGene_gn=CHD6;refGene_tx=XM_019826498.2;refGene_gn=CHD6;refGene_tx=XM_019826499.2;refGene_gn=CHD6;refGene_tx=XM_023251244.1;refGene_gn=CHD6;refGene_tx=XM_023251245.1;refGene_gn=CHD6;refGene_tx=XM_023251246.1;refGene_gn=CHD6;refGene_tx=XM_023251247.1;refGene_gn=CHD6;refGene_tx=XM_023251248.1
A3 18965858 18965976
refGene_gn=CHD6;refGene_tx=XM_019826498.2;refGene_gn=CHD6;refGene_tx=XM_019826499.2;refGene_gn=CHD6;refGene_tx=XM_023251244.1;refGene_gn=CHD6;refGene_tx=XM_023251245.1;refGene_gn=CHD6;refGene_tx=XM_023251246.1;refGene_gn=CHD6;refGene_tx=XM_023251247.1;refGene_gn=CHD6;refGene_tx=XM_023251248.1
A3 18970974 18971061
refGene_gn=CHD6;refGene_tx=XM_019826498.2;refGene_gn=CHD6;refGene_tx=XM_019826499.2;refGene_gn=CHD6;refGene_tx=XM_023251244.1;refGene_gn=CHD6;refGene_tx=XM_023251245.1;refGene_gn=CHD6;refGene_tx=XM_023251246.1;refGene_gn=CHD6;refGene_tx=XM_023251247.1;refGene_gn=CHD6;refGene_tx=XM_023251248.1
A3 18971246 18971381
refGene_gn=CHD6;refGene_tx=XM_019826498.2;refGene_gn=CHD6;refGene_tx=XM_019826499.2;refGene_gn=CHD6;refGene_tx=XM_023251244.1;refGene_gn=CHD6;refGene_tx=XM_023251245.1;refGene_gn=CHD6;refGene_tx=XM_023251246.1;refGene_gn=CHD6;refGene_tx=XM_023251247.1;refGene_gn=CHD6;refGene_tx=XM_023251248.1
A3 18976575 18976697
refGene_gn=CHD6;refGene_tx=XM_019826498.2;refGene_gn=CHD6;refGene_tx=XM_019826499.2;refGene_gn=CHD6;refGene_tx=XM_023251244.1;refGene_gn=CHD6;refGene_tx=XM_023251245.1;refGene_gn=CHD6;refGene_tx=XM_023251246.1;refGene_gn=CHD6;refGene_tx=XM_023251247.1;refGene_gn=CHD6;refGene_tx=XM_023251248.1
A3 18978127 18978371
refGene_gn=CHD6;refGene_tx=XM_019826498.2;refGene_gn=CHD6;refGene_tx=X

M_019826499.2;refGene_gn=CHD6;refGene_tx=XM_019826502.2;refGene_gn=CHD6;refGene_tx=XM_023251244.1;refGene_gn=CHD6;refGene_tx=XM_023251245.1;refGene_gn=CHD6;refGene_tx=XM_023251246.1;refGene_gn=CHD6;refGene_tx=XM_023251247.1;refGene_gn=CHD6;refGene_tx=XM_023251248.1
A3 18979587 18979764
refGene_gn=CHD6;refGene_tx=XM_019826498.2;refGene_gn=CHD6;refGene_tx=XM_019826499.2;refGene_gn=CHD6;refGene_tx=XM_019826502.2;refGene_gn=CHD6;refGene_tx=XM_023251244.1;refGene_gn=CHD6;refGene_tx=XM_023251245.1;refGene_gn=CHD6;refGene_tx=XM_023251246.1;refGene_gn=CHD6;refGene_tx=XM_023251247.1;refGene_gn=CHD6;refGene_tx=XM_023251248.1
A3 18980360 18980504
refGene_gn=CHD6;refGene_tx=XM_019826498.2;refGene_gn=CHD6;refGene_tx=XM_019826499.2;refGene_gn=CHD6;refGene_tx=XM_019826502.2;refGene_gn=CHD6;refGene_tx=XM_023251244.1;refGene_gn=CHD6;refGene_tx=XM_023251245.1;refGene_gn=CHD6;refGene_tx=XM_023251246.1;refGene_gn=CHD6;refGene_tx=XM_023251247.1;refGene_gn=CHD6;refGene_tx=XM_023251248.1
A3 18983442 18983698
refGene_gn=CHD6;refGene_tx=XM_019826498.2;refGene_gn=CHD6;refGene_tx=XM_019826499.2;refGene_gn=CHD6;refGene_tx=XM_019826502.2;refGene_gn=CHD6;refGene_tx=XM_023251244.1;refGene_gn=CHD6;refGene_tx=XM_023251245.1;refGene_gn=CHD6;refGene_tx=XM_023251246.1;refGene_gn=CHD6;refGene_tx=XM_023251247.1;refGene_gn=CHD6;refGene_tx=XM_023251248.1
A3 18984508 18984719
refGene_gn=CHD6;refGene_tx=XM_019826498.2;refGene_gn=CHD6;refGene_tx=XM_019826499.2;refGene_gn=CHD6;refGene_tx=XM_019826502.2;refGene_gn=CHD6;refGene_tx=XM_023251244.1;refGene_gn=CHD6;refGene_tx=XM_023251245.1;refGene_gn=CHD6;refGene_tx=XM_023251246.1;refGene_gn=CHD6;refGene_tx=XM_023251247.1;refGene_gn=CHD6;refGene_tx=XM_023251248.1
A3 18993324 18993520
refGene_gn=CHD6;refGene_tx=XM_019826498.2;refGene_gn=CHD6;refGene_tx=XM_019826499.2;refGene_gn=CHD6;refGene_tx=XM_019826502.2;refGene_gn=CHD6;refGene_tx=XM_023251244.1;refGene_gn=CHD6;refGene_tx=XM_023251245.1;refGene_gn=CHD6;refGene_tx=XM_023251246.1;refGene_gn=CHD6;refGene_tx=XM_023251247.1;refGene_gn=CHD6;refGene_tx=XM_023251248.1
A3 19006589 19006754
refGene_gn=CHD6;refGene_tx=XM_019826498.2;refGene_gn=CHD6;refGene_tx=XM_019826499.2;refGene_gn=CHD6;refGene_tx=XM_019826502.2;refGene_gn=CHD6;refGene_tx=XM_023251244.1;refGene_gn=CHD6;refGene_tx=XM_023251245.1;refGene_gn=CHD6;refGene_tx=XM_023251246.1;refGene_gn=CHD6;refGene_tx=XM_023251247.1;refGene_gn=CHD6;refGene_tx=XM_023251248.1
A3 19008059 19008239
refGene_gn=CHD6;refGene_tx=XM_019826498.2;refGene_gn=CHD6;refGene_tx=XM_019826499.2;refGene_gn=CHD6;refGene_tx=XM_019826502.2;refGene_gn=CHD6;refGene_tx=XM_023251244.1;refGene_gn=CHD6;refGene_tx=XM_023251245.1;refGene_gn=CHD6;refGene_tx=XM_023251246.1;refGene_gn=CHD6;refGene_tx=XM_023251247.1;refGene_gn=CHD6;refGene_tx=XM_023251248.1
A3 19009303 19009414
refGene_gn=CHD6;refGene_tx=XM_019826498.2;refGene_gn=CHD6;refGene_tx=XM_019826499.2;refGene_gn=CHD6;refGene_tx=XM_019826502.2;refGene_gn=CHD6;refGene_tx=XM_023251244.1;refGene_gn=CHD6;refGene_tx=XM_023251245.1;

refGene_gn=CHD6;refGene_tx=XM_023251246.1;refGene_gn=CHD6;refGene_tx=XM_023251247.1;refGene_gn=CHD6;refGene_tx=XM_023251248.1
A3 19011384 19011587
refGene_gn=CHD6;refGene_tx=XM_019826498.2;refGene_gn=CHD6;refGene_tx=XM_019826499.2;refGene_gn=CHD6;refGene_tx=XM_019826502.2;refGene_gn=CHD6;refGene_tx=XM_023251244.1;refGene_gn=CHD6;refGene_tx=XM_023251245.1;refGene_gn=CHD6;refGene_tx=XM_023251246.1;refGene_gn=CHD6;refGene_tx=XM_023251247.1;refGene_gn=CHD6;refGene_tx=XM_023251248.1
A3 19012262 19012462
refGene_gn=CHD6;refGene_tx=XM_019826498.2;refGene_gn=CHD6;refGene_tx=XM_019826499.2;refGene_gn=CHD6;refGene_tx=XM_019826502.2;refGene_gn=CHD6;refGene_tx=XM_023251244.1;refGene_gn=CHD6;refGene_tx=XM_023251245.1;refGene_gn=CHD6;refGene_tx=XM_023251246.1;refGene_gn=CHD6;refGene_tx=XM_023251247.1;refGene_gn=CHD6;refGene_tx=XM_023251248.1
A3 19013157 19013317
refGene_gn=CHD6;refGene_tx=XM_019826498.2;refGene_gn=CHD6;refGene_tx=XM_019826499.2;refGene_gn=CHD6;refGene_tx=XM_019826502.2;refGene_gn=CHD6;refGene_tx=XM_023251244.1;refGene_gn=CHD6;refGene_tx=XM_023251245.1;refGene_gn=CHD6;refGene_tx=XM_023251246.1;refGene_gn=CHD6;refGene_tx=XM_023251247.1;refGene_gn=CHD6;refGene_tx=XM_023251248.1
A3 19017901 19017991
refGene_gn=CHD6;refGene_tx=XM_019826498.2;refGene_gn=CHD6;refGene_tx=XM_019826499.2;refGene_gn=CHD6;refGene_tx=XM_019826502.2;refGene_gn=CHD6;refGene_tx=XM_023251244.1;refGene_gn=CHD6;refGene_tx=XM_023251245.1;refGene_gn=CHD6;refGene_tx=XM_023251246.1;refGene_gn=CHD6;refGene_tx=XM_023251247.1;refGene_gn=CHD6;refGene_tx=XM_023251248.1
A3 19018666 19018770
refGene_gn=CHD6;refGene_tx=XM_019826498.2;refGene_gn=CHD6;refGene_tx=XM_019826499.2;refGene_gn=CHD6;refGene_tx=XM_019826502.2;refGene_gn=CHD6;refGene_tx=XM_023251244.1;refGene_gn=CHD6;refGene_tx=XM_023251245.1;refGene_gn=CHD6;refGene_tx=XM_023251246.1;refGene_gn=CHD6;refGene_tx=XM_023251247.1;refGene_gn=CHD6;refGene_tx=XM_023251248.1
A3 19023983 19024113
refGene_gn=CHD6;refGene_tx=XM_019826498.2;refGene_gn=CHD6;refGene_tx=XM_019826499.2;refGene_gn=CHD6;refGene_tx=XM_019826502.2;refGene_gn=CHD6;refGene_tx=XM_023251244.1;refGene_gn=CHD6;refGene_tx=XM_023251245.1;refGene_gn=CHD6;refGene_tx=XM_023251246.1;refGene_gn=CHD6;refGene_tx=XM_023251247.1;refGene_gn=CHD6;refGene_tx=XM_023251248.1
A3 19025745 19025806
refGene_gn=CHD6;refGene_tx=XM_019826498.2;refGene_gn=CHD6;refGene_tx=XM_019826499.2;refGene_gn=CHD6;refGene_tx=XM_019826502.2;refGene_gn=CHD6;refGene_tx=XM_023251244.1;refGene_gn=CHD6;refGene_tx=XM_023251245.1;refGene_gn=CHD6;refGene_tx=XM_023251246.1;refGene_gn=CHD6;refGene_tx=XM_023251247.1;refGene_gn=CHD6;refGene_tx=XM_023251248.1
A3 19040376 19040437
refGene_gn=CHD6;refGene_tx=XM_019826498.2;refGene_gn=CHD6;refGene_tx=XM_019826499.2;refGene_gn=CHD6;refGene_tx=XM_019826502.2;refGene_gn=CHD6;refGene_tx=XM_023251244.1;refGene_gn=CHD6;refGene_tx=XM_023251245.1;refGene_gn=CHD6;refGene_tx=XM_023251246.1;refGene_gn=CHD6;refGene_tx=XM_023251247.1;refGene_gn=CHD6;refGene_tx=XM_023251248.1

A3 19041104 19041321
refGene_gn=CHD6;refGene_tx=XM_019826498.2;refGene_gn=CHD6;refGene_tx=XM_019826499.2;refGene_gn=CHD6;refGene_tx=XM_019826502.2;refGene_gn=CHD6;refGene_tx=XM_023251244.1;refGene_gn=CHD6;refGene_tx=XM_023251245.1;refGene_gn=CHD6;refGene_tx=XM_023251246.1;refGene_gn=CHD6;refGene_tx=XM_023251247.1;refGene_gn=CHD6;refGene_tx=XM_023251248.1
A3 19042998 19043207
refGene_gn=CHD6;refGene_tx=XM_019826498.2;refGene_gn=CHD6;refGene_tx=XM_019826499.2;refGene_gn=CHD6;refGene_tx=XM_019826502.2;refGene_gn=CHD6;refGene_tx=XM_023251244.1;refGene_gn=CHD6;refGene_tx=XM_023251245.1;refGene_gn=CHD6;refGene_tx=XM_023251246.1;refGene_gn=CHD6;refGene_tx=XM_023251247.1;refGene_gn=CHD6;refGene_tx=XM_023251248.1
A3 19044326 19045904
refGene_gn=CHD6;refGene_tx=XM_019826498.2;refGene_gn=CHD6;refGene_tx=XM_019826499.2;refGene_gn=CHD6;refGene_tx=XM_019826502.2;refGene_gn=CHD6;refGene_tx=XM_023251244.1;refGene_gn=CHD6;refGene_tx=XM_023251245.1;refGene_gn=CHD6;refGene_tx=XM_023251246.1;refGene_gn=CHD6;refGene_tx=XM_023251247.1;refGene_gn=CHD6;refGene_tx=XM_023251248.1
A3 19049022 19049174
refGene_gn=CHD6;refGene_tx=XM_019826498.2;refGene_gn=CHD6;refGene_tx=XM_019826499.2;refGene_gn=CHD6;refGene_tx=XM_019826502.2;refGene_gn=CHD6;refGene_tx=XM_023251244.1;refGene_gn=CHD6;refGene_tx=XM_023251245.1;refGene_gn=CHD6;refGene_tx=XM_023251246.1;refGene_gn=CHD6;refGene_tx=XM_023251247.1;refGene_gn=CHD6;refGene_tx=XM_023251248.1
A3 19049546 19049753
refGene_gn=CHD6;refGene_tx=XM_019826498.2;refGene_gn=CHD6;refGene_tx=XM_019826499.2;refGene_gn=CHD6;refGene_tx=XM_019826502.2;refGene_gn=CHD6;refGene_tx=XM_023251244.1;refGene_gn=CHD6;refGene_tx=XM_023251245.1;refGene_gn=CHD6;refGene_tx=XM_023251246.1;refGene_gn=CHD6;refGene_tx=XM_023251247.1;refGene_gn=CHD6;refGene_tx=XM_023251248.1
A3 19050758 19051211
refGene_gn=CHD6;refGene_tx=XM_019826498.2;refGene_gn=CHD6;refGene_tx=XM_019826499.2;refGene_gn=CHD6;refGene_tx=XM_019826502.2;refGene_gn=CHD6;refGene_tx=XM_023251244.1;refGene_gn=CHD6;refGene_tx=XM_023251245.1;refGene_gn=CHD6;refGene_tx=XM_023251246.1;refGene_gn=CHD6;refGene_tx=XM_023251247.1;refGene_gn=CHD6;refGene_tx=XM_023251248.1
A3 19053128 19053320
refGene_gn=CHD6;refGene_tx=XM_019826498.2;refGene_gn=CHD6;refGene_tx=XM_019826499.2;refGene_gn=CHD6;refGene_tx=XM_019826502.2;refGene_gn=CHD6;refGene_tx=XM_023251244.1;refGene_gn=CHD6;refGene_tx=XM_023251245.1;refGene_gn=CHD6;refGene_tx=XM_023251246.1;refGene_gn=CHD6;refGene_tx=XM_023251247.1;refGene_gn=CHD6;refGene_tx=XM_023251248.1
A3 19054307 19054427
refGene_gn=CHD6;refGene_tx=XM_019826498.2;refGene_gn=CHD6;refGene_tx=XM_019826499.2;refGene_gn=CHD6;refGene_tx=XM_019826502.2;refGene_gn=CHD6;refGene_tx=XM_023251244.1;refGene_gn=CHD6;refGene_tx=XM_023251245.1;refGene_gn=CHD6;refGene_tx=XM_023251246.1;refGene_gn=CHD6;refGene_tx=XM_023251247.1
A3 19061128 19062025
refGene_gn=CHD6;refGene_tx=XM_019826498.2;refGene_gn=CHD6;refGene_tx=X

M_019826499.2;refGene_gn=CHD6;refGene_tx=XM_019826502.2;refGene_gn=CHD6;refGene_tx=XM_023251244.1;refGene_gn=CHD6;refGene_tx=XM_023251245.1;refGene_gn=CHD6;refGene_tx=XM_023251246.1;refGene_gn=CHD6;refGene_tx=XM_023251247.1;refGene_gn=CHD6;refGene_tx=XM_023251248.1
A3 19095422 19095586 refGene_gn=EMILIN3;refGene_tx=XM_023251249.1
A3 19096876 19096999 refGene_gn=EMILIN3;refGene_tx=XM_023251249.1
A3 19098148 19098372 refGene_gn=EMILIN3;refGene_tx=XM_023251249.1
A3 19098643 19100397 refGene_gn=EMILIN3;refGene_tx=XM_023251249.1
A3 19102466 19102611
refGene_gn=LPIN3;refGene_tx=XM_006929748.4;refGene_gn=LPIN3;refGene_tx=XM_019826503.2;refGene_gn=LPIN3;refGene_tx=XM_019826504.2;refGene_gn=LPIN3;refGene_tx=XM_019826505.2
A3 19102786 19102890
refGene_gn=LPIN3;refGene_tx=XM_006929748.4;refGene_gn=LPIN3;refGene_tx=XM_019826503.2;refGene_gn=LPIN3;refGene_tx=XM_019826504.2;refGene_gn=LPIN3;refGene_tx=XM_019826505.2
A3 19102975 19103090
refGene_gn=LPIN3;refGene_tx=XM_006929748.4;refGene_gn=LPIN3;refGene_tx=XM_019826503.2;refGene_gn=LPIN3;refGene_tx=XM_019826504.2;refGene_gn=LPIN3;refGene_tx=XM_019826505.2
A3 19103247 19103400
refGene_gn=LPIN3;refGene_tx=XM_006929748.4;refGene_gn=LPIN3;refGene_tx=XM_019826503.2;refGene_gn=LPIN3;refGene_tx=XM_019826504.2;refGene_gn=LPIN3;refGene_tx=XM_019826505.2
A3 19103883 19103970
refGene_gn=LPIN3;refGene_tx=XM_006929748.4;refGene_gn=LPIN3;refGene_tx=XM_019826503.2;refGene_gn=LPIN3;refGene_tx=XM_019826504.2;refGene_gn=LPIN3;refGene_tx=XM_019826505.2
A3 19104150 19104299
refGene_gn=LPIN3;refGene_tx=XM_006929748.4;refGene_gn=LPIN3;refGene_tx=XM_019826503.2;refGene_gn=LPIN3;refGene_tx=XM_019826504.2;refGene_gn=LPIN3;refGene_tx=XM_019826505.2
A3 19105249 19105388
refGene_gn=LPIN3;refGene_tx=XM_006929748.4;refGene_gn=LPIN3;refGene_tx=XM_019826503.2;refGene_gn=LPIN3;refGene_tx=XM_019826504.2;refGene_gn=LPIN3;refGene_tx=XM_019826505.2
A3 19106111 19106155
refGene_gn=LPIN3;refGene_tx=XM_006929748.4;refGene_gn=LPIN3;refGene_tx=XM_019826503.2;refGene_gn=LPIN3;refGene_tx=XM_019826504.2;refGene_gn=LPIN3;refGene_tx=XM_019826505.2
A3 19106230 19106323
refGene_gn=LPIN3;refGene_tx=XM_006929748.4;refGene_gn=LPIN3;refGene_tx=XM_019826503.2;refGene_gn=LPIN3;refGene_tx=XM_019826504.2;refGene_gn=LPIN3;refGene_tx=XM_019826505.2
A3 19107985 19108055
refGene_gn=LPIN3;refGene_tx=XM_006929748.4;refGene_gn=LPIN3;refGene_tx=XM_019826503.2;refGene_gn=LPIN3;refGene_tx=XM_019826504.2;refGene_gn=LPIN3;refGene_tx=XM_019826505.2
A3 19108153 19108247
refGene_gn=LPIN3;refGene_tx=XM_006929748.4;refGene_gn=LPIN3;refGene_tx

=XM_019826503.2;refGene_gn=LPIN3;refGene_tx=XM_019826504.2;refGene_gn=LPIN3;refGene_tx=XM_019826505.2
A3 19108655 19108807
refGene_gn=LPIN3;refGene_tx=XM_006929748.4;refGene_gn=LPIN3;refGene_tx=XM_019826503.2;refGene_gn=LPIN3;refGene_tx=XM_019826504.2;refGene_gn=LPIN3;refGene_tx=XM_019826505.2
A3 19108945 19109045
refGene_gn=LPIN3;refGene_tx=XM_006929748.4;refGene_gn=LPIN3;refGene_tx=XM_019826503.2;refGene_gn=LPIN3;refGene_tx=XM_019826504.2;refGene_gn=LPIN3;refGene_tx=XM_019826505.2
A3 19109832 19110175
refGene_gn=LPIN3;refGene_tx=XM_006929748.4;refGene_gn=LPIN3;refGene_tx=XM_019826503.2;refGene_gn=LPIN3;refGene_tx=XM_019826504.2;refGene_gn=LPIN3;refGene_tx=XM_019826505.2
A3 19110333 19110457
refGene_gn=LPIN3;refGene_tx=XM_006929748.4;refGene_gn=LPIN3;refGene_tx=XM_019826503.2;refGene_gn=LPIN3;refGene_tx=XM_019826504.2;refGene_gn=LPIN3;refGene_tx=XM_019826505.2
A3 19110799 19110883
refGene_gn=LPIN3;refGene_tx=XM_006929748.4;refGene_gn=LPIN3;refGene_tx=XM_019826503.2;refGene_gn=LPIN3;refGene_tx=XM_019826504.2;refGene_gn=LPIN3;refGene_tx=XM_019826505.2
A3 19111089 19111355
refGene_gn=LPIN3;refGene_tx=XM_006929748.4;refGene_gn=LPIN3;refGene_tx=XM_019826503.2;refGene_gn=LPIN3;refGene_tx=XM_019826504.2;refGene_gn=LPIN3;refGene_tx=XM_019826505.2
A3 19112376 19112472
refGene_gn=LPIN3;refGene_tx=XM_006929748.4;refGene_gn=LPIN3;refGene_tx=XM_019826503.2;refGene_gn=LPIN3;refGene_tx=XM_019826504.2;refGene_gn=LPIN3;refGene_tx=XM_019826505.2
A3 19113681 19113881
refGene_gn=LPIN3;refGene_tx=XM_006929748.4;refGene_gn=LPIN3;refGene_tx=XM_019826503.2;refGene_gn=LPIN3;refGene_tx=XM_019826504.2;refGene_gn=LPIN3;refGene_tx=XM_019826505.2
A3 19113974 19114127
refGene_gn=LPIN3;refGene_tx=XM_019826503.2;refGene_gn=LPIN3;refGene_tx=XM_019826504.2
A3 19117346 19117449
refGene_gn=LPIN3;refGene_tx=XM_019826503.2;refGene_gn=LPIN3;refGene_tx=XM_019826504.2
A3 19236977 19239837 refGene_gn=ZHX3;refGene_tx=XM_019826507.2
A3 19241221 19241232 refGene_gn=ZHX3;refGene_tx=XM_019826507.2
A3 19253316 19253359
refGene_gn=PLCG1;refGene_tx=XM_023251250.1;refGene_gn=PLCG1;refGene_tx=XM_023251251.1;refGene_gn=PLCG1;refGene_tx=XM_023251252.1
A3 19253555 19253743
refGene_gn=PLCG1;refGene_tx=XM_023251250.1;refGene_gn=PLCG1;refGene_tx=XM_023251251.1;refGene_gn=PLCG1;refGene_tx=XM_023251252.1
A3 19253838 19253927
refGene_gn=PLCG1;refGene_tx=XM_023251250.1;refGene_gn=PLCG1;refGene_tx

=XM_023251251.1;refGene_gn=PLCG1;refGene_tx=XM_023251252.1
A3 19254044 19254206
refGene_gn=PLCG1;refGene_tx=XM_023251250.1;refGene_gn=PLCG1;refGene_tx
=XM_023251251.1;refGene_gn=PLCG1;refGene_tx=XM_023251252.1
A3 19254292 19254407
refGene_gn=PLCG1;refGene_tx=XM_023251250.1;refGene_gn=PLCG1;refGene_tx
=XM_023251251.1;refGene_gn=PLCG1;refGene_tx=XM_023251252.1
A3 19254883 19255032
refGene_gn=PLCG1;refGene_tx=XM_023251250.1;refGene_gn=PLCG1;refGene_tx
=XM_023251251.1;refGene_gn=PLCG1;refGene_tx=XM_023251252.1
A3 19255103 19255328
refGene_gn=PLCG1;refGene_tx=XM_023251250.1;refGene_gn=PLCG1;refGene_tx
=XM_023251251.1;refGene_gn=PLCG1;refGene_tx=XM_023251252.1
A3 19255468 19255565
refGene_gn=PLCG1;refGene_tx=XM_023251250.1;refGene_gn=PLCG1;refGene_tx
=XM_023251251.1;refGene_gn=PLCG1;refGene_tx=XM_023251252.1
A3 19256911 19257069
refGene_gn=PLCG1;refGene_tx=XM_023251250.1;refGene_gn=PLCG1;refGene_tx
=XM_023251251.1;refGene_gn=PLCG1;refGene_tx=XM_023251252.1
A3 19257546 19257616
refGene_gn=PLCG1;refGene_tx=XM_023251250.1;refGene_gn=PLCG1;refGene_tx
=XM_023251251.1;refGene_gn=PLCG1;refGene_tx=XM_023251252.1
A3 19257900 19257997
refGene_gn=PLCG1;refGene_tx=XM_023251250.1;refGene_gn=PLCG1;refGene_tx
=XM_023251251.1;refGene_gn=PLCG1;refGene_tx=XM_023251252.1
A3 19258223 19258327
refGene_gn=PLCG1;refGene_tx=XM_023251250.1;refGene_gn=PLCG1;refGene_tx
=XM_023251251.1;refGene_gn=PLCG1;refGene_tx=XM_023251252.1
A3 19259103 19259181
refGene_gn=PLCG1;refGene_tx=XM_023251250.1;refGene_gn=PLCG1;refGene_tx
=XM_023251251.1;refGene_gn=PLCG1;refGene_tx=XM_023251252.1
A3 19260066 19260247
refGene_gn=PLCG1;refGene_tx=XM_023251250.1;refGene_gn=PLCG1;refGene_tx
=XM_023251251.1;refGene_gn=PLCG1;refGene_tx=XM_023251252.1
A3 19260335 19260455
refGene_gn=PLCG1;refGene_tx=XM_023251250.1;refGene_gn=PLCG1;refGene_tx
=XM_023251251.1;refGene_gn=PLCG1;refGene_tx=XM_023251252.1
A3 19260529 19260730
refGene_gn=PLCG1;refGene_tx=XM_023251250.1;refGene_gn=PLCG1;refGene_tx
=XM_023251251.1;refGene_gn=PLCG1;refGene_tx=XM_023251252.1
A3 19260974 19261162
refGene_gn=PLCG1;refGene_tx=XM_023251250.1;refGene_gn=PLCG1;refGene_tx
=XM_023251251.1;refGene_gn=PLCG1;refGene_tx=XM_023251252.1
A3 19261241 19261343
refGene_gn=PLCG1;refGene_tx=XM_023251250.1;refGene_gn=PLCG1;refGene_tx
=XM_023251251.1;refGene_gn=PLCG1;refGene_tx=XM_023251252.1
A3 19261425 19261548
refGene_gn=PLCG1;refGene_tx=XM_023251250.1;refGene_gn=PLCG1;refGene_tx
=XM_023251251.1;refGene_gn=PLCG1;refGene_tx=XM_023251252.1
A3 19261663 19261832

refGene_gn=PLCG1;refGene_tx=XM_023251250.1;refGene_gn=PLCG1;refGene_tx=XM_023251251.1;refGene_gn=PLCG1;refGene_tx=XM_023251252.1
A3 19262574 19262695
refGene_gn=PLCG1;refGene_tx=XM_023251250.1;refGene_gn=PLCG1;refGene_tx=XM_023251251.1;refGene_gn=PLCG1;refGene_tx=XM_023251252.1
A3 19262769 19262855
refGene_gn=PLCG1;refGene_tx=XM_023251250.1;refGene_gn=PLCG1;refGene_tx=XM_023251251.1;refGene_gn=PLCG1;refGene_tx=XM_023251252.1
A3 19262941 19263060
refGene_gn=PLCG1;refGene_tx=XM_023251250.1;refGene_gn=PLCG1;refGene_tx=XM_023251251.1;refGene_gn=PLCG1;refGene_tx=XM_023251252.1
A3 19263317 19263419
refGene_gn=PLCG1;refGene_tx=XM_023251250.1;refGene_gn=PLCG1;refGene_tx=XM_023251251.1;refGene_gn=PLCG1;refGene_tx=XM_023251252.1
A3 19263522 19263595
refGene_gn=PLCG1;refGene_tx=XM_023251250.1;refGene_gn=PLCG1;refGene_tx=XM_023251251.1;refGene_gn=PLCG1;refGene_tx=XM_023251252.1
A3 19263797 19263832
refGene_gn=PLCG1;refGene_tx=XM_023251250.1;refGene_gn=PLCG1;refGene_tx=XM_023251251.1;refGene_gn=PLCG1;refGene_tx=XM_023251252.1
A3 19264065 19264149
refGene_gn=PLCG1;refGene_tx=XM_023251250.1;refGene_gn=PLCG1;refGene_tx=XM_023251251.1;refGene_gn=PLCG1;refGene_tx=XM_023251252.1
A3 19264245 19264330
refGene_gn=PLCG1;refGene_tx=XM_023251250.1;refGene_gn=PLCG1;refGene_tx=XM_023251251.1;refGene_gn=PLCG1;refGene_tx=XM_023251252.1
A3 19266846 19266894
refGene_gn=PLCG1;refGene_tx=XM_023251250.1;refGene_gn=PLCG1;refGene_tx=XM_023251251.1;refGene_gn=PLCG1;refGene_tx=XM_023251252.1
A3 19267052 19267146
refGene_gn=PLCG1;refGene_tx=XM_023251250.1;refGene_gn=PLCG1;refGene_tx=XM_023251251.1;refGene_gn=PLCG1;refGene_tx=XM_023251252.1
A3 19267261 19267414
refGene_gn=PLCG1;refGene_tx=XM_023251250.1;refGene_gn=PLCG1;refGene_tx=XM_023251251.1;refGene_gn=PLCG1;refGene_tx=XM_023251252.1
A3 19287967 19288184
refGene_gn=PLCG1;refGene_tx=XM_023251250.1;refGene_gn=PLCG1;refGene_tx=XM_023251251.1
A3 19299943 19300046 refGene_gn=TOP1;refGene_tx=XM_023251253.1
A3 19301040 19301190 refGene_gn=TOP1;refGene_tx=XM_023251253.1
A3 19301419 19301514 refGene_gn=TOP1;refGene_tx=XM_023251253.1
A3 19305304 19305432 refGene_gn=TOP1;refGene_tx=XM_023251253.1
A3 19306380 19306495 refGene_gn=TOP1;refGene_tx=XM_023251253.1
A3 19307047 19307116 refGene_gn=TOP1;refGene_tx=XM_023251253.1
A3 19308122 19308308 refGene_gn=TOP1;refGene_tx=XM_023251253.1
A3 19309765 19309909 refGene_gn=TOP1;refGene_tx=XM_023251253.1
A3 19323016 19323161 refGene_gn=TOP1;refGene_tx=XM_023251253.1
A3 19323441 19323629 refGene_gn=TOP1;refGene_tx=XM_023251253.1
A3 19325792 19325915 refGene_gn=TOP1;refGene_tx=XM_023251253.1
A3 19326827 19326949 refGene_gn=TOP1;refGene_tx=XM_023251253.1

A3 19330829 19330945 refGene_gn=TOP1;refGene_tx=XM_023251253.1
A3 19339090 19339197 refGene_gn=TOP1;refGene_tx=XM_023251253.1
A3 19342634 19342710 refGene_gn=TOP1;refGene_tx=XM_023251253.1
A3 19343767 19343863 refGene_gn=TOP1;refGene_tx=XM_023251253.1
A3 19347126 19347182 refGene_gn=TOP1;refGene_tx=XM_023251253.1
A3 19348129 19348253 refGene_gn=TOP1;refGene_tx=XM_023251253.1
A3 19361750 19361853 refGene_gn=TOP1;refGene_tx=XM_023251253.1
A3 19393448 19393473 refGene_gn=TOP1;refGene_tx=XM_023251253.1
A3 19393803 19393836 refGene_gn=TOP1;refGene_tx=XM_023251253.1
A3 19693221 19694193 refGene_gn=MAFB;refGene_tx=XM_023251254.1
A3 19694968 19695026
refGene_gn=LOC109498000;refGene_tx=XM_023251466.1
A3 19698127 19699026
refGene_gn=LOC109498000;refGene_tx=XM_023251466.1
A3 19699638 19699914
refGene_gn=LOC109498000;refGene_tx=XM_023251466.1
A3 21066306 21066351
refGene_gn=DHX35;refGene_tx=XM_003983466.5;refGene_gn=DHX35;refGene_tx=XM_006929758.4;refGene_gn=DHX35;refGene_tx=XM_019826510.2;refGene_gn=DHX35;refGene_tx=XM_019826511.2
A3 21069859 21069971
refGene_gn=DHX35;refGene_tx=XM_003983466.5;refGene_gn=DHX35;refGene_tx=XM_006929758.4;refGene_gn=DHX35;refGene_tx=XM_019826510.2;refGene_gn=DHX35;refGene_tx=XM_019826511.2
A3 21072603 21072675
refGene_gn=DHX35;refGene_tx=XM_003983466.5;refGene_gn=DHX35;refGene_tx=XM_006929758.4;refGene_gn=DHX35;refGene_tx=XM_019826510.2;refGene_gn=DHX35;refGene_tx=XM_019826511.2
A3 21074534 21074616
refGene_gn=DHX35;refGene_tx=XM_003983466.5;refGene_gn=DHX35;refGene_tx=XM_006929758.4;refGene_gn=DHX35;refGene_tx=XM_019826510.2;refGene_gn=DHX35;refGene_tx=XM_019826511.2
A3 21076777 21076907
refGene_gn=DHX35;refGene_tx=XM_003983466.5;refGene_gn=DHX35;refGene_tx=XM_006929758.4;refGene_gn=DHX35;refGene_tx=XM_019826510.2;refGene_gn=DHX35;refGene_tx=XM_019826511.2
A3 21078409 21078487
refGene_gn=DHX35;refGene_tx=XM_003983466.5;refGene_gn=DHX35;refGene_tx=XM_006929758.4;refGene_gn=DHX35;refGene_tx=XM_019826510.2;refGene_gn=DHX35;refGene_tx=XM_019826511.2
A3 21080395 21080490
refGene_gn=DHX35;refGene_tx=XM_003983466.5;refGene_gn=DHX35;refGene_tx=XM_006929758.4;refGene_gn=DHX35;refGene_tx=XM_019826510.2;refGene_gn=DHX35;refGene_tx=XM_019826511.2
A3 21082323 21082419
refGene_gn=DHX35;refGene_tx=XM_003983466.5;refGene_gn=DHX35;refGene_tx=XM_006929758.4;refGene_gn=DHX35;refGene_tx=XM_019826510.2;refGene_gn=DHX35;refGene_tx=XM_019826511.2
A3 21085591 21085646
refGene_gn=DHX35;refGene_tx=XM_003983466.5;refGene_gn=DHX35;refGene_tx

=XM_006929758.4;refGene_gn=DHX35;refGene_tx=XM_019826510.2;refGene_gn=DHX35;refGene_tx=XM_019826511.2
A3 21088362 21088487
refGene_gn=DHX35;refGene_tx=XM_003983466.5;refGene_gn=DHX35;refGene_tx=XM_006929758.4;refGene_gn=DHX35;refGene_tx=XM_019826510.2;refGene_gn=DHX35;refGene_tx=XM_019826511.2
A3 21092773 21092984
refGene_gn=DHX35;refGene_tx=XM_003983466.5;refGene_gn=DHX35;refGene_tx=XM_006929758.4;refGene_gn=DHX35;refGene_tx=XM_019826510.2;refGene_gn=DHX35;refGene_tx=XM_019826511.2
A3 21094870 21095029
refGene_gn=DHX35;refGene_tx=XM_003983466.5;refGene_gn=DHX35;refGene_tx=XM_006929758.4;refGene_gn=DHX35;refGene_tx=XM_019826511.2
A3 21095897 21095994
refGene_gn=DHX35;refGene_tx=XM_003983466.5;refGene_gn=DHX35;refGene_tx=XM_006929758.4;refGene_gn=DHX35;refGene_tx=XM_019826510.2;refGene_gn=DHX35;refGene_tx=XM_019826511.2
A3 21097188 21097301
refGene_gn=DHX35;refGene_tx=XM_003983466.5;refGene_gn=DHX35;refGene_tx=XM_006929758.4;refGene_gn=DHX35;refGene_tx=XM_019826510.2;refGene_gn=DHX35;refGene_tx=XM_019826511.2
A3 21103599 21103659
refGene_gn=DHX35;refGene_tx=XM_003983466.5;refGene_gn=DHX35;refGene_tx=XM_019826510.2;refGene_gn=DHX35;refGene_tx=XM_019826511.2
A3 21105611 21105681
refGene_gn=DHX35;refGene_tx=XM_003983466.5;refGene_gn=DHX35;refGene_tx=XM_019826510.2
A3 21106523 21106585
refGene_gn=DHX35;refGene_tx=XM_003983466.5;refGene_gn=DHX35;refGene_tx=XM_019826510.2
A3 21108942 21109047
refGene_gn=DHX35;refGene_tx=XM_003983466.5;refGene_gn=DHX35;refGene_tx=XM_019826510.2
A3 21114091 21114169
refGene_gn=DHX35;refGene_tx=XM_003983466.5;refGene_gn=DHX35;refGene_tx=XM_006929758.4;refGene_gn=DHX35;refGene_tx=XM_019826510.2;refGene_gn=DHX35;refGene_tx=XM_019826511.2
A3 21121127 21121220
refGene_gn=DHX35;refGene_tx=XM_003983466.5;refGene_gn=DHX35;refGene_tx=XM_006929758.4;refGene_gn=DHX35;refGene_tx=XM_019826510.2;refGene_gn=DHX35;refGene_tx=XM_019826511.2
A3 21126023 21126157
refGene_gn=DHX35;refGene_tx=XM_003983466.5;refGene_gn=DHX35;refGene_tx=XM_006929758.4;refGene_gn=DHX35;refGene_tx=XM_019826510.2;refGene_gn=DHX35;refGene_tx=XM_019826511.2
A3 21131153 21131193
refGene_gn=DHX35;refGene_tx=XM_003983466.5;refGene_gn=DHX35;refGene_tx=XM_006929758.4;refGene_gn=DHX35;refGene_tx=XM_019826510.2;refGene_gn=DHX35;refGene_tx=XM_019826511.2
A3 21138437 21139428 refGene_gn=FAM83D;refGene_tx=XM_023251255.1

A3 21142537 21142662 refGene_gn=FAM83D;refGene_tx=XM_023251255.1
A3 21146889 21147057 refGene_gn=FAM83D;refGene_tx=XM_023251255.1
A3 21158479 21158956 refGene_gn=FAM83D;refGene_tx=XM_023251255.1
A3 21165317 21165827 refGene_gn=PPP1R16B;refGene_tx=XM_023251256.1
A3 21174166 21174335 refGene_gn=PPP1R16B;refGene_tx=XM_023251256.1
A3 21174426 21174556 refGene_gn=PPP1R16B;refGene_tx=XM_023251256.1
A3 21175806 21175882 refGene_gn=PPP1R16B;refGene_tx=XM_023251256.1
A3 21176415 21176541 refGene_gn=PPP1R16B;refGene_tx=XM_023251256.1
A3 21179392 21179517 refGene_gn=PPP1R16B;refGene_tx=XM_023251256.1
A3 21181416 21181520 refGene_gn=PPP1R16B;refGene_tx=XM_023251256.1
A3 21183494 21183640 refGene_gn=PPP1R16B;refGene_tx=XM_023251256.1
A3 21187620 21187691 refGene_gn=PPP1R16B;refGene_tx=XM_023251256.1
A3 21235142 21235392 refGene_gn=PPP1R16B;refGene_tx=XM_023251256.1
A3 21289697 21289955 refGene_gn=ACTR5;refGene_tx=XM_023251257.1
A3 21296227 21296360 refGene_gn=ACTR5;refGene_tx=XM_023251257.1
A3 21296934 21297074 refGene_gn=ACTR5;refGene_tx=XM_023251257.1
A3 21298277 21298394 refGene_gn=ACTR5;refGene_tx=XM_023251257.1
A3 21306736 21306919 refGene_gn=ACTR5;refGene_tx=XM_023251257.1
A3 21307976 21308194 refGene_gn=ACTR5;refGene_tx=XM_023251257.1
A3 21314496 21314666 refGene_gn=ACTR5;refGene_tx=XM_023251257.1
A3 21316795 21317025 refGene_gn=ACTR5;refGene_tx=XM_023251257.1
A3 21318222 21318621 refGene_gn=ACTR5;refGene_tx=XM_023251257.1
A3 21326406 21327594 refGene_gn=SLC32A1;refGene_tx=XM_011280886.3
A3 21329714 21330107 refGene_gn=SLC32A1;refGene_tx=XM_011280886.3
A3 21393596 21393649 refGene_gn=ARHGAP40;refGene_tx=XM_023251545.1
A3 21394466 21394508 refGene_gn=ARHGAP40;refGene_tx=XM_023251545.1
A3 21394510 21394552 refGene_gn=ARHGAP40;refGene_tx=XM_023251545.1
A3 21394554 21394603 refGene_gn=ARHGAP40;refGene_tx=XM_023251545.1
A3 21395291 21395392 refGene_gn=ARHGAP40;refGene_tx=XM_023251545.1
A3 21395394 21395463 refGene_gn=ARHGAP40;refGene_tx=XM_023251545.1
A3 21395935 21396056 refGene_gn=ARHGAP40;refGene_tx=XM_023251545.1
A3 21397186 21397254 refGene_gn=ARHGAP40;refGene_tx=XM_023251545.1
A3 21397737 21397751 refGene_gn=ARHGAP40;refGene_tx=XM_023251545.1
A3 21397868 21397994 refGene_gn=ARHGAP40;refGene_tx=XM_023251545.1
A3 21399373 21399676 refGene_gn=ARHGAP40;refGene_tx=XM_023251545.1
A3 21401291 21401352 refGene_gn=ARHGAP40;refGene_tx=XM_023251545.1
A3 21401970 21402125 refGene_gn=ARHGAP40;refGene_tx=XM_023251545.1
A3 21402368 21402498 refGene_gn=ARHGAP40;refGene_tx=XM_023251545.1
A3 21402843 21403062 refGene_gn=ARHGAP40;refGene_tx=XM_023251545.1
A3 21405675 21405853 refGene_gn=ARHGAP40;refGene_tx=XM_023251545.1
A3 21408193 21408267 refGene_gn=ARHGAP40;refGene_tx=XM_023251545.1
A3 21410860 21410918 refGene_gn=ARHGAP40;refGene_tx=XM_023251545.1
A3 21424316 21424435 refGene_gn=ADIG;refGene_tx=XM_003983469.4
A3 21427713 21427840 refGene_gn=ADIG;refGene_tx=XM_003983469.4
A3 21434840 21435034
refGene_gn=RALGAPB;refGene_tx=XM_003983470.5;refGene_gn=RALGAPB;refGene_tx=XM_006929759.4;refGene_gn=RALGAPB;refGene_tx=XM_019826517.2;refGene_gn=RALGAPB;refGene_tx=XM_019826518.2;refGene_gn=RALGAPB;refGene_tx=XM_019826519.2;refGene_gn=RALGAPB;refGene_tx=XM_019826520.2;refGene_gn=RALGAPB;refGene_tx=XM_019826521.2

A3 21435524 21435673
refGene_gn=RALGAPB;refGene_tx=XM_003983470.5;refGene_gn=RALGAPB;refGene_tx=XM_006929759.4;refGene_gn=RALGAPB;refGene_tx=XM_019826517.2;refGene_gn=RALGAPB;refGene_tx=XM_019826518.2;refGene_gn=RALGAPB;refGene_tx=XM_019826519.2;refGene_gn=RALGAPB;refGene_tx=XM_019826520.2;refGene_gn=RALGAPB;refGene_tx=XM_019826521.2
A3 21438593 21438672
refGene_gn=RALGAPB;refGene_tx=XM_003983470.5;refGene_gn=RALGAPB;refGene_tx=XM_006929759.4;refGene_gn=RALGAPB;refGene_tx=XM_019826517.2;refGene_gn=RALGAPB;refGene_tx=XM_019826518.2;refGene_gn=RALGAPB;refGene_tx=XM_019826519.2;refGene_gn=RALGAPB;refGene_tx=XM_019826520.2;refGene_gn=RALGAPB;refGene_tx=XM_019826521.2
A3 21439500 21439609
refGene_gn=RALGAPB;refGene_tx=XM_003983470.5;refGene_gn=RALGAPB;refGene_tx=XM_006929759.4;refGene_gn=RALGAPB;refGene_tx=XM_019826517.2;refGene_gn=RALGAPB;refGene_tx=XM_019826518.2;refGene_gn=RALGAPB;refGene_tx=XM_019826519.2;refGene_gn=RALGAPB;refGene_tx=XM_019826520.2;refGene_gn=RALGAPB;refGene_tx=XM_019826521.2
A3 21443794 21443931
refGene_gn=RALGAPB;refGene_tx=XM_003983470.5;refGene_gn=RALGAPB;refGene_tx=XM_006929759.4;refGene_gn=RALGAPB;refGene_tx=XM_019826517.2;refGene_gn=RALGAPB;refGene_tx=XM_019826518.2;refGene_gn=RALGAPB;refGene_tx=XM_019826519.2;refGene_gn=RALGAPB;refGene_tx=XM_019826520.2;refGene_gn=RALGAPB;refGene_tx=XM_019826521.2
A3 21445490 21445613
refGene_gn=RALGAPB;refGene_tx=XM_003983470.5;refGene_gn=RALGAPB;refGene_tx=XM_006929759.4;refGene_gn=RALGAPB;refGene_tx=XM_019826517.2;refGene_gn=RALGAPB;refGene_tx=XM_019826518.2;refGene_gn=RALGAPB;refGene_tx=XM_019826519.2;refGene_gn=RALGAPB;refGene_tx=XM_019826520.2;refGene_gn=RALGAPB;refGene_tx=XM_019826521.2
A3 21446929 21447095
refGene_gn=RALGAPB;refGene_tx=XM_003983470.5;refGene_gn=RALGAPB;refGene_tx=XM_006929759.4;refGene_gn=RALGAPB;refGene_tx=XM_019826517.2;refGene_gn=RALGAPB;refGene_tx=XM_019826518.2;refGene_gn=RALGAPB;refGene_tx=XM_019826519.2;refGene_gn=RALGAPB;refGene_tx=XM_019826520.2;refGene_gn=RALGAPB;refGene_tx=XM_019826521.2
A3 21451159 21451318
refGene_gn=RALGAPB;refGene_tx=XM_003983470.5;refGene_gn=RALGAPB;refGene_tx=XM_006929759.4;refGene_gn=RALGAPB;refGene_tx=XM_019826517.2;refGene_gn=RALGAPB;refGene_tx=XM_019826518.2;refGene_gn=RALGAPB;refGene_tx=XM_019826519.2;refGene_gn=RALGAPB;refGene_tx=XM_019826520.2;refGene_gn=RALGAPB;refGene_tx=XM_019826521.2
A3 21455403 21455613
refGene_gn=RALGAPB;refGene_tx=XM_003983470.5;refGene_gn=RALGAPB;refGene_tx=XM_006929759.4;refGene_gn=RALGAPB;refGene_tx=XM_019826517.2;refGene_gn=RALGAPB;refGene_tx=XM_019826518.2;refGene_gn=RALGAPB;refGene_tx=XM_019826519.2;refGene_gn=RALGAPB;refGene_tx=XM_019826520.2;refGene_gn=RALGAPB;refGene_tx=XM_019826521.2
A3 21457860 21458013
refGene_gn=RALGAPB;refGene_tx=XM_003983470.5;refGene_gn=RALGAPB;refGen

e_tx=XM_006929759.4;refGene_gn=RALGAPB;refGene_tx=XM_019826517.2;refGene_gn=RALGAPB;refGene_tx=XM_019826518.2;refGene_gn=RALGAPB;refGene_tx=XM_019826519.2;refGene_gn=RALGAPB;refGene_tx=XM_019826520.2;refGene_gn=RALGAPB;refGene_tx=XM_019826521.2
A3 21459596 21459703
refGene_gn=RALGAPB;refGene_tx=XM_003983470.5;refGene_gn=RALGAPB;refGene_tx=XM_006929759.4;refGene_gn=RALGAPB;refGene_tx=XM_019826517.2;refGene_gn=RALGAPB;refGene_tx=XM_019826518.2;refGene_gn=RALGAPB;refGene_tx=XM_019826519.2;refGene_gn=RALGAPB;refGene_tx=XM_019826520.2;refGene_gn=RALGAPB;refGene_tx=XM_019826521.2
A3 21462235 21462423
refGene_gn=RALGAPB;refGene_tx=XM_003983470.5;refGene_gn=RALGAPB;refGene_tx=XM_006929759.4;refGene_gn=RALGAPB;refGene_tx=XM_019826517.2;refGene_gn=RALGAPB;refGene_tx=XM_019826518.2;refGene_gn=RALGAPB;refGene_tx=XM_019826519.2;refGene_gn=RALGAPB;refGene_tx=XM_019826520.2;refGene_gn=RALGAPB;refGene_tx=XM_019826521.2
A3 21467867 21468019
refGene_gn=RALGAPB;refGene_tx=XM_003983470.5;refGene_gn=RALGAPB;refGene_tx=XM_006929759.4;refGene_gn=RALGAPB;refGene_tx=XM_019826517.2;refGene_gn=RALGAPB;refGene_tx=XM_019826518.2;refGene_gn=RALGAPB;refGene_tx=XM_019826519.2;refGene_gn=RALGAPB;refGene_tx=XM_019826520.2;refGene_gn=RALGAPB;refGene_tx=XM_019826521.2
A3 21471814 21471997
refGene_gn=RALGAPB;refGene_tx=XM_003983470.5;refGene_gn=RALGAPB;refGene_tx=XM_006929759.4;refGene_gn=RALGAPB;refGene_tx=XM_019826517.2;refGene_gn=RALGAPB;refGene_tx=XM_019826518.2;refGene_gn=RALGAPB;refGene_tx=XM_019826519.2;refGene_gn=RALGAPB;refGene_tx=XM_019826520.2;refGene_gn=RALGAPB;refGene_tx=XM_019826521.2
A3 21473074 21473208
refGene_gn=RALGAPB;refGene_tx=XM_003983470.5;refGene_gn=RALGAPB;refGene_tx=XM_006929759.4;refGene_gn=RALGAPB;refGene_tx=XM_019826517.2;refGene_gn=RALGAPB;refGene_tx=XM_019826518.2;refGene_gn=RALGAPB;refGene_tx=XM_019826519.2;refGene_gn=RALGAPB;refGene_tx=XM_019826520.2;refGene_gn=RALGAPB;refGene_tx=XM_019826521.2
A3 21474090 21474099
refGene_gn=RALGAPB;refGene_tx=XM_019826517.2;refGene_gn=RALGAPB;refGene_tx=XM_019826519.2;refGene_gn=RALGAPB;refGene_tx=XM_019826521.2
A3 21475411 21475541
refGene_gn=RALGAPB;refGene_tx=XM_003983470.5;refGene_gn=RALGAPB;refGene_tx=XM_006929759.4;refGene_gn=RALGAPB;refGene_tx=XM_019826517.2;refGene_gn=RALGAPB;refGene_tx=XM_019826518.2;refGene_gn=RALGAPB;refGene_tx=XM_019826519.2;refGene_gn=RALGAPB;refGene_tx=XM_019826520.2;refGene_gn=RALGAPB;refGene_tx=XM_019826521.2
A3 21476947 21477012
refGene_gn=RALGAPB;refGene_tx=XM_003983470.5;refGene_gn=RALGAPB;refGene_tx=XM_006929759.4;refGene_gn=RALGAPB;refGene_tx=XM_019826517.2;refGene_gn=RALGAPB;refGene_tx=XM_019826518.2;refGene_gn=RALGAPB;refGene_tx=XM_019826519.2;refGene_gn=RALGAPB;refGene_tx=XM_019826520.2;refGene_gn=RALGAPB;refGene_tx=XM_019826521.2
A3 21479173 21479321

refGene_gn=RALGAPB;refGene_tx=XM_003983470.5;refGene_gn=RALGAPB;refGene_tx=XM_006929759.4;refGene_gn=RALGAPB;refGene_tx=XM_019826517.2;refGene_gn=RALGAPB;refGene_tx=XM_019826518.2;refGene_gn=RALGAPB;refGene_tx=XM_019826519.2;refGene_gn=RALGAPB;refGene_tx=XM_019826520.2;refGene_gn=RALGAPB;refGene_tx=XM_019826521.2
A3 21479829 21479944
refGene_gn=RALGAPB;refGene_tx=XM_003983470.5;refGene_gn=RALGAPB;refGene_tx=XM_006929759.4;refGene_gn=RALGAPB;refGene_tx=XM_019826517.2;refGene_gn=RALGAPB;refGene_tx=XM_019826518.2;refGene_gn=RALGAPB;refGene_tx=XM_019826519.2;refGene_gn=RALGAPB;refGene_tx=XM_019826520.2;refGene_gn=RALGAPB;refGene_tx=XM_019826521.2
A3 21480476 21480644
refGene_gn=RALGAPB;refGene_tx=XM_003983470.5;refGene_gn=RALGAPB;refGene_tx=XM_006929759.4;refGene_gn=RALGAPB;refGene_tx=XM_019826517.2;refGene_gn=RALGAPB;refGene_tx=XM_019826518.2;refGene_gn=RALGAPB;refGene_tx=XM_019826519.2;refGene_gn=RALGAPB;refGene_tx=XM_019826520.2;refGene_gn=RALGAPB;refGene_tx=XM_019826521.2
A3 21483715 21483917
refGene_gn=RALGAPB;refGene_tx=XM_003983470.5;refGene_gn=RALGAPB;refGene_tx=XM_006929759.4;refGene_gn=RALGAPB;refGene_tx=XM_019826517.2;refGene_gn=RALGAPB;refGene_tx=XM_019826518.2;refGene_gn=RALGAPB;refGene_tx=XM_019826519.2;refGene_gn=RALGAPB;refGene_tx=XM_019826520.2;refGene_gn=RALGAPB;refGene_tx=XM_019826521.2
A3 21485137 21485173
refGene_gn=RALGAPB;refGene_tx=XM_019826517.2;refGene_gn=RALGAPB;refGene_tx=XM_019826518.2;refGene_gn=RALGAPB;refGene_tx=XM_019826519.2;refGene_gn=RALGAPB;refGene_tx=XM_019826520.2
A3 21487398 21487615
refGene_gn=RALGAPB;refGene_tx=XM_003983470.5;refGene_gn=RALGAPB;refGene_tx=XM_006929759.4;refGene_gn=RALGAPB;refGene_tx=XM_019826517.2;refGene_gn=RALGAPB;refGene_tx=XM_019826518.2;refGene_gn=RALGAPB;refGene_tx=XM_019826519.2;refGene_gn=RALGAPB;refGene_tx=XM_019826520.2;refGene_gn=RALGAPB;refGene_tx=XM_019826521.2
A3 21487744 21487893
refGene_gn=RALGAPB;refGene_tx=XM_003983470.5;refGene_gn=RALGAPB;refGene_tx=XM_006929759.4;refGene_gn=RALGAPB;refGene_tx=XM_019826517.2;refGene_gn=RALGAPB;refGene_tx=XM_019826518.2;refGene_gn=RALGAPB;refGene_tx=XM_019826519.2;refGene_gn=RALGAPB;refGene_tx=XM_019826520.2;refGene_gn=RALGAPB;refGene_tx=XM_019826521.2
A3 21488955 21489134
refGene_gn=RALGAPB;refGene_tx=XM_003983470.5;refGene_gn=RALGAPB;refGene_tx=XM_006929759.4;refGene_gn=RALGAPB;refGene_tx=XM_019826517.2;refGene_gn=RALGAPB;refGene_tx=XM_019826518.2;refGene_gn=RALGAPB;refGene_tx=XM_019826519.2;refGene_gn=RALGAPB;refGene_tx=XM_019826520.2;refGene_gn=RALGAPB;refGene_tx=XM_019826521.2
A3 21489950 21490082
refGene_gn=RALGAPB;refGene_tx=XM_003983470.5;refGene_gn=RALGAPB;refGene_tx=XM_006929759.4;refGene_gn=RALGAPB;refGene_tx=XM_019826517.2;refGene_gn=RALGAPB;refGene_tx=XM_019826518.2;refGene_gn=RALGAPB;refGene_tx=XM_019826519.2;refGene_gn=RALGAPB;refGene_tx=XM_019826520.2;refGene_gn

=RALGAPB;refGene_tx=XM_019826521.2
A3 21495427 21495614
refGene_gn=RALGAPB;refGene_tx=XM_003983470.5;refGene_gn=RALGAPB;refGene_tx=XM_006929759.4;refGene_gn=RALGAPB;refGene_tx=XM_019826517.2;refGene_gn=RALGAPB;refGene_tx=XM_019826518.2;refGene_gn=RALGAPB;refGene_tx=XM_019826519.2;refGene_gn=RALGAPB;refGene_tx=XM_019826520.2;refGene_gn=RALGAPB;refGene_tx=XM_019826521.2
A3 21497455 21497619
refGene_gn=RALGAPB;refGene_tx=XM_003983470.5;refGene_gn=RALGAPB;refGene_tx=XM_006929759.4;refGene_gn=RALGAPB;refGene_tx=XM_019826517.2;refGene_gn=RALGAPB;refGene_tx=XM_019826518.2;refGene_gn=RALGAPB;refGene_tx=XM_019826519.2;refGene_gn=RALGAPB;refGene_tx=XM_019826520.2;refGene_gn=RALGAPB;refGene_tx=XM_019826521.2
A3 21502185 21502388
refGene_gn=RALGAPB;refGene_tx=XM_003983470.5;refGene_gn=RALGAPB;refGene_tx=XM_006929759.4;refGene_gn=RALGAPB;refGene_tx=XM_019826517.2;refGene_gn=RALGAPB;refGene_tx=XM_019826518.2;refGene_gn=RALGAPB;refGene_tx=XM_019826519.2;refGene_gn=RALGAPB;refGene_tx=XM_019826520.2;refGene_gn=RALGAPB;refGene_tx=XM_019826521.2
A3 21505799 21505985
refGene_gn=RALGAPB;refGene_tx=XM_003983470.5;refGene_gn=RALGAPB;refGene_tx=XM_006929759.4;refGene_gn=RALGAPB;refGene_tx=XM_019826517.2;refGene_gn=RALGAPB;refGene_tx=XM_019826518.2;refGene_gn=RALGAPB;refGene_tx=XM_019826519.2;refGene_gn=RALGAPB;refGene_tx=XM_019826520.2;refGene_gn=RALGAPB;refGene_tx=XM_019826521.2
A3 21566380 21566425
refGene_gn=LOC109498011;refGene_tx=XM_023251258.1
A3 21567557 21567634
refGene_gn=LOC109498011;refGene_tx=XM_023251258.1
A3 21568484 21568548
refGene_gn=LOC109498011;refGene_tx=XM_023251258.1
A3 21569447 21569493
refGene_gn=LOC109498011;refGene_tx=XM_023251258.1
A3 21571057 21571125
refGene_gn=LOC109498011;refGene_tx=XM_023251258.1
A3 21573933 21574104
refGene_gn=LOC109498011;refGene_tx=XM_023251258.1
A3 21575250 21575310
refGene_gn=LOC109498011;refGene_tx=XM_023251258.1
A3 21576593 21576770
refGene_gn=LOC109498011;refGene_tx=XM_023251258.1
A3 21577502 21577594
refGene_gn=LOC109498011;refGene_tx=XM_023251258.1
A3 21578718 21578782
refGene_gn=LOC109498011;refGene_tx=XM_023251258.1
A3 21581756 21581820
refGene_gn=LOC109498011;refGene_tx=XM_023251258.1
A3 21586080 21586236
refGene_gn=LOC109498011;refGene_tx=XM_023251258.1
A3 21586655 21586696

refGene_gn=LOC109498011;refGene_tx=XM_023251258.1
A3 21623411 21623456
refGene_gn=LOC101097963;refGene_tx=XM_003983473.3
A3 21625046 21625123
refGene_gn=LOC101097963;refGene_tx=XM_003983473.3
A3 21625927 21625991
refGene_gn=LOC101097963;refGene_tx=XM_003983473.3
A3 21627989 21628032
refGene_gn=LOC101097963;refGene_tx=XM_003983473.3
A3 21628824 21628892
refGene_gn=LOC101097963;refGene_tx=XM_003983473.3
A3 21630185 21630353
refGene_gn=LOC101097963;refGene_tx=XM_003983473.3
A3 21632318 21632378
refGene_gn=LOC101097963;refGene_tx=XM_003983473.3
A3 21633942 21634119
refGene_gn=LOC101097963;refGene_tx=XM_003983473.3
A3 21634629 21634721
refGene_gn=LOC101097963;refGene_tx=XM_003983473.3
A3 21635320 21635384
refGene_gn=LOC101097963;refGene_tx=XM_003983473.3
A3 21639496 21639560
refGene_gn=LOC101097963;refGene_tx=XM_003983473.3
A3 21640148 21640304
refGene_gn=LOC101097963;refGene_tx=XM_003983473.3
A3 21644072 21644201
refGene_gn=LOC101097963;refGene_tx=XM_003983473.3
A3 21645321 21645436
refGene_gn=LOC101097963;refGene_tx=XM_003983473.3
A3 21647341 21647465
refGene_gn=LOC101097963;refGene_tx=XM_003983473.3
A3 21656913 21656952 refGene_gn=BPI;refGene_tx=XM_003983542.4
A3 21657727 21658057 refGene_gn=BPI;refGene_tx=XM_019826524.1
A3 21658529 21658606
refGene_gn=BPI;refGene_tx=XM_003983542.4;refGene_gn=BPI;refGene_tx=XM_
019826524.1
A3 21659682 21659746
refGene_gn=BPI;refGene_tx=XM_003983542.4;refGene_gn=BPI;refGene_tx=XM_
019826524.1
A3 21662200 21662243
refGene_gn=BPI;refGene_tx=XM_003983542.4;refGene_gn=BPI;refGene_tx=XM_
019826524.1
A3 21664416 21664484
refGene_gn=BPI;refGene_tx=XM_003983542.4;refGene_gn=BPI;refGene_tx=XM_
019826524.1
A3 21665485 21665653
refGene_gn=BPI;refGene_tx=XM_003983542.4;refGene_gn=BPI;refGene_tx=XM_
019826524.1
A3 21667460 21667520
refGene_gn=BPI;refGene_tx=XM_003983542.4;refGene_gn=BPI;refGene_tx=XM_

019826524.1
A3 21668396 21668573
refGene_gn=BPI;refGene_tx=XM_003983542.4;refGene_gn=BPI;refGene_tx=XM_
019826524.1
A3 21672042 21672134
refGene_gn=BPI;refGene_tx=XM_003983542.4;refGene_gn=BPI;refGene_tx=XM_
019826524.1
A3 21673539 21673603
refGene_gn=BPI;refGene_tx=XM_003983542.4;refGene_gn=BPI;refGene_tx=XM_
019826524.1
A3 21675733 21675797
refGene_gn=BPI;refGene_tx=XM_003983542.4;refGene_gn=BPI;refGene_tx=XM_
019826524.1
A3 21676999 21677161
refGene_gn=BPI;refGene_tx=XM_003983542.4;refGene_gn=BPI;refGene_tx=XM_
019826524.1
A3 21678542 21678671
refGene_gn=BPI;refGene_tx=XM_003983542.4;refGene_gn=BPI;refGene_tx=XM_
019826524.1
A3 21679898 21680013
refGene_gn=BPI;refGene_tx=XM_003983542.4;refGene_gn=BPI;refGene_tx=XM_
019826524.1
A3 21682667 21682797
refGene_gn=BPI;refGene_tx=XM_003983542.4;refGene_gn=BPI;refGene_tx=XM_
019826524.1
A3 21792167 21792177 refGene_gn=TGM2;refGene_tx=XM_003983474.3
A3 21795740 21795920 refGene_gn=TGM2;refGene_tx=XM_003983474.3
A3 21800525 21800768 refGene_gn=TGM2;refGene_tx=XM_003983474.3
A3 21804059 21804178 refGene_gn=TGM2;refGene_tx=XM_003983474.3
A3 21806668 21806797 refGene_gn=TGM2;refGene_tx=XM_003983474.3
A3 21807864 21808042 refGene_gn=TGM2;refGene_tx=XM_003983474.3
A3 21811641 21811777 refGene_gn=TGM2;refGene_tx=XM_003983474.3
A3 21812472 21812576 refGene_gn=TGM2;refGene_tx=XM_003983474.3
A3 21814133 21814376 refGene_gn=TGM2;refGene_tx=XM_003983474.3
A3 21816406 21816679 refGene_gn=TGM2;refGene_tx=XM_003983474.3
A3 21820144 21820305 refGene_gn=TGM2;refGene_tx=XM_003983474.3
A3 21820768 21820905 refGene_gn=TGM2;refGene_tx=XM_003983474.3
A3 21821618 21821769 refGene_gn=TGM2;refGene_tx=XM_003983474.3
A3 21853713 21853863
refGene_gn=RPRD1B;refGene_tx=XM_003983478.4;refGene_gn=RPRD1B;refGene_
tx=XM_006929760.3
A3 21872892 21873068
refGene_gn=RPRD1B;refGene_tx=XM_003983478.4;refGene_gn=RPRD1B;refGene_
tx=XM_006929760.3
A3 21879557 21879684
refGene_gn=RPRD1B;refGene_tx=XM_003983478.4;refGene_gn=RPRD1B;refGene_
tx=XM_006929760.3
A3 21881484 21881597
refGene_gn=RPRD1B;refGene_tx=XM_003983478.4;refGene_gn=RPRD1B;refGene_
tx=XM_006929760.3

A3 21888978 21889112
refGene_gn=RPRD1B;refGene_tx=XM_003983478.4;refGene_gn=RPRD1B;refGene_
tx=XM_006929760.3
A3 21896497 21896627
refGene_gn=RPRD1B;refGene_tx=XM_003983478.4;refGene_gn=RPRD1B;refGene_
tx=XM_006929760.3
A3 21901440 21901591
refGene_gn=RPRD1B;refGene_tx=XM_003983478.4;refGene_gn=RPRD1B;refGene_
tx=XM_006929760.3
A3 21919010 21921312 refGene_gn=TTI1;refGene_tx=XM_003983479.4
A3 21925702 21925842 refGene_gn=TTI1;refGene_tx=XM_003983479.4
A3 21925843 21925913 refGene_gn=TTI1;refGene_tx=XM_003983479.4
A3 21928864 21929013 refGene_gn=TTI1;refGene_tx=XM_003983479.4
A3 21931612 21931753 refGene_gn=TTI1;refGene_tx=XM_003983479.4
A3 21933741 21933946 refGene_gn=TTI1;refGene_tx=XM_003983479.4
A3 21934220 21934308 refGene_gn=TTI1;refGene_tx=XM_003983479.4
A3 21946582 21946766 refGene_gn=TTI1;refGene_tx=XM_003983479.4
A3 21971972 21972245
refGene_gn=VSTM2L;refGene_tx=XM_003983480.5;refGene_gn=VSTM2L;refGene_
tx=XM_019826526.2
A3 21972546 21972687 refGene_gn=VSTM2L;refGene_tx=XM_019826527.2
A3 21982396 21982447
refGene_gn=VSTM2L;refGene_tx=XM_003983480.5;refGene_gn=VSTM2L;refGene_
tx=XM_019826526.2;refGene_gn=VSTM2L;refGene_tx=XM_019826527.2
A3 21983772 21983942
refGene_gn=VSTM2L;refGene_tx=XM_003983480.5;refGene_gn=VSTM2L;refGene_
tx=XM_019826526.2;refGene_gn=VSTM2L;refGene_tx=XM_019826527.2
A3 21999436 21999535
refGene_gn=VSTM2L;refGene_tx=XM_019826526.2;refGene_gn=VSTM2L;refGene_
tx=XM_019826527.2
A3 22008233 22008354
refGene_gn=VSTM2L;refGene_tx=XM_003983480.5;refGene_gn=VSTM2L;refGene_
tx=XM_019826526.2;refGene_gn=VSTM2L;refGene_tx=XM_019826527.2
A3 22033378 22033467 refGene_gn=CTNNBL1;refGene_tx=XM_003983481.4
A3 22043553 22043626 refGene_gn=CTNNBL1;refGene_tx=XM_003983481.4
A3 22043873 22044011 refGene_gn=CTNNBL1;refGene_tx=XM_003983481.4
A3 22059709 22059790 refGene_gn=CTNNBL1;refGene_tx=XM_003983481.4
A3 22061797 22061895 refGene_gn=CTNNBL1;refGene_tx=XM_003983481.4
A3 22095355 22095537 refGene_gn=CTNNBL1;refGene_tx=XM_003983481.4
A3 22117815 22117964 refGene_gn=CTNNBL1;refGene_tx=XM_003983481.4
A3 22119453 22119512 refGene_gn=CTNNBL1;refGene_tx=XM_003983481.4
A3 22119750 22119823 refGene_gn=CTNNBL1;refGene_tx=XM_003983481.4
A3 22129222 22129314 refGene_gn=CTNNBL1;refGene_tx=XM_003983481.4
A3 22132457 22132551 refGene_gn=CTNNBL1;refGene_tx=XM_003983481.4
A3 22140239 22140337 refGene_gn=CTNNBL1;refGene_tx=XM_003983481.4
A3 22149721 22149861 refGene_gn=CTNNBL1;refGene_tx=XM_003983481.4
A3 22159687 22159794 refGene_gn=CTNNBL1;refGene_tx=XM_003983481.4
A3 22163858 22164047 refGene_gn=CTNNBL1;refGene_tx=XM_003983481.4
A3 22194789 22194819 refGene_gn=CTNNBL1;refGene_tx=XM_003983481.4
A3 22326472 22326702

refGene_gn=NNAT;refGene_tx=XM_003983482.5;refGene_gn=NNAT;refGene_tx=XM_003983483.5;refGene_gn=NNAT;refGene_tx=XM_019826528.2
A3 22326928 22327009
refGene_gn=NNAT;refGene_tx=XM_003983482.5;refGene_gn=NNAT;refGene_tx=XM_019826528.2
A3 22327939 22328011
refGene_gn=NNAT;refGene_tx=XM_003983482.5;refGene_gn=NNAT;refGene_tx=XM_003983483.5;refGene_gn=NNAT;refGene_tx=XM_019826528.2
A3 22330101 22330365
refGene_gn=BLCAP;refGene_tx=NM_001009852.1;refGene_gn=BLCAP;refGene_tx=XM_019826171.2
A3 22426689 22426898
refGene_gn=SRC;refGene_tx=XM_003983484.5;refGene_gn=SRC;refGene_tx=XM_019826529.2;refGene_gn=SRC;refGene_tx=XM_019826530.2
A3 22427192 22427324
refGene_gn=SRC;refGene_tx=XM_003983484.5;refGene_gn=SRC;refGene_tx=XM_019826529.2;refGene_gn=SRC;refGene_tx=XM_019826530.2
A3 22427416 22427570
refGene_gn=SRC;refGene_tx=XM_003983484.5;refGene_gn=SRC;refGene_tx=XM_019826529.2;refGene_gn=SRC;refGene_tx=XM_019826530.2
A3 22428288 22428365
refGene_gn=SRC;refGene_tx=XM_003983484.5;refGene_gn=SRC;refGene_tx=XM_019826529.2;refGene_gn=SRC;refGene_tx=XM_019826530.2
A3 22429944 22430124
refGene_gn=SRC;refGene_tx=XM_003983484.5;refGene_gn=SRC;refGene_tx=XM_019826529.2;refGene_gn=SRC;refGene_tx=XM_019826530.2
A3 22431546 22431702
refGene_gn=SRC;refGene_tx=XM_003983484.5;refGene_gn=SRC;refGene_tx=XM_019826529.2;refGene_gn=SRC;refGene_tx=XM_019826530.2
A3 22433153 22433303
refGene_gn=SRC;refGene_tx=XM_003983484.5;refGene_gn=SRC;refGene_tx=XM_019826529.2;refGene_gn=SRC;refGene_tx=XM_019826530.2
A3 22435127 22435231
refGene_gn=SRC;refGene_tx=XM_003983484.5;refGene_gn=SRC;refGene_tx=XM_019826529.2;refGene_gn=SRC;refGene_tx=XM_019826530.2
A3 22435400 22435499
refGene_gn=SRC;refGene_tx=XM_003983484.5;refGene_gn=SRC;refGene_tx=XM_019826529.2;refGene_gn=SRC;refGene_tx=XM_019826530.2
A3 22436354 22436387 refGene_gn=SRC;refGene_tx=XM_019826529.2
A3 22441782 22441800
refGene_gn=SRC;refGene_tx=XM_019826529.2;refGene_gn=SRC;refGene_tx=XM_019826530.2
A3 22442764 22442864
refGene_gn=SRC;refGene_tx=XM_003983484.5;refGene_gn=SRC;refGene_tx=XM_019826529.2;refGene_gn=SRC;refGene_tx=XM_019826530.2
A3 22444583 22444833
refGene_gn=SRC;refGene_tx=XM_003983484.5;refGene_gn=SRC;refGene_tx=XM_019826529.2;refGene_gn=SRC;refGene_tx=XM_019826530.2
A3 22496424 22496532
refGene_gn=MANBAL;refGene_tx=XM_003983486.5;refGene_gn=MANBAL;refGene_

tx=XM_006929761.2;refGene_gn=MANBAL;refGene_tx=XM_023251260.1
A3	22511441	22511654
refGene_gn=MANBAL;refGene_tx=XM_003983486.5;refGene_gn=MANBAL;refGene_
tx=XM_006929761.2;refGene_gn=MANBAL;refGene_tx=XM_023251260.1
A3	22514333	22514408	refGene_gn=MANBAL;refGene_tx=XM_003983486.5
A3	22522083	22522293	refGene_gn=MANBAL;refGene_tx=XM_023251260.1
A3	22547481	22547561
refGene_gn=GHRH;refGene_tx=XM_019826531.1;refGene_gn=GHRH;refGene_tx=X
M_019826533.1
A3	22547890	22547995
refGene_gn=GHRH;refGene_tx=XM_019826531.1;refGene_gn=GHRH;refGene_tx=X
M_019826533.1
A3	22549408	22549528
refGene_gn=GHRH;refGene_tx=XM_019826531.1;refGene_gn=GHRH;refGene_tx=X
M_019826533.1
A3	22551473	22551489
refGene_gn=GHRH;refGene_tx=XM_019826531.1;refGene_gn=GHRH;refGene_tx=X
M_019826533.1
A3	22558314	22558327
refGene_gn=RPN2;refGene_tx=XM_003983488.4;refGene_gn=RPN2;refGene_tx=X
M_006929762.3
A3	22561040	22561088	refGene_gn=RPN2;refGene_tx=XM_006929762.3
A3	22562618	22562748
refGene_gn=RPN2;refGene_tx=XM_003983488.4;refGene_gn=RPN2;refGene_tx=X
M_006929762.3
A3	22564268	22564344
refGene_gn=RPN2;refGene_tx=XM_003983488.4;refGene_gn=RPN2;refGene_tx=X
M_006929762.3
A3	22565965	22566061
refGene_gn=RPN2;refGene_tx=XM_003983488.4;refGene_gn=RPN2;refGene_tx=X
M_006929762.3
A3	22568050	22568137
refGene_gn=RPN2;refGene_tx=XM_003983488.4;refGene_gn=RPN2;refGene_tx=X
M_006929762.3
A3	22569330	22569525
refGene_gn=RPN2;refGene_tx=XM_003983488.4;refGene_gn=RPN2;refGene_tx=X
M_006929762.3
A3	22572274	22572389
refGene_gn=RPN2;refGene_tx=XM_003983488.4;refGene_gn=RPN2;refGene_tx=X
M_006929762.3
A3	22574020	22574112
refGene_gn=RPN2;refGene_tx=XM_003983488.4;refGene_gn=RPN2;refGene_tx=X
M_006929762.3
A3	22587474	22587580
refGene_gn=RPN2;refGene_tx=XM_003983488.4;refGene_gn=RPN2;refGene_tx=X
M_006929762.3
A3	22589628	22589747
refGene_gn=RPN2;refGene_tx=XM_003983488.4;refGene_gn=RPN2;refGene_tx=X
M_006929762.3
A3	22591314	22591491

refGene_gn=RPN2;refGene_tx=XM_003983488.4;refGene_gn=RPN2;refGene_tx=XM_006929762.3
A3 22593951 22594086
refGene_gn=RPN2;refGene_tx=XM_003983488.4;refGene_gn=RPN2;refGene_tx=XM_006929762.3
A3 22595076 22595152
refGene_gn=RPN2;refGene_tx=XM_003983488.4;refGene_gn=RPN2;refGene_tx=XM_006929762.3
A3 22600120 22600296
refGene_gn=RPN2;refGene_tx=XM_003983488.4;refGene_gn=RPN2;refGene_tx=XM_006929762.3
A3 22600843 22600939
refGene_gn=RPN2;refGene_tx=XM_003983488.4;refGene_gn=RPN2;refGene_tx=XM_006929762.3
A3 22613876 22614070
refGene_gn=RPN2;refGene_tx=XM_003983488.4;refGene_gn=RPN2;refGene_tx=XM_006929762.3
A3 22618123 22618569
refGene_gn=MROH8;refGene_tx=XM_011280885.3;refGene_gn=MROH8;refGene_tx=XM_019826538.2;refGene_gn=MROH8;refGene_tx=XM_019826540.2;refGene_gn=MROH8;refGene_tx=XM_019826542.2;refGene_gn=RPN2;refGene_tx=XM_003983488.4;refGene_gn=RPN2;refGene_tx=XM_006929762.3
A3 22621691 22621813
refGene_gn=MROH8;refGene_tx=XM_011280885.3;refGene_gn=MROH8;refGene_tx=XM_019826538.2;refGene_gn=MROH8;refGene_tx=XM_019826540.2;refGene_gn=MROH8;refGene_tx=XM_019826542.2
A3 22624115 22624261
refGene_gn=MROH8;refGene_tx=XM_011280885.3;refGene_gn=MROH8;refGene_tx=XM_019826538.2;refGene_gn=MROH8;refGene_tx=XM_019826540.2;refGene_gn=MROH8;refGene_tx=XM_019826542.2
A3 22628282 22628405
refGene_gn=MROH8;refGene_tx=XM_011280885.3;refGene_gn=MROH8;refGene_tx=XM_019826538.2;refGene_gn=MROH8;refGene_tx=XM_019826540.2;refGene_gn=MROH8;refGene_tx=XM_019826542.2
A3 22634712 22634792
refGene_gn=MROH8;refGene_tx=XM_011280885.3;refGene_gn=MROH8;refGene_tx=XM_019826538.2;refGene_gn=MROH8;refGene_tx=XM_019826540.2
A3 22635769 22635914
refGene_gn=MROH8;refGene_tx=XM_011280885.3;refGene_gn=MROH8;refGene_tx=XM_019826538.2;refGene_gn=MROH8;refGene_tx=XM_019826540.2
A3 22641957 22642053
refGene_gn=MROH8;refGene_tx=XM_011280885.3;refGene_gn=MROH8;refGene_tx=XM_019826538.2;refGene_gn=MROH8;refGene_tx=XM_019826540.2;refGene_gn=MROH8;refGene_tx=XM_019826542.2
A3 22643541 22643646
refGene_gn=MROH8;refGene_tx=XM_011280885.3;refGene_gn=MROH8;refGene_tx=XM_019826538.2;refGene_gn=MROH8;refGene_tx=XM_019826540.2;refGene_gn=MROH8;refGene_tx=XM_019826542.2
A3 22647795 22647931
refGene_gn=MROH8;refGene_tx=XM_011280885.3;refGene_gn=MROH8;refGene_tx

=XM_019826538.2;refGene_gn=MROH8;refGene_tx=XM_019826540.2;refGene_gn=MROH8;refGene_tx=XM_019826542.2
A3 22649454 22649573
refGene_gn=MROH8;refGene_tx=XM_011280885.3;refGene_gn=MROH8;refGene_tx=XM_019826538.2;refGene_gn=MROH8;refGene_tx=XM_019826540.2;refGene_gn=MROH8;refGene_tx=XM_019826542.2
A3 22650884 22651037
refGene_gn=MROH8;refGene_tx=XM_011280885.3;refGene_gn=MROH8;refGene_tx=XM_019826538.2;refGene_gn=MROH8;refGene_tx=XM_019826540.2;refGene_gn=MROH8;refGene_tx=XM_019826542.2
A3 22652653 22652814
refGene_gn=MROH8;refGene_tx=XM_011280885.3;refGene_gn=MROH8;refGene_tx=XM_019826538.2;refGene_gn=MROH8;refGene_tx=XM_019826540.2;refGene_gn=MROH8;refGene_tx=XM_019826542.2
A3 22656627 22656763
refGene_gn=MROH8;refGene_tx=XM_011280885.3;refGene_gn=MROH8;refGene_tx=XM_019826538.2;refGene_gn=MROH8;refGene_tx=XM_019826540.2;refGene_gn=MROH8;refGene_tx=XM_019826542.2
A3 22659396 22659587
refGene_gn=MROH8;refGene_tx=XM_011280885.3;refGene_gn=MROH8;refGene_tx=XM_019826538.2;refGene_gn=MROH8;refGene_tx=XM_019826540.2;refGene_gn=MROH8;refGene_tx=XM_019826542.2
A3 22660925 22661063
refGene_gn=MROH8;refGene_tx=XM_011280885.3;refGene_gn=MROH8;refGene_tx=XM_019826538.2;refGene_gn=MROH8;refGene_tx=XM_019826540.2;refGene_gn=MROH8;refGene_tx=XM_019826542.2
A3 22661486 22661601
refGene_gn=MROH8;refGene_tx=XM_011280885.3;refGene_gn=MROH8;refGene_tx=XM_019826538.2;refGene_gn=MROH8;refGene_tx=XM_019826540.2;refGene_gn=MROH8;refGene_tx=XM_019826542.2
A3 22662132 22662273
refGene_gn=MROH8;refGene_tx=XM_011280885.3;refGene_gn=MROH8;refGene_tx=XM_019826538.2;refGene_gn=MROH8;refGene_tx=XM_019826540.2;refGene_gn=MROH8;refGene_tx=XM_019826542.2
A3 22663358 22663523
refGene_gn=MROH8;refGene_tx=XM_011280885.3;refGene_gn=MROH8;refGene_tx=XM_019826538.2;refGene_gn=MROH8;refGene_tx=XM_019826542.2
A3 22664179 22664335
refGene_gn=MROH8;refGene_tx=XM_011280885.3;refGene_gn=MROH8;refGene_tx=XM_019826538.2;refGene_gn=MROH8;refGene_tx=XM_019826540.2;refGene_gn=MROH8;refGene_tx=XM_019826542.2
A3 22666481 22666604
refGene_gn=MROH8;refGene_tx=XM_011280885.3;refGene_gn=MROH8;refGene_tx=XM_019826538.2;refGene_gn=MROH8;refGene_tx=XM_019826540.2;refGene_gn=MROH8;refGene_tx=XM_019826542.2
A3 22668084 22668184
refGene_gn=MROH8;refGene_tx=XM_011280885.3;refGene_gn=MROH8;refGene_tx=XM_019826538.2;refGene_gn=MROH8;refGene_tx=XM_019826540.2;refGene_gn=MROH8;refGene_tx=XM_019826542.2
A3 22668435 22668501

refGene_gn=MROH8;refGene_tx=XM_011280885.3;refGene_gn=MROH8;refGene_tx=XM_019826540.2;refGene_gn=MROH8;refGene_tx=XM_019826542.2
A3 22673748 22673897
refGene_gn=MROH8;refGene_tx=XM_011280885.3;refGene_gn=MROH8;refGene_tx=XM_019826538.2;refGene_gn=MROH8;refGene_tx=XM_019826540.2;refGene_gn=MROH8;refGene_tx=XM_019826542.2
A3 22689217 22689373
refGene_gn=RBL1;refGene_tx=XM_003983544.5;refGene_gn=RBL1;refGene_tx=XM_023251262.1;refGene_gn=RBL1;refGene_tx=XM_023251264.1
A3 22693585 22693719
refGene_gn=RBL1;refGene_tx=XM_003983544.5;refGene_gn=RBL1;refGene_tx=XM_023251262.1;refGene_gn=RBL1;refGene_tx=XM_023251264.1
A3 22701014 22701215
refGene_gn=RBL1;refGene_tx=XM_003983544.5;refGene_gn=RBL1;refGene_tx=XM_023251262.1;refGene_gn=RBL1;refGene_tx=XM_023251263.1;refGene_gn=RBL1;refGene_tx=XM_023251264.1
A3 22701580 22701645
refGene_gn=RBL1;refGene_tx=XM_003983544.5;refGene_gn=RBL1;refGene_tx=XM_023251262.1;refGene_gn=RBL1;refGene_tx=XM_023251263.1;refGene_gn=RBL1;refGene_tx=XM_023251264.1
A3 22701747 22701876
refGene_gn=RBL1;refGene_tx=XM_003983544.5;refGene_gn=RBL1;refGene_tx=XM_023251262.1;refGene_gn=RBL1;refGene_tx=XM_023251263.1;refGene_gn=RBL1;refGene_tx=XM_023251264.1
A3 22701987 22702148
refGene_gn=RBL1;refGene_tx=XM_003983544.5;refGene_gn=RBL1;refGene_tx=XM_023251262.1;refGene_gn=RBL1;refGene_tx=XM_023251263.1;refGene_gn=RBL1;refGene_tx=XM_023251264.1
A3 22703940 22703990
refGene_gn=RBL1;refGene_tx=XM_003983544.5;refGene_gn=RBL1;refGene_tx=XM_023251262.1;refGene_gn=RBL1;refGene_tx=XM_023251263.1;refGene_gn=RBL1;refGene_tx=XM_023251264.1
A3 22705401 22705588
refGene_gn=RBL1;refGene_tx=XM_003983544.5;refGene_gn=RBL1;refGene_tx=XM_023251262.1;refGene_gn=RBL1;refGene_tx=XM_023251263.1;refGene_gn=RBL1;refGene_tx=XM_023251264.1
A3 22706681 22706848
refGene_gn=RBL1;refGene_tx=XM_003983544.5;refGene_gn=RBL1;refGene_tx=XM_023251262.1;refGene_gn=RBL1;refGene_tx=XM_023251263.1;refGene_gn=RBL1;refGene_tx=XM_023251264.1
A3 22711280 22711393
refGene_gn=RBL1;refGene_tx=XM_003983544.5;refGene_gn=RBL1;refGene_tx=XM_023251262.1;refGene_gn=RBL1;refGene_tx=XM_023251263.1;refGene_gn=RBL1;refGene_tx=XM_023251264.1
A3 22711582 22711686
refGene_gn=RBL1;refGene_tx=XM_003983544.5;refGene_gn=RBL1;refGene_tx=XM_023251262.1;refGene_gn=RBL1;refGene_tx=XM_023251263.1;refGene_gn=RBL1;refGene_tx=XM_023251264.1
A3 22716599 22716737
refGene_gn=RBL1;refGene_tx=XM_003983544.5;refGene_gn=RBL1;refGene_tx=X

M_023251262.1;refGene_gn=RBL1;refGene_tx=XM_023251263.1;refGene_gn=RBL1;refGene_tx=XM_023251264.1
A3	22718228	22718393	refGene_gn=RBL1;refGene_tx=XM_003983544.5;refGene_gn=RBL1;refGene_tx=XM_023251262.1;refGene_gn=RBL1;refGene_tx=XM_023251263.1;refGene_gn=RBL1;refGene_tx=XM_023251264.1
A3	22720909	22721057	refGene_gn=RBL1;refGene_tx=XM_003983544.5;refGene_gn=RBL1;refGene_tx=XM_023251262.1;refGene_gn=RBL1;refGene_tx=XM_023251263.1;refGene_gn=RBL1;refGene_tx=XM_023251264.1
A3	22727439	22727706	refGene_gn=RBL1;refGene_tx=XM_003983544.5;refGene_gn=RBL1;refGene_tx=XM_023251262.1;refGene_gn=RBL1;refGene_tx=XM_023251263.1;refGene_gn=RBL1;refGene_tx=XM_023251264.1
A3	22728518	22728730	refGene_gn=RBL1;refGene_tx=XM_003983544.5;refGene_gn=RBL1;refGene_tx=XM_023251262.1;refGene_gn=RBL1;refGene_tx=XM_023251263.1;refGene_gn=RBL1;refGene_tx=XM_023251264.1
A3	22735581	22735758	refGene_gn=RBL1;refGene_tx=XM_003983544.5;refGene_gn=RBL1;refGene_tx=XM_023251262.1;refGene_gn=RBL1;refGene_tx=XM_023251263.1;refGene_gn=RBL1;refGene_tx=XM_023251264.1
A3	22737209	22737281	refGene_gn=RBL1;refGene_tx=XM_003983544.5;refGene_gn=RBL1;refGene_tx=XM_023251262.1;refGene_gn=RBL1;refGene_tx=XM_023251263.1;refGene_gn=RBL1;refGene_tx=XM_023251264.1
A3	22739188	22739279	refGene_gn=RBL1;refGene_tx=XM_003983544.5;refGene_gn=RBL1;refGene_tx=XM_023251262.1;refGene_gn=RBL1;refGene_tx=XM_023251263.1;refGene_gn=RBL1;refGene_tx=XM_023251264.1
A3	22744640	22744789	refGene_gn=RBL1;refGene_tx=XM_003983544.5;refGene_gn=RBL1;refGene_tx=XM_023251262.1;refGene_gn=RBL1;refGene_tx=XM_023251263.1;refGene_gn=RBL1;refGene_tx=XM_023251264.1
A3	22746188	22746353	refGene_gn=RBL1;refGene_tx=XM_003983544.5;refGene_gn=RBL1;refGene_tx=XM_023251262.1;refGene_gn=RBL1;refGene_tx=XM_023251263.1;refGene_gn=RBL1;refGene_tx=XM_023251264.1
A3	22748533	22748704	refGene_gn=RBL1;refGene_tx=XM_003983544.5;refGene_gn=RBL1;refGene_tx=XM_023251262.1;refGene_gn=RBL1;refGene_tx=XM_023251263.1
A3	22764801	22765009	refGene_gn=SAMHD1;refGene_tx=XM_003983547.4
A3	22768578	22768645	refGene_gn=SAMHD1;refGene_tx=XM_003983547.4
A3	22774154	22774227	refGene_gn=SAMHD1;refGene_tx=XM_003983547.4
A3	22780206	22780367	refGene_gn=SAMHD1;refGene_tx=XM_003983547.4
A3	22783228	22783344	refGene_gn=SAMHD1;refGene_tx=XM_003983547.4
A3	22785522	22785593	refGene_gn=SAMHD1;refGene_tx=XM_003983547.4
A3	22791667	22791823	refGene_gn=SAMHD1;refGene_tx=XM_003983547.4
A3	22794307	22794408	refGene_gn=SAMHD1;refGene_tx=XM_003983547.4
A3	22794584	22794693	refGene_gn=SAMHD1;refGene_tx=XM_003983547.4

A3	22795533	22795625	refGene_gn=SAMHD1;refGene_tx=XM_003983547.4
A3	22796710	22796826	refGene_gn=SAMHD1;refGene_tx=XM_003983547.4
A3	22802309	22802449	refGene_gn=SAMHD1;refGene_tx=XM_003983547.4
A3	22803225	22803321	refGene_gn=SAMHD1;refGene_tx=XM_003983547.4
A3	22805269	22805374	refGene_gn=SAMHD1;refGene_tx=XM_003983547.4
A3	22806558	22806696	refGene_gn=SAMHD1;refGene_tx=XM_003983547.4
A3	22809600	22809735	refGene_gn=SAMHD1;refGene_tx=XM_003983547.4
A3	22813435	22813625
refGene_gn=TLDC2;refGene_tx=XM_003983548.3;refGene_gn=TLDC2;refGene_tx=XM_019826544.1
A3	22814633	22814707
refGene_gn=TLDC2;refGene_tx=XM_003983548.3;refGene_gn=TLDC2;refGene_tx=XM_019826544.1
A3	22819948	22820044
refGene_gn=TLDC2;refGene_tx=XM_003983548.3;refGene_gn=TLDC2;refGene_tx=XM_019826544.1
A3	22823073	22823226
refGene_gn=TLDC2;refGene_tx=XM_003983548.3;refGene_gn=TLDC2;refGene_tx=XM_019826544.1
A3	22823706	22823856
refGene_gn=TLDC2;refGene_tx=XM_003983548.3;refGene_gn=TLDC2;refGene_tx=XM_019826544.1
A3	22825096	22825129
refGene_gn=TLDC2;refGene_tx=XM_003983548.3;refGene_gn=TLDC2;refGene_tx=XM_019826544.1
A3	22834568	22835243
refGene_gn=SOGA1;refGene_tx=XM_006929781.3;refGene_gn=SOGA1;refGene_tx=XM_006929782.3;refGene_gn=SOGA1;refGene_tx=XM_019826546.2;refGene_gn=SOGA1;refGene_tx=XM_019826548.2
A3	22853948	22854173
refGene_gn=SOGA1;refGene_tx=XM_006929781.3;refGene_gn=SOGA1;refGene_tx=XM_006929782.3;refGene_gn=SOGA1;refGene_tx=XM_019826546.2;refGene_gn=SOGA1;refGene_tx=XM_019826548.2
A3	22860258	22860417
refGene_gn=SOGA1;refGene_tx=XM_006929781.3;refGene_gn=SOGA1;refGene_tx=XM_006929782.3;refGene_gn=SOGA1;refGene_tx=XM_019826546.2;refGene_gn=SOGA1;refGene_tx=XM_019826548.2
A3	22872160	22872223
refGene_gn=SOGA1;refGene_tx=XM_006929781.3;refGene_gn=SOGA1;refGene_tx=XM_006929782.3;refGene_gn=SOGA1;refGene_tx=XM_019826546.2;refGene_gn=SOGA1;refGene_tx=XM_019826548.2
A3	22872905	22874087
refGene_gn=SOGA1;refGene_tx=XM_006929781.3;refGene_gn=SOGA1;refGene_tx=XM_006929782.3;refGene_gn=SOGA1;refGene_tx=XM_019826546.2;refGene_gn=SOGA1;refGene_tx=XM_019826548.2
A3	22876187	22876328
refGene_gn=SOGA1;refGene_tx=XM_006929781.3;refGene_gn=SOGA1;refGene_tx=XM_019826546.2;refGene_gn=SOGA1;refGene_tx=XM_019826548.2
A3	22879567	22879693
refGene_gn=SOGA1;refGene_tx=XM_006929781.3;refGene_gn=SOGA1;refGene_tx

=XM_006929782.3;refGene_gn=SOGA1;refGene_tx=XM_019826546.2;refGene_gn=SOGA1;refGene_tx=XM_019826548.2
A3 22880900 22881131
refGene_gn=SOGA1;refGene_tx=XM_006929781.3;refGene_gn=SOGA1;refGene_tx=XM_006929782.3;refGene_gn=SOGA1;refGene_tx=XM_019826546.2;refGene_gn=SOGA1;refGene_tx=XM_019826548.2
A3 22887323 22887428
refGene_gn=SOGA1;refGene_tx=XM_006929781.3;refGene_gn=SOGA1;refGene_tx=XM_006929782.3;refGene_gn=SOGA1;refGene_tx=XM_019826546.2;refGene_gn=SOGA1;refGene_tx=XM_019826548.2
A3 22888371 22888539
refGene_gn=SOGA1;refGene_tx=XM_006929781.3;refGene_gn=SOGA1;refGene_tx=XM_006929782.3;refGene_gn=SOGA1;refGene_tx=XM_019826546.2;refGene_gn=SOGA1;refGene_tx=XM_019826548.2
A3 22890052 22890304
refGene_gn=SOGA1;refGene_tx=XM_006929781.3;refGene_gn=SOGA1;refGene_tx=XM_006929782.3;refGene_gn=SOGA1;refGene_tx=XM_019826546.2;refGene_gn=SOGA1;refGene_tx=XM_019826548.2
A3 22891524 22891596
refGene_gn=SOGA1;refGene_tx=XM_006929781.3;refGene_gn=SOGA1;refGene_tx=XM_006929782.3;refGene_gn=SOGA1;refGene_tx=XM_019826546.2;refGene_gn=SOGA1;refGene_tx=XM_019826548.2
A3 22892858 22892929
refGene_gn=SOGA1;refGene_tx=XM_006929781.3;refGene_gn=SOGA1;refGene_tx=XM_006929782.3;refGene_gn=SOGA1;refGene_tx=XM_019826546.2;refGene_gn=SOGA1;refGene_tx=XM_019826548.2
A3 22894280 22895631
refGene_gn=SOGA1;refGene_tx=XM_006929781.3;refGene_gn=SOGA1;refGene_tx=XM_006929782.3;refGene_gn=SOGA1;refGene_tx=XM_019826546.2;refGene_gn=SOGA1;refGene_tx=XM_019826548.2
A3 22899439 22899592
refGene_gn=SOGA1;refGene_tx=XM_006929781.3;refGene_gn=SOGA1;refGene_tx=XM_006929782.3;refGene_gn=SOGA1;refGene_tx=XM_019826546.2;refGene_gn=SOGA1;refGene_tx=XM_019826548.2
A3 22910758 22910792
refGene_gn=DSN1;refGene_tx=XM_003983549.5;refGene_gn=DSN1;refGene_tx=XM_011280884.3
A3 22911016 22911343 refGene_gn=DSN1;refGene_tx=XM_003983549.5
A3 22913389 22913463
refGene_gn=DSN1;refGene_tx=XM_003983549.5;refGene_gn=DSN1;refGene_tx=XM_011280884.3
A3 22914335 22914408
refGene_gn=DSN1;refGene_tx=XM_003983549.5;refGene_gn=DSN1;refGene_tx=XM_011280884.3
A3 22916512 22916600
refGene_gn=DSN1;refGene_tx=XM_003983549.5;refGene_gn=DSN1;refGene_tx=XM_011280884.3
A3 22918261 22918321
refGene_gn=DSN1;refGene_tx=XM_003983549.5;refGene_gn=DSN1;refGene_tx=XM_011280884.3

A3 22918717 22918792
refGene_gn=DSN1;refGene_tx=XM_003983549.5;refGene_gn=DSN1;refGene_tx=XM_011280884.3
A3 22919955 22920103
refGene_gn=DSN1;refGene_tx=XM_003983549.5;refGene_gn=DSN1;refGene_tx=XM_011280884.3
A3 22921228 22921316
refGene_gn=DSN1;refGene_tx=XM_003983549.5;refGene_gn=DSN1;refGene_tx=XM_011280884.3
A3 22921889 22921972
refGene_gn=DSN1;refGene_tx=XM_003983549.5;refGene_gn=DSN1;refGene_tx=XM_011280884.3
A3 22945923 22945980
refGene_gn=NDRG3;refGene_tx=XM_011280881.3;refGene_gn=NDRG3;refGene_tx=XM_019826549.2;refGene_gn=NDRG3;refGene_tx=XM_019826550.2;refGene_gn=NDRG3;refGene_tx=XM_019826551.2;refGene_gn=NDRG3;refGene_tx=XM_023251266.1
A3 22955583 22955619
refGene_gn=NDRG3;refGene_tx=XM_011280881.3;refGene_gn=NDRG3;refGene_tx=XM_019826549.2;refGene_gn=NDRG3;refGene_tx=XM_019826550.2
A3 22963884 22963890 refGene_gn=NDRG3;refGene_tx=XM_019826553.2
A3 22967122 22967228
refGene_gn=NDRG3;refGene_tx=XM_011280881.3;refGene_gn=NDRG3;refGene_tx=XM_019826549.2;refGene_gn=NDRG3;refGene_tx=XM_019826550.2;refGene_gn=NDRG3;refGene_tx=XM_019826551.2;refGene_gn=NDRG3;refGene_tx=XM_019826553.2;refGene_gn=NDRG3;refGene_tx=XM_023251266.1
A3 22968431 22968552
refGene_gn=NDRG3;refGene_tx=XM_011280881.3;refGene_gn=NDRG3;refGene_tx=XM_019826549.2;refGene_gn=NDRG3;refGene_tx=XM_019826550.2;refGene_gn=NDRG3;refGene_tx=XM_019826551.2;refGene_gn=NDRG3;refGene_tx=XM_019826553.2;refGene_gn=NDRG3;refGene_tx=XM_023251266.1
A3 22970425 22970488
refGene_gn=NDRG3;refGene_tx=XM_011280881.3;refGene_gn=NDRG3;refGene_tx=XM_019826549.2;refGene_gn=NDRG3;refGene_tx=XM_019826550.2;refGene_gn=NDRG3;refGene_tx=XM_019826551.2;refGene_gn=NDRG3;refGene_tx=XM_019826553.2;refGene_gn=NDRG3;refGene_tx=XM_023251266.1
A3 22971832 22971893
refGene_gn=NDRG3;refGene_tx=XM_011280881.3;refGene_gn=NDRG3;refGene_tx=XM_019826549.2;refGene_gn=NDRG3;refGene_tx=XM_019826550.2;refGene_gn=NDRG3;refGene_tx=XM_019826551.2;refGene_gn=NDRG3;refGene_tx=XM_019826553.2;refGene_gn=NDRG3;refGene_tx=XM_023251266.1
A3 22973619 22973706
refGene_gn=NDRG3;refGene_tx=XM_011280881.3;refGene_gn=NDRG3;refGene_tx=XM_019826549.2;refGene_gn=NDRG3;refGene_tx=XM_019826550.2;refGene_gn=NDRG3;refGene_tx=XM_019826551.2;refGene_gn=NDRG3;refGene_tx=XM_019826553.2;refGene_gn=NDRG3;refGene_tx=XM_023251266.1
A3 22978116 22978173
refGene_gn=NDRG3;refGene_tx=XM_011280881.3;refGene_gn=NDRG3;refGene_tx=XM_019826549.2;refGene_gn=NDRG3;refGene_tx=XM_019826550.2;refGene_gn=NDRG3;refGene_tx=XM_019826551.2;refGene_gn=NDRG3;refGene_tx=XM_0198265

53.2;refGene_gn=NDRG3;refGene_tx=XM_023251266.1
A3	22985354 22985458
refGene_gn=NDRG3;refGene_tx=XM_011280881.3;refGene_gn=NDRG3;refGene_tx=XM_019826549.2;refGene_gn=NDRG3;refGene_tx=XM_019826550.2;refGene_gn=NDRG3;refGene_tx=XM_019826551.2;refGene_gn=NDRG3;refGene_tx=XM_019826553.2;refGene_gn=NDRG3;refGene_tx=XM_023251266.1
A3	22986647 22986713
refGene_gn=NDRG3;refGene_tx=XM_011280881.3;refGene_gn=NDRG3;refGene_tx=XM_019826549.2;refGene_gn=NDRG3;refGene_tx=XM_019826550.2;refGene_gn=NDRG3;refGene_tx=XM_019826551.2;refGene_gn=NDRG3;refGene_tx=XM_019826553.2;refGene_gn=NDRG3;refGene_tx=XM_023251266.1
A3	22986835 22986887
refGene_gn=NDRG3;refGene_tx=XM_011280881.3;refGene_gn=NDRG3;refGene_tx=XM_019826549.2;refGene_gn=NDRG3;refGene_tx=XM_019826550.2;refGene_gn=NDRG3;refGene_tx=XM_019826551.2;refGene_gn=NDRG3;refGene_tx=XM_019826553.2;refGene_gn=NDRG3;refGene_tx=XM_023251266.1
A3	22990762 22990810
refGene_gn=NDRG3;refGene_tx=XM_011280881.3;refGene_gn=NDRG3;refGene_tx=XM_019826549.2;refGene_gn=NDRG3;refGene_tx=XM_019826550.2;refGene_gn=NDRG3;refGene_tx=XM_019826551.2;refGene_gn=NDRG3;refGene_tx=XM_019826553.2;refGene_gn=NDRG3;refGene_tx=XM_023251266.1
A3	22993180 22993216
refGene_gn=NDRG3;refGene_tx=XM_011280881.3;refGene_gn=NDRG3;refGene_tx=XM_019826549.2;refGene_gn=NDRG3;refGene_tx=XM_019826550.2;refGene_gn=NDRG3;refGene_tx=XM_019826551.2;refGene_gn=NDRG3;refGene_tx=XM_019826553.2;refGene_gn=NDRG3;refGene_tx=XM_023251266.1
A3	22993304 22993356
refGene_gn=NDRG3;refGene_tx=XM_011280881.3;refGene_gn=NDRG3;refGene_tx=XM_019826549.2;refGene_gn=NDRG3;refGene_tx=XM_019826550.2;refGene_gn=NDRG3;refGene_tx=XM_019826551.2;refGene_gn=NDRG3;refGene_tx=XM_019826553.2;refGene_gn=NDRG3;refGene_tx=XM_023251266.1
A3	22994565 22994604
refGene_gn=NDRG3;refGene_tx=XM_019826549.2;refGene_gn=NDRG3;refGene_tx=XM_019826550.2;refGene_gn=NDRG3;refGene_tx=XM_019826551.2;refGene_gn=NDRG3;refGene_tx=XM_019826553.2
A3	22995716 22995898
refGene_gn=NDRG3;refGene_tx=XM_011280881.3;refGene_gn=NDRG3;refGene_tx=XM_019826549.2;refGene_gn=NDRG3;refGene_tx=XM_019826550.2;refGene_gn=NDRG3;refGene_tx=XM_019826551.2;refGene_gn=NDRG3;refGene_tx=XM_019826553.2;refGene_gn=NDRG3;refGene_tx=XM_023251266.1
A3	23004308 23004399
refGene_gn=SLA2;refGene_tx=XM_011280880.3;refGene_gn=SLA2;refGene_tx=XM_019826555.2
A3	23008682 23008782
refGene_gn=SLA2;refGene_tx=XM_011280880.3;refGene_gn=SLA2;refGene_tx=XM_019826555.2
A3	23009142 23009229
refGene_gn=SLA2;refGene_tx=XM_011280880.3;refGene_gn=SLA2;refGene_tx=XM_019826555.2
A3	23011287 23011391

refGene_gn=SLA2;refGene_tx=XM_011280880.3;refGene_gn=SLA2;refGene_tx=XM_019826555.2
A3 23030463 23030613
refGene_gn=SLA2;refGene_tx=XM_011280880.3;refGene_gn=SLA2;refGene_tx=XM_019826555.2
A3 23031476 23031609
refGene_gn=SLA2;refGene_tx=XM_011280880.3;refGene_gn=SLA2;refGene_tx=XM_019826555.2
A3 23032011 23032177
refGene_gn=SLA2;refGene_tx=XM_011280880.3;refGene_gn=SLA2;refGene_tx=XM_019826555.2
A3 23033399 23033613
refGene_gn=LOC102899611;refGene_tx=XM_006929790.2;refGene_gn=LOC102899611;refGene_tx=XM_006929791.3
A3 23034916 23035046
refGene_gn=LOC102899611;refGene_tx=XM_006929790.2
A3 23039659 23039763
refGene_gn=LOC102899611;refGene_tx=XM_006929790.2;refGene_gn=LOC102899611;refGene_tx=XM_006929791.3
A3 23041346 23041460
refGene_gn=LOC102899611;refGene_tx=XM_006929790.2;refGene_gn=LOC102899611;refGene_tx=XM_006929791.3
A3 23053741 23054263
refGene_gn=TGIF2;refGene_tx=XM_003983553.5;refGene_gn=TGIF2;refGene_tx=XM_006929785.3;refGene_gn=TGIF2;refGene_tx=XM_006929786.4;refGene_gn=TGIF2;refGene_tx=XM_006929788.2;refGene_gn=TGIF2;refGene_tx=XM_023251267.1
A3 23062450 23062642
refGene_gn=TGIF2;refGene_tx=XM_003983553.5;refGene_gn=TGIF2;refGene_tx=XM_006929785.3;refGene_gn=TGIF2;refGene_tx=XM_006929786.4;refGene_gn=TGIF2;refGene_tx=XM_006929788.2;refGene_gn=TGIF2;refGene_tx=XM_023251267.1
A3 23065444 23065520
refGene_gn=LOC109498511;refGene_tx=XM_023251639.1
A3 23066316 23066749
refGene_gn=LOC109498511;refGene_tx=XM_023251639.1
A3 23068377 23068492
refGene_gn=LOC109498511;refGene_tx=XM_023251639.1
A3 23098408 23098581
refGene_gn=MYL9;refGene_tx=XM_003983555.3;refGene_gn=MYL9;refGene_tx=XM_003983556.3
A3 23099434 23099596 refGene_gn=MYL9;refGene_tx=XM_003983555.3
A3 23101389 23101573
refGene_gn=MYL9;refGene_tx=XM_003983555.3;refGene_gn=MYL9;refGene_tx=XM_003983556.3
A3 23116159 23116378
refGene_gn=DLGAP4;refGene_tx=XM_023251268.1;refGene_gn=DLGAP4;refGene_tx=XM_023251269.1;refGene_gn=DLGAP4;refGene_tx=XM_023251270.1;refGene_gn=DLGAP4;refGene_tx=XM_023251271.1;refGene_gn=DLGAP4;refGene_tx=XM_023251272.1;refGene_gn=DLGAP4;refGene_tx=XM_023251273.1;refGene_gn=DLGAP

4;refGene_tx=XM_023251275.1;refGene_gn=DLGAP4;refGene_tx=XM_023251276.1;refGene_gn=DLGAP4;refGene_tx=XM_023251277.1;refGene_gn=DLGAP4;refGene_tx=XM_023251278.1
A3 23116380 23116389 refGene_gn=DLGAP4;refGene_tx=XM_023251274.1
A3 23117158 23117314 refGene_gn=DLGAP4;refGene_tx=XM_023251268.1;refGene_gn=DLGAP4;refGene_tx=XM_023251269.1;refGene_gn=DLGAP4;refGene_tx=XM_023251270.1;refGene_gn=DLGAP4;refGene_tx=XM_023251271.1;refGene_gn=DLGAP4;refGene_tx=XM_023251272.1;refGene_gn=DLGAP4;refGene_tx=XM_023251273.1;refGene_gn=DLGAP4;refGene_tx=XM_023251274.1;refGene_gn=DLGAP4;refGene_tx=XM_023251275.1;refGene_gn=DLGAP4;refGene_tx=XM_023251276.1;refGene_gn=DLGAP4;refGene_tx=XM_023251277.1;refGene_gn=DLGAP4;refGene_tx=XM_023251278.1
A3 23118217 23118309 refGene_gn=DLGAP4;refGene_tx=XM_023251268.1;refGene_gn=DLGAP4;refGene_tx=XM_023251269.1;refGene_gn=DLGAP4;refGene_tx=XM_023251270.1;refGene_gn=DLGAP4;refGene_tx=XM_023251271.1;refGene_gn=DLGAP4;refGene_tx=XM_023251272.1;refGene_gn=DLGAP4;refGene_tx=XM_023251273.1;refGene_gn=DLGAP4;refGene_tx=XM_023251274.1;refGene_gn=DLGAP4;refGene_tx=XM_023251275.1;refGene_gn=DLGAP4;refGene_tx=XM_023251276.1;refGene_gn=DLGAP4;refGene_tx=XM_023251277.1;refGene_gn=DLGAP4;refGene_tx=XM_023251278.1
A3 23144780 23145193 refGene_gn=DLGAP4;refGene_tx=XM_023251268.1;refGene_gn=DLGAP4;refGene_tx=XM_023251269.1;refGene_gn=DLGAP4;refGene_tx=XM_023251270.1;refGene_gn=DLGAP4;refGene_tx=XM_023251271.1;refGene_gn=DLGAP4;refGene_tx=XM_023251272.1;refGene_gn=DLGAP4;refGene_tx=XM_023251273.1;refGene_gn=DLGAP4;refGene_tx=XM_023251274.1;refGene_gn=DLGAP4;refGene_tx=XM_023251275.1;refGene_gn=DLGAP4;refGene_tx=XM_023251276.1;refGene_gn=DLGAP4;refGene_tx=XM_023251277.1;refGene_gn=DLGAP4;refGene_tx=XM_023251278.1
A3 23145716 23145805 refGene_gn=DLGAP4;refGene_tx=XM_023251268.1;refGene_gn=DLGAP4;refGene_tx=XM_023251269.1;refGene_gn=DLGAP4;refGene_tx=XM_023251271.1;refGene_gn=DLGAP4;refGene_tx=XM_023251273.1;refGene_gn=DLGAP4;refGene_tx=XM_023251274.1;refGene_gn=DLGAP4;refGene_tx=XM_023251275.1;refGene_gn=DLGAP4;refGene_tx=XM_023251276.1;refGene_gn=DLGAP4;refGene_tx=XM_023251277.1
A3 23146050 23146130 refGene_gn=DLGAP4;refGene_tx=XM_023251270.1;refGene_gn=DLGAP4;refGene_tx=XM_023251272.1
A3 23148303 23148665 refGene_gn=DLGAP4;refGene_tx=XM_023251268.1;refGene_gn=DLGAP4;refGene_tx=XM_023251269.1;refGene_gn=DLGAP4;refGene_tx=XM_023251270.1;refGene_gn=DLGAP4;refGene_tx=XM_023251271.1;refGene_gn=DLGAP4;refGene_tx=XM_023251272.1;refGene_gn=DLGAP4;refGene_tx=XM_023251273.1;refGene_gn=DLGAP4;refGene_tx=XM_023251274.1;refGene_gn=DLGAP4;refGene_tx=XM_023251275.1;refGene_gn=DLGAP4;refGene_tx=XM_023251276.1;refGene_gn=DLGAP4;refGene_tx=XM_023251277.1
A3 23164090 23164151 refGene_gn=DLGAP4;refGene_tx=XM_023251276.1
A3 23175334 23175548 refGene_gn=DLGAP4;refGene_tx=XM_023251275.1
A3 23179401 23179432 refGene_gn=DLGAP4;refGene_tx=XM_023251277.1
A3 23188205 23188446

refGene_gn=DLGAP4;refGene_tx=XM_023251268.1;refGene_gn=DLGAP4;refGene_tx=XM_023251269.1;refGene_gn=DLGAP4;refGene_tx=XM_023251270.1;refGene_gn=DLGAP4;refGene_tx=XM_023251271.1;refGene_gn=DLGAP4;refGene_tx=XM_023251272.1;refGene_gn=DLGAP4;refGene_tx=XM_023251273.1;refGene_gn=DLGAP4;refGene_tx=XM_023251274.1
A3 23191193 23191244
refGene_gn=DLGAP4;refGene_tx=XM_023251268.1;refGene_gn=DLGAP4;refGene_tx=XM_023251269.1;refGene_gn=DLGAP4;refGene_tx=XM_023251270.1;refGene_gn=DLGAP4;refGene_tx=XM_023251271.1;refGene_gn=DLGAP4;refGene_tx=XM_023251272.1;refGene_gn=DLGAP4;refGene_tx=XM_023251274.1
A3 23191789 23191936
refGene_gn=DLGAP4;refGene_tx=XM_023251268.1;refGene_gn=DLGAP4;refGene_tx=XM_023251269.1;refGene_gn=DLGAP4;refGene_tx=XM_023251270.1;refGene_gn=DLGAP4;refGene_tx=XM_023251274.1
A3 23193904 23194019
refGene_gn=DLGAP4;refGene_tx=XM_023251268.1;refGene_gn=DLGAP4;refGene_tx=XM_023251269.1;refGene_gn=DLGAP4;refGene_tx=XM_023251270.1;refGene_gn=DLGAP4;refGene_tx=XM_023251271.1;refGene_gn=DLGAP4;refGene_tx=XM_023251272.1;refGene_gn=DLGAP4;refGene_tx=XM_023251273.1;refGene_gn=DLGAP4;refGene_tx=XM_023251274.1
A3 23196333 23196575
refGene_gn=DLGAP4;refGene_tx=XM_023251268.1;refGene_gn=DLGAP4;refGene_tx=XM_023251269.1;refGene_gn=DLGAP4;refGene_tx=XM_023251270.1;refGene_gn=DLGAP4;refGene_tx=XM_023251271.1;refGene_gn=DLGAP4;refGene_tx=XM_023251272.1;refGene_gn=DLGAP4;refGene_tx=XM_023251273.1;refGene_gn=DLGAP4;refGene_tx=XM_023251274.1
A3 23199264 23200263
refGene_gn=DLGAP4;refGene_tx=XM_023251268.1;refGene_gn=DLGAP4;refGene_tx=XM_023251269.1;refGene_gn=DLGAP4;refGene_tx=XM_023251270.1;refGene_gn=DLGAP4;refGene_tx=XM_023251271.1;refGene_gn=DLGAP4;refGene_tx=XM_023251272.1;refGene_gn=DLGAP4;refGene_tx=XM_023251273.1;refGene_gn=DLGAP4;refGene_tx=XM_023251274.1
A3 23358491 23358659
refGene_gn=AAR2;refGene_tx=XM_003983558.5;refGene_gn=AAR2;refGene_tx=XM_006929792.2;refGene_gn=AAR2;refGene_tx=XM_006929793.2;refGene_gn=AAR2;refGene_tx=XM_006929794.1;refGene_gn=AAR2;refGene_tx=XM_023251279.1
A3 23368006 23368236
refGene_gn=AAR2;refGene_tx=XM_003983558.5;refGene_gn=AAR2;refGene_tx=XM_006929792.2;refGene_gn=AAR2;refGene_tx=XM_006929793.2;refGene_gn=AAR2;refGene_tx=XM_006929794.1;refGene_gn=AAR2;refGene_tx=XM_023251279.1
A3 23371580 23372337
refGene_gn=AAR2;refGene_tx=XM_003983558.5;refGene_gn=AAR2;refGene_tx=XM_006929792.2;refGene_gn=AAR2;refGene_tx=XM_006929793.2;refGene_gn=AAR2;refGene_tx=XM_006929794.1;refGene_gn=AAR2;refGene_tx=XM_023251279.1
A3 23381233 23381242
refGene_gn=EPB41L1;refGene_tx=XM_019826561.2;refGene_gn=EPB41L1;refGene_tx=XM_019826562.2;refGene_gn=EPB41L1;refGene_tx=XM_019826563.2;refGene_gn=EPB41L1;refGene_tx=XM_019826564.2;refGene_gn=EPB41L1;refGene_tx=XM_019826565.2;refGene_gn=EPB41L1;refGene_tx=XM_019826566.2;refGene_gn=EPB41L1;refGene_tx=XM_019826567.2;refGene_gn=EPB41L1;refGene_tx=XM_01

9826568.2;refGene_gn=EPB41L1;refGene_tx=XM_019826569.2;refGene_gn=EPB41L1;refGene_tx=XM_019826570.2;refGene_gn=EPB41L1;refGene_tx=XM_019826571.2;refGene_gn=EPB41L1;refGene_tx=XM_019826572.2;refGene_gn=EPB41L1;refGene_tx=XM_019826573.2;refGene_gn=EPB41L1;refGene_tx=XM_019826574.2;refGene_gn=EPB41L1;refGene_tx=XM_023251280.1
A3 23386771 23386888
refGene_gn=EPB41L1;refGene_tx=XM_019826561.2;refGene_gn=EPB41L1;refGene_tx=XM_019826562.2;refGene_gn=EPB41L1;refGene_tx=XM_019826563.2;refGene_gn=EPB41L1;refGene_tx=XM_019826564.2;refGene_gn=EPB41L1;refGene_tx=XM_019826565.2;refGene_gn=EPB41L1;refGene_tx=XM_019826566.2;refGene_gn=EPB41L1;refGene_tx=XM_019826567.2;refGene_gn=EPB41L1;refGene_tx=XM_019826568.2;refGene_gn=EPB41L1;refGene_tx=XM_019826569.2;refGene_gn=EPB41L1;refGene_tx=XM_019826570.2;refGene_gn=EPB41L1;refGene_tx=XM_019826571.2;refGene_gn=EPB41L1;refGene_tx=XM_019826572.2;refGene_gn=EPB41L1;refGene_tx=XM_019826573.2;refGene_gn=EPB41L1;refGene_tx=XM_019826574.2;refGene_gn=EPB41L1;refGene_tx=XM_023251280.1
A3 23387215 23387296
refGene_gn=EPB41L1;refGene_tx=XM_019826561.2;refGene_gn=EPB41L1;refGene_tx=XM_019826562.2;refGene_gn=EPB41L1;refGene_tx=XM_019826563.2;refGene_gn=EPB41L1;refGene_tx=XM_019826564.2;refGene_gn=EPB41L1;refGene_tx=XM_019826565.2;refGene_gn=EPB41L1;refGene_tx=XM_019826566.2;refGene_gn=EPB41L1;refGene_tx=XM_019826567.2;refGene_gn=EPB41L1;refGene_tx=XM_019826568.2;refGene_gn=EPB41L1;refGene_tx=XM_019826569.2;refGene_gn=EPB41L1;refGene_tx=XM_019826570.2;refGene_gn=EPB41L1;refGene_tx=XM_019826571.2;refGene_gn=EPB41L1;refGene_tx=XM_019826572.2;refGene_gn=EPB41L1;refGene_tx=XM_019826573.2;refGene_gn=EPB41L1;refGene_tx=XM_019826574.2;refGene_gn=EPB41L1;refGene_tx=XM_023251280.1
A3 23389303 23389387
refGene_gn=EPB41L1;refGene_tx=XM_019826561.2;refGene_gn=EPB41L1;refGene_tx=XM_019826562.2;refGene_gn=EPB41L1;refGene_tx=XM_019826563.2;refGene_gn=EPB41L1;refGene_tx=XM_019826564.2;refGene_gn=EPB41L1;refGene_tx=XM_019826565.2;refGene_gn=EPB41L1;refGene_tx=XM_019826566.2;refGene_gn=EPB41L1;refGene_tx=XM_019826569.2;refGene_gn=EPB41L1;refGene_tx=XM_019826570.2;refGene_gn=EPB41L1;refGene_tx=XM_019826571.2;refGene_gn=EPB41L1;refGene_tx=XM_019826572.2;refGene_gn=EPB41L1;refGene_tx=XM_019826573.2;refGene_gn=EPB41L1;refGene_tx=XM_019826574.2;refGene_gn=EPB41L1;refGene_tx=XM_023251280.1
A3 23390190 23390277
refGene_gn=EPB41L1;refGene_tx=XM_019826561.2;refGene_gn=EPB41L1;refGene_tx=XM_019826562.2;refGene_gn=EPB41L1;refGene_tx=XM_019826563.2;refGene_gn=EPB41L1;refGene_tx=XM_019826564.2;refGene_gn=EPB41L1;refGene_tx=XM_019826565.2;refGene_gn=EPB41L1;refGene_tx=XM_019826566.2;refGene_gn=EPB41L1;refGene_tx=XM_019826569.2;refGene_gn=EPB41L1;refGene_tx=XM_019826570.2;refGene_gn=EPB41L1;refGene_tx=XM_019826571.2;refGene_gn=EPB41L1;refGene_tx=XM_019826572.2;refGene_gn=EPB41L1;refGene_tx=XM_019826573.2;refGene_gn=EPB41L1;refGene_tx=XM_019826574.2;refGene_gn=EPB41L1;refGene_tx=XM_023251280.1
A3 23396237 23396321
refGene_gn=EPB41L1;refGene_tx=XM_019826561.2;refGene_gn=EPB41L1;refGene_tx=XM_019826562.2;refGene_gn=EPB41L1;refGene_tx=XM_019826563.2;refGe

ne_gn=EPB41L1;refGene_tx=XM_019826564.2;refGene_gn=EPB41L1;refGene_tx=XM_019826565.2;refGene_gn=EPB41L1;refGene_tx=XM_019826566.2;refGene_gn=EPB41L1;refGene_tx=XM_019826567.2;refGene_gn=EPB41L1;refGene_tx=XM_019826568.2;refGene_gn=EPB41L1;refGene_tx=XM_019826569.2;refGene_gn=EPB41L1;refGene_tx=XM_019826570.2;refGene_gn=EPB41L1;refGene_tx=XM_019826571.2;refGene_gn=EPB41L1;refGene_tx=XM_019826572.2;refGene_gn=EPB41L1;refGene_tx=XM_019826573.2;refGene_gn=EPB41L1;refGene_tx=XM_019826574.2;refGene_gn=EPB41L1;refGene_tx=XM_023251280.1
A3 23398229 23398334
refGene_gn=EPB41L1;refGene_tx=XM_019826561.2;refGene_gn=EPB41L1;refGene_tx=XM_019826562.2;refGene_gn=EPB41L1;refGene_tx=XM_019826563.2;refGene_gn=EPB41L1;refGene_tx=XM_019826564.2;refGene_gn=EPB41L1;refGene_tx=XM_019826565.2;refGene_gn=EPB41L1;refGene_tx=XM_019826566.2;refGene_gn=EPB41L1;refGene_tx=XM_019826567.2;refGene_gn=EPB41L1;refGene_tx=XM_019826568.2;refGene_gn=EPB41L1;refGene_tx=XM_019826569.2;refGene_gn=EPB41L1;refGene_tx=XM_019826570.2;refGene_gn=EPB41L1;refGene_tx=XM_019826571.2;refGene_gn=EPB41L1;refGene_tx=XM_019826572.2;refGene_gn=EPB41L1;refGene_tx=XM_019826573.2;refGene_gn=EPB41L1;refGene_tx=XM_019826574.2;refGene_gn=EPB41L1;refGene_tx=XM_023251280.1
A3 23400174 23400585
refGene_gn=EPB41L1;refGene_tx=XM_019826561.2;refGene_gn=EPB41L1;refGene_tx=XM_019826562.2;refGene_gn=EPB41L1;refGene_tx=XM_019826563.2;refGene_gn=EPB41L1;refGene_tx=XM_019826564.2;refGene_gn=EPB41L1;refGene_tx=XM_019826565.2;refGene_gn=EPB41L1;refGene_tx=XM_019826566.2;refGene_gn=EPB41L1;refGene_tx=XM_019826567.2;refGene_gn=EPB41L1;refGene_tx=XM_019826568.2;refGene_gn=EPB41L1;refGene_tx=XM_019826569.2;refGene_gn=EPB41L1;refGene_tx=XM_019826570.2;refGene_gn=EPB41L1;refGene_tx=XM_019826571.2;refGene_gn=EPB41L1;refGene_tx=XM_019826572.2;refGene_gn=EPB41L1;refGene_tx=XM_023251280.1
A3 23401645 23401753
refGene_gn=EPB41L1;refGene_tx=XM_019826561.2;refGene_gn=EPB41L1;refGene_tx=XM_019826562.2;refGene_gn=EPB41L1;refGene_tx=XM_019826563.2;refGene_gn=EPB41L1;refGene_tx=XM_019826564.2;refGene_gn=EPB41L1;refGene_tx=XM_019826565.2;refGene_gn=EPB41L1;refGene_tx=XM_019826567.2;refGene_gn=EPB41L1;refGene_tx=XM_019826568.2;refGene_gn=EPB41L1;refGene_tx=XM_023251280.1
A3 23402264 23402348
refGene_gn=EPB41L1;refGene_tx=XM_019826561.2;refGene_gn=EPB41L1;refGene_tx=XM_019826562.2;refGene_gn=EPB41L1;refGene_tx=XM_019826563.2;refGene_gn=EPB41L1;refGene_tx=XM_019826564.2;refGene_gn=EPB41L1;refGene_tx=XM_019826567.2;refGene_gn=EPB41L1;refGene_tx=XM_019826568.2;refGene_gn=EPB41L1;refGene_tx=XM_023251280.1
A3 23408428 23408533 refGene_gn=EPB41L1;refGene_tx=XM_019826566.2
A3 23411338 23411521
refGene_gn=EPB41L1;refGene_tx=XM_019826561.2;refGene_gn=EPB41L1;refGene_tx=XM_019826562.2;refGene_gn=EPB41L1;refGene_tx=XM_019826563.2;refGene_gn=EPB41L1;refGene_tx=XM_019826564.2;refGene_gn=EPB41L1;refGene_tx=XM_019826565.2;refGene_gn=EPB41L1;refGene_tx=XM_019826566.2;refGene_gn=EPB41L1;refGene_tx=XM_019826567.2;refGene_gn=EPB41L1;refGene_tx=XM_019826568.2;refGene_gn=EPB41L1;refGene_tx=XM_019826569.2;refGene_gn=EPB4

1L1;refGene_tx=XM_019826570.2;refGene_gn=EPB41L1;refGene_tx=XM_0198265
71.2;refGene_gn=EPB41L1;refGene_tx=XM_019826572.2;refGene_gn=EPB41L1;r
efGene_tx=XM_019826573.2;refGene_gn=EPB41L1;refGene_tx=XM_019826574.2;
refGene_gn=EPB41L1;refGene_tx=XM_023251280.1
A3 23413939 23413975
refGene_gn=EPB41L1;refGene_tx=XM_019826561.2;refGene_gn=EPB41L1;refGen
e_tx=XM_019826562.2;refGene_gn=EPB41L1;refGene_tx=XM_019826563.2;refGe
ne_gn=EPB41L1;refGene_tx=XM_019826564.2;refGene_gn=EPB41L1;refGene_tx=
XM_019826565.2;refGene_gn=EPB41L1;refGene_tx=XM_019826566.2;refGene_gn
=EPB41L1;refGene_tx=XM_019826567.2;refGene_gn=EPB41L1;refGene_tx=XM_01
9826568.2;refGene_gn=EPB41L1;refGene_tx=XM_019826569.2;refGene_gn=EPB4
1L1;refGene_tx=XM_019826570.2;refGene_gn=EPB41L1;refGene_tx=XM_0198265
73.2;refGene_gn=EPB41L1;refGene_tx=XM_023251280.1
A3 23414877 23415026
refGene_gn=EPB41L1;refGene_tx=XM_019826561.2;refGene_gn=EPB41L1;refGen
e_tx=XM_019826562.2;refGene_gn=EPB41L1;refGene_tx=XM_019826563.2;refGe
ne_gn=EPB41L1;refGene_tx=XM_019826564.2;refGene_gn=EPB41L1;refGene_tx=
XM_019826565.2;refGene_gn=EPB41L1;refGene_tx=XM_019826566.2;refGene_gn
=EPB41L1;refGene_tx=XM_019826567.2;refGene_gn=EPB41L1;refGene_tx=XM_01
9826568.2;refGene_gn=EPB41L1;refGene_tx=XM_019826569.2;refGene_gn=EPB4
1L1;refGene_tx=XM_019826570.2;refGene_gn=EPB41L1;refGene_tx=XM_0198265
71.2;refGene_gn=EPB41L1;refGene_tx=XM_019826572.2;refGene_gn=EPB41L1;r
efGene_tx=XM_019826573.2;refGene_gn=EPB41L1;refGene_tx=XM_019826574.2;
refGene_gn=EPB41L1;refGene_tx=XM_023251280.1
A3 23417721 23417897
refGene_gn=EPB41L1;refGene_tx=XM_019826561.2;refGene_gn=EPB41L1;refGen
e_tx=XM_019826562.2;refGene_gn=EPB41L1;refGene_tx=XM_019826563.2;refGe
ne_gn=EPB41L1;refGene_tx=XM_019826564.2;refGene_gn=EPB41L1;refGene_tx=
XM_019826565.2;refGene_gn=EPB41L1;refGene_tx=XM_019826566.2;refGene_gn
=EPB41L1;refGene_tx=XM_019826567.2;refGene_gn=EPB41L1;refGene_tx=XM_01
9826568.2;refGene_gn=EPB41L1;refGene_tx=XM_019826569.2;refGene_gn=EPB4
1L1;refGene_tx=XM_019826570.2;refGene_gn=EPB41L1;refGene_tx=XM_0198265
71.2;refGene_gn=EPB41L1;refGene_tx=XM_019826572.2;refGene_gn=EPB41L1;r
efGene_tx=XM_019826573.2;refGene_gn=EPB41L1;refGene_tx=XM_019826574.2;
refGene_gn=EPB41L1;refGene_tx=XM_023251280.1
A3 23418115 23418213
refGene_gn=EPB41L1;refGene_tx=XM_019826561.2;refGene_gn=EPB41L1;refGen
e_tx=XM_019826562.2;refGene_gn=EPB41L1;refGene_tx=XM_019826563.2;refGe
ne_gn=EPB41L1;refGene_tx=XM_019826564.2;refGene_gn=EPB41L1;refGene_tx=
XM_019826565.2;refGene_gn=EPB41L1;refGene_tx=XM_019826566.2;refGene_gn
=EPB41L1;refGene_tx=XM_019826567.2;refGene_gn=EPB41L1;refGene_tx=XM_01
9826568.2;refGene_gn=EPB41L1;refGene_tx=XM_019826569.2;refGene_gn=EPB4
1L1;refGene_tx=XM_019826570.2;refGene_gn=EPB41L1;refGene_tx=XM_0198265
71.2;refGene_gn=EPB41L1;refGene_tx=XM_019826572.2;refGene_gn=EPB41L1;r
efGene_tx=XM_019826573.2;refGene_gn=EPB41L1;refGene_tx=XM_019826574.2;
refGene_gn=EPB41L1;refGene_tx=XM_023251280.1
A3 23419580 23419733
refGene_gn=EPB41L1;refGene_tx=XM_019826561.2;refGene_gn=EPB41L1;refGen
e_tx=XM_019826562.2;refGene_gn=EPB41L1;refGene_tx=XM_019826563.2;refGe
ne_gn=EPB41L1;refGene_tx=XM_019826564.2;refGene_gn=EPB41L1;refGene_tx=

XM_019826565.2;refGene_gn=EPB41L1;refGene_tx=XM_019826566.2;refGene_gn=EPB41L1;refGene_tx=XM_019826567.2;refGene_gn=EPB41L1;refGene_tx=XM_019826568.2;refGene_gn=EPB41L1;refGene_tx=XM_019826569.2;refGene_gn=EPB41L1;refGene_tx=XM_019826570.2;refGene_gn=EPB41L1;refGene_tx=XM_019826571.2;refGene_gn=EPB41L1;refGene_tx=XM_019826572.2;refGene_gn=EPB41L1;refGene_tx=XM_019826573.2;refGene_gn=EPB41L1;refGene_tx=XM_019826574.2;refGene_gn=EPB41L1;refGene_tx=XM_023251280.1
A3 23420233 23420321
refGene_gn=EPB41L1;refGene_tx=XM_019826561.2;refGene_gn=EPB41L1;refGene_tx=XM_019826562.2;refGene_gn=EPB41L1;refGene_tx=XM_019826563.2;refGene_gn=EPB41L1;refGene_tx=XM_019826564.2;refGene_gn=EPB41L1;refGene_tx=XM_019826565.2;refGene_gn=EPB41L1;refGene_tx=XM_019826566.2;refGene_gn=EPB41L1;refGene_tx=XM_019826567.2;refGene_gn=EPB41L1;refGene_tx=XM_019826568.2;refGene_gn=EPB41L1;refGene_tx=XM_019826569.2;refGene_gn=EPB41L1;refGene_tx=XM_019826570.2;refGene_gn=EPB41L1;refGene_tx=XM_019826571.2;refGene_gn=EPB41L1;refGene_tx=XM_019826572.2;refGene_gn=EPB41L1;refGene_tx=XM_019826573.2;refGene_gn=EPB41L1;refGene_tx=XM_019826574.2;refGene_gn=EPB41L1;refGene_tx=XM_023251280.1
A3 23422675 23422894
refGene_gn=EPB41L1;refGene_tx=XM_019826561.2;refGene_gn=EPB41L1;refGene_tx=XM_019826562.2;refGene_gn=EPB41L1;refGene_tx=XM_019826563.2;refGene_gn=EPB41L1;refGene_tx=XM_019826564.2;refGene_gn=EPB41L1;refGene_tx=XM_019826565.2;refGene_gn=EPB41L1;refGene_tx=XM_019826566.2;refGene_gn=EPB41L1;refGene_tx=XM_019826567.2;refGene_gn=EPB41L1;refGene_tx=XM_019826568.2;refGene_gn=EPB41L1;refGene_tx=XM_019826569.2;refGene_gn=EPB41L1;refGene_tx=XM_019826570.2;refGene_gn=EPB41L1;refGene_tx=XM_019826571.2;refGene_gn=EPB41L1;refGene_tx=XM_019826572.2;refGene_gn=EPB41L1;refGene_tx=XM_019826573.2;refGene_gn=EPB41L1;refGene_tx=XM_019826574.2;refGene_gn=EPB41L1;refGene_tx=XM_023251280.1
A3 23425708 23425784
refGene_gn=EPB41L1;refGene_tx=XM_019826561.2;refGene_gn=EPB41L1;refGene_tx=XM_019826562.2;refGene_gn=EPB41L1;refGene_tx=XM_019826563.2;refGene_gn=EPB41L1;refGene_tx=XM_019826564.2;refGene_gn=EPB41L1;refGene_tx=XM_019826565.2;refGene_gn=EPB41L1;refGene_tx=XM_019826566.2;refGene_gn=EPB41L1;refGene_tx=XM_019826567.2;refGene_gn=EPB41L1;refGene_tx=XM_019826568.2;refGene_gn=EPB41L1;refGene_tx=XM_019826569.2;refGene_gn=EPB41L1;refGene_tx=XM_019826570.2;refGene_gn=EPB41L1;refGene_tx=XM_019826571.2;refGene_gn=EPB41L1;refGene_tx=XM_019826572.2;refGene_gn=EPB41L1;refGene_tx=XM_019826573.2;refGene_gn=EPB41L1;refGene_tx=XM_019826574.2;refGene_gn=EPB41L1;refGene_tx=XM_023251280.1
A3 23429313 23429356
refGene_gn=EPB41L1;refGene_tx=XM_019826561.2;refGene_gn=EPB41L1;refGene_tx=XM_019826562.2;refGene_gn=EPB41L1;refGene_tx=XM_019826563.2;refGene_gn=EPB41L1;refGene_tx=XM_019826564.2;refGene_gn=EPB41L1;refGene_tx=XM_019826565.2;refGene_gn=EPB41L1;refGene_tx=XM_019826566.2;refGene_gn=EPB41L1;refGene_tx=XM_019826567.2;refGene_gn=EPB41L1;refGene_tx=XM_019826568.2;refGene_gn=EPB41L1;refGene_tx=XM_019826569.2;refGene_gn=EPB41L1;refGene_tx=XM_019826570.2;refGene_gn=EPB41L1;refGene_tx=XM_019826571.2;refGene_gn=EPB41L1;refGene_tx=XM_019826572.2;refGene_gn=EPB41L1;refGene_tx=XM_019826573.2;refGene_gn=EPB41L1;refGene_tx=XM_019826574.2;

refGene_gn=EPB41L1;refGene_tx=XM_023251280.1
A3 23429953 23430058
refGene_gn=EPB41L1;refGene_tx=XM_019826561.2;refGene_gn=EPB41L1;refGene_tx=XM_019826562.2;refGene_gn=EPB41L1;refGene_tx=XM_019826563.2;refGene_gn=EPB41L1;refGene_tx=XM_019826564.2;refGene_gn=EPB41L1;refGene_tx=XM_019826565.2;refGene_gn=EPB41L1;refGene_tx=XM_019826566.2;refGene_gn=EPB41L1;refGene_tx=XM_019826567.2;refGene_gn=EPB41L1;refGene_tx=XM_019826568.2;refGene_gn=EPB41L1;refGene_tx=XM_019826569.2;refGene_gn=EPB41L1;refGene_tx=XM_019826570.2;refGene_gn=EPB41L1;refGene_tx=XM_019826571.2;refGene_gn=EPB41L1;refGene_tx=XM_019826572.2;refGene_gn=EPB41L1;refGene_tx=XM_019826573.2;refGene_gn=EPB41L1;refGene_tx=XM_019826574.2;refGene_gn=EPB41L1;refGene_tx=XM_023251280.1
A3 23432002 23432167
refGene_gn=EPB41L1;refGene_tx=XM_019826561.2;refGene_gn=EPB41L1;refGene_tx=XM_019826562.2;refGene_gn=EPB41L1;refGene_tx=XM_019826563.2;refGene_gn=EPB41L1;refGene_tx=XM_019826564.2;refGene_gn=EPB41L1;refGene_tx=XM_019826565.2;refGene_gn=EPB41L1;refGene_tx=XM_019826566.2;refGene_gn=EPB41L1;refGene_tx=XM_019826567.2;refGene_gn=EPB41L1;refGene_tx=XM_019826568.2;refGene_gn=EPB41L1;refGene_tx=XM_019826569.2;refGene_gn=EPB41L1;refGene_tx=XM_019826570.2;refGene_gn=EPB41L1;refGene_tx=XM_019826571.2;refGene_gn=EPB41L1;refGene_tx=XM_019826572.2;refGene_gn=EPB41L1;refGene_tx=XM_019826573.2;refGene_gn=EPB41L1;refGene_tx=XM_019826574.2;refGene_gn=EPB41L1;refGene_tx=XM_023251280.1
A3 23433385 23433562
refGene_gn=EPB41L1;refGene_tx=XM_019826561.2;refGene_gn=EPB41L1;refGene_tx=XM_019826562.2;refGene_gn=EPB41L1;refGene_tx=XM_019826563.2;refGene_gn=EPB41L1;refGene_tx=XM_019826564.2;refGene_gn=EPB41L1;refGene_tx=XM_019826565.2;refGene_gn=EPB41L1;refGene_tx=XM_019826566.2;refGene_gn=EPB41L1;refGene_tx=XM_019826567.2;refGene_gn=EPB41L1;refGene_tx=XM_019826568.2;refGene_gn=EPB41L1;refGene_tx=XM_019826569.2;refGene_gn=EPB41L1;refGene_tx=XM_019826570.2;refGene_gn=EPB41L1;refGene_tx=XM_019826571.2;refGene_gn=EPB41L1;refGene_tx=XM_019826572.2;refGene_gn=EPB41L1;refGene_tx=XM_019826573.2;refGene_gn=EPB41L1;refGene_tx=XM_019826574.2;refGene_gn=EPB41L1;refGene_tx=XM_023251280.1
A3 23548623 23548942
refGene_gn=CNBD2;refGene_tx=XM_006929807.3;refGene_gn=CNBD2;refGene_tx=XM_019826577.2;refGene_gn=CNBD2;refGene_tx=XM_019826581.2;refGene_gn=CNBD2;refGene_tx=XM_019826582.2;refGene_gn=CNBD2;refGene_tx=XM_019826583.2;refGene_gn=CNBD2;refGene_tx=XM_019826584.2;refGene_gn=CNBD2;refGene_tx=XM_023251285.1;refGene_gn=CNBD2;refGene_tx=XM_023251286.1
A3 23555601 23555765
refGene_gn=CNBD2;refGene_tx=XM_006929807.3;refGene_gn=CNBD2;refGene_tx=XM_019826577.2;refGene_gn=CNBD2;refGene_tx=XM_019826581.2;refGene_gn=CNBD2;refGene_tx=XM_019826583.2;refGene_gn=CNBD2;refGene_tx=XM_019826584.2;refGene_gn=CNBD2;refGene_tx=XM_023251285.1;refGene_gn=CNBD2;refGene_tx=XM_023251286.1
A3 23561801 23561922
refGene_gn=CNBD2;refGene_tx=XM_006929807.3;refGene_gn=CNBD2;refGene_tx=XM_019826577.2;refGene_gn=CNBD2;refGene_tx=XM_019826581.2;refGene_gn=CNBD2;refGene_tx=XM_019826582.2;refGene_gn=CNBD2;refGene_tx=XM_0198265

83.2;refGene_gn=CNBD2;refGene_tx=XM_019826584.2;refGene_gn=CNBD2;refGene_tx=XM_023251285.1;refGene_gn=CNBD2;refGene_tx=XM_023251286.1
A3 23564149 23564327
refGene_gn=CNBD2;refGene_tx=XM_006929807.3;refGene_gn=CNBD2;refGene_tx=XM_019826577.2;refGene_gn=CNBD2;refGene_tx=XM_019826581.2;refGene_gn=CNBD2;refGene_tx=XM_019826582.2;refGene_gn=CNBD2;refGene_tx=XM_019826583.2;refGene_gn=CNBD2;refGene_tx=XM_019826584.2;refGene_gn=CNBD2;refGene_tx=XM_023251285.1;refGene_gn=CNBD2;refGene_tx=XM_023251286.1
A3 23569476 23569591
refGene_gn=CNBD2;refGene_tx=XM_006929807.3;refGene_gn=CNBD2;refGene_tx=XM_019826577.2;refGene_gn=CNBD2;refGene_tx=XM_019826581.2;refGene_gn=CNBD2;refGene_tx=XM_019826582.2;refGene_gn=CNBD2;refGene_tx=XM_019826583.2;refGene_gn=CNBD2;refGene_tx=XM_019826584.2;refGene_gn=CNBD2;refGene_tx=XM_023251285.1;refGene_gn=CNBD2;refGene_tx=XM_023251286.1
A3 23578248 23578387
refGene_gn=CNBD2;refGene_tx=XM_006929807.3;refGene_gn=CNBD2;refGene_tx=XM_019826577.2;refGene_gn=CNBD2;refGene_tx=XM_019826581.2;refGene_gn=CNBD2;refGene_tx=XM_019826582.2;refGene_gn=CNBD2;refGene_tx=XM_019826583.2;refGene_gn=CNBD2;refGene_tx=XM_019826584.2;refGene_gn=CNBD2;refGene_tx=XM_023251285.1;refGene_gn=CNBD2;refGene_tx=XM_023251286.1
A3 23580970 23581122
refGene_gn=CNBD2;refGene_tx=XM_006929807.3;refGene_gn=CNBD2;refGene_tx=XM_019826577.2;refGene_gn=CNBD2;refGene_tx=XM_019826581.2;refGene_gn=CNBD2;refGene_tx=XM_019826582.2;refGene_gn=CNBD2;refGene_tx=XM_019826583.2;refGene_gn=CNBD2;refGene_tx=XM_019826584.2;refGene_gn=CNBD2;refGene_tx=XM_023251285.1;refGene_gn=CNBD2;refGene_tx=XM_023251286.1
A3 23581620 23581777
refGene_gn=CNBD2;refGene_tx=XM_006929807.3;refGene_gn=CNBD2;refGene_tx=XM_019826577.2;refGene_gn=CNBD2;refGene_tx=XM_019826581.2;refGene_gn=CNBD2;refGene_tx=XM_019826582.2;refGene_gn=CNBD2;refGene_tx=XM_019826583.2;refGene_gn=CNBD2;refGene_tx=XM_019826584.2;refGene_gn=CNBD2;refGene_tx=XM_023251285.1;refGene_gn=CNBD2;refGene_tx=XM_023251286.1
A3 23587156 23587320
refGene_gn=CNBD2;refGene_tx=XM_006929807.3;refGene_gn=CNBD2;refGene_tx=XM_019826577.2;refGene_gn=CNBD2;refGene_tx=XM_019826581.2;refGene_gn=CNBD2;refGene_tx=XM_019826582.2;refGene_gn=CNBD2;refGene_tx=XM_019826583.2;refGene_gn=CNBD2;refGene_tx=XM_019826584.2;refGene_gn=CNBD2;refGene_tx=XM_023251285.1;refGene_gn=CNBD2;refGene_tx=XM_023251286.1
A3 23591717 23591771
refGene_gn=CNBD2;refGene_tx=XM_006929807.3;refGene_gn=CNBD2;refGene_tx=XM_019826577.2;refGene_gn=CNBD2;refGene_tx=XM_019826581.2;refGene_gn=CNBD2;refGene_tx=XM_019826582.2;refGene_gn=CNBD2;refGene_tx=XM_019826583.2;refGene_gn=CNBD2;refGene_tx=XM_019826584.2;refGene_gn=CNBD2;refGene_tx=XM_023251285.1;refGene_gn=CNBD2;refGene_tx=XM_023251286.1
A3 23594209 23594347
refGene_gn=CNBD2;refGene_tx=XM_006929807.3;refGene_gn=CNBD2;refGene_tx=XM_019826577.2;refGene_gn=CNBD2;refGene_tx=XM_019826581.2;refGene_gn=CNBD2;refGene_tx=XM_019826582.2;refGene_gn=CNBD2;refGene_tx=XM_023251285.1;refGene_gn=CNBD2;refGene_tx=XM_023251286.1
A3 23598537 23598612

refGene_gn=CNBD2;refGene_tx=XM_006929807.3;refGene_gn=CNBD2;refGene_tx=XM_019826581.2
A3 23610701 23610755
refGene_gn=CNBD2;refGene_tx=XM_019826577.2;refGene_gn=CNBD2;refGene_tx=XM_019826582.2
A3 23613050 23613311
refGene_gn=CNBD2;refGene_tx=XM_019826577.2;refGene_gn=CNBD2;refGene_tx=XM_019826582.2
A3 23613530 23614070 refGene_gn=SCAND1;refGene_tx=XM_004001388.5
A3 23617745 23617888
refGene_gn=PHF20;refGene_tx=XM_006929808.4;refGene_gn=PHF20;refGene_tx=XM_006929809.4;refGene_gn=PHF20;refGene_tx=XM_019826575.2;refGene_gn=PHF20;refGene_tx=XM_019826576.2;refGene_gn=PHF20;refGene_tx=XM_023251281.1;refGene_gn=PHF20;refGene_tx=XM_023251283.1;refGene_gn=PHF20;refGene_tx=XM_023251284.1
A3 23623294 23623478
refGene_gn=PHF20;refGene_tx=XM_006929808.4;refGene_gn=PHF20;refGene_tx=XM_006929809.4;refGene_gn=PHF20;refGene_tx=XM_019826575.2;refGene_gn=PHF20;refGene_tx=XM_019826576.2;refGene_gn=PHF20;refGene_tx=XM_023251281.1;refGene_gn=PHF20;refGene_tx=XM_023251283.1;refGene_gn=PHF20;refGene_tx=XM_023251284.1
A3 23625460 23625866
refGene_gn=PHF20;refGene_tx=XM_006929808.4;refGene_gn=PHF20;refGene_tx=XM_006929809.4;refGene_gn=PHF20;refGene_tx=XM_019826575.2;refGene_gn=PHF20;refGene_tx=XM_019826576.2;refGene_gn=PHF20;refGene_tx=XM_023251281.1;refGene_gn=PHF20;refGene_tx=XM_023251283.1;refGene_gn=PHF20;refGene_tx=XM_023251284.1
A3 23631779 23631975
refGene_gn=PHF20;refGene_tx=XM_006929808.4;refGene_gn=PHF20;refGene_tx=XM_006929809.4;refGene_gn=PHF20;refGene_tx=XM_019826575.2;refGene_gn=PHF20;refGene_tx=XM_019826576.2;refGene_gn=PHF20;refGene_tx=XM_023251281.1;refGene_gn=PHF20;refGene_tx=XM_023251283.1;refGene_gn=PHF20;refGene_tx=XM_023251284.1
A3 23634865 23634965
refGene_gn=PHF20;refGene_tx=XM_006929808.4;refGene_gn=PHF20;refGene_tx=XM_006929809.4;refGene_gn=PHF20;refGene_tx=XM_019826575.2;refGene_gn=PHF20;refGene_tx=XM_019826576.2;refGene_gn=PHF20;refGene_tx=XM_023251281.1;refGene_gn=PHF20;refGene_tx=XM_023251283.1;refGene_gn=PHF20;refGene_tx=XM_023251284.1
A3 23642796 23642975
refGene_gn=PHF20;refGene_tx=XM_006929808.4;refGene_gn=PHF20;refGene_tx=XM_006929809.4;refGene_gn=PHF20;refGene_tx=XM_019826575.2;refGene_gn=PHF20;refGene_tx=XM_019826576.2;refGene_gn=PHF20;refGene_tx=XM_023251281.1;refGene_gn=PHF20;refGene_tx=XM_023251283.1;refGene_gn=PHF20;refGene_tx=XM_023251284.1
A3 23643941 23644106
refGene_gn=PHF20;refGene_tx=XM_006929808.4;refGene_gn=PHF20;refGene_tx=XM_006929809.4;refGene_gn=PHF20;refGene_tx=XM_019826575.2;refGene_gn=PHF20;refGene_tx=XM_019826576.2;refGene_gn=PHF20;refGene_tx=XM_023251281.1;refGene_gn=PHF20;refGene_tx=XM_023251283.1;refGene_gn=PHF20;refGe

ne_tx=XM_023251284.1
A3	23645919	23646018
refGene_gn=PHF20;refGene_tx=XM_006929808.4;refGene_gn=PHF20;refGene_tx=XM_006929809.4;refGene_gn=PHF20;refGene_tx=XM_019826575.2;refGene_gn=PHF20;refGene_tx=XM_023251281.1
A3	23662429	23662708
refGene_gn=PHF20;refGene_tx=XM_006929808.4;refGene_gn=PHF20;refGene_tx=XM_006929809.4;refGene_gn=PHF20;refGene_tx=XM_019826575.2;refGene_gn=PHF20;refGene_tx=XM_019826576.2;refGene_gn=PHF20;refGene_tx=XM_023251281.1;refGene_gn=PHF20;refGene_tx=XM_023251283.1;refGene_gn=PHF20;refGene_tx=XM_023251284.1
A3	23689635	23689815
refGene_gn=PHF20;refGene_tx=XM_006929808.4;refGene_gn=PHF20;refGene_tx=XM_006929809.4;refGene_gn=PHF20;refGene_tx=XM_019826575.2;refGene_gn=PHF20;refGene_tx=XM_019826576.2;refGene_gn=PHF20;refGene_tx=XM_023251281.1;refGene_gn=PHF20;refGene_tx=XM_023251283.1;refGene_gn=PHF20;refGene_tx=XM_023251284.1
A3	23690341	23690530
refGene_gn=PHF20;refGene_tx=XM_006929808.4;refGene_gn=PHF20;refGene_tx=XM_006929809.4;refGene_gn=PHF20;refGene_tx=XM_019826575.2;refGene_gn=PHF20;refGene_tx=XM_019826576.2;refGene_gn=PHF20;refGene_tx=XM_023251281.1;refGene_gn=PHF20;refGene_tx=XM_023251283.1;refGene_gn=PHF20;refGene_tx=XM_023251284.1
A3	23691665	23691779
refGene_gn=PHF20;refGene_tx=XM_006929808.4;refGene_gn=PHF20;refGene_tx=XM_006929809.4;refGene_gn=PHF20;refGene_tx=XM_019826575.2;refGene_gn=PHF20;refGene_tx=XM_019826576.2;refGene_gn=PHF20;refGene_tx=XM_023251281.1;refGene_gn=PHF20;refGene_tx=XM_023251283.1;refGene_gn=PHF20;refGene_tx=XM_023251284.1
A3	23695956	23696344
refGene_gn=PHF20;refGene_tx=XM_006929808.4;refGene_gn=PHF20;refGene_tx=XM_006929809.4;refGene_gn=PHF20;refGene_tx=XM_019826575.2;refGene_gn=PHF20;refGene_tx=XM_019826576.2;refGene_gn=PHF20;refGene_tx=XM_023251281.1;refGene_gn=PHF20;refGene_tx=XM_023251283.1;refGene_gn=PHF20;refGene_tx=XM_023251284.1
A3	23700457	23700537
refGene_gn=PHF20;refGene_tx=XM_006929808.4;refGene_gn=PHF20;refGene_tx=XM_006929809.4;refGene_gn=PHF20;refGene_tx=XM_019826575.2;refGene_gn=PHF20;refGene_tx=XM_023251281.1;refGene_gn=PHF20;refGene_tx=XM_023251283.1
A3	23709820	23709905
refGene_gn=PHF20;refGene_tx=XM_006929808.4;refGene_gn=PHF20;refGene_tx=XM_006929809.4;refGene_gn=PHF20;refGene_tx=XM_019826575.2;refGene_gn=PHF20;refGene_tx=XM_019826576.2;refGene_gn=PHF20;refGene_tx=XM_023251281.1;refGene_gn=PHF20;refGene_tx=XM_023251283.1
A3	23711898	23712070
refGene_gn=PHF20;refGene_tx=XM_006929808.4;refGene_gn=PHF20;refGene_tx=XM_006929809.4;refGene_gn=PHF20;refGene_tx=XM_019826575.2;refGene_gn=PHF20;refGene_tx=XM_023251281.1;refGene_gn=PHF20;refGene_tx=XM_023251283.1;refGene_gn=PHF20;refGene_tx=XM_023251284.1

A3 23742269 23742352
refGene_gn=PHF20;refGene_tx=XM_006929808.4;refGene_gn=PHF20;refGene_tx=XM_006929809.4;refGene_gn=PHF20;refGene_tx=XM_019826575.2;refGene_gn=PHF20;refGene_tx=XM_023251281.1;refGene_gn=PHF20;refGene_tx=XM_023251283.1;refGene_gn=PHF20;refGene_tx=XM_023251284.1
A3 23789098 23789149
refGene_gn=RBM39;refGene_tx=XM_011280872.3;refGene_gn=RBM39;refGene_tx=XM_011280873.3;refGene_gn=RBM39;refGene_tx=XM_011280874.3;refGene_gn=RBM39;refGene_tx=XM_011280875.3
A3 23790927 23790977
refGene_gn=RBM39;refGene_tx=XM_011280872.3;refGene_gn=RBM39;refGene_tx=XM_011280873.3;refGene_gn=RBM39;refGene_tx=XM_011280874.3;refGene_gn=RBM39;refGene_tx=XM_011280875.3
A3 23795545 23795740
refGene_gn=RBM39;refGene_tx=XM_011280872.3;refGene_gn=RBM39;refGene_tx=XM_011280873.3;refGene_gn=RBM39;refGene_tx=XM_011280874.3;refGene_gn=RBM39;refGene_tx=XM_011280875.3
A3 23798075 23798141
refGene_gn=RBM39;refGene_tx=XM_011280872.3;refGene_gn=RBM39;refGene_tx=XM_011280873.3
A3 23798237 23798291
refGene_gn=RBM39;refGene_tx=XM_011280872.3;refGene_gn=RBM39;refGene_tx=XM_011280873.3;refGene_gn=RBM39;refGene_tx=XM_011280874.3;refGene_gn=RBM39;refGene_tx=XM_011280875.3
A3 23800296 23800414
refGene_gn=RBM39;refGene_tx=XM_011280872.3;refGene_gn=RBM39;refGene_tx=XM_011280873.3;refGene_gn=RBM39;refGene_tx=XM_011280874.3;refGene_gn=RBM39;refGene_tx=XM_011280875.3
A3 23801768 23801921
refGene_gn=RBM39;refGene_tx=XM_011280872.3;refGene_gn=RBM39;refGene_tx=XM_011280873.3;refGene_gn=RBM39;refGene_tx=XM_011280874.3;refGene_gn=RBM39;refGene_tx=XM_011280875.3
A3 23804379 23804517
refGene_gn=RBM39;refGene_tx=XM_011280872.3;refGene_gn=RBM39;refGene_tx=XM_011280873.3;refGene_gn=RBM39;refGene_tx=XM_011280874.3;refGene_gn=RBM39;refGene_tx=XM_011280875.3
A3 23805636 23805702
refGene_gn=RBM39;refGene_tx=XM_011280872.3;refGene_gn=RBM39;refGene_tx=XM_011280873.3;refGene_gn=RBM39;refGene_tx=XM_011280874.3;refGene_gn=RBM39;refGene_tx=XM_011280875.3
A3 23807948 23808153
refGene_gn=RBM39;refGene_tx=XM_011280872.3;refGene_gn=RBM39;refGene_tx=XM_011280873.3;refGene_gn=RBM39;refGene_tx=XM_011280874.3;refGene_gn=RBM39;refGene_tx=XM_011280875.3
A3 23809175 23809253
refGene_gn=RBM39;refGene_tx=XM_011280872.3;refGene_gn=RBM39;refGene_tx=XM_011280873.3;refGene_gn=RBM39;refGene_tx=XM_011280874.3;refGene_gn=RBM39;refGene_tx=XM_011280875.3
A3 23812946 23812997
refGene_gn=RBM39;refGene_tx=XM_011280872.3;refGene_gn=RBM39;refGene_tx

=XM_011280873.3;refGene_gn=RBM39;refGene_tx=XM_011280874.3;refGene_gn=RBM39;refGene_tx=XM_011280875.3
A3 23815300 23815382
refGene_gn=RBM39;refGene_tx=XM_011280872.3;refGene_gn=RBM39;refGene_tx=XM_011280873.3;refGene_gn=RBM39;refGene_tx=XM_011280874.3;refGene_gn=RBM39;refGene_tx=XM_011280875.3
A3 23817220 23817326
refGene_gn=RBM39;refGene_tx=XM_011280872.3;refGene_gn=RBM39;refGene_tx=XM_011280873.3;refGene_gn=RBM39;refGene_tx=XM_011280874.3;refGene_gn=RBM39;refGene_tx=XM_011280875.3
A3 23817838 23817917
refGene_gn=RBM39;refGene_tx=XM_011280872.3;refGene_gn=RBM39;refGene_tx=XM_011280873.3;refGene_gn=RBM39;refGene_tx=XM_011280874.3;refGene_gn=RBM39;refGene_tx=XM_011280875.3
A3 23818006 23818107
refGene_gn=RBM39;refGene_tx=XM_011280872.3;refGene_gn=RBM39;refGene_tx=XM_011280873.3;refGene_gn=RBM39;refGene_tx=XM_011280874.3;refGene_gn=RBM39;refGene_tx=XM_011280875.3
A3 23821090 23821199 refGene_gn=ROMO1;refGene_tx=XM_003983564.5
A3 23822180 23822311 refGene_gn=ROMO1;refGene_tx=XM_003983564.5
A3 23822661 23822758
refGene_gn=NFS1;refGene_tx=XM_003983566.3;refGene_gn=NFS1;refGene_tx=XM_011280871.2
A3 23823405 23823515
refGene_gn=NFS1;refGene_tx=XM_003983566.3;refGene_gn=NFS1;refGene_tx=XM_011280871.2;refGene_gn=NFS1;refGene_tx=XM_019826585.1
A3 23824239 23824356
refGene_gn=NFS1;refGene_tx=XM_003983566.3;refGene_gn=NFS1;refGene_tx=XM_011280871.2;refGene_gn=NFS1;refGene_tx=XM_019826585.1
A3 23825165 23825249
refGene_gn=NFS1;refGene_tx=XM_003983566.3;refGene_gn=NFS1;refGene_tx=XM_011280871.2;refGene_gn=NFS1;refGene_tx=XM_019826585.1
A3 23830162 23830315
refGene_gn=NFS1;refGene_tx=XM_011280871.2;refGene_gn=NFS1;refGene_tx=XM_019826585.1
A3 23834230 23834324
refGene_gn=NFS1;refGene_tx=XM_003983566.3;refGene_gn=NFS1;refGene_tx=XM_011280871.2;refGene_gn=NFS1;refGene_tx=XM_019826585.1
A3 23834998 23835133
refGene_gn=NFS1;refGene_tx=XM_003983566.3;refGene_gn=NFS1;refGene_tx=XM_011280871.2;refGene_gn=NFS1;refGene_tx=XM_019826585.1
A3 23839102 23839260
refGene_gn=NFS1;refGene_tx=XM_003983566.3;refGene_gn=NFS1;refGene_tx=XM_011280871.2;refGene_gn=NFS1;refGene_tx=XM_019826585.1
A3 23839685 23839791
refGene_gn=NFS1;refGene_tx=XM_003983566.3;refGene_gn=NFS1;refGene_tx=XM_011280871.2;refGene_gn=NFS1;refGene_tx=XM_019826585.1
A3 23839870 23839952
refGene_gn=NFS1;refGene_tx=XM_003983566.3;refGene_gn=NFS1;refGene_tx=XM_011280871.2;refGene_gn=NFS1;refGene_tx=XM_019826585.1

A3 23840580 23840664
refGene_gn=NFS1;refGene_tx=XM_003983566.3;refGene_gn=NFS1;refGene_tx=XM_011280871.2;refGene_gn=NFS1;refGene_tx=XM_019826585.1
A3 23841217 23841307
refGene_gn=NFS1;refGene_tx=XM_003983566.3;refGene_gn=NFS1;refGene_tx=XM_011280871.2;refGene_gn=NFS1;refGene_tx=XM_019826585.1
A3 23843950 23844014
refGene_gn=NFS1;refGene_tx=XM_003983566.3;refGene_gn=NFS1;refGene_tx=XM_011280871.2;refGene_gn=NFS1;refGene_tx=XM_019826585.1
A3 23856160 23859007
refGene_gn=LOC101100631;refGene_tx=XM_006929812.3;refGene_gn=LOC101100631;refGene_tx=XM_019826593.2;refGene_gn=LOC101100631;refGene_tx=XM_019826594.2;refGene_gn=LOC101100631;refGene_tx=XM_023251288.1;refGene_gn=LOC101100631;refGene_tx=XM_023251289.1
A3 23877003 23877132
refGene_gn=LOC101100631;refGene_tx=XM_003983572.4;refGene_gn=LOC101100631;refGene_tx=XM_003983573.5;refGene_gn=LOC101100631;refGene_tx=XM_006929810.4;refGene_gn=LOC101100631;refGene_tx=XM_006929811.3;refGene_gn=LOC101100631;refGene_tx=XM_019826595.2;refGene_gn=LOC101100631;refGene_tx=XM_023251289.1
A3 23877231 23877411
refGene_gn=LOC101100631;refGene_tx=XM_003983572.4;refGene_gn=LOC101100631;refGene_tx=XM_003983573.5;refGene_gn=LOC101100631;refGene_tx=XM_006929810.4;refGene_gn=LOC101100631;refGene_tx=XM_006929811.3;refGene_gn=LOC101100631;refGene_tx=XM_019826595.2;refGene_gn=LOC101100631;refGene_tx=XM_023251289.1
A3 23877567 23877642
refGene_gn=LOC101100631;refGene_tx=XM_003983572.4;refGene_gn=LOC101100631;refGene_tx=XM_003983573.5;refGene_gn=LOC101100631;refGene_tx=XM_006929810.4;refGene_gn=LOC101100631;refGene_tx=XM_006929811.3;refGene_gn=LOC101100631;refGene_tx=XM_019826595.2;refGene_gn=LOC101100631;refGene_tx=XM_023251289.1
A3 23877723 23877795
refGene_gn=LOC101100631;refGene_tx=XM_003983572.4;refGene_gn=LOC101100631;refGene_tx=XM_003983573.5;refGene_gn=LOC101100631;refGene_tx=XM_006929810.4;refGene_gn=LOC101100631;refGene_tx=XM_006929811.3;refGene_gn=LOC101100631;refGene_tx=XM_019826595.2;refGene_gn=LOC101100631;refGene_tx=XM_023251289.1
A3 23877929 23878010
refGene_gn=LOC101100631;refGene_tx=XM_003983572.4;refGene_gn=LOC101100631;refGene_tx=XM_003983573.5;refGene_gn=LOC101100631;refGene_tx=XM_006929810.4;refGene_gn=LOC101100631;refGene_tx=XM_006929811.3;refGene_gn=LOC101100631;refGene_tx=XM_019826595.2;refGene_gn=LOC101100631;refGene_tx=XM_023251289.1
A3 23878179 23878269
refGene_gn=LOC101100631;refGene_tx=XM_003983572.4;refGene_gn=LOC101100631;refGene_tx=XM_003983573.5;refGene_gn=LOC101100631;refGene_tx=XM_006929810.4;refGene_gn=LOC101100631;refGene_tx=XM_006929811.3;refGene_gn=LOC101100631;refGene_tx=XM_019826595.2;refGene_gn=LOC101100631;refGene_tx=XM_023251289.1

A3 23878376 23878463
refGene_gn=LOC101100631;refGene_tx=XM_003983572.4;refGene_gn=LOC101100631;refGene_tx=XM_003983573.5;refGene_gn=LOC101100631;refGene_tx=XM_006929810.4;refGene_gn=LOC101100631;refGene_tx=XM_006929811.3;refGene_gn=LOC101100631;refGene_tx=XM_019826595.2;refGene_gn=LOC101100631;refGene_tx=XM_023251289.1
A3 23878604 23878691
refGene_gn=LOC101100631;refGene_tx=XM_003983572.4;refGene_gn=LOC101100631;refGene_tx=XM_003983573.5;refGene_gn=LOC101100631;refGene_tx=XM_006929810.4;refGene_gn=LOC101100631;refGene_tx=XM_006929811.3;refGene_gn=LOC101100631;refGene_tx=XM_019826595.2;refGene_gn=LOC101100631;refGene_tx=XM_023251289.1
A3 23878784 23878844
refGene_gn=LOC101100631;refGene_tx=XM_003983572.4;refGene_gn=LOC101100631;refGene_tx=XM_003983573.5;refGene_gn=LOC101100631;refGene_tx=XM_006929810.4;refGene_gn=LOC101100631;refGene_tx=XM_006929811.3;refGene_gn=LOC101100631;refGene_tx=XM_019826595.2;refGene_gn=LOC101100631;refGene_tx=XM_023251289.1
A3 23878923 23879057
refGene_gn=LOC101100631;refGene_tx=XM_003983572.4;refGene_gn=LOC101100631;refGene_tx=XM_003983573.5;refGene_gn=LOC101100631;refGene_tx=XM_006929810.4;refGene_gn=LOC101100631;refGene_tx=XM_006929811.3;refGene_gn=LOC101100631;refGene_tx=XM_019826595.2;refGene_gn=LOC101100631;refGene_tx=XM_023251289.1
A3 23879161 23879216
refGene_gn=LOC101100631;refGene_tx=XM_003983572.4;refGene_gn=LOC101100631;refGene_tx=XM_003983573.5;refGene_gn=LOC101100631;refGene_tx=XM_006929810.4;refGene_gn=LOC101100631;refGene_tx=XM_006929811.3;refGene_gn=LOC101100631;refGene_tx=XM_019826595.2;refGene_gn=LOC101100631;refGene_tx=XM_023251289.1
A3 23879462 23879514
refGene_gn=LOC101100631;refGene_tx=XM_003983572.4;refGene_gn=LOC101100631;refGene_tx=XM_003983573.5;refGene_gn=LOC101100631;refGene_tx=XM_006929810.4;refGene_gn=LOC101100631;refGene_tx=XM_006929811.3;refGene_gn=LOC101100631;refGene_tx=XM_019826595.2;refGene_gn=LOC101100631;refGene_tx=XM_023251289.1
A3 23881111 23881245
refGene_gn=LOC101100631;refGene_tx=XM_003983572.4;refGene_gn=LOC101100631;refGene_tx=XM_003983573.5;refGene_gn=LOC101100631;refGene_tx=XM_006929810.4;refGene_gn=LOC101100631;refGene_tx=XM_006929811.3;refGene_gn=LOC101100631;refGene_tx=XM_019826595.2;refGene_gn=LOC101100631;refGene_tx=XM_023251289.1
A3 23881396 23881633
refGene_gn=LOC101100631;refGene_tx=XM_003983572.4;refGene_gn=LOC101100631;refGene_tx=XM_003983573.5;refGene_gn=LOC101100631;refGene_tx=XM_006929810.4;refGene_gn=LOC101100631;refGene_tx=XM_006929811.3;refGene_gn=LOC101100631;refGene_tx=XM_019826595.2;refGene_gn=LOC101100631;refGene_tx=XM_023251289.1
A3 23881832 23881973
refGene_gn=LOC101100631;refGene_tx=XM_003983572.4;refGene_gn=LOC101100

631;refGene_tx=XM_003983573.5;refGene_gn=LOC101100631;refGene_tx=XM_006929810.4;refGene_gn=LOC101100631;refGene_tx=XM_006929811.3;refGene_gn=LOC101100631;refGene_tx=XM_019826595.2;refGene_gn=LOC101100631;refGene_tx=XM_023251289.1
A3 23885485 23885635
refGene_gn=SPAG4;refGene_tx=XM_003983574.5;refGene_gn=SPAG4;refGene_tx=XM_019826596.2;refGene_gn=SPAG4;refGene_tx=XM_019826600.2;refGene_gn=SPAG4;refGene_tx=XM_023251290.1;refGene_gn=SPAG4;refGene_tx=XM_023251291.1;refGene_gn=SPAG4;refGene_tx=XM_023251292.1
A3 23885747 23885837
refGene_gn=SPAG4;refGene_tx=XM_003983574.5;refGene_gn=SPAG4;refGene_tx=XM_019826596.2;refGene_gn=SPAG4;refGene_tx=XM_019826600.2;refGene_gn=SPAG4;refGene_tx=XM_023251290.1;refGene_gn=SPAG4;refGene_tx=XM_023251291.1;refGene_gn=SPAG4;refGene_tx=XM_023251292.1
A3 23886355 23886523
refGene_gn=SPAG4;refGene_tx=XM_003983574.5;refGene_gn=SPAG4;refGene_tx=XM_019826596.2;refGene_gn=SPAG4;refGene_tx=XM_023251290.1;refGene_gn=SPAG4;refGene_tx=XM_023251291.1;refGene_gn=SPAG4;refGene_tx=XM_023251292.1
A3 23886861 23886977
refGene_gn=SPAG4;refGene_tx=XM_003983574.5;refGene_gn=SPAG4;refGene_tx=XM_019826596.2;refGene_gn=SPAG4;refGene_tx=XM_019826600.2;refGene_gn=SPAG4;refGene_tx=XM_023251290.1;refGene_gn=SPAG4;refGene_tx=XM_023251291.1;refGene_gn=SPAG4;refGene_tx=XM_023251292.1
A3 23887144 23887220
refGene_gn=SPAG4;refGene_tx=XM_003983574.5;refGene_gn=SPAG4;refGene_tx=XM_019826596.2;refGene_gn=SPAG4;refGene_tx=XM_019826600.2;refGene_gn=SPAG4;refGene_tx=XM_023251290.1;refGene_gn=SPAG4;refGene_tx=XM_023251291.1;refGene_gn=SPAG4;refGene_tx=XM_023251292.1
A3 23887478 23887587
refGene_gn=SPAG4;refGene_tx=XM_003983574.5;refGene_gn=SPAG4;refGene_tx=XM_019826596.2;refGene_gn=SPAG4;refGene_tx=XM_019826600.2;refGene_gn=SPAG4;refGene_tx=XM_023251290.1;refGene_gn=SPAG4;refGene_tx=XM_023251291.1;refGene_gn=SPAG4;refGene_tx=XM_023251292.1
A3 23887708 23887734
refGene_gn=SPAG4;refGene_tx=XM_003983574.5;refGene_gn=SPAG4;refGene_tx=XM_019826596.2;refGene_gn=SPAG4;refGene_tx=XM_019826600.2;refGene_gn=SPAG4;refGene_tx=XM_023251290.1;refGene_gn=SPAG4;refGene_tx=XM_023251291.1;refGene_gn=SPAG4;refGene_tx=XM_023251292.1
A3 23888051 23888370
refGene_gn=SPAG4;refGene_tx=XM_003983574.5;refGene_gn=SPAG4;refGene_tx=XM_019826596.2;refGene_gn=SPAG4;refGene_tx=XM_019826600.2;refGene_gn=SPAG4;refGene_tx=XM_023251290.1;refGene_gn=SPAG4;refGene_tx=XM_023251291.1;refGene_gn=SPAG4;refGene_tx=XM_023251292.1
A3 23888587 23888654
refGene_gn=SPAG4;refGene_tx=XM_003983574.5;refGene_gn=SPAG4;refGene_tx=XM_019826596.2;refGene_gn=SPAG4;refGene_tx=XM_019826600.2;refGene_gn=SPAG4;refGene_tx=XM_023251290.1;refGene_gn=SPAG4;refGene_tx=XM_023251291.1;refGene_gn=SPAG4;refGene_tx=XM_023251292.1
A3 23888941 23889161

refGene_gn=SPAG4;refGene_tx=XM_003983574.5;refGene_gn=SPAG4;refGene_tx=XM_019826596.2;refGene_gn=SPAG4;refGene_tx=XM_019826600.2;refGene_gn=SPAG4;refGene_tx=XM_023251290.1;refGene_gn=SPAG4;refGene_tx=XM_023251291.1;refGene_gn=SPAG4;refGene_tx=XM_023251292.1
A3 23889553 23889863
refGene_gn=SPAG4;refGene_tx=XM_003983574.5;refGene_gn=SPAG4;refGene_tx=XM_019826596.2;refGene_gn=SPAG4;refGene_tx=XM_019826600.2;refGene_gn=SPAG4;refGene_tx=XM_023251291.1;refGene_gn=SPAG4;refGene_tx=XM_023251292.1
A3 23893122 23893201 refGene_gn=FER1L4;refGene_tx=XM_023251302.1
A3 23893463 23893519 refGene_gn=FER1L4;refGene_tx=XM_023251302.1
A3 23893893 23893982 refGene_gn=FER1L4;refGene_tx=XM_023251302.1
A3 23894082 23894182 refGene_gn=FER1L4;refGene_tx=XM_023251302.1
A3 23895317 23895409 refGene_gn=FER1L4;refGene_tx=XM_023251302.1
A3 23895604 23895646 refGene_gn=FER1L4;refGene_tx=XM_023251302.1
A3 23895942 23896107 refGene_gn=FER1L4;refGene_tx=XM_023251302.1
A3 23896610 23896671 refGene_gn=FER1L4;refGene_tx=XM_023251302.1
A3 23896797 23896929 refGene_gn=FER1L4;refGene_tx=XM_023251302.1
A3 23897191 23897254 refGene_gn=FER1L4;refGene_tx=XM_023251302.1
A3 23897334 23897419 refGene_gn=FER1L4;refGene_tx=XM_023251302.1
A3 23897646 23897806 refGene_gn=FER1L4;refGene_tx=XM_023251302.1
A3 23897926 23898110 refGene_gn=FER1L4;refGene_tx=XM_023251302.1
A3 23898187 23898374 refGene_gn=FER1L4;refGene_tx=XM_023251302.1
A3 23898980 23899339 refGene_gn=FER1L4;refGene_tx=XM_023251302.1
A3 23899653 23899762 refGene_gn=FER1L4;refGene_tx=XM_023251302.1
A3 23899953 23900128 refGene_gn=FER1L4;refGene_tx=XM_023251302.1
A3 23900245 23900366 refGene_gn=FER1L4;refGene_tx=XM_023251302.1
A3 23900783 23900884 refGene_gn=FER1L4;refGene_tx=XM_023251302.1
A3 23905678 23905766 refGene_gn=FER1L4;refGene_tx=XM_023251302.1
A3 23905839 23905956 refGene_gn=FER1L4;refGene_tx=XM_023251302.1
A3 23906322 23906475 refGene_gn=FER1L4;refGene_tx=XM_023251302.1
A3 23907594 23907748 refGene_gn=FER1L4;refGene_tx=XM_023251302.1
A3 23907873 23907998 refGene_gn=FER1L4;refGene_tx=XM_023251302.1
A3 23908413 23908548 refGene_gn=FER1L4;refGene_tx=XM_023251302.1
A3 23908674 23908848 refGene_gn=FER1L4;refGene_tx=XM_023251302.1
A3 23909299 23909416 refGene_gn=FER1L4;refGene_tx=XM_023251302.1
A3 23909543 23909705 refGene_gn=FER1L4;refGene_tx=XM_023251302.1
A3 23916058 23916257 refGene_gn=FER1L4;refGene_tx=XM_023251302.1
A3 23916399 23916458 refGene_gn=FER1L4;refGene_tx=XM_023251302.1
A3 23916534 23916666 refGene_gn=FER1L4;refGene_tx=XM_023251302.1
A3 23917268 23917311 refGene_gn=FER1L4;refGene_tx=XM_023251302.1
A3 23917404 23917511 refGene_gn=FER1L4;refGene_tx=XM_023251302.1
A3 23917587 23917725 refGene_gn=FER1L4;refGene_tx=XM_023251302.1
A3 23917820 23917937 refGene_gn=FER1L4;refGene_tx=XM_023251302.1
A3 23921970 23922098 refGene_gn=FER1L4;refGene_tx=XM_023251302.1
A3 23922297 23922468 refGene_gn=FER1L4;refGene_tx=XM_023251302.1
A3 23922888 23923043 refGene_gn=FER1L4;refGene_tx=XM_023251302.1
A3 23923622 23923783 refGene_gn=FER1L4;refGene_tx=XM_023251302.1
A3 23924343 23924429 refGene_gn=FER1L4;refGene_tx=XM_023251302.1
A3 23925593 23925692 refGene_gn=FER1L4;refGene_tx=XM_023251302.1

A3 23925934 23926176 refGene_gn=FER1L4;refGene_tx=XM_023251302.1
A3 23926618 23926896 refGene_gn=FER1L4;refGene_tx=XM_023251302.1
A3 23926992 23927093 refGene_gn=FER1L4;refGene_tx=XM_023251302.1
A3 23927189 23927394 refGene_gn=FER1L4;refGene_tx=XM_023251302.1
A3 23929226 23929306
refGene_gn=ERGIC3;refGene_tx=XM_003983575.4;refGene_gn=ERGIC3;refGene_
tx=XM_019826603.2
A3 23929506 23929562
refGene_gn=ERGIC3;refGene_tx=XM_003983575.4;refGene_gn=ERGIC3;refGene_
tx=XM_019826603.2
A3 23929642 23929779
refGene_gn=ERGIC3;refGene_tx=XM_003983575.4;refGene_gn=ERGIC3;refGene_
tx=XM_019826603.2
A3 23930380 23930445
refGene_gn=ERGIC3;refGene_tx=XM_003983575.4;refGene_gn=ERGIC3;refGene_
tx=XM_019826603.2
A3 23930522 23930619
refGene_gn=ERGIC3;refGene_tx=XM_003983575.4;refGene_gn=ERGIC3;refGene_
tx=XM_019826603.2
A3 23931428 23931460
refGene_gn=ERGIC3;refGene_tx=XM_003983575.4;refGene_gn=ERGIC3;refGene_
tx=XM_019826603.2
A3 23932083 23932098 refGene_gn=ERGIC3;refGene_tx=XM_019826603.2
A3 23935925 23935983
refGene_gn=ERGIC3;refGene_tx=XM_003983575.4;refGene_gn=ERGIC3;refGene_
tx=XM_019826603.2
A3 23936104 23936270
refGene_gn=ERGIC3;refGene_tx=XM_003983575.4;refGene_gn=ERGIC3;refGene_
tx=XM_019826603.2
A3 23937092 23937186
refGene_gn=ERGIC3;refGene_tx=XM_003983575.4;refGene_gn=ERGIC3;refGene_
tx=XM_019826603.2
A3 23942150 23942270
refGene_gn=ERGIC3;refGene_tx=XM_003983575.4;refGene_gn=ERGIC3;refGene_
tx=XM_019826603.2
A3 23942480 23942568
refGene_gn=ERGIC3;refGene_tx=XM_003983575.4;refGene_gn=ERGIC3;refGene_
tx=XM_019826603.2
A3 23942678 23942749
refGene_gn=ERGIC3;refGene_tx=XM_003983575.4;refGene_gn=ERGIC3;refGene_
tx=XM_019826603.2
A3 23942887 23942975
refGene_gn=ERGIC3;refGene_tx=XM_003983575.4;refGene_gn=ERGIC3;refGene_
tx=XM_019826603.2
A3 23947964 23948140
refGene_gn=CA3H20orf173;refGene_tx=XM_023251301.1
A3 23948220 23948398
refGene_gn=CA3H20orf173;refGene_tx=XM_023251301.1
A3 23949283 23949340
refGene_gn=CA3H20orf173;refGene_tx=XM_023251301.1

A3	23974048 23974306
refGene_gn=CEP250;refGene_tx=XM_006929814.2;refGene_gn=CEP250;refGene_tx=XM_023251293.1;refGene_gn=CEP250;refGene_tx=XM_023251294.1;refGene_gn=CEP250;refGene_tx=XM_023251295.1;refGene_gn=CEP250;refGene_tx=XM_023251296.1;refGene_gn=CEP250;refGene_tx=XM_023251297.1;refGene_gn=CEP250;refGene_tx=XM_023251299.1;refGene_gn=CEP250;refGene_tx=XM_023251300.1
A3	23976177 23976240
refGene_gn=CEP250;refGene_tx=XM_006929814.2;refGene_gn=CEP250;refGene_tx=XM_023251293.1;refGene_gn=CEP250;refGene_tx=XM_023251294.1;refGene_gn=CEP250;refGene_tx=XM_023251295.1;refGene_gn=CEP250;refGene_tx=XM_023251296.1;refGene_gn=CEP250;refGene_tx=XM_023251297.1;refGene_gn=CEP250;refGene_tx=XM_023251299.1;refGene_gn=CEP250;refGene_tx=XM_023251300.1
A3	23977207 23977294
refGene_gn=CEP250;refGene_tx=XM_006929814.2;refGene_gn=CEP250;refGene_tx=XM_023251293.1;refGene_gn=CEP250;refGene_tx=XM_023251294.1;refGene_gn=CEP250;refGene_tx=XM_023251295.1;refGene_gn=CEP250;refGene_tx=XM_023251296.1;refGene_gn=CEP250;refGene_tx=XM_023251297.1;refGene_gn=CEP250;refGene_tx=XM_023251299.1;refGene_gn=CEP250;refGene_tx=XM_023251300.1
A3	23978132 23978301
refGene_gn=CEP250;refGene_tx=XM_006929814.2;refGene_gn=CEP250;refGene_tx=XM_023251293.1;refGene_gn=CEP250;refGene_tx=XM_023251294.1;refGene_gn=CEP250;refGene_tx=XM_023251295.1;refGene_gn=CEP250;refGene_tx=XM_023251296.1;refGene_gn=CEP250;refGene_tx=XM_023251297.1;refGene_gn=CEP250;refGene_tx=XM_023251298.1;refGene_gn=CEP250;refGene_tx=XM_023251299.1;refGene_gn=CEP250;refGene_tx=XM_023251300.1
A3	23978494 23978587
refGene_gn=CEP250;refGene_tx=XM_006929814.2;refGene_gn=CEP250;refGene_tx=XM_023251293.1;refGene_gn=CEP250;refGene_tx=XM_023251294.1;refGene_gn=CEP250;refGene_tx=XM_023251295.1;refGene_gn=CEP250;refGene_tx=XM_023251296.1;refGene_gn=CEP250;refGene_tx=XM_023251297.1;refGene_gn=CEP250;refGene_tx=XM_023251298.1;refGene_gn=CEP250;refGene_tx=XM_023251299.1;refGene_gn=CEP250;refGene_tx=XM_023251300.1
A3	23981128 23983738
refGene_gn=CEP250;refGene_tx=XM_006929814.2;refGene_gn=CEP250;refGene_tx=XM_023251293.1;refGene_gn=CEP250;refGene_tx=XM_023251294.1;refGene_gn=CEP250;refGene_tx=XM_023251295.1;refGene_gn=CEP250;refGene_tx=XM_023251296.1;refGene_gn=CEP250;refGene_tx=XM_023251297.1;refGene_gn=CEP250;refGene_tx=XM_023251298.1;refGene_gn=CEP250;refGene_tx=XM_023251299.1;refGene_gn=CEP250;refGene_tx=XM_023251300.1
A3	23984242 23984364
refGene_gn=CEP250;refGene_tx=XM_006929814.2;refGene_gn=CEP250;refGene_tx=XM_023251293.1;refGene_gn=CEP250;refGene_tx=XM_023251294.1;refGene_gn=CEP250;refGene_tx=XM_023251295.1;refGene_gn=CEP250;refGene_tx=XM_023251296.1;refGene_gn=CEP250;refGene_tx=XM_023251297.1;refGene_gn=CEP250;refGene_tx=XM_023251298.1;refGene_gn=CEP250;refGene_tx=XM_023251299.1;refGene_gn=CEP250;refGene_tx=XM_023251300.1
A3	23985406 23985527

refGene_gn=CEP250;refGene_tx=XM_006929814.2;refGene_gn=CEP250;refGene_
tx=XM_023251293.1;refGene_gn=CEP250;refGene_tx=XM_023251294.1;refGene_
gn=CEP250;refGene_tx=XM_023251295.1;refGene_gn=CEP250;refGene_tx=XM_02
3251296.1;refGene_gn=CEP250;refGene_tx=XM_023251297.1;refGene_gn=CEP25
0;refGene_tx=XM_023251298.1;refGene_gn=CEP250;refGene_tx=XM_023251299.
1;refGene_gn=CEP250;refGene_tx=XM_023251300.1
A3 23986471 23986593
refGene_gn=CEP250;refGene_tx=XM_006929814.2;refGene_gn=CEP250;refGene_
tx=XM_023251293.1;refGene_gn=CEP250;refGene_tx=XM_023251294.1;refGene_
gn=CEP250;refGene_tx=XM_023251295.1;refGene_gn=CEP250;refGene_tx=XM_02
3251296.1;refGene_gn=CEP250;refGene_tx=XM_023251297.1;refGene_gn=CEP25
0;refGene_tx=XM_023251298.1;refGene_gn=CEP250;refGene_tx=XM_023251299.
1;refGene_gn=CEP250;refGene_tx=XM_023251300.1
A3 23987823 23988163
refGene_gn=CEP250;refGene_tx=XM_006929814.2;refGene_gn=CEP250;refGene_
tx=XM_023251293.1;refGene_gn=CEP250;refGene_tx=XM_023251294.1;refGene_
gn=CEP250;refGene_tx=XM_023251295.1;refGene_gn=CEP250;refGene_tx=XM_02
3251296.1;refGene_gn=CEP250;refGene_tx=XM_023251297.1;refGene_gn=CEP25
0;refGene_tx=XM_023251298.1;refGene_gn=CEP250;refGene_tx=XM_023251299.
1;refGene_gn=CEP250;refGene_tx=XM_023251300.1
A3 23989138 23989277
refGene_gn=CEP250;refGene_tx=XM_006929814.2;refGene_gn=CEP250;refGene_
tx=XM_023251293.1;refGene_gn=CEP250;refGene_tx=XM_023251294.1;refGene_
gn=CEP250;refGene_tx=XM_023251295.1;refGene_gn=CEP250;refGene_tx=XM_02
3251296.1;refGene_gn=CEP250;refGene_tx=XM_023251297.1;refGene_gn=CEP25
0;refGene_tx=XM_023251298.1;refGene_gn=CEP250;refGene_tx=XM_023251299.
1;refGene_gn=CEP250;refGene_tx=XM_023251300.1
A3 23991115 23991249
refGene_gn=CEP250;refGene_tx=XM_006929814.2;refGene_gn=CEP250;refGene_
tx=XM_023251293.1;refGene_gn=CEP250;refGene_tx=XM_023251294.1;refGene_
gn=CEP250;refGene_tx=XM_023251295.1;refGene_gn=CEP250;refGene_tx=XM_02
3251296.1;refGene_gn=CEP250;refGene_tx=XM_023251297.1;refGene_gn=CEP25
0;refGene_tx=XM_023251298.1;refGene_gn=CEP250;refGene_tx=XM_023251299.
1;refGene_gn=CEP250;refGene_tx=XM_023251300.1
A3 23992204 23992348
refGene_gn=CEP250;refGene_tx=XM_006929814.2;refGene_gn=CEP250;refGene_
tx=XM_023251293.1;refGene_gn=CEP250;refGene_tx=XM_023251294.1;refGene_
gn=CEP250;refGene_tx=XM_023251295.1;refGene_gn=CEP250;refGene_tx=XM_02
3251296.1;refGene_gn=CEP250;refGene_tx=XM_023251297.1;refGene_gn=CEP25
0;refGene_tx=XM_023251298.1;refGene_gn=CEP250;refGene_tx=XM_023251299.
1;refGene_gn=CEP250;refGene_tx=XM_023251300.1
A3 23993945 23994080
refGene_gn=CEP250;refGene_tx=XM_006929814.2;refGene_gn=CEP250;refGene_
tx=XM_023251293.1;refGene_gn=CEP250;refGene_tx=XM_023251294.1;refGene_
gn=CEP250;refGene_tx=XM_023251295.1;refGene_gn=CEP250;refGene_tx=XM_02
3251296.1;refGene_gn=CEP250;refGene_tx=XM_023251297.1;refGene_gn=CEP25
0;refGene_tx=XM_023251298.1;refGene_gn=CEP250;refGene_tx=XM_023251299.
1;refGene_gn=CEP250;refGene_tx=XM_023251300.1
A3 23994478 23994646
refGene_gn=CEP250;refGene_tx=XM_006929814.2;refGene_gn=CEP250;refGene_

tx=XM_023251293.1;refGene_gn=CEP250;refGene_tx=XM_023251294.1;refGene_gn=CEP250;refGene_tx=XM_023251295.1;refGene_gn=CEP250;refGene_tx=XM_023251297.1;refGene_gn=CEP250;refGene_tx=XM_023251298.1;refGene_gn=CEP250;refGene_tx=XM_023251299.1;refGene_gn=CEP250;refGene_tx=XM_023251300.1
A3 24000837 24001007
refGene_gn=CEP250;refGene_tx=XM_006929814.2;refGene_gn=CEP250;refGene_tx=XM_023251293.1;refGene_gn=CEP250;refGene_tx=XM_023251294.1;refGene_gn=CEP250;refGene_tx=XM_023251295.1;refGene_gn=CEP250;refGene_tx=XM_023251296.1;refGene_gn=CEP250;refGene_tx=XM_023251297.1;refGene_gn=CEP250;refGene_tx=XM_023251298.1;refGene_gn=CEP250;refGene_tx=XM_023251299.1;refGene_gn=CEP250;refGene_tx=XM_023251300.1
A3 24001240 24001368
refGene_gn=CEP250;refGene_tx=XM_006929814.2;refGene_gn=CEP250;refGene_tx=XM_023251293.1;refGene_gn=CEP250;refGene_tx=XM_023251294.1;refGene_gn=CEP250;refGene_tx=XM_023251295.1;refGene_gn=CEP250;refGene_tx=XM_023251296.1;refGene_gn=CEP250;refGene_tx=XM_023251297.1;refGene_gn=CEP250;refGene_tx=XM_023251298.1;refGene_gn=CEP250;refGene_tx=XM_023251299.1;refGene_gn=CEP250;refGene_tx=XM_023251300.1
A3 24001590 24001784
refGene_gn=CEP250;refGene_tx=XM_006929814.2;refGene_gn=CEP250;refGene_tx=XM_023251293.1;refGene_gn=CEP250;refGene_tx=XM_023251294.1;refGene_gn=CEP250;refGene_tx=XM_023251295.1;refGene_gn=CEP250;refGene_tx=XM_023251296.1;refGene_gn=CEP250;refGene_tx=XM_023251297.1;refGene_gn=CEP250;refGene_tx=XM_023251298.1;refGene_gn=CEP250;refGene_tx=XM_023251299.1;refGene_gn=CEP250;refGene_tx=XM_023251300.1
A3 24003076 24003307
refGene_gn=CEP250;refGene_tx=XM_006929814.2;refGene_gn=CEP250;refGene_tx=XM_023251293.1;refGene_gn=CEP250;refGene_tx=XM_023251294.1;refGene_gn=CEP250;refGene_tx=XM_023251295.1;refGene_gn=CEP250;refGene_tx=XM_023251296.1;refGene_gn=CEP250;refGene_tx=XM_023251297.1;refGene_gn=CEP250;refGene_tx=XM_023251298.1;refGene_gn=CEP250;refGene_tx=XM_023251299.1;refGene_gn=CEP250;refGene_tx=XM_023251300.1
A3 24004018 24004165
refGene_gn=CEP250;refGene_tx=XM_006929814.2;refGene_gn=CEP250;refGene_tx=XM_023251293.1;refGene_gn=CEP250;refGene_tx=XM_023251294.1;refGene_gn=CEP250;refGene_tx=XM_023251295.1;refGene_gn=CEP250;refGene_tx=XM_023251296.1;refGene_gn=CEP250;refGene_tx=XM_023251297.1;refGene_gn=CEP250;refGene_tx=XM_023251298.1;refGene_gn=CEP250;refGene_tx=XM_023251299.1;refGene_gn=CEP250;refGene_tx=XM_023251300.1
A3 24004989 24005134
refGene_gn=CEP250;refGene_tx=XM_006929814.2;refGene_gn=CEP250;refGene_tx=XM_023251293.1;refGene_gn=CEP250;refGene_tx=XM_023251294.1;refGene_gn=CEP250;refGene_tx=XM_023251295.1;refGene_gn=CEP250;refGene_tx=XM_023251296.1;refGene_gn=CEP250;refGene_tx=XM_023251297.1;refGene_gn=CEP250;refGene_tx=XM_023251298.1;refGene_gn=CEP250;refGene_tx=XM_023251299.1;refGene_gn=CEP250;refGene_tx=XM_023251300.1
A3 24006884 24007067
refGene_gn=CEP250;refGene_tx=XM_006929814.2;refGene_gn=CEP250;refGene_tx=XM_023251293.1;refGene_gn=CEP250;refGene_tx=XM_023251294.1;refGene_

gn=CEP250;refGene_tx=XM_023251295.1;refGene_gn=CEP250;refGene_tx=XM_02
3251296.1;refGene_gn=CEP250;refGene_tx=XM_023251297.1;refGene_gn=CEP25
0;refGene_tx=XM_023251298.1;refGene_gn=CEP250;refGene_tx=XM_023251299.
1;refGene_gn=CEP250;refGene_tx=XM_023251300.1
A3 24007550 24007729
refGene_gn=CEP250;refGene_tx=XM_006929814.2;refGene_gn=CEP250;refGene_
tx=XM_023251293.1;refGene_gn=CEP250;refGene_tx=XM_023251294.1;refGene_
gn=CEP250;refGene_tx=XM_023251295.1;refGene_gn=CEP250;refGene_tx=XM_02
3251296.1;refGene_gn=CEP250;refGene_tx=XM_023251297.1;refGene_gn=CEP25
0;refGene_tx=XM_023251298.1;refGene_gn=CEP250;refGene_tx=XM_023251299.
1;refGene_gn=CEP250;refGene_tx=XM_023251300.1
A3 24008646 24008808
refGene_gn=CEP250;refGene_tx=XM_006929814.2;refGene_gn=CEP250;refGene_
tx=XM_023251293.1;refGene_gn=CEP250;refGene_tx=XM_023251294.1;refGene_
gn=CEP250;refGene_tx=XM_023251295.1;refGene_gn=CEP250;refGene_tx=XM_02
3251296.1;refGene_gn=CEP250;refGene_tx=XM_023251297.1;refGene_gn=CEP25
0;refGene_tx=XM_023251298.1;refGene_gn=CEP250;refGene_tx=XM_023251299.
1;refGene_gn=CEP250;refGene_tx=XM_023251300.1
A3 24009325 24009433
refGene_gn=CEP250;refGene_tx=XM_006929814.2;refGene_gn=CEP250;refGene_
tx=XM_023251293.1;refGene_gn=CEP250;refGene_tx=XM_023251294.1;refGene_
gn=CEP250;refGene_tx=XM_023251295.1;refGene_gn=CEP250;refGene_tx=XM_02
3251296.1;refGene_gn=CEP250;refGene_tx=XM_023251297.1;refGene_gn=CEP25
0;refGene_tx=XM_023251298.1;refGene_gn=CEP250;refGene_tx=XM_023251299.
1;refGene_gn=CEP250;refGene_tx=XM_023251300.1
A3 24011352 24011449
refGene_gn=CEP250;refGene_tx=XM_006929814.2;refGene_gn=CEP250;refGene_
tx=XM_023251293.1;refGene_gn=CEP250;refGene_tx=XM_023251294.1;refGene_
gn=CEP250;refGene_tx=XM_023251295.1;refGene_gn=CEP250;refGene_tx=XM_02
3251296.1;refGene_gn=CEP250;refGene_tx=XM_023251297.1;refGene_gn=CEP25
0;refGene_tx=XM_023251298.1;refGene_gn=CEP250;refGene_tx=XM_023251299.
1;refGene_gn=CEP250;refGene_tx=XM_023251300.1
A3 24011674 24011793 refGene_gn=CEP250;refGene_tx=XM_023251300.1
A3 24013074 24013326
refGene_gn=CEP250;refGene_tx=XM_006929814.2;refGene_gn=CEP250;refGene_
tx=XM_023251293.1;refGene_gn=CEP250;refGene_tx=XM_023251294.1;refGene_
gn=CEP250;refGene_tx=XM_023251295.1;refGene_gn=CEP250;refGene_tx=XM_02
3251296.1;refGene_gn=CEP250;refGene_tx=XM_023251297.1;refGene_gn=CEP25
0;refGene_tx=XM_023251298.1;refGene_gn=CEP250;refGene_tx=XM_023251299.
1
A3 24013564 24013671
refGene_gn=CEP250;refGene_tx=XM_006929814.2;refGene_gn=CEP250;refGene_
tx=XM_023251293.1;refGene_gn=CEP250;refGene_tx=XM_023251294.1;refGene_
gn=CEP250;refGene_tx=XM_023251295.1;refGene_gn=CEP250;refGene_tx=XM_02
3251296.1;refGene_gn=CEP250;refGene_tx=XM_023251297.1;refGene_gn=CEP25
0;refGene_tx=XM_023251298.1;refGene_gn=CEP250;refGene_tx=XM_023251299.
1
A3 24014463 24014629
refGene_gn=CEP250;refGene_tx=XM_006929814.2;refGene_gn=CEP250;refGene_
tx=XM_023251293.1;refGene_gn=CEP250;refGene_tx=XM_023251294.1;refGene_

gn=CEP250;refGene_tx=XM_023251295.1;refGene_gn=CEP250;refGene_tx=XM_02
3251296.1;refGene_gn=CEP250;refGene_tx=XM_023251297.1;refGene_gn=CEP25
0;refGene_tx=XM_023251298.1;refGene_gn=CEP250;refGene_tx=XM_023251299.
1
A3 24014859 24014942
refGene_gn=CEP250;refGene_tx=XM_006929814.2;refGene_gn=CEP250;refGene_
tx=XM_023251293.1;refGene_gn=CEP250;refGene_tx=XM_023251294.1;refGene_
gn=CEP250;refGene_tx=XM_023251295.1;refGene_gn=CEP250;refGene_tx=XM_02
3251296.1;refGene_gn=CEP250;refGene_tx=XM_023251298.1
A3 24015899 24015956
refGene_gn=CEP250;refGene_tx=XM_006929814.2;refGene_gn=CEP250;refGene_
tx=XM_023251293.1;refGene_gn=CEP250;refGene_tx=XM_023251294.1;refGene_
gn=CEP250;refGene_tx=XM_023251295.1;refGene_gn=CEP250;refGene_tx=XM_02
3251296.1;refGene_gn=CEP250;refGene_tx=XM_023251298.1
A3 24016630 24016816
refGene_gn=CEP250;refGene_tx=XM_006929814.2;refGene_gn=CEP250;refGene_
tx=XM_023251293.1;refGene_gn=CEP250;refGene_tx=XM_023251294.1;refGene_
gn=CEP250;refGene_tx=XM_023251295.1;refGene_gn=CEP250;refGene_tx=XM_02
3251296.1;refGene_gn=CEP250;refGene_tx=XM_023251298.1
A3 24038506 24039131
refGene_gn=GDF5;refGene_tx=XM_003983579.4;refGene_gn=GDF5;refGene_tx=X
M_006929819.3
A3 24041723 24042598
refGene_gn=GDF5;refGene_tx=XM_003983579.4;refGene_gn=GDF5;refGene_tx=X
M_006929819.3
A3 24057495 24057519
refGene_gn=LOC101081793;refGene_tx=XM_003983580.4;refGene_gn=LOC101081
793;refGene_tx=XM_006929820.3;refGene_gn=LOC101081793;refGene_tx=XM_01
9826606.2;refGene_gn=LOC101081793;refGene_tx=XM_019826607.2
A3 24072190 24072289
refGene_gn=LOC101081793;refGene_tx=XM_003983580.4;refGene_gn=LOC101081
793;refGene_tx=XM_006929820.3;refGene_gn=LOC101081793;refGene_tx=XM_01
9826606.2;refGene_gn=LOC101081793;refGene_tx=XM_019826607.2
A3 24081568 24081664
refGene_gn=LOC101081793;refGene_tx=XM_003983580.4;refGene_gn=LOC101081
793;refGene_tx=XM_006929820.3;refGene_gn=LOC101081793;refGene_tx=XM_01
9826606.2
A3 24083130 24083238
refGene_gn=LOC101081793;refGene_tx=XM_003983580.4;refGene_gn=LOC101081
793;refGene_tx=XM_006929820.3
A3 24086034 24086175
refGene_gn=LOC101081793;refGene_tx=XM_006929820.3
A3 24091349 24091422
refGene_gn=LOC101081793;refGene_tx=XM_003983580.4;refGene_gn=LOC101081
793;refGene_tx=XM_006929820.3;refGene_gn=LOC101081793;refGene_tx=XM_01
9826606.2;refGene_gn=LOC101081793;refGene_tx=XM_019826607.2
A3 24102760 24102818
refGene_gn=LOC101081793;refGene_tx=XM_003983580.4;refGene_gn=LOC101081
793;refGene_tx=XM_006929820.3;refGene_gn=LOC101081793;refGene_tx=XM_01
9826606.2;refGene_gn=LOC101081793;refGene_tx=XM_019826607.2

A3 24117533 24117642
refGene_gn=LOC101081793;refGene_tx=XM_003983580.4;refGene_gn=LOC101081793;refGene_tx=XM_006929820.3;refGene_gn=LOC101081793;refGene_tx=XM_019826606.2;refGene_gn=LOC101081793;refGene_tx=XM_019826607.2
A3 24151133 24151211
refGene_gn=LOC101081793;refGene_tx=XM_003983580.4;refGene_gn=LOC101081793;refGene_tx=XM_006929820.3;refGene_gn=LOC101081793;refGene_tx=XM_019826606.2;refGene_gn=LOC101081793;refGene_tx=XM_019826607.2
A3 24157697 24157811
refGene_gn=LOC101081793;refGene_tx=XM_003983580.4;refGene_gn=LOC101081793;refGene_tx=XM_006929820.3;refGene_gn=LOC101081793;refGene_tx=XM_019826606.2;refGene_gn=LOC101081793;refGene_tx=XM_019826607.2
A3 24160747 24160882
refGene_gn=LOC101081793;refGene_tx=XM_003983580.4;refGene_gn=LOC101081793;refGene_tx=XM_006929820.3;refGene_gn=LOC101081793;refGene_tx=XM_019826606.2;refGene_gn=LOC101081793;refGene_tx=XM_019826607.2
A3 24173768 24174281 refGene_gn=FAM83C;refGene_tx=XM_003983583.3
A3 24176464 24176632 refGene_gn=FAM83C;refGene_tx=XM_003983583.3
A3 24176843 24176968 refGene_gn=FAM83C;refGene_tx=XM_003983583.3
A3 24177475 24178916 refGene_gn=FAM83C;refGene_tx=XM_003983583.3
A3 24181461 24181568 refGene_gn=EIF6;refGene_tx=XM_006929821.3
A3 24181685 24181771 refGene_gn=EIF6;refGene_tx=XM_006929821.3
A3 24183338 24183514 refGene_gn=EIF6;refGene_tx=XM_006929821.3
A3 24184065 24184242 refGene_gn=EIF6;refGene_tx=XM_006929821.3
A3 24184440 24184622 refGene_gn=EIF6;refGene_tx=XM_006929821.3
A3 24187513 24187523 refGene_gn=EIF6;refGene_tx=XM_006929821.3
A3 24188616 24188784
refGene_gn=LOC109498025;refGene_tx=XM_023251306.1
A3 24191845 24192183 refGene_gn=MMP24;refGene_tx=XM_023251672.1
A3 24194480 24194747 refGene_gn=MMP24;refGene_tx=XM_023251672.1
A3 24196209 24196348 refGene_gn=MMP24;refGene_tx=XM_023251672.1
A3 24198830 24199045 refGene_gn=MMP24;refGene_tx=XM_023251672.1
A3 24200832 24200994 refGene_gn=MMP24;refGene_tx=XM_023251672.1
A3 24210146 24210451 refGene_gn=MMP24;refGene_tx=XM_023251672.1
A3 24211966 24212083 refGene_gn=MMP24;refGene_tx=XM_023251672.1
A3 24216103 24216252 refGene_gn=MMP24;refGene_tx=XM_023251672.1
A3 24234335 24234908 refGene_gn=MMP24;refGene_tx=XM_023251672.1
A3 24239375 24239453 refGene_gn=MMP24;refGene_tx=XM_023251672.1
A3 24263658 24263774 refGene_gn=PROCR;refGene_tx=XM_003983585.4
A3 24264021 24264300 refGene_gn=PROCR;refGene_tx=XM_003983585.4
A3 24264739 24264991 refGene_gn=PROCR;refGene_tx=XM_003983585.4
A3 24267127 24267206 refGene_gn=PROCR;refGene_tx=XM_003983585.4
A3 24285395 24285502
refGene_gn=EDEM2;refGene_tx=XM_019826610.2;refGene_gn=EDEM2;refGene_tx=XM_023251303.1;refGene_gn=EDEM2;refGene_tx=XM_023251304.1;refGene_gn=EDEM2;refGene_tx=XM_023251305.1
A3 24285714 24285825
refGene_gn=EDEM2;refGene_tx=XM_019826610.2;refGene_gn=EDEM2;refGene_tx=XM_023251303.1;refGene_gn=EDEM2;refGene_tx=XM_023251304.1;refGene_gn=EDEM2;refGene_tx=XM_023251305.1

A3	24287644	24287684
refGene_gn=EDEM2;refGene_tx=XM_019826610.2;refGene_gn=EDEM2;refGene_tx=XM_023251303.1;refGene_gn=EDEM2;refGene_tx=XM_023251304.1;refGene_gn=EDEM2;refGene_tx=XM_023251305.1
A3	24291059	24291165
refGene_gn=EDEM2;refGene_tx=XM_019826610.2;refGene_gn=EDEM2;refGene_tx=XM_023251303.1;refGene_gn=EDEM2;refGene_tx=XM_023251304.1;refGene_gn=EDEM2;refGene_tx=XM_023251305.1
A3	24295516	24295561
refGene_gn=EDEM2;refGene_tx=XM_019826610.2;refGene_gn=EDEM2;refGene_tx=XM_023251303.1
A3	24296185	24296311
refGene_gn=EDEM2;refGene_tx=XM_019826610.2;refGene_gn=EDEM2;refGene_tx=XM_023251303.1;refGene_gn=EDEM2;refGene_tx=XM_023251304.1
A3	24299125	24299337
refGene_gn=EDEM2;refGene_tx=XM_019826610.2;refGene_gn=EDEM2;refGene_tx=XM_023251303.1;refGene_gn=EDEM2;refGene_tx=XM_023251304.1;refGene_gn=EDEM2;refGene_tx=XM_023251305.1
A3	24301136	24301278
refGene_gn=EDEM2;refGene_tx=XM_019826610.2;refGene_gn=EDEM2;refGene_tx=XM_023251303.1;refGene_gn=EDEM2;refGene_tx=XM_023251304.1;refGene_gn=EDEM2;refGene_tx=XM_023251305.1
A3	24302804	24302929
refGene_gn=EDEM2;refGene_tx=XM_019826610.2;refGene_gn=EDEM2;refGene_tx=XM_023251303.1;refGene_gn=EDEM2;refGene_tx=XM_023251304.1;refGene_gn=EDEM2;refGene_tx=XM_023251305.1
A3	24309863	24310008
refGene_gn=EDEM2;refGene_tx=XM_019826610.2;refGene_gn=EDEM2;refGene_tx=XM_023251303.1;refGene_gn=EDEM2;refGene_tx=XM_023251304.1;refGene_gn=EDEM2;refGene_tx=XM_023251305.1
A3	24313536	24313658
refGene_gn=EDEM2;refGene_tx=XM_019826610.2;refGene_gn=EDEM2;refGene_tx=XM_023251303.1;refGene_gn=EDEM2;refGene_tx=XM_023251304.1;refGene_gn=EDEM2;refGene_tx=XM_023251305.1
A3	24315817	24316315
refGene_gn=EDEM2;refGene_tx=XM_019826610.2;refGene_gn=EDEM2;refGene_tx=XM_023251303.1;refGene_gn=EDEM2;refGene_tx=XM_023251304.1;refGene_gn=EDEM2;refGene_tx=XM_023251305.1
A3	24348901	24349069	refGene_gn=TRPC4AP;refGene_tx=XM_023251307.1
A3	24362452	24362581	refGene_gn=TRPC4AP;refGene_tx=XM_023251307.1
A3	24365384	24365501	refGene_gn=TRPC4AP;refGene_tx=XM_023251307.1
A3	24372021	24372079	refGene_gn=TRPC4AP;refGene_tx=XM_023251307.1
A3	24374441	24374497	refGene_gn=TRPC4AP;refGene_tx=XM_023251307.1
A3	24377359	24377488	refGene_gn=TRPC4AP;refGene_tx=XM_023251307.1
A3	24383230	24383438	refGene_gn=TRPC4AP;refGene_tx=XM_023251307.1
A3	24391981	24392004	refGene_gn=TRPC4AP;refGene_tx=XM_023251307.1
A3	24392005	24392168	refGene_gn=TRPC4AP;refGene_tx=XM_023251307.1
A3	24402010	24402153	refGene_gn=TRPC4AP;refGene_tx=XM_023251307.1
A3	24406897	24407029	refGene_gn=TRPC4AP;refGene_tx=XM_023251307.1
A3	24408711	24408770	refGene_gn=TRPC4AP;refGene_tx=XM_023251307.1

A3 24411033 24411135 refGene_gn=TRPC4AP;refGene_tx=XM_023251307.1
A3 24412250 24412334 refGene_gn=TRPC4AP;refGene_tx=XM_023251307.1
A3 24413418 24413509 refGene_gn=TRPC4AP;refGene_tx=XM_023251307.1
A3 24414548 24414689 refGene_gn=TRPC4AP;refGene_tx=XM_023251307.1
A3 24415290 24415399 refGene_gn=TRPC4AP;refGene_tx=XM_023251307.1
A3 24416008 24416121 refGene_gn=TRPC4AP;refGene_tx=XM_023251307.1
A3 24416951 24417158 refGene_gn=TRPC4AP;refGene_tx=XM_023251307.1
A3 24417336 24417474 refGene_gn=TRPC4AP;refGene_tx=XM_023251307.1
A3 24418413 24418425
refGene_gn=MYH7B;refGene_tx=XM_003983604.5;refGene_gn=MYH7B;refGene_tx
=XM_006929822.4;refGene_gn=MYH7B;refGene_tx=XM_019826614.2
A3 24418516 24418654
refGene_gn=MYH7B;refGene_tx=XM_003983604.5;refGene_gn=MYH7B;refGene_tx
=XM_006929822.4;refGene_gn=MYH7B;refGene_tx=XM_019826614.2
A3 24419114 24419210
refGene_gn=MYH7B;refGene_tx=XM_003983604.5;refGene_gn=MYH7B;refGene_tx
=XM_006929822.4;refGene_gn=MYH7B;refGene_tx=XM_019826614.2
A3 24419303 24419408
refGene_gn=MYH7B;refGene_tx=XM_003983604.5;refGene_gn=MYH7B;refGene_tx
=XM_006929822.4;refGene_gn=MYH7B;refGene_tx=XM_019826614.2
A3 24419499 24419670
refGene_gn=MYH7B;refGene_tx=XM_003983604.5;refGene_gn=MYH7B;refGene_tx
=XM_006929822.4;refGene_gn=MYH7B;refGene_tx=XM_019826614.2
A3 24419775 24419901
refGene_gn=MYH7B;refGene_tx=XM_003983604.5;refGene_gn=MYH7B;refGene_tx
=XM_006929822.4;refGene_gn=MYH7B;refGene_tx=XM_019826614.2
A3 24419975 24420372
refGene_gn=MYH7B;refGene_tx=XM_003983604.5;refGene_gn=MYH7B;refGene_tx
=XM_006929822.4;refGene_gn=MYH7B;refGene_tx=XM_019826614.2
A3 24420849 24420965
refGene_gn=MYH7B;refGene_tx=XM_003983604.5;refGene_gn=MYH7B;refGene_tx
=XM_006929822.4;refGene_gn=MYH7B;refGene_tx=XM_019826614.2
A3 24421067 24421192
refGene_gn=MYH7B;refGene_tx=XM_003983604.5;refGene_gn=MYH7B;refGene_tx
=XM_006929822.4;refGene_gn=MYH7B;refGene_tx=XM_019826614.2
A3 24421278 24421628
refGene_gn=MYH7B;refGene_tx=XM_003983604.5;refGene_gn=MYH7B;refGene_tx
=XM_006929822.4;refGene_gn=MYH7B;refGene_tx=XM_019826614.2
A3 24421770 24421967
refGene_gn=MYH7B;refGene_tx=XM_003983604.5;refGene_gn=MYH7B;refGene_tx
=XM_006929822.4;refGene_gn=MYH7B;refGene_tx=XM_019826614.2
A3 24422058 24422177
refGene_gn=MYH7B;refGene_tx=XM_003983604.5;refGene_gn=MYH7B;refGene_tx
=XM_006929822.4;refGene_gn=MYH7B;refGene_tx=XM_019826614.2
A3 24422264 24422391
refGene_gn=MYH7B;refGene_tx=XM_003983604.5;refGene_gn=MYH7B;refGene_tx
=XM_006929822.4;refGene_gn=MYH7B;refGene_tx=XM_019826614.2
A3 24423221 24423611
refGene_gn=MYH7B;refGene_tx=XM_003983604.5;refGene_gn=MYH7B;refGene_tx
=XM_006929822.4;refGene_gn=MYH7B;refGene_tx=XM_019826614.2

A3	24423698	24423789
refGene_gn=MYH7B;refGene_tx=XM_003983604.5;refGene_gn=MYH7B;refGene_tx=XM_006929822.4;refGene_gn=MYH7B;refGene_tx=XM_019826614.2
A3	24424137	24424283
refGene_gn=MYH7B;refGene_tx=XM_003983604.5;refGene_gn=MYH7B;refGene_tx=XM_006929822.4;refGene_gn=MYH7B;refGene_tx=XM_019826614.2
A3	24424385	24424562
refGene_gn=MYH7B;refGene_tx=XM_003983604.5;refGene_gn=MYH7B;refGene_tx=XM_006929822.4;refGene_gn=MYH7B;refGene_tx=XM_019826614.2
A3	24425198	24425441
refGene_gn=MYH7B;refGene_tx=XM_003983604.5;refGene_gn=MYH7B;refGene_tx=XM_006929822.4;refGene_gn=MYH7B;refGene_tx=XM_019826614.2
A3	24426375	24426631
refGene_gn=MYH7B;refGene_tx=XM_003983604.5;refGene_gn=MYH7B;refGene_tx=XM_006929822.4;refGene_gn=MYH7B;refGene_tx=XM_019826614.2
A3	24427380	24427517
refGene_gn=MYH7B;refGene_tx=XM_003983604.5;refGene_gn=MYH7B;refGene_tx=XM_006929822.4;refGene_gn=MYH7B;refGene_tx=XM_019826614.2
A3	24427624	24427748
refGene_gn=MYH7B;refGene_tx=XM_003983604.5;refGene_gn=MYH7B;refGene_tx=XM_006929822.4;refGene_gn=MYH7B;refGene_tx=XM_019826614.2
A3	24429568	24429686
refGene_gn=MYH7B;refGene_tx=XM_003983604.5;refGene_gn=MYH7B;refGene_tx=XM_006929822.4;refGene_gn=MYH7B;refGene_tx=XM_019826614.2
A3	24429886	24429974
refGene_gn=MYH7B;refGene_tx=XM_003983604.5;refGene_gn=MYH7B;refGene_tx=XM_006929822.4;refGene_gn=MYH7B;refGene_tx=XM_019826614.2
A3	24430414	24430491
refGene_gn=MYH7B;refGene_tx=XM_003983604.5;refGene_gn=MYH7B;refGene_tx=XM_006929822.4;refGene_gn=MYH7B;refGene_tx=XM_019826614.2
A3	24430576	24430709
refGene_gn=MYH7B;refGene_tx=XM_003983604.5;refGene_gn=MYH7B;refGene_tx=XM_006929822.4;refGene_gn=MYH7B;refGene_tx=XM_019826614.2
A3	24430800	24430980
refGene_gn=MYH7B;refGene_tx=XM_003983604.5;refGene_gn=MYH7B;refGene_tx=XM_006929822.4;refGene_gn=MYH7B;refGene_tx=XM_019826614.2
A3	24432177	24432348
refGene_gn=MYH7B;refGene_tx=XM_003983604.5;refGene_gn=MYH7B;refGene_tx=XM_006929822.4;refGene_gn=MYH7B;refGene_tx=XM_019826614.2
A3	24432522	24432672
refGene_gn=MYH7B;refGene_tx=XM_003983604.5;refGene_gn=MYH7B;refGene_tx=XM_006929822.4;refGene_gn=MYH7B;refGene_tx=XM_019826614.2
A3	24432763	24432882
refGene_gn=MYH7B;refGene_tx=XM_003983604.5;refGene_gn=MYH7B;refGene_tx=XM_006929822.4;refGene_gn=MYH7B;refGene_tx=XM_019826614.2
A3	24433188	24433327
refGene_gn=MYH7B;refGene_tx=XM_003983604.5;refGene_gn=MYH7B;refGene_tx=XM_006929822.4;refGene_gn=MYH7B;refGene_tx=XM_019826614.2
A3	24433480	24433584
refGene_gn=MYH7B;refGene_tx=XM_003983604.5;refGene_gn=MYH7B;refGene_tx

=XM_006929822.4;refGene_gn=MYH7B;refGene_tx=XM_019826614.2
A3 24434088 24434187
refGene_gn=MYH7B;refGene_tx=XM_003983604.5;refGene_gn=MYH7B;refGene_tx
=XM_006929822.4;refGene_gn=MYH7B;refGene_tx=XM_019826614.2
A3 24435378 24435442
refGene_gn=MYH7B;refGene_tx=XM_003983604.5;refGene_gn=MYH7B;refGene_tx
=XM_006929822.4;refGene_gn=MYH7B;refGene_tx=XM_019826614.2
A3 24435527 24435620
refGene_gn=MYH7B;refGene_tx=XM_003983604.5;refGene_gn=MYH7B;refGene_tx
=XM_006929822.4;refGene_gn=MYH7B;refGene_tx=XM_019826614.2
A3 24435795 24435819
refGene_gn=MYH7B;refGene_tx=XM_003983604.5;refGene_gn=MYH7B;refGene_tx
=XM_006929822.4
A3 24438033 24438130
refGene_gn=MYH7B;refGene_tx=XM_003983604.5;refGene_gn=MYH7B;refGene_tx
=XM_006929822.4
A3 24439193 24439221
refGene_gn=MYH7B;refGene_tx=XM_003983604.5;refGene_gn=MYH7B;refGene_tx
=XM_006929822.4
A3 24439531 24439688
refGene_gn=MYH7B;refGene_tx=XM_003983604.5;refGene_gn=MYH7B;refGene_tx
=XM_006929822.4
A3 24440121 24440265
refGene_gn=MYH7B;refGene_tx=XM_003983604.5;refGene_gn=MYH7B;refGene_tx
=XM_006929822.4
A3 24440451 24440558 refGene_gn=MYH7B;refGene_tx=XM_003983604.5
A3 24442059 24442160
refGene_gn=MYH7B;refGene_tx=XM_003983604.5;refGene_gn=MYH7B;refGene_tx
=XM_006929822.4
A3 24443043 24443117 refGene_gn=MYH7B;refGene_tx=XM_003983604.5
A3 24465023 24465152
refGene_gn=GSS;refGene_tx=XM_006929824.2;refGene_gn=GSS;refGene_tx=XM_
019826615.1
A3 24470765 24470911
refGene_gn=GSS;refGene_tx=XM_006929824.2;refGene_gn=GSS;refGene_tx=XM_
006929825.2;refGene_gn=GSS;refGene_tx=XM_019826615.1
A3 24473764 24473840
refGene_gn=GSS;refGene_tx=XM_006929824.2;refGene_gn=GSS;refGene_tx=XM_
006929825.2;refGene_gn=GSS;refGene_tx=XM_019826615.1
A3 24474131 24474271
refGene_gn=GSS;refGene_tx=XM_006929824.2;refGene_gn=GSS;refGene_tx=XM_
006929825.2;refGene_gn=GSS;refGene_tx=XM_019826615.1
A3 24474951 24475068
refGene_gn=GSS;refGene_tx=XM_006929824.2;refGene_gn=GSS;refGene_tx=XM_
006929825.2;refGene_gn=GSS;refGene_tx=XM_019826615.1
A3 24479791 24479872
refGene_gn=GSS;refGene_tx=XM_006929824.2;refGene_gn=GSS;refGene_tx=XM_
006929825.2;refGene_gn=GSS;refGene_tx=XM_019826615.1
A3 24479974 24480052
refGene_gn=GSS;refGene_tx=XM_006929824.2;refGene_gn=GSS;refGene_tx=XM_

006929825.2;refGene_gn=GSS;refGene_tx=XM_019826615.1
A3	24482645	24482712
refGene_gn=GSS;refGene_tx=XM_006929824.2;refGene_gn=GSS;refGene_tx=XM_006929825.2;refGene_gn=GSS;refGene_tx=XM_019826615.1
A3	24485922	24486117
refGene_gn=GSS;refGene_tx=XM_006929824.2;refGene_gn=GSS;refGene_tx=XM_006929825.2;refGene_gn=GSS;refGene_tx=XM_019826615.1
A3	24486521	24486603
refGene_gn=GSS;refGene_tx=XM_006929824.2;refGene_gn=GSS;refGene_tx=XM_006929825.2;refGene_gn=GSS;refGene_tx=XM_019826615.1
A3	24488019	24488209
refGene_gn=GSS;refGene_tx=XM_006929824.2;refGene_gn=GSS;refGene_tx=XM_006929825.2;refGene_gn=GSS;refGene_tx=XM_019826615.1
A3	24488782	24488882	refGene_gn=GSS;refGene_tx=XM_019826615.1
A3	24489190	24489314
refGene_gn=GSS;refGene_tx=XM_006929824.2;refGene_gn=GSS;refGene_tx=XM_006929825.2
A3	24490946	24491074
refGene_gn=ACSS2;refGene_tx=XM_003983588.5;refGene_gn=ACSS2;refGene_tx=XM_003983590.5
A3	24491217	24491292
refGene_gn=ACSS2;refGene_tx=XM_003983588.5;refGene_gn=ACSS2;refGene_tx=XM_003983590.5
A3	24492033	24492210
refGene_gn=ACSS2;refGene_tx=XM_003983588.5;refGene_gn=ACSS2;refGene_tx=XM_003983590.5
A3	24492547	24492616
refGene_gn=ACSS2;refGene_tx=XM_003983588.5;refGene_gn=ACSS2;refGene_tx=XM_003983590.5
A3	24494192	24494301
refGene_gn=ACSS2;refGene_tx=XM_003983588.5;refGene_gn=ACSS2;refGene_tx=XM_003983590.5
A3	24495904	24495985
refGene_gn=ACSS2;refGene_tx=XM_003983588.5;refGene_gn=ACSS2;refGene_tx=XM_003983590.5
A3	24496180	24496237
refGene_gn=ACSS2;refGene_tx=XM_003983588.5;refGene_gn=ACSS2;refGene_tx=XM_003983590.5
A3	24496318	24496451
refGene_gn=ACSS2;refGene_tx=XM_003983588.5;refGene_gn=ACSS2;refGene_tx=XM_003983590.5
A3	24496646	24496780
refGene_gn=ACSS2;refGene_tx=XM_003983588.5;refGene_gn=ACSS2;refGene_tx=XM_003983590.5
A3	24497089	24497260
refGene_gn=ACSS2;refGene_tx=XM_003983588.5;refGene_gn=ACSS2;refGene_tx=XM_003983590.5
A3	24498706	24498844
refGene_gn=ACSS2;refGene_tx=XM_003983588.5;refGene_gn=ACSS2;refGene_tx=XM_003983590.5

A3 24504951 24504990 refGene_gn=ACSS2;refGene_tx=XM_003983590.5
A3 24505902 24506017
refGene_gn=ACSS2;refGene_tx=XM_003983588.5;refGene_gn=ACSS2;refGene_tx=XM_003983590.5
A3 24506171 24506247
refGene_gn=ACSS2;refGene_tx=XM_003983588.5;refGene_gn=ACSS2;refGene_tx=XM_003983590.5
A3 24506528 24506601
refGene_gn=ACSS2;refGene_tx=XM_003983588.5;refGene_gn=ACSS2;refGene_tx=XM_003983590.5
A3 24506856 24506960
refGene_gn=ACSS2;refGene_tx=XM_003983588.5;refGene_gn=ACSS2;refGene_tx=XM_003983590.5
A3 24507163 24507255
refGene_gn=ACSS2;refGene_tx=XM_003983588.5;refGene_gn=ACSS2;refGene_tx=XM_003983590.5
A3 24536508 24536704
refGene_gn=ACSS2;refGene_tx=XM_003983588.5;refGene_gn=ACSS2;refGene_tx=XM_003983590.5
A3 24542304 24542494
refGene_gn=ACSS2;refGene_tx=XM_003983588.5;refGene_gn=ACSS2;refGene_tx=XM_003983590.5
A3 24545497 24545666
refGene_gn=GGT7;refGene_tx=XM_003983592.4;refGene_gn=GGT7;refGene_tx=XM_019826616.1;refGene_gn=GGT7;refGene_tx=XM_019826617.1;refGene_gn=GGT7;refGene_tx=XM_023251308.1
A3 24551904 24552140
refGene_gn=GGT7;refGene_tx=XM_003983592.4;refGene_gn=GGT7;refGene_tx=XM_019826616.1;refGene_gn=GGT7;refGene_tx=XM_019826617.1;refGene_gn=GGT7;refGene_tx=XM_023251308.1
A3 24552505 24552657
refGene_gn=GGT7;refGene_tx=XM_003983592.4;refGene_gn=GGT7;refGene_tx=XM_019826616.1;refGene_gn=GGT7;refGene_tx=XM_019826617.1;refGene_gn=GGT7;refGene_tx=XM_023251308.1
A3 24554573 24554691
refGene_gn=GGT7;refGene_tx=XM_003983592.4;refGene_gn=GGT7;refGene_tx=XM_019826616.1;refGene_gn=GGT7;refGene_tx=XM_019826617.1;refGene_gn=GGT7;refGene_tx=XM_023251308.1
A3 24555541 24555609
refGene_gn=GGT7;refGene_tx=XM_003983592.4;refGene_gn=GGT7;refGene_tx=XM_019826617.1;refGene_gn=GGT7;refGene_tx=XM_023251308.1
A3 24555830 24555939
refGene_gn=GGT7;refGene_tx=XM_003983592.4;refGene_gn=GGT7;refGene_tx=XM_019826616.1;refGene_gn=GGT7;refGene_tx=XM_019826617.1;refGene_gn=GGT7;refGene_tx=XM_023251308.1
A3 24556254 24556451
refGene_gn=GGT7;refGene_tx=XM_003983592.4;refGene_gn=GGT7;refGene_tx=XM_019826616.1;refGene_gn=GGT7;refGene_tx=XM_019826617.1;refGene_gn=GGT7;refGene_tx=XM_023251308.1
A3 24558463 24558551

refGene_gn=GGT7;refGene_tx=XM_003983592.4;refGene_gn=GGT7;refGene_tx=XM_019826616.1;refGene_gn=GGT7;refGene_tx=XM_019826617.1;refGene_gn=GGT7;refGene_tx=XM_023251308.1
A3 24560044 24560172
refGene_gn=GGT7;refGene_tx=XM_003983592.4;refGene_gn=GGT7;refGene_tx=XM_019826616.1;refGene_gn=GGT7;refGene_tx=XM_019826617.1;refGene_gn=GGT7;refGene_tx=XM_023251308.1
A3 24560344 24560433
refGene_gn=GGT7;refGene_tx=XM_003983592.4;refGene_gn=GGT7;refGene_tx=XM_019826616.1;refGene_gn=GGT7;refGene_tx=XM_019826617.1;refGene_gn=GGT7;refGene_tx=XM_023251308.1
A3 24562095 24562245
refGene_gn=GGT7;refGene_tx=XM_003983592.4;refGene_gn=GGT7;refGene_tx=XM_019826616.1;refGene_gn=GGT7;refGene_tx=XM_019826617.1;refGene_gn=GGT7;refGene_tx=XM_023251308.1
A3 24562362 24562480
refGene_gn=GGT7;refGene_tx=XM_003983592.4;refGene_gn=GGT7;refGene_tx=XM_019826616.1;refGene_gn=GGT7;refGene_tx=XM_019826617.1;refGene_gn=GGT7;refGene_tx=XM_023251308.1
A3 24563173 24563311
refGene_gn=GGT7;refGene_tx=XM_003983592.4;refGene_gn=GGT7;refGene_tx=XM_019826616.1;refGene_gn=GGT7;refGene_tx=XM_019826617.1;refGene_gn=GGT7;refGene_tx=XM_023251308.1
A3 24564460 24564560
refGene_gn=GGT7;refGene_tx=XM_003983592.4;refGene_gn=GGT7;refGene_tx=XM_019826616.1;refGene_gn=GGT7;refGene_tx=XM_019826617.1;refGene_gn=GGT7;refGene_tx=XM_023251308.1
A3 24567299 24567322 refGene_gn=GGT7;refGene_tx=XM_019826617.1
A3 24568241 24568405
refGene_gn=GGT7;refGene_tx=XM_003983592.4;refGene_gn=GGT7;refGene_tx=XM_019826616.1;refGene_gn=GGT7;refGene_tx=XM_023251308.1
A3 24627572 24627807
refGene_gn=NCOA6;refGene_tx=XM_011280858.3;refGene_gn=NCOA6;refGene_tx=XM_011280860.3;refGene_gn=NCOA6;refGene_tx=XM_011280862.3;refGene_gn=NCOA6;refGene_tx=XM_011280865.3
A3 24634738 24634894
refGene_gn=NCOA6;refGene_tx=XM_011280858.3;refGene_gn=NCOA6;refGene_tx=XM_011280860.3;refGene_gn=NCOA6;refGene_tx=XM_011280862.3;refGene_gn=NCOA6;refGene_tx=XM_011280865.3
A3 24641338 24641461
refGene_gn=NCOA6;refGene_tx=XM_011280858.3;refGene_gn=NCOA6;refGene_tx=XM_011280860.3;refGene_gn=NCOA6;refGene_tx=XM_011280862.3;refGene_gn=NCOA6;refGene_tx=XM_011280865.3
A3 24650545 24650674
refGene_gn=NCOA6;refGene_tx=XM_011280858.3;refGene_gn=NCOA6;refGene_tx=XM_011280860.3;refGene_gn=NCOA6;refGene_tx=XM_011280862.3;refGene_gn=NCOA6;refGene_tx=XM_011280865.3
A3 24651379 24652288
refGene_gn=NCOA6;refGene_tx=XM_011280858.3;refGene_gn=NCOA6;refGene_tx=XM_011280860.3;refGene_gn=NCOA6;refGene_tx=XM_011280862.3;refGene_gn=

NCOA6;refGene_tx=XM_011280865.3
A3 24654331 24654478
refGene_gn=NCOA6;refGene_tx=XM_011280858.3;refGene_gn=NCOA6;refGene_tx=XM_011280860.3;refGene_gn=NCOA6;refGene_tx=XM_011280862.3;refGene_gn=NCOA6;refGene_tx=XM_011280865.3
A3 24657333 24658450
refGene_gn=NCOA6;refGene_tx=XM_011280858.3;refGene_gn=NCOA6;refGene_tx=XM_011280860.3;refGene_gn=NCOA6;refGene_tx=XM_011280862.3;refGene_gn=NCOA6;refGene_tx=XM_011280865.3
A3 24661103 24661225
refGene_gn=NCOA6;refGene_tx=XM_011280858.3;refGene_gn=NCOA6;refGene_tx=XM_011280860.3;refGene_gn=NCOA6;refGene_tx=XM_011280862.3;refGene_gn=NCOA6;refGene_tx=XM_011280865.3
A3 24664885 24667891
refGene_gn=NCOA6;refGene_tx=XM_011280858.3;refGene_gn=NCOA6;refGene_tx=XM_011280860.3;refGene_gn=NCOA6;refGene_tx=XM_011280862.3
A3 24670674 24670743
refGene_gn=NCOA6;refGene_tx=XM_011280858.3;refGene_gn=NCOA6;refGene_tx=XM_011280860.3;refGene_gn=NCOA6;refGene_tx=XM_011280862.3;refGene_gn=NCOA6;refGene_tx=XM_011280865.3
A3 24673511 24673548
refGene_gn=NCOA6;refGene_tx=XM_011280858.3;refGene_gn=NCOA6;refGene_tx=XM_011280860.3;refGene_gn=NCOA6;refGene_tx=XM_011280862.3;refGene_gn=NCOA6;refGene_tx=XM_011280865.3
A3 24676994 24677143
refGene_gn=NCOA6;refGene_tx=XM_011280858.3;refGene_gn=NCOA6;refGene_tx=XM_011280860.3;refGene_gn=NCOA6;refGene_tx=XM_011280862.3;refGene_gn=NCOA6;refGene_tx=XM_011280865.3
A3 24684795 24684839
refGene_gn=NCOA6;refGene_tx=XM_011280858.3;refGene_gn=NCOA6;refGene_tx=XM_011280860.3;refGene_gn=NCOA6;refGene_tx=XM_011280862.3;refGene_gn=NCOA6;refGene_tx=XM_011280865.3
A3 24689801 24690054
refGene_gn=TP53INP2;refGene_tx=XM_023251309.1;refGene_gn=TP53INP2;refGene_tx=XM_023251310.1;refGene_gn=TP53INP2;refGene_tx=XM_023251311.1;refGene_gn=TP53INP2;refGene_tx=XM_023251312.1
A3 24690642 24690931
refGene_gn=TP53INP2;refGene_tx=XM_023251309.1;refGene_gn=TP53INP2;refGene_tx=XM_023251310.1;refGene_gn=TP53INP2;refGene_tx=XM_023251311.1;refGene_gn=TP53INP2;refGene_tx=XM_023251312.1
A3 24691333 24691457
refGene_gn=TP53INP2;refGene_tx=XM_023251309.1;refGene_gn=TP53INP2;refGene_tx=XM_023251310.1;refGene_gn=TP53INP2;refGene_tx=XM_023251311.1;refGene_gn=TP53INP2;refGene_tx=XM_023251312.1
A3 24721344 24721474 refGene_gn=PIGU;refGene_tx=XM_003983593.5
A3 24732321 24732386 refGene_gn=PIGU;refGene_tx=XM_003983593.5
A3 24744007 24744067
refGene_gn=PIGU;refGene_tx=XM_003983593.5;refGene_gn=PIGU;refGene_tx=XM_019826620.2
A3 24744908 24744971

refGene_gn=PIGU;refGene_tx=XM_003983593.5;refGene_gn=PIGU;refGene_tx=XM_019826620.2
A3 24751925 24752035
refGene_gn=PIGU;refGene_tx=XM_003983593.5;refGene_gn=PIGU;refGene_tx=XM_019826620.2
A3 24754241 24754342
refGene_gn=PIGU;refGene_tx=XM_003983593.5;refGene_gn=PIGU;refGene_tx=XM_019826620.2
A3 24771864 24771962
refGene_gn=PIGU;refGene_tx=XM_003983593.5;refGene_gn=PIGU;refGene_tx=XM_019826620.2
A3 24779152 24779307
refGene_gn=PIGU;refGene_tx=XM_003983593.5;refGene_gn=PIGU;refGene_tx=XM_019826620.2
A3 24780938 24781082
refGene_gn=PIGU;refGene_tx=XM_003983593.5;refGene_gn=PIGU;refGene_tx=XM_019826620.2
A3 24783769 24783894
refGene_gn=PIGU;refGene_tx=XM_003983593.5;refGene_gn=PIGU;refGene_tx=XM_019826620.2
A3 24790774 24790917
refGene_gn=PIGU;refGene_tx=XM_003983593.5;refGene_gn=PIGU;refGene_tx=XM_019826620.2
A3 24805309 24805423
refGene_gn=PIGU;refGene_tx=XM_003983593.5;refGene_gn=PIGU;refGene_tx=XM_019826620.2
A3 24806426 24806589 refGene_gn=MAP1LC3A;refGene_tx=XM_003983595.4
A3 24806853 24806960 refGene_gn=MAP1LC3A;refGene_tx=XM_003983595.4
A3 24807036 24807092 refGene_gn=MAP1LC3A;refGene_tx=XM_003983595.4
A3 24807389 24807429 refGene_gn=MAP1LC3A;refGene_tx=XM_003983595.4
A3 24826484 24826528
refGene_gn=DYNLRB1;refGene_tx=XM_003983596.5;refGene_gn=DYNLRB1;refGene_tx=XM_011280855.2;refGene_gn=DYNLRB1;refGene_tx=XM_011280856.3;refGene_gn=DYNLRB1;refGene_tx=XM_023251313.1
A3 24831370 24831538
refGene_gn=DYNLRB1;refGene_tx=XM_003983596.5;refGene_gn=DYNLRB1;refGene_tx=XM_011280855.2;refGene_gn=DYNLRB1;refGene_tx=XM_011280856.3;refGene_gn=DYNLRB1;refGene_tx=XM_023251313.1
A3 24837997 24838073
refGene_gn=DYNLRB1;refGene_tx=XM_003983596.5;refGene_gn=DYNLRB1;refGene_tx=XM_011280855.2
A3 24842761 24842809 refGene_gn=DYNLRB1;refGene_tx=XM_011280855.2
A3 24846020 24846023 refGene_gn=DYNLRB1;refGene_tx=XM_003983596.5
A3 24852038 24852138
refGene_gn=ITCH;refGene_tx=XM_003983597.5;refGene_gn=ITCH;refGene_tx=XM_006929828.4;refGene_gn=ITCH;refGene_tx=XM_011280854.3;refGene_gn=ITCH;refGene_tx=XM_019826622.1
A3 24852811 24852884
refGene_gn=ITCH;refGene_tx=XM_003983597.5;refGene_gn=ITCH;refGene_tx=XM_006929828.4;refGene_gn=ITCH;refGene_tx=XM_011280854.3;refGene_gn=ITC

H;refGene_tx=XM_019826622.1
A3 24856641 24856738
refGene_gn=ITCH;refGene_tx=XM_003983597.5;refGene_gn=ITCH;refGene_tx=XM_006929828.4;refGene_gn=ITCH;refGene_tx=XM_011280854.3;refGene_gn=ITCH;refGene_tx=XM_019826622.1
A3 24859529 24859634
refGene_gn=ITCH;refGene_tx=XM_003983597.5;refGene_gn=ITCH;refGene_tx=XM_006929828.4;refGene_gn=ITCH;refGene_tx=XM_011280854.3;refGene_gn=ITCH;refGene_tx=XM_019826622.1
A3 24860078 24860199
refGene_gn=ITCH;refGene_tx=XM_003983597.5;refGene_gn=ITCH;refGene_tx=XM_006929828.4;refGene_gn=ITCH;refGene_tx=XM_011280854.3;refGene_gn=ITCH;refGene_tx=XM_019826622.1
A3 24865539 24865680
refGene_gn=ITCH;refGene_tx=XM_003983597.5;refGene_gn=ITCH;refGene_tx=XM_006929828.4;refGene_gn=ITCH;refGene_tx=XM_011280854.3;refGene_gn=ITCH;refGene_tx=XM_019826622.1
A3 24866010 24866144
refGene_gn=ITCH;refGene_tx=XM_003983597.5;refGene_gn=ITCH;refGene_tx=XM_006929828.4;refGene_gn=ITCH;refGene_tx=XM_011280854.3;refGene_gn=ITCH;refGene_tx=XM_019826622.1
A3 24866379 24866539
refGene_gn=ITCH;refGene_tx=XM_003983597.5;refGene_gn=ITCH;refGene_tx=XM_006929828.4;refGene_gn=ITCH;refGene_tx=XM_011280854.3;refGene_gn=ITCH;refGene_tx=XM_019826622.1
A3 24868494 24868583
refGene_gn=ITCH;refGene_tx=XM_003983597.5;refGene_gn=ITCH;refGene_tx=XM_006929828.4;refGene_gn=ITCH;refGene_tx=XM_011280854.3;refGene_gn=ITCH;refGene_tx=XM_019826622.1
A3 24873109 24873181
refGene_gn=ITCH;refGene_tx=XM_003983597.5;refGene_gn=ITCH;refGene_tx=XM_006929828.4;refGene_gn=ITCH;refGene_tx=XM_011280854.3;refGene_gn=ITCH;refGene_tx=XM_019826622.1
A3 24874057 24874130
refGene_gn=ITCH;refGene_tx=XM_003983597.5;refGene_gn=ITCH;refGene_tx=XM_006929828.4;refGene_gn=ITCH;refGene_tx=XM_011280854.3;refGene_gn=ITCH;refGene_tx=XM_019826622.1
A3 24880678 24880807
refGene_gn=ITCH;refGene_tx=XM_003983597.5;refGene_gn=ITCH;refGene_tx=XM_006929828.4;refGene_gn=ITCH;refGene_tx=XM_011280854.3;refGene_gn=ITCH;refGene_tx=XM_019826622.1
A3 24885138 24885223
refGene_gn=ITCH;refGene_tx=XM_003983597.5;refGene_gn=ITCH;refGene_tx=XM_006929828.4;refGene_gn=ITCH;refGene_tx=XM_011280854.3;refGene_gn=ITCH;refGene_tx=XM_019826622.1
A3 24890701 24890771
refGene_gn=ITCH;refGene_tx=XM_003983597.5;refGene_gn=ITCH;refGene_tx=XM_006929828.4;refGene_gn=ITCH;refGene_tx=XM_011280854.3;refGene_gn=ITCH;refGene_tx=XM_019826622.1
A3 24895364 24895539

refGene_gn=ITCH;refGene_tx=XM_003983597.5;refGene_gn=ITCH;refGene_tx=XM_006929828.4;refGene_gn=ITCH;refGene_tx=XM_011280854.3;refGene_gn=ITCH;refGene_tx=XM_019826622.1
A3 24896773 24896869
refGene_gn=ITCH;refGene_tx=XM_003983597.5;refGene_gn=ITCH;refGene_tx=XM_006929828.4;refGene_gn=ITCH;refGene_tx=XM_011280854.3;refGene_gn=ITCH;refGene_tx=XM_019826622.1
A3 24899172 24899362
refGene_gn=ITCH;refGene_tx=XM_003983597.5;refGene_gn=ITCH;refGene_tx=XM_006929828.4;refGene_gn=ITCH;refGene_tx=XM_011280854.3;refGene_gn=ITCH;refGene_tx=XM_019826622.1
A3 24900710 24900868
refGene_gn=ITCH;refGene_tx=XM_003983597.5;refGene_gn=ITCH;refGene_tx=XM_006929828.4;refGene_gn=ITCH;refGene_tx=XM_011280854.3;refGene_gn=ITCH;refGene_tx=XM_019826622.1
A3 24916199 24916245
refGene_gn=ITCH;refGene_tx=XM_003983597.5;refGene_gn=ITCH;refGene_tx=XM_006929828.4;refGene_gn=ITCH;refGene_tx=XM_011280854.3;refGene_gn=ITCH;refGene_tx=XM_019826622.1
A3 24925858 24925996
refGene_gn=ITCH;refGene_tx=XM_003983597.5;refGene_gn=ITCH;refGene_tx=XM_006929828.4;refGene_gn=ITCH;refGene_tx=XM_011280854.3;refGene_gn=ITCH;refGene_tx=XM_019826622.1
A3 24927576 24927701
refGene_gn=ITCH;refGene_tx=XM_003983597.5;refGene_gn=ITCH;refGene_tx=XM_006929828.4;refGene_gn=ITCH;refGene_tx=XM_011280854.3;refGene_gn=ITCH;refGene_tx=XM_019826622.1
A3 24932371 24932513
refGene_gn=ITCH;refGene_tx=XM_003983597.5;refGene_gn=ITCH;refGene_tx=XM_006929828.4;refGene_gn=ITCH;refGene_tx=XM_011280854.3
A3 24949533 24949603
refGene_gn=ITCH;refGene_tx=XM_003983597.5;refGene_gn=ITCH;refGene_tx=XM_006929828.4;refGene_gn=ITCH;refGene_tx=XM_011280854.3
A3 25053070 25053098 refGene_gn=AHCY;refGene_tx=XM_003983598.4
A3 25058717 25058908
refGene_gn=AHCY;refGene_tx=XM_003983598.4;refGene_gn=AHCY;refGene_tx=XM_006929829.4
A3 25060036 25060112
refGene_gn=AHCY;refGene_tx=XM_003983598.4;refGene_gn=AHCY;refGene_tx=XM_006929829.4
A3 25061574 25061724
refGene_gn=AHCY;refGene_tx=XM_003983598.4;refGene_gn=AHCY;refGene_tx=XM_006929829.4
A3 25062629 25062742
refGene_gn=AHCY;refGene_tx=XM_003983598.4;refGene_gn=AHCY;refGene_tx=XM_006929829.4
A3 25063174 25063382
refGene_gn=AHCY;refGene_tx=XM_003983598.4;refGene_gn=AHCY;refGene_tx=XM_006929829.4
A3 25063489 25063577

refGene_gn=AHCY;refGene_tx=XM_003983598.4;refGene_gn=AHCY;refGene_tx=XM_006929829.4
A3 25063670 25063788
refGene_gn=AHCY;refGene_tx=XM_003983598.4;refGene_gn=AHCY;refGene_tx=XM_006929829.4
A3 25064827 25065022
refGene_gn=AHCY;refGene_tx=XM_003983598.4;refGene_gn=AHCY;refGene_tx=XM_006929829.4
A3 25066049 25066181
refGene_gn=AHCY;refGene_tx=XM_003983598.4;refGene_gn=AHCY;refGene_tx=XM_006929829.4
A3 25082292 25082478
refGene_gn=ASIP;refGene_tx=NM_001009190.1;refGene_gn=ASIP;refGene_tx=XM_006929783.3;refGene_gn=ASIP;refGene_tx=XM_019826162.2;refGene_gn=ASIP;refGene_tx=XM_019826165.2
A3 25084806 25084868
refGene_gn=ASIP;refGene_tx=NM_001009190.1;refGene_gn=ASIP;refGene_tx=XM_006929783.3;refGene_gn=ASIP;refGene_tx=XM_019826162.2;refGene_gn=ASIP;refGene_tx=XM_019826165.2
A3 25086529 25086699
refGene_gn=ASIP;refGene_tx=NM_001009190.1;refGene_gn=ASIP;refGene_tx=XM_006929783.3;refGene_gn=ASIP;refGene_tx=XM_019826162.2;refGene_gn=ASIP;refGene_tx=XM_019826165.2
A3 25149418 25149501 refGene_gn=ASIP;refGene_tx=XM_019826162.2
A3 25149904 25149952 refGene_gn=ASIP;refGene_tx=XM_019826162.2
A3 25246436 25246451 refGene_gn=EIF2S2;refGene_tx=XM_011280853.3
A3 25246743 25247001 refGene_gn=EIF2S2;refGene_tx=XM_023251314.1
A3 25252005 25252183
refGene_gn=EIF2S2;refGene_tx=XM_011280853.3;refGene_gn=EIF2S2;refGene_tx=XM_023251314.1
A3 25253342 25253446
refGene_gn=EIF2S2;refGene_tx=XM_011280853.3;refGene_gn=EIF2S2;refGene_tx=XM_023251314.1
A3 25257325 25257461
refGene_gn=EIF2S2;refGene_tx=XM_011280853.3;refGene_gn=EIF2S2;refGene_tx=XM_023251314.1
A3 25258111 25258212
refGene_gn=EIF2S2;refGene_tx=XM_011280853.3;refGene_gn=EIF2S2;refGene_tx=XM_023251314.1
A3 25258774 25258923
refGene_gn=EIF2S2;refGene_tx=XM_011280853.3;refGene_gn=EIF2S2;refGene_tx=XM_023251314.1
A3 25260914 25260971
refGene_gn=EIF2S2;refGene_tx=XM_011280853.3;refGene_gn=EIF2S2;refGene_tx=XM_023251314.1
A3 25263138 25263224
refGene_gn=EIF2S2;refGene_tx=XM_011280853.3;refGene_gn=EIF2S2;refGene_tx=XM_023251314.1
A3 25263811 25263987
refGene_gn=EIF2S2;refGene_tx=XM_011280853.3;refGene_gn=EIF2S2;refGene_

tx=XM_023251314.1
A3	25274705 25275089
refGene_gn=RALY;refGene_tx=XM_019826623.2;refGene_gn=RALY;refGene_tx=XM_019826624.2;refGene_gn=RALY;refGene_tx=XM_019826625.1;refGene_gn=RALY;refGene_tx=XM_019826628.2;refGene_gn=RALY;refGene_tx=XM_019826629.2;refGene_gn=RALY;refGene_tx=XM_023251316.1
A3	25276278 25276323
refGene_gn=RALY;refGene_tx=XM_003983608.5;refGene_gn=RALY;refGene_tx=XM_006929833.4
A3	25277513 25277704
refGene_gn=RALY;refGene_tx=XM_003983608.5;refGene_gn=RALY;refGene_tx=XM_006929833.4;refGene_gn=RALY;refGene_tx=XM_019826623.2;refGene_gn=RALY;refGene_tx=XM_019826624.2;refGene_gn=RALY;refGene_tx=XM_019826625.1;refGene_gn=RALY;refGene_tx=XM_019826628.2;refGene_gn=RALY;refGene_tx=XM_019826629.2;refGene_gn=RALY;refGene_tx=XM_023251316.1
A3	25277911 25278025
refGene_gn=RALY;refGene_tx=XM_003983608.5;refGene_gn=RALY;refGene_tx=XM_006929833.4;refGene_gn=RALY;refGene_tx=XM_019826623.2;refGene_gn=RALY;refGene_tx=XM_019826624.2;refGene_gn=RALY;refGene_tx=XM_019826625.1;refGene_gn=RALY;refGene_tx=XM_019826628.2;refGene_gn=RALY;refGene_tx=XM_019826629.2;refGene_gn=RALY;refGene_tx=XM_023251316.1
A3	25278747 25278914
refGene_gn=RALY;refGene_tx=XM_003983608.5;refGene_gn=RALY;refGene_tx=XM_006929833.4;refGene_gn=RALY;refGene_tx=XM_019826623.2;refGene_gn=RALY;refGene_tx=XM_019826624.2;refGene_gn=RALY;refGene_tx=XM_019826625.1;refGene_gn=RALY;refGene_tx=XM_019826628.2;refGene_gn=RALY;refGene_tx=XM_019826629.2;refGene_gn=RALY;refGene_tx=XM_023251316.1
A3	25280976 25281024
refGene_gn=RALY;refGene_tx=XM_006929833.4;refGene_gn=RALY;refGene_tx=XM_019826623.2;refGene_gn=RALY;refGene_tx=XM_019826624.2;refGene_gn=RALY;refGene_tx=XM_019826625.1;refGene_gn=RALY;refGene_tx=XM_019826628.2;refGene_gn=RALY;refGene_tx=XM_023251316.1
A3	25281223 25281296
refGene_gn=RALY;refGene_tx=XM_003983608.5;refGene_gn=RALY;refGene_tx=XM_006929833.4;refGene_gn=RALY;refGene_tx=XM_019826623.2;refGene_gn=RALY;refGene_tx=XM_019826624.2;refGene_gn=RALY;refGene_tx=XM_019826625.1;refGene_gn=RALY;refGene_tx=XM_019826628.2;refGene_gn=RALY;refGene_tx=XM_019826629.2;refGene_gn=RALY;refGene_tx=XM_023251316.1
A3	25282703 25282959
refGene_gn=RALY;refGene_tx=XM_003983608.5;refGene_gn=RALY;refGene_tx=XM_006929833.4;refGene_gn=RALY;refGene_tx=XM_019826623.2;refGene_gn=RALY;refGene_tx=XM_019826624.2;refGene_gn=RALY;refGene_tx=XM_019826625.1;refGene_gn=RALY;refGene_tx=XM_019826628.2;refGene_gn=RALY;refGene_tx=XM_019826629.2;refGene_gn=RALY;refGene_tx=XM_023251316.1
A3	25472552 25472617 refGene_gn=CHMP4B;refGene_tx=XM_006929835.3
A3	25473822 25473949 refGene_gn=CHMP4B;refGene_tx=XM_006929835.3
A3	25475030 25475145 refGene_gn=CHMP4B;refGene_tx=XM_006929835.3
A3	25477369 25477547 refGene_gn=CHMP4B;refGene_tx=XM_006929835.3
A3	25525994 25526184 refGene_gn=CHMP4B;refGene_tx=XM_006929835.3
A3	25538764 25539294

refGene_gn=ZNF341;refGene_tx=XM_023251317.1;refGene_gn=ZNF341;refGene_tx=XM_023251318.1;refGene_gn=ZNF341;refGene_tx=XM_023251319.1
A3 25541432 25541503
refGene_gn=ZNF341;refGene_tx=XM_023251317.1;refGene_gn=ZNF341;refGene_tx=XM_023251318.1;refGene_gn=ZNF341;refGene_tx=XM_023251319.1
A3 25541759 25541871
refGene_gn=ZNF341;refGene_tx=XM_023251317.1;refGene_gn=ZNF341;refGene_tx=XM_023251318.1;refGene_gn=ZNF341;refGene_tx=XM_023251319.1
A3 25545095 25545228
refGene_gn=ZNF341;refGene_tx=XM_023251317.1;refGene_gn=ZNF341;refGene_tx=XM_023251318.1;refGene_gn=ZNF341;refGene_tx=XM_023251319.1
A3 25548262 25548359
refGene_gn=ZNF341;refGene_tx=XM_023251317.1;refGene_gn=ZNF341;refGene_tx=XM_023251318.1;refGene_gn=ZNF341;refGene_tx=XM_023251319.1
A3 25556114 25556323
refGene_gn=ZNF341;refGene_tx=XM_023251317.1;refGene_gn=ZNF341;refGene_tx=XM_023251318.1;refGene_gn=ZNF341;refGene_tx=XM_023251319.1
A3 25560908 25561099
refGene_gn=ZNF341;refGene_tx=XM_023251317.1;refGene_gn=ZNF341;refGene_tx=XM_023251318.1;refGene_gn=ZNF341;refGene_tx=XM_023251319.1
A3 25561335 25561529
refGene_gn=ZNF341;refGene_tx=XM_023251317.1;refGene_gn=ZNF341;refGene_tx=XM_023251318.1;refGene_gn=ZNF341;refGene_tx=XM_023251319.1
A3 25563461 25563552
refGene_gn=ZNF341;refGene_tx=XM_023251317.1;refGene_gn=ZNF341;refGene_tx=XM_023251318.1;refGene_gn=ZNF341;refGene_tx=XM_023251319.1
A3 25564913 25565109
refGene_gn=ZNF341;refGene_tx=XM_023251317.1;refGene_gn=ZNF341;refGene_tx=XM_023251318.1;refGene_gn=ZNF341;refGene_tx=XM_023251319.1
A3 25568887 25569139
refGene_gn=ZNF341;refGene_tx=XM_023251317.1;refGene_gn=ZNF341;refGene_tx=XM_023251318.1
A3 25572536 25572686
refGene_gn=ZNF341;refGene_tx=XM_023251317.1;refGene_gn=ZNF341;refGene_tx=XM_023251318.1;refGene_gn=ZNF341;refGene_tx=XM_023251319.1
A3 25575857 25576054 refGene_gn=ZNF341;refGene_tx=XM_023251318.1
A3 25577689 25577800
refGene_gn=ZNF341;refGene_tx=XM_023251317.1;refGene_gn=ZNF341;refGene_tx=XM_023251318.1;refGene_gn=ZNF341;refGene_tx=XM_023251319.1
A3 25583735 25583766 refGene_gn=ZNF341;refGene_tx=XM_023251318.1
A3 25589498 25589611
refGene_gn=PXMP4;refGene_tx=XM_003983645.4;refGene_gn=PXMP4;refGene_tx=XM_006929836.3;refGene_gn=PXMP4;refGene_tx=XM_006929837.4;refGene_gn=PXMP4;refGene_tx=XM_006929840.3;refGene_gn=PXMP4;refGene_tx=XM_011280849.2
A3 25595704 25595767
refGene_gn=PXMP4;refGene_tx=XM_003983645.4;refGene_gn=PXMP4;refGene_tx=XM_006929836.3;refGene_gn=PXMP4;refGene_tx=XM_006929837.4;refGene_gn=PXMP4;refGene_tx=XM_006929840.3;refGene_gn=PXMP4;refGene_tx=XM_011280849.2

A3 25598577 25598776
refGene_gn=PXMP4;refGene_tx=XM_003983645.4;refGene_gn=PXMP4;refGene_tx
=XM_006929836.3;refGene_gn=PXMP4;refGene_tx=XM_011280849.2
A3 25602709 25602973
refGene_gn=PXMP4;refGene_tx=XM_003983645.4;refGene_gn=PXMP4;refGene_tx
=XM_006929836.3;refGene_gn=PXMP4;refGene_tx=XM_006929837.4;refGene_gn=
PXMP4;refGene_tx=XM_006929840.3;refGene_gn=PXMP4;refGene_tx=XM_0112808
49.2
A3 25607316 25607396
refGene_gn=PXMP4;refGene_tx=XM_003983645.4;refGene_gn=PXMP4;refGene_tx
=XM_006929840.3
A3 25610104 25610316
refGene_gn=PXMP4;refGene_tx=XM_006929836.3;refGene_gn=PXMP4;refGene_tx
=XM_006929837.4
A3 25615581 25615842 refGene_gn=E2F1;refGene_tx=XM_023251320.1
A3 25621172 25621263 refGene_gn=E2F1;refGene_tx=XM_023251320.1
A3 25621637 25621857 refGene_gn=E2F1;refGene_tx=XM_023251320.1
A3 25622684 25622837 refGene_gn=E2F1;refGene_tx=XM_023251320.1
A3 25623745 25623860 refGene_gn=E2F1;refGene_tx=XM_023251320.1
A3 25623954 25624177 refGene_gn=E2F1;refGene_tx=XM_023251320.1
A3 25624307 25624555 refGene_gn=E2F1;refGene_tx=XM_023251320.1
A3 25627081 25627198
refGene_gn=NECAB3;refGene_tx=XM_011280897.3;refGene_gn=NECAB3;refGene_
tx=XM_023251322.1
A3 25629161 25629186
refGene_gn=NECAB3;refGene_tx=XM_011280897.3;refGene_gn=NECAB3;refGene_
tx=XM_023251322.1
A3 25630687 25630796
refGene_gn=NECAB3;refGene_tx=XM_011280897.3;refGene_gn=NECAB3;refGene_
tx=XM_023251322.1
A3 25631791 25631817
refGene_gn=NECAB3;refGene_tx=XM_011280897.3;refGene_gn=NECAB3;refGene_
tx=XM_023251322.1
A3 25632036 25632134
refGene_gn=NECAB3;refGene_tx=XM_011280897.3;refGene_gn=NECAB3;refGene_
tx=XM_023251322.1
A3 25633343 25634627 refGene_gn=ACTL10;refGene_tx=XM_023250921.1
A3 25635010 25635081 refGene_gn=ACTL10;refGene_tx=XM_023250921.1
A3 25635343 25635395 refGene_gn=ACTL10;refGene_tx=XM_023250921.1
A3 25637222 25637633
refGene_gn=CA3H20orf144;refGene_tx=XM_006929843.4
A3 25638354 25638600
refGene_gn=CA3H20orf144;refGene_tx=XM_006929843.4
A3 25640032 25640169
refGene_gn=NECAB3;refGene_tx=XM_011280897.3;refGene_gn=NECAB3;refGene_
tx=XM_023251322.1
A3 25640429 25640548
refGene_gn=NECAB3;refGene_tx=XM_011280897.3;refGene_gn=NECAB3;refGene_
tx=XM_023251322.1
A3 25640653 25640733

refGene_gn=NECAB3;refGene_tx=XM_011280897.3;refGene_gn=NECAB3;refGene_tx=XM_023251322.1
A3 25640837 25640891
refGene_gn=NECAB3;refGene_tx=XM_011280897.3;refGene_gn=NECAB3;refGene_tx=XM_023251322.1
A3 25641405 25641518
refGene_gn=NECAB3;refGene_tx=XM_011280897.3;refGene_gn=NECAB3;refGene_tx=XM_023251322.1
A3 25641687 25641765 refGene_gn=NECAB3;refGene_tx=XM_011280897.3
A3 25642274 25642366
refGene_gn=NECAB3;refGene_tx=XM_011280897.3;refGene_gn=NECAB3;refGene_tx=XM_023251322.1
A3 25642455 25642484
refGene_gn=NECAB3;refGene_tx=XM_011280897.3;refGene_gn=NECAB3;refGene_tx=XM_023251322.1
A3 25654055 25654355
refGene_gn=CBFA2T2;refGene_tx=XM_003983612.4;refGene_gn=CBFA2T2;refGene_tx=XM_006929844.3;refGene_gn=CBFA2T2;refGene_tx=XM_006929845.3;refGene_gn=CBFA2T2;refGene_tx=XM_006929846.3;refGene_gn=CBFA2T2;refGene_tx=XM_019826635.2
A3 25659439 25659630
refGene_gn=CBFA2T2;refGene_tx=XM_003983612.4;refGene_gn=CBFA2T2;refGene_tx=XM_006929844.3;refGene_gn=CBFA2T2;refGene_tx=XM_006929845.3;refGene_gn=CBFA2T2;refGene_tx=XM_006929846.3;refGene_gn=CBFA2T2;refGene_tx=XM_019826635.2
A3 25662914 25662983
refGene_gn=CBFA2T2;refGene_tx=XM_003983612.4;refGene_gn=CBFA2T2;refGene_tx=XM_006929844.3;refGene_gn=CBFA2T2;refGene_tx=XM_006929845.3;refGene_gn=CBFA2T2;refGene_tx=XM_019826635.2
A3 25672344 25672540
refGene_gn=CBFA2T2;refGene_tx=XM_003983612.4;refGene_gn=CBFA2T2;refGene_tx=XM_006929844.3;refGene_gn=CBFA2T2;refGene_tx=XM_006929845.3;refGene_gn=CBFA2T2;refGene_tx=XM_006929846.3;refGene_gn=CBFA2T2;refGene_tx=XM_019826635.2
A3 25673984 25674070
refGene_gn=CBFA2T2;refGene_tx=XM_003983612.4;refGene_gn=CBFA2T2;refGene_tx=XM_006929844.3;refGene_gn=CBFA2T2;refGene_tx=XM_006929845.3;refGene_gn=CBFA2T2;refGene_tx=XM_006929846.3;refGene_gn=CBFA2T2;refGene_tx=XM_019826635.2
A3 25674836 25675090
refGene_gn=CBFA2T2;refGene_tx=XM_003983612.4;refGene_gn=CBFA2T2;refGene_tx=XM_006929844.3;refGene_gn=CBFA2T2;refGene_tx=XM_006929845.3;refGene_gn=CBFA2T2;refGene_tx=XM_006929846.3;refGene_gn=CBFA2T2;refGene_tx=XM_019826635.2
A3 25675750 25675932
refGene_gn=CBFA2T2;refGene_tx=XM_003983612.4;refGene_gn=CBFA2T2;refGene_tx=XM_006929844.3;refGene_gn=CBFA2T2;refGene_tx=XM_006929845.3;refGene_gn=CBFA2T2;refGene_tx=XM_006929846.3;refGene_gn=CBFA2T2;refGene_tx=XM_019826635.2
A3 25680480 25680570

refGene_gn=CBFA2T2;refGene_tx=XM_003983612.4;refGene_gn=CBFA2T2;refGene_tx=XM_006929844.3;refGene_gn=CBFA2T2;refGene_tx=XM_006929845.3;refGene_gn=CBFA2T2;refGene_tx=XM_006929846.3;refGene_gn=CBFA2T2;refGene_tx=XM_019826635.2
A3 25688324 25688566
refGene_gn=CBFA2T2;refGene_tx=XM_003983612.4;refGene_gn=CBFA2T2;refGene_tx=XM_006929844.3;refGene_gn=CBFA2T2;refGene_tx=XM_006929845.3;refGene_gn=CBFA2T2;refGene_tx=XM_006929846.3;refGene_gn=CBFA2T2;refGene_tx=XM_019826635.2
A3 25692844 25692988
refGene_gn=CBFA2T2;refGene_tx=XM_003983612.4;refGene_gn=CBFA2T2;refGene_tx=XM_006929844.3;refGene_gn=CBFA2T2;refGene_tx=XM_006929845.3;refGene_gn=CBFA2T2;refGene_tx=XM_006929846.3;refGene_gn=CBFA2T2;refGene_tx=XM_019826635.2
A3 25791614 25791648
refGene_gn=CBFA2T2;refGene_tx=XM_003983612.4;refGene_gn=CBFA2T2;refGene_tx=XM_006929844.3;refGene_gn=CBFA2T2;refGene_tx=XM_006929846.3
A3 25835947 25836257 refGene_gn=SNTA1;refGene_tx=XM_011280896.3
A3 25840273 25840459 refGene_gn=SNTA1;refGene_tx=XM_011280896.3
A3 25855987 25856192 refGene_gn=SNTA1;refGene_tx=XM_011280896.3
A3 25859059 25859267 refGene_gn=SNTA1;refGene_tx=XM_011280896.3
A3 25859410 25859541 refGene_gn=SNTA1;refGene_tx=XM_011280896.3
A3 25861512 25861709 refGene_gn=SNTA1;refGene_tx=XM_011280896.3
A3 25862659 25862847 refGene_gn=SNTA1;refGene_tx=XM_011280896.3
A3 25862931 25863024 refGene_gn=SNTA1;refGene_tx=XM_011280896.3
A3 25871402 25871694
refGene_gn=CDK5RAP1;refGene_tx=XM_011280848.3;refGene_gn=CDK5RAP1;refGene_tx=XM_023251324.1
A3 25874713 25874817
refGene_gn=CDK5RAP1;refGene_tx=XM_011280848.3;refGene_gn=CDK5RAP1;refGene_tx=XM_023251324.1
A3 25875571 25875606
refGene_gn=CDK5RAP1;refGene_tx=XM_011280848.3;refGene_gn=CDK5RAP1;refGene_tx=XM_023251324.1
A3 25876606 25876707
refGene_gn=CDK5RAP1;refGene_tx=XM_011280848.3;refGene_gn=CDK5RAP1;refGene_tx=XM_023251324.1
A3 25879238 25879449
refGene_gn=CDK5RAP1;refGene_tx=XM_011280848.3;refGene_gn=CDK5RAP1;refGene_tx=XM_023251324.1
A3 25881150 25881271
refGene_gn=CDK5RAP1;refGene_tx=XM_011280848.3;refGene_gn=CDK5RAP1;refGene_tx=XM_023251324.1
A3 25885627 25885858
refGene_gn=CDK5RAP1;refGene_tx=XM_011280848.3;refGene_gn=CDK5RAP1;refGene_tx=XM_023251324.1
A3 25887499 25887597
refGene_gn=CDK5RAP1;refGene_tx=XM_011280848.3;refGene_gn=CDK5RAP1;refGene_tx=XM_023251324.1
A3 25889046 25889102

refGene_gn=CDK5RAP1;refGene_tx=XM_011280848.3;refGene_gn=CDK5RAP1;refGene_tx=XM_023251324.1
A3 25890932 25891063
refGene_gn=CDK5RAP1;refGene_tx=XM_011280848.3;refGene_gn=CDK5RAP1;refGene_tx=XM_023251324.1
A3 25893756 25893906
refGene_gn=CDK5RAP1;refGene_tx=XM_011280848.3;refGene_gn=CDK5RAP1;refGene_tx=XM_023251324.1
A3 25902812 25902953
refGene_gn=CDK5RAP1;refGene_tx=XM_011280848.3;refGene_gn=CDK5RAP1;refGene_tx=XM_023251324.1
A3 25905834 25905915
refGene_gn=CDK5RAP1;refGene_tx=XM_011280848.3;refGene_gn=CDK5RAP1;refGene_tx=XM_023251324.1
A3 25918172 25918231
refGene_gn=LOC105260421;refGene_tx=XM_019826637.1;refGene_gn=LOC105260421;refGene_tx=XM_019826638.1
A3 25919504 25919584
refGene_gn=LOC105260421;refGene_tx=XM_019826637.1
A3 25920737 25920798
refGene_gn=LOC105260421;refGene_tx=XM_019826637.1;refGene_gn=LOC105260421;refGene_tx=XM_019826638.1
A3 25922379 25922410
refGene_gn=LOC105260421;refGene_tx=XM_019826637.1;refGene_gn=LOC105260421;refGene_tx=XM_019826638.1
A3 25922694 25922762
refGene_gn=LOC105260421;refGene_tx=XM_019826637.1;refGene_gn=LOC105260421;refGene_tx=XM_019826638.1
A3 25923321 25923444
refGene_gn=LOC105260421;refGene_tx=XM_019826637.1;refGene_gn=LOC105260421;refGene_tx=XM_019826638.1
A3 25924303 25924450
refGene_gn=LOC105260421;refGene_tx=XM_019826637.1;refGene_gn=LOC105260421;refGene_tx=XM_019826638.1
A3 25924891 25924962
refGene_gn=LOC105260421;refGene_tx=XM_019826637.1;refGene_gn=LOC105260421;refGene_tx=XM_019826638.1
A3 25925663 25925721
refGene_gn=LOC105260421;refGene_tx=XM_019826637.1;refGene_gn=LOC105260421;refGene_tx=XM_019826638.1
A3 25926054 25926133
refGene_gn=LOC105260421;refGene_tx=XM_019826637.1;refGene_gn=LOC105260421;refGene_tx=XM_019826638.1
A3 25927042 25927198
refGene_gn=LOC105260421;refGene_tx=XM_019826637.1;refGene_gn=LOC105260421;refGene_tx=XM_019826638.1
A3 25928676 25928778
refGene_gn=LOC105260421;refGene_tx=XM_019826637.1;refGene_gn=LOC105260421;refGene_tx=XM_019826638.1
A3 25929104 25929588

refGene_gn=LOC105260421;refGene_tx=XM_019826637.1;refGene_gn=LOC105260421;refGene_tx=XM_019826638.1
A3 25931890 25932152 refGene_gn=LOC105260421;refGene_tx=XM_019826637.1;refGene_gn=LOC105260421;refGene_tx=XM_019826638.1
A3 25944136 25944169 refGene_gn=LOC101099199;refGene_tx=XM_003983649.2
A3 25945137 25945217 refGene_gn=LOC101099199;refGene_tx=XM_003983649.2
A3 25947220 25947284 refGene_gn=LOC101099199;refGene_tx=XM_003983649.2
A3 25949259 25949305 refGene_gn=LOC101099199;refGene_tx=XM_003983649.2
A3 25949540 25949608 refGene_gn=LOC101099199;refGene_tx=XM_003983649.2
A3 25950637 25950790 refGene_gn=LOC101099199;refGene_tx=XM_003983649.2
A3 25951776 25951830 refGene_gn=LOC101099199;refGene_tx=XM_003983649.2
A3 25954605 25954788 refGene_gn=LOC101099199;refGene_tx=XM_003983649.2
A3 25955585 25955671 refGene_gn=LOC101099199;refGene_tx=XM_003983649.2
A3 25956103 25956167 refGene_gn=LOC101099199;refGene_tx=XM_003983649.2
A3 25956264 25956349 refGene_gn=LOC101099199;refGene_tx=XM_003983649.2
A3 25957093 25957225 refGene_gn=LOC101099199;refGene_tx=XM_003983649.2
A3 25958551 25958656 refGene_gn=LOC101099199;refGene_tx=XM_003983649.2
A3 25959759 25959976 refGene_gn=LOC101099199;refGene_tx=XM_003983649.2
A3 25961519 25961628 refGene_gn=LOC101099199;refGene_tx=XM_003983649.2
A3 25971215 25971382 refGene_gn=BPIFB1;refGene_tx=XM_023251326.1
A3 25972311 25972375 refGene_gn=BPIFB1;refGene_tx=XM_023251326.1
A3 25973260 25973306 refGene_gn=BPIFB1;refGene_tx=XM_023251326.1
A3 25973816 25973884 refGene_gn=BPIFB1;refGene_tx=XM_023251326.1
A3 25975702 25975861 refGene_gn=BPIFB1;refGene_tx=XM_023251326.1
A3 25976373 25976427 refGene_gn=BPIFB1;refGene_tx=XM_023251326.1
A3 25977356 25977536 refGene_gn=BPIFB1;refGene_tx=XM_023251326.1
A3 25978530 25978616 refGene_gn=BPIFB1;refGene_tx=XM_023251326.1
A3 25980389 25980453 refGene_gn=BPIFB1;refGene_tx=XM_023251326.1
A3 25983397 25983479 refGene_gn=BPIFB1;refGene_tx=XM_023251326.1
A3 25984233 25984383 refGene_gn=BPIFB1;refGene_tx=XM_023251326.1
A3 25985282 25985390 refGene_gn=BPIFB1;refGene_tx=XM_023251326.1
A3 25985996 25986138 refGene_gn=BPIFB1;refGene_tx=XM_023251326.1
A3 25987191 25987306 refGene_gn=BPIFB1;refGene_tx=XM_023251326.1
A3 26016731 26016828 refGene_gn=BPIFA1;refGene_tx=XM_023251327.1

A3 26017692 26017842 refGene_gn=BPIFA1;refGene_tx=XM_023251327.1
A3 26017984 26018092 refGene_gn=BPIFA1;refGene_tx=XM_023251327.1
A3 26019568 26019728 refGene_gn=BPIFA1;refGene_tx=XM_023251327.1
A3 26019902 26020062 refGene_gn=BPIFA1;refGene_tx=XM_023251327.1
A3 26028693 26028728 refGene_gn=BPIFA3;refGene_tx=XM_003983615.4
A3 26029228 26029292 refGene_gn=BPIFA3;refGene_tx=XM_003983615.4
A3 26029708 26029793 refGene_gn=BPIFA3;refGene_tx=XM_003983615.4
A3 26031035 26031185 refGene_gn=BPIFA3;refGene_tx=XM_003983615.4
A3 26031819 26031927 refGene_gn=BPIFA3;refGene_tx=XM_003983615.4
A3 26033782 26033933 refGene_gn=BPIFA3;refGene_tx=XM_003983615.4
A3 26036885 26037000 refGene_gn=BPIFA3;refGene_tx=XM_003983615.4
A3 26050610 26050666
refGene_gn=LOC100652387;refGene_tx=NM_001243083.1
A3 26051805 26051869
refGene_gn=LOC100652387;refGene_tx=NM_001243083.1
A3 26052498 26052580
refGene_gn=LOC100652387;refGene_tx=NM_001243083.1
A3 26054712 26054862
refGene_gn=LOC100652387;refGene_tx=NM_001243083.1
A3 26055759 26055864
refGene_gn=LOC100652387;refGene_tx=NM_001243083.1
A3 26057464 26057594
refGene_gn=LOC100652387;refGene_tx=NM_001243083.1
A3 26058779 26058879
refGene_gn=LOC100652387;refGene_tx=NM_001243083.1
A3 26072665 26072706 refGene_gn=BPIFA2;refGene_tx=XM_023251328.1
A3 26073535 26073599 refGene_gn=BPIFA2;refGene_tx=XM_023251328.1
A3 26074675 26074757 refGene_gn=BPIFA2;refGene_tx=XM_023251328.1
A3 26077101 26077254 refGene_gn=BPIFA2;refGene_tx=XM_023251328.1
A3 26078421 26078529 refGene_gn=BPIFA2;refGene_tx=XM_023251328.1
A3 26079190 26079335 refGene_gn=BPIFA2;refGene_tx=XM_023251328.1
A3 26081024 26081181 refGene_gn=BPIFA2;refGene_tx=XM_023251328.1
A3 26122305 26122329 refGene_gn=BPIFB4;refGene_tx=XM_023250913.1
A3 26126736 26126813 refGene_gn=BPIFB4;refGene_tx=XM_023250913.1
A3 26129464 26129528 refGene_gn=BPIFB4;refGene_tx=XM_023250913.1
A3 26131452 26131495 refGene_gn=BPIFB4;refGene_tx=XM_023250913.1
A3 26133730 26133798 refGene_gn=BPIFB4;refGene_tx=XM_023250913.1
A3 26134877 26135048 refGene_gn=BPIFB4;refGene_tx=XM_023250913.1
A3 26138129 26138183 refGene_gn=BPIFB4;refGene_tx=XM_023250913.1
A3 26139944 26140148 refGene_gn=BPIFB4;refGene_tx=XM_023250913.1
A3 26141507 26141599 refGene_gn=BPIFB4;refGene_tx=XM_023250913.1
A3 26142715 26142776 refGene_gn=BPIFB4;refGene_tx=XM_023250913.1
A3 26143233 26143297 refGene_gn=BPIFB4;refGene_tx=XM_023250913.1
A3 26146489 26146633 refGene_gn=BPIFB4;refGene_tx=XM_023250913.1
A3 26147445 26147550 refGene_gn=BPIFB4;refGene_tx=XM_023250913.1
A3 26148518 26149038 refGene_gn=BPIFB4;refGene_tx=XM_023250913.1
A3 26149702 26149765 refGene_gn=BPIFB4;refGene_tx=XM_023250913.1
A3 26150552 26150658 refGene_gn=BPIFB4;refGene_tx=XM_023250913.1
A3 26156737 26156767 refGene_gn=BPIFB3;refGene_tx=XM_003983617.4
A3 26157464 26157541 refGene_gn=BPIFB3;refGene_tx=XM_003983617.4

| | | | |
|---|---|---|---|
| A3 | 26158252 | 26158316 | refGene_gn=BPIFB3;refGene_tx=XM_003983617.4 |
| A3 | 26159085 | 26159128 | refGene_gn=BPIFB3;refGene_tx=XM_003983617.4 |
| A3 | 26160303 | 26160371 | refGene_gn=BPIFB3;refGene_tx=XM_003983617.4 |
| A3 | 26161246 | 26161417 | refGene_gn=BPIFB3;refGene_tx=XM_003983617.4 |
| A3 | 26163306 | 26163360 | refGene_gn=BPIFB3;refGene_tx=XM_003983617.4 |
| A3 | 26164998 | 26165178 | refGene_gn=BPIFB3;refGene_tx=XM_003983617.4 |
| A3 | 26165296 | 26165388 | refGene_gn=BPIFB3;refGene_tx=XM_003983617.4 |
| A3 | 26166052 | 26166113 | refGene_gn=BPIFB3;refGene_tx=XM_003983617.4 |
| A3 | 26168404 | 26168468 | refGene_gn=BPIFB3;refGene_tx=XM_003983617.4 |
| A3 | 26169950 | 26170091 | refGene_gn=BPIFB3;refGene_tx=XM_003983617.4 |
| A3 | 26170553 | 26170658 | refGene_gn=BPIFB3;refGene_tx=XM_003983617.4 |
| A3 | 26172216 | 26172373 | refGene_gn=BPIFB3;refGene_tx=XM_003983617.4 |
| A3 | 26173079 | 26173194 | refGene_gn=BPIFB3;refGene_tx=XM_003983617.4 |
| A3 | 26179996 | 26180035 | refGene_gn=BPIFB6;refGene_tx=XM_019828127.2 |
| A3 | 26180641 | 26180718 | refGene_gn=BPIFB6;refGene_tx=XM_019828127.2 |
| A3 | 26181377 | 26181441 | refGene_gn=BPIFB6;refGene_tx=XM_019828127.2 |
| A3 | 26182306 | 26182352 | refGene_gn=BPIFB6;refGene_tx=XM_019828127.2 |
| A3 | 26183751 | 26183819 | refGene_gn=BPIFB6;refGene_tx=XM_019828127.2 |
| A3 | 26184589 | 26184763 | refGene_gn=BPIFB6;refGene_tx=XM_019828127.2 |
| A3 | 26185391 | 26185445 | refGene_gn=BPIFB6;refGene_tx=XM_019828127.2 |
| A3 | 26186492 | 26186669 | refGene_gn=BPIFB6;refGene_tx=XM_019828127.2 |
| A3 | 26187460 | 26187552 | refGene_gn=BPIFB6;refGene_tx=XM_019828127.2 |
| A3 | 26188349 | 26188410 | refGene_gn=BPIFB6;refGene_tx=XM_019828127.2 |
| A3 | 26188848 | 26188912 | refGene_gn=BPIFB6;refGene_tx=XM_019828127.2 |
| A3 | 26189070 | 26189220 | refGene_gn=BPIFB6;refGene_tx=XM_019828127.2 |
| A3 | 26189691 | 26189796 | refGene_gn=BPIFB6;refGene_tx=XM_019828127.2 |
| A3 | 26190831 | 26190931 | refGene_gn=BPIFB6;refGene_tx=XM_019828127.2 |
| A3 | 26192257 | 26192354 | refGene_gn=BPIFB6;refGene_tx=XM_019828127.2 |
| A3 | 26199864 | 26199906 | refGene_gn=BPIFB2;refGene_tx=XM_023251329.1 |
| A3 | 26201463 | 26201540 | refGene_gn=BPIFB2;refGene_tx=XM_023251329.1 |
| A3 | 26201852 | 26201916 | refGene_gn=BPIFB2;refGene_tx=XM_023251329.1 |
| A3 | 26202671 | 26202717 | refGene_gn=BPIFB2;refGene_tx=XM_023251329.1 |
| A3 | 26202901 | 26202969 | refGene_gn=BPIFB2;refGene_tx=XM_023251329.1 |
| A3 | 26203795 | 26203966 | refGene_gn=BPIFB2;refGene_tx=XM_023251329.1 |
| A3 | 26204479 | 26204533 | refGene_gn=BPIFB2;refGene_tx=XM_023251329.1 |
| A3 | 26204755 | 26204941 | refGene_gn=BPIFB2;refGene_tx=XM_023251329.1 |
| A3 | 26205187 | 26205279 | refGene_gn=BPIFB2;refGene_tx=XM_023251329.1 |
| A3 | 26209238 | 26209299 | refGene_gn=BPIFB2;refGene_tx=XM_023251329.1 |
| A3 | 26210633 | 26210694 | refGene_gn=BPIFB2;refGene_tx=XM_023251329.1 |
| A3 | 26212495 | 26212642 | refGene_gn=BPIFB2;refGene_tx=XM_023251329.1 |
| A3 | 26213447 | 26213552 | refGene_gn=BPIFB2;refGene_tx=XM_023251329.1 |
| A3 | 26215382 | 26215476 | refGene_gn=BPIFB2;refGene_tx=XM_023251329.1 |
| A3 | 26217937 | 26218046 | refGene_gn=BPIFB2;refGene_tx=XM_023251329.1 |
| A3 | 26221635 | 26221700 | refGene_gn=SUN5;refGene_tx=XM_011280846.3 |
| A3 | 26223251 | 26223310 | refGene_gn=SUN5;refGene_tx=XM_011280846.3 |
| A3 | 26223502 | 26223577 | refGene_gn=SUN5;refGene_tx=XM_011280846.3 |
| A3 | 26224473 | 26224540 | refGene_gn=SUN5;refGene_tx=XM_011280846.3 |
| A3 | 26225802 | 26225864 | refGene_gn=SUN5;refGene_tx=XM_011280846.3 |
| A3 | 26229578 | 26229628 | refGene_gn=SUN5;refGene_tx=XM_011280846.3 |
| A3 | 26230997 | 26231032 | refGene_gn=SUN5;refGene_tx=XM_011280846.3 |

A3 26231592 26231701 refGene_gn=SUN5;refGene_tx=XM_011280846.3
A3 26236311 26236390 refGene_gn=SUN5;refGene_tx=XM_011280846.3
A3 26237248 26237364 refGene_gn=SUN5;refGene_tx=XM_011280846.3
A3 26237871 26238039 refGene_gn=SUN5;refGene_tx=XM_011280846.3
A3 26238535 26238622 refGene_gn=SUN5;refGene_tx=XM_011280846.3
A3 26239676 26239802 refGene_gn=SUN5;refGene_tx=XM_011280846.3
A3 26262561 26262987 refGene_gn=EFCAB8;refGene_tx=XM_019826643.2
A3 26263931 26264030 refGene_gn=EFCAB8;refGene_tx=XM_019826643.2
A3 26264114 26264319 refGene_gn=EFCAB8;refGene_tx=XM_019826643.2
A3 26265490 26265623 refGene_gn=EFCAB8;refGene_tx=XM_019826643.2
A3 26271021 26271190 refGene_gn=EFCAB8;refGene_tx=XM_019826643.2
A3 26277160 26277298 refGene_gn=EFCAB8;refGene_tx=XM_019826643.2
A3 26277860 26278082 refGene_gn=EFCAB8;refGene_tx=XM_019826643.2
A3 26287354 26287489 refGene_gn=EFCAB8;refGene_tx=XM_019826643.2
A3 26288612 26288825 refGene_gn=EFCAB8;refGene_tx=XM_019826643.2
A3 26289690 26289895 refGene_gn=EFCAB8;refGene_tx=XM_019826643.2
A3 26290882 26290953 refGene_gn=EFCAB8;refGene_tx=XM_019826643.2
A3 26291726 26291954 refGene_gn=EFCAB8;refGene_tx=XM_019826643.2
A3 26293138 26293252 refGene_gn=EFCAB8;refGene_tx=XM_019826643.2
A3 26293706 26293844 refGene_gn=EFCAB8;refGene_tx=XM_019826643.2
A3 26295255 26295407 refGene_gn=EFCAB8;refGene_tx=XM_019826643.2
A3 26295729 26295797 refGene_gn=EFCAB8;refGene_tx=XM_019826643.2
A3 26302387 26302518 refGene_gn=EFCAB8;refGene_tx=XM_019826643.2
A3 26305628 26305702 refGene_gn=EFCAB8;refGene_tx=XM_019826643.2
A3 26308859 26308984 refGene_gn=EFCAB8;refGene_tx=XM_019826643.2
A3 26309522 26309607 refGene_gn=EFCAB8;refGene_tx=XM_019826643.2
A3 26309826 26309932 refGene_gn=EFCAB8;refGene_tx=XM_019826643.2
A3 26312916 26313052 refGene_gn=EFCAB8;refGene_tx=XM_019826643.2
A3 26315374 26315478 refGene_gn=EFCAB8;refGene_tx=XM_019826643.2
A3 26316175 26316213 refGene_gn=EFCAB8;refGene_tx=XM_019826643.2
A3 26316215 26316293 refGene_gn=EFCAB8;refGene_tx=XM_019826643.2
A3 26323117 26323271 refGene_gn=EFCAB8;refGene_tx=XM_019826643.2
A3 26325585 26325637 refGene_gn=EFCAB8;refGene_tx=XM_019826643.2
A3 26329716 26329805 refGene_gn=EFCAB8;refGene_tx=XM_019826643.2
A3 26338910 26338967 refGene_gn=MAPRE1;refGene_tx=XM_023251331.1
A3 26340566 26340719 refGene_gn=MAPRE1;refGene_tx=XM_023251331.1
A3 26344866 26344988 refGene_gn=MAPRE1;refGene_tx=XM_023251331.1
A3 26347509 26347717 refGene_gn=MAPRE1;refGene_tx=XM_023251331.1
A3 26350064 26350210 refGene_gn=MAPRE1;refGene_tx=XM_023251331.1
A3 26361996 26362117 refGene_gn=MAPRE1;refGene_tx=XM_023251331.1
A3 26373678 26373820
refGene_gn=DNMT3B;refGene_tx=XM_006929850.4;refGene_gn=DNMT3B;refGene_tx=XM_019826654.2;refGene_gn=DNMT3B;refGene_tx=XM_019826656.2;refGene_gn=DNMT3B;refGene_tx=XM_019826659.1;refGene_gn=DNMT3B;refGene_tx=XM_023251332.1;refGene_gn=DNMT3B;refGene_tx=XM_023251333.1;refGene_gn=DNMT3B;refGene_tx=XM_023251334.1;refGene_gn=DNMT3B;refGene_tx=XM_023251335.1;refGene_gn=DNMT3B;refGene_tx=XM_023251338.1
A3 26375413 26375532
refGene_gn=DNMT3B;refGene_tx=XM_006929850.4;refGene_gn=DNMT3B;refGene_tx=XM_019826654.2;refGene_gn=DNMT3B;refGene_tx=XM_019826656.2;refGene_

gn=DNMT3B;refGene_tx=XM_019826659.1;refGene_gn=DNMT3B;refGene_tx=XM_02
3251332.1;refGene_gn=DNMT3B;refGene_tx=XM_023251333.1;refGene_gn=DNMT3
B;refGene_tx=XM_023251337.1
A3 26376299 26376369
refGene_gn=DNMT3B;refGene_tx=XM_006929850.4;refGene_gn=DNMT3B;refGene_
tx=XM_019826654.2;refGene_gn=DNMT3B;refGene_tx=XM_019826656.2;refGene_
gn=DNMT3B;refGene_tx=XM_019826659.1;refGene_gn=DNMT3B;refGene_tx=XM_02
3251332.1;refGene_gn=DNMT3B;refGene_tx=XM_023251333.1
A3 26379186 26379272
refGene_gn=DNMT3B;refGene_tx=XM_006929850.4;refGene_gn=DNMT3B;refGene_
tx=XM_019826654.2;refGene_gn=DNMT3B;refGene_tx=XM_019826656.2;refGene_
gn=DNMT3B;refGene_tx=XM_019826659.1;refGene_gn=DNMT3B;refGene_tx=XM_02
3251332.1;refGene_gn=DNMT3B;refGene_tx=XM_023251333.1;refGene_gn=DNMT3
B;refGene_tx=XM_023251334.1;refGene_gn=DNMT3B;refGene_tx=XM_023251335.
1;refGene_gn=DNMT3B;refGene_tx=XM_023251337.1;refGene_gn=DNMT3B;refGen
e_tx=XM_023251338.1
A3 26380349 26380498
refGene_gn=DNMT3B;refGene_tx=XM_006929850.4;refGene_gn=DNMT3B;refGene_
tx=XM_019826654.2;refGene_gn=DNMT3B;refGene_tx=XM_019826656.2;refGene_
gn=DNMT3B;refGene_tx=XM_019826659.1;refGene_gn=DNMT3B;refGene_tx=XM_02
3251332.1;refGene_gn=DNMT3B;refGene_tx=XM_023251333.1;refGene_gn=DNMT3
B;refGene_tx=XM_023251334.1;refGene_gn=DNMT3B;refGene_tx=XM_023251335.
1;refGene_gn=DNMT3B;refGene_tx=XM_023251337.1;refGene_gn=DNMT3B;refGen
e_tx=XM_023251338.1
A3 26380808 26380899
refGene_gn=DNMT3B;refGene_tx=XM_006929850.4;refGene_gn=DNMT3B;refGene_
tx=XM_019826654.2;refGene_gn=DNMT3B;refGene_tx=XM_019826656.2;refGene_
gn=DNMT3B;refGene_tx=XM_019826659.1;refGene_gn=DNMT3B;refGene_tx=XM_02
3251332.1;refGene_gn=DNMT3B;refGene_tx=XM_023251333.1;refGene_gn=DNMT3
B;refGene_tx=XM_023251334.1;refGene_gn=DNMT3B;refGene_tx=XM_023251335.
1;refGene_gn=DNMT3B;refGene_tx=XM_023251337.1;refGene_gn=DNMT3B;refGen
e_tx=XM_023251338.1
A3 26381876 26382022
refGene_gn=DNMT3B;refGene_tx=XM_006929850.4;refGene_gn=DNMT3B;refGene_
tx=XM_019826654.2;refGene_gn=DNMT3B;refGene_tx=XM_019826656.2;refGene_
gn=DNMT3B;refGene_tx=XM_019826659.1;refGene_gn=DNMT3B;refGene_tx=XM_02
3251332.1;refGene_gn=DNMT3B;refGene_tx=XM_023251333.1;refGene_gn=DNMT3
B;refGene_tx=XM_023251334.1;refGene_gn=DNMT3B;refGene_tx=XM_023251335.
1;refGene_gn=DNMT3B;refGene_tx=XM_023251337.1;refGene_gn=DNMT3B;refGen
e_tx=XM_023251338.1
A3 26383017 26383102
refGene_gn=DNMT3B;refGene_tx=XM_006929850.4;refGene_gn=DNMT3B;refGene_
tx=XM_019826654.2;refGene_gn=DNMT3B;refGene_tx=XM_019826656.2;refGene_
gn=DNMT3B;refGene_tx=XM_019826659.1;refGene_gn=DNMT3B;refGene_tx=XM_02
3251332.1;refGene_gn=DNMT3B;refGene_tx=XM_023251333.1;refGene_gn=DNMT3
B;refGene_tx=XM_023251334.1;refGene_gn=DNMT3B;refGene_tx=XM_023251335.
1;refGene_gn=DNMT3B;refGene_tx=XM_023251337.1;refGene_gn=DNMT3B;refGen
e_tx=XM_023251338.1
A3 26383661 26383845
refGene_gn=DNMT3B;refGene_tx=XM_006929850.4;refGene_gn=DNMT3B;refGene_

tx=XM_019826654.2;refGene_gn=DNMT3B;refGene_tx=XM_019826656.2;refGene_
gn=DNMT3B;refGene_tx=XM_019826659.1;refGene_gn=DNMT3B;refGene_tx=XM_02
3251332.1;refGene_gn=DNMT3B;refGene_tx=XM_023251333.1;refGene_gn=DNMT3
B;refGene_tx=XM_023251334.1;refGene_gn=DNMT3B;refGene_tx=XM_023251335.
1;refGene_gn=DNMT3B;refGene_tx=XM_023251337.1;refGene_gn=DNMT3B;refGen
e_tx=XM_023251338.1
A3 26384473 26384586
refGene_gn=DNMT3B;refGene_tx=XM_006929850.4;refGene_gn=DNMT3B;refGene_
tx=XM_019826654.2;refGene_gn=DNMT3B;refGene_tx=XM_019826656.2;refGene_
gn=DNMT3B;refGene_tx=XM_019826659.1;refGene_gn=DNMT3B;refGene_tx=XM_02
3251332.1;refGene_gn=DNMT3B;refGene_tx=XM_023251333.1;refGene_gn=DNMT3
B;refGene_tx=XM_023251334.1;refGene_gn=DNMT3B;refGene_tx=XM_023251335.
1;refGene_gn=DNMT3B;refGene_tx=XM_023251337.1;refGene_gn=DNMT3B;refGen
e_tx=XM_023251338.1
A3 26384893 26384973
refGene_gn=DNMT3B;refGene_tx=XM_006929850.4;refGene_gn=DNMT3B;refGene_
tx=XM_019826654.2;refGene_gn=DNMT3B;refGene_tx=XM_019826656.2;refGene_
gn=DNMT3B;refGene_tx=XM_019826659.1;refGene_gn=DNMT3B;refGene_tx=XM_02
3251332.1;refGene_gn=DNMT3B;refGene_tx=XM_023251333.1;refGene_gn=DNMT3
B;refGene_tx=XM_023251334.1;refGene_gn=DNMT3B;refGene_tx=XM_023251335.
1;refGene_gn=DNMT3B;refGene_tx=XM_023251337.1;refGene_gn=DNMT3B;refGen
e_tx=XM_023251338.1
A3 26386341 26386386
refGene_gn=DNMT3B;refGene_tx=XM_006929850.4;refGene_gn=DNMT3B;refGene_
tx=XM_019826654.2;refGene_gn=DNMT3B;refGene_tx=XM_019826656.2;refGene_
gn=DNMT3B;refGene_tx=XM_019826659.1;refGene_gn=DNMT3B;refGene_tx=XM_02
3251332.1;refGene_gn=DNMT3B;refGene_tx=XM_023251333.1;refGene_gn=DNMT3
B;refGene_tx=XM_023251334.1;refGene_gn=DNMT3B;refGene_tx=XM_023251335.
1;refGene_gn=DNMT3B;refGene_tx=XM_023251337.1;refGene_gn=DNMT3B;refGen
e_tx=XM_023251338.1
A3 26386546 26386672
refGene_gn=DNMT3B;refGene_tx=XM_006929850.4;refGene_gn=DNMT3B;refGene_
tx=XM_019826654.2;refGene_gn=DNMT3B;refGene_tx=XM_019826656.2;refGene_
gn=DNMT3B;refGene_tx=XM_019826659.1;refGene_gn=DNMT3B;refGene_tx=XM_02
3251332.1;refGene_gn=DNMT3B;refGene_tx=XM_023251333.1;refGene_gn=DNMT3
B;refGene_tx=XM_023251334.1;refGene_gn=DNMT3B;refGene_tx=XM_023251335.
1;refGene_gn=DNMT3B;refGene_tx=XM_023251337.1;refGene_gn=DNMT3B;refGen
e_tx=XM_023251338.1
A3 26387849 26387909
refGene_gn=DNMT3B;refGene_tx=XM_019826654.2;refGene_gn=DNMT3B;refGene_
tx=XM_019826656.2;refGene_gn=DNMT3B;refGene_tx=XM_019826659.1;refGene_
gn=DNMT3B;refGene_tx=XM_023251333.1;refGene_gn=DNMT3B;refGene_tx=XM_02
3251334.1;refGene_gn=DNMT3B;refGene_tx=XM_023251337.1
A3 26388716 26388861
refGene_gn=DNMT3B;refGene_tx=XM_006929850.4;refGene_gn=DNMT3B;refGene_
tx=XM_019826654.2;refGene_gn=DNMT3B;refGene_tx=XM_019826656.2;refGene_
gn=DNMT3B;refGene_tx=XM_019826659.1;refGene_gn=DNMT3B;refGene_tx=XM_02
3251332.1;refGene_gn=DNMT3B;refGene_tx=XM_023251333.1;refGene_gn=DNMT3
B;refGene_tx=XM_023251334.1;refGene_gn=DNMT3B;refGene_tx=XM_023251335.
1;refGene_gn=DNMT3B;refGene_tx=XM_023251337.1;refGene_gn=DNMT3B;refGen

e_tx=XM_023251338.1
A3 26389740 26389848
refGene_gn=DNMT3B;refGene_tx=XM_006929850.4;refGene_gn=DNMT3B;refGene_tx=XM_019826654.2;refGene_gn=DNMT3B;refGene_tx=XM_019826656.2;refGene_gn=DNMT3B;refGene_tx=XM_019826659.1;refGene_gn=DNMT3B;refGene_tx=XM_023251332.1;refGene_gn=DNMT3B;refGene_tx=XM_023251333.1;refGene_gn=DNMT3B;refGene_tx=XM_023251334.1;refGene_gn=DNMT3B;refGene_tx=XM_023251335.1;refGene_gn=DNMT3B;refGene_tx=XM_023251337.1;refGene_gn=DNMT3B;refGene_tx=XM_023251338.1
A3 26392354 26392513
refGene_gn=DNMT3B;refGene_tx=XM_006929850.4;refGene_gn=DNMT3B;refGene_tx=XM_019826654.2;refGene_gn=DNMT3B;refGene_tx=XM_019826656.2;refGene_gn=DNMT3B;refGene_tx=XM_019826659.1;refGene_gn=DNMT3B;refGene_tx=XM_023251332.1;refGene_gn=DNMT3B;refGene_tx=XM_023251333.1;refGene_gn=DNMT3B;refGene_tx=XM_023251334.1;refGene_gn=DNMT3B;refGene_tx=XM_023251335.1;refGene_gn=DNMT3B;refGene_tx=XM_023251337.1;refGene_gn=DNMT3B;refGene_tx=XM_023251338.1
A3 26393666 26393876
refGene_gn=DNMT3B;refGene_tx=XM_006929850.4;refGene_gn=DNMT3B;refGene_tx=XM_019826654.2;refGene_gn=DNMT3B;refGene_tx=XM_019826656.2;refGene_gn=DNMT3B;refGene_tx=XM_019826659.1;refGene_gn=DNMT3B;refGene_tx=XM_023251332.1;refGene_gn=DNMT3B;refGene_tx=XM_023251333.1;refGene_gn=DNMT3B;refGene_tx=XM_023251334.1;refGene_gn=DNMT3B;refGene_tx=XM_023251335.1;refGene_gn=DNMT3B;refGene_tx=XM_023251337.1;refGene_gn=DNMT3B;refGene_tx=XM_023251338.1
A3 26394535 26394709
refGene_gn=DNMT3B;refGene_tx=XM_006929850.4;refGene_gn=DNMT3B;refGene_tx=XM_019826654.2;refGene_gn=DNMT3B;refGene_tx=XM_019826656.2;refGene_gn=DNMT3B;refGene_tx=XM_019826659.1;refGene_gn=DNMT3B;refGene_tx=XM_023251332.1;refGene_gn=DNMT3B;refGene_tx=XM_023251333.1;refGene_gn=DNMT3B;refGene_tx=XM_023251334.1;refGene_gn=DNMT3B;refGene_tx=XM_023251335.1;refGene_gn=DNMT3B;refGene_tx=XM_023251337.1
A3 26395630 26395738
refGene_gn=DNMT3B;refGene_tx=XM_006929850.4;refGene_gn=DNMT3B;refGene_tx=XM_019826654.2;refGene_gn=DNMT3B;refGene_tx=XM_019826656.2;refGene_gn=DNMT3B;refGene_tx=XM_023251332.1;refGene_gn=DNMT3B;refGene_tx=XM_023251333.1;refGene_gn=DNMT3B;refGene_tx=XM_023251334.1;refGene_gn=DNMT3B;refGene_tx=XM_023251335.1;refGene_gn=DNMT3B;refGene_tx=XM_023251337.1;refGene_gn=DNMT3B;refGene_tx=XM_023251338.1
A3 26397723 26397785
refGene_gn=DNMT3B;refGene_tx=XM_006929850.4;refGene_gn=DNMT3B;refGene_tx=XM_019826654.2;refGene_gn=DNMT3B;refGene_tx=XM_019826656.2;refGene_gn=DNMT3B;refGene_tx=XM_023251332.1;refGene_gn=DNMT3B;refGene_tx=XM_023251333.1;refGene_gn=DNMT3B;refGene_tx=XM_023251334.1;refGene_gn=DNMT3B;refGene_tx=XM_023251335.1;refGene_gn=DNMT3B;refGene_tx=XM_023251337.1;refGene_gn=DNMT3B;refGene_tx=XM_023251338.1
A3 26398765 26398928
refGene_gn=DNMT3B;refGene_tx=XM_006929850.4;refGene_gn=DNMT3B;refGene_tx=XM_019826654.2;refGene_gn=DNMT3B;refGene_tx=XM_019826656.2;refGene_gn=DNMT3B;refGene_tx=XM_023251332.1;refGene_gn=DNMT3B;refGene_tx=XM_02

3251333.1;refGene_gn=DNMT3B;refGene_tx=XM_023251334.1;refGene_gn=DNMT3B;refGene_tx=XM_023251335.1;refGene_gn=DNMT3B;refGene_tx=XM_023251337.1;refGene_gn=DNMT3B;refGene_tx=XM_023251338.1
A3 26399121 26399151
refGene_gn=DNMT3B;refGene_tx=XM_006929850.4;refGene_gn=DNMT3B;refGene_tx=XM_019826654.2
A3 26407057 26407159
refGene_gn=DNMT3B;refGene_tx=XM_023251332.1;refGene_gn=DNMT3B;refGene_tx=XM_023251334.1;refGene_gn=DNMT3B;refGene_tx=XM_023251335.1;refGene_gn=DNMT3B;refGene_tx=XM_023251337.1;refGene_gn=DNMT3B;refGene_tx=XM_023251338.1
A3 26432107 26432191 refGene_gn=COMMD7;refGene_tx=XM_023251339.1
A3 26439597 26439651 refGene_gn=COMMD7;refGene_tx=XM_023251339.1
A3 26439744 26439847 refGene_gn=COMMD7;refGene_tx=XM_023251339.1
A3 26448692 26448749 refGene_gn=COMMD7;refGene_tx=XM_023251339.1
A3 26448834 26448872 refGene_gn=COMMD7;refGene_tx=XM_023251339.1
A3 26449513 26449604 refGene_gn=COMMD7;refGene_tx=XM_023251339.1
A3 26449914 26449964 refGene_gn=COMMD7;refGene_tx=XM_023251339.1
A3 26450342 26450391 refGene_gn=COMMD7;refGene_tx=XM_023251339.1
A3 26450914 26450991 refGene_gn=COMMD7;refGene_tx=XM_023251339.1
A3 26490489 26490558
refGene_gn=LOC109498496;refGene_tx=XM_019828003.1
A3 26490559 26490606
refGene_gn=LOC109498496;refGene_tx=XM_019828003.1
A3 26490607 26490718
refGene_gn=LOC109498496;refGene_tx=XM_019828003.1
A3 26490719 26490828
refGene_gn=LOC109498496;refGene_tx=XM_019828003.1
A3 26541722 26542043 refGene_gn=NOL4L;refGene_tx=XM_023251347.1
A3 26589131 26589194 refGene_gn=NOL4L;refGene_tx=XM_006929852.3
A3 26591349 26591505
refGene_gn=NOL4L;refGene_tx=XM_006929852.3;refGene_gn=NOL4L;refGene_tx=XM_023251347.1;refGene_gn=NOL4L;refGene_tx=XM_023251348.1
A3 26597224 26597336
refGene_gn=NOL4L;refGene_tx=XM_006929852.3;refGene_gn=NOL4L;refGene_tx=XM_023251347.1;refGene_gn=NOL4L;refGene_tx=XM_023251348.1
A3 26605525 26605635
refGene_gn=NOL4L;refGene_tx=XM_006929852.3;refGene_gn=NOL4L;refGene_tx=XM_023251347.1;refGene_gn=NOL4L;refGene_tx=XM_023251348.1
A3 26637414 26637556
refGene_gn=NOL4L;refGene_tx=XM_003983626.4;refGene_gn=NOL4L;refGene_tx=XM_006929852.3;refGene_gn=NOL4L;refGene_tx=XM_006929854.4;refGene_gn=NOL4L;refGene_tx=XM_023251347.1;refGene_gn=NOL4L;refGene_tx=XM_023251348.1
A3 26653459 26653737
refGene_gn=NOL4L;refGene_tx=XM_003983626.4;refGene_gn=NOL4L;refGene_tx=XM_006929852.3;refGene_gn=NOL4L;refGene_tx=XM_006929854.4;refGene_gn=NOL4L;refGene_tx=XM_023251347.1;refGene_gn=NOL4L;refGene_tx=XM_023251348.1
A3 26655757 26655943

refGene_gn=NOL4L;refGene_tx=XM_003983626.4;refGene_gn=NOL4L;refGene_tx=XM_006929852.3;refGene_gn=NOL4L;refGene_tx=XM_006929854.4;refGene_gn=NOL4L;refGene_tx=XM_023251347.1;refGene_gn=NOL4L;refGene_tx=XM_023251348.1
A3 26656024 26656216
refGene_gn=NOL4L;refGene_tx=XM_003983626.4;refGene_gn=NOL4L;refGene_tx=XM_006929852.3;refGene_gn=NOL4L;refGene_tx=XM_006929854.4;refGene_gn=NOL4L;refGene_tx=XM_023251347.1;refGene_gn=NOL4L;refGene_tx=XM_023251348.1
A3 26656496 26656619
refGene_gn=NOL4L;refGene_tx=XM_003983626.4;refGene_gn=NOL4L;refGene_tx=XM_006929852.3;refGene_gn=NOL4L;refGene_tx=XM_006929854.4;refGene_gn=NOL4L;refGene_tx=XM_023251347.1;refGene_gn=NOL4L;refGene_tx=XM_023251348.1
A3 26656923 26657125
refGene_gn=NOL4L;refGene_tx=XM_003983626.4;refGene_gn=NOL4L;refGene_tx=XM_006929852.3;refGene_gn=NOL4L;refGene_tx=XM_006929854.4;refGene_gn=NOL4L;refGene_tx=XM_023251347.1;refGene_gn=NOL4L;refGene_tx=XM_023251348.1
A3 26661513 26661734
refGene_gn=NOL4L;refGene_tx=XM_003983626.4;refGene_gn=NOL4L;refGene_tx=XM_006929852.3;refGene_gn=NOL4L;refGene_tx=XM_006929854.4;refGene_gn=NOL4L;refGene_tx=XM_023251347.1;refGene_gn=NOL4L;refGene_tx=XM_023251348.1
A3 26671685 26674604
refGene_gn=ASXL1;refGene_tx=XM_023251340.1;refGene_gn=ASXL1;refGene_tx=XM_023251341.1;refGene_gn=ASXL1;refGene_tx=XM_023251342.1;refGene_gn=ASXL1;refGene_tx=XM_023251344.1;refGene_gn=ASXL1;refGene_tx=XM_023251345.1;refGene_gn=ASXL1;refGene_tx=XM_023251346.1
A3 26675131 26675765
refGene_gn=ASXL1;refGene_tx=XM_023251340.1;refGene_gn=ASXL1;refGene_tx=XM_023251341.1;refGene_gn=ASXL1;refGene_tx=XM_023251342.1;refGene_gn=ASXL1;refGene_tx=XM_023251344.1;refGene_gn=ASXL1;refGene_tx=XM_023251345.1;refGene_gn=ASXL1;refGene_tx=XM_023251346.1
A3 26676049 26676155
refGene_gn=ASXL1;refGene_tx=XM_023251340.1;refGene_gn=ASXL1;refGene_tx=XM_023251341.1;refGene_gn=ASXL1;refGene_tx=XM_023251342.1;refGene_gn=ASXL1;refGene_tx=XM_023251344.1;refGene_gn=ASXL1;refGene_tx=XM_023251345.1;refGene_gn=ASXL1;refGene_tx=XM_023251346.1
A3 26677505 26677602
refGene_gn=ASXL1;refGene_tx=XM_023251340.1;refGene_gn=ASXL1;refGene_tx=XM_023251341.1;refGene_gn=ASXL1;refGene_tx=XM_023251342.1;refGene_gn=ASXL1;refGene_tx=XM_023251344.1;refGene_gn=ASXL1;refGene_tx=XM_023251345.1;refGene_gn=ASXL1;refGene_tx=XM_023251346.1
A3 26677697 26677861
refGene_gn=ASXL1;refGene_tx=XM_023251340.1;refGene_gn=ASXL1;refGene_tx=XM_023251341.1;refGene_gn=ASXL1;refGene_tx=XM_023251342.1;refGene_gn=ASXL1;refGene_tx=XM_023251344.1;refGene_gn=ASXL1;refGene_tx=XM_023251345.1;refGene_gn=ASXL1;refGene_tx=XM_023251346.1
A3 26679114 26679267

refGene_gn=ASXL1;refGene_tx=XM_023251340.1;refGene_gn=ASXL1;refGene_tx=XM_023251341.1;refGene_gn=ASXL1;refGene_tx=XM_023251342.1;refGene_gn=ASXL1;refGene_tx=XM_023251344.1;refGene_gn=ASXL1;refGene_tx=XM_023251345.1
A3 26679728 26679822
refGene_gn=ASXL1;refGene_tx=XM_023251340.1;refGene_gn=ASXL1;refGene_tx=XM_023251341.1;refGene_gn=ASXL1;refGene_tx=XM_023251342.1;refGene_gn=ASXL1;refGene_tx=XM_023251344.1;refGene_gn=ASXL1;refGene_tx=XM_023251345.1;refGene_gn=ASXL1;refGene_tx=XM_023251346.1
A3 26680410 26680508
refGene_gn=ASXL1;refGene_tx=XM_023251340.1;refGene_gn=ASXL1;refGene_tx=XM_023251341.1;refGene_gn=ASXL1;refGene_tx=XM_023251342.1;refGene_gn=ASXL1;refGene_tx=XM_023251344.1;refGene_gn=ASXL1;refGene_tx=XM_023251345.1;refGene_gn=ASXL1;refGene_tx=XM_023251346.1
A3 26680585 26680706
refGene_gn=ASXL1;refGene_tx=XM_023251340.1;refGene_gn=ASXL1;refGene_tx=XM_023251341.1;refGene_gn=ASXL1;refGene_tx=XM_023251342.1;refGene_gn=ASXL1;refGene_tx=XM_023251344.1;refGene_gn=ASXL1;refGene_tx=XM_023251345.1;refGene_gn=ASXL1;refGene_tx=XM_023251346.1
A3 26740155 26740264
refGene_gn=ASXL1;refGene_tx=XM_023251340.1;refGene_gn=ASXL1;refGene_tx=XM_023251341.1;refGene_gn=ASXL1;refGene_tx=XM_023251342.1;refGene_gn=ASXL1;refGene_tx=XM_023251344.1;refGene_gn=ASXL1;refGene_tx=XM_023251345.1;refGene_gn=ASXL1;refGene_tx=XM_023251346.1
A3 26741616 26741619
refGene_gn=ASXL1;refGene_tx=XM_023251340.1;refGene_gn=ASXL1;refGene_tx=XM_023251342.1;refGene_gn=ASXL1;refGene_tx=XM_023251344.1;refGene_gn=ASXL1;refGene_tx=XM_023251346.1
A3 26742679 26742762
refGene_gn=ASXL1;refGene_tx=XM_023251340.1;refGene_gn=ASXL1;refGene_tx=XM_023251341.1;refGene_gn=ASXL1;refGene_tx=XM_023251342.1;refGene_gn=ASXL1;refGene_tx=XM_023251344.1;refGene_gn=ASXL1;refGene_tx=XM_023251345.1;refGene_gn=ASXL1;refGene_tx=XM_023251346.1
A3 26748519 26748576
refGene_gn=ASXL1;refGene_tx=XM_023251340.1;refGene_gn=ASXL1;refGene_tx=XM_023251341.1;refGene_gn=ASXL1;refGene_tx=XM_023251346.1
A3 26769831 26769928
refGene_gn=KIF3B;refGene_tx=XM_003983627.5;refGene_gn=KIF3B;refGene_tx=XM_006929856.4;refGene_gn=KIF3B;refGene_tx=XM_019826666.1
A3 26770814 26770993
refGene_gn=KIF3B;refGene_tx=XM_003983627.5;refGene_gn=KIF3B;refGene_tx=XM_006929856.4;refGene_gn=KIF3B;refGene_tx=XM_019826666.1
A3 26773166 26773272
refGene_gn=KIF3B;refGene_tx=XM_003983627.5;refGene_gn=KIF3B;refGene_tx=XM_006929856.4;refGene_gn=KIF3B;refGene_tx=XM_019826666.1
A3 26773950 26774064
refGene_gn=KIF3B;refGene_tx=XM_003983627.5;refGene_gn=KIF3B;refGene_tx=XM_006929856.4;refGene_gn=KIF3B;refGene_tx=XM_019826666.1
A3 26779932 26780051
refGene_gn=KIF3B;refGene_tx=XM_003983627.5;refGene_gn=KIF3B;refGene_tx

=XM_006929856.4;refGene_gn=KIF3B;refGene_tx=XM_019826666.1
A3	26780169 26780292
refGene_gn=KIF3B;refGene_tx=XM_003983627.5;refGene_gn=KIF3B;refGene_tx=XM_006929856.4;refGene_gn=KIF3B;refGene_tx=XM_019826666.1
A3	26780506 26780608
refGene_gn=KIF3B;refGene_tx=XM_003983627.5;refGene_gn=KIF3B;refGene_tx=XM_006929856.4;refGene_gn=KIF3B;refGene_tx=XM_019826666.1
A3	26783614 26785018
refGene_gn=KIF3B;refGene_tx=XM_003983627.5;refGene_gn=KIF3B;refGene_tx=XM_006929856.4;refGene_gn=KIF3B;refGene_tx=XM_019826666.1
A3	26835279 26835468 refGene_gn=POFUT1;refGene_tx=XM_003983628.4
A3	26838285 26838528 refGene_gn=POFUT1;refGene_tx=XM_003983628.4
A3	26840347 26840540 refGene_gn=POFUT1;refGene_tx=XM_003983628.4
A3	26850825 26850938 refGene_gn=POFUT1;refGene_tx=XM_003983628.4
A3	26852315 26852498 refGene_gn=POFUT1;refGene_tx=XM_003983628.4
A3	26856777 26856899 refGene_gn=POFUT1;refGene_tx=XM_003983628.4
A3	26858551 26858684 refGene_gn=POFUT1;refGene_tx=XM_003983628.4
A3	26864525 26864785 refGene_gn=PLAGL2;refGene_tx=XM_023251350.1
A3	26867363 26868594 refGene_gn=PLAGL2;refGene_tx=XM_023251350.1
A3	26874871 26875384
refGene_gn=LOC111559887;refGene_tx=XM_023252173.1
A3	26875385 26875489
refGene_gn=LOC111559887;refGene_tx=XM_023252173.1
A3	26875490 26875725
refGene_gn=LOC111559887;refGene_tx=XM_023252173.1
A3	26894961 26895111
refGene_gn=TM9SF4;refGene_tx=XM_003983630.5;refGene_gn=TM9SF4;refGene_tx=XM_011280843.2;refGene_gn=TM9SF4;refGene_tx=XM_019826667.1
A3	26898075 26898165
refGene_gn=TM9SF4;refGene_tx=XM_003983630.5;refGene_gn=TM9SF4;refGene_tx=XM_011280843.2;refGene_gn=TM9SF4;refGene_tx=XM_019826667.1
A3	26899009 26899129
refGene_gn=TM9SF4;refGene_tx=XM_003983630.5;refGene_gn=TM9SF4;refGene_tx=XM_011280843.2;refGene_gn=TM9SF4;refGene_tx=XM_019826667.1
A3	26900420 26900484
refGene_gn=TM9SF4;refGene_tx=XM_003983630.5;refGene_gn=TM9SF4;refGene_tx=XM_011280843.2;refGene_gn=TM9SF4;refGene_tx=XM_019826667.1
A3	26900998 26901174
refGene_gn=TM9SF4;refGene_tx=XM_003983630.5;refGene_gn=TM9SF4;refGene_tx=XM_011280843.2;refGene_gn=TM9SF4;refGene_tx=XM_019826667.1
A3	26902720 26902804
refGene_gn=TM9SF4;refGene_tx=XM_003983630.5;refGene_gn=TM9SF4;refGene_tx=XM_011280843.2;refGene_gn=TM9SF4;refGene_tx=XM_019826667.1
A3	26905619 26905695
refGene_gn=TM9SF4;refGene_tx=XM_003983630.5;refGene_gn=TM9SF4;refGene_tx=XM_011280843.2;refGene_gn=TM9SF4;refGene_tx=XM_019826667.1
A3	26905790 26905872
refGene_gn=TM9SF4;refGene_tx=XM_003983630.5;refGene_gn=TM9SF4;refGene_tx=XM_011280843.2;refGene_gn=TM9SF4;refGene_tx=XM_019826667.1
A3	26906917 26907050

refGene_gn=TM9SF4;refGene_tx=XM_003983630.5;refGene_gn=TM9SF4;refGene_tx=XM_011280843.2;refGene_gn=TM9SF4;refGene_tx=XM_019826667.1
A3 26909538 26909609 refGene_gn=TM9SF4;refGene_tx=XM_003983630.5;refGene_gn=TM9SF4;refGene_tx=XM_011280843.2;refGene_gn=TM9SF4;refGene_tx=XM_019826667.1
A3 26910957 26911069 refGene_gn=TM9SF4;refGene_tx=XM_003983630.5;refGene_gn=TM9SF4;refGene_tx=XM_011280843.2;refGene_gn=TM9SF4;refGene_tx=XM_019826667.1
A3 26911198 26911317 refGene_gn=TM9SF4;refGene_tx=XM_003983630.5;refGene_gn=TM9SF4;refGene_tx=XM_011280843.2;refGene_gn=TM9SF4;refGene_tx=XM_019826667.1
A3 26912632 26912756 refGene_gn=TM9SF4;refGene_tx=XM_003983630.5;refGene_gn=TM9SF4;refGene_tx=XM_011280843.2;refGene_gn=TM9SF4;refGene_tx=XM_019826667.1
A3 26913897 26914027 refGene_gn=TM9SF4;refGene_tx=XM_003983630.5;refGene_gn=TM9SF4;refGene_tx=XM_011280843.2;refGene_gn=TM9SF4;refGene_tx=XM_019826667.1
A3 26914127 26914296 refGene_gn=TM9SF4;refGene_tx=XM_003983630.5;refGene_gn=TM9SF4;refGene_tx=XM_011280843.2;refGene_gn=TM9SF4;refGene_tx=XM_019826667.1
A3 26919103 26919203 refGene_gn=TM9SF4;refGene_tx=XM_003983630.5;refGene_gn=TM9SF4;refGene_tx=XM_019826667.1
A3 26924424 26924538 refGene_gn=TM9SF4;refGene_tx=XM_003983630.5;refGene_gn=TM9SF4;refGene_tx=XM_019826667.1
A3 26924634 26924649 refGene_gn=TM9SF4;refGene_tx=XM_019826667.1
A3 26945697 26945712 refGene_gn=TM9SF4;refGene_tx=XM_003983630.5
A3 26950595 26950795 refGene_gn=HCK;refGene_tx=XM_006929858.4
A3 26953986 26954118 refGene_gn=HCK;refGene_tx=XM_006929858.4
A3 26954946 26955100 refGene_gn=HCK;refGene_tx=XM_006929858.4
A3 26961117 26961194 refGene_gn=HCK;refGene_tx=XM_006929858.4
A3 26961989 26962169 refGene_gn=HCK;refGene_tx=XM_006929858.4
A3 26963641 26963794 refGene_gn=HCK;refGene_tx=XM_006929858.4
A3 26964328 26964478 refGene_gn=HCK;refGene_tx=XM_006929858.4
A3 26967393 26967497 refGene_gn=HCK;refGene_tx=XM_006929858.4
A3 26970912 26971011 refGene_gn=HCK;refGene_tx=XM_006929858.4
A3 26972262 26972362 refGene_gn=HCK;refGene_tx=XM_006929858.4
A3 26972711 26972754 refGene_gn=HCK;refGene_tx=XM_006929858.4
A3 26975805 26975919 refGene_gn=HCK;refGene_tx=XM_006929858.4
A3 27011204 27011515 refGene_gn=CCM2L;refGene_tx=XM_023251352.1
A3 27012434 27012573 refGene_gn=CCM2L;refGene_tx=XM_023251352.1
A3 27013147 27013277 refGene_gn=CCM2L;refGene_tx=XM_023251352.1
A3 27015831 27015895 refGene_gn=CCM2L;refGene_tx=XM_023251352.1
A3 27018155 27018291 refGene_gn=CCM2L;refGene_tx=XM_023251352.1
A3 27021137 27021601 refGene_gn=CCM2L;refGene_tx=XM_023251352.1
A3 27022390 27022574 refGene_gn=CCM2L;refGene_tx=XM_023251352.1
A3 27022672 27022756 refGene_gn=CCM2L;refGene_tx=XM_023251352.1
A3 27024289 27024457 refGene_gn=CCM2L;refGene_tx=XM_023251352.1
A3 27027252 27027282 refGene_gn=CCM2L;refGene_tx=XM_023251352.1

A3 27039046 27040002 refGene_gn=XKR7;refGene_tx=XM_023251353.1
A3 27041280 27041483 refGene_gn=XKR7;refGene_tx=XM_023251353.1
A3 27062423 27063010 refGene_gn=XKR7;refGene_tx=XM_023251353.1
A3 27077315 27077402 refGene_gn=PDRG1;refGene_tx=XM_003983632.4
A3 27079014 27079090 refGene_gn=PDRG1;refGene_tx=XM_003983632.4
A3 27080047 27080122 refGene_gn=PDRG1;refGene_tx=XM_003983632.4
A3 27082037 27082118 refGene_gn=PDRG1;refGene_tx=XM_003983632.4
A3 27082716 27082799 refGene_gn=PDRG1;refGene_tx=XM_003983632.4
A3 27085688 27085765
refGene_gn=TTLL9;refGene_tx=XM_006929859.3;refGene_gn=TTLL9;refGene_tx=XM_019826670.1;refGene_gn=TTLL9;refGene_tx=XM_019826671.1;refGene_gn=TTLL9;refGene_tx=XM_019826672.1
A3 27089887 27090012
refGene_gn=TTLL9;refGene_tx=XM_006929859.3;refGene_gn=TTLL9;refGene_tx=XM_019826670.1;refGene_gn=TTLL9;refGene_tx=XM_019826671.1;refGene_gn=TTLL9;refGene_tx=XM_019826672.1
A3 27091421 27091535
refGene_gn=TTLL9;refGene_tx=XM_006929859.3;refGene_gn=TTLL9;refGene_tx=XM_019826670.1;refGene_gn=TTLL9;refGene_tx=XM_019826671.1;refGene_gn=TTLL9;refGene_tx=XM_019826672.1
A3 27093262 27093459
refGene_gn=TTLL9;refGene_tx=XM_006929859.3;refGene_gn=TTLL9;refGene_tx=XM_019826670.1;refGene_gn=TTLL9;refGene_tx=XM_019826671.1;refGene_gn=TTLL9;refGene_tx=XM_019826672.1
A3 27094526 27094585
refGene_gn=TTLL9;refGene_tx=XM_006929859.3;refGene_gn=TTLL9;refGene_tx=XM_019826670.1;refGene_gn=TTLL9;refGene_tx=XM_019826671.1;refGene_gn=TTLL9;refGene_tx=XM_019826672.1
A3 27099938 27100026
refGene_gn=TTLL9;refGene_tx=XM_006929859.3;refGene_gn=TTLL9;refGene_tx=XM_019826670.1;refGene_gn=TTLL9;refGene_tx=XM_019826671.1;refGene_gn=TTLL9;refGene_tx=XM_019826672.1
A3 27100880 27100921
refGene_gn=TTLL9;refGene_tx=XM_006929859.3;refGene_gn=TTLL9;refGene_tx=XM_019826670.1;refGene_gn=TTLL9;refGene_tx=XM_019826671.1;refGene_gn=TTLL9;refGene_tx=XM_019826672.1
A3 27101841 27101932
refGene_gn=TTLL9;refGene_tx=XM_006929859.3;refGene_gn=TTLL9;refGene_tx=XM_019826670.1;refGene_gn=TTLL9;refGene_tx=XM_019826671.1;refGene_gn=TTLL9;refGene_tx=XM_019826672.1
A3 27103089 27103164
refGene_gn=TTLL9;refGene_tx=XM_006929859.3;refGene_gn=TTLL9;refGene_tx=XM_019826670.1;refGene_gn=TTLL9;refGene_tx=XM_019826671.1;refGene_gn=TTLL9;refGene_tx=XM_019826672.1
A3 27103761 27103830
refGene_gn=TTLL9;refGene_tx=XM_006929859.3;refGene_gn=TTLL9;refGene_tx=XM_019826670.1;refGene_gn=TTLL9;refGene_tx=XM_019826671.1;refGene_gn=TTLL9;refGene_tx=XM_019826672.1
A3 27113029 27113215
refGene_gn=TTLL9;refGene_tx=XM_006929859.3;refGene_gn=TTLL9;refGene_tx

=XM_019826670.1;refGene_gn=TTLL9;refGene_tx=XM_019826671.1;refGene_gn=TTLL9;refGene_tx=XM_019826672.1
A3 27113986 27114098
refGene_gn=TTLL9;refGene_tx=XM_006929859.3;refGene_gn=TTLL9;refGene_tx=XM_019826670.1;refGene_gn=TTLL9;refGene_tx=XM_019826671.1;refGene_gn=TTLL9;refGene_tx=XM_019826672.1
A3 27114282 27114366
refGene_gn=TTLL9;refGene_tx=XM_006929859.3;refGene_gn=TTLL9;refGene_tx=XM_019826670.1;refGene_gn=TTLL9;refGene_tx=XM_019826671.1
A3 27116189 27116282
refGene_gn=TTLL9;refGene_tx=XM_006929859.3;refGene_gn=TTLL9;refGene_tx=XM_019826670.1;refGene_gn=TTLL9;refGene_tx=XM_019826671.1;refGene_gn=TTLL9;refGene_tx=XM_019826672.1
A3 27123344 27123388
refGene_gn=TTLL9;refGene_tx=XM_006929859.3;refGene_gn=TTLL9;refGene_tx=XM_019826670.1;refGene_gn=TTLL9;refGene_tx=XM_019826671.1;refGene_gn=TTLL9;refGene_tx=XM_019826672.1
A3 27124391 27124421
refGene_gn=TTLL9;refGene_tx=XM_019826670.1;refGene_gn=TTLL9;refGene_tx=XM_019826671.1;refGene_gn=TTLL9;refGene_tx=XM_019826672.1
A3 27126762 27126777 refGene_gn=TTLL9;refGene_tx=XM_006929859.3
A3 27134239 27134251 refGene_gn=DUSP15;refGene_tx=XM_023251357.1
A3 27134489 27134510 refGene_gn=DUSP15;refGene_tx=XM_023251356.1
A3 27135243 27135277
refGene_gn=DUSP15;refGene_tx=XM_023251356.1;refGene_gn=DUSP15;refGene_tx=XM_023251357.1
A3 27136893 27136976
refGene_gn=DUSP15;refGene_tx=XM_023251356.1;refGene_gn=DUSP15;refGene_tx=XM_023251357.1
A3 27139272 27139322
refGene_gn=DUSP15;refGene_tx=XM_023251356.1;refGene_gn=DUSP15;refGene_tx=XM_023251357.1
A3 27140061 27140136
refGene_gn=DUSP15;refGene_tx=XM_023251356.1;refGene_gn=DUSP15;refGene_tx=XM_023251357.1
A3 27141290 27141462
refGene_gn=DUSP15;refGene_tx=XM_023251356.1;refGene_gn=DUSP15;refGene_tx=XM_023251357.1
A3 27142315 27142588
refGene_gn=DUSP15;refGene_tx=XM_023251356.1;refGene_gn=DUSP15;refGene_tx=XM_023251357.1
A3 27145339 27145513
refGene_gn=LOC101080774;refGene_tx=XM_023251355.1
A3 27149927 27150117
refGene_gn=LOC101080774;refGene_tx=XM_023251355.1
A3 27150430 27150531
refGene_gn=LOC101080774;refGene_tx=XM_023251355.1
A3 27151452 27151558
refGene_gn=LOC101080774;refGene_tx=XM_023251355.1
A3 27151676 27151742

refGene_gn=LOC101080774;refGene_tx=XM_023251355.1
A3 27152558 27152927
refGene_gn=LOC101080774;refGene_tx=XM_023251355.1
A3 27153520 27153597
refGene_gn=LOC101080774;refGene_tx=XM_023251355.1
A3 27153598 27153640
refGene_gn=LOC101080774;refGene_tx=XM_023251355.1
A3 27154125 27154260
refGene_gn=LOC101080774;refGene_tx=XM_023251355.1
A3 27156899 27157892 refGene_gn=FOXS1;refGene_tx=XM_003983633.5
A3 27167798 27167879 refGene_gn=MYLK2;refGene_tx=XM_006929860.3
A3 27168801 27168934 refGene_gn=MYLK2;refGene_tx=XM_006929860.3
A3 27169116 27169269 refGene_gn=MYLK2;refGene_tx=XM_006929860.3
A3 27170060 27170189 refGene_gn=MYLK2;refGene_tx=XM_006929860.3
A3 27170328 27170399 refGene_gn=MYLK2;refGene_tx=XM_006929860.3
A3 27172925 27173067 refGene_gn=MYLK2;refGene_tx=XM_006929860.3
A3 27173151 27173261 refGene_gn=MYLK2;refGene_tx=XM_006929860.3
A3 27175486 27175580 refGene_gn=MYLK2;refGene_tx=XM_006929860.3
A3 27176180 27176286 refGene_gn=MYLK2;refGene_tx=XM_006929860.3
A3 27177837 27178139 refGene_gn=MYLK2;refGene_tx=XM_006929860.3
A3 27179049 27179578 refGene_gn=MYLK2;refGene_tx=XM_006929860.3
A3 27180048 27180100 refGene_gn=MYLK2;refGene_tx=XM_006929860.3
A3 27196352 27196463
refGene_gn=TPX2;refGene_tx=XM_011280841.3;refGene_gn=TPX2;refGene_tx=XM_011280842.3;refGene_gn=TPX2;refGene_tx=XM_019826677.2;refGene_gn=TPX2;refGene_tx=XM_019826678.2;refGene_gn=TPX2;refGene_tx=XM_019826679.2;refGene_gn=TPX2;refGene_tx=XM_019826680.2;refGene_gn=TPX2;refGene_tx=XM_019826681.2;refGene_gn=TPX2;refGene_tx=XM_023251359.1
A3 27198023 27198211
refGene_gn=TPX2;refGene_tx=XM_011280841.3;refGene_gn=TPX2;refGene_tx=XM_011280842.3;refGene_gn=TPX2;refGene_tx=XM_019826677.2;refGene_gn=TPX2;refGene_tx=XM_019826678.2;refGene_gn=TPX2;refGene_tx=XM_019826679.2;refGene_gn=TPX2;refGene_tx=XM_019826680.2;refGene_gn=TPX2;refGene_tx=XM_019826681.2;refGene_gn=TPX2;refGene_tx=XM_023251359.1
A3 27198915 27199027
refGene_gn=TPX2;refGene_tx=XM_011280841.3;refGene_gn=TPX2;refGene_tx=XM_011280842.3;refGene_gn=TPX2;refGene_tx=XM_019826677.2;refGene_gn=TPX2;refGene_tx=XM_019826678.2;refGene_gn=TPX2;refGene_tx=XM_019826679.2;refGene_gn=TPX2;refGene_tx=XM_019826680.2;refGene_gn=TPX2;refGene_tx=XM_019826681.2;refGene_gn=TPX2;refGene_tx=XM_023251359.1
A3 27201559 27201706
refGene_gn=TPX2;refGene_tx=XM_011280841.3;refGene_gn=TPX2;refGene_tx=XM_011280842.3;refGene_gn=TPX2;refGene_tx=XM_019826677.2;refGene_gn=TPX2;refGene_tx=XM_019826678.2;refGene_gn=TPX2;refGene_tx=XM_019826679.2;refGene_gn=TPX2;refGene_tx=XM_019826680.2;refGene_gn=TPX2;refGene_tx=XM_019826681.2;refGene_gn=TPX2;refGene_tx=XM_023251359.1
A3 27202139 27202316
refGene_gn=TPX2;refGene_tx=XM_011280841.3;refGene_gn=TPX2;refGene_tx=XM_011280842.3;refGene_gn=TPX2;refGene_tx=XM_019826677.2;refGene_gn=TPX2;refGene_tx=XM_019826678.2;refGene_gn=TPX2;refGene_tx=XM_019826679.2;

refGene_gn=TPX2;refGene_tx=XM_019826680.2;refGene_gn=TPX2;refGene_tx=XM_019826681.2;refGene_gn=TPX2;refGene_tx=XM_023251359.1
A3 27203581 27203677
refGene_gn=TPX2;refGene_tx=XM_011280841.3;refGene_gn=TPX2;refGene_tx=XM_011280842.3;refGene_gn=TPX2;refGene_tx=XM_019826677.2;refGene_gn=TPX2;refGene_tx=XM_019826678.2;refGene_gn=TPX2;refGene_tx=XM_019826679.2;refGene_gn=TPX2;refGene_tx=XM_019826680.2;refGene_gn=TPX2;refGene_tx=XM_019826681.2;refGene_gn=TPX2;refGene_tx=XM_023251359.1
A3 27212561 27212778
refGene_gn=TPX2;refGene_tx=XM_011280841.3;refGene_gn=TPX2;refGene_tx=XM_011280842.3;refGene_gn=TPX2;refGene_tx=XM_019826677.2;refGene_gn=TPX2;refGene_tx=XM_019826678.2;refGene_gn=TPX2;refGene_tx=XM_019826679.2;refGene_gn=TPX2;refGene_tx=XM_019826680.2;refGene_gn=TPX2;refGene_tx=XM_019826681.2;refGene_gn=TPX2;refGene_tx=XM_023251359.1
A3 27214072 27214214
refGene_gn=TPX2;refGene_tx=XM_011280841.3;refGene_gn=TPX2;refGene_tx=XM_011280842.3;refGene_gn=TPX2;refGene_tx=XM_019826677.2;refGene_gn=TPX2;refGene_tx=XM_019826678.2;refGene_gn=TPX2;refGene_tx=XM_019826679.2;refGene_gn=TPX2;refGene_tx=XM_019826680.2;refGene_gn=TPX2;refGene_tx=XM_019826681.2;refGene_gn=TPX2;refGene_tx=XM_023251359.1
A3 27217282 27217451
refGene_gn=TPX2;refGene_tx=XM_011280841.3;refGene_gn=TPX2;refGene_tx=XM_011280842.3;refGene_gn=TPX2;refGene_tx=XM_019826677.2;refGene_gn=TPX2;refGene_tx=XM_019826678.2;refGene_gn=TPX2;refGene_tx=XM_019826679.2;refGene_gn=TPX2;refGene_tx=XM_019826680.2;refGene_gn=TPX2;refGene_tx=XM_019826681.2;refGene_gn=TPX2;refGene_tx=XM_023251359.1
A3 27219666 27219818
refGene_gn=TPX2;refGene_tx=XM_011280841.3;refGene_gn=TPX2;refGene_tx=XM_011280842.3;refGene_gn=TPX2;refGene_tx=XM_019826677.2;refGene_gn=TPX2;refGene_tx=XM_019826678.2;refGene_gn=TPX2;refGene_tx=XM_019826679.2;refGene_gn=TPX2;refGene_tx=XM_019826680.2;refGene_gn=TPX2;refGene_tx=XM_019826681.2;refGene_gn=TPX2;refGene_tx=XM_023251359.1
A3 27220924 27221046
refGene_gn=TPX2;refGene_tx=XM_011280841.3;refGene_gn=TPX2;refGene_tx=XM_011280842.3;refGene_gn=TPX2;refGene_tx=XM_019826677.2;refGene_gn=TPX2;refGene_tx=XM_019826678.2;refGene_gn=TPX2;refGene_tx=XM_019826679.2;refGene_gn=TPX2;refGene_tx=XM_019826680.2;refGene_gn=TPX2;refGene_tx=XM_019826681.2;refGene_gn=TPX2;refGene_tx=XM_023251359.1
A3 27222561 27222690
refGene_gn=TPX2;refGene_tx=XM_011280841.3;refGene_gn=TPX2;refGene_tx=XM_011280842.3;refGene_gn=TPX2;refGene_tx=XM_019826677.2;refGene_gn=TPX2;refGene_tx=XM_019826678.2;refGene_gn=TPX2;refGene_tx=XM_019826679.2;refGene_gn=TPX2;refGene_tx=XM_019826680.2;refGene_gn=TPX2;refGene_tx=XM_019826681.2;refGene_gn=TPX2;refGene_tx=XM_023251359.1
A3 27223791 27223920
refGene_gn=TPX2;refGene_tx=XM_011280841.3;refGene_gn=TPX2;refGene_tx=XM_011280842.3;refGene_gn=TPX2;refGene_tx=XM_019826677.2;refGene_gn=TPX2;refGene_tx=XM_019826678.2;refGene_gn=TPX2;refGene_tx=XM_019826679.2;refGene_gn=TPX2;refGene_tx=XM_019826680.2;refGene_gn=TPX2;refGene_tx=XM_019826681.2;refGene_gn=TPX2;refGene_tx=XM_023251359.1

A3 27231078 27231205
refGene_gn=TPX2;refGene_tx=XM_011280841.3;refGene_gn=TPX2;refGene_tx=XM_011280842.3;refGene_gn=TPX2;refGene_tx=XM_019826677.2;refGene_gn=TPX2;refGene_tx=XM_019826678.2;refGene_gn=TPX2;refGene_tx=XM_019826679.2;refGene_gn=TPX2;refGene_tx=XM_019826680.2;refGene_gn=TPX2;refGene_tx=XM_019826681.2;refGene_gn=TPX2;refGene_tx=XM_023251359.1
A3 27237447 27237570
refGene_gn=TPX2;refGene_tx=XM_011280841.3;refGene_gn=TPX2;refGene_tx=XM_011280842.3;refGene_gn=TPX2;refGene_tx=XM_019826677.2;refGene_gn=TPX2;refGene_tx=XM_019826678.2;refGene_gn=TPX2;refGene_tx=XM_019826679.2;refGene_gn=TPX2;refGene_tx=XM_019826680.2;refGene_gn=TPX2;refGene_tx=XM_019826681.2;refGene_gn=TPX2;refGene_tx=XM_023251359.1
A3 27239428 27239534
refGene_gn=TPX2;refGene_tx=XM_011280841.3;refGene_gn=TPX2;refGene_tx=XM_011280842.3;refGene_gn=TPX2;refGene_tx=XM_019826677.2;refGene_gn=TPX2;refGene_tx=XM_019826678.2;refGene_gn=TPX2;refGene_tx=XM_019826679.2;refGene_gn=TPX2;refGene_tx=XM_019826680.2;refGene_gn=TPX2;refGene_tx=XM_019826681.2;refGene_gn=TPX2;refGene_tx=XM_023251359.1
A3 27268082 27268646 refGene_gn=BCL2L1;refGene_tx=NM_001009228.1
A3 27313136 27313274 refGene_gn=BCL2L1;refGene_tx=NM_001009228.1
A3 27332536 27332673
refGene_gn=LOC101096643;refGene_tx=XM_003983639.3;refGene_gn=LOC101096643;refGene_tx=XM_019826684.2;refGene_gn=LOC101096643;refGene_tx=XM_019826686.2;refGene_gn=LOC101096643;refGene_tx=XM_019826688.2
A3 27333429 27333455
refGene_gn=LOC101096643;refGene_tx=XM_019826685.2
A3 27333944 27334076
refGene_gn=LOC101096643;refGene_tx=XM_003983639.3;refGene_gn=LOC101096643;refGene_tx=XM_019826684.2;refGene_gn=LOC101096643;refGene_tx=XM_019826685.2;refGene_gn=LOC101096643;refGene_tx=XM_019826686.2;refGene_gn=LOC101096643;refGene_tx=XM_019826688.2
A3 27334869 27335034
refGene_gn=LOC101096643;refGene_tx=XM_003983639.3;refGene_gn=LOC101096643;refGene_tx=XM_019826684.2;refGene_gn=LOC101096643;refGene_tx=XM_019826685.2;refGene_gn=LOC101096643;refGene_tx=XM_019826686.2;refGene_gn=LOC101096643;refGene_tx=XM_019826688.2
A3 27336613 27336695
refGene_gn=LOC101096643;refGene_tx=XM_003983639.3;refGene_gn=LOC101096643;refGene_tx=XM_019826684.2;refGene_gn=LOC101096643;refGene_tx=XM_019826685.2
A3 27337728 27337761
refGene_gn=LOC101096643;refGene_tx=XM_019826684.2;refGene_gn=LOC101096643;refGene_tx=XM_019826685.2
A3 27362431 27362473 refGene_gn=ID1;refGene_tx=XM_003983640.4
A3 27362698 27363121 refGene_gn=ID1;refGene_tx=XM_003983640.4
A3 27386399 27387006
refGene_gn=LOC105260422;refGene_tx=XM_023251361.1
A3 27390148 27390251
refGene_gn=HM13;refGene_tx=XM_003983641.4;refGene_gn=HM13;refGene_tx=XM_003983642.4;refGene_gn=HM13;refGene_tx=XM_019826689.2

A3 27391150 27391297 refGene_gn=HM13;refGene_tx=XM_003983642.4
A3 27393026 27393112
refGene_gn=HM13;refGene_tx=XM_003983641.4;refGene_gn=HM13;refGene_tx=XM_003983642.4;refGene_gn=HM13;refGene_tx=XM_019826689.2
A3 27395772 27395912
refGene_gn=LOC105260422;refGene_tx=XM_023251361.1
A3 27397585 27397688
refGene_gn=HM13;refGene_tx=XM_003983641.4;refGene_gn=HM13;refGene_tx=XM_003983642.4;refGene_gn=HM13;refGene_tx=XM_019826689.2
A3 27400270 27400307
refGene_gn=HM13;refGene_tx=XM_003983641.4;refGene_gn=HM13;refGene_tx=XM_003983642.4;refGene_gn=HM13;refGene_tx=XM_019826689.2
A3 27403421 27403505
refGene_gn=HM13;refGene_tx=XM_003983641.4;refGene_gn=HM13;refGene_tx=XM_003983642.4;refGene_gn=HM13;refGene_tx=XM_019826689.2
A3 27406675 27406733
refGene_gn=HM13;refGene_tx=XM_003983641.4;refGene_gn=HM13;refGene_tx=XM_003983642.4;refGene_gn=HM13;refGene_tx=XM_019826689.2
A3 27407540 27407666
refGene_gn=HM13;refGene_tx=XM_003983641.4;refGene_gn=HM13;refGene_tx=XM_003983642.4;refGene_gn=HM13;refGene_tx=XM_019826689.2
A3 27407744 27407884
refGene_gn=HM13;refGene_tx=XM_003983641.4;refGene_gn=HM13;refGene_tx=XM_003983642.4;refGene_gn=HM13;refGene_tx=XM_019826689.2
A3 27409729 27409818
refGene_gn=HM13;refGene_tx=XM_003983641.4;refGene_gn=HM13;refGene_tx=XM_003983642.4;refGene_gn=HM13;refGene_tx=XM_019826689.2
A3 27414876 27414959
refGene_gn=HM13;refGene_tx=XM_003983641.4;refGene_gn=HM13;refGene_tx=XM_003983642.4;refGene_gn=HM13;refGene_tx=XM_019826689.2
A3 27419927 27420056 refGene_gn=HM13;refGene_tx=XM_019826689.2
A3 27420431 27420530
refGene_gn=HM13;refGene_tx=XM_003983641.4;refGene_gn=HM13;refGene_tx=XM_003983642.4;refGene_gn=HM13;refGene_tx=XM_019826689.2
A3 27428811 27428994
refGene_gn=HM13;refGene_tx=XM_003983641.4;refGene_gn=HM13;refGene_tx=XM_003983642.4;refGene_gn=HM13;refGene_tx=XM_019826689.2
A3 27454951 27455220 refGene_gn=REM1;refGene_tx=XM_011280932.2
A3 27456422 27456624 refGene_gn=REM1;refGene_tx=XM_011280932.2
A3 27459319 27459402 refGene_gn=REM1;refGene_tx=XM_011280932.2
A3 27460463 27460803 refGene_gn=REM1;refGene_tx=XM_011280932.2
A3 27464839 27464897 refGene_gn=DEFB124;refGene_tx=XM_019826691.1
A3 27467321 27467476 refGene_gn=DEFB124;refGene_tx=XM_019826691.1
A3 27480058 27480252 refGene_gn=DEFB123;refGene_tx=XM_003983689.5
A3 27488499 27488557 refGene_gn=DEFB123;refGene_tx=XM_003983689.5
A3 27494684 27494742
refGene_gn=LOC101087728;refGene_tx=XM_019828049.2
A3 27496737 27496874
refGene_gn=LOC101087728;refGene_tx=XM_019828049.2
A3 27502500 27502581

refGene_gn=LOC101087728;refGene_tx=XM_019828049.2
A3	27512213 27512271 refGene_gn=DEFB121;refGene_tx=XM_023251362.1
A3	27513426 27513584 refGene_gn=DEFB121;refGene_tx=XM_023251362.1
A3	27522740 27522795
refGene_gn=LOC101081718;refGene_tx=XM_011280931.3
A3	27531789 27531983
refGene_gn=LOC101081718;refGene_tx=XM_011280931.3
A3	27536574 27536700
refGene_gn=LOC109498512;refGene_tx=XM_019828050.1
A3	27536980 27537047
refGene_gn=LOC109498512;refGene_tx=XM_019828050.1
A3	27537048 27537079
refGene_gn=LOC109498512;refGene_tx=XM_019828050.1
A3	27545928 27546000
refGene_gn=LOC109498512;refGene_tx=XM_019828050.1
A3	27546714 27546781
refGene_gn=LOC109498512;refGene_tx=XM_019828050.1
A3	27609905 27609972 refGene_gn=DEFB116;refGene_tx=XM_003983665.3
A3	27614811 27615050 refGene_gn=DEFB116;refGene_tx=XM_003983665.3
A3	27654209 27654448
refGene_gn=LOC109498039;refGene_tx=XM_019826695.2
A3	27657044 27657138
refGene_gn=LOC109498039;refGene_tx=XM_019826695.2
A3	27672029 27672087 refGene_gn=DEFB125;refGene_tx=XM_019826694.2
A3	27678614 27678938 refGene_gn=DEFB125;refGene_tx=XM_019826694.2
A3	27679035 27679130 refGene_gn=DEFB125;refGene_tx=XM_019826694.2
A3	27692043 27692092 refGene_gn=DEFB127;refGene_tx=XM_003983666.4
A3	27693628 27693882 refGene_gn=DEFB127;refGene_tx=XM_003983666.4
A3	27709946 27710182 refGene_gn=DEFB128;refGene_tx=XM_023251363.1
A3	27711452 27711504 refGene_gn=DEFB128;refGene_tx=XM_023251363.1
A3	27747260 27747318 refGene_gn=DEFB129;refGene_tx=XM_003983667.4
A3	27749451 27749897 refGene_gn=DEFB129;refGene_tx=XM_003983667.4
A3	27781248 27781438 refGene_gn=DEFB132;refGene_tx=XM_023252174.1
A3	27782600 27782742 refGene_gn=DEFB132;refGene_tx=XM_023252174.1
A3	27782845 27782894 refGene_gn=DEFB132;refGene_tx=XM_023252174.1
A3	27810545 27810606
refGene_gn=CA3H20orf96;refGene_tx=XM_023251364.1;refGene_gn=CA3H20orf9
6;refGene_tx=XM_023251365.1;refGene_gn=CA3H20orf96;refGene_tx=XM_02325
1366.1
A3	27811953 27812072
refGene_gn=CA3H20orf96;refGene_tx=XM_023251364.1;refGene_gn=CA3H20orf9
6;refGene_tx=XM_023251365.1;refGene_gn=CA3H20orf96;refGene_tx=XM_02325
1366.1
A3	27812743 27812830
refGene_gn=CA3H20orf96;refGene_tx=XM_023251364.1;refGene_gn=CA3H20orf9
6;refGene_tx=XM_023251365.1;refGene_gn=CA3H20orf96;refGene_tx=XM_02325
1366.1
A3	27812989 27813091
refGene_gn=CA3H20orf96;refGene_tx=XM_023251364.1;refGene_gn=CA3H20orf9
6;refGene_tx=XM_023251365.1;refGene_gn=CA3H20orf96;refGene_tx=XM_02325

1366.1
A3 27813176 27813334
refGene_gn=CA3H20orf96;refGene_tx=XM_023251364.1;refGene_gn=CA3H20orf9
6;refGene_tx=XM_023251365.1;refGene_gn=CA3H20orf96;refGene_tx=XM_02325
1366.1
A3 27814032 27814132
refGene_gn=CA3H20orf96;refGene_tx=XM_023251364.1;refGene_gn=CA3H20orf9
6;refGene_tx=XM_023251365.1;refGene_gn=CA3H20orf96;refGene_tx=XM_02325
1366.1
A3 27817328 27817487
refGene_gn=CA3H20orf96;refGene_tx=XM_023251364.1;refGene_gn=CA3H20orf9
6;refGene_tx=XM_023251365.1;refGene_gn=CA3H20orf96;refGene_tx=XM_02325
1366.1
A3 27822787 27822900
refGene_gn=CA3H20orf96;refGene_tx=XM_023251364.1;refGene_gn=CA3H20orf9
6;refGene_tx=XM_023251365.1;refGene_gn=CA3H20orf96;refGene_tx=XM_02325
1366.1
A3 27834247 27834365
refGene_gn=CA3H20orf96;refGene_tx=XM_023251364.1
A3 27834942 27835024
refGene_gn=CA3H20orf96;refGene_tx=XM_023251364.1;refGene_gn=CA3H20orf9
6;refGene_tx=XM_023251365.1
A3 27835258 27835278
refGene_gn=CA3H20orf96;refGene_tx=XM_023251364.1
A3 27840697 27841921 refGene_gn=ZCCHC3;refGene_tx=XM_023251367.1
A3 27864010 27864955 refGene_gn=SOX12;refGene_tx=XM_023251368.1
A3 27881157 27881346
refGene_gn=NRSN2;refGene_tx=XM_003983669.3;refGene_gn=NRSN2;refGene_tx
=XM_019826697.2
A3 27883820 27884246
refGene_gn=NRSN2;refGene_tx=XM_003983669.3;refGene_gn=NRSN2;refGene_tx
=XM_019826697.2
A3 27911561 27911852 refGene_gn=TRIB3;refGene_tx=XM_003983670.4
A3 27913638 27913928 refGene_gn=TRIB3;refGene_tx=XM_003983670.4
A3 27915872 27916365 refGene_gn=TRIB3;refGene_tx=XM_003983670.4
A3 27924771 27924793 refGene_gn=RBCK1;refGene_tx=XM_006929871.2
A3 27925876 27926021 refGene_gn=RBCK1;refGene_tx=XM_006929871.2
A3 27930801 27930895 refGene_gn=RBCK1;refGene_tx=XM_006929871.2
A3 27931037 27931236
refGene_gn=RBCK1;refGene_tx=XM_006929871.2;refGene_gn=RBCK1;refGene_tx
=XM_019826704.1
A3 27932418 27932540
refGene_gn=RBCK1;refGene_tx=XM_006929871.2;refGene_gn=RBCK1;refGene_tx
=XM_019826704.1
A3 27932661 27932835
refGene_gn=RBCK1;refGene_tx=XM_006929871.2;refGene_gn=RBCK1;refGene_tx
=XM_019826704.1
A3 27933984 27934145 refGene_gn=RBCK1;refGene_tx=XM_006929871.2
A3 27935292 27935404
refGene_gn=RBCK1;refGene_tx=XM_006929871.2;refGene_gn=RBCK1;refGene_tx

=XM_019826704.1
A3	27938156	27938336
refGene_gn=RBCK1;refGene_tx=XM_006929871.2;refGene_gn=RBCK1;refGene_tx=XM_019826704.1
A3	27938831	27938930
refGene_gn=RBCK1;refGene_tx=XM_006929871.2;refGene_gn=RBCK1;refGene_tx=XM_019826704.1
A3	27939278	27939422
refGene_gn=RBCK1;refGene_tx=XM_006929871.2;refGene_gn=RBCK1;refGene_tx=XM_019826704.1
A3	27940407	27940488
refGene_gn=RBCK1;refGene_tx=XM_006929871.2;refGene_gn=RBCK1;refGene_tx=XM_019826704.1
A3	27946210	27946466
refGene_gn=TBC1D20;refGene_tx=XM_003983673.5;refGene_gn=TBC1D20;refGene_tx=XM_006929875.4;refGene_gn=TBC1D20;refGene_tx=XM_019826705.2
A3	27946741	27946929
refGene_gn=TBC1D20;refGene_tx=XM_003983673.5;refGene_gn=TBC1D20;refGene_tx=XM_006929875.4;refGene_gn=TBC1D20;refGene_tx=XM_019826705.2
A3	27947740	27947882
refGene_gn=TBC1D20;refGene_tx=XM_003983673.5;refGene_gn=TBC1D20;refGene_tx=XM_006929875.4;refGene_gn=TBC1D20;refGene_tx=XM_019826705.2
A3	27948811	27948913
refGene_gn=TBC1D20;refGene_tx=XM_003983673.5;refGene_gn=TBC1D20;refGene_tx=XM_006929875.4;refGene_gn=TBC1D20;refGene_tx=XM_019826705.2
A3	27949103	27949290
refGene_gn=TBC1D20;refGene_tx=XM_003983673.5;refGene_gn=TBC1D20;refGene_tx=XM_006929875.4;refGene_gn=TBC1D20;refGene_tx=XM_019826705.2
A3	27953058	27953139
refGene_gn=TBC1D20;refGene_tx=XM_003983673.5;refGene_gn=TBC1D20;refGene_tx=XM_006929875.4;refGene_gn=TBC1D20;refGene_tx=XM_019826705.2
A3	27954805	27954991
refGene_gn=TBC1D20;refGene_tx=XM_003983673.5;refGene_gn=TBC1D20;refGene_tx=XM_006929875.4;refGene_gn=TBC1D20;refGene_tx=XM_019826705.2
A3	27962951	27963021	refGene_gn=TBC1D20;refGene_tx=XM_003983673.5
A3	27975224	27975340	refGene_gn=CSNK2A1;refGene_tx=XM_003983674.5
A3	27977342	27977429	refGene_gn=CSNK2A1;refGene_tx=XM_003983674.5
A3	27978518	27978667	refGene_gn=CSNK2A1;refGene_tx=XM_003983674.5
A3	27979434	27979535	refGene_gn=CSNK2A1;refGene_tx=XM_003983674.5
A3	27981166	27981268	refGene_gn=CSNK2A1;refGene_tx=XM_003983674.5
A3	27982191	27982302	refGene_gn=CSNK2A1;refGene_tx=XM_003983674.5
A3	27986527	27986611	refGene_gn=CSNK2A1;refGene_tx=XM_003983674.5
A3	27988091	27988151	refGene_gn=CSNK2A1;refGene_tx=XM_003983674.5
A3	27989503	27989554	refGene_gn=CSNK2A1;refGene_tx=XM_003983674.5
A3	27990067	27990169	refGene_gn=CSNK2A1;refGene_tx=XM_003983674.5
A3	27996185	27996297	refGene_gn=CSNK2A1;refGene_tx=XM_003983674.5
A3	27998300	27998401	refGene_gn=CSNK2A1;refGene_tx=XM_003983674.5
A3	28078987	28079062	refGene_gn=TCF15;refGene_tx=XM_023251369.1
A3	28084479	28084998	refGene_gn=TCF15;refGene_tx=XM_023251369.1
A3	28120715	28120919	refGene_gn=SRXN1;refGene_tx=XM_006929901.4

A3 28124750 28124978 refGene_gn=SRXN1;refGene_tx=XM_006929901.4
A3 28136984 28137512 refGene_gn=SCRT2;refGene_tx=XM_023251370.1
A3 28137513 28137776 refGene_gn=SCRT2;refGene_tx=XM_023251370.1
A3 28148570 28148703 refGene_gn=SCRT2;refGene_tx=XM_023251370.1
A3 28217629 28217842
refGene_gn=SLC52A3;refGene_tx=XM_011280930.3;refGene_gn=SLC52A3;refGene_tx=XM_019826706.2;refGene_gn=SLC52A3;refGene_tx=XM_019826707.2
A3 28218349 28218473
refGene_gn=SLC52A3;refGene_tx=XM_011280930.3;refGene_gn=SLC52A3;refGene_tx=XM_019826706.2;refGene_gn=SLC52A3;refGene_tx=XM_019826707.2
A3 28220143 28220649
refGene_gn=SLC52A3;refGene_tx=XM_011280930.3;refGene_gn=SLC52A3;refGene_tx=XM_019826706.2;refGene_gn=SLC52A3;refGene_tx=XM_019826707.2
A3 28221675 28222242
refGene_gn=SLC52A3;refGene_tx=XM_011280930.3;refGene_gn=SLC52A3;refGene_tx=XM_019826706.2;refGene_gn=SLC52A3;refGene_tx=XM_019826707.2
A3 28255749 28256643
refGene_gn=FAM110A;refGene_tx=XM_003983676.3;refGene_gn=FAM110A;refGene_tx=XM_019826712.2;refGene_gn=FAM110A;refGene_tx=XM_019826713.1;refGene_gn=FAM110A;refGene_tx=XM_019826715.2;refGene_gn=FAM110A;refGene_tx=XM_019826716.2
A3 28289135 28289296
refGene_gn=ANGPT4;refGene_tx=XM_003983700.5;refGene_gn=ANGPT4;refGene_tx=XM_019826717.2
A3 28290119 28290250
refGene_gn=ANGPT4;refGene_tx=XM_003983700.5;refGene_gn=ANGPT4;refGene_tx=XM_019826717.2
A3 28294680 28294847
refGene_gn=ANGPT4;refGene_tx=XM_003983700.5;refGene_gn=ANGPT4;refGene_tx=XM_019826717.2
A3 28296372 28296474
refGene_gn=ANGPT4;refGene_tx=XM_003983700.5;refGene_gn=ANGPT4;refGene_tx=XM_019826717.2
A3 28299079 28299195
refGene_gn=ANGPT4;refGene_tx=XM_003983700.5;refGene_gn=ANGPT4;refGene_tx=XM_019826717.2
A3 28301256 28301489
refGene_gn=ANGPT4;refGene_tx=XM_003983700.5;refGene_gn=ANGPT4;refGene_tx=XM_019826717.2
A3 28304182 28304304
refGene_gn=ANGPT4;refGene_tx=XM_003983700.5;refGene_gn=ANGPT4;refGene_tx=XM_019826717.2
A3 28305496 28305652 refGene_gn=ANGPT4;refGene_tx=XM_003983700.5
A3 28328095 28328407
refGene_gn=ANGPT4;refGene_tx=XM_003983700.5;refGene_gn=ANGPT4;refGene_tx=XM_019826717.2
A3 28359854 28359958 refGene_gn=RSPO4;refGene_tx=XM_011280951.3
A3 28362359 28362545 refGene_gn=RSPO4;refGene_tx=XM_011280951.3
A3 28364336 28364477 refGene_gn=RSPO4;refGene_tx=XM_011280951.3
A3 28365001 28365190 refGene_gn=RSPO4;refGene_tx=XM_011280951.3

A3 28395592 28395671 refGene_gn=RSPO4;refGene_tx=XM_011280951.3
A3 28486184 28486190 refGene_gn=PSMF1;refGene_tx=XM_006929926.4
A3 28490617 28490761
refGene_gn=PSMF1;refGene_tx=XM_006929926.4;refGene_gn=PSMF1;refGene_tx=XM_006929927.3
A3 28497097 28497250
refGene_gn=PSMF1;refGene_tx=XM_006929926.4;refGene_gn=PSMF1;refGene_tx=XM_006929927.3
A3 28498641 28498724
refGene_gn=PSMF1;refGene_tx=XM_006929926.4;refGene_gn=PSMF1;refGene_tx=XM_006929927.3
A3 28509851 28510037
refGene_gn=PSMF1;refGene_tx=XM_006929926.4;refGene_gn=PSMF1;refGene_tx=XM_006929927.3
A3 28530815 28530869
refGene_gn=PSMF1;refGene_tx=XM_006929926.4;refGene_gn=PSMF1;refGene_tx=XM_006929927.3
A3 28532008 28532167
refGene_gn=PSMF1;refGene_tx=XM_006929926.4;refGene_gn=PSMF1;refGene_tx=XM_006929927.3
A3 28532697 28532749
refGene_gn=PSMF1;refGene_tx=XM_006929926.4;refGene_gn=PSMF1;refGene_tx=XM_006929927.3
A3 28547320 28548019
refGene_gn=TMEM74B;refGene_tx=XM_011280929.3;refGene_gn=TMEM74B;refGene_tx=XM_019826723.1;refGene_gn=TMEM74B;refGene_tx=XM_023251372.1;refGene_gn=TMEM74B;refGene_tx=XM_023251373.1
A3 28560149 28560320
refGene_gn=CA3H20orf202;refGene_tx=XM_023252175.1
A3 28563957 28564191
refGene_gn=CA3H20orf202;refGene_tx=XM_023252175.1
A3 28579758 28579902 refGene_gn=RAD21L1;refGene_tx=XM_023251374.1
A3 28581499 28581629 refGene_gn=RAD21L1;refGene_tx=XM_023251374.1
A3 28583570 28583664 refGene_gn=RAD21L1;refGene_tx=XM_023251374.1
A3 28585918 28586025 refGene_gn=RAD21L1;refGene_tx=XM_023251374.1
A3 28589689 28589860 refGene_gn=RAD21L1;refGene_tx=XM_023251374.1
A3 28591301 28591397 refGene_gn=RAD21L1;refGene_tx=XM_023251374.1
A3 28593303 28593414 refGene_gn=RAD21L1;refGene_tx=XM_023251374.1
A3 28594985 28595212 refGene_gn=RAD21L1;refGene_tx=XM_023251374.1
A3 28595450 28595550 refGene_gn=RAD21L1;refGene_tx=XM_023251374.1
A3 28596397 28596522 refGene_gn=RAD21L1;refGene_tx=XM_023251374.1
A3 28598916 28599012 refGene_gn=RAD21L1;refGene_tx=XM_023251374.1
A3 28601314 28601392 refGene_gn=RAD21L1;refGene_tx=XM_023251374.1
A3 28605162 28605351 refGene_gn=RAD21L1;refGene_tx=XM_023251374.1
A3 28639592 28639774
refGene_gn=SNPH;refGene_tx=XM_023251375.1;refGene_gn=SNPH;refGene_tx=XM_023251376.1;refGene_gn=SNPH;refGene_tx=XM_023251377.1;refGene_gn=SNPH;refGene_tx=XM_023251378.1
A3 28640374 28640581
refGene_gn=SNPH;refGene_tx=XM_023251375.1;refGene_gn=SNPH;refGene_tx=X

M_023251376.1;refGene_gn=SNPH;refGene_tx=XM_023251377.1;refGene_gn=SNPH;refGene_tx=XM_023251378.1
A3 28643971 28644121
refGene_gn=SNPH;refGene_tx=XM_023251375.1;refGene_gn=SNPH;refGene_tx=XM_023251376.1;refGene_gn=SNPH;refGene_tx=XM_023251377.1;refGene_gn=SNPH;refGene_tx=XM_023251378.1
A3 28647150 28648321
refGene_gn=SNPH;refGene_tx=XM_023251375.1;refGene_gn=SNPH;refGene_tx=XM_023251376.1;refGene_gn=SNPH;refGene_tx=XM_023251377.1;refGene_gn=SNPH;refGene_tx=XM_023251378.1
A3 28652554 28652609
refGene_gn=SDCBP2;refGene_tx=XM_003983734.5;refGene_gn=SDCBP2;refGene_tx=XM_006929922.4;refGene_gn=SDCBP2;refGene_tx=XM_019826725.2;refGene_gn=SDCBP2;refGene_tx=XM_019826726.2;refGene_gn=SDCBP2;refGene_tx=XM_019826727.1
A3 28652848 28652940
refGene_gn=SDCBP2;refGene_tx=XM_003983734.5;refGene_gn=SDCBP2;refGene_tx=XM_006929922.4;refGene_gn=SDCBP2;refGene_tx=XM_019826725.2;refGene_gn=SDCBP2;refGene_tx=XM_019826726.2;refGene_gn=SDCBP2;refGene_tx=XM_019826727.1
A3 28655055 28655227
refGene_gn=SDCBP2;refGene_tx=XM_003983734.5;refGene_gn=SDCBP2;refGene_tx=XM_006929922.4;refGene_gn=SDCBP2;refGene_tx=XM_019826725.2;refGene_gn=SDCBP2;refGene_tx=XM_019826726.2;refGene_gn=SDCBP2;refGene_tx=XM_019826727.1
A3 28655300 28655476
refGene_gn=SDCBP2;refGene_tx=XM_003983734.5;refGene_gn=SDCBP2;refGene_tx=XM_006929922.4;refGene_gn=SDCBP2;refGene_tx=XM_019826725.2;refGene_gn=SDCBP2;refGene_tx=XM_019826726.2;refGene_gn=SDCBP2;refGene_tx=XM_019826727.1
A3 28656063 28656222
refGene_gn=SDCBP2;refGene_tx=XM_003983734.5;refGene_gn=SDCBP2;refGene_tx=XM_006929922.4;refGene_gn=SDCBP2;refGene_tx=XM_019826725.2;refGene_gn=SDCBP2;refGene_tx=XM_019826726.2;refGene_gn=SDCBP2;refGene_tx=XM_019826727.1
A3 28660829 28660930
refGene_gn=SDCBP2;refGene_tx=XM_003983734.5;refGene_gn=SDCBP2;refGene_tx=XM_006929922.4;refGene_gn=SDCBP2;refGene_tx=XM_019826725.2;refGene_gn=SDCBP2;refGene_tx=XM_019826726.2
A3 28661616 28661671
refGene_gn=SDCBP2;refGene_tx=XM_003983734.5;refGene_gn=SDCBP2;refGene_tx=XM_006929922.4;refGene_gn=SDCBP2;refGene_tx=XM_019826725.2;refGene_gn=SDCBP2;refGene_tx=XM_019826726.2
A3 28662328 28662382
refGene_gn=SDCBP2;refGene_tx=XM_003983734.5;refGene_gn=SDCBP2;refGene_tx=XM_006929922.4;refGene_gn=SDCBP2;refGene_tx=XM_019826725.2;refGene_gn=SDCBP2;refGene_tx=XM_019826726.2
A3 28699466 28699595 refGene_gn=FKBP1A;refGene_tx=XM_006929921.4
A3 28703023 28703136 refGene_gn=FKBP1A;refGene_tx=XM_006929921.4
A3 28722658 28722706 refGene_gn=FKBP1A;refGene_tx=XM_006929921.4

A3 28722793 28722830 refGene_gn=FKBP1A;refGene_tx=XM_006929921.4
A3 28740577 28740740
refGene_gn=NSFL1C;refGene_tx=XM_023251379.1;refGene_gn=NSFL1C;refGene_tx=XM_023251380.1
A3 28741768 28741933
refGene_gn=NSFL1C;refGene_tx=XM_023251379.1;refGene_gn=NSFL1C;refGene_tx=XM_023251380.1
A3 28748312 28748450
refGene_gn=NSFL1C;refGene_tx=XM_023251379.1;refGene_gn=NSFL1C;refGene_tx=XM_023251380.1
A3 28749294 28749404
refGene_gn=NSFL1C;refGene_tx=XM_023251379.1;refGene_gn=NSFL1C;refGene_tx=XM_023251380.1
A3 28750746 28750839
refGene_gn=NSFL1C;refGene_tx=XM_023251379.1;refGene_gn=NSFL1C;refGene_tx=XM_023251380.1
A3 28751464 28751630
refGene_gn=NSFL1C;refGene_tx=XM_023251379.1;refGene_gn=NSFL1C;refGene_tx=XM_023251380.1
A3 28752239 28752245 refGene_gn=NSFL1C;refGene_tx=XM_023251379.1
A3 28754110 28754185
refGene_gn=NSFL1C;refGene_tx=XM_023251379.1;refGene_gn=NSFL1C;refGene_tx=XM_023251380.1
A3 28760237 28760335
refGene_gn=NSFL1C;refGene_tx=XM_023251379.1;refGene_gn=NSFL1C;refGene_tx=XM_023251380.1
A3 28762863 28762968
refGene_gn=NSFL1C;refGene_tx=XM_023251379.1;refGene_gn=NSFL1C;refGene_tx=XM_023251380.1
A3 28770810 28770941 refGene_gn=SIRPB2;refGene_tx=XM_006929949.3
A3 28772214 28772283 refGene_gn=SIRPB2;refGene_tx=XM_006929949.3
A3 28772810 28773152 refGene_gn=SIRPB2;refGene_tx=XM_006929949.3
A3 28774361 28774727 refGene_gn=SIRPB2;refGene_tx=XM_006929949.3
A3 28782059 28782255 refGene_gn=SIRPB2;refGene_tx=XM_006929949.3
A3 28837387 28837710
refGene_gn=LOC111559785;refGene_tx=XM_023251381.1
A3 28850673 28851046
refGene_gn=LOC111559785;refGene_tx=XM_023251381.1
A3 28877026 28877112
refGene_gn=LOC101086051;refGene_tx=XM_019826731.2;refGene_gn=LOC101086051;refGene_tx=XM_019826732.2;refGene_gn=LOC101086051;refGene_tx=XM_019826733.2
A3 28877506 28877857
refGene_gn=LOC101086051;refGene_tx=XM_019826731.2;refGene_gn=LOC101086051;refGene_tx=XM_019826732.2;refGene_gn=LOC101086051;refGene_tx=XM_019826733.2
A3 28879774 28879817
refGene_gn=LOC101086051;refGene_tx=XM_019826733.2
A3 28884773 28884846
refGene_gn=LOC101086051;refGene_tx=XM_019826731.2;refGene_gn=LOC101086

051;refGene_tx=XM_019826732.2
A3 28891263 28891442
refGene_gn=LOC101085793;refGene_tx=XM_023252176.1
A3 28898898 28899237
refGene_gn=LOC101085793;refGene_tx=XM_023252176.1
A3 28899805 28900126
refGene_gn=LOC101085793;refGene_tx=XM_023252176.1
A3 28900816 28900946
refGene_gn=LOC101085793;refGene_tx=XM_023252176.1
A3 28925278 28925652
refGene_gn=LOC105259814;refGene_tx=XM_023251382.1
A3 28926221 28926542
refGene_gn=LOC105259814;refGene_tx=XM_023251382.1
A3 28927235 28927592
refGene_gn=LOC105259814;refGene_tx=XM_023251382.1
A3 28939346 28939419
refGene_gn=LOC105259814;refGene_tx=XM_023251382.1
A3 28957828 28957911
refGene_gn=LOC109498046;refGene_tx=XM_023252177.1
A3 28958304 28958643
refGene_gn=LOC109498046;refGene_tx=XM_023252177.1
A3 28959222 28959543
refGene_gn=LOC109498046;refGene_tx=XM_023252177.1
A3 28959629 28959759
refGene_gn=LOC109498046;refGene_tx=XM_023252177.1
A3 28960717 28960773
refGene_gn=LOC109498046;refGene_tx=XM_023252177.1
A3 28961478 28961628
refGene_gn=LOC109498046;refGene_tx=XM_023252177.1
A3 28961692 28961836
refGene_gn=LOC109498046;refGene_tx=XM_023252177.1
A3 28973950 28974020
refGene_gn=LOC109498046;refGene_tx=XM_023252177.1
A3 28984116 28984223
refGene_gn=LOC109498497;refGene_tx=XM_019828004.2
A3 28984338 28984665
refGene_gn=LOC109498497;refGene_tx=XM_019828004.2
A3 28990733 28990899
refGene_gn=LOC109498497;refGene_tx=XM_019828004.2
A3 28995747 28995767
refGene_gn=LOC101085539;refGene_tx=XM_019826729.1
A3 28996239 28996377
refGene_gn=LOC101085539;refGene_tx=XM_019826729.1
A3 28996553 28996580
refGene_gn=LOC101085539;refGene_tx=XM_019826729.1
A3 28999167 28999521
refGene_gn=LOC101085539;refGene_tx=XM_019826729.1
A3 29006717 29006883
refGene_gn=LOC101085539;refGene_tx=XM_019826729.1
A3 29009093 29024890 refGene_gn=LOC101085033

A3 29124970 29125049 refGene_gn=SIRPA;refGene_tx=XM_023251383.1
A3 29143579 29143936 refGene_gn=SIRPA;refGene_tx=XM_023251383.1
A3 29148230 29148551 refGene_gn=SIRPA;refGene_tx=XM_023251383.1
A3 29149114 29149469 refGene_gn=SIRPA;refGene_tx=XM_023251383.1
A3 29150753 29150759 refGene_gn=SIRPA;refGene_tx=XM_023251383.1
A3 29151335 29151466 refGene_gn=SIRPA;refGene_tx=XM_023251383.1
A3 29154181 29154206 refGene_gn=SIRPA;refGene_tx=XM_023251383.1
A3 29160929 29160969 refGene_gn=SIRPA;refGene_tx=XM_023251383.1
A3 29163274 29163529 refGene_gn=SIRPA;refGene_tx=XM_023251383.1
A3 29204572 29205211
refGene_gn=PDYN;refGene_tx=XM_003983754.3;refGene_gn=PDYN;refGene_tx=XM_019826735.1
A3 29206518 29206647
refGene_gn=PDYN;refGene_tx=XM_003983754.3;refGene_gn=PDYN;refGene_tx=XM_019826735.1
A3 29308557 29308863
refGene_gn=STK35;refGene_tx=XM_019826736.2;refGene_gn=STK35;refGene_tx=XM_023251384.1
A3 29309453 29310051
refGene_gn=STK35;refGene_tx=XM_019826736.2;refGene_gn=STK35;refGene_tx=XM_023251384.1;refGene_gn=STK35;refGene_tx=XM_023251385.1
A3 29322531 29323244
refGene_gn=STK35;refGene_tx=XM_019826736.2;refGene_gn=STK35;refGene_tx=XM_023251384.1;refGene_gn=STK35;refGene_tx=XM_023251385.1
A3 29481193 29481200 refGene_gn=TGM3;refGene_tx=XM_003983751.3
A3 29491872 29492046 refGene_gn=TGM3;refGene_tx=XM_003983751.3
A3 29492389 29492632 refGene_gn=TGM3;refGene_tx=XM_003983751.3
A3 29493260 29493379 refGene_gn=TGM3;refGene_tx=XM_003983751.3
A3 29494607 29494736 refGene_gn=TGM3;refGene_tx=XM_003983751.3
A3 29497460 29497638 refGene_gn=TGM3;refGene_tx=XM_003983751.3
A3 29497728 29497864 refGene_gn=TGM3;refGene_tx=XM_003983751.3
A3 29506539 29506643 refGene_gn=TGM3;refGene_tx=XM_003983751.3
A3 29507786 29508032 refGene_gn=TGM3;refGene_tx=XM_003983751.3
A3 29511667 29511976 refGene_gn=TGM3;refGene_tx=XM_003983751.3
A3 29514369 29514527 refGene_gn=TGM3;refGene_tx=XM_003983751.3
A3 29517641 29517775 refGene_gn=TGM3;refGene_tx=XM_003983751.3
A3 29518115 29518263 refGene_gn=TGM3;refGene_tx=XM_003983751.3
A3 29551969 29551976 refGene_gn=TGM6;refGene_tx=XM_019826737.1
A3 29565168 29565342 refGene_gn=TGM6;refGene_tx=XM_019826737.1
A3 29565866 29566109 refGene_gn=TGM6;refGene_tx=XM_019826737.1
A3 29566870 29566989 refGene_gn=TGM6;refGene_tx=XM_019826737.1
A3 29568493 29568622 refGene_gn=TGM6;refGene_tx=XM_019826737.1
A3 29569934 29570112 refGene_gn=TGM6;refGene_tx=XM_019826737.1
A3 29571137 29571276 refGene_gn=TGM6;refGene_tx=XM_019826737.1
A3 29573450 29573554 refGene_gn=TGM6;refGene_tx=XM_019826737.1
A3 29573631 29573874 refGene_gn=TGM6;refGene_tx=XM_019826737.1
A3 29583467 29583809 refGene_gn=TGM6;refGene_tx=XM_019826737.1
A3 29586771 29586926 refGene_gn=TGM6;refGene_tx=XM_019826737.1
A3 29587199 29587333 refGene_gn=TGM6;refGene_tx=XM_019826737.1
A3 29588383 29588537 refGene_gn=TGM6;refGene_tx=XM_019826737.1

A3	29604707	29604745	refGene_gn=SNRPB;refGene_tx=XM_019826739.1
A3	29604866	29604877	refGene_gn=SNRPB;refGene_tx=XM_003983731.5
A3	29605611	29605737
refGene_gn=SNRPB;refGene_tx=XM_003983731.5;refGene_gn=SNRPB;refGene_tx
=XM_019826739.1
A3	29606055	29606194
refGene_gn=SNRPB;refGene_tx=XM_003983731.5;refGene_gn=SNRPB;refGene_tx
=XM_019826739.1
A3	29606737	29606890
refGene_gn=SNRPB;refGene_tx=XM_003983731.5;refGene_gn=SNRPB;refGene_tx
=XM_019826739.1
A3	29608946	29609058
refGene_gn=SNRPB;refGene_tx=XM_003983731.5;refGene_gn=SNRPB;refGene_tx
=XM_019826739.1
A3	29611958	29612110
refGene_gn=SNRPB;refGene_tx=XM_003983731.5;refGene_gn=SNRPB;refGene_tx
=XM_019826739.1
A3	29614910	29614913
refGene_gn=SNRPB;refGene_tx=XM_003983731.5;refGene_gn=SNRPB;refGene_tx
=XM_019826739.1
A3	29663392	29663426	refGene_gn=TMC2;refGene_tx=XM_023252178.1
A3	29665678	29665997	refGene_gn=TMC2;refGene_tx=XM_023252178.1
A3	29669988	29670141	refGene_gn=TMC2;refGene_tx=XM_023252178.1
A3	29680543	29680634	refGene_gn=TMC2;refGene_tx=XM_023252178.1
A3	29684076	29684158	refGene_gn=TMC2;refGene_tx=XM_023252178.1
A3	29684522	29684629	refGene_gn=TMC2;refGene_tx=XM_023252178.1
A3	29692926	29693069	refGene_gn=TMC2;refGene_tx=XM_023252178.1
A3	29694395	29694543	refGene_gn=TMC2;refGene_tx=XM_023252178.1
A3	29696994	29697183	refGene_gn=TMC2;refGene_tx=XM_023252178.1
A3	29699836	29700016	refGene_gn=TMC2;refGene_tx=XM_023252178.1
A3	29701497	29701647	refGene_gn=TMC2;refGene_tx=XM_023252178.1
A3	29703649	29703778	refGene_gn=TMC2;refGene_tx=XM_023252178.1
A3	29707061	29707129	refGene_gn=TMC2;refGene_tx=XM_023252178.1
A3	29707974	29708214	refGene_gn=TMC2;refGene_tx=XM_023252178.1
A3	29714381	29714507	refGene_gn=TMC2;refGene_tx=XM_023252178.1
A3	29718469	29718548	refGene_gn=TMC2;refGene_tx=XM_023252178.1
A3	29719384	29719502	refGene_gn=TMC2;refGene_tx=XM_023252178.1
A3	29720655	29720873	refGene_gn=TMC2;refGene_tx=XM_023252178.1
A3	29731510	29731513	refGene_gn=NOP56;refGene_tx=XM_003983729.4
A3	29731666	29731756	refGene_gn=NOP56;refGene_tx=XM_003983729.4
A3	29732153	29732268	refGene_gn=NOP56;refGene_tx=XM_003983729.4
A3	29732824	29732986	refGene_gn=NOP56;refGene_tx=XM_003983729.4
A3	29733161	29733360	refGene_gn=NOP56;refGene_tx=XM_003983729.4
A3	29733727	29733915	refGene_gn=NOP56;refGene_tx=XM_003983729.4
A3	29733992	29734144	refGene_gn=NOP56;refGene_tx=XM_003983729.4
A3	29734321	29734422	refGene_gn=NOP56;refGene_tx=XM_003983729.4
A3	29734790	29734939	refGene_gn=NOP56;refGene_tx=XM_003983729.4
A3	29735168	29735290	refGene_gn=NOP56;refGene_tx=XM_003983729.4
A3	29735478	29735616	refGene_gn=NOP56;refGene_tx=XM_003983729.4
A3	29736325	29736700	refGene_gn=NOP56;refGene_tx=XM_003983729.4

A3 29736838 29736919 refGene_gn=IDH3B;refGene_tx=XM_003983728.3
A3 29737146 29737233 refGene_gn=IDH3B;refGene_tx=XM_003983727.4
A3 29737894 29737955
refGene_gn=IDH3B;refGene_tx=XM_003983727.4;refGene_gn=IDH3B;refGene_tx=XM_003983728.3
A3 29738069 29738164
refGene_gn=IDH3B;refGene_tx=XM_003983727.4;refGene_gn=IDH3B;refGene_tx=XM_003983728.3
A3 29738410 29738557
refGene_gn=IDH3B;refGene_tx=XM_003983727.4;refGene_gn=IDH3B;refGene_tx=XM_003983728.3
A3 29738643 29738746
refGene_gn=IDH3B;refGene_tx=XM_003983727.4;refGene_gn=IDH3B;refGene_tx=XM_003983728.3
A3 29738840 29738974
refGene_gn=IDH3B;refGene_tx=XM_003983727.4;refGene_gn=IDH3B;refGene_tx=XM_003983728.3
A3 29739077 29739210
refGene_gn=IDH3B;refGene_tx=XM_003983727.4;refGene_gn=IDH3B;refGene_tx=XM_003983728.3
A3 29739286 29739347
refGene_gn=IDH3B;refGene_tx=XM_003983727.4;refGene_gn=IDH3B;refGene_tx=XM_003983728.3
A3 29740830 29740951
refGene_gn=IDH3B;refGene_tx=XM_003983727.4;refGene_gn=IDH3B;refGene_tx=XM_003983728.3
A3 29741050 29741149
refGene_gn=IDH3B;refGene_tx=XM_003983727.4;refGene_gn=IDH3B;refGene_tx=XM_003983728.3
A3 29741257 29741338
refGene_gn=IDH3B;refGene_tx=XM_003983727.4;refGene_gn=IDH3B;refGene_tx=XM_003983728.3
A3 29741466 29741502
refGene_gn=IDH3B;refGene_tx=XM_003983727.4;refGene_gn=IDH3B;refGene_tx=XM_003983728.3
A3 29767647 29767650 refGene_gn=EBF4;refGene_tx=XM_023251386.1
A3 29768145 29768282
refGene_gn=EBF4;refGene_tx=XM_023251386.1;refGene_gn=EBF4;refGene_tx=XM_023251387.1
A3 29777839 29777996
refGene_gn=EBF4;refGene_tx=XM_023251386.1;refGene_gn=EBF4;refGene_tx=XM_023251387.1
A3 29778222 29778286
refGene_gn=EBF4;refGene_tx=XM_023251386.1;refGene_gn=EBF4;refGene_tx=XM_023251387.1
A3 29778437 29778493
refGene_gn=EBF4;refGene_tx=XM_023251386.1;refGene_gn=EBF4;refGene_tx=XM_023251387.1
A3 29780100 29780174
refGene_gn=EBF4;refGene_tx=XM_023251386.1;refGene_gn=EBF4;refGene_tx=X

M_023251387.1
A3 29781282 29781351
refGene_gn=EBF4;refGene_tx=XM_023251386.1;refGene_gn=EBF4;refGene_tx=XM_023251387.1
A3 29815392 29815474
refGene_gn=EBF4;refGene_tx=XM_023251386.1;refGene_gn=EBF4;refGene_tx=XM_023251387.1
A3 29816295 29816413
refGene_gn=EBF4;refGene_tx=XM_023251386.1;refGene_gn=EBF4;refGene_tx=XM_023251387.1
A3 29816514 29816648
refGene_gn=EBF4;refGene_tx=XM_023251386.1;refGene_gn=EBF4;refGene_tx=XM_023251387.1
A3 29816743 29816870
refGene_gn=EBF4;refGene_tx=XM_023251386.1;refGene_gn=EBF4;refGene_tx=XM_023251387.1
A3 29818459 29818548
refGene_gn=EBF4;refGene_tx=XM_023251386.1;refGene_gn=EBF4;refGene_tx=XM_023251387.1
A3 29818695 29818761
refGene_gn=EBF4;refGene_tx=XM_023251386.1;refGene_gn=EBF4;refGene_tx=XM_023251387.1
A3 29818862 29819040
refGene_gn=EBF4;refGene_tx=XM_023251386.1;refGene_gn=EBF4;refGene_tx=XM_023251387.1
A3 29819113 29819302
refGene_gn=EBF4;refGene_tx=XM_023251386.1;refGene_gn=EBF4;refGene_tx=XM_023251387.1
A3 29822031 29822229
refGene_gn=EBF4;refGene_tx=XM_023251386.1;refGene_gn=EBF4;refGene_tx=XM_023251387.1
A3 29823945 29824016
refGene_gn=EBF4;refGene_tx=XM_023251386.1;refGene_gn=EBF4;refGene_tx=XM_023251387.1
A3 29835549 29835791
refGene_gn=CPXM1;refGene_tx=XM_003983725.4;refGene_gn=CPXM1;refGene_tx=XM_006929918.3
A3 29835889 29835992
refGene_gn=CPXM1;refGene_tx=XM_003983725.4;refGene_gn=CPXM1;refGene_tx=XM_006929918.3
A3 29837594 29837734
refGene_gn=CPXM1;refGene_tx=XM_003983725.4;refGene_gn=CPXM1;refGene_tx=XM_006929918.3
A3 29837916 29838214
refGene_gn=CPXM1;refGene_tx=XM_003983725.4;refGene_gn=CPXM1;refGene_tx=XM_006929918.3
A3 29838297 29838477
refGene_gn=CPXM1;refGene_tx=XM_003983725.4;refGene_gn=CPXM1;refGene_tx=XM_006929918.3
A3 29838557 29838754

refGene_gn=CPXM1;refGene_tx=XM_003983725.4;refGene_gn=CPXM1;refGene_tx=XM_006929918.3
A3 29838837 29838961
refGene_gn=CPXM1;refGene_tx=XM_003983725.4;refGene_gn=CPXM1;refGene_tx=XM_006929918.3
A3 29839320 29839409
refGene_gn=CPXM1;refGene_tx=XM_003983725.4;refGene_gn=CPXM1;refGene_tx=XM_006929918.3
A3 29839494 29839645
refGene_gn=CPXM1;refGene_tx=XM_003983725.4;refGene_gn=CPXM1;refGene_tx=XM_006929918.3
A3 29840265 29840356
refGene_gn=CPXM1;refGene_tx=XM_003983725.4;refGene_gn=CPXM1;refGene_tx=XM_006929918.3
A3 29840458 29840598
refGene_gn=CPXM1;refGene_tx=XM_003983725.4;refGene_gn=CPXM1;refGene_tx=XM_006929918.3
A3 29840722 29840832
refGene_gn=CPXM1;refGene_tx=XM_003983725.4;refGene_gn=CPXM1;refGene_tx=XM_006929918.3
A3 29841032 29841194
refGene_gn=CPXM1;refGene_tx=XM_003983725.4;refGene_gn=CPXM1;refGene_tx=XM_006929918.3
A3 29842563 29842735
refGene_gn=CPXM1;refGene_tx=XM_003983725.4;refGene_gn=CPXM1;refGene_tx=XM_006929918.3
A3 29849747 29850010
refGene_gn=CA3H20orf141;refGene_tx=XM_003983747.5
A3 29850086 29850327
refGene_gn=CA3H20orf141;refGene_tx=XM_003983747.5
A3 29850837 29850905 refGene_gn=TMEM239;refGene_tx=XM_019826743.1
A3 29850988 29851523
refGene_gn=TMEM239;refGene_tx=XM_003983724.5;refGene_gn=TMEM239;refGene_tx=XM_006929917.3;refGene_gn=TMEM239;refGene_tx=XM_019826743.1
A3 29865572 29865820
refGene_gn=PCED1A;refGene_tx=XM_023251388.1;refGene_gn=PCED1A;refGene_tx=XM_023251389.1
A3 29866162 29866438
refGene_gn=PCED1A;refGene_tx=XM_023251388.1;refGene_gn=PCED1A;refGene_tx=XM_023251389.1
A3 29868100 29868347
refGene_gn=PCED1A;refGene_tx=XM_023251388.1;refGene_gn=PCED1A;refGene_tx=XM_023251389.1
A3 29868465 29868616
refGene_gn=PCED1A;refGene_tx=XM_023251388.1;refGene_gn=PCED1A;refGene_tx=XM_023251389.1
A3 29868718 29868957
refGene_gn=PCED1A;refGene_tx=XM_023251388.1;refGene_gn=PCED1A;refGene_tx=XM_023251389.1
A3 29869067 29869147

refGene_gn=PCED1A;refGene_tx=XM_023251388.1;refGene_gn=PCED1A;refGene_tx=XM_023251389.1
A3 29869669 29869793
refGene_gn=PCED1A;refGene_tx=XM_023251388.1;refGene_gn=PCED1A;refGene_tx=XM_023251389.1
A3 29870669 29870722
refGene_gn=VPS16;refGene_tx=XM_011280925.3;refGene_gn=VPS16;refGene_tx=XM_019826744.2
A3 29885526 29885615
refGene_gn=VPS16;refGene_tx=XM_011280925.3;refGene_gn=VPS16;refGene_tx=XM_019826744.2
A3 29885841 29885939
refGene_gn=VPS16;refGene_tx=XM_011280925.3;refGene_gn=VPS16;refGene_tx=XM_019826744.2
A3 29886020 29886149
refGene_gn=VPS16;refGene_tx=XM_011280925.3;refGene_gn=VPS16;refGene_tx=XM_019826744.2
A3 29886228 29886373
refGene_gn=VPS16;refGene_tx=XM_011280925.3;refGene_gn=VPS16;refGene_tx=XM_019826744.2
A3 29886498 29886614
refGene_gn=VPS16;refGene_tx=XM_011280925.3;refGene_gn=VPS16;refGene_tx=XM_019826744.2
A3 29886711 29886834
refGene_gn=VPS16;refGene_tx=XM_011280925.3;refGene_gn=VPS16;refGene_tx=XM_019826744.2
A3 29886963 29887019
refGene_gn=VPS16;refGene_tx=XM_011280925.3;refGene_gn=VPS16;refGene_tx=XM_019826744.2
A3 29887371 29887461
refGene_gn=VPS16;refGene_tx=XM_011280925.3;refGene_gn=VPS16;refGene_tx=XM_019826744.2
A3 29887567 29887662
refGene_gn=VPS16;refGene_tx=XM_011280925.3;refGene_gn=VPS16;refGene_tx=XM_019826744.2
A3 29887763 29887840
refGene_gn=VPS16;refGene_tx=XM_011280925.3;refGene_gn=VPS16;refGene_tx=XM_019826744.2
A3 29888293 29888425
refGene_gn=VPS16;refGene_tx=XM_011280925.3;refGene_gn=VPS16;refGene_tx=XM_019826744.2
A3 29888558 29888686
refGene_gn=VPS16;refGene_tx=XM_011280925.3;refGene_gn=VPS16;refGene_tx=XM_019826744.2
A3 29888822 29888858
refGene_gn=VPS16;refGene_tx=XM_011280925.3;refGene_gn=VPS16;refGene_tx=XM_019826744.2
A3 29889047 29889156
refGene_gn=VPS16;refGene_tx=XM_011280925.3;refGene_gn=VPS16;refGene_tx=XM_019826744.2

A3 29889324 29889459
refGene_gn=VPS16;refGene_tx=XM_011280925.3;refGene_gn=VPS16;refGene_tx
=XM_019826744.2
A3 29889554 29889663
refGene_gn=VPS16;refGene_tx=XM_011280925.3;refGene_gn=VPS16;refGene_tx
=XM_019826744.2
A3 29889754 29889852
refGene_gn=VPS16;refGene_tx=XM_011280925.3;refGene_gn=VPS16;refGene_tx
=XM_019826744.2
A3 29889935 29890043
refGene_gn=VPS16;refGene_tx=XM_011280925.3;refGene_gn=VPS16;refGene_tx
=XM_019826744.2
A3 29890359 29890437
refGene_gn=VPS16;refGene_tx=XM_011280925.3;refGene_gn=VPS16;refGene_tx
=XM_019826744.2
A3 29890974 29891144
refGene_gn=VPS16;refGene_tx=XM_011280925.3;refGene_gn=VPS16;refGene_tx
=XM_019826744.2
A3 29891226 29891323
refGene_gn=VPS16;refGene_tx=XM_011280925.3;refGene_gn=VPS16;refGene_tx
=XM_019826744.2
A3 29891995 29892112
refGene_gn=VPS16;refGene_tx=XM_011280925.3;refGene_gn=VPS16;refGene_tx
=XM_019826744.2
A3 29892215 29892360 refGene_gn=VPS16;refGene_tx=XM_011280925.3
A3 29945929 29945956
refGene_gn=PTPRA;refGene_tx=XM_023251390.1;refGene_gn=PTPRA;refGene_tx
=XM_023251391.1;refGene_gn=PTPRA;refGene_tx=XM_023251392.1
A3 29985439 29985518
refGene_gn=PTPRA;refGene_tx=XM_019826746.2;refGene_gn=PTPRA;refGene_tx
=XM_023251390.1;refGene_gn=PTPRA;refGene_tx=XM_023251391.1;refGene_gn=
PTPRA;refGene_tx=XM_023251392.1;refGene_gn=PTPRA;refGene_tx=XM_0232513
93.1
A3 29986088 29986430
refGene_gn=PTPRA;refGene_tx=XM_019826746.2;refGene_gn=PTPRA;refGene_tx
=XM_023251390.1;refGene_gn=PTPRA;refGene_tx=XM_023251391.1;refGene_gn=
PTPRA;refGene_tx=XM_023251392.1;refGene_gn=PTPRA;refGene_tx=XM_0232513
93.1
A3 29994507 29994534
refGene_gn=PTPRA;refGene_tx=XM_019826746.2;refGene_gn=PTPRA;refGene_tx
=XM_023251390.1;refGene_gn=PTPRA;refGene_tx=XM_023251391.1;refGene_gn=
PTPRA;refGene_tx=XM_023251393.1
A3 30009909 30009994
refGene_gn=PTPRA;refGene_tx=XM_019826746.2;refGene_gn=PTPRA;refGene_tx
=XM_023251390.1;refGene_gn=PTPRA;refGene_tx=XM_023251391.1;refGene_gn=
PTPRA;refGene_tx=XM_023251392.1;refGene_gn=PTPRA;refGene_tx=XM_0232513
93.1
A3 30011195 30011269
refGene_gn=PTPRA;refGene_tx=XM_019826746.2;refGene_gn=PTPRA;refGene_tx
=XM_019826747.1;refGene_gn=PTPRA;refGene_tx=XM_023251390.1;refGene_gn=

PTPRA;refGene_tx=XM_023251391.1;refGene_gn=PTPRA;refGene_tx=XM_023251392.1;refGene_gn=PTPRA;refGene_tx=XM_023251393.1
A3 30011485 30011622
refGene_gn=PTPRA;refGene_tx=XM_019826746.2;refGene_gn=PTPRA;refGene_tx=XM_019826747.1;refGene_gn=PTPRA;refGene_tx=XM_023251390.1;refGene_gn=PTPRA;refGene_tx=XM_023251391.1;refGene_gn=PTPRA;refGene_tx=XM_023251392.1;refGene_gn=PTPRA;refGene_tx=XM_023251393.1
A3 30020640 30020734 refGene_gn=PTPRA;refGene_tx=XM_019826747.1
A3 30025090 30025181
refGene_gn=PTPRA;refGene_tx=XM_019826746.2;refGene_gn=PTPRA;refGene_tx=XM_019826747.1;refGene_gn=PTPRA;refGene_tx=XM_023251390.1;refGene_gn=PTPRA;refGene_tx=XM_023251391.1;refGene_gn=PTPRA;refGene_tx=XM_023251392.1;refGene_gn=PTPRA;refGene_tx=XM_023251393.1
A3 30026098 30026175
refGene_gn=PTPRA;refGene_tx=XM_019826746.2;refGene_gn=PTPRA;refGene_tx=XM_019826747.1;refGene_gn=PTPRA;refGene_tx=XM_023251390.1;refGene_gn=PTPRA;refGene_tx=XM_023251391.1;refGene_gn=PTPRA;refGene_tx=XM_023251392.1;refGene_gn=PTPRA;refGene_tx=XM_023251393.1
A3 30034030 30034369
refGene_gn=LOC109498061;refGene_tx=XM_023251394.1
A3 30034370 30034437
refGene_gn=LOC109498061;refGene_tx=XM_023251394.1
A3 30034439 30034549
refGene_gn=LOC109498061;refGene_tx=XM_023251394.1
A3 30035678 30035715
refGene_gn=PTPRA;refGene_tx=XM_019826746.2;refGene_gn=PTPRA;refGene_tx=XM_019826747.1;refGene_gn=PTPRA;refGene_tx=XM_023251390.1;refGene_gn=PTPRA;refGene_tx=XM_023251391.1;refGene_gn=PTPRA;refGene_tx=XM_023251392.1;refGene_gn=PTPRA;refGene_tx=XM_023251393.1
A3 30039482 30039580
refGene_gn=PTPRA;refGene_tx=XM_019826746.2;refGene_gn=PTPRA;refGene_tx=XM_019826747.1;refGene_gn=PTPRA;refGene_tx=XM_023251390.1;refGene_gn=PTPRA;refGene_tx=XM_023251391.1;refGene_gn=PTPRA;refGene_tx=XM_023251392.1;refGene_gn=PTPRA;refGene_tx=XM_023251393.1
A3 30040703 30040823
refGene_gn=PTPRA;refGene_tx=XM_019826746.2;refGene_gn=PTPRA;refGene_tx=XM_019826747.1;refGene_gn=PTPRA;refGene_tx=XM_023251390.1;refGene_gn=PTPRA;refGene_tx=XM_023251391.1;refGene_gn=PTPRA;refGene_tx=XM_023251392.1;refGene_gn=PTPRA;refGene_tx=XM_023251393.1
A3 30041579 30041746
refGene_gn=PTPRA;refGene_tx=XM_019826746.2;refGene_gn=PTPRA;refGene_tx=XM_019826747.1;refGene_gn=PTPRA;refGene_tx=XM_023251390.1;refGene_gn=PTPRA;refGene_tx=XM_023251391.1;refGene_gn=PTPRA;refGene_tx=XM_023251392.1;refGene_gn=PTPRA;refGene_tx=XM_023251393.1
A3 30042246 30042382
refGene_gn=PTPRA;refGene_tx=XM_019826746.2;refGene_gn=PTPRA;refGene_tx=XM_019826747.1;refGene_gn=PTPRA;refGene_tx=XM_023251390.1;refGene_gn=PTPRA;refGene_tx=XM_023251391.1;refGene_gn=PTPRA;refGene_tx=XM_023251392.1;refGene_gn=PTPRA;refGene_tx=XM_023251393.1
A3 30043090 30043240

refGene_gn=PTPRA;refGene_tx=XM_019826746.2;refGene_gn=PTPRA;refGene_tx=XM_019826747.1;refGene_gn=PTPRA;refGene_tx=XM_023251390.1;refGene_gn=PTPRA;refGene_tx=XM_023251391.1;refGene_gn=PTPRA;refGene_tx=XM_023251392.1;refGene_gn=PTPRA;refGene_tx=XM_023251393.1
A3 30044400 30044494
refGene_gn=PTPRA;refGene_tx=XM_019826746.2;refGene_gn=PTPRA;refGene_tx=XM_019826747.1;refGene_gn=PTPRA;refGene_tx=XM_023251390.1;refGene_gn=PTPRA;refGene_tx=XM_023251391.1;refGene_gn=PTPRA;refGene_tx=XM_023251392.1;refGene_gn=PTPRA;refGene_tx=XM_023251393.1
A3 30044793 30044870
refGene_gn=PTPRA;refGene_tx=XM_019826746.2;refGene_gn=PTPRA;refGene_tx=XM_019826747.1;refGene_gn=PTPRA;refGene_tx=XM_023251390.1;refGene_gn=PTPRA;refGene_tx=XM_023251391.1;refGene_gn=PTPRA;refGene_tx=XM_023251392.1;refGene_gn=PTPRA;refGene_tx=XM_023251393.1
A3 30045345 30045480
refGene_gn=PTPRA;refGene_tx=XM_019826746.2;refGene_gn=PTPRA;refGene_tx=XM_019826747.1;refGene_gn=PTPRA;refGene_tx=XM_023251390.1;refGene_gn=PTPRA;refGene_tx=XM_023251391.1;refGene_gn=PTPRA;refGene_tx=XM_023251392.1;refGene_gn=PTPRA;refGene_tx=XM_023251393.1
A3 30053995 30054121
refGene_gn=PTPRA;refGene_tx=XM_019826746.2;refGene_gn=PTPRA;refGene_tx=XM_019826747.1;refGene_gn=PTPRA;refGene_tx=XM_023251390.1;refGene_gn=PTPRA;refGene_tx=XM_023251391.1;refGene_gn=PTPRA;refGene_tx=XM_023251392.1;refGene_gn=PTPRA;refGene_tx=XM_023251393.1
A3 30054200 30054352
refGene_gn=PTPRA;refGene_tx=XM_019826746.2;refGene_gn=PTPRA;refGene_tx=XM_019826747.1;refGene_gn=PTPRA;refGene_tx=XM_023251390.1;refGene_gn=PTPRA;refGene_tx=XM_023251391.1;refGene_gn=PTPRA;refGene_tx=XM_023251392.1;refGene_gn=PTPRA;refGene_tx=XM_023251393.1
A3 30060253 30060389
refGene_gn=PTPRA;refGene_tx=XM_019826746.2;refGene_gn=PTPRA;refGene_tx=XM_019826747.1;refGene_gn=PTPRA;refGene_tx=XM_023251390.1;refGene_gn=PTPRA;refGene_tx=XM_023251391.1;refGene_gn=PTPRA;refGene_tx=XM_023251392.1;refGene_gn=PTPRA;refGene_tx=XM_023251393.1
A3 30061120 30061195
refGene_gn=PTPRA;refGene_tx=XM_019826746.2;refGene_gn=PTPRA;refGene_tx=XM_019826747.1;refGene_gn=PTPRA;refGene_tx=XM_023251390.1;refGene_gn=PTPRA;refGene_tx=XM_023251391.1;refGene_gn=PTPRA;refGene_tx=XM_023251392.1;refGene_gn=PTPRA;refGene_tx=XM_023251393.1
A3 30066956 30067110 refGene_gn=GNRH2;refGene_tx=XM_023251395.1
A3 30067229 30067304 refGene_gn=GNRH2;refGene_tx=XM_023251395.1
A3 30067306 30067596 refGene_gn=GNRH2;refGene_tx=XM_023251395.1
A3 30068554 30068766 refGene_gn=MRPS26;refGene_tx=XM_006929915.3
A3 30068860 30069007 refGene_gn=MRPS26;refGene_tx=XM_006929915.3
A3 30069096 30069220 refGene_gn=MRPS26;refGene_tx=XM_006929915.3
A3 30069956 30070091 refGene_gn=MRPS26;refGene_tx=XM_006929915.3
A3 30091933 30092053 refGene_gn=OXT;refGene_tx=XM_023250914.1
A3 30092211 30092413 refGene_gn=OXT;refGene_tx=XM_023250914.1
A3 30092690 30092746 refGene_gn=OXT;refGene_tx=XM_023250914.1
A3 30103791 30103970 refGene_gn=AVP;refGene_tx=XM_023251396.1

A3 30104207 30104409 refGene_gn=AVP;refGene_tx=XM_023251396.1
A3 30105583 30105703 refGene_gn=AVP;refGene_tx=XM_023251396.1
A3 30119524 30119733
refGene_gn=UBOX5;refGene_tx=XM_003983712.4;refGene_gn=UBOX5;refGene_tx=XM_006929912.3;refGene_gn=UBOX5;refGene_tx=XM_006929913.3;refGene_gn=UBOX5;refGene_tx=XM_019826750.1
A3 30122150 30122312
refGene_gn=UBOX5;refGene_tx=XM_003983712.4;refGene_gn=UBOX5;refGene_tx=XM_006929912.3;refGene_gn=UBOX5;refGene_tx=XM_006929913.3;refGene_gn=UBOX5;refGene_tx=XM_019826750.1
A3 30124657 30125858
refGene_gn=UBOX5;refGene_tx=XM_003983712.4;refGene_gn=UBOX5;refGene_tx=XM_006929912.3;refGene_gn=UBOX5;refGene_tx=XM_006929913.3;refGene_gn=UBOX5;refGene_tx=XM_019826750.1
A3 30127504 30127558
refGene_gn=UBOX5;refGene_tx=XM_003983712.4;refGene_gn=UBOX5;refGene_tx=XM_006929912.3;refGene_gn=UBOX5;refGene_tx=XM_006929913.3;refGene_gn=UBOX5;refGene_tx=XM_019826750.1
A3 30147680 30150125
refGene_gn=FASTKD5;refGene_tx=XM_006929910.3;refGene_gn=FASTKD5;refGene_tx=XM_006929911.3;refGene_gn=FASTKD5;refGene_tx=XM_019826749.1
A3 30161639 30162338
refGene_gn=LZTS3;refGene_tx=XM_023251398.1;refGene_gn=LZTS3;refGene_tx=XM_023251399.1;refGene_gn=LZTS3;refGene_tx=XM_023251400.1;refGene_gn=LZTS3;refGene_tx=XM_023251401.1;refGene_gn=LZTS3;refGene_tx=XM_023251402.1;refGene_gn=LZTS3;refGene_tx=XM_023251403.1;refGene_gn=LZTS3;refGene_tx=XM_023251405.1
A3 30162678 30163542
refGene_gn=LZTS3;refGene_tx=XM_023251398.1;refGene_gn=LZTS3;refGene_tx=XM_023251399.1;refGene_gn=LZTS3;refGene_tx=XM_023251400.1;refGene_gn=LZTS3;refGene_tx=XM_023251401.1;refGene_gn=LZTS3;refGene_tx=XM_023251402.1;refGene_gn=LZTS3;refGene_tx=XM_023251403.1;refGene_gn=LZTS3;refGene_tx=XM_023251405.1
A3 30163860 30164337
refGene_gn=LZTS3;refGene_tx=XM_023251398.1;refGene_gn=LZTS3;refGene_tx=XM_023251399.1;refGene_gn=LZTS3;refGene_tx=XM_023251400.1;refGene_gn=LZTS3;refGene_tx=XM_023251401.1;refGene_gn=LZTS3;refGene_tx=XM_023251402.1;refGene_gn=LZTS3;refGene_tx=XM_023251403.1;refGene_gn=LZTS3;refGene_tx=XM_023251405.1
A3 30164918 30164990
refGene_gn=LZTS3;refGene_tx=XM_023251398.1;refGene_gn=LZTS3;refGene_tx=XM_023251399.1;refGene_gn=LZTS3;refGene_tx=XM_023251400.1;refGene_gn=LZTS3;refGene_tx=XM_023251401.1
A3 30181863 30182030 refGene_gn=DDRGK1;refGene_tx=XM_003983711.5
A3 30182408 30182457 refGene_gn=DDRGK1;refGene_tx=XM_003983711.5
A3 30182971 30183028 refGene_gn=DDRGK1;refGene_tx=XM_003983711.5
A3 30185581 30185620 refGene_gn=DDRGK1;refGene_tx=XM_003983711.5
A3 30185903 30186026 refGene_gn=DDRGK1;refGene_tx=XM_003983711.5
A3 30188980 30189082 refGene_gn=DDRGK1;refGene_tx=XM_003983711.5
A3 30189312 30189425 refGene_gn=DDRGK1;refGene_tx=XM_003983711.5

A3 30190652 30190859 refGene_gn=DDRGK1;refGene_tx=XM_003983711.5
A3 30192022 30192188 refGene_gn=DDRGK1;refGene_tx=XM_003983711.5
A3 30197612 30197694
refGene_gn=ITPA;refGene_tx=XM_003983710.5;refGene_gn=ITPA;refGene_tx=XM_019826751.2
A3 30198569 30198581 refGene_gn=ITPA;refGene_tx=XM_019826752.1
A3 30199928 30199993 refGene_gn=ITPA;refGene_tx=XM_006929909.4
A3 30200108 30200166
refGene_gn=ITPA;refGene_tx=XM_003983710.5;refGene_gn=ITPA;refGene_tx=XM_006929909.4;refGene_gn=ITPA;refGene_tx=XM_019826752.1
A3 30200258 30200323
refGene_gn=ITPA;refGene_tx=XM_003983710.5;refGene_gn=ITPA;refGene_tx=XM_019826751.2;refGene_gn=ITPA;refGene_tx=XM_019826752.1
A3 30200942 30201016
refGene_gn=ITPA;refGene_tx=XM_003983710.5;refGene_gn=ITPA;refGene_tx=XM_006929909.4;refGene_gn=ITPA;refGene_tx=XM_019826751.2;refGene_gn=ITPA;refGene_tx=XM_019826752.1
A3 30201829 30201861
refGene_gn=ITPA;refGene_tx=XM_003983710.5;refGene_gn=ITPA;refGene_tx=XM_006929909.4;refGene_gn=ITPA;refGene_tx=XM_019826751.2;refGene_gn=ITPA;refGene_tx=XM_019826752.1
A3 30202491 30202607
refGene_gn=ITPA;refGene_tx=XM_003983710.5;refGene_gn=ITPA;refGene_tx=XM_006929909.4;refGene_gn=ITPA;refGene_tx=XM_019826751.2;refGene_gn=ITPA;refGene_tx=XM_019826752.1
A3 30205910 30205987
refGene_gn=ITPA;refGene_tx=XM_003983710.5;refGene_gn=ITPA;refGene_tx=XM_006929909.4;refGene_gn=ITPA;refGene_tx=XM_019826751.2;refGene_gn=ITPA;refGene_tx=XM_019826752.1
A3 30207127 30207269
refGene_gn=ITPA;refGene_tx=XM_003983710.5;refGene_gn=ITPA;refGene_tx=XM_006929909.4;refGene_gn=ITPA;refGene_tx=XM_019826751.2;refGene_gn=ITPA;refGene_tx=XM_019826752.1
A3 30210216 30210314
refGene_gn=SLC4A11;refGene_tx=XM_006929906.3;refGene_gn=SLC4A11;refGene_tx=XM_006929908.4;refGene_gn=SLC4A11;refGene_tx=XM_019826756.2;refGene_gn=SLC4A11;refGene_tx=XM_019826757.2;refGene_gn=SLC4A11;refGene_tx=XM_019826758.2;refGene_gn=SLC4A11;refGene_tx=XM_019826760.2;refGene_gn=SLC4A11;refGene_tx=XM_019826761.2;refGene_gn=SLC4A11;refGene_tx=XM_019826763.2;refGene_gn=SLC4A11;refGene_tx=XM_019826764.2;refGene_gn=SLC4A11;refGene_tx=XM_019826765.2;refGene_gn=SLC4A11;refGene_tx=XM_019826766.2;refGene_gn=SLC4A11;refGene_tx=XM_019826769.1;refGene_gn=SLC4A11;refGene_tx=XM_023251407.1;refGene_gn=SLC4A11;refGene_tx=XM_023251408.1;refGene_gn=SLC4A11;refGene_tx=XM_023251410.1;refGene_gn=SLC4A11;refGene_tx=XM_023251411.1
A3 30210455 30210625
refGene_gn=SLC4A11;refGene_tx=XM_006929906.3;refGene_gn=SLC4A11;refGene_tx=XM_006929908.4;refGene_gn=SLC4A11;refGene_tx=XM_019826756.2;refGene_gn=SLC4A11;refGene_tx=XM_019826757.2;refGene_gn=SLC4A11;refGene_tx=XM_019826758.2;refGene_gn=SLC4A11;refGene_tx=XM_019826760.2;refGene_gn

=SLC4A11;refGene_tx=XM_019826761.2;refGene_gn=SLC4A11;refGene_tx=XM_019826763.2;refGene_gn=SLC4A11;refGene_tx=XM_019826764.2;refGene_gn=SLC4A11;refGene_tx=XM_019826765.2;refGene_gn=SLC4A11;refGene_tx=XM_019826766.2;refGene_gn=SLC4A11;refGene_tx=XM_019826769.1;refGene_gn=SLC4A11;refGene_tx=XM_023251407.1;refGene_gn=SLC4A11;refGene_tx=XM_023251408.1;refGene_gn=SLC4A11;refGene_tx=XM_023251410.1;refGene_gn=SLC4A11;refGene_tx=XM_023251411.1
A3 30210710 30210906
refGene_gn=SLC4A11;refGene_tx=XM_006929906.3;refGene_gn=SLC4A11;refGene_tx=XM_006929908.4;refGene_gn=SLC4A11;refGene_tx=XM_019826756.2;refGene_gn=SLC4A11;refGene_tx=XM_019826757.2;refGene_gn=SLC4A11;refGene_tx=XM_019826758.2;refGene_gn=SLC4A11;refGene_tx=XM_019826760.2;refGene_gn=SLC4A11;refGene_tx=XM_019826761.2;refGene_gn=SLC4A11;refGene_tx=XM_019826763.2;refGene_gn=SLC4A11;refGene_tx=XM_019826764.2;refGene_gn=SLC4A11;refGene_tx=XM_019826765.2;refGene_gn=SLC4A11;refGene_tx=XM_019826766.2;refGene_gn=SLC4A11;refGene_tx=XM_019826769.1;refGene_gn=SLC4A11;refGene_tx=XM_023251407.1;refGene_gn=SLC4A11;refGene_tx=XM_023251408.1;refGene_gn=SLC4A11;refGene_tx=XM_023251410.1;refGene_gn=SLC4A11;refGene_tx=XM_023251411.1
A3 30211137 30211311
refGene_gn=SLC4A11;refGene_tx=XM_006929906.3;refGene_gn=SLC4A11;refGene_tx=XM_006929908.4;refGene_gn=SLC4A11;refGene_tx=XM_019826756.2;refGene_gn=SLC4A11;refGene_tx=XM_019826757.2;refGene_gn=SLC4A11;refGene_tx=XM_019826758.2;refGene_gn=SLC4A11;refGene_tx=XM_019826760.2;refGene_gn=SLC4A11;refGene_tx=XM_019826761.2;refGene_gn=SLC4A11;refGene_tx=XM_019826763.2;refGene_gn=SLC4A11;refGene_tx=XM_019826764.2;refGene_gn=SLC4A11;refGene_tx=XM_019826765.2;refGene_gn=SLC4A11;refGene_tx=XM_019826766.2;refGene_gn=SLC4A11;refGene_tx=XM_019826769.1;refGene_gn=SLC4A11;refGene_tx=XM_023251407.1;refGene_gn=SLC4A11;refGene_tx=XM_023251408.1;refGene_gn=SLC4A11;refGene_tx=XM_023251410.1;refGene_gn=SLC4A11;refGene_tx=XM_023251411.1
A3 30211402 30211571
refGene_gn=SLC4A11;refGene_tx=XM_006929906.3;refGene_gn=SLC4A11;refGene_tx=XM_006929908.4;refGene_gn=SLC4A11;refGene_tx=XM_019826756.2;refGene_gn=SLC4A11;refGene_tx=XM_019826757.2;refGene_gn=SLC4A11;refGene_tx=XM_019826758.2;refGene_gn=SLC4A11;refGene_tx=XM_019826760.2;refGene_gn=SLC4A11;refGene_tx=XM_019826761.2;refGene_gn=SLC4A11;refGene_tx=XM_019826763.2;refGene_gn=SLC4A11;refGene_tx=XM_019826764.2;refGene_gn=SLC4A11;refGene_tx=XM_019826765.2;refGene_gn=SLC4A11;refGene_tx=XM_019826766.2;refGene_gn=SLC4A11;refGene_tx=XM_019826769.1;refGene_gn=SLC4A11;refGene_tx=XM_023251407.1;refGene_gn=SLC4A11;refGene_tx=XM_023251408.1;refGene_gn=SLC4A11;refGene_tx=XM_023251410.1;refGene_gn=SLC4A11;refGene_tx=XM_023251411.1
A3 30211658 30211765
refGene_gn=SLC4A11;refGene_tx=XM_006929906.3;refGene_gn=SLC4A11;refGene_tx=XM_006929908.4;refGene_gn=SLC4A11;refGene_tx=XM_019826756.2;refGene_gn=SLC4A11;refGene_tx=XM_019826757.2;refGene_gn=SLC4A11;refGene_tx=XM_019826758.2;refGene_gn=SLC4A11;refGene_tx=XM_019826760.2;refGene_gn=SLC4A11;refGene_tx=XM_019826761.2;refGene_gn=SLC4A11;refGene_tx=XM_019826763.2;refGene_gn=SLC4A11;refGene_tx=XM_019826764.2;refGene_gn=SLC4

A11;refGene_tx=XM_019826765.2;refGene_gn=SLC4A11;refGene_tx=XM_019826766.2;refGene_gn=SLC4A11;refGene_tx=XM_019826769.1;refGene_gn=SLC4A11;refGene_tx=XM_023251407.1;refGene_gn=SLC4A11;refGene_tx=XM_023251408.1;refGene_gn=SLC4A11;refGene_tx=XM_023251410.1;refGene_gn=SLC4A11;refGene_tx=XM_023251411.1
A3 30211841 30212103
refGene_gn=SLC4A11;refGene_tx=XM_006929906.3;refGene_gn=SLC4A11;refGene_tx=XM_006929908.4;refGene_gn=SLC4A11;refGene_tx=XM_019826756.2;refGene_gn=SLC4A11;refGene_tx=XM_019826757.2;refGene_gn=SLC4A11;refGene_tx=XM_019826758.2;refGene_gn=SLC4A11;refGene_tx=XM_019826760.2;refGene_gn=SLC4A11;refGene_tx=XM_019826761.2;refGene_gn=SLC4A11;refGene_tx=XM_019826763.2;refGene_gn=SLC4A11;refGene_tx=XM_019826764.2;refGene_gn=SLC4A11;refGene_tx=XM_019826765.2;refGene_gn=SLC4A11;refGene_tx=XM_019826766.2;refGene_gn=SLC4A11;refGene_tx=XM_019826769.1;refGene_gn=SLC4A11;refGene_tx=XM_023251407.1;refGene_gn=SLC4A11;refGene_tx=XM_023251408.1;refGene_gn=SLC4A11;refGene_tx=XM_023251410.1;refGene_gn=SLC4A11;refGene_tx=XM_023251411.1
A3 30212520 30212594
refGene_gn=SLC4A11;refGene_tx=XM_006929906.3;refGene_gn=SLC4A11;refGene_tx=XM_006929908.4;refGene_gn=SLC4A11;refGene_tx=XM_019826756.2;refGene_gn=SLC4A11;refGene_tx=XM_019826757.2;refGene_gn=SLC4A11;refGene_tx=XM_019826758.2;refGene_gn=SLC4A11;refGene_tx=XM_019826760.2;refGene_gn=SLC4A11;refGene_tx=XM_019826761.2;refGene_gn=SLC4A11;refGene_tx=XM_019826763.2;refGene_gn=SLC4A11;refGene_tx=XM_019826764.2;refGene_gn=SLC4A11;refGene_tx=XM_019826765.2;refGene_gn=SLC4A11;refGene_tx=XM_019826766.2;refGene_gn=SLC4A11;refGene_tx=XM_019826769.1;refGene_gn=SLC4A11;refGene_tx=XM_023251407.1;refGene_gn=SLC4A11;refGene_tx=XM_023251408.1;refGene_gn=SLC4A11;refGene_tx=XM_023251410.1;refGene_gn=SLC4A11;refGene_tx=XM_023251411.1
A3 30212864 30212997
refGene_gn=SLC4A11;refGene_tx=XM_006929906.3;refGene_gn=SLC4A11;refGene_tx=XM_006929908.4;refGene_gn=SLC4A11;refGene_tx=XM_019826756.2;refGene_gn=SLC4A11;refGene_tx=XM_019826757.2;refGene_gn=SLC4A11;refGene_tx=XM_019826758.2;refGene_gn=SLC4A11;refGene_tx=XM_019826760.2;refGene_gn=SLC4A11;refGene_tx=XM_019826761.2;refGene_gn=SLC4A11;refGene_tx=XM_019826763.2;refGene_gn=SLC4A11;refGene_tx=XM_019826764.2;refGene_gn=SLC4A11;refGene_tx=XM_019826765.2;refGene_gn=SLC4A11;refGene_tx=XM_019826766.2;refGene_gn=SLC4A11;refGene_tx=XM_019826769.1;refGene_gn=SLC4A11;refGene_tx=XM_023251407.1;refGene_gn=SLC4A11;refGene_tx=XM_023251408.1;refGene_gn=SLC4A11;refGene_tx=XM_023251410.1;refGene_gn=SLC4A11;refGene_tx=XM_023251411.1
A3 30213087 30213201
refGene_gn=SLC4A11;refGene_tx=XM_006929906.3;refGene_gn=SLC4A11;refGene_tx=XM_006929908.4;refGene_gn=SLC4A11;refGene_tx=XM_019826756.2;refGene_gn=SLC4A11;refGene_tx=XM_019826757.2;refGene_gn=SLC4A11;refGene_tx=XM_019826758.2;refGene_gn=SLC4A11;refGene_tx=XM_019826760.2;refGene_gn=SLC4A11;refGene_tx=XM_019826761.2;refGene_gn=SLC4A11;refGene_tx=XM_019826763.2;refGene_gn=SLC4A11;refGene_tx=XM_019826764.2;refGene_gn=SLC4A11;refGene_tx=XM_019826765.2;refGene_gn=SLC4A11;refGene_tx=XM_019826766.2;refGene_gn=SLC4A11;refGene_tx=XM_019826769.1;refGene_gn=SLC4A11;r

efGene_tx=XM_023251407.1;refGene_gn=SLC4A11;refGene_tx=XM_023251408.1;
refGene_gn=SLC4A11;refGene_tx=XM_023251410.1;refGene_gn=SLC4A11;refGen
e_tx=XM_023251411.1
A3 30213285 30213411
refGene_gn=SLC4A11;refGene_tx=XM_006929906.3;refGene_gn=SLC4A11;refGen
e_tx=XM_006929908.4;refGene_gn=SLC4A11;refGene_tx=XM_019826756.2;refGe
ne_gn=SLC4A11;refGene_tx=XM_019826757.2;refGene_gn=SLC4A11;refGene_tx=
XM_019826758.2;refGene_gn=SLC4A11;refGene_tx=XM_019826760.2;refGene_gn
=SLC4A11;refGene_tx=XM_019826761.2;refGene_gn=SLC4A11;refGene_tx=XM_01
9826763.2;refGene_gn=SLC4A11;refGene_tx=XM_019826764.2;refGene_gn=SLC4
A11;refGene_tx=XM_019826765.2;refGene_gn=SLC4A11;refGene_tx=XM_0198267
66.2;refGene_gn=SLC4A11;refGene_tx=XM_019826769.1;refGene_gn=SLC4A11;r
efGene_tx=XM_023251407.1;refGene_gn=SLC4A11;refGene_tx=XM_023251408.1;
refGene_gn=SLC4A11;refGene_tx=XM_023251410.1;refGene_gn=SLC4A11;refGen
e_tx=XM_023251411.1
A3 30213522 30213616
refGene_gn=SLC4A11;refGene_tx=XM_006929906.3;refGene_gn=SLC4A11;refGen
e_tx=XM_006929908.4;refGene_gn=SLC4A11;refGene_tx=XM_019826756.2;refGe
ne_gn=SLC4A11;refGene_tx=XM_019826757.2;refGene_gn=SLC4A11;refGene_tx=
XM_019826758.2;refGene_gn=SLC4A11;refGene_tx=XM_019826760.2;refGene_gn
=SLC4A11;refGene_tx=XM_019826761.2;refGene_gn=SLC4A11;refGene_tx=XM_01
9826763.2;refGene_gn=SLC4A11;refGene_tx=XM_019826764.2;refGene_gn=SLC4
A11;refGene_tx=XM_019826765.2;refGene_gn=SLC4A11;refGene_tx=XM_0198267
66.2;refGene_gn=SLC4A11;refGene_tx=XM_019826769.1;refGene_gn=SLC4A11;r
efGene_tx=XM_023251407.1;refGene_gn=SLC4A11;refGene_tx=XM_023251408.1;
refGene_gn=SLC4A11;refGene_tx=XM_023251410.1;refGene_gn=SLC4A11;refGen
e_tx=XM_023251411.1
A3 30213702 30213915
refGene_gn=SLC4A11;refGene_tx=XM_006929906.3;refGene_gn=SLC4A11;refGen
e_tx=XM_006929908.4;refGene_gn=SLC4A11;refGene_tx=XM_019826756.2;refGe
ne_gn=SLC4A11;refGene_tx=XM_019826757.2;refGene_gn=SLC4A11;refGene_tx=
XM_019826758.2;refGene_gn=SLC4A11;refGene_tx=XM_019826760.2;refGene_gn
=SLC4A11;refGene_tx=XM_019826761.2;refGene_gn=SLC4A11;refGene_tx=XM_01
9826763.2;refGene_gn=SLC4A11;refGene_tx=XM_019826764.2;refGene_gn=SLC4
A11;refGene_tx=XM_019826765.2;refGene_gn=SLC4A11;refGene_tx=XM_0198267
66.2;refGene_gn=SLC4A11;refGene_tx=XM_019826769.1;refGene_gn=SLC4A11;r
efGene_tx=XM_023251407.1;refGene_gn=SLC4A11;refGene_tx=XM_023251408.1;
refGene_gn=SLC4A11;refGene_tx=XM_023251410.1;refGene_gn=SLC4A11;refGen
e_tx=XM_023251411.1
A3 30217202 30217326
refGene_gn=SLC4A11;refGene_tx=XM_006929906.3;refGene_gn=SLC4A11;refGen
e_tx=XM_006929908.4;refGene_gn=SLC4A11;refGene_tx=XM_019826756.2;refGe
ne_gn=SLC4A11;refGene_tx=XM_019826757.2;refGene_gn=SLC4A11;refGene_tx=
XM_019826758.2;refGene_gn=SLC4A11;refGene_tx=XM_019826760.2;refGene_gn
=SLC4A11;refGene_tx=XM_019826761.2;refGene_gn=SLC4A11;refGene_tx=XM_01
9826763.2;refGene_gn=SLC4A11;refGene_tx=XM_019826764.2;refGene_gn=SLC4
A11;refGene_tx=XM_019826765.2;refGene_gn=SLC4A11;refGene_tx=XM_0198267
66.2;refGene_gn=SLC4A11;refGene_tx=XM_019826769.1;refGene_gn=SLC4A11;r
efGene_tx=XM_023251407.1;refGene_gn=SLC4A11;refGene_tx=XM_023251408.1;
refGene_gn=SLC4A11;refGene_tx=XM_023251410.1;refGene_gn=SLC4A11;refGen

e_tx=XM_023251411.1
A3	30217540	30217622
refGene_gn=SLC4A11;refGene_tx=XM_006929906.3;refGene_gn=SLC4A11;refGene_tx=XM_006929908.4;refGene_gn=SLC4A11;refGene_tx=XM_019826756.2;refGene_gn=SLC4A11;refGene_tx=XM_019826757.2;refGene_gn=SLC4A11;refGene_tx=XM_019826758.2;refGene_gn=SLC4A11;refGene_tx=XM_019826760.2;refGene_gn=SLC4A11;refGene_tx=XM_019826761.2;refGene_gn=SLC4A11;refGene_tx=XM_019826763.2;refGene_gn=SLC4A11;refGene_tx=XM_019826764.2;refGene_gn=SLC4A11;refGene_tx=XM_019826765.2;refGene_gn=SLC4A11;refGene_tx=XM_019826766.2;refGene_gn=SLC4A11;refGene_tx=XM_019826769.1;refGene_gn=SLC4A11;refGene_tx=XM_023251407.1;refGene_gn=SLC4A11;refGene_tx=XM_023251408.1;refGene_gn=SLC4A11;refGene_tx=XM_023251410.1;refGene_gn=SLC4A11;refGene_tx=XM_023251411.1
A3	30217705	30217937
refGene_gn=SLC4A11;refGene_tx=XM_006929906.3;refGene_gn=SLC4A11;refGene_tx=XM_006929908.4;refGene_gn=SLC4A11;refGene_tx=XM_019826756.2;refGene_gn=SLC4A11;refGene_tx=XM_019826757.2;refGene_gn=SLC4A11;refGene_tx=XM_019826758.2;refGene_gn=SLC4A11;refGene_tx=XM_019826760.2;refGene_gn=SLC4A11;refGene_tx=XM_019826761.2;refGene_gn=SLC4A11;refGene_tx=XM_019826763.2;refGene_gn=SLC4A11;refGene_tx=XM_019826764.2;refGene_gn=SLC4A11;refGene_tx=XM_019826765.2;refGene_gn=SLC4A11;refGene_tx=XM_019826766.2;refGene_gn=SLC4A11;refGene_tx=XM_019826769.1;refGene_gn=SLC4A11;refGene_tx=XM_023251407.1;refGene_gn=SLC4A11;refGene_tx=XM_023251408.1;refGene_gn=SLC4A11;refGene_tx=XM_023251410.1;refGene_gn=SLC4A11;refGene_tx=XM_023251411.1
A3	30218224	30218274
refGene_gn=SLC4A11;refGene_tx=XM_006929906.3;refGene_gn=SLC4A11;refGene_tx=XM_006929908.4;refGene_gn=SLC4A11;refGene_tx=XM_019826756.2;refGene_gn=SLC4A11;refGene_tx=XM_019826757.2;refGene_gn=SLC4A11;refGene_tx=XM_019826758.2;refGene_gn=SLC4A11;refGene_tx=XM_019826760.2;refGene_gn=SLC4A11;refGene_tx=XM_019826769.1;refGene_gn=SLC4A11;refGene_tx=XM_023251407.1;refGene_gn=SLC4A11;refGene_tx=XM_023251408.1;refGene_gn=SLC4A11;refGene_tx=XM_023251410.1
A3	30218402	30218555
refGene_gn=SLC4A11;refGene_tx=XM_006929906.3;refGene_gn=SLC4A11;refGene_tx=XM_006929908.4;refGene_gn=SLC4A11;refGene_tx=XM_019826756.2;refGene_gn=SLC4A11;refGene_tx=XM_019826757.2;refGene_gn=SLC4A11;refGene_tx=XM_019826758.2;refGene_gn=SLC4A11;refGene_tx=XM_019826761.2;refGene_gn=SLC4A11;refGene_tx=XM_019826763.2;refGene_gn=SLC4A11;refGene_tx=XM_019826764.2;refGene_gn=SLC4A11;refGene_tx=XM_019826765.2;refGene_gn=SLC4A11;refGene_tx=XM_019826766.2;refGene_gn=SLC4A11;refGene_tx=XM_019826769.1;refGene_gn=SLC4A11;refGene_tx=XM_023251407.1;refGene_gn=SLC4A11;refGene_tx=XM_023251410.1;refGene_gn=SLC4A11;refGene_tx=XM_023251411.1
A3	30220848	30220896
refGene_gn=SLC4A11;refGene_tx=XM_006929906.3;refGene_gn=SLC4A11;refGene_tx=XM_006929908.4;refGene_gn=SLC4A11;refGene_tx=XM_019826756.2;refGene_gn=SLC4A11;refGene_tx=XM_019826757.2;refGene_gn=SLC4A11;refGene_tx=XM_019826758.2;refGene_gn=SLC4A11;refGene_tx=XM_019826760.2;refGene_gn=SLC4A11;refGene_tx=XM_019826769.1;refGene_gn=SLC4A11;refGene_tx=XM_023251407.1;refGene_gn=SLC4A11;refGene_tx=XM_023251408.1;refGene_gn=SLC4

A11;refGene_tx=XM_023251410.1
A3	30221179 30221447
refGene_gn=SLC4A11;refGene_tx=XM_006929906.3;refGene_gn=SLC4A11;refGene_tx=XM_006929908.4;refGene_gn=SLC4A11;refGene_tx=XM_019826760.2;refGene_gn=SLC4A11;refGene_tx=XM_023251407.1;refGene_gn=SLC4A11;refGene_tx=XM_023251408.1
A3	30222485 30222528 refGene_gn=SLC4A11;refGene_tx=XM_019826757.2
A3	30232972 30233085
refGene_gn=CA3H20orf194;refGene_tx=XM_011280923.3;refGene_gn=CA3H20orf194;refGene_tx=XM_011280924.3;refGene_gn=CA3H20orf194;refGene_tx=XM_019826770.2;refGene_gn=CA3H20orf194;refGene_tx=XM_019826771.2
A3	30233637 30233731
refGene_gn=CA3H20orf194;refGene_tx=XM_011280923.3;refGene_gn=CA3H20orf194;refGene_tx=XM_011280924.3;refGene_gn=CA3H20orf194;refGene_tx=XM_019826770.2;refGene_gn=CA3H20orf194;refGene_tx=XM_019826771.2
A3	30235045 30235111
refGene_gn=CA3H20orf194;refGene_tx=XM_011280923.3;refGene_gn=CA3H20orf194;refGene_tx=XM_011280924.3;refGene_gn=CA3H20orf194;refGene_tx=XM_019826770.2;refGene_gn=CA3H20orf194;refGene_tx=XM_019826771.2
A3	30236223 30236429
refGene_gn=CA3H20orf194;refGene_tx=XM_011280923.3;refGene_gn=CA3H20orf194;refGene_tx=XM_011280924.3;refGene_gn=CA3H20orf194;refGene_tx=XM_019826770.2;refGene_gn=CA3H20orf194;refGene_tx=XM_019826771.2
A3	30238155 30238230
refGene_gn=CA3H20orf194;refGene_tx=XM_011280923.3;refGene_gn=CA3H20orf194;refGene_tx=XM_011280924.3;refGene_gn=CA3H20orf194;refGene_tx=XM_019826770.2;refGene_gn=CA3H20orf194;refGene_tx=XM_019826771.2
A3	30238556 30238663
refGene_gn=CA3H20orf194;refGene_tx=XM_011280923.3;refGene_gn=CA3H20orf194;refGene_tx=XM_011280924.3;refGene_gn=CA3H20orf194;refGene_tx=XM_019826770.2;refGene_gn=CA3H20orf194;refGene_tx=XM_019826771.2
A3	30243032 30243119
refGene_gn=CA3H20orf194;refGene_tx=XM_011280923.3;refGene_gn=CA3H20orf194;refGene_tx=XM_011280924.3;refGene_gn=CA3H20orf194;refGene_tx=XM_019826770.2;refGene_gn=CA3H20orf194;refGene_tx=XM_019826771.2
A3	30245634 30245770
refGene_gn=CA3H20orf194;refGene_tx=XM_011280923.3;refGene_gn=CA3H20orf194;refGene_tx=XM_011280924.3;refGene_gn=CA3H20orf194;refGene_tx=XM_019826770.2;refGene_gn=CA3H20orf194;refGene_tx=XM_019826771.2
A3	30255132 30255170
refGene_gn=CA3H20orf194;refGene_tx=XM_011280923.3;refGene_gn=CA3H20orf194;refGene_tx=XM_011280924.3;refGene_gn=CA3H20orf194;refGene_tx=XM_019826770.2;refGene_gn=CA3H20orf194;refGene_tx=XM_019826771.2
A3	30265130 30265256
refGene_gn=CA3H20orf194;refGene_tx=XM_011280923.3;refGene_gn=CA3H20orf194;refGene_tx=XM_011280924.3;refGene_gn=CA3H20orf194;refGene_tx=XM_019826770.2;refGene_gn=CA3H20orf194;refGene_tx=XM_019826771.2
A3	30270670 30270829
refGene_gn=CA3H20orf194;refGene_tx=XM_011280923.3;refGene_gn=CA3H20orf194;refGene_tx=XM_011280924.3;refGene_gn=CA3H20orf194;refGene_tx=XM_01

9826770.2;refGene_gn=CA3H20orf194;refGene_tx=XM_019826771.2
A3 30272115 30272204
refGene_gn=CA3H20orf194;refGene_tx=XM_011280923.3;refGene_gn=CA3H20orf194;refGene_tx=XM_011280924.3;refGene_gn=CA3H20orf194;refGene_tx=XM_019826770.2;refGene_gn=CA3H20orf194;refGene_tx=XM_019826771.2
A3 30274087 30274205
refGene_gn=CA3H20orf194;refGene_tx=XM_011280923.3;refGene_gn=CA3H20orf194;refGene_tx=XM_011280924.3;refGene_gn=CA3H20orf194;refGene_tx=XM_019826770.2;refGene_gn=CA3H20orf194;refGene_tx=XM_019826771.2
A3 30274710 30274812
refGene_gn=CA3H20orf194;refGene_tx=XM_011280923.3;refGene_gn=CA3H20orf194;refGene_tx=XM_011280924.3;refGene_gn=CA3H20orf194;refGene_tx=XM_019826770.2;refGene_gn=CA3H20orf194;refGene_tx=XM_019826771.2
A3 30275226 30275315
refGene_gn=CA3H20orf194;refGene_tx=XM_011280923.3;refGene_gn=CA3H20orf194;refGene_tx=XM_011280924.3;refGene_gn=CA3H20orf194;refGene_tx=XM_019826770.2;refGene_gn=CA3H20orf194;refGene_tx=XM_019826771.2
A3 30276390 30276537
refGene_gn=CA3H20orf194;refGene_tx=XM_011280923.3;refGene_gn=CA3H20orf194;refGene_tx=XM_011280924.3;refGene_gn=CA3H20orf194;refGene_tx=XM_019826770.2;refGene_gn=CA3H20orf194;refGene_tx=XM_019826771.2
A3 30280139 30280243
refGene_gn=CA3H20orf194;refGene_tx=XM_011280923.3;refGene_gn=CA3H20orf194;refGene_tx=XM_011280924.3;refGene_gn=CA3H20orf194;refGene_tx=XM_019826770.2;refGene_gn=CA3H20orf194;refGene_tx=XM_019826771.2
A3 30291163 30291251
refGene_gn=CA3H20orf194;refGene_tx=XM_011280923.3;refGene_gn=CA3H20orf194;refGene_tx=XM_011280924.3;refGene_gn=CA3H20orf194;refGene_tx=XM_019826770.2;refGene_gn=CA3H20orf194;refGene_tx=XM_019826771.2
A3 30292386 30292437
refGene_gn=CA3H20orf194;refGene_tx=XM_011280923.3;refGene_gn=CA3H20orf194;refGene_tx=XM_011280924.3;refGene_gn=CA3H20orf194;refGene_tx=XM_019826770.2;refGene_gn=CA3H20orf194;refGene_tx=XM_019826771.2
A3 30293372 30293443
refGene_gn=CA3H20orf194;refGene_tx=XM_011280923.3;refGene_gn=CA3H20orf194;refGene_tx=XM_011280924.3;refGene_gn=CA3H20orf194;refGene_tx=XM_019826770.2;refGene_gn=CA3H20orf194;refGene_tx=XM_019826771.2
A3 30294007 30294119
refGene_gn=CA3H20orf194;refGene_tx=XM_011280923.3;refGene_gn=CA3H20orf194;refGene_tx=XM_011280924.3;refGene_gn=CA3H20orf194;refGene_tx=XM_019826770.2;refGene_gn=CA3H20orf194;refGene_tx=XM_019826771.2
A3 30297559 30297605
refGene_gn=CA3H20orf194;refGene_tx=XM_011280923.3;refGene_gn=CA3H20orf194;refGene_tx=XM_011280924.3;refGene_gn=CA3H20orf194;refGene_tx=XM_019826770.2;refGene_gn=CA3H20orf194;refGene_tx=XM_019826771.2
A3 30297978 30298023
refGene_gn=CA3H20orf194;refGene_tx=XM_011280923.3;refGene_gn=CA3H20orf194;refGene_tx=XM_011280924.3;refGene_gn=CA3H20orf194;refGene_tx=XM_019826770.2;refGene_gn=CA3H20orf194;refGene_tx=XM_019826771.2
A3 30299766 30299843

refGene_gn=CA3H20orf194;refGene_tx=XM_011280923.3;refGene_gn=CA3H20orf194;refGene_tx=XM_011280924.3;refGene_gn=CA3H20orf194;refGene_tx=XM_019826770.2;refGene_gn=CA3H20orf194;refGene_tx=XM_019826771.2
A3 30301232 30301320
refGene_gn=CA3H20orf194;refGene_tx=XM_011280923.3;refGene_gn=CA3H20orf194;refGene_tx=XM_011280924.3;refGene_gn=CA3H20orf194;refGene_tx=XM_019826770.2;refGene_gn=CA3H20orf194;refGene_tx=XM_019826771.2
A3 30305377 30305414
refGene_gn=CA3H20orf194;refGene_tx=XM_011280923.3;refGene_gn=CA3H20orf194;refGene_tx=XM_011280924.3;refGene_gn=CA3H20orf194;refGene_tx=XM_019826770.2;refGene_gn=CA3H20orf194;refGene_tx=XM_019826771.2
A3 30306394 30306476
refGene_gn=CA3H20orf194;refGene_tx=XM_011280923.3;refGene_gn=CA3H20orf194;refGene_tx=XM_011280924.3;refGene_gn=CA3H20orf194;refGene_tx=XM_019826770.2;refGene_gn=CA3H20orf194;refGene_tx=XM_019826771.2
A3 30310195 30310331
refGene_gn=CA3H20orf194;refGene_tx=XM_011280923.3;refGene_gn=CA3H20orf194;refGene_tx=XM_011280924.3;refGene_gn=CA3H20orf194;refGene_tx=XM_019826770.2;refGene_gn=CA3H20orf194;refGene_tx=XM_019826771.2
A3 30313512 30313568
refGene_gn=CA3H20orf194;refGene_tx=XM_011280923.3;refGene_gn=CA3H20orf194;refGene_tx=XM_011280924.3;refGene_gn=CA3H20orf194;refGene_tx=XM_019826770.2;refGene_gn=CA3H20orf194;refGene_tx=XM_019826771.2
A3 30317313 30317412
refGene_gn=CA3H20orf194;refGene_tx=XM_011280923.3;refGene_gn=CA3H20orf194;refGene_tx=XM_011280924.3;refGene_gn=CA3H20orf194;refGene_tx=XM_019826770.2;refGene_gn=CA3H20orf194;refGene_tx=XM_019826771.2
A3 30328185 30328263
refGene_gn=CA3H20orf194;refGene_tx=XM_011280923.3;refGene_gn=CA3H20orf194;refGene_tx=XM_011280924.3;refGene_gn=CA3H20orf194;refGene_tx=XM_019826770.2;refGene_gn=CA3H20orf194;refGene_tx=XM_019826771.2
A3 30335436 30335543
refGene_gn=CA3H20orf194;refGene_tx=XM_011280923.3;refGene_gn=CA3H20orf194;refGene_tx=XM_011280924.3;refGene_gn=CA3H20orf194;refGene_tx=XM_019826770.2;refGene_gn=CA3H20orf194;refGene_tx=XM_019826771.2
A3 30336410 30336507
refGene_gn=CA3H20orf194;refGene_tx=XM_011280923.3;refGene_gn=CA3H20orf194;refGene_tx=XM_011280924.3;refGene_gn=CA3H20orf194;refGene_tx=XM_019826770.2;refGene_gn=CA3H20orf194;refGene_tx=XM_019826771.2
A3 30337389 30337514
refGene_gn=CA3H20orf194;refGene_tx=XM_011280923.3;refGene_gn=CA3H20orf194;refGene_tx=XM_011280924.3;refGene_gn=CA3H20orf194;refGene_tx=XM_019826770.2;refGene_gn=CA3H20orf194;refGene_tx=XM_019826771.2
A3 30337690 30337721
refGene_gn=CA3H20orf194;refGene_tx=XM_019826771.2
A3 30341157 30341277
refGene_gn=CA3H20orf194;refGene_tx=XM_011280923.3;refGene_gn=CA3H20orf194;refGene_tx=XM_011280924.3;refGene_gn=CA3H20orf194;refGene_tx=XM_019826770.2
A3 30344243 30344323

refGene_gn=CA3H20orf194;refGene_tx=XM_011280923.3;refGene_gn=CA3H20orf194;refGene_tx=XM_011280924.3;refGene_gn=CA3H20orf194;refGene_tx=XM_019826770.2
A3 30362038 30362121
refGene_gn=CA3H20orf194;refGene_tx=XM_011280923.3;refGene_gn=CA3H20orf194;refGene_tx=XM_011280924.3;refGene_gn=CA3H20orf194;refGene_tx=XM_019826770.2
A3 30408602 30408979
refGene_gn=ATRN;refGene_tx=XM_019826774.2;refGene_gn=ATRN;refGene_tx=XM_019826775.2;refGene_gn=ATRN;refGene_tx=XM_019826776.2;refGene_gn=ATRN;refGene_tx=XM_019826777.2;refGene_gn=ATRN;refGene_tx=XM_019826778.2
A3 30444851 30444935
refGene_gn=ATRN;refGene_tx=XM_019826774.2;refGene_gn=ATRN;refGene_tx=XM_019826775.2;refGene_gn=ATRN;refGene_tx=XM_019826776.2;refGene_gn=ATRN;refGene_tx=XM_019826777.2;refGene_gn=ATRN;refGene_tx=XM_019826778.2
A3 30448697 30448811
refGene_gn=ATRN;refGene_tx=XM_019826774.2;refGene_gn=ATRN;refGene_tx=XM_019826775.2;refGene_gn=ATRN;refGene_tx=XM_019826776.2;refGene_gn=ATRN;refGene_tx=XM_019826777.2;refGene_gn=ATRN;refGene_tx=XM_019826778.2
A3 30455589 30455718
refGene_gn=ATRN;refGene_tx=XM_019826774.2;refGene_gn=ATRN;refGene_tx=XM_019826775.2;refGene_gn=ATRN;refGene_tx=XM_019826776.2;refGene_gn=ATRN;refGene_tx=XM_019826777.2;refGene_gn=ATRN;refGene_tx=XM_019826778.2
A3 30460726 30460932
refGene_gn=ATRN;refGene_tx=XM_019826774.2;refGene_gn=ATRN;refGene_tx=XM_019826775.2;refGene_gn=ATRN;refGene_tx=XM_019826776.2;refGene_gn=ATRN;refGene_tx=XM_019826777.2;refGene_gn=ATRN;refGene_tx=XM_019826778.2
A3 30462037 30462206
refGene_gn=ATRN;refGene_tx=XM_019826774.2;refGene_gn=ATRN;refGene_tx=XM_019826775.2;refGene_gn=ATRN;refGene_tx=XM_019826776.2;refGene_gn=ATRN;refGene_tx=XM_019826777.2;refGene_gn=ATRN;refGene_tx=XM_019826778.2
A3 30473208 30473299
refGene_gn=ATRN;refGene_tx=XM_019826774.2;refGene_gn=ATRN;refGene_tx=XM_019826775.2;refGene_gn=ATRN;refGene_tx=XM_019826776.2;refGene_gn=ATRN;refGene_tx=XM_019826777.2;refGene_gn=ATRN;refGene_tx=XM_019826778.2
A3 30474239 30474483
refGene_gn=ATRN;refGene_tx=XM_019826774.2;refGene_gn=ATRN;refGene_tx=XM_019826775.2;refGene_gn=ATRN;refGene_tx=XM_019826776.2;refGene_gn=ATRN;refGene_tx=XM_019826777.2;refGene_gn=ATRN;refGene_tx=XM_019826778.2
A3 30475228 30475412
refGene_gn=ATRN;refGene_tx=XM_019826774.2;refGene_gn=ATRN;refGene_tx=XM_019826775.2;refGene_gn=ATRN;refGene_tx=XM_019826776.2;refGene_gn=ATRN;refGene_tx=XM_019826777.2;refGene_gn=ATRN;refGene_tx=XM_019826778.2
A3 30476450 30476605
refGene_gn=ATRN;refGene_tx=XM_019826774.2;refGene_gn=ATRN;refGene_tx=XM_019826775.2;refGene_gn=ATRN;refGene_tx=XM_019826776.2;refGene_gn=ATRN;refGene_tx=XM_019826777.2;refGene_gn=ATRN;refGene_tx=XM_019826778.2
A3 30479452 30479537
refGene_gn=ATRN;refGene_tx=XM_019826774.2;refGene_gn=ATRN;refGene_tx=XM_019826775.2;refGene_gn=ATRN;refGene_tx=XM_019826776.2;refGene_gn=ATR

N;refGene_tx=XM_019826777.2;refGene_gn=ATRN;refGene_tx=XM_019826778.2
A3 30487981 30488202
refGene_gn=ATRN;refGene_tx=XM_019826774.2;refGene_gn=ATRN;refGene_tx=XM_019826775.2;refGene_gn=ATRN;refGene_tx=XM_019826776.2;refGene_gn=ATRN;refGene_tx=XM_019826777.2;refGene_gn=ATRN;refGene_tx=XM_019826778.2
A3 30490390 30490512
refGene_gn=ATRN;refGene_tx=XM_019826774.2;refGene_gn=ATRN;refGene_tx=XM_019826775.2;refGene_gn=ATRN;refGene_tx=XM_019826776.2;refGene_gn=ATRN;refGene_tx=XM_019826777.2;refGene_gn=ATRN;refGene_tx=XM_019826778.2
A3 30491314 30491453
refGene_gn=ATRN;refGene_tx=XM_019826774.2;refGene_gn=ATRN;refGene_tx=XM_019826775.2;refGene_gn=ATRN;refGene_tx=XM_019826776.2;refGene_gn=ATRN;refGene_tx=XM_019826777.2;refGene_gn=ATRN;refGene_tx=XM_019826778.2
A3 30493319 30493510
refGene_gn=ATRN;refGene_tx=XM_019826774.2;refGene_gn=ATRN;refGene_tx=XM_019826775.2;refGene_gn=ATRN;refGene_tx=XM_019826776.2;refGene_gn=ATRN;refGene_tx=XM_019826777.2;refGene_gn=ATRN;refGene_tx=XM_019826778.2
A3 30495949 30496169
refGene_gn=ATRN;refGene_tx=XM_019826774.2;refGene_gn=ATRN;refGene_tx=XM_019826775.2;refGene_gn=ATRN;refGene_tx=XM_019826776.2;refGene_gn=ATRN;refGene_tx=XM_019826777.2;refGene_gn=ATRN;refGene_tx=XM_019826778.2
A3 30497414 30497600
refGene_gn=ATRN;refGene_tx=XM_019826774.2;refGene_gn=ATRN;refGene_tx=XM_019826775.2;refGene_gn=ATRN;refGene_tx=XM_019826776.2;refGene_gn=ATRN;refGene_tx=XM_019826777.2;refGene_gn=ATRN;refGene_tx=XM_019826778.2
A3 30498100 30498334
refGene_gn=ATRN;refGene_tx=XM_019826774.2;refGene_gn=ATRN;refGene_tx=XM_019826775.2;refGene_gn=ATRN;refGene_tx=XM_019826776.2;refGene_gn=ATRN;refGene_tx=XM_019826777.2;refGene_gn=ATRN;refGene_tx=XM_019826778.2
A3 30504504 30504642
refGene_gn=ATRN;refGene_tx=XM_019826774.2;refGene_gn=ATRN;refGene_tx=XM_019826775.2;refGene_gn=ATRN;refGene_tx=XM_019826776.2;refGene_gn=ATRN;refGene_tx=XM_019826777.2;refGene_gn=ATRN;refGene_tx=XM_019826778.2
A3 30507382 30507476
refGene_gn=ATRN;refGene_tx=XM_019826774.2;refGene_gn=ATRN;refGene_tx=XM_019826775.2;refGene_gn=ATRN;refGene_tx=XM_019826776.2;refGene_gn=ATRN;refGene_tx=XM_019826777.2;refGene_gn=ATRN;refGene_tx=XM_019826778.2
A3 30509597 30509650
refGene_gn=ATRN;refGene_tx=XM_019826774.2;refGene_gn=ATRN;refGene_tx=XM_019826775.2;refGene_gn=ATRN;refGene_tx=XM_019826776.2;refGene_gn=ATRN;refGene_tx=XM_019826777.2;refGene_gn=ATRN;refGene_tx=XM_019826778.2
A3 30512540 30512635
refGene_gn=ATRN;refGene_tx=XM_019826774.2;refGene_gn=ATRN;refGene_tx=XM_019826775.2;refGene_gn=ATRN;refGene_tx=XM_019826776.2;refGene_gn=ATRN;refGene_tx=XM_019826777.2;refGene_gn=ATRN;refGene_tx=XM_019826778.2
A3 30515251 30515330
refGene_gn=ATRN;refGene_tx=XM_019826774.2;refGene_gn=ATRN;refGene_tx=XM_019826775.2;refGene_gn=ATRN;refGene_tx=XM_019826776.2;refGene_gn=ATRN;refGene_tx=XM_019826777.2;refGene_gn=ATRN;refGene_tx=XM_019826778.2
A3 30517393 30517551

refGene_gn=ATRN;refGene_tx=XM_019826774.2;refGene_gn=ATRN;refGene_tx=XM_019826775.2;refGene_gn=ATRN;refGene_tx=XM_019826776.2;refGene_gn=ATRN;refGene_tx=XM_019826777.2;refGene_gn=ATRN;refGene_tx=XM_019826778.2
A3 30538802 30538864
refGene_gn=ATRN;refGene_tx=XM_019826774.2;refGene_gn=ATRN;refGene_tx=XM_019826775.2;refGene_gn=ATRN;refGene_tx=XM_019826776.2;refGene_gn=ATRN;refGene_tx=XM_019826777.2;refGene_gn=ATRN;refGene_tx=XM_019826778.2
A3 30542425 30542459 refGene_gn=ATRN;refGene_tx=XM_019826777.2
A3 30545117 30545262 refGene_gn=ATRN;refGene_tx=XM_019826776.2
A3 30556062 30556141
refGene_gn=ATRN;refGene_tx=XM_019826774.2;refGene_gn=ATRN;refGene_tx=XM_019826775.2;refGene_gn=ATRN;refGene_tx=XM_019826778.2
A3 30558371 30558479
refGene_gn=ATRN;refGene_tx=XM_019826774.2;refGene_gn=ATRN;refGene_tx=XM_019826775.2;refGene_gn=ATRN;refGene_tx=XM_019826778.2
A3 30563649 30563764
refGene_gn=ATRN;refGene_tx=XM_019826774.2;refGene_gn=ATRN;refGene_tx=XM_019826775.2;refGene_gn=ATRN;refGene_tx=XM_019826778.2
A3 30563916 30563922 refGene_gn=ATRN;refGene_tx=XM_019826778.2
A3 30566064 30566413
refGene_gn=ATRN;refGene_tx=XM_019826774.2;refGene_gn=ATRN;refGene_tx=XM_019826775.2;refGene_gn=ATRN;refGene_tx=XM_019826778.2
A3 30578691 30578772 refGene_gn=GFRA4;refGene_tx=XM_023252179.1
A3 30578904 30578996 refGene_gn=GFRA4;refGene_tx=XM_023252179.1
A3 30579676 30579811 refGene_gn=GFRA4;refGene_tx=XM_023252179.1
A3 30579904 30580014 refGene_gn=GFRA4;refGene_tx=XM_023252179.1
A3 30580196 30580580 refGene_gn=GFRA4;refGene_tx=XM_023252179.1
A3 30580767 30580888 refGene_gn=GFRA4;refGene_tx=XM_023252179.1
A3 30582427 30582470 refGene_gn=GFRA4;refGene_tx=XM_023252179.1
A3 30587723 30587809
refGene_gn=ADAM33;refGene_tx=XM_006929892.4;refGene_gn=ADAM33;refGene_tx=XM_006929893.4;refGene_gn=ADAM33;refGene_tx=XM_006929894.4;refGene_gn=ADAM33;refGene_tx=XM_006929895.4
A3 30587941 30588033
refGene_gn=ADAM33;refGene_tx=XM_006929892.4;refGene_gn=ADAM33;refGene_tx=XM_006929893.4;refGene_gn=ADAM33;refGene_tx=XM_006929894.4;refGene_gn=ADAM33;refGene_tx=XM_006929895.4
A3 30589343 30589491
refGene_gn=ADAM33;refGene_tx=XM_006929892.4;refGene_gn=ADAM33;refGene_tx=XM_006929893.4;refGene_gn=ADAM33;refGene_tx=XM_006929894.4;refGene_gn=ADAM33;refGene_tx=XM_006929895.4
A3 30589574 30589683
refGene_gn=ADAM33;refGene_tx=XM_006929892.4;refGene_gn=ADAM33;refGene_tx=XM_006929893.4;refGene_gn=ADAM33;refGene_tx=XM_006929894.4;refGene_gn=ADAM33;refGene_tx=XM_006929895.4
A3 30589765 30589843
refGene_gn=ADAM33;refGene_tx=XM_006929892.4;refGene_gn=ADAM33;refGene_tx=XM_006929893.4;refGene_gn=ADAM33;refGene_tx=XM_006929894.4;refGene_gn=ADAM33;refGene_tx=XM_006929895.4
A3 30589946 30590145

refGene_gn=ADAM33;refGene_tx=XM_006929892.4;refGene_gn=ADAM33;refGene_tx=XM_006929893.4;refGene_gn=ADAM33;refGene_tx=XM_006929894.4;refGene_gn=ADAM33;refGene_tx=XM_006929895.4
A3 30590239 30590348
refGene_gn=ADAM33;refGene_tx=XM_006929892.4;refGene_gn=ADAM33;refGene_tx=XM_006929893.4;refGene_gn=ADAM33;refGene_tx=XM_006929894.4;refGene_gn=ADAM33;refGene_tx=XM_006929895.4
A3 30590467 30590663
refGene_gn=ADAM33;refGene_tx=XM_006929892.4;refGene_gn=ADAM33;refGene_tx=XM_006929893.4;refGene_gn=ADAM33;refGene_tx=XM_006929894.4;refGene_gn=ADAM33;refGene_tx=XM_006929895.4
A3 30590906 30590996
refGene_gn=ADAM33;refGene_tx=XM_006929892.4;refGene_gn=ADAM33;refGene_tx=XM_006929893.4;refGene_gn=ADAM33;refGene_tx=XM_006929894.4;refGene_gn=ADAM33;refGene_tx=XM_006929895.4
A3 30591111 30591289
refGene_gn=ADAM33;refGene_tx=XM_006929892.4;refGene_gn=ADAM33;refGene_tx=XM_006929893.4;refGene_gn=ADAM33;refGene_tx=XM_006929894.4;refGene_gn=ADAM33;refGene_tx=XM_006929895.4
A3 30591826 30591969
refGene_gn=ADAM33;refGene_tx=XM_006929892.4;refGene_gn=ADAM33;refGene_tx=XM_006929893.4;refGene_gn=ADAM33;refGene_tx=XM_006929894.4;refGene_gn=ADAM33;refGene_tx=XM_006929895.4
A3 30592069 30592154
refGene_gn=ADAM33;refGene_tx=XM_006929892.4;refGene_gn=ADAM33;refGene_tx=XM_006929893.4;refGene_gn=ADAM33;refGene_tx=XM_006929894.4;refGene_gn=ADAM33;refGene_tx=XM_006929895.4
A3 30592241 30592408
refGene_gn=ADAM33;refGene_tx=XM_006929892.4;refGene_gn=ADAM33;refGene_tx=XM_006929893.4;refGene_gn=ADAM33;refGene_tx=XM_006929894.4;refGene_gn=ADAM33;refGene_tx=XM_006929895.4
A3 30592594 30592666
refGene_gn=ADAM33;refGene_tx=XM_006929892.4;refGene_gn=ADAM33;refGene_tx=XM_006929893.4;refGene_gn=ADAM33;refGene_tx=XM_006929894.4;refGene_gn=ADAM33;refGene_tx=XM_006929895.4
A3 30592747 30592813
refGene_gn=ADAM33;refGene_tx=XM_006929892.4;refGene_gn=ADAM33;refGene_tx=XM_006929893.4;refGene_gn=ADAM33;refGene_tx=XM_006929894.4;refGene_gn=ADAM33;refGene_tx=XM_006929895.4
A3 30593033 30593223
refGene_gn=ADAM33;refGene_tx=XM_006929892.4;refGene_gn=ADAM33;refGene_tx=XM_006929893.4;refGene_gn=ADAM33;refGene_tx=XM_006929894.4;refGene_gn=ADAM33;refGene_tx=XM_006929895.4
A3 30593308 30593385
refGene_gn=ADAM33;refGene_tx=XM_006929892.4;refGene_gn=ADAM33;refGene_tx=XM_006929893.4;refGene_gn=ADAM33;refGene_tx=XM_006929894.4;refGene_gn=ADAM33;refGene_tx=XM_006929895.4
A3 30593584 30593663
refGene_gn=ADAM33;refGene_tx=XM_006929892.4;refGene_gn=ADAM33;refGene_tx=XM_006929893.4;refGene_gn=ADAM33;refGene_tx=XM_006929894.4;refGene_

gn=ADAM33;refGene_tx=XM_006929895.4
A3 30595560 30595637
refGene_gn=ADAM33;refGene_tx=XM_006929892.4;refGene_gn=ADAM33;refGene_tx=XM_006929893.4;refGene_gn=ADAM33;refGene_tx=XM_006929894.4;refGene_gn=ADAM33;refGene_tx=XM_006929895.4
A3 30597802 30597882
refGene_gn=ADAM33;refGene_tx=XM_006929892.4;refGene_gn=ADAM33;refGene_tx=XM_006929893.4;refGene_gn=ADAM33;refGene_tx=XM_006929894.4;refGene_gn=ADAM33;refGene_tx=XM_006929895.4
A3 30600085 30600761
refGene_gn=ADAM33;refGene_tx=XM_006929892.4;refGene_gn=ADAM33;refGene_tx=XM_006929893.4;refGene_gn=ADAM33;refGene_tx=XM_006929894.4;refGene_gn=ADAM33;refGene_tx=XM_006929895.4
A3 30606455 30606563
refGene_gn=SIGLEC1;refGene_tx=XM_023251413.1;refGene_gn=SIGLEC1;refGene_tx=XM_023251414.1
A3 30607218 30607330
refGene_gn=SIGLEC1;refGene_tx=XM_023251413.1;refGene_gn=SIGLEC1;refGene_tx=XM_023251414.1;refGene_gn=SIGLEC1;refGene_tx=XM_023251415.1
A3 30607704 30607807
refGene_gn=SIGLEC1;refGene_tx=XM_023251413.1;refGene_gn=SIGLEC1;refGene_tx=XM_023251414.1;refGene_gn=SIGLEC1;refGene_tx=XM_023251415.1
A3 30608079 30608382
refGene_gn=SIGLEC1;refGene_tx=XM_023251413.1;refGene_gn=SIGLEC1;refGene_tx=XM_023251414.1;refGene_gn=SIGLEC1;refGene_tx=XM_023251415.1
A3 30609222 30609483
refGene_gn=SIGLEC1;refGene_tx=XM_023251413.1;refGene_gn=SIGLEC1;refGene_tx=XM_023251414.1;refGene_gn=SIGLEC1;refGene_tx=XM_023251415.1
A3 30610145 30610448
refGene_gn=SIGLEC1;refGene_tx=XM_023251413.1;refGene_gn=SIGLEC1;refGene_tx=XM_023251414.1;refGene_gn=SIGLEC1;refGene_tx=XM_023251415.1
A3 30610769 30611021
refGene_gn=SIGLEC1;refGene_tx=XM_023251413.1;refGene_gn=SIGLEC1;refGene_tx=XM_023251414.1;refGene_gn=SIGLEC1;refGene_tx=XM_023251415.1
A3 30611108 30611378
refGene_gn=SIGLEC1;refGene_tx=XM_023251413.1;refGene_gn=SIGLEC1;refGene_tx=XM_023251414.1;refGene_gn=SIGLEC1;refGene_tx=XM_023251415.1
A3 30611670 30611922
refGene_gn=SIGLEC1;refGene_tx=XM_023251413.1;refGene_gn=SIGLEC1;refGene_tx=XM_023251414.1;refGene_gn=SIGLEC1;refGene_tx=XM_023251415.1
A3 30612858 30613170
refGene_gn=SIGLEC1;refGene_tx=XM_023251413.1;refGene_gn=SIGLEC1;refGene_tx=XM_023251414.1;refGene_gn=SIGLEC1;refGene_tx=XM_023251415.1
A3 30613279 30613540
refGene_gn=SIGLEC1;refGene_tx=XM_023251413.1;refGene_gn=SIGLEC1;refGene_tx=XM_023251414.1;refGene_gn=SIGLEC1;refGene_tx=XM_023251415.1
A3 30614856 30615159
refGene_gn=SIGLEC1;refGene_tx=XM_023251413.1;refGene_gn=SIGLEC1;refGene_tx=XM_023251414.1;refGene_gn=SIGLEC1;refGene_tx=XM_023251415.1
A3 30615376 30615631

refGene_gn=SIGLEC1;refGene_tx=XM_023251413.1;refGene_gn=SIGLEC1;refGene_tx=XM_023251414.1;refGene_gn=SIGLEC1;refGene_tx=XM_023251415.1
A3 30615891 30616215
refGene_gn=SIGLEC1;refGene_tx=XM_023251413.1;refGene_gn=SIGLEC1;refGene_tx=XM_023251414.1;refGene_gn=SIGLEC1;refGene_tx=XM_023251415.1
A3 30616964 30617222
refGene_gn=SIGLEC1;refGene_tx=XM_023251413.1;refGene_gn=SIGLEC1;refGene_tx=XM_023251414.1;refGene_gn=SIGLEC1;refGene_tx=XM_023251415.1
A3 30617916 30618216
refGene_gn=SIGLEC1;refGene_tx=XM_023251413.1;refGene_gn=SIGLEC1;refGene_tx=XM_023251414.1;refGene_gn=SIGLEC1;refGene_tx=XM_023251415.1
A3 30620582 30620837
refGene_gn=SIGLEC1;refGene_tx=XM_023251413.1;refGene_gn=SIGLEC1;refGene_tx=XM_023251414.1;refGene_gn=SIGLEC1;refGene_tx=XM_023251415.1
A3 30621173 30621440
refGene_gn=SIGLEC1;refGene_tx=XM_023251413.1;refGene_gn=SIGLEC1;refGene_tx=XM_023251414.1;refGene_gn=SIGLEC1;refGene_tx=XM_023251415.1
A3 30622651 30622948
refGene_gn=SIGLEC1;refGene_tx=XM_023251413.1;refGene_gn=SIGLEC1;refGene_tx=XM_023251414.1;refGene_gn=SIGLEC1;refGene_tx=XM_023251415.1
A3 30623225 30623585
refGene_gn=SIGLEC1;refGene_tx=XM_023251413.1;refGene_gn=SIGLEC1;refGene_tx=XM_023251414.1;refGene_gn=SIGLEC1;refGene_tx=XM_023251415.1
A3 30623882 30623931
refGene_gn=SIGLEC1;refGene_tx=XM_023251413.1;refGene_gn=SIGLEC1;refGene_tx=XM_023251414.1;refGene_gn=SIGLEC1;refGene_tx=XM_023251415.1
A3 30650353 30650396 refGene_gn=HSPA12B;refGene_tx=XM_023251416.1
A3 30652228 30652326 refGene_gn=HSPA12B;refGene_tx=XM_023251416.1
A3 30653296 30653421 refGene_gn=HSPA12B;refGene_tx=XM_023251416.1
A3 30656902 30657089 refGene_gn=HSPA12B;refGene_tx=XM_023251416.1
A3 30657435 30657540 refGene_gn=HSPA12B;refGene_tx=XM_023251416.1
A3 30657823 30657940 refGene_gn=HSPA12B;refGene_tx=XM_023251416.1
A3 30659859 30660034 refGene_gn=HSPA12B;refGene_tx=XM_023251416.1
A3 30660944 30661031 refGene_gn=HSPA12B;refGene_tx=XM_023251416.1
A3 30661470 30661575 refGene_gn=HSPA12B;refGene_tx=XM_023251416.1
A3 30661681 30661940 refGene_gn=HSPA12B;refGene_tx=XM_023251416.1
A3 30662405 30662509 refGene_gn=HSPA12B;refGene_tx=XM_023251416.1
A3 30663128 30663766 refGene_gn=HSPA12B;refGene_tx=XM_023251416.1
A3 30665675 30665776
refGene_gn=CA3H20orf27;refGene_tx=XM_003983684.5;refGene_gn=CA3H20orf27;refGene_tx=XM_023251417.1;refGene_gn=CA3H20orf27;refGene_tx=XM_023251418.1
A3 30666007 30666130
refGene_gn=CA3H20orf27;refGene_tx=XM_003983684.5;refGene_gn=CA3H20orf27;refGene_tx=XM_023251417.1;refGene_gn=CA3H20orf27;refGene_tx=XM_023251418.1
A3 30667014 30667153
refGene_gn=CA3H20orf27;refGene_tx=XM_003983684.5;refGene_gn=CA3H20orf27;refGene_tx=XM_023251417.1;refGene_gn=CA3H20orf27;refGene_tx=XM_023251418.1

A3 30669537 30669680
refGene_gn=CA3H20orf27;refGene_tx=XM_003983684.5;refGene_gn=CA3H20orf27;refGene_tx=XM_023251417.1;refGene_gn=CA3H20orf27;refGene_tx=XM_023251418.1
A3 30671167 30671186
refGene_gn=CA3H20orf27;refGene_tx=XM_003983684.5;refGene_gn=CA3H20orf27;refGene_tx=XM_023251417.1;refGene_gn=CA3H20orf27;refGene_tx=XM_023251418.1
A3 30686557 30686882 refGene_gn=SPEF1;refGene_tx=XM_019826780.1
A3 30687136 30687244
refGene_gn=SPEF1;refGene_tx=XM_003983683.5;refGene_gn=SPEF1;refGene_tx=XM_019826780.1
A3 30687359 30687483
refGene_gn=SPEF1;refGene_tx=XM_003983683.5;refGene_gn=SPEF1;refGene_tx=XM_019826780.1
A3 30687687 30687748
refGene_gn=SPEF1;refGene_tx=XM_003983683.5;refGene_gn=SPEF1;refGene_tx=XM_019826780.1
A3 30687903 30687943
refGene_gn=SPEF1;refGene_tx=XM_003983683.5;refGene_gn=SPEF1;refGene_tx=XM_019826780.1
A3 30688135 30688292
refGene_gn=SPEF1;refGene_tx=XM_003983683.5;refGene_gn=SPEF1;refGene_tx=XM_019826780.1
A3 30688584 30688696
refGene_gn=SPEF1;refGene_tx=XM_003983683.5;refGene_gn=SPEF1;refGene_tx=XM_019826780.1
A3 30689826 30689935
refGene_gn=SPEF1;refGene_tx=XM_003983683.5;refGene_gn=SPEF1;refGene_tx=XM_019826780.1
A3 30693262 30695083 refGene_gn=CENPB;refGene_tx=XM_019828060.2
A3 30703682 30703882 refGene_gn=CDC25B;refGene_tx=XM_019826783.2
A3 30704696 30704806 refGene_gn=CDC25B;refGene_tx=XM_019826783.2
A3 30705469 30705521 refGene_gn=CDC25B;refGene_tx=XM_019826783.2
A3 30707501 30707543 refGene_gn=CDC25B;refGene_tx=XM_019826783.2
A3 30707791 30707828 refGene_gn=CDC25B;refGene_tx=XM_019826783.2
A3 30708084 30708292 refGene_gn=CDC25B;refGene_tx=XM_019826783.2
A3 30708294 30708434 refGene_gn=CDC25B;refGene_tx=XM_019826783.2
A3 30708621 30708756 refGene_gn=CDC25B;refGene_tx=XM_019826783.2
A3 30709136 30709217 refGene_gn=CDC25B;refGene_tx=XM_019826783.2
A3 30709332 30709509 refGene_gn=CDC25B;refGene_tx=XM_019826783.2
A3 30709680 30709776 refGene_gn=CDC25B;refGene_tx=XM_019826783.2
A3 30710351 30710414 refGene_gn=CDC25B;refGene_tx=XM_019826783.2
A3 30710562 30710661 refGene_gn=CDC25B;refGene_tx=XM_019826783.2
A3 30710901 30711035 refGene_gn=CDC25B;refGene_tx=XM_019826783.2
A3 30711751 30711863 refGene_gn=CDC25B;refGene_tx=XM_019826783.2
A3 30711981 30712122 refGene_gn=CDC25B;refGene_tx=XM_019826783.2
A3 30722513 30722689
refGene_gn=AP5S1;refGene_tx=XM_003983682.4;refGene_gn=AP5S1;refGene_tx=XM_006929886.4;refGene_gn=AP5S1;refGene_tx=XM_019826784.2

A3 30723958 30724385
refGene_gn=AP5S1;refGene_tx=XM_003983682.4;refGene_gn=AP5S1;refGene_tx
=XM_006929886.4;refGene_gn=AP5S1;refGene_tx=XM_019826784.2
A3 30744923 30744982 refGene_gn=MAVS;refGene_tx=XM_019826785.2
A3 30745748 30745896
refGene_gn=MAVS;refGene_tx=XM_003983681.5;refGene_gn=MAVS;refGene_tx=X
M_006929884.4;refGene_gn=MAVS;refGene_tx=XM_006929885.4;refGene_gn=MAV
S;refGene_tx=XM_019826785.2;refGene_gn=MAVS;refGene_tx=XM_019826786.2
A3 30747040 30747215
refGene_gn=MAVS;refGene_tx=XM_003983681.5;refGene_gn=MAVS;refGene_tx=X
M_006929884.4;refGene_gn=MAVS;refGene_tx=XM_006929885.4;refGene_gn=MAV
S;refGene_tx=XM_019826785.2;refGene_gn=MAVS;refGene_tx=XM_019826786.2
A3 30748417 30748593
refGene_gn=MAVS;refGene_tx=XM_003983681.5;refGene_gn=MAVS;refGene_tx=X
M_006929884.4;refGene_gn=MAVS;refGene_tx=XM_006929885.4;refGene_gn=MAV
S;refGene_tx=XM_019826785.2;refGene_gn=MAVS;refGene_tx=XM_019826786.2
A3 30749289 30749446
refGene_gn=MAVS;refGene_tx=XM_003983681.5;refGene_gn=MAVS;refGene_tx=X
M_006929884.4;refGene_gn=MAVS;refGene_tx=XM_006929885.4;refGene_gn=MAV
S;refGene_tx=XM_019826785.2;refGene_gn=MAVS;refGene_tx=XM_019826786.2
A3 30751150 30751659
refGene_gn=MAVS;refGene_tx=XM_003983681.5;refGene_gn=MAVS;refGene_tx=X
M_006929884.4;refGene_gn=MAVS;refGene_tx=XM_006929885.4;refGene_gn=MAV
S;refGene_tx=XM_019826785.2;refGene_gn=MAVS;refGene_tx=XM_019826786.2
A3 30752572 30753010
refGene_gn=MAVS;refGene_tx=XM_003983681.5;refGene_gn=MAVS;refGene_tx=X
M_006929884.4;refGene_gn=MAVS;refGene_tx=XM_006929885.4;refGene_gn=MAV
S;refGene_tx=XM_019826785.2;refGene_gn=MAVS;refGene_tx=XM_019826786.2
A3 30780100 30780368
refGene_gn=LOC109498514;refGene_tx=XM_019828061.1
A3 30797253 30797366
refGene_gn=LOC101092510;refGene_tx=XM_019826787.1
A3 30803430 30803607
refGene_gn=LOC101092510;refGene_tx=XM_019826787.1
A3 30805660 30805784
refGene_gn=LOC101092510;refGene_tx=XM_019826787.1
A3 30807636 30807762
refGene_gn=LOC101092510;refGene_tx=XM_019826787.1
A3 30811222 30811273
refGene_gn=LOC101092510;refGene_tx=XM_019826787.1
A3 30820609 30820748
refGene_gn=RNF24;refGene_tx=XM_006929880.4;refGene_gn=RNF24;refGene_tx
=XM_019826788.2;refGene_gn=RNF24;refGene_tx=XM_019826789.2;refGene_gn=
RNF24;refGene_tx=XM_023251420.1
A3 30821721 30821801
refGene_gn=RNF24;refGene_tx=XM_006929880.4;refGene_gn=RNF24;refGene_tx
=XM_019826788.2;refGene_gn=RNF24;refGene_tx=XM_019826789.2;refGene_gn=
RNF24;refGene_tx=XM_023251420.1
A3 30837239 30837281
refGene_gn=RNF24;refGene_tx=XM_006929880.4;refGene_gn=RNF24;refGene_tx

=XM_019826788.2;refGene_gn=RNF24;refGene_tx=XM_019826789.2;refGene_gn=RNF24;refGene_tx=XM_023251420.1
A3 30839497 30839540
refGene_gn=RNF24;refGene_tx=XM_006929880.4;refGene_gn=RNF24;refGene_tx=XM_019826788.2;refGene_gn=RNF24;refGene_tx=XM_019826789.2;refGene_gn=RNF24;refGene_tx=XM_023251420.1
A3 30845885 30846028
refGene_gn=RNF24;refGene_tx=XM_006929880.4;refGene_gn=RNF24;refGene_tx=XM_019826788.2;refGene_gn=RNF24;refGene_tx=XM_019826789.2;refGene_gn=RNF24;refGene_tx=XM_023251420.1
A3 31008846 31009054
refGene_gn=SMOX;refGene_tx=XM_023251421.1;refGene_gn=SMOX;refGene_tx=XM_023251422.1;refGene_gn=SMOX;refGene_tx=XM_023251423.1;refGene_gn=SMOX;refGene_tx=XM_023251424.1;refGene_gn=SMOX;refGene_tx=XM_023251425.1;refGene_gn=SMOX;refGene_tx=XM_023251426.1
A3 31012825 31013052
refGene_gn=SMOX;refGene_tx=XM_023251421.1;refGene_gn=SMOX;refGene_tx=XM_023251422.1;refGene_gn=SMOX;refGene_tx=XM_023251423.1;refGene_gn=SMOX;refGene_tx=XM_023251424.1;refGene_gn=SMOX;refGene_tx=XM_023251425.1;refGene_gn=SMOX;refGene_tx=XM_023251426.1
A3 31017763 31017937
refGene_gn=SMOX;refGene_tx=XM_023251421.1;refGene_gn=SMOX;refGene_tx=XM_023251422.1;refGene_gn=SMOX;refGene_tx=XM_023251423.1;refGene_gn=SMOX;refGene_tx=XM_023251425.1;refGene_gn=SMOX;refGene_tx=XM_023251426.1
A3 31018100 31018860
refGene_gn=SMOX;refGene_tx=XM_023251421.1;refGene_gn=SMOX;refGene_tx=XM_023251422.1;refGene_gn=SMOX;refGene_tx=XM_023251423.1;refGene_gn=SMOX;refGene_tx=XM_023251424.1;refGene_gn=SMOX;refGene_tx=XM_023251425.1;refGene_gn=SMOX;refGene_tx=XM_023251426.1
A3 31019439 31019600
refGene_gn=SMOX;refGene_tx=XM_023251421.1;refGene_gn=SMOX;refGene_tx=XM_023251422.1;refGene_gn=SMOX;refGene_tx=XM_023251423.1;refGene_gn=SMOX;refGene_tx=XM_023251424.1;refGene_gn=SMOX;refGene_tx=XM_023251425.1;refGene_gn=SMOX;refGene_tx=XM_023251426.1
A3 31021467 31021557
refGene_gn=SMOX;refGene_tx=XM_023251421.1;refGene_gn=SMOX;refGene_tx=XM_023251422.1;refGene_gn=SMOX;refGene_tx=XM_023251424.1
A3 31022007 31022145
refGene_gn=SMOX;refGene_tx=XM_023251421.1;refGene_gn=SMOX;refGene_tx=XM_023251422.1;refGene_gn=SMOX;refGene_tx=XM_023251423.1;refGene_gn=SMOX;refGene_tx=XM_023251424.1
A3 31032438 31032450 refGene_gn=SMOX;refGene_tx=XM_023251426.1
A3 31037230 31037281 refGene_gn=SMOX;refGene_tx=XM_023251425.1
A3 31041970 31042575 refGene_gn=ADRA1D;refGene_tx=XM_023251427.1
A3 31060320 31061452 refGene_gn=ADRA1D;refGene_tx=XM_023251427.1
A3 31442238 31443021
refGene_gn=PRNP;refGene_tx=XM_019826795.2;refGene_gn=PRNP;refGene_tx=XM_023251428.1
A3 31462255 31462792 refGene_gn=PRND;refGene_tx=XM_006929928.3
A3 31503784 31503854

refGene_gn=RASSF2;refGene_tx=XM_003983737.5;refGene_gn=RASSF2;refGene_tx=XM_023251430.1;refGene_gn=RASSF2;refGene_tx=XM_023251431.1
A3 31505328 31505426
refGene_gn=RASSF2;refGene_tx=XM_003983737.5;refGene_gn=RASSF2;refGene_tx=XM_023251430.1;refGene_gn=RASSF2;refGene_tx=XM_023251431.1
A3 31506638 31506760
refGene_gn=RASSF2;refGene_tx=XM_003983737.5;refGene_gn=RASSF2;refGene_tx=XM_023251430.1;refGene_gn=RASSF2;refGene_tx=XM_023251431.1
A3 31507229 31507281
refGene_gn=RASSF2;refGene_tx=XM_003983737.5;refGene_gn=RASSF2;refGene_tx=XM_023251430.1;refGene_gn=RASSF2;refGene_tx=XM_023251431.1
A3 31508623 31508725
refGene_gn=RASSF2;refGene_tx=XM_003983737.5;refGene_gn=RASSF2;refGene_tx=XM_023251430.1;refGene_gn=RASSF2;refGene_tx=XM_023251431.1
A3 31509548 31509709
refGene_gn=RASSF2;refGene_tx=XM_003983737.5;refGene_gn=RASSF2;refGene_tx=XM_023251430.1;refGene_gn=RASSF2;refGene_tx=XM_023251431.1
A3 31511174 31511263
refGene_gn=RASSF2;refGene_tx=XM_003983737.5;refGene_gn=RASSF2;refGene_tx=XM_023251430.1;refGene_gn=RASSF2;refGene_tx=XM_023251431.1
A3 31513514 31513666
refGene_gn=RASSF2;refGene_tx=XM_003983737.5;refGene_gn=RASSF2;refGene_tx=XM_023251430.1;refGene_gn=RASSF2;refGene_tx=XM_023251431.1
A3 31515947 31516023
refGene_gn=RASSF2;refGene_tx=XM_003983737.5;refGene_gn=RASSF2;refGene_tx=XM_023251430.1;refGene_gn=RASSF2;refGene_tx=XM_023251431.1
A3 31518680 31518739
refGene_gn=RASSF2;refGene_tx=XM_003983737.5;refGene_gn=RASSF2;refGene_tx=XM_023251430.1;refGene_gn=RASSF2;refGene_tx=XM_023251431.1
A3 31581029 31581262
refGene_gn=SLC23A2;refGene_tx=XM_019826798.2;refGene_gn=SLC23A2;refGene_tx=XM_019826800.2;refGene_gn=SLC23A2;refGene_tx=XM_023251432.1;refGene_gn=SLC23A2;refGene_tx=XM_023251433.1;refGene_gn=SLC23A2;refGene_tx=XM_023251434.1;refGene_gn=SLC23A2;refGene_tx=XM_023251435.1;refGene_gn=SLC23A2;refGene_tx=XM_023251436.1;refGene_gn=SLC23A2;refGene_tx=XM_023251438.1
A3 31583027 31583123
refGene_gn=SLC23A2;refGene_tx=XM_019826798.2;refGene_gn=SLC23A2;refGene_tx=XM_019826800.2;refGene_gn=SLC23A2;refGene_tx=XM_023251432.1;refGene_gn=SLC23A2;refGene_tx=XM_023251433.1;refGene_gn=SLC23A2;refGene_tx=XM_023251434.1;refGene_gn=SLC23A2;refGene_tx=XM_023251435.1;refGene_gn=SLC23A2;refGene_tx=XM_023251436.1;refGene_gn=SLC23A2;refGene_tx=XM_023251438.1
A3 31585929 31586067
refGene_gn=SLC23A2;refGene_tx=XM_019826798.2;refGene_gn=SLC23A2;refGene_tx=XM_019826800.2;refGene_gn=SLC23A2;refGene_tx=XM_023251432.1;refGene_gn=SLC23A2;refGene_tx=XM_023251433.1;refGene_gn=SLC23A2;refGene_tx=XM_023251434.1;refGene_gn=SLC23A2;refGene_tx=XM_023251435.1;refGene_gn=SLC23A2;refGene_tx=XM_023251436.1;refGene_gn=SLC23A2;refGene_tx=XM_023251438.1

A3 31586720 31586850
refGene_gn=SLC23A2;refGene_tx=XM_019826798.2;refGene_gn=SLC23A2;refGene_tx=XM_019826800.2;refGene_gn=SLC23A2;refGene_tx=XM_023251432.1;refGene_gn=SLC23A2;refGene_tx=XM_023251433.1;refGene_gn=SLC23A2;refGene_tx=XM_023251434.1;refGene_gn=SLC23A2;refGene_tx=XM_023251435.1;refGene_gn=SLC23A2;refGene_tx=XM_023251436.1;refGene_gn=SLC23A2;refGene_tx=XM_023251438.1
A3 31592125 31592231
refGene_gn=SLC23A2;refGene_tx=XM_019826798.2;refGene_gn=SLC23A2;refGene_tx=XM_019826800.2;refGene_gn=SLC23A2;refGene_tx=XM_023251432.1;refGene_gn=SLC23A2;refGene_tx=XM_023251433.1;refGene_gn=SLC23A2;refGene_tx=XM_023251434.1;refGene_gn=SLC23A2;refGene_tx=XM_023251435.1;refGene_gn=SLC23A2;refGene_tx=XM_023251436.1;refGene_gn=SLC23A2;refGene_tx=XM_023251438.1
A3 31593379 31593527
refGene_gn=SLC23A2;refGene_tx=XM_019826798.2;refGene_gn=SLC23A2;refGene_tx=XM_019826800.2;refGene_gn=SLC23A2;refGene_tx=XM_023251432.1;refGene_gn=SLC23A2;refGene_tx=XM_023251433.1;refGene_gn=SLC23A2;refGene_tx=XM_023251434.1;refGene_gn=SLC23A2;refGene_tx=XM_023251435.1;refGene_gn=SLC23A2;refGene_tx=XM_023251436.1;refGene_gn=SLC23A2;refGene_tx=XM_023251438.1
A3 31597788 31597945
refGene_gn=SLC23A2;refGene_tx=XM_019826798.2;refGene_gn=SLC23A2;refGene_tx=XM_019826800.2;refGene_gn=SLC23A2;refGene_tx=XM_023251432.1;refGene_gn=SLC23A2;refGene_tx=XM_023251433.1;refGene_gn=SLC23A2;refGene_tx=XM_023251434.1;refGene_gn=SLC23A2;refGene_tx=XM_023251435.1;refGene_gn=SLC23A2;refGene_tx=XM_023251436.1;refGene_gn=SLC23A2;refGene_tx=XM_023251438.1
A3 31598446 31598567
refGene_gn=SLC23A2;refGene_tx=XM_019826798.2;refGene_gn=SLC23A2;refGene_tx=XM_019826800.2;refGene_gn=SLC23A2;refGene_tx=XM_023251432.1;refGene_gn=SLC23A2;refGene_tx=XM_023251433.1;refGene_gn=SLC23A2;refGene_tx=XM_023251434.1;refGene_gn=SLC23A2;refGene_tx=XM_023251435.1;refGene_gn=SLC23A2;refGene_tx=XM_023251436.1;refGene_gn=SLC23A2;refGene_tx=XM_023251438.1
A3 31605892 31606074
refGene_gn=SLC23A2;refGene_tx=XM_019826798.2;refGene_gn=SLC23A2;refGene_tx=XM_019826800.2;refGene_gn=SLC23A2;refGene_tx=XM_023251432.1;refGene_gn=SLC23A2;refGene_tx=XM_023251433.1;refGene_gn=SLC23A2;refGene_tx=XM_023251434.1;refGene_gn=SLC23A2;refGene_tx=XM_023251435.1;refGene_gn=SLC23A2;refGene_tx=XM_023251436.1;refGene_gn=SLC23A2;refGene_tx=XM_023251438.1
A3 31607075 31607146
refGene_gn=SLC23A2;refGene_tx=XM_019826798.2;refGene_gn=SLC23A2;refGene_tx=XM_019826800.2;refGene_gn=SLC23A2;refGene_tx=XM_023251432.1;refGene_gn=SLC23A2;refGene_tx=XM_023251433.1;refGene_gn=SLC23A2;refGene_tx=XM_023251434.1;refGene_gn=SLC23A2;refGene_tx=XM_023251435.1;refGene_gn=SLC23A2;refGene_tx=XM_023251436.1;refGene_gn=SLC23A2;refGene_tx=XM_023251438.1
A3 31608156 31608245

refGene_gn=SLC23A2;refGene_tx=XM_019826798.2;refGene_gn=SLC23A2;refGene_tx=XM_019826800.2;refGene_gn=SLC23A2;refGene_tx=XM_023251432.1;refGene_gn=SLC23A2;refGene_tx=XM_023251433.1;refGene_gn=SLC23A2;refGene_tx=XM_023251434.1;refGene_gn=SLC23A2;refGene_tx=XM_023251435.1;refGene_gn=SLC23A2;refGene_tx=XM_023251436.1;refGene_gn=SLC23A2;refGene_tx=XM_023251438.1
A3 31623092 31623250
refGene_gn=SLC23A2;refGene_tx=XM_019826798.2;refGene_gn=SLC23A2;refGene_tx=XM_019826800.2;refGene_gn=SLC23A2;refGene_tx=XM_023251432.1;refGene_gn=SLC23A2;refGene_tx=XM_023251433.1;refGene_gn=SLC23A2;refGene_tx=XM_023251434.1;refGene_gn=SLC23A2;refGene_tx=XM_023251435.1;refGene_gn=SLC23A2;refGene_tx=XM_023251436.1;refGene_gn=SLC23A2;refGene_tx=XM_023251438.1
A3 31625531 31625648
refGene_gn=SLC23A2;refGene_tx=XM_019826798.2;refGene_gn=SLC23A2;refGene_tx=XM_019826800.2;refGene_gn=SLC23A2;refGene_tx=XM_023251432.1;refGene_gn=SLC23A2;refGene_tx=XM_023251433.1;refGene_gn=SLC23A2;refGene_tx=XM_023251434.1;refGene_gn=SLC23A2;refGene_tx=XM_023251435.1;refGene_gn=SLC23A2;refGene_tx=XM_023251436.1;refGene_gn=SLC23A2;refGene_tx=XM_023251438.1
A3 31634415 31634514
refGene_gn=SLC23A2;refGene_tx=XM_019826798.2;refGene_gn=SLC23A2;refGene_tx=XM_019826800.2;refGene_gn=SLC23A2;refGene_tx=XM_023251432.1;refGene_gn=SLC23A2;refGene_tx=XM_023251433.1;refGene_gn=SLC23A2;refGene_tx=XM_023251434.1;refGene_gn=SLC23A2;refGene_tx=XM_023251435.1;refGene_gn=SLC23A2;refGene_tx=XM_023251438.1
A3 31648209 31648308 refGene_gn=SLC23A2;refGene_tx=XM_019826800.2
A3 31654586 31654694
refGene_gn=SLC23A2;refGene_tx=XM_019826798.2;refGene_gn=SLC23A2;refGene_tx=XM_023251432.1;refGene_gn=SLC23A2;refGene_tx=XM_023251433.1;refGene_gn=SLC23A2;refGene_tx=XM_023251434.1;refGene_gn=SLC23A2;refGene_tx=XM_023251435.1;refGene_gn=SLC23A2;refGene_tx=XM_023251436.1
A3 31790277 31790418
refGene_gn=TMEM230;refGene_tx=XM_006929932.3;refGene_gn=TMEM230;refGene_tx=XM_006929933.2
A3 31796156 31796279
refGene_gn=TMEM230;refGene_tx=XM_006929932.3;refGene_gn=TMEM230;refGene_tx=XM_006929933.2
A3 31797150 31797229 refGene_gn=TMEM230;refGene_tx=XM_006929933.2
A3 31797601 31797700
refGene_gn=TMEM230;refGene_tx=XM_006929932.3;refGene_gn=TMEM230;refGene_tx=XM_006929933.2
A3 31801726 31801806 refGene_gn=PCNA;refGene_tx=XM_003983740.4
A3 31801907 31802031 refGene_gn=PCNA;refGene_tx=XM_003983740.4
A3 31804363 31804558 refGene_gn=PCNA;refGene_tx=XM_003983740.4
A3 31805620 31805688 refGene_gn=PCNA;refGene_tx=XM_003983740.4
A3 31805839 31805937 refGene_gn=PCNA;refGene_tx=XM_003983740.4
A3 31806772 31806993 refGene_gn=PCNA;refGene_tx=XM_003983740.4
A3 31817051 31817108 refGene_gn=CDS2;refGene_tx=XM_011280916.3
A3 31856233 31856370 refGene_gn=CDS2;refGene_tx=XM_011280916.3

A3 31858237 31858334 refGene_gn=CDS2;refGene_tx=XM_011280916.3
A3 31859301 31859399 refGene_gn=CDS2;refGene_tx=XM_011280916.3
A3 31860409 31860549 refGene_gn=CDS2;refGene_tx=XM_011280916.3
A3 31862338 31862397 refGene_gn=CDS2;refGene_tx=XM_011280916.3
A3 31863049 31863132 refGene_gn=CDS2;refGene_tx=XM_011280916.3
A3 31865598 31865686 refGene_gn=CDS2;refGene_tx=XM_011280916.3
A3 31866202 31866271 refGene_gn=CDS2;refGene_tx=XM_011280916.3
A3 31867199 31867352 refGene_gn=CDS2;refGene_tx=XM_011280916.3
A3 31869098 31869218 refGene_gn=CDS2;refGene_tx=XM_011280916.3
A3 31869942 31870046 refGene_gn=CDS2;refGene_tx=XM_011280916.3
A3 31870306 31870439 refGene_gn=CDS2;refGene_tx=XM_011280916.3
A3 31930697 31931394 refGene_gn=PROKR2;refGene_tx=XM_023251439.1
A3 31939299 31939757 refGene_gn=PROKR2;refGene_tx=XM_023251439.1
A3 32112944 32113134 refGene_gn=GPCPD1;refGene_tx=XM_023251441.1
A3 32118555 32118716 refGene_gn=GPCPD1;refGene_tx=XM_023251441.1
A3 32119150 32119286 refGene_gn=GPCPD1;refGene_tx=XM_023251441.1
A3 32119786 32119923 refGene_gn=GPCPD1;refGene_tx=XM_023251441.1
A3 32121439 32121505 refGene_gn=GPCPD1;refGene_tx=XM_023251441.1
A3 32123210 32123272 refGene_gn=GPCPD1;refGene_tx=XM_023251441.1
A3 32124664 32124704 refGene_gn=GPCPD1;refGene_tx=XM_023251441.1
A3 32125432 32125510 refGene_gn=GPCPD1;refGene_tx=XM_023251441.1
A3 32127963 32128056 refGene_gn=GPCPD1;refGene_tx=XM_023251441.1
A3 32130482 32130537 refGene_gn=GPCPD1;refGene_tx=XM_023251441.1
A3 32131265 32131398 refGene_gn=GPCPD1;refGene_tx=XM_023251441.1
A3 32131704 32131867 refGene_gn=GPCPD1;refGene_tx=XM_023251441.1
A3 32134001 32134233 refGene_gn=GPCPD1;refGene_tx=XM_023251441.1
A3 32138046 32138170 refGene_gn=GPCPD1;refGene_tx=XM_023251441.1
A3 32140045 32140087 refGene_gn=GPCPD1;refGene_tx=XM_023251441.1
A3 32141976 32142052 refGene_gn=GPCPD1;refGene_tx=XM_023251441.1
A3 32147841 32147926 refGene_gn=GPCPD1;refGene_tx=XM_023251441.1
A3 32152184 32152281 refGene_gn=GPCPD1;refGene_tx=XM_023251441.1
A3 32154784 32154833 refGene_gn=GPCPD1;refGene_tx=XM_023251441.1
A3 32285234 32285412
refGene_gn=CA3H20orf196;refGene_tx=XM_003983743.4;refGene_gn=CA3H20orf196;refGene_tx=XM_006929935.3;refGene_gn=CA3H20orf196;refGene_tx=XM_006929936.3;refGene_gn=CA3H20orf196;refGene_tx=XM_006929940.3;refGene_gn=CA3H20orf196;refGene_tx=XM_011280913.2;refGene_gn=CA3H20orf196;refGene_tx=XM_011280914.2;refGene_gn=CA3H20orf196;refGene_tx=XM_011280915.2;refGene_gn=CA3H20orf196;refGene_tx=XM_019826803.1;refGene_gn=CA3H20orf196;refGene_tx=XM_019826805.2;refGene_gn=CA3H20orf196;refGene_tx=XM_019826806.2;refGene_gn=CA3H20orf196;refGene_tx=XM_019826807.2;refGene_gn=CA3H20orf196;refGene_tx=XM_019826810.2;refGene_gn=CA3H20orf196;refGene_tx=XM_023251442.1
A3 32285506 32285517
refGene_gn=CA3H20orf196;refGene_tx=XM_011280914.2
A3 32295159 32295305
refGene_gn=CA3H20orf196;refGene_tx=XM_019826810.2
A3 32303050 32303064
refGene_gn=CA3H20orf196;refGene_tx=XM_011280913.2
A3 32349489 32349500

refGene_gn=CA3H20orf196;refGene_tx=XM_011280915.2
A3 32361014 32361457
refGene_gn=CA3H20orf196;refGene_tx=XM_003983743.4;refGene_gn=CA3H20orf196;refGene_tx=XM_006929935.3;refGene_gn=CA3H20orf196;refGene_tx=XM_006929936.3;refGene_gn=CA3H20orf196;refGene_tx=XM_006929940.3;refGene_gn=CA3H20orf196;refGene_tx=XM_019826803.1;refGene_gn=CA3H20orf196;refGene_tx=XM_019826805.2;refGene_gn=CA3H20orf196;refGene_tx=XM_019826806.2;refGene_gn=CA3H20orf196;refGene_tx=XM_019826807.2;refGene_gn=CA3H20orf196;refGene_tx=XM_023251442.1
A3 32391141 32391190
refGene_gn=CHGB;refGene_tx=XM_011280910.2;refGene_gn=CHGB;refGene_tx=XM_011280912.2
A3 32395361 32395408
refGene_gn=CHGB;refGene_tx=XM_011280910.2;refGene_gn=CHGB;refGene_tx=XM_011280912.2
A3 32395931 32396025
refGene_gn=CHGB;refGene_tx=XM_011280910.2;refGene_gn=CHGB;refGene_tx=XM_011280912.2
A3 32400748 32402484
refGene_gn=CHGB;refGene_tx=XM_011280910.2;refGene_gn=CHGB;refGene_tx=XM_011280912.2
A3 32403578 32403662
refGene_gn=CHGB;refGene_tx=XM_011280910.2;refGene_gn=CHGB;refGene_tx=XM_011280912.2
A3 32416813 32417005
refGene_gn=TRMT6;refGene_tx=XM_003983744.5;refGene_gn=TRMT6;refGene_tx=XM_006929942.3;refGene_gn=TRMT6;refGene_tx=XM_019826818.2
A3 32418368 32418483
refGene_gn=TRMT6;refGene_tx=XM_003983744.5;refGene_gn=TRMT6;refGene_tx=XM_006929942.3;refGene_gn=TRMT6;refGene_tx=XM_019826817.2;refGene_gn=TRMT6;refGene_tx=XM_019826818.2
A3 32418553 32418656
refGene_gn=TRMT6;refGene_tx=XM_003983744.5;refGene_gn=TRMT6;refGene_tx=XM_006929942.3;refGene_gn=TRMT6;refGene_tx=XM_019826817.2;refGene_gn=TRMT6;refGene_tx=XM_019826818.2
A3 32419509 32419595
refGene_gn=TRMT6;refGene_tx=XM_003983744.5;refGene_gn=TRMT6;refGene_tx=XM_006929942.3;refGene_gn=TRMT6;refGene_tx=XM_019826817.2;refGene_gn=TRMT6;refGene_tx=XM_019826818.2
A3 32420051 32420410
refGene_gn=TRMT6;refGene_tx=XM_003983744.5;refGene_gn=TRMT6;refGene_tx=XM_006929942.3;refGene_gn=TRMT6;refGene_tx=XM_019826817.2;refGene_gn=TRMT6;refGene_tx=XM_019826818.2
A3 32421318 32421443
refGene_gn=TRMT6;refGene_tx=XM_003983744.5;refGene_gn=TRMT6;refGene_tx=XM_006929942.3;refGene_gn=TRMT6;refGene_tx=XM_019826817.2;refGene_gn=TRMT6;refGene_tx=XM_019826818.2
A3 32421716 32421800
refGene_gn=TRMT6;refGene_tx=XM_003983744.5;refGene_gn=TRMT6;refGene_tx=XM_006929942.3;refGene_gn=TRMT6;refGene_tx=XM_019826817.2;refGene_gn=

TRMT6;refGene_tx=XM_019826818.2
A3 32421923 32422015
refGene_gn=TRMT6;refGene_tx=XM_003983744.5;refGene_gn=TRMT6;refGene_tx=XM_019826817.2;refGene_gn=TRMT6;refGene_tx=XM_019826818.2
A3 32422550 32422660
refGene_gn=TRMT6;refGene_tx=XM_003983744.5;refGene_gn=TRMT6;refGene_tx=XM_019826817.2;refGene_gn=TRMT6;refGene_tx=XM_019826818.2
A3 32424088 32424216
refGene_gn=TRMT6;refGene_tx=XM_003983744.5;refGene_gn=TRMT6;refGene_tx=XM_019826817.2
A3 32427691 32427819
refGene_gn=TRMT6;refGene_tx=XM_003983744.5;refGene_gn=TRMT6;refGene_tx=XM_019826817.2;refGene_gn=TRMT6;refGene_tx=XM_019826818.2
A3 32428429 32428469 refGene_gn=MCM8;refGene_tx=XM_023251443.1
A3 32433639 32433792 refGene_gn=MCM8;refGene_tx=XM_023251443.1
A3 32434103 32434187 refGene_gn=MCM8;refGene_tx=XM_023251443.1
A3 32436361 32436444 refGene_gn=MCM8;refGene_tx=XM_023251443.1
A3 32436857 32437007 refGene_gn=MCM8;refGene_tx=XM_023251443.1
A3 32438892 32438996 refGene_gn=MCM8;refGene_tx=XM_023251443.1
A3 32439767 32439966 refGene_gn=MCM8;refGene_tx=XM_023251443.1
A3 32443723 32443809 refGene_gn=MCM8;refGene_tx=XM_023251443.1
A3 32451700 32451852 refGene_gn=MCM8;refGene_tx=XM_023251443.1
A3 32452097 32452293 refGene_gn=MCM8;refGene_tx=XM_023251443.1
A3 32455194 32455225 refGene_gn=MCM8;refGene_tx=XM_023251443.1
A3 32455737 32455878 refGene_gn=MCM8;refGene_tx=XM_023251443.1
A3 32459098 32459240 refGene_gn=MCM8;refGene_tx=XM_023251443.1
A3 32462610 32462806 refGene_gn=MCM8;refGene_tx=XM_023251443.1
A3 32464518 32464738 refGene_gn=MCM8;refGene_tx=XM_023251443.1
A3 32465029 32465239 refGene_gn=MCM8;refGene_tx=XM_023251443.1
A3 32466507 32466584 refGene_gn=MCM8;refGene_tx=XM_023251443.1
A3 32470222 32470352 refGene_gn=MCM8;refGene_tx=XM_023251443.1
A3 32470353 32470413 refGene_gn=MCM8;refGene_tx=XM_023251443.1
A3 32472648 32472741 refGene_gn=MCM8;refGene_tx=XM_023251443.1
A3 32488514 32488820 refGene_gn=CRLS1;refGene_tx=XM_023251446.1
A3 32489680 32489689 refGene_gn=CRLS1;refGene_tx=XM_003983745.5
A3 32492014 32492152
refGene_gn=CRLS1;refGene_tx=XM_003983745.5;refGene_gn=CRLS1;refGene_tx=XM_011280909.3;refGene_gn=CRLS1;refGene_tx=XM_023251446.1;refGene_gn=CRLS1;refGene_tx=XM_023251447.1
A3 32496956 32497086
refGene_gn=CRLS1;refGene_tx=XM_003983745.5;refGene_gn=CRLS1;refGene_tx=XM_011280909.3;refGene_gn=CRLS1;refGene_tx=XM_023251446.1;refGene_gn=CRLS1;refGene_tx=XM_023251447.1
A3 32507032 32507118
refGene_gn=CRLS1;refGene_tx=XM_003983745.5;refGene_gn=CRLS1;refGene_tx=XM_011280909.3;refGene_gn=CRLS1;refGene_tx=XM_023251446.1;refGene_gn=CRLS1;refGene_tx=XM_023251447.1
A3 32508487 32508556
refGene_gn=CRLS1;refGene_tx=XM_003983745.5;refGene_gn=CRLS1;refGene_tx=XM_011280909.3;refGene_gn=CRLS1;refGene_tx=XM_023251446.1;refGene_gn=

CRLS1;refGene_tx=XM_023251447.1
A3 32510312 32510404
refGene_gn=CRLS1;refGene_tx=XM_003983745.5;refGene_gn=CRLS1;refGene_tx=XM_011280909.3;refGene_gn=CRLS1;refGene_tx=XM_023251446.1;refGene_gn=CRLS1;refGene_tx=XM_023251447.1
A3 32512478 32512566
refGene_gn=CRLS1;refGene_tx=XM_003983745.5;refGene_gn=CRLS1;refGene_tx=XM_011280909.3;refGene_gn=CRLS1;refGene_tx=XM_023251446.1;refGene_gn=CRLS1;refGene_tx=XM_023251447.1
A3 32517313 32518559 refGene_gn=LRRN4;refGene_tx=XM_019828065.2
A3 32520121 32520259 refGene_gn=LRRN4;refGene_tx=XM_019828065.2
A3 32524053 32524258 refGene_gn=LRRN4;refGene_tx=XM_019828065.2
A3 32525335 32525969 refGene_gn=LRRN4;refGene_tx=XM_019828065.2
A3 32545112 32545286
refGene_gn=FERMT1;refGene_tx=XM_023251444.1;refGene_gn=FERMT1;refGene_tx=XM_023251445.1
A3 32546951 32547093
refGene_gn=FERMT1;refGene_tx=XM_023251444.1;refGene_gn=FERMT1;refGene_tx=XM_023251445.1
A3 32548207 32548332
refGene_gn=FERMT1;refGene_tx=XM_023251444.1;refGene_gn=FERMT1;refGene_tx=XM_023251445.1
A3 32548956 32549178
refGene_gn=FERMT1;refGene_tx=XM_023251444.1;refGene_gn=FERMT1;refGene_tx=XM_023251445.1
A3 32550365 32550472
refGene_gn=FERMT1;refGene_tx=XM_023251444.1;refGene_gn=FERMT1;refGene_tx=XM_023251445.1
A3 32551421 32551546
refGene_gn=FERMT1;refGene_tx=XM_023251444.1;refGene_gn=FERMT1;refGene_tx=XM_023251445.1
A3 32559290 32559340
refGene_gn=FERMT1;refGene_tx=XM_023251444.1;refGene_gn=FERMT1;refGene_tx=XM_023251445.1
A3 32560835 32560967
refGene_gn=FERMT1;refGene_tx=XM_023251444.1;refGene_gn=FERMT1;refGene_tx=XM_023251445.1
A3 32561398 32561506
refGene_gn=FERMT1;refGene_tx=XM_023251444.1;refGene_gn=FERMT1;refGene_tx=XM_023251445.1
A3 32566464 32566567
refGene_gn=FERMT1;refGene_tx=XM_023251444.1;refGene_gn=FERMT1;refGene_tx=XM_023251445.1
A3 32568307 32568521
refGene_gn=FERMT1;refGene_tx=XM_023251444.1;refGene_gn=FERMT1;refGene_tx=XM_023251445.1
A3 32574429 32574663
refGene_gn=FERMT1;refGene_tx=XM_023251444.1;refGene_gn=FERMT1;refGene_tx=XM_023251445.1
A3 32575306 32575457 refGene_gn=FERMT1;refGene_tx=XM_023251444.1

A3 33154510 33154853 refGene_gn=BMP2;refGene_tx=XM_003983769.5
A3 33162498 33163343 refGene_gn=BMP2;refGene_tx=XM_003983769.5
A3 34143967 34144038 refGene_gn=HAO1;refGene_tx=XM_003983770.4
A3 34145893 34145963 refGene_gn=HAO1;refGene_tx=XM_003983770.4
A3 34146074 34146233 refGene_gn=HAO1;refGene_tx=XM_003983770.4
A3 34152444 34152536 refGene_gn=HAO1;refGene_tx=XM_003983770.4
A3 34158636 34158812 refGene_gn=HAO1;refGene_tx=XM_003983770.4
A3 34166001 34166257 refGene_gn=HAO1;refGene_tx=XM_003983770.4
A3 34183052 34183204 refGene_gn=HAO1;refGene_tx=XM_003983770.4
A3 34189635 34189772 refGene_gn=HAO1;refGene_tx=XM_003983770.4
A3 34228454 34228813 refGene_gn=TMX4;refGene_tx=XM_023251449.1
A3 34229977 34230041 refGene_gn=TMX4;refGene_tx=XM_023251449.1
A3 34234915 34235017 refGene_gn=TMX4;refGene_tx=XM_023251449.1
A3 34244478 34244524 refGene_gn=TMX4;refGene_tx=XM_023251449.1
A3 34247084 34247213 refGene_gn=TMX4;refGene_tx=XM_023251449.1
A3 34248777 34248823 refGene_gn=TMX4;refGene_tx=XM_023251449.1
A3 34257847 34257963 refGene_gn=TMX4;refGene_tx=XM_023251449.1
A3 34269683 34269856 refGene_gn=TMX4;refGene_tx=XM_023251449.1
A3 34386223 34386322
refGene_gn=PLCB1;refGene_tx=XM_023251450.1;refGene_gn=PLCB1;refGene_tx
=XM_023251451.1;refGene_gn=PLCB1;refGene_tx=XM_023251452.1
A3 34403538 34403616
refGene_gn=PLCB1;refGene_tx=XM_023251450.1;refGene_gn=PLCB1;refGene_tx
=XM_023251451.1;refGene_gn=PLCB1;refGene_tx=XM_023251452.1
A3 34619635 34619704
refGene_gn=PLCB1;refGene_tx=XM_023251450.1;refGene_gn=PLCB1;refGene_tx
=XM_023251451.1;refGene_gn=PLCB1;refGene_tx=XM_023251452.1
A3 34838309 34838447
refGene_gn=PLCB1;refGene_tx=XM_023251450.1;refGene_gn=PLCB1;refGene_tx
=XM_023251451.1;refGene_gn=PLCB1;refGene_tx=XM_023251452.1
A3 34864060 34864140
refGene_gn=PLCB1;refGene_tx=XM_023251450.1;refGene_gn=PLCB1;refGene_tx
=XM_023251451.1;refGene_gn=PLCB1;refGene_tx=XM_023251452.1
A3 34865952 34866006
refGene_gn=PLCB1;refGene_tx=XM_023251450.1;refGene_gn=PLCB1;refGene_tx
=XM_023251451.1;refGene_gn=PLCB1;refGene_tx=XM_023251452.1
A3 34867119 34867195
refGene_gn=PLCB1;refGene_tx=XM_023251450.1;refGene_gn=PLCB1;refGene_tx
=XM_023251451.1;refGene_gn=PLCB1;refGene_tx=XM_023251452.1
A3 34874893 34874994
refGene_gn=PLCB1;refGene_tx=XM_023251450.1;refGene_gn=PLCB1;refGene_tx
=XM_023251451.1;refGene_gn=PLCB1;refGene_tx=XM_023251452.1
A3 34876421 34876588
refGene_gn=PLCB1;refGene_tx=XM_023251450.1;refGene_gn=PLCB1;refGene_tx
=XM_023251451.1;refGene_gn=PLCB1;refGene_tx=XM_023251452.1
A3 34899108 34899255
refGene_gn=PLCB1;refGene_tx=XM_023251450.1;refGene_gn=PLCB1;refGene_tx
=XM_023251451.1;refGene_gn=PLCB1;refGene_tx=XM_023251452.1
A3 34910469 34910627
refGene_gn=PLCB1;refGene_tx=XM_023251450.1;refGene_gn=PLCB1;refGene_tx

=XM_023251451.1;refGene_gn=PLCB1;refGene_tx=XM_023251452.1
A3	34920667	34920750
refGene_gn=PLCB1;refGene_tx=XM_023251450.1;refGene_gn=PLCB1;refGene_tx=XM_023251451.1;refGene_gn=PLCB1;refGene_tx=XM_023251452.1
A3	34927031	34927116
refGene_gn=PLCB1;refGene_tx=XM_023251450.1;refGene_gn=PLCB1;refGene_tx=XM_023251451.1;refGene_gn=PLCB1;refGene_tx=XM_023251452.1
A3	34928447	34928625
refGene_gn=PLCB1;refGene_tx=XM_023251450.1;refGene_gn=PLCB1;refGene_tx=XM_023251451.1;refGene_gn=PLCB1;refGene_tx=XM_023251452.1
A3	34933183	34933251
refGene_gn=PLCB1;refGene_tx=XM_023251450.1;refGene_gn=PLCB1;refGene_tx=XM_023251451.1;refGene_gn=PLCB1;refGene_tx=XM_023251452.1
A3	34935200	34935297
refGene_gn=PLCB1;refGene_tx=XM_023251450.1;refGene_gn=PLCB1;refGene_tx=XM_023251451.1;refGene_gn=PLCB1;refGene_tx=XM_023251452.1
A3	34937523	34937608
refGene_gn=PLCB1;refGene_tx=XM_023251450.1;refGene_gn=PLCB1;refGene_tx=XM_023251451.1;refGene_gn=PLCB1;refGene_tx=XM_023251452.1
A3	34939295	34939420
refGene_gn=PLCB1;refGene_tx=XM_023251450.1;refGene_gn=PLCB1;refGene_tx=XM_023251451.1;refGene_gn=PLCB1;refGene_tx=XM_023251452.1
A3	34944372	34944527
refGene_gn=PLCB1;refGene_tx=XM_023251450.1;refGene_gn=PLCB1;refGene_tx=XM_023251451.1;refGene_gn=PLCB1;refGene_tx=XM_023251452.1
A3	34947018	34947183
refGene_gn=PLCB1;refGene_tx=XM_023251450.1;refGene_gn=PLCB1;refGene_tx=XM_023251451.1;refGene_gn=PLCB1;refGene_tx=XM_023251452.1
A3	34949198	34949298
refGene_gn=PLCB1;refGene_tx=XM_023251450.1;refGene_gn=PLCB1;refGene_tx=XM_023251451.1;refGene_gn=PLCB1;refGene_tx=XM_023251452.1
A3	34949978	34950083
refGene_gn=PLCB1;refGene_tx=XM_023251450.1;refGene_gn=PLCB1;refGene_tx=XM_023251451.1;refGene_gn=PLCB1;refGene_tx=XM_023251452.1
A3	34951126	34951236
refGene_gn=PLCB1;refGene_tx=XM_023251450.1;refGene_gn=PLCB1;refGene_tx=XM_023251451.1;refGene_gn=PLCB1;refGene_tx=XM_023251452.1
A3	34966010	34966143
refGene_gn=PLCB1;refGene_tx=XM_023251450.1;refGene_gn=PLCB1;refGene_tx=XM_023251451.1;refGene_gn=PLCB1;refGene_tx=XM_023251452.1
A3	34968969	34969023
refGene_gn=PLCB1;refGene_tx=XM_023251450.1;refGene_gn=PLCB1;refGene_tx=XM_023251451.1;refGene_gn=PLCB1;refGene_tx=XM_023251452.1
A3	34972862	34973082
refGene_gn=PLCB1;refGene_tx=XM_023251450.1;refGene_gn=PLCB1;refGene_tx=XM_023251451.1;refGene_gn=PLCB1;refGene_tx=XM_023251452.1
A3	34976813	34976994
refGene_gn=PLCB1;refGene_tx=XM_023251450.1;refGene_gn=PLCB1;refGene_tx=XM_023251451.1;refGene_gn=PLCB1;refGene_tx=XM_023251452.1
A3	34995857	34995934

refGene_gn=PLCB1;refGene_tx=XM_023251450.1;refGene_gn=PLCB1;refGene_tx=XM_023251451.1;refGene_gn=PLCB1;refGene_tx=XM_023251452.1
A3 34996040 34996130
refGene_gn=PLCB1;refGene_tx=XM_023251450.1;refGene_gn=PLCB1;refGene_tx=XM_023251451.1;refGene_gn=PLCB1;refGene_tx=XM_023251452.1
A3 34996600 34996658
refGene_gn=PLCB1;refGene_tx=XM_023251450.1;refGene_gn=PLCB1;refGene_tx=XM_023251451.1;refGene_gn=PLCB1;refGene_tx=XM_023251452.1
A3 34997263 34997350
refGene_gn=PLCB1;refGene_tx=XM_023251450.1;refGene_gn=PLCB1;refGene_tx=XM_023251451.1;refGene_gn=PLCB1;refGene_tx=XM_023251452.1
A3 35009082 35009181
refGene_gn=PLCB1;refGene_tx=XM_023251451.1;refGene_gn=PLCB1;refGene_tx=XM_023251452.1
A3 35079898 35080126 refGene_gn=PLCB1;refGene_tx=XM_023251450.1
A3 35513700 35513784
refGene_gn=PLCB4;refGene_tx=XM_019826823.2;refGene_gn=PLCB4;refGene_tx=XM_023251453.1;refGene_gn=PLCB4;refGene_tx=XM_023251454.1;refGene_gn=PLCB4;refGene_tx=XM_023251455.1;refGene_gn=PLCB4;refGene_tx=XM_023251456.1;refGene_gn=PLCB4;refGene_tx=XM_023251457.1;refGene_gn=PLCB4;refGene_tx=XM_023251458.1;refGene_gn=PLCB4;refGene_tx=XM_023251459.1;refGene_gn=PLCB4;refGene_tx=XM_023251460.1
A3 35536518 35536599
refGene_gn=PLCB4;refGene_tx=XM_019826823.2;refGene_gn=PLCB4;refGene_tx=XM_023251453.1;refGene_gn=PLCB4;refGene_tx=XM_023251454.1;refGene_gn=PLCB4;refGene_tx=XM_023251455.1;refGene_gn=PLCB4;refGene_tx=XM_023251456.1;refGene_gn=PLCB4;refGene_tx=XM_023251457.1;refGene_gn=PLCB4;refGene_tx=XM_023251458.1;refGene_gn=PLCB4;refGene_tx=XM_023251459.1;refGene_gn=PLCB4;refGene_tx=XM_023251460.1
A3 35537470 35537530
refGene_gn=PLCB4;refGene_tx=XM_019826823.2;refGene_gn=PLCB4;refGene_tx=XM_023251453.1;refGene_gn=PLCB4;refGene_tx=XM_023251454.1;refGene_gn=PLCB4;refGene_tx=XM_023251455.1;refGene_gn=PLCB4;refGene_tx=XM_023251456.1;refGene_gn=PLCB4;refGene_tx=XM_023251457.1;refGene_gn=PLCB4;refGene_tx=XM_023251458.1;refGene_gn=PLCB4;refGene_tx=XM_023251459.1;refGene_gn=PLCB4;refGene_tx=XM_023251460.1
A3 35539063 35539207
refGene_gn=PLCB4;refGene_tx=XM_019826823.2;refGene_gn=PLCB4;refGene_tx=XM_023251453.1;refGene_gn=PLCB4;refGene_tx=XM_023251454.1;refGene_gn=PLCB4;refGene_tx=XM_023251455.1;refGene_gn=PLCB4;refGene_tx=XM_023251456.1;refGene_gn=PLCB4;refGene_tx=XM_023251457.1;refGene_gn=PLCB4;refGene_tx=XM_023251458.1;refGene_gn=PLCB4;refGene_tx=XM_023251459.1;refGene_gn=PLCB4;refGene_tx=XM_023251460.1
A3 35563734 35563814
refGene_gn=PLCB4;refGene_tx=XM_019826823.2;refGene_gn=PLCB4;refGene_tx=XM_023251453.1;refGene_gn=PLCB4;refGene_tx=XM_023251454.1;refGene_gn=PLCB4;refGene_tx=XM_023251455.1;refGene_gn=PLCB4;refGene_tx=XM_023251456.1;refGene_gn=PLCB4;refGene_tx=XM_023251457.1;refGene_gn=PLCB4;refGene_tx=XM_023251458.1;refGene_gn=PLCB4;refGene_tx=XM_023251459.1;refGene_gn=PLCB4;refGene_tx=XM_023251460.1

A3 35566532 35566586
refGene_gn=PLCB4;refGene_tx=XM_019826823.2;refGene_gn=PLCB4;refGene_tx=XM_023251453.1;refGene_gn=PLCB4;refGene_tx=XM_023251454.1;refGene_gn=PLCB4;refGene_tx=XM_023251455.1;refGene_gn=PLCB4;refGene_tx=XM_023251456.1;refGene_gn=PLCB4;refGene_tx=XM_023251457.1;refGene_gn=PLCB4;refGene_tx=XM_023251458.1;refGene_gn=PLCB4;refGene_tx=XM_023251459.1;refGene_gn=PLCB4;refGene_tx=XM_023251460.1
A3 35572500 35572582
refGene_gn=PLCB4;refGene_tx=XM_019826823.2;refGene_gn=PLCB4;refGene_tx=XM_023251453.1;refGene_gn=PLCB4;refGene_tx=XM_023251454.1;refGene_gn=PLCB4;refGene_tx=XM_023251455.1;refGene_gn=PLCB4;refGene_tx=XM_023251456.1;refGene_gn=PLCB4;refGene_tx=XM_023251457.1;refGene_gn=PLCB4;refGene_tx=XM_023251458.1;refGene_gn=PLCB4;refGene_tx=XM_023251459.1;refGene_gn=PLCB4;refGene_tx=XM_023251460.1
A3 35573530 35573631
refGene_gn=PLCB4;refGene_tx=XM_019826823.2;refGene_gn=PLCB4;refGene_tx=XM_023251453.1;refGene_gn=PLCB4;refGene_tx=XM_023251454.1;refGene_gn=PLCB4;refGene_tx=XM_023251455.1;refGene_gn=PLCB4;refGene_tx=XM_023251456.1;refGene_gn=PLCB4;refGene_tx=XM_023251457.1;refGene_gn=PLCB4;refGene_tx=XM_023251458.1;refGene_gn=PLCB4;refGene_tx=XM_023251459.1;refGene_gn=PLCB4;refGene_tx=XM_023251460.1
A3 35574251 35574309
refGene_gn=PLCB4;refGene_tx=XM_019826823.2;refGene_gn=PLCB4;refGene_tx=XM_023251453.1;refGene_gn=PLCB4;refGene_tx=XM_023251454.1;refGene_gn=PLCB4;refGene_tx=XM_023251455.1;refGene_gn=PLCB4;refGene_tx=XM_023251456.1;refGene_gn=PLCB4;refGene_tx=XM_023251457.1;refGene_gn=PLCB4;refGene_tx=XM_023251458.1;refGene_gn=PLCB4;refGene_tx=XM_023251459.1;refGene_gn=PLCB4;refGene_tx=XM_023251460.1
A3 35581205 35581314
refGene_gn=PLCB4;refGene_tx=XM_019826823.2;refGene_gn=PLCB4;refGene_tx=XM_023251453.1;refGene_gn=PLCB4;refGene_tx=XM_023251454.1;refGene_gn=PLCB4;refGene_tx=XM_023251455.1;refGene_gn=PLCB4;refGene_tx=XM_023251456.1;refGene_gn=PLCB4;refGene_tx=XM_023251457.1;refGene_gn=PLCB4;refGene_tx=XM_023251458.1;refGene_gn=PLCB4;refGene_tx=XM_023251459.1;refGene_gn=PLCB4;refGene_tx=XM_023251460.1
A3 35585902 35586113
refGene_gn=PLCB4;refGene_tx=XM_019826823.2;refGene_gn=PLCB4;refGene_tx=XM_023251453.1;refGene_gn=PLCB4;refGene_tx=XM_023251454.1;refGene_gn=PLCB4;refGene_tx=XM_023251455.1;refGene_gn=PLCB4;refGene_tx=XM_023251456.1;refGene_gn=PLCB4;refGene_tx=XM_023251457.1;refGene_gn=PLCB4;refGene_tx=XM_023251458.1;refGene_gn=PLCB4;refGene_tx=XM_023251459.1;refGene_gn=PLCB4;refGene_tx=XM_023251460.1
A3 35587522 35587616
refGene_gn=PLCB4;refGene_tx=XM_019826823.2;refGene_gn=PLCB4;refGene_tx=XM_023251453.1;refGene_gn=PLCB4;refGene_tx=XM_023251454.1;refGene_gn=PLCB4;refGene_tx=XM_023251455.1;refGene_gn=PLCB4;refGene_tx=XM_023251456.1;refGene_gn=PLCB4;refGene_tx=XM_023251457.1;refGene_gn=PLCB4;refGene_tx=XM_023251458.1;refGene_gn=PLCB4;refGene_tx=XM_023251459.1;refGene_gn=PLCB4;refGene_tx=XM_023251460.1
A3 35590500 35590580

refGene_gn=PLCB4;refGene_tx=XM_019826823.2;refGene_gn=PLCB4;refGene_tx=XM_023251453.1;refGene_gn=PLCB4;refGene_tx=XM_023251454.1;refGene_gn=PLCB4;refGene_tx=XM_023251455.1;refGene_gn=PLCB4;refGene_tx=XM_023251456.1;refGene_gn=PLCB4;refGene_tx=XM_023251457.1;refGene_gn=PLCB4;refGene_tx=XM_023251458.1;refGene_gn=PLCB4;refGene_tx=XM_023251459.1;refGene_gn=PLCB4;refGene_tx=XM_023251460.1
A3 35591028 35591113
refGene_gn=PLCB4;refGene_tx=XM_019826823.2;refGene_gn=PLCB4;refGene_tx=XM_023251453.1;refGene_gn=PLCB4;refGene_tx=XM_023251454.1;refGene_gn=PLCB4;refGene_tx=XM_023251455.1;refGene_gn=PLCB4;refGene_tx=XM_023251456.1;refGene_gn=PLCB4;refGene_tx=XM_023251457.1;refGene_gn=PLCB4;refGene_tx=XM_023251458.1;refGene_gn=PLCB4;refGene_tx=XM_023251459.1;refGene_gn=PLCB4;refGene_tx=XM_023251460.1
A3 35593858 35593949
refGene_gn=PLCB4;refGene_tx=XM_019826823.2;refGene_gn=PLCB4;refGene_tx=XM_023251453.1;refGene_gn=PLCB4;refGene_tx=XM_023251454.1;refGene_gn=PLCB4;refGene_tx=XM_023251455.1;refGene_gn=PLCB4;refGene_tx=XM_023251456.1;refGene_gn=PLCB4;refGene_tx=XM_023251457.1;refGene_gn=PLCB4;refGene_tx=XM_023251458.1;refGene_gn=PLCB4;refGene_tx=XM_023251459.1;refGene_gn=PLCB4;refGene_tx=XM_023251460.1
A3 35595664 35595760
refGene_gn=PLCB4;refGene_tx=XM_019826823.2;refGene_gn=PLCB4;refGene_tx=XM_023251453.1;refGene_gn=PLCB4;refGene_tx=XM_023251454.1;refGene_gn=PLCB4;refGene_tx=XM_023251455.1;refGene_gn=PLCB4;refGene_tx=XM_023251456.1;refGene_gn=PLCB4;refGene_tx=XM_023251457.1;refGene_gn=PLCB4;refGene_tx=XM_023251458.1;refGene_gn=PLCB4;refGene_tx=XM_023251459.1;refGene_gn=PLCB4;refGene_tx=XM_023251460.1
A3 35601907 35602008
refGene_gn=PLCB4;refGene_tx=XM_019826823.2;refGene_gn=PLCB4;refGene_tx=XM_023251453.1;refGene_gn=PLCB4;refGene_tx=XM_023251454.1;refGene_gn=PLCB4;refGene_tx=XM_023251455.1;refGene_gn=PLCB4;refGene_tx=XM_023251456.1;refGene_gn=PLCB4;refGene_tx=XM_023251457.1;refGene_gn=PLCB4;refGene_tx=XM_023251458.1;refGene_gn=PLCB4;refGene_tx=XM_023251459.1;refGene_gn=PLCB4;refGene_tx=XM_023251460.1
A3 35605239 35605275
refGene_gn=PLCB4;refGene_tx=XM_019826823.2;refGene_gn=PLCB4;refGene_tx=XM_023251453.1;refGene_gn=PLCB4;refGene_tx=XM_023251454.1;refGene_gn=PLCB4;refGene_tx=XM_023251455.1;refGene_gn=PLCB4;refGene_tx=XM_023251456.1;refGene_gn=PLCB4;refGene_tx=XM_023251457.1;refGene_gn=PLCB4;refGene_tx=XM_023251458.1;refGene_gn=PLCB4;refGene_tx=XM_023251459.1;refGene_gn=PLCB4;refGene_tx=XM_023251460.1
A3 35607438 35607580
refGene_gn=PLCB4;refGene_tx=XM_019826823.2;refGene_gn=PLCB4;refGene_tx=XM_023251453.1;refGene_gn=PLCB4;refGene_tx=XM_023251454.1;refGene_gn=PLCB4;refGene_tx=XM_023251455.1;refGene_gn=PLCB4;refGene_tx=XM_023251456.1;refGene_gn=PLCB4;refGene_tx=XM_023251457.1;refGene_gn=PLCB4;refGene_tx=XM_023251458.1;refGene_gn=PLCB4;refGene_tx=XM_023251459.1;refGene_gn=PLCB4;refGene_tx=XM_023251460.1
A3 35608195 35608280
refGene_gn=PLCB4;refGene_tx=XM_019826823.2;refGene_gn=PLCB4;refGene_tx

=XM_023251453.1;refGene_gn=PLCB4;refGene_tx=XM_023251454.1;refGene_gn=PLCB4;refGene_tx=XM_023251455.1;refGene_gn=PLCB4;refGene_tx=XM_023251456.1;refGene_gn=PLCB4;refGene_tx=XM_023251457.1;refGene_gn=PLCB4;refGene_tx=XM_023251458.1;refGene_gn=PLCB4;refGene_tx=XM_023251459.1;refGene_gn=PLCB4;refGene_tx=XM_023251460.1
A3 35608621 35608746
refGene_gn=PLCB4;refGene_tx=XM_019826823.2;refGene_gn=PLCB4;refGene_tx=XM_023251453.1;refGene_gn=PLCB4;refGene_tx=XM_023251454.1;refGene_gn=PLCB4;refGene_tx=XM_023251455.1;refGene_gn=PLCB4;refGene_tx=XM_023251456.1;refGene_gn=PLCB4;refGene_tx=XM_023251457.1;refGene_gn=PLCB4;refGene_tx=XM_023251458.1;refGene_gn=PLCB4;refGene_tx=XM_023251459.1;refGene_gn=PLCB4;refGene_tx=XM_023251460.1
A3 35610903 35610955
refGene_gn=PLCB4;refGene_tx=XM_019826823.2;refGene_gn=PLCB4;refGene_tx=XM_023251453.1;refGene_gn=PLCB4;refGene_tx=XM_023251454.1;refGene_gn=PLCB4;refGene_tx=XM_023251455.1;refGene_gn=PLCB4;refGene_tx=XM_023251456.1;refGene_gn=PLCB4;refGene_tx=XM_023251457.1;refGene_gn=PLCB4;refGene_tx=XM_023251458.1;refGene_gn=PLCB4;refGene_tx=XM_023251459.1;refGene_gn=PLCB4;refGene_tx=XM_023251460.1
A3 35625365 35625468
refGene_gn=PLCB4;refGene_tx=XM_019826823.2;refGene_gn=PLCB4;refGene_tx=XM_023251453.1;refGene_gn=PLCB4;refGene_tx=XM_023251454.1;refGene_gn=PLCB4;refGene_tx=XM_023251455.1;refGene_gn=PLCB4;refGene_tx=XM_023251456.1;refGene_gn=PLCB4;refGene_tx=XM_023251457.1;refGene_gn=PLCB4;refGene_tx=XM_023251458.1;refGene_gn=PLCB4;refGene_tx=XM_023251459.1;refGene_gn=PLCB4;refGene_tx=XM_023251460.1
A3 35626778 35626943
refGene_gn=PLCB4;refGene_tx=XM_019826823.2;refGene_gn=PLCB4;refGene_tx=XM_023251453.1;refGene_gn=PLCB4;refGene_tx=XM_023251454.1;refGene_gn=PLCB4;refGene_tx=XM_023251455.1;refGene_gn=PLCB4;refGene_tx=XM_023251456.1;refGene_gn=PLCB4;refGene_tx=XM_023251457.1;refGene_gn=PLCB4;refGene_tx=XM_023251458.1;refGene_gn=PLCB4;refGene_tx=XM_023251459.1;refGene_gn=PLCB4;refGene_tx=XM_023251460.1
A3 35629460 35629665
refGene_gn=PLCB4;refGene_tx=XM_019826823.2;refGene_gn=PLCB4;refGene_tx=XM_023251453.1;refGene_gn=PLCB4;refGene_tx=XM_023251454.1;refGene_gn=PLCB4;refGene_tx=XM_023251455.1;refGene_gn=PLCB4;refGene_tx=XM_023251456.1;refGene_gn=PLCB4;refGene_tx=XM_023251457.1;refGene_gn=PLCB4;refGene_tx=XM_023251458.1;refGene_gn=PLCB4;refGene_tx=XM_023251459.1;refGene_gn=PLCB4;refGene_tx=XM_023251460.1
A3 35643455 35643544
refGene_gn=PLCB4;refGene_tx=XM_019826823.2;refGene_gn=PLCB4;refGene_tx=XM_023251453.1;refGene_gn=PLCB4;refGene_tx=XM_023251454.1;refGene_gn=PLCB4;refGene_tx=XM_023251455.1;refGene_gn=PLCB4;refGene_tx=XM_023251456.1;refGene_gn=PLCB4;refGene_tx=XM_023251457.1;refGene_gn=PLCB4;refGene_tx=XM_023251458.1;refGene_gn=PLCB4;refGene_tx=XM_023251459.1;refGene_gn=PLCB4;refGene_tx=XM_023251460.1
A3 35644092 35644243
refGene_gn=PLCB4;refGene_tx=XM_019826823.2;refGene_gn=PLCB4;refGene_tx=XM_023251453.1;refGene_gn=PLCB4;refGene_tx=XM_023251454.1;refGene_gn=

PLCB4;refGene_tx=XM_023251455.1;refGene_gn=PLCB4;refGene_tx=XM_0232514
56.1;refGene_gn=PLCB4;refGene_tx=XM_023251457.1;refGene_gn=PLCB4;refGe
ne_tx=XM_023251458.1;refGene_gn=PLCB4;refGene_tx=XM_023251459.1;refGen
e_gn=PLCB4;refGene_tx=XM_023251460.1
A3 35649246 35649296
refGene_gn=PLCB4;refGene_tx=XM_019826823.2;refGene_gn=PLCB4;refGene_tx
=XM_023251453.1;refGene_gn=PLCB4;refGene_tx=XM_023251454.1;refGene_gn=
PLCB4;refGene_tx=XM_023251455.1;refGene_gn=PLCB4;refGene_tx=XM_0232514
56.1;refGene_gn=PLCB4;refGene_tx=XM_023251457.1;refGene_gn=PLCB4;refGe
ne_tx=XM_023251458.1;refGene_gn=PLCB4;refGene_tx=XM_023251459.1;refGen
e_gn=PLCB4;refGene_tx=XM_023251460.1
A3 35649429 35649495
refGene_gn=PLCB4;refGene_tx=XM_019826823.2;refGene_gn=PLCB4;refGene_tx
=XM_023251453.1;refGene_gn=PLCB4;refGene_tx=XM_023251454.1;refGene_gn=
PLCB4;refGene_tx=XM_023251455.1;refGene_gn=PLCB4;refGene_tx=XM_0232514
56.1;refGene_gn=PLCB4;refGene_tx=XM_023251457.1;refGene_gn=PLCB4;refGe
ne_tx=XM_023251458.1;refGene_gn=PLCB4;refGene_tx=XM_023251459.1;refGen
e_gn=PLCB4;refGene_tx=XM_023251460.1
A3 35659843 35659959
refGene_gn=PLCB4;refGene_tx=XM_019826823.2;refGene_gn=PLCB4;refGene_tx
=XM_023251453.1;refGene_gn=PLCB4;refGene_tx=XM_023251454.1;refGene_gn=
PLCB4;refGene_tx=XM_023251455.1;refGene_gn=PLCB4;refGene_tx=XM_0232514
56.1;refGene_gn=PLCB4;refGene_tx=XM_023251457.1;refGene_gn=PLCB4;refGe
ne_tx=XM_023251458.1;refGene_gn=PLCB4;refGene_tx=XM_023251459.1;refGen
e_gn=PLCB4;refGene_tx=XM_023251460.1
A3 35661251 35661327
refGene_gn=PLCB4;refGene_tx=XM_019826823.2;refGene_gn=PLCB4;refGene_tx
=XM_023251453.1;refGene_gn=PLCB4;refGene_tx=XM_023251454.1;refGene_gn=
PLCB4;refGene_tx=XM_023251455.1;refGene_gn=PLCB4;refGene_tx=XM_0232514
56.1;refGene_gn=PLCB4;refGene_tx=XM_023251457.1;refGene_gn=PLCB4;refGe
ne_tx=XM_023251458.1;refGene_gn=PLCB4;refGene_tx=XM_023251459.1;refGen
e_gn=PLCB4;refGene_tx=XM_023251460.1
A3 35663102 35663278
refGene_gn=PLCB4;refGene_tx=XM_019826823.2;refGene_gn=PLCB4;refGene_tx
=XM_023251453.1;refGene_gn=PLCB4;refGene_tx=XM_023251454.1;refGene_gn=
PLCB4;refGene_tx=XM_023251455.1;refGene_gn=PLCB4;refGene_tx=XM_0232514
56.1;refGene_gn=PLCB4;refGene_tx=XM_023251457.1;refGene_gn=PLCB4;refGe
ne_tx=XM_023251458.1;refGene_gn=PLCB4;refGene_tx=XM_023251459.1;refGen
e_gn=PLCB4;refGene_tx=XM_023251460.1
A3 35666568 35666670
refGene_gn=PLCB4;refGene_tx=XM_019826823.2;refGene_gn=PLCB4;refGene_tx
=XM_023251453.1;refGene_gn=PLCB4;refGene_tx=XM_023251454.1;refGene_gn=
PLCB4;refGene_tx=XM_023251455.1;refGene_gn=PLCB4;refGene_tx=XM_0232514
56.1;refGene_gn=PLCB4;refGene_tx=XM_023251457.1;refGene_gn=PLCB4;refGe
ne_tx=XM_023251458.1;refGene_gn=PLCB4;refGene_tx=XM_023251459.1;refGen
e_gn=PLCB4;refGene_tx=XM_023251460.1
A3 35671579 35671637
refGene_gn=PLCB4;refGene_tx=XM_019826823.2;refGene_gn=PLCB4;refGene_tx
=XM_023251453.1;refGene_gn=PLCB4;refGene_tx=XM_023251454.1;refGene_gn=
PLCB4;refGene_tx=XM_023251455.1;refGene_gn=PLCB4;refGene_tx=XM_0232514

56.1;refGene_gn=PLCB4;refGene_tx=XM_023251457.1;refGene_gn=PLCB4;refGene_tx=XM_023251458.1;refGene_gn=PLCB4;refGene_tx=XM_023251459.1;refGene_gn=PLCB4;refGene_tx=XM_023251460.1
A3 35672037 35672124
refGene_gn=PLCB4;refGene_tx=XM_019826823.2;refGene_gn=PLCB4;refGene_tx=XM_023251453.1;refGene_gn=PLCB4;refGene_tx=XM_023251454.1;refGene_gn=PLCB4;refGene_tx=XM_023251455.1;refGene_gn=PLCB4;refGene_tx=XM_023251456.1;refGene_gn=PLCB4;refGene_tx=XM_023251457.1;refGene_gn=PLCB4;refGene_tx=XM_023251458.1;refGene_gn=PLCB4;refGene_tx=XM_023251459.1;refGene_gn=PLCB4;refGene_tx=XM_023251460.1
A3 35675098 35675135
refGene_gn=PLCB4;refGene_tx=XM_019826823.2;refGene_gn=PLCB4;refGene_tx=XM_023251453.1;refGene_gn=PLCB4;refGene_tx=XM_023251454.1;refGene_gn=PLCB4;refGene_tx=XM_023251455.1;refGene_gn=PLCB4;refGene_tx=XM_023251456.1;refGene_gn=PLCB4;refGene_tx=XM_023251457.1;refGene_gn=PLCB4;refGene_tx=XM_023251458.1;refGene_gn=PLCB4;refGene_tx=XM_023251459.1
A3 35677234 35677323
refGene_gn=PLCB4;refGene_tx=XM_019826823.2;refGene_gn=PLCB4;refGene_tx=XM_023251453.1;refGene_gn=PLCB4;refGene_tx=XM_023251454.1;refGene_gn=PLCB4;refGene_tx=XM_023251455.1;refGene_gn=PLCB4;refGene_tx=XM_023251456.1;refGene_gn=PLCB4;refGene_tx=XM_023251457.1;refGene_gn=PLCB4;refGene_tx=XM_023251458.1;refGene_gn=PLCB4;refGene_tx=XM_023251459.1;refGene_gn=PLCB4;refGene_tx=XM_023251460.1
A3 35730324 35730388 refGene_gn=LAMP5;refGene_tx=XM_003983774.4
A3 35730930 35731103 refGene_gn=LAMP5;refGene_tx=XM_003983774.4
A3 35731480 35731612 refGene_gn=LAMP5;refGene_tx=XM_003983774.4
A3 35731739 35731845 refGene_gn=LAMP5;refGene_tx=XM_003983774.4
A3 35733221 35733410 refGene_gn=LAMP5;refGene_tx=XM_003983774.4
A3 35745464 35745643 refGene_gn=LAMP5;refGene_tx=XM_003983774.4
A3 35755354 35755510
refGene_gn=PAK5;refGene_tx=XM_006929957.4;refGene_gn=PAK5;refGene_tx=XM_019826827.2;refGene_gn=PAK5;refGene_tx=XM_019826828.2;refGene_gn=PAK5;refGene_tx=XM_023251461.1;refGene_gn=PAK5;refGene_tx=XM_023251462.1;refGene_gn=PAK5;refGene_tx=XM_023251463.1;refGene_gn=PAK5;refGene_tx=XM_023251464.1;refGene_gn=PAK5;refGene_tx=XM_023251465.1
A3 35757947 35758082
refGene_gn=PAK5;refGene_tx=XM_006929957.4;refGene_gn=PAK5;refGene_tx=XM_019826827.2;refGene_gn=PAK5;refGene_tx=XM_019826828.2;refGene_gn=PAK5;refGene_tx=XM_023251461.1;refGene_gn=PAK5;refGene_tx=XM_023251462.1;refGene_gn=PAK5;refGene_tx=XM_023251463.1;refGene_gn=PAK5;refGene_tx=XM_023251464.1;refGene_gn=PAK5;refGene_tx=XM_023251465.1
A3 35759648 35759774
refGene_gn=PAK5;refGene_tx=XM_006929957.4;refGene_gn=PAK5;refGene_tx=XM_019826827.2;refGene_gn=PAK5;refGene_tx=XM_019826828.2;refGene_gn=PAK5;refGene_tx=XM_023251461.1;refGene_gn=PAK5;refGene_tx=XM_023251462.1;refGene_gn=PAK5;refGene_tx=XM_023251463.1;refGene_gn=PAK5;refGene_tx=XM_023251464.1;refGene_gn=PAK5;refGene_tx=XM_023251465.1
A3 35772203 35772330
refGene_gn=PAK5;refGene_tx=XM_006929957.4;refGene_gn=PAK5;refGene_tx=XM_019826827.2;refGene_gn=PAK5;refGene_tx=XM_019826828.2;refGene_gn=PAK

5;refGene_tx=XM_023251461.1;refGene_gn=PAK5;refGene_tx=XM_023251462.1;
refGene_gn=PAK5;refGene_tx=XM_023251463.1;refGene_gn=PAK5;refGene_tx=X
M_023251464.1;refGene_gn=PAK5;refGene_tx=XM_023251465.1
A3 35776362 35776496
refGene_gn=PAK5;refGene_tx=XM_006929957.4;refGene_gn=PAK5;refGene_tx=X
M_019826827.2;refGene_gn=PAK5;refGene_tx=XM_019826828.2;refGene_gn=PAK
5;refGene_tx=XM_023251461.1;refGene_gn=PAK5;refGene_tx=XM_023251462.1;
refGene_gn=PAK5;refGene_tx=XM_023251463.1;refGene_gn=PAK5;refGene_tx=X
M_023251464.1;refGene_gn=PAK5;refGene_tx=XM_023251465.1
A3 35778865 35779357
refGene_gn=PAK5;refGene_tx=XM_006929957.4;refGene_gn=PAK5;refGene_tx=X
M_019826827.2;refGene_gn=PAK5;refGene_tx=XM_019826828.2;refGene_gn=PAK
5;refGene_tx=XM_023251461.1;refGene_gn=PAK5;refGene_tx=XM_023251462.1;
refGene_gn=PAK5;refGene_tx=XM_023251463.1;refGene_gn=PAK5;refGene_tx=X
M_023251464.1;refGene_gn=PAK5;refGene_tx=XM_023251465.1
A3 35795646 35796432
refGene_gn=PAK5;refGene_tx=XM_006929957.4;refGene_gn=PAK5;refGene_tx=X
M_019826827.2;refGene_gn=PAK5;refGene_tx=XM_019826828.2;refGene_gn=PAK
5;refGene_tx=XM_023251461.1;refGene_gn=PAK5;refGene_tx=XM_023251462.1;
refGene_gn=PAK5;refGene_tx=XM_023251463.1;refGene_gn=PAK5;refGene_tx=X
M_023251464.1;refGene_gn=PAK5;refGene_tx=XM_023251465.1
A3 35857297 35857501
refGene_gn=PAK5;refGene_tx=XM_006929957.4;refGene_gn=PAK5;refGene_tx=X
M_019826827.2;refGene_gn=PAK5;refGene_tx=XM_019826828.2;refGene_gn=PAK
5;refGene_tx=XM_023251461.1;refGene_gn=PAK5;refGene_tx=XM_023251462.1;
refGene_gn=PAK5;refGene_tx=XM_023251463.1;refGene_gn=PAK5;refGene_tx=X
M_023251464.1;refGene_gn=PAK5;refGene_tx=XM_023251465.1
A3 36202221 36202567 refGene_gn=ANKEF1;refGene_tx=XM_011280903.3
A3 36207539 36207739
refGene_gn=ANKEF1;refGene_tx=XM_011280903.3;refGene_gn=ANKEF1;refGene_
tx=XM_011280904.3
A3 36208492 36208642
refGene_gn=ANKEF1;refGene_tx=XM_011280903.3;refGene_gn=ANKEF1;refGene_
tx=XM_011280904.3;refGene_gn=ANKEF1;refGene_tx=XM_011280905.3
A3 36209661 36209785
refGene_gn=ANKEF1;refGene_tx=XM_011280903.3;refGene_gn=ANKEF1;refGene_
tx=XM_011280904.3;refGene_gn=ANKEF1;refGene_tx=XM_011280905.3
A3 36213471 36214294
refGene_gn=ANKEF1;refGene_tx=XM_011280903.3;refGene_gn=ANKEF1;refGene_
tx=XM_011280904.3;refGene_gn=ANKEF1;refGene_tx=XM_011280905.3
A3 36216063 36216290
refGene_gn=ANKEF1;refGene_tx=XM_011280903.3;refGene_gn=ANKEF1;refGene_
tx=XM_011280904.3;refGene_gn=ANKEF1;refGene_tx=XM_011280905.3
A3 36217663 36217827
refGene_gn=ANKEF1;refGene_tx=XM_011280903.3;refGene_gn=ANKEF1;refGene_
tx=XM_011280904.3;refGene_gn=ANKEF1;refGene_tx=XM_011280905.3
A3 36219713 36219848
refGene_gn=ANKEF1;refGene_tx=XM_011280903.3;refGene_gn=ANKEF1;refGene_
tx=XM_011280904.3;refGene_gn=ANKEF1;refGene_tx=XM_011280905.3
A3 36220692 36220851

refGene_gn=ANKEF1;refGene_tx=XM_011280903.3;refGene_gn=ANKEF1;refGene_tx=XM_011280904.3;refGene_gn=ANKEF1;refGene_tx=XM_011280905.3
A3 36422744 36422816
refGene_gn=SNAP25;refGene_tx=XM_003983780.4;refGene_gn=SNAP25;refGene_tx=XM_003983781.4;refGene_gn=SNAP25;refGene_tx=XM_019826829.1;refGene_gn=SNAP25;refGene_tx=XM_019826831.2;refGene_gn=SNAP25;refGene_tx=XM_019826832.2
A3 36424704 36424746
refGene_gn=SNAP25;refGene_tx=XM_003983780.4;refGene_gn=SNAP25;refGene_tx=XM_003983781.4;refGene_gn=SNAP25;refGene_tx=XM_019826829.1;refGene_gn=SNAP25;refGene_tx=XM_019826831.2;refGene_gn=SNAP25;refGene_tx=XM_019826832.2
A3 36431793 36431842
refGene_gn=SNAP25;refGene_tx=XM_003983780.4;refGene_gn=SNAP25;refGene_tx=XM_003983781.4;refGene_gn=SNAP25;refGene_tx=XM_019826829.1;refGene_gn=SNAP25;refGene_tx=XM_019826831.2;refGene_gn=SNAP25;refGene_tx=XM_019826832.2
A3 36438575 36438693 refGene_gn=SNAP25;refGene_tx=XM_003983781.4
A3 36438863 36438981
refGene_gn=SNAP25;refGene_tx=XM_003983780.4;refGene_gn=SNAP25;refGene_tx=XM_019826829.1;refGene_gn=SNAP25;refGene_tx=XM_019826831.2;refGene_gn=SNAP25;refGene_tx=XM_019826832.2
A3 36442639 36442765
refGene_gn=SNAP25;refGene_tx=XM_003983780.4;refGene_gn=SNAP25;refGene_tx=XM_003983781.4;refGene_gn=SNAP25;refGene_tx=XM_019826829.1;refGene_gn=SNAP25;refGene_tx=XM_019826831.2;refGene_gn=SNAP25;refGene_tx=XM_019826832.2
A3 36445503 36445648
refGene_gn=SNAP25;refGene_tx=XM_003983780.4;refGene_gn=SNAP25;refGene_tx=XM_003983781.4;refGene_gn=SNAP25;refGene_tx=XM_019826829.1;refGene_gn=SNAP25;refGene_tx=XM_019826831.2;refGene_gn=SNAP25;refGene_tx=XM_019826832.2
A3 36451641 36451710
refGene_gn=SNAP25;refGene_tx=XM_003983780.4;refGene_gn=SNAP25;refGene_tx=XM_003983781.4;refGene_gn=SNAP25;refGene_tx=XM_019826829.1;refGene_gn=SNAP25;refGene_tx=XM_019826831.2;refGene_gn=SNAP25;refGene_tx=XM_019826832.2
A3 36545694 36546135
refGene_gn=MKKS;refGene_tx=XM_003983782.4;refGene_gn=MKKS;refGene_tx=XM_019826834.1
A3 36547980 36548091
refGene_gn=MKKS;refGene_tx=XM_003983782.4;refGene_gn=MKKS;refGene_tx=XM_019826834.1
A3 36549018 36549194
refGene_gn=MKKS;refGene_tx=XM_003983782.4;refGene_gn=MKKS;refGene_tx=XM_019826834.1
A3 36551246 36552231
refGene_gn=MKKS;refGene_tx=XM_003983782.4;refGene_gn=MKKS;refGene_tx=XM_019826834.1
A3 36593008 36593035 refGene_gn=SLX4IP;refGene_tx=XM_003983783.4

A3 36684890 36684980 refGene_gn=SLX4IP;refGene_tx=XM_003983783.4
A3 36688927 36689048 refGene_gn=SLX4IP;refGene_tx=XM_003983783.4
A3 36726510 36726588 refGene_gn=SLX4IP;refGene_tx=XM_003983783.4
A3 36729714 36729803 refGene_gn=SLX4IP;refGene_tx=XM_003983783.4
A3 36746874 36746975 refGene_gn=SLX4IP;refGene_tx=XM_003983783.4
A3 36748065 36748780 refGene_gn=SLX4IP;refGene_tx=XM_003983783.4
A3 36764691 36765149 refGene_gn=JAG1;refGene_tx=XM_003983784.5
A3 36765962 36766113 refGene_gn=JAG1;refGene_tx=XM_003983784.5
A3 36766287 36766419 refGene_gn=JAG1;refGene_tx=XM_003983784.5
A3 36766597 36766831 refGene_gn=JAG1;refGene_tx=XM_003983784.5
A3 36766951 36767061 refGene_gn=JAG1;refGene_tx=XM_003983784.5
A3 36767665 36767779 refGene_gn=JAG1;refGene_tx=XM_003983784.5
A3 36769176 36769262 refGene_gn=JAG1;refGene_tx=XM_003983784.5
A3 36769728 36769756 refGene_gn=JAG1;refGene_tx=XM_003983784.5
A3 36770252 36770369 refGene_gn=JAG1;refGene_tx=XM_003983784.5
A3 36770505 36770619 refGene_gn=JAG1;refGene_tx=XM_003983784.5
A3 36770724 36770838 refGene_gn=JAG1;refGene_tx=XM_003983784.5
A3 36771365 36771479 refGene_gn=JAG1;refGene_tx=XM_003983784.5
A3 36771881 36772046 refGene_gn=JAG1;refGene_tx=XM_003983784.5
A3 36772825 36772976 refGene_gn=JAG1;refGene_tx=XM_003983784.5
A3 36773605 36773779 refGene_gn=JAG1;refGene_tx=XM_003983784.5
A3 36774119 36774166 refGene_gn=JAG1;refGene_tx=XM_003983784.5
A3 36774583 36774697 refGene_gn=JAG1;refGene_tx=XM_003983784.5
A3 36775288 36775402 refGene_gn=JAG1;refGene_tx=XM_003983784.5
A3 36776620 36776734 refGene_gn=JAG1;refGene_tx=XM_003983784.5
A3 36777163 36777283 refGene_gn=JAG1;refGene_tx=XM_003983784.5
A3 36777787 36777918 refGene_gn=JAG1;refGene_tx=XM_003983784.5
A3 36781762 36781823 refGene_gn=JAG1;refGene_tx=XM_003983784.5
A3 36783534 36783789 refGene_gn=JAG1;refGene_tx=XM_003983784.5
A3 36789357 36789409 refGene_gn=JAG1;refGene_tx=XM_003983784.5
A3 36798012 36798318 refGene_gn=JAG1;refGene_tx=XM_003983784.5
A3 36798744 36798825 refGene_gn=JAG1;refGene_tx=XM_003983784.5
A3 37980348 37980695
refGene_gn=BTBD3;refGene_tx=XM_003983785.5;refGene_gn=BTBD3;refGene_tx
=XM_023251467.1;refGene_gn=BTBD3;refGene_tx=XM_023251468.1
A3 37981186 37981277
refGene_gn=BTBD3;refGene_tx=XM_003983785.5;refGene_gn=BTBD3;refGene_tx
=XM_023251467.1
A3 37981868 37981987
refGene_gn=BTBD3;refGene_tx=XM_003983785.5;refGene_gn=BTBD3;refGene_tx
=XM_019826836.2;refGene_gn=BTBD3;refGene_tx=XM_023251467.1
A3 37985029 37986062
refGene_gn=BTBD3;refGene_tx=XM_003983785.5;refGene_gn=BTBD3;refGene_tx
=XM_019826836.2;refGene_gn=BTBD3;refGene_tx=XM_023251467.1;refGene_gn=
BTBD3;refGene_tx=XM_023251468.1
A3 38865142 38865259 refGene_gn=SPTLC3;refGene_tx=XM_011280956.3
A3 38894462 38894648 refGene_gn=SPTLC3;refGene_tx=XM_011280956.3
A3 38907413 38907568 refGene_gn=SPTLC3;refGene_tx=XM_011280956.3
A3 38908205 38908354 refGene_gn=SPTLC3;refGene_tx=XM_011280956.3
A3 38925989 38926114 refGene_gn=SPTLC3;refGene_tx=XM_011280956.3

A3	38928533	38928627	refGene_gn=SPTLC3;refGene_tx=XM_011280956.3
A3	38942953	38943059	refGene_gn=SPTLC3;refGene_tx=XM_011280956.3
A3	38952452	38952672	refGene_gn=SPTLC3;refGene_tx=XM_011280956.3
A3	38961216	38961343	refGene_gn=SPTLC3;refGene_tx=XM_011280956.3
A3	38984035	38984171	refGene_gn=SPTLC3;refGene_tx=XM_011280956.3
A3	38991196	38991326	refGene_gn=SPTLC3;refGene_tx=XM_011280956.3
A3	38995923	38996037	refGene_gn=SPTLC3;refGene_tx=XM_011280956.3
A3	39044087	39044225	refGene_gn=ISM1;refGene_tx=XM_023251469.1
A3	39091488	39091728	refGene_gn=ISM1;refGene_tx=XM_023251469.1
A3	39100666	39100931	refGene_gn=ISM1;refGene_tx=XM_023251469.1
A3	39109680	39109824	refGene_gn=ISM1;refGene_tx=XM_023251469.1
A3	39113289	39113379	refGene_gn=ISM1;refGene_tx=XM_023251469.1
A3	39117233	39117751	refGene_gn=ISM1;refGene_tx=XM_023251469.1
A3	39182256	39182349
refGene_gn=TASP1;refGene_tx=XM_003983788.4;refGene_gn=TASP1;refGene_tx=XM_019826841.1;refGene_gn=TASP1;refGene_tx=XM_019826846.2
A3	39219869	39219943
refGene_gn=TASP1;refGene_tx=XM_003983788.4;refGene_gn=TASP1;refGene_tx=XM_019826841.1;refGene_gn=TASP1;refGene_tx=XM_019826846.2
A3	39242406	39242517
refGene_gn=TASP1;refGene_tx=XM_003983788.4;refGene_gn=TASP1;refGene_tx=XM_019826841.1;refGene_gn=TASP1;refGene_tx=XM_019826846.2;refGene_gn=TASP1;refGene_tx=XM_023251470.1
A3	39299486	39299597
refGene_gn=TASP1;refGene_tx=XM_003983788.4;refGene_gn=TASP1;refGene_tx=XM_019826841.1;refGene_gn=TASP1;refGene_tx=XM_019826846.2;refGene_gn=TASP1;refGene_tx=XM_023251470.1
A3	39340090	39340169
refGene_gn=TASP1;refGene_tx=XM_003983788.4;refGene_gn=TASP1;refGene_tx=XM_019826841.1;refGene_gn=TASP1;refGene_tx=XM_019826846.2;refGene_gn=TASP1;refGene_tx=XM_023251470.1
A3	39345074	39345194
refGene_gn=TASP1;refGene_tx=XM_003983788.4;refGene_gn=TASP1;refGene_tx=XM_019826846.2;refGene_gn=TASP1;refGene_tx=XM_023251470.1
A3	39372167	39372274
refGene_gn=TASP1;refGene_tx=XM_003983788.4;refGene_gn=TASP1;refGene_tx=XM_019826841.1;refGene_gn=TASP1;refGene_tx=XM_019826846.2;refGene_gn=TASP1;refGene_tx=XM_023251470.1
A3	39390827	39390907
refGene_gn=TASP1;refGene_tx=XM_003983788.4;refGene_gn=TASP1;refGene_tx=XM_019826841.1;refGene_gn=TASP1;refGene_tx=XM_019826846.2;refGene_gn=TASP1;refGene_tx=XM_023251470.1
A3	39402314	39402399
refGene_gn=TASP1;refGene_tx=XM_003983788.4;refGene_gn=TASP1;refGene_tx=XM_019826841.1;refGene_gn=TASP1;refGene_tx=XM_019826846.2;refGene_gn=TASP1;refGene_tx=XM_023251470.1
A3	39411171	39411292
refGene_gn=TASP1;refGene_tx=XM_003983788.4;refGene_gn=TASP1;refGene_tx=XM_019826841.1;refGene_gn=TASP1;refGene_tx=XM_019826846.2;refGene_gn=TASP1;refGene_tx=XM_023251470.1

A3	39435947	39436016
refGene_gn=TASP1;refGene_tx=XM_003983788.4;refGene_gn=TASP1;refGene_tx=XM_019826841.1;refGene_gn=TASP1;refGene_tx=XM_023251470.1
A3	39438295	39438363
refGene_gn=TASP1;refGene_tx=XM_003983788.4;refGene_gn=TASP1;refGene_tx=XM_019826841.1;refGene_gn=TASP1;refGene_tx=XM_023251470.1
A3	39442087	39442232
refGene_gn=TASP1;refGene_tx=XM_003983788.4;refGene_gn=TASP1;refGene_tx=XM_019826841.1;refGene_gn=TASP1;refGene_tx=XM_023251470.1
A3	39512263	39512557
refGene_gn=ESF1;refGene_tx=XM_011280957.3;refGene_gn=ESF1;refGene_tx=XM_011280958.3
A3	39513276	39513423
refGene_gn=ESF1;refGene_tx=XM_011280957.3;refGene_gn=ESF1;refGene_tx=XM_011280958.3
A3	39515624	39515707
refGene_gn=ESF1;refGene_tx=XM_011280957.3;refGene_gn=ESF1;refGene_tx=XM_011280958.3
A3	39523946	39524034
refGene_gn=ESF1;refGene_tx=XM_011280957.3;refGene_gn=ESF1;refGene_tx=XM_011280958.3
A3	39527820	39527942
refGene_gn=ESF1;refGene_tx=XM_011280957.3;refGene_gn=ESF1;refGene_tx=XM_011280958.3
A3	39554329	39554491
refGene_gn=ESF1;refGene_tx=XM_011280957.3;refGene_gn=ESF1;refGene_tx=XM_011280958.3
A3	39560599	39560747
refGene_gn=ESF1;refGene_tx=XM_011280957.3;refGene_gn=ESF1;refGene_tx=XM_011280958.3
A3	39562407	39562522
refGene_gn=ESF1;refGene_tx=XM_011280957.3;refGene_gn=ESF1;refGene_tx=XM_011280958.3
A3	39564660	39564813
refGene_gn=ESF1;refGene_tx=XM_011280957.3;refGene_gn=ESF1;refGene_tx=XM_011280958.3
A3	39565833	39565934
refGene_gn=ESF1;refGene_tx=XM_011280957.3;refGene_gn=ESF1;refGene_tx=XM_011280958.3
A3	39568604	39568718
refGene_gn=ESF1;refGene_tx=XM_011280957.3;refGene_gn=ESF1;refGene_tx=XM_011280958.3
A3	39569531	39569935
refGene_gn=ESF1;refGene_tx=XM_011280957.3;refGene_gn=ESF1;refGene_tx=XM_011280958.3
A3	39575261	39575895
refGene_gn=ESF1;refGene_tx=XM_011280957.3;refGene_gn=ESF1;refGene_tx=XM_011280958.3
A3	39577404	39577626
refGene_gn=NDUFAF5;refGene_tx=XM_011280959.3;refGene_gn=NDUFAF5;refGen

e_tx=XM_011280960.3;refGene_gn=NDUFAF5;refGene_tx=XM_019826847.2;refGene_gn=NDUFAF5;refGene_tx=XM_023251471.1
A3 39580299 39580340
refGene_gn=NDUFAF5;refGene_tx=XM_011280959.3;refGene_gn=NDUFAF5;refGene_tx=XM_011280960.3;refGene_gn=NDUFAF5;refGene_tx=XM_019826847.2;refGene_gn=NDUFAF5;refGene_tx=XM_023251471.1
A3 39582407 39582471
refGene_gn=NDUFAF5;refGene_tx=XM_011280959.3;refGene_gn=NDUFAF5;refGene_tx=XM_011280960.3;refGene_gn=NDUFAF5;refGene_tx=XM_019826847.2;refGene_gn=NDUFAF5;refGene_tx=XM_023251471.1
A3 39585068 39585116
refGene_gn=NDUFAF5;refGene_tx=XM_011280959.3;refGene_gn=NDUFAF5;refGene_tx=XM_011280960.3;refGene_gn=NDUFAF5;refGene_tx=XM_019826847.2;refGene_gn=NDUFAF5;refGene_tx=XM_023251471.1;refGene_gn=NDUFAF5;refGene_tx=XM_023251474.1;refGene_gn=NDUFAF5;refGene_tx=XM_023251475.1
A3 39586435 39586539
refGene_gn=NDUFAF5;refGene_tx=XM_011280959.3;refGene_gn=NDUFAF5;refGene_tx=XM_011280960.3;refGene_gn=NDUFAF5;refGene_tx=XM_023251471.1;refGene_gn=NDUFAF5;refGene_tx=XM_023251472.1;refGene_gn=NDUFAF5;refGene_tx=XM_023251473.1
A3 39588426 39588466
refGene_gn=NDUFAF5;refGene_tx=XM_011280959.3;refGene_gn=NDUFAF5;refGene_tx=XM_011280960.3;refGene_gn=NDUFAF5;refGene_tx=XM_023251471.1;refGene_gn=NDUFAF5;refGene_tx=XM_023251472.1;refGene_gn=NDUFAF5;refGene_tx=XM_023251473.1;refGene_gn=NDUFAF5;refGene_tx=XM_023251474.1;refGene_gn=NDUFAF5;refGene_tx=XM_023251475.1
A3 39591713 39591911
refGene_gn=NDUFAF5;refGene_tx=XM_011280959.3;refGene_gn=NDUFAF5;refGene_tx=XM_019826847.2;refGene_gn=NDUFAF5;refGene_tx=XM_023251471.1;refGene_gn=NDUFAF5;refGene_tx=XM_023251472.1;refGene_gn=NDUFAF5;refGene_tx=XM_023251473.1;refGene_gn=NDUFAF5;refGene_tx=XM_023251474.1;refGene_gn=NDUFAF5;refGene_tx=XM_023251475.1;refGene_gn=NDUFAF5;refGene_tx=XM_023251476.1
A3 39596782 39596843
refGene_gn=NDUFAF5;refGene_tx=XM_011280959.3;refGene_gn=NDUFAF5;refGene_tx=XM_011280960.3;refGene_gn=NDUFAF5;refGene_tx=XM_019826847.2;refGene_gn=NDUFAF5;refGene_tx=XM_023251471.1;refGene_gn=NDUFAF5;refGene_tx=XM_023251472.1;refGene_gn=NDUFAF5;refGene_tx=XM_023251473.1;refGene_gn=NDUFAF5;refGene_tx=XM_023251474.1;refGene_gn=NDUFAF5;refGene_tx=XM_023251475.1;refGene_gn=NDUFAF5;refGene_tx=XM_023251476.1
A3 39601749 39601772 refGene_gn=NDUFAF5;refGene_tx=XM_023251471.1
A3 39604824 39604908
refGene_gn=NDUFAF5;refGene_tx=XM_011280959.3;refGene_gn=NDUFAF5;refGene_tx=XM_011280960.3;refGene_gn=NDUFAF5;refGene_tx=XM_019826847.2;refGene_gn=NDUFAF5;refGene_tx=XM_023251472.1;refGene_gn=NDUFAF5;refGene_tx=XM_023251473.1;refGene_gn=NDUFAF5;refGene_tx=XM_023251474.1;refGene_gn=NDUFAF5;refGene_tx=XM_023251475.1;refGene_gn=NDUFAF5;refGene_tx=XM_023251476.1
A3 39605260 39605343
refGene_gn=NDUFAF5;refGene_tx=XM_011280959.3;refGene_gn=NDUFAF5;refGen

e_tx=XM_011280960.3;refGene_gn=NDUFAF5;refGene_tx=XM_019826847.2;refGene_gn=NDUFAF5;refGene_tx=XM_023251472.1;refGene_gn=NDUFAF5;refGene_tx=XM_023251473.1;refGene_gn=NDUFAF5;refGene_tx=XM_023251474.1;refGene_gn=NDUFAF5;refGene_tx=XM_023251475.1;refGene_gn=NDUFAF5;refGene_tx=XM_023251476.1
A3 39605517 39605610
refGene_gn=NDUFAF5;refGene_tx=XM_011280959.3;refGene_gn=NDUFAF5;refGene_tx=XM_011280960.3;refGene_gn=NDUFAF5;refGene_tx=XM_019826847.2;refGene_gn=NDUFAF5;refGene_tx=XM_023251472.1;refGene_gn=NDUFAF5;refGene_tx=XM_023251473.1;refGene_gn=NDUFAF5;refGene_tx=XM_023251474.1;refGene_gn=NDUFAF5;refGene_tx=XM_023251475.1;refGene_gn=NDUFAF5;refGene_tx=XM_023251476.1
A3 39630597 39630720
refGene_gn=SEL1L2;refGene_tx=XM_006929968.3;refGene_gn=SEL1L2;refGene_tx=XM_019826851.2;refGene_gn=SEL1L2;refGene_tx=XM_023251477.1;refGene_gn=SEL1L2;refGene_tx=XM_023251478.1;refGene_gn=SEL1L2;refGene_tx=XM_023251479.1;refGene_gn=SEL1L2;refGene_tx=XM_023251480.1;refGene_gn=SEL1L2;refGene_tx=XM_023251482.1;refGene_gn=SEL1L2;refGene_tx=XM_023251483.1
A3 39630854 39630983
refGene_gn=SEL1L2;refGene_tx=XM_006929968.3;refGene_gn=SEL1L2;refGene_tx=XM_019826851.2;refGene_gn=SEL1L2;refGene_tx=XM_023251477.1;refGene_gn=SEL1L2;refGene_tx=XM_023251478.1;refGene_gn=SEL1L2;refGene_tx=XM_023251479.1;refGene_gn=SEL1L2;refGene_tx=XM_023251480.1;refGene_gn=SEL1L2;refGene_tx=XM_023251482.1;refGene_gn=SEL1L2;refGene_tx=XM_023251483.1
A3 39637123 39637369 refGene_gn=SEL1L2;refGene_tx=XM_023251481.1
A3 39639040 39639213
refGene_gn=SEL1L2;refGene_tx=XM_006929968.3;refGene_gn=SEL1L2;refGene_tx=XM_019826851.2;refGene_gn=SEL1L2;refGene_tx=XM_023251477.1;refGene_gn=SEL1L2;refGene_tx=XM_023251478.1;refGene_gn=SEL1L2;refGene_tx=XM_023251479.1;refGene_gn=SEL1L2;refGene_tx=XM_023251480.1;refGene_gn=SEL1L2;refGene_tx=XM_023251481.1;refGene_gn=SEL1L2;refGene_tx=XM_023251482.1
A3 39644079 39644154
refGene_gn=SEL1L2;refGene_tx=XM_006929968.3;refGene_gn=SEL1L2;refGene_tx=XM_019826851.2;refGene_gn=SEL1L2;refGene_tx=XM_023251477.1;refGene_gn=SEL1L2;refGene_tx=XM_023251478.1;refGene_gn=SEL1L2;refGene_tx=XM_023251479.1;refGene_gn=SEL1L2;refGene_tx=XM_023251480.1;refGene_gn=SEL1L2;refGene_tx=XM_023251481.1;refGene_gn=SEL1L2;refGene_tx=XM_023251482.1;refGene_gn=SEL1L2;refGene_tx=XM_023251483.1
A3 39644261 39644427
refGene_gn=SEL1L2;refGene_tx=XM_006929968.3;refGene_gn=SEL1L2;refGene_tx=XM_019826851.2;refGene_gn=SEL1L2;refGene_tx=XM_023251477.1;refGene_gn=SEL1L2;refGene_tx=XM_023251478.1;refGene_gn=SEL1L2;refGene_tx=XM_023251479.1;refGene_gn=SEL1L2;refGene_tx=XM_023251480.1;refGene_gn=SEL1L2;refGene_tx=XM_023251481.1;refGene_gn=SEL1L2;refGene_tx=XM_023251482.1
A3 39645556 39645705
refGene_gn=SEL1L2;refGene_tx=XM_006929968.3;refGene_gn=SEL1L2;refGene_

tx=XM_019826851.2;refGene_gn=SEL1L2;refGene_tx=XM_023251477.1;refGene_
gn=SEL1L2;refGene_tx=XM_023251478.1;refGene_gn=SEL1L2;refGene_tx=XM_02
3251479.1;refGene_gn=SEL1L2;refGene_tx=XM_023251480.1;refGene_gn=SEL1L
2;refGene_tx=XM_023251481.1;refGene_gn=SEL1L2;refGene_tx=XM_023251483.
1
A3 39649665 39649753
refGene_gn=SEL1L2;refGene_tx=XM_006929968.3;refGene_gn=SEL1L2;refGene_
tx=XM_019826851.2;refGene_gn=SEL1L2;refGene_tx=XM_023251477.1;refGene_
gn=SEL1L2;refGene_tx=XM_023251478.1;refGene_gn=SEL1L2;refGene_tx=XM_02
3251479.1;refGene_gn=SEL1L2;refGene_tx=XM_023251480.1;refGene_gn=SEL1L
2;refGene_tx=XM_023251481.1;refGene_gn=SEL1L2;refGene_tx=XM_023251483.
1
A3 39650324 39650387
refGene_gn=SEL1L2;refGene_tx=XM_006929968.3;refGene_gn=SEL1L2;refGene_
tx=XM_019826851.2;refGene_gn=SEL1L2;refGene_tx=XM_023251477.1;refGene_
gn=SEL1L2;refGene_tx=XM_023251478.1;refGene_gn=SEL1L2;refGene_tx=XM_02
3251479.1;refGene_gn=SEL1L2;refGene_tx=XM_023251480.1;refGene_gn=SEL1L
2;refGene_tx=XM_023251481.1;refGene_gn=SEL1L2;refGene_tx=XM_023251482.
1;refGene_gn=SEL1L2;refGene_tx=XM_023251483.1
A3 39654578 39654656
refGene_gn=SEL1L2;refGene_tx=XM_006929968.3;refGene_gn=SEL1L2;refGene_
tx=XM_019826851.2;refGene_gn=SEL1L2;refGene_tx=XM_023251477.1;refGene_
gn=SEL1L2;refGene_tx=XM_023251478.1;refGene_gn=SEL1L2;refGene_tx=XM_02
3251479.1;refGene_gn=SEL1L2;refGene_tx=XM_023251480.1;refGene_gn=SEL1L
2;refGene_tx=XM_023251481.1;refGene_gn=SEL1L2;refGene_tx=XM_023251482.
1;refGene_gn=SEL1L2;refGene_tx=XM_023251483.1
A3 39655557 39655626
refGene_gn=SEL1L2;refGene_tx=XM_006929968.3;refGene_gn=SEL1L2;refGene_
tx=XM_019826851.2;refGene_gn=SEL1L2;refGene_tx=XM_023251477.1;refGene_
gn=SEL1L2;refGene_tx=XM_023251478.1;refGene_gn=SEL1L2;refGene_tx=XM_02
3251479.1;refGene_gn=SEL1L2;refGene_tx=XM_023251480.1;refGene_gn=SEL1L
2;refGene_tx=XM_023251481.1;refGene_gn=SEL1L2;refGene_tx=XM_023251482.
1;refGene_gn=SEL1L2;refGene_tx=XM_023251483.1
A3 39663544 39663601
refGene_gn=SEL1L2;refGene_tx=XM_006929968.3;refGene_gn=SEL1L2;refGene_
tx=XM_019826851.2;refGene_gn=SEL1L2;refGene_tx=XM_023251477.1;refGene_
gn=SEL1L2;refGene_tx=XM_023251478.1;refGene_gn=SEL1L2;refGene_tx=XM_02
3251479.1;refGene_gn=SEL1L2;refGene_tx=XM_023251480.1;refGene_gn=SEL1L
2;refGene_tx=XM_023251481.1;refGene_gn=SEL1L2;refGene_tx=XM_023251482.
1;refGene_gn=SEL1L2;refGene_tx=XM_023251483.1
A3 39664300 39664455
refGene_gn=SEL1L2;refGene_tx=XM_006929968.3;refGene_gn=SEL1L2;refGene_
tx=XM_019826851.2;refGene_gn=SEL1L2;refGene_tx=XM_023251477.1;refGene_
gn=SEL1L2;refGene_tx=XM_023251478.1;refGene_gn=SEL1L2;refGene_tx=XM_02
3251479.1;refGene_gn=SEL1L2;refGene_tx=XM_023251480.1;refGene_gn=SEL1L
2;refGene_tx=XM_023251481.1;refGene_gn=SEL1L2;refGene_tx=XM_023251482.
1;refGene_gn=SEL1L2;refGene_tx=XM_023251483.1
A3 39665855 39665937
refGene_gn=SEL1L2;refGene_tx=XM_006929968.3;refGene_gn=SEL1L2;refGene_
tx=XM_019826851.2;refGene_gn=SEL1L2;refGene_tx=XM_023251477.1;refGene_

gn=SEL1L2;refGene_tx=XM_023251478.1;refGene_gn=SEL1L2;refGene_tx=XM_02
3251479.1;refGene_gn=SEL1L2;refGene_tx=XM_023251480.1;refGene_gn=SEL1L
2;refGene_tx=XM_023251481.1;refGene_gn=SEL1L2;refGene_tx=XM_023251482.
1;refGene_gn=SEL1L2;refGene_tx=XM_023251483.1
A3 39666027 39666087
refGene_gn=SEL1L2;refGene_tx=XM_006929968.3;refGene_gn=SEL1L2;refGene_
tx=XM_019826851.2;refGene_gn=SEL1L2;refGene_tx=XM_023251477.1;refGene_
gn=SEL1L2;refGene_tx=XM_023251478.1;refGene_gn=SEL1L2;refGene_tx=XM_02
3251479.1;refGene_gn=SEL1L2;refGene_tx=XM_023251480.1;refGene_gn=SEL1L
2;refGene_tx=XM_023251481.1;refGene_gn=SEL1L2;refGene_tx=XM_023251482.
1;refGene_gn=SEL1L2;refGene_tx=XM_023251483.1
A3 39666754 39666808
refGene_gn=SEL1L2;refGene_tx=XM_006929968.3;refGene_gn=SEL1L2;refGene_
tx=XM_019826851.2;refGene_gn=SEL1L2;refGene_tx=XM_023251477.1;refGene_
gn=SEL1L2;refGene_tx=XM_023251478.1;refGene_gn=SEL1L2;refGene_tx=XM_02
3251479.1;refGene_gn=SEL1L2;refGene_tx=XM_023251480.1;refGene_gn=SEL1L
2;refGene_tx=XM_023251481.1;refGene_gn=SEL1L2;refGene_tx=XM_023251482.
1;refGene_gn=SEL1L2;refGene_tx=XM_023251483.1
A3 39689032 39689195
refGene_gn=SEL1L2;refGene_tx=XM_006929968.3;refGene_gn=SEL1L2;refGene_
tx=XM_019826851.2;refGene_gn=SEL1L2;refGene_tx=XM_023251477.1;refGene_
gn=SEL1L2;refGene_tx=XM_023251478.1;refGene_gn=SEL1L2;refGene_tx=XM_02
3251479.1;refGene_gn=SEL1L2;refGene_tx=XM_023251480.1;refGene_gn=SEL1L
2;refGene_tx=XM_023251481.1;refGene_gn=SEL1L2;refGene_tx=XM_023251482.
1;refGene_gn=SEL1L2;refGene_tx=XM_023251483.1
A3 39693619 39693718
refGene_gn=SEL1L2;refGene_tx=XM_006929968.3;refGene_gn=SEL1L2;refGene_
tx=XM_019826851.2;refGene_gn=SEL1L2;refGene_tx=XM_023251477.1;refGene_
gn=SEL1L2;refGene_tx=XM_023251478.1;refGene_gn=SEL1L2;refGene_tx=XM_02
3251479.1;refGene_gn=SEL1L2;refGene_tx=XM_023251480.1;refGene_gn=SEL1L
2;refGene_tx=XM_023251481.1;refGene_gn=SEL1L2;refGene_tx=XM_023251482.
1;refGene_gn=SEL1L2;refGene_tx=XM_023251483.1
A3 39705689 39705692 refGene_gn=SEL1L2;refGene_tx=XM_023251483.1
A3 39709619 39709789 refGene_gn=SEL1L2;refGene_tx=XM_023251483.1
A3 39728546 39728602 refGene_gn=SEL1L2;refGene_tx=XM_023251483.1
A3 39746605 39746663 refGene_gn=SEL1L2;refGene_tx=XM_023251483.1
A3 39753763 39753809 refGene_gn=MACROD2;refGene_tx=XM_023251484.1
A3 39759248 39759365 refGene_gn=MACROD2;refGene_tx=XM_023251484.1
A3 39823212 39823320 refGene_gn=MACROD2;refGene_tx=XM_023251484.1
A3 40111490 40113440
refGene_gn=FLRT3;refGene_tx=XM_006929960.4;refGene_gn=FLRT3;refGene_tx
=XM_019826854.2;refGene_gn=FLRT3;refGene_tx=XM_019826855.2
A3 40275251 40275281 refGene_gn=MACROD2;refGene_tx=XM_023251484.1
A3 40489354 40489471 refGene_gn=MACROD2;refGene_tx=XM_023251484.1
A3 41013587 41013709 refGene_gn=MACROD2;refGene_tx=XM_023251484.1
A3 41216636 41216667 refGene_gn=MACROD2;refGene_tx=XM_023251484.1
A3 41285095 41285169 refGene_gn=MACROD2;refGene_tx=XM_023251484.1
A3 41608147 41608229 refGene_gn=MACROD2;refGene_tx=XM_023251484.1
A3 41624974 41625016 refGene_gn=MACROD2;refGene_tx=XM_023251484.1
A3 41632746 41632830 refGene_gn=MACROD2;refGene_tx=XM_023251484.1

A3 41672495 41672558 refGene_gn=MACROD2;refGene_tx=XM_023251484.1
A3 41677137 41677206 refGene_gn=MACROD2;refGene_tx=XM_023251484.1
A3 41706535 41706613 refGene_gn=MACROD2;refGene_tx=XM_023251484.1
A3 41718005 41718080 refGene_gn=MACROD2;refGene_tx=XM_023251484.1
A3 41718333 41718426 refGene_gn=MACROD2;refGene_tx=XM_023251484.1
A3 41768922 41769000 refGene_gn=MACROD2;refGene_tx=XM_023251484.1
A3 41771867 41771929 refGene_gn=MACROD2;refGene_tx=XM_023251484.1
A3 41962852 41963011
refGene_gn=KIF16B;refGene_tx=XM_006929963.4;refGene_gn=KIF16B;refGene_tx=XM_006929964.4;refGene_gn=KIF16B;refGene_tx=XM_006929965.4;refGene_gn=KIF16B;refGene_tx=XM_019826860.2
A3 41972039 41972132 refGene_gn=KIF16B;refGene_tx=XM_006929963.4
A3 41995666 41995750
refGene_gn=KIF16B;refGene_tx=XM_006929963.4;refGene_gn=KIF16B;refGene_tx=XM_006929964.4;refGene_gn=KIF16B;refGene_tx=XM_006929965.4
A3 42015482 42015572
refGene_gn=KIF16B;refGene_tx=XM_006929963.4;refGene_gn=KIF16B;refGene_tx=XM_006929964.4;refGene_gn=KIF16B;refGene_tx=XM_006929965.4;refGene_gn=KIF16B;refGene_tx=XM_019826860.2
A3 42038564 42038687
refGene_gn=KIF16B;refGene_tx=XM_006929963.4;refGene_gn=KIF16B;refGene_tx=XM_006929964.4;refGene_gn=KIF16B;refGene_tx=XM_006929965.4;refGene_gn=KIF16B;refGene_tx=XM_019826860.2
A3 42054496 42054547
refGene_gn=KIF16B;refGene_tx=XM_006929963.4;refGene_gn=KIF16B;refGene_tx=XM_006929964.4;refGene_gn=KIF16B;refGene_tx=XM_006929965.4;refGene_gn=KIF16B;refGene_tx=XM_019826860.2
A3 42055540 42055637
refGene_gn=KIF16B;refGene_tx=XM_006929963.4;refGene_gn=KIF16B;refGene_tx=XM_006929964.4;refGene_gn=KIF16B;refGene_tx=XM_006929965.4;refGene_gn=KIF16B;refGene_tx=XM_019826860.2
A3 42057521 42057674
refGene_gn=KIF16B;refGene_tx=XM_006929963.4;refGene_gn=KIF16B;refGene_tx=XM_006929964.4;refGene_gn=KIF16B;refGene_tx=XM_006929965.4;refGene_gn=KIF16B;refGene_tx=XM_019826860.2
A3 42061352 42062711
refGene_gn=KIF16B;refGene_tx=XM_006929963.4;refGene_gn=KIF16B;refGene_tx=XM_006929964.4;refGene_gn=KIF16B;refGene_tx=XM_006929965.4;refGene_gn=KIF16B;refGene_tx=XM_019826860.2
A3 42064072 42064126
refGene_gn=KIF16B;refGene_tx=XM_006929963.4;refGene_gn=KIF16B;refGene_tx=XM_006929964.4;refGene_gn=KIF16B;refGene_tx=XM_006929965.4;refGene_gn=KIF16B;refGene_tx=XM_019826860.2
A3 42064473 42064506
refGene_gn=KIF16B;refGene_tx=XM_006929963.4;refGene_gn=KIF16B;refGene_tx=XM_006929964.4;refGene_gn=KIF16B;refGene_tx=XM_019826860.2
A3 42088862 42088951
refGene_gn=KIF16B;refGene_tx=XM_006929963.4;refGene_gn=KIF16B;refGene_tx=XM_006929964.4;refGene_gn=KIF16B;refGene_tx=XM_006929965.4;refGene_gn=KIF16B;refGene_tx=XM_019826860.2

A3	42090381	42090464
refGene_gn=KIF16B;refGene_tx=XM_006929963.4;refGene_gn=KIF16B;refGene_tx=XM_006929964.4;refGene_gn=KIF16B;refGene_tx=XM_006929965.4;refGene_gn=KIF16B;refGene_tx=XM_019826860.2
A3	42105223	42105361
refGene_gn=KIF16B;refGene_tx=XM_006929963.4;refGene_gn=KIF16B;refGene_tx=XM_006929964.4;refGene_gn=KIF16B;refGene_tx=XM_006929965.4;refGene_gn=KIF16B;refGene_tx=XM_019826860.2
A3	42107025	42107077
refGene_gn=KIF16B;refGene_tx=XM_006929963.4;refGene_gn=KIF16B;refGene_tx=XM_006929964.4;refGene_gn=KIF16B;refGene_tx=XM_006929965.4;refGene_gn=KIF16B;refGene_tx=XM_019826860.2
A3	42107957	42108077
refGene_gn=KIF16B;refGene_tx=XM_006929963.4;refGene_gn=KIF16B;refGene_tx=XM_006929964.4;refGene_gn=KIF16B;refGene_tx=XM_006929965.4;refGene_gn=KIF16B;refGene_tx=XM_019826860.2
A3	42183830	42183890
refGene_gn=KIF16B;refGene_tx=XM_006929963.4;refGene_gn=KIF16B;refGene_tx=XM_006929964.4;refGene_gn=KIF16B;refGene_tx=XM_006929965.4;refGene_gn=KIF16B;refGene_tx=XM_019826860.2
A3	42186887	42186953
refGene_gn=KIF16B;refGene_tx=XM_006929963.4;refGene_gn=KIF16B;refGene_tx=XM_006929964.4;refGene_gn=KIF16B;refGene_tx=XM_006929965.4;refGene_gn=KIF16B;refGene_tx=XM_019826860.2
A3	42194750	42194926
refGene_gn=KIF16B;refGene_tx=XM_006929963.4;refGene_gn=KIF16B;refGene_tx=XM_006929964.4;refGene_gn=KIF16B;refGene_tx=XM_006929965.4;refGene_gn=KIF16B;refGene_tx=XM_019826860.2
A3	42195886	42196018
refGene_gn=KIF16B;refGene_tx=XM_006929963.4;refGene_gn=KIF16B;refGene_tx=XM_006929964.4;refGene_gn=KIF16B;refGene_tx=XM_006929965.4;refGene_gn=KIF16B;refGene_tx=XM_019826860.2
A3	42196199	42196368
refGene_gn=KIF16B;refGene_tx=XM_006929963.4;refGene_gn=KIF16B;refGene_tx=XM_006929964.4;refGene_gn=KIF16B;refGene_tx=XM_006929965.4;refGene_gn=KIF16B;refGene_tx=XM_019826860.2
A3	42197847	42197990
refGene_gn=KIF16B;refGene_tx=XM_006929963.4;refGene_gn=KIF16B;refGene_tx=XM_006929964.4;refGene_gn=KIF16B;refGene_tx=XM_006929965.4;refGene_gn=KIF16B;refGene_tx=XM_019826860.2
A3	42200611	42200721
refGene_gn=KIF16B;refGene_tx=XM_006929963.4;refGene_gn=KIF16B;refGene_tx=XM_006929964.4;refGene_gn=KIF16B;refGene_tx=XM_006929965.4;refGene_gn=KIF16B;refGene_tx=XM_019826860.2
A3	42203102	42203200
refGene_gn=KIF16B;refGene_tx=XM_006929963.4;refGene_gn=KIF16B;refGene_tx=XM_006929964.4;refGene_gn=KIF16B;refGene_tx=XM_006929965.4;refGene_gn=KIF16B;refGene_tx=XM_019826860.2
A3	42205504	42205621
refGene_gn=KIF16B;refGene_tx=XM_006929963.4;refGene_gn=KIF16B;refGene_

tx=XM_006929964.4;refGene_gn=KIF16B;refGene_tx=XM_006929965.4;refGene_gn=KIF16B;refGene_tx=XM_019826860.2
A3 42211597 42211711
refGene_gn=KIF16B;refGene_tx=XM_006929963.4;refGene_gn=KIF16B;refGene_tx=XM_006929964.4;refGene_gn=KIF16B;refGene_tx=XM_006929965.4;refGene_gn=KIF16B;refGene_tx=XM_019826860.2
A3 42214332 42214402
refGene_gn=KIF16B;refGene_tx=XM_006929963.4;refGene_gn=KIF16B;refGene_tx=XM_006929964.4;refGene_gn=KIF16B;refGene_tx=XM_006929965.4;refGene_gn=KIF16B;refGene_tx=XM_019826860.2
A3 42252728 42252775
refGene_gn=KIF16B;refGene_tx=XM_006929963.4;refGene_gn=KIF16B;refGene_tx=XM_006929964.4;refGene_gn=KIF16B;refGene_tx=XM_006929965.4;refGene_gn=KIF16B;refGene_tx=XM_019826860.2
A3 42413739 42413803
refGene_gn=SNRPB2;refGene_tx=XM_003983797.4;refGene_gn=SNRPB2;refGene_tx=XM_006929969.4
A3 42414128 42414301
refGene_gn=SNRPB2;refGene_tx=XM_003983797.4;refGene_gn=SNRPB2;refGene_tx=XM_006929969.4
A3 42419097 42419238
refGene_gn=SNRPB2;refGene_tx=XM_003983797.4;refGene_gn=SNRPB2;refGene_tx=XM_006929969.4
A3 42421674 42421725
refGene_gn=SNRPB2;refGene_tx=XM_003983797.4;refGene_gn=SNRPB2;refGene_tx=XM_006929969.4
A3 42423141 42423230
refGene_gn=SNRPB2;refGene_tx=XM_003983797.4;refGene_gn=SNRPB2;refGene_tx=XM_006929969.4
A3 42423679 42423839
refGene_gn=SNRPB2;refGene_tx=XM_003983797.4;refGene_gn=SNRPB2;refGene_tx=XM_006929969.4
A3 42430973 42431088 refGene_gn=OTOR;refGene_tx=XM_003983798.3
A3 42431448 42431588 refGene_gn=OTOR;refGene_tx=XM_003983798.3
A3 42432475 42432586 refGene_gn=OTOR;refGene_tx=XM_003983798.3
A3 42433469 42433493 refGene_gn=OTOR;refGene_tx=XM_003983798.3
A3 42809619 42809796 refGene_gn=PCSK2;refGene_tx=XM_003983799.5
A3 42848311 42848416 refGene_gn=PCSK2;refGene_tx=XM_003983799.5
A3 42954789 42954903 refGene_gn=PCSK2;refGene_tx=XM_003983799.5
A3 42957097 42957206 refGene_gn=PCSK2;refGene_tx=XM_003983799.5
A3 42965849 42965887 refGene_gn=PCSK2;refGene_tx=XM_003983799.5
A3 43009475 43009552 refGene_gn=PCSK2;refGene_tx=XM_003983799.5
A3 43028698 43028787 refGene_gn=PCSK2;refGene_tx=XM_003983799.5
A3 43039611 43039787 refGene_gn=PCSK2;refGene_tx=XM_003983799.5
A3 43061201 43061417 refGene_gn=PCSK2;refGene_tx=XM_003983799.5
A3 43063658 43063759 refGene_gn=PCSK2;refGene_tx=XM_003983799.5
A3 43070466 43070694 refGene_gn=PCSK2;refGene_tx=XM_003983799.5
A3 43080584 43081071 refGene_gn=PCSK2;refGene_tx=XM_003983799.5
A3 43091715 43092701
refGene_gn=BFSP1;refGene_tx=XM_019826862.2;refGene_gn=BFSP1;refGene_tx

=XM_023251485.1;refGene_gn=BFSP1;refGene_tx=XM_023251486.1;refGene_gn=BFSP1;refGene_tx=XM_023251487.1;refGene_gn=BFSP1;refGene_tx=XM_023251488.1;refGene_gn=BFSP1;refGene_tx=XM_023251489.1
A3 43094692 43094778
refGene_gn=BFSP1;refGene_tx=XM_019826862.2;refGene_gn=BFSP1;refGene_tx=XM_023251485.1;refGene_gn=BFSP1;refGene_tx=XM_023251486.1;refGene_gn=BFSP1;refGene_tx=XM_023251487.1;refGene_gn=BFSP1;refGene_tx=XM_023251488.1;refGene_gn=BFSP1;refGene_tx=XM_023251489.1
A3 43097176 43097397
refGene_gn=BFSP1;refGene_tx=XM_019826862.2;refGene_gn=BFSP1;refGene_tx=XM_023251485.1;refGene_gn=BFSP1;refGene_tx=XM_023251486.1;refGene_gn=BFSP1;refGene_tx=XM_023251487.1;refGene_gn=BFSP1;refGene_tx=XM_023251488.1;refGene_gn=BFSP1;refGene_tx=XM_023251489.1
A3 43107166 43107274
refGene_gn=BFSP1;refGene_tx=XM_019826862.2;refGene_gn=BFSP1;refGene_tx=XM_023251485.1;refGene_gn=BFSP1;refGene_tx=XM_023251486.1;refGene_gn=BFSP1;refGene_tx=XM_023251487.1;refGene_gn=BFSP1;refGene_tx=XM_023251488.1;refGene_gn=BFSP1;refGene_tx=XM_023251489.1
A3 43111489 43111582
refGene_gn=BFSP1;refGene_tx=XM_019826862.2;refGene_gn=BFSP1;refGene_tx=XM_023251485.1;refGene_gn=BFSP1;refGene_tx=XM_023251486.1;refGene_gn=BFSP1;refGene_tx=XM_023251487.1;refGene_gn=BFSP1;refGene_tx=XM_023251488.1;refGene_gn=BFSP1;refGene_tx=XM_023251489.1
A3 43114360 43114456
refGene_gn=BFSP1;refGene_tx=XM_019826862.2;refGene_gn=BFSP1;refGene_tx=XM_023251485.1;refGene_gn=BFSP1;refGene_tx=XM_023251486.1;refGene_gn=BFSP1;refGene_tx=XM_023251487.1;refGene_gn=BFSP1;refGene_tx=XM_023251488.1;refGene_gn=BFSP1;refGene_tx=XM_023251489.1
A3 43124875 43124936
refGene_gn=BFSP1;refGene_tx=XM_019826862.2;refGene_gn=BFSP1;refGene_tx=XM_023251485.1;refGene_gn=BFSP1;refGene_tx=XM_023251486.1;refGene_gn=BFSP1;refGene_tx=XM_023251487.1;refGene_gn=BFSP1;refGene_tx=XM_023251488.1;refGene_gn=BFSP1;refGene_tx=XM_023251489.1
A3 43132390 43132764
refGene_gn=BFSP1;refGene_tx=XM_023251485.1;refGene_gn=BFSP1;refGene_tx=XM_023251486.1;refGene_gn=BFSP1;refGene_tx=XM_023251487.1;refGene_gn=BFSP1;refGene_tx=XM_023251488.1;refGene_gn=BFSP1;refGene_tx=XM_023251489.1
A3 43145566 43145568 refGene_gn=BFSP1;refGene_tx=XM_019826862.2
A3 43162947 43162950 refGene_gn=DSTN;refGene_tx=XM_003983801.5
A3 43190610 43190918 refGene_gn=DSTN;refGene_tx=XM_003983801.5
A3 43194181 43194258 refGene_gn=DSTN;refGene_tx=XM_003983801.5
A3 43196473 43196583 refGene_gn=DSTN;refGene_tx=XM_003983801.5
A3 43203486 43203525
refGene_gn=RRBP1;refGene_tx=XM_023251490.1;refGene_gn=RRBP1;refGene_tx=XM_023251491.1;refGene_gn=RRBP1;refGene_tx=XM_023251492.1;refGene_gn=RRBP1;refGene_tx=XM_023251493.1
A3 43204084 43204228
refGene_gn=RRBP1;refGene_tx=XM_023251490.1;refGene_gn=RRBP1;refGene_tx=XM_023251491.1;refGene_gn=RRBP1;refGene_tx=XM_023251492.1;refGene_gn=

RRBP1;refGene_tx=XM_023251493.1
A3	43204707	43204806
refGene_gn=RRBP1;refGene_tx=XM_023251490.1;refGene_gn=RRBP1;refGene_tx=XM_023251491.1;refGene_gn=RRBP1;refGene_tx=XM_023251492.1;refGene_gn=RRBP1;refGene_tx=XM_023251493.1
A3	43205076	43205160
refGene_gn=RRBP1;refGene_tx=XM_023251490.1;refGene_gn=RRBP1;refGene_tx=XM_023251491.1;refGene_gn=RRBP1;refGene_tx=XM_023251492.1;refGene_gn=RRBP1;refGene_tx=XM_023251493.1
A3	43205895	43206003
refGene_gn=RRBP1;refGene_tx=XM_023251490.1;refGene_gn=RRBP1;refGene_tx=XM_023251491.1;refGene_gn=RRBP1;refGene_tx=XM_023251492.1
A3	43207753	43207837
refGene_gn=RRBP1;refGene_tx=XM_023251490.1;refGene_gn=RRBP1;refGene_tx=XM_023251491.1;refGene_gn=RRBP1;refGene_tx=XM_023251492.1;refGene_gn=RRBP1;refGene_tx=XM_023251493.1
A3	43208404	43208500
refGene_gn=RRBP1;refGene_tx=XM_023251490.1;refGene_gn=RRBP1;refGene_tx=XM_023251491.1;refGene_gn=RRBP1;refGene_tx=XM_023251492.1;refGene_gn=RRBP1;refGene_tx=XM_023251493.1
A3	43209063	43209135
refGene_gn=RRBP1;refGene_tx=XM_023251490.1;refGene_gn=RRBP1;refGene_tx=XM_023251491.1;refGene_gn=RRBP1;refGene_tx=XM_023251492.1;refGene_gn=RRBP1;refGene_tx=XM_023251493.1
A3	43209570	43209663
refGene_gn=RRBP1;refGene_tx=XM_023251490.1;refGene_gn=RRBP1;refGene_tx=XM_023251491.1;refGene_gn=RRBP1;refGene_tx=XM_023251492.1;refGene_gn=RRBP1;refGene_tx=XM_023251493.1
A3	43210367	43210457
refGene_gn=RRBP1;refGene_tx=XM_023251490.1;refGene_gn=RRBP1;refGene_tx=XM_023251491.1;refGene_gn=RRBP1;refGene_tx=XM_023251492.1;refGene_gn=RRBP1;refGene_tx=XM_023251493.1
A3	43210611	43210695
refGene_gn=RRBP1;refGene_tx=XM_023251490.1;refGene_gn=RRBP1;refGene_tx=XM_023251491.1;refGene_gn=RRBP1;refGene_tx=XM_023251492.1;refGene_gn=RRBP1;refGene_tx=XM_023251493.1
A3	43210776	43210869
refGene_gn=RRBP1;refGene_tx=XM_023251490.1;refGene_gn=RRBP1;refGene_tx=XM_023251491.1;refGene_gn=RRBP1;refGene_tx=XM_023251492.1;refGene_gn=RRBP1;refGene_tx=XM_023251493.1
A3	43212916	43213009
refGene_gn=RRBP1;refGene_tx=XM_023251490.1;refGene_gn=RRBP1;refGene_tx=XM_023251491.1;refGene_gn=RRBP1;refGene_tx=XM_023251492.1;refGene_gn=RRBP1;refGene_tx=XM_023251493.1
A3	43213859	43213950
refGene_gn=RRBP1;refGene_tx=XM_023251490.1;refGene_gn=RRBP1;refGene_tx=XM_023251491.1;refGene_gn=RRBP1;refGene_tx=XM_023251492.1;refGene_gn=RRBP1;refGene_tx=XM_023251493.1
A3	43216693	43216719
refGene_gn=RRBP1;refGene_tx=XM_023251490.1;refGene_gn=RRBP1;refGene_tx

=XM_023251491.1;refGene_gn=RRBP1;refGene_tx=XM_023251492.1;refGene_gn=RRBP1;refGene_tx=XM_023251493.1
A3 43216865 43217044
refGene_gn=RRBP1;refGene_tx=XM_023251490.1;refGene_gn=RRBP1;refGene_tx=XM_023251491.1;refGene_gn=RRBP1;refGene_tx=XM_023251492.1;refGene_gn=RRBP1;refGene_tx=XM_023251493.1
A3 43218867 43219006
refGene_gn=RRBP1;refGene_tx=XM_023251490.1;refGene_gn=RRBP1;refGene_tx=XM_023251491.1;refGene_gn=RRBP1;refGene_tx=XM_023251492.1;refGene_gn=RRBP1;refGene_tx=XM_023251493.1
A3 43221260 43221414
refGene_gn=RRBP1;refGene_tx=XM_023251490.1;refGene_gn=RRBP1;refGene_tx=XM_023251491.1;refGene_gn=RRBP1;refGene_tx=XM_023251492.1;refGene_gn=RRBP1;refGene_tx=XM_023251493.1
A3 43223776 43223895
refGene_gn=RRBP1;refGene_tx=XM_023251490.1;refGene_gn=RRBP1;refGene_tx=XM_023251491.1;refGene_gn=RRBP1;refGene_tx=XM_023251492.1;refGene_gn=RRBP1;refGene_tx=XM_023251493.1
A3 43224618 43224771
refGene_gn=RRBP1;refGene_tx=XM_023251490.1;refGene_gn=RRBP1;refGene_tx=XM_023251491.1;refGene_gn=RRBP1;refGene_tx=XM_023251492.1;refGene_gn=RRBP1;refGene_tx=XM_023251493.1
A3 43228012 43228135
refGene_gn=RRBP1;refGene_tx=XM_023251490.1;refGene_gn=RRBP1;refGene_tx=XM_023251491.1;refGene_gn=RRBP1;refGene_tx=XM_023251492.1;refGene_gn=RRBP1;refGene_tx=XM_023251493.1
A3 43229344 43229493
refGene_gn=RRBP1;refGene_tx=XM_023251490.1;refGene_gn=RRBP1;refGene_tx=XM_023251491.1;refGene_gn=RRBP1;refGene_tx=XM_023251492.1;refGene_gn=RRBP1;refGene_tx=XM_023251493.1
A3 43242404 43244349
refGene_gn=RRBP1;refGene_tx=XM_023251490.1;refGene_gn=RRBP1;refGene_tx=XM_023251491.1;refGene_gn=RRBP1;refGene_tx=XM_023251492.1;refGene_gn=RRBP1;refGene_tx=XM_023251493.1
A3 43313002 43313128
refGene_gn=BANF2;refGene_tx=XM_006929973.3;refGene_gn=BANF2;refGene_tx=XM_023251494.1
A3 43321994 43322141
refGene_gn=BANF2;refGene_tx=XM_006929973.3;refGene_gn=BANF2;refGene_tx=XM_023251494.1
A3 43494467 43494518
refGene_gn=SNX5;refGene_tx=XM_003983818.5;refGene_gn=SNX5;refGene_tx=XM_011280964.2
A3 43496070 43496156
refGene_gn=SNX5;refGene_tx=XM_003983818.5;refGene_gn=SNX5;refGene_tx=XM_011280964.2
A3 43499160 43499320
refGene_gn=SNX5;refGene_tx=XM_003983818.5;refGene_gn=SNX5;refGene_tx=XM_011280964.2
A3 43500675 43500762

refGene_gn=SNX5;refGene_tx=XM_003983818.5;refGene_gn=SNX5;refGene_tx=XM_011280964.2
A3	43500849	43500889
refGene_gn=SNX5;refGene_tx=XM_003983818.5;refGene_gn=SNX5;refGene_tx=XM_011280964.2
A3	43501846	43501922
refGene_gn=SNX5;refGene_tx=XM_003983818.5;refGene_gn=SNX5;refGene_tx=XM_011280964.2
A3	43502009	43502115
refGene_gn=SNX5;refGene_tx=XM_003983818.5;refGene_gn=SNX5;refGene_tx=XM_011280964.2
A3	43503308	43503404
refGene_gn=SNX5;refGene_tx=XM_003983818.5;refGene_gn=SNX5;refGene_tx=XM_011280964.2
A3	43504512	43504636
refGene_gn=SNX5;refGene_tx=XM_003983818.5;refGene_gn=SNX5;refGene_tx=XM_011280964.2
A3	43505804	43505926
refGene_gn=SNX5;refGene_tx=XM_003983818.5;refGene_gn=SNX5;refGene_tx=XM_011280964.2
A3	43506713	43506824	refGene_gn=SNX5;refGene_tx=XM_011280964.2
A3	43507961	43508066	refGene_gn=SNX5;refGene_tx=XM_011280964.2
A3	43519518	43519566	refGene_gn=SNX5;refGene_tx=XM_011280964.2
A3	43520976	43521490
refGene_gn=MGME1;refGene_tx=XM_003983804.3;refGene_gn=MGME1;refGene_tx=XM_006929974.2;refGene_gn=MGME1;refGene_tx=XM_006929975.4;refGene_gn=MGME1;refGene_tx=XM_006929976.4
A3	43525769	43525986
refGene_gn=MGME1;refGene_tx=XM_003983804.3;refGene_gn=MGME1;refGene_tx=XM_006929974.2;refGene_gn=MGME1;refGene_tx=XM_006929975.4;refGene_gn=MGME1;refGene_tx=XM_006929976.4
A3	43536499	43536632
refGene_gn=MGME1;refGene_tx=XM_003983804.3;refGene_gn=MGME1;refGene_tx=XM_006929974.2;refGene_gn=MGME1;refGene_tx=XM_006929975.4;refGene_gn=MGME1;refGene_tx=XM_006929976.4
A3	43537022	43537184
refGene_gn=MGME1;refGene_tx=XM_003983804.3;refGene_gn=MGME1;refGene_tx=XM_006929974.2;refGene_gn=MGME1;refGene_tx=XM_006929975.4;refGene_gn=MGME1;refGene_tx=XM_006929976.4
A3	43557978	43558292
refGene_gn=OVOL2;refGene_tx=XM_011280965.3;refGene_gn=OVOL2;refGene_tx=XM_019826869.2;refGene_gn=OVOL2;refGene_tx=XM_019826871.2;refGene_gn=OVOL2;refGene_tx=XM_019826872.2;refGene_gn=OVOL2;refGene_tx=XM_019826873.2;refGene_gn=OVOL2;refGene_tx=XM_019826874.2
A3	43573179	43573369
refGene_gn=OVOL2;refGene_tx=XM_011280965.3;refGene_gn=OVOL2;refGene_tx=XM_019826869.2;refGene_gn=OVOL2;refGene_tx=XM_019826871.2;refGene_gn=OVOL2;refGene_tx=XM_019826872.2;refGene_gn=OVOL2;refGene_tx=XM_019826873.2;refGene_gn=OVOL2;refGene_tx=XM_019826874.2
A3	43586247	43586471

refGene_gn=OVOL2;refGene_tx=XM_011280965.3;refGene_gn=OVOL2;refGene_tx=XM_019826869.2;refGene_gn=OVOL2;refGene_tx=XM_019826872.2;refGene_gn=OVOL2;refGene_tx=XM_019826873.2
A3 43587132 43587232 refGene_gn=OVOL2;refGene_tx=XM_011280965.3
A3 43588685 43589024
refGene_gn=OVOL2;refGene_tx=XM_019826869.2;refGene_gn=OVOL2;refGene_tx=XM_019826871.2
A3 43640225 43640321 refGene_gn=PET117;refGene_tx=XM_003983806.4
A3 43643560 43643710 refGene_gn=PET117;refGene_tx=XM_003983806.4
A3 43644024 43644283 refGene_gn=KAT14;refGene_tx=XM_023251495.1
A3 43646164 43646283 refGene_gn=KAT14;refGene_tx=XM_023251495.1
A3 43650319 43650441 refGene_gn=KAT14;refGene_tx=XM_023251495.1
A3 43658109 43658291 refGene_gn=KAT14;refGene_tx=XM_023251495.1
A3 43661107 43661524 refGene_gn=KAT14;refGene_tx=XM_023251495.1
A3 43661655 43662113 refGene_gn=KAT14;refGene_tx=XM_023251495.1
A3 43662114 43662225 refGene_gn=KAT14;refGene_tx=XM_023251495.1
A3 43675054 43675191 refGene_gn=KAT14;refGene_tx=XM_023251495.1
A3 43676043 43676219 refGene_gn=KAT14;refGene_tx=XM_023251495.1
A3 43677997 43678188 refGene_gn=KAT14;refGene_tx=XM_023251495.1
A3 43681605 43681779 refGene_gn=KAT14;refGene_tx=XM_023251495.1
A3 43777625 43777658
refGene_gn=ZNF133;refGene_tx=XM_003983809.5;refGene_gn=ZNF133;refGene_tx=XM_006929977.4;refGene_gn=ZNF133;refGene_tx=XM_006929978.4
A3 43784662 43784789
refGene_gn=ZNF133;refGene_tx=XM_003983809.5;refGene_gn=ZNF133;refGene_tx=XM_006929977.4;refGene_gn=ZNF133;refGene_tx=XM_006929978.4;refGene_gn=ZNF133;refGene_tx=XM_006929979.4;refGene_gn=ZNF133;refGene_tx=XM_006929980.4
A3 43784928 43785024
refGene_gn=ZNF133;refGene_tx=XM_003983809.5;refGene_gn=ZNF133;refGene_tx=XM_006929977.4;refGene_gn=ZNF133;refGene_tx=XM_006929978.4;refGene_gn=ZNF133;refGene_tx=XM_006929979.4;refGene_gn=ZNF133;refGene_tx=XM_006929980.4
A3 43787298 43789046
refGene_gn=ZNF133;refGene_tx=XM_003983809.5;refGene_gn=ZNF133;refGene_tx=XM_006929977.4;refGene_gn=ZNF133;refGene_tx=XM_006929978.4;refGene_gn=ZNF133;refGene_tx=XM_006929979.4;refGene_gn=ZNF133;refGene_tx=XM_006929980.4
A3 43817669 43817793 refGene_gn=DZANK1;refGene_tx=XM_023251496.1
A3 43818232 43818307 refGene_gn=DZANK1;refGene_tx=XM_023251496.1
A3 43822204 43822332 refGene_gn=DZANK1;refGene_tx=XM_023251496.1
A3 43822830 43822836 refGene_gn=DZANK1;refGene_tx=XM_023251496.1
A3 43822837 43822912 refGene_gn=DZANK1;refGene_tx=XM_023251496.1
A3 43826558 43826656 refGene_gn=DZANK1;refGene_tx=XM_023251496.1
A3 43827108 43827205 refGene_gn=DZANK1;refGene_tx=XM_023251496.1
A3 43829045 43829120 refGene_gn=DZANK1;refGene_tx=XM_023251496.1
A3 43831062 43831166 refGene_gn=DZANK1;refGene_tx=XM_023251496.1
A3 43835067 43835275 refGene_gn=DZANK1;refGene_tx=XM_023251496.1
A3 43836624 43836789 refGene_gn=DZANK1;refGene_tx=XM_023251496.1
A3 43837590 43837713 refGene_gn=DZANK1;refGene_tx=XM_023251496.1

A3 43846487 43846580 refGene_gn=DZANK1;refGene_tx=XM_023251496.1
A3 43854306 43854420 refGene_gn=DZANK1;refGene_tx=XM_023251496.1
A3 43858997 43859115 refGene_gn=DZANK1;refGene_tx=XM_023251496.1
A3 43866447 43866533 refGene_gn=DZANK1;refGene_tx=XM_023251496.1
A3 43870402 43870527 refGene_gn=DZANK1;refGene_tx=XM_023251496.1
A3 43871566 43871684 refGene_gn=DZANK1;refGene_tx=XM_023251496.1
A3 43873083 43873198 refGene_gn=DZANK1;refGene_tx=XM_023251496.1
A3 43881870 43882024 refGene_gn=DZANK1;refGene_tx=XM_023251496.1
A3 43885394 43885503 refGene_gn=DZANK1;refGene_tx=XM_023251496.1
A3 43888590 43888652
refGene_gn=POLR3F;refGene_tx=XM_003983811.4;refGene_gn=POLR3F;refGene_tx=XM_006929982.3
A3 43890352 43890470
refGene_gn=POLR3F;refGene_tx=XM_003983811.4;refGene_gn=POLR3F;refGene_tx=XM_006929982.3
A3 43893248 43893316
refGene_gn=POLR3F;refGene_tx=XM_003983811.4;refGene_gn=POLR3F;refGene_tx=XM_006929982.3
A3 43893682 43893703 refGene_gn=POLR3F;refGene_tx=XM_006929982.3
A3 43893827 43893895
refGene_gn=POLR3F;refGene_tx=XM_003983811.4;refGene_gn=POLR3F;refGene_tx=XM_006929982.3
A3 43895613 43895726
refGene_gn=POLR3F;refGene_tx=XM_003983811.4;refGene_gn=POLR3F;refGene_tx=XM_006929982.3
A3 43897730 43897874
refGene_gn=POLR3F;refGene_tx=XM_003983811.4;refGene_gn=POLR3F;refGene_tx=XM_006929982.3
A3 43898059 43898167
refGene_gn=POLR3F;refGene_tx=XM_003983811.4;refGene_gn=POLR3F;refGene_tx=XM_006929982.3
A3 43899713 43899905
refGene_gn=POLR3F;refGene_tx=XM_003983811.4;refGene_gn=POLR3F;refGene_tx=XM_006929982.3
A3 43900899 43900977
refGene_gn=POLR3F;refGene_tx=XM_003983811.4;refGene_gn=POLR3F;refGene_tx=XM_006929982.3
A3 43905348 43905575 refGene_gn=RBBP9;refGene_tx=XM_003983820.5
A3 43905970 43906056 refGene_gn=RBBP9;refGene_tx=XM_003983820.5
A3 43909340 43909446 refGene_gn=RBBP9;refGene_tx=XM_003983820.5
A3 43910450 43910493 refGene_gn=RBBP9;refGene_tx=XM_003983820.5
A3 43911707 43911806 refGene_gn=RBBP9;refGene_tx=XM_003983820.5
A3 43927052 43927273
refGene_gn=SEC23B;refGene_tx=XM_003983812.5;refGene_gn=SEC23B;refGene_tx=XM_006929984.4
A3 43928644 43928702
refGene_gn=SEC23B;refGene_tx=XM_003983812.5;refGene_gn=SEC23B;refGene_tx=XM_006929984.4
A3 43933035 43933122
refGene_gn=SEC23B;refGene_tx=XM_003983812.5;refGene_gn=SEC23B;refGene_

tx=XM_006929984.4
A3 43934712 43934949
refGene_gn=SEC23B;refGene_tx=XM_003983812.5;refGene_gn=SEC23B;refGene_
tx=XM_006929984.4
A3 43935053 43935139
refGene_gn=SEC23B;refGene_tx=XM_003983812.5;refGene_gn=SEC23B;refGene_
tx=XM_006929984.4
A3 43936593 43936738
refGene_gn=SEC23B;refGene_tx=XM_003983812.5;refGene_gn=SEC23B;refGene_
tx=XM_006929984.4
A3 43937135 43937294
refGene_gn=SEC23B;refGene_tx=XM_003983812.5;refGene_gn=SEC23B;refGene_
tx=XM_006929984.4
A3 43937654 43937770
refGene_gn=SEC23B;refGene_tx=XM_003983812.5;refGene_gn=SEC23B;refGene_
tx=XM_006929984.4
A3 43940827 43940951
refGene_gn=SEC23B;refGene_tx=XM_003983812.5;refGene_gn=SEC23B;refGene_
tx=XM_006929984.4
A3 43942249 43942330
refGene_gn=SEC23B;refGene_tx=XM_003983812.5;refGene_gn=SEC23B;refGene_
tx=XM_006929984.4
A3 43947661 43947751
refGene_gn=SEC23B;refGene_tx=XM_003983812.5;refGene_gn=SEC23B;refGene_
tx=XM_006929984.4
A3 43948658 43948765
refGene_gn=SEC23B;refGene_tx=XM_003983812.5;refGene_gn=SEC23B;refGene_
tx=XM_006929984.4
A3 43950199 43950353
refGene_gn=SEC23B;refGene_tx=XM_003983812.5;refGene_gn=SEC23B;refGene_
tx=XM_006929984.4
A3 43954026 43954104
refGene_gn=SEC23B;refGene_tx=XM_003983812.5;refGene_gn=SEC23B;refGene_
tx=XM_006929984.4
A3 43957114 43957276
refGene_gn=SEC23B;refGene_tx=XM_003983812.5;refGene_gn=SEC23B;refGene_
tx=XM_006929984.4
A3 43958621 43958708
refGene_gn=SEC23B;refGene_tx=XM_003983812.5;refGene_gn=SEC23B;refGene_
tx=XM_006929984.4
A3 43961345 43961501
refGene_gn=SEC23B;refGene_tx=XM_003983812.5;refGene_gn=SEC23B;refGene_
tx=XM_006929984.4
A3 43961876 43961942
refGene_gn=SEC23B;refGene_tx=XM_003983812.5;refGene_gn=SEC23B;refGene_
tx=XM_006929984.4
A3 43969788 43969878
refGene_gn=SEC23B;refGene_tx=XM_003983812.5;refGene_gn=SEC23B;refGene_
tx=XM_006929984.4
A3 43975798 43975916

refGene_gn=LOC111559796;refGene_tx=XM_023251497.1
A3 43976755 43976922
refGene_gn=LOC111559796;refGene_tx=XM_023251497.1
A3 43989395 43989438
refGene_gn=DTD1;refGene_tx=XM_023251498.1;refGene_gn=DTD1;refGene_tx=XM_023251499.1;refGene_gn=DTD1;refGene_tx=XM_023251500.1
A3 43993943 43994034
refGene_gn=DTD1;refGene_tx=XM_023251498.1;refGene_gn=DTD1;refGene_tx=XM_023251499.1;refGene_gn=DTD1;refGene_tx=XM_023251500.1
A3 43994949 43995185
refGene_gn=DTD1;refGene_tx=XM_023251498.1;refGene_gn=DTD1;refGene_tx=XM_023251499.1;refGene_gn=DTD1;refGene_tx=XM_023251500.1
A3 44020664 44020771
refGene_gn=DTD1;refGene_tx=XM_023251498.1;refGene_gn=DTD1;refGene_tx=XM_023251499.1;refGene_gn=DTD1;refGene_tx=XM_023251500.1
A3 44021479 44021481 refGene_gn=DTD1;refGene_tx=XM_023251500.1
A3 44022719 44022725 refGene_gn=DTD1;refGene_tx=XM_023251500.1
A3 44029377 44029383 refGene_gn=DTD1;refGene_tx=XM_023251500.1
A3 44031959 44032038 refGene_gn=DTD1;refGene_tx=XM_023251500.1
A3 44182803 44182956
refGene_gn=DTD1;refGene_tx=XM_023251498.1;refGene_gn=DTD1;refGene_tx=XM_023251499.1
A3 44241381 44241852 refGene_gn=SCP2D1;refGene_tx=XM_003983814.5
A3 44552167 44552609
refGene_gn=SLC24A3;refGene_tx=XM_023251501.1;refGene_gn=SLC24A3;refGene_tx=XM_023251502.1
A3 44615157 44615286
refGene_gn=SLC24A3;refGene_tx=XM_023251501.1;refGene_gn=SLC24A3;refGene_tx=XM_023251502.1
A3 44835526 44835603
refGene_gn=SLC24A3;refGene_tx=XM_023251501.1;refGene_gn=SLC24A3;refGene_tx=XM_023251502.1
A3 44895456 44895531
refGene_gn=SLC24A3;refGene_tx=XM_023251501.1;refGene_gn=SLC24A3;refGene_tx=XM_023251502.1
A3 44898953 44899038
refGene_gn=SLC24A3;refGene_tx=XM_023251501.1;refGene_gn=SLC24A3;refGene_tx=XM_023251502.1
A3 44899413 44899517
refGene_gn=SLC24A3;refGene_tx=XM_023251501.1;refGene_gn=SLC24A3;refGene_tx=XM_023251502.1
A3 44968310 44968385
refGene_gn=SLC24A3;refGene_tx=XM_023251501.1;refGene_gn=SLC24A3;refGene_tx=XM_023251502.1
A3 44978691 44978717
refGene_gn=SLC24A3;refGene_tx=XM_023251501.1;refGene_gn=SLC24A3;refGene_tx=XM_023251502.1
A3 44986877 44986931
refGene_gn=SLC24A3;refGene_tx=XM_023251501.1;refGene_gn=SLC24A3;refGene_tx=XM_023251502.1

A3 44992730 44992864
refGene_gn=SLC24A3;refGene_tx=XM_023251501.1;refGene_gn=SLC24A3;refGene_tx=XM_023251502.1
A3 44994749 44994910
refGene_gn=SLC24A3;refGene_tx=XM_023251501.1;refGene_gn=SLC24A3;refGene_tx=XM_023251502.1
A3 44995696 44995967
refGene_gn=SLC24A3;refGene_tx=XM_023251501.1;refGene_gn=SLC24A3;refGene_tx=XM_023251502.1
A3 45002884 45003051
refGene_gn=SLC24A3;refGene_tx=XM_023251501.1;refGene_gn=SLC24A3;refGene_tx=XM_023251502.1
A3 45006229 45006344
refGene_gn=SLC24A3;refGene_tx=XM_023251501.1;refGene_gn=SLC24A3;refGene_tx=XM_023251502.1
A3 45007969 45008082 refGene_gn=SLC24A3;refGene_tx=XM_023251502.1
A3 45028864 45028930
refGene_gn=SLC24A3;refGene_tx=XM_023251501.1;refGene_gn=SLC24A3;refGene_tx=XM_023251502.1
A3 45033404 45033796
refGene_gn=SLC24A3;refGene_tx=XM_023251501.1;refGene_gn=SLC24A3;refGene_tx=XM_023251502.1
A3 45178926 45178983
refGene_gn=RIN2;refGene_tx=XM_006929986.4;refGene_gn=RIN2;refGene_tx=XM_019826878.1;refGene_gn=RIN2;refGene_tx=XM_019826879.2
A3 45213616 45213733 refGene_gn=RIN2;refGene_tx=XM_019826877.2
A3 45222569 45222670
refGene_gn=RIN2;refGene_tx=XM_006929986.4;refGene_gn=RIN2;refGene_tx=XM_019826877.2;refGene_gn=RIN2;refGene_tx=XM_019826878.1;refGene_gn=RIN2;refGene_tx=XM_019826879.2
A3 45223045 45223068 refGene_gn=RIN2;refGene_tx=XM_006929987.4
A3 45243893 45244086
refGene_gn=RIN2;refGene_tx=XM_006929986.4;refGene_gn=RIN2;refGene_tx=XM_006929987.4;refGene_gn=RIN2;refGene_tx=XM_006929989.4;refGene_gn=RIN2;refGene_tx=XM_011280989.3;refGene_gn=RIN2;refGene_tx=XM_019826877.2;refGene_gn=RIN2;refGene_tx=XM_019826878.1;refGene_gn=RIN2;refGene_tx=XM_019826879.2
A3 45248079 45248191
refGene_gn=RIN2;refGene_tx=XM_006929986.4;refGene_gn=RIN2;refGene_tx=XM_006929987.4;refGene_gn=RIN2;refGene_tx=XM_006929989.4;refGene_gn=RIN2;refGene_tx=XM_011280989.3;refGene_gn=RIN2;refGene_tx=XM_019826877.2;refGene_gn=RIN2;refGene_tx=XM_019826878.1;refGene_gn=RIN2;refGene_tx=XM_019826879.2;refGene_gn=RIN2;refGene_tx=XM_019826880.2
A3 45252164 45252237
refGene_gn=RIN2;refGene_tx=XM_006929986.4;refGene_gn=RIN2;refGene_tx=XM_006929987.4;refGene_gn=RIN2;refGene_tx=XM_006929989.4;refGene_gn=RIN2;refGene_tx=XM_011280989.3;refGene_gn=RIN2;refGene_tx=XM_019826877.2;refGene_gn=RIN2;refGene_tx=XM_019826878.1;refGene_gn=RIN2;refGene_tx=XM_019826879.2;refGene_gn=RIN2;refGene_tx=XM_019826880.2
A3 45257117 45257209

refGene_gn=RIN2;refGene_tx=XM_006929986.4;refGene_gn=RIN2;refGene_tx=XM_006929987.4;refGene_gn=RIN2;refGene_tx=XM_006929989.4;refGene_gn=RIN2;refGene_tx=XM_011280989.3;refGene_gn=RIN2;refGene_tx=XM_019826877.2;refGene_gn=RIN2;refGene_tx=XM_019826878.1;refGene_gn=RIN2;refGene_tx=XM_019826879.2;refGene_gn=RIN2;refGene_tx=XM_019826880.2
A3 45260293 45261433
refGene_gn=RIN2;refGene_tx=XM_006929986.4;refGene_gn=RIN2;refGene_tx=XM_006929987.4;refGene_gn=RIN2;refGene_tx=XM_006929989.4;refGene_gn=RIN2;refGene_tx=XM_011280989.3;refGene_gn=RIN2;refGene_tx=XM_019826877.2;refGene_gn=RIN2;refGene_tx=XM_019826878.1;refGene_gn=RIN2;refGene_tx=XM_019826879.2;refGene_gn=RIN2;refGene_tx=XM_019826880.2
A3 45275860 45276166
refGene_gn=RIN2;refGene_tx=XM_006929986.4;refGene_gn=RIN2;refGene_tx=XM_006929987.4;refGene_gn=RIN2;refGene_tx=XM_006929989.4;refGene_gn=RIN2;refGene_tx=XM_011280989.3;refGene_gn=RIN2;refGene_tx=XM_019826877.2;refGene_gn=RIN2;refGene_tx=XM_019826878.1;refGene_gn=RIN2;refGene_tx=XM_019826879.2;refGene_gn=RIN2;refGene_tx=XM_019826880.2
A3 45278010 45278142
refGene_gn=RIN2;refGene_tx=XM_006929986.4;refGene_gn=RIN2;refGene_tx=XM_006929987.4;refGene_gn=RIN2;refGene_tx=XM_006929989.4;refGene_gn=RIN2;refGene_tx=XM_011280989.3;refGene_gn=RIN2;refGene_tx=XM_019826877.2;refGene_gn=RIN2;refGene_tx=XM_019826878.1;refGene_gn=RIN2;refGene_tx=XM_019826879.2;refGene_gn=RIN2;refGene_tx=XM_019826880.2
A3 45282952 45283116
refGene_gn=RIN2;refGene_tx=XM_006929986.4;refGene_gn=RIN2;refGene_tx=XM_006929987.4;refGene_gn=RIN2;refGene_tx=XM_006929989.4;refGene_gn=RIN2;refGene_tx=XM_011280989.3;refGene_gn=RIN2;refGene_tx=XM_019826877.2;refGene_gn=RIN2;refGene_tx=XM_019826878.1;refGene_gn=RIN2;refGene_tx=XM_019826879.2;refGene_gn=RIN2;refGene_tx=XM_019826880.2
A3 45286650 45286974
refGene_gn=RIN2;refGene_tx=XM_006929986.4;refGene_gn=RIN2;refGene_tx=XM_006929987.4;refGene_gn=RIN2;refGene_tx=XM_006929989.4;refGene_gn=RIN2;refGene_tx=XM_011280989.3;refGene_gn=RIN2;refGene_tx=XM_019826877.2;refGene_gn=RIN2;refGene_tx=XM_019826878.1;refGene_gn=RIN2;refGene_tx=XM_019826879.2;refGene_gn=RIN2;refGene_tx=XM_019826880.2
A3 45299303 45299356
refGene_gn=NAA20;refGene_tx=XM_006929992.4;refGene_gn=NAA20;refGene_tx=XM_011280985.2;refGene_gn=NAA20;refGene_tx=XM_011280986.2;refGene_gn=NAA20;refGene_tx=XM_011280987.2;refGene_gn=NAA20;refGene_tx=XM_019826889.2
A3 45299719 45299759
refGene_gn=NAA20;refGene_tx=XM_006929992.4;refGene_gn=NAA20;refGene_tx=XM_011280986.2;refGene_gn=NAA20;refGene_tx=XM_019826889.2
A3 45304041 45304066
refGene_gn=NAA20;refGene_tx=XM_006929992.4;refGene_gn=NAA20;refGene_tx=XM_011280985.2;refGene_gn=NAA20;refGene_tx=XM_011280987.2
A3 45304481 45304565 refGene_gn=NAA20;refGene_tx=XM_019826889.2
A3 45305285 45305291 refGene_gn=NAA20;refGene_tx=XM_006929994.4
A3 45307325 45307416
refGene_gn=NAA20;refGene_tx=XM_006929992.4;refGene_gn=NAA20;refGene_tx

=XM_006929994.4;refGene_gn=NAA20;refGene_tx=XM_011280985.2;refGene_gn=NAA20;refGene_tx=XM_011280986.2;refGene_gn=NAA20;refGene_tx=XM_011280987.2;refGene_gn=NAA20;refGene_tx=XM_019826889.2
A3 45308108 45308244
refGene_gn=NAA20;refGene_tx=XM_006929992.4;refGene_gn=NAA20;refGene_tx=XM_006929994.4;refGene_gn=NAA20;refGene_tx=XM_011280985.2;refGene_gn=NAA20;refGene_tx=XM_011280986.2;refGene_gn=NAA20;refGene_tx=XM_011280987.2;refGene_gn=NAA20;refGene_tx=XM_019826889.2
A3 45313803 45313949
refGene_gn=NAA20;refGene_tx=XM_006929992.4;refGene_gn=NAA20;refGene_tx=XM_006929994.4;refGene_gn=NAA20;refGene_tx=XM_011280985.2;refGene_gn=NAA20;refGene_tx=XM_011280986.2;refGene_gn=NAA20;refGene_tx=XM_019826889.2
A3 45314382 45314468
refGene_gn=NAA20;refGene_tx=XM_006929992.4;refGene_gn=NAA20;refGene_tx=XM_006929994.4;refGene_gn=NAA20;refGene_tx=XM_011280985.2;refGene_gn=NAA20;refGene_tx=XM_011280986.2;refGene_gn=NAA20;refGene_tx=XM_011280987.2;refGene_gn=NAA20;refGene_tx=XM_019826889.2
A3 45317709 45317877 refGene_gn=CRNKL1;refGene_tx=XM_011280988.3
A3 45318829 45319078 refGene_gn=CRNKL1;refGene_tx=XM_011280988.3
A3 45319520 45319622 refGene_gn=CRNKL1;refGene_tx=XM_011280988.3
A3 45320785 45321025 refGene_gn=CRNKL1;refGene_tx=XM_011280988.3
A3 45321803 45321884 refGene_gn=CRNKL1;refGene_tx=XM_011280988.3
A3 45322556 45322616 refGene_gn=CRNKL1;refGene_tx=XM_011280988.3
A3 45323331 45323523 refGene_gn=CRNKL1;refGene_tx=XM_011280988.3
A3 45324979 45325150 refGene_gn=CRNKL1;refGene_tx=XM_011280988.3
A3 45326291 45326470 refGene_gn=CRNKL1;refGene_tx=XM_011280988.3
A3 45328222 45328389 refGene_gn=CRNKL1;refGene_tx=XM_011280988.3
A3 45328859 45329018 refGene_gn=CRNKL1;refGene_tx=XM_011280988.3
A3 45329839 45329931 refGene_gn=CRNKL1;refGene_tx=XM_011280988.3
A3 45330967 45331120 refGene_gn=CRNKL1;refGene_tx=XM_011280988.3
A3 45332924 45332975 refGene_gn=CRNKL1;refGene_tx=XM_011280988.3
A3 45341290 45341433
refGene_gn=CFAP61;refGene_tx=XM_011280983.3;refGene_gn=CFAP61;refGene_tx=XM_019826881.2;refGene_gn=CFAP61;refGene_tx=XM_019826883.2;refGene_gn=CFAP61;refGene_tx=XM_019826885.2;refGene_gn=CFAP61;refGene_tx=XM_019826888.2;refGene_gn=CFAP61;refGene_tx=XM_023251503.1
A3 45349871 45350022
refGene_gn=CFAP61;refGene_tx=XM_011280983.3;refGene_gn=CFAP61;refGene_tx=XM_019826881.2;refGene_gn=CFAP61;refGene_tx=XM_019826882.2;refGene_gn=CFAP61;refGene_tx=XM_019826883.2;refGene_gn=CFAP61;refGene_tx=XM_019826885.2;refGene_gn=CFAP61;refGene_tx=XM_019826888.2;refGene_gn=CFAP61;refGene_tx=XM_023251503.1
A3 45354043 45354120
refGene_gn=CFAP61;refGene_tx=XM_011280983.3;refGene_gn=CFAP61;refGene_tx=XM_019826881.2;refGene_gn=CFAP61;refGene_tx=XM_019826882.2;refGene_gn=CFAP61;refGene_tx=XM_019826883.2;refGene_gn=CFAP61;refGene_tx=XM_019826885.2;refGene_gn=CFAP61;refGene_tx=XM_019826888.2;refGene_gn=CFAP61;refGene_tx=XM_023251503.1
A3 45354980 45355048

refGene_gn=CFAP61;refGene_tx=XM_011280983.3;refGene_gn=CFAP61;refGene_tx=XM_019826881.2;refGene_gn=CFAP61;refGene_tx=XM_019826882.2;refGene_gn=CFAP61;refGene_tx=XM_019826883.2;refGene_gn=CFAP61;refGene_tx=XM_019826885.2;refGene_gn=CFAP61;refGene_tx=XM_019826888.2;refGene_gn=CFAP61;refGene_tx=XM_023251503.1
A3 45355512 45355639
refGene_gn=CFAP61;refGene_tx=XM_011280983.3;refGene_gn=CFAP61;refGene_tx=XM_019826881.2;refGene_gn=CFAP61;refGene_tx=XM_019826882.2;refGene_gn=CFAP61;refGene_tx=XM_019826883.2;refGene_gn=CFAP61;refGene_tx=XM_019826885.2;refGene_gn=CFAP61;refGene_tx=XM_019826888.2;refGene_gn=CFAP61;refGene_tx=XM_023251503.1
A3 45369913 45370046
refGene_gn=CFAP61;refGene_tx=XM_011280983.3;refGene_gn=CFAP61;refGene_tx=XM_019826881.2;refGene_gn=CFAP61;refGene_tx=XM_019826882.2;refGene_gn=CFAP61;refGene_tx=XM_019826883.2;refGene_gn=CFAP61;refGene_tx=XM_019826885.2;refGene_gn=CFAP61;refGene_tx=XM_019826888.2;refGene_gn=CFAP61;refGene_tx=XM_023251503.1
A3 45389263 45389423
refGene_gn=CFAP61;refGene_tx=XM_011280983.3;refGene_gn=CFAP61;refGene_tx=XM_019826881.2;refGene_gn=CFAP61;refGene_tx=XM_019826882.2;refGene_gn=CFAP61;refGene_tx=XM_019826883.2;refGene_gn=CFAP61;refGene_tx=XM_019826885.2;refGene_gn=CFAP61;refGene_tx=XM_019826888.2;refGene_gn=CFAP61;refGene_tx=XM_023251503.1
A3 45416326 45416418
refGene_gn=CFAP61;refGene_tx=XM_011280983.3;refGene_gn=CFAP61;refGene_tx=XM_019826881.2;refGene_gn=CFAP61;refGene_tx=XM_019826882.2;refGene_gn=CFAP61;refGene_tx=XM_019826883.2;refGene_gn=CFAP61;refGene_tx=XM_019826885.2;refGene_gn=CFAP61;refGene_tx=XM_019826888.2;refGene_gn=CFAP61;refGene_tx=XM_023251503.1
A3 45422879 45422954
refGene_gn=CFAP61;refGene_tx=XM_011280983.3;refGene_gn=CFAP61;refGene_tx=XM_019826881.2;refGene_gn=CFAP61;refGene_tx=XM_019826882.2;refGene_gn=CFAP61;refGene_tx=XM_019826883.2;refGene_gn=CFAP61;refGene_tx=XM_019826885.2;refGene_gn=CFAP61;refGene_tx=XM_019826888.2;refGene_gn=CFAP61;refGene_tx=XM_023251503.1
A3 45427829 45428008
refGene_gn=CFAP61;refGene_tx=XM_011280983.3;refGene_gn=CFAP61;refGene_tx=XM_019826881.2;refGene_gn=CFAP61;refGene_tx=XM_019826882.2;refGene_gn=CFAP61;refGene_tx=XM_019826885.2;refGene_gn=CFAP61;refGene_tx=XM_019826888.2;refGene_gn=CFAP61;refGene_tx=XM_023251503.1
A3 45429783 45429823
refGene_gn=CFAP61;refGene_tx=XM_011280983.3;refGene_gn=CFAP61;refGene_tx=XM_019826881.2;refGene_gn=CFAP61;refGene_tx=XM_019826882.2;refGene_gn=CFAP61;refGene_tx=XM_019826885.2;refGene_gn=CFAP61;refGene_tx=XM_019826888.2;refGene_gn=CFAP61;refGene_tx=XM_023251503.1
A3 45433563 45433703
refGene_gn=CFAP61;refGene_tx=XM_011280983.3;refGene_gn=CFAP61;refGene_tx=XM_019826881.2;refGene_gn=CFAP61;refGene_tx=XM_019826882.2;refGene_gn=CFAP61;refGene_tx=XM_019826883.2;refGene_gn=CFAP61;refGene_tx=XM_019826885.2;refGene_gn=CFAP61;refGene_tx=XM_019826888.2;refGene_gn=CFAP6

1;refGene_tx=XM_023251503.1
A3 45455973 45456100
refGene_gn=CFAP61;refGene_tx=XM_011280983.3;refGene_gn=CFAP61;refGene_tx=XM_019826881.2;refGene_gn=CFAP61;refGene_tx=XM_019826882.2;refGene_gn=CFAP61;refGene_tx=XM_019826883.2;refGene_gn=CFAP61;refGene_tx=XM_019826885.2;refGene_gn=CFAP61;refGene_tx=XM_023251503.1
A3 45460570 45460648
refGene_gn=CFAP61;refGene_tx=XM_011280983.3;refGene_gn=CFAP61;refGene_tx=XM_019826882.2;refGene_gn=CFAP61;refGene_tx=XM_019826883.2;refGene_gn=CFAP61;refGene_tx=XM_019826885.2;refGene_gn=CFAP61;refGene_tx=XM_023251503.1
A3 45466576 45466783
refGene_gn=CFAP61;refGene_tx=XM_011280983.3;refGene_gn=CFAP61;refGene_tx=XM_019826881.2;refGene_gn=CFAP61;refGene_tx=XM_019826882.2;refGene_gn=CFAP61;refGene_tx=XM_019826883.2;refGene_gn=CFAP61;refGene_tx=XM_019826885.2;refGene_gn=CFAP61;refGene_tx=XM_019826888.2;refGene_gn=CFAP61;refGene_tx=XM_023251503.1
A3 45469543 45469678
refGene_gn=CFAP61;refGene_tx=XM_011280983.3;refGene_gn=CFAP61;refGene_tx=XM_019826881.2;refGene_gn=CFAP61;refGene_tx=XM_019826882.2;refGene_gn=CFAP61;refGene_tx=XM_019826883.2;refGene_gn=CFAP61;refGene_tx=XM_019826885.2;refGene_gn=CFAP61;refGene_tx=XM_023251503.1
A3 45482341 45482494 refGene_gn=CFAP61;refGene_tx=XM_019826885.2
A3 45501207 45501335
refGene_gn=CFAP61;refGene_tx=XM_011280983.3;refGene_gn=CFAP61;refGene_tx=XM_019826881.2;refGene_gn=CFAP61;refGene_tx=XM_019826882.2;refGene_gn=CFAP61;refGene_tx=XM_019826883.2;refGene_gn=CFAP61;refGene_tx=XM_023251503.1
A3 45516400 45516499
refGene_gn=CFAP61;refGene_tx=XM_011280983.3;refGene_gn=CFAP61;refGene_tx=XM_019826881.2;refGene_gn=CFAP61;refGene_tx=XM_019826882.2;refGene_gn=CFAP61;refGene_tx=XM_019826883.2;refGene_gn=CFAP61;refGene_tx=XM_023251503.1
A3 45522913 45523082
refGene_gn=CFAP61;refGene_tx=XM_011280983.3;refGene_gn=CFAP61;refGene_tx=XM_019826881.2;refGene_gn=CFAP61;refGene_tx=XM_019826882.2;refGene_gn=CFAP61;refGene_tx=XM_019826883.2;refGene_gn=CFAP61;refGene_tx=XM_023251503.1
A3 45532139 45532314
refGene_gn=CFAP61;refGene_tx=XM_011280983.3;refGene_gn=CFAP61;refGene_tx=XM_019826881.2;refGene_gn=CFAP61;refGene_tx=XM_019826882.2;refGene_gn=CFAP61;refGene_tx=XM_019826883.2;refGene_gn=CFAP61;refGene_tx=XM_023251503.1
A3 45543955 45544248
refGene_gn=CFAP61;refGene_tx=XM_011280983.3;refGene_gn=CFAP61;refGene_tx=XM_019826881.2;refGene_gn=CFAP61;refGene_tx=XM_019826882.2;refGene_gn=CFAP61;refGene_tx=XM_019826883.2;refGene_gn=CFAP61;refGene_tx=XM_023251503.1
A3 45551220 45551548
refGene_gn=CFAP61;refGene_tx=XM_011280983.3;refGene_gn=CFAP61;refGene_

tx=XM_019826881.2;refGene_gn=CFAP61;refGene_tx=XM_019826882.2;refGene_gn=CFAP61;refGene_tx=XM_019826883.2;refGene_gn=CFAP61;refGene_tx=XM_023251503.1
A3 45552909 45553001 refGene_gn=CFAP61;refGene_tx=XM_011280983.3;refGene_gn=CFAP61;refGene_tx=XM_019826881.2;refGene_gn=CFAP61;refGene_tx=XM_019826882.2;refGene_gn=CFAP61;refGene_tx=XM_019826883.2;refGene_gn=CFAP61;refGene_tx=XM_023251503.1
A3 45560760 45560966 refGene_gn=CFAP61;refGene_tx=XM_011280983.3;refGene_gn=CFAP61;refGene_tx=XM_019826881.2;refGene_gn=CFAP61;refGene_tx=XM_019826882.2;refGene_gn=CFAP61;refGene_tx=XM_019826883.2;refGene_gn=CFAP61;refGene_tx=XM_023251503.1
A3 45607687 45607778 refGene_gn=CFAP61;refGene_tx=XM_011280983.3;refGene_gn=CFAP61;refGene_tx=XM_019826881.2;refGene_gn=CFAP61;refGene_tx=XM_019826882.2;refGene_gn=CFAP61;refGene_tx=XM_019826883.2;refGene_gn=CFAP61;refGene_tx=XM_023251503.1
A3 45616991 45617192 refGene_gn=CFAP61;refGene_tx=XM_011280983.3;refGene_gn=CFAP61;refGene_tx=XM_019826881.2;refGene_gn=CFAP61;refGene_tx=XM_019826882.2;refGene_gn=CFAP61;refGene_tx=XM_019826883.2;refGene_gn=CFAP61;refGene_tx=XM_023251503.1
A3 45624892 45626419 refGene_gn=INSM1;refGene_tx=XM_023251504.1
A3 45650386 45650391 refGene_gn=RALGAPA2;refGene_tx=XM_006929996.4;refGene_gn=RALGAPA2;refGene_tx=XM_019826890.2;refGene_gn=RALGAPA2;refGene_tx=XM_019826891.2;refGene_gn=RALGAPA2;refGene_tx=XM_019826892.2;refGene_gn=RALGAPA2;refGene_tx=XM_019826893.2
A3 45666495 45666617 refGene_gn=RALGAPA2;refGene_tx=XM_006929996.4;refGene_gn=RALGAPA2;refGene_tx=XM_019826890.2;refGene_gn=RALGAPA2;refGene_tx=XM_019826891.2;refGene_gn=RALGAPA2;refGene_tx=XM_019826892.2;refGene_gn=RALGAPA2;refGene_tx=XM_019826893.2
A3 45730368 45730496 refGene_gn=RALGAPA2;refGene_tx=XM_006929996.4;refGene_gn=RALGAPA2;refGene_tx=XM_019826890.2;refGene_gn=RALGAPA2;refGene_tx=XM_019826891.2;refGene_gn=RALGAPA2;refGene_tx=XM_019826892.2;refGene_gn=RALGAPA2;refGene_tx=XM_019826893.2
A3 45749710 45749869 refGene_gn=RALGAPA2;refGene_tx=XM_006929996.4;refGene_gn=RALGAPA2;refGene_tx=XM_019826890.2;refGene_gn=RALGAPA2;refGene_tx=XM_019826891.2;refGene_gn=RALGAPA2;refGene_tx=XM_019826892.2;refGene_gn=RALGAPA2;refGene_tx=XM_019826893.2
A3 45755904 45756060 refGene_gn=RALGAPA2;refGene_tx=XM_006929996.4;refGene_gn=RALGAPA2;refGene_tx=XM_019826890.2;refGene_gn=RALGAPA2;refGene_tx=XM_019826891.2;refGene_gn=RALGAPA2;refGene_tx=XM_019826892.2;refGene_gn=RALGAPA2;refGene_tx=XM_019826893.2
A3 45757979 45758103

refGene_gn=RALGAPA2;refGene_tx=XM_006929996.4;refGene_gn=RALGAPA2;refGene_tx=XM_019826890.2;refGene_gn=RALGAPA2;refGene_tx=XM_019826891.2;refGene_gn=RALGAPA2;refGene_tx=XM_019826892.2;refGene_gn=RALGAPA2;refGene_tx=XM_019826893.2
A3 45760765 45760817 refGene_gn=RALGAPA2;refGene_tx=XM_019826894.2
A3 45762461 45762533
refGene_gn=RALGAPA2;refGene_tx=XM_006929996.4;refGene_gn=RALGAPA2;refGene_tx=XM_019826890.2;refGene_gn=RALGAPA2;refGene_tx=XM_019826891.2;refGene_gn=RALGAPA2;refGene_tx=XM_019826892.2;refGene_gn=RALGAPA2;refGene_tx=XM_019826893.2;refGene_gn=RALGAPA2;refGene_tx=XM_019826894.2
A3 45763392 45764164
refGene_gn=RALGAPA2;refGene_tx=XM_006929996.4;refGene_gn=RALGAPA2;refGene_tx=XM_019826890.2;refGene_gn=RALGAPA2;refGene_tx=XM_019826891.2;refGene_gn=RALGAPA2;refGene_tx=XM_019826892.2;refGene_gn=RALGAPA2;refGene_tx=XM_019826893.2;refGene_gn=RALGAPA2;refGene_tx=XM_019826894.2
A3 45769329 45769513
refGene_gn=RALGAPA2;refGene_tx=XM_006929996.4;refGene_gn=RALGAPA2;refGene_tx=XM_019826890.2;refGene_gn=RALGAPA2;refGene_tx=XM_019826891.2;refGene_gn=RALGAPA2;refGene_tx=XM_019826892.2;refGene_gn=RALGAPA2;refGene_tx=XM_019826893.2;refGene_gn=RALGAPA2;refGene_tx=XM_019826894.2
A3 45771522 45771660
refGene_gn=RALGAPA2;refGene_tx=XM_006929996.4;refGene_gn=RALGAPA2;refGene_tx=XM_019826890.2;refGene_gn=RALGAPA2;refGene_tx=XM_019826891.2;refGene_gn=RALGAPA2;refGene_tx=XM_019826892.2;refGene_gn=RALGAPA2;refGene_tx=XM_019826893.2;refGene_gn=RALGAPA2;refGene_tx=XM_019826894.2
A3 45771939 45772008
refGene_gn=RALGAPA2;refGene_tx=XM_006929996.4;refGene_gn=RALGAPA2;refGene_tx=XM_019826890.2;refGene_gn=RALGAPA2;refGene_tx=XM_019826891.2;refGene_gn=RALGAPA2;refGene_tx=XM_019826892.2;refGene_gn=RALGAPA2;refGene_tx=XM_019826893.2;refGene_gn=RALGAPA2;refGene_tx=XM_019826894.2
A3 45773893 45774004
refGene_gn=RALGAPA2;refGene_tx=XM_006929996.4;refGene_gn=RALGAPA2;refGene_tx=XM_019826890.2;refGene_gn=RALGAPA2;refGene_tx=XM_019826891.2;refGene_gn=RALGAPA2;refGene_tx=XM_019826892.2;refGene_gn=RALGAPA2;refGene_tx=XM_019826893.2;refGene_gn=RALGAPA2;refGene_tx=XM_019826894.2
A3 45780058 45780167
refGene_gn=RALGAPA2;refGene_tx=XM_006929996.4;refGene_gn=RALGAPA2;refGene_tx=XM_019826890.2;refGene_gn=RALGAPA2;refGene_tx=XM_019826891.2;refGene_gn=RALGAPA2;refGene_tx=XM_019826892.2;refGene_gn=RALGAPA2;refGene_tx=XM_019826893.2;refGene_gn=RALGAPA2;refGene_tx=XM_019826894.2
A3 45787472 45787531
refGene_gn=RALGAPA2;refGene_tx=XM_006929996.4;refGene_gn=RALGAPA2;refGene_tx=XM_019826890.2;refGene_gn=RALGAPA2;refGene_tx=XM_019826891.2;refGene_gn=RALGAPA2;refGene_tx=XM_019826892.2;refGene_gn=RALGAPA2;refGene_tx=XM_019826893.2;refGene_gn=RALGAPA2;refGene_tx=XM_019826894.2
A3 45788371 45788500
refGene_gn=RALGAPA2;refGene_tx=XM_006929996.4;refGene_gn=RALGAPA2;refGene_tx=XM_019826890.2;refGene_gn=RALGAPA2;refGene_tx=XM_019826891.2;refGene_gn=RALGAPA2;refGene_tx=XM_019826892.2;refGene_gn=RALGAPA2;refGene_tx=XM_019826893.2;refGene_gn=RALGAPA2;refGene_tx=XM_019826894.2

A3	45794413 45794542
refGene_gn=RALGAPA2;refGene_tx=XM_006929996.4;refGene_gn=RALGAPA2;refGene_tx=XM_019826890.2;refGene_gn=RALGAPA2;refGene_tx=XM_019826891.2;refGene_gn=RALGAPA2;refGene_tx=XM_019826892.2;refGene_gn=RALGAPA2;refGene_tx=XM_019826893.2;refGene_gn=RALGAPA2;refGene_tx=XM_019826894.2
A3	45819102 45819258
refGene_gn=RALGAPA2;refGene_tx=XM_006929996.4;refGene_gn=RALGAPA2;refGene_tx=XM_019826890.2;refGene_gn=RALGAPA2;refGene_tx=XM_019826891.2;refGene_gn=RALGAPA2;refGene_tx=XM_019826892.2;refGene_gn=RALGAPA2;refGene_tx=XM_019826893.2;refGene_gn=RALGAPA2;refGene_tx=XM_019826894.2
A3	45819496 45819595
refGene_gn=RALGAPA2;refGene_tx=XM_006929996.4;refGene_gn=RALGAPA2;refGene_tx=XM_019826890.2;refGene_gn=RALGAPA2;refGene_tx=XM_019826891.2;refGene_gn=RALGAPA2;refGene_tx=XM_019826892.2;refGene_gn=RALGAPA2;refGene_tx=XM_019826893.2;refGene_gn=RALGAPA2;refGene_tx=XM_019826894.2
A3	45820451 45820645
refGene_gn=RALGAPA2;refGene_tx=XM_006929996.4;refGene_gn=RALGAPA2;refGene_tx=XM_019826890.2;refGene_gn=RALGAPA2;refGene_tx=XM_019826891.2;refGene_gn=RALGAPA2;refGene_tx=XM_019826892.2;refGene_gn=RALGAPA2;refGene_tx=XM_019826893.2;refGene_gn=RALGAPA2;refGene_tx=XM_019826894.2
A3	45830877 45831078
refGene_gn=RALGAPA2;refGene_tx=XM_006929996.4;refGene_gn=RALGAPA2;refGene_tx=XM_019826890.2;refGene_gn=RALGAPA2;refGene_tx=XM_019826891.2;refGene_gn=RALGAPA2;refGene_tx=XM_019826892.2;refGene_gn=RALGAPA2;refGene_tx=XM_019826893.2;refGene_gn=RALGAPA2;refGene_tx=XM_019826894.2
A3	45832471 45832562
refGene_gn=RALGAPA2;refGene_tx=XM_006929996.4;refGene_gn=RALGAPA2;refGene_tx=XM_019826890.2;refGene_gn=RALGAPA2;refGene_tx=XM_019826891.2;refGene_gn=RALGAPA2;refGene_tx=XM_019826892.2;refGene_gn=RALGAPA2;refGene_tx=XM_019826893.2;refGene_gn=RALGAPA2;refGene_tx=XM_019826894.2
A3	45837410 45837508
refGene_gn=RALGAPA2;refGene_tx=XM_006929996.4;refGene_gn=RALGAPA2;refGene_tx=XM_019826890.2;refGene_gn=RALGAPA2;refGene_tx=XM_019826891.2;refGene_gn=RALGAPA2;refGene_tx=XM_019826892.2;refGene_gn=RALGAPA2;refGene_tx=XM_019826893.2;refGene_gn=RALGAPA2;refGene_tx=XM_019826894.2
A3	45839261 45839393
refGene_gn=RALGAPA2;refGene_tx=XM_006929996.4;refGene_gn=RALGAPA2;refGene_tx=XM_019826890.2;refGene_gn=RALGAPA2;refGene_tx=XM_019826891.2;refGene_gn=RALGAPA2;refGene_tx=XM_019826892.2;refGene_gn=RALGAPA2;refGene_tx=XM_019826893.2;refGene_gn=RALGAPA2;refGene_tx=XM_019826894.2
A3	45845359 45845500
refGene_gn=RALGAPA2;refGene_tx=XM_019826890.2;refGene_gn=RALGAPA2;refGene_tx=XM_019826891.2;refGene_gn=RALGAPA2;refGene_tx=XM_019826892.2;refGene_gn=RALGAPA2;refGene_tx=XM_019826894.2
A3	45849297 45849462
refGene_gn=RALGAPA2;refGene_tx=XM_006929996.4;refGene_gn=RALGAPA2;refGene_tx=XM_019826890.2;refGene_gn=RALGAPA2;refGene_tx=XM_019826891.2;refGene_gn=RALGAPA2;refGene_tx=XM_019826892.2;refGene_gn=RALGAPA2;refGene_tx=XM_019826893.2;refGene_gn=RALGAPA2;refGene_tx=XM_019826894.2
A3	45851892 45852130

refGene_gn=RALGAPA2;refGene_tx=XM_006929996.4;refGene_gn=RALGAPA2;refGene_tx=XM_019826890.2;refGene_gn=RALGAPA2;refGene_tx=XM_019826891.2;refGene_gn=RALGAPA2;refGene_tx=XM_019826892.2;refGene_gn=RALGAPA2;refGene_tx=XM_019826893.2;refGene_gn=RALGAPA2;refGene_tx=XM_019826894.2
A3 45858446 45858558
refGene_gn=RALGAPA2;refGene_tx=XM_006929996.4;refGene_gn=RALGAPA2;refGene_tx=XM_019826890.2;refGene_gn=RALGAPA2;refGene_tx=XM_019826891.2;refGene_gn=RALGAPA2;refGene_tx=XM_019826892.2;refGene_gn=RALGAPA2;refGene_tx=XM_019826893.2;refGene_gn=RALGAPA2;refGene_tx=XM_019826894.2
A3 45862625 45862774
refGene_gn=RALGAPA2;refGene_tx=XM_006929996.4;refGene_gn=RALGAPA2;refGene_tx=XM_019826890.2;refGene_gn=RALGAPA2;refGene_tx=XM_019826891.2;refGene_gn=RALGAPA2;refGene_tx=XM_019826892.2;refGene_gn=RALGAPA2;refGene_tx=XM_019826893.2;refGene_gn=RALGAPA2;refGene_tx=XM_019826894.2
A3 45865929 45866067
refGene_gn=RALGAPA2;refGene_tx=XM_006929996.4;refGene_gn=RALGAPA2;refGene_tx=XM_019826890.2;refGene_gn=RALGAPA2;refGene_tx=XM_019826891.2;refGene_gn=RALGAPA2;refGene_tx=XM_019826892.2;refGene_gn=RALGAPA2;refGene_tx=XM_019826893.2;refGene_gn=RALGAPA2;refGene_tx=XM_019826894.2
A3 45867389 45867557
refGene_gn=RALGAPA2;refGene_tx=XM_006929996.4;refGene_gn=RALGAPA2;refGene_tx=XM_019826890.2;refGene_gn=RALGAPA2;refGene_tx=XM_019826891.2;refGene_gn=RALGAPA2;refGene_tx=XM_019826892.2;refGene_gn=RALGAPA2;refGene_tx=XM_019826893.2;refGene_gn=RALGAPA2;refGene_tx=XM_019826894.2
A3 45882935 45883163
refGene_gn=RALGAPA2;refGene_tx=XM_006929996.4;refGene_gn=RALGAPA2;refGene_tx=XM_019826890.2;refGene_gn=RALGAPA2;refGene_tx=XM_019826891.2;refGene_gn=RALGAPA2;refGene_tx=XM_019826892.2;refGene_gn=RALGAPA2;refGene_tx=XM_019826893.2;refGene_gn=RALGAPA2;refGene_tx=XM_019826894.2
A3 45891670 45891870
refGene_gn=RALGAPA2;refGene_tx=XM_006929996.4;refGene_gn=RALGAPA2;refGene_tx=XM_019826890.2;refGene_gn=RALGAPA2;refGene_tx=XM_019826891.2;refGene_gn=RALGAPA2;refGene_tx=XM_019826892.2;refGene_gn=RALGAPA2;refGene_tx=XM_019826893.2;refGene_gn=RALGAPA2;refGene_tx=XM_019826894.2
A3 45894007 45894146
refGene_gn=RALGAPA2;refGene_tx=XM_006929996.4;refGene_gn=RALGAPA2;refGene_tx=XM_019826890.2;refGene_gn=RALGAPA2;refGene_tx=XM_019826891.2;refGene_gn=RALGAPA2;refGene_tx=XM_019826892.2;refGene_gn=RALGAPA2;refGene_tx=XM_019826893.2;refGene_gn=RALGAPA2;refGene_tx=XM_019826894.2
A3 45897415 45897531
refGene_gn=RALGAPA2;refGene_tx=XM_006929996.4;refGene_gn=RALGAPA2;refGene_tx=XM_019826890.2;refGene_gn=RALGAPA2;refGene_tx=XM_019826891.2;refGene_gn=RALGAPA2;refGene_tx=XM_019826892.2;refGene_gn=RALGAPA2;refGene_tx=XM_019826893.2;refGene_gn=RALGAPA2;refGene_tx=XM_019826894.2
A3 45898335 45898513
refGene_gn=RALGAPA2;refGene_tx=XM_006929996.4;refGene_gn=RALGAPA2;refGene_tx=XM_019826890.2;refGene_gn=RALGAPA2;refGene_tx=XM_019826891.2;refGene_gn=RALGAPA2;refGene_tx=XM_019826892.2;refGene_gn=RALGAPA2;refGene_tx=XM_019826893.2;refGene_gn=RALGAPA2;refGene_tx=XM_019826894.2
A3 45901036 45901080

refGene_gn=RALGAPA2;refGene_tx=XM_006929996.4;refGene_gn=RALGAPA2;refGene_tx=XM_019826890.2;refGene_gn=RALGAPA2;refGene_tx=XM_019826891.2;refGene_gn=RALGAPA2;refGene_tx=XM_019826892.2;refGene_gn=RALGAPA2;refGene_tx=XM_019826893.2;refGene_gn=RALGAPA2;refGene_tx=XM_019826894.2
A3 45910926 45910984
refGene_gn=RALGAPA2;refGene_tx=XM_006929996.4;refGene_gn=RALGAPA2;refGene_tx=XM_019826890.2;refGene_gn=RALGAPA2;refGene_tx=XM_019826891.2;refGene_gn=RALGAPA2;refGene_tx=XM_019826892.2;refGene_gn=RALGAPA2;refGene_tx=XM_019826893.2;refGene_gn=RALGAPA2;refGene_tx=XM_019826894.2
A3 45933719 45933772
refGene_gn=RALGAPA2;refGene_tx=XM_006929996.4;refGene_gn=RALGAPA2;refGene_tx=XM_019826890.2;refGene_gn=RALGAPA2;refGene_tx=XM_019826891.2;refGene_gn=RALGAPA2;refGene_tx=XM_019826892.2;refGene_gn=RALGAPA2;refGene_tx=XM_019826893.2;refGene_gn=RALGAPA2;refGene_tx=XM_019826894.2
A3 45938980 45939091
refGene_gn=RALGAPA2;refGene_tx=XM_006929996.4;refGene_gn=RALGAPA2;refGene_tx=XM_019826890.2;refGene_gn=RALGAPA2;refGene_tx=XM_019826891.2;refGene_gn=RALGAPA2;refGene_tx=XM_019826892.2;refGene_gn=RALGAPA2;refGene_tx=XM_019826893.2;refGene_gn=RALGAPA2;refGene_tx=XM_019826894.2
A3 45968454 45968560
refGene_gn=RALGAPA2;refGene_tx=XM_006929996.4;refGene_gn=RALGAPA2;refGene_tx=XM_019826890.2;refGene_gn=RALGAPA2;refGene_tx=XM_019826891.2;refGene_gn=RALGAPA2;refGene_tx=XM_019826892.2;refGene_gn=RALGAPA2;refGene_tx=XM_019826893.2;refGene_gn=RALGAPA2;refGene_tx=XM_019826894.2
A3 46417491 46417647
refGene_gn=KIZ;refGene_tx=XM_023251505.1;refGene_gn=KIZ;refGene_tx=XM_023251506.1
A3 46426484 46426577
refGene_gn=KIZ;refGene_tx=XM_023251505.1;refGene_gn=KIZ;refGene_tx=XM_023251506.1
A3 46463302 46463948
refGene_gn=KIZ;refGene_tx=XM_023251505.1;refGene_gn=KIZ;refGene_tx=XM_023251506.1
A3 46464334 46464653
refGene_gn=KIZ;refGene_tx=XM_023251505.1;refGene_gn=KIZ;refGene_tx=XM_023251506.1
A3 46505667 46505788
refGene_gn=KIZ;refGene_tx=XM_023251505.1;refGene_gn=KIZ;refGene_tx=XM_023251506.1
A3 46517344 46517510
refGene_gn=KIZ;refGene_tx=XM_023251505.1;refGene_gn=KIZ;refGene_tx=XM_023251506.1
A3 46518384 46518450
refGene_gn=KIZ;refGene_tx=XM_023251505.1;refGene_gn=KIZ;refGene_tx=XM_023251506.1
A3 46532474 46532579
refGene_gn=KIZ;refGene_tx=XM_023251505.1;refGene_gn=KIZ;refGene_tx=XM_023251506.1
A3 46535472 46535569
refGene_gn=KIZ;refGene_tx=XM_023251505.1;refGene_gn=KIZ;refGene_tx=XM_

023251506.1
A3	46545800	46545844
refGene_gn=KIZ;refGene_tx=XM_023251505.1;refGene_gn=KIZ;refGene_tx=XM_
023251506.1
A3	46548216	46548320
refGene_gn=KIZ;refGene_tx=XM_023251505.1;refGene_gn=KIZ;refGene_tx=XM_
023251506.1
A3	46597989	46598064	refGene_gn=XRN2;refGene_tx=XM_011280982.3
A3	46612331	46612459	refGene_gn=XRN2;refGene_tx=XM_011280982.3
A3	46612541	46612653	refGene_gn=XRN2;refGene_tx=XM_011280982.3
A3	46615019	46615131	refGene_gn=XRN2;refGene_tx=XM_011280982.3
A3	46616662	46616721	refGene_gn=XRN2;refGene_tx=XM_011280982.3
A3	46616797	46616887	refGene_gn=XRN2;refGene_tx=XM_011280982.3
A3	46617739	46617812	refGene_gn=XRN2;refGene_tx=XM_011280982.3
A3	46617940	46617991	refGene_gn=XRN2;refGene_tx=XM_011280982.3
A3	46618450	46618608	refGene_gn=XRN2;refGene_tx=XM_011280982.3
A3	46619620	46619695	refGene_gn=XRN2;refGene_tx=XM_011280982.3
A3	46619781	46619915	refGene_gn=XRN2;refGene_tx=XM_011280982.3
A3	46620014	46620072	refGene_gn=XRN2;refGene_tx=XM_011280982.3
A3	46620155	46620263	refGene_gn=XRN2;refGene_tx=XM_011280982.3
A3	46624387	46624432	refGene_gn=XRN2;refGene_tx=XM_011280982.3
A3	46626053	46626185	refGene_gn=XRN2;refGene_tx=XM_011280982.3
A3	46629481	46629600	refGene_gn=XRN2;refGene_tx=XM_011280982.3
A3	46631681	46631817	refGene_gn=XRN2;refGene_tx=XM_011280982.3
A3	46633639	46633747	refGene_gn=XRN2;refGene_tx=XM_011280982.3
A3	46633840	46633930	refGene_gn=XRN2;refGene_tx=XM_011280982.3
A3	46634707	46634780	refGene_gn=XRN2;refGene_tx=XM_011280982.3
A3	46643785	46643869	refGene_gn=XRN2;refGene_tx=XM_011280982.3
A3	46644969	46645067	refGene_gn=XRN2;refGene_tx=XM_011280982.3
A3	46645673	46645753	refGene_gn=XRN2;refGene_tx=XM_011280982.3
A3	46646957	46647014	refGene_gn=XRN2;refGene_tx=XM_011280982.3
A3	46654988	46655057	refGene_gn=XRN2;refGene_tx=XM_011280982.3
A3	46655154	46655286	refGene_gn=XRN2;refGene_tx=XM_011280982.3
A3	46656314	46656442	refGene_gn=XRN2;refGene_tx=XM_011280982.3
A3	46668079	46668143	refGene_gn=XRN2;refGene_tx=XM_011280982.3
A3	46673196	46673335	refGene_gn=XRN2;refGene_tx=XM_011280982.3
A3	46676024	46676090	refGene_gn=XRN2;refGene_tx=XM_011280982.3
A3	46682891	46683508	refGene_gn=NKX2-4;refGene_tx=XM_023250915.1
A3	46683901	46684352	refGene_gn=NKX2-4;refGene_tx=XM_023250915.1
A3	46782678	46783241
refGene_gn=NKX2-2;refGene_tx=XM_023251508.1;refGene_gn=NKX2-2;refGene_
tx=XM_023251509.1
A3	46784192	46784451
refGene_gn=NKX2-2;refGene_tx=XM_023251508.1;refGene_gn=NKX2-2;refGene_
tx=XM_023251509.1
A3	46957128	46957381	refGene_gn=PAX1;refGene_tx=XM_003983829.4
A3	46957842	46958472	refGene_gn=PAX1;refGene_tx=XM_003983829.4
A3	46960083	46960226	refGene_gn=PAX1;refGene_tx=XM_003983829.4
A3	46960751	46960974	refGene_gn=PAX1;refGene_tx=XM_003983829.4
A3	46965835	46965927	refGene_gn=PAX1;refGene_tx=XM_003983829.4

A3 47719767 47721072
refGene_gn=FOXA2;refGene_tx=XM_003983832.4;refGene_gn=FOXA2;refGene_tx=XM_019826897.2
A3 47722074 47722161
refGene_gn=FOXA2;refGene_tx=XM_003983832.4;refGene_gn=FOXA2;refGene_tx=XM_019826897.2
A3 48110496 48111660 refGene_gn=SSTR4;refGene_tx=XM_003983838.5
A3 48125270 48127289 refGene_gn=THBD;refGene_tx=XM_006929999.4
A3 48136480 48136526 refGene_gn=CD93;refGene_tx=XM_011280979.3
A3 48162648 48162673 refGene_gn=CD93;refGene_tx=XM_011280981.3
A3 48162922 48164844
refGene_gn=CD93;refGene_tx=XM_011280979.3;refGene_gn=CD93;refGene_tx=XM_011280981.3
A3 48436804 48437227 refGene_gn=NXT1;refGene_tx=XM_003983834.5
A3 48460225 48461601
refGene_gn=GZF1;refGene_tx=XM_023251511.1;refGene_gn=GZF1;refGene_tx=XM_023251512.1
A3 48462961 48463056
refGene_gn=GZF1;refGene_tx=XM_023251511.1;refGene_gn=GZF1;refGene_tx=XM_023251512.1
A3 48464643 48464811
refGene_gn=GZF1;refGene_tx=XM_023251511.1;refGene_gn=GZF1;refGene_tx=XM_023251512.1
A3 48465563 48465721
refGene_gn=GZF1;refGene_tx=XM_023251511.1;refGene_gn=GZF1;refGene_tx=XM_023251512.1
A3 48466268 48466619
refGene_gn=GZF1;refGene_tx=XM_023251511.1;refGene_gn=GZF1;refGene_tx=XM_023251512.1
A3 48473106 48473217
refGene_gn=NAPB;refGene_tx=XM_003983836.5;refGene_gn=NAPB;refGene_tx=XM_011280978.3;refGene_gn=NAPB;refGene_tx=XM_019826898.2;refGene_gn=NAPB;refGene_tx=XM_023251513.1;refGene_gn=NAPB;refGene_tx=XM_023251514.1
A3 48474211 48474262
refGene_gn=NAPB;refGene_tx=XM_003983836.5;refGene_gn=NAPB;refGene_tx=XM_011280978.3;refGene_gn=NAPB;refGene_tx=XM_019826898.2;refGene_gn=NAPB;refGene_tx=XM_023251513.1;refGene_gn=NAPB;refGene_tx=XM_023251514.1;refGene_gn=NAPB;refGene_tx=XM_023251515.1
A3 48474632 48474701
refGene_gn=NAPB;refGene_tx=XM_003983836.5;refGene_gn=NAPB;refGene_tx=XM_011280978.3;refGene_gn=NAPB;refGene_tx=XM_019826898.2;refGene_gn=NAPB;refGene_tx=XM_023251513.1;refGene_gn=NAPB;refGene_tx=XM_023251514.1;refGene_gn=NAPB;refGene_tx=XM_023251515.1
A3 48475678 48475783
refGene_gn=NAPB;refGene_tx=XM_003983836.5;refGene_gn=NAPB;refGene_tx=XM_011280978.3;refGene_gn=NAPB;refGene_tx=XM_019826898.2;refGene_gn=NAPB;refGene_tx=XM_023251513.1;refGene_gn=NAPB;refGene_tx=XM_023251515.1
A3 48482971 48483056
refGene_gn=NAPB;refGene_tx=XM_003983836.5;refGene_gn=NAPB;refGene_tx=XM_011280978.3;refGene_gn=NAPB;refGene_tx=XM_019826898.2;refGene_gn=NAP

B;refGene_tx=XM_023251513.1;refGene_gn=NAPB;refGene_tx=XM_023251514.1;
refGene_gn=NAPB;refGene_tx=XM_023251515.1
A3 48483228 48483284
refGene_gn=NAPB;refGene_tx=XM_003983836.5;refGene_gn=NAPB;refGene_tx=XM_011280978.3;refGene_gn=NAPB;refGene_tx=XM_019826898.2;refGene_gn=NAPB;refGene_tx=XM_023251513.1;refGene_gn=NAPB;refGene_tx=XM_023251514.1;refGene_gn=NAPB;refGene_tx=XM_023251515.1
A3 48490582 48490660
refGene_gn=NAPB;refGene_tx=XM_003983836.5;refGene_gn=NAPB;refGene_tx=XM_011280978.3;refGene_gn=NAPB;refGene_tx=XM_019826898.2;refGene_gn=NAPB;refGene_tx=XM_023251513.1;refGene_gn=NAPB;refGene_tx=XM_023251514.1;refGene_gn=NAPB;refGene_tx=XM_023251515.1
A3 48490810 48490857
refGene_gn=NAPB;refGene_tx=XM_003983836.5;refGene_gn=NAPB;refGene_tx=XM_019826898.2;refGene_gn=NAPB;refGene_tx=XM_023251513.1;refGene_gn=NAPB;refGene_tx=XM_023251514.1;refGene_gn=NAPB;refGene_tx=XM_023251515.1
A3 48492689 48492806
refGene_gn=NAPB;refGene_tx=XM_003983836.5;refGene_gn=NAPB;refGene_tx=XM_011280978.3;refGene_gn=NAPB;refGene_tx=XM_019826898.2;refGene_gn=NAPB;refGene_tx=XM_023251513.1;refGene_gn=NAPB;refGene_tx=XM_023251514.1;refGene_gn=NAPB;refGene_tx=XM_023251515.1
A3 48495457 48495537
refGene_gn=NAPB;refGene_tx=XM_003983836.5;refGene_gn=NAPB;refGene_tx=XM_019826898.2;refGene_gn=NAPB;refGene_tx=XM_023251513.1;refGene_gn=NAPB;refGene_tx=XM_023251514.1;refGene_gn=NAPB;refGene_tx=XM_023251515.1
A3 48500106 48500258
refGene_gn=NAPB;refGene_tx=XM_023251513.1;refGene_gn=NAPB;refGene_tx=XM_023251514.1;refGene_gn=NAPB;refGene_tx=XM_023251515.1
A3 48505385 48505423 refGene_gn=NAPB;refGene_tx=XM_019826898.2
A3 48511555 48511653 refGene_gn=NAPB;refGene_tx=XM_003983836.5
A3 48527459 48527678
refGene_gn=CSTL1;refGene_tx=XM_003983840.4;refGene_gn=CSTL1;refGene_tx=XM_019826900.2
A3 48530757 48530871
refGene_gn=CSTL1;refGene_tx=XM_003983840.4;refGene_gn=CSTL1;refGene_tx=XM_019826900.2
A3 48531600 48531684
refGene_gn=CSTL1;refGene_tx=XM_003983840.4;refGene_gn=CSTL1;refGene_tx=XM_019826900.2
A3 48539412 48539496
refGene_gn=CST11;refGene_tx=XM_003983841.3;refGene_gn=CST11;refGene_tx=XM_019826901.2
A3 48541040 48541145 refGene_gn=CST11;refGene_tx=XM_003983841.3
A3 48541770 48541998
refGene_gn=CST11;refGene_tx=XM_003983841.3;refGene_gn=CST11;refGene_tx=XM_019826901.2
A3 48589021 48589252 refGene_gn=CST8;refGene_tx=XM_006930002.4
A3 48592710 48592824 refGene_gn=CST8;refGene_tx=XM_006930002.4
A3 48599236 48599320 refGene_gn=CST8;refGene_tx=XM_006930002.4
A3 48632300 48632528

refGene_gn=LOC101088321;refGene_tx=XM_003983843.5;refGene_gn=LOC101088321;refGene_tx=XM_019826902.2
A3 48638134 48638248
refGene_gn=LOC101088321;refGene_tx=XM_003983843.5;refGene_gn=LOC101088321;refGene_tx=XM_019826902.2
A3 48641980 48642064
refGene_gn=LOC101088321;refGene_tx=XM_003983843.5;refGene_gn=LOC101088321;refGene_tx=XM_019826902.2
A3 48650611 48650704
refGene_gn=LOC101089343;refGene_tx=XM_019828130.2
A3 48651588 48651708
refGene_gn=LOC101089343;refGene_tx=XM_019828130.2
A3 48653446 48653686
refGene_gn=LOC101089343;refGene_tx=XM_019828130.2
A3 48703677 48703761 refGene_gn=CST3;refGene_tx=NM_001184972.1
A3 48704588 48704702 refGene_gn=CST3;refGene_tx=NM_001184972.1
A3 48707299 48707545 refGene_gn=CST3;refGene_tx=NM_001184972.1
A3 48786201 48786285
refGene_gn=LOC101088567;refGene_tx=XM_003983844.3
A3 48787160 48787274
refGene_gn=LOC101088567;refGene_tx=XM_003983844.3
A3 48789914 48790136
refGene_gn=LOC101088567;refGene_tx=XM_003983844.3
A3 49474887 49475367
refGene_gn=SYNDIG1;refGene_tx=XM_019826904.1;refGene_gn=SYNDIG1;refGene_tx=XM_019826907.2;refGene_gn=SYNDIG1;refGene_tx=XM_019826909.2;refGene_gn=SYNDIG1;refGene_tx=XM_019826911.2;refGene_gn=SYNDIG1;refGene_tx=XM_023251516.1;refGene_gn=SYNDIG1;refGene_tx=XM_023251517.1;refGene_gn=SYNDIG1;refGene_tx=XM_023251518.1;refGene_gn=SYNDIG1;refGene_tx=XM_023251519.1;refGene_gn=SYNDIG1;refGene_tx=XM_023251520.1;refGene_gn=SYNDIG1;refGene_tx=XM_023251521.1;refGene_gn=SYNDIG1;refGene_tx=XM_023251522.1;refGene_gn=SYNDIG1;refGene_tx=XM_023251523.1
A3 49506658 49506796
refGene_gn=SYNDIG1;refGene_tx=XM_019826904.1;refGene_gn=SYNDIG1;refGene_tx=XM_019826907.2;refGene_gn=SYNDIG1;refGene_tx=XM_019826909.2;refGene_gn=SYNDIG1;refGene_tx=XM_019826911.2;refGene_gn=SYNDIG1;refGene_tx=XM_023251516.1;refGene_gn=SYNDIG1;refGene_tx=XM_023251517.1;refGene_gn=SYNDIG1;refGene_tx=XM_023251518.1;refGene_gn=SYNDIG1;refGene_tx=XM_023251519.1;refGene_gn=SYNDIG1;refGene_tx=XM_023251520.1;refGene_gn=SYNDIG1;refGene_tx=XM_023251521.1;refGene_gn=SYNDIG1;refGene_tx=XM_023251523.1
A3 49547378 49547505
refGene_gn=SYNDIG1;refGene_tx=XM_019826904.1;refGene_gn=SYNDIG1;refGene_tx=XM_019826907.2;refGene_gn=SYNDIG1;refGene_tx=XM_019826909.2;refGene_gn=SYNDIG1;refGene_tx=XM_019826911.2;refGene_gn=SYNDIG1;refGene_tx=XM_023251516.1;refGene_gn=SYNDIG1;refGene_tx=XM_023251517.1;refGene_gn=SYNDIG1;refGene_tx=XM_023251518.1;refGene_gn=SYNDIG1;refGene_tx=XM_023251519.1;refGene_gn=SYNDIG1;refGene_tx=XM_023251520.1
A3 49562877 49562883 refGene_gn=SYNDIG1;refGene_tx=XM_023251523.1
A3 49574110 49574269

refGene_gn=SYNDIG1;refGene_tx=XM_019826904.1;refGene_gn=SYNDIG1;refGene_tx=XM_019826907.2;refGene_gn=SYNDIG1;refGene_tx=XM_019826909.2;refGene_gn=SYNDIG1;refGene_tx=XM_019826911.2;refGene_gn=SYNDIG1;refGene_tx=XM_023251516.1;refGene_gn=SYNDIG1;refGene_tx=XM_023251517.1;refGene_gn=SYNDIG1;refGene_tx=XM_023251518.1;refGene_gn=SYNDIG1;refGene_tx=XM_023251519.1;refGene_gn=SYNDIG1;refGene_tx=XM_023251520.1;refGene_gn=SYNDIG1;refGene_tx=XM_023251521.1;refGene_gn=SYNDIG1;refGene_tx=XM_023251522.1
A3 49891662 49891891 refGene_gn=CST7;refGene_tx=XM_003983848.5;refGene_gn=CST7;refGene_tx=XM_006930004.4;refGene_gn=CST7;refGene_tx=XM_019826917.2;refGene_gn=CST7;refGene_tx=XM_019826919.2;refGene_gn=CST7;refGene_tx=XM_019826920.2
A3 49891994 49892190 refGene_gn=CST7;refGene_tx=XM_019826918.2
A3 49901300 49901494 refGene_gn=CST7;refGene_tx=XM_003983848.5;refGene_gn=CST7;refGene_tx=XM_006930004.4;refGene_gn=CST7;refGene_tx=XM_019826917.2;refGene_gn=CST7;refGene_tx=XM_019826918.2;refGene_gn=CST7;refGene_tx=XM_019826919.2;refGene_gn=CST7;refGene_tx=XM_019826920.2
A3 49903160 49903427 refGene_gn=CST7;refGene_tx=XM_003983848.5;refGene_gn=CST7;refGene_tx=XM_006930004.4;refGene_gn=CST7;refGene_tx=XM_019826917.2;refGene_gn=CST7;refGene_tx=XM_019826918.2;refGene_gn=CST7;refGene_tx=XM_019826919.2;refGene_gn=CST7;refGene_tx=XM_019826920.2
A3 49903667 49903673 refGene_gn=CST7;refGene_tx=XM_019826920.2
A3 49903854 49903932 refGene_gn=CST7;refGene_tx=XM_003983848.5;refGene_gn=CST7;refGene_tx=XM_019826919.2
A3 49909708 49909918 refGene_gn=APMAP;refGene_tx=XM_006930005.4
A3 49915510 49915703 refGene_gn=APMAP;refGene_tx=XM_006930005.4
A3 49916146 49916281 refGene_gn=APMAP;refGene_tx=XM_006930005.4
A3 49916881 49917056 refGene_gn=APMAP;refGene_tx=XM_006930005.4
A3 49918267 49918384 refGene_gn=APMAP;refGene_tx=XM_006930005.4
A3 49921439 49921532 refGene_gn=APMAP;refGene_tx=XM_006930005.4
A3 49927958 49928074 refGene_gn=APMAP;refGene_tx=XM_006930005.4
A3 49933001 49933118 refGene_gn=APMAP;refGene_tx=XM_006930005.4
A3 49944086 49944122 refGene_gn=APMAP;refGene_tx=XM_006930005.4
A3 49944123 49944179 refGene_gn=APMAP;refGene_tx=XM_006930005.4
A3 49958587 49958767 refGene_gn=ACSS1;refGene_tx=XM_003983846.5;refGene_gn=ACSS1;refGene_tx=XM_023251525.1
A3 49965534 49965653 refGene_gn=ACSS1;refGene_tx=XM_003983846.5;refGene_gn=ACSS1;refGene_tx=XM_023251525.1
A3 49968004 49968068 refGene_gn=ACSS1;refGene_tx=XM_003983846.5;refGene_gn=ACSS1;refGene_tx=XM_023251525.1
A3 49968183 49968311 refGene_gn=ACSS1;refGene_tx=XM_003983846.5;refGene_gn=ACSS1;refGene_tx=XM_023251525.1
A3 49969055 49969182

refGene_gn=ACSS1;refGene_tx=XM_003983846.5;refGene_gn=ACSS1;refGene_tx=XM_023251525.1
A3 49969571 49969684 refGene_gn=ACSS1;refGene_tx=XM_003983846.5;refGene_gn=ACSS1;refGene_tx=XM_023251525.1
A3 49970840 49970933 refGene_gn=ACSS1;refGene_tx=XM_003983846.5;refGene_gn=ACSS1;refGene_tx=XM_023251525.1
A3 49974922 49975060 refGene_gn=ACSS1;refGene_tx=XM_003983846.5;refGene_gn=ACSS1;refGene_tx=XM_023251525.1
A3 49976351 49976499 refGene_gn=ACSS1;refGene_tx=XM_003983846.5;refGene_gn=ACSS1;refGene_tx=XM_023251525.1
A3 49980999 49981152 refGene_gn=ACSS1;refGene_tx=XM_003983846.5;refGene_gn=ACSS1;refGene_tx=XM_023251525.1
A3 49981548 49981724 refGene_gn=ACSS1;refGene_tx=XM_003983846.5;refGene_gn=ACSS1;refGene_tx=XM_023251525.1
A3 49988713 49988913 refGene_gn=ACSS1;refGene_tx=XM_003983846.5;refGene_gn=ACSS1;refGene_tx=XM_023251525.1
A3 50007284 50007381 refGene_gn=ACSS1;refGene_tx=XM_003983846.5;refGene_gn=ACSS1;refGene_tx=XM_023251525.1
A3 50017653 50018176 refGene_gn=ACSS1;refGene_tx=XM_003983846.5
A3 50018348 50018568 refGene_gn=ACSS1;refGene_tx=XM_023251525.1
A3 50041630 50041917 refGene_gn=VSX1;refGene_tx=XM_019826923.2
A3 50043346 50043527 refGene_gn=VSX1;refGene_tx=XM_019826923.2
A3 50044632 50044756 refGene_gn=VSX1;refGene_tx=XM_019826923.2
A3 50045197 50045276 refGene_gn=VSX1;refGene_tx=XM_019826923.2
A3 50047030 50047454 refGene_gn=VSX1;refGene_tx=XM_019826923.2
A3 50195981 50196273 refGene_gn=ENTPD6;refGene_tx=XM_003983852.5;refGene_gn=ENTPD6;refGene_tx=XM_006930007.4;refGene_gn=ENTPD6;refGene_tx=XM_006930008.4;refGene_gn=ENTPD6;refGene_tx=XM_006930009.4;refGene_gn=ENTPD6;refGene_tx=XM_011280971.3;refGene_gn=ENTPD6;refGene_tx=XM_019826924.2
A3 50197876 50197953 refGene_gn=ENTPD6;refGene_tx=XM_003983852.5;refGene_gn=ENTPD6;refGene_tx=XM_006930007.4;refGene_gn=ENTPD6;refGene_tx=XM_006930008.4;refGene_gn=ENTPD6;refGene_tx=XM_006930009.4;refGene_gn=ENTPD6;refGene_tx=XM_011280971.3;refGene_gn=ENTPD6;refGene_tx=XM_019826924.2
A3 50200021 50200165 refGene_gn=ENTPD6;refGene_tx=XM_003983852.5;refGene_gn=ENTPD6;refGene_tx=XM_006930007.4;refGene_gn=ENTPD6;refGene_tx=XM_006930008.4;refGene_gn=ENTPD6;refGene_tx=XM_006930009.4;refGene_gn=ENTPD6;refGene_tx=XM_011280971.3;refGene_gn=ENTPD6;refGene_tx=XM_019826924.2
A3 50201326 50201402 refGene_gn=ENTPD6;refGene_tx=XM_003983852.5;refGene_gn=ENTPD6;refGene_

tx=XM_006930007.4;refGene_gn=ENTPD6;refGene_tx=XM_006930008.4;refGene_gn=ENTPD6;refGene_tx=XM_006930009.4;refGene_gn=ENTPD6;refGene_tx=XM_011280971.3;refGene_gn=ENTPD6;refGene_tx=XM_019826924.2
A3 50202061 50202097
refGene_gn=ENTPD6;refGene_tx=XM_003983852.5;refGene_gn=ENTPD6;refGene_tx=XM_006930007.4;refGene_gn=ENTPD6;refGene_tx=XM_006930008.4;refGene_gn=ENTPD6;refGene_tx=XM_006930009.4;refGene_gn=ENTPD6;refGene_tx=XM_011280971.3
A3 50202993 50203082
refGene_gn=ENTPD6;refGene_tx=XM_003983852.5;refGene_gn=ENTPD6;refGene_tx=XM_006930007.4;refGene_gn=ENTPD6;refGene_tx=XM_006930008.4;refGene_gn=ENTPD6;refGene_tx=XM_006930009.4;refGene_gn=ENTPD6;refGene_tx=XM_011280971.3;refGene_gn=ENTPD6;refGene_tx=XM_019826924.2
A3 50203884 50203964
refGene_gn=ENTPD6;refGene_tx=XM_003983852.5;refGene_gn=ENTPD6;refGene_tx=XM_006930007.4;refGene_gn=ENTPD6;refGene_tx=XM_006930008.4;refGene_gn=ENTPD6;refGene_tx=XM_006930009.4;refGene_gn=ENTPD6;refGene_tx=XM_011280971.3;refGene_gn=ENTPD6;refGene_tx=XM_019826924.2
A3 50204616 50204681
refGene_gn=ENTPD6;refGene_tx=XM_003983852.5;refGene_gn=ENTPD6;refGene_tx=XM_006930007.4;refGene_gn=ENTPD6;refGene_tx=XM_006930008.4;refGene_gn=ENTPD6;refGene_tx=XM_006930009.4;refGene_gn=ENTPD6;refGene_tx=XM_011280971.3;refGene_gn=ENTPD6;refGene_tx=XM_019826924.2
A3 50215659 50215761
refGene_gn=ENTPD6;refGene_tx=XM_003983852.5;refGene_gn=ENTPD6;refGene_tx=XM_006930007.4;refGene_gn=ENTPD6;refGene_tx=XM_006930008.4;refGene_gn=ENTPD6;refGene_tx=XM_006930009.4;refGene_gn=ENTPD6;refGene_tx=XM_011280971.3;refGene_gn=ENTPD6;refGene_tx=XM_019826924.2
A3 50218279 50218420
refGene_gn=ENTPD6;refGene_tx=XM_003983852.5;refGene_gn=ENTPD6;refGene_tx=XM_006930007.4;refGene_gn=ENTPD6;refGene_tx=XM_006930008.4;refGene_gn=ENTPD6;refGene_tx=XM_006930009.4;refGene_gn=ENTPD6;refGene_tx=XM_011280971.3;refGene_gn=ENTPD6;refGene_tx=XM_019826924.2
A3 50220059 50220116
refGene_gn=ENTPD6;refGene_tx=XM_003983852.5;refGene_gn=ENTPD6;refGene_tx=XM_006930007.4;refGene_gn=ENTPD6;refGene_tx=XM_006930008.4;refGene_gn=ENTPD6;refGene_tx=XM_006930009.4;refGene_gn=ENTPD6;refGene_tx=XM_011280971.3;refGene_gn=ENTPD6;refGene_tx=XM_019826924.2
A3 50221295 50221408
refGene_gn=ENTPD6;refGene_tx=XM_003983852.5;refGene_gn=ENTPD6;refGene_tx=XM_006930007.4;refGene_gn=ENTPD6;refGene_tx=XM_006930008.4;refGene_gn=ENTPD6;refGene_tx=XM_006930009.4;refGene_gn=ENTPD6;refGene_tx=XM_011280971.3;refGene_gn=ENTPD6;refGene_tx=XM_019826924.2
A3 50221518 50221617
refGene_gn=ENTPD6;refGene_tx=XM_003983852.5;refGene_gn=ENTPD6;refGene_tx=XM_006930007.4;refGene_gn=ENTPD6;refGene_tx=XM_006930008.4;refGene_gn=ENTPD6;refGene_tx=XM_006930009.4;refGene_gn=ENTPD6;refGene_tx=XM_011280971.3;refGene_gn=ENTPD6;refGene_tx=XM_019826924.2
A3 50240534 50240777 refGene_gn=PYGB;refGene_tx=XM_006930010.4
A3 50254667 50254769 refGene_gn=PYGB;refGene_tx=XM_006930010.4

A3	50269467 50269546 refGene_gn=PYGB;refGene_tx=XM_006930010.4
A3	50272062 50272166 refGene_gn=PYGB;refGene_tx=XM_006930010.4
A3	50276412 50276544 refGene_gn=PYGB;refGene_tx=XM_006930010.4
A3	50278407 50278519 refGene_gn=PYGB;refGene_tx=XM_006930010.4
A3	50278929 50279012 refGene_gn=PYGB;refGene_tx=XM_006930010.4
A3	50280202 50280346 refGene_gn=PYGB;refGene_tx=XM_006930010.4
A3	50280913 50281006 refGene_gn=PYGB;refGene_tx=XM_006930010.4
A3	50282111 50282258 refGene_gn=PYGB;refGene_tx=XM_006930010.4
A3	50283116 50283280 refGene_gn=PYGB;refGene_tx=XM_006930010.4
A3	50284214 50284329 refGene_gn=PYGB;refGene_tx=XM_006930010.4
A3	50285738 50285840 refGene_gn=PYGB;refGene_tx=XM_006930010.4
A3	50286585 50286733 refGene_gn=PYGB;refGene_tx=XM_006930010.4
A3	50289651 50289710 refGene_gn=PYGB;refGene_tx=XM_006930010.4
A3	50291353 50291495 refGene_gn=PYGB;refGene_tx=XM_006930010.4
A3	50293281 50293489 refGene_gn=PYGB;refGene_tx=XM_006930010.4
A3	50294877 50295012 refGene_gn=PYGB;refGene_tx=XM_006930010.4
A3	50296281 50296348 refGene_gn=PYGB;refGene_tx=XM_006930010.4
A3	50297702 50297855 refGene_gn=PYGB;refGene_tx=XM_006930010.4
A3	50302230 50302270
refGene_gn=ABHD12;refGene_tx=XM_003983854.5;refGene_gn=ABHD12;refGene_
tx=XM_023251526.1
A3	50303434 50303562
refGene_gn=ABHD12;refGene_tx=XM_003983854.5;refGene_gn=ABHD12;refGene_
tx=XM_023251526.1
A3	50304745 50304824
refGene_gn=ABHD12;refGene_tx=XM_003983854.5;refGene_gn=ABHD12;refGene_
tx=XM_023251526.1
A3	50307149 50307232
refGene_gn=ABHD12;refGene_tx=XM_003983854.5;refGene_gn=ABHD12;refGene_
tx=XM_023251526.1
A3	50308245 50308325
refGene_gn=ABHD12;refGene_tx=XM_003983854.5;refGene_gn=ABHD12;refGene_
tx=XM_023251526.1
A3	50308794 50308832
refGene_gn=ABHD12;refGene_tx=XM_003983854.5;refGene_gn=ABHD12;refGene_
tx=XM_023251526.1
A3	50309906 50310036
refGene_gn=ABHD12;refGene_tx=XM_003983854.5;refGene_gn=ABHD12;refGene_
tx=XM_023251526.1
A3	50312345 50312391
refGene_gn=ABHD12;refGene_tx=XM_003983854.5;refGene_gn=ABHD12;refGene_
tx=XM_023251526.1
A3	50315202 50315233
refGene_gn=ABHD12;refGene_tx=XM_003983854.5;refGene_gn=ABHD12;refGene_
tx=XM_023251526.1
A3	50319083 50319203
refGene_gn=ABHD12;refGene_tx=XM_003983854.5;refGene_gn=ABHD12;refGene_
tx=XM_023251526.1
A3	50321766 50321872
refGene_gn=ABHD12;refGene_tx=XM_003983854.5;refGene_gn=ABHD12;refGene_

tx=XM_023251526.1
A3 50329805 50329930
refGene_gn=ABHD12;refGene_tx=XM_003983854.5;refGene_gn=ABHD12;refGene_tx=XM_023251526.1
A3 50388232 50388246 refGene_gn=ABHD12;refGene_tx=XM_023251526.1
A3 50389097 50389288 refGene_gn=ABHD12;refGene_tx=XM_003983854.5
A3 50395301 50395376
refGene_gn=GINS1;refGene_tx=XM_003983855.5;refGene_gn=GINS1;refGene_tx=XM_019826926.2
A3 50399688 50399753
refGene_gn=GINS1;refGene_tx=XM_003983855.5;refGene_gn=GINS1;refGene_tx=XM_019826926.2
A3 50404003 50404102
refGene_gn=GINS1;refGene_tx=XM_003983855.5;refGene_gn=GINS1;refGene_tx=XM_019826926.2
A3 50404632 50404723
refGene_gn=GINS1;refGene_tx=XM_003983855.5;refGene_gn=GINS1;refGene_tx=XM_019826926.2
A3 50408017 50408134
refGene_gn=GINS1;refGene_tx=XM_003983855.5;refGene_gn=GINS1;refGene_tx=XM_019826926.2
A3 50411127 50411220 refGene_gn=GINS1;refGene_tx=XM_019826926.2
A3 50414226 50414301 refGene_gn=GINS1;refGene_tx=XM_003983855.5
A3 50417517 50417586 refGene_gn=GINS1;refGene_tx=XM_003983855.5
A3 50439948 50440140 refGene_gn=NINL;refGene_tx=XM_023251527.1
A3 50443945 50444059 refGene_gn=NINL;refGene_tx=XM_023251527.1
A3 50446161 50446371 refGene_gn=NINL;refGene_tx=XM_023251527.1
A3 50450815 50450929 refGene_gn=NINL;refGene_tx=XM_023251527.1
A3 50451875 50452034 refGene_gn=NINL;refGene_tx=XM_023251527.1
A3 50473168 50473238 refGene_gn=NINL;refGene_tx=XM_023251527.1
A3 50474574 50474679 refGene_gn=NINL;refGene_tx=XM_023251527.1
A3 50482465 50483632 refGene_gn=NINL;refGene_tx=XM_023251527.1
A3 50485128 50485167 refGene_gn=NINL;refGene_tx=XM_023251527.1
A3 50485461 50485790 refGene_gn=NINL;refGene_tx=XM_023251527.1
A3 50486762 50486872 refGene_gn=NINL;refGene_tx=XM_023251527.1
A3 50489652 50489782 refGene_gn=NINL;refGene_tx=XM_023251527.1
A3 50498838 50498919 refGene_gn=NINL;refGene_tx=XM_023251527.1
A3 50499519 50499630 refGene_gn=NINL;refGene_tx=XM_023251527.1
A3 50500701 50500876 refGene_gn=NINL;refGene_tx=XM_023251527.1
A3 50506363 50506504 refGene_gn=NINL;refGene_tx=XM_023251527.1
A3 50507939 50508076 refGene_gn=NINL;refGene_tx=XM_023251527.1
A3 50512104 50512275 refGene_gn=NINL;refGene_tx=XM_023251527.1
A3 50514668 50514821 refGene_gn=NINL;refGene_tx=XM_023251527.1
A3 50515529 50515720 refGene_gn=NINL;refGene_tx=XM_023251527.1
A3 50517705 50517772 refGene_gn=NINL;refGene_tx=XM_023251527.1
A3 50520228 50520407 refGene_gn=NINL;refGene_tx=XM_023251527.1
A3 50524289 50524386 refGene_gn=NINL;refGene_tx=XM_023251527.1
A3 50532711 50532905 refGene_gn=NINL;refGene_tx=XM_023251527.1
A3 50569227 50569234 refGene_gn=NINL;refGene_tx=XM_023251527.1
A3 50643280 50643937

refGene_gn=NANP;refGene_tx=XM_003983857.5;refGene_gn=NANP;refGene_tx=XM_006930012.4
A3 50659083 50659230
refGene_gn=NANP;refGene_tx=XM_003983857.5;refGene_gn=NANP;refGene_tx=XM_006930012.4
A3 50844261 50844416
refGene_gn=LOC105260261;refGene_tx=XM_023251528.1;refGene_gn=LOC105260261;refGene_tx=XM_023251529.1
A3 50845759 50845953
refGene_gn=LOC105260261;refGene_tx=XM_023251528.1;refGene_gn=LOC105260261;refGene_tx=XM_023251529.1
A3 50846101 50846478
refGene_gn=LOC105260261;refGene_tx=XM_023251528.1;refGene_gn=LOC105260261;refGene_tx=XM_023251529.1
A3 50847268 50847484
refGene_gn=LOC105260261;refGene_tx=XM_023251528.1;refGene_gn=LOC105260261;refGene_tx=XM_023251529.1
A3 50847865 50847969
refGene_gn=LOC105260261;refGene_tx=XM_023251528.1;refGene_gn=LOC105260261;refGene_tx=XM_023251529.1
A3 50848220 50848260
refGene_gn=LOC105260261;refGene_tx=XM_023251529.1
A3 50848471 50848541
refGene_gn=LOC105260261;refGene_tx=XM_023251528.1
A3 50849192 50849282
refGene_gn=LOC105260261;refGene_tx=XM_023251528.1
A3 53213762 53213852
refGene_gn=LOC102899245;refGene_tx=XM_023252180.1
A3 53214473 53214543
refGene_gn=LOC102899245;refGene_tx=XM_023252180.1
A3 53215045 53215149
refGene_gn=LOC102899245;refGene_tx=XM_023252180.1
A3 53215530 53215746
refGene_gn=LOC102899245;refGene_tx=XM_023252180.1
A3 53216536 53216913
refGene_gn=LOC102899245;refGene_tx=XM_023252180.1
A3 53217061 53217255
refGene_gn=LOC102899245;refGene_tx=XM_023252180.1
A3 53218599 53218787
refGene_gn=LOC102899245;refGene_tx=XM_023252180.1
A3 53220250 53220415
refGene_gn=LOC102899245;refGene_tx=XM_023252180.1
A3 53456250 53457900 refGene_gn=SOWAHC;refGene_tx=XM_006930014.4
A3 53458704 53458731 refGene_gn=SEPT10;refGene_tx=XM_019826929.2
A3 53484923 53484992
refGene_gn=SEPT10;refGene_tx=XM_019826929.2;refGene_gn=SEPT10;refGene_tx=XM_019826932.2;refGene_gn=SEPT10;refGene_tx=XM_019826933.2;refGene_gn=SEPT10;refGene_tx=XM_023251530.1;refGene_gn=SEPT10;refGene_tx=XM_023251531.1
A3 53490715 53490836

refGene_gn=SEPT10;refGene_tx=XM_019826929.2;refGene_gn=SEPT10;refGene_tx=XM_019826932.2;refGene_gn=SEPT10;refGene_tx=XM_019826933.2;refGene_gn=SEPT10;refGene_tx=XM_023251530.1;refGene_gn=SEPT10;refGene_tx=XM_023251531.1
A3 53491240 53491436
refGene_gn=SEPT10;refGene_tx=XM_019826929.2;refGene_gn=SEPT10;refGene_tx=XM_019826932.2;refGene_gn=SEPT10;refGene_tx=XM_019826933.2;refGene_gn=SEPT10;refGene_tx=XM_023251530.1;refGene_gn=SEPT10;refGene_tx=XM_023251531.1
A3 53496011 53496198
refGene_gn=SEPT10;refGene_tx=XM_019826929.2;refGene_gn=SEPT10;refGene_tx=XM_019826932.2;refGene_gn=SEPT10;refGene_tx=XM_019826933.2;refGene_gn=SEPT10;refGene_tx=XM_023251530.1;refGene_gn=SEPT10;refGene_tx=XM_023251531.1
A3 53499267 53499429
refGene_gn=SEPT10;refGene_tx=XM_019826929.2;refGene_gn=SEPT10;refGene_tx=XM_019826932.2;refGene_gn=SEPT10;refGene_tx=XM_019826933.2;refGene_gn=SEPT10;refGene_tx=XM_023251530.1;refGene_gn=SEPT10;refGene_tx=XM_023251531.1
A3 53501017 53501114
refGene_gn=SEPT10;refGene_tx=XM_019826929.2;refGene_gn=SEPT10;refGene_tx=XM_019826932.2;refGene_gn=SEPT10;refGene_tx=XM_019826933.2;refGene_gn=SEPT10;refGene_tx=XM_023251530.1;refGene_gn=SEPT10;refGene_tx=XM_023251531.1
A3 53502302 53502471
refGene_gn=SEPT10;refGene_tx=XM_019826929.2;refGene_gn=SEPT10;refGene_tx=XM_019826932.2;refGene_gn=SEPT10;refGene_tx=XM_019826933.2;refGene_gn=SEPT10;refGene_tx=XM_023251530.1;refGene_gn=SEPT10;refGene_tx=XM_023251531.1
A3 53517252 53517385
refGene_gn=SEPT10;refGene_tx=XM_019826929.2;refGene_gn=SEPT10;refGene_tx=XM_019826932.2;refGene_gn=SEPT10;refGene_tx=XM_019826933.2;refGene_gn=SEPT10;refGene_tx=XM_023251530.1;refGene_gn=SEPT10;refGene_tx=XM_023251531.1
A3 53522707 53522895
refGene_gn=SEPT10;refGene_tx=XM_019826929.2;refGene_gn=SEPT10;refGene_tx=XM_019826932.2;refGene_gn=SEPT10;refGene_tx=XM_019826933.2;refGene_gn=SEPT10;refGene_tx=XM_023251530.1;refGene_gn=SEPT10;refGene_tx=XM_023251531.1
A3 53523410 53523528
refGene_gn=SEPT10;refGene_tx=XM_019826929.2;refGene_gn=SEPT10;refGene_tx=XM_019826932.2;refGene_gn=SEPT10;refGene_tx=XM_019826933.2;refGene_gn=SEPT10;refGene_tx=XM_023251530.1
A3 53524654 53524818
refGene_gn=SEPT10;refGene_tx=XM_019826929.2;refGene_gn=SEPT10;refGene_tx=XM_019826932.2;refGene_gn=SEPT10;refGene_tx=XM_019826933.2;refGene_gn=SEPT10;refGene_tx=XM_023251531.1
A3 53595955 53596124 refGene_gn=SH3RF3;refGene_tx=XM_011281104.3
A3 53605265 53605600 refGene_gn=SH3RF3;refGene_tx=XM_011281104.3
A3 53651061 53651396 refGene_gn=SH3RF3;refGene_tx=XM_011281104.3

A3 53665933 53666181 refGene_gn=SH3RF3;refGene_tx=XM_011281104.3
A3 53668253 53668424 refGene_gn=SH3RF3;refGene_tx=XM_011281104.3
A3 53676137 53676217 refGene_gn=SH3RF3;refGene_tx=XM_011281104.3
A3 53695026 53695380 refGene_gn=SH3RF3;refGene_tx=XM_011281104.3
A3 53729076 53729172 refGene_gn=SH3RF3;refGene_tx=XM_011281104.3
A3 53747904 53748180 refGene_gn=SH3RF3;refGene_tx=XM_011281104.3
A3 53958228 53958801 refGene_gn=SH3RF3;refGene_tx=XM_011281104.3
A3 54153582 54153633
refGene_gn=EDAR;refGene_tx=XM_003983868.4;refGene_gn=EDAR;refGene_tx=XM_019826939.2;refGene_gn=EDAR;refGene_tx=XM_019826940.2
A3 54154357 54154480
refGene_gn=EDAR;refGene_tx=XM_003983868.4;refGene_gn=EDAR;refGene_tx=XM_019826939.2;refGene_gn=EDAR;refGene_tx=XM_019826940.2
A3 54155239 54155421
refGene_gn=EDAR;refGene_tx=XM_003983868.4;refGene_gn=EDAR;refGene_tx=XM_019826939.2;refGene_gn=EDAR;refGene_tx=XM_019826940.2
A3 54162427 54162513
refGene_gn=EDAR;refGene_tx=XM_003983868.4;refGene_gn=EDAR;refGene_tx=XM_019826939.2;refGene_gn=EDAR;refGene_tx=XM_019826940.2
A3 54169960 54170047
refGene_gn=EDAR;refGene_tx=XM_003983868.4;refGene_gn=EDAR;refGene_tx=XM_019826939.2;refGene_gn=EDAR;refGene_tx=XM_019826940.2
A3 54171635 54171764
refGene_gn=EDAR;refGene_tx=XM_003983868.4;refGene_gn=EDAR;refGene_tx=XM_019826939.2
A3 54171880 54171955
refGene_gn=EDAR;refGene_tx=XM_003983868.4;refGene_gn=EDAR;refGene_tx=XM_019826939.2
A3 54172239 54172312
refGene_gn=EDAR;refGene_tx=XM_003983868.4;refGene_gn=EDAR;refGene_tx=XM_019826939.2;refGene_gn=EDAR;refGene_tx=XM_019826940.2
A3 54174870 54175027
refGene_gn=EDAR;refGene_tx=XM_003983868.4;refGene_gn=EDAR;refGene_tx=XM_019826939.2;refGene_gn=EDAR;refGene_tx=XM_019826940.2
A3 54176132 54176193
refGene_gn=EDAR;refGene_tx=XM_003983868.4;refGene_gn=EDAR;refGene_tx=XM_019826939.2;refGene_gn=EDAR;refGene_tx=XM_019826940.2
A3 54181163 54181486
refGene_gn=EDAR;refGene_tx=XM_003983868.4;refGene_gn=EDAR;refGene_tx=XM_019826939.2;refGene_gn=EDAR;refGene_tx=XM_019826940.2
A3 54224631 54224797
refGene_gn=CCDC138;refGene_tx=XM_003983858.4;refGene_gn=CCDC138;refGene_tx=XM_011281075.3;refGene_gn=CCDC138;refGene_tx=XM_011281076.3;refGene_gn=CCDC138;refGene_tx=XM_019826936.2
A3 54226523 54226662
refGene_gn=CCDC138;refGene_tx=XM_003983858.4;refGene_gn=CCDC138;refGene_tx=XM_011281075.3;refGene_gn=CCDC138;refGene_tx=XM_011281076.3;refGene_gn=CCDC138;refGene_tx=XM_019826936.2
A3 54268335 54268512
refGene_gn=CCDC138;refGene_tx=XM_003983858.4;refGene_gn=CCDC138;refGen

e_tx=XM_011281075.3;refGene_gn=CCDC138;refGene_tx=XM_011281076.3;refGene_gn=CCDC138;refGene_tx=XM_019826936.2
A3	54289532 54289725
refGene_gn=CCDC138;refGene_tx=XM_003983858.4;refGene_gn=CCDC138;refGene_tx=XM_011281075.3;refGene_gn=CCDC138;refGene_tx=XM_011281076.3;refGene_gn=CCDC138;refGene_tx=XM_019826936.2
A3	54304610 54304727
refGene_gn=CCDC138;refGene_tx=XM_003983858.4;refGene_gn=CCDC138;refGene_tx=XM_011281075.3;refGene_gn=CCDC138;refGene_tx=XM_011281076.3;refGene_gn=CCDC138;refGene_tx=XM_019826936.2
A3	54318012 54318177
refGene_gn=CCDC138;refGene_tx=XM_003983858.4;refGene_gn=CCDC138;refGene_tx=XM_011281075.3;refGene_gn=CCDC138;refGene_tx=XM_011281076.3;refGene_gn=CCDC138;refGene_tx=XM_019826936.2
A3	54321292 54321400
refGene_gn=CCDC138;refGene_tx=XM_003983858.4;refGene_gn=CCDC138;refGene_tx=XM_011281075.3;refGene_gn=CCDC138;refGene_tx=XM_011281076.3
A3	54321497 54321575
refGene_gn=CCDC138;refGene_tx=XM_003983858.4;refGene_gn=CCDC138;refGene_tx=XM_011281075.3;refGene_gn=CCDC138;refGene_tx=XM_011281076.3;refGene_gn=CCDC138;refGene_tx=XM_019826936.2
A3	54333457 54333577
refGene_gn=CCDC138;refGene_tx=XM_003983858.4;refGene_gn=CCDC138;refGene_tx=XM_011281075.3;refGene_gn=CCDC138;refGene_tx=XM_011281076.3;refGene_gn=CCDC138;refGene_tx=XM_019826936.2
A3	54340255 54340414
refGene_gn=CCDC138;refGene_tx=XM_003983858.4;refGene_gn=CCDC138;refGene_tx=XM_011281075.3;refGene_gn=CCDC138;refGene_tx=XM_011281076.3;refGene_gn=CCDC138;refGene_tx=XM_019826936.2
A3	54345954 54346139
refGene_gn=CCDC138;refGene_tx=XM_003983858.4;refGene_gn=CCDC138;refGene_tx=XM_011281075.3;refGene_gn=CCDC138;refGene_tx=XM_011281076.3;refGene_gn=CCDC138;refGene_tx=XM_019826936.2
A3	54355039 54355167
refGene_gn=CCDC138;refGene_tx=XM_003983858.4;refGene_gn=CCDC138;refGene_tx=XM_011281075.3;refGene_gn=CCDC138;refGene_tx=XM_011281076.3;refGene_gn=CCDC138;refGene_tx=XM_019826936.2
A3	54356917 54357038
refGene_gn=CCDC138;refGene_tx=XM_003983858.4;refGene_gn=CCDC138;refGene_tx=XM_011281075.3;refGene_gn=CCDC138;refGene_tx=XM_011281076.3;refGene_gn=CCDC138;refGene_tx=XM_019826936.2
A3	54358485 54358543
refGene_gn=CCDC138;refGene_tx=XM_003983858.4;refGene_gn=CCDC138;refGene_tx=XM_011281075.3;refGene_gn=CCDC138;refGene_tx=XM_011281076.3;refGene_gn=CCDC138;refGene_tx=XM_019826936.2
A3	54359935 54360028
refGene_gn=CCDC138;refGene_tx=XM_003983858.4;refGene_gn=CCDC138;refGene_tx=XM_011281075.3;refGene_gn=CCDC138;refGene_tx=XM_011281076.3;refGene_gn=CCDC138;refGene_tx=XM_019826936.2
A3	54362824 54363130

refGene_gn=LOC101094012;refGene_tx=XM_023251533.1
A3 54363522 54363857
refGene_gn=LOC101094012;refGene_tx=XM_023251533.1
A3 54364199 54364473
refGene_gn=LOC101094012;refGene_tx=XM_023251533.1
A3 54365145 54365291
refGene_gn=LOC101094012;refGene_tx=XM_023251533.1
A3 54368786 54368888
refGene_gn=LOC101094012;refGene_tx=XM_023251533.1
A3 54370775 54370980
refGene_gn=LOC101094012;refGene_tx=XM_023251533.1
A3 54377491 54377670
refGene_gn=LOC101094012;refGene_tx=XM_023251533.1
A3 54378183 54378276
refGene_gn=LOC101094012;refGene_tx=XM_023251533.1
A3 54378924 54379095
refGene_gn=LOC101094012;refGene_tx=XM_023251533.1
A3 54381908 54381986
refGene_gn=LOC101094012;refGene_tx=XM_023251533.1
A3 54382622 54386403
refGene_gn=LOC101094012;refGene_tx=XM_023251533.1
A3 54386404 54387247
refGene_gn=LOC101094012;refGene_tx=XM_023251533.1
A3 54388211 54388306
refGene_gn=LOC101094012;refGene_tx=XM_023251533.1
A3 54391874 54392010
refGene_gn=LOC101094012;refGene_tx=XM_023251533.1
A3 54396102 54396186
refGene_gn=LOC101094012;refGene_tx=XM_023251533.1
A3 54396269 54396449
refGene_gn=LOC101094012;refGene_tx=XM_023251533.1
A3 54398630 54398777
refGene_gn=LOC101094012;refGene_tx=XM_023251533.1
A3 54398856 54398994
refGene_gn=LOC101094012;refGene_tx=XM_023251533.1
A3 54399263 54399392
refGene_gn=LOC101094012;refGene_tx=XM_023251533.1
A3 54399393 54399426
refGene_gn=LOC101094012;refGene_tx=XM_023251533.1
A3 54400335 54400459
refGene_gn=LOC101094012;refGene_tx=XM_023251533.1
A3 54400658 54400834
refGene_gn=LOC101094012;refGene_tx=XM_023251533.1
A3 54400921 54401103
refGene_gn=LOC101094012;refGene_tx=XM_023251533.1
A3 54402717 54402927
refGene_gn=LOC101094012;refGene_tx=XM_023251533.1
A3 54405476 54405564
refGene_gn=LOC101094012;refGene_tx=XM_023251533.1
A3 54409603 54409796

refGene_gn=LOC101094012;refGene_tx=XM_023251533.1
A3	54411854 54412000
refGene_gn=LOC101094012;refGene_tx=XM_023251533.1
A3	54412342 54412573
refGene_gn=LOC101094012;refGene_tx=XM_023251533.1
A3	54417600 54417753
refGene_gn=LOC101094012;refGene_tx=XM_023251533.1
A3	54418240 54418352
refGene_gn=LOC101094012;refGene_tx=XM_023251533.1
A3	54419747 54419815
refGene_gn=LOC101094012;refGene_tx=XM_023251533.1
A3	54445594 54445666
refGene_gn=LOC101094012;refGene_tx=XM_023251533.1
A3	54472885 54473000
refGene_gn=LIMS1;refGene_tx=XM_003983861.5;refGene_gn=LIMS1;refGene_tx=XM_003983862.5
A3	54474983 54475131
refGene_gn=LIMS1;refGene_tx=XM_006930019.3;refGene_gn=LIMS1;refGene_tx=XM_006930020.3;refGene_gn=LIMS1;refGene_tx=XM_006930021.4;refGene_gn=LIMS1;refGene_tx=XM_019826946.2;refGene_gn=LIMS1;refGene_tx=XM_019826947.2;refGene_gn=LIMS1;refGene_tx=XM_023251534.1;refGene_gn=LIMS1;refGene_tx=XM_023251535.1
A3	54475319 54475395
refGene_gn=LIMS1;refGene_tx=XM_003983861.5;refGene_gn=LIMS1;refGene_tx=XM_003983862.5;refGene_gn=LIMS1;refGene_tx=XM_006930019.3;refGene_gn=LIMS1;refGene_tx=XM_006930020.3;refGene_gn=LIMS1;refGene_tx=XM_006930021.4;refGene_gn=LIMS1;refGene_tx=XM_019826946.2;refGene_gn=LIMS1;refGene_tx=XM_019826947.2;refGene_gn=LIMS1;refGene_tx=XM_023251534.1;refGene_gn=LIMS1;refGene_tx=XM_023251535.1
A3	54476950 54476999
refGene_gn=LIMS1;refGene_tx=XM_003983861.5;refGene_gn=LIMS1;refGene_tx=XM_003983862.5;refGene_gn=LIMS1;refGene_tx=XM_006930019.3;refGene_gn=LIMS1;refGene_tx=XM_006930020.3;refGene_gn=LIMS1;refGene_tx=XM_006930021.4;refGene_gn=LIMS1;refGene_tx=XM_019826946.2;refGene_gn=LIMS1;refGene_tx=XM_019826947.2;refGene_gn=LIMS1;refGene_tx=XM_023251534.1;refGene_gn=LIMS1;refGene_tx=XM_023251535.1
A3	54477777 54477870
refGene_gn=LIMS1;refGene_tx=XM_003983861.5;refGene_gn=LIMS1;refGene_tx=XM_003983862.5;refGene_gn=LIMS1;refGene_tx=XM_006930019.3;refGene_gn=LIMS1;refGene_tx=XM_006930020.3;refGene_gn=LIMS1;refGene_tx=XM_006930021.4;refGene_gn=LIMS1;refGene_tx=XM_019826946.2;refGene_gn=LIMS1;refGene_tx=XM_019826947.2;refGene_gn=LIMS1;refGene_tx=XM_023251534.1;refGene_gn=LIMS1;refGene_tx=XM_023251535.1
A3	54478387 54478538
refGene_gn=LIMS1;refGene_tx=XM_003983861.5;refGene_gn=LIMS1;refGene_tx=XM_003983862.5;refGene_gn=LIMS1;refGene_tx=XM_006930019.3;refGene_gn=LIMS1;refGene_tx=XM_006930020.3;refGene_gn=LIMS1;refGene_tx=XM_006930021.4;refGene_gn=LIMS1;refGene_tx=XM_019826946.2;refGene_gn=LIMS1;refGene_tx=XM_019826947.2;refGene_gn=LIMS1;refGene_tx=XM_023251534.1;refGene_gn=LIMS1;refGene_tx=XM_023251535.1

A3 54479925 54480075
refGene_gn=LIMS1;refGene_tx=XM_003983861.5;refGene_gn=LIMS1;refGene_tx=XM_003983862.5;refGene_gn=LIMS1;refGene_tx=XM_006930019.3;refGene_gn=LIMS1;refGene_tx=XM_006930020.3;refGene_gn=LIMS1;refGene_tx=XM_006930021.4;refGene_gn=LIMS1;refGene_tx=XM_019826946.2;refGene_gn=LIMS1;refGene_tx=XM_019826947.2;refGene_gn=LIMS1;refGene_tx=XM_023251534.1;refGene_gn=LIMS1;refGene_tx=XM_023251535.1
A3 54480491 54480612
refGene_gn=LIMS1;refGene_tx=XM_003983861.5;refGene_gn=LIMS1;refGene_tx=XM_003983862.5;refGene_gn=LIMS1;refGene_tx=XM_006930019.3;refGene_gn=LIMS1;refGene_tx=XM_006930020.3;refGene_gn=LIMS1;refGene_tx=XM_006930021.4;refGene_gn=LIMS1;refGene_tx=XM_019826946.2;refGene_gn=LIMS1;refGene_tx=XM_019826947.2;refGene_gn=LIMS1;refGene_tx=XM_023251534.1;refGene_gn=LIMS1;refGene_tx=XM_023251535.1
A3 54481415 54481482
refGene_gn=LIMS1;refGene_tx=XM_003983861.5;refGene_gn=LIMS1;refGene_tx=XM_003983862.5;refGene_gn=LIMS1;refGene_tx=XM_006930019.3;refGene_gn=LIMS1;refGene_tx=XM_006930020.3;refGene_gn=LIMS1;refGene_tx=XM_006930021.4;refGene_gn=LIMS1;refGene_tx=XM_019826946.2;refGene_gn=LIMS1;refGene_tx=XM_019826947.2;refGene_gn=LIMS1;refGene_tx=XM_023251534.1;refGene_gn=LIMS1;refGene_tx=XM_023251535.1
A3 54496962 54497122
refGene_gn=LIMS1;refGene_tx=XM_003983861.5;refGene_gn=LIMS1;refGene_tx=XM_003983862.5;refGene_gn=LIMS1;refGene_tx=XM_006930019.3;refGene_gn=LIMS1;refGene_tx=XM_006930020.3;refGene_gn=LIMS1;refGene_tx=XM_006930021.4;refGene_gn=LIMS1;refGene_tx=XM_019826946.2;refGene_gn=LIMS1;refGene_tx=XM_019826947.2;refGene_gn=LIMS1;refGene_tx=XM_023251534.1;refGene_gn=LIMS1;refGene_tx=XM_023251535.1
A3 54502072 54502254
refGene_gn=LIMS1;refGene_tx=XM_003983861.5;refGene_gn=LIMS1;refGene_tx=XM_006930019.3;refGene_gn=LIMS1;refGene_tx=XM_006930020.3
A3 54537785 54537892
refGene_gn=LIMS1;refGene_tx=XM_003983862.5;refGene_gn=LIMS1;refGene_tx=XM_019826946.2
A3 54620776 54620808 refGene_gn=LIMS1;refGene_tx=XM_023251534.1
A3 54678522 54678593
refGene_gn=GCC2;refGene_tx=XM_003983863.5;refGene_gn=GCC2;refGene_tx=XM_019826950.2
A3 54685043 54685245
refGene_gn=GCC2;refGene_tx=XM_003983863.5;refGene_gn=GCC2;refGene_tx=XM_019826950.2
A3 54693990 54694130
refGene_gn=GCC2;refGene_tx=XM_003983863.5;refGene_gn=GCC2;refGene_tx=XM_019826950.2
A3 54695137 54695332
refGene_gn=GCC2;refGene_tx=XM_003983863.5;refGene_gn=GCC2;refGene_tx=XM_019826950.2
A3 54697018 54697236
refGene_gn=GCC2;refGene_tx=XM_003983863.5;refGene_gn=GCC2;refGene_tx=XM_019826950.2

A3 54699934 54700111
refGene_gn=GCC2;refGene_tx=XM_003983863.5;refGene_gn=GCC2;refGene_tx=XM_019826950.2
A3 54701879 54702001
refGene_gn=GCC2;refGene_tx=XM_003983863.5;refGene_gn=GCC2;refGene_tx=XM_019826950.2
A3 54702906 54703044
refGene_gn=GCC2;refGene_tx=XM_003983863.5;refGene_gn=GCC2;refGene_tx=XM_019826950.2
A3 54703722 54703800
refGene_gn=GCC2;refGene_tx=XM_003983863.5;refGene_gn=GCC2;refGene_tx=XM_019826950.2
A3 54703887 54703988
refGene_gn=GCC2;refGene_tx=XM_003983863.5;refGene_gn=GCC2;refGene_tx=XM_019826950.2
A3 54719873 54720036
refGene_gn=GCC2;refGene_tx=XM_003983863.5;refGene_gn=GCC2;refGene_tx=XM_019826950.2
A3 54720912 54721017
refGene_gn=GCC2;refGene_tx=XM_003983863.5;refGene_gn=GCC2;refGene_tx=XM_019826950.2
A3 54721316 54721481
refGene_gn=GCC2;refGene_tx=XM_003983863.5;refGene_gn=GCC2;refGene_tx=XM_019826950.2
A3 54724236 54724356
refGene_gn=GCC2;refGene_tx=XM_003983863.5;refGene_gn=GCC2;refGene_tx=XM_019826950.2
A3 54730919 54731018
refGene_gn=GCC2;refGene_tx=XM_003983863.5;refGene_gn=GCC2;refGene_tx=XM_019826950.2
A3 54731243 54731344
refGene_gn=GCC2;refGene_tx=XM_003983863.5;refGene_gn=GCC2;refGene_tx=XM_019826950.2
A3 54733823 54733893
refGene_gn=GCC2;refGene_tx=XM_003983863.5;refGene_gn=GCC2;refGene_tx=XM_019826950.2
A3 54734583 54737049
refGene_gn=GCC2;refGene_tx=XM_003983863.5;refGene_gn=GCC2;refGene_tx=XM_019826950.2
A3 54737749 54737854
refGene_gn=GCC2;refGene_tx=XM_003983863.5;refGene_gn=GCC2;refGene_tx=XM_019826950.2
A3 54740822 54740890
refGene_gn=GCC2;refGene_tx=XM_003983863.5;refGene_gn=GCC2;refGene_tx=XM_019826950.2
A3 54742008 54742093
refGene_gn=GCC2;refGene_tx=XM_003983863.5;refGene_gn=GCC2;refGene_tx=XM_019826950.2
A3 54743078 54743135
refGene_gn=GCC2;refGene_tx=XM_003983863.5;refGene_gn=GCC2;refGene_tx=X

M_019826950.2
A3	54743408	54743510
refGene_gn=GCC2;refGene_tx=XM_003983863.5;refGene_gn=GCC2;refGene_tx=X
M_019826950.2
A3	54791252	54791368	refGene_gn=SULT1C4;refGene_tx=XM_003983869.5
A3	54792316	54792497	refGene_gn=SULT1C4;refGene_tx=XM_003983869.5
A3	54795820	54795915	refGene_gn=SULT1C4;refGene_tx=XM_003983869.5
A3	54796132	54796259	refGene_gn=SULT1C4;refGene_tx=XM_003983869.5
A3	54796538	54796636	refGene_gn=SULT1C4;refGene_tx=XM_003983869.5
A3	54797214	54797340	refGene_gn=SULT1C4;refGene_tx=XM_003983869.5
A3	54800397	54800566	refGene_gn=SULT1C4;refGene_tx=XM_003983869.5
A3	54911562	54911675
refGene_gn=LOC101097138;refGene_tx=XM_011281074.3
A3	54913472	54913501
refGene_gn=LOC101097138;refGene_tx=XM_019828131.2
A3	54914102	54914283
refGene_gn=LOC101097138;refGene_tx=XM_011281074.3;refGene_gn=LOC101097
138;refGene_tx=XM_019828131.2
A3	54914413	54914508
refGene_gn=LOC101097138;refGene_tx=XM_011281074.3;refGene_gn=LOC101097
138;refGene_tx=XM_019828131.2
A3	54914941	54915068
refGene_gn=LOC101097138;refGene_tx=XM_011281074.3;refGene_gn=LOC101097
138;refGene_tx=XM_019828131.2
A3	54918713	54918811
refGene_gn=LOC101097138;refGene_tx=XM_011281074.3;refGene_gn=LOC101097
138;refGene_tx=XM_019828131.2
A3	54924916	54925042
refGene_gn=LOC101097138;refGene_tx=XM_011281074.3;refGene_gn=LOC101097
138;refGene_tx=XM_019828131.2
A3	54925473	54925624
refGene_gn=LOC101097138;refGene_tx=XM_011281074.3;refGene_gn=LOC101097
138;refGene_tx=XM_019828131.2
A3	54953457	54953570
refGene_gn=LOC101095165;refGene_tx=XM_003983864.5;refGene_gn=LOC101095
165;refGene_tx=XM_019826951.2
A3	54953805	54953986
refGene_gn=LOC101095165;refGene_tx=XM_003983864.5;refGene_gn=LOC101095
165;refGene_tx=XM_019826951.2
A3	54955165	54955260
refGene_gn=LOC101095165;refGene_tx=XM_003983864.5;refGene_gn=LOC101095
165;refGene_tx=XM_019826951.2
A3	54956206	54956333
refGene_gn=LOC101095165;refGene_tx=XM_003983864.5;refGene_gn=LOC101095
165;refGene_tx=XM_019826951.2
A3	54959186	54959284
refGene_gn=LOC101095165;refGene_tx=XM_003983864.5;refGene_gn=LOC101095
165;refGene_tx=XM_019826951.2
A3	54984143	54984272
refGene_gn=LOC101095165;refGene_tx=XM_003983864.5;refGene_gn=LOC101095

165;refGene_tx=XM_019826951.2
A3 54986484 54986656
refGene_gn=LOC101095165;refGene_tx=XM_003983864.5;refGene_gn=LOC101095165;refGene_tx=XM_019826951.2
A3 55272064 55272691
refGene_gn=SLC5A7;refGene_tx=XM_006930026.3;refGene_gn=SLC5A7;refGene_tx=XM_019826952.2
A3 55273979 55274197
refGene_gn=SLC5A7;refGene_tx=XM_006930026.3;refGene_gn=SLC5A7;refGene_tx=XM_019826952.2
A3 55277954 55278108
refGene_gn=SLC5A7;refGene_tx=XM_006930026.3;refGene_gn=SLC5A7;refGene_tx=XM_019826952.2
A3 55281502 55281646
refGene_gn=SLC5A7;refGene_tx=XM_006930026.3;refGene_gn=SLC5A7;refGene_tx=XM_019826952.2
A3 55285045 55285194
refGene_gn=SLC5A7;refGene_tx=XM_006930026.3;refGene_gn=SLC5A7;refGene_tx=XM_019826952.2
A3 55289375 55289531
refGene_gn=SLC5A7;refGene_tx=XM_006930026.3;refGene_gn=SLC5A7;refGene_tx=XM_019826952.2
A3 55290276 55290390
refGene_gn=SLC5A7;refGene_tx=XM_006930026.3;refGene_gn=SLC5A7;refGene_tx=XM_019826952.2
A3 55294692 55294870
refGene_gn=SLC5A7;refGene_tx=XM_006930026.3;refGene_gn=SLC5A7;refGene_tx=XM_019826952.2
A3 56282882 56283784
refGene_gn=LOC111559893;refGene_tx=XM_023252181.1
A3 56283966 56284573
refGene_gn=LOC111559893;refGene_tx=XM_023252181.1
A3 56288358 56288475
refGene_gn=LOC111559893;refGene_tx=XM_023252181.1
A3 56300898 56301009
refGene_gn=LOC111559893;refGene_tx=XM_023252181.1
A3 56326188 56327134
refGene_gn=ST6GAL2;refGene_tx=XM_019826953.2;refGene_gn=ST6GAL2;refGene_tx=XM_023251537.1;refGene_gn=ST6GAL2;refGene_tx=XM_023251538.1;refGene_gn=ST6GAL2;refGene_tx=XM_023251539.1
A3 56336921 56337019
refGene_gn=ST6GAL2;refGene_tx=XM_019826953.2;refGene_gn=ST6GAL2;refGene_tx=XM_023251537.1;refGene_gn=ST6GAL2;refGene_tx=XM_023251538.1;refGene_gn=ST6GAL2;refGene_tx=XM_023251539.1
A3 56338726 56338828
refGene_gn=ST6GAL2;refGene_tx=XM_019826953.2;refGene_gn=ST6GAL2;refGene_tx=XM_023251537.1;refGene_gn=ST6GAL2;refGene_tx=XM_023251538.1;refGene_gn=ST6GAL2;refGene_tx=XM_023251539.1
A3 56341598 56341773
refGene_gn=ST6GAL2;refGene_tx=XM_019826953.2;refGene_gn=ST6GAL2;refGen

e_tx=XM_023251537.1;refGene_gn=ST6GAL2;refGene_tx=XM_023251538.1;refGene_gn=ST6GAL2;refGene_tx=XM_023251539.1
A3 56364845 56365117
refGene_gn=ST6GAL2;refGene_tx=XM_019826953.2;refGene_gn=ST6GAL2;refGene_tx=XM_023251537.1;refGene_gn=ST6GAL2;refGene_tx=XM_023251538.1;refGene_gn=ST6GAL2;refGene_tx=XM_023251539.1
A3 56720144 56720238
refGene_gn=UXS1;refGene_tx=XM_023251540.1;refGene_gn=UXS1;refGene_tx=XM_023251541.1
A3 56749623 56749651
refGene_gn=UXS1;refGene_tx=XM_023251540.1;refGene_gn=UXS1;refGene_tx=XM_023251541.1
A3 56750920 56750984
refGene_gn=UXS1;refGene_tx=XM_023251540.1;refGene_gn=UXS1;refGene_tx=XM_023251541.1
A3 56752059 56752103
refGene_gn=UXS1;refGene_tx=XM_023251540.1;refGene_gn=UXS1;refGene_tx=XM_023251541.1
A3 56757148 56757209
refGene_gn=UXS1;refGene_tx=XM_023251540.1;refGene_gn=UXS1;refGene_tx=XM_023251541.1
A3 56768654 56768835
refGene_gn=UXS1;refGene_tx=XM_023251540.1;refGene_gn=UXS1;refGene_tx=XM_023251541.1
A3 56782836 56782941
refGene_gn=UXS1;refGene_tx=XM_023251540.1;refGene_gn=UXS1;refGene_tx=XM_023251541.1
A3 56785947 56786007
refGene_gn=UXS1;refGene_tx=XM_023251540.1;refGene_gn=UXS1;refGene_tx=XM_023251541.1
A3 56788812 56788934
refGene_gn=UXS1;refGene_tx=XM_023251540.1;refGene_gn=UXS1;refGene_tx=XM_023251541.1
A3 56799311 56799431
refGene_gn=UXS1;refGene_tx=XM_023251540.1;refGene_gn=UXS1;refGene_tx=XM_023251541.1
A3 56806880 56806924
refGene_gn=UXS1;refGene_tx=XM_023251540.1;refGene_gn=UXS1;refGene_tx=XM_023251541.1
A3 56810488 56810549
refGene_gn=UXS1;refGene_tx=XM_023251540.1;refGene_gn=UXS1;refGene_tx=XM_023251541.1
A3 56813655 56813713
refGene_gn=UXS1;refGene_tx=XM_023251540.1;refGene_gn=UXS1;refGene_tx=XM_023251541.1
A3 56815624 56815728
refGene_gn=UXS1;refGene_tx=XM_023251540.1;refGene_gn=UXS1;refGene_tx=XM_023251541.1
A3 56816085 56816217
refGene_gn=UXS1;refGene_tx=XM_023251540.1;refGene_gn=UXS1;refGene_tx=X

M_023251541.1
A3 56833168 56833318
refGene_gn=CA3H2orf40;refGene_tx=XM_003983874.5
A3 56837368 56837526
refGene_gn=CA3H2orf40;refGene_tx=XM_003983874.5
A3 56839525 56839573
refGene_gn=CA3H2orf40;refGene_tx=XM_003983874.5
A3 56844360 56844439
refGene_gn=CA3H2orf40;refGene_tx=XM_003983874.5
A3 57012098 57012293
refGene_gn=NCK2;refGene_tx=XM_023251542.1;refGene_gn=NCK2;refGene_tx=XM_023251543.1;refGene_gn=NCK2;refGene_tx=XM_023251544.1
A3 57032609 57033331
refGene_gn=NCK2;refGene_tx=XM_023251542.1;refGene_gn=NCK2;refGene_tx=XM_023251543.1;refGene_gn=NCK2;refGene_tx=XM_023251544.1
A3 57055070 57055296
refGene_gn=NCK2;refGene_tx=XM_023251542.1;refGene_gn=NCK2;refGene_tx=XM_023251543.1;refGene_gn=NCK2;refGene_tx=XM_023251544.1
A3 57485575 57485734
refGene_gn=FHL2;refGene_tx=XM_003983876.5;refGene_gn=FHL2;refGene_tx=XM_019826957.2;refGene_gn=FHL2;refGene_tx=XM_023251546.1
A3 57494603 57494778
refGene_gn=FHL2;refGene_tx=XM_003983876.5;refGene_gn=FHL2;refGene_tx=XM_019826957.2;refGene_gn=FHL2;refGene_tx=XM_023251546.1
A3 57500242 57500412
refGene_gn=FHL2;refGene_tx=XM_003983876.5;refGene_gn=FHL2;refGene_tx=XM_019826957.2;refGene_gn=FHL2;refGene_tx=XM_023251546.1
A3 57503346 57503533
refGene_gn=FHL2;refGene_tx=XM_003983876.5;refGene_gn=FHL2;refGene_tx=XM_019826957.2;refGene_gn=FHL2;refGene_tx=XM_023251546.1
A3 57508140 57508292
refGene_gn=FHL2;refGene_tx=XM_003983876.5;refGene_gn=FHL2;refGene_tx=XM_019826957.2;refGene_gn=FHL2;refGene_tx=XM_023251546.1
A3 57524869 57524926
refGene_gn=CA3H2orf49;refGene_tx=XM_006930029.4
A3 57528598 57528721
refGene_gn=CA3H2orf49;refGene_tx=XM_006930029.4
A3 57528847 57528974
refGene_gn=CA3H2orf49;refGene_tx=XM_006930029.4
A3 57531757 57531867
refGene_gn=CA3H2orf49;refGene_tx=XM_006930029.4
A3 57531868 57531925
refGene_gn=CA3H2orf49;refGene_tx=XM_006930029.4
A3 57534407 57534506
refGene_gn=CA3H2orf49;refGene_tx=XM_006930029.4
A3 57560703 57561391
refGene_gn=TGFBRAP1;refGene_tx=XM_003983878.5;refGene_gn=TGFBRAP1;refGene_tx=XM_023251547.1
A3 57571596 57571791
refGene_gn=TGFBRAP1;refGene_tx=XM_003983878.5;refGene_gn=TGFBRAP1;refG

ene_tx=XM_023251547.1
A3 57574104 57574259
refGene_gn=TGFBRAP1;refGene_tx=XM_003983878.5;refGene_gn=TGFBRAP1;refGene_tx=XM_023251547.1
A3 57585438 57585521
refGene_gn=TGFBRAP1;refGene_tx=XM_003983878.5;refGene_gn=TGFBRAP1;refGene_tx=XM_023251547.1
A3 57588586 57588928
refGene_gn=TGFBRAP1;refGene_tx=XM_003983878.5;refGene_gn=TGFBRAP1;refGene_tx=XM_023251547.1
A3 57590652 57590710
refGene_gn=TGFBRAP1;refGene_tx=XM_003983878.5;refGene_gn=TGFBRAP1;refGene_tx=XM_023251547.1
A3 57592549 57592693
refGene_gn=TGFBRAP1;refGene_tx=XM_003983878.5;refGene_gn=TGFBRAP1;refGene_tx=XM_023251547.1
A3 57594628 57594772
refGene_gn=TGFBRAP1;refGene_tx=XM_003983878.5;refGene_gn=TGFBRAP1;refGene_tx=XM_023251547.1
A3 57595355 57595515
refGene_gn=TGFBRAP1;refGene_tx=XM_003983878.5;refGene_gn=TGFBRAP1;refGene_tx=XM_023251547.1
A3 57598582 57599016
refGene_gn=TGFBRAP1;refGene_tx=XM_003983878.5;refGene_gn=TGFBRAP1;refGene_tx=XM_023251547.1
A3 57600681 57600858
refGene_gn=TGFBRAP1;refGene_tx=XM_003983878.5;refGene_gn=TGFBRAP1;refGene_tx=XM_023251547.1
A3 57617811 57619098 refGene_gn=GPR45;refGene_tx=XM_023250916.1
A3 57755995 57756087 refGene_gn=MRPS9;refGene_tx=XM_006930030.4
A3 57758108 57758278 refGene_gn=MRPS9;refGene_tx=XM_006930030.4
A3 57762601 57762710 refGene_gn=MRPS9;refGene_tx=XM_006930030.4
A3 57763215 57763382 refGene_gn=MRPS9;refGene_tx=XM_023251548.1
A3 57763703 57763872
refGene_gn=MRPS9;refGene_tx=XM_006930030.4;refGene_gn=MRPS9;refGene_tx=XM_023251548.1
A3 57766428 57766504
refGene_gn=MRPS9;refGene_tx=XM_006930030.4;refGene_gn=MRPS9;refGene_tx=XM_023251548.1
A3 57767564 57767650
refGene_gn=MRPS9;refGene_tx=XM_006930030.4;refGene_gn=MRPS9;refGene_tx=XM_023251548.1
A3 57776575 57776655
refGene_gn=MRPS9;refGene_tx=XM_006930030.4;refGene_gn=MRPS9;refGene_tx=XM_023251548.1
A3 57788290 57788321
refGene_gn=MRPS9;refGene_tx=XM_006930030.4;refGene_gn=MRPS9;refGene_tx=XM_023251548.1
A3 57788406 57788469
refGene_gn=MRPS9;refGene_tx=XM_006930030.4;refGene_gn=MRPS9;refGene_tx

=XM_023251548.1
A3	57809437	57809617	refGene_gn=MRPS9;refGene_tx=XM_006930030.4;refGene_gn=MRPS9;refGene_tx=XM_023251548.1
A3	57820717	57820834	refGene_gn=MRPS9;refGene_tx=XM_006930030.4;refGene_gn=MRPS9;refGene_tx=XM_023251548.1
A3	57976297	57977806	refGene_gn=POU3F3;refGene_tx=XM_023251549.1
A3	59820900	59821121	refGene_gn=TMEM182;refGene_tx=XM_003983881.4;refGene_gn=TMEM182;refGene_tx=XM_019826964.2;refGene_gn=TMEM182;refGene_tx=XM_019826965.2
A3	59836685	59836823	refGene_gn=TMEM182;refGene_tx=XM_003983881.4;refGene_gn=TMEM182;refGene_tx=XM_019826964.2;refGene_gn=TMEM182;refGene_tx=XM_019826965.2
A3	59869931	59870030	refGene_gn=TMEM182;refGene_tx=XM_003983881.4;refGene_gn=TMEM182;refGene_tx=XM_019826964.2;refGene_gn=TMEM182;refGene_tx=XM_019826965.2
A3	59871897	59871997	refGene_gn=TMEM182;refGene_tx=XM_003983881.4;refGene_gn=TMEM182;refGene_tx=XM_019826964.2;refGene_gn=TMEM182;refGene_tx=XM_019826965.2
A3	59872235	59872367	refGene_gn=TMEM182;refGene_tx=XM_003983881.4
A3	59879655	59879853	refGene_gn=TMEM182;refGene_tx=XM_019826964.2;refGene_gn=TMEM182;refGene_tx=XM_019826965.2
A3	59896889	59897054	refGene_gn=MFSD9;refGene_tx=XM_023251550.1;refGene_gn=MFSD9;refGene_tx=XM_023251551.1
A3	59902073	59902161	refGene_gn=MFSD9;refGene_tx=XM_023251550.1;refGene_gn=MFSD9;refGene_tx=XM_023251551.1;refGene_gn=MFSD9;refGene_tx=XM_023251552.1
A3	59903367	59903411	refGene_gn=MFSD9;refGene_tx=XM_023251550.1;refGene_gn=MFSD9;refGene_tx=XM_023251552.1
A3	59903729	59903859	refGene_gn=MFSD9;refGene_tx=XM_023251550.1;refGene_gn=MFSD9;refGene_tx=XM_023251552.1
A3	59906411	59906621	refGene_gn=MFSD9;refGene_tx=XM_023251550.1;refGene_gn=MFSD9;refGene_tx=XM_023251551.1;refGene_gn=MFSD9;refGene_tx=XM_023251552.1
A3	59909808	59910599	refGene_gn=MFSD9;refGene_tx=XM_023251550.1;refGene_gn=MFSD9;refGene_tx=XM_023251551.1;refGene_gn=MFSD9;refGene_tx=XM_023251552.1
A3	59921096	59921467	refGene_gn=SLC9A2;refGene_tx=XM_011281070.3
A3	59922789	59922880	refGene_gn=SLC9A2;refGene_tx=XM_011281070.3
A3	59924393	59924525	refGene_gn=SLC9A2;refGene_tx=XM_011281070.3
A3	59926619	59926716	refGene_gn=SLC9A2;refGene_tx=XM_011281070.3
A3	59934217	59934379	refGene_gn=SLC9A2;refGene_tx=XM_011281070.3
A3	59940944	59941015	refGene_gn=SLC9A2;refGene_tx=XM_011281070.3
A3	59941579	59941669	refGene_gn=SLC9A2;refGene_tx=XM_011281070.3
A3	59952148	59952351	refGene_gn=SLC9A2;refGene_tx=XM_011281070.3

A3	59952750 59952968 refGene_gn=SLC9A2;refGene_tx=XM_011281070.3
A3	59972265 59972516 refGene_gn=SLC9A2;refGene_tx=XM_011281070.3
A3	59979752 59980216 refGene_gn=SLC9A2;refGene_tx=XM_011281070.3
A3	60019822 60020120 refGene_gn=SLC9A2;refGene_tx=XM_011281070.3
A3	60081276 60081632 refGene_gn=SLC9A4;refGene_tx=XM_003983882.3
A3	60088859 60088947 refGene_gn=SLC9A4;refGene_tx=XM_003983882.3
A3	60090410 60090542 refGene_gn=SLC9A4;refGene_tx=XM_003983882.3
A3	60095836 60095933 refGene_gn=SLC9A4;refGene_tx=XM_003983882.3
A3	60101455 60101617 refGene_gn=SLC9A4;refGene_tx=XM_003983882.3
A3	60105639 60105710 refGene_gn=SLC9A4;refGene_tx=XM_003983882.3
A3	60109020 60109107 refGene_gn=SLC9A4;refGene_tx=XM_003983882.3
A3	60109715 60109918 refGene_gn=SLC9A4;refGene_tx=XM_003983882.3
A3	60113111 60113329 refGene_gn=SLC9A4;refGene_tx=XM_003983882.3
A3	60114506 60114766 refGene_gn=SLC9A4;refGene_tx=XM_003983882.3
A3	60142306 60142770 refGene_gn=SLC9A4;refGene_tx=XM_003983882.3
A3	60148203 60148489 refGene_gn=SLC9A4;refGene_tx=XM_003983882.3
A3	60173353 60173727
refGene_gn=IL18RAP;refGene_tx=XM_003983898.5;refGene_gn=IL18RAP;refGen
e_tx=XM_019826970.2
A3	60175305 60175479
refGene_gn=IL18RAP;refGene_tx=XM_003983898.5;refGene_gn=IL18RAP;refGen
e_tx=XM_019826970.2
A3	60178088 60178226 refGene_gn=IL18RAP;refGene_tx=XM_003983898.5
A3	60179363 60179515
refGene_gn=IL18RAP;refGene_tx=XM_003983898.5;refGene_gn=IL18RAP;refGen
e_tx=XM_019826970.2
A3	60181999 60182123
refGene_gn=IL18RAP;refGene_tx=XM_003983898.5;refGene_gn=IL18RAP;refGen
e_tx=XM_019826970.2
A3	60183584 60183650
refGene_gn=IL18RAP;refGene_tx=XM_003983898.5;refGene_gn=IL18RAP;refGen
e_tx=XM_019826970.2
A3	60187952 60188103
refGene_gn=IL18RAP;refGene_tx=XM_003983898.5;refGene_gn=IL18RAP;refGen
e_tx=XM_019826970.2
A3	60195303 60195493
refGene_gn=IL18RAP;refGene_tx=XM_003983898.5;refGene_gn=IL18RAP;refGen
e_tx=XM_019826970.2
A3	60195835 60196154
refGene_gn=IL18RAP;refGene_tx=XM_003983898.5;refGene_gn=IL18RAP;refGen
e_tx=XM_019826970.2
A3	60198194 60198267
refGene_gn=IL18RAP;refGene_tx=XM_003983898.5;refGene_gn=IL18RAP;refGen
e_tx=XM_019826970.2
A3	60216112 60216468
refGene_gn=IL18R1;refGene_tx=NM_001009863.2;refGene_gn=IL18R1;refGene_
tx=XM_023250909.1;refGene_gn=IL18R1;refGene_tx=XM_023250910.1
A3	60218290 60218449
refGene_gn=IL18R1;refGene_tx=NM_001009863.2;refGene_gn=IL18R1;refGene_
tx=XM_023250909.1;refGene_gn=IL18R1;refGene_tx=XM_023250910.1

A3 60223928 60224090
refGene_gn=IL18R1;refGene_tx=NM_001009863.2;refGene_gn=IL18R1;refGene_tx=XM_023250909.1;refGene_gn=IL18R1;refGene_tx=XM_023250910.1
A3 60225250 60225390
refGene_gn=IL18R1;refGene_tx=NM_001009863.2;refGene_gn=IL18R1;refGene_tx=XM_023250909.1;refGene_gn=IL18R1;refGene_tx=XM_023250910.1
A3 60227105 60227226
refGene_gn=IL18R1;refGene_tx=NM_001009863.2;refGene_gn=IL18R1;refGene_tx=XM_023250909.1;refGene_gn=IL18R1;refGene_tx=XM_023250910.1
A3 60230845 60230908
refGene_gn=IL18R1;refGene_tx=NM_001009863.2;refGene_gn=IL18R1;refGene_tx=XM_023250909.1;refGene_gn=IL18R1;refGene_tx=XM_023250910.1
A3 60233085 60233236
refGene_gn=IL18R1;refGene_tx=NM_001009863.2;refGene_gn=IL18R1;refGene_tx=XM_023250909.1;refGene_gn=IL18R1;refGene_tx=XM_023250910.1
A3 60236592 60236758
refGene_gn=IL18R1;refGene_tx=NM_001009863.2;refGene_gn=IL18R1;refGene_tx=XM_023250909.1;refGene_gn=IL18R1;refGene_tx=XM_023250910.1
A3 60240681 60240925
refGene_gn=IL18R1;refGene_tx=NM_001009863.2;refGene_gn=IL18R1;refGene_tx=XM_023250909.1;refGene_gn=IL18R1;refGene_tx=XM_023250910.1
A3 60244868 60244926
refGene_gn=IL18R1;refGene_tx=NM_001009863.2;refGene_gn=IL18R1;refGene_tx=XM_023250909.1;refGene_gn=IL18R1;refGene_tx=XM_023250910.1
A3 60256893 60257279
refGene_gn=IL1RL1;refGene_tx=XM_003983883.3;refGene_gn=IL1RL1;refGene_tx=XM_019826972.1;refGene_gn=IL1RL1;refGene_tx=XM_019826973.2;refGene_gn=IL1RL1;refGene_tx=XM_019826976.1
A3 60260144 60260312
refGene_gn=IL1RL1;refGene_tx=XM_003983883.3;refGene_gn=IL1RL1;refGene_tx=XM_019826972.1;refGene_gn=IL1RL1;refGene_tx=XM_019826973.2;refGene_gn=IL1RL1;refGene_tx=XM_019826976.1
A3 60261297 60261444
refGene_gn=IL1RL1;refGene_tx=XM_003983883.3;refGene_gn=IL1RL1;refGene_tx=XM_019826972.1;refGene_gn=IL1RL1;refGene_tx=XM_019826973.2
A3 60267120 60267266
refGene_gn=IL1RL1;refGene_tx=XM_003983883.3;refGene_gn=IL1RL1;refGene_tx=XM_019826972.1;refGene_gn=IL1RL1;refGene_tx=XM_019826973.2;refGene_gn=IL1RL1;refGene_tx=XM_019826976.1
A3 60267359 60267501
refGene_gn=IL1RL1;refGene_tx=XM_003983883.3;refGene_gn=IL1RL1;refGene_tx=XM_019826972.1;refGene_gn=IL1RL1;refGene_tx=XM_019826973.2;refGene_gn=IL1RL1;refGene_tx=XM_019826976.1
A3 60268907 60268979
refGene_gn=IL1RL1;refGene_tx=XM_003983883.3;refGene_gn=IL1RL1;refGene_tx=XM_019826972.1;refGene_gn=IL1RL1;refGene_tx=XM_019826973.2;refGene_gn=IL1RL1;refGene_tx=XM_019826976.1
A3 60271768 60271931
refGene_gn=IL1RL1;refGene_tx=XM_003983883.3;refGene_gn=IL1RL1;refGene_tx=XM_019826972.1;refGene_gn=IL1RL1;refGene_tx=XM_019826973.2;refGene_

gn=IL1RL1;refGene_tx=XM_019826976.1
A3 60272046 60272221
refGene_gn=IL1RL1;refGene_tx=XM_003983883.3;refGene_gn=IL1RL1;refGene_tx=XM_019826972.1;refGene_gn=IL1RL1;refGene_tx=XM_019826973.2;refGene_gn=IL1RL1;refGene_tx=XM_019826976.1
A3 60273250 60273461
refGene_gn=IL1RL1;refGene_tx=XM_003983883.3;refGene_gn=IL1RL1;refGene_tx=XM_019826972.1;refGene_gn=IL1RL1;refGene_tx=XM_019826973.2;refGene_gn=IL1RL1;refGene_tx=XM_019826976.1
A3 60274003 60274064
refGene_gn=IL1RL1;refGene_tx=XM_003983883.3;refGene_gn=IL1RL1;refGene_tx=XM_019826972.1;refGene_gn=IL1RL1;refGene_tx=XM_019826973.2;refGene_gn=IL1RL1;refGene_tx=XM_019826976.1
A3 60381870 60382238
refGene_gn=IL1RL2;refGene_tx=XM_011281068.3;refGene_gn=IL1RL2;refGene_tx=XM_019826977.2;refGene_gn=IL1RL2;refGene_tx=XM_019826978.2;refGene_gn=IL1RL2;refGene_tx=XM_019826979.2;refGene_gn=IL1RL2;refGene_tx=XM_019826983.2;refGene_gn=IL1RL2;refGene_tx=XM_023251556.1;refGene_gn=IL1RL2;refGene_tx=XM_023251557.1
A3 60384107 60384269
refGene_gn=IL1RL2;refGene_tx=XM_011281068.3;refGene_gn=IL1RL2;refGene_tx=XM_019826977.2;refGene_gn=IL1RL2;refGene_tx=XM_019826978.2;refGene_gn=IL1RL2;refGene_tx=XM_019826979.2;refGene_gn=IL1RL2;refGene_tx=XM_019826983.2;refGene_gn=IL1RL2;refGene_tx=XM_023251556.1;refGene_gn=IL1RL2;refGene_tx=XM_023251557.1
A3 60389203 60389259 refGene_gn=IL1RL2;refGene_tx=XM_019826984.2
A3 60393988 60394092 refGene_gn=IL1RL2;refGene_tx=XM_019826982.2
A3 60394183 60394327
refGene_gn=IL1RL2;refGene_tx=XM_011281068.3;refGene_gn=IL1RL2;refGene_tx=XM_019826977.2;refGene_gn=IL1RL2;refGene_tx=XM_019826978.2;refGene_gn=IL1RL2;refGene_tx=XM_019826979.2;refGene_gn=IL1RL2;refGene_tx=XM_019826982.2;refGene_gn=IL1RL2;refGene_tx=XM_019826983.2;refGene_gn=IL1RL2;refGene_tx=XM_019826984.2;refGene_gn=IL1RL2;refGene_tx=XM_023251556.1;refGene_gn=IL1RL2;refGene_tx=XM_023251557.1
A3 60396095 60396232
refGene_gn=IL1RL2;refGene_tx=XM_011281068.3;refGene_gn=IL1RL2;refGene_tx=XM_019826977.2;refGene_gn=IL1RL2;refGene_tx=XM_019826978.2;refGene_gn=IL1RL2;refGene_tx=XM_019826979.2;refGene_gn=IL1RL2;refGene_tx=XM_019826982.2;refGene_gn=IL1RL2;refGene_tx=XM_019826983.2;refGene_gn=IL1RL2;refGene_tx=XM_019826984.2;refGene_gn=IL1RL2;refGene_tx=XM_023251556.1;refGene_gn=IL1RL2;refGene_tx=XM_023251557.1
A3 60397230 60397360
refGene_gn=IL1RL2;refGene_tx=XM_011281068.3;refGene_gn=IL1RL2;refGene_tx=XM_019826977.2;refGene_gn=IL1RL2;refGene_tx=XM_019826978.2;refGene_gn=IL1RL2;refGene_tx=XM_019826979.2;refGene_gn=IL1RL2;refGene_tx=XM_019826982.2;refGene_gn=IL1RL2;refGene_tx=XM_019826983.2;refGene_gn=IL1RL2;refGene_tx=XM_019826984.2;refGene_gn=IL1RL2;refGene_tx=XM_023251556.1;refGene_gn=IL1RL2;refGene_tx=XM_023251557.1
A3 60404761 60404836
refGene_gn=IL1RL2;refGene_tx=XM_011281068.3;refGene_gn=IL1RL2;refGene_

tx=XM_019826977.2;refGene_gn=IL1RL2;refGene_tx=XM_019826978.2;refGene_gn=IL1RL2;refGene_tx=XM_019826979.2;refGene_gn=IL1RL2;refGene_tx=XM_019826982.2;refGene_gn=IL1RL2;refGene_tx=XM_019826983.2;refGene_gn=IL1RL2;refGene_tx=XM_019826984.2;refGene_gn=IL1RL2;refGene_tx=XM_023251556.1;refGene_gn=IL1RL2;refGene_tx=XM_023251557.1
A3 60406433 60406593
refGene_gn=IL1RL2;refGene_tx=XM_011281068.3;refGene_gn=IL1RL2;refGene_tx=XM_019826977.2;refGene_gn=IL1RL2;refGene_tx=XM_019826978.2;refGene_gn=IL1RL2;refGene_tx=XM_019826979.2;refGene_gn=IL1RL2;refGene_tx=XM_019826982.2;refGene_gn=IL1RL2;refGene_tx=XM_019826983.2;refGene_gn=IL1RL2;refGene_tx=XM_019826984.2;refGene_gn=IL1RL2;refGene_tx=XM_023251556.1;refGene_gn=IL1RL2;refGene_tx=XM_023251557.1
A3 60414751 60414947
refGene_gn=IL1RL2;refGene_tx=XM_011281068.3;refGene_gn=IL1RL2;refGene_tx=XM_019826977.2;refGene_gn=IL1RL2;refGene_tx=XM_019826978.2;refGene_gn=IL1RL2;refGene_tx=XM_019826979.2;refGene_gn=IL1RL2;refGene_tx=XM_019826982.2;refGene_gn=IL1RL2;refGene_tx=XM_019826983.2;refGene_gn=IL1RL2;refGene_tx=XM_019826984.2;refGene_gn=IL1RL2;refGene_tx=XM_023251556.1
A3 60417967 60418202
refGene_gn=IL1RL2;refGene_tx=XM_011281068.3;refGene_gn=IL1RL2;refGene_tx=XM_019826977.2;refGene_gn=IL1RL2;refGene_tx=XM_019826978.2;refGene_gn=IL1RL2;refGene_tx=XM_019826979.2;refGene_gn=IL1RL2;refGene_tx=XM_019826982.2;refGene_gn=IL1RL2;refGene_tx=XM_019826983.2;refGene_gn=IL1RL2;refGene_tx=XM_019826984.2;refGene_gn=IL1RL2;refGene_tx=XM_023251556.1;refGene_gn=IL1RL2;refGene_tx=XM_023251557.1
A3 60418976 60419034 refGene_gn=IL1RL2;refGene_tx=XM_011281068.3
A3 60419130 60419266 refGene_gn=IL1RL2;refGene_tx=XM_023251556.1
A3 60419500 60419948
refGene_gn=IL1RL2;refGene_tx=XM_019826977.2;refGene_gn=IL1RL2;refGene_tx=XM_019826978.2;refGene_gn=IL1RL2;refGene_tx=XM_019826979.2;refGene_gn=IL1RL2;refGene_tx=XM_019826982.2;refGene_gn=IL1RL2;refGene_tx=XM_019826984.2
A3 60432972 60433391
refGene_gn=IL1R1;refGene_tx=XM_003983884.5;refGene_gn=IL1R1;refGene_tx=XM_006930031.4;refGene_gn=IL1R1;refGene_tx=XM_006930032.4;refGene_gn=IL1R1;refGene_tx=XM_019826985.2;refGene_gn=IL1R1;refGene_tx=XM_019826986.2;refGene_gn=IL1R1;refGene_tx=XM_023251558.1;refGene_gn=IL1R1;refGene_tx=XM_023251559.1
A3 60433920 60434088
refGene_gn=IL1R1;refGene_tx=XM_003983884.5;refGene_gn=IL1R1;refGene_tx=XM_006930031.4;refGene_gn=IL1R1;refGene_tx=XM_006930032.4;refGene_gn=IL1R1;refGene_tx=XM_019826985.2;refGene_gn=IL1R1;refGene_tx=XM_019826986.2;refGene_gn=IL1R1;refGene_tx=XM_023251558.1;refGene_gn=IL1R1;refGene_tx=XM_023251559.1
A3 60434856 60435000
refGene_gn=IL1R1;refGene_tx=XM_003983884.5;refGene_gn=IL1R1;refGene_tx=XM_006930031.4;refGene_gn=IL1R1;refGene_tx=XM_006930032.4;refGene_gn=IL1R1;refGene_tx=XM_019826985.2;refGene_gn=IL1R1;refGene_tx=XM_019826986.2;refGene_gn=IL1R1;refGene_tx=XM_019826987.1;refGene_gn=IL1R1;refGe

ne_tx=XM_023251558.1;refGene_gn=IL1R1;refGene_tx=XM_023251559.1
A3 60437248 60437318 refGene_gn=IL1R1;refGene_tx=XM_019826987.1
A3 60437460 60437612
refGene_gn=IL1R1;refGene_tx=XM_003983884.5;refGene_gn=IL1R1;refGene_tx=XM_006930031.4;refGene_gn=IL1R1;refGene_tx=XM_006930032.4;refGene_gn=IL1R1;refGene_tx=XM_019826985.2;refGene_gn=IL1R1;refGene_tx=XM_019826986.2;refGene_gn=IL1R1;refGene_tx=XM_019826987.1;refGene_gn=IL1R1;refGene_tx=XM_023251558.1;refGene_gn=IL1R1;refGene_tx=XM_023251559.1
A3 60438372 60438490
refGene_gn=IL1R1;refGene_tx=XM_003983884.5;refGene_gn=IL1R1;refGene_tx=XM_006930031.4;refGene_gn=IL1R1;refGene_tx=XM_006930032.4;refGene_gn=IL1R1;refGene_tx=XM_019826985.2;refGene_gn=IL1R1;refGene_tx=XM_019826986.2;refGene_gn=IL1R1;refGene_tx=XM_019826987.1;refGene_gn=IL1R1;refGene_tx=XM_023251558.1;refGene_gn=IL1R1;refGene_tx=XM_023251559.1
A3 60441202 60441348 refGene_gn=IL1R1;refGene_tx=XM_019826988.2
A3 60443766 60443832
refGene_gn=IL1R1;refGene_tx=XM_003983884.5;refGene_gn=IL1R1;refGene_tx=XM_006930031.4;refGene_gn=IL1R1;refGene_tx=XM_006930032.4;refGene_gn=IL1R1;refGene_tx=XM_019826985.2;refGene_gn=IL1R1;refGene_tx=XM_019826986.2;refGene_gn=IL1R1;refGene_tx=XM_019826987.1;refGene_gn=IL1R1;refGene_tx=XM_019826988.2;refGene_gn=IL1R1;refGene_tx=XM_023251558.1;refGene_gn=IL1R1;refGene_tx=XM_023251559.1
A3 60445822 60445991
refGene_gn=IL1R1;refGene_tx=XM_003983884.5;refGene_gn=IL1R1;refGene_tx=XM_006930031.4;refGene_gn=IL1R1;refGene_tx=XM_006930032.4;refGene_gn=IL1R1;refGene_tx=XM_019826985.2;refGene_gn=IL1R1;refGene_tx=XM_019826987.1;refGene_gn=IL1R1;refGene_tx=XM_019826988.2;refGene_gn=IL1R1;refGene_tx=XM_023251558.1;refGene_gn=IL1R1;refGene_tx=XM_023251559.1
A3 60447016 60447206
refGene_gn=IL1R1;refGene_tx=XM_003983884.5;refGene_gn=IL1R1;refGene_tx=XM_006930031.4;refGene_gn=IL1R1;refGene_tx=XM_006930032.4;refGene_gn=IL1R1;refGene_tx=XM_019826985.2;refGene_gn=IL1R1;refGene_tx=XM_019826987.1;refGene_gn=IL1R1;refGene_tx=XM_019826988.2;refGene_gn=IL1R1;refGene_tx=XM_023251558.1;refGene_gn=IL1R1;refGene_tx=XM_023251559.1
A3 60447315 60447550
refGene_gn=IL1R1;refGene_tx=XM_003983884.5;refGene_gn=IL1R1;refGene_tx=XM_006930031.4;refGene_gn=IL1R1;refGene_tx=XM_006930032.4;refGene_gn=IL1R1;refGene_tx=XM_019826985.2;refGene_gn=IL1R1;refGene_tx=XM_019826987.1;refGene_gn=IL1R1;refGene_tx=XM_019826988.2;refGene_gn=IL1R1;refGene_tx=XM_023251558.1;refGene_gn=IL1R1;refGene_tx=XM_023251559.1
A3 60456787 60456845
refGene_gn=IL1R1;refGene_tx=XM_003983884.5;refGene_gn=IL1R1;refGene_tx=XM_006930031.4;refGene_gn=IL1R1;refGene_tx=XM_006930032.4;refGene_gn=IL1R1;refGene_tx=XM_019826985.2;refGene_gn=IL1R1;refGene_tx=XM_019826986.2;refGene_gn=IL1R1;refGene_tx=XM_019826987.1;refGene_gn=IL1R1;refGene_tx=XM_019826988.2;refGene_gn=IL1R1;refGene_tx=XM_023251558.1;refGene_gn=IL1R1;refGene_tx=XM_023251559.1
A3 60592051 60592191
refGene_gn=IL1R2;refGene_tx=XM_003983885.4;refGene_gn=IL1R2;refGene_tx=XM_019826989.2

A3 60594260 60594403
refGene_gn=IL1R2;refGene_tx=XM_003983885.4;refGene_gn=IL1R2;refGene_tx=XM_019826989.2
A3 60595943 60596079
refGene_gn=IL1R2;refGene_tx=XM_003983885.4;refGene_gn=IL1R2;refGene_tx=XM_019826989.2
A3 60597125 60597188
refGene_gn=IL1R2;refGene_tx=XM_003983885.4;refGene_gn=IL1R2;refGene_tx=XM_019826989.2
A3 60599315 60599490
refGene_gn=IL1R2;refGene_tx=XM_003983885.4;refGene_gn=IL1R2;refGene_tx=XM_019826989.2
A3 60602247 60602428
refGene_gn=IL1R2;refGene_tx=XM_003983885.4;refGene_gn=IL1R2;refGene_tx=XM_019826989.2
A3 60608328 60608593
refGene_gn=IL1R2;refGene_tx=XM_003983885.4;refGene_gn=IL1R2;refGene_tx=XM_019826989.2
A3 60609410 60609535
refGene_gn=IL1R2;refGene_tx=XM_003983885.4;refGene_gn=IL1R2;refGene_tx=XM_019826989.2
A3 60610542 60610571 refGene_gn=IL1R2;refGene_tx=XM_019826989.2
A3 60626008 60626022 refGene_gn=IL1R2;refGene_tx=XM_003983885.4
A3 60700322 60700406 refGene_gn=MAP4K4;refGene_tx=XM_023251560.1
A3 60702102 60702242 refGene_gn=MAP4K4;refGene_tx=XM_023251560.1
A3 60702998 60703158 refGene_gn=MAP4K4;refGene_tx=XM_023251560.1
A3 60703715 60703865 refGene_gn=MAP4K4;refGene_tx=XM_023251560.1
A3 60705881 60705982 refGene_gn=MAP4K4;refGene_tx=XM_023251560.1
A3 60708306 60708441 refGene_gn=MAP4K4;refGene_tx=XM_023251560.1
A3 60714562 60714706 refGene_gn=MAP4K4;refGene_tx=XM_023251560.1
A3 60721999 60722060 refGene_gn=MAP4K4;refGene_tx=XM_023251560.1
A3 60722061 60722171 refGene_gn=MAP4K4;refGene_tx=XM_023251560.1
A3 60722493 60722611 refGene_gn=MAP4K4;refGene_tx=XM_023251560.1
A3 60724688 60724880 refGene_gn=MAP4K4;refGene_tx=XM_023251560.1
A3 60725941 60726062 refGene_gn=MAP4K4;refGene_tx=XM_023251560.1
A3 60726855 60727031 refGene_gn=MAP4K4;refGene_tx=XM_023251560.1
A3 60729063 60729161 refGene_gn=MAP4K4;refGene_tx=XM_023251560.1
A3 60729860 60730012 refGene_gn=MAP4K4;refGene_tx=XM_023251560.1
A3 60731439 60731546 refGene_gn=MAP4K4;refGene_tx=XM_023251560.1
A3 60735207 60735369 refGene_gn=MAP4K4;refGene_tx=XM_023251560.1
A3 60736340 60736460 refGene_gn=MAP4K4;refGene_tx=XM_023251560.1
A3 60737132 60737219 refGene_gn=MAP4K4;refGene_tx=XM_023251560.1
A3 60741034 60741196 refGene_gn=MAP4K4;refGene_tx=XM_023251560.1
A3 60755142 60755353 refGene_gn=MAP4K4;refGene_tx=XM_023251560.1
A3 60756864 60756937 refGene_gn=MAP4K4;refGene_tx=XM_023251560.1
A3 60759273 60759449 refGene_gn=MAP4K4;refGene_tx=XM_023251560.1
A3 60762784 60762863 refGene_gn=MAP4K4;refGene_tx=XM_023251560.1
A3 60764326 60764381 refGene_gn=MAP4K4;refGene_tx=XM_023251560.1
A3 60766953 60767084 refGene_gn=MAP4K4;refGene_tx=XM_023251560.1
A3 60769619 60769710 refGene_gn=MAP4K4;refGene_tx=XM_023251560.1

A3 60773653 60773764 refGene_gn=MAP4K4;refGene_tx=XM_023251560.1
A3 60775014 60775140 refGene_gn=MAP4K4;refGene_tx=XM_023251560.1
A3 60807721 60807778 refGene_gn=MAP4K4;refGene_tx=XM_023251560.1
A3 60889274 60889340 refGene_gn=MAP4K4;refGene_tx=XM_023251560.1
A3 60889682 60889739 refGene_gn=MAP4K4;refGene_tx=XM_023251560.1
A3 60984911 60984983 refGene_gn=RFX8;refGene_tx=XM_019827009.2
A3 61088427 61088501
refGene_gn=RFX8;refGene_tx=XM_023251561.1;refGene_gn=RFX8;refGene_tx=XM_023251564.1
A3 61095770 61095894
refGene_gn=RFX8;refGene_tx=XM_006930033.3;refGene_gn=RFX8;refGene_tx=XM_019827006.2;refGene_gn=RFX8;refGene_tx=XM_019827007.1;refGene_gn=RFX8;refGene_tx=XM_019827008.2;refGene_gn=RFX8;refGene_tx=XM_023251561.1;refGene_gn=RFX8;refGene_tx=XM_023251562.1;refGene_gn=RFX8;refGene_tx=XM_023251563.1;refGene_gn=RFX8;refGene_tx=XM_023251564.1
A3 61106505 61106567
refGene_gn=RFX8;refGene_tx=XM_006930033.3;refGene_gn=RFX8;refGene_tx=XM_019827006.2;refGene_gn=RFX8;refGene_tx=XM_019827007.1;refGene_gn=RFX8;refGene_tx=XM_019827008.2;refGene_gn=RFX8;refGene_tx=XM_019827009.2;refGene_gn=RFX8;refGene_tx=XM_023251561.1;refGene_gn=RFX8;refGene_tx=XM_023251562.1;refGene_gn=RFX8;refGene_tx=XM_023251563.1;refGene_gn=RFX8;refGene_tx=XM_023251564.1
A3 61107663 61107750
refGene_gn=RFX8;refGene_tx=XM_006930033.3;refGene_gn=RFX8;refGene_tx=XM_019827006.2;refGene_gn=RFX8;refGene_tx=XM_019827007.1;refGene_gn=RFX8;refGene_tx=XM_019827008.2;refGene_gn=RFX8;refGene_tx=XM_019827009.2;refGene_gn=RFX8;refGene_tx=XM_023251561.1;refGene_gn=RFX8;refGene_tx=XM_023251562.1;refGene_gn=RFX8;refGene_tx=XM_023251563.1;refGene_gn=RFX8;refGene_tx=XM_023251564.1
A3 61126841 61126905
refGene_gn=RFX8;refGene_tx=XM_006930033.3;refGene_gn=RFX8;refGene_tx=XM_019827006.2;refGene_gn=RFX8;refGene_tx=XM_019827007.1;refGene_gn=RFX8;refGene_tx=XM_019827008.2;refGene_gn=RFX8;refGene_tx=XM_019827009.2;refGene_gn=RFX8;refGene_tx=XM_023251561.1;refGene_gn=RFX8;refGene_tx=XM_023251562.1;refGene_gn=RFX8;refGene_tx=XM_023251563.1;refGene_gn=RFX8;refGene_tx=XM_023251564.1
A3 61130867 61130978
refGene_gn=RFX8;refGene_tx=XM_006930033.3;refGene_gn=RFX8;refGene_tx=XM_019827006.2;refGene_gn=RFX8;refGene_tx=XM_019827007.1;refGene_gn=RFX8;refGene_tx=XM_019827008.2;refGene_gn=RFX8;refGene_tx=XM_019827009.2;refGene_gn=RFX8;refGene_tx=XM_023251561.1;refGene_gn=RFX8;refGene_tx=XM_023251562.1;refGene_gn=RFX8;refGene_tx=XM_023251563.1;refGene_gn=RFX8;refGene_tx=XM_023251564.1
A3 61131565 61131643
refGene_gn=RFX8;refGene_tx=XM_006930033.3;refGene_gn=RFX8;refGene_tx=XM_019827006.2;refGene_gn=RFX8;refGene_tx=XM_019827007.1;refGene_gn=RFX8;refGene_tx=XM_019827008.2;refGene_gn=RFX8;refGene_tx=XM_019827009.2;refGene_gn=RFX8;refGene_tx=XM_023251561.1;refGene_gn=RFX8;refGene_tx=XM_023251562.1;refGene_gn=RFX8;refGene_tx=XM_023251563.1;refGene_gn=RFX8;refGene_tx=XM_023251564.1

A3 61134234 61134348
refGene_gn=RFX8;refGene_tx=XM_006930033.3;refGene_gn=RFX8;refGene_tx=XM_019827006.2;refGene_gn=RFX8;refGene_tx=XM_019827007.1;refGene_gn=RFX8;refGene_tx=XM_019827008.2;refGene_gn=RFX8;refGene_tx=XM_019827009.2;refGene_gn=RFX8;refGene_tx=XM_023251561.1;refGene_gn=RFX8;refGene_tx=XM_023251562.1;refGene_gn=RFX8;refGene_tx=XM_023251563.1;refGene_gn=RFX8;refGene_tx=XM_023251564.1
A3 61135208 61135359
refGene_gn=RFX8;refGene_tx=XM_006930033.3;refGene_gn=RFX8;refGene_tx=XM_019827006.2;refGene_gn=RFX8;refGene_tx=XM_019827007.1;refGene_gn=RFX8;refGene_tx=XM_019827008.2;refGene_gn=RFX8;refGene_tx=XM_019827009.2;refGene_gn=RFX8;refGene_tx=XM_023251561.1;refGene_gn=RFX8;refGene_tx=XM_023251562.1;refGene_gn=RFX8;refGene_tx=XM_023251563.1
A3 61137608 61137667
refGene_gn=RFX8;refGene_tx=XM_006930033.3;refGene_gn=RFX8;refGene_tx=XM_019827006.2;refGene_gn=RFX8;refGene_tx=XM_019827007.1;refGene_gn=RFX8;refGene_tx=XM_019827008.2;refGene_gn=RFX8;refGene_tx=XM_019827009.2;refGene_gn=RFX8;refGene_tx=XM_023251561.1;refGene_gn=RFX8;refGene_tx=XM_023251562.1;refGene_gn=RFX8;refGene_tx=XM_023251563.1
A3 61139302 61139459
refGene_gn=RFX8;refGene_tx=XM_006930033.3;refGene_gn=RFX8;refGene_tx=XM_019827006.2;refGene_gn=RFX8;refGene_tx=XM_019827007.1;refGene_gn=RFX8;refGene_tx=XM_019827008.2;refGene_gn=RFX8;refGene_tx=XM_019827009.2;refGene_gn=RFX8;refGene_tx=XM_023251561.1;refGene_gn=RFX8;refGene_tx=XM_023251562.1;refGene_gn=RFX8;refGene_tx=XM_023251563.1;refGene_gn=RFX8;refGene_tx=XM_023251564.1
A3 61141875 61141970
refGene_gn=RFX8;refGene_tx=XM_006930033.3;refGene_gn=RFX8;refGene_tx=XM_019827006.2;refGene_gn=RFX8;refGene_tx=XM_019827007.1;refGene_gn=RFX8;refGene_tx=XM_019827008.2;refGene_gn=RFX8;refGene_tx=XM_019827009.2;refGene_gn=RFX8;refGene_tx=XM_023251561.1;refGene_gn=RFX8;refGene_tx=XM_023251562.1;refGene_gn=RFX8;refGene_tx=XM_023251563.1;refGene_gn=RFX8;refGene_tx=XM_023251564.1
A3 61145113 61145228
refGene_gn=RFX8;refGene_tx=XM_006930033.3;refGene_gn=RFX8;refGene_tx=XM_019827006.2;refGene_gn=RFX8;refGene_tx=XM_019827007.1;refGene_gn=RFX8;refGene_tx=XM_019827008.2;refGene_gn=RFX8;refGene_tx=XM_019827009.2;refGene_gn=RFX8;refGene_tx=XM_023251561.1;refGene_gn=RFX8;refGene_tx=XM_023251562.1;refGene_gn=RFX8;refGene_tx=XM_023251563.1;refGene_gn=RFX8;refGene_tx=XM_023251564.1
A3 61148074 61148391
refGene_gn=RFX8;refGene_tx=XM_006930033.3;refGene_gn=RFX8;refGene_tx=XM_019827006.2;refGene_gn=RFX8;refGene_tx=XM_019827007.1;refGene_gn=RFX8;refGene_tx=XM_019827008.2;refGene_gn=RFX8;refGene_tx=XM_019827009.2;refGene_gn=RFX8;refGene_tx=XM_023251561.1;refGene_gn=RFX8;refGene_tx=XM_023251562.1;refGene_gn=RFX8;refGene_tx=XM_023251563.1;refGene_gn=RFX8;refGene_tx=XM_023251564.1
A3 61152710 61152887
refGene_gn=RFX8;refGene_tx=XM_006930033.3;refGene_gn=RFX8;refGene_tx=XM_019827006.2;refGene_gn=RFX8;refGene_tx=XM_019827007.1;refGene_gn=RFX

8;refGene_tx=XM_019827008.2;refGene_gn=RFX8;refGene_tx=XM_019827009.2;
refGene_gn=RFX8;refGene_tx=XM_023251561.1;refGene_gn=RFX8;refGene_tx=X
M_023251562.1;refGene_gn=RFX8;refGene_tx=XM_023251563.1;refGene_gn=RFX
8;refGene_tx=XM_023251564.1
A3 61167335 61167752 refGene_gn=CREG2;refGene_tx=XM_023251565.1
A3 61170401 61170571 refGene_gn=CREG2;refGene_tx=XM_023251565.1
A3 61195419 61195533 refGene_gn=CREG2;refGene_tx=XM_023251565.1
A3 61198659 61198807 refGene_gn=CREG2;refGene_tx=XM_023251565.1
A3 61248686 61249143 refGene_gn=RNF149;refGene_tx=XM_023251568.1
A3 61260366 61260617 refGene_gn=RNF149;refGene_tx=XM_023251568.1
A3 61261552 61261621 refGene_gn=RNF149;refGene_tx=XM_023251568.1
A3 61267273 61267356 refGene_gn=RNF149;refGene_tx=XM_023251568.1
A3 61269513 61269610 refGene_gn=RNF149;refGene_tx=XM_023251568.1
A3 61272932 61273131 refGene_gn=RNF149;refGene_tx=XM_023251568.1
A3 61276781 61276825 refGene_gn=RNF149;refGene_tx=XM_023251568.1
A3 61283051 61283249
refGene_gn=CNOT11;refGene_tx=XM_023251566.1;refGene_gn=CNOT11;refGene_
tx=XM_023251567.1
A3 61283329 61283426
refGene_gn=CNOT11;refGene_tx=XM_023251566.1;refGene_gn=CNOT11;refGene_
tx=XM_023251567.1
A3 61284816 61285019
refGene_gn=CNOT11;refGene_tx=XM_023251566.1;refGene_gn=CNOT11;refGene_
tx=XM_023251567.1
A3 61286254 61286457
refGene_gn=CNOT11;refGene_tx=XM_023251566.1;refGene_gn=CNOT11;refGene_
tx=XM_023251567.1
A3 61288118 61288271 refGene_gn=CNOT11;refGene_tx=XM_023251566.1
A3 61296575 61296740
refGene_gn=CNOT11;refGene_tx=XM_023251566.1;refGene_gn=CNOT11;refGene_
tx=XM_023251567.1
A3 61299600 61300123
refGene_gn=CNOT11;refGene_tx=XM_023251566.1;refGene_gn=CNOT11;refGene_
tx=XM_023251567.1
A3 61323588 61323730
refGene_gn=LOC109498518;refGene_tx=XM_023251569.1;refGene_gn=LOC109498
518;refGene_tx=XM_023251570.1;refGene_gn=LOC109498518;refGene_tx=XM_02
3251571.1
A3 61354848 61354899
refGene_gn=LOC109498518;refGene_tx=XM_023251569.1;refGene_gn=LOC109498
518;refGene_tx=XM_023251571.1
A3 61360530 61360691
refGene_gn=LOC109498518;refGene_tx=XM_023251569.1;refGene_gn=LOC109498
518;refGene_tx=XM_023251570.1;refGene_gn=LOC109498518;refGene_tx=XM_02
3251571.1
A3 61362216 61362606
refGene_gn=LOC109498518;refGene_tx=XM_023251569.1;refGene_gn=LOC109498
518;refGene_tx=XM_023251570.1;refGene_gn=LOC109498518;refGene_tx=XM_02
3251571.1
A3 61363795 61363922

refGene_gn=TBC1D8;refGene_tx=XM_023251572.1;refGene_gn=TBC1D8;refGene_tx=XM_023251573.1;refGene_gn=TBC1D8;refGene_tx=XM_023251574.1;refGene_gn=TBC1D8;refGene_tx=XM_023251575.1;refGene_gn=TBC1D8;refGene_tx=XM_023251576.1
A3 61404515 61404671
refGene_gn=TBC1D8;refGene_tx=XM_023251572.1;refGene_gn=TBC1D8;refGene_tx=XM_023251573.1;refGene_gn=TBC1D8;refGene_tx=XM_023251574.1;refGene_gn=TBC1D8;refGene_tx=XM_023251575.1
A3 61423896 61424015
refGene_gn=TBC1D8;refGene_tx=XM_023251572.1;refGene_gn=TBC1D8;refGene_tx=XM_023251573.1;refGene_gn=TBC1D8;refGene_tx=XM_023251574.1;refGene_gn=TBC1D8;refGene_tx=XM_023251575.1;refGene_gn=TBC1D8;refGene_tx=XM_023251576.1
A3 61428401 61428630
refGene_gn=TBC1D8;refGene_tx=XM_023251572.1;refGene_gn=TBC1D8;refGene_tx=XM_023251573.1;refGene_gn=TBC1D8;refGene_tx=XM_023251574.1;refGene_gn=TBC1D8;refGene_tx=XM_023251575.1;refGene_gn=TBC1D8;refGene_tx=XM_023251576.1
A3 61431394 61431635
refGene_gn=TBC1D8;refGene_tx=XM_023251572.1;refGene_gn=TBC1D8;refGene_tx=XM_023251573.1;refGene_gn=TBC1D8;refGene_tx=XM_023251574.1;refGene_gn=TBC1D8;refGene_tx=XM_023251575.1;refGene_gn=TBC1D8;refGene_tx=XM_023251576.1
A3 61442369 61442577
refGene_gn=TBC1D8;refGene_tx=XM_023251572.1;refGene_gn=TBC1D8;refGene_tx=XM_023251573.1;refGene_gn=TBC1D8;refGene_tx=XM_023251574.1;refGene_gn=TBC1D8;refGene_tx=XM_023251575.1;refGene_gn=TBC1D8;refGene_tx=XM_023251576.1
A3 61444189 61444384
refGene_gn=TBC1D8;refGene_tx=XM_023251572.1;refGene_gn=TBC1D8;refGene_tx=XM_023251573.1;refGene_gn=TBC1D8;refGene_tx=XM_023251574.1;refGene_gn=TBC1D8;refGene_tx=XM_023251575.1;refGene_gn=TBC1D8;refGene_tx=XM_023251576.1
A3 61445327 61445507
refGene_gn=TBC1D8;refGene_tx=XM_023251572.1;refGene_gn=TBC1D8;refGene_tx=XM_023251573.1;refGene_gn=TBC1D8;refGene_tx=XM_023251574.1;refGene_gn=TBC1D8;refGene_tx=XM_023251575.1;refGene_gn=TBC1D8;refGene_tx=XM_023251576.1
A3 61446904 61447055
refGene_gn=TBC1D8;refGene_tx=XM_023251572.1;refGene_gn=TBC1D8;refGene_tx=XM_023251573.1;refGene_gn=TBC1D8;refGene_tx=XM_023251574.1;refGene_gn=TBC1D8;refGene_tx=XM_023251575.1;refGene_gn=TBC1D8;refGene_tx=XM_023251576.1
A3 61449562 61449777
refGene_gn=TBC1D8;refGene_tx=XM_023251572.1;refGene_gn=TBC1D8;refGene_tx=XM_023251573.1;refGene_gn=TBC1D8;refGene_tx=XM_023251574.1;refGene_gn=TBC1D8;refGene_tx=XM_023251575.1;refGene_gn=TBC1D8;refGene_tx=XM_023251576.1
A3 61450694 61450812
refGene_gn=TBC1D8;refGene_tx=XM_023251572.1;refGene_gn=TBC1D8;refGene_

tx=XM_023251573.1;refGene_gn=TBC1D8;refGene_tx=XM_023251574.1;refGene_gn=TBC1D8;refGene_tx=XM_023251575.1;refGene_gn=TBC1D8;refGene_tx=XM_023251576.1
A3 61453151 61453437
refGene_gn=TBC1D8;refGene_tx=XM_023251572.1;refGene_gn=TBC1D8;refGene_tx=XM_023251573.1;refGene_gn=TBC1D8;refGene_tx=XM_023251574.1;refGene_gn=TBC1D8;refGene_tx=XM_023251575.1;refGene_gn=TBC1D8;refGene_tx=XM_023251576.1
A3 61454185 61454315
refGene_gn=TBC1D8;refGene_tx=XM_023251572.1;refGene_gn=TBC1D8;refGene_tx=XM_023251573.1;refGene_gn=TBC1D8;refGene_tx=XM_023251574.1;refGene_gn=TBC1D8;refGene_tx=XM_023251575.1;refGene_gn=TBC1D8;refGene_tx=XM_023251576.1
A3 61454465 61454564
refGene_gn=TBC1D8;refGene_tx=XM_023251572.1;refGene_gn=TBC1D8;refGene_tx=XM_023251573.1;refGene_gn=TBC1D8;refGene_tx=XM_023251574.1;refGene_gn=TBC1D8;refGene_tx=XM_023251575.1;refGene_gn=TBC1D8;refGene_tx=XM_023251576.1
A3 61455259 61455328
refGene_gn=TBC1D8;refGene_tx=XM_023251572.1;refGene_gn=TBC1D8;refGene_tx=XM_023251573.1;refGene_gn=TBC1D8;refGene_tx=XM_023251574.1;refGene_gn=TBC1D8;refGene_tx=XM_023251575.1;refGene_gn=TBC1D8;refGene_tx=XM_023251576.1
A3 61460680 61460921
refGene_gn=TBC1D8;refGene_tx=XM_023251572.1;refGene_gn=TBC1D8;refGene_tx=XM_023251573.1;refGene_gn=TBC1D8;refGene_tx=XM_023251574.1;refGene_gn=TBC1D8;refGene_tx=XM_023251575.1;refGene_gn=TBC1D8;refGene_tx=XM_023251576.1
A3 61461467 61461533
refGene_gn=TBC1D8;refGene_tx=XM_023251572.1;refGene_gn=TBC1D8;refGene_tx=XM_023251573.1;refGene_gn=TBC1D8;refGene_tx=XM_023251574.1;refGene_gn=TBC1D8;refGene_tx=XM_023251575.1;refGene_gn=TBC1D8;refGene_tx=XM_023251576.1
A3 61472888 61472978
refGene_gn=TBC1D8;refGene_tx=XM_023251572.1;refGene_gn=TBC1D8;refGene_tx=XM_023251573.1;refGene_gn=TBC1D8;refGene_tx=XM_023251574.1;refGene_gn=TBC1D8;refGene_tx=XM_023251575.1;refGene_gn=TBC1D8;refGene_tx=XM_023251576.1
A3 61473366 61473464
refGene_gn=TBC1D8;refGene_tx=XM_023251572.1;refGene_gn=TBC1D8;refGene_tx=XM_023251573.1;refGene_gn=TBC1D8;refGene_tx=XM_023251574.1;refGene_gn=TBC1D8;refGene_tx=XM_023251575.1;refGene_gn=TBC1D8;refGene_tx=XM_023251576.1
A3 61474779 61475241
refGene_gn=TBC1D8;refGene_tx=XM_023251572.1;refGene_gn=TBC1D8;refGene_tx=XM_023251573.1;refGene_gn=TBC1D8;refGene_tx=XM_023251574.1;refGene_gn=TBC1D8;refGene_tx=XM_023251575.1;refGene_gn=TBC1D8;refGene_tx=XM_023251576.1
A3 61477098 61477130
refGene_gn=RPL31;refGene_tx=XM_003983889.3;refGene_gn=RPL31;refGene_tx

=XM_023251577.1
A3	61477454	61477567
refGene_gn=RPL31;refGene_tx=XM_003983889.3;refGene_gn=RPL31;refGene_tx
=XM_023251577.1
A3	61478887	61479013
refGene_gn=RPL31;refGene_tx=XM_003983889.3;refGene_gn=RPL31;refGene_tx
=XM_023251577.1
A3	61484977	61485048	refGene_gn=RPL31;refGene_tx=XM_023251577.1
A3	61485054	61485161	refGene_gn=RPL31;refGene_tx=XM_003983889.3
A3	61491370	61491578
refGene_gn=NPAS2;refGene_tx=XM_003983890.4;refGene_gn=NPAS2;refGene_tx
=XM_019827020.2;refGene_gn=NPAS2;refGene_tx=XM_019827021.2;refGene_gn=
NPAS2;refGene_tx=XM_023251578.1;refGene_gn=NPAS2;refGene_tx=XM_0232515
79.1;refGene_gn=NPAS2;refGene_tx=XM_023251580.1;refGene_gn=NPAS2;refGe
ne_tx=XM_023251581.1;refGene_gn=NPAS2;refGene_tx=XM_023251582.1;refGen
e_gn=NPAS2;refGene_tx=XM_023251583.1
A3	61493465	61493646
refGene_gn=NPAS2;refGene_tx=XM_003983890.4;refGene_gn=NPAS2;refGene_tx
=XM_019827020.2;refGene_gn=NPAS2;refGene_tx=XM_019827021.2;refGene_gn=
NPAS2;refGene_tx=XM_023251578.1;refGene_gn=NPAS2;refGene_tx=XM_0232515
79.1;refGene_gn=NPAS2;refGene_tx=XM_023251580.1;refGene_gn=NPAS2;refGe
ne_tx=XM_023251582.1;refGene_gn=NPAS2;refGene_tx=XM_023251583.1
A3	61496075	61496168
refGene_gn=NPAS2;refGene_tx=XM_003983890.4;refGene_gn=NPAS2;refGene_tx
=XM_019827020.2;refGene_gn=NPAS2;refGene_tx=XM_019827021.2;refGene_gn=
NPAS2;refGene_tx=XM_023251578.1;refGene_gn=NPAS2;refGene_tx=XM_0232515
79.1;refGene_gn=NPAS2;refGene_tx=XM_023251581.1;refGene_gn=NPAS2;refGe
ne_tx=XM_023251582.1
A3	61496452	61496643
refGene_gn=NPAS2;refGene_tx=XM_003983890.4;refGene_gn=NPAS2;refGene_tx
=XM_019827020.2;refGene_gn=NPAS2;refGene_tx=XM_019827021.2;refGene_gn=
NPAS2;refGene_tx=XM_023251578.1;refGene_gn=NPAS2;refGene_tx=XM_0232515
79.1;refGene_gn=NPAS2;refGene_tx=XM_023251580.1;refGene_gn=NPAS2;refGe
ne_tx=XM_023251581.1;refGene_gn=NPAS2;refGene_tx=XM_023251582.1
A3	61498630	61498828
refGene_gn=NPAS2;refGene_tx=XM_003983890.4;refGene_gn=NPAS2;refGene_tx
=XM_019827020.2;refGene_gn=NPAS2;refGene_tx=XM_019827021.2;refGene_gn=
NPAS2;refGene_tx=XM_023251578.1;refGene_gn=NPAS2;refGene_tx=XM_0232515
79.1;refGene_gn=NPAS2;refGene_tx=XM_023251580.1;refGene_gn=NPAS2;refGe
ne_tx=XM_023251581.1;refGene_gn=NPAS2;refGene_tx=XM_023251582.1;refGen
e_gn=NPAS2;refGene_tx=XM_023251583.1
A3	61504353	61504500
refGene_gn=NPAS2;refGene_tx=XM_003983890.4;refGene_gn=NPAS2;refGene_tx
=XM_019827020.2;refGene_gn=NPAS2;refGene_tx=XM_019827021.2;refGene_gn=
NPAS2;refGene_tx=XM_023251578.1;refGene_gn=NPAS2;refGene_tx=XM_0232515
79.1;refGene_gn=NPAS2;refGene_tx=XM_023251580.1;refGene_gn=NPAS2;refGe
ne_tx=XM_023251581.1;refGene_gn=NPAS2;refGene_tx=XM_023251582.1;refGen
e_gn=NPAS2;refGene_tx=XM_023251583.1
A3	61508262	61508352
refGene_gn=NPAS2;refGene_tx=XM_003983890.4;refGene_gn=NPAS2;refGene_tx

=XM_019827020.2;refGene_gn=NPAS2;refGene_tx=XM_019827021.2;refGene_gn=NPAS2;refGene_tx=XM_023251578.1;refGene_gn=NPAS2;refGene_tx=XM_023251579.1;refGene_gn=NPAS2;refGene_tx=XM_023251580.1;refGene_gn=NPAS2;refGene_tx=XM_023251581.1;refGene_gn=NPAS2;refGene_tx=XM_023251582.1;refGene_gn=NPAS2;refGene_tx=XM_023251583.1
A3 61510556 61510666
refGene_gn=NPAS2;refGene_tx=XM_003983890.4;refGene_gn=NPAS2;refGene_tx=XM_019827020.2;refGene_gn=NPAS2;refGene_tx=XM_019827021.2;refGene_gn=NPAS2;refGene_tx=XM_023251578.1;refGene_gn=NPAS2;refGene_tx=XM_023251579.1;refGene_gn=NPAS2;refGene_tx=XM_023251580.1;refGene_gn=NPAS2;refGene_tx=XM_023251581.1;refGene_gn=NPAS2;refGene_tx=XM_023251582.1;refGene_gn=NPAS2;refGene_tx=XM_023251583.1
A3 61511192 61511334
refGene_gn=NPAS2;refGene_tx=XM_003983890.4;refGene_gn=NPAS2;refGene_tx=XM_019827020.2;refGene_gn=NPAS2;refGene_tx=XM_019827021.2;refGene_gn=NPAS2;refGene_tx=XM_023251578.1;refGene_gn=NPAS2;refGene_tx=XM_023251579.1;refGene_gn=NPAS2;refGene_tx=XM_023251580.1;refGene_gn=NPAS2;refGene_tx=XM_023251581.1;refGene_gn=NPAS2;refGene_tx=XM_023251582.1;refGene_gn=NPAS2;refGene_tx=XM_023251583.1
A3 61514084 61514169
refGene_gn=NPAS2;refGene_tx=XM_003983890.4;refGene_gn=NPAS2;refGene_tx=XM_019827020.2;refGene_gn=NPAS2;refGene_tx=XM_019827021.2;refGene_gn=NPAS2;refGene_tx=XM_023251578.1;refGene_gn=NPAS2;refGene_tx=XM_023251579.1;refGene_gn=NPAS2;refGene_tx=XM_023251580.1;refGene_gn=NPAS2;refGene_tx=XM_023251581.1;refGene_gn=NPAS2;refGene_tx=XM_023251582.1;refGene_gn=NPAS2;refGene_tx=XM_023251583.1
A3 61516310 61516458
refGene_gn=NPAS2;refGene_tx=XM_003983890.4;refGene_gn=NPAS2;refGene_tx=XM_019827020.2;refGene_gn=NPAS2;refGene_tx=XM_019827021.2;refGene_gn=NPAS2;refGene_tx=XM_023251578.1;refGene_gn=NPAS2;refGene_tx=XM_023251579.1;refGene_gn=NPAS2;refGene_tx=XM_023251580.1;refGene_gn=NPAS2;refGene_tx=XM_023251581.1;refGene_gn=NPAS2;refGene_tx=XM_023251582.1;refGene_gn=NPAS2;refGene_tx=XM_023251583.1
A3 61518885 61518992
refGene_gn=NPAS2;refGene_tx=XM_003983890.4;refGene_gn=NPAS2;refGene_tx=XM_019827020.2;refGene_gn=NPAS2;refGene_tx=XM_019827021.2;refGene_gn=NPAS2;refGene_tx=XM_023251578.1;refGene_gn=NPAS2;refGene_tx=XM_023251579.1;refGene_gn=NPAS2;refGene_tx=XM_023251580.1;refGene_gn=NPAS2;refGene_tx=XM_023251581.1;refGene_gn=NPAS2;refGene_tx=XM_023251582.1;refGene_gn=NPAS2;refGene_tx=XM_023251583.1
A3 61519742 61519825
refGene_gn=NPAS2;refGene_tx=XM_003983890.4;refGene_gn=NPAS2;refGene_tx=XM_019827020.2;refGene_gn=NPAS2;refGene_tx=XM_019827021.2;refGene_gn=NPAS2;refGene_tx=XM_023251578.1;refGene_gn=NPAS2;refGene_tx=XM_023251579.1;refGene_gn=NPAS2;refGene_tx=XM_023251580.1;refGene_gn=NPAS2;refGene_tx=XM_023251581.1;refGene_gn=NPAS2;refGene_tx=XM_023251582.1;refGene_gn=NPAS2;refGene_tx=XM_023251583.1
A3 61520517 61520636
refGene_gn=NPAS2;refGene_tx=XM_003983890.4;refGene_gn=NPAS2;refGene_tx=XM_019827020.2;refGene_gn=NPAS2;refGene_tx=XM_019827021.2;refGene_gn=

NPAS2;refGene_tx=XM_023251578.1;refGene_gn=NPAS2;refGene_tx=XM_0232515
79.1;refGene_gn=NPAS2;refGene_tx=XM_023251580.1;refGene_gn=NPAS2;refGe
ne_tx=XM_023251581.1;refGene_gn=NPAS2;refGene_tx=XM_023251582.1;refGen
e_gn=NPAS2;refGene_tx=XM_023251583.1
A3 61531372 61531486
refGene_gn=NPAS2;refGene_tx=XM_003983890.4;refGene_gn=NPAS2;refGene_tx
=XM_019827020.2;refGene_gn=NPAS2;refGene_tx=XM_019827021.2;refGene_gn=
NPAS2;refGene_tx=XM_023251578.1;refGene_gn=NPAS2;refGene_tx=XM_0232515
79.1;refGene_gn=NPAS2;refGene_tx=XM_023251580.1;refGene_gn=NPAS2;refGe
ne_tx=XM_023251581.1;refGene_gn=NPAS2;refGene_tx=XM_023251582.1;refGen
e_gn=NPAS2;refGene_tx=XM_023251583.1
A3 61532511 61532632
refGene_gn=NPAS2;refGene_tx=XM_003983890.4;refGene_gn=NPAS2;refGene_tx
=XM_019827020.2;refGene_gn=NPAS2;refGene_tx=XM_019827021.2;refGene_gn=
NPAS2;refGene_tx=XM_023251578.1;refGene_gn=NPAS2;refGene_tx=XM_0232515
79.1;refGene_gn=NPAS2;refGene_tx=XM_023251580.1;refGene_gn=NPAS2;refGe
ne_tx=XM_023251581.1;refGene_gn=NPAS2;refGene_tx=XM_023251582.1;refGen
e_gn=NPAS2;refGene_tx=XM_023251583.1
A3 61542705 61542795
refGene_gn=NPAS2;refGene_tx=XM_003983890.4;refGene_gn=NPAS2;refGene_tx
=XM_019827020.2;refGene_gn=NPAS2;refGene_tx=XM_023251578.1;refGene_gn=
NPAS2;refGene_tx=XM_023251579.1;refGene_gn=NPAS2;refGene_tx=XM_0232515
80.1;refGene_gn=NPAS2;refGene_tx=XM_023251581.1;refGene_gn=NPAS2;refGe
ne_tx=XM_023251582.1;refGene_gn=NPAS2;refGene_tx=XM_023251583.1
A3 61547460 61547552
refGene_gn=NPAS2;refGene_tx=XM_003983890.4;refGene_gn=NPAS2;refGene_tx
=XM_019827020.2;refGene_gn=NPAS2;refGene_tx=XM_023251578.1;refGene_gn=
NPAS2;refGene_tx=XM_023251579.1;refGene_gn=NPAS2;refGene_tx=XM_0232515
80.1;refGene_gn=NPAS2;refGene_tx=XM_023251581.1;refGene_gn=NPAS2;refGe
ne_tx=XM_023251582.1;refGene_gn=NPAS2;refGene_tx=XM_023251583.1
A3 61554534 61554683
refGene_gn=NPAS2;refGene_tx=XM_003983890.4;refGene_gn=NPAS2;refGene_tx
=XM_023251578.1;refGene_gn=NPAS2;refGene_tx=XM_023251579.1;refGene_gn=
NPAS2;refGene_tx=XM_023251580.1;refGene_gn=NPAS2;refGene_tx=XM_0232515
81.1;refGene_gn=NPAS2;refGene_tx=XM_023251582.1;refGene_gn=NPAS2;refGe
ne_tx=XM_023251583.1
A3 61574519 61574551
refGene_gn=NPAS2;refGene_tx=XM_003983890.4;refGene_gn=NPAS2;refGene_tx
=XM_023251578.1;refGene_gn=NPAS2;refGene_tx=XM_023251579.1;refGene_gn=
NPAS2;refGene_tx=XM_023251580.1;refGene_gn=NPAS2;refGene_tx=XM_0232515
81.1;refGene_gn=NPAS2;refGene_tx=XM_023251582.1;refGene_gn=NPAS2;refGe
ne_tx=XM_023251583.1
A3 61859137 61859280
refGene_gn=PDCL3;refGene_tx=XM_003983891.4;refGene_gn=PDCL3;refGene_tx
=XM_019827023.2
A3 61862684 61862893
refGene_gn=PDCL3;refGene_tx=XM_003983891.4;refGene_gn=PDCL3;refGene_tx
=XM_019827023.2
A3 61863674 61863818
refGene_gn=PDCL3;refGene_tx=XM_003983891.4;refGene_gn=PDCL3;refGene_tx

=XM_019827023.2
A3	61864083	61864174
refGene_gn=PDCL3;refGene_tx=XM_003983891.4;refGene_gn=PDCL3;refGene_tx=XM_019827023.2
A3	61866286	61866416
refGene_gn=PDCL3;refGene_tx=XM_003983891.4;refGene_gn=PDCL3;refGene_tx=XM_019827023.2
A3	61867746	61867752
refGene_gn=PDCL3;refGene_tx=XM_003983891.4;refGene_gn=PDCL3;refGene_tx=XM_019827023.2
A3	61902369	61902402
refGene_gn=NMS;refGene_tx=XM_006930035.4;refGene_gn=NMS;refGene_tx=XM_019827032.2
A3	61903308	61903347
refGene_gn=NMS;refGene_tx=XM_006930034.4;refGene_gn=NMS;refGene_tx=XM_019827029.2;refGene_gn=NMS;refGene_tx=XM_019827030.2
A3	61904418	61904460
refGene_gn=NMS;refGene_tx=XM_006930034.4;refGene_gn=NMS;refGene_tx=XM_006930035.4;refGene_gn=NMS;refGene_tx=XM_019827029.2;refGene_gn=NMS;refGene_tx=XM_019827030.2;refGene_gn=NMS;refGene_tx=XM_019827032.2
A3	61905025	61905061
refGene_gn=NMS;refGene_tx=XM_006930034.4;refGene_gn=NMS;refGene_tx=XM_006930035.4;refGene_gn=NMS;refGene_tx=XM_019827029.2;refGene_gn=NMS;refGene_tx=XM_019827030.2;refGene_gn=NMS;refGene_tx=XM_019827032.2
A3	61906499	61906577
refGene_gn=NMS;refGene_tx=XM_006930034.4;refGene_gn=NMS;refGene_tx=XM_006930035.4
A3	61908077	61908101
refGene_gn=NMS;refGene_tx=XM_006930034.4;refGene_gn=NMS;refGene_tx=XM_006930035.4;refGene_gn=NMS;refGene_tx=XM_019827029.2
A3	61911325	61911364
refGene_gn=NMS;refGene_tx=XM_006930034.4;refGene_gn=NMS;refGene_tx=XM_006930035.4;refGene_gn=NMS;refGene_tx=XM_019827029.2;refGene_gn=NMS;refGene_tx=XM_019827030.2;refGene_gn=NMS;refGene_tx=XM_019827032.2
A3	61911693	61911749
refGene_gn=NMS;refGene_tx=XM_006930034.4;refGene_gn=NMS;refGene_tx=XM_006930035.4;refGene_gn=NMS;refGene_tx=XM_019827029.2;refGene_gn=NMS;refGene_tx=XM_019827030.2;refGene_gn=NMS;refGene_tx=XM_019827032.2
A3	61913638	61913972
refGene_gn=NMS;refGene_tx=XM_006930034.4;refGene_gn=NMS;refGene_tx=XM_006930035.4;refGene_gn=NMS;refGene_tx=XM_019827029.2;refGene_gn=NMS;refGene_tx=XM_019827030.2;refGene_gn=NMS;refGene_tx=XM_019827032.2
A3	61959708	61959751
refGene_gn=CHST10;refGene_tx=XM_019827033.2;refGene_gn=CHST10;refGene_tx=XM_019827035.2;refGene_gn=CHST10;refGene_tx=XM_023251584.1
A3	61960302	61960434
refGene_gn=CHST10;refGene_tx=XM_019827033.2;refGene_gn=CHST10;refGene_tx=XM_019827035.2;refGene_gn=CHST10;refGene_tx=XM_023251584.1
A3	61967435	61967527
refGene_gn=CHST10;refGene_tx=XM_019827033.2;refGene_gn=CHST10;refGene_

tx=XM_019827035.2;refGene_gn=CHST10;refGene_tx=XM_023251584.1
A3 61967884 61967926 refGene_gn=CHST10;refGene_tx=XM_019827036.2
A3 61972969 61973204
refGene_gn=CHST10;refGene_tx=XM_019827033.2;refGene_gn=CHST10;refGene_tx=XM_019827035.2;refGene_gn=CHST10;refGene_tx=XM_019827036.2;refGene_gn=CHST10;refGene_tx=XM_023251584.1
A3 61975357 61975463
refGene_gn=CHST10;refGene_tx=XM_019827033.2;refGene_gn=CHST10;refGene_tx=XM_019827035.2;refGene_gn=CHST10;refGene_tx=XM_019827036.2;refGene_gn=CHST10;refGene_tx=XM_023251584.1
A3 61977155 61977693
refGene_gn=CHST10;refGene_tx=XM_019827033.2;refGene_gn=CHST10;refGene_tx=XM_019827035.2;refGene_gn=CHST10;refGene_tx=XM_019827036.2;refGene_gn=CHST10;refGene_tx=XM_023251584.1
A3 62031377 62032059 refGene_gn=LONRF2;refGene_tx=XM_023251585.1
A3 62040336 62040455 refGene_gn=LONRF2;refGene_tx=XM_023251585.1
A3 62059486 62059609 refGene_gn=LONRF2;refGene_tx=XM_023251585.1
A3 62061732 62061891 refGene_gn=LONRF2;refGene_tx=XM_023251585.1
A3 62062765 62062967 refGene_gn=LONRF2;refGene_tx=XM_023251585.1
A3 62063600 62063700 refGene_gn=LONRF2;refGene_tx=XM_023251585.1
A3 62063973 62064088 refGene_gn=LONRF2;refGene_tx=XM_023251585.1
A3 62066957 62067079 refGene_gn=LONRF2;refGene_tx=XM_023251585.1
A3 62068761 62068920 refGene_gn=LONRF2;refGene_tx=XM_023251585.1
A3 62072339 62072502 refGene_gn=LONRF2;refGene_tx=XM_023251585.1
A3 62075675 62075825 refGene_gn=LONRF2;refGene_tx=XM_023251585.1
A3 62077464 62077659 refGene_gn=LONRF2;refGene_tx=XM_023251585.1
A3 62270818 62270871 refGene_gn=AFF3;refGene_tx=XM_023251586.1
A3 62357202 62357277 refGene_gn=AFF3;refGene_tx=XM_023251586.1
A3 62359579 62359700 refGene_gn=AFF3;refGene_tx=XM_023251586.1
A3 62360950 62361263 refGene_gn=AFF3;refGene_tx=XM_023251586.1
A3 62361392 62361772 refGene_gn=AFF3;refGene_tx=XM_023251586.1
A3 62519875 62519923 refGene_gn=AFF3;refGene_tx=XM_023251586.1
A3 62599173 62599254 refGene_gn=AFF3;refGene_tx=XM_023251586.1
A3 62614966 62615003 refGene_gn=AFF3;refGene_tx=XM_023251586.1
A3 62626158 62626210 refGene_gn=AFF3;refGene_tx=XM_023251586.1
A3 62680107 62680159 refGene_gn=AFF3;refGene_tx=XM_023251586.1
A3 62702923 62702964 refGene_gn=AFF3;refGene_tx=XM_023251586.1
A3 62751131 62751318 refGene_gn=AFF3;refGene_tx=XM_023251586.1
A3 62758694 62759798 refGene_gn=AFF3;refGene_tx=XM_023251586.1
A3 62764976 62765101 refGene_gn=AFF3;refGene_tx=XM_023251586.1
A3 62768321 62768523 refGene_gn=AFF3;refGene_tx=XM_023251586.1
A3 62771148 62771270 refGene_gn=AFF3;refGene_tx=XM_023251586.1
A3 62780582 62780646 refGene_gn=AFF3;refGene_tx=XM_023251586.1
A3 62782976 62783113 refGene_gn=AFF3;refGene_tx=XM_023251586.1
A3 62787924 62787996 refGene_gn=AFF3;refGene_tx=XM_023251586.1
A3 62789615 62789709 refGene_gn=AFF3;refGene_tx=XM_023251586.1
A3 62793215 62793265 refGene_gn=AFF3;refGene_tx=XM_023251586.1
A3 62793418 62793642 refGene_gn=AFF3;refGene_tx=XM_023251586.1
A3 62796293 62796415 refGene_gn=AFF3;refGene_tx=XM_023251586.1
A3 62868846 62868900

refGene_gn=REV1;refGene_tx=XM_023251587.1;refGene_gn=REV1;refGene_tx=XM_023251588.1
A3 62870285 62870412
refGene_gn=REV1;refGene_tx=XM_023251587.1;refGene_gn=REV1;refGene_tx=XM_023251588.1
A3 62886465 62886634
refGene_gn=REV1;refGene_tx=XM_011281065.3;refGene_gn=REV1;refGene_tx=XM_019827050.2;refGene_gn=REV1;refGene_tx=XM_019827051.2;refGene_gn=REV1;refGene_tx=XM_019827052.2;refGene_gn=REV1;refGene_tx=XM_023251587.1;refGene_gn=REV1;refGene_tx=XM_023251588.1
A3 62891100 62891253
refGene_gn=REV1;refGene_tx=XM_011281065.3;refGene_gn=REV1;refGene_tx=XM_019827050.2;refGene_gn=REV1;refGene_tx=XM_019827051.2;refGene_gn=REV1;refGene_tx=XM_019827052.2;refGene_gn=REV1;refGene_tx=XM_023251587.1;refGene_gn=REV1;refGene_tx=XM_023251588.1
A3 62894128 62894835
refGene_gn=REV1;refGene_tx=XM_011281065.3;refGene_gn=REV1;refGene_tx=XM_019827050.2;refGene_gn=REV1;refGene_tx=XM_019827051.2;refGene_gn=REV1;refGene_tx=XM_019827052.2;refGene_gn=REV1;refGene_tx=XM_023251587.1;refGene_gn=REV1;refGene_tx=XM_023251588.1
A3 62897575 62897683
refGene_gn=REV1;refGene_tx=XM_011281065.3;refGene_gn=REV1;refGene_tx=XM_019827050.2;refGene_gn=REV1;refGene_tx=XM_019827051.2;refGene_gn=REV1;refGene_tx=XM_019827052.2;refGene_gn=REV1;refGene_tx=XM_023251587.1;refGene_gn=REV1;refGene_tx=XM_023251588.1
A3 62899066 62899183
refGene_gn=REV1;refGene_tx=XM_011281065.3;refGene_gn=REV1;refGene_tx=XM_019827050.2;refGene_gn=REV1;refGene_tx=XM_019827051.2;refGene_gn=REV1;refGene_tx=XM_019827052.2;refGene_gn=REV1;refGene_tx=XM_023251587.1;refGene_gn=REV1;refGene_tx=XM_023251588.1
A3 62903616 62903725
refGene_gn=REV1;refGene_tx=XM_011281065.3;refGene_gn=REV1;refGene_tx=XM_019827050.2;refGene_gn=REV1;refGene_tx=XM_019827051.2;refGene_gn=REV1;refGene_tx=XM_019827052.2;refGene_gn=REV1;refGene_tx=XM_019827053.2;refGene_gn=REV1;refGene_tx=XM_023251587.1;refGene_gn=REV1;refGene_tx=XM_023251588.1
A3 62908049 62908178
refGene_gn=REV1;refGene_tx=XM_011281065.3;refGene_gn=REV1;refGene_tx=XM_019827050.2;refGene_gn=REV1;refGene_tx=XM_019827051.2;refGene_gn=REV1;refGene_tx=XM_019827052.2;refGene_gn=REV1;refGene_tx=XM_019827053.2;refGene_gn=REV1;refGene_tx=XM_019827054.2;refGene_gn=REV1;refGene_tx=XM_023251587.1;refGene_gn=REV1;refGene_tx=XM_023251588.1
A3 62910344 62910499
refGene_gn=REV1;refGene_tx=XM_011281065.3;refGene_gn=REV1;refGene_tx=XM_019827050.2;refGene_gn=REV1;refGene_tx=XM_019827051.2;refGene_gn=REV1;refGene_tx=XM_019827052.2;refGene_gn=REV1;refGene_tx=XM_019827053.2;refGene_gn=REV1;refGene_tx=XM_019827054.2;refGene_gn=REV1;refGene_tx=XM_023251587.1;refGene_gn=REV1;refGene_tx=XM_023251588.1
A3 62913132 62913252
refGene_gn=REV1;refGene_tx=XM_011281065.3;refGene_gn=REV1;refGene_tx=X

M_019827050.2;refGene_gn=REV1;refGene_tx=XM_019827051.2;refGene_gn=REV
1;refGene_tx=XM_019827052.2;refGene_gn=REV1;refGene_tx=XM_019827053.2;
refGene_gn=REV1;refGene_tx=XM_019827054.2;refGene_gn=REV1;refGene_tx=X
M_023251587.1
A3 62918031 62918252
refGene_gn=REV1;refGene_tx=XM_011281065.3;refGene_gn=REV1;refGene_tx=X
M_019827050.2;refGene_gn=REV1;refGene_tx=XM_019827051.2;refGene_gn=REV
1;refGene_tx=XM_019827052.2;refGene_gn=REV1;refGene_tx=XM_019827053.2;
refGene_gn=REV1;refGene_tx=XM_019827054.2;refGene_gn=REV1;refGene_tx=X
M_023251587.1;refGene_gn=REV1;refGene_tx=XM_023251588.1
A3 62918516 62918689
refGene_gn=REV1;refGene_tx=XM_011281065.3;refGene_gn=REV1;refGene_tx=X
M_019827050.2;refGene_gn=REV1;refGene_tx=XM_019827051.2;refGene_gn=REV
1;refGene_tx=XM_019827052.2;refGene_gn=REV1;refGene_tx=XM_019827053.2;
refGene_gn=REV1;refGene_tx=XM_019827054.2;refGene_gn=REV1;refGene_tx=X
M_023251587.1;refGene_gn=REV1;refGene_tx=XM_023251588.1
A3 62924295 62924398
refGene_gn=REV1;refGene_tx=XM_011281065.3;refGene_gn=REV1;refGene_tx=X
M_019827050.2;refGene_gn=REV1;refGene_tx=XM_019827051.2;refGene_gn=REV
1;refGene_tx=XM_019827052.2;refGene_gn=REV1;refGene_tx=XM_019827053.2;
refGene_gn=REV1;refGene_tx=XM_019827054.2;refGene_gn=REV1;refGene_tx=X
M_023251587.1;refGene_gn=REV1;refGene_tx=XM_023251588.1
A3 62925535 62925689
refGene_gn=REV1;refGene_tx=XM_011281065.3;refGene_gn=REV1;refGene_tx=X
M_019827050.2;refGene_gn=REV1;refGene_tx=XM_019827051.2;refGene_gn=REV
1;refGene_tx=XM_019827052.2;refGene_gn=REV1;refGene_tx=XM_019827053.2;
refGene_gn=REV1;refGene_tx=XM_019827054.2;refGene_gn=REV1;refGene_tx=X
M_023251587.1;refGene_gn=REV1;refGene_tx=XM_023251588.1
A3 62925886 62926089
refGene_gn=REV1;refGene_tx=XM_011281065.3;refGene_gn=REV1;refGene_tx=X
M_019827050.2;refGene_gn=REV1;refGene_tx=XM_019827051.2;refGene_gn=REV
1;refGene_tx=XM_019827052.2;refGene_gn=REV1;refGene_tx=XM_019827053.2;
refGene_gn=REV1;refGene_tx=XM_019827054.2;refGene_gn=REV1;refGene_tx=X
M_023251587.1;refGene_gn=REV1;refGene_tx=XM_023251588.1
A3 62927332 62927566
refGene_gn=REV1;refGene_tx=XM_011281065.3;refGene_gn=REV1;refGene_tx=X
M_019827050.2;refGene_gn=REV1;refGene_tx=XM_019827051.2;refGene_gn=REV
1;refGene_tx=XM_019827052.2;refGene_gn=REV1;refGene_tx=XM_019827053.2;
refGene_gn=REV1;refGene_tx=XM_019827054.2;refGene_gn=REV1;refGene_tx=X
M_023251587.1;refGene_gn=REV1;refGene_tx=XM_023251588.1
A3 62928612 62928733
refGene_gn=REV1;refGene_tx=XM_011281065.3;refGene_gn=REV1;refGene_tx=X
M_019827050.2;refGene_gn=REV1;refGene_tx=XM_019827051.2;refGene_gn=REV
1;refGene_tx=XM_019827052.2;refGene_gn=REV1;refGene_tx=XM_019827053.2;
refGene_gn=REV1;refGene_tx=XM_019827054.2;refGene_gn=REV1;refGene_tx=X
M_023251587.1;refGene_gn=REV1;refGene_tx=XM_023251588.1
A3 62931583 62931804
refGene_gn=REV1;refGene_tx=XM_011281065.3;refGene_gn=REV1;refGene_tx=X
M_019827050.2;refGene_gn=REV1;refGene_tx=XM_019827051.2;refGene_gn=REV
1;refGene_tx=XM_019827052.2;refGene_gn=REV1;refGene_tx=XM_019827053.2;

refGene_gn=REV1;refGene_tx=XM_019827054.2;refGene_gn=REV1;refGene_tx=XM_023251587.1;refGene_gn=REV1;refGene_tx=XM_023251588.1
A3 62931904 62932061
refGene_gn=REV1;refGene_tx=XM_011281065.3;refGene_gn=REV1;refGene_tx=XM_019827050.2;refGene_gn=REV1;refGene_tx=XM_019827051.2;refGene_gn=REV1;refGene_tx=XM_019827052.2;refGene_gn=REV1;refGene_tx=XM_019827053.2;refGene_gn=REV1;refGene_tx=XM_019827054.2;refGene_gn=REV1;refGene_tx=XM_023251587.1;refGene_gn=REV1;refGene_tx=XM_023251588.1
A3 62932311 62932414
refGene_gn=REV1;refGene_tx=XM_011281065.3;refGene_gn=REV1;refGene_tx=XM_019827050.2;refGene_gn=REV1;refGene_tx=XM_019827051.2;refGene_gn=REV1;refGene_tx=XM_019827052.2;refGene_gn=REV1;refGene_tx=XM_019827053.2;refGene_gn=REV1;refGene_tx=XM_019827054.2;refGene_gn=REV1;refGene_tx=XM_023251587.1;refGene_gn=REV1;refGene_tx=XM_023251588.1
A3 62932757 62932869
refGene_gn=REV1;refGene_tx=XM_011281065.3;refGene_gn=REV1;refGene_tx=XM_019827050.2;refGene_gn=REV1;refGene_tx=XM_019827051.2;refGene_gn=REV1;refGene_tx=XM_019827052.2;refGene_gn=REV1;refGene_tx=XM_019827053.2;refGene_gn=REV1;refGene_tx=XM_019827054.2;refGene_gn=REV1;refGene_tx=XM_023251587.1;refGene_gn=REV1;refGene_tx=XM_023251588.1
A3 62934653 62934761 refGene_gn=EIF5B;refGene_tx=XM_023251589.1
A3 62935133 62935295 refGene_gn=EIF5B;refGene_tx=XM_023251589.1
A3 62937309 62937448 refGene_gn=EIF5B;refGene_tx=XM_023251589.1
A3 62939284 62939449 refGene_gn=EIF5B;refGene_tx=XM_023251589.1
A3 62939563 62939640 refGene_gn=EIF5B;refGene_tx=XM_023251589.1
A3 62939747 62939879 refGene_gn=EIF5B;refGene_tx=XM_023251589.1
A3 62941100 62941232 refGene_gn=EIF5B;refGene_tx=XM_023251589.1
A3 62942608 62942770 refGene_gn=EIF5B;refGene_tx=XM_023251589.1
A3 62942905 62943088 refGene_gn=EIF5B;refGene_tx=XM_023251589.1
A3 62943501 62943633 refGene_gn=EIF5B;refGene_tx=XM_023251589.1
A3 62956510 62956652 refGene_gn=EIF5B;refGene_tx=XM_023251589.1
A3 62957193 62957261 refGene_gn=EIF5B;refGene_tx=XM_023251589.1
A3 62960081 62960192 refGene_gn=EIF5B;refGene_tx=XM_023251589.1
A3 62960364 62960472 refGene_gn=EIF5B;refGene_tx=XM_023251589.1
A3 62969340 62969373 refGene_gn=EIF5B;refGene_tx=XM_023251589.1
A3 62969374 62969628 refGene_gn=EIF5B;refGene_tx=XM_023251589.1
A3 62974430 62974505 refGene_gn=EIF5B;refGene_tx=XM_023251589.1
A3 62976522 62976612 refGene_gn=EIF5B;refGene_tx=XM_023251589.1
A3 62977303 62977402 refGene_gn=EIF5B;refGene_tx=XM_023251589.1
A3 62987114 62987265 refGene_gn=EIF5B;refGene_tx=XM_023251589.1
A3 62987838 62988056 refGene_gn=EIF5B;refGene_tx=XM_023251589.1
A3 62990225 62990898 refGene_gn=EIF5B;refGene_tx=XM_023251589.1
A3 62991548 62991633 refGene_gn=EIF5B;refGene_tx=XM_023251589.1
A3 62991732 62991858 refGene_gn=EIF5B;refGene_tx=XM_023251589.1
A3 62994303 62994335 refGene_gn=EIF5B;refGene_tx=XM_023251589.1
A3 63030197 63030386
refGene_gn=TXNDC9;refGene_tx=XM_003983904.5;refGene_gn=TXNDC9;refGene_tx=XM_006930046.4;refGene_gn=TXNDC9;refGene_tx=XM_006930047.4;refGene_gn=TXNDC9;refGene_tx=XM_023251590.1
A3 63038015 63038134

refGene_gn=TXNDC9;refGene_tx=XM_003983904.5;refGene_gn=TXNDC9;refGene_tx=XM_006930046.4;refGene_gn=TXNDC9;refGene_tx=XM_006930047.4;refGene_gn=TXNDC9;refGene_tx=XM_023251590.1
A3 63042610 63042865
refGene_gn=TXNDC9;refGene_tx=XM_003983904.5;refGene_gn=TXNDC9;refGene_tx=XM_006930046.4;refGene_gn=TXNDC9;refGene_tx=XM_006930047.4;refGene_gn=TXNDC9;refGene_tx=XM_023251590.1
A3 63045116 63045234
refGene_gn=TXNDC9;refGene_tx=XM_003983904.5;refGene_gn=TXNDC9;refGene_tx=XM_006930046.4;refGene_gn=TXNDC9;refGene_tx=XM_006930047.4;refGene_gn=TXNDC9;refGene_tx=XM_023251590.1
A3 63112365 63112408 refGene_gn=LYG1;refGene_tx=XM_023250918.1
A3 63114017 63114122 refGene_gn=LYG1;refGene_tx=XM_023250918.1
A3 63115046 63115240 refGene_gn=LYG1;refGene_tx=XM_023250918.1
A3 63116747 63116880 refGene_gn=LYG1;refGene_tx=XM_023250918.1
A3 63117072 63117191 refGene_gn=LYG1;refGene_tx=XM_023250918.1
A3 63157769 63157812 refGene_gn=LYG2;refGene_tx=XM_003983906.4
A3 63161950 63162091 refGene_gn=LYG2;refGene_tx=XM_003983906.4
A3 63162865 63163062 refGene_gn=LYG2;refGene_tx=XM_003983906.4
A3 63163851 63163990 refGene_gn=LYG2;refGene_tx=XM_003983906.4
A3 63165324 63165443 refGene_gn=LYG2;refGene_tx=XM_003983906.4
A3 63195668 63195801
refGene_gn=MRPL30;refGene_tx=XM_006930048.4;refGene_gn=MRPL30;refGene_tx=XM_019827056.2
A3 63195879 63195953
refGene_gn=MRPL30;refGene_tx=XM_006930048.4;refGene_gn=MRPL30;refGene_tx=XM_019827056.2
A3 63196183 63196330
refGene_gn=MRPL30;refGene_tx=XM_006930048.4;refGene_gn=MRPL30;refGene_tx=XM_019827056.2
A3 63200225 63200306
refGene_gn=MRPL30;refGene_tx=XM_006930048.4;refGene_gn=MRPL30;refGene_tx=XM_019827056.2
A3 63201960 63202038
refGene_gn=MRPL30;refGene_tx=XM_006930048.4;refGene_gn=MRPL30;refGene_tx=XM_019827056.2
A3 63206233 63206305
refGene_gn=MRPL30;refGene_tx=XM_006930048.4;refGene_gn=MRPL30;refGene_tx=XM_019827056.2
A3 63206474 63206625 refGene_gn=MITD1;refGene_tx=XM_003983908.4
A3 63212731 63212833
refGene_gn=MITD1;refGene_tx=XM_003983908.4;refGene_gn=MITD1;refGene_tx=XM_019827055.2
A3 63214702 63214839
refGene_gn=MITD1;refGene_tx=XM_003983908.4;refGene_gn=MITD1;refGene_tx=XM_019827055.2
A3 63214919 63215006
refGene_gn=MITD1;refGene_tx=XM_003983908.4;refGene_gn=MITD1;refGene_tx=XM_019827055.2
A3 63215706 63215822

refGene_gn=MITD1;refGene_tx=XM_003983908.4;refGene_gn=MITD1;refGene_tx=XM_019827055.2
A3 63216713 63216774
refGene_gn=MITD1;refGene_tx=XM_003983908.4;refGene_gn=MITD1;refGene_tx=XM_019827055.2
A3 63216887 63216983
refGene_gn=MITD1;refGene_tx=XM_003983908.4;refGene_gn=MITD1;refGene_tx=XM_019827055.2
A3 63228619 63229741
refGene_gn=LIPT1;refGene_tx=XM_011281062.3;refGene_gn=LIPT1;refGene_tx=XM_019827057.2;refGene_gn=LIPT1;refGene_tx=XM_023251595.1
A3 63236254 63236544
refGene_gn=CA3H2orf15;refGene_tx=XM_019827066.2
A3 63236546 63236599
refGene_gn=CA3H2orf15;refGene_tx=XM_019827066.2
A3 63250555 63250624
refGene_gn=CA3H2orf15;refGene_tx=XM_019827066.2
A3 63251432 63251503
refGene_gn=CA3H2orf15;refGene_tx=XM_019827066.2
A3 63270261 63270372
refGene_gn=TSGA10;refGene_tx=XM_019827063.2;refGene_gn=TSGA10;refGene_tx=XM_023251591.1;refGene_gn=TSGA10;refGene_tx=XM_023251592.1;refGene_gn=TSGA10;refGene_tx=XM_023251594.1
A3 63280849 63280915
refGene_gn=TSGA10;refGene_tx=XM_019827063.2;refGene_gn=TSGA10;refGene_tx=XM_023251591.1;refGene_gn=TSGA10;refGene_tx=XM_023251592.1;refGene_gn=TSGA10;refGene_tx=XM_023251594.1
A3 63281922 63282090
refGene_gn=TSGA10;refGene_tx=XM_019827063.2;refGene_gn=TSGA10;refGene_tx=XM_023251591.1;refGene_gn=TSGA10;refGene_tx=XM_023251592.1;refGene_gn=TSGA10;refGene_tx=XM_023251593.1;refGene_gn=TSGA10;refGene_tx=XM_023251594.1
A3 63282475 63282631
refGene_gn=TSGA10;refGene_tx=XM_019827063.2;refGene_gn=TSGA10;refGene_tx=XM_023251591.1;refGene_gn=TSGA10;refGene_tx=XM_023251592.1;refGene_gn=TSGA10;refGene_tx=XM_023251593.1;refGene_gn=TSGA10;refGene_tx=XM_023251594.1
A3 63286674 63286845
refGene_gn=TSGA10;refGene_tx=XM_019827063.2;refGene_gn=TSGA10;refGene_tx=XM_023251591.1;refGene_gn=TSGA10;refGene_tx=XM_023251592.1;refGene_gn=TSGA10;refGene_tx=XM_023251593.1;refGene_gn=TSGA10;refGene_tx=XM_023251594.1
A3 63286926 63287004
refGene_gn=TSGA10;refGene_tx=XM_019827063.2;refGene_gn=TSGA10;refGene_tx=XM_023251591.1;refGene_gn=TSGA10;refGene_tx=XM_023251592.1;refGene_gn=TSGA10;refGene_tx=XM_023251593.1;refGene_gn=TSGA10;refGene_tx=XM_023251594.1
A3 63287945 63288097
refGene_gn=TSGA10;refGene_tx=XM_019827063.2;refGene_gn=TSGA10;refGene_tx=XM_023251591.1;refGene_gn=TSGA10;refGene_tx=XM_023251592.1;refGene_

gn=TSGA10;refGene_tx=XM_023251593.1;refGene_gn=TSGA10;refGene_tx=XM_023251594.1
A3	63306762 63306878 refGene_gn=TSGA10;refGene_tx=XM_023251594.1
A3	63309692 63309851 refGene_gn=TSGA10;refGene_tx=XM_023251594.1
A3	63314119 63314171 refGene_gn=TSGA10;refGene_tx=XM_023251594.1
A3	63315736 63315905
refGene_gn=TSGA10;refGene_tx=XM_019827063.2;refGene_gn=TSGA10;refGene_tx=XM_023251591.1;refGene_gn=TSGA10;refGene_tx=XM_023251592.1;refGene_gn=TSGA10;refGene_tx=XM_023251593.1;refGene_gn=TSGA10;refGene_tx=XM_023251594.1
A3	63318245 63318356
refGene_gn=TSGA10;refGene_tx=XM_019827063.2;refGene_gn=TSGA10;refGene_tx=XM_023251591.1;refGene_gn=TSGA10;refGene_tx=XM_023251592.1;refGene_gn=TSGA10;refGene_tx=XM_023251593.1;refGene_gn=TSGA10;refGene_tx=XM_023251594.1
A3	63323640 63323826
refGene_gn=TSGA10;refGene_tx=XM_019827063.2;refGene_gn=TSGA10;refGene_tx=XM_023251591.1;refGene_gn=TSGA10;refGene_tx=XM_023251592.1;refGene_gn=TSGA10;refGene_tx=XM_023251593.1;refGene_gn=TSGA10;refGene_tx=XM_023251594.1
A3	63360425 63361331
refGene_gn=LOC101087915;refGene_tx=XM_011281059.3
A3	63385623 63385833
refGene_gn=TSGA10;refGene_tx=XM_019827063.2;refGene_gn=TSGA10;refGene_tx=XM_023251591.1;refGene_gn=TSGA10;refGene_tx=XM_023251592.1;refGene_gn=TSGA10;refGene_tx=XM_023251593.1;refGene_gn=TSGA10;refGene_tx=XM_023251594.1
A3	63406175 63406378
refGene_gn=TSGA10;refGene_tx=XM_019827063.2;refGene_gn=TSGA10;refGene_tx=XM_023251591.1;refGene_gn=TSGA10;refGene_tx=XM_023251592.1;refGene_gn=TSGA10;refGene_tx=XM_023251593.1;refGene_gn=TSGA10;refGene_tx=XM_023251594.1
A3	63408275 63408380
refGene_gn=TSGA10;refGene_tx=XM_019827063.2;refGene_gn=TSGA10;refGene_tx=XM_023251591.1;refGene_gn=TSGA10;refGene_tx=XM_023251592.1;refGene_gn=TSGA10;refGene_tx=XM_023251593.1;refGene_gn=TSGA10;refGene_tx=XM_023251594.1
A3	63408551 63408701
refGene_gn=TSGA10;refGene_tx=XM_019827063.2;refGene_gn=TSGA10;refGene_tx=XM_023251591.1;refGene_gn=TSGA10;refGene_tx=XM_023251592.1;refGene_gn=TSGA10;refGene_tx=XM_023251593.1;refGene_gn=TSGA10;refGene_tx=XM_023251594.1
A3	63415143 63415168
refGene_gn=TSGA10;refGene_tx=XM_019827063.2;refGene_gn=TSGA10;refGene_tx=XM_023251591.1;refGene_gn=TSGA10;refGene_tx=XM_023251592.1;refGene_gn=TSGA10;refGene_tx=XM_023251593.1;refGene_gn=TSGA10;refGene_tx=XM_023251594.1
A3	63538895 63538965
refGene_gn=KIAA1211L;refGene_tx=XM_023251596.1;refGene_gn=KIAA1211L;refGene_tx=XM_023251597.1;refGene_gn=KIAA1211L;refGene_tx=XM_023251598.1

A3 63551401 63551570
refGene_gn=KIAA1211L;refGene_tx=XM_023251596.1;refGene_gn=KIAA1211L;refGene_tx=XM_023251597.1;refGene_gn=KIAA1211L;refGene_tx=XM_023251598.1
A3 63558311 63558447
refGene_gn=KIAA1211L;refGene_tx=XM_023251596.1;refGene_gn=KIAA1211L;refGene_tx=XM_023251597.1;refGene_gn=KIAA1211L;refGene_tx=XM_023251598.1
A3 63558797 63558959
refGene_gn=KIAA1211L;refGene_tx=XM_023251596.1;refGene_gn=KIAA1211L;refGene_tx=XM_023251597.1;refGene_gn=KIAA1211L;refGene_tx=XM_023251598.1;refGene_gn=KIAA1211L;refGene_tx=XM_023251599.1
A3 63565146 63565374
refGene_gn=KIAA1211L;refGene_tx=XM_023251596.1;refGene_gn=KIAA1211L;refGene_tx=XM_023251597.1;refGene_gn=KIAA1211L;refGene_tx=XM_023251598.1;refGene_gn=KIAA1211L;refGene_tx=XM_023251599.1
A3 63569323 63570881
refGene_gn=KIAA1211L;refGene_tx=XM_023251596.1;refGene_gn=KIAA1211L;refGene_tx=XM_023251597.1;refGene_gn=KIAA1211L;refGene_tx=XM_023251598.1;refGene_gn=KIAA1211L;refGene_tx=XM_023251599.1
A3 63596263 63596436
refGene_gn=KIAA1211L;refGene_tx=XM_023251596.1;refGene_gn=KIAA1211L;refGene_tx=XM_023251597.1;refGene_gn=KIAA1211L;refGene_tx=XM_023251598.1;refGene_gn=KIAA1211L;refGene_tx=XM_023251599.1
A3 63597444 63597589
refGene_gn=KIAA1211L;refGene_tx=XM_023251596.1;refGene_gn=KIAA1211L;refGene_tx=XM_023251597.1;refGene_gn=KIAA1211L;refGene_tx=XM_023251598.1;refGene_gn=KIAA1211L;refGene_tx=XM_023251599.1
A3 63598677 63598820
refGene_gn=KIAA1211L;refGene_tx=XM_023251596.1;refGene_gn=KIAA1211L;refGene_tx=XM_023251597.1;refGene_gn=KIAA1211L;refGene_tx=XM_023251598.1;refGene_gn=KIAA1211L;refGene_tx=XM_023251599.1
A3 63678080 63678174
refGene_gn=MGAT4A;refGene_tx=XM_023251617.1;refGene_gn=MGAT4A;refGene_tx=XM_023251618.1;refGene_gn=MGAT4A;refGene_tx=XM_023251619.1;refGene_gn=MGAT4A;refGene_tx=XM_023251620.1
A3 63715927 63716095
refGene_gn=MGAT4A;refGene_tx=XM_023251617.1;refGene_gn=MGAT4A;refGene_tx=XM_023251618.1;refGene_gn=MGAT4A;refGene_tx=XM_023251619.1;refGene_gn=MGAT4A;refGene_tx=XM_023251620.1
A3 63718491 63718632
refGene_gn=MGAT4A;refGene_tx=XM_023251617.1;refGene_gn=MGAT4A;refGene_tx=XM_023251618.1;refGene_gn=MGAT4A;refGene_tx=XM_023251619.1;refGene_gn=MGAT4A;refGene_tx=XM_023251620.1
A3 63740118 63740252
refGene_gn=MGAT4A;refGene_tx=XM_023251617.1;refGene_gn=MGAT4A;refGene_tx=XM_023251618.1;refGene_gn=MGAT4A;refGene_tx=XM_023251619.1;refGene_gn=MGAT4A;refGene_tx=XM_023251620.1
A3 63761305 63761352
refGene_gn=MGAT4A;refGene_tx=XM_023251617.1;refGene_gn=MGAT4A;refGene_tx=XM_023251618.1;refGene_gn=MGAT4A;refGene_tx=XM_023251619.1;refGene_gn=MGAT4A;refGene_tx=XM_023251620.1

A3	63763098	63763212
refGene_gn=MGAT4A;refGene_tx=XM_023251617.1;refGene_gn=MGAT4A;refGene_tx=XM_023251618.1;refGene_gn=MGAT4A;refGene_tx=XM_023251619.1;refGene_gn=MGAT4A;refGene_tx=XM_023251620.1
A3	63764836	63764912
refGene_gn=MGAT4A;refGene_tx=XM_023251617.1;refGene_gn=MGAT4A;refGene_tx=XM_023251618.1;refGene_gn=MGAT4A;refGene_tx=XM_023251619.1;refGene_gn=MGAT4A;refGene_tx=XM_023251620.1
A3	63774015	63774130
refGene_gn=MGAT4A;refGene_tx=XM_023251617.1;refGene_gn=MGAT4A;refGene_tx=XM_023251618.1;refGene_gn=MGAT4A;refGene_tx=XM_023251619.1;refGene_gn=MGAT4A;refGene_tx=XM_023251620.1
A3	63775205	63775336
refGene_gn=MGAT4A;refGene_tx=XM_023251617.1;refGene_gn=MGAT4A;refGene_tx=XM_023251618.1;refGene_gn=MGAT4A;refGene_tx=XM_023251619.1;refGene_gn=MGAT4A;refGene_tx=XM_023251620.1
A3	63778219	63778327
refGene_gn=MGAT4A;refGene_tx=XM_023251617.1;refGene_gn=MGAT4A;refGene_tx=XM_023251618.1;refGene_gn=MGAT4A;refGene_tx=XM_023251619.1;refGene_gn=MGAT4A;refGene_tx=XM_023251620.1
A3	63778450	63778644
refGene_gn=MGAT4A;refGene_tx=XM_023251617.1;refGene_gn=MGAT4A;refGene_tx=XM_023251618.1;refGene_gn=MGAT4A;refGene_tx=XM_023251619.1;refGene_gn=MGAT4A;refGene_tx=XM_023251620.1
A3	63783057	63783136
refGene_gn=MGAT4A;refGene_tx=XM_023251617.1;refGene_gn=MGAT4A;refGene_tx=XM_023251618.1;refGene_gn=MGAT4A;refGene_tx=XM_023251619.1;refGene_gn=MGAT4A;refGene_tx=XM_023251620.1
A3	63783790	63783857
refGene_gn=MGAT4A;refGene_tx=XM_023251617.1;refGene_gn=MGAT4A;refGene_tx=XM_023251618.1;refGene_gn=MGAT4A;refGene_tx=XM_023251619.1;refGene_gn=MGAT4A;refGene_tx=XM_023251620.1
A3	63790159	63790272
refGene_gn=MGAT4A;refGene_tx=XM_023251617.1;refGene_gn=MGAT4A;refGene_tx=XM_023251618.1;refGene_gn=MGAT4A;refGene_tx=XM_023251619.1;refGene_gn=MGAT4A;refGene_tx=XM_023251620.1
A3	63790379	63790406
refGene_gn=MGAT4A;refGene_tx=XM_023251617.1;refGene_gn=MGAT4A;refGene_tx=XM_023251618.1;refGene_gn=MGAT4A;refGene_tx=XM_023251619.1;refGene_gn=MGAT4A;refGene_tx=XM_023251620.1
A3	63796930	63797067
refGene_gn=UNC50;refGene_tx=XM_003983911.5;refGene_gn=UNC50;refGene_tx=XM_006930051.2;refGene_gn=UNC50;refGene_tx=XM_019827067.1
A3	63798521	63798623
refGene_gn=UNC50;refGene_tx=XM_003983911.5;refGene_gn=UNC50;refGene_tx=XM_006930051.2;refGene_gn=UNC50;refGene_tx=XM_019827067.1
A3	63798708	63798848
refGene_gn=UNC50;refGene_tx=XM_003983911.5;refGene_gn=UNC50;refGene_tx=XM_006930051.2;refGene_gn=UNC50;refGene_tx=XM_019827067.1
A3	63812120	63812241

refGene_gn=UNC50;refGene_tx=XM_003983911.5;refGene_gn=UNC50;refGene_tx=XM_006930051.2;refGene_gn=UNC50;refGene_tx=XM_019827067.1
A3 63813028 63813308
refGene_gn=UNC50;refGene_tx=XM_003983911.5;refGene_gn=UNC50;refGene_tx=XM_006930051.2;refGene_gn=UNC50;refGene_tx=XM_019827067.1
A3 63814948 63815047
refGene_gn=LOC101088658;refGene_tx=XM_003983912.5
A3 63821007 63821091
refGene_gn=LOC101088658;refGene_tx=XM_003983912.5
A3 63822685 63822727
refGene_gn=LOC101088658;refGene_tx=XM_003983912.5
A3 63837626 63837759
refGene_gn=INPP4A;refGene_tx=XM_023251606.1;refGene_gn=INPP4A;refGene_tx=XM_023251607.1;refGene_gn=INPP4A;refGene_tx=XM_023251608.1;refGene_gn=INPP4A;refGene_tx=XM_023251612.1;refGene_gn=INPP4A;refGene_tx=XM_023251613.1;refGene_gn=INPP4A;refGene_tx=XM_023251614.1;refGene_gn=INPP4A;refGene_tx=XM_023251615.1
A3 63842583 63842764
refGene_gn=INPP4A;refGene_tx=XM_023251600.1;refGene_gn=INPP4A;refGene_tx=XM_023251601.1;refGene_gn=INPP4A;refGene_tx=XM_023251602.1;refGene_gn=INPP4A;refGene_tx=XM_023251603.1;refGene_gn=INPP4A;refGene_tx=XM_023251604.1;refGene_gn=INPP4A;refGene_tx=XM_023251609.1;refGene_gn=INPP4A;refGene_tx=XM_023251610.1;refGene_gn=INPP4A;refGene_tx=XM_023251611.1;refGene_gn=INPP4A;refGene_tx=XM_023251616.1
A3 63846205 63846350 refGene_gn=INPP4A;refGene_tx=XM_023251605.1
A3 63846974 63847129
refGene_gn=INPP4A;refGene_tx=XM_023251600.1;refGene_gn=INPP4A;refGene_tx=XM_023251601.1;refGene_gn=INPP4A;refGene_tx=XM_023251602.1;refGene_gn=INPP4A;refGene_tx=XM_023251603.1;refGene_gn=INPP4A;refGene_tx=XM_023251604.1;refGene_gn=INPP4A;refGene_tx=XM_023251605.1;refGene_gn=INPP4A;refGene_tx=XM_023251606.1;refGene_gn=INPP4A;refGene_tx=XM_023251607.1;refGene_gn=INPP4A;refGene_tx=XM_023251608.1;refGene_gn=INPP4A;refGene_tx=XM_023251609.1;refGene_gn=INPP4A;refGene_tx=XM_023251610.1;refGene_gn=INPP4A;refGene_tx=XM_023251611.1;refGene_gn=INPP4A;refGene_tx=XM_023251612.1;refGene_gn=INPP4A;refGene_tx=XM_023251613.1;refGene_gn=INPP4A;refGene_tx=XM_023251614.1;refGene_gn=INPP4A;refGene_tx=XM_023251615.1;refGene_gn=INPP4A;refGene_tx=XM_023251616.1
A3 63850751 63850864
refGene_gn=INPP4A;refGene_tx=XM_023251600.1;refGene_gn=INPP4A;refGene_tx=XM_023251601.1;refGene_gn=INPP4A;refGene_tx=XM_023251602.1;refGene_gn=INPP4A;refGene_tx=XM_023251603.1;refGene_gn=INPP4A;refGene_tx=XM_023251604.1;refGene_gn=INPP4A;refGene_tx=XM_023251605.1;refGene_gn=INPP4A;refGene_tx=XM_023251606.1;refGene_gn=INPP4A;refGene_tx=XM_023251607.1;refGene_gn=INPP4A;refGene_tx=XM_023251608.1;refGene_gn=INPP4A;refGene_tx=XM_023251609.1;refGene_gn=INPP4A;refGene_tx=XM_023251610.1;refGene_gn=INPP4A;refGene_tx=XM_023251611.1;refGene_gn=INPP4A;refGene_tx=XM_023251612.1;refGene_gn=INPP4A;refGene_tx=XM_023251613.1;refGene_gn=INPP4A;refGene_tx=XM_023251614.1;refGene_gn=INPP4A;refGene_tx=XM_023251615.1;refGene_gn=INPP4A;refGene_tx=XM_023251616.1
A3 63855379 63855477

refGene_gn=INPP4A;refGene_tx=XM_023251600.1;refGene_gn=INPP4A;refGene_tx=XM_023251601.1;refGene_gn=INPP4A;refGene_tx=XM_023251602.1;refGene_gn=INPP4A;refGene_tx=XM_023251603.1;refGene_gn=INPP4A;refGene_tx=XM_023251604.1;refGene_gn=INPP4A;refGene_tx=XM_023251605.1;refGene_gn=INPP4A;refGene_tx=XM_023251606.1;refGene_gn=INPP4A;refGene_tx=XM_023251607.1;refGene_gn=INPP4A;refGene_tx=XM_023251608.1;refGene_gn=INPP4A;refGene_tx=XM_023251609.1;refGene_gn=INPP4A;refGene_tx=XM_023251610.1;refGene_gn=INPP4A;refGene_tx=XM_023251611.1;refGene_gn=INPP4A;refGene_tx=XM_023251612.1;refGene_gn=INPP4A;refGene_tx=XM_023251613.1;refGene_gn=INPP4A;refGene_tx=XM_023251614.1;refGene_gn=INPP4A;refGene_tx=XM_023251615.1;refGene_gn=INPP4A;refGene_tx=XM_023251616.1
A3 63857875 63858016
refGene_gn=INPP4A;refGene_tx=XM_023251600.1;refGene_gn=INPP4A;refGene_tx=XM_023251601.1;refGene_gn=INPP4A;refGene_tx=XM_023251602.1;refGene_gn=INPP4A;refGene_tx=XM_023251603.1;refGene_gn=INPP4A;refGene_tx=XM_023251604.1;refGene_gn=INPP4A;refGene_tx=XM_023251605.1;refGene_gn=INPP4A;refGene_tx=XM_023251606.1;refGene_gn=INPP4A;refGene_tx=XM_023251607.1;refGene_gn=INPP4A;refGene_tx=XM_023251608.1;refGene_gn=INPP4A;refGene_tx=XM_023251609.1;refGene_gn=INPP4A;refGene_tx=XM_023251610.1;refGene_gn=INPP4A;refGene_tx=XM_023251611.1;refGene_gn=INPP4A;refGene_tx=XM_023251612.1;refGene_gn=INPP4A;refGene_tx=XM_023251613.1;refGene_gn=INPP4A;refGene_tx=XM_023251614.1;refGene_gn=INPP4A;refGene_tx=XM_023251615.1;refGene_gn=INPP4A;refGene_tx=XM_023251616.1
A3 63858326 63858453
refGene_gn=INPP4A;refGene_tx=XM_023251600.1;refGene_gn=INPP4A;refGene_tx=XM_023251601.1;refGene_gn=INPP4A;refGene_tx=XM_023251602.1;refGene_gn=INPP4A;refGene_tx=XM_023251603.1;refGene_gn=INPP4A;refGene_tx=XM_023251604.1;refGene_gn=INPP4A;refGene_tx=XM_023251605.1;refGene_gn=INPP4A;refGene_tx=XM_023251606.1;refGene_gn=INPP4A;refGene_tx=XM_023251607.1;refGene_gn=INPP4A;refGene_tx=XM_023251608.1;refGene_gn=INPP4A;refGene_tx=XM_023251609.1;refGene_gn=INPP4A;refGene_tx=XM_023251610.1;refGene_gn=INPP4A;refGene_tx=XM_023251611.1;refGene_gn=INPP4A;refGene_tx=XM_023251612.1;refGene_gn=INPP4A;refGene_tx=XM_023251613.1;refGene_gn=INPP4A;refGene_tx=XM_023251614.1;refGene_gn=INPP4A;refGene_tx=XM_023251615.1;refGene_gn=INPP4A;refGene_tx=XM_023251616.1
A3 63859637 63859761
refGene_gn=INPP4A;refGene_tx=XM_023251600.1;refGene_gn=INPP4A;refGene_tx=XM_023251601.1;refGene_gn=INPP4A;refGene_tx=XM_023251602.1;refGene_gn=INPP4A;refGene_tx=XM_023251603.1;refGene_gn=INPP4A;refGene_tx=XM_023251604.1;refGene_gn=INPP4A;refGene_tx=XM_023251605.1;refGene_gn=INPP4A;refGene_tx=XM_023251606.1;refGene_gn=INPP4A;refGene_tx=XM_023251607.1;refGene_gn=INPP4A;refGene_tx=XM_023251608.1;refGene_gn=INPP4A;refGene_tx=XM_023251609.1;refGene_gn=INPP4A;refGene_tx=XM_023251610.1;refGene_gn=INPP4A;refGene_tx=XM_023251611.1;refGene_gn=INPP4A;refGene_tx=XM_023251612.1;refGene_gn=INPP4A;refGene_tx=XM_023251613.1;refGene_gn=INPP4A;refGene_tx=XM_023251614.1;refGene_gn=INPP4A;refGene_tx=XM_023251615.1;refGene_gn=INPP4A;refGene_tx=XM_023251616.1
A3 63860770 63860943
refGene_gn=INPP4A;refGene_tx=XM_023251600.1;refGene_gn=INPP4A;refGene_tx=XM_023251601.1;refGene_gn=INPP4A;refGene_tx=XM_023251602.1;refGene_

gn=INPP4A;refGene_tx=XM_023251603.1;refGene_gn=INPP4A;refGene_tx=XM_02
3251604.1;refGene_gn=INPP4A;refGene_tx=XM_023251605.1;refGene_gn=INPP4
A;refGene_tx=XM_023251606.1;refGene_gn=INPP4A;refGene_tx=XM_023251607.
1;refGene_gn=INPP4A;refGene_tx=XM_023251608.1;refGene_gn=INPP4A;refGen
e_tx=XM_023251609.1;refGene_gn=INPP4A;refGene_tx=XM_023251610.1;refGen
e_gn=INPP4A;refGene_tx=XM_023251611.1;refGene_gn=INPP4A;refGene_tx=XM_
023251612.1;refGene_gn=INPP4A;refGene_tx=XM_023251613.1;refGene_gn=INP
P4A;refGene_tx=XM_023251614.1;refGene_gn=INPP4A;refGene_tx=XM_02325161
5.1;refGene_gn=INPP4A;refGene_tx=XM_023251616.1
A3 63865014 63865047
refGene_gn=INPP4A;refGene_tx=XM_023251600.1;refGene_gn=INPP4A;refGene_
tx=XM_023251601.1;refGene_gn=INPP4A;refGene_tx=XM_023251602.1;refGene_
gn=INPP4A;refGene_tx=XM_023251603.1;refGene_gn=INPP4A;refGene_tx=XM_02
3251605.1;refGene_gn=INPP4A;refGene_tx=XM_023251606.1;refGene_gn=INPP4
A;refGene_tx=XM_023251607.1;refGene_gn=INPP4A;refGene_tx=XM_023251608.
1;refGene_gn=INPP4A;refGene_tx=XM_023251609.1;refGene_gn=INPP4A;refGen
e_tx=XM_023251610.1;refGene_gn=INPP4A;refGene_tx=XM_023251612.1;refGen
e_gn=INPP4A;refGene_tx=XM_023251613.1;refGene_gn=INPP4A;refGene_tx=XM_
023251614.1;refGene_gn=INPP4A;refGene_tx=XM_023251616.1
A3 63868702 63868958
refGene_gn=INPP4A;refGene_tx=XM_023251600.1;refGene_gn=INPP4A;refGene_
tx=XM_023251601.1;refGene_gn=INPP4A;refGene_tx=XM_023251602.1;refGene_
gn=INPP4A;refGene_tx=XM_023251603.1;refGene_gn=INPP4A;refGene_tx=XM_02
3251604.1;refGene_gn=INPP4A;refGene_tx=XM_023251605.1;refGene_gn=INPP4
A;refGene_tx=XM_023251606.1;refGene_gn=INPP4A;refGene_tx=XM_023251607.
1;refGene_gn=INPP4A;refGene_tx=XM_023251608.1;refGene_gn=INPP4A;refGen
e_tx=XM_023251609.1;refGene_gn=INPP4A;refGene_tx=XM_023251610.1;refGen
e_gn=INPP4A;refGene_tx=XM_023251611.1;refGene_gn=INPP4A;refGene_tx=XM_
023251612.1;refGene_gn=INPP4A;refGene_tx=XM_023251613.1;refGene_gn=INP
P4A;refGene_tx=XM_023251614.1;refGene_gn=INPP4A;refGene_tx=XM_02325161
5.1;refGene_gn=INPP4A;refGene_tx=XM_023251616.1
A3 63870022 63870241
refGene_gn=INPP4A;refGene_tx=XM_023251600.1;refGene_gn=INPP4A;refGene_
tx=XM_023251601.1;refGene_gn=INPP4A;refGene_tx=XM_023251602.1;refGene_
gn=INPP4A;refGene_tx=XM_023251603.1;refGene_gn=INPP4A;refGene_tx=XM_02
3251604.1;refGene_gn=INPP4A;refGene_tx=XM_023251605.1;refGene_gn=INPP4
A;refGene_tx=XM_023251606.1;refGene_gn=INPP4A;refGene_tx=XM_023251607.
1;refGene_gn=INPP4A;refGene_tx=XM_023251608.1;refGene_gn=INPP4A;refGen
e_tx=XM_023251609.1;refGene_gn=INPP4A;refGene_tx=XM_023251610.1;refGen
e_gn=INPP4A;refGene_tx=XM_023251611.1;refGene_gn=INPP4A;refGene_tx=XM_
023251612.1;refGene_gn=INPP4A;refGene_tx=XM_023251613.1;refGene_gn=INP
P4A;refGene_tx=XM_023251614.1;refGene_gn=INPP4A;refGene_tx=XM_02325161
5.1;refGene_gn=INPP4A;refGene_tx=XM_023251616.1
A3 63871522 63871706
refGene_gn=INPP4A;refGene_tx=XM_023251600.1;refGene_gn=INPP4A;refGene_
tx=XM_023251601.1;refGene_gn=INPP4A;refGene_tx=XM_023251602.1;refGene_
gn=INPP4A;refGene_tx=XM_023251603.1;refGene_gn=INPP4A;refGene_tx=XM_02
3251604.1;refGene_gn=INPP4A;refGene_tx=XM_023251605.1;refGene_gn=INPP4
A;refGene_tx=XM_023251606.1;refGene_gn=INPP4A;refGene_tx=XM_023251607.
1;refGene_gn=INPP4A;refGene_tx=XM_023251608.1;refGene_gn=INPP4A;refGen

e_tx=XM_023251609.1;refGene_gn=INPP4A;refGene_tx=XM_023251610.1;refGen
e_gn=INPP4A;refGene_tx=XM_023251611.1;refGene_gn=INPP4A;refGene_tx=XM_
023251612.1;refGene_gn=INPP4A;refGene_tx=XM_023251613.1;refGene_gn=INP
P4A;refGene_tx=XM_023251614.1;refGene_gn=INPP4A;refGene_tx=XM_02325161
5.1;refGene_gn=INPP4A;refGene_tx=XM_023251616.1
A3 63874267 63874285
refGene_gn=INPP4A;refGene_tx=XM_023251600.1;refGene_gn=INPP4A;refGene_
tx=XM_023251601.1;refGene_gn=INPP4A;refGene_tx=XM_023251602.1;refGene_
gn=INPP4A;refGene_tx=XM_023251604.1;refGene_gn=INPP4A;refGene_tx=XM_02
3251605.1;refGene_gn=INPP4A;refGene_tx=XM_023251606.1;refGene_gn=INPP4
A;refGene_tx=XM_023251607.1;refGene_gn=INPP4A;refGene_tx=XM_023251609.
1;refGene_gn=INPP4A;refGene_tx=XM_023251610.1;refGene_gn=INPP4A;refGen
e_tx=XM_023251611.1;refGene_gn=INPP4A;refGene_tx=XM_023251612.1;refGen
e_gn=INPP4A;refGene_tx=XM_023251613.1;refGene_gn=INPP4A;refGene_tx=XM_
023251616.1
A3 63876566 63876675
refGene_gn=INPP4A;refGene_tx=XM_023251600.1;refGene_gn=INPP4A;refGene_
tx=XM_023251601.1;refGene_gn=INPP4A;refGene_tx=XM_023251602.1;refGene_
gn=INPP4A;refGene_tx=XM_023251603.1;refGene_gn=INPP4A;refGene_tx=XM_02
3251604.1;refGene_gn=INPP4A;refGene_tx=XM_023251605.1;refGene_gn=INPP4
A;refGene_tx=XM_023251606.1;refGene_gn=INPP4A;refGene_tx=XM_023251607.
1;refGene_gn=INPP4A;refGene_tx=XM_023251608.1;refGene_gn=INPP4A;refGen
e_tx=XM_023251609.1;refGene_gn=INPP4A;refGene_tx=XM_023251610.1;refGen
e_gn=INPP4A;refGene_tx=XM_023251611.1;refGene_gn=INPP4A;refGene_tx=XM_
023251612.1;refGene_gn=INPP4A;refGene_tx=XM_023251613.1;refGene_gn=INP
P4A;refGene_tx=XM_023251614.1;refGene_gn=INPP4A;refGene_tx=XM_02325161
5.1;refGene_gn=INPP4A;refGene_tx=XM_023251616.1
A3 63877203 63877308
refGene_gn=INPP4A;refGene_tx=XM_023251600.1;refGene_gn=INPP4A;refGene_
tx=XM_023251601.1;refGene_gn=INPP4A;refGene_tx=XM_023251602.1;refGene_
gn=INPP4A;refGene_tx=XM_023251603.1;refGene_gn=INPP4A;refGene_tx=XM_02
3251604.1;refGene_gn=INPP4A;refGene_tx=XM_023251605.1;refGene_gn=INPP4
A;refGene_tx=XM_023251606.1;refGene_gn=INPP4A;refGene_tx=XM_023251607.
1;refGene_gn=INPP4A;refGene_tx=XM_023251608.1;refGene_gn=INPP4A;refGen
e_tx=XM_023251609.1;refGene_gn=INPP4A;refGene_tx=XM_023251610.1;refGen
e_gn=INPP4A;refGene_tx=XM_023251611.1;refGene_gn=INPP4A;refGene_tx=XM_
023251612.1;refGene_gn=INPP4A;refGene_tx=XM_023251613.1;refGene_gn=INP
P4A;refGene_tx=XM_023251614.1;refGene_gn=INPP4A;refGene_tx=XM_02325161
5.1;refGene_gn=INPP4A;refGene_tx=XM_023251616.1
A3 63879141 63879272
refGene_gn=INPP4A;refGene_tx=XM_023251600.1;refGene_gn=INPP4A;refGene_
tx=XM_023251601.1;refGene_gn=INPP4A;refGene_tx=XM_023251602.1;refGene_
gn=INPP4A;refGene_tx=XM_023251603.1;refGene_gn=INPP4A;refGene_tx=XM_02
3251604.1;refGene_gn=INPP4A;refGene_tx=XM_023251605.1;refGene_gn=INPP4
A;refGene_tx=XM_023251606.1;refGene_gn=INPP4A;refGene_tx=XM_023251607.
1;refGene_gn=INPP4A;refGene_tx=XM_023251608.1;refGene_gn=INPP4A;refGen
e_tx=XM_023251609.1;refGene_gn=INPP4A;refGene_tx=XM_023251610.1;refGen
e_gn=INPP4A;refGene_tx=XM_023251611.1;refGene_gn=INPP4A;refGene_tx=XM_
023251612.1;refGene_gn=INPP4A;refGene_tx=XM_023251613.1;refGene_gn=INP
P4A;refGene_tx=XM_023251614.1;refGene_gn=INPP4A;refGene_tx=XM_02325161

5.1;refGene_gn=INPP4A;refGene_tx=XM_023251616.1
A3 63882159 63882307
refGene_gn=INPP4A;refGene_tx=XM_023251600.1;refGene_gn=INPP4A;refGene_tx=XM_023251601.1;refGene_gn=INPP4A;refGene_tx=XM_023251602.1;refGene_gn=INPP4A;refGene_tx=XM_023251603.1;refGene_gn=INPP4A;refGene_tx=XM_023251604.1;refGene_gn=INPP4A;refGene_tx=XM_023251605.1;refGene_gn=INPP4A;refGene_tx=XM_023251606.1;refGene_gn=INPP4A;refGene_tx=XM_023251607.1;refGene_gn=INPP4A;refGene_tx=XM_023251608.1;refGene_gn=INPP4A;refGene_tx=XM_023251609.1;refGene_gn=INPP4A;refGene_tx=XM_023251610.1;refGene_gn=INPP4A;refGene_tx=XM_023251611.1;refGene_gn=INPP4A;refGene_tx=XM_023251612.1;refGene_gn=INPP4A;refGene_tx=XM_023251613.1;refGene_gn=INPP4A;refGene_tx=XM_023251614.1;refGene_gn=INPP4A;refGene_tx=XM_023251615.1;refGene_gn=INPP4A;refGene_tx=XM_023251616.1
A3 63882786 63882877
refGene_gn=INPP4A;refGene_tx=XM_023251600.1;refGene_gn=INPP4A;refGene_tx=XM_023251601.1;refGene_gn=INPP4A;refGene_tx=XM_023251602.1;refGene_gn=INPP4A;refGene_tx=XM_023251603.1;refGene_gn=INPP4A;refGene_tx=XM_023251604.1;refGene_gn=INPP4A;refGene_tx=XM_023251605.1;refGene_gn=INPP4A;refGene_tx=XM_023251606.1;refGene_gn=INPP4A;refGene_tx=XM_023251607.1;refGene_gn=INPP4A;refGene_tx=XM_023251608.1;refGene_gn=INPP4A;refGene_tx=XM_023251609.1;refGene_gn=INPP4A;refGene_tx=XM_023251610.1;refGene_gn=INPP4A;refGene_tx=XM_023251611.1;refGene_gn=INPP4A;refGene_tx=XM_023251612.1;refGene_gn=INPP4A;refGene_tx=XM_023251613.1;refGene_gn=INPP4A;refGene_tx=XM_023251614.1;refGene_gn=INPP4A;refGene_tx=XM_023251615.1;refGene_gn=INPP4A;refGene_tx=XM_023251616.1
A3 63883681 63883793
refGene_gn=INPP4A;refGene_tx=XM_023251600.1;refGene_gn=INPP4A;refGene_tx=XM_023251601.1;refGene_gn=INPP4A;refGene_tx=XM_023251602.1;refGene_gn=INPP4A;refGene_tx=XM_023251603.1;refGene_gn=INPP4A;refGene_tx=XM_023251604.1;refGene_gn=INPP4A;refGene_tx=XM_023251605.1;refGene_gn=INPP4A;refGene_tx=XM_023251606.1;refGene_gn=INPP4A;refGene_tx=XM_023251607.1;refGene_gn=INPP4A;refGene_tx=XM_023251608.1;refGene_gn=INPP4A;refGene_tx=XM_023251609.1;refGene_gn=INPP4A;refGene_tx=XM_023251610.1;refGene_gn=INPP4A;refGene_tx=XM_023251611.1;refGene_gn=INPP4A;refGene_tx=XM_023251612.1;refGene_gn=INPP4A;refGene_tx=XM_023251613.1;refGene_gn=INPP4A;refGene_tx=XM_023251614.1;refGene_gn=INPP4A;refGene_tx=XM_023251615.1;refGene_gn=INPP4A;refGene_tx=XM_023251616.1
A3 63885365 63885445
refGene_gn=INPP4A;refGene_tx=XM_023251600.1;refGene_gn=INPP4A;refGene_tx=XM_023251601.1;refGene_gn=INPP4A;refGene_tx=XM_023251602.1;refGene_gn=INPP4A;refGene_tx=XM_023251603.1;refGene_gn=INPP4A;refGene_tx=XM_023251604.1;refGene_gn=INPP4A;refGene_tx=XM_023251605.1;refGene_gn=INPP4A;refGene_tx=XM_023251606.1;refGene_gn=INPP4A;refGene_tx=XM_023251607.1;refGene_gn=INPP4A;refGene_tx=XM_023251608.1;refGene_gn=INPP4A;refGene_tx=XM_023251609.1;refGene_gn=INPP4A;refGene_tx=XM_023251610.1;refGene_gn=INPP4A;refGene_tx=XM_023251611.1;refGene_gn=INPP4A;refGene_tx=XM_023251612.1;refGene_gn=INPP4A;refGene_tx=XM_023251613.1;refGene_gn=INPP4A;refGene_tx=XM_023251614.1;refGene_gn=INPP4A;refGene_tx=XM_023251615.1;refGene_gn=INPP4A;refGene_tx=XM_023251616.1
A3 63885734 63885851

refGene_gn=INPP4A;refGene_tx=XM_023251600.1;refGene_gn=INPP4A;refGene_tx=XM_023251601.1;refGene_gn=INPP4A;refGene_tx=XM_023251602.1;refGene_gn=INPP4A;refGene_tx=XM_023251603.1;refGene_gn=INPP4A;refGene_tx=XM_023251604.1;refGene_gn=INPP4A;refGene_tx=XM_023251605.1;refGene_gn=INPP4A;refGene_tx=XM_023251606.1;refGene_gn=INPP4A;refGene_tx=XM_023251607.1;refGene_gn=INPP4A;refGene_tx=XM_023251608.1;refGene_gn=INPP4A;refGene_tx=XM_023251609.1;refGene_gn=INPP4A;refGene_tx=XM_023251610.1;refGene_gn=INPP4A;refGene_tx=XM_023251611.1;refGene_gn=INPP4A;refGene_tx=XM_023251612.1;refGene_gn=INPP4A;refGene_tx=XM_023251613.1;refGene_gn=INPP4A;refGene_tx=XM_023251614.1;refGene_gn=INPP4A;refGene_tx=XM_023251615.1;refGene_gn=INPP4A;refGene_tx=XM_023251616.1
A3 63888126 63888245
refGene_gn=INPP4A;refGene_tx=XM_023251600.1;refGene_gn=INPP4A;refGene_tx=XM_023251601.1;refGene_gn=INPP4A;refGene_tx=XM_023251602.1;refGene_gn=INPP4A;refGene_tx=XM_023251603.1;refGene_gn=INPP4A;refGene_tx=XM_023251604.1;refGene_gn=INPP4A;refGene_tx=XM_023251605.1;refGene_gn=INPP4A;refGene_tx=XM_023251606.1;refGene_gn=INPP4A;refGene_tx=XM_023251607.1;refGene_gn=INPP4A;refGene_tx=XM_023251608.1;refGene_gn=INPP4A;refGene_tx=XM_023251609.1;refGene_gn=INPP4A;refGene_tx=XM_023251610.1;refGene_gn=INPP4A;refGene_tx=XM_023251611.1;refGene_gn=INPP4A;refGene_tx=XM_023251612.1;refGene_gn=INPP4A;refGene_tx=XM_023251613.1;refGene_gn=INPP4A;refGene_tx=XM_023251614.1;refGene_gn=INPP4A;refGene_tx=XM_023251615.1
A3 63897417 63897462
refGene_gn=INPP4A;refGene_tx=XM_023251600.1;refGene_gn=INPP4A;refGene_tx=XM_023251601.1;refGene_gn=INPP4A;refGene_tx=XM_023251602.1;refGene_gn=INPP4A;refGene_tx=XM_023251603.1;refGene_gn=INPP4A;refGene_tx=XM_023251604.1;refGene_gn=INPP4A;refGene_tx=XM_023251605.1;refGene_gn=INPP4A;refGene_tx=XM_023251606.1;refGene_gn=INPP4A;refGene_tx=XM_023251607.1;refGene_gn=INPP4A;refGene_tx=XM_023251608.1;refGene_gn=INPP4A;refGene_tx=XM_023251609.1;refGene_gn=INPP4A;refGene_tx=XM_023251610.1;refGene_gn=INPP4A;refGene_tx=XM_023251611.1;refGene_gn=INPP4A;refGene_tx=XM_023251612.1;refGene_gn=INPP4A;refGene_tx=XM_023251613.1;refGene_gn=INPP4A;refGene_tx=XM_023251614.1;refGene_gn=INPP4A;refGene_tx=XM_023251615.1
A3 63898009 63898115
refGene_gn=INPP4A;refGene_tx=XM_023251600.1;refGene_gn=INPP4A;refGene_tx=XM_023251601.1;refGene_gn=INPP4A;refGene_tx=XM_023251602.1;refGene_gn=INPP4A;refGene_tx=XM_023251603.1;refGene_gn=INPP4A;refGene_tx=XM_023251604.1;refGene_gn=INPP4A;refGene_tx=XM_023251605.1;refGene_gn=INPP4A;refGene_tx=XM_023251606.1;refGene_gn=INPP4A;refGene_tx=XM_023251607.1;refGene_gn=INPP4A;refGene_tx=XM_023251608.1;refGene_gn=INPP4A;refGene_tx=XM_023251609.1;refGene_gn=INPP4A;refGene_tx=XM_023251610.1;refGene_gn=INPP4A;refGene_tx=XM_023251611.1;refGene_gn=INPP4A;refGene_tx=XM_023251612.1;refGene_gn=INPP4A;refGene_tx=XM_023251613.1;refGene_gn=INPP4A;refGene_tx=XM_023251614.1;refGene_gn=INPP4A;refGene_tx=XM_023251615.1
A3 63983852 63985264 refGene_gn=CNGA3;refGene_tx=XM_019827070.2
A3 63988220 63988327 refGene_gn=CNGA3;refGene_tx=XM_019827070.2
A3 63990593 63990701 refGene_gn=CNGA3;refGene_tx=XM_019827070.2

| | | | |
|---|---|---|---|
| A3 | 63992043 | 63992115 | refGene_gn=CNGA3;refGene_tx=XM_019827070.2 |
| A3 | 63999909 | 63999963 | refGene_gn=CNGA3;refGene_tx=XM_019827070.2 |
| A3 | 64003091 | 64003271 | refGene_gn=CNGA3;refGene_tx=XM_019827070.2 |
| A3 | 64005751 | 64005865 | refGene_gn=CNGA3;refGene_tx=XM_019827070.2 |
| A3 | 64015401 | 64015487 | refGene_gn=CNGA3;refGene_tx=XM_019827070.2 |
| A3 | 64058783 | 64058833 | refGene_gn=VWA3B;refGene_tx=XM_019827072.2 |
| A3 | 64058835 | 64058944 | refGene_gn=VWA3B;refGene_tx=XM_019827072.2 |
| A3 | 64059077 | 64059294 | refGene_gn=VWA3B;refGene_tx=XM_019827072.2 |
| A3 | 64069935 | 64070036 | refGene_gn=VWA3B;refGene_tx=XM_019827072.2 |
| A3 | 64072378 | 64072516 | refGene_gn=VWA3B;refGene_tx=XM_019827072.2 |
| A3 | 64073963 | 64074088 | refGene_gn=VWA3B;refGene_tx=XM_019827072.2 |
| A3 | 64079049 | 64079161 | refGene_gn=VWA3B;refGene_tx=XM_019827072.2 |
| A3 | 64095375 | 64095577 | refGene_gn=VWA3B;refGene_tx=XM_019827072.2 |
| A3 | 64102268 | 64102319 | refGene_gn=VWA3B;refGene_tx=XM_019827072.2 |
| A3 | 64107954 | 64108073 | refGene_gn=VWA3B;refGene_tx=XM_019827072.2 |
| A3 | 64120278 | 64120426 | refGene_gn=VWA3B;refGene_tx=XM_019827072.2 |
| A3 | 64120517 | 64120605 | refGene_gn=VWA3B;refGene_tx=XM_019827072.2 |
| A3 | 64121404 | 64121521 | refGene_gn=VWA3B;refGene_tx=XM_019827072.2 |
| A3 | 64130177 | 64130335 | refGene_gn=VWA3B;refGene_tx=XM_019827072.2 |
| A3 | 64132895 | 64133026 | refGene_gn=VWA3B;refGene_tx=XM_019827072.2 |
| A3 | 64138789 | 64138972 | refGene_gn=VWA3B;refGene_tx=XM_019827072.2 |
| A3 | 64146492 | 64146591 | refGene_gn=VWA3B;refGene_tx=XM_019827072.2 |
| A3 | 64156734 | 64156866 | refGene_gn=VWA3B;refGene_tx=XM_019827072.2 |
| A3 | 64157944 | 64158083 | refGene_gn=VWA3B;refGene_tx=XM_019827072.2 |
| A3 | 64165850 | 64166005 | refGene_gn=VWA3B;refGene_tx=XM_019827072.2 |
| A3 | 64172980 | 64173177 | refGene_gn=VWA3B;refGene_tx=XM_019827072.2 |
| A3 | 64193297 | 64193423 | refGene_gn=VWA3B;refGene_tx=XM_019827072.2 |
| A3 | 64224173 | 64224289 | refGene_gn=VWA3B;refGene_tx=XM_019827072.2 |
| A3 | 64230387 | 64230557 | refGene_gn=VWA3B;refGene_tx=XM_019827072.2 |
| A3 | 64237801 | 64237961 | refGene_gn=VWA3B;refGene_tx=XM_019827072.2 |
| A3 | 64240158 | 64240409 | refGene_gn=VWA3B;refGene_tx=XM_019827072.2 |
| A3 | 64242747 | 64242842 | refGene_gn=VWA3B;refGene_tx=XM_019827072.2 |
| A3 | 64255689 | 64255885 | refGene_gn=VWA3B;refGene_tx=XM_019827072.2 |
| A3 | 64369260 | 64369441 | refGene_gn=TMEM131;refGene_tx=XM_003983920.5 |
| A3 | 64429138 | 64429200 | refGene_gn=TMEM131;refGene_tx=XM_003983920.5 |
| A3 | 64463391 | 64463432 | refGene_gn=TMEM131;refGene_tx=XM_003983920.5 |
| A3 | 64483207 | 64483276 | refGene_gn=TMEM131;refGene_tx=XM_003983920.5 |
| A3 | 64506105 | 64506229 | refGene_gn=TMEM131;refGene_tx=XM_003983920.5 |
| A3 | 64524179 | 64524296 | refGene_gn=TMEM131;refGene_tx=XM_003983920.5 |
| A3 | 64526561 | 64526684 | refGene_gn=TMEM131;refGene_tx=XM_003983920.5 |
| A3 | 64531988 | 64532069 | refGene_gn=TMEM131;refGene_tx=XM_003983920.5 |
| A3 | 64534298 | 64534449 | refGene_gn=TMEM131;refGene_tx=XM_003983920.5 |
| A3 | 64534553 | 64534610 | refGene_gn=TMEM131;refGene_tx=XM_003983920.5 |
| A3 | 64535545 | 64535607 | refGene_gn=TMEM131;refGene_tx=XM_003983920.5 |
| A3 | 64545736 | 64545845 | refGene_gn=TMEM131;refGene_tx=XM_003983920.5 |
| A3 | 64549286 | 64549395 | refGene_gn=TMEM131;refGene_tx=XM_003983920.5 |
| A3 | 64550198 | 64550352 | refGene_gn=TMEM131;refGene_tx=XM_003983920.5 |
| A3 | 64550445 | 64550616 | refGene_gn=TMEM131;refGene_tx=XM_003983920.5 |
| A3 | 64551688 | 64551799 | refGene_gn=TMEM131;refGene_tx=XM_003983920.5 |
| A3 | 64551896 | 64552031 | refGene_gn=TMEM131;refGene_tx=XM_003983920.5 |

A3 64555920 64556025 refGene_gn=TMEM131;refGene_tx=XM_003983920.5
A3 64557389 64557476 refGene_gn=TMEM131;refGene_tx=XM_003983920.5
A3 64561140 64561293 refGene_gn=TMEM131;refGene_tx=XM_003983920.5
A3 64561393 64561469 refGene_gn=TMEM131;refGene_tx=XM_003983920.5
A3 64561638 64561756 refGene_gn=TMEM131;refGene_tx=XM_003983920.5
A3 64564342 64564481 refGene_gn=TMEM131;refGene_tx=XM_003983920.5
A3 64564596 64564706 refGene_gn=TMEM131;refGene_tx=XM_003983920.5
A3 64565173 64565240 refGene_gn=TMEM131;refGene_tx=XM_003983920.5
A3 64570194 64570346 refGene_gn=TMEM131;refGene_tx=XM_003983920.5
A3 64570950 64571093 refGene_gn=TMEM131;refGene_tx=XM_003983920.5
A3 64571510 64571697 refGene_gn=TMEM131;refGene_tx=XM_003983920.5
A3 64572794 64572980 refGene_gn=TMEM131;refGene_tx=XM_003983920.5
A3 64573785 64573944 refGene_gn=TMEM131;refGene_tx=XM_003983920.5
A3 64574363 64574947 refGene_gn=TMEM131;refGene_tx=XM_003983920.5
A3 64588597 64588773 refGene_gn=TMEM131;refGene_tx=XM_003983920.5
A3 64592279 64592407 refGene_gn=TMEM131;refGene_tx=XM_003983920.5
A3 64601206 64601331 refGene_gn=TMEM131;refGene_tx=XM_003983920.5
A3 64601534 64601684 refGene_gn=TMEM131;refGene_tx=XM_003983920.5
A3 64605592 64605758 refGene_gn=TMEM131;refGene_tx=XM_003983920.5
A3 64606913 64607035 refGene_gn=TMEM131;refGene_tx=XM_003983920.5
A3 64607123 64607220 refGene_gn=TMEM131;refGene_tx=XM_003983920.5
A3 64608032 64608130 refGene_gn=TMEM131;refGene_tx=XM_003983920.5
A3 64608900 64609061 refGene_gn=TMEM131;refGene_tx=XM_003983920.5
A3 64610112 64610397 refGene_gn=TMEM131;refGene_tx=XM_003983920.5
A3 64648810 64648934
refGene_gn=LOC101097059;refGene_tx=XM_011281054.3;refGene_gn=LOC101097
059;refGene_tx=XM_023251621.1
A3 64650672 64650785
refGene_gn=LOC101097059;refGene_tx=XM_011281054.3;refGene_gn=LOC101097
059;refGene_tx=XM_023251621.1
A3 64650888 64651029
refGene_gn=LOC101097059;refGene_tx=XM_011281054.3;refGene_gn=LOC101097
059;refGene_tx=XM_023251621.1
A3 64651118 64651311
refGene_gn=LOC101097059;refGene_tx=XM_011281054.3;refGene_gn=LOC101097
059;refGene_tx=XM_023251621.1
A3 64653457 64653664
refGene_gn=LOC101097059;refGene_tx=XM_011281054.3;refGene_gn=LOC101097
059;refGene_tx=XM_023251621.1
A3 64654224 64654417
refGene_gn=LOC101097059;refGene_tx=XM_011281054.3;refGene_gn=LOC101097
059;refGene_tx=XM_023251621.1
A3 64655456 64655508
refGene_gn=LOC101097059;refGene_tx=XM_011281054.3;refGene_gn=LOC101097
059;refGene_tx=XM_023251621.1
A3 64655734 64655781
refGene_gn=LOC101097059;refGene_tx=XM_011281054.3;refGene_gn=LOC101097
059;refGene_tx=XM_023251621.1
A3 64655863 64655951
refGene_gn=LOC101097059;refGene_tx=XM_011281054.3;refGene_gn=LOC101097

059;refGene_tx=XM_023251621.1
A3 64656052 64656191
refGene_gn=LOC101097059;refGene_tx=XM_011281054.3;refGene_gn=LOC101097
059;refGene_tx=XM_023251621.1
A3 64660668 64660829
refGene_gn=LOC101097059;refGene_tx=XM_011281054.3;refGene_gn=LOC101097
059;refGene_tx=XM_023251621.1
A3 64661498 64661900
refGene_gn=LOC101097059;refGene_tx=XM_023251621.1
A3 64677107 64677240
refGene_gn=LOC102902283;refGene_tx=XM_023251622.1;refGene_gn=LOC102902
283;refGene_tx=XM_023251623.1
A3 64679186 64679337
refGene_gn=LOC102902283;refGene_tx=XM_023251622.1;refGene_gn=LOC102902
283;refGene_tx=XM_023251623.1
A3 64680697 64680805
refGene_gn=LOC102902283;refGene_tx=XM_023251622.1;refGene_gn=LOC102902
283;refGene_tx=XM_023251623.1
A3 64687532 64687604
refGene_gn=LOC102902283;refGene_tx=XM_023251622.1;refGene_gn=LOC102902
283;refGene_tx=XM_023251623.1
A3 64688541 64688631
refGene_gn=LOC102902283;refGene_tx=XM_023251622.1;refGene_gn=LOC102902
283;refGene_tx=XM_023251623.1
A3 64700207 64700258
refGene_gn=LOC102902283;refGene_tx=XM_023251622.1
A3 64700922 64701003
refGene_gn=LOC102902283;refGene_tx=XM_023251622.1;refGene_gn=LOC102902
283;refGene_tx=XM_023251623.1
A3 64702932 64702989
refGene_gn=LOC102902283;refGene_tx=XM_023251622.1;refGene_gn=LOC102902
283;refGene_tx=XM_023251623.1
A3 64712140 64712203
refGene_gn=LOC102902283;refGene_tx=XM_023251622.1;refGene_gn=LOC102902
283;refGene_tx=XM_023251623.1
A3 64715059 64715120
refGene_gn=LOC102902283;refGene_tx=XM_023251622.1;refGene_gn=LOC102902
283;refGene_tx=XM_023251623.1
A3 64725554 64725602 refGene_gn=ACTR1B;refGene_tx=XM_003983921.4
A3 64727655 64727720 refGene_gn=ACTR1B;refGene_tx=XM_003983921.4
A3 64728986 64729062 refGene_gn=ACTR1B;refGene_tx=XM_003983921.4
A3 64730223 64730349 refGene_gn=ACTR1B;refGene_tx=XM_003983921.4
A3 64730702 64730827 refGene_gn=ACTR1B;refGene_tx=XM_003983921.4
A3 64731059 64731276 refGene_gn=ACTR1B;refGene_tx=XM_003983921.4
A3 64731411 64731504 refGene_gn=ACTR1B;refGene_tx=XM_003983921.4
A3 64731595 64731770 refGene_gn=ACTR1B;refGene_tx=XM_003983921.4
A3 64733069 64733131 refGene_gn=ACTR1B;refGene_tx=XM_003983921.4
A3 64733373 64733414 refGene_gn=ACTR1B;refGene_tx=XM_003983921.4
A3 64733577 64733680 refGene_gn=ACTR1B;refGene_tx=XM_003983921.4
A3 64742531 64742644

refGene_gn=LOC101090761;refGene_tx=XM_003983922.5
A3 64743207 64743307
refGene_gn=LOC101090761;refGene_tx=XM_003983922.5
A3 64743472 64743546
refGene_gn=LOC101090761;refGene_tx=XM_003983922.5
A3 64744099 64744202
refGene_gn=LOC101090761;refGene_tx=XM_003983922.5
A3 64755518 64755560
refGene_gn=FAM178B;refGene_tx=XM_006930055.3;refGene_gn=FAM178B;refGene_tx=XM_006930056.3;refGene_gn=FAM178B;refGene_tx=XM_019827077.2;refGene_gn=FAM178B;refGene_tx=XM_019827083.2;refGene_gn=FAM178B;refGene_tx=XM_019827086.2;refGene_gn=FAM178B;refGene_tx=XM_023251624.1;refGene_gn=FAM178B;refGene_tx=XM_023251625.1
A3 64780547 64780617
refGene_gn=FAM178B;refGene_tx=XM_006930055.3;refGene_gn=FAM178B;refGene_tx=XM_006930056.3;refGene_gn=FAM178B;refGene_tx=XM_019827077.2;refGene_gn=FAM178B;refGene_tx=XM_019827083.2;refGene_gn=FAM178B;refGene_tx=XM_019827086.2;refGene_gn=FAM178B;refGene_tx=XM_023251624.1;refGene_gn=FAM178B;refGene_tx=XM_023251625.1
A3 64780819 64781259
refGene_gn=FAM178B;refGene_tx=XM_006930055.3;refGene_gn=FAM178B;refGene_tx=XM_006930056.3;refGene_gn=FAM178B;refGene_tx=XM_019827077.2;refGene_gn=FAM178B;refGene_tx=XM_019827083.2;refGene_gn=FAM178B;refGene_tx=XM_019827086.2;refGene_gn=FAM178B;refGene_tx=XM_023251624.1;refGene_gn=FAM178B;refGene_tx=XM_023251625.1
A3 64782766 64782828
refGene_gn=FAM178B;refGene_tx=XM_006930055.3;refGene_gn=FAM178B;refGene_tx=XM_006930056.3;refGene_gn=FAM178B;refGene_tx=XM_019827077.2;refGene_gn=FAM178B;refGene_tx=XM_019827083.2;refGene_gn=FAM178B;refGene_tx=XM_019827086.2;refGene_gn=FAM178B;refGene_tx=XM_023251624.1;refGene_gn=FAM178B;refGene_tx=XM_023251625.1
A3 64784670 64784778
refGene_gn=FAM178B;refGene_tx=XM_006930055.3;refGene_gn=FAM178B;refGene_tx=XM_006930056.3;refGene_gn=FAM178B;refGene_tx=XM_019827077.2;refGene_gn=FAM178B;refGene_tx=XM_019827083.2;refGene_gn=FAM178B;refGene_tx=XM_019827086.2;refGene_gn=FAM178B;refGene_tx=XM_023251624.1;refGene_gn=FAM178B;refGene_tx=XM_023251625.1
A3 64790095 64790248
refGene_gn=FAM178B;refGene_tx=XM_006930055.3;refGene_gn=FAM178B;refGene_tx=XM_006930056.3;refGene_gn=FAM178B;refGene_tx=XM_019827077.2;refGene_gn=FAM178B;refGene_tx=XM_019827083.2;refGene_gn=FAM178B;refGene_tx=XM_019827086.2;refGene_gn=FAM178B;refGene_tx=XM_023251624.1;refGene_gn=FAM178B;refGene_tx=XM_023251625.1
A3 64797167 64797273
refGene_gn=FAM178B;refGene_tx=XM_006930055.3;refGene_gn=FAM178B;refGene_tx=XM_006930056.3;refGene_gn=FAM178B;refGene_tx=XM_019827077.2;refGene_gn=FAM178B;refGene_tx=XM_019827083.2;refGene_gn=FAM178B;refGene_tx=XM_019827086.2;refGene_gn=FAM178B;refGene_tx=XM_023251624.1;refGene_gn=FAM178B;refGene_tx=XM_023251625.1
A3 64801019 64801104

refGene_gn=FAM178B;refGene_tx=XM_006930055.3;refGene_gn=FAM178B;refGene_tx=XM_006930056.3;refGene_gn=FAM178B;refGene_tx=XM_019827077.2;refGene_gn=FAM178B;refGene_tx=XM_019827083.2;refGene_gn=FAM178B;refGene_tx=XM_019827086.2;refGene_gn=FAM178B;refGene_tx=XM_023251624.1;refGene_gn=FAM178B;refGene_tx=XM_023251625.1
A3 64825801 64825916
refGene_gn=FAM178B;refGene_tx=XM_006930055.3;refGene_gn=FAM178B;refGene_tx=XM_006930056.3;refGene_gn=FAM178B;refGene_tx=XM_019827077.2;refGene_gn=FAM178B;refGene_tx=XM_019827083.2;refGene_gn=FAM178B;refGene_tx=XM_019827086.2;refGene_gn=FAM178B;refGene_tx=XM_023251624.1;refGene_gn=FAM178B;refGene_tx=XM_023251625.1
A3 64832391 64832485
refGene_gn=FAM178B;refGene_tx=XM_006930055.3;refGene_gn=FAM178B;refGene_tx=XM_006930056.3;refGene_gn=FAM178B;refGene_tx=XM_019827077.2;refGene_gn=FAM178B;refGene_tx=XM_019827083.2;refGene_gn=FAM178B;refGene_tx=XM_019827086.2;refGene_gn=FAM178B;refGene_tx=XM_023251624.1;refGene_gn=FAM178B;refGene_tx=XM_023251625.1
A3 64832766 64832943
refGene_gn=FAM178B;refGene_tx=XM_006930055.3;refGene_gn=FAM178B;refGene_tx=XM_006930056.3;refGene_gn=FAM178B;refGene_tx=XM_019827077.2;refGene_gn=FAM178B;refGene_tx=XM_019827083.2;refGene_gn=FAM178B;refGene_tx=XM_019827086.2;refGene_gn=FAM178B;refGene_tx=XM_023251624.1;refGene_gn=FAM178B;refGene_tx=XM_023251625.1
A3 64833215 64833313
refGene_gn=FAM178B;refGene_tx=XM_006930055.3;refGene_gn=FAM178B;refGene_tx=XM_006930056.3;refGene_gn=FAM178B;refGene_tx=XM_019827077.2;refGene_gn=FAM178B;refGene_tx=XM_019827083.2;refGene_gn=FAM178B;refGene_tx=XM_019827086.2;refGene_gn=FAM178B;refGene_tx=XM_023251624.1;refGene_gn=FAM178B;refGene_tx=XM_023251625.1
A3 64846758 64846846
refGene_gn=FAM178B;refGene_tx=XM_006930055.3;refGene_gn=FAM178B;refGene_tx=XM_006930056.3;refGene_gn=FAM178B;refGene_tx=XM_019827077.2;refGene_gn=FAM178B;refGene_tx=XM_019827083.2;refGene_gn=FAM178B;refGene_tx=XM_019827086.2;refGene_gn=FAM178B;refGene_tx=XM_023251624.1;refGene_gn=FAM178B;refGene_tx=XM_023251625.1
A3 64853835 64853961
refGene_gn=FAM178B;refGene_tx=XM_006930055.3;refGene_gn=FAM178B;refGene_tx=XM_006930056.3;refGene_gn=FAM178B;refGene_tx=XM_019827077.2;refGene_gn=FAM178B;refGene_tx=XM_019827083.2;refGene_gn=FAM178B;refGene_tx=XM_019827086.2;refGene_gn=FAM178B;refGene_tx=XM_023251624.1;refGene_gn=FAM178B;refGene_tx=XM_023251625.1
A3 64855374 64855437
refGene_gn=FAM178B;refGene_tx=XM_019827077.2;refGene_gn=FAM178B;refGene_tx=XM_023251624.1;refGene_gn=FAM178B;refGene_tx=XM_023251625.1
A3 64867277 64867337 refGene_gn=FAM178B;refGene_tx=XM_019827086.2
A3 64870155 64870233
refGene_gn=FAM178B;refGene_tx=XM_006930055.3;refGene_gn=FAM178B;refGene_tx=XM_019827077.2;refGene_gn=FAM178B;refGene_tx=XM_019827083.2;refGene_gn=FAM178B;refGene_tx=XM_023251624.1;refGene_gn=FAM178B;refGene_tx=XM_023251625.1

A3 64870768 64871026
refGene_gn=FAM178B;refGene_tx=XM_006930055.3;refGene_gn=FAM178B;refGene_tx=XM_019827077.2;refGene_gn=FAM178B;refGene_tx=XM_019827083.2;refGene_gn=FAM178B;refGene_tx=XM_023251624.1;refGene_gn=FAM178B;refGene_tx=XM_023251625.1
A3 64871550 64871556 refGene_gn=FAM178B;refGene_tx=XM_023251625.1
A3 64872555 64872585
refGene_gn=FAM178B;refGene_tx=XM_006930055.3;refGene_gn=FAM178B;refGene_tx=XM_006930056.3;refGene_gn=FAM178B;refGene_tx=XM_023251624.1
A3 64880067 64880176
refGene_gn=SEMA4C;refGene_tx=XM_023251626.1;refGene_gn=SEMA4C;refGene_tx=XM_023251627.1
A3 64881634 64881783
refGene_gn=SEMA4C;refGene_tx=XM_023251626.1;refGene_gn=SEMA4C;refGene_tx=XM_023251627.1
A3 64882142 64882205
refGene_gn=SEMA4C;refGene_tx=XM_023251626.1;refGene_gn=SEMA4C;refGene_tx=XM_023251627.1
A3 64882378 64882477
refGene_gn=SEMA4C;refGene_tx=XM_023251626.1;refGene_gn=SEMA4C;refGene_tx=XM_023251627.1
A3 64882579 64882676
refGene_gn=SEMA4C;refGene_tx=XM_023251626.1;refGene_gn=SEMA4C;refGene_tx=XM_023251627.1
A3 64882798 64882915
refGene_gn=SEMA4C;refGene_tx=XM_023251626.1;refGene_gn=SEMA4C;refGene_tx=XM_023251627.1
A3 64883020 64883172
refGene_gn=SEMA4C;refGene_tx=XM_023251626.1;refGene_gn=SEMA4C;refGene_tx=XM_023251627.1
A3 64883255 64883431
refGene_gn=SEMA4C;refGene_tx=XM_023251626.1;refGene_gn=SEMA4C;refGene_tx=XM_023251627.1
A3 64883730 64883875
refGene_gn=SEMA4C;refGene_tx=XM_023251626.1;refGene_gn=SEMA4C;refGene_tx=XM_023251627.1
A3 64883966 64884189
refGene_gn=SEMA4C;refGene_tx=XM_023251626.1;refGene_gn=SEMA4C;refGene_tx=XM_023251627.1
A3 64884323 64884436
refGene_gn=SEMA4C;refGene_tx=XM_023251626.1;refGene_gn=SEMA4C;refGene_tx=XM_023251627.1
A3 64885365 64885523
refGene_gn=SEMA4C;refGene_tx=XM_023251626.1;refGene_gn=SEMA4C;refGene_tx=XM_023251627.1
A3 64885612 64885683
refGene_gn=SEMA4C;refGene_tx=XM_023251626.1;refGene_gn=SEMA4C;refGene_tx=XM_023251627.1
A3 64885809 64886639
refGene_gn=SEMA4C;refGene_tx=XM_023251626.1;refGene_gn=SEMA4C;refGene_

tx=XM_023251627.1
A3 64889011 64889111 refGene_gn=ANKRD39;refGene_tx=XM_003983946.5
A3 64893796 64893900 refGene_gn=ANKRD39;refGene_tx=XM_003983946.5
A3 64894314 64894518 refGene_gn=ANKRD39;refGene_tx=XM_003983946.5
A3 64902272 64902416 refGene_gn=ANKRD39;refGene_tx=XM_003983946.5
A3 64904974 64905001 refGene_gn=ANKRD23;refGene_tx=XM_003983947.3
A3 64906331 64906478 refGene_gn=ANKRD23;refGene_tx=XM_003983947.3
A3 64906651 64906777 refGene_gn=ANKRD23;refGene_tx=XM_003983947.3
A3 64907802 64907928 refGene_gn=ANKRD23;refGene_tx=XM_003983947.3
A3 64908196 64908295 refGene_gn=ANKRD23;refGene_tx=XM_003983947.3
A3 64908379 64908478 refGene_gn=ANKRD23;refGene_tx=XM_003983947.3
A3 64908633 64908732 refGene_gn=ANKRD23;refGene_tx=XM_003983947.3
A3 64908874 64908973 refGene_gn=ANKRD23;refGene_tx=XM_003983947.3
A3 64909063 64909159 refGene_gn=ANKRD23;refGene_tx=XM_003983947.3
A3 64916018 64916083 refGene_gn=CNNM3;refGene_tx=XM_023251636.1
A3 64916860 64916988 refGene_gn=CNNM3;refGene_tx=XM_023251635.1
A3 64919299 64919438
refGene_gn=CNNM3;refGene_tx=XM_023251635.1;refGene_gn=CNNM3;refGene_tx
=XM_023251636.1
A3 64919876 64920010
refGene_gn=CNNM3;refGene_tx=XM_023251635.1;refGene_gn=CNNM3;refGene_tx
=XM_023251636.1
A3 64920598 64920695
refGene_gn=CNNM3;refGene_tx=XM_023251635.1;refGene_gn=CNNM3;refGene_tx
=XM_023251636.1
A3 64920899 64921069
refGene_gn=CNNM3;refGene_tx=XM_023251635.1;refGene_gn=CNNM3;refGene_tx
=XM_023251636.1
A3 64921437 64921587
refGene_gn=CNNM3;refGene_tx=XM_023251635.1;refGene_gn=CNNM3;refGene_tx
=XM_023251636.1
A3 64922558 64922702
refGene_gn=CNNM3;refGene_tx=XM_023251635.1;refGene_gn=CNNM3;refGene_tx
=XM_023251636.1
A3 64930405 64931735
refGene_gn=CNNM3;refGene_tx=XM_023251635.1;refGene_gn=CNNM3;refGene_tx
=XM_023251636.1
A3 64937434 64937632
refGene_gn=CNNM4;refGene_tx=XM_023251633.1;refGene_gn=CNNM4;refGene_tx
=XM_023251634.1
A3 64937940 64938122
refGene_gn=CNNM4;refGene_tx=XM_023251633.1;refGene_gn=CNNM4;refGene_tx
=XM_023251634.1
A3 64941361 64941424 refGene_gn=CNNM4;refGene_tx=XM_023251633.1
A3 64943287 64943384
refGene_gn=CNNM4;refGene_tx=XM_023251633.1;refGene_gn=CNNM4;refGene_tx
=XM_023251634.1
A3 64943692 64943862
refGene_gn=CNNM4;refGene_tx=XM_023251633.1;refGene_gn=CNNM4;refGene_tx
=XM_023251634.1

A3	64946318	64946453
refGene_gn=CNNM4;refGene_tx=XM_023251633.1;refGene_gn=CNNM4;refGene_tx
=XM_023251634.1
A3	64946730	64946874
refGene_gn=CNNM4;refGene_tx=XM_023251633.1;refGene_gn=CNNM4;refGene_tx
=XM_023251634.1
A3	64974780	64976182
refGene_gn=CNNM4;refGene_tx=XM_023251633.1;refGene_gn=CNNM4;refGene_tx
=XM_023251634.1
A3	64992381	64992568
refGene_gn=LMAN2L;refGene_tx=XM_003983926.4;refGene_gn=LMAN2L;refGene_
tx=XM_003983927.4;refGene_gn=LMAN2L;refGene_tx=XM_019827108.2
A3	64993729	64993848
refGene_gn=LMAN2L;refGene_tx=XM_003983926.4;refGene_gn=LMAN2L;refGene_
tx=XM_003983927.4;refGene_gn=LMAN2L;refGene_tx=XM_019827108.2
A3	64996689	64996807
refGene_gn=LMAN2L;refGene_tx=XM_003983926.4;refGene_gn=LMAN2L;refGene_
tx=XM_003983927.4;refGene_gn=LMAN2L;refGene_tx=XM_019827108.2;refGene_
gn=LMAN2L;refGene_tx=XM_019827109.2;refGene_gn=LMAN2L;refGene_tx=XM_01
9827110.2;refGene_gn=LMAN2L;refGene_tx=XM_019827112.2;refGene_gn=LMAN2
L;refGene_tx=XM_023251637.1;refGene_gn=LMAN2L;refGene_tx=XM_023251638.
1
A3	64997600	64997683
refGene_gn=LMAN2L;refGene_tx=XM_003983926.4;refGene_gn=LMAN2L;refGene_
tx=XM_003983927.4;refGene_gn=LMAN2L;refGene_tx=XM_019827108.2
A3	65014390	65014423
refGene_gn=LMAN2L;refGene_tx=XM_003983927.4;refGene_gn=LMAN2L;refGene_
tx=XM_019827108.2;refGene_gn=LMAN2L;refGene_tx=XM_019827109.2;refGene_
gn=LMAN2L;refGene_tx=XM_019827110.2;refGene_gn=LMAN2L;refGene_tx=XM_02
3251637.1;refGene_gn=LMAN2L;refGene_tx=XM_023251638.1
A3	65015494	65015656
refGene_gn=LMAN2L;refGene_tx=XM_003983926.4;refGene_gn=LMAN2L;refGene_
tx=XM_003983927.4;refGene_gn=LMAN2L;refGene_tx=XM_019827108.2;refGene_
gn=LMAN2L;refGene_tx=XM_019827109.2;refGene_gn=LMAN2L;refGene_tx=XM_01
9827110.2;refGene_gn=LMAN2L;refGene_tx=XM_019827112.2;refGene_gn=LMAN2
L;refGene_tx=XM_023251637.1;refGene_gn=LMAN2L;refGene_tx=XM_023251638.
1
A3	65015745	65015860
refGene_gn=LMAN2L;refGene_tx=XM_003983926.4;refGene_gn=LMAN2L;refGene_
tx=XM_003983927.4;refGene_gn=LMAN2L;refGene_tx=XM_019827108.2;refGene_
gn=LMAN2L;refGene_tx=XM_019827109.2;refGene_gn=LMAN2L;refGene_tx=XM_01
9827110.2;refGene_gn=LMAN2L;refGene_tx=XM_019827112.2;refGene_gn=LMAN2
L;refGene_tx=XM_023251637.1;refGene_gn=LMAN2L;refGene_tx=XM_023251638.
1
A3	65019064	65019184
refGene_gn=LMAN2L;refGene_tx=XM_003983926.4;refGene_gn=LMAN2L;refGene_
tx=XM_003983927.4;refGene_gn=LMAN2L;refGene_tx=XM_019827108.2;refGene_
gn=LMAN2L;refGene_tx=XM_019827109.2;refGene_gn=LMAN2L;refGene_tx=XM_01
9827110.2;refGene_gn=LMAN2L;refGene_tx=XM_019827112.2;refGene_gn=LMAN2
L;refGene_tx=XM_023251637.1;refGene_gn=LMAN2L;refGene_tx=XM_023251638.

1
A3 65020000 65020143
refGene_gn=LMAN2L;refGene_tx=XM_003983926.4;refGene_gn=LMAN2L;refGene_tx=XM_003983927.4;refGene_gn=LMAN2L;refGene_tx=XM_019827109.2;refGene_gn=LMAN2L;refGene_tx=XM_019827110.2;refGene_gn=LMAN2L;refGene_tx=XM_019827112.2;refGene_gn=LMAN2L;refGene_tx=XM_023251637.1;refGene_gn=LMAN2L;refGene_tx=XM_023251638.1
A3 65022642 65022870
refGene_gn=FER1L5;refGene_tx=XM_006930060.3;refGene_gn=FER1L5;refGene_tx=XM_019827088.2;refGene_gn=FER1L5;refGene_tx=XM_019827089.2;refGene_gn=FER1L5;refGene_tx=XM_019827091.2;refGene_gn=FER1L5;refGene_tx=XM_019827092.2;refGene_gn=FER1L5;refGene_tx=XM_019827094.2;refGene_gn=FER1L5;refGene_tx=XM_019827096.2;refGene_gn=FER1L5;refGene_tx=XM_023251628.1
A3 65022981 65023126
refGene_gn=FER1L5;refGene_tx=XM_006930060.3;refGene_gn=FER1L5;refGene_tx=XM_019827088.2;refGene_gn=FER1L5;refGene_tx=XM_019827089.2;refGene_gn=FER1L5;refGene_tx=XM_019827091.2;refGene_gn=FER1L5;refGene_tx=XM_019827092.2;refGene_gn=FER1L5;refGene_tx=XM_019827094.2;refGene_gn=FER1L5;refGene_tx=XM_019827095.2;refGene_gn=FER1L5;refGene_tx=XM_019827096.2;refGene_gn=FER1L5;refGene_tx=XM_023251628.1
A3 65023440 65023550
refGene_gn=FER1L5;refGene_tx=XM_006930060.3;refGene_gn=FER1L5;refGene_tx=XM_019827088.2;refGene_gn=FER1L5;refGene_tx=XM_019827089.2;refGene_gn=FER1L5;refGene_tx=XM_019827091.2;refGene_gn=FER1L5;refGene_tx=XM_019827092.2;refGene_gn=FER1L5;refGene_tx=XM_019827094.2;refGene_gn=FER1L5;refGene_tx=XM_019827096.2;refGene_gn=FER1L5;refGene_tx=XM_023251628.1
A3 65023724 65023918
refGene_gn=FER1L5;refGene_tx=XM_006930060.3;refGene_gn=FER1L5;refGene_tx=XM_019827088.2;refGene_gn=FER1L5;refGene_tx=XM_019827089.2;refGene_gn=FER1L5;refGene_tx=XM_019827091.2;refGene_gn=FER1L5;refGene_tx=XM_019827092.2;refGene_gn=FER1L5;refGene_tx=XM_019827094.2;refGene_gn=FER1L5;refGene_tx=XM_019827095.2;refGene_gn=FER1L5;refGene_tx=XM_019827096.2;refGene_gn=FER1L5;refGene_tx=XM_023251628.1
A3 65023992 65024092
refGene_gn=FER1L5;refGene_tx=XM_006930060.3;refGene_gn=FER1L5;refGene_tx=XM_019827088.2;refGene_gn=FER1L5;refGene_tx=XM_019827089.2;refGene_gn=FER1L5;refGene_tx=XM_019827091.2;refGene_gn=FER1L5;refGene_tx=XM_019827092.2;refGene_gn=FER1L5;refGene_tx=XM_019827094.2;refGene_gn=FER1L5;refGene_tx=XM_019827095.2;refGene_gn=FER1L5;refGene_tx=XM_019827096.2;refGene_gn=FER1L5;refGene_tx=XM_023251628.1
A3 65024291 65024433
refGene_gn=FER1L5;refGene_tx=XM_006930060.3;refGene_gn=FER1L5;refGene_tx=XM_019827088.2;refGene_gn=FER1L5;refGene_tx=XM_019827089.2;refGene_gn=FER1L5;refGene_tx=XM_019827091.2;refGene_gn=FER1L5;refGene_tx=XM_019827092.2;refGene_gn=FER1L5;refGene_tx=XM_019827094.2;refGene_gn=FER1L5;refGene_tx=XM_019827095.2;refGene_gn=FER1L5;refGene_tx=XM_019827096.2;refGene_gn=FER1L5;refGene_tx=XM_023251628.1
A3 65024632 65024722

refGene_gn=FER1L5;refGene_tx=XM_006930060.3;refGene_gn=FER1L5;refGene_tx=XM_019827088.2;refGene_gn=FER1L5;refGene_tx=XM_019827089.2;refGene_gn=FER1L5;refGene_tx=XM_019827091.2;refGene_gn=FER1L5;refGene_tx=XM_019827092.2;refGene_gn=FER1L5;refGene_tx=XM_019827094.2;refGene_gn=FER1L5;refGene_tx=XM_019827095.2;refGene_gn=FER1L5;refGene_tx=XM_019827096.2;refGene_gn=FER1L5;refGene_tx=XM_023251628.1
A3 65024963 65025052
refGene_gn=FER1L5;refGene_tx=XM_006930060.3;refGene_gn=FER1L5;refGene_tx=XM_019827088.2;refGene_gn=FER1L5;refGene_tx=XM_019827089.2;refGene_gn=FER1L5;refGene_tx=XM_019827091.2;refGene_gn=FER1L5;refGene_tx=XM_019827092.2;refGene_gn=FER1L5;refGene_tx=XM_019827094.2;refGene_gn=FER1L5;refGene_tx=XM_019827095.2;refGene_gn=FER1L5;refGene_tx=XM_019827096.2;refGene_gn=FER1L5;refGene_tx=XM_023251628.1
A3 65025709 65025874 refGene_gn=FER1L5;refGene_tx=XM_019827097.2
A3 65026226 65026366
refGene_gn=FER1L5;refGene_tx=XM_006930060.3;refGene_gn=FER1L5;refGene_tx=XM_019827088.2;refGene_gn=FER1L5;refGene_tx=XM_019827089.2;refGene_gn=FER1L5;refGene_tx=XM_019827091.2;refGene_gn=FER1L5;refGene_tx=XM_019827092.2;refGene_gn=FER1L5;refGene_tx=XM_019827094.2;refGene_gn=FER1L5;refGene_tx=XM_019827095.2;refGene_gn=FER1L5;refGene_tx=XM_019827096.2;refGene_gn=FER1L5;refGene_tx=XM_019827097.2;refGene_gn=FER1L5;refGene_tx=XM_023251628.1
A3 65026568 65026717
refGene_gn=FER1L5;refGene_tx=XM_006930060.3;refGene_gn=FER1L5;refGene_tx=XM_019827088.2;refGene_gn=FER1L5;refGene_tx=XM_019827089.2;refGene_gn=FER1L5;refGene_tx=XM_019827091.2;refGene_gn=FER1L5;refGene_tx=XM_019827092.2;refGene_gn=FER1L5;refGene_tx=XM_019827094.2;refGene_gn=FER1L5;refGene_tx=XM_019827095.2;refGene_gn=FER1L5;refGene_tx=XM_019827096.2;refGene_gn=FER1L5;refGene_tx=XM_019827097.2;refGene_gn=FER1L5;refGene_tx=XM_023251628.1
A3 65026927 65027098
refGene_gn=FER1L5;refGene_tx=XM_006930060.3;refGene_gn=FER1L5;refGene_tx=XM_019827088.2;refGene_gn=FER1L5;refGene_tx=XM_019827089.2;refGene_gn=FER1L5;refGene_tx=XM_019827091.2;refGene_gn=FER1L5;refGene_tx=XM_019827092.2;refGene_gn=FER1L5;refGene_tx=XM_019827094.2;refGene_gn=FER1L5;refGene_tx=XM_019827095.2;refGene_gn=FER1L5;refGene_tx=XM_019827096.2;refGene_gn=FER1L5;refGene_tx=XM_019827097.2;refGene_gn=FER1L5;refGene_tx=XM_023251628.1
A3 65027470 65027562
refGene_gn=FER1L5;refGene_tx=XM_006930060.3;refGene_gn=FER1L5;refGene_tx=XM_019827088.2;refGene_gn=FER1L5;refGene_tx=XM_019827089.2;refGene_gn=FER1L5;refGene_tx=XM_019827091.2;refGene_gn=FER1L5;refGene_tx=XM_019827092.2;refGene_gn=FER1L5;refGene_tx=XM_019827094.2;refGene_gn=FER1L5;refGene_tx=XM_019827095.2;refGene_gn=FER1L5;refGene_tx=XM_019827096.2;refGene_gn=FER1L5;refGene_tx=XM_019827097.2;refGene_gn=FER1L5;refGene_tx=XM_023251628.1
A3 65027793 65027955
refGene_gn=FER1L5;refGene_tx=XM_006930060.3;refGene_gn=FER1L5;refGene_tx=XM_019827088.2;refGene_gn=FER1L5;refGene_tx=XM_019827089.2;refGene_gn=FER1L5;refGene_tx=XM_019827091.2;refGene_gn=FER1L5;refGene_tx=XM_01

9827092.2;refGene_gn=FER1L5;refGene_tx=XM_019827094.2;refGene_gn=FER1L5;refGene_tx=XM_019827095.2;refGene_gn=FER1L5;refGene_tx=XM_019827096.2;refGene_gn=FER1L5;refGene_tx=XM_019827097.2;refGene_gn=FER1L5;refGene_tx=XM_023251628.1
A3 65028498 65028618
refGene_gn=FER1L5;refGene_tx=XM_006930060.3;refGene_gn=FER1L5;refGene_tx=XM_019827088.2;refGene_gn=FER1L5;refGene_tx=XM_019827089.2;refGene_gn=FER1L5;refGene_tx=XM_019827091.2;refGene_gn=FER1L5;refGene_tx=XM_019827092.2;refGene_gn=FER1L5;refGene_tx=XM_019827094.2;refGene_gn=FER1L5;refGene_tx=XM_019827095.2;refGene_gn=FER1L5;refGene_tx=XM_019827096.2;refGene_gn=FER1L5;refGene_tx=XM_019827097.2;refGene_gn=FER1L5;refGene_tx=XM_023251628.1
A3 65028982 65029084
refGene_gn=FER1L5;refGene_tx=XM_006930060.3;refGene_gn=FER1L5;refGene_tx=XM_019827088.2;refGene_gn=FER1L5;refGene_tx=XM_019827089.2;refGene_gn=FER1L5;refGene_tx=XM_019827091.2;refGene_gn=FER1L5;refGene_tx=XM_019827092.2;refGene_gn=FER1L5;refGene_tx=XM_019827094.2;refGene_gn=FER1L5;refGene_tx=XM_019827095.2;refGene_gn=FER1L5;refGene_tx=XM_019827096.2;refGene_gn=FER1L5;refGene_tx=XM_019827097.2;refGene_gn=FER1L5;refGene_tx=XM_023251628.1
A3 65029217 65029268
refGene_gn=FER1L5;refGene_tx=XM_006930060.3;refGene_gn=FER1L5;refGene_tx=XM_019827088.2;refGene_gn=FER1L5;refGene_tx=XM_019827089.2;refGene_gn=FER1L5;refGene_tx=XM_019827091.2;refGene_gn=FER1L5;refGene_tx=XM_019827092.2;refGene_gn=FER1L5;refGene_tx=XM_019827094.2;refGene_gn=FER1L5;refGene_tx=XM_019827095.2;refGene_gn=FER1L5;refGene_tx=XM_019827096.2;refGene_gn=FER1L5;refGene_tx=XM_019827097.2;refGene_gn=FER1L5;refGene_tx=XM_023251628.1
A3 65029914 65029940
refGene_gn=FER1L5;refGene_tx=XM_006930060.3;refGene_gn=FER1L5;refGene_tx=XM_019827088.2;refGene_gn=FER1L5;refGene_tx=XM_019827089.2;refGene_gn=FER1L5;refGene_tx=XM_019827091.2;refGene_gn=FER1L5;refGene_tx=XM_019827092.2;refGene_gn=FER1L5;refGene_tx=XM_019827094.2;refGene_gn=FER1L5;refGene_tx=XM_019827095.2;refGene_gn=FER1L5;refGene_tx=XM_019827096.2;refGene_gn=FER1L5;refGene_tx=XM_019827097.2;refGene_gn=FER1L5;refGene_tx=XM_023251628.1
A3 65030086 65030249
refGene_gn=FER1L5;refGene_tx=XM_006930060.3;refGene_gn=FER1L5;refGene_tx=XM_019827088.2;refGene_gn=FER1L5;refGene_tx=XM_019827089.2;refGene_gn=FER1L5;refGene_tx=XM_019827091.2;refGene_gn=FER1L5;refGene_tx=XM_019827092.2;refGene_gn=FER1L5;refGene_tx=XM_019827094.2;refGene_gn=FER1L5;refGene_tx=XM_019827095.2;refGene_gn=FER1L5;refGene_tx=XM_019827096.2;refGene_gn=FER1L5;refGene_tx=XM_019827097.2;refGene_gn=FER1L5;refGene_tx=XM_023251628.1
A3 65030334 65030481
refGene_gn=FER1L5;refGene_tx=XM_006930060.3;refGene_gn=FER1L5;refGene_tx=XM_019827088.2;refGene_gn=FER1L5;refGene_tx=XM_019827089.2;refGene_gn=FER1L5;refGene_tx=XM_019827091.2;refGene_gn=FER1L5;refGene_tx=XM_019827092.2;refGene_gn=FER1L5;refGene_tx=XM_019827094.2;refGene_gn=FER1L5;refGene_tx=XM_019827095.2;refGene_gn=FER1L5;refGene_tx=XM_019827096.

2;refGene_gn=FER1L5;refGene_tx=XM_019827097.2;refGene_gn=FER1L5;refGene_tx=XM_023251628.1
A3 65031510 65031612
refGene_gn=FER1L5;refGene_tx=XM_006930060.3;refGene_gn=FER1L5;refGene_tx=XM_019827088.2;refGene_gn=FER1L5;refGene_tx=XM_019827089.2;refGene_gn=FER1L5;refGene_tx=XM_019827091.2;refGene_gn=FER1L5;refGene_tx=XM_019827094.2;refGene_gn=FER1L5;refGene_tx=XM_019827095.2;refGene_gn=FER1L5;refGene_tx=XM_019827096.2;refGene_gn=FER1L5;refGene_tx=XM_019827097.2;refGene_gn=FER1L5;refGene_tx=XM_023251628.1
A3 65031923 65032085
refGene_gn=FER1L5;refGene_tx=XM_006930060.3;refGene_gn=FER1L5;refGene_tx=XM_019827088.2;refGene_gn=FER1L5;refGene_tx=XM_019827089.2;refGene_gn=FER1L5;refGene_tx=XM_019827091.2;refGene_gn=FER1L5;refGene_tx=XM_019827092.2;refGene_gn=FER1L5;refGene_tx=XM_019827094.2;refGene_gn=FER1L5;refGene_tx=XM_019827095.2;refGene_gn=FER1L5;refGene_tx=XM_019827096.2;refGene_gn=FER1L5;refGene_tx=XM_019827097.2;refGene_gn=FER1L5;refGene_tx=XM_023251628.1
A3 65032293 65032475
refGene_gn=FER1L5;refGene_tx=XM_006930060.3;refGene_gn=FER1L5;refGene_tx=XM_019827088.2;refGene_gn=FER1L5;refGene_tx=XM_019827089.2;refGene_gn=FER1L5;refGene_tx=XM_019827091.2;refGene_gn=FER1L5;refGene_tx=XM_019827092.2;refGene_gn=FER1L5;refGene_tx=XM_019827094.2;refGene_gn=FER1L5;refGene_tx=XM_019827095.2;refGene_gn=FER1L5;refGene_tx=XM_019827096.2;refGene_gn=FER1L5;refGene_tx=XM_019827097.2;refGene_gn=FER1L5;refGene_tx=XM_023251628.1
A3 65033301 65033379
refGene_gn=FER1L5;refGene_tx=XM_006930060.3;refGene_gn=FER1L5;refGene_tx=XM_019827088.2;refGene_gn=FER1L5;refGene_tx=XM_019827089.2;refGene_gn=FER1L5;refGene_tx=XM_019827092.2;refGene_gn=FER1L5;refGene_tx=XM_019827094.2;refGene_gn=FER1L5;refGene_tx=XM_019827095.2;refGene_gn=FER1L5;refGene_tx=XM_019827096.2;refGene_gn=FER1L5;refGene_tx=XM_019827097.2;refGene_gn=FER1L5;refGene_tx=XM_023251628.1
A3 65033510 65033655
refGene_gn=FER1L5;refGene_tx=XM_006930060.3;refGene_gn=FER1L5;refGene_tx=XM_019827088.2;refGene_gn=FER1L5;refGene_tx=XM_019827089.2;refGene_gn=FER1L5;refGene_tx=XM_019827091.2;refGene_gn=FER1L5;refGene_tx=XM_019827092.2;refGene_gn=FER1L5;refGene_tx=XM_019827094.2;refGene_gn=FER1L5;refGene_tx=XM_019827095.2;refGene_gn=FER1L5;refGene_tx=XM_019827096.2;refGene_gn=FER1L5;refGene_tx=XM_019827097.2;refGene_gn=FER1L5;refGene_tx=XM_023251628.1
A3 65033900 65034074
refGene_gn=FER1L5;refGene_tx=XM_006930060.3;refGene_gn=FER1L5;refGene_tx=XM_019827088.2;refGene_gn=FER1L5;refGene_tx=XM_019827089.2;refGene_gn=FER1L5;refGene_tx=XM_019827091.2;refGene_gn=FER1L5;refGene_tx=XM_019827092.2;refGene_gn=FER1L5;refGene_tx=XM_019827094.2;refGene_gn=FER1L5;refGene_tx=XM_019827095.2;refGene_gn=FER1L5;refGene_tx=XM_019827096.2;refGene_gn=FER1L5;refGene_tx=XM_019827097.2;refGene_gn=FER1L5;refGene_tx=XM_023251628.1
A3 65034152 65034316
refGene_gn=FER1L5;refGene_tx=XM_006930060.3;refGene_gn=FER1L5;refGene_

tx=XM_019827088.2;refGene_gn=FER1L5;refGene_tx=XM_019827089.2;refGene_gn=FER1L5;refGene_tx=XM_019827091.2;refGene_gn=FER1L5;refGene_tx=XM_019827092.2;refGene_gn=FER1L5;refGene_tx=XM_019827094.2;refGene_gn=FER1L5;refGene_tx=XM_019827095.2;refGene_gn=FER1L5;refGene_tx=XM_019827096.2;refGene_gn=FER1L5;refGene_tx=XM_019827097.2;refGene_gn=FER1L5;refGene_tx=XM_023251628.1
A3 65034895 65034998
refGene_gn=FER1L5;refGene_tx=XM_006930060.3;refGene_gn=FER1L5;refGene_tx=XM_019827088.2;refGene_gn=FER1L5;refGene_tx=XM_019827089.2;refGene_gn=FER1L5;refGene_tx=XM_019827091.2;refGene_gn=FER1L5;refGene_tx=XM_019827092.2;refGene_gn=FER1L5;refGene_tx=XM_019827094.2;refGene_gn=FER1L5;refGene_tx=XM_019827095.2;refGene_gn=FER1L5;refGene_tx=XM_019827096.2;refGene_gn=FER1L5;refGene_tx=XM_019827097.2;refGene_gn=FER1L5;refGene_tx=XM_023251628.1
A3 65036166 65036281
refGene_gn=FER1L5;refGene_tx=XM_006930060.3;refGene_gn=FER1L5;refGene_tx=XM_019827088.2;refGene_gn=FER1L5;refGene_tx=XM_019827089.2;refGene_gn=FER1L5;refGene_tx=XM_019827091.2;refGene_gn=FER1L5;refGene_tx=XM_019827092.2;refGene_gn=FER1L5;refGene_tx=XM_019827094.2;refGene_gn=FER1L5;refGene_tx=XM_019827095.2;refGene_gn=FER1L5;refGene_tx=XM_019827096.2;refGene_gn=FER1L5;refGene_tx=XM_019827097.2;refGene_gn=FER1L5;refGene_tx=XM_023251628.1
A3 65036565 65036729
refGene_gn=FER1L5;refGene_tx=XM_006930060.3;refGene_gn=FER1L5;refGene_tx=XM_019827088.2;refGene_gn=FER1L5;refGene_tx=XM_019827089.2;refGene_gn=FER1L5;refGene_tx=XM_019827091.2;refGene_gn=FER1L5;refGene_tx=XM_019827092.2;refGene_gn=FER1L5;refGene_tx=XM_019827094.2;refGene_gn=FER1L5;refGene_tx=XM_019827095.2;refGene_gn=FER1L5;refGene_tx=XM_019827096.2;refGene_gn=FER1L5;refGene_tx=XM_019827097.2;refGene_gn=FER1L5;refGene_tx=XM_023251628.1
A3 65037921 65038053
refGene_gn=FER1L5;refGene_tx=XM_006930060.3;refGene_gn=FER1L5;refGene_tx=XM_019827088.2;refGene_gn=FER1L5;refGene_tx=XM_019827089.2;refGene_gn=FER1L5;refGene_tx=XM_019827091.2;refGene_gn=FER1L5;refGene_tx=XM_019827092.2;refGene_gn=FER1L5;refGene_tx=XM_019827094.2;refGene_gn=FER1L5;refGene_tx=XM_019827095.2;refGene_gn=FER1L5;refGene_tx=XM_019827096.2;refGene_gn=FER1L5;refGene_tx=XM_019827097.2;refGene_gn=FER1L5;refGene_tx=XM_023251628.1
A3 65039066 65039222
refGene_gn=FER1L5;refGene_tx=XM_006930060.3;refGene_gn=FER1L5;refGene_tx=XM_019827088.2;refGene_gn=FER1L5;refGene_tx=XM_019827089.2;refGene_gn=FER1L5;refGene_tx=XM_019827091.2;refGene_gn=FER1L5;refGene_tx=XM_019827092.2;refGene_gn=FER1L5;refGene_tx=XM_019827094.2;refGene_gn=FER1L5;refGene_tx=XM_019827095.2;refGene_gn=FER1L5;refGene_tx=XM_019827096.2;refGene_gn=FER1L5;refGene_tx=XM_019827097.2;refGene_gn=FER1L5;refGene_tx=XM_023251628.1
A3 65039298 65039476
refGene_gn=FER1L5;refGene_tx=XM_006930060.3;refGene_gn=FER1L5;refGene_tx=XM_019827088.2;refGene_gn=FER1L5;refGene_tx=XM_019827089.2;refGene_gn=FER1L5;refGene_tx=XM_019827091.2;refGene_gn=FER1L5;refGene_tx=XM_01

9827092.2;refGene_gn=FER1L5;refGene_tx=XM_019827094.2;refGene_gn=FER1L5;refGene_tx=XM_019827095.2;refGene_gn=FER1L5;refGene_tx=XM_019827096.2;refGene_gn=FER1L5;refGene_tx=XM_019827097.2;refGene_gn=FER1L5;refGene_tx=XM_023251628.1
A3 65040115 65040216
refGene_gn=FER1L5;refGene_tx=XM_006930060.3;refGene_gn=FER1L5;refGene_tx=XM_019827088.2;refGene_gn=FER1L5;refGene_tx=XM_019827089.2;refGene_gn=FER1L5;refGene_tx=XM_019827091.2;refGene_gn=FER1L5;refGene_tx=XM_019827092.2;refGene_gn=FER1L5;refGene_tx=XM_019827094.2;refGene_gn=FER1L5;refGene_tx=XM_019827095.2;refGene_gn=FER1L5;refGene_tx=XM_019827096.2;refGene_gn=FER1L5;refGene_tx=XM_019827097.2;refGene_gn=FER1L5;refGene_tx=XM_023251628.1
A3 65041062 65041187
refGene_gn=FER1L5;refGene_tx=XM_006930060.3;refGene_gn=FER1L5;refGene_tx=XM_019827088.2;refGene_gn=FER1L5;refGene_tx=XM_019827089.2;refGene_gn=FER1L5;refGene_tx=XM_019827091.2;refGene_gn=FER1L5;refGene_tx=XM_019827092.2;refGene_gn=FER1L5;refGene_tx=XM_019827094.2;refGene_gn=FER1L5;refGene_tx=XM_019827095.2;refGene_gn=FER1L5;refGene_tx=XM_019827096.2;refGene_gn=FER1L5;refGene_tx=XM_019827097.2;refGene_gn=FER1L5;refGene_tx=XM_023251628.1
A3 65053017 65053195
refGene_gn=FER1L5;refGene_tx=XM_006930060.3;refGene_gn=FER1L5;refGene_tx=XM_019827088.2;refGene_gn=FER1L5;refGene_tx=XM_019827089.2;refGene_gn=FER1L5;refGene_tx=XM_019827091.2;refGene_gn=FER1L5;refGene_tx=XM_019827092.2;refGene_gn=FER1L5;refGene_tx=XM_019827094.2;refGene_gn=FER1L5;refGene_tx=XM_019827095.2;refGene_gn=FER1L5;refGene_tx=XM_019827096.2;refGene_gn=FER1L5;refGene_tx=XM_019827097.2;refGene_gn=FER1L5;refGene_tx=XM_023251628.1
A3 65054707 65054833
refGene_gn=FER1L5;refGene_tx=XM_006930060.3;refGene_gn=FER1L5;refGene_tx=XM_019827088.2;refGene_gn=FER1L5;refGene_tx=XM_019827089.2;refGene_gn=FER1L5;refGene_tx=XM_019827091.2;refGene_gn=FER1L5;refGene_tx=XM_019827092.2;refGene_gn=FER1L5;refGene_tx=XM_019827094.2;refGene_gn=FER1L5;refGene_tx=XM_019827095.2;refGene_gn=FER1L5;refGene_tx=XM_019827096.2;refGene_gn=FER1L5;refGene_tx=XM_019827097.2;refGene_gn=FER1L5;refGene_tx=XM_023251628.1
A3 65055803 65055898
refGene_gn=FER1L5;refGene_tx=XM_006930060.3;refGene_gn=FER1L5;refGene_tx=XM_019827088.2;refGene_gn=FER1L5;refGene_tx=XM_019827089.2;refGene_gn=FER1L5;refGene_tx=XM_019827091.2;refGene_gn=FER1L5;refGene_tx=XM_019827092.2;refGene_gn=FER1L5;refGene_tx=XM_019827094.2;refGene_gn=FER1L5;refGene_tx=XM_019827095.2;refGene_gn=FER1L5;refGene_tx=XM_019827096.2;refGene_gn=FER1L5;refGene_tx=XM_019827097.2;refGene_gn=FER1L5;refGene_tx=XM_023251628.1
A3 65055974 65056057
refGene_gn=FER1L5;refGene_tx=XM_006930060.3;refGene_gn=FER1L5;refGene_tx=XM_019827088.2;refGene_gn=FER1L5;refGene_tx=XM_019827089.2;refGene_gn=FER1L5;refGene_tx=XM_019827091.2;refGene_gn=FER1L5;refGene_tx=XM_019827092.2;refGene_gn=FER1L5;refGene_tx=XM_019827094.2;refGene_gn=FER1L5;refGene_tx=XM_019827095.2;refGene_gn=FER1L5;refGene_tx=XM_019827096.

2;refGene_gn=FER1L5;refGene_tx=XM_019827097.2;refGene_gn=FER1L5;refGene_tx=XM_023251628.1
A3 65056131 65056194
refGene_gn=FER1L5;refGene_tx=XM_006930060.3;refGene_gn=FER1L5;refGene_tx=XM_019827088.2;refGene_gn=FER1L5;refGene_tx=XM_019827089.2;refGene_gn=FER1L5;refGene_tx=XM_019827091.2;refGene_gn=FER1L5;refGene_tx=XM_019827092.2;refGene_gn=FER1L5;refGene_tx=XM_019827094.2;refGene_gn=FER1L5;refGene_tx=XM_019827095.2;refGene_gn=FER1L5;refGene_tx=XM_019827096.2;refGene_gn=FER1L5;refGene_tx=XM_019827097.2;refGene_gn=FER1L5;refGene_tx=XM_023251628.1;refGene_gn=LMAN2L;refGene_tx=XM_019827108.2
A3 65059040 65059109
refGene_gn=FER1L5;refGene_tx=XM_006930060.3;refGene_gn=FER1L5;refGene_tx=XM_019827088.2;refGene_gn=FER1L5;refGene_tx=XM_019827089.2;refGene_gn=FER1L5;refGene_tx=XM_019827091.2;refGene_gn=FER1L5;refGene_tx=XM_019827092.2;refGene_gn=FER1L5;refGene_tx=XM_019827094.2;refGene_gn=FER1L5;refGene_tx=XM_019827095.2;refGene_gn=FER1L5;refGene_tx=XM_019827096.2;refGene_gn=FER1L5;refGene_tx=XM_019827097.2;refGene_gn=FER1L5;refGene_tx=XM_023251628.1
A3 65060945 65060998
refGene_gn=FER1L5;refGene_tx=XM_006930060.3;refGene_gn=FER1L5;refGene_tx=XM_019827088.2;refGene_gn=FER1L5;refGene_tx=XM_019827089.2;refGene_gn=FER1L5;refGene_tx=XM_019827091.2;refGene_gn=FER1L5;refGene_tx=XM_019827092.2;refGene_gn=FER1L5;refGene_tx=XM_019827094.2;refGene_gn=FER1L5;refGene_tx=XM_019827095.2;refGene_gn=FER1L5;refGene_tx=XM_019827096.2;refGene_gn=FER1L5;refGene_tx=XM_019827097.2;refGene_gn=FER1L5;refGene_tx=XM_023251628.1
A3 65061438 65061562
refGene_gn=FER1L5;refGene_tx=XM_006930060.3;refGene_gn=FER1L5;refGene_tx=XM_019827088.2;refGene_gn=FER1L5;refGene_tx=XM_019827089.2;refGene_gn=FER1L5;refGene_tx=XM_019827091.2;refGene_gn=FER1L5;refGene_tx=XM_019827092.2;refGene_gn=FER1L5;refGene_tx=XM_019827095.2;refGene_gn=FER1L5;refGene_tx=XM_019827096.2;refGene_gn=FER1L5;refGene_tx=XM_019827097.2;refGene_gn=FER1L5;refGene_tx=XM_023251628.1
A3 65061794 65061910
refGene_gn=FER1L5;refGene_tx=XM_006930060.3;refGene_gn=FER1L5;refGene_tx=XM_019827088.2;refGene_gn=FER1L5;refGene_tx=XM_019827089.2;refGene_gn=FER1L5;refGene_tx=XM_019827091.2;refGene_gn=FER1L5;refGene_tx=XM_019827092.2;refGene_gn=FER1L5;refGene_tx=XM_019827095.2;refGene_gn=FER1L5;refGene_tx=XM_019827096.2;refGene_gn=FER1L5;refGene_tx=XM_019827097.2;refGene_gn=FER1L5;refGene_tx=XM_023251628.1
A3 65062518 65062549
refGene_gn=FER1L5;refGene_tx=XM_006930060.3;refGene_gn=FER1L5;refGene_tx=XM_019827088.2;refGene_gn=FER1L5;refGene_tx=XM_019827089.2;refGene_gn=FER1L5;refGene_tx=XM_019827091.2;refGene_gn=FER1L5;refGene_tx=XM_019827092.2;refGene_gn=FER1L5;refGene_tx=XM_019827094.2;refGene_gn=FER1L5;refGene_tx=XM_019827095.2;refGene_gn=FER1L5;refGene_tx=XM_019827096.2;refGene_gn=FER1L5;refGene_tx=XM_019827097.2;refGene_gn=FER1L5;refGene_tx=XM_023251628.1
A3 65066207 65066258
refGene_gn=FER1L5;refGene_tx=XM_006930060.3;refGene_gn=FER1L5;refGene_

tx=XM_019827088.2;refGene_gn=FER1L5;refGene_tx=XM_019827089.2;refGene_gn=FER1L5;refGene_tx=XM_019827091.2;refGene_gn=FER1L5;refGene_tx=XM_019827092.2;refGene_gn=FER1L5;refGene_tx=XM_019827094.2;refGene_gn=FER1L5;refGene_tx=XM_019827095.2;refGene_gn=FER1L5;refGene_tx=XM_019827096.2;refGene_gn=FER1L5;refGene_tx=XM_019827097.2;refGene_gn=FER1L5;refGene_tx=XM_023251628.1
A3 65067026 65067089
refGene_gn=FER1L5;refGene_tx=XM_006930060.3;refGene_gn=FER1L5;refGene_tx=XM_019827088.2;refGene_gn=FER1L5;refGene_tx=XM_019827089.2;refGene_gn=FER1L5;refGene_tx=XM_019827091.2;refGene_gn=FER1L5;refGene_tx=XM_019827092.2;refGene_gn=FER1L5;refGene_tx=XM_019827094.2;refGene_gn=FER1L5;refGene_tx=XM_019827095.2;refGene_gn=FER1L5;refGene_tx=XM_019827096.2;refGene_gn=FER1L5;refGene_tx=XM_019827097.2;refGene_gn=FER1L5;refGene_tx=XM_023251628.1
A3 65068143 65068272
refGene_gn=FER1L5;refGene_tx=XM_006930060.3;refGene_gn=FER1L5;refGene_tx=XM_019827088.2;refGene_gn=FER1L5;refGene_tx=XM_019827089.2;refGene_gn=FER1L5;refGene_tx=XM_019827091.2;refGene_gn=FER1L5;refGene_tx=XM_019827092.2;refGene_gn=FER1L5;refGene_tx=XM_019827094.2;refGene_gn=FER1L5;refGene_tx=XM_019827095.2;refGene_gn=FER1L5;refGene_tx=XM_019827096.2;refGene_gn=FER1L5;refGene_tx=XM_019827097.2;refGene_gn=FER1L5;refGene_tx=XM_023251628.1
A3 65069394 65069504
refGene_gn=FER1L5;refGene_tx=XM_006930060.3;refGene_gn=FER1L5;refGene_tx=XM_019827088.2;refGene_gn=FER1L5;refGene_tx=XM_019827089.2;refGene_gn=FER1L5;refGene_tx=XM_019827091.2;refGene_gn=FER1L5;refGene_tx=XM_019827092.2;refGene_gn=FER1L5;refGene_tx=XM_019827094.2;refGene_gn=FER1L5;refGene_tx=XM_019827095.2;refGene_gn=FER1L5;refGene_tx=XM_019827096.2;refGene_gn=FER1L5;refGene_tx=XM_019827097.2
A3 65069972 65070027
refGene_gn=FER1L5;refGene_tx=XM_006930060.3;refGene_gn=FER1L5;refGene_tx=XM_019827088.2;refGene_gn=FER1L5;refGene_tx=XM_019827089.2;refGene_gn=FER1L5;refGene_tx=XM_019827091.2;refGene_gn=FER1L5;refGene_tx=XM_019827092.2;refGene_gn=FER1L5;refGene_tx=XM_019827094.2;refGene_gn=FER1L5;refGene_tx=XM_019827095.2;refGene_gn=FER1L5;refGene_tx=XM_019827097.2
A3 65071753 65071862
refGene_gn=FER1L5;refGene_tx=XM_006930060.3;refGene_gn=FER1L5;refGene_tx=XM_019827088.2;refGene_gn=FER1L5;refGene_tx=XM_019827089.2;refGene_gn=FER1L5;refGene_tx=XM_019827091.2;refGene_gn=FER1L5;refGene_tx=XM_019827092.2;refGene_gn=FER1L5;refGene_tx=XM_019827094.2;refGene_gn=FER1L5;refGene_tx=XM_019827095.2;refGene_gn=FER1L5;refGene_tx=XM_019827097.2
A3 65072562 65072654
refGene_gn=FER1L5;refGene_tx=XM_006930060.3;refGene_gn=FER1L5;refGene_tx=XM_019827088.2;refGene_gn=FER1L5;refGene_tx=XM_019827089.2;refGene_gn=FER1L5;refGene_tx=XM_019827091.2;refGene_gn=FER1L5;refGene_tx=XM_019827092.2;refGene_gn=FER1L5;refGene_tx=XM_019827094.2;refGene_gn=FER1L5;refGene_tx=XM_019827095.2;refGene_gn=FER1L5;refGene_tx=XM_019827097.2

A3 65073269 65073322
refGene_gn=FER1L5;refGene_tx=XM_006930060.3;refGene_gn=FER1L5;refGene_tx=XM_019827088.2;refGene_gn=FER1L5;refGene_tx=XM_019827089.2;refGene_gn=FER1L5;refGene_tx=XM_019827091.2;refGene_gn=FER1L5;refGene_tx=XM_019827092.2;refGene_gn=FER1L5;refGene_tx=XM_019827094.2;refGene_gn=FER1L5;refGene_tx=XM_019827095.2;refGene_gn=FER1L5;refGene_tx=XM_019827097.2
A3 65075220 65075305
refGene_gn=FER1L5;refGene_tx=XM_006930060.3;refGene_gn=FER1L5;refGene_tx=XM_019827088.2;refGene_gn=FER1L5;refGene_tx=XM_019827089.2;refGene_gn=FER1L5;refGene_tx=XM_019827091.2;refGene_gn=FER1L5;refGene_tx=XM_019827092.2;refGene_gn=FER1L5;refGene_tx=XM_019827094.2;refGene_gn=FER1L5;refGene_tx=XM_019827095.2;refGene_gn=FER1L5;refGene_tx=XM_019827097.2
A3 65083824 65084039
refGene_gn=KANSL3;refGene_tx=XM_019827098.2;refGene_gn=KANSL3;refGene_tx=XM_023251629.1;refGene_gn=KANSL3;refGene_tx=XM_023251630.1;refGene_gn=KANSL3;refGene_tx=XM_023251631.1;refGene_gn=KANSL3;refGene_tx=XM_023251632.1
A3 65086673 65086844
refGene_gn=KANSL3;refGene_tx=XM_019827098.2;refGene_gn=KANSL3;refGene_tx=XM_019827100.2;refGene_gn=KANSL3;refGene_tx=XM_019827101.2;refGene_gn=KANSL3;refGene_tx=XM_019827102.2;refGene_gn=KANSL3;refGene_tx=XM_023251629.1;refGene_gn=KANSL3;refGene_tx=XM_023251630.1;refGene_gn=KANSL3;refGene_tx=XM_023251631.1;refGene_gn=KANSL3;refGene_tx=XM_023251632.1
A3 65090525 65090581 refGene_gn=KANSL3;refGene_tx=XM_019827103.2
A3 65097001 65097092
refGene_gn=KANSL3;refGene_tx=XM_019827098.2;refGene_gn=KANSL3;refGene_tx=XM_019827100.2;refGene_gn=KANSL3;refGene_tx=XM_019827101.2;refGene_gn=KANSL3;refGene_tx=XM_019827102.2;refGene_gn=KANSL3;refGene_tx=XM_019827103.2;refGene_gn=KANSL3;refGene_tx=XM_019827104.2;refGene_gn=KANSL3;refGene_tx=XM_019827106.2;refGene_gn=KANSL3;refGene_tx=XM_023251629.1;refGene_gn=KANSL3;refGene_tx=XM_023251630.1;refGene_gn=KANSL3;refGene_tx=XM_023251631.1;refGene_gn=KANSL3;refGene_tx=XM_023251632.1
A3 65097218 65097404
refGene_gn=KANSL3;refGene_tx=XM_019827098.2;refGene_gn=KANSL3;refGene_tx=XM_019827100.2;refGene_gn=KANSL3;refGene_tx=XM_019827101.2;refGene_gn=KANSL3;refGene_tx=XM_019827102.2;refGene_gn=KANSL3;refGene_tx=XM_019827103.2;refGene_gn=KANSL3;refGene_tx=XM_019827104.2;refGene_gn=KANSL3;refGene_tx=XM_019827106.2;refGene_gn=KANSL3;refGene_tx=XM_023251629.1;refGene_gn=KANSL3;refGene_tx=XM_023251630.1;refGene_gn=KANSL3;refGene_tx=XM_023251631.1;refGene_gn=KANSL3;refGene_tx=XM_023251632.1
A3 65103522 65103654
refGene_gn=KANSL3;refGene_tx=XM_019827098.2;refGene_gn=KANSL3;refGene_tx=XM_019827100.2;refGene_gn=KANSL3;refGene_tx=XM_019827101.2;refGene_gn=KANSL3;refGene_tx=XM_019827102.2;refGene_gn=KANSL3;refGene_tx=XM_019827103.2;refGene_gn=KANSL3;refGene_tx=XM_019827104.2;refGene_gn=KANSL3;refGene_tx=XM_019827106.2;refGene_gn=KANSL3;refGene_tx=XM_023251629.1;refGene_gn=KANSL3;refGene_tx=XM_023251630.1;refGene_gn=KANSL3;refGen

e_tx=XM_023251631.1;refGene_gn=KANSL3;refGene_tx=XM_023251632.1
A3 65103904 65104021
refGene_gn=KANSL3;refGene_tx=XM_019827098.2;refGene_gn=KANSL3;refGene_tx=XM_019827100.2;refGene_gn=KANSL3;refGene_tx=XM_019827101.2;refGene_gn=KANSL3;refGene_tx=XM_019827102.2;refGene_gn=KANSL3;refGene_tx=XM_019827103.2;refGene_gn=KANSL3;refGene_tx=XM_019827104.2;refGene_gn=KANSL3;refGene_tx=XM_019827106.2;refGene_gn=KANSL3;refGene_tx=XM_023251629.1;refGene_gn=KANSL3;refGene_tx=XM_023251630.1;refGene_gn=KANSL3;refGene_tx=XM_023251631.1;refGene_gn=KANSL3;refGene_tx=XM_023251632.1
A3 65104268 65104370
refGene_gn=KANSL3;refGene_tx=XM_019827098.2;refGene_gn=KANSL3;refGene_tx=XM_019827100.2;refGene_gn=KANSL3;refGene_tx=XM_019827101.2;refGene_gn=KANSL3;refGene_tx=XM_019827102.2;refGene_gn=KANSL3;refGene_tx=XM_019827103.2;refGene_gn=KANSL3;refGene_tx=XM_019827104.2;refGene_gn=KANSL3;refGene_tx=XM_019827106.2;refGene_gn=KANSL3;refGene_tx=XM_023251629.1;refGene_gn=KANSL3;refGene_tx=XM_023251630.1;refGene_gn=KANSL3;refGene_tx=XM_023251631.1;refGene_gn=KANSL3;refGene_tx=XM_023251632.1
A3 65104473 65104545
refGene_gn=KANSL3;refGene_tx=XM_019827098.2;refGene_gn=KANSL3;refGene_tx=XM_019827100.2;refGene_gn=KANSL3;refGene_tx=XM_019827101.2;refGene_gn=KANSL3;refGene_tx=XM_019827102.2;refGene_gn=KANSL3;refGene_tx=XM_019827103.2;refGene_gn=KANSL3;refGene_tx=XM_019827104.2;refGene_gn=KANSL3;refGene_tx=XM_019827106.2;refGene_gn=KANSL3;refGene_tx=XM_023251629.1;refGene_gn=KANSL3;refGene_tx=XM_023251630.1;refGene_gn=KANSL3;refGene_tx=XM_023251631.1;refGene_gn=KANSL3;refGene_tx=XM_023251632.1
A3 65105681 65105756
refGene_gn=KANSL3;refGene_tx=XM_019827098.2;refGene_gn=KANSL3;refGene_tx=XM_019827100.2;refGene_gn=KANSL3;refGene_tx=XM_019827101.2;refGene_gn=KANSL3;refGene_tx=XM_019827102.2;refGene_gn=KANSL3;refGene_tx=XM_019827103.2;refGene_gn=KANSL3;refGene_tx=XM_019827104.2;refGene_gn=KANSL3;refGene_tx=XM_019827106.2;refGene_gn=KANSL3;refGene_tx=XM_023251629.1;refGene_gn=KANSL3;refGene_tx=XM_023251630.1;refGene_gn=KANSL3;refGene_tx=XM_023251631.1;refGene_gn=KANSL3;refGene_tx=XM_023251632.1
A3 65105932 65106090
refGene_gn=KANSL3;refGene_tx=XM_019827098.2;refGene_gn=KANSL3;refGene_tx=XM_019827100.2;refGene_gn=KANSL3;refGene_tx=XM_019827101.2;refGene_gn=KANSL3;refGene_tx=XM_019827102.2;refGene_gn=KANSL3;refGene_tx=XM_019827103.2;refGene_gn=KANSL3;refGene_tx=XM_019827104.2;refGene_gn=KANSL3;refGene_tx=XM_019827106.2;refGene_gn=KANSL3;refGene_tx=XM_023251629.1;refGene_gn=KANSL3;refGene_tx=XM_023251630.1;refGene_gn=KANSL3;refGene_tx=XM_023251631.1;refGene_gn=KANSL3;refGene_tx=XM_023251632.1
A3 65108353 65108417
refGene_gn=KANSL3;refGene_tx=XM_019827098.2;refGene_gn=KANSL3;refGene_tx=XM_019827100.2;refGene_gn=KANSL3;refGene_tx=XM_019827101.2;refGene_gn=KANSL3;refGene_tx=XM_019827102.2;refGene_gn=KANSL3;refGene_tx=XM_019827103.2;refGene_gn=KANSL3;refGene_tx=XM_019827104.2;refGene_gn=KANSL3;refGene_tx=XM_019827106.2;refGene_gn=KANSL3;refGene_tx=XM_023251629.1;refGene_gn=KANSL3;refGene_tx=XM_023251630.1;refGene_gn=KANSL3;refGene_tx=XM_023251631.1;refGene_gn=KANSL3;refGene_tx=XM_023251632.1
A3 65108857 65109058

refGene_gn=KANSL3;refGene_tx=XM_019827098.2;refGene_gn=KANSL3;refGene_tx=XM_019827100.2;refGene_gn=KANSL3;refGene_tx=XM_019827101.2;refGene_gn=KANSL3;refGene_tx=XM_019827102.2;refGene_gn=KANSL3;refGene_tx=XM_019827103.2;refGene_gn=KANSL3;refGene_tx=XM_019827104.2;refGene_gn=KANSL3;refGene_tx=XM_019827106.2;refGene_gn=KANSL3;refGene_tx=XM_023251629.1;refGene_gn=KANSL3;refGene_tx=XM_023251630.1;refGene_gn=KANSL3;refGene_tx=XM_023251631.1;refGene_gn=KANSL3;refGene_tx=XM_023251632.1
A3 65109264 65109421
refGene_gn=KANSL3;refGene_tx=XM_019827098.2;refGene_gn=KANSL3;refGene_tx=XM_019827100.2;refGene_gn=KANSL3;refGene_tx=XM_019827101.2;refGene_gn=KANSL3;refGene_tx=XM_019827102.2;refGene_gn=KANSL3;refGene_tx=XM_019827103.2;refGene_gn=KANSL3;refGene_tx=XM_019827104.2;refGene_gn=KANSL3;refGene_tx=XM_019827106.2;refGene_gn=KANSL3;refGene_tx=XM_023251629.1;refGene_gn=KANSL3;refGene_tx=XM_023251630.1;refGene_gn=KANSL3;refGene_tx=XM_023251631.1;refGene_gn=KANSL3;refGene_tx=XM_023251632.1
A3 65110954 65111032
refGene_gn=KANSL3;refGene_tx=XM_019827098.2;refGene_gn=KANSL3;refGene_tx=XM_019827100.2;refGene_gn=KANSL3;refGene_tx=XM_019827101.2;refGene_gn=KANSL3;refGene_tx=XM_019827102.2;refGene_gn=KANSL3;refGene_tx=XM_019827103.2;refGene_gn=KANSL3;refGene_tx=XM_019827104.2;refGene_gn=KANSL3;refGene_tx=XM_019827106.2;refGene_gn=KANSL3;refGene_tx=XM_023251629.1;refGene_gn=KANSL3;refGene_tx=XM_023251630.1
A3 65112635 65112827
refGene_gn=KANSL3;refGene_tx=XM_019827098.2;refGene_gn=KANSL3;refGene_tx=XM_019827100.2;refGene_gn=KANSL3;refGene_tx=XM_019827101.2;refGene_gn=KANSL3;refGene_tx=XM_019827102.2;refGene_gn=KANSL3;refGene_tx=XM_019827103.2;refGene_gn=KANSL3;refGene_tx=XM_019827104.2;refGene_gn=KANSL3;refGene_tx=XM_019827106.2;refGene_gn=KANSL3;refGene_tx=XM_023251629.1;refGene_gn=KANSL3;refGene_tx=XM_023251630.1;refGene_gn=KANSL3;refGene_tx=XM_023251631.1;refGene_gn=KANSL3;refGene_tx=XM_023251632.1
A3 65113403 65113488
refGene_gn=KANSL3;refGene_tx=XM_019827098.2;refGene_gn=KANSL3;refGene_tx=XM_019827100.2;refGene_gn=KANSL3;refGene_tx=XM_019827101.2;refGene_gn=KANSL3;refGene_tx=XM_019827102.2;refGene_gn=KANSL3;refGene_tx=XM_019827103.2;refGene_gn=KANSL3;refGene_tx=XM_019827104.2;refGene_gn=KANSL3;refGene_tx=XM_019827106.2;refGene_gn=KANSL3;refGene_tx=XM_023251629.1;refGene_gn=KANSL3;refGene_tx=XM_023251630.1;refGene_gn=KANSL3;refGene_tx=XM_023251631.1;refGene_gn=KANSL3;refGene_tx=XM_023251632.1
A3 65113914 65114042
refGene_gn=KANSL3;refGene_tx=XM_019827098.2;refGene_gn=KANSL3;refGene_tx=XM_019827100.2;refGene_gn=KANSL3;refGene_tx=XM_019827101.2;refGene_gn=KANSL3;refGene_tx=XM_019827102.2;refGene_gn=KANSL3;refGene_tx=XM_019827103.2;refGene_gn=KANSL3;refGene_tx=XM_019827104.2;refGene_gn=KANSL3;refGene_tx=XM_019827106.2;refGene_gn=KANSL3;refGene_tx=XM_023251629.1;refGene_gn=KANSL3;refGene_tx=XM_023251630.1;refGene_gn=KANSL3;refGene_tx=XM_023251631.1;refGene_gn=KANSL3;refGene_tx=XM_023251632.1
A3 65114918 65115028
refGene_gn=KANSL3;refGene_tx=XM_019827098.2;refGene_gn=KANSL3;refGene_tx=XM_019827100.2;refGene_gn=KANSL3;refGene_tx=XM_019827101.2;refGene_gn=KANSL3;refGene_tx=XM_019827102.2;refGene_gn=KANSL3;refGene_tx=XM_01

9827103.2;refGene_gn=KANSL3;refGene_tx=XM_019827104.2;refGene_gn=KANSL3;refGene_tx=XM_019827106.2;refGene_gn=KANSL3;refGene_tx=XM_023251629.1;refGene_gn=KANSL3;refGene_tx=XM_023251630.1;refGene_gn=KANSL3;refGene_tx=XM_023251631.1;refGene_gn=KANSL3;refGene_tx=XM_023251632.1
A3 65115435 65115658
refGene_gn=KANSL3;refGene_tx=XM_019827098.2;refGene_gn=KANSL3;refGene_tx=XM_019827100.2;refGene_gn=KANSL3;refGene_tx=XM_019827101.2;refGene_gn=KANSL3;refGene_tx=XM_019827102.2;refGene_gn=KANSL3;refGene_tx=XM_019827103.2;refGene_gn=KANSL3;refGene_tx=XM_019827104.2;refGene_gn=KANSL3;refGene_tx=XM_019827106.2;refGene_gn=KANSL3;refGene_tx=XM_023251629.1;refGene_gn=KANSL3;refGene_tx=XM_023251630.1;refGene_gn=KANSL3;refGene_tx=XM_023251631.1;refGene_gn=KANSL3;refGene_tx=XM_023251632.1
A3 65115993 65116127
refGene_gn=KANSL3;refGene_tx=XM_019827098.2;refGene_gn=KANSL3;refGene_tx=XM_019827100.2;refGene_gn=KANSL3;refGene_tx=XM_019827101.2;refGene_gn=KANSL3;refGene_tx=XM_019827102.2;refGene_gn=KANSL3;refGene_tx=XM_019827103.2;refGene_gn=KANSL3;refGene_tx=XM_019827104.2;refGene_gn=KANSL3;refGene_tx=XM_019827106.2;refGene_gn=KANSL3;refGene_tx=XM_023251629.1;refGene_gn=KANSL3;refGene_tx=XM_023251630.1;refGene_gn=KANSL3;refGene_tx=XM_023251631.1;refGene_gn=KANSL3;refGene_tx=XM_023251632.1
A3 65123645 65123692
refGene_gn=KANSL3;refGene_tx=XM_019827098.2;refGene_gn=KANSL3;refGene_tx=XM_019827100.2;refGene_gn=KANSL3;refGene_tx=XM_019827101.2;refGene_gn=KANSL3;refGene_tx=XM_019827102.2;refGene_gn=KANSL3;refGene_tx=XM_019827103.2;refGene_gn=KANSL3;refGene_tx=XM_019827104.2;refGene_gn=KANSL3;refGene_tx=XM_019827106.2;refGene_gn=KANSL3;refGene_tx=XM_023251629.1;refGene_gn=KANSL3;refGene_tx=XM_023251631.1
A3 65123869 65123903
refGene_gn=KANSL3;refGene_tx=XM_019827098.2;refGene_gn=KANSL3;refGene_tx=XM_019827100.2;refGene_gn=KANSL3;refGene_tx=XM_019827101.2;refGene_gn=KANSL3;refGene_tx=XM_019827102.2;refGene_gn=KANSL3;refGene_tx=XM_019827103.2;refGene_gn=KANSL3;refGene_tx=XM_019827104.2;refGene_gn=KANSL3;refGene_tx=XM_019827106.2;refGene_gn=KANSL3;refGene_tx=XM_023251629.1;refGene_gn=KANSL3;refGene_tx=XM_023251630.1;refGene_gn=KANSL3;refGene_tx=XM_023251631.1;refGene_gn=KANSL3;refGene_tx=XM_023251632.1
A3 65131199 65132339 refGene_gn=OXER1;refGene_tx=XM_011281088.3
A3 65134366 65134445 refGene_gn=HAAO;refGene_tx=XM_003983932.5
A3 65134548 65134631 refGene_gn=HAAO;refGene_tx=XM_003983932.5
A3 65134817 65134886 refGene_gn=HAAO;refGene_tx=XM_003983932.5
A3 65136706 65136852 refGene_gn=HAAO;refGene_tx=XM_003983932.5
A3 65137120 65137164 refGene_gn=HAAO;refGene_tx=XM_003983932.5
A3 65137494 65137584 refGene_gn=HAAO;refGene_tx=XM_003983932.5
A3 65140702 65140809 refGene_gn=HAAO;refGene_tx=XM_003983932.5
A3 65141122 65141206 refGene_gn=HAAO;refGene_tx=XM_003983932.5
A3 65146154 65146233 refGene_gn=HAAO;refGene_tx=XM_003983932.5
A3 65148712 65148792 refGene_gn=HAAO;refGene_tx=XM_003983932.5
A3 65524020 65525472 refGene_gn=ZFP36L2;refGene_tx=XM_011281046.2
A3 65525990 65526041 refGene_gn=ZFP36L2;refGene_tx=XM_011281046.2
A3 65530384 65530777 refGene_gn=THADA;refGene_tx=XM_006930069.4
A3 65537226 65537396 refGene_gn=THADA;refGene_tx=XM_006930069.4

A3 65579322 65579454 refGene_gn=THADA;refGene_tx=XM_006930069.4
A3 65582069 65582223 refGene_gn=THADA;refGene_tx=XM_006930069.4
A3 65585371 65585444 refGene_gn=THADA;refGene_tx=XM_006930069.4
A3 65585777 65585869 refGene_gn=THADA;refGene_tx=XM_006930069.4
A3 65585870 65585897 refGene_gn=THADA;refGene_tx=XM_006930069.4
A3 65586461 65586841 refGene_gn=THADA;refGene_tx=XM_006930069.4
A3 65609371 65609466 refGene_gn=THADA;refGene_tx=XM_006930069.4
A3 65629078 65629194 refGene_gn=THADA;refGene_tx=XM_006930069.4
A3 65681446 65681615 refGene_gn=THADA;refGene_tx=XM_006930069.4
A3 65713496 65713628 refGene_gn=THADA;refGene_tx=XM_006930069.4
A3 65715153 65715243 refGene_gn=THADA;refGene_tx=XM_006930069.4
A3 65757564 65757656 refGene_gn=THADA;refGene_tx=XM_006930069.4
A3 65769169 65769292 refGene_gn=THADA;refGene_tx=XM_006930069.4
A3 65774659 65774773 refGene_gn=THADA;refGene_tx=XM_006930069.4
A3 65778156 65778289 refGene_gn=THADA;refGene_tx=XM_006930069.4
A3 65797486 65797596 refGene_gn=THADA;refGene_tx=XM_006930069.4
A3 65808887 65809045 refGene_gn=THADA;refGene_tx=XM_006930069.4
A3 65810494 65810653 refGene_gn=THADA;refGene_tx=XM_006930069.4
A3 65814198 65814335 refGene_gn=THADA;refGene_tx=XM_006930069.4
A3 65814609 65814745 refGene_gn=THADA;refGene_tx=XM_006930069.4
A3 65819065 65819279 refGene_gn=THADA;refGene_tx=XM_006930069.4
A3 65821766 65821918 refGene_gn=THADA;refGene_tx=XM_006930069.4
A3 65830234 65830358 refGene_gn=THADA;refGene_tx=XM_006930069.4
A3 65836091 65836214 refGene_gn=THADA;refGene_tx=XM_006930069.4
A3 65836944 65837100 refGene_gn=THADA;refGene_tx=XM_006930069.4
A3 65840329 65840481 refGene_gn=THADA;refGene_tx=XM_006930069.4
A3 65842796 65843503 refGene_gn=THADA;refGene_tx=XM_006930069.4
A3 65845578 65845799 refGene_gn=THADA;refGene_tx=XM_006930069.4
A3 65846752 65846847 refGene_gn=THADA;refGene_tx=XM_006930069.4
A3 65849636 65849824 refGene_gn=THADA;refGene_tx=XM_006930069.4
A3 65853539 65853588 refGene_gn=THADA;refGene_tx=XM_006930069.4
A3 65854094 65854127 refGene_gn=THADA;refGene_tx=XM_006930069.4
A3 65854246 65854395 refGene_gn=THADA;refGene_tx=XM_006930069.4
A3 65859080 65859211 refGene_gn=THADA;refGene_tx=XM_006930069.4
A3 65859928 65860023 refGene_gn=THADA;refGene_tx=XM_006930069.4
A3 65860308 65860384 refGene_gn=THADA;refGene_tx=XM_006930069.4
A3 65906127 65906250 refGene_gn=PLEKHH2;refGene_tx=XM_011281045.3
A3 65925563 65925622
refGene_gn=C1GALT1C1L;refGene_tx=XM_019828005.1
A3 65925624 65925765
refGene_gn=C1GALT1C1L;refGene_tx=XM_019828005.1
A3 65925767 65925929
refGene_gn=C1GALT1C1L;refGene_tx=XM_019828005.1
A3 65925931 65925981
refGene_gn=C1GALT1C1L;refGene_tx=XM_019828005.1
A3 65925982 65926137
refGene_gn=C1GALT1C1L;refGene_tx=XM_019828005.1
A3 65928650 65928713 refGene_gn=PLEKHH2;refGene_tx=XM_011281045.3
A3 65938307 65938457 refGene_gn=PLEKHH2;refGene_tx=XM_011281045.3
A3 65940092 65940176 refGene_gn=PLEKHH2;refGene_tx=XM_011281045.3

A3 65940993 65941075 refGene_gn=PLEKHH2;refGene_tx=XM_011281045.3
A3 65945806 65945992 refGene_gn=PLEKHH2;refGene_tx=XM_011281045.3
A3 65948443 65949405 refGene_gn=PLEKHH2;refGene_tx=XM_011281045.3
A3 65952652 65952728 refGene_gn=PLEKHH2;refGene_tx=XM_011281045.3
A3 65954216 65954311 refGene_gn=PLEKHH2;refGene_tx=XM_011281045.3
A3 65954896 65955041 refGene_gn=PLEKHH2;refGene_tx=XM_011281045.3
A3 65957590 65957727 refGene_gn=PLEKHH2;refGene_tx=XM_011281045.3
A3 65957827 65957938 refGene_gn=PLEKHH2;refGene_tx=XM_011281045.3
A3 65958095 65958182 refGene_gn=PLEKHH2;refGene_tx=XM_011281045.3
A3 65960527 65960683 refGene_gn=PLEKHH2;refGene_tx=XM_011281045.3
A3 65968638 65968719 refGene_gn=PLEKHH2;refGene_tx=XM_011281045.3
A3 65972579 65972759 refGene_gn=PLEKHH2;refGene_tx=XM_011281045.3
A3 65976623 65976732 refGene_gn=PLEKHH2;refGene_tx=XM_011281045.3
A3 65977666 65977779 refGene_gn=PLEKHH2;refGene_tx=XM_011281045.3
A3 65982184 65982364 refGene_gn=PLEKHH2;refGene_tx=XM_011281045.3
A3 65984423 65984521 refGene_gn=PLEKHH2;refGene_tx=XM_011281045.3
A3 65986458 65986636 refGene_gn=PLEKHH2;refGene_tx=XM_011281045.3
A3 65987768 65987924 refGene_gn=PLEKHH2;refGene_tx=XM_011281045.3
A3 65989763 65989861 refGene_gn=PLEKHH2;refGene_tx=XM_011281045.3
A3 65997739 65997881 refGene_gn=PLEKHH2;refGene_tx=XM_011281045.3
A3 66002907 66003053 refGene_gn=PLEKHH2;refGene_tx=XM_011281045.3
A3 66004104 66004234 refGene_gn=PLEKHH2;refGene_tx=XM_011281045.3
A3 66008369 66008456 refGene_gn=PLEKHH2;refGene_tx=XM_011281045.3
A3 66010139 66010277 refGene_gn=PLEKHH2;refGene_tx=XM_011281045.3
A3 66011240 66011426 refGene_gn=PLEKHH2;refGene_tx=XM_011281045.3
A3 66019018 66019026
refGene_gn=DYNC2LI1;refGene_tx=XM_006930070.3;refGene_gn=DYNC2LI1;refGene_tx=XM_019827128.2
A3 66021258 66021376
refGene_gn=DYNC2LI1;refGene_tx=XM_006930070.3;refGene_gn=DYNC2LI1;refGene_tx=XM_019827128.2
A3 66032626 66032661
refGene_gn=DYNC2LI1;refGene_tx=XM_006930070.3;refGene_gn=DYNC2LI1;refGene_tx=XM_019827128.2
A3 66035432 66035502
refGene_gn=DYNC2LI1;refGene_tx=XM_006930070.3;refGene_gn=DYNC2LI1;refGene_tx=XM_019827128.2
A3 66037246 66037335
refGene_gn=DYNC2LI1;refGene_tx=XM_006930070.3;refGene_gn=DYNC2LI1;refGene_tx=XM_019827128.2
A3 66042962 66043149
refGene_gn=DYNC2LI1;refGene_tx=XM_006930070.3;refGene_gn=DYNC2LI1;refGene_tx=XM_019827128.2
A3 66044078 66044147
refGene_gn=DYNC2LI1;refGene_tx=XM_006930070.3;refGene_gn=DYNC2LI1;refGene_tx=XM_019827128.2
A3 66044751 66044829
refGene_gn=DYNC2LI1;refGene_tx=XM_006930070.3;refGene_gn=DYNC2LI1;refGene_tx=XM_019827128.2
A3 66048997 66049074

refGene_gn=DYNC2LI1;refGene_tx=XM_006930070.3;refGene_gn=DYNC2LI1;refGene_tx=XM_019827128.2
A3 66049683 66049754
refGene_gn=DYNC2LI1;refGene_tx=XM_006930070.3;refGene_gn=DYNC2LI1;refGene_tx=XM_019827128.2
A3 66052919 66053017
refGene_gn=DYNC2LI1;refGene_tx=XM_006930070.3;refGene_gn=DYNC2LI1;refGene_tx=XM_019827128.2
A3 66053366 66053459
refGene_gn=DYNC2LI1;refGene_tx=XM_006930070.3;refGene_gn=DYNC2LI1;refGene_tx=XM_019827128.2
A3 66057200 66057263
refGene_gn=DYNC2LI1;refGene_tx=XM_006930070.3;refGene_gn=DYNC2LI1;refGene_tx=XM_019827128.2
A3 66060417 66060611
refGene_gn=ABCG5;refGene_tx=XM_003983934.5;refGene_gn=ABCG5;refGene_tx=XM_023251640.1
A3 66061753 66061866
refGene_gn=ABCG5;refGene_tx=XM_003983934.5;refGene_gn=ABCG5;refGene_tx=XM_023251640.1
A3 66068485 66068671
refGene_gn=ABCG5;refGene_tx=XM_003983934.5;refGene_gn=ABCG5;refGene_tx=XM_023251640.1
A3 66074225 66074364
refGene_gn=ABCG5;refGene_tx=XM_003983934.5;refGene_gn=ABCG5;refGene_tx=XM_023251640.1
A3 66075157 66075363
refGene_gn=ABCG5;refGene_tx=XM_003983934.5;refGene_gn=ABCG5;refGene_tx=XM_023251640.1
A3 66075445 66075659
refGene_gn=ABCG5;refGene_tx=XM_003983934.5;refGene_gn=ABCG5;refGene_tx=XM_023251640.1
A3 66076122 66076252
refGene_gn=ABCG5;refGene_tx=XM_003983934.5;refGene_gn=ABCG5;refGene_tx=XM_023251640.1
A3 66076993 66077133
refGene_gn=ABCG5;refGene_tx=XM_003983934.5;refGene_gn=ABCG5;refGene_tx=XM_023251640.1
A3 66078294 66078427
refGene_gn=ABCG5;refGene_tx=XM_003983934.5;refGene_gn=ABCG5;refGene_tx=XM_023251640.1
A3 66083591 66083690
refGene_gn=ABCG5;refGene_tx=XM_003983934.5;refGene_gn=ABCG5;refGene_tx=XM_023251640.1
A3 66083771 66083908
refGene_gn=ABCG5;refGene_tx=XM_003983934.5;refGene_gn=ABCG5;refGene_tx=XM_023251640.1
A3 66090516 66090638 refGene_gn=ABCG5;refGene_tx=XM_003983934.5
A3 66091320 66091466 refGene_gn=ABCG5;refGene_tx=XM_003983934.5
A3 66091851 66091914 refGene_gn=ABCG8;refGene_tx=XM_006930071.3

| | | | |
|---|---|---|---|
| A3 | 66097483 | 66097585 | refGene_gn=ABCG8;refGene_tx=XM_006930071.3 |
| A3 | 66098782 | 66098933 | refGene_gn=ABCG8;refGene_tx=XM_006930071.3 |
| A3 | 66101194 | 66101433 | refGene_gn=ABCG8;refGene_tx=XM_006930071.3 |
| A3 | 66101954 | 66102087 | refGene_gn=ABCG8;refGene_tx=XM_006930071.3 |
| A3 | 66102187 | 66102457 | refGene_gn=ABCG8;refGene_tx=XM_006930071.3 |
| A3 | 66104853 | 66105016 | refGene_gn=ABCG8;refGene_tx=XM_006930071.3 |
| A3 | 66105105 | 66105189 | refGene_gn=ABCG8;refGene_tx=XM_006930071.3 |
| A3 | 66106080 | 66106280 | refGene_gn=ABCG8;refGene_tx=XM_006930071.3 |
| A3 | 66106723 | 66106800 | refGene_gn=ABCG8;refGene_tx=XM_006930071.3 |
| A3 | 66107509 | 66107777 | refGene_gn=ABCG8;refGene_tx=XM_006930071.3 |
| A3 | 66108752 | 66108880 | refGene_gn=ABCG8;refGene_tx=XM_006930071.3 |
| A3 | 66108965 | 66109103 | refGene_gn=ABCG8;refGene_tx=XM_006930071.3 |
| A3 | 66117905 | 66117962 | refGene_gn=LRPPRC;refGene_tx=XM_011281043.3 |
| A3 | 66119822 | 66119965 | refGene_gn=LRPPRC;refGene_tx=XM_011281043.3 |
| A3 | 66122162 | 66122247 | refGene_gn=LRPPRC;refGene_tx=XM_011281043.3 |
| A3 | 66124118 | 66124193 | refGene_gn=LRPPRC;refGene_tx=XM_011281043.3 |
| A3 | 66126701 | 66126817 | refGene_gn=LRPPRC;refGene_tx=XM_011281043.3 |
| A3 | 66126949 | 66127089 | refGene_gn=LRPPRC;refGene_tx=XM_011281043.3 |
| A3 | 66128837 | 66129042 | refGene_gn=LRPPRC;refGene_tx=XM_011281043.3 |
| A3 | 66132500 | 66132589 | refGene_gn=LRPPRC;refGene_tx=XM_011281043.3 |
| A3 | 66138581 | 66138708 | refGene_gn=LRPPRC;refGene_tx=XM_011281043.3 |
| A3 | 66141847 | 66141956 | refGene_gn=LRPPRC;refGene_tx=XM_011281043.3 |
| A3 | 66142092 | 66142235 | refGene_gn=LRPPRC;refGene_tx=XM_011281043.3 |
| A3 | 66149720 | 66149811 | refGene_gn=LRPPRC;refGene_tx=XM_011281043.3 |
| A3 | 66150607 | 66150676 | refGene_gn=LRPPRC;refGene_tx=XM_011281043.3 |
| A3 | 66160198 | 66160305 | refGene_gn=LRPPRC;refGene_tx=XM_011281043.3 |
| A3 | 66160774 | 66160899 | refGene_gn=LRPPRC;refGene_tx=XM_011281043.3 |
| A3 | 66170656 | 66170864 | refGene_gn=LRPPRC;refGene_tx=XM_011281043.3 |
| A3 | 66172283 | 66172369 | refGene_gn=LRPPRC;refGene_tx=XM_011281043.3 |
| A3 | 66173075 | 66173206 | refGene_gn=LRPPRC;refGene_tx=XM_011281043.3 |
| A3 | 66174234 | 66174348 | refGene_gn=LRPPRC;refGene_tx=XM_011281043.3 |
| A3 | 66174699 | 66174744 | refGene_gn=LRPPRC;refGene_tx=XM_011281043.3 |
| A3 | 66175079 | 66175157 | refGene_gn=LRPPRC;refGene_tx=XM_011281043.3 |
| A3 | 66175378 | 66175485 | refGene_gn=LRPPRC;refGene_tx=XM_011281043.3 |
| A3 | 66176604 | 66176662 | refGene_gn=LRPPRC;refGene_tx=XM_011281043.3 |
| A3 | 66178678 | 66178706 | refGene_gn=LRPPRC;refGene_tx=XM_011281043.3 |
| A3 | 66191586 | 66191653 | refGene_gn=LRPPRC;refGene_tx=XM_011281043.3 |
| A3 | 66194659 | 66194753 | refGene_gn=LRPPRC;refGene_tx=XM_011281043.3 |
| A3 | 66196241 | 66196360 | refGene_gn=LRPPRC;refGene_tx=XM_011281043.3 |
| A3 | 66205171 | 66205279 | refGene_gn=LRPPRC;refGene_tx=XM_011281043.3 |
| A3 | 66205360 | 66205466 | refGene_gn=LRPPRC;refGene_tx=XM_011281043.3 |
| A3 | 66205720 | 66205866 | refGene_gn=LRPPRC;refGene_tx=XM_011281043.3 |
| A3 | 66206183 | 66206328 | refGene_gn=LRPPRC;refGene_tx=XM_011281043.3 |
| A3 | 66206678 | 66206805 | refGene_gn=LRPPRC;refGene_tx=XM_011281043.3 |
| A3 | 66207647 | 66207734 | refGene_gn=LRPPRC;refGene_tx=XM_011281043.3 |
| A3 | 66208433 | 66208492 | refGene_gn=LRPPRC;refGene_tx=XM_011281043.3 |
| A3 | 66208599 | 66208721 | refGene_gn=LRPPRC;refGene_tx=XM_011281043.3 |
| A3 | 66211652 | 66211775 | refGene_gn=LRPPRC;refGene_tx=XM_011281043.3 |
| A3 | 66213100 | 66213297 | refGene_gn=LRPPRC;refGene_tx=XM_011281043.3 |
| A3 | 66228846 | 66228995 | refGene_gn=LRPPRC;refGene_tx=XM_011281043.3 |

A3 66446162 66447008
refGene_gn=PPM1B;refGene_tx=XM_003983935.4;refGene_gn=PPM1B;refGene_tx=XM_003983936.5;refGene_gn=PPM1B;refGene_tx=XM_019827129.2;refGene_gn=PPM1B;refGene_tx=XM_019827130.2
A3 66455808 66455926
refGene_gn=PPM1B;refGene_tx=XM_003983935.4;refGene_gn=PPM1B;refGene_tx=XM_003983936.5;refGene_gn=PPM1B;refGene_tx=XM_019827129.2;refGene_gn=PPM1B;refGene_tx=XM_019827130.2
A3 66459186 66459298
refGene_gn=PPM1B;refGene_tx=XM_003983935.4;refGene_gn=PPM1B;refGene_tx=XM_003983936.5;refGene_gn=PPM1B;refGene_tx=XM_019827129.2;refGene_gn=PPM1B;refGene_tx=XM_019827130.2
A3 66459687 66459745
refGene_gn=PPM1B;refGene_tx=XM_003983935.4;refGene_gn=PPM1B;refGene_tx=XM_003983936.5;refGene_gn=PPM1B;refGene_tx=XM_019827129.2;refGene_gn=PPM1B;refGene_tx=XM_019827130.2
A3 66460513 66460540 refGene_gn=PPM1B;refGene_tx=XM_019827130.2
A3 66469831 66470137 refGene_gn=PPM1B;refGene_tx=XM_003983935.4
A3 66471753 66471783 refGene_gn=PPM1B;refGene_tx=XM_003983936.5
A3 66481667 66481712 refGene_gn=PPM1B;refGene_tx=XM_019827129.2
A3 66509155 66509573
refGene_gn=SLC3A1;refGene_tx=XM_003983937.5;refGene_gn=SLC3A1;refGene_tx=XM_019827133.2
A3 66514340 66514520
refGene_gn=SLC3A1;refGene_tx=XM_003983937.5;refGene_gn=SLC3A1;refGene_tx=XM_019827133.2
A3 66514672 66514827
refGene_gn=SLC3A1;refGene_tx=XM_003983937.5;refGene_gn=SLC3A1;refGene_tx=XM_019827133.2
A3 66520439 66520565 refGene_gn=SLC3A1;refGene_tx=XM_003983937.5
A3 66531644 66531764
refGene_gn=SLC3A1;refGene_tx=XM_003983937.5;refGene_gn=SLC3A1;refGene_tx=XM_019827133.2
A3 66534538 66534663
refGene_gn=SLC3A1;refGene_tx=XM_003983937.5;refGene_gn=SLC3A1;refGene_tx=XM_019827133.2
A3 66536783 66536979
refGene_gn=SLC3A1;refGene_tx=XM_003983937.5;refGene_gn=SLC3A1;refGene_tx=XM_019827133.2
A3 66538835 66538901 refGene_gn=PREPL;refGene_tx=XM_019827131.2
A3 66539587 66539755
refGene_gn=SLC3A1;refGene_tx=XM_003983937.5;refGene_gn=SLC3A1;refGene_tx=XM_019827133.2
A3 66540652 66540769
refGene_gn=SLC3A1;refGene_tx=XM_003983937.5;refGene_gn=SLC3A1;refGene_tx=XM_019827133.2
A3 66545529 66545970
refGene_gn=SLC3A1;refGene_tx=XM_003983937.5;refGene_gn=SLC3A1;refGene_tx=XM_019827133.2
A3 66546642 66546732

refGene_gn=PREPL;refGene_tx=XM_011281039.3;refGene_gn=PREPL;refGene_tx=XM_011281040.3;refGene_gn=PREPL;refGene_tx=XM_011281041.3;refGene_gn=PREPL;refGene_tx=XM_011281042.3;refGene_gn=PREPL;refGene_tx=XM_019827132.2
A3 66546905 66546908 refGene_gn=PREPL;refGene_tx=XM_019827131.2
A3 66547105 66547179 refGene_gn=PREPL;refGene_tx=XM_011281039.3;refGene_gn=PREPL;refGene_tx=XM_011281040.3;refGene_gn=PREPL;refGene_tx=XM_011281041.3;refGene_gn=PREPL;refGene_tx=XM_011281042.3;refGene_gn=PREPL;refGene_tx=XM_019827131.2;refGene_gn=PREPL;refGene_tx=XM_019827132.2
A3 66547989 66548113 refGene_gn=PREPL;refGene_tx=XM_011281039.3;refGene_gn=PREPL;refGene_tx=XM_011281040.3;refGene_gn=PREPL;refGene_tx=XM_011281041.3;refGene_gn=PREPL;refGene_tx=XM_011281042.3;refGene_gn=PREPL;refGene_tx=XM_019827131.2;refGene_gn=PREPL;refGene_tx=XM_019827132.2
A3 66548506 66548656 refGene_gn=PREPL;refGene_tx=XM_011281039.3;refGene_gn=PREPL;refGene_tx=XM_011281040.3;refGene_gn=PREPL;refGene_tx=XM_011281041.3;refGene_gn=PREPL;refGene_tx=XM_011281042.3;refGene_gn=PREPL;refGene_tx=XM_019827131.2;refGene_gn=PREPL;refGene_tx=XM_019827132.2
A3 66551941 66552158 refGene_gn=PREPL;refGene_tx=XM_011281039.3;refGene_gn=PREPL;refGene_tx=XM_011281040.3;refGene_gn=PREPL;refGene_tx=XM_011281041.3;refGene_gn=PREPL;refGene_tx=XM_011281042.3;refGene_gn=PREPL;refGene_tx=XM_019827131.2;refGene_gn=PREPL;refGene_tx=XM_019827132.2
A3 66555183 66555359 refGene_gn=PREPL;refGene_tx=XM_011281039.3;refGene_gn=PREPL;refGene_tx=XM_011281040.3;refGene_gn=PREPL;refGene_tx=XM_011281041.3;refGene_gn=PREPL;refGene_tx=XM_011281042.3;refGene_gn=PREPL;refGene_tx=XM_019827131.2;refGene_gn=PREPL;refGene_tx=XM_019827132.2
A3 66557438 66557636 refGene_gn=PREPL;refGene_tx=XM_011281039.3;refGene_gn=PREPL;refGene_tx=XM_011281040.3;refGene_gn=PREPL;refGene_tx=XM_011281041.3;refGene_gn=PREPL;refGene_tx=XM_011281042.3;refGene_gn=PREPL;refGene_tx=XM_019827131.2;refGene_gn=PREPL;refGene_tx=XM_019827132.2
A3 66565779 66565965 refGene_gn=PREPL;refGene_tx=XM_011281039.3;refGene_gn=PREPL;refGene_tx=XM_011281040.3;refGene_gn=PREPL;refGene_tx=XM_011281041.3;refGene_gn=PREPL;refGene_tx=XM_011281042.3;refGene_gn=PREPL;refGene_tx=XM_019827131.2;refGene_gn=PREPL;refGene_tx=XM_019827132.2
A3 66566541 66566758 refGene_gn=PREPL;refGene_tx=XM_011281039.3;refGene_gn=PREPL;refGene_tx=XM_011281040.3;refGene_gn=PREPL;refGene_tx=XM_011281041.3;refGene_gn=PREPL;refGene_tx=XM_011281042.3;refGene_gn=PREPL;refGene_tx=XM_019827131.2;refGene_gn=PREPL;refGene_tx=XM_019827132.2
A3 66583228 66583364 refGene_gn=PREPL;refGene_tx=XM_011281039.3;refGene_gn=PREPL;refGene_tx=XM_011281040.3;refGene_gn=PREPL;refGene_tx=XM_011281041.3;refGene_gn=PREPL;refGene_tx=XM_011281042.3;refGene_gn=PREPL;refGene_tx=XM_019827131.2;refGene_gn=PREPL;refGene_tx=XM_019827132.2

A3 66584782 66584989
refGene_gn=PREPL;refGene_tx=XM_011281039.3;refGene_gn=PREPL;refGene_tx=XM_011281040.3;refGene_gn=PREPL;refGene_tx=XM_011281041.3;refGene_gn=PREPL;refGene_tx=XM_011281042.3;refGene_gn=PREPL;refGene_tx=XM_019827131.2
A3 66585529 66585596
refGene_gn=PREPL;refGene_tx=XM_011281039.3;refGene_gn=PREPL;refGene_tx=XM_011281040.3;refGene_gn=PREPL;refGene_tx=XM_011281041.3;refGene_gn=PREPL;refGene_tx=XM_011281042.3;refGene_gn=PREPL;refGene_tx=XM_019827131.2;refGene_gn=PREPL;refGene_tx=XM_019827132.2
A3 66586222 66586345
refGene_gn=PREPL;refGene_tx=XM_011281039.3;refGene_gn=PREPL;refGene_tx=XM_011281040.3;refGene_gn=PREPL;refGene_tx=XM_011281041.3;refGene_gn=PREPL;refGene_tx=XM_011281042.3;refGene_gn=PREPL;refGene_tx=XM_019827131.2;refGene_gn=PREPL;refGene_tx=XM_019827132.2
A3 66599258 66599477
refGene_gn=PREPL;refGene_tx=XM_011281039.3;refGene_gn=PREPL;refGene_tx=XM_011281040.3;refGene_gn=PREPL;refGene_tx=XM_019827131.2;refGene_gn=PREPL;refGene_tx=XM_019827132.2
A3 66601592 66601862
refGene_gn=CAMKMT;refGene_tx=XM_003983956.4;refGene_gn=CAMKMT;refGene_tx=XM_011281035.3;refGene_gn=CAMKMT;refGene_tx=XM_011281037.3;refGene_gn=CAMKMT;refGene_tx=XM_011281038.3;refGene_gn=CAMKMT;refGene_tx=XM_019827134.2;refGene_gn=CAMKMT;refGene_tx=XM_019827135.2
A3 66617681 66617854
refGene_gn=CAMKMT;refGene_tx=XM_003983956.4;refGene_gn=CAMKMT;refGene_tx=XM_011281035.3;refGene_gn=CAMKMT;refGene_tx=XM_011281037.3;refGene_gn=CAMKMT;refGene_tx=XM_011281038.3;refGene_gn=CAMKMT;refGene_tx=XM_019827134.2;refGene_gn=CAMKMT;refGene_tx=XM_019827135.2
A3 66624547 66624612
refGene_gn=CAMKMT;refGene_tx=XM_003983956.4;refGene_gn=CAMKMT;refGene_tx=XM_011281035.3;refGene_gn=CAMKMT;refGene_tx=XM_011281037.3;refGene_gn=CAMKMT;refGene_tx=XM_011281038.3;refGene_gn=CAMKMT;refGene_tx=XM_019827134.2;refGene_gn=CAMKMT;refGene_tx=XM_019827135.2
A3 66949536 66949597
refGene_gn=CAMKMT;refGene_tx=XM_003983956.4;refGene_gn=CAMKMT;refGene_tx=XM_011281035.3;refGene_gn=CAMKMT;refGene_tx=XM_011281037.3;refGene_gn=CAMKMT;refGene_tx=XM_011281038.3;refGene_gn=CAMKMT;refGene_tx=XM_019827134.2;refGene_gn=CAMKMT;refGene_tx=XM_019827135.2
A3 66951683 66951738
refGene_gn=CAMKMT;refGene_tx=XM_003983956.4;refGene_gn=CAMKMT;refGene_tx=XM_011281035.3;refGene_gn=CAMKMT;refGene_tx=XM_011281037.3;refGene_gn=CAMKMT;refGene_tx=XM_019827134.2;refGene_gn=CAMKMT;refGene_tx=XM_019827135.2
A3 66952799 66952863
refGene_gn=CAMKMT;refGene_tx=XM_003983956.4;refGene_gn=CAMKMT;refGene_tx=XM_011281035.3;refGene_gn=CAMKMT;refGene_tx=XM_011281037.3;refGene_gn=CAMKMT;refGene_tx=XM_019827134.2;refGene_gn=CAMKMT;refGene_tx=XM_019827135.2
A3 66961055 66961122

refGene_gn=CAMKMT;refGene_tx=XM_003983956.4;refGene_gn=CAMKMT;refGene_tx=XM_011281035.3;refGene_gn=CAMKMT;refGene_tx=XM_011281037.3;refGene_gn=CAMKMT;refGene_tx=XM_019827134.2;refGene_gn=CAMKMT;refGene_tx=XM_019827135.2
A3 66989191 66989266
refGene_gn=CAMKMT;refGene_tx=XM_003983956.4;refGene_gn=CAMKMT;refGene_tx=XM_011281035.3;refGene_gn=CAMKMT;refGene_tx=XM_011281038.3;refGene_gn=CAMKMT;refGene_tx=XM_019827134.2
A3 66996768 66996832
refGene_gn=CAMKMT;refGene_tx=XM_003983956.4;refGene_gn=CAMKMT;refGene_tx=XM_011281035.3;refGene_gn=CAMKMT;refGene_tx=XM_011281037.3;refGene_gn=CAMKMT;refGene_tx=XM_011281038.3;refGene_gn=CAMKMT;refGene_tx=XM_019827134.2;refGene_gn=CAMKMT;refGene_tx=XM_019827135.2
A3 67008331 67008463
refGene_gn=CAMKMT;refGene_tx=XM_003983956.4;refGene_gn=CAMKMT;refGene_tx=XM_011281035.3;refGene_gn=CAMKMT;refGene_tx=XM_011281037.3;refGene_gn=CAMKMT;refGene_tx=XM_011281038.3
A3 67009197 67009224 refGene_gn=CAMKMT;refGene_tx=XM_003983956.4
A3 67014359 67014437
refGene_gn=CAMKMT;refGene_tx=XM_011281035.3;refGene_gn=CAMKMT;refGene_tx=XM_011281037.3;refGene_gn=CAMKMT;refGene_tx=XM_011281038.3;refGene_gn=CAMKMT;refGene_tx=XM_019827134.2;refGene_gn=CAMKMT;refGene_tx=XM_019827135.2
A3 67165082 67165137
refGene_gn=SIX3;refGene_tx=XM_011281034.3;refGene_gn=SIX3;refGene_tx=XM_023251642.1;refGene_gn=SIX3;refGene_tx=XM_023251643.1;refGene_gn=SIX3;refGene_tx=XM_023251644.1
A3 67172773 67173584
refGene_gn=SIX3;refGene_tx=XM_011281034.3;refGene_gn=SIX3;refGene_tx=XM_023251642.1;refGene_gn=SIX3;refGene_tx=XM_023251643.1;refGene_gn=SIX3;refGene_tx=XM_023251644.1
A3 67175417 67175610
refGene_gn=SIX3;refGene_tx=XM_011281034.3;refGene_gn=SIX3;refGene_tx=XM_023251642.1;refGene_gn=SIX3;refGene_tx=XM_023251643.1;refGene_gn=SIX3;refGene_tx=XM_023251644.1
A3 67219959 67219968 refGene_gn=SIX3;refGene_tx=XM_023251644.1
A3 67234333 67234664
refGene_gn=SIX2;refGene_tx=XM_023251645.1;refGene_gn=SIX2;refGene_tx=XM_023251646.1
A3 67236810 67237376
refGene_gn=SIX2;refGene_tx=XM_023251645.1;refGene_gn=SIX2;refGene_tx=XM_023251646.1
A3 67259050 67259071 refGene_gn=SIX3;refGene_tx=XM_023251643.1
A3 67266753 67266843 refGene_gn=SIX3;refGene_tx=XM_023251642.1
A3 67572791 67572846
refGene_gn=SRBD1;refGene_tx=XM_019827137.2;refGene_gn=SRBD1;refGene_tx=XM_023251647.1;refGene_gn=SRBD1;refGene_tx=XM_023251648.1
A3 67575463 67575530 refGene_gn=SRBD1;refGene_tx=XM_019827136.2
A3 67580229 67580519
refGene_gn=SRBD1;refGene_tx=XM_003983938.5;refGene_gn=SRBD1;refGene_tx

=XM_006930073.4;refGene_gn=SRBD1;refGene_tx=XM_019827136.2;refGene_gn=SRBD1;refGene_tx=XM_019827137.2;refGene_gn=SRBD1;refGene_tx=XM_023251647.1;refGene_gn=SRBD1;refGene_tx=XM_023251648.1
A3 67584012 67584197
refGene_gn=SRBD1;refGene_tx=XM_003983938.5;refGene_gn=SRBD1;refGene_tx=XM_006930073.4;refGene_gn=SRBD1;refGene_tx=XM_019827136.2;refGene_gn=SRBD1;refGene_tx=XM_019827137.2;refGene_gn=SRBD1;refGene_tx=XM_023251647.1;refGene_gn=SRBD1;refGene_tx=XM_023251648.1
A3 67603137 67603320
refGene_gn=SRBD1;refGene_tx=XM_003983938.5;refGene_gn=SRBD1;refGene_tx=XM_006930073.4;refGene_gn=SRBD1;refGene_tx=XM_019827136.2;refGene_gn=SRBD1;refGene_tx=XM_019827137.2;refGene_gn=SRBD1;refGene_tx=XM_023251647.1;refGene_gn=SRBD1;refGene_tx=XM_023251648.1
A3 67607157 67607334
refGene_gn=SRBD1;refGene_tx=XM_003983938.5;refGene_gn=SRBD1;refGene_tx=XM_006930073.4;refGene_gn=SRBD1;refGene_tx=XM_019827136.2;refGene_gn=SRBD1;refGene_tx=XM_019827137.2;refGene_gn=SRBD1;refGene_tx=XM_023251647.1;refGene_gn=SRBD1;refGene_tx=XM_023251648.1
A3 67608616 67608723
refGene_gn=SRBD1;refGene_tx=XM_003983938.5;refGene_gn=SRBD1;refGene_tx=XM_006930073.4;refGene_gn=SRBD1;refGene_tx=XM_019827136.2;refGene_gn=SRBD1;refGene_tx=XM_019827137.2;refGene_gn=SRBD1;refGene_tx=XM_023251647.1;refGene_gn=SRBD1;refGene_tx=XM_023251648.1
A3 67664610 67664693
refGene_gn=SRBD1;refGene_tx=XM_003983938.5;refGene_gn=SRBD1;refGene_tx=XM_006930073.4;refGene_gn=SRBD1;refGene_tx=XM_019827136.2;refGene_gn=SRBD1;refGene_tx=XM_019827137.2;refGene_gn=SRBD1;refGene_tx=XM_023251647.1;refGene_gn=SRBD1;refGene_tx=XM_023251648.1
A3 67673845 67673937
refGene_gn=SRBD1;refGene_tx=XM_003983938.5;refGene_gn=SRBD1;refGene_tx=XM_006930073.4;refGene_gn=SRBD1;refGene_tx=XM_019827136.2;refGene_gn=SRBD1;refGene_tx=XM_019827137.2;refGene_gn=SRBD1;refGene_tx=XM_023251647.1;refGene_gn=SRBD1;refGene_tx=XM_023251648.1
A3 67719461 67719569
refGene_gn=SRBD1;refGene_tx=XM_003983938.5;refGene_gn=SRBD1;refGene_tx=XM_006930073.4;refGene_gn=SRBD1;refGene_tx=XM_019827136.2;refGene_gn=SRBD1;refGene_tx=XM_019827137.2;refGene_gn=SRBD1;refGene_tx=XM_023251647.1;refGene_gn=SRBD1;refGene_tx=XM_023251648.1
A3 67720308 67720399
refGene_gn=SRBD1;refGene_tx=XM_003983938.5;refGene_gn=SRBD1;refGene_tx=XM_006930073.4;refGene_gn=SRBD1;refGene_tx=XM_019827136.2;refGene_gn=SRBD1;refGene_tx=XM_019827137.2;refGene_gn=SRBD1;refGene_tx=XM_023251647.1;refGene_gn=SRBD1;refGene_tx=XM_023251648.1
A3 67723587 67723745
refGene_gn=SRBD1;refGene_tx=XM_003983938.5;refGene_gn=SRBD1;refGene_tx=XM_006930073.4;refGene_gn=SRBD1;refGene_tx=XM_019827136.2;refGene_gn=SRBD1;refGene_tx=XM_019827137.2;refGene_gn=SRBD1;refGene_tx=XM_023251647.1;refGene_gn=SRBD1;refGene_tx=XM_023251648.1
A3 67728339 67728447
refGene_gn=SRBD1;refGene_tx=XM_003983938.5;refGene_gn=SRBD1;refGene_tx

=XM_006930073.4;refGene_gn=SRBD1;refGene_tx=XM_019827136.2;refGene_gn=SRBD1;refGene_tx=XM_019827137.2;refGene_gn=SRBD1;refGene_tx=XM_023251647.1;refGene_gn=SRBD1;refGene_tx=XM_023251648.1
A3 67733163 67733267
refGene_gn=SRBD1;refGene_tx=XM_003983938.5;refGene_gn=SRBD1;refGene_tx=XM_006930073.4;refGene_gn=SRBD1;refGene_tx=XM_019827136.2;refGene_gn=SRBD1;refGene_tx=XM_019827137.2;refGene_gn=SRBD1;refGene_tx=XM_023251647.1;refGene_gn=SRBD1;refGene_tx=XM_023251648.1
A3 67741516 67741652
refGene_gn=SRBD1;refGene_tx=XM_003983938.5;refGene_gn=SRBD1;refGene_tx=XM_006930073.4;refGene_gn=SRBD1;refGene_tx=XM_019827136.2;refGene_gn=SRBD1;refGene_tx=XM_019827137.2;refGene_gn=SRBD1;refGene_tx=XM_023251647.1;refGene_gn=SRBD1;refGene_tx=XM_023251648.1
A3 67742932 67743029
refGene_gn=SRBD1;refGene_tx=XM_003983938.5;refGene_gn=SRBD1;refGene_tx=XM_006930073.4;refGene_gn=SRBD1;refGene_tx=XM_019827136.2;refGene_gn=SRBD1;refGene_tx=XM_019827137.2;refGene_gn=SRBD1;refGene_tx=XM_023251647.1;refGene_gn=SRBD1;refGene_tx=XM_023251648.1
A3 67747570 67747709
refGene_gn=SRBD1;refGene_tx=XM_003983938.5;refGene_gn=SRBD1;refGene_tx=XM_006930073.4;refGene_gn=SRBD1;refGene_tx=XM_019827136.2;refGene_gn=SRBD1;refGene_tx=XM_019827137.2;refGene_gn=SRBD1;refGene_tx=XM_023251647.1;refGene_gn=SRBD1;refGene_tx=XM_023251648.1
A3 67749017 67749135
refGene_gn=SRBD1;refGene_tx=XM_003983938.5;refGene_gn=SRBD1;refGene_tx=XM_006930073.4;refGene_gn=SRBD1;refGene_tx=XM_019827136.2;refGene_gn=SRBD1;refGene_tx=XM_019827137.2;refGene_gn=SRBD1;refGene_tx=XM_023251647.1;refGene_gn=SRBD1;refGene_tx=XM_023251648.1
A3 67752718 67752885
refGene_gn=SRBD1;refGene_tx=XM_003983938.5;refGene_gn=SRBD1;refGene_tx=XM_006930073.4;refGene_gn=SRBD1;refGene_tx=XM_019827136.2;refGene_gn=SRBD1;refGene_tx=XM_019827137.2;refGene_gn=SRBD1;refGene_tx=XM_023251647.1;refGene_gn=SRBD1;refGene_tx=XM_023251648.1
A3 67760730 67761117
refGene_gn=SRBD1;refGene_tx=XM_003983938.5;refGene_gn=SRBD1;refGene_tx=XM_006930073.4;refGene_gn=SRBD1;refGene_tx=XM_019827136.2;refGene_gn=SRBD1;refGene_tx=XM_019827137.2;refGene_gn=SRBD1;refGene_tx=XM_023251647.1;refGene_gn=SRBD1;refGene_tx=XM_023251648.1
A3 67767857 67768038
refGene_gn=SRBD1;refGene_tx=XM_003983938.5;refGene_gn=SRBD1;refGene_tx=XM_006930073.4;refGene_gn=SRBD1;refGene_tx=XM_019827136.2;refGene_gn=SRBD1;refGene_tx=XM_019827137.2;refGene_gn=SRBD1;refGene_tx=XM_023251647.1;refGene_gn=SRBD1;refGene_tx=XM_023251648.1
A3 67770500 67770580
refGene_gn=SRBD1;refGene_tx=XM_003983938.5;refGene_gn=SRBD1;refGene_tx=XM_006930073.4;refGene_gn=SRBD1;refGene_tx=XM_019827136.2;refGene_gn=SRBD1;refGene_tx=XM_019827137.2;refGene_gn=SRBD1;refGene_tx=XM_023251647.1;refGene_gn=SRBD1;refGene_tx=XM_023251648.1
A3 67817006 67817354
refGene_gn=PRKCE;refGene_tx=XM_011281033.2;refGene_gn=PRKCE;refGene_tx

=XM_019827139.2
A3	68003834	68003898
refGene_gn=PRKCE;refGene_tx=XM_011281033.2;refGene_gn=PRKCE;refGene_tx=XM_019827139.2
A3	68133134	68133294
refGene_gn=PRKCE;refGene_tx=XM_011281033.2;refGene_gn=PRKCE;refGene_tx=XM_019827139.2
A3	68135382	68135417
refGene_gn=PRKCE;refGene_tx=XM_011281033.2;refGene_gn=PRKCE;refGene_tx=XM_019827139.2
A3	68136617	68136703
refGene_gn=PRKCE;refGene_tx=XM_011281033.2;refGene_gn=PRKCE;refGene_tx=XM_019827139.2
A3	68141006	68141136
refGene_gn=PRKCE;refGene_tx=XM_011281033.2;refGene_gn=PRKCE;refGene_tx=XM_019827139.2
A3	68158137	68158280
refGene_gn=PRKCE;refGene_tx=XM_011281033.2;refGene_gn=PRKCE;refGene_tx=XM_019827139.2
A3	68160845	68160942
refGene_gn=PRKCE;refGene_tx=XM_011281033.2;refGene_gn=PRKCE;refGene_tx=XM_019827139.2
A3	68163773	68163973
refGene_gn=PRKCE;refGene_tx=XM_011281033.2;refGene_gn=PRKCE;refGene_tx=XM_019827139.2
A3	68166941	68167115
refGene_gn=PRKCE;refGene_tx=XM_011281033.2;refGene_gn=PRKCE;refGene_tx=XM_019827139.2
A3	68231844	68231999
refGene_gn=PRKCE;refGene_tx=XM_011281033.2;refGene_gn=PRKCE;refGene_tx=XM_019827139.2
A3	68284030	68284169
refGene_gn=PRKCE;refGene_tx=XM_011281033.2;refGene_gn=PRKCE;refGene_tx=XM_019827139.2
A3	68289593	68289782
refGene_gn=PRKCE;refGene_tx=XM_011281033.2;refGene_gn=PRKCE;refGene_tx=XM_019827139.2
A3	68294495	68294642
refGene_gn=PRKCE;refGene_tx=XM_011281033.2;refGene_gn=PRKCE;refGene_tx=XM_019827139.2
A3	68299954	68299969	refGene_gn=PRKCE;refGene_tx=XM_019827139.2
A3	68318253	68318400	refGene_gn=PRKCE;refGene_tx=XM_011281033.2
A3	68406075	68406101
refGene_gn=EPAS1;refGene_tx=XM_023251650.1;refGene_gn=EPAS1;refGene_tx=XM_023251651.1
A3	68453484	68453675
refGene_gn=EPAS1;refGene_tx=XM_023251650.1;refGene_gn=EPAS1;refGene_tx=XM_023251651.1
A3	68462002	68462154
refGene_gn=EPAS1;refGene_tx=XM_023251650.1;refGene_gn=EPAS1;refGene_tx

=XM_023251651.1
A3 68462575 68462660
refGene_gn=EPAS1;refGene_tx=XM_023251650.1;refGene_gn=EPAS1;refGene_tx=XM_023251651.1
A3 68466052 68466171
refGene_gn=EPAS1;refGene_tx=XM_023251650.1;refGene_gn=EPAS1;refGene_tx=XM_023251651.1
A3 68466265 68466471
refGene_gn=EPAS1;refGene_tx=XM_023251650.1;refGene_gn=EPAS1;refGene_tx=XM_023251651.1
A3 68475435 68475542
refGene_gn=EPAS1;refGene_tx=XM_023251650.1;refGene_gn=EPAS1;refGene_tx=XM_023251651.1
A3 68480728 68480876
refGene_gn=EPAS1;refGene_tx=XM_023251650.1;refGene_gn=EPAS1;refGene_tx=XM_023251651.1
A3 68481536 68481751
refGene_gn=EPAS1;refGene_tx=XM_023251650.1;refGene_gn=EPAS1;refGene_tx=XM_023251651.1
A3 68482873 68483064
refGene_gn=EPAS1;refGene_tx=XM_023251650.1;refGene_gn=EPAS1;refGene_tx=XM_023251651.1
A3 68483578 68483689
refGene_gn=EPAS1;refGene_tx=XM_023251650.1;refGene_gn=EPAS1;refGene_tx=XM_023251651.1
A3 68484680 68485174
refGene_gn=EPAS1;refGene_tx=XM_023251650.1;refGene_gn=EPAS1;refGene_tx=XM_023251651.1
A3 68486035 68486171
refGene_gn=EPAS1;refGene_tx=XM_023251650.1;refGene_gn=EPAS1;refGene_tx=XM_023251651.1
A3 68486356 68486480
refGene_gn=EPAS1;refGene_tx=XM_023251650.1;refGene_gn=EPAS1;refGene_tx=XM_023251651.1
A3 68486803 68486965
refGene_gn=EPAS1;refGene_tx=XM_023251650.1;refGene_gn=EPAS1;refGene_tx=XM_023251651.1
A3 68487523 68487675
refGene_gn=EPAS1;refGene_tx=XM_023251650.1;refGene_gn=EPAS1;refGene_tx=XM_023251651.1
A3 68564164 68564280 refGene_gn=TMEM247;refGene_tx=XM_023252182.1
A3 68564731 68564799 refGene_gn=TMEM247;refGene_tx=XM_023252182.1
A3 68564886 68564903 refGene_gn=TMEM247;refGene_tx=XM_023252182.1
A3 68564949 68565288 refGene_gn=TMEM247;refGene_tx=XM_023252182.1
A3 68568351 68568540 refGene_gn=TMEM247;refGene_tx=XM_023252182.1
A3 68595576 68596257
refGene_gn=ATP6V1E2;refGene_tx=XM_003983959.4;refGene_gn=ATP6V1E2;refGene_tx=XM_006930089.3
A3 68619680 68619849
refGene_gn=RHOQ;refGene_tx=XM_023251654.1;refGene_gn=RHOQ;refGene_tx=X

M_023251655.1
A3 68620410 68620469
refGene_gn=RHOQ;refGene_tx=XM_023251654.1;refGene_gn=RHOQ;refGene_tx=XM_023251655.1
A3 68652742 68652907
refGene_gn=RHOQ;refGene_tx=XM_023251654.1;refGene_gn=RHOQ;refGene_tx=XM_023251655.1
A3 68653225 68653321
refGene_gn=RHOQ;refGene_tx=XM_023251654.1;refGene_gn=RHOQ;refGene_tx=XM_023251655.1
A3 68654284 68654391 refGene_gn=RHOQ;refGene_tx=XM_023251655.1
A3 68654615 68654771
refGene_gn=RHOQ;refGene_tx=XM_023251654.1;refGene_gn=RHOQ;refGene_tx=XM_023251655.1
A3 68655203 68655317
refGene_gn=PIGF;refGene_tx=XM_003983960.5;refGene_gn=PIGF;refGene_tx=XM_006930090.4;refGene_gn=PIGF;refGene_tx=XM_006930091.4;refGene_gn=PIGF;refGene_tx=XM_019827145.2;refGene_gn=PIGF;refGene_tx=XM_019827146.2;refGene_gn=PIGF;refGene_tx=XM_019827147.2
A3 68670423 68670532
refGene_gn=PIGF;refGene_tx=XM_003983960.5;refGene_gn=PIGF;refGene_tx=XM_006930090.4;refGene_gn=PIGF;refGene_tx=XM_006930091.4;refGene_gn=PIGF;refGene_tx=XM_019827145.2;refGene_gn=PIGF;refGene_tx=XM_019827147.2
A3 68684074 68684191
refGene_gn=PIGF;refGene_tx=XM_003983960.5;refGene_gn=PIGF;refGene_tx=XM_006930090.4;refGene_gn=PIGF;refGene_tx=XM_006930091.4;refGene_gn=PIGF;refGene_tx=XM_019827145.2;refGene_gn=PIGF;refGene_tx=XM_019827146.2;refGene_gn=PIGF;refGene_tx=XM_019827147.2
A3 68685652 68685744
refGene_gn=PIGF;refGene_tx=XM_003983960.5;refGene_gn=PIGF;refGene_tx=XM_006930090.4;refGene_gn=PIGF;refGene_tx=XM_006930091.4;refGene_gn=PIGF;refGene_tx=XM_019827145.2;refGene_gn=PIGF;refGene_tx=XM_019827146.2;refGene_gn=PIGF;refGene_tx=XM_019827147.2
A3 68686955 68687183
refGene_gn=PIGF;refGene_tx=XM_003983960.5;refGene_gn=PIGF;refGene_tx=XM_006930090.4;refGene_gn=PIGF;refGene_tx=XM_006930091.4;refGene_gn=PIGF;refGene_tx=XM_019827145.2;refGene_gn=PIGF;refGene_tx=XM_019827146.2
A3 68688391 68688492
refGene_gn=PIGF;refGene_tx=XM_006930090.4;refGene_gn=PIGF;refGene_tx=XM_019827146.2
A3 68689276 68689292 refGene_gn=CRIPT;refGene_tx=XM_003983961.4
A3 68694219 68694285 refGene_gn=CRIPT;refGene_tx=XM_003983961.4
A3 68695027 68695082 refGene_gn=CRIPT;refGene_tx=XM_003983961.4
A3 68699152 68699256 refGene_gn=CRIPT;refGene_tx=XM_003983961.4
A3 68699589 68699654 refGene_gn=CRIPT;refGene_tx=XM_003983961.4
A3 68750053 68750074 refGene_gn=PIGF;refGene_tx=XM_019827147.2
A3 68822470 68824081
refGene_gn=SOCS5;refGene_tx=XM_003983962.5;refGene_gn=SOCS5;refGene_tx=XM_019827142.2;refGene_gn=SOCS5;refGene_tx=XM_019827143.2;refGene_gn=SOCS5;refGene_tx=XM_019827144.2;refGene_gn=SOCS5;refGene_tx=XM_0232516

52.1;refGene_gn=SOCS5;refGene_tx=XM_023251653.1
A3 68882495 68882632
refGene_gn=MCFD2;refGene_tx=XM_023251656.1;refGene_gn=MCFD2;refGene_tx=XM_023251657.1;refGene_gn=MCFD2;refGene_tx=XM_023251658.1
A3 68889023 68889048
refGene_gn=MCFD2;refGene_tx=XM_023251656.1;refGene_gn=MCFD2;refGene_tx=XM_023251657.1;refGene_gn=MCFD2;refGene_tx=XM_023251658.1
A3 68898381 68898427 refGene_gn=MCFD2;refGene_tx=XM_023251659.1
A3 68898887 68898970 refGene_gn=MCFD2;refGene_tx=XM_023251659.1
A3 68911891 68912002
refGene_gn=MCFD2;refGene_tx=XM_023251656.1;refGene_gn=MCFD2;refGene_tx=XM_023251657.1;refGene_gn=MCFD2;refGene_tx=XM_023251658.1;refGene_gn=MCFD2;refGene_tx=XM_023251659.1
A3 68933945 68934008 refGene_gn=MCFD2;refGene_tx=XM_023251661.1
A3 68937538 68937670 refGene_gn=MCFD2;refGene_tx=XM_006930094.3
A3 68938837 68938906
refGene_gn=MCFD2;refGene_tx=XM_023251660.1;refGene_gn=MCFD2;refGene_tx=XM_023251662.1
A3 68939078 68939238
refGene_gn=MCFD2;refGene_tx=XM_006930094.3;refGene_gn=MCFD2;refGene_tx=XM_023251656.1;refGene_gn=MCFD2;refGene_tx=XM_023251657.1;refGene_gn=MCFD2;refGene_tx=XM_023251658.1;refGene_gn=MCFD2;refGene_tx=XM_023251659.1;refGene_gn=MCFD2;refGene_tx=XM_023251660.1;refGene_gn=MCFD2;refGene_tx=XM_023251661.1;refGene_gn=MCFD2;refGene_tx=XM_023251662.1
A3 68940021 68940170
refGene_gn=MCFD2;refGene_tx=XM_006930094.3;refGene_gn=MCFD2;refGene_tx=XM_023251656.1;refGene_gn=MCFD2;refGene_tx=XM_023251657.1;refGene_gn=MCFD2;refGene_tx=XM_023251658.1;refGene_gn=MCFD2;refGene_tx=XM_023251659.1;refGene_gn=MCFD2;refGene_tx=XM_023251660.1;refGene_gn=MCFD2;refGene_tx=XM_023251661.1;refGene_gn=MCFD2;refGene_tx=XM_023251662.1
A3 68976415 68976599 refGene_gn=TTC7A;refGene_tx=XM_003983963.5
A3 68985760 68985924 refGene_gn=TTC7A;refGene_tx=XM_003983963.5
A3 68990957 68991126 refGene_gn=TTC7A;refGene_tx=XM_003983963.5
A3 69007224 69007355 refGene_gn=TTC7A;refGene_tx=XM_003983963.5
A3 69009843 69009959 refGene_gn=TTC7A;refGene_tx=XM_003983963.5
A3 69019548 69019622 refGene_gn=TTC7A;refGene_tx=XM_006930095.4
A3 69020715 69020794
refGene_gn=TTC7A;refGene_tx=XM_003983963.5;refGene_gn=TTC7A;refGene_tx=XM_006930095.4;refGene_gn=TTC7A;refGene_tx=XM_019827149.2
A3 69021565 69021723
refGene_gn=TTC7A;refGene_tx=XM_003983963.5;refGene_gn=TTC7A;refGene_tx=XM_006930095.4;refGene_gn=TTC7A;refGene_tx=XM_019827149.2
A3 69022292 69022356
refGene_gn=TTC7A;refGene_tx=XM_003983963.5;refGene_gn=TTC7A;refGene_tx=XM_006930095.4;refGene_gn=TTC7A;refGene_tx=XM_019827149.2
A3 69032072 69032210
refGene_gn=TTC7A;refGene_tx=XM_003983963.5;refGene_gn=TTC7A;refGene_tx=XM_006930095.4;refGene_gn=TTC7A;refGene_tx=XM_019827149.2
A3 69033770 69033854
refGene_gn=TTC7A;refGene_tx=XM_003983963.5;refGene_gn=TTC7A;refGene_tx

=XM_006930095.4;refGene_gn=TTC7A;refGene_tx=XM_019827149.2
A3	69038091	69038196
refGene_gn=TTC7A;refGene_tx=XM_003983963.5;refGene_gn=TTC7A;refGene_tx
=XM_006930095.4;refGene_gn=TTC7A;refGene_tx=XM_019827149.2
A3	69047438	69047556
refGene_gn=TTC7A;refGene_tx=XM_003983963.5;refGene_gn=TTC7A;refGene_tx
=XM_006930095.4;refGene_gn=TTC7A;refGene_tx=XM_019827149.2
A3	69048846	69048904
refGene_gn=TTC7A;refGene_tx=XM_003983963.5;refGene_gn=TTC7A;refGene_tx
=XM_006930095.4;refGene_gn=TTC7A;refGene_tx=XM_019827149.2
A3	69050038	69050111
refGene_gn=TTC7A;refGene_tx=XM_003983963.5;refGene_gn=TTC7A;refGene_tx
=XM_006930095.4;refGene_gn=TTC7A;refGene_tx=XM_019827149.2
A3	69053119	69053280
refGene_gn=TTC7A;refGene_tx=XM_003983963.5;refGene_gn=TTC7A;refGene_tx
=XM_006930095.4;refGene_gn=TTC7A;refGene_tx=XM_019827149.2
A3	69068501	69068618
refGene_gn=TTC7A;refGene_tx=XM_003983963.5;refGene_gn=TTC7A;refGene_tx
=XM_006930095.4;refGene_gn=TTC7A;refGene_tx=XM_019827149.2
A3	69072149	69072247
refGene_gn=TTC7A;refGene_tx=XM_003983963.5;refGene_gn=TTC7A;refGene_tx
=XM_006930095.4;refGene_gn=TTC7A;refGene_tx=XM_019827149.2
A3	69074088	69074223
refGene_gn=TTC7A;refGene_tx=XM_003983963.5;refGene_gn=TTC7A;refGene_tx
=XM_006930095.4;refGene_gn=TTC7A;refGene_tx=XM_019827149.2
A3	69082066	69082269
refGene_gn=TTC7A;refGene_tx=XM_003983963.5;refGene_gn=TTC7A;refGene_tx
=XM_006930095.4;refGene_gn=TTC7A;refGene_tx=XM_019827149.2
A3	69093984	69094206
refGene_gn=TTC7A;refGene_tx=XM_003983963.5;refGene_gn=TTC7A;refGene_tx
=XM_006930095.4;refGene_gn=TTC7A;refGene_tx=XM_019827149.2
A3	69159322	69159351	refGene_gn=CALM2;refGene_tx=XM_003983965.5
A3	69160085	69160221	refGene_gn=CALM2;refGene_tx=XM_003983965.5
A3	69160455	69160562	refGene_gn=CALM2;refGene_tx=XM_003983965.5
A3	69160659	69160803	refGene_gn=CALM2;refGene_tx=XM_003983965.5
A3	69167804	69167835	refGene_gn=CALM2;refGene_tx=XM_003983965.5
A3	69173689	69173692	refGene_gn=CALM2;refGene_tx=XM_003983965.5
A3	69241190	69241285
refGene_gn=LOC109498520;refGene_tx=XM_023252183.1
A3	69246545	69246693
refGene_gn=LOC109498520;refGene_tx=XM_023252183.1
A3	69252944	69253306
refGene_gn=LOC109498520;refGene_tx=XM_023252183.1
A3	69281742	69281837
refGene_gn=LOC109498520;refGene_tx=XM_023252183.1
A3	69283079	69283219
refGene_gn=LOC109498520;refGene_tx=XM_023252183.1
A3	69336418	69336494	refGene_gn=EPCAM;refGene_tx=XM_019827150.2
A3	69338684	69338792	refGene_gn=EPCAM;refGene_tx=XM_019827150.2
A3	69339103	69339344	refGene_gn=EPCAM;refGene_tx=XM_019827150.2

A3 69340217 69340283 refGene_gn=EPCAM;refGene_tx=XM_019827150.2
A3 69341494 69341558 refGene_gn=EPCAM;refGene_tx=XM_019827150.2
A3 69342986 69343088 refGene_gn=EPCAM;refGene_tx=XM_019827150.2
A3 69344784 69344985 refGene_gn=EPCAM;refGene_tx=XM_019827150.2
A3 69347270 69347315 refGene_gn=EPCAM;refGene_tx=XM_019827150.2
A3 69348690 69348732 refGene_gn=EPCAM;refGene_tx=XM_019827150.2
A3 69362221 69362432 refGene_gn=MSH2;refGene_tx=XM_023251663.1
A3 69374960 69375115 refGene_gn=MSH2;refGene_tx=XM_023251663.1
A3 69378692 69378971 refGene_gn=MSH2;refGene_tx=XM_023251663.1
A3 69379915 69380062 refGene_gn=MSH2;refGene_tx=XM_023251663.1
A3 69382303 69382453 refGene_gn=MSH2;refGene_tx=XM_023251663.1
A3 69384831 69384965 refGene_gn=MSH2;refGene_tx=XM_023251663.1
A3 69397526 69397726 refGene_gn=MSH2;refGene_tx=XM_023251663.1
A3 69412965 69413075 refGene_gn=MSH2;refGene_tx=XM_023251663.1
A3 69431077 69431201 refGene_gn=MSH2;refGene_tx=XM_023251663.1
A3 69432674 69432825 refGene_gn=MSH2;refGene_tx=XM_023251663.1
A3 69435642 69435740 refGene_gn=MSH2;refGene_tx=XM_023251663.1
A3 69439377 69439623 refGene_gn=MSH2;refGene_tx=XM_023251663.1
A3 69440815 69441020 refGene_gn=MSH2;refGene_tx=XM_023251663.1
A3 69442214 69442462 refGene_gn=MSH2;refGene_tx=XM_023251663.1
A3 69444386 69444562 refGene_gn=MSH2;refGene_tx=XM_023251663.1
A3 69447107 69447278 refGene_gn=MSH2;refGene_tx=XM_023251663.1
A3 69474359 69475261 refGene_gn=KCNK12;refGene_tx=XM_023251665.1
A3 69523323 69523714 refGene_gn=KCNK12;refGene_tx=XM_023251665.1
A3 69657003 69657257
refGene_gn=MSH6;refGene_tx=XM_003983968.4;refGene_gn=MSH6;refGene_tx=X
M_019827151.2
A3 69667319 69667519
refGene_gn=MSH6;refGene_tx=XM_003983968.4;refGene_gn=MSH6;refGene_tx=X
M_019827151.2
A3 69669799 69669969
refGene_gn=MSH6;refGene_tx=XM_003983968.4;refGene_gn=MSH6;refGene_tx=X
M_019827151.2
A3 69671217 69673759
refGene_gn=MSH6;refGene_tx=XM_003983968.4;refGene_gn=MSH6;refGene_tx=X
M_019827151.2
A3 69675588 69675857
refGene_gn=MSH6;refGene_tx=XM_003983968.4;refGene_gn=MSH6;refGene_tx=X
M_019827151.2
A3 69676938 69677056
refGene_gn=MSH6;refGene_tx=XM_003983968.4;refGene_gn=MSH6;refGene_tx=X
M_019827151.2
A3 69677366 69677456
refGene_gn=MSH6;refGene_tx=XM_003983968.4;refGene_gn=MSH6;refGene_tx=X
M_019827151.2
A3 69677918 69678073
refGene_gn=MSH6;refGene_tx=XM_003983968.4;refGene_gn=MSH6;refGene_tx=X
M_019827151.2
A3 69678176 69678376
refGene_gn=MSH6;refGene_tx=XM_003983968.4;refGene_gn=MSH6;refGene_tx=X

M_019827151.2
A3	69678482	69678564
refGene_gn=MSH6;refGene_tx=XM_003983968.4;refGene_gn=MSH6;refGene_tx=X
M_019827151.2
A3	69679815	69679945	refGene_gn=FBXO11;refGene_tx=XM_023251664.1
A3	69680027	69680126	refGene_gn=FBXO11;refGene_tx=XM_023251664.1
A3	69680748	69680857	refGene_gn=FBXO11;refGene_tx=XM_023251664.1
A3	69681204	69681312	refGene_gn=FBXO11;refGene_tx=XM_023251664.1
A3	69681978	69682089	refGene_gn=FBXO11;refGene_tx=XM_023251664.1
A3	69684402	69684546	refGene_gn=FBXO11;refGene_tx=XM_023251664.1
A3	69684635	69684712	refGene_gn=FBXO11;refGene_tx=XM_023251664.1
A3	69685697	69685783	refGene_gn=FBXO11;refGene_tx=XM_023251664.1
A3	69685871	69685994	refGene_gn=FBXO11;refGene_tx=XM_023251664.1
A3	69686970	69687065	refGene_gn=FBXO11;refGene_tx=XM_023251664.1
A3	69687921	69688007	refGene_gn=FBXO11;refGene_tx=XM_023251664.1
A3	69689195	69689413	refGene_gn=FBXO11;refGene_tx=XM_023251664.1
A3	69697616	69697754	refGene_gn=FBXO11;refGene_tx=XM_023251664.1
A3	69697840	69697947	refGene_gn=FBXO11;refGene_tx=XM_023251664.1
A3	69698272	69698384	refGene_gn=FBXO11;refGene_tx=XM_023251664.1
A3	69698469	69698576	refGene_gn=FBXO11;refGene_tx=XM_023251664.1
A3	69700106	69700239	refGene_gn=FBXO11;refGene_tx=XM_023251664.1
A3	69700319	69700403	refGene_gn=FBXO11;refGene_tx=XM_023251664.1
A3	69701120	69701250	refGene_gn=FBXO11;refGene_tx=XM_023251664.1
A3	69703333	69703478	refGene_gn=FBXO11;refGene_tx=XM_023251664.1
A3	69703899	69703981	refGene_gn=FBXO11;refGene_tx=XM_023251664.1
A3	69704359	69704487	refGene_gn=FBXO11;refGene_tx=XM_023251664.1
A3	69763086	69763306	refGene_gn=FBXO11;refGene_tx=XM_023251664.1
A3	70130654	70131191
refGene_gn=FOXN2;refGene_tx=XM_003983970.5;refGene_gn=FOXN2;refGene_tx
=XM_019827153.2
A3	70143536	70143661
refGene_gn=FOXN2;refGene_tx=XM_003983970.5;refGene_gn=FOXN2;refGene_tx
=XM_019827153.2
A3	70146086	70146151
refGene_gn=FOXN2;refGene_tx=XM_003983970.5;refGene_gn=FOXN2;refGene_tx
=XM_019827153.2
A3	70154097	70154166
refGene_gn=FOXN2;refGene_tx=XM_003983970.5;refGene_gn=FOXN2;refGene_tx
=XM_019827153.2
A3	70156987	70157511
refGene_gn=FOXN2;refGene_tx=XM_003983970.5;refGene_gn=FOXN2;refGene_tx
=XM_019827153.2
A3	70213255	70213312
refGene_gn=PPP1R21;refGene_tx=XM_003983971.5;refGene_gn=PPP1R21;refGen
e_tx=XM_019827155.2
A3	70222605	70222674
refGene_gn=PPP1R21;refGene_tx=XM_003983971.5;refGene_gn=PPP1R21;refGen
e_tx=XM_019827155.2
A3	70225921	70226068
refGene_gn=PPP1R21;refGene_tx=XM_003983971.5;refGene_gn=PPP1R21;refGen

e_tx=XM_019827155.2
A3 70228085 70228187
refGene_gn=PPP1R21;refGene_tx=XM_003983971.5;refGene_gn=PPP1R21;refGen
e_tx=XM_019827155.2
A3 70229532 70229697
refGene_gn=PPP1R21;refGene_tx=XM_003983971.5;refGene_gn=PPP1R21;refGen
e_tx=XM_019827155.2
A3 70229874 70229933
refGene_gn=PPP1R21;refGene_tx=XM_003983971.5;refGene_gn=PPP1R21;refGen
e_tx=XM_019827155.2
A3 70230940 70231035
refGene_gn=PPP1R21;refGene_tx=XM_003983971.5;refGene_gn=PPP1R21;refGen
e_tx=XM_019827155.2
A3 70234681 70234734
refGene_gn=PPP1R21;refGene_tx=XM_003983971.5;refGene_gn=PPP1R21;refGen
e_tx=XM_019827155.2
A3 70235221 70235371
refGene_gn=PPP1R21;refGene_tx=XM_003983971.5;refGene_gn=PPP1R21;refGen
e_tx=XM_019827155.2
A3 70239425 70239527
refGene_gn=PPP1R21;refGene_tx=XM_003983971.5;refGene_gn=PPP1R21;refGen
e_tx=XM_019827155.2
A3 70239615 70239704
refGene_gn=PPP1R21;refGene_tx=XM_003983971.5;refGene_gn=PPP1R21;refGen
e_tx=XM_019827155.2
A3 70242268 70242405
refGene_gn=PPP1R21;refGene_tx=XM_003983971.5;refGene_gn=PPP1R21;refGen
e_tx=XM_019827155.2
A3 70247253 70247346 refGene_gn=PPP1R21;refGene_tx=XM_003983971.5
A3 70251947 70252075
refGene_gn=PPP1R21;refGene_tx=XM_003983971.5;refGene_gn=PPP1R21;refGen
e_tx=XM_019827155.2
A3 70254633 70254786
refGene_gn=PPP1R21;refGene_tx=XM_003983971.5;refGene_gn=PPP1R21;refGen
e_tx=XM_019827155.2
A3 70258949 70259042
refGene_gn=PPP1R21;refGene_tx=XM_003983971.5;refGene_gn=PPP1R21;refGen
e_tx=XM_019827155.2
A3 70262459 70262702
refGene_gn=PPP1R21;refGene_tx=XM_003983971.5;refGene_gn=PPP1R21;refGen
e_tx=XM_019827155.2
A3 70268106 70268139
refGene_gn=PPP1R21;refGene_tx=XM_003983971.5;refGene_gn=PPP1R21;refGen
e_tx=XM_019827155.2
A3 70269291 70269408
refGene_gn=PPP1R21;refGene_tx=XM_003983971.5;refGene_gn=PPP1R21;refGen
e_tx=XM_019827155.2
A3 70272539 70272638
refGene_gn=PPP1R21;refGene_tx=XM_003983971.5;refGene_gn=PPP1R21;refGen
e_tx=XM_019827155.2

A3	70273589	70273718
refGene_gn=PPP1R21;refGene_tx=XM_003983971.5;refGene_gn=PPP1R21;refGene_tx=XM_019827155.2
A3	70276208	70276238
refGene_gn=PPP1R21;refGene_tx=XM_003983971.5;refGene_gn=PPP1R21;refGene_tx=XM_019827155.2
A3	70318422	70320352
refGene_gn=STON1;refGene_tx=XM_003983974.5;refGene_gn=STON1;refGene_tx=XM_019827156.2;refGene_gn=STON1;refGene_tx=XM_019827157.2;refGene_gn=STON1;refGene_tx=XM_019827159.2;refGene_gn=STON1;refGene_tx=XM_019827160.2
A3	70327999	70328202
refGene_gn=STON1;refGene_tx=XM_003983974.5;refGene_gn=STON1;refGene_tx=XM_019827156.2;refGene_gn=STON1;refGene_tx=XM_019827157.2;refGene_gn=STON1;refGene_tx=XM_019827159.2;refGene_gn=STON1;refGene_tx=XM_019827160.2
A3	70331060	70331135
refGene_gn=STON1;refGene_tx=XM_003983974.5;refGene_gn=STON1;refGene_tx=XM_019827156.2;refGene_gn=STON1;refGene_tx=XM_019827157.2;refGene_gn=STON1;refGene_tx=XM_019827159.2;refGene_gn=STON1;refGene_tx=XM_019827160.2
A3	70338966	70339024	refGene_gn=LHCGR;refGene_tx=XM_023251668.1
A3	70346236	70346243	refGene_gn=LHCGR;refGene_tx=XM_023251670.1
A3	70350712	70350733	refGene_gn=GTF2A1L;refGene_tx=XM_011281028.2
A3	70353032	70353134	refGene_gn=GTF2A1L;refGene_tx=XM_011281028.2
A3	70353526	70353650	refGene_gn=GTF2A1L;refGene_tx=XM_011281028.2
A3	70361673	70361729	refGene_gn=GTF2A1L;refGene_tx=XM_011281028.2
A3	70363988	70364073	refGene_gn=GTF2A1L;refGene_tx=XM_011281028.2
A3	70365946	70366533	refGene_gn=GTF2A1L;refGene_tx=XM_011281028.2
A3	70384771	70385032	refGene_gn=GTF2A1L;refGene_tx=XM_011281028.2
A3	70385747	70385837	refGene_gn=GTF2A1L;refGene_tx=XM_011281028.2
A3	70392455	70392563	refGene_gn=GTF2A1L;refGene_tx=XM_011281028.2
A3	70395455	70395471	refGene_gn=LHCGR;refGene_tx=XM_023251669.1
A3	70398833	70398837	refGene_gn=LHCGR;refGene_tx=XM_023251671.1
A3	70400924	70402077
refGene_gn=LHCGR;refGene_tx=XM_023251666.1;refGene_gn=LHCGR;refGene_tx=XM_023251667.1
A3	70407229	70407310
refGene_gn=LHCGR;refGene_tx=XM_023251666.1;refGene_gn=LHCGR;refGene_tx=XM_023251667.1;refGene_gn=LHCGR;refGene_tx=XM_023251668.1;refGene_gn=LHCGR;refGene_tx=XM_023251669.1;refGene_gn=LHCGR;refGene_tx=XM_023251670.1;refGene_gn=LHCGR;refGene_tx=XM_023251671.1
A3	70410548	70410734
refGene_gn=LHCGR;refGene_tx=XM_023251666.1;refGene_gn=LHCGR;refGene_tx=XM_023251667.1;refGene_gn=LHCGR;refGene_tx=XM_023251668.1;refGene_gn=LHCGR;refGene_tx=XM_023251669.1;refGene_gn=LHCGR;refGene_tx=XM_023251670.1;refGene_gn=LHCGR;refGene_tx=XM_023251671.1
A3	70414536	70414611
refGene_gn=LHCGR;refGene_tx=XM_023251666.1;refGene_gn=LHCGR;refGene_tx=XM_023251668.1;refGene_gn=LHCGR;refGene_tx=XM_023251669.1;refGene_gn=

LHCGR;refGene_tx=XM_023251670.1;refGene_gn=LHCGR;refGene_tx=XM_0232516
71.1
A3 70419647 70419716
refGene_gn=LHCGR;refGene_tx=XM_023251666.1;refGene_gn=LHCGR;refGene_tx
=XM_023251667.1;refGene_gn=LHCGR;refGene_tx=XM_023251668.1;refGene_gn=
LHCGR;refGene_tx=XM_023251669.1;refGene_gn=LHCGR;refGene_tx=XM_0232516
70.1;refGene_gn=LHCGR;refGene_tx=XM_023251671.1
A3 70429171 70429249
refGene_gn=LHCGR;refGene_tx=XM_023251666.1;refGene_gn=LHCGR;refGene_tx
=XM_023251667.1;refGene_gn=LHCGR;refGene_tx=XM_023251668.1;refGene_gn=
LHCGR;refGene_tx=XM_023251669.1;refGene_gn=LHCGR;refGene_tx=XM_0232516
70.1;refGene_gn=LHCGR;refGene_tx=XM_023251671.1
A3 70429337 70429412
refGene_gn=LHCGR;refGene_tx=XM_023251666.1;refGene_gn=LHCGR;refGene_tx
=XM_023251667.1;refGene_gn=LHCGR;refGene_tx=XM_023251668.1;refGene_gn=
LHCGR;refGene_tx=XM_023251669.1;refGene_gn=LHCGR;refGene_tx=XM_0232516
70.1;refGene_gn=LHCGR;refGene_tx=XM_023251671.1
A3 70431502 70431577
refGene_gn=LHCGR;refGene_tx=XM_023251666.1;refGene_gn=LHCGR;refGene_tx
=XM_023251667.1;refGene_gn=LHCGR;refGene_tx=XM_023251668.1;refGene_gn=
LHCGR;refGene_tx=XM_023251669.1;refGene_gn=LHCGR;refGene_tx=XM_0232516
70.1;refGene_gn=LHCGR;refGene_tx=XM_023251671.1
A3 70434524 70434599
refGene_gn=LHCGR;refGene_tx=XM_023251666.1;refGene_gn=LHCGR;refGene_tx
=XM_023251667.1;refGene_gn=LHCGR;refGene_tx=XM_023251668.1;refGene_gn=
LHCGR;refGene_tx=XM_023251669.1;refGene_gn=LHCGR;refGene_tx=XM_0232516
70.1;refGene_gn=LHCGR;refGene_tx=XM_023251671.1
A3 70436529 70436601
refGene_gn=LHCGR;refGene_tx=XM_023251666.1;refGene_gn=LHCGR;refGene_tx
=XM_023251667.1;refGene_gn=LHCGR;refGene_tx=XM_023251668.1;refGene_gn=
LHCGR;refGene_tx=XM_023251669.1;refGene_gn=LHCGR;refGene_tx=XM_0232516
70.1;refGene_gn=LHCGR;refGene_tx=XM_023251671.1
A3 70455188 70455352
refGene_gn=LHCGR;refGene_tx=XM_023251666.1;refGene_gn=LHCGR;refGene_tx
=XM_023251667.1;refGene_gn=LHCGR;refGene_tx=XM_023251668.1;refGene_gn=
LHCGR;refGene_tx=XM_023251669.1;refGene_gn=LHCGR;refGene_tx=XM_0232516
70.1;refGene_gn=LHCGR;refGene_tx=XM_023251671.1
A3 70631857 70633082
refGene_gn=FSHR;refGene_tx=NM_001048014.1;refGene_gn=FSHR;refGene_tx=X
M_006930085.3;refGene_gn=FSHR;refGene_tx=XM_019826174.2
A3 70637952 70638138
refGene_gn=FSHR;refGene_tx=NM_001048014.1;refGene_gn=FSHR;refGene_tx=X
M_006930085.3;refGene_gn=FSHR;refGene_tx=XM_019826174.2
A3 70650732 70650807
refGene_gn=FSHR;refGene_tx=NM_001048014.1;refGene_gn=FSHR;refGene_tx=X
M_006930085.3;refGene_gn=FSHR;refGene_tx=XM_019826174.2
A3 70650913 70650982
refGene_gn=FSHR;refGene_tx=NM_001048014.1;refGene_gn=FSHR;refGene_tx=X
M_006930085.3;refGene_gn=FSHR;refGene_tx=XM_019826174.2
A3 70656447 70656525

refGene_gn=FSHR;refGene_tx=NM_001048014.1;refGene_gn=FSHR;refGene_tx=XM_006930085.3;refGene_gn=FSHR;refGene_tx=XM_019826174.2
A3 70657908 70657980
refGene_gn=FSHR;refGene_tx=NM_001048014.1;refGene_gn=FSHR;refGene_tx=XM_006930085.3;refGene_gn=FSHR;refGene_tx=XM_019826174.2
A3 70679196 70679271
refGene_gn=FSHR;refGene_tx=NM_001048014.1;refGene_gn=FSHR;refGene_tx=XM_006930085.3;refGene_gn=FSHR;refGene_tx=XM_019826174.2
A3 70681701 70681776
refGene_gn=FSHR;refGene_tx=NM_001048014.1;refGene_gn=FSHR;refGene_tx=XM_019826174.2
A3 70724300 70724372
refGene_gn=FSHR;refGene_tx=NM_001048014.1;refGene_gn=FSHR;refGene_tx=XM_006930085.3;refGene_gn=FSHR;refGene_tx=XM_019826174.2
A3 70802119 70802271
refGene_gn=FSHR;refGene_tx=NM_001048014.1;refGene_gn=FSHR;refGene_tx=XM_006930085.3;refGene_gn=FSHR;refGene_tx=XM_019826174.2
A3 71455579 71455887
refGene_gn=NRXN1;refGene_tx=XM_003983983.5;refGene_gn=NRXN1;refGene_tx=XM_011281021.3;refGene_gn=NRXN1;refGene_tx=XM_011281023.3;refGene_gn=NRXN1;refGene_tx=XM_011281024.3;refGene_gn=NRXN1;refGene_tx=XM_011281027.3;refGene_gn=NRXN1;refGene_tx=XM_019827164.2;refGene_gn=NRXN1;refGene_tx=XM_019827165.2;refGene_gn=NRXN1;refGene_tx=XM_019827166.2;refGene_gn=NRXN1;refGene_tx=XM_019827168.2;refGene_gn=NRXN1;refGene_tx=XM_019827169.2;refGene_gn=NRXN1;refGene_tx=XM_019827170.2;refGene_gn=NRXN1;refGene_tx=XM_019827171.2;refGene_gn=NRXN1;refGene_tx=XM_019827172.2;refGene_gn=NRXN1;refGene_tx=XM_019827173.2;refGene_gn=NRXN1;refGene_tx=XM_019827174.2;refGene_gn=NRXN1;refGene_tx=XM_019827175.2;refGene_gn=NRXN1;refGene_tx=XM_019827176.2;refGene_gn=NRXN1;refGene_tx=XM_019827177.2;refGene_gn=NRXN1;refGene_tx=XM_019827178.2;refGene_gn=NRXN1;refGene_tx=XM_019827179.2;refGene_gn=NRXN1;refGene_tx=XM_019827180.2;refGene_gn=NRXN1;refGene_tx=XM_019827181.2;refGene_gn=NRXN1;refGene_tx=XM_019827182.2;refGene_gn=NRXN1;refGene_tx=XM_019827183.2;refGene_gn=NRXN1;refGene_tx=XM_019827184.2;refGene_gn=NRXN1;refGene_tx=XM_019827185.2;refGene_gn=NRXN1;refGene_tx=XM_019827186.2;refGene_gn=NRXN1;refGene_tx=XM_019827187.2;refGene_gn=NRXN1;refGene_tx=XM_019827188.2;refGene_gn=NRXN1;refGene_tx=XM_019827189.2
A3 71477336 71477424
refGene_gn=NRXN1;refGene_tx=XM_003983983.5;refGene_gn=NRXN1;refGene_tx=XM_011281021.3;refGene_gn=NRXN1;refGene_tx=XM_011281023.3;refGene_gn=NRXN1;refGene_tx=XM_011281024.3;refGene_gn=NRXN1;refGene_tx=XM_011281027.3;refGene_gn=NRXN1;refGene_tx=XM_019827164.2;refGene_gn=NRXN1;refGene_tx=XM_019827165.2;refGene_gn=NRXN1;refGene_tx=XM_019827166.2;refGene_gn=NRXN1;refGene_tx=XM_019827168.2;refGene_gn=NRXN1;refGene_tx=XM_019827169.2;refGene_gn=NRXN1;refGene_tx=XM_019827170.2;refGene_gn=NRXN1;refGene_tx=XM_019827171.2;refGene_gn=NRXN1;refGene_tx=XM_019827172.2;refGene_gn=NRXN1;refGene_tx=XM_019827173.2;refGene_gn=NRXN1;refGene_tx=XM_019827174.2;refGene_gn=NRXN1;refGene_tx=XM_019827175.2;refGene_gn=NRXN1;refGene_tx=XM_019827176.2;refGene_gn=NRXN1;refGene_tx=XM_019827177.2;refGene_gn=NRXN1;refGene_tx=XM_019827178.2;refGene_gn=NRXN1;refGen

e_tx=XM_019827179.2;refGene_gn=NRXN1;refGene_tx=XM_019827180.2;refGene_gn=NRXN1;refGene_tx=XM_019827181.2;refGene_gn=NRXN1;refGene_tx=XM_019827182.2;refGene_gn=NRXN1;refGene_tx=XM_019827183.2;refGene_gn=NRXN1;refGene_tx=XM_019827184.2;refGene_gn=NRXN1;refGene_tx=XM_019827185.2;refGene_gn=NRXN1;refGene_tx=XM_019827186.2;refGene_gn=NRXN1;refGene_tx=XM_019827187.2;refGene_gn=NRXN1;refGene_tx=XM_019827188.2;refGene_gn=NRXN1;refGene_tx=XM_019827189.2
A3 71507399 71507432
refGene_gn=NRXN1;refGene_tx=XM_019827186.2;refGene_gn=NRXN1;refGene_tx=XM_019827187.2
A3 71588243 71588563
refGene_gn=NRXN1;refGene_tx=XM_003983983.5;refGene_gn=NRXN1;refGene_tx=XM_011281021.3;refGene_gn=NRXN1;refGene_tx=XM_011281023.3;refGene_gn=NRXN1;refGene_tx=XM_011281024.3;refGene_gn=NRXN1;refGene_tx=XM_011281027.3;refGene_gn=NRXN1;refGene_tx=XM_019827164.2;refGene_gn=NRXN1;refGene_tx=XM_019827165.2;refGene_gn=NRXN1;refGene_tx=XM_019827166.2;refGene_gn=NRXN1;refGene_tx=XM_019827168.2;refGene_gn=NRXN1;refGene_tx=XM_019827169.2;refGene_gn=NRXN1;refGene_tx=XM_019827170.2;refGene_gn=NRXN1;refGene_tx=XM_019827171.2;refGene_gn=NRXN1;refGene_tx=XM_019827172.2;refGene_gn=NRXN1;refGene_tx=XM_019827173.2;refGene_gn=NRXN1;refGene_tx=XM_019827174.2;refGene_gn=NRXN1;refGene_tx=XM_019827175.2;refGene_gn=NRXN1;refGene_tx=XM_019827176.2;refGene_gn=NRXN1;refGene_tx=XM_019827177.2;refGene_gn=NRXN1;refGene_tx=XM_019827178.2;refGene_gn=NRXN1;refGene_tx=XM_019827179.2;refGene_gn=NRXN1;refGene_tx=XM_019827180.2;refGene_gn=NRXN1;refGene_tx=XM_019827181.2;refGene_gn=NRXN1;refGene_tx=XM_019827182.2;refGene_gn=NRXN1;refGene_tx=XM_019827183.2;refGene_gn=NRXN1;refGene_tx=XM_019827184.2;refGene_gn=NRXN1;refGene_tx=XM_019827185.2;refGene_gn=NRXN1;refGene_tx=XM_019827188.2;refGene_gn=NRXN1;refGene_tx=XM_019827189.2
A3 71589965 71590055
refGene_gn=NRXN1;refGene_tx=XM_011281021.3;refGene_gn=NRXN1;refGene_tx=XM_011281023.3;refGene_gn=NRXN1;refGene_tx=XM_011281024.3;refGene_gn=NRXN1;refGene_tx=XM_019827164.2;refGene_gn=NRXN1;refGene_tx=XM_019827165.2;refGene_gn=NRXN1;refGene_tx=XM_019827166.2;refGene_gn=NRXN1;refGene_tx=XM_019827168.2;refGene_gn=NRXN1;refGene_tx=XM_019827169.2;refGene_gn=NRXN1;refGene_tx=XM_019827170.2;refGene_gn=NRXN1;refGene_tx=XM_019827171.2;refGene_gn=NRXN1;refGene_tx=XM_019827172.2;refGene_gn=NRXN1;refGene_tx=XM_019827173.2;refGene_gn=NRXN1;refGene_tx=XM_019827174.2;refGene_gn=NRXN1;refGene_tx=XM_019827175.2;refGene_gn=NRXN1;refGene_tx=XM_019827176.2;refGene_gn=NRXN1;refGene_tx=XM_019827183.2;refGene_gn=NRXN1;refGene_tx=XM_019827184.2;refGene_gn=NRXN1;refGene_tx=XM_019827188.2
A3 71627114 71627286
refGene_gn=NRXN1;refGene_tx=XM_003983983.5;refGene_gn=NRXN1;refGene_tx=XM_011281021.3;refGene_gn=NRXN1;refGene_tx=XM_011281023.3;refGene_gn=NRXN1;refGene_tx=XM_011281024.3;refGene_gn=NRXN1;refGene_tx=XM_011281027.3;refGene_gn=NRXN1;refGene_tx=XM_019827164.2;refGene_gn=NRXN1;refGene_tx=XM_019827165.2;refGene_gn=NRXN1;refGene_tx=XM_019827166.2;refGene_gn=NRXN1;refGene_tx=XM_019827168.2;refGene_gn=NRXN1;refGene_tx=XM_019827169.2;refGene_gn=NRXN1;refGene_tx=XM_019827170.2;refGene_gn=NRXN1;

refGene_tx=XM_019827171.2;refGene_gn=NRXN1;refGene_tx=XM_019827172.2;refGene_gn=NRXN1;refGene_tx=XM_019827173.2;refGene_gn=NRXN1;refGene_tx=XM_019827174.2;refGene_gn=NRXN1;refGene_tx=XM_019827175.2;refGene_gn=NRXN1;refGene_tx=XM_019827176.2;refGene_gn=NRXN1;refGene_tx=XM_019827177.2;refGene_gn=NRXN1;refGene_tx=XM_019827178.2;refGene_gn=NRXN1;refGene_tx=XM_019827179.2;refGene_gn=NRXN1;refGene_tx=XM_019827180.2;refGene_gn=NRXN1;refGene_tx=XM_019827181.2;refGene_gn=NRXN1;refGene_tx=XM_019827182.2;refGene_gn=NRXN1;refGene_tx=XM_019827183.2;refGene_gn=NRXN1;refGene_tx=XM_019827184.2;refGene_gn=NRXN1;refGene_tx=XM_019827185.2;refGene_gn=NRXN1;refGene_tx=XM_019827188.2;refGene_gn=NRXN1;refGene_tx=XM_019827189.2
A3 71768015 71768197
refGene_gn=NRXN1;refGene_tx=XM_003983983.5;refGene_gn=NRXN1;refGene_tx=XM_011281021.3;refGene_gn=NRXN1;refGene_tx=XM_011281023.3;refGene_gn=NRXN1;refGene_tx=XM_011281024.3;refGene_gn=NRXN1;refGene_tx=XM_011281027.3;refGene_gn=NRXN1;refGene_tx=XM_019827164.2;refGene_gn=NRXN1;refGene_tx=XM_019827165.2;refGene_gn=NRXN1;refGene_tx=XM_019827166.2;refGene_gn=NRXN1;refGene_tx=XM_019827168.2;refGene_gn=NRXN1;refGene_tx=XM_019827169.2;refGene_gn=NRXN1;refGene_tx=XM_019827170.2;refGene_gn=NRXN1;refGene_tx=XM_019827171.2;refGene_gn=NRXN1;refGene_tx=XM_019827172.2;refGene_gn=NRXN1;refGene_tx=XM_019827173.2;refGene_gn=NRXN1;refGene_tx=XM_019827174.2;refGene_gn=NRXN1;refGene_tx=XM_019827175.2;refGene_gn=NRXN1;refGene_tx=XM_019827176.2;refGene_gn=NRXN1;refGene_tx=XM_019827177.2;refGene_gn=NRXN1;refGene_tx=XM_019827178.2;refGene_gn=NRXN1;refGene_tx=XM_019827179.2;refGene_gn=NRXN1;refGene_tx=XM_019827180.2;refGene_gn=NRXN1;refGene_tx=XM_019827181.2;refGene_gn=NRXN1;refGene_tx=XM_019827182.2;refGene_gn=NRXN1;refGene_tx=XM_019827183.2;refGene_gn=NRXN1;refGene_tx=XM_019827184.2;refGene_gn=NRXN1;refGene_tx=XM_019827185.2;refGene_gn=NRXN1;refGene_tx=XM_019827188.2;refGene_gn=NRXN1;refGene_tx=XM_019827189.2
A3 71882877 71883127
refGene_gn=NRXN1;refGene_tx=XM_011281027.3;refGene_gn=NRXN1;refGene_tx=XM_019827183.2;refGene_gn=NRXN1;refGene_tx=XM_019827184.2;refGene_gn=NRXN1;refGene_tx=XM_019827185.2
A3 72004729 72004849
refGene_gn=NRXN1;refGene_tx=XM_003983983.5;refGene_gn=NRXN1;refGene_tx=XM_011281021.3;refGene_gn=NRXN1;refGene_tx=XM_011281023.3;refGene_gn=NRXN1;refGene_tx=XM_011281024.3;refGene_gn=NRXN1;refGene_tx=XM_019827164.2;refGene_gn=NRXN1;refGene_tx=XM_019827165.2;refGene_gn=NRXN1;refGene_tx=XM_019827166.2;refGene_gn=NRXN1;refGene_tx=XM_019827168.2;refGene_gn=NRXN1;refGene_tx=XM_019827169.2;refGene_gn=NRXN1;refGene_tx=XM_019827170.2;refGene_gn=NRXN1;refGene_tx=XM_019827171.2;refGene_gn=NRXN1;refGene_tx=XM_019827172.2;refGene_gn=NRXN1;refGene_tx=XM_019827173.2;refGene_gn=NRXN1;refGene_tx=XM_019827174.2;refGene_gn=NRXN1;refGene_tx=XM_019827175.2;refGene_gn=NRXN1;refGene_tx=XM_019827176.2;refGene_gn=NRXN1;refGene_tx=XM_019827177.2;refGene_gn=NRXN1;refGene_tx=XM_019827178.2;refGene_gn=NRXN1;refGene_tx=XM_019827179.2;refGene_gn=NRXN1;refGene_tx=XM_019827180.2;refGene_gn=NRXN1;refGene_tx=XM_019827181.2;refGene_gn=NRXN1;refGene_tx=XM_019827182.2;refGene_gn=NRXN1;refGene_tx=XM_019827188.2;refGene_gn=NRXN1;refGene_tx=XM_019827189.2

A3	72005770	72005797
refGene_gn=NRXN1;refGene_tx=XM_019827164.2;refGene_gn=NRXN1;refGene_tx=XM_019827165.2;refGene_gn=NRXN1;refGene_tx=XM_019827166.2;refGene_gn=NRXN1;refGene_tx=XM_019827168.2;refGene_gn=NRXN1;refGene_tx=XM_019827169.2;refGene_gn=NRXN1;refGene_tx=XM_019827170.2;refGene_gn=NRXN1;refGene_tx=XM_019827175.2;refGene_gn=NRXN1;refGene_tx=XM_019827177.2
A3	72011102	72011276
refGene_gn=NRXN1;refGene_tx=XM_003983983.5;refGene_gn=NRXN1;refGene_tx=XM_011281021.3;refGene_gn=NRXN1;refGene_tx=XM_011281023.3;refGene_gn=NRXN1;refGene_tx=XM_011281024.3;refGene_gn=NRXN1;refGene_tx=XM_019827164.2;refGene_gn=NRXN1;refGene_tx=XM_019827165.2;refGene_gn=NRXN1;refGene_tx=XM_019827166.2;refGene_gn=NRXN1;refGene_tx=XM_019827168.2;refGene_gn=NRXN1;refGene_tx=XM_019827169.2;refGene_gn=NRXN1;refGene_tx=XM_019827170.2;refGene_gn=NRXN1;refGene_tx=XM_019827171.2;refGene_gn=NRXN1;refGene_tx=XM_019827172.2;refGene_gn=NRXN1;refGene_tx=XM_019827173.2;refGene_gn=NRXN1;refGene_tx=XM_019827174.2;refGene_gn=NRXN1;refGene_tx=XM_019827175.2;refGene_gn=NRXN1;refGene_tx=XM_019827176.2;refGene_gn=NRXN1;refGene_tx=XM_019827177.2;refGene_gn=NRXN1;refGene_tx=XM_019827178.2;refGene_gn=NRXN1;refGene_tx=XM_019827179.2;refGene_gn=NRXN1;refGene_tx=XM_019827180.2;refGene_gn=NRXN1;refGene_tx=XM_019827181.2;refGene_gn=NRXN1;refGene_tx=XM_019827182.2;refGene_gn=NRXN1;refGene_tx=XM_019827188.2;refGene_gn=NRXN1;refGene_tx=XM_019827189.2
A3	72032847	72033038
refGene_gn=NRXN1;refGene_tx=XM_003983983.5;refGene_gn=NRXN1;refGene_tx=XM_011281021.3;refGene_gn=NRXN1;refGene_tx=XM_011281023.3;refGene_gn=NRXN1;refGene_tx=XM_011281024.3;refGene_gn=NRXN1;refGene_tx=XM_019827164.2;refGene_gn=NRXN1;refGene_tx=XM_019827165.2;refGene_gn=NRXN1;refGene_tx=XM_019827166.2;refGene_gn=NRXN1;refGene_tx=XM_019827168.2;refGene_gn=NRXN1;refGene_tx=XM_019827169.2;refGene_gn=NRXN1;refGene_tx=XM_019827170.2;refGene_gn=NRXN1;refGene_tx=XM_019827171.2;refGene_gn=NRXN1;refGene_tx=XM_019827172.2;refGene_gn=NRXN1;refGene_tx=XM_019827173.2;refGene_gn=NRXN1;refGene_tx=XM_019827174.2;refGene_gn=NRXN1;refGene_tx=XM_019827175.2;refGene_gn=NRXN1;refGene_tx=XM_019827176.2;refGene_gn=NRXN1;refGene_tx=XM_019827177.2;refGene_gn=NRXN1;refGene_tx=XM_019827178.2;refGene_gn=NRXN1;refGene_tx=XM_019827179.2;refGene_gn=NRXN1;refGene_tx=XM_019827180.2;refGene_gn=NRXN1;refGene_tx=XM_019827181.2;refGene_gn=NRXN1;refGene_tx=XM_019827182.2;refGene_gn=NRXN1;refGene_tx=XM_019827188.2;refGene_gn=NRXN1;refGene_tx=XM_019827189.2
A3	72034299	72034681
refGene_gn=NRXN1;refGene_tx=XM_003983983.5;refGene_gn=NRXN1;refGene_tx=XM_011281021.3;refGene_gn=NRXN1;refGene_tx=XM_011281023.3;refGene_gn=NRXN1;refGene_tx=XM_011281024.3;refGene_gn=NRXN1;refGene_tx=XM_019827164.2;refGene_gn=NRXN1;refGene_tx=XM_019827165.2;refGene_gn=NRXN1;refGene_tx=XM_019827166.2;refGene_gn=NRXN1;refGene_tx=XM_019827168.2;refGene_gn=NRXN1;refGene_tx=XM_019827169.2;refGene_gn=NRXN1;refGene_tx=XM_019827170.2;refGene_gn=NRXN1;refGene_tx=XM_019827171.2;refGene_gn=NRXN1;refGene_tx=XM_019827172.2;refGene_gn=NRXN1;refGene_tx=XM_019827173.2;refGene_gn=NRXN1;refGene_tx=XM_019827174.2;refGene_gn=NRXN1;refGene_tx=XM_019827175.2;refGene_gn=NRXN1;refGene_tx=XM_019827176.2;refGene_gn=NRXN1;refGene_tx=XM_019827177.2;refGene_gn=NRXN1;refGene_tx=XM_01982717

8.2;refGene_gn=NRXN1;refGene_tx=XM_019827179.2;refGene_gn=NRXN1;refGene_tx=XM_019827180.2;refGene_gn=NRXN1;refGene_tx=XM_019827181.2;refGene_gn=NRXN1;refGene_tx=XM_019827182.2;refGene_gn=NRXN1;refGene_tx=XM_019827188.2;refGene_gn=NRXN1;refGene_tx=XM_019827189.2
A3 72047125 72047248
refGene_gn=NRXN1;refGene_tx=XM_003983983.5;refGene_gn=NRXN1;refGene_tx=XM_011281021.3;refGene_gn=NRXN1;refGene_tx=XM_011281023.3;refGene_gn=NRXN1;refGene_tx=XM_011281024.3;refGene_gn=NRXN1;refGene_tx=XM_019827164.2;refGene_gn=NRXN1;refGene_tx=XM_019827165.2;refGene_gn=NRXN1;refGene_tx=XM_019827166.2;refGene_gn=NRXN1;refGene_tx=XM_019827168.2;refGene_gn=NRXN1;refGene_tx=XM_019827169.2;refGene_gn=NRXN1;refGene_tx=XM_019827170.2;refGene_gn=NRXN1;refGene_tx=XM_019827171.2;refGene_gn=NRXN1;refGene_tx=XM_019827172.2;refGene_gn=NRXN1;refGene_tx=XM_019827173.2;refGene_gn=NRXN1;refGene_tx=XM_019827174.2;refGene_gn=NRXN1;refGene_tx=XM_019827175.2;refGene_gn=NRXN1;refGene_tx=XM_019827176.2;refGene_gn=NRXN1;refGene_tx=XM_019827177.2;refGene_gn=NRXN1;refGene_tx=XM_019827178.2;refGene_gn=NRXN1;refGene_tx=XM_019827179.2;refGene_gn=NRXN1;refGene_tx=XM_019827180.2;refGene_gn=NRXN1;refGene_tx=XM_019827181.2;refGene_gn=NRXN1;refGene_tx=XM_019827182.2;refGene_gn=NRXN1;refGene_tx=XM_019827188.2;refGene_gn=NRXN1;refGene_tx=XM_019827189.2
A3 72071432 72071459
refGene_gn=NRXN1;refGene_tx=XM_003983983.5;refGene_gn=NRXN1;refGene_tx=XM_011281021.3;refGene_gn=NRXN1;refGene_tx=XM_011281023.3;refGene_gn=NRXN1;refGene_tx=XM_011281024.3;refGene_gn=NRXN1;refGene_tx=XM_019827164.2;refGene_gn=NRXN1;refGene_tx=XM_019827165.2;refGene_gn=NRXN1;refGene_tx=XM_019827166.2;refGene_gn=NRXN1;refGene_tx=XM_019827168.2;refGene_gn=NRXN1;refGene_tx=XM_019827169.2;refGene_gn=NRXN1;refGene_tx=XM_019827171.2;refGene_gn=NRXN1;refGene_tx=XM_019827172.2;refGene_gn=NRXN1;refGene_tx=XM_019827173.2;refGene_gn=NRXN1;refGene_tx=XM_019827175.2;refGene_gn=NRXN1;refGene_tx=XM_019827176.2;refGene_gn=NRXN1;refGene_tx=XM_019827177.2
A3 72074020 72074224
refGene_gn=NRXN1;refGene_tx=XM_003983983.5;refGene_gn=NRXN1;refGene_tx=XM_011281021.3;refGene_gn=NRXN1;refGene_tx=XM_011281023.3;refGene_gn=NRXN1;refGene_tx=XM_011281024.3;refGene_gn=NRXN1;refGene_tx=XM_019827164.2;refGene_gn=NRXN1;refGene_tx=XM_019827165.2;refGene_gn=NRXN1;refGene_tx=XM_019827166.2;refGene_gn=NRXN1;refGene_tx=XM_019827168.2;refGene_gn=NRXN1;refGene_tx=XM_019827169.2;refGene_gn=NRXN1;refGene_tx=XM_019827170.2;refGene_gn=NRXN1;refGene_tx=XM_019827171.2;refGene_gn=NRXN1;refGene_tx=XM_019827172.2;refGene_gn=NRXN1;refGene_tx=XM_019827173.2;refGene_gn=NRXN1;refGene_tx=XM_019827174.2;refGene_gn=NRXN1;refGene_tx=XM_019827175.2;refGene_gn=NRXN1;refGene_tx=XM_019827176.2;refGene_gn=NRXN1;refGene_tx=XM_019827177.2;refGene_gn=NRXN1;refGene_tx=XM_019827178.2;refGene_gn=NRXN1;refGene_tx=XM_019827179.2;refGene_gn=NRXN1;refGene_tx=XM_019827180.2;refGene_gn=NRXN1;refGene_tx=XM_019827181.2;refGene_gn=NRXN1;refGene_tx=XM_019827182.2;refGene_gn=NRXN1;refGene_tx=XM_019827188.2;refGene_gn=NRXN1;refGene_tx=XM_019827189.2
A3 72080762 72081146
refGene_gn=NRXN1;refGene_tx=XM_003983983.5;refGene_gn=NRXN1;refGene_tx=XM_011281021.3;refGene_gn=NRXN1;refGene_tx=XM_011281023.3;refGene_gn=

NRXN1;refGene_tx=XM_011281024.3;refGene_gn=NRXN1;refGene_tx=XM_0198271
64.2;refGene_gn=NRXN1;refGene_tx=XM_019827165.2;refGene_gn=NRXN1;refGe
ne_tx=XM_019827166.2;refGene_gn=NRXN1;refGene_tx=XM_019827168.2;refGen
e_gn=NRXN1;refGene_tx=XM_019827169.2;refGene_gn=NRXN1;refGene_tx=XM_01
9827170.2;refGene_gn=NRXN1;refGene_tx=XM_019827171.2;refGene_gn=NRXN1;
refGene_tx=XM_019827172.2;refGene_gn=NRXN1;refGene_tx=XM_019827173.2;r
efGene_gn=NRXN1;refGene_tx=XM_019827174.2;refGene_gn=NRXN1;refGene_tx=
XM_019827175.2;refGene_gn=NRXN1;refGene_tx=XM_019827176.2;refGene_gn=N
RXN1;refGene_tx=XM_019827177.2;refGene_gn=NRXN1;refGene_tx=XM_01982717
8.2;refGene_gn=NRXN1;refGene_tx=XM_019827179.2;refGene_gn=NRXN1;refGen
e_tx=XM_019827180.2;refGene_gn=NRXN1;refGene_tx=XM_019827181.2;refGene
_gn=NRXN1;refGene_tx=XM_019827182.2;refGene_gn=NRXN1;refGene_tx=XM_019
827188.2;refGene_gn=NRXN1;refGene_tx=XM_019827189.2
A3 72094133 72094572
refGene_gn=NRXN1;refGene_tx=XM_003983983.5;refGene_gn=NRXN1;refGene_tx
=XM_011281021.3;refGene_gn=NRXN1;refGene_tx=XM_011281023.3;refGene_gn=
NRXN1;refGene_tx=XM_011281024.3;refGene_gn=NRXN1;refGene_tx=XM_0198271
64.2;refGene_gn=NRXN1;refGene_tx=XM_019827165.2;refGene_gn=NRXN1;refGe
ne_tx=XM_019827166.2;refGene_gn=NRXN1;refGene_tx=XM_019827168.2;refGen
e_gn=NRXN1;refGene_tx=XM_019827169.2;refGene_gn=NRXN1;refGene_tx=XM_01
9827170.2;refGene_gn=NRXN1;refGene_tx=XM_019827171.2;refGene_gn=NRXN1;
refGene_tx=XM_019827172.2;refGene_gn=NRXN1;refGene_tx=XM_019827173.2;r
efGene_gn=NRXN1;refGene_tx=XM_019827174.2;refGene_gn=NRXN1;refGene_tx=
XM_019827175.2;refGene_gn=NRXN1;refGene_tx=XM_019827176.2;refGene_gn=N
RXN1;refGene_tx=XM_019827177.2;refGene_gn=NRXN1;refGene_tx=XM_01982717
8.2;refGene_gn=NRXN1;refGene_tx=XM_019827179.2;refGene_gn=NRXN1;refGen
e_tx=XM_019827180.2;refGene_gn=NRXN1;refGene_tx=XM_019827181.2;refGene
_gn=NRXN1;refGene_tx=XM_019827182.2;refGene_gn=NRXN1;refGene_tx=XM_019
827188.2;refGene_gn=NRXN1;refGene_tx=XM_019827189.2
A3 72157789 72157951
refGene_gn=NRXN1;refGene_tx=XM_003983983.5;refGene_gn=NRXN1;refGene_tx
=XM_011281021.3;refGene_gn=NRXN1;refGene_tx=XM_011281023.3;refGene_gn=
NRXN1;refGene_tx=XM_011281024.3;refGene_gn=NRXN1;refGene_tx=XM_0198271
64.2;refGene_gn=NRXN1;refGene_tx=XM_019827165.2;refGene_gn=NRXN1;refGe
ne_tx=XM_019827166.2;refGene_gn=NRXN1;refGene_tx=XM_019827168.2;refGen
e_gn=NRXN1;refGene_tx=XM_019827169.2;refGene_gn=NRXN1;refGene_tx=XM_01
9827170.2;refGene_gn=NRXN1;refGene_tx=XM_019827171.2;refGene_gn=NRXN1;
refGene_tx=XM_019827172.2;refGene_gn=NRXN1;refGene_tx=XM_019827173.2;r
efGene_gn=NRXN1;refGene_tx=XM_019827174.2;refGene_gn=NRXN1;refGene_tx=
XM_019827175.2;refGene_gn=NRXN1;refGene_tx=XM_019827176.2;refGene_gn=N
RXN1;refGene_tx=XM_019827177.2;refGene_gn=NRXN1;refGene_tx=XM_01982717
8.2;refGene_gn=NRXN1;refGene_tx=XM_019827179.2;refGene_gn=NRXN1;refGen
e_tx=XM_019827180.2;refGene_gn=NRXN1;refGene_tx=XM_019827181.2;refGene
_gn=NRXN1;refGene_tx=XM_019827182.2;refGene_gn=NRXN1;refGene_tx=XM_019
827188.2;refGene_gn=NRXN1;refGene_tx=XM_019827189.2
A3 72159002 72159026
refGene_gn=NRXN1;refGene_tx=XM_003983983.5;refGene_gn=NRXN1;refGene_tx
=XM_011281021.3;refGene_gn=NRXN1;refGene_tx=XM_019827164.2;refGene_gn=
NRXN1;refGene_tx=XM_019827165.2;refGene_gn=NRXN1;refGene_tx=XM_0198271
66.2;refGene_gn=NRXN1;refGene_tx=XM_019827168.2;refGene_gn=NRXN1;refGe

ne_tx=XM_019827170.2;refGene_gn=NRXN1;refGene_tx=XM_019827171.2;refGen
e_gn=NRXN1;refGene_tx=XM_019827172.2;refGene_gn=NRXN1;refGene_tx=XM_01
9827173.2;refGene_gn=NRXN1;refGene_tx=XM_019827174.2;refGene_gn=NRXN1;
refGene_tx=XM_019827175.2;refGene_gn=NRXN1;refGene_tx=XM_019827177.2;r
efGene_gn=NRXN1;refGene_tx=XM_019827178.2;refGene_gn=NRXN1;refGene_tx=
XM_019827188.2;refGene_gn=NRXN1;refGene_tx=XM_019827189.2
A3 72161133 72161435
refGene_gn=NRXN1;refGene_tx=XM_003983983.5;refGene_gn=NRXN1;refGene_tx
=XM_011281021.3;refGene_gn=NRXN1;refGene_tx=XM_011281023.3;refGene_gn=
NRXN1;refGene_tx=XM_011281024.3;refGene_gn=NRXN1;refGene_tx=XM_0198271
64.2;refGene_gn=NRXN1;refGene_tx=XM_019827165.2;refGene_gn=NRXN1;refGe
ne_tx=XM_019827166.2;refGene_gn=NRXN1;refGene_tx=XM_019827168.2;refGen
e_gn=NRXN1;refGene_tx=XM_019827169.2;refGene_gn=NRXN1;refGene_tx=XM_01
9827170.2;refGene_gn=NRXN1;refGene_tx=XM_019827171.2;refGene_gn=NRXN1;
refGene_tx=XM_019827172.2;refGene_gn=NRXN1;refGene_tx=XM_019827173.2;r
efGene_gn=NRXN1;refGene_tx=XM_019827174.2;refGene_gn=NRXN1;refGene_tx=
XM_019827175.2;refGene_gn=NRXN1;refGene_tx=XM_019827176.2;refGene_gn=N
RXN1;refGene_tx=XM_019827177.2;refGene_gn=NRXN1;refGene_tx=XM_01982717
8.2;refGene_gn=NRXN1;refGene_tx=XM_019827179.2;refGene_gn=NRXN1;refGen
e_tx=XM_019827180.2;refGene_gn=NRXN1;refGene_tx=XM_019827181.2;refGene
_gn=NRXN1;refGene_tx=XM_019827182.2;refGene_gn=NRXN1;refGene_tx=XM_019
827188.2;refGene_gn=NRXN1;refGene_tx=XM_019827189.2
A3 72481192 72481204
refGene_gn=NRXN1;refGene_tx=XM_003983983.5;refGene_gn=NRXN1;refGene_tx
=XM_011281021.3;refGene_gn=NRXN1;refGene_tx=XM_011281023.3;refGene_gn=
NRXN1;refGene_tx=XM_011281024.3;refGene_gn=NRXN1;refGene_tx=XM_0198271
64.2;refGene_gn=NRXN1;refGene_tx=XM_019827165.2;refGene_gn=NRXN1;refGe
ne_tx=XM_019827166.2;refGene_gn=NRXN1;refGene_tx=XM_019827169.2;refGen
e_gn=NRXN1;refGene_tx=XM_019827170.2;refGene_gn=NRXN1;refGene_tx=XM_01
9827171.2;refGene_gn=NRXN1;refGene_tx=XM_019827172.2;refGene_gn=NRXN1;
refGene_tx=XM_019827174.2;refGene_gn=NRXN1;refGene_tx=XM_019827177.2;r
efGene_gn=NRXN1;refGene_tx=XM_019827178.2;refGene_gn=NRXN1;refGene_tx=
XM_019827179.2
A3 72481979 72482009
refGene_gn=NRXN1;refGene_tx=XM_003983983.5;refGene_gn=NRXN1;refGene_tx
=XM_011281021.3;refGene_gn=NRXN1;refGene_tx=XM_011281023.3;refGene_gn=
NRXN1;refGene_tx=XM_011281024.3;refGene_gn=NRXN1;refGene_tx=XM_0198271
64.2;refGene_gn=NRXN1;refGene_tx=XM_019827165.2;refGene_gn=NRXN1;refGe
ne_tx=XM_019827166.2;refGene_gn=NRXN1;refGene_tx=XM_019827168.2;refGen
e_gn=NRXN1;refGene_tx=XM_019827169.2;refGene_gn=NRXN1;refGene_tx=XM_01
9827170.2;refGene_gn=NRXN1;refGene_tx=XM_019827171.2;refGene_gn=NRXN1;
refGene_tx=XM_019827172.2;refGene_gn=NRXN1;refGene_tx=XM_019827173.2;r
efGene_gn=NRXN1;refGene_tx=XM_019827174.2;refGene_gn=NRXN1;refGene_tx=
XM_019827176.2;refGene_gn=NRXN1;refGene_tx=XM_019827177.2;refGene_gn=N
RXN1;refGene_tx=XM_019827178.2;refGene_gn=NRXN1;refGene_tx=XM_01982717
9.2;refGene_gn=NRXN1;refGene_tx=XM_019827180.2
A3 72485234 72485252
refGene_gn=NRXN1;refGene_tx=XM_003983983.5;refGene_gn=NRXN1;refGene_tx
=XM_011281021.3;refGene_gn=NRXN1;refGene_tx=XM_011281023.3;refGene_gn=
NRXN1;refGene_tx=XM_011281024.3;refGene_gn=NRXN1;refGene_tx=XM_0198271

64.2;refGene_gn=NRXN1;refGene_tx=XM_019827165.2;refGene_gn=NRXN1;refGene_tx=XM_019827166.2;refGene_gn=NRXN1;refGene_tx=XM_019827168.2;refGene_gn=NRXN1;refGene_tx=XM_019827169.2;refGene_gn=NRXN1;refGene_tx=XM_019827170.2;refGene_gn=NRXN1;refGene_tx=XM_019827171.2;refGene_gn=NRXN1;refGene_tx=XM_019827172.2;refGene_gn=NRXN1;refGene_tx=XM_019827173.2;refGene_gn=NRXN1;refGene_tx=XM_019827174.2;refGene_gn=NRXN1;refGene_tx=XM_019827176.2;refGene_gn=NRXN1;refGene_tx=XM_019827177.2;refGene_gn=NRXN1;refGene_tx=XM_019827178.2;refGene_gn=NRXN1;refGene_tx=XM_019827179.2;refGene_gn=NRXN1;refGene_tx=XM_019827180.2
A3 72586280 72587052
refGene_gn=NRXN1;refGene_tx=XM_003983983.5;refGene_gn=NRXN1;refGene_tx=XM_011281021.3;refGene_gn=NRXN1;refGene_tx=XM_011281023.3;refGene_gn=NRXN1;refGene_tx=XM_011281024.3;refGene_gn=NRXN1;refGene_tx=XM_019827164.2;refGene_gn=NRXN1;refGene_tx=XM_019827165.2;refGene_gn=NRXN1;refGene_tx=XM_019827166.2;refGene_gn=NRXN1;refGene_tx=XM_019827168.2;refGene_gn=NRXN1;refGene_tx=XM_019827169.2;refGene_gn=NRXN1;refGene_tx=XM_019827170.2;refGene_gn=NRXN1;refGene_tx=XM_019827171.2;refGene_gn=NRXN1;refGene_tx=XM_019827172.2;refGene_gn=NRXN1;refGene_tx=XM_019827173.2;refGene_gn=NRXN1;refGene_tx=XM_019827174.2;refGene_gn=NRXN1;refGene_tx=XM_019827175.2;refGene_gn=NRXN1;refGene_tx=XM_019827176.2;refGene_gn=NRXN1;refGene_tx=XM_019827177.2;refGene_gn=NRXN1;refGene_tx=XM_019827178.2;refGene_gn=NRXN1;refGene_tx=XM_019827179.2;refGene_gn=NRXN1;refGene_tx=XM_019827180.2;refGene_gn=NRXN1;refGene_tx=XM_019827181.2;refGene_gn=NRXN1;refGene_tx=XM_019827182.2;refGene_gn=NRXN1;refGene_tx=XM_019827188.2;refGene_gn=NRXN1;refGene_tx=XM_019827189.2
A3 74881169 74881375
refGene_gn=ASB3;refGene_tx=XM_011281016.3;refGene_gn=ASB3;refGene_tx=XM_011281017.3;refGene_gn=ASB3;refGene_tx=XM_011281018.3;refGene_gn=ASB3;refGene_tx=XM_019827190.2;refGene_gn=ASB3;refGene_tx=XM_019827191.2
A3 74898432 74898566
refGene_gn=ASB3;refGene_tx=XM_011281016.3;refGene_gn=ASB3;refGene_tx=XM_011281017.3;refGene_gn=ASB3;refGene_tx=XM_011281018.3;refGene_gn=ASB3;refGene_tx=XM_019827190.2;refGene_gn=ASB3;refGene_tx=XM_019827191.2
A3 74904100 74904361
refGene_gn=ASB3;refGene_tx=XM_011281016.3;refGene_gn=ASB3;refGene_tx=XM_011281017.3;refGene_gn=ASB3;refGene_tx=XM_011281018.3;refGene_gn=ASB3;refGene_tx=XM_019827190.2
A3 74915768 74915966
refGene_gn=ASB3;refGene_tx=XM_011281016.3;refGene_gn=ASB3;refGene_tx=XM_011281017.3;refGene_gn=ASB3;refGene_tx=XM_011281018.3;refGene_gn=ASB3;refGene_tx=XM_019827190.2;refGene_gn=ASB3;refGene_tx=XM_019827191.2
A3 74918848 74919026
refGene_gn=ASB3;refGene_tx=XM_011281016.3;refGene_gn=ASB3;refGene_tx=XM_011281017.3;refGene_gn=ASB3;refGene_tx=XM_011281018.3;refGene_gn=ASB3;refGene_tx=XM_019827190.2;refGene_gn=ASB3;refGene_tx=XM_019827191.2
A3 74921813 74921949
refGene_gn=ASB3;refGene_tx=XM_011281016.3;refGene_gn=ASB3;refGene_tx=XM_011281017.3;refGene_gn=ASB3;refGene_tx=XM_011281018.3;refGene_gn=ASB3;refGene_tx=XM_019827190.2;refGene_gn=ASB3;refGene_tx=XM_019827191.2
A3 74922539 74922652

refGene_gn=ASB3;refGene_tx=XM_011281016.3;refGene_gn=ASB3;refGene_tx=XM_011281017.3;refGene_gn=ASB3;refGene_tx=XM_011281018.3;refGene_gn=ASB3;refGene_tx=XM_019827190.2;refGene_gn=ASB3;refGene_tx=XM_019827191.2
A3 74949190 74949349
refGene_gn=ASB3;refGene_tx=XM_011281016.3;refGene_gn=ASB3;refGene_tx=XM_011281017.3;refGene_gn=ASB3;refGene_tx=XM_011281018.3;refGene_gn=ASB3;refGene_tx=XM_019827190.2;refGene_gn=ASB3;refGene_tx=XM_019827191.2
A3 74958924 74959120
refGene_gn=ASB3;refGene_tx=XM_011281016.3;refGene_gn=ASB3;refGene_tx=XM_011281017.3;refGene_gn=ASB3;refGene_tx=XM_011281018.3;refGene_gn=ASB3;refGene_tx=XM_019827190.2;refGene_gn=ASB3;refGene_tx=XM_019827191.2
A3 74971317 74971452
refGene_gn=CHAC2;refGene_tx=XM_003983992.4;refGene_gn=CHAC2;refGene_tx=XM_019827193.2
A3 74976120 74976156 refGene_gn=CHAC2;refGene_tx=XM_003983992.4
A3 74978189 74978573
refGene_gn=CHAC2;refGene_tx=XM_003983992.4;refGene_gn=CHAC2;refGene_tx=XM_019827193.2
A3 74997125 74997287
refGene_gn=ERLEC1;refGene_tx=XM_003983993.4;refGene_gn=ERLEC1;refGene_tx=XM_006930101.3;refGene_gn=ERLEC1;refGene_tx=XM_019827192.2
A3 75003010 75003115
refGene_gn=ERLEC1;refGene_tx=XM_003983993.4;refGene_gn=ERLEC1;refGene_tx=XM_006930101.3;refGene_gn=ERLEC1;refGene_tx=XM_019827192.2
A3 75004750 75004831
refGene_gn=ERLEC1;refGene_tx=XM_003983993.4;refGene_gn=ERLEC1;refGene_tx=XM_006930101.3;refGene_gn=ERLEC1;refGene_tx=XM_019827192.2
A3 75005799 75005877
refGene_gn=ERLEC1;refGene_tx=XM_003983993.4;refGene_gn=ERLEC1;refGene_tx=XM_006930101.3;refGene_gn=ERLEC1;refGene_tx=XM_019827192.2
A3 75006012 75006076
refGene_gn=ERLEC1;refGene_tx=XM_003983993.4;refGene_gn=ERLEC1;refGene_tx=XM_006930101.3;refGene_gn=ERLEC1;refGene_tx=XM_019827192.2
A3 75006581 75006616
refGene_gn=ERLEC1;refGene_tx=XM_003983993.4;refGene_gn=ERLEC1;refGene_tx=XM_006930101.3;refGene_gn=ERLEC1;refGene_tx=XM_019827192.2
A3 75008883 75009107
refGene_gn=ERLEC1;refGene_tx=XM_003983993.4;refGene_gn=ERLEC1;refGene_tx=XM_019827192.2
A3 75009199 75009329
refGene_gn=ERLEC1;refGene_tx=XM_003983993.4;refGene_gn=ERLEC1;refGene_tx=XM_019827192.2
A3 75016794 75016956
refGene_gn=ERLEC1;refGene_tx=XM_003983993.4;refGene_gn=ERLEC1;refGene_tx=XM_006930101.3
A3 75017814 75017874
refGene_gn=ERLEC1;refGene_tx=XM_003983993.4;refGene_gn=ERLEC1;refGene_tx=XM_006930101.3;refGene_gn=ERLEC1;refGene_tx=XM_019827192.2
A3 75021989 75022114
refGene_gn=ERLEC1;refGene_tx=XM_003983993.4;refGene_gn=ERLEC1;refGene_

tx=XM_006930101.3;refGene_gn=ERLEC1;refGene_tx=XM_019827192.2
A3	75023854 75023932
refGene_gn=ERLEC1;refGene_tx=XM_003983993.4;refGene_gn=ERLEC1;refGene_tx=XM_006930101.3;refGene_gn=ERLEC1;refGene_tx=XM_019827192.2
A3	75024874 75024950
refGene_gn=ERLEC1;refGene_tx=XM_003983993.4;refGene_gn=ERLEC1;refGene_tx=XM_006930101.3;refGene_gn=ERLEC1;refGene_tx=XM_019827192.2
A3	75027207 75027279
refGene_gn=ERLEC1;refGene_tx=XM_003983993.4;refGene_gn=ERLEC1;refGene_tx=XM_006930101.3;refGene_gn=ERLEC1;refGene_tx=XM_019827192.2
A3	75041831 75043454 refGene_gn=GPR75;refGene_tx=XM_006930102.4
A3	75059033 75059168
refGene_gn=PSME4;refGene_tx=XM_023251673.1;refGene_gn=PSME4;refGene_tx=XM_023251674.1;refGene_gn=PSME4;refGene_tx=XM_023251675.1;refGene_gn=PSME4;refGene_tx=XM_023251676.1
A3	75059915 75060049
refGene_gn=PSME4;refGene_tx=XM_023251673.1;refGene_gn=PSME4;refGene_tx=XM_023251674.1;refGene_gn=PSME4;refGene_tx=XM_023251675.1;refGene_gn=PSME4;refGene_tx=XM_023251676.1
A3	75061850 75062013
refGene_gn=PSME4;refGene_tx=XM_023251673.1;refGene_gn=PSME4;refGene_tx=XM_023251674.1;refGene_gn=PSME4;refGene_tx=XM_023251675.1;refGene_gn=PSME4;refGene_tx=XM_023251676.1
A3	75064548 75064704
refGene_gn=PSME4;refGene_tx=XM_023251673.1;refGene_gn=PSME4;refGene_tx=XM_023251674.1;refGene_gn=PSME4;refGene_tx=XM_023251675.1;refGene_gn=PSME4;refGene_tx=XM_023251676.1
A3	75065392 75065521
refGene_gn=PSME4;refGene_tx=XM_023251673.1;refGene_gn=PSME4;refGene_tx=XM_023251674.1;refGene_gn=PSME4;refGene_tx=XM_023251675.1;refGene_gn=PSME4;refGene_tx=XM_023251676.1
A3	75074504 75074590
refGene_gn=PSME4;refGene_tx=XM_023251673.1;refGene_gn=PSME4;refGene_tx=XM_023251674.1;refGene_gn=PSME4;refGene_tx=XM_023251675.1;refGene_gn=PSME4;refGene_tx=XM_023251676.1
A3	75076494 75076703
refGene_gn=PSME4;refGene_tx=XM_023251673.1;refGene_gn=PSME4;refGene_tx=XM_023251674.1;refGene_gn=PSME4;refGene_tx=XM_023251675.1;refGene_gn=PSME4;refGene_tx=XM_023251676.1
A3	75077059 75077191
refGene_gn=PSME4;refGene_tx=XM_023251673.1;refGene_gn=PSME4;refGene_tx=XM_023251674.1;refGene_gn=PSME4;refGene_tx=XM_023251675.1;refGene_gn=PSME4;refGene_tx=XM_023251676.1
A3	75077961 75078053
refGene_gn=PSME4;refGene_tx=XM_023251673.1;refGene_gn=PSME4;refGene_tx=XM_023251674.1;refGene_gn=PSME4;refGene_tx=XM_023251675.1;refGene_gn=PSME4;refGene_tx=XM_023251676.1
A3	75078847 75078952
refGene_gn=PSME4;refGene_tx=XM_023251673.1;refGene_gn=PSME4;refGene_tx=XM_023251674.1;refGene_gn=PSME4;refGene_tx=XM_023251675.1;refGene_gn=

PSME4;refGene_tx=XM_023251676.1
A3 75080676 75080829
refGene_gn=PSME4;refGene_tx=XM_023251673.1;refGene_gn=PSME4;refGene_tx=XM_023251674.1;refGene_gn=PSME4;refGene_tx=XM_023251675.1;refGene_gn=PSME4;refGene_tx=XM_023251676.1
A3 75081386 75081512
refGene_gn=PSME4;refGene_tx=XM_023251673.1;refGene_gn=PSME4;refGene_tx=XM_023251674.1;refGene_gn=PSME4;refGene_tx=XM_023251676.1
A3 75082517 75082587
refGene_gn=PSME4;refGene_tx=XM_023251673.1;refGene_gn=PSME4;refGene_tx=XM_023251674.1;refGene_gn=PSME4;refGene_tx=XM_023251675.1;refGene_gn=PSME4;refGene_tx=XM_023251676.1
A3 75083127 75083281
refGene_gn=PSME4;refGene_tx=XM_023251673.1;refGene_gn=PSME4;refGene_tx=XM_023251674.1;refGene_gn=PSME4;refGene_tx=XM_023251675.1;refGene_gn=PSME4;refGene_tx=XM_023251676.1
A3 75084476 75084558
refGene_gn=PSME4;refGene_tx=XM_023251673.1;refGene_gn=PSME4;refGene_tx=XM_023251674.1;refGene_gn=PSME4;refGene_tx=XM_023251675.1;refGene_gn=PSME4;refGene_tx=XM_023251676.1
A3 75085460 75085590
refGene_gn=PSME4;refGene_tx=XM_023251673.1;refGene_gn=PSME4;refGene_tx=XM_023251674.1;refGene_gn=PSME4;refGene_tx=XM_023251675.1;refGene_gn=PSME4;refGene_tx=XM_023251676.1
A3 75086326 75086380
refGene_gn=PSME4;refGene_tx=XM_023251673.1;refGene_gn=PSME4;refGene_tx=XM_023251674.1;refGene_gn=PSME4;refGene_tx=XM_023251675.1;refGene_gn=PSME4;refGene_tx=XM_023251676.1
A3 75087076 75087213
refGene_gn=PSME4;refGene_tx=XM_023251673.1;refGene_gn=PSME4;refGene_tx=XM_023251674.1;refGene_gn=PSME4;refGene_tx=XM_023251675.1;refGene_gn=PSME4;refGene_tx=XM_023251676.1
A3 75088281 75088491
refGene_gn=PSME4;refGene_tx=XM_023251673.1;refGene_gn=PSME4;refGene_tx=XM_023251674.1;refGene_gn=PSME4;refGene_tx=XM_023251675.1;refGene_gn=PSME4;refGene_tx=XM_023251676.1
A3 75090537 75090669
refGene_gn=PSME4;refGene_tx=XM_023251673.1;refGene_gn=PSME4;refGene_tx=XM_023251674.1;refGene_gn=PSME4;refGene_tx=XM_023251675.1;refGene_gn=PSME4;refGene_tx=XM_023251676.1
A3 75091640 75091736
refGene_gn=PSME4;refGene_tx=XM_023251673.1;refGene_gn=PSME4;refGene_tx=XM_023251674.1;refGene_gn=PSME4;refGene_tx=XM_023251675.1;refGene_gn=PSME4;refGene_tx=XM_023251676.1
A3 75091883 75091946
refGene_gn=PSME4;refGene_tx=XM_023251673.1;refGene_gn=PSME4;refGene_tx=XM_023251674.1;refGene_gn=PSME4;refGene_tx=XM_023251675.1;refGene_gn=PSME4;refGene_tx=XM_023251676.1
A3 75093709 75093808
refGene_gn=PSME4;refGene_tx=XM_023251673.1;refGene_gn=PSME4;refGene_tx

=XM_023251674.1;refGene_gn=PSME4;refGene_tx=XM_023251675.1;refGene_gn=PSME4;refGene_tx=XM_023251676.1
A3	75093901	75093957
refGene_gn=PSME4;refGene_tx=XM_023251673.1;refGene_gn=PSME4;refGene_tx=XM_023251674.1;refGene_gn=PSME4;refGene_tx=XM_023251675.1;refGene_gn=PSME4;refGene_tx=XM_023251676.1
A3	75094159	75094216
refGene_gn=PSME4;refGene_tx=XM_023251673.1;refGene_gn=PSME4;refGene_tx=XM_023251674.1;refGene_gn=PSME4;refGene_tx=XM_023251675.1;refGene_gn=PSME4;refGene_tx=XM_023251676.1
A3	75095241	75095297
refGene_gn=PSME4;refGene_tx=XM_023251673.1;refGene_gn=PSME4;refGene_tx=XM_023251674.1;refGene_gn=PSME4;refGene_tx=XM_023251675.1;refGene_gn=PSME4;refGene_tx=XM_023251676.1
A3	75102658	75102754
refGene_gn=PSME4;refGene_tx=XM_023251673.1;refGene_gn=PSME4;refGene_tx=XM_023251674.1;refGene_gn=PSME4;refGene_tx=XM_023251675.1;refGene_gn=PSME4;refGene_tx=XM_023251676.1
A3	75103702	75103860
refGene_gn=PSME4;refGene_tx=XM_023251673.1;refGene_gn=PSME4;refGene_tx=XM_023251674.1;refGene_gn=PSME4;refGene_tx=XM_023251675.1;refGene_gn=PSME4;refGene_tx=XM_023251676.1
A3	75104358	75104574
refGene_gn=PSME4;refGene_tx=XM_023251673.1;refGene_gn=PSME4;refGene_tx=XM_023251674.1;refGene_gn=PSME4;refGene_tx=XM_023251675.1;refGene_gn=PSME4;refGene_tx=XM_023251676.1
A3	75106656	75106724
refGene_gn=PSME4;refGene_tx=XM_023251673.1;refGene_gn=PSME4;refGene_tx=XM_023251674.1;refGene_gn=PSME4;refGene_tx=XM_023251675.1;refGene_gn=PSME4;refGene_tx=XM_023251676.1
A3	75107217	75107287
refGene_gn=PSME4;refGene_tx=XM_023251673.1;refGene_gn=PSME4;refGene_tx=XM_023251674.1;refGene_gn=PSME4;refGene_tx=XM_023251675.1;refGene_gn=PSME4;refGene_tx=XM_023251676.1
A3	75107508	75107607
refGene_gn=PSME4;refGene_tx=XM_023251673.1;refGene_gn=PSME4;refGene_tx=XM_023251674.1;refGene_gn=PSME4;refGene_tx=XM_023251675.1;refGene_gn=PSME4;refGene_tx=XM_023251676.1
A3	75109665	75109816
refGene_gn=PSME4;refGene_tx=XM_023251673.1;refGene_gn=PSME4;refGene_tx=XM_023251674.1;refGene_gn=PSME4;refGene_tx=XM_023251675.1;refGene_gn=PSME4;refGene_tx=XM_023251676.1
A3	75110076	75110141
refGene_gn=PSME4;refGene_tx=XM_023251673.1;refGene_gn=PSME4;refGene_tx=XM_023251674.1;refGene_gn=PSME4;refGene_tx=XM_023251675.1;refGene_gn=PSME4;refGene_tx=XM_023251676.1
A3	75111749	75111839
refGene_gn=PSME4;refGene_tx=XM_023251673.1;refGene_gn=PSME4;refGene_tx=XM_023251674.1;refGene_gn=PSME4;refGene_tx=XM_023251675.1;refGene_gn=PSME4;refGene_tx=XM_023251676.1

A3 75112604 75112791
refGene_gn=PSME4;refGene_tx=XM_023251673.1;refGene_gn=PSME4;refGene_tx=XM_023251674.1;refGene_gn=PSME4;refGene_tx=XM_023251675.1;refGene_gn=PSME4;refGene_tx=XM_023251676.1
A3 75115443 75115709
refGene_gn=PSME4;refGene_tx=XM_023251673.1;refGene_gn=PSME4;refGene_tx=XM_023251674.1;refGene_gn=PSME4;refGene_tx=XM_023251675.1;refGene_gn=PSME4;refGene_tx=XM_023251676.1
A3 75116459 75116552
refGene_gn=PSME4;refGene_tx=XM_023251673.1;refGene_gn=PSME4;refGene_tx=XM_023251674.1;refGene_gn=PSME4;refGene_tx=XM_023251675.1;refGene_gn=PSME4;refGene_tx=XM_023251676.1
A3 75118804 75118927
refGene_gn=PSME4;refGene_tx=XM_023251673.1;refGene_gn=PSME4;refGene_tx=XM_023251674.1;refGene_gn=PSME4;refGene_tx=XM_023251675.1;refGene_gn=PSME4;refGene_tx=XM_023251676.1
A3 75120046 75120121
refGene_gn=PSME4;refGene_tx=XM_023251673.1;refGene_gn=PSME4;refGene_tx=XM_023251674.1;refGene_gn=PSME4;refGene_tx=XM_023251675.1;refGene_gn=PSME4;refGene_tx=XM_023251676.1
A3 75120650 75120714
refGene_gn=PSME4;refGene_tx=XM_023251673.1;refGene_gn=PSME4;refGene_tx=XM_023251674.1;refGene_gn=PSME4;refGene_tx=XM_023251675.1;refGene_gn=PSME4;refGene_tx=XM_023251676.1
A3 75122215 75122365
refGene_gn=PSME4;refGene_tx=XM_023251673.1;refGene_gn=PSME4;refGene_tx=XM_023251674.1;refGene_gn=PSME4;refGene_tx=XM_023251675.1;refGene_gn=PSME4;refGene_tx=XM_023251676.1
A3 75124655 75124700
refGene_gn=PSME4;refGene_tx=XM_023251673.1;refGene_gn=PSME4;refGene_tx=XM_023251674.1;refGene_gn=PSME4;refGene_tx=XM_023251675.1;refGene_gn=PSME4;refGene_tx=XM_023251676.1
A3 75126680 75126746
refGene_gn=PSME4;refGene_tx=XM_023251673.1;refGene_gn=PSME4;refGene_tx=XM_023251675.1;refGene_gn=PSME4;refGene_tx=XM_023251676.1
A3 75135611 75135728
refGene_gn=PSME4;refGene_tx=XM_023251673.1;refGene_gn=PSME4;refGene_tx=XM_023251674.1;refGene_gn=PSME4;refGene_tx=XM_023251675.1;refGene_gn=PSME4;refGene_tx=XM_023251676.1
A3 75136367 75136508
refGene_gn=PSME4;refGene_tx=XM_023251673.1;refGene_gn=PSME4;refGene_tx=XM_023251674.1;refGene_gn=PSME4;refGene_tx=XM_023251675.1;refGene_gn=PSME4;refGene_tx=XM_023251676.1
A3 75155715 75155957
refGene_gn=PSME4;refGene_tx=XM_023251673.1;refGene_gn=PSME4;refGene_tx=XM_023251674.1;refGene_gn=PSME4;refGene_tx=XM_023251675.1
A3 75258496 75258554 refGene_gn=ACYP2;refGene_tx=XM_006930104.4
A3 75277905 75277922
refGene_gn=ACYP2;refGene_tx=XM_006930104.4;refGene_gn=ACYP2;refGene_tx=XM_019827199.2

A3 75279327 75279437
refGene_gn=ACYP2;refGene_tx=XM_006930104.4;refGene_gn=ACYP2;refGene_tx=XM_019827199.2
A3 75383167 75383486 refGene_gn=TSPYL6;refGene_tx=XM_023250922.1
A3 75383487 75383995 refGene_gn=TSPYL6;refGene_tx=XM_023250922.1
A3 75384346 75384383 refGene_gn=TSPYL6;refGene_tx=XM_023250922.1
A3 75431066 75431181
refGene_gn=ACYP2;refGene_tx=XM_006930104.4;refGene_gn=ACYP2;refGene_tx=XM_019827199.2
A3 75455532 75455558
refGene_gn=CA3H2orf73;refGene_tx=XM_019827198.2
A3 75459863 75460077
refGene_gn=CA3H2orf73;refGene_tx=XM_019827198.2
A3 75466816 75466970
refGene_gn=CA3H2orf73;refGene_tx=XM_019827198.2
A3 75480024 75480140
refGene_gn=CA3H2orf73;refGene_tx=XM_019827198.2
A3 75480928 75481297
refGene_gn=CA3H2orf73;refGene_tx=XM_019827198.2
A3 75638379 75638527
refGene_gn=SPTBN1;refGene_tx=XM_023251677.1;refGene_gn=SPTBN1;refGene_tx=XM_023251678.1
A3 75670550 75670659
refGene_gn=SPTBN1;refGene_tx=XM_003983998.5;refGene_gn=SPTBN1;refGene_tx=XM_023251679.1
A3 75709337 75709489
refGene_gn=SPTBN1;refGene_tx=XM_003983998.5;refGene_gn=SPTBN1;refGene_tx=XM_023251677.1;refGene_gn=SPTBN1;refGene_tx=XM_023251678.1;refGene_gn=SPTBN1;refGene_tx=XM_023251679.1
A3 75719926 75720100
refGene_gn=SPTBN1;refGene_tx=XM_003983998.5;refGene_gn=SPTBN1;refGene_tx=XM_023251677.1;refGene_gn=SPTBN1;refGene_tx=XM_023251678.1;refGene_gn=SPTBN1;refGene_tx=XM_023251679.1
A3 75723896 75723988
refGene_gn=SPTBN1;refGene_tx=XM_003983998.5;refGene_gn=SPTBN1;refGene_tx=XM_023251677.1;refGene_gn=SPTBN1;refGene_tx=XM_023251678.1;refGene_gn=SPTBN1;refGene_tx=XM_023251679.1
A3 75725204 75725285
refGene_gn=SPTBN1;refGene_tx=XM_003983998.5;refGene_gn=SPTBN1;refGene_tx=XM_023251677.1;refGene_gn=SPTBN1;refGene_tx=XM_023251678.1;refGene_gn=SPTBN1;refGene_tx=XM_023251679.1
A3 75725674 75725790
refGene_gn=SPTBN1;refGene_tx=XM_003983998.5;refGene_gn=SPTBN1;refGene_tx=XM_023251677.1;refGene_gn=SPTBN1;refGene_tx=XM_023251678.1;refGene_gn=SPTBN1;refGene_tx=XM_023251679.1
A3 75729052 75729165
refGene_gn=SPTBN1;refGene_tx=XM_003983998.5;refGene_gn=SPTBN1;refGene_tx=XM_023251677.1;refGene_gn=SPTBN1;refGene_tx=XM_023251678.1;refGene_gn=SPTBN1;refGene_tx=XM_023251679.1
A3 75729922 75730110

refGene_gn=SPTBN1;refGene_tx=XM_003983998.5;refGene_gn=SPTBN1;refGene_tx=XM_023251677.1;refGene_gn=SPTBN1;refGene_tx=XM_023251678.1;refGene_gn=SPTBN1;refGene_tx=XM_023251679.1
A3 75731104 75731222
refGene_gn=SPTBN1;refGene_tx=XM_003983998.5;refGene_gn=SPTBN1;refGene_tx=XM_023251677.1;refGene_gn=SPTBN1;refGene_tx=XM_023251678.1;refGene_gn=SPTBN1;refGene_tx=XM_023251679.1
A3 75732843 75733002
refGene_gn=SPTBN1;refGene_tx=XM_003983998.5;refGene_gn=SPTBN1;refGene_tx=XM_023251677.1;refGene_gn=SPTBN1;refGene_tx=XM_023251678.1;refGene_gn=SPTBN1;refGene_tx=XM_023251679.1
A3 75733668 75733971
refGene_gn=SPTBN1;refGene_tx=XM_003983998.5;refGene_gn=SPTBN1;refGene_tx=XM_023251677.1;refGene_gn=SPTBN1;refGene_tx=XM_023251678.1;refGene_gn=SPTBN1;refGene_tx=XM_023251679.1
A3 75735712 75735866
refGene_gn=SPTBN1;refGene_tx=XM_003983998.5;refGene_gn=SPTBN1;refGene_tx=XM_023251677.1;refGene_gn=SPTBN1;refGene_tx=XM_023251678.1;refGene_gn=SPTBN1;refGene_tx=XM_023251679.1
A3 75736542 75737413
refGene_gn=SPTBN1;refGene_tx=XM_003983998.5;refGene_gn=SPTBN1;refGene_tx=XM_023251677.1;refGene_gn=SPTBN1;refGene_tx=XM_023251678.1;refGene_gn=SPTBN1;refGene_tx=XM_023251679.1
A3 75737512 75737650
refGene_gn=SPTBN1;refGene_tx=XM_003983998.5;refGene_gn=SPTBN1;refGene_tx=XM_023251677.1;refGene_gn=SPTBN1;refGene_tx=XM_023251678.1;refGene_gn=SPTBN1;refGene_tx=XM_023251679.1
A3 75738453 75739210
refGene_gn=SPTBN1;refGene_tx=XM_003983998.5;refGene_gn=SPTBN1;refGene_tx=XM_023251677.1;refGene_gn=SPTBN1;refGene_tx=XM_023251678.1;refGene_gn=SPTBN1;refGene_tx=XM_023251679.1
A3 75740063 75740266
refGene_gn=SPTBN1;refGene_tx=XM_003983998.5;refGene_gn=SPTBN1;refGene_tx=XM_023251677.1;refGene_gn=SPTBN1;refGene_tx=XM_023251678.1;refGene_gn=SPTBN1;refGene_tx=XM_023251679.1
A3 75745251 75745342
refGene_gn=SPTBN1;refGene_tx=XM_003983998.5;refGene_gn=SPTBN1;refGene_tx=XM_023251677.1;refGene_gn=SPTBN1;refGene_tx=XM_023251678.1;refGene_gn=SPTBN1;refGene_tx=XM_023251679.1
A3 75750543 75750690
refGene_gn=SPTBN1;refGene_tx=XM_003983998.5;refGene_gn=SPTBN1;refGene_tx=XM_023251677.1;refGene_gn=SPTBN1;refGene_tx=XM_023251678.1;refGene_gn=SPTBN1;refGene_tx=XM_023251679.1
A3 75751540 75751804
refGene_gn=SPTBN1;refGene_tx=XM_003983998.5;refGene_gn=SPTBN1;refGene_tx=XM_023251677.1;refGene_gn=SPTBN1;refGene_tx=XM_023251678.1;refGene_gn=SPTBN1;refGene_tx=XM_023251679.1
A3 75753170 75753395
refGene_gn=SPTBN1;refGene_tx=XM_003983998.5;refGene_gn=SPTBN1;refGene_tx=XM_023251677.1;refGene_gn=SPTBN1;refGene_tx=XM_023251678.1;refGene_

gn=SPTBN1;refGene_tx=XM_023251679.1
A3 75753867 75753957
refGene_gn=SPTBN1;refGene_tx=XM_003983998.5;refGene_gn=SPTBN1;refGene_tx=XM_023251677.1;refGene_gn=SPTBN1;refGene_tx=XM_023251678.1;refGene_gn=SPTBN1;refGene_tx=XM_023251679.1
A3 75754124 75754406
refGene_gn=SPTBN1;refGene_tx=XM_003983998.5;refGene_gn=SPTBN1;refGene_tx=XM_023251677.1;refGene_gn=SPTBN1;refGene_tx=XM_023251678.1;refGene_gn=SPTBN1;refGene_tx=XM_023251679.1
A3 75755052 75755183
refGene_gn=SPTBN1;refGene_tx=XM_003983998.5;refGene_gn=SPTBN1;refGene_tx=XM_023251677.1;refGene_gn=SPTBN1;refGene_tx=XM_023251678.1;refGene_gn=SPTBN1;refGene_tx=XM_023251679.1
A3 75756451 75756656
refGene_gn=SPTBN1;refGene_tx=XM_003983998.5;refGene_gn=SPTBN1;refGene_tx=XM_023251677.1;refGene_gn=SPTBN1;refGene_tx=XM_023251678.1;refGene_gn=SPTBN1;refGene_tx=XM_023251679.1
A3 75757038 75757413
refGene_gn=SPTBN1;refGene_tx=XM_003983998.5;refGene_gn=SPTBN1;refGene_tx=XM_023251677.1;refGene_gn=SPTBN1;refGene_tx=XM_023251678.1;refGene_gn=SPTBN1;refGene_tx=XM_023251679.1
A3 75761427 75761672
refGene_gn=SPTBN1;refGene_tx=XM_003983998.5;refGene_gn=SPTBN1;refGene_tx=XM_023251677.1;refGene_gn=SPTBN1;refGene_tx=XM_023251678.1;refGene_gn=SPTBN1;refGene_tx=XM_023251679.1
A3 75762904 75763043
refGene_gn=SPTBN1;refGene_tx=XM_003983998.5;refGene_gn=SPTBN1;refGene_tx=XM_023251677.1;refGene_gn=SPTBN1;refGene_tx=XM_023251678.1;refGene_gn=SPTBN1;refGene_tx=XM_023251679.1
A3 75763840 75763925
refGene_gn=SPTBN1;refGene_tx=XM_003983998.5;refGene_gn=SPTBN1;refGene_tx=XM_023251677.1;refGene_gn=SPTBN1;refGene_tx=XM_023251678.1;refGene_gn=SPTBN1;refGene_tx=XM_023251679.1
A3 75765476 75765673
refGene_gn=SPTBN1;refGene_tx=XM_003983998.5;refGene_gn=SPTBN1;refGene_tx=XM_023251677.1;refGene_gn=SPTBN1;refGene_tx=XM_023251678.1;refGene_gn=SPTBN1;refGene_tx=XM_023251679.1
A3 75766792 75766905
refGene_gn=SPTBN1;refGene_tx=XM_003983998.5;refGene_gn=SPTBN1;refGene_tx=XM_023251677.1;refGene_gn=SPTBN1;refGene_tx=XM_023251678.1;refGene_gn=SPTBN1;refGene_tx=XM_023251679.1
A3 75767591 75767742
refGene_gn=SPTBN1;refGene_tx=XM_003983998.5;refGene_gn=SPTBN1;refGene_tx=XM_023251677.1;refGene_gn=SPTBN1;refGene_tx=XM_023251678.1;refGene_gn=SPTBN1;refGene_tx=XM_023251679.1
A3 75772045 75772284
refGene_gn=SPTBN1;refGene_tx=XM_023251677.1;refGene_gn=SPTBN1;refGene_tx=XM_023251678.1;refGene_gn=SPTBN1;refGene_tx=XM_023251679.1
A3 75773506 75773680
refGene_gn=SPTBN1;refGene_tx=XM_023251677.1;refGene_gn=SPTBN1;refGene_

tx=XM_023251678.1;refGene_gn=SPTBN1;refGene_tx=XM_023251679.1
A3 75780431 75780474
refGene_gn=SPTBN1;refGene_tx=XM_023251677.1;refGene_gn=SPTBN1;refGene_tx=XM_023251678.1;refGene_gn=SPTBN1;refGene_tx=XM_023251679.1
A3 75781336 75781555
refGene_gn=SPTBN1;refGene_tx=XM_023251677.1;refGene_gn=SPTBN1;refGene_tx=XM_023251678.1;refGene_gn=SPTBN1;refGene_tx=XM_023251679.1
A3 75828413 75828610
refGene_gn=EML6;refGene_tx=XM_003983999.5;refGene_gn=EML6;refGene_tx=XM_019827201.2;refGene_gn=EML6;refGene_tx=XM_019827202.2;refGene_gn=EML6;refGene_tx=XM_019827203.2;refGene_gn=EML6;refGene_tx=XM_019827204.2;refGene_gn=EML6;refGene_tx=XM_019827205.2
A3 75939323 75939483
refGene_gn=EML6;refGene_tx=XM_003983999.5;refGene_gn=EML6;refGene_tx=XM_019827201.2;refGene_gn=EML6;refGene_tx=XM_019827202.2;refGene_gn=EML6;refGene_tx=XM_019827203.2;refGene_gn=EML6;refGene_tx=XM_019827204.2;refGene_gn=EML6;refGene_tx=XM_019827205.2
A3 75943158 75943257
refGene_gn=EML6;refGene_tx=XM_003983999.5;refGene_gn=EML6;refGene_tx=XM_019827201.2;refGene_gn=EML6;refGene_tx=XM_019827202.2;refGene_gn=EML6;refGene_tx=XM_019827203.2;refGene_gn=EML6;refGene_tx=XM_019827204.2;refGene_gn=EML6;refGene_tx=XM_019827205.2
A3 75946088 75946157
refGene_gn=EML6;refGene_tx=XM_003983999.5;refGene_gn=EML6;refGene_tx=XM_019827201.2;refGene_gn=EML6;refGene_tx=XM_019827202.2;refGene_gn=EML6;refGene_tx=XM_019827203.2;refGene_gn=EML6;refGene_tx=XM_019827204.2;refGene_gn=EML6;refGene_tx=XM_019827205.2
A3 75954002 75954188
refGene_gn=EML6;refGene_tx=XM_003983999.5;refGene_gn=EML6;refGene_tx=XM_019827201.2;refGene_gn=EML6;refGene_tx=XM_019827202.2;refGene_gn=EML6;refGene_tx=XM_019827203.2;refGene_gn=EML6;refGene_tx=XM_019827204.2;refGene_gn=EML6;refGene_tx=XM_019827205.2
A3 75956023 75956159
refGene_gn=EML6;refGene_tx=XM_003983999.5;refGene_gn=EML6;refGene_tx=XM_019827201.2;refGene_gn=EML6;refGene_tx=XM_019827202.2;refGene_gn=EML6;refGene_tx=XM_019827203.2;refGene_gn=EML6;refGene_tx=XM_019827204.2;refGene_gn=EML6;refGene_tx=XM_019827205.2
A3 75985023 75985225
refGene_gn=EML6;refGene_tx=XM_003983999.5;refGene_gn=EML6;refGene_tx=XM_019827201.2;refGene_gn=EML6;refGene_tx=XM_019827202.2;refGene_gn=EML6;refGene_tx=XM_019827203.2;refGene_gn=EML6;refGene_tx=XM_019827204.2;refGene_gn=EML6;refGene_tx=XM_019827205.2
A3 75987781 75987919
refGene_gn=EML6;refGene_tx=XM_003983999.5;refGene_gn=EML6;refGene_tx=XM_019827201.2;refGene_gn=EML6;refGene_tx=XM_019827202.2;refGene_gn=EML6;refGene_tx=XM_019827203.2;refGene_gn=EML6;refGene_tx=XM_019827204.2;refGene_gn=EML6;refGene_tx=XM_019827205.2
A3 75989652 75989909
refGene_gn=EML6;refGene_tx=XM_003983999.5;refGene_gn=EML6;refGene_tx=XM_019827201.2;refGene_gn=EML6;refGene_tx=XM_019827202.2;refGene_gn=EML

6;refGene_tx=XM_019827203.2;refGene_gn=EML6;refGene_tx=XM_019827204.2;
refGene_gn=EML6;refGene_tx=XM_019827205.2
A3 75992141 75992354
refGene_gn=EML6;refGene_tx=XM_003983999.5;refGene_gn=EML6;refGene_tx=XM_019827201.2;refGene_gn=EML6;refGene_tx=XM_019827202.2;refGene_gn=EML6;refGene_tx=XM_019827203.2;refGene_gn=EML6;refGene_tx=XM_019827204.2;refGene_gn=EML6;refGene_tx=XM_019827205.2
A3 75996681 75996849
refGene_gn=EML6;refGene_tx=XM_003983999.5;refGene_gn=EML6;refGene_tx=XM_019827201.2;refGene_gn=EML6;refGene_tx=XM_019827202.2;refGene_gn=EML6;refGene_tx=XM_019827203.2;refGene_gn=EML6;refGene_tx=XM_019827204.2;refGene_gn=EML6;refGene_tx=XM_019827205.2
A3 75997950 75998057
refGene_gn=EML6;refGene_tx=XM_003983999.5;refGene_gn=EML6;refGene_tx=XM_019827201.2;refGene_gn=EML6;refGene_tx=XM_019827202.2;refGene_gn=EML6;refGene_tx=XM_019827203.2;refGene_gn=EML6;refGene_tx=XM_019827204.2;refGene_gn=EML6;refGene_tx=XM_019827205.2
A3 76001766 76001885
refGene_gn=EML6;refGene_tx=XM_003983999.5;refGene_gn=EML6;refGene_tx=XM_019827201.2;refGene_gn=EML6;refGene_tx=XM_019827202.2;refGene_gn=EML6;refGene_tx=XM_019827203.2;refGene_gn=EML6;refGene_tx=XM_019827204.2;refGene_gn=EML6;refGene_tx=XM_019827205.2
A3 76004335 76004522
refGene_gn=EML6;refGene_tx=XM_003983999.5;refGene_gn=EML6;refGene_tx=XM_019827201.2;refGene_gn=EML6;refGene_tx=XM_019827202.2;refGene_gn=EML6;refGene_tx=XM_019827203.2;refGene_gn=EML6;refGene_tx=XM_019827204.2;refGene_gn=EML6;refGene_tx=XM_019827205.2
A3 76005795 76005901
refGene_gn=EML6;refGene_tx=XM_003983999.5;refGene_gn=EML6;refGene_tx=XM_019827201.2;refGene_gn=EML6;refGene_tx=XM_019827202.2;refGene_gn=EML6;refGene_tx=XM_019827203.2;refGene_gn=EML6;refGene_tx=XM_019827204.2;refGene_gn=EML6;refGene_tx=XM_019827205.2
A3 76012785 76012879
refGene_gn=EML6;refGene_tx=XM_003983999.5;refGene_gn=EML6;refGene_tx=XM_019827201.2;refGene_gn=EML6;refGene_tx=XM_019827202.2;refGene_gn=EML6;refGene_tx=XM_019827203.2;refGene_gn=EML6;refGene_tx=XM_019827204.2;refGene_gn=EML6;refGene_tx=XM_019827205.2
A3 76022712 76022813
refGene_gn=EML6;refGene_tx=XM_003983999.5;refGene_gn=EML6;refGene_tx=XM_019827201.2;refGene_gn=EML6;refGene_tx=XM_019827202.2;refGene_gn=EML6;refGene_tx=XM_019827203.2;refGene_gn=EML6;refGene_tx=XM_019827204.2;refGene_gn=EML6;refGene_tx=XM_019827205.2
A3 76024105 76024308
refGene_gn=EML6;refGene_tx=XM_003983999.5;refGene_gn=EML6;refGene_tx=XM_019827201.2;refGene_gn=EML6;refGene_tx=XM_019827202.2;refGene_gn=EML6;refGene_tx=XM_019827203.2;refGene_gn=EML6;refGene_tx=XM_019827204.2;refGene_gn=EML6;refGene_tx=XM_019827205.2
A3 76026964 76027076
refGene_gn=EML6;refGene_tx=XM_003983999.5;refGene_gn=EML6;refGene_tx=XM_019827201.2;refGene_gn=EML6;refGene_tx=XM_019827202.2;refGene_gn=EML

6;refGene_tx=XM_019827203.2;refGene_gn=EML6;refGene_tx=XM_019827204.2;
refGene_gn=EML6;refGene_tx=XM_019827205.2
A3 76027336 76027464
refGene_gn=EML6;refGene_tx=XM_003983999.5;refGene_gn=EML6;refGene_tx=X
M_019827201.2;refGene_gn=EML6;refGene_tx=XM_019827202.2;refGene_gn=EML
6;refGene_tx=XM_019827203.2;refGene_gn=EML6;refGene_tx=XM_019827204.2;
refGene_gn=EML6;refGene_tx=XM_019827205.2
A3 76031840 76031982
refGene_gn=EML6;refGene_tx=XM_003983999.5;refGene_gn=EML6;refGene_tx=X
M_019827201.2;refGene_gn=EML6;refGene_tx=XM_019827202.2;refGene_gn=EML
6;refGene_tx=XM_019827203.2;refGene_gn=EML6;refGene_tx=XM_019827204.2;
refGene_gn=EML6;refGene_tx=XM_019827205.2
A3 76035178 76035331
refGene_gn=EML6;refGene_tx=XM_003983999.5;refGene_gn=EML6;refGene_tx=X
M_019827201.2;refGene_gn=EML6;refGene_tx=XM_019827202.2;refGene_gn=EML
6;refGene_tx=XM_019827203.2;refGene_gn=EML6;refGene_tx=XM_019827204.2;
refGene_gn=EML6;refGene_tx=XM_019827205.2
A3 76035494 76035626
refGene_gn=EML6;refGene_tx=XM_003983999.5;refGene_gn=EML6;refGene_tx=X
M_019827201.2;refGene_gn=EML6;refGene_tx=XM_019827202.2;refGene_gn=EML
6;refGene_tx=XM_019827203.2;refGene_gn=EML6;refGene_tx=XM_019827204.2;
refGene_gn=EML6;refGene_tx=XM_019827205.2
A3 76042244 76042333
refGene_gn=EML6;refGene_tx=XM_003983999.5;refGene_gn=EML6;refGene_tx=X
M_019827201.2;refGene_gn=EML6;refGene_tx=XM_019827202.2;refGene_gn=EML
6;refGene_tx=XM_019827203.2;refGene_gn=EML6;refGene_tx=XM_019827204.2;
refGene_gn=EML6;refGene_tx=XM_019827205.2
A3 76046900 76047077
refGene_gn=EML6;refGene_tx=XM_003983999.5;refGene_gn=EML6;refGene_tx=X
M_019827201.2;refGene_gn=EML6;refGene_tx=XM_019827202.2;refGene_gn=EML
6;refGene_tx=XM_019827203.2;refGene_gn=EML6;refGene_tx=XM_019827204.2;
refGene_gn=EML6;refGene_tx=XM_019827205.2
A3 76055193 76055395
refGene_gn=EML6;refGene_tx=XM_003983999.5;refGene_gn=EML6;refGene_tx=X
M_019827201.2;refGene_gn=EML6;refGene_tx=XM_019827202.2;refGene_gn=EML
6;refGene_tx=XM_019827203.2;refGene_gn=EML6;refGene_tx=XM_019827204.2;
refGene_gn=EML6;refGene_tx=XM_019827205.2
A3 76055502 76055629
refGene_gn=EML6;refGene_tx=XM_003983999.5;refGene_gn=EML6;refGene_tx=X
M_019827201.2;refGene_gn=EML6;refGene_tx=XM_019827202.2;refGene_gn=EML
6;refGene_tx=XM_019827203.2;refGene_gn=EML6;refGene_tx=XM_019827204.2;
refGene_gn=EML6;refGene_tx=XM_019827205.2
A3 76076688 76076703
refGene_gn=EML6;refGene_tx=XM_019827201.2;refGene_gn=EML6;refGene_tx=X
M_019827203.2;refGene_gn=EML6;refGene_tx=XM_019827205.2
A3 76078027 76078036
refGene_gn=EML6;refGene_tx=XM_019827201.2;refGene_gn=EML6;refGene_tx=X
M_019827202.2;refGene_gn=EML6;refGene_tx=XM_019827203.2;refGene_gn=EML
6;refGene_tx=XM_019827205.2
A3 76078922 76079001

refGene_gn=EML6;refGene_tx=XM_003983999.5;refGene_gn=EML6;refGene_tx=XM_019827201.2;refGene_gn=EML6;refGene_tx=XM_019827202.2;refGene_gn=EML6;refGene_tx=XM_019827203.2;refGene_gn=EML6;refGene_tx=XM_019827204.2;refGene_gn=EML6;refGene_tx=XM_019827205.2
A3 76081432 76081562
refGene_gn=EML6;refGene_tx=XM_003983999.5;refGene_gn=EML6;refGene_tx=XM_019827201.2;refGene_gn=EML6;refGene_tx=XM_019827202.2;refGene_gn=EML6;refGene_tx=XM_019827203.2;refGene_gn=EML6;refGene_tx=XM_019827204.2;refGene_gn=EML6;refGene_tx=XM_019827205.2
A3 76083032 76083131
refGene_gn=EML6;refGene_tx=XM_003983999.5;refGene_gn=EML6;refGene_tx=XM_019827201.2;refGene_gn=EML6;refGene_tx=XM_019827202.2;refGene_gn=EML6;refGene_tx=XM_019827203.2;refGene_gn=EML6;refGene_tx=XM_019827204.2;refGene_gn=EML6;refGene_tx=XM_019827205.2
A3 76083762 76083783
refGene_gn=EML6;refGene_tx=XM_019827201.2;refGene_gn=EML6;refGene_tx=XM_019827202.2;refGene_gn=EML6;refGene_tx=XM_019827204.2;refGene_gn=EML6;refGene_tx=XM_019827205.2
A3 76084146 76084320
refGene_gn=EML6;refGene_tx=XM_003983999.5;refGene_gn=EML6;refGene_tx=XM_019827201.2;refGene_gn=EML6;refGene_tx=XM_019827202.2;refGene_gn=EML6;refGene_tx=XM_019827203.2;refGene_gn=EML6;refGene_tx=XM_019827204.2;refGene_gn=EML6;refGene_tx=XM_019827205.2
A3 76088185 76088394
refGene_gn=EML6;refGene_tx=XM_003983999.5;refGene_gn=EML6;refGene_tx=XM_019827201.2;refGene_gn=EML6;refGene_tx=XM_019827202.2;refGene_gn=EML6;refGene_tx=XM_019827203.2;refGene_gn=EML6;refGene_tx=XM_019827204.2;refGene_gn=EML6;refGene_tx=XM_019827205.2
A3 76088689 76088847
refGene_gn=EML6;refGene_tx=XM_003983999.5;refGene_gn=EML6;refGene_tx=XM_019827201.2;refGene_gn=EML6;refGene_tx=XM_019827202.2;refGene_gn=EML6;refGene_tx=XM_019827203.2;refGene_gn=EML6;refGene_tx=XM_019827204.2;refGene_gn=EML6;refGene_tx=XM_019827205.2
A3 76089479 76089594
refGene_gn=EML6;refGene_tx=XM_003983999.5;refGene_gn=EML6;refGene_tx=XM_019827201.2;refGene_gn=EML6;refGene_tx=XM_019827202.2;refGene_gn=EML6;refGene_tx=XM_019827203.2;refGene_gn=EML6;refGene_tx=XM_019827204.2;refGene_gn=EML6;refGene_tx=XM_019827205.2
A3 76091194 76091383
refGene_gn=EML6;refGene_tx=XM_003983999.5;refGene_gn=EML6;refGene_tx=XM_019827201.2;refGene_gn=EML6;refGene_tx=XM_019827202.2;refGene_gn=EML6;refGene_tx=XM_019827203.2;refGene_gn=EML6;refGene_tx=XM_019827204.2;refGene_gn=EML6;refGene_tx=XM_019827205.2
A3 76093067 76093240
refGene_gn=EML6;refGene_tx=XM_003983999.5;refGene_gn=EML6;refGene_tx=XM_019827201.2;refGene_gn=EML6;refGene_tx=XM_019827202.2;refGene_gn=EML6;refGene_tx=XM_019827203.2;refGene_gn=EML6;refGene_tx=XM_019827204.2;refGene_gn=EML6;refGene_tx=XM_019827205.2
A3 76093657 76093820
refGene_gn=EML6;refGene_tx=XM_003983999.5;refGene_gn=EML6;refGene_tx=X

M_019827201.2;refGene_gn=EML6;refGene_tx=XM_019827202.2;refGene_gn=EML6;refGene_tx=XM_019827203.2;refGene_gn=EML6;refGene_tx=XM_019827204.2;refGene_gn=EML6;refGene_tx=XM_019827205.2
A3 76095435 76095539
refGene_gn=EML6;refGene_tx=XM_003983999.5;refGene_gn=EML6;refGene_tx=XM_019827201.2;refGene_gn=EML6;refGene_tx=XM_019827202.2;refGene_gn=EML6;refGene_tx=XM_019827203.2;refGene_gn=EML6;refGene_tx=XM_019827204.2;refGene_gn=EML6;refGene_tx=XM_019827205.2
A3 76096448 76096602
refGene_gn=EML6;refGene_tx=XM_003983999.5;refGene_gn=EML6;refGene_tx=XM_019827201.2;refGene_gn=EML6;refGene_tx=XM_019827202.2;refGene_gn=EML6;refGene_tx=XM_019827203.2;refGene_gn=EML6;refGene_tx=XM_019827204.2
A3 76097004 76097105
refGene_gn=EML6;refGene_tx=XM_003983999.5;refGene_gn=EML6;refGene_tx=XM_019827201.2;refGene_gn=EML6;refGene_tx=XM_019827202.2;refGene_gn=EML6;refGene_tx=XM_019827203.2;refGene_gn=EML6;refGene_tx=XM_019827204.2
A3 76099550 76099575
refGene_gn=EML6;refGene_tx=XM_003983999.5;refGene_gn=EML6;refGene_tx=XM_019827201.2;refGene_gn=EML6;refGene_tx=XM_019827202.2;refGene_gn=EML6;refGene_tx=XM_019827203.2;refGene_gn=EML6;refGene_tx=XM_019827204.2;refGene_gn=EML6;refGene_tx=XM_019827205.2
A3 76102571 76102614
refGene_gn=RTN4;refGene_tx=XM_019827207.2;refGene_gn=RTN4;refGene_tx=XM_019827208.2;refGene_gn=RTN4;refGene_tx=XM_019827209.2;refGene_gn=RTN4;refGene_tx=XM_019827210.2
A3 76102993 76103052
refGene_gn=RTN4;refGene_tx=XM_019827207.2;refGene_gn=RTN4;refGene_tx=XM_019827208.2;refGene_gn=RTN4;refGene_tx=XM_019827209.2;refGene_gn=RTN4;refGene_tx=XM_019827210.2
A3 76103260 76103307
refGene_gn=RTN4;refGene_tx=XM_019827207.2;refGene_gn=RTN4;refGene_tx=XM_019827208.2;refGene_gn=RTN4;refGene_tx=XM_019827209.2;refGene_gn=RTN4;refGene_tx=XM_019827210.2
A3 76104038 76104108
refGene_gn=RTN4;refGene_tx=XM_019827207.2;refGene_gn=RTN4;refGene_tx=XM_019827208.2;refGene_gn=RTN4;refGene_tx=XM_019827209.2;refGene_gn=RTN4;refGene_tx=XM_019827210.2
A3 76112882 76113021
refGene_gn=RTN4;refGene_tx=XM_019827207.2;refGene_gn=RTN4;refGene_tx=XM_019827208.2;refGene_gn=RTN4;refGene_tx=XM_019827209.2;refGene_gn=RTN4;refGene_tx=XM_019827210.2
A3 76117494 76117702
refGene_gn=RTN4;refGene_tx=XM_019827207.2;refGene_gn=RTN4;refGene_tx=XM_019827208.2;refGene_gn=RTN4;refGene_tx=XM_019827209.2;refGene_gn=RTN4;refGene_tx=XM_019827210.2
A3 76137182 76137216 refGene_gn=RTN4;refGene_tx=XM_019827210.2
A3 76148410 76150801 refGene_gn=RTN4;refGene_tx=XM_019827207.2
A3 76151472 76151529
refGene_gn=RTN4;refGene_tx=XM_019827207.2;refGene_gn=RTN4;refGene_tx=XM_019827208.2

A3 76173794 76174353
refGene_gn=RTN4;refGene_tx=XM_019827207.2;refGene_gn=RTN4;refGene_tx=XM_019827208.2;refGene_gn=RTN4;refGene_tx=XM_019827209.2
A3 76337295 76337492
refGene_gn=CLHC1;refGene_tx=XM_003984007.3;refGene_gn=CLHC1;refGene_tx=XM_019827218.2;refGene_gn=CLHC1;refGene_tx=XM_019827221.1;refGene_gn=CLHC1;refGene_tx=XM_019827222.1;refGene_gn=CLHC1;refGene_tx=XM_019827226.1;refGene_gn=CLHC1;refGene_tx=XM_019827228.1;refGene_gn=CLHC1;refGene_tx=XM_019827229.2
A3 76337945 76338031 refGene_gn=CLHC1;refGene_tx=XM_019827223.1
A3 76339267 76339447
refGene_gn=CLHC1;refGene_tx=XM_003984007.3;refGene_gn=CLHC1;refGene_tx=XM_019827218.2;refGene_gn=CLHC1;refGene_tx=XM_019827221.1;refGene_gn=CLHC1;refGene_tx=XM_019827222.1;refGene_gn=CLHC1;refGene_tx=XM_019827223.1;refGene_gn=CLHC1;refGene_tx=XM_019827226.1;refGene_gn=CLHC1;refGene_tx=XM_019827228.1;refGene_gn=CLHC1;refGene_tx=XM_019827229.2
A3 76345908 76346111
refGene_gn=CLHC1;refGene_tx=XM_003984007.3;refGene_gn=CLHC1;refGene_tx=XM_019827218.2;refGene_gn=CLHC1;refGene_tx=XM_019827221.1;refGene_gn=CLHC1;refGene_tx=XM_019827222.1;refGene_gn=CLHC1;refGene_tx=XM_019827223.1;refGene_gn=CLHC1;refGene_tx=XM_019827226.1;refGene_gn=CLHC1;refGene_tx=XM_019827228.1;refGene_gn=CLHC1;refGene_tx=XM_019827229.2
A3 76352875 76353050
refGene_gn=CLHC1;refGene_tx=XM_003984007.3;refGene_gn=CLHC1;refGene_tx=XM_019827218.2;refGene_gn=CLHC1;refGene_tx=XM_019827221.1;refGene_gn=CLHC1;refGene_tx=XM_019827222.1;refGene_gn=CLHC1;refGene_tx=XM_019827223.1;refGene_gn=CLHC1;refGene_tx=XM_019827226.1;refGene_gn=CLHC1;refGene_tx=XM_019827228.1;refGene_gn=CLHC1;refGene_tx=XM_019827229.2
A3 76359937 76360044
refGene_gn=CLHC1;refGene_tx=XM_003984007.3;refGene_gn=CLHC1;refGene_tx=XM_019827218.2;refGene_gn=CLHC1;refGene_tx=XM_019827221.1;refGene_gn=CLHC1;refGene_tx=XM_019827222.1;refGene_gn=CLHC1;refGene_tx=XM_019827223.1;refGene_gn=CLHC1;refGene_tx=XM_019827226.1;refGene_gn=CLHC1;refGene_tx=XM_019827228.1;refGene_gn=CLHC1;refGene_tx=XM_019827229.2
A3 76361552 76361637
refGene_gn=CLHC1;refGene_tx=XM_003984007.3;refGene_gn=CLHC1;refGene_tx=XM_019827218.2;refGene_gn=CLHC1;refGene_tx=XM_019827221.1;refGene_gn=CLHC1;refGene_tx=XM_019827222.1;refGene_gn=CLHC1;refGene_tx=XM_019827223.1;refGene_gn=CLHC1;refGene_tx=XM_019827226.1;refGene_gn=CLHC1;refGene_tx=XM_019827228.1;refGene_gn=CLHC1;refGene_tx=XM_019827229.2
A3 76362731 76362844
refGene_gn=CLHC1;refGene_tx=XM_003984007.3;refGene_gn=CLHC1;refGene_tx=XM_019827218.2;refGene_gn=CLHC1;refGene_tx=XM_019827221.1;refGene_gn=CLHC1;refGene_tx=XM_019827222.1;refGene_gn=CLHC1;refGene_tx=XM_019827223.1;refGene_gn=CLHC1;refGene_tx=XM_019827226.1;refGene_gn=CLHC1;refGene_tx=XM_019827228.1;refGene_gn=CLHC1;refGene_tx=XM_019827229.2
A3 76362954 76363156
refGene_gn=CLHC1;refGene_tx=XM_003984007.3;refGene_gn=CLHC1;refGene_tx=XM_019827218.2;refGene_gn=CLHC1;refGene_tx=XM_019827221.1;refGene_gn=CLHC1;refGene_tx=XM_019827222.1;refGene_gn=CLHC1;refGene_tx=XM_0198272

23.1;refGene_gn=CLHC1;refGene_tx=XM_019827226.1;refGene_gn=CLHC1;refGene_tx=XM_019827228.1;refGene_gn=CLHC1;refGene_tx=XM_019827229.2
A3	76364584	76364718
refGene_gn=CLHC1;refGene_tx=XM_003984007.3;refGene_gn=CLHC1;refGene_tx=XM_019827218.2;refGene_gn=CLHC1;refGene_tx=XM_019827221.1;refGene_gn=CLHC1;refGene_tx=XM_019827222.1;refGene_gn=CLHC1;refGene_tx=XM_019827223.1;refGene_gn=CLHC1;refGene_tx=XM_019827226.1;refGene_gn=CLHC1;refGene_tx=XM_019827228.1;refGene_gn=CLHC1;refGene_tx=XM_019827229.2
A3	76368514	76368702
refGene_gn=CLHC1;refGene_tx=XM_003984007.3;refGene_gn=CLHC1;refGene_tx=XM_019827218.2;refGene_gn=CLHC1;refGene_tx=XM_019827221.1;refGene_gn=CLHC1;refGene_tx=XM_019827222.1;refGene_gn=CLHC1;refGene_tx=XM_019827223.1
A3	76370590	76370722
refGene_gn=CLHC1;refGene_tx=XM_019827221.1;refGene_gn=CLHC1;refGene_tx=XM_019827222.1
A3	76372840	76373017
refGene_gn=CLHC1;refGene_tx=XM_003984007.3;refGene_gn=CLHC1;refGene_tx=XM_019827218.2;refGene_gn=CLHC1;refGene_tx=XM_019827223.1;refGene_gn=CLHC1;refGene_tx=XM_019827226.1
A3	76378402	76378450	refGene_gn=RPS27A;refGene_tx=XM_003984008.5
A3	76379106	76379161	refGene_gn=RPS27A;refGene_tx=XM_003984008.5
A3	76379555	76379641	refGene_gn=RPS27A;refGene_tx=XM_003984008.5
A3	76380006	76380138	refGene_gn=RPS27A;refGene_tx=XM_003984008.5
A3	76380705	76380855	refGene_gn=RPS27A;refGene_tx=XM_003984008.5
A3	76381921	76382094
refGene_gn=MTIF2;refGene_tx=XM_006930105.3;refGene_gn=MTIF2;refGene_tx=XM_019827211.1;refGene_gn=MTIF2;refGene_tx=XM_019827212.1;refGene_gn=MTIF2;refGene_tx=XM_019827213.1;refGene_gn=MTIF2;refGene_tx=XM_019827214.2;refGene_gn=MTIF2;refGene_tx=XM_019827215.2;refGene_gn=MTIF2;refGene_tx=XM_019827216.2
A3	76382735	76382876
refGene_gn=MTIF2;refGene_tx=XM_006930105.3;refGene_gn=MTIF2;refGene_tx=XM_019827211.1;refGene_gn=MTIF2;refGene_tx=XM_019827212.1;refGene_gn=MTIF2;refGene_tx=XM_019827213.1;refGene_gn=MTIF2;refGene_tx=XM_019827214.2;refGene_gn=MTIF2;refGene_tx=XM_019827215.2;refGene_gn=MTIF2;refGene_tx=XM_019827216.2
A3	76384241	76384406
refGene_gn=MTIF2;refGene_tx=XM_006930105.3;refGene_gn=MTIF2;refGene_tx=XM_019827211.1;refGene_gn=MTIF2;refGene_tx=XM_019827212.1;refGene_gn=MTIF2;refGene_tx=XM_019827213.1;refGene_gn=MTIF2;refGene_tx=XM_019827214.2;refGene_gn=MTIF2;refGene_tx=XM_019827215.2;refGene_gn=MTIF2;refGene_tx=XM_019827216.2
A3	76386556	76386697
refGene_gn=MTIF2;refGene_tx=XM_006930105.3;refGene_gn=MTIF2;refGene_tx=XM_019827211.1;refGene_gn=MTIF2;refGene_tx=XM_019827212.1;refGene_gn=MTIF2;refGene_tx=XM_019827213.1;refGene_gn=MTIF2;refGene_tx=XM_019827214.2;refGene_gn=MTIF2;refGene_tx=XM_019827215.2;refGene_gn=MTIF2;refGene_tx=XM_019827216.2
A3	76387499	76387752

refGene_gn=MTIF2;refGene_tx=XM_006930105.3;refGene_gn=MTIF2;refGene_tx=XM_019827211.1;refGene_gn=MTIF2;refGene_tx=XM_019827212.1;refGene_gn=MTIF2;refGene_tx=XM_019827213.1;refGene_gn=MTIF2;refGene_tx=XM_019827214.2;refGene_gn=MTIF2;refGene_tx=XM_019827215.2;refGene_gn=MTIF2;refGene_tx=XM_019827216.2
A3 76392877 76393082
refGene_gn=MTIF2;refGene_tx=XM_006930105.3;refGene_gn=MTIF2;refGene_tx=XM_019827211.1;refGene_gn=MTIF2;refGene_tx=XM_019827212.1;refGene_gn=MTIF2;refGene_tx=XM_019827213.1;refGene_gn=MTIF2;refGene_tx=XM_019827214.2;refGene_gn=MTIF2;refGene_tx=XM_019827215.2;refGene_gn=MTIF2;refGene_tx=XM_019827216.2
A3 76394815 76394940
refGene_gn=MTIF2;refGene_tx=XM_006930105.3;refGene_gn=MTIF2;refGene_tx=XM_019827211.1;refGene_gn=MTIF2;refGene_tx=XM_019827212.1;refGene_gn=MTIF2;refGene_tx=XM_019827213.1;refGene_gn=MTIF2;refGene_tx=XM_019827214.2;refGene_gn=MTIF2;refGene_tx=XM_019827215.2;refGene_gn=MTIF2;refGene_tx=XM_019827216.2
A3 76397872 76398012
refGene_gn=MTIF2;refGene_tx=XM_006930105.3;refGene_gn=MTIF2;refGene_tx=XM_019827211.1;refGene_gn=MTIF2;refGene_tx=XM_019827212.1;refGene_gn=MTIF2;refGene_tx=XM_019827213.1;refGene_gn=MTIF2;refGene_tx=XM_019827214.2;refGene_gn=MTIF2;refGene_tx=XM_019827215.2;refGene_gn=MTIF2;refGene_tx=XM_019827216.2
A3 76399815 76399992
refGene_gn=MTIF2;refGene_tx=XM_006930105.3;refGene_gn=MTIF2;refGene_tx=XM_019827211.1;refGene_gn=MTIF2;refGene_tx=XM_019827212.1;refGene_gn=MTIF2;refGene_tx=XM_019827213.1;refGene_gn=MTIF2;refGene_tx=XM_019827214.2;refGene_gn=MTIF2;refGene_tx=XM_019827215.2;refGene_gn=MTIF2;refGene_tx=XM_019827216.2
A3 76401128 76401289
refGene_gn=MTIF2;refGene_tx=XM_006930105.3;refGene_gn=MTIF2;refGene_tx=XM_019827211.1;refGene_gn=MTIF2;refGene_tx=XM_019827212.1;refGene_gn=MTIF2;refGene_tx=XM_019827213.1;refGene_gn=MTIF2;refGene_tx=XM_019827214.2;refGene_gn=MTIF2;refGene_tx=XM_019827215.2;refGene_gn=MTIF2;refGene_tx=XM_019827216.2
A3 76401749 76401921
refGene_gn=MTIF2;refGene_tx=XM_006930105.3;refGene_gn=MTIF2;refGene_tx=XM_019827211.1;refGene_gn=MTIF2;refGene_tx=XM_019827212.1;refGene_gn=MTIF2;refGene_tx=XM_019827213.1;refGene_gn=MTIF2;refGene_tx=XM_019827214.2;refGene_gn=MTIF2;refGene_tx=XM_019827215.2;refGene_gn=MTIF2;refGene_tx=XM_019827216.2
A3 76407599 76407711
refGene_gn=MTIF2;refGene_tx=XM_006930105.3;refGene_gn=MTIF2;refGene_tx=XM_019827211.1;refGene_gn=MTIF2;refGene_tx=XM_019827212.1;refGene_gn=MTIF2;refGene_tx=XM_019827213.1;refGene_gn=MTIF2;refGene_tx=XM_019827214.2;refGene_gn=MTIF2;refGene_tx=XM_019827215.2;refGene_gn=MTIF2;refGene_tx=XM_019827216.2
A3 76408078 76408297
refGene_gn=MTIF2;refGene_tx=XM_006930105.3;refGene_gn=MTIF2;refGene_tx=XM_019827211.1;refGene_gn=MTIF2;refGene_tx=XM_019827212.1;refGene_gn=

MTIF2;refGene_tx=XM_019827213.1;refGene_gn=MTIF2;refGene_tx=XM_019827214.2;refGene_gn=MTIF2;refGene_tx=XM_019827215.2;refGene_gn=MTIF2;refGene_tx=XM_019827216.2
A3 76424328 76424412
refGene_gn=LOC101093429;refGene_tx=XM_003984010.5
A3 76425467 76425899
refGene_gn=LOC101093429;refGene_tx=XM_003984010.5
A3 76431473 76431538
refGene_gn=CCDC88A;refGene_tx=XM_011281012.3;refGene_gn=CCDC88A;refGene_tx=XM_019827231.2;refGene_gn=CCDC88A;refGene_tx=XM_019827232.2;refGene_gn=CCDC88A;refGene_tx=XM_019827233.2;refGene_gn=CCDC88A;refGene_tx=XM_019827234.2;refGene_gn=CCDC88A;refGene_tx=XM_019827235.2
A3 76435018 76435481
refGene_gn=CCDC88A;refGene_tx=XM_011281012.3;refGene_gn=CCDC88A;refGene_tx=XM_019827231.2;refGene_gn=CCDC88A;refGene_tx=XM_019827232.2;refGene_gn=CCDC88A;refGene_tx=XM_019827233.2;refGene_gn=CCDC88A;refGene_tx=XM_019827234.2;refGene_gn=CCDC88A;refGene_tx=XM_019827235.2
A3 76435668 76435937
refGene_gn=CCDC88A;refGene_tx=XM_011281012.3;refGene_gn=CCDC88A;refGene_tx=XM_019827231.2;refGene_gn=CCDC88A;refGene_tx=XM_019827232.2;refGene_gn=CCDC88A;refGene_tx=XM_019827233.2;refGene_gn=CCDC88A;refGene_tx=XM_019827234.2;refGene_gn=CCDC88A;refGene_tx=XM_019827235.2
A3 76438328 76438409
refGene_gn=CCDC88A;refGene_tx=XM_011281012.3;refGene_gn=CCDC88A;refGene_tx=XM_019827231.2;refGene_gn=CCDC88A;refGene_tx=XM_019827232.2;refGene_gn=CCDC88A;refGene_tx=XM_019827234.2;refGene_gn=CCDC88A;refGene_tx=XM_019827235.2
A3 76439390 76439462
refGene_gn=CCDC88A;refGene_tx=XM_011281012.3;refGene_gn=CCDC88A;refGene_tx=XM_019827231.2;refGene_gn=CCDC88A;refGene_tx=XM_019827232.2;refGene_gn=CCDC88A;refGene_tx=XM_019827233.2;refGene_gn=CCDC88A;refGene_tx=XM_019827234.2;refGene_gn=CCDC88A;refGene_tx=XM_019827235.2
A3 76440088 76440292
refGene_gn=CCDC88A;refGene_tx=XM_011281012.3;refGene_gn=CCDC88A;refGene_tx=XM_019827231.2;refGene_gn=CCDC88A;refGene_tx=XM_019827232.2;refGene_gn=CCDC88A;refGene_tx=XM_019827233.2;refGene_gn=CCDC88A;refGene_tx=XM_019827234.2
A3 76441282 76441366
refGene_gn=CCDC88A;refGene_tx=XM_011281012.3;refGene_gn=CCDC88A;refGene_tx=XM_019827231.2;refGene_gn=CCDC88A;refGene_tx=XM_019827232.2;refGene_gn=CCDC88A;refGene_tx=XM_019827233.2
A3 76444856 76445071
refGene_gn=CCDC88A;refGene_tx=XM_011281012.3;refGene_gn=CCDC88A;refGene_tx=XM_019827231.2;refGene_gn=CCDC88A;refGene_tx=XM_019827232.2;refGene_gn=CCDC88A;refGene_tx=XM_019827233.2;refGene_gn=CCDC88A;refGene_tx=XM_019827234.2;refGene_gn=CCDC88A;refGene_tx=XM_019827235.2
A3 76445193 76445286
refGene_gn=CCDC88A;refGene_tx=XM_011281012.3;refGene_gn=CCDC88A;refGene_tx=XM_019827231.2;refGene_gn=CCDC88A;refGene_tx=XM_019827232.2;refGene_gn=CCDC88A;refGene_tx=XM_019827233.2;refGene_gn=CCDC88A;refGene_tx=

XM_019827234.2;refGene_gn=CCDC88A;refGene_tx=XM_019827235.2
A3 76448370 76448516
refGene_gn=CCDC88A;refGene_tx=XM_011281012.3;refGene_gn=CCDC88A;refGene_tx=XM_019827231.2;refGene_gn=CCDC88A;refGene_tx=XM_019827232.2;refGene_gn=CCDC88A;refGene_tx=XM_019827233.2;refGene_gn=CCDC88A;refGene_tx=XM_019827234.2;refGene_gn=CCDC88A;refGene_tx=XM_019827235.2
A3 76452804 76452991
refGene_gn=CCDC88A;refGene_tx=XM_011281012.3;refGene_gn=CCDC88A;refGene_tx=XM_019827231.2;refGene_gn=CCDC88A;refGene_tx=XM_019827232.2;refGene_gn=CCDC88A;refGene_tx=XM_019827233.2;refGene_gn=CCDC88A;refGene_tx=XM_019827234.2;refGene_gn=CCDC88A;refGene_tx=XM_019827235.2
A3 76456130 76456274
refGene_gn=CCDC88A;refGene_tx=XM_011281012.3;refGene_gn=CCDC88A;refGene_tx=XM_019827231.2;refGene_gn=CCDC88A;refGene_tx=XM_019827232.2;refGene_gn=CCDC88A;refGene_tx=XM_019827233.2;refGene_gn=CCDC88A;refGene_tx=XM_019827234.2;refGene_gn=CCDC88A;refGene_tx=XM_019827235.2
A3 76456458 76456736
refGene_gn=CCDC88A;refGene_tx=XM_011281012.3;refGene_gn=CCDC88A;refGene_tx=XM_019827231.2;refGene_gn=CCDC88A;refGene_tx=XM_019827232.2;refGene_gn=CCDC88A;refGene_tx=XM_019827233.2;refGene_gn=CCDC88A;refGene_tx=XM_019827234.2;refGene_gn=CCDC88A;refGene_tx=XM_019827235.2
A3 76458039 76458201
refGene_gn=CCDC88A;refGene_tx=XM_011281012.3;refGene_gn=CCDC88A;refGene_tx=XM_019827231.2;refGene_gn=CCDC88A;refGene_tx=XM_019827232.2;refGene_gn=CCDC88A;refGene_tx=XM_019827233.2;refGene_gn=CCDC88A;refGene_tx=XM_019827234.2;refGene_gn=CCDC88A;refGene_tx=XM_019827235.2
A3 76463699 76463864
refGene_gn=CCDC88A;refGene_tx=XM_011281012.3;refGene_gn=CCDC88A;refGene_tx=XM_019827231.2;refGene_gn=CCDC88A;refGene_tx=XM_019827232.2;refGene_gn=CCDC88A;refGene_tx=XM_019827233.2;refGene_gn=CCDC88A;refGene_tx=XM_019827234.2;refGene_gn=CCDC88A;refGene_tx=XM_019827235.2
A3 76467054 76467196
refGene_gn=CCDC88A;refGene_tx=XM_011281012.3;refGene_gn=CCDC88A;refGene_tx=XM_019827231.2;refGene_gn=CCDC88A;refGene_tx=XM_019827232.2;refGene_gn=CCDC88A;refGene_tx=XM_019827233.2;refGene_gn=CCDC88A;refGene_tx=XM_019827234.2;refGene_gn=CCDC88A;refGene_tx=XM_019827235.2
A3 76472648 76472776
refGene_gn=CCDC88A;refGene_tx=XM_011281012.3;refGene_gn=CCDC88A;refGene_tx=XM_019827231.2;refGene_gn=CCDC88A;refGene_tx=XM_019827232.2;refGene_gn=CCDC88A;refGene_tx=XM_019827233.2;refGene_gn=CCDC88A;refGene_tx=XM_019827234.2;refGene_gn=CCDC88A;refGene_tx=XM_019827235.2
A3 76474184 76475255
refGene_gn=CCDC88A;refGene_tx=XM_011281012.3;refGene_gn=CCDC88A;refGene_tx=XM_019827231.2;refGene_gn=CCDC88A;refGene_tx=XM_019827232.2;refGene_gn=CCDC88A;refGene_tx=XM_019827233.2;refGene_gn=CCDC88A;refGene_tx=XM_019827234.2;refGene_gn=CCDC88A;refGene_tx=XM_019827235.2
A3 76476424 76476562
refGene_gn=CCDC88A;refGene_tx=XM_011281012.3;refGene_gn=CCDC88A;refGene_tx=XM_019827231.2;refGene_gn=CCDC88A;refGene_tx=XM_019827232.2;refGene_gn=CCDC88A;refGene_tx=XM_019827233.2;refGene_gn=CCDC88A;refGene_tx=

XM_019827234.2;refGene_gn=CCDC88A;refGene_tx=XM_019827235.2
A3 76481078 76481263
refGene_gn=CCDC88A;refGene_tx=XM_011281012.3;refGene_gn=CCDC88A;refGene_tx=XM_019827231.2;refGene_gn=CCDC88A;refGene_tx=XM_019827232.2;refGene_gn=CCDC88A;refGene_tx=XM_019827233.2;refGene_gn=CCDC88A;refGene_tx=XM_019827234.2;refGene_gn=CCDC88A;refGene_tx=XM_019827235.2
A3 76484789 76484934
refGene_gn=CCDC88A;refGene_tx=XM_011281012.3;refGene_gn=CCDC88A;refGene_tx=XM_019827231.2;refGene_gn=CCDC88A;refGene_tx=XM_019827232.2;refGene_gn=CCDC88A;refGene_tx=XM_019827233.2;refGene_gn=CCDC88A;refGene_tx=XM_019827234.2;refGene_gn=CCDC88A;refGene_tx=XM_019827235.2
A3 76485519 76485666
refGene_gn=CCDC88A;refGene_tx=XM_011281012.3;refGene_gn=CCDC88A;refGene_tx=XM_019827231.2;refGene_gn=CCDC88A;refGene_tx=XM_019827232.2;refGene_gn=CCDC88A;refGene_tx=XM_019827233.2;refGene_gn=CCDC88A;refGene_tx=XM_019827234.2;refGene_gn=CCDC88A;refGene_tx=XM_019827235.2
A3 76487353 76487512
refGene_gn=CCDC88A;refGene_tx=XM_011281012.3;refGene_gn=CCDC88A;refGene_tx=XM_019827231.2;refGene_gn=CCDC88A;refGene_tx=XM_019827232.2;refGene_gn=CCDC88A;refGene_tx=XM_019827233.2;refGene_gn=CCDC88A;refGene_tx=XM_019827234.2;refGene_gn=CCDC88A;refGene_tx=XM_019827235.2
A3 76490379 76490461
refGene_gn=CCDC88A;refGene_tx=XM_011281012.3;refGene_gn=CCDC88A;refGene_tx=XM_019827231.2;refGene_gn=CCDC88A;refGene_tx=XM_019827232.2;refGene_gn=CCDC88A;refGene_tx=XM_019827233.2;refGene_gn=CCDC88A;refGene_tx=XM_019827234.2;refGene_gn=CCDC88A;refGene_tx=XM_019827235.2
A3 76505030 76505203
refGene_gn=CCDC88A;refGene_tx=XM_011281012.3;refGene_gn=CCDC88A;refGene_tx=XM_019827231.2;refGene_gn=CCDC88A;refGene_tx=XM_019827232.2;refGene_gn=CCDC88A;refGene_tx=XM_019827233.2;refGene_gn=CCDC88A;refGene_tx=XM_019827234.2;refGene_gn=CCDC88A;refGene_tx=XM_019827235.2
A3 76512805 76512946
refGene_gn=CCDC88A;refGene_tx=XM_011281012.3;refGene_gn=CCDC88A;refGene_tx=XM_019827231.2;refGene_gn=CCDC88A;refGene_tx=XM_019827232.2;refGene_gn=CCDC88A;refGene_tx=XM_019827233.2;refGene_gn=CCDC88A;refGene_tx=XM_019827234.2;refGene_gn=CCDC88A;refGene_tx=XM_019827235.2
A3 76514506 76514590
refGene_gn=CCDC88A;refGene_tx=XM_011281012.3;refGene_gn=CCDC88A;refGene_tx=XM_019827231.2;refGene_gn=CCDC88A;refGene_tx=XM_019827232.2;refGene_gn=CCDC88A;refGene_tx=XM_019827233.2;refGene_gn=CCDC88A;refGene_tx=XM_019827234.2;refGene_gn=CCDC88A;refGene_tx=XM_019827235.2
A3 76521978 76522037
refGene_gn=CCDC88A;refGene_tx=XM_011281012.3;refGene_gn=CCDC88A;refGene_tx=XM_019827231.2;refGene_gn=CCDC88A;refGene_tx=XM_019827232.2;refGene_gn=CCDC88A;refGene_tx=XM_019827233.2;refGene_gn=CCDC88A;refGene_tx=XM_019827234.2;refGene_gn=CCDC88A;refGene_tx=XM_019827235.2
A3 76524000 76524070
refGene_gn=CCDC88A;refGene_tx=XM_011281012.3;refGene_gn=CCDC88A;refGene_tx=XM_019827231.2;refGene_gn=CCDC88A;refGene_tx=XM_019827232.2;refGene_gn=CCDC88A;refGene_tx=XM_019827233.2;refGene_gn=CCDC88A;refGene_tx=

XM_019827234.2;refGene_gn=CCDC88A;refGene_tx=XM_019827235.2
A3 76537442 76537551
refGene_gn=CCDC88A;refGene_tx=XM_011281012.3;refGene_gn=CCDC88A;refGene_tx=XM_019827231.2;refGene_gn=CCDC88A;refGene_tx=XM_019827232.2;refGene_gn=CCDC88A;refGene_tx=XM_019827233.2;refGene_gn=CCDC88A;refGene_tx=XM_019827234.2;refGene_gn=CCDC88A;refGene_tx=XM_019827235.2
A3 76570219 76570320
refGene_gn=CCDC88A;refGene_tx=XM_011281012.3;refGene_gn=CCDC88A;refGene_tx=XM_019827231.2;refGene_gn=CCDC88A;refGene_tx=XM_019827232.2;refGene_gn=CCDC88A;refGene_tx=XM_019827233.2;refGene_gn=CCDC88A;refGene_tx=XM_019827234.2;refGene_gn=CCDC88A;refGene_tx=XM_019827235.2
A3 76570416 76570479
refGene_gn=CCDC88A;refGene_tx=XM_011281012.3;refGene_gn=CCDC88A;refGene_tx=XM_019827231.2;refGene_gn=CCDC88A;refGene_tx=XM_019827232.2;refGene_gn=CCDC88A;refGene_tx=XM_019827233.2;refGene_gn=CCDC88A;refGene_tx=XM_019827234.2;refGene_gn=CCDC88A;refGene_tx=XM_019827235.2
A3 76647066 76647181
refGene_gn=CFAP36;refGene_tx=XM_003984011.5;refGene_gn=CFAP36;refGene_tx=XM_006930106.4
A3 76650248 76650313
refGene_gn=CFAP36;refGene_tx=XM_003984011.5;refGene_gn=CFAP36;refGene_tx=XM_006930106.4
A3 76651283 76651385
refGene_gn=CFAP36;refGene_tx=XM_003984011.5;refGene_gn=CFAP36;refGene_tx=XM_006930106.4
A3 76664439 76664554
refGene_gn=CFAP36;refGene_tx=XM_003984011.5;refGene_gn=CFAP36;refGene_tx=XM_006930106.4
A3 76668192 76668280
refGene_gn=CFAP36;refGene_tx=XM_003984011.5;refGene_gn=CFAP36;refGene_tx=XM_006930106.4
A3 76670012 76670064
refGene_gn=CFAP36;refGene_tx=XM_003984011.5;refGene_gn=CFAP36;refGene_tx=XM_006930106.4
A3 76671836 76671936
refGene_gn=CFAP36;refGene_tx=XM_003984011.5;refGene_gn=CFAP36;refGene_tx=XM_006930106.4
A3 76675539 76675679
refGene_gn=CFAP36;refGene_tx=XM_003984011.5;refGene_gn=CFAP36;refGene_tx=XM_006930106.4
A3 76675873 76676020
refGene_gn=CFAP36;refGene_tx=XM_003984011.5;refGene_gn=CFAP36;refGene_tx=XM_006930106.4
A3 76676571 76676673
refGene_gn=CFAP36;refGene_tx=XM_003984011.5;refGene_gn=CFAP36;refGene_tx=XM_006930106.4
A3 76680429 76680525
refGene_gn=PPP4R3B;refGene_tx=XM_003984012.5;refGene_gn=PPP4R3B;refGene_tx=XM_006930107.4;refGene_gn=PPP4R3B;refGene_tx=XM_019827236.2;refGene_gn=PPP4R3B;refGene_tx=XM_019827237.2

A3 76687105 76687308
refGene_gn=PPP4R3B;refGene_tx=XM_003984012.5;refGene_gn=PPP4R3B;refGene_tx=XM_006930107.4;refGene_gn=PPP4R3B;refGene_tx=XM_019827236.2;refGene_gn=PPP4R3B;refGene_tx=XM_019827237.2
A3 76689134 76689319
refGene_gn=PPP4R3B;refGene_tx=XM_003984012.5;refGene_gn=PPP4R3B;refGene_tx=XM_006930107.4;refGene_gn=PPP4R3B;refGene_tx=XM_019827236.2;refGene_gn=PPP4R3B;refGene_tx=XM_019827237.2
A3 76689719 76689859
refGene_gn=PPP4R3B;refGene_tx=XM_003984012.5;refGene_gn=PPP4R3B;refGene_tx=XM_006930107.4;refGene_gn=PPP4R3B;refGene_tx=XM_019827236.2;refGene_gn=PPP4R3B;refGene_tx=XM_019827237.2
A3 76692144 76692314
refGene_gn=PPP4R3B;refGene_tx=XM_003984012.5;refGene_gn=PPP4R3B;refGene_tx=XM_006930107.4;refGene_gn=PPP4R3B;refGene_tx=XM_019827236.2;refGene_gn=PPP4R3B;refGene_tx=XM_019827237.2
A3 76695504 76695663
refGene_gn=PPP4R3B;refGene_tx=XM_003984012.5;refGene_gn=PPP4R3B;refGene_tx=XM_006930107.4;refGene_gn=PPP4R3B;refGene_tx=XM_019827236.2;refGene_gn=PPP4R3B;refGene_tx=XM_019827237.2
A3 76697709 76697751
refGene_gn=PPP4R3B;refGene_tx=XM_003984012.5;refGene_gn=PPP4R3B;refGene_tx=XM_006930107.4;refGene_gn=PPP4R3B;refGene_tx=XM_019827236.2;refGene_gn=PPP4R3B;refGene_tx=XM_019827237.2
A3 76701253 76701356
refGene_gn=PPP4R3B;refGene_tx=XM_003984012.5;refGene_gn=PPP4R3B;refGene_tx=XM_006930107.4;refGene_gn=PPP4R3B;refGene_tx=XM_019827236.2;refGene_gn=PPP4R3B;refGene_tx=XM_019827237.2
A3 76702655 76702787
refGene_gn=PPP4R3B;refGene_tx=XM_003984012.5;refGene_gn=PPP4R3B;refGene_tx=XM_006930107.4;refGene_gn=PPP4R3B;refGene_tx=XM_019827236.2;refGene_gn=PPP4R3B;refGene_tx=XM_019827237.2
A3 76714157 76714274
refGene_gn=PPP4R3B;refGene_tx=XM_003984012.5;refGene_gn=PPP4R3B;refGene_tx=XM_006930107.4
A3 76715666 76715783
refGene_gn=PPP4R3B;refGene_tx=XM_003984012.5;refGene_gn=PPP4R3B;refGene_tx=XM_006930107.4;refGene_gn=PPP4R3B;refGene_tx=XM_019827236.2
A3 76717078 76717156
refGene_gn=PPP4R3B;refGene_tx=XM_003984012.5;refGene_gn=PPP4R3B;refGene_tx=XM_006930107.4;refGene_gn=PPP4R3B;refGene_tx=XM_019827236.2;refGene_gn=PPP4R3B;refGene_tx=XM_019827237.2
A3 76723408 76724032
refGene_gn=PPP4R3B;refGene_tx=XM_003984012.5;refGene_gn=PPP4R3B;refGene_tx=XM_006930107.4;refGene_gn=PPP4R3B;refGene_tx=XM_019827236.2;refGene_gn=PPP4R3B;refGene_tx=XM_019827237.2
A3 76732440 76732539
refGene_gn=PPP4R3B;refGene_tx=XM_003984012.5;refGene_gn=PPP4R3B;refGene_tx=XM_006930107.4;refGene_gn=PPP4R3B;refGene_tx=XM_019827236.2;refGene_gn=PPP4R3B;refGene_tx=XM_019827237.2

A3 76742764 76742820
refGene_gn=PPP4R3B;refGene_tx=XM_003984012.5;refGene_gn=PPP4R3B;refGene_tx=XM_006930107.4;refGene_gn=PPP4R3B;refGene_tx=XM_019827236.2;refGene_gn=PPP4R3B;refGene_tx=XM_019827237.2
A3 76744174 76744316
refGene_gn=PPP4R3B;refGene_tx=XM_003984012.5;refGene_gn=PPP4R3B;refGene_tx=XM_006930107.4;refGene_gn=PPP4R3B;refGene_tx=XM_019827236.2;refGene_gn=PPP4R3B;refGene_tx=XM_019827237.2
A3 76767947 76768106
refGene_gn=PNPT1;refGene_tx=XM_003984014.5;refGene_gn=PNPT1;refGene_tx=XM_019827239.1
A3 76769217 76769265
refGene_gn=PNPT1;refGene_tx=XM_003984014.5;refGene_gn=PNPT1;refGene_tx=XM_019827239.1
A3 76773244 76773323
refGene_gn=PNPT1;refGene_tx=XM_003984014.5;refGene_gn=PNPT1;refGene_tx=XM_019827239.1
A3 76774263 76774319
refGene_gn=PNPT1;refGene_tx=XM_003984014.5;refGene_gn=PNPT1;refGene_tx=XM_019827239.1
A3 76774395 76774502
refGene_gn=PNPT1;refGene_tx=XM_003984014.5;refGene_gn=PNPT1;refGene_tx=XM_019827239.1
A3 76775534 76775618
refGene_gn=PNPT1;refGene_tx=XM_003984014.5;refGene_gn=PNPT1;refGene_tx=XM_019827239.1
A3 76775938 76776022
refGene_gn=PNPT1;refGene_tx=XM_003984014.5;refGene_gn=PNPT1;refGene_tx=XM_019827239.1
A3 76777516 76777580
refGene_gn=PNPT1;refGene_tx=XM_003984014.5;refGene_gn=PNPT1;refGene_tx=XM_019827239.1
A3 76777671 76777743
refGene_gn=PNPT1;refGene_tx=XM_003984014.5;refGene_gn=PNPT1;refGene_tx=XM_019827239.1
A3 76778248 76778355
refGene_gn=PNPT1;refGene_tx=XM_003984014.5;refGene_gn=PNPT1;refGene_tx=XM_019827239.1
A3 76782904 76782958
refGene_gn=PNPT1;refGene_tx=XM_003984014.5;refGene_gn=PNPT1;refGene_tx=XM_019827239.1
A3 76784478 76784568
refGene_gn=PNPT1;refGene_tx=XM_003984014.5;refGene_gn=PNPT1;refGene_tx=XM_019827239.1
A3 76784650 76784717
refGene_gn=PNPT1;refGene_tx=XM_003984014.5;refGene_gn=PNPT1;refGene_tx=XM_019827239.1
A3 76787560 76787597
refGene_gn=PNPT1;refGene_tx=XM_003984014.5;refGene_gn=PNPT1;refGene_tx=XM_019827239.1

A3 76789575 76789646
refGene_gn=PNPT1;refGene_tx=XM_003984014.5;refGene_gn=PNPT1;refGene_tx=XM_019827239.1
A3 76793162 76793265
refGene_gn=PNPT1;refGene_tx=XM_003984014.5;refGene_gn=PNPT1;refGene_tx=XM_019827239.1
A3 76793735 76793832
refGene_gn=PNPT1;refGene_tx=XM_003984014.5;refGene_gn=PNPT1;refGene_tx=XM_019827239.1
A3 76795479 76795537
refGene_gn=PNPT1;refGene_tx=XM_003984014.5;refGene_gn=PNPT1;refGene_tx=XM_019827239.1
A3 76795897 76795949
refGene_gn=PNPT1;refGene_tx=XM_003984014.5;refGene_gn=PNPT1;refGene_tx=XM_019827239.1
A3 76797058 76797245
refGene_gn=PNPT1;refGene_tx=XM_003984014.5;refGene_gn=PNPT1;refGene_tx=XM_019827239.1
A3 76804215 76804329
refGene_gn=PNPT1;refGene_tx=XM_003984014.5;refGene_gn=PNPT1;refGene_tx=XM_019827239.1
A3 76805465 76805513
refGene_gn=PNPT1;refGene_tx=XM_003984014.5;refGene_gn=PNPT1;refGene_tx=XM_019827239.1
A3 76805609 76805673
refGene_gn=PNPT1;refGene_tx=XM_003984014.5;refGene_gn=PNPT1;refGene_tx=XM_019827239.1
A3 76806675 76806725
refGene_gn=PNPT1;refGene_tx=XM_003984014.5;refGene_gn=PNPT1;refGene_tx=XM_019827239.1
A3 76807248 76807354
refGene_gn=PNPT1;refGene_tx=XM_003984014.5;refGene_gn=PNPT1;refGene_tx=XM_019827239.1
A3 76808048 76808132
refGene_gn=PNPT1;refGene_tx=XM_003984014.5;refGene_gn=PNPT1;refGene_tx=XM_019827239.1
A3 76808712 76808773 refGene_gn=PNPT1;refGene_tx=XM_003984014.5
A3 76815686 76815847 refGene_gn=PNPT1;refGene_tx=XM_003984014.5
A3 76939524 76939686
refGene_gn=EFEMP1;refGene_tx=XM_003984015.4;refGene_gn=EFEMP1;refGene_tx=XM_006930109.3
A3 76944193 76944389
refGene_gn=EFEMP1;refGene_tx=XM_003984015.4;refGene_gn=EFEMP1;refGene_tx=XM_006930109.3
A3 76944473 76944597
refGene_gn=EFEMP1;refGene_tx=XM_003984015.4;refGene_gn=EFEMP1;refGene_tx=XM_006930109.3
A3 76949204 76949324
refGene_gn=EFEMP1;refGene_tx=XM_003984015.4;refGene_gn=EFEMP1;refGene_tx=XM_006930109.3

A3 76950658 76950778
refGene_gn=EFEMP1;refGene_tx=XM_003984015.4;refGene_gn=EFEMP1;refGene_tx=XM_006930109.3
A3 76951762 76951882
refGene_gn=EFEMP1;refGene_tx=XM_003984015.4;refGene_gn=EFEMP1;refGene_tx=XM_006930109.3
A3 76955881 76956004
refGene_gn=EFEMP1;refGene_tx=XM_003984015.4;refGene_gn=EFEMP1;refGene_tx=XM_006930109.3
A3 76992586 76993003
refGene_gn=EFEMP1;refGene_tx=XM_003984015.4;refGene_gn=EFEMP1;refGene_tx=XM_006930109.3
A3 76993167 76993216
refGene_gn=EFEMP1;refGene_tx=XM_003984015.4;refGene_gn=EFEMP1;refGene_tx=XM_006930109.3
A3 76997391 76997475
refGene_gn=EFEMP1;refGene_tx=XM_003984015.4;refGene_gn=EFEMP1;refGene_tx=XM_006930109.3
A3 77265372 77265639
refGene_gn=CCDC85A;refGene_tx=XM_003984016.4;refGene_gn=CCDC85A;refGene_tx=XM_006930110.4;refGene_gn=CCDC85A;refGene_tx=XM_006930111.4;refGene_gn=CCDC85A;refGene_tx=XM_019827240.2;refGene_gn=CCDC85A;refGene_tx=XM_019827241.2;refGene_gn=CCDC85A;refGene_tx=XM_023251680.1
A3 77272308 77273257
refGene_gn=CCDC85A;refGene_tx=XM_003984016.4;refGene_gn=CCDC85A;refGene_tx=XM_006930110.4;refGene_gn=CCDC85A;refGene_tx=XM_006930111.4;refGene_gn=CCDC85A;refGene_tx=XM_019827240.2;refGene_gn=CCDC85A;refGene_tx=XM_019827241.2;refGene_gn=CCDC85A;refGene_tx=XM_023251680.1
A3 77429026 77429103
refGene_gn=CCDC85A;refGene_tx=XM_003984016.4;refGene_gn=CCDC85A;refGene_tx=XM_006930110.4;refGene_gn=CCDC85A;refGene_tx=XM_006930111.4;refGene_gn=CCDC85A;refGene_tx=XM_019827240.2;refGene_gn=CCDC85A;refGene_tx=XM_019827241.2
A3 77455388 77455523
refGene_gn=CCDC85A;refGene_tx=XM_003984016.4;refGene_gn=CCDC85A;refGene_tx=XM_019827240.2
A3 77458576 77458696
refGene_gn=CCDC85A;refGene_tx=XM_003984016.4;refGene_gn=CCDC85A;refGene_tx=XM_006930110.4;refGene_gn=CCDC85A;refGene_tx=XM_019827240.2;refGene_gn=CCDC85A;refGene_tx=XM_019827241.2
A3 77462272 77462371
refGene_gn=CCDC85A;refGene_tx=XM_019827240.2;refGene_gn=CCDC85A;refGene_tx=XM_019827241.2
A3 77466372 77466462
refGene_gn=CCDC85A;refGene_tx=XM_003984016.4;refGene_gn=CCDC85A;refGene_tx=XM_006930110.4;refGene_gn=CCDC85A;refGene_tx=XM_006930111.4;refGene_gn=CCDC85A;refGene_tx=XM_019827240.2;refGene_gn=CCDC85A;refGene_tx=XM_019827241.2;refGene_gn=CCDC85A;refGene_tx=XM_023251680.1
A3 78828240 78828376
refGene_gn=VRK2;refGene_tx=XM_023251681.1;refGene_gn=VRK2;refGene_tx=X

M_023251682.1;refGene_gn=VRK2;refGene_tx=XM_023251683.1;refGene_gn=VRK
2;refGene_tx=XM_023251685.1
A3 78831523 78831554 refGene_gn=VRK2;refGene_tx=XM_019827243.1
A3 78850587 78850657 refGene_gn=VRK2;refGene_tx=XM_023251684.1
A3 78854772 78854822
refGene_gn=VRK2;refGene_tx=XM_019827243.1;refGene_gn=VRK2;refGene_tx=X
M_023251681.1;refGene_gn=VRK2;refGene_tx=XM_023251682.1;refGene_gn=VRK
2;refGene_tx=XM_023251683.1;refGene_gn=VRK2;refGene_tx=XM_023251684.1;
refGene_gn=VRK2;refGene_tx=XM_023251685.1
A3 78855536 78855606
refGene_gn=VRK2;refGene_tx=XM_019827243.1;refGene_gn=VRK2;refGene_tx=X
M_023251681.1;refGene_gn=VRK2;refGene_tx=XM_023251682.1;refGene_gn=VRK
2;refGene_tx=XM_023251683.1;refGene_gn=VRK2;refGene_tx=XM_023251684.1;
refGene_gn=VRK2;refGene_tx=XM_023251685.1
A3 78857055 78857143
refGene_gn=VRK2;refGene_tx=XM_019827243.1;refGene_gn=VRK2;refGene_tx=X
M_019827244.2;refGene_gn=VRK2;refGene_tx=XM_023251681.1;refGene_gn=VRK
2;refGene_tx=XM_023251682.1;refGene_gn=VRK2;refGene_tx=XM_023251683.1;
refGene_gn=VRK2;refGene_tx=XM_023251684.1;refGene_gn=VRK2;refGene_tx=X
M_023251685.1
A3 78859507 78859613
refGene_gn=VRK2;refGene_tx=XM_019827243.1;refGene_gn=VRK2;refGene_tx=X
M_019827244.2;refGene_gn=VRK2;refGene_tx=XM_023251681.1;refGene_gn=VRK
2;refGene_tx=XM_023251682.1;refGene_gn=VRK2;refGene_tx=XM_023251683.1;
refGene_gn=VRK2;refGene_tx=XM_023251684.1;refGene_gn=VRK2;refGene_tx=X
M_023251685.1
A3 78860820 78860913
refGene_gn=VRK2;refGene_tx=XM_019827243.1;refGene_gn=VRK2;refGene_tx=X
M_019827244.2;refGene_gn=VRK2;refGene_tx=XM_023251681.1;refGene_gn=VRK
2;refGene_tx=XM_023251682.1;refGene_gn=VRK2;refGene_tx=XM_023251683.1;
refGene_gn=VRK2;refGene_tx=XM_023251684.1;refGene_gn=VRK2;refGene_tx=X
M_023251685.1
A3 78876879 78877014
refGene_gn=VRK2;refGene_tx=XM_019827243.1;refGene_gn=VRK2;refGene_tx=X
M_019827244.2;refGene_gn=VRK2;refGene_tx=XM_023251681.1;refGene_gn=VRK
2;refGene_tx=XM_023251682.1;refGene_gn=VRK2;refGene_tx=XM_023251683.1;
refGene_gn=VRK2;refGene_tx=XM_023251684.1
A3 78892322 78892455
refGene_gn=VRK2;refGene_tx=XM_019827243.1;refGene_gn=VRK2;refGene_tx=X
M_019827244.2;refGene_gn=VRK2;refGene_tx=XM_023251681.1;refGene_gn=VRK
2;refGene_tx=XM_023251682.1;refGene_gn=VRK2;refGene_tx=XM_023251683.1;
refGene_gn=VRK2;refGene_tx=XM_023251684.1;refGene_gn=VRK2;refGene_tx=X
M_023251685.1
A3 78902439 78902560
refGene_gn=VRK2;refGene_tx=XM_019827243.1;refGene_gn=VRK2;refGene_tx=X
M_019827244.2;refGene_gn=VRK2;refGene_tx=XM_023251681.1;refGene_gn=VRK
2;refGene_tx=XM_023251682.1;refGene_gn=VRK2;refGene_tx=XM_023251683.1;
refGene_gn=VRK2;refGene_tx=XM_023251684.1;refGene_gn=VRK2;refGene_tx=X
M_023251685.1
A3 78904853 78904912

refGene_gn=VRK2;refGene_tx=XM_019827243.1;refGene_gn=VRK2;refGene_tx=XM_019827244.2;refGene_gn=VRK2;refGene_tx=XM_023251681.1;refGene_gn=VRK2;refGene_tx=XM_023251682.1;refGene_gn=VRK2;refGene_tx=XM_023251683.1;refGene_gn=VRK2;refGene_tx=XM_023251684.1;refGene_gn=VRK2;refGene_tx=XM_023251685.1
A3 78909403 78909570
refGene_gn=VRK2;refGene_tx=XM_019827243.1;refGene_gn=VRK2;refGene_tx=XM_019827244.2;refGene_gn=VRK2;refGene_tx=XM_023251681.1;refGene_gn=VRK2;refGene_tx=XM_023251682.1;refGene_gn=VRK2;refGene_tx=XM_023251683.1;refGene_gn=VRK2;refGene_tx=XM_023251684.1;refGene_gn=VRK2;refGene_tx=XM_023251685.1
A3 78914682 78914844
refGene_gn=VRK2;refGene_tx=XM_019827243.1;refGene_gn=VRK2;refGene_tx=XM_019827244.2;refGene_gn=VRK2;refGene_tx=XM_023251681.1;refGene_gn=VRK2;refGene_tx=XM_023251682.1;refGene_gn=VRK2;refGene_tx=XM_023251683.1;refGene_gn=VRK2;refGene_tx=XM_023251684.1;refGene_gn=VRK2;refGene_tx=XM_023251685.1
A3 78929646 78929949
refGene_gn=VRK2;refGene_tx=XM_019827243.1;refGene_gn=VRK2;refGene_tx=XM_019827244.2;refGene_gn=VRK2;refGene_tx=XM_023251681.1;refGene_gn=VRK2;refGene_tx=XM_023251682.1;refGene_gn=VRK2;refGene_tx=XM_023251683.1;refGene_gn=VRK2;refGene_tx=XM_023251684.1;refGene_gn=VRK2;refGene_tx=XM_023251685.1
A3 78930020 78930056 refGene_gn=FANCL;refGene_tx=XM_023251686.1
A3 78930363 78930435 refGene_gn=FANCL;refGene_tx=XM_023251686.1
A3 78931763 78931880 refGene_gn=FANCL;refGene_tx=XM_023251686.1
A3 78932721 78932803 refGene_gn=FANCL;refGene_tx=XM_023251686.1
A3 78932881 78932927 refGene_gn=FANCL;refGene_tx=XM_023251686.1
A3 78933411 78933495 refGene_gn=FANCL;refGene_tx=XM_023251686.1
A3 78937719 78937870 refGene_gn=FANCL;refGene_tx=XM_023251686.1
A3 78969339 78969462 refGene_gn=FANCL;refGene_tx=XM_023251686.1
A3 78973439 78973508 refGene_gn=FANCL;refGene_tx=XM_023251686.1
A3 78978493 78978590 refGene_gn=FANCL;refGene_tx=XM_023251686.1
A3 78992773 78992874 refGene_gn=FANCL;refGene_tx=XM_023251686.1
A3 78992973 78993024 refGene_gn=FANCL;refGene_tx=XM_023251686.1
A3 78997549 78997606 refGene_gn=FANCL;refGene_tx=XM_023251686.1
A3 79000090 79000151 refGene_gn=FANCL;refGene_tx=XM_023251686.1
A3 79002282 79002341 refGene_gn=FANCL;refGene_tx=XM_023251686.1
A3 79013856 79013943 refGene_gn=FANCL;refGene_tx=XM_023251686.1
A3 81018669 81018771
refGene_gn=BCL11A;refGene_tx=XM_003984025.5;refGene_gn=BCL11A;refGene_tx=XM_011281010.3;refGene_gn=BCL11A;refGene_tx=XM_019827251.2
A3 81026608 81028629
refGene_gn=BCL11A;refGene_tx=XM_003984025.5;refGene_gn=BCL11A;refGene_tx=XM_011281010.3;refGene_gn=BCL11A;refGene_tx=XM_019827251.2;refGene_gn=BCL11A;refGene_tx=XM_019827252.2;refGene_gn=BCL11A;refGene_tx=XM_019827253.2;refGene_gn=BCL11A;refGene_tx=XM_023251687.1;refGene_gn=BCL11A;refGene_tx=XM_023251688.1
A3 81033979 81034081
refGene_gn=BCL11A;refGene_tx=XM_003984025.5;refGene_gn=BCL11A;refGene_

tx=XM_011281010.3;refGene_gn=BCL11A;refGene_tx=XM_019827251.2;refGene_gn=BCL11A;refGene_tx=XM_019827253.2;refGene_gn=BCL11A;refGene_tx=XM_023251687.1;refGene_gn=BCL11A;refGene_tx=XM_023251688.1
A3 81094682 81094755 refGene_gn=BCL11A;refGene_tx=XM_023251688.1
A3 81111481 81111811
refGene_gn=BCL11A;refGene_tx=XM_003984025.5;refGene_gn=BCL11A;refGene_tx=XM_011281010.3;refGene_gn=BCL11A;refGene_tx=XM_019827251.2;refGene_gn=BCL11A;refGene_tx=XM_019827252.2;refGene_gn=BCL11A;refGene_tx=XM_019827253.2;refGene_gn=BCL11A;refGene_tx=XM_023251687.1
A3 81118750 81118805
refGene_gn=BCL11A;refGene_tx=XM_003984025.5;refGene_gn=BCL11A;refGene_tx=XM_011281010.3;refGene_gn=BCL11A;refGene_tx=XM_019827251.2;refGene_gn=BCL11A;refGene_tx=XM_019827252.2;refGene_gn=BCL11A;refGene_tx=XM_023251687.1
A3 81300667 81300684
refGene_gn=PAPOLG;refGene_tx=XM_011281009.3;refGene_gn=PAPOLG;refGene_tx=XM_019827255.2;refGene_gn=PAPOLG;refGene_tx=XM_019827256.2
A3 81303728 81303890
refGene_gn=PAPOLG;refGene_tx=XM_011281009.3;refGene_gn=PAPOLG;refGene_tx=XM_019827255.2;refGene_gn=PAPOLG;refGene_tx=XM_019827256.2
A3 81305362 81305429
refGene_gn=PAPOLG;refGene_tx=XM_011281009.3;refGene_gn=PAPOLG;refGene_tx=XM_019827255.2;refGene_gn=PAPOLG;refGene_tx=XM_019827256.2
A3 81309156 81309238
refGene_gn=PAPOLG;refGene_tx=XM_011281009.3;refGene_gn=PAPOLG;refGene_tx=XM_019827255.2;refGene_gn=PAPOLG;refGene_tx=XM_019827256.2
A3 81309465 81309575
refGene_gn=PAPOLG;refGene_tx=XM_011281009.3;refGene_gn=PAPOLG;refGene_tx=XM_019827255.2;refGene_gn=PAPOLG;refGene_tx=XM_019827256.2
A3 81311085 81311139
refGene_gn=PAPOLG;refGene_tx=XM_011281009.3;refGene_gn=PAPOLG;refGene_tx=XM_019827255.2;refGene_gn=PAPOLG;refGene_tx=XM_019827256.2
A3 81312935 81313047
refGene_gn=PAPOLG;refGene_tx=XM_011281009.3;refGene_gn=PAPOLG;refGene_tx=XM_019827255.2;refGene_gn=PAPOLG;refGene_tx=XM_019827256.2
A3 81315071 81315161
refGene_gn=PAPOLG;refGene_tx=XM_011281009.3;refGene_gn=PAPOLG;refGene_tx=XM_019827255.2;refGene_gn=PAPOLG;refGene_tx=XM_019827256.2
A3 81317605 81317744
refGene_gn=PAPOLG;refGene_tx=XM_011281009.3;refGene_gn=PAPOLG;refGene_tx=XM_019827255.2;refGene_gn=PAPOLG;refGene_tx=XM_019827256.2
A3 81318459 81318532
refGene_gn=PAPOLG;refGene_tx=XM_011281009.3;refGene_gn=PAPOLG;refGene_tx=XM_019827255.2;refGene_gn=PAPOLG;refGene_tx=XM_019827256.2
A3 81319294 81319415
refGene_gn=PAPOLG;refGene_tx=XM_011281009.3;refGene_gn=PAPOLG;refGene_tx=XM_019827255.2;refGene_gn=PAPOLG;refGene_tx=XM_019827256.2
A3 81319752 81319837
refGene_gn=PAPOLG;refGene_tx=XM_011281009.3;refGene_gn=PAPOLG;refGene_tx=XM_019827255.2;refGene_gn=PAPOLG;refGene_tx=XM_019827256.2

A3 81320228 81320282
refGene_gn=PAPOLG;refGene_tx=XM_011281009.3;refGene_gn=PAPOLG;refGene_tx=XM_019827255.2;refGene_gn=PAPOLG;refGene_tx=XM_019827256.2
A3 81321840 81321960
refGene_gn=PAPOLG;refGene_tx=XM_011281009.3;refGene_gn=PAPOLG;refGene_tx=XM_019827255.2;refGene_gn=PAPOLG;refGene_tx=XM_019827256.2
A3 81322873 81322983
refGene_gn=PAPOLG;refGene_tx=XM_011281009.3;refGene_gn=PAPOLG;refGene_tx=XM_019827255.2;refGene_gn=PAPOLG;refGene_tx=XM_019827256.2
A3 81323683 81323754
refGene_gn=PAPOLG;refGene_tx=XM_019827255.2;refGene_gn=PAPOLG;refGene_tx=XM_019827256.2
A3 81324605 81324727 refGene_gn=PAPOLG;refGene_tx=XM_011281009.3
A3 81324967 81325143 refGene_gn=PAPOLG;refGene_tx=XM_011281009.3
A3 81325527 81325616 refGene_gn=PAPOLG;refGene_tx=XM_011281009.3
A3 81325696 81325917 refGene_gn=PAPOLG;refGene_tx=XM_011281009.3
A3 81326406 81326472 refGene_gn=PAPOLG;refGene_tx=XM_011281009.3
A3 81326657 81326714 refGene_gn=PAPOLG;refGene_tx=XM_011281009.3
A3 81328752 81328851 refGene_gn=PAPOLG;refGene_tx=XM_011281009.3
A3 81419918 81419928 refGene_gn=REL;refGene_tx=XM_003984028.5
A3 81430337 81430480 refGene_gn=REL;refGene_tx=XM_003984028.5
A3 81430969 81431118 refGene_gn=REL;refGene_tx=XM_003984028.5
A3 81436378 81436470 refGene_gn=REL;refGene_tx=XM_003984028.5
A3 81449111 81449252 refGene_gn=REL;refGene_tx=XM_003984028.5
A3 81450363 81450468 refGene_gn=REL;refGene_tx=XM_003984028.5
A3 81451170 81451383 refGene_gn=REL;refGene_tx=XM_003984028.5
A3 81452185 81452254 refGene_gn=REL;refGene_tx=XM_003984028.5
A3 81452382 81452448 refGene_gn=REL;refGene_tx=XM_003984028.5
A3 81453652 81454416 refGene_gn=REL;refGene_tx=XM_003984028.5
A3 81467246 81467285
refGene_gn=PUS10;refGene_tx=XM_006930121.3;refGene_gn=PUS10;refGene_tx=XM_019827257.2
A3 81469352 81469452
refGene_gn=PUS10;refGene_tx=XM_006930121.3;refGene_gn=PUS10;refGene_tx=XM_019827257.2
A3 81472722 81472865
refGene_gn=PUS10;refGene_tx=XM_006930121.3;refGene_gn=PUS10;refGene_tx=XM_019827257.2
A3 81480897 81481015
refGene_gn=PUS10;refGene_tx=XM_006930121.3;refGene_gn=PUS10;refGene_tx=XM_019827257.2
A3 81482066 81482122
refGene_gn=PUS10;refGene_tx=XM_006930121.3;refGene_gn=PUS10;refGene_tx=XM_019827257.2
A3 81482215 81482292
refGene_gn=PUS10;refGene_tx=XM_006930121.3;refGene_gn=PUS10;refGene_tx=XM_019827257.2
A3 81483140 81483197
refGene_gn=PUS10;refGene_tx=XM_006930121.3;refGene_gn=PUS10;refGene_tx=XM_019827257.2

A3	81489511	81489637
refGene_gn=PUS10;refGene_tx=XM_006930121.3;refGene_gn=PUS10;refGene_tx=XM_019827257.2
A3	81490712	81490798
refGene_gn=PUS10;refGene_tx=XM_006930121.3;refGene_gn=PUS10;refGene_tx=XM_019827257.2
A3	81492150	81492215
refGene_gn=PUS10;refGene_tx=XM_006930121.3;refGene_gn=PUS10;refGene_tx=XM_019827257.2
A3	81494619	81494665
refGene_gn=PUS10;refGene_tx=XM_006930121.3;refGene_gn=PUS10;refGene_tx=XM_019827257.2
A3	81494910	81494972
refGene_gn=PUS10;refGene_tx=XM_006930121.3;refGene_gn=PUS10;refGene_tx=XM_019827257.2
A3	81497082	81497194
refGene_gn=PUS10;refGene_tx=XM_006930121.3;refGene_gn=PUS10;refGene_tx=XM_019827257.2
A3	81501169	81501204
refGene_gn=PUS10;refGene_tx=XM_006930121.3;refGene_gn=PUS10;refGene_tx=XM_019827257.2
A3	81531144	81531231
refGene_gn=PUS10;refGene_tx=XM_006930121.3;refGene_gn=PUS10;refGene_tx=XM_019827257.2
A3	81531963	81532227
refGene_gn=PUS10;refGene_tx=XM_006930121.3;refGene_gn=PUS10;refGene_tx=XM_019827257.2
A3	81535297	81535423
refGene_gn=PUS10;refGene_tx=XM_006930121.3;refGene_gn=PUS10;refGene_tx=XM_019827257.2
A3	81539822	81539914	refGene_gn=PEX13;refGene_tx=XM_003984030.5
A3	81553166	81553861	refGene_gn=PEX13;refGene_tx=XM_003984030.5
A3	81554393	81554519	refGene_gn=PEX13;refGene_tx=XM_003984030.5
A3	81556842	81557141	refGene_gn=PEX13;refGene_tx=XM_003984030.5
A3	81575219	81575369
refGene_gn=KIAA1841;refGene_tx=XM_023251689.1;refGene_gn=KIAA1841;refGene_tx=XM_023251690.1;refGene_gn=KIAA1841;refGene_tx=XM_023251691.1;refGene_gn=KIAA1841;refGene_tx=XM_023251692.1
A3	81585527	81585714
refGene_gn=KIAA1841;refGene_tx=XM_023251689.1;refGene_gn=KIAA1841;refGene_tx=XM_023251690.1;refGene_gn=KIAA1841;refGene_tx=XM_023251691.1;refGene_gn=KIAA1841;refGene_tx=XM_023251692.1
A3	81586524	81586618
refGene_gn=KIAA1841;refGene_tx=XM_023251689.1;refGene_gn=KIAA1841;refGene_tx=XM_023251690.1;refGene_gn=KIAA1841;refGene_tx=XM_023251691.1;refGene_gn=KIAA1841;refGene_tx=XM_023251692.1
A3	81588633	81588872
refGene_gn=KIAA1841;refGene_tx=XM_023251689.1;refGene_gn=KIAA1841;refGene_tx=XM_023251690.1;refGene_gn=KIAA1841;refGene_tx=XM_023251691.1;refGene_gn=KIAA1841;refGene_tx=XM_023251692.1;refGene_gn=KIAA1841;refGen

e_tx=XM_023251693.1;refGene_gn=KIAA1841;refGene_tx=XM_023251694.1
A3 81592238 81592297
refGene_gn=KIAA1841;refGene_tx=XM_023251689.1;refGene_gn=KIAA1841;refGene_tx=XM_023251690.1;refGene_gn=KIAA1841;refGene_tx=XM_023251691.1;refGene_gn=KIAA1841;refGene_tx=XM_023251692.1;refGene_gn=KIAA1841;refGene_tx=XM_023251693.1;refGene_gn=KIAA1841;refGene_tx=XM_023251694.1
A3 81595058 81595219
refGene_gn=KIAA1841;refGene_tx=XM_023251689.1;refGene_gn=KIAA1841;refGene_tx=XM_023251690.1;refGene_gn=KIAA1841;refGene_tx=XM_023251691.1;refGene_gn=KIAA1841;refGene_tx=XM_023251692.1;refGene_gn=KIAA1841;refGene_tx=XM_023251693.1;refGene_gn=KIAA1841;refGene_tx=XM_023251694.1
A3 81599671 81599758
refGene_gn=KIAA1841;refGene_tx=XM_023251689.1;refGene_gn=KIAA1841;refGene_tx=XM_023251690.1;refGene_gn=KIAA1841;refGene_tx=XM_023251691.1;refGene_gn=KIAA1841;refGene_tx=XM_023251692.1;refGene_gn=KIAA1841;refGene_tx=XM_023251693.1;refGene_gn=KIAA1841;refGene_tx=XM_023251694.1
A3 81599870 81599981
refGene_gn=KIAA1841;refGene_tx=XM_023251689.1;refGene_gn=KIAA1841;refGene_tx=XM_023251690.1;refGene_gn=KIAA1841;refGene_tx=XM_023251691.1;refGene_gn=KIAA1841;refGene_tx=XM_023251692.1;refGene_gn=KIAA1841;refGene_tx=XM_023251693.1;refGene_gn=KIAA1841;refGene_tx=XM_023251694.1
A3 81602501 81602625
refGene_gn=KIAA1841;refGene_tx=XM_023251689.1;refGene_gn=KIAA1841;refGene_tx=XM_023251690.1;refGene_gn=KIAA1841;refGene_tx=XM_023251691.1;refGene_gn=KIAA1841;refGene_tx=XM_023251692.1;refGene_gn=KIAA1841;refGene_tx=XM_023251693.1;refGene_gn=KIAA1841;refGene_tx=XM_023251694.1
A3 81606098 81606251
refGene_gn=KIAA1841;refGene_tx=XM_023251689.1;refGene_gn=KIAA1841;refGene_tx=XM_023251690.1;refGene_gn=KIAA1841;refGene_tx=XM_023251691.1;refGene_gn=KIAA1841;refGene_tx=XM_023251692.1;refGene_gn=KIAA1841;refGene_tx=XM_023251693.1;refGene_gn=KIAA1841;refGene_tx=XM_023251694.1
A3 81614421 81614567
refGene_gn=KIAA1841;refGene_tx=XM_023251689.1;refGene_gn=KIAA1841;refGene_tx=XM_023251690.1;refGene_gn=KIAA1841;refGene_tx=XM_023251691.1;refGene_gn=KIAA1841;refGene_tx=XM_023251692.1;refGene_gn=KIAA1841;refGene_tx=XM_023251693.1;refGene_gn=KIAA1841;refGene_tx=XM_023251694.1
A3 81615302 81615402
refGene_gn=KIAA1841;refGene_tx=XM_023251689.1;refGene_gn=KIAA1841;refGene_tx=XM_023251690.1;refGene_gn=KIAA1841;refGene_tx=XM_023251691.1;refGene_gn=KIAA1841;refGene_tx=XM_023251692.1;refGene_gn=KIAA1841;refGene_tx=XM_023251693.1;refGene_gn=KIAA1841;refGene_tx=XM_023251694.1
A3 81616866 81616899
refGene_gn=KIAA1841;refGene_tx=XM_023251689.1;refGene_gn=KIAA1841;refGene_tx=XM_023251690.1;refGene_gn=KIAA1841;refGene_tx=XM_023251691.1;refGene_gn=KIAA1841;refGene_tx=XM_023251692.1;refGene_gn=KIAA1841;refGene_tx=XM_023251693.1;refGene_gn=KIAA1841;refGene_tx=XM_023251694.1
A3 81617250 81617293 refGene_gn=KIAA1841;refGene_tx=XM_023251692.1
A3 81617731 81617831
refGene_gn=KIAA1841;refGene_tx=XM_023251689.1;refGene_gn=KIAA1841;refGene_tx=XM_023251690.1;refGene_gn=KIAA1841;refGene_tx=XM_023251691.1;re

fGene_gn=KIAA1841;refGene_tx=XM_023251692.1;refGene_gn=KIAA1841;refGene_tx=XM_023251693.1;refGene_gn=KIAA1841;refGene_tx=XM_023251694.1
A3 81629378 81629470
refGene_gn=KIAA1841;refGene_tx=XM_023251689.1;refGene_gn=KIAA1841;refGene_tx=XM_023251690.1;refGene_gn=KIAA1841;refGene_tx=XM_023251691.1;refGene_gn=KIAA1841;refGene_tx=XM_023251693.1;refGene_gn=KIAA1841;refGene_tx=XM_023251694.1
A3 81630349 81630378
refGene_gn=KIAA1841;refGene_tx=XM_023251689.1;refGene_gn=KIAA1841;refGene_tx=XM_023251690.1;refGene_gn=KIAA1841;refGene_tx=XM_023251691.1;refGene_gn=KIAA1841;refGene_tx=XM_023251693.1;refGene_gn=KIAA1841;refGene_tx=XM_023251694.1
A3 81630455 81630529
refGene_gn=KIAA1841;refGene_tx=XM_023251689.1;refGene_gn=KIAA1841;refGene_tx=XM_023251690.1;refGene_gn=KIAA1841;refGene_tx=XM_023251691.1;refGene_gn=KIAA1841;refGene_tx=XM_023251693.1;refGene_gn=KIAA1841;refGene_tx=XM_023251694.1
A3 81631342 81631431
refGene_gn=KIAA1841;refGene_tx=XM_023251689.1;refGene_gn=KIAA1841;refGene_tx=XM_023251690.1;refGene_gn=KIAA1841;refGene_tx=XM_023251691.1;refGene_gn=KIAA1841;refGene_tx=XM_023251693.1;refGene_gn=KIAA1841;refGene_tx=XM_023251694.1
A3 81631574 81631622 refGene_gn=KIAA1841;refGene_tx=XM_023251691.1
A3 81633125 81633217
refGene_gn=KIAA1841;refGene_tx=XM_023251689.1;refGene_gn=KIAA1841;refGene_tx=XM_023251690.1;refGene_gn=KIAA1841;refGene_tx=XM_023251693.1;refGene_gn=KIAA1841;refGene_tx=XM_023251694.1
A3 81634116 81634153
refGene_gn=KIAA1841;refGene_tx=XM_023251689.1;refGene_gn=KIAA1841;refGene_tx=XM_023251690.1;refGene_gn=KIAA1841;refGene_tx=XM_023251693.1;refGene_gn=KIAA1841;refGene_tx=XM_023251694.1
A3 81702282 81702398
refGene_gn=CA3H2orf74;refGene_tx=XM_006930122.3
A3 81702629 81702743
refGene_gn=CA3H2orf74;refGene_tx=XM_006930122.3
A3 81702844 81703025
refGene_gn=CA3H2orf74;refGene_tx=XM_006930122.3
A3 81703801 81703999
refGene_gn=CA3H2orf74;refGene_tx=XM_006930122.3
A3 81712606 81712921
refGene_gn=LOC109498186;refGene_tx=XM_019827258.2
A3 81714199 81714261 refGene_gn=AHSA2P;refGene_tx=XM_023252184.1
A3 81720323 81721290 refGene_gn=AHSA2P;refGene_tx=XM_023252184.1
A3 81722278 81722353 refGene_gn=AHSA2P;refGene_tx=XM_023252184.1
A3 81741626 81742234 refGene_gn=USP34;refGene_tx=XM_023251695.1
A3 81742436 81742596 refGene_gn=USP34;refGene_tx=XM_023251695.1
A3 81743263 81743407 refGene_gn=USP34;refGene_tx=XM_023251695.1
A3 81743513 81743654 refGene_gn=USP34;refGene_tx=XM_023251695.1
A3 81745773 81745853 refGene_gn=USP34;refGene_tx=XM_023251695.1
A3 81754032 81754156 refGene_gn=USP34;refGene_tx=XM_023251695.1

| | | | |
|---|---|---|---|
| A3 | 81755146 | 81755271 | refGene_gn=USP34;refGene_tx=XM_023251695.1 |
| A3 | 81755387 | 81755492 | refGene_gn=USP34;refGene_tx=XM_023251695.1 |
| A3 | 81756470 | 81756578 | refGene_gn=USP34;refGene_tx=XM_023251695.1 |
| A3 | 81757383 | 81757510 | refGene_gn=USP34;refGene_tx=XM_023251695.1 |
| A3 | 81759048 | 81759127 | refGene_gn=USP34;refGene_tx=XM_023251695.1 |
| A3 | 81760558 | 81760716 | refGene_gn=USP34;refGene_tx=XM_023251695.1 |
| A3 | 81761699 | 81762334 | refGene_gn=USP34;refGene_tx=XM_023251695.1 |
| A3 | 81767466 | 81767614 | refGene_gn=USP34;refGene_tx=XM_023251695.1 |
| A3 | 81768643 | 81768748 | refGene_gn=USP34;refGene_tx=XM_023251695.1 |
| A3 | 81769743 | 81769788 | refGene_gn=USP34;refGene_tx=XM_023251695.1 |
| A3 | 81769867 | 81769972 | refGene_gn=USP34;refGene_tx=XM_023251695.1 |
| A3 | 81770052 | 81770101 | refGene_gn=USP34;refGene_tx=XM_023251695.1 |
| A3 | 81773602 | 81773754 | refGene_gn=USP34;refGene_tx=XM_023251695.1 |
| A3 | 81774867 | 81774942 | refGene_gn=USP34;refGene_tx=XM_023251695.1 |
| A3 | 81775046 | 81775215 | refGene_gn=USP34;refGene_tx=XM_023251695.1 |
| A3 | 81775817 | 81775903 | refGene_gn=USP34;refGene_tx=XM_023251695.1 |
| A3 | 81777630 | 81777711 | refGene_gn=USP34;refGene_tx=XM_023251695.1 |
| A3 | 81780313 | 81780379 | refGene_gn=USP34;refGene_tx=XM_023251695.1 |
| A3 | 81780493 | 81780541 | refGene_gn=USP34;refGene_tx=XM_023251695.1 |
| A3 | 81780623 | 81780699 | refGene_gn=USP34;refGene_tx=XM_023251695.1 |
| A3 | 81780794 | 81780859 | refGene_gn=USP34;refGene_tx=XM_023251695.1 |
| A3 | 81784668 | 81784764 | refGene_gn=USP34;refGene_tx=XM_023251695.1 |
| A3 | 81784898 | 81784952 | refGene_gn=USP34;refGene_tx=XM_023251695.1 |
| A3 | 81788545 | 81788624 | refGene_gn=USP34;refGene_tx=XM_023251695.1 |
| A3 | 81789714 | 81789868 | refGene_gn=USP34;refGene_tx=XM_023251695.1 |
| A3 | 81793615 | 81793788 | refGene_gn=USP34;refGene_tx=XM_023251695.1 |
| A3 | 81796654 | 81796749 | refGene_gn=USP34;refGene_tx=XM_023251695.1 |
| A3 | 81797395 | 81797424 | refGene_gn=USP34;refGene_tx=XM_023251695.1 |
| A3 | 81797426 | 81797472 | refGene_gn=USP34;refGene_tx=XM_023251695.1 |
| A3 | 81797563 | 81797620 | refGene_gn=USP34;refGene_tx=XM_023251695.1 |
| A3 | 81797703 | 81797850 | refGene_gn=USP34;refGene_tx=XM_023251695.1 |
| A3 | 81798667 | 81798733 | refGene_gn=USP34;refGene_tx=XM_023251695.1 |
| A3 | 81802839 | 81802864 | refGene_gn=USP34;refGene_tx=XM_023251695.1 |
| A3 | 81802865 | 81802885 | refGene_gn=USP34;refGene_tx=XM_023251695.1 |
| A3 | 81802886 | 81802967 | refGene_gn=USP34;refGene_tx=XM_023251695.1 |
| A3 | 81802969 | 81802986 | refGene_gn=USP34;refGene_tx=XM_023251695.1 |
| A3 | 81803468 | 81803673 | refGene_gn=USP34;refGene_tx=XM_023251695.1 |
| A3 | 81813983 | 81814104 | refGene_gn=USP34;refGene_tx=XM_023251695.1 |
| A3 | 81814206 | 81814262 | refGene_gn=USP34;refGene_tx=XM_023251695.1 |
| A3 | 81815546 | 81815651 | refGene_gn=USP34;refGene_tx=XM_023251695.1 |
| A3 | 81816822 | 81816975 | refGene_gn=USP34;refGene_tx=XM_023251695.1 |
| A3 | 81817967 | 81818092 | refGene_gn=USP34;refGene_tx=XM_023251695.1 |
| A3 | 81818222 | 81818263 | refGene_gn=USP34;refGene_tx=XM_023251695.1 |
| A3 | 81821471 | 81821554 | refGene_gn=USP34;refGene_tx=XM_023251695.1 |
| A3 | 81824420 | 81824621 | refGene_gn=USP34;refGene_tx=XM_023251695.1 |
| A3 | 81834582 | 81834669 | refGene_gn=USP34;refGene_tx=XM_023251695.1 |
| A3 | 81835156 | 81835240 | refGene_gn=USP34;refGene_tx=XM_023251695.1 |
| A3 | 81836032 | 81836155 | refGene_gn=USP34;refGene_tx=XM_023251695.1 |
| A3 | 81837196 | 81837322 | refGene_gn=USP34;refGene_tx=XM_023251695.1 |
| A3 | 81841022 | 81841232 | refGene_gn=USP34;refGene_tx=XM_023251695.1 |

A3 81841640 81841741 refGene_gn=USP34;refGene_tx=XM_023251695.1
A3 81847372 81847520 refGene_gn=USP34;refGene_tx=XM_023251695.1
A3 81847616 81847743 refGene_gn=USP34;refGene_tx=XM_023251695.1
A3 81850457 81850617 refGene_gn=USP34;refGene_tx=XM_023251695.1
A3 81850721 81850821 refGene_gn=USP34;refGene_tx=XM_023251695.1
A3 81852614 81852728 refGene_gn=USP34;refGene_tx=XM_023251695.1
A3 81853703 81853858 refGene_gn=USP34;refGene_tx=XM_023251695.1
A3 81856311 81856394 refGene_gn=USP34;refGene_tx=XM_023251695.1
A3 81861671 81861767 refGene_gn=USP34;refGene_tx=XM_023251695.1
A3 81864048 81864138 refGene_gn=USP34;refGene_tx=XM_023251695.1
A3 81872724 81872852 refGene_gn=USP34;refGene_tx=XM_023251695.1
A3 81872941 81873057 refGene_gn=USP34;refGene_tx=XM_023251695.1
A3 81877228 81877443 refGene_gn=USP34;refGene_tx=XM_023251695.1
A3 81879640 81880254 refGene_gn=USP34;refGene_tx=XM_023251695.1
A3 81881084 81881215 refGene_gn=USP34;refGene_tx=XM_023251695.1
A3 81881549 81881585 refGene_gn=USP34;refGene_tx=XM_023251695.1
A3 81882236 81882366 refGene_gn=USP34;refGene_tx=XM_023251695.1
A3 81882533 81882659 refGene_gn=USP34;refGene_tx=XM_023251695.1
A3 81895869 81895962 refGene_gn=USP34;refGene_tx=XM_023251695.1
A3 81896051 81896133 refGene_gn=USP34;refGene_tx=XM_023251695.1
A3 81905652 81905714 refGene_gn=USP34;refGene_tx=XM_023251695.1
A3 81906692 81906885 refGene_gn=USP34;refGene_tx=XM_023251695.1
A3 81910192 81910260 refGene_gn=USP34;refGene_tx=XM_023251695.1
A3 81926026 81926176 refGene_gn=USP34;refGene_tx=XM_023251695.1
A3 81926337 81926388 refGene_gn=USP34;refGene_tx=XM_023251695.1
A3 81933602 81934023 refGene_gn=USP34;refGene_tx=XM_023251695.1
A3 81947931 81948019 refGene_gn=USP34;refGene_tx=XM_023251695.1
A3 81991115 81991158 refGene_gn=USP34;refGene_tx=XM_023251695.1
A3 81991318 81991330
refGene_gn=LOC102902567;refGene_tx=XM_023252185.1
A3 81991364 81991843
refGene_gn=LOC102902567;refGene_tx=XM_023252185.1
A3 81992944 81993053
refGene_gn=LOC102902567;refGene_tx=XM_023252185.1
A3 82027575 82027722
refGene_gn=XPO1;refGene_tx=XM_019827268.2;refGene_gn=XPO1;refGene_tx=XM_023251696.1;refGene_gn=XPO1;refGene_tx=XM_023251697.1;refGene_gn=XPO1;refGene_tx=XM_023251698.1
A3 82028821 82028918
refGene_gn=XPO1;refGene_tx=XM_019827268.2;refGene_gn=XPO1;refGene_tx=XM_023251696.1;refGene_gn=XPO1;refGene_tx=XM_023251697.1;refGene_gn=XPO1;refGene_tx=XM_023251698.1
A3 82029661 82029821
refGene_gn=XPO1;refGene_tx=XM_019827268.2;refGene_gn=XPO1;refGene_tx=XM_023251696.1;refGene_gn=XPO1;refGene_tx=XM_023251697.1;refGene_gn=XPO1;refGene_tx=XM_023251698.1
A3 82029903 82030038
refGene_gn=XPO1;refGene_tx=XM_019827268.2;refGene_gn=XPO1;refGene_tx=XM_023251696.1;refGene_gn=XPO1;refGene_tx=XM_023251697.1;refGene_gn=XPO1;refGene_tx=XM_023251698.1

A3 82030713 82030882
refGene_gn=XPO1;refGene_tx=XM_019827268.2;refGene_gn=XPO1;refGene_tx=XM_023251696.1;refGene_gn=XPO1;refGene_tx=XM_023251697.1;refGene_gn=XPO1;refGene_tx=XM_023251698.1
A3 82031565 82031760
refGene_gn=XPO1;refGene_tx=XM_019827268.2;refGene_gn=XPO1;refGene_tx=XM_023251696.1;refGene_gn=XPO1;refGene_tx=XM_023251697.1;refGene_gn=XPO1;refGene_tx=XM_023251698.1
A3 82033398 82033505
refGene_gn=XPO1;refGene_tx=XM_019827268.2;refGene_gn=XPO1;refGene_tx=XM_023251696.1;refGene_gn=XPO1;refGene_tx=XM_023251697.1;refGene_gn=XPO1;refGene_tx=XM_023251698.1
A3 82033804 82033988
refGene_gn=XPO1;refGene_tx=XM_019827268.2;refGene_gn=XPO1;refGene_tx=XM_023251696.1;refGene_gn=XPO1;refGene_tx=XM_023251697.1;refGene_gn=XPO1;refGene_tx=XM_023251698.1
A3 82034337 82034472
refGene_gn=XPO1;refGene_tx=XM_019827268.2;refGene_gn=XPO1;refGene_tx=XM_023251696.1;refGene_gn=XPO1;refGene_tx=XM_023251697.1;refGene_gn=XPO1;refGene_tx=XM_023251698.1
A3 82035040 82035204
refGene_gn=XPO1;refGene_tx=XM_019827268.2;refGene_gn=XPO1;refGene_tx=XM_023251696.1;refGene_gn=XPO1;refGene_tx=XM_023251697.1;refGene_gn=XPO1;refGene_tx=XM_023251698.1
A3 82035304 82035461
refGene_gn=XPO1;refGene_tx=XM_019827268.2;refGene_gn=XPO1;refGene_tx=XM_023251696.1;refGene_gn=XPO1;refGene_tx=XM_023251697.1;refGene_gn=XPO1;refGene_tx=XM_023251698.1
A3 82035539 82035721
refGene_gn=XPO1;refGene_tx=XM_019827268.2;refGene_gn=XPO1;refGene_tx=XM_023251696.1;refGene_gn=XPO1;refGene_tx=XM_023251697.1;refGene_gn=XPO1;refGene_tx=XM_023251698.1
A3 82035912 82036051
refGene_gn=XPO1;refGene_tx=XM_019827268.2;refGene_gn=XPO1;refGene_tx=XM_023251696.1;refGene_gn=XPO1;refGene_tx=XM_023251697.1;refGene_gn=XPO1;refGene_tx=XM_023251698.1
A3 82036820 82037018
refGene_gn=XPO1;refGene_tx=XM_019827268.2;refGene_gn=XPO1;refGene_tx=XM_023251696.1;refGene_gn=XPO1;refGene_tx=XM_023251697.1;refGene_gn=XPO1;refGene_tx=XM_023251698.1
A3 82038347 82038506
refGene_gn=XPO1;refGene_tx=XM_019827268.2;refGene_gn=XPO1;refGene_tx=XM_023251696.1;refGene_gn=XPO1;refGene_tx=XM_023251697.1;refGene_gn=XPO1;refGene_tx=XM_023251698.1
A3 82039167 82039296
refGene_gn=XPO1;refGene_tx=XM_019827268.2;refGene_gn=XPO1;refGene_tx=XM_023251696.1;refGene_gn=XPO1;refGene_tx=XM_023251697.1;refGene_gn=XPO1;refGene_tx=XM_023251698.1
A3 82040418 82040538
refGene_gn=XPO1;refGene_tx=XM_019827268.2;refGene_gn=XPO1;refGene_tx=X

M_023251696.1;refGene_gn=XPO1;refGene_tx=XM_023251697.1;refGene_gn=XPO1;refGene_tx=XM_023251698.1
A3 82040619 82040668 refGene_gn=XPO1;refGene_tx=XM_019827268.2;refGene_gn=XPO1;refGene_tx=XM_023251696.1;refGene_gn=XPO1;refGene_tx=XM_023251697.1;refGene_gn=XPO1;refGene_tx=XM_023251698.1
A3 82041887 82042069 refGene_gn=XPO1;refGene_tx=XM_019827268.2;refGene_gn=XPO1;refGene_tx=XM_023251696.1;refGene_gn=XPO1;refGene_tx=XM_023251697.1;refGene_gn=XPO1;refGene_tx=XM_023251698.1
A3 82043351 82043396 refGene_gn=XPO1;refGene_tx=XM_019827268.2;refGene_gn=XPO1;refGene_tx=XM_023251696.1;refGene_gn=XPO1;refGene_tx=XM_023251697.1;refGene_gn=XPO1;refGene_tx=XM_023251698.1
A3 82043608 82043670 refGene_gn=XPO1;refGene_tx=XM_023251696.1;refGene_gn=XPO1;refGene_tx=XM_023251697.1
A3 82058056 82058129 refGene_gn=XPO1;refGene_tx=XM_023251696.1;refGene_gn=XPO1;refGene_tx=XM_023251697.1
A3 82061564 82061666 refGene_gn=XPO1;refGene_tx=XM_023251696.1;refGene_gn=XPO1;refGene_tx=XM_023251697.1
A3 82066808 82066934 refGene_gn=XPO1;refGene_tx=XM_023251696.1;refGene_gn=XPO1;refGene_tx=XM_023251697.1
A3 82298434 82298573 refGene_gn=FAM161A;refGene_tx=XM_003984032.5;refGene_gn=FAM161A;refGene_tx=XM_003984033.5
A3 82299012 82299167 refGene_gn=FAM161A;refGene_tx=XM_003984032.5;refGene_gn=FAM161A;refGene_tx=XM_003984033.5
A3 82306899 82306999 refGene_gn=FAM161A;refGene_tx=XM_003984032.5;refGene_gn=FAM161A;refGene_tx=XM_003984033.5
A3 82308194 82308285 refGene_gn=FAM161A;refGene_tx=XM_006930123.4
A3 82309035 82309203 refGene_gn=FAM161A;refGene_tx=XM_003984033.5;refGene_gn=FAM161A;refGene_tx=XM_006930123.4
A3 82309558 82310719 refGene_gn=FAM161A;refGene_tx=XM_003984032.5;refGene_gn=FAM161A;refGene_tx=XM_003984033.5;refGene_gn=FAM161A;refGene_tx=XM_006930123.4
A3 82312976 82313215 refGene_gn=FAM161A;refGene_tx=XM_003984032.5;refGene_gn=FAM161A;refGene_tx=XM_003984033.5;refGene_gn=FAM161A;refGene_tx=XM_006930123.4
A3 82323980 82324166 refGene_gn=FAM161A;refGene_tx=XM_003984032.5;refGene_gn=FAM161A;refGene_tx=XM_003984033.5;refGene_gn=FAM161A;refGene_tx=XM_006930123.4
A3 82335741 82335756 refGene_gn=CCT4;refGene_tx=XM_003984034.5
A3 82335991 82336105 refGene_gn=CCT4;refGene_tx=XM_003984034.5

A3 82337528 82337763 refGene_gn=CCT4;refGene_tx=XM_003984034.5
A3 82337896 82338027 refGene_gn=CCT4;refGene_tx=XM_003984034.5
A3 82338118 82338229 refGene_gn=CCT4;refGene_tx=XM_003984034.5
A3 82338307 82338404 refGene_gn=CCT4;refGene_tx=XM_003984034.5
A3 82340419 82340559 refGene_gn=CCT4;refGene_tx=XM_003984034.5
A3 82341774 82341907 refGene_gn=CCT4;refGene_tx=XM_003984034.5
A3 82342592 82342714 refGene_gn=CCT4;refGene_tx=XM_003984034.5
A3 82344126 82344269 refGene_gn=CCT4;refGene_tx=XM_003984034.5
A3 82345377 82345486 refGene_gn=CCT4;refGene_tx=XM_003984034.5
A3 82346395 82346485 refGene_gn=CCT4;refGene_tx=XM_003984034.5
A3 82347259 82347312 refGene_gn=CCT4;refGene_tx=XM_003984034.5
A3 82350024 82350151 refGene_gn=CCT4;refGene_tx=XM_003984034.5
A3 82360581 82360758
refGene_gn=COMMD1;refGene_tx=XM_003984035.4;refGene_gn=COMMD1;refGene_tx=XM_019827269.2;refGene_gn=COMMD1;refGene_tx=XM_019827272.2
A3 82444649 82444931
refGene_gn=COMMD1;refGene_tx=XM_003984035.4;refGene_gn=COMMD1;refGene_tx=XM_019827269.2;refGene_gn=COMMD1;refGene_tx=XM_019827272.2
A3 82538924 82538987 refGene_gn=COMMD1;refGene_tx=XM_019827269.2
A3 82541887 82541923 refGene_gn=COMMD1;refGene_tx=XM_019827272.2
A3 82549758 82549866
refGene_gn=COMMD1;refGene_tx=XM_003984035.4;refGene_gn=COMMD1;refGene_tx=XM_019827269.2
A3 82641544 82642738
refGene_gn=B3GNT2;refGene_tx=XM_006930126.4;refGene_gn=B3GNT2;refGene_tx=XM_006930127.3;refGene_gn=B3GNT2;refGene_tx=XM_023251700.1;refGene_gn=B3GNT2;refGene_tx=XM_023251701.1;refGene_gn=B3GNT2;refGene_tx=XM_023251702.1
A3 82858995 82859274 refGene_gn=TMEM17;refGene_tx=XM_003984037.5
A3 82860167 82860281 refGene_gn=TMEM17;refGene_tx=XM_003984037.5
A3 82860453 82860557 refGene_gn=TMEM17;refGene_tx=XM_003984037.5
A3 82863971 82864071 refGene_gn=TMEM17;refGene_tx=XM_003984037.5
A3 83004754 83004756 refGene_gn=EHBP1;refGene_tx=XM_019827277.2
A3 83036306 83036410
refGene_gn=EHBP1;refGene_tx=XM_003984038.4;refGene_gn=EHBP1;refGene_tx=XM_019827273.2;refGene_gn=EHBP1;refGene_tx=XM_019827274.2;refGene_gn=EHBP1;refGene_tx=XM_019827275.2;refGene_gn=EHBP1;refGene_tx=XM_019827276.2;refGene_gn=EHBP1;refGene_tx=XM_019827278.2;refGene_gn=EHBP1;refGene_tx=XM_019827279.2;refGene_gn=EHBP1;refGene_tx=XM_023251703.1;refGene_gn=EHBP1;refGene_tx=XM_023251704.1;refGene_gn=EHBP1;refGene_tx=XM_023251705.1;refGene_gn=EHBP1;refGene_tx=XM_023251706.1;refGene_gn=EHBP1;refGene_tx=XM_023251707.1
A3 83101543 83101601
refGene_gn=EHBP1;refGene_tx=XM_003984038.4;refGene_gn=EHBP1;refGene_tx=XM_019827273.2;refGene_gn=EHBP1;refGene_tx=XM_019827274.2;refGene_gn=EHBP1;refGene_tx=XM_019827275.2;refGene_gn=EHBP1;refGene_tx=XM_019827276.2;refGene_gn=EHBP1;refGene_tx=XM_019827277.2;refGene_gn=EHBP1;refGene_tx=XM_019827278.2;refGene_gn=EHBP1;refGene_tx=XM_019827279.2;refGene_gn=EHBP1;refGene_tx=XM_023251703.1;refGene_gn=EHBP1;refGene_tx=XM_023251704.1;refGene_gn=EHBP1;refGene_tx=XM_023251705.1;refGene_gn=EHBP1;

refGene_tx=XM_023251706.1;refGene_gn=EHBP1;refGene_tx=XM_023251707.1
A3 83110850 83110946
refGene_gn=EHBP1;refGene_tx=XM_003984038.4;refGene_gn=EHBP1;refGene_tx
=XM_019827273.2;refGene_gn=EHBP1;refGene_tx=XM_019827274.2;refGene_gn=
EHBP1;refGene_tx=XM_019827275.2;refGene_gn=EHBP1;refGene_tx=XM_0198272
76.2;refGene_gn=EHBP1;refGene_tx=XM_019827277.2;refGene_gn=EHBP1;refGe
ne_tx=XM_019827278.2;refGene_gn=EHBP1;refGene_tx=XM_019827279.2;refGen
e_gn=EHBP1;refGene_tx=XM_023251703.1;refGene_gn=EHBP1;refGene_tx=XM_02
3251704.1;refGene_gn=EHBP1;refGene_tx=XM_023251705.1;refGene_gn=EHBP1;
refGene_tx=XM_023251706.1;refGene_gn=EHBP1;refGene_tx=XM_023251707.1
A3 83117472 83117526
refGene_gn=EHBP1;refGene_tx=XM_003984038.4;refGene_gn=EHBP1;refGene_tx
=XM_019827273.2;refGene_gn=EHBP1;refGene_tx=XM_019827274.2;refGene_gn=
EHBP1;refGene_tx=XM_019827275.2;refGene_gn=EHBP1;refGene_tx=XM_0198272
76.2;refGene_gn=EHBP1;refGene_tx=XM_019827277.2;refGene_gn=EHBP1;refGe
ne_tx=XM_019827278.2;refGene_gn=EHBP1;refGene_tx=XM_019827279.2;refGen
e_gn=EHBP1;refGene_tx=XM_023251703.1;refGene_gn=EHBP1;refGene_tx=XM_02
3251704.1;refGene_gn=EHBP1;refGene_tx=XM_023251705.1;refGene_gn=EHBP1;
refGene_tx=XM_023251706.1;refGene_gn=EHBP1;refGene_tx=XM_023251707.1
A3 83167214 83167396
refGene_gn=EHBP1;refGene_tx=XM_003984038.4;refGene_gn=EHBP1;refGene_tx
=XM_019827273.2;refGene_gn=EHBP1;refGene_tx=XM_019827274.2;refGene_gn=
EHBP1;refGene_tx=XM_019827275.2;refGene_gn=EHBP1;refGene_tx=XM_0198272
76.2;refGene_gn=EHBP1;refGene_tx=XM_019827277.2;refGene_gn=EHBP1;refGe
ne_tx=XM_019827278.2;refGene_gn=EHBP1;refGene_tx=XM_019827279.2;refGen
e_gn=EHBP1;refGene_tx=XM_023251703.1;refGene_gn=EHBP1;refGene_tx=XM_02
3251704.1;refGene_gn=EHBP1;refGene_tx=XM_023251705.1;refGene_gn=EHBP1;
refGene_tx=XM_023251706.1;refGene_gn=EHBP1;refGene_tx=XM_023251707.1
A3 83168498 83168587 refGene_gn=EHBP1;refGene_tx=XM_019827280.2
A3 83170089 83170229
refGene_gn=EHBP1;refGene_tx=XM_003984038.4;refGene_gn=EHBP1;refGene_tx
=XM_019827273.2;refGene_gn=EHBP1;refGene_tx=XM_019827274.2;refGene_gn=
EHBP1;refGene_tx=XM_019827275.2;refGene_gn=EHBP1;refGene_tx=XM_0198272
76.2;refGene_gn=EHBP1;refGene_tx=XM_019827277.2;refGene_gn=EHBP1;refGe
ne_tx=XM_019827278.2;refGene_gn=EHBP1;refGene_tx=XM_019827279.2;refGen
e_gn=EHBP1;refGene_tx=XM_019827280.2;refGene_gn=EHBP1;refGene_tx=XM_02
3251703.1;refGene_gn=EHBP1;refGene_tx=XM_023251704.1;refGene_gn=EHBP1;
refGene_tx=XM_023251705.1;refGene_gn=EHBP1;refGene_tx=XM_023251706.1;r
efGene_gn=EHBP1;refGene_tx=XM_023251707.1
A3 83183396 83183501
refGene_gn=EHBP1;refGene_tx=XM_019827273.2;refGene_gn=EHBP1;refGene_tx
=XM_019827274.2;refGene_gn=EHBP1;refGene_tx=XM_019827275.2;refGene_gn=
EHBP1;refGene_tx=XM_019827276.2;refGene_gn=EHBP1;refGene_tx=XM_0198272
77.2;refGene_gn=EHBP1;refGene_tx=XM_019827278.2;refGene_gn=EHBP1;refGe
ne_tx=XM_019827280.2;refGene_gn=EHBP1;refGene_tx=XM_023251703.1;refGen
e_gn=EHBP1;refGene_tx=XM_023251704.1;refGene_gn=EHBP1;refGene_tx=XM_02
3251705.1;refGene_gn=EHBP1;refGene_tx=XM_023251706.1
A3 83184136 83184259
refGene_gn=EHBP1;refGene_tx=XM_003984038.4;refGene_gn=EHBP1;refGene_tx
=XM_019827273.2;refGene_gn=EHBP1;refGene_tx=XM_019827274.2;refGene_gn=

EHBP1;refGene_tx=XM_019827275.2;refGene_gn=EHBP1;refGene_tx=XM_0198272
76.2;refGene_gn=EHBP1;refGene_tx=XM_019827277.2;refGene_gn=EHBP1;refGe
ne_tx=XM_019827278.2;refGene_gn=EHBP1;refGene_tx=XM_019827279.2;refGen
e_gn=EHBP1;refGene_tx=XM_019827280.2;refGene_gn=EHBP1;refGene_tx=XM_02
3251703.1;refGene_gn=EHBP1;refGene_tx=XM_023251704.1;refGene_gn=EHBP1;
refGene_tx=XM_023251705.1;refGene_gn=EHBP1;refGene_tx=XM_023251706.1;r
efGene_gn=EHBP1;refGene_tx=XM_023251707.1
A3 83188030 83188271
refGene_gn=EHBP1;refGene_tx=XM_003984038.4;refGene_gn=EHBP1;refGene_tx
=XM_019827273.2;refGene_gn=EHBP1;refGene_tx=XM_019827274.2;refGene_gn=
EHBP1;refGene_tx=XM_019827275.2;refGene_gn=EHBP1;refGene_tx=XM_0198272
76.2;refGene_gn=EHBP1;refGene_tx=XM_019827277.2;refGene_gn=EHBP1;refGe
ne_tx=XM_019827278.2;refGene_gn=EHBP1;refGene_tx=XM_019827279.2;refGen
e_gn=EHBP1;refGene_tx=XM_019827280.2;refGene_gn=EHBP1;refGene_tx=XM_02
3251703.1;refGene_gn=EHBP1;refGene_tx=XM_023251704.1;refGene_gn=EHBP1;
refGene_tx=XM_023251705.1;refGene_gn=EHBP1;refGene_tx=XM_023251706.1;r
efGene_gn=EHBP1;refGene_tx=XM_023251707.1
A3 83195901 83196091
refGene_gn=EHBP1;refGene_tx=XM_003984038.4;refGene_gn=EHBP1;refGene_tx
=XM_019827273.2;refGene_gn=EHBP1;refGene_tx=XM_019827274.2;refGene_gn=
EHBP1;refGene_tx=XM_019827275.2;refGene_gn=EHBP1;refGene_tx=XM_0198272
76.2;refGene_gn=EHBP1;refGene_tx=XM_019827277.2;refGene_gn=EHBP1;refGe
ne_tx=XM_019827278.2;refGene_gn=EHBP1;refGene_tx=XM_019827279.2;refGen
e_gn=EHBP1;refGene_tx=XM_019827280.2;refGene_gn=EHBP1;refGene_tx=XM_02
3251703.1;refGene_gn=EHBP1;refGene_tx=XM_023251704.1;refGene_gn=EHBP1;
refGene_tx=XM_023251705.1;refGene_gn=EHBP1;refGene_tx=XM_023251706.1;r
efGene_gn=EHBP1;refGene_tx=XM_023251707.1
A3 83263671 83263850
refGene_gn=EHBP1;refGene_tx=XM_003984038.4;refGene_gn=EHBP1;refGene_tx
=XM_019827273.2;refGene_gn=EHBP1;refGene_tx=XM_019827274.2;refGene_gn=
EHBP1;refGene_tx=XM_019827275.2;refGene_gn=EHBP1;refGene_tx=XM_0198272
76.2;refGene_gn=EHBP1;refGene_tx=XM_019827277.2;refGene_gn=EHBP1;refGe
ne_tx=XM_019827278.2;refGene_gn=EHBP1;refGene_tx=XM_019827279.2;refGen
e_gn=EHBP1;refGene_tx=XM_019827280.2;refGene_gn=EHBP1;refGene_tx=XM_02
3251703.1;refGene_gn=EHBP1;refGene_tx=XM_023251704.1;refGene_gn=EHBP1;
refGene_tx=XM_023251705.1;refGene_gn=EHBP1;refGene_tx=XM_023251706.1;r
efGene_gn=EHBP1;refGene_tx=XM_023251707.1
A3 83264912 83264961
refGene_gn=EHBP1;refGene_tx=XM_003984038.4;refGene_gn=EHBP1;refGene_tx
=XM_019827273.2;refGene_gn=EHBP1;refGene_tx=XM_019827274.2;refGene_gn=
EHBP1;refGene_tx=XM_019827275.2;refGene_gn=EHBP1;refGene_tx=XM_0198272
76.2;refGene_gn=EHBP1;refGene_tx=XM_019827277.2;refGene_gn=EHBP1;refGe
ne_tx=XM_019827278.2;refGene_gn=EHBP1;refGene_tx=XM_019827279.2;refGen
e_gn=EHBP1;refGene_tx=XM_019827280.2;refGene_gn=EHBP1;refGene_tx=XM_02
3251703.1;refGene_gn=EHBP1;refGene_tx=XM_023251704.1;refGene_gn=EHBP1;
refGene_tx=XM_023251705.1;refGene_gn=EHBP1;refGene_tx=XM_023251706.1;r
efGene_gn=EHBP1;refGene_tx=XM_023251707.1
A3 83268982 83269885
refGene_gn=EHBP1;refGene_tx=XM_003984038.4;refGene_gn=EHBP1;refGene_tx
=XM_019827273.2;refGene_gn=EHBP1;refGene_tx=XM_019827274.2;refGene_gn=

EHBP1;refGene_tx=XM_019827275.2;refGene_gn=EHBP1;refGene_tx=XM_019827276.2;refGene_gn=EHBP1;refGene_tx=XM_019827277.2;refGene_gn=EHBP1;refGene_tx=XM_019827278.2;refGene_gn=EHBP1;refGene_tx=XM_019827279.2;refGene_gn=EHBP1;refGene_tx=XM_019827280.2;refGene_gn=EHBP1;refGene_tx=XM_023251703.1;refGene_gn=EHBP1;refGene_tx=XM_023251704.1;refGene_gn=EHBP1;refGene_tx=XM_023251705.1;refGene_gn=EHBP1;refGene_tx=XM_023251706.1;refGene_gn=EHBP1;refGene_tx=XM_023251707.1
A3 83278484 83278628
refGene_gn=EHBP1;refGene_tx=XM_003984038.4;refGene_gn=EHBP1;refGene_tx=XM_019827273.2;refGene_gn=EHBP1;refGene_tx=XM_019827274.2;refGene_gn=EHBP1;refGene_tx=XM_019827275.2;refGene_gn=EHBP1;refGene_tx=XM_019827276.2;refGene_gn=EHBP1;refGene_tx=XM_019827277.2;refGene_gn=EHBP1;refGene_tx=XM_019827278.2;refGene_gn=EHBP1;refGene_tx=XM_019827279.2;refGene_gn=EHBP1;refGene_tx=XM_019827280.2;refGene_gn=EHBP1;refGene_tx=XM_023251703.1;refGene_gn=EHBP1;refGene_tx=XM_023251704.1;refGene_gn=EHBP1;refGene_tx=XM_023251705.1;refGene_gn=EHBP1;refGene_tx=XM_023251706.1;refGene_gn=EHBP1;refGene_tx=XM_023251707.1
A3 83305157 83305305
refGene_gn=EHBP1;refGene_tx=XM_003984038.4;refGene_gn=EHBP1;refGene_tx=XM_019827273.2;refGene_gn=EHBP1;refGene_tx=XM_019827274.2;refGene_gn=EHBP1;refGene_tx=XM_019827275.2;refGene_gn=EHBP1;refGene_tx=XM_019827276.2;refGene_gn=EHBP1;refGene_tx=XM_019827277.2;refGene_gn=EHBP1;refGene_tx=XM_019827278.2;refGene_gn=EHBP1;refGene_tx=XM_019827279.2;refGene_gn=EHBP1;refGene_tx=XM_019827280.2;refGene_gn=EHBP1;refGene_tx=XM_023251703.1;refGene_gn=EHBP1;refGene_tx=XM_023251704.1;refGene_gn=EHBP1;refGene_tx=XM_023251705.1;refGene_gn=EHBP1;refGene_tx=XM_023251706.1;refGene_gn=EHBP1;refGene_tx=XM_023251707.1
A3 83313651 83313759
refGene_gn=EHBP1;refGene_tx=XM_003984038.4;refGene_gn=EHBP1;refGene_tx=XM_019827273.2;refGene_gn=EHBP1;refGene_tx=XM_019827274.2;refGene_gn=EHBP1;refGene_tx=XM_019827275.2;refGene_gn=EHBP1;refGene_tx=XM_019827276.2;refGene_gn=EHBP1;refGene_tx=XM_019827277.2;refGene_gn=EHBP1;refGene_tx=XM_019827280.2;refGene_gn=EHBP1;refGene_tx=XM_023251703.1;refGene_gn=EHBP1;refGene_tx=XM_023251704.1;refGene_gn=EHBP1;refGene_tx=XM_023251705.1;refGene_gn=EHBP1;refGene_tx=XM_023251706.1
A3 83316990 83317115
refGene_gn=EHBP1;refGene_tx=XM_003984038.4;refGene_gn=EHBP1;refGene_tx=XM_019827273.2;refGene_gn=EHBP1;refGene_tx=XM_019827274.2;refGene_gn=EHBP1;refGene_tx=XM_019827275.2;refGene_gn=EHBP1;refGene_tx=XM_019827276.2;refGene_gn=EHBP1;refGene_tx=XM_019827277.2;refGene_gn=EHBP1;refGene_tx=XM_019827278.2;refGene_gn=EHBP1;refGene_tx=XM_019827279.2;refGene_gn=EHBP1;refGene_tx=XM_019827280.2;refGene_gn=EHBP1;refGene_tx=XM_023251703.1;refGene_gn=EHBP1;refGene_tx=XM_023251704.1;refGene_gn=EHBP1;refGene_tx=XM_023251705.1;refGene_gn=EHBP1;refGene_tx=XM_023251706.1;refGene_gn=EHBP1;refGene_tx=XM_023251707.1
A3 83319472 83319611
refGene_gn=EHBP1;refGene_tx=XM_003984038.4;refGene_gn=EHBP1;refGene_tx=XM_019827273.2;refGene_gn=EHBP1;refGene_tx=XM_019827274.2;refGene_gn=EHBP1;refGene_tx=XM_019827275.2;refGene_gn=EHBP1;refGene_tx=XM_019827276.2;refGene_gn=EHBP1;refGene_tx=XM_019827277.2;refGene_gn=EHBP1;refGe

ne_tx=XM_019827278.2;refGene_gn=EHBP1;refGene_tx=XM_019827279.2;refGene_gn=EHBP1;refGene_tx=XM_019827280.2;refGene_gn=EHBP1;refGene_tx=XM_023251703.1;refGene_gn=EHBP1;refGene_tx=XM_023251704.1;refGene_gn=EHBP1;refGene_tx=XM_023251705.1;refGene_gn=EHBP1;refGene_tx=XM_023251706.1;refGene_gn=EHBP1;refGene_tx=XM_023251707.1
A3 83319823 83319930
refGene_gn=EHBP1;refGene_tx=XM_003984038.4;refGene_gn=EHBP1;refGene_tx=XM_019827273.2;refGene_gn=EHBP1;refGene_tx=XM_019827274.2;refGene_gn=EHBP1;refGene_tx=XM_019827275.2;refGene_gn=EHBP1;refGene_tx=XM_019827276.2;refGene_gn=EHBP1;refGene_tx=XM_019827277.2;refGene_gn=EHBP1;refGene_tx=XM_019827278.2;refGene_gn=EHBP1;refGene_tx=XM_019827279.2;refGene_gn=EHBP1;refGene_tx=XM_019827280.2;refGene_gn=EHBP1;refGene_tx=XM_023251703.1;refGene_gn=EHBP1;refGene_tx=XM_023251704.1;refGene_gn=EHBP1;refGene_tx=XM_023251705.1;refGene_gn=EHBP1;refGene_tx=XM_023251706.1;refGene_gn=EHBP1;refGene_tx=XM_023251707.1
A3 83322571 83322695
refGene_gn=EHBP1;refGene_tx=XM_003984038.4;refGene_gn=EHBP1;refGene_tx=XM_019827273.2;refGene_gn=EHBP1;refGene_tx=XM_019827274.2;refGene_gn=EHBP1;refGene_tx=XM_019827275.2;refGene_gn=EHBP1;refGene_tx=XM_019827276.2;refGene_gn=EHBP1;refGene_tx=XM_019827277.2;refGene_gn=EHBP1;refGene_tx=XM_019827278.2;refGene_gn=EHBP1;refGene_tx=XM_019827279.2;refGene_gn=EHBP1;refGene_tx=XM_019827280.2;refGene_gn=EHBP1;refGene_tx=XM_023251703.1;refGene_gn=EHBP1;refGene_tx=XM_023251704.1;refGene_gn=EHBP1;refGene_tx=XM_023251705.1;refGene_gn=EHBP1;refGene_tx=XM_023251706.1;refGene_gn=EHBP1;refGene_tx=XM_023251707.1
A3 83357618 83357718
refGene_gn=EHBP1;refGene_tx=XM_003984038.4;refGene_gn=EHBP1;refGene_tx=XM_019827273.2;refGene_gn=EHBP1;refGene_tx=XM_019827274.2;refGene_gn=EHBP1;refGene_tx=XM_019827275.2;refGene_gn=EHBP1;refGene_tx=XM_019827276.2;refGene_gn=EHBP1;refGene_tx=XM_019827277.2;refGene_gn=EHBP1;refGene_tx=XM_019827278.2;refGene_gn=EHBP1;refGene_tx=XM_019827279.2;refGene_gn=EHBP1;refGene_tx=XM_019827280.2;refGene_gn=EHBP1;refGene_tx=XM_023251703.1;refGene_gn=EHBP1;refGene_tx=XM_023251704.1;refGene_gn=EHBP1;refGene_tx=XM_023251705.1;refGene_gn=EHBP1;refGene_tx=XM_023251706.1;refGene_gn=EHBP1;refGene_tx=XM_023251707.1
A3 83358847 83358921
refGene_gn=EHBP1;refGene_tx=XM_003984038.4;refGene_gn=EHBP1;refGene_tx=XM_019827273.2;refGene_gn=EHBP1;refGene_tx=XM_019827274.2;refGene_gn=EHBP1;refGene_tx=XM_019827275.2;refGene_gn=EHBP1;refGene_tx=XM_019827276.2;refGene_gn=EHBP1;refGene_tx=XM_019827277.2;refGene_gn=EHBP1;refGene_tx=XM_019827278.2;refGene_gn=EHBP1;refGene_tx=XM_019827279.2;refGene_gn=EHBP1;refGene_tx=XM_019827280.2;refGene_gn=EHBP1;refGene_tx=XM_023251703.1;refGene_gn=EHBP1;refGene_tx=XM_023251704.1;refGene_gn=EHBP1;refGene_tx=XM_023251705.1;refGene_gn=EHBP1;refGene_tx=XM_023251706.1;refGene_gn=EHBP1;refGene_tx=XM_023251707.1
A3 83365208 83365323
refGene_gn=EHBP1;refGene_tx=XM_003984038.4;refGene_gn=EHBP1;refGene_tx=XM_019827273.2;refGene_gn=EHBP1;refGene_tx=XM_019827274.2;refGene_gn=EHBP1;refGene_tx=XM_019827275.2;refGene_gn=EHBP1;refGene_tx=XM_019827276.2;refGene_gn=EHBP1;refGene_tx=XM_019827277.2;refGene_gn=EHBP1;refGe

ne_tx=XM_019827278.2;refGene_gn=EHBP1;refGene_tx=XM_019827279.2;refGene_gn=EHBP1;refGene_tx=XM_019827280.2;refGene_gn=EHBP1;refGene_tx=XM_023251703.1;refGene_gn=EHBP1;refGene_tx=XM_023251704.1;refGene_gn=EHBP1;refGene_tx=XM_023251705.1;refGene_gn=EHBP1;refGene_tx=XM_023251706.1;refGene_gn=EHBP1;refGene_tx=XM_023251707.1
A3 83365551 83365642
refGene_gn=EHBP1;refGene_tx=XM_003984038.4;refGene_gn=EHBP1;refGene_tx=XM_019827273.2;refGene_gn=EHBP1;refGene_tx=XM_019827274.2;refGene_gn=EHBP1;refGene_tx=XM_019827275.2;refGene_gn=EHBP1;refGene_tx=XM_019827276.2;refGene_gn=EHBP1;refGene_tx=XM_019827277.2;refGene_gn=EHBP1;refGene_tx=XM_019827278.2;refGene_gn=EHBP1;refGene_tx=XM_019827279.2;refGene_gn=EHBP1;refGene_tx=XM_019827280.2;refGene_gn=EHBP1;refGene_tx=XM_023251703.1;refGene_gn=EHBP1;refGene_tx=XM_023251704.1;refGene_gn=EHBP1;refGene_tx=XM_023251705.1;refGene_gn=EHBP1;refGene_tx=XM_023251706.1;refGene_gn=EHBP1;refGene_tx=XM_023251707.1
A3 83369934 83370166
refGene_gn=LOC109498522;refGene_tx=XM_019828087.1
A3 83370440 83370949
refGene_gn=LOC109498522;refGene_tx=XM_019828087.1
A3 83371249 83371342
refGene_gn=LOC109498522;refGene_tx=XM_019828087.1
A3 83371426 83371534
refGene_gn=LOC109498522;refGene_tx=XM_019828087.1
A3 83371617 83371824
refGene_gn=LOC109498522;refGene_tx=XM_019828087.1
A3 83373250 83373347
refGene_gn=OTX1;refGene_tx=XM_011281004.3;refGene_gn=OTX1;refGene_tx=XM_023251708.1;refGene_gn=OTX1;refGene_tx=XM_023251709.1;refGene_gn=OTX1;refGene_tx=XM_023251710.1
A3 83374355 83374507
refGene_gn=OTX1;refGene_tx=XM_011281004.3;refGene_gn=OTX1;refGene_tx=XM_023251708.1;refGene_gn=OTX1;refGene_tx=XM_023251709.1;refGene_gn=OTX1;refGene_tx=XM_023251710.1
A3 83374987 83375161 refGene_gn=OTX1;refGene_tx=XM_023251711.1
A3 83375771 83376590
refGene_gn=OTX1;refGene_tx=XM_011281004.3;refGene_gn=OTX1;refGene_tx=XM_023251708.1;refGene_gn=OTX1;refGene_tx=XM_023251709.1;refGene_gn=OTX1;refGene_tx=XM_023251710.1;refGene_gn=OTX1;refGene_tx=XM_023251711.1
A3 83440511 83440562 refGene_gn=WDPCP;refGene_tx=XM_019827284.1
A3 83448409 83448438 refGene_gn=WDPCP;refGene_tx=XM_019827284.1
A3 83472482 83472562 refGene_gn=WDPCP;refGene_tx=XM_019827284.1
A3 83497208 83497368 refGene_gn=WDPCP;refGene_tx=XM_019827284.1
A3 83577110 83577213 refGene_gn=WDPCP;refGene_tx=XM_019827284.1
A3 83626829 83626893 refGene_gn=WDPCP;refGene_tx=XM_019827284.1
A3 83664662 83664786 refGene_gn=WDPCP;refGene_tx=XM_019827284.1
A3 83668025 83668214 refGene_gn=WDPCP;refGene_tx=XM_019827284.1
A3 83680437 83681047 refGene_gn=WDPCP;refGene_tx=XM_019827284.1
A3 83692882 83693074 refGene_gn=WDPCP;refGene_tx=XM_019827284.1
A3 83696643 83696777 refGene_gn=WDPCP;refGene_tx=XM_019827284.1
A3 83726265 83726290 refGene_gn=WDPCP;refGene_tx=XM_019827284.1

A3 83826791 83826794 refGene_gn=MDH1;refGene_tx=NM_001009329.1
A3 83832573 83832672 refGene_gn=MDH1;refGene_tx=NM_001009329.1
A3 83833512 83833609 refGene_gn=MDH1;refGene_tx=NM_001009329.1
A3 83836212 83836388 refGene_gn=MDH1;refGene_tx=NM_001009329.1
A3 83838518 83838641 refGene_gn=MDH1;refGene_tx=NM_001009329.1
A3 83844818 83844995 refGene_gn=MDH1;refGene_tx=NM_001009329.1
A3 83845505 83845619 refGene_gn=MDH1;refGene_tx=NM_001009329.1
A3 83846164 83846254 refGene_gn=MDH1;refGene_tx=NM_001009329.1
A3 83846794 83846920 refGene_gn=MDH1;refGene_tx=NM_001009329.1
A3 84070124 84070143 refGene_gn=UGP2;refGene_tx=XM_003984040.5
A3 84087650 84087851
refGene_gn=UGP2;refGene_tx=XM_003984040.5;refGene_gn=UGP2;refGene_tx=XM_006930132.4;refGene_gn=UGP2;refGene_tx=XM_019827285.1;refGene_gn=VPS54;refGene_tx=XM_019827282.1
A3 84088594 84088702
refGene_gn=UGP2;refGene_tx=XM_003984040.5;refGene_gn=UGP2;refGene_tx=XM_006930132.4;refGene_gn=UGP2;refGene_tx=XM_019827285.1
A3 84123814 84124000
refGene_gn=UGP2;refGene_tx=XM_003984040.5;refGene_gn=UGP2;refGene_tx=XM_006930132.4;refGene_gn=UGP2;refGene_tx=XM_019827285.1
A3 84125162 84125296
refGene_gn=UGP2;refGene_tx=XM_003984040.5;refGene_gn=UGP2;refGene_tx=XM_006930132.4;refGene_gn=UGP2;refGene_tx=XM_019827285.1
A3 84127128 84127426
refGene_gn=UGP2;refGene_tx=XM_003984040.5;refGene_gn=UGP2;refGene_tx=XM_006930132.4;refGene_gn=UGP2;refGene_tx=XM_019827285.1
A3 84128572 84128770
refGene_gn=UGP2;refGene_tx=XM_003984040.5;refGene_gn=UGP2;refGene_tx=XM_006930132.4;refGene_gn=UGP2;refGene_tx=XM_019827285.1
A3 84129348 84129591
refGene_gn=UGP2;refGene_tx=XM_003984040.5;refGene_gn=UGP2;refGene_tx=XM_006930132.4;refGene_gn=UGP2;refGene_tx=XM_019827285.1
A3 84133211 84133316
refGene_gn=UGP2;refGene_tx=XM_003984040.5;refGene_gn=UGP2;refGene_tx=XM_006930132.4;refGene_gn=UGP2;refGene_tx=XM_019827285.1
A3 84134855 84134963
refGene_gn=UGP2;refGene_tx=XM_003984040.5;refGene_gn=UGP2;refGene_tx=XM_006930132.4;refGene_gn=UGP2;refGene_tx=XM_019827285.1
A3 84137182 84137288
refGene_gn=VPS54;refGene_tx=XM_011281000.3;refGene_gn=VPS54;refGene_tx=XM_011281002.3;refGene_gn=VPS54;refGene_tx=XM_019827281.2
A3 84141146 84141241
refGene_gn=VPS54;refGene_tx=XM_011281000.3;refGene_gn=VPS54;refGene_tx=XM_011281002.3;refGene_gn=VPS54;refGene_tx=XM_019827281.2;refGene_gn=VPS54;refGene_tx=XM_019827282.1
A3 84143786 84143894
refGene_gn=VPS54;refGene_tx=XM_011281000.3;refGene_gn=VPS54;refGene_tx=XM_011281002.3;refGene_gn=VPS54;refGene_tx=XM_019827281.2;refGene_gn=VPS54;refGene_tx=XM_019827282.1
A3 84153956 84154037

refGene_gn=VPS54;refGene_tx=XM_011281000.3;refGene_gn=VPS54;refGene_tx=XM_011281002.3;refGene_gn=VPS54;refGene_tx=XM_019827281.2;refGene_gn=VPS54;refGene_tx=XM_019827282.1
A3 84154143 84154265
refGene_gn=VPS54;refGene_tx=XM_011281000.3;refGene_gn=VPS54;refGene_tx=XM_011281002.3;refGene_gn=VPS54;refGene_tx=XM_019827281.2;refGene_gn=VPS54;refGene_tx=XM_019827282.1
A3 84154816 84154904
refGene_gn=VPS54;refGene_tx=XM_011281000.3;refGene_gn=VPS54;refGene_tx=XM_011281002.3;refGene_gn=VPS54;refGene_tx=XM_019827281.2;refGene_gn=VPS54;refGene_tx=XM_019827282.1
A3 84155850 84155956
refGene_gn=VPS54;refGene_tx=XM_011281000.3;refGene_gn=VPS54;refGene_tx=XM_011281002.3;refGene_gn=VPS54;refGene_tx=XM_019827281.2;refGene_gn=VPS54;refGene_tx=XM_019827282.1
A3 84158522 84158586
refGene_gn=VPS54;refGene_tx=XM_011281000.3;refGene_gn=VPS54;refGene_tx=XM_011281002.3;refGene_gn=VPS54;refGene_tx=XM_019827281.2;refGene_gn=VPS54;refGene_tx=XM_019827282.1
A3 84162230 84162343
refGene_gn=VPS54;refGene_tx=XM_011281000.3;refGene_gn=VPS54;refGene_tx=XM_011281002.3;refGene_gn=VPS54;refGene_tx=XM_019827281.2;refGene_gn=VPS54;refGene_tx=XM_019827282.1
A3 84163034 84163216
refGene_gn=VPS54;refGene_tx=XM_011281000.3;refGene_gn=VPS54;refGene_tx=XM_011281002.3;refGene_gn=VPS54;refGene_tx=XM_019827281.2;refGene_gn=VPS54;refGene_tx=XM_019827282.1
A3 84164162 84164292
refGene_gn=VPS54;refGene_tx=XM_011281000.3;refGene_gn=VPS54;refGene_tx=XM_011281002.3;refGene_gn=VPS54;refGene_tx=XM_019827281.2;refGene_gn=VPS54;refGene_tx=XM_019827282.1
A3 84169700 84170041
refGene_gn=VPS54;refGene_tx=XM_011281000.3;refGene_gn=VPS54;refGene_tx=XM_011281002.3;refGene_gn=VPS54;refGene_tx=XM_019827281.2;refGene_gn=VPS54;refGene_tx=XM_019827282.1
A3 84173528 84173625
refGene_gn=VPS54;refGene_tx=XM_011281000.3;refGene_gn=VPS54;refGene_tx=XM_011281002.3;refGene_gn=VPS54;refGene_tx=XM_019827281.2;refGene_gn=VPS54;refGene_tx=XM_019827282.1
A3 84174611 84174667
refGene_gn=VPS54;refGene_tx=XM_011281000.3;refGene_gn=VPS54;refGene_tx=XM_011281002.3;refGene_gn=VPS54;refGene_tx=XM_019827281.2;refGene_gn=VPS54;refGene_tx=XM_019827282.1
A3 84177111 84177219
refGene_gn=VPS54;refGene_tx=XM_011281000.3;refGene_gn=VPS54;refGene_tx=XM_011281002.3;refGene_gn=VPS54;refGene_tx=XM_019827281.2;refGene_gn=VPS54;refGene_tx=XM_019827282.1
A3 84178109 84178236
refGene_gn=VPS54;refGene_tx=XM_011281000.3;refGene_gn=VPS54;refGene_tx=XM_011281002.3;refGene_gn=VPS54;refGene_tx=XM_019827281.2;refGene_gn=

VPS54;refGene_tx=XM_019827282.1
A3 84188324 84188710
refGene_gn=VPS54;refGene_tx=XM_011281000.3;refGene_gn=VPS54;refGene_tx=XM_011281002.3;refGene_gn=VPS54;refGene_tx=XM_019827281.2;refGene_gn=VPS54;refGene_tx=XM_019827282.1
A3 84192049 84192181
refGene_gn=VPS54;refGene_tx=XM_011281000.3;refGene_gn=VPS54;refGene_tx=XM_011281002.3;refGene_gn=VPS54;refGene_tx=XM_019827281.2;refGene_gn=VPS54;refGene_tx=XM_019827282.1
A3 84194624 84194659
refGene_gn=VPS54;refGene_tx=XM_011281000.3;refGene_gn=VPS54;refGene_tx=XM_011281002.3;refGene_gn=VPS54;refGene_tx=XM_019827281.2;refGene_gn=VPS54;refGene_tx=XM_019827282.1
A3 84196597 84196676
refGene_gn=VPS54;refGene_tx=XM_011281000.3;refGene_gn=VPS54;refGene_tx=XM_011281002.3;refGene_gn=VPS54;refGene_tx=XM_019827281.2;refGene_gn=VPS54;refGene_tx=XM_019827282.1
A3 84205259 84205501
refGene_gn=VPS54;refGene_tx=XM_011281000.3;refGene_gn=VPS54;refGene_tx=XM_011281002.3;refGene_gn=VPS54;refGene_tx=XM_019827281.2;refGene_gn=VPS54;refGene_tx=XM_019827282.1
A3 84207236 84207372
refGene_gn=VPS54;refGene_tx=XM_011281000.3;refGene_gn=VPS54;refGene_tx=XM_011281002.3;refGene_gn=VPS54;refGene_tx=XM_019827281.2;refGene_gn=VPS54;refGene_tx=XM_019827282.1
A3 84216038 84216165
refGene_gn=LOC109498189;refGene_tx=XM_023252186.1
A3 84223524 84223721
refGene_gn=LOC109498189;refGene_tx=XM_023252186.1
A3 84223805 84224531
refGene_gn=LOC109498189;refGene_tx=XM_023252186.1
A3 84284084 84284651
refGene_gn=PELI1;refGene_tx=XM_019827286.2;refGene_gn=PELI1;refGene_tx=XM_023251712.1
A3 84285434 84285623
refGene_gn=PELI1;refGene_tx=XM_019827286.2;refGene_gn=PELI1;refGene_tx=XM_023251712.1
A3 84285719 84285917
refGene_gn=PELI1;refGene_tx=XM_019827286.2;refGene_gn=PELI1;refGene_tx=XM_023251712.1
A3 84290082 84290184
refGene_gn=PELI1;refGene_tx=XM_019827286.2;refGene_gn=PELI1;refGene_tx=XM_023251712.1
A3 84293196 84293326
refGene_gn=PELI1;refGene_tx=XM_019827286.2;refGene_gn=PELI1;refGene_tx=XM_023251712.1
A3 84296194 84296265
refGene_gn=PELI1;refGene_tx=XM_019827286.2;refGene_gn=PELI1;refGene_tx=XM_023251712.1
A3 84607223 84607259 refGene_gn=LGALSL;refGene_tx=XM_023251713.1

A3 84608072 84608144 refGene_gn=LGALSL;refGene_tx=XM_023251713.1
A3 84608294 84608383 refGene_gn=LGALSL;refGene_tx=XM_023251713.1
A3 84609024 84609202 refGene_gn=LGALSL;refGene_tx=XM_023251713.1
A3 84611372 84611516 refGene_gn=LGALSL;refGene_tx=XM_023251713.1
A3 84687110 84689036
refGene_gn=AFTPH;refGene_tx=XM_011280996.3;refGene_gn=AFTPH;refGene_tx=XM_011280997.3;refGene_gn=AFTPH;refGene_tx=XM_019827287.2;refGene_gn=AFTPH;refGene_tx=XM_019827288.2
A3 84700723 84700875
refGene_gn=AFTPH;refGene_tx=XM_011280996.3;refGene_gn=AFTPH;refGene_tx=XM_011280997.3;refGene_gn=AFTPH;refGene_tx=XM_019827287.2
A3 84703204 84703331
refGene_gn=AFTPH;refGene_tx=XM_011280996.3;refGene_gn=AFTPH;refGene_tx=XM_011280997.3;refGene_gn=AFTPH;refGene_tx=XM_019827287.2
A3 84703736 84703793
refGene_gn=AFTPH;refGene_tx=XM_011280996.3;refGene_gn=AFTPH;refGene_tx=XM_011280997.3;refGene_gn=AFTPH;refGene_tx=XM_019827287.2
A3 84707490 84707613
refGene_gn=AFTPH;refGene_tx=XM_011280996.3;refGene_gn=AFTPH;refGene_tx=XM_011280997.3;refGene_gn=AFTPH;refGene_tx=XM_019827287.2;refGene_gn=AFTPH;refGene_tx=XM_019827288.2
A3 84713867 84713928
refGene_gn=AFTPH;refGene_tx=XM_011280996.3;refGene_gn=AFTPH;refGene_tx=XM_011280997.3;refGene_gn=AFTPH;refGene_tx=XM_019827287.2;refGene_gn=AFTPH;refGene_tx=XM_019827288.2
A3 84715448 84715532
refGene_gn=AFTPH;refGene_tx=XM_011280996.3;refGene_gn=AFTPH;refGene_tx=XM_019827287.2;refGene_gn=AFTPH;refGene_tx=XM_019827288.2
A3 84719419 84719543
refGene_gn=AFTPH;refGene_tx=XM_011280996.3;refGene_gn=AFTPH;refGene_tx=XM_011280997.3;refGene_gn=AFTPH;refGene_tx=XM_019827287.2;refGene_gn=AFTPH;refGene_tx=XM_019827288.2
A3 84726278 84726423
refGene_gn=AFTPH;refGene_tx=XM_011280996.3;refGene_gn=AFTPH;refGene_tx=XM_011280997.3;refGene_gn=AFTPH;refGene_tx=XM_019827287.2;refGene_gn=AFTPH;refGene_tx=XM_019827288.2
A3 84764836 84765806
refGene_gn=SERTAD2;refGene_tx=XM_019827289.2;refGene_gn=SERTAD2;refGene_tx=XM_023251714.1;refGene_gn=SERTAD2;refGene_tx=XM_023251715.1;refGene_gn=SERTAD2;refGene_tx=XM_023251716.1;refGene_gn=SERTAD2;refGene_tx=XM_023251717.1;refGene_gn=SERTAD2;refGene_tx=XM_023251718.1;refGene_gn=SERTAD2;refGene_tx=XM_023251719.1;refGene_gn=SERTAD2;refGene_tx=XM_023251720.1
A3 84778949 84779080 refGene_gn=SERTAD2;refGene_tx=XM_019827289.2
A3 85051732 85052259 refGene_gn=SLC1A4;refGene_tx=NM_001278844.1
A3 85060269 85060312 refGene_gn=SLC1A4;refGene_tx=NM_001278844.1
A3 85062507 85062570 refGene_gn=SLC1A4;refGene_tx=NM_001278844.1
A3 85063935 85064102 refGene_gn=SLC1A4;refGene_tx=NM_001278844.1
A3 85066757 85066991 refGene_gn=SLC1A4;refGene_tx=NM_001278844.1
A3 85068189 85068384 refGene_gn=SLC1A4;refGene_tx=NM_001278844.1

A3	85068691	85068826	refGene_gn=SLC1A4;refGene_tx=NM_001278844.1
A3	85070789	85071024	refGene_gn=SLC1A4;refGene_tx=NM_001278844.1
A3	85117089	85117452
refGene_gn=CEP68;refGene_tx=XM_006930142.4;refGene_gn=CEP68;refGene_tx
=XM_019827291.2
A3	85119071	85120598
refGene_gn=CEP68;refGene_tx=XM_006930142.4;refGene_gn=CEP68;refGene_tx
=XM_006930143.4;refGene_gn=CEP68;refGene_tx=XM_019827291.2
A3	85122246	85122369
refGene_gn=CEP68;refGene_tx=XM_006930142.4;refGene_gn=CEP68;refGene_tx
=XM_006930143.4;refGene_gn=CEP68;refGene_tx=XM_019827291.2
A3	85124765	85124862
refGene_gn=CEP68;refGene_tx=XM_006930142.4;refGene_gn=CEP68;refGene_tx
=XM_006930143.4;refGene_gn=CEP68;refGene_tx=XM_019827291.2
A3	85128204	85128377
refGene_gn=CEP68;refGene_tx=XM_006930142.4;refGene_gn=CEP68;refGene_tx
=XM_006930143.4;refGene_gn=CEP68;refGene_tx=XM_019827291.2
A3	85129268	85129292
refGene_gn=CEP68;refGene_tx=XM_006930142.4;refGene_gn=CEP68;refGene_tx
=XM_006930143.4;refGene_gn=CEP68;refGene_tx=XM_019827291.2
A3	85133881	85134079	refGene_gn=RAB1A;refGene_tx=XM_003984057.5
A3	85134560	85134692	refGene_gn=RAB1A;refGene_tx=XM_003984057.5
A3	85136230	85136326	refGene_gn=RAB1A;refGene_tx=XM_003984057.5
A3	85140110	85140206	refGene_gn=RAB1A;refGene_tx=XM_003984057.5
A3	85144678	85144751	refGene_gn=RAB1A;refGene_tx=XM_003984057.5
A3	85162826	85162849	refGene_gn=RAB1A;refGene_tx=XM_003984057.5
A3	85272811	85272859
refGene_gn=ACTR2;refGene_tx=XM_003984059.5;refGene_gn=ACTR2;refGene_tx
=XM_003984060.5
A3	85280800	85280911
refGene_gn=ACTR2;refGene_tx=XM_003984059.5;refGene_gn=ACTR2;refGene_tx
=XM_003984060.5
A3	85282980	85282995	refGene_gn=ACTR2;refGene_tx=XM_003984060.5
A3	85286179	85286395
refGene_gn=ACTR2;refGene_tx=XM_003984059.5;refGene_gn=ACTR2;refGene_tx
=XM_003984060.5
A3	85290440	85290513
refGene_gn=ACTR2;refGene_tx=XM_003984059.5;refGene_gn=ACTR2;refGene_tx
=XM_003984060.5
A3	85294256	85294393
refGene_gn=ACTR2;refGene_tx=XM_003984059.5;refGene_gn=ACTR2;refGene_tx
=XM_003984060.5
A3	85296214	85296364
refGene_gn=ACTR2;refGene_tx=XM_003984059.5;refGene_gn=ACTR2;refGene_tx
=XM_003984060.5
A3	85300211	85300357
refGene_gn=ACTR2;refGene_tx=XM_003984059.5;refGene_gn=ACTR2;refGene_tx
=XM_003984060.5
A3	85303750	85303883
refGene_gn=ACTR2;refGene_tx=XM_003984059.5;refGene_gn=ACTR2;refGene_tx

=XM_003984060.5
A3 85307918 85308089
refGene_gn=ACTR2;refGene_tx=XM_003984059.5;refGene_gn=ACTR2;refGene_tx
=XM_003984060.5
A3 85350548 85351214
refGene_gn=SPRED2;refGene_tx=XM_011281129.2;refGene_gn=SPRED2;refGene_
tx=XM_019827292.2
A3 85353469 85353619
refGene_gn=SPRED2;refGene_tx=XM_011281129.2;refGene_gn=SPRED2;refGene_
tx=XM_019827292.2
A3 85369938 85370003
refGene_gn=SPRED2;refGene_tx=XM_011281129.2;refGene_gn=SPRED2;refGene_
tx=XM_019827292.2
A3 85372155 85372324
refGene_gn=SPRED2;refGene_tx=XM_011281129.2;refGene_gn=SPRED2;refGene_
tx=XM_019827292.2
A3 85381249 85381427
refGene_gn=SPRED2;refGene_tx=XM_011281129.2;refGene_gn=SPRED2;refGene_
tx=XM_019827292.2
A3 85401710 85401727 refGene_gn=SPRED2;refGene_tx=XM_019827292.2
A3 85462370 85462396 refGene_gn=SPRED2;refGene_tx=XM_011281129.2
A3 85991981 85992317
refGene_gn=LOC101092677;refGene_tx=XM_003984167.1
A3 86387810 86387819 refGene_gn=MEIS1;refGene_tx=XM_023251722.1
A3 86388604 86388616
refGene_gn=MEIS1;refGene_tx=XM_006930144.4;refGene_gn=MEIS1;refGene_tx
=XM_006930145.4;refGene_gn=MEIS1;refGene_tx=XM_019827300.2;refGene_gn=
MEIS1;refGene_tx=XM_023251723.1
A3 86390487 86390714
refGene_gn=MEIS1;refGene_tx=XM_006930144.4;refGene_gn=MEIS1;refGene_tx
=XM_006930145.4;refGene_gn=MEIS1;refGene_tx=XM_019827300.2;refGene_gn=
MEIS1;refGene_tx=XM_023251722.1;refGene_gn=MEIS1;refGene_tx=XM_0232517
23.1
A3 86392566 86392708
refGene_gn=MEIS1;refGene_tx=XM_006930144.4;refGene_gn=MEIS1;refGene_tx
=XM_006930145.4;refGene_gn=MEIS1;refGene_tx=XM_019827300.2;refGene_gn=
MEIS1;refGene_tx=XM_023251722.1;refGene_gn=MEIS1;refGene_tx=XM_0232517
23.1
A3 86393225 86393276
refGene_gn=MEIS1;refGene_tx=XM_006930144.4;refGene_gn=MEIS1;refGene_tx
=XM_006930145.4;refGene_gn=MEIS1;refGene_tx=XM_019827300.2;refGene_gn=
MEIS1;refGene_tx=XM_023251722.1;refGene_gn=MEIS1;refGene_tx=XM_0232517
23.1
A3 86394070 86394121
refGene_gn=MEIS1;refGene_tx=XM_006930144.4;refGene_gn=MEIS1;refGene_tx
=XM_006930145.4;refGene_gn=MEIS1;refGene_tx=XM_019827300.2;refGene_gn=
MEIS1;refGene_tx=XM_023251722.1;refGene_gn=MEIS1;refGene_tx=XM_0232517
23.1
A3 86395651 86395798
refGene_gn=MEIS1;refGene_tx=XM_006930144.4;refGene_gn=MEIS1;refGene_tx

=XM_006930145.4;refGene_gn=MEIS1;refGene_tx=XM_019827300.2;refGene_gn=MEIS1;refGene_tx=XM_023251722.1;refGene_gn=MEIS1;refGene_tx=XM_023251723.1
A3 86417488 86417600
refGene_gn=MEIS1;refGene_tx=XM_006930144.4;refGene_gn=MEIS1;refGene_tx=XM_006930145.4;refGene_gn=MEIS1;refGene_tx=XM_019827300.2;refGene_gn=MEIS1;refGene_tx=XM_023251722.1;refGene_gn=MEIS1;refGene_tx=XM_023251723.1
A3 86465057 86465203
refGene_gn=MEIS1;refGene_tx=XM_006930144.4;refGene_gn=MEIS1;refGene_tx=XM_006930145.4;refGene_gn=MEIS1;refGene_tx=XM_019827300.2;refGene_gn=MEIS1;refGene_tx=XM_023251722.1;refGene_gn=MEIS1;refGene_tx=XM_023251723.1
A3 86501899 86501976
refGene_gn=MEIS1;refGene_tx=XM_006930144.4;refGene_gn=MEIS1;refGene_tx=XM_006930145.4;refGene_gn=MEIS1;refGene_tx=XM_019827300.2;refGene_gn=MEIS1;refGene_tx=XM_023251722.1;refGene_gn=MEIS1;refGene_tx=XM_023251723.1
A3 86521076 86521216
refGene_gn=MEIS1;refGene_tx=XM_006930144.4;refGene_gn=MEIS1;refGene_tx=XM_006930145.4;refGene_gn=MEIS1;refGene_tx=XM_019827300.2;refGene_gn=MEIS1;refGene_tx=XM_023251722.1;refGene_gn=MEIS1;refGene_tx=XM_023251723.1
A3 86522278 86522368
refGene_gn=MEIS1;refGene_tx=XM_006930144.4;refGene_gn=MEIS1;refGene_tx=XM_006930145.4;refGene_gn=MEIS1;refGene_tx=XM_019827300.2;refGene_gn=MEIS1;refGene_tx=XM_023251722.1;refGene_gn=MEIS1;refGene_tx=XM_023251723.1
A3 86522668 86522727
refGene_gn=MEIS1;refGene_tx=XM_006930145.4;refGene_gn=MEIS1;refGene_tx=XM_023251723.1
A3 86524877 86525161
refGene_gn=MEIS1;refGene_tx=XM_006930144.4;refGene_gn=MEIS1;refGene_tx=XM_019827300.2;refGene_gn=MEIS1;refGene_tx=XM_023251722.1
A3 87298542 87298765
refGene_gn=ETAA1;refGene_tx=XM_003984065.5;refGene_gn=ETAA1;refGene_tx=XM_019827293.2
A3 87299198 87299291
refGene_gn=ETAA1;refGene_tx=XM_019827293.2;refGene_gn=ETAA1;refGene_tx=XM_019827295.2
A3 87299915 87300044
refGene_gn=ETAA1;refGene_tx=XM_003984065.5;refGene_gn=ETAA1;refGene_tx=XM_006930147.4;refGene_gn=ETAA1;refGene_tx=XM_019827293.2;refGene_gn=ETAA1;refGene_tx=XM_019827295.2
A3 87300312 87300389
refGene_gn=ETAA1;refGene_tx=XM_003984065.5;refGene_gn=ETAA1;refGene_tx=XM_006930147.4;refGene_gn=ETAA1;refGene_tx=XM_019827293.2;refGene_gn=ETAA1;refGene_tx=XM_019827295.2
A3 87304316 87304429
refGene_gn=ETAA1;refGene_tx=XM_003984065.5;refGene_gn=ETAA1;refGene_tx

=XM_006930147.4;refGene_gn=ETAA1;refGene_tx=XM_019827293.2;refGene_gn=ETAA1;refGene_tx=XM_019827295.2
A3 87304682 87306718
refGene_gn=ETAA1;refGene_tx=XM_003984065.5;refGene_gn=ETAA1;refGene_tx=XM_006930147.4;refGene_gn=ETAA1;refGene_tx=XM_019827293.2;refGene_gn=ETAA1;refGene_tx=XM_019827295.2
A3 87312576 87312695
refGene_gn=ETAA1;refGene_tx=XM_003984065.5;refGene_gn=ETAA1;refGene_tx=XM_006930147.4;refGene_gn=ETAA1;refGene_tx=XM_019827293.2;refGene_gn=ETAA1;refGene_tx=XM_019827295.2
A3 87352717 87353755
refGene_gn=LOC111556217;refGene_tx=XM_006930148.4;refGene_gn=LOC111556217;refGene_tx=XM_011281128.3
A3 87874227 87874392
refGene_gn=C1D;refGene_tx=XM_003984067.5;refGene_gn=C1D;refGene_tx=XM_006930150.4;refGene_gn=C1D;refGene_tx=XM_006930151.4
A3 87877055 87877111
refGene_gn=C1D;refGene_tx=XM_003984067.5;refGene_gn=C1D;refGene_tx=XM_006930150.4;refGene_gn=C1D;refGene_tx=XM_006930151.4
A3 87877421 87877488
refGene_gn=C1D;refGene_tx=XM_003984067.5;refGene_gn=C1D;refGene_tx=XM_006930150.4;refGene_gn=C1D;refGene_tx=XM_006930151.4
A3 87878214 87878352
refGene_gn=C1D;refGene_tx=XM_003984067.5;refGene_gn=C1D;refGene_tx=XM_006930150.4;refGene_gn=C1D;refGene_tx=XM_006930151.4
A3 87934550 87934758 refGene_gn=WDR92;refGene_tx=XM_003984068.4
A3 87937776 87937874 refGene_gn=WDR92;refGene_tx=XM_003984068.4
A3 87940185 87940320 refGene_gn=WDR92;refGene_tx=XM_003984068.4
A3 87941510 87941626 refGene_gn=WDR92;refGene_tx=XM_003984068.4
A3 87943777 87943879 refGene_gn=WDR92;refGene_tx=XM_003984068.4
A3 87946491 87946622 refGene_gn=WDR92;refGene_tx=XM_003984068.4
A3 87949673 87949774 refGene_gn=WDR92;refGene_tx=XM_003984068.4
A3 87958764 87958947 refGene_gn=WDR92;refGene_tx=XM_003984068.4
A3 87959458 87959665 refGene_gn=PNO1;refGene_tx=XM_003984070.4
A3 87959903 87960053 refGene_gn=PNO1;refGene_tx=XM_003984070.4
A3 87963817 87963901 refGene_gn=PNO1;refGene_tx=XM_003984070.4
A3 87964315 87964376 refGene_gn=PNO1;refGene_tx=XM_003984070.4
A3 87964617 87964735 refGene_gn=PNO1;refGene_tx=XM_003984070.4
A3 87970030 87970101 refGene_gn=PNO1;refGene_tx=XM_003984070.4
A3 87970755 87970823 refGene_gn=PNO1;refGene_tx=XM_003984070.4
A3 87974248 87974296
refGene_gn=PPP3R1;refGene_tx=XM_019827307.2;refGene_gn=PPP3R1;refGene_tx=XM_023251727.1
A3 87978661 87978846
refGene_gn=PPP3R1;refGene_tx=XM_019827307.2;refGene_gn=PPP3R1;refGene_tx=XM_023251727.1
A3 87979993 87980053
refGene_gn=PPP3R1;refGene_tx=XM_019827307.2;refGene_gn=PPP3R1;refGene_tx=XM_023251727.1
A3 87981521 87981698

refGene_gn=PPP3R1;refGene_tx=XM_019827307.2;refGene_gn=PPP3R1;refGene_tx=XM_023251727.1
A3 87981864 87981889 refGene_gn=PPP3R1;refGene_tx=XM_019827307.2
A3 88004717 88004757 refGene_gn=PPP3R1;refGene_tx=XM_023251727.1
A3 88006761 88006920
refGene_gn=LOC102900168;refGene_tx=XM_023251726.1
A3 88006921 88007362
refGene_gn=LOC102900168;refGene_tx=XM_023251726.1
A3 88039659 88039662 refGene_gn=PPP3R1;refGene_tx=XM_023251727.1
A3 88072342 88072507 refGene_gn=CNRIP1;refGene_tx=XM_003984072.5
A3 88086728 88086879 refGene_gn=CNRIP1;refGene_tx=XM_003984072.5
A3 88088899 88089078 refGene_gn=CNRIP1;refGene_tx=XM_003984072.5
A3 88122561 88122603 refGene_gn=PLEK;refGene_tx=XM_003984073.5
A3 88132304 88132460 refGene_gn=PLEK;refGene_tx=XM_003984073.5
A3 88132713 88132895 refGene_gn=PLEK;refGene_tx=XM_003984073.5
A3 88134495 88134587 refGene_gn=PLEK;refGene_tx=XM_003984073.5
A3 88137499 88137684 refGene_gn=PLEK;refGene_tx=XM_003984073.5
A3 88139443 88139548 refGene_gn=PLEK;refGene_tx=XM_003984073.5
A3 88144265 88144349 refGene_gn=PLEK;refGene_tx=XM_003984073.5
A3 88145004 88145074 refGene_gn=PLEK;refGene_tx=XM_003984073.5
A3 88146469 88146606 refGene_gn=PLEK;refGene_tx=XM_003984073.5
A3 88191196 88191358
refGene_gn=FBXO48;refGene_tx=XM_006930152.3;refGene_gn=FBXO48;refGene_tx=XM_006930153.3;refGene_gn=FBXO48;refGene_tx=XM_019827310.2
A3 88191846 88192152
refGene_gn=FBXO48;refGene_tx=XM_006930152.3;refGene_gn=FBXO48;refGene_tx=XM_006930153.3;refGene_gn=FBXO48;refGene_tx=XM_019827310.2
A3 88194777 88194873
refGene_gn=APLF;refGene_tx=XM_011281125.3;refGene_gn=APLF;refGene_tx=XM_019827311.2;refGene_gn=APLF;refGene_tx=XM_019827312.2
A3 88209149 88209221
refGene_gn=APLF;refGene_tx=XM_011281125.3;refGene_gn=APLF;refGene_tx=XM_019827312.2
A3 88219606 88219779
refGene_gn=APLF;refGene_tx=XM_011281125.3;refGene_gn=APLF;refGene_tx=XM_019827311.2;refGene_gn=APLF;refGene_tx=XM_019827312.2
A3 88228935 88229083
refGene_gn=APLF;refGene_tx=XM_011281125.3;refGene_gn=APLF;refGene_tx=XM_019827311.2;refGene_gn=APLF;refGene_tx=XM_019827312.2
A3 88229643 88229776
refGene_gn=APLF;refGene_tx=XM_011281125.3;refGene_gn=APLF;refGene_tx=XM_019827311.2;refGene_gn=APLF;refGene_tx=XM_019827312.2
A3 88237734 88237880
refGene_gn=APLF;refGene_tx=XM_011281125.3;refGene_gn=APLF;refGene_tx=XM_019827311.2;refGene_gn=APLF;refGene_tx=XM_019827312.2
A3 88249267 88249623
refGene_gn=APLF;refGene_tx=XM_011281125.3;refGene_gn=APLF;refGene_tx=XM_019827311.2;refGene_gn=APLF;refGene_tx=XM_019827312.2
A3 88250273 88250399
refGene_gn=APLF;refGene_tx=XM_011281125.3;refGene_gn=APLF;refGene_tx=X

M_019827311.2;refGene_gn=APLF;refGene_tx=XM_019827312.2
A3 88268731 88268778
refGene_gn=APLF;refGene_tx=XM_011281125.3;refGene_gn=APLF;refGene_tx=XM_019827311.2;refGene_gn=APLF;refGene_tx=XM_019827312.2
A3 88278684 88278890
refGene_gn=APLF;refGene_tx=XM_011281125.3;refGene_gn=APLF;refGene_tx=XM_019827311.2;refGene_gn=APLF;refGene_tx=XM_019827312.2
A3 88311380 88311865
refGene_gn=PROKR1;refGene_tx=XM_003984076.5;refGene_gn=PROKR1;refGene_tx=XM_019827318.2;refGene_gn=PROKR1;refGene_tx=XM_023251729.1
A3 88320977 88321674
refGene_gn=PROKR1;refGene_tx=XM_003984076.5;refGene_gn=PROKR1;refGene_tx=XM_019827318.2;refGene_gn=PROKR1;refGene_tx=XM_023251729.1
A3 88345007 88345295
refGene_gn=LOC101092021;refGene_tx=XM_011281124.3
A3 88345296 88345916
refGene_gn=LOC101092021;refGene_tx=XM_011281124.3
A3 88345917 88345978
refGene_gn=LOC101092021;refGene_tx=XM_011281124.3
A3 88379486 88379547
refGene_gn=ARHGAP25;refGene_tx=XM_003984077.5;refGene_gn=ARHGAP25;refGene_tx=XM_003984078.5
A3 88398772 88398797 refGene_gn=ARHGAP25;refGene_tx=XM_019827314.2
A3 88420944 88421144
refGene_gn=ARHGAP25;refGene_tx=XM_003984077.5;refGene_gn=ARHGAP25;refGene_tx=XM_003984078.5;refGene_gn=ARHGAP25;refGene_tx=XM_019827314.2
A3 88430822 88430910
refGene_gn=ARHGAP25;refGene_tx=XM_003984077.5;refGene_gn=ARHGAP25;refGene_tx=XM_003984078.5;refGene_gn=ARHGAP25;refGene_tx=XM_019827314.2
A3 88436150 88436267
refGene_gn=ARHGAP25;refGene_tx=XM_003984077.5;refGene_gn=ARHGAP25;refGene_tx=XM_019827314.2
A3 88442651 88442763 refGene_gn=ARHGAP25;refGene_tx=XM_019827315.2
A3 88449273 88449481
refGene_gn=ARHGAP25;refGene_tx=XM_003984077.5;refGene_gn=ARHGAP25;refGene_tx=XM_003984078.5;refGene_gn=ARHGAP25;refGene_tx=XM_019827314.2;refGene_gn=ARHGAP25;refGene_tx=XM_019827315.2
A3 88454448 88454581
refGene_gn=ARHGAP25;refGene_tx=XM_003984077.5;refGene_gn=ARHGAP25;refGene_tx=XM_003984078.5;refGene_gn=ARHGAP25;refGene_tx=XM_019827314.2;refGene_gn=ARHGAP25;refGene_tx=XM_019827315.2
A3 88459701 88459775
refGene_gn=ARHGAP25;refGene_tx=XM_003984077.5;refGene_gn=ARHGAP25;refGene_tx=XM_003984078.5;refGene_gn=ARHGAP25;refGene_tx=XM_019827314.2;refGene_gn=ARHGAP25;refGene_tx=XM_019827315.2
A3 88461277 88461399
refGene_gn=ARHGAP25;refGene_tx=XM_003984077.5;refGene_gn=ARHGAP25;refGene_tx=XM_003984078.5;refGene_gn=ARHGAP25;refGene_tx=XM_019827314.2;refGene_gn=ARHGAP25;refGene_tx=XM_019827315.2
A3 88462224 88462421

refGene_gn=ARHGAP25;refGene_tx=XM_003984077.5;refGene_gn=ARHGAP25;refGene_tx=XM_003984078.5;refGene_gn=ARHGAP25;refGene_tx=XM_019827314.2;refGene_gn=ARHGAP25;refGene_tx=XM_019827315.2
A3 88465233 88465766
refGene_gn=ARHGAP25;refGene_tx=XM_003984077.5;refGene_gn=ARHGAP25;refGene_tx=XM_003984078.5;refGene_gn=ARHGAP25;refGene_tx=XM_019827314.2;refGene_gn=ARHGAP25;refGene_tx=XM_019827315.2
A3 88468551 88468759
refGene_gn=ARHGAP25;refGene_tx=XM_003984077.5;refGene_gn=ARHGAP25;refGene_tx=XM_003984078.5;refGene_gn=ARHGAP25;refGene_tx=XM_019827314.2;refGene_gn=ARHGAP25;refGene_tx=XM_019827315.2
A3 88504247 88505188 refGene_gn=BMP10;refGene_tx=XM_023251730.1
A3 88509791 88510125 refGene_gn=BMP10;refGene_tx=XM_023251730.1
A3 88560562 88560645 refGene_gn=GKN2;refGene_tx=XM_019827321.2
A3 88561772 88561929 refGene_gn=GKN2;refGene_tx=XM_019827321.2
A3 88562790 88562901 refGene_gn=GKN2;refGene_tx=XM_019827321.2
A3 88565156 88565294 refGene_gn=GKN2;refGene_tx=XM_019827321.2
A3 88565724 88565775 refGene_gn=GKN2;refGene_tx=XM_019827321.2
A3 88586111 88586123 refGene_gn=GKN1;refGene_tx=XM_011281123.3
A3 88588091 88588145 refGene_gn=GKN1;refGene_tx=XM_011281123.3
A3 88588235 88588373 refGene_gn=GKN1;refGene_tx=XM_011281123.3
A3 88589320 88589431 refGene_gn=GKN1;refGene_tx=XM_011281123.3
A3 88590337 88590485 refGene_gn=GKN1;refGene_tx=XM_011281123.3
A3 88591127 88591219 refGene_gn=GKN1;refGene_tx=XM_011281123.3
A3 88610220 88610328 refGene_gn=GKN2;refGene_tx=XM_019827321.2
A3 88622451 88622603
refGene_gn=ANTXR1;refGene_tx=XM_003984080.5;refGene_gn=ANTXR1;refGene_tx=XM_006930154.4
A3 88626261 88626420 refGene_gn=GKN2;refGene_tx=XM_019827321.2
A3 88647995 88648067
refGene_gn=ANTXR1;refGene_tx=XM_003984080.5;refGene_gn=ANTXR1;refGene_tx=XM_006930154.4
A3 88653003 88653075
refGene_gn=ANTXR1;refGene_tx=XM_003984080.5;refGene_gn=ANTXR1;refGene_tx=XM_006930154.4
A3 88675092 88675174
refGene_gn=ANTXR1;refGene_tx=XM_003984080.5;refGene_gn=ANTXR1;refGene_tx=XM_006930154.4
A3 88675750 88675784
refGene_gn=ANTXR1;refGene_tx=XM_003984080.5;refGene_gn=ANTXR1;refGene_tx=XM_006930154.4
A3 88677002 88677082
refGene_gn=ANTXR1;refGene_tx=XM_003984080.5;refGene_gn=ANTXR1;refGene_tx=XM_006930154.4
A3 88679966 88680035
refGene_gn=ANTXR1;refGene_tx=XM_003984080.5;refGene_gn=ANTXR1;refGene_tx=XM_006930154.4
A3 88681812 88681893
refGene_gn=ANTXR1;refGene_tx=XM_003984080.5;refGene_gn=ANTXR1;refGene_tx=XM_006930154.4

A3	88695503	88695564
refGene_gn=ANTXR1;refGene_tx=XM_003984080.5;refGene_gn=ANTXR1;refGene_tx=XM_006930154.4
A3	88705978	88706077
refGene_gn=ANTXR1;refGene_tx=XM_003984080.5;refGene_gn=ANTXR1;refGene_tx=XM_006930154.4
A3	88724662	88724732
refGene_gn=ANTXR1;refGene_tx=XM_003984080.5;refGene_gn=ANTXR1;refGene_tx=XM_006930154.4
A3	88726226	88726305
refGene_gn=ANTXR1;refGene_tx=XM_003984080.5;refGene_gn=ANTXR1;refGene_tx=XM_006930154.4
A3	88754344	88754440
refGene_gn=ANTXR1;refGene_tx=XM_003984080.5;refGene_gn=ANTXR1;refGene_tx=XM_006930154.4
A3	88771747	88771789
refGene_gn=ANTXR1;refGene_tx=XM_003984080.5;refGene_gn=ANTXR1;refGene_tx=XM_006930154.4
A3	88779829	88779925
refGene_gn=ANTXR1;refGene_tx=XM_003984080.5;refGene_gn=ANTXR1;refGene_tx=XM_006930154.4
A3	88780446	88780614
refGene_gn=ANTXR1;refGene_tx=XM_003984080.5;refGene_gn=ANTXR1;refGene_tx=XM_006930154.4
A3	88791536	88791617
refGene_gn=ANTXR1;refGene_tx=XM_003984080.5;refGene_gn=ANTXR1;refGene_tx=XM_006930154.4
A3	88794485	88794578	refGene_gn=ANTXR1;refGene_tx=XM_006930154.4
A3	88839060	88839321	refGene_gn=ANTXR1;refGene_tx=XM_003984080.5
A3	88884092	88884629
refGene_gn=LOC102900641;refGene_tx=XM_023252187.1
A3	88887886	88888015
refGene_gn=LOC102900641;refGene_tx=XM_023252187.1
A3	88903594	88903639
refGene_gn=GFPT1;refGene_tx=XM_006930156.4;refGene_gn=GFPT1;refGene_tx=XM_023251731.1
A3	88904292	88904454
refGene_gn=GFPT1;refGene_tx=XM_006930156.4;refGene_gn=GFPT1;refGene_tx=XM_023251731.1
A3	88905384	88905552
refGene_gn=GFPT1;refGene_tx=XM_006930156.4;refGene_gn=GFPT1;refGene_tx=XM_023251731.1
A3	88906115	88906243
refGene_gn=GFPT1;refGene_tx=XM_006930156.4;refGene_gn=GFPT1;refGene_tx=XM_023251731.1
A3	88906502	88906617
refGene_gn=GFPT1;refGene_tx=XM_006930156.4;refGene_gn=GFPT1;refGene_tx=XM_023251731.1
A3	88914371	88914529
refGene_gn=GFPT1;refGene_tx=XM_006930156.4;refGene_gn=GFPT1;refGene_tx

=XM_023251731.1
A3	88914923	88915044
refGene_gn=GFPT1;refGene_tx=XM_006930156.4;refGene_gn=GFPT1;refGene_tx
=XM_023251731.1
A3	88919289	88919387
refGene_gn=GFPT1;refGene_tx=XM_006930156.4;refGene_gn=GFPT1;refGene_tx
=XM_023251731.1
A3	88924004	88924100
refGene_gn=GFPT1;refGene_tx=XM_006930156.4;refGene_gn=GFPT1;refGene_tx
=XM_023251731.1
A3	88924996	88925160
refGene_gn=GFPT1;refGene_tx=XM_006930156.4;refGene_gn=GFPT1;refGene_tx
=XM_023251731.1
A3	88927063	88927169
refGene_gn=GFPT1;refGene_tx=XM_006930156.4;refGene_gn=GFPT1;refGene_tx
=XM_023251731.1
A3	88930412	88930466 refGene_gn=GFPT1;refGene_tx=XM_006930156.4
A3	88930670	88930750
refGene_gn=GFPT1;refGene_tx=XM_006930156.4;refGene_gn=GFPT1;refGene_tx
=XM_023251731.1
A3	88931341	88931403
refGene_gn=GFPT1;refGene_tx=XM_006930156.4;refGene_gn=GFPT1;refGene_tx
=XM_023251731.1
A3	88934603	88934738
refGene_gn=GFPT1;refGene_tx=XM_006930156.4;refGene_gn=GFPT1;refGene_tx
=XM_023251731.1
A3	88936508	88936567
refGene_gn=GFPT1;refGene_tx=XM_006930156.4;refGene_gn=GFPT1;refGene_tx
=XM_023251731.1
A3	88939813	88939939
refGene_gn=GFPT1;refGene_tx=XM_006930156.4;refGene_gn=GFPT1;refGene_tx
=XM_023251731.1
A3	88943923	88944031
refGene_gn=GFPT1;refGene_tx=XM_006930156.4;refGene_gn=GFPT1;refGene_tx
=XM_023251731.1
A3	88946144	88946252
refGene_gn=GFPT1;refGene_tx=XM_006930156.4;refGene_gn=GFPT1;refGene_tx
=XM_023251731.1
A3	88956809	88956816
refGene_gn=GFPT1;refGene_tx=XM_006930156.4;refGene_gn=GFPT1;refGene_tx
=XM_023251731.1
A3	88968699	88968744
refGene_gn=NFU1;refGene_tx=XM_003984083.5;refGene_gn=NFU1;refGene_tx=X
M_011281119.3;refGene_gn=NFU1;refGene_tx=XM_011281120.3;refGene_gn=NFU
1;refGene_tx=XM_011281121.3;refGene_gn=NFU1;refGene_tx=XM_023251732.1
A3	88970343	88970518
refGene_gn=NFU1;refGene_tx=XM_003984083.5;refGene_gn=NFU1;refGene_tx=X
M_011281119.3;refGene_gn=NFU1;refGene_tx=XM_011281120.3;refGene_gn=NFU
1;refGene_tx=XM_011281121.3;refGene_gn=NFU1;refGene_tx=XM_023251732.1
A3	88971996	88972057

refGene_gn=NFU1;refGene_tx=XM_003984083.5;refGene_gn=NFU1;refGene_tx=XM_011281119.3;refGene_gn=NFU1;refGene_tx=XM_011281120.3;refGene_gn=NFU1;refGene_tx=XM_011281121.3;refGene_gn=NFU1;refGene_tx=XM_023251732.1
A3 88976058 88976173
refGene_gn=NFU1;refGene_tx=XM_003984083.5;refGene_gn=NFU1;refGene_tx=XM_011281119.3;refGene_gn=NFU1;refGene_tx=XM_011281120.3;refGene_gn=NFU1;refGene_tx=XM_011281121.3;refGene_gn=NFU1;refGene_tx=XM_023251732.1
A3 88980276 88980343
refGene_gn=NFU1;refGene_tx=XM_003984083.5;refGene_gn=NFU1;refGene_tx=XM_011281119.3;refGene_gn=NFU1;refGene_tx=XM_011281120.3;refGene_gn=NFU1;refGene_tx=XM_023251732.1
A3 88985989 88986125
refGene_gn=NFU1;refGene_tx=XM_003984083.5;refGene_gn=NFU1;refGene_tx=XM_011281119.3;refGene_gn=NFU1;refGene_tx=XM_011281120.3;refGene_gn=NFU1;refGene_tx=XM_023251732.1
A3 88992737 88992838
refGene_gn=NFU1;refGene_tx=XM_003984083.5;refGene_gn=NFU1;refGene_tx=XM_011281119.3;refGene_gn=NFU1;refGene_tx=XM_011281120.3;refGene_gn=NFU1;refGene_tx=XM_023251732.1
A3 88995662 88995724
refGene_gn=NFU1;refGene_tx=XM_003984083.5;refGene_gn=NFU1;refGene_tx=XM_011281120.3
A3 89017218 89018348
refGene_gn=AAK1;refGene_tx=XM_011281115.3;refGene_gn=AAK1;refGene_tx=XM_019827331.2;refGene_gn=AAK1;refGene_tx=XM_019827332.2;refGene_gn=AAK1;refGene_tx=XM_019827334.2;refGene_gn=AAK1;refGene_tx=XM_019827335.2;refGene_gn=AAK1;refGene_tx=XM_019827336.2;refGene_gn=AAK1;refGene_tx=XM_019827337.2;refGene_gn=AAK1;refGene_tx=XM_019827338.2;refGene_gn=AAK1;refGene_tx=XM_019827339.2;refGene_gn=AAK1;refGene_tx=XM_019827340.2
A3 89027288 89027383
refGene_gn=AAK1;refGene_tx=XM_011281118.3;refGene_gn=AAK1;refGene_tx=XM_019827341.2;refGene_gn=AAK1;refGene_tx=XM_019827342.2;refGene_gn=AAK1;refGene_tx=XM_019827343.2
A3 89028304 89028415
refGene_gn=AAK1;refGene_tx=XM_011281115.3;refGene_gn=AAK1;refGene_tx=XM_011281118.3;refGene_gn=AAK1;refGene_tx=XM_019827331.2;refGene_gn=AAK1;refGene_tx=XM_019827332.2;refGene_gn=AAK1;refGene_tx=XM_019827334.2;refGene_gn=AAK1;refGene_tx=XM_019827335.2;refGene_gn=AAK1;refGene_tx=XM_019827336.2;refGene_gn=AAK1;refGene_tx=XM_019827338.2;refGene_gn=AAK1;refGene_tx=XM_019827341.2;refGene_gn=AAK1;refGene_tx=XM_019827342.2;refGene_gn=AAK1;refGene_tx=XM_019827343.2
A3 89030152 89030263
refGene_gn=AAK1;refGene_tx=XM_011281115.3;refGene_gn=AAK1;refGene_tx=XM_011281118.3;refGene_gn=AAK1;refGene_tx=XM_019827331.2;refGene_gn=AAK1;refGene_tx=XM_019827332.2;refGene_gn=AAK1;refGene_tx=XM_019827334.2;refGene_gn=AAK1;refGene_tx=XM_019827335.2;refGene_gn=AAK1;refGene_tx=XM_019827338.2;refGene_gn=AAK1;refGene_tx=XM_019827341.2;refGene_gn=AAK1;refGene_tx=XM_019827342.2;refGene_gn=AAK1;refGene_tx=XM_019827343.2
A3 89031661 89031763
refGene_gn=AAK1;refGene_tx=XM_011281115.3;refGene_gn=AAK1;refGene_tx=X

M_011281118.3;refGene_gn=AAK1;refGene_tx=XM_019827331.2;refGene_gn=AAK1;refGene_tx=XM_019827332.2;refGene_gn=AAK1;refGene_tx=XM_019827334.2;refGene_gn=AAK1;refGene_tx=XM_019827335.2;refGene_gn=AAK1;refGene_tx=XM_019827336.2;refGene_gn=AAK1;refGene_tx=XM_019827337.2;refGene_gn=AAK1;refGene_tx=XM_019827338.2;refGene_gn=AAK1;refGene_tx=XM_019827341.2;refGene_gn=AAK1;refGene_tx=XM_019827342.2;refGene_gn=AAK1;refGene_tx=XM_019827343.2

A3 89032783 89032885

refGene_gn=AAK1;refGene_tx=XM_011281115.3;refGene_gn=AAK1;refGene_tx=XM_011281118.3;refGene_gn=AAK1;refGene_tx=XM_019827331.2;refGene_gn=AAK1;refGene_tx=XM_019827332.2;refGene_gn=AAK1;refGene_tx=XM_019827334.2;refGene_gn=AAK1;refGene_tx=XM_019827335.2;refGene_gn=AAK1;refGene_tx=XM_019827336.2;refGene_gn=AAK1;refGene_tx=XM_019827337.2;refGene_gn=AAK1;refGene_tx=XM_019827338.2;refGene_gn=AAK1;refGene_tx=XM_019827341.2;refGene_gn=AAK1;refGene_tx=XM_019827342.2;refGene_gn=AAK1;refGene_tx=XM_019827343.2

A3 89045047 89045143

refGene_gn=AAK1;refGene_tx=XM_011281115.3;refGene_gn=AAK1;refGene_tx=XM_011281118.3;refGene_gn=AAK1;refGene_tx=XM_019827331.2;refGene_gn=AAK1;refGene_tx=XM_019827332.2;refGene_gn=AAK1;refGene_tx=XM_019827334.2;refGene_gn=AAK1;refGene_tx=XM_019827336.2;refGene_gn=AAK1;refGene_tx=XM_019827337.2;refGene_gn=AAK1;refGene_tx=XM_019827338.2;refGene_gn=AAK1;refGene_tx=XM_019827339.2;refGene_gn=AAK1;refGene_tx=XM_019827341.2;refGene_gn=AAK1;refGene_tx=XM_019827342.2;refGene_gn=AAK1;refGene_tx=XM_019827343.2

A3 89051373 89051478

refGene_gn=AAK1;refGene_tx=XM_011281115.3;refGene_gn=AAK1;refGene_tx=XM_011281118.3;refGene_gn=AAK1;refGene_tx=XM_019827331.2;refGene_gn=AAK1;refGene_tx=XM_019827332.2;refGene_gn=AAK1;refGene_tx=XM_019827334.2;refGene_gn=AAK1;refGene_tx=XM_019827335.2;refGene_gn=AAK1;refGene_tx=XM_019827336.2;refGene_gn=AAK1;refGene_tx=XM_019827337.2;refGene_gn=AAK1;refGene_tx=XM_019827338.2;refGene_gn=AAK1;refGene_tx=XM_019827339.2;refGene_gn=AAK1;refGene_tx=XM_019827340.2;refGene_gn=AAK1;refGene_tx=XM_019827341.2;refGene_gn=AAK1;refGene_tx=XM_019827342.2;refGene_gn=AAK1;refGene_tx=XM_019827343.2

A3 89053453 89053629

refGene_gn=AAK1;refGene_tx=XM_011281115.3;refGene_gn=AAK1;refGene_tx=XM_011281118.3;refGene_gn=AAK1;refGene_tx=XM_019827331.2;refGene_gn=AAK1;refGene_tx=XM_019827332.2;refGene_gn=AAK1;refGene_tx=XM_019827334.2;refGene_gn=AAK1;refGene_tx=XM_019827335.2;refGene_gn=AAK1;refGene_tx=XM_019827336.2;refGene_gn=AAK1;refGene_tx=XM_019827337.2;refGene_gn=AAK1;refGene_tx=XM_019827338.2;refGene_gn=AAK1;refGene_tx=XM_019827339.2;refGene_gn=AAK1;refGene_tx=XM_019827340.2;refGene_gn=AAK1;refGene_tx=XM_019827341.2;refGene_gn=AAK1;refGene_tx=XM_019827342.2;refGene_gn=AAK1;refGene_tx=XM_019827343.2

A3 89054891 89055124

refGene_gn=AAK1;refGene_tx=XM_011281115.3;refGene_gn=AAK1;refGene_tx=XM_011281118.3;refGene_gn=AAK1;refGene_tx=XM_019827331.2;refGene_gn=AAK1;refGene_tx=XM_019827332.2;refGene_gn=AAK1;refGene_tx=XM_019827334.2;refGene_gn=AAK1;refGene_tx=XM_019827335.2;refGene_gn=AAK1;refGene_tx=X

M_019827336.2;refGene_gn=AAK1;refGene_tx=XM_019827337.2;refGene_gn=AAK
1;refGene_tx=XM_019827338.2;refGene_gn=AAK1;refGene_tx=XM_019827339.2;
refGene_gn=AAK1;refGene_tx=XM_019827340.2;refGene_gn=AAK1;refGene_tx=X
M_019827341.2;refGene_gn=AAK1;refGene_tx=XM_019827342.2;refGene_gn=AAK
1;refGene_tx=XM_019827343.2
A3 89059361 89059649
refGene_gn=AAK1;refGene_tx=XM_011281115.3;refGene_gn=AAK1;refGene_tx=X
M_011281118.3;refGene_gn=AAK1;refGene_tx=XM_019827331.2;refGene_gn=AAK
1;refGene_tx=XM_019827332.2;refGene_gn=AAK1;refGene_tx=XM_019827334.2;
refGene_gn=AAK1;refGene_tx=XM_019827335.2;refGene_gn=AAK1;refGene_tx=X
M_019827336.2;refGene_gn=AAK1;refGene_tx=XM_019827337.2;refGene_gn=AAK
1;refGene_tx=XM_019827339.2;refGene_gn=AAK1;refGene_tx=XM_019827340.2;
refGene_gn=AAK1;refGene_tx=XM_019827341.2;refGene_gn=AAK1;refGene_tx=X
M_019827342.2
A3 89062880 89063197
refGene_gn=AAK1;refGene_tx=XM_011281115.3;refGene_gn=AAK1;refGene_tx=X
M_011281118.3;refGene_gn=AAK1;refGene_tx=XM_019827331.2;refGene_gn=AAK
1;refGene_tx=XM_019827332.2;refGene_gn=AAK1;refGene_tx=XM_019827334.2;
refGene_gn=AAK1;refGene_tx=XM_019827335.2;refGene_gn=AAK1;refGene_tx=X
M_019827336.2;refGene_gn=AAK1;refGene_tx=XM_019827337.2;refGene_gn=AAK
1;refGene_tx=XM_019827338.2;refGene_gn=AAK1;refGene_tx=XM_019827339.2;
refGene_gn=AAK1;refGene_tx=XM_019827340.2;refGene_gn=AAK1;refGene_tx=X
M_019827341.2;refGene_gn=AAK1;refGene_tx=XM_019827342.2;refGene_gn=AAK
1;refGene_tx=XM_019827343.2
A3 89064036 89064191
refGene_gn=AAK1;refGene_tx=XM_011281115.3;refGene_gn=AAK1;refGene_tx=X
M_011281118.3;refGene_gn=AAK1;refGene_tx=XM_019827331.2;refGene_gn=AAK
1;refGene_tx=XM_019827332.2;refGene_gn=AAK1;refGene_tx=XM_019827334.2;
refGene_gn=AAK1;refGene_tx=XM_019827335.2;refGene_gn=AAK1;refGene_tx=X
M_019827336.2;refGene_gn=AAK1;refGene_tx=XM_019827337.2;refGene_gn=AAK
1;refGene_tx=XM_019827338.2;refGene_gn=AAK1;refGene_tx=XM_019827339.2;
refGene_gn=AAK1;refGene_tx=XM_019827340.2;refGene_gn=AAK1;refGene_tx=X
M_019827341.2;refGene_gn=AAK1;refGene_tx=XM_019827342.2;refGene_gn=AAK
1;refGene_tx=XM_019827343.2
A3 89068360 89068440
refGene_gn=AAK1;refGene_tx=XM_011281115.3;refGene_gn=AAK1;refGene_tx=X
M_011281118.3;refGene_gn=AAK1;refGene_tx=XM_019827331.2;refGene_gn=AAK
1;refGene_tx=XM_019827332.2;refGene_gn=AAK1;refGene_tx=XM_019827334.2;
refGene_gn=AAK1;refGene_tx=XM_019827335.2;refGene_gn=AAK1;refGene_tx=X
M_019827336.2;refGene_gn=AAK1;refGene_tx=XM_019827337.2;refGene_gn=AAK
1;refGene_tx=XM_019827338.2;refGene_gn=AAK1;refGene_tx=XM_019827339.2;
refGene_gn=AAK1;refGene_tx=XM_019827340.2;refGene_gn=AAK1;refGene_tx=X
M_019827341.2;refGene_gn=AAK1;refGene_tx=XM_019827342.2;refGene_gn=AAK
1;refGene_tx=XM_019827343.2
A3 89070632 89070736
refGene_gn=AAK1;refGene_tx=XM_011281115.3;refGene_gn=AAK1;refGene_tx=X
M_011281118.3;refGene_gn=AAK1;refGene_tx=XM_019827331.2;refGene_gn=AAK
1;refGene_tx=XM_019827332.2;refGene_gn=AAK1;refGene_tx=XM_019827334.2;
refGene_gn=AAK1;refGene_tx=XM_019827335.2;refGene_gn=AAK1;refGene_tx=X
M_019827336.2;refGene_gn=AAK1;refGene_tx=XM_019827337.2;refGene_gn=AAK

1;refGene_tx=XM_019827338.2;refGene_gn=AAK1;refGene_tx=XM_019827339.2;
refGene_gn=AAK1;refGene_tx=XM_019827340.2;refGene_gn=AAK1;refGene_tx=X
M_019827341.2;refGene_gn=AAK1;refGene_tx=XM_019827342.2;refGene_gn=AAK
1;refGene_tx=XM_019827343.2
A3 89075070 89075203
refGene_gn=AAK1;refGene_tx=XM_011281115.3;refGene_gn=AAK1;refGene_tx=X
M_011281118.3;refGene_gn=AAK1;refGene_tx=XM_019827331.2;refGene_gn=AAK
1;refGene_tx=XM_019827332.2;refGene_gn=AAK1;refGene_tx=XM_019827334.2;
refGene_gn=AAK1;refGene_tx=XM_019827335.2;refGene_gn=AAK1;refGene_tx=X
M_019827336.2;refGene_gn=AAK1;refGene_tx=XM_019827337.2;refGene_gn=AAK
1;refGene_tx=XM_019827338.2;refGene_gn=AAK1;refGene_tx=XM_019827339.2;
refGene_gn=AAK1;refGene_tx=XM_019827340.2;refGene_gn=AAK1;refGene_tx=X
M_019827341.2;refGene_gn=AAK1;refGene_tx=XM_019827342.2;refGene_gn=AAK
1;refGene_tx=XM_019827343.2
A3 89075706 89075788
refGene_gn=AAK1;refGene_tx=XM_011281115.3;refGene_gn=AAK1;refGene_tx=X
M_011281118.3;refGene_gn=AAK1;refGene_tx=XM_019827331.2;refGene_gn=AAK
1;refGene_tx=XM_019827332.2;refGene_gn=AAK1;refGene_tx=XM_019827334.2;
refGene_gn=AAK1;refGene_tx=XM_019827335.2;refGene_gn=AAK1;refGene_tx=X
M_019827336.2;refGene_gn=AAK1;refGene_tx=XM_019827337.2;refGene_gn=AAK
1;refGene_tx=XM_019827338.2;refGene_gn=AAK1;refGene_tx=XM_019827339.2;
refGene_gn=AAK1;refGene_tx=XM_019827340.2;refGene_gn=AAK1;refGene_tx=X
M_019827341.2;refGene_gn=AAK1;refGene_tx=XM_019827342.2;refGene_gn=AAK
1;refGene_tx=XM_019827343.2
A3 89077118 89077240
refGene_gn=AAK1;refGene_tx=XM_011281115.3;refGene_gn=AAK1;refGene_tx=X
M_011281118.3;refGene_gn=AAK1;refGene_tx=XM_019827331.2;refGene_gn=AAK
1;refGene_tx=XM_019827332.2;refGene_gn=AAK1;refGene_tx=XM_019827334.2;
refGene_gn=AAK1;refGene_tx=XM_019827335.2;refGene_gn=AAK1;refGene_tx=X
M_019827336.2;refGene_gn=AAK1;refGene_tx=XM_019827337.2;refGene_gn=AAK
1;refGene_tx=XM_019827338.2;refGene_gn=AAK1;refGene_tx=XM_019827339.2;
refGene_gn=AAK1;refGene_tx=XM_019827340.2;refGene_gn=AAK1;refGene_tx=X
M_019827341.2;refGene_gn=AAK1;refGene_tx=XM_019827342.2;refGene_gn=AAK
1;refGene_tx=XM_019827343.2
A3 89086416 89086559
refGene_gn=AAK1;refGene_tx=XM_011281115.3;refGene_gn=AAK1;refGene_tx=X
M_011281118.3;refGene_gn=AAK1;refGene_tx=XM_019827331.2;refGene_gn=AAK
1;refGene_tx=XM_019827332.2;refGene_gn=AAK1;refGene_tx=XM_019827334.2;
refGene_gn=AAK1;refGene_tx=XM_019827335.2;refGene_gn=AAK1;refGene_tx=X
M_019827336.2;refGene_gn=AAK1;refGene_tx=XM_019827337.2;refGene_gn=AAK
1;refGene_tx=XM_019827338.2;refGene_gn=AAK1;refGene_tx=XM_019827339.2;
refGene_gn=AAK1;refGene_tx=XM_019827340.2;refGene_gn=AAK1;refGene_tx=X
M_019827341.2;refGene_gn=AAK1;refGene_tx=XM_019827342.2;refGene_gn=AAK
1;refGene_tx=XM_019827343.2
A3 89088313 89088422
refGene_gn=AAK1;refGene_tx=XM_011281115.3;refGene_gn=AAK1;refGene_tx=X
M_011281118.3;refGene_gn=AAK1;refGene_tx=XM_019827331.2;refGene_gn=AAK
1;refGene_tx=XM_019827332.2;refGene_gn=AAK1;refGene_tx=XM_019827334.2;
refGene_gn=AAK1;refGene_tx=XM_019827335.2;refGene_gn=AAK1;refGene_tx=X
M_019827336.2;refGene_gn=AAK1;refGene_tx=XM_019827337.2;refGene_gn=AAK

1;refGene_tx=XM_019827338.2;refGene_gn=AAK1;refGene_tx=XM_019827339.2;
refGene_gn=AAK1;refGene_tx=XM_019827340.2;refGene_gn=AAK1;refGene_tx=X
M_019827341.2;refGene_gn=AAK1;refGene_tx=XM_019827342.2;refGene_gn=AAK
1;refGene_tx=XM_019827343.2
A3 89103746 89103865
refGene_gn=AAK1;refGene_tx=XM_011281115.3;refGene_gn=AAK1;refGene_tx=X
M_011281118.3;refGene_gn=AAK1;refGene_tx=XM_019827331.2;refGene_gn=AAK
1;refGene_tx=XM_019827332.2;refGene_gn=AAK1;refGene_tx=XM_019827334.2;
refGene_gn=AAK1;refGene_tx=XM_019827335.2;refGene_gn=AAK1;refGene_tx=X
M_019827336.2;refGene_gn=AAK1;refGene_tx=XM_019827337.2;refGene_gn=AAK
1;refGene_tx=XM_019827338.2;refGene_gn=AAK1;refGene_tx=XM_019827339.2;
refGene_gn=AAK1;refGene_tx=XM_019827340.2;refGene_gn=AAK1;refGene_tx=X
M_019827341.2;refGene_gn=AAK1;refGene_tx=XM_019827342.2;refGene_gn=AAK
1;refGene_tx=XM_019827343.2
A3 89182487 89182650
refGene_gn=AAK1;refGene_tx=XM_011281115.3;refGene_gn=AAK1;refGene_tx=X
M_011281118.3;refGene_gn=AAK1;refGene_tx=XM_019827331.2;refGene_gn=AAK
1;refGene_tx=XM_019827332.2;refGene_gn=AAK1;refGene_tx=XM_019827334.2;
refGene_gn=AAK1;refGene_tx=XM_019827335.2;refGene_gn=AAK1;refGene_tx=X
M_019827336.2;refGene_gn=AAK1;refGene_tx=XM_019827337.2;refGene_gn=AAK
1;refGene_tx=XM_019827338.2;refGene_gn=AAK1;refGene_tx=XM_019827339.2;
refGene_gn=AAK1;refGene_tx=XM_019827340.2;refGene_gn=AAK1;refGene_tx=X
M_019827341.2;refGene_gn=AAK1;refGene_tx=XM_019827342.2;refGene_gn=AAK
1;refGene_tx=XM_019827343.2
A3 89310076 89310079
refGene_gn=ANXA4;refGene_tx=XM_011281114.3;refGene_gn=ANXA4;refGene_tx
=XM_019827344.2;refGene_gn=ANXA4;refGene_tx=XM_023251733.1
A3 89318752 89318840
refGene_gn=ANXA4;refGene_tx=XM_011281114.3;refGene_gn=ANXA4;refGene_tx
=XM_019827344.2;refGene_gn=ANXA4;refGene_tx=XM_023251733.1
A3 89323393 89323488
refGene_gn=ANXA4;refGene_tx=XM_011281114.3;refGene_gn=ANXA4;refGene_tx
=XM_019827344.2;refGene_gn=ANXA4;refGene_tx=XM_023251733.1
A3 89325639 89325753
refGene_gn=ANXA4;refGene_tx=XM_011281114.3;refGene_gn=ANXA4;refGene_tx
=XM_019827344.2;refGene_gn=ANXA4;refGene_tx=XM_023251733.1
A3 89327001 89327092
refGene_gn=ANXA4;refGene_tx=XM_011281114.3;refGene_gn=ANXA4;refGene_tx
=XM_019827344.2;refGene_gn=ANXA4;refGene_tx=XM_023251733.1
A3 89329057 89329137
refGene_gn=ANXA4;refGene_tx=XM_011281114.3;refGene_gn=ANXA4;refGene_tx
=XM_019827344.2;refGene_gn=ANXA4;refGene_tx=XM_023251733.1
A3 89331271 89331328
refGene_gn=ANXA4;refGene_tx=XM_011281114.3;refGene_gn=ANXA4;refGene_tx
=XM_019827344.2;refGene_gn=ANXA4;refGene_tx=XM_023251733.1
A3 89336246 89336340
refGene_gn=ANXA4;refGene_tx=XM_011281114.3;refGene_gn=ANXA4;refGene_tx
=XM_019827344.2;refGene_gn=ANXA4;refGene_tx=XM_023251733.1
A3 89340358 89340454
refGene_gn=ANXA4;refGene_tx=XM_011281114.3;refGene_gn=ANXA4;refGene_tx

=XM_019827344.2;refGene_gn=ANXA4;refGene_tx=XM_023251733.1
A3 89340984 89341043
refGene_gn=ANXA4;refGene_tx=XM_011281114.3;refGene_gn=ANXA4;refGene_tx=XM_019827344.2;refGene_gn=ANXA4;refGene_tx=XM_023251733.1
A3 89342227 89342350
refGene_gn=ANXA4;refGene_tx=XM_011281114.3;refGene_gn=ANXA4;refGene_tx=XM_019827344.2;refGene_gn=ANXA4;refGene_tx=XM_023251733.1
A3 89344665 89344725
refGene_gn=ANXA4;refGene_tx=XM_011281114.3;refGene_gn=ANXA4;refGene_tx=XM_019827344.2;refGene_gn=ANXA4;refGene_tx=XM_023251733.1
A3 89350145 89350405 refGene_gn=GMCL1;refGene_tx=XM_003984088.5
A3 89356719 89356843 refGene_gn=GMCL1;refGene_tx=XM_003984088.5
A3 89358636 89358733 refGene_gn=GMCL1;refGene_tx=XM_003984088.5
A3 89360086 89360184 refGene_gn=GMCL1;refGene_tx=XM_003984088.5
A3 89362763 89362876 refGene_gn=GMCL1;refGene_tx=XM_003984088.5
A3 89363523 89363589 refGene_gn=GMCL1;refGene_tx=XM_003984088.5
A3 89365454 89365539 refGene_gn=GMCL1;refGene_tx=XM_003984088.5
A3 89367708 89367799 refGene_gn=GMCL1;refGene_tx=XM_003984088.5
A3 89368896 89369034 refGene_gn=GMCL1;refGene_tx=XM_003984088.5
A3 89371351 89371421 refGene_gn=GMCL1;refGene_tx=XM_003984088.5
A3 89374779 89374855 refGene_gn=GMCL1;refGene_tx=XM_003984088.5
A3 89379396 89379542 refGene_gn=GMCL1;refGene_tx=XM_003984088.5
A3 89381489 89381577 refGene_gn=GMCL1;refGene_tx=XM_003984088.5
A3 89385729 89385858 refGene_gn=GMCL1;refGene_tx=XM_003984088.5
A3 89393202 89393236 refGene_gn=SNRNP27;refGene_tx=XM_003984089.4
A3 89394235 89394356 refGene_gn=SNRNP27;refGene_tx=XM_003984089.4
A3 89395845 89395958 refGene_gn=SNRNP27;refGene_tx=XM_003984089.4
A3 89396757 89396837 refGene_gn=SNRNP27;refGene_tx=XM_003984089.4
A3 89401642 89401707 refGene_gn=SNRNP27;refGene_tx=XM_003984089.4
A3 89402719 89402774 refGene_gn=SNRNP27;refGene_tx=XM_003984089.4
A3 89414538 89414611
refGene_gn=MXD1;refGene_tx=XM_003984090.5;refGene_gn=MXD1;refGene_tx=XM_006930161.4
A3 89415356 89415456
refGene_gn=MXD1;refGene_tx=XM_003984090.5;refGene_gn=MXD1;refGene_tx=XM_006930161.4
A3 89420250 89420280
refGene_gn=MXD1;refGene_tx=XM_003984090.5;refGene_gn=MXD1;refGene_tx=XM_006930161.4
A3 89435192 89435307
refGene_gn=MXD1;refGene_tx=XM_003984090.5;refGene_gn=MXD1;refGene_tx=XM_006930161.4
A3 89436924 89437084
refGene_gn=MXD1;refGene_tx=XM_003984090.5;refGene_gn=MXD1;refGene_tx=XM_006930161.4
A3 89437317 89437505
refGene_gn=MXD1;refGene_tx=XM_003984090.5;refGene_gn=MXD1;refGene_tx=XM_006930161.4
A3 89457310 89458468 refGene_gn=ASPRV1;refGene_tx=XM_004001507.5
A3 89566962 89568033 refGene_gn=PCBP1;refGene_tx=XM_011281113.3

A3 89618740 89618949
refGene_gn=CA3H2orf42;refGene_tx=XM_003984093.4
A3 89621869 89622032
refGene_gn=CA3H2orf42;refGene_tx=XM_003984093.4
A3 89623409 89623510
refGene_gn=CA3H2orf42;refGene_tx=XM_003984093.4
A3 89624790 89624898
refGene_gn=CA3H2orf42;refGene_tx=XM_003984093.4
A3 89629695 89629800
refGene_gn=CA3H2orf42;refGene_tx=XM_003984093.4
A3 89631452 89631557
refGene_gn=CA3H2orf42;refGene_tx=XM_003984093.4
A3 89633103 89633214
refGene_gn=CA3H2orf42;refGene_tx=XM_003984093.4
A3 89634147 89634750
refGene_gn=CA3H2orf42;refGene_tx=XM_003984093.4
A3 89634752 89634969
refGene_gn=CA3H2orf42;refGene_tx=XM_003984093.4
A3 89664841 89664968
refGene_gn=TIA1;refGene_tx=XM_003984094.5;refGene_gn=TIA1;refGene_tx=XM_003984095.5;refGene_gn=TIA1;refGene_tx=XM_006930164.4;refGene_gn=TIA1;refGene_tx=XM_006930168.4;refGene_gn=TIA1;refGene_tx=XM_011281107.3;refGene_gn=TIA1;refGene_tx=XM_011281108.3;refGene_gn=TIA1;refGene_tx=XM_011281109.3;refGene_gn=TIA1;refGene_tx=XM_019827349.2;refGene_gn=TIA1;refGene_tx=XM_023251734.1;refGene_gn=TIA1;refGene_tx=XM_023251735.1;refGene_gn=TIA1;refGene_tx=XM_023251736.1;refGene_gn=TIA1;refGene_tx=XM_023251737.1;refGene_gn=TIA1;refGene_tx=XM_023251738.1
A3 89666251 89666397
refGene_gn=TIA1;refGene_tx=XM_003984094.5;refGene_gn=TIA1;refGene_tx=XM_003984095.5;refGene_gn=TIA1;refGene_tx=XM_006930164.4;refGene_gn=TIA1;refGene_tx=XM_006930168.4;refGene_gn=TIA1;refGene_tx=XM_011281107.3;refGene_gn=TIA1;refGene_tx=XM_011281108.3;refGene_gn=TIA1;refGene_tx=XM_011281109.3;refGene_gn=TIA1;refGene_tx=XM_019827349.2;refGene_gn=TIA1;refGene_tx=XM_023251734.1;refGene_gn=TIA1;refGene_tx=XM_023251735.1;refGene_gn=TIA1;refGene_tx=XM_023251736.1;refGene_gn=TIA1;refGene_tx=XM_023251737.1;refGene_gn=TIA1;refGene_tx=XM_023251738.1
A3 89667560 89667684
refGene_gn=TIA1;refGene_tx=XM_003984094.5;refGene_gn=TIA1;refGene_tx=XM_003984095.5;refGene_gn=TIA1;refGene_tx=XM_006930164.4;refGene_gn=TIA1;refGene_tx=XM_006930168.4;refGene_gn=TIA1;refGene_tx=XM_011281107.3;refGene_gn=TIA1;refGene_tx=XM_011281108.3;refGene_gn=TIA1;refGene_tx=XM_011281109.3;refGene_gn=TIA1;refGene_tx=XM_019827349.2;refGene_gn=TIA1;refGene_tx=XM_023251734.1;refGene_gn=TIA1;refGene_tx=XM_023251735.1;refGene_gn=TIA1;refGene_tx=XM_023251736.1;refGene_gn=TIA1;refGene_tx=XM_023251737.1;refGene_gn=TIA1;refGene_tx=XM_023251738.1
A3 89668053 89668138
refGene_gn=TIA1;refGene_tx=XM_003984094.5;refGene_gn=TIA1;refGene_tx=XM_003984095.5;refGene_gn=TIA1;refGene_tx=XM_006930164.4;refGene_gn=TIA1;refGene_tx=XM_006930168.4;refGene_gn=TIA1;refGene_tx=XM_011281107.3;refGene_gn=TIA1;refGene_tx=XM_011281108.3;refGene_gn=TIA1;refGene_tx=X

M_011281109.3;refGene_gn=TIA1;refGene_tx=XM_019827349.2;refGene_gn=TIA1;refGene_tx=XM_023251734.1;refGene_gn=TIA1;refGene_tx=XM_023251735.1;refGene_gn=TIA1;refGene_tx=XM_023251736.1;refGene_gn=TIA1;refGene_tx=XM_023251737.1;refGene_gn=TIA1;refGene_tx=XM_023251738.1
A3 89668254 89668350
refGene_gn=TIA1;refGene_tx=XM_003984094.5;refGene_gn=TIA1;refGene_tx=XM_003984095.5;refGene_gn=TIA1;refGene_tx=XM_006930164.4;refGene_gn=TIA1;refGene_tx=XM_006930168.4;refGene_gn=TIA1;refGene_tx=XM_011281107.3;refGene_gn=TIA1;refGene_tx=XM_011281108.3;refGene_gn=TIA1;refGene_tx=XM_011281109.3;refGene_gn=TIA1;refGene_tx=XM_019827349.2;refGene_gn=TIA1;refGene_tx=XM_023251734.1;refGene_gn=TIA1;refGene_tx=XM_023251735.1;refGene_gn=TIA1;refGene_tx=XM_023251736.1;refGene_gn=TIA1;refGene_tx=XM_023251737.1;refGene_gn=TIA1;refGene_tx=XM_023251738.1
A3 89668746 89668855
refGene_gn=TIA1;refGene_tx=XM_003984094.5;refGene_gn=TIA1;refGene_tx=XM_003984095.5;refGene_gn=TIA1;refGene_tx=XM_006930164.4;refGene_gn=TIA1;refGene_tx=XM_006930168.4;refGene_gn=TIA1;refGene_tx=XM_011281107.3;refGene_gn=TIA1;refGene_tx=XM_011281108.3;refGene_gn=TIA1;refGene_tx=XM_011281109.3;refGene_gn=TIA1;refGene_tx=XM_019827349.2;refGene_gn=TIA1;refGene_tx=XM_023251734.1;refGene_gn=TIA1;refGene_tx=XM_023251735.1;refGene_gn=TIA1;refGene_tx=XM_023251736.1;refGene_gn=TIA1;refGene_tx=XM_023251737.1;refGene_gn=TIA1;refGene_tx=XM_023251738.1
A3 89676822 89676898
refGene_gn=TIA1;refGene_tx=XM_003984094.5;refGene_gn=TIA1;refGene_tx=XM_003984095.5;refGene_gn=TIA1;refGene_tx=XM_006930164.4;refGene_gn=TIA1;refGene_tx=XM_006930168.4;refGene_gn=TIA1;refGene_tx=XM_011281107.3;refGene_gn=TIA1;refGene_tx=XM_011281108.3;refGene_gn=TIA1;refGene_tx=XM_011281109.3;refGene_gn=TIA1;refGene_tx=XM_019827349.2;refGene_gn=TIA1;refGene_tx=XM_023251734.1;refGene_gn=TIA1;refGene_tx=XM_023251735.1;refGene_gn=TIA1;refGene_tx=XM_023251736.1;refGene_gn=TIA1;refGene_tx=XM_023251737.1;refGene_gn=TIA1;refGene_tx=XM_023251738.1
A3 89677635 89677672 refGene_gn=TIA1;refGene_tx=XM_023251742.1
A3 89680085 89680173
refGene_gn=TIA1;refGene_tx=XM_003984094.5;refGene_gn=TIA1;refGene_tx=XM_003984095.5;refGene_gn=TIA1;refGene_tx=XM_006930164.4;refGene_gn=TIA1;refGene_tx=XM_006930168.4;refGene_gn=TIA1;refGene_tx=XM_011281107.3;refGene_gn=TIA1;refGene_tx=XM_011281108.3;refGene_gn=TIA1;refGene_tx=XM_011281109.3;refGene_gn=TIA1;refGene_tx=XM_019827349.2;refGene_gn=TIA1;refGene_tx=XM_023251734.1;refGene_gn=TIA1;refGene_tx=XM_023251735.1;refGene_gn=TIA1;refGene_tx=XM_023251736.1;refGene_gn=TIA1;refGene_tx=XM_023251739.1;refGene_gn=TIA1;refGene_tx=XM_023251740.1;refGene_gn=TIA1;refGene_tx=XM_023251741.1;refGene_gn=TIA1;refGene_tx=XM_023251742.1
A3 89680671 89680790
refGene_gn=TIA1;refGene_tx=XM_023251735.1;refGene_gn=TIA1;refGene_tx=XM_023251736.1;refGene_gn=TIA1;refGene_tx=XM_023251739.1;refGene_gn=TIA1;refGene_tx=XM_023251740.1;refGene_gn=TIA1;refGene_tx=XM_023251741.1
A3 89681401 89681434
refGene_gn=TIA1;refGene_tx=XM_003984095.5;refGene_gn=TIA1;refGene_tx=XM_006930168.4;refGene_gn=TIA1;refGene_tx=XM_011281107.3;refGene_gn=TIA1;refGene_tx=XM_011281108.3;refGene_gn=TIA1;refGene_tx=XM_011281109.3;

refGene_gn=TIA1;refGene_tx=XM_019827349.2;refGene_gn=TIA1;refGene_tx=XM_023251739.1;refGene_gn=TIA1;refGene_tx=XM_023251740.1;refGene_gn=TIA1;refGene_tx=XM_023251741.1;refGene_gn=TIA1;refGene_tx=XM_023251742.1
A3 89681606 89681661
refGene_gn=TIA1;refGene_tx=XM_003984094.5;refGene_gn=TIA1;refGene_tx=XM_003984095.5;refGene_gn=TIA1;refGene_tx=XM_006930164.4;refGene_gn=TIA1;refGene_tx=XM_006930168.4;refGene_gn=TIA1;refGene_tx=XM_011281107.3;refGene_gn=TIA1;refGene_tx=XM_011281108.3;refGene_gn=TIA1;refGene_tx=XM_011281109.3;refGene_gn=TIA1;refGene_tx=XM_019827349.2;refGene_gn=TIA1;refGene_tx=XM_023251734.1;refGene_gn=TIA1;refGene_tx=XM_023251739.1;refGene_gn=TIA1;refGene_tx=XM_023251740.1;refGene_gn=TIA1;refGene_tx=XM_023251741.1;refGene_gn=TIA1;refGene_tx=XM_023251742.1
A3 89682460 89682559
refGene_gn=TIA1;refGene_tx=XM_003984094.5;refGene_gn=TIA1;refGene_tx=XM_003984095.5;refGene_gn=TIA1;refGene_tx=XM_006930164.4;refGene_gn=TIA1;refGene_tx=XM_006930168.4;refGene_gn=TIA1;refGene_tx=XM_011281107.3;refGene_gn=TIA1;refGene_tx=XM_011281108.3;refGene_gn=TIA1;refGene_tx=XM_011281109.3;refGene_gn=TIA1;refGene_tx=XM_019827349.2;refGene_gn=TIA1;refGene_tx=XM_023251734.1;refGene_gn=TIA1;refGene_tx=XM_023251739.1;refGene_gn=TIA1;refGene_tx=XM_023251740.1;refGene_gn=TIA1;refGene_tx=XM_023251741.1;refGene_gn=TIA1;refGene_tx=XM_023251742.1
A3 89686157 89686260
refGene_gn=TIA1;refGene_tx=XM_003984094.5;refGene_gn=TIA1;refGene_tx=XM_003984095.5;refGene_gn=TIA1;refGene_tx=XM_006930164.4;refGene_gn=TIA1;refGene_tx=XM_006930168.4;refGene_gn=TIA1;refGene_tx=XM_011281107.3;refGene_gn=TIA1;refGene_tx=XM_011281108.3;refGene_gn=TIA1;refGene_tx=XM_011281109.3;refGene_gn=TIA1;refGene_tx=XM_019827349.2;refGene_gn=TIA1;refGene_tx=XM_023251734.1;refGene_gn=TIA1;refGene_tx=XM_023251739.1;refGene_gn=TIA1;refGene_tx=XM_023251740.1;refGene_gn=TIA1;refGene_tx=XM_023251741.1;refGene_gn=TIA1;refGene_tx=XM_023251742.1
A3 89694928 89694954
refGene_gn=TIA1;refGene_tx=XM_003984094.5;refGene_gn=TIA1;refGene_tx=XM_003984095.5;refGene_gn=TIA1;refGene_tx=XM_006930164.4;refGene_gn=TIA1;refGene_tx=XM_006930168.4;refGene_gn=TIA1;refGene_tx=XM_011281107.3;refGene_gn=TIA1;refGene_tx=XM_011281108.3;refGene_gn=TIA1;refGene_tx=XM_023251734.1;refGene_gn=TIA1;refGene_tx=XM_023251739.1;refGene_gn=TIA1;refGene_tx=XM_023251740.1;refGene_gn=TIA1;refGene_tx=XM_023251741.1;refGene_gn=TIA1;refGene_tx=XM_023251742.1
A3 89700767 89700879 refGene_gn=PCYOX1;refGene_tx=XM_003984096.4
A3 89701969 89702176 refGene_gn=PCYOX1;refGene_tx=XM_003984096.4
A3 89703227 89703402 refGene_gn=PCYOX1;refGene_tx=XM_003984096.4
A3 89708326 89708538 refGene_gn=PCYOX1;refGene_tx=XM_003984096.4
A3 89708867 89709020 refGene_gn=PCYOX1;refGene_tx=XM_003984096.4
A3 89709436 89710095 refGene_gn=PCYOX1;refGene_tx=XM_003984096.4
A3 89712991 89713042 refGene_gn=SNRPG;refGene_tx=XM_003984098.3
A3 89718350 89718475 refGene_gn=SNRPG;refGene_tx=XM_003984098.3
A3 89719746 89719769 refGene_gn=SNRPG;refGene_tx=XM_003984098.3
A3 89723104 89723136 refGene_gn=SNRPG;refGene_tx=XM_003984098.3
A3 89725942 89726131 refGene_gn=FAM136A;refGene_tx=XM_006930169.4
A3 89728138 89728279 refGene_gn=FAM136A;refGene_tx=XM_006930169.4

A3 89729192 89729279 refGene_gn=FAM136A;refGene_tx=XM_006930169.4
A3 89827136 89827150
refGene_gn=TGFA;refGene_tx=XM_023251743.1;refGene_gn=TGFA;refGene_tx=XM_023251744.1
A3 89836980 89836988 refGene_gn=TGFA;refGene_tx=XM_023251745.1
A3 89839150 89839260
refGene_gn=TGFA;refGene_tx=XM_023251743.1;refGene_gn=TGFA;refGene_tx=XM_023251744.1;refGene_gn=TGFA;refGene_tx=XM_023251745.1
A3 89842206 89842356
refGene_gn=TGFA;refGene_tx=XM_023251743.1;refGene_gn=TGFA;refGene_tx=XM_023251744.1;refGene_gn=TGFA;refGene_tx=XM_023251745.1
A3 89855736 89855857
refGene_gn=TGFA;refGene_tx=XM_023251743.1;refGene_gn=TGFA;refGene_tx=XM_023251744.1;refGene_gn=TGFA;refGene_tx=XM_023251745.1
A3 89906558 89906612
refGene_gn=TGFA;refGene_tx=XM_023251743.1;refGene_gn=TGFA;refGene_tx=XM_023251744.1;refGene_gn=TGFA;refGene_tx=XM_023251745.1
A3 89943149 89943189
refGene_gn=TGFA;refGene_tx=XM_023251743.1;refGene_gn=TGFA;refGene_tx=XM_023251744.1;refGene_gn=TGFA;refGene_tx=XM_023251745.1
A3 90022295 90022600 refGene_gn=ADD2;refGene_tx=XM_023251746.1
A3 90027799 90027925 refGene_gn=ADD2;refGene_tx=XM_023251746.1
A3 90029558 90029574 refGene_gn=ADD2;refGene_tx=XM_023251746.1
A3 90029575 90029707 refGene_gn=ADD2;refGene_tx=XM_023251746.1
A3 90031613 90031703 refGene_gn=ADD2;refGene_tx=XM_023251746.1
A3 90032591 90032711 refGene_gn=ADD2;refGene_tx=XM_023251746.1
A3 90033765 90034023 refGene_gn=ADD2;refGene_tx=XM_023251746.1
A3 90037801 90037978 refGene_gn=ADD2;refGene_tx=XM_023251746.1
A3 90042475 90042574 refGene_gn=ADD2;refGene_tx=XM_023251746.1
A3 90044497 90044641 refGene_gn=ADD2;refGene_tx=XM_023251746.1
A3 90046059 90046209 refGene_gn=ADD2;refGene_tx=XM_023251746.1
A3 90048663 90048744 refGene_gn=ADD2;refGene_tx=XM_023251746.1
A3 90049143 90049295 refGene_gn=ADD2;refGene_tx=XM_023251746.1
A3 90056235 90056374 refGene_gn=ADD2;refGene_tx=XM_023251746.1
A3 90058125 90058308 refGene_gn=ADD2;refGene_tx=XM_023251746.1
A3 90063962 90064010 refGene_gn=FIGLA;refGene_tx=XM_023251753.1
A3 90137661 90137886 refGene_gn=FIGLA;refGene_tx=XM_023251753.1
A3 90139491 90139644 refGene_gn=FIGLA;refGene_tx=XM_023251753.1
A3 90142236 90142482 refGene_gn=FIGLA;refGene_tx=XM_023251753.1
A3 90144093 90144115 refGene_gn=CLEC4F;refGene_tx=XM_023251751.1
A3 90150664 90150815
refGene_gn=CLEC4F;refGene_tx=XM_006930232.4;refGene_gn=CLEC4F;refGene_tx=XM_019827358.2;refGene_gn=CLEC4F;refGene_tx=XM_019827360.2;refGene_gn=CLEC4F;refGene_tx=XM_023251747.1;refGene_gn=CLEC4F;refGene_tx=XM_023251749.1;refGene_gn=CLEC4F;refGene_tx=XM_023251750.1
A3 90151505 90151624
refGene_gn=CLEC4F;refGene_tx=XM_006930232.4;refGene_gn=CLEC4F;refGene_tx=XM_019827358.2;refGene_gn=CLEC4F;refGene_tx=XM_019827360.2;refGene_gn=CLEC4F;refGene_tx=XM_023251747.1;refGene_gn=CLEC4F;refGene_tx=XM_023251749.1;refGene_gn=CLEC4F;refGene_tx=XM_023251750.1;refGene_gn=CLEC4

F;refGene_tx=XM_023251751.1
A3	90152505	90152657
refGene_gn=CLEC4F;refGene_tx=XM_006930232.4;refGene_gn=CLEC4F;refGene_tx=XM_019827358.2;refGene_gn=CLEC4F;refGene_tx=XM_019827360.2;refGene_gn=CLEC4F;refGene_tx=XM_023251747.1;refGene_gn=CLEC4F;refGene_tx=XM_023251749.1;refGene_gn=CLEC4F;refGene_tx=XM_023251750.1;refGene_gn=CLEC4F;refGene_tx=XM_023251751.1
A3	90157384	90158503
refGene_gn=CLEC4F;refGene_tx=XM_006930232.4;refGene_gn=CLEC4F;refGene_tx=XM_019827358.2;refGene_gn=CLEC4F;refGene_tx=XM_019827360.2;refGene_gn=CLEC4F;refGene_tx=XM_023251747.1;refGene_gn=CLEC4F;refGene_tx=XM_023251749.1;refGene_gn=CLEC4F;refGene_tx=XM_023251750.1;refGene_gn=CLEC4F;refGene_tx=XM_023251751.1
A3	90160429	90160540
refGene_gn=CLEC4F;refGene_tx=XM_006930232.4;refGene_gn=CLEC4F;refGene_tx=XM_019827358.2;refGene_gn=CLEC4F;refGene_tx=XM_019827360.2;refGene_gn=CLEC4F;refGene_tx=XM_023251747.1;refGene_gn=CLEC4F;refGene_tx=XM_023251749.1;refGene_gn=CLEC4F;refGene_tx=XM_023251750.1;refGene_gn=CLEC4F;refGene_tx=XM_023251751.1
A3	90160859	90160976
refGene_gn=CLEC4F;refGene_tx=XM_006930232.4;refGene_gn=CLEC4F;refGene_tx=XM_019827358.2;refGene_gn=CLEC4F;refGene_tx=XM_019827360.2;refGene_gn=CLEC4F;refGene_tx=XM_023251747.1;refGene_gn=CLEC4F;refGene_tx=XM_023251749.1;refGene_gn=CLEC4F;refGene_tx=XM_023251751.1
A3	90161825	90161886
refGene_gn=CLEC4F;refGene_tx=XM_006930232.4;refGene_gn=CLEC4F;refGene_tx=XM_019827358.2;refGene_gn=CLEC4F;refGene_tx=XM_019827360.2;refGene_gn=CLEC4F;refGene_tx=XM_023251747.1;refGene_gn=CLEC4F;refGene_tx=XM_023251749.1;refGene_gn=CLEC4F;refGene_tx=XM_023251750.1;refGene_gn=CLEC4F;refGene_tx=XM_023251751.1
A3	90181139	90181290
refGene_gn=CD207;refGene_tx=XM_019827370.2;refGene_gn=CD207;refGene_tx=XM_023251752.1
A3	90181763	90181882
refGene_gn=CD207;refGene_tx=XM_019827370.2;refGene_gn=CD207;refGene_tx=XM_023251752.1
A3	90182912	90183064
refGene_gn=CD207;refGene_tx=XM_019827370.2;refGene_gn=CD207;refGene_tx=XM_023251752.1
A3	90183665	90184040
refGene_gn=CD207;refGene_tx=XM_019827370.2;refGene_gn=CD207;refGene_tx=XM_023251752.1
A3	90185404	90185521
refGene_gn=CD207;refGene_tx=XM_019827370.2;refGene_gn=CD207;refGene_tx=XM_023251752.1
A3	90208793	90208937
refGene_gn=CD207;refGene_tx=XM_019827370.2;refGene_gn=CD207;refGene_tx=XM_023251752.1
A3	90219165	90219256	refGene_gn=CD207;refGene_tx=XM_019827370.2
A3	90230233	90230504	refGene_gn=CD207;refGene_tx=XM_023251752.1

A3 90230704 90230954 refGene_gn=VAX2;refGene_tx=XM_011281258.3
A3 90248898 90249086 refGene_gn=VAX2;refGene_tx=XM_011281258.3
A3 90258423 90258870 refGene_gn=VAX2;refGene_tx=XM_011281258.3
A3 90261543 90261661 refGene_gn=ATP6V1B1;refGene_tx=XM_003984103.4
A3 90267452 90267508 refGene_gn=ATP6V1B1;refGene_tx=XM_003984103.4
A3 90280929 90281028 refGene_gn=ATP6V1B1;refGene_tx=XM_003984103.4
A3 90281226 90281320 refGene_gn=ATP6V1B1;refGene_tx=XM_003984103.4
A3 90281892 90281970 refGene_gn=ATP6V1B1;refGene_tx=XM_003984103.4
A3 90282856 90282996 refGene_gn=ATP6V1B1;refGene_tx=XM_003984103.4
A3 90283404 90283506 refGene_gn=ATP6V1B1;refGene_tx=XM_003984103.4
A3 90284090 90284188 refGene_gn=ATP6V1B1;refGene_tx=XM_003984103.4
A3 90285585 90285709 refGene_gn=ATP6V1B1;refGene_tx=XM_003984103.4
A3 90285889 90286040 refGene_gn=ATP6V1B1;refGene_tx=XM_003984103.4
A3 90286286 90286369 refGene_gn=ATP6V1B1;refGene_tx=XM_003984103.4
A3 90287412 90287517 refGene_gn=ATP6V1B1;refGene_tx=XM_003984103.4
A3 90287715 90287845 refGene_gn=ATP6V1B1;refGene_tx=XM_003984103.4
A3 90287932 90288096 refGene_gn=ATP6V1B1;refGene_tx=XM_003984103.4
A3 90290445 90290510 refGene_gn=ANKRD53;refGene_tx=XM_003984177.5
A3 90290760 90290965 refGene_gn=ANKRD53;refGene_tx=XM_003984177.5
A3 90291296 90291496 refGene_gn=ANKRD53;refGene_tx=XM_003984177.5
A3 90292777 90292942 refGene_gn=ANKRD53;refGene_tx=XM_003984177.5
A3 90293346 90293467 refGene_gn=ANKRD53;refGene_tx=XM_003984177.5
A3 90295454 90296174 refGene_gn=ANKRD53;refGene_tx=XM_003984177.5
A3 90299349 90299465
refGene_gn=TEX261;refGene_tx=XM_003984104.4;refGene_gn=TEX261;refGene_
tx=XM_011281228.3;refGene_gn=TEX261;refGene_tx=XM_019827380.2
A3 90299632 90299735
refGene_gn=TEX261;refGene_tx=XM_003984104.4;refGene_gn=TEX261;refGene_
tx=XM_011281228.3;refGene_gn=TEX261;refGene_tx=XM_019827380.2
A3 90300464 90300532
refGene_gn=TEX261;refGene_tx=XM_003984104.4;refGene_gn=TEX261;refGene_
tx=XM_011281228.3;refGene_gn=TEX261;refGene_tx=XM_019827380.2
A3 90302411 90302565
refGene_gn=TEX261;refGene_tx=XM_003984104.4;refGene_gn=TEX261;refGene_
tx=XM_011281228.3;refGene_gn=TEX261;refGene_tx=XM_019827380.2
A3 90303695 90303775
refGene_gn=TEX261;refGene_tx=XM_003984104.4;refGene_gn=TEX261;refGene_
tx=XM_011281228.3;refGene_gn=TEX261;refGene_tx=XM_019827380.2
A3 90304713 90304783
refGene_gn=TEX261;refGene_tx=XM_003984104.4;refGene_gn=TEX261;refGene_
tx=XM_019827380.2
A3 90305417 90305436 refGene_gn=TEX261;refGene_tx=XM_011281228.3
A3 90327765 90327794
refGene_gn=NAGK;refGene_tx=XM_003984105.5;refGene_gn=NAGK;refGene_tx=X
M_019827372.2
A3 90329542 90329627
refGene_gn=NAGK;refGene_tx=XM_003984105.5;refGene_gn=NAGK;refGene_tx=X
M_019827372.2
A3 90329780 90329879
refGene_gn=NAGK;refGene_tx=XM_003984105.5;refGene_gn=NAGK;refGene_tx=X

M_019827371.2;refGene_gn=NAGK;refGene_tx=XM_019827372.2;refGene_gn=NAGK;refGene_tx=XM_023251755.1
A3 90330453 90330595
refGene_gn=NAGK;refGene_tx=XM_003984105.5;refGene_gn=NAGK;refGene_tx=XM_019827371.2;refGene_gn=NAGK;refGene_tx=XM_019827372.2;refGene_gn=NAGK;refGene_tx=XM_023251755.1
A3 90331396 90331507
refGene_gn=NAGK;refGene_tx=XM_003984105.5;refGene_gn=NAGK;refGene_tx=XM_019827371.2;refGene_gn=NAGK;refGene_tx=XM_019827372.2;refGene_gn=NAGK;refGene_tx=XM_023251755.1
A3 90332207 90332320
refGene_gn=NAGK;refGene_tx=XM_003984105.5;refGene_gn=NAGK;refGene_tx=XM_019827371.2;refGene_gn=NAGK;refGene_tx=XM_019827372.2;refGene_gn=NAGK;refGene_tx=XM_023251755.1
A3 90333770 90333858
refGene_gn=NAGK;refGene_tx=XM_003984105.5;refGene_gn=NAGK;refGene_tx=XM_019827371.2;refGene_gn=NAGK;refGene_tx=XM_019827372.2;refGene_gn=NAGK;refGene_tx=XM_023251755.1
A3 90334845 90334943
refGene_gn=NAGK;refGene_tx=XM_003984105.5;refGene_gn=NAGK;refGene_tx=XM_019827371.2;refGene_gn=NAGK;refGene_tx=XM_019827372.2;refGene_gn=NAGK;refGene_tx=XM_023251755.1
A3 90335436 90335515
refGene_gn=NAGK;refGene_tx=XM_003984105.5;refGene_gn=NAGK;refGene_tx=XM_019827371.2;refGene_gn=NAGK;refGene_tx=XM_019827372.2;refGene_gn=NAGK;refGene_tx=XM_023251755.1
A3 90336359 90336550
refGene_gn=NAGK;refGene_tx=XM_003984105.5;refGene_gn=NAGK;refGene_tx=XM_019827371.2;refGene_gn=NAGK;refGene_tx=XM_023251755.1
A3 90338120 90338137 refGene_gn=NAGK;refGene_tx=XM_019827372.2
A3 90344505 90344562
refGene_gn=PAIP2B;refGene_tx=XM_003984106.5;refGene_gn=PAIP2B;refGene_tx=XM_006930172.4;refGene_gn=PAIP2B;refGene_tx=XM_006930173.4;refGene_gn=PAIP2B;refGene_tx=XM_019827382.2
A3 90345471 90345648
refGene_gn=PAIP2B;refGene_tx=XM_003984106.5;refGene_gn=PAIP2B;refGene_tx=XM_006930172.4;refGene_gn=PAIP2B;refGene_tx=XM_006930173.4;refGene_gn=PAIP2B;refGene_tx=XM_019827382.2
A3 90346075 90346142 refGene_gn=PAIP2B;refGene_tx=XM_006930172.4
A3 90347605 90347743
refGene_gn=PAIP2B;refGene_tx=XM_003984106.5;refGene_gn=PAIP2B;refGene_tx=XM_006930172.4;refGene_gn=PAIP2B;refGene_tx=XM_006930173.4;refGene_gn=PAIP2B;refGene_tx=XM_019827382.2
A3 90407410 90407526
refGene_gn=ZNF638;refGene_tx=XM_011281226.3;refGene_gn=ZNF638;refGene_tx=XM_011281227.3;refGene_gn=ZNF638;refGene_tx=XM_019827383.2;refGene_gn=ZNF638;refGene_tx=XM_019827384.2;refGene_gn=ZNF638;refGene_tx=XM_019827387.2;refGene_gn=ZNF638;refGene_tx=XM_019827388.2;refGene_gn=ZNF638;refGene_tx=XM_019827389.2
A3 90474064 90475577

refGene_gn=ZNF638;refGene_tx=XM_011281226.3;refGene_gn=ZNF638;refGene_tx=XM_011281227.3;refGene_gn=ZNF638;refGene_tx=XM_019827383.2;refGene_gn=ZNF638;refGene_tx=XM_019827384.2;refGene_gn=ZNF638;refGene_tx=XM_019827387.2;refGene_gn=ZNF638;refGene_tx=XM_019827388.2;refGene_gn=ZNF638;refGene_tx=XM_019827389.2;refGene_gn=ZNF638;refGene_tx=XM_019827390.1;refGene_gn=ZNF638;refGene_tx=XM_019827391.1
A3 90480665 90480727
refGene_gn=ZNF638;refGene_tx=XM_011281226.3;refGene_gn=ZNF638;refGene_tx=XM_011281227.3;refGene_gn=ZNF638;refGene_tx=XM_019827383.2;refGene_gn=ZNF638;refGene_tx=XM_019827384.2;refGene_gn=ZNF638;refGene_tx=XM_019827387.2;refGene_gn=ZNF638;refGene_tx=XM_019827388.2;refGene_gn=ZNF638;refGene_tx=XM_019827389.2;refGene_gn=ZNF638;refGene_tx=XM_019827390.1;refGene_gn=ZNF638;refGene_tx=XM_019827391.1
A3 90485916 90485955
refGene_gn=ZNF638;refGene_tx=XM_011281226.3;refGene_gn=ZNF638;refGene_tx=XM_011281227.3;refGene_gn=ZNF638;refGene_tx=XM_019827383.2;refGene_gn=ZNF638;refGene_tx=XM_019827384.2;refGene_gn=ZNF638;refGene_tx=XM_019827388.2;refGene_gn=ZNF638;refGene_tx=XM_019827389.2;refGene_gn=ZNF638;refGene_tx=XM_019827390.1;refGene_gn=ZNF638;refGene_tx=XM_019827391.1
A3 90486929 90487222
refGene_gn=ZNF638;refGene_tx=XM_011281226.3;refGene_gn=ZNF638;refGene_tx=XM_011281227.3;refGene_gn=ZNF638;refGene_tx=XM_019827383.2;refGene_gn=ZNF638;refGene_tx=XM_019827384.2;refGene_gn=ZNF638;refGene_tx=XM_019827387.2;refGene_gn=ZNF638;refGene_tx=XM_019827388.2;refGene_gn=ZNF638;refGene_tx=XM_019827389.2;refGene_gn=ZNF638;refGene_tx=XM_019827390.1;refGene_gn=ZNF638;refGene_tx=XM_019827391.1
A3 90488263 90488541
refGene_gn=ZNF638;refGene_tx=XM_011281226.3;refGene_gn=ZNF638;refGene_tx=XM_011281227.3;refGene_gn=ZNF638;refGene_tx=XM_019827383.2;refGene_gn=ZNF638;refGene_tx=XM_019827384.2;refGene_gn=ZNF638;refGene_tx=XM_019827387.2;refGene_gn=ZNF638;refGene_tx=XM_019827388.2;refGene_gn=ZNF638;refGene_tx=XM_019827389.2;refGene_gn=ZNF638;refGene_tx=XM_019827390.1;refGene_gn=ZNF638;refGene_tx=XM_019827391.1
A3 90490401 90490548
refGene_gn=ZNF638;refGene_tx=XM_011281226.3;refGene_gn=ZNF638;refGene_tx=XM_011281227.3;refGene_gn=ZNF638;refGene_tx=XM_019827383.2;refGene_gn=ZNF638;refGene_tx=XM_019827384.2;refGene_gn=ZNF638;refGene_tx=XM_019827387.2;refGene_gn=ZNF638;refGene_tx=XM_019827388.2;refGene_gn=ZNF638;refGene_tx=XM_019827389.2;refGene_gn=ZNF638;refGene_tx=XM_019827390.1;refGene_gn=ZNF638;refGene_tx=XM_019827391.1
A3 90491677 90491800
refGene_gn=ZNF638;refGene_tx=XM_011281226.3;refGene_gn=ZNF638;refGene_tx=XM_011281227.3;refGene_gn=ZNF638;refGene_tx=XM_019827383.2;refGene_gn=ZNF638;refGene_tx=XM_019827384.2;refGene_gn=ZNF638;refGene_tx=XM_019827387.2;refGene_gn=ZNF638;refGene_tx=XM_019827388.2;refGene_gn=ZNF638;refGene_tx=XM_019827389.2;refGene_gn=ZNF638;refGene_tx=XM_019827390.1;refGene_gn=ZNF638;refGene_tx=XM_019827391.1
A3 90497182 90497241
refGene_gn=ZNF638;refGene_tx=XM_011281226.3;refGene_gn=ZNF638;refGene_

tx=XM_011281227.3;refGene_gn=ZNF638;refGene_tx=XM_019827383.2;refGene_gn=ZNF638;refGene_tx=XM_019827384.2;refGene_gn=ZNF638;refGene_tx=XM_019827387.2;refGene_gn=ZNF638;refGene_tx=XM_019827388.2;refGene_gn=ZNF638;refGene_tx=XM_019827389.2;refGene_gn=ZNF638;refGene_tx=XM_019827390.1;refGene_gn=ZNF638;refGene_tx=XM_019827391.1
A3 90497604 90497657
refGene_gn=ZNF638;refGene_tx=XM_011281226.3;refGene_gn=ZNF638;refGene_tx=XM_011281227.3;refGene_gn=ZNF638;refGene_tx=XM_019827383.2;refGene_gn=ZNF638;refGene_tx=XM_019827384.2;refGene_gn=ZNF638;refGene_tx=XM_019827387.2;refGene_gn=ZNF638;refGene_tx=XM_019827388.2;refGene_gn=ZNF638;refGene_tx=XM_019827389.2;refGene_gn=ZNF638;refGene_tx=XM_019827390.1;refGene_gn=ZNF638;refGene_tx=XM_019827391.1
A3 90506309 90506360
refGene_gn=ZNF638;refGene_tx=XM_011281226.3;refGene_gn=ZNF638;refGene_tx=XM_011281227.3;refGene_gn=ZNF638;refGene_tx=XM_019827383.2;refGene_gn=ZNF638;refGene_tx=XM_019827384.2;refGene_gn=ZNF638;refGene_tx=XM_019827387.2;refGene_gn=ZNF638;refGene_tx=XM_019827388.2;refGene_gn=ZNF638;refGene_tx=XM_019827389.2;refGene_gn=ZNF638;refGene_tx=XM_019827390.1;refGene_gn=ZNF638;refGene_tx=XM_019827391.1
A3 90509109 90509181
refGene_gn=ZNF638;refGene_tx=XM_011281226.3;refGene_gn=ZNF638;refGene_tx=XM_011281227.3;refGene_gn=ZNF638;refGene_tx=XM_019827383.2;refGene_gn=ZNF638;refGene_tx=XM_019827384.2;refGene_gn=ZNF638;refGene_tx=XM_019827387.2;refGene_gn=ZNF638;refGene_tx=XM_019827388.2;refGene_gn=ZNF638;refGene_tx=XM_019827389.2;refGene_gn=ZNF638;refGene_tx=XM_019827390.1;refGene_gn=ZNF638;refGene_tx=XM_019827391.1
A3 90510619 90510706
refGene_gn=ZNF638;refGene_tx=XM_011281226.3;refGene_gn=ZNF638;refGene_tx=XM_011281227.3;refGene_gn=ZNF638;refGene_tx=XM_019827383.2;refGene_gn=ZNF638;refGene_tx=XM_019827384.2;refGene_gn=ZNF638;refGene_tx=XM_019827387.2;refGene_gn=ZNF638;refGene_tx=XM_019827388.2;refGene_gn=ZNF638;refGene_tx=XM_019827389.2;refGene_gn=ZNF638;refGene_tx=XM_019827390.1;refGene_gn=ZNF638;refGene_tx=XM_019827391.1
A3 90510963 90511032
refGene_gn=ZNF638;refGene_tx=XM_011281226.3;refGene_gn=ZNF638;refGene_tx=XM_011281227.3;refGene_gn=ZNF638;refGene_tx=XM_019827383.2;refGene_gn=ZNF638;refGene_tx=XM_019827384.2;refGene_gn=ZNF638;refGene_tx=XM_019827387.2;refGene_gn=ZNF638;refGene_tx=XM_019827388.2;refGene_gn=ZNF638;refGene_tx=XM_019827389.2;refGene_gn=ZNF638;refGene_tx=XM_019827390.1;refGene_gn=ZNF638;refGene_tx=XM_019827391.1
A3 90511417 90511458
refGene_gn=ZNF638;refGene_tx=XM_011281226.3;refGene_gn=ZNF638;refGene_tx=XM_011281227.3;refGene_gn=ZNF638;refGene_tx=XM_019827383.2;refGene_gn=ZNF638;refGene_tx=XM_019827384.2;refGene_gn=ZNF638;refGene_tx=XM_019827387.2;refGene_gn=ZNF638;refGene_tx=XM_019827388.2;refGene_gn=ZNF638;refGene_tx=XM_019827389.2;refGene_gn=ZNF638;refGene_tx=XM_019827390.1;refGene_gn=ZNF638;refGene_tx=XM_019827391.1
A3 90513823 90513955
refGene_gn=ZNF638;refGene_tx=XM_011281226.3;refGene_gn=ZNF638;refGene_tx=XM_011281227.3;refGene_gn=ZNF638;refGene_tx=XM_019827383.2;refGene_

gn=ZNF638;refGene_tx=XM_019827384.2;refGene_gn=ZNF638;refGene_tx=XM_01
9827387.2;refGene_gn=ZNF638;refGene_tx=XM_019827390.1;refGene_gn=ZNF63
8;refGene_tx=XM_019827391.1
A3 90516229 90516358
refGene_gn=ZNF638;refGene_tx=XM_011281226.3;refGene_gn=ZNF638;refGene_
tx=XM_011281227.3;refGene_gn=ZNF638;refGene_tx=XM_019827383.2;refGene_
gn=ZNF638;refGene_tx=XM_019827384.2;refGene_gn=ZNF638;refGene_tx=XM_01
9827387.2;refGene_gn=ZNF638;refGene_tx=XM_019827388.2;refGene_gn=ZNF63
8;refGene_tx=XM_019827389.2;refGene_gn=ZNF638;refGene_tx=XM_019827390.
1;refGene_gn=ZNF638;refGene_tx=XM_019827391.1
A3 90518522 90518564
refGene_gn=ZNF638;refGene_tx=XM_011281226.3;refGene_gn=ZNF638;refGene_
tx=XM_011281227.3;refGene_gn=ZNF638;refGene_tx=XM_019827383.2;refGene_
gn=ZNF638;refGene_tx=XM_019827384.2;refGene_gn=ZNF638;refGene_tx=XM_01
9827387.2;refGene_gn=ZNF638;refGene_tx=XM_019827388.2;refGene_gn=ZNF63
8;refGene_tx=XM_019827389.2;refGene_gn=ZNF638;refGene_tx=XM_019827390.
1;refGene_gn=ZNF638;refGene_tx=XM_019827391.1
A3 90519028 90519163
refGene_gn=ZNF638;refGene_tx=XM_011281226.3;refGene_gn=ZNF638;refGene_
tx=XM_011281227.3;refGene_gn=ZNF638;refGene_tx=XM_019827383.2;refGene_
gn=ZNF638;refGene_tx=XM_019827384.2;refGene_gn=ZNF638;refGene_tx=XM_01
9827387.2;refGene_gn=ZNF638;refGene_tx=XM_019827388.2;refGene_gn=ZNF63
8;refGene_tx=XM_019827389.2;refGene_gn=ZNF638;refGene_tx=XM_019827390.
1;refGene_gn=ZNF638;refGene_tx=XM_019827391.1
A3 90521512 90521638
refGene_gn=ZNF638;refGene_tx=XM_011281226.3;refGene_gn=ZNF638;refGene_
tx=XM_011281227.3;refGene_gn=ZNF638;refGene_tx=XM_019827383.2;refGene_
gn=ZNF638;refGene_tx=XM_019827384.2;refGene_gn=ZNF638;refGene_tx=XM_01
9827387.2;refGene_gn=ZNF638;refGene_tx=XM_019827388.2;refGene_gn=ZNF63
8;refGene_tx=XM_019827389.2;refGene_gn=ZNF638;refGene_tx=XM_019827390.
1;refGene_gn=ZNF638;refGene_tx=XM_019827391.1
A3 90526956 90526994
refGene_gn=ZNF638;refGene_tx=XM_011281226.3;refGene_gn=ZNF638;refGene_
tx=XM_011281227.3;refGene_gn=ZNF638;refGene_tx=XM_019827383.2;refGene_
gn=ZNF638;refGene_tx=XM_019827384.2;refGene_gn=ZNF638;refGene_tx=XM_01
9827387.2;refGene_gn=ZNF638;refGene_tx=XM_019827388.2;refGene_gn=ZNF63
8;refGene_tx=XM_019827389.2;refGene_gn=ZNF638;refGene_tx=XM_019827390.
1;refGene_gn=ZNF638;refGene_tx=XM_019827391.1
A3 90531634 90532859
refGene_gn=ZNF638;refGene_tx=XM_011281226.3;refGene_gn=ZNF638;refGene_
tx=XM_011281227.3;refGene_gn=ZNF638;refGene_tx=XM_019827383.2;refGene_
gn=ZNF638;refGene_tx=XM_019827384.2;refGene_gn=ZNF638;refGene_tx=XM_01
9827387.2;refGene_gn=ZNF638;refGene_tx=XM_019827388.2;refGene_gn=ZNF63
8;refGene_tx=XM_019827389.2;refGene_gn=ZNF638;refGene_tx=XM_019827390.
1;refGene_gn=ZNF638;refGene_tx=XM_019827391.1
A3 90533457 90533523
refGene_gn=ZNF638;refGene_tx=XM_011281226.3;refGene_gn=ZNF638;refGene_
tx=XM_011281227.3;refGene_gn=ZNF638;refGene_tx=XM_019827383.2;refGene_
gn=ZNF638;refGene_tx=XM_019827384.2;refGene_gn=ZNF638;refGene_tx=XM_01
9827387.2;refGene_gn=ZNF638;refGene_tx=XM_019827388.2;refGene_gn=ZNF63

8;refGene_tx=XM_019827389.2;refGene_gn=ZNF638;refGene_tx=XM_019827390.
1;refGene_gn=ZNF638;refGene_tx=XM_019827391.1
A3 90535132 90536087
refGene_gn=ZNF638;refGene_tx=XM_011281226.3;refGene_gn=ZNF638;refGene_
tx=XM_011281227.3;refGene_gn=ZNF638;refGene_tx=XM_019827383.2;refGene_
gn=ZNF638;refGene_tx=XM_019827384.2;refGene_gn=ZNF638;refGene_tx=XM_01
9827387.2;refGene_gn=ZNF638;refGene_tx=XM_019827388.2;refGene_gn=ZNF63
8;refGene_tx=XM_019827389.2;refGene_gn=ZNF638;refGene_tx=XM_019827390.
1;refGene_gn=ZNF638;refGene_tx=XM_019827391.1
A3 90536597 90536717
refGene_gn=ZNF638;refGene_tx=XM_011281226.3;refGene_gn=ZNF638;refGene_
tx=XM_019827383.2;refGene_gn=ZNF638;refGene_tx=XM_019827384.2;refGene_
gn=ZNF638;refGene_tx=XM_019827387.2;refGene_gn=ZNF638;refGene_tx=XM_01
9827388.2;refGene_gn=ZNF638;refGene_tx=XM_019827389.2;refGene_gn=ZNF63
8;refGene_tx=XM_019827390.1;refGene_gn=ZNF638;refGene_tx=XM_019827391.
1
A3 90539385 90539484
refGene_gn=ZNF638;refGene_tx=XM_011281226.3;refGene_gn=ZNF638;refGene_
tx=XM_011281227.3;refGene_gn=ZNF638;refGene_tx=XM_019827383.2;refGene_
gn=ZNF638;refGene_tx=XM_019827384.2;refGene_gn=ZNF638;refGene_tx=XM_01
9827387.2;refGene_gn=ZNF638;refGene_tx=XM_019827388.2;refGene_gn=ZNF63
8;refGene_tx=XM_019827389.2;refGene_gn=ZNF638;refGene_tx=XM_019827390.
1;refGene_gn=ZNF638;refGene_tx=XM_019827391.1
A3 90540205 90540324
refGene_gn=ZNF638;refGene_tx=XM_011281226.3;refGene_gn=ZNF638;refGene_
tx=XM_011281227.3;refGene_gn=ZNF638;refGene_tx=XM_019827383.2;refGene_
gn=ZNF638;refGene_tx=XM_019827384.2;refGene_gn=ZNF638;refGene_tx=XM_01
9827387.2;refGene_gn=ZNF638;refGene_tx=XM_019827388.2;refGene_gn=ZNF63
8;refGene_tx=XM_019827389.2;refGene_gn=ZNF638;refGene_tx=XM_019827390.
1;refGene_gn=ZNF638;refGene_tx=XM_019827391.1
A3 90541892 90541958
refGene_gn=ZNF638;refGene_tx=XM_011281226.3;refGene_gn=ZNF638;refGene_
tx=XM_011281227.3;refGene_gn=ZNF638;refGene_tx=XM_019827383.2;refGene_
gn=ZNF638;refGene_tx=XM_019827384.2;refGene_gn=ZNF638;refGene_tx=XM_01
9827387.2;refGene_gn=ZNF638;refGene_tx=XM_019827388.2;refGene_gn=ZNF63
8;refGene_tx=XM_019827389.2;refGene_gn=ZNF638;refGene_tx=XM_019827390.
1;refGene_gn=ZNF638;refGene_tx=XM_019827391.1
A3 90566093 90566181 refGene_gn=DYSF;refGene_tx=XM_003984110.5
A3 90597437 90597493 refGene_gn=DYSF;refGene_tx=XM_003984110.5
A3 90598417 90598509 refGene_gn=DYSF;refGene_tx=XM_003984110.5
A3 90616577 90616683 refGene_gn=DYSF;refGene_tx=XM_003984110.5
A3 90627748 90627863 refGene_gn=DYSF;refGene_tx=XM_003984110.5
A3 90629105 90629198 refGene_gn=DYSF;refGene_tx=XM_003984110.5
A3 90629558 90629770 refGene_gn=DYSF;refGene_tx=XM_003984110.5
A3 90631886 90632015 refGene_gn=DYSF;refGene_tx=XM_003984110.5
A3 90632467 90632530 refGene_gn=DYSF;refGene_tx=XM_003984110.5
A3 90633279 90633330 refGene_gn=DYSF;refGene_tx=XM_003984110.5
A3 90634820 90634851 refGene_gn=DYSF;refGene_tx=XM_003984110.5
A3 90635431 90635547 refGene_gn=DYSF;refGene_tx=XM_003984110.5
A3 90640981 90641108 refGene_gn=DYSF;refGene_tx=XM_003984110.5

A3	90642658	90642762	refGene_gn=DYSF;refGene_tx=XM_003984110.5
A3	90652962	90653031	refGene_gn=DYSF;refGene_tx=XM_003984110.5
A3	90653574	90653618	refGene_gn=DYSF;refGene_tx=XM_003984110.5
A3	90657077	90657160	refGene_gn=DYSF;refGene_tx=XM_003984110.5
A3	90664646	90664688	refGene_gn=DYSF;refGene_tx=XM_003984110.5
A3	90666322	90666438	refGene_gn=DYSF;refGene_tx=XM_003984110.5
A3	90666958	90667072	refGene_gn=DYSF;refGene_tx=XM_003984110.5
A3	90668527	90668705	refGene_gn=DYSF;refGene_tx=XM_003984110.5
A3	90669267	90669392	refGene_gn=DYSF;refGene_tx=XM_003984110.5
A3	90671253	90671360	refGene_gn=DYSF;refGene_tx=XM_003984110.5
A3	90676269	90676462	refGene_gn=DYSF;refGene_tx=XM_003984110.5
A3	90678572	90678728	refGene_gn=DYSF;refGene_tx=XM_003984110.5
A3	90682431	90682563	refGene_gn=DYSF;refGene_tx=XM_003984110.5
A3	90682649	90682816	refGene_gn=DYSF;refGene_tx=XM_003984110.5
A3	90683928	90684043	refGene_gn=DYSF;refGene_tx=XM_003984110.5
A3	90684322	90684428	refGene_gn=DYSF;refGene_tx=XM_003984110.5
A3	90684698	90684841	refGene_gn=DYSF;refGene_tx=XM_003984110.5
A3	90686597	90686771	refGene_gn=DYSF;refGene_tx=XM_003984110.5
A3	90700364	90700458	refGene_gn=DYSF;refGene_tx=XM_003984110.5
A3	90701001	90701079	refGene_gn=DYSF;refGene_tx=XM_003984110.5
A3	90709096	90709278	refGene_gn=DYSF;refGene_tx=XM_003984110.5
A3	90711961	90712102	refGene_gn=DYSF;refGene_tx=XM_003984110.5
A3	90712909	90712939	refGene_gn=DYSF;refGene_tx=XM_003984110.5
A3	90715185	90715215	refGene_gn=DYSF;refGene_tx=XM_003984110.5
A3	90723856	90723958	refGene_gn=DYSF;refGene_tx=XM_003984110.5
A3	90724068	90724230	refGene_gn=DYSF;refGene_tx=XM_003984110.5
A3	90725289	90725455	refGene_gn=DYSF;refGene_tx=XM_003984110.5
A3	90725946	90726023	refGene_gn=DYSF;refGene_tx=XM_003984110.5
A3	90757557	90757656	refGene_gn=DYSF;refGene_tx=XM_003984110.5
A3	90765199	90765328	refGene_gn=DYSF;refGene_tx=XM_003984110.5
A3	90765646	90765802	refGene_gn=DYSF;refGene_tx=XM_003984110.5
A3	90767344	90767436	refGene_gn=DYSF;refGene_tx=XM_003984110.5
A3	90770038	90770209	refGene_gn=DYSF;refGene_tx=XM_003984110.5
A3	90772365	90772508	refGene_gn=DYSF;refGene_tx=XM_003984110.5
A3	90774390	90774464	refGene_gn=DYSF;refGene_tx=XM_003984110.5
A3	90774465	90774531	refGene_gn=DYSF;refGene_tx=XM_003984110.5
A3	90775791	90775880	refGene_gn=DYSF;refGene_tx=XM_003984110.5
A3	90776147	90776243	refGene_gn=DYSF;refGene_tx=XM_003984110.5
A3	90776591	90776733	refGene_gn=DYSF;refGene_tx=XM_003984110.5
A3	90780544	90780644	refGene_gn=DYSF;refGene_tx=XM_003984110.5
A3	90785275	90785454	refGene_gn=DYSF;refGene_tx=XM_003984110.5
A3	90787001	90787111	refGene_gn=DYSF;refGene_tx=XM_003984110.5
A3	90788445	90788593	refGene_gn=DYSF;refGene_tx=XM_003984110.5
A3	90792461	90792500	refGene_gn=DYSF;refGene_tx=XM_003984110.5
A3	91171578	91171971
refGene_gn=LOC101100712;refGene_tx=XM_003984115.5;refGene_gn=LOC101100
712;refGene_tx=XM_023251759.1
A3	91172328	91172613
refGene_gn=LOC101100712;refGene_tx=XM_003984115.5;refGene_gn=LOC101100
712;refGene_tx=XM_023251759.1

A3	91174130	91174286
refGene_gn=LOC101100712;refGene_tx=XM_003984115.5;refGene_gn=LOC101100712;refGene_tx=XM_023251759.1
A3	91174548	91174824
refGene_gn=LOC101100712;refGene_tx=XM_003984115.5;refGene_gn=LOC101100712;refGene_tx=XM_023251759.1
A3	91183894	91184119
refGene_gn=LOC101100712;refGene_tx=XM_003984115.5;refGene_gn=LOC101100712;refGene_tx=XM_023251759.1
A3	91187636	91187840
refGene_gn=LOC101100712;refGene_tx=XM_003984115.5;refGene_gn=LOC101100712;refGene_tx=XM_023251759.1
A3	91212128	91212515
refGene_gn=LOC102899149;refGene_tx=XM_006930182.3
A3	91213170	91213473
refGene_gn=LOC102899149;refGene_tx=XM_006930182.3
A3	91217585	91217712
refGene_gn=EXOC6B;refGene_tx=XM_023251760.1;refGene_gn=EXOC6B;refGene_tx=XM_023251761.1;refGene_gn=EXOC6B;refGene_tx=XM_023251762.1;refGene_gn=EXOC6B;refGene_tx=XM_023251766.1
A3	91220704	91220716
refGene_gn=EXOC6B;refGene_tx=XM_023251760.1;refGene_gn=EXOC6B;refGene_tx=XM_023251761.1;refGene_gn=EXOC6B;refGene_tx=XM_023251766.1
A3	91221794	91221907
refGene_gn=EXOC6B;refGene_tx=XM_023251760.1;refGene_gn=EXOC6B;refGene_tx=XM_023251761.1;refGene_gn=EXOC6B;refGene_tx=XM_023251762.1;refGene_gn=EXOC6B;refGene_tx=XM_023251766.1
A3	91366275	91366293	refGene_gn=EXOC6B;refGene_tx=XM_023251765.1
A3	91367696	91367729	refGene_gn=EXOC6B;refGene_tx=XM_023251764.1
A3	91371696	91371770
refGene_gn=EXOC6B;refGene_tx=XM_023251760.1;refGene_gn=EXOC6B;refGene_tx=XM_023251761.1;refGene_gn=EXOC6B;refGene_tx=XM_023251762.1;refGene_gn=EXOC6B;refGene_tx=XM_023251764.1;refGene_gn=EXOC6B;refGene_tx=XM_023251765.1;refGene_gn=EXOC6B;refGene_tx=XM_023251766.1
A3	91398588	91398730
refGene_gn=EXOC6B;refGene_tx=XM_023251760.1;refGene_gn=EXOC6B;refGene_tx=XM_023251761.1;refGene_gn=EXOC6B;refGene_tx=XM_023251762.1;refGene_gn=EXOC6B;refGene_tx=XM_023251764.1;refGene_gn=EXOC6B;refGene_tx=XM_023251765.1;refGene_gn=EXOC6B;refGene_tx=XM_023251766.1
A3	91467205	91467385
refGene_gn=EXOC6B;refGene_tx=XM_023251760.1;refGene_gn=EXOC6B;refGene_tx=XM_023251761.1;refGene_gn=EXOC6B;refGene_tx=XM_023251762.1;refGene_gn=EXOC6B;refGene_tx=XM_023251764.1;refGene_gn=EXOC6B;refGene_tx=XM_023251765.1;refGene_gn=EXOC6B;refGene_tx=XM_023251766.1
A3	91475820	91475955
refGene_gn=EXOC6B;refGene_tx=XM_023251760.1;refGene_gn=EXOC6B;refGene_tx=XM_023251761.1;refGene_gn=EXOC6B;refGene_tx=XM_023251762.1;refGene_gn=EXOC6B;refGene_tx=XM_023251764.1;refGene_gn=EXOC6B;refGene_tx=XM_023251765.1;refGene_gn=EXOC6B;refGene_tx=XM_023251766.1
A3	91487449	91487561

refGene_gn=EXOC6B;refGene_tx=XM_023251760.1;refGene_gn=EXOC6B;refGene_tx=XM_023251761.1;refGene_gn=EXOC6B;refGene_tx=XM_023251762.1;refGene_gn=EXOC6B;refGene_tx=XM_023251764.1;refGene_gn=EXOC6B;refGene_tx=XM_023251765.1;refGene_gn=EXOC6B;refGene_tx=XM_023251766.1
A3 91500140 91500250
refGene_gn=EXOC6B;refGene_tx=XM_023251760.1;refGene_gn=EXOC6B;refGene_tx=XM_023251761.1;refGene_gn=EXOC6B;refGene_tx=XM_023251762.1;refGene_gn=EXOC6B;refGene_tx=XM_023251764.1;refGene_gn=EXOC6B;refGene_tx=XM_023251765.1;refGene_gn=EXOC6B;refGene_tx=XM_023251766.1
A3 91501394 91501500
refGene_gn=EXOC6B;refGene_tx=XM_023251760.1;refGene_gn=EXOC6B;refGene_tx=XM_023251761.1;refGene_gn=EXOC6B;refGene_tx=XM_023251762.1;refGene_gn=EXOC6B;refGene_tx=XM_023251764.1;refGene_gn=EXOC6B;refGene_tx=XM_023251765.1;refGene_gn=EXOC6B;refGene_tx=XM_023251766.1
A3 91502957 91503055
refGene_gn=EXOC6B;refGene_tx=XM_023251760.1;refGene_gn=EXOC6B;refGene_tx=XM_023251761.1;refGene_gn=EXOC6B;refGene_tx=XM_023251762.1;refGene_gn=EXOC6B;refGene_tx=XM_023251764.1;refGene_gn=EXOC6B;refGene_tx=XM_023251765.1;refGene_gn=EXOC6B;refGene_tx=XM_023251766.1
A3 91503835 91503907
refGene_gn=EXOC6B;refGene_tx=XM_023251760.1;refGene_gn=EXOC6B;refGene_tx=XM_023251761.1;refGene_gn=EXOC6B;refGene_tx=XM_023251762.1;refGene_gn=EXOC6B;refGene_tx=XM_023251764.1;refGene_gn=EXOC6B;refGene_tx=XM_023251765.1;refGene_gn=EXOC6B;refGene_tx=XM_023251766.1
A3 91513275 91513396
refGene_gn=EXOC6B;refGene_tx=XM_023251760.1;refGene_gn=EXOC6B;refGene_tx=XM_023251761.1;refGene_gn=EXOC6B;refGene_tx=XM_023251762.1;refGene_gn=EXOC6B;refGene_tx=XM_023251764.1;refGene_gn=EXOC6B;refGene_tx=XM_023251765.1;refGene_gn=EXOC6B;refGene_tx=XM_023251766.1
A3 91514657 91514704
refGene_gn=EXOC6B;refGene_tx=XM_023251760.1;refGene_gn=EXOC6B;refGene_tx=XM_023251761.1;refGene_gn=EXOC6B;refGene_tx=XM_023251762.1;refGene_gn=EXOC6B;refGene_tx=XM_023251764.1;refGene_gn=EXOC6B;refGene_tx=XM_023251765.1;refGene_gn=EXOC6B;refGene_tx=XM_023251766.1
A3 91515101 91515185
refGene_gn=EXOC6B;refGene_tx=XM_023251760.1;refGene_gn=EXOC6B;refGene_tx=XM_023251761.1;refGene_gn=EXOC6B;refGene_tx=XM_023251762.1;refGene_gn=EXOC6B;refGene_tx=XM_023251764.1;refGene_gn=EXOC6B;refGene_tx=XM_023251765.1;refGene_gn=EXOC6B;refGene_tx=XM_023251766.1
A3 91558825 91558894
refGene_gn=EXOC6B;refGene_tx=XM_023251760.1;refGene_gn=EXOC6B;refGene_tx=XM_023251761.1;refGene_gn=EXOC6B;refGene_tx=XM_023251762.1;refGene_gn=EXOC6B;refGene_tx=XM_023251764.1;refGene_gn=EXOC6B;refGene_tx=XM_023251765.1;refGene_gn=EXOC6B;refGene_tx=XM_023251766.1
A3 91575867 91576044
refGene_gn=EXOC6B;refGene_tx=XM_023251760.1;refGene_gn=EXOC6B;refGene_tx=XM_023251761.1;refGene_gn=EXOC6B;refGene_tx=XM_023251762.1;refGene_gn=EXOC6B;refGene_tx=XM_023251764.1;refGene_gn=EXOC6B;refGene_tx=XM_023251765.1;refGene_gn=EXOC6B;refGene_tx=XM_023251766.1
A3 91589418 91589442 refGene_gn=EXOC6B;refGene_tx=XM_023251766.1

A3 91713991 91714196
refGene_gn=EXOC6B;refGene_tx=XM_023251760.1;refGene_gn=EXOC6B;refGene_tx=XM_023251761.1;refGene_gn=EXOC6B;refGene_tx=XM_023251762.1;refGene_gn=EXOC6B;refGene_tx=XM_023251764.1;refGene_gn=EXOC6B;refGene_tx=XM_023251765.1
A3 91726562 91726608
refGene_gn=EXOC6B;refGene_tx=XM_023251760.1;refGene_gn=EXOC6B;refGene_tx=XM_023251761.1;refGene_gn=EXOC6B;refGene_tx=XM_023251762.1;refGene_gn=EXOC6B;refGene_tx=XM_023251764.1;refGene_gn=EXOC6B;refGene_tx=XM_023251765.1
A3 91726719 91726798
refGene_gn=EXOC6B;refGene_tx=XM_023251760.1;refGene_gn=EXOC6B;refGene_tx=XM_023251761.1;refGene_gn=EXOC6B;refGene_tx=XM_023251762.1;refGene_gn=EXOC6B;refGene_tx=XM_023251764.1;refGene_gn=EXOC6B;refGene_tx=XM_023251765.1
A3 91867855 91868177 refGene_gn=SPR;refGene_tx=XM_003984117.5
A3 91868775 91869066 refGene_gn=SPR;refGene_tx=XM_003984117.5
A3 91869902 91870093 refGene_gn=SPR;refGene_tx=XM_003984117.5
A3 91896515 91897047 refGene_gn=EMX1;refGene_tx=XM_023251768.1
A3 91902820 91903005 refGene_gn=EMX1;refGene_tx=XM_023251768.1
A3 91912348 91912516 refGene_gn=EMX1;refGene_tx=XM_023251768.1
A3 91923802 91923880
refGene_gn=SFXN5;refGene_tx=XM_003984118.5;refGene_gn=SFXN5;refGene_tx=XM_006930185.4;refGene_gn=SFXN5;refGene_tx=XM_019827405.2;refGene_gn=SFXN5;refGene_tx=XM_019827406.2
A3 91939960 91940078
refGene_gn=SFXN5;refGene_tx=XM_003984118.5;refGene_gn=SFXN5;refGene_tx=XM_006930185.4;refGene_gn=SFXN5;refGene_tx=XM_019827405.2;refGene_gn=SFXN5;refGene_tx=XM_019827406.2
A3 91945979 91946065
refGene_gn=SFXN5;refGene_tx=XM_003984118.5;refGene_gn=SFXN5;refGene_tx=XM_006930185.4;refGene_gn=SFXN5;refGene_tx=XM_019827406.2
A3 91948060 91948176
refGene_gn=SFXN5;refGene_tx=XM_003984118.5;refGene_gn=SFXN5;refGene_tx=XM_006930185.4;refGene_gn=SFXN5;refGene_tx=XM_019827406.2
A3 91962548 91962639
refGene_gn=SFXN5;refGene_tx=XM_003984118.5;refGene_gn=SFXN5;refGene_tx=XM_006930185.4;refGene_gn=SFXN5;refGene_tx=XM_019827405.2;refGene_gn=SFXN5;refGene_tx=XM_019827406.2
A3 91973786 91973852
refGene_gn=SFXN5;refGene_tx=XM_003984118.5;refGene_gn=SFXN5;refGene_tx=XM_006930185.4;refGene_gn=SFXN5;refGene_tx=XM_019827405.2;refGene_gn=SFXN5;refGene_tx=XM_019827406.2
A3 91975226 91975283
refGene_gn=SFXN5;refGene_tx=XM_003984118.5;refGene_gn=SFXN5;refGene_tx=XM_006930185.4;refGene_gn=SFXN5;refGene_tx=XM_019827405.2;refGene_gn=SFXN5;refGene_tx=XM_019827406.2
A3 91976291 91976345
refGene_gn=SFXN5;refGene_tx=XM_003984118.5;refGene_gn=SFXN5;refGene_tx=XM_006930185.4;refGene_gn=SFXN5;refGene_tx=XM_019827405.2;refGene_gn=

SFXN5;refGene_tx=XM_019827406.2
A3 91995439 91995465
refGene_gn=SFXN5;refGene_tx=XM_003984118.5;refGene_gn=SFXN5;refGene_tx=XM_006930185.4;refGene_gn=SFXN5;refGene_tx=XM_019827405.2;refGene_gn=SFXN5;refGene_tx=XM_019827406.2
A3 91997445 91997500
refGene_gn=SFXN5;refGene_tx=XM_003984118.5;refGene_gn=SFXN5;refGene_tx=XM_006930185.4;refGene_gn=SFXN5;refGene_tx=XM_019827405.2;refGene_gn=SFXN5;refGene_tx=XM_019827406.2
A3 91997973 91998000
refGene_gn=SFXN5;refGene_tx=XM_003984118.5;refGene_gn=SFXN5;refGene_tx=XM_006930185.4;refGene_gn=SFXN5;refGene_tx=XM_019827405.2
A3 92010510 92010588
refGene_gn=SFXN5;refGene_tx=XM_003984118.5;refGene_gn=SFXN5;refGene_tx=XM_006930185.4;refGene_gn=SFXN5;refGene_tx=XM_019827405.2
A3 92024074 92024143
refGene_gn=SFXN5;refGene_tx=XM_003984118.5;refGene_gn=SFXN5;refGene_tx=XM_019827405.2
A3 92037005 92037107
refGene_gn=SFXN5;refGene_tx=XM_003984118.5;refGene_gn=SFXN5;refGene_tx=XM_006930185.4;refGene_gn=SFXN5;refGene_tx=XM_019827405.2
A3 92040811 92041015
refGene_gn=RAB11FIP5;refGene_tx=XM_011281223.2;refGene_gn=RAB11FIP5;refGene_tx=XM_011281224.2;refGene_gn=RAB11FIP5;refGene_tx=XM_019827408.2
A3 92041289 92041479
refGene_gn=RAB11FIP5;refGene_tx=XM_011281223.2;refGene_gn=RAB11FIP5;refGene_tx=XM_011281224.2;refGene_gn=RAB11FIP5;refGene_tx=XM_019827408.2
A3 92044782 92046765
refGene_gn=RAB11FIP5;refGene_tx=XM_011281223.2;refGene_gn=RAB11FIP5;refGene_tx=XM_019827408.2;refGene_gn=RAB11FIP5;refGene_tx=XM_019827409.2
A3 92054206 92054906
refGene_gn=RAB11FIP5;refGene_tx=XM_011281223.2;refGene_gn=RAB11FIP5;refGene_tx=XM_011281224.2;refGene_gn=RAB11FIP5;refGene_tx=XM_019827408.2;refGene_gn=RAB11FIP5;refGene_tx=XM_019827409.2
A3 92055044 92055481
refGene_gn=RAB11FIP5;refGene_tx=XM_011281223.2;refGene_gn=RAB11FIP5;refGene_tx=XM_011281224.2;refGene_gn=RAB11FIP5;refGene_tx=XM_019827408.2;refGene_gn=RAB11FIP5;refGene_tx=XM_019827409.2
A3 92076298 92076729
refGene_gn=RAB11FIP5;refGene_tx=XM_011281223.2;refGene_gn=RAB11FIP5;refGene_tx=XM_011281224.2;refGene_gn=RAB11FIP5;refGene_tx=XM_019827408.2;refGene_gn=RAB11FIP5;refGene_tx=XM_019827409.2
A3 92150517 92150869 refGene_gn=NOTO;refGene_tx=XM_023251769.1
A3 92152435 92152650 refGene_gn=NOTO;refGene_tx=XM_023251769.1
A3 92155055 92155214 refGene_gn=NOTO;refGene_tx=XM_023251769.1
A3 92157269 92157365 refGene_gn=SMYD5;refGene_tx=XM_003984119.5
A3 92161208 92161317 refGene_gn=SMYD5;refGene_tx=XM_003984119.5
A3 92162042 92162182 refGene_gn=SMYD5;refGene_tx=XM_003984119.5
A3 92162650 92162772 refGene_gn=SMYD5;refGene_tx=XM_003984119.5
A3 92163160 92163230 refGene_gn=SMYD5;refGene_tx=XM_003984119.5

| | | | |
|---|---|---|---|
| A3 | 92163851 | 92163956 | refGene_gn=SMYD5;refGene_tx=XM_003984119.5 |
| A3 | 92164732 | 92164795 | refGene_gn=SMYD5;refGene_tx=XM_003984119.5 |
| A3 | 92165010 | 92165081 | refGene_gn=SMYD5;refGene_tx=XM_003984119.5 |
| A3 | 92165384 | 92165491 | refGene_gn=SMYD5;refGene_tx=XM_003984119.5 |
| A3 | 92165881 | 92165938 | refGene_gn=SMYD5;refGene_tx=XM_003984119.5 |
| A3 | 92166793 | 92166888 | refGene_gn=SMYD5;refGene_tx=XM_003984119.5 |
| A3 | 92167545 | 92167616 | refGene_gn=SMYD5;refGene_tx=XM_003984119.5 |
| A3 | 92167712 | 92167857 | refGene_gn=SMYD5;refGene_tx=XM_003984119.5 |
| A3 | 92170052 | 92170173 | refGene_gn=PRADC1;refGene_tx=XM_003984120.5 |
| A3 | 92170423 | 92170591 | refGene_gn=PRADC1;refGene_tx=XM_003984120.5 |
| A3 | 92170872 | 92170982 | refGene_gn=PRADC1;refGene_tx=XM_003984120.5 |
| A3 | 92171508 | 92171609 | refGene_gn=PRADC1;refGene_tx=XM_003984120.5 |
| A3 | 92173529 | 92173596 | refGene_gn=PRADC1;refGene_tx=XM_003984120.5 |
| A3 | 92174841 | 92174847 | refGene_gn=CCT7;refGene_tx=XM_003984121.4 |
| A3 | 92180082 | 92180236 | refGene_gn=CCT7;refGene_tx=XM_003984121.4 |
| A3 | 92180865 | 92180972 | refGene_gn=CCT7;refGene_tx=XM_003984121.4 |
| A3 | 92183165 | 92183291 | refGene_gn=CCT7;refGene_tx=XM_003984121.4 |
| A3 | 92184082 | 92184135 | refGene_gn=CCT7;refGene_tx=XM_003984121.4 |
| A3 | 92184488 | 92184660 | refGene_gn=CCT7;refGene_tx=XM_003984121.4 |
| A3 | 92188575 | 92188740 | refGene_gn=CCT7;refGene_tx=XM_003984121.4 |
| A3 | 92189585 | 92189774 | refGene_gn=CCT7;refGene_tx=XM_003984121.4 |
| A3 | 92190573 | 92190671 | refGene_gn=CCT7;refGene_tx=XM_003984121.4 |
| A3 | 92191052 | 92191185 | refGene_gn=CCT7;refGene_tx=XM_003984121.4 |
| A3 | 92191641 | 92191848 | refGene_gn=CCT7;refGene_tx=XM_003984121.4 |
| A3 | 92192781 | 92193003 | refGene_gn=CCT7;refGene_tx=XM_003984121.4 |
| A3 | 92198865 | 92198928 | refGene_gn=FBXO41;refGene_tx=XM_023251770.1 |
| A3 | 92199060 | 92199176 | refGene_gn=FBXO41;refGene_tx=XM_023251770.1 |
| A3 | 92200614 | 92200773 | refGene_gn=FBXO41;refGene_tx=XM_023251770.1 |
| A3 | 92200952 | 92201071 | refGene_gn=FBXO41;refGene_tx=XM_023251770.1 |
| A3 | 92202415 | 92202511 | refGene_gn=FBXO41;refGene_tx=XM_023251770.1 |
| A3 | 92202691 | 92202844 | refGene_gn=FBXO41;refGene_tx=XM_023251770.1 |
| A3 | 92202966 | 92203082 | refGene_gn=FBXO41;refGene_tx=XM_023251770.1 |
| A3 | 92203296 | 92203538 | refGene_gn=FBXO41;refGene_tx=XM_023251770.1 |
| A3 | 92204368 | 92204727 | refGene_gn=FBXO41;refGene_tx=XM_023251770.1 |
| A3 | 92204997 | 92205071 | refGene_gn=FBXO41;refGene_tx=XM_023251770.1 |
| A3 | 92205555 | 92205781 | refGene_gn=FBXO41;refGene_tx=XM_023251770.1 |
| A3 | 92207263 | 92208156 | refGene_gn=FBXO41;refGene_tx=XM_023251770.1 |
| A3 | 92228656 | 92229975 | refGene_gn=EGR4;refGene_tx=XM_006930186.4 |
| A3 | 92230376 | 92230818 | refGene_gn=EGR4;refGene_tx=XM_006930186.4 |
| A3 | 92291314 | 92291623 | refGene_gn=ALMS1;refGene_tx=XM_023252188.1 |
| A3 | 92309890 | 92310016 | refGene_gn=ALMS1;refGene_tx=XM_023252188.1 |
| A3 | 92321234 | 92321430 | refGene_gn=ALMS1;refGene_tx=XM_023252188.1 |
| A3 | 92326693 | 92326811 | refGene_gn=ALMS1;refGene_tx=XM_023252188.1 |
| A3 | 92328038 | 92329813 | refGene_gn=ALMS1;refGene_tx=XM_023252188.1 |
| A3 | 92330074 | 92330193 | refGene_gn=ALMS1;refGene_tx=XM_023252188.1 |
| A3 | 92330298 | 92330448 | refGene_gn=ALMS1;refGene_tx=XM_023252188.1 |
| A3 | 92335497 | 92335657 | refGene_gn=ALMS1;refGene_tx=XM_023252188.1 |
| A3 | 92352868 | 92354632 | refGene_gn=ALMS1;refGene_tx=XM_023252188.1 |
| A3 | 92354771 | 92356846 | refGene_gn=ALMS1;refGene_tx=XM_023252188.1 |
| A3 | 92356887 | 92359990 | refGene_gn=ALMS1;refGene_tx=XM_023252188.1 |

A3 92361263 92361397 refGene_gn=ALMS1;refGene_tx=XM_023252188.1
A3 92393384 92393457 refGene_gn=ALMS1;refGene_tx=XM_023252188.1
A3 92393458 92393958 refGene_gn=ALMS1;refGene_tx=XM_023252188.1
A3 92394834 92395727 refGene_gn=ALMS1;refGene_tx=XM_023252188.1
A3 92422782 92423024 refGene_gn=ALMS1;refGene_tx=XM_023252188.1
A3 92426634 92426760 refGene_gn=ALMS1;refGene_tx=XM_023252188.1
A3 92439061 92439232 refGene_gn=ALMS1;refGene_tx=XM_023252188.1
A3 92441817 92441955 refGene_gn=ALMS1;refGene_tx=XM_023252188.1
A3 92445556 92445727 refGene_gn=ALMS1;refGene_tx=XM_023252188.1
A3 92470083 92471231 refGene_gn=ALMS1;refGene_tx=XM_023252188.1
A3 92511715 92511836 refGene_gn=ALMS1;refGene_tx=XM_023252188.1
A3 92512910 92513114 refGene_gn=ALMS1;refGene_tx=XM_023252188.1
A3 92513414 92513644 refGene_gn=ALMS1;refGene_tx=XM_023252188.1
A3 92514376 92514560 refGene_gn=ALMS1;refGene_tx=XM_023252188.1
A3 92515472 92515536 refGene_gn=ALMS1;refGene_tx=XM_023252188.1
A3 92519087 92519187 refGene_gn=ALMS1;refGene_tx=XM_023252188.1
A3 92520238 92520283 refGene_gn=ALMS1;refGene_tx=XM_023252188.1
A3 92536718 92537494 refGene_gn=NAT8;refGene_tx=XM_003984184.4
A3 92538129 92538343 refGene_gn=NAT8;refGene_tx=XM_003984184.4
A3 92540426 92540561
refGene_gn=TPRKB;refGene_tx=XM_019827418.2;refGene_gn=TPRKB;refGene_tx=XM_019827419.2;refGene_gn=TPRKB;refGene_tx=XM_019827420.2;refGene_gn=TPRKB;refGene_tx=XM_019827421.2;refGene_gn=TPRKB;refGene_tx=XM_019827422.2
A3 92545476 92545563 refGene_gn=TPRKB;refGene_tx=XM_003984127.3
A3 92546794 92546971
refGene_gn=TPRKB;refGene_tx=XM_003984127.3;refGene_gn=TPRKB;refGene_tx=XM_019827418.2;refGene_gn=TPRKB;refGene_tx=XM_019827419.2;refGene_gn=TPRKB;refGene_tx=XM_019827420.2;refGene_gn=TPRKB;refGene_tx=XM_019827421.2;refGene_gn=TPRKB;refGene_tx=XM_019827422.2
A3 92548302 92548425
refGene_gn=TPRKB;refGene_tx=XM_003984127.3;refGene_gn=TPRKB;refGene_tx=XM_019827418.2;refGene_gn=TPRKB;refGene_tx=XM_019827419.2;refGene_gn=TPRKB;refGene_tx=XM_019827420.2;refGene_gn=TPRKB;refGene_tx=XM_019827421.2
A3 92549778 92549919
refGene_gn=TPRKB;refGene_tx=XM_003984127.3;refGene_gn=TPRKB;refGene_tx=XM_019827418.2;refGene_gn=TPRKB;refGene_tx=XM_019827419.2;refGene_gn=TPRKB;refGene_tx=XM_019827420.2;refGene_gn=TPRKB;refGene_tx=XM_019827421.2;refGene_gn=TPRKB;refGene_tx=XM_019827422.2
A3 92563585 92563781
refGene_gn=DUSP11;refGene_tx=XM_006930191.3;refGene_gn=DUSP11;refGene_tx=XM_023251771.1
A3 92564670 92564856
refGene_gn=DUSP11;refGene_tx=XM_006930191.3;refGene_gn=DUSP11;refGene_tx=XM_023251771.1
A3 92565320 92565396
refGene_gn=DUSP11;refGene_tx=XM_006930191.3;refGene_gn=DUSP11;refGene_tx=XM_023251771.1
A3 92565672 92565719

refGene_gn=DUSP11;refGene_tx=XM_006930191.3;refGene_gn=DUSP11;refGene_tx=XM_023251771.1
A3 92568223 92568284
refGene_gn=DUSP11;refGene_tx=XM_006930191.3;refGene_gn=DUSP11;refGene_tx=XM_023251771.1
A3 92572089 92572213
refGene_gn=DUSP11;refGene_tx=XM_006930191.3;refGene_gn=DUSP11;refGene_tx=XM_023251771.1
A3 92576121 92576253
refGene_gn=DUSP11;refGene_tx=XM_006930191.3;refGene_gn=DUSP11;refGene_tx=XM_023251771.1
A3 92579149 92579225
refGene_gn=DUSP11;refGene_tx=XM_006930191.3;refGene_gn=DUSP11;refGene_tx=XM_023251771.1
A3 92580900 92581001 refGene_gn=DUSP11;refGene_tx=XM_006930191.3
A3 92584394 92584447 refGene_gn=DUSP11;refGene_tx=XM_023251771.1
A3 92586277 92586374
refGene_gn=CA3H2orf78;refGene_tx=XM_003984186.4
A3 92589367 92590129
refGene_gn=CA3H2orf78;refGene_tx=XM_003984186.4
A3 92590983 92592902
refGene_gn=CA3H2orf78;refGene_tx=XM_003984186.4
A3 92605878 92605959 refGene_gn=STAMBP;refGene_tx=XM_003984128.5
A3 92607789 92608007
refGene_gn=STAMBP;refGene_tx=XM_003984128.5;refGene_gn=STAMBP;refGene_tx=XM_006930193.4
A3 92618382 92618458
refGene_gn=STAMBP;refGene_tx=XM_003984128.5;refGene_gn=STAMBP;refGene_tx=XM_006930193.4
A3 92618744 92618840
refGene_gn=STAMBP;refGene_tx=XM_003984128.5;refGene_gn=STAMBP;refGene_tx=XM_006930193.4
A3 92619912 92620279
refGene_gn=STAMBP;refGene_tx=XM_003984128.5;refGene_gn=STAMBP;refGene_tx=XM_006930193.4
A3 92622613 92622738
refGene_gn=STAMBP;refGene_tx=XM_003984128.5;refGene_gn=STAMBP;refGene_tx=XM_006930193.4
A3 92623616 92623754
refGene_gn=STAMBP;refGene_tx=XM_003984128.5;refGene_gn=STAMBP;refGene_tx=XM_006930193.4
A3 92628774 92628887
refGene_gn=STAMBP;refGene_tx=XM_003984128.5;refGene_gn=STAMBP;refGene_tx=XM_006930193.4
A3 92629426 92629526
refGene_gn=STAMBP;refGene_tx=XM_003984128.5;refGene_gn=STAMBP;refGene_tx=XM_006930193.4
A3 92630889 92630946
refGene_gn=STAMBP;refGene_tx=XM_003984128.5;refGene_gn=STAMBP;refGene_tx=XM_006930193.4

A3	92669370	92669496	refGene_gn=ACTG2;refGene_tx=XM_003984129.5
A3	92670338	92670467	refGene_gn=ACTG2;refGene_tx=XM_003984129.5
A3	92675655	92675766	refGene_gn=ACTG2;refGene_tx=XM_003984129.5
A3	92676013	92676098	refGene_gn=ACTG2;refGene_tx=XM_003984129.5
A3	92677612	92677774	refGene_gn=ACTG2;refGene_tx=XM_003984129.5
A3	92678551	92678743	refGene_gn=ACTG2;refGene_tx=XM_003984129.5
A3	92680995	92681177	refGene_gn=ACTG2;refGene_tx=XM_003984129.5
A3	92683565	92683709	refGene_gn=ACTG2;refGene_tx=XM_003984129.5
A3	92689609	92689751
refGene_gn=DGUOK;refGene_tx=XM_003984131.5;refGene_gn=DGUOK;refGene_tx=XM_006930194.4;refGene_gn=DGUOK;refGene_tx=XM_019827423.2;refGene_gn=DGUOK;refGene_tx=XM_019827424.2;refGene_gn=DGUOK;refGene_tx=XM_023251772.1;refGene_gn=DGUOK;refGene_tx=XM_023251773.1;refGene_gn=DGUOK;refGene_tx=XM_023251774.1;refGene_gn=DGUOK;refGene_tx=XM_023251775.1
A3	92700515	92700628
refGene_gn=DGUOK;refGene_tx=XM_003984131.5;refGene_gn=DGUOK;refGene_tx=XM_019827423.2;refGene_gn=DGUOK;refGene_tx=XM_019827424.2;refGene_gn=DGUOK;refGene_tx=XM_023251772.1;refGene_gn=DGUOK;refGene_tx=XM_023251773.1;refGene_gn=DGUOK;refGene_tx=XM_023251774.1;refGene_gn=DGUOK;refGene_tx=XM_023251775.1
A3	92702816	92702914	refGene_gn=DGUOK;refGene_tx=XM_006930194.4
A3	92708236	92708424
refGene_gn=DGUOK;refGene_tx=XM_003984131.5;refGene_gn=DGUOK;refGene_tx=XM_006930194.4;refGene_gn=DGUOK;refGene_tx=XM_019827423.2;refGene_gn=DGUOK;refGene_tx=XM_019827424.2;refGene_gn=DGUOK;refGene_tx=XM_019827425.2;refGene_gn=DGUOK;refGene_tx=XM_023251772.1;refGene_gn=DGUOK;refGene_tx=XM_023251773.1;refGene_gn=DGUOK;refGene_tx=XM_023251774.1;refGene_gn=DGUOK;refGene_tx=XM_023251775.1;refGene_gn=DGUOK;refGene_tx=XM_023251776.1
A3	92711885	92712033
refGene_gn=DGUOK;refGene_tx=XM_003984131.5;refGene_gn=DGUOK;refGene_tx=XM_006930194.4;refGene_gn=DGUOK;refGene_tx=XM_019827423.2;refGene_gn=DGUOK;refGene_tx=XM_019827425.2;refGene_gn=DGUOK;refGene_tx=XM_023251772.1;refGene_gn=DGUOK;refGene_tx=XM_023251773.1;refGene_gn=DGUOK;refGene_tx=XM_023251774.1;refGene_gn=DGUOK;refGene_tx=XM_023251775.1;refGene_gn=DGUOK;refGene_tx=XM_023251776.1
A3	92717002	92717267
refGene_gn=DGUOK;refGene_tx=XM_003984131.5;refGene_gn=DGUOK;refGene_tx=XM_006930194.4;refGene_gn=DGUOK;refGene_tx=XM_019827423.2;refGene_gn=DGUOK;refGene_tx=XM_019827424.2;refGene_gn=DGUOK;refGene_tx=XM_019827425.2;refGene_gn=DGUOK;refGene_tx=XM_023251772.1;refGene_gn=DGUOK;refGene_tx=XM_023251773.1;refGene_gn=DGUOK;refGene_tx=XM_023251774.1;refGene_gn=DGUOK;refGene_tx=XM_023251775.1;refGene_gn=DGUOK;refGene_tx=XM_023251776.1
A3	92717725	92717825
refGene_gn=DGUOK;refGene_tx=XM_003984131.5;refGene_gn=DGUOK;refGene_tx=XM_006930194.4;refGene_gn=DGUOK;refGene_tx=XM_019827425.2;refGene_gn=DGUOK;refGene_tx=XM_023251772.1;refGene_gn=DGUOK;refGene_tx=XM_023251773.1;refGene_gn=DGUOK;refGene_tx=XM_023251774.1;refGene_gn=DGUOK;refGene_tx=XM_023251775.1;refGene_gn=DGUOK;refGene_tx=XM_023251776.1

A3 92718265 92718292
refGene_gn=DGUOK;refGene_tx=XM_003984131.5;refGene_gn=DGUOK;refGene_tx=XM_006930194.4;refGene_gn=DGUOK;refGene_tx=XM_019827425.2;refGene_gn=DGUOK;refGene_tx=XM_023251772.1;refGene_gn=DGUOK;refGene_tx=XM_023251773.1;refGene_gn=DGUOK;refGene_tx=XM_023251774.1;refGene_gn=DGUOK;refGene_tx=XM_023251775.1;refGene_gn=DGUOK;refGene_tx=XM_023251776.1
A3 92743341 92743644
refGene_gn=TET3;refGene_tx=XM_019827426.2;refGene_gn=TET3;refGene_tx=XM_019827427.2;refGene_gn=TET3;refGene_tx=XM_019827429.2;refGene_gn=TET3;refGene_tx=XM_023251777.1
A3 92757702 92757759
refGene_gn=TET3;refGene_tx=XM_019827426.2;refGene_gn=TET3;refGene_tx=XM_019827427.2;refGene_gn=TET3;refGene_tx=XM_019827429.2;refGene_gn=TET3;refGene_tx=XM_023251777.1
A3 92794464 92796598
refGene_gn=TET3;refGene_tx=XM_019827426.2;refGene_gn=TET3;refGene_tx=XM_019827427.2;refGene_gn=TET3;refGene_tx=XM_019827429.2;refGene_gn=TET3;refGene_tx=XM_023251777.1
A3 92815449 92815540
refGene_gn=TET3;refGene_tx=XM_019827426.2;refGene_gn=TET3;refGene_tx=XM_019827427.2;refGene_gn=TET3;refGene_tx=XM_019827429.2;refGene_gn=TET3;refGene_tx=XM_023251777.1
A3 92821755 92821849
refGene_gn=TET3;refGene_tx=XM_019827426.2;refGene_gn=TET3;refGene_tx=XM_019827427.2;refGene_gn=TET3;refGene_tx=XM_019827429.2;refGene_gn=TET3;refGene_tx=XM_023251777.1
A3 92828455 92828664
refGene_gn=TET3;refGene_tx=XM_019827426.2;refGene_gn=TET3;refGene_tx=XM_019827427.2;refGene_gn=TET3;refGene_tx=XM_019827429.2;refGene_gn=TET3;refGene_tx=XM_023251777.1
A3 92830685 92830836
refGene_gn=TET3;refGene_tx=XM_019827426.2;refGene_gn=TET3;refGene_tx=XM_019827427.2;refGene_gn=TET3;refGene_tx=XM_019827429.2;refGene_gn=TET3;refGene_tx=XM_023251777.1
A3 92833526 92833616
refGene_gn=TET3;refGene_tx=XM_019827426.2;refGene_gn=TET3;refGene_tx=XM_019827427.2;refGene_gn=TET3;refGene_tx=XM_019827429.2;refGene_gn=TET3;refGene_tx=XM_023251777.1
A3 92834127 92834265
refGene_gn=TET3;refGene_tx=XM_019827426.2;refGene_gn=TET3;refGene_tx=XM_019827427.2;refGene_gn=TET3;refGene_tx=XM_019827429.2;refGene_gn=TET3;refGene_tx=XM_023251777.1
A3 92838727 92839064
refGene_gn=TET3;refGene_tx=XM_019827426.2;refGene_gn=TET3;refGene_tx=XM_019827427.2;refGene_gn=TET3;refGene_tx=XM_019827429.2;refGene_gn=TET3;refGene_tx=XM_023251777.1
A3 92839647 92841425
refGene_gn=TET3;refGene_tx=XM_019827426.2;refGene_gn=TET3;refGene_tx=XM_019827427.2;refGene_gn=TET3;refGene_tx=XM_019827429.2;refGene_gn=TET3;refGene_tx=XM_023251777.1

A3 92887184 92887250
refGene_gn=BOLA3;refGene_tx=XM_023251778.1;refGene_gn=BOLA3;refGene_tx=XM_023251779.1
A3 92890596 92890685
refGene_gn=BOLA3;refGene_tx=XM_023251778.1;refGene_gn=BOLA3;refGene_tx=XM_023251779.1
A3 92892553 92892677
refGene_gn=BOLA3;refGene_tx=XM_023251778.1;refGene_gn=BOLA3;refGene_tx=XM_023251779.1
A3 92894575 92894578 refGene_gn=BOLA3;refGene_tx=XM_023251779.1
A3 92894852 92894906 refGene_gn=BOLA3;refGene_tx=XM_023251778.1
A3 92902930 92903008
refGene_gn=MOB1A;refGene_tx=XM_011281218.3;refGene_gn=MOB1A;refGene_tx=XM_011281219.3
A3 92904602 92904766
refGene_gn=MOB1A;refGene_tx=XM_011281218.3;refGene_gn=MOB1A;refGene_tx=XM_011281219.3
A3 92909216 92909350
refGene_gn=MOB1A;refGene_tx=XM_011281218.3;refGene_gn=MOB1A;refGene_tx=XM_011281219.3
A3 92910720 92910814
refGene_gn=MOB1A;refGene_tx=XM_011281218.3;refGene_gn=MOB1A;refGene_tx=XM_011281219.3
A3 92915976 92916143
refGene_gn=MOB1A;refGene_tx=XM_011281218.3;refGene_gn=MOB1A;refGene_tx=XM_011281219.3
A3 92919939 92919953
refGene_gn=MOB1A;refGene_tx=XM_011281218.3;refGene_gn=MOB1A;refGene_tx=XM_011281219.3
A3 92934628 92934729 refGene_gn=MTHFD2;refGene_tx=XM_003984134.4
A3 92939380 92939565 refGene_gn=MTHFD2;refGene_tx=XM_003984134.4
A3 92941373 92941496 refGene_gn=MTHFD2;refGene_tx=XM_003984134.4
A3 92942410 92942563 refGene_gn=MTHFD2;refGene_tx=XM_003984134.4
A3 92943037 92943145 refGene_gn=MTHFD2;refGene_tx=XM_003984134.4
A3 92944003 92944096 refGene_gn=MTHFD2;refGene_tx=XM_003984134.4
A3 92944473 92944599 refGene_gn=MTHFD2;refGene_tx=XM_003984134.4
A3 92945821 92945985 refGene_gn=MTHFD2;refGene_tx=XM_003984134.4
A3 92949626 92949792
refGene_gn=SLC4A5;refGene_tx=XM_019827439.2;refGene_gn=SLC4A5;refGene_tx=XM_019827444.2;refGene_gn=SLC4A5;refGene_tx=XM_023251781.1;refGene_gn=SLC4A5;refGene_tx=XM_023251782.1
A3 92951824 92951892
refGene_gn=SLC4A5;refGene_tx=XM_019827439.2;refGene_gn=SLC4A5;refGene_tx=XM_019827441.2;refGene_gn=SLC4A5;refGene_tx=XM_019827444.2;refGene_gn=SLC4A5;refGene_tx=XM_023251781.1;refGene_gn=SLC4A5;refGene_tx=XM_023251782.1;refGene_gn=SLC4A5;refGene_tx=XM_023251783.1
A3 92952958 92953043
refGene_gn=SLC4A5;refGene_tx=XM_019827441.2;refGene_gn=SLC4A5;refGene_tx=XM_023251783.1
A3 92954644 92954800

refGene_gn=SLC4A5;refGene_tx=XM_019827439.2;refGene_gn=SLC4A5;refGene_tx=XM_019827441.2;refGene_gn=SLC4A5;refGene_tx=XM_019827444.2;refGene_gn=SLC4A5;refGene_tx=XM_023251781.1;refGene_gn=SLC4A5;refGene_tx=XM_023251782.1
A3 92960348 92960522
refGene_gn=SLC4A5;refGene_tx=XM_019827439.2;refGene_gn=SLC4A5;refGene_tx=XM_019827441.2;refGene_gn=SLC4A5;refGene_tx=XM_019827444.2;refGene_gn=SLC4A5;refGene_tx=XM_023251781.1;refGene_gn=SLC4A5;refGene_tx=XM_023251782.1;refGene_gn=SLC4A5;refGene_tx=XM_023251783.1
A3 92961140 92961209
refGene_gn=SLC4A5;refGene_tx=XM_019827439.2;refGene_gn=SLC4A5;refGene_tx=XM_019827441.2;refGene_gn=SLC4A5;refGene_tx=XM_019827444.2;refGene_gn=SLC4A5;refGene_tx=XM_023251781.1;refGene_gn=SLC4A5;refGene_tx=XM_023251782.1;refGene_gn=SLC4A5;refGene_tx=XM_023251783.1
A3 92965557 92965630
refGene_gn=SLC4A5;refGene_tx=XM_019827439.2;refGene_gn=SLC4A5;refGene_tx=XM_019827441.2;refGene_gn=SLC4A5;refGene_tx=XM_019827444.2;refGene_gn=SLC4A5;refGene_tx=XM_023251781.1;refGene_gn=SLC4A5;refGene_tx=XM_023251782.1;refGene_gn=SLC4A5;refGene_tx=XM_023251783.1
A3 92966902 92967081
refGene_gn=SLC4A5;refGene_tx=XM_019827439.2;refGene_gn=SLC4A5;refGene_tx=XM_019827441.2;refGene_gn=SLC4A5;refGene_tx=XM_019827444.2;refGene_gn=SLC4A5;refGene_tx=XM_023251781.1;refGene_gn=SLC4A5;refGene_tx=XM_023251782.1;refGene_gn=SLC4A5;refGene_tx=XM_023251783.1
A3 92967836 92967998
refGene_gn=SLC4A5;refGene_tx=XM_019827439.2;refGene_gn=SLC4A5;refGene_tx=XM_019827441.2;refGene_gn=SLC4A5;refGene_tx=XM_019827444.2;refGene_gn=SLC4A5;refGene_tx=XM_023251781.1;refGene_gn=SLC4A5;refGene_tx=XM_023251782.1;refGene_gn=SLC4A5;refGene_tx=XM_023251783.1
A3 92969417 92969531
refGene_gn=SLC4A5;refGene_tx=XM_019827439.2;refGene_gn=SLC4A5;refGene_tx=XM_019827441.2;refGene_gn=SLC4A5;refGene_tx=XM_019827444.2;refGene_gn=SLC4A5;refGene_tx=XM_023251781.1;refGene_gn=SLC4A5;refGene_tx=XM_023251782.1;refGene_gn=SLC4A5;refGene_tx=XM_023251783.1
A3 92974268 92974469
refGene_gn=SLC4A5;refGene_tx=XM_019827439.2;refGene_gn=SLC4A5;refGene_tx=XM_019827441.2;refGene_gn=SLC4A5;refGene_tx=XM_019827444.2;refGene_gn=SLC4A5;refGene_tx=XM_023251781.1;refGene_gn=SLC4A5;refGene_tx=XM_023251782.1;refGene_gn=SLC4A5;refGene_tx=XM_023251783.1
A3 92975760 92975819
refGene_gn=SLC4A5;refGene_tx=XM_019827439.2;refGene_gn=SLC4A5;refGene_tx=XM_019827441.2;refGene_gn=SLC4A5;refGene_tx=XM_019827444.2;refGene_gn=SLC4A5;refGene_tx=XM_023251781.1;refGene_gn=SLC4A5;refGene_tx=XM_023251782.1;refGene_gn=SLC4A5;refGene_tx=XM_023251783.1
A3 92979552 92979824
refGene_gn=SLC4A5;refGene_tx=XM_019827439.2;refGene_gn=SLC4A5;refGene_tx=XM_019827441.2;refGene_gn=SLC4A5;refGene_tx=XM_019827444.2;refGene_gn=SLC4A5;refGene_tx=XM_023251781.1;refGene_gn=SLC4A5;refGene_tx=XM_023251782.1;refGene_gn=SLC4A5;refGene_tx=XM_023251783.1
A3 92980512 92980646

refGene_gn=SLC4A5;refGene_tx=XM_019827439.2;refGene_gn=SLC4A5;refGene_tx=XM_019827441.2;refGene_gn=SLC4A5;refGene_tx=XM_019827444.2;refGene_gn=SLC4A5;refGene_tx=XM_023251781.1;refGene_gn=SLC4A5;refGene_tx=XM_023251782.1;refGene_gn=SLC4A5;refGene_tx=XM_023251783.1
A3 92982887 92983062
refGene_gn=SLC4A5;refGene_tx=XM_019827439.2;refGene_gn=SLC4A5;refGene_tx=XM_019827441.2;refGene_gn=SLC4A5;refGene_tx=XM_019827444.2;refGene_gn=SLC4A5;refGene_tx=XM_023251781.1;refGene_gn=SLC4A5;refGene_tx=XM_023251782.1;refGene_gn=SLC4A5;refGene_tx=XM_023251783.1
A3 92984733 92984919
refGene_gn=SLC4A5;refGene_tx=XM_019827439.2;refGene_gn=SLC4A5;refGene_tx=XM_019827441.2;refGene_gn=SLC4A5;refGene_tx=XM_019827444.2;refGene_gn=SLC4A5;refGene_tx=XM_023251781.1;refGene_gn=SLC4A5;refGene_tx=XM_023251782.1;refGene_gn=SLC4A5;refGene_tx=XM_023251783.1
A3 92985292 92985447
refGene_gn=SLC4A5;refGene_tx=XM_019827439.2;refGene_gn=SLC4A5;refGene_tx=XM_019827441.2;refGene_gn=SLC4A5;refGene_tx=XM_019827444.2;refGene_gn=SLC4A5;refGene_tx=XM_023251781.1;refGene_gn=SLC4A5;refGene_tx=XM_023251782.1;refGene_gn=SLC4A5;refGene_tx=XM_023251783.1
A3 92986611 92986699
refGene_gn=SLC4A5;refGene_tx=XM_019827439.2;refGene_gn=SLC4A5;refGene_tx=XM_019827441.2;refGene_gn=SLC4A5;refGene_tx=XM_019827444.2;refGene_gn=SLC4A5;refGene_tx=XM_023251781.1;refGene_gn=SLC4A5;refGene_tx=XM_023251782.1;refGene_gn=SLC4A5;refGene_tx=XM_023251783.1
A3 92987151 92987309
refGene_gn=SLC4A5;refGene_tx=XM_019827439.2;refGene_gn=SLC4A5;refGene_tx=XM_019827441.2;refGene_gn=SLC4A5;refGene_tx=XM_019827444.2;refGene_gn=SLC4A5;refGene_tx=XM_023251781.1;refGene_gn=SLC4A5;refGene_tx=XM_023251782.1;refGene_gn=SLC4A5;refGene_tx=XM_023251783.1
A3 92990538 92990594
refGene_gn=SLC4A5;refGene_tx=XM_019827439.2;refGene_gn=SLC4A5;refGene_tx=XM_019827441.2;refGene_gn=SLC4A5;refGene_tx=XM_019827444.2;refGene_gn=SLC4A5;refGene_tx=XM_023251781.1;refGene_gn=SLC4A5;refGene_tx=XM_023251782.1;refGene_gn=SLC4A5;refGene_tx=XM_023251783.1
A3 92993137 92993233
refGene_gn=SLC4A5;refGene_tx=XM_019827439.2;refGene_gn=SLC4A5;refGene_tx=XM_019827441.2;refGene_gn=SLC4A5;refGene_tx=XM_019827444.2;refGene_gn=SLC4A5;refGene_tx=XM_023251781.1;refGene_gn=SLC4A5;refGene_tx=XM_023251782.1;refGene_gn=SLC4A5;refGene_tx=XM_023251783.1
A3 92994510 92994666
refGene_gn=SLC4A5;refGene_tx=XM_019827439.2;refGene_gn=SLC4A5;refGene_tx=XM_019827441.2;refGene_gn=SLC4A5;refGene_tx=XM_019827444.2;refGene_gn=SLC4A5;refGene_tx=XM_023251781.1;refGene_gn=SLC4A5;refGene_tx=XM_023251782.1;refGene_gn=SLC4A5;refGene_tx=XM_023251783.1
A3 92995398 92995559
refGene_gn=SLC4A5;refGene_tx=XM_019827439.2;refGene_gn=SLC4A5;refGene_tx=XM_019827441.2;refGene_gn=SLC4A5;refGene_tx=XM_019827444.2;refGene_gn=SLC4A5;refGene_tx=XM_023251781.1;refGene_gn=SLC4A5;refGene_tx=XM_023251782.1;refGene_gn=SLC4A5;refGene_tx=XM_023251783.1
A3 93014301 93014431

refGene_gn=SLC4A5;refGene_tx=XM_019827439.2;refGene_gn=SLC4A5;refGene_tx=XM_019827441.2;refGene_gn=SLC4A5;refGene_tx=XM_019827444.2;refGene_gn=SLC4A5;refGene_tx=XM_023251781.1;refGene_gn=SLC4A5;refGene_tx=XM_023251782.1;refGene_gn=SLC4A5;refGene_tx=XM_023251783.1
A3 93027221 93027416
refGene_gn=SLC4A5;refGene_tx=XM_019827439.2;refGene_gn=SLC4A5;refGene_tx=XM_019827441.2;refGene_gn=SLC4A5;refGene_tx=XM_023251781.1;refGene_gn=SLC4A5;refGene_tx=XM_023251782.1;refGene_gn=SLC4A5;refGene_tx=XM_023251783.1
A3 93033814 93033895
refGene_gn=SLC4A5;refGene_tx=XM_019827439.2;refGene_gn=SLC4A5;refGene_tx=XM_019827441.2;refGene_gn=SLC4A5;refGene_tx=XM_023251781.1;refGene_gn=SLC4A5;refGene_tx=XM_023251782.1;refGene_gn=SLC4A5;refGene_tx=XM_023251783.1
A3 93046337 93046377
refGene_gn=SLC4A5;refGene_tx=XM_019827439.2;refGene_gn=SLC4A5;refGene_tx=XM_019827441.2;refGene_gn=SLC4A5;refGene_tx=XM_023251781.1;refGene_gn=SLC4A5;refGene_tx=XM_023251783.1
A3 93075965 93076103
refGene_gn=DCTN1;refGene_tx=XM_006930201.4;refGene_gn=DCTN1;refGene_tx=XM_011281215.2;refGene_gn=DCTN1;refGene_tx=XM_019827431.2;refGene_gn=DCTN1;refGene_tx=XM_019827432.2;refGene_gn=DCTN1;refGene_tx=XM_019827433.2;refGene_gn=DCTN1;refGene_tx=XM_019827434.2;refGene_gn=DCTN1;refGene_tx=XM_019827435.2;refGene_gn=DCTN1;refGene_tx=XM_019827436.2;refGene_gn=DCTN1;refGene_tx=XM_019827437.2;refGene_gn=DCTN1;refGene_tx=XM_019827438.2
A3 93076500 93076590
refGene_gn=DCTN1;refGene_tx=XM_006930201.4;refGene_gn=DCTN1;refGene_tx=XM_011281215.2;refGene_gn=DCTN1;refGene_tx=XM_019827431.2;refGene_gn=DCTN1;refGene_tx=XM_019827432.2;refGene_gn=DCTN1;refGene_tx=XM_019827433.2;refGene_gn=DCTN1;refGene_tx=XM_019827434.2;refGene_gn=DCTN1;refGene_tx=XM_019827435.2;refGene_gn=DCTN1;refGene_tx=XM_019827436.2;refGene_gn=DCTN1;refGene_tx=XM_019827437.2;refGene_gn=DCTN1;refGene_tx=XM_019827438.2
A3 93077107 93077187
refGene_gn=DCTN1;refGene_tx=XM_006930201.4;refGene_gn=DCTN1;refGene_tx=XM_011281215.2;refGene_gn=DCTN1;refGene_tx=XM_019827431.2;refGene_gn=DCTN1;refGene_tx=XM_019827432.2;refGene_gn=DCTN1;refGene_tx=XM_019827433.2;refGene_gn=DCTN1;refGene_tx=XM_019827434.2;refGene_gn=DCTN1;refGene_tx=XM_019827435.2;refGene_gn=DCTN1;refGene_tx=XM_019827436.2;refGene_gn=DCTN1;refGene_tx=XM_019827437.2;refGene_gn=DCTN1;refGene_tx=XM_019827438.2
A3 93077444 93077631
refGene_gn=DCTN1;refGene_tx=XM_006930201.4;refGene_gn=DCTN1;refGene_tx=XM_011281215.2;refGene_gn=DCTN1;refGene_tx=XM_019827431.2;refGene_gn=DCTN1;refGene_tx=XM_019827432.2;refGene_gn=DCTN1;refGene_tx=XM_019827433.2;refGene_gn=DCTN1;refGene_tx=XM_019827434.2;refGene_gn=DCTN1;refGene_tx=XM_019827435.2;refGene_gn=DCTN1;refGene_tx=XM_019827436.2;refGene_gn=DCTN1;refGene_tx=XM_019827437.2;refGene_gn=DCTN1;refGene_tx=XM_019827438.2

A3 93077746 93077880
refGene_gn=DCTN1;refGene_tx=XM_006930201.4;refGene_gn=DCTN1;refGene_tx=XM_011281215.2;refGene_gn=DCTN1;refGene_tx=XM_019827431.2;refGene_gn=DCTN1;refGene_tx=XM_019827432.2;refGene_gn=DCTN1;refGene_tx=XM_019827433.2;refGene_gn=DCTN1;refGene_tx=XM_019827434.2;refGene_gn=DCTN1;refGene_tx=XM_019827435.2;refGene_gn=DCTN1;refGene_tx=XM_019827436.2;refGene_gn=DCTN1;refGene_tx=XM_019827437.2;refGene_gn=DCTN1;refGene_tx=XM_019827438.2
A3 93078089 93078104
refGene_gn=DCTN1;refGene_tx=XM_011281215.2;refGene_gn=DCTN1;refGene_tx=XM_019827431.2;refGene_gn=DCTN1;refGene_tx=XM_019827433.2;refGene_gn=DCTN1;refGene_tx=XM_019827434.2;refGene_gn=DCTN1;refGene_tx=XM_019827435.2;refGene_gn=DCTN1;refGene_tx=XM_019827437.2
A3 93079502 93079669
refGene_gn=DCTN1;refGene_tx=XM_006930201.4;refGene_gn=DCTN1;refGene_tx=XM_011281215.2;refGene_gn=DCTN1;refGene_tx=XM_019827431.2;refGene_gn=DCTN1;refGene_tx=XM_019827432.2;refGene_gn=DCTN1;refGene_tx=XM_019827433.2;refGene_gn=DCTN1;refGene_tx=XM_019827434.2;refGene_gn=DCTN1;refGene_tx=XM_019827435.2;refGene_gn=DCTN1;refGene_tx=XM_019827436.2;refGene_gn=DCTN1;refGene_tx=XM_019827437.2;refGene_gn=DCTN1;refGene_tx=XM_019827438.2
A3 93079957 93080100
refGene_gn=DCTN1;refGene_tx=XM_006930201.4;refGene_gn=DCTN1;refGene_tx=XM_011281215.2;refGene_gn=DCTN1;refGene_tx=XM_019827431.2;refGene_gn=DCTN1;refGene_tx=XM_019827432.2;refGene_gn=DCTN1;refGene_tx=XM_019827433.2;refGene_gn=DCTN1;refGene_tx=XM_019827434.2;refGene_gn=DCTN1;refGene_tx=XM_019827435.2;refGene_gn=DCTN1;refGene_tx=XM_019827436.2;refGene_gn=DCTN1;refGene_tx=XM_019827437.2;refGene_gn=DCTN1;refGene_tx=XM_019827438.2
A3 93080353 93080479
refGene_gn=DCTN1;refGene_tx=XM_006930201.4;refGene_gn=DCTN1;refGene_tx=XM_011281215.2;refGene_gn=DCTN1;refGene_tx=XM_019827431.2;refGene_gn=DCTN1;refGene_tx=XM_019827432.2;refGene_gn=DCTN1;refGene_tx=XM_019827433.2;refGene_gn=DCTN1;refGene_tx=XM_019827434.2;refGene_gn=DCTN1;refGene_tx=XM_019827435.2;refGene_gn=DCTN1;refGene_tx=XM_019827436.2;refGene_gn=DCTN1;refGene_tx=XM_019827437.2;refGene_gn=DCTN1;refGene_tx=XM_019827438.2
A3 93080699 93080831
refGene_gn=DCTN1;refGene_tx=XM_006930201.4;refGene_gn=DCTN1;refGene_tx=XM_011281215.2;refGene_gn=DCTN1;refGene_tx=XM_019827431.2;refGene_gn=DCTN1;refGene_tx=XM_019827432.2;refGene_gn=DCTN1;refGene_tx=XM_019827433.2;refGene_gn=DCTN1;refGene_tx=XM_019827434.2;refGene_gn=DCTN1;refGene_tx=XM_019827435.2;refGene_gn=DCTN1;refGene_tx=XM_019827436.2;refGene_gn=DCTN1;refGene_tx=XM_019827437.2;refGene_gn=DCTN1;refGene_tx=XM_019827438.2
A3 93080914 93081076
refGene_gn=DCTN1;refGene_tx=XM_006930201.4;refGene_gn=DCTN1;refGene_tx=XM_011281215.2;refGene_gn=DCTN1;refGene_tx=XM_019827431.2;refGene_gn=DCTN1;refGene_tx=XM_019827432.2;refGene_gn=DCTN1;refGene_tx=XM_019827433.2;refGene_gn=DCTN1;refGene_tx=XM_019827434.2;refGene_gn=DCTN1;refGe

ne_tx=XM_019827435.2;refGene_gn=DCTN1;refGene_tx=XM_019827436.2;refGene_gn=DCTN1;refGene_tx=XM_019827437.2;refGene_gn=DCTN1;refGene_tx=XM_019827438.2
A3 93081240 93081390
refGene_gn=DCTN1;refGene_tx=XM_006930201.4;refGene_gn=DCTN1;refGene_tx=XM_011281215.2;refGene_gn=DCTN1;refGene_tx=XM_019827431.2;refGene_gn=DCTN1;refGene_tx=XM_019827432.2;refGene_gn=DCTN1;refGene_tx=XM_019827433.2;refGene_gn=DCTN1;refGene_tx=XM_019827434.2;refGene_gn=DCTN1;refGene_tx=XM_019827435.2;refGene_gn=DCTN1;refGene_tx=XM_019827436.2;refGene_gn=DCTN1;refGene_tx=XM_019827437.2;refGene_gn=DCTN1;refGene_tx=XM_019827438.2
A3 93081502 93081565
refGene_gn=DCTN1;refGene_tx=XM_006930201.4;refGene_gn=DCTN1;refGene_tx=XM_011281215.2;refGene_gn=DCTN1;refGene_tx=XM_019827431.2;refGene_gn=DCTN1;refGene_tx=XM_019827432.2;refGene_gn=DCTN1;refGene_tx=XM_019827433.2;refGene_gn=DCTN1;refGene_tx=XM_019827434.2;refGene_gn=DCTN1;refGene_tx=XM_019827435.2;refGene_gn=DCTN1;refGene_tx=XM_019827436.2;refGene_gn=DCTN1;refGene_tx=XM_019827437.2;refGene_gn=DCTN1;refGene_tx=XM_019827438.2
A3 93081831 93081900
refGene_gn=DCTN1;refGene_tx=XM_006930201.4;refGene_gn=DCTN1;refGene_tx=XM_011281215.2;refGene_gn=DCTN1;refGene_tx=XM_019827431.2;refGene_gn=DCTN1;refGene_tx=XM_019827432.2;refGene_gn=DCTN1;refGene_tx=XM_019827433.2;refGene_gn=DCTN1;refGene_tx=XM_019827434.2;refGene_gn=DCTN1;refGene_tx=XM_019827435.2;refGene_gn=DCTN1;refGene_tx=XM_019827436.2;refGene_gn=DCTN1;refGene_tx=XM_019827437.2;refGene_gn=DCTN1;refGene_tx=XM_019827438.2
A3 93082177 93082346
refGene_gn=DCTN1;refGene_tx=XM_006930201.4;refGene_gn=DCTN1;refGene_tx=XM_011281215.2;refGene_gn=DCTN1;refGene_tx=XM_019827431.2;refGene_gn=DCTN1;refGene_tx=XM_019827432.2;refGene_gn=DCTN1;refGene_tx=XM_019827433.2;refGene_gn=DCTN1;refGene_tx=XM_019827434.2;refGene_gn=DCTN1;refGene_tx=XM_019827435.2;refGene_gn=DCTN1;refGene_tx=XM_019827436.2;refGene_gn=DCTN1;refGene_tx=XM_019827437.2;refGene_gn=DCTN1;refGene_tx=XM_019827438.2
A3 93082449 93082610
refGene_gn=DCTN1;refGene_tx=XM_006930201.4;refGene_gn=DCTN1;refGene_tx=XM_011281215.2;refGene_gn=DCTN1;refGene_tx=XM_019827431.2;refGene_gn=DCTN1;refGene_tx=XM_019827432.2;refGene_gn=DCTN1;refGene_tx=XM_019827433.2;refGene_gn=DCTN1;refGene_tx=XM_019827434.2;refGene_gn=DCTN1;refGene_tx=XM_019827435.2;refGene_gn=DCTN1;refGene_tx=XM_019827436.2;refGene_gn=DCTN1;refGene_tx=XM_019827437.2;refGene_gn=DCTN1;refGene_tx=XM_019827438.2
A3 93083206 93083359
refGene_gn=DCTN1;refGene_tx=XM_006930201.4;refGene_gn=DCTN1;refGene_tx=XM_011281215.2;refGene_gn=DCTN1;refGene_tx=XM_019827431.2;refGene_gn=DCTN1;refGene_tx=XM_019827432.2;refGene_gn=DCTN1;refGene_tx=XM_019827433.2;refGene_gn=DCTN1;refGene_tx=XM_019827434.2;refGene_gn=DCTN1;refGene_tx=XM_019827435.2;refGene_gn=DCTN1;refGene_tx=XM_019827436.2;refGene_gn=DCTN1;refGene_tx=XM_019827437.2;refGene_gn=DCTN1;refGene_tx=XM_01

9827438.2
A3	93083588	93083705
refGene_gn=DCTN1;refGene_tx=XM_006930201.4;refGene_gn=DCTN1;refGene_tx=XM_011281215.2;refGene_gn=DCTN1;refGene_tx=XM_019827431.2;refGene_gn=DCTN1;refGene_tx=XM_019827432.2;refGene_gn=DCTN1;refGene_tx=XM_019827433.2;refGene_gn=DCTN1;refGene_tx=XM_019827434.2;refGene_gn=DCTN1;refGene_tx=XM_019827435.2;refGene_gn=DCTN1;refGene_tx=XM_019827436.2;refGene_gn=DCTN1;refGene_tx=XM_019827437.2;refGene_gn=DCTN1;refGene_tx=XM_019827438.2
A3	93083793	93083985
refGene_gn=DCTN1;refGene_tx=XM_006930201.4;refGene_gn=DCTN1;refGene_tx=XM_011281215.2;refGene_gn=DCTN1;refGene_tx=XM_019827431.2;refGene_gn=DCTN1;refGene_tx=XM_019827432.2;refGene_gn=DCTN1;refGene_tx=XM_019827433.2;refGene_gn=DCTN1;refGene_tx=XM_019827434.2;refGene_gn=DCTN1;refGene_tx=XM_019827435.2;refGene_gn=DCTN1;refGene_tx=XM_019827436.2;refGene_gn=DCTN1;refGene_tx=XM_019827437.2;refGene_gn=DCTN1;refGene_tx=XM_019827438.2
A3	93084159	93084264
refGene_gn=DCTN1;refGene_tx=XM_006930201.4;refGene_gn=DCTN1;refGene_tx=XM_011281215.2;refGene_gn=DCTN1;refGene_tx=XM_019827431.2;refGene_gn=DCTN1;refGene_tx=XM_019827432.2;refGene_gn=DCTN1;refGene_tx=XM_019827433.2;refGene_gn=DCTN1;refGene_tx=XM_019827434.2;refGene_gn=DCTN1;refGene_tx=XM_019827435.2;refGene_gn=DCTN1;refGene_tx=XM_019827436.2;refGene_gn=DCTN1;refGene_tx=XM_019827437.2;refGene_gn=DCTN1;refGene_tx=XM_019827438.2
A3	93084373	93084533
refGene_gn=DCTN1;refGene_tx=XM_006930201.4;refGene_gn=DCTN1;refGene_tx=XM_011281215.2;refGene_gn=DCTN1;refGene_tx=XM_019827431.2;refGene_gn=DCTN1;refGene_tx=XM_019827432.2;refGene_gn=DCTN1;refGene_tx=XM_019827433.2;refGene_gn=DCTN1;refGene_tx=XM_019827434.2;refGene_gn=DCTN1;refGene_tx=XM_019827435.2;refGene_gn=DCTN1;refGene_tx=XM_019827436.2;refGene_gn=DCTN1;refGene_tx=XM_019827437.2;refGene_gn=DCTN1;refGene_tx=XM_019827438.2
A3	93084654	93084733
refGene_gn=DCTN1;refGene_tx=XM_006930201.4;refGene_gn=DCTN1;refGene_tx=XM_011281215.2;refGene_gn=DCTN1;refGene_tx=XM_019827431.2;refGene_gn=DCTN1;refGene_tx=XM_019827432.2;refGene_gn=DCTN1;refGene_tx=XM_019827433.2;refGene_gn=DCTN1;refGene_tx=XM_019827434.2;refGene_gn=DCTN1;refGene_tx=XM_019827435.2;refGene_gn=DCTN1;refGene_tx=XM_019827436.2;refGene_gn=DCTN1;refGene_tx=XM_019827437.2;refGene_gn=DCTN1;refGene_tx=XM_019827438.2
A3	93084809	93085014
refGene_gn=DCTN1;refGene_tx=XM_006930201.4;refGene_gn=DCTN1;refGene_tx=XM_011281215.2;refGene_gn=DCTN1;refGene_tx=XM_019827431.2;refGene_gn=DCTN1;refGene_tx=XM_019827432.2;refGene_gn=DCTN1;refGene_tx=XM_019827433.2;refGene_gn=DCTN1;refGene_tx=XM_019827434.2;refGene_gn=DCTN1;refGene_tx=XM_019827435.2;refGene_gn=DCTN1;refGene_tx=XM_019827436.2;refGene_gn=DCTN1;refGene_tx=XM_019827437.2;refGene_gn=DCTN1;refGene_tx=XM_019827438.2
A3	93085163	93085361

refGene_gn=DCTN1;refGene_tx=XM_006930201.4;refGene_gn=DCTN1;refGene_tx=XM_011281215.2;refGene_gn=DCTN1;refGene_tx=XM_019827431.2;refGene_gn=DCTN1;refGene_tx=XM_019827432.2;refGene_gn=DCTN1;refGene_tx=XM_019827433.2;refGene_gn=DCTN1;refGene_tx=XM_019827434.2;refGene_gn=DCTN1;refGene_tx=XM_019827435.2;refGene_gn=DCTN1;refGene_tx=XM_019827436.2;refGene_gn=DCTN1;refGene_tx=XM_019827437.2;refGene_gn=DCTN1;refGene_tx=XM_019827438.2
A3 93085710 93085902
refGene_gn=DCTN1;refGene_tx=XM_006930201.4;refGene_gn=DCTN1;refGene_tx=XM_011281215.2;refGene_gn=DCTN1;refGene_tx=XM_019827431.2;refGene_gn=DCTN1;refGene_tx=XM_019827432.2;refGene_gn=DCTN1;refGene_tx=XM_019827433.2;refGene_gn=DCTN1;refGene_tx=XM_019827434.2;refGene_gn=DCTN1;refGene_tx=XM_019827435.2;refGene_gn=DCTN1;refGene_tx=XM_019827436.2;refGene_gn=DCTN1;refGene_tx=XM_019827437.2;refGene_gn=DCTN1;refGene_tx=XM_019827438.2
A3 93087146 93087167
refGene_gn=DCTN1;refGene_tx=XM_006930201.4;refGene_gn=DCTN1;refGene_tx=XM_011281215.2;refGene_gn=DCTN1;refGene_tx=XM_019827431.2;refGene_gn=DCTN1;refGene_tx=XM_019827432.2;refGene_gn=DCTN1;refGene_tx=XM_019827433.2;refGene_gn=DCTN1;refGene_tx=XM_019827434.2;refGene_gn=DCTN1;refGene_tx=XM_019827435.2;refGene_gn=DCTN1;refGene_tx=XM_019827436.2
A3 93088550 93088580
refGene_gn=DCTN1;refGene_tx=XM_006930201.4;refGene_gn=DCTN1;refGene_tx=XM_011281215.2;refGene_gn=DCTN1;refGene_tx=XM_019827431.2;refGene_gn=DCTN1;refGene_tx=XM_019827432.2;refGene_gn=DCTN1;refGene_tx=XM_019827433.2;refGene_gn=DCTN1;refGene_tx=XM_019827434.2;refGene_gn=DCTN1;refGene_tx=XM_019827436.2
A3 93090908 93090929
refGene_gn=DCTN1;refGene_tx=XM_011281215.2;refGene_gn=DCTN1;refGene_tx=XM_019827431.2;refGene_gn=DCTN1;refGene_tx=XM_019827432.2;refGene_gn=DCTN1;refGene_tx=XM_019827434.2;refGene_gn=DCTN1;refGene_tx=XM_019827436.2
A3 93091631 93091666
refGene_gn=DCTN1;refGene_tx=XM_011281215.2;refGene_gn=DCTN1;refGene_tx=XM_019827431.2;refGene_gn=DCTN1;refGene_tx=XM_019827432.2;refGene_gn=DCTN1;refGene_tx=XM_019827433.2;refGene_gn=DCTN1;refGene_tx=XM_019827434.2;refGene_gn=DCTN1;refGene_tx=XM_019827435.2;refGene_gn=DCTN1;refGene_tx=XM_019827436.2;refGene_gn=DCTN1;refGene_tx=XM_019827437.2;refGene_gn=DCTN1;refGene_tx=XM_019827438.2
A3 93091866 93091945
refGene_gn=DCTN1;refGene_tx=XM_011281215.2;refGene_gn=DCTN1;refGene_tx=XM_019827431.2;refGene_gn=DCTN1;refGene_tx=XM_019827432.2;refGene_gn=DCTN1;refGene_tx=XM_019827433.2;refGene_gn=DCTN1;refGene_tx=XM_019827434.2;refGene_gn=DCTN1;refGene_tx=XM_019827435.2;refGene_gn=DCTN1;refGene_tx=XM_019827436.2;refGene_gn=DCTN1;refGene_tx=XM_019827437.2;refGene_gn=DCTN1;refGene_tx=XM_019827438.2
A3 93092200 93092446
refGene_gn=DCTN1;refGene_tx=XM_011281215.2;refGene_gn=DCTN1;refGene_tx=XM_019827431.2;refGene_gn=DCTN1;refGene_tx=XM_019827432.2;refGene_gn=DCTN1;refGene_tx=XM_019827433.2;refGene_gn=DCTN1;refGene_tx=XM_0198274

34.2;refGene_gn=DCTN1;refGene_tx=XM_019827435.2;refGene_gn=DCTN1;refGene_tx=XM_019827436.2;refGene_gn=DCTN1;refGene_tx=XM_019827437.2;refGene_gn=DCTN1;refGene_tx=XM_019827438.2
A3 93094125 93094158
refGene_gn=DCTN1;refGene_tx=XM_019827431.2;refGene_gn=DCTN1;refGene_tx=XM_019827432.2;refGene_gn=DCTN1;refGene_tx=XM_019827433.2;refGene_gn=DCTN1;refGene_tx=XM_019827434.2;refGene_gn=DCTN1;refGene_tx=XM_019827435.2;refGene_gn=DCTN1;refGene_tx=XM_019827436.2;refGene_gn=DCTN1;refGene_tx=XM_019827437.2;refGene_gn=DCTN1;refGene_tx=XM_019827438.2
A3 93113623 93113812
refGene_gn=LOC109498222;refGene_tx=XM_019827451.2;refGene_gn=LOC109498222;refGene_tx=XM_019827452.2
A3 93113931 93114038
refGene_gn=LOC109498222;refGene_tx=XM_019827451.2;refGene_gn=LOC109498222;refGene_tx=XM_019827452.2
A3 93114506 93114695
refGene_gn=LOC109498222;refGene_tx=XM_019827451.2;refGene_gn=LOC109498222;refGene_tx=XM_019827452.2
A3 93117441 93117531
refGene_gn=LOC109498222;refGene_tx=XM_019827451.2
A3 93117794 93118034
refGene_gn=LOC109498222;refGene_tx=XM_019827451.2;refGene_gn=LOC109498222;refGene_tx=XM_019827452.2
A3 93118173 93118351
refGene_gn=LOC109498222;refGene_tx=XM_019827451.2;refGene_gn=LOC109498222;refGene_tx=XM_019827452.2
A3 93119066 93120395
refGene_gn=CA3H2orf81;refGene_tx=XM_019827453.2;refGene_gn=CA3H2orf81;refGene_tx=XM_023251784.1
A3 93120573 93120795
refGene_gn=CA3H2orf81;refGene_tx=XM_019827453.2;refGene_gn=CA3H2orf81;refGene_tx=XM_023251784.1
A3 93120875 93121097
refGene_gn=CA3H2orf81;refGene_tx=XM_019827453.2;refGene_gn=CA3H2orf81;refGene_tx=XM_023251784.1
A3 93123743 93123761
refGene_gn=CA3H2orf81;refGene_tx=XM_019827453.2;refGene_gn=CA3H2orf81;refGene_tx=XM_023251784.1
A3 93123983 93124027 refGene_gn=WDR54;refGene_tx=XM_006930204.3
A3 93124407 93124630
refGene_gn=WDR54;refGene_tx=XM_003984140.4;refGene_gn=WDR54;refGene_tx=XM_006930204.3;refGene_gn=WDR54;refGene_tx=XM_006930205.3;refGene_gn=WDR54;refGene_tx=XM_006930206.3
A3 93124782 93124845
refGene_gn=WDR54;refGene_tx=XM_003984140.4;refGene_gn=WDR54;refGene_tx=XM_006930204.3;refGene_gn=WDR54;refGene_tx=XM_006930205.3;refGene_gn=WDR54;refGene_tx=XM_006930206.3
A3 93125238 93125305
refGene_gn=WDR54;refGene_tx=XM_003984140.4;refGene_gn=WDR54;refGene_tx=XM_006930204.3;refGene_gn=WDR54;refGene_tx=XM_006930205.3;refGene_gn=

WDR54;refGene_tx=XM_006930206.3
A3 93125395 93125449
refGene_gn=WDR54;refGene_tx=XM_003984140.4;refGene_gn=WDR54;refGene_tx
=XM_006930204.3;refGene_gn=WDR54;refGene_tx=XM_006930205.3;refGene_gn=
WDR54;refGene_tx=XM_006930206.3
A3 93125776 93125904
refGene_gn=WDR54;refGene_tx=XM_003984140.4;refGene_gn=WDR54;refGene_tx
=XM_006930204.3;refGene_gn=WDR54;refGene_tx=XM_006930205.3;refGene_gn=
WDR54;refGene_tx=XM_006930206.3
A3 93126794 93126895
refGene_gn=WDR54;refGene_tx=XM_003984140.4;refGene_gn=WDR54;refGene_tx
=XM_006930204.3;refGene_gn=WDR54;refGene_tx=XM_006930205.3;refGene_gn=
WDR54;refGene_tx=XM_006930206.3
A3 93126969 93127132
refGene_gn=WDR54;refGene_tx=XM_003984140.4;refGene_gn=WDR54;refGene_tx
=XM_006930204.3;refGene_gn=WDR54;refGene_tx=XM_006930205.3;refGene_gn=
WDR54;refGene_tx=XM_006930206.3
A3 93127317 93127392
refGene_gn=WDR54;refGene_tx=XM_003984140.4;refGene_gn=WDR54;refGene_tx
=XM_006930204.3;refGene_gn=WDR54;refGene_tx=XM_006930205.3;refGene_gn=
WDR54;refGene_tx=XM_006930206.3
A3 93127469 93127601
refGene_gn=WDR54;refGene_tx=XM_003984140.4;refGene_gn=WDR54;refGene_tx
=XM_006930204.3;refGene_gn=WDR54;refGene_tx=XM_006930205.3;refGene_gn=
WDR54;refGene_tx=XM_006930206.3
A3 93127963 93128483
refGene_gn=RTKN;refGene_tx=XM_003984142.5;refGene_gn=RTKN;refGene_tx=X
M_006930211.4;refGene_gn=RTKN;refGene_tx=XM_006930213.4;refGene_gn=RTK
N;refGene_tx=XM_011281212.3;refGene_gn=RTKN;refGene_tx=XM_019827445.2;
refGene_gn=RTKN;refGene_tx=XM_019827446.2;refGene_gn=RTKN;refGene_tx=X
M_019827447.2;refGene_gn=RTKN;refGene_tx=XM_019827448.2;refGene_gn=RTK
N;refGene_tx=XM_019827449.2
A3 93128557 93128670 refGene_gn=RTKN;refGene_tx=XM_019827450.2
A3 93129067 93129172
refGene_gn=RTKN;refGene_tx=XM_003984142.5;refGene_gn=RTKN;refGene_tx=X
M_006930211.4;refGene_gn=RTKN;refGene_tx=XM_006930213.4;refGene_gn=RTK
N;refGene_tx=XM_011281212.3;refGene_gn=RTKN;refGene_tx=XM_019827445.2;
refGene_gn=RTKN;refGene_tx=XM_019827446.2;refGene_gn=RTKN;refGene_tx=X
M_019827447.2;refGene_gn=RTKN;refGene_tx=XM_019827448.2;refGene_gn=RTK
N;refGene_tx=XM_019827449.2;refGene_gn=RTKN;refGene_tx=XM_019827450.2
A3 93129335 93129504
refGene_gn=RTKN;refGene_tx=XM_003984142.5;refGene_gn=RTKN;refGene_tx=X
M_006930211.4;refGene_gn=RTKN;refGene_tx=XM_006930213.4;refGene_gn=RTK
N;refGene_tx=XM_011281212.3;refGene_gn=RTKN;refGene_tx=XM_019827445.2;
refGene_gn=RTKN;refGene_tx=XM_019827446.2;refGene_gn=RTKN;refGene_tx=X
M_019827447.2;refGene_gn=RTKN;refGene_tx=XM_019827448.2;refGene_gn=RTK
N;refGene_tx=XM_019827449.2;refGene_gn=RTKN;refGene_tx=XM_019827450.2
A3 93130175 93130304
refGene_gn=RTKN;refGene_tx=XM_003984142.5;refGene_gn=RTKN;refGene_tx=X
M_006930211.4;refGene_gn=RTKN;refGene_tx=XM_006930213.4;refGene_gn=RTK

N;refGene_tx=XM_011281212.3;refGene_gn=RTKN;refGene_tx=XM_019827445.2;
refGene_gn=RTKN;refGene_tx=XM_019827446.2;refGene_gn=RTKN;refGene_tx=X
M_019827447.2;refGene_gn=RTKN;refGene_tx=XM_019827448.2;refGene_gn=RTK
N;refGene_tx=XM_019827449.2;refGene_gn=RTKN;refGene_tx=XM_019827450.2
A3 93130537 93130644
refGene_gn=RTKN;refGene_tx=XM_003984142.5;refGene_gn=RTKN;refGene_tx=X
M_006930211.4;refGene_gn=RTKN;refGene_tx=XM_006930213.4;refGene_gn=RTK
N;refGene_tx=XM_011281212.3;refGene_gn=RTKN;refGene_tx=XM_019827445.2;
refGene_gn=RTKN;refGene_tx=XM_019827446.2;refGene_gn=RTKN;refGene_tx=X
M_019827447.2;refGene_gn=RTKN;refGene_tx=XM_019827448.2;refGene_gn=RTK
N;refGene_tx=XM_019827449.2;refGene_gn=RTKN;refGene_tx=XM_019827450.2
A3 93130750 93130845
refGene_gn=RTKN;refGene_tx=XM_003984142.5;refGene_gn=RTKN;refGene_tx=X
M_006930211.4;refGene_gn=RTKN;refGene_tx=XM_006930213.4;refGene_gn=RTK
N;refGene_tx=XM_011281212.3;refGene_gn=RTKN;refGene_tx=XM_019827445.2;
refGene_gn=RTKN;refGene_tx=XM_019827446.2;refGene_gn=RTKN;refGene_tx=X
M_019827447.2;refGene_gn=RTKN;refGene_tx=XM_019827448.2;refGene_gn=RTK
N;refGene_tx=XM_019827449.2;refGene_gn=RTKN;refGene_tx=XM_019827450.2
A3 93131595 93131805
refGene_gn=RTKN;refGene_tx=XM_003984142.5;refGene_gn=RTKN;refGene_tx=X
M_006930211.4;refGene_gn=RTKN;refGene_tx=XM_006930213.4;refGene_gn=RTK
N;refGene_tx=XM_011281212.3;refGene_gn=RTKN;refGene_tx=XM_019827445.2;
refGene_gn=RTKN;refGene_tx=XM_019827446.2;refGene_gn=RTKN;refGene_tx=X
M_019827447.2;refGene_gn=RTKN;refGene_tx=XM_019827448.2;refGene_gn=RTK
N;refGene_tx=XM_019827449.2;refGene_gn=RTKN;refGene_tx=XM_019827450.2
A3 93132025 93132143
refGene_gn=RTKN;refGene_tx=XM_003984142.5;refGene_gn=RTKN;refGene_tx=X
M_006930211.4;refGene_gn=RTKN;refGene_tx=XM_006930213.4;refGene_gn=RTK
N;refGene_tx=XM_011281212.3;refGene_gn=RTKN;refGene_tx=XM_019827445.2;
refGene_gn=RTKN;refGene_tx=XM_019827446.2;refGene_gn=RTKN;refGene_tx=X
M_019827447.2;refGene_gn=RTKN;refGene_tx=XM_019827448.2;refGene_gn=RTK
N;refGene_tx=XM_019827449.2;refGene_gn=RTKN;refGene_tx=XM_019827450.2
A3 93132238 93132292
refGene_gn=RTKN;refGene_tx=XM_003984142.5;refGene_gn=RTKN;refGene_tx=X
M_006930211.4;refGene_gn=RTKN;refGene_tx=XM_006930213.4;refGene_gn=RTK
N;refGene_tx=XM_011281212.3;refGene_gn=RTKN;refGene_tx=XM_019827445.2;
refGene_gn=RTKN;refGene_tx=XM_019827446.2;refGene_gn=RTKN;refGene_tx=X
M_019827447.2;refGene_gn=RTKN;refGene_tx=XM_019827448.2;refGene_gn=RTK
N;refGene_tx=XM_019827449.2;refGene_gn=RTKN;refGene_tx=XM_019827450.2
A3 93132389 93132451
refGene_gn=RTKN;refGene_tx=XM_003984142.5;refGene_gn=RTKN;refGene_tx=X
M_006930211.4;refGene_gn=RTKN;refGene_tx=XM_006930213.4;refGene_gn=RTK
N;refGene_tx=XM_011281212.3;refGene_gn=RTKN;refGene_tx=XM_019827445.2;
refGene_gn=RTKN;refGene_tx=XM_019827446.2;refGene_gn=RTKN;refGene_tx=X
M_019827447.2;refGene_gn=RTKN;refGene_tx=XM_019827448.2;refGene_gn=RTK
N;refGene_tx=XM_019827449.2;refGene_gn=RTKN;refGene_tx=XM_019827450.2
A3 93134178 93134378
refGene_gn=RTKN;refGene_tx=XM_003984142.5;refGene_gn=RTKN;refGene_tx=X
M_006930211.4;refGene_gn=RTKN;refGene_tx=XM_006930213.4;refGene_gn=RTK
N;refGene_tx=XM_011281212.3;refGene_gn=RTKN;refGene_tx=XM_019827445.2;

refGene_gn=RTKN;refGene_tx=XM_019827446.2;refGene_gn=RTKN;refGene_tx=XM_019827447.2;refGene_gn=RTKN;refGene_tx=XM_019827448.2;refGene_gn=RTKN;refGene_tx=XM_019827449.2;refGene_gn=RTKN;refGene_tx=XM_019827450.2
A3 93136843 93136888 refGene_gn=RTKN;refGene_tx=XM_019827448.2
A3 93136995 93137019 refGene_gn=RTKN;refGene_tx=XM_019827449.2
A3 93140562 93140634
refGene_gn=RTKN;refGene_tx=XM_003984142.5;refGene_gn=RTKN;refGene_tx=XM_006930211.4;refGene_gn=RTKN;refGene_tx=XM_011281212.3;refGene_gn=RTKN;refGene_tx=XM_019827445.2;refGene_gn=RTKN;refGene_tx=XM_019827447.2;refGene_gn=RTKN;refGene_tx=XM_019827450.2
A3 93142649 93142760 refGene_gn=RTKN;refGene_tx=XM_019827446.2
A3 93151746 93151804 refGene_gn=INO80B;refGene_tx=XM_011281213.2
A3 93152036 93152229 refGene_gn=INO80B;refGene_tx=XM_011281213.2
A3 93152445 93152564 refGene_gn=INO80B;refGene_tx=XM_011281213.2
A3 93152734 93152904 refGene_gn=INO80B;refGene_tx=XM_011281213.2
A3 93153899 93154430 refGene_gn=INO80B;refGene_tx=XM_011281213.2
A3 93155174 93155243
refGene_gn=WBP1;refGene_tx=XM_003984144.4;refGene_gn=WBP1;refGene_tx=XM_019827454.2
A3 93155570 93155672 refGene_gn=WBP1;refGene_tx=XM_019827454.2
A3 93156210 93156313
refGene_gn=WBP1;refGene_tx=XM_003984144.4;refGene_gn=WBP1;refGene_tx=XM_019827454.2
A3 93156437 93156614
refGene_gn=WBP1;refGene_tx=XM_003984144.4;refGene_gn=WBP1;refGene_tx=XM_019827454.2
A3 93156787 93157242
refGene_gn=WBP1;refGene_tx=XM_003984144.4;refGene_gn=WBP1;refGene_tx=XM_019827454.2
A3 93157836 93159580 refGene_gn=MOGS;refGene_tx=XM_003984145.4
A3 93159819 93160016 refGene_gn=MOGS;refGene_tx=XM_003984145.4
A3 93160483 93160710 refGene_gn=MOGS;refGene_tx=XM_003984145.4
A3 93160902 93161245 refGene_gn=MOGS;refGene_tx=XM_003984145.4
A3 93164780 93164914 refGene_gn=MRPL53;refGene_tx=XM_003984146.5
A3 93165021 93165134 refGene_gn=MRPL53;refGene_tx=XM_003984146.5
A3 93165236 93165328 refGene_gn=MRPL53;refGene_tx=XM_003984146.5
A3 93166572 93166829
refGene_gn=CCDC142;refGene_tx=XM_006930214.4;refGene_gn=CCDC142;refGene_tx=XM_019827455.2;refGene_gn=CCDC142;refGene_tx=XM_019827456.2;refGene_gn=CCDC142;refGene_tx=XM_019827457.2
A3 93166985 93167185
refGene_gn=CCDC142;refGene_tx=XM_006930214.4;refGene_gn=CCDC142;refGene_tx=XM_019827455.2;refGene_gn=CCDC142;refGene_tx=XM_019827456.2;refGene_gn=CCDC142;refGene_tx=XM_019827457.2
A3 93167310 93167488
refGene_gn=CCDC142;refGene_tx=XM_006930214.4;refGene_gn=CCDC142;refGene_tx=XM_019827455.2;refGene_gn=CCDC142;refGene_tx=XM_019827456.2;refGene_gn=CCDC142;refGene_tx=XM_019827457.2
A3 93167698 93167813
refGene_gn=CCDC142;refGene_tx=XM_006930214.4;refGene_gn=CCDC142;refGen

e_tx=XM_019827455.2;refGene_gn=CCDC142;refGene_tx=XM_019827456.2;refGene_gn=CCDC142;refGene_tx=XM_019827457.2
A3 93171361 93171475
refGene_gn=CCDC142;refGene_tx=XM_006930214.4;refGene_gn=CCDC142;refGene_tx=XM_019827455.2;refGene_gn=CCDC142;refGene_tx=XM_019827456.2;refGene_gn=CCDC142;refGene_tx=XM_019827457.2
A3 93171547 93171678
refGene_gn=CCDC142;refGene_tx=XM_006930214.4;refGene_gn=CCDC142;refGene_tx=XM_019827455.2;refGene_gn=CCDC142;refGene_tx=XM_019827456.2;refGene_gn=CCDC142;refGene_tx=XM_019827457.2
A3 93171770 93171962
refGene_gn=CCDC142;refGene_tx=XM_006930214.4;refGene_gn=CCDC142;refGene_tx=XM_019827455.2;refGene_gn=CCDC142;refGene_tx=XM_019827456.2;refGene_gn=CCDC142;refGene_tx=XM_019827457.2
A3 93172039 93172126
refGene_gn=CCDC142;refGene_tx=XM_006930214.4;refGene_gn=CCDC142;refGene_tx=XM_019827455.2;refGene_gn=CCDC142;refGene_tx=XM_019827457.2
A3 93172422 93173497
refGene_gn=CCDC142;refGene_tx=XM_006930214.4;refGene_gn=CCDC142;refGene_tx=XM_019827455.2;refGene_gn=CCDC142;refGene_tx=XM_019827456.2;refGene_gn=CCDC142;refGene_tx=XM_019827457.2
A3 93173757 93173797
refGene_gn=TTC31;refGene_tx=XM_023251789.1;refGene_gn=TTC31;refGene_tx=XM_023251790.1;refGene_gn=TTC31;refGene_tx=XM_023251791.1
A3 93173954 93174043
refGene_gn=TTC31;refGene_tx=XM_023251789.1;refGene_gn=TTC31;refGene_tx=XM_023251790.1
A3 93177002 93177105
refGene_gn=TTC31;refGene_tx=XM_023251789.1;refGene_gn=TTC31;refGene_tx=XM_023251790.1
A3 93177225 93177464
refGene_gn=TTC31;refGene_tx=XM_023251789.1;refGene_gn=TTC31;refGene_tx=XM_023251790.1;refGene_gn=TTC31;refGene_tx=XM_023251791.1
A3 93177636 93177720
refGene_gn=TTC31;refGene_tx=XM_023251789.1;refGene_gn=TTC31;refGene_tx=XM_023251790.1;refGene_gn=TTC31;refGene_tx=XM_023251791.1
A3 93178478 93178535
refGene_gn=TTC31;refGene_tx=XM_023251789.1;refGene_gn=TTC31;refGene_tx=XM_023251790.1;refGene_gn=TTC31;refGene_tx=XM_023251791.1
A3 93178635 93178716
refGene_gn=TTC31;refGene_tx=XM_023251789.1;refGene_gn=TTC31;refGene_tx=XM_023251790.1;refGene_gn=TTC31;refGene_tx=XM_023251791.1
A3 93178829 93179024
refGene_gn=TTC31;refGene_tx=XM_023251789.1;refGene_gn=TTC31;refGene_tx=XM_023251790.1;refGene_gn=TTC31;refGene_tx=XM_023251791.1
A3 93179270 93179322
refGene_gn=TTC31;refGene_tx=XM_023251789.1;refGene_gn=TTC31;refGene_tx=XM_023251790.1;refGene_gn=TTC31;refGene_tx=XM_023251791.1
A3 93179411 93179502
refGene_gn=TTC31;refGene_tx=XM_023251789.1;refGene_gn=TTC31;refGene_tx

=XM_023251790.1;refGene_gn=TTC31;refGene_tx=XM_023251791.1
A3 93179590 93179732
refGene_gn=TTC31;refGene_tx=XM_023251789.1;refGene_gn=TTC31;refGene_tx=XM_023251790.1;refGene_gn=TTC31;refGene_tx=XM_023251791.1
A3 93179947 93180049
refGene_gn=TTC31;refGene_tx=XM_023251789.1;refGene_gn=TTC31;refGene_tx=XM_023251790.1;refGene_gn=TTC31;refGene_tx=XM_023251791.1
A3 93180315 93180609
refGene_gn=TTC31;refGene_tx=XM_023251789.1;refGene_gn=TTC31;refGene_tx=XM_023251790.1;refGene_gn=TTC31;refGene_tx=XM_023251791.1
A3 93186393 93186770 refGene_gn=LBX2;refGene_tx=XM_003984148.4
A3 93188031 93188530 refGene_gn=LBX2;refGene_tx=XM_003984148.4
A3 93194435 93194483 refGene_gn=PCGF1;refGene_tx=XM_003984149.4
A3 93194586 93194667 refGene_gn=PCGF1;refGene_tx=XM_003984149.4
A3 93194802 93194889 refGene_gn=PCGF1;refGene_tx=XM_003984149.4
A3 93194986 93195020 refGene_gn=PCGF1;refGene_tx=XM_003984149.4
A3 93195203 93195309 refGene_gn=PCGF1;refGene_tx=XM_003984149.4
A3 93195432 93195504 refGene_gn=PCGF1;refGene_tx=XM_003984149.4
A3 93195710 93195863 refGene_gn=PCGF1;refGene_tx=XM_003984149.4
A3 93196019 93196125 refGene_gn=PCGF1;refGene_tx=XM_003984149.4
A3 93196552 93196645 refGene_gn=PCGF1;refGene_tx=XM_003984149.4
A3 93202836 93203236 refGene_gn=TLX2;refGene_tx=XM_011281205.3
A3 93203630 93203868 refGene_gn=TLX2;refGene_tx=XM_011281205.3
A3 93204013 93204230 refGene_gn=TLX2;refGene_tx=XM_011281205.3
A3 93206232 93206389
refGene_gn=DQX1;refGene_tx=XM_006930219.4;refGene_gn=DQX1;refGene_tx=XM_019827468.2
A3 93206634 93206825
refGene_gn=DQX1;refGene_tx=XM_006930219.4;refGene_gn=DQX1;refGene_tx=XM_019827468.2;refGene_gn=DQX1;refGene_tx=XM_019827470.2;refGene_gn=DQX1;refGene_tx=XM_019827471.2
A3 93207149 93207340
refGene_gn=DQX1;refGene_tx=XM_006930219.4;refGene_gn=DQX1;refGene_tx=XM_019827468.2;refGene_gn=DQX1;refGene_tx=XM_019827470.2
A3 93207512 93207632
refGene_gn=DQX1;refGene_tx=XM_006930219.4;refGene_gn=DQX1;refGene_tx=XM_019827468.2;refGene_gn=DQX1;refGene_tx=XM_019827470.2;refGene_gn=DQX1;refGene_tx=XM_019827471.2
A3 93209213 93209405
refGene_gn=DQX1;refGene_tx=XM_006930219.4;refGene_gn=DQX1;refGene_tx=XM_019827468.2;refGene_gn=DQX1;refGene_tx=XM_019827470.2;refGene_gn=DQX1;refGene_tx=XM_019827471.2
A3 93209507 93209868
refGene_gn=DQX1;refGene_tx=XM_006930219.4;refGene_gn=DQX1;refGene_tx=XM_019827468.2;refGene_gn=DQX1;refGene_tx=XM_019827470.2;refGene_gn=DQX1;refGene_tx=XM_019827471.2
A3 93209958 93210192
refGene_gn=DQX1;refGene_tx=XM_006930219.4;refGene_gn=DQX1;refGene_tx=XM_019827468.2;refGene_gn=DQX1;refGene_tx=XM_019827470.2;refGene_gn=DQX1;refGene_tx=XM_019827471.2

A3 93210526 93210911
refGene_gn=DQX1;refGene_tx=XM_006930219.4;refGene_gn=DQX1;refGene_tx=XM_019827468.2;refGene_gn=DQX1;refGene_tx=XM_019827470.2;refGene_gn=DQX1;refGene_tx=XM_019827471.2
A3 93211296 93211490
refGene_gn=DQX1;refGene_tx=XM_006930219.4;refGene_gn=DQX1;refGene_tx=XM_019827468.2;refGene_gn=DQX1;refGene_tx=XM_019827470.2;refGene_gn=DQX1;refGene_tx=XM_019827471.2
A3 93211736 93211973
refGene_gn=DQX1;refGene_tx=XM_006930219.4;refGene_gn=DQX1;refGene_tx=XM_019827468.2;refGene_gn=DQX1;refGene_tx=XM_019827470.2;refGene_gn=DQX1;refGene_tx=XM_019827471.2
A3 93213017 93213054 refGene_gn=AUP1;refGene_tx=XM_003984151.4
A3 93213163 93213282 refGene_gn=AUP1;refGene_tx=XM_003984151.4
A3 93213467 93213583 refGene_gn=AUP1;refGene_tx=XM_003984151.4
A3 93213695 93213815 refGene_gn=AUP1;refGene_tx=XM_003984151.4
A3 93213946 93214049 refGene_gn=AUP1;refGene_tx=XM_003984151.4
A3 93214150 93214217 refGene_gn=AUP1;refGene_tx=XM_003984151.4
A3 93214470 93214544 refGene_gn=AUP1;refGene_tx=XM_003984151.4
A3 93214639 93214712 refGene_gn=AUP1;refGene_tx=XM_003984151.4
A3 93214949 93215134 refGene_gn=AUP1;refGene_tx=XM_003984151.4
A3 93215332 93215483 refGene_gn=AUP1;refGene_tx=XM_003984151.4
A3 93215562 93215700 refGene_gn=AUP1;refGene_tx=XM_003984151.4
A3 93215781 93215876 refGene_gn=AUP1;refGene_tx=XM_003984151.4
A3 93215878 93215917 refGene_gn=AUP1;refGene_tx=XM_003984151.4
A3 93216204 93216710
refGene_gn=HTRA2;refGene_tx=XM_003984153.4;refGene_gn=HTRA2;refGene_tx=XM_019827476.1
A3 93216810 93217015
refGene_gn=HTRA2;refGene_tx=XM_003984153.4;refGene_gn=HTRA2;refGene_tx=XM_019827476.1
A3 93217104 93217299
refGene_gn=HTRA2;refGene_tx=XM_003984153.4;refGene_gn=HTRA2;refGene_tx=XM_019827476.1
A3 93217539 93217572
refGene_gn=HTRA2;refGene_tx=XM_003984153.4;refGene_gn=HTRA2;refGene_tx=XM_019827476.1
A3 93217806 93217912
refGene_gn=HTRA2;refGene_tx=XM_003984153.4;refGene_gn=HTRA2;refGene_tx=XM_019827476.1
A3 93218090 93218160
refGene_gn=HTRA2;refGene_tx=XM_003984153.4;refGene_gn=HTRA2;refGene_tx=XM_019827476.1
A3 93218263 93218267 refGene_gn=HTRA2;refGene_tx=XM_019827476.1
A3 93218846 93218942 refGene_gn=HTRA2;refGene_tx=XM_003984153.4
A3 93219049 93219215 refGene_gn=HTRA2;refGene_tx=XM_003984153.4
A3 93219843 93219917
refGene_gn=LOXL3;refGene_tx=XM_006930220.3;refGene_gn=LOXL3;refGene_tx=XM_019827460.2;refGene_gn=LOXL3;refGene_tx=XM_019827461.2;refGene_gn=LOXL3;refGene_tx=XM_019827462.2;refGene_gn=LOXL3;refGene_tx=XM_0198274

64.2;refGene_gn=LOXL3;refGene_tx=XM_019827465.1;refGene_gn=LOXL3;refGene_tx=XM_019827466.1
A3	93220101	93220213
refGene_gn=LOXL3;refGene_tx=XM_006930220.3;refGene_gn=LOXL3;refGene_tx=XM_019827460.2;refGene_gn=LOXL3;refGene_tx=XM_019827461.2;refGene_gn=LOXL3;refGene_tx=XM_019827462.2;refGene_gn=LOXL3;refGene_tx=XM_019827464.2;refGene_gn=LOXL3;refGene_tx=XM_019827465.1;refGene_gn=LOXL3;refGene_tx=XM_019827466.1
A3	93220304	93220441
refGene_gn=LOXL3;refGene_tx=XM_006930220.3;refGene_gn=LOXL3;refGene_tx=XM_019827460.2;refGene_gn=LOXL3;refGene_tx=XM_019827461.2;refGene_gn=LOXL3;refGene_tx=XM_019827462.2;refGene_gn=LOXL3;refGene_tx=XM_019827464.2;refGene_gn=LOXL3;refGene_tx=XM_019827465.1;refGene_gn=LOXL3;refGene_tx=XM_019827466.1
A3	93220520	93220636
refGene_gn=LOXL3;refGene_tx=XM_006930220.3;refGene_gn=LOXL3;refGene_tx=XM_019827460.2;refGene_gn=LOXL3;refGene_tx=XM_019827461.2;refGene_gn=LOXL3;refGene_tx=XM_019827462.2;refGene_gn=LOXL3;refGene_tx=XM_019827464.2;refGene_gn=LOXL3;refGene_tx=XM_019827465.1;refGene_gn=LOXL3;refGene_tx=XM_019827466.1
A3	93220728	93220972
refGene_gn=LOXL3;refGene_tx=XM_006930220.3;refGene_gn=LOXL3;refGene_tx=XM_019827460.2;refGene_gn=LOXL3;refGene_tx=XM_019827461.2;refGene_gn=LOXL3;refGene_tx=XM_019827462.2;refGene_gn=LOXL3;refGene_tx=XM_019827464.2;refGene_gn=LOXL3;refGene_tx=XM_019827465.1;refGene_gn=LOXL3;refGene_tx=XM_019827466.1
A3	93221168	93221339
refGene_gn=LOXL3;refGene_tx=XM_006930220.3;refGene_gn=LOXL3;refGene_tx=XM_019827460.2;refGene_gn=LOXL3;refGene_tx=XM_019827461.2;refGene_gn=LOXL3;refGene_tx=XM_019827462.2;refGene_gn=LOXL3;refGene_tx=XM_019827463.2;refGene_gn=LOXL3;refGene_tx=XM_019827464.2;refGene_gn=LOXL3;refGene_tx=XM_019827465.1
A3	93221476	93221644
refGene_gn=LOXL3;refGene_tx=XM_006930220.3;refGene_gn=LOXL3;refGene_tx=XM_019827460.2;refGene_gn=LOXL3;refGene_tx=XM_019827461.2;refGene_gn=LOXL3;refGene_tx=XM_019827462.2;refGene_gn=LOXL3;refGene_tx=XM_019827463.2;refGene_gn=LOXL3;refGene_tx=XM_019827464.2;refGene_gn=LOXL3;refGene_tx=XM_019827465.1
A3	93221884	93222039
refGene_gn=LOXL3;refGene_tx=XM_006930220.3;refGene_gn=LOXL3;refGene_tx=XM_019827460.2;refGene_gn=LOXL3;refGene_tx=XM_019827461.2;refGene_gn=LOXL3;refGene_tx=XM_019827462.2;refGene_gn=LOXL3;refGene_tx=XM_019827463.2;refGene_gn=LOXL3;refGene_tx=XM_019827464.2;refGene_gn=LOXL3;refGene_tx=XM_019827465.1
A3	93222183	93222364
refGene_gn=LOXL3;refGene_tx=XM_006930220.3;refGene_gn=LOXL3;refGene_tx=XM_019827460.2;refGene_gn=LOXL3;refGene_tx=XM_019827461.2;refGene_gn=LOXL3;refGene_tx=XM_019827462.2;refGene_gn=LOXL3;refGene_tx=XM_019827463.2;refGene_gn=LOXL3;refGene_tx=XM_019827465.1
A3	93222606	93222826

refGene_gn=LOXL3;refGene_tx=XM_006930220.3;refGene_gn=LOXL3;refGene_tx=XM_019827460.2;refGene_gn=LOXL3;refGene_tx=XM_019827461.2;refGene_gn=LOXL3;refGene_tx=XM_019827462.2;refGene_gn=LOXL3;refGene_tx=XM_019827463.2
A3 93224993 93225053 refGene_gn=LOXL3;refGene_tx=XM_006930220.3;refGene_gn=LOXL3;refGene_tx=XM_019827460.2;refGene_gn=LOXL3;refGene_tx=XM_019827461.2;refGene_gn=LOXL3;refGene_tx=XM_019827463.2
A3 93225695 93225752 refGene_gn=LOXL3;refGene_tx=XM_006930220.3;refGene_gn=LOXL3;refGene_tx=XM_019827460.2;refGene_gn=LOXL3;refGene_tx=XM_019827463.2
A3 93233451 93233666 refGene_gn=LOXL3;refGene_tx=XM_006930220.3;refGene_gn=LOXL3;refGene_tx=XM_019827460.2;refGene_gn=LOXL3;refGene_tx=XM_019827461.2;refGene_gn=LOXL3;refGene_tx=XM_019827462.2;refGene_gn=LOXL3;refGene_tx=XM_019827463.2
A3 93234354 93234518 refGene_gn=LOXL3;refGene_tx=XM_006930220.3;refGene_gn=LOXL3;refGene_tx=XM_019827460.2;refGene_gn=LOXL3;refGene_tx=XM_019827461.2;refGene_gn=LOXL3;refGene_tx=XM_019827462.2;refGene_gn=LOXL3;refGene_tx=XM_019827463.2
A3 93236335 93237311 refGene_gn=LOXL3;refGene_tx=XM_006930220.3;refGene_gn=LOXL3;refGene_tx=XM_019827460.2;refGene_gn=LOXL3;refGene_tx=XM_019827461.2;refGene_gn=LOXL3;refGene_tx=XM_019827462.2;refGene_gn=LOXL3;refGene_tx=XM_019827463.2
A3 93238993 93239053 refGene_gn=DOK1;refGene_tx=XM_003984155.3;refGene_gn=DOK1;refGene_tx=XM_019827474.1
A3 93239412 93239712 refGene_gn=DOK1;refGene_tx=XM_003984155.3;refGene_gn=DOK1;refGene_tx=XM_019827474.1
A3 93239862 93239956 refGene_gn=DOK1;refGene_tx=XM_003984155.3;refGene_gn=DOK1;refGene_tx=XM_019827474.1
A3 93240174 93240359 refGene_gn=DOK1;refGene_tx=XM_003984155.3
A3 93240597 93241440 refGene_gn=DOK1;refGene_tx=XM_003984155.3;refGene_gn=DOK1;refGene_tx=XM_019827472.1;refGene_gn=DOK1;refGene_tx=XM_019827473.1;refGene_gn=DOK1;refGene_tx=XM_019827474.1
A3 93243155 93243338 refGene_gn=M1AP;refGene_tx=XM_003984193.4;refGene_gn=M1AP;refGene_tx=XM_019827482.2
A3 93243843 93243855 refGene_gn=M1AP;refGene_tx=XM_003984193.4
A3 93244317 93244458 refGene_gn=M1AP;refGene_tx=XM_003984193.4;refGene_gn=M1AP;refGene_tx=XM_019827482.2
A3 93245581 93245788 refGene_gn=M1AP;refGene_tx=XM_003984193.4;refGene_gn=M1AP;refGene_tx=XM_019827482.2

A3 93260505 93260647
refGene_gn=M1AP;refGene_tx=XM_003984193.4;refGene_gn=M1AP;refGene_tx=XM_019827482.2
A3 93261611 93261774
refGene_gn=M1AP;refGene_tx=XM_003984193.4;refGene_gn=M1AP;refGene_tx=XM_019827482.2
A3 93265556 93265730
refGene_gn=M1AP;refGene_tx=XM_003984193.4;refGene_gn=M1AP;refGene_tx=XM_019827482.2
A3 93288097 93288266
refGene_gn=M1AP;refGene_tx=XM_003984193.4;refGene_gn=M1AP;refGene_tx=XM_019827482.2
A3 93289743 93289929
refGene_gn=M1AP;refGene_tx=XM_003984193.4;refGene_gn=M1AP;refGene_tx=XM_019827482.2
A3 93306943 93307183
refGene_gn=M1AP;refGene_tx=XM_003984193.4;refGene_gn=M1AP;refGene_tx=XM_019827482.2
A3 93317698 93317843
refGene_gn=SEMA4F;refGene_tx=XM_003984157.4;refGene_gn=SEMA4F;refGene_tx=XM_006930222.3
A3 93320122 93320274
refGene_gn=SEMA4F;refGene_tx=XM_003984157.4;refGene_gn=SEMA4F;refGene_tx=XM_006930222.3
A3 93320935 93320995
refGene_gn=SEMA4F;refGene_tx=XM_003984157.4;refGene_gn=SEMA4F;refGene_tx=XM_006930222.3;refGene_gn=SEMA4F;refGene_tx=XM_019827483.2
A3 93321228 93321327
refGene_gn=SEMA4F;refGene_tx=XM_003984157.4;refGene_gn=SEMA4F;refGene_tx=XM_019827483.2
A3 93326239 93326333
refGene_gn=SEMA4F;refGene_tx=XM_003984157.4;refGene_gn=SEMA4F;refGene_tx=XM_006930222.3;refGene_gn=SEMA4F;refGene_tx=XM_019827483.2
A3 93336430 93336550
refGene_gn=SEMA4F;refGene_tx=XM_003984157.4;refGene_gn=SEMA4F;refGene_tx=XM_006930222.3;refGene_gn=SEMA4F;refGene_tx=XM_019827483.2
A3 93336645 93336797
refGene_gn=SEMA4F;refGene_tx=XM_003984157.4;refGene_gn=SEMA4F;refGene_tx=XM_006930222.3;refGene_gn=SEMA4F;refGene_tx=XM_019827483.2
A3 93337474 93337653
refGene_gn=SEMA4F;refGene_tx=XM_003984157.4;refGene_gn=SEMA4F;refGene_tx=XM_006930222.3;refGene_gn=SEMA4F;refGene_tx=XM_019827483.2
A3 93337863 93338011
refGene_gn=SEMA4F;refGene_tx=XM_003984157.4;refGene_gn=SEMA4F;refGene_tx=XM_006930222.3;refGene_gn=SEMA4F;refGene_tx=XM_019827483.2
A3 93338136 93338359
refGene_gn=SEMA4F;refGene_tx=XM_003984157.4;refGene_gn=SEMA4F;refGene_tx=XM_006930222.3;refGene_gn=SEMA4F;refGene_tx=XM_019827483.2
A3 93338497 93338607
refGene_gn=SEMA4F;refGene_tx=XM_003984157.4;refGene_gn=SEMA4F;refGene_

tx=XM_006930222.3;refGene_gn=SEMA4F;refGene_tx=XM_019827483.2
A3 93338721 93338882
refGene_gn=SEMA4F;refGene_tx=XM_003984157.4;refGene_gn=SEMA4F;refGene_tx=XM_006930222.3;refGene_gn=SEMA4F;refGene_tx=XM_019827483.2
A3 93342194 93342253
refGene_gn=SEMA4F;refGene_tx=XM_003984157.4;refGene_gn=SEMA4F;refGene_tx=XM_006930222.3;refGene_gn=SEMA4F;refGene_tx=XM_019827483.2
A3 93342481 93343092
refGene_gn=SEMA4F;refGene_tx=XM_003984157.4;refGene_gn=SEMA4F;refGene_tx=XM_006930222.3;refGene_gn=SEMA4F;refGene_tx=XM_019827483.2
A3 93491424 93491487
refGene_gn=HK2;refGene_tx=XM_003984158.5;refGene_gn=HK2;refGene_tx=XM_019827485.2
A3 93510574 93510737
refGene_gn=HK2;refGene_tx=XM_003984158.5;refGene_gn=HK2;refGene_tx=XM_019827484.2;refGene_gn=HK2;refGene_tx=XM_019827485.2
A3 93526540 93526689
refGene_gn=HK2;refGene_tx=XM_003984158.5;refGene_gn=HK2;refGene_tx=XM_019827484.2;refGene_gn=HK2;refGene_tx=XM_019827485.2
A3 93531457 93531577
refGene_gn=HK2;refGene_tx=XM_003984158.5;refGene_gn=HK2;refGene_tx=XM_019827484.2;refGene_gn=HK2;refGene_tx=XM_019827485.2
A3 93532190 93532286
refGene_gn=HK2;refGene_tx=XM_003984158.5;refGene_gn=HK2;refGene_tx=XM_019827484.2;refGene_gn=HK2;refGene_tx=XM_019827485.2
A3 93533096 93533196
refGene_gn=HK2;refGene_tx=XM_003984158.5;refGene_gn=HK2;refGene_tx=XM_019827484.2;refGene_gn=HK2;refGene_tx=XM_019827485.2
A3 93533482 93533666
refGene_gn=HK2;refGene_tx=XM_003984158.5;refGene_gn=HK2;refGene_tx=XM_019827484.2;refGene_gn=HK2;refGene_tx=XM_019827485.2
A3 93536469 93536625
refGene_gn=HK2;refGene_tx=XM_003984158.5;refGene_gn=HK2;refGene_tx=XM_019827484.2;refGene_gn=HK2;refGene_tx=XM_019827485.2
A3 93537894 93538128
refGene_gn=HK2;refGene_tx=XM_003984158.5;refGene_gn=HK2;refGene_tx=XM_019827484.2;refGene_gn=HK2;refGene_tx=XM_019827485.2
A3 93539115 93539420
refGene_gn=HK2;refGene_tx=XM_003984158.5;refGene_gn=HK2;refGene_tx=XM_019827484.2;refGene_gn=HK2;refGene_tx=XM_019827485.2
A3 93540758 93540907
refGene_gn=HK2;refGene_tx=XM_003984158.5;refGene_gn=HK2;refGene_tx=XM_019827484.2;refGene_gn=HK2;refGene_tx=XM_019827485.2
A3 93541137 93541257
refGene_gn=HK2;refGene_tx=XM_003984158.5;refGene_gn=HK2;refGene_tx=XM_019827484.2
A3 93545015 93545111
refGene_gn=HK2;refGene_tx=XM_003984158.5;refGene_gn=HK2;refGene_tx=XM_019827484.2;refGene_gn=HK2;refGene_tx=XM_019827485.2
A3 93545565 93545665

refGene_gn=HK2;refGene_tx=XM_003984158.5;refGene_gn=HK2;refGene_tx=XM_019827484.2;refGene_gn=HK2;refGene_tx=XM_019827485.2
A3 93545762 93545946
refGene_gn=HK2;refGene_tx=XM_003984158.5;refGene_gn=HK2;refGene_tx=XM_019827484.2;refGene_gn=HK2;refGene_tx=XM_019827485.2
A3 93547196 93547352
refGene_gn=HK2;refGene_tx=XM_003984158.5;refGene_gn=HK2;refGene_tx=XM_019827484.2;refGene_gn=HK2;refGene_tx=XM_019827485.2
A3 93548589 93548823
refGene_gn=HK2;refGene_tx=XM_003984158.5;refGene_gn=HK2;refGene_tx=XM_019827484.2;refGene_gn=HK2;refGene_tx=XM_019827485.2
A3 93551200 93551345
refGene_gn=HK2;refGene_tx=XM_003984158.5;refGene_gn=HK2;refGene_tx=XM_019827484.2;refGene_gn=HK2;refGene_tx=XM_019827485.2
A3 93586632 93586842
refGene_gn=POLE4;refGene_tx=XM_006930225.3;refGene_gn=POLE4;refGene_tx=XM_019827486.2;refGene_gn=POLE4;refGene_tx=XM_019827487.2
A3 93587423 93587508
refGene_gn=POLE4;refGene_tx=XM_006930225.3;refGene_gn=POLE4;refGene_tx=XM_019827486.2;refGene_gn=POLE4;refGene_tx=XM_019827487.2
A3 93588133 93588175
refGene_gn=POLE4;refGene_tx=XM_006930225.3;refGene_gn=POLE4;refGene_tx=XM_019827486.2;refGene_gn=POLE4;refGene_tx=XM_019827487.2
A3 93598481 93598495 refGene_gn=POLE4;refGene_tx=XM_006930225.3
A3 93694403 93694695 refGene_gn=TACR1;refGene_tx=XM_003984160.4
A3 93695799 93695996 refGene_gn=TACR1;refGene_tx=XM_003984160.4
A3 93697076 93697227 refGene_gn=TACR1;refGene_tx=XM_003984160.4
A3 93760649 93760844 refGene_gn=TACR1;refGene_tx=XM_003984160.4
A3 93833466 93833855 refGene_gn=TACR1;refGene_tx=XM_003984160.4
A3 94062299 94062673
refGene_gn=EVA1A;refGene_tx=XM_023251793.1;refGene_gn=EVA1A;refGene_tx=XM_023251794.1;refGene_gn=EVA1A;refGene_tx=XM_023251795.1;refGene_gn=EVA1A;refGene_tx=XM_023251796.1;refGene_gn=EVA1A;refGene_tx=XM_023251797.1;refGene_gn=EVA1A;refGene_tx=XM_023251798.1
A3 94077849 94077895
refGene_gn=EVA1A;refGene_tx=XM_023251793.1;refGene_gn=EVA1A;refGene_tx=XM_023251794.1;refGene_gn=EVA1A;refGene_tx=XM_023251795.1;refGene_gn=EVA1A;refGene_tx=XM_023251796.1;refGene_gn=EVA1A;refGene_tx=XM_023251797.1;refGene_gn=EVA1A;refGene_tx=XM_023251798.1
A3 94187323 94187426 refGene_gn=MRPL19;refGene_tx=XM_003984162.4
A3 94187600 94187718 refGene_gn=MRPL19;refGene_tx=XM_003984162.4
A3 94192799 94192918 refGene_gn=MRPL19;refGene_tx=XM_003984162.4
A3 94193132 94193267 refGene_gn=MRPL19;refGene_tx=XM_003984162.4
A3 94197056 94197238 refGene_gn=MRPL19;refGene_tx=XM_003984162.4
A3 94197405 94197627 refGene_gn=MRPL19;refGene_tx=XM_003984162.4
A3 94218929 94219047
refGene_gn=GCFC2;refGene_tx=XM_023251800.1;refGene_gn=GCFC2;refGene_tx=XM_023251801.1;refGene_gn=GCFC2;refGene_tx=XM_023251802.1
A3 94220370 94220495
refGene_gn=GCFC2;refGene_tx=XM_023251800.1;refGene_gn=GCFC2;refGene_tx

=XM_023251801.1;refGene_gn=GCFC2;refGene_tx=XM_023251802.1
A3 94223330 94223477
refGene_gn=GCFC2;refGene_tx=XM_023251800.1;refGene_gn=GCFC2;refGene_tx=XM_023251801.1;refGene_gn=GCFC2;refGene_tx=XM_023251802.1
A3 94225932 94225999
refGene_gn=GCFC2;refGene_tx=XM_023251800.1;refGene_gn=GCFC2;refGene_tx=XM_023251801.1;refGene_gn=GCFC2;refGene_tx=XM_023251802.1
A3 94227774 94227851
refGene_gn=GCFC2;refGene_tx=XM_023251800.1;refGene_gn=GCFC2;refGene_tx=XM_023251801.1;refGene_gn=GCFC2;refGene_tx=XM_023251802.1
A3 94234758 94234880
refGene_gn=GCFC2;refGene_tx=XM_023251800.1;refGene_gn=GCFC2;refGene_tx=XM_023251801.1;refGene_gn=GCFC2;refGene_tx=XM_023251802.1;refGene_gn=GCFC2;refGene_tx=XM_023251803.1
A3 94235127 94235138 refGene_gn=GCFC2;refGene_tx=XM_023251804.1
A3 94241379 94241530
refGene_gn=GCFC2;refGene_tx=XM_023251800.1;refGene_gn=GCFC2;refGene_tx=XM_023251802.1;refGene_gn=GCFC2;refGene_tx=XM_023251803.1;refGene_gn=GCFC2;refGene_tx=XM_023251804.1
A3 94244242 94244442
refGene_gn=GCFC2;refGene_tx=XM_023251800.1;refGene_gn=GCFC2;refGene_tx=XM_023251802.1;refGene_gn=GCFC2;refGene_tx=XM_023251803.1;refGene_gn=GCFC2;refGene_tx=XM_023251804.1
A3 94245661 94245774
refGene_gn=GCFC2;refGene_tx=XM_023251800.1;refGene_gn=GCFC2;refGene_tx=XM_023251801.1;refGene_gn=GCFC2;refGene_tx=XM_023251802.1;refGene_gn=GCFC2;refGene_tx=XM_023251803.1;refGene_gn=GCFC2;refGene_tx=XM_023251804.1
A3 94246289 94246371
refGene_gn=GCFC2;refGene_tx=XM_023251800.1;refGene_gn=GCFC2;refGene_tx=XM_023251801.1;refGene_gn=GCFC2;refGene_tx=XM_023251802.1;refGene_gn=GCFC2;refGene_tx=XM_023251803.1;refGene_gn=GCFC2;refGene_tx=XM_023251804.1
A3 94247320 94247444
refGene_gn=GCFC2;refGene_tx=XM_023251800.1;refGene_gn=GCFC2;refGene_tx=XM_023251801.1;refGene_gn=GCFC2;refGene_tx=XM_023251802.1;refGene_gn=GCFC2;refGene_tx=XM_023251803.1;refGene_gn=GCFC2;refGene_tx=XM_023251804.1
A3 94249416 94249603
refGene_gn=GCFC2;refGene_tx=XM_023251800.1;refGene_gn=GCFC2;refGene_tx=XM_023251801.1;refGene_gn=GCFC2;refGene_tx=XM_023251802.1;refGene_gn=GCFC2;refGene_tx=XM_023251803.1;refGene_gn=GCFC2;refGene_tx=XM_023251804.1
A3 94254470 94254528 refGene_gn=GCFC2;refGene_tx=XM_023251805.1
A3 94261610 94261623 refGene_gn=GCFC2;refGene_tx=XM_023251806.1
A3 94261980 94262096
refGene_gn=GCFC2;refGene_tx=XM_023251800.1;refGene_gn=GCFC2;refGene_tx=XM_023251801.1;refGene_gn=GCFC2;refGene_tx=XM_023251802.1;refGene_gn=GCFC2;refGene_tx=XM_023251803.1;refGene_gn=GCFC2;refGene_tx=XM_023251804.1;refGene_gn=GCFC2;refGene_tx=XM_023251805.1;refGene_gn=GCFC2;refGe

ne_tx=XM_023251806.1
A3 94265080 94265178
refGene_gn=GCFC2;refGene_tx=XM_023251800.1;refGene_gn=GCFC2;refGene_tx=XM_023251801.1;refGene_gn=GCFC2;refGene_tx=XM_023251802.1;refGene_gn=GCFC2;refGene_tx=XM_023251803.1;refGene_gn=GCFC2;refGene_tx=XM_023251804.1;refGene_gn=GCFC2;refGene_tx=XM_023251805.1;refGene_gn=GCFC2;refGene_tx=XM_023251806.1
A3 94266526 94266751
refGene_gn=GCFC2;refGene_tx=XM_023251800.1;refGene_gn=GCFC2;refGene_tx=XM_023251801.1;refGene_gn=GCFC2;refGene_tx=XM_023251802.1;refGene_gn=GCFC2;refGene_tx=XM_023251803.1;refGene_gn=GCFC2;refGene_tx=XM_023251804.1;refGene_gn=GCFC2;refGene_tx=XM_023251805.1;refGene_gn=GCFC2;refGene_tx=XM_023251806.1
A3 94271002 94271128
refGene_gn=GCFC2;refGene_tx=XM_023251800.1;refGene_gn=GCFC2;refGene_tx=XM_023251801.1;refGene_gn=GCFC2;refGene_tx=XM_023251803.1;refGene_gn=GCFC2;refGene_tx=XM_023251804.1;refGene_gn=GCFC2;refGene_tx=XM_023251805.1;refGene_gn=GCFC2;refGene_tx=XM_023251806.1
A3 94274231 94274502
refGene_gn=GCFC2;refGene_tx=XM_023251800.1;refGene_gn=GCFC2;refGene_tx=XM_023251801.1;refGene_gn=GCFC2;refGene_tx=XM_023251803.1;refGene_gn=GCFC2;refGene_tx=XM_023251804.1;refGene_gn=GCFC2;refGene_tx=XM_023251805.1;refGene_gn=GCFC2;refGene_tx=XM_023251806.1
A3 95166826 95167048
refGene_gn=LRRTM4;refGene_tx=XM_006930227.4;refGene_gn=LRRTM4;refGene_tx=XM_019827488.2
A3 95866070 95867623
refGene_gn=LRRTM4;refGene_tx=XM_006930227.4;refGene_gn=LRRTM4;refGene_tx=XM_019827488.2;refGene_gn=LRRTM4;refGene_tx=XM_019827493.2
A3 95869456 95869460 refGene_gn=LRRTM4;refGene_tx=XM_006930227.4
A3 95869892 95869899
refGene_gn=LRRTM4;refGene_tx=XM_019827488.2;refGene_gn=LRRTM4;refGene_tx=XM_019827493.2
A3 97118500 97118579
refGene_gn=LOC101081885;refGene_tx=XM_003984197.2
A3 97119134 97119253
refGene_gn=LOC101081885;refGene_tx=XM_003984197.2
A3 97119466 97119604
refGene_gn=LOC101081885;refGene_tx=XM_003984197.2
A3 97120203 97120330
refGene_gn=LOC101081885;refGene_tx=XM_003984197.2
A3 97120613 97120681
refGene_gn=LOC101081885;refGene_tx=XM_003984197.2
A3 97155593 97155661
refGene_gn=LOC101082144;refGene_tx=XM_003984198.3
A3 97155944 97156071
refGene_gn=LOC101082144;refGene_tx=XM_003984198.3
A3 97156670 97156808
refGene_gn=LOC101082144;refGene_tx=XM_003984198.3
A3 97157020 97157139

refGene_gn=LOC101082144;refGene_tx=XM_003984198.3
A3 97157715 97157794
refGene_gn=LOC101082144;refGene_tx=XM_003984198.3
A3 97604920 97605022
refGene_gn=CTNNA2;refGene_tx=XM_023251808.1;refGene_gn=CTNNA2;refGene_tx=XM_023251809.1;refGene_gn=CTNNA2;refGene_tx=XM_023251810.1;refGene_gn=CTNNA2;refGene_tx=XM_023251811.1;refGene_gn=CTNNA2;refGene_tx=XM_023251812.1
A3 97691659 97691855
refGene_gn=CTNNA2;refGene_tx=XM_023251808.1;refGene_gn=CTNNA2;refGene_tx=XM_023251809.1;refGene_gn=CTNNA2;refGene_tx=XM_023251810.1;refGene_gn=CTNNA2;refGene_tx=XM_023251811.1;refGene_gn=CTNNA2;refGene_tx=XM_023251812.1
A3 97817778 97817945
refGene_gn=CTNNA2;refGene_tx=XM_023251808.1;refGene_gn=CTNNA2;refGene_tx=XM_023251809.1;refGene_gn=CTNNA2;refGene_tx=XM_023251810.1;refGene_gn=CTNNA2;refGene_tx=XM_023251811.1;refGene_gn=CTNNA2;refGene_tx=XM_023251812.1
A3 97831825 97831945
refGene_gn=CTNNA2;refGene_tx=XM_023251808.1;refGene_gn=CTNNA2;refGene_tx=XM_023251809.1;refGene_gn=CTNNA2;refGene_tx=XM_023251810.1;refGene_gn=CTNNA2;refGene_tx=XM_023251811.1;refGene_gn=CTNNA2;refGene_tx=XM_023251812.1
A3 97835484 97835751
refGene_gn=CTNNA2;refGene_tx=XM_023251808.1;refGene_gn=CTNNA2;refGene_tx=XM_023251809.1;refGene_gn=CTNNA2;refGene_tx=XM_023251810.1;refGene_gn=CTNNA2;refGene_tx=XM_023251811.1;refGene_gn=CTNNA2;refGene_tx=XM_023251812.1
A3 97873843 97874047
refGene_gn=CTNNA2;refGene_tx=XM_023251808.1;refGene_gn=CTNNA2;refGene_tx=XM_023251809.1;refGene_gn=CTNNA2;refGene_tx=XM_023251810.1;refGene_gn=CTNNA2;refGene_tx=XM_023251811.1;refGene_gn=CTNNA2;refGene_tx=XM_023251812.1
A3 98230920 98232489
refGene_gn=LRRTM1;refGene_tx=XM_003984200.5;refGene_gn=LRRTM1;refGene_tx=XM_006930241.4;refGene_gn=LRRTM1;refGene_tx=XM_023251813.1
A3 98333716 98333797
refGene_gn=CTNNA2;refGene_tx=XM_023251808.1;refGene_gn=CTNNA2;refGene_tx=XM_023251809.1;refGene_gn=CTNNA2;refGene_tx=XM_023251810.1;refGene_gn=CTNNA2;refGene_tx=XM_023251811.1;refGene_gn=CTNNA2;refGene_tx=XM_023251812.1
A3 98367938 98368091
refGene_gn=CTNNA2;refGene_tx=XM_023251808.1;refGene_gn=CTNNA2;refGene_tx=XM_023251809.1;refGene_gn=CTNNA2;refGene_tx=XM_023251810.1;refGene_gn=CTNNA2;refGene_tx=XM_023251811.1;refGene_gn=CTNNA2;refGene_tx=XM_023251812.1
A3 98491196 98491289
refGene_gn=CTNNA2;refGene_tx=XM_023251808.1;refGene_gn=CTNNA2;refGene_tx=XM_023251809.1;refGene_gn=CTNNA2;refGene_tx=XM_023251810.1;refGene_gn=CTNNA2;refGene_tx=XM_023251811.1;refGene_gn=CTNNA2;refGene_tx=XM_02

3251812.1
A3	98492130	98492287
refGene_gn=CTNNA2;refGene_tx=XM_023251808.1;refGene_gn=CTNNA2;refGene_tx=XM_023251809.1;refGene_gn=CTNNA2;refGene_tx=XM_023251810.1;refGene_gn=CTNNA2;refGene_tx=XM_023251811.1;refGene_gn=CTNNA2;refGene_tx=XM_023251812.1
A3	98498155	98498356
refGene_gn=CTNNA2;refGene_tx=XM_023251808.1;refGene_gn=CTNNA2;refGene_tx=XM_023251809.1;refGene_gn=CTNNA2;refGene_tx=XM_023251810.1;refGene_gn=CTNNA2;refGene_tx=XM_023251811.1;refGene_gn=CTNNA2;refGene_tx=XM_023251812.1
A3	98517034	98517186
refGene_gn=CTNNA2;refGene_tx=XM_023251808.1;refGene_gn=CTNNA2;refGene_tx=XM_023251809.1;refGene_gn=CTNNA2;refGene_tx=XM_023251810.1;refGene_gn=CTNNA2;refGene_tx=XM_023251811.1;refGene_gn=CTNNA2;refGene_tx=XM_023251812.1
A3	98526755	98526869
refGene_gn=CTNNA2;refGene_tx=XM_023251808.1;refGene_gn=CTNNA2;refGene_tx=XM_023251809.1;refGene_gn=CTNNA2;refGene_tx=XM_023251810.1;refGene_gn=CTNNA2;refGene_tx=XM_023251811.1;refGene_gn=CTNNA2;refGene_tx=XM_023251812.1
A3	98534896	98535078
refGene_gn=CTNNA2;refGene_tx=XM_023251808.1;refGene_gn=CTNNA2;refGene_tx=XM_023251809.1;refGene_gn=CTNNA2;refGene_tx=XM_023251810.1;refGene_gn=CTNNA2;refGene_tx=XM_023251811.1;refGene_gn=CTNNA2;refGene_tx=XM_023251812.1
A3	98560160	98560266
refGene_gn=CTNNA2;refGene_tx=XM_023251808.1;refGene_gn=CTNNA2;refGene_tx=XM_023251809.1;refGene_gn=CTNNA2;refGene_tx=XM_023251810.1;refGene_gn=CTNNA2;refGene_tx=XM_023251811.1;refGene_gn=CTNNA2;refGene_tx=XM_023251812.1
A3	98564560	98564695
refGene_gn=CTNNA2;refGene_tx=XM_023251808.1;refGene_gn=CTNNA2;refGene_tx=XM_023251809.1;refGene_gn=CTNNA2;refGene_tx=XM_023251810.1;refGene_gn=CTNNA2;refGene_tx=XM_023251811.1;refGene_gn=CTNNA2;refGene_tx=XM_023251812.1
A3	98575545	98575689
refGene_gn=CTNNA2;refGene_tx=XM_023251808.1;refGene_gn=CTNNA2;refGene_tx=XM_023251809.1;refGene_gn=CTNNA2;refGene_tx=XM_023251812.1
A3	98583756	98583771	refGene_gn=CTNNA2;refGene_tx=XM_023251812.1
A3	98603275	98603392
refGene_gn=CTNNA2;refGene_tx=XM_023251808.1;refGene_gn=CTNNA2;refGene_tx=XM_023251810.1
A3	98605991	98606279
refGene_gn=CTNNA2;refGene_tx=XM_023251808.1;refGene_gn=CTNNA2;refGene_tx=XM_023251809.1;refGene_gn=CTNNA2;refGene_tx=XM_023251810.1;refGene_gn=CTNNA2;refGene_tx=XM_023251811.1
A3	100793201	100793522
refGene_gn=LOC101089093;refGene_tx=XM_011281195.2
A3	101774344	101774371

refGene_gn=SUCLG1;refGene_tx=XM_011281193.2;refGene_gn=SUCLG1;refGene_tx=XM_011281194.3;refGene_gn=SUCLG1;refGene_tx=XM_019827510.1
A3 101775951 101776140
refGene_gn=SUCLG1;refGene_tx=XM_011281193.2;refGene_gn=SUCLG1;refGene_tx=XM_011281194.3;refGene_gn=SUCLG1;refGene_tx=XM_019827510.1
A3 101782362 101782514
refGene_gn=SUCLG1;refGene_tx=XM_011281193.2;refGene_gn=SUCLG1;refGene_tx=XM_011281194.3;refGene_gn=SUCLG1;refGene_tx=XM_019827510.1
A3 101784349 101784433
refGene_gn=SUCLG1;refGene_tx=XM_011281193.2;refGene_gn=SUCLG1;refGene_tx=XM_011281194.3;refGene_gn=SUCLG1;refGene_tx=XM_019827510.1
A3 101794851 101794909
refGene_gn=SUCLG1;refGene_tx=XM_011281193.2;refGene_gn=SUCLG1;refGene_tx=XM_011281194.3;refGene_gn=SUCLG1;refGene_tx=XM_019827510.1
A3 101795056 101795269
refGene_gn=SUCLG1;refGene_tx=XM_011281193.2;refGene_gn=SUCLG1;refGene_tx=XM_011281194.3;refGene_gn=SUCLG1;refGene_tx=XM_019827510.1
A3 101798573 101798690
refGene_gn=SUCLG1;refGene_tx=XM_011281193.2;refGene_gn=SUCLG1;refGene_tx=XM_011281194.3;refGene_gn=SUCLG1;refGene_tx=XM_019827510.1
A3 101806448 101806552
refGene_gn=SUCLG1;refGene_tx=XM_011281193.2;refGene_gn=SUCLG1;refGene_tx=XM_011281194.3;refGene_gn=SUCLG1;refGene_tx=XM_019827510.1
A3 101811968 101812122
refGene_gn=SUCLG1;refGene_tx=XM_019827510.1
A3 101813372 101813390
refGene_gn=SUCLG1;refGene_tx=XM_019827510.1
A3 101816400 101816449
refGene_gn=SUCLG1;refGene_tx=XM_011281194.3
A3 101816745 101816842
refGene_gn=SUCLG1;refGene_tx=XM_011281193.2
A3 101846529 101846754
refGene_gn=DNAH6;refGene_tx=XM_023251818.1
A3 101857714 101857891
refGene_gn=DNAH6;refGene_tx=XM_023251818.1
A3 101861343 101861600
refGene_gn=DNAH6;refGene_tx=XM_023251818.1
A3 101879658 101879926
refGene_gn=DNAH6;refGene_tx=XM_023251818.1
A3 101883951 101884086
refGene_gn=DNAH6;refGene_tx=XM_023251818.1
A3 101884174 101884295
refGene_gn=DNAH6;refGene_tx=XM_023251818.1
A3 101885940 101886058
refGene_gn=DNAH6;refGene_tx=XM_023251818.1
A3 101887629 101887798
refGene_gn=DNAH6;refGene_tx=XM_023251818.1
A3 101889984 101890101
refGene_gn=DNAH6;refGene_tx=XM_023251818.1
A3 101891829 101892030

refGene_gn=DNAH6;refGene_tx=XM_023251818.1
A3 101894577 101894698
refGene_gn=DNAH6;refGene_tx=XM_023251818.1
A3 101897813 101897965
refGene_gn=DNAH6;refGene_tx=XM_023251818.1
A3 101898778 101898931
refGene_gn=DNAH6;refGene_tx=XM_023251818.1
A3 101902838 101903090
refGene_gn=DNAH6;refGene_tx=XM_023251818.1
A3 101906414 101906543
refGene_gn=DNAH6;refGene_tx=XM_023251818.1
A3 101910642 101910756
refGene_gn=DNAH6;refGene_tx=XM_023251818.1
A3 101912726 101912870
refGene_gn=DNAH6;refGene_tx=XM_023251818.1
A3 101919119 101919332
refGene_gn=DNAH6;refGene_tx=XM_023251818.1
A3 101920160 101920253
refGene_gn=DNAH6;refGene_tx=XM_023251818.1
A3 101921715 101921835
refGene_gn=DNAH6;refGene_tx=XM_023251818.1
A3 101924544 101924725
refGene_gn=DNAH6;refGene_tx=XM_023251818.1
A3 101926796 101926893
refGene_gn=DNAH6;refGene_tx=XM_023251818.1
A3 101929264 101929484
refGene_gn=DNAH6;refGene_tx=XM_023251818.1
A3 101930913 101931078
refGene_gn=DNAH6;refGene_tx=XM_023251818.1
A3 101931163 101931277
refGene_gn=DNAH6;refGene_tx=XM_023251818.1
A3 101932851 101932977
refGene_gn=DNAH6;refGene_tx=XM_023251818.1
A3 101933060 101933216
refGene_gn=DNAH6;refGene_tx=XM_023251818.1
A3 101933500 101933662
refGene_gn=DNAH6;refGene_tx=XM_023251818.1
A3 101937666 101937804
refGene_gn=DNAH6;refGene_tx=XM_023251818.1
A3 101939021 101939189
refGene_gn=DNAH6;refGene_tx=XM_023251818.1
A3 101942165 101942314
refGene_gn=DNAH6;refGene_tx=XM_023251818.1
A3 101943653 101943761
refGene_gn=DNAH6;refGene_tx=XM_023251818.1
A3 101947608 101948164
refGene_gn=DNAH6;refGene_tx=XM_023251818.1
A3 101948955 101949078
refGene_gn=DNAH6;refGene_tx=XM_023251818.1
A3 101953401 101953584

refGene_gn=DNAH6;refGene_tx=XM_023251818.1
A3 101954151 101954295
refGene_gn=DNAH6;refGene_tx=XM_023251818.1
A3 101958587 101958809
refGene_gn=DNAH6;refGene_tx=XM_023251818.1
A3 101959630 101959778
refGene_gn=DNAH6;refGene_tx=XM_023251818.1
A3 101961836 101961994
refGene_gn=DNAH6;refGene_tx=XM_023251818.1
A3 101963433 101963565
refGene_gn=DNAH6;refGene_tx=XM_023251818.1
A3 101967936 101968108
refGene_gn=DNAH6;refGene_tx=XM_023251818.1
A3 101969221 101969368
refGene_gn=DNAH6;refGene_tx=XM_023251818.1
A3 101970178 101970252
refGene_gn=DNAH6;refGene_tx=XM_023251818.1
A3 101971925 101972080
refGene_gn=DNAH6;refGene_tx=XM_023251818.1
A3 101981453 101981685
refGene_gn=DNAH6;refGene_tx=XM_023251818.1
A3 101989778 101989931
refGene_gn=DNAH6;refGene_tx=XM_023251818.1
A3 101991194 101991335
refGene_gn=DNAH6;refGene_tx=XM_023251818.1
A3 101992673 101992916
refGene_gn=DNAH6;refGene_tx=XM_023251818.1
A3 101995600 101995768
refGene_gn=DNAH6;refGene_tx=XM_023251818.1
A3 101996468 101996704
refGene_gn=DNAH6;refGene_tx=XM_023251818.1
A3 101997821 101998083
refGene_gn=DNAH6;refGene_tx=XM_023251818.1
A3 101999195 101999319
refGene_gn=DNAH6;refGene_tx=XM_023251818.1
A3 102000049 102000246
refGene_gn=DNAH6;refGene_tx=XM_023251818.1
A3 102001215 102001419
refGene_gn=DNAH6;refGene_tx=XM_023251818.1
A3 102002156 102002282
refGene_gn=DNAH6;refGene_tx=XM_023251818.1
A3 102004231 102004396
refGene_gn=DNAH6;refGene_tx=XM_023251818.1
A3 102006606 102006674
refGene_gn=DNAH6;refGene_tx=XM_023251818.1
A3 102008855 102009036
refGene_gn=DNAH6;refGene_tx=XM_023251818.1
A3 102013441 102013621
refGene_gn=DNAH6;refGene_tx=XM_023251818.1
A3 102015372 102015606

refGene_gn=DNAH6;refGene_tx=XM_023251818.1
A3 102027315 102027451
refGene_gn=DNAH6;refGene_tx=XM_023251818.1
A3 102031879 102032049
refGene_gn=DNAH6;refGene_tx=XM_023251818.1
A3 102037338 102037529
refGene_gn=DNAH6;refGene_tx=XM_023251818.1
A3 102039529 102039690
refGene_gn=DNAH6;refGene_tx=XM_023251818.1
A3 102043458 102043523
refGene_gn=DNAH6;refGene_tx=XM_023251818.1
A3 102043524 102043548
refGene_gn=DNAH6;refGene_tx=XM_023251818.1
A3 102044298 102044445
refGene_gn=DNAH6;refGene_tx=XM_023251818.1
A3 102045919 102046058
refGene_gn=DNAH6;refGene_tx=XM_023251818.1
A3 102051118 102051238
refGene_gn=DNAH6;refGene_tx=XM_023251818.1
A3 102051782 102051904
refGene_gn=DNAH6;refGene_tx=XM_023251818.1
A3 102059039 102059169
refGene_gn=DNAH6;refGene_tx=XM_023251818.1
A3 102061633 102061761
refGene_gn=DNAH6;refGene_tx=XM_023251818.1
A3 102064911 102065097
refGene_gn=DNAH6;refGene_tx=XM_023251818.1
A3 102065858 102065931
refGene_gn=DNAH6;refGene_tx=XM_023251818.1
A3 102066866 102067018
refGene_gn=DNAH6;refGene_tx=XM_023251818.1
A3 102069570 102069793
refGene_gn=DNAH6;refGene_tx=XM_023251818.1
A3 102074265 102074369
refGene_gn=DNAH6;refGene_tx=XM_023251818.1
A3 102076759 102076946
refGene_gn=TRABD2A;refGene_tx=XM_006930243.4;refGene_gn=TRABD2A;refGene_tx=XM_019827497.2;refGene_gn=TRABD2A;refGene_tx=XM_019827503.2;refGene_gn=TRABD2A;refGene_tx=XM_019827504.2;refGene_gn=TRABD2A;refGene_tx=XM_023251814.1
A3 102079106 102079364
refGene_gn=TRABD2A;refGene_tx=XM_006930243.4;refGene_gn=TRABD2A;refGene_tx=XM_019827497.2;refGene_gn=TRABD2A;refGene_tx=XM_019827503.2;refGene_gn=TRABD2A;refGene_tx=XM_019827504.2;refGene_gn=TRABD2A;refGene_tx=XM_019827509.2;refGene_gn=TRABD2A;refGene_tx=XM_023251814.1
A3 102079512 102079576
refGene_gn=TRABD2A;refGene_tx=XM_019827506.2
A3 102085661 102085752
refGene_gn=TRABD2A;refGene_tx=XM_006930243.4;refGene_gn=TRABD2A;refGene_tx=XM_019827497.2;refGene_gn=TRABD2A;refGene_tx=XM_019827503.2;refGe

ne_gn=TRABD2A;refGene_tx=XM_019827506.2;refGene_gn=TRABD2A;refGene_tx=XM_019827509.2;refGene_gn=TRABD2A;refGene_tx=XM_023251814.1
A3	102090958	102091133
refGene_gn=TRABD2A;refGene_tx=XM_006930243.4;refGene_gn=TRABD2A;refGene_tx=XM_019827497.2;refGene_gn=TRABD2A;refGene_tx=XM_019827503.2;refGene_gn=TRABD2A;refGene_tx=XM_019827504.2;refGene_gn=TRABD2A;refGene_tx=XM_019827506.2;refGene_gn=TRABD2A;refGene_tx=XM_019827509.2;refGene_gn=TRABD2A;refGene_tx=XM_023251814.1
A3	102093640	102093787
refGene_gn=TRABD2A;refGene_tx=XM_006930243.4;refGene_gn=TRABD2A;refGene_tx=XM_019827497.2;refGene_gn=TRABD2A;refGene_tx=XM_019827503.2;refGene_gn=TRABD2A;refGene_tx=XM_019827504.2;refGene_gn=TRABD2A;refGene_tx=XM_019827506.2;refGene_gn=TRABD2A;refGene_tx=XM_019827509.2;refGene_gn=TRABD2A;refGene_tx=XM_023251814.1
A3	102119977	102120538
refGene_gn=TRABD2A;refGene_tx=XM_006930243.4;refGene_gn=TRABD2A;refGene_tx=XM_019827497.2;refGene_gn=TRABD2A;refGene_tx=XM_019827503.2;refGene_gn=TRABD2A;refGene_tx=XM_019827504.2;refGene_gn=TRABD2A;refGene_tx=XM_019827506.2;refGene_gn=TRABD2A;refGene_tx=XM_019827509.2;refGene_gn=TRABD2A;refGene_tx=XM_023251814.1
A3	102129550	102129658
refGene_gn=TRABD2A;refGene_tx=XM_006930243.4;refGene_gn=TRABD2A;refGene_tx=XM_019827497.2;refGene_gn=TRABD2A;refGene_tx=XM_019827503.2;refGene_gn=TRABD2A;refGene_tx=XM_019827504.2;refGene_gn=TRABD2A;refGene_tx=XM_019827506.2;refGene_gn=TRABD2A;refGene_tx=XM_019827509.2
A3	102135433	102135505
refGene_gn=TRABD2A;refGene_tx=XM_023251814.1
A3	102157435	102158052
refGene_gn=TMSB10;refGene_tx=XM_023251816.1
A3	102158328	102158443
refGene_gn=TMSB10;refGene_tx=XM_023251816.1;refGene_gn=TMSB10;refGene_tx=XM_023251817.1
A3	102158701	102158736
refGene_gn=TMSB10;refGene_tx=XM_023251817.1
A3	102209437	102209453
refGene_gn=KCMF1;refGene_tx=XM_023251819.1
A3	102257238	102257406
refGene_gn=KCMF1;refGene_tx=XM_023251819.1
A3	102261926	102262066
refGene_gn=KCMF1;refGene_tx=XM_023251819.1
A3	102277147	102277249
refGene_gn=KCMF1;refGene_tx=XM_023251819.1
A3	102280419	102280594
refGene_gn=KCMF1;refGene_tx=XM_023251819.1
A3	102282694	102282977
refGene_gn=KCMF1;refGene_tx=XM_023251819.1
A3	102286946	102287208
refGene_gn=KCMF1;refGene_tx=XM_023251819.1
A3	102343909	102344152
refGene_gn=TCF7L1;refGene_tx=XM_023251821.1;refGene_gn=TCF7L1;refGene_

tx=XM_023251822.1
A3	102344235	102344299
refGene_gn=TCF7L1;refGene_tx=XM_023251821.1;refGene_gn=TCF7L1;refGene_tx=XM_023251822.1
A3	102344514	102344642
refGene_gn=TCF7L1;refGene_tx=XM_023251821.1;refGene_gn=TCF7L1;refGene_tx=XM_023251822.1
A3	102481304	102481388
refGene_gn=TCF7L1;refGene_tx=XM_023251821.1;refGene_gn=TCF7L1;refGene_tx=XM_023251822.1
A3	102494968	102495101
refGene_gn=TCF7L1;refGene_tx=XM_023251821.1;refGene_gn=TCF7L1;refGene_tx=XM_023251822.1
A3	102496186	102496292
refGene_gn=TCF7L1;refGene_tx=XM_023251821.1;refGene_gn=TCF7L1;refGene_tx=XM_023251822.1
A3	102496594	102496678
refGene_gn=TCF7L1;refGene_tx=XM_023251821.1;refGene_gn=TCF7L1;refGene_tx=XM_023251822.1
A3	102497422	102497566
refGene_gn=TCF7L1;refGene_tx=XM_023251821.1;refGene_gn=TCF7L1;refGene_tx=XM_023251822.1
A3	102498280	102498440
refGene_gn=TCF7L1;refGene_tx=XM_023251821.1;refGene_gn=TCF7L1;refGene_tx=XM_023251822.1
A3	102498526	102498634
refGene_gn=TCF7L1;refGene_tx=XM_023251821.1;refGene_gn=TCF7L1;refGene_tx=XM_023251822.1
A3	102499554	102499630
refGene_gn=TCF7L1;refGene_tx=XM_023251821.1;refGene_gn=TCF7L1;refGene_tx=XM_023251822.1
A3	102500764	102501198
refGene_gn=TCF7L1;refGene_tx=XM_023251821.1;refGene_gn=TCF7L1;refGene_tx=XM_023251822.1
A3	102507277	102507323
refGene_gn=LOC101090095;refGene_tx=XM_023251826.1;refGene_gn=LOC101090095;refGene_tx=XM_023251827.1
A3	102507552	102508034
refGene_gn=LOC101090095;refGene_tx=XM_023251826.1;refGene_gn=LOC101090095;refGene_tx=XM_023251827.1
A3	102508872	102509646
refGene_gn=TGOLN2;refGene_tx=XM_023252189.1
A3	102511050	102511134
refGene_gn=TGOLN2;refGene_tx=XM_023252189.1
A3	102512413	102512419
refGene_gn=TGOLN2;refGene_tx=XM_023252189.1
A3	102518004	102518138
refGene_gn=RETSAT;refGene_tx=XM_003984224.5
A3	102518384	102518544
refGene_gn=RETSAT;refGene_tx=XM_003984224.5

A3 102518754 102518921
refGene_gn=RETSAT;refGene_tx=XM_003984224.5
A3 102519001 102519111
refGene_gn=RETSAT;refGene_tx=XM_003984224.5
A3 102519335 102519474
refGene_gn=RETSAT;refGene_tx=XM_003984224.5
A3 102523139 102523259
refGene_gn=RETSAT;refGene_tx=XM_003984224.5
A3 102525239 102525437
refGene_gn=RETSAT;refGene_tx=XM_003984224.5
A3 102525905 102526107
refGene_gn=RETSAT;refGene_tx=XM_003984224.5
A3 102526638 102526880
refGene_gn=RETSAT;refGene_tx=XM_003984224.5
A3 102527218 102527401
refGene_gn=RETSAT;refGene_tx=XM_003984224.5
A3 102530402 102530580
refGene_gn=RETSAT;refGene_tx=XM_003984224.5
A3 102533094 102533145
refGene_gn=ELMOD3;refGene_tx=XM_023251823.1;refGene_gn=ELMOD3;refGene_
tx=XM_023251824.1
A3 102537479 102537557
refGene_gn=ELMOD3;refGene_tx=XM_023251823.1;refGene_gn=ELMOD3;refGene_
tx=XM_023251824.1
A3 102538003 102538073
refGene_gn=ELMOD3;refGene_tx=XM_023251823.1;refGene_gn=ELMOD3;refGene_
tx=XM_023251824.1
A3 102542229 102542298
refGene_gn=ELMOD3;refGene_tx=XM_023251823.1;refGene_gn=ELMOD3;refGene_
tx=XM_023251824.1
A3 102543302 102543391
refGene_gn=ELMOD3;refGene_tx=XM_023251823.1;refGene_gn=ELMOD3;refGene_
tx=XM_023251824.1
A3 102544845 102544969
refGene_gn=ELMOD3;refGene_tx=XM_023251823.1;refGene_gn=ELMOD3;refGene_
tx=XM_023251824.1
A3 102545220 102545343
refGene_gn=ELMOD3;refGene_tx=XM_023251823.1;refGene_gn=ELMOD3;refGene_
tx=XM_023251824.1
A3 102546942 102547073
refGene_gn=ELMOD3;refGene_tx=XM_023251823.1;refGene_gn=ELMOD3;refGene_
tx=XM_023251824.1
A3 102555423 102555500
refGene_gn=ELMOD3;refGene_tx=XM_023251823.1;refGene_gn=ELMOD3;refGene_
tx=XM_023251824.1
A3 102555768 102555896
refGene_gn=ELMOD3;refGene_tx=XM_023251823.1;refGene_gn=ELMOD3;refGene_
tx=XM_023251824.1
A3 102556370 102556573
refGene_gn=ELMOD3;refGene_tx=XM_023251823.1;refGene_gn=ELMOD3;refGene_

tx=XM_023251824.1
A3	102559486	102559552
refGene_gn=CAPG;refGene_tx=XM_003984204.3;refGene_gn=CAPG;refGene_tx=XM_006930250.3;refGene_gn=CAPG;refGene_tx=XM_006930251.4;refGene_gn=CAPG;refGene_tx=XM_006930252.3;refGene_gn=CAPG;refGene_tx=XM_006930253.4;refGene_gn=CAPG;refGene_tx=XM_011281188.2;refGene_gn=CAPG;refGene_tx=XM_011281189.2
A3	102560108	102560197
refGene_gn=CAPG;refGene_tx=XM_003984204.3;refGene_gn=CAPG;refGene_tx=XM_006930250.3;refGene_gn=CAPG;refGene_tx=XM_006930251.4;refGene_gn=CAPG;refGene_tx=XM_006930252.3;refGene_gn=CAPG;refGene_tx=XM_006930253.4;refGene_gn=CAPG;refGene_tx=XM_011281188.2;refGene_gn=CAPG;refGene_tx=XM_011281189.2
A3	102561871	102562004
refGene_gn=CAPG;refGene_tx=XM_003984204.3;refGene_gn=CAPG;refGene_tx=XM_006930250.3;refGene_gn=CAPG;refGene_tx=XM_006930251.4;refGene_gn=CAPG;refGene_tx=XM_006930252.3;refGene_gn=CAPG;refGene_tx=XM_006930253.4;refGene_gn=CAPG;refGene_tx=XM_011281188.2;refGene_gn=CAPG;refGene_tx=XM_011281189.2
A3	102562451	102562544
refGene_gn=CAPG;refGene_tx=XM_003984204.3;refGene_gn=CAPG;refGene_tx=XM_006930250.3;refGene_gn=CAPG;refGene_tx=XM_006930251.4;refGene_gn=CAPG;refGene_tx=XM_006930252.3;refGene_gn=CAPG;refGene_tx=XM_006930253.4;refGene_gn=CAPG;refGene_tx=XM_011281188.2;refGene_gn=CAPG;refGene_tx=XM_011281189.2
A3	102562863	102563013
refGene_gn=CAPG;refGene_tx=XM_003984204.3;refGene_gn=CAPG;refGene_tx=XM_006930250.3;refGene_gn=CAPG;refGene_tx=XM_006930251.4;refGene_gn=CAPG;refGene_tx=XM_006930252.3;refGene_gn=CAPG;refGene_tx=XM_006930253.4;refGene_gn=CAPG;refGene_tx=XM_011281188.2;refGene_gn=CAPG;refGene_tx=XM_011281189.2
A3	102564200	102564368
refGene_gn=CAPG;refGene_tx=XM_003984204.3;refGene_gn=CAPG;refGene_tx=XM_006930250.3;refGene_gn=CAPG;refGene_tx=XM_006930251.4;refGene_gn=CAPG;refGene_tx=XM_006930252.3;refGene_gn=CAPG;refGene_tx=XM_006930253.4;refGene_gn=CAPG;refGene_tx=XM_011281188.2;refGene_gn=CAPG;refGene_tx=XM_011281189.2
A3	102564574	102564729
refGene_gn=CAPG;refGene_tx=XM_003984204.3;refGene_gn=CAPG;refGene_tx=XM_006930250.3;refGene_gn=CAPG;refGene_tx=XM_006930251.4;refGene_gn=CAPG;refGene_tx=XM_006930252.3;refGene_gn=CAPG;refGene_tx=XM_006930253.4;refGene_gn=CAPG;refGene_tx=XM_011281188.2;refGene_gn=CAPG;refGene_tx=XM_011281189.2
A3	102564824	102564997
refGene_gn=CAPG;refGene_tx=XM_003984204.3;refGene_gn=CAPG;refGene_tx=XM_006930250.3;refGene_gn=CAPG;refGene_tx=XM_006930251.4;refGene_gn=CAPG;refGene_tx=XM_006930252.3;refGene_gn=CAPG;refGene_tx=XM_006930253.4;refGene_gn=CAPG;refGene_tx=XM_011281188.2;refGene_gn=CAPG;refGene_tx=XM_011281189.2
A3	102565174	102565197

refGene_gn=CAPG;refGene_tx=XM_003984204.3;refGene_gn=CAPG;refGene_tx=XM_006930250.3;refGene_gn=CAPG;refGene_tx=XM_006930251.4;refGene_gn=CAPG;refGene_tx=XM_006930252.3;refGene_gn=CAPG;refGene_tx=XM_006930253.4;refGene_gn=CAPG;refGene_tx=XM_011281188.2;refGene_gn=CAPG;refGene_tx=XM_011281189.2
A3 102594847 102594850
refGene_gn=SH2D6;refGene_tx=XM_023251829.1
A3 102595295 102595356
refGene_gn=SH2D6;refGene_tx=XM_023251829.1
A3 102595725 102595804
refGene_gn=SH2D6;refGene_tx=XM_023251829.1
A3 102595982 102596013
refGene_gn=SH2D6;refGene_tx=XM_023251829.1
A3 102596138 102596163
refGene_gn=SH2D6;refGene_tx=XM_023251829.1
A3 102596313 102596391
refGene_gn=SH2D6;refGene_tx=XM_023251829.1
A3 102596566 102596625
refGene_gn=SH2D6;refGene_tx=XM_023251829.1
A3 102596901 102596985
refGene_gn=SH2D6;refGene_tx=XM_023251829.1
A3 102597154 102597313
refGene_gn=SH2D6;refGene_tx=XM_023251829.1
A3 102597935 102598052
refGene_gn=SH2D6;refGene_tx=XM_023251829.1
A3 102677622 102677713
refGene_gn=MAT2A;refGene_tx=XM_003984206.5
A3 102679651 102679729
refGene_gn=MAT2A;refGene_tx=XM_003984206.5
A3 102679826 102679949
refGene_gn=MAT2A;refGene_tx=XM_003984206.5
A3 102680354 102680467
refGene_gn=MAT2A;refGene_tx=XM_003984206.5
A3 102680554 102680698
refGene_gn=MAT2A;refGene_tx=XM_003984206.5
A3 102680866 102681085
refGene_gn=MAT2A;refGene_tx=XM_003984206.5
A3 102681309 102681492
refGene_gn=MAT2A;refGene_tx=XM_003984206.5
A3 102681639 102681773
refGene_gn=MAT2A;refGene_tx=XM_003984206.5
A3 102682470 102682573
refGene_gn=MAT2A;refGene_tx=XM_003984206.5
A3 102687239 102687432
refGene_gn=GGCX;refGene_tx=XM_011281186.3;refGene_gn=GGCX;refGene_tx=XM_011281187.3;refGene_gn=GGCX;refGene_tx=XM_019827517.2;refGene_gn=GGCX;refGene_tx=XM_023251830.1
A3 102687866 102688062
refGene_gn=GGCX;refGene_tx=XM_011281186.3;refGene_gn=GGCX;refGene_tx=XM_011281187.3;refGene_gn=GGCX;refGene_tx=XM_019827517.2;refGene_gn=GGC

X;refGene_tx=XM_023251830.1
A3 102688228 102688376
refGene_gn=GGCX;refGene_tx=XM_011281186.3;refGene_gn=GGCX;refGene_tx=XM_011281187.3;refGene_gn=GGCX;refGene_tx=XM_019827517.2;refGene_gn=GGCX;refGene_tx=XM_023251830.1
A3 102688657 102688788
refGene_gn=GGCX;refGene_tx=XM_011281186.3;refGene_gn=GGCX;refGene_tx=XM_011281187.3;refGene_gn=GGCX;refGene_tx=XM_019827517.2;refGene_gn=GGCX;refGene_tx=XM_023251830.1
A3 102688979 102689149
refGene_gn=GGCX;refGene_tx=XM_011281186.3;refGene_gn=GGCX;refGene_tx=XM_011281187.3;refGene_gn=GGCX;refGene_tx=XM_019827517.2;refGene_gn=GGCX;refGene_tx=XM_023251830.1
A3 102689606 102689758
refGene_gn=GGCX;refGene_tx=XM_011281186.3;refGene_gn=GGCX;refGene_tx=XM_011281187.3;refGene_gn=GGCX;refGene_tx=XM_019827517.2;refGene_gn=GGCX;refGene_tx=XM_023251830.1
A3 102690145 102690277
refGene_gn=GGCX;refGene_tx=XM_011281186.3;refGene_gn=GGCX;refGene_tx=XM_011281187.3;refGene_gn=GGCX;refGene_tx=XM_019827517.2;refGene_gn=GGCX;refGene_tx=XM_023251830.1
A3 102690406 102690672
refGene_gn=GGCX;refGene_tx=XM_011281186.3;refGene_gn=GGCX;refGene_tx=XM_011281187.3;refGene_gn=GGCX;refGene_tx=XM_019827517.2;refGene_gn=GGCX;refGene_tx=XM_023251830.1
A3 102691027 102691191
refGene_gn=GGCX;refGene_tx=XM_011281186.3;refGene_gn=GGCX;refGene_tx=XM_011281187.3;refGene_gn=GGCX;refGene_tx=XM_019827517.2;refGene_gn=GGCX;refGene_tx=XM_023251830.1
A3 102692979 102693086
refGene_gn=GGCX;refGene_tx=XM_011281186.3;refGene_gn=GGCX;refGene_tx=XM_011281187.3;refGene_gn=GGCX;refGene_tx=XM_019827517.2;refGene_gn=GGCX;refGene_tx=XM_023251830.1
A3 102693521 102693600
refGene_gn=GGCX;refGene_tx=XM_011281186.3;refGene_gn=GGCX;refGene_tx=XM_011281187.3;refGene_gn=GGCX;refGene_tx=XM_019827517.2;refGene_gn=GGCX;refGene_tx=XM_023251830.1
A3 102694757 102695047
refGene_gn=GGCX;refGene_tx=XM_011281187.3;refGene_gn=GGCX;refGene_tx=XM_019827517.2
A3 102695240 102695406
refGene_gn=GGCX;refGene_tx=XM_011281186.3;refGene_gn=GGCX;refGene_tx=XM_011281187.3;refGene_gn=GGCX;refGene_tx=XM_023251830.1
A3 102695733 102695892
refGene_gn=GGCX;refGene_tx=XM_011281186.3;refGene_gn=GGCX;refGene_tx=XM_023251830.1
A3 102697008 102697179
refGene_gn=GGCX;refGene_tx=XM_011281186.3;refGene_gn=GGCX;refGene_tx=XM_023251830.1
A3 102697560 102697603

refGene_gn=GGCX;refGene_tx=XM_011281186.3
A3 102707604 102707607
refGene_gn=VAMP8;refGene_tx=XM_006930255.4
A3 102708954 102709113
refGene_gn=VAMP8;refGene_tx=XM_006930255.4
A3 102710362 102710506
refGene_gn=VAMP8;refGene_tx=XM_006930255.4
A3 102712856 102712859
refGene_gn=VAMP5;refGene_tx=XM_019827521.2;refGene_gn=VAMP5;refGene_tx=XM_019827522.2
A3 102717519 102717651
refGene_gn=VAMP5;refGene_tx=XM_019827520.2;refGene_gn=VAMP5;refGene_tx=XM_019827521.2
A3 102718373 102718511
refGene_gn=VAMP5;refGene_tx=XM_019827520.2;refGene_gn=VAMP5;refGene_tx=XM_019827521.2;refGene_gn=VAMP5;refGene_tx=XM_019827522.2
A3 102718977 102719187
refGene_gn=VAMP5;refGene_tx=XM_019827520.2;refGene_gn=VAMP5;refGene_tx=XM_019827521.2;refGene_gn=VAMP5;refGene_tx=XM_019827522.2
A3 102721692 102721778
refGene_gn=RNF181;refGene_tx=XM_003984209.4
A3 102722401 102722532
refGene_gn=RNF181;refGene_tx=XM_003984209.4
A3 102722711 102722821
refGene_gn=RNF181;refGene_tx=XM_003984209.4
A3 102723006 102723081
refGene_gn=RNF181;refGene_tx=XM_003984209.4
A3 102723348 102723408
refGene_gn=RNF181;refGene_tx=XM_003984209.4
A3 102724750 102724992
refGene_gn=TMEM150A;refGene_tx=XM_003984210.5;refGene_gn=TMEM150A;refGene_tx=XM_019827518.2;refGene_gn=TMEM150A;refGene_tx=XM_023251832.1;refGene_gn=TMEM150A;refGene_tx=XM_023251833.1
A3 102725256 102725434
refGene_gn=TMEM150A;refGene_tx=XM_003984210.5;refGene_gn=TMEM150A;refGene_tx=XM_019827518.2;refGene_gn=TMEM150A;refGene_tx=XM_023251832.1;refGene_gn=TMEM150A;refGene_tx=XM_023251833.1
A3 102725618 102725746
refGene_gn=TMEM150A;refGene_tx=XM_003984210.5;refGene_gn=TMEM150A;refGene_tx=XM_019827518.2;refGene_gn=TMEM150A;refGene_tx=XM_023251832.1;refGene_gn=TMEM150A;refGene_tx=XM_023251833.1
A3 102726104 102726172
refGene_gn=TMEM150A;refGene_tx=XM_003984210.5;refGene_gn=TMEM150A;refGene_tx=XM_019827518.2;refGene_gn=TMEM150A;refGene_tx=XM_023251832.1;refGene_gn=TMEM150A;refGene_tx=XM_023251833.1
A3 102726775 102726862
refGene_gn=TMEM150A;refGene_tx=XM_003984210.5;refGene_gn=TMEM150A;refGene_tx=XM_019827518.2;refGene_gn=TMEM150A;refGene_tx=XM_023251832.1;refGene_gn=TMEM150A;refGene_tx=XM_023251833.1
A3 102727199 102727247

refGene_gn=TMEM150A;refGene_tx=XM_003984210.5;refGene_gn=TMEM150A;refGene_tx=XM_019827518.2;refGene_gn=TMEM150A;refGene_tx=XM_023251832.1;refGene_gn=TMEM150A;refGene_tx=XM_023251833.1
A3 102727693 102727758
refGene_gn=TMEM150A;refGene_tx=XM_003984210.5;refGene_gn=TMEM150A;refGene_tx=XM_023251832.1
A3 102734120 102734243
refGene_gn=CA3H2orf68;refGene_tx=XM_003984211.5
A3 102734608 102734760
refGene_gn=CA3H2orf68;refGene_tx=XM_003984211.5
A3 102735725 102735844
refGene_gn=CA3H2orf68;refGene_tx=XM_003984211.5
A3 102735922 102736029
refGene_gn=CA3H2orf68;refGene_tx=XM_003984211.5
A3 102737162 102737430
refGene_gn=USP39;refGene_tx=XM_003984212.4
A3 102741997 102742067
refGene_gn=USP39;refGene_tx=XM_003984212.4
A3 102743829 102743924
refGene_gn=USP39;refGene_tx=XM_003984212.4
A3 102744953 102745090
refGene_gn=USP39;refGene_tx=XM_003984212.4
A3 102746320 102746473
refGene_gn=USP39;refGene_tx=XM_003984212.4
A3 102752538 102752764
refGene_gn=USP39;refGene_tx=XM_003984212.4
A3 102754562 102754640
refGene_gn=USP39;refGene_tx=XM_003984212.4
A3 102755850 102755918
refGene_gn=USP39;refGene_tx=XM_003984212.4
A3 102756636 102756825
refGene_gn=USP39;refGene_tx=XM_003984212.4
A3 102760722 102760865
refGene_gn=USP39;refGene_tx=XM_003984212.4
A3 102766612 102766748
refGene_gn=USP39;refGene_tx=XM_003984212.4
A3 102769186 102769273
refGene_gn=USP39;refGene_tx=XM_003984212.4
A3 102770347 102770395
refGene_gn=USP39;refGene_tx=XM_003984212.4
A3 102780701 102780764
refGene_gn=SFTPB;refGene_tx=XM_011281183.3;refGene_gn=SFTPB;refGene_tx=XM_019827523.2;refGene_gn=SFTPB;refGene_tx=XM_019827524.2;refGene_gn=SFTPB;refGene_tx=XM_019827525.2
A3 102781284 102781365
refGene_gn=SFTPB;refGene_tx=XM_011281183.3;refGene_gn=SFTPB;refGene_tx=XM_019827523.2;refGene_gn=SFTPB;refGene_tx=XM_019827524.2;refGene_gn=SFTPB;refGene_tx=XM_019827525.2
A3 102782248 102782409
refGene_gn=SFTPB;refGene_tx=XM_011281183.3;refGene_gn=SFTPB;refGene_tx

=XM_019827523.2;refGene_gn=SFTPB;refGene_tx=XM_019827524.2;refGene_gn=SFTPB;refGene_tx=XM_019827525.2
A3 102782566 102782750
refGene_gn=SFTPB;refGene_tx=XM_011281183.3;refGene_gn=SFTPB;refGene_tx=XM_019827523.2;refGene_gn=SFTPB;refGene_tx=XM_019827524.2;refGene_gn=SFTPB;refGene_tx=XM_019827525.2
A3 102785315 102785405
refGene_gn=SFTPB;refGene_tx=XM_011281183.3;refGene_gn=SFTPB;refGene_tx=XM_019827523.2;refGene_gn=SFTPB;refGene_tx=XM_019827524.2;refGene_gn=SFTPB;refGene_tx=XM_019827525.2
A3 102785652 102785805
refGene_gn=SFTPB;refGene_tx=XM_011281183.3;refGene_gn=SFTPB;refGene_tx=XM_019827523.2;refGene_gn=SFTPB;refGene_tx=XM_019827524.2;refGene_gn=SFTPB;refGene_tx=XM_019827525.2
A3 102786382 102786508
refGene_gn=SFTPB;refGene_tx=XM_011281183.3;refGene_gn=SFTPB;refGene_tx=XM_019827523.2;refGene_gn=SFTPB;refGene_tx=XM_019827524.2;refGene_gn=SFTPB;refGene_tx=XM_019827525.2
A3 102786897 102786969
refGene_gn=SFTPB;refGene_tx=XM_011281183.3;refGene_gn=SFTPB;refGene_tx=XM_019827523.2;refGene_gn=SFTPB;refGene_tx=XM_019827524.2;refGene_gn=SFTPB;refGene_tx=XM_019827525.2
A3 102787318 102787446
refGene_gn=SFTPB;refGene_tx=XM_011281183.3;refGene_gn=SFTPB;refGene_tx=XM_019827523.2;refGene_gn=SFTPB;refGene_tx=XM_019827524.2;refGene_gn=SFTPB;refGene_tx=XM_019827525.2
A3 102787771 102787835
refGene_gn=SFTPB;refGene_tx=XM_011281183.3;refGene_gn=SFTPB;refGene_tx=XM_019827523.2;refGene_gn=SFTPB;refGene_tx=XM_019827524.2;refGene_gn=SFTPB;refGene_tx=XM_019827525.2
A3 102906496 102907279
refGene_gn=ATOH8;refGene_tx=XM_011281182.2
A3 102916187 102916379
refGene_gn=ATOH8;refGene_tx=XM_011281182.2
A3 102937415 102937421
refGene_gn=ATOH8;refGene_tx=XM_011281182.2
A3 102983532 102983781
refGene_gn=ST3GAL5;refGene_tx=XM_019827531.2;refGene_gn=ST3GAL5;refGene_tx=XM_023251836.1;refGene_gn=ST3GAL5;refGene_tx=XM_023251837.1;refGene_gn=ST3GAL5;refGene_tx=XM_023251838.1;refGene_gn=ST3GAL5;refGene_tx=XM_023251839.1
A3 102988162 102988321
refGene_gn=ST3GAL5;refGene_tx=XM_019827531.2;refGene_gn=ST3GAL5;refGene_tx=XM_023251836.1;refGene_gn=ST3GAL5;refGene_tx=XM_023251837.1;refGene_gn=ST3GAL5;refGene_tx=XM_023251838.1
A3 102990047 102990234
refGene_gn=ST3GAL5;refGene_tx=XM_019827531.2;refGene_gn=ST3GAL5;refGene_tx=XM_023251836.1;refGene_gn=ST3GAL5;refGene_tx=XM_023251837.1;refGene_gn=ST3GAL5;refGene_tx=XM_023251838.1;refGene_gn=ST3GAL5;refGene_tx=XM_023251839.1

A3	102993988	102994353
refGene_gn=ST3GAL5;refGene_tx=XM_019827531.2;refGene_gn=ST3GAL5;refGene_tx=XM_023251836.1;refGene_gn=ST3GAL5;refGene_tx=XM_023251837.1;refGene_gn=ST3GAL5;refGene_tx=XM_023251838.1;refGene_gn=ST3GAL5;refGene_tx=XM_023251839.1
A3	102998835	102998940
refGene_gn=ST3GAL5;refGene_tx=XM_019827531.2;refGene_gn=ST3GAL5;refGene_tx=XM_023251836.1;refGene_gn=ST3GAL5;refGene_tx=XM_023251839.1
A3	103003985	103004097
refGene_gn=ST3GAL5;refGene_tx=XM_019827531.2;refGene_gn=ST3GAL5;refGene_tx=XM_023251836.1;refGene_gn=ST3GAL5;refGene_tx=XM_023251837.1;refGene_gn=ST3GAL5;refGene_tx=XM_023251838.1;refGene_gn=ST3GAL5;refGene_tx=XM_023251839.1
A3	103006877	103007001
refGene_gn=ST3GAL5;refGene_tx=XM_019827531.2;refGene_gn=ST3GAL5;refGene_tx=XM_023251836.1;refGene_gn=ST3GAL5;refGene_tx=XM_023251837.1;refGene_gn=ST3GAL5;refGene_tx=XM_023251838.1;refGene_gn=ST3GAL5;refGene_tx=XM_023251839.1
A3	103032259	103032347
refGene_gn=ST3GAL5;refGene_tx=XM_023251836.1;refGene_gn=ST3GAL5;refGene_tx=XM_023251837.1;refGene_gn=ST3GAL5;refGene_tx=XM_023251838.1;refGene_gn=ST3GAL5;refGene_tx=XM_023251839.1
A3	103123899	103124000
refGene_gn=POLR1A;refGene_tx=XM_023251840.1;refGene_gn=POLR1A;refGene_tx=XM_023251841.1;refGene_gn=POLR1A;refGene_tx=XM_023251842.1;refGene_gn=POLR1A;refGene_tx=XM_023251843.1;refGene_gn=POLR1A;refGene_tx=XM_023251844.1;refGene_gn=POLR1A;refGene_tx=XM_023251845.1
A3	103124374	103124539
refGene_gn=POLR1A;refGene_tx=XM_023251840.1;refGene_gn=POLR1A;refGene_tx=XM_023251841.1;refGene_gn=POLR1A;refGene_tx=XM_023251842.1;refGene_gn=POLR1A;refGene_tx=XM_023251843.1;refGene_gn=POLR1A;refGene_tx=XM_023251844.1;refGene_gn=POLR1A;refGene_tx=XM_023251845.1
A3	103125032	103125150
refGene_gn=POLR1A;refGene_tx=XM_023251840.1;refGene_gn=POLR1A;refGene_tx=XM_023251841.1;refGene_gn=POLR1A;refGene_tx=XM_023251842.1;refGene_gn=POLR1A;refGene_tx=XM_023251843.1;refGene_gn=POLR1A;refGene_tx=XM_023251844.1;refGene_gn=POLR1A;refGene_tx=XM_023251845.1
A3	103126285	103126486
refGene_gn=POLR1A;refGene_tx=XM_023251840.1;refGene_gn=POLR1A;refGene_tx=XM_023251841.1;refGene_gn=POLR1A;refGene_tx=XM_023251842.1;refGene_gn=POLR1A;refGene_tx=XM_023251843.1;refGene_gn=POLR1A;refGene_tx=XM_023251844.1;refGene_gn=POLR1A;refGene_tx=XM_023251845.1
A3	103127326	103127626
refGene_gn=POLR1A;refGene_tx=XM_023251840.1;refGene_gn=POLR1A;refGene_tx=XM_023251841.1;refGene_gn=POLR1A;refGene_tx=XM_023251842.1;refGene_gn=POLR1A;refGene_tx=XM_023251843.1;refGene_gn=POLR1A;refGene_tx=XM_023251844.1;refGene_gn=POLR1A;refGene_tx=XM_023251845.1
A3	103128208	103128319
refGene_gn=POLR1A;refGene_tx=XM_023251840.1;refGene_gn=POLR1A;refGene_tx=XM_023251841.1;refGene_gn=POLR1A;refGene_tx=XM_023251842.1;refGene_

gn=POLR1A;refGene_tx=XM_023251843.1;refGene_gn=POLR1A;refGene_tx=XM_023251844.1;refGene_gn=POLR1A;refGene_tx=XM_023251845.1
A3 103129871 103129998
refGene_gn=POLR1A;refGene_tx=XM_023251840.1;refGene_gn=POLR1A;refGene_tx=XM_023251841.1;refGene_gn=POLR1A;refGene_tx=XM_023251842.1;refGene_gn=POLR1A;refGene_tx=XM_023251843.1;refGene_gn=POLR1A;refGene_tx=XM_023251844.1;refGene_gn=POLR1A;refGene_tx=XM_023251845.1
A3 103134521 103134679
refGene_gn=POLR1A;refGene_tx=XM_023251840.1;refGene_gn=POLR1A;refGene_tx=XM_023251841.1;refGene_gn=POLR1A;refGene_tx=XM_023251842.1;refGene_gn=POLR1A;refGene_tx=XM_023251843.1;refGene_gn=POLR1A;refGene_tx=XM_023251844.1;refGene_gn=POLR1A;refGene_tx=XM_023251845.1
A3 103135160 103135296
refGene_gn=POLR1A;refGene_tx=XM_023251840.1;refGene_gn=POLR1A;refGene_tx=XM_023251841.1;refGene_gn=POLR1A;refGene_tx=XM_023251842.1;refGene_gn=POLR1A;refGene_tx=XM_023251843.1;refGene_gn=POLR1A;refGene_tx=XM_023251844.1;refGene_gn=POLR1A;refGene_tx=XM_023251845.1
A3 103136220 103136388
refGene_gn=POLR1A;refGene_tx=XM_023251840.1;refGene_gn=POLR1A;refGene_tx=XM_023251841.1;refGene_gn=POLR1A;refGene_tx=XM_023251842.1;refGene_gn=POLR1A;refGene_tx=XM_023251843.1;refGene_gn=POLR1A;refGene_tx=XM_023251844.1;refGene_gn=POLR1A;refGene_tx=XM_023251845.1
A3 103138341 103138556
refGene_gn=POLR1A;refGene_tx=XM_023251840.1;refGene_gn=POLR1A;refGene_tx=XM_023251841.1;refGene_gn=POLR1A;refGene_tx=XM_023251842.1;refGene_gn=POLR1A;refGene_tx=XM_023251843.1;refGene_gn=POLR1A;refGene_tx=XM_023251844.1;refGene_gn=POLR1A;refGene_tx=XM_023251845.1
A3 103139424 103139646
refGene_gn=POLR1A;refGene_tx=XM_023251840.1;refGene_gn=POLR1A;refGene_tx=XM_023251841.1;refGene_gn=POLR1A;refGene_tx=XM_023251842.1;refGene_gn=POLR1A;refGene_tx=XM_023251843.1;refGene_gn=POLR1A;refGene_tx=XM_023251844.1;refGene_gn=POLR1A;refGene_tx=XM_023251845.1
A3 103140562 103140728
refGene_gn=POLR1A;refGene_tx=XM_023251840.1;refGene_gn=POLR1A;refGene_tx=XM_023251841.1;refGene_gn=POLR1A;refGene_tx=XM_023251842.1;refGene_gn=POLR1A;refGene_tx=XM_023251843.1;refGene_gn=POLR1A;refGene_tx=XM_023251844.1;refGene_gn=POLR1A;refGene_tx=XM_023251845.1
A3 103142700 103142783
refGene_gn=POLR1A;refGene_tx=XM_023251840.1;refGene_gn=POLR1A;refGene_tx=XM_023251841.1;refGene_gn=POLR1A;refGene_tx=XM_023251842.1;refGene_gn=POLR1A;refGene_tx=XM_023251843.1;refGene_gn=POLR1A;refGene_tx=XM_023251844.1;refGene_gn=POLR1A;refGene_tx=XM_023251845.1
A3 103144100 103144253
refGene_gn=POLR1A;refGene_tx=XM_023251840.1;refGene_gn=POLR1A;refGene_tx=XM_023251841.1;refGene_gn=POLR1A;refGene_tx=XM_023251842.1;refGene_gn=POLR1A;refGene_tx=XM_023251843.1;refGene_gn=POLR1A;refGene_tx=XM_023251844.1;refGene_gn=POLR1A;refGene_tx=XM_023251845.1
A3 103145119 103145218
refGene_gn=POLR1A;refGene_tx=XM_023251840.1;refGene_gn=POLR1A;refGene_tx=XM_023251841.1;refGene_gn=POLR1A;refGene_tx=XM_023251842.1;refGene_

gn=POLR1A;refGene_tx=XM_023251843.1;refGene_gn=POLR1A;refGene_tx=XM_023251844.1;refGene_gn=POLR1A;refGene_tx=XM_023251845.1
A3 103146862 103147021
refGene_gn=POLR1A;refGene_tx=XM_023251840.1;refGene_gn=POLR1A;refGene_tx=XM_023251841.1;refGene_gn=POLR1A;refGene_tx=XM_023251842.1;refGene_gn=POLR1A;refGene_tx=XM_023251843.1;refGene_gn=POLR1A;refGene_tx=XM_023251844.1;refGene_gn=POLR1A;refGene_tx=XM_023251845.1
A3 103147137 103147220
refGene_gn=POLR1A;refGene_tx=XM_023251840.1;refGene_gn=POLR1A;refGene_tx=XM_023251841.1;refGene_gn=POLR1A;refGene_tx=XM_023251842.1;refGene_gn=POLR1A;refGene_tx=XM_023251843.1;refGene_gn=POLR1A;refGene_tx=XM_023251844.1;refGene_gn=POLR1A;refGene_tx=XM_023251845.1
A3 103149779 103149963
refGene_gn=POLR1A;refGene_tx=XM_023251840.1;refGene_gn=POLR1A;refGene_tx=XM_023251841.1;refGene_gn=POLR1A;refGene_tx=XM_023251842.1;refGene_gn=POLR1A;refGene_tx=XM_023251843.1;refGene_gn=POLR1A;refGene_tx=XM_023251844.1;refGene_gn=POLR1A;refGene_tx=XM_023251845.1
A3 103151280 103151430
refGene_gn=POLR1A;refGene_tx=XM_023251840.1;refGene_gn=POLR1A;refGene_tx=XM_023251841.1;refGene_gn=POLR1A;refGene_tx=XM_023251842.1;refGene_gn=POLR1A;refGene_tx=XM_023251843.1;refGene_gn=POLR1A;refGene_tx=XM_023251844.1;refGene_gn=POLR1A;refGene_tx=XM_023251845.1
A3 103163870 103164062
refGene_gn=POLR1A;refGene_tx=XM_023251840.1;refGene_gn=POLR1A;refGene_tx=XM_023251841.1;refGene_gn=POLR1A;refGene_tx=XM_023251842.1;refGene_gn=POLR1A;refGene_tx=XM_023251843.1;refGene_gn=POLR1A;refGene_tx=XM_023251844.1;refGene_gn=POLR1A;refGene_tx=XM_023251845.1
A3 103168668 103168923
refGene_gn=POLR1A;refGene_tx=XM_023251840.1;refGene_gn=POLR1A;refGene_tx=XM_023251841.1;refGene_gn=POLR1A;refGene_tx=XM_023251842.1;refGene_gn=POLR1A;refGene_tx=XM_023251843.1;refGene_gn=POLR1A;refGene_tx=XM_023251844.1;refGene_gn=POLR1A;refGene_tx=XM_023251845.1
A3 103175816 103176047
refGene_gn=POLR1A;refGene_tx=XM_023251840.1;refGene_gn=POLR1A;refGene_tx=XM_023251841.1;refGene_gn=POLR1A;refGene_tx=XM_023251842.1;refGene_gn=POLR1A;refGene_tx=XM_023251843.1;refGene_gn=POLR1A;refGene_tx=XM_023251844.1;refGene_gn=POLR1A;refGene_tx=XM_023251845.1
A3 103178306 103178429
refGene_gn=POLR1A;refGene_tx=XM_023251840.1;refGene_gn=POLR1A;refGene_tx=XM_023251841.1;refGene_gn=POLR1A;refGene_tx=XM_023251842.1;refGene_gn=POLR1A;refGene_tx=XM_023251843.1;refGene_gn=POLR1A;refGene_tx=XM_023251844.1;refGene_gn=POLR1A;refGene_tx=XM_023251845.1
A3 103178567 103178738
refGene_gn=POLR1A;refGene_tx=XM_023251840.1;refGene_gn=POLR1A;refGene_tx=XM_023251841.1;refGene_gn=POLR1A;refGene_tx=XM_023251842.1;refGene_gn=POLR1A;refGene_tx=XM_023251843.1;refGene_gn=POLR1A;refGene_tx=XM_023251844.1;refGene_gn=POLR1A;refGene_tx=XM_023251845.1
A3 103181405 103181568
refGene_gn=POLR1A;refGene_tx=XM_023251840.1;refGene_gn=POLR1A;refGene_tx=XM_023251841.1;refGene_gn=POLR1A;refGene_tx=XM_023251842.1;refGene_

gn=POLR1A;refGene_tx=XM_023251843.1;refGene_gn=POLR1A;refGene_tx=XM_023251844.1;refGene_gn=POLR1A;refGene_tx=XM_023251845.1
A3 103181851 103181957
refGene_gn=POLR1A;refGene_tx=XM_023251840.1;refGene_gn=POLR1A;refGene_tx=XM_023251841.1;refGene_gn=POLR1A;refGene_tx=XM_023251842.1;refGene_gn=POLR1A;refGene_tx=XM_023251843.1;refGene_gn=POLR1A;refGene_tx=XM_023251844.1;refGene_gn=POLR1A;refGene_tx=XM_023251845.1
A3 103183529 103183616
refGene_gn=POLR1A;refGene_tx=XM_023251840.1;refGene_gn=POLR1A;refGene_tx=XM_023251841.1;refGene_gn=POLR1A;refGene_tx=XM_023251842.1;refGene_gn=POLR1A;refGene_tx=XM_023251843.1;refGene_gn=POLR1A;refGene_tx=XM_023251844.1;refGene_gn=POLR1A;refGene_tx=XM_023251845.1
A3 103189587 103189691
refGene_gn=POLR1A;refGene_tx=XM_023251840.1;refGene_gn=POLR1A;refGene_tx=XM_023251841.1;refGene_gn=POLR1A;refGene_tx=XM_023251842.1;refGene_gn=POLR1A;refGene_tx=XM_023251843.1;refGene_gn=POLR1A;refGene_tx=XM_023251844.1;refGene_gn=POLR1A;refGene_tx=XM_023251845.1
A3 103189811 103189897
refGene_gn=POLR1A;refGene_tx=XM_023251840.1;refGene_gn=POLR1A;refGene_tx=XM_023251841.1;refGene_gn=POLR1A;refGene_tx=XM_023251842.1;refGene_gn=POLR1A;refGene_tx=XM_023251843.1;refGene_gn=POLR1A;refGene_tx=XM_023251844.1;refGene_gn=POLR1A;refGene_tx=XM_023251845.1
A3 103190938 103191046
refGene_gn=POLR1A;refGene_tx=XM_023251840.1;refGene_gn=POLR1A;refGene_tx=XM_023251841.1;refGene_gn=POLR1A;refGene_tx=XM_023251842.1;refGene_gn=POLR1A;refGene_tx=XM_023251843.1;refGene_gn=POLR1A;refGene_tx=XM_023251844.1;refGene_gn=POLR1A;refGene_tx=XM_023251845.1
A3 103194038 103194188
refGene_gn=POLR1A;refGene_tx=XM_023251840.1;refGene_gn=POLR1A;refGene_tx=XM_023251841.1;refGene_gn=POLR1A;refGene_tx=XM_023251842.1;refGene_gn=POLR1A;refGene_tx=XM_023251843.1;refGene_gn=POLR1A;refGene_tx=XM_023251844.1;refGene_gn=POLR1A;refGene_tx=XM_023251845.1
A3 103195299 103195504
refGene_gn=POLR1A;refGene_tx=XM_023251840.1;refGene_gn=POLR1A;refGene_tx=XM_023251841.1;refGene_gn=POLR1A;refGene_tx=XM_023251842.1;refGene_gn=POLR1A;refGene_tx=XM_023251843.1;refGene_gn=POLR1A;refGene_tx=XM_023251844.1;refGene_gn=POLR1A;refGene_tx=XM_023251845.1
A3 103198144 103198221
refGene_gn=POLR1A;refGene_tx=XM_023251840.1;refGene_gn=POLR1A;refGene_tx=XM_023251841.1;refGene_gn=POLR1A;refGene_tx=XM_023251842.1;refGene_gn=POLR1A;refGene_tx=XM_023251843.1;refGene_gn=POLR1A;refGene_tx=XM_023251844.1
A3 103198426 103198443
refGene_gn=POLR1A;refGene_tx=XM_023251845.1
A3 103198655 103198780
refGene_gn=PTCD3;refGene_tx=XM_023251846.1;refGene_gn=PTCD3;refGene_tx=XM_023251847.1
A3 103200950 103201003
refGene_gn=PTCD3;refGene_tx=XM_023251846.1;refGene_gn=PTCD3;refGene_tx=XM_023251847.1

A3	103201121	103201158
refGene_gn=PTCD3;refGene_tx=XM_023251846.1;refGene_gn=PTCD3;refGene_tx
=XM_023251847.1
A3	103203312	103203358
refGene_gn=PTCD3;refGene_tx=XM_023251846.1;refGene_gn=PTCD3;refGene_tx
=XM_023251847.1
A3	103207160	103207229
refGene_gn=PTCD3;refGene_tx=XM_023251846.1;refGene_gn=PTCD3;refGene_tx
=XM_023251847.1
A3	103207704	103207809
refGene_gn=PTCD3;refGene_tx=XM_023251846.1;refGene_gn=PTCD3;refGene_tx
=XM_023251847.1
A3	103209763	103209887
refGene_gn=PTCD3;refGene_tx=XM_023251846.1;refGene_gn=PTCD3;refGene_tx
=XM_023251847.1
A3	103211687	103211803
refGene_gn=PTCD3;refGene_tx=XM_023251846.1;refGene_gn=PTCD3;refGene_tx
=XM_023251847.1
A3	103213270	103213338
refGene_gn=PTCD3;refGene_tx=XM_023251846.1;refGene_gn=PTCD3;refGene_tx
=XM_023251847.1
A3	103214712	103214800
refGene_gn=PTCD3;refGene_tx=XM_023251846.1;refGene_gn=PTCD3;refGene_tx
=XM_023251847.1
A3	103215198	103215259
refGene_gn=PTCD3;refGene_tx=XM_023251846.1;refGene_gn=PTCD3;refGene_tx
=XM_023251847.1
A3	103215555	103215641
refGene_gn=PTCD3;refGene_tx=XM_023251846.1;refGene_gn=PTCD3;refGene_tx
=XM_023251847.1
A3	103216315	103216460
refGene_gn=PTCD3;refGene_tx=XM_023251846.1;refGene_gn=PTCD3;refGene_tx
=XM_023251847.1
A3	103217372	103217423
refGene_gn=PTCD3;refGene_tx=XM_023251846.1;refGene_gn=PTCD3;refGene_tx
=XM_023251847.1
A3	103219189	103219279
refGene_gn=PTCD3;refGene_tx=XM_023251846.1;refGene_gn=PTCD3;refGene_tx
=XM_023251847.1
A3	103219743	103219772
refGene_gn=PTCD3;refGene_tx=XM_023251846.1;refGene_gn=PTCD3;refGene_tx
=XM_023251847.1
A3	103220846	103220953
refGene_gn=PTCD3;refGene_tx=XM_023251846.1;refGene_gn=PTCD3;refGene_tx
=XM_023251847.1
A3	103221717	103221796
refGene_gn=PTCD3;refGene_tx=XM_023251846.1;refGene_gn=PTCD3;refGene_tx
=XM_023251847.1
A3	103221889	103221980
refGene_gn=PTCD3;refGene_tx=XM_023251846.1;refGene_gn=PTCD3;refGene_tx

=XM_023251847.1
A3	103222853	103222939
refGene_gn=PTCD3;refGene_tx=XM_023251846.1;refGene_gn=PTCD3;refGene_tx=XM_023251847.1
A3	103223456	103223605
refGene_gn=PTCD3;refGene_tx=XM_023251846.1;refGene_gn=PTCD3;refGene_tx=XM_023251847.1
A3	103225027	103225069
refGene_gn=PTCD3;refGene_tx=XM_023251846.1
A3	103225475	103225634
refGene_gn=PTCD3;refGene_tx=XM_023251846.1;refGene_gn=PTCD3;refGene_tx=XM_023251847.1
A3	103225910	103225983
refGene_gn=PTCD3;refGene_tx=XM_023251846.1
A3	103231954	103232568
refGene_gn=IMMT;refGene_tx=XM_006930260.4;refGene_gn=IMMT;refGene_tx=XM_006930262.4;refGene_gn=IMMT;refGene_tx=XM_006930264.4;refGene_gn=IMMT;refGene_tx=XM_011281181.3;refGene_gn=IMMT;refGene_tx=XM_019827536.2;refGene_gn=IMMT;refGene_tx=XM_019827537.2
A3	103233565	103233695
refGene_gn=IMMT;refGene_tx=XM_006930260.4;refGene_gn=IMMT;refGene_tx=XM_006930262.4;refGene_gn=IMMT;refGene_tx=XM_006930264.4;refGene_gn=IMMT;refGene_tx=XM_011281181.3;refGene_gn=IMMT;refGene_tx=XM_019827536.2;refGene_gn=IMMT;refGene_tx=XM_019827537.2
A3	103234407	103234539
refGene_gn=IMMT;refGene_tx=XM_006930260.4;refGene_gn=IMMT;refGene_tx=XM_006930262.4;refGene_gn=IMMT;refGene_tx=XM_006930264.4;refGene_gn=IMMT;refGene_tx=XM_011281181.3;refGene_gn=IMMT;refGene_tx=XM_019827536.2;refGene_gn=IMMT;refGene_tx=XM_019827537.2
A3	103236184	103236408
refGene_gn=IMMT;refGene_tx=XM_006930260.4;refGene_gn=IMMT;refGene_tx=XM_006930262.4;refGene_gn=IMMT;refGene_tx=XM_006930264.4;refGene_gn=IMMT;refGene_tx=XM_011281181.3;refGene_gn=IMMT;refGene_tx=XM_019827536.2;refGene_gn=IMMT;refGene_tx=XM_019827537.2
A3	103237733	103237748
refGene_gn=IMMT;refGene_tx=XM_006930260.4;refGene_gn=IMMT;refGene_tx=XM_006930262.4;refGene_gn=IMMT;refGene_tx=XM_006930264.4;refGene_gn=IMMT;refGene_tx=XM_011281181.3;refGene_gn=IMMT;refGene_tx=XM_019827536.2;refGene_gn=IMMT;refGene_tx=XM_019827537.2
A3	103242570	103242700
refGene_gn=IMMT;refGene_tx=XM_006930260.4;refGene_gn=IMMT;refGene_tx=XM_006930262.4;refGene_gn=IMMT;refGene_tx=XM_006930264.4;refGene_gn=IMMT;refGene_tx=XM_011281181.3;refGene_gn=IMMT;refGene_tx=XM_019827536.2;refGene_gn=IMMT;refGene_tx=XM_019827537.2
A3	103243191	103243327
refGene_gn=IMMT;refGene_tx=XM_006930260.4;refGene_gn=IMMT;refGene_tx=XM_006930262.4;refGene_gn=IMMT;refGene_tx=XM_006930264.4;refGene_gn=IMMT;refGene_tx=XM_011281181.3;refGene_gn=IMMT;refGene_tx=XM_019827536.2;refGene_gn=IMMT;refGene_tx=XM_019827537.2
A3	103244573	103244677

refGene_gn=IMMT;refGene_tx=XM_006930260.4;refGene_gn=IMMT;refGene_tx=XM_006930262.4;refGene_gn=IMMT;refGene_tx=XM_006930264.4;refGene_gn=IMMT;refGene_tx=XM_011281181.3;refGene_gn=IMMT;refGene_tx=XM_019827536.2;refGene_gn=IMMT;refGene_tx=XM_019827537.2
A3 103246911 103247048
refGene_gn=IMMT;refGene_tx=XM_006930260.4;refGene_gn=IMMT;refGene_tx=XM_006930262.4;refGene_gn=IMMT;refGene_tx=XM_006930264.4;refGene_gn=IMMT;refGene_tx=XM_011281181.3;refGene_gn=IMMT;refGene_tx=XM_019827536.2;refGene_gn=IMMT;refGene_tx=XM_019827537.2
A3 103249690 103249786
refGene_gn=IMMT;refGene_tx=XM_006930260.4;refGene_gn=IMMT;refGene_tx=XM_011281181.3;refGene_gn=IMMT;refGene_tx=XM_019827536.2
A3 103250136 103250271
refGene_gn=IMMT;refGene_tx=XM_006930260.4;refGene_gn=IMMT;refGene_tx=XM_006930262.4;refGene_gn=IMMT;refGene_tx=XM_011281181.3;refGene_gn=IMMT;refGene_tx=XM_019827536.2;refGene_gn=IMMT;refGene_tx=XM_019827537.2
A3 103251968 103252080
refGene_gn=IMMT;refGene_tx=XM_006930260.4;refGene_gn=IMMT;refGene_tx=XM_006930262.4;refGene_gn=IMMT;refGene_tx=XM_006930264.4;refGene_gn=IMMT;refGene_tx=XM_011281181.3;refGene_gn=IMMT;refGene_tx=XM_019827536.2;refGene_gn=IMMT;refGene_tx=XM_019827537.2
A3 103256086 103256276
refGene_gn=IMMT;refGene_tx=XM_006930260.4;refGene_gn=IMMT;refGene_tx=XM_006930262.4;refGene_gn=IMMT;refGene_tx=XM_006930264.4;refGene_gn=IMMT;refGene_tx=XM_011281181.3;refGene_gn=IMMT;refGene_tx=XM_019827536.2;refGene_gn=IMMT;refGene_tx=XM_019827537.2
A3 103257017 103257091
refGene_gn=IMMT;refGene_tx=XM_006930260.4;refGene_gn=IMMT;refGene_tx=XM_006930262.4;refGene_gn=IMMT;refGene_tx=XM_006930264.4;refGene_gn=IMMT;refGene_tx=XM_011281181.3;refGene_gn=IMMT;refGene_tx=XM_019827536.2;refGene_gn=IMMT;refGene_tx=XM_019827537.2
A3 103271695 103271740
refGene_gn=IMMT;refGene_tx=XM_006930260.4;refGene_gn=IMMT;refGene_tx=XM_006930262.4;refGene_gn=IMMT;refGene_tx=XM_006930264.4;refGene_gn=IMMT;refGene_tx=XM_011281181.3;refGene_gn=IMMT;refGene_tx=XM_019827536.2;refGene_gn=IMMT;refGene_tx=XM_019827537.2
A3 103275475 103275518
refGene_gn=MRPL35;refGene_tx=XM_003984217.5
A3 103279180 103279217
refGene_gn=MRPL35;refGene_tx=XM_019827539.2
A3 103280941 103281131
refGene_gn=MRPL35;refGene_tx=XM_003984217.5;refGene_gn=MRPL35;refGene_tx=XM_019827539.2
A3 103282194 103282339
refGene_gn=MRPL35;refGene_tx=XM_003984217.5;refGene_gn=MRPL35;refGene_tx=XM_019827539.2
A3 103285051 103285240
refGene_gn=MRPL35;refGene_tx=XM_003984217.5;refGene_gn=MRPL35;refGene_tx=XM_019827539.2
A3 103292690 103292762

refGene_gn=REEP1;refGene_tx=XM_023251848.1;refGene_gn=REEP1;refGene_tx=XM_023251849.1;refGene_gn=REEP1;refGene_tx=XM_023251850.1;refGene_gn=REEP1;refGene_tx=XM_023251851.1;refGene_gn=REEP1;refGene_tx=XM_023251852.1;refGene_gn=REEP1;refGene_tx=XM_023251853.1
A3 103295752 103295904
refGene_gn=REEP1;refGene_tx=XM_023251848.1;refGene_gn=REEP1;refGene_tx=XM_023251849.1;refGene_gn=REEP1;refGene_tx=XM_023251850.1;refGene_gn=REEP1;refGene_tx=XM_023251851.1
A3 103301427 103301463
refGene_gn=REEP1;refGene_tx=XM_023251848.1;refGene_gn=REEP1;refGene_tx=XM_023251849.1;refGene_gn=REEP1;refGene_tx=XM_023251850.1;refGene_gn=REEP1;refGene_tx=XM_023251851.1;refGene_gn=REEP1;refGene_tx=XM_023251852.1
A3 103305437 103305615
refGene_gn=REEP1;refGene_tx=XM_023251848.1;refGene_gn=REEP1;refGene_tx=XM_023251849.1;refGene_gn=REEP1;refGene_tx=XM_023251850.1;refGene_gn=REEP1;refGene_tx=XM_023251851.1;refGene_gn=REEP1;refGene_tx=XM_023251852.1;refGene_gn=REEP1;refGene_tx=XM_023251853.1
A3 103313627 103313780
refGene_gn=REEP1;refGene_tx=XM_023251848.1;refGene_gn=REEP1;refGene_tx=XM_023251849.1;refGene_gn=REEP1;refGene_tx=XM_023251850.1;refGene_gn=REEP1;refGene_tx=XM_023251852.1;refGene_gn=REEP1;refGene_tx=XM_023251853.1
A3 103323092 103323206
refGene_gn=REEP1;refGene_tx=XM_023251848.1;refGene_gn=REEP1;refGene_tx=XM_023251849.1;refGene_gn=REEP1;refGene_tx=XM_023251850.1;refGene_gn=REEP1;refGene_tx=XM_023251851.1;refGene_gn=REEP1;refGene_tx=XM_023251852.1;refGene_gn=REEP1;refGene_tx=XM_023251853.1
A3 103326254 103326375
refGene_gn=REEP1;refGene_tx=XM_023251848.1;refGene_gn=REEP1;refGene_tx=XM_023251849.1;refGene_gn=REEP1;refGene_tx=XM_023251850.1;refGene_gn=REEP1;refGene_tx=XM_023251851.1;refGene_gn=REEP1;refGene_tx=XM_023251852.1;refGene_gn=REEP1;refGene_tx=XM_023251853.1
A3 103334812 103334889
refGene_gn=REEP1;refGene_tx=XM_023251848.1;refGene_gn=REEP1;refGene_tx=XM_023251849.1;refGene_gn=REEP1;refGene_tx=XM_023251850.1;refGene_gn=REEP1;refGene_tx=XM_023251851.1;refGene_gn=REEP1;refGene_tx=XM_023251852.1;refGene_gn=REEP1;refGene_tx=XM_023251853.1
A3 103348505 103348578
refGene_gn=REEP1;refGene_tx=XM_023251848.1;refGene_gn=REEP1;refGene_tx=XM_023251849.1;refGene_gn=REEP1;refGene_tx=XM_023251851.1;refGene_gn=REEP1;refGene_tx=XM_023251852.1;refGene_gn=REEP1;refGene_tx=XM_023251853.1
A3 103403490 103403522
refGene_gn=REEP1;refGene_tx=XM_023251848.1;refGene_gn=REEP1;refGene_tx=XM_023251849.1;refGene_gn=REEP1;refGene_tx=XM_023251851.1;refGene_gn=REEP1;refGene_tx=XM_023251852.1;refGene_gn=REEP1;refGene_tx=XM_023251853.1
A3 103483011 103483197
refGene_gn=KDM3A;refGene_tx=XM_023251854.1

A3 103489747 103489903
refGene_gn=KDM3A;refGene_tx=XM_023251854.1
A3 103491018 103491129
refGene_gn=KDM3A;refGene_tx=XM_023251854.1
A3 103494324 103494427
refGene_gn=KDM3A;refGene_tx=XM_023251854.1
A3 103495128 103495253
refGene_gn=KDM3A;refGene_tx=XM_023251854.1
A3 103495511 103495584
refGene_gn=KDM3A;refGene_tx=XM_023251854.1
A3 103495694 103495783
refGene_gn=KDM3A;refGene_tx=XM_023251854.1
A3 103501525 103501689
refGene_gn=KDM3A;refGene_tx=XM_023251854.1
A3 103503387 103503908
refGene_gn=KDM3A;refGene_tx=XM_023251854.1
A3 103509170 103509375
refGene_gn=KDM3A;refGene_tx=XM_023251854.1
A3 103512802 103513017
refGene_gn=KDM3A;refGene_tx=XM_023251854.1
A3 103516466 103516480
refGene_gn=KDM3A;refGene_tx=XM_023251854.1
A3 103516481 103516620
refGene_gn=KDM3A;refGene_tx=XM_023251854.1
A3 103516758 103516854
refGene_gn=KDM3A;refGene_tx=XM_023251854.1
A3 103517257 103517385
refGene_gn=KDM3A;refGene_tx=XM_023251854.1
A3 103518867 103519063
refGene_gn=KDM3A;refGene_tx=XM_023251854.1
A3 103521123 103521296
refGene_gn=KDM3A;refGene_tx=XM_023251854.1
A3 103521672 103521909
refGene_gn=KDM3A;refGene_tx=XM_023251854.1
A3 103523322 103523494
refGene_gn=KDM3A;refGene_tx=XM_023251854.1
A3 103525852 103525940
refGene_gn=KDM3A;refGene_tx=XM_023251854.1
A3 103526616 103526747
refGene_gn=KDM3A;refGene_tx=XM_023251854.1
A3 103531036 103531156
refGene_gn=KDM3A;refGene_tx=XM_023251854.1
A3 103531238 103531378
refGene_gn=KDM3A;refGene_tx=XM_023251854.1
A3 103533110 103533287
refGene_gn=KDM3A;refGene_tx=XM_023251854.1
A3 103533370 103533505
refGene_gn=KDM3A;refGene_tx=XM_023251854.1
A3 103534468 103534549
refGene_gn=KDM3A;refGene_tx=XM_023251854.1

A3	103561349	103561492
refGene_gn=LOC101094500;refGene_tx=XM_003984233.5;refGene_gn=LOC101094
500;refGene_tx=XM_006930275.3
A3	103562869	103562984
refGene_gn=LOC101094500;refGene_tx=XM_003984233.5;refGene_gn=LOC101094
500;refGene_tx=XM_006930275.3
A3	103566126	103566248
refGene_gn=LOC101094500;refGene_tx=XM_003984233.5;refGene_gn=LOC101094
500;refGene_tx=XM_006930275.3
A3	103576629	103576809
refGene_gn=LOC101094500;refGene_tx=XM_003984233.5;refGene_gn=LOC101094
500;refGene_tx=XM_006930275.3
A3	103593877	103593938
refGene_gn=LOC101094500;refGene_tx=XM_003984233.5;refGene_gn=LOC101094
500;refGene_tx=XM_006930275.3
A3	103618405	103618450
refGene_gn=LOC101094500;refGene_tx=XM_003984233.5
A3	103648641	103650217
refGene_gn=LOC101094500;refGene_tx=XM_006930272.4;refGene_gn=LOC101094
500;refGene_tx=XM_006930273.4;refGene_gn=LOC101094500;refGene_tx=XM_01
9827542.2
A3	103657215	103657331
refGene_gn=LOC101094500;refGene_tx=XM_006930272.4;refGene_gn=LOC101094
500;refGene_tx=XM_019827542.2
A3	103667683	103667823
refGene_gn=LOC101094500;refGene_tx=XM_006930272.4;refGene_gn=LOC101094
500;refGene_tx=XM_006930273.4;refGene_gn=LOC101094500;refGene_tx=XM_00
6930275.3
A3	103670027	103670253
refGene_gn=LOC101094500;refGene_tx=XM_006930272.4;refGene_gn=LOC101094
500;refGene_tx=XM_006930273.4;refGene_gn=LOC101094500;refGene_tx=XM_00
6930275.3
A3	103753228	103753370
refGene_gn=RMND5A;refGene_tx=XM_023251855.1;refGene_gn=RMND5A;refGene_
tx=XM_023251856.1;refGene_gn=RMND5A;refGene_tx=XM_023251857.1;refGene_
gn=RMND5A;refGene_tx=XM_023251858.1;refGene_gn=RMND5A;refGene_tx=XM_02
3251859.1
A3	103773661	103773804
refGene_gn=RMND5A;refGene_tx=XM_023251855.1;refGene_gn=RMND5A;refGene_
tx=XM_023251856.1;refGene_gn=RMND5A;refGene_tx=XM_023251857.1;refGene_
gn=RMND5A;refGene_tx=XM_023251858.1
A3	103782253	103782358
refGene_gn=RMND5A;refGene_tx=XM_023251855.1;refGene_gn=RMND5A;refGene_
tx=XM_023251856.1;refGene_gn=RMND5A;refGene_tx=XM_023251857.1
A3	103790068	103790203
refGene_gn=RMND5A;refGene_tx=XM_023251855.1;refGene_gn=RMND5A;refGene_
tx=XM_023251856.1;refGene_gn=RMND5A;refGene_tx=XM_023251857.1;refGene_
gn=RMND5A;refGene_tx=XM_023251858.1;refGene_gn=RMND5A;refGene_tx=XM_02
3251859.1
A3	103792002	103792103

refGene_gn=RMND5A;refGene_tx=XM_023251855.1;refGene_gn=RMND5A;refGene_tx=XM_023251856.1;refGene_gn=RMND5A;refGene_tx=XM_023251857.1;refGene_gn=RMND5A;refGene_tx=XM_023251858.1;refGene_gn=RMND5A;refGene_tx=XM_023251859.1
A3 103803072 103803239
refGene_gn=RMND5A;refGene_tx=XM_023251855.1;refGene_gn=RMND5A;refGene_tx=XM_023251856.1;refGene_gn=RMND5A;refGene_tx=XM_023251857.1;refGene_gn=RMND5A;refGene_tx=XM_023251858.1;refGene_gn=RMND5A;refGene_tx=XM_023251859.1
A3 103803897 103804063
refGene_gn=RMND5A;refGene_tx=XM_023251855.1;refGene_gn=RMND5A;refGene_tx=XM_023251856.1;refGene_gn=RMND5A;refGene_tx=XM_023251857.1;refGene_gn=RMND5A;refGene_tx=XM_023251858.1;refGene_gn=RMND5A;refGene_tx=XM_023251859.1
A3 103807125 103807228
refGene_gn=RMND5A;refGene_tx=XM_023251855.1;refGene_gn=RMND5A;refGene_tx=XM_023251856.1;refGene_gn=RMND5A;refGene_tx=XM_023251857.1;refGene_gn=RMND5A;refGene_tx=XM_023251858.1;refGene_gn=RMND5A;refGene_tx=XM_023251859.1
A3 103809155 103809310
refGene_gn=RMND5A;refGene_tx=XM_023251855.1;refGene_gn=RMND5A;refGene_tx=XM_023251856.1;refGene_gn=RMND5A;refGene_tx=XM_023251857.1;refGene_gn=RMND5A;refGene_tx=XM_023251858.1;refGene_gn=RMND5A;refGene_tx=XM_023251859.1
A3 103810552 103810616
refGene_gn=RMND5A;refGene_tx=XM_023251855.1;refGene_gn=RMND5A;refGene_tx=XM_023251856.1;refGene_gn=RMND5A;refGene_tx=XM_023251857.1;refGene_gn=RMND5A;refGene_tx=XM_023251858.1;refGene_gn=RMND5A;refGene_tx=XM_023251859.1
A3 103823472 103823524
refGene_gn=CD8A;refGene_tx=NM_001009843.2
A3 103826189 103826220
refGene_gn=CD8A;refGene_tx=NM_001009843.2
A3 103826859 103826970
refGene_gn=CD8A;refGene_tx=NM_001009843.2
A3 103827190 103827301
refGene_gn=CD8A;refGene_tx=NM_001009843.2
A3 103827917 103828283
refGene_gn=CD8A;refGene_tx=NM_001009843.2
A3 103828371 103828420
refGene_gn=CD8A;refGene_tx=NM_001009843.2
A3 103869266 103869279
refGene_gn=CD8B;refGene_tx=NM_001009867.1
A3 103871356 103871393
refGene_gn=CD8B;refGene_tx=NM_001009867.1
A3 103872874 103872964
refGene_gn=CD8B;refGene_tx=NM_001009867.1
A3 103876604 103876694
refGene_gn=CD8B;refGene_tx=NM_001009867.1
A3 103880357 103880717

refGene_gn=CD8B;refGene_tx=NM_001009867.1
A3	103883860	103883903
refGene_gn=CD8B;refGene_tx=NM_001009867.1
A3	103889605	103890244
refGene_gn=LOC101098787;refGene_tx=XM_023252190.1
A3	103903288	103904062
refGene_gn=KRCC1;refGene_tx=XM_011281176.2;refGene_gn=KRCC1;refGene_tx
=XM_011281177.2
A3	103928152	103928289
refGene_gn=SMYD1;refGene_tx=XM_003984235.4
A3	103942751	103942928
refGene_gn=SMYD1;refGene_tx=XM_003984235.4
A3	103946395	103946609
refGene_gn=SMYD1;refGene_tx=XM_003984235.4
A3	103951923	103952054
refGene_gn=SMYD1;refGene_tx=XM_003984235.4
A3	103954443	103954482
refGene_gn=SMYD1;refGene_tx=XM_003984235.4
A3	103957392	103957540
refGene_gn=SMYD1;refGene_tx=XM_003984235.4
A3	103957542	103957584
refGene_gn=SMYD1;refGene_tx=XM_003984235.4
A3	103962372	103962465
refGene_gn=SMYD1;refGene_tx=XM_003984235.4
A3	103964592	103964756
refGene_gn=SMYD1;refGene_tx=XM_003984235.4
A3	103966050	103966219
refGene_gn=SMYD1;refGene_tx=XM_003984235.4
A3	103967505	103967664
refGene_gn=SMYD1;refGene_tx=XM_003984235.4
A3	103976518	103976569
refGene_gn=FABP1;refGene_tx=XM_003984236.4
A3	103978444	103978537
refGene_gn=FABP1;refGene_tx=XM_003984236.4
A3	103980640	103980813
refGene_gn=FABP1;refGene_tx=XM_003984236.4
A3	103982416	103982483
refGene_gn=FABP1;refGene_tx=XM_003984236.4
A3	104014197	104014212
refGene_gn=THNSL2;refGene_tx=XM_006930284.4
A3	104015172	104015395
refGene_gn=THNSL2;refGene_tx=XM_006930280.4;refGene_gn=THNSL2;refGene_
tx=XM_006930281.4;refGene_gn=THNSL2;refGene_tx=XM_006930282.4;refGene_
gn=THNSL2;refGene_tx=XM_006930283.4;refGene_gn=THNSL2;refGene_tx=XM_00
6930284.4;refGene_gn=THNSL2;refGene_tx=XM_006930285.4;refGene_gn=THNSL
2;refGene_tx=XM_006930286.4;refGene_gn=THNSL2;refGene_tx=XM_019827545.
2;refGene_gn=THNSL2;refGene_tx=XM_019827546.2;refGene_gn=THNSL2;refGen
e_tx=XM_023251860.1
A3	104016908	104017103
refGene_gn=THNSL2;refGene_tx=XM_006930280.4;refGene_gn=THNSL2;refGene_

tx=XM_006930281.4;refGene_gn=THNSL2;refGene_tx=XM_006930282.4;refGene_gn=THNSL2;refGene_tx=XM_006930283.4;refGene_gn=THNSL2;refGene_tx=XM_006930284.4;refGene_gn=THNSL2;refGene_tx=XM_006930285.4;refGene_gn=THNSL2;refGene_tx=XM_006930286.4;refGene_gn=THNSL2;refGene_tx=XM_019827545.2;refGene_gn=THNSL2;refGene_tx=XM_019827546.2;refGene_gn=THNSL2;refGene_tx=XM_023251860.1
A3 104017510 104017663
refGene_gn=THNSL2;refGene_tx=XM_006930280.4;refGene_gn=THNSL2;refGene_tx=XM_006930281.4;refGene_gn=THNSL2;refGene_tx=XM_006930282.4;refGene_gn=THNSL2;refGene_tx=XM_006930283.4;refGene_gn=THNSL2;refGene_tx=XM_006930284.4;refGene_gn=THNSL2;refGene_tx=XM_006930285.4;refGene_gn=THNSL2;refGene_tx=XM_006930286.4;refGene_gn=THNSL2;refGene_tx=XM_019827545.2;refGene_gn=THNSL2;refGene_tx=XM_019827546.2;refGene_gn=THNSL2;refGene_tx=XM_023251860.1
A3 104021452 104021683
refGene_gn=THNSL2;refGene_tx=XM_006930280.4;refGene_gn=THNSL2;refGene_tx=XM_006930281.4;refGene_gn=THNSL2;refGene_tx=XM_006930282.4;refGene_gn=THNSL2;refGene_tx=XM_006930283.4;refGene_gn=THNSL2;refGene_tx=XM_006930284.4;refGene_gn=THNSL2;refGene_tx=XM_006930285.4;refGene_gn=THNSL2;refGene_tx=XM_006930286.4;refGene_gn=THNSL2;refGene_tx=XM_019827546.2;refGene_gn=THNSL2;refGene_tx=XM_023251860.1
A3 104025062 104025211
refGene_gn=THNSL2;refGene_tx=XM_006930280.4;refGene_gn=THNSL2;refGene_tx=XM_006930281.4;refGene_gn=THNSL2;refGene_tx=XM_006930282.4;refGene_gn=THNSL2;refGene_tx=XM_006930283.4;refGene_gn=THNSL2;refGene_tx=XM_006930284.4;refGene_gn=THNSL2;refGene_tx=XM_006930285.4;refGene_gn=THNSL2;refGene_tx=XM_006930286.4;refGene_gn=THNSL2;refGene_tx=XM_019827545.2;refGene_gn=THNSL2;refGene_tx=XM_019827546.2;refGene_gn=THNSL2;refGene_tx=XM_023251860.1
A3 104025313 104025439
refGene_gn=THNSL2;refGene_tx=XM_006930280.4;refGene_gn=THNSL2;refGene_tx=XM_006930281.4;refGene_gn=THNSL2;refGene_tx=XM_006930282.4;refGene_gn=THNSL2;refGene_tx=XM_006930283.4;refGene_gn=THNSL2;refGene_tx=XM_006930284.4;refGene_gn=THNSL2;refGene_tx=XM_006930285.4;refGene_gn=THNSL2;refGene_tx=XM_006930286.4;refGene_gn=THNSL2;refGene_tx=XM_019827545.2;refGene_gn=THNSL2;refGene_tx=XM_019827546.2;refGene_gn=THNSL2;refGene_tx=XM_023251860.1
A3 104026345 104026598
refGene_gn=THNSL2;refGene_tx=XM_006930285.4;refGene_gn=THNSL2;refGene_tx=XM_006930286.4
A3 104027648 104027800
refGene_gn=THNSL2;refGene_tx=XM_006930280.4;refGene_gn=THNSL2;refGene_tx=XM_006930281.4;refGene_gn=THNSL2;refGene_tx=XM_006930282.4;refGene_gn=THNSL2;refGene_tx=XM_006930283.4;refGene_gn=THNSL2;refGene_tx=XM_006930284.4;refGene_gn=THNSL2;refGene_tx=XM_019827545.2;refGene_gn=THNSL2;refGene_tx=XM_023251860.1
A3 104028199 104028425
refGene_gn=THNSL2;refGene_tx=XM_006930280.4;refGene_gn=THNSL2;refGene_tx=XM_006930281.4;refGene_gn=THNSL2;refGene_tx=XM_006930282.4;refGene_gn=THNSL2;refGene_tx=XM_006930283.4;refGene_gn=THNSL2;refGene_tx=XM_00

6930284.4;refGene_gn=THNSL2;refGene_tx=XM_019827545.2;refGene_gn=THNSL2;refGene_tx=XM_019827546.2;refGene_gn=THNSL2;refGene_tx=XM_023251860.1
A3 104197159 104197779
refGene_gn=FOXI3;refGene_tx=XM_011281245.3
A3 104200408 104200916
refGene_gn=FOXI3;refGene_tx=XM_011281245.3
A3 104200917 104201049
refGene_gn=FOXI3;refGene_tx=XM_011281245.3
A3 104219934 104219987
refGene_gn=TEX37;refGene_tx=XM_019827547.2;refGene_gn=TEX37;refGene_tx=XM_019827550.2;refGene_gn=TEX37;refGene_tx=XM_023251862.1
A3 104261626 104261655
refGene_gn=TEX37;refGene_tx=XM_003984251.4;refGene_gn=TEX37;refGene_tx=XM_006930288.3
A3 104262073 104262110
refGene_gn=TEX37;refGene_tx=XM_003984251.4;refGene_gn=TEX37;refGene_tx=XM_006930287.4;refGene_gn=TEX37;refGene_tx=XM_006930288.3;refGene_gn=TEX37;refGene_tx=XM_019827547.2;refGene_gn=TEX37;refGene_tx=XM_019827549.2;refGene_gn=TEX37;refGene_tx=XM_019827550.2;refGene_gn=TEX37;refGene_tx=XM_023251862.1;refGene_gn=TEX37;refGene_tx=XM_023251863.1;refGene_gn=TEX37;refGene_tx=XM_023251864.1
A3 104262843 104262936
refGene_gn=TEX37;refGene_tx=XM_003984251.4;refGene_gn=TEX37;refGene_tx=XM_006930287.4;refGene_gn=TEX37;refGene_tx=XM_019827547.2;refGene_gn=TEX37;refGene_tx=XM_019827549.2;refGene_gn=TEX37;refGene_tx=XM_023251862.1;refGene_gn=TEX37;refGene_tx=XM_023251863.1;refGene_gn=TEX37;refGene_tx=XM_023251864.1
A3 104265509 104265938
refGene_gn=TEX37;refGene_tx=XM_003984251.4;refGene_gn=TEX37;refGene_tx=XM_006930287.4;refGene_gn=TEX37;refGene_tx=XM_006930288.3;refGene_gn=TEX37;refGene_tx=XM_019827547.2;refGene_gn=TEX37;refGene_tx=XM_019827549.2;refGene_gn=TEX37;refGene_tx=XM_019827550.2;refGene_gn=TEX37;refGene_tx=XM_023251862.1;refGene_gn=TEX37;refGene_tx=XM_023251863.1;refGene_gn=TEX37;refGene_tx=XM_023251864.1
A3 104287766 104287967
refGene_gn=EIF2AK3;refGene_tx=XM_019827552.2
A3 104291527 104291590
refGene_gn=EIF2AK3;refGene_tx=XM_019827552.2
A3 104295660 104295762
refGene_gn=EIF2AK3;refGene_tx=XM_019827552.2
A3 104301334 104301502
refGene_gn=EIF2AK3;refGene_tx=XM_019827552.2
A3 104305277 104306055
refGene_gn=EIF2AK3;refGene_tx=XM_019827552.2
A3 104307287 104307437
refGene_gn=EIF2AK3;refGene_tx=XM_019827552.2
A3 104311508 104311631
refGene_gn=EIF2AK3;refGene_tx=XM_019827552.2
A3 104315036 104315149

refGene_gn=EIF2AK3;refGene_tx=XM_019827552.2
A3 104315492 104315713
refGene_gn=EIF2AK3;refGene_tx=XM_019827552.2
A3 104317039 104317162
refGene_gn=EIF2AK3;refGene_tx=XM_019827552.2
A3 104317853 104317994
refGene_gn=EIF2AK3;refGene_tx=XM_019827552.2
A3 104319327 104319490
refGene_gn=EIF2AK3;refGene_tx=XM_019827552.2
A3 104320661 104320896
refGene_gn=EIF2AK3;refGene_tx=XM_019827552.2
A3 104322877 104323008
refGene_gn=EIF2AK3;refGene_tx=XM_019827552.2
A3 104324730 104324925
refGene_gn=EIF2AK3;refGene_tx=XM_019827552.2
A3 104343641 104343771
refGene_gn=EIF2AK3;refGene_tx=XM_019827552.2
A3 104355685 104355993
refGene_gn=EIF2AK3;refGene_tx=XM_019827552.2
A3 104403886 104404171
refGene_gn=RPIA;refGene_tx=XM_003984239.5;refGene_gn=RPIA;refGene_tx=XM_019827555.2
A3 104412808 104412869
refGene_gn=RPIA;refGene_tx=XM_003984239.5;refGene_gn=RPIA;refGene_tx=XM_019827555.2
A3 104414268 104414324
refGene_gn=RPIA;refGene_tx=XM_003984239.5;refGene_gn=RPIA;refGene_tx=XM_019827555.2
A3 104422035 104422095
refGene_gn=RPIA;refGene_tx=XM_003984239.5;refGene_gn=RPIA;refGene_tx=XM_019827555.2
A3 104424435 104424500
refGene_gn=RPIA;refGene_tx=XM_003984239.5;refGene_gn=RPIA;refGene_tx=XM_019827555.2
A3 104425939 104426008
refGene_gn=RPIA;refGene_tx=XM_003984239.5
A3 104426740 104426882
refGene_gn=RPIA;refGene_tx=XM_003984239.5;refGene_gn=RPIA;refGene_tx=XM_019827555.2
A3 104429093 104429193
refGene_gn=RPIA;refGene_tx=XM_003984239.5;refGene_gn=RPIA;refGene_tx=XM_019827555.2
A3 104434480 104434578
refGene_gn=RPIA;refGene_tx=XM_003984239.5;refGene_gn=RPIA;refGene_tx=XM_019827555.2
A3 104466611 104555199
refGene_gn=LOC101100802;refGene_gn=LOC109498252
A3 104599771 104600411 refGene_gn=LOC111559716
A3 104607428 104607973 refGene_gn=LOC111559844
A3 104615504 104619070 refGene_gn=LOC111559896

A3	104730951	104730976
refGene_gn=PAX8;refGene_tx=XM_019827558.2;refGene_gn=PAX8;refGene_tx=XM_023251865.1;refGene_gn=PAX8;refGene_tx=XM_023251866.1;refGene_gn=PAX8;refGene_tx=XM_023251867.1;refGene_gn=PAX8;refGene_tx=XM_023251868.1;refGene_gn=PAX8;refGene_tx=XM_023251869.1
A3	104763639	104763805
refGene_gn=PAX8;refGene_tx=XM_019827558.2;refGene_gn=PAX8;refGene_tx=XM_023251865.1;refGene_gn=PAX8;refGene_tx=XM_023251866.1;refGene_gn=PAX8;refGene_tx=XM_023251867.1;refGene_gn=PAX8;refGene_tx=XM_023251868.1;refGene_gn=PAX8;refGene_tx=XM_023251869.1
A3	104766217	104766418
refGene_gn=PAX8;refGene_tx=XM_019827558.2;refGene_gn=PAX8;refGene_tx=XM_023251865.1;refGene_gn=PAX8;refGene_tx=XM_023251866.1;refGene_gn=PAX8;refGene_tx=XM_023251867.1;refGene_gn=PAX8;refGene_tx=XM_023251868.1;refGene_gn=PAX8;refGene_tx=XM_023251869.1
A3	104767781	104767870
refGene_gn=PAX8;refGene_tx=XM_019827558.2;refGene_gn=PAX8;refGene_tx=XM_023251865.1;refGene_gn=PAX8;refGene_tx=XM_023251866.1;refGene_gn=PAX8;refGene_tx=XM_023251867.1;refGene_gn=PAX8;refGene_tx=XM_023251868.1;refGene_gn=PAX8;refGene_tx=XM_023251869.1
A3	104768188	104768314
refGene_gn=PAX8;refGene_tx=XM_019827558.2;refGene_gn=PAX8;refGene_tx=XM_023251865.1;refGene_gn=PAX8;refGene_tx=XM_023251866.1;refGene_gn=PAX8;refGene_tx=XM_023251867.1;refGene_gn=PAX8;refGene_tx=XM_023251868.1;refGene_gn=PAX8;refGene_tx=XM_023251869.1
A3	104768574	104768750
refGene_gn=PAX8;refGene_tx=XM_019827558.2;refGene_gn=PAX8;refGene_tx=XM_023251865.1;refGene_gn=PAX8;refGene_tx=XM_023251866.1;refGene_gn=PAX8;refGene_tx=XM_023251867.1;refGene_gn=PAX8;refGene_tx=XM_023251868.1;refGene_gn=PAX8;refGene_tx=XM_023251869.1
A3	104772831	104772952
refGene_gn=PAX8;refGene_tx=XM_019827558.2;refGene_gn=PAX8;refGene_tx=XM_023251865.1;refGene_gn=PAX8;refGene_tx=XM_023251866.1;refGene_gn=PAX8;refGene_tx=XM_023251867.1;refGene_gn=PAX8;refGene_tx=XM_023251868.1;refGene_gn=PAX8;refGene_tx=XM_023251869.1
A3	104773423	104773630
refGene_gn=PAX8;refGene_tx=XM_019827558.2;refGene_gn=PAX8;refGene_tx=XM_023251865.1;refGene_gn=PAX8;refGene_tx=XM_023251866.1;refGene_gn=PAX8;refGene_tx=XM_023251867.1
A3	104781107	104781209
refGene_gn=PAX8;refGene_tx=XM_019827558.2;refGene_gn=PAX8;refGene_tx=XM_023251865.1;refGene_gn=PAX8;refGene_tx=XM_023251866.1;refGene_gn=PAX8;refGene_tx=XM_023251867.1;refGene_gn=PAX8;refGene_tx=XM_023251868.1;refGene_gn=PAX8;refGene_tx=XM_023251869.1
A3	104787449	104787536
refGene_gn=PAX8;refGene_tx=XM_019827558.2;refGene_gn=PAX8;refGene_tx=XM_023251865.1;refGene_gn=PAX8;refGene_tx=XM_023251866.1;refGene_gn=PAX8;refGene_tx=XM_023251867.1;refGene_gn=PAX8;refGene_tx=XM_023251868.1;refGene_gn=PAX8;refGene_tx=XM_023251869.1
A3	104788969	104789046

refGene_gn=PAX8;refGene_tx=XM_019827558.2;refGene_gn=PAX8;refGene_tx=XM_023251865.1;refGene_gn=PAX8;refGene_tx=XM_023251868.1;refGene_gn=PAX8;refGene_tx=XM_023251869.1
A3 104802681 104802939
refGene_gn=PSD4;refGene_tx=XM_006930290.4;refGene_gn=PSD4;refGene_tx=XM_019827563.2;refGene_gn=PSD4;refGene_tx=XM_019827564.2;refGene_gn=PSD4;refGene_tx=XM_023251870.1;refGene_gn=PSD4;refGene_tx=XM_023251871.1;refGene_gn=PSD4;refGene_tx=XM_023251872.1
A3 104804321 104804465
refGene_gn=PSD4;refGene_tx=XM_006930290.4;refGene_gn=PSD4;refGene_tx=XM_019827563.2;refGene_gn=PSD4;refGene_tx=XM_019827564.2;refGene_gn=PSD4;refGene_tx=XM_023251870.1;refGene_gn=PSD4;refGene_tx=XM_023251871.1;refGene_gn=PSD4;refGene_tx=XM_023251872.1
A3 104804659 104804804
refGene_gn=PSD4;refGene_tx=XM_006930290.4;refGene_gn=PSD4;refGene_tx=XM_019827563.2;refGene_gn=PSD4;refGene_tx=XM_019827564.2;refGene_gn=PSD4;refGene_tx=XM_023251870.1;refGene_gn=PSD4;refGene_tx=XM_023251871.1;refGene_gn=PSD4;refGene_tx=XM_023251872.1
A3 104805324 104805488
refGene_gn=PSD4;refGene_tx=XM_006930290.4;refGene_gn=PSD4;refGene_tx=XM_019827563.2;refGene_gn=PSD4;refGene_tx=XM_019827564.2;refGene_gn=PSD4;refGene_tx=XM_023251870.1;refGene_gn=PSD4;refGene_tx=XM_023251871.1;refGene_gn=PSD4;refGene_tx=XM_023251872.1
A3 104805608 104805679
refGene_gn=PSD4;refGene_tx=XM_006930290.4;refGene_gn=PSD4;refGene_tx=XM_019827563.2;refGene_gn=PSD4;refGene_tx=XM_019827564.2;refGene_gn=PSD4;refGene_tx=XM_023251870.1;refGene_gn=PSD4;refGene_tx=XM_023251871.1;refGene_gn=PSD4;refGene_tx=XM_023251872.1
A3 104806609 104806770
refGene_gn=PSD4;refGene_tx=XM_006930290.4;refGene_gn=PSD4;refGene_tx=XM_019827563.2;refGene_gn=PSD4;refGene_tx=XM_019827564.2;refGene_gn=PSD4;refGene_tx=XM_023251870.1;refGene_gn=PSD4;refGene_tx=XM_023251871.1;refGene_gn=PSD4;refGene_tx=XM_023251872.1
A3 104807114 104807158
refGene_gn=PSD4;refGene_tx=XM_006930290.4;refGene_gn=PSD4;refGene_tx=XM_019827563.2;refGene_gn=PSD4;refGene_tx=XM_019827564.2;refGene_gn=PSD4;refGene_tx=XM_023251870.1;refGene_gn=PSD4;refGene_tx=XM_023251871.1;refGene_gn=PSD4;refGene_tx=XM_023251872.1
A3 104808195 104808285
refGene_gn=PSD4;refGene_tx=XM_006930290.4;refGene_gn=PSD4;refGene_tx=XM_019827563.2;refGene_gn=PSD4;refGene_tx=XM_019827564.2;refGene_gn=PSD4;refGene_tx=XM_023251870.1;refGene_gn=PSD4;refGene_tx=XM_023251871.1;refGene_gn=PSD4;refGene_tx=XM_023251872.1
A3 104808469 104808528
refGene_gn=PSD4;refGene_tx=XM_006930290.4;refGene_gn=PSD4;refGene_tx=XM_019827563.2;refGene_gn=PSD4;refGene_tx=XM_019827564.2;refGene_gn=PSD4;refGene_tx=XM_023251870.1;refGene_gn=PSD4;refGene_tx=XM_023251871.1;refGene_gn=PSD4;refGene_tx=XM_023251872.1
A3 104808769 104808882
refGene_gn=PSD4;refGene_tx=XM_006930290.4;refGene_gn=PSD4;refGene_tx=X

M_019827563.2;refGene_gn=PSD4;refGene_tx=XM_019827564.2;refGene_gn=PSD
4;refGene_tx=XM_023251870.1;refGene_gn=PSD4;refGene_tx=XM_023251871.1;
refGene_gn=PSD4;refGene_tx=XM_023251872.1
A3 104809029 104809110
refGene_gn=PSD4;refGene_tx=XM_006930290.4;refGene_gn=PSD4;refGene_tx=X
M_019827563.2;refGene_gn=PSD4;refGene_tx=XM_019827564.2;refGene_gn=PSD
4;refGene_tx=XM_023251870.1;refGene_gn=PSD4;refGene_tx=XM_023251871.1;
refGene_gn=PSD4;refGene_tx=XM_023251872.1
A3 104809530 104809707
refGene_gn=PSD4;refGene_tx=XM_006930290.4;refGene_gn=PSD4;refGene_tx=X
M_019827563.2;refGene_gn=PSD4;refGene_tx=XM_019827564.2;refGene_gn=PSD
4;refGene_tx=XM_023251870.1;refGene_gn=PSD4;refGene_tx=XM_023251871.1;
refGene_gn=PSD4;refGene_tx=XM_023251872.1
A3 104812644 104813029
refGene_gn=PSD4;refGene_tx=XM_006930290.4;refGene_gn=PSD4;refGene_tx=X
M_019827563.2;refGene_gn=PSD4;refGene_tx=XM_019827564.2;refGene_gn=PSD
4;refGene_tx=XM_023251870.1;refGene_gn=PSD4;refGene_tx=XM_023251871.1;
refGene_gn=PSD4;refGene_tx=XM_023251872.1
A3 104813534 104813613
refGene_gn=PSD4;refGene_tx=XM_006930290.4;refGene_gn=PSD4;refGene_tx=X
M_019827563.2;refGene_gn=PSD4;refGene_tx=XM_019827564.2;refGene_gn=PSD
4;refGene_tx=XM_023251870.1;refGene_gn=PSD4;refGene_tx=XM_023251871.1;
refGene_gn=PSD4;refGene_tx=XM_023251872.1
A3 104813899 104814004
refGene_gn=PSD4;refGene_tx=XM_006930290.4;refGene_gn=PSD4;refGene_tx=X
M_019827563.2;refGene_gn=PSD4;refGene_tx=XM_019827564.2;refGene_gn=PSD
4;refGene_tx=XM_023251870.1;refGene_gn=PSD4;refGene_tx=XM_023251871.1;
refGene_gn=PSD4;refGene_tx=XM_023251872.1
A3 104815121 104816090
refGene_gn=PSD4;refGene_tx=XM_006930290.4;refGene_gn=PSD4;refGene_tx=X
M_019827563.2;refGene_gn=PSD4;refGene_tx=XM_019827564.2;refGene_gn=PSD
4;refGene_tx=XM_023251870.1;refGene_gn=PSD4;refGene_tx=XM_023251871.1;
refGene_gn=PSD4;refGene_tx=XM_023251872.1
A3 104848924 104849140
refGene_gn=LOC101096719;refGene_tx=XM_003984241.5;refGene_gn=LOC101096
719;refGene_tx=XM_006930291.4;refGene_gn=LOC101096719;refGene_tx=XM_00
6930292.4;refGene_gn=LOC101096719;refGene_tx=XM_019827565.2;refGene_gn
=LOC101096719;refGene_tx=XM_019827566.2
A3 104850205 104850318
refGene_gn=LOC101096719;refGene_tx=XM_003984241.5;refGene_gn=LOC101096
719;refGene_tx=XM_006930291.4;refGene_gn=LOC101096719;refGene_tx=XM_00
6930292.4;refGene_gn=LOC101096719;refGene_tx=XM_019827565.2;refGene_gn
=LOC101096719;refGene_tx=XM_019827566.2
A3 104851692 104851781
refGene_gn=LOC101096719;refGene_tx=XM_003984241.5;refGene_gn=LOC101096
719;refGene_tx=XM_006930291.4;refGene_gn=LOC101096719;refGene_tx=XM_00
6930292.4;refGene_gn=LOC101096719;refGene_tx=XM_019827565.2;refGene_gn
=LOC101096719;refGene_tx=XM_019827566.2
A3 104853719 104853835
refGene_gn=LOC101096719;refGene_tx=XM_003984241.5;refGene_gn=LOC101096

719;refGene_tx=XM_006930291.4;refGene_gn=LOC101096719;refGene_tx=XM_006930292.4;refGene_gn=LOC101096719;refGene_tx=XM_019827565.2;refGene_gn=LOC101096719;refGene_tx=XM_019827566.2
A3 104862015 104862094
refGene_gn=LOC101096719;refGene_tx=XM_019827565.2
A3 104863337 104863347
refGene_gn=LOC101096719;refGene_tx=XM_006930291.4
A3 104898941 104899154
refGene_gn=IL1F10;refGene_tx=XM_003984242.5;refGene_gn=IL1F10;refGene_tx=XM_019827568.2
A3 104899433 104899561
refGene_gn=IL1F10;refGene_tx=XM_003984242.5;refGene_gn=IL1F10;refGene_tx=XM_019827568.2
A3 104899841 104899927
refGene_gn=IL1F10;refGene_tx=XM_003984242.5;refGene_gn=IL1F10;refGene_tx=XM_019827568.2
A3 104901719 104901751
refGene_gn=IL1F10;refGene_tx=XM_003984242.5;refGene_gn=IL1F10;refGene_tx=XM_019827568.2
A3 104911595 104911820
refGene_gn=LOC101097230;refGene_tx=XM_006930293.3;refGene_gn=LOC101097230;refGene_tx=XM_019827567.2
A3 104912022 104912150
refGene_gn=LOC101097230;refGene_tx=XM_006930293.3;refGene_gn=LOC101097230;refGene_tx=XM_019827567.2
A3 104913186 104913272
refGene_gn=LOC101097230;refGene_tx=XM_006930293.3;refGene_gn=LOC101097230;refGene_tx=XM_019827567.2
A3 104914616 104914645
refGene_gn=LOC101097230;refGene_tx=XM_006930293.3;refGene_gn=LOC101097230;refGene_tx=XM_019827567.2
A3 104936409 104936622
refGene_gn=IL36G;refGene_tx=XM_019827573.2;refGene_gn=IL36G;refGene_tx=XM_019827574.1;refGene_gn=IL36G;refGene_tx=XM_023251874.1
A3 104937415 104937555
refGene_gn=IL36G;refGene_tx=XM_019827573.2;refGene_gn=IL36G;refGene_tx=XM_019827574.1;refGene_gn=IL36G;refGene_tx=XM_023251874.1
A3 104940168 104940276
refGene_gn=IL36G;refGene_tx=XM_019827573.2;refGene_gn=IL36G;refGene_tx=XM_019827574.1;refGene_gn=IL36G;refGene_tx=XM_023251874.1
A3 104940832 104940920
refGene_gn=IL36G;refGene_tx=XM_019827573.2;refGene_gn=IL36G;refGene_tx=XM_019827574.1;refGene_gn=IL36G;refGene_tx=XM_023251874.1
A3 104951395 104951605
refGene_gn=IL36G;refGene_tx=XM_023251875.1
A3 104954497 104954637
refGene_gn=IL36G;refGene_tx=XM_019827573.2;refGene_gn=IL36G;refGene_tx=XM_019827574.1;refGene_gn=IL36G;refGene_tx=XM_023251874.1;refGene_gn=IL36G;refGene_tx=XM_023251875.1
A3 104955246 104955351

refGene_gn=IL36G;refGene_tx=XM_019827573.2;refGene_gn=IL36G;refGene_tx=XM_019827574.1;refGene_gn=IL36G;refGene_tx=XM_023251874.1;refGene_gn=IL36G;refGene_tx=XM_023251875.1
A3 104955805 104955873
refGene_gn=IL36G;refGene_tx=XM_019827574.1;refGene_gn=IL36G;refGene_tx=XM_023251874.1;refGene_gn=IL36G;refGene_tx=XM_023251875.1
A3 104956293 104956325
refGene_gn=IL36G;refGene_tx=XM_019827574.1
A3 104956370 104956465
refGene_gn=IL36G;refGene_tx=XM_023251874.1;refGene_gn=IL36G;refGene_tx=XM_023251875.1
A3 104988032 104988095
refGene_gn=IL36G;refGene_tx=XM_019827573.2
A3 104989544 104989623
refGene_gn=IL36G;refGene_tx=XM_019827573.2
A3 105050647 105050694
refGene_gn=IL1B;refGene_tx=NM_001077414.1
A3 105051231 105051280
refGene_gn=IL1B;refGene_tx=NM_001077414.1
A3 105052907 105053112
refGene_gn=IL1B;refGene_tx=NM_001077414.1
A3 105053654 105053816
refGene_gn=IL1B;refGene_tx=NM_001077414.1
A3 105055062 105055193
refGene_gn=IL1B;refGene_tx=NM_001077414.1
A3 105056164 105056374
refGene_gn=IL1B;refGene_tx=NM_001077414.1
A3 105113200 105113247
refGene_gn=IL1A;refGene_tx=NM_001009351.1;refGene_gn=IL1A;refGene_tx=XM_006930269.4;refGene_gn=IL1A;refGene_tx=XM_006930270.4;refGene_gn=IL1A;refGene_tx=XM_019826172.2;refGene_gn=IL1A;refGene_tx=XM_023250911.1
A3 105114214 105114263
refGene_gn=IL1A;refGene_tx=NM_001009351.1;refGene_gn=IL1A;refGene_tx=XM_006930269.4;refGene_gn=IL1A;refGene_tx=XM_006930270.4;refGene_gn=IL1A;refGene_tx=XM_019826172.2;refGene_gn=IL1A;refGene_tx=XM_023250911.1
A3 105115052 105115275
refGene_gn=IL1A;refGene_tx=NM_001009351.1;refGene_gn=IL1A;refGene_tx=XM_006930269.4;refGene_gn=IL1A;refGene_tx=XM_006930270.4;refGene_gn=IL1A;refGene_tx=XM_019826172.2;refGene_gn=IL1A;refGene_tx=XM_023250911.1
A3 105117058 105117232
refGene_gn=IL1A;refGene_tx=NM_001009351.1;refGene_gn=IL1A;refGene_tx=XM_006930269.4;refGene_gn=IL1A;refGene_tx=XM_006930270.4;refGene_gn=IL1A;refGene_tx=XM_023250911.1
A3 105118593 105118715
refGene_gn=IL1A;refGene_tx=NM_001009351.1;refGene_gn=IL1A;refGene_tx=XM_006930269.4;refGene_gn=IL1A;refGene_tx=XM_006930270.4;refGene_gn=IL1A;refGene_tx=XM_019826172.2;refGene_gn=IL1A;refGene_tx=XM_023250911.1
A3 105120920 105121118
refGene_gn=IL1A;refGene_tx=NM_001009351.1;refGene_gn=IL1A;refGene_tx=XM_006930269.4;refGene_gn=IL1A;refGene_tx=XM_006930270.4;refGene_gn=IL1

A;refGene_tx=XM_019826172.2;refGene_gn=IL1A;refGene_tx=XM_023250911.1
A3 105138422 105138453
refGene_gn=CKAP2L;refGene_tx=XM_003984244.4
A3 105140893 105140960
refGene_gn=CKAP2L;refGene_tx=XM_003984244.4
A3 105142596 105142648
refGene_gn=CKAP2L;refGene_tx=XM_003984244.4
A3 105150675 105151931
refGene_gn=CKAP2L;refGene_tx=XM_003984244.4;refGene_gn=CKAP2L;refGene_tx=XM_019827571.2;refGene_gn=CKAP2L;refGene_tx=XM_019827572.2
A3 105155996 105156204
refGene_gn=CKAP2L;refGene_tx=XM_003984244.4;refGene_gn=CKAP2L;refGene_tx=XM_019827571.2;refGene_gn=CKAP2L;refGene_tx=XM_019827572.2
A3 105160895 105161051
refGene_gn=CKAP2L;refGene_tx=XM_003984244.4;refGene_gn=CKAP2L;refGene_tx=XM_019827571.2;refGene_gn=CKAP2L;refGene_tx=XM_019827572.2
A3 105165849 105165913
refGene_gn=CKAP2L;refGene_tx=XM_003984244.4;refGene_gn=CKAP2L;refGene_tx=XM_019827571.2;refGene_gn=CKAP2L;refGene_tx=XM_019827572.2
A3 105167037 105167227
refGene_gn=CKAP2L;refGene_tx=XM_003984244.4;refGene_gn=CKAP2L;refGene_tx=XM_019827571.2;refGene_gn=CKAP2L;refGene_tx=XM_019827572.2
A3 105168621 105168847
refGene_gn=CKAP2L;refGene_tx=XM_003984244.4;refGene_gn=CKAP2L;refGene_tx=XM_019827571.2;refGene_gn=CKAP2L;refGene_tx=XM_019827572.2
A3 105177668 105177724
refGene_gn=NT5DC4;refGene_tx=XM_023252191.1
A3 105178559 105178838
refGene_gn=NT5DC4;refGene_tx=XM_023252191.1
A3 105178964 105179031
refGene_gn=NT5DC4;refGene_tx=XM_023252191.1
A3 105179274 105179382
refGene_gn=NT5DC4;refGene_tx=XM_023252191.1
A3 105180114 105180147
refGene_gn=NT5DC4;refGene_tx=XM_023252191.1
A3 105180748 105180786
refGene_gn=NT5DC4;refGene_tx=XM_023252191.1
A3 105180788 105180831
refGene_gn=NT5DC4;refGene_tx=XM_023252191.1
A3 105181069 105181123
refGene_gn=NT5DC4;refGene_tx=XM_023252191.1
A3 105181411 105181505
refGene_gn=NT5DC4;refGene_tx=XM_023252191.1
A3 105181840 105181898
refGene_gn=NT5DC4;refGene_tx=XM_023252191.1
A3 105182007 105182099
refGene_gn=NT5DC4;refGene_tx=XM_023252191.1
A3 105182335 105182435
refGene_gn=NT5DC4;refGene_tx=XM_023252191.1
A3 105182519 105182633

refGene_gn=NT5DC4;refGene_tx=XM_023252191.1
A3 105182717 105182791
refGene_gn=NT5DC4;refGene_tx=XM_023252191.1
A3 105190514 105190661
refGene_gn=NT5DC4;refGene_tx=XM_023252191.1
A3 105190860 105191156
refGene_gn=NT5DC4;refGene_tx=XM_023252191.1
A3 105191563 105191965
refGene_gn=NT5DC4;refGene_tx=XM_023252191.1
A3 105236937 105237052
refGene_gn=SLC20A1;refGene_tx=NM_001009840.1
A3 105237053 105237094
refGene_gn=SLC20A1;refGene_tx=NM_001009840.1
A3 105238098 105238183
refGene_gn=SLC20A1;refGene_tx=NM_001009840.1
A3 105239168 105239354
refGene_gn=SLC20A1;refGene_tx=NM_001009840.1
A3 105241072 105241631
refGene_gn=SLC20A1;refGene_tx=NM_001009840.1
A3 105241751 105242021
refGene_gn=SLC20A1;refGene_tx=NM_001009840.1
A3 105243794 105243914
refGene_gn=SLC20A1;refGene_tx=NM_001009840.1
A3 105246614 105246711
refGene_gn=SLC20A1;refGene_tx=NM_001009840.1
A3 105251346 105251432
refGene_gn=SLC20A1;refGene_tx=NM_001009840.1
A3 105251618 105251759
refGene_gn=SLC20A1;refGene_tx=NM_001009840.1
A3 105251905 105252251
refGene_gn=SLC20A1;refGene_tx=NM_001009840.1
A3 105309091 105309115
refGene_gn=CHCHD5;refGene_tx=XM_011281175.3
A3 105311971 105312137
refGene_gn=CHCHD5;refGene_tx=XM_011281175.3
A3 105312237 105312389
refGene_gn=CHCHD5;refGene_tx=XM_011281175.3
A3 105322080 105322963
refGene_gn=POLR1B;refGene_tx=XM_003984245.4
A3 105323607 105323861
refGene_gn=POLR1B;refGene_tx=XM_003984245.4
A3 105325014 105325211
refGene_gn=POLR1B;refGene_tx=XM_003984245.4
A3 105328080 105328237
refGene_gn=POLR1B;refGene_tx=XM_003984245.4
A3 105328846 105329017
refGene_gn=POLR1B;refGene_tx=XM_003984245.4
A3 105333774 105333908
refGene_gn=POLR1B;refGene_tx=XM_003984245.4
A3 105334817 105335099

refGene_gn=POLR1B;refGene_tx=XM_003984245.4
A3	105336176	105336348
refGene_gn=POLR1B;refGene_tx=XM_003984245.4
A3	105338715	105338887
refGene_gn=POLR1B;refGene_tx=XM_003984245.4
A3	105339631	105339855
refGene_gn=POLR1B;refGene_tx=XM_003984245.4
A3	105340628	105340765
refGene_gn=POLR1B;refGene_tx=XM_003984245.4
A3	105341863	105341996
refGene_gn=POLR1B;refGene_tx=XM_003984245.4
A3	105343680	105343827
refGene_gn=POLR1B;refGene_tx=XM_003984245.4
A3	105344079	105344247
refGene_gn=POLR1B;refGene_tx=XM_003984245.4
A3	105348150	105348327
refGene_gn=POLR1B;refGene_tx=XM_003984245.4
A3	105359825	105359940
refGene_gn=TTL;refGene_tx=XM_011281173.2;refGene_gn=TTL;refGene_tx=XM_019827581.2
A3	105361510	105361580
refGene_gn=TTL;refGene_tx=XM_011281174.3
A3	105362867	105363011
refGene_gn=TTL;refGene_tx=XM_011281173.2;refGene_gn=TTL;refGene_tx=XM_011281174.3
A3	105372384	105372654
refGene_gn=TTL;refGene_tx=XM_011281173.2;refGene_gn=TTL;refGene_tx=XM_011281174.3;refGene_gn=TTL;refGene_tx=XM_019827581.2
A3	105373578	105373714
refGene_gn=TTL;refGene_tx=XM_011281173.2;refGene_gn=TTL;refGene_tx=XM_011281174.3;refGene_gn=TTL;refGene_tx=XM_019827581.2
A3	105382154	105382387
refGene_gn=TTL;refGene_tx=XM_011281173.2;refGene_gn=TTL;refGene_tx=XM_011281174.3;refGene_gn=TTL;refGene_tx=XM_019827581.2
A3	105393389	105393468
refGene_gn=TTL;refGene_tx=XM_011281173.2;refGene_gn=TTL;refGene_tx=XM_011281174.3;refGene_gn=TTL;refGene_tx=XM_019827581.2
A3	105396413	105396570
refGene_gn=TTL;refGene_tx=XM_011281173.2;refGene_gn=TTL;refGene_tx=XM_011281174.3;refGene_gn=TTL;refGene_tx=XM_019827581.2
A3	105406853	105407059
refGene_gn=LOC102901615;refGene_tx=XM_019827580.2;refGene_gn=LOC102901615;refGene_tx=XM_023251877.1;refGene_gn=LOC102901615;refGene_tx=XM_023251878.1;refGene_gn=LOC102901615;refGene_tx=XM_023251879.1;refGene_gn=LOC102901615;refGene_tx=XM_023251880.1;refGene_gn=LOC102901615;refGene_tx=XM_023251881.1;refGene_gn=LOC102901615;refGene_tx=XM_023251882.1
A3	105418160	105418344
refGene_gn=LOC102901615;refGene_tx=XM_019827580.2;refGene_gn=LOC102901615;refGene_tx=XM_023251877.1;refGene_gn=LOC102901615;refGene_tx=XM_023251878.1;refGene_gn=LOC102901615;refGene_tx=XM_023251879.1;refGene_gn

=LOC102901615;refGene_tx=XM_023251880.1;refGene_gn=LOC102901615;refGene_tx=XM_023251881.1;refGene_gn=LOC102901615;refGene_tx=XM_023251882.1
A3 105419832 105423120
refGene_gn=LOC102901615;refGene_tx=XM_019827580.2;refGene_gn=LOC102901615;refGene_tx=XM_023251877.1;refGene_gn=LOC102901615;refGene_tx=XM_023251878.1;refGene_gn=LOC102901615;refGene_tx=XM_023251879.1;refGene_gn=LOC102901615;refGene_tx=XM_023251880.1;refGene_gn=LOC102901615;refGene_tx=XM_023251881.1;refGene_gn=LOC102901615;refGene_tx=XM_023251882.1
A3 105425429 105425615
refGene_gn=LOC102901615;refGene_tx=XM_019827580.2;refGene_gn=LOC102901615;refGene_tx=XM_023251877.1;refGene_gn=LOC102901615;refGene_tx=XM_023251878.1;refGene_gn=LOC102901615;refGene_tx=XM_023251879.1;refGene_gn=LOC102901615;refGene_tx=XM_023251880.1;refGene_gn=LOC102901615;refGene_tx=XM_023251882.1
A3 105427939 105428112
refGene_gn=LOC102901615;refGene_tx=XM_019827580.2;refGene_gn=LOC102901615;refGene_tx=XM_023251877.1;refGene_gn=LOC102901615;refGene_tx=XM_023251878.1;refGene_gn=LOC102901615;refGene_tx=XM_023251879.1;refGene_gn=LOC102901615;refGene_tx=XM_023251880.1;refGene_gn=LOC102901615;refGene_tx=XM_023251881.1;refGene_gn=LOC102901615;refGene_tx=XM_023251882.1
A3 105430924 105431042
refGene_gn=LOC102901615;refGene_tx=XM_019827580.2;refGene_gn=LOC102901615;refGene_tx=XM_023251877.1;refGene_gn=LOC102901615;refGene_tx=XM_023251878.1;refGene_gn=LOC102901615;refGene_tx=XM_023251879.1;refGene_gn=LOC102901615;refGene_tx=XM_023251880.1;refGene_gn=LOC102901615;refGene_tx=XM_023251881.1
A3 105432614 105432803
refGene_gn=LOC102901615;refGene_tx=XM_019827580.2;refGene_gn=LOC102901615;refGene_tx=XM_023251877.1;refGene_gn=LOC102901615;refGene_tx=XM_023251878.1;refGene_gn=LOC102901615;refGene_tx=XM_023251879.1;refGene_gn=LOC102901615;refGene_tx=XM_023251880.1;refGene_gn=LOC102901615;refGene_tx=XM_023251881.1;refGene_gn=LOC102901615;refGene_tx=XM_023251882.1
A3 105444884 105446353
refGene_gn=ZC3H6;refGene_tx=XM_003984258.5;refGene_gn=ZC3H6;refGene_tx=XM_019827582.2
A3 105450665 105450908
refGene_gn=ZC3H6;refGene_tx=XM_003984258.5;refGene_gn=ZC3H6;refGene_tx=XM_019827582.2
A3 105451466 105451963
refGene_gn=ZC3H6;refGene_tx=XM_003984258.5;refGene_gn=ZC3H6;refGene_tx=XM_019827582.2
A3 105453252 105453506
refGene_gn=ZC3H6;refGene_tx=XM_003984258.5;refGene_gn=ZC3H6;refGene_tx=XM_019827582.2
A3 105454389 105454499
refGene_gn=ZC3H6;refGene_tx=XM_003984258.5;refGene_gn=ZC3H6;refGene_tx=XM_019827582.2
A3 105458441 105458553
refGene_gn=ZC3H6;refGene_tx=XM_003984258.5;refGene_gn=ZC3H6;refGene_tx=XM_019827582.2

A3	105459389	105459506
refGene_gn=ZC3H6;refGene_tx=XM_003984258.5;refGene_gn=ZC3H6;refGene_tx=XM_019827582.2
A3	105465845	105465979
refGene_gn=ZC3H6;refGene_tx=XM_003984258.5;refGene_gn=ZC3H6;refGene_tx=XM_019827582.2
A3	105468300	105468547
refGene_gn=ZC3H6;refGene_tx=XM_003984258.5;refGene_gn=ZC3H6;refGene_tx=XM_019827582.2
A3	105482563	105482686
refGene_gn=ZC3H6;refGene_tx=XM_003984258.5;refGene_gn=ZC3H6;refGene_tx=XM_019827582.2
A3	105490972	105491153
refGene_gn=ZC3H6;refGene_tx=XM_003984258.5;refGene_gn=ZC3H6;refGene_tx=XM_019827582.2
A3	105515315	105515347
refGene_gn=ZC3H6;refGene_tx=XM_003984258.5;refGene_gn=ZC3H6;refGene_tx=XM_019827582.2
A3	105520738	105520815
refGene_gn=ZC3H8;refGene_tx=XM_003984259.5;refGene_gn=ZC3H8;refGene_tx=XM_006930300.4;refGene_gn=ZC3H8;refGene_tx=XM_006930301.3;refGene_gn=ZC3H8;refGene_tx=XM_006930302.4;refGene_gn=ZC3H8;refGene_tx=XM_019827583.2;refGene_gn=ZC3H8;refGene_tx=XM_019827584.2;refGene_gn=ZC3H8;refGene_tx=XM_019827585.2;refGene_gn=ZC3H8;refGene_tx=XM_019827586.2
A3	105524997	105525094
refGene_gn=ZC3H8;refGene_tx=XM_003984259.5;refGene_gn=ZC3H8;refGene_tx=XM_006930300.4;refGene_gn=ZC3H8;refGene_tx=XM_006930301.3;refGene_gn=ZC3H8;refGene_tx=XM_006930302.4;refGene_gn=ZC3H8;refGene_tx=XM_019827583.2;refGene_gn=ZC3H8;refGene_tx=XM_019827584.2;refGene_gn=ZC3H8;refGene_tx=XM_019827586.2
A3	105533318	105533550
refGene_gn=ZC3H8;refGene_tx=XM_003984259.5;refGene_gn=ZC3H8;refGene_tx=XM_006930300.4;refGene_gn=ZC3H8;refGene_tx=XM_006930301.3;refGene_gn=ZC3H8;refGene_tx=XM_006930302.4;refGene_gn=ZC3H8;refGene_tx=XM_019827583.2;refGene_gn=ZC3H8;refGene_tx=XM_019827584.2;refGene_gn=ZC3H8;refGene_tx=XM_019827585.2;refGene_gn=ZC3H8;refGene_tx=XM_019827586.2
A3	105535588	105535722
refGene_gn=ZC3H8;refGene_tx=XM_003984259.5;refGene_gn=ZC3H8;refGene_tx=XM_006930300.4;refGene_gn=ZC3H8;refGene_tx=XM_006930301.3;refGene_gn=ZC3H8;refGene_tx=XM_006930302.4;refGene_gn=ZC3H8;refGene_tx=XM_019827583.2;refGene_gn=ZC3H8;refGene_tx=XM_019827584.2;refGene_gn=ZC3H8;refGene_tx=XM_019827585.2;refGene_gn=ZC3H8;refGene_tx=XM_019827586.2
A3	105537577	105537700
refGene_gn=ZC3H8;refGene_tx=XM_003984259.5;refGene_gn=ZC3H8;refGene_tx=XM_006930300.4;refGene_gn=ZC3H8;refGene_tx=XM_006930301.3;refGene_gn=ZC3H8;refGene_tx=XM_006930302.4;refGene_gn=ZC3H8;refGene_tx=XM_019827583.2;refGene_gn=ZC3H8;refGene_tx=XM_019827584.2;refGene_gn=ZC3H8;refGene_tx=XM_019827585.2
A3	105538298	105538410
refGene_gn=ZC3H8;refGene_tx=XM_003984259.5;refGene_gn=ZC3H8;refGene_tx

=XM_006930300.4;refGene_gn=ZC3H8;refGene_tx=XM_006930301.3;refGene_gn=ZC3H8;refGene_tx=XM_006930302.4;refGene_gn=ZC3H8;refGene_tx=XM_019827583.2;refGene_gn=ZC3H8;refGene_tx=XM_019827584.2;refGene_gn=ZC3H8;refGene_tx=XM_019827585.2;refGene_gn=ZC3H8;refGene_tx=XM_019827586.2
A3 105539785 105539895
refGene_gn=ZC3H8;refGene_tx=XM_003984259.5;refGene_gn=ZC3H8;refGene_tx=XM_006930300.4;refGene_gn=ZC3H8;refGene_tx=XM_006930301.3;refGene_gn=ZC3H8;refGene_tx=XM_006930302.4;refGene_gn=ZC3H8;refGene_tx=XM_019827583.2;refGene_gn=ZC3H8;refGene_tx=XM_019827584.2;refGene_gn=ZC3H8;refGene_tx=XM_019827585.2;refGene_gn=ZC3H8;refGene_tx=XM_019827586.2
A3 105541002 105541032
refGene_gn=ZC3H8;refGene_tx=XM_003984259.5;refGene_gn=ZC3H8;refGene_tx=XM_006930300.4;refGene_gn=ZC3H8;refGene_tx=XM_006930301.3;refGene_gn=ZC3H8;refGene_tx=XM_006930302.4;refGene_gn=ZC3H8;refGene_tx=XM_019827583.2;refGene_gn=ZC3H8;refGene_tx=XM_019827584.2;refGene_gn=ZC3H8;refGene_tx=XM_019827585.2;refGene_gn=ZC3H8;refGene_tx=XM_019827586.2
A3 105565447 105565820
refGene_gn=FBLN7;refGene_tx=XM_023251884.1
A3 105567194 105567333
refGene_gn=FBLN7;refGene_tx=XM_023251884.1
A3 105568274 105568412
refGene_gn=FBLN7;refGene_tx=XM_023251884.1
A3 105569364 105569505
refGene_gn=FBLN7;refGene_tx=XM_023251884.1
A3 105572349 105572475
refGene_gn=FBLN7;refGene_tx=XM_023251884.1
A3 105584689 105584860
refGene_gn=FBLN7;refGene_tx=XM_023251884.1
A3 105590232 105590392
refGene_gn=FBLN7;refGene_tx=XM_023251884.1
A3 105614798 105614876
refGene_gn=FBLN7;refGene_tx=XM_023251884.1
A3 105627606 105627666
refGene_gn=TMEM87B;refGene_tx=XM_011281171.2;refGene_gn=TMEM87B;refGene_tx=XM_019827588.2
A3 105628814 105628845
refGene_gn=TMEM87B;refGene_tx=XM_011281171.2;refGene_gn=TMEM87B;refGene_tx=XM_019827588.2
A3 105631753 105631806
refGene_gn=TMEM87B;refGene_tx=XM_011281171.2;refGene_gn=TMEM87B;refGene_tx=XM_019827588.2
A3 105633723 105633797
refGene_gn=TMEM87B;refGene_tx=XM_011281171.2;refGene_gn=TMEM87B;refGene_tx=XM_019827588.2
A3 105637137 105637211
refGene_gn=TMEM87B;refGene_tx=XM_011281171.2;refGene_gn=TMEM87B;refGene_tx=XM_019827588.2
A3 105638571 105638675
refGene_gn=TMEM87B;refGene_tx=XM_011281171.2;refGene_gn=TMEM87B;refGene_tx=XM_019827588.2

A3 105640958 105641017
refGene_gn=TMEM87B;refGene_tx=XM_011281171.2;refGene_gn=TMEM87B;refGene_tx=XM_019827588.2
A3 105641103 105641212
refGene_gn=TMEM87B;refGene_tx=XM_011281171.2;refGene_gn=TMEM87B;refGene_tx=XM_019827588.2
A3 105642886 105642958
refGene_gn=TMEM87B;refGene_tx=XM_019827588.2
A3 105646829 105646901
refGene_gn=TMEM87B;refGene_tx=XM_011281171.2
A3 105648525 105648619
refGene_gn=TMEM87B;refGene_tx=XM_011281171.2;refGene_gn=TMEM87B;refGene_tx=XM_019827588.2
A3 105652992 105653092
refGene_gn=TMEM87B;refGene_tx=XM_011281171.2;refGene_gn=TMEM87B;refGene_tx=XM_019827588.2
A3 105657267 105657451
refGene_gn=TMEM87B;refGene_tx=XM_011281171.2;refGene_gn=TMEM87B;refGene_tx=XM_019827588.2
A3 105657922 105657984
refGene_gn=TMEM87B;refGene_tx=XM_011281171.2;refGene_gn=TMEM87B;refGene_tx=XM_019827588.2
A3 105659368 105659459
refGene_gn=TMEM87B;refGene_tx=XM_011281171.2;refGene_gn=TMEM87B;refGene_tx=XM_019827588.2
A3 105661276 105661330
refGene_gn=TMEM87B;refGene_tx=XM_011281171.2;refGene_gn=TMEM87B;refGene_tx=XM_019827588.2
A3 105662783 105662921
refGene_gn=TMEM87B;refGene_tx=XM_011281171.2;refGene_gn=TMEM87B;refGene_tx=XM_019827588.2
A3 105665634 105665726
refGene_gn=TMEM87B;refGene_tx=XM_011281171.2;refGene_gn=TMEM87B;refGene_tx=XM_019827588.2
A3 105670841 105670902
refGene_gn=TMEM87B;refGene_tx=XM_011281171.2;refGene_gn=TMEM87B;refGene_tx=XM_019827588.2
A3 105674494 105674689
refGene_gn=TMEM87B;refGene_tx=XM_011281171.2;refGene_gn=TMEM87B;refGene_tx=XM_019827588.2
A3 105676473 105676507
refGene_gn=MERTK;refGene_tx=XM_023251887.1
A3 105688557 105689071
refGene_gn=MERTK;refGene_tx=XM_023251885.1;refGene_gn=MERTK;refGene_tx=XM_023251886.1
A3 105696631 105696768
refGene_gn=MERTK;refGene_tx=XM_023251885.1;refGene_gn=MERTK;refGene_tx=XM_023251886.1;refGene_gn=MERTK;refGene_tx=XM_023251887.1
A3 105697678 105697838
refGene_gn=MERTK;refGene_tx=XM_023251885.1;refGene_gn=MERTK;refGene_tx

=XM_023251886.1;refGene_gn=MERTK;refGene_tx=XM_023251887.1
A3 105699303 105699413
refGene_gn=MERTK;refGene_tx=XM_023251885.1;refGene_gn=MERTK;refGene_tx
=XM_023251886.1;refGene_gn=MERTK;refGene_tx=XM_023251887.1
A3 105705463 105705582
refGene_gn=MERTK;refGene_tx=XM_023251885.1;refGene_gn=MERTK;refGene_tx
=XM_023251886.1;refGene_gn=MERTK;refGene_tx=XM_023251887.1
A3 105708071 105708164
refGene_gn=MERTK;refGene_tx=XM_023251885.1;refGene_gn=MERTK;refGene_tx
=XM_023251886.1;refGene_gn=MERTK;refGene_tx=XM_023251887.1
A3 105712181 105712262
refGene_gn=MERTK;refGene_tx=XM_023251885.1;refGene_gn=MERTK;refGene_tx
=XM_023251886.1;refGene_gn=MERTK;refGene_tx=XM_023251887.1
A3 105713398 105713494
refGene_gn=MERTK;refGene_tx=XM_023251885.1;refGene_gn=MERTK;refGene_tx
=XM_023251886.1;refGene_gn=MERTK;refGene_tx=XM_023251887.1
A3 105714998 105715084
refGene_gn=MERTK;refGene_tx=XM_023251885.1;refGene_gn=MERTK;refGene_tx
=XM_023251886.1;refGene_gn=MERTK;refGene_tx=XM_023251887.1
A3 105717768 105717922
refGene_gn=MERTK;refGene_tx=XM_023251885.1;refGene_gn=MERTK;refGene_tx
=XM_023251886.1;refGene_gn=MERTK;refGene_tx=XM_023251887.1
A3 105720206 105720360
refGene_gn=MERTK;refGene_tx=XM_023251885.1;refGene_gn=MERTK;refGene_tx
=XM_023251886.1;refGene_gn=MERTK;refGene_tx=XM_023251887.1
A3 105728223 105728375
refGene_gn=MERTK;refGene_tx=XM_023251885.1;refGene_gn=MERTK;refGene_tx
=XM_023251886.1;refGene_gn=MERTK;refGene_tx=XM_023251887.1
A3 105732720 105732904
refGene_gn=MERTK;refGene_tx=XM_023251885.1;refGene_gn=MERTK;refGene_tx
=XM_023251886.1;refGene_gn=MERTK;refGene_tx=XM_023251887.1
A3 105740409 105740525
refGene_gn=MERTK;refGene_tx=XM_023251885.1;refGene_gn=MERTK;refGene_tx
=XM_023251886.1;refGene_gn=MERTK;refGene_tx=XM_023251887.1
A3 105743646 105743733
refGene_gn=MERTK;refGene_tx=XM_023251885.1;refGene_gn=MERTK;refGene_tx
=XM_023251886.1;refGene_gn=MERTK;refGene_tx=XM_023251887.1
A3 105761047 105761221
refGene_gn=MERTK;refGene_tx=XM_023251885.1;refGene_gn=MERTK;refGene_tx
=XM_023251886.1;refGene_gn=MERTK;refGene_tx=XM_023251887.1
A3 105763204 105763305
refGene_gn=MERTK;refGene_tx=XM_023251885.1;refGene_gn=MERTK;refGene_tx
=XM_023251886.1;refGene_gn=MERTK;refGene_tx=XM_023251887.1
A3 105773771 105774177
refGene_gn=MERTK;refGene_tx=XM_023251885.1;refGene_gn=MERTK;refGene_tx
=XM_023251886.1;refGene_gn=MERTK;refGene_tx=XM_023251887.1
A3 105802031 105802092
refGene_gn=MERTK;refGene_tx=XM_023251885.1;refGene_gn=MERTK;refGene_tx
=XM_023251887.1
A3 105823758 105823792

refGene_gn=MERTK;refGene_tx=XM_023251886.1
A3 105825987 105826200
refGene_gn=ANAPC1;refGene_tx=XM_003984262.5
A3 105827786 105827948
refGene_gn=ANAPC1;refGene_tx=XM_003984262.5
A3 105834169 105834221
refGene_gn=ANAPC1;refGene_tx=XM_003984262.5
A3 105834446 105834547
refGene_gn=ANAPC1;refGene_tx=XM_003984262.5
A3 105835202 105835285
refGene_gn=ANAPC1;refGene_tx=XM_003984262.5
A3 105836598 105836672
refGene_gn=ANAPC1;refGene_tx=XM_003984262.5
A3 105838969 105839115
refGene_gn=ANAPC1;refGene_tx=XM_003984262.5
A3 105839338 105839559
refGene_gn=ANAPC1;refGene_tx=XM_003984262.5
A3 105840249 105840459
refGene_gn=ANAPC1;refGene_tx=XM_003984262.5
A3 105843251 105843347
refGene_gn=ANAPC1;refGene_tx=XM_003984262.5
A3 105844773 105844864
refGene_gn=ANAPC1;refGene_tx=XM_003984262.5
A3 105844962 105845028
refGene_gn=ANAPC1;refGene_tx=XM_003984262.5
A3 105848747 105848882
refGene_gn=ANAPC1;refGene_tx=XM_003984262.5
A3 105853055 105853196
refGene_gn=ANAPC1;refGene_tx=XM_003984262.5
A3 105854623 105854684
refGene_gn=ANAPC1;refGene_tx=XM_003984262.5
A3 105856803 105856991
refGene_gn=ANAPC1;refGene_tx=XM_003984262.5
A3 105857631 105857706
refGene_gn=ANAPC1;refGene_tx=XM_003984262.5
A3 105860326 105860595
refGene_gn=ANAPC1;refGene_tx=XM_003984262.5
A3 105861824 105861916
refGene_gn=ANAPC1;refGene_tx=XM_003984262.5
A3 105862686 105862835
refGene_gn=ANAPC1;refGene_tx=XM_003984262.5
A3 105866007 105866086
refGene_gn=ANAPC1;refGene_tx=XM_003984262.5
A3 105866970 105867007
refGene_gn=ANAPC1;refGene_tx=XM_003984262.5
A3 105867101 105867172
refGene_gn=ANAPC1;refGene_tx=XM_003984262.5
A3 105869653 105869832
refGene_gn=ANAPC1;refGene_tx=XM_003984262.5
A3 105873211 105873239

refGene_gn=ANAPC1;refGene_tx=XM_003984262.5
A3	105873241	105873425
refGene_gn=ANAPC1;refGene_tx=XM_003984262.5
A3	105875695	105875814
refGene_gn=ANAPC1;refGene_tx=XM_003984262.5
A3	105880397	105880669
refGene_gn=ANAPC1;refGene_tx=XM_003984262.5
A3	105883015	105883250
refGene_gn=ANAPC1;refGene_tx=XM_003984262.5
A3	105884758	105884850
refGene_gn=ANAPC1;refGene_tx=XM_003984262.5
A3	105886420	105886555
refGene_gn=ANAPC1;refGene_tx=XM_003984262.5
A3	105886743	105886826
refGene_gn=ANAPC1;refGene_tx=XM_003984262.5
A3	105887733	105887812
refGene_gn=ANAPC1;refGene_tx=XM_003984262.5
A3	105888530	105888605
refGene_gn=ANAPC1;refGene_tx=XM_003984262.5
A3	105891052	105891129
refGene_gn=ANAPC1;refGene_tx=XM_003984262.5
A3	105891621	105891712
refGene_gn=ANAPC1;refGene_tx=XM_003984262.5
A3	105891807	105891861
refGene_gn=ANAPC1;refGene_tx=XM_003984262.5
A3	105892765	105892959
refGene_gn=ANAPC1;refGene_tx=XM_003984262.5
A3	105896549	105896636
refGene_gn=ANAPC1;refGene_tx=XM_003984262.5
A3	105899338	105899456
refGene_gn=ANAPC1;refGene_tx=XM_003984262.5
A3	105900405	105900483
refGene_gn=ANAPC1;refGene_tx=XM_003984262.5
A3	105901105	105901173
refGene_gn=ANAPC1;refGene_tx=XM_003984262.5
A3	105902191	105902330
refGene_gn=ANAPC1;refGene_tx=XM_003984262.5
A3	105904455	105904542
refGene_gn=ANAPC1;refGene_tx=XM_003984262.5
A3	105906264	105906360
refGene_gn=ANAPC1;refGene_tx=XM_003984262.5
A3	105908044	105908243
refGene_gn=ANAPC1;refGene_tx=XM_003984262.5
A3	105913965	105914100
refGene_gn=ANAPC1;refGene_tx=XM_003984262.5
A3	105920151	105920267
refGene_gn=ANAPC1;refGene_tx=XM_003984262.5
A3	105928841	105928901
refGene_gn=LOC102901812;refGene_tx=XM_011281241.3;refGene_gn=LOC102901
812;refGene_tx=XM_019827595.2;refGene_gn=LOC102901812;refGene_tx=XM_01

9827596.2
A3	105928992	105929167
refGene_gn=LOC102901812;refGene_tx=XM_011281241.3;refGene_gn=LOC102901812;refGene_tx=XM_019827595.2;refGene_gn=LOC102901812;refGene_tx=XM_019827596.2
A3	105929691	105929777
refGene_gn=LOC102901812;refGene_tx=XM_011281241.3;refGene_gn=LOC102901812;refGene_tx=XM_019827595.2;refGene_gn=LOC102901812;refGene_tx=XM_019827596.2
A3	105930127	105930186
refGene_gn=LOC102901812;refGene_tx=XM_011281241.3;refGene_gn=LOC102901812;refGene_tx=XM_019827595.2;refGene_gn=LOC102901812;refGene_tx=XM_019827596.2
A3	105931760	105931955
refGene_gn=LOC102901812;refGene_tx=XM_011281241.3;refGene_gn=LOC102901812;refGene_tx=XM_019827595.2;refGene_gn=LOC102901812;refGene_tx=XM_019827596.2
A3	105932141	105932258
refGene_gn=LOC102901812;refGene_tx=XM_011281241.3;refGene_gn=LOC102901812;refGene_tx=XM_019827595.2;refGene_gn=LOC102901812;refGene_tx=XM_019827596.2
A3	105932725	105932838
refGene_gn=LOC102901812;refGene_tx=XM_011281241.3;refGene_gn=LOC102901812;refGene_tx=XM_019827595.2;refGene_gn=LOC102901812;refGene_tx=XM_019827596.2
A3	105933205	105933213
refGene_gn=LOC102901812;refGene_tx=XM_011281241.3
A3	106472996	106473095
refGene_gn=BCL2L11;refGene_tx=XM_003984263.5;refGene_gn=BCL2L11;refGene_tx=XM_006930307.4;refGene_gn=BCL2L11;refGene_tx=XM_006930308.3;refGene_gn=BCL2L11;refGene_tx=XM_006930311.3;refGene_gn=BCL2L11;refGene_tx=XM_019827598.2;refGene_gn=BCL2L11;refGene_tx=XM_019827600.2
A3	106485605	106485715
refGene_gn=BCL2L11;refGene_tx=XM_003984263.5;refGene_gn=BCL2L11;refGene_tx=XM_006930307.4;refGene_gn=BCL2L11;refGene_tx=XM_006930308.3;refGene_gn=BCL2L11;refGene_tx=XM_006930311.3;refGene_gn=BCL2L11;refGene_tx=XM_019827598.2
A3	106514519	106514901
refGene_gn=BCL2L11;refGene_tx=XM_003984263.5;refGene_gn=BCL2L11;refGene_tx=XM_006930307.4;refGene_gn=BCL2L11;refGene_tx=XM_006930308.3;refGene_gn=BCL2L11;refGene_tx=XM_006930311.3;refGene_gn=BCL2L11;refGene_tx=XM_019827598.2;refGene_gn=BCL2L11;refGene_tx=XM_019827600.2
A3	106516962	106517901
refGene_gn=LOC111559849;refGene_tx=XM_023251889.1;refGene_gn=LOC111559849;refGene_tx=XM_023251890.1;refGene_gn=LOC111559849;refGene_tx=XM_023251891.1;refGene_gn=LOC111559849;refGene_tx=XM_023251892.1;refGene_gn=LOC111559849;refGene_tx=XM_023251893.1;refGene_gn=LOC111559849;refGene_tx=XM_023251894.1
A3	106518146	106518509
refGene_gn=LOC111559849;refGene_tx=XM_023251889.1;refGene_gn=LOC111559

849;refGene_tx=XM_023251890.1;refGene_gn=LOC111559849;refGene_tx=XM_023251891.1;refGene_gn=LOC111559849;refGene_tx=XM_023251892.1;refGene_gn=LOC111559849;refGene_tx=XM_023251894.1
A3 106640218 106640692
refGene_gn=LOC111559897;refGene_tx=XM_023252192.1
A3 106667161 106667251
refGene_gn=LOC111559897;refGene_tx=XM_023252192.1
A3 106691040 106691347
refGene_gn=LOC111559897;refGene_tx=XM_023252192.1
A3 106707822 106707939
refGene_gn=LOC111559897;refGene_tx=XM_023252192.1
A3 106709164 106709300
refGene_gn=LOC111559897;refGene_tx=XM_023252192.1
A3 106710656 106710750
refGene_gn=LOC111559897;refGene_tx=XM_023252192.1
A3 106809184 106809332
refGene_gn=LOC101088661;refGene_tx=XM_023252193.1
A3 106813466 106813539
refGene_gn=LOC101088661;refGene_tx=XM_023252193.1
A3 106815330 106815417
refGene_gn=LOC101088661;refGene_tx=XM_023252193.1
A3 106816866 106816981
refGene_gn=LOC101088661;refGene_tx=XM_023252193.1
A3 106821471 106821552
refGene_gn=LOC101088661;refGene_tx=XM_023252193.1
A3 106894585 106894611
refGene_gn=BUB1;refGene_tx=XM_011281169.3
A3 106898040 106898100
refGene_gn=BUB1;refGene_tx=XM_011281169.3
A3 106899061 106899200
refGene_gn=BUB1;refGene_tx=XM_011281169.3
A3 106900476 106900673
refGene_gn=BUB1;refGene_tx=XM_011281169.3
A3 106902759 106902803
refGene_gn=BUB1;refGene_tx=XM_011281169.3
A3 106903428 106903529
refGene_gn=BUB1;refGene_tx=XM_011281169.3
A3 106905499 106905552
refGene_gn=BUB1;refGene_tx=XM_011281169.3
A3 106905645 106905830
refGene_gn=BUB1;refGene_tx=XM_011281169.3
A3 106907123 106907275
refGene_gn=BUB1;refGene_tx=XM_011281169.3
A3 106912403 106912663
refGene_gn=BUB1;refGene_tx=XM_011281169.3
A3 106914468 106914527
refGene_gn=BUB1;refGene_tx=XM_011281169.3
A3 106915819 106915948
refGene_gn=BUB1;refGene_tx=XM_011281169.3
A3 106916043 106916154

refGene_gn=BUB1;refGene_tx=XM_011281169.3
A3 106916908 106917008
refGene_gn=BUB1;refGene_tx=XM_011281169.3
A3 106918059 106918141
refGene_gn=BUB1;refGene_tx=XM_011281169.3
A3 106921154 106921329
refGene_gn=BUB1;refGene_tx=XM_011281169.3
A3 106923314 106923402
refGene_gn=BUB1;refGene_tx=XM_011281169.3
A3 106927476 106927715
refGene_gn=BUB1;refGene_tx=XM_011281169.3
A3 106928877 106929021
refGene_gn=BUB1;refGene_tx=XM_011281169.3
A3 106937613 106937729
refGene_gn=BUB1;refGene_tx=XM_011281169.3
A3 106938048 106938210
refGene_gn=BUB1;refGene_tx=XM_011281169.3
A3 106938366 106938524
refGene_gn=BUB1;refGene_tx=XM_011281169.3
A3 106938616 106938791
refGene_gn=BUB1;refGene_tx=XM_011281169.3
A3 106939356 106939463
refGene_gn=BUB1;refGene_tx=XM_011281169.3
A3 106941309 106941505
refGene_gn=BUB1;refGene_tx=XM_011281169.3
A3 106949332 106949478
refGene_gn=LOC101089184;refGene_tx=XM_023251895.1;refGene_gn=LOC101089184;refGene_tx=XM_023251896.1;refGene_gn=LOC101089184;refGene_tx=XM_023251897.1;refGene_gn=LOC101089184;refGene_tx=XM_023251898.1
A3 106950698 106950823
refGene_gn=LOC101089184;refGene_tx=XM_023251895.1;refGene_gn=LOC101089184;refGene_tx=XM_023251896.1;refGene_gn=LOC101089184;refGene_tx=XM_023251897.1;refGene_gn=LOC101089184;refGene_tx=XM_023251898.1
A3 106951057 106951171
refGene_gn=LOC101089184;refGene_tx=XM_023251895.1;refGene_gn=LOC101089184;refGene_tx=XM_023251896.1;refGene_gn=LOC101089184;refGene_tx=XM_023251897.1;refGene_gn=LOC101089184;refGene_tx=XM_023251898.1
A3 106953006 106953097
refGene_gn=LOC101089184;refGene_tx=XM_023251895.1;refGene_gn=LOC101089184;refGene_tx=XM_023251896.1;refGene_gn=LOC101089184;refGene_tx=XM_023251897.1;refGene_gn=LOC101089184;refGene_tx=XM_023251898.1
A3 106953250 106953470
refGene_gn=LOC101089184;refGene_tx=XM_023251895.1;refGene_gn=LOC101089184;refGene_tx=XM_023251896.1;refGene_gn=LOC101089184;refGene_tx=XM_023251897.1;refGene_gn=LOC101089184;refGene_tx=XM_023251898.1
A3 106955498 106955568
refGene_gn=LOC101089184;refGene_tx=XM_023251895.1;refGene_gn=LOC101089184;refGene_tx=XM_023251896.1;refGene_gn=LOC101089184;refGene_tx=XM_023251897.1;refGene_gn=LOC101089184;refGene_tx=XM_023251898.1
A3 106956120 106956208

refGene_gn=LOC101089184;refGene_tx=XM_023251895.1;refGene_gn=LOC101089184;refGene_tx=XM_023251896.1;refGene_gn=LOC101089184;refGene_tx=XM_023251897.1;refGene_gn=LOC101089184;refGene_tx=XM_023251898.1
A3 106957357 106957422
refGene_gn=LOC101089184;refGene_tx=XM_023251895.1;refGene_gn=LOC101089184;refGene_tx=XM_023251896.1;refGene_gn=LOC101089184;refGene_tx=XM_023251897.1
A3 106958682 106958868
refGene_gn=LOC101089184;refGene_tx=XM_023251895.1;refGene_gn=LOC101089184;refGene_tx=XM_023251896.1;refGene_gn=LOC101089184;refGene_tx=XM_023251897.1
A3 106960301 106960446
refGene_gn=LOC101089184;refGene_tx=XM_023251895.1;refGene_gn=LOC101089184;refGene_tx=XM_023251896.1;refGene_gn=LOC101089184;refGene_tx=XM_023251897.1
A3 106969195 106969397
refGene_gn=LOC101089184;refGene_tx=XM_023251895.1;refGene_gn=LOC101089184;refGene_tx=XM_023251896.1;refGene_gn=LOC101089184;refGene_tx=XM_023251897.1;refGene_gn=LOC101089184;refGene_tx=XM_023251898.1
A3 106971409 106971536
refGene_gn=LOC101089184;refGene_tx=XM_023251895.1;refGene_gn=LOC101089184;refGene_tx=XM_023251896.1;refGene_gn=LOC101089184;refGene_tx=XM_023251897.1;refGene_gn=LOC101089184;refGene_tx=XM_023251898.1
A3 106972960 106973062
refGene_gn=LOC101089184;refGene_tx=XM_023251895.1;refGene_gn=LOC101089184;refGene_tx=XM_023251896.1;refGene_gn=LOC101089184;refGene_tx=XM_023251897.1;refGene_gn=LOC101089184;refGene_tx=XM_023251898.1
A3 106973290 106973409
refGene_gn=LOC101089184;refGene_tx=XM_023251895.1;refGene_gn=LOC101089184;refGene_tx=XM_023251896.1;refGene_gn=LOC101089184;refGene_tx=XM_023251897.1;refGene_gn=LOC101089184;refGene_tx=XM_023251898.1
A3 106973509 106973626
refGene_gn=LOC101089184;refGene_tx=XM_023251895.1;refGene_gn=LOC101089184;refGene_tx=XM_023251896.1;refGene_gn=LOC101089184;refGene_tx=XM_023251897.1;refGene_gn=LOC101089184;refGene_tx=XM_023251898.1
A3 106977979 106978108
refGene_gn=LOC101089184;refGene_tx=XM_023251895.1;refGene_gn=LOC101089184;refGene_tx=XM_023251896.1;refGene_gn=LOC101089184;refGene_tx=XM_023251897.1;refGene_gn=LOC101089184;refGene_tx=XM_023251898.1
A3 106980906 106981021
refGene_gn=LOC101089184;refGene_tx=XM_023251895.1;refGene_gn=LOC101089184;refGene_tx=XM_023251896.1;refGene_gn=LOC101089184;refGene_tx=XM_023251897.1;refGene_gn=LOC101089184;refGene_tx=XM_023251898.1
A3 106982053 106982145
refGene_gn=LOC101089184;refGene_tx=XM_023251895.1;refGene_gn=LOC101089184;refGene_tx=XM_023251898.1
A3 106990940 106991015
refGene_gn=LOC101089184;refGene_tx=XM_023251895.1;refGene_gn=LOC101089184;refGene_tx=XM_023251898.1
A3 106991131 106991363

refGene_gn=LOC111559898;refGene_tx=XM_023252194.1
A3 107006221 107006373
refGene_gn=LOC111559898;refGene_tx=XM_023252194.1
A3 107009719 107009743
refGene_gn=LOC102902265;refGene_tx=XM_023252195.1
A3 107010876 107011140
refGene_gn=LOC102902265;refGene_tx=XM_023252195.1
A3 107017791 107017860
refGene_gn=NPHP1;refGene_tx=XM_011281167.3;refGene_gn=NPHP1;refGene_tx=XM_019827602.2;refGene_gn=NPHP1;refGene_tx=XM_019827603.2;refGene_gn=NPHP1;refGene_tx=XM_019827604.2;refGene_gn=NPHP1;refGene_tx=XM_023251899.1
A3 107020997 107021071
refGene_gn=NPHP1;refGene_tx=XM_011281167.3;refGene_gn=NPHP1;refGene_tx=XM_019827602.2;refGene_gn=NPHP1;refGene_tx=XM_019827603.2;refGene_gn=NPHP1;refGene_tx=XM_019827604.2;refGene_gn=NPHP1;refGene_tx=XM_023251899.1
A3 107026984 107027045
refGene_gn=NPHP1;refGene_tx=XM_011281167.3;refGene_gn=NPHP1;refGene_tx=XM_019827602.2;refGene_gn=NPHP1;refGene_tx=XM_019827603.2;refGene_gn=NPHP1;refGene_tx=XM_019827604.2;refGene_gn=NPHP1;refGene_tx=XM_023251899.1
A3 107028660 107028785
refGene_gn=NPHP1;refGene_tx=XM_011281167.3;refGene_gn=NPHP1;refGene_tx=XM_019827602.2;refGene_gn=NPHP1;refGene_tx=XM_019827603.2;refGene_gn=NPHP1;refGene_tx=XM_019827604.2;refGene_gn=NPHP1;refGene_tx=XM_023251899.1
A3 107031033 107031262
refGene_gn=NPHP1;refGene_tx=XM_011281167.3;refGene_gn=NPHP1;refGene_tx=XM_019827602.2;refGene_gn=NPHP1;refGene_tx=XM_019827603.2;refGene_gn=NPHP1;refGene_tx=XM_019827604.2;refGene_gn=NPHP1;refGene_tx=XM_023251899.1
A3 107031563 107031665
refGene_gn=NPHP1;refGene_tx=XM_011281167.3;refGene_gn=NPHP1;refGene_tx=XM_019827602.2;refGene_gn=NPHP1;refGene_tx=XM_019827603.2;refGene_gn=NPHP1;refGene_tx=XM_019827604.2;refGene_gn=NPHP1;refGene_tx=XM_023251899.1
A3 107035456 107035557
refGene_gn=NPHP1;refGene_tx=XM_011281167.3;refGene_gn=NPHP1;refGene_tx=XM_019827602.2;refGene_gn=NPHP1;refGene_tx=XM_019827603.2;refGene_gn=NPHP1;refGene_tx=XM_019827604.2;refGene_gn=NPHP1;refGene_tx=XM_023251899.1
A3 107035877 107035920
refGene_gn=NPHP1;refGene_tx=XM_011281167.3;refGene_gn=NPHP1;refGene_tx=XM_019827602.2;refGene_gn=NPHP1;refGene_tx=XM_019827603.2;refGene_gn=NPHP1;refGene_tx=XM_019827604.2;refGene_gn=NPHP1;refGene_tx=XM_023251899.1
A3 107037221 107037309
refGene_gn=NPHP1;refGene_tx=XM_011281167.3;refGene_gn=NPHP1;refGene_tx=XM_019827602.2;refGene_gn=NPHP1;refGene_tx=XM_019827603.2;refGene_gn=

NPHP1;refGene_tx=XM_019827604.2;refGene_gn=NPHP1;refGene_tx=XM_0232518
99.1
A3 107038876 107038971
refGene_gn=NPHP1;refGene_tx=XM_011281167.3;refGene_gn=NPHP1;refGene_tx
=XM_019827602.2;refGene_gn=NPHP1;refGene_tx=XM_019827603.2;refGene_gn=
NPHP1;refGene_tx=XM_019827604.2;refGene_gn=NPHP1;refGene_tx=XM_0232518
99.1
A3 107041022 107041151
refGene_gn=NPHP1;refGene_tx=XM_011281167.3;refGene_gn=NPHP1;refGene_tx
=XM_019827602.2;refGene_gn=NPHP1;refGene_tx=XM_019827603.2;refGene_gn=
NPHP1;refGene_tx=XM_019827604.2;refGene_gn=NPHP1;refGene_tx=XM_0232518
99.1
A3 107054288 107054363
refGene_gn=NPHP1;refGene_tx=XM_011281167.3;refGene_gn=NPHP1;refGene_tx
=XM_019827602.2;refGene_gn=NPHP1;refGene_tx=XM_019827603.2;refGene_gn=
NPHP1;refGene_tx=XM_019827604.2;refGene_gn=NPHP1;refGene_tx=XM_0232518
99.1
A3 107056123 107056234
refGene_gn=NPHP1;refGene_tx=XM_011281167.3;refGene_gn=NPHP1;refGene_tx
=XM_019827602.2;refGene_gn=NPHP1;refGene_tx=XM_019827603.2;refGene_gn=
NPHP1;refGene_tx=XM_019827604.2;refGene_gn=NPHP1;refGene_tx=XM_0232518
99.1
A3 107057512 107057595
refGene_gn=NPHP1;refGene_tx=XM_011281167.3;refGene_gn=NPHP1;refGene_tx
=XM_019827602.2;refGene_gn=NPHP1;refGene_tx=XM_019827603.2;refGene_gn=
NPHP1;refGene_tx=XM_019827604.2;refGene_gn=NPHP1;refGene_tx=XM_0232518
99.1
A3 107060513 107060590
refGene_gn=NPHP1;refGene_tx=XM_011281167.3;refGene_gn=NPHP1;refGene_tx
=XM_019827602.2;refGene_gn=NPHP1;refGene_tx=XM_019827603.2;refGene_gn=
NPHP1;refGene_tx=XM_023251899.1
A3 107061534 107061634
refGene_gn=NPHP1;refGene_tx=XM_011281167.3;refGene_gn=NPHP1;refGene_tx
=XM_019827602.2;refGene_gn=NPHP1;refGene_tx=XM_019827603.2;refGene_gn=
NPHP1;refGene_tx=XM_023251899.1
A3 107064654 107064778
refGene_gn=NPHP1;refGene_tx=XM_023251899.1
A3 107070353 107070466
refGene_gn=NPHP1;refGene_tx=XM_011281167.3;refGene_gn=NPHP1;refGene_tx
=XM_019827602.2;refGene_gn=NPHP1;refGene_tx=XM_019827603.2;refGene_gn=
NPHP1;refGene_tx=XM_019827604.2
A3 107073717 107073791
refGene_gn=NPHP1;refGene_tx=XM_011281167.3;refGene_gn=NPHP1;refGene_tx
=XM_019827602.2;refGene_gn=NPHP1;refGene_tx=XM_019827603.2;refGene_gn=
NPHP1;refGene_tx=XM_019827604.2
A3 107077209 107077254
refGene_gn=NPHP1;refGene_tx=XM_011281167.3;refGene_gn=NPHP1;refGene_tx
=XM_019827602.2;refGene_gn=NPHP1;refGene_tx=XM_019827603.2;refGene_gn=
NPHP1;refGene_tx=XM_019827604.2
A3 107078980 107079253

refGene_gn=NPHP1;refGene_tx=XM_011281167.3;refGene_gn=NPHP1;refGene_tx=XM_019827602.2;refGene_gn=NPHP1;refGene_tx=XM_019827603.2;refGene_gn=NPHP1;refGene_tx=XM_019827604.2
A3 107092998 107093106
refGene_gn=MALL;refGene_tx=XM_019827606.2;refGene_gn=MALL;refGene_tx=XM_019827607.2;refGene_gn=MALL;refGene_tx=XM_019827608.2
A3 107113392 107113560
refGene_gn=MALL;refGene_tx=XM_019827606.2;refGene_gn=MALL;refGene_tx=XM_019827607.2
A3 107115975 107116098
refGene_gn=MALL;refGene_tx=XM_019827606.2;refGene_gn=MALL;refGene_tx=XM_019827607.2;refGene_gn=MALL;refGene_tx=XM_019827608.2
A3 107118160 107118378
refGene_gn=MALL;refGene_tx=XM_019827606.2;refGene_gn=MALL;refGene_tx=XM_019827607.2;refGene_gn=MALL;refGene_tx=XM_019827608.2
A3 107151203 107151296
refGene_gn=MAL;refGene_tx=XM_003984268.5;refGene_gn=MAL;refGene_tx=XM_019827609.2
A3 107172411 107172579
refGene_gn=MAL;refGene_tx=XM_003984268.5
A3 107173792 107173918
refGene_gn=MAL;refGene_tx=XM_003984268.5
A3 107178296 107178371
refGene_gn=MAL;refGene_tx=XM_003984268.5;refGene_gn=MAL;refGene_tx=XM_019827609.2
A3 107206739 107206964
refGene_gn=MRPS5;refGene_tx=XM_003984272.5;refGene_gn=MRPS5;refGene_tx=XM_019827610.2
A3 107210101 107210238
refGene_gn=MRPS5;refGene_tx=XM_003984272.5;refGene_gn=MRPS5;refGene_tx=XM_019827610.2
A3 107215608 107215671
refGene_gn=MRPS5;refGene_tx=XM_003984272.5;refGene_gn=MRPS5;refGene_tx=XM_019827610.2
A3 107215955 107216013
refGene_gn=MRPS5;refGene_tx=XM_003984272.5;refGene_gn=MRPS5;refGene_tx=XM_019827610.2
A3 107216710 107216757
refGene_gn=MRPS5;refGene_tx=XM_003984272.5;refGene_gn=MRPS5;refGene_tx=XM_019827610.2
A3 107218790 107218881
refGene_gn=MRPS5;refGene_tx=XM_003984272.5;refGene_gn=MRPS5;refGene_tx=XM_019827610.2
A3 107220488 107220523
refGene_gn=MRPS5;refGene_tx=XM_003984272.5;refGene_gn=MRPS5;refGene_tx=XM_019827610.2
A3 107223495 107223729
refGene_gn=MRPS5;refGene_tx=XM_003984272.5;refGene_gn=MRPS5;refGene_tx=XM_019827610.2
A3 107225379 107225505

refGene_gn=MRPS5;refGene_tx=XM_003984272.5
A3 107230049 107230187
refGene_gn=MRPS5;refGene_tx=XM_003984272.5
A3 107232188 107232269
refGene_gn=MRPS5;refGene_tx=XM_003984272.5
A3 107236836 107236894
refGene_gn=MRPS5;refGene_tx=XM_003984272.5
A3 107248900 107249970
refGene_gn=ZNF514;refGene_tx=XM_006930315.4;refGene_gn=ZNF514;refGene_tx=XM_019827611.2;refGene_gn=ZNF514;refGene_tx=XM_019827612.2;refGene_gn=ZNF514;refGene_tx=XM_019827613.2;refGene_gn=ZNF514;refGene_tx=XM_019827614.2;refGene_gn=ZNF514;refGene_tx=XM_019827615.2;refGene_gn=ZNF514;refGene_tx=XM_019827618.2
A3 107252563 107252647
refGene_gn=ZNF514;refGene_tx=XM_006930315.4;refGene_gn=ZNF514;refGene_tx=XM_019827611.2;refGene_gn=ZNF514;refGene_tx=XM_019827612.2;refGene_gn=ZNF514;refGene_tx=XM_019827613.2;refGene_gn=ZNF514;refGene_tx=XM_019827614.2;refGene_gn=ZNF514;refGene_tx=XM_019827615.2;refGene_gn=ZNF514;refGene_tx=XM_019827618.2
A3 107252988 107253115
refGene_gn=ZNF514;refGene_tx=XM_006930315.4;refGene_gn=ZNF514;refGene_tx=XM_019827611.2;refGene_gn=ZNF514;refGene_tx=XM_019827612.2;refGene_gn=ZNF514;refGene_tx=XM_019827613.2;refGene_gn=ZNF514;refGene_tx=XM_019827614.2;refGene_gn=ZNF514;refGene_tx=XM_019827615.2;refGene_gn=ZNF514;refGene_tx=XM_019827618.2
A3 107258122 107258211
refGene_gn=ZNF514;refGene_tx=XM_006930315.4;refGene_gn=ZNF514;refGene_tx=XM_019827611.2;refGene_gn=ZNF514;refGene_tx=XM_019827612.2;refGene_gn=ZNF514;refGene_tx=XM_019827613.2;refGene_gn=ZNF514;refGene_tx=XM_019827618.2
A3 107259679 107259698
refGene_gn=ZNF514;refGene_tx=XM_006930315.4;refGene_gn=ZNF514;refGene_tx=XM_019827611.2;refGene_gn=ZNF514;refGene_tx=XM_019827612.2;refGene_gn=ZNF514;refGene_tx=XM_019827613.2;refGene_gn=ZNF514;refGene_tx=XM_019827618.2
A3 107276353 107276386
refGene_gn=ZNF2;refGene_tx=XM_023251907.1;refGene_gn=ZNF2;refGene_tx=XM_023251908.1;refGene_gn=ZNF2;refGene_tx=XM_023251909.1
A3 107277826 107277953
refGene_gn=ZNF2;refGene_tx=XM_023251907.1;refGene_gn=ZNF2;refGene_tx=XM_023251908.1;refGene_gn=ZNF2;refGene_tx=XM_023251909.1
A3 107280718 107280832
refGene_gn=ZNF2;refGene_tx=XM_023251907.1;refGene_gn=ZNF2;refGene_tx=XM_023251909.1
A3 107281361 107282368
refGene_gn=ZNF2;refGene_tx=XM_023251907.1;refGene_gn=ZNF2;refGene_tx=XM_023251908.1;refGene_gn=ZNF2;refGene_tx=XM_023251909.1
A3 107298668 107299302
refGene_gn=ZNF2;refGene_tx=XM_023251909.1
A3 107305621 107305640

refGene_gn=ZNF2;refGene_tx=XM_023251901.1
A3	107306911	107307000
refGene_gn=ZNF2;refGene_tx=XM_023251901.1;refGene_gn=ZNF2;refGene_tx=XM_023251902.1;refGene_gn=ZNF2;refGene_tx=XM_023251903.1;refGene_gn=ZNF2;refGene_tx=XM_023251904.1;refGene_gn=ZNF2;refGene_tx=XM_023251906.1
A3	107310386	107310492
refGene_gn=ZNF2;refGene_tx=XM_023251901.1;refGene_gn=ZNF2;refGene_tx=XM_023251902.1;refGene_gn=ZNF2;refGene_tx=XM_023251903.1;refGene_gn=ZNF2;refGene_tx=XM_023251904.1;refGene_gn=ZNF2;refGene_tx=XM_023251906.1
A3	107310983	107311070
refGene_gn=ZNF2;refGene_tx=XM_023251901.1;refGene_gn=ZNF2;refGene_tx=XM_023251902.1;refGene_gn=ZNF2;refGene_tx=XM_023251903.1;refGene_gn=ZNF2;refGene_tx=XM_023251904.1;refGene_gn=ZNF2;refGene_tx=XM_023251906.1
A3	107315747	107317018
refGene_gn=ZNF2;refGene_tx=XM_023251901.1;refGene_gn=ZNF2;refGene_tx=XM_023251902.1;refGene_gn=ZNF2;refGene_tx=XM_023251903.1;refGene_gn=ZNF2;refGene_tx=XM_023251904.1;refGene_gn=ZNF2;refGene_tx=XM_023251906.1;refGene_gn=ZNF2;refGene_tx=XM_023251909.1
A3	107343751	107344007
refGene_gn=PROM2;refGene_tx=XM_003984283.4
A3	107344648	107344698
refGene_gn=PROM2;refGene_tx=XM_003984283.4
A3	107345125	107345328
refGene_gn=PROM2;refGene_tx=XM_003984283.4
A3	107345423	107345544
refGene_gn=PROM2;refGene_tx=XM_003984283.4
A3	107345824	107345888
refGene_gn=PROM2;refGene_tx=XM_003984283.4
A3	107346175	107346265
refGene_gn=PROM2;refGene_tx=XM_003984283.4
A3	107346554	107346754
refGene_gn=PROM2;refGene_tx=XM_003984283.4
A3	107347104	107347179
refGene_gn=PROM2;refGene_tx=XM_003984283.4
A3	107347977	107348041
refGene_gn=PROM2;refGene_tx=XM_003984283.4
A3	107348221	107348381
refGene_gn=PROM2;refGene_tx=XM_003984283.4
A3	107348667	107348820
refGene_gn=PROM2;refGene_tx=XM_003984283.4
A3	107349695	107349819
refGene_gn=PROM2;refGene_tx=XM_003984283.4
A3	107350332	107350424
refGene_gn=PROM2;refGene_tx=XM_003984283.4
A3	107350539	107350624
refGene_gn=PROM2;refGene_tx=XM_003984283.4
A3	107352975	107353122
refGene_gn=PROM2;refGene_tx=XM_003984283.4
A3	107353630	107353702
refGene_gn=PROM2;refGene_tx=XM_003984283.4

A3	107354518	107354611
refGene_gn=PROM2;refGene_tx=XM_003984283.4
A3	107354747	107354801
refGene_gn=PROM2;refGene_tx=XM_003984283.4
A3	107355140	107355221
refGene_gn=PROM2;refGene_tx=XM_003984283.4
A3	107355441	107355510
refGene_gn=PROM2;refGene_tx=XM_003984283.4
A3	107356206	107356296
refGene_gn=PROM2;refGene_tx=XM_003984283.4
A3	107356525	107356632
refGene_gn=PROM2;refGene_tx=XM_003984283.4
A3	107356987	107357051
refGene_gn=PROM2;refGene_tx=XM_003984283.4
A3	107364914	107365013
refGene_gn=KCNIP3;refGene_tx=XM_003984274.5
A3	107377038	107377204
refGene_gn=KCNIP3;refGene_tx=XM_003984274.5
A3	107422923	107423026
refGene_gn=KCNIP3;refGene_tx=XM_003984275.5
A3	107439744	107439869
refGene_gn=KCNIP3;refGene_tx=XM_003984274.5;refGene_gn=KCNIP3;refGene_tx=XM_003984275.5
A3	107440367	107440437
refGene_gn=KCNIP3;refGene_tx=XM_003984274.5;refGene_gn=KCNIP3;refGene_tx=XM_003984275.5
A3	107440653	107440724
refGene_gn=KCNIP3;refGene_tx=XM_003984274.5;refGene_gn=KCNIP3;refGene_tx=XM_003984275.5
A3	107446712	107446820
refGene_gn=KCNIP3;refGene_tx=XM_003984274.5;refGene_gn=KCNIP3;refGene_tx=XM_003984275.5
A3	107447526	107447631
refGene_gn=KCNIP3;refGene_tx=XM_003984274.5;refGene_gn=KCNIP3;refGene_tx=XM_003984275.5
A3	107448291	107448354
refGene_gn=KCNIP3;refGene_tx=XM_003984274.5;refGene_gn=KCNIP3;refGene_tx=XM_003984275.5
A3	107449015	107449063
refGene_gn=KCNIP3;refGene_tx=XM_003984274.5;refGene_gn=KCNIP3;refGene_tx=XM_003984275.5
A3	107473821	107473868
refGene_gn=LOC101087731;refGene_tx=XM_019827628.2
A3	107474173	107474415
refGene_gn=LOC101087731;refGene_tx=XM_003984276.5;refGene_gn=LOC101087731;refGene_tx=XM_006930325.4;refGene_gn=LOC101087731;refGene_tx=XM_006930326.4;refGene_gn=LOC101087731;refGene_tx=XM_006930327.4;refGene_gn=LOC101087731;refGene_tx=XM_019827628.2
A3	107476457	107476674
refGene_gn=LOC101087731;refGene_tx=XM_003984276.5;refGene_gn=LOC101087

731;refGene_tx=XM_006930325.4;refGene_gn=LOC101087731;refGene_tx=XM_006930326.4;refGene_gn=LOC101087731;refGene_tx=XM_006930327.4;refGene_gn=LOC101087731;refGene_tx=XM_019827628.2
A3 107478806 107478866
refGene_gn=LOC101087731;refGene_tx=XM_003984276.5;refGene_gn=LOC101087731;refGene_tx=XM_006930325.4;refGene_gn=LOC101087731;refGene_tx=XM_006930326.4;refGene_gn=LOC101087731;refGene_tx=XM_006930327.4;refGene_gn=LOC101087731;refGene_tx=XM_019827628.2
A3 107479058 107479221
refGene_gn=LOC101087731;refGene_tx=XM_003984276.5;refGene_gn=LOC101087731;refGene_tx=XM_006930325.4;refGene_gn=LOC101087731;refGene_tx=XM_006930326.4;refGene_gn=LOC101087731;refGene_tx=XM_006930327.4;refGene_gn=LOC101087731;refGene_tx=XM_019827628.2
A3 107480113 107480222
refGene_gn=LOC101087731;refGene_tx=XM_003984276.5;refGene_gn=LOC101087731;refGene_tx=XM_006930325.4;refGene_gn=LOC101087731;refGene_tx=XM_006930326.4;refGene_gn=LOC101087731;refGene_tx=XM_006930327.4;refGene_gn=LOC101087731;refGene_tx=XM_019827628.2
A3 107480332 107480420
refGene_gn=LOC101087731;refGene_tx=XM_003984276.5;refGene_gn=LOC101087731;refGene_tx=XM_006930325.4;refGene_gn=LOC101087731;refGene_tx=XM_006930326.4;refGene_gn=LOC101087731;refGene_tx=XM_006930327.4;refGene_gn=LOC101087731;refGene_tx=XM_019827628.2
A3 107480532 107480595
refGene_gn=LOC101087731;refGene_tx=XM_003984276.5;refGene_gn=LOC101087731;refGene_tx=XM_006930325.4;refGene_gn=LOC101087731;refGene_tx=XM_006930326.4;refGene_gn=LOC101087731;refGene_tx=XM_006930327.4;refGene_gn=LOC101087731;refGene_tx=XM_019827628.2
A3 107481028 107481145
refGene_gn=GPAT2;refGene_tx=XM_003984278.5;refGene_gn=GPAT2;refGene_tx=XM_019827619.2;refGene_gn=GPAT2;refGene_tx=XM_019827621.2;refGene_gn=GPAT2;refGene_tx=XM_019827623.2;refGene_gn=GPAT2;refGene_tx=XM_019827624.2;refGene_gn=GPAT2;refGene_tx=XM_019827626.2;refGene_gn=GPAT2;refGene_tx=XM_019827627.2
A3 107481423 107481479
refGene_gn=GPAT2;refGene_tx=XM_003984278.5;refGene_gn=GPAT2;refGene_tx=XM_019827619.2;refGene_gn=GPAT2;refGene_tx=XM_019827621.2;refGene_gn=GPAT2;refGene_tx=XM_019827623.2;refGene_gn=GPAT2;refGene_tx=XM_019827624.2;refGene_gn=GPAT2;refGene_tx=XM_019827626.2;refGene_gn=GPAT2;refGene_tx=XM_019827627.2
A3 107481720 107481786
refGene_gn=GPAT2;refGene_tx=XM_003984278.5;refGene_gn=GPAT2;refGene_tx=XM_019827619.2;refGene_gn=GPAT2;refGene_tx=XM_019827621.2;refGene_gn=GPAT2;refGene_tx=XM_019827623.2;refGene_gn=GPAT2;refGene_tx=XM_019827624.2;refGene_gn=GPAT2;refGene_tx=XM_019827626.2;refGene_gn=GPAT2;refGene_tx=XM_019827627.2
A3 107481883 107482004
refGene_gn=GPAT2;refGene_tx=XM_003984278.5;refGene_gn=GPAT2;refGene_tx=XM_019827619.2;refGene_gn=GPAT2;refGene_tx=XM_019827621.2;refGene_gn=GPAT2;refGene_tx=XM_019827623.2;refGene_gn=GPAT2;refGene_tx=XM_0198276

24.2;refGene_gn=GPAT2;refGene_tx=XM_019827626.2;refGene_gn=GPAT2;refGene_tx=XM_019827627.2
A3 107482084 107482216
refGene_gn=GPAT2;refGene_tx=XM_003984278.5;refGene_gn=GPAT2;refGene_tx=XM_019827619.2;refGene_gn=GPAT2;refGene_tx=XM_019827621.2;refGene_gn=GPAT2;refGene_tx=XM_019827623.2;refGene_gn=GPAT2;refGene_tx=XM_019827624.2;refGene_gn=GPAT2;refGene_tx=XM_019827626.2;refGene_gn=GPAT2;refGene_tx=XM_019827627.2
A3 107482689 107482767
refGene_gn=GPAT2;refGene_tx=XM_003984278.5;refGene_gn=GPAT2;refGene_tx=XM_019827619.2;refGene_gn=GPAT2;refGene_tx=XM_019827621.2;refGene_gn=GPAT2;refGene_tx=XM_019827623.2;refGene_gn=GPAT2;refGene_tx=XM_019827624.2;refGene_gn=GPAT2;refGene_tx=XM_019827626.2;refGene_gn=GPAT2;refGene_tx=XM_019827627.2
A3 107482944 107483093
refGene_gn=GPAT2;refGene_tx=XM_003984278.5;refGene_gn=GPAT2;refGene_tx=XM_019827619.2;refGene_gn=GPAT2;refGene_tx=XM_019827621.2;refGene_gn=GPAT2;refGene_tx=XM_019827623.2;refGene_gn=GPAT2;refGene_tx=XM_019827624.2;refGene_gn=GPAT2;refGene_tx=XM_019827626.2;refGene_gn=GPAT2;refGene_tx=XM_019827627.2
A3 107483181 107483440
refGene_gn=GPAT2;refGene_tx=XM_003984278.5;refGene_gn=GPAT2;refGene_tx=XM_019827619.2;refGene_gn=GPAT2;refGene_tx=XM_019827621.2;refGene_gn=GPAT2;refGene_tx=XM_019827623.2;refGene_gn=GPAT2;refGene_tx=XM_019827624.2;refGene_gn=GPAT2;refGene_tx=XM_019827626.2;refGene_gn=GPAT2;refGene_tx=XM_019827627.2
A3 107483535 107483606
refGene_gn=GPAT2;refGene_tx=XM_003984278.5;refGene_gn=GPAT2;refGene_tx=XM_019827619.2;refGene_gn=GPAT2;refGene_tx=XM_019827621.2;refGene_gn=GPAT2;refGene_tx=XM_019827623.2;refGene_gn=GPAT2;refGene_tx=XM_019827624.2;refGene_gn=GPAT2;refGene_tx=XM_019827626.2;refGene_gn=GPAT2;refGene_tx=XM_019827627.2
A3 107484329 107484448
refGene_gn=GPAT2;refGene_tx=XM_003984278.5;refGene_gn=GPAT2;refGene_tx=XM_019827619.2;refGene_gn=GPAT2;refGene_tx=XM_019827621.2;refGene_gn=GPAT2;refGene_tx=XM_019827623.2;refGene_gn=GPAT2;refGene_tx=XM_019827624.2;refGene_gn=GPAT2;refGene_tx=XM_019827626.2;refGene_gn=GPAT2;refGene_tx=XM_019827627.2
A3 107484799 107484882
refGene_gn=GPAT2;refGene_tx=XM_003984278.5;refGene_gn=GPAT2;refGene_tx=XM_019827619.2;refGene_gn=GPAT2;refGene_tx=XM_019827621.2;refGene_gn=GPAT2;refGene_tx=XM_019827623.2;refGene_gn=GPAT2;refGene_tx=XM_019827624.2;refGene_gn=GPAT2;refGene_tx=XM_019827626.2;refGene_gn=GPAT2;refGene_tx=XM_019827627.2
A3 107485051 107485171
refGene_gn=GPAT2;refGene_tx=XM_003984278.5;refGene_gn=GPAT2;refGene_tx=XM_019827619.2;refGene_gn=GPAT2;refGene_tx=XM_019827621.2;refGene_gn=GPAT2;refGene_tx=XM_019827623.2;refGene_gn=GPAT2;refGene_tx=XM_019827624.2;refGene_gn=GPAT2;refGene_tx=XM_019827626.2;refGene_gn=GPAT2;refGene_tx=XM_019827627.2

A3 107485743 107485956
refGene_gn=GPAT2;refGene_tx=XM_003984278.5;refGene_gn=GPAT2;refGene_tx
=XM_019827619.2;refGene_gn=GPAT2;refGene_tx=XM_019827621.2;refGene_gn=
GPAT2;refGene_tx=XM_019827623.2;refGene_gn=GPAT2;refGene_tx=XM_0198276
24.2;refGene_gn=GPAT2;refGene_tx=XM_019827626.2;refGene_gn=GPAT2;refGe
ne_tx=XM_019827627.2
A3 107486052 107486152
refGene_gn=GPAT2;refGene_tx=XM_003984278.5;refGene_gn=GPAT2;refGene_tx
=XM_019827619.2;refGene_gn=GPAT2;refGene_tx=XM_019827621.2;refGene_gn=
GPAT2;refGene_tx=XM_019827623.2;refGene_gn=GPAT2;refGene_tx=XM_0198276
24.2;refGene_gn=GPAT2;refGene_tx=XM_019827626.2;refGene_gn=GPAT2;refGe
ne_tx=XM_019827627.2
A3 107486458 107486595
refGene_gn=GPAT2;refGene_tx=XM_003984278.5;refGene_gn=GPAT2;refGene_tx
=XM_019827619.2;refGene_gn=GPAT2;refGene_tx=XM_019827621.2;refGene_gn=
GPAT2;refGene_tx=XM_019827623.2;refGene_gn=GPAT2;refGene_tx=XM_0198276
24.2;refGene_gn=GPAT2;refGene_tx=XM_019827626.2;refGene_gn=GPAT2;refGe
ne_tx=XM_019827627.2
A3 107486976 107487076
refGene_gn=GPAT2;refGene_tx=XM_003984278.5;refGene_gn=GPAT2;refGene_tx
=XM_019827619.2;refGene_gn=GPAT2;refGene_tx=XM_019827621.2;refGene_gn=
GPAT2;refGene_tx=XM_019827623.2;refGene_gn=GPAT2;refGene_tx=XM_0198276
24.2;refGene_gn=GPAT2;refGene_tx=XM_019827626.2;refGene_gn=GPAT2;refGe
ne_tx=XM_019827627.2
A3 107489334 107489463
refGene_gn=GPAT2;refGene_tx=XM_003984278.5;refGene_gn=GPAT2;refGene_tx
=XM_019827619.2;refGene_gn=GPAT2;refGene_tx=XM_019827621.2;refGene_gn=
GPAT2;refGene_tx=XM_019827623.2;refGene_gn=GPAT2;refGene_tx=XM_0198276
24.2;refGene_gn=GPAT2;refGene_tx=XM_019827626.2;refGene_gn=GPAT2;refGe
ne_tx=XM_019827627.2
A3 107489707 107489812
refGene_gn=GPAT2;refGene_tx=XM_003984278.5;refGene_gn=GPAT2;refGene_tx
=XM_019827619.2;refGene_gn=GPAT2;refGene_tx=XM_019827621.2;refGene_gn=
GPAT2;refGene_tx=XM_019827623.2;refGene_gn=GPAT2;refGene_tx=XM_0198276
24.2;refGene_gn=GPAT2;refGene_tx=XM_019827626.2;refGene_gn=GPAT2;refGe
ne_tx=XM_019827627.2
A3 107490242 107490316
refGene_gn=GPAT2;refGene_tx=XM_003984278.5;refGene_gn=GPAT2;refGene_tx
=XM_019827619.2;refGene_gn=GPAT2;refGene_tx=XM_019827621.2;refGene_gn=
GPAT2;refGene_tx=XM_019827623.2;refGene_gn=GPAT2;refGene_tx=XM_0198276
24.2;refGene_gn=GPAT2;refGene_tx=XM_019827626.2;refGene_gn=GPAT2;refGe
ne_tx=XM_019827627.2
A3 107490992 107491106
refGene_gn=GPAT2;refGene_tx=XM_003984278.5;refGene_gn=GPAT2;refGene_tx
=XM_019827619.2;refGene_gn=GPAT2;refGene_tx=XM_019827621.2;refGene_gn=
GPAT2;refGene_tx=XM_019827623.2;refGene_gn=GPAT2;refGene_tx=XM_0198276
24.2;refGene_gn=GPAT2;refGene_tx=XM_019827626.2;refGene_gn=GPAT2;refGe
ne_tx=XM_019827627.2
A3 107491236 107491365
refGene_gn=GPAT2;refGene_tx=XM_003984278.5;refGene_gn=GPAT2;refGene_tx

=XM_019827619.2;refGene_gn=GPAT2;refGene_tx=XM_019827621.2;refGene_gn=GPAT2;refGene_tx=XM_019827623.2;refGene_gn=GPAT2;refGene_tx=XM_019827624.2;refGene_gn=GPAT2;refGene_tx=XM_019827626.2;refGene_gn=GPAT2;refGene_tx=XM_019827627.2
A3 107494969 107495107
refGene_gn=GPAT2;refGene_tx=XM_003984278.5;refGene_gn=GPAT2;refGene_tx=XM_019827619.2;refGene_gn=GPAT2;refGene_tx=XM_019827621.2;refGene_gn=GPAT2;refGene_tx=XM_019827623.2;refGene_gn=GPAT2;refGene_tx=XM_019827624.2;refGene_gn=GPAT2;refGene_tx=XM_019827626.2
A3 107554432 107554442
refGene_gn=ADRA2B;refGene_tx=XM_006930331.3
A3 107562230 107563555
refGene_gn=ADRA2B;refGene_tx=XM_006930331.3
A3 107571136 107571537
refGene_gn=ASTL;refGene_tx=XM_011281240.3
A3 107575996 107576151
refGene_gn=ASTL;refGene_tx=XM_011281240.3
A3 107576249 107576331
refGene_gn=ASTL;refGene_tx=XM_011281240.3
A3 107580689 107580871
refGene_gn=ASTL;refGene_tx=XM_011281240.3
A3 107581634 107581752
refGene_gn=ASTL;refGene_tx=XM_011281240.3
A3 107582687 107582778
refGene_gn=ASTL;refGene_tx=XM_011281240.3
A3 107584114 107584176
refGene_gn=ASTL;refGene_tx=XM_011281240.3
A3 107586271 107586397
refGene_gn=ASTL;refGene_tx=XM_011281240.3
A3 107587029 107587075
refGene_gn=ASTL;refGene_tx=XM_011281240.3
A3 107592863 107593078
refGene_gn=DUSP2;refGene_tx=XM_003984285.5
A3 107593180 107593400
refGene_gn=DUSP2;refGene_tx=XM_003984285.5
A3 107593835 107593957
refGene_gn=DUSP2;refGene_tx=XM_003984285.5
A3 107594054 107594442
refGene_gn=DUSP2;refGene_tx=XM_003984285.5
A3 107627031 107627216
refGene_gn=STARD7;refGene_tx=XM_003984286.4
A3 107628066 107628151
refGene_gn=STARD7;refGene_tx=XM_003984286.4
A3 107637031 107637131
refGene_gn=STARD7;refGene_tx=XM_003984286.4
A3 107637573 107637656
refGene_gn=STARD7;refGene_tx=XM_003984286.4
A3 107637736 107637847
refGene_gn=STARD7;refGene_tx=XM_003984286.4
A3 107639350 107639400

refGene_gn=STARD7;refGene_tx=XM_003984286.4
A3	107639967	107640176
refGene_gn=STARD7;refGene_tx=XM_003984286.4
A3	107650064	107650384
refGene_gn=STARD7;refGene_tx=XM_003984286.4
A3	107661387	107661695
refGene_gn=TMEM127;refGene_tx=XM_003984287.4
A3	107662473	107662638
refGene_gn=TMEM127;refGene_tx=XM_003984287.4
A3	107671357	107671601
refGene_gn=TMEM127;refGene_tx=XM_003984287.4
A3	107672568	107672707
refGene_gn=CIAO1;refGene_tx=XM_003984288.5
A3	107673288	107673437
refGene_gn=CIAO1;refGene_tx=XM_003984288.5
A3	107673591	107673703
refGene_gn=CIAO1;refGene_tx=XM_003984288.5
A3	107673805	107673894
refGene_gn=CIAO1;refGene_tx=XM_003984288.5
A3	107674140	107674342
refGene_gn=CIAO1;refGene_tx=XM_003984288.5
A3	107674973	107675061
refGene_gn=CIAO1;refGene_tx=XM_003984288.5
A3	107675937	107676178
refGene_gn=CIAO1;refGene_tx=XM_003984288.5
A3	107679525	107679669
refGene_gn=SNRNP200;refGene_tx=XM_003984289.5
A3	107679772	107679865
refGene_gn=SNRNP200;refGene_tx=XM_003984289.5
A3	107680935	107681017
refGene_gn=SNRNP200;refGene_tx=XM_003984289.5
A3	107681132	107681293
refGene_gn=SNRNP200;refGene_tx=XM_003984289.5
A3	107681600	107681777
refGene_gn=SNRNP200;refGene_tx=XM_003984289.5
A3	107681875	107682019
refGene_gn=SNRNP200;refGene_tx=XM_003984289.5
A3	107682240	107682362
refGene_gn=SNRNP200;refGene_tx=XM_003984289.5
A3	107682586	107682751
refGene_gn=SNRNP200;refGene_tx=XM_003984289.5
A3	107682845	107683035
refGene_gn=SNRNP200;refGene_tx=XM_003984289.5
A3	107683530	107683639
refGene_gn=SNRNP200;refGene_tx=XM_003984289.5
A3	107684682	107684791
refGene_gn=SNRNP200;refGene_tx=XM_003984289.5
A3	107685642	107685794
refGene_gn=SNRNP200;refGene_tx=XM_003984289.5
A3	107685978	107686157

refGene_gn=SNRNP200;refGene_tx=XM_003984289.5
A3	107686246	107686438
refGene_gn=SNRNP200;refGene_tx=XM_003984289.5
A3	107686792	107687020
refGene_gn=SNRNP200;refGene_tx=XM_003984289.5
A3	107687333	107687494
refGene_gn=SNRNP200;refGene_tx=XM_003984289.5
A3	107688900	107689074
refGene_gn=SNRNP200;refGene_tx=XM_003984289.5
A3	107689281	107689471
refGene_gn=SNRNP200;refGene_tx=XM_003984289.5
A3	107689583	107689738
refGene_gn=SNRNP200;refGene_tx=XM_003984289.5
A3	107690030	107690149
refGene_gn=SNRNP200;refGene_tx=XM_003984289.5
A3	107690484	107690591
refGene_gn=SNRNP200;refGene_tx=XM_003984289.5
A3	107691460	107691544
refGene_gn=SNRNP200;refGene_tx=XM_003984289.5
A3	107691817	107691898
refGene_gn=SNRNP200;refGene_tx=XM_003984289.5
A3	107692011	107692164
refGene_gn=SNRNP200;refGene_tx=XM_003984289.5
A3	107692567	107692765
refGene_gn=SNRNP200;refGene_tx=XM_003984289.5
A3	107693564	107693753
refGene_gn=SNRNP200;refGene_tx=XM_003984289.5
A3	107693934	107694066
refGene_gn=SNRNP200;refGene_tx=XM_003984289.5
A3	107694611	107694722
refGene_gn=SNRNP200;refGene_tx=XM_003984289.5
A3	107694964	107695114
refGene_gn=SNRNP200;refGene_tx=XM_003984289.5
A3	107696333	107696457
refGene_gn=SNRNP200;refGene_tx=XM_003984289.5
A3	107696659	107696853
refGene_gn=SNRNP200;refGene_tx=XM_003984289.5
A3	107698351	107698522
refGene_gn=SNRNP200;refGene_tx=XM_003984289.5
A3	107699050	107699206
refGene_gn=SNRNP200;refGene_tx=XM_003984289.5
A3	107699416	107699554
refGene_gn=SNRNP200;refGene_tx=XM_003984289.5
A3	107699852	107700026
refGene_gn=SNRNP200;refGene_tx=XM_003984289.5
A3	107700138	107700222
refGene_gn=SNRNP200;refGene_tx=XM_003984289.5
A3	107700811	107700948
refGene_gn=SNRNP200;refGene_tx=XM_003984289.5
A3	107701133	107701233

refGene_gn=SNRNP200;refGene_tx=XM_003984289.5
A3 107701342 107701495
refGene_gn=SNRNP200;refGene_tx=XM_003984289.5
A3 107701867 107701966
refGene_gn=SNRNP200;refGene_tx=XM_003984289.5
A3 107703571 107703627
refGene_gn=SNRNP200;refGene_tx=XM_003984289.5
A3 107704019 107704212
refGene_gn=SNRNP200;refGene_tx=XM_003984289.5
A3 107704926 107705098
refGene_gn=SNRNP200;refGene_tx=XM_003984289.5
A3 107707206 107707370
refGene_gn=SNRNP200;refGene_tx=XM_003984289.5
A3 107707882 107707927
refGene_gn=SNRNP200;refGene_tx=XM_003984289.5
A3 107718513 107718573
refGene_gn=NCAPH;refGene_tx=XM_006930332.3;refGene_gn=NCAPH;refGene_tx
=XM_006930333.3;refGene_gn=NCAPH;refGene_tx=XM_006930334.3
A3 107721785 107721861
refGene_gn=NCAPH;refGene_tx=XM_006930332.3;refGene_gn=NCAPH;refGene_tx
=XM_006930333.3;refGene_gn=NCAPH;refGene_tx=XM_006930334.3
A3 107722198 107722290
refGene_gn=NCAPH;refGene_tx=XM_006930332.3;refGene_gn=NCAPH;refGene_tx
=XM_006930333.3;refGene_gn=NCAPH;refGene_tx=XM_006930334.3
A3 107724482 107724596
refGene_gn=NCAPH;refGene_tx=XM_006930332.3;refGene_gn=NCAPH;refGene_tx
=XM_006930333.3;refGene_gn=NCAPH;refGene_tx=XM_006930334.3
A3 107725264 107725447
refGene_gn=NCAPH;refGene_tx=XM_006930332.3;refGene_gn=NCAPH;refGene_tx
=XM_006930333.3;refGene_gn=NCAPH;refGene_tx=XM_006930334.3
A3 107726642 107726753
refGene_gn=NCAPH;refGene_tx=XM_006930332.3;refGene_gn=NCAPH;refGene_tx
=XM_006930333.3;refGene_gn=NCAPH;refGene_tx=XM_006930334.3
A3 107729348 107729471
refGene_gn=NCAPH;refGene_tx=XM_006930332.3;refGene_gn=NCAPH;refGene_tx
=XM_006930333.3;refGene_gn=NCAPH;refGene_tx=XM_006930334.3
A3 107731287 107731394
refGene_gn=NCAPH;refGene_tx=XM_006930332.3;refGene_gn=NCAPH;refGene_tx
=XM_006930333.3;refGene_gn=NCAPH;refGene_tx=XM_006930334.3
A3 107731745 107731894
refGene_gn=NCAPH;refGene_tx=XM_006930332.3;refGene_gn=NCAPH;refGene_tx
=XM_006930333.3;refGene_gn=NCAPH;refGene_tx=XM_006930334.3
A3 107735585 107735791
refGene_gn=NCAPH;refGene_tx=XM_006930332.3;refGene_gn=NCAPH;refGene_tx
=XM_006930333.3;refGene_gn=NCAPH;refGene_tx=XM_006930334.3
A3 107736503 107736595
refGene_gn=NCAPH;refGene_tx=XM_006930332.3;refGene_gn=NCAPH;refGene_tx
=XM_006930333.3;refGene_gn=NCAPH;refGene_tx=XM_006930334.3
A3 107738216 107738406
refGene_gn=NCAPH;refGene_tx=XM_006930332.3;refGene_gn=NCAPH;refGene_tx

=XM_006930333.3;refGene_gn=NCAPH;refGene_tx=XM_006930334.3
A3 107745025 107745150
refGene_gn=NCAPH;refGene_tx=XM_006930332.3;refGene_gn=NCAPH;refGene_tx
=XM_006930333.3;refGene_gn=NCAPH;refGene_tx=XM_006930334.3
A3 107745818 107745954
refGene_gn=NCAPH;refGene_tx=XM_006930332.3;refGene_gn=NCAPH;refGene_tx
=XM_006930333.3;refGene_gn=NCAPH;refGene_tx=XM_006930334.3
A3 107746279 107746372
refGene_gn=NCAPH;refGene_tx=XM_006930332.3;refGene_gn=NCAPH;refGene_tx
=XM_006930333.3;refGene_gn=NCAPH;refGene_tx=XM_006930334.3
A3 107747290 107747381
refGene_gn=NCAPH;refGene_tx=XM_006930332.3;refGene_gn=NCAPH;refGene_tx
=XM_006930333.3;refGene_gn=NCAPH;refGene_tx=XM_006930334.3
A3 107747541 107747761
refGene_gn=NCAPH;refGene_tx=XM_006930332.3;refGene_gn=NCAPH;refGene_tx
=XM_006930333.3;refGene_gn=NCAPH;refGene_tx=XM_006930334.3
A3 107752338 107752357
refGene_gn=NCAPH;refGene_tx=XM_006930332.3;refGene_gn=NCAPH;refGene_tx
=XM_006930333.3
A3 107757750 107759394
refGene_gn=ITPRIPL1;refGene_tx=XM_003984291.3;refGene_gn=ITPRIPL1;refG
ene_tx=XM_006930337.4;refGene_gn=ITPRIPL1;refGene_tx=XM_019827630.2
A3 107800233 107800454
refGene_gn=NEURL3;refGene_tx=XM_023251914.1
A3 107801103 107801172
refGene_gn=NEURL3;refGene_tx=XM_023251914.1
A3 107802105 107802591
refGene_gn=NEURL3;refGene_tx=XM_023251914.1
A3 107805301 107805329
refGene_gn=NEURL3;refGene_tx=XM_023251914.1
A3 107838341 107838345
refGene_gn=ARID5A;refGene_tx=XM_006930339.4
A3 107847134 107847250
refGene_gn=ARID5A;refGene_tx=XM_006930339.4
A3 107848727 107848866
refGene_gn=ARID5A;refGene_tx=XM_006930339.4
A3 107849209 107849262
refGene_gn=ARID5A;refGene_tx=XM_006930339.4;refGene_gn=ARID5A;refGene_
tx=XM_006930340.4
A3 107849746 107849844
refGene_gn=ARID5A;refGene_tx=XM_006930339.4;refGene_gn=ARID5A;refGene_
tx=XM_006930340.4
A3 107850147 107850307
refGene_gn=ARID5A;refGene_tx=XM_006930339.4;refGene_gn=ARID5A;refGene_
tx=XM_006930340.4
A3 107850726 107851965
refGene_gn=ARID5A;refGene_tx=XM_006930339.4;refGene_gn=ARID5A;refGene_
tx=XM_006930340.4
A3 107878807 107878833
refGene_gn=MTA3;refGene_tx=XM_023251915.1

A3	107895063	107895210
refGene_gn=MTA3;refGene_tx=XM_023251915.1
A3	107897938	107898025
refGene_gn=MTA3;refGene_tx=XM_023251915.1
A3	107907211	107907434
refGene_gn=MTA3;refGene_tx=XM_023251915.1
A3	107908383	107908535
refGene_gn=MTA3;refGene_tx=XM_023251915.1
A3	107912216	107912341
refGene_gn=MTA3;refGene_tx=XM_023251915.1
A3	107917644	107917703
refGene_gn=MTA3;refGene_tx=XM_023251915.1
A3	107919679	107919754
refGene_gn=MTA3;refGene_tx=XM_023251915.1
A3	107928604	107928664
refGene_gn=MTA3;refGene_tx=XM_023251915.1
A3	107928665	107928794
refGene_gn=MTA3;refGene_tx=XM_023251915.1
A3	107947732	107947832
refGene_gn=MTA3;refGene_tx=XM_023251915.1
A3	107951766	107951869
refGene_gn=MTA3;refGene_tx=XM_023251915.1
A3	107965334	107965452
refGene_gn=MTA3;refGene_tx=XM_023251915.1
A3	107968288	107968352
refGene_gn=MTA3;refGene_tx=XM_023251915.1
A3	107992530	107992657
refGene_gn=MTA3;refGene_tx=XM_023251915.1
A3	108042186	108042280
refGene_gn=MTA3;refGene_tx=XM_023251915.1
A3	108052120	108052188
refGene_gn=MTA3;refGene_tx=XM_023251915.1
A3	108053863	108053891
refGene_gn=MTA3;refGene_tx=XM_023251915.1
A3	108099210	108100791
refGene_gn=KCNG3;refGene_tx=XM_023251916.1;refGene_gn=KCNG3;refGene_tx=XM_023251917.1;refGene_gn=KCNG3;refGene_tx=XM_023251918.1;refGene_gn=KCNG3;refGene_tx=XM_023251919.1;refGene_gn=KCNG3;refGene_tx=XM_023251920.1;refGene_gn=KCNG3;refGene_tx=XM_023251921.1;refGene_gn=KCNG3;refGene_tx=XM_023251922.1;refGene_gn=KCNG3;refGene_tx=XM_023251923.1;refGene_gn=KCNG3;refGene_tx=XM_023251924.1;refGene_gn=KCNG3;refGene_tx=XM_023251925.1;refGene_gn=KCNG3;refGene_tx=XM_023251926.1;refGene_gn=KCNG3;refGene_tx=XM_023251927.1;refGene_gn=KCNG3;refGene_tx=XM_023251928.1;refGene_gn=LOC111559899;refGene_tx=XM_023252196.1
A3	108143910	108144556
refGene_gn=KCNG3;refGene_tx=XM_023251916.1;refGene_gn=KCNG3;refGene_tx=XM_023251917.1;refGene_gn=KCNG3;refGene_tx=XM_023251918.1;refGene_gn=KCNG3;refGene_tx=XM_023251919.1;refGene_gn=KCNG3;refGene_tx=XM_023251920.1;refGene_gn=KCNG3;refGene_tx=XM_023251921.1;refGene_gn=KCNG3;refGene_tx=XM_023251922.1;refGene_gn=KCNG3;refGene_tx=XM_023251923.1;refGen

e_gn=KCNG3;refGene_tx=XM_023251924.1;refGene_gn=KCNG3;refGene_tx=XM_023251925.1;refGene_gn=KCNG3;refGene_tx=XM_023251926.1;refGene_gn=KCNG3;refGene_tx=XM_023251927.1;refGene_gn=KCNG3;refGene_tx=XM_023251928.1
A3 108157762 108157838
refGene_gn=KCNG3;refGene_tx=XM_023251916.1;refGene_gn=KCNG3;refGene_tx=XM_023251917.1;refGene_gn=KCNG3;refGene_tx=XM_023251919.1;refGene_gn=KCNG3;refGene_tx=XM_023251920.1;refGene_gn=KCNG3;refGene_tx=XM_023251922.1;refGene_gn=KCNG3;refGene_tx=XM_023251923.1;refGene_gn=KCNG3;refGene_tx=XM_023251924.1;refGene_gn=KCNG3;refGene_tx=XM_023251925.1;refGene_gn=KCNG3;refGene_tx=XM_023251926.1;refGene_gn=KCNG3;refGene_tx=XM_023251927.1;refGene_gn=KCNG3;refGene_tx=XM_023251928.1
A3 108158899 108159076
refGene_gn=KCNG3;refGene_tx=XM_023251916.1;refGene_gn=KCNG3;refGene_tx=XM_023251917.1;refGene_gn=KCNG3;refGene_tx=XM_023251926.1
A3 108181866 108181949
refGene_gn=KCNG3;refGene_tx=XM_023251922.1;refGene_gn=KCNG3;refGene_tx=XM_023251925.1;refGene_gn=KCNG3;refGene_tx=XM_023251928.1
A3 108186077 108186202
refGene_gn=KCNG3;refGene_tx=XM_023251919.1;refGene_gn=KCNG3;refGene_tx=XM_023251920.1;refGene_gn=KCNG3;refGene_tx=XM_023251923.1;refGene_gn=KCNG3;refGene_tx=XM_023251924.1;refGene_gn=KCNG3;refGene_tx=XM_023251927.1
A3 108187259 108187275
refGene_gn=KCNG3;refGene_tx=XM_023251919.1;refGene_gn=KCNG3;refGene_tx=XM_023251927.1
A3 108200702 108200709
refGene_gn=KCNG3;refGene_tx=XM_023251920.1
A3 108204026 108204098
refGene_gn=LOC101092350;refGene_tx=XM_003984292.5
A3 108223536 108223668
refGene_gn=LOC101092350;refGene_tx=XM_003984292.5
A3 108224733 108224874
refGene_gn=LOC101092350;refGene_tx=XM_003984292.5
A3 108242041 108242515
refGene_gn=EML4;refGene_tx=XM_019827633.2;refGene_gn=EML4;refGene_tx=XM_019827634.2;refGene_gn=EML4;refGene_tx=XM_019827635.2;refGene_gn=EML4;refGene_tx=XM_019827636.2;refGene_gn=EML4;refGene_tx=XM_019827637.2;refGene_gn=EML4;refGene_tx=XM_019827638.2;refGene_gn=EML4;refGene_tx=XM_019827639.2;refGene_gn=EML4;refGene_tx=XM_019827640.2;refGene_gn=EML4;refGene_tx=XM_019827641.2
A3 108243376 108243507
refGene_gn=EML4;refGene_tx=XM_019827633.2;refGene_gn=EML4;refGene_tx=XM_019827634.2;refGene_gn=EML4;refGene_tx=XM_019827635.2;refGene_gn=EML4;refGene_tx=XM_019827636.2;refGene_gn=EML4;refGene_tx=XM_019827637.2;refGene_gn=EML4;refGene_tx=XM_019827638.2;refGene_gn=EML4;refGene_tx=XM_019827639.2;refGene_gn=EML4;refGene_tx=XM_019827640.2;refGene_gn=EML4;refGene_tx=XM_019827641.2
A3 108246380 108246479
refGene_gn=EML4;refGene_tx=XM_019827633.2;refGene_gn=EML4;refGene_tx=XM_019827634.2;refGene_gn=EML4;refGene_tx=XM_019827635.2;refGene_gn=EML

4;refGene_tx=XM_019827636.2;refGene_gn=EML4;refGene_tx=XM_019827637.2;
refGene_gn=EML4;refGene_tx=XM_019827638.2;refGene_gn=EML4;refGene_tx=X
M_019827639.2;refGene_gn=EML4;refGene_tx=XM_019827640.2;refGene_gn=EML
4;refGene_tx=XM_019827641.2
A3 108246987 108247075
refGene_gn=EML4;refGene_tx=XM_019827633.2;refGene_gn=EML4;refGene_tx=X
M_019827634.2;refGene_gn=EML4;refGene_tx=XM_019827635.2;refGene_gn=EML
4;refGene_tx=XM_019827636.2;refGene_gn=EML4;refGene_tx=XM_019827637.2;
refGene_gn=EML4;refGene_tx=XM_019827638.2;refGene_gn=EML4;refGene_tx=X
M_019827639.2;refGene_gn=EML4;refGene_tx=XM_019827640.2;refGene_gn=EML
4;refGene_tx=XM_019827641.2
A3 108257011 108257109
refGene_gn=EML4;refGene_tx=XM_019827633.2;refGene_gn=EML4;refGene_tx=X
M_019827634.2;refGene_gn=EML4;refGene_tx=XM_019827635.2;refGene_gn=EML
4;refGene_tx=XM_019827636.2;refGene_gn=EML4;refGene_tx=XM_019827637.2;
refGene_gn=EML4;refGene_tx=XM_019827638.2;refGene_gn=EML4;refGene_tx=X
M_019827639.2;refGene_gn=EML4;refGene_tx=XM_019827640.2;refGene_gn=EML
4;refGene_tx=XM_019827641.2
A3 108258439 108258528
refGene_gn=EML4;refGene_tx=XM_019827633.2;refGene_gn=EML4;refGene_tx=X
M_019827634.2;refGene_gn=EML4;refGene_tx=XM_019827635.2;refGene_gn=EML
4;refGene_tx=XM_019827636.2;refGene_gn=EML4;refGene_tx=XM_019827637.2;
refGene_gn=EML4;refGene_tx=XM_019827638.2;refGene_gn=EML4;refGene_tx=X
M_019827639.2;refGene_gn=EML4;refGene_tx=XM_019827640.2;refGene_gn=EML
4;refGene_tx=XM_019827641.2
A3 108268930 108268998
refGene_gn=EML4;refGene_tx=XM_019827633.2;refGene_gn=EML4;refGene_tx=X
M_019827634.2;refGene_gn=EML4;refGene_tx=XM_019827635.2;refGene_gn=EML
4;refGene_tx=XM_019827636.2;refGene_gn=EML4;refGene_tx=XM_019827637.2;
refGene_gn=EML4;refGene_tx=XM_019827638.2;refGene_gn=EML4;refGene_tx=X
M_019827639.2;refGene_gn=EML4;refGene_tx=XM_019827640.2;refGene_gn=EML
4;refGene_tx=XM_019827641.2
A3 108269703 108269835
refGene_gn=EML4;refGene_tx=XM_019827633.2;refGene_gn=EML4;refGene_tx=X
M_019827634.2;refGene_gn=EML4;refGene_tx=XM_019827635.2;refGene_gn=EML
4;refGene_tx=XM_019827636.2;refGene_gn=EML4;refGene_tx=XM_019827637.2;
refGene_gn=EML4;refGene_tx=XM_019827638.2;refGene_gn=EML4;refGene_tx=X
M_019827639.2;refGene_gn=EML4;refGene_tx=XM_019827640.2;refGene_gn=EML
4;refGene_tx=XM_019827641.2
A3 108270141 108270267
refGene_gn=EML4;refGene_tx=XM_019827633.2;refGene_gn=EML4;refGene_tx=X
M_019827634.2;refGene_gn=EML4;refGene_tx=XM_019827635.2;refGene_gn=EML
4;refGene_tx=XM_019827636.2;refGene_gn=EML4;refGene_tx=XM_019827637.2;
refGene_gn=EML4;refGene_tx=XM_019827638.2;refGene_gn=EML4;refGene_tx=X
M_019827639.2;refGene_gn=EML4;refGene_tx=XM_019827640.2;refGene_gn=EML
4;refGene_tx=XM_019827641.2
A3 108272207 108272359
refGene_gn=EML4;refGene_tx=XM_019827633.2;refGene_gn=EML4;refGene_tx=X
M_019827634.2;refGene_gn=EML4;refGene_tx=XM_019827635.2;refGene_gn=EML
4;refGene_tx=XM_019827636.2;refGene_gn=EML4;refGene_tx=XM_019827637.2;

refGene_gn=EML4;refGene_tx=XM_019827638.2;refGene_gn=EML4;refGene_tx=XM_019827639.2;refGene_gn=EML4;refGene_tx=XM_019827640.2;refGene_gn=EML4;refGene_tx=XM_019827641.2
A3 108283600 108283736
refGene_gn=EML4;refGene_tx=XM_019827633.2;refGene_gn=EML4;refGene_tx=XM_019827634.2;refGene_gn=EML4;refGene_tx=XM_019827635.2;refGene_gn=EML4;refGene_tx=XM_019827636.2;refGene_gn=EML4;refGene_tx=XM_019827637.2;refGene_gn=EML4;refGene_tx=XM_019827638.2;refGene_gn=EML4;refGene_tx=XM_019827639.2;refGene_gn=EML4;refGene_tx=XM_019827640.2;refGene_gn=EML4;refGene_tx=XM_019827641.2
A3 108283847 108283982
refGene_gn=EML4;refGene_tx=XM_019827633.2;refGene_gn=EML4;refGene_tx=XM_019827634.2;refGene_gn=EML4;refGene_tx=XM_019827635.2;refGene_gn=EML4;refGene_tx=XM_019827636.2;refGene_gn=EML4;refGene_tx=XM_019827637.2;refGene_gn=EML4;refGene_tx=XM_019827638.2;refGene_gn=EML4;refGene_tx=XM_019827639.2;refGene_gn=EML4;refGene_tx=XM_019827640.2;refGene_gn=EML4;refGene_tx=XM_019827641.2
A3 108289072 108289168
refGene_gn=EML4;refGene_tx=XM_019827633.2;refGene_gn=EML4;refGene_tx=XM_019827634.2;refGene_gn=EML4;refGene_tx=XM_019827635.2;refGene_gn=EML4;refGene_tx=XM_019827636.2;refGene_gn=EML4;refGene_tx=XM_019827637.2;refGene_gn=EML4;refGene_tx=XM_019827638.2;refGene_gn=EML4;refGene_tx=XM_019827639.2;refGene_gn=EML4;refGene_tx=XM_019827640.2;refGene_gn=EML4;refGene_tx=XM_019827641.2
A3 108289893 108290004
refGene_gn=EML4;refGene_tx=XM_019827633.2;refGene_gn=EML4;refGene_tx=XM_019827634.2;refGene_gn=EML4;refGene_tx=XM_019827635.2;refGene_gn=EML4;refGene_tx=XM_019827636.2;refGene_gn=EML4;refGene_tx=XM_019827637.2;refGene_gn=EML4;refGene_tx=XM_019827638.2;refGene_gn=EML4;refGene_tx=XM_019827639.2;refGene_gn=EML4;refGene_tx=XM_019827640.2;refGene_gn=EML4;refGene_tx=XM_019827641.2
A3 108291111 108291181
refGene_gn=EML4;refGene_tx=XM_019827633.2;refGene_gn=EML4;refGene_tx=XM_019827634.2;refGene_gn=EML4;refGene_tx=XM_019827635.2;refGene_gn=EML4;refGene_tx=XM_019827636.2;refGene_gn=EML4;refGene_tx=XM_019827637.2;refGene_gn=EML4;refGene_tx=XM_019827638.2;refGene_gn=EML4;refGene_tx=XM_019827639.2;refGene_gn=EML4;refGene_tx=XM_019827640.2;refGene_gn=EML4;refGene_tx=XM_019827641.2
A3 108292122 108292272
refGene_gn=EML4;refGene_tx=XM_019827633.2;refGene_gn=EML4;refGene_tx=XM_019827634.2;refGene_gn=EML4;refGene_tx=XM_019827635.2;refGene_gn=EML4;refGene_tx=XM_019827636.2;refGene_gn=EML4;refGene_tx=XM_019827637.2;refGene_gn=EML4;refGene_tx=XM_019827638.2;refGene_gn=EML4;refGene_tx=XM_019827639.2;refGene_gn=EML4;refGene_tx=XM_019827640.2;refGene_gn=EML4;refGene_tx=XM_019827641.2
A3 108294232 108294356
refGene_gn=EML4;refGene_tx=XM_019827633.2;refGene_gn=EML4;refGene_tx=XM_019827634.2;refGene_gn=EML4;refGene_tx=XM_019827635.2;refGene_gn=EML4;refGene_tx=XM_019827636.2;refGene_gn=EML4;refGene_tx=XM_019827637.2;refGene_gn=EML4;refGene_tx=XM_019827638.2;refGene_gn=EML4;refGene_tx=X

M_019827639.2;refGene_gn=EML4;refGene_tx=XM_019827640.2;refGene_gn=EML
4;refGene_tx=XM_019827641.2
A3 108307144 108307177
refGene_gn=EML4;refGene_tx=XM_019827633.2;refGene_gn=EML4;refGene_tx=X
M_019827634.2;refGene_gn=EML4;refGene_tx=XM_019827636.2;refGene_gn=EML
4;refGene_tx=XM_019827638.2;refGene_gn=EML4;refGene_tx=XM_019827639.2;
refGene_gn=EML4;refGene_tx=XM_019827641.2
A3 108307381 108307407
refGene_gn=EML4;refGene_tx=XM_019827633.2;refGene_gn=EML4;refGene_tx=X
M_019827634.2;refGene_gn=EML4;refGene_tx=XM_019827635.2;refGene_gn=EML
4;refGene_tx=XM_019827636.2;refGene_gn=EML4;refGene_tx=XM_019827637.2;
refGene_gn=EML4;refGene_tx=XM_019827638.2;refGene_gn=EML4;refGene_tx=X
M_019827639.2;refGene_gn=EML4;refGene_tx=XM_019827640.2;refGene_gn=EML
4;refGene_tx=XM_019827641.2
A3 108307901 108308030
refGene_gn=EML4;refGene_tx=XM_019827633.2;refGene_gn=EML4;refGene_tx=X
M_019827634.2;refGene_gn=EML4;refGene_tx=XM_019827635.2;refGene_gn=EML
4;refGene_tx=XM_019827636.2;refGene_gn=EML4;refGene_tx=XM_019827637.2;
refGene_gn=EML4;refGene_tx=XM_019827638.2;refGene_gn=EML4;refGene_tx=X
M_019827639.2;refGene_gn=EML4;refGene_tx=XM_019827640.2;refGene_gn=EML
4;refGene_tx=XM_019827641.2
A3 108310147 108310321
refGene_gn=EML4;refGene_tx=XM_019827633.2;refGene_gn=EML4;refGene_tx=X
M_019827634.2;refGene_gn=EML4;refGene_tx=XM_019827635.2;refGene_gn=EML
4;refGene_tx=XM_019827636.2;refGene_gn=EML4;refGene_tx=XM_019827637.2;
refGene_gn=EML4;refGene_tx=XM_019827641.2
A3 108314150 108314280
refGene_gn=EML4;refGene_tx=XM_019827633.2;refGene_gn=EML4;refGene_tx=X
M_019827634.2;refGene_gn=EML4;refGene_tx=XM_019827635.2;refGene_gn=EML
4;refGene_tx=XM_019827636.2;refGene_gn=EML4;refGene_tx=XM_019827637.2;
refGene_gn=EML4;refGene_tx=XM_019827638.2;refGene_gn=EML4;refGene_tx=X
M_019827639.2;refGene_gn=EML4;refGene_tx=XM_019827640.2;refGene_gn=EML
4;refGene_tx=XM_019827641.2
A3 108331174 108331357
refGene_gn=EML4;refGene_tx=XM_019827633.2;refGene_gn=EML4;refGene_tx=X
M_019827634.2;refGene_gn=EML4;refGene_tx=XM_019827635.2;refGene_gn=EML
4;refGene_tx=XM_019827636.2;refGene_gn=EML4;refGene_tx=XM_019827637.2;
refGene_gn=EML4;refGene_tx=XM_019827638.2;refGene_gn=EML4;refGene_tx=X
M_019827639.2;refGene_gn=EML4;refGene_tx=XM_019827640.2
A3 108340876 108340943
refGene_gn=EML4;refGene_tx=XM_019827633.2;refGene_gn=EML4;refGene_tx=X
M_019827634.2;refGene_gn=EML4;refGene_tx=XM_019827635.2;refGene_gn=EML
4;refGene_tx=XM_019827637.2;refGene_gn=EML4;refGene_tx=XM_019827638.2;
refGene_gn=EML4;refGene_tx=XM_019827639.2;refGene_gn=EML4;refGene_tx=X
M_019827640.2
A3 108401205 108401230
refGene_gn=EML4;refGene_tx=XM_019827636.2
A3 108506086 108506106
refGene_gn=PKDCC;refGene_tx=XM_023251930.1
A3 108512867 108512956

refGene_gn=PKDCC;refGene_tx=XM_023251929.1
A3 108513176 108513350
refGene_gn=PKDCC;refGene_tx=XM_023251929.1;refGene_gn=PKDCC;refGene_tx=XM_023251930.1
A3 108514951 108515059
refGene_gn=PKDCC;refGene_tx=XM_023251929.1;refGene_gn=PKDCC;refGene_tx=XM_023251930.1
A3 108515315 108515395
refGene_gn=PKDCC;refGene_tx=XM_023251929.1;refGene_gn=PKDCC;refGene_tx=XM_023251930.1
A3 108516048 108516320
refGene_gn=PKDCC;refGene_tx=XM_023251929.1;refGene_gn=PKDCC;refGene_tx=XM_023251930.1
A3 108516921 108517044
refGene_gn=PKDCC;refGene_tx=XM_023251929.1;refGene_gn=PKDCC;refGene_tx=XM_023251930.1
A3 108521291 108521930
refGene_gn=PKDCC;refGene_tx=XM_023251929.1;refGene_gn=PKDCC;refGene_tx=XM_023251930.1
A3 108600753 108600889
refGene_gn=CA3H2orf91;refGene_tx=XM_006944354.1
A3 108600891 108600950
refGene_gn=CA3H2orf91;refGene_tx=XM_006944354.1
A3 108600951 108601049
refGene_gn=CA3H2orf91;refGene_tx=XM_006944354.1
A3 108606357 108606466
refGene_gn=CA3H2orf91;refGene_tx=XM_006944354.1
A3 109937102 109938901
refGene_gn=SLC8A1;refGene_tx=NM_001009848.1
A3 110181807 110181914
refGene_gn=SLC8A1;refGene_tx=NM_001009848.1
A3 110185044 110185065
refGene_gn=SLC8A1;refGene_tx=NM_001009848.1
A3 110185502 110185520
refGene_gn=SLC8A1;refGene_tx=NM_001009848.1
A3 110185631 110185646
refGene_gn=SLC8A1;refGene_tx=NM_001009848.1
A3 110189512 110189581
refGene_gn=SLC8A1;refGene_tx=NM_001009848.1
A3 110194168 110194299
refGene_gn=SLC8A1;refGene_tx=NM_001009848.1
A3 110198226 110198326
refGene_gn=SLC8A1;refGene_tx=NM_001009848.1
A3 110221684 110221960
refGene_gn=SLC8A1;refGene_tx=NM_001009848.1
A3 110244259 110244636
refGene_gn=SLC8A1;refGene_tx=NM_001009848.1
A3 110499360 110499486
refGene_gn=THUMPD2;refGene_tx=XM_003984301.5;refGene_gn=THUMPD2;refGene_tx=XM_006930344.4;refGene_gn=THUMPD2;refGene_tx=XM_006930345.4;refGe

ne_gn=THUMPD2;refGene_tx=XM_019827646.2
A3	110507950	110508086
refGene_gn=THUMPD2;refGene_tx=XM_003984301.5;refGene_gn=THUMPD2;refGene_tx=XM_006930344.4;refGene_gn=THUMPD2;refGene_tx=XM_006930345.4;refGene_gn=THUMPD2;refGene_tx=XM_019827646.2
A3	110509721	110510149
refGene_gn=THUMPD2;refGene_tx=XM_003984301.5;refGene_gn=THUMPD2;refGene_tx=XM_006930344.4;refGene_gn=THUMPD2;refGene_tx=XM_006930345.4;refGene_gn=THUMPD2;refGene_tx=XM_019827646.2
A3	110511330	110511408
refGene_gn=THUMPD2;refGene_tx=XM_003984301.5;refGene_gn=THUMPD2;refGene_tx=XM_006930344.4;refGene_gn=THUMPD2;refGene_tx=XM_019827646.2
A3	110513719	110513772
refGene_gn=THUMPD2;refGene_tx=XM_003984301.5;refGene_gn=THUMPD2;refGene_tx=XM_006930344.4;refGene_gn=THUMPD2;refGene_tx=XM_006930345.4;refGene_gn=THUMPD2;refGene_tx=XM_019827646.2
A3	110515536	110515624
refGene_gn=THUMPD2;refGene_tx=XM_003984301.5;refGene_gn=THUMPD2;refGene_tx=XM_006930344.4;refGene_gn=THUMPD2;refGene_tx=XM_006930345.4;refGene_gn=THUMPD2;refGene_tx=XM_019827646.2
A3	110522413	110522485
refGene_gn=THUMPD2;refGene_tx=XM_003984301.5;refGene_gn=THUMPD2;refGene_tx=XM_006930344.4;refGene_gn=THUMPD2;refGene_tx=XM_006930345.4;refGene_gn=THUMPD2;refGene_tx=XM_019827646.2
A3	110522999	110523114
refGene_gn=THUMPD2;refGene_tx=XM_003984301.5;refGene_gn=THUMPD2;refGene_tx=XM_006930344.4;refGene_gn=THUMPD2;refGene_tx=XM_006930345.4
A3	110530466	110530575
refGene_gn=THUMPD2;refGene_tx=XM_003984301.5;refGene_gn=THUMPD2;refGene_tx=XM_006930344.4;refGene_gn=THUMPD2;refGene_tx=XM_006930345.4;refGene_gn=THUMPD2;refGene_tx=XM_019827646.2
A3	110533544	110533869
refGene_gn=THUMPD2;refGene_tx=XM_003984301.5;refGene_gn=THUMPD2;refGene_tx=XM_006930344.4;refGene_gn=THUMPD2;refGene_tx=XM_006930345.4
A3	110556774	110557016
refGene_gn=TMEM178A;refGene_tx=XM_019827648.2;refGene_gn=TMEM178A;refGene_tx=XM_023251931.1;refGene_gn=TMEM178A;refGene_tx=XM_023251932.1
A3	110567162	110567300
refGene_gn=TMEM178A;refGene_tx=XM_019827648.2;refGene_gn=TMEM178A;refGene_tx=XM_023251931.1
A3	110570357	110570471
refGene_gn=TMEM178A;refGene_tx=XM_019827648.2;refGene_gn=TMEM178A;refGene_tx=XM_023251931.1;refGene_gn=TMEM178A;refGene_tx=XM_023251932.1
A3	110585074	110585270
refGene_gn=TMEM178A;refGene_tx=XM_019827648.2
A3	110607812	110608212
refGene_gn=TMEM178A;refGene_tx=XM_023251931.1;refGene_gn=TMEM178A;refGene_tx=XM_023251932.1
A3	110804509	110804605
refGene_gn=MAP4K3;refGene_tx=XM_023251934.1;refGene_gn=MAP4K3;refGene_

tx=XM_023251935.1
A3 110870428 110870486
refGene_gn=MAP4K3;refGene_tx=XM_023251934.1;refGene_gn=MAP4K3;refGene_
tx=XM_023251935.1
A3 110912099 110912190
refGene_gn=MAP4K3;refGene_tx=XM_023251934.1;refGene_gn=MAP4K3;refGene_
tx=XM_023251935.1
A3 110924713 110924778
refGene_gn=MAP4K3;refGene_tx=XM_023251934.1;refGene_gn=MAP4K3;refGene_
tx=XM_023251935.1
A3 110931325 110931381
refGene_gn=MAP4K3;refGene_tx=XM_023251934.1;refGene_gn=MAP4K3;refGene_
tx=XM_023251935.1
A3 110931977 110932025
refGene_gn=MAP4K3;refGene_tx=XM_023251934.1;refGene_gn=MAP4K3;refGene_
tx=XM_023251935.1
A3 110936217 110936260
refGene_gn=MAP4K3;refGene_tx=XM_023251934.1;refGene_gn=MAP4K3;refGene_
tx=XM_023251935.1
A3 110937990 110938063
refGene_gn=MAP4K3;refGene_tx=XM_023251934.1;refGene_gn=MAP4K3;refGene_
tx=XM_023251935.1
A3 110940900 110941032
refGene_gn=MAP4K3;refGene_tx=XM_023251934.1;refGene_gn=MAP4K3;refGene_
tx=XM_023251935.1
A3 110941241 110941304
refGene_gn=MAP4K3;refGene_tx=XM_023251934.1;refGene_gn=MAP4K3;refGene_
tx=XM_023251935.1
A3 110941389 110941471
refGene_gn=MAP4K3;refGene_tx=XM_023251934.1;refGene_gn=MAP4K3;refGene_
tx=XM_023251935.1
A3 110941558 110941669
refGene_gn=MAP4K3;refGene_tx=XM_023251934.1;refGene_gn=MAP4K3;refGene_
tx=XM_023251935.1
A3 110945949 110946028
refGene_gn=MAP4K3;refGene_tx=XM_023251934.1;refGene_gn=MAP4K3;refGene_
tx=XM_023251935.1
A3 110950748 110950807
refGene_gn=MAP4K3;refGene_tx=XM_023251934.1;refGene_gn=MAP4K3;refGene_
tx=XM_023251935.1
A3 110952060 110952123
refGene_gn=MAP4K3;refGene_tx=XM_023251934.1
A3 110966424 110966483
refGene_gn=MAP4K3;refGene_tx=XM_023251934.1;refGene_gn=MAP4K3;refGene_
tx=XM_023251935.1
A3 110972271 110972310
refGene_gn=MAP4K3;refGene_tx=XM_023251934.1;refGene_gn=MAP4K3;refGene_
tx=XM_023251935.1
A3 110972542 110972596
refGene_gn=MAP4K3;refGene_tx=XM_023251934.1;refGene_gn=MAP4K3;refGene_

tx=XM_023251935.1
A3	110974570	110974613
refGene_gn=MAP4K3;refGene_tx=XM_023251934.1;refGene_gn=MAP4K3;refGene_tx=XM_023251935.1
A3	110976512	110976672
refGene_gn=MAP4K3;refGene_tx=XM_023251934.1;refGene_gn=MAP4K3;refGene_tx=XM_023251935.1
A3	110977941	110978054
refGene_gn=MAP4K3;refGene_tx=XM_023251934.1;refGene_gn=MAP4K3;refGene_tx=XM_023251935.1
A3	110982074	110982116
refGene_gn=MAP4K3;refGene_tx=XM_023251934.1;refGene_gn=MAP4K3;refGene_tx=XM_023251935.1
A3	110983935	110984020
refGene_gn=MAP4K3;refGene_tx=XM_023251934.1;refGene_gn=MAP4K3;refGene_tx=XM_023251935.1
A3	110985991	110986071
refGene_gn=MAP4K3;refGene_tx=XM_023251934.1;refGene_gn=MAP4K3;refGene_tx=XM_023251935.1
A3	110987189	110987250
refGene_gn=MAP4K3;refGene_tx=XM_023251934.1;refGene_gn=MAP4K3;refGene_tx=XM_023251935.1
A3	110987570	110987688
refGene_gn=MAP4K3;refGene_tx=XM_023251934.1;refGene_gn=MAP4K3;refGene_tx=XM_023251935.1
A3	110992523	110992582
refGene_gn=MAP4K3;refGene_tx=XM_023251934.1;refGene_gn=MAP4K3;refGene_tx=XM_023251935.1
A3	110993583	110993687
refGene_gn=MAP4K3;refGene_tx=XM_023251934.1;refGene_gn=MAP4K3;refGene_tx=XM_023251935.1
A3	110998201	110998373
refGene_gn=MAP4K3;refGene_tx=XM_023251934.1;refGene_gn=MAP4K3;refGene_tx=XM_023251935.1
A3	111001932	111002001
refGene_gn=MAP4K3;refGene_tx=XM_023251934.1;refGene_gn=MAP4K3;refGene_tx=XM_023251935.1
A3	111002085	111002178
refGene_gn=MAP4K3;refGene_tx=XM_023251934.1;refGene_gn=MAP4K3;refGene_tx=XM_023251935.1
A3	111007390	111007461
refGene_gn=MAP4K3;refGene_tx=XM_023251934.1;refGene_gn=MAP4K3;refGene_tx=XM_023251935.1
A3	111009982	111010038
refGene_gn=MAP4K3;refGene_tx=XM_023251934.1;refGene_gn=MAP4K3;refGene_tx=XM_023251935.1
A3	111010712	111010800
refGene_gn=MAP4K3;refGene_tx=XM_023251934.1;refGene_gn=MAP4K3;refGene_tx=XM_023251935.1
A3	111029965	111030133

refGene_gn=CDKL4;refGene_tx=XM_003984302.5;refGene_gn=CDKL4;refGene_tx=XM_006930347.4;refGene_gn=CDKL4;refGene_tx=XM_019827652.2
A3 111033376 111033498
refGene_gn=CDKL4;refGene_tx=XM_003984302.5;refGene_gn=CDKL4;refGene_tx=XM_006930347.4;refGene_gn=CDKL4;refGene_tx=XM_019827652.2
A3 111044747 111044820
refGene_gn=CDKL4;refGene_tx=XM_003984302.5;refGene_gn=CDKL4;refGene_tx=XM_006930347.4;refGene_gn=CDKL4;refGene_tx=XM_019827652.2
A3 111050904 111050995
refGene_gn=CDKL4;refGene_tx=XM_003984302.5;refGene_gn=CDKL4;refGene_tx=XM_006930347.4;refGene_gn=CDKL4;refGene_tx=XM_019827652.2
A3 111052139 111052337
refGene_gn=CDKL4;refGene_tx=XM_003984302.5;refGene_gn=CDKL4;refGene_tx=XM_006930347.4;refGene_gn=CDKL4;refGene_tx=XM_019827652.2
A3 111054454 111054537
refGene_gn=CDKL4;refGene_tx=XM_003984302.5;refGene_gn=CDKL4;refGene_tx=XM_006930347.4;refGene_gn=CDKL4;refGene_tx=XM_019827652.2
A3 111056656 111056713
refGene_gn=CDKL4;refGene_tx=XM_003984302.5;refGene_gn=CDKL4;refGene_tx=XM_006930347.4;refGene_gn=CDKL4;refGene_tx=XM_019827652.2
A3 111061566 111061701
refGene_gn=CDKL4;refGene_tx=XM_003984302.5;refGene_gn=CDKL4;refGene_tx=XM_006930347.4;refGene_gn=CDKL4;refGene_tx=XM_019827652.2
A3 111064325 111064427
refGene_gn=CDKL4;refGene_tx=XM_003984302.5;refGene_gn=CDKL4;refGene_tx=XM_006930347.4;refGene_gn=CDKL4;refGene_tx=XM_019827652.2
A3 111120430 111120517
refGene_gn=SOS1;refGene_tx=XM_011281161.3;refGene_gn=SOS1;refGene_tx=XM_011281162.3;refGene_gn=SOS1;refGene_tx=XM_011281163.3
A3 111192617 111192743
refGene_gn=SOS1;refGene_tx=XM_011281161.3;refGene_gn=SOS1;refGene_tx=XM_011281162.3;refGene_gn=SOS1;refGene_tx=XM_011281163.3
A3 111200326 111200458
refGene_gn=SOS1;refGene_tx=XM_011281161.3;refGene_gn=SOS1;refGene_tx=XM_011281162.3;refGene_gn=SOS1;refGene_tx=XM_011281163.3
A3 111202363 111202528
refGene_gn=SOS1;refGene_tx=XM_011281161.3;refGene_gn=SOS1;refGene_tx=XM_011281162.3;refGene_gn=SOS1;refGene_tx=XM_011281163.3
A3 111204449 111204659
refGene_gn=SOS1;refGene_tx=XM_011281161.3;refGene_gn=SOS1;refGene_tx=XM_011281162.3;refGene_gn=SOS1;refGene_tx=XM_011281163.3
A3 111208907 111209051
refGene_gn=SOS1;refGene_tx=XM_011281161.3;refGene_gn=SOS1;refGene_tx=XM_011281162.3;refGene_gn=SOS1;refGene_tx=XM_011281163.3
A3 111215511 111215622
refGene_gn=SOS1;refGene_tx=XM_011281161.3;refGene_gn=SOS1;refGene_tx=XM_011281162.3;refGene_gn=SOS1;refGene_tx=XM_011281163.3
A3 111215699 111215798
refGene_gn=SOS1;refGene_tx=XM_011281161.3;refGene_gn=SOS1;refGene_tx=XM_011281162.3;refGene_gn=SOS1;refGene_tx=XM_011281163.3

A3	111229279	111229407
refGene_gn=SOS1;refGene_tx=XM_011281161.3;refGene_gn=SOS1;refGene_tx=XM_011281162.3;refGene_gn=SOS1;refGene_tx=XM_011281163.3
A3	111230170	111230826
refGene_gn=SOS1;refGene_tx=XM_011281161.3;refGene_gn=SOS1;refGene_tx=XM_011281162.3;refGene_gn=SOS1;refGene_tx=XM_011281163.3
A3	111240006	111240088
refGene_gn=SOS1;refGene_tx=XM_011281161.3;refGene_gn=SOS1;refGene_tx=XM_011281162.3;refGene_gn=SOS1;refGene_tx=XM_011281163.3
A3	111240933	111241056
refGene_gn=SOS1;refGene_tx=XM_011281161.3;refGene_gn=SOS1;refGene_tx=XM_011281162.3;refGene_gn=SOS1;refGene_tx=XM_011281163.3
A3	111241350	111241454
refGene_gn=SOS1;refGene_tx=XM_011281161.3;refGene_gn=SOS1;refGene_tx=XM_011281162.3;refGene_gn=SOS1;refGene_tx=XM_011281163.3
A3	111242632	111242855
refGene_gn=SOS1;refGene_tx=XM_011281161.3;refGene_gn=SOS1;refGene_tx=XM_011281162.3;refGene_gn=SOS1;refGene_tx=XM_011281163.3
A3	111243964	111244084
refGene_gn=SOS1;refGene_tx=XM_011281161.3;refGene_gn=SOS1;refGene_tx=XM_011281162.3;refGene_gn=SOS1;refGene_tx=XM_011281163.3
A3	111249438	111249601
refGene_gn=SOS1;refGene_tx=XM_011281161.3;refGene_gn=SOS1;refGene_tx=XM_011281162.3;refGene_gn=SOS1;refGene_tx=XM_011281163.3
A3	111250102	111250220
refGene_gn=SOS1;refGene_tx=XM_011281161.3;refGene_gn=SOS1;refGene_tx=XM_011281162.3;refGene_gn=SOS1;refGene_tx=XM_011281163.3
A3	111258659	111258832
refGene_gn=SOS1;refGene_tx=XM_011281161.3;refGene_gn=SOS1;refGene_tx=XM_011281162.3;refGene_gn=SOS1;refGene_tx=XM_011281163.3
A3	111259046	111259163
refGene_gn=SOS1;refGene_tx=XM_011281161.3;refGene_gn=SOS1;refGene_tx=XM_011281162.3;refGene_gn=SOS1;refGene_tx=XM_011281163.3
A3	111260661	111260926
refGene_gn=SOS1;refGene_tx=XM_011281161.3;refGene_gn=SOS1;refGene_tx=XM_011281162.3;refGene_gn=SOS1;refGene_tx=XM_011281163.3
A3	111266672	111266717
refGene_gn=SOS1;refGene_tx=XM_011281161.3;refGene_gn=SOS1;refGene_tx=XM_011281163.3
A3	111268417	111268536
refGene_gn=SOS1;refGene_tx=XM_011281161.3;refGene_gn=SOS1;refGene_tx=XM_011281162.3;refGene_gn=SOS1;refGene_tx=XM_011281163.3
A3	111269411	111269903
refGene_gn=SOS1;refGene_tx=XM_011281161.3;refGene_gn=SOS1;refGene_tx=XM_011281162.3;refGene_gn=SOS1;refGene_tx=XM_011281163.3
A3	111282933	111283063
refGene_gn=ARHGEF33;refGene_tx=XM_023251936.1;refGene_gn=ARHGEF33;refGene_tx=XM_023251937.1
A3	111284855	111284907
refGene_gn=ARHGEF33;refGene_tx=XM_023251938.1;refGene_gn=ARHGEF33;refG

ene_tx=XM_023251939.1;refGene_gn=ARHGEF33;refGene_tx=XM_023251940.1;refGene_gn=ARHGEF33;refGene_tx=XM_023251941.1
A3 111289945 111290085
refGene_gn=ARHGEF33;refGene_tx=XM_023251936.1;refGene_gn=ARHGEF33;refGene_tx=XM_023251937.1;refGene_gn=ARHGEF33;refGene_tx=XM_023251938.1;refGene_gn=ARHGEF33;refGene_tx=XM_023251939.1;refGene_gn=ARHGEF33;refGene_tx=XM_023251940.1;refGene_gn=ARHGEF33;refGene_tx=XM_023251941.1
A3 111295168 111295985
refGene_gn=ARHGEF33;refGene_tx=XM_023251936.1;refGene_gn=ARHGEF33;refGene_tx=XM_023251937.1;refGene_gn=ARHGEF33;refGene_tx=XM_023251938.1;refGene_gn=ARHGEF33;refGene_tx=XM_023251939.1;refGene_gn=ARHGEF33;refGene_tx=XM_023251940.1;refGene_gn=ARHGEF33;refGene_tx=XM_023251941.1
A3 111298728 111298893
refGene_gn=ARHGEF33;refGene_tx=XM_023251936.1;refGene_gn=ARHGEF33;refGene_tx=XM_023251937.1;refGene_gn=ARHGEF33;refGene_tx=XM_023251938.1;refGene_gn=ARHGEF33;refGene_tx=XM_023251939.1;refGene_gn=ARHGEF33;refGene_tx=XM_023251940.1;refGene_gn=ARHGEF33;refGene_tx=XM_023251941.1
A3 111299888 111300037
refGene_gn=ARHGEF33;refGene_tx=XM_023251936.1;refGene_gn=ARHGEF33;refGene_tx=XM_023251937.1;refGene_gn=ARHGEF33;refGene_tx=XM_023251938.1;refGene_gn=ARHGEF33;refGene_tx=XM_023251939.1;refGene_gn=ARHGEF33;refGene_tx=XM_023251940.1;refGene_gn=ARHGEF33;refGene_tx=XM_023251941.1
A3 111301707 111301791
refGene_gn=ARHGEF33;refGene_tx=XM_023251936.1;refGene_gn=ARHGEF33;refGene_tx=XM_023251937.1;refGene_gn=ARHGEF33;refGene_tx=XM_023251938.1;refGene_gn=ARHGEF33;refGene_tx=XM_023251939.1;refGene_gn=ARHGEF33;refGene_tx=XM_023251940.1;refGene_gn=ARHGEF33;refGene_tx=XM_023251941.1
A3 111302962 111303046
refGene_gn=ARHGEF33;refGene_tx=XM_023251936.1;refGene_gn=ARHGEF33;refGene_tx=XM_023251937.1;refGene_gn=ARHGEF33;refGene_tx=XM_023251938.1;refGene_gn=ARHGEF33;refGene_tx=XM_023251939.1;refGene_gn=ARHGEF33;refGene_tx=XM_023251940.1;refGene_gn=ARHGEF33;refGene_tx=XM_023251941.1
A3 111305005 111305138
refGene_gn=ARHGEF33;refGene_tx=XM_023251936.1;refGene_gn=ARHGEF33;refGene_tx=XM_023251937.1;refGene_gn=ARHGEF33;refGene_tx=XM_023251938.1;refGene_gn=ARHGEF33;refGene_tx=XM_023251939.1;refGene_gn=ARHGEF33;refGene_tx=XM_023251940.1;refGene_gn=ARHGEF33;refGene_tx=XM_023251941.1
A3 111310367 111310497
refGene_gn=ARHGEF33;refGene_tx=XM_023251936.1;refGene_gn=ARHGEF33;refGene_tx=XM_023251937.1;refGene_gn=ARHGEF33;refGene_tx=XM_023251938.1;refGene_gn=ARHGEF33;refGene_tx=XM_023251939.1;refGene_gn=ARHGEF33;refGene_tx=XM_023251940.1;refGene_gn=ARHGEF33;refGene_tx=XM_023251941.1
A3 111315087 111315312
refGene_gn=ARHGEF33;refGene_tx=XM_023251936.1;refGene_gn=ARHGEF33;refGene_tx=XM_023251937.1;refGene_gn=ARHGEF33;refGene_tx=XM_023251938.1;refGene_gn=ARHGEF33;refGene_tx=XM_023251939.1;refGene_gn=ARHGEF33;refGene_tx=XM_023251940.1;refGene_gn=ARHGEF33;refGene_tx=XM_023251941.1
A3 111316353 111316413
refGene_gn=ARHGEF33;refGene_tx=XM_023251936.1;refGene_gn=ARHGEF33;refGene_tx=XM_023251937.1;refGene_gn=ARHGEF33;refGene_tx=XM_023251938.1;re

fGene_gn=ARHGEF33;refGene_tx=XM_023251939.1;refGene_gn=ARHGEF33;refGene_tx=XM_023251940.1;refGene_gn=ARHGEF33;refGene_tx=XM_023251941.1
A3 111319156 111319299
refGene_gn=ARHGEF33;refGene_tx=XM_023251936.1;refGene_gn=ARHGEF33;refGene_tx=XM_023251937.1;refGene_gn=ARHGEF33;refGene_tx=XM_023251938.1;refGene_gn=ARHGEF33;refGene_tx=XM_023251939.1;refGene_gn=ARHGEF33;refGene_tx=XM_023251940.1;refGene_gn=ARHGEF33;refGene_tx=XM_023251941.1
A3 111320445 111320567
refGene_gn=ARHGEF33;refGene_tx=XM_023251936.1;refGene_gn=ARHGEF33;refGene_tx=XM_023251937.1;refGene_gn=ARHGEF33;refGene_tx=XM_023251938.1;refGene_gn=ARHGEF33;refGene_tx=XM_023251939.1;refGene_gn=ARHGEF33;refGene_tx=XM_023251940.1;refGene_gn=ARHGEF33;refGene_tx=XM_023251941.1
A3 111320886 111321051
refGene_gn=ARHGEF33;refGene_tx=XM_023251936.1;refGene_gn=ARHGEF33;refGene_tx=XM_023251937.1;refGene_gn=ARHGEF33;refGene_tx=XM_023251938.1;refGene_gn=ARHGEF33;refGene_tx=XM_023251939.1;refGene_gn=ARHGEF33;refGene_tx=XM_023251940.1;refGene_gn=ARHGEF33;refGene_tx=XM_023251941.1
A3 111328292 111328342
refGene_gn=ARHGEF33;refGene_tx=XM_023251936.1;refGene_gn=ARHGEF33;refGene_tx=XM_023251937.1;refGene_gn=ARHGEF33;refGene_tx=XM_023251938.1;refGene_gn=ARHGEF33;refGene_tx=XM_023251939.1;refGene_gn=ARHGEF33;refGene_tx=XM_023251940.1;refGene_gn=ARHGEF33;refGene_tx=XM_023251941.1
A3 111329983 111330008
refGene_gn=ARHGEF33;refGene_tx=XM_023251936.1;refGene_gn=ARHGEF33;refGene_tx=XM_023251937.1;refGene_gn=ARHGEF33;refGene_tx=XM_023251938.1;refGene_gn=ARHGEF33;refGene_tx=XM_023251939.1;refGene_gn=ARHGEF33;refGene_tx=XM_023251940.1;refGene_gn=ARHGEF33;refGene_tx=XM_023251941.1
A3 111359819 111359922
refGene_gn=MORN2;refGene_tx=XM_003984303.4
A3 111361651 111361788
refGene_gn=MORN2;refGene_tx=XM_003984303.4
A3 111374680 111374892
refGene_gn=DHX57;refGene_tx=XM_006930350.2;refGene_gn=DHX57;refGene_tx=XM_019827653.2;refGene_gn=DHX57;refGene_tx=XM_019827654.1
A3 111376645 111376798
refGene_gn=DHX57;refGene_tx=XM_006930350.2;refGene_gn=DHX57;refGene_tx=XM_019827653.2;refGene_gn=DHX57;refGene_tx=XM_019827654.1
A3 111377839 111378025
refGene_gn=DHX57;refGene_tx=XM_006930350.2;refGene_gn=DHX57;refGene_tx=XM_019827653.2;refGene_gn=DHX57;refGene_tx=XM_019827654.1
A3 111378329 111379168
refGene_gn=DHX57;refGene_tx=XM_006930350.2;refGene_gn=DHX57;refGene_tx=XM_019827653.2;refGene_gn=DHX57;refGene_tx=XM_019827654.1
A3 111380890 111381138
refGene_gn=DHX57;refGene_tx=XM_006930350.2;refGene_gn=DHX57;refGene_tx=XM_019827653.2;refGene_gn=DHX57;refGene_tx=XM_019827654.1
A3 111382605 111382727
refGene_gn=DHX57;refGene_tx=XM_006930350.2;refGene_gn=DHX57;refGene_tx=XM_019827653.2;refGene_gn=DHX57;refGene_tx=XM_019827654.1
A3 111384449 111384645

refGene_gn=DHX57;refGene_tx=XM_006930350.2;refGene_gn=DHX57;refGene_tx=XM_019827653.2;refGene_gn=DHX57;refGene_tx=XM_019827654.1
A3 111385581 111385706
refGene_gn=DHX57;refGene_tx=XM_006930350.2;refGene_gn=DHX57;refGene_tx=XM_019827653.2;refGene_gn=DHX57;refGene_tx=XM_019827654.1
A3 111390047 111390181
refGene_gn=DHX57;refGene_tx=XM_006930350.2;refGene_gn=DHX57;refGene_tx=XM_019827653.2;refGene_gn=DHX57;refGene_tx=XM_019827654.1
A3 111391037 111391092
refGene_gn=DHX57;refGene_tx=XM_006930350.2;refGene_gn=DHX57;refGene_tx=XM_019827653.2;refGene_gn=DHX57;refGene_tx=XM_019827654.1
A3 111392908 111393114
refGene_gn=DHX57;refGene_tx=XM_006930350.2;refGene_gn=DHX57;refGene_tx=XM_019827653.2;refGene_gn=DHX57;refGene_tx=XM_019827654.1
A3 111397857 111397974
refGene_gn=DHX57;refGene_tx=XM_006930350.2;refGene_gn=DHX57;refGene_tx=XM_019827653.2;refGene_gn=DHX57;refGene_tx=XM_019827654.1
A3 111401272 111401369
refGene_gn=DHX57;refGene_tx=XM_006930350.2;refGene_gn=DHX57;refGene_tx=XM_019827653.2;refGene_gn=DHX57;refGene_tx=XM_019827654.1
A3 111402424 111402598
refGene_gn=DHX57;refGene_tx=XM_006930350.2;refGene_gn=DHX57;refGene_tx=XM_019827653.2;refGene_gn=DHX57;refGene_tx=XM_019827654.1
A3 111403427 111403628
refGene_gn=DHX57;refGene_tx=XM_006930350.2;refGene_gn=DHX57;refGene_tx=XM_019827653.2;refGene_gn=DHX57;refGene_tx=XM_019827654.1
A3 111405251 111405528
refGene_gn=DHX57;refGene_tx=XM_006930350.2;refGene_gn=DHX57;refGene_tx=XM_019827653.2;refGene_gn=DHX57;refGene_tx=XM_019827654.1
A3 111409617 111409713
refGene_gn=DHX57;refGene_tx=XM_006930350.2;refGene_gn=DHX57;refGene_tx=XM_019827653.2;refGene_gn=DHX57;refGene_tx=XM_019827654.1
A3 111409801 111409885
refGene_gn=DHX57;refGene_tx=XM_006930350.2;refGene_gn=DHX57;refGene_tx=XM_019827653.2;refGene_gn=DHX57;refGene_tx=XM_019827654.1
A3 111412972 111413107
refGene_gn=DHX57;refGene_tx=XM_006930350.2;refGene_gn=DHX57;refGene_tx=XM_019827653.2;refGene_gn=DHX57;refGene_tx=XM_019827654.1
A3 111414033 111414108
refGene_gn=DHX57;refGene_tx=XM_006930350.2;refGene_gn=DHX57;refGene_tx=XM_019827653.2;refGene_gn=DHX57;refGene_tx=XM_019827654.1
A3 111414779 111414914
refGene_gn=DHX57;refGene_tx=XM_006930350.2;refGene_gn=DHX57;refGene_tx=XM_019827653.2;refGene_gn=DHX57;refGene_tx=XM_019827654.1
A3 111417851 111418052
refGene_gn=DHX57;refGene_tx=XM_006930350.2;refGene_gn=DHX57;refGene_tx=XM_019827653.2;refGene_gn=DHX57;refGene_tx=XM_019827654.1
A3 111419879 111420023
refGene_gn=DHX57;refGene_tx=XM_006930350.2;refGene_gn=DHX57;refGene_tx=XM_019827653.2;refGene_gn=DHX57;refGene_tx=XM_019827654.1

A3 111427192 111427568
refGene_gn=GEMIN6;refGene_tx=XM_003984305.3;refGene_gn=GEMIN6;refGene_tx=XM_006930349.1
A3 111430998 111431136
refGene_gn=GEMIN6;refGene_tx=XM_003984305.3;refGene_gn=GEMIN6;refGene_tx=XM_006930349.1
A3 111431700 111431786
refGene_gn=GEMIN6;refGene_tx=XM_003984305.3
A3 111439686 111441453
refGene_gn=LOC101096974;refGene_tx=XM_011281159.3;refGene_gn=LOC101096974;refGene_tx=XM_011281160.3;refGene_gn=LOC101096974;refGene_tx=XM_019827655.2;refGene_gn=LOC101096974;refGene_tx=XM_019827656.2;refGene_gn=LOC101096974;refGene_tx=XM_019827657.2
A3 111454550 111454578
refGene_gn=SRSF7;refGene_tx=XM_003984306.3;refGene_gn=SRSF7;refGene_tx=XM_003984307.4;refGene_gn=SRSF7;refGene_tx=XM_006930351.3;refGene_gn=SRSF7;refGene_tx=XM_006930352.3
A3 111455597 111455778
refGene_gn=SRSF7;refGene_tx=XM_003984306.3;refGene_gn=SRSF7;refGene_tx=XM_003984307.4;refGene_gn=SRSF7;refGene_tx=XM_006930351.3;refGene_gn=SRSF7;refGene_tx=XM_006930352.3
A3 111456016 111456193
refGene_gn=SRSF7;refGene_tx=XM_003984306.3;refGene_gn=SRSF7;refGene_tx=XM_003984307.4;refGene_gn=SRSF7;refGene_tx=XM_006930351.3;refGene_gn=SRSF7;refGene_tx=XM_006930352.3
A3 111457059 111457134
refGene_gn=SRSF7;refGene_tx=XM_003984306.3;refGene_gn=SRSF7;refGene_tx=XM_003984307.4;refGene_gn=SRSF7;refGene_tx=XM_006930351.3;refGene_gn=SRSF7;refGene_tx=XM_006930352.3
A3 111457217 111457328
refGene_gn=SRSF7;refGene_tx=XM_003984306.3;refGene_gn=SRSF7;refGene_tx=XM_003984307.4;refGene_gn=SRSF7;refGene_tx=XM_006930351.3;refGene_gn=SRSF7;refGene_tx=XM_006930352.3
A3 111458965 111459019
refGene_gn=SRSF7;refGene_tx=XM_003984306.3;refGene_gn=SRSF7;refGene_tx=XM_003984307.4;refGene_gn=SRSF7;refGene_tx=XM_006930351.3;refGene_gn=SRSF7;refGene_tx=XM_006930352.3
A3 111459560 111459596
refGene_gn=SRSF7;refGene_tx=XM_003984306.3;refGene_gn=SRSF7;refGene_tx=XM_006930351.3
A3 111460248 111460303
refGene_gn=SRSF7;refGene_tx=XM_003984306.3;refGene_gn=SRSF7;refGene_tx=XM_003984307.4;refGene_gn=SRSF7;refGene_tx=XM_006930351.3;refGene_gn=SRSF7;refGene_tx=XM_006930352.3
A3 111474856 111474934
refGene_gn=GALM;refGene_tx=XM_003984308.5;refGene_gn=GALM;refGene_tx=XM_019827659.2
A3 111475716 111475891
refGene_gn=GALM;refGene_tx=XM_003984308.5;refGene_gn=GALM;refGene_tx=XM_019827659.2

A3 111477260 111477402
refGene_gn=GALM;refGene_tx=XM_003984308.5;refGene_gn=GALM;refGene_tx=XM_019827659.2
A3 111501293 111501532
refGene_gn=GALM;refGene_tx=XM_019827658.2
A3 111506601 111506683
refGene_gn=GALM;refGene_tx=XM_003984308.5;refGene_gn=GALM;refGene_tx=XM_019827658.2;refGene_gn=GALM;refGene_tx=XM_019827659.2
A3 111512471 111512678
refGene_gn=GALM;refGene_tx=XM_003984308.5;refGene_gn=GALM;refGene_tx=XM_019827658.2;refGene_gn=GALM;refGene_tx=XM_019827659.2
A3 111517972 111518127
refGene_gn=GALM;refGene_tx=XM_003984308.5;refGene_gn=GALM;refGene_tx=XM_019827658.2;refGene_gn=GALM;refGene_tx=XM_019827659.2
A3 111527022 111527212
refGene_gn=GALM;refGene_tx=XM_003984308.5;refGene_gn=GALM;refGene_tx=XM_019827658.2
A3 111587761 111587956
refGene_gn=HNRNPLL;refGene_tx=XM_006930353.4;refGene_gn=HNRNPLL;refGene_tx=XM_006930354.4
A3 111588133 111588146
refGene_gn=GALM;refGene_tx=XM_019827659.2
A3 111597759 111597878
refGene_gn=HNRNPLL;refGene_tx=XM_006930353.4;refGene_gn=HNRNPLL;refGene_tx=XM_006930354.4
A3 111602624 111602862
refGene_gn=HNRNPLL;refGene_tx=XM_006930353.4;refGene_gn=HNRNPLL;refGene_tx=XM_006930354.4
A3 111604732 111604818
refGene_gn=HNRNPLL;refGene_tx=XM_006930353.4;refGene_gn=HNRNPLL;refGene_tx=XM_006930354.4
A3 111606397 111606494
refGene_gn=HNRNPLL;refGene_tx=XM_006930353.4;refGene_gn=HNRNPLL;refGene_tx=XM_006930354.4
A3 111606579 111606652
refGene_gn=HNRNPLL;refGene_tx=XM_006930353.4;refGene_gn=HNRNPLL;refGene_tx=XM_006930354.4
A3 111611120 111611192
refGene_gn=HNRNPLL;refGene_tx=XM_006930353.4
A3 111616204 111616422
refGene_gn=HNRNPLL;refGene_tx=XM_006930353.4;refGene_gn=HNRNPLL;refGene_tx=XM_006930354.4
A3 111619280 111619402
refGene_gn=HNRNPLL;refGene_tx=XM_006930353.4;refGene_gn=HNRNPLL;refGene_tx=XM_006930354.4
A3 111619878 111620080
refGene_gn=HNRNPLL;refGene_tx=XM_006930353.4;refGene_gn=HNRNPLL;refGene_tx=XM_006930354.4
A3 111620730 111620788
refGene_gn=HNRNPLL;refGene_tx=XM_006930353.4;refGene_gn=HNRNPLL;refGen

e_tx=XM_006930354.4
A3 111620878 111620977
refGene_gn=HNRNPLL;refGene_tx=XM_006930353.4;refGene_gn=HNRNPLL;refGene_tx=XM_006930354.4
A3 111624450 111624506
refGene_gn=HNRNPLL;refGene_tx=XM_006930353.4;refGene_gn=HNRNPLL;refGene_tx=XM_006930354.4
A3 111782223 111782341
refGene_gn=ATL2;refGene_tx=XM_023251942.1;refGene_gn=ATL2;refGene_tx=XM_023251943.1;refGene_gn=ATL2;refGene_tx=XM_023251944.1;refGene_gn=ATL2;refGene_tx=XM_023251945.1;refGene_gn=ATL2;refGene_tx=XM_023251946.1
A3 111782825 111782886
refGene_gn=ATL2;refGene_tx=XM_019827660.2
A3 111807513 111807758
refGene_gn=ATL2;refGene_tx=XM_019827660.2;refGene_gn=ATL2;refGene_tx=XM_023251942.1;refGene_gn=ATL2;refGene_tx=XM_023251943.1;refGene_gn=ATL2;refGene_tx=XM_023251944.1;refGene_gn=ATL2;refGene_tx=XM_023251945.1;refGene_gn=ATL2;refGene_tx=XM_023251946.1
A3 111825086 111825221
refGene_gn=ATL2;refGene_tx=XM_019827660.2;refGene_gn=ATL2;refGene_tx=XM_023251942.1;refGene_gn=ATL2;refGene_tx=XM_023251943.1;refGene_gn=ATL2;refGene_tx=XM_023251944.1;refGene_gn=ATL2;refGene_tx=XM_023251945.1;refGene_gn=ATL2;refGene_tx=XM_023251946.1
A3 111826515 111826620
refGene_gn=ATL2;refGene_tx=XM_019827660.2;refGene_gn=ATL2;refGene_tx=XM_023251942.1;refGene_gn=ATL2;refGene_tx=XM_023251943.1;refGene_gn=ATL2;refGene_tx=XM_023251944.1;refGene_gn=ATL2;refGene_tx=XM_023251945.1;refGene_gn=ATL2;refGene_tx=XM_023251946.1
A3 111830265 111830316
refGene_gn=ATL2;refGene_tx=XM_019827660.2;refGene_gn=ATL2;refGene_tx=XM_023251942.1;refGene_gn=ATL2;refGene_tx=XM_023251943.1;refGene_gn=ATL2;refGene_tx=XM_023251944.1;refGene_gn=ATL2;refGene_tx=XM_023251945.1;refGene_gn=ATL2;refGene_tx=XM_023251946.1
A3 111830610 111830667
refGene_gn=ATL2;refGene_tx=XM_019827660.2;refGene_gn=ATL2;refGene_tx=XM_023251942.1;refGene_gn=ATL2;refGene_tx=XM_023251943.1;refGene_gn=ATL2;refGene_tx=XM_023251944.1;refGene_gn=ATL2;refGene_tx=XM_023251945.1;refGene_gn=ATL2;refGene_tx=XM_023251946.1
A3 111832114 111832207
refGene_gn=ATL2;refGene_tx=XM_019827660.2;refGene_gn=ATL2;refGene_tx=XM_023251942.1;refGene_gn=ATL2;refGene_tx=XM_023251943.1;refGene_gn=ATL2;refGene_tx=XM_023251944.1;refGene_gn=ATL2;refGene_tx=XM_023251945.1;refGene_gn=ATL2;refGene_tx=XM_023251946.1
A3 111834722 111834861
refGene_gn=ATL2;refGene_tx=XM_019827660.2;refGene_gn=ATL2;refGene_tx=XM_023251942.1;refGene_gn=ATL2;refGene_tx=XM_023251943.1;refGene_gn=ATL2;refGene_tx=XM_023251944.1;refGene_gn=ATL2;refGene_tx=XM_023251945.1;refGene_gn=ATL2;refGene_tx=XM_023251946.1
A3 111835651 111835779
refGene_gn=ATL2;refGene_tx=XM_019827660.2;refGene_gn=ATL2;refGene_tx=X

M_023251942.1;refGene_gn=ATL2;refGene_tx=XM_023251943.1;refGene_gn=ATL2;refGene_tx=XM_023251944.1;refGene_gn=ATL2;refGene_tx=XM_023251945.1;refGene_gn=ATL2;refGene_tx=XM_023251946.1
A3 111836812 111836869
refGene_gn=ATL2;refGene_tx=XM_019827660.2;refGene_gn=ATL2;refGene_tx=XM_023251942.1;refGene_gn=ATL2;refGene_tx=XM_023251943.1;refGene_gn=ATL2;refGene_tx=XM_023251944.1;refGene_gn=ATL2;refGene_tx=XM_023251945.1;refGene_gn=ATL2;refGene_tx=XM_023251946.1
A3 111838349 111838421
refGene_gn=ATL2;refGene_tx=XM_019827660.2;refGene_gn=ATL2;refGene_tx=XM_023251942.1;refGene_gn=ATL2;refGene_tx=XM_023251943.1;refGene_gn=ATL2;refGene_tx=XM_023251944.1;refGene_gn=ATL2;refGene_tx=XM_023251945.1;refGene_gn=ATL2;refGene_tx=XM_023251946.1
A3 111839040 111839472
refGene_gn=ATL2;refGene_tx=XM_019827660.2;refGene_gn=ATL2;refGene_tx=XM_023251942.1;refGene_gn=ATL2;refGene_tx=XM_023251943.1;refGene_gn=ATL2;refGene_tx=XM_023251944.1;refGene_gn=ATL2;refGene_tx=XM_023251945.1;refGene_gn=ATL2;refGene_tx=XM_023251946.1
A3 111840956 111840971
refGene_gn=ATL2;refGene_tx=XM_019827660.2;refGene_gn=ATL2;refGene_tx=XM_023251942.1;refGene_gn=ATL2;refGene_tx=XM_023251943.1;refGene_gn=ATL2;refGene_tx=XM_023251944.1
A3 111841066 111841174
refGene_gn=ATL2;refGene_tx=XM_023251943.1;refGene_gn=ATL2;refGene_tx=XM_023251944.1;refGene_gn=ATL2;refGene_tx=XM_023251946.1
A3 111841557 111841677
refGene_gn=ATL2;refGene_tx=XM_019827660.2;refGene_gn=ATL2;refGene_tx=XM_023251942.1;refGene_gn=ATL2;refGene_tx=XM_023251945.1
A3 112022870 112023916
refGene_gn=LOC101099873;refGene_tx=XM_023251947.1;refGene_gn=LOC101099873;refGene_tx=XM_023251948.1
A3 112026714 112027330
refGene_gn=LOC101099873;refGene_tx=XM_023251947.1;refGene_gn=LOC101099873;refGene_tx=XM_023251948.1
A3 112063202 112063256
refGene_gn=RMDN2;refGene_tx=XM_023251949.1;refGene_gn=RMDN2;refGene_tx=XM_023251950.1;refGene_gn=RMDN2;refGene_tx=XM_023251951.1;refGene_gn=RMDN2;refGene_tx=XM_023251952.1;refGene_gn=RMDN2;refGene_tx=XM_023251954.1
A3 112073871 112073952
refGene_gn=RMDN2;refGene_tx=XM_023251949.1;refGene_gn=RMDN2;refGene_tx=XM_023251950.1;refGene_gn=RMDN2;refGene_tx=XM_023251951.1;refGene_gn=RMDN2;refGene_tx=XM_023251952.1;refGene_gn=RMDN2;refGene_tx=XM_023251954.1
A3 112074040 112074094
refGene_gn=RMDN2;refGene_tx=XM_023251949.1;refGene_gn=RMDN2;refGene_tx=XM_023251950.1;refGene_gn=RMDN2;refGene_tx=XM_023251951.1;refGene_gn=RMDN2;refGene_tx=XM_023251952.1;refGene_gn=RMDN2;refGene_tx=XM_023251954.1
A3 112082425 112082524

refGene_gn=RMDN2;refGene_tx=XM_023251949.1;refGene_gn=RMDN2;refGene_tx=XM_023251950.1;refGene_gn=RMDN2;refGene_tx=XM_023251951.1;refGene_gn=RMDN2;refGene_tx=XM_023251952.1;refGene_gn=RMDN2;refGene_tx=XM_023251954.1
A3 112093641 112093719
refGene_gn=RMDN2;refGene_tx=XM_023251949.1;refGene_gn=RMDN2;refGene_tx=XM_023251951.1;refGene_gn=RMDN2;refGene_tx=XM_023251952.1;refGene_gn=RMDN2;refGene_tx=XM_023251954.1
A3 112095132 112095208
refGene_gn=RMDN2;refGene_tx=XM_023251949.1;refGene_gn=RMDN2;refGene_tx=XM_023251950.1;refGene_gn=RMDN2;refGene_tx=XM_023251951.1;refGene_gn=RMDN2;refGene_tx=XM_023251952.1;refGene_gn=RMDN2;refGene_tx=XM_023251954.1
A3 112106954 112107015
refGene_gn=RMDN2;refGene_tx=XM_023251949.1;refGene_gn=RMDN2;refGene_tx=XM_023251950.1;refGene_gn=RMDN2;refGene_tx=XM_023251951.1;refGene_gn=RMDN2;refGene_tx=XM_023251952.1;refGene_gn=RMDN2;refGene_tx=XM_023251954.1
A3 112107687 112107695
refGene_gn=RMDN2;refGene_tx=XM_023251953.1
A3 112110960 112111063
refGene_gn=RMDN2;refGene_tx=XM_023251949.1;refGene_gn=RMDN2;refGene_tx=XM_023251950.1;refGene_gn=RMDN2;refGene_tx=XM_023251951.1;refGene_gn=RMDN2;refGene_tx=XM_023251952.1;refGene_gn=RMDN2;refGene_tx=XM_023251953.1;refGene_gn=RMDN2;refGene_tx=XM_023251954.1
A3 112112192 112112367
refGene_gn=RMDN2;refGene_tx=XM_023251949.1;refGene_gn=RMDN2;refGene_tx=XM_023251950.1;refGene_gn=RMDN2;refGene_tx=XM_023251951.1;refGene_gn=RMDN2;refGene_tx=XM_023251952.1;refGene_gn=RMDN2;refGene_tx=XM_023251953.1;refGene_gn=RMDN2;refGene_tx=XM_023251954.1
A3 112128348 112129059
refGene_gn=RMDN2;refGene_tx=XM_023251949.1;refGene_gn=RMDN2;refGene_tx=XM_023251950.1;refGene_gn=RMDN2;refGene_tx=XM_023251953.1
A3 112129681 112129698
refGene_gn=RMDN2;refGene_tx=XM_023251949.1;refGene_gn=RMDN2;refGene_tx=XM_023251950.1;refGene_gn=RMDN2;refGene_tx=XM_023251953.1;refGene_gn=RMDN2;refGene_tx=XM_023251954.1
A3 112136993 112137445
refGene_gn=RMDN2;refGene_tx=XM_023251951.1;refGene_gn=RMDN2;refGene_tx=XM_023251952.1
A3 112373137 112373902
refGene_gn=CDC42EP3;refGene_tx=XM_003984317.5
A3 112594450 112594596
refGene_gn=QPCT;refGene_tx=XM_003984318.4
A3 112595848 112595965
refGene_gn=QPCT;refGene_tx=XM_003984318.4
A3 112596794 112596868
refGene_gn=QPCT;refGene_tx=XM_019827673.1
A3 112597340 112597440
refGene_gn=QPCT;refGene_tx=XM_003984318.4;refGene_gn=QPCT;refGene_tx=X

M_019827673.1
A3	112599440	112599617
refGene_gn=QPCT;refGene_tx=XM_003984318.4;refGene_gn=QPCT;refGene_tx=X
M_019827673.1
A3	112604581	112604860
refGene_gn=QPCT;refGene_tx=XM_003984318.4;refGene_gn=QPCT;refGene_tx=X
M_019827673.1
A3	112610667	112610814
refGene_gn=QPCT;refGene_tx=XM_003984318.4;refGene_gn=QPCT;refGene_tx=X
M_019827673.1
A3	112618704	112618824
refGene_gn=QPCT;refGene_tx=XM_003984318.4;refGene_gn=QPCT;refGene_tx=X
M_019827673.1
A3	112647486	112647774
refGene_gn=PRKD3;refGene_tx=XM_003984319.5;refGene_gn=PRKD3;refGene_tx
=XM_023251955.1
A3	112681152	112681291
refGene_gn=PRKD3;refGene_tx=XM_003984319.5;refGene_gn=PRKD3;refGene_tx
=XM_023251955.1
A3	112682830	112682962
refGene_gn=PRKD3;refGene_tx=XM_003984319.5;refGene_gn=PRKD3;refGene_tx
=XM_023251955.1
A3	112683935	112684093
refGene_gn=PRKD3;refGene_tx=XM_003984319.5;refGene_gn=PRKD3;refGene_tx
=XM_023251955.1
A3	112689249	112689442
refGene_gn=PRKD3;refGene_tx=XM_003984319.5;refGene_gn=PRKD3;refGene_tx
=XM_023251955.1
A3	112695328	112695406
refGene_gn=PRKD3;refGene_tx=XM_003984319.5;refGene_gn=PRKD3;refGene_tx
=XM_023251955.1
A3	112698126	112698310
refGene_gn=PRKD3;refGene_tx=XM_003984319.5;refGene_gn=PRKD3;refGene_tx
=XM_023251955.1
A3	112699777	112699901
refGene_gn=PRKD3;refGene_tx=XM_003984319.5;refGene_gn=PRKD3;refGene_tx
=XM_023251955.1
A3	112701574	112701652
refGene_gn=PRKD3;refGene_tx=XM_003984319.5;refGene_gn=PRKD3;refGene_tx
=XM_023251955.1
A3	112702720	112702997
refGene_gn=PRKD3;refGene_tx=XM_003984319.5;refGene_gn=PRKD3;refGene_tx
=XM_023251955.1
A3	112705854	112705907
refGene_gn=PRKD3;refGene_tx=XM_003984319.5;refGene_gn=PRKD3;refGene_tx
=XM_023251955.1
A3	112710692	112710765
refGene_gn=PRKD3;refGene_tx=XM_003984319.5;refGene_gn=PRKD3;refGene_tx
=XM_023251955.1
A3	112711825	112711932

refGene_gn=PRKD3;refGene_tx=XM_003984319.5;refGene_gn=PRKD3;refGene_tx=XM_023251955.1
A3 112718552 112718714
refGene_gn=PRKD3;refGene_tx=XM_003984319.5;refGene_gn=PRKD3;refGene_tx=XM_023251955.1
A3 112718933 112719032
refGene_gn=PRKD3;refGene_tx=XM_003984319.5;refGene_gn=PRKD3;refGene_tx=XM_023251955.1
A3 112721733 112722001
refGene_gn=PRKD3;refGene_tx=XM_003984319.5;refGene_gn=PRKD3;refGene_tx=XM_023251955.1
A3 112724112 112724198
refGene_gn=PRKD3;refGene_tx=XM_003984319.5;refGene_gn=PRKD3;refGene_tx=XM_023251955.1
A3 112725317 112725491
refGene_gn=PRKD3;refGene_tx=XM_003984319.5;refGene_gn=PRKD3;refGene_tx=XM_023251955.1
A3 112730001 112730214
refGene_gn=NDUFAF7;refGene_tx=XM_011281155.2;refGene_gn=NDUFAF7;refGene_tx=XM_019827677.1;refGene_gn=NDUFAF7;refGene_tx=XM_019827678.1
A3 112730948 112731122
refGene_gn=NDUFAF7;refGene_tx=XM_011281155.2;refGene_gn=NDUFAF7;refGene_tx=XM_019827677.1;refGene_gn=NDUFAF7;refGene_tx=XM_019827678.1
A3 112731529 112731673
refGene_gn=NDUFAF7;refGene_tx=XM_011281155.2;refGene_gn=NDUFAF7;refGene_tx=XM_019827677.1;refGene_gn=NDUFAF7;refGene_tx=XM_019827678.1
A3 112734016 112734127
refGene_gn=NDUFAF7;refGene_tx=XM_011281155.2;refGene_gn=NDUFAF7;refGene_tx=XM_019827677.1;refGene_gn=NDUFAF7;refGene_tx=XM_019827678.1
A3 112735037 112735096
refGene_gn=NDUFAF7;refGene_tx=XM_011281155.2;refGene_gn=NDUFAF7;refGene_tx=XM_019827677.1;refGene_gn=NDUFAF7;refGene_tx=XM_019827678.1
A3 112735935 112736149
refGene_gn=NDUFAF7;refGene_tx=XM_011281155.2;refGene_gn=NDUFAF7;refGene_tx=XM_019827677.1;refGene_gn=NDUFAF7;refGene_tx=XM_019827678.1
A3 112739209 112739320
refGene_gn=NDUFAF7;refGene_tx=XM_011281155.2;refGene_gn=NDUFAF7;refGene_tx=XM_019827677.1;refGene_gn=NDUFAF7;refGene_tx=XM_019827678.1
A3 112740311 112740392
refGene_gn=NDUFAF7;refGene_tx=XM_011281155.2;refGene_gn=NDUFAF7;refGene_tx=XM_019827677.1;refGene_gn=NDUFAF7;refGene_tx=XM_019827678.1
A3 112753673 112753834
refGene_gn=NDUFAF7;refGene_tx=XM_011281155.2;refGene_gn=NDUFAF7;refGene_tx=XM_019827677.1
A3 112754126 112754233
refGene_gn=NDUFAF7;refGene_tx=XM_011281155.2;refGene_gn=NDUFAF7;refGene_tx=XM_019827677.1
A3 112754321 112754477
refGene_gn=CEBPZ;refGene_tx=XM_023252197.1
A3 112756935 112758434

refGene_gn=CEBPZ;refGene_tx=XM_023252197.1
A3 112761475 112761707
refGene_gn=CEBPZ;refGene_tx=XM_023252197.1
A3 112762282 112762463
refGene_gn=CEBPZ;refGene_tx=XM_023252197.1
A3 112763693 112763782
refGene_gn=CEBPZ;refGene_tx=XM_023252197.1
A3 112766860 112766914
refGene_gn=CEBPZ;refGene_tx=XM_023252197.1
A3 112767477 112767580
refGene_gn=CEBPZ;refGene_tx=XM_023252197.1
A3 112767683 112767752
refGene_gn=CEBPZ;refGene_tx=XM_023252197.1
A3 112769166 112769233
refGene_gn=CEBPZ;refGene_tx=XM_023252197.1
A3 112770537 112770638
refGene_gn=CEBPZ;refGene_tx=XM_023252197.1
A3 112771439 112771497
refGene_gn=CEBPZ;refGene_tx=XM_023252197.1
A3 112771777 112771965
refGene_gn=CEBPZ;refGene_tx=XM_023252197.1
A3 112777428 112777490
refGene_gn=CEBPZ;refGene_tx=XM_023252197.1
A3 112777687 112777769
refGene_gn=CEBPZ;refGene_tx=XM_023252197.1
A3 112778557 112778697
refGene_gn=CEBPZ;refGene_tx=XM_023252197.1
A3 112778738 112778821
refGene_gn=CEBPZOS;refGene_tx=XM_019827679.1;refGene_gn=CEBPZOS;refGene_tx=XM_019827680.1;refGene_gn=CEBPZOS;refGene_tx=XM_019827681.2;refGene_gn=CEBPZOS;refGene_tx=XM_019827682.2;refGene_gn=CEBPZOS;refGene_tx=XM_019827683.2;refGene_gn=CEBPZOS;refGene_tx=XM_019827684.2
A3 112779401 112779446
refGene_gn=CEBPZOS;refGene_tx=XM_019827679.1;refGene_gn=CEBPZOS;refGene_tx=XM_019827680.1;refGene_gn=CEBPZOS;refGene_tx=XM_019827681.2;refGene_gn=CEBPZOS;refGene_tx=XM_019827682.2;refGene_gn=CEBPZOS;refGene_tx=XM_019827683.2;refGene_gn=CEBPZOS;refGene_tx=XM_019827684.2
A3 112780532 112780647
refGene_gn=CEBPZOS;refGene_tx=XM_019827679.1;refGene_gn=CEBPZOS;refGene_tx=XM_019827680.1;refGene_gn=CEBPZOS;refGene_tx=XM_019827681.2;refGene_gn=CEBPZOS;refGene_tx=XM_019827682.2;refGene_gn=CEBPZOS;refGene_tx=XM_019827683.2;refGene_gn=CEBPZOS;refGene_tx=XM_019827684.2
A3 112789132 112789217
refGene_gn=SULT6B1;refGene_tx=XM_023251958.1
A3 112790645 112790758
refGene_gn=SULT6B1;refGene_tx=XM_023251958.1
A3 112795128 112795218
refGene_gn=SULT6B1;refGene_tx=XM_023251958.1
A3 112796364 112796491
refGene_gn=SULT6B1;refGene_tx=XM_023251958.1

A3 112801784 112801879
refGene_gn=SULT6B1;refGene_tx=XM_023251958.1
A3 112804989 112805146
refGene_gn=SULT6B1;refGene_tx=XM_023251958.1
A3 112807525 112807656
refGene_gn=SULT6B1;refGene_tx=XM_023251958.1
A3 112815806 112815854
refGene_gn=LOC109498301;refGene_tx=XM_023252198.1
A3 112816092 112816717
refGene_gn=LOC109498301;refGene_tx=XM_023252198.1
A3 112817046 112817108
refGene_gn=LOC109498301;refGene_tx=XM_023252198.1
A3 112822541 112822660
refGene_gn=EIF2AK2;refGene_tx=XM_019827674.2;refGene_gn=EIF2AK2;refGene_tx=XM_023251956.1;refGene_gn=EIF2AK2;refGene_tx=XM_023251957.1
A3 112823531 112823652
refGene_gn=EIF2AK2;refGene_tx=XM_019827674.2;refGene_gn=EIF2AK2;refGene_tx=XM_023251956.1;refGene_gn=EIF2AK2;refGene_tx=XM_023251957.1
A3 112826589 112826747
refGene_gn=EIF2AK2;refGene_tx=XM_019827674.2;refGene_gn=EIF2AK2;refGene_tx=XM_023251956.1;refGene_gn=EIF2AK2;refGene_tx=XM_023251957.1
A3 112828908 112829035
refGene_gn=EIF2AK2;refGene_tx=XM_019827674.2;refGene_gn=EIF2AK2;refGene_tx=XM_023251956.1;refGene_gn=EIF2AK2;refGene_tx=XM_023251957.1
A3 112830261 112830341
refGene_gn=EIF2AK2;refGene_tx=XM_019827674.2;refGene_gn=EIF2AK2;refGene_tx=XM_023251956.1;refGene_gn=EIF2AK2;refGene_tx=XM_023251957.1
A3 112830501 112830595
refGene_gn=EIF2AK2;refGene_tx=XM_019827674.2;refGene_gn=EIF2AK2;refGene_tx=XM_023251956.1;refGene_gn=EIF2AK2;refGene_tx=XM_023251957.1
A3 112833244 112833279
refGene_gn=EIF2AK2;refGene_tx=XM_019827674.2;refGene_gn=EIF2AK2;refGene_tx=XM_023251956.1;refGene_gn=EIF2AK2;refGene_tx=XM_023251957.1
A3 112834838 112834901
refGene_gn=EIF2AK2;refGene_tx=XM_019827674.2;refGene_gn=EIF2AK2;refGene_tx=XM_023251956.1;refGene_gn=EIF2AK2;refGene_tx=XM_023251957.1
A3 112836695 112836818
refGene_gn=EIF2AK2;refGene_tx=XM_019827674.2;refGene_gn=EIF2AK2;refGene_tx=XM_023251956.1;refGene_gn=EIF2AK2;refGene_tx=XM_023251957.1
A3 112839741 112839861
refGene_gn=EIF2AK2;refGene_tx=XM_019827674.2;refGene_gn=EIF2AK2;refGene_tx=XM_023251956.1;refGene_gn=EIF2AK2;refGene_tx=XM_023251957.1
A3 112842090 112842271
refGene_gn=EIF2AK2;refGene_tx=XM_019827674.2;refGene_gn=EIF2AK2;refGene_tx=XM_023251956.1;refGene_gn=EIF2AK2;refGene_tx=XM_023251957.1
A3 112844828 112844957
refGene_gn=EIF2AK2;refGene_tx=XM_019827674.2;refGene_gn=EIF2AK2;refGene_tx=XM_023251956.1;refGene_gn=EIF2AK2;refGene_tx=XM_023251957.1
A3 112847307 112847409
refGene_gn=EIF2AK2;refGene_tx=XM_019827674.2;refGene_gn=EIF2AK2;refGen

e_tx=XM_023251956.1;refGene_gn=EIF2AK2;refGene_tx=XM_023251957.1
A3 112848443 112848497
refGene_gn=EIF2AK2;refGene_tx=XM_019827674.2;refGene_gn=EIF2AK2;refGene_tx=XM_023251956.1;refGene_gn=EIF2AK2;refGene_tx=XM_023251957.1
A3 112848573 112848696
refGene_gn=EIF2AK2;refGene_tx=XM_019827674.2;refGene_gn=EIF2AK2;refGene_tx=XM_023251956.1;refGene_gn=EIF2AK2;refGene_tx=XM_023251957.1
A3 112858040 112858096
refGene_gn=GPATCH11;refGene_tx=XM_003984324.4;refGene_gn=GPATCH11;refGene_tx=XM_006930360.2
A3 112858794 112858876
refGene_gn=GPATCH11;refGene_tx=XM_003984324.4;refGene_gn=GPATCH11;refGene_tx=XM_006930360.2
A3 112859747 112859903
refGene_gn=GPATCH11;refGene_tx=XM_003984324.4;refGene_gn=GPATCH11;refGene_tx=XM_006930360.2;refGene_gn=GPATCH11;refGene_tx=XM_019827685.2
A3 112860971 112861062
refGene_gn=GPATCH11;refGene_tx=XM_003984324.4;refGene_gn=GPATCH11;refGene_tx=XM_006930360.2;refGene_gn=GPATCH11;refGene_tx=XM_019827685.2
A3 112861190 112861311
refGene_gn=GPATCH11;refGene_tx=XM_003984324.4;refGene_gn=GPATCH11;refGene_tx=XM_006930360.2;refGene_gn=GPATCH11;refGene_tx=XM_019827685.2
A3 112862360 112862402
refGene_gn=GPATCH11;refGene_tx=XM_003984324.4;refGene_gn=GPATCH11;refGene_tx=XM_006930360.2;refGene_gn=GPATCH11;refGene_tx=XM_019827685.2
A3 112863188 112863415
refGene_gn=GPATCH11;refGene_tx=XM_003984324.4;refGene_gn=GPATCH11;refGene_tx=XM_006930360.2;refGene_gn=GPATCH11;refGene_tx=XM_019827685.2
A3 112863972 112864031
refGene_gn=GPATCH11;refGene_tx=XM_003984324.4;refGene_gn=GPATCH11;refGene_tx=XM_006930360.2;refGene_gn=GPATCH11;refGene_tx=XM_019827685.2
A3 112869553 112869679
refGene_gn=HEATR5B;refGene_tx=XM_006930362.4
A3 112872817 112873029
refGene_gn=HEATR5B;refGene_tx=XM_006930362.4
A3 112875406 112875515
refGene_gn=HEATR5B;refGene_tx=XM_006930362.4
A3 112877075 112877225
refGene_gn=HEATR5B;refGene_tx=XM_006930362.4
A3 112882403 112882575
refGene_gn=HEATR5B;refGene_tx=XM_006930362.4
A3 112884218 112884376
refGene_gn=HEATR5B;refGene_tx=XM_006930362.4
A3 112885624 112885874
refGene_gn=HEATR5B;refGene_tx=XM_006930362.4
A3 112889384 112889540
refGene_gn=HEATR5B;refGene_tx=XM_006930362.4
A3 112890034 112890285
refGene_gn=HEATR5B;refGene_tx=XM_006930362.4
A3 112891869 112891981

refGene_gn=HEATR5B;refGene_tx=XM_006930362.4
A3 112893229 112893382
refGene_gn=HEATR5B;refGene_tx=XM_006930362.4
A3 112894561 112894661
refGene_gn=HEATR5B;refGene_tx=XM_006930362.4
A3 112894973 112895083
refGene_gn=HEATR5B;refGene_tx=XM_006930362.4
A3 112897018 112897182
refGene_gn=HEATR5B;refGene_tx=XM_006930362.4
A3 112897738 112897914
refGene_gn=HEATR5B;refGene_tx=XM_006930362.4
A3 112901528 112901634
refGene_gn=HEATR5B;refGene_tx=XM_006930362.4
A3 112904215 112904406
refGene_gn=HEATR5B;refGene_tx=XM_006930362.4
A3 112908594 112908754
refGene_gn=HEATR5B;refGene_tx=XM_006930362.4
A3 112910373 112910563
refGene_gn=HEATR5B;refGene_tx=XM_006930362.4
A3 112912655 112912825
refGene_gn=HEATR5B;refGene_tx=XM_006930362.4
A3 112920392 112920537
refGene_gn=HEATR5B;refGene_tx=XM_006930362.4
A3 112925373 112925613
refGene_gn=HEATR5B;refGene_tx=XM_006930362.4
A3 112926109 112926361
refGene_gn=HEATR5B;refGene_tx=XM_006930362.4
A3 112931553 112931735
refGene_gn=HEATR5B;refGene_tx=XM_006930362.4
A3 112934227 112934296
refGene_gn=HEATR5B;refGene_tx=XM_006930362.4
A3 112941349 112941529
refGene_gn=HEATR5B;refGene_tx=XM_006930362.4
A3 112946207 112946445
refGene_gn=HEATR5B;refGene_tx=XM_006930362.4
A3 112947644 112947899
refGene_gn=HEATR5B;refGene_tx=XM_006930362.4
A3 112949842 112949970
refGene_gn=HEATR5B;refGene_tx=XM_006930362.4
A3 112953157 112953302
refGene_gn=HEATR5B;refGene_tx=XM_006930362.4
A3 112954484 112954751
refGene_gn=HEATR5B;refGene_tx=XM_006930362.4
A3 112955814 112956039
refGene_gn=HEATR5B;refGene_tx=XM_006930362.4
A3 112962021 112962173
refGene_gn=HEATR5B;refGene_tx=XM_006930362.4
A3 112963377 112963591
refGene_gn=HEATR5B;refGene_tx=XM_006930362.4
A3 112967719 112968024

refGene_gn=HEATR5B;refGene_tx=XM_006930362.4
A3 112984139 112984373
refGene_gn=STRN;refGene_tx=XM_023251959.1;refGene_gn=STRN;refGene_tx=XM_023251960.1
A3 113015037 113015141
refGene_gn=STRN;refGene_tx=XM_023251959.1;refGene_gn=STRN;refGene_tx=XM_023251960.1
A3 113023700 113023774
refGene_gn=STRN;refGene_tx=XM_023251959.1;refGene_gn=STRN;refGene_tx=XM_023251960.1
A3 113031924 113032003
refGene_gn=STRN;refGene_tx=XM_023251959.1;refGene_gn=STRN;refGene_tx=XM_023251960.1
A3 113034862 113035030
refGene_gn=STRN;refGene_tx=XM_023251959.1;refGene_gn=STRN;refGene_tx=XM_023251960.1
A3 113036903 113037039
refGene_gn=STRN;refGene_tx=XM_023251959.1;refGene_gn=STRN;refGene_tx=XM_023251960.1
A3 113042193 113042329
refGene_gn=STRN;refGene_tx=XM_023251959.1;refGene_gn=STRN;refGene_tx=XM_023251960.1
A3 113045338 113045449
refGene_gn=STRN;refGene_tx=XM_023251959.1;refGene_gn=STRN;refGene_tx=XM_023251960.1
A3 113047507 113047594
refGene_gn=STRN;refGene_tx=XM_023251959.1
A3 113048058 113048202
refGene_gn=STRN;refGene_tx=XM_023251959.1;refGene_gn=STRN;refGene_tx=XM_023251960.1
A3 113052484 113052621
refGene_gn=STRN;refGene_tx=XM_023251959.1;refGene_gn=STRN;refGene_tx=XM_023251960.1
A3 113057739 113057915
refGene_gn=STRN;refGene_tx=XM_023251959.1;refGene_gn=STRN;refGene_tx=XM_023251960.1
A3 113060027 113060075
refGene_gn=STRN;refGene_tx=XM_023251959.1;refGene_gn=STRN;refGene_tx=XM_023251960.1
A3 113060863 113060985
refGene_gn=STRN;refGene_tx=XM_023251959.1;refGene_gn=STRN;refGene_tx=XM_023251960.1
A3 113064050 113064218
refGene_gn=STRN;refGene_tx=XM_023251959.1;refGene_gn=STRN;refGene_tx=XM_023251960.1
A3 113065916 113066057
refGene_gn=STRN;refGene_tx=XM_023251959.1;refGene_gn=STRN;refGene_tx=XM_023251960.1
A3 113070505 113070610
refGene_gn=STRN;refGene_tx=XM_023251959.1;refGene_gn=STRN;refGene_tx=X

M_023251960.1
A3 113072222 113072309
refGene_gn=STRN;refGene_tx=XM_023251959.1;refGene_gn=STRN;refGene_tx=XM_023251960.1
A3 113072429 113072599
refGene_gn=STRN;refGene_tx=XM_023251959.1;refGene_gn=STRN;refGene_tx=XM_023251960.1
A3 113103410 113103589
refGene_gn=VIT;refGene_tx=XM_006930363.2;refGene_gn=VIT;refGene_tx=XM_006930364.2;refGene_gn=VIT;refGene_tx=XM_006930365.3
A3 113107306 113107820
refGene_gn=VIT;refGene_tx=XM_006930363.2;refGene_gn=VIT;refGene_tx=XM_006930364.2;refGene_gn=VIT;refGene_tx=XM_006930365.3
A3 113110289 113110516
refGene_gn=VIT;refGene_tx=XM_006930363.2;refGene_gn=VIT;refGene_tx=XM_006930364.2;refGene_gn=VIT;refGene_tx=XM_006930365.3
A3 113114164 113114268
refGene_gn=VIT;refGene_tx=XM_006930363.2;refGene_gn=VIT;refGene_tx=XM_006930364.2;refGene_gn=VIT;refGene_tx=XM_006930365.3
A3 113127372 113127520
refGene_gn=VIT;refGene_tx=XM_006930363.2;refGene_gn=VIT;refGene_tx=XM_006930364.2;refGene_gn=VIT;refGene_tx=XM_006930365.3
A3 113131919 113131982
refGene_gn=VIT;refGene_tx=XM_006930363.2;refGene_gn=VIT;refGene_tx=XM_006930364.2;refGene_gn=VIT;refGene_tx=XM_006930365.3
A3 113133464 113133509
refGene_gn=VIT;refGene_tx=XM_006930363.2;refGene_gn=VIT;refGene_tx=XM_006930365.3
A3 113140262 113140319
refGene_gn=VIT;refGene_tx=XM_006930363.2;refGene_gn=VIT;refGene_tx=XM_006930364.2;refGene_gn=VIT;refGene_tx=XM_006930365.3
A3 113145328 113145520
refGene_gn=VIT;refGene_tx=XM_006930363.2;refGene_gn=VIT;refGene_tx=XM_006930364.2;refGene_gn=VIT;refGene_tx=XM_006930365.3
A3 113152565 113152643
refGene_gn=VIT;refGene_tx=XM_006930363.2;refGene_gn=VIT;refGene_tx=XM_006930364.2;refGene_gn=VIT;refGene_tx=XM_006930365.3
A3 113157954 113158088
refGene_gn=VIT;refGene_tx=XM_006930363.2;refGene_gn=VIT;refGene_tx=XM_006930364.2;refGene_gn=VIT;refGene_tx=XM_006930365.3
A3 113164314 113164471
refGene_gn=VIT;refGene_tx=XM_006930363.2;refGene_gn=VIT;refGene_tx=XM_006930364.2;refGene_gn=VIT;refGene_tx=XM_006930365.3
A3 113177999 113178065
refGene_gn=VIT;refGene_tx=XM_006930363.2;refGene_gn=VIT;refGene_tx=XM_006930364.2;refGene_gn=VIT;refGene_tx=XM_006930365.3
A3 113185236 113185306
refGene_gn=VIT;refGene_tx=XM_006930363.2;refGene_gn=VIT;refGene_tx=XM_006930364.2;refGene_gn=VIT;refGene_tx=XM_006930365.3
A3 113188493 113188583

refGene_gn=VIT;refGene_tx=XM_006930363.2;refGene_gn=VIT;refGene_tx=XM_006930364.2
A3 113267515 113267769
refGene_gn=FEZ2;refGene_tx=XM_023251966.1;refGene_gn=FEZ2;refGene_tx=XM_023251967.1
A3 113272604 113272713
refGene_gn=FEZ2;refGene_tx=XM_023251966.1;refGene_gn=FEZ2;refGene_tx=XM_023251967.1
A3 113278557 113278674
refGene_gn=FEZ2;refGene_tx=XM_023251966.1;refGene_gn=FEZ2;refGene_tx=XM_023251967.1
A3 113280560 113280702
refGene_gn=FEZ2;refGene_tx=XM_023251966.1;refGene_gn=FEZ2;refGene_tx=XM_023251967.1
A3 113281758 113282027
refGene_gn=FEZ2;refGene_tx=XM_023251966.1;refGene_gn=FEZ2;refGene_tx=XM_023251967.1
A3 113302106 113302187
refGene_gn=FEZ2;refGene_tx=XM_023251966.1
A3 113304344 113304420
refGene_gn=FEZ2;refGene_tx=XM_023251966.1;refGene_gn=FEZ2;refGene_tx=XM_023251967.1
A3 113307116 113307182
refGene_gn=FEZ2;refGene_tx=XM_023251966.1;refGene_gn=FEZ2;refGene_tx=XM_023251967.1
A3 113309598 113309615
refGene_gn=FEZ2;refGene_tx=XM_023251966.1;refGene_gn=FEZ2;refGene_tx=XM_023251967.1
A3 113314104 113314281
refGene_gn=CRIM1;refGene_tx=XM_023251961.1;refGene_gn=CRIM1;refGene_tx=XM_023251962.1;refGene_gn=CRIM1;refGene_tx=XM_023251963.1;refGene_gn=CRIM1;refGene_tx=XM_023251964.1;refGene_gn=CRIM1;refGene_tx=XM_023251965.1
A3 113315616 113315807
refGene_gn=CRIM1;refGene_tx=XM_023251961.1;refGene_gn=CRIM1;refGene_tx=XM_023251962.1;refGene_gn=CRIM1;refGene_tx=XM_023251963.1;refGene_gn=CRIM1;refGene_tx=XM_023251964.1;refGene_gn=CRIM1;refGene_tx=XM_023251965.1
A3 113318516 113318639
refGene_gn=CRIM1;refGene_tx=XM_023251961.1;refGene_gn=CRIM1;refGene_tx=XM_023251962.1;refGene_gn=CRIM1;refGene_tx=XM_023251963.1;refGene_gn=CRIM1;refGene_tx=XM_023251964.1;refGene_gn=CRIM1;refGene_tx=XM_023251965.1
A3 113326229 113326424
refGene_gn=CRIM1;refGene_tx=XM_023251961.1;refGene_gn=CRIM1;refGene_tx=XM_023251962.1;refGene_gn=CRIM1;refGene_tx=XM_023251963.1;refGene_gn=CRIM1;refGene_tx=XM_023251964.1;refGene_gn=CRIM1;refGene_tx=XM_023251965.1
A3 113343409 113343631
refGene_gn=CRIM1;refGene_tx=XM_023251961.1;refGene_gn=CRIM1;refGene_tx

=XM_023251962.1;refGene_gn=CRIM1;refGene_tx=XM_023251964.1;refGene_gn=CRIM1;refGene_tx=XM_023251965.1
A3	113347662	113347878
refGene_gn=CRIM1;refGene_tx=XM_023251961.1;refGene_gn=CRIM1;refGene_tx=XM_023251962.1;refGene_gn=CRIM1;refGene_tx=XM_023251963.1;refGene_gn=CRIM1;refGene_tx=XM_023251964.1;refGene_gn=CRIM1;refGene_tx=XM_023251965.1
A3	113351649	113351859
refGene_gn=CRIM1;refGene_tx=XM_023251961.1;refGene_gn=CRIM1;refGene_tx=XM_023251963.1;refGene_gn=CRIM1;refGene_tx=XM_023251964.1;refGene_gn=CRIM1;refGene_tx=XM_023251965.1
A3	113353013	113353135
refGene_gn=CRIM1;refGene_tx=XM_023251961.1;refGene_gn=CRIM1;refGene_tx=XM_023251962.1;refGene_gn=CRIM1;refGene_tx=XM_023251963.1;refGene_gn=CRIM1;refGene_tx=XM_023251964.1;refGene_gn=CRIM1;refGene_tx=XM_023251965.1
A3	113356102	113356259
refGene_gn=CRIM1;refGene_tx=XM_023251961.1;refGene_gn=CRIM1;refGene_tx=XM_023251962.1;refGene_gn=CRIM1;refGene_tx=XM_023251963.1;refGene_gn=CRIM1;refGene_tx=XM_023251964.1;refGene_gn=CRIM1;refGene_tx=XM_023251965.1
A3	113364926	113365055
refGene_gn=CRIM1;refGene_tx=XM_023251961.1;refGene_gn=CRIM1;refGene_tx=XM_023251962.1;refGene_gn=CRIM1;refGene_tx=XM_023251963.1;refGene_gn=CRIM1;refGene_tx=XM_023251964.1;refGene_gn=CRIM1;refGene_tx=XM_023251965.1
A3	113384220	113384418
refGene_gn=CRIM1;refGene_tx=XM_023251961.1;refGene_gn=CRIM1;refGene_tx=XM_023251962.1;refGene_gn=CRIM1;refGene_tx=XM_023251963.1;refGene_gn=CRIM1;refGene_tx=XM_023251964.1;refGene_gn=CRIM1;refGene_tx=XM_023251965.1
A3	113387078	113387261
refGene_gn=CRIM1;refGene_tx=XM_023251961.1;refGene_gn=CRIM1;refGene_tx=XM_023251962.1;refGene_gn=CRIM1;refGene_tx=XM_023251963.1;refGene_gn=CRIM1;refGene_tx=XM_023251964.1;refGene_gn=CRIM1;refGene_tx=XM_023251965.1
A3	113394248	113394370
refGene_gn=CRIM1;refGene_tx=XM_023251961.1;refGene_gn=CRIM1;refGene_tx=XM_023251962.1;refGene_gn=CRIM1;refGene_tx=XM_023251963.1;refGene_gn=CRIM1;refGene_tx=XM_023251964.1
A3	113416117	113416238
refGene_gn=CRIM1;refGene_tx=XM_023251961.1;refGene_gn=CRIM1;refGene_tx=XM_023251962.1;refGene_gn=CRIM1;refGene_tx=XM_023251963.1;refGene_gn=CRIM1;refGene_tx=XM_023251964.1
A3	113417301	113417544
refGene_gn=CRIM1;refGene_tx=XM_023251961.1;refGene_gn=CRIM1;refGene_tx=XM_023251962.1;refGene_gn=CRIM1;refGene_tx=XM_023251963.1
A3	113463166	113463340
refGene_gn=CRIM1;refGene_tx=XM_023251961.1;refGene_gn=CRIM1;refGene_tx=XM_023251962.1;refGene_gn=CRIM1;refGene_tx=XM_023251963.1;refGene_gn=

CRIM1;refGene_tx=XM_023251964.1;refGene_gn=CRIM1;refGene_tx=XM_0232519
65.1
A3	113503369	113503700
refGene_gn=CRIM1;refGene_tx=XM_023251961.1;refGene_gn=CRIM1;refGene_tx
=XM_023251962.1;refGene_gn=CRIM1;refGene_tx=XM_023251963.1;refGene_gn=
CRIM1;refGene_tx=XM_023251964.1;refGene_gn=CRIM1;refGene_tx=XM_0232519
65.1
A3	115674342	115674395
refGene_gn=FAM98A;refGene_tx=XM_003984328.4
A3	115677577	115677726
refGene_gn=FAM98A;refGene_tx=XM_003984328.4
A3	115679803	115679938
refGene_gn=FAM98A;refGene_tx=XM_003984328.4
A3	115683764	115683949
refGene_gn=FAM98A;refGene_tx=XM_003984328.4
A3	115684960	115685041
refGene_gn=FAM98A;refGene_tx=XM_003984328.4
A3	115685597	115685714
refGene_gn=FAM98A;refGene_tx=XM_003984328.4
A3	115688384	115688552
refGene_gn=FAM98A;refGene_tx=XM_003984328.4
A3	115688641	115689310
refGene_gn=FAM98A;refGene_tx=XM_003984328.4
A3	115704908	115704917
refGene_gn=RASGRP3;refGene_tx=XM_011281141.3;refGene_gn=RASGRP3;refGen
e_tx=XM_011281144.2;refGene_gn=RASGRP3;refGene_tx=XM_011281145.3;refGe
ne_gn=RASGRP3;refGene_tx=XM_011281146.3;refGene_gn=RASGRP3;refGene_tx=
XM_011281147.2;refGene_gn=RASGRP3;refGene_tx=XM_019827692.2;refGene_gn
=RASGRP3;refGene_tx=XM_019827693.1;refGene_gn=RASGRP3;refGene_tx=XM_01
9827694.2
A3	115708274	115708639
refGene_gn=RASGRP3;refGene_tx=XM_011281141.3;refGene_gn=RASGRP3;refGen
e_tx=XM_011281144.2;refGene_gn=RASGRP3;refGene_tx=XM_011281145.3;refGe
ne_gn=RASGRP3;refGene_tx=XM_011281146.3;refGene_gn=RASGRP3;refGene_tx=
XM_011281147.2;refGene_gn=RASGRP3;refGene_tx=XM_019827692.2;refGene_gn
=RASGRP3;refGene_tx=XM_019827693.1;refGene_gn=RASGRP3;refGene_tx=XM_01
9827694.2
A3	115708964	115709090
refGene_gn=RASGRP3;refGene_tx=XM_011281141.3;refGene_gn=RASGRP3;refGen
e_tx=XM_011281144.2;refGene_gn=RASGRP3;refGene_tx=XM_011281145.3;refGe
ne_gn=RASGRP3;refGene_tx=XM_011281146.3;refGene_gn=RASGRP3;refGene_tx=
XM_011281147.2;refGene_gn=RASGRP3;refGene_tx=XM_019827692.2;refGene_gn
=RASGRP3;refGene_tx=XM_019827693.1;refGene_gn=RASGRP3;refGene_tx=XM_01
9827694.2
A3	115712060	115712097
refGene_gn=RASGRP3;refGene_tx=XM_011281141.3;refGene_gn=RASGRP3;refGen
e_tx=XM_011281144.2;refGene_gn=RASGRP3;refGene_tx=XM_011281145.3;refGe
ne_gn=RASGRP3;refGene_tx=XM_011281146.3;refGene_gn=RASGRP3;refGene_tx=
XM_011281147.2;refGene_gn=RASGRP3;refGene_tx=XM_019827692.2;refGene_gn
=RASGRP3;refGene_tx=XM_019827693.1;refGene_gn=RASGRP3;refGene_tx=XM_01

9827694.2
A3 115715074 115715222
refGene_gn=RASGRP3;refGene_tx=XM_011281141.3;refGene_gn=RASGRP3;refGene_tx=XM_011281144.2;refGene_gn=RASGRP3;refGene_tx=XM_011281145.3;refGene_gn=RASGRP3;refGene_tx=XM_011281146.3;refGene_gn=RASGRP3;refGene_tx=XM_011281147.2;refGene_gn=RASGRP3;refGene_tx=XM_019827692.2;refGene_gn=RASGRP3;refGene_tx=XM_019827693.1;refGene_gn=RASGRP3;refGene_tx=XM_019827694.2
A3 115718297 115718413
refGene_gn=RASGRP3;refGene_tx=XM_011281141.3;refGene_gn=RASGRP3;refGene_tx=XM_011281144.2;refGene_gn=RASGRP3;refGene_tx=XM_011281145.3;refGene_gn=RASGRP3;refGene_tx=XM_011281146.3;refGene_gn=RASGRP3;refGene_tx=XM_011281147.2;refGene_gn=RASGRP3;refGene_tx=XM_019827692.2;refGene_gn=RASGRP3;refGene_tx=XM_019827693.1;refGene_gn=RASGRP3;refGene_tx=XM_019827694.2
A3 115722736 115722853
refGene_gn=RASGRP3;refGene_tx=XM_011281141.3;refGene_gn=RASGRP3;refGene_tx=XM_011281144.2;refGene_gn=RASGRP3;refGene_tx=XM_011281145.3;refGene_gn=RASGRP3;refGene_tx=XM_011281146.3;refGene_gn=RASGRP3;refGene_tx=XM_011281147.2;refGene_gn=RASGRP3;refGene_tx=XM_019827692.2;refGene_gn=RASGRP3;refGene_tx=XM_019827693.1;refGene_gn=RASGRP3;refGene_tx=XM_019827694.2
A3 115726652 115726730
refGene_gn=RASGRP3;refGene_tx=XM_011281141.3;refGene_gn=RASGRP3;refGene_tx=XM_011281144.2;refGene_gn=RASGRP3;refGene_tx=XM_011281145.3;refGene_gn=RASGRP3;refGene_tx=XM_011281146.3;refGene_gn=RASGRP3;refGene_tx=XM_011281147.2;refGene_gn=RASGRP3;refGene_tx=XM_019827692.2;refGene_gn=RASGRP3;refGene_tx=XM_019827693.1;refGene_gn=RASGRP3;refGene_tx=XM_019827694.2
A3 115733055 115733331
refGene_gn=RASGRP3;refGene_tx=XM_011281141.3;refGene_gn=RASGRP3;refGene_tx=XM_011281144.2;refGene_gn=RASGRP3;refGene_tx=XM_011281145.3;refGene_gn=RASGRP3;refGene_tx=XM_011281146.3;refGene_gn=RASGRP3;refGene_tx=XM_011281147.2;refGene_gn=RASGRP3;refGene_tx=XM_019827692.2;refGene_gn=RASGRP3;refGene_tx=XM_019827693.1;refGene_gn=RASGRP3;refGene_tx=XM_019827694.2
A3 115735592 115735709
refGene_gn=RASGRP3;refGene_tx=XM_011281141.3;refGene_gn=RASGRP3;refGene_tx=XM_011281144.2;refGene_gn=RASGRP3;refGene_tx=XM_011281145.3;refGene_gn=RASGRP3;refGene_tx=XM_011281146.3;refGene_gn=RASGRP3;refGene_tx=XM_011281147.2;refGene_gn=RASGRP3;refGene_tx=XM_019827692.2;refGene_gn=RASGRP3;refGene_tx=XM_019827693.1;refGene_gn=RASGRP3;refGene_tx=XM_019827694.2
A3 115736088 115736262
refGene_gn=RASGRP3;refGene_tx=XM_011281141.3;refGene_gn=RASGRP3;refGene_tx=XM_011281144.2;refGene_gn=RASGRP3;refGene_tx=XM_011281145.3;refGene_gn=RASGRP3;refGene_tx=XM_011281146.3;refGene_gn=RASGRP3;refGene_tx=XM_011281147.2;refGene_gn=RASGRP3;refGene_tx=XM_019827692.2;refGene_gn=RASGRP3;refGene_tx=XM_019827693.1;refGene_gn=RASGRP3;refGene_tx=XM_019827694.2

A3	115737707	115737855
refGene_gn=RASGRP3;refGene_tx=XM_011281141.3;refGene_gn=RASGRP3;refGene_tx=XM_011281144.2;refGene_gn=RASGRP3;refGene_tx=XM_011281145.3;refGene_gn=RASGRP3;refGene_tx=XM_011281146.3;refGene_gn=RASGRP3;refGene_tx=XM_011281147.2;refGene_gn=RASGRP3;refGene_tx=XM_019827692.2;refGene_gn=RASGRP3;refGene_tx=XM_019827693.1;refGene_gn=RASGRP3;refGene_tx=XM_019827694.2
A3	115739107	115739239
refGene_gn=RASGRP3;refGene_tx=XM_011281141.3;refGene_gn=RASGRP3;refGene_tx=XM_011281144.2;refGene_gn=RASGRP3;refGene_tx=XM_011281145.3;refGene_gn=RASGRP3;refGene_tx=XM_011281146.3;refGene_gn=RASGRP3;refGene_tx=XM_011281147.2;refGene_gn=RASGRP3;refGene_tx=XM_019827692.2;refGene_gn=RASGRP3;refGene_tx=XM_019827693.1;refGene_gn=RASGRP3;refGene_tx=XM_019827694.2
A3	115739803	115739866
refGene_gn=RASGRP3;refGene_tx=XM_011281141.3;refGene_gn=RASGRP3;refGene_tx=XM_011281144.2;refGene_gn=RASGRP3;refGene_tx=XM_011281145.3;refGene_gn=RASGRP3;refGene_tx=XM_011281146.3;refGene_gn=RASGRP3;refGene_tx=XM_011281147.2;refGene_gn=RASGRP3;refGene_tx=XM_019827692.2;refGene_gn=RASGRP3;refGene_tx=XM_019827693.1;refGene_gn=RASGRP3;refGene_tx=XM_019827694.2
A3	115741084	115741187
refGene_gn=RASGRP3;refGene_tx=XM_011281141.3;refGene_gn=RASGRP3;refGene_tx=XM_011281144.2;refGene_gn=RASGRP3;refGene_tx=XM_011281145.3;refGene_gn=RASGRP3;refGene_tx=XM_011281146.3;refGene_gn=RASGRP3;refGene_tx=XM_011281147.2;refGene_gn=RASGRP3;refGene_tx=XM_019827692.2;refGene_gn=RASGRP3;refGene_tx=XM_019827693.1;refGene_gn=RASGRP3;refGene_tx=XM_019827694.2
A3	115742166	115742236
refGene_gn=RASGRP3;refGene_tx=XM_011281141.3;refGene_gn=RASGRP3;refGene_tx=XM_011281144.2;refGene_gn=RASGRP3;refGene_tx=XM_011281145.3;refGene_gn=RASGRP3;refGene_tx=XM_011281146.3;refGene_gn=RASGRP3;refGene_tx=XM_011281147.2;refGene_gn=RASGRP3;refGene_tx=XM_019827692.2;refGene_gn=RASGRP3;refGene_tx=XM_019827693.1;refGene_gn=RASGRP3;refGene_tx=XM_019827694.2
A3	115847273	115847455
refGene_gn=LTBP1;refGene_tx=XM_006930373.4;refGene_gn=LTBP1;refGene_tx=XM_006930374.4;refGene_gn=LTBP1;refGene_tx=XM_006930376.4;refGene_gn=LTBP1;refGene_tx=XM_023251968.1;refGene_gn=LTBP1;refGene_tx=XM_023251969.1;refGene_gn=LTBP1;refGene_tx=XM_023251970.1;refGene_gn=LTBP1;refGene_tx=XM_023251971.1;refGene_gn=LTBP1;refGene_tx=XM_023251972.1;refGene_gn=LTBP1;refGene_tx=XM_023251973.1
A3	115848542	115848692
refGene_gn=LTBP1;refGene_tx=XM_006930373.4;refGene_gn=LTBP1;refGene_tx=XM_006930374.4;refGene_gn=LTBP1;refGene_tx=XM_006930376.4;refGene_gn=LTBP1;refGene_tx=XM_023251968.1;refGene_gn=LTBP1;refGene_tx=XM_023251969.1;refGene_gn=LTBP1;refGene_tx=XM_023251970.1;refGene_gn=LTBP1;refGene_tx=XM_023251971.1;refGene_gn=LTBP1;refGene_tx=XM_023251972.1;refGene_gn=LTBP1;refGene_tx=XM_023251973.1
A3	115852639	115852762

refGene_gn=LTBP1;refGene_tx=XM_006930373.4;refGene_gn=LTBP1;refGene_tx=XM_006930374.4;refGene_gn=LTBP1;refGene_tx=XM_006930376.4;refGene_gn=LTBP1;refGene_tx=XM_023251968.1;refGene_gn=LTBP1;refGene_tx=XM_023251969.1;refGene_gn=LTBP1;refGene_tx=XM_023251970.1;refGene_gn=LTBP1;refGene_tx=XM_023251971.1;refGene_gn=LTBP1;refGene_tx=XM_023251972.1;refGene_gn=LTBP1;refGene_tx=XM_023251973.1
A3 115872868 115873039
refGene_gn=LTBP1;refGene_tx=XM_006930373.4;refGene_gn=LTBP1;refGene_tx=XM_006930374.4;refGene_gn=LTBP1;refGene_tx=XM_006930376.4;refGene_gn=LTBP1;refGene_tx=XM_023251968.1;refGene_gn=LTBP1;refGene_tx=XM_023251969.1;refGene_gn=LTBP1;refGene_tx=XM_023251970.1;refGene_gn=LTBP1;refGene_tx=XM_023251971.1;refGene_gn=LTBP1;refGene_tx=XM_023251972.1;refGene_gn=LTBP1;refGene_tx=XM_023251973.1
A3 115874249 115874390
refGene_gn=LTBP1;refGene_tx=XM_006930373.4;refGene_gn=LTBP1;refGene_tx=XM_006930374.4;refGene_gn=LTBP1;refGene_tx=XM_006930376.4;refGene_gn=LTBP1;refGene_tx=XM_023251968.1;refGene_gn=LTBP1;refGene_tx=XM_023251969.1;refGene_gn=LTBP1;refGene_tx=XM_023251970.1;refGene_gn=LTBP1;refGene_tx=XM_023251971.1;refGene_gn=LTBP1;refGene_tx=XM_023251972.1;refGene_gn=LTBP1;refGene_tx=XM_023251973.1
A3 115875072 115875201
refGene_gn=LTBP1;refGene_tx=XM_006930373.4;refGene_gn=LTBP1;refGene_tx=XM_006930374.4;refGene_gn=LTBP1;refGene_tx=XM_006930376.4;refGene_gn=LTBP1;refGene_tx=XM_023251968.1;refGene_gn=LTBP1;refGene_tx=XM_023251969.1;refGene_gn=LTBP1;refGene_tx=XM_023251970.1;refGene_gn=LTBP1;refGene_tx=XM_023251971.1;refGene_gn=LTBP1;refGene_tx=XM_023251972.1;refGene_gn=LTBP1;refGene_tx=XM_023251973.1
A3 115877102 115877189
refGene_gn=LTBP1;refGene_tx=XM_006930373.4;refGene_gn=LTBP1;refGene_tx=XM_006930374.4;refGene_gn=LTBP1;refGene_tx=XM_006930376.4;refGene_gn=LTBP1;refGene_tx=XM_023251968.1;refGene_gn=LTBP1;refGene_tx=XM_023251969.1;refGene_gn=LTBP1;refGene_tx=XM_023251970.1;refGene_gn=LTBP1;refGene_tx=XM_023251971.1;refGene_gn=LTBP1;refGene_tx=XM_023251972.1;refGene_gn=LTBP1;refGene_tx=XM_023251973.1
A3 115877945 115878122
refGene_gn=LTBP1;refGene_tx=XM_006930373.4;refGene_gn=LTBP1;refGene_tx=XM_006930374.4;refGene_gn=LTBP1;refGene_tx=XM_006930376.4;refGene_gn=LTBP1;refGene_tx=XM_023251968.1;refGene_gn=LTBP1;refGene_tx=XM_023251969.1;refGene_gn=LTBP1;refGene_tx=XM_023251970.1;refGene_gn=LTBP1;refGene_tx=XM_023251971.1;refGene_gn=LTBP1;refGene_tx=XM_023251972.1;refGene_gn=LTBP1;refGene_tx=XM_023251973.1
A3 115891153 115891297
refGene_gn=LTBP1;refGene_tx=XM_006930373.4;refGene_gn=LTBP1;refGene_tx=XM_006930374.4;refGene_gn=LTBP1;refGene_tx=XM_006930376.4;refGene_gn=LTBP1;refGene_tx=XM_023251968.1;refGene_gn=LTBP1;refGene_tx=XM_023251969.1;refGene_gn=LTBP1;refGene_tx=XM_023251970.1;refGene_gn=LTBP1;refGene_tx=XM_023251971.1;refGene_gn=LTBP1;refGene_tx=XM_023251972.1
A3 115894813 115894939
refGene_gn=LTBP1;refGene_tx=XM_006930374.4;refGene_gn=LTBP1;refGene_tx=XM_023251968.1;refGene_gn=LTBP1;refGene_tx=XM_023251969.1;refGene_gn=

LTBP1;refGene_tx=XM_023251971.1;refGene_gn=LTBP1;refGene_tx=XM_0232519
72.1
A3 115922844 115922970
refGene_gn=LTBP1;refGene_tx=XM_006930373.4;refGene_gn=LTBP1;refGene_tx
=XM_006930374.4;refGene_gn=LTBP1;refGene_tx=XM_006930376.4;refGene_gn=
LTBP1;refGene_tx=XM_023251968.1;refGene_gn=LTBP1;refGene_tx=XM_0232519
69.1;refGene_gn=LTBP1;refGene_tx=XM_023251970.1;refGene_gn=LTBP1;refGe
ne_tx=XM_023251971.1;refGene_gn=LTBP1;refGene_tx=XM_023251972.1;refGen
e_gn=LTBP1;refGene_tx=XM_023251973.1
A3 115928830 115928953
refGene_gn=LTBP1;refGene_tx=XM_006930373.4;refGene_gn=LTBP1;refGene_tx
=XM_006930374.4;refGene_gn=LTBP1;refGene_tx=XM_006930376.4;refGene_gn=
LTBP1;refGene_tx=XM_023251968.1;refGene_gn=LTBP1;refGene_tx=XM_0232519
69.1;refGene_gn=LTBP1;refGene_tx=XM_023251970.1;refGene_gn=LTBP1;refGe
ne_tx=XM_023251971.1;refGene_gn=LTBP1;refGene_tx=XM_023251972.1;refGen
e_gn=LTBP1;refGene_tx=XM_023251973.1
A3 115935135 115935258
refGene_gn=LTBP1;refGene_tx=XM_006930373.4;refGene_gn=LTBP1;refGene_tx
=XM_006930374.4;refGene_gn=LTBP1;refGene_tx=XM_006930376.4;refGene_gn=
LTBP1;refGene_tx=XM_023251968.1;refGene_gn=LTBP1;refGene_tx=XM_0232519
69.1;refGene_gn=LTBP1;refGene_tx=XM_023251970.1;refGene_gn=LTBP1;refGe
ne_tx=XM_023251971.1;refGene_gn=LTBP1;refGene_tx=XM_023251972.1;refGen
e_gn=LTBP1;refGene_tx=XM_023251973.1
A3 115937616 115937739
refGene_gn=LTBP1;refGene_tx=XM_006930373.4;refGene_gn=LTBP1;refGene_tx
=XM_006930374.4;refGene_gn=LTBP1;refGene_tx=XM_006930376.4;refGene_gn=
LTBP1;refGene_tx=XM_023251968.1;refGene_gn=LTBP1;refGene_tx=XM_0232519
69.1;refGene_gn=LTBP1;refGene_tx=XM_023251970.1;refGene_gn=LTBP1;refGe
ne_tx=XM_023251971.1;refGene_gn=LTBP1;refGene_tx=XM_023251972.1;refGen
e_gn=LTBP1;refGene_tx=XM_023251973.1
A3 115941452 115941575
refGene_gn=LTBP1;refGene_tx=XM_006930373.4;refGene_gn=LTBP1;refGene_tx
=XM_006930374.4;refGene_gn=LTBP1;refGene_tx=XM_006930376.4;refGene_gn=
LTBP1;refGene_tx=XM_023251968.1;refGene_gn=LTBP1;refGene_tx=XM_0232519
69.1;refGene_gn=LTBP1;refGene_tx=XM_023251970.1;refGene_gn=LTBP1;refGe
ne_tx=XM_023251971.1;refGene_gn=LTBP1;refGene_tx=XM_023251972.1;refGen
e_gn=LTBP1;refGene_tx=XM_023251973.1
A3 115953429 115953549
refGene_gn=LTBP1;refGene_tx=XM_006930373.4;refGene_gn=LTBP1;refGene_tx
=XM_006930374.4;refGene_gn=LTBP1;refGene_tx=XM_006930376.4;refGene_gn=
LTBP1;refGene_tx=XM_023251968.1;refGene_gn=LTBP1;refGene_tx=XM_0232519
69.1;refGene_gn=LTBP1;refGene_tx=XM_023251970.1;refGene_gn=LTBP1;refGe
ne_tx=XM_023251971.1;refGene_gn=LTBP1;refGene_tx=XM_023251972.1;refGen
e_gn=LTBP1;refGene_tx=XM_023251973.1
A3 115957255 115957378
refGene_gn=LTBP1;refGene_tx=XM_006930373.4;refGene_gn=LTBP1;refGene_tx
=XM_006930374.4;refGene_gn=LTBP1;refGene_tx=XM_006930376.4;refGene_gn=
LTBP1;refGene_tx=XM_023251968.1;refGene_gn=LTBP1;refGene_tx=XM_0232519
69.1;refGene_gn=LTBP1;refGene_tx=XM_023251970.1;refGene_gn=LTBP1;refGe
ne_tx=XM_023251971.1;refGene_gn=LTBP1;refGene_tx=XM_023251972.1;refGen

e_gn=LTBP1;refGene_tx=XM_023251973.1
A3 115957784 115957910
refGene_gn=LTBP1;refGene_tx=XM_006930373.4;refGene_gn=LTBP1;refGene_tx=XM_006930374.4;refGene_gn=LTBP1;refGene_tx=XM_006930376.4;refGene_gn=LTBP1;refGene_tx=XM_023251968.1;refGene_gn=LTBP1;refGene_tx=XM_023251969.1;refGene_gn=LTBP1;refGene_tx=XM_023251970.1;refGene_gn=LTBP1;refGene_tx=XM_023251971.1;refGene_gn=LTBP1;refGene_tx=XM_023251972.1;refGene_gn=LTBP1;refGene_tx=XM_023251973.1
A3 115959131 115959257
refGene_gn=LTBP1;refGene_tx=XM_006930373.4;refGene_gn=LTBP1;refGene_tx=XM_006930374.4;refGene_gn=LTBP1;refGene_tx=XM_006930376.4;refGene_gn=LTBP1;refGene_tx=XM_023251968.1;refGene_gn=LTBP1;refGene_tx=XM_023251969.1;refGene_gn=LTBP1;refGene_tx=XM_023251970.1;refGene_gn=LTBP1;refGene_tx=XM_023251971.1;refGene_gn=LTBP1;refGene_tx=XM_023251972.1;refGene_gn=LTBP1;refGene_tx=XM_023251973.1
A3 115969810 115969909
refGene_gn=LTBP1;refGene_tx=XM_006930373.4;refGene_gn=LTBP1;refGene_tx=XM_006930374.4;refGene_gn=LTBP1;refGene_tx=XM_006930376.4;refGene_gn=LTBP1;refGene_tx=XM_023251968.1;refGene_gn=LTBP1;refGene_tx=XM_023251969.1;refGene_gn=LTBP1;refGene_tx=XM_023251970.1;refGene_gn=LTBP1;refGene_tx=XM_023251971.1;refGene_gn=LTBP1;refGene_tx=XM_023251972.1;refGene_gn=LTBP1;refGene_tx=XM_023251973.1
A3 115970407 115970507
refGene_gn=LTBP1;refGene_tx=XM_006930373.4;refGene_gn=LTBP1;refGene_tx=XM_006930374.4;refGene_gn=LTBP1;refGene_tx=XM_006930376.4;refGene_gn=LTBP1;refGene_tx=XM_023251968.1;refGene_gn=LTBP1;refGene_tx=XM_023251969.1;refGene_gn=LTBP1;refGene_tx=XM_023251970.1;refGene_gn=LTBP1;refGene_tx=XM_023251971.1;refGene_gn=LTBP1;refGene_tx=XM_023251972.1;refGene_gn=LTBP1;refGene_tx=XM_023251973.1
A3 115973783 115973806
refGene_gn=LTBP1;refGene_tx=XM_006930373.4;refGene_gn=LTBP1;refGene_tx=XM_006930374.4;refGene_gn=LTBP1;refGene_tx=XM_006930376.4;refGene_gn=LTBP1;refGene_tx=XM_023251968.1;refGene_gn=LTBP1;refGene_tx=XM_023251969.1;refGene_gn=LTBP1;refGene_tx=XM_023251970.1;refGene_gn=LTBP1;refGene_tx=XM_023251971.1;refGene_gn=LTBP1;refGene_tx=XM_023251972.1;refGene_gn=LTBP1;refGene_tx=XM_023251973.1
A3 115975855 115976083
refGene_gn=LTBP1;refGene_tx=XM_006930373.4;refGene_gn=LTBP1;refGene_tx=XM_006930374.4;refGene_gn=LTBP1;refGene_tx=XM_006930376.4;refGene_gn=LTBP1;refGene_tx=XM_023251968.1;refGene_gn=LTBP1;refGene_tx=XM_023251969.1;refGene_gn=LTBP1;refGene_tx=XM_023251970.1;refGene_gn=LTBP1;refGene_tx=XM_023251971.1;refGene_gn=LTBP1;refGene_tx=XM_023251972.1;refGene_gn=LTBP1;refGene_tx=XM_023251973.1
A3 115978830 115978998
refGene_gn=LTBP1;refGene_tx=XM_006930373.4;refGene_gn=LTBP1;refGene_tx=XM_006930374.4;refGene_gn=LTBP1;refGene_tx=XM_006930376.4;refGene_gn=LTBP1;refGene_tx=XM_023251968.1;refGene_gn=LTBP1;refGene_tx=XM_023251969.1;refGene_gn=LTBP1;refGene_tx=XM_023251970.1;refGene_gn=LTBP1;refGene_tx=XM_023251971.1;refGene_gn=LTBP1;refGene_tx=XM_023251972.1;refGene_gn=LTBP1;refGene_tx=XM_023251973.1

A3 115989196 115989319
refGene_gn=LTBP1;refGene_tx=XM_006930373.4;refGene_gn=LTBP1;refGene_tx=XM_006930374.4;refGene_gn=LTBP1;refGene_tx=XM_006930376.4;refGene_gn=LTBP1;refGene_tx=XM_023251968.1;refGene_gn=LTBP1;refGene_tx=XM_023251969.1;refGene_gn=LTBP1;refGene_tx=XM_023251970.1;refGene_gn=LTBP1;refGene_tx=XM_023251971.1;refGene_gn=LTBP1;refGene_tx=XM_023251972.1;refGene_gn=LTBP1;refGene_tx=XM_023251973.1
A3 116011148 116011220
refGene_gn=LTBP1;refGene_tx=XM_006930373.4;refGene_gn=LTBP1;refGene_tx=XM_006930374.4;refGene_gn=LTBP1;refGene_tx=XM_006930376.4;refGene_gn=LTBP1;refGene_tx=XM_023251968.1;refGene_gn=LTBP1;refGene_tx=XM_023251969.1;refGene_gn=LTBP1;refGene_tx=XM_023251970.1;refGene_gn=LTBP1;refGene_tx=XM_023251971.1;refGene_gn=LTBP1;refGene_tx=XM_023251972.1;refGene_gn=LTBP1;refGene_tx=XM_023251973.1
A3 116016318 116016421
refGene_gn=LTBP1;refGene_tx=XM_006930373.4;refGene_gn=LTBP1;refGene_tx=XM_006930374.4;refGene_gn=LTBP1;refGene_tx=XM_006930376.4;refGene_gn=LTBP1;refGene_tx=XM_023251968.1;refGene_gn=LTBP1;refGene_tx=XM_023251969.1;refGene_gn=LTBP1;refGene_tx=XM_023251970.1;refGene_gn=LTBP1;refGene_tx=XM_023251971.1;refGene_gn=LTBP1;refGene_tx=XM_023251972.1;refGene_gn=LTBP1;refGene_tx=XM_023251973.1
A3 116037885 116038160
refGene_gn=LTBP1;refGene_tx=XM_006930373.4;refGene_gn=LTBP1;refGene_tx=XM_006930374.4;refGene_gn=LTBP1;refGene_tx=XM_006930376.4;refGene_gn=LTBP1;refGene_tx=XM_023251968.1;refGene_gn=LTBP1;refGene_tx=XM_023251969.1;refGene_gn=LTBP1;refGene_tx=XM_023251970.1;refGene_gn=LTBP1;refGene_tx=XM_023251971.1;refGene_gn=LTBP1;refGene_tx=XM_023251972.1;refGene_gn=LTBP1;refGene_tx=XM_023251973.1
A3 116039535 116039760
refGene_gn=LTBP1;refGene_tx=XM_006930373.4;refGene_gn=LTBP1;refGene_tx=XM_006930374.4;refGene_gn=LTBP1;refGene_tx=XM_006930376.4;refGene_gn=LTBP1;refGene_tx=XM_023251968.1;refGene_gn=LTBP1;refGene_tx=XM_023251969.1;refGene_gn=LTBP1;refGene_tx=XM_023251970.1;refGene_gn=LTBP1;refGene_tx=XM_023251971.1;refGene_gn=LTBP1;refGene_tx=XM_023251972.1;refGene_gn=LTBP1;refGene_tx=XM_023251973.1
A3 116081353 116081576
refGene_gn=LTBP1;refGene_tx=XM_006930373.4;refGene_gn=LTBP1;refGene_tx=XM_006930374.4;refGene_gn=LTBP1;refGene_tx=XM_006930376.4;refGene_gn=LTBP1;refGene_tx=XM_023251968.1;refGene_gn=LTBP1;refGene_tx=XM_023251969.1;refGene_gn=LTBP1;refGene_tx=XM_023251970.1;refGene_gn=LTBP1;refGene_tx=XM_023251971.1;refGene_gn=LTBP1;refGene_tx=XM_023251973.1
A3 116108349 116108519
refGene_gn=LTBP1;refGene_tx=XM_023251968.1;refGene_gn=LTBP1;refGene_tx=XM_023251969.1;refGene_gn=LTBP1;refGene_tx=XM_023251970.1;refGene_gn=LTBP1;refGene_tx=XM_023251971.1;refGene_gn=LTBP1;refGene_tx=XM_023251972.1;refGene_gn=LTBP1;refGene_tx=XM_023251973.1
A3 116201816 116202114
refGene_gn=LTBP1;refGene_tx=XM_023251968.1;refGene_gn=LTBP1;refGene_tx=XM_023251969.1;refGene_gn=LTBP1;refGene_tx=XM_023251970.1;refGene_gn=LTBP1;refGene_tx=XM_023251971.1;refGene_gn=LTBP1;refGene_tx=XM_0232519

72.1;refGene_gn=LTBP1;refGene_tx=XM_023251973.1
A3 116246351 116246422
refGene_gn=LTBP1;refGene_tx=XM_023251968.1;refGene_gn=LTBP1;refGene_tx=XM_023251969.1;refGene_gn=LTBP1;refGene_tx=XM_023251970.1;refGene_gn=LTBP1;refGene_tx=XM_023251971.1;refGene_gn=LTBP1;refGene_tx=XM_023251972.1;refGene_gn=LTBP1;refGene_tx=XM_023251973.1
A3 116247466 116247954
refGene_gn=LTBP1;refGene_tx=XM_023251968.1;refGene_gn=LTBP1;refGene_tx=XM_023251969.1;refGene_gn=LTBP1;refGene_tx=XM_023251970.1;refGene_gn=LTBP1;refGene_tx=XM_023251971.1;refGene_gn=LTBP1;refGene_tx=XM_023251972.1;refGene_gn=LTBP1;refGene_tx=XM_023251973.1
A3 116356302 116356425
refGene_gn=TTC27;refGene_tx=XM_023251974.1
A3 116361437 116361538
refGene_gn=TTC27;refGene_tx=XM_023251974.1
A3 116366039 116366151
refGene_gn=TTC27;refGene_tx=XM_023251974.1
A3 116367441 116367639
refGene_gn=TTC27;refGene_tx=XM_023251974.1
A3 116387355 116387521
refGene_gn=TTC27;refGene_tx=XM_023251974.1
A3 116394111 116394164
refGene_gn=TTC27;refGene_tx=XM_023251974.1
A3 116398272 116398371
refGene_gn=TTC27;refGene_tx=XM_023251974.1
A3 116419513 116419741
refGene_gn=TTC27;refGene_tx=XM_023251974.1
A3 116440070 116440201
refGene_gn=TTC27;refGene_tx=XM_023251974.1
A3 116445002 116445090
refGene_gn=TTC27;refGene_tx=XM_023251974.1
A3 116473554 116473668
refGene_gn=TTC27;refGene_tx=XM_023251974.1
A3 116503943 116504010
refGene_gn=TTC27;refGene_tx=XM_023251974.1
A3 116507567 116507680
refGene_gn=TTC27;refGene_tx=XM_023251974.1
A3 116515622 116515756
refGene_gn=TTC27;refGene_tx=XM_023251974.1
A3 116519261 116519426
refGene_gn=TTC27;refGene_tx=XM_023251974.1
A3 116531064 116531167
refGene_gn=TTC27;refGene_tx=XM_023251974.1
A3 116538877 116539018
refGene_gn=TTC27;refGene_tx=XM_023251974.1
A3 116545240 116545370
refGene_gn=TTC27;refGene_tx=XM_023251974.1
A3 116545772 116545950
refGene_gn=TTC27;refGene_tx=XM_023251974.1
A3 116547337 116547437

refGene_gn=TTC27;refGene_tx=XM_023251974.1
A3 116564555 116564735
refGene_gn=BIRC6;refGene_tx=XM_019827703.2
A3 116565866 116566001
refGene_gn=BIRC6;refGene_tx=XM_019827703.2
A3 116568986 116569175
refGene_gn=BIRC6;refGene_tx=XM_019827703.2
A3 116575361 116575439
refGene_gn=BIRC6;refGene_tx=XM_019827703.2
A3 116579059 116579221
refGene_gn=BIRC6;refGene_tx=XM_019827703.2
A3 116581148 116581366
refGene_gn=BIRC6;refGene_tx=XM_019827703.2
A3 116588451 116588562
refGene_gn=BIRC6;refGene_tx=XM_019827703.2
A3 116589193 116589339
refGene_gn=BIRC6;refGene_tx=XM_019827703.2
A3 116603884 116604095
refGene_gn=BIRC6;refGene_tx=XM_019827703.2
A3 116623312 116623481
refGene_gn=BIRC6;refGene_tx=XM_019827703.2
A3 116624363 116624528
refGene_gn=BIRC6;refGene_tx=XM_019827703.2
A3 116628661 116628879
refGene_gn=BIRC6;refGene_tx=XM_019827703.2
A3 116635753 116636054
refGene_gn=BIRC6;refGene_tx=XM_019827703.2
A3 116637556 116637753
refGene_gn=BIRC6;refGene_tx=XM_019827703.2
A3 116639106 116639280
refGene_gn=BIRC6;refGene_tx=XM_019827703.2
A3 116640308 116640476
refGene_gn=BIRC6;refGene_tx=XM_019827703.2
A3 116641131 116641263
refGene_gn=BIRC6;refGene_tx=XM_019827703.2
A3 116646373 116646503
refGene_gn=BIRC6;refGene_tx=XM_019827703.2
A3 116647456 116647600
refGene_gn=BIRC6;refGene_tx=XM_019827703.2
A3 116648706 116649490
refGene_gn=BIRC6;refGene_tx=XM_019827703.2
A3 116650916 116651138
refGene_gn=BIRC6;refGene_tx=XM_019827703.2
A3 116655098 116655207
refGene_gn=BIRC6;refGene_tx=XM_019827703.2
A3 116655471 116655728
refGene_gn=BIRC6;refGene_tx=XM_019827703.2
A3 116657605 116657885
refGene_gn=BIRC6;refGene_tx=XM_019827703.2
A3 116659473 116659674

refGene_gn=BIRC6;refGene_tx=XM_019827703.2
A3 116660318 116660513
refGene_gn=BIRC6;refGene_tx=XM_019827703.2
A3 116660646 116660695
refGene_gn=BIRC6;refGene_tx=XM_019827703.2
A3 116660696 116660744
refGene_gn=BIRC6;refGene_tx=XM_019827703.2
A3 116661721 116661897
refGene_gn=BIRC6;refGene_tx=XM_019827703.2
A3 116662868 116663431
refGene_gn=BIRC6;refGene_tx=XM_019827703.2
A3 116668642 116668770
refGene_gn=BIRC6;refGene_tx=XM_019827703.2
A3 116671591 116671725
refGene_gn=BIRC6;refGene_tx=XM_019827703.2
A3 116672967 116673078
refGene_gn=BIRC6;refGene_tx=XM_019827703.2
A3 116676062 116676189
refGene_gn=BIRC6;refGene_tx=XM_019827703.2
A3 116676677 116676832
refGene_gn=BIRC6;refGene_tx=XM_019827703.2
A3 116679206 116679323
refGene_gn=BIRC6;refGene_tx=XM_019827703.2
A3 116681893 116682047
refGene_gn=BIRC6;refGene_tx=XM_019827703.2
A3 116683156 116683290
refGene_gn=BIRC6;refGene_tx=XM_019827703.2
A3 116687248 116687404
refGene_gn=BIRC6;refGene_tx=XM_019827703.2
A3 116688030 116688214
refGene_gn=BIRC6;refGene_tx=XM_019827703.2
A3 116688786 116689002
refGene_gn=BIRC6;refGene_tx=XM_019827703.2
A3 116691044 116691176
refGene_gn=BIRC6;refGene_tx=XM_019827703.2
A3 116694582 116694710
refGene_gn=BIRC6;refGene_tx=XM_019827703.2
A3 116696504 116696615
refGene_gn=BIRC6;refGene_tx=XM_019827703.2
A3 116697560 116697694
refGene_gn=BIRC6;refGene_tx=XM_019827703.2
A3 116698177 116698397
refGene_gn=BIRC6;refGene_tx=XM_019827703.2
A3 116699010 116699357
refGene_gn=BIRC6;refGene_tx=XM_019827703.2
A3 116699800 116700009
refGene_gn=BIRC6;refGene_tx=XM_019827703.2
A3 116700196 116700411
refGene_gn=BIRC6;refGene_tx=XM_019827703.2
A3 116702794 116702894

refGene_gn=BIRC6;refGene_tx=XM_019827703.2
A3	116703131	116703455
refGene_gn=BIRC6;refGene_tx=XM_019827703.2
A3	116704244	116704432
refGene_gn=BIRC6;refGene_tx=XM_019827703.2
A3	116713013	116713148
refGene_gn=BIRC6;refGene_tx=XM_019827703.2
A3	116716060	116716194
refGene_gn=BIRC6;refGene_tx=XM_019827703.2
A3	116717037	116717185
refGene_gn=BIRC6;refGene_tx=XM_019827703.2
A3	116718587	116718685
refGene_gn=BIRC6;refGene_tx=XM_019827703.2
A3	116719788	116719920
refGene_gn=BIRC6;refGene_tx=XM_019827703.2
A3	116720017	116720179
refGene_gn=BIRC6;refGene_tx=XM_019827703.2
A3	116720956	116721090
refGene_gn=BIRC6;refGene_tx=XM_019827703.2
A3	116722085	116722264
refGene_gn=BIRC6;refGene_tx=XM_019827703.2
A3	116724260	116724392
refGene_gn=BIRC6;refGene_tx=XM_019827703.2
A3	116724886	116724976
refGene_gn=BIRC6;refGene_tx=XM_019827703.2
A3	116726088	116726249
refGene_gn=BIRC6;refGene_tx=XM_019827703.2
A3	116727040	116727266
refGene_gn=BIRC6;refGene_tx=XM_019827703.2
A3	116729230	116729380
refGene_gn=BIRC6;refGene_tx=XM_019827703.2
A3	116734115	116735510
refGene_gn=BIRC6;refGene_tx=XM_019827703.2
A3	116740864	116740923
refGene_gn=BIRC6;refGene_tx=XM_019827703.2
A3	116745665	116745826
refGene_gn=BIRC6;refGene_tx=XM_019827703.2
A3	116745905	116746128
refGene_gn=BIRC6;refGene_tx=XM_019827703.2
A3	116751291	116751374
refGene_gn=BIRC6;refGene_tx=XM_019827703.2
A3	116752219	116752331
refGene_gn=BIRC6;refGene_tx=XM_019827703.2
A3	116755081	116755275
refGene_gn=BIRC6;refGene_tx=XM_019827703.2
A3	116760897	116761035
refGene_gn=BIRC6;refGene_tx=XM_019827703.2
A3	116763711	116763893
refGene_gn=BIRC6;refGene_tx=XM_019827703.2
A3	116789141	116789466

refGene_gn=BIRC6;refGene_tx=XM_019827703.2
A3 116830248 116830386
refGene_gn=YIPF4;refGene_tx=XM_003984341.5
A3 116834592 116834706
refGene_gn=YIPF4;refGene_tx=XM_003984341.5
A3 116838689 116838767
refGene_gn=YIPF4;refGene_tx=XM_003984341.5
A3 116841982 116842154
refGene_gn=YIPF4;refGene_tx=XM_003984341.5
A3 116844273 116844427
refGene_gn=YIPF4;refGene_tx=XM_003984341.5
A3 116858432 116858511
refGene_gn=YIPF4;refGene_tx=XM_003984341.5
A3 116873525 116873847
refGene_gn=NLRC4;refGene_tx=XM_019827705.2
A3 116874673 116876665
refGene_gn=NLRC4;refGene_tx=XM_019827705.2
A3 116881952 116882045
refGene_gn=NLRC4;refGene_tx=XM_019827705.2
A3 116883750 116883921
refGene_gn=NLRC4;refGene_tx=XM_019827705.2
A3 116886161 116886254
refGene_gn=NLRC4;refGene_tx=XM_019827705.2
A3 116887413 116887581
refGene_gn=NLRC4;refGene_tx=XM_019827705.2
A3 116900000 116900290
refGene_gn=NLRC4;refGene_tx=XM_019827705.2
A3 116904053 116904557
refGene_gn=SLC30A6;refGene_tx=XM_011281136.3;refGene_gn=SLC30A6;refGen
e_tx=XM_011281137.3;refGene_gn=SLC30A6;refGene_tx=XM_019827704.2
A3 116911857 116911926
refGene_gn=SLC30A6;refGene_tx=XM_011281136.3;refGene_gn=SLC30A6;refGen
e_tx=XM_011281137.3;refGene_gn=SLC30A6;refGene_tx=XM_019827704.2
A3 116912906 116912954
refGene_gn=SLC30A6;refGene_tx=XM_011281136.3;refGene_gn=SLC30A6;refGen
e_tx=XM_011281137.3;refGene_gn=SLC30A6;refGene_tx=XM_019827704.2
A3 116913829 116913932
refGene_gn=SLC30A6;refGene_tx=XM_011281136.3;refGene_gn=SLC30A6;refGen
e_tx=XM_011281137.3;refGene_gn=SLC30A6;refGene_tx=XM_019827704.2
A3 116919303 116919423
refGene_gn=SLC30A6;refGene_tx=XM_011281136.3;refGene_gn=SLC30A6;refGen
e_tx=XM_011281137.3;refGene_gn=SLC30A6;refGene_tx=XM_019827704.2
A3 116919727 116919776
refGene_gn=SLC30A6;refGene_tx=XM_011281136.3;refGene_gn=SLC30A6;refGen
e_tx=XM_011281137.3;refGene_gn=SLC30A6;refGene_tx=XM_019827704.2
A3 116921715 116921810
refGene_gn=SLC30A6;refGene_tx=XM_011281136.3;refGene_gn=SLC30A6;refGen
e_tx=XM_011281137.3;refGene_gn=SLC30A6;refGene_tx=XM_019827704.2
A3 116922727 116922763
refGene_gn=SLC30A6;refGene_tx=XM_011281136.3;refGene_gn=SLC30A6;refGen

e_tx=XM_011281137.3
A3	116923351	116923432
refGene_gn=SLC30A6;refGene_tx=XM_011281136.3;refGene_gn=SLC30A6;refGene_tx=XM_011281137.3
A3	116941732	116941798
refGene_gn=SLC30A6;refGene_tx=XM_011281136.3
A3	116942790	116942833
refGene_gn=SLC30A6;refGene_tx=XM_011281136.3;refGene_gn=SLC30A6;refGene_tx=XM_011281137.3
A3	116944350	116944435
refGene_gn=SLC30A6;refGene_tx=XM_011281136.3;refGene_gn=SLC30A6;refGene_tx=XM_011281137.3
A3	116949478	116949565
refGene_gn=SLC30A6;refGene_tx=XM_011281136.3;refGene_gn=SLC30A6;refGene_tx=XM_011281137.3
A3	116953728	116953731
refGene_gn=SLC30A6;refGene_tx=XM_011281136.3;refGene_gn=SLC30A6;refGene_tx=XM_011281137.3
A3	116962566	116962689
refGene_gn=SPAST;refGene_tx=XM_011281134.3;refGene_gn=SPAST;refGene_tx=XM_011281135.3;refGene_gn=SPAST;refGene_tx=XM_019827706.2
A3	116965831	116965872
refGene_gn=SPAST;refGene_tx=XM_011281134.3;refGene_gn=SPAST;refGene_tx=XM_011281135.3;refGene_gn=SPAST;refGene_tx=XM_019827706.2
A3	116967542	116967613
refGene_gn=SPAST;refGene_tx=XM_011281134.3;refGene_gn=SPAST;refGene_tx=XM_011281135.3;refGene_gn=SPAST;refGene_tx=XM_019827706.2
A3	116969002	116969082
refGene_gn=SPAST;refGene_tx=XM_011281134.3;refGene_gn=SPAST;refGene_tx=XM_011281135.3;refGene_gn=SPAST;refGene_tx=XM_019827706.2
A3	116970025	116970068
refGene_gn=SPAST;refGene_tx=XM_011281134.3;refGene_gn=SPAST;refGene_tx=XM_011281135.3;refGene_gn=SPAST;refGene_tx=XM_019827706.2
A3	116972508	116972588
refGene_gn=SPAST;refGene_tx=XM_011281134.3;refGene_gn=SPAST;refGene_tx=XM_011281135.3;refGene_gn=SPAST;refGene_tx=XM_019827706.2
A3	116972734	116972826
refGene_gn=SPAST;refGene_tx=XM_011281134.3;refGene_gn=SPAST;refGene_tx=XM_011281135.3;refGene_gn=SPAST;refGene_tx=XM_019827706.2
A3	116973066	116973142
refGene_gn=SPAST;refGene_tx=XM_011281134.3;refGene_gn=SPAST;refGene_tx=XM_011281135.3;refGene_gn=SPAST;refGene_tx=XM_019827706.2
A3	116974478	116974550
refGene_gn=SPAST;refGene_tx=XM_011281134.3;refGene_gn=SPAST;refGene_tx=XM_011281135.3;refGene_gn=SPAST;refGene_tx=XM_019827706.2
A3	116976106	116976181
refGene_gn=SPAST;refGene_tx=XM_011281134.3;refGene_gn=SPAST;refGene_tx=XM_011281135.3;refGene_gn=SPAST;refGene_tx=XM_019827706.2
A3	116980937	116981031
refGene_gn=SPAST;refGene_tx=XM_011281134.3;refGene_gn=SPAST;refGene_tx

=XM_011281135.3;refGene_gn=SPAST;refGene_tx=XM_019827706.2
A3 116981510 116981644
refGene_gn=SPAST;refGene_tx=XM_011281134.3;refGene_gn=SPAST;refGene_tx
=XM_011281135.3;refGene_gn=SPAST;refGene_tx=XM_019827706.2
A3 116982220 116982408
refGene_gn=SPAST;refGene_tx=XM_011281134.3;refGene_gn=SPAST;refGene_tx
=XM_011281135.3;refGene_gn=SPAST;refGene_tx=XM_019827706.2
A3 116987964 116988060
refGene_gn=SPAST;refGene_tx=XM_011281134.3;refGene_gn=SPAST;refGene_tx
=XM_011281135.3
A3 116994655 116994739
refGene_gn=SPAST;refGene_tx=XM_011281134.3;refGene_gn=SPAST;refGene_tx
=XM_011281135.3;refGene_gn=SPAST;refGene_tx=XM_019827706.2
A3 116996912 116996999
refGene_gn=SPAST;refGene_tx=XM_011281134.3;refGene_gn=SPAST;refGene_tx
=XM_011281135.3;refGene_gn=SPAST;refGene_tx=XM_019827706.2
A3 117012722 117013137
refGene_gn=SPAST;refGene_tx=XM_011281134.3;refGene_gn=SPAST;refGene_tx
=XM_011281135.3;refGene_gn=SPAST;refGene_tx=XM_019827706.2
A3 117026998 117027034
refGene_gn=DPY30;refGene_tx=XM_003984342.3;refGene_gn=DPY30;refGene_tx
=XM_003984343.4;refGene_gn=DPY30;refGene_tx=XM_019827707.2
A3 117027119 117027167
refGene_gn=DPY30;refGene_tx=XM_003984342.3;refGene_gn=DPY30;refGene_tx
=XM_003984343.4;refGene_gn=DPY30;refGene_tx=XM_019827707.2
A3 117033066 117033209
refGene_gn=DPY30;refGene_tx=XM_003984342.3;refGene_gn=DPY30;refGene_tx
=XM_003984343.4;refGene_gn=DPY30;refGene_tx=XM_019827707.2
A3 117035410 117035483
refGene_gn=DPY30;refGene_tx=XM_003984342.3;refGene_gn=DPY30;refGene_tx
=XM_003984343.4;refGene_gn=DPY30;refGene_tx=XM_019827707.2
A3 117043242 117043548 refGene_gn=DPP9-
AS1;refGene_tx=XM_023252199.1
A3 117043586 117043622 refGene_gn=DPP9-
AS1;refGene_tx=XM_023252199.1
A3 117043654 117043715
refGene_gn=MEMO1;refGene_tx=XM_023251975.1
A3 117106097 117106179
refGene_gn=MEMO1;refGene_tx=XM_023251975.1
A3 117122726 117122795
refGene_gn=MEMO1;refGene_tx=XM_023251975.1
A3 117130283 117130396
refGene_gn=MEMO1;refGene_tx=XM_023251975.1
A3 117132328 117132440
refGene_gn=MEMO1;refGene_tx=XM_023251975.1
A3 117149592 117149735
refGene_gn=MEMO1;refGene_tx=XM_023251975.1
A3 117158079 117158156
refGene_gn=MEMO1;refGene_tx=XM_023251975.1
A3 117168075 117168180

refGene_gn=MEMO1;refGene_tx=XM_023251975.1
A3 117169527 117169659
refGene_gn=MEMO1;refGene_tx=XM_023251975.1
A3 117391178 117391459
refGene_gn=SRD5A2;refGene_tx=XM_006930379.3
A3 117415678 117415842
refGene_gn=SRD5A2;refGene_tx=XM_006930379.3
A3 117420386 117420488
refGene_gn=SRD5A2;refGene_tx=XM_006930379.3
A3 117423887 117424038
refGene_gn=SRD5A2;refGene_tx=XM_006930379.3
A3 117427048 117427115
refGene_gn=SRD5A2;refGene_tx=XM_006930379.3
A3 117507345 117507387
refGene_gn=XDH;refGene_tx=NM_001009217.1;refGene_gn=XDH;refGene_tx=XM_023250907.1
A3 117513725 117513783
refGene_gn=XDH;refGene_tx=NM_001009217.1;refGene_gn=XDH;refGene_tx=XM_023250907.1
A3 117520306 117521058
refGene_gn=LOC101092676;refGene_tx=XM_003984079.4
A3 117521060 117521246
refGene_gn=LOC101092676;refGene_tx=XM_003984079.4
A3 117525849 117526183
refGene_gn=LOC101092676;refGene_tx=XM_003984079.4
A3 117551523 117551620
refGene_gn=XDH;refGene_tx=NM_001009217.1;refGene_gn=XDH;refGene_tx=XM_023250907.1
A3 117553037 117553146
refGene_gn=XDH;refGene_tx=NM_001009217.1;refGene_gn=XDH;refGene_tx=XM_023250907.1
A3 117555023 117555150
refGene_gn=XDH;refGene_tx=NM_001009217.1;refGene_gn=XDH;refGene_tx=XM_023250907.1
A3 117556096 117556158
refGene_gn=XDH;refGene_tx=NM_001009217.1;refGene_gn=XDH;refGene_tx=XM_023250907.1
A3 117559241 117559310
refGene_gn=XDH;refGene_tx=NM_001009217.1;refGene_gn=XDH;refGene_tx=XM_023250907.1
A3 117559647 117559737
refGene_gn=XDH;refGene_tx=NM_001009217.1;refGene_gn=XDH;refGene_tx=XM_023250907.1
A3 117560870 117561012
refGene_gn=XDH;refGene_tx=NM_001009217.1;refGene_gn=XDH;refGene_tx=XM_023250907.1
A3 117563363 117563456
refGene_gn=XDH;refGene_tx=NM_001009217.1;refGene_gn=XDH;refGene_tx=XM_023250907.1
A3 117564061 117564213

refGene_gn=XDH;refGene_tx=NM_001009217.1;refGene_gn=XDH;refGene_tx=XM_023250907.1
A3 117565258 117565352
refGene_gn=XDH;refGene_tx=NM_001009217.1;refGene_gn=XDH;refGene_tx=XM_023250907.1
A3 117566705 117566815
refGene_gn=XDH;refGene_tx=NM_001009217.1;refGene_gn=XDH;refGene_tx=XM_023250907.1
A3 117569141 117569326
refGene_gn=XDH;refGene_tx=NM_001009217.1;refGene_gn=XDH;refGene_tx=XM_023250907.1
A3 117571358 117571533
refGene_gn=XDH;refGene_tx=NM_001009217.1;refGene_gn=XDH;refGene_tx=XM_023250907.1
A3 117573266 117573350
refGene_gn=XDH;refGene_tx=NM_001009217.1;refGene_gn=XDH;refGene_tx=XM_023250907.1
A3 117574800 117574970
refGene_gn=XDH;refGene_tx=NM_001009217.1;refGene_gn=XDH;refGene_tx=XM_023250907.1
A3 117582998 117583122
refGene_gn=XDH;refGene_tx=NM_001009217.1;refGene_gn=XDH;refGene_tx=XM_023250907.1
A3 117584731 117584851
refGene_gn=XDH;refGene_tx=NM_001009217.1;refGene_gn=XDH;refGene_tx=XM_023250907.1
A3 117585318 117585415
refGene_gn=XDH;refGene_tx=NM_001009217.1;refGene_gn=XDH;refGene_tx=XM_023250907.1
A3 117586408 117586533
refGene_gn=XDH;refGene_tx=NM_001009217.1;refGene_gn=XDH;refGene_tx=XM_023250907.1
A3 117587316 117587450
refGene_gn=XDH;refGene_tx=NM_001009217.1;refGene_gn=XDH;refGene_tx=XM_023250907.1
A3 117587868 117587956
refGene_gn=XDH;refGene_tx=NM_001009217.1;refGene_gn=XDH;refGene_tx=XM_023250907.1
A3 117589075 117589162
refGene_gn=XDH;refGene_tx=NM_001009217.1;refGene_gn=XDH;refGene_tx=XM_023250907.1
A3 117593439 117593631
refGene_gn=XDH;refGene_tx=NM_001009217.1;refGene_gn=XDH;refGene_tx=XM_023250907.1
A3 117593800 117593946
refGene_gn=XDH;refGene_tx=NM_001009217.1;refGene_gn=XDH;refGene_tx=XM_023250907.1
A3 117594746 117594828
refGene_gn=XDH;refGene_tx=NM_001009217.1;refGene_gn=XDH;refGene_tx=XM_023250907.1

A3	117595378	117595474
refGene_gn=XDH;refGene_tx=NM_001009217.1;refGene_gn=XDH;refGene_tx=XM_023250907.1
A3	117596765	117596894
refGene_gn=XDH;refGene_tx=NM_001009217.1;refGene_gn=XDH;refGene_tx=XM_023250907.1
A3	117597562	117597637
refGene_gn=XDH;refGene_tx=NM_001009217.1;refGene_gn=XDH;refGene_tx=XM_023250907.1
A3	117599138	117599191
refGene_gn=XDH;refGene_tx=NM_001009217.1;refGene_gn=XDH;refGene_tx=XM_023250907.1
A3	117600898	117601013
refGene_gn=XDH;refGene_tx=NM_001009217.1;refGene_gn=XDH;refGene_tx=XM_023250907.1
A3	117602034	117602100
refGene_gn=XDH;refGene_tx=NM_001009217.1;refGene_gn=XDH;refGene_tx=XM_023250907.1
A3	117602363	117602552
refGene_gn=XDH;refGene_tx=NM_001009217.1;refGene_gn=XDH;refGene_tx=XM_023250907.1
A3	117603845	117604022
refGene_gn=XDH;refGene_tx=NM_001009217.1;refGene_gn=XDH;refGene_tx=XM_023250907.1
A3	117605651	117605702
refGene_gn=XDH;refGene_tx=NM_001009217.1;refGene_gn=XDH;refGene_tx=XM_023250907.1
A3	117658724	117659252
refGene_gn=EHD3;refGene_tx=XM_003984350.4
A3	117662295	117662460
refGene_gn=EHD3;refGene_tx=XM_003984350.4
A3	117663102	117663515
refGene_gn=EHD3;refGene_tx=XM_003984350.4
A3	117671019	117671117
refGene_gn=EHD3;refGene_tx=XM_003984350.4
A3	117674327	117674504
refGene_gn=EHD3;refGene_tx=XM_003984350.4
A3	117683171	117683398
refGene_gn=EHD3;refGene_tx=XM_003984350.4
A3	117708761	117708986
refGene_gn=CAPN14;refGene_tx=XM_003984351.3
A3	117710366	117710436
refGene_gn=CAPN14;refGene_tx=XM_003984351.3
A3	117711256	117711375
refGene_gn=CAPN14;refGene_tx=XM_003984351.3
A3	117711511	117711648
refGene_gn=CAPN14;refGene_tx=XM_003984351.3
A3	117712133	117712308
refGene_gn=CAPN14;refGene_tx=XM_003984351.3
A3	117712907	117712973

refGene_gn=CAPN14;refGene_tx=XM_003984351.3
A3 117715853 117715939
refGene_gn=CAPN14;refGene_tx=XM_003984351.3
A3 117717204 117717279
refGene_gn=CAPN14;refGene_tx=XM_003984351.3
A3 117718337 117718501
refGene_gn=CAPN14;refGene_tx=XM_003984351.3
A3 117719700 117719864
refGene_gn=CAPN14;refGene_tx=XM_003984351.3
A3 117720387 117720396
refGene_gn=CAPN14;refGene_tx=XM_003984351.3
A3 117722963 117723169
refGene_gn=CAPN14;refGene_tx=XM_003984351.3
A3 117724145 117724182
refGene_gn=CAPN14;refGene_tx=XM_003984351.3
A3 117724932 117724989
refGene_gn=CAPN14;refGene_tx=XM_003984351.3
A3 117726458 117726516
refGene_gn=CAPN14;refGene_tx=XM_003984351.3
A3 117729087 117729152
refGene_gn=CAPN14;refGene_tx=XM_003984351.3
A3 117730452 117730521
refGene_gn=CAPN14;refGene_tx=XM_003984351.3
A3 117731405 117731484
refGene_gn=CAPN14;refGene_tx=XM_003984351.3
A3 117732089 117732206
refGene_gn=CAPN14;refGene_tx=XM_003984351.3
A3 117732558 117732614
refGene_gn=CAPN14;refGene_tx=XM_003984351.3
A3 117734956 117734983
refGene_gn=CAPN14;refGene_tx=XM_003984351.3
A3 117778549 117778678
refGene_gn=GALNT14;refGene_tx=XM_019827708.2;refGene_gn=GALNT14;refGene_tx=XM_019827709.2;refGene_gn=GALNT14;refGene_tx=XM_019827712.2;refGene_gn=GALNT14;refGene_tx=XM_019827714.2;refGene_gn=GALNT14;refGene_tx=XM_019827715.2
A3 117910985 117911155
refGene_gn=GALNT14;refGene_tx=XM_019827708.2;refGene_gn=GALNT14;refGene_tx=XM_019827709.2;refGene_gn=GALNT14;refGene_tx=XM_019827712.2;refGene_gn=GALNT14;refGene_tx=XM_019827714.2;refGene_gn=GALNT14;refGene_tx=XM_019827715.2
A3 117937657 117937756
refGene_gn=GALNT14;refGene_tx=XM_019827708.2;refGene_gn=GALNT14;refGene_tx=XM_019827709.2;refGene_gn=GALNT14;refGene_tx=XM_019827712.2;refGene_gn=GALNT14;refGene_tx=XM_019827714.2;refGene_gn=GALNT14;refGene_tx=XM_019827715.2
A3 117945241 117945309
refGene_gn=GALNT14;refGene_tx=XM_019827708.2;refGene_gn=GALNT14;refGene_tx=XM_019827709.2;refGene_gn=GALNT14;refGene_tx=XM_019827712.2;refGene_gn=GALNT14;refGene_tx=XM_019827714.2;refGene_gn=GALNT14;refGene_tx=

XM_019827715.2
A3 117947640 117947706
refGene_gn=GALNT14;refGene_tx=XM_019827708.2;refGene_gn=GALNT14;refGene_tx=XM_019827709.2;refGene_gn=GALNT14;refGene_tx=XM_019827712.2;refGene_gn=GALNT14;refGene_tx=XM_019827714.2;refGene_gn=GALNT14;refGene_tx=XM_019827715.2
A3 117947877 117947999
refGene_gn=GALNT14;refGene_tx=XM_019827708.2;refGene_gn=GALNT14;refGene_tx=XM_019827709.2;refGene_gn=GALNT14;refGene_tx=XM_019827712.2;refGene_gn=GALNT14;refGene_tx=XM_019827714.2;refGene_gn=GALNT14;refGene_tx=XM_019827715.2
A3 117955658 117955746
refGene_gn=GALNT14;refGene_tx=XM_019827708.2;refGene_gn=GALNT14;refGene_tx=XM_019827709.2;refGene_gn=GALNT14;refGene_tx=XM_019827712.2;refGene_gn=GALNT14;refGene_tx=XM_019827714.2;refGene_gn=GALNT14;refGene_tx=XM_019827715.2
A3 117956473 117956558
refGene_gn=GALNT14;refGene_tx=XM_019827708.2;refGene_gn=GALNT14;refGene_tx=XM_019827709.2;refGene_gn=GALNT14;refGene_tx=XM_019827712.2;refGene_gn=GALNT14;refGene_tx=XM_019827714.2;refGene_gn=GALNT14;refGene_tx=XM_019827715.2
A3 117960218 117960322
refGene_gn=GALNT14;refGene_tx=XM_019827708.2;refGene_gn=GALNT14;refGene_tx=XM_019827709.2;refGene_gn=GALNT14;refGene_tx=XM_019827712.2;refGene_gn=GALNT14;refGene_tx=XM_019827714.2;refGene_gn=GALNT14;refGene_tx=XM_019827715.2
A3 117968563 117968690
refGene_gn=GALNT14;refGene_tx=XM_019827708.2;refGene_gn=GALNT14;refGene_tx=XM_019827709.2;refGene_gn=GALNT14;refGene_tx=XM_019827712.2;refGene_gn=GALNT14;refGene_tx=XM_019827714.2;refGene_gn=GALNT14;refGene_tx=XM_019827715.2
A3 117970839 117970932
refGene_gn=GALNT14;refGene_tx=XM_019827708.2;refGene_gn=GALNT14;refGene_tx=XM_019827709.2;refGene_gn=GALNT14;refGene_tx=XM_019827712.2;refGene_gn=GALNT14;refGene_tx=XM_019827714.2;refGene_gn=GALNT14;refGene_tx=XM_019827715.2
A3 117972349 117972632
refGene_gn=GALNT14;refGene_tx=XM_019827714.2
A3 117974998 117975082
refGene_gn=GALNT14;refGene_tx=XM_019827708.2;refGene_gn=GALNT14;refGene_tx=XM_019827709.2;refGene_gn=GALNT14;refGene_tx=XM_019827712.2
A3 117975556 117975701
refGene_gn=GALNT14;refGene_tx=XM_019827708.2;refGene_gn=GALNT14;refGene_tx=XM_019827709.2;refGene_gn=GALNT14;refGene_tx=XM_019827712.2
A3 117979600 117979669
refGene_gn=GALNT14;refGene_tx=XM_019827712.2
A3 117986090 117986210
refGene_gn=GALNT14;refGene_tx=XM_019827708.2;refGene_gn=GALNT14;refGene_tx=XM_019827709.2;refGene_gn=GALNT14;refGene_tx=XM_019827715.2
A3 117987403 117987562

refGene_gn=GALNT14;refGene_tx=XM_019827708.2;refGene_gn=GALNT14;refGene_tx=XM_019827709.2;refGene_gn=GALNT14;refGene_tx=XM_019827715.2
A3 118087505 118087703
refGene_gn=CAPN13;refGene_tx=XM_023252200.1
A3 118097133 118097212
refGene_gn=CAPN13;refGene_tx=XM_023252200.1
A3 118098231 118098347
refGene_gn=CAPN13;refGene_tx=XM_023252200.1
A3 118103702 118103839
refGene_gn=CAPN13;refGene_tx=XM_023252200.1
A3 118108993 118109168
refGene_gn=CAPN13;refGene_tx=XM_023252200.1
A3 118110105 118110180
refGene_gn=CAPN13;refGene_tx=XM_023252200.1
A3 118114755 118114841
refGene_gn=CAPN13;refGene_tx=XM_023252200.1
A3 118118494 118118569
refGene_gn=CAPN13;refGene_tx=XM_023252200.1
A3 118119606 118119752
refGene_gn=CAPN13;refGene_tx=XM_023252200.1
A3 118121364 118121507
refGene_gn=CAPN13;refGene_tx=XM_023252200.1
A3 118125421 118125433
refGene_gn=CAPN13;refGene_tx=XM_023252200.1
A3 118127602 118127802
refGene_gn=CAPN13;refGene_tx=XM_023252200.1
A3 118128826 118128863
refGene_gn=CAPN13;refGene_tx=XM_023252200.1
A3 118129221 118129278
refGene_gn=CAPN13;refGene_tx=XM_023252200.1
A3 118132385 118132443
refGene_gn=CAPN13;refGene_tx=XM_023252200.1
A3 118132555 118132614
refGene_gn=CAPN13;refGene_tx=XM_023252200.1
A3 118134215 118134284
refGene_gn=CAPN13;refGene_tx=XM_023252200.1
A3 118136132 118136208
refGene_gn=CAPN13;refGene_tx=XM_023252200.1
A3 118138451 118138580
refGene_gn=CAPN13;refGene_tx=XM_023252200.1
A3 118139592 118139648
refGene_gn=CAPN13;refGene_tx=XM_023252200.1
A3 118140223 118140250
refGene_gn=CAPN13;refGene_tx=XM_023252200.1
A3 118211545 118212045
refGene_gn=LCLAT1;refGene_tx=XM_003984353.5;refGene_gn=LCLAT1;refGene_tx=XM_011281295.3;refGene_gn=LCLAT1;refGene_tx=XM_019827716.2
A3 118278978 118279095
refGene_gn=LCLAT1;refGene_tx=XM_003984353.5;refGene_gn=LCLAT1;refGene_tx=XM_011281295.3;refGene_gn=LCLAT1;refGene_tx=XM_019827716.2

A3	118285162	118285309
refGene_gn=LCLAT1;refGene_tx=XM_003984353.5;refGene_gn=LCLAT1;refGene_tx=XM_011281295.3;refGene_gn=LCLAT1;refGene_tx=XM_019827716.2
A3	118317926	118318125
refGene_gn=LCLAT1;refGene_tx=XM_003984353.5;refGene_gn=LCLAT1;refGene_tx=XM_011281295.3;refGene_gn=LCLAT1;refGene_tx=XM_019827716.2
A3	118328082	118328247
refGene_gn=LCLAT1;refGene_tx=XM_003984353.5;refGene_gn=LCLAT1;refGene_tx=XM_011281295.3;refGene_gn=LCLAT1;refGene_tx=XM_019827716.2
A3	118559830	118560019
refGene_gn=LBH;refGene_tx=XM_003984354.4;refGene_gn=LBH;refGene_tx=XM_019827719.1
A3	118570237	118570411
refGene_gn=LBH;refGene_tx=XM_019827720.1
A3	118581401	118581504
refGene_gn=LBH;refGene_tx=XM_003984354.4;refGene_gn=LBH;refGene_tx=XM_019827719.1;refGene_gn=LBH;refGene_tx=XM_019827720.1
A3	118581829	118581849
refGene_gn=LBH;refGene_tx=XM_019827719.1
A3	118584123	118584149
refGene_gn=LBH;refGene_tx=XM_003984354.4;refGene_gn=LBH;refGene_tx=XM_019827720.1
A3	118654458	118654683
refGene_gn=YPEL5;refGene_tx=XM_003984355.5;refGene_gn=YPEL5;refGene_tx=XM_011281293.2;refGene_gn=YPEL5;refGene_tx=XM_019827721.2;refGene_gn=YPEL5;refGene_tx=XM_023251977.1
A3	118656202	118656343
refGene_gn=YPEL5;refGene_tx=XM_003984355.5;refGene_gn=YPEL5;refGene_tx=XM_011281293.2;refGene_gn=YPEL5;refGene_tx=XM_019827721.2;refGene_gn=YPEL5;refGene_tx=XM_023251977.1
A3	118665987	118666005
refGene_gn=LOC109498313;refGene_tx=XM_023252201.1
A3	118668238	118668395
refGene_gn=LOC109498313;refGene_tx=XM_023252201.1
A3	118668487	118669365
refGene_gn=LOC109498313;refGene_tx=XM_023252201.1
A3	118883093	118883787
refGene_gn=ALK;refGene_tx=XM_003984359.5
A3	119080948	119081068
refGene_gn=ALK;refGene_tx=XM_003984359.5
A3	119101507	119101672
refGene_gn=ALK;refGene_tx=XM_003984359.5
A3	119243425	119243627
refGene_gn=ALK;refGene_tx=XM_003984359.5
A3	119374509	119374637
refGene_gn=ALK;refGene_tx=XM_003984359.5
A3	119432366	119432498
refGene_gn=ALK;refGene_tx=XM_003984359.5
A3	119439956	119440088
refGene_gn=ALK;refGene_tx=XM_003984359.5

A3 119442392 119442493
refGene_gn=ALK;refGene_tx=XM_003984359.5
A3 119461297 119461467
refGene_gn=ALK;refGene_tx=XM_003984359.5
A3 119476472 119476567
refGene_gn=ALK;refGene_tx=XM_003984359.5
A3 119476738 119476867
refGene_gn=ALK;refGene_tx=XM_003984359.5
A3 119502150 119502313
refGene_gn=ALK;refGene_tx=XM_003984359.5
A3 119512761 119512912
refGene_gn=ALK;refGene_tx=XM_003984359.5
A3 119519081 119519213
refGene_gn=ALK;refGene_tx=XM_003984359.5
A3 119519957 119520102
refGene_gn=ALK;refGene_tx=XM_003984359.5
A3 119522659 119522842
refGene_gn=ALK;refGene_tx=XM_003984359.5
A3 119524012 119524111
refGene_gn=ALK;refGene_tx=XM_003984359.5;refGene_gn=ALK;refGene_tx=XM_023251978.1;refGene_gn=ALK;refGene_tx=XM_023251980.1
A3 119524626 119524779
refGene_gn=ALK;refGene_tx=XM_003984359.5;refGene_gn=ALK;refGene_tx=XM_023251978.1;refGene_gn=ALK;refGene_tx=XM_023251980.1
A3 119525836 119525941
refGene_gn=ALK;refGene_tx=XM_003984359.5;refGene_gn=ALK;refGene_tx=XM_023251978.1;refGene_gn=ALK;refGene_tx=XM_023251980.1
A3 119526837 119527024
refGene_gn=ALK;refGene_tx=XM_003984359.5;refGene_gn=ALK;refGene_tx=XM_023251978.1;refGene_gn=ALK;refGene_tx=XM_023251980.1
A3 119527729 119527820
refGene_gn=ALK;refGene_tx=XM_003984359.5;refGene_gn=ALK;refGene_tx=XM_023251978.1;refGene_gn=ALK;refGene_tx=XM_023251980.1
A3 119527952 119528017
refGene_gn=ALK;refGene_tx=XM_003984359.5;refGene_gn=ALK;refGene_tx=XM_023251978.1;refGene_gn=ALK;refGene_tx=XM_023251980.1
A3 119529746 119529876
refGene_gn=ALK;refGene_tx=XM_003984359.5;refGene_gn=ALK;refGene_tx=XM_023251978.1;refGene_gn=ALK;refGene_tx=XM_023251980.1
A3 119535403 119535501
refGene_gn=ALK;refGene_tx=XM_003984359.5;refGene_gn=ALK;refGene_tx=XM_023251978.1;refGene_gn=ALK;refGene_tx=XM_023251980.1
A3 119539132 119539225
refGene_gn=ALK;refGene_tx=XM_003984359.5;refGene_gn=ALK;refGene_tx=XM_023251978.1;refGene_gn=ALK;refGene_tx=XM_023251980.1
A3 119541598 119541700
refGene_gn=ALK;refGene_tx=XM_003984359.5;refGene_gn=ALK;refGene_tx=XM_023251978.1;refGene_gn=ALK;refGene_tx=XM_023251980.1
A3 119551063 119551198
refGene_gn=ALK;refGene_tx=XM_003984359.5;refGene_gn=ALK;refGene_tx=XM_

023251978.1;refGene_gn=ALK;refGene_tx=XM_023251980.1
A3 119551898 119551989
refGene_gn=ALK;refGene_tx=XM_003984359.5;refGene_gn=ALK;refGene_tx=XM_
023251978.1;refGene_gn=ALK;refGene_tx=XM_023251980.1
A3 119554960 119555647
refGene_gn=ALK;refGene_tx=XM_003984359.5;refGene_gn=ALK;refGene_tx=XM_
023251978.1;refGene_gn=ALK;refGene_tx=XM_023251980.1
A3 119565821 119566143
refGene_gn=CLIP4;refGene_tx=XM_006930385.4;refGene_gn=CLIP4;refGene_tx
=XM_011281292.3;refGene_gn=CLIP4;refGene_tx=XM_023251981.1;refGene_gn=
CLIP4;refGene_tx=XM_023251982.1;refGene_gn=CLIP4;refGene_tx=XM_0232519
83.1
A3 119571502 119571579
refGene_gn=CLIP4;refGene_tx=XM_006930385.4;refGene_gn=CLIP4;refGene_tx
=XM_011281292.3;refGene_gn=CLIP4;refGene_tx=XM_023251981.1;refGene_gn=
CLIP4;refGene_tx=XM_023251982.1;refGene_gn=CLIP4;refGene_tx=XM_0232519
83.1;refGene_gn=CLIP4;refGene_tx=XM_023251985.1
A3 119578185 119578250
refGene_gn=CLIP4;refGene_tx=XM_006930385.4;refGene_gn=CLIP4;refGene_tx
=XM_011281292.3;refGene_gn=CLIP4;refGene_tx=XM_023251981.1;refGene_gn=
CLIP4;refGene_tx=XM_023251982.1;refGene_gn=CLIP4;refGene_tx=XM_0232519
83.1;refGene_gn=CLIP4;refGene_tx=XM_023251985.1
A3 119582272 119582396
refGene_gn=CLIP4;refGene_tx=XM_006930385.4;refGene_gn=CLIP4;refGene_tx
=XM_011281292.3;refGene_gn=CLIP4;refGene_tx=XM_023251981.1;refGene_gn=
CLIP4;refGene_tx=XM_023251982.1;refGene_gn=CLIP4;refGene_tx=XM_0232519
83.1;refGene_gn=CLIP4;refGene_tx=XM_023251985.1
A3 119585844 119585979
refGene_gn=CLIP4;refGene_tx=XM_006930385.4;refGene_gn=CLIP4;refGene_tx
=XM_011281292.3;refGene_gn=CLIP4;refGene_tx=XM_023251981.1;refGene_gn=
CLIP4;refGene_tx=XM_023251982.1;refGene_gn=CLIP4;refGene_tx=XM_0232519
83.1;refGene_gn=CLIP4;refGene_tx=XM_023251985.1
A3 119589588 119589732
refGene_gn=CLIP4;refGene_tx=XM_006930385.4;refGene_gn=CLIP4;refGene_tx
=XM_011281292.3;refGene_gn=CLIP4;refGene_tx=XM_023251981.1;refGene_gn=
CLIP4;refGene_tx=XM_023251982.1;refGene_gn=CLIP4;refGene_tx=XM_0232519
83.1;refGene_gn=CLIP4;refGene_tx=XM_023251985.1
A3 119590512 119590602
refGene_gn=CLIP4;refGene_tx=XM_006930385.4;refGene_gn=CLIP4;refGene_tx
=XM_011281292.3;refGene_gn=CLIP4;refGene_tx=XM_023251981.1;refGene_gn=
CLIP4;refGene_tx=XM_023251982.1;refGene_gn=CLIP4;refGene_tx=XM_0232519
83.1;refGene_gn=CLIP4;refGene_tx=XM_023251985.1
A3 119594833 119594971
refGene_gn=CLIP4;refGene_tx=XM_006930385.4;refGene_gn=CLIP4;refGene_tx
=XM_011281292.3;refGene_gn=CLIP4;refGene_tx=XM_023251981.1;refGene_gn=
CLIP4;refGene_tx=XM_023251982.1;refGene_gn=CLIP4;refGene_tx=XM_0232519
83.1;refGene_gn=CLIP4;refGene_tx=XM_023251985.1
A3 119601893 119602029
refGene_gn=CLIP4;refGene_tx=XM_006930385.4;refGene_gn=CLIP4;refGene_tx
=XM_011281292.3;refGene_gn=CLIP4;refGene_tx=XM_023251981.1;refGene_gn=

CLIP4;refGene_tx=XM_023251982.1;refGene_gn=CLIP4;refGene_tx=XM_0232519
83.1;refGene_gn=CLIP4;refGene_tx=XM_023251985.1
A3 119606137 119606374
refGene_gn=CLIP4;refGene_tx=XM_006930385.4;refGene_gn=CLIP4;refGene_tx
=XM_011281292.3;refGene_gn=CLIP4;refGene_tx=XM_023251981.1;refGene_gn=
CLIP4;refGene_tx=XM_023251982.1;refGene_gn=CLIP4;refGene_tx=XM_0232519
83.1;refGene_gn=CLIP4;refGene_tx=XM_023251985.1
A3 119609536 119609655
refGene_gn=CLIP4;refGene_tx=XM_006930385.4;refGene_gn=CLIP4;refGene_tx
=XM_011281292.3;refGene_gn=CLIP4;refGene_tx=XM_023251981.1;refGene_gn=
CLIP4;refGene_tx=XM_023251982.1;refGene_gn=CLIP4;refGene_tx=XM_0232519
83.1;refGene_gn=CLIP4;refGene_tx=XM_023251985.1
A3 119611429 119611591
refGene_gn=CLIP4;refGene_tx=XM_006930385.4;refGene_gn=CLIP4;refGene_tx
=XM_011281292.3;refGene_gn=CLIP4;refGene_tx=XM_023251981.1;refGene_gn=
CLIP4;refGene_tx=XM_023251982.1;refGene_gn=CLIP4;refGene_tx=XM_0232519
83.1;refGene_gn=CLIP4;refGene_tx=XM_023251985.1
A3 119613135 119613229
refGene_gn=CLIP4;refGene_tx=XM_006930385.4;refGene_gn=CLIP4;refGene_tx
=XM_011281292.3;refGene_gn=CLIP4;refGene_tx=XM_023251981.1;refGene_gn=
CLIP4;refGene_tx=XM_023251982.1;refGene_gn=CLIP4;refGene_tx=XM_0232519
83.1;refGene_gn=CLIP4;refGene_tx=XM_023251985.1
A3 119614026 119614166
refGene_gn=CLIP4;refGene_tx=XM_006930385.4;refGene_gn=CLIP4;refGene_tx
=XM_023251981.1;refGene_gn=CLIP4;refGene_tx=XM_023251982.1;refGene_gn=
CLIP4;refGene_tx=XM_023251983.1;refGene_gn=CLIP4;refGene_tx=XM_0232519
85.1
A3 119622141 119622274
refGene_gn=CLIP4;refGene_tx=XM_006930385.4;refGene_gn=CLIP4;refGene_tx
=XM_023251981.1;refGene_gn=CLIP4;refGene_tx=XM_023251982.1;refGene_gn=
CLIP4;refGene_tx=XM_023251983.1;refGene_gn=CLIP4;refGene_tx=XM_0232519
85.1
A3 119628841 119628850
refGene_gn=LOC111559902;refGene_tx=XM_023252202.1
A3 119629697 119630441
refGene_gn=LOC111559902;refGene_tx=XM_023252202.1
A3 119655007 119658738
refGene_gn=CA3H2orf71;refGene_tx=XM_003984440.4;refGene_gn=CA3H2orf71;
refGene_tx=XM_019827723.2
A3 119663818 119664002
refGene_gn=CA3H2orf71;refGene_tx=XM_003984440.4;refGene_gn=CA3H2orf71;
refGene_tx=XM_019827723.2
A3 119751559 119751751
refGene_gn=WDR43;refGene_tx=XM_023251986.1
A3 119751855 119751902
refGene_gn=WDR43;refGene_tx=XM_023251986.1
A3 119754164 119754234
refGene_gn=WDR43;refGene_tx=XM_023251986.1
A3 119754696 119754810
refGene_gn=WDR43;refGene_tx=XM_023251986.1

A3 119757349 119757413
refGene_gn=WDR43;refGene_tx=XM_023251986.1
A3 119758414 119758446
refGene_gn=WDR43;refGene_tx=XM_023251986.1
A3 119759062 119759149
refGene_gn=WDR43;refGene_tx=XM_023251986.1
A3 119763991 119764123
refGene_gn=WDR43;refGene_tx=XM_023251986.1
A3 119765582 119765714
refGene_gn=WDR43;refGene_tx=XM_023251986.1
A3 119766818 119766905
refGene_gn=WDR43;refGene_tx=XM_023251986.1
A3 119768426 119768598
refGene_gn=WDR43;refGene_tx=XM_023251986.1
A3 119771545 119771610
refGene_gn=WDR43;refGene_tx=XM_023251986.1
A3 119772378 119772481
refGene_gn=WDR43;refGene_tx=XM_023251986.1
A3 119780090 119780230
refGene_gn=WDR43;refGene_tx=XM_023251986.1
A3 119781254 119781375
refGene_gn=WDR43;refGene_tx=XM_023251986.1
A3 119786209 119786331
refGene_gn=WDR43;refGene_tx=XM_023251986.1
A3 119791333 119791471
refGene_gn=WDR43;refGene_tx=XM_023251986.1
A3 119798266 119798491
refGene_gn=WDR43;refGene_tx=XM_023251986.1
A3 119807925 119808924
refGene_gn=TRMT61B;refGene_tx=XM_019827725.2;refGene_gn=TRMT61B;refGene_tx=XM_023251987.1
A3 119810059 119810162
refGene_gn=TRMT61B;refGene_tx=XM_019827725.2;refGene_gn=TRMT61B;refGene_tx=XM_023251987.1
A3 119812851 119812863
refGene_gn=TRMT61B;refGene_tx=XM_023251987.1
A3 119813071 119813250
refGene_gn=TRMT61B;refGene_tx=XM_023251987.1
A3 119817105 119817209
refGene_gn=TRMT61B;refGene_tx=XM_019827725.2
A3 119818182 119818274
refGene_gn=TRMT61B;refGene_tx=XM_019827725.2;refGene_gn=TRMT61B;refGene_tx=XM_023251987.1
A3 119818942 119819169
refGene_gn=TRMT61B;refGene_tx=XM_019827725.2;refGene_gn=TRMT61B;refGene_tx=XM_023251987.1
A3 119819874 119819952
refGene_gn=TRMT61B;refGene_tx=XM_019827725.2;refGene_gn=TRMT61B;refGene_tx=XM_023251987.1
A3 119820036 119820080

refGene_gn=TRMT61B;refGene_tx=XM_019827725.2;refGene_gn=TRMT61B;refGene_tx=XM_023251987.1
A3 119820332 119820421
refGene_gn=SPDYA;refGene_tx=XM_003984363.5;refGene_gn=SPDYA;refGene_tx=XM_023251988.1;refGene_gn=SPDYA;refGene_tx=XM_023251989.1;refGene_gn=SPDYA;refGene_tx=XM_023251990.1;refGene_gn=SPDYA;refGene_tx=XM_023251991.1
A3 119824555 119824575
refGene_gn=SPDYA;refGene_tx=XM_019827726.2
A3 119827341 119827630
refGene_gn=SPDYA;refGene_tx=XM_003984363.5;refGene_gn=SPDYA;refGene_tx=XM_019827726.2;refGene_gn=SPDYA;refGene_tx=XM_023251988.1;refGene_gn=SPDYA;refGene_tx=XM_023251989.1;refGene_gn=SPDYA;refGene_tx=XM_023251990.1;refGene_gn=SPDYA;refGene_tx=XM_023251991.1
A3 119841097 119841269
refGene_gn=SPDYA;refGene_tx=XM_003984363.5;refGene_gn=SPDYA;refGene_tx=XM_019827726.2;refGene_gn=SPDYA;refGene_tx=XM_023251988.1;refGene_gn=SPDYA;refGene_tx=XM_023251989.1;refGene_gn=SPDYA;refGene_tx=XM_023251990.1;refGene_gn=SPDYA;refGene_tx=XM_023251991.1
A3 119846845 119846931
refGene_gn=SPDYA;refGene_tx=XM_003984363.5;refGene_gn=SPDYA;refGene_tx=XM_019827726.2;refGene_gn=SPDYA;refGene_tx=XM_023251988.1;refGene_gn=SPDYA;refGene_tx=XM_023251989.1;refGene_gn=SPDYA;refGene_tx=XM_023251990.1;refGene_gn=SPDYA;refGene_tx=XM_023251991.1
A3 119849051 119849110
refGene_gn=SPDYA;refGene_tx=XM_003984363.5;refGene_gn=SPDYA;refGene_tx=XM_019827726.2;refGene_gn=SPDYA;refGene_tx=XM_023251988.1;refGene_gn=SPDYA;refGene_tx=XM_023251989.1;refGene_gn=SPDYA;refGene_tx=XM_023251990.1;refGene_gn=SPDYA;refGene_tx=XM_023251991.1
A3 119852201 119852454
refGene_gn=SPDYA;refGene_tx=XM_003984363.5;refGene_gn=SPDYA;refGene_tx=XM_019827726.2;refGene_gn=SPDYA;refGene_tx=XM_023251988.1;refGene_gn=SPDYA;refGene_tx=XM_023251989.1;refGene_gn=SPDYA;refGene_tx=XM_023251990.1;refGene_gn=SPDYA;refGene_tx=XM_023251991.1
A3 119853537 119853555
refGene_gn=SPDYA;refGene_tx=XM_003984363.5;refGene_gn=SPDYA;refGene_tx=XM_019827726.2
A3 119868138 119868243
refGene_gn=PPP1CB;refGene_tx=XM_006930392.4;refGene_gn=PPP1CB;refGene_tx=XM_019827728.2
A3 119871022 119871157
refGene_gn=PPP1CB;refGene_tx=XM_006930392.4;refGene_gn=PPP1CB;refGene_tx=XM_019827728.2
A3 119874013 119874165
refGene_gn=PPP1CB;refGene_tx=XM_006930392.4;refGene_gn=PPP1CB;refGene_tx=XM_019827728.2
A3 119879397 119879469
refGene_gn=PPP1CB;refGene_tx=XM_006930392.4;refGene_gn=PPP1CB;refGene_tx=XM_019827728.2
A3 119881813 119881918

refGene_gn=PPP1CB;refGene_tx=XM_006930392.4;refGene_gn=PPP1CB;refGene_tx=XM_019827728.2
A3 119884029 119884260
refGene_gn=PPP1CB;refGene_tx=XM_006930392.4;refGene_gn=PPP1CB;refGene_tx=XM_019827728.2
A3 119886140 119886272
refGene_gn=PPP1CB;refGene_tx=XM_006930392.4;refGene_gn=PPP1CB;refGene_tx=XM_019827728.2
A3 119909616 119909668
refGene_gn=PPP1CB;refGene_tx=XM_006930392.4;refGene_gn=PPP1CB;refGene_tx=XM_019827728.2
A3 120014645 120014849
refGene_gn=PLB1;refGene_tx=XM_019827729.2;refGene_gn=PLB1;refGene_tx=XM_019827733.2;refGene_gn=PLB1;refGene_tx=XM_023251993.1
A3 120016242 120016317
refGene_gn=PLB1;refGene_tx=XM_019827729.2;refGene_gn=PLB1;refGene_tx=XM_019827733.2;refGene_gn=PLB1;refGene_tx=XM_023251993.1
A3 120022254 120022350
refGene_gn=PLB1;refGene_tx=XM_019827729.2;refGene_gn=PLB1;refGene_tx=XM_019827733.2;refGene_gn=PLB1;refGene_tx=XM_023251993.1
A3 120022819 120022924
refGene_gn=PLB1;refGene_tx=XM_019827729.2;refGene_gn=PLB1;refGene_tx=XM_019827733.2;refGene_gn=PLB1;refGene_tx=XM_023251993.1
A3 120024212 120024294
refGene_gn=PLB1;refGene_tx=XM_019827729.2;refGene_gn=PLB1;refGene_tx=XM_019827733.2;refGene_gn=PLB1;refGene_tx=XM_023251993.1
A3 120025477 120025572
refGene_gn=PLB1;refGene_tx=XM_019827729.2;refGene_gn=PLB1;refGene_tx=XM_019827733.2;refGene_gn=PLB1;refGene_tx=XM_023251993.1
A3 120026103 120026169
refGene_gn=PLB1;refGene_tx=XM_019827729.2;refGene_gn=PLB1;refGene_tx=XM_019827733.2;refGene_gn=PLB1;refGene_tx=XM_023251993.1
A3 120027530 120027611
refGene_gn=PLB1;refGene_tx=XM_019827729.2;refGene_gn=PLB1;refGene_tx=XM_019827733.2;refGene_gn=PLB1;refGene_tx=XM_023251993.1
A3 120028564 120028616
refGene_gn=PLB1;refGene_tx=XM_019827729.2;refGene_gn=PLB1;refGene_tx=XM_019827733.2;refGene_gn=PLB1;refGene_tx=XM_023251993.1
A3 120032250 120032350
refGene_gn=PLB1;refGene_tx=XM_019827729.2;refGene_gn=PLB1;refGene_tx=XM_019827733.2;refGene_gn=PLB1;refGene_tx=XM_023251993.1
A3 120032562 120032606
refGene_gn=PLB1;refGene_tx=XM_019827729.2;refGene_gn=PLB1;refGene_tx=XM_019827733.2;refGene_gn=PLB1;refGene_tx=XM_023251993.1
A3 120032839 120032907
refGene_gn=PLB1;refGene_tx=XM_019827729.2;refGene_gn=PLB1;refGene_tx=XM_019827733.2;refGene_gn=PLB1;refGene_tx=XM_023251993.1
A3 120034841 120034900
refGene_gn=PLB1;refGene_tx=XM_019827729.2;refGene_gn=PLB1;refGene_tx=XM_019827733.2;refGene_gn=PLB1;refGene_tx=XM_023251993.1

A3	120035658	120035719
refGene_gn=PLB1;refGene_tx=XM_019827729.2;refGene_gn=PLB1;refGene_tx=XM_019827733.2;refGene_gn=PLB1;refGene_tx=XM_023251993.1
A3	120038884	120038950
refGene_gn=PLB1;refGene_tx=XM_019827729.2;refGene_gn=PLB1;refGene_tx=XM_019827733.2;refGene_gn=PLB1;refGene_tx=XM_023251993.1
A3	120044878	120044950
refGene_gn=PLB1;refGene_tx=XM_019827729.2;refGene_gn=PLB1;refGene_tx=XM_019827733.2;refGene_gn=PLB1;refGene_tx=XM_023251993.1
A3	120045484	120045580
refGene_gn=PLB1;refGene_tx=XM_019827729.2;refGene_gn=PLB1;refGene_tx=XM_019827733.2;refGene_gn=PLB1;refGene_tx=XM_023251993.1
A3	120046688	120046793
refGene_gn=PLB1;refGene_tx=XM_019827729.2;refGene_gn=PLB1;refGene_tx=XM_019827733.2;refGene_gn=PLB1;refGene_tx=XM_023251993.1
A3	120047457	120047539
refGene_gn=PLB1;refGene_tx=XM_019827729.2;refGene_gn=PLB1;refGene_tx=XM_019827733.2;refGene_gn=PLB1;refGene_tx=XM_023251993.1
A3	120048580	120048681
refGene_gn=PLB1;refGene_tx=XM_019827729.2;refGene_gn=PLB1;refGene_tx=XM_019827733.2;refGene_gn=PLB1;refGene_tx=XM_023251993.1
A3	120049503	120049569
refGene_gn=PLB1;refGene_tx=XM_019827729.2;refGene_gn=PLB1;refGene_tx=XM_019827733.2;refGene_gn=PLB1;refGene_tx=XM_023251993.1
A3	120050124	120050205
refGene_gn=PLB1;refGene_tx=XM_019827729.2;refGene_gn=PLB1;refGene_tx=XM_019827733.2;refGene_gn=PLB1;refGene_tx=XM_023251993.1
A3	120050593	120050645
refGene_gn=PLB1;refGene_tx=XM_019827729.2;refGene_gn=PLB1;refGene_tx=XM_019827733.2;refGene_gn=PLB1;refGene_tx=XM_023251993.1
A3	120052094	120052203
refGene_gn=PLB1;refGene_tx=XM_019827729.2;refGene_gn=PLB1;refGene_tx=XM_019827733.2;refGene_gn=PLB1;refGene_tx=XM_023251993.1
A3	120052851	120052895
refGene_gn=PLB1;refGene_tx=XM_019827729.2;refGene_gn=PLB1;refGene_tx=XM_019827733.2;refGene_gn=PLB1;refGene_tx=XM_023251993.1
A3	120057943	120058017
refGene_gn=PLB1;refGene_tx=XM_019827729.2;refGene_gn=PLB1;refGene_tx=XM_019827733.2;refGene_gn=PLB1;refGene_tx=XM_023251993.1
A3	120058895	120058954
refGene_gn=PLB1;refGene_tx=XM_019827729.2;refGene_gn=PLB1;refGene_tx=XM_019827733.2;refGene_gn=PLB1;refGene_tx=XM_023251993.1
A3	120059832	120059893
refGene_gn=PLB1;refGene_tx=XM_019827729.2;refGene_gn=PLB1;refGene_tx=XM_019827733.2;refGene_gn=PLB1;refGene_tx=XM_023251993.1
A3	120060429	120060468
refGene_gn=PLB1;refGene_tx=XM_019827729.2;refGene_gn=PLB1;refGene_tx=XM_019827733.2;refGene_gn=PLB1;refGene_tx=XM_023251993.1
A3	120061432	120061504
refGene_gn=PLB1;refGene_tx=XM_019827729.2;refGene_gn=PLB1;refGene_tx=X

M_019827733.2;refGene_gn=PLB1;refGene_tx=XM_023251993.1
A3 120061723 120061819
refGene_gn=PLB1;refGene_tx=XM_019827729.2;refGene_gn=PLB1;refGene_tx=XM_019827733.2;refGene_gn=PLB1;refGene_tx=XM_023251993.1
A3 120061941 120062046
refGene_gn=PLB1;refGene_tx=XM_019827729.2;refGene_gn=PLB1;refGene_tx=XM_019827733.2;refGene_gn=PLB1;refGene_tx=XM_023251993.1
A3 120064693 120064775
refGene_gn=PLB1;refGene_tx=XM_019827729.2;refGene_gn=PLB1;refGene_tx=XM_019827733.2;refGene_gn=PLB1;refGene_tx=XM_023251993.1
A3 120066769 120066870
refGene_gn=PLB1;refGene_tx=XM_019827729.2;refGene_gn=PLB1;refGene_tx=XM_019827733.2;refGene_gn=PLB1;refGene_tx=XM_023251993.1
A3 120067157 120067223
refGene_gn=PLB1;refGene_tx=XM_019827729.2;refGene_gn=PLB1;refGene_tx=XM_019827733.2;refGene_gn=PLB1;refGene_tx=XM_023251993.1
A3 120070115 120070196
refGene_gn=PLB1;refGene_tx=XM_019827729.2;refGene_gn=PLB1;refGene_tx=XM_019827733.2;refGene_gn=PLB1;refGene_tx=XM_023251993.1
A3 120070533 120070585
refGene_gn=PLB1;refGene_tx=XM_019827729.2;refGene_gn=PLB1;refGene_tx=XM_019827733.2;refGene_gn=PLB1;refGene_tx=XM_023251993.1
A3 120073761 120073870
refGene_gn=PLB1;refGene_tx=XM_019827729.2;refGene_gn=PLB1;refGene_tx=XM_019827733.2;refGene_gn=PLB1;refGene_tx=XM_023251993.1
A3 120078283 120078327
refGene_gn=PLB1;refGene_tx=XM_019827729.2;refGene_gn=PLB1;refGene_tx=XM_019827733.2;refGene_gn=PLB1;refGene_tx=XM_023251993.1
A3 120079832 120079906
refGene_gn=PLB1;refGene_tx=XM_019827729.2;refGene_gn=PLB1;refGene_tx=XM_019827733.2;refGene_gn=PLB1;refGene_tx=XM_023251993.1
A3 120081884 120081943
refGene_gn=PLB1;refGene_tx=XM_019827729.2;refGene_gn=PLB1;refGene_tx=XM_019827733.2;refGene_gn=PLB1;refGene_tx=XM_023251993.1
A3 120087635 120087699
refGene_gn=PLB1;refGene_tx=XM_019827729.2;refGene_gn=PLB1;refGene_tx=XM_019827733.2;refGene_gn=PLB1;refGene_tx=XM_023251993.1
A3 120089842 120089917
refGene_gn=PLB1;refGene_tx=XM_019827729.2;refGene_gn=PLB1;refGene_tx=XM_019827733.2;refGene_gn=PLB1;refGene_tx=XM_023251993.1
A3 120090775 120090847
refGene_gn=PLB1;refGene_tx=XM_019827729.2;refGene_gn=PLB1;refGene_tx=XM_019827733.2;refGene_gn=PLB1;refGene_tx=XM_023251993.1
A3 120096090 120096147
refGene_gn=PLB1;refGene_tx=XM_019827729.2;refGene_gn=PLB1;refGene_tx=XM_019827733.2;refGene_gn=PLB1;refGene_tx=XM_023251993.1
A3 120097222 120097318
refGene_gn=PLB1;refGene_tx=XM_019827729.2;refGene_gn=PLB1;refGene_tx=XM_019827733.2;refGene_gn=PLB1;refGene_tx=XM_023251993.1
A3 120098701 120098777

refGene_gn=PLB1;refGene_tx=XM_019827729.2;refGene_gn=PLB1;refGene_tx=XM_019827733.2;refGene_gn=PLB1;refGene_tx=XM_023251993.1
A3 120099062 120099142
refGene_gn=PLB1;refGene_tx=XM_019827729.2;refGene_gn=PLB1;refGene_tx=XM_019827733.2;refGene_gn=PLB1;refGene_tx=XM_023251993.1
A3 120099893 120099956
refGene_gn=PLB1;refGene_tx=XM_019827729.2;refGene_gn=PLB1;refGene_tx=XM_019827733.2;refGene_gn=PLB1;refGene_tx=XM_023251993.1
A3 120102853 120102940
refGene_gn=PLB1;refGene_tx=XM_019827729.2;refGene_gn=PLB1;refGene_tx=XM_019827733.2;refGene_gn=PLB1;refGene_tx=XM_023251993.1
A3 120104797 120104849
refGene_gn=PLB1;refGene_tx=XM_019827729.2;refGene_gn=PLB1;refGene_tx=XM_019827733.2;refGene_gn=PLB1;refGene_tx=XM_023251993.1
A3 120105191 120105282
refGene_gn=PLB1;refGene_tx=XM_019827729.2
A3 120108117 120108161
refGene_gn=PLB1;refGene_tx=XM_019827729.2;refGene_gn=PLB1;refGene_tx=XM_023251993.1
A3 120108734 120108775
refGene_gn=PLB1;refGene_tx=XM_019827729.2;refGene_gn=PLB1;refGene_tx=XM_023251993.1
A3 120112802 120112861
refGene_gn=PLB1;refGene_tx=XM_019827729.2
A3 120114000 120114067
refGene_gn=PLB1;refGene_tx=XM_019827729.2
A3 120115194 120115256
refGene_gn=PLB1;refGene_tx=XM_019827729.2
A3 120136251 120136306
refGene_gn=PLB1;refGene_tx=XM_019827729.2
A3 120223005 120223524
refGene_gn=FOSL2;refGene_tx=XM_003984367.4;refGene_gn=FOSL2;refGene_tx=XM_019827735.2;refGene_gn=FOSL2;refGene_tx=XM_019827736.2
A3 120226204 120226312
refGene_gn=FOSL2;refGene_tx=XM_003984367.4;refGene_gn=FOSL2;refGene_tx=XM_019827735.2;refGene_gn=FOSL2;refGene_tx=XM_019827736.2
A3 120231786 120232038
refGene_gn=FOSL2;refGene_tx=XM_003984367.4;refGene_gn=FOSL2;refGene_tx=XM_019827735.2;refGene_gn=FOSL2;refGene_tx=XM_019827736.2
A3 120241833 120241932
refGene_gn=FOSL2;refGene_tx=XM_003984367.4
A3 120294768 120294832
refGene_gn=BABAM2;refGene_tx=XM_011281289.2
A3 120295085 120295194
refGene_gn=BABAM2;refGene_tx=XM_019827747.2
A3 120297214 120297353
refGene_gn=BABAM2;refGene_tx=XM_019827744.2;refGene_gn=BABAM2;refGene_tx=XM_019827745.2;refGene_gn=BABAM2;refGene_tx=XM_019827746.2;refGene_gn=BABAM2;refGene_tx=XM_023251995.1;refGene_gn=BABAM2;refGene_tx=XM_023251996.1

A3 120328370 120328524
refGene_gn=BABAM2;refGene_tx=XM_011281289.2;refGene_gn=BABAM2;refGene_tx=XM_019827744.2;refGene_gn=BABAM2;refGene_tx=XM_019827745.2;refGene_gn=BABAM2;refGene_tx=XM_019827746.2;refGene_gn=BABAM2;refGene_tx=XM_019827747.2;refGene_gn=BABAM2;refGene_tx=XM_023251995.1;refGene_gn=BABAM2;refGene_tx=XM_023251996.1
A3 120376059 120376142
refGene_gn=BABAM2;refGene_tx=XM_011281289.2;refGene_gn=BABAM2;refGene_tx=XM_019827744.2;refGene_gn=BABAM2;refGene_tx=XM_019827745.2;refGene_gn=BABAM2;refGene_tx=XM_019827746.2;refGene_gn=BABAM2;refGene_tx=XM_019827747.2;refGene_gn=BABAM2;refGene_tx=XM_023251995.1;refGene_gn=BABAM2;refGene_tx=XM_023251996.1
A3 120378699 120378770
refGene_gn=BABAM2;refGene_tx=XM_011281289.2;refGene_gn=BABAM2;refGene_tx=XM_019827744.2;refGene_gn=BABAM2;refGene_tx=XM_019827745.2;refGene_gn=BABAM2;refGene_tx=XM_019827746.2;refGene_gn=BABAM2;refGene_tx=XM_019827747.2;refGene_gn=BABAM2;refGene_tx=XM_023251995.1;refGene_gn=BABAM2;refGene_tx=XM_023251996.1
A3 120382073 120382173
refGene_gn=BABAM2;refGene_tx=XM_011281289.2;refGene_gn=BABAM2;refGene_tx=XM_019827744.2;refGene_gn=BABAM2;refGene_tx=XM_019827745.2;refGene_gn=BABAM2;refGene_tx=XM_019827746.2;refGene_gn=BABAM2;refGene_tx=XM_019827747.2;refGene_gn=BABAM2;refGene_tx=XM_023251995.1;refGene_gn=BABAM2;refGene_tx=XM_023251996.1
A3 120477001 120477111
refGene_gn=BABAM2;refGene_tx=XM_011281289.2;refGene_gn=BABAM2;refGene_tx=XM_019827744.2;refGene_gn=BABAM2;refGene_tx=XM_019827745.2;refGene_gn=BABAM2;refGene_tx=XM_019827746.2;refGene_gn=BABAM2;refGene_tx=XM_019827747.2;refGene_gn=BABAM2;refGene_tx=XM_023251995.1;refGene_gn=BABAM2;refGene_tx=XM_023251996.1
A3 120551804 120551879
refGene_gn=BABAM2;refGene_tx=XM_011281289.2;refGene_gn=BABAM2;refGene_tx=XM_019827744.2;refGene_gn=BABAM2;refGene_tx=XM_019827745.2;refGene_gn=BABAM2;refGene_tx=XM_019827746.2;refGene_gn=BABAM2;refGene_tx=XM_019827747.2;refGene_gn=BABAM2;refGene_tx=XM_023251995.1;refGene_gn=BABAM2;refGene_tx=XM_023251996.1
A3 120571963 120572158
refGene_gn=BABAM2;refGene_tx=XM_011281289.2;refGene_gn=BABAM2;refGene_tx=XM_019827744.2;refGene_gn=BABAM2;refGene_tx=XM_019827745.2;refGene_gn=BABAM2;refGene_tx=XM_019827746.2;refGene_gn=BABAM2;refGene_tx=XM_019827747.2;refGene_gn=BABAM2;refGene_tx=XM_023251995.1;refGene_gn=BABAM2;refGene_tx=XM_023251996.1
A3 120605695 120605790
refGene_gn=BABAM2;refGene_tx=XM_011281289.2;refGene_gn=BABAM2;refGene_tx=XM_019827744.2;refGene_gn=BABAM2;refGene_tx=XM_019827745.2;refGene_gn=BABAM2;refGene_tx=XM_019827746.2;refGene_gn=BABAM2;refGene_tx=XM_019827747.2;refGene_gn=BABAM2;refGene_tx=XM_023251995.1;refGene_gn=BABAM2;refGene_tx=XM_023251996.1
A3 120653106 120653183
refGene_gn=BABAM2;refGene_tx=XM_011281289.2;refGene_gn=BABAM2;refGene_

tx=XM_019827744.2;refGene_gn=BABAM2;refGene_tx=XM_019827745.2;refGene_gn=BABAM2;refGene_tx=XM_019827746.2;refGene_gn=BABAM2;refGene_tx=XM_019827747.2;refGene_gn=BABAM2;refGene_tx=XM_023251995.1;refGene_gn=BABAM2;refGene_tx=XM_023251996.1
A3 120691070 120691198
refGene_gn=BABAM2;refGene_tx=XM_011281289.2;refGene_gn=BABAM2;refGene_tx=XM_019827744.2;refGene_gn=BABAM2;refGene_tx=XM_019827745.2;refGene_gn=BABAM2;refGene_tx=XM_019827746.2;refGene_gn=BABAM2;refGene_tx=XM_019827747.2;refGene_gn=BABAM2;refGene_tx=XM_023251995.1;refGene_gn=BABAM2;refGene_tx=XM_023251996.1
A3 120695030 120695119
refGene_gn=RBKS;refGene_tx=XM_003984371.4;refGene_gn=RBKS;refGene_tx=XM_023251998.1;refGene_gn=RBKS;refGene_tx=XM_023251999.1;refGene_gn=RBKS;refGene_tx=XM_023252000.1
A3 120711038 120711171
refGene_gn=RBKS;refGene_tx=XM_003984371.4;refGene_gn=RBKS;refGene_tx=XM_023251998.1;refGene_gn=RBKS;refGene_tx=XM_023251999.1;refGene_gn=RBKS;refGene_tx=XM_023252000.1
A3 120723350 120723414
refGene_gn=RBKS;refGene_tx=XM_003984371.4;refGene_gn=RBKS;refGene_tx=XM_023251998.1;refGene_gn=RBKS;refGene_tx=XM_023251999.1;refGene_gn=RBKS;refGene_tx=XM_023252000.1
A3 120724355 120724418
refGene_gn=RBKS;refGene_tx=XM_003984371.4;refGene_gn=RBKS;refGene_tx=XM_023251998.1;refGene_gn=RBKS;refGene_tx=XM_023251999.1;refGene_gn=RBKS;refGene_tx=XM_023252000.1
A3 120727756 120727921
refGene_gn=RBKS;refGene_tx=XM_003984371.4;refGene_gn=RBKS;refGene_tx=XM_023251998.1;refGene_gn=RBKS;refGene_tx=XM_023251999.1;refGene_gn=RBKS;refGene_tx=XM_023252000.1
A3 120735660 120735752
refGene_gn=RBKS;refGene_tx=XM_003984371.4;refGene_gn=RBKS;refGene_tx=XM_023251998.1;refGene_gn=RBKS;refGene_tx=XM_023251999.1;refGene_gn=RBKS;refGene_tx=XM_023252000.1
A3 120742902 120743091
refGene_gn=RBKS;refGene_tx=XM_003984371.4;refGene_gn=RBKS;refGene_tx=XM_023251998.1;refGene_gn=RBKS;refGene_tx=XM_023251999.1;refGene_gn=RBKS;refGene_tx=XM_023252000.1
A3 120744363 120744414
refGene_gn=RBKS;refGene_tx=XM_023251999.1
A3 120788850 120789024
refGene_gn=RBKS;refGene_tx=XM_003984371.4
A3 120791759 120791844
refGene_gn=MRPL33;refGene_tx=XM_003984372.3;refGene_gn=MRPL33;refGene_tx=XM_019827749.1;refGene_gn=MRPL33;refGene_tx=XM_023252001.1
A3 120796367 120796484
refGene_gn=RBKS;refGene_tx=XM_023251998.1;refGene_gn=RBKS;refGene_tx=XM_023252000.1
A3 120796781 120796942
refGene_gn=MRPL33;refGene_tx=XM_019827749.1

A3 120798224 120798331
refGene_gn=MRPL33;refGene_tx=XM_003984372.3;refGene_gn=MRPL33;refGene_tx=XM_019827749.1
A3 120800534 120800553
refGene_gn=MRPL33;refGene_tx=XM_003984372.3;refGene_gn=MRPL33;refGene_tx=XM_019827749.1;refGene_gn=MRPL33;refGene_tx=XM_023252001.1
A3 120801412 120801508
refGene_gn=MRPL33;refGene_tx=XM_003984372.3;refGene_gn=MRPL33;refGene_tx=XM_019827749.1;refGene_gn=MRPL33;refGene_tx=XM_023252001.1
A3 120803487 120803512
refGene_gn=MRPL33;refGene_tx=XM_023252001.1
A3 120868130 120868180
refGene_gn=SLC4A1AP;refGene_tx=XM_011281288.3
A3 120870267 120870337
refGene_gn=SLC4A1AP;refGene_tx=XM_011281288.3
A3 120871982 120872050
refGene_gn=SLC4A1AP;refGene_tx=XM_011281288.3
A3 120873272 120873359
refGene_gn=SLC4A1AP;refGene_tx=XM_011281288.3
A3 120876071 120876312
refGene_gn=SLC4A1AP;refGene_tx=XM_011281288.3
A3 120878507 120878619
refGene_gn=SLC4A1AP;refGene_tx=XM_011281288.3
A3 120883962 120884149
refGene_gn=SLC4A1AP;refGene_tx=XM_011281288.3
A3 120884516 120884586
refGene_gn=SLC4A1AP;refGene_tx=XM_011281288.3
A3 120885384 120885545
refGene_gn=SLC4A1AP;refGene_tx=XM_011281288.3
A3 120893497 120893637
refGene_gn=SLC4A1AP;refGene_tx=XM_011281288.3
A3 120893968 120894029
refGene_gn=SLC4A1AP;refGene_tx=XM_011281288.3
A3 120895771 120895894
refGene_gn=SLC4A1AP;refGene_tx=XM_011281288.3
A3 120898135 120898331
refGene_gn=SLC4A1AP;refGene_tx=XM_011281288.3
A3 120898784 120899447
refGene_gn=SLC4A1AP;refGene_tx=XM_011281288.3
A3 120901701 120902114
refGene_gn=SUPT7L;refGene_tx=XM_006930396.3;refGene_gn=SUPT7L;refGene_tx=XM_006930397.3;refGene_gn=SUPT7L;refGene_tx=XM_006930398.3;refGene_gn=SUPT7L;refGene_tx=XM_019827748.2;refGene_gn=SUPT7L;refGene_tx=XM_023252003.1
A3 120909032 120909357
refGene_gn=SUPT7L;refGene_tx=XM_006930396.3;refGene_gn=SUPT7L;refGene_tx=XM_006930397.3;refGene_gn=SUPT7L;refGene_tx=XM_006930398.3;refGene_gn=SUPT7L;refGene_tx=XM_019827748.2;refGene_gn=SUPT7L;refGene_tx=XM_023252003.1
A3 120910389 120910627

refGene_gn=SUPT7L;refGene_tx=XM_006930396.3;refGene_gn=SUPT7L;refGene_tx=XM_006930397.3;refGene_gn=SUPT7L;refGene_tx=XM_006930398.3;refGene_gn=SUPT7L;refGene_tx=XM_019827748.2;refGene_gn=SUPT7L;refGene_tx=XM_023252003.1
A3 120912300 120912563
refGene_gn=SUPT7L;refGene_tx=XM_006930396.3;refGene_gn=SUPT7L;refGene_tx=XM_006930397.3;refGene_gn=SUPT7L;refGene_tx=XM_006930398.3;refGene_gn=SUPT7L;refGene_tx=XM_019827748.2;refGene_gn=SUPT7L;refGene_tx=XM_023252003.1
A3 120916463 120916549
refGene_gn=GPN1;refGene_tx=XM_003984375.4;refGene_gn=GPN1;refGene_tx=XM_023252004.1
A3 120918301 120918409
refGene_gn=GPN1;refGene_tx=XM_003984375.4;refGene_gn=GPN1;refGene_tx=XM_023252004.1
A3 120922696 120922787
refGene_gn=GPN1;refGene_tx=XM_003984375.4;refGene_gn=GPN1;refGene_tx=XM_023252004.1
A3 120923438 120923478
refGene_gn=GPN1;refGene_tx=XM_003984375.4;refGene_gn=GPN1;refGene_tx=XM_023252004.1
A3 120924973 120925056
refGene_gn=GPN1;refGene_tx=XM_003984375.4;refGene_gn=GPN1;refGene_tx=XM_023252004.1
A3 120926380 120926527
refGene_gn=GPN1;refGene_tx=XM_003984375.4;refGene_gn=GPN1;refGene_tx=XM_023252004.1
A3 120927166 120927212
refGene_gn=GPN1;refGene_tx=XM_003984375.4;refGene_gn=GPN1;refGene_tx=XM_023252004.1
A3 120928156 120928251
refGene_gn=GPN1;refGene_tx=XM_003984375.4;refGene_gn=GPN1;refGene_tx=XM_023252004.1
A3 120928460 120928539
refGene_gn=GPN1;refGene_tx=XM_003984375.4;refGene_gn=GPN1;refGene_tx=XM_023252004.1
A3 120931214 120931252
refGene_gn=GPN1;refGene_tx=XM_003984375.4;refGene_gn=GPN1;refGene_tx=XM_023252004.1
A3 120932133 120932200
refGene_gn=GPN1;refGene_tx=XM_023252004.1
A3 120933663 120933703
refGene_gn=GPN1;refGene_tx=XM_023252004.1
A3 120935883 120935977
refGene_gn=GPN1;refGene_tx=XM_003984375.4;refGene_gn=GPN1;refGene_tx=XM_023252004.1
A3 120936789 120936801
refGene_gn=GPN1;refGene_tx=XM_023252004.1
A3 120936991 120937461
refGene_gn=CCDC121;refGene_tx=XM_003984444.5

A3 120938153 120939120
refGene_gn=CCDC121;refGene_tx=XM_003984444.5
A3 120945031 120945340
refGene_gn=ZNF512;refGene_tx=XM_006930400.4;refGene_gn=ZNF512;refGene_tx=XM_006930401.4
A3 120949519 120949618
refGene_gn=ZNF512;refGene_tx=XM_006930400.4;refGene_gn=ZNF512;refGene_tx=XM_006930401.4
A3 120950779 120950842
refGene_gn=ZNF512;refGene_tx=XM_006930400.4;refGene_gn=ZNF512;refGene_tx=XM_006930401.4
A3 120951764 120951866
refGene_gn=ZNF512;refGene_tx=XM_006930400.4;refGene_gn=ZNF512;refGene_tx=XM_006930401.4
A3 120959258 120959453
refGene_gn=ZNF512;refGene_tx=XM_006930400.4;refGene_gn=ZNF512;refGene_tx=XM_006930401.4
A3 120962856 120963024
refGene_gn=ZNF512;refGene_tx=XM_006930400.4;refGene_gn=ZNF512;refGene_tx=XM_006930401.4
A3 120963636 120963735
refGene_gn=ZNF512;refGene_tx=XM_006930400.4;refGene_gn=ZNF512;refGene_tx=XM_006930401.4
A3 120964150 120964237
refGene_gn=ZNF512;refGene_tx=XM_006930400.4;refGene_gn=ZNF512;refGene_tx=XM_006930401.4
A3 120964759 120964884
refGene_gn=ZNF512;refGene_tx=XM_006930400.4;refGene_gn=ZNF512;refGene_tx=XM_006930401.4
A3 120965539 120965623
refGene_gn=ZNF512;refGene_tx=XM_006930400.4;refGene_gn=ZNF512;refGene_tx=XM_006930401.4
A3 120966302 120966398
refGene_gn=ZNF512;refGene_tx=XM_006930400.4;refGene_gn=ZNF512;refGene_tx=XM_006930401.4
A3 120967599 120967784
refGene_gn=ZNF512;refGene_tx=XM_006930400.4;refGene_gn=ZNF512;refGene_tx=XM_006930401.4
A3 120976987 120977046
refGene_gn=ZNF512;refGene_tx=XM_006930400.4;refGene_gn=ZNF512;refGene_tx=XM_006930401.4
A3 120977568 120977598
refGene_gn=ZNF512;refGene_tx=XM_006930400.4;refGene_gn=ZNF512;refGene_tx=XM_006930401.4
A3 120980580 120997030
refGene_gn=LOC109498324;refGene_tx=XM_019827751.2
A3 120997593 120997689
refGene_gn=LOC109498324;refGene_tx=XM_019827751.2
A3 120997898 120997956
refGene_gn=LOC109498324;refGene_tx=XM_019827751.2

A3 120998612 120998655
refGene_gn=LOC109498324;refGene_tx=XM_019827751.2
A3 121015634 121015778
refGene_gn=LOC109498324;refGene_tx=XM_019827751.2
A3 121032007 121032121
refGene_gn=GCKR;refGene_tx=XM_023250920.1
A3 121032220 121032267
refGene_gn=GCKR;refGene_tx=XM_023250920.1
A3 121032269 121032314
refGene_gn=GCKR;refGene_tx=XM_023250920.1
A3 121032506 121032603
refGene_gn=GCKR;refGene_tx=XM_023250920.1
A3 121032976 121033053
refGene_gn=GCKR;refGene_tx=XM_023250920.1
A3 121033441 121033539
refGene_gn=GCKR;refGene_tx=XM_023250920.1
A3 121033780 121033891
refGene_gn=GCKR;refGene_tx=XM_023250920.1
A3 121034533 121034639
refGene_gn=GCKR;refGene_tx=XM_023250920.1
A3 121034747 121034748
refGene_gn=GCKR;refGene_tx=XM_023250920.1
A3 121035845 121035951
refGene_gn=GCKR;refGene_tx=XM_023250920.1
A3 121038230 121038325
refGene_gn=GCKR;refGene_tx=XM_023250920.1
A3 121039302 121039356
refGene_gn=GCKR;refGene_tx=XM_023250920.1
A3 121039535 121039602
refGene_gn=GCKR;refGene_tx=XM_023250920.1
A3 121039957 121040010
refGene_gn=GCKR;refGene_tx=XM_023250920.1
A3 121040329 121040398
refGene_gn=GCKR;refGene_tx=XM_023250920.1
A3 121040399 121040408
refGene_gn=GCKR;refGene_tx=XM_023250920.1
A3 121041051 121041120
refGene_gn=GCKR;refGene_tx=XM_023250920.1
A3 121041310 121041445
refGene_gn=GCKR;refGene_tx=XM_023250920.1
A3 121041446 121041468
refGene_gn=GCKR;refGene_tx=XM_023250920.1
A3 121041764 121041823
refGene_gn=GCKR;refGene_tx=XM_023250920.1
A3 121044064 121044185
refGene_gn=FNDC4;refGene_tx=XM_003984380.5
A3 121044262 121044378
refGene_gn=FNDC4;refGene_tx=XM_003984380.5
A3 121044607 121044812
refGene_gn=FNDC4;refGene_tx=XM_003984380.5

A3 121045263 121045353
refGene_gn=FNDC4;refGene_tx=XM_003984380.5
A3 121045645 121045770
refGene_gn=FNDC4;refGene_tx=XM_003984380.5
A3 121045937 121045973
refGene_gn=FNDC4;refGene_tx=XM_003984380.5
A3 121048523 121048562
refGene_gn=IFT172;refGene_tx=XM_003984381.5
A3 121051531 121051675
refGene_gn=IFT172;refGene_tx=XM_003984381.5;refGene_gn=IFT172;refGene_tx=XM_019827754.2
A3 121051940 121052053
refGene_gn=IFT172;refGene_tx=XM_003984381.5;refGene_gn=IFT172;refGene_tx=XM_019827754.2
A3 121052493 121052533
refGene_gn=IFT172;refGene_tx=XM_003984381.5;refGene_gn=IFT172;refGene_tx=XM_019827754.2
A3 121052825 121052891
refGene_gn=IFT172;refGene_tx=XM_003984381.5;refGene_gn=IFT172;refGene_tx=XM_019827754.2
A3 121053099 121053179
refGene_gn=IFT172;refGene_tx=XM_003984381.5;refGene_gn=IFT172;refGene_tx=XM_019827754.2
A3 121053391 121053479
refGene_gn=IFT172;refGene_tx=XM_003984381.5;refGene_gn=IFT172;refGene_tx=XM_019827754.2;refGene_gn=IFT172;refGene_tx=XM_019827755.1
A3 121054382 121054597
refGene_gn=IFT172;refGene_tx=XM_003984381.5;refGene_gn=IFT172;refGene_tx=XM_019827754.2;refGene_gn=IFT172;refGene_tx=XM_019827755.1
A3 121055514 121055638
refGene_gn=IFT172;refGene_tx=XM_003984381.5;refGene_gn=IFT172;refGene_tx=XM_019827754.2;refGene_gn=IFT172;refGene_tx=XM_019827755.1
A3 121056038 121056134
refGene_gn=IFT172;refGene_tx=XM_003984381.5;refGene_gn=IFT172;refGene_tx=XM_019827754.2;refGene_gn=IFT172;refGene_tx=XM_019827755.1
A3 121057024 121057186
refGene_gn=IFT172;refGene_tx=XM_003984381.5;refGene_gn=IFT172;refGene_tx=XM_019827754.2;refGene_gn=IFT172;refGene_tx=XM_019827755.1
A3 121057559 121057613
refGene_gn=IFT172;refGene_tx=XM_003984381.5;refGene_gn=IFT172;refGene_tx=XM_019827754.2;refGene_gn=IFT172;refGene_tx=XM_019827755.1
A3 121057855 121057959
refGene_gn=IFT172;refGene_tx=XM_003984381.5;refGene_gn=IFT172;refGene_tx=XM_019827754.2;refGene_gn=IFT172;refGene_tx=XM_019827755.1
A3 121058134 121058220
refGene_gn=IFT172;refGene_tx=XM_003984381.5;refGene_gn=IFT172;refGene_tx=XM_019827754.2;refGene_gn=IFT172;refGene_tx=XM_019827755.1
A3 121062311 121062424
refGene_gn=IFT172;refGene_tx=XM_003984381.5;refGene_gn=IFT172;refGene_tx=XM_019827754.2;refGene_gn=IFT172;refGene_tx=XM_019827755.1

A3	121063261	121063429
refGene_gn=IFT172;refGene_tx=XM_003984381.5;refGene_gn=IFT172;refGene_tx=XM_019827754.2;refGene_gn=IFT172;refGene_tx=XM_019827755.1
A3	121067503	121067640
refGene_gn=IFT172;refGene_tx=XM_003984381.5;refGene_gn=IFT172;refGene_tx=XM_019827754.2;refGene_gn=IFT172;refGene_tx=XM_019827755.1
A3	121067875	121067983
refGene_gn=IFT172;refGene_tx=XM_003984381.5;refGene_gn=IFT172;refGene_tx=XM_019827754.2;refGene_gn=IFT172;refGene_tx=XM_019827755.1
A3	121069337	121069422
refGene_gn=IFT172;refGene_tx=XM_003984381.5;refGene_gn=IFT172;refGene_tx=XM_019827754.2;refGene_gn=IFT172;refGene_tx=XM_019827755.1
A3	121069725	121069818
refGene_gn=IFT172;refGene_tx=XM_003984381.5;refGene_gn=IFT172;refGene_tx=XM_019827754.2;refGene_gn=IFT172;refGene_tx=XM_019827755.1
A3	121070314	121070392
refGene_gn=IFT172;refGene_tx=XM_003984381.5;refGene_gn=IFT172;refGene_tx=XM_019827754.2;refGene_gn=IFT172;refGene_tx=XM_019827755.1
A3	121070615	121070864
refGene_gn=IFT172;refGene_tx=XM_003984381.5;refGene_gn=IFT172;refGene_tx=XM_019827754.2;refGene_gn=IFT172;refGene_tx=XM_019827755.1
A3	121071043	121071122
refGene_gn=IFT172;refGene_tx=XM_003984381.5;refGene_gn=IFT172;refGene_tx=XM_019827754.2;refGene_gn=IFT172;refGene_tx=XM_019827755.1
A3	121071319	121071440
refGene_gn=IFT172;refGene_tx=XM_003984381.5;refGene_gn=IFT172;refGene_tx=XM_019827754.2;refGene_gn=IFT172;refGene_tx=XM_019827755.1
A3	121071596	121071741
refGene_gn=IFT172;refGene_tx=XM_003984381.5;refGene_gn=IFT172;refGene_tx=XM_019827754.2;refGene_gn=IFT172;refGene_tx=XM_019827755.1
A3	121072466	121072556
refGene_gn=IFT172;refGene_tx=XM_003984381.5;refGene_gn=IFT172;refGene_tx=XM_019827754.2;refGene_gn=IFT172;refGene_tx=XM_019827755.1
A3	121073145	121073243
refGene_gn=IFT172;refGene_tx=XM_003984381.5;refGene_gn=IFT172;refGene_tx=XM_019827754.2;refGene_gn=IFT172;refGene_tx=XM_019827755.1
A3	121073413	121073549
refGene_gn=IFT172;refGene_tx=XM_003984381.5;refGene_gn=IFT172;refGene_tx=XM_019827754.2;refGene_gn=IFT172;refGene_tx=XM_019827755.1
A3	121073634	121073751
refGene_gn=IFT172;refGene_tx=XM_003984381.5;refGene_gn=IFT172;refGene_tx=XM_019827754.2;refGene_gn=IFT172;refGene_tx=XM_019827755.1
A3	121074072	121074215
refGene_gn=IFT172;refGene_tx=XM_003984381.5;refGene_gn=IFT172;refGene_tx=XM_019827754.2;refGene_gn=IFT172;refGene_tx=XM_019827755.1
A3	121075028	121075122
refGene_gn=IFT172;refGene_tx=XM_003984381.5;refGene_gn=IFT172;refGene_tx=XM_019827754.2;refGene_gn=IFT172;refGene_tx=XM_019827755.1
A3	121075235	121075300
refGene_gn=IFT172;refGene_tx=XM_003984381.5;refGene_gn=IFT172;refGene_

tx=XM_019827754.2;refGene_gn=IFT172;refGene_tx=XM_019827755.1
A3 121075495 121075676
refGene_gn=IFT172;refGene_tx=XM_003984381.5;refGene_gn=IFT172;refGene_tx=XM_019827754.2;refGene_gn=IFT172;refGene_tx=XM_019827755.1
A3 121075910 121076020
refGene_gn=IFT172;refGene_tx=XM_003984381.5;refGene_gn=IFT172;refGene_tx=XM_019827754.2;refGene_gn=IFT172;refGene_tx=XM_019827755.1
A3 121076129 121076259
refGene_gn=IFT172;refGene_tx=XM_003984381.5;refGene_gn=IFT172;refGene_tx=XM_019827754.2;refGene_gn=IFT172;refGene_tx=XM_019827755.1
A3 121078734 121078833
refGene_gn=IFT172;refGene_tx=XM_003984381.5;refGene_gn=IFT172;refGene_tx=XM_019827754.2;refGene_gn=IFT172;refGene_tx=XM_019827755.1
A3 121078999 121079109
refGene_gn=IFT172;refGene_tx=XM_003984381.5;refGene_gn=IFT172;refGene_tx=XM_019827754.2;refGene_gn=IFT172;refGene_tx=XM_019827755.1
A3 121079232 121079296
refGene_gn=IFT172;refGene_tx=XM_003984381.5;refGene_gn=IFT172;refGene_tx=XM_019827754.2;refGene_gn=IFT172;refGene_tx=XM_019827755.1
A3 121079415 121079502
refGene_gn=IFT172;refGene_tx=XM_003984381.5;refGene_gn=IFT172;refGene_tx=XM_019827754.2;refGene_gn=IFT172;refGene_tx=XM_019827755.1
A3 121079792 121079909
refGene_gn=IFT172;refGene_tx=XM_003984381.5;refGene_gn=IFT172;refGene_tx=XM_019827754.2;refGene_gn=IFT172;refGene_tx=XM_019827755.1
A3 121080748 121080859
refGene_gn=IFT172;refGene_tx=XM_003984381.5;refGene_gn=IFT172;refGene_tx=XM_019827754.2;refGene_gn=IFT172;refGene_tx=XM_019827755.1
A3 121081261 121081381
refGene_gn=IFT172;refGene_tx=XM_003984381.5;refGene_gn=IFT172;refGene_tx=XM_019827754.2;refGene_gn=IFT172;refGene_tx=XM_019827755.1
A3 121081890 121081986
refGene_gn=IFT172;refGene_tx=XM_003984381.5;refGene_gn=IFT172;refGene_tx=XM_019827754.2;refGene_gn=IFT172;refGene_tx=XM_019827755.1
A3 121082207 121082267
refGene_gn=IFT172;refGene_tx=XM_003984381.5;refGene_gn=IFT172;refGene_tx=XM_019827754.2;refGene_gn=IFT172;refGene_tx=XM_019827755.1
A3 121082351 121082450
refGene_gn=IFT172;refGene_tx=XM_003984381.5;refGene_gn=IFT172;refGene_tx=XM_019827754.2;refGene_gn=IFT172;refGene_tx=XM_019827755.1
A3 121082772 121082926
refGene_gn=IFT172;refGene_tx=XM_003984381.5;refGene_gn=IFT172;refGene_tx=XM_019827754.2;refGene_gn=IFT172;refGene_tx=XM_019827755.1
A3 121083464 121083556
refGene_gn=IFT172;refGene_tx=XM_003984381.5;refGene_gn=IFT172;refGene_tx=XM_019827754.2;refGene_gn=IFT172;refGene_tx=XM_019827755.1
A3 121083763 121083853
refGene_gn=IFT172;refGene_tx=XM_003984381.5;refGene_gn=IFT172;refGene_tx=XM_019827754.2;refGene_gn=IFT172;refGene_tx=XM_019827755.1
A3 121084209 121084317

refGene_gn=KRTCAP3;refGene_tx=XM_003984382.3;refGene_gn=KRTCAP3;refGene_tx=XM_023252008.1
A3	121084399	121084534
refGene_gn=KRTCAP3;refGene_tx=XM_003984382.3;refGene_gn=KRTCAP3;refGene_tx=XM_023252008.1
A3	121084661	121084868
refGene_gn=KRTCAP3;refGene_tx=XM_003984382.3;refGene_gn=KRTCAP3;refGene_tx=XM_023252008.1
A3	121085038	121085098
refGene_gn=KRTCAP3;refGene_tx=XM_003984382.3;refGene_gn=KRTCAP3;refGene_tx=XM_023252008.1
A3	121085185	121085545
refGene_gn=KRTCAP3;refGene_tx=XM_003984382.3;refGene_gn=KRTCAP3;refGene_tx=XM_023252008.1
A3	121086157	121086262
refGene_gn=NRBP1;refGene_tx=XM_011281287.3;refGene_gn=NRBP1;refGene_tx=XM_019827756.2
A3	121086346	121086402
refGene_gn=NRBP1;refGene_tx=XM_011281287.3;refGene_gn=NRBP1;refGene_tx=XM_019827756.2
A3	121086667	121086731
refGene_gn=NRBP1;refGene_tx=XM_011281287.3;refGene_gn=NRBP1;refGene_tx=XM_019827756.2
A3	121086815	121086869
refGene_gn=NRBP1;refGene_tx=XM_011281287.3;refGene_gn=NRBP1;refGene_tx=XM_019827756.2
A3	121087068	121087204
refGene_gn=NRBP1;refGene_tx=XM_011281287.3;refGene_gn=NRBP1;refGene_tx=XM_019827756.2
A3	121087292	121087343
refGene_gn=NRBP1;refGene_tx=XM_011281287.3;refGene_gn=NRBP1;refGene_tx=XM_019827756.2
A3	121087490	121087596
refGene_gn=NRBP1;refGene_tx=XM_011281287.3;refGene_gn=NRBP1;refGene_tx=XM_019827756.2
A3	121087777	121087910
refGene_gn=NRBP1;refGene_tx=XM_011281287.3;refGene_gn=NRBP1;refGene_tx=XM_019827756.2
A3	121089124	121089223
refGene_gn=NRBP1;refGene_tx=XM_011281287.3;refGene_gn=NRBP1;refGene_tx=XM_019827756.2
A3	121089377	121089436
refGene_gn=NRBP1;refGene_tx=XM_011281287.3;refGene_gn=NRBP1;refGene_tx=XM_019827756.2
A3	121089625	121089709
refGene_gn=NRBP1;refGene_tx=XM_011281287.3;refGene_gn=NRBP1;refGene_tx=XM_019827756.2
A3	121090711	121090735
refGene_gn=NRBP1;refGene_tx=XM_019827756.2
A3	121091212	121091307

refGene_gn=NRBP1;refGene_tx=XM_011281287.3;refGene_gn=NRBP1;refGene_tx=XM_019827756.2
A3 121091673 121091714
refGene_gn=NRBP1;refGene_tx=XM_011281287.3;refGene_gn=NRBP1;refGene_tx=XM_019827756.2
A3 121091845 121091935
refGene_gn=NRBP1;refGene_tx=XM_011281287.3;refGene_gn=NRBP1;refGene_tx=XM_019827756.2
A3 121092315 121092417
refGene_gn=NRBP1;refGene_tx=XM_011281287.3;refGene_gn=NRBP1;refGene_tx=XM_019827756.2
A3 121092607 121092730
refGene_gn=NRBP1;refGene_tx=XM_011281287.3;refGene_gn=NRBP1;refGene_tx=XM_019827756.2
A3 121092893 121093103
refGene_gn=NRBP1;refGene_tx=XM_011281287.3;refGene_gn=NRBP1;refGene_tx=XM_019827756.2
A3 121111893 121112013
refGene_gn=PPM1G;refGene_tx=XM_003984384.3
A3 121125121 121125191
refGene_gn=PPM1G;refGene_tx=XM_003984384.3
A3 121125704 121125790
refGene_gn=PPM1G;refGene_tx=XM_003984384.3
A3 121126363 121126496
refGene_gn=PPM1G;refGene_tx=XM_003984384.3
A3 121127147 121127563
refGene_gn=PPM1G;refGene_tx=XM_003984384.3
A3 121128514 121128649
refGene_gn=PPM1G;refGene_tx=XM_003984384.3
A3 121128994 121129229
refGene_gn=PPM1G;refGene_tx=XM_003984384.3
A3 121129612 121129742
refGene_gn=PPM1G;refGene_tx=XM_003984384.3
A3 121130039 121130142
refGene_gn=PPM1G;refGene_tx=XM_003984384.3
A3 121130463 121130670
refGene_gn=PPM1G;refGene_tx=XM_003984384.3
A3 121131880 121131935
refGene_gn=ZNF513;refGene_tx=XM_003984392.4
A3 121132178 121132340
refGene_gn=ZNF513;refGene_tx=XM_003984392.4
A3 121133321 121133909
refGene_gn=ZNF513;refGene_tx=XM_003984392.4
A3 121134004 121134831
refGene_gn=ZNF513;refGene_tx=XM_003984392.4
A3 121135663 121135777
refGene_gn=SNX17;refGene_tx=XM_003984385.4
A3 121135870 121135912
refGene_gn=SNX17;refGene_tx=XM_003984385.4
A3 121136003 121136078

refGene_gn=SNX17;refGene_tx=XM_003984385.4
A3 121136206 121136284
refGene_gn=SNX17;refGene_tx=XM_003984385.4
A3 121136447 121136573
refGene_gn=SNX17;refGene_tx=XM_003984385.4
A3 121136695 121136899
refGene_gn=SNX17;refGene_tx=XM_003984385.4
A3 121136994 121137087
refGene_gn=SNX17;refGene_tx=XM_003984385.4
A3 121137371 121137441
refGene_gn=SNX17;refGene_tx=XM_003984385.4
A3 121137681 121137769
refGene_gn=SNX17;refGene_tx=XM_003984385.4
A3 121137933 121138024
refGene_gn=SNX17;refGene_tx=XM_003984385.4
A3 121138130 121138241
refGene_gn=SNX17;refGene_tx=XM_003984385.4
A3 121138519 121138584
refGene_gn=SNX17;refGene_tx=XM_003984385.4
A3 121139108 121139226
refGene_gn=SNX17;refGene_tx=XM_003984385.4
A3 121140239 121140314
refGene_gn=SNX17;refGene_tx=XM_003984385.4
A3 121140782 121140845
refGene_gn=SNX17;refGene_tx=XM_003984385.4
A3 121141236 121141267
refGene_gn=EIF2B4;refGene_tx=XM_003984393.5;refGene_gn=EIF2B4;refGene_tx=XM_006930407.4
A3 121141554 121141692
refGene_gn=EIF2B4;refGene_tx=XM_003984393.5;refGene_gn=EIF2B4;refGene_tx=XM_006930406.4;refGene_gn=EIF2B4;refGene_tx=XM_006930407.4;refGene_gn=EIF2B4;refGene_tx=XM_019827757.2
A3 121142011 121142147
refGene_gn=EIF2B4;refGene_tx=XM_003984393.5;refGene_gn=EIF2B4;refGene_tx=XM_006930406.4;refGene_gn=EIF2B4;refGene_tx=XM_006930407.4;refGene_gn=EIF2B4;refGene_tx=XM_019827757.2
A3 121142343 121142550
refGene_gn=EIF2B4;refGene_tx=XM_003984393.5;refGene_gn=EIF2B4;refGene_tx=XM_006930406.4;refGene_gn=EIF2B4;refGene_tx=XM_006930407.4;refGene_gn=EIF2B4;refGene_tx=XM_019827757.2
A3 121142823 121142906
refGene_gn=EIF2B4;refGene_tx=XM_003984393.5;refGene_gn=EIF2B4;refGene_tx=XM_006930406.4;refGene_gn=EIF2B4;refGene_tx=XM_006930407.4;refGene_gn=EIF2B4;refGene_tx=XM_019827757.2
A3 121143072 121143164
refGene_gn=EIF2B4;refGene_tx=XM_003984393.5;refGene_gn=EIF2B4;refGene_tx=XM_006930406.4;refGene_gn=EIF2B4;refGene_tx=XM_006930407.4;refGene_gn=EIF2B4;refGene_tx=XM_019827757.2
A3 121143399 121143514
refGene_gn=EIF2B4;refGene_tx=XM_003984393.5;refGene_gn=EIF2B4;refGene_

tx=XM_006930406.4;refGene_gn=EIF2B4;refGene_tx=XM_006930407.4;refGene_gn=EIF2B4;refGene_tx=XM_019827757.2;refGene_gn=EIF2B4;refGene_tx=XM_019827758.2;refGene_gn=EIF2B4;refGene_tx=XM_019827759.1
A3 121143720 121143797
refGene_gn=EIF2B4;refGene_tx=XM_003984393.5;refGene_gn=EIF2B4;refGene_tx=XM_006930406.4;refGene_gn=EIF2B4;refGene_tx=XM_006930407.4;refGene_gn=EIF2B4;refGene_tx=XM_019827757.2;refGene_gn=EIF2B4;refGene_tx=XM_019827758.2;refGene_gn=EIF2B4;refGene_tx=XM_019827759.1
A3 121144985 121145088
refGene_gn=EIF2B4;refGene_tx=XM_003984393.5;refGene_gn=EIF2B4;refGene_tx=XM_006930406.4;refGene_gn=EIF2B4;refGene_tx=XM_006930407.4;refGene_gn=EIF2B4;refGene_tx=XM_019827757.2;refGene_gn=EIF2B4;refGene_tx=XM_019827758.2;refGene_gn=EIF2B4;refGene_tx=XM_019827759.1
A3 121145330 121145458
refGene_gn=EIF2B4;refGene_tx=XM_003984393.5;refGene_gn=EIF2B4;refGene_tx=XM_006930406.4;refGene_gn=EIF2B4;refGene_tx=XM_006930407.4;refGene_gn=EIF2B4;refGene_tx=XM_019827757.2;refGene_gn=EIF2B4;refGene_tx=XM_019827758.2;refGene_gn=EIF2B4;refGene_tx=XM_019827759.1
A3 121145596 121145774
refGene_gn=EIF2B4;refGene_tx=XM_003984393.5;refGene_gn=EIF2B4;refGene_tx=XM_006930406.4;refGene_gn=EIF2B4;refGene_tx=XM_006930407.4;refGene_gn=EIF2B4;refGene_tx=XM_019827757.2;refGene_gn=EIF2B4;refGene_tx=XM_019827758.2;refGene_gn=EIF2B4;refGene_tx=XM_019827759.1
A3 121146180 121146361
refGene_gn=EIF2B4;refGene_tx=XM_003984393.5;refGene_gn=EIF2B4;refGene_tx=XM_006930406.4;refGene_gn=EIF2B4;refGene_tx=XM_006930407.4;refGene_gn=EIF2B4;refGene_tx=XM_019827757.2;refGene_gn=EIF2B4;refGene_tx=XM_019827758.2;refGene_gn=EIF2B4;refGene_tx=XM_019827759.1
A3 121146482 121146682
refGene_gn=EIF2B4;refGene_tx=XM_003984393.5;refGene_gn=EIF2B4;refGene_tx=XM_006930406.4;refGene_gn=EIF2B4;refGene_tx=XM_006930407.4;refGene_gn=EIF2B4;refGene_tx=XM_019827757.2;refGene_gn=EIF2B4;refGene_tx=XM_019827758.2;refGene_gn=EIF2B4;refGene_tx=XM_019827759.1
A3 121159989 121160236
refGene_gn=GTF3C2;refGene_tx=XM_006930408.4
A3 121160399 121160721
refGene_gn=GTF3C2;refGene_tx=XM_006930408.4
A3 121161309 121161595
refGene_gn=GTF3C2;refGene_tx=XM_006930408.4
A3 121164233 121164328
refGene_gn=GTF3C2;refGene_tx=XM_006930408.4
A3 121164741 121164819
refGene_gn=GTF3C2;refGene_tx=XM_006930408.4
A3 121164941 121165040
refGene_gn=GTF3C2;refGene_tx=XM_006930408.4
A3 121166113 121166341
refGene_gn=GTF3C2;refGene_tx=XM_006930408.4
A3 121166514 121166626
refGene_gn=GTF3C2;refGene_tx=XM_006930408.4
A3 121166826 121166935

refGene_gn=GTF3C2;refGene_tx=XM_006930408.4
A3 121168031 121168057
refGene_gn=GTF3C2;refGene_tx=XM_006930408.4
A3 121168244 121168374
refGene_gn=GTF3C2;refGene_tx=XM_006930408.4
A3 121170506 121170651
refGene_gn=GTF3C2;refGene_tx=XM_006930408.4
A3 121170756 121170918
refGene_gn=GTF3C2;refGene_tx=XM_006930408.4
A3 121171119 121171207
refGene_gn=GTF3C2;refGene_tx=XM_006930408.4
A3 121171457 121171586
refGene_gn=GTF3C2;refGene_tx=XM_006930408.4
A3 121172088 121172238
refGene_gn=GTF3C2;refGene_tx=XM_006930408.4
A3 121172388 121172496
refGene_gn=GTF3C2;refGene_tx=XM_006930408.4
A3 121172793 121173012
refGene_gn=GTF3C2;refGene_tx=XM_006930408.4
A3 121177119 121177189
refGene_gn=MPV17;refGene_tx=XM_003984396.4;refGene_gn=MPV17;refGene_tx=XM_006930409.3;refGene_gn=MPV17;refGene_tx=XM_019827760.1
A3 121183918 121184034
refGene_gn=MPV17;refGene_tx=XM_003984396.4;refGene_gn=MPV17;refGene_tx=XM_006930409.3;refGene_gn=MPV17;refGene_tx=XM_019827760.1
A3 121184254 121184347
refGene_gn=MPV17;refGene_tx=XM_003984396.4;refGene_gn=MPV17;refGene_tx=XM_006930409.3;refGene_gn=MPV17;refGene_tx=XM_019827760.1
A3 121184441 121184537
refGene_gn=MPV17;refGene_tx=XM_003984396.4;refGene_gn=MPV17;refGene_tx=XM_006930409.3;refGene_gn=MPV17;refGene_tx=XM_019827760.1
A3 121184794 121184827
refGene_gn=MPV17;refGene_tx=XM_003984396.4;refGene_gn=MPV17;refGene_tx=XM_006930409.3;refGene_gn=MPV17;refGene_tx=XM_019827760.1
A3 121185065 121185118
refGene_gn=MPV17;refGene_tx=XM_003984396.4;refGene_gn=MPV17;refGene_tx=XM_006930409.3;refGene_gn=MPV17;refGene_tx=XM_019827760.1
A3 121186110 121186180
refGene_gn=MPV17;refGene_tx=XM_003984396.4;refGene_gn=MPV17;refGene_tx=XM_006930409.3;refGene_gn=MPV17;refGene_tx=XM_019827760.1
A3 121188212 121188593
refGene_gn=UCN;refGene_tx=XM_023252010.1
A3 121189240 121189257
refGene_gn=TRIM54;refGene_tx=XM_003984445.5
A3 121189551 121189644
refGene_gn=TRIM54;refGene_tx=XM_003984445.5
A3 121189779 121189904
refGene_gn=TRIM54;refGene_tx=XM_003984445.5
A3 121190016 121190039
refGene_gn=TRIM54;refGene_tx=XM_003984445.5

A3	121190313	121190547
refGene_gn=TRIM54;refGene_tx=XM_003984445.5
A3	121190986	121191082
refGene_gn=TRIM54;refGene_tx=XM_003984445.5
A3	121193937	121194109
refGene_gn=TRIM54;refGene_tx=XM_003984445.5
A3	121194631	121194804
refGene_gn=TRIM54;refGene_tx=XM_003984445.5
A3	121211381	121211549
refGene_gn=TRIM54;refGene_tx=XM_003984445.5
A3	121214052	121214105
refGene_gn=DNAJC5G;refGene_tx=XM_023252009.1
A3	121214604	121214746
refGene_gn=DNAJC5G;refGene_tx=XM_023252009.1
A3	121214943	121215157
refGene_gn=DNAJC5G;refGene_tx=XM_023252009.1
A3	121215465	121215538
refGene_gn=DNAJC5G;refGene_tx=XM_023252009.1
A3	121215540	121215577
refGene_gn=DNAJC5G;refGene_tx=XM_023252009.1
A3	121226287	121226382
refGene_gn=SLC30A3;refGene_tx=XM_003984397.5;refGene_gn=SLC30A3;refGene_tx=XM_006930410.3
A3	121230282	121230464
refGene_gn=SLC30A3;refGene_tx=XM_003984397.5;refGene_gn=SLC30A3;refGene_tx=XM_006930410.3;refGene_gn=SLC30A3;refGene_tx=XM_023252012.1
A3	121230938	121231085
refGene_gn=SLC30A3;refGene_tx=XM_003984397.5;refGene_gn=SLC30A3;refGene_tx=XM_006930410.3;refGene_gn=SLC30A3;refGene_tx=XM_023252012.1
A3	121231180	121231334
refGene_gn=SLC30A3;refGene_tx=XM_003984397.5
A3	121231929	121232128
refGene_gn=SLC30A3;refGene_tx=XM_003984397.5;refGene_gn=SLC30A3;refGene_tx=XM_006930410.3;refGene_gn=SLC30A3;refGene_tx=XM_023252012.1
A3	121232359	121232465
refGene_gn=SLC30A3;refGene_tx=XM_003984397.5;refGene_gn=SLC30A3;refGene_tx=XM_006930410.3;refGene_gn=SLC30A3;refGene_tx=XM_023252012.1
A3	121232722	121232857
refGene_gn=SLC30A3;refGene_tx=XM_003984397.5;refGene_gn=SLC30A3;refGene_tx=XM_006930410.3;refGene_gn=SLC30A3;refGene_tx=XM_023252012.1
A3	121234026	121234404
refGene_gn=SLC30A3;refGene_tx=XM_003984397.5;refGene_gn=SLC30A3;refGene_tx=XM_006930410.3;refGene_gn=SLC30A3;refGene_tx=XM_023252012.1
A3	121243780	121243883
refGene_gn=CAD;refGene_tx=XM_006930444.3;refGene_gn=CAD;refGene_tx=XM_011281284.2
A3	121243980	121244075
refGene_gn=CAD;refGene_tx=XM_006930444.3;refGene_gn=CAD;refGene_tx=XM_011281284.2
A3	121244284	121244386

refGene_gn=CAD;refGene_tx=XM_006930444.3;refGene_gn=CAD;refGene_tx=XM_011281284.2
A3 121244473 121244629
refGene_gn=CAD;refGene_tx=XM_006930444.3;refGene_gn=CAD;refGene_tx=XM_011281284.2
A3 121244825 121244951
refGene_gn=CAD;refGene_tx=XM_006930444.3;refGene_gn=CAD;refGene_tx=XM_011281284.2
A3 121245127 121245340
refGene_gn=CAD;refGene_tx=XM_006930444.3;refGene_gn=CAD;refGene_tx=XM_011281284.2
A3 121245861 121245936
refGene_gn=CAD;refGene_tx=XM_006930444.3;refGene_gn=CAD;refGene_tx=XM_011281284.2
A3 121246046 121246216
refGene_gn=CAD;refGene_tx=XM_006930444.3;refGene_gn=CAD;refGene_tx=XM_011281284.2
A3 121246320 121246365
refGene_gn=CAD;refGene_tx=XM_006930444.3;refGene_gn=CAD;refGene_tx=XM_011281284.2
A3 121246653 121246704
refGene_gn=CAD;refGene_tx=XM_006930444.3
A3 121246988 121247085
refGene_gn=CAD;refGene_tx=XM_006930444.3;refGene_gn=CAD;refGene_tx=XM_011281284.2
A3 121247534 121247636
refGene_gn=CAD;refGene_tx=XM_006930444.3;refGene_gn=CAD;refGene_tx=XM_011281284.2
A3 121247862 121248003
refGene_gn=CAD;refGene_tx=XM_006930444.3;refGene_gn=CAD;refGene_tx=XM_011281284.2
A3 121248104 121248295
refGene_gn=CAD;refGene_tx=XM_006930444.3;refGene_gn=CAD;refGene_tx=XM_011281284.2
A3 121248655 121248857
refGene_gn=CAD;refGene_tx=XM_006930444.3;refGene_gn=CAD;refGene_tx=XM_011281284.2
A3 121249067 121249199
refGene_gn=CAD;refGene_tx=XM_006930444.3;refGene_gn=CAD;refGene_tx=XM_011281284.2
A3 121249371 121249536
refGene_gn=CAD;refGene_tx=XM_006930444.3;refGene_gn=CAD;refGene_tx=XM_011281284.2
A3 121249703 121249870
refGene_gn=CAD;refGene_tx=XM_006930444.3;refGene_gn=CAD;refGene_tx=XM_011281284.2
A3 121250158 121250240
refGene_gn=CAD;refGene_tx=XM_006930444.3;refGene_gn=CAD;refGene_tx=XM_011281284.2
A3 121250470 121250710

refGene_gn=CAD;refGene_tx=XM_006930444.3;refGene_gn=CAD;refGene_tx=XM_011281284.2
A3 121251135 121251240
refGene_gn=CAD;refGene_tx=XM_006930444.3;refGene_gn=CAD;refGene_tx=XM_011281284.2
A3 121251368 121251551
refGene_gn=CAD;refGene_tx=XM_006930444.3;refGene_gn=CAD;refGene_tx=XM_011281284.2
A3 121252147 121252315
refGene_gn=CAD;refGene_tx=XM_006930444.3;refGene_gn=CAD;refGene_tx=XM_011281284.2
A3 121252629 121252848
refGene_gn=CAD;refGene_tx=XM_006930444.3;refGene_gn=CAD;refGene_tx=XM_011281284.2
A3 121253051 121253234
refGene_gn=CAD;refGene_tx=XM_006930444.3;refGene_gn=CAD;refGene_tx=XM_011281284.2
A3 121253314 121253539
refGene_gn=CAD;refGene_tx=XM_006930444.3;refGene_gn=CAD;refGene_tx=XM_011281284.2
A3 121253726 121253825
refGene_gn=CAD;refGene_tx=XM_006930444.3;refGene_gn=CAD;refGene_tx=XM_011281284.2
A3 121254186 121254433
refGene_gn=CAD;refGene_tx=XM_006930444.3;refGene_gn=CAD;refGene_tx=XM_011281284.2
A3 121254644 121254889
refGene_gn=CAD;refGene_tx=XM_006930444.3;refGene_gn=CAD;refGene_tx=XM_011281284.2
A3 121255588 121255701
refGene_gn=CAD;refGene_tx=XM_006930444.3;refGene_gn=CAD;refGene_tx=XM_011281284.2
A3 121258457 121258588
refGene_gn=CAD;refGene_tx=XM_006930444.3;refGene_gn=CAD;refGene_tx=XM_011281284.2
A3 121258762 121258887
refGene_gn=CAD;refGene_tx=XM_006930444.3;refGene_gn=CAD;refGene_tx=XM_011281284.2
A3 121259158 121259347
refGene_gn=CAD;refGene_tx=XM_006930444.3;refGene_gn=CAD;refGene_tx=XM_011281284.2
A3 121259551 121259773
refGene_gn=CAD;refGene_tx=XM_006930444.3;refGene_gn=CAD;refGene_tx=XM_011281284.2
A3 121259906 121260140
refGene_gn=CAD;refGene_tx=XM_006930444.3;refGene_gn=CAD;refGene_tx=XM_011281284.2
A3 121260279 121260411
refGene_gn=CAD;refGene_tx=XM_006930444.3;refGene_gn=CAD;refGene_tx=XM_011281284.2

A3 121260674 121260820
refGene_gn=CAD;refGene_tx=XM_006930444.3;refGene_gn=CAD;refGene_tx=XM_011281284.2
A3 121261156 121261269
refGene_gn=CAD;refGene_tx=XM_006930444.3;refGene_gn=CAD;refGene_tx=XM_011281284.2
A3 121261452 121261638
refGene_gn=CAD;refGene_tx=XM_006930444.3;refGene_gn=CAD;refGene_tx=XM_011281284.2
A3 121261841 121262013
refGene_gn=CAD;refGene_tx=XM_006930444.3;refGene_gn=CAD;refGene_tx=XM_011281284.2
A3 121262216 121262358
refGene_gn=CAD;refGene_tx=XM_006930444.3;refGene_gn=CAD;refGene_tx=XM_011281284.2
A3 121262529 121262672
refGene_gn=CAD;refGene_tx=XM_006930444.3;refGene_gn=CAD;refGene_tx=XM_011281284.2
A3 121263782 121263912
refGene_gn=CAD;refGene_tx=XM_006930444.3;refGene_gn=CAD;refGene_tx=XM_011281284.2
A3 121265767 121265907
refGene_gn=CAD;refGene_tx=XM_006930444.3;refGene_gn=CAD;refGene_tx=XM_011281284.2
A3 121266160 121266242
refGene_gn=CAD;refGene_tx=XM_006930444.3;refGene_gn=CAD;refGene_tx=XM_011281284.2
A3 121266909 121267014
refGene_gn=ATRAID;refGene_tx=XM_003984398.5
A3 121267236 121267334
refGene_gn=ATRAID;refGene_tx=XM_003984398.5
A3 121267634 121267756
refGene_gn=ATRAID;refGene_tx=XM_003984398.5
A3 121267856 121267928
refGene_gn=ATRAID;refGene_tx=XM_003984398.5
A3 121268139 121268211
refGene_gn=ATRAID;refGene_tx=XM_003984398.5
A3 121270404 121270526
refGene_gn=ATRAID;refGene_tx=XM_003984398.5
A3 121271218 121271317
refGene_gn=ATRAID;refGene_tx=XM_003984398.5
A3 121275482 121275875
refGene_gn=SLC5A6;refGene_tx=XM_003984400.4;refGene_gn=SLC5A6;refGene_tx=XM_006930411.4;refGene_gn=SLC5A6;refGene_tx=XM_006930412.4;refGene_gn=SLC5A6;refGene_tx=XM_006930413.4;refGene_gn=SLC5A6;refGene_tx=XM_006930414.4;refGene_gn=SLC5A6;refGene_tx=XM_006930415.3;refGene_gn=SLC5A6;refGene_tx=XM_006930416.4;refGene_gn=SLC5A6;refGene_tx=XM_011281282.3;refGene_gn=SLC5A6;refGene_tx=XM_019827764.2;refGene_gn=SLC5A6;refGene_tx=XM_019827765.2;refGene_gn=SLC5A6;refGene_tx=XM_019827766.2
A3 121276225 121276291

refGene_gn=SLC5A6;refGene_tx=XM_003984400.4;refGene_gn=SLC5A6;refGene_tx=XM_006930411.4;refGene_gn=SLC5A6;refGene_tx=XM_006930412.4;refGene_gn=SLC5A6;refGene_tx=XM_006930413.4;refGene_gn=SLC5A6;refGene_tx=XM_006930414.4;refGene_gn=SLC5A6;refGene_tx=XM_006930415.3;refGene_gn=SLC5A6;refGene_tx=XM_006930416.4;refGene_gn=SLC5A6;refGene_tx=XM_011281282.3;refGene_gn=SLC5A6;refGene_tx=XM_019827764.2;refGene_gn=SLC5A6;refGene_tx=XM_019827765.2;refGene_gn=SLC5A6;refGene_tx=XM_019827766.2
A3 121276626 121276678
refGene_gn=SLC5A6;refGene_tx=XM_003984400.4;refGene_gn=SLC5A6;refGene_tx=XM_006930411.4;refGene_gn=SLC5A6;refGene_tx=XM_006930412.4;refGene_gn=SLC5A6;refGene_tx=XM_006930413.4;refGene_gn=SLC5A6;refGene_tx=XM_006930414.4;refGene_gn=SLC5A6;refGene_tx=XM_006930415.3;refGene_gn=SLC5A6;refGene_tx=XM_006930416.4;refGene_gn=SLC5A6;refGene_tx=XM_011281282.3;refGene_gn=SLC5A6;refGene_tx=XM_019827764.2;refGene_gn=SLC5A6;refGene_tx=XM_019827765.2;refGene_gn=SLC5A6;refGene_tx=XM_019827766.2
A3 121277054 121277122
refGene_gn=SLC5A6;refGene_tx=XM_003984400.4;refGene_gn=SLC5A6;refGene_tx=XM_006930411.4;refGene_gn=SLC5A6;refGene_tx=XM_006930412.4;refGene_gn=SLC5A6;refGene_tx=XM_006930413.4;refGene_gn=SLC5A6;refGene_tx=XM_006930414.4;refGene_gn=SLC5A6;refGene_tx=XM_006930415.3;refGene_gn=SLC5A6;refGene_tx=XM_006930416.4;refGene_gn=SLC5A6;refGene_tx=XM_011281282.3;refGene_gn=SLC5A6;refGene_tx=XM_019827764.2;refGene_gn=SLC5A6;refGene_tx=XM_019827765.2;refGene_gn=SLC5A6;refGene_tx=XM_019827766.2
A3 121277440 121277595
refGene_gn=SLC5A6;refGene_tx=XM_003984400.4;refGene_gn=SLC5A6;refGene_tx=XM_006930411.4;refGene_gn=SLC5A6;refGene_tx=XM_006930412.4;refGene_gn=SLC5A6;refGene_tx=XM_006930413.4;refGene_gn=SLC5A6;refGene_tx=XM_006930414.4;refGene_gn=SLC5A6;refGene_tx=XM_006930415.3;refGene_gn=SLC5A6;refGene_tx=XM_006930416.4;refGene_gn=SLC5A6;refGene_tx=XM_011281282.3;refGene_gn=SLC5A6;refGene_tx=XM_019827764.2;refGene_gn=SLC5A6;refGene_tx=XM_019827765.2;refGene_gn=SLC5A6;refGene_tx=XM_019827766.2
A3 121278016 121278157
refGene_gn=SLC5A6;refGene_tx=XM_003984400.4;refGene_gn=SLC5A6;refGene_tx=XM_006930411.4;refGene_gn=SLC5A6;refGene_tx=XM_006930412.4;refGene_gn=SLC5A6;refGene_tx=XM_006930413.4;refGene_gn=SLC5A6;refGene_tx=XM_006930414.4;refGene_gn=SLC5A6;refGene_tx=XM_006930415.3;refGene_gn=SLC5A6;refGene_tx=XM_006930416.4;refGene_gn=SLC5A6;refGene_tx=XM_011281282.3;refGene_gn=SLC5A6;refGene_tx=XM_019827764.2;refGene_gn=SLC5A6;refGene_tx=XM_019827765.2;refGene_gn=SLC5A6;refGene_tx=XM_019827766.2
A3 121278362 121278492
refGene_gn=SLC5A6;refGene_tx=XM_003984400.4;refGene_gn=SLC5A6;refGene_tx=XM_006930411.4;refGene_gn=SLC5A6;refGene_tx=XM_006930412.4;refGene_gn=SLC5A6;refGene_tx=XM_006930413.4;refGene_gn=SLC5A6;refGene_tx=XM_006930414.4;refGene_gn=SLC5A6;refGene_tx=XM_006930415.3;refGene_gn=SLC5A6;refGene_tx=XM_006930416.4;refGene_gn=SLC5A6;refGene_tx=XM_011281282.3;refGene_gn=SLC5A6;refGene_tx=XM_019827764.2;refGene_gn=SLC5A6;refGene_tx=XM_019827765.2;refGene_gn=SLC5A6;refGene_tx=XM_019827766.2
A3 121279058 121279147
refGene_gn=SLC5A6;refGene_tx=XM_003984400.4;refGene_gn=SLC5A6;refGene_tx=XM_006930411.4;refGene_gn=SLC5A6;refGene_tx=XM_006930412.4;refGene_

gn=SLC5A6;refGene_tx=XM_006930413.4;refGene_gn=SLC5A6;refGene_tx=XM_00
6930414.4;refGene_gn=SLC5A6;refGene_tx=XM_006930415.3;refGene_gn=SLC5A
6;refGene_tx=XM_006930416.4;refGene_gn=SLC5A6;refGene_tx=XM_011281282.
3;refGene_gn=SLC5A6;refGene_tx=XM_019827764.2;refGene_gn=SLC5A6;refGen
e_tx=XM_019827765.2;refGene_gn=SLC5A6;refGene_tx=XM_019827766.2
A3 121279575 121279688
refGene_gn=SLC5A6;refGene_tx=XM_003984400.4;refGene_gn=SLC5A6;refGene_
tx=XM_006930411.4;refGene_gn=SLC5A6;refGene_tx=XM_006930412.4;refGene_
gn=SLC5A6;refGene_tx=XM_006930413.4;refGene_gn=SLC5A6;refGene_tx=XM_00
6930414.4;refGene_gn=SLC5A6;refGene_tx=XM_006930415.3;refGene_gn=SLC5A
6;refGene_tx=XM_006930416.4;refGene_gn=SLC5A6;refGene_tx=XM_011281282.
3;refGene_gn=SLC5A6;refGene_tx=XM_019827764.2;refGene_gn=SLC5A6;refGen
e_tx=XM_019827765.2;refGene_gn=SLC5A6;refGene_tx=XM_019827766.2
A3 121280033 121280101
refGene_gn=SLC5A6;refGene_tx=XM_003984400.4;refGene_gn=SLC5A6;refGene_
tx=XM_006930411.4;refGene_gn=SLC5A6;refGene_tx=XM_006930412.4;refGene_
gn=SLC5A6;refGene_tx=XM_006930413.4;refGene_gn=SLC5A6;refGene_tx=XM_00
6930414.4;refGene_gn=SLC5A6;refGene_tx=XM_006930415.3;refGene_gn=SLC5A
6;refGene_tx=XM_006930416.4;refGene_gn=SLC5A6;refGene_tx=XM_011281282.
3;refGene_gn=SLC5A6;refGene_tx=XM_019827764.2;refGene_gn=SLC5A6;refGen
e_tx=XM_019827765.2;refGene_gn=SLC5A6;refGene_tx=XM_019827766.2
A3 121280512 121280599
refGene_gn=SLC5A6;refGene_tx=XM_003984400.4;refGene_gn=SLC5A6;refGene_
tx=XM_006930411.4;refGene_gn=SLC5A6;refGene_tx=XM_006930412.4;refGene_
gn=SLC5A6;refGene_tx=XM_006930413.4;refGene_gn=SLC5A6;refGene_tx=XM_00
6930414.4;refGene_gn=SLC5A6;refGene_tx=XM_006930415.3;refGene_gn=SLC5A
6;refGene_tx=XM_006930416.4;refGene_gn=SLC5A6;refGene_tx=XM_011281282.
3;refGene_gn=SLC5A6;refGene_tx=XM_019827764.2;refGene_gn=SLC5A6;refGen
e_tx=XM_019827765.2;refGene_gn=SLC5A6;refGene_tx=XM_019827766.2
A3 121280746 121280928
refGene_gn=SLC5A6;refGene_tx=XM_003984400.4;refGene_gn=SLC5A6;refGene_
tx=XM_006930411.4;refGene_gn=SLC5A6;refGene_tx=XM_006930412.4;refGene_
gn=SLC5A6;refGene_tx=XM_006930413.4;refGene_gn=SLC5A6;refGene_tx=XM_00
6930414.4;refGene_gn=SLC5A6;refGene_tx=XM_006930415.3;refGene_gn=SLC5A
6;refGene_tx=XM_006930416.4;refGene_gn=SLC5A6;refGene_tx=XM_011281282.
3;refGene_gn=SLC5A6;refGene_tx=XM_019827764.2;refGene_gn=SLC5A6;refGen
e_tx=XM_019827765.2;refGene_gn=SLC5A6;refGene_tx=XM_019827766.2
A3 121281143 121281247
refGene_gn=SLC5A6;refGene_tx=XM_003984400.4;refGene_gn=SLC5A6;refGene_
tx=XM_006930411.4;refGene_gn=SLC5A6;refGene_tx=XM_006930412.4;refGene_
gn=SLC5A6;refGene_tx=XM_006930413.4;refGene_gn=SLC5A6;refGene_tx=XM_00
6930414.4;refGene_gn=SLC5A6;refGene_tx=XM_006930415.3;refGene_gn=SLC5A
6;refGene_tx=XM_006930416.4;refGene_gn=SLC5A6;refGene_tx=XM_011281282.
3;refGene_gn=SLC5A6;refGene_tx=XM_019827764.2;refGene_gn=SLC5A6;refGen
e_tx=XM_019827765.2;refGene_gn=SLC5A6;refGene_tx=XM_019827766.2
A3 121281496 121281615
refGene_gn=SLC5A6;refGene_tx=XM_003984400.4;refGene_gn=SLC5A6;refGene_
tx=XM_006930411.4;refGene_gn=SLC5A6;refGene_tx=XM_006930412.4;refGene_
gn=SLC5A6;refGene_tx=XM_006930413.4;refGene_gn=SLC5A6;refGene_tx=XM_00
6930414.4;refGene_gn=SLC5A6;refGene_tx=XM_006930415.3;refGene_gn=SLC5A

6;refGene_tx=XM_006930416.4;refGene_gn=SLC5A6;refGene_tx=XM_011281282.
3;refGene_gn=SLC5A6;refGene_tx=XM_019827764.2;refGene_gn=SLC5A6;refGen
e_tx=XM_019827765.2;refGene_gn=SLC5A6;refGene_tx=XM_019827766.2
A3 121281963 121282116
refGene_gn=SLC5A6;refGene_tx=XM_003984400.4;refGene_gn=SLC5A6;refGene_
tx=XM_006930411.4;refGene_gn=SLC5A6;refGene_tx=XM_006930412.4;refGene_
gn=SLC5A6;refGene_tx=XM_006930413.4;refGene_gn=SLC5A6;refGene_tx=XM_00
6930414.4;refGene_gn=SLC5A6;refGene_tx=XM_006930415.3;refGene_gn=SLC5A
6;refGene_tx=XM_006930416.4;refGene_gn=SLC5A6;refGene_tx=XM_011281282.
3;refGene_gn=SLC5A6;refGene_tx=XM_019827764.2;refGene_gn=SLC5A6;refGen
e_tx=XM_019827765.2;refGene_gn=SLC5A6;refGene_tx=XM_019827766.2
A3 121336286 121336466
refGene_gn=TCF23;refGene_tx=XM_003984401.5
A3 121338261 121338504
refGene_gn=TCF23;refGene_tx=XM_003984401.5
A3 121339093 121339303
refGene_gn=TCF23;refGene_tx=XM_003984401.5
A3 121351942 121353142
refGene_gn=PRR30;refGene_tx=XM_011281277.3;refGene_gn=PRR30;refGene_tx
=XM_011281279.3;refGene_gn=PRR30;refGene_tx=XM_019827767.2
A3 121359423 121359558
refGene_gn=PREB;refGene_tx=XM_003984402.4;refGene_gn=PREB;refGene_tx=X
M_019827771.1
A3 121360156 121360349
refGene_gn=PREB;refGene_tx=XM_003984402.4;refGene_gn=PREB;refGene_tx=X
M_019827771.1
A3 121360535 121360756
refGene_gn=PREB;refGene_tx=XM_003984402.4;refGene_gn=PREB;refGene_tx=X
M_019827771.1
A3 121360937 121361018
refGene_gn=PREB;refGene_tx=XM_003984402.4;refGene_gn=PREB;refGene_tx=X
M_019827771.1
A3 121361117 121361242
refGene_gn=PREB;refGene_tx=XM_003984402.4;refGene_gn=PREB;refGene_tx=X
M_019827771.1
A3 121361441 121361615
refGene_gn=PREB;refGene_tx=XM_003984402.4;refGene_gn=PREB;refGene_tx=X
M_019827771.1
A3 121361757 121361830
refGene_gn=PREB;refGene_tx=XM_003984402.4;refGene_gn=PREB;refGene_tx=X
M_019827771.1
A3 121362019 121362179
refGene_gn=PREB;refGene_tx=XM_003984402.4
A3 121362351 121362446
refGene_gn=PREB;refGene_tx=XM_003984402.4;refGene_gn=PREB;refGene_tx=X
M_019827771.1
A3 121363071 121363283
refGene_gn=ABHD1;refGene_tx=XM_006930445.4
A3 121363407 121363573
refGene_gn=ABHD1;refGene_tx=XM_006930445.4

A3 121363670 121363719
refGene_gn=ABHD1;refGene_tx=XM_006930445.4
A3 121363880 121364054
refGene_gn=ABHD1;refGene_tx=XM_006930445.4
A3 121364179 121364292
refGene_gn=ABHD1;refGene_tx=XM_006930445.4
A3 121364464 121364510
refGene_gn=ABHD1;refGene_tx=XM_006930445.4
A3 121364707 121364890
refGene_gn=ABHD1;refGene_tx=XM_006930445.4
A3 121365231 121365392
refGene_gn=ABHD1;refGene_tx=XM_006930445.4
A3 121368551 121368665
refGene_gn=ABHD1;refGene_tx=XM_006930445.4
A3 121376070 121376172
refGene_gn=KHK;refGene_tx=XM_019827773.2
A3 121385320 121385397
refGene_gn=CGREF1;refGene_tx=XM_003984406.5;refGene_gn=CGREF1;refGene_tx=XM_023252013.1;refGene_gn=CGREF1;refGene_tx=XM_023252014.1;refGene_gn=CGREF1;refGene_tx=XM_023252015.1
A3 121386500 121386566
refGene_gn=CGREF1;refGene_tx=XM_003984406.5;refGene_gn=CGREF1;refGene_tx=XM_023252013.1;refGene_gn=CGREF1;refGene_tx=XM_023252014.1;refGene_gn=CGREF1;refGene_tx=XM_023252015.1
A3 121386644 121386715
refGene_gn=CGREF1;refGene_tx=XM_003984406.5;refGene_gn=CGREF1;refGene_tx=XM_023252013.1;refGene_gn=CGREF1;refGene_tx=XM_023252014.1;refGene_gn=CGREF1;refGene_tx=XM_023252015.1
A3 121386855 121386980
refGene_gn=CGREF1;refGene_tx=XM_003984406.5;refGene_gn=CGREF1;refGene_tx=XM_023252013.1;refGene_gn=CGREF1;refGene_tx=XM_023252014.1;refGene_gn=CGREF1;refGene_tx=XM_023252015.1
A3 121387198 121387645
refGene_gn=CGREF1;refGene_tx=XM_003984406.5;refGene_gn=CGREF1;refGene_tx=XM_023252013.1;refGene_gn=CGREF1;refGene_tx=XM_023252014.1;refGene_gn=CGREF1;refGene_tx=XM_023252015.1
A3 121389292 121389378
refGene_gn=KHK;refGene_tx=XM_003984404.3;refGene_gn=KHK;refGene_tx=XM_003984405.3;refGene_gn=KHK;refGene_tx=XM_006930418.2;refGene_gn=KHK;refGene_tx=XM_019827772.1
A3 121389469 121389627
refGene_gn=KHK;refGene_tx=XM_003984404.3;refGene_gn=KHK;refGene_tx=XM_003984405.3;refGene_gn=KHK;refGene_tx=XM_006930418.2;refGene_gn=KHK;refGene_tx=XM_019827772.1
A3 121389760 121389853
refGene_gn=KHK;refGene_tx=XM_003984404.3;refGene_gn=KHK;refGene_tx=XM_003984405.3;refGene_gn=KHK;refGene_tx=XM_006930418.2;refGene_gn=KHK;refGene_tx=XM_019827772.1;refGene_gn=KHK;refGene_tx=XM_019827774.2;refGene_gn=KHK;refGene_tx=XM_019827775.2
A3 121390165 121390240

refGene_gn=KHK;refGene_tx=XM_019827776.2
A3	121390901	121391048
refGene_gn=KHK;refGene_tx=XM_003984404.3;refGene_gn=KHK;refGene_tx=XM_003984405.3;refGene_gn=KHK;refGene_tx=XM_006930418.2;refGene_gn=KHK;refGene_tx=XM_019827772.1;refGene_gn=KHK;refGene_tx=XM_019827773.2;refGene_gn=KHK;refGene_tx=XM_019827774.2;refGene_gn=KHK;refGene_tx=XM_019827775.2;refGene_gn=KHK;refGene_tx=XM_019827776.2
A3	121391623	121391696
refGene_gn=KHK;refGene_tx=XM_003984404.3;refGene_gn=KHK;refGene_tx=XM_003984405.3;refGene_gn=KHK;refGene_tx=XM_006930418.2;refGene_gn=KHK;refGene_tx=XM_019827772.1;refGene_gn=KHK;refGene_tx=XM_019827773.2;refGene_gn=KHK;refGene_tx=XM_019827774.2;refGene_gn=KHK;refGene_tx=XM_019827775.2;refGene_gn=KHK;refGene_tx=XM_019827776.2
A3	121393529	121393664
refGene_gn=KHK;refGene_tx=XM_003984404.3;refGene_gn=KHK;refGene_tx=XM_019827772.1;refGene_gn=KHK;refGene_tx=XM_019827773.2;refGene_gn=KHK;refGene_tx=XM_019827774.2;refGene_gn=KHK;refGene_tx=XM_019827775.2;refGene_gn=KHK;refGene_tx=XM_019827776.2
A3	121393853	121393988
refGene_gn=KHK;refGene_tx=XM_003984405.3;refGene_gn=KHK;refGene_tx=XM_019827772.1;refGene_gn=KHK;refGene_tx=XM_019827773.2;refGene_gn=KHK;refGene_tx=XM_019827774.2;refGene_gn=KHK;refGene_tx=XM_019827775.2;refGene_gn=KHK;refGene_tx=XM_019827776.2
A3	121396011	121396128
refGene_gn=KHK;refGene_tx=XM_003984404.3;refGene_gn=KHK;refGene_tx=XM_003984405.3;refGene_gn=KHK;refGene_tx=XM_006930418.2;refGene_gn=KHK;refGene_tx=XM_019827772.1;refGene_gn=KHK;refGene_tx=XM_019827773.2;refGene_gn=KHK;refGene_tx=XM_019827774.2;refGene_gn=KHK;refGene_tx=XM_019827775.2;refGene_gn=KHK;refGene_tx=XM_019827776.2
A3	121400048	121400140
refGene_gn=KHK;refGene_tx=XM_003984404.3;refGene_gn=KHK;refGene_tx=XM_003984405.3;refGene_gn=KHK;refGene_tx=XM_006930418.2;refGene_gn=KHK;refGene_tx=XM_019827772.1;refGene_gn=KHK;refGene_tx=XM_019827773.2;refGene_gn=KHK;refGene_tx=XM_019827774.2;refGene_gn=KHK;refGene_tx=XM_019827775.2;refGene_gn=KHK;refGene_tx=XM_019827776.2
A3	121401392	121401730
refGene_gn=EMILIN1;refGene_tx=XM_023252016.1
A3	121402151	121402289
refGene_gn=EMILIN1;refGene_tx=XM_023252016.1
A3	121402456	121402474
refGene_gn=EMILIN1;refGene_tx=XM_023252016.1
A3	121402887	121403004
refGene_gn=EMILIN1;refGene_tx=XM_023252016.1
A3	121403381	121405331
refGene_gn=EMILIN1;refGene_tx=XM_023252016.1
A3	121406423	121406644
refGene_gn=EMILIN1;refGene_tx=XM_023252016.1
A3	121407111	121407237
refGene_gn=EMILIN1;refGene_tx=XM_023252016.1
A3	121408180	121408350

refGene_gn=EMILIN1;refGene_tx=XM_023252016.1
A3	121414685	121414799
refGene_gn=OST4;refGene_tx=XM_003984409.3
A3	121415930	121416102
refGene_gn=AGBL5;refGene_tx=XM_003984408.5;refGene_gn=AGBL5;refGene_tx=XM_019827778.2;refGene_gn=AGBL5;refGene_tx=XM_019827779.1
A3	121416458	121416589
refGene_gn=AGBL5;refGene_tx=XM_003984408.5;refGene_gn=AGBL5;refGene_tx=XM_019827778.2;refGene_gn=AGBL5;refGene_tx=XM_019827779.1;refGene_gn=AGBL5;refGene_tx=XM_019827780.1
A3	121417084	121417131
refGene_gn=AGBL5;refGene_tx=XM_019827780.1
A3	121417423	121417536
refGene_gn=AGBL5;refGene_tx=XM_003984408.5;refGene_gn=AGBL5;refGene_tx=XM_019827778.2;refGene_gn=AGBL5;refGene_tx=XM_019827779.1;refGene_gn=AGBL5;refGene_tx=XM_019827780.1
A3	121418615	121418771
refGene_gn=AGBL5;refGene_tx=XM_003984408.5;refGene_gn=AGBL5;refGene_tx=XM_019827778.2;refGene_gn=AGBL5;refGene_tx=XM_019827779.1;refGene_gn=AGBL5;refGene_tx=XM_019827780.1
A3	121424897	121424998
refGene_gn=AGBL5;refGene_tx=XM_019827781.2
A3	121426379	121426594
refGene_gn=AGBL5;refGene_tx=XM_003984408.5;refGene_gn=AGBL5;refGene_tx=XM_019827778.2;refGene_gn=AGBL5;refGene_tx=XM_019827779.1;refGene_gn=AGBL5;refGene_tx=XM_019827780.1;refGene_gn=AGBL5;refGene_tx=XM_019827781.2
A3	121427221	121427424
refGene_gn=AGBL5;refGene_tx=XM_003984408.5;refGene_gn=AGBL5;refGene_tx=XM_019827778.2;refGene_gn=AGBL5;refGene_tx=XM_019827779.1;refGene_gn=AGBL5;refGene_tx=XM_019827780.1;refGene_gn=AGBL5;refGene_tx=XM_019827781.2
A3	121428060	121428196
refGene_gn=AGBL5;refGene_tx=XM_003984408.5;refGene_gn=AGBL5;refGene_tx=XM_019827778.2;refGene_gn=AGBL5;refGene_tx=XM_019827779.1;refGene_gn=AGBL5;refGene_tx=XM_019827780.1;refGene_gn=AGBL5;refGene_tx=XM_019827781.2
A3	121428528	121428698
refGene_gn=AGBL5;refGene_tx=XM_003984408.5;refGene_gn=AGBL5;refGene_tx=XM_019827778.2;refGene_gn=AGBL5;refGene_tx=XM_019827779.1;refGene_gn=AGBL5;refGene_tx=XM_019827780.1;refGene_gn=AGBL5;refGene_tx=XM_019827781.2
A3	121429179	121429636
refGene_gn=AGBL5;refGene_tx=XM_003984408.5;refGene_gn=AGBL5;refGene_tx=XM_019827778.2;refGene_gn=AGBL5;refGene_tx=XM_019827779.1;refGene_gn=AGBL5;refGene_tx=XM_019827780.1;refGene_gn=AGBL5;refGene_tx=XM_019827781.2
A3	121430064	121430243
refGene_gn=AGBL5;refGene_tx=XM_003984408.5;refGene_gn=AGBL5;refGene_tx=XM_019827778.2;refGene_gn=AGBL5;refGene_tx=XM_019827779.1;refGene_gn=

AGBL5;refGene_tx=XM_019827780.1;refGene_gn=AGBL5;refGene_tx=XM_0198277
81.2
A3 121430488 121430666
refGene_gn=AGBL5;refGene_tx=XM_003984408.5;refGene_gn=AGBL5;refGene_tx
=XM_019827778.2;refGene_gn=AGBL5;refGene_tx=XM_019827779.1;refGene_gn=
AGBL5;refGene_tx=XM_019827780.1;refGene_gn=AGBL5;refGene_tx=XM_0198277
81.2
A3 121431229 121431393
refGene_gn=AGBL5;refGene_tx=XM_003984408.5;refGene_gn=AGBL5;refGene_tx
=XM_019827778.2;refGene_gn=AGBL5;refGene_tx=XM_019827779.1;refGene_gn=
AGBL5;refGene_tx=XM_019827780.1;refGene_gn=AGBL5;refGene_tx=XM_0198277
81.2
A3 121431712 121431884
refGene_gn=AGBL5;refGene_tx=XM_003984408.5;refGene_gn=AGBL5;refGene_tx
=XM_019827778.2;refGene_gn=AGBL5;refGene_tx=XM_019827779.1;refGene_gn=
AGBL5;refGene_tx=XM_019827780.1;refGene_gn=AGBL5;refGene_tx=XM_0198277
81.2
A3 121432132 121432347
refGene_gn=AGBL5;refGene_tx=XM_003984408.5;refGene_gn=AGBL5;refGene_tx
=XM_019827778.2;refGene_gn=AGBL5;refGene_tx=XM_019827779.1;refGene_gn=
AGBL5;refGene_tx=XM_019827780.1;refGene_gn=AGBL5;refGene_tx=XM_0198277
81.2
A3 121439355 121439482
refGene_gn=TMEM214;refGene_tx=XM_023252017.1
A3 121439691 121439843
refGene_gn=TMEM214;refGene_tx=XM_023252017.1
A3 121439972 121440141
refGene_gn=TMEM214;refGene_tx=XM_023252017.1;refGene_gn=TMEM214;refGen
e_tx=XM_023252018.1
A3 121440331 121440453
refGene_gn=TMEM214;refGene_tx=XM_023252017.1;refGene_gn=TMEM214;refGen
e_tx=XM_023252018.1
A3 121440998 121441116
refGene_gn=TMEM214;refGene_tx=XM_023252017.1;refGene_gn=TMEM214;refGen
e_tx=XM_023252018.1
A3 121441287 121441401
refGene_gn=TMEM214;refGene_tx=XM_023252017.1;refGene_gn=TMEM214;refGen
e_tx=XM_023252018.1
A3 121441586 121441635
refGene_gn=TMEM214;refGene_tx=XM_023252017.1;refGene_gn=TMEM214;refGen
e_tx=XM_023252018.1
A3 121441902 121441994
refGene_gn=TMEM214;refGene_tx=XM_023252017.1;refGene_gn=TMEM214;refGen
e_tx=XM_023252018.1
A3 121442380 121442522
refGene_gn=TMEM214;refGene_tx=XM_023252017.1;refGene_gn=TMEM214;refGen
e_tx=XM_023252018.1
A3 121442882 121442984
refGene_gn=TMEM214;refGene_tx=XM_023252017.1;refGene_gn=TMEM214;refGen
e_tx=XM_023252018.1

A3 121443495 121443577
refGene_gn=TMEM214;refGene_tx=XM_023252017.1;refGene_gn=TMEM214;refGene_tx=XM_023252018.1
A3 121443686 121443792
refGene_gn=TMEM214;refGene_tx=XM_023252017.1;refGene_gn=TMEM214;refGene_tx=XM_023252018.1
A3 121444235 121444318
refGene_gn=TMEM214;refGene_tx=XM_023252017.1;refGene_gn=TMEM214;refGene_tx=XM_023252018.1
A3 121444577 121444712
refGene_gn=TMEM214;refGene_tx=XM_023252017.1;refGene_gn=TMEM214;refGene_tx=XM_023252018.1
A3 121445009 121445160
refGene_gn=TMEM214;refGene_tx=XM_023252017.1;refGene_gn=TMEM214;refGene_tx=XM_023252018.1
A3 121445716 121445916
refGene_gn=TMEM214;refGene_tx=XM_023252017.1;refGene_gn=TMEM214;refGene_tx=XM_023252018.1
A3 121446826 121446971
refGene_gn=TMEM214;refGene_tx=XM_023252017.1;refGene_gn=TMEM214;refGene_tx=XM_023252018.1
A3 121452129 121452198
refGene_gn=MAPRE3;refGene_tx=XM_006930422.4;refGene_gn=MAPRE3;refGene_tx=XM_019827785.2;refGene_gn=MAPRE3;refGene_tx=XM_019827786.2;refGene_gn=MAPRE3;refGene_tx=XM_023252019.1
A3 121452441 121452594
refGene_gn=MAPRE3;refGene_tx=XM_006930422.4;refGene_gn=MAPRE3;refGene_tx=XM_019827785.2;refGene_gn=MAPRE3;refGene_tx=XM_019827786.2;refGene_gn=MAPRE3;refGene_tx=XM_023252019.1
A3 121452711 121452866
refGene_gn=MAPRE3;refGene_tx=XM_006930422.4;refGene_gn=MAPRE3;refGene_tx=XM_019827785.2;refGene_gn=MAPRE3;refGene_tx=XM_019827786.2;refGene_gn=MAPRE3;refGene_tx=XM_023252019.1
A3 121454172 121454374
refGene_gn=MAPRE3;refGene_tx=XM_006930422.4;refGene_gn=MAPRE3;refGene_tx=XM_019827785.2;refGene_gn=MAPRE3;refGene_tx=XM_019827786.2;refGene_gn=MAPRE3;refGene_tx=XM_023252019.1
A3 121454992 121455138
refGene_gn=MAPRE3;refGene_tx=XM_006930422.4;refGene_gn=MAPRE3;refGene_tx=XM_019827785.2;refGene_gn=MAPRE3;refGene_tx=XM_019827786.2;refGene_gn=MAPRE3;refGene_tx=XM_023252019.1
A3 121455772 121455893
refGene_gn=MAPRE3;refGene_tx=XM_006930422.4;refGene_gn=MAPRE3;refGene_tx=XM_019827785.2;refGene_gn=MAPRE3;refGene_tx=XM_019827786.2;refGene_gn=MAPRE3;refGene_tx=XM_023252019.1
A3 121527626 121527712
refGene_gn=DPYSL5;refGene_tx=XM_011281274.3;refGene_gn=DPYSL5;refGene_tx=XM_019827783.2;refGene_gn=DPYSL5;refGene_tx=XM_019827784.2
A3 121529590 121529759
refGene_gn=DPYSL5;refGene_tx=XM_011281274.3;refGene_gn=DPYSL5;refGene_

tx=XM_019827783.2;refGene_gn=DPYSL5;refGene_tx=XM_019827784.2
A3 121534583 121534791
refGene_gn=DPYSL5;refGene_tx=XM_011281274.3;refGene_gn=DPYSL5;refGene_tx=XM_019827783.2;refGene_gn=DPYSL5;refGene_tx=XM_019827784.2
A3 121536171 121536314
refGene_gn=DPYSL5;refGene_tx=XM_011281274.3;refGene_gn=DPYSL5;refGene_tx=XM_019827783.2;refGene_gn=DPYSL5;refGene_tx=XM_019827784.2
A3 121537942 121538084
refGene_gn=DPYSL5;refGene_tx=XM_011281274.3;refGene_gn=DPYSL5;refGene_tx=XM_019827783.2;refGene_gn=DPYSL5;refGene_tx=XM_019827784.2
A3 121544783 121544940
refGene_gn=DPYSL5;refGene_tx=XM_011281274.3;refGene_gn=DPYSL5;refGene_tx=XM_019827783.2;refGene_gn=DPYSL5;refGene_tx=XM_019827784.2
A3 121545886 121545962
refGene_gn=DPYSL5;refGene_tx=XM_011281274.3;refGene_gn=DPYSL5;refGene_tx=XM_019827783.2;refGene_gn=DPYSL5;refGene_tx=XM_019827784.2
A3 121547760 121547805
refGene_gn=DPYSL5;refGene_tx=XM_011281274.3;refGene_gn=DPYSL5;refGene_tx=XM_019827783.2;refGene_gn=DPYSL5;refGene_tx=XM_019827784.2
A3 121550037 121550106
refGene_gn=DPYSL5;refGene_tx=XM_011281274.3;refGene_gn=DPYSL5;refGene_tx=XM_019827783.2;refGene_gn=DPYSL5;refGene_tx=XM_019827784.2
A3 121550960 121551140
refGene_gn=DPYSL5;refGene_tx=XM_011281274.3;refGene_gn=DPYSL5;refGene_tx=XM_019827783.2;refGene_gn=DPYSL5;refGene_tx=XM_019827784.2
A3 121553775 121553934
refGene_gn=DPYSL5;refGene_tx=XM_011281274.3;refGene_gn=DPYSL5;refGene_tx=XM_019827783.2;refGene_gn=DPYSL5;refGene_tx=XM_019827784.2
A3 121581147 121581408
refGene_gn=DPYSL5;refGene_tx=XM_011281274.3;refGene_gn=DPYSL5;refGene_tx=XM_019827783.2;refGene_gn=DPYSL5;refGene_tx=XM_019827784.2
A3 121656472 121656607
refGene_gn=CENPA;refGene_tx=XM_003984412.5
A3 121656924 121657002
refGene_gn=CENPA;refGene_tx=XM_003984412.5
A3 121657520 121657630
refGene_gn=CENPA;refGene_tx=XM_003984412.5
A3 121665635 121665729
refGene_gn=CENPA;refGene_tx=XM_003984412.5
A3 121671097 121671567
refGene_gn=SLC35F6;refGene_tx=XM_011281273.3
A3 121673120 121673231
refGene_gn=SLC35F6;refGene_tx=XM_011281273.3
A3 121674041 121674254
refGene_gn=SLC35F6;refGene_tx=XM_011281273.3
A3 121674496 121674668
refGene_gn=SLC35F6;refGene_tx=XM_011281273.3
A3 121675425 121675498
refGene_gn=SLC35F6;refGene_tx=XM_011281273.3
A3 121684270 121684347

refGene_gn=SLC35F6;refGene_tx=XM_011281273.3
A3 121711794 121712717
refGene_gn=KCNK3;refGene_tx=XM_003984414.5
A3 121746809 121747092
refGene_gn=KCNK3;refGene_tx=XM_003984414.5
A3 121750955 121751165
refGene_gn=CIB4;refGene_tx=XM_023252021.1
A3 121797647 121797785
refGene_gn=CIB4;refGene_tx=XM_003984415.5;refGene_gn=CIB4;refGene_tx=XM_019827788.2;refGene_gn=CIB4;refGene_tx=XM_023252020.1
A3 121798194 121798229
refGene_gn=CIB4;refGene_tx=XM_003984415.5;refGene_gn=CIB4;refGene_tx=XM_019827788.2;refGene_gn=CIB4;refGene_tx=XM_023252020.1
A3 121805256 121805353
refGene_gn=CIB4;refGene_tx=XM_003984415.5;refGene_gn=CIB4;refGene_tx=XM_019827788.2;refGene_gn=CIB4;refGene_tx=XM_023252020.1
A3 121835956 121836098
refGene_gn=CIB4;refGene_tx=XM_003984415.5;refGene_gn=CIB4;refGene_tx=XM_019827788.2;refGene_gn=CIB4;refGene_tx=XM_023252020.1;refGene_gn=CIB4;refGene_tx=XM_023252021.1
A3 121845869 121845979
refGene_gn=CIB4;refGene_tx=XM_003984415.5;refGene_gn=CIB4;refGene_tx=XM_019827788.2;refGene_gn=CIB4;refGene_tx=XM_023252020.1;refGene_gn=CIB4;refGene_tx=XM_023252021.1
A3 121846119 121846287
refGene_gn=CIB4;refGene_tx=XM_019827788.2;refGene_gn=CIB4;refGene_tx=XM_023252021.1
A3 121846842 121846931
refGene_gn=CIB4;refGene_tx=XM_003984415.5;refGene_gn=CIB4;refGene_tx=XM_023252020.1
A3 121847660 121847706
refGene_gn=CIB4;refGene_tx=XM_023252020.1
A3 121848707 121848738
refGene_gn=CIB4;refGene_tx=XM_003984415.5
A3 121850625 121850770
refGene_gn=CA3H2orf70;refGene_tx=XM_003984416.5;refGene_gn=CA3H2orf70;refGene_tx=XM_023252022.1
A3 121851878 121851991
refGene_gn=CA3H2orf70;refGene_tx=XM_003984416.5;refGene_gn=CA3H2orf70;refGene_tx=XM_023252022.1
A3 121853205 121853479
refGene_gn=CA3H2orf70;refGene_tx=XM_003984416.5;refGene_gn=CA3H2orf70;refGene_tx=XM_023252022.1
A3 121856383 121856424
refGene_gn=CA3H2orf70;refGene_tx=XM_023252022.1
A3 121863606 121863680
refGene_gn=CA3H2orf70;refGene_tx=XM_003984416.5
A3 121867701 121867780
refGene_gn=OTOF;refGene_tx=XM_019827790.2;refGene_gn=OTOF;refGene_tx=XM_019827793.2;refGene_gn=OTOF;refGene_tx=XM_023252024.1

A3	121882869	121882928
refGene_gn=OTOF;refGene_tx=XM_019827790.2;refGene_gn=OTOF;refGene_tx=XM_019827793.2;refGene_gn=OTOF;refGene_tx=XM_023252024.1
A3	121896757	121896846
refGene_gn=OTOF;refGene_tx=XM_019827790.2;refGene_gn=OTOF;refGene_tx=XM_019827793.2;refGene_gn=OTOF;refGene_tx=XM_023252024.1
A3	121903623	121903723
refGene_gn=OTOF;refGene_tx=XM_019827790.2;refGene_gn=OTOF;refGene_tx=XM_019827793.2;refGene_gn=OTOF;refGene_tx=XM_023252023.1;refGene_gn=OTOF;refGene_tx=XM_023252024.1
A3	121906175	121906357
refGene_gn=OTOF;refGene_tx=XM_019827790.2;refGene_gn=OTOF;refGene_tx=XM_019827793.2;refGene_gn=OTOF;refGene_tx=XM_023252023.1;refGene_gn=OTOF;refGene_tx=XM_023252024.1
A3	121918787	121918861
refGene_gn=OTOF;refGene_tx=XM_019827790.2;refGene_gn=OTOF;refGene_tx=XM_019827793.2;refGene_gn=OTOF;refGene_tx=XM_023252023.1;refGene_gn=OTOF;refGene_tx=XM_023252024.1
A3	121920471	121920598
refGene_gn=OTOF;refGene_tx=XM_019827790.2;refGene_gn=OTOF;refGene_tx=XM_019827793.2;refGene_gn=OTOF;refGene_tx=XM_023252023.1;refGene_gn=OTOF;refGene_tx=XM_023252024.1
A3	121921067	121921122
refGene_gn=OTOF;refGene_tx=XM_019827790.2;refGene_gn=OTOF;refGene_tx=XM_019827793.2;refGene_gn=OTOF;refGene_tx=XM_023252023.1;refGene_gn=OTOF;refGene_tx=XM_023252024.1
A3	121927428	121927560
refGene_gn=OTOF;refGene_tx=XM_019827790.2;refGene_gn=OTOF;refGene_tx=XM_019827793.2;refGene_gn=OTOF;refGene_tx=XM_023252023.1;refGene_gn=OTOF;refGene_tx=XM_023252024.1
A3	121931072	121931135
refGene_gn=OTOF;refGene_tx=XM_019827790.2;refGene_gn=OTOF;refGene_tx=XM_019827793.2;refGene_gn=OTOF;refGene_tx=XM_023252023.1;refGene_gn=OTOF;refGene_tx=XM_023252024.1
A3	121931522	121931607
refGene_gn=OTOF;refGene_tx=XM_019827790.2;refGene_gn=OTOF;refGene_tx=XM_019827793.2;refGene_gn=OTOF;refGene_tx=XM_023252023.1;refGene_gn=OTOF;refGene_tx=XM_023252024.1
A3	121934187	121934347
refGene_gn=OTOF;refGene_tx=XM_019827790.2;refGene_gn=OTOF;refGene_tx=XM_019827793.2;refGene_gn=OTOF;refGene_tx=XM_023252023.1;refGene_gn=OTOF;refGene_tx=XM_023252024.1
A3	121934705	121934892
refGene_gn=OTOF;refGene_tx=XM_019827790.2;refGene_gn=OTOF;refGene_tx=XM_019827793.2;refGene_gn=OTOF;refGene_tx=XM_023252023.1;refGene_gn=OTOF;refGene_tx=XM_023252024.1
A3	121937039	121937226
refGene_gn=OTOF;refGene_tx=XM_019827790.2;refGene_gn=OTOF;refGene_tx=XM_019827793.2;refGene_gn=OTOF;refGene_tx=XM_023252023.1;refGene_gn=OTOF;refGene_tx=XM_023252024.1

A3	121938986	121939210
refGene_gn=OTOF;refGene_tx=XM_019827790.2;refGene_gn=OTOF;refGene_tx=XM_019827793.2;refGene_gn=OTOF;refGene_tx=XM_023252023.1;refGene_gn=OTOF;refGene_tx=XM_023252024.1
A3	121939665	121939774
refGene_gn=OTOF;refGene_tx=XM_019827790.2;refGene_gn=OTOF;refGene_tx=XM_019827793.2;refGene_gn=OTOF;refGene_tx=XM_023252023.1;refGene_gn=OTOF;refGene_tx=XM_023252024.1
A3	121940288	121940469
refGene_gn=OTOF;refGene_tx=XM_019827790.2;refGene_gn=OTOF;refGene_tx=XM_019827793.2;refGene_gn=OTOF;refGene_tx=XM_023252023.1;refGene_gn=OTOF;refGene_tx=XM_023252024.1
A3	121940587	121940708
refGene_gn=OTOF;refGene_tx=XM_019827790.2;refGene_gn=OTOF;refGene_tx=XM_019827793.2;refGene_gn=OTOF;refGene_tx=XM_023252023.1;refGene_gn=OTOF;refGene_tx=XM_023252024.1
A3	121941295	121941396
refGene_gn=OTOF;refGene_tx=XM_019827790.2;refGene_gn=OTOF;refGene_tx=XM_019827793.2;refGene_gn=OTOF;refGene_tx=XM_023252023.1;refGene_gn=OTOF;refGene_tx=XM_023252024.1
A3	121941516	121941607
refGene_gn=OTOF;refGene_tx=XM_019827790.2;refGene_gn=OTOF;refGene_tx=XM_019827793.2;refGene_gn=OTOF;refGene_tx=XM_023252023.1;refGene_gn=OTOF;refGene_tx=XM_023252024.1
A3	121941750	121941867
refGene_gn=OTOF;refGene_tx=XM_019827790.2;refGene_gn=OTOF;refGene_tx=XM_019827793.2;refGene_gn=OTOF;refGene_tx=XM_023252023.1;refGene_gn=OTOF;refGene_tx=XM_023252024.1
A3	121942205	121942358
refGene_gn=OTOF;refGene_tx=XM_019827790.2;refGene_gn=OTOF;refGene_tx=XM_019827793.2;refGene_gn=OTOF;refGene_tx=XM_023252023.1;refGene_gn=OTOF;refGene_tx=XM_023252024.1
A3	121942719	121942909
refGene_gn=OTOF;refGene_tx=XM_019827790.2;refGene_gn=OTOF;refGene_tx=XM_019827793.2;refGene_gn=OTOF;refGene_tx=XM_023252023.1;refGene_gn=OTOF;refGene_tx=XM_023252024.1
A3	121942989	121943114
refGene_gn=OTOF;refGene_tx=XM_019827790.2;refGene_gn=OTOF;refGene_tx=XM_019827793.2;refGene_gn=OTOF;refGene_tx=XM_023252023.1;refGene_gn=OTOF;refGene_tx=XM_023252024.1
A3	121943458	121943593
refGene_gn=OTOF;refGene_tx=XM_019827790.2;refGene_gn=OTOF;refGene_tx=XM_019827793.2;refGene_gn=OTOF;refGene_tx=XM_023252023.1;refGene_gn=OTOF;refGene_tx=XM_023252024.1
A3	121944287	121944449
refGene_gn=OTOF;refGene_tx=XM_019827790.2;refGene_gn=OTOF;refGene_tx=XM_019827793.2;refGene_gn=OTOF;refGene_tx=XM_023252023.1;refGene_gn=OTOF;refGene_tx=XM_023252024.1
A3	121944629	121944749
refGene_gn=OTOF;refGene_tx=XM_019827790.2;refGene_gn=OTOF;refGene_tx=X

M_019827793.2;refGene_gn=OTOF;refGene_tx=XM_023252023.1;refGene_gn=OTOF;refGene_tx=XM_023252024.1
A3 121945206 121945368
refGene_gn=OTOF;refGene_tx=XM_019827790.2;refGene_gn=OTOF;refGene_tx=XM_019827793.2;refGene_gn=OTOF;refGene_tx=XM_023252023.1;refGene_gn=OTOF;refGene_tx=XM_023252024.1
A3 121945469 121945632
refGene_gn=OTOF;refGene_tx=XM_019827790.2;refGene_gn=OTOF;refGene_tx=XM_019827793.2;refGene_gn=OTOF;refGene_tx=XM_023252023.1;refGene_gn=OTOF;refGene_tx=XM_023252024.1
A3 121946136 121946267
refGene_gn=OTOF;refGene_tx=XM_019827790.2;refGene_gn=OTOF;refGene_tx=XM_019827793.2;refGene_gn=OTOF;refGene_tx=XM_023252023.1;refGene_gn=OTOF;refGene_tx=XM_023252024.1
A3 121947804 121947834
refGene_gn=OTOF;refGene_tx=XM_019827790.2;refGene_gn=OTOF;refGene_tx=XM_019827793.2;refGene_gn=OTOF;refGene_tx=XM_023252023.1;refGene_gn=OTOF;refGene_tx=XM_023252024.1
A3 121948231 121948363
refGene_gn=OTOF;refGene_tx=XM_019827790.2;refGene_gn=OTOF;refGene_tx=XM_019827793.2;refGene_gn=OTOF;refGene_tx=XM_023252023.1;refGene_gn=OTOF;refGene_tx=XM_023252024.1
A3 121950365 121950432
refGene_gn=OTOF;refGene_tx=XM_019827790.2;refGene_gn=OTOF;refGene_tx=XM_019827793.2;refGene_gn=OTOF;refGene_tx=XM_023252023.1;refGene_gn=OTOF;refGene_tx=XM_023252024.1
A3 121951292 121951429
refGene_gn=OTOF;refGene_tx=XM_019827790.2;refGene_gn=OTOF;refGene_tx=XM_019827793.2;refGene_gn=OTOF;refGene_tx=XM_023252023.1;refGene_gn=OTOF;refGene_tx=XM_023252024.1
A3 121951556 121951691
refGene_gn=OTOF;refGene_tx=XM_019827790.2;refGene_gn=OTOF;refGene_tx=XM_019827793.2;refGene_gn=OTOF;refGene_tx=XM_023252023.1;refGene_gn=OTOF;refGene_tx=XM_023252024.1
A3 121952005 121952143
refGene_gn=OTOF;refGene_tx=XM_019827790.2;refGene_gn=OTOF;refGene_tx=XM_019827793.2;refGene_gn=OTOF;refGene_tx=XM_023252023.1;refGene_gn=OTOF;refGene_tx=XM_023252024.1
A3 121952478 121952606
refGene_gn=OTOF;refGene_tx=XM_019827790.2;refGene_gn=OTOF;refGene_tx=XM_019827793.2;refGene_gn=OTOF;refGene_tx=XM_023252023.1;refGene_gn=OTOF;refGene_tx=XM_023252024.1
A3 121952704 121952875
refGene_gn=OTOF;refGene_tx=XM_019827790.2;refGene_gn=OTOF;refGene_tx=XM_019827793.2;refGene_gn=OTOF;refGene_tx=XM_023252023.1;refGene_gn=OTOF;refGene_tx=XM_023252024.1
A3 121953511 121953672
refGene_gn=OTOF;refGene_tx=XM_019827790.2;refGene_gn=OTOF;refGene_tx=XM_019827793.2;refGene_gn=OTOF;refGene_tx=XM_023252023.1;refGene_gn=OTOF;refGene_tx=XM_023252024.1

A3	121954344	121954487
refGene_gn=OTOF;refGene_tx=XM_019827790.2;refGene_gn=OTOF;refGene_tx=XM_019827793.2;refGene_gn=OTOF;refGene_tx=XM_023252023.1;refGene_gn=OTOF;refGene_tx=XM_023252024.1
A3	121954870	121954959
refGene_gn=OTOF;refGene_tx=XM_019827790.2;refGene_gn=OTOF;refGene_tx=XM_019827793.2;refGene_gn=OTOF;refGene_tx=XM_023252023.1;refGene_gn=OTOF;refGene_tx=XM_023252024.1
A3	121956579	121956678
refGene_gn=OTOF;refGene_tx=XM_019827790.2;refGene_gn=OTOF;refGene_tx=XM_019827793.2;refGene_gn=OTOF;refGene_tx=XM_023252023.1;refGene_gn=OTOF;refGene_tx=XM_023252024.1
A3	121956809	121957051
refGene_gn=OTOF;refGene_tx=XM_019827790.2;refGene_gn=OTOF;refGene_tx=XM_019827793.2;refGene_gn=OTOF;refGene_tx=XM_023252023.1;refGene_gn=OTOF;refGene_tx=XM_023252024.1
A3	121957166	121957345
refGene_gn=OTOF;refGene_tx=XM_019827790.2;refGene_gn=OTOF;refGene_tx=XM_019827793.2;refGene_gn=OTOF;refGene_tx=XM_023252023.1;refGene_gn=OTOF;refGene_tx=XM_023252024.1
A3	121957436	121957537
refGene_gn=OTOF;refGene_tx=XM_019827790.2;refGene_gn=OTOF;refGene_tx=XM_019827793.2;refGene_gn=OTOF;refGene_tx=XM_023252023.1;refGene_gn=OTOF;refGene_tx=XM_023252024.1
A3	121958011	121958122
refGene_gn=OTOF;refGene_tx=XM_023252024.1
A3	121959614	121959795
refGene_gn=OTOF;refGene_tx=XM_019827790.2;refGene_gn=OTOF;refGene_tx=XM_019827793.2;refGene_gn=OTOF;refGene_tx=XM_023252023.1;refGene_gn=OTOF;refGene_tx=XM_023252024.1
A3	121961365	121961422
refGene_gn=DRC1;refGene_tx=XM_023252025.1;refGene_gn=DRC1;refGene_tx=XM_023252026.1
A3	121962630	121962733
refGene_gn=DRC1;refGene_tx=XM_023252025.1;refGene_gn=DRC1;refGene_tx=XM_023252026.1
A3	121963075	121963219
refGene_gn=DRC1;refGene_tx=XM_023252025.1;refGene_gn=DRC1;refGene_tx=XM_023252026.1
A3	121964255	121964440
refGene_gn=DRC1;refGene_tx=XM_023252025.1;refGene_gn=DRC1;refGene_tx=XM_023252026.1
A3	121966978	121967065
refGene_gn=DRC1;refGene_tx=XM_023252025.1;refGene_gn=DRC1;refGene_tx=XM_023252026.1
A3	121967609	121967699
refGene_gn=DRC1;refGene_tx=XM_023252025.1;refGene_gn=DRC1;refGene_tx=XM_023252026.1
A3	121968933	121969046
refGene_gn=DRC1;refGene_tx=XM_023252025.1;refGene_gn=DRC1;refGene_tx=X

M_023252026.1
A3 121971077 121971310
refGene_gn=DRC1;refGene_tx=XM_023252025.1;refGene_gn=DRC1;refGene_tx=XM_023252026.1
A3 121971584 121971719
refGene_gn=DRC1;refGene_tx=XM_023252025.1;refGene_gn=DRC1;refGene_tx=XM_023252026.1
A3 121974884 121975024
refGene_gn=DRC1;refGene_tx=XM_023252025.1;refGene_gn=DRC1;refGene_tx=XM_023252026.1
A3 121982951 121983074
refGene_gn=DRC1;refGene_tx=XM_023252025.1;refGene_gn=DRC1;refGene_tx=XM_023252026.1
A3 121983949 121984036
refGene_gn=DRC1;refGene_tx=XM_023252025.1;refGene_gn=DRC1;refGene_tx=XM_023252026.1
A3 121984973 121985111
refGene_gn=DRC1;refGene_tx=XM_023252025.1;refGene_gn=DRC1;refGene_tx=XM_023252026.1
A3 121985712 121985896
refGene_gn=DRC1;refGene_tx=XM_023252025.1;refGene_gn=DRC1;refGene_tx=XM_023252026.1
A3 121987823 121987936
refGene_gn=DRC1;refGene_tx=XM_023252025.1;refGene_gn=DRC1;refGene_tx=XM_023252026.1
A3 121988026 121988065
refGene_gn=DRC1;refGene_tx=XM_023252026.1
A3 121992147 121992235
refGene_gn=DRC1;refGene_tx=XM_023252025.1
A3 122003378 122003530
refGene_gn=DRC1;refGene_tx=XM_023252025.1
A3 122014607 122014706
refGene_gn=SELENOI;refGene_tx=XM_011281271.3
A3 122017322 122017505
refGene_gn=SELENOI;refGene_tx=XM_011281271.3
A3 122019746 122019927
refGene_gn=SELENOI;refGene_tx=XM_011281271.3;refGene_gn=SELENOI;refGene_tx=XM_019827799.2
A3 122021305 122021354
refGene_gn=SELENOI;refGene_tx=XM_011281271.3;refGene_gn=SELENOI;refGene_tx=XM_019827799.2
A3 122027249 122027358
refGene_gn=SELENOI;refGene_tx=XM_011281271.3;refGene_gn=SELENOI;refGene_tx=XM_019827799.2
A3 122028311 122028574
refGene_gn=SELENOI;refGene_tx=XM_011281271.3;refGene_gn=SELENOI;refGene_tx=XM_019827799.2
A3 122034455 122034530
refGene_gn=SELENOI;refGene_tx=XM_011281271.3;refGene_gn=SELENOI;refGene_tx=XM_019827799.2

A3	122036635	122036744
refGene_gn=SELENOI;refGene_tx=XM_011281271.3;refGene_gn=SELENOI;refGen
e_tx=XM_019827799.2
A3	122037201	122037270
refGene_gn=SELENOI;refGene_tx=XM_011281271.3;refGene_gn=SELENOI;refGen
e_tx=XM_019827799.2
A3	122054266	122054323
refGene_gn=SELENOI;refGene_tx=XM_011281271.3;refGene_gn=SELENOI;refGen
e_tx=XM_019827799.2
A3	122080649	122080733
refGene_gn=ADGRF3;refGene_tx=XM_006930448.4
A3	122082037	122082200
refGene_gn=ADGRF3;refGene_tx=XM_006930448.4;refGene_gn=ADGRF3;refGene_
tx=XM_019827800.2
A3	122082604	122082745
refGene_gn=ADGRF3;refGene_tx=XM_006930448.4;refGene_gn=ADGRF3;refGene_
tx=XM_019827800.2
A3	122083216	122083390
refGene_gn=ADGRF3;refGene_tx=XM_006930448.4;refGene_gn=ADGRF3;refGene_
tx=XM_019827800.2
A3	122083870	122084116
refGene_gn=ADGRF3;refGene_tx=XM_006930448.4;refGene_gn=ADGRF3;refGene_
tx=XM_019827800.2
A3	122084848	122085058
refGene_gn=ADGRF3;refGene_tx=XM_006930448.4;refGene_gn=ADGRF3;refGene_
tx=XM_019827800.2
A3	122085563	122085707
refGene_gn=ADGRF3;refGene_tx=XM_006930448.4;refGene_gn=ADGRF3;refGene_
tx=XM_019827800.2
A3	122085878	122086075
refGene_gn=ADGRF3;refGene_tx=XM_006930448.4;refGene_gn=ADGRF3;refGene_
tx=XM_019827800.2
A3	122086469	122086649
refGene_gn=ADGRF3;refGene_tx=XM_006930448.4;refGene_gn=ADGRF3;refGene_
tx=XM_019827800.2
A3	122087575	122088955
refGene_gn=ADGRF3;refGene_tx=XM_006930448.4;refGene_gn=ADGRF3;refGene_
tx=XM_019827800.2
A3	122089425	122089467
refGene_gn=ADGRF3;refGene_tx=XM_006930448.4;refGene_gn=ADGRF3;refGene_
tx=XM_019827800.2
A3	122089555	122089717
refGene_gn=ADGRF3;refGene_tx=XM_006930448.4;refGene_gn=ADGRF3;refGene_
tx=XM_019827800.2
A3	122101112	122101148
refGene_gn=HADHB;refGene_tx=XM_003984419.5;refGene_gn=HADHB;refGene_tx
=XM_019827803.2
A3	122102831	122102996
refGene_gn=HADHB;refGene_tx=XM_003984419.5;refGene_gn=HADHB;refGene_tx
=XM_019827803.2

A3	122103402	122103477
refGene_gn=HADHB;refGene_tx=XM_003984419.5;refGene_gn=HADHB;refGene_tx=XM_019827803.2
A3	122103801	122103889
refGene_gn=HADHB;refGene_tx=XM_003984419.5;refGene_gn=HADHB;refGene_tx=XM_019827803.2
A3	122104478	122104526
refGene_gn=HADHB;refGene_tx=XM_003984419.5;refGene_gn=HADHB;refGene_tx=XM_019827803.2
A3	122104606	122104686
refGene_gn=HADHB;refGene_tx=XM_003984419.5;refGene_gn=HADHB;refGene_tx=XM_019827803.2
A3	122106971	122107093
refGene_gn=HADHB;refGene_tx=XM_003984419.5;refGene_gn=HADHB;refGene_tx=XM_019827803.2
A3	122108458	122108639
refGene_gn=HADHB;refGene_tx=XM_003984419.5;refGene_gn=HADHB;refGene_tx=XM_019827803.2
A3	122108945	122109133
refGene_gn=HADHB;refGene_tx=XM_003984419.5;refGene_gn=HADHB;refGene_tx=XM_019827803.2
A3	122110690	122110778
refGene_gn=HADHB;refGene_tx=XM_003984419.5;refGene_gn=HADHB;refGene_tx=XM_019827803.2
A3	122114417	122114517
refGene_gn=HADHB;refGene_tx=XM_003984419.5;refGene_gn=HADHB;refGene_tx=XM_019827803.2
A3	122119922	122119967
refGene_gn=HADHB;refGene_tx=XM_003984419.5;refGene_gn=HADHB;refGene_tx=XM_019827803.2
A3	122123940	122124040
refGene_gn=HADHB;refGene_tx=XM_003984419.5;refGene_gn=HADHB;refGene_tx=XM_019827803.2
A3	122132048	122132093
refGene_gn=HADHB;refGene_tx=XM_003984419.5;refGene_gn=HADHB;refGene_tx=XM_019827803.2
A3	122132199	122132266
refGene_gn=HADHB;refGene_tx=XM_003984419.5;refGene_gn=HADHB;refGene_tx=XM_019827803.2
A3	122140952	122141019
refGene_gn=HADHA;refGene_tx=XM_003984421.5
A3	122149978	122150020
refGene_gn=HADHA;refGene_tx=XM_003984421.5
A3	122150113	122150184
refGene_gn=HADHA;refGene_tx=XM_003984421.5
A3	122151495	122151629
refGene_gn=HADHA;refGene_tx=XM_003984421.5
A3	122154732	122154871
refGene_gn=HADHA;refGene_tx=XM_003984421.5
A3	122155798	122155918

refGene_gn=HADHA;refGene_tx=XM_003984421.5
A3	122158447	122158550
refGene_gn=HADHA;refGene_tx=XM_003984421.5
A3	122167714	122167837
refGene_gn=HADHA;refGene_tx=XM_003984421.5
A3	122168339	122168458
refGene_gn=HADHA;refGene_tx=XM_003984421.5
A3	122170691	122170748
refGene_gn=HADHA;refGene_tx=XM_003984421.5
A3	122172616	122172726
refGene_gn=HADHA;refGene_tx=XM_003984421.5
A3	122176730	122176865
refGene_gn=HADHA;refGene_tx=XM_003984421.5
A3	122179126	122179298
refGene_gn=HADHA;refGene_tx=XM_003984421.5
A3	122181597	122181684
refGene_gn=HADHA;refGene_tx=XM_003984421.5
A3	122185150	122185291
refGene_gn=HADHA;refGene_tx=XM_003984421.5
A3	122185728	122185797
refGene_gn=HADHA;refGene_tx=XM_003984421.5
A3	122186587	122186783
refGene_gn=HADHA;refGene_tx=XM_003984421.5
A3	122187471	122187586
refGene_gn=HADHA;refGene_tx=XM_003984421.5
A3	122188110	122188256
refGene_gn=HADHA;refGene_tx=XM_003984421.5
A3	122188344	122188490
refGene_gn=HADHA;refGene_tx=XM_003984421.5
A3	122191504	122192522
refGene_gn=GAREM2;refGene_tx=XM_023252027.1
A3	122193004	122193174
refGene_gn=GAREM2;refGene_tx=XM_023252027.1
A3	122194155	122195193
refGene_gn=GAREM2;refGene_tx=XM_023252027.1
A3	122195888	122196019
refGene_gn=GAREM2;refGene_tx=XM_023252027.1
A3	122201981	122202122
refGene_gn=GAREM2;refGene_tx=XM_023252027.1
A3	122204943	122205055
refGene_gn=GAREM2;refGene_tx=XM_023252027.1
A3	122235150	122235234
refGene_gn=RAB10;refGene_tx=XM_003984422.5
A3	122240263	122240365
refGene_gn=RAB10;refGene_tx=XM_003984422.5
A3	122240861	122240951
refGene_gn=RAB10;refGene_tx=XM_003984422.5
A3	122246154	122246293
refGene_gn=RAB10;refGene_tx=XM_003984422.5
A3	122261531	122261592

refGene_gn=RAB10;refGene_tx=XM_003984422.5
A3 122309760 122309887
refGene_gn=RAB10;refGene_tx=XM_003984422.5
A3 122360986 122362537
refGene_gn=KIF3C;refGene_tx=XM_003984423.5
A3 122382094 122382196
refGene_gn=KIF3C;refGene_tx=XM_003984423.5
A3 122382840 122382963
refGene_gn=KIF3C;refGene_tx=XM_003984423.5
A3 122384102 122384221
refGene_gn=KIF3C;refGene_tx=XM_003984423.5
A3 122384714 122384831
refGene_gn=KIF3C;refGene_tx=XM_003984423.5
A3 122395691 122395800
refGene_gn=KIF3C;refGene_tx=XM_003984423.5
A3 122395977 122396150
refGene_gn=KIF3C;refGene_tx=XM_003984423.5
A3 122396393 122396487
refGene_gn=KIF3C;refGene_tx=XM_003984423.5
A3 122442663 122442720
refGene_gn=ASXL2;refGene_tx=XM_006930426.4;refGene_gn=ASXL2;refGene_tx=XM_019827805.2
A3 122491180 122491263
refGene_gn=ASXL2;refGene_tx=XM_006930426.4;refGene_gn=ASXL2;refGene_tx=XM_019827805.2
A3 122500543 122500546
refGene_gn=ASXL2;refGene_tx=XM_006930426.4
A3 122518728 122518837
refGene_gn=ASXL2;refGene_tx=XM_006930426.4;refGene_gn=ASXL2;refGene_tx=XM_019827805.2
A3 122522846 122522997
refGene_gn=ASXL2;refGene_tx=XM_006930426.4;refGene_gn=ASXL2;refGene_tx=XM_019827805.2
A3 122525835 122525872
refGene_gn=ASXL2;refGene_tx=XM_019827806.2
A3 122539588 122539704
refGene_gn=ASXL2;refGene_tx=XM_006930426.4;refGene_gn=ASXL2;refGene_tx=XM_019827805.2;refGene_gn=ASXL2;refGene_tx=XM_019827806.2
A3 122544724 122544854
refGene_gn=ASXL2;refGene_tx=XM_006930426.4;refGene_gn=ASXL2;refGene_tx=XM_019827805.2;refGene_gn=ASXL2;refGene_tx=XM_019827806.2;refGene_gn=ASXL2;refGene_tx=XM_019827807.2;refGene_gn=ASXL2;refGene_tx=XM_019827808.2
A3 122545640 122545784
refGene_gn=ASXL2;refGene_tx=XM_006930426.4;refGene_gn=ASXL2;refGene_tx=XM_019827805.2;refGene_gn=ASXL2;refGene_tx=XM_019827806.2;refGene_gn=ASXL2;refGene_tx=XM_019827807.2;refGene_gn=ASXL2;refGene_tx=XM_019827808.2
A3 122553130 122553294
refGene_gn=ASXL2;refGene_tx=XM_006930426.4;refGene_gn=ASXL2;refGene_tx

=XM_019827805.2;refGene_gn=ASXL2;refGene_tx=XM_019827806.2;refGene_gn=ASXL2;refGene_tx=XM_019827807.2;refGene_gn=ASXL2;refGene_tx=XM_019827808.2
A3 122558309 122558406
refGene_gn=ASXL2;refGene_tx=XM_006930426.4;refGene_gn=ASXL2;refGene_tx=XM_019827805.2;refGene_gn=ASXL2;refGene_tx=XM_019827806.2;refGene_gn=ASXL2;refGene_tx=XM_019827807.2;refGene_gn=ASXL2;refGene_tx=XM_019827808.2
A3 122560991 122561097
refGene_gn=ASXL2;refGene_tx=XM_006930426.4;refGene_gn=ASXL2;refGene_tx=XM_019827805.2;refGene_gn=ASXL2;refGene_tx=XM_019827806.2;refGene_gn=ASXL2;refGene_tx=XM_019827807.2;refGene_gn=ASXL2;refGene_tx=XM_019827808.2
A3 122562989 122563707
refGene_gn=ASXL2;refGene_tx=XM_006930426.4;refGene_gn=ASXL2;refGene_tx=XM_019827805.2;refGene_gn=ASXL2;refGene_tx=XM_019827806.2;refGene_gn=ASXL2;refGene_tx=XM_019827807.2;refGene_gn=ASXL2;refGene_tx=XM_019827808.2
A3 122566770 122569239
refGene_gn=ASXL2;refGene_tx=XM_006930426.4;refGene_gn=ASXL2;refGene_tx=XM_019827805.2;refGene_gn=ASXL2;refGene_tx=XM_019827806.2;refGene_gn=ASXL2;refGene_tx=XM_019827807.2;refGene_gn=ASXL2;refGene_tx=XM_019827808.2
A3 122653667 122653734
refGene_gn=DTNB;refGene_tx=XM_019827809.1;refGene_gn=DTNB;refGene_tx=XM_019827811.1;refGene_gn=DTNB;refGene_tx=XM_019827813.1;refGene_gn=DTNB;refGene_tx=XM_019827814.1;refGene_gn=DTNB;refGene_tx=XM_019827817.1;refGene_gn=DTNB;refGene_tx=XM_019827822.1
A3 122663187 122663268
refGene_gn=DTNB;refGene_tx=XM_019827809.1;refGene_gn=DTNB;refGene_tx=XM_019827811.1;refGene_gn=DTNB;refGene_tx=XM_019827813.1;refGene_gn=DTNB;refGene_tx=XM_019827814.1;refGene_gn=DTNB;refGene_tx=XM_019827817.1;refGene_gn=DTNB;refGene_tx=XM_019827822.1
A3 122669021 122669235
refGene_gn=DTNB;refGene_tx=XM_019827809.1;refGene_gn=DTNB;refGene_tx=XM_019827811.1;refGene_gn=DTNB;refGene_tx=XM_019827813.1;refGene_gn=DTNB;refGene_tx=XM_019827814.1;refGene_gn=DTNB;refGene_tx=XM_019827817.1;refGene_gn=DTNB;refGene_tx=XM_019827822.1
A3 122684158 122684244
refGene_gn=DTNB;refGene_tx=XM_019827809.1;refGene_gn=DTNB;refGene_tx=XM_019827811.1;refGene_gn=DTNB;refGene_tx=XM_019827813.1;refGene_gn=DTNB;refGene_tx=XM_019827814.1;refGene_gn=DTNB;refGene_tx=XM_019827817.1;refGene_gn=DTNB;refGene_tx=XM_019827822.1
A3 122695735 122695890
refGene_gn=DTNB;refGene_tx=XM_019827809.1;refGene_gn=DTNB;refGene_tx=XM_019827811.1;refGene_gn=DTNB;refGene_tx=XM_019827813.1;refGene_gn=DTNB;refGene_tx=XM_019827814.1;refGene_gn=DTNB;refGene_tx=XM_019827817.1;refGene_gn=DTNB;refGene_tx=XM_019827822.1
A3 122707460 122707566
refGene_gn=DTNB;refGene_tx=XM_019827809.1;refGene_gn=DTNB;refGene_tx=X

M_019827811.1;refGene_gn=DTNB;refGene_tx=XM_019827813.1;refGene_gn=DTN
B;refGene_tx=XM_019827814.1;refGene_gn=DTNB;refGene_tx=XM_019827817.1;
refGene_gn=DTNB;refGene_tx=XM_019827822.1
A3 122711623 122711790
refGene_gn=DTNB;refGene_tx=XM_019827809.1;refGene_gn=DTNB;refGene_tx=X
M_019827811.1;refGene_gn=DTNB;refGene_tx=XM_019827813.1;refGene_gn=DTN
B;refGene_tx=XM_019827814.1;refGene_gn=DTNB;refGene_tx=XM_019827817.1;
refGene_gn=DTNB;refGene_tx=XM_019827822.1
A3 122751169 122751294
refGene_gn=DTNB;refGene_tx=XM_019827809.1;refGene_gn=DTNB;refGene_tx=X
M_019827811.1;refGene_gn=DTNB;refGene_tx=XM_019827813.1;refGene_gn=DTN
B;refGene_tx=XM_019827814.1;refGene_gn=DTNB;refGene_tx=XM_019827817.1;
refGene_gn=DTNB;refGene_tx=XM_019827822.1
A3 122791387 122791465
refGene_gn=DTNB;refGene_tx=XM_019827809.1;refGene_gn=DTNB;refGene_tx=X
M_019827811.1;refGene_gn=DTNB;refGene_tx=XM_019827813.1;refGene_gn=DTN
B;refGene_tx=XM_019827814.1;refGene_gn=DTNB;refGene_tx=XM_019827817.1;
refGene_gn=DTNB;refGene_tx=XM_019827822.1
A3 122807341 122807431
refGene_gn=DTNB;refGene_tx=XM_019827809.1;refGene_gn=DTNB;refGene_tx=X
M_019827811.1;refGene_gn=DTNB;refGene_tx=XM_019827813.1;refGene_gn=DTN
B;refGene_tx=XM_019827814.1;refGene_gn=DTNB;refGene_tx=XM_019827817.1;
refGene_gn=DTNB;refGene_tx=XM_019827822.1
A3 122810608 122810696
refGene_gn=DTNB;refGene_tx=XM_019827809.1;refGene_gn=DTNB;refGene_tx=X
M_019827811.1;refGene_gn=DTNB;refGene_tx=XM_019827813.1;refGene_gn=DTN
B;refGene_tx=XM_019827814.1;refGene_gn=DTNB;refGene_tx=XM_019827817.1;
refGene_gn=DTNB;refGene_tx=XM_019827822.1
A3 122819776 122819818
refGene_gn=DTNB;refGene_tx=XM_019827809.1;refGene_gn=DTNB;refGene_tx=X
M_019827811.1;refGene_gn=DTNB;refGene_tx=XM_019827814.1;refGene_gn=DTN
B;refGene_tx=XM_019827817.1
A3 122826278 122826364
refGene_gn=DTNB;refGene_tx=XM_019827809.1;refGene_gn=DTNB;refGene_tx=X
M_019827811.1;refGene_gn=DTNB;refGene_tx=XM_019827813.1;refGene_gn=DTN
B;refGene_tx=XM_019827814.1;refGene_gn=DTNB;refGene_tx=XM_019827817.1;
refGene_gn=DTNB;refGene_tx=XM_019827822.1
A3 122827210 122827324
refGene_gn=DTNB;refGene_tx=XM_019827809.1;refGene_gn=DTNB;refGene_tx=X
M_019827811.1;refGene_gn=DTNB;refGene_tx=XM_019827813.1;refGene_gn=DTN
B;refGene_tx=XM_019827814.1;refGene_gn=DTNB;refGene_tx=XM_019827817.1;
refGene_gn=DTNB;refGene_tx=XM_019827822.1
A3 122832003 122832100
refGene_gn=DTNB;refGene_tx=XM_019827809.1;refGene_gn=DTNB;refGene_tx=X
M_019827811.1;refGene_gn=DTNB;refGene_tx=XM_019827813.1;refGene_gn=DTN
B;refGene_tx=XM_019827814.1;refGene_gn=DTNB;refGene_tx=XM_019827817.1;
refGene_gn=DTNB;refGene_tx=XM_019827822.1
A3 122838014 122838035
refGene_gn=DTNB;refGene_tx=XM_019827809.1;refGene_gn=DTNB;refGene_tx=X
M_019827811.1;refGene_gn=DTNB;refGene_tx=XM_019827813.1;refGene_gn=DTN

B;refGene_tx=XM_019827814.1;refGene_gn=DTNB;refGene_tx=XM_019827817.1
A3 122859432 122859592
refGene_gn=DTNB;refGene_tx=XM_019827809.1;refGene_gn=DTNB;refGene_tx=XM_019827811.1;refGene_gn=DTNB;refGene_tx=XM_019827813.1;refGene_gn=DTNB;refGene_tx=XM_019827814.1;refGene_gn=DTNB;refGene_tx=XM_019827817.1;refGene_gn=DTNB;refGene_tx=XM_019827822.1
A3 122860424 122860514
refGene_gn=DTNB;refGene_tx=XM_019827809.1;refGene_gn=DTNB;refGene_tx=XM_019827811.1;refGene_gn=DTNB;refGene_tx=XM_019827813.1;refGene_gn=DTNB;refGene_tx=XM_019827814.1;refGene_gn=DTNB;refGene_tx=XM_019827822.1
A3 122860651 122860698
refGene_gn=DTNB;refGene_tx=XM_019827811.1;refGene_gn=DTNB;refGene_tx=XM_019827813.1
A3 122864354 122864408
refGene_gn=DTNB;refGene_tx=XM_019827809.1
A3 122867890 122867895
refGene_gn=DTNB;refGene_tx=XM_019827809.1;refGene_gn=DTNB;refGene_tx=XM_019827814.1;refGene_gn=DTNB;refGene_tx=XM_019827817.1
A3 122869789 122870115
refGene_gn=DTNB;refGene_tx=XM_019827822.1
A3 122926844 122926907
refGene_gn=DNMT3A;refGene_tx=XM_019827823.2;refGene_gn=DNMT3A;refGene_tx=XM_023252030.1;refGene_gn=DNMT3A;refGene_tx=XM_023252031.1;refGene_gn=DNMT3A;refGene_tx=XM_023252032.1;refGene_gn=DNMT3A;refGene_tx=XM_023252033.1;refGene_gn=DNMT3A;refGene_tx=XM_023252034.1
A3 122939105 122939210
refGene_gn=DNMT3A;refGene_tx=XM_019827823.2;refGene_gn=DNMT3A;refGene_tx=XM_023252030.1;refGene_gn=DNMT3A;refGene_tx=XM_023252031.1;refGene_gn=DNMT3A;refGene_tx=XM_023252032.1;refGene_gn=DNMT3A;refGene_tx=XM_023252033.1;refGene_gn=DNMT3A;refGene_tx=XM_023252034.1
A3 122955135 122955406
refGene_gn=DNMT3A;refGene_tx=XM_019827823.2;refGene_gn=DNMT3A;refGene_tx=XM_023252030.1;refGene_gn=DNMT3A;refGene_tx=XM_023252031.1;refGene_gn=DNMT3A;refGene_tx=XM_023252032.1;refGene_gn=DNMT3A;refGene_tx=XM_023252033.1;refGene_gn=DNMT3A;refGene_tx=XM_023252034.1
A3 122961539 122961583
refGene_gn=DNMT3A;refGene_tx=XM_019827823.2;refGene_gn=DNMT3A;refGene_tx=XM_023252030.1;refGene_gn=DNMT3A;refGene_tx=XM_023252031.1;refGene_gn=DNMT3A;refGene_tx=XM_023252032.1;refGene_gn=DNMT3A;refGene_tx=XM_023252033.1;refGene_gn=DNMT3A;refGene_tx=XM_023252034.1
A3 122962037 122962184
refGene_gn=DNMT3A;refGene_tx=XM_019827823.2;refGene_gn=DNMT3A;refGene_tx=XM_023252030.1;refGene_gn=DNMT3A;refGene_tx=XM_023252031.1;refGene_gn=DNMT3A;refGene_tx=XM_023252032.1;refGene_gn=DNMT3A;refGene_tx=XM_023252033.1;refGene_gn=DNMT3A;refGene_tx=XM_023252034.1
A3 122985734 122985950
refGene_gn=DNMT3A;refGene_tx=XM_019827823.2;refGene_gn=DNMT3A;refGene_tx=XM_023252030.1;refGene_gn=DNMT3A;refGene_tx=XM_023252031.1;refGene_gn=DNMT3A;refGene_tx=XM_023252032.1;refGene_gn=DNMT3A;refGene_tx=XM_023252033.1;refGene_gn=DNMT3A;refGene_tx=XM_023252034.1

A3 122986214 122986373
refGene_gn=DNMT3A;refGene_tx=XM_019827823.2;refGene_gn=DNMT3A;refGene_tx=XM_023252030.1;refGene_gn=DNMT3A;refGene_tx=XM_023252031.1;refGene_gn=DNMT3A;refGene_tx=XM_023252032.1;refGene_gn=DNMT3A;refGene_tx=XM_023252033.1;refGene_gn=DNMT3A;refGene_tx=XM_023252034.1
A3 122986743 122986851
refGene_gn=DNMT3A;refGene_tx=XM_019827823.2;refGene_gn=DNMT3A;refGene_tx=XM_023252030.1;refGene_gn=DNMT3A;refGene_tx=XM_023252031.1;refGene_gn=DNMT3A;refGene_tx=XM_023252032.1;refGene_gn=DNMT3A;refGene_tx=XM_023252033.1;refGene_gn=DNMT3A;refGene_tx=XM_023252034.1
A3 122987073 122987230
refGene_gn=DNMT3A;refGene_tx=XM_019827823.2;refGene_gn=DNMT3A;refGene_tx=XM_023252030.1;refGene_gn=DNMT3A;refGene_tx=XM_023252031.1;refGene_gn=DNMT3A;refGene_tx=XM_023252032.1;refGene_gn=DNMT3A;refGene_tx=XM_023252033.1;refGene_gn=DNMT3A;refGene_tx=XM_023252034.1
A3 122987560 122987710
refGene_gn=DNMT3A;refGene_tx=XM_019827823.2;refGene_gn=DNMT3A;refGene_tx=XM_023252030.1;refGene_gn=DNMT3A;refGene_tx=XM_023252031.1;refGene_gn=DNMT3A;refGene_tx=XM_023252032.1;refGene_gn=DNMT3A;refGene_tx=XM_023252033.1;refGene_gn=DNMT3A;refGene_tx=XM_023252034.1
A3 122987817 122987862
refGene_gn=DNMT3A;refGene_tx=XM_019827823.2;refGene_gn=DNMT3A;refGene_tx=XM_023252030.1;refGene_gn=DNMT3A;refGene_tx=XM_023252031.1;refGene_gn=DNMT3A;refGene_tx=XM_023252032.1;refGene_gn=DNMT3A;refGene_tx=XM_023252033.1;refGene_gn=DNMT3A;refGene_tx=XM_023252034.1
A3 122988606 122988686
refGene_gn=DNMT3A;refGene_tx=XM_019827823.2;refGene_gn=DNMT3A;refGene_tx=XM_023252030.1;refGene_gn=DNMT3A;refGene_tx=XM_023252031.1;refGene_gn=DNMT3A;refGene_tx=XM_023252032.1;refGene_gn=DNMT3A;refGene_tx=XM_023252033.1;refGene_gn=DNMT3A;refGene_tx=XM_023252034.1
A3 122989199 122989312
refGene_gn=DNMT3A;refGene_tx=XM_019827823.2;refGene_gn=DNMT3A;refGene_tx=XM_023252030.1;refGene_gn=DNMT3A;refGene_tx=XM_023252031.1;refGene_gn=DNMT3A;refGene_tx=XM_023252032.1;refGene_gn=DNMT3A;refGene_tx=XM_023252033.1;refGene_gn=DNMT3A;refGene_tx=XM_023252034.1
A3 122989468 122989652
refGene_gn=DNMT3A;refGene_tx=XM_019827823.2;refGene_gn=DNMT3A;refGene_tx=XM_023252030.1;refGene_gn=DNMT3A;refGene_tx=XM_023252031.1;refGene_gn=DNMT3A;refGene_tx=XM_023252032.1;refGene_gn=DNMT3A;refGene_tx=XM_023252033.1;refGene_gn=DNMT3A;refGene_tx=XM_023252034.1
A3 122989868 122989953
refGene_gn=DNMT3A;refGene_tx=XM_019827823.2;refGene_gn=DNMT3A;refGene_tx=XM_023252030.1;refGene_gn=DNMT3A;refGene_tx=XM_023252031.1;refGene_gn=DNMT3A;refGene_tx=XM_023252032.1;refGene_gn=DNMT3A;refGene_tx=XM_023252033.1;refGene_gn=DNMT3A;refGene_tx=XM_023252034.1
A3 122992194 122992340
refGene_gn=DNMT3A;refGene_tx=XM_019827823.2;refGene_gn=DNMT3A;refGene_tx=XM_023252030.1;refGene_gn=DNMT3A;refGene_tx=XM_023252031.1;refGene_gn=DNMT3A;refGene_tx=XM_023252032.1;refGene_gn=DNMT3A;refGene_tx=XM_023252033.1;refGene_gn=DNMT3A;refGene_tx=XM_023252034.1

A3 122993095 122993186
refGene_gn=DNMT3A;refGene_tx=XM_019827823.2;refGene_gn=DNMT3A;refGene_tx=XM_023252030.1;refGene_gn=DNMT3A;refGene_tx=XM_023252031.1;refGene_gn=DNMT3A;refGene_tx=XM_023252032.1;refGene_gn=DNMT3A;refGene_tx=XM_023252033.1;refGene_gn=DNMT3A;refGene_tx=XM_023252034.1
A3 122993375 122993524
refGene_gn=DNMT3A;refGene_tx=XM_019827823.2;refGene_gn=DNMT3A;refGene_tx=XM_023252030.1;refGene_gn=DNMT3A;refGene_tx=XM_023252031.1;refGene_gn=DNMT3A;refGene_tx=XM_023252032.1;refGene_gn=DNMT3A;refGene_tx=XM_023252033.1;refGene_gn=DNMT3A;refGene_tx=XM_023252034.1
A3 122994547 122994633
refGene_gn=DNMT3A;refGene_tx=XM_019827823.2;refGene_gn=DNMT3A;refGene_tx=XM_023252030.1;refGene_gn=DNMT3A;refGene_tx=XM_023252031.1;refGene_gn=DNMT3A;refGene_tx=XM_023252032.1;refGene_gn=DNMT3A;refGene_tx=XM_023252033.1;refGene_gn=DNMT3A;refGene_tx=XM_023252034.1
A3 122996457 122996527
refGene_gn=DNMT3A;refGene_tx=XM_019827823.2;refGene_gn=DNMT3A;refGene_tx=XM_023252030.1;refGene_gn=DNMT3A;refGene_tx=XM_023252031.1;refGene_gn=DNMT3A;refGene_tx=XM_023252032.1;refGene_gn=DNMT3A;refGene_tx=XM_023252033.1;refGene_gn=DNMT3A;refGene_tx=XM_023252034.1
A3 122997362 122997481
refGene_gn=DNMT3A;refGene_tx=XM_019827823.2;refGene_gn=DNMT3A;refGene_tx=XM_023252030.1;refGene_gn=DNMT3A;refGene_tx=XM_023252031.1;refGene_gn=DNMT3A;refGene_tx=XM_023252032.1;refGene_gn=DNMT3A;refGene_tx=XM_023252033.1;refGene_gn=DNMT3A;refGene_tx=XM_023252034.1
A3 122998559 122998701
refGene_gn=DNMT3A;refGene_tx=XM_019827823.2;refGene_gn=DNMT3A;refGene_tx=XM_023252030.1;refGene_gn=DNMT3A;refGene_tx=XM_023252031.1;refGene_gn=DNMT3A;refGene_tx=XM_023252032.1;refGene_gn=DNMT3A;refGene_tx=XM_023252033.1;refGene_gn=DNMT3A;refGene_tx=XM_023252034.1
A3 123052553 123053369
refGene_gn=POMC;refGene_tx=XM_023252035.1
A3 123060734 123060840
refGene_gn=EFR3B;refGene_tx=XM_023252037.1;refGene_gn=EFR3B;refGene_tx=XM_023252038.1;refGene_gn=EFR3B;refGene_tx=XM_023252039.1
A3 123061238 123061288
refGene_gn=EFR3B;refGene_tx=XM_023252037.1;refGene_gn=EFR3B;refGene_tx=XM_023252038.1;refGene_gn=EFR3B;refGene_tx=XM_023252039.1
A3 123062607 123062714
refGene_gn=EFR3B;refGene_tx=XM_023252037.1;refGene_gn=EFR3B;refGene_tx=XM_023252038.1;refGene_gn=EFR3B;refGene_tx=XM_023252039.1
A3 123064001 123064050
refGene_gn=EFR3B;refGene_tx=XM_023252037.1;refGene_gn=EFR3B;refGene_tx=XM_023252038.1;refGene_gn=EFR3B;refGene_tx=XM_023252039.1
A3 123066988 123067080
refGene_gn=EFR3B;refGene_tx=XM_023252037.1;refGene_gn=EFR3B;refGene_tx=XM_023252038.1;refGene_gn=EFR3B;refGene_tx=XM_023252039.1
A3 123067844 123067972
refGene_gn=EFR3B;refGene_tx=XM_023252037.1;refGene_gn=EFR3B;refGene_tx=XM_023252038.1;refGene_gn=EFR3B;refGene_tx=XM_023252039.1

A3 123068714 123068782
refGene_gn=EFR3B;refGene_tx=XM_023252037.1;refGene_gn=EFR3B;refGene_tx=XM_023252038.1;refGene_gn=EFR3B;refGene_tx=XM_023252039.1
A3 123071183 123071315
refGene_gn=EFR3B;refGene_tx=XM_023252037.1;refGene_gn=EFR3B;refGene_tx=XM_023252038.1;refGene_gn=EFR3B;refGene_tx=XM_023252039.1
A3 123072518 123072680
refGene_gn=EFR3B;refGene_tx=XM_023252037.1;refGene_gn=EFR3B;refGene_tx=XM_023252038.1;refGene_gn=EFR3B;refGene_tx=XM_023252039.1
A3 123073314 123073390
refGene_gn=EFR3B;refGene_tx=XM_023252037.1;refGene_gn=EFR3B;refGene_tx=XM_023252038.1;refGene_gn=EFR3B;refGene_tx=XM_023252039.1
A3 123073505 123073678
refGene_gn=EFR3B;refGene_tx=XM_023252037.1;refGene_gn=EFR3B;refGene_tx=XM_023252038.1;refGene_gn=EFR3B;refGene_tx=XM_023252039.1
A3 123075660 123075712
refGene_gn=EFR3B;refGene_tx=XM_023252037.1;refGene_gn=EFR3B;refGene_tx=XM_023252038.1;refGene_gn=EFR3B;refGene_tx=XM_023252039.1
A3 123076103 123076215
refGene_gn=EFR3B;refGene_tx=XM_023252037.1;refGene_gn=EFR3B;refGene_tx=XM_023252038.1;refGene_gn=EFR3B;refGene_tx=XM_023252039.1
A3 123076973 123077135
refGene_gn=EFR3B;refGene_tx=XM_023252037.1;refGene_gn=EFR3B;refGene_tx=XM_023252038.1;refGene_gn=EFR3B;refGene_tx=XM_023252039.1
A3 123077396 123077532
refGene_gn=EFR3B;refGene_tx=XM_023252037.1;refGene_gn=EFR3B;refGene_tx=XM_023252038.1;refGene_gn=EFR3B;refGene_tx=XM_023252039.1
A3 123078254 123078333
refGene_gn=EFR3B;refGene_tx=XM_023252037.1;refGene_gn=EFR3B;refGene_tx=XM_023252038.1;refGene_gn=EFR3B;refGene_tx=XM_023252039.1
A3 123078814 123078949
refGene_gn=EFR3B;refGene_tx=XM_023252037.1;refGene_gn=EFR3B;refGene_tx=XM_023252038.1;refGene_gn=EFR3B;refGene_tx=XM_023252039.1
A3 123080208 123080358
refGene_gn=EFR3B;refGene_tx=XM_023252037.1;refGene_gn=EFR3B;refGene_tx=XM_023252038.1;refGene_gn=EFR3B;refGene_tx=XM_023252039.1
A3 123081170 123081292
refGene_gn=EFR3B;refGene_tx=XM_023252037.1;refGene_gn=EFR3B;refGene_tx=XM_023252038.1;refGene_gn=EFR3B;refGene_tx=XM_023252039.1
A3 123096010 123096161
refGene_gn=EFR3B;refGene_tx=XM_023252037.1;refGene_gn=EFR3B;refGene_tx=XM_023252038.1;refGene_gn=EFR3B;refGene_tx=XM_023252039.1
A3 123103434 123103562
refGene_gn=EFR3B;refGene_tx=XM_023252037.1;refGene_gn=EFR3B;refGene_tx=XM_023252038.1;refGene_gn=EFR3B;refGene_tx=XM_023252039.1
A3 123104560 123104637
refGene_gn=EFR3B;refGene_tx=XM_023252037.1;refGene_gn=EFR3B;refGene_tx=XM_023252038.1;refGene_gn=EFR3B;refGene_tx=XM_023252039.1
A3 123140261 123140346
refGene_gn=EFR3B;refGene_tx=XM_023252037.1;refGene_gn=EFR3B;refGene_tx

=XM_023252038.1;refGene_gn=EFR3B;refGene_tx=XM_023252039.1
A3 123182419 123182506
refGene_gn=DNAJC27;refGene_tx=XM_003984435.5;refGene_gn=DNAJC27;refGene_tx=XM_006930438.4;refGene_gn=DNAJC27;refGene_tx=XM_019827827.2;refGene_gn=DNAJC27;refGene_tx=XM_019827828.2
A3 123187951 123188034
refGene_gn=DNAJC27;refGene_tx=XM_003984435.5;refGene_gn=DNAJC27;refGene_tx=XM_006930438.4;refGene_gn=DNAJC27;refGene_tx=XM_019827827.2;refGene_gn=DNAJC27;refGene_tx=XM_019827828.2
A3 123192118 123192188
refGene_gn=DNAJC27;refGene_tx=XM_003984435.5;refGene_gn=DNAJC27;refGene_tx=XM_006930438.4;refGene_gn=DNAJC27;refGene_tx=XM_019827827.2;refGene_gn=DNAJC27;refGene_tx=XM_019827828.2
A3 123205237 123205402
refGene_gn=DNAJC27;refGene_tx=XM_003984435.5;refGene_gn=DNAJC27;refGene_tx=XM_006930438.4;refGene_gn=DNAJC27;refGene_tx=XM_019827827.2;refGene_gn=DNAJC27;refGene_tx=XM_019827828.2
A3 123205912 123206035
refGene_gn=DNAJC27;refGene_tx=XM_003984435.5;refGene_gn=DNAJC27;refGene_tx=XM_006930438.4;refGene_gn=DNAJC27;refGene_tx=XM_019827827.2;refGene_gn=DNAJC27;refGene_tx=XM_019827828.2
A3 123206946 123206997
refGene_gn=DNAJC27;refGene_tx=XM_019827827.2;refGene_gn=DNAJC27;refGene_tx=XM_019827828.2
A3 123209265 123209426
refGene_gn=DNAJC27;refGene_tx=XM_003984435.5;refGene_gn=DNAJC27;refGene_tx=XM_006930438.4
A3 123212524 123212657
refGene_gn=DNAJC27;refGene_tx=XM_003984435.5;refGene_gn=DNAJC27;refGene_tx=XM_006930438.4
A3 123242102 123242777
refGene_gn=ADCY3;refGene_tx=XM_011281268.3;refGene_gn=ADCY3;refGene_tx=XM_011281269.3;refGene_gn=ADCY3;refGene_tx=XM_019827829.2
A3 123287800 123287950
refGene_gn=ADCY3;refGene_tx=XM_011281268.3;refGene_gn=ADCY3;refGene_tx=XM_011281269.3;refGene_gn=ADCY3;refGene_tx=XM_019827829.2
A3 123310871 123311002
refGene_gn=ADCY3;refGene_tx=XM_011281268.3;refGene_gn=ADCY3;refGene_tx=XM_011281269.3;refGene_gn=ADCY3;refGene_tx=XM_019827829.2
A3 123311538 123311650
refGene_gn=ADCY3;refGene_tx=XM_011281268.3;refGene_gn=ADCY3;refGene_tx=XM_011281269.3;refGene_gn=ADCY3;refGene_tx=XM_019827829.2
A3 123311802 123311930
refGene_gn=ADCY3;refGene_tx=XM_011281268.3;refGene_gn=ADCY3;refGene_tx=XM_011281269.3;refGene_gn=ADCY3;refGene_tx=XM_019827829.2
A3 123313030 123313189
refGene_gn=ADCY3;refGene_tx=XM_011281268.3;refGene_gn=ADCY3;refGene_tx=XM_011281269.3;refGene_gn=ADCY3;refGene_tx=XM_019827829.2
A3 123313931 123314109
refGene_gn=ADCY3;refGene_tx=XM_011281268.3;refGene_gn=ADCY3;refGene_tx

=XM_011281269.3;refGene_gn=ADCY3;refGene_tx=XM_019827829.2
A3 123318360 123318489
refGene_gn=ADCY3;refGene_tx=XM_011281268.3;refGene_gn=ADCY3;refGene_tx=XM_011281269.3;refGene_gn=ADCY3;refGene_tx=XM_019827829.2
A3 123320092 123320235
refGene_gn=ADCY3;refGene_tx=XM_011281268.3;refGene_gn=ADCY3;refGene_tx=XM_011281269.3;refGene_gn=ADCY3;refGene_tx=XM_019827829.2
A3 123320371 123320533
refGene_gn=ADCY3;refGene_tx=XM_011281268.3;refGene_gn=ADCY3;refGene_tx=XM_011281269.3;refGene_gn=ADCY3;refGene_tx=XM_019827829.2
A3 123322151 123322239
refGene_gn=ADCY3;refGene_tx=XM_011281268.3;refGene_gn=ADCY3;refGene_tx=XM_011281269.3;refGene_gn=ADCY3;refGene_tx=XM_019827829.2
A3 123323013 123323130
refGene_gn=ADCY3;refGene_tx=XM_011281268.3;refGene_gn=ADCY3;refGene_tx=XM_011281269.3;refGene_gn=ADCY3;refGene_tx=XM_019827829.2
A3 123324877 123325137
refGene_gn=ADCY3;refGene_tx=XM_011281268.3;refGene_gn=ADCY3;refGene_tx=XM_011281269.3;refGene_gn=ADCY3;refGene_tx=XM_019827829.2
A3 123325394 123325460
refGene_gn=ADCY3;refGene_tx=XM_011281268.3;refGene_gn=ADCY3;refGene_tx=XM_011281269.3;refGene_gn=ADCY3;refGene_tx=XM_019827829.2
A3 123326900 123326982
refGene_gn=ADCY3;refGene_tx=XM_011281268.3;refGene_gn=ADCY3;refGene_tx=XM_011281269.3;refGene_gn=ADCY3;refGene_tx=XM_019827829.2
A3 123327824 123327983
refGene_gn=ADCY3;refGene_tx=XM_011281268.3;refGene_gn=ADCY3;refGene_tx=XM_011281269.3;refGene_gn=ADCY3;refGene_tx=XM_019827829.2
A3 123328498 123328645
refGene_gn=ADCY3;refGene_tx=XM_011281268.3;refGene_gn=ADCY3;refGene_tx=XM_011281269.3;refGene_gn=ADCY3;refGene_tx=XM_019827829.2
A3 123329182 123329302
refGene_gn=ADCY3;refGene_tx=XM_011281268.3;refGene_gn=ADCY3;refGene_tx=XM_011281269.3;refGene_gn=ADCY3;refGene_tx=XM_019827829.2
A3 123330286 123330410
refGene_gn=ADCY3;refGene_tx=XM_011281268.3;refGene_gn=ADCY3;refGene_tx=XM_011281269.3;refGene_gn=ADCY3;refGene_tx=XM_019827829.2
A3 123331073 123331402
refGene_gn=ADCY3;refGene_tx=XM_011281268.3;refGene_gn=ADCY3;refGene_tx=XM_011281269.3;refGene_gn=ADCY3;refGene_tx=XM_019827829.2
A3 123331743 123331926
refGene_gn=ADCY3;refGene_tx=XM_011281269.3
A3 123332537 123332541
refGene_gn=CENPO;refGene_tx=XM_006930440.3
A3 123333660 123333797
refGene_gn=CENPO;refGene_tx=XM_006930439.3;refGene_gn=CENPO;refGene_tx=XM_006930440.3;refGene_gn=CENPO;refGene_tx=XM_019827830.1
A3 123334733 123334905
refGene_gn=CENPO;refGene_tx=XM_006930439.3;refGene_gn=CENPO;refGene_tx=XM_006930440.3;refGene_gn=CENPO;refGene_tx=XM_019827830.1

A3 123336651 123336911
refGene_gn=CENPO;refGene_tx=XM_006930439.3;refGene_gn=CENPO;refGene_tx=XM_006930440.3;refGene_gn=CENPO;refGene_tx=XM_019827830.1
A3 123337593 123337711
refGene_gn=CENPO;refGene_tx=XM_006930439.3;refGene_gn=CENPO;refGene_tx=XM_006930440.3;refGene_gn=CENPO;refGene_tx=XM_019827830.1
A3 123342128 123342167
refGene_gn=CENPO;refGene_tx=XM_019827830.1
A3 123343727 123343894
refGene_gn=CENPO;refGene_tx=XM_006930439.3;refGene_gn=CENPO;refGene_tx=XM_006930440.3
A3 123345335 123345366
refGene_gn=CENPO;refGene_tx=XM_006930439.3;refGene_gn=CENPO;refGene_tx=XM_006930440.3
A3 123345813 123346065
refGene_gn=PTRHD1;refGene_tx=XM_003984439.5
A3 123348515 123348686
refGene_gn=PTRHD1;refGene_tx=XM_003984439.5
A3 123362005 123362176
refGene_gn=NCOA1;refGene_tx=XM_023252041.1;refGene_gn=NCOA1;refGene_tx=XM_023252042.1;refGene_gn=NCOA1;refGene_tx=XM_023252043.1;refGene_gn=NCOA1;refGene_tx=XM_023252044.1;refGene_gn=NCOA1;refGene_tx=XM_023252045.1;refGene_gn=NCOA1;refGene_tx=XM_023252046.1
A3 123362273 123362318
refGene_gn=NCOA1;refGene_tx=XM_023252047.1;refGene_gn=NCOA1;refGene_tx=XM_023252048.1
A3 123366021 123366111
refGene_gn=NCOA1;refGene_tx=XM_023252041.1;refGene_gn=NCOA1;refGene_tx=XM_023252042.1;refGene_gn=NCOA1;refGene_tx=XM_023252043.1;refGene_gn=NCOA1;refGene_tx=XM_023252044.1;refGene_gn=NCOA1;refGene_tx=XM_023252045.1;refGene_gn=NCOA1;refGene_tx=XM_023252046.1;refGene_gn=NCOA1;refGene_tx=XM_023252047.1;refGene_gn=NCOA1;refGene_tx=XM_023252048.1
A3 123369957 123370141
refGene_gn=NCOA1;refGene_tx=XM_023252041.1;refGene_gn=NCOA1;refGene_tx=XM_023252042.1;refGene_gn=NCOA1;refGene_tx=XM_023252043.1;refGene_gn=NCOA1;refGene_tx=XM_023252044.1;refGene_gn=NCOA1;refGene_tx=XM_023252045.1;refGene_gn=NCOA1;refGene_tx=XM_023252046.1;refGene_gn=NCOA1;refGene_tx=XM_023252047.1;refGene_gn=NCOA1;refGene_tx=XM_023252048.1
A3 123375508 123375683
refGene_gn=NCOA1;refGene_tx=XM_023252041.1;refGene_gn=NCOA1;refGene_tx=XM_023252042.1;refGene_gn=NCOA1;refGene_tx=XM_023252043.1;refGene_gn=NCOA1;refGene_tx=XM_023252044.1;refGene_gn=NCOA1;refGene_tx=XM_023252045.1;refGene_gn=NCOA1;refGene_tx=XM_023252046.1;refGene_gn=NCOA1;refGene_tx=XM_023252047.1;refGene_gn=NCOA1;refGene_tx=XM_023252048.1
A3 123382690 123383093
refGene_gn=NCOA1;refGene_tx=XM_023252041.1;refGene_gn=NCOA1;refGene_tx=XM_023252042.1;refGene_gn=NCOA1;refGene_tx=XM_023252043.1;refGene_gn=NCOA1;refGene_tx=XM_023252044.1;refGene_gn=NCOA1;refGene_tx=XM_023252045.1;refGene_gn=NCOA1;refGene_tx=XM_023252046.1;refGene_gn=NCOA1;refGene_tx=XM_023252047.1;refGene_gn=NCOA1;refGene_tx=XM_023252048.1

A3	123384879	123384981
refGene_gn=NCOA1;refGene_tx=XM_023252041.1;refGene_gn=NCOA1;refGene_tx=XM_023252042.1;refGene_gn=NCOA1;refGene_tx=XM_023252043.1;refGene_gn=NCOA1;refGene_tx=XM_023252044.1;refGene_gn=NCOA1;refGene_tx=XM_023252045.1;refGene_gn=NCOA1;refGene_tx=XM_023252046.1;refGene_gn=NCOA1;refGene_tx=XM_023252047.1;refGene_gn=NCOA1;refGene_tx=XM_023252048.1
A3	123393409	123393724
refGene_gn=NCOA1;refGene_tx=XM_023252041.1;refGene_gn=NCOA1;refGene_tx=XM_023252042.1;refGene_gn=NCOA1;refGene_tx=XM_023252043.1;refGene_gn=NCOA1;refGene_tx=XM_023252044.1;refGene_gn=NCOA1;refGene_tx=XM_023252045.1;refGene_gn=NCOA1;refGene_tx=XM_023252046.1;refGene_gn=NCOA1;refGene_tx=XM_023252047.1;refGene_gn=NCOA1;refGene_tx=XM_023252048.1
A3	123394741	123394910
refGene_gn=NCOA1;refGene_tx=XM_023252041.1;refGene_gn=NCOA1;refGene_tx=XM_023252042.1;refGene_gn=NCOA1;refGene_tx=XM_023252043.1;refGene_gn=NCOA1;refGene_tx=XM_023252044.1;refGene_gn=NCOA1;refGene_tx=XM_023252045.1;refGene_gn=NCOA1;refGene_tx=XM_023252046.1;refGene_gn=NCOA1;refGene_tx=XM_023252047.1;refGene_gn=NCOA1;refGene_tx=XM_023252048.1
A3	123396394	123396512
refGene_gn=NCOA1;refGene_tx=XM_023252041.1;refGene_gn=NCOA1;refGene_tx=XM_023252042.1;refGene_gn=NCOA1;refGene_tx=XM_023252043.1;refGene_gn=NCOA1;refGene_tx=XM_023252044.1;refGene_gn=NCOA1;refGene_tx=XM_023252045.1;refGene_gn=NCOA1;refGene_tx=XM_023252046.1;refGene_gn=NCOA1;refGene_tx=XM_023252047.1;refGene_gn=NCOA1;refGene_tx=XM_023252048.1
A3	123404920	123405101
refGene_gn=NCOA1;refGene_tx=XM_023252041.1;refGene_gn=NCOA1;refGene_tx=XM_023252042.1;refGene_gn=NCOA1;refGene_tx=XM_023252043.1;refGene_gn=NCOA1;refGene_tx=XM_023252044.1;refGene_gn=NCOA1;refGene_tx=XM_023252045.1;refGene_gn=NCOA1;refGene_tx=XM_023252046.1;refGene_gn=NCOA1;refGene_tx=XM_023252047.1;refGene_gn=NCOA1;refGene_tx=XM_023252048.1
A3	123408380	123409701
refGene_gn=NCOA1;refGene_tx=XM_023252041.1;refGene_gn=NCOA1;refGene_tx=XM_023252042.1;refGene_gn=NCOA1;refGene_tx=XM_023252043.1;refGene_gn=NCOA1;refGene_tx=XM_023252044.1;refGene_gn=NCOA1;refGene_tx=XM_023252045.1;refGene_gn=NCOA1;refGene_tx=XM_023252046.1;refGene_gn=NCOA1;refGene_tx=XM_023252047.1;refGene_gn=NCOA1;refGene_tx=XM_023252048.1
A3	123411298	123411446
refGene_gn=NCOA1;refGene_tx=XM_023252041.1;refGene_gn=NCOA1;refGene_tx=XM_023252042.1;refGene_gn=NCOA1;refGene_tx=XM_023252043.1;refGene_gn=NCOA1;refGene_tx=XM_023252044.1;refGene_gn=NCOA1;refGene_tx=XM_023252045.1;refGene_gn=NCOA1;refGene_tx=XM_023252046.1;refGene_gn=NCOA1;refGene_tx=XM_023252047.1;refGene_gn=NCOA1;refGene_tx=XM_023252048.1
A3	123419701	123419842
refGene_gn=NCOA1;refGene_tx=XM_023252041.1;refGene_gn=NCOA1;refGene_tx=XM_023252042.1;refGene_gn=NCOA1;refGene_tx=XM_023252043.1;refGene_gn=NCOA1;refGene_tx=XM_023252044.1;refGene_gn=NCOA1;refGene_tx=XM_023252045.1;refGene_gn=NCOA1;refGene_tx=XM_023252046.1;refGene_gn=NCOA1;refGene_tx=XM_023252047.1;refGene_gn=NCOA1;refGene_tx=XM_023252048.1
A3	123424257	123424353
refGene_gn=NCOA1;refGene_tx=XM_023252041.1;refGene_gn=NCOA1;refGene_tx

=XM_023252042.1;refGene_gn=NCOA1;refGene_tx=XM_023252043.1;refGene_gn=NCOA1;refGene_tx=XM_023252044.1;refGene_gn=NCOA1;refGene_tx=XM_023252045.1;refGene_gn=NCOA1;refGene_tx=XM_023252046.1;refGene_gn=NCOA1;refGene_tx=XM_023252047.1;refGene_gn=NCOA1;refGene_tx=XM_023252048.1
A3 123425646 123425826
refGene_gn=NCOA1;refGene_tx=XM_023252041.1;refGene_gn=NCOA1;refGene_tx=XM_023252042.1;refGene_gn=NCOA1;refGene_tx=XM_023252043.1;refGene_gn=NCOA1;refGene_tx=XM_023252044.1;refGene_gn=NCOA1;refGene_tx=XM_023252045.1;refGene_gn=NCOA1;refGene_tx=XM_023252046.1;refGene_gn=NCOA1;refGene_tx=XM_023252047.1;refGene_gn=NCOA1;refGene_tx=XM_023252048.1
A3 123427973 123428151
refGene_gn=NCOA1;refGene_tx=XM_023252041.1;refGene_gn=NCOA1;refGene_tx=XM_023252042.1;refGene_gn=NCOA1;refGene_tx=XM_023252043.1;refGene_gn=NCOA1;refGene_tx=XM_023252044.1;refGene_gn=NCOA1;refGene_tx=XM_023252045.1;refGene_gn=NCOA1;refGene_tx=XM_023252046.1;refGene_gn=NCOA1;refGene_tx=XM_023252047.1;refGene_gn=NCOA1;refGene_tx=XM_023252048.1
A3 123436666 123436764
refGene_gn=NCOA1;refGene_tx=XM_023252041.1;refGene_gn=NCOA1;refGene_tx=XM_023252042.1;refGene_gn=NCOA1;refGene_tx=XM_023252043.1;refGene_gn=NCOA1;refGene_tx=XM_023252044.1;refGene_gn=NCOA1;refGene_tx=XM_023252045.1;refGene_gn=NCOA1;refGene_tx=XM_023252046.1;refGene_gn=NCOA1;refGene_tx=XM_023252047.1;refGene_gn=NCOA1;refGene_tx=XM_023252048.1
A3 123443249 123443416
refGene_gn=NCOA1;refGene_tx=XM_023252041.1;refGene_gn=NCOA1;refGene_tx=XM_023252042.1;refGene_gn=NCOA1;refGene_tx=XM_023252043.1;refGene_gn=NCOA1;refGene_tx=XM_023252044.1;refGene_gn=NCOA1;refGene_tx=XM_023252045.1;refGene_gn=NCOA1;refGene_tx=XM_023252046.1;refGene_gn=NCOA1;refGene_tx=XM_023252047.1;refGene_gn=NCOA1;refGene_tx=XM_023252048.1
A3 123450706 123450795
refGene_gn=NCOA1;refGene_tx=XM_023252041.1;refGene_gn=NCOA1;refGene_tx=XM_023252042.1;refGene_gn=NCOA1;refGene_tx=XM_023252043.1;refGene_gn=NCOA1;refGene_tx=XM_023252044.1;refGene_gn=NCOA1;refGene_tx=XM_023252045.1;refGene_gn=NCOA1;refGene_tx=XM_023252046.1;refGene_gn=NCOA1;refGene_tx=XM_023252047.1;refGene_gn=NCOA1;refGene_tx=XM_023252048.1
A3 123697794 123697882
refGene_gn=LOC102902347;refGene_tx=XM_023252203.1
A3 123727258 123727327
refGene_gn=LOC102902347;refGene_tx=XM_023252203.1
A3 123738701 123739525
refGene_gn=LOC102902347;refGene_tx=XM_023252203.1
A3 123743945 123744044
refGene_gn=LOC102902347;refGene_tx=XM_023252203.1
A3 123773723 123773754
refGene_gn=ITSN2;refGene_tx=XM_019827831.2;refGene_gn=ITSN2;refGene_tx=XM_019827835.2;refGene_gn=ITSN2;refGene_tx=XM_019827837.2;refGene_gn=ITSN2;refGene_tx=XM_023252049.1;refGene_gn=ITSN2;refGene_tx=XM_023252050.1;refGene_gn=ITSN2;refGene_tx=XM_023252051.1;refGene_gn=ITSN2;refGene_tx=XM_023252052.1;refGene_gn=ITSN2;refGene_tx=XM_023252053.1;refGene_gn=ITSN2;refGene_tx=XM_023252054.1;refGene_gn=ITSN2;refGene_tx=XM_023252055.1;refGene_gn=ITSN2;refGene_tx=XM_023252056.1;refGene_gn=ITSN2;

refGene_tx=XM_023252057.1;refGene_gn=ITSN2;refGene_tx=XM_023252058.1;r
efGene_gn=ITSN2;refGene_tx=XM_023252059.1
A3 123775875 123775977
refGene_gn=ITSN2;refGene_tx=XM_019827831.2;refGene_gn=ITSN2;refGene_tx
=XM_023252049.1;refGene_gn=ITSN2;refGene_tx=XM_023252050.1;refGene_gn=
ITSN2;refGene_tx=XM_023252051.1;refGene_gn=ITSN2;refGene_tx=XM_0232520
52.1;refGene_gn=ITSN2;refGene_tx=XM_023252053.1;refGene_gn=ITSN2;refGe
ne_tx=XM_023252054.1;refGene_gn=ITSN2;refGene_tx=XM_023252055.1
A3 123780907 123781000
refGene_gn=ITSN2;refGene_tx=XM_019827831.2;refGene_gn=ITSN2;refGene_tx
=XM_019827835.2;refGene_gn=ITSN2;refGene_tx=XM_019827837.2;refGene_gn=
ITSN2;refGene_tx=XM_019827838.2;refGene_gn=ITSN2;refGene_tx=XM_0232520
49.1;refGene_gn=ITSN2;refGene_tx=XM_023252050.1;refGene_gn=ITSN2;refGe
ne_tx=XM_023252051.1;refGene_gn=ITSN2;refGene_tx=XM_023252052.1;refGen
e_gn=ITSN2;refGene_tx=XM_023252053.1;refGene_gn=ITSN2;refGene_tx=XM_02
3252054.1;refGene_gn=ITSN2;refGene_tx=XM_023252055.1;refGene_gn=ITSN2;
refGene_tx=XM_023252056.1;refGene_gn=ITSN2;refGene_tx=XM_023252057.1;r
efGene_gn=ITSN2;refGene_tx=XM_023252058.1;refGene_gn=ITSN2;refGene_tx=
XM_023252059.1
A3 123783181 123783245
refGene_gn=ITSN2;refGene_tx=XM_019827831.2;refGene_gn=ITSN2;refGene_tx
=XM_019827835.2;refGene_gn=ITSN2;refGene_tx=XM_019827837.2;refGene_gn=
ITSN2;refGene_tx=XM_019827838.2;refGene_gn=ITSN2;refGene_tx=XM_0232520
49.1;refGene_gn=ITSN2;refGene_tx=XM_023252050.1;refGene_gn=ITSN2;refGe
ne_tx=XM_023252051.1;refGene_gn=ITSN2;refGene_tx=XM_023252052.1;refGen
e_gn=ITSN2;refGene_tx=XM_023252053.1;refGene_gn=ITSN2;refGene_tx=XM_02
3252054.1;refGene_gn=ITSN2;refGene_tx=XM_023252055.1;refGene_gn=ITSN2;
refGene_tx=XM_023252056.1;refGene_gn=ITSN2;refGene_tx=XM_023252057.1;r
efGene_gn=ITSN2;refGene_tx=XM_023252058.1;refGene_gn=ITSN2;refGene_tx=
XM_023252059.1
A3 123784078 123784242
refGene_gn=ITSN2;refGene_tx=XM_019827831.2;refGene_gn=ITSN2;refGene_tx
=XM_019827835.2;refGene_gn=ITSN2;refGene_tx=XM_019827837.2;refGene_gn=
ITSN2;refGene_tx=XM_019827838.2;refGene_gn=ITSN2;refGene_tx=XM_0232520
49.1;refGene_gn=ITSN2;refGene_tx=XM_023252050.1;refGene_gn=ITSN2;refGe
ne_tx=XM_023252051.1;refGene_gn=ITSN2;refGene_tx=XM_023252052.1;refGen
e_gn=ITSN2;refGene_tx=XM_023252053.1;refGene_gn=ITSN2;refGene_tx=XM_02
3252054.1;refGene_gn=ITSN2;refGene_tx=XM_023252055.1;refGene_gn=ITSN2;
refGene_tx=XM_023252056.1;refGene_gn=ITSN2;refGene_tx=XM_023252057.1;r
efGene_gn=ITSN2;refGene_tx=XM_023252058.1;refGene_gn=ITSN2;refGene_tx=
XM_023252059.1
A3 123785992 123786199
refGene_gn=ITSN2;refGene_tx=XM_019827831.2;refGene_gn=ITSN2;refGene_tx
=XM_019827835.2;refGene_gn=ITSN2;refGene_tx=XM_019827837.2;refGene_gn=
ITSN2;refGene_tx=XM_019827838.2;refGene_gn=ITSN2;refGene_tx=XM_0232520
49.1;refGene_gn=ITSN2;refGene_tx=XM_023252050.1;refGene_gn=ITSN2;refGe
ne_tx=XM_023252051.1;refGene_gn=ITSN2;refGene_tx=XM_023252052.1;refGen
e_gn=ITSN2;refGene_tx=XM_023252053.1;refGene_gn=ITSN2;refGene_tx=XM_02
3252054.1;refGene_gn=ITSN2;refGene_tx=XM_023252055.1;refGene_gn=ITSN2;
refGene_tx=XM_023252056.1;refGene_gn=ITSN2;refGene_tx=XM_023252057.1;r

efGene_gn=ITSN2;refGene_tx=XM_023252058.1;refGene_gn=ITSN2;refGene_tx=XM_023252059.1
A3 123786343 123786440
refGene_gn=ITSN2;refGene_tx=XM_019827831.2;refGene_gn=ITSN2;refGene_tx=XM_019827835.2;refGene_gn=ITSN2;refGene_tx=XM_019827837.2;refGene_gn=ITSN2;refGene_tx=XM_019827838.2;refGene_gn=ITSN2;refGene_tx=XM_023252049.1;refGene_gn=ITSN2;refGene_tx=XM_023252050.1;refGene_gn=ITSN2;refGene_tx=XM_023252051.1;refGene_gn=ITSN2;refGene_tx=XM_023252052.1;refGene_gn=ITSN2;refGene_tx=XM_023252053.1;refGene_gn=ITSN2;refGene_tx=XM_023252054.1;refGene_gn=ITSN2;refGene_tx=XM_023252055.1;refGene_gn=ITSN2;refGene_tx=XM_023252056.1;refGene_gn=ITSN2;refGene_tx=XM_023252057.1;refGene_gn=ITSN2;refGene_tx=XM_023252058.1;refGene_gn=ITSN2;refGene_tx=XM_023252059.1
A3 123787747 123787887
refGene_gn=ITSN2;refGene_tx=XM_019827831.2;refGene_gn=ITSN2;refGene_tx=XM_019827835.2;refGene_gn=ITSN2;refGene_tx=XM_019827837.2;refGene_gn=ITSN2;refGene_tx=XM_019827838.2;refGene_gn=ITSN2;refGene_tx=XM_023252049.1;refGene_gn=ITSN2;refGene_tx=XM_023252050.1;refGene_gn=ITSN2;refGene_tx=XM_023252051.1;refGene_gn=ITSN2;refGene_tx=XM_023252052.1;refGene_gn=ITSN2;refGene_tx=XM_023252053.1;refGene_gn=ITSN2;refGene_tx=XM_023252054.1;refGene_gn=ITSN2;refGene_tx=XM_023252055.1;refGene_gn=ITSN2;refGene_tx=XM_023252056.1;refGene_gn=ITSN2;refGene_tx=XM_023252057.1;refGene_gn=ITSN2;refGene_tx=XM_023252058.1;refGene_gn=ITSN2;refGene_tx=XM_023252059.1
A3 123792343 123792407
refGene_gn=ITSN2;refGene_tx=XM_019827831.2;refGene_gn=ITSN2;refGene_tx=XM_019827835.2;refGene_gn=ITSN2;refGene_tx=XM_019827837.2;refGene_gn=ITSN2;refGene_tx=XM_019827838.2;refGene_gn=ITSN2;refGene_tx=XM_023252049.1;refGene_gn=ITSN2;refGene_tx=XM_023252050.1;refGene_gn=ITSN2;refGene_tx=XM_023252051.1;refGene_gn=ITSN2;refGene_tx=XM_023252052.1;refGene_gn=ITSN2;refGene_tx=XM_023252053.1;refGene_gn=ITSN2;refGene_tx=XM_023252054.1;refGene_gn=ITSN2;refGene_tx=XM_023252055.1;refGene_gn=ITSN2;refGene_tx=XM_023252056.1;refGene_gn=ITSN2;refGene_tx=XM_023252057.1;refGene_gn=ITSN2;refGene_tx=XM_023252058.1;refGene_gn=ITSN2;refGene_tx=XM_023252059.1
A3 123792934 123793072
refGene_gn=ITSN2;refGene_tx=XM_019827831.2;refGene_gn=ITSN2;refGene_tx=XM_019827835.2;refGene_gn=ITSN2;refGene_tx=XM_019827837.2;refGene_gn=ITSN2;refGene_tx=XM_019827838.2;refGene_gn=ITSN2;refGene_tx=XM_023252049.1;refGene_gn=ITSN2;refGene_tx=XM_023252050.1;refGene_gn=ITSN2;refGene_tx=XM_023252051.1;refGene_gn=ITSN2;refGene_tx=XM_023252052.1;refGene_gn=ITSN2;refGene_tx=XM_023252053.1;refGene_gn=ITSN2;refGene_tx=XM_023252054.1;refGene_gn=ITSN2;refGene_tx=XM_023252055.1;refGene_gn=ITSN2;refGene_tx=XM_023252056.1;refGene_gn=ITSN2;refGene_tx=XM_023252057.1;refGene_gn=ITSN2;refGene_tx=XM_023252058.1;refGene_gn=ITSN2;refGene_tx=XM_023252059.1
A3 123793517 123793603
refGene_gn=ITSN2;refGene_tx=XM_019827831.2;refGene_gn=ITSN2;refGene_tx=XM_019827835.2;refGene_gn=ITSN2;refGene_tx=XM_019827837.2;refGene_gn=ITSN2;refGene_tx=XM_019827838.2;refGene_gn=ITSN2;refGene_tx=XM_0232520

49.1;refGene_gn=ITSN2;refGene_tx=XM_023252050.1;refGene_gn=ITSN2;refGene_tx=XM_023252051.1;refGene_gn=ITSN2;refGene_tx=XM_023252052.1;refGene_gn=ITSN2;refGene_tx=XM_023252053.1;refGene_gn=ITSN2;refGene_tx=XM_023252054.1;refGene_gn=ITSN2;refGene_tx=XM_023252055.1;refGene_gn=ITSN2;refGene_tx=XM_023252056.1;refGene_gn=ITSN2;refGene_tx=XM_023252057.1;refGene_gn=ITSN2;refGene_tx=XM_023252058.1;refGene_gn=ITSN2;refGene_tx=XM_023252059.1
A3 123794523 123794786
refGene_gn=ITSN2;refGene_tx=XM_019827831.2;refGene_gn=ITSN2;refGene_tx=XM_019827835.2;refGene_gn=ITSN2;refGene_tx=XM_019827837.2;refGene_gn=ITSN2;refGene_tx=XM_019827838.2;refGene_gn=ITSN2;refGene_tx=XM_023252049.1;refGene_gn=ITSN2;refGene_tx=XM_023252050.1;refGene_gn=ITSN2;refGene_tx=XM_023252051.1;refGene_gn=ITSN2;refGene_tx=XM_023252052.1;refGene_gn=ITSN2;refGene_tx=XM_023252053.1;refGene_gn=ITSN2;refGene_tx=XM_023252054.1;refGene_gn=ITSN2;refGene_tx=XM_023252055.1;refGene_gn=ITSN2;refGene_tx=XM_023252056.1;refGene_gn=ITSN2;refGene_tx=XM_023252057.1;refGene_gn=ITSN2;refGene_tx=XM_023252058.1;refGene_gn=ITSN2;refGene_tx=XM_023252059.1
A3 123795677 123795827
refGene_gn=ITSN2;refGene_tx=XM_019827831.2;refGene_gn=ITSN2;refGene_tx=XM_019827835.2;refGene_gn=ITSN2;refGene_tx=XM_019827837.2;refGene_gn=ITSN2;refGene_tx=XM_019827838.2;refGene_gn=ITSN2;refGene_tx=XM_023252049.1;refGene_gn=ITSN2;refGene_tx=XM_023252050.1;refGene_gn=ITSN2;refGene_tx=XM_023252051.1;refGene_gn=ITSN2;refGene_tx=XM_023252052.1;refGene_gn=ITSN2;refGene_tx=XM_023252053.1;refGene_gn=ITSN2;refGene_tx=XM_023252054.1;refGene_gn=ITSN2;refGene_tx=XM_023252055.1;refGene_gn=ITSN2;refGene_tx=XM_023252056.1;refGene_gn=ITSN2;refGene_tx=XM_023252057.1;refGene_gn=ITSN2;refGene_tx=XM_023252058.1;refGene_gn=ITSN2;refGene_tx=XM_023252059.1
A3 123798757 123798898
refGene_gn=ITSN2;refGene_tx=XM_019827831.2;refGene_gn=ITSN2;refGene_tx=XM_019827835.2;refGene_gn=ITSN2;refGene_tx=XM_019827837.2;refGene_gn=ITSN2;refGene_tx=XM_019827838.2;refGene_gn=ITSN2;refGene_tx=XM_023252049.1;refGene_gn=ITSN2;refGene_tx=XM_023252050.1;refGene_gn=ITSN2;refGene_tx=XM_023252051.1;refGene_gn=ITSN2;refGene_tx=XM_023252052.1;refGene_gn=ITSN2;refGene_tx=XM_023252053.1;refGene_gn=ITSN2;refGene_tx=XM_023252054.1;refGene_gn=ITSN2;refGene_tx=XM_023252055.1;refGene_gn=ITSN2;refGene_tx=XM_023252056.1;refGene_gn=ITSN2;refGene_tx=XM_023252057.1;refGene_gn=ITSN2;refGene_tx=XM_023252058.1;refGene_gn=ITSN2;refGene_tx=XM_023252059.1
A3 123799892 123799980
refGene_gn=ITSN2;refGene_tx=XM_019827831.2;refGene_gn=ITSN2;refGene_tx=XM_019827835.2;refGene_gn=ITSN2;refGene_tx=XM_019827837.2;refGene_gn=ITSN2;refGene_tx=XM_019827838.2;refGene_gn=ITSN2;refGene_tx=XM_023252049.1;refGene_gn=ITSN2;refGene_tx=XM_023252050.1;refGene_gn=ITSN2;refGene_tx=XM_023252051.1;refGene_gn=ITSN2;refGene_tx=XM_023252052.1;refGene_gn=ITSN2;refGene_tx=XM_023252053.1;refGene_gn=ITSN2;refGene_tx=XM_023252054.1;refGene_gn=ITSN2;refGene_tx=XM_023252055.1;refGene_gn=ITSN2;refGene_tx=XM_023252056.1;refGene_gn=ITSN2;refGene_tx=XM_023252057.1;refGene_gn=ITSN2;refGene_tx=XM_023252058.1;refGene_gn=ITSN2;refGene_tx=

XM_023252059.1
A3	123803958	123804098
refGene_gn=ITSN2;refGene_tx=XM_019827831.2;refGene_gn=ITSN2;refGene_tx=XM_019827835.2;refGene_gn=ITSN2;refGene_tx=XM_019827837.2;refGene_gn=ITSN2;refGene_tx=XM_019827838.2;refGene_gn=ITSN2;refGene_tx=XM_023252049.1;refGene_gn=ITSN2;refGene_tx=XM_023252050.1;refGene_gn=ITSN2;refGene_tx=XM_023252051.1;refGene_gn=ITSN2;refGene_tx=XM_023252052.1;refGene_gn=ITSN2;refGene_tx=XM_023252053.1;refGene_gn=ITSN2;refGene_tx=XM_023252054.1;refGene_gn=ITSN2;refGene_tx=XM_023252055.1;refGene_gn=ITSN2;refGene_tx=XM_023252056.1;refGene_gn=ITSN2;refGene_tx=XM_023252057.1;refGene_gn=ITSN2;refGene_tx=XM_023252058.1;refGene_gn=ITSN2;refGene_tx=XM_023252059.1
A3	123805141	123805222
refGene_gn=ITSN2;refGene_tx=XM_019827831.2;refGene_gn=ITSN2;refGene_tx=XM_019827835.2;refGene_gn=ITSN2;refGene_tx=XM_019827838.2;refGene_gn=ITSN2;refGene_tx=XM_023252049.1;refGene_gn=ITSN2;refGene_tx=XM_023252050.1;refGene_gn=ITSN2;refGene_tx=XM_023252051.1;refGene_gn=ITSN2;refGene_tx=XM_023252052.1;refGene_gn=ITSN2;refGene_tx=XM_023252053.1;refGene_gn=ITSN2;refGene_tx=XM_023252055.1;refGene_gn=ITSN2;refGene_tx=XM_023252056.1;refGene_gn=ITSN2;refGene_tx=XM_023252057.1;refGene_gn=ITSN2;refGene_tx=XM_023252058.1
A3	123816876	123817013
refGene_gn=ITSN2;refGene_tx=XM_019827831.2;refGene_gn=ITSN2;refGene_tx=XM_019827835.2;refGene_gn=ITSN2;refGene_tx=XM_019827837.2;refGene_gn=ITSN2;refGene_tx=XM_019827838.2;refGene_gn=ITSN2;refGene_tx=XM_023252049.1;refGene_gn=ITSN2;refGene_tx=XM_023252050.1;refGene_gn=ITSN2;refGene_tx=XM_023252051.1;refGene_gn=ITSN2;refGene_tx=XM_023252052.1;refGene_gn=ITSN2;refGene_tx=XM_023252053.1;refGene_gn=ITSN2;refGene_tx=XM_023252054.1;refGene_gn=ITSN2;refGene_tx=XM_023252055.1;refGene_gn=ITSN2;refGene_tx=XM_023252056.1;refGene_gn=ITSN2;refGene_tx=XM_023252057.1;refGene_gn=ITSN2;refGene_tx=XM_023252058.1;refGene_gn=ITSN2;refGene_tx=XM_023252059.1
A3	123819954	123820097
refGene_gn=ITSN2;refGene_tx=XM_019827831.2;refGene_gn=ITSN2;refGene_tx=XM_019827835.2;refGene_gn=ITSN2;refGene_tx=XM_019827837.2;refGene_gn=ITSN2;refGene_tx=XM_019827838.2;refGene_gn=ITSN2;refGene_tx=XM_023252049.1;refGene_gn=ITSN2;refGene_tx=XM_023252050.1;refGene_gn=ITSN2;refGene_tx=XM_023252051.1;refGene_gn=ITSN2;refGene_tx=XM_023252052.1;refGene_gn=ITSN2;refGene_tx=XM_023252053.1;refGene_gn=ITSN2;refGene_tx=XM_023252054.1;refGene_gn=ITSN2;refGene_tx=XM_023252055.1;refGene_gn=ITSN2;refGene_tx=XM_023252056.1;refGene_gn=ITSN2;refGene_tx=XM_023252057.1;refGene_gn=ITSN2;refGene_tx=XM_023252058.1;refGene_gn=ITSN2;refGene_tx=XM_023252059.1
A3	123821356	123821454
refGene_gn=ITSN2;refGene_tx=XM_019827831.2;refGene_gn=ITSN2;refGene_tx=XM_019827835.2;refGene_gn=ITSN2;refGene_tx=XM_019827837.2;refGene_gn=ITSN2;refGene_tx=XM_019827838.2;refGene_gn=ITSN2;refGene_tx=XM_023252049.1;refGene_gn=ITSN2;refGene_tx=XM_023252050.1;refGene_gn=ITSN2;refGene_tx=XM_023252051.1;refGene_gn=ITSN2;refGene_tx=XM_023252052.1;refGene_gn=ITSN2;refGene_tx=XM_023252053.1;refGene_gn=ITSN2;refGene_tx=XM_02

3252054.1;refGene_gn=ITSN2;refGene_tx=XM_023252055.1;refGene_gn=ITSN2;
refGene_tx=XM_023252056.1;refGene_gn=ITSN2;refGene_tx=XM_023252057.1;r
efGene_gn=ITSN2;refGene_tx=XM_023252058.1;refGene_gn=ITSN2;refGene_tx=
XM_023252059.1
A3 123826972 123827148
refGene_gn=ITSN2;refGene_tx=XM_019827831.2;refGene_gn=ITSN2;refGene_tx
=XM_019827835.2;refGene_gn=ITSN2;refGene_tx=XM_019827837.2;refGene_gn=
ITSN2;refGene_tx=XM_019827838.2;refGene_gn=ITSN2;refGene_tx=XM_0232520
49.1;refGene_gn=ITSN2;refGene_tx=XM_023252050.1;refGene_gn=ITSN2;refGe
ne_tx=XM_023252051.1;refGene_gn=ITSN2;refGene_tx=XM_023252052.1;refGen
e_gn=ITSN2;refGene_tx=XM_023252053.1;refGene_gn=ITSN2;refGene_tx=XM_02
3252054.1;refGene_gn=ITSN2;refGene_tx=XM_023252055.1;refGene_gn=ITSN2;
refGene_tx=XM_023252056.1;refGene_gn=ITSN2;refGene_tx=XM_023252057.1;r
efGene_gn=ITSN2;refGene_tx=XM_023252058.1;refGene_gn=ITSN2;refGene_tx=
XM_023252059.1
A3 123827439 123827584
refGene_gn=ITSN2;refGene_tx=XM_019827831.2;refGene_gn=ITSN2;refGene_tx
=XM_019827835.2;refGene_gn=ITSN2;refGene_tx=XM_019827837.2;refGene_gn=
ITSN2;refGene_tx=XM_019827838.2;refGene_gn=ITSN2;refGene_tx=XM_0232520
49.1;refGene_gn=ITSN2;refGene_tx=XM_023252050.1;refGene_gn=ITSN2;refGe
ne_tx=XM_023252051.1;refGene_gn=ITSN2;refGene_tx=XM_023252052.1;refGen
e_gn=ITSN2;refGene_tx=XM_023252053.1;refGene_gn=ITSN2;refGene_tx=XM_02
3252054.1;refGene_gn=ITSN2;refGene_tx=XM_023252055.1;refGene_gn=ITSN2;
refGene_tx=XM_023252056.1;refGene_gn=ITSN2;refGene_tx=XM_023252057.1;r
efGene_gn=ITSN2;refGene_tx=XM_023252058.1;refGene_gn=ITSN2;refGene_tx=
XM_023252059.1
A3 123829622 123829828
refGene_gn=ITSN2;refGene_tx=XM_019827831.2;refGene_gn=ITSN2;refGene_tx
=XM_019827835.2;refGene_gn=ITSN2;refGene_tx=XM_019827837.2;refGene_gn=
ITSN2;refGene_tx=XM_019827838.2;refGene_gn=ITSN2;refGene_tx=XM_0232520
49.1;refGene_gn=ITSN2;refGene_tx=XM_023252050.1;refGene_gn=ITSN2;refGe
ne_tx=XM_023252051.1;refGene_gn=ITSN2;refGene_tx=XM_023252052.1;refGen
e_gn=ITSN2;refGene_tx=XM_023252053.1;refGene_gn=ITSN2;refGene_tx=XM_02
3252054.1;refGene_gn=ITSN2;refGene_tx=XM_023252055.1;refGene_gn=ITSN2;
refGene_tx=XM_023252056.1;refGene_gn=ITSN2;refGene_tx=XM_023252057.1;r
efGene_gn=ITSN2;refGene_tx=XM_023252058.1;refGene_gn=ITSN2;refGene_tx=
XM_023252059.1
A3 123832200 123832265
refGene_gn=ITSN2;refGene_tx=XM_019827831.2;refGene_gn=ITSN2;refGene_tx
=XM_019827835.2;refGene_gn=ITSN2;refGene_tx=XM_019827837.2;refGene_gn=
ITSN2;refGene_tx=XM_019827838.2;refGene_gn=ITSN2;refGene_tx=XM_0232520
49.1;refGene_gn=ITSN2;refGene_tx=XM_023252050.1;refGene_gn=ITSN2;refGe
ne_tx=XM_023252051.1;refGene_gn=ITSN2;refGene_tx=XM_023252052.1;refGen
e_gn=ITSN2;refGene_tx=XM_023252053.1;refGene_gn=ITSN2;refGene_tx=XM_02
3252054.1;refGene_gn=ITSN2;refGene_tx=XM_023252055.1;refGene_gn=ITSN2;
refGene_tx=XM_023252056.1;refGene_gn=ITSN2;refGene_tx=XM_023252057.1;r
efGene_gn=ITSN2;refGene_tx=XM_023252058.1;refGene_gn=ITSN2;refGene_tx=
XM_023252059.1
A3 123834080 123834247
refGene_gn=ITSN2;refGene_tx=XM_019827831.2;refGene_gn=ITSN2;refGene_tx

=XM_019827835.2;refGene_gn=ITSN2;refGene_tx=XM_019827837.2;refGene_gn=ITSN2;refGene_tx=XM_019827838.2;refGene_gn=ITSN2;refGene_tx=XM_023252049.1;refGene_gn=ITSN2;refGene_tx=XM_023252050.1;refGene_gn=ITSN2;refGene_tx=XM_023252051.1;refGene_gn=ITSN2;refGene_tx=XM_023252052.1;refGene_gn=ITSN2;refGene_tx=XM_023252053.1;refGene_gn=ITSN2;refGene_tx=XM_023252054.1;refGene_gn=ITSN2;refGene_tx=XM_023252055.1;refGene_gn=ITSN2;refGene_tx=XM_023252056.1;refGene_gn=ITSN2;refGene_tx=XM_023252057.1;refGene_gn=ITSN2;refGene_tx=XM_023252058.1;refGene_gn=ITSN2;refGene_tx=XM_023252059.1

A3 123837974 123838020
refGene_gn=ITSN2;refGene_tx=XM_019827831.2;refGene_gn=ITSN2;refGene_tx=XM_019827835.2;refGene_gn=ITSN2;refGene_tx=XM_019827837.2;refGene_gn=ITSN2;refGene_tx=XM_019827838.2;refGene_gn=ITSN2;refGene_tx=XM_023252049.1;refGene_gn=ITSN2;refGene_tx=XM_023252050.1;refGene_gn=ITSN2;refGene_tx=XM_023252051.1;refGene_gn=ITSN2;refGene_tx=XM_023252052.1;refGene_gn=ITSN2;refGene_tx=XM_023252053.1;refGene_gn=ITSN2;refGene_tx=XM_023252054.1;refGene_gn=ITSN2;refGene_tx=XM_023252055.1;refGene_gn=ITSN2;refGene_tx=XM_023252056.1;refGene_gn=ITSN2;refGene_tx=XM_023252057.1;refGene_gn=ITSN2;refGene_tx=XM_023252058.1;refGene_gn=ITSN2;refGene_tx=XM_023252059.1

A3 123838102 123838224
refGene_gn=ITSN2;refGene_tx=XM_019827831.2;refGene_gn=ITSN2;refGene_tx=XM_019827835.2;refGene_gn=ITSN2;refGene_tx=XM_019827837.2;refGene_gn=ITSN2;refGene_tx=XM_019827838.2;refGene_gn=ITSN2;refGene_tx=XM_023252049.1;refGene_gn=ITSN2;refGene_tx=XM_023252050.1;refGene_gn=ITSN2;refGene_tx=XM_023252051.1;refGene_gn=ITSN2;refGene_tx=XM_023252052.1;refGene_gn=ITSN2;refGene_tx=XM_023252053.1;refGene_gn=ITSN2;refGene_tx=XM_023252054.1;refGene_gn=ITSN2;refGene_tx=XM_023252055.1;refGene_gn=ITSN2;refGene_tx=XM_023252056.1;refGene_gn=ITSN2;refGene_tx=XM_023252057.1;refGene_gn=ITSN2;refGene_tx=XM_023252058.1;refGene_gn=ITSN2;refGene_tx=XM_023252059.1

A3 123839963 123840060
refGene_gn=ITSN2;refGene_tx=XM_019827831.2;refGene_gn=ITSN2;refGene_tx=XM_019827835.2;refGene_gn=ITSN2;refGene_tx=XM_019827837.2;refGene_gn=ITSN2;refGene_tx=XM_019827838.2;refGene_gn=ITSN2;refGene_tx=XM_023252049.1;refGene_gn=ITSN2;refGene_tx=XM_023252050.1;refGene_gn=ITSN2;refGene_tx=XM_023252051.1;refGene_gn=ITSN2;refGene_tx=XM_023252052.1;refGene_gn=ITSN2;refGene_tx=XM_023252053.1;refGene_gn=ITSN2;refGene_tx=XM_023252054.1;refGene_gn=ITSN2;refGene_tx=XM_023252055.1;refGene_gn=ITSN2;refGene_tx=XM_023252056.1;refGene_gn=ITSN2;refGene_tx=XM_023252057.1;refGene_gn=ITSN2;refGene_tx=XM_023252058.1;refGene_gn=ITSN2;refGene_tx=XM_023252059.1

A3 123840742 123840934
refGene_gn=ITSN2;refGene_tx=XM_019827831.2;refGene_gn=ITSN2;refGene_tx=XM_019827835.2;refGene_gn=ITSN2;refGene_tx=XM_019827837.2;refGene_gn=ITSN2;refGene_tx=XM_019827838.2;refGene_gn=ITSN2;refGene_tx=XM_023252049.1;refGene_gn=ITSN2;refGene_tx=XM_023252050.1;refGene_gn=ITSN2;refGene_tx=XM_023252051.1;refGene_gn=ITSN2;refGene_tx=XM_023252052.1;refGene_gn=ITSN2;refGene_tx=XM_023252053.1;refGene_gn=ITSN2;refGene_tx=XM_023252054.1;refGene_gn=ITSN2;refGene_tx=XM_023252055.1;refGene_gn=ITSN2;

refGene_tx=XM_023252056.1;refGene_gn=ITSN2;refGene_tx=XM_023252057.1;r
efGene_gn=ITSN2;refGene_tx=XM_023252058.1;refGene_gn=ITSN2;refGene_tx=
XM_023252059.1
A3 123868250 123868372
refGene_gn=ITSN2;refGene_tx=XM_019827831.2;refGene_gn=ITSN2;refGene_tx
=XM_019827835.2;refGene_gn=ITSN2;refGene_tx=XM_019827837.2;refGene_gn=
ITSN2;refGene_tx=XM_019827838.2;refGene_gn=ITSN2;refGene_tx=XM_0232520
49.1;refGene_gn=ITSN2;refGene_tx=XM_023252050.1;refGene_gn=ITSN2;refGe
ne_tx=XM_023252051.1;refGene_gn=ITSN2;refGene_tx=XM_023252052.1;refGen
e_gn=ITSN2;refGene_tx=XM_023252053.1;refGene_gn=ITSN2;refGene_tx=XM_02
3252054.1;refGene_gn=ITSN2;refGene_tx=XM_023252055.1;refGene_gn=ITSN2;
refGene_tx=XM_023252056.1;refGene_gn=ITSN2;refGene_tx=XM_023252057.1;r
efGene_gn=ITSN2;refGene_tx=XM_023252058.1;refGene_gn=ITSN2;refGene_tx=
XM_023252059.1
A3 123871905 123872012
refGene_gn=ITSN2;refGene_tx=XM_019827831.2;refGene_gn=ITSN2;refGene_tx
=XM_019827835.2;refGene_gn=ITSN2;refGene_tx=XM_019827837.2;refGene_gn=
ITSN2;refGene_tx=XM_019827838.2;refGene_gn=ITSN2;refGene_tx=XM_0232520
49.1;refGene_gn=ITSN2;refGene_tx=XM_023252050.1;refGene_gn=ITSN2;refGe
ne_tx=XM_023252051.1;refGene_gn=ITSN2;refGene_tx=XM_023252052.1;refGen
e_gn=ITSN2;refGene_tx=XM_023252053.1;refGene_gn=ITSN2;refGene_tx=XM_02
3252054.1;refGene_gn=ITSN2;refGene_tx=XM_023252055.1;refGene_gn=ITSN2;
refGene_tx=XM_023252056.1;refGene_gn=ITSN2;refGene_tx=XM_023252057.1;r
efGene_gn=ITSN2;refGene_tx=XM_023252058.1;refGene_gn=ITSN2;refGene_tx=
XM_023252059.1
A3 123872867 123873051
refGene_gn=ITSN2;refGene_tx=XM_019827831.2;refGene_gn=ITSN2;refGene_tx
=XM_019827835.2;refGene_gn=ITSN2;refGene_tx=XM_019827837.2;refGene_gn=
ITSN2;refGene_tx=XM_019827838.2;refGene_gn=ITSN2;refGene_tx=XM_0232520
49.1;refGene_gn=ITSN2;refGene_tx=XM_023252050.1;refGene_gn=ITSN2;refGe
ne_tx=XM_023252051.1;refGene_gn=ITSN2;refGene_tx=XM_023252052.1;refGen
e_gn=ITSN2;refGene_tx=XM_023252053.1;refGene_gn=ITSN2;refGene_tx=XM_02
3252054.1;refGene_gn=ITSN2;refGene_tx=XM_023252055.1;refGene_gn=ITSN2;
refGene_tx=XM_023252056.1;refGene_gn=ITSN2;refGene_tx=XM_023252057.1;r
efGene_gn=ITSN2;refGene_tx=XM_023252058.1;refGene_gn=ITSN2;refGene_tx=
XM_023252059.1
A3 123877264 123877363
refGene_gn=ITSN2;refGene_tx=XM_019827831.2;refGene_gn=ITSN2;refGene_tx
=XM_019827835.2;refGene_gn=ITSN2;refGene_tx=XM_019827837.2;refGene_gn=
ITSN2;refGene_tx=XM_019827838.2;refGene_gn=ITSN2;refGene_tx=XM_0232520
49.1;refGene_gn=ITSN2;refGene_tx=XM_023252050.1;refGene_gn=ITSN2;refGe
ne_tx=XM_023252051.1;refGene_gn=ITSN2;refGene_tx=XM_023252052.1;refGen
e_gn=ITSN2;refGene_tx=XM_023252053.1;refGene_gn=ITSN2;refGene_tx=XM_02
3252054.1;refGene_gn=ITSN2;refGene_tx=XM_023252056.1;refGene_gn=ITSN2;
refGene_tx=XM_023252057.1;refGene_gn=ITSN2;refGene_tx=XM_023252058.1;r
efGene_gn=ITSN2;refGene_tx=XM_023252059.1
A3 123880242 123880410
refGene_gn=ITSN2;refGene_tx=XM_019827831.2;refGene_gn=ITSN2;refGene_tx
=XM_019827835.2;refGene_gn=ITSN2;refGene_tx=XM_019827837.2;refGene_gn=
ITSN2;refGene_tx=XM_019827838.2;refGene_gn=ITSN2;refGene_tx=XM_0232520

49.1;refGene_gn=ITSN2;refGene_tx=XM_023252050.1;refGene_gn=ITSN2;refGene_tx=XM_023252051.1;refGene_gn=ITSN2;refGene_tx=XM_023252052.1;refGene_gn=ITSN2;refGene_tx=XM_023252053.1;refGene_gn=ITSN2;refGene_tx=XM_023252054.1;refGene_gn=ITSN2;refGene_tx=XM_023252055.1;refGene_gn=ITSN2;refGene_tx=XM_023252056.1;refGene_gn=ITSN2;refGene_tx=XM_023252057.1;refGene_gn=ITSN2;refGene_tx=XM_023252058.1;refGene_gn=ITSN2;refGene_tx=XM_023252059.1
A3	123880764	123880980
refGene_gn=ITSN2;refGene_tx=XM_019827831.2;refGene_gn=ITSN2;refGene_tx=XM_019827835.2;refGene_gn=ITSN2;refGene_tx=XM_019827837.2;refGene_gn=ITSN2;refGene_tx=XM_019827838.2;refGene_gn=ITSN2;refGene_tx=XM_023252049.1;refGene_gn=ITSN2;refGene_tx=XM_023252050.1;refGene_gn=ITSN2;refGene_tx=XM_023252051.1;refGene_gn=ITSN2;refGene_tx=XM_023252052.1;refGene_gn=ITSN2;refGene_tx=XM_023252053.1;refGene_gn=ITSN2;refGene_tx=XM_023252054.1;refGene_gn=ITSN2;refGene_tx=XM_023252055.1;refGene_gn=ITSN2;refGene_tx=XM_023252056.1;refGene_gn=ITSN2;refGene_tx=XM_023252057.1;refGene_gn=ITSN2;refGene_tx=XM_023252058.1;refGene_gn=ITSN2;refGene_tx=XM_023252059.1
A3	123881523	123881645
refGene_gn=ITSN2;refGene_tx=XM_019827831.2;refGene_gn=ITSN2;refGene_tx=XM_019827835.2;refGene_gn=ITSN2;refGene_tx=XM_019827837.2;refGene_gn=ITSN2;refGene_tx=XM_019827838.2;refGene_gn=ITSN2;refGene_tx=XM_023252049.1;refGene_gn=ITSN2;refGene_tx=XM_023252050.1;refGene_gn=ITSN2;refGene_tx=XM_023252051.1;refGene_gn=ITSN2;refGene_tx=XM_023252052.1;refGene_gn=ITSN2;refGene_tx=XM_023252053.1;refGene_gn=ITSN2;refGene_tx=XM_023252054.1;refGene_gn=ITSN2;refGene_tx=XM_023252055.1;refGene_gn=ITSN2;refGene_tx=XM_023252056.1;refGene_gn=ITSN2;refGene_tx=XM_023252057.1;refGene_gn=ITSN2;refGene_tx=XM_023252058.1;refGene_gn=ITSN2;refGene_tx=XM_023252059.1
A3	123882441	123882524
refGene_gn=ITSN2;refGene_tx=XM_019827831.2;refGene_gn=ITSN2;refGene_tx=XM_019827835.2;refGene_gn=ITSN2;refGene_tx=XM_019827837.2;refGene_gn=ITSN2;refGene_tx=XM_019827838.2;refGene_gn=ITSN2;refGene_tx=XM_023252049.1;refGene_gn=ITSN2;refGene_tx=XM_023252050.1;refGene_gn=ITSN2;refGene_tx=XM_023252051.1;refGene_gn=ITSN2;refGene_tx=XM_023252052.1;refGene_gn=ITSN2;refGene_tx=XM_023252053.1;refGene_gn=ITSN2;refGene_tx=XM_023252054.1;refGene_gn=ITSN2;refGene_tx=XM_023252055.1;refGene_gn=ITSN2;refGene_tx=XM_023252056.1;refGene_gn=ITSN2;refGene_tx=XM_023252057.1;refGene_gn=ITSN2;refGene_tx=XM_023252058.1;refGene_gn=ITSN2;refGene_tx=XM_023252059.1
A3	123885083	123885167
refGene_gn=ITSN2;refGene_tx=XM_019827831.2;refGene_gn=ITSN2;refGene_tx=XM_019827835.2;refGene_gn=ITSN2;refGene_tx=XM_019827837.2;refGene_gn=ITSN2;refGene_tx=XM_019827838.2;refGene_gn=ITSN2;refGene_tx=XM_023252049.1;refGene_gn=ITSN2;refGene_tx=XM_023252050.1;refGene_gn=ITSN2;refGene_tx=XM_023252051.1;refGene_gn=ITSN2;refGene_tx=XM_023252052.1;refGene_gn=ITSN2;refGene_tx=XM_023252053.1;refGene_gn=ITSN2;refGene_tx=XM_023252054.1;refGene_gn=ITSN2;refGene_tx=XM_023252055.1;refGene_gn=ITSN2;refGene_tx=XM_023252056.1;refGene_gn=ITSN2;refGene_tx=XM_023252057.1;refGene_gn=ITSN2;refGene_tx=XM_023252058.1;refGene_gn=ITSN2;refGene_tx=

XM_023252059.1
A3	123885569	123885743
refGene_gn=ITSN2;refGene_tx=XM_019827831.2;refGene_gn=ITSN2;refGene_tx=XM_019827835.2;refGene_gn=ITSN2;refGene_tx=XM_019827837.2;refGene_gn=ITSN2;refGene_tx=XM_019827838.2;refGene_gn=ITSN2;refGene_tx=XM_023252049.1;refGene_gn=ITSN2;refGene_tx=XM_023252050.1;refGene_gn=ITSN2;refGene_tx=XM_023252051.1;refGene_gn=ITSN2;refGene_tx=XM_023252052.1;refGene_gn=ITSN2;refGene_tx=XM_023252053.1;refGene_gn=ITSN2;refGene_tx=XM_023252054.1;refGene_gn=ITSN2;refGene_tx=XM_023252055.1;refGene_gn=ITSN2;refGene_tx=XM_023252056.1;refGene_gn=ITSN2;refGene_tx=XM_023252057.1;refGene_gn=ITSN2;refGene_tx=XM_023252058.1;refGene_gn=ITSN2;refGene_tx=XM_023252059.1
A3	123886205	123886363
refGene_gn=ITSN2;refGene_tx=XM_019827831.2;refGene_gn=ITSN2;refGene_tx=XM_019827835.2;refGene_gn=ITSN2;refGene_tx=XM_019827837.2;refGene_gn=ITSN2;refGene_tx=XM_019827838.2;refGene_gn=ITSN2;refGene_tx=XM_023252049.1;refGene_gn=ITSN2;refGene_tx=XM_023252050.1;refGene_gn=ITSN2;refGene_tx=XM_023252051.1;refGene_gn=ITSN2;refGene_tx=XM_023252052.1;refGene_gn=ITSN2;refGene_tx=XM_023252053.1;refGene_gn=ITSN2;refGene_tx=XM_023252054.1;refGene_gn=ITSN2;refGene_tx=XM_023252055.1;refGene_gn=ITSN2;refGene_tx=XM_023252056.1;refGene_gn=ITSN2;refGene_tx=XM_023252057.1;refGene_gn=ITSN2;refGene_tx=XM_023252058.1;refGene_gn=ITSN2;refGene_tx=XM_023252059.1
A3	123910046	123910172
refGene_gn=FAM228B;refGene_tx=XM_023252204.1
A3	123910768	123910864
refGene_gn=FAM228B;refGene_tx=XM_023252204.1
A3	123910865	123910914
refGene_gn=FAM228B;refGene_tx=XM_023252204.1
A3	123913691	123913793
refGene_gn=FAM228B;refGene_tx=XM_023252204.1
A3	123919611	123919768
refGene_gn=FAM228B;refGene_tx=XM_023252204.1
A3	123919862	123919950
refGene_gn=FAM228B;refGene_tx=XM_023252204.1
A3	123924480	123924561
refGene_gn=FAM228B;refGene_tx=XM_023252204.1
A3	123925182	123925374
refGene_gn=FAM228B;refGene_tx=XM_023252204.1
A3	123927889	123927958
refGene_gn=FAM228B;refGene_tx=XM_023252204.1
A3	123930424	123930523
refGene_gn=FAM228B;refGene_tx=XM_023252204.1
A3	123933729	123933846
refGene_gn=PFN4;refGene_tx=XM_019827847.2;refGene_gn=PFN4;refGene_tx=XM_019827849.2
A3	123934209	123934347
refGene_gn=PFN4;refGene_tx=XM_019827847.2;refGene_gn=PFN4;refGene_tx=XM_019827849.2
A3	123935160	123935266

refGene_gn=PFN4;refGene_tx=XM_019827847.2;refGene_gn=PFN4;refGene_tx=XM_019827849.2
A3 123937959 123937988
refGene_gn=PFN4;refGene_tx=XM_019827847.2;refGene_gn=PFN4;refGene_tx=XM_019827849.2
A3 123948847 123948848
refGene_gn=TP53I3;refGene_tx=XM_003984458.5;refGene_gn=TP53I3;refGene_tx=XM_006930452.4
A3 123949183 123949335
refGene_gn=TP53I3;refGene_tx=XM_003984458.5;refGene_gn=TP53I3;refGene_tx=XM_006930452.4
A3 123950423 123950691
refGene_gn=TP53I3;refGene_tx=XM_003984458.5;refGene_gn=TP53I3;refGene_tx=XM_006930452.4
A3 123951894 123952107
refGene_gn=TP53I3;refGene_tx=XM_003984458.5;refGene_gn=TP53I3;refGene_tx=XM_006930452.4
A3 123952983 123953180
refGene_gn=TP53I3;refGene_tx=XM_003984458.5;refGene_gn=TP53I3;refGene_tx=XM_006930452.4
A3 123953956 123954139
refGene_gn=TP53I3;refGene_tx=XM_003984458.5;refGene_gn=TP53I3;refGene_tx=XM_006930452.4
A3 123954937 123954967
refGene_gn=SF3B6;refGene_tx=XM_003984459.5
A3 123957914 123958033
refGene_gn=SF3B6;refGene_tx=XM_003984459.5
A3 123962593 123962752
refGene_gn=FKBP1B;refGene_tx=XM_019827844.2;refGene_gn=FKBP1B;refGene_tx=XM_023252061.1
A3 123964043 123964182
refGene_gn=SF3B6;refGene_tx=XM_003984459.5
A3 123964351 123964441
refGene_gn=SF3B6;refGene_tx=XM_003984459.5
A3 123969237 123969411
refGene_gn=FKBP1B;refGene_tx=XM_023252062.1;refGene_gn=FKBP1B;refGene_tx=XM_023252064.1
A3 123970454 123970567
refGene_gn=FKBP1B;refGene_tx=XM_019827844.2;refGene_gn=FKBP1B;refGene_tx=XM_023252061.1;refGene_gn=FKBP1B;refGene_tx=XM_023252062.1;refGene_gn=FKBP1B;refGene_tx=XM_023252064.1
A3 123975243 123975291
refGene_gn=FKBP1B;refGene_tx=XM_023252061.1;refGene_gn=FKBP1B;refGene_tx=XM_023252062.1;refGene_gn=FKBP1B;refGene_tx=XM_023252064.1
A3 123979790 123979827
refGene_gn=FKBP1B;refGene_tx=XM_023252061.1;refGene_gn=FKBP1B;refGene_tx=XM_023252062.1;refGene_gn=FKBP1B;refGene_tx=XM_023252064.1
A3 123991604 123993434
refGene_gn=WDCP;refGene_tx=XM_023252060.1
A3 123996292 123996410

refGene_gn=WDCP;refGene_tx=XM_023252060.1
A3 123997715 123997843
refGene_gn=WDCP;refGene_tx=XM_023252060.1
A3 124006533 124006558
refGene_gn=MFSD2B;refGene_tx=XM_023252065.1
A3 124007683 124007860
refGene_gn=MFSD2B;refGene_tx=XM_023252065.1
A3 124008269 124008413
refGene_gn=MFSD2B;refGene_tx=XM_023252065.1
A3 124008794 124008904
refGene_gn=MFSD2B;refGene_tx=XM_023252065.1
A3 124009138 124009219
refGene_gn=MFSD2B;refGene_tx=XM_023252065.1
A3 124009528 124009612
refGene_gn=MFSD2B;refGene_tx=XM_023252065.1
A3 124010052 124010174
refGene_gn=MFSD2B;refGene_tx=XM_023252065.1
A3 124010287 124010378
refGene_gn=MFSD2B;refGene_tx=XM_023252065.1
A3 124012543 124012674
refGene_gn=MFSD2B;refGene_tx=XM_023252065.1
A3 124012756 124012835
refGene_gn=MFSD2B;refGene_tx=XM_023252065.1
A3 124013125 124013249
refGene_gn=MFSD2B;refGene_tx=XM_023252065.1
A3 124013721 124013846
refGene_gn=MFSD2B;refGene_tx=XM_023252065.1
A3 124014978 124015104
refGene_gn=MFSD2B;refGene_tx=XM_023252065.1
A3 124018241 124018325
refGene_gn=MFSD2B;refGene_tx=XM_023252065.1
A3 124024188 124024381
refGene_gn=UBXN2A;refGene_tx=XM_003984460.4;refGene_gn=UBXN2A;refGene_tx=XM_006930454.3
A3 124035279 124035435
refGene_gn=UBXN2A;refGene_tx=XM_003984460.4;refGene_gn=UBXN2A;refGene_tx=XM_006930454.3
A3 124036960 124037098
refGene_gn=UBXN2A;refGene_tx=XM_003984460.4;refGene_gn=UBXN2A;refGene_tx=XM_006930454.3
A3 124040268 124040375
refGene_gn=UBXN2A;refGene_tx=XM_003984460.4;refGene_gn=UBXN2A;refGene_tx=XM_006930454.3
A3 124045155 124045294
refGene_gn=UBXN2A;refGene_tx=XM_003984460.4;refGene_gn=UBXN2A;refGene_tx=XM_006930454.3
A3 124061053 124061094
refGene_gn=UBXN2A;refGene_tx=XM_003984460.4;refGene_gn=UBXN2A;refGene_tx=XM_006930454.3
A3 124085204 124085420

refGene_gn=ATAD2B;refGene_tx=XM_023252066.1;refGene_gn=ATAD2B;refGene_tx=XM_023252067.1;refGene_gn=ATAD2B;refGene_tx=XM_023252068.1
A3 124108212 124108364
refGene_gn=ATAD2B;refGene_tx=XM_023252066.1;refGene_gn=ATAD2B;refGene_tx=XM_023252067.1;refGene_gn=ATAD2B;refGene_tx=XM_023252068.1
A3 124113338 124113388
refGene_gn=ATAD2B;refGene_tx=XM_023252066.1;refGene_gn=ATAD2B;refGene_tx=XM_023252067.1;refGene_gn=ATAD2B;refGene_tx=XM_023252068.1
A3 124113750 124113904
refGene_gn=ATAD2B;refGene_tx=XM_023252066.1;refGene_gn=ATAD2B;refGene_tx=XM_023252067.1;refGene_gn=ATAD2B;refGene_tx=XM_023252068.1
A3 124115096 124115199
refGene_gn=ATAD2B;refGene_tx=XM_023252066.1;refGene_gn=ATAD2B;refGene_tx=XM_023252067.1;refGene_gn=ATAD2B;refGene_tx=XM_023252068.1
A3 124117425 124117534
refGene_gn=ATAD2B;refGene_tx=XM_023252066.1;refGene_gn=ATAD2B;refGene_tx=XM_023252067.1;refGene_gn=ATAD2B;refGene_tx=XM_023252068.1
A3 124119723 124119840
refGene_gn=ATAD2B;refGene_tx=XM_023252066.1;refGene_gn=ATAD2B;refGene_tx=XM_023252067.1;refGene_gn=ATAD2B;refGene_tx=XM_023252068.1
A3 124122128 124122204
refGene_gn=ATAD2B;refGene_tx=XM_023252066.1;refGene_gn=ATAD2B;refGene_tx=XM_023252067.1;refGene_gn=ATAD2B;refGene_tx=XM_023252068.1
A3 124142423 124142522
refGene_gn=ATAD2B;refGene_tx=XM_023252066.1;refGene_gn=ATAD2B;refGene_tx=XM_023252067.1;refGene_gn=ATAD2B;refGene_tx=XM_023252068.1
A3 124145023 124145135
refGene_gn=ATAD2B;refGene_tx=XM_023252066.1;refGene_gn=ATAD2B;refGene_tx=XM_023252067.1;refGene_gn=ATAD2B;refGene_tx=XM_023252068.1
A3 124148162 124148278
refGene_gn=ATAD2B;refGene_tx=XM_023252066.1;refGene_gn=ATAD2B;refGene_tx=XM_023252067.1;refGene_gn=ATAD2B;refGene_tx=XM_023252068.1
A3 124149851 124150026
refGene_gn=ATAD2B;refGene_tx=XM_023252066.1;refGene_gn=ATAD2B;refGene_tx=XM_023252067.1;refGene_gn=ATAD2B;refGene_tx=XM_023252068.1
A3 124154709 124154798
refGene_gn=ATAD2B;refGene_tx=XM_023252066.1;refGene_gn=ATAD2B;refGene_tx=XM_023252067.1;refGene_gn=ATAD2B;refGene_tx=XM_023252068.1
A3 124170133 124170293
refGene_gn=ATAD2B;refGene_tx=XM_023252066.1;refGene_gn=ATAD2B;refGene_tx=XM_023252067.1;refGene_gn=ATAD2B;refGene_tx=XM_023252068.1
A3 124177056 124177147
refGene_gn=ATAD2B;refGene_tx=XM_023252066.1;refGene_gn=ATAD2B;refGene_tx=XM_023252067.1;refGene_gn=ATAD2B;refGene_tx=XM_023252068.1
A3 124182639 124182951
refGene_gn=ATAD2B;refGene_tx=XM_023252066.1;refGene_gn=ATAD2B;refGene_tx=XM_023252067.1;refGene_gn=ATAD2B;refGene_tx=XM_023252068.1
A3 124185262 124185398
refGene_gn=ATAD2B;refGene_tx=XM_023252066.1;refGene_gn=ATAD2B;refGene_tx=XM_023252067.1;refGene_gn=ATAD2B;refGene_tx=XM_023252068.1

A3 124193694 124193881
refGene_gn=ATAD2B;refGene_tx=XM_023252066.1;refGene_gn=ATAD2B;refGene_tx=XM_023252067.1;refGene_gn=ATAD2B;refGene_tx=XM_023252068.1
A3 124204587 124204773
refGene_gn=ATAD2B;refGene_tx=XM_023252066.1;refGene_gn=ATAD2B;refGene_tx=XM_023252067.1;refGene_gn=ATAD2B;refGene_tx=XM_023252068.1
A3 124217695 124217831
refGene_gn=ATAD2B;refGene_tx=XM_023252066.1;refGene_gn=ATAD2B;refGene_tx=XM_023252067.1;refGene_gn=ATAD2B;refGene_tx=XM_023252068.1
A3 124219792 124219989
refGene_gn=ATAD2B;refGene_tx=XM_023252066.1;refGene_gn=ATAD2B;refGene_tx=XM_023252067.1;refGene_gn=ATAD2B;refGene_tx=XM_023252068.1
A3 124223035 124223195
refGene_gn=ATAD2B;refGene_tx=XM_023252066.1;refGene_gn=ATAD2B;refGene_tx=XM_023252067.1;refGene_gn=ATAD2B;refGene_tx=XM_023252068.1
A3 124235608 124235731
refGene_gn=ATAD2B;refGene_tx=XM_023252066.1;refGene_gn=ATAD2B;refGene_tx=XM_023252067.1;refGene_gn=ATAD2B;refGene_tx=XM_023252068.1
A3 124238887 124239025
refGene_gn=ATAD2B;refGene_tx=XM_023252066.1;refGene_gn=ATAD2B;refGene_tx=XM_023252067.1;refGene_gn=ATAD2B;refGene_tx=XM_023252068.1
A3 124242127 124242811
refGene_gn=ATAD2B;refGene_tx=XM_023252066.1;refGene_gn=ATAD2B;refGene_tx=XM_023252067.1;refGene_gn=ATAD2B;refGene_tx=XM_023252068.1
A3 124245629 124245757
refGene_gn=ATAD2B;refGene_tx=XM_023252066.1;refGene_gn=ATAD2B;refGene_tx=XM_023252067.1;refGene_gn=ATAD2B;refGene_tx=XM_023252068.1
A3 124246216 124246450
refGene_gn=ATAD2B;refGene_tx=XM_023252066.1;refGene_gn=ATAD2B;refGene_tx=XM_023252067.1;refGene_gn=ATAD2B;refGene_tx=XM_023252068.1
A3 124248216 124248258
refGene_gn=ATAD2B;refGene_tx=XM_023252067.1;refGene_gn=ATAD2B;refGene_tx=XM_023252068.1
A3 124279554 124279738
refGene_gn=KLHL29;refGene_tx=XM_023252069.1;refGene_gn=KLHL29;refGene_tx=XM_023252070.1;refGene_gn=KLHL29;refGene_tx=XM_023252071.1
A3 124281793 124281938
refGene_gn=KLHL29;refGene_tx=XM_023252069.1;refGene_gn=KLHL29;refGene_tx=XM_023252070.1
A3 124282298 124282492
refGene_gn=KLHL29;refGene_tx=XM_023252069.1;refGene_gn=KLHL29;refGene_tx=XM_023252070.1;refGene_gn=KLHL29;refGene_tx=XM_023252071.1
A3 124287595 124287776
refGene_gn=KLHL29;refGene_tx=XM_023252069.1;refGene_gn=KLHL29;refGene_tx=XM_023252070.1;refGene_gn=KLHL29;refGene_tx=XM_023252071.1
A3 124287983 124288166
refGene_gn=KLHL29;refGene_tx=XM_023252069.1;refGene_gn=KLHL29;refGene_tx=XM_023252070.1;refGene_gn=KLHL29;refGene_tx=XM_023252071.1
A3 124288309 124288508
refGene_gn=KLHL29;refGene_tx=XM_023252069.1;refGene_gn=KLHL29;refGene_

tx=XM_023252070.1;refGene_gn=KLHL29;refGene_tx=XM_023252071.1
A3 124290928 124291188
refGene_gn=KLHL29;refGene_tx=XM_023252069.1;refGene_gn=KLHL29;refGene_
tx=XM_023252070.1;refGene_gn=KLHL29;refGene_tx=XM_023252071.1
A3 124291909 124292112
refGene_gn=KLHL29;refGene_tx=XM_023252069.1;refGene_gn=KLHL29;refGene_
tx=XM_023252070.1;refGene_gn=KLHL29;refGene_tx=XM_023252071.1
A3 124300831 124300970
refGene_gn=KLHL29;refGene_tx=XM_023252069.1;refGene_gn=KLHL29;refGene_
tx=XM_023252070.1;refGene_gn=KLHL29;refGene_tx=XM_023252071.1
A3 124337132 124337645
refGene_gn=KLHL29;refGene_tx=XM_023252069.1;refGene_gn=KLHL29;refGene_
tx=XM_023252070.1;refGene_gn=KLHL29;refGene_tx=XM_023252071.1
A3 124340146 124340288
refGene_gn=KLHL29;refGene_tx=XM_023252069.1;refGene_gn=KLHL29;refGene_
tx=XM_023252070.1;refGene_gn=KLHL29;refGene_tx=XM_023252071.1
A3 124422730 124423015
refGene_gn=KLHL29;refGene_tx=XM_023252069.1;refGene_gn=KLHL29;refGene_
tx=XM_023252070.1;refGene_gn=KLHL29;refGene_tx=XM_023252071.1
A3 126606950 126607009
refGene_gn=TDRD15;refGene_tx=XM_019827852.2
A3 126611335 126617558
refGene_gn=TDRD15;refGene_tx=XM_019827852.2
A3 126693089 126693156
refGene_gn=APOB;refGene_tx=XM_003984468.5
A3 126693431 126693470
refGene_gn=APOB;refGene_tx=XM_003984468.5
A3 126694636 126694752
refGene_gn=APOB;refGene_tx=XM_003984468.5
A3 126695952 126696098
refGene_gn=APOB;refGene_tx=XM_003984468.5
A3 126698127 126698281
refGene_gn=APOB;refGene_tx=XM_003984468.5
A3 126699118 126699274
refGene_gn=APOB;refGene_tx=XM_003984468.5
A3 126700787 126700912
refGene_gn=APOB;refGene_tx=XM_003984468.5
A3 126701558 126701644
refGene_gn=APOB;refGene_tx=XM_003984468.5
A3 126702806 126703026
refGene_gn=APOB;refGene_tx=XM_003984468.5
A3 126703810 126704038
refGene_gn=APOB;refGene_tx=XM_003984468.5
A3 126704253 126704371
refGene_gn=APOB;refGene_tx=XM_003984468.5
A3 126704478 126704625
refGene_gn=APOB;refGene_tx=XM_003984468.5
A3 126705325 126705537
refGene_gn=APOB;refGene_tx=XM_003984468.5
A3 126705804 126706042

refGene_gn=APOB;refGene_tx=XM_003984468.5
A3	126706616	126706793
refGene_gn=APOB;refGene_tx=XM_003984468.5
A3	126708170	126708362
refGene_gn=APOB;refGene_tx=XM_003984468.5
A3	126709287	126709455
refGene_gn=APOB;refGene_tx=XM_003984468.5
A3	126710148	126710360
refGene_gn=APOB;refGene_tx=XM_003984468.5
A3	126712880	126713063
refGene_gn=APOB;refGene_tx=XM_003984468.5
A3	126713877	126713999
refGene_gn=APOB;refGene_tx=XM_003984468.5
A3	126715669	126715880
refGene_gn=APOB;refGene_tx=XM_003984468.5
A3	126716430	126716606
refGene_gn=APOB;refGene_tx=XM_003984468.5
A3	126716708	126716896
refGene_gn=APOB;refGene_tx=XM_003984468.5
A3	126717377	126717523
refGene_gn=APOB;refGene_tx=XM_003984468.5
A3	126719471	126719845
refGene_gn=APOB;refGene_tx=XM_003984468.5
A3	126720350	126727547
refGene_gn=APOB;refGene_tx=XM_003984468.5
A3	126727938	126728053
refGene_gn=APOB;refGene_tx=XM_003984468.5
A3	126728154	126728338
refGene_gn=APOB;refGene_tx=XM_003984468.5
A3	126729499	126731128
refGene_gn=APOB;refGene_tx=XM_003984468.5
A3	126885391	126885545
refGene_gn=LDAH;refGene_tx=XM_023252072.1;refGene_gn=LDAH;refGene_tx=X
M_023252073.1
A3	126901991	126902135
refGene_gn=LDAH;refGene_tx=XM_023252072.1;refGene_gn=LDAH;refGene_tx=X
M_023252073.1
A3	126916189	126916359
refGene_gn=LDAH;refGene_tx=XM_023252072.1;refGene_gn=LDAH;refGene_tx=X
M_023252073.1
A3	126949552	126949787
refGene_gn=LDAH;refGene_tx=XM_023252072.1;refGene_gn=LDAH;refGene_tx=X
M_023252073.1
A3	126964934	126965017
refGene_gn=LDAH;refGene_tx=XM_023252072.1;refGene_gn=LDAH;refGene_tx=X
M_023252073.1
A3	126982169	126982394
refGene_gn=LDAH;refGene_tx=XM_023252072.1;refGene_gn=LDAH;refGene_tx=X
M_023252073.1
A3	126998246	126999214

refGene_gn=GDF7;refGene_tx=XM_023252074.1
A3 127002059 127002459
refGene_gn=GDF7;refGene_tx=XM_023252074.1
A3 127018367 127018399
refGene_gn=HS1BP3;refGene_tx=XM_003984469.5
A3 127023101 127023267
refGene_gn=HS1BP3;refGene_tx=XM_003984469.5
A3 127026975 127027183
refGene_gn=HS1BP3;refGene_tx=XM_003984469.5
A3 127029824 127030044
refGene_gn=HS1BP3;refGene_tx=XM_003984469.5
A3 127041367 127041531
refGene_gn=HS1BP3;refGene_tx=XM_003984469.5
A3 127042401 127042516
refGene_gn=HS1BP3;refGene_tx=XM_003984469.5
A3 127047238 127047491
refGene_gn=HS1BP3;refGene_tx=XM_003984469.5
A3 127191473 127192064
refGene_gn=RHOB;refGene_tx=XM_003984470.3
A3 127220500 127220734
refGene_gn=LOC101101211;refGene_tx=XM_003984477.5
A3 127221243 127221331
refGene_gn=LOC101101211;refGene_tx=XM_003984477.5
A3 127222989 127223114
refGene_gn=LOC101101211;refGene_tx=XM_003984477.5
A3 127224484 127224614
refGene_gn=LOC101101211;refGene_tx=XM_003984477.5
A3 127224788 127224922
refGene_gn=LOC101101211;refGene_tx=XM_003984477.5
A3 127229483 127229627
refGene_gn=LOC101101211;refGene_tx=XM_003984477.5
A3 127229628 127230237
refGene_gn=LOC101101211;refGene_tx=XM_003984477.5
A3 127297188 127297239
refGene_gn=PUM2;refGene_tx=XM_023252075.1;refGene_gn=PUM2;refGene_tx=XM_023252076.1;refGene_gn=PUM2;refGene_tx=XM_023252077.1;refGene_gn=PUM2;refGene_tx=XM_023252078.1
A3 127305117 127305226
refGene_gn=PUM2;refGene_tx=XM_023252075.1;refGene_gn=PUM2;refGene_tx=XM_023252076.1;refGene_gn=PUM2;refGene_tx=XM_023252077.1;refGene_gn=PUM2;refGene_tx=XM_023252078.1
A3 127310646 127310834
refGene_gn=PUM2;refGene_tx=XM_011281306.2;refGene_gn=PUM2;refGene_tx=XM_023252075.1;refGene_gn=PUM2;refGene_tx=XM_023252076.1;refGene_gn=PUM2;refGene_tx=XM_023252077.1;refGene_gn=PUM2;refGene_tx=XM_023252078.1
A3 127311398 127311568
refGene_gn=PUM2;refGene_tx=XM_011281306.2;refGene_gn=PUM2;refGene_tx=XM_023252075.1;refGene_gn=PUM2;refGene_tx=XM_023252076.1;refGene_gn=PUM2;refGene_tx=XM_023252077.1;refGene_gn=PUM2;refGene_tx=XM_023252078.1
A3 127314721 127314992

refGene_gn=PUM2;refGene_tx=XM_011281306.2;refGene_gn=PUM2;refGene_tx=XM_023252075.1;refGene_gn=PUM2;refGene_tx=XM_023252076.1;refGene_gn=PUM2;refGene_tx=XM_023252077.1;refGene_gn=PUM2;refGene_tx=XM_023252078.1
A3 127315227 127315321
refGene_gn=PUM2;refGene_tx=XM_011281306.2;refGene_gn=PUM2;refGene_tx=XM_023252075.1;refGene_gn=PUM2;refGene_tx=XM_023252076.1;refGene_gn=PUM2;refGene_tx=XM_023252077.1;refGene_gn=PUM2;refGene_tx=XM_023252078.1
A3 127323001 127323127
refGene_gn=PUM2;refGene_tx=XM_011281306.2;refGene_gn=PUM2;refGene_tx=XM_023252075.1;refGene_gn=PUM2;refGene_tx=XM_023252076.1;refGene_gn=PUM2;refGene_tx=XM_023252077.1;refGene_gn=PUM2;refGene_tx=XM_023252078.1
A3 127325926 127326069
refGene_gn=PUM2;refGene_tx=XM_011281306.2;refGene_gn=PUM2;refGene_tx=XM_023252075.1;refGene_gn=PUM2;refGene_tx=XM_023252076.1;refGene_gn=PUM2;refGene_tx=XM_023252077.1;refGene_gn=PUM2;refGene_tx=XM_023252078.1
A3 127327261 127327391
refGene_gn=PUM2;refGene_tx=XM_011281306.2;refGene_gn=PUM2;refGene_tx=XM_023252075.1;refGene_gn=PUM2;refGene_tx=XM_023252076.1;refGene_gn=PUM2;refGene_tx=XM_023252077.1;refGene_gn=PUM2;refGene_tx=XM_023252078.1
A3 127338309 127338453
refGene_gn=PUM2;refGene_tx=XM_011281306.2;refGene_gn=PUM2;refGene_tx=XM_023252075.1;refGene_gn=PUM2;refGene_tx=XM_023252076.1;refGene_gn=PUM2;refGene_tx=XM_023252077.1;refGene_gn=PUM2;refGene_tx=XM_023252078.1
A3 127338569 127338860
refGene_gn=PUM2;refGene_tx=XM_011281306.2;refGene_gn=PUM2;refGene_tx=XM_023252075.1;refGene_gn=PUM2;refGene_tx=XM_023252076.1;refGene_gn=PUM2;refGene_tx=XM_023252077.1;refGene_gn=PUM2;refGene_tx=XM_023252078.1
A3 127343038 127343275
refGene_gn=PUM2;refGene_tx=XM_011281306.2;refGene_gn=PUM2;refGene_tx=XM_023252075.1;refGene_gn=PUM2;refGene_tx=XM_023252076.1
A3 127359274 127359542
refGene_gn=PUM2;refGene_tx=XM_011281306.2;refGene_gn=PUM2;refGene_tx=XM_023252075.1;refGene_gn=PUM2;refGene_tx=XM_023252076.1;refGene_gn=PUM2;refGene_tx=XM_023252077.1;refGene_gn=PUM2;refGene_tx=XM_023252078.1
A3 127360592 127360722
refGene_gn=PUM2;refGene_tx=XM_011281306.2;refGene_gn=PUM2;refGene_tx=XM_023252075.1;refGene_gn=PUM2;refGene_tx=XM_023252076.1;refGene_gn=PUM2;refGene_tx=XM_023252077.1;refGene_gn=PUM2;refGene_tx=XM_023252078.1
A3 127363121 127363256
refGene_gn=PUM2;refGene_tx=XM_011281306.2;refGene_gn=PUM2;refGene_tx=XM_023252075.1;refGene_gn=PUM2;refGene_tx=XM_023252076.1;refGene_gn=PUM2;refGene_tx=XM_023252077.1;refGene_gn=PUM2;refGene_tx=XM_023252078.1
A3 127367377 127367515
refGene_gn=PUM2;refGene_tx=XM_011281306.2;refGene_gn=PUM2;refGene_tx=XM_023252075.1;refGene_gn=PUM2;refGene_tx=XM_023252076.1;refGene_gn=PUM2;refGene_tx=XM_023252077.1;refGene_gn=PUM2;refGene_tx=XM_023252078.1
A3 127368320 127368446
refGene_gn=PUM2;refGene_tx=XM_011281306.2;refGene_gn=PUM2;refGene_tx=XM_023252075.1;refGene_gn=PUM2;refGene_tx=XM_023252076.1;refGene_gn=PUM2;refGene_tx=XM_023252077.1;refGene_gn=PUM2;refGene_tx=XM_023252078.1

A3 127368685 127368807
refGene_gn=PUM2;refGene_tx=XM_011281306.2;refGene_gn=PUM2;refGene_tx=XM_023252075.1;refGene_gn=PUM2;refGene_tx=XM_023252076.1;refGene_gn=PUM2;refGene_tx=XM_023252077.1;refGene_gn=PUM2;refGene_tx=XM_023252078.1
A3 127369341 127369534
refGene_gn=PUM2;refGene_tx=XM_011281306.2;refGene_gn=PUM2;refGene_tx=XM_023252075.1;refGene_gn=PUM2;refGene_tx=XM_023252076.1;refGene_gn=PUM2;refGene_tx=XM_023252077.1;refGene_gn=PUM2;refGene_tx=XM_023252078.1
A3 127371161 127371293
refGene_gn=PUM2;refGene_tx=XM_011281306.2;refGene_gn=PUM2;refGene_tx=XM_023252075.1;refGene_gn=PUM2;refGene_tx=XM_023252076.1;refGene_gn=PUM2;refGene_tx=XM_023252077.1;refGene_gn=PUM2;refGene_tx=XM_023252078.1
A3 127388343 127388409
refGene_gn=SDC1;refGene_tx=XM_003984478.4
A3 127406896 127406978
refGene_gn=SDC1;refGene_tx=XM_003984478.4
A3 127407964 127408443
refGene_gn=SDC1;refGene_tx=XM_003984478.4
A3 127409029 127409165
refGene_gn=SDC1;refGene_tx=XM_003984478.4
A3 127409324 127409494
refGene_gn=SDC1;refGene_tx=XM_003984478.4
A3 127442877 127443083
refGene_gn=LOC109498531;refGene_tx=XM_019828118.2
A3 127444757 127444896
refGene_gn=LOC109498531;refGene_tx=XM_019828118.2
A3 127445516 127445913
refGene_gn=LOC109498531;refGene_tx=XM_019828118.2
A3 127454765 127454809
refGene_gn=LOC109498531;refGene_tx=XM_019828118.2
A3 127553128 127553239
refGene_gn=LAPTM4A;refGene_tx=XM_003984474.5
A3 127561482 127561603
refGene_gn=LAPTM4A;refGene_tx=XM_003984474.5
A3 127565409 127565486
refGene_gn=LAPTM4A;refGene_tx=XM_003984474.5
A3 127565586 127565709
refGene_gn=LAPTM4A;refGene_tx=XM_003984474.5
A3 127568656 127568752
refGene_gn=LAPTM4A;refGene_tx=XM_003984474.5
A3 127569279 127569378
refGene_gn=LAPTM4A;refGene_tx=XM_003984474.5
A3 127570611 127570686
refGene_gn=LAPTM4A;refGene_tx=XM_003984474.5
A3 127582654 127582871
refGene_gn=MATN3;refGene_tx=XM_023252079.1
A3 127587455 127588022
refGene_gn=MATN3;refGene_tx=XM_023252079.1
A3 127591069 127591195
refGene_gn=MATN3;refGene_tx=XM_023252079.1

A3 127591993 127592119
refGene_gn=MATN3;refGene_tx=XM_023252079.1
A3 127593770 127593896
refGene_gn=MATN3;refGene_tx=XM_023252079.1
A3 127597535 127597661
refGene_gn=MATN3;refGene_tx=XM_023252079.1
A3 127601502 127601613
refGene_gn=MATN3;refGene_tx=XM_023252079.1
A3 127603685 127603741
refGene_gn=MATN3;refGene_tx=XM_023252079.1
A3 127606809 127606833
refGene_gn=WDR35;refGene_tx=XM_011281307.3;refGene_gn=WDR35;refGene_tx=XM_011281308.3;refGene_gn=WDR35;refGene_tx=XM_019827872.1
A3 127607525 127607643
refGene_gn=WDR35;refGene_tx=XM_011281307.3;refGene_gn=WDR35;refGene_tx=XM_011281308.3;refGene_gn=WDR35;refGene_tx=XM_019827872.1
A3 127609363 127609435
refGene_gn=WDR35;refGene_tx=XM_011281307.3;refGene_gn=WDR35;refGene_tx=XM_011281308.3;refGene_gn=WDR35;refGene_tx=XM_019827872.1
A3 127610457 127610550
refGene_gn=WDR35;refGene_tx=XM_011281307.3;refGene_gn=WDR35;refGene_tx=XM_011281308.3;refGene_gn=WDR35;refGene_tx=XM_019827872.1
A3 127612258 127612387
refGene_gn=WDR35;refGene_tx=XM_011281307.3;refGene_gn=WDR35;refGene_tx=XM_011281308.3;refGene_gn=WDR35;refGene_tx=XM_019827871.2;refGene_gn=WDR35;refGene_tx=XM_019827872.1
A3 127615738 127615872
refGene_gn=WDR35;refGene_tx=XM_011281307.3;refGene_gn=WDR35;refGene_tx=XM_011281308.3;refGene_gn=WDR35;refGene_tx=XM_019827871.2;refGene_gn=WDR35;refGene_tx=XM_019827872.1
A3 127617018 127617184
refGene_gn=WDR35;refGene_tx=XM_011281307.3;refGene_gn=WDR35;refGene_tx=XM_011281308.3;refGene_gn=WDR35;refGene_tx=XM_019827871.2;refGene_gn=WDR35;refGene_tx=XM_019827872.1
A3 127617314 127617460
refGene_gn=WDR35;refGene_tx=XM_011281307.3;refGene_gn=WDR35;refGene_tx=XM_011281308.3;refGene_gn=WDR35;refGene_tx=XM_019827871.2;refGene_gn=WDR35;refGene_tx=XM_019827872.1
A3 127621563 127621689
refGene_gn=WDR35;refGene_tx=XM_011281307.3;refGene_gn=WDR35;refGene_tx=XM_011281308.3;refGene_gn=WDR35;refGene_tx=XM_019827871.2;refGene_gn=WDR35;refGene_tx=XM_019827872.1
A3 127624230 127624416
refGene_gn=WDR35;refGene_tx=XM_011281307.3;refGene_gn=WDR35;refGene_tx=XM_011281308.3;refGene_gn=WDR35;refGene_tx=XM_019827871.2;refGene_gn=WDR35;refGene_tx=XM_019827872.1
A3 127628538 127628571
refGene_gn=WDR35;refGene_tx=XM_011281308.3;refGene_gn=WDR35;refGene_tx=XM_019827871.2;refGene_gn=WDR35;refGene_tx=XM_019827872.1
A3 127628932 127628993

refGene_gn=WDR35;refGene_tx=XM_011281307.3;refGene_gn=WDR35;refGene_tx=XM_011281308.3;refGene_gn=WDR35;refGene_tx=XM_019827871.2;refGene_gn=WDR35;refGene_tx=XM_019827872.1
A3 127635934 127636079
refGene_gn=WDR35;refGene_tx=XM_011281307.3;refGene_gn=WDR35;refGene_tx=XM_011281308.3;refGene_gn=WDR35;refGene_tx=XM_019827871.2;refGene_gn=WDR35;refGene_tx=XM_019827872.1
A3 127639936 127640006
refGene_gn=WDR35;refGene_tx=XM_011281307.3;refGene_gn=WDR35;refGene_tx=XM_011281308.3;refGene_gn=WDR35;refGene_tx=XM_019827871.2;refGene_gn=WDR35;refGene_tx=XM_019827872.1
A3 127641268 127641322
refGene_gn=WDR35;refGene_tx=XM_011281307.3;refGene_gn=WDR35;refGene_tx=XM_011281308.3;refGene_gn=WDR35;refGene_tx=XM_019827871.2;refGene_gn=WDR35;refGene_tx=XM_019827872.1
A3 127645709 127645819
refGene_gn=WDR35;refGene_tx=XM_011281307.3;refGene_gn=WDR35;refGene_tx=XM_011281308.3;refGene_gn=WDR35;refGene_tx=XM_019827871.2;refGene_gn=WDR35;refGene_tx=XM_019827872.1
A3 127646539 127646750
refGene_gn=WDR35;refGene_tx=XM_011281307.3;refGene_gn=WDR35;refGene_tx=XM_011281308.3;refGene_gn=WDR35;refGene_tx=XM_019827871.2;refGene_gn=WDR35;refGene_tx=XM_019827872.1
A3 127651897 127651978
refGene_gn=WDR35;refGene_tx=XM_011281307.3;refGene_gn=WDR35;refGene_tx=XM_011281308.3;refGene_gn=WDR35;refGene_tx=XM_019827871.2;refGene_gn=WDR35;refGene_tx=XM_019827872.1
A3 127654453 127654590
refGene_gn=WDR35;refGene_tx=XM_011281307.3;refGene_gn=WDR35;refGene_tx=XM_011281308.3;refGene_gn=WDR35;refGene_tx=XM_019827871.2;refGene_gn=WDR35;refGene_tx=XM_019827872.1
A3 127655141 127655345
refGene_gn=WDR35;refGene_tx=XM_011281307.3;refGene_gn=WDR35;refGene_tx=XM_011281308.3;refGene_gn=WDR35;refGene_tx=XM_019827871.2;refGene_gn=WDR35;refGene_tx=XM_019827872.1
A3 127656468 127656615
refGene_gn=WDR35;refGene_tx=XM_011281307.3;refGene_gn=WDR35;refGene_tx=XM_011281308.3;refGene_gn=WDR35;refGene_tx=XM_019827871.2;refGene_gn=WDR35;refGene_tx=XM_019827872.1
A3 127657471 127657604
refGene_gn=WDR35;refGene_tx=XM_011281307.3;refGene_gn=WDR35;refGene_tx=XM_011281308.3;refGene_gn=WDR35;refGene_tx=XM_019827871.2;refGene_gn=WDR35;refGene_tx=XM_019827872.1
A3 127660118 127660229
refGene_gn=WDR35;refGene_tx=XM_011281307.3;refGene_gn=WDR35;refGene_tx=XM_011281308.3;refGene_gn=WDR35;refGene_tx=XM_019827871.2;refGene_gn=WDR35;refGene_tx=XM_019827872.1
A3 127661187 127661352
refGene_gn=WDR35;refGene_tx=XM_011281307.3;refGene_gn=WDR35;refGene_tx=XM_011281308.3;refGene_gn=WDR35;refGene_tx=XM_019827871.2;refGene_gn=

WDR35;refGene_tx=XM_019827872.1
A3 127662458 127662599
refGene_gn=WDR35;refGene_tx=XM_011281307.3;refGene_gn=WDR35;refGene_tx
=XM_011281308.3;refGene_gn=WDR35;refGene_tx=XM_019827871.2;refGene_gn=
WDR35;refGene_tx=XM_019827872.1
A3 127663257 127663414
refGene_gn=WDR35;refGene_tx=XM_011281307.3;refGene_gn=WDR35;refGene_tx
=XM_011281308.3;refGene_gn=WDR35;refGene_tx=XM_019827871.2
A3 127664999 127665240
refGene_gn=WDR35;refGene_tx=XM_011281307.3;refGene_gn=WDR35;refGene_tx
=XM_011281308.3;refGene_gn=WDR35;refGene_tx=XM_019827871.2
A3 127665539 127665690
refGene_gn=WDR35;refGene_tx=XM_011281307.3;refGene_gn=WDR35;refGene_tx
=XM_011281308.3;refGene_gn=WDR35;refGene_tx=XM_019827871.2
A3 127670511 127670663
refGene_gn=TTC32;refGene_tx=XM_011281309.3
A3 127674329 127674496
refGene_gn=TTC32;refGene_tx=XM_011281309.3
A3 127675460 127675600
refGene_gn=TTC32;refGene_tx=XM_011281309.3
A3 128127933 128128601
refGene_gn=OSR1;refGene_tx=XM_011281310.3
A3 128129361 128129497
refGene_gn=OSR1;refGene_tx=XM_011281310.3
A3 128785523 128785556
refGene_gn=NT5C1B;refGene_tx=XM_003984482.5;refGene_gn=NT5C1B;refGene_
tx=XM_011281312.3;refGene_gn=NT5C1B;refGene_tx=XM_011281313.3
A3 128787336 128787426
refGene_gn=NT5C1B;refGene_tx=XM_003984482.5;refGene_gn=NT5C1B;refGene_
tx=XM_011281312.3;refGene_gn=NT5C1B;refGene_tx=XM_011281313.3
A3 128787763 128787946
refGene_gn=NT5C1B;refGene_tx=XM_003984482.5;refGene_gn=NT5C1B;refGene_
tx=XM_011281313.3
A3 128788557 128788695
refGene_gn=NT5C1B;refGene_tx=XM_003984482.5;refGene_gn=NT5C1B;refGene_
tx=XM_011281312.3;refGene_gn=NT5C1B;refGene_tx=XM_011281313.3
A3 128790032 128790464
refGene_gn=NT5C1B;refGene_tx=XM_003984482.5;refGene_gn=NT5C1B;refGene_
tx=XM_011281312.3;refGene_gn=NT5C1B;refGene_tx=XM_011281313.3
A3 128790804 128790972
refGene_gn=NT5C1B;refGene_tx=XM_003984482.5;refGene_gn=NT5C1B;refGene_
tx=XM_011281312.3;refGene_gn=NT5C1B;refGene_tx=XM_011281313.3
A3 128792076 128792206
refGene_gn=NT5C1B;refGene_tx=XM_003984482.5;refGene_gn=NT5C1B;refGene_
tx=XM_011281312.3;refGene_gn=NT5C1B;refGene_tx=XM_011281313.3
A3 128794620 128794743
refGene_gn=NT5C1B;refGene_tx=XM_003984482.5;refGene_gn=NT5C1B;refGene_
tx=XM_011281312.3;refGene_gn=NT5C1B;refGene_tx=XM_011281313.3
A3 128795134 128795319
refGene_gn=NT5C1B;refGene_tx=XM_003984482.5;refGene_gn=NT5C1B;refGene_

tx=XM_011281312.3;refGene_gn=NT5C1B;refGene_tx=XM_011281313.3
A3 128803590 128803602
refGene_gn=NT5C1B;refGene_tx=XM_011281313.3
A3 128804132 128804465
refGene_gn=NT5C1B;refGene_tx=XM_003984482.5;refGene_gn=NT5C1B;refGene_tx=XM_011281312.3
A3 128807238 128807955
refGene_gn=RDH14;refGene_tx=XM_003984483.4;refGene_gn=RDH14;refGene_tx=XM_011281311.3
A3 128811250 128811868
refGene_gn=RDH14;refGene_tx=XM_003984483.4;refGene_gn=RDH14;refGene_tx=XM_011281311.3
A3 129321487 129322963
refGene_gn=KCNS3;refGene_tx=XM_011281314.2;refGene_gn=KCNS3;refGene_tx=XM_023252080.1;refGene_gn=KCNS3;refGene_tx=XM_023252081.1
A3 129393671 129394253
refGene_gn=MSGN1;refGene_tx=XM_003984485.1
A3 129425719 129425748
refGene_gn=GEN1;refGene_tx=XM_023252084.1
A3 129426105 129427421
refGene_gn=GEN1;refGene_tx=XM_011281315.3;refGene_gn=GEN1;refGene_tx=XM_023252083.1;refGene_gn=GEN1;refGene_tx=XM_023252084.1
A3 129428430 129428571
refGene_gn=GEN1;refGene_tx=XM_011281315.3;refGene_gn=GEN1;refGene_tx=XM_023252083.1;refGene_gn=GEN1;refGene_tx=XM_023252084.1
A3 129429942 129430004
refGene_gn=GEN1;refGene_tx=XM_011281315.3;refGene_gn=GEN1;refGene_tx=XM_023252083.1;refGene_gn=GEN1;refGene_tx=XM_023252084.1
A3 129431576 129431707
refGene_gn=GEN1;refGene_tx=XM_011281315.3;refGene_gn=GEN1;refGene_tx=XM_023252083.1;refGene_gn=GEN1;refGene_tx=XM_023252084.1
A3 129433093 129433174
refGene_gn=GEN1;refGene_tx=XM_011281315.3;refGene_gn=GEN1;refGene_tx=XM_023252083.1;refGene_gn=GEN1;refGene_tx=XM_023252084.1
A3 129433265 129433302
refGene_gn=GEN1;refGene_tx=XM_011281315.3;refGene_gn=GEN1;refGene_tx=XM_023252083.1;refGene_gn=GEN1;refGene_tx=XM_023252084.1
A3 129433606 129433757
refGene_gn=GEN1;refGene_tx=XM_011281315.3;refGene_gn=GEN1;refGene_tx=XM_023252083.1;refGene_gn=GEN1;refGene_tx=XM_023252084.1
A3 129435657 129435749
refGene_gn=GEN1;refGene_tx=XM_011281315.3;refGene_gn=GEN1;refGene_tx=XM_023252083.1;refGene_gn=GEN1;refGene_tx=XM_023252084.1
A3 129437964 129438038
refGene_gn=GEN1;refGene_tx=XM_011281315.3;refGene_gn=GEN1;refGene_tx=XM_023252083.1;refGene_gn=GEN1;refGene_tx=XM_023252084.1
A3 129440845 129440956
refGene_gn=GEN1;refGene_tx=XM_011281315.3;refGene_gn=GEN1;refGene_tx=XM_023252083.1;refGene_gn=GEN1;refGene_tx=XM_023252084.1
A3 129442339 129442516

refGene_gn=GEN1;refGene_tx=XM_011281315.3;refGene_gn=GEN1;refGene_tx=XM_023252083.1;refGene_gn=GEN1;refGene_tx=XM_023252084.1
A3 129446306 129446493
refGene_gn=GEN1;refGene_tx=XM_011281315.3;refGene_gn=GEN1;refGene_tx=XM_023252083.1;refGene_gn=GEN1;refGene_tx=XM_023252084.1
A3 129447174 129447335
refGene_gn=GEN1;refGene_tx=XM_011281315.3;refGene_gn=GEN1;refGene_tx=XM_023252083.1;refGene_gn=GEN1;refGene_tx=XM_023252084.1
A3 129463975 129464113
refGene_gn=SMC6;refGene_tx=XM_003984487.5;refGene_gn=SMC6;refGene_tx=XM_023252082.1
A3 129466025 129466143
refGene_gn=SMC6;refGene_tx=XM_003984487.5;refGene_gn=SMC6;refGene_tx=XM_023252082.1
A3 129468265 129468371
refGene_gn=SMC6;refGene_tx=XM_003984487.5;refGene_gn=SMC6;refGene_tx=XM_023252082.1
A3 129474267 129474404
refGene_gn=SMC6;refGene_tx=XM_003984487.5;refGene_gn=SMC6;refGene_tx=XM_023252082.1
A3 129475032 129475094
refGene_gn=SMC6;refGene_tx=XM_003984487.5;refGene_gn=SMC6;refGene_tx=XM_023252082.1
A3 129478944 129479040
refGene_gn=SMC6;refGene_tx=XM_003984487.5;refGene_gn=SMC6;refGene_tx=XM_023252082.1
A3 129480004 129480106
refGene_gn=SMC6;refGene_tx=XM_003984487.5;refGene_gn=SMC6;refGene_tx=XM_023252082.1
A3 129482905 129483025
refGene_gn=SMC6;refGene_tx=XM_003984487.5;refGene_gn=SMC6;refGene_tx=XM_023252082.1
A3 129483127 129483226
refGene_gn=SMC6;refGene_tx=XM_003984487.5;refGene_gn=SMC6;refGene_tx=XM_023252082.1
A3 129486011 129486158
refGene_gn=SMC6;refGene_tx=XM_003984487.5;refGene_gn=SMC6;refGene_tx=XM_023252082.1
A3 129486426 129486515
refGene_gn=SMC6;refGene_tx=XM_003984487.5;refGene_gn=SMC6;refGene_tx=XM_023252082.1
A3 129486694 129486859
refGene_gn=SMC6;refGene_tx=XM_003984487.5;refGene_gn=SMC6;refGene_tx=XM_023252082.1
A3 129487187 129487366
refGene_gn=SMC6;refGene_tx=XM_003984487.5;refGene_gn=SMC6;refGene_tx=XM_023252082.1
A3 129488445 129488650
refGene_gn=SMC6;refGene_tx=XM_003984487.5;refGene_gn=SMC6;refGene_tx=XM_023252082.1

A3 129491953 129492068
refGene_gn=SMC6;refGene_tx=XM_003984487.5;refGene_gn=SMC6;refGene_tx=XM_023252082.1
A3 129493978 129494139
refGene_gn=SMC6;refGene_tx=XM_003984487.5;refGene_gn=SMC6;refGene_tx=XM_023252082.1
A3 129500047 129500183
refGene_gn=SMC6;refGene_tx=XM_003984487.5;refGene_gn=SMC6;refGene_tx=XM_023252082.1
A3 129501691 129501772
refGene_gn=SMC6;refGene_tx=XM_003984487.5;refGene_gn=SMC6;refGene_tx=XM_023252082.1
A3 129503109 129503280
refGene_gn=SMC6;refGene_tx=XM_003984487.5;refGene_gn=SMC6;refGene_tx=XM_023252082.1
A3 129517465 129517603
refGene_gn=SMC6;refGene_tx=XM_003984487.5;refGene_gn=SMC6;refGene_tx=XM_023252082.1
A3 129518598 129518744
refGene_gn=SMC6;refGene_tx=XM_003984487.5;refGene_gn=SMC6;refGene_tx=XM_023252082.1
A3 129522288 129522414
refGene_gn=SMC6;refGene_tx=XM_003984487.5;refGene_gn=SMC6;refGene_tx=XM_023252082.1
A3 129527076 129527182
refGene_gn=SMC6;refGene_tx=XM_003984487.5;refGene_gn=SMC6;refGene_tx=XM_023252082.1
A3 129529888 129530041
refGene_gn=SMC6;refGene_tx=XM_003984487.5;refGene_gn=SMC6;refGene_tx=XM_023252082.1
A3 129533743 129533841
refGene_gn=SMC6;refGene_tx=XM_003984487.5;refGene_gn=SMC6;refGene_tx=XM_023252082.1
A3 129534635 129534750
refGene_gn=SMC6;refGene_tx=XM_003984487.5;refGene_gn=SMC6;refGene_tx=XM_023252082.1
A3 129544174 129544372
refGene_gn=VSNL1;refGene_tx=XM_003984488.5
A3 129546177 129546393
refGene_gn=VSNL1;refGene_tx=XM_003984488.5
A3 129601874 129602036
refGene_gn=VSNL1;refGene_tx=XM_003984488.5
A3 129667842 129671119
refGene_gn=RAD51AP2;refGene_tx=XM_023252085.1
A3 129671760 129671844
refGene_gn=RAD51AP2;refGene_tx=XM_023252085.1
A3 129675007 129675122
refGene_gn=RAD51AP2;refGene_tx=XM_023252085.1
A3 129675123 129675163
refGene_gn=RAD51AP2;refGene_tx=XM_023252085.1

A3 130255887 130255966
refGene_gn=LOC111559873;refGene_tx=XM_023252086.1
A3 130457325 130457395
refGene_gn=FAM49A;refGene_tx=XM_019827884.2;refGene_gn=FAM49A;refGene_tx=XM_019827886.2;refGene_gn=FAM49A;refGene_tx=XM_023252087.1;refGene_gn=FAM49A;refGene_tx=XM_023252088.1
A3 130472908 130473030
refGene_gn=FAM49A;refGene_tx=XM_019827884.2;refGene_gn=FAM49A;refGene_tx=XM_019827886.2;refGene_gn=FAM49A;refGene_tx=XM_023252087.1;refGene_gn=FAM49A;refGene_tx=XM_023252088.1
A3 130474273 130474379
refGene_gn=FAM49A;refGene_tx=XM_019827884.2;refGene_gn=FAM49A;refGene_tx=XM_019827886.2;refGene_gn=FAM49A;refGene_tx=XM_023252087.1;refGene_gn=FAM49A;refGene_tx=XM_023252088.1
A3 130476172 130476309
refGene_gn=FAM49A;refGene_tx=XM_019827884.2;refGene_gn=FAM49A;refGene_tx=XM_019827886.2;refGene_gn=FAM49A;refGene_tx=XM_023252087.1;refGene_gn=FAM49A;refGene_tx=XM_023252088.1
A3 130476991 130477069
refGene_gn=FAM49A;refGene_tx=XM_019827884.2;refGene_gn=FAM49A;refGene_tx=XM_019827886.2;refGene_gn=FAM49A;refGene_tx=XM_023252087.1;refGene_gn=FAM49A;refGene_tx=XM_023252088.1
A3 130477250 130477367
refGene_gn=FAM49A;refGene_tx=XM_019827884.2;refGene_gn=FAM49A;refGene_tx=XM_019827886.2;refGene_gn=FAM49A;refGene_tx=XM_023252087.1;refGene_gn=FAM49A;refGene_tx=XM_023252088.1
A3 130477457 130477537
refGene_gn=FAM49A;refGene_tx=XM_019827884.2;refGene_gn=FAM49A;refGene_tx=XM_019827886.2;refGene_gn=FAM49A;refGene_tx=XM_023252087.1;refGene_gn=FAM49A;refGene_tx=XM_023252088.1
A3 130478984 130479111
refGene_gn=FAM49A;refGene_tx=XM_019827884.2;refGene_gn=FAM49A;refGene_tx=XM_019827886.2;refGene_gn=FAM49A;refGene_tx=XM_023252087.1;refGene_gn=FAM49A;refGene_tx=XM_023252088.1
A3 130483520 130483591
refGene_gn=FAM49A;refGene_tx=XM_019827884.2;refGene_gn=FAM49A;refGene_tx=XM_019827886.2;refGene_gn=FAM49A;refGene_tx=XM_023252087.1;refGene_gn=FAM49A;refGene_tx=XM_023252088.1
A3 130485864 130485928
refGene_gn=FAM49A;refGene_tx=XM_019827884.2;refGene_gn=FAM49A;refGene_tx=XM_019827886.2;refGene_gn=FAM49A;refGene_tx=XM_023252087.1;refGene_gn=FAM49A;refGene_tx=XM_023252088.1
A3 130929412 130929629
refGene_gn=LOC111559876;refGene_tx=XM_023252089.1;refGene_gn=LOC111559876;refGene_tx=XM_023252090.1
A3 130929839 130929960
refGene_gn=LOC111559876;refGene_tx=XM_023252089.1
A3 130930372 130930522
refGene_gn=LOC111559876;refGene_tx=XM_023252089.1;refGene_gn=LOC111559876;refGene_tx=XM_023252090.1

A3 130931559 130931831
refGene_gn=LOC111559876;refGene_tx=XM_023252089.1;refGene_gn=LOC111559
876;refGene_tx=XM_023252090.1
A3 130963981 130964586
refGene_gn=MYCN;refGene_tx=XM_023252091.1
A3 130966834 130967633
refGene_gn=MYCN;refGene_tx=XM_023252091.1
A3 131226919 131227050
refGene_gn=DDX1;refGene_tx=XM_011281319.3
A3 131227723 131227844
refGene_gn=DDX1;refGene_tx=XM_011281319.3
A3 131228127 131228223
refGene_gn=DDX1;refGene_tx=XM_011281319.3
A3 131229204 131229247
refGene_gn=DDX1;refGene_tx=XM_011281319.3
A3 131229330 131229403
refGene_gn=DDX1;refGene_tx=XM_011281319.3
A3 131229555 131229628
refGene_gn=DDX1;refGene_tx=XM_011281319.3
A3 131231150 131231242
refGene_gn=DDX1;refGene_tx=XM_011281319.3
A3 131234665 131234812
refGene_gn=DDX1;refGene_tx=XM_011281319.3
A3 131236766 131236818
refGene_gn=DDX1;refGene_tx=XM_011281319.3
A3 131238529 131238718
refGene_gn=DDX1;refGene_tx=XM_011281319.3
A3 131239684 131239774
refGene_gn=DDX1;refGene_tx=XM_011281319.3
A3 131240602 131240701
refGene_gn=DDX1;refGene_tx=XM_011281319.3
A3 131244059 131244120
refGene_gn=DDX1;refGene_tx=XM_011281319.3
A3 131249388 131249527
refGene_gn=DDX1;refGene_tx=XM_011281319.3
A3 131250226 131250341
refGene_gn=DDX1;refGene_tx=XM_011281319.3
A3 131250449 131250526
refGene_gn=DDX1;refGene_tx=XM_011281319.3
A3 131252419 131252492
refGene_gn=DDX1;refGene_tx=XM_011281319.3
A3 131253185 131253262
refGene_gn=DDX1;refGene_tx=XM_011281319.3
A3 131253630 131253714
refGene_gn=DDX1;refGene_tx=XM_011281319.3
A3 131253942 131254026
refGene_gn=DDX1;refGene_tx=XM_011281319.3
A3 131255783 131255831
refGene_gn=DDX1;refGene_tx=XM_011281319.3
A3 131258030 131258127

refGene_gn=DDX1;refGene_tx=XM_011281319.3
A3 131259089 131259119
refGene_gn=DDX1;refGene_tx=XM_011281319.3
A3 131260008 131260072
refGene_gn=DDX1;refGene_tx=XM_011281319.3
A3 131260357 131260409
refGene_gn=DDX1;refGene_tx=XM_011281319.3
A3 131263270 131263286
refGene_gn=DDX1;refGene_tx=XM_011281319.3
A3 131264916 131265147
refGene_gn=NBAS;refGene_tx=XM_019827889.2;refGene_gn=NBAS;refGene_tx=XM_019827890.2
A3 131288356 131288473
refGene_gn=NBAS;refGene_tx=XM_006930475.4
A3 131290783 131290838
refGene_gn=NBAS;refGene_tx=XM_006930475.4;refGene_gn=NBAS;refGene_tx=XM_019827889.2;refGene_gn=NBAS;refGene_tx=XM_019827890.2
A3 131293022 131293059
refGene_gn=NBAS;refGene_tx=XM_006930475.4;refGene_gn=NBAS;refGene_tx=XM_019827889.2;refGene_gn=NBAS;refGene_tx=XM_019827890.2
A3 131295779 131295857
refGene_gn=NBAS;refGene_tx=XM_006930475.4;refGene_gn=NBAS;refGene_tx=XM_019827889.2;refGene_gn=NBAS;refGene_tx=XM_019827890.2
A3 131296210 131296258
refGene_gn=NBAS;refGene_tx=XM_006930475.4;refGene_gn=NBAS;refGene_tx=XM_019827889.2;refGene_gn=NBAS;refGene_tx=XM_019827890.2
A3 131301429 131301473
refGene_gn=NBAS;refGene_tx=XM_006930475.4;refGene_gn=NBAS;refGene_tx=XM_019827889.2;refGene_gn=NBAS;refGene_tx=XM_019827890.2
A3 131311195 131311329
refGene_gn=NBAS;refGene_tx=XM_006930475.4;refGene_gn=NBAS;refGene_tx=XM_019827889.2;refGene_gn=NBAS;refGene_tx=XM_019827890.2
A3 131313733 131313867
refGene_gn=NBAS;refGene_tx=XM_006930475.4;refGene_gn=NBAS;refGene_tx=XM_019827889.2;refGene_gn=NBAS;refGene_tx=XM_019827890.2
A3 131315623 131315722
refGene_gn=NBAS;refGene_tx=XM_006930475.4;refGene_gn=NBAS;refGene_tx=XM_019827889.2;refGene_gn=NBAS;refGene_tx=XM_019827890.2
A3 131340645 131340784
refGene_gn=NBAS;refGene_tx=XM_006930475.4;refGene_gn=NBAS;refGene_tx=XM_019827889.2;refGene_gn=NBAS;refGene_tx=XM_019827890.2
A3 131347103 131347172
refGene_gn=NBAS;refGene_tx=XM_006930475.4;refGene_gn=NBAS;refGene_tx=XM_019827889.2;refGene_gn=NBAS;refGene_tx=XM_019827890.2;refGene_gn=NBAS;refGene_tx=XM_019827891.2
A3 131356337 131356466
refGene_gn=NBAS;refGene_tx=XM_006930475.4;refGene_gn=NBAS;refGene_tx=XM_019827889.2;refGene_gn=NBAS;refGene_tx=XM_019827890.2;refGene_gn=NBAS;refGene_tx=XM_019827891.2
A3 131366355 131366419

refGene_gn=NBAS;refGene_tx=XM_006930475.4;refGene_gn=NBAS;refGene_tx=XM_019827889.2;refGene_gn=NBAS;refGene_tx=XM_019827890.2;refGene_gn=NBAS;refGene_tx=XM_019827891.2
A3 131367442 131367636
refGene_gn=NBAS;refGene_tx=XM_006930475.4;refGene_gn=NBAS;refGene_tx=XM_019827889.2;refGene_gn=NBAS;refGene_tx=XM_019827890.2;refGene_gn=NBAS;refGene_tx=XM_019827891.2
A3 131368815 131369073
refGene_gn=NBAS;refGene_tx=XM_006930475.4;refGene_gn=NBAS;refGene_tx=XM_019827889.2;refGene_gn=NBAS;refGene_tx=XM_019827890.2;refGene_gn=NBAS;refGene_tx=XM_019827891.2
A3 131369797 131369923
refGene_gn=NBAS;refGene_tx=XM_006930475.4;refGene_gn=NBAS;refGene_tx=XM_019827889.2;refGene_gn=NBAS;refGene_tx=XM_019827890.2;refGene_gn=NBAS;refGene_tx=XM_019827891.2
A3 131373568 131373720
refGene_gn=NBAS;refGene_tx=XM_006930475.4;refGene_gn=NBAS;refGene_tx=XM_019827889.2;refGene_gn=NBAS;refGene_tx=XM_019827890.2;refGene_gn=NBAS;refGene_tx=XM_019827891.2
A3 131374608 131374749
refGene_gn=NBAS;refGene_tx=XM_006930475.4;refGene_gn=NBAS;refGene_tx=XM_019827889.2;refGene_gn=NBAS;refGene_tx=XM_019827890.2;refGene_gn=NBAS;refGene_tx=XM_019827891.2
A3 131375008 131375087
refGene_gn=NBAS;refGene_tx=XM_006930475.4;refGene_gn=NBAS;refGene_tx=XM_019827889.2;refGene_gn=NBAS;refGene_tx=XM_019827890.2;refGene_gn=NBAS;refGene_tx=XM_019827891.2
A3 131380803 131380908
refGene_gn=NBAS;refGene_tx=XM_006930475.4;refGene_gn=NBAS;refGene_tx=XM_019827889.2;refGene_gn=NBAS;refGene_tx=XM_019827890.2;refGene_gn=NBAS;refGene_tx=XM_019827891.2
A3 131381534 131381671
refGene_gn=NBAS;refGene_tx=XM_006930475.4;refGene_gn=NBAS;refGene_tx=XM_019827889.2;refGene_gn=NBAS;refGene_tx=XM_019827890.2;refGene_gn=NBAS;refGene_tx=XM_019827891.2
A3 131409969 131410053
refGene_gn=NBAS;refGene_tx=XM_006930475.4;refGene_gn=NBAS;refGene_tx=XM_019827889.2;refGene_gn=NBAS;refGene_tx=XM_019827890.2;refGene_gn=NBAS;refGene_tx=XM_019827891.2
A3 131416449 131416603
refGene_gn=NBAS;refGene_tx=XM_006930475.4;refGene_gn=NBAS;refGene_tx=XM_019827889.2;refGene_gn=NBAS;refGene_tx=XM_019827890.2;refGene_gn=NBAS;refGene_tx=XM_019827891.2
A3 131421703 131421889
refGene_gn=NBAS;refGene_tx=XM_006930475.4;refGene_gn=NBAS;refGene_tx=XM_019827889.2;refGene_gn=NBAS;refGene_tx=XM_019827890.2;refGene_gn=NBAS;refGene_tx=XM_019827891.2
A3 131423581 131423755
refGene_gn=NBAS;refGene_tx=XM_006930475.4;refGene_gn=NBAS;refGene_tx=XM_019827889.2;refGene_gn=NBAS;refGene_tx=XM_019827890.2;refGene_gn=NBA

S;refGene_tx=XM_019827891.2
A3	131433783	131433917
refGene_gn=NBAS;refGene_tx=XM_006930475.4;refGene_gn=NBAS;refGene_tx=XM_019827889.2;refGene_gn=NBAS;refGene_tx=XM_019827890.2;refGene_gn=NBAS;refGene_tx=XM_019827891.2
A3	131439831	131439894
refGene_gn=NBAS;refGene_tx=XM_006930475.4;refGene_gn=NBAS;refGene_tx=XM_019827889.2;refGene_gn=NBAS;refGene_tx=XM_019827890.2;refGene_gn=NBAS;refGene_tx=XM_019827891.2
A3	131442375	131442498
refGene_gn=NBAS;refGene_tx=XM_006930475.4;refGene_gn=NBAS;refGene_tx=XM_019827889.2;refGene_gn=NBAS;refGene_tx=XM_019827890.2;refGene_gn=NBAS;refGene_tx=XM_019827891.2
A3	131446609	131446712
refGene_gn=NBAS;refGene_tx=XM_006930475.4;refGene_gn=NBAS;refGene_tx=XM_019827889.2;refGene_gn=NBAS;refGene_tx=XM_019827890.2;refGene_gn=NBAS;refGene_tx=XM_019827891.2
A3	131450386	131450616
refGene_gn=NBAS;refGene_tx=XM_006930475.4;refGene_gn=NBAS;refGene_tx=XM_019827889.2;refGene_gn=NBAS;refGene_tx=XM_019827890.2;refGene_gn=NBAS;refGene_tx=XM_019827891.2
A3	131453723	131453836
refGene_gn=NBAS;refGene_tx=XM_006930475.4;refGene_gn=NBAS;refGene_tx=XM_019827889.2;refGene_gn=NBAS;refGene_tx=XM_019827890.2;refGene_gn=NBAS;refGene_tx=XM_019827891.2
A3	131460411	131460525
refGene_gn=NBAS;refGene_tx=XM_006930475.4;refGene_gn=NBAS;refGene_tx=XM_019827889.2;refGene_gn=NBAS;refGene_tx=XM_019827890.2;refGene_gn=NBAS;refGene_tx=XM_019827891.2
A3	131465589	131465703
refGene_gn=NBAS;refGene_tx=XM_006930475.4;refGene_gn=NBAS;refGene_tx=XM_019827889.2;refGene_gn=NBAS;refGene_tx=XM_019827890.2;refGene_gn=NBAS;refGene_tx=XM_019827891.2
A3	131470236	131470394
refGene_gn=NBAS;refGene_tx=XM_006930475.4;refGene_gn=NBAS;refGene_tx=XM_019827889.2;refGene_gn=NBAS;refGene_tx=XM_019827890.2;refGene_gn=NBAS;refGene_tx=XM_019827891.2
A3	131471393	131471483
refGene_gn=NBAS;refGene_tx=XM_006930475.4;refGene_gn=NBAS;refGene_tx=XM_019827889.2;refGene_gn=NBAS;refGene_tx=XM_019827890.2;refGene_gn=NBAS;refGene_tx=XM_019827891.2
A3	131493916	131494081
refGene_gn=NBAS;refGene_tx=XM_006930475.4;refGene_gn=NBAS;refGene_tx=XM_019827889.2;refGene_gn=NBAS;refGene_tx=XM_019827890.2;refGene_gn=NBAS;refGene_tx=XM_019827891.2
A3	131497138	131497252
refGene_gn=NBAS;refGene_tx=XM_006930475.4;refGene_gn=NBAS;refGene_tx=XM_019827889.2;refGene_gn=NBAS;refGene_tx=XM_019827890.2;refGene_gn=NBAS;refGene_tx=XM_019827891.2
A3	131497576	131497700

refGene_gn=NBAS;refGene_tx=XM_006930475.4;refGene_gn=NBAS;refGene_tx=XM_019827889.2;refGene_gn=NBAS;refGene_tx=XM_019827890.2;refGene_gn=NBAS;refGene_tx=XM_019827891.2
A3 131508309 131508386
refGene_gn=NBAS;refGene_tx=XM_006930475.4;refGene_gn=NBAS;refGene_tx=XM_019827889.2;refGene_gn=NBAS;refGene_tx=XM_019827890.2;refGene_gn=NBAS;refGene_tx=XM_019827891.2
A3 131508769 131508907
refGene_gn=NBAS;refGene_tx=XM_006930475.4;refGene_gn=NBAS;refGene_tx=XM_019827889.2;refGene_gn=NBAS;refGene_tx=XM_019827890.2;refGene_gn=NBAS;refGene_tx=XM_019827891.2
A3 131521157 131521387
refGene_gn=NBAS;refGene_tx=XM_006930475.4;refGene_gn=NBAS;refGene_tx=XM_019827889.2;refGene_gn=NBAS;refGene_tx=XM_019827890.2;refGene_gn=NBAS;refGene_tx=XM_019827891.2
A3 131525474 131525585
refGene_gn=NBAS;refGene_tx=XM_006930475.4;refGene_gn=NBAS;refGene_tx=XM_019827889.2;refGene_gn=NBAS;refGene_tx=XM_019827890.2;refGene_gn=NBAS;refGene_tx=XM_019827891.2
A3 131533381 131533632
refGene_gn=NBAS;refGene_tx=XM_006930475.4;refGene_gn=NBAS;refGene_tx=XM_019827889.2;refGene_gn=NBAS;refGene_tx=XM_019827890.2;refGene_gn=NBAS;refGene_tx=XM_019827891.2
A3 131534938 131535273
refGene_gn=NBAS;refGene_tx=XM_006930475.4;refGene_gn=NBAS;refGene_tx=XM_019827889.2;refGene_gn=NBAS;refGene_tx=XM_019827890.2;refGene_gn=NBAS;refGene_tx=XM_019827891.2
A3 131572327 131572546
refGene_gn=NBAS;refGene_tx=XM_006930475.4;refGene_gn=NBAS;refGene_tx=XM_019827889.2;refGene_gn=NBAS;refGene_tx=XM_019827890.2;refGene_gn=NBAS;refGene_tx=XM_019827891.2
A3 131575696 131575899
refGene_gn=NBAS;refGene_tx=XM_006930475.4;refGene_gn=NBAS;refGene_tx=XM_019827889.2;refGene_gn=NBAS;refGene_tx=XM_019827890.2;refGene_gn=NBAS;refGene_tx=XM_019827891.2
A3 131578213 131578303
refGene_gn=NBAS;refGene_tx=XM_006930475.4;refGene_gn=NBAS;refGene_tx=XM_019827889.2;refGene_gn=NBAS;refGene_tx=XM_019827890.2;refGene_gn=NBAS;refGene_tx=XM_019827891.2
A3 131588885 131589081
refGene_gn=NBAS;refGene_tx=XM_006930475.4;refGene_gn=NBAS;refGene_tx=XM_019827889.2;refGene_gn=NBAS;refGene_tx=XM_019827890.2;refGene_gn=NBAS;refGene_tx=XM_019827891.2
A3 131602462 131602602
refGene_gn=NBAS;refGene_tx=XM_006930475.4;refGene_gn=NBAS;refGene_tx=XM_019827889.2;refGene_gn=NBAS;refGene_tx=XM_019827890.2;refGene_gn=NBAS;refGene_tx=XM_019827891.2
A3 131606533 131606678
refGene_gn=NBAS;refGene_tx=XM_006930475.4;refGene_gn=NBAS;refGene_tx=XM_019827889.2;refGene_gn=NBAS;refGene_tx=XM_019827890.2;refGene_gn=NBA

S;refGene_tx=XM_019827891.2
A3 131613210 131613339
refGene_gn=NBAS;refGene_tx=XM_006930475.4;refGene_gn=NBAS;refGene_tx=XM_019827889.2;refGene_gn=NBAS;refGene_tx=XM_019827890.2;refGene_gn=NBAS;refGene_tx=XM_019827891.2
A3 131624095 131624371
refGene_gn=NBAS;refGene_tx=XM_006930475.4;refGene_gn=NBAS;refGene_tx=XM_019827889.2;refGene_gn=NBAS;refGene_tx=XM_019827890.2;refGene_gn=NBAS;refGene_tx=XM_019827891.2
A3 132069319 132070198
refGene_gn=FAM84A;refGene_tx=XM_023252092.1
A3 133531626 133532095
refGene_gn=TRIB2;refGene_tx=XM_003984494.5
A3 133549472 133549765
refGene_gn=TRIB2;refGene_tx=XM_003984494.5
A3 133554520 133554790
refGene_gn=TRIB2;refGene_tx=XM_003984494.5
A3 134305130 134305290
refGene_gn=LPIN1;refGene_tx=XM_003984496.5;refGene_gn=LPIN1;refGene_tx=XM_011281321.3;refGene_gn=LPIN1;refGene_tx=XM_011281322.1;refGene_gn=LPIN1;refGene_tx=XM_011281323.2;refGene_gn=LPIN1;refGene_tx=XM_019827894.1;refGene_gn=LPIN1;refGene_tx=XM_019827896.2;refGene_gn=LPIN1;refGene_tx=XM_023252093.1;refGene_gn=LPIN1;refGene_tx=XM_023252094.1
A3 134308241 134308345
refGene_gn=LPIN1;refGene_tx=XM_003984496.5;refGene_gn=LPIN1;refGene_tx=XM_011281321.3;refGene_gn=LPIN1;refGene_tx=XM_011281322.1;refGene_gn=LPIN1;refGene_tx=XM_011281323.2;refGene_gn=LPIN1;refGene_tx=XM_019827894.1;refGene_gn=LPIN1;refGene_tx=XM_019827896.2;refGene_gn=LPIN1;refGene_tx=XM_023252093.1;refGene_gn=LPIN1;refGene_tx=XM_023252094.1
A3 134309109 134309224
refGene_gn=LPIN1;refGene_tx=XM_003984496.5;refGene_gn=LPIN1;refGene_tx=XM_011281321.3;refGene_gn=LPIN1;refGene_tx=XM_011281322.1;refGene_gn=LPIN1;refGene_tx=XM_011281323.2;refGene_gn=LPIN1;refGene_tx=XM_019827894.1;refGene_gn=LPIN1;refGene_tx=XM_019827896.2;refGene_gn=LPIN1;refGene_tx=XM_023252093.1;refGene_gn=LPIN1;refGene_tx=XM_023252094.1
A3 134315458 134315611
refGene_gn=LPIN1;refGene_tx=XM_003984496.5;refGene_gn=LPIN1;refGene_tx=XM_011281321.3;refGene_gn=LPIN1;refGene_tx=XM_011281322.1;refGene_gn=LPIN1;refGene_tx=XM_011281323.2;refGene_gn=LPIN1;refGene_tx=XM_019827894.1;refGene_gn=LPIN1;refGene_tx=XM_019827896.2;refGene_gn=LPIN1;refGene_tx=XM_023252093.1;refGene_gn=LPIN1;refGene_tx=XM_023252094.1
A3 134325185 134325272
refGene_gn=LPIN1;refGene_tx=XM_003984496.5;refGene_gn=LPIN1;refGene_tx=XM_011281321.3;refGene_gn=LPIN1;refGene_tx=XM_011281322.1;refGene_gn=LPIN1;refGene_tx=XM_011281323.2;refGene_gn=LPIN1;refGene_tx=XM_019827894.1;refGene_gn=LPIN1;refGene_tx=XM_019827896.2;refGene_gn=LPIN1;refGene_tx=XM_023252093.1;refGene_gn=LPIN1;refGene_tx=XM_023252094.1
A3 134325753 134325902
refGene_gn=LPIN1;refGene_tx=XM_003984496.5;refGene_gn=LPIN1;refGene_tx=XM_011281321.3;refGene_gn=LPIN1;refGene_tx=XM_011281322.1;refGene_gn=

LPIN1;refGene_tx=XM_011281323.2;refGene_gn=LPIN1;refGene_tx=XM_019827894.1;refGene_gn=LPIN1;refGene_tx=XM_019827896.2;refGene_gn=LPIN1;refGene_tx=XM_023252093.1;refGene_gn=LPIN1;refGene_tx=XM_023252094.1
A3 134326878 134327005
refGene_gn=LPIN1;refGene_tx=XM_003984496.5;refGene_gn=LPIN1;refGene_tx=XM_011281321.3;refGene_gn=LPIN1;refGene_tx=XM_011281322.1;refGene_gn=LPIN1;refGene_tx=XM_011281323.2;refGene_gn=LPIN1;refGene_tx=XM_019827894.1;refGene_gn=LPIN1;refGene_tx=XM_019827896.2;refGene_gn=LPIN1;refGene_tx=XM_023252093.1;refGene_gn=LPIN1;refGene_tx=XM_023252094.1
A3 134331708 134331788
refGene_gn=LPIN1;refGene_tx=XM_003984496.5;refGene_gn=LPIN1;refGene_tx=XM_011281321.3;refGene_gn=LPIN1;refGene_tx=XM_011281322.1;refGene_gn=LPIN1;refGene_tx=XM_011281323.2;refGene_gn=LPIN1;refGene_tx=XM_019827894.1;refGene_gn=LPIN1;refGene_tx=XM_019827896.2;refGene_gn=LPIN1;refGene_tx=XM_023252093.1;refGene_gn=LPIN1;refGene_tx=XM_023252094.1
A3 134334829 134334922
refGene_gn=LPIN1;refGene_tx=XM_003984496.5;refGene_gn=LPIN1;refGene_tx=XM_011281321.3;refGene_gn=LPIN1;refGene_tx=XM_011281322.1;refGene_gn=LPIN1;refGene_tx=XM_011281323.2;refGene_gn=LPIN1;refGene_tx=XM_019827894.1;refGene_gn=LPIN1;refGene_tx=XM_019827896.2;refGene_gn=LPIN1;refGene_tx=XM_023252093.1;refGene_gn=LPIN1;refGene_tx=XM_023252094.1
A3 134338263 134338333
refGene_gn=LPIN1;refGene_tx=XM_003984496.5;refGene_gn=LPIN1;refGene_tx=XM_011281321.3;refGene_gn=LPIN1;refGene_tx=XM_011281322.1;refGene_gn=LPIN1;refGene_tx=XM_011281323.2;refGene_gn=LPIN1;refGene_tx=XM_019827894.1;refGene_gn=LPIN1;refGene_tx=XM_019827896.2;refGene_gn=LPIN1;refGene_tx=XM_023252093.1;refGene_gn=LPIN1;refGene_tx=XM_023252094.1
A3 134338980 134339074
refGene_gn=LPIN1;refGene_tx=XM_003984496.5;refGene_gn=LPIN1;refGene_tx=XM_011281321.3;refGene_gn=LPIN1;refGene_tx=XM_011281322.1;refGene_gn=LPIN1;refGene_tx=XM_011281323.2;refGene_gn=LPIN1;refGene_tx=XM_019827894.1;refGene_gn=LPIN1;refGene_tx=XM_019827896.2;refGene_gn=LPIN1;refGene_tx=XM_023252093.1;refGene_gn=LPIN1;refGene_tx=XM_023252094.1
A3 134340768 134340959
refGene_gn=LPIN1;refGene_tx=XM_003984496.5;refGene_gn=LPIN1;refGene_tx=XM_011281321.3;refGene_gn=LPIN1;refGene_tx=XM_011281322.1;refGene_gn=LPIN1;refGene_tx=XM_011281323.2;refGene_gn=LPIN1;refGene_tx=XM_019827894.1;refGene_gn=LPIN1;refGene_tx=XM_019827896.2;refGene_gn=LPIN1;refGene_tx=XM_023252093.1;refGene_gn=LPIN1;refGene_tx=XM_023252094.1
A3 134341593 134341687
refGene_gn=LPIN1;refGene_tx=XM_003984496.5;refGene_gn=LPIN1;refGene_tx=XM_011281321.3;refGene_gn=LPIN1;refGene_tx=XM_011281322.1;refGene_gn=LPIN1;refGene_tx=XM_011281323.2;refGene_gn=LPIN1;refGene_tx=XM_019827894.1;refGene_gn=LPIN1;refGene_tx=XM_019827896.2;refGene_gn=LPIN1;refGene_tx=XM_023252093.1;refGene_gn=LPIN1;refGene_tx=XM_023252094.1
A3 134342802 134343112
refGene_gn=LPIN1;refGene_tx=XM_003984496.5;refGene_gn=LPIN1;refGene_tx=XM_011281321.3;refGene_gn=LPIN1;refGene_tx=XM_011281322.1;refGene_gn=LPIN1;refGene_tx=XM_011281323.2;refGene_gn=LPIN1;refGene_tx=XM_019827894.1;refGene_gn=LPIN1;refGene_tx=XM_019827896.2;refGene_gn=LPIN1;refGe

ne_tx=XM_023252093.1;refGene_gn=LPIN1;refGene_tx=XM_023252094.1
A3	134344421	134344551
refGene_gn=LPIN1;refGene_tx=XM_003984496.5;refGene_gn=LPIN1;refGene_tx=XM_011281321.3;refGene_gn=LPIN1;refGene_tx=XM_011281322.1;refGene_gn=LPIN1;refGene_tx=XM_011281323.2;refGene_gn=LPIN1;refGene_tx=XM_019827894.1;refGene_gn=LPIN1;refGene_tx=XM_019827896.2;refGene_gn=LPIN1;refGene_tx=XM_023252093.1;refGene_gn=LPIN1;refGene_tx=XM_023252094.1
A3	134347830	134347938
refGene_gn=LPIN1;refGene_tx=XM_003984496.5;refGene_gn=LPIN1;refGene_tx=XM_011281321.3;refGene_gn=LPIN1;refGene_tx=XM_011281323.2;refGene_gn=LPIN1;refGene_tx=XM_019827894.1;refGene_gn=LPIN1;refGene_tx=XM_019827896.2;refGene_gn=LPIN1;refGene_tx=XM_023252093.1
A3	134350258	134350384
refGene_gn=LPIN1;refGene_tx=XM_003984496.5;refGene_gn=LPIN1;refGene_tx=XM_011281321.3;refGene_gn=LPIN1;refGene_tx=XM_011281322.1;refGene_gn=LPIN1;refGene_tx=XM_011281323.2;refGene_gn=LPIN1;refGene_tx=XM_019827894.1;refGene_gn=LPIN1;refGene_tx=XM_019827896.2;refGene_gn=LPIN1;refGene_tx=XM_023252093.1;refGene_gn=LPIN1;refGene_tx=XM_023252094.1
A3	134352465	134352773
refGene_gn=LPIN1;refGene_tx=XM_003984496.5;refGene_gn=LPIN1;refGene_tx=XM_011281321.3;refGene_gn=LPIN1;refGene_tx=XM_011281322.1;refGene_gn=LPIN1;refGene_tx=XM_011281323.2;refGene_gn=LPIN1;refGene_tx=XM_019827894.1;refGene_gn=LPIN1;refGene_tx=XM_019827896.2;refGene_gn=LPIN1;refGene_tx=XM_023252093.1;refGene_gn=LPIN1;refGene_tx=XM_023252094.1
A3	134354279	134354342
refGene_gn=LPIN1;refGene_tx=XM_011281321.3;refGene_gn=LPIN1;refGene_tx=XM_011281322.1;refGene_gn=LPIN1;refGene_tx=XM_011281323.2;refGene_gn=LPIN1;refGene_tx=XM_019827894.1;refGene_gn=LPIN1;refGene_tx=XM_019827896.2
A3	134356403	134356499
refGene_gn=LPIN1;refGene_tx=XM_003984496.5;refGene_gn=LPIN1;refGene_tx=XM_011281321.3;refGene_gn=LPIN1;refGene_tx=XM_011281322.1;refGene_gn=LPIN1;refGene_tx=XM_011281323.2;refGene_gn=LPIN1;refGene_tx=XM_019827894.1;refGene_gn=LPIN1;refGene_tx=XM_019827896.2;refGene_gn=LPIN1;refGene_tx=XM_023252093.1;refGene_gn=LPIN1;refGene_tx=XM_023252094.1
A3	134357249	134357450
refGene_gn=LPIN1;refGene_tx=XM_003984496.5;refGene_gn=LPIN1;refGene_tx=XM_011281321.3;refGene_gn=LPIN1;refGene_tx=XM_011281322.1;refGene_gn=LPIN1;refGene_tx=XM_011281323.2;refGene_gn=LPIN1;refGene_tx=XM_019827894.1;refGene_gn=LPIN1;refGene_tx=XM_019827896.2;refGene_gn=LPIN1;refGene_tx=XM_023252093.1;refGene_gn=LPIN1;refGene_tx=XM_023252094.1
A3	134397923	134397980
refGene_gn=LPIN1;refGene_tx=XM_003984496.5;refGene_gn=LPIN1;refGene_tx=XM_011281321.3;refGene_gn=LPIN1;refGene_tx=XM_023252094.1
A3	134432765	134432843
refGene_gn=LPIN1;refGene_tx=XM_003984496.5;refGene_gn=LPIN1;refGene_tx=XM_011281321.3;refGene_gn=LPIN1;refGene_tx=XM_023252094.1
A3	134441100	134441275
refGene_gn=NTSR2;refGene_tx=XM_019828119.1
A3	134441277	134441541

refGene_gn=NTSR2;refGene_tx=XM_019828119.1
A3	134441543	134441671
refGene_gn=NTSR2;refGene_tx=XM_019828119.1
A3	134445850	134445961
refGene_gn=NTSR2;refGene_tx=XM_019828119.1
A3	134445963	134446102
refGene_gn=NTSR2;refGene_tx=XM_019828119.1
A3	134448004	134448095
refGene_gn=NTSR2;refGene_tx=XM_019828119.1
A3	134449067	134449320
refGene_gn=NTSR2;refGene_tx=XM_019828119.1
A3	134457003	134457167
refGene_gn=GREB1;refGene_tx=XM_023252095.1;refGene_gn=GREB1;refGene_tx=XM_023252096.1;refGene_gn=GREB1;refGene_tx=XM_023252097.1;refGene_gn=GREB1;refGene_tx=XM_023252098.1;refGene_gn=GREB1;refGene_tx=XM_023252099.1;refGene_gn=GREB1;refGene_tx=XM_023252100.1;refGene_gn=GREB1;refGene_tx=XM_023252101.1
A3	134458325	134458464
refGene_gn=GREB1;refGene_tx=XM_023252095.1;refGene_gn=GREB1;refGene_tx=XM_023252096.1;refGene_gn=GREB1;refGene_tx=XM_023252097.1;refGene_gn=GREB1;refGene_tx=XM_023252098.1;refGene_gn=GREB1;refGene_tx=XM_023252099.1;refGene_gn=GREB1;refGene_tx=XM_023252100.1;refGene_gn=GREB1;refGene_tx=XM_023252101.1
A3	134458932	134459133
refGene_gn=GREB1;refGene_tx=XM_023252095.1;refGene_gn=GREB1;refGene_tx=XM_023252096.1;refGene_gn=GREB1;refGene_tx=XM_023252097.1;refGene_gn=GREB1;refGene_tx=XM_023252098.1;refGene_gn=GREB1;refGene_tx=XM_023252099.1;refGene_gn=GREB1;refGene_tx=XM_023252100.1;refGene_gn=GREB1;refGene_tx=XM_023252101.1
A3	134461165	134461301
refGene_gn=GREB1;refGene_tx=XM_023252095.1;refGene_gn=GREB1;refGene_tx=XM_023252096.1;refGene_gn=GREB1;refGene_tx=XM_023252097.1;refGene_gn=GREB1;refGene_tx=XM_023252098.1;refGene_gn=GREB1;refGene_tx=XM_023252099.1;refGene_gn=GREB1;refGene_tx=XM_023252100.1;refGene_gn=GREB1;refGene_tx=XM_023252101.1
A3	134463172	134463391
refGene_gn=GREB1;refGene_tx=XM_023252095.1;refGene_gn=GREB1;refGene_tx=XM_023252096.1;refGene_gn=GREB1;refGene_tx=XM_023252097.1;refGene_gn=GREB1;refGene_tx=XM_023252098.1;refGene_gn=GREB1;refGene_tx=XM_023252099.1;refGene_gn=GREB1;refGene_tx=XM_023252100.1;refGene_gn=GREB1;refGene_tx=XM_023252101.1
A3	134465393	134465568
refGene_gn=GREB1;refGene_tx=XM_023252095.1;refGene_gn=GREB1;refGene_tx=XM_023252096.1;refGene_gn=GREB1;refGene_tx=XM_023252097.1;refGene_gn=GREB1;refGene_tx=XM_023252098.1;refGene_gn=GREB1;refGene_tx=XM_023252099.1;refGene_gn=GREB1;refGene_tx=XM_023252100.1;refGene_gn=GREB1;refGene_tx=XM_023252101.1
A3	134465947	134466152
refGene_gn=GREB1;refGene_tx=XM_023252095.1;refGene_gn=GREB1;refGene_tx=XM_023252096.1;refGene_gn=GREB1;refGene_tx=XM_023252097.1;refGene_gn=

GREB1;refGene_tx=XM_023252098.1;refGene_gn=GREB1;refGene_tx=XM_0232520
99.1;refGene_gn=GREB1;refGene_tx=XM_023252100.1;refGene_gn=GREB1;refGe
ne_tx=XM_023252101.1
A3 134469737 134469899
refGene_gn=GREB1;refGene_tx=XM_023252095.1;refGene_gn=GREB1;refGene_tx
=XM_023252096.1;refGene_gn=GREB1;refGene_tx=XM_023252097.1;refGene_gn=
GREB1;refGene_tx=XM_023252098.1;refGene_gn=GREB1;refGene_tx=XM_0232520
99.1;refGene_gn=GREB1;refGene_tx=XM_023252100.1;refGene_gn=GREB1;refGe
ne_tx=XM_023252101.1
A3 134473347 134473490
refGene_gn=GREB1;refGene_tx=XM_023252095.1;refGene_gn=GREB1;refGene_tx
=XM_023252096.1;refGene_gn=GREB1;refGene_tx=XM_023252097.1;refGene_gn=
GREB1;refGene_tx=XM_023252098.1;refGene_gn=GREB1;refGene_tx=XM_0232520
99.1;refGene_gn=GREB1;refGene_tx=XM_023252100.1;refGene_gn=GREB1;refGe
ne_tx=XM_023252101.1
A3 134474450 134474609
refGene_gn=GREB1;refGene_tx=XM_023252095.1;refGene_gn=GREB1;refGene_tx
=XM_023252096.1;refGene_gn=GREB1;refGene_tx=XM_023252097.1;refGene_gn=
GREB1;refGene_tx=XM_023252098.1;refGene_gn=GREB1;refGene_tx=XM_0232520
99.1;refGene_gn=GREB1;refGene_tx=XM_023252100.1;refGene_gn=GREB1;refGe
ne_tx=XM_023252101.1
A3 134477136 134477239
refGene_gn=GREB1;refGene_tx=XM_023252095.1;refGene_gn=GREB1;refGene_tx
=XM_023252096.1;refGene_gn=GREB1;refGene_tx=XM_023252097.1;refGene_gn=
GREB1;refGene_tx=XM_023252098.1;refGene_gn=GREB1;refGene_tx=XM_0232520
99.1;refGene_gn=GREB1;refGene_tx=XM_023252100.1;refGene_gn=GREB1;refGe
ne_tx=XM_023252101.1
A3 134479742 134480356
refGene_gn=GREB1;refGene_tx=XM_023252095.1;refGene_gn=GREB1;refGene_tx
=XM_023252096.1;refGene_gn=GREB1;refGene_tx=XM_023252097.1;refGene_gn=
GREB1;refGene_tx=XM_023252098.1;refGene_gn=GREB1;refGene_tx=XM_0232520
99.1;refGene_gn=GREB1;refGene_tx=XM_023252100.1;refGene_gn=GREB1;refGe
ne_tx=XM_023252101.1
A3 134483218 134483308
refGene_gn=GREB1;refGene_tx=XM_023252095.1;refGene_gn=GREB1;refGene_tx
=XM_023252096.1;refGene_gn=GREB1;refGene_tx=XM_023252097.1;refGene_gn=
GREB1;refGene_tx=XM_023252098.1;refGene_gn=GREB1;refGene_tx=XM_0232520
99.1;refGene_gn=GREB1;refGene_tx=XM_023252100.1;refGene_gn=GREB1;refGe
ne_tx=XM_023252101.1
A3 134484623 134484823
refGene_gn=GREB1;refGene_tx=XM_023252095.1;refGene_gn=GREB1;refGene_tx
=XM_023252096.1;refGene_gn=GREB1;refGene_tx=XM_023252097.1;refGene_gn=
GREB1;refGene_tx=XM_023252098.1;refGene_gn=GREB1;refGene_tx=XM_0232520
99.1;refGene_gn=GREB1;refGene_tx=XM_023252100.1;refGene_gn=GREB1;refGe
ne_tx=XM_023252101.1
A3 134486723 134486839
refGene_gn=GREB1;refGene_tx=XM_023252095.1;refGene_gn=GREB1;refGene_tx
=XM_023252096.1;refGene_gn=GREB1;refGene_tx=XM_023252097.1;refGene_gn=
GREB1;refGene_tx=XM_023252098.1;refGene_gn=GREB1;refGene_tx=XM_0232520
99.1;refGene_gn=GREB1;refGene_tx=XM_023252100.1;refGene_gn=GREB1;refGe

ne_tx=XM_023252101.1
A3 134488195 134488535
refGene_gn=GREB1;refGene_tx=XM_023252095.1;refGene_gn=GREB1;refGene_tx=XM_023252096.1;refGene_gn=GREB1;refGene_tx=XM_023252097.1;refGene_gn=GREB1;refGene_tx=XM_023252098.1;refGene_gn=GREB1;refGene_tx=XM_023252099.1;refGene_gn=GREB1;refGene_tx=XM_023252100.1;refGene_gn=GREB1;refGene_tx=XM_023252101.1;refGene_gn=GREB1;refGene_tx=XM_023252102.1
A3 134492238 134492375
refGene_gn=GREB1;refGene_tx=XM_023252095.1;refGene_gn=GREB1;refGene_tx=XM_023252096.1;refGene_gn=GREB1;refGene_tx=XM_023252097.1;refGene_gn=GREB1;refGene_tx=XM_023252098.1;refGene_gn=GREB1;refGene_tx=XM_023252099.1;refGene_gn=GREB1;refGene_tx=XM_023252100.1;refGene_gn=GREB1;refGene_tx=XM_023252101.1;refGene_gn=GREB1;refGene_tx=XM_023252102.1
A3 134493297 134493493
refGene_gn=GREB1;refGene_tx=XM_023252095.1;refGene_gn=GREB1;refGene_tx=XM_023252096.1;refGene_gn=GREB1;refGene_tx=XM_023252097.1;refGene_gn=GREB1;refGene_tx=XM_023252098.1;refGene_gn=GREB1;refGene_tx=XM_023252099.1;refGene_gn=GREB1;refGene_tx=XM_023252100.1;refGene_gn=GREB1;refGene_tx=XM_023252101.1;refGene_gn=GREB1;refGene_tx=XM_023252102.1
A3 134494449 134494630
refGene_gn=GREB1;refGene_tx=XM_023252095.1;refGene_gn=GREB1;refGene_tx=XM_023252096.1;refGene_gn=GREB1;refGene_tx=XM_023252097.1;refGene_gn=GREB1;refGene_tx=XM_023252098.1;refGene_gn=GREB1;refGene_tx=XM_023252099.1;refGene_gn=GREB1;refGene_tx=XM_023252100.1;refGene_gn=GREB1;refGene_tx=XM_023252101.1;refGene_gn=GREB1;refGene_tx=XM_023252102.1
A3 134495410 134495608
refGene_gn=GREB1;refGene_tx=XM_023252095.1;refGene_gn=GREB1;refGene_tx=XM_023252096.1;refGene_gn=GREB1;refGene_tx=XM_023252097.1;refGene_gn=GREB1;refGene_tx=XM_023252098.1;refGene_gn=GREB1;refGene_tx=XM_023252099.1;refGene_gn=GREB1;refGene_tx=XM_023252100.1;refGene_gn=GREB1;refGene_tx=XM_023252101.1;refGene_gn=GREB1;refGene_tx=XM_023252102.1
A3 134496212 134496332
refGene_gn=GREB1;refGene_tx=XM_023252095.1;refGene_gn=GREB1;refGene_tx=XM_023252096.1;refGene_gn=GREB1;refGene_tx=XM_023252097.1;refGene_gn=GREB1;refGene_tx=XM_023252098.1;refGene_gn=GREB1;refGene_tx=XM_023252099.1;refGene_gn=GREB1;refGene_tx=XM_023252100.1;refGene_gn=GREB1;refGene_tx=XM_023252101.1;refGene_gn=GREB1;refGene_tx=XM_023252102.1
A3 134497046 134497175
refGene_gn=GREB1;refGene_tx=XM_023252095.1;refGene_gn=GREB1;refGene_tx=XM_023252096.1;refGene_gn=GREB1;refGene_tx=XM_023252097.1;refGene_gn=GREB1;refGene_tx=XM_023252098.1;refGene_gn=GREB1;refGene_tx=XM_023252099.1;refGene_gn=GREB1;refGene_tx=XM_023252100.1;refGene_gn=GREB1;refGene_tx=XM_023252101.1;refGene_gn=GREB1;refGene_tx=XM_023252102.1
A3 134500986 134501337
refGene_gn=GREB1;refGene_tx=XM_023252095.1;refGene_gn=GREB1;refGene_tx=XM_023252096.1;refGene_gn=GREB1;refGene_tx=XM_023252097.1;refGene_gn=GREB1;refGene_tx=XM_023252098.1;refGene_gn=GREB1;refGene_tx=XM_023252099.1;refGene_gn=GREB1;refGene_tx=XM_023252100.1;refGene_gn=GREB1;refGene_tx=XM_023252101.1;refGene_gn=GREB1;refGene_tx=XM_023252102.1
A3 134503297 134503483

refGene_gn=GREB1;refGene_tx=XM_023252095.1;refGene_gn=GREB1;refGene_tx=XM_023252096.1;refGene_gn=GREB1;refGene_tx=XM_023252097.1;refGene_gn=GREB1;refGene_tx=XM_023252098.1;refGene_gn=GREB1;refGene_tx=XM_023252099.1;refGene_gn=GREB1;refGene_tx=XM_023252100.1;refGene_gn=GREB1;refGene_tx=XM_023252101.1;refGene_gn=GREB1;refGene_tx=XM_023252102.1
A3 134506633 134506777
refGene_gn=GREB1;refGene_tx=XM_023252095.1;refGene_gn=GREB1;refGene_tx=XM_023252096.1;refGene_gn=GREB1;refGene_tx=XM_023252097.1;refGene_gn=GREB1;refGene_tx=XM_023252098.1;refGene_gn=GREB1;refGene_tx=XM_023252099.1;refGene_gn=GREB1;refGene_tx=XM_023252100.1;refGene_gn=GREB1;refGene_tx=XM_023252101.1;refGene_gn=GREB1;refGene_tx=XM_023252102.1
A3 134507264 134507378
refGene_gn=GREB1;refGene_tx=XM_023252095.1;refGene_gn=GREB1;refGene_tx=XM_023252096.1;refGene_gn=GREB1;refGene_tx=XM_023252097.1;refGene_gn=GREB1;refGene_tx=XM_023252098.1;refGene_gn=GREB1;refGene_tx=XM_023252099.1;refGene_gn=GREB1;refGene_tx=XM_023252100.1;refGene_gn=GREB1;refGene_tx=XM_023252101.1;refGene_gn=GREB1;refGene_tx=XM_023252102.1
A3 134510886 134511015
refGene_gn=GREB1;refGene_tx=XM_023252095.1;refGene_gn=GREB1;refGene_tx=XM_023252096.1;refGene_gn=GREB1;refGene_tx=XM_023252097.1;refGene_gn=GREB1;refGene_tx=XM_023252098.1;refGene_gn=GREB1;refGene_tx=XM_023252099.1;refGene_gn=GREB1;refGene_tx=XM_023252100.1;refGene_gn=GREB1;refGene_tx=XM_023252101.1;refGene_gn=GREB1;refGene_tx=XM_023252102.1
A3 134513174 134513318
refGene_gn=GREB1;refGene_tx=XM_023252095.1;refGene_gn=GREB1;refGene_tx=XM_023252096.1;refGene_gn=GREB1;refGene_tx=XM_023252097.1;refGene_gn=GREB1;refGene_tx=XM_023252098.1;refGene_gn=GREB1;refGene_tx=XM_023252099.1;refGene_gn=GREB1;refGene_tx=XM_023252100.1;refGene_gn=GREB1;refGene_tx=XM_023252101.1;refGene_gn=GREB1;refGene_tx=XM_023252102.1
A3 134514706 134514889
refGene_gn=GREB1;refGene_tx=XM_023252095.1;refGene_gn=GREB1;refGene_tx=XM_023252096.1;refGene_gn=GREB1;refGene_tx=XM_023252097.1;refGene_gn=GREB1;refGene_tx=XM_023252098.1;refGene_gn=GREB1;refGene_tx=XM_023252099.1;refGene_gn=GREB1;refGene_tx=XM_023252100.1;refGene_gn=GREB1;refGene_tx=XM_023252101.1;refGene_gn=GREB1;refGene_tx=XM_023252102.1
A3 134522515 134522692
refGene_gn=GREB1;refGene_tx=XM_023252095.1;refGene_gn=GREB1;refGene_tx=XM_023252096.1;refGene_gn=GREB1;refGene_tx=XM_023252097.1;refGene_gn=GREB1;refGene_tx=XM_023252098.1;refGene_gn=GREB1;refGene_tx=XM_023252099.1;refGene_gn=GREB1;refGene_tx=XM_023252100.1;refGene_gn=GREB1;refGene_tx=XM_023252101.1;refGene_gn=GREB1;refGene_tx=XM_023252102.1
A3 134525381 134525501
refGene_gn=GREB1;refGene_tx=XM_023252095.1;refGene_gn=GREB1;refGene_tx=XM_023252096.1;refGene_gn=GREB1;refGene_tx=XM_023252097.1;refGene_gn=GREB1;refGene_tx=XM_023252098.1;refGene_gn=GREB1;refGene_tx=XM_023252099.1;refGene_gn=GREB1;refGene_tx=XM_023252100.1;refGene_gn=GREB1;refGene_tx=XM_023252101.1;refGene_gn=GREB1;refGene_tx=XM_023252102.1
A3 134531651 134531808
refGene_gn=GREB1;refGene_tx=XM_023252095.1;refGene_gn=GREB1;refGene_tx=XM_023252096.1;refGene_gn=GREB1;refGene_tx=XM_023252097.1;refGene_gn=

GREB1;refGene_tx=XM_023252098.1;refGene_gn=GREB1;refGene_tx=XM_0232520
99.1;refGene_gn=GREB1;refGene_tx=XM_023252100.1;refGene_gn=GREB1;refGe
ne_tx=XM_023252101.1;refGene_gn=GREB1;refGene_tx=XM_023252102.1
A3	134616368	134616476
refGene_gn=E2F6;refGene_tx=XM_023252103.1
A3	134624990	134625045
refGene_gn=E2F6;refGene_tx=XM_023252103.1
A3	134628819	134629036
refGene_gn=E2F6;refGene_tx=XM_023252103.1
A3	134630759	134630915
refGene_gn=E2F6;refGene_tx=XM_023252103.1
A3	134632518	134632633
refGene_gn=E2F6;refGene_tx=XM_023252103.1
A3	134635560	134635708
refGene_gn=E2F6;refGene_tx=XM_023252103.1
A3	134636307	134636357
refGene_gn=E2F6;refGene_tx=XM_023252103.1
A3	134726578	134726719
refGene_gn=ROCK2;refGene_tx=XM_006930517.4;refGene_gn=ROCK2;refGene_tx
=XM_019827905.2;refGene_gn=ROCK2;refGene_tx=XM_023252104.1;refGene_gn=
ROCK2;refGene_tx=XM_023252105.1;refGene_gn=ROCK2;refGene_tx=XM_0232521
06.1;refGene_gn=ROCK2;refGene_tx=XM_023252107.1
A3	134784377	134784459
refGene_gn=ROCK2;refGene_tx=XM_006930517.4;refGene_gn=ROCK2;refGene_tx
=XM_019827905.2;refGene_gn=ROCK2;refGene_tx=XM_023252104.1;refGene_gn=
ROCK2;refGene_tx=XM_023252105.1;refGene_gn=ROCK2;refGene_tx=XM_0232521
06.1;refGene_gn=ROCK2;refGene_tx=XM_023252107.1
A3	134785415	134785516
refGene_gn=ROCK2;refGene_tx=XM_006930517.4;refGene_gn=ROCK2;refGene_tx
=XM_019827905.2;refGene_gn=ROCK2;refGene_tx=XM_023252104.1;refGene_gn=
ROCK2;refGene_tx=XM_023252105.1;refGene_gn=ROCK2;refGene_tx=XM_0232521
06.1;refGene_gn=ROCK2;refGene_tx=XM_023252107.1
A3	134798832	134798970
refGene_gn=ROCK2;refGene_tx=XM_006930517.4;refGene_gn=ROCK2;refGene_tx
=XM_019827905.2;refGene_gn=ROCK2;refGene_tx=XM_023252104.1;refGene_gn=
ROCK2;refGene_tx=XM_023252105.1;refGene_gn=ROCK2;refGene_tx=XM_0232521
06.1;refGene_gn=ROCK2;refGene_tx=XM_023252107.1
A3	134810312	134810573
refGene_gn=ROCK2;refGene_tx=XM_006930517.4;refGene_gn=ROCK2;refGene_tx
=XM_019827905.2;refGene_gn=ROCK2;refGene_tx=XM_023252104.1;refGene_gn=
ROCK2;refGene_tx=XM_023252105.1;refGene_gn=ROCK2;refGene_tx=XM_0232521
06.1;refGene_gn=ROCK2;refGene_tx=XM_023252107.1
A3	134819547	134819692
refGene_gn=ROCK2;refGene_tx=XM_006930517.4;refGene_gn=ROCK2;refGene_tx
=XM_019827905.2;refGene_gn=ROCK2;refGene_tx=XM_023252104.1;refGene_gn=
ROCK2;refGene_tx=XM_023252105.1;refGene_gn=ROCK2;refGene_tx=XM_0232521
06.1;refGene_gn=ROCK2;refGene_tx=XM_023252107.1
A3	134821540	134821679
refGene_gn=ROCK2;refGene_tx=XM_006930517.4;refGene_gn=ROCK2;refGene_tx
=XM_019827905.2;refGene_gn=ROCK2;refGene_tx=XM_023252104.1;refGene_gn=

ROCK2;refGene_tx=XM_023252105.1;refGene_gn=ROCK2;refGene_tx=XM_023252106.1;refGene_gn=ROCK2;refGene_tx=XM_023252107.1
A3 134825396 134825488
refGene_gn=ROCK2;refGene_tx=XM_006930517.4;refGene_gn=ROCK2;refGene_tx=XM_019827905.2;refGene_gn=ROCK2;refGene_tx=XM_023252104.1;refGene_gn=ROCK2;refGene_tx=XM_023252105.1;refGene_gn=ROCK2;refGene_tx=XM_023252106.1;refGene_gn=ROCK2;refGene_tx=XM_023252107.1
A3 134826277 134826437
refGene_gn=ROCK2;refGene_tx=XM_006930517.4;refGene_gn=ROCK2;refGene_tx=XM_019827905.2;refGene_gn=ROCK2;refGene_tx=XM_023252104.1;refGene_gn=ROCK2;refGene_tx=XM_023252105.1;refGene_gn=ROCK2;refGene_tx=XM_023252106.1;refGene_gn=ROCK2;refGene_tx=XM_023252107.1
A3 134828367 134828428
refGene_gn=ROCK2;refGene_tx=XM_006930517.4;refGene_gn=ROCK2;refGene_tx=XM_019827905.2;refGene_gn=ROCK2;refGene_tx=XM_023252104.1;refGene_gn=ROCK2;refGene_tx=XM_023252105.1;refGene_gn=ROCK2;refGene_tx=XM_023252106.1;refGene_gn=ROCK2;refGene_tx=XM_023252107.1
A3 134829025 134829037
refGene_gn=ROCK2;refGene_tx=XM_006930517.4;refGene_gn=ROCK2;refGene_tx=XM_019827905.2;refGene_gn=ROCK2;refGene_tx=XM_023252104.1;refGene_gn=ROCK2;refGene_tx=XM_023252105.1;refGene_gn=ROCK2;refGene_tx=XM_023252106.1;refGene_gn=ROCK2;refGene_tx=XM_023252107.1
A3 134830344 134830424
refGene_gn=ROCK2;refGene_tx=XM_006930517.4;refGene_gn=ROCK2;refGene_tx=XM_019827905.2;refGene_gn=ROCK2;refGene_tx=XM_023252104.1;refGene_gn=ROCK2;refGene_tx=XM_023252105.1;refGene_gn=ROCK2;refGene_tx=XM_023252106.1;refGene_gn=ROCK2;refGene_tx=XM_023252107.1
A3 134830575 134830624
refGene_gn=ROCK2;refGene_tx=XM_006930517.4;refGene_gn=ROCK2;refGene_tx=XM_019827905.2;refGene_gn=ROCK2;refGene_tx=XM_023252104.1;refGene_gn=ROCK2;refGene_tx=XM_023252105.1;refGene_gn=ROCK2;refGene_tx=XM_023252106.1;refGene_gn=ROCK2;refGene_tx=XM_023252107.1
A3 134831219 134831355
refGene_gn=ROCK2;refGene_tx=XM_006930517.4;refGene_gn=ROCK2;refGene_tx=XM_019827905.2;refGene_gn=ROCK2;refGene_tx=XM_023252104.1;refGene_gn=ROCK2;refGene_tx=XM_023252105.1;refGene_gn=ROCK2;refGene_tx=XM_023252106.1;refGene_gn=ROCK2;refGene_tx=XM_023252107.1
A3 134831460 134831552
refGene_gn=ROCK2;refGene_tx=XM_006930517.4;refGene_gn=ROCK2;refGene_tx=XM_019827905.2;refGene_gn=ROCK2;refGene_tx=XM_023252104.1;refGene_gn=ROCK2;refGene_tx=XM_023252105.1;refGene_gn=ROCK2;refGene_tx=XM_023252106.1;refGene_gn=ROCK2;refGene_tx=XM_023252107.1
A3 134831785 134832032
refGene_gn=ROCK2;refGene_tx=XM_006930517.4;refGene_gn=ROCK2;refGene_tx=XM_019827905.2;refGene_gn=ROCK2;refGene_tx=XM_023252104.1;refGene_gn=ROCK2;refGene_tx=XM_023252105.1;refGene_gn=ROCK2;refGene_tx=XM_023252106.1;refGene_gn=ROCK2;refGene_tx=XM_023252107.1
A3 134832390 134832497
refGene_gn=ROCK2;refGene_tx=XM_006930517.4;refGene_gn=ROCK2;refGene_tx=XM_019827905.2;refGene_gn=ROCK2;refGene_tx=XM_023252104.1;refGene_gn=

ROCK2;refGene_tx=XM_023252105.1;refGene_gn=ROCK2;refGene_tx=XM_023252106.1;refGene_gn=ROCK2;refGene_tx=XM_023252107.1
A3 134835164 134835324
refGene_gn=ROCK2;refGene_tx=XM_006930517.4;refGene_gn=ROCK2;refGene_tx=XM_019827905.2;refGene_gn=ROCK2;refGene_tx=XM_023252104.1;refGene_gn=ROCK2;refGene_tx=XM_023252105.1;refGene_gn=ROCK2;refGene_tx=XM_023252106.1;refGene_gn=ROCK2;refGene_tx=XM_023252107.1
A3 134837978 134838139
refGene_gn=ROCK2;refGene_tx=XM_006930517.4;refGene_gn=ROCK2;refGene_tx=XM_019827905.2;refGene_gn=ROCK2;refGene_tx=XM_023252104.1;refGene_gn=ROCK2;refGene_tx=XM_023252105.1;refGene_gn=ROCK2;refGene_tx=XM_023252106.1;refGene_gn=ROCK2;refGene_tx=XM_023252107.1
A3 134838696 134838881
refGene_gn=ROCK2;refGene_tx=XM_006930517.4;refGene_gn=ROCK2;refGene_tx=XM_019827905.2;refGene_gn=ROCK2;refGene_tx=XM_023252104.1;refGene_gn=ROCK2;refGene_tx=XM_023252105.1;refGene_gn=ROCK2;refGene_tx=XM_023252106.1;refGene_gn=ROCK2;refGene_tx=XM_023252107.1
A3 134842522 134842592
refGene_gn=ROCK2;refGene_tx=XM_006930517.4;refGene_gn=ROCK2;refGene_tx=XM_019827905.2;refGene_gn=ROCK2;refGene_tx=XM_023252104.1;refGene_gn=ROCK2;refGene_tx=XM_023252105.1;refGene_gn=ROCK2;refGene_tx=XM_023252106.1;refGene_gn=ROCK2;refGene_tx=XM_023252107.1
A3 134843512 134843616
refGene_gn=ROCK2;refGene_tx=XM_006930517.4;refGene_gn=ROCK2;refGene_tx=XM_019827905.2;refGene_gn=ROCK2;refGene_tx=XM_023252104.1;refGene_gn=ROCK2;refGene_tx=XM_023252105.1;refGene_gn=ROCK2;refGene_tx=XM_023252106.1;refGene_gn=ROCK2;refGene_tx=XM_023252107.1
A3 134843777 134843964
refGene_gn=ROCK2;refGene_tx=XM_006930517.4;refGene_gn=ROCK2;refGene_tx=XM_019827905.2;refGene_gn=ROCK2;refGene_tx=XM_023252104.1;refGene_gn=ROCK2;refGene_tx=XM_023252105.1;refGene_gn=ROCK2;refGene_tx=XM_023252106.1;refGene_gn=ROCK2;refGene_tx=XM_023252107.1
A3 134846152 134846246
refGene_gn=ROCK2;refGene_tx=XM_006930517.4;refGene_gn=ROCK2;refGene_tx=XM_019827905.2;refGene_gn=ROCK2;refGene_tx=XM_023252104.1;refGene_gn=ROCK2;refGene_tx=XM_023252105.1;refGene_gn=ROCK2;refGene_tx=XM_023252106.1;refGene_gn=ROCK2;refGene_tx=XM_023252107.1
A3 134846348 134846443
refGene_gn=ROCK2;refGene_tx=XM_006930517.4;refGene_gn=ROCK2;refGene_tx=XM_019827905.2;refGene_gn=ROCK2;refGene_tx=XM_023252104.1;refGene_gn=ROCK2;refGene_tx=XM_023252105.1;refGene_gn=ROCK2;refGene_tx=XM_023252106.1;refGene_gn=ROCK2;refGene_tx=XM_023252107.1
A3 134847194 134847374
refGene_gn=ROCK2;refGene_tx=XM_006930517.4;refGene_gn=ROCK2;refGene_tx=XM_019827905.2;refGene_gn=ROCK2;refGene_tx=XM_023252104.1;refGene_gn=ROCK2;refGene_tx=XM_023252105.1;refGene_gn=ROCK2;refGene_tx=XM_023252106.1;refGene_gn=ROCK2;refGene_tx=XM_023252107.1
A3 134847553 134847722
refGene_gn=ROCK2;refGene_tx=XM_006930517.4;refGene_gn=ROCK2;refGene_tx=XM_019827905.2;refGene_gn=ROCK2;refGene_tx=XM_023252104.1;refGene_gn=

ROCK2;refGene_tx=XM_023252105.1;refGene_gn=ROCK2;refGene_tx=XM_0232521
06.1;refGene_gn=ROCK2;refGene_tx=XM_023252107.1
A3 134848070 134848247
refGene_gn=ROCK2;refGene_tx=XM_006930517.4;refGene_gn=ROCK2;refGene_tx
=XM_023252104.1;refGene_gn=ROCK2;refGene_tx=XM_023252105.1;refGene_gn=
ROCK2;refGene_tx=XM_023252106.1;refGene_gn=ROCK2;refGene_tx=XM_0232521
07.1
A3 134850056 134850127
refGene_gn=ROCK2;refGene_tx=XM_006930517.4;refGene_gn=ROCK2;refGene_tx
=XM_019827905.2;refGene_gn=ROCK2;refGene_tx=XM_023252104.1;refGene_gn=
ROCK2;refGene_tx=XM_023252105.1;refGene_gn=ROCK2;refGene_tx=XM_0232521
06.1;refGene_gn=ROCK2;refGene_tx=XM_023252107.1
A3 134851075 134851164
refGene_gn=ROCK2;refGene_tx=XM_006930517.4;refGene_gn=ROCK2;refGene_tx
=XM_019827905.2;refGene_gn=ROCK2;refGene_tx=XM_023252104.1;refGene_gn=
ROCK2;refGene_tx=XM_023252105.1;refGene_gn=ROCK2;refGene_tx=XM_0232521
06.1;refGene_gn=ROCK2;refGene_tx=XM_023252107.1
A3 134851560 134851639
refGene_gn=ROCK2;refGene_tx=XM_006930517.4;refGene_gn=ROCK2;refGene_tx
=XM_019827905.2;refGene_gn=ROCK2;refGene_tx=XM_023252104.1;refGene_gn=
ROCK2;refGene_tx=XM_023252105.1;refGene_gn=ROCK2;refGene_tx=XM_0232521
06.1;refGene_gn=ROCK2;refGene_tx=XM_023252107.1
A3 134852773 134853035
refGene_gn=ROCK2;refGene_tx=XM_006930517.4;refGene_gn=ROCK2;refGene_tx
=XM_019827905.2;refGene_gn=ROCK2;refGene_tx=XM_023252104.1;refGene_gn=
ROCK2;refGene_tx=XM_023252105.1;refGene_gn=ROCK2;refGene_tx=XM_0232521
06.1;refGene_gn=ROCK2;refGene_tx=XM_023252107.1
A3 134853122 134853336
refGene_gn=ROCK2;refGene_tx=XM_006930517.4;refGene_gn=ROCK2;refGene_tx
=XM_019827905.2;refGene_gn=ROCK2;refGene_tx=XM_023252104.1;refGene_gn=
ROCK2;refGene_tx=XM_023252105.1;refGene_gn=ROCK2;refGene_tx=XM_0232521
06.1;refGene_gn=ROCK2;refGene_tx=XM_023252107.1
A3 134854921 134854955
refGene_gn=ROCK2;refGene_tx=XM_023252105.1;refGene_gn=ROCK2;refGene_tx
=XM_023252106.1
A3 134860799 134860887
refGene_gn=ROCK2;refGene_tx=XM_006930517.4;refGene_gn=ROCK2;refGene_tx
=XM_019827905.2;refGene_gn=ROCK2;refGene_tx=XM_023252104.1
A3 134862178 134862182
refGene_gn=ROCK2;refGene_tx=XM_023252107.1
A3 134875948 134876075
refGene_gn=PQLC3;refGene_tx=XM_023252108.1
A3 134878071 134878141
refGene_gn=PQLC3;refGene_tx=XM_023252108.1
A3 134878226 134878309
refGene_gn=PQLC3;refGene_tx=XM_023252108.1
A3 134883618 134883761
refGene_gn=PQLC3;refGene_tx=XM_023252108.1
A3 134895760 134895952
refGene_gn=CA3H2orf50;refGene_tx=XM_003984497.3

A3 134897961 134898001
refGene_gn=CA3H2orf50;refGene_tx=XM_003984497.3
A3 134901618 134902081
refGene_gn=CA3H2orf50;refGene_tx=XM_003984497.3
A3 134903894 134904024
refGene_gn=CA3H2orf50;refGene_tx=XM_003984497.3
A3 135044306 135045791
refGene_gn=KCNF1;refGene_tx=XM_004001599.5
A3 135124889 135124908
refGene_gn=PDIA6;refGene_tx=XM_023252110.1
A3 135130730 135130872
refGene_gn=PDIA6;refGene_tx=XM_023252110.1
A3 135135076 135135134
refGene_gn=PDIA6;refGene_tx=XM_023252110.1
A3 135135585 135135712
refGene_gn=PDIA6;refGene_tx=XM_023252110.1
A3 135137439 135137546
refGene_gn=PDIA6;refGene_tx=XM_023252110.1
A3 135138755 135138886
refGene_gn=PDIA6;refGene_tx=XM_023252110.1
A3 135139270 135139385
refGene_gn=PDIA6;refGene_tx=XM_023252110.1
A3 135139916 135140057
refGene_gn=PDIA6;refGene_tx=XM_023252110.1
A3 135140814 135140899
refGene_gn=PDIA6;refGene_tx=XM_023252110.1
A3 135141026 135141099
refGene_gn=PDIA6;refGene_tx=XM_023252110.1
A3 135142208 135142367
refGene_gn=PDIA6;refGene_tx=XM_023252110.1
A3 135145973 135146070
refGene_gn=PDIA6;refGene_tx=XM_023252110.1
A3 135146666 135146735
refGene_gn=PDIA6;refGene_tx=XM_023252110.1
A3 135147765 135147855
refGene_gn=ATP6V1C2;refGene_tx=XM_023252109.1
A3 135148155 135148288
refGene_gn=ATP6V1C2;refGene_tx=XM_023252109.1
A3 135152349 135152447
refGene_gn=ATP6V1C2;refGene_tx=XM_023252109.1
A3 135153287 135153303
refGene_gn=ATP6V1C2;refGene_tx=XM_023252109.1
A3 135153304 135153423
refGene_gn=ATP6V1C2;refGene_tx=XM_023252109.1
A3 135155761 135155855
refGene_gn=ATP6V1C2;refGene_tx=XM_023252109.1
A3 135155965 135156058
refGene_gn=ATP6V1C2;refGene_tx=XM_023252109.1
A3 135158267 135158336
refGene_gn=ATP6V1C2;refGene_tx=XM_023252109.1

A3 135158906 135159005
refGene_gn=ATP6V1C2;refGene_tx=XM_023252109.1
A3 135161626 135161718
refGene_gn=ATP6V1C2;refGene_tx=XM_023252109.1
A3 135166384 135166479
refGene_gn=ATP6V1C2;refGene_tx=XM_023252109.1
A3 135173506 135173592
refGene_gn=ATP6V1C2;refGene_tx=XM_023252109.1
A3 135197486 135197554
refGene_gn=ATP6V1C2;refGene_tx=XM_023252109.1
A3 135199957 135200086
refGene_gn=ATP6V1C2;refGene_tx=XM_023252109.1
A3 135223543 135223609
refGene_gn=NOL10;refGene_tx=XM_003984501.5
A3 135226332 135226378
refGene_gn=NOL10;refGene_tx=XM_003984501.5
A3 135228846 135228945
refGene_gn=NOL10;refGene_tx=XM_003984501.5
A3 135231987 135232065
refGene_gn=NOL10;refGene_tx=XM_003984501.5
A3 135233425 135233463
refGene_gn=NOL10;refGene_tx=XM_003984501.5
A3 135234664 135234801
refGene_gn=NOL10;refGene_tx=XM_003984501.5
A3 135235575 135235641
refGene_gn=NOL10;refGene_tx=XM_003984501.5
A3 135237258 135237319
refGene_gn=NOL10;refGene_tx=XM_003984501.5
A3 135241756 135241842
refGene_gn=NOL10;refGene_tx=XM_003984501.5
A3 135244321 135244400
refGene_gn=NOL10;refGene_tx=XM_003984501.5
A3 135246109 135246259
refGene_gn=NOL10;refGene_tx=XM_003984501.5
A3 135248225 135248292
refGene_gn=NOL10;refGene_tx=XM_003984501.5
A3 135255201 135255254
refGene_gn=NOL10;refGene_tx=XM_003984501.5
A3 135286153 135286280
refGene_gn=NOL10;refGene_tx=XM_003984501.5
A3 135288704 135288784
refGene_gn=NOL10;refGene_tx=XM_003984501.5
A3 135288964 135289063
refGene_gn=NOL10;refGene_tx=XM_003984501.5
A3 135291187 135291277
refGene_gn=NOL10;refGene_tx=XM_003984501.5
A3 135299873 135300047
refGene_gn=NOL10;refGene_tx=XM_003984501.5
A3 135300300 135300548
refGene_gn=NOL10;refGene_tx=XM_003984501.5

A3	135309413	135309516
refGene_gn=NOL10;refGene_tx=XM_003984501.5
A3	135312882	135313002
refGene_gn=NOL10;refGene_tx=XM_003984501.5
A3	135433559	135433661
refGene_gn=ODC1;refGene_tx=XM_023252111.1;refGene_gn=ODC1;refGene_tx=XM_023252112.1;refGene_gn=ODC1;refGene_tx=XM_023252113.1
A3	135433944	135434118
refGene_gn=ODC1;refGene_tx=XM_023252111.1;refGene_gn=ODC1;refGene_tx=XM_023252112.1;refGene_gn=ODC1;refGene_tx=XM_023252113.1
A3	135434294	135434467
refGene_gn=ODC1;refGene_tx=XM_023252111.1;refGene_gn=ODC1;refGene_tx=XM_023252112.1;refGene_gn=ODC1;refGene_tx=XM_023252113.1
A3	135434727	135434862
refGene_gn=ODC1;refGene_tx=XM_023252111.1;refGene_gn=ODC1;refGene_tx=XM_023252112.1;refGene_gn=ODC1;refGene_tx=XM_023252113.1
A3	135434979	135435061
refGene_gn=ODC1;refGene_tx=XM_023252111.1;refGene_gn=ODC1;refGene_tx=XM_023252112.1;refGene_gn=ODC1;refGene_tx=XM_023252113.1
A3	135435245	135435329
refGene_gn=ODC1;refGene_tx=XM_023252111.1;refGene_gn=ODC1;refGene_tx=XM_023252112.1;refGene_gn=ODC1;refGene_tx=XM_023252113.1
A3	135436106	135436269
refGene_gn=ODC1;refGene_tx=XM_023252111.1;refGene_gn=ODC1;refGene_tx=XM_023252112.1;refGene_gn=ODC1;refGene_tx=XM_023252113.1
A3	135436351	135436464
refGene_gn=ODC1;refGene_tx=XM_023252111.1;refGene_gn=ODC1;refGene_tx=XM_023252112.1;refGene_gn=ODC1;refGene_tx=XM_023252113.1
A3	135436757	135436972
refGene_gn=ODC1;refGene_tx=XM_023252111.1;refGene_gn=ODC1;refGene_tx=XM_023252112.1;refGene_gn=ODC1;refGene_tx=XM_023252113.1
A3	135437467	135437612
refGene_gn=ODC1;refGene_tx=XM_023252111.1;refGene_gn=ODC1;refGene_tx=XM_023252112.1;refGene_gn=ODC1;refGene_tx=XM_023252113.1
A3	135448588	135448686
refGene_gn=HPCAL1;refGene_tx=XM_003984505.4
A3	135451355	135451461
refGene_gn=HPCAL1;refGene_tx=XM_003984505.4
A3	135453865	135454243
refGene_gn=HPCAL1;refGene_tx=XM_003984505.4
A3	135674285	135674516
refGene_gn=RRM2;refGene_tx=XM_023252115.1
A3	135685216	135685369
refGene_gn=RRM2;refGene_tx=XM_023252116.1
A3	135685447	135685561
refGene_gn=RRM2;refGene_tx=XM_023252115.1;refGene_gn=RRM2;refGene_tx=XM_023252116.1
A3	135685652	135685757
refGene_gn=RRM2;refGene_tx=XM_023252115.1;refGene_gn=RRM2;refGene_tx=XM_023252116.1

A3 135686729 135686863
refGene_gn=RRM2;refGene_tx=XM_023252115.1;refGene_gn=RRM2;refGene_tx=XM_023252116.1
A3 135686991 135687086
refGene_gn=RRM2;refGene_tx=XM_023252115.1;refGene_gn=RRM2;refGene_tx=XM_023252116.1
A3 135688904 135689038
refGene_gn=RRM2;refGene_tx=XM_023252115.1;refGene_gn=RRM2;refGene_tx=XM_023252116.1
A3 135689570 135689687
refGene_gn=RRM2;refGene_tx=XM_023252115.1;refGene_gn=RRM2;refGene_tx=XM_023252116.1
A3 135689988 135690132
refGene_gn=RRM2;refGene_tx=XM_023252115.1;refGene_gn=RRM2;refGene_tx=XM_023252116.1
A3 135690506 135690581
refGene_gn=RRM2;refGene_tx=XM_023252115.1;refGene_gn=RRM2;refGene_tx=XM_023252116.1
A3 135690662 135690773
refGene_gn=RRM2;refGene_tx=XM_023252115.1;refGene_gn=RRM2;refGene_tx=XM_023252116.1
A3 135727057 135727396
refGene_gn=CYS1;refGene_tx=XM_023252117.1;refGene_gn=CYS1;refGene_tx=XM_023252118.1
A3 135743159 135743212
refGene_gn=CYS1;refGene_tx=XM_023252117.1;refGene_gn=CYS1;refGene_tx=XM_023252118.1
A3 135743820 135743938
refGene_gn=CYS1;refGene_tx=XM_023252117.1
A3 135747334 135747449
refGene_gn=CYS1;refGene_tx=XM_023252118.1
A3 135758155 135758436
refGene_gn=KLF11;refGene_tx=XM_006930484.4;refGene_gn=KLF11;refGene_tx=XM_023252119.1
A3 135760191 135761170
refGene_gn=KLF11;refGene_tx=XM_006930484.4;refGene_gn=KLF11;refGene_tx=XM_023252119.1
A3 135762637 135762907
refGene_gn=KLF11;refGene_tx=XM_006930484.4;refGene_gn=KLF11;refGene_tx=XM_023252119.1
A3 135764847 135764889
refGene_gn=KLF11;refGene_tx=XM_023252119.1
A3 135792558 135792673
refGene_gn=GRHL1;refGene_tx=XM_023252120.1
A3 135793786 135793851
refGene_gn=GRHL1;refGene_tx=XM_023252120.1
A3 135797276 135797362
refGene_gn=GRHL1;refGene_tx=XM_023252120.1
A3 135797700 135797792
refGene_gn=GRHL1;refGene_tx=XM_023252120.1

A3 135800082 135800120
refGene_gn=GRHL1;refGene_tx=XM_023252120.1
A3 135801198 135801207
refGene_gn=GRHL1;refGene_tx=XM_023252120.1
A3 135801208 135801339
refGene_gn=GRHL1;refGene_tx=XM_023252120.1
A3 135803474 135803526
refGene_gn=GRHL1;refGene_tx=XM_023252120.1
A3 135812309 135812468
refGene_gn=GRHL1;refGene_tx=XM_023252120.1
A3 135835793 135835888
refGene_gn=GRHL1;refGene_tx=XM_023252120.1
A3 135836889 135837001
refGene_gn=GRHL1;refGene_tx=XM_023252120.1
A3 135837448 135837605
refGene_gn=GRHL1;refGene_tx=XM_023252120.1
A3 135839055 135839132
refGene_gn=GRHL1;refGene_tx=XM_023252120.1
A3 135840173 135840564
refGene_gn=GRHL1;refGene_tx=XM_023252120.1
A3 135842442 135842513
refGene_gn=GRHL1;refGene_tx=XM_023252120.1
A3 135844878 135845065
refGene_gn=GRHL1;refGene_tx=XM_023252120.1
A3 135848324 135848344
refGene_gn=GRHL1;refGene_tx=XM_023252120.1
A3 135866024 135866229
refGene_gn=TAF1B;refGene_tx=XM_023252121.1
A3 135872880 135873103
refGene_gn=TAF1B;refGene_tx=XM_023252121.1
A3 135873774 135873845
refGene_gn=TAF1B;refGene_tx=XM_023252121.1
A3 135878380 135878471
refGene_gn=TAF1B;refGene_tx=XM_023252121.1
A3 135881160 135881207
refGene_gn=TAF1B;refGene_tx=XM_023252121.1
A3 135881769 135881947
refGene_gn=TAF1B;refGene_tx=XM_023252121.1
A3 135883736 135883884
refGene_gn=TAF1B;refGene_tx=XM_023252121.1
A3 135904485 135904585
refGene_gn=TAF1B;refGene_tx=XM_023252121.1
A3 135910546 135910703
refGene_gn=TAF1B;refGene_tx=XM_023252121.1
A3 135917553 135917689
refGene_gn=TAF1B;refGene_tx=XM_023252121.1
A3 135922498 135922580
refGene_gn=TAF1B;refGene_tx=XM_023252121.1
A3 135923886 135923894
refGene_gn=TAF1B;refGene_tx=XM_023252121.1

A3 135925382 135925480
refGene_gn=TAF1B;refGene_tx=XM_023252121.1
A3 135928710 135928798
refGene_gn=TAF1B;refGene_tx=XM_023252121.1
A3 135931804 135931903
refGene_gn=TAF1B;refGene_tx=XM_023252121.1
A3 135933733 135933751
refGene_gn=TAF1B;refGene_tx=XM_023252121.1
A3 136126045 136126339
refGene_gn=YWHAQ;refGene_tx=XM_003984508.5
A3 136157929 136158053
refGene_gn=YWHAQ;refGene_tx=XM_003984508.5
A3 136160735 136160899
refGene_gn=YWHAQ;refGene_tx=XM_003984508.5
A3 136161099 136161195
refGene_gn=YWHAQ;refGene_tx=XM_003984508.5
A3 136163182 136163242
refGene_gn=YWHAQ;refGene_tx=XM_003984508.5
A3 136186158 136186255
refGene_gn=ADAM17;refGene_tx=XM_003984509.4
A3 136196482 136196615
refGene_gn=ADAM17;refGene_tx=XM_003984509.4
A3 136203057 136203188
refGene_gn=ADAM17;refGene_tx=XM_003984509.4
A3 136203995 136204084
refGene_gn=ADAM17;refGene_tx=XM_003984509.4
A3 136208773 136208942
refGene_gn=ADAM17;refGene_tx=XM_003984509.4
A3 136210410 136210544
refGene_gn=ADAM17;refGene_tx=XM_003984509.4
A3 136213514 136213604
refGene_gn=ADAM17;refGene_tx=XM_003984509.4
A3 136215633 136215747
refGene_gn=ADAM17;refGene_tx=XM_003984509.4
A3 136217759 136217904
refGene_gn=ADAM17;refGene_tx=XM_003984509.4
A3 136218011 136218100
refGene_gn=ADAM17;refGene_tx=XM_003984509.4
A3 136219447 136219600
refGene_gn=ADAM17;refGene_tx=XM_003984509.4
A3 136225694 136225894
refGene_gn=ADAM17;refGene_tx=XM_003984509.4
A3 136226949 136227053
refGene_gn=ADAM17;refGene_tx=XM_003984509.4
A3 136232304 136232439
refGene_gn=ADAM17;refGene_tx=XM_003984509.4
A3 136234036 136234167
refGene_gn=ADAM17;refGene_tx=XM_003984509.4
A3 136234486 136234565
refGene_gn=ADAM17;refGene_tx=XM_003984509.4

A3 136235488 136235577
refGene_gn=ADAM17;refGene_tx=XM_003984509.4
A3 136237205 136237256
refGene_gn=ADAM17;refGene_tx=XM_003984509.4
A3 136237673 136238027
refGene_gn=ADAM17;refGene_tx=XM_003984509.4
A3 136239644 136239827
refGene_gn=IAH1;refGene_tx=XM_011281327.3
A3 136241008 136241130
refGene_gn=IAH1;refGene_tx=XM_011281327.3
A3 136243040 136243202
refGene_gn=IAH1;refGene_tx=XM_011281327.3
A3 136245971 136246120
refGene_gn=IAH1;refGene_tx=XM_011281327.3
A3 136251287 136251340
refGene_gn=IAH1;refGene_tx=XM_011281327.3
A3 136252932 136253004
refGene_gn=IAH1;refGene_tx=XM_011281327.3
A3 136255912 136256014
refGene_gn=CPSF3;refGene_tx=XM_011281328.3;refGene_gn=CPSF3;refGene_tx=XM_019827943.2
A3 136258452 136258549
refGene_gn=CPSF3;refGene_tx=XM_011281328.3;refGene_gn=CPSF3;refGene_tx=XM_019827943.2
A3 136259708 136259778
refGene_gn=CPSF3;refGene_tx=XM_011281328.3;refGene_gn=CPSF3;refGene_tx=XM_019827943.2
A3 136264412 136264500
refGene_gn=CPSF3;refGene_tx=XM_011281328.3;refGene_gn=CPSF3;refGene_tx=XM_019827943.2
A3 136266002 136266097
refGene_gn=CPSF3;refGene_tx=XM_011281328.3;refGene_gn=CPSF3;refGene_tx=XM_019827943.2
A3 136267460 136267559
refGene_gn=CPSF3;refGene_tx=XM_011281328.3;refGene_gn=CPSF3;refGene_tx=XM_019827943.2
A3 136268761 136268870
refGene_gn=CPSF3;refGene_tx=XM_011281328.3;refGene_gn=CPSF3;refGene_tx=XM_019827943.2
A3 136272254 136272407
refGene_gn=CPSF3;refGene_tx=XM_011281328.3;refGene_gn=CPSF3;refGene_tx=XM_019827943.2
A3 136275521 136275668
refGene_gn=CPSF3;refGene_tx=XM_011281328.3;refGene_gn=CPSF3;refGene_tx=XM_019827943.2
A3 136276575 136276734
refGene_gn=CPSF3;refGene_tx=XM_011281328.3;refGene_gn=CPSF3;refGene_tx=XM_019827943.2
A3 136277986 136278162
refGene_gn=CPSF3;refGene_tx=XM_011281328.3;refGene_gn=CPSF3;refGene_tx

=XM_019827943.2
A3	136284470	136284621
refGene_gn=CPSF3;refGene_tx=XM_011281328.3;refGene_gn=CPSF3;refGene_tx=XM_019827943.2
A3	136285075	136285165
refGene_gn=CPSF3;refGene_tx=XM_011281328.3;refGene_gn=CPSF3;refGene_tx=XM_019827943.2
A3	136286323	136286501
refGene_gn=CPSF3;refGene_tx=XM_011281328.3;refGene_gn=CPSF3;refGene_tx=XM_019827943.2
A3	136287724	136287853
refGene_gn=CPSF3;refGene_tx=XM_011281328.3;refGene_gn=CPSF3;refGene_tx=XM_019827943.2
A3	136288949	136289047
refGene_gn=CPSF3;refGene_tx=XM_011281328.3;refGene_gn=CPSF3;refGene_tx=XM_019827943.2
A3	136290121	136290185
refGene_gn=CPSF3;refGene_tx=XM_011281328.3;refGene_gn=CPSF3;refGene_tx=XM_019827943.2
A3	136294607	136294657
refGene_gn=CPSF3;refGene_tx=XM_011281328.3
A3	136299502	136299574
refGene_gn=ITGB1BP1;refGene_tx=XM_003984512.4;refGene_gn=ITGB1BP1;refGene_tx=XM_006930489.4;refGene_gn=ITGB1BP1;refGene_tx=XM_006930491.3;refGene_gn=ITGB1BP1;refGene_tx=XM_019827944.2;refGene_gn=ITGB1BP1;refGene_tx=XM_019827945.2;refGene_gn=ITGB1BP1;refGene_tx=XM_019827946.2;refGene_gn=ITGB1BP1;refGene_tx=XM_023252132.1
A3	136301703	136301782
refGene_gn=ITGB1BP1;refGene_tx=XM_003984512.4;refGene_gn=ITGB1BP1;refGene_tx=XM_006930489.4;refGene_gn=ITGB1BP1;refGene_tx=XM_006930491.3;refGene_gn=ITGB1BP1;refGene_tx=XM_019827944.2;refGene_gn=ITGB1BP1;refGene_tx=XM_019827945.2;refGene_gn=ITGB1BP1;refGene_tx=XM_019827946.2;refGene_gn=ITGB1BP1;refGene_tx=XM_023252132.1
A3	136303125	136303262
refGene_gn=ITGB1BP1;refGene_tx=XM_003984512.4;refGene_gn=ITGB1BP1;refGene_tx=XM_006930489.4;refGene_gn=ITGB1BP1;refGene_tx=XM_006930491.3;refGene_gn=ITGB1BP1;refGene_tx=XM_019827944.2;refGene_gn=ITGB1BP1;refGene_tx=XM_019827945.2;refGene_gn=ITGB1BP1;refGene_tx=XM_019827946.2;refGene_gn=ITGB1BP1;refGene_tx=XM_023252132.1
A3	136305906	136305999
refGene_gn=ITGB1BP1;refGene_tx=XM_003984512.4;refGene_gn=ITGB1BP1;refGene_tx=XM_006930489.4;refGene_gn=ITGB1BP1;refGene_tx=XM_006930491.3;refGene_gn=ITGB1BP1;refGene_tx=XM_019827944.2;refGene_gn=ITGB1BP1;refGene_tx=XM_019827945.2;refGene_gn=ITGB1BP1;refGene_tx=XM_019827946.2;refGene_gn=ITGB1BP1;refGene_tx=XM_023252132.1
A3	136306521	136306671
refGene_gn=ITGB1BP1;refGene_tx=XM_003984512.4;refGene_gn=ITGB1BP1;refGene_tx=XM_006930489.4;refGene_gn=ITGB1BP1;refGene_tx=XM_006930491.3;refGene_gn=ITGB1BP1;refGene_tx=XM_019827944.2;refGene_gn=ITGB1BP1;refGene_tx=XM_019827945.2;refGene_gn=ITGB1BP1;refGene_tx=XM_019827946.2;refG

ene_gn=ITGB1BP1;refGene_tx=XM_023252132.1
A3	136307151	136307223
refGene_gn=ITGB1BP1;refGene_tx=XM_003984512.4;refGene_gn=ITGB1BP1;refGene_tx=XM_006930489.4;refGene_gn=ITGB1BP1;refGene_tx=XM_006930491.3;refGene_gn=ITGB1BP1;refGene_tx=XM_019827944.2;refGene_gn=ITGB1BP1;refGene_tx=XM_019827945.2;refGene_gn=ITGB1BP1;refGene_tx=XM_019827946.2;refGene_gn=ITGB1BP1;refGene_tx=XM_023252132.1
A3	136310264	136310339
refGene_gn=ASAP2;refGene_tx=XM_023252122.1;refGene_gn=ASAP2;refGene_tx=XM_023252123.1;refGene_gn=ASAP2;refGene_tx=XM_023252124.1;refGene_gn=ASAP2;refGene_tx=XM_023252125.1;refGene_gn=ASAP2;refGene_tx=XM_023252127.1;refGene_gn=ASAP2;refGene_tx=XM_023252128.1;refGene_gn=ASAP2;refGene_tx=XM_023252129.1;refGene_gn=ASAP2;refGene_tx=XM_023252130.1
A3	136312000	136312123
refGene_gn=ASAP2;refGene_tx=XM_023252122.1;refGene_gn=ASAP2;refGene_tx=XM_023252123.1;refGene_gn=ASAP2;refGene_tx=XM_023252124.1;refGene_gn=ASAP2;refGene_tx=XM_023252125.1;refGene_gn=ASAP2;refGene_tx=XM_023252127.1;refGene_gn=ASAP2;refGene_tx=XM_023252128.1;refGene_gn=ASAP2;refGene_tx=XM_023252129.1;refGene_gn=ASAP2;refGene_tx=XM_023252130.1
A3	136312550	136312642
refGene_gn=ASAP2;refGene_tx=XM_023252122.1;refGene_gn=ASAP2;refGene_tx=XM_023252123.1;refGene_gn=ASAP2;refGene_tx=XM_023252124.1;refGene_gn=ASAP2;refGene_tx=XM_023252125.1;refGene_gn=ASAP2;refGene_tx=XM_023252127.1;refGene_gn=ASAP2;refGene_tx=XM_023252128.1;refGene_gn=ASAP2;refGene_tx=XM_023252129.1;refGene_gn=ASAP2;refGene_tx=XM_023252130.1
A3	136313077	136313127
refGene_gn=ASAP2;refGene_tx=XM_023252122.1;refGene_gn=ASAP2;refGene_tx=XM_023252123.1;refGene_gn=ASAP2;refGene_tx=XM_023252124.1;refGene_gn=ASAP2;refGene_tx=XM_023252125.1;refGene_gn=ASAP2;refGene_tx=XM_023252127.1;refGene_gn=ASAP2;refGene_tx=XM_023252128.1;refGene_gn=ASAP2;refGene_tx=XM_023252129.1;refGene_gn=ASAP2;refGene_tx=XM_023252130.1
A3	136319888	136320060
refGene_gn=ASAP2;refGene_tx=XM_023252122.1;refGene_gn=ASAP2;refGene_tx=XM_023252123.1;refGene_gn=ASAP2;refGene_tx=XM_023252124.1;refGene_gn=ASAP2;refGene_tx=XM_023252125.1;refGene_gn=ASAP2;refGene_tx=XM_023252127.1;refGene_gn=ASAP2;refGene_tx=XM_023252128.1;refGene_gn=ASAP2;refGene_tx=XM_023252129.1;refGene_gn=ASAP2;refGene_tx=XM_023252130.1
A3	136321351	136321486
refGene_gn=ASAP2;refGene_tx=XM_023252122.1;refGene_gn=ASAP2;refGene_tx=XM_023252123.1;refGene_gn=ASAP2;refGene_tx=XM_023252124.1;refGene_gn=ASAP2;refGene_tx=XM_023252125.1;refGene_gn=ASAP2;refGene_tx=XM_023252130.1
A3	136323572	136323646
refGene_gn=ASAP2;refGene_tx=XM_023252131.1
A3	136324106	136324359
refGene_gn=ASAP2;refGene_tx=XM_023252122.1;refGene_gn=ASAP2;refGene_tx=XM_023252123.1;refGene_gn=ASAP2;refGene_tx=XM_023252124.1;refGene_gn=ASAP2;refGene_tx=XM_023252125.1;refGene_gn=ASAP2;refGene_tx=XM_023252127.1;refGene_gn=ASAP2;refGene_tx=XM_023252128.1;refGene_gn=ASAP2;refGene_tx=XM_023252129.1;refGene_gn=ASAP2;refGene_tx=XM_023252130.1;refGen

e_gn=ASAP2;refGene_tx=XM_023252131.1
A3 136326586 136326700
refGene_gn=ASAP2;refGene_tx=XM_023252122.1;refGene_gn=ASAP2;refGene_tx=XM_023252123.1;refGene_gn=ASAP2;refGene_tx=XM_023252124.1;refGene_gn=ASAP2;refGene_tx=XM_023252125.1;refGene_gn=ASAP2;refGene_tx=XM_023252127.1;refGene_gn=ASAP2;refGene_tx=XM_023252128.1;refGene_gn=ASAP2;refGene_tx=XM_023252129.1;refGene_gn=ASAP2;refGene_tx=XM_023252130.1;refGene_gn=ASAP2;refGene_tx=XM_023252131.1
A3 136327929 136327997
refGene_gn=ASAP2;refGene_tx=XM_023252122.1;refGene_gn=ASAP2;refGene_tx=XM_023252123.1;refGene_gn=ASAP2;refGene_tx=XM_023252124.1;refGene_gn=ASAP2;refGene_tx=XM_023252125.1;refGene_gn=ASAP2;refGene_tx=XM_023252127.1;refGene_gn=ASAP2;refGene_tx=XM_023252128.1;refGene_gn=ASAP2;refGene_tx=XM_023252129.1;refGene_gn=ASAP2;refGene_tx=XM_023252130.1;refGene_gn=ASAP2;refGene_tx=XM_023252131.1
A3 136329053 136329169
refGene_gn=ASAP2;refGene_tx=XM_023252122.1;refGene_gn=ASAP2;refGene_tx=XM_023252123.1;refGene_gn=ASAP2;refGene_tx=XM_023252124.1;refGene_gn=ASAP2;refGene_tx=XM_023252125.1;refGene_gn=ASAP2;refGene_tx=XM_023252127.1;refGene_gn=ASAP2;refGene_tx=XM_023252128.1;refGene_gn=ASAP2;refGene_tx=XM_023252129.1;refGene_gn=ASAP2;refGene_tx=XM_023252130.1;refGene_gn=ASAP2;refGene_tx=XM_023252131.1
A3 136330735 136330821
refGene_gn=ASAP2;refGene_tx=XM_023252122.1;refGene_gn=ASAP2;refGene_tx=XM_023252123.1;refGene_gn=ASAP2;refGene_tx=XM_023252124.1;refGene_gn=ASAP2;refGene_tx=XM_023252125.1;refGene_gn=ASAP2;refGene_tx=XM_023252127.1;refGene_gn=ASAP2;refGene_tx=XM_023252128.1;refGene_gn=ASAP2;refGene_tx=XM_023252129.1;refGene_gn=ASAP2;refGene_tx=XM_023252130.1;refGene_gn=ASAP2;refGene_tx=XM_023252131.1
A3 136332216 136332406
refGene_gn=ASAP2;refGene_tx=XM_023252122.1;refGene_gn=ASAP2;refGene_tx=XM_023252123.1;refGene_gn=ASAP2;refGene_tx=XM_023252124.1;refGene_gn=ASAP2;refGene_tx=XM_023252125.1;refGene_gn=ASAP2;refGene_tx=XM_023252127.1;refGene_gn=ASAP2;refGene_tx=XM_023252128.1;refGene_gn=ASAP2;refGene_tx=XM_023252129.1;refGene_gn=ASAP2;refGene_tx=XM_023252130.1;refGene_gn=ASAP2;refGene_tx=XM_023252131.1
A3 136338557 136338652
refGene_gn=ASAP2;refGene_tx=XM_023252122.1;refGene_gn=ASAP2;refGene_tx=XM_023252123.1;refGene_gn=ASAP2;refGene_tx=XM_023252124.1;refGene_gn=ASAP2;refGene_tx=XM_023252125.1;refGene_gn=ASAP2;refGene_tx=XM_023252127.1;refGene_gn=ASAP2;refGene_tx=XM_023252128.1;refGene_gn=ASAP2;refGene_tx=XM_023252129.1;refGene_gn=ASAP2;refGene_tx=XM_023252130.1;refGene_gn=ASAP2;refGene_tx=XM_023252131.1
A3 136345624 136345758
refGene_gn=ASAP2;refGene_tx=XM_023252122.1;refGene_gn=ASAP2;refGene_tx=XM_023252123.1;refGene_gn=ASAP2;refGene_tx=XM_023252124.1;refGene_gn=ASAP2;refGene_tx=XM_023252125.1;refGene_gn=ASAP2;refGene_tx=XM_023252127.1;refGene_gn=ASAP2;refGene_tx=XM_023252128.1;refGene_gn=ASAP2;refGene_tx=XM_023252129.1;refGene_gn=ASAP2;refGene_tx=XM_023252130.1;refGene_gn=ASAP2;refGene_tx=XM_023252131.1

A3	136350067	136350234
refGene_gn=ASAP2;refGene_tx=XM_023252122.1;refGene_gn=ASAP2;refGene_tx=XM_023252123.1;refGene_gn=ASAP2;refGene_tx=XM_023252124.1;refGene_gn=ASAP2;refGene_tx=XM_023252125.1;refGene_gn=ASAP2;refGene_tx=XM_023252127.1;refGene_gn=ASAP2;refGene_tx=XM_023252128.1;refGene_gn=ASAP2;refGene_tx=XM_023252129.1;refGene_gn=ASAP2;refGene_tx=XM_023252130.1;refGene_gn=ASAP2;refGene_tx=XM_023252131.1
A3	136350315	136350364
refGene_gn=ASAP2;refGene_tx=XM_023252122.1;refGene_gn=ASAP2;refGene_tx=XM_023252123.1;refGene_gn=ASAP2;refGene_tx=XM_023252124.1;refGene_gn=ASAP2;refGene_tx=XM_023252125.1;refGene_gn=ASAP2;refGene_tx=XM_023252127.1;refGene_gn=ASAP2;refGene_tx=XM_023252128.1;refGene_gn=ASAP2;refGene_tx=XM_023252129.1;refGene_gn=ASAP2;refGene_tx=XM_023252130.1;refGene_gn=ASAP2;refGene_tx=XM_023252131.1
A3	136354590	136354678
refGene_gn=ASAP2;refGene_tx=XM_023252122.1;refGene_gn=ASAP2;refGene_tx=XM_023252123.1;refGene_gn=ASAP2;refGene_tx=XM_023252124.1;refGene_gn=ASAP2;refGene_tx=XM_023252125.1;refGene_gn=ASAP2;refGene_tx=XM_023252127.1;refGene_gn=ASAP2;refGene_tx=XM_023252128.1;refGene_gn=ASAP2;refGene_tx=XM_023252129.1;refGene_gn=ASAP2;refGene_tx=XM_023252130.1;refGene_gn=ASAP2;refGene_tx=XM_023252131.1
A3	136360963	136361033
refGene_gn=ASAP2;refGene_tx=XM_023252122.1;refGene_gn=ASAP2;refGene_tx=XM_023252123.1;refGene_gn=ASAP2;refGene_tx=XM_023252124.1;refGene_gn=ASAP2;refGene_tx=XM_023252125.1;refGene_gn=ASAP2;refGene_tx=XM_023252127.1;refGene_gn=ASAP2;refGene_tx=XM_023252128.1;refGene_gn=ASAP2;refGene_tx=XM_023252129.1;refGene_gn=ASAP2;refGene_tx=XM_023252131.1
A3	136361133	136361237
refGene_gn=ASAP2;refGene_tx=XM_023252122.1;refGene_gn=ASAP2;refGene_tx=XM_023252123.1;refGene_gn=ASAP2;refGene_tx=XM_023252124.1;refGene_gn=ASAP2;refGene_tx=XM_023252125.1;refGene_gn=ASAP2;refGene_tx=XM_023252127.1;refGene_gn=ASAP2;refGene_tx=XM_023252128.1;refGene_gn=ASAP2;refGene_tx=XM_023252129.1;refGene_gn=ASAP2;refGene_tx=XM_023252131.1
A3	136368863	136368872
refGene_gn=ASAP2;refGene_tx=XM_023252122.1;refGene_gn=ASAP2;refGene_tx=XM_023252123.1;refGene_gn=ASAP2;refGene_tx=XM_023252127.1;refGene_gn=ASAP2;refGene_tx=XM_023252131.1
A3	136369755	136369842
refGene_gn=ASAP2;refGene_tx=XM_023252122.1;refGene_gn=ASAP2;refGene_tx=XM_023252123.1;refGene_gn=ASAP2;refGene_tx=XM_023252124.1;refGene_gn=ASAP2;refGene_tx=XM_023252125.1;refGene_gn=ASAP2;refGene_tx=XM_023252127.1;refGene_gn=ASAP2;refGene_tx=XM_023252128.1;refGene_gn=ASAP2;refGene_tx=XM_023252129.1;refGene_gn=ASAP2;refGene_tx=XM_023252130.1;refGene_gn=ASAP2;refGene_tx=XM_023252131.1
A3	136370136	136370212
refGene_gn=ASAP2;refGene_tx=XM_023252122.1;refGene_gn=ASAP2;refGene_tx=XM_023252123.1;refGene_gn=ASAP2;refGene_tx=XM_023252124.1;refGene_gn=ASAP2;refGene_tx=XM_023252125.1;refGene_gn=ASAP2;refGene_tx=XM_023252127.1;refGene_gn=ASAP2;refGene_tx=XM_023252128.1;refGene_gn=ASAP2;refGene_tx=XM_023252129.1;refGene_gn=ASAP2;refGene_tx=XM_023252130.1;refGen

e_gn=ASAP2;refGene_tx=XM_023252131.1
A3 136376161 136376247
refGene_gn=ASAP2;refGene_tx=XM_023252122.1;refGene_gn=ASAP2;refGene_tx=XM_023252123.1;refGene_gn=ASAP2;refGene_tx=XM_023252124.1;refGene_gn=ASAP2;refGene_tx=XM_023252125.1;refGene_gn=ASAP2;refGene_tx=XM_023252127.1;refGene_gn=ASAP2;refGene_tx=XM_023252128.1;refGene_gn=ASAP2;refGene_tx=XM_023252129.1;refGene_gn=ASAP2;refGene_tx=XM_023252130.1;refGene_gn=ASAP2;refGene_tx=XM_023252131.1
A3 136381684 136381814
refGene_gn=ASAP2;refGene_tx=XM_023252122.1;refGene_gn=ASAP2;refGene_tx=XM_023252123.1;refGene_gn=ASAP2;refGene_tx=XM_023252124.1;refGene_gn=ASAP2;refGene_tx=XM_023252125.1;refGene_gn=ASAP2;refGene_tx=XM_023252127.1;refGene_gn=ASAP2;refGene_tx=XM_023252128.1;refGene_gn=ASAP2;refGene_tx=XM_023252129.1;refGene_gn=ASAP2;refGene_tx=XM_023252130.1;refGene_gn=ASAP2;refGene_tx=XM_023252131.1
A3 136383905 136383955
refGene_gn=ASAP2;refGene_tx=XM_023252122.1;refGene_gn=ASAP2;refGene_tx=XM_023252123.1;refGene_gn=ASAP2;refGene_tx=XM_023252124.1;refGene_gn=ASAP2;refGene_tx=XM_023252125.1;refGene_gn=ASAP2;refGene_tx=XM_023252127.1;refGene_gn=ASAP2;refGene_tx=XM_023252128.1;refGene_gn=ASAP2;refGene_tx=XM_023252129.1;refGene_gn=ASAP2;refGene_tx=XM_023252130.1;refGene_gn=ASAP2;refGene_tx=XM_023252131.1
A3 136385723 136385798
refGene_gn=ASAP2;refGene_tx=XM_023252122.1;refGene_gn=ASAP2;refGene_tx=XM_023252123.1;refGene_gn=ASAP2;refGene_tx=XM_023252124.1;refGene_gn=ASAP2;refGene_tx=XM_023252125.1;refGene_gn=ASAP2;refGene_tx=XM_023252127.1;refGene_gn=ASAP2;refGene_tx=XM_023252128.1;refGene_gn=ASAP2;refGene_tx=XM_023252129.1;refGene_gn=ASAP2;refGene_tx=XM_023252130.1;refGene_gn=ASAP2;refGene_tx=XM_023252131.1
A3 136403309 136403455
refGene_gn=ASAP2;refGene_tx=XM_023252122.1;refGene_gn=ASAP2;refGene_tx=XM_023252123.1;refGene_gn=ASAP2;refGene_tx=XM_023252124.1;refGene_gn=ASAP2;refGene_tx=XM_023252125.1;refGene_gn=ASAP2;refGene_tx=XM_023252127.1;refGene_gn=ASAP2;refGene_tx=XM_023252128.1;refGene_gn=ASAP2;refGene_tx=XM_023252129.1;refGene_gn=ASAP2;refGene_tx=XM_023252130.1;refGene_gn=ASAP2;refGene_tx=XM_023252131.1
A3 136422713 136422786
refGene_gn=ASAP2;refGene_tx=XM_023252122.1;refGene_gn=ASAP2;refGene_tx=XM_023252123.1;refGene_gn=ASAP2;refGene_tx=XM_023252124.1;refGene_gn=ASAP2;refGene_tx=XM_023252125.1;refGene_gn=ASAP2;refGene_tx=XM_023252127.1;refGene_gn=ASAP2;refGene_tx=XM_023252128.1;refGene_gn=ASAP2;refGene_tx=XM_023252129.1;refGene_gn=ASAP2;refGene_tx=XM_023252130.1;refGene_gn=ASAP2;refGene_tx=XM_023252131.1
A3 136486313 136486439
refGene_gn=ASAP2;refGene_tx=XM_023252122.1;refGene_gn=ASAP2;refGene_tx=XM_023252123.1;refGene_gn=ASAP2;refGene_tx=XM_023252124.1;refGene_gn=ASAP2;refGene_tx=XM_023252125.1;refGene_gn=ASAP2;refGene_tx=XM_023252127.1;refGene_gn=ASAP2;refGene_tx=XM_023252128.1;refGene_gn=ASAP2;refGene_tx=XM_023252129.1;refGene_gn=ASAP2;refGene_tx=XM_023252130.1;refGene_gn=ASAP2;refGene_tx=XM_023252131.1

A3	136639748	136639823
refGene_gn=MBOAT2;refGene_tx=XM_023252133.1;refGene_gn=MBOAT2;refGene_tx=XM_023252134.1;refGene_gn=MBOAT2;refGene_tx=XM_023252135.1
A3	136689369	136689515
refGene_gn=MBOAT2;refGene_tx=XM_023252133.1;refGene_gn=MBOAT2;refGene_tx=XM_023252134.1;refGene_gn=MBOAT2;refGene_tx=XM_023252135.1
A3	136694996	136695074
refGene_gn=MBOAT2;refGene_tx=XM_023252133.1;refGene_gn=MBOAT2;refGene_tx=XM_023252134.1;refGene_gn=MBOAT2;refGene_tx=XM_023252135.1;refGene_gn=MBOAT2;refGene_tx=XM_023252136.1
A3	136727238	136727334
refGene_gn=MBOAT2;refGene_tx=XM_023252133.1;refGene_gn=MBOAT2;refGene_tx=XM_023252135.1;refGene_gn=MBOAT2;refGene_tx=XM_023252136.1
A3	136739618	136739674
refGene_gn=MBOAT2;refGene_tx=XM_023252133.1;refGene_gn=MBOAT2;refGene_tx=XM_023252134.1;refGene_gn=MBOAT2;refGene_tx=XM_023252135.1;refGene_gn=MBOAT2;refGene_tx=XM_023252136.1
A3	136741221	136741276
refGene_gn=MBOAT2;refGene_tx=XM_023252133.1;refGene_gn=MBOAT2;refGene_tx=XM_023252134.1;refGene_gn=MBOAT2;refGene_tx=XM_023252135.1;refGene_gn=MBOAT2;refGene_tx=XM_023252136.1
A3	136746816	136747000
refGene_gn=MBOAT2;refGene_tx=XM_023252133.1;refGene_gn=MBOAT2;refGene_tx=XM_023252134.1;refGene_gn=MBOAT2;refGene_tx=XM_023252135.1;refGene_gn=MBOAT2;refGene_tx=XM_023252136.1
A3	136751609	136751802
refGene_gn=MBOAT2;refGene_tx=XM_023252133.1;refGene_gn=MBOAT2;refGene_tx=XM_023252134.1;refGene_gn=MBOAT2;refGene_tx=XM_023252135.1;refGene_gn=MBOAT2;refGene_tx=XM_023252136.1
A3	136752169	136752273
refGene_gn=MBOAT2;refGene_tx=XM_023252133.1;refGene_gn=MBOAT2;refGene_tx=XM_023252134.1;refGene_gn=MBOAT2;refGene_tx=XM_023252135.1;refGene_gn=MBOAT2;refGene_tx=XM_023252136.1
A3	136755882	136755947
refGene_gn=MBOAT2;refGene_tx=XM_023252133.1;refGene_gn=MBOAT2;refGene_tx=XM_023252134.1;refGene_gn=MBOAT2;refGene_tx=XM_023252135.1;refGene_gn=MBOAT2;refGene_tx=XM_023252136.1
A3	136757561	136757694
refGene_gn=MBOAT2;refGene_tx=XM_023252133.1;refGene_gn=MBOAT2;refGene_tx=XM_023252134.1;refGene_gn=MBOAT2;refGene_tx=XM_023252135.1;refGene_gn=MBOAT2;refGene_tx=XM_023252136.1
A3	136759860	136760012
refGene_gn=MBOAT2;refGene_tx=XM_023252133.1;refGene_gn=MBOAT2;refGene_tx=XM_023252134.1;refGene_gn=MBOAT2;refGene_tx=XM_023252136.1
A3	136761974	136762200
refGene_gn=MBOAT2;refGene_tx=XM_023252133.1;refGene_gn=MBOAT2;refGene_tx=XM_023252134.1;refGene_gn=MBOAT2;refGene_tx=XM_023252135.1;refGene_gn=MBOAT2;refGene_tx=XM_023252136.1
A3	136788401	136788509
refGene_gn=KIDINS220;refGene_tx=XM_011281334.3

A3	136796880	136796933
refGene_gn=KIDINS220;refGene_tx=XM_011281334.3
A3	136796934	136796977
refGene_gn=KIDINS220;refGene_tx=XM_011281334.3
A3	136797702	136797801
refGene_gn=KIDINS220;refGene_tx=XM_011281334.3
A3	136802709	136802808
refGene_gn=KIDINS220;refGene_tx=XM_011281334.3
A3	136803755	136803857
refGene_gn=KIDINS220;refGene_tx=XM_011281334.3
A3	136806571	136806670
refGene_gn=KIDINS220;refGene_tx=XM_011281334.3
A3	136808372	136808570
refGene_gn=KIDINS220;refGene_tx=XM_011281334.3
A3	136810871	136810970
refGene_gn=KIDINS220;refGene_tx=XM_011281334.3
A3	136812610	136812709
refGene_gn=KIDINS220;refGene_tx=XM_011281334.3
A3	136813986	136814085
refGene_gn=KIDINS220;refGene_tx=XM_011281334.3
A3	136817279	136817457
refGene_gn=KIDINS220;refGene_tx=XM_011281334.3
A3	136821116	136821281
refGene_gn=KIDINS220;refGene_tx=XM_011281334.3
A3	136822545	136822725
refGene_gn=KIDINS220;refGene_tx=XM_011281334.3
A3	136823546	136823712
refGene_gn=KIDINS220;refGene_tx=XM_011281334.3
A3	136825627	136825779
refGene_gn=KIDINS220;refGene_tx=XM_011281334.3
A3	136825959	136826249
refGene_gn=KIDINS220;refGene_tx=XM_011281334.3
A3	136827464	136827605
refGene_gn=KIDINS220;refGene_tx=XM_011281334.3
A3	136828269	136828513
refGene_gn=KIDINS220;refGene_tx=XM_011281334.3
A3	136828678	136828767
refGene_gn=KIDINS220;refGene_tx=XM_011281334.3
A3	136830215	136830360
refGene_gn=KIDINS220;refGene_tx=XM_011281334.3
A3	136834881	136835044
refGene_gn=KIDINS220;refGene_tx=XM_011281334.3
A3	136854398	136854577
refGene_gn=KIDINS220;refGene_tx=XM_011281334.3
A3	136855900	136856127
refGene_gn=KIDINS220;refGene_tx=XM_011281334.3
A3	136858494	136858608
refGene_gn=KIDINS220;refGene_tx=XM_011281334.3
A3	136859363	136859420
refGene_gn=KIDINS220;refGene_tx=XM_011281334.3

A3	136863431	136863467
refGene_gn=KIDINS220;refGene_tx=XM_011281334.3
A3	136865058	136865091
refGene_gn=KIDINS220;refGene_tx=XM_011281334.3
A3	136869698	136869830
refGene_gn=KIDINS220;refGene_tx=XM_011281334.3
A3	136872057	136872156
refGene_gn=KIDINS220;refGene_tx=XM_011281334.3
A3	136873157	136873394
refGene_gn=KIDINS220;refGene_tx=XM_011281334.3
A3	136876024	136877287
refGene_gn=KIDINS220;refGene_tx=XM_011281334.3
A3	136909421	136909478
refGene_gn=ID2;refGene_tx=XM_003984514.4
A3	136909824	136910172
refGene_gn=ID2;refGene_tx=XM_003984514.4
A3	138294145	138294277
refGene_gn=RNF144A;refGene_tx=XM_023252141.1
A3	138306377	138306674
refGene_gn=RNF144A;refGene_tx=XM_023252138.1;refGene_gn=RNF144A;refGene_tx=XM_023252139.1;refGene_gn=RNF144A;refGene_tx=XM_023252142.1;refGene_gn=RNF144A;refGene_tx=XM_023252143.1;refGene_gn=RNF144A;refGene_tx=XM_023252144.1;refGene_gn=RNF144A;refGene_tx=XM_023252145.1;refGene_gn=RNF144A;refGene_tx=XM_023252146.1;refGene_gn=RNF144A;refGene_tx=XM_023252147.1;refGene_gn=RNF144A;refGene_tx=XM_023252148.1
A3	138307406	138307496
refGene_gn=RNF144A;refGene_tx=XM_023252138.1;refGene_gn=RNF144A;refGene_tx=XM_023252139.1;refGene_gn=RNF144A;refGene_tx=XM_023252141.1;refGene_gn=RNF144A;refGene_tx=XM_023252142.1;refGene_gn=RNF144A;refGene_tx=XM_023252143.1;refGene_gn=RNF144A;refGene_tx=XM_023252144.1;refGene_gn=RNF144A;refGene_tx=XM_023252145.1;refGene_gn=RNF144A;refGene_tx=XM_023252146.1;refGene_gn=RNF144A;refGene_tx=XM_023252147.1;refGene_gn=RNF144A;refGene_tx=XM_023252148.1
A3	138313365	138313513
refGene_gn=RNF144A;refGene_tx=XM_023252138.1;refGene_gn=RNF144A;refGene_tx=XM_023252139.1;refGene_gn=RNF144A;refGene_tx=XM_023252141.1;refGene_gn=RNF144A;refGene_tx=XM_023252142.1;refGene_gn=RNF144A;refGene_tx=XM_023252143.1;refGene_gn=RNF144A;refGene_tx=XM_023252144.1;refGene_gn=RNF144A;refGene_tx=XM_023252145.1;refGene_gn=RNF144A;refGene_tx=XM_023252146.1;refGene_gn=RNF144A;refGene_tx=XM_023252147.1;refGene_gn=RNF144A;refGene_tx=XM_023252148.1
A3	138319695	138319903
refGene_gn=RNF144A;refGene_tx=XM_023252138.1;refGene_gn=RNF144A;refGene_tx=XM_023252139.1;refGene_gn=RNF144A;refGene_tx=XM_023252141.1;refGene_gn=RNF144A;refGene_tx=XM_023252142.1;refGene_gn=RNF144A;refGene_tx=XM_023252143.1;refGene_gn=RNF144A;refGene_tx=XM_023252144.1;refGene_gn=RNF144A;refGene_tx=XM_023252145.1;refGene_gn=RNF144A;refGene_tx=XM_023252146.1;refGene_gn=RNF144A;refGene_tx=XM_023252147.1;refGene_gn=RNF144A;refGene_tx=XM_023252148.1
A3	138327415	138327476

refGene_gn=RNF144A;refGene_tx=XM_023252138.1;refGene_gn=RNF144A;refGen
e_tx=XM_023252139.1;refGene_gn=RNF144A;refGene_tx=XM_023252141.1;refGe
ne_gn=RNF144A;refGene_tx=XM_023252142.1;refGene_gn=RNF144A;refGene_tx=
XM_023252143.1;refGene_gn=RNF144A;refGene_tx=XM_023252144.1;refGene_gn
=RNF144A;refGene_tx=XM_023252145.1;refGene_gn=RNF144A;refGene_tx=XM_02
3252146.1;refGene_gn=RNF144A;refGene_tx=XM_023252147.1;refGene_gn=RNF1
44A;refGene_tx=XM_023252148.1
A3 138327634 138327739
refGene_gn=RNF144A;refGene_tx=XM_023252138.1;refGene_gn=RNF144A;refGen
e_tx=XM_023252139.1;refGene_gn=RNF144A;refGene_tx=XM_023252141.1;refGe
ne_gn=RNF144A;refGene_tx=XM_023252142.1;refGene_gn=RNF144A;refGene_tx=
XM_023252143.1;refGene_gn=RNF144A;refGene_tx=XM_023252144.1;refGene_gn
=RNF144A;refGene_tx=XM_023252145.1;refGene_gn=RNF144A;refGene_tx=XM_02
3252146.1;refGene_gn=RNF144A;refGene_tx=XM_023252147.1;refGene_gn=RNF1
44A;refGene_tx=XM_023252148.1
A3 138338745 138338891
refGene_gn=RNF144A;refGene_tx=XM_023252138.1;refGene_gn=RNF144A;refGen
e_tx=XM_023252139.1;refGene_gn=RNF144A;refGene_tx=XM_023252141.1;refGe
ne_gn=RNF144A;refGene_tx=XM_023252142.1;refGene_gn=RNF144A;refGene_tx=
XM_023252143.1;refGene_gn=RNF144A;refGene_tx=XM_023252144.1;refGene_gn
=RNF144A;refGene_tx=XM_023252145.1;refGene_gn=RNF144A;refGene_tx=XM_02
3252146.1;refGene_gn=RNF144A;refGene_tx=XM_023252147.1;refGene_gn=RNF1
44A;refGene_tx=XM_023252148.1
A3 138342217 138342351
refGene_gn=RNF144A;refGene_tx=XM_023252138.1;refGene_gn=RNF144A;refGen
e_tx=XM_023252139.1
A3 138343948 138343977
refGene_gn=RNF144A;refGene_tx=XM_023252139.1
A3 138355090 138355275
refGene_gn=RNF144A;refGene_tx=XM_023252138.1;refGene_gn=RNF144A;refGen
e_tx=XM_023252142.1
A3 138429614 138429779
refGene_gn=RSAD2;refGene_tx=XM_003984516.4;refGene_gn=RSAD2;refGene_tx
=XM_019827951.2
A3 138431884 138431917
refGene_gn=RSAD2;refGene_tx=XM_003984516.4;refGene_gn=RSAD2;refGene_tx
=XM_019827951.2
A3 138432730 138432880
refGene_gn=RSAD2;refGene_tx=XM_003984516.4;refGene_gn=RSAD2;refGene_tx
=XM_019827951.2
A3 138435499 138435729
refGene_gn=RSAD2;refGene_tx=XM_003984516.4;refGene_gn=RSAD2;refGene_tx
=XM_019827951.2
A3 138439428 138439590
refGene_gn=RSAD2;refGene_tx=XM_003984516.4
A3 138445394 138445746
refGene_gn=RSAD2;refGene_tx=XM_003984516.4;refGene_gn=RSAD2;refGene_tx
=XM_019827951.2
A3 138452916 138453576
refGene_gn=CMPK2;refGene_tx=XM_023252150.1;refGene_gn=CMPK2;refGene_tx

=XM_023252151.1
A3	138455192	138455307
refGene_gn=CMPK2;refGene_tx=XM_023252150.1;refGene_gn=CMPK2;refGene_tx=XM_023252151.1
A3	138456230	138456432
refGene_gn=CMPK2;refGene_tx=XM_023252150.1;refGene_gn=CMPK2;refGene_tx=XM_023252151.1
A3	138462187	138462421
refGene_gn=CMPK2;refGene_tx=XM_023252150.1;refGene_gn=CMPK2;refGene_tx=XM_023252151.1
A3	138463249	138463253
refGene_gn=CMPK2;refGene_tx=XM_023252151.1
A3	138463875	138463993
refGene_gn=CMPK2;refGene_tx=XM_023252150.1
A3	139386934	139388338
refGene_gn=SOX11;refGene_tx=XM_023252152.1
A3	140723456	140723480
refGene_gn=DCDC2C;refGene_tx=XM_023252154.1;refGene_gn=DCDC2C;refGene_tx=XM_023252156.1;refGene_gn=DCDC2C;refGene_tx=XM_023252157.1;refGene_gn=DCDC2C;refGene_tx=XM_023252160.1
A3	140736447	140736462
refGene_gn=DCDC2C;refGene_tx=XM_023252155.1;refGene_gn=DCDC2C;refGene_tx=XM_023252158.1
A3	140754559	140754601
refGene_gn=DCDC2C;refGene_tx=XM_023252154.1;refGene_gn=DCDC2C;refGene_tx=XM_023252155.1;refGene_gn=DCDC2C;refGene_tx=XM_023252156.1;refGene_gn=DCDC2C;refGene_tx=XM_023252160.1
A3	140756635	140756704
refGene_gn=DCDC2C;refGene_tx=XM_023252154.1;refGene_gn=DCDC2C;refGene_tx=XM_023252155.1;refGene_gn=DCDC2C;refGene_tx=XM_023252156.1;refGene_gn=DCDC2C;refGene_tx=XM_023252157.1;refGene_gn=DCDC2C;refGene_tx=XM_023252158.1;refGene_gn=DCDC2C;refGene_tx=XM_023252160.1
A3	140764004	140764105
refGene_gn=DCDC2C;refGene_tx=XM_023252154.1;refGene_gn=DCDC2C;refGene_tx=XM_023252155.1;refGene_gn=DCDC2C;refGene_tx=XM_023252156.1;refGene_gn=DCDC2C;refGene_tx=XM_023252157.1;refGene_gn=DCDC2C;refGene_tx=XM_023252158.1;refGene_gn=DCDC2C;refGene_tx=XM_023252160.1
A3	140766516	140766643
refGene_gn=DCDC2C;refGene_tx=XM_023252154.1;refGene_gn=DCDC2C;refGene_tx=XM_023252155.1;refGene_gn=DCDC2C;refGene_tx=XM_023252156.1;refGene_gn=DCDC2C;refGene_tx=XM_023252157.1;refGene_gn=DCDC2C;refGene_tx=XM_023252158.1;refGene_gn=DCDC2C;refGene_tx=XM_023252160.1
A3	140773016	140773059
refGene_gn=DCDC2C;refGene_tx=XM_023252154.1;refGene_gn=DCDC2C;refGene_tx=XM_023252155.1;refGene_gn=DCDC2C;refGene_tx=XM_023252156.1;refGene_gn=DCDC2C;refGene_tx=XM_023252157.1;refGene_gn=DCDC2C;refGene_tx=XM_023252158.1;refGene_gn=DCDC2C;refGene_tx=XM_023252160.1
A3	140776104	140776242
refGene_gn=DCDC2C;refGene_tx=XM_023252154.1;refGene_gn=DCDC2C;refGene_tx=XM_023252155.1;refGene_gn=DCDC2C;refGene_tx=XM_023252156.1;refGene_

gn=DCDC2C;refGene_tx=XM_023252157.1;refGene_gn=DCDC2C;refGene_tx=XM_023252158.1;refGene_gn=DCDC2C;refGene_tx=XM_023252160.1
A3 140780533 140780662
refGene_gn=DCDC2C;refGene_tx=XM_023252154.1;refGene_gn=DCDC2C;refGene_tx=XM_023252155.1;refGene_gn=DCDC2C;refGene_tx=XM_023252156.1;refGene_gn=DCDC2C;refGene_tx=XM_023252157.1;refGene_gn=DCDC2C;refGene_tx=XM_023252158.1
A3 140785577 140785654
refGene_gn=DCDC2C;refGene_tx=XM_023252154.1;refGene_gn=DCDC2C;refGene_tx=XM_023252155.1;refGene_gn=DCDC2C;refGene_tx=XM_023252156.1;refGene_gn=DCDC2C;refGene_tx=XM_023252157.1;refGene_gn=DCDC2C;refGene_tx=XM_023252158.1;refGene_gn=DCDC2C;refGene_tx=XM_023252160.1
A3 140797028 140797080
refGene_gn=DCDC2C;refGene_tx=XM_023252154.1;refGene_gn=DCDC2C;refGene_tx=XM_023252155.1;refGene_gn=DCDC2C;refGene_tx=XM_023252156.1;refGene_gn=DCDC2C;refGene_tx=XM_023252157.1;refGene_gn=DCDC2C;refGene_tx=XM_023252158.1;refGene_gn=DCDC2C;refGene_tx=XM_023252160.1
A3 140805127 140805414
refGene_gn=DCDC2C;refGene_tx=XM_023252154.1;refGene_gn=DCDC2C;refGene_tx=XM_023252155.1;refGene_gn=DCDC2C;refGene_tx=XM_023252156.1;refGene_gn=DCDC2C;refGene_tx=XM_023252157.1;refGene_gn=DCDC2C;refGene_tx=XM_023252158.1;refGene_gn=DCDC2C;refGene_tx=XM_023252160.1
A3 140806509 140806716
refGene_gn=ALLC;refGene_tx=XM_003984539.5;refGene_gn=ALLC;refGene_tx=XM_006930498.4;refGene_gn=ALLC;refGene_tx=XM_019827963.2
A3 140807408 140807533
refGene_gn=ALLC;refGene_tx=XM_003984539.5;refGene_gn=ALLC;refGene_tx=XM_006930498.4;refGene_gn=ALLC;refGene_tx=XM_019827963.2
A3 140809807 140809916
refGene_gn=ALLC;refGene_tx=XM_003984539.5;refGene_gn=ALLC;refGene_tx=XM_006930498.4;refGene_gn=ALLC;refGene_tx=XM_019827963.2
A3 140810897 140810971
refGene_gn=ALLC;refGene_tx=XM_003984539.5;refGene_gn=ALLC;refGene_tx=XM_006930498.4;refGene_gn=ALLC;refGene_tx=XM_019827963.2
A3 140811393 140811549
refGene_gn=ALLC;refGene_tx=XM_003984539.5;refGene_gn=ALLC;refGene_tx=XM_006930498.4;refGene_gn=ALLC;refGene_tx=XM_019827963.2
A3 140814306 140814439
refGene_gn=ALLC;refGene_tx=XM_003984539.5;refGene_gn=ALLC;refGene_tx=XM_006930498.4;refGene_gn=ALLC;refGene_tx=XM_019827963.2
A3 140815265 140815345
refGene_gn=ALLC;refGene_tx=XM_003984539.5;refGene_gn=ALLC;refGene_tx=XM_006930498.4;refGene_gn=ALLC;refGene_tx=XM_019827963.2
A3 140816426 140816552
refGene_gn=ALLC;refGene_tx=XM_003984539.5;refGene_gn=ALLC;refGene_tx=XM_006930498.4;refGene_gn=ALLC;refGene_tx=XM_019827963.2
A3 140817654 140817742
refGene_gn=ALLC;refGene_tx=XM_003984539.5;refGene_gn=ALLC;refGene_tx=XM_006930498.4;refGene_gn=ALLC;refGene_tx=XM_019827963.2
A3 140819165 140819216

refGene_gn=ALLC;refGene_tx=XM_003984539.5;refGene_gn=ALLC;refGene_tx=XM_006930498.4
A3 140821382 140821472
refGene_gn=ALLC;refGene_tx=XM_003984539.5;refGene_gn=ALLC;refGene_tx=XM_006930498.4
A3 140837912 140838322
refGene_gn=COLEC11;refGene_tx=XM_003984518.5;refGene_gn=COLEC11;refGene_tx=XM_006930502.4;refGene_gn=COLEC11;refGene_tx=XM_011281340.3;refGene_gn=COLEC11;refGene_tx=XM_011281341.3;refGene_gn=COLEC11;refGene_tx=XM_019827964.2;refGene_gn=COLEC11;refGene_tx=XM_019827965.2;refGene_gn=COLEC11;refGene_tx=XM_019827966.2
A3 140838464 140838560
refGene_gn=COLEC11;refGene_tx=XM_003984518.5;refGene_gn=COLEC11;refGene_tx=XM_006930502.4;refGene_gn=COLEC11;refGene_tx=XM_011281340.3;refGene_gn=COLEC11;refGene_tx=XM_011281341.3;refGene_gn=COLEC11;refGene_tx=XM_019827964.2;refGene_gn=COLEC11;refGene_tx=XM_019827965.2;refGene_gn=COLEC11;refGene_tx=XM_019827966.2
A3 140839649 140839703
refGene_gn=COLEC11;refGene_tx=XM_003984518.5;refGene_gn=COLEC11;refGene_tx=XM_006930502.4;refGene_gn=COLEC11;refGene_tx=XM_011281340.3;refGene_gn=COLEC11;refGene_tx=XM_011281341.3;refGene_gn=COLEC11;refGene_tx=XM_019827964.2;refGene_gn=COLEC11;refGene_tx=XM_019827965.2;refGene_gn=COLEC11;refGene_tx=XM_019827966.2
A3 140842488 140842560
refGene_gn=COLEC11;refGene_tx=XM_006930502.4;refGene_gn=COLEC11;refGene_tx=XM_011281340.3;refGene_gn=COLEC11;refGene_tx=XM_019827964.2;refGene_gn=COLEC11;refGene_tx=XM_019827965.2
A3 140854791 140854863
refGene_gn=COLEC11;refGene_tx=XM_003984518.5;refGene_gn=COLEC11;refGene_tx=XM_006930502.4;refGene_gn=COLEC11;refGene_tx=XM_011281340.3;refGene_gn=COLEC11;refGene_tx=XM_011281341.3;refGene_gn=COLEC11;refGene_tx=XM_019827964.2;refGene_gn=COLEC11;refGene_tx=XM_019827965.2;refGene_gn=COLEC11;refGene_tx=XM_019827966.2
A3 140856218 140856357
refGene_gn=COLEC11;refGene_tx=XM_019827964.2;refGene_gn=COLEC11;refGene_tx=XM_019827966.2
A3 140861948 140862066
refGene_gn=COLEC11;refGene_tx=XM_003984518.5;refGene_gn=COLEC11;refGene_tx=XM_006930502.4;refGene_gn=COLEC11;refGene_tx=XM_011281340.3;refGene_gn=COLEC11;refGene_tx=XM_011281341.3;refGene_gn=COLEC11;refGene_tx=XM_019827965.2
A3 140876575 140876653
refGene_gn=RPS7;refGene_tx=NM_001009832.1;refGene_gn=RPS7;refGene_tx=XM_023250908.1
A3 140877252 140877403
refGene_gn=RPS7;refGene_tx=NM_001009832.1;refGene_gn=RPS7;refGene_tx=XM_023250908.1
A3 140880081 140880146
refGene_gn=RPS7;refGene_tx=NM_001009832.1;refGene_gn=RPS7;refGene_tx=XM_023250908.1

A3 140880860 140881004
refGene_gn=RPS7;refGene_tx=NM_001009832.1;refGene_gn=RPS7;refGene_tx=XM_023250908.1
A3 140881730 140881802
refGene_gn=RPS7;refGene_tx=NM_001009832.1;refGene_gn=RPS7;refGene_tx=XM_023250908.1
A3 140882054 140882129
refGene_gn=RPS7;refGene_tx=NM_001009832.1;refGene_gn=RPS7;refGene_tx=XM_023250908.1
A3 140894675 140894806
refGene_gn=RNASEH1;refGene_tx=XM_011281342.3;refGene_gn=RNASEH1;refGene_tx=XM_011281343.3;refGene_gn=RNASEH1;refGene_tx=XM_019827968.2
A3 140895686 140895802
refGene_gn=RNASEH1;refGene_tx=XM_011281342.3;refGene_gn=RNASEH1;refGene_tx=XM_011281343.3;refGene_gn=RNASEH1;refGene_tx=XM_019827968.2
A3 140900890 140901049
refGene_gn=RNASEH1;refGene_tx=XM_011281342.3;refGene_gn=RNASEH1;refGene_tx=XM_011281343.3;refGene_gn=RNASEH1;refGene_tx=XM_019827968.2
A3 140902840 140902940
refGene_gn=RNASEH1;refGene_tx=XM_011281342.3;refGene_gn=RNASEH1;refGene_tx=XM_011281343.3;refGene_gn=RNASEH1;refGene_tx=XM_019827968.2
A3 140903788 140903843
refGene_gn=RNASEH1;refGene_tx=XM_011281342.3;refGene_gn=RNASEH1;refGene_tx=XM_011281343.3;refGene_gn=RNASEH1;refGene_tx=XM_019827968.2
A3 140904175 140904260
refGene_gn=RNASEH1;refGene_tx=XM_011281342.3;refGene_gn=RNASEH1;refGene_tx=XM_011281343.3;refGene_gn=RNASEH1;refGene_tx=XM_019827968.2
A3 140905329 140905454
refGene_gn=RNASEH1;refGene_tx=XM_011281342.3;refGene_gn=RNASEH1;refGene_tx=XM_011281343.3;refGene_gn=RNASEH1;refGene_tx=XM_019827968.2
A3 140906974 140907061
refGene_gn=RNASEH1;refGene_tx=XM_011281342.3;refGene_gn=RNASEH1;refGene_tx=XM_011281343.3;refGene_gn=RNASEH1;refGene_tx=XM_019827968.2
A3 140928607 140928727
refGene_gn=ADI1;refGene_tx=XM_023252162.1
A3 140932941 140933061
refGene_gn=ADI1;refGene_tx=XM_023252162.1
A3 140934094 140934227
refGene_gn=ADI1;refGene_tx=XM_023252162.1
A3 140934229 140934273
refGene_gn=ADI1;refGene_tx=XM_023252162.1
A3 140935507 140935627
refGene_gn=ADI1;refGene_tx=XM_023252162.1
A3 140961045 140961288
refGene_gn=TRAPPC12;refGene_tx=XM_003984542.4
A3 140961430 140961518
refGene_gn=TRAPPC12;refGene_tx=XM_003984542.4
A3 140961866 140961967
refGene_gn=TRAPPC12;refGene_tx=XM_003984542.4
A3 140975299 140975398

refGene_gn=TRAPPC12;refGene_tx=XM_003984542.4
A3 140980489 140980563
refGene_gn=TRAPPC12;refGene_tx=XM_003984542.4
A3 140982753 140982826
refGene_gn=TRAPPC12;refGene_tx=XM_003984542.4
A3 140997601 140997714
refGene_gn=TRAPPC12;refGene_tx=XM_003984542.4
A3 141012858 141012997
refGene_gn=TRAPPC12;refGene_tx=XM_003984542.4
A3 141013575 141013689
refGene_gn=TRAPPC12;refGene_tx=XM_003984542.4
A3 141025156 141025273
refGene_gn=TRAPPC12;refGene_tx=XM_003984542.4
A3 141035329 141036514
refGene_gn=TRAPPC12;refGene_tx=XM_003984542.4
A3 141047614 141047656
refGene_gn=EIPR1;refGene_tx=XM_011281345.3;refGene_gn=EIPR1;refGene_tx
=XM_011281346.3
A3 141061163 141061247
refGene_gn=EIPR1;refGene_tx=XM_011281345.3
A3 141073074 141073207
refGene_gn=EIPR1;refGene_tx=XM_011281345.3;refGene_gn=EIPR1;refGene_tx
=XM_011281346.3
A3 141100784 141100833
refGene_gn=EIPR1;refGene_tx=XM_019827970.2
A3 141133770 141133927
refGene_gn=EIPR1;refGene_tx=XM_011281345.3;refGene_gn=EIPR1;refGene_tx
=XM_011281346.3;refGene_gn=EIPR1;refGene_tx=XM_019827970.2;refGene_gn=
EIPR1;refGene_tx=XM_019827971.2
A3 141142645 141142680
refGene_gn=EIPR1;refGene_tx=XM_019827972.2
A3 141152874 141152974
refGene_gn=EIPR1;refGene_tx=XM_011281345.3;refGene_gn=EIPR1;refGene_tx
=XM_011281346.3;refGene_gn=EIPR1;refGene_tx=XM_019827970.2;refGene_gn=
EIPR1;refGene_tx=XM_019827971.2;refGene_gn=EIPR1;refGene_tx=XM_0198279
72.2
A3 141163946 141164083
refGene_gn=EIPR1;refGene_tx=XM_011281345.3;refGene_gn=EIPR1;refGene_tx
=XM_011281346.3;refGene_gn=EIPR1;refGene_tx=XM_019827970.2;refGene_gn=
EIPR1;refGene_tx=XM_019827971.2;refGene_gn=EIPR1;refGene_tx=XM_0198279
72.2
A3 141167278 141167446
refGene_gn=EIPR1;refGene_tx=XM_011281345.3;refGene_gn=EIPR1;refGene_tx
=XM_011281346.3;refGene_gn=EIPR1;refGene_tx=XM_019827970.2;refGene_gn=
EIPR1;refGene_tx=XM_019827971.2;refGene_gn=EIPR1;refGene_tx=XM_0198279
72.2
A3 141168717 141168885
refGene_gn=EIPR1;refGene_tx=XM_011281345.3;refGene_gn=EIPR1;refGene_tx
=XM_011281346.3;refGene_gn=EIPR1;refGene_tx=XM_019827970.2;refGene_gn=
EIPR1;refGene_tx=XM_019827971.2;refGene_gn=EIPR1;refGene_tx=XM_0198279

72.2
A3	141170892	141171067
refGene_gn=EIPR1;refGene_tx=XM_011281345.3;refGene_gn=EIPR1;refGene_tx=XM_011281346.3;refGene_gn=EIPR1;refGene_tx=XM_019827970.2;refGene_gn=EIPR1;refGene_tx=XM_019827971.2;refGene_gn=EIPR1;refGene_tx=XM_019827972.2
A3	142048821	142048876
refGene_gn=MYT1L;refGene_tx=XM_011281347.3;refGene_gn=MYT1L;refGene_tx=XM_019827976.2;refGene_gn=MYT1L;refGene_tx=XM_019827977.2;refGene_gn=MYT1L;refGene_tx=XM_019827978.2;refGene_gn=MYT1L;refGene_tx=XM_019827979.2;refGene_gn=MYT1L;refGene_tx=XM_019827980.2;refGene_gn=MYT1L;refGene_tx=XM_019827981.2;refGene_gn=MYT1L;refGene_tx=XM_019827982.2;refGene_gn=MYT1L;refGene_tx=XM_019827983.2;refGene_gn=MYT1L;refGene_tx=XM_019827984.2;refGene_gn=MYT1L;refGene_tx=XM_019827985.2;refGene_gn=MYT1L;refGene_tx=XM_019827986.2;refGene_gn=MYT1L;refGene_tx=XM_019827987.2;refGene_gn=MYT1L;refGene_tx=XM_019827988.2;refGene_gn=MYT1L;refGene_tx=XM_019827989.2;refGene_gn=MYT1L;refGene_tx=XM_019827990.2;refGene_gn=MYT1L;refGene_tx=XM_019827991.2;refGene_gn=MYT1L;refGene_tx=XM_019827992.2
A3	142049047	142049081
refGene_gn=MYT1L;refGene_tx=XM_011281347.3;refGene_gn=MYT1L;refGene_tx=XM_019827976.2;refGene_gn=MYT1L;refGene_tx=XM_019827977.2;refGene_gn=MYT1L;refGene_tx=XM_019827978.2;refGene_gn=MYT1L;refGene_tx=XM_019827979.2;refGene_gn=MYT1L;refGene_tx=XM_019827980.2;refGene_gn=MYT1L;refGene_tx=XM_019827981.2;refGene_gn=MYT1L;refGene_tx=XM_019827982.2;refGene_gn=MYT1L;refGene_tx=XM_019827983.2;refGene_gn=MYT1L;refGene_tx=XM_019827984.2;refGene_gn=MYT1L;refGene_tx=XM_019827985.2;refGene_gn=MYT1L;refGene_tx=XM_019827986.2;refGene_gn=MYT1L;refGene_tx=XM_019827987.2;refGene_gn=MYT1L;refGene_tx=XM_019827988.2;refGene_gn=MYT1L;refGene_tx=XM_019827989.2;refGene_gn=MYT1L;refGene_tx=XM_019827990.2;refGene_gn=MYT1L;refGene_tx=XM_019827991.2;refGene_gn=MYT1L;refGene_tx=XM_019827992.2
A3	142049375	142049438
refGene_gn=MYT1L;refGene_tx=XM_011281347.3;refGene_gn=MYT1L;refGene_tx=XM_019827976.2;refGene_gn=MYT1L;refGene_tx=XM_019827977.2;refGene_gn=MYT1L;refGene_tx=XM_019827978.2;refGene_gn=MYT1L;refGene_tx=XM_019827979.2;refGene_gn=MYT1L;refGene_tx=XM_019827980.2;refGene_gn=MYT1L;refGene_tx=XM_019827981.2;refGene_gn=MYT1L;refGene_tx=XM_019827982.2;refGene_gn=MYT1L;refGene_tx=XM_019827983.2;refGene_gn=MYT1L;refGene_tx=XM_019827984.2;refGene_gn=MYT1L;refGene_tx=XM_019827985.2;refGene_gn=MYT1L;refGene_tx=XM_019827986.2;refGene_gn=MYT1L;refGene_tx=XM_019827987.2;refGene_gn=MYT1L;refGene_tx=XM_019827988.2;refGene_gn=MYT1L;refGene_tx=XM_019827989.2;refGene_gn=MYT1L;refGene_tx=XM_019827990.2;refGene_gn=MYT1L;refGene_tx=XM_019827991.2;refGene_gn=MYT1L;refGene_tx=XM_019827992.2
A3	142080160	142080513
refGene_gn=MYT1L;refGene_tx=XM_011281347.3;refGene_gn=MYT1L;refGene_tx=XM_019827976.2;refGene_gn=MYT1L;refGene_tx=XM_019827977.2;refGene_gn=MYT1L;refGene_tx=XM_019827978.2;refGene_gn=MYT1L;refGene_tx=XM_019827979.2;refGene_gn=MYT1L;refGene_tx=XM_019827980.2;refGene_gn=MYT1L;refGe

ne_tx=XM_019827981.2;refGene_gn=MYT1L;refGene_tx=XM_019827982.2;refGen
e_gn=MYT1L;refGene_tx=XM_019827983.2;refGene_gn=MYT1L;refGene_tx=XM_01
9827984.2;refGene_gn=MYT1L;refGene_tx=XM_019827985.2;refGene_gn=MYT1L;
refGene_tx=XM_019827986.2;refGene_gn=MYT1L;refGene_tx=XM_019827987.2;r
efGene_gn=MYT1L;refGene_tx=XM_019827988.2;refGene_gn=MYT1L;refGene_tx=
XM_019827989.2;refGene_gn=MYT1L;refGene_tx=XM_019827990.2;refGene_gn=M
YT1L;refGene_tx=XM_019827991.2;refGene_gn=MYT1L;refGene_tx=XM_01982799
2.2
A3 142097414 142098392
refGene_gn=MYT1L;refGene_tx=XM_011281347.3;refGene_gn=MYT1L;refGene_tx
=XM_019827976.2;refGene_gn=MYT1L;refGene_tx=XM_019827977.2;refGene_gn=
MYT1L;refGene_tx=XM_019827978.2;refGene_gn=MYT1L;refGene_tx=XM_0198279
79.2;refGene_gn=MYT1L;refGene_tx=XM_019827980.2;refGene_gn=MYT1L;refGe
ne_tx=XM_019827981.2;refGene_gn=MYT1L;refGene_tx=XM_019827982.2;refGen
e_gn=MYT1L;refGene_tx=XM_019827983.2;refGene_gn=MYT1L;refGene_tx=XM_01
9827984.2;refGene_gn=MYT1L;refGene_tx=XM_019827985.2;refGene_gn=MYT1L;
refGene_tx=XM_019827986.2;refGene_gn=MYT1L;refGene_tx=XM_019827987.2;r
efGene_gn=MYT1L;refGene_tx=XM_019827988.2;refGene_gn=MYT1L;refGene_tx=
XM_019827989.2;refGene_gn=MYT1L;refGene_tx=XM_019827990.2;refGene_gn=M
YT1L;refGene_tx=XM_019827991.2;refGene_gn=MYT1L;refGene_tx=XM_01982799
2.2
A3 142102749 142102884
refGene_gn=MYT1L;refGene_tx=XM_011281347.3;refGene_gn=MYT1L;refGene_tx
=XM_019827976.2;refGene_gn=MYT1L;refGene_tx=XM_019827977.2;refGene_gn=
MYT1L;refGene_tx=XM_019827978.2;refGene_gn=MYT1L;refGene_tx=XM_0198279
79.2;refGene_gn=MYT1L;refGene_tx=XM_019827980.2;refGene_gn=MYT1L;refGe
ne_tx=XM_019827981.2;refGene_gn=MYT1L;refGene_tx=XM_019827982.2;refGen
e_gn=MYT1L;refGene_tx=XM_019827983.2;refGene_gn=MYT1L;refGene_tx=XM_01
9827984.2;refGene_gn=MYT1L;refGene_tx=XM_019827985.2;refGene_gn=MYT1L;
refGene_tx=XM_019827986.2;refGene_gn=MYT1L;refGene_tx=XM_019827987.2;r
efGene_gn=MYT1L;refGene_tx=XM_019827988.2;refGene_gn=MYT1L;refGene_tx=
XM_019827989.2;refGene_gn=MYT1L;refGene_tx=XM_019827990.2;refGene_gn=M
YT1L;refGene_tx=XM_019827991.2;refGene_gn=MYT1L;refGene_tx=XM_01982799
2.2
A3 142107806 142107897
refGene_gn=MYT1L;refGene_tx=XM_011281347.3;refGene_gn=MYT1L;refGene_tx
=XM_019827976.2;refGene_gn=MYT1L;refGene_tx=XM_019827977.2;refGene_gn=
MYT1L;refGene_tx=XM_019827978.2;refGene_gn=MYT1L;refGene_tx=XM_0198279
79.2;refGene_gn=MYT1L;refGene_tx=XM_019827980.2;refGene_gn=MYT1L;refGe
ne_tx=XM_019827981.2;refGene_gn=MYT1L;refGene_tx=XM_019827982.2;refGen
e_gn=MYT1L;refGene_tx=XM_019827983.2;refGene_gn=MYT1L;refGene_tx=XM_01
9827984.2;refGene_gn=MYT1L;refGene_tx=XM_019827985.2;refGene_gn=MYT1L;
refGene_tx=XM_019827986.2;refGene_gn=MYT1L;refGene_tx=XM_019827987.2;r
efGene_gn=MYT1L;refGene_tx=XM_019827988.2;refGene_gn=MYT1L;refGene_tx=
XM_019827989.2;refGene_gn=MYT1L;refGene_tx=XM_019827990.2;refGene_gn=M
YT1L;refGene_tx=XM_019827991.2;refGene_gn=MYT1L;refGene_tx=XM_01982799
2.2
A3 142109529 142109637
refGene_gn=MYT1L;refGene_tx=XM_011281347.3;refGene_gn=MYT1L;refGene_tx
=XM_019827976.2;refGene_gn=MYT1L;refGene_tx=XM_019827977.2;refGene_gn=

MYT1L;refGene_tx=XM_019827978.2;refGene_gn=MYT1L;refGene_tx=XM_0198279
79.2;refGene_gn=MYT1L;refGene_tx=XM_019827980.2;refGene_gn=MYT1L;refGe
ne_tx=XM_019827981.2;refGene_gn=MYT1L;refGene_tx=XM_019827982.2;refGen
e_gn=MYT1L;refGene_tx=XM_019827983.2;refGene_gn=MYT1L;refGene_tx=XM_01
9827984.2;refGene_gn=MYT1L;refGene_tx=XM_019827985.2;refGene_gn=MYT1L;
refGene_tx=XM_019827986.2;refGene_gn=MYT1L;refGene_tx=XM_019827987.2;r
efGene_gn=MYT1L;refGene_tx=XM_019827988.2;refGene_gn=MYT1L;refGene_tx=
XM_019827989.2;refGene_gn=MYT1L;refGene_tx=XM_019827990.2;refGene_gn=M
YT1L;refGene_tx=XM_019827991.2;refGene_gn=MYT1L;refGene_tx=XM_01982799
2.2
A3 142113611 142113823
refGene_gn=MYT1L;refGene_tx=XM_011281347.3;refGene_gn=MYT1L;refGene_tx
=XM_019827976.2;refGene_gn=MYT1L;refGene_tx=XM_019827977.2;refGene_gn=
MYT1L;refGene_tx=XM_019827978.2;refGene_gn=MYT1L;refGene_tx=XM_0198279
79.2;refGene_gn=MYT1L;refGene_tx=XM_019827980.2;refGene_gn=MYT1L;refGe
ne_tx=XM_019827981.2;refGene_gn=MYT1L;refGene_tx=XM_019827982.2;refGen
e_gn=MYT1L;refGene_tx=XM_019827983.2;refGene_gn=MYT1L;refGene_tx=XM_01
9827984.2;refGene_gn=MYT1L;refGene_tx=XM_019827985.2;refGene_gn=MYT1L;
refGene_tx=XM_019827986.2;refGene_gn=MYT1L;refGene_tx=XM_019827987.2;r
efGene_gn=MYT1L;refGene_tx=XM_019827988.2;refGene_gn=MYT1L;refGene_tx=
XM_019827989.2;refGene_gn=MYT1L;refGene_tx=XM_019827990.2;refGene_gn=M
YT1L;refGene_tx=XM_019827991.2;refGene_gn=MYT1L;refGene_tx=XM_01982799
2.2
A3 142120981 142121232
refGene_gn=MYT1L;refGene_tx=XM_011281347.3;refGene_gn=MYT1L;refGene_tx
=XM_019827976.2;refGene_gn=MYT1L;refGene_tx=XM_019827977.2;refGene_gn=
MYT1L;refGene_tx=XM_019827978.2;refGene_gn=MYT1L;refGene_tx=XM_0198279
79.2;refGene_gn=MYT1L;refGene_tx=XM_019827980.2;refGene_gn=MYT1L;refGe
ne_tx=XM_019827981.2;refGene_gn=MYT1L;refGene_tx=XM_019827982.2;refGen
e_gn=MYT1L;refGene_tx=XM_019827983.2;refGene_gn=MYT1L;refGene_tx=XM_01
9827984.2;refGene_gn=MYT1L;refGene_tx=XM_019827985.2;refGene_gn=MYT1L;
refGene_tx=XM_019827986.2;refGene_gn=MYT1L;refGene_tx=XM_019827987.2;r
efGene_gn=MYT1L;refGene_tx=XM_019827988.2;refGene_gn=MYT1L;refGene_tx=
XM_019827989.2;refGene_gn=MYT1L;refGene_tx=XM_019827990.2;refGene_gn=M
YT1L;refGene_tx=XM_019827991.2;refGene_gn=MYT1L;refGene_tx=XM_01982799
2.2
A3 142123526 142123763
refGene_gn=MYT1L;refGene_tx=XM_011281347.3;refGene_gn=MYT1L;refGene_tx
=XM_019827976.2;refGene_gn=MYT1L;refGene_tx=XM_019827977.2;refGene_gn=
MYT1L;refGene_tx=XM_019827978.2;refGene_gn=MYT1L;refGene_tx=XM_0198279
79.2;refGene_gn=MYT1L;refGene_tx=XM_019827980.2;refGene_gn=MYT1L;refGe
ne_tx=XM_019827981.2;refGene_gn=MYT1L;refGene_tx=XM_019827982.2;refGen
e_gn=MYT1L;refGene_tx=XM_019827983.2;refGene_gn=MYT1L;refGene_tx=XM_01
9827984.2;refGene_gn=MYT1L;refGene_tx=XM_019827985.2;refGene_gn=MYT1L;
refGene_tx=XM_019827986.2;refGene_gn=MYT1L;refGene_tx=XM_019827987.2;r
efGene_gn=MYT1L;refGene_tx=XM_019827988.2;refGene_gn=MYT1L;refGene_tx=
XM_019827989.2;refGene_gn=MYT1L;refGene_tx=XM_019827990.2;refGene_gn=M
YT1L;refGene_tx=XM_019827991.2;refGene_gn=MYT1L;refGene_tx=XM_01982799
2.2
A3 142125977 142126099

refGene_gn=MYT1L;refGene_tx=XM_011281347.3;refGene_gn=MYT1L;refGene_tx
=XM_019827976.2;refGene_gn=MYT1L;refGene_tx=XM_019827977.2;refGene_gn=
MYT1L;refGene_tx=XM_019827978.2;refGene_gn=MYT1L;refGene_tx=XM_0198279
79.2;refGene_gn=MYT1L;refGene_tx=XM_019827980.2;refGene_gn=MYT1L;refGe
ne_tx=XM_019827981.2;refGene_gn=MYT1L;refGene_tx=XM_019827982.2;refGen
e_gn=MYT1L;refGene_tx=XM_019827983.2;refGene_gn=MYT1L;refGene_tx=XM_01
9827984.2;refGene_gn=MYT1L;refGene_tx=XM_019827985.2;refGene_gn=MYT1L;
refGene_tx=XM_019827986.2;refGene_gn=MYT1L;refGene_tx=XM_019827987.2;r
efGene_gn=MYT1L;refGene_tx=XM_019827988.2;refGene_gn=MYT1L;refGene_tx=
XM_019827989.2;refGene_gn=MYT1L;refGene_tx=XM_019827990.2;refGene_gn=M
YT1L;refGene_tx=XM_019827991.2;refGene_gn=MYT1L;refGene_tx=XM_01982799
2.2
A3 142126460 142126523
refGene_gn=MYT1L;refGene_tx=XM_019827976.2;refGene_gn=MYT1L;refGene_tx
=XM_019827977.2;refGene_gn=MYT1L;refGene_tx=XM_019827978.2;refGene_gn=
MYT1L;refGene_tx=XM_019827979.2;refGene_gn=MYT1L;refGene_tx=XM_0198279
80.2;refGene_gn=MYT1L;refGene_tx=XM_019827981.2;refGene_gn=MYT1L;refGe
ne_tx=XM_019827982.2;refGene_gn=MYT1L;refGene_tx=XM_019827983.2;refGen
e_gn=MYT1L;refGene_tx=XM_019827984.2;refGene_gn=MYT1L;refGene_tx=XM_01
9827985.2;refGene_gn=MYT1L;refGene_tx=XM_019827989.2;refGene_gn=MYT1L;
refGene_tx=XM_019827990.2
A3 142126986 142127055
refGene_gn=MYT1L;refGene_tx=XM_011281347.3;refGene_gn=MYT1L;refGene_tx
=XM_019827976.2;refGene_gn=MYT1L;refGene_tx=XM_019827977.2;refGene_gn=
MYT1L;refGene_tx=XM_019827978.2;refGene_gn=MYT1L;refGene_tx=XM_0198279
79.2;refGene_gn=MYT1L;refGene_tx=XM_019827980.2;refGene_gn=MYT1L;refGe
ne_tx=XM_019827981.2;refGene_gn=MYT1L;refGene_tx=XM_019827982.2;refGen
e_gn=MYT1L;refGene_tx=XM_019827983.2;refGene_gn=MYT1L;refGene_tx=XM_01
9827984.2;refGene_gn=MYT1L;refGene_tx=XM_019827985.2;refGene_gn=MYT1L;
refGene_tx=XM_019827986.2;refGene_gn=MYT1L;refGene_tx=XM_019827987.2;r
efGene_gn=MYT1L;refGene_tx=XM_019827988.2;refGene_gn=MYT1L;refGene_tx=
XM_019827989.2;refGene_gn=MYT1L;refGene_tx=XM_019827990.2;refGene_gn=M
YT1L;refGene_tx=XM_019827991.2;refGene_gn=MYT1L;refGene_tx=XM_01982799
2.2
A3 142129821 142130011
refGene_gn=MYT1L;refGene_tx=XM_019827989.2;refGene_gn=MYT1L;refGene_tx
=XM_019827990.2;refGene_gn=MYT1L;refGene_tx=XM_019827991.2;refGene_gn=
MYT1L;refGene_tx=XM_019827992.2
A3 142136529 142137528
refGene_gn=LOC109498475;refGene_tx=XM_019827994.2
A3 142155853 142155916
refGene_gn=MYT1L;refGene_tx=XM_011281347.3;refGene_gn=MYT1L;refGene_tx
=XM_019827976.2;refGene_gn=MYT1L;refGene_tx=XM_019827977.2;refGene_gn=
MYT1L;refGene_tx=XM_019827978.2;refGene_gn=MYT1L;refGene_tx=XM_0198279
79.2;refGene_gn=MYT1L;refGene_tx=XM_019827980.2;refGene_gn=MYT1L;refGe
ne_tx=XM_019827981.2;refGene_gn=MYT1L;refGene_tx=XM_019827982.2;refGen
e_gn=MYT1L;refGene_tx=XM_019827983.2;refGene_gn=MYT1L;refGene_tx=XM_01
9827984.2;refGene_gn=MYT1L;refGene_tx=XM_019827985.2;refGene_gn=MYT1L;
refGene_tx=XM_019827986.2;refGene_gn=MYT1L;refGene_tx=XM_019827987.2;r
efGene_gn=MYT1L;refGene_tx=XM_019827988.2

A3	142161439	142161523
refGene_gn=MYT1L;refGene_tx=XM_011281347.3;refGene_gn=MYT1L;refGene_tx=XM_019827976.2;refGene_gn=MYT1L;refGene_tx=XM_019827977.2;refGene_gn=MYT1L;refGene_tx=XM_019827978.2;refGene_gn=MYT1L;refGene_tx=XM_019827979.2;refGene_gn=MYT1L;refGene_tx=XM_019827980.2;refGene_gn=MYT1L;refGene_tx=XM_019827981.2;refGene_gn=MYT1L;refGene_tx=XM_019827982.2;refGene_gn=MYT1L;refGene_tx=XM_019827983.2;refGene_gn=MYT1L;refGene_tx=XM_019827984.2;refGene_gn=MYT1L;refGene_tx=XM_019827985.2;refGene_gn=MYT1L;refGene_tx=XM_019827986.2;refGene_gn=MYT1L;refGene_tx=XM_019827987.2;refGene_gn=MYT1L;refGene_tx=XM_019827988.2
A3	142162873	142163095
refGene_gn=MYT1L;refGene_tx=XM_011281347.3;refGene_gn=MYT1L;refGene_tx=XM_019827976.2;refGene_gn=MYT1L;refGene_tx=XM_019827977.2;refGene_gn=MYT1L;refGene_tx=XM_019827978.2;refGene_gn=MYT1L;refGene_tx=XM_019827979.2;refGene_gn=MYT1L;refGene_tx=XM_019827980.2;refGene_gn=MYT1L;refGene_tx=XM_019827981.2;refGene_gn=MYT1L;refGene_tx=XM_019827982.2;refGene_gn=MYT1L;refGene_tx=XM_019827983.2;refGene_gn=MYT1L;refGene_tx=XM_019827984.2;refGene_gn=MYT1L;refGene_tx=XM_019827985.2;refGene_gn=MYT1L;refGene_tx=XM_019827986.2;refGene_gn=MYT1L;refGene_tx=XM_019827987.2;refGene_gn=MYT1L;refGene_tx=XM_019827988.2
A3	142176716	142176808
refGene_gn=MYT1L;refGene_tx=XM_011281347.3;refGene_gn=MYT1L;refGene_tx=XM_019827976.2;refGene_gn=MYT1L;refGene_tx=XM_019827977.2;refGene_gn=MYT1L;refGene_tx=XM_019827978.2;refGene_gn=MYT1L;refGene_tx=XM_019827979.2;refGene_gn=MYT1L;refGene_tx=XM_019827980.2;refGene_gn=MYT1L;refGene_tx=XM_019827981.2;refGene_gn=MYT1L;refGene_tx=XM_019827982.2;refGene_gn=MYT1L;refGene_tx=XM_019827983.2;refGene_gn=MYT1L;refGene_tx=XM_019827984.2;refGene_gn=MYT1L;refGene_tx=XM_019827985.2;refGene_gn=MYT1L;refGene_tx=XM_019827986.2;refGene_gn=MYT1L;refGene_tx=XM_019827987.2;refGene_gn=MYT1L;refGene_tx=XM_019827988.2
A3	142183654	142183764
refGene_gn=MYT1L;refGene_tx=XM_011281347.3;refGene_gn=MYT1L;refGene_tx=XM_019827976.2;refGene_gn=MYT1L;refGene_tx=XM_019827977.2;refGene_gn=MYT1L;refGene_tx=XM_019827978.2;refGene_gn=MYT1L;refGene_tx=XM_019827979.2;refGene_gn=MYT1L;refGene_tx=XM_019827980.2;refGene_gn=MYT1L;refGene_tx=XM_019827981.2;refGene_gn=MYT1L;refGene_tx=XM_019827982.2;refGene_gn=MYT1L;refGene_tx=XM_019827983.2;refGene_gn=MYT1L;refGene_tx=XM_019827984.2;refGene_gn=MYT1L;refGene_tx=XM_019827985.2;refGene_gn=MYT1L;refGene_tx=XM_019827986.2;refGene_gn=MYT1L;refGene_tx=XM_019827987.2;refGene_gn=MYT1L;refGene_tx=XM_019827988.2
A3	142188372	142188516
refGene_gn=MYT1L;refGene_tx=XM_011281347.3;refGene_gn=MYT1L;refGene_tx=XM_019827976.2;refGene_gn=MYT1L;refGene_tx=XM_019827977.2;refGene_gn=MYT1L;refGene_tx=XM_019827978.2;refGene_gn=MYT1L;refGene_tx=XM_019827979.2;refGene_gn=MYT1L;refGene_tx=XM_019827980.2;refGene_gn=MYT1L;refGene_tx=XM_019827981.2;refGene_gn=MYT1L;refGene_tx=XM_019827982.2;refGene_gn=MYT1L;refGene_tx=XM_019827983.2;refGene_gn=MYT1L;refGene_tx=XM_019827984.2;refGene_gn=MYT1L;refGene_tx=XM_019827985.2;refGene_gn=MYT1L;refGene_tx=XM_019827986.2;refGene_gn=MYT1L;refGene_tx=XM_019827987.2;refGene_gn=MYT1L;refGene_tx=XM_019827988.2

A3	142188810	142188951
refGene_gn=MYT1L;refGene_tx=XM_011281347.3;refGene_gn=MYT1L;refGene_tx=XM_019827976.2;refGene_gn=MYT1L;refGene_tx=XM_019827977.2;refGene_gn=MYT1L;refGene_tx=XM_019827978.2;refGene_gn=MYT1L;refGene_tx=XM_019827979.2;refGene_gn=MYT1L;refGene_tx=XM_019827980.2;refGene_gn=MYT1L;refGene_tx=XM_019827981.2;refGene_gn=MYT1L;refGene_tx=XM_019827982.2;refGene_gn=MYT1L;refGene_tx=XM_019827983.2;refGene_gn=MYT1L;refGene_tx=XM_019827984.2;refGene_gn=MYT1L;refGene_tx=XM_019827985.2;refGene_gn=MYT1L;refGene_tx=XM_019827986.2;refGene_gn=MYT1L;refGene_tx=XM_019827987.2;refGene_gn=MYT1L;refGene_tx=XM_019827988.2
A3	142207801	142207804
refGene_gn=PXDN;refGene_tx=XM_006930507.3
A3	142208421	142208525
refGene_gn=PXDN;refGene_tx=XM_006930507.3
A3	142237479	142237880
refGene_gn=PXDN;refGene_tx=XM_023252163.1;refGene_gn=PXDN;refGene_tx=XM_023252164.1
A3	142244451	142244468
refGene_gn=PXDN;refGene_tx=XM_019827995.2
A3	142250385	142250457
refGene_gn=PXDN;refGene_tx=XM_006930507.3;refGene_gn=PXDN;refGene_tx=XM_019827995.2;refGene_gn=PXDN;refGene_tx=XM_023252163.1;refGene_gn=PXDN;refGene_tx=XM_023252164.1
A3	142251234	142251306
refGene_gn=PXDN;refGene_tx=XM_006930507.3;refGene_gn=PXDN;refGene_tx=XM_019827995.2;refGene_gn=PXDN;refGene_tx=XM_023252163.1;refGene_gn=PXDN;refGene_tx=XM_023252164.1
A3	142255139	142255211
refGene_gn=PXDN;refGene_tx=XM_006930507.3;refGene_gn=PXDN;refGene_tx=XM_019827995.2;refGene_gn=PXDN;refGene_tx=XM_023252163.1
A3	142258081	142258153
refGene_gn=PXDN;refGene_tx=XM_006930507.3;refGene_gn=PXDN;refGene_tx=XM_019827995.2;refGene_gn=PXDN;refGene_tx=XM_023252163.1;refGene_gn=PXDN;refGene_tx=XM_023252164.1
A3	142258516	142258588
refGene_gn=PXDN;refGene_tx=XM_006930507.3;refGene_gn=PXDN;refGene_tx=XM_019827995.2;refGene_gn=PXDN;refGene_tx=XM_023252163.1;refGene_gn=PXDN;refGene_tx=XM_023252164.1
A3	142259472	142259642
refGene_gn=PXDN;refGene_tx=XM_006930507.3;refGene_gn=PXDN;refGene_tx=XM_019827995.2;refGene_gn=PXDN;refGene_tx=XM_023252163.1;refGene_gn=PXDN;refGene_tx=XM_023252164.1
A3	142261054	142261172
refGene_gn=PXDN;refGene_tx=XM_006930507.3;refGene_gn=PXDN;refGene_tx=XM_019827995.2;refGene_gn=PXDN;refGene_tx=XM_023252163.1;refGene_gn=PXDN;refGene_tx=XM_023252164.1
A3	142263106	142263276
refGene_gn=PXDN;refGene_tx=XM_006930507.3;refGene_gn=PXDN;refGene_tx=XM_019827995.2;refGene_gn=PXDN;refGene_tx=XM_023252163.1;refGene_gn=PXDN;refGene_tx=XM_023252164.1

A3	142266244	142266517
refGene_gn=PXDN;refGene_tx=XM_006930507.3;refGene_gn=PXDN;refGene_tx=XM_019827995.2;refGene_gn=PXDN;refGene_tx=XM_023252163.1;refGene_gn=PXDN;refGene_tx=XM_023252164.1
A3	142267720	142267837
refGene_gn=PXDN;refGene_tx=XM_006930507.3;refGene_gn=PXDN;refGene_tx=XM_019827995.2;refGene_gn=PXDN;refGene_tx=XM_023252163.1;refGene_gn=PXDN;refGene_tx=XM_023252164.1
A3	142268867	142269026
refGene_gn=PXDN;refGene_tx=XM_006930507.3;refGene_gn=PXDN;refGene_tx=XM_019827995.2;refGene_gn=PXDN;refGene_tx=XM_023252163.1;refGene_gn=PXDN;refGene_tx=XM_023252164.1
A3	142270056	142270169
refGene_gn=PXDN;refGene_tx=XM_006930507.3;refGene_gn=PXDN;refGene_tx=XM_019827995.2;refGene_gn=PXDN;refGene_tx=XM_023252163.1;refGene_gn=PXDN;refGene_tx=XM_023252164.1
A3	142270900	142271057
refGene_gn=PXDN;refGene_tx=XM_006930507.3;refGene_gn=PXDN;refGene_tx=XM_019827995.2;refGene_gn=PXDN;refGene_tx=XM_023252163.1;refGene_gn=PXDN;refGene_tx=XM_023252164.1
A3	142274700	142274809
refGene_gn=PXDN;refGene_tx=XM_006930507.3;refGene_gn=PXDN;refGene_tx=XM_019827995.2;refGene_gn=PXDN;refGene_tx=XM_023252163.1;refGene_gn=PXDN;refGene_tx=XM_023252164.1
A3	142275351	142275509
refGene_gn=PXDN;refGene_tx=XM_006930507.3;refGene_gn=PXDN;refGene_tx=XM_019827995.2;refGene_gn=PXDN;refGene_tx=XM_023252163.1;refGene_gn=PXDN;refGene_tx=XM_023252164.1
A3	142279115	142280619
refGene_gn=PXDN;refGene_tx=XM_006930507.3;refGene_gn=PXDN;refGene_tx=XM_019827995.2;refGene_gn=PXDN;refGene_tx=XM_023252163.1;refGene_gn=PXDN;refGene_tx=XM_023252164.1
A3	142282934	142283069
refGene_gn=PXDN;refGene_tx=XM_006930507.3;refGene_gn=PXDN;refGene_tx=XM_019827995.2;refGene_gn=PXDN;refGene_tx=XM_023252163.1;refGene_gn=PXDN;refGene_tx=XM_023252164.1
A3	142283686	142283895
refGene_gn=PXDN;refGene_tx=XM_006930507.3;refGene_gn=PXDN;refGene_tx=XM_019827995.2;refGene_gn=PXDN;refGene_tx=XM_023252163.1;refGene_gn=PXDN;refGene_tx=XM_023252164.1
A3	142287691	142287812
refGene_gn=PXDN;refGene_tx=XM_006930507.3;refGene_gn=PXDN;refGene_tx=XM_019827995.2;refGene_gn=PXDN;refGene_tx=XM_023252163.1;refGene_gn=PXDN;refGene_tx=XM_023252164.1
A3	142287914	142288047
refGene_gn=PXDN;refGene_tx=XM_006930507.3;refGene_gn=PXDN;refGene_tx=XM_019827995.2;refGene_gn=PXDN;refGene_tx=XM_023252163.1;refGene_gn=PXDN;refGene_tx=XM_023252164.1
A3	142290316	142290430
refGene_gn=PXDN;refGene_tx=XM_006930507.3;refGene_gn=PXDN;refGene_tx=X

M_019827995.2;refGene_gn=PXDN;refGene_tx=XM_023252163.1;refGene_gn=PXDN;refGene_tx=XM_023252164.1
A3 142291084 142291204
refGene_gn=PXDN;refGene_tx=XM_006930507.3;refGene_gn=PXDN;refGene_tx=XM_019827995.2;refGene_gn=PXDN;refGene_tx=XM_023252163.1;refGene_gn=PXDN;refGene_tx=XM_023252164.1
A3 142316145 142316184
refGene_gn=TPO;refGene_tx=XM_006930524.4
A3 142317119 142317243
refGene_gn=TPO;refGene_tx=XM_006930524.4
A3 142322362 142322462
refGene_gn=TPO;refGene_tx=XM_006930524.4
A3 142326087 142326219
refGene_gn=TPO;refGene_tx=XM_006930524.4
A3 142329509 142329680
refGene_gn=TPO;refGene_tx=XM_006930524.4
A3 142330317 142330526
refGene_gn=TPO;refGene_tx=XM_006930524.4
A3 142330918 142331156
refGene_gn=TPO;refGene_tx=XM_006930524.4
A3 142332217 142332388
refGene_gn=TPO;refGene_tx=XM_006930524.4
A3 142332979 142333238
refGene_gn=TPO;refGene_tx=XM_006930524.4
A3 142335320 142335839
refGene_gn=TPO;refGene_tx=XM_006930524.4
A3 142336901 142337108
refGene_gn=TPO;refGene_tx=XM_006930524.4
A3 142338731 142338861
refGene_gn=TPO;refGene_tx=XM_006930524.4
A3 142345439 142345572
refGene_gn=TPO;refGene_tx=XM_006930524.4
A3 142347595 142347765
refGene_gn=TPO;refGene_tx=XM_006930524.4
A3 142355245 142355330
refGene_gn=TPO;refGene_tx=XM_006930524.4
A3 142360020 142360150
refGene_gn=TPO;refGene_tx=XM_006930524.4
A3 142382858 142382993
refGene_gn=SNTG2;refGene_tx=XM_003984546.5;refGene_gn=SNTG2;refGene_tx=XM_023252165.1
A3 142398680 142398791
refGene_gn=SNTG2;refGene_tx=XM_003984546.5;refGene_gn=SNTG2;refGene_tx=XM_023252165.1
A3 142400872 142400965
refGene_gn=SNTG2;refGene_tx=XM_003984546.5;refGene_gn=SNTG2;refGene_tx=XM_023252165.1
A3 142410913 142411114
refGene_gn=SNTG2;refGene_tx=XM_003984546.5;refGene_gn=SNTG2;refGene_tx=XM_023252165.1

A3 142419688 142419760
refGene_gn=SNTG2;refGene_tx=XM_003984546.5;refGene_gn=SNTG2;refGene_tx=XM_023252165.1
A3 142427645 142427762
refGene_gn=SNTG2;refGene_tx=XM_003984546.5;refGene_gn=SNTG2;refGene_tx=XM_023252165.1
A3 142432288 142432327
refGene_gn=SNTG2;refGene_tx=XM_003984546.5;refGene_gn=SNTG2;refGene_tx=XM_023252165.1
A3 142433974 142434104
refGene_gn=SNTG2;refGene_tx=XM_003984546.5;refGene_gn=SNTG2;refGene_tx=XM_023252165.1
A3 142447831 142447959
refGene_gn=SNTG2;refGene_tx=XM_003984546.5;refGene_gn=SNTG2;refGene_tx=XM_023252165.1
A3 142472504 142472596
refGene_gn=SNTG2;refGene_tx=XM_003984546.5;refGene_gn=SNTG2;refGene_tx=XM_023252165.1
A3 142479604 142479692
refGene_gn=SNTG2;refGene_tx=XM_003984546.5;refGene_gn=SNTG2;refGene_tx=XM_023252165.1
A3 142489295 142489337
refGene_gn=SNTG2;refGene_tx=XM_003984546.5;refGene_gn=SNTG2;refGene_tx=XM_023252165.1
A3 142489441 142489485
refGene_gn=SNTG2;refGene_tx=XM_003984546.5;refGene_gn=SNTG2;refGene_tx=XM_023252165.1
A3 142502125 142502183
refGene_gn=SNTG2;refGene_tx=XM_003984546.5;refGene_gn=SNTG2;refGene_tx=XM_023252165.1
A3 142502271 142502328
refGene_gn=SNTG2;refGene_tx=XM_003984546.5;refGene_gn=SNTG2;refGene_tx=XM_023252165.1
A3 142510292 142510556
refGene_gn=SNTG2;refGene_tx=XM_003984546.5;refGene_gn=SNTG2;refGene_tx=XM_023252165.1
A3 142559919 142559991
refGene_gn=SNTG2;refGene_tx=XM_023252165.1
A3 142597596 142597907
refGene_gn=LOC109498652;refGene_tx=XM_019828135.2
A3 142598894 142598972
refGene_gn=LOC109498652;refGene_tx=XM_019828135.2
A3 142607242 142607462
refGene_gn=LOC109498652;refGene_tx=XM_019828135.2
A3 142680805 142681031
refGene_gn=TMEM18;refGene_tx=XM_003984521.5;refGene_gn=TMEM18;refGene_tx=XM_006930509.4;refGene_gn=TMEM18;refGene_tx=XM_023252166.1
A3 142682088 142682209
refGene_gn=TMEM18;refGene_tx=XM_003984521.5;refGene_gn=TMEM18;refGene_tx=XM_006930509.4

A3 142683802 142683857
refGene_gn=TMEM18;refGene_tx=XM_003984521.5;refGene_gn=TMEM18;refGene_tx=XM_006930509.4;refGene_gn=TMEM18;refGene_tx=XM_023252166.1
A3 142686672 142686766
refGene_gn=TMEM18;refGene_tx=XM_003984521.5;refGene_gn=TMEM18;refGene_tx=XM_006930509.4;refGene_gn=TMEM18;refGene_tx=XM_023252166.1
A3 142686855 142686945
refGene_gn=TMEM18;refGene_tx=XM_003984521.5;refGene_gn=TMEM18;refGene_tx=XM_006930509.4;refGene_gn=TMEM18;refGene_tx=XM_023252166.1
A3 142982968 142983200
refGene_gn=ALKAL2;refGene_tx=XM_023252168.1;refGene_gn=ALKAL2;refGene_tx=XM_023252169.1;refGene_gn=ALKAL2;refGene_tx=XM_023252170.1
A3 142984538 142984592
refGene_gn=ALKAL2;refGene_tx=XM_023252168.1;refGene_gn=ALKAL2;refGene_tx=XM_023252169.1;refGene_gn=ALKAL2;refGene_tx=XM_023252170.1
A3 142984673 142984754
refGene_gn=ALKAL2;refGene_tx=XM_023252168.1;refGene_gn=ALKAL2;refGene_tx=XM_023252169.1
A3 142987142 142987297
refGene_gn=ALKAL2;refGene_tx=XM_023252168.1
A3 142987500 142987565
refGene_gn=ALKAL2;refGene_tx=XM_023252169.1;refGene_gn=ALKAL2;refGene_tx=XM_023252170.1
A3 142993393 142993471
refGene_gn=ACP1;refGene_tx=XM_003984522.5;refGene_gn=ACP1;refGene_tx=XM_006930511.4
A3 142993646 142993752
refGene_gn=ACP1;refGene_tx=XM_003984522.5;refGene_gn=ACP1;refGene_tx=XM_006930511.4
A3 142995794 142995856
refGene_gn=ACP1;refGene_tx=XM_003984522.5;refGene_gn=ACP1;refGene_tx=XM_006930511.4
A3 143001748 143002023
refGene_gn=ACP1;refGene_tx=XM_003984522.5;refGene_gn=ACP1;refGene_tx=XM_006930511.4;refGene_gn=ACP1;refGene_tx=XM_023252167.1
A3 143002110 143002184
refGene_gn=ACP1;refGene_tx=XM_003984522.5;refGene_gn=ACP1;refGene_tx=XM_006930511.4;refGene_gn=ACP1;refGene_tx=XM_023252167.1
A3 143006313 143006522
refGene_gn=ACP1;refGene_tx=XM_003984522.5;refGene_gn=ACP1;refGene_tx=XM_006930511.4;refGene_gn=ACP1;refGene_tx=XM_023252167.1
A3 143006673 143006765
refGene_gn=ACP1;refGene_tx=XM_023252167.1
A3 143006791 143006882
refGene_gn=SH3YL1;refGene_tx=XM_011281348.3;refGene_gn=SH3YL1;refGene_tx=XM_011281349.3
A3 143007404 143007527
refGene_gn=SH3YL1;refGene_tx=XM_011281348.3
A3 143007970 143008031
refGene_gn=SH3YL1;refGene_tx=XM_019827999.2

A3 143020370 143020481
refGene_gn=SH3YL1;refGene_tx=XM_011281348.3;refGene_gn=SH3YL1;refGene_tx=XM_011281349.3;refGene_gn=SH3YL1;refGene_tx=XM_019827999.2
A3 143023294 143023408
refGene_gn=SH3YL1;refGene_tx=XM_011281348.3;refGene_gn=SH3YL1;refGene_tx=XM_011281349.3;refGene_gn=SH3YL1;refGene_tx=XM_019827999.2
A3 143025134 143025199
refGene_gn=SH3YL1;refGene_tx=XM_011281348.3;refGene_gn=SH3YL1;refGene_tx=XM_011281349.3;refGene_gn=SH3YL1;refGene_tx=XM_019827999.2
A3 143030262 143030375
refGene_gn=SH3YL1;refGene_tx=XM_011281348.3;refGene_gn=SH3YL1;refGene_tx=XM_011281349.3;refGene_gn=SH3YL1;refGene_tx=XM_019827999.2
A3 143031285 143031414
refGene_gn=SH3YL1;refGene_tx=XM_011281348.3;refGene_gn=SH3YL1;refGene_tx=XM_011281349.3;refGene_gn=SH3YL1;refGene_tx=XM_019827999.2
A3 143033246 143033415
refGene_gn=SH3YL1;refGene_tx=XM_011281348.3;refGene_gn=SH3YL1;refGene_tx=XM_011281349.3;refGene_gn=SH3YL1;refGene_tx=XM_019827999.2
A3 143034075 143034154
refGene_gn=SH3YL1;refGene_tx=XM_011281348.3;refGene_gn=SH3YL1;refGene_tx=XM_011281349.3;refGene_gn=SH3YL1;refGene_tx=XM_019827999.2
A3 143040211 143040268
refGene_gn=SH3YL1;refGene_tx=XM_011281348.3;refGene_gn=SH3YL1;refGene_tx=XM_011281349.3;refGene_gn=SH3YL1;refGene_tx=XM_019827999.2
A3 143046683 143046874
refGene_gn=SH3YL1;refGene_tx=XM_011281348.3;refGene_gn=SH3YL1;refGene_tx=XM_011281349.3;refGene_gn=SH3YL1;refGene_tx=XM_019827999.2
A3 143165975 143166941
refGene_gn=FAM110C;refGene_tx=XM_023252171.1;refGene_gn=FAM110C;refGene_tx=XM_023252172.1
A3 143171150 143171152
refGene_gn=FAM110C;refGene_tx=XM_023252172.1
B1 299869 300007
refGene_gn=LOC102899133;refGene_tx=XM_023252247.1;refGene_gn=LOC102899133;refGene_tx=XM_023252248.1;refGene_gn=LOC102899133;refGene_tx=XM_023252249.1;refGene_gn=LOC102899133;refGene_tx=XM_023252250.1;refGene_gn=LOC102899133;refGene_tx=XM_023252251.1;refGene_gn=LOC102899133;refGene_tx=XM_023252252.1;refGene_gn=LOC102899133;refGene_tx=XM_023252253.1;refGene_gn=LOC102899133;refGene_tx=XM_023252254.1
B1 300630 300757
refGene_gn=LOC102899133;refGene_tx=XM_023252247.1;refGene_gn=LOC102899133;refGene_tx=XM_023252248.1;refGene_gn=LOC102899133;refGene_tx=XM_023252249.1;refGene_gn=LOC102899133;refGene_tx=XM_023252250.1;refGene_gn=LOC102899133;refGene_tx=XM_023252251.1;refGene_gn=LOC102899133;refGene_tx=XM_023252252.1;refGene_gn=LOC102899133;refGene_tx=XM_023252253.1;refGene_gn=LOC102899133;refGene_tx=XM_023252254.1
B1 301469 301565
refGene_gn=LOC102899133;refGene_tx=XM_023252247.1;refGene_gn=LOC102899133;refGene_tx=XM_023252248.1;refGene_gn=LOC102899133;refGene_tx=XM_023252249.1;refGene_gn=LOC102899133;refGene_tx=XM_023252250.1;refGene_gn

=LOC102899133;refGene_tx=XM_023252251.1;refGene_gn=LOC102899133;refGene_tx=XM_023252252.1;refGene_gn=LOC102899133;refGene_tx=XM_023252253.1
B1 302087 302170
refGene_gn=LOC102899133;refGene_tx=XM_019828160.1;refGene_gn=LOC102899133;refGene_tx=XM_023252247.1;refGene_gn=LOC102899133;refGene_tx=XM_023252248.1;refGene_gn=LOC102899133;refGene_tx=XM_023252249.1;refGene_gn=LOC102899133;refGene_tx=XM_023252250.1;refGene_gn=LOC102899133;refGene_tx=XM_023252251.1;refGene_gn=LOC102899133;refGene_tx=XM_023252252.1;refGene_gn=LOC102899133;refGene_tx=XM_023252253.1;refGene_gn=LOC102899133;refGene_tx=XM_023252254.1
B1 302609 304733
refGene_gn=LOC102899133;refGene_tx=XM_019828160.1;refGene_gn=LOC102899133;refGene_tx=XM_023252247.1;refGene_gn=LOC102899133;refGene_tx=XM_023252248.1;refGene_gn=LOC102899133;refGene_tx=XM_023252249.1;refGene_gn=LOC102899133;refGene_tx=XM_023252250.1;refGene_gn=LOC102899133;refGene_tx=XM_023252251.1;refGene_gn=LOC102899133;refGene_tx=XM_023252252.1;refGene_gn=LOC102899133;refGene_tx=XM_023252253.1;refGene_gn=LOC102899133;refGene_tx=XM_023252254.1
B1 334800 334938
refGene_gn=LOC105259656;refGene_tx=XM_023252258.1;refGene_gn=LOC105259656;refGene_tx=XM_023252259.1;refGene_gn=LOC105259656;refGene_tx=XM_023252260.1;refGene_gn=LOC105259656;refGene_tx=XM_023252261.1;refGene_gn=LOC105259656;refGene_tx=XM_023252262.1
B1 335664 335791
refGene_gn=LOC105259656;refGene_tx=XM_023252258.1;refGene_gn=LOC105259656;refGene_tx=XM_023252259.1;refGene_gn=LOC105259656;refGene_tx=XM_023252260.1;refGene_gn=LOC105259656;refGene_tx=XM_023252261.1;refGene_gn=LOC105259656;refGene_tx=XM_023252262.1
B1 336515 336611
refGene_gn=LOC105259656;refGene_tx=XM_023252258.1;refGene_gn=LOC105259656;refGene_tx=XM_023252259.1;refGene_gn=LOC105259656;refGene_tx=XM_023252260.1;refGene_gn=LOC105259656;refGene_tx=XM_023252261.1;refGene_gn=LOC105259656;refGene_tx=XM_023252262.1
B1 337131 337214
refGene_gn=LOC105259656;refGene_tx=XM_023252258.1;refGene_gn=LOC105259656;refGene_tx=XM_023252259.1;refGene_gn=LOC105259656;refGene_tx=XM_023252260.1;refGene_gn=LOC105259656;refGene_tx=XM_023252261.1;refGene_gn=LOC105259656;refGene_tx=XM_023252262.1
B1 337654 337768
refGene_gn=LOC105259656;refGene_tx=XM_023252258.1;refGene_gn=LOC105259656;refGene_tx=XM_023252259.1;refGene_gn=LOC105259656;refGene_tx=XM_023252260.1;refGene_gn=LOC105259656;refGene_tx=XM_023252261.1;refGene_gn=LOC105259656;refGene_tx=XM_023252262.1
B1 380332 380368
refGene_gn=ZNF596;refGene_tx=XM_011281357.3;refGene_gn=ZNF596;refGene_tx=XM_011281358.3;refGene_gn=ZNF596;refGene_tx=XM_023252255.1;refGene_gn=ZNF596;refGene_tx=XM_023252256.1;refGene_gn=ZNF596;refGene_tx=XM_023252257.1
B1 382422 382549
refGene_gn=ZNF596;refGene_tx=XM_011281357.3;refGene_gn=ZNF596;refGene_

tx=XM_011281358.3;refGene_gn=ZNF596;refGene_tx=XM_011281359.3;refGene_gn=ZNF596;refGene_tx=XM_023252255.1;refGene_gn=ZNF596;refGene_tx=XM_023252256.1;refGene_gn=ZNF596;refGene_tx=XM_023252257.1
B1 383276 383372
refGene_gn=ZNF596;refGene_tx=XM_011281357.3;refGene_gn=ZNF596;refGene_tx=XM_011281358.3;refGene_gn=ZNF596;refGene_tx=XM_011281359.3;refGene_gn=ZNF596;refGene_tx=XM_023252255.1;refGene_gn=ZNF596;refGene_tx=XM_023252256.1;refGene_gn=ZNF596;refGene_tx=XM_023252257.1
B1 383844 383927
refGene_gn=ZNF596;refGene_tx=XM_011281357.3;refGene_gn=ZNF596;refGene_tx=XM_011281358.3;refGene_gn=ZNF596;refGene_tx=XM_011281359.3
B1 384361 385606
refGene_gn=ZNF596;refGene_tx=XM_011281357.3;refGene_gn=ZNF596;refGene_tx=XM_011281358.3;refGene_gn=ZNF596;refGene_tx=XM_011281359.3;refGene_gn=ZNF596;refGene_tx=XM_023252257.1
B1 388229 388363
refGene_gn=ZNF596;refGene_tx=XM_023252255.1;refGene_gn=ZNF596;refGene_tx=XM_023252256.1
B1 389798 389828 refGene_gn=ZNF596;refGene_tx=XM_023252256.1
B1 409666 410994
refGene_gn=LOC111560063;refGene_tx=XM_023252244.1
B1 411271 411305
refGene_gn=LOC111560063;refGene_tx=XM_023252244.1
B1 435080 436712
refGene_gn=LOC101089843;refGene_tx=XM_011280045.3
B1 453092 455054
refGene_gn=LOC101090710;refGene_tx=XM_019828163.2;refGene_gn=LOC101090710;refGene_tx=XM_019828165.2;refGene_gn=LOC101090710;refGene_tx=XM_019828166.2;refGene_gn=LOC101090710;refGene_tx=XM_019828167.2;refGene_gn=LOC101090710;refGene_tx=XM_019828168.2;refGene_gn=LOC101090710;refGene_tx=XM_019828169.2;refGene_gn=LOC101090710;refGene_tx=XM_019828171.2;refGene_gn=LOC101090710;refGene_tx=XM_019828172.2;refGene_gn=LOC101090710;refGene_tx=XM_019828173.2;refGene_gn=LOC101090710;refGene_tx=XM_019828174.2;refGene_gn=LOC101090710;refGene_tx=XM_019828176.2;refGene_gn=LOC101090710;refGene_tx=XM_019828177.2;refGene_gn=LOC101090710;refGene_tx=XM_023252265.1
B1 455509 455592
refGene_gn=LOC101090710;refGene_tx=XM_019828163.2;refGene_gn=LOC101090710;refGene_tx=XM_019828165.2;refGene_gn=LOC101090710;refGene_tx=XM_019828166.2;refGene_gn=LOC101090710;refGene_tx=XM_019828167.2;refGene_gn=LOC101090710;refGene_tx=XM_019828168.2;refGene_gn=LOC101090710;refGene_tx=XM_019828169.2;refGene_gn=LOC101090710;refGene_tx=XM_019828171.2;refGene_gn=LOC101090710;refGene_tx=XM_019828172.2;refGene_gn=LOC101090710;refGene_tx=XM_019828173.2;refGene_gn=LOC101090710;refGene_tx=XM_019828174.2;refGene_gn=LOC101090710;refGene_tx=XM_019828176.2;refGene_gn=LOC101090710;refGene_tx=XM_019828177.2;refGene_gn=LOC101090710;refGene_tx=XM_023252265.1
B1 456095 456191
refGene_gn=LOC101090710;refGene_tx=XM_019828163.2;refGene_gn=LOC101090710;refGene_tx=XM_019828165.2;refGene_gn=LOC101090710;refGene_tx=XM_01

9828166.2;refGene_gn=LOC101090710;refGene_tx=XM_019828167.2;refGene_gn=LOC101090710;refGene_tx=XM_019828168.2;refGene_gn=LOC101090710;refGene_tx=XM_019828169.2;refGene_gn=LOC101090710;refGene_tx=XM_019828171.2;refGene_gn=LOC101090710;refGene_tx=XM_019828172.2;refGene_gn=LOC101090710;refGene_tx=XM_019828173.2;refGene_gn=LOC101090710;refGene_tx=XM_019828174.2;refGene_gn=LOC101090710;refGene_tx=XM_023252265.1
B1 456914 457041
refGene_gn=LOC101090710;refGene_tx=XM_019828163.2;refGene_gn=LOC101090710;refGene_tx=XM_019828165.2;refGene_gn=LOC101090710;refGene_tx=XM_019828166.2;refGene_gn=LOC101090710;refGene_tx=XM_019828167.2;refGene_gn=LOC101090710;refGene_tx=XM_019828168.2;refGene_gn=LOC101090710;refGene_tx=XM_019828169.2;refGene_gn=LOC101090710;refGene_tx=XM_019828171.2;refGene_gn=LOC101090710;refGene_tx=XM_019828172.2;refGene_gn=LOC101090710;refGene_tx=XM_019828173.2;refGene_gn=LOC101090710;refGene_tx=XM_019828174.2;refGene_gn=LOC101090710;refGene_tx=XM_019828176.2;refGene_gn=LOC101090710;refGene_tx=XM_019828177.2;refGene_gn=LOC101090710;refGene_tx=XM_023252265.1
B1 457767 457881
refGene_gn=LOC101090710;refGene_tx=XM_019828163.2;refGene_gn=LOC101090710;refGene_tx=XM_019828165.2;refGene_gn=LOC101090710;refGene_tx=XM_019828166.2;refGene_gn=LOC101090710;refGene_tx=XM_019828167.2;refGene_gn=LOC101090710;refGene_tx=XM_019828168.2;refGene_gn=LOC101090710;refGene_tx=XM_019828169.2;refGene_gn=LOC101090710;refGene_tx=XM_019828171.2;refGene_gn=LOC101090710;refGene_tx=XM_019828172.2;refGene_gn=LOC101090710;refGene_tx=XM_019828173.2;refGene_gn=LOC101090710;refGene_tx=XM_019828174.2;refGene_gn=LOC101090710;refGene_tx=XM_019828176.2;refGene_gn=LOC101090710;refGene_tx=XM_019828177.2;refGene_gn=LOC101090710;refGene_tx=XM_023252265.1
B1 565749 565883
refGene_gn=FBXO25;refGene_tx=XM_003984548.4;refGene_gn=FBXO25;refGene_tx=XM_003984549.4;refGene_gn=FBXO25;refGene_tx=XM_006930530.3;refGene_gn=FBXO25;refGene_tx=XM_006930531.3;refGene_gn=FBXO25;refGene_tx=XM_006930532.3;refGene_gn=FBXO25;refGene_tx=XM_006930534.3;refGene_gn=FBXO25;refGene_tx=XM_019828179.2;refGene_gn=FBXO25;refGene_tx=XM_019828180.2;refGene_gn=FBXO25;refGene_tx=XM_023252266.1;refGene_gn=FBXO25;refGene_tx=XM_023252268.1
B1 583794 583898
refGene_gn=FBXO25;refGene_tx=XM_003984548.4;refGene_gn=FBXO25;refGene_tx=XM_003984549.4;refGene_gn=FBXO25;refGene_tx=XM_006930530.3;refGene_gn=FBXO25;refGene_tx=XM_006930531.3;refGene_gn=FBXO25;refGene_tx=XM_006930532.3;refGene_gn=FBXO25;refGene_tx=XM_006930534.3;refGene_gn=FBXO25;refGene_tx=XM_019828179.2;refGene_gn=FBXO25;refGene_tx=XM_019828180.2;refGene_gn=FBXO25;refGene_tx=XM_023252266.1;refGene_gn=FBXO25;refGene_tx=XM_023252268.1
B1 584812 584862
refGene_gn=FBXO25;refGene_tx=XM_003984548.4;refGene_gn=FBXO25;refGene_tx=XM_003984549.4;refGene_gn=FBXO25;refGene_tx=XM_006930530.3;refGene_gn=FBXO25;refGene_tx=XM_006930531.3;refGene_gn=FBXO25;refGene_tx=XM_006930532.3;refGene_gn=FBXO25;refGene_tx=XM_006930534.3;refGene_gn=FBXO25;refGene_tx=XM_019828179.2;refGene_gn=FBXO25;refGene_tx=XM_019828180.

2;refGene_gn=FBXO25;refGene_tx=XM_023252266.1;refGene_gn=FBXO25;refGene_tx=XM_023252268.1
B1 587319 587433
refGene_gn=FBXO25;refGene_tx=XM_003984548.4;refGene_gn=FBXO25;refGene_tx=XM_003984549.4;refGene_gn=FBXO25;refGene_tx=XM_006930530.3;refGene_gn=FBXO25;refGene_tx=XM_006930531.3;refGene_gn=FBXO25;refGene_tx=XM_006930532.3;refGene_gn=FBXO25;refGene_tx=XM_019828179.2;refGene_gn=FBXO25;refGene_tx=XM_019828180.2;refGene_gn=FBXO25;refGene_tx=XM_023252266.1;refGene_gn=FBXO25;refGene_tx=XM_023252268.1
B1 595752 595846
refGene_gn=FBXO25;refGene_tx=XM_003984548.4;refGene_gn=FBXO25;refGene_tx=XM_003984549.4;refGene_gn=FBXO25;refGene_tx=XM_006930530.3;refGene_gn=FBXO25;refGene_tx=XM_006930531.3;refGene_gn=FBXO25;refGene_tx=XM_006930532.3;refGene_gn=FBXO25;refGene_tx=XM_006930534.3;refGene_gn=FBXO25;refGene_tx=XM_019828179.2;refGene_gn=FBXO25;refGene_tx=XM_019828180.2;refGene_gn=FBXO25;refGene_tx=XM_023252266.1;refGene_gn=FBXO25;refGene_tx=XM_023252268.1
B1 597883 598068
refGene_gn=FBXO25;refGene_tx=XM_003984548.4;refGene_gn=FBXO25;refGene_tx=XM_003984549.4;refGene_gn=FBXO25;refGene_tx=XM_006930530.3;refGene_gn=FBXO25;refGene_tx=XM_006930531.3;refGene_gn=FBXO25;refGene_tx=XM_006930532.3;refGene_gn=FBXO25;refGene_tx=XM_006930534.3;refGene_gn=FBXO25;refGene_tx=XM_019828179.2;refGene_gn=FBXO25;refGene_tx=XM_019828180.2;refGene_gn=FBXO25;refGene_tx=XM_023252266.1;refGene_gn=FBXO25;refGene_tx=XM_023252268.1
B1 601751 601934
refGene_gn=FBXO25;refGene_tx=XM_003984548.4;refGene_gn=FBXO25;refGene_tx=XM_003984549.4;refGene_gn=FBXO25;refGene_tx=XM_006930530.3;refGene_gn=FBXO25;refGene_tx=XM_006930531.3;refGene_gn=FBXO25;refGene_tx=XM_006930532.3;refGene_gn=FBXO25;refGene_tx=XM_006930534.3;refGene_gn=FBXO25;refGene_tx=XM_019828179.2;refGene_gn=FBXO25;refGene_tx=XM_019828180.2;refGene_gn=FBXO25;refGene_tx=XM_023252266.1;refGene_gn=FBXO25;refGene_tx=XM_023252268.1
B1 603214 603358
refGene_gn=FBXO25;refGene_tx=XM_003984548.4;refGene_gn=FBXO25;refGene_tx=XM_003984549.4;refGene_gn=FBXO25;refGene_tx=XM_006930530.3;refGene_gn=FBXO25;refGene_tx=XM_006930531.3;refGene_gn=FBXO25;refGene_tx=XM_006930532.3;refGene_gn=FBXO25;refGene_tx=XM_006930534.3;refGene_gn=FBXO25;refGene_tx=XM_019828179.2;refGene_gn=FBXO25;refGene_tx=XM_019828180.2;refGene_gn=FBXO25;refGene_tx=XM_023252266.1;refGene_gn=FBXO25;refGene_tx=XM_023252268.1
B1 605706 605733
refGene_gn=FBXO25;refGene_tx=XM_003984548.4;refGene_gn=FBXO25;refGene_tx=XM_006930530.3;refGene_gn=FBXO25;refGene_tx=XM_006930534.3;refGene_gn=FBXO25;refGene_tx=XM_019828179.2;refGene_gn=FBXO25;refGene_tx=XM_023252266.1;refGene_gn=FBXO25;refGene_tx=XM_023252268.1
B1 606272 606362
refGene_gn=FBXO25;refGene_tx=XM_003984548.4;refGene_gn=FBXO25;refGene_tx=XM_003984549.4;refGene_gn=FBXO25;refGene_tx=XM_006930530.3;refGene_gn=FBXO25;refGene_tx=XM_006930531.3;refGene_gn=FBXO25;refGene_tx=XM_00

6930532.3;refGene_gn=FBXO25;refGene_tx=XM_006930534.3;refGene_gn=FBXO2
5;refGene_tx=XM_019828179.2;refGene_gn=FBXO25;refGene_tx=XM_019828180.
2;refGene_gn=FBXO25;refGene_tx=XM_023252266.1;refGene_gn=FBXO25;refGen
e_tx=XM_023252268.1
B1 621639 621985 refGene_gn=TDRP;refGene_tx=XM_023252269.1
B1 623128 623232 refGene_gn=TDRP;refGene_tx=XM_023252269.1
B1 656830 656938 refGene_gn=TDRP;refGene_tx=XM_023252269.1
B1 747473 747481 refGene_gn=ERICH1;refGene_tx=XM_019828186.2
B1 755803 755871
refGene_gn=ERICH1;refGene_tx=XM_006930536.4;refGene_gn=ERICH1;refGene_
tx=XM_006930537.3
B1 760254 760449
refGene_gn=ERICH1;refGene_tx=XM_006930536.4;refGene_gn=ERICH1;refGene_
tx=XM_006930537.3;refGene_gn=ERICH1;refGene_tx=XM_019828186.2
B1 762653 763244
refGene_gn=ERICH1;refGene_tx=XM_006930536.4;refGene_gn=ERICH1;refGene_
tx=XM_019828186.2
B1 781245 781398
refGene_gn=ERICH1;refGene_tx=XM_006930536.4;refGene_gn=ERICH1;refGene_
tx=XM_006930537.3;refGene_gn=ERICH1;refGene_tx=XM_019828186.2
B1 789011 789158
refGene_gn=ERICH1;refGene_tx=XM_006930536.4;refGene_gn=ERICH1;refGene_
tx=XM_006930537.3;refGene_gn=ERICH1;refGene_tx=XM_019828186.2
B1 807560 807582
refGene_gn=ERICH1;refGene_tx=XM_006930536.4;refGene_gn=ERICH1;refGene_
tx=XM_006930537.3;refGene_gn=ERICH1;refGene_tx=XM_019828186.2
B1 813094 813112 refGene_gn=DLGAP2;refGene_tx=XM_023252245.1
B1 998979 999034 refGene_gn=DLGAP2;refGene_tx=XM_023252245.1
B1 1249064 1249097 refGene_gn=DLGAP2;refGene_tx=XM_023252245.1
B1 1429421 1429487 refGene_gn=DLGAP2;refGene_tx=XM_023252245.1
B1 1483131 1484201 refGene_gn=DLGAP2;refGene_tx=XM_023252245.1
B1 1494431 1494643 refGene_gn=DLGAP2;refGene_tx=XM_023252245.1
B1 1533033 1533181 refGene_gn=DLGAP2;refGene_tx=XM_023252245.1
B1 1543526 1543746 refGene_gn=DLGAP2;refGene_tx=XM_023252245.1
B1 1571040 1571390 refGene_gn=DLGAP2;refGene_tx=XM_023252245.1
B1 1572371 1572442 refGene_gn=DLGAP2;refGene_tx=XM_023252245.1
B1 1573619 1573689 refGene_gn=DLGAP2;refGene_tx=XM_023252245.1
B1 1573842 1573995 refGene_gn=DLGAP2;refGene_tx=XM_023252245.1
B1 1582276 1582853 refGene_gn=DLGAP2;refGene_tx=XM_023252245.1
B1 1592445 1592537 refGene_gn=DLGAP2;refGene_tx=XM_023252245.1
B1 1602278 1602431 refGene_gn=DLGAP2;refGene_tx=XM_023252245.1
B1 1605733 1605952 refGene_gn=DLGAP2;refGene_tx=XM_023252245.1
B1 1639312 1639855 refGene_gn=CLN8;refGene_tx=XM_006930538.4
B1 1646861 1647185 refGene_gn=CLN8;refGene_tx=XM_006930538.4
B1 1684872 1685042
refGene_gn=ARHGEF10;refGene_tx=XM_023252270.1;refGene_gn=ARHGEF10;refG
ene_tx=XM_023252271.1;refGene_gn=ARHGEF10;refGene_tx=XM_023252272.1
B1 1687991 1688014 refGene_gn=ARHGEF10;refGene_tx=XM_023252273.1
B1 1698273 1698341
refGene_gn=ARHGEF10;refGene_tx=XM_023252270.1;refGene_gn=ARHGEF10;refG

ene_tx=XM_023252271.1;refGene_gn=ARHGEF10;refGene_tx=XM_023252272.1;refGene_gn=ARHGEF10;refGene_tx=XM_023252273.1;refGene_gn=ARHGEF10;refGene_tx=XM_023252274.1;refGene_gn=ARHGEF10;refGene_tx=XM_023252275.1;refGene_gn=ARHGEF10;refGene_tx=XM_023252276.1;refGene_gn=ARHGEF10;refGene_tx=XM_023252277.1
B1 1706003 1706153
refGene_gn=ARHGEF10;refGene_tx=XM_023252270.1;refGene_gn=ARHGEF10;refGene_tx=XM_023252271.1;refGene_gn=ARHGEF10;refGene_tx=XM_023252272.1;refGene_gn=ARHGEF10;refGene_tx=XM_023252273.1;refGene_gn=ARHGEF10;refGene_tx=XM_023252274.1;refGene_gn=ARHGEF10;refGene_tx=XM_023252275.1;refGene_gn=ARHGEF10;refGene_tx=XM_023252276.1;refGene_gn=ARHGEF10;refGene_tx=XM_023252277.1
B1 1706747 1707035
refGene_gn=ARHGEF10;refGene_tx=XM_023252270.1;refGene_gn=ARHGEF10;refGene_tx=XM_023252271.1;refGene_gn=ARHGEF10;refGene_tx=XM_023252272.1;refGene_gn=ARHGEF10;refGene_tx=XM_023252273.1;refGene_gn=ARHGEF10;refGene_tx=XM_023252274.1;refGene_gn=ARHGEF10;refGene_tx=XM_023252275.1;refGene_gn=ARHGEF10;refGene_tx=XM_023252276.1;refGene_gn=ARHGEF10;refGene_tx=XM_023252277.1
B1 1709391 1709455
refGene_gn=ARHGEF10;refGene_tx=XM_023252270.1;refGene_gn=ARHGEF10;refGene_tx=XM_023252271.1;refGene_gn=ARHGEF10;refGene_tx=XM_023252272.1;refGene_gn=ARHGEF10;refGene_tx=XM_023252273.1;refGene_gn=ARHGEF10;refGene_tx=XM_023252274.1;refGene_gn=ARHGEF10;refGene_tx=XM_023252275.1;refGene_gn=ARHGEF10;refGene_tx=XM_023252276.1;refGene_gn=ARHGEF10;refGene_tx=XM_023252277.1
B1 1713965 1714042
refGene_gn=ARHGEF10;refGene_tx=XM_023252270.1;refGene_gn=ARHGEF10;refGene_tx=XM_023252271.1;refGene_gn=ARHGEF10;refGene_tx=XM_023252272.1;refGene_gn=ARHGEF10;refGene_tx=XM_023252273.1;refGene_gn=ARHGEF10;refGene_tx=XM_023252274.1;refGene_gn=ARHGEF10;refGene_tx=XM_023252275.1;refGene_gn=ARHGEF10;refGene_tx=XM_023252276.1;refGene_gn=ARHGEF10;refGene_tx=XM_023252277.1
B1 1716115 1716172
refGene_gn=ARHGEF10;refGene_tx=XM_023252270.1;refGene_gn=ARHGEF10;refGene_tx=XM_023252271.1;refGene_gn=ARHGEF10;refGene_tx=XM_023252272.1;refGene_gn=ARHGEF10;refGene_tx=XM_023252273.1;refGene_gn=ARHGEF10;refGene_tx=XM_023252274.1;refGene_gn=ARHGEF10;refGene_tx=XM_023252275.1;refGene_gn=ARHGEF10;refGene_tx=XM_023252276.1;refGene_gn=ARHGEF10;refGene_tx=XM_023252277.1
B1 1723142 1723306
refGene_gn=ARHGEF10;refGene_tx=XM_023252270.1;refGene_gn=ARHGEF10;refGene_tx=XM_023252271.1;refGene_gn=ARHGEF10;refGene_tx=XM_023252272.1;refGene_gn=ARHGEF10;refGene_tx=XM_023252273.1;refGene_gn=ARHGEF10;refGene_tx=XM_023252274.1;refGene_gn=ARHGEF10;refGene_tx=XM_023252275.1;refGene_gn=ARHGEF10;refGene_tx=XM_023252276.1;refGene_gn=ARHGEF10;refGene_tx=XM_023252277.1
B1 1725700 1725817
refGene_gn=ARHGEF10;refGene_tx=XM_023252270.1;refGene_gn=ARHGEF10;refGene_tx=XM_023252271.1;refGene_gn=ARHGEF10;refGene_tx=XM_023252273.1;re

fGene_gn=ARHGEF10;refGene_tx=XM_023252274.1;refGene_gn=ARHGEF10;refGene_tx=XM_023252275.1;refGene_gn=ARHGEF10;refGene_tx=XM_023252276.1;refGene_gn=ARHGEF10;refGene_tx=XM_023252277.1
B1 1728303 1728418
refGene_gn=ARHGEF10;refGene_tx=XM_023252270.1;refGene_gn=ARHGEF10;refGene_tx=XM_023252271.1;refGene_gn=ARHGEF10;refGene_tx=XM_023252272.1;refGene_gn=ARHGEF10;refGene_tx=XM_023252273.1;refGene_gn=ARHGEF10;refGene_tx=XM_023252274.1;refGene_gn=ARHGEF10;refGene_tx=XM_023252275.1;refGene_gn=ARHGEF10;refGene_tx=XM_023252276.1;refGene_gn=ARHGEF10;refGene_tx=XM_023252277.1
B1 1731637 1731744
refGene_gn=ARHGEF10;refGene_tx=XM_023252270.1;refGene_gn=ARHGEF10;refGene_tx=XM_023252271.1;refGene_gn=ARHGEF10;refGene_tx=XM_023252272.1;refGene_gn=ARHGEF10;refGene_tx=XM_023252273.1;refGene_gn=ARHGEF10;refGene_tx=XM_023252274.1;refGene_gn=ARHGEF10;refGene_tx=XM_023252275.1;refGene_gn=ARHGEF10;refGene_tx=XM_023252276.1;refGene_gn=ARHGEF10;refGene_tx=XM_023252277.1
B1 1735735 1735813
refGene_gn=ARHGEF10;refGene_tx=XM_023252270.1;refGene_gn=ARHGEF10;refGene_tx=XM_023252271.1;refGene_gn=ARHGEF10;refGene_tx=XM_023252272.1;refGene_gn=ARHGEF10;refGene_tx=XM_023252273.1;refGene_gn=ARHGEF10;refGene_tx=XM_023252274.1;refGene_gn=ARHGEF10;refGene_tx=XM_023252275.1;refGene_gn=ARHGEF10;refGene_tx=XM_023252276.1;refGene_gn=ARHGEF10;refGene_tx=XM_023252277.1
B1 1736049 1736229
refGene_gn=ARHGEF10;refGene_tx=XM_023252270.1;refGene_gn=ARHGEF10;refGene_tx=XM_023252271.1;refGene_gn=ARHGEF10;refGene_tx=XM_023252272.1;refGene_gn=ARHGEF10;refGene_tx=XM_023252273.1;refGene_gn=ARHGEF10;refGene_tx=XM_023252274.1;refGene_gn=ARHGEF10;refGene_tx=XM_023252275.1;refGene_gn=ARHGEF10;refGene_tx=XM_023252276.1;refGene_gn=ARHGEF10;refGene_tx=XM_023252277.1
B1 1737891 1738008
refGene_gn=ARHGEF10;refGene_tx=XM_023252270.1;refGene_gn=ARHGEF10;refGene_tx=XM_023252271.1;refGene_gn=ARHGEF10;refGene_tx=XM_023252272.1;refGene_gn=ARHGEF10;refGene_tx=XM_023252273.1;refGene_gn=ARHGEF10;refGene_tx=XM_023252274.1;refGene_gn=ARHGEF10;refGene_tx=XM_023252275.1;refGene_gn=ARHGEF10;refGene_tx=XM_023252276.1;refGene_gn=ARHGEF10;refGene_tx=XM_023252277.1
B1 1739931 1740024
refGene_gn=ARHGEF10;refGene_tx=XM_023252270.1;refGene_gn=ARHGEF10;refGene_tx=XM_023252271.1;refGene_gn=ARHGEF10;refGene_tx=XM_023252272.1;refGene_gn=ARHGEF10;refGene_tx=XM_023252273.1;refGene_gn=ARHGEF10;refGene_tx=XM_023252274.1;refGene_gn=ARHGEF10;refGene_tx=XM_023252275.1;refGene_gn=ARHGEF10;refGene_tx=XM_023252276.1;refGene_gn=ARHGEF10;refGene_tx=XM_023252277.1
B1 1743190 1743361
refGene_gn=ARHGEF10;refGene_tx=XM_023252270.1;refGene_gn=ARHGEF10;refGene_tx=XM_023252271.1;refGene_gn=ARHGEF10;refGene_tx=XM_023252272.1;refGene_gn=ARHGEF10;refGene_tx=XM_023252273.1;refGene_gn=ARHGEF10;refGene_tx=XM_023252274.1;refGene_gn=ARHGEF10;refGene_tx=XM_023252275.1;refG

ene_gn=ARHGEF10;refGene_tx=XM_023252276.1;refGene_gn=ARHGEF10;refGene_tx=XM_023252277.1
B1	1744988	1745134
refGene_gn=ARHGEF10;refGene_tx=XM_023252270.1;refGene_gn=ARHGEF10;refGene_tx=XM_023252271.1;refGene_gn=ARHGEF10;refGene_tx=XM_023252272.1;refGene_gn=ARHGEF10;refGene_tx=XM_023252273.1;refGene_gn=ARHGEF10;refGene_tx=XM_023252274.1;refGene_gn=ARHGEF10;refGene_tx=XM_023252275.1;refGene_gn=ARHGEF10;refGene_tx=XM_023252276.1;refGene_gn=ARHGEF10;refGene_tx=XM_023252277.1
B1	1749463	1749636
refGene_gn=ARHGEF10;refGene_tx=XM_023252270.1;refGene_gn=ARHGEF10;refGene_tx=XM_023252271.1;refGene_gn=ARHGEF10;refGene_tx=XM_023252272.1;refGene_gn=ARHGEF10;refGene_tx=XM_023252273.1;refGene_gn=ARHGEF10;refGene_tx=XM_023252274.1;refGene_gn=ARHGEF10;refGene_tx=XM_023252275.1;refGene_gn=ARHGEF10;refGene_tx=XM_023252276.1;refGene_gn=ARHGEF10;refGene_tx=XM_023252277.1
B1	1754423	1754539
refGene_gn=ARHGEF10;refGene_tx=XM_023252270.1;refGene_gn=ARHGEF10;refGene_tx=XM_023252271.1;refGene_gn=ARHGEF10;refGene_tx=XM_023252272.1;refGene_gn=ARHGEF10;refGene_tx=XM_023252273.1;refGene_gn=ARHGEF10;refGene_tx=XM_023252274.1;refGene_gn=ARHGEF10;refGene_tx=XM_023252275.1;refGene_gn=ARHGEF10;refGene_tx=XM_023252276.1;refGene_gn=ARHGEF10;refGene_tx=XM_023252277.1
B1	1754924	1755052
refGene_gn=ARHGEF10;refGene_tx=XM_023252270.1;refGene_gn=ARHGEF10;refGene_tx=XM_023252271.1;refGene_gn=ARHGEF10;refGene_tx=XM_023252272.1;refGene_gn=ARHGEF10;refGene_tx=XM_023252273.1;refGene_gn=ARHGEF10;refGene_tx=XM_023252274.1;refGene_gn=ARHGEF10;refGene_tx=XM_023252275.1;refGene_gn=ARHGEF10;refGene_tx=XM_023252276.1;refGene_gn=ARHGEF10;refGene_tx=XM_023252277.1
B1	1755213	1755314
refGene_gn=ARHGEF10;refGene_tx=XM_023252270.1;refGene_gn=ARHGEF10;refGene_tx=XM_023252271.1;refGene_gn=ARHGEF10;refGene_tx=XM_023252272.1;refGene_gn=ARHGEF10;refGene_tx=XM_023252273.1;refGene_gn=ARHGEF10;refGene_tx=XM_023252274.1;refGene_gn=ARHGEF10;refGene_tx=XM_023252275.1;refGene_gn=ARHGEF10;refGene_tx=XM_023252276.1;refGene_gn=ARHGEF10;refGene_tx=XM_023252277.1
B1	1757029	1757151
refGene_gn=ARHGEF10;refGene_tx=XM_023252270.1;refGene_gn=ARHGEF10;refGene_tx=XM_023252271.1;refGene_gn=ARHGEF10;refGene_tx=XM_023252272.1;refGene_gn=ARHGEF10;refGene_tx=XM_023252273.1;refGene_gn=ARHGEF10;refGene_tx=XM_023252274.1;refGene_gn=ARHGEF10;refGene_tx=XM_023252275.1;refGene_gn=ARHGEF10;refGene_tx=XM_023252276.1;refGene_gn=ARHGEF10;refGene_tx=XM_023252277.1
B1	1759066	1759153
refGene_gn=ARHGEF10;refGene_tx=XM_023252270.1;refGene_gn=ARHGEF10;refGene_tx=XM_023252271.1;refGene_gn=ARHGEF10;refGene_tx=XM_023252272.1;refGene_gn=ARHGEF10;refGene_tx=XM_023252273.1;refGene_gn=ARHGEF10;refGene_tx=XM_023252274.1;refGene_gn=ARHGEF10;refGene_tx=XM_023252275.1;refGene_gn=ARHGEF10;refGene_tx=XM_023252276.1;refGene_gn=ARHGEF10;refGene_

tx=XM_023252277.1
B1	1761409	1761600
refGene_gn=ARHGEF10;refGene_tx=XM_023252270.1;refGene_gn=ARHGEF10;refGene_tx=XM_023252271.1;refGene_gn=ARHGEF10;refGene_tx=XM_023252272.1;refGene_gn=ARHGEF10;refGene_tx=XM_023252273.1;refGene_gn=ARHGEF10;refGene_tx=XM_023252274.1;refGene_gn=ARHGEF10;refGene_tx=XM_023252275.1;refGene_gn=ARHGEF10;refGene_tx=XM_023252276.1;refGene_gn=ARHGEF10;refGene_tx=XM_023252277.1
B1	1762102	1762260
refGene_gn=ARHGEF10;refGene_tx=XM_023252270.1;refGene_gn=ARHGEF10;refGene_tx=XM_023252271.1;refGene_gn=ARHGEF10;refGene_tx=XM_023252272.1;refGene_gn=ARHGEF10;refGene_tx=XM_023252273.1;refGene_gn=ARHGEF10;refGene_tx=XM_023252274.1;refGene_gn=ARHGEF10;refGene_tx=XM_023252275.1;refGene_gn=ARHGEF10;refGene_tx=XM_023252276.1;refGene_gn=ARHGEF10;refGene_tx=XM_023252277.1
B1	1766570	1766713
refGene_gn=ARHGEF10;refGene_tx=XM_023252270.1;refGene_gn=ARHGEF10;refGene_tx=XM_023252271.1;refGene_gn=ARHGEF10;refGene_tx=XM_023252272.1;refGene_gn=ARHGEF10;refGene_tx=XM_023252273.1;refGene_gn=ARHGEF10;refGene_tx=XM_023252274.1;refGene_gn=ARHGEF10;refGene_tx=XM_023252275.1;refGene_gn=ARHGEF10;refGene_tx=XM_023252276.1;refGene_gn=ARHGEF10;refGene_tx=XM_023252277.1
B1	1770867	1771042
refGene_gn=ARHGEF10;refGene_tx=XM_023252270.1;refGene_gn=ARHGEF10;refGene_tx=XM_023252271.1;refGene_gn=ARHGEF10;refGene_tx=XM_023252272.1;refGene_gn=ARHGEF10;refGene_tx=XM_023252273.1;refGene_gn=ARHGEF10;refGene_tx=XM_023252274.1;refGene_gn=ARHGEF10;refGene_tx=XM_023252275.1;refGene_gn=ARHGEF10;refGene_tx=XM_023252276.1;refGene_gn=ARHGEF10;refGene_tx=XM_023252277.1
B1	1778329	1778452
refGene_gn=ARHGEF10;refGene_tx=XM_023252270.1;refGene_gn=ARHGEF10;refGene_tx=XM_023252271.1;refGene_gn=ARHGEF10;refGene_tx=XM_023252272.1;refGene_gn=ARHGEF10;refGene_tx=XM_023252273.1;refGene_gn=ARHGEF10;refGene_tx=XM_023252274.1;refGene_gn=ARHGEF10;refGene_tx=XM_023252275.1;refGene_gn=ARHGEF10;refGene_tx=XM_023252276.1;refGene_gn=ARHGEF10;refGene_tx=XM_023252277.1
B1	1781249	1781761
refGene_gn=ARHGEF10;refGene_tx=XM_023252270.1;refGene_gn=ARHGEF10;refGene_tx=XM_023252271.1;refGene_gn=ARHGEF10;refGene_tx=XM_023252272.1;refGene_gn=ARHGEF10;refGene_tx=XM_023252273.1;refGene_gn=ARHGEF10;refGene_tx=XM_023252274.1;refGene_gn=ARHGEF10;refGene_tx=XM_023252275.1;refGene_gn=ARHGEF10;refGene_tx=XM_023252276.1;refGene_gn=ARHGEF10;refGene_tx=XM_023252277.1
B1	1818541	1820596
refGene_gn=KBTBD11;refGene_tx=XM_023252278.1;refGene_gn=KBTBD11;refGene_tx=XM_023252279.1;refGene_gn=KBTBD11;refGene_tx=XM_023252280.1;refGene_gn=KBTBD11;refGene_tx=XM_023252281.1;refGene_gn=KBTBD11;refGene_tx=XM_023252282.1;refGene_gn=KBTBD11;refGene_tx=XM_023252283.1;refGene_gn=KBTBD11;refGene_tx=XM_023252284.1;refGene_gn=KBTBD11;refGene_tx=XM_023252285.1;refGene_gn=KBTBD11;refGene_tx=XM_023252286.1;refGene_gn=KBTB

D11;refGene_tx=XM_023252287.1;refGene_gn=KBTBD11;refGene_tx=XM_0232522
88.1;refGene_gn=KBTBD11;refGene_tx=XM_023252289.1;refGene_gn=KBTBD11;r
efGene_tx=XM_023252290.1
B1 1860328 1860451 refGene_gn=MYOM2;refGene_tx=XM_003984560.4
B1 1865516 1865635
refGene_gn=MYOM2;refGene_tx=XM_003984560.4;refGene_gn=MYOM2;refGene_tx
=XM_006930539.3;refGene_gn=MYOM2;refGene_tx=XM_011281363.3
B1 1867936 1868077
refGene_gn=MYOM2;refGene_tx=XM_003984560.4;refGene_gn=MYOM2;refGene_tx
=XM_006930539.3;refGene_gn=MYOM2;refGene_tx=XM_011281363.3
B1 1871546 1871685
refGene_gn=MYOM2;refGene_tx=XM_003984560.4;refGene_gn=MYOM2;refGene_tx
=XM_006930539.3;refGene_gn=MYOM2;refGene_tx=XM_011281363.3
B1 1871802 1871960
refGene_gn=MYOM2;refGene_tx=XM_003984560.4;refGene_gn=MYOM2;refGene_tx
=XM_006930539.3;refGene_gn=MYOM2;refGene_tx=XM_011281363.3
B1 1873142 1873235
refGene_gn=MYOM2;refGene_tx=XM_003984560.4;refGene_gn=MYOM2;refGene_tx
=XM_006930539.3;refGene_gn=MYOM2;refGene_tx=XM_011281363.3
B1 1880156 1880245
refGene_gn=MYOM2;refGene_tx=XM_003984560.4;refGene_gn=MYOM2;refGene_tx
=XM_006930539.3;refGene_gn=MYOM2;refGene_tx=XM_011281363.3
B1 1880321 1880372
refGene_gn=MYOM2;refGene_tx=XM_003984560.4;refGene_gn=MYOM2;refGene_tx
=XM_006930539.3;refGene_gn=MYOM2;refGene_tx=XM_011281363.3
B1 1882962 1883127
refGene_gn=MYOM2;refGene_tx=XM_003984560.4;refGene_gn=MYOM2;refGene_tx
=XM_006930539.3;refGene_gn=MYOM2;refGene_tx=XM_011281363.3
B1 1885674 1885836
refGene_gn=MYOM2;refGene_tx=XM_003984560.4;refGene_gn=MYOM2;refGene_tx
=XM_006930539.3;refGene_gn=MYOM2;refGene_tx=XM_011281363.3
B1 1888205 1888347
refGene_gn=MYOM2;refGene_tx=XM_003984560.4;refGene_gn=MYOM2;refGene_tx
=XM_006930539.3;refGene_gn=MYOM2;refGene_tx=XM_011281363.3
B1 1890350 1890550
refGene_gn=MYOM2;refGene_tx=XM_003984560.4;refGene_gn=MYOM2;refGene_tx
=XM_006930539.3;refGene_gn=MYOM2;refGene_tx=XM_011281363.3
B1 1891197 1891251
refGene_gn=MYOM2;refGene_tx=XM_003984560.4;refGene_gn=MYOM2;refGene_tx
=XM_006930539.3;refGene_gn=MYOM2;refGene_tx=XM_011281363.3
B1 1895179 1895307
refGene_gn=MYOM2;refGene_tx=XM_003984560.4;refGene_gn=MYOM2;refGene_tx
=XM_006930539.3;refGene_gn=MYOM2;refGene_tx=XM_011281363.3
B1 1898934 1899118
refGene_gn=MYOM2;refGene_tx=XM_003984560.4;refGene_gn=MYOM2;refGene_tx
=XM_006930539.3;refGene_gn=MYOM2;refGene_tx=XM_011281363.3
B1 1900327 1900502
refGene_gn=MYOM2;refGene_tx=XM_003984560.4;refGene_gn=MYOM2;refGene_tx
=XM_006930539.3;refGene_gn=MYOM2;refGene_tx=XM_011281363.3
B1 1902038 1902160

refGene_gn=MYOM2;refGene_tx=XM_003984560.4;refGene_gn=MYOM2;refGene_tx=XM_006930539.3;refGene_gn=MYOM2;refGene_tx=XM_011281363.3
B1 1903018 1903206
refGene_gn=MYOM2;refGene_tx=XM_003984560.4;refGene_gn=MYOM2;refGene_tx=XM_006930539.3;refGene_gn=MYOM2;refGene_tx=XM_011281363.3
B1 1904799 1904926
refGene_gn=MYOM2;refGene_tx=XM_003984560.4;refGene_gn=MYOM2;refGene_tx=XM_006930539.3;refGene_gn=MYOM2;refGene_tx=XM_011281363.3
B1 1905346 1905525
refGene_gn=MYOM2;refGene_tx=XM_003984560.4;refGene_gn=MYOM2;refGene_tx=XM_006930539.3;refGene_gn=MYOM2;refGene_tx=XM_011281363.3
B1 1906547 1906662
refGene_gn=MYOM2;refGene_tx=XM_003984560.4;refGene_gn=MYOM2;refGene_tx=XM_006930539.3;refGene_gn=MYOM2;refGene_tx=XM_011281363.3
B1 1907606 1907763
refGene_gn=MYOM2;refGene_tx=XM_003984560.4;refGene_gn=MYOM2;refGene_tx=XM_006930539.3;refGene_gn=MYOM2;refGene_tx=XM_011281363.3
B1 1907845 1907952
refGene_gn=MYOM2;refGene_tx=XM_003984560.4;refGene_gn=MYOM2;refGene_tx=XM_006930539.3;refGene_gn=MYOM2;refGene_tx=XM_011281363.3
B1 1910406 1910451
refGene_gn=MYOM2;refGene_tx=XM_003984560.4;refGene_gn=MYOM2;refGene_tx=XM_006930539.3;refGene_gn=MYOM2;refGene_tx=XM_011281363.3
B1 1911006 1911143
refGene_gn=MYOM2;refGene_tx=XM_003984560.4;refGene_gn=MYOM2;refGene_tx=XM_006930539.3;refGene_gn=MYOM2;refGene_tx=XM_011281363.3
B1 1915711 1915856
refGene_gn=MYOM2;refGene_tx=XM_003984560.4;refGene_gn=MYOM2;refGene_tx=XM_006930539.3;refGene_gn=MYOM2;refGene_tx=XM_011281363.3
B1 1915966 1916026
refGene_gn=MYOM2;refGene_tx=XM_003984560.4;refGene_gn=MYOM2;refGene_tx=XM_006930539.3;refGene_gn=MYOM2;refGene_tx=XM_011281363.3
B1 1917172 1917240
refGene_gn=MYOM2;refGene_tx=XM_003984560.4;refGene_gn=MYOM2;refGene_tx=XM_006930539.3;refGene_gn=MYOM2;refGene_tx=XM_011281363.3
B1 1922073 1922187
refGene_gn=MYOM2;refGene_tx=XM_003984560.4;refGene_gn=MYOM2;refGene_tx=XM_006930539.3;refGene_gn=MYOM2;refGene_tx=XM_011281363.3
B1 1922375 1922463
refGene_gn=MYOM2;refGene_tx=XM_003984560.4;refGene_gn=MYOM2;refGene_tx=XM_006930539.3;refGene_gn=MYOM2;refGene_tx=XM_011281363.3
B1 1922980 1923019
refGene_gn=MYOM2;refGene_tx=XM_003984560.4;refGene_gn=MYOM2;refGene_tx=XM_006930539.3;refGene_gn=MYOM2;refGene_tx=XM_011281363.3
B1 1925625 1925731
refGene_gn=MYOM2;refGene_tx=XM_003984560.4;refGene_gn=MYOM2;refGene_tx=XM_006930539.3;refGene_gn=MYOM2;refGene_tx=XM_011281363.3
B1 1926800 1926964
refGene_gn=MYOM2;refGene_tx=XM_003984560.4;refGene_gn=MYOM2;refGene_tx=XM_006930539.3;refGene_gn=MYOM2;refGene_tx=XM_011281363.3

B1	1927169	1927206
refGene_gn=MYOM2;refGene_tx=XM_003984560.4;refGene_gn=MYOM2;refGene_tx=XM_006930539.3;refGene_gn=MYOM2;refGene_tx=XM_011281363.3
B1	1928484	1928507
refGene_gn=MYOM2;refGene_tx=XM_003984560.4;refGene_gn=MYOM2;refGene_tx=XM_006930539.3;refGene_gn=MYOM2;refGene_tx=XM_011281363.3
B1	1930282	1930338
refGene_gn=MYOM2;refGene_tx=XM_003984560.4;refGene_gn=MYOM2;refGene_tx=XM_006930539.3;refGene_gn=MYOM2;refGene_tx=XM_011281363.3
B1	1931456	1931771
refGene_gn=MYOM2;refGene_tx=XM_003984560.4;refGene_gn=MYOM2;refGene_tx=XM_006930539.3;refGene_gn=MYOM2;refGene_tx=XM_011281363.3
B1	2780645	2780805	refGene_gn=CSMD1;refGene_tx=XM_023252291.1
B1	2784300	2784433	refGene_gn=CSMD1;refGene_tx=XM_023252291.1
B1	2791250	2791338	refGene_gn=CSMD1;refGene_tx=XM_023252291.1
B1	2791918	2792031	refGene_gn=CSMD1;refGene_tx=XM_023252291.1
B1	2794481	2794643	refGene_gn=CSMD1;refGene_tx=XM_023252291.1
B1	2797169	2797214	refGene_gn=CSMD1;refGene_tx=XM_023252291.1
B1	2798491	2798671	refGene_gn=CSMD1;refGene_tx=XM_023252291.1
B1	2801067	2801179	refGene_gn=CSMD1;refGene_tx=XM_023252291.1
B1	2804084	2804158	refGene_gn=CSMD1;refGene_tx=XM_023252291.1
B1	2806124	2806298	refGene_gn=CSMD1;refGene_tx=XM_023252291.1
B1	2806587	2806761	refGene_gn=CSMD1;refGene_tx=XM_023252291.1
B1	2809099	2809279	refGene_gn=CSMD1;refGene_tx=XM_023252291.1
B1	2810359	2810536	refGene_gn=CSMD1;refGene_tx=XM_023252291.1
B1	2818000	2818183	refGene_gn=CSMD1;refGene_tx=XM_023252291.1
B1	2819127	2819301	refGene_gn=CSMD1;refGene_tx=XM_023252291.1
B1	2836410	2836584	refGene_gn=CSMD1;refGene_tx=XM_023252291.1
B1	2856604	2856778	refGene_gn=CSMD1;refGene_tx=XM_023252291.1
B1	2861355	2861550	refGene_gn=CSMD1;refGene_tx=XM_023252291.1
B1	2878223	2878409	refGene_gn=CSMD1;refGene_tx=XM_023252291.1
B1	2911508	2911697	refGene_gn=CSMD1;refGene_tx=XM_023252291.1
B1	2916658	2916805	refGene_gn=CSMD1;refGene_tx=XM_023252291.1
B1	2924244	2924433	refGene_gn=CSMD1;refGene_tx=XM_023252291.1
B1	2931757	2931871	refGene_gn=CSMD1;refGene_tx=XM_023252291.1
B1	2932875	2932956	refGene_gn=CSMD1;refGene_tx=XM_023252291.1
B1	2933749	2933895	refGene_gn=CSMD1;refGene_tx=XM_023252291.1
B1	2936022	2936200	refGene_gn=CSMD1;refGene_tx=XM_023252291.1
B1	2944414	2944603	refGene_gn=CSMD1;refGene_tx=XM_023252291.1
B1	2954742	2954952	refGene_gn=CSMD1;refGene_tx=XM_023252291.1
B1	2963942	2964059	refGene_gn=CSMD1;refGene_tx=XM_023252291.1
B1	2970038	2970108	refGene_gn=CSMD1;refGene_tx=XM_023252291.1
B1	2974196	2974315	refGene_gn=CSMD1;refGene_tx=XM_023252291.1
B1	2997794	2997899	refGene_gn=CSMD1;refGene_tx=XM_023252291.1
B1	3011130	3011227	refGene_gn=CSMD1;refGene_tx=XM_023252291.1
B1	3011586	3011711	refGene_gn=CSMD1;refGene_tx=XM_023252291.1
B1	3012945	3013149	refGene_gn=CSMD1;refGene_tx=XM_023252291.1
B1	3020091	3020187	refGene_gn=CSMD1;refGene_tx=XM_023252291.1
B1	3021899	3022013	refGene_gn=CSMD1;refGene_tx=XM_023252291.1
B1	3026472	3026589	refGene_gn=CSMD1;refGene_tx=XM_023252291.1

B1 3034037 3034232 refGene_gn=CSMD1;refGene_tx=XM_023252291.1
B1 3039070 3039258 refGene_gn=CSMD1;refGene_tx=XM_023252291.1
B1 3043041 3043180 refGene_gn=CSMD1;refGene_tx=XM_023252291.1
B1 3048359 3048551 refGene_gn=CSMD1;refGene_tx=XM_023252291.1
B1 3080024 3080227 refGene_gn=CSMD1;refGene_tx=XM_023252291.1
B1 3096380 3096507 refGene_gn=CSMD1;refGene_tx=XM_023252291.1
B1 3096973 3097165 refGene_gn=CSMD1;refGene_tx=XM_023252291.1
B1 3121718 3121875 refGene_gn=CSMD1;refGene_tx=XM_023252291.1
B1 3126220 3126390 refGene_gn=CSMD1;refGene_tx=XM_023252291.1
B1 3137034 3137223 refGene_gn=CSMD1;refGene_tx=XM_023252291.1
B1 3139953 3140169 refGene_gn=CSMD1;refGene_tx=XM_023252291.1
B1 3141617 3141734 refGene_gn=CSMD1;refGene_tx=XM_023252291.1
B1 3150961 3151150 refGene_gn=CSMD1;refGene_tx=XM_023252291.1
B1 3159779 3159967 refGene_gn=CSMD1;refGene_tx=XM_023252291.1
B1 3163667 3163806 refGene_gn=CSMD1;refGene_tx=XM_023252291.1
B1 3171552 3171747 refGene_gn=CSMD1;refGene_tx=XM_023252291.1
B1 3173031 3173358 refGene_gn=CSMD1;refGene_tx=XM_023252291.1
B1 3174486 3174669 refGene_gn=CSMD1;refGene_tx=XM_023252291.1
B1 3234699 3234812 refGene_gn=CSMD1;refGene_tx=XM_023252291.1
B1 3256623 3256727 refGene_gn=CSMD1;refGene_tx=XM_023252291.1
B1 3330608 3330730 refGene_gn=CSMD1;refGene_tx=XM_023252291.1
B1 3343687 3343812 refGene_gn=CSMD1;refGene_tx=XM_023252291.1
B1 3367545 3367633 refGene_gn=CSMD1;refGene_tx=XM_023252291.1
B1 3452636 3452714 refGene_gn=CSMD1;refGene_tx=XM_023252291.1
B1 3511150 3511263 refGene_gn=CSMD1;refGene_tx=XM_023252291.1
B1 3785340 3785548 refGene_gn=CSMD1;refGene_tx=XM_023252291.1
B1 3812831 3813026 refGene_gn=CSMD1;refGene_tx=XM_023252291.1
B1 4189137 4189250 refGene_gn=CSMD1;refGene_tx=XM_023252291.1
B1 4430839 4431056 refGene_gn=CSMD1;refGene_tx=XM_023252291.1
B1 4810554 4810639 refGene_gn=CSMD1;refGene_tx=XM_023252291.1
B1 6234836 6234876 refGene_gn=MCPH1;refGene_tx=NM_001009348.1
B1 6236315 6236407 refGene_gn=MCPH1;refGene_tx=NM_001009348.1
B1 6245059 6245178 refGene_gn=MCPH1;refGene_tx=NM_001009348.1
B1 6272810 6272898 refGene_gn=MCPH1;refGene_tx=NM_001009348.1
B1 6275979 6276094 refGene_gn=MCPH1;refGene_tx=NM_001009348.1
B1 6281757 6281901 refGene_gn=MCPH1;refGene_tx=NM_001009348.1
B1 6285561 6285648 refGene_gn=MCPH1;refGene_tx=NM_001009348.1
B1 6288186 6288616 refGene_gn=MCPH1;refGene_tx=NM_001009348.1
B1 6288617 6289450 refGene_gn=MCPH1;refGene_tx=NM_001009348.1
B1 6299894 6300004 refGene_gn=MCPH1;refGene_tx=NM_001009348.1
B1 6337169 6337207 refGene_gn=MCPH1;refGene_tx=NM_001009348.1
B1 6339515 6339678 refGene_gn=MCPH1;refGene_tx=NM_001009348.1
B1 6363679 6363757 refGene_gn=MCPH1;refGene_tx=NM_001009348.1
B1 6366678 6366839
refGene_gn=ANGPT2;refGene_tx=XM_006944160.3;refGene_gn=ANGPT2;refGene_
tx=XM_006944161.3
B1 6370603 6370734
refGene_gn=ANGPT2;refGene_tx=XM_006944160.3;refGene_gn=ANGPT2;refGene_
tx=XM_006944161.3
B1 6375032 6375199

refGene_gn=ANGPT2;refGene_tx=XM_006944160.3;refGene_gn=ANGPT2;refGene_tx=XM_006944161.3
B1	6375763	6375865
refGene_gn=ANGPT2;refGene_tx=XM_006944160.3;refGene_gn=ANGPT2;refGene_tx=XM_006944161.3
B1	6382076	6382207
refGene_gn=ANGPT2;refGene_tx=XM_006944160.3;refGene_gn=ANGPT2;refGene_tx=XM_006944161.3
B1	6384633	6384863
refGene_gn=ANGPT2;refGene_tx=XM_006944160.3;refGene_gn=ANGPT2;refGene_tx=XM_006944161.3
B1	6389626	6389748
refGene_gn=ANGPT2;refGene_tx=XM_006944160.3;refGene_gn=ANGPT2;refGene_tx=XM_006944161.3
B1	6393127	6393283
refGene_gn=ANGPT2;refGene_tx=XM_006944160.3;refGene_gn=ANGPT2;refGene_tx=XM_006944161.3
B1	6422823	6423108
refGene_gn=ANGPT2;refGene_tx=XM_006944160.3;refGene_gn=ANGPT2;refGene_tx=XM_006944161.3
B1	6490555	6490793	refGene_gn=MCPH1;refGene_tx=NM_001009348.1
B1	6503407	6503463	refGene_gn=MCPH1;refGene_tx=NM_001009348.1
B1	6723115	6723334	refGene_gn=AGPAT5;refGene_tx=XM_011281365.3
B1	6738332	6738402	refGene_gn=AGPAT5;refGene_tx=XM_011281365.3
B1	6743283	6743399	refGene_gn=AGPAT5;refGene_tx=XM_011281365.3
B1	6746894	6746984	refGene_gn=AGPAT5;refGene_tx=XM_011281365.3
B1	6758373	6758464	refGene_gn=AGPAT5;refGene_tx=XM_011281365.3
B1	6771685	6771847	refGene_gn=AGPAT5;refGene_tx=XM_011281365.3
B1	6778151	6778275	refGene_gn=AGPAT5;refGene_tx=XM_011281365.3
B1	6780488	6780714	refGene_gn=AGPAT5;refGene_tx=XM_011281365.3
B1	6825871	6826983	refGene_gn=XKR5;refGene_tx=XM_023252292.1
B1	6829299	6829411	refGene_gn=XKR5;refGene_tx=XM_023252292.1
B1	6835401	6835571	refGene_gn=XKR5;refGene_tx=XM_023252292.1
B1	6837436	6837646	refGene_gn=XKR5;refGene_tx=XM_023252292.1
B1	6838706	6838891	refGene_gn=XKR5;refGene_tx=XM_023252292.1
B1	6847981	6848165	refGene_gn=XKR5;refGene_tx=XM_023252292.1
B1	6850061	6850119	refGene_gn=XKR5;refGene_tx=XM_023252292.1
B1	6855367	6855481
refGene_gn=LOC102902196;refGene_tx=XM_019828195.2;refGene_gn=LOC102902196;refGene_tx=XM_023252293.1
B1	6862006	6862155	refGene_gn=DEFB1;refGene_tx=XM_006930540.3
B1	6867764	6867910
refGene_gn=LOC102902196;refGene_tx=XM_019828195.2;refGene_gn=LOC102902196;refGene_tx=XM_023252293.1
B1	6873968	6874026	refGene_gn=DEFB1;refGene_tx=XM_006930540.3
B1	6891297	6891450
refGene_gn=LOC102902196;refGene_tx=XM_019828195.2;refGene_gn=LOC102902196;refGene_tx=XM_023252293.1
B1	6910610	6910746
refGene_gn=LOC102902196;refGene_tx=XM_019828195.2;refGene_gn=LOC102902

196;refGene_tx=XM_023252293.1
B1	6936038	6936123
refGene_gn=LOC102902196;refGene_tx=XM_019828195.2;refGene_gn=LOC102902196;refGene_tx=XM_023252293.1
B1	6936821	6936988
refGene_gn=LOC102902196;refGene_tx=XM_019828195.2;refGene_gn=LOC102902196;refGene_tx=XM_023252293.1
B1	6973533	6973679
refGene_gn=LOC101101212;refGene_tx=XM_003984552.2
B1	6974641	6974780
refGene_gn=LOC101101212;refGene_tx=XM_003984552.2
B1	6994632	6994823
refGene_gn=LOC102899407;refGene_tx=XM_006930554.3
B1	6996273	6996328
refGene_gn=LOC102899407;refGene_tx=XM_006930554.3
B1	7010349	7010483
refGene_gn=LOC101083583;refGene_tx=XM_003984563.2
B1	7013673	7013823
refGene_gn=LOC101083583;refGene_tx=XM_003984563.2
B1	7014996	7015054
refGene_gn=LOC101083583;refGene_tx=XM_003984563.2
B1	7022837	7022989
refGene_gn=LOC111560075;refGene_tx=XM_023252294.1
B1	7025279	7025328
refGene_gn=LOC111560075;refGene_tx=XM_023252294.1
B1	7030697	7030767
refGene_gn=LOC105260478;refGene_tx=XM_019828197.2
B1	7031127	7031169
refGene_gn=LOC105260478;refGene_tx=XM_019828197.2
B1	7034138	7034266
refGene_gn=LOC105260478;refGene_tx=XM_019828197.2
B1	7056613	7056662
refGene_gn=LOC109499336;refGene_tx=XM_023252246.1
B1	7059936	7061120
refGene_gn=LOC109499336;refGene_tx=XM_023252246.1
B1	7072607	7074215
refGene_gn=LOC111560019;refGene_tx=XM_023252217.1
B1	7090164	7090245
refGene_gn=LOC111556279;refGene_tx=XM_023252295.1
B1	7090726	7091611
refGene_gn=LOC111556279;refGene_tx=XM_023252295.1
B1	7116549	7118157
refGene_gn=LOC105260280;refGene_tx=XM_011280069.2
B1	7154447	7154459
refGene_gn=LOC102902320;refGene_tx=XM_006930541.2;refGene_gn=LOC102902320;refGene_tx=XM_019828204.1;refGene_gn=LOC102902320;refGene_tx=XM_023252296.1;refGene_gn=LOC102902320;refGene_tx=XM_023252297.1;refGene_gn=LOC102902320;refGene_tx=XM_023252298.1;refGene_gn=LOC102902320;refGene_tx=XM_023252299.1
B1	7155776	7155903

refGene_gn=LOC102902320;refGene_tx=XM_006930541.2;refGene_gn=LOC102902320;refGene_tx=XM_019828204.1;refGene_gn=LOC102902320;refGene_tx=XM_023252296.1;refGene_gn=LOC102902320;refGene_tx=XM_023252297.1;refGene_gn=LOC102902320;refGene_tx=XM_023252298.1;refGene_gn=LOC102902320;refGene_tx=XM_023252299.1
B1 7157078 7157174
refGene_gn=LOC102902320;refGene_tx=XM_006930541.2;refGene_gn=LOC102902320;refGene_tx=XM_019828204.1;refGene_gn=LOC102902320;refGene_tx=XM_023252296.1;refGene_gn=LOC102902320;refGene_tx=XM_023252297.1;refGene_gn=LOC102902320;refGene_tx=XM_023252298.1;refGene_gn=LOC102902320;refGene_tx=XM_023252299.1
B1 7157449 7157517
refGene_gn=LOC102902320;refGene_tx=XM_006930541.2
B1 7168126 7168233
refGene_gn=LOC102902320;refGene_tx=XM_006930541.2;refGene_gn=LOC102902320;refGene_tx=XM_019828204.1;refGene_gn=LOC102902320;refGene_tx=XM_023252296.1;refGene_gn=LOC102902320;refGene_tx=XM_023252297.1;refGene_gn=LOC102902320;refGene_tx=XM_023252298.1;refGene_gn=LOC102902320;refGene_tx=XM_023252299.1
B1 7190927 7191575
refGene_gn=LOC101084354;refGene_tx=XM_006930542.3;refGene_gn=LOC101084354;refGene_tx=XM_006930543.3;refGene_gn=LOC101084354;refGene_tx=XM_019828205.1;refGene_gn=LOC101084354;refGene_tx=XM_019828206.1;refGene_gn=LOC101084354;refGene_tx=XM_019828207.1;refGene_gn=LOC101084354;refGene_tx=XM_019828208.1;refGene_gn=LOC101084354;refGene_tx=XM_019828209.2;refGene_gn=LOC101084354;refGene_tx=XM_019828210.1;refGene_gn=LOC101084354;refGene_tx=XM_023252300.1;refGene_gn=LOC101084354;refGene_tx=XM_023252301.1;refGene_gn=LOC101084354;refGene_tx=XM_023252302.1
B1 7192015 7192098
refGene_gn=LOC101084354;refGene_tx=XM_006930542.3;refGene_gn=LOC101084354;refGene_tx=XM_006930543.3;refGene_gn=LOC101084354;refGene_tx=XM_019828205.1;refGene_gn=LOC101084354;refGene_tx=XM_019828206.1;refGene_gn=LOC101084354;refGene_tx=XM_019828207.1;refGene_gn=LOC101084354;refGene_tx=XM_019828208.1;refGene_gn=LOC101084354;refGene_tx=XM_019828209.2;refGene_gn=LOC101084354;refGene_tx=XM_019828210.1;refGene_gn=LOC101084354;refGene_tx=XM_023252300.1;refGene_gn=LOC101084354;refGene_tx=XM_023252301.1;refGene_gn=LOC101084354;refGene_tx=XM_023252302.1
B1 7192565 7192661
refGene_gn=LOC101084354;refGene_tx=XM_006930542.3;refGene_gn=LOC101084354;refGene_tx=XM_006930543.3;refGene_gn=LOC101084354;refGene_tx=XM_019828205.1;refGene_gn=LOC101084354;refGene_tx=XM_019828206.1;refGene_gn=LOC101084354;refGene_tx=XM_019828207.1;refGene_gn=LOC101084354;refGene_tx=XM_019828208.1;refGene_gn=LOC101084354;refGene_tx=XM_019828209.2;refGene_gn=LOC101084354;refGene_tx=XM_019828210.1;refGene_gn=LOC101084354;refGene_tx=XM_023252300.1;refGene_gn=LOC101084354;refGene_tx=XM_023252301.1;refGene_gn=LOC101084354;refGene_tx=XM_023252302.1
B1 7193381 7193508
refGene_gn=LOC101084354;refGene_tx=XM_006930542.3;refGene_gn=LOC101084354;refGene_tx=XM_006930543.3;refGene_gn=LOC101084354;refGene_tx=XM_019828205.1;refGene_gn=LOC101084354;refGene_tx=XM_019828206.1;refGene_gn

=LOC101084354;refGene_tx=XM_019828207.1;refGene_gn=LOC101084354;refGene_tx=XM_019828208.1;refGene_gn=LOC101084354;refGene_tx=XM_019828209.2;refGene_gn=LOC101084354;refGene_tx=XM_019828210.1;refGene_gn=LOC101084354;refGene_tx=XM_023252300.1;refGene_gn=LOC101084354;refGene_tx=XM_023252301.1;refGene_gn=LOC101084354;refGene_tx=XM_023252302.1
B1 7195256 7195268
refGene_gn=LOC101084354;refGene_tx=XM_006930542.3;refGene_gn=LOC101084354;refGene_tx=XM_006930543.3;refGene_gn=LOC101084354;refGene_tx=XM_019828205.1;refGene_gn=LOC101084354;refGene_tx=XM_019828206.1;refGene_gn=LOC101084354;refGene_tx=XM_019828207.1;refGene_gn=LOC101084354;refGene_tx=XM_019828208.1;refGene_gn=LOC101084354;refGene_tx=XM_019828209.2;refGene_gn=LOC101084354;refGene_tx=XM_019828210.1;refGene_gn=LOC101084354;refGene_tx=XM_023252300.1;refGene_gn=LOC101084354;refGene_tx=XM_023252301.1;refGene_gn=LOC101084354;refGene_tx=XM_023252302.1
B1 7233829 7233959
refGene_gn=LOC109499338;refGene_tx=XM_019829895.2
B1 7237127 7237288
refGene_gn=LOC109499338;refGene_tx=XM_019829895.2
B1 7263006 7263201
refGene_gn=LOC111560020;refGene_tx=XM_023252218.1
B1 7278852 7278910
refGene_gn=LOC109498699;refGene_tx=XM_019828211.2
B1 7285586 7285768
refGene_gn=LOC109498699;refGene_tx=XM_019828211.2
B1 7337370 7337523
refGene_gn=GPM6A;refGene_tx=XM_011281366.3;refGene_gn=GPM6A;refGene_tx=XM_019828212.1;refGene_gn=GPM6A;refGene_tx=XM_023252303.1
B1 7342300 7342366
refGene_gn=GPM6A;refGene_tx=XM_011281366.3;refGene_gn=GPM6A;refGene_tx=XM_019828212.1;refGene_gn=GPM6A;refGene_tx=XM_023252303.1
B1 7342921 7342998
refGene_gn=GPM6A;refGene_tx=XM_011281366.3;refGene_gn=GPM6A;refGene_tx=XM_019828212.1;refGene_gn=GPM6A;refGene_tx=XM_023252303.1
B1 7349450 7349604
refGene_gn=GPM6A;refGene_tx=XM_011281366.3;refGene_gn=GPM6A;refGene_tx=XM_019828212.1;refGene_gn=GPM6A;refGene_tx=XM_023252303.1
B1 7360831 7360988
refGene_gn=GPM6A;refGene_tx=XM_011281366.3;refGene_gn=GPM6A;refGene_tx=XM_019828212.1;refGene_gn=GPM6A;refGene_tx=XM_023252303.1
B1 7379863 7380056
refGene_gn=GPM6A;refGene_tx=XM_011281366.3;refGene_gn=GPM6A;refGene_tx=XM_019828212.1;refGene_gn=GPM6A;refGene_tx=XM_023252303.1
B1 7497164 7497201 refGene_gn=GPM6A;refGene_tx=XM_011281366.3
B1 7498484 7498491 refGene_gn=GPM6A;refGene_tx=XM_019828212.1
B1 7692070 7692074 refGene_gn=GPM6A;refGene_tx=XM_023252303.1
B1 7757472 7757545 refGene_gn=WDR17;refGene_tx=XM_023252308.1
B1 7759914 7759980
refGene_gn=WDR17;refGene_tx=XM_019828217.2;refGene_gn=WDR17;refGene_tx=XM_019828225.2;refGene_gn=WDR17;refGene_tx=XM_023252304.1;refGene_gn=WDR17;refGene_tx=XM_023252305.1;refGene_gn=WDR17;refGene_tx=XM_0232523

06.1;refGene_gn=WDR17;refGene_tx=XM_023252309.1
B1 7771721 7771850
refGene_gn=WDR17;refGene_tx=XM_019828217.2;refGene_gn=WDR17;refGene_tx=XM_019828219.1;refGene_gn=WDR17;refGene_tx=XM_019828224.1;refGene_gn=WDR17;refGene_tx=XM_019828225.2;refGene_gn=WDR17;refGene_tx=XM_023252304.1;refGene_gn=WDR17;refGene_tx=XM_023252305.1;refGene_gn=WDR17;refGene_tx=XM_023252306.1;refGene_gn=WDR17;refGene_tx=XM_023252307.1;refGene_gn=WDR17;refGene_tx=XM_023252308.1;refGene_gn=WDR17;refGene_tx=XM_023252309.1
B1 7779719 7779903
refGene_gn=WDR17;refGene_tx=XM_019828217.2;refGene_gn=WDR17;refGene_tx=XM_019828219.1;refGene_gn=WDR17;refGene_tx=XM_019828225.2;refGene_gn=WDR17;refGene_tx=XM_023252304.1;refGene_gn=WDR17;refGene_tx=XM_023252305.1;refGene_gn=WDR17;refGene_tx=XM_023252306.1;refGene_gn=WDR17;refGene_tx=XM_023252307.1;refGene_gn=WDR17;refGene_tx=XM_023252309.1
B1 7783817 7784048
refGene_gn=WDR17;refGene_tx=XM_019828217.2;refGene_gn=WDR17;refGene_tx=XM_019828219.1;refGene_gn=WDR17;refGene_tx=XM_019828224.1;refGene_gn=WDR17;refGene_tx=XM_019828225.2;refGene_gn=WDR17;refGene_tx=XM_023252304.1;refGene_gn=WDR17;refGene_tx=XM_023252305.1;refGene_gn=WDR17;refGene_tx=XM_023252306.1;refGene_gn=WDR17;refGene_tx=XM_023252307.1;refGene_gn=WDR17;refGene_tx=XM_023252308.1;refGene_gn=WDR17;refGene_tx=XM_023252309.1
B1 7789160 7789412
refGene_gn=WDR17;refGene_tx=XM_019828217.2;refGene_gn=WDR17;refGene_tx=XM_019828219.1;refGene_gn=WDR17;refGene_tx=XM_019828224.1;refGene_gn=WDR17;refGene_tx=XM_019828225.2;refGene_gn=WDR17;refGene_tx=XM_023252304.1;refGene_gn=WDR17;refGene_tx=XM_023252305.1;refGene_gn=WDR17;refGene_tx=XM_023252306.1;refGene_gn=WDR17;refGene_tx=XM_023252307.1;refGene_gn=WDR17;refGene_tx=XM_023252308.1;refGene_gn=WDR17;refGene_tx=XM_023252309.1
B1 7791376 7791499
refGene_gn=WDR17;refGene_tx=XM_019828217.2;refGene_gn=WDR17;refGene_tx=XM_019828219.1;refGene_gn=WDR17;refGene_tx=XM_019828224.1;refGene_gn=WDR17;refGene_tx=XM_019828225.2;refGene_gn=WDR17;refGene_tx=XM_023252304.1;refGene_gn=WDR17;refGene_tx=XM_023252305.1;refGene_gn=WDR17;refGene_tx=XM_023252306.1;refGene_gn=WDR17;refGene_tx=XM_023252307.1;refGene_gn=WDR17;refGene_tx=XM_023252308.1;refGene_gn=WDR17;refGene_tx=XM_023252309.1
B1 7794126 7794311
refGene_gn=WDR17;refGene_tx=XM_019828217.2;refGene_gn=WDR17;refGene_tx=XM_019828219.1;refGene_gn=WDR17;refGene_tx=XM_019828224.1;refGene_gn=WDR17;refGene_tx=XM_019828225.2;refGene_gn=WDR17;refGene_tx=XM_023252304.1;refGene_gn=WDR17;refGene_tx=XM_023252305.1;refGene_gn=WDR17;refGene_tx=XM_023252306.1;refGene_gn=WDR17;refGene_tx=XM_023252307.1;refGene_gn=WDR17;refGene_tx=XM_023252308.1;refGene_gn=WDR17;refGene_tx=XM_023252309.1
B1 7796588 7796757
refGene_gn=WDR17;refGene_tx=XM_019828217.2;refGene_gn=WDR17;refGene_tx=XM_019828219.1;refGene_gn=WDR17;refGene_tx=XM_019828224.1;refGene_gn=

WDR17;refGene_tx=XM_019828225.2;refGene_gn=WDR17;refGene_tx=XM_0232523
04.1;refGene_gn=WDR17;refGene_tx=XM_023252305.1;refGene_gn=WDR17;refGe
ne_tx=XM_023252306.1;refGene_gn=WDR17;refGene_tx=XM_023252307.1;refGen
e_gn=WDR17;refGene_tx=XM_023252308.1;refGene_gn=WDR17;refGene_tx=XM_02
3252309.1
B1 7799340 7799432
refGene_gn=WDR17;refGene_tx=XM_019828217.2;refGene_gn=WDR17;refGene_tx
=XM_019828219.1;refGene_gn=WDR17;refGene_tx=XM_019828224.1;refGene_gn=
WDR17;refGene_tx=XM_019828225.2;refGene_gn=WDR17;refGene_tx=XM_0232523
04.1;refGene_gn=WDR17;refGene_tx=XM_023252305.1;refGene_gn=WDR17;refGe
ne_tx=XM_023252306.1;refGene_gn=WDR17;refGene_tx=XM_023252307.1;refGen
e_gn=WDR17;refGene_tx=XM_023252308.1;refGene_gn=WDR17;refGene_tx=XM_02
3252309.1
B1 7802078 7802161
refGene_gn=WDR17;refGene_tx=XM_019828217.2;refGene_gn=WDR17;refGene_tx
=XM_019828219.1;refGene_gn=WDR17;refGene_tx=XM_019828224.1;refGene_gn=
WDR17;refGene_tx=XM_019828225.2;refGene_gn=WDR17;refGene_tx=XM_0232523
04.1;refGene_gn=WDR17;refGene_tx=XM_023252305.1;refGene_gn=WDR17;refGe
ne_tx=XM_023252306.1;refGene_gn=WDR17;refGene_tx=XM_023252307.1;refGen
e_gn=WDR17;refGene_tx=XM_023252308.1;refGene_gn=WDR17;refGene_tx=XM_02
3252309.1
B1 7803790 7803877
refGene_gn=WDR17;refGene_tx=XM_019828217.2;refGene_gn=WDR17;refGene_tx
=XM_019828219.1;refGene_gn=WDR17;refGene_tx=XM_019828224.1;refGene_gn=
WDR17;refGene_tx=XM_019828225.2;refGene_gn=WDR17;refGene_tx=XM_0232523
04.1;refGene_gn=WDR17;refGene_tx=XM_023252305.1;refGene_gn=WDR17;refGe
ne_tx=XM_023252306.1;refGene_gn=WDR17;refGene_tx=XM_023252307.1;refGen
e_gn=WDR17;refGene_tx=XM_023252308.1;refGene_gn=WDR17;refGene_tx=XM_02
3252309.1
B1 7809141 7809306
refGene_gn=WDR17;refGene_tx=XM_019828217.2;refGene_gn=WDR17;refGene_tx
=XM_019828219.1;refGene_gn=WDR17;refGene_tx=XM_019828224.1;refGene_gn=
WDR17;refGene_tx=XM_019828225.2;refGene_gn=WDR17;refGene_tx=XM_0232523
04.1;refGene_gn=WDR17;refGene_tx=XM_023252305.1;refGene_gn=WDR17;refGe
ne_tx=XM_023252306.1;refGene_gn=WDR17;refGene_tx=XM_023252307.1;refGen
e_gn=WDR17;refGene_tx=XM_023252308.1;refGene_gn=WDR17;refGene_tx=XM_02
3252309.1
B1 7811224 7811427
refGene_gn=WDR17;refGene_tx=XM_019828217.2;refGene_gn=WDR17;refGene_tx
=XM_019828219.1;refGene_gn=WDR17;refGene_tx=XM_019828224.1;refGene_gn=
WDR17;refGene_tx=XM_019828225.2;refGene_gn=WDR17;refGene_tx=XM_0232523
04.1;refGene_gn=WDR17;refGene_tx=XM_023252305.1;refGene_gn=WDR17;refGe
ne_tx=XM_023252306.1;refGene_gn=WDR17;refGene_tx=XM_023252307.1;refGen
e_gn=WDR17;refGene_tx=XM_023252308.1;refGene_gn=WDR17;refGene_tx=XM_02
3252309.1
B1 7812411 7812561
refGene_gn=WDR17;refGene_tx=XM_019828217.2;refGene_gn=WDR17;refGene_tx
=XM_019828219.1;refGene_gn=WDR17;refGene_tx=XM_019828224.1;refGene_gn=
WDR17;refGene_tx=XM_019828225.2;refGene_gn=WDR17;refGene_tx=XM_0232523
04.1;refGene_gn=WDR17;refGene_tx=XM_023252305.1;refGene_gn=WDR17;refGe

ne_tx=XM_023252306.1;refGene_gn=WDR17;refGene_tx=XM_023252307.1;refGene_gn=WDR17;refGene_tx=XM_023252308.1;refGene_gn=WDR17;refGene_tx=XM_023252309.1
B1	7812648	7812779
refGene_gn=WDR17;refGene_tx=XM_019828217.2;refGene_gn=WDR17;refGene_tx=XM_019828219.1;refGene_gn=WDR17;refGene_tx=XM_019828224.1;refGene_gn=WDR17;refGene_tx=XM_019828225.2;refGene_gn=WDR17;refGene_tx=XM_023252304.1;refGene_gn=WDR17;refGene_tx=XM_023252305.1;refGene_gn=WDR17;refGene_tx=XM_023252306.1;refGene_gn=WDR17;refGene_tx=XM_023252307.1;refGene_gn=WDR17;refGene_tx=XM_023252308.1;refGene_gn=WDR17;refGene_tx=XM_023252309.1
B1	7813081	7813207
refGene_gn=WDR17;refGene_tx=XM_019828217.2;refGene_gn=WDR17;refGene_tx=XM_019828219.1;refGene_gn=WDR17;refGene_tx=XM_019828224.1;refGene_gn=WDR17;refGene_tx=XM_019828225.2;refGene_gn=WDR17;refGene_tx=XM_023252304.1;refGene_gn=WDR17;refGene_tx=XM_023252305.1;refGene_gn=WDR17;refGene_tx=XM_023252306.1;refGene_gn=WDR17;refGene_tx=XM_023252307.1;refGene_gn=WDR17;refGene_tx=XM_023252308.1;refGene_gn=WDR17;refGene_tx=XM_023252309.1
B1	7814226	7814382
refGene_gn=WDR17;refGene_tx=XM_019828217.2;refGene_gn=WDR17;refGene_tx=XM_019828219.1;refGene_gn=WDR17;refGene_tx=XM_019828224.1;refGene_gn=WDR17;refGene_tx=XM_019828225.2;refGene_gn=WDR17;refGene_tx=XM_023252304.1;refGene_gn=WDR17;refGene_tx=XM_023252305.1;refGene_gn=WDR17;refGene_tx=XM_023252306.1;refGene_gn=WDR17;refGene_tx=XM_023252307.1;refGene_gn=WDR17;refGene_tx=XM_023252308.1;refGene_gn=WDR17;refGene_tx=XM_023252309.1
B1	7817193	7817258
refGene_gn=WDR17;refGene_tx=XM_019828217.2;refGene_gn=WDR17;refGene_tx=XM_019828219.1;refGene_gn=WDR17;refGene_tx=XM_019828224.1;refGene_gn=WDR17;refGene_tx=XM_019828225.2;refGene_gn=WDR17;refGene_tx=XM_023252304.1;refGene_gn=WDR17;refGene_tx=XM_023252305.1;refGene_gn=WDR17;refGene_tx=XM_023252306.1;refGene_gn=WDR17;refGene_tx=XM_023252307.1;refGene_gn=WDR17;refGene_tx=XM_023252308.1;refGene_gn=WDR17;refGene_tx=XM_023252309.1
B1	7820215	7820348
refGene_gn=WDR17;refGene_tx=XM_019828217.2;refGene_gn=WDR17;refGene_tx=XM_019828219.1;refGene_gn=WDR17;refGene_tx=XM_019828224.1;refGene_gn=WDR17;refGene_tx=XM_019828225.2;refGene_gn=WDR17;refGene_tx=XM_023252304.1;refGene_gn=WDR17;refGene_tx=XM_023252305.1;refGene_gn=WDR17;refGene_tx=XM_023252306.1;refGene_gn=WDR17;refGene_tx=XM_023252307.1;refGene_gn=WDR17;refGene_tx=XM_023252308.1;refGene_gn=WDR17;refGene_tx=XM_023252309.1
B1	7821127	7821219
refGene_gn=WDR17;refGene_tx=XM_019828217.2;refGene_gn=WDR17;refGene_tx=XM_019828219.1;refGene_gn=WDR17;refGene_tx=XM_019828224.1;refGene_gn=WDR17;refGene_tx=XM_019828225.2;refGene_gn=WDR17;refGene_tx=XM_023252304.1;refGene_gn=WDR17;refGene_tx=XM_023252305.1;refGene_gn=WDR17;refGene_tx=XM_023252306.1;refGene_gn=WDR17;refGene_tx=XM_023252307.1;refGene_gn=WDR17;refGene_tx=XM_023252308.1;refGene_gn=WDR17;refGene_tx=XM_02

3252309.1
B1	7822273	7822373
refGene_gn=WDR17;refGene_tx=XM_019828217.2;refGene_gn=WDR17;refGene_tx=XM_019828219.1;refGene_gn=WDR17;refGene_tx=XM_019828224.1;refGene_gn=WDR17;refGene_tx=XM_019828225.2;refGene_gn=WDR17;refGene_tx=XM_023252304.1;refGene_gn=WDR17;refGene_tx=XM_023252305.1;refGene_gn=WDR17;refGene_tx=XM_023252306.1;refGene_gn=WDR17;refGene_tx=XM_023252307.1;refGene_gn=WDR17;refGene_tx=XM_023252308.1;refGene_gn=WDR17;refGene_tx=XM_023252309.1
B1	7823191	7823331
refGene_gn=WDR17;refGene_tx=XM_019828217.2;refGene_gn=WDR17;refGene_tx=XM_019828219.1;refGene_gn=WDR17;refGene_tx=XM_019828224.1;refGene_gn=WDR17;refGene_tx=XM_019828225.2;refGene_gn=WDR17;refGene_tx=XM_023252304.1;refGene_gn=WDR17;refGene_tx=XM_023252305.1;refGene_gn=WDR17;refGene_tx=XM_023252306.1;refGene_gn=WDR17;refGene_tx=XM_023252307.1;refGene_gn=WDR17;refGene_tx=XM_023252308.1;refGene_gn=WDR17;refGene_tx=XM_023252309.1
B1	7826031	7826052
refGene_gn=WDR17;refGene_tx=XM_019828217.2;refGene_gn=WDR17;refGene_tx=XM_019828219.1;refGene_gn=WDR17;refGene_tx=XM_019828224.1;refGene_gn=WDR17;refGene_tx=XM_019828225.2;refGene_gn=WDR17;refGene_tx=XM_023252306.1;refGene_gn=WDR17;refGene_tx=XM_023252307.1;refGene_gn=WDR17;refGene_tx=XM_023252308.1;refGene_gn=WDR17;refGene_tx=XM_023252309.1
B1	7828183	7828295
refGene_gn=WDR17;refGene_tx=XM_019828217.2;refGene_gn=WDR17;refGene_tx=XM_019828219.1;refGene_gn=WDR17;refGene_tx=XM_019828224.1;refGene_gn=WDR17;refGene_tx=XM_019828225.2;refGene_gn=WDR17;refGene_tx=XM_023252304.1;refGene_gn=WDR17;refGene_tx=XM_023252305.1;refGene_gn=WDR17;refGene_tx=XM_023252306.1;refGene_gn=WDR17;refGene_tx=XM_023252307.1;refGene_gn=WDR17;refGene_tx=XM_023252308.1;refGene_gn=WDR17;refGene_tx=XM_023252309.1
B1	7831438	7831580
refGene_gn=WDR17;refGene_tx=XM_019828217.2;refGene_gn=WDR17;refGene_tx=XM_019828219.1;refGene_gn=WDR17;refGene_tx=XM_019828224.1;refGene_gn=WDR17;refGene_tx=XM_019828225.2;refGene_gn=WDR17;refGene_tx=XM_023252304.1;refGene_gn=WDR17;refGene_tx=XM_023252305.1;refGene_gn=WDR17;refGene_tx=XM_023252306.1;refGene_gn=WDR17;refGene_tx=XM_023252307.1;refGene_gn=WDR17;refGene_tx=XM_023252308.1;refGene_gn=WDR17;refGene_tx=XM_023252309.1
B1	7831857	7831960
refGene_gn=WDR17;refGene_tx=XM_019828217.2;refGene_gn=WDR17;refGene_tx=XM_019828219.1;refGene_gn=WDR17;refGene_tx=XM_019828224.1;refGene_gn=WDR17;refGene_tx=XM_019828225.2;refGene_gn=WDR17;refGene_tx=XM_023252304.1;refGene_gn=WDR17;refGene_tx=XM_023252305.1;refGene_gn=WDR17;refGene_tx=XM_023252306.1;refGene_gn=WDR17;refGene_tx=XM_023252307.1;refGene_gn=WDR17;refGene_tx=XM_023252308.1;refGene_gn=WDR17;refGene_tx=XM_023252309.1
B1	7833017	7833119
refGene_gn=WDR17;refGene_tx=XM_019828217.2;refGene_gn=WDR17;refGene_tx=XM_019828219.1;refGene_gn=WDR17;refGene_tx=XM_019828224.1;refGene_gn=

WDR17;refGene_tx=XM_019828225.2;refGene_gn=WDR17;refGene_tx=XM_0232523
04.1;refGene_gn=WDR17;refGene_tx=XM_023252305.1;refGene_gn=WDR17;refGe
ne_tx=XM_023252306.1;refGene_gn=WDR17;refGene_tx=XM_023252307.1;refGen
e_gn=WDR17;refGene_tx=XM_023252308.1;refGene_gn=WDR17;refGene_tx=XM_02
3252309.1
B1 7836993 7837092
refGene_gn=WDR17;refGene_tx=XM_019828217.2;refGene_gn=WDR17;refGene_tx
=XM_019828219.1;refGene_gn=WDR17;refGene_tx=XM_019828224.1;refGene_gn=
WDR17;refGene_tx=XM_019828225.2;refGene_gn=WDR17;refGene_tx=XM_0232523
04.1;refGene_gn=WDR17;refGene_tx=XM_023252305.1;refGene_gn=WDR17;refGe
ne_tx=XM_023252306.1;refGene_gn=WDR17;refGene_tx=XM_023252307.1;refGen
e_gn=WDR17;refGene_tx=XM_023252308.1
B1 7837411 7837607
refGene_gn=WDR17;refGene_tx=XM_019828217.2;refGene_gn=WDR17;refGene_tx
=XM_019828219.1;refGene_gn=WDR17;refGene_tx=XM_019828224.1;refGene_gn=
WDR17;refGene_tx=XM_019828225.2;refGene_gn=WDR17;refGene_tx=XM_0232523
04.1;refGene_gn=WDR17;refGene_tx=XM_023252305.1;refGene_gn=WDR17;refGe
ne_tx=XM_023252306.1;refGene_gn=WDR17;refGene_tx=XM_023252307.1;refGen
e_gn=WDR17;refGene_tx=XM_023252308.1
B1 7839740 7839860
refGene_gn=WDR17;refGene_tx=XM_019828217.2;refGene_gn=WDR17;refGene_tx
=XM_019828219.1;refGene_gn=WDR17;refGene_tx=XM_019828224.1;refGene_gn=
WDR17;refGene_tx=XM_019828225.2;refGene_gn=WDR17;refGene_tx=XM_0232523
04.1;refGene_gn=WDR17;refGene_tx=XM_023252305.1;refGene_gn=WDR17;refGe
ne_tx=XM_023252306.1;refGene_gn=WDR17;refGene_tx=XM_023252307.1;refGen
e_gn=WDR17;refGene_tx=XM_023252308.1
B1 7844822 7844929
refGene_gn=SPATA4;refGene_tx=XM_006930546.4;refGene_gn=SPATA4;refGene_
tx=XM_011281368.3;refGene_gn=SPATA4;refGene_tx=XM_011281370.3
B1 7846091 7846114
refGene_gn=SPATA4;refGene_tx=XM_011281369.3;refGene_gn=SPATA4;refGene_
tx=XM_019828226.2
B1 7846432 7846522
refGene_gn=SPATA4;refGene_tx=XM_006930546.4;refGene_gn=SPATA4;refGene_
tx=XM_011281368.3;refGene_gn=SPATA4;refGene_tx=XM_011281369.3;refGene_
gn=SPATA4;refGene_tx=XM_019828226.2
B1 7847812 7847929
refGene_gn=SPATA4;refGene_tx=XM_006930546.4;refGene_gn=SPATA4;refGene_
tx=XM_011281368.3;refGene_gn=SPATA4;refGene_tx=XM_011281369.3;refGene_
gn=SPATA4;refGene_tx=XM_011281370.3;refGene_gn=SPATA4;refGene_tx=XM_01
9828226.2
B1 7855245 7855466
refGene_gn=SPATA4;refGene_tx=XM_006930546.4;refGene_gn=SPATA4;refGene_
tx=XM_011281368.3;refGene_gn=SPATA4;refGene_tx=XM_011281369.3;refGene_
gn=SPATA4;refGene_tx=XM_011281370.3;refGene_gn=SPATA4;refGene_tx=XM_01
9828226.2
B1 7855571 7855690
refGene_gn=SPATA4;refGene_tx=XM_006930546.4;refGene_gn=SPATA4;refGene_
tx=XM_011281368.3;refGene_gn=SPATA4;refGene_tx=XM_011281369.3;refGene_
gn=SPATA4;refGene_tx=XM_011281370.3;refGene_gn=SPATA4;refGene_tx=XM_01

9828226.2
B1	7856291	7856421
refGene_gn=SPATA4;refGene_tx=XM_006930546.4;refGene_gn=SPATA4;refGene_tx=XM_011281368.3;refGene_gn=SPATA4;refGene_tx=XM_011281369.3;refGene_gn=SPATA4;refGene_tx=XM_011281370.3;refGene_gn=SPATA4;refGene_tx=XM_019828226.2
B1	7857261	7857479
refGene_gn=SPATA4;refGene_tx=XM_006930546.4;refGene_gn=SPATA4;refGene_tx=XM_011281368.3;refGene_gn=SPATA4;refGene_tx=XM_011281369.3;refGene_gn=SPATA4;refGene_tx=XM_011281370.3;refGene_gn=SPATA4;refGene_tx=XM_019828226.2
B1	7865128	7865256
refGene_gn=ASB5;refGene_tx=XM_006930548.4;refGene_gn=ASB5;refGene_tx=XM_006930549.4;refGene_gn=ASB5;refGene_tx=XM_019828227.2;refGene_gn=ASB5;refGene_tx=XM_019828228.2
B1	7866499	7866691
refGene_gn=ASB5;refGene_tx=XM_006930548.4;refGene_gn=ASB5;refGene_tx=XM_006930549.4;refGene_gn=ASB5;refGene_tx=XM_019828227.2;refGene_gn=ASB5;refGene_tx=XM_019828228.2
B1	7868933	7869068
refGene_gn=ASB5;refGene_tx=XM_006930548.4;refGene_gn=ASB5;refGene_tx=XM_006930549.4;refGene_gn=ASB5;refGene_tx=XM_019828227.2;refGene_gn=ASB5;refGene_tx=XM_019828228.2
B1	7869230	7869381
refGene_gn=ASB5;refGene_tx=XM_006930548.4;refGene_gn=ASB5;refGene_tx=XM_006930549.4;refGene_gn=ASB5;refGene_tx=XM_019828227.2;refGene_gn=ASB5;refGene_tx=XM_019828228.2
B1	7870685	7870793
refGene_gn=ASB5;refGene_tx=XM_006930548.4;refGene_gn=ASB5;refGene_tx=XM_006930549.4;refGene_gn=ASB5;refGene_tx=XM_019828227.2;refGene_gn=ASB5;refGene_tx=XM_019828228.2
B1	7872970	7873050
refGene_gn=ASB5;refGene_tx=XM_006930548.4;refGene_gn=ASB5;refGene_tx=XM_006930549.4;refGene_gn=ASB5;refGene_tx=XM_019828227.2;refGene_gn=ASB5;refGene_tx=XM_019828228.2
B1	7878376	7878437	refGene_gn=ASB5;refGene_tx=XM_019828228.2
B1	7886783	7886907	refGene_gn=ASB5;refGene_tx=XM_006930548.4
B1	7891009	7891046	refGene_gn=ASB5;refGene_tx=XM_006930549.4
B1	7916355	7916551	refGene_gn=ASB5;refGene_tx=XM_019828227.2
B1	7952646	7952789	refGene_gn=SPCS3;refGene_tx=XM_023252310.1
B1	7954253	7954327	refGene_gn=SPCS3;refGene_tx=XM_023252310.1
B1	7956709	7956786	refGene_gn=SPCS3;refGene_tx=XM_023252310.1
B1	7960320	7960436	refGene_gn=SPCS3;refGene_tx=XM_023252310.1
B1	7961375	7961508	refGene_gn=SPCS3;refGene_tx=XM_023252310.1
B1	8260105	8260220	refGene_gn=VEGFC;refGene_tx=XM_023252311.1
B1	8263313	8263647	refGene_gn=VEGFC;refGene_tx=XM_023252311.1
B1	8263952	8264059	refGene_gn=VEGFC;refGene_tx=XM_023252311.1
B1	8279429	8279581	refGene_gn=VEGFC;refGene_tx=XM_023252311.1
B1	8294396	8294587	refGene_gn=VEGFC;refGene_tx=XM_023252311.1
B1	8296023	8296237	refGene_gn=VEGFC;refGene_tx=XM_023252311.1

| | | | |
|---|---|---|---|
| B1 | 8370173 | 8370320 | refGene_gn=VEGFC;refGene_tx=XM_023252311.1 |
| B1 | 8858630 | 8858753 | refGene_gn=NEIL3;refGene_tx=XM_019828229.2 |
| B1 | 8873641 | 8873769 | refGene_gn=NEIL3;refGene_tx=XM_019828229.2 |
| B1 | 8882791 | 8882926 | refGene_gn=NEIL3;refGene_tx=XM_019828229.2 |
| B1 | 8883206 | 8883420 | refGene_gn=NEIL3;refGene_tx=XM_019828229.2 |
| B1 | 8885960 | 8886035 | refGene_gn=NEIL3;refGene_tx=XM_019828229.2 |
| B1 | 8888094 | 8888261 | refGene_gn=NEIL3;refGene_tx=XM_019828229.2 |
| B1 | 8896367 | 8896525 | refGene_gn=NEIL3;refGene_tx=XM_019828229.2 |
| B1 | 8898651 | 8899072 | refGene_gn=NEIL3;refGene_tx=XM_019828229.2 |
| B1 | 8906297 | 8906472 | refGene_gn=NEIL3;refGene_tx=XM_019828229.2 |
| B1 | 8907924 | 8908107 | refGene_gn=NEIL3;refGene_tx=XM_019828229.2 |
| B1 | 8974393 | 8974494 | refGene_gn=AGA;refGene_tx=XM_023252312.1 |
| B1 | 8975961 | 8976095 | refGene_gn=AGA;refGene_tx=XM_023252312.1 |
| B1 | 8976804 | 8976912 | refGene_gn=AGA;refGene_tx=XM_023252312.1 |
| B1 | 8977755 | 8977831 | refGene_gn=AGA;refGene_tx=XM_023252312.1 |
| B1 | 8978772 | 8978887 | refGene_gn=AGA;refGene_tx=XM_023252312.1 |
| B1 | 8980320 | 8980433 | refGene_gn=AGA;refGene_tx=XM_023252312.1 |
| B1 | 8981736 | 8981849 | refGene_gn=AGA;refGene_tx=XM_023252312.1 |
| B1 | 8982620 | 8982774 | refGene_gn=AGA;refGene_tx=XM_023252312.1 |
| B1 | 8984607 | 8984734 | refGene_gn=AGA;refGene_tx=XM_023252312.1 |
| B1 | 13633767 | 13633986 | refGene_gn=TENM3;refGene_tx=XM_023252313.1 |
| B1 | 13633987 | 13634000 | refGene_gn=TENM3;refGene_tx=XM_023252313.1 |
| B1 | 13655315 | 13655594 | refGene_gn=TENM3;refGene_tx=XM_023252313.1 |
| B1 | 13895504 | 13895742 | refGene_gn=TENM3;refGene_tx=XM_023252313.1 |
| B1 | 13922362 | 13922601 | refGene_gn=TENM3;refGene_tx=XM_023252313.1 |
| B1 | 13950324 | 13950447 | refGene_gn=TENM3;refGene_tx=XM_023252313.1 |
| B1 | 13969603 | 13969818 | refGene_gn=TENM3;refGene_tx=XM_023252313.1 |
| B1 | 13975652 | 13975863 | refGene_gn=TENM3;refGene_tx=XM_023252313.1 |
| B1 | 13976357 | 13976459 | refGene_gn=TENM3;refGene_tx=XM_023252313.1 |
| B1 | 13976648 | 13976843 | refGene_gn=TENM3;refGene_tx=XM_023252313.1 |
| B1 | 13978056 | 13978257 | refGene_gn=TENM3;refGene_tx=XM_023252313.1 |
| B1 | 13984932 | 13985118 | refGene_gn=TENM3;refGene_tx=XM_023252313.1 |
| B1 | 14006922 | 14006949 | refGene_gn=TENM3;refGene_tx=XM_023252313.1 |
| B1 | 14008517 | 14008538 | refGene_gn=TENM3;refGene_tx=XM_023252313.1 |
| B1 | 14009455 | 14009602 | refGene_gn=TENM3;refGene_tx=XM_023252313.1 |
| B1 | 14023399 | 14023616 | refGene_gn=TENM3;refGene_tx=XM_023252313.1 |
| B1 | 14024631 | 14024751 | refGene_gn=TENM3;refGene_tx=XM_023252313.1 |
| B1 | 14025555 | 14025817 | refGene_gn=TENM3;refGene_tx=XM_023252313.1 |
| B1 | 14031430 | 14031698 | refGene_gn=TENM3;refGene_tx=XM_023252313.1 |
| B1 | 14033248 | 14033392 | refGene_gn=TENM3;refGene_tx=XM_023252313.1 |
| B1 | 14037698 | 14037948 | refGene_gn=TENM3;refGene_tx=XM_023252313.1 |
| B1 | 14047811 | 14048044 | refGene_gn=TENM3;refGene_tx=XM_023252313.1 |
| B1 | 14049125 | 14049280 | refGene_gn=TENM3;refGene_tx=XM_023252313.1 |
| B1 | 14050275 | 14051150 | refGene_gn=TENM3;refGene_tx=XM_023252313.1 |
| B1 | 14068898 | 14069074 | refGene_gn=TENM3;refGene_tx=XM_023252313.1 |
| B1 | 14070846 | 14071082 | refGene_gn=TENM3;refGene_tx=XM_023252313.1 |
| B1 | 14078730 | 14079027 | refGene_gn=TENM3;refGene_tx=XM_023252313.1 |
| B1 | 14082100 | 14083712 | refGene_gn=TENM3;refGene_tx=XM_023252313.1 |
| B1 | 14086401 | 14086532 | refGene_gn=TENM3;refGene_tx=XM_023252313.1 |
| B1 | 14089386 | 14090142 | refGene_gn=TENM3;refGene_tx=XM_023252313.1 |

B1 14159562 14159699
refGene_gn=DCTD;refGene_tx=XM_023252314.1;refGene_gn=DCTD;refGene_tx=XM_023252315.1
B1 14161024 14161121 refGene_gn=DCTD;refGene_tx=XM_023252314.1
B1 14162402 14162519
refGene_gn=DCTD;refGene_tx=XM_023252314.1;refGene_gn=DCTD;refGene_tx=XM_023252315.1
B1 14182540 14182676
refGene_gn=DCTD;refGene_tx=XM_023252314.1;refGene_gn=DCTD;refGene_tx=XM_023252315.1
B1 14183070 14183178
refGene_gn=DCTD;refGene_tx=XM_023252314.1;refGene_gn=DCTD;refGene_tx=XM_023252315.1
B1 14283160 14283291 refGene_gn=WWC2;refGene_tx=XM_023252316.1
B1 14293100 14293102 refGene_gn=WWC2;refGene_tx=XM_023252317.1
B1 14383781 14383891
refGene_gn=WWC2;refGene_tx=XM_023252316.1;refGene_gn=WWC2;refGene_tx=XM_023252317.1
B1 14395760 14395964
refGene_gn=WWC2;refGene_tx=XM_023252316.1;refGene_gn=WWC2;refGene_tx=XM_023252317.1
B1 14396777 14396854
refGene_gn=WWC2;refGene_tx=XM_023252316.1;refGene_gn=WWC2;refGene_tx=XM_023252317.1
B1 14416970 14417050
refGene_gn=WWC2;refGene_tx=XM_019828241.2;refGene_gn=WWC2;refGene_tx=XM_023252316.1;refGene_gn=WWC2;refGene_tx=XM_023252317.1
B1 14422726 14422856
refGene_gn=WWC2;refGene_tx=XM_019828241.2;refGene_gn=WWC2;refGene_tx=XM_023252316.1;refGene_gn=WWC2;refGene_tx=XM_023252317.1
B1 14426275 14426422
refGene_gn=WWC2;refGene_tx=XM_019828241.2;refGene_gn=WWC2;refGene_tx=XM_023252316.1;refGene_gn=WWC2;refGene_tx=XM_023252317.1
B1 14427449 14427523
refGene_gn=WWC2;refGene_tx=XM_019828241.2;refGene_gn=WWC2;refGene_tx=XM_023252316.1;refGene_gn=WWC2;refGene_tx=XM_023252317.1
B1 14431223 14431466
refGene_gn=WWC2;refGene_tx=XM_019828241.2;refGene_gn=WWC2;refGene_tx=XM_023252316.1;refGene_gn=WWC2;refGene_tx=XM_023252317.1
B1 14437085 14437175
refGene_gn=WWC2;refGene_tx=XM_019828241.2;refGene_gn=WWC2;refGene_tx=XM_023252316.1;refGene_gn=WWC2;refGene_tx=XM_023252317.1
B1 14438222 14438842
refGene_gn=WWC2;refGene_tx=XM_019828241.2;refGene_gn=WWC2;refGene_tx=XM_023252316.1;refGene_gn=WWC2;refGene_tx=XM_023252317.1
B1 14441903 14442033
refGene_gn=WWC2;refGene_tx=XM_019828241.2;refGene_gn=WWC2;refGene_tx=XM_023252316.1;refGene_gn=WWC2;refGene_tx=XM_023252317.1
B1 14442863 14442944
refGene_gn=WWC2;refGene_tx=XM_019828241.2;refGene_gn=WWC2;refGene_tx=X

M_023252316.1;refGene_gn=WWC2;refGene_tx=XM_023252317.1
B1 14443042 14443129
refGene_gn=WWC2;refGene_tx=XM_019828241.2;refGene_gn=WWC2;refGene_tx=XM_023252316.1;refGene_gn=WWC2;refGene_tx=XM_023252317.1
B1 14445764 14445957
refGene_gn=WWC2;refGene_tx=XM_019828241.2;refGene_gn=WWC2;refGene_tx=XM_023252316.1;refGene_gn=WWC2;refGene_tx=XM_023252317.1
B1 14447590 14447755
refGene_gn=WWC2;refGene_tx=XM_019828241.2;refGene_gn=WWC2;refGene_tx=XM_023252316.1;refGene_gn=WWC2;refGene_tx=XM_023252317.1
B1 14460355 14460486
refGene_gn=WWC2;refGene_tx=XM_019828241.2;refGene_gn=WWC2;refGene_tx=XM_023252316.1;refGene_gn=WWC2;refGene_tx=XM_023252317.1
B1 14462655 14462854
refGene_gn=WWC2;refGene_tx=XM_019828241.2;refGene_gn=WWC2;refGene_tx=XM_023252316.1;refGene_gn=WWC2;refGene_tx=XM_023252317.1
B1 14464082 14464247
refGene_gn=WWC2;refGene_tx=XM_019828241.2;refGene_gn=WWC2;refGene_tx=XM_023252316.1;refGene_gn=WWC2;refGene_tx=XM_023252317.1
B1 14465628 14465721
refGene_gn=WWC2;refGene_tx=XM_019828241.2;refGene_gn=WWC2;refGene_tx=XM_023252316.1;refGene_gn=WWC2;refGene_tx=XM_023252317.1
B1 14468329 14468572
refGene_gn=WWC2;refGene_tx=XM_019828241.2;refGene_gn=WWC2;refGene_tx=XM_023252316.1;refGene_gn=WWC2;refGene_tx=XM_023252317.1
B1 14487567 14487695
refGene_gn=WWC2;refGene_tx=XM_019828241.2;refGene_gn=WWC2;refGene_tx=XM_023252316.1;refGene_gn=WWC2;refGene_tx=XM_023252317.1
B1 14490209 14490276
refGene_gn=WWC2;refGene_tx=XM_019828241.2;refGene_gn=WWC2;refGene_tx=XM_023252316.1;refGene_gn=WWC2;refGene_tx=XM_023252317.1
B1 14494110 14494773
refGene_gn=LOC101095096;refGene_tx=XM_003984603.2
B1 14496332 14497106
refGene_gn=LOC101087918;refGene_tx=XM_019828242.2
B1 14591743 14592015
refGene_gn=CDKN2AIP;refGene_tx=XM_011281373.3;refGene_gn=CDKN2AIP;refGene_tx=XM_011281374.3
B1 14592475 14592606
refGene_gn=CDKN2AIP;refGene_tx=XM_011281373.3;refGene_gn=CDKN2AIP;refGene_tx=XM_011281374.3
B1 14593038 14594387
refGene_gn=CDKN2AIP;refGene_tx=XM_011281373.3;refGene_gn=CDKN2AIP;refGene_tx=XM_011281374.3
B1 14642429 14642601 refGene_gn=ING2;refGene_tx=XM_003984576.5
B1 14647309 14647574 refGene_gn=ING2;refGene_tx=XM_003984576.5
B1 14647575 14647981 refGene_gn=ING2;refGene_tx=XM_003984576.5
B1 14744003 14744054 refGene_gn=RWDD4;refGene_tx=XM_019828248.2
B1 14745594 14745627
refGene_gn=RWDD4;refGene_tx=XM_011281375.2;refGene_gn=RWDD4;refGene_tx

=XM_019828249.2
B1	14748173 14748176 refGene_gn=RWDD4;refGene_tx=XM_011281375.2
B1	14748311 14748361
refGene_gn=RWDD4;refGene_tx=XM_011281375.2;refGene_gn=RWDD4;refGene_tx=XM_019828248.2;refGene_gn=RWDD4;refGene_tx=XM_019828249.2
B1	14750614 14750732
refGene_gn=RWDD4;refGene_tx=XM_011281375.2;refGene_gn=RWDD4;refGene_tx=XM_019828248.2;refGene_gn=RWDD4;refGene_tx=XM_019828249.2
B1	14752565 14752713
refGene_gn=RWDD4;refGene_tx=XM_011281375.2;refGene_gn=RWDD4;refGene_tx=XM_019828248.2;refGene_gn=RWDD4;refGene_tx=XM_019828249.2
B1	14752797 14752907
refGene_gn=RWDD4;refGene_tx=XM_011281375.2;refGene_gn=RWDD4;refGene_tx=XM_019828248.2;refGene_gn=RWDD4;refGene_tx=XM_019828249.2
B1	14757602 14757683
refGene_gn=RWDD4;refGene_tx=XM_011281375.2;refGene_gn=RWDD4;refGene_tx=XM_019828248.2;refGene_gn=RWDD4;refGene_tx=XM_019828249.2
B1	14760117 14760141
refGene_gn=RWDD4;refGene_tx=XM_011281375.2;refGene_gn=RWDD4;refGene_tx=XM_019828248.2;refGene_gn=RWDD4;refGene_tx=XM_019828249.2
B1	14763695 14763899
refGene_gn=TRAPPC11;refGene_tx=XM_011281376.3;refGene_gn=TRAPPC11;refGene_tx=XM_019828244.2
B1	14765444 14765614
refGene_gn=TRAPPC11;refGene_tx=XM_011281376.3;refGene_gn=TRAPPC11;refGene_tx=XM_019828244.2
B1	14766260 14766331
refGene_gn=TRAPPC11;refGene_tx=XM_011281376.3;refGene_gn=TRAPPC11;refGene_tx=XM_019828244.2
B1	14767168 14767283
refGene_gn=TRAPPC11;refGene_tx=XM_011281376.3;refGene_gn=TRAPPC11;refGene_tx=XM_019828244.2
B1	14770700 14770800
refGene_gn=TRAPPC11;refGene_tx=XM_011281376.3;refGene_gn=TRAPPC11;refGene_tx=XM_019828244.2
B1	14771135 14771209
refGene_gn=TRAPPC11;refGene_tx=XM_011281376.3;refGene_gn=TRAPPC11;refGene_tx=XM_019828244.2
B1	14772957 14773054
refGene_gn=TRAPPC11;refGene_tx=XM_011281376.3;refGene_gn=TRAPPC11;refGene_tx=XM_019828244.2
B1	14775116 14775250
refGene_gn=TRAPPC11;refGene_tx=XM_011281376.3;refGene_gn=TRAPPC11;refGene_tx=XM_019828244.2
B1	14775905 14776053
refGene_gn=TRAPPC11;refGene_tx=XM_011281376.3;refGene_gn=TRAPPC11;refGene_tx=XM_019828244.2
B1	14778027 14778121
refGene_gn=TRAPPC11;refGene_tx=XM_011281376.3;refGene_gn=TRAPPC11;refGene_tx=XM_019828244.2

B1	14779093 14779173
refGene_gn=TRAPPC11;refGene_tx=XM_011281376.3;refGene_gn=TRAPPC11;refGene_tx=XM_019828244.2
B1	14779275 14779354
refGene_gn=TRAPPC11;refGene_tx=XM_011281376.3;refGene_gn=TRAPPC11;refGene_tx=XM_019828244.2
B1	14779437 14779492
refGene_gn=TRAPPC11;refGene_tx=XM_011281376.3;refGene_gn=TRAPPC11;refGene_tx=XM_019828244.2
B1	14779831 14779977
refGene_gn=TRAPPC11;refGene_tx=XM_011281376.3;refGene_gn=TRAPPC11;refGene_tx=XM_019828244.2
B1	14780419 14780481
refGene_gn=TRAPPC11;refGene_tx=XM_011281376.3;refGene_gn=TRAPPC11;refGene_tx=XM_019828244.2
B1	14780591 14780724
refGene_gn=TRAPPC11;refGene_tx=XM_011281376.3;refGene_gn=TRAPPC11;refGene_tx=XM_019828244.2
B1	14781842 14781973
refGene_gn=TRAPPC11;refGene_tx=XM_011281376.3;refGene_gn=TRAPPC11;refGene_tx=XM_019828244.2
B1	14785644 14785800
refGene_gn=TRAPPC11;refGene_tx=XM_011281376.3;refGene_gn=TRAPPC11;refGene_tx=XM_019828244.2
B1	14787162 14787350
refGene_gn=TRAPPC11;refGene_tx=XM_011281376.3;refGene_gn=TRAPPC11;refGene_tx=XM_019828244.2
B1	14787485 14787634
refGene_gn=TRAPPC11;refGene_tx=XM_011281376.3;refGene_gn=TRAPPC11;refGene_tx=XM_019828244.2
B1	14787815 14787937
refGene_gn=TRAPPC11;refGene_tx=XM_011281376.3;refGene_gn=TRAPPC11;refGene_tx=XM_019828244.2
B1	14788467 14788587
refGene_gn=TRAPPC11;refGene_tx=XM_011281376.3;refGene_gn=TRAPPC11;refGene_tx=XM_019828244.2
B1	14792106 14792172
refGene_gn=TRAPPC11;refGene_tx=XM_011281376.3;refGene_gn=TRAPPC11;refGene_tx=XM_019828244.2
B1	14792282 14792439
refGene_gn=TRAPPC11;refGene_tx=XM_011281376.3;refGene_gn=TRAPPC11;refGene_tx=XM_019828244.2
B1	14797001 14797113
refGene_gn=TRAPPC11;refGene_tx=XM_011281376.3;refGene_gn=TRAPPC11;refGene_tx=XM_019828244.2
B1	14800230 14800322
refGene_gn=TRAPPC11;refGene_tx=XM_011281376.3;refGene_gn=TRAPPC11;refGene_tx=XM_019828244.2
B1	14801303 14801437
refGene_gn=TRAPPC11;refGene_tx=XM_011281376.3;refGene_gn=TRAPPC11;refG

ene_tx=XM_019828244.2
B1	14803288 14803456
refGene_gn=TRAPPC11;refGene_tx=XM_011281376.3;refGene_gn=TRAPPC11;refG
ene_tx=XM_019828244.2
B1	14807470 14807515 refGene_gn=TRAPPC11;refGene_tx=XM_011281376.3
B1	14811295 14811300 refGene_gn=TRAPPC11;refGene_tx=XM_019828244.2
B1	14817603 14817658 refGene_gn=TRAPPC11;refGene_tx=XM_019828244.2
B1	14974826 14974992 refGene_gn=STOX2;refGene_tx=XM_011281377.3
B1	15053292 15053317
refGene_gn=STOX2;refGene_tx=XM_023252318.1;refGene_gn=STOX2;refGene_tx
=XM_023252319.1
B1	15067550 15067703
refGene_gn=STOX2;refGene_tx=XM_011281377.3;refGene_gn=STOX2;refGene_tx
=XM_019828246.2;refGene_gn=STOX2;refGene_tx=XM_023252318.1;refGene_gn=
STOX2;refGene_tx=XM_023252319.1
B1	15076058 15078324
refGene_gn=STOX2;refGene_tx=XM_011281377.3;refGene_gn=STOX2;refGene_tx
=XM_019828246.2;refGene_gn=STOX2;refGene_tx=XM_023252318.1;refGene_gn=
STOX2;refGene_tx=XM_023252319.1
B1	15083833 15084029
refGene_gn=STOX2;refGene_tx=XM_011281377.3;refGene_gn=STOX2;refGene_tx
=XM_019828246.2;refGene_gn=STOX2;refGene_tx=XM_023252318.1;refGene_gn=
STOX2;refGene_tx=XM_023252319.1
B1	15157016 15157222 refGene_gn=ENPP6;refGene_tx=XM_003984581.5
B1	15161801 15161925 refGene_gn=ENPP6;refGene_tx=XM_003984581.5
B1	15175486 15175624 refGene_gn=ENPP6;refGene_tx=XM_003984581.5
B1	15178521 15178539 refGene_gn=ENPP6;refGene_tx=XM_003984581.5
B1	15178540 15178702 refGene_gn=ENPP6;refGene_tx=XM_003984581.5
B1	15179621 15179763 refGene_gn=ENPP6;refGene_tx=XM_003984581.5
B1	15186274 15186386 refGene_gn=ENPP6;refGene_tx=XM_003984581.5
B1	15210083 15210263 refGene_gn=ENPP6;refGene_tx=XM_003984581.5
B1	15264744 15264985 refGene_gn=ENPP6;refGene_tx=XM_003984581.5
B1	15384394 15384703
refGene_gn=IRF2;refGene_tx=XM_006930564.4;refGene_gn=IRF2;refGene_tx=X
M_006930565.4;refGene_gn=IRF2;refGene_tx=XM_011281380.3;refGene_gn=IRF
2;refGene_tx=XM_019828250.2;refGene_gn=IRF2;refGene_tx=XM_019828251.2
B1	15386024 15386071
refGene_gn=IRF2;refGene_tx=XM_006930564.4;refGene_gn=IRF2;refGene_tx=X
M_006930565.4;refGene_gn=IRF2;refGene_tx=XM_011281380.3;refGene_gn=IRF
2;refGene_tx=XM_019828250.2;refGene_gn=IRF2;refGene_tx=XM_019828251.2
B1	15393639 15393804
refGene_gn=IRF2;refGene_tx=XM_006930564.4;refGene_gn=IRF2;refGene_tx=X
M_006930565.4;refGene_gn=IRF2;refGene_tx=XM_011281380.3;refGene_gn=IRF
2;refGene_tx=XM_019828250.2;refGene_gn=IRF2;refGene_tx=XM_019828251.2
B1	15405450 15405568
refGene_gn=IRF2;refGene_tx=XM_006930564.4;refGene_gn=IRF2;refGene_tx=X
M_006930565.4;refGene_gn=IRF2;refGene_tx=XM_011281380.3;refGene_gn=IRF
2;refGene_tx=XM_019828250.2;refGene_gn=IRF2;refGene_tx=XM_019828251.2
B1	15414786 15414833
refGene_gn=IRF2;refGene_tx=XM_006930564.4;refGene_gn=IRF2;refGene_tx=X

M_006930565.4;refGene_gn=IRF2;refGene_tx=XM_011281380.3;refGene_gn=IRF2;refGene_tx=XM_019828250.2;refGene_gn=IRF2;refGene_tx=XM_019828251.2
B1 15415155 15415332 refGene_gn=IRF2;refGene_tx=XM_006930564.4;refGene_gn=IRF2;refGene_tx=XM_006930565.4;refGene_gn=IRF2;refGene_tx=XM_011281380.3;refGene_gn=IRF2;refGene_tx=XM_019828250.2;refGene_gn=IRF2;refGene_tx=XM_019828251.2
B1 15416947 15417047 refGene_gn=IRF2;refGene_tx=XM_006930564.4;refGene_gn=IRF2;refGene_tx=XM_006930565.4;refGene_gn=IRF2;refGene_tx=XM_011281380.3;refGene_gn=IRF2;refGene_tx=XM_019828250.2;refGene_gn=IRF2;refGene_tx=XM_019828251.2
B1 15425496 15425592 refGene_gn=IRF2;refGene_tx=XM_006930564.4;refGene_gn=IRF2;refGene_tx=XM_006930565.4;refGene_gn=IRF2;refGene_tx=XM_011281380.3;refGene_gn=IRF2;refGene_tx=XM_019828250.2;refGene_gn=IRF2;refGene_tx=XM_019828251.2
B1 15468757 15468814 refGene_gn=IRF2;refGene_tx=XM_019828250.2
B1 15586445 15586675 refGene_gn=CASP3;refGene_tx=NM_001009338.1
B1 15588068 15588189 refGene_gn=CASP3;refGene_tx=NM_001009338.1
B1 15588973 15589149 refGene_gn=CASP3;refGene_tx=NM_001009338.1
B1 15589468 15589597 refGene_gn=CASP3;refGene_tx=NM_001009338.1
B1 15592613 15592738 refGene_gn=CASP3;refGene_tx=NM_001009338.1
B1 15595076 15595129 refGene_gn=CASP3;refGene_tx=NM_001009338.1
B1 15615163 15615343 refGene_gn=PRIMPOL;refGene_tx=XM_011281381.3;refGene_gn=PRIMPOL;refGene_tx=XM_019828252.2;refGene_gn=PRIMPOL;refGene_tx=XM_019828253.2;refGene_gn=PRIMPOL;refGene_tx=XM_019828254.2;refGene_gn=PRIMPOL;refGene_tx=XM_019828255.2;refGene_gn=PRIMPOL;refGene_tx=XM_019828256.2
B1 15621316 15621414 refGene_gn=PRIMPOL;refGene_tx=XM_011281381.3;refGene_gn=PRIMPOL;refGene_tx=XM_019828252.2;refGene_gn=PRIMPOL;refGene_tx=XM_019828253.2;refGene_gn=PRIMPOL;refGene_tx=XM_019828254.2;refGene_gn=PRIMPOL;refGene_tx=XM_019828255.2;refGene_gn=PRIMPOL;refGene_tx=XM_019828256.2
B1 15622310 15622440 refGene_gn=PRIMPOL;refGene_tx=XM_011281381.3;refGene_gn=PRIMPOL;refGene_tx=XM_019828252.2;refGene_gn=PRIMPOL;refGene_tx=XM_019828253.2;refGene_gn=PRIMPOL;refGene_tx=XM_019828254.2;refGene_gn=PRIMPOL;refGene_tx=XM_019828255.2;refGene_gn=PRIMPOL;refGene_tx=XM_019828256.2
B1 15631393 15631541 refGene_gn=PRIMPOL;refGene_tx=XM_011281381.3;refGene_gn=PRIMPOL;refGene_tx=XM_019828252.2;refGene_gn=PRIMPOL;refGene_tx=XM_019828253.2;refGene_gn=PRIMPOL;refGene_tx=XM_019828254.2;refGene_gn=PRIMPOL;refGene_tx=XM_019828255.2;refGene_gn=PRIMPOL;refGene_tx=XM_019828256.2
B1 15636761 15637046 refGene_gn=PRIMPOL;refGene_tx=XM_011281381.3;refGene_gn=PRIMPOL;refGene_tx=XM_019828252.2;refGene_gn=PRIMPOL;refGene_tx=XM_019828253.2;refGene_gn=PRIMPOL;refGene_tx=XM_019828254.2;refGene_gn=PRIMPOL;refGene_tx=XM_019828255.2;refGene_gn=PRIMPOL;refGene_tx=XM_019828256.2
B1 15653669 15653832 refGene_gn=PRIMPOL;refGene_tx=XM_011281381.3;refGene_gn=PRIMPOL;refGene_tx=XM_019828252.2;refGene_gn=PRIMPOL;refGene_tx=XM_019828253.2;refGene_gn=PRIMPOL;refGene_tx=XM_019828254.2;refGene_gn=PRIMPOL;refGene_tx=

XM_019828255.2;refGene_gn=PRIMPOL;refGene_tx=XM_019828256.2
B1 15655475 15655564
refGene_gn=PRIMPOL;refGene_tx=XM_011281381.3;refGene_gn=PRIMPOL;refGene_tx=XM_019828252.2;refGene_gn=PRIMPOL;refGene_tx=XM_019828253.2;refGene_gn=PRIMPOL;refGene_tx=XM_019828254.2;refGene_gn=PRIMPOL;refGene_tx=XM_019828255.2;refGene_gn=PRIMPOL;refGene_tx=XM_019828256.2
B1 15658292 15658382
refGene_gn=PRIMPOL;refGene_tx=XM_011281381.3;refGene_gn=PRIMPOL;refGene_tx=XM_019828252.2;refGene_gn=PRIMPOL;refGene_tx=XM_019828253.2;refGene_gn=PRIMPOL;refGene_tx=XM_019828254.2;refGene_gn=PRIMPOL;refGene_tx=XM_019828255.2;refGene_gn=PRIMPOL;refGene_tx=XM_019828256.2
B1 15658459 15658568
refGene_gn=PRIMPOL;refGene_tx=XM_011281381.3;refGene_gn=PRIMPOL;refGene_tx=XM_019828252.2;refGene_gn=PRIMPOL;refGene_tx=XM_019828253.2;refGene_gn=PRIMPOL;refGene_tx=XM_019828254.2;refGene_gn=PRIMPOL;refGene_tx=XM_019828255.2;refGene_gn=PRIMPOL;refGene_tx=XM_019828256.2
B1 15663300 15663383
refGene_gn=PRIMPOL;refGene_tx=XM_011281381.3;refGene_gn=PRIMPOL;refGene_tx=XM_019828252.2;refGene_gn=PRIMPOL;refGene_tx=XM_019828253.2;refGene_gn=PRIMPOL;refGene_tx=XM_019828254.2;refGene_gn=PRIMPOL;refGene_tx=XM_019828255.2;refGene_gn=PRIMPOL;refGene_tx=XM_019828256.2
B1 15663463 15663510
refGene_gn=PRIMPOL;refGene_tx=XM_011281381.3;refGene_gn=PRIMPOL;refGene_tx=XM_019828252.2;refGene_gn=PRIMPOL;refGene_tx=XM_019828253.2;refGene_gn=PRIMPOL;refGene_tx=XM_019828254.2;refGene_gn=PRIMPOL;refGene_tx=XM_019828255.2;refGene_gn=PRIMPOL;refGene_tx=XM_019828256.2
B1 15666261 15666510
refGene_gn=PRIMPOL;refGene_tx=XM_011281381.3;refGene_gn=PRIMPOL;refGene_tx=XM_019828252.2;refGene_gn=PRIMPOL;refGene_tx=XM_019828253.2;refGene_gn=PRIMPOL;refGene_tx=XM_019828254.2;refGene_gn=PRIMPOL;refGene_tx=XM_019828255.2;refGene_gn=PRIMPOL;refGene_tx=XM_019828256.2
B1 15672214 15672328 refGene_gn=CENPU;refGene_tx=XM_023252320.1
B1 15673588 15673745 refGene_gn=CENPU;refGene_tx=XM_023252320.1
B1 15675651 15675713 refGene_gn=CENPU;refGene_tx=XM_023252320.1
B1 15676832 15676880 refGene_gn=CENPU;refGene_tx=XM_023252320.1
B1 15677098 15677177 refGene_gn=CENPU;refGene_tx=XM_023252320.1
B1 15689102 15689211 refGene_gn=CENPU;refGene_tx=XM_023252320.1
B1 15690290 15690341 refGene_gn=CENPU;refGene_tx=XM_023252320.1
B1 15697048 15697291 refGene_gn=CENPU;refGene_tx=XM_023252320.1
B1 15697735 15697796 refGene_gn=CENPU;refGene_tx=XM_023252320.1
B1 15705540 15705646 refGene_gn=CENPU;refGene_tx=XM_023252320.1
B1 15709136 15709254 refGene_gn=CENPU;refGene_tx=XM_023252320.1
B1 15710590 15710639 refGene_gn=CENPU;refGene_tx=XM_023252320.1
B1 15713886 15713927 refGene_gn=CENPU;refGene_tx=XM_023252320.1
B1 15729581 15729722
refGene_gn=ACSL1;refGene_tx=XM_003984584.5;refGene_gn=ACSL1;refGene_tx=XM_019828257.2;refGene_gn=ACSL1;refGene_tx=XM_019828258.2;refGene_gn=ACSL1;refGene_tx=XM_023252321.1;refGene_gn=ACSL1;refGene_tx=XM_023252322.1
B1 15730086 15730158

refGene_gn=ACSL1;refGene_tx=XM_003984584.5;refGene_gn=ACSL1;refGene_tx=XM_019828257.2;refGene_gn=ACSL1;refGene_tx=XM_019828258.2;refGene_gn=ACSL1;refGene_tx=XM_023252321.1;refGene_gn=ACSL1;refGene_tx=XM_023252322.1
B1 15730264 15730366
refGene_gn=ACSL1;refGene_tx=XM_003984584.5;refGene_gn=ACSL1;refGene_tx=XM_019828257.2;refGene_gn=ACSL1;refGene_tx=XM_019828258.2;refGene_gn=ACSL1;refGene_tx=XM_023252321.1;refGene_gn=ACSL1;refGene_tx=XM_023252322.1
B1 15731853 15731997
refGene_gn=ACSL1;refGene_tx=XM_003984584.5;refGene_gn=ACSL1;refGene_tx=XM_019828257.2;refGene_gn=ACSL1;refGene_tx=XM_019828258.2;refGene_gn=ACSL1;refGene_tx=XM_023252321.1;refGene_gn=ACSL1;refGene_tx=XM_023252322.1
B1 15733565 15733682
refGene_gn=ACSL1;refGene_tx=XM_003984584.5;refGene_gn=ACSL1;refGene_tx=XM_019828257.2;refGene_gn=ACSL1;refGene_tx=XM_019828258.2;refGene_gn=ACSL1;refGene_tx=XM_023252321.1;refGene_gn=ACSL1;refGene_tx=XM_023252322.1
B1 15735165 15735254
refGene_gn=ACSL1;refGene_tx=XM_003984584.5;refGene_gn=ACSL1;refGene_tx=XM_019828257.2;refGene_gn=ACSL1;refGene_tx=XM_019828258.2;refGene_gn=ACSL1;refGene_tx=XM_023252321.1;refGene_gn=ACSL1;refGene_tx=XM_023252322.1
B1 15737002 15737075
refGene_gn=ACSL1;refGene_tx=XM_003984584.5;refGene_gn=ACSL1;refGene_tx=XM_019828257.2;refGene_gn=ACSL1;refGene_tx=XM_019828258.2;refGene_gn=ACSL1;refGene_tx=XM_023252321.1;refGene_gn=ACSL1;refGene_tx=XM_023252322.1
B1 15738030 15738126
refGene_gn=ACSL1;refGene_tx=XM_003984584.5;refGene_gn=ACSL1;refGene_tx=XM_019828257.2;refGene_gn=ACSL1;refGene_tx=XM_019828258.2;refGene_gn=ACSL1;refGene_tx=XM_023252321.1;refGene_gn=ACSL1;refGene_tx=XM_023252322.1
B1 15739277 15739412
refGene_gn=ACSL1;refGene_tx=XM_003984584.5;refGene_gn=ACSL1;refGene_tx=XM_019828257.2;refGene_gn=ACSL1;refGene_tx=XM_019828258.2;refGene_gn=ACSL1;refGene_tx=XM_023252321.1;refGene_gn=ACSL1;refGene_tx=XM_023252322.1
B1 15740208 15740379
refGene_gn=LOC101090856;refGene_tx=XM_011281382.3
B1 15742523 15742658
refGene_gn=ACSL1;refGene_tx=XM_003984584.5;refGene_gn=ACSL1;refGene_tx=XM_019828257.2;refGene_gn=ACSL1;refGene_tx=XM_019828258.2;refGene_gn=ACSL1;refGene_tx=XM_023252321.1;refGene_gn=ACSL1;refGene_tx=XM_023252322.1
B1 15744429 15744507 refGene_gn=ACSL1;refGene_tx=XM_023252322.1
B1 15744577 15744655
refGene_gn=ACSL1;refGene_tx=XM_003984584.5;refGene_gn=ACSL1;refGene_tx=XM_019828257.2;refGene_gn=ACSL1;refGene_tx=XM_019828258.2;refGene_gn=

ACSL1;refGene_tx=XM_023252321.1
B1 15747039 15747113
refGene_gn=ACSL1;refGene_tx=XM_003984584.5;refGene_gn=ACSL1;refGene_tx
=XM_019828257.2;refGene_gn=ACSL1;refGene_tx=XM_019828258.2;refGene_gn=
ACSL1;refGene_tx=XM_023252321.1;refGene_gn=ACSL1;refGene_tx=XM_0232523
22.1
B1 15747966 15748018
refGene_gn=ACSL1;refGene_tx=XM_003984584.5;refGene_gn=ACSL1;refGene_tx
=XM_019828257.2;refGene_gn=ACSL1;refGene_tx=XM_019828258.2;refGene_gn=
ACSL1;refGene_tx=XM_023252321.1;refGene_gn=ACSL1;refGene_tx=XM_0232523
22.1
B1 15748183 15748216
refGene_gn=ACSL1;refGene_tx=XM_003984584.5;refGene_gn=ACSL1;refGene_tx
=XM_019828257.2;refGene_gn=ACSL1;refGene_tx=XM_019828258.2;refGene_gn=
ACSL1;refGene_tx=XM_023252321.1;refGene_gn=ACSL1;refGene_tx=XM_0232523
22.1
B1 15753703 15753882
refGene_gn=ACSL1;refGene_tx=XM_003984584.5;refGene_gn=ACSL1;refGene_tx
=XM_019828257.2;refGene_gn=ACSL1;refGene_tx=XM_019828258.2;refGene_gn=
ACSL1;refGene_tx=XM_023252321.1;refGene_gn=ACSL1;refGene_tx=XM_0232523
22.1
B1 15754103 15754203
refGene_gn=ACSL1;refGene_tx=XM_003984584.5;refGene_gn=ACSL1;refGene_tx
=XM_019828257.2;refGene_gn=ACSL1;refGene_tx=XM_019828258.2;refGene_gn=
ACSL1;refGene_tx=XM_023252321.1;refGene_gn=ACSL1;refGene_tx=XM_0232523
22.1
B1 15756511 15756613
refGene_gn=ACSL1;refGene_tx=XM_003984584.5;refGene_gn=ACSL1;refGene_tx
=XM_019828257.2;refGene_gn=ACSL1;refGene_tx=XM_019828258.2;refGene_gn=
ACSL1;refGene_tx=XM_023252321.1;refGene_gn=ACSL1;refGene_tx=XM_0232523
22.1
B1 15759175 15759240
refGene_gn=ACSL1;refGene_tx=XM_003984584.5;refGene_gn=ACSL1;refGene_tx
=XM_019828257.2;refGene_gn=ACSL1;refGene_tx=XM_019828258.2;refGene_gn=
ACSL1;refGene_tx=XM_023252321.1;refGene_gn=ACSL1;refGene_tx=XM_0232523
22.1
B1 15763360 15763475
refGene_gn=ACSL1;refGene_tx=XM_003984584.5;refGene_gn=ACSL1;refGene_tx
=XM_019828257.2;refGene_gn=ACSL1;refGene_tx=XM_019828258.2;refGene_gn=
ACSL1;refGene_tx=XM_023252321.1;refGene_gn=ACSL1;refGene_tx=XM_0232523
22.1
B1 15777713 15777908
refGene_gn=ACSL1;refGene_tx=XM_003984584.5;refGene_gn=ACSL1;refGene_tx
=XM_019828257.2;refGene_gn=ACSL1;refGene_tx=XM_019828258.2;refGene_gn=
ACSL1;refGene_tx=XM_023252321.1;refGene_gn=ACSL1;refGene_tx=XM_0232523
22.1
B1 15987520 15987815 refGene_gn=HELT;refGene_tx=XM_023253385.1
B1 15987922 15988077 refGene_gn=HELT;refGene_tx=XM_023253385.1
B1 15988343 15988440 refGene_gn=HELT;refGene_tx=XM_023253385.1
B1 15988855 15989355 refGene_gn=HELT;refGene_tx=XM_023253385.1

B1 16104865 16104976 refGene_gn=SLC25A4;refGene_tx=XM_023252323.1
B1 16106289 16106776 refGene_gn=SLC25A4;refGene_tx=XM_023252323.1
B1 16107301 16107442 refGene_gn=SLC25A4;refGene_tx=XM_023252323.1
B1 16108279 16108437 refGene_gn=SLC25A4;refGene_tx=XM_023252323.1
B1 16168454 16168862 refGene_gn=SNX25;refGene_tx=XM_023252358.1
B1 16196639 16196724
refGene_gn=SNX25;refGene_tx=XM_003984589.5;refGene_gn=SNX25;refGene_tx=XM_023252358.1
B1 16208239 16208456
refGene_gn=SNX25;refGene_tx=XM_003984589.5;refGene_gn=SNX25;refGene_tx=XM_023252358.1
B1 16216033 16216206
refGene_gn=SNX25;refGene_tx=XM_003984589.5;refGene_gn=SNX25;refGene_tx=XM_023252358.1
B1 16217335 16217522
refGene_gn=SNX25;refGene_tx=XM_003984589.5;refGene_gn=SNX25;refGene_tx=XM_023252358.1
B1 16237484 16237555
refGene_gn=SNX25;refGene_tx=XM_003984589.5;refGene_gn=SNX25;refGene_tx=XM_023252358.1
B1 16255781 16255963
refGene_gn=SNX25;refGene_tx=XM_003984589.5;refGene_gn=SNX25;refGene_tx=XM_023252358.1
B1 16268015 16268147
refGene_gn=SNX25;refGene_tx=XM_003984589.5;refGene_gn=SNX25;refGene_tx=XM_023252358.1
B1 16270120 16270393
refGene_gn=SNX25;refGene_tx=XM_003984589.5;refGene_gn=SNX25;refGene_tx=XM_023252358.1
B1 16273413 16273578
refGene_gn=SNX25;refGene_tx=XM_003984589.5;refGene_gn=SNX25;refGene_tx=XM_023252358.1
B1 16276514 16276646
refGene_gn=SNX25;refGene_tx=XM_003984589.5;refGene_gn=SNX25;refGene_tx=XM_023252358.1
B1 16279399 16279540
refGene_gn=SNX25;refGene_tx=XM_003984589.5;refGene_gn=SNX25;refGene_tx=XM_023252358.1
B1 16283314 16283428
refGene_gn=SNX25;refGene_tx=XM_003984589.5;refGene_gn=SNX25;refGene_tx=XM_023252358.1
B1 16288480 16288645
refGene_gn=SNX25;refGene_tx=XM_003984589.5;refGene_gn=SNX25;refGene_tx=XM_023252358.1
B1 16291254 16291372
refGene_gn=SNX25;refGene_tx=XM_003984589.5;refGene_gn=SNX25;refGene_tx=XM_023252358.1
B1 16295301 16295368
refGene_gn=SNX25;refGene_tx=XM_003984589.5;refGene_gn=SNX25;refGene_tx=XM_023252358.1

B1 16298266 16298448
refGene_gn=SNX25;refGene_tx=XM_003984589.5;refGene_gn=SNX25;refGene_tx
=XM_023252358.1
B1 16298967 16299068
refGene_gn=SNX25;refGene_tx=XM_003984589.5;refGene_gn=SNX25;refGene_tx
=XM_023252358.1
B1 16299599 16299680
refGene_gn=SNX25;refGene_tx=XM_003984589.5;refGene_gn=SNX25;refGene_tx
=XM_023252358.1
B1 16303898 16303964
refGene_gn=LRP2BP;refGene_tx=XM_023252359.1;refGene_gn=LRP2BP;refGene_
tx=XM_023252360.1;refGene_gn=LRP2BP;refGene_tx=XM_023252361.1;refGene_
gn=LRP2BP;refGene_tx=XM_023252362.1;refGene_gn=LRP2BP;refGene_tx=XM_02
3252363.1;refGene_gn=LRP2BP;refGene_tx=XM_023252364.1;refGene_gn=LRP2B
P;refGene_tx=XM_023252365.1;refGene_gn=LRP2BP;refGene_tx=XM_023252366.
1;refGene_gn=LRP2BP;refGene_tx=XM_023252367.1
B1 16306636 16306811
refGene_gn=LRP2BP;refGene_tx=XM_023252359.1;refGene_gn=LRP2BP;refGene_
tx=XM_023252360.1;refGene_gn=LRP2BP;refGene_tx=XM_023252361.1;refGene_
gn=LRP2BP;refGene_tx=XM_023252362.1;refGene_gn=LRP2BP;refGene_tx=XM_02
3252363.1;refGene_gn=LRP2BP;refGene_tx=XM_023252364.1;refGene_gn=LRP2B
P;refGene_tx=XM_023252365.1;refGene_gn=LRP2BP;refGene_tx=XM_023252366.
1;refGene_gn=LRP2BP;refGene_tx=XM_023252367.1
B1 16308444 16308668
refGene_gn=LRP2BP;refGene_tx=XM_023252359.1;refGene_gn=LRP2BP;refGene_
tx=XM_023252360.1;refGene_gn=LRP2BP;refGene_tx=XM_023252361.1;refGene_
gn=LRP2BP;refGene_tx=XM_023252362.1;refGene_gn=LRP2BP;refGene_tx=XM_02
3252363.1;refGene_gn=LRP2BP;refGene_tx=XM_023252364.1;refGene_gn=LRP2B
P;refGene_tx=XM_023252365.1;refGene_gn=LRP2BP;refGene_tx=XM_023252366.
1;refGene_gn=LRP2BP;refGene_tx=XM_023252367.1
B1 16309723 16309829
refGene_gn=LRP2BP;refGene_tx=XM_023252359.1;refGene_gn=LRP2BP;refGene_
tx=XM_023252360.1;refGene_gn=LRP2BP;refGene_tx=XM_023252361.1;refGene_
gn=LRP2BP;refGene_tx=XM_023252362.1;refGene_gn=LRP2BP;refGene_tx=XM_02
3252363.1;refGene_gn=LRP2BP;refGene_tx=XM_023252364.1;refGene_gn=LRP2B
P;refGene_tx=XM_023252365.1;refGene_gn=LRP2BP;refGene_tx=XM_023252366.
1;refGene_gn=LRP2BP;refGene_tx=XM_023252367.1
B1 16309916 16310059
refGene_gn=LRP2BP;refGene_tx=XM_023252359.1;refGene_gn=LRP2BP;refGene_
tx=XM_023252360.1;refGene_gn=LRP2BP;refGene_tx=XM_023252361.1;refGene_
gn=LRP2BP;refGene_tx=XM_023252362.1;refGene_gn=LRP2BP;refGene_tx=XM_02
3252363.1;refGene_gn=LRP2BP;refGene_tx=XM_023252364.1;refGene_gn=LRP2B
P;refGene_tx=XM_023252365.1;refGene_gn=LRP2BP;refGene_tx=XM_023252366.
1;refGene_gn=LRP2BP;refGene_tx=XM_023252367.1
B1 16324525 16324639
refGene_gn=LRP2BP;refGene_tx=XM_023252359.1;refGene_gn=LRP2BP;refGene_
tx=XM_023252360.1;refGene_gn=LRP2BP;refGene_tx=XM_023252361.1;refGene_
gn=LRP2BP;refGene_tx=XM_023252362.1;refGene_gn=LRP2BP;refGene_tx=XM_02
3252363.1;refGene_gn=LRP2BP;refGene_tx=XM_023252364.1;refGene_gn=LRP2B
P;refGene_tx=XM_023252365.1;refGene_gn=LRP2BP;refGene_tx=XM_023252366.

1;refGene_gn=LRP2BP;refGene_tx=XM_023252367.1
B1 16326324 16326437
refGene_gn=LRP2BP;refGene_tx=XM_023252359.1;refGene_gn=LRP2BP;refGene_tx=XM_023252360.1;refGene_gn=LRP2BP;refGene_tx=XM_023252361.1;refGene_gn=LRP2BP;refGene_tx=XM_023252362.1;refGene_gn=LRP2BP;refGene_tx=XM_023252363.1;refGene_gn=LRP2BP;refGene_tx=XM_023252364.1;refGene_gn=LRP2BP;refGene_tx=XM_023252365.1;refGene_gn=LRP2BP;refGene_tx=XM_023252366.1;refGene_gn=LRP2BP;refGene_tx=XM_023252367.1
B1 16327151 16327257
refGene_gn=LRP2BP;refGene_tx=XM_023252359.1;refGene_gn=LRP2BP;refGene_tx=XM_023252360.1;refGene_gn=LRP2BP;refGene_tx=XM_023252361.1;refGene_gn=LRP2BP;refGene_tx=XM_023252362.1;refGene_gn=LRP2BP;refGene_tx=XM_023252363.1;refGene_gn=LRP2BP;refGene_tx=XM_023252364.1;refGene_gn=LRP2BP;refGene_tx=XM_023252365.1;refGene_gn=LRP2BP;refGene_tx=XM_023252366.1;refGene_gn=LRP2BP;refGene_tx=XM_023252367.1
B1 16338711 16338738 refGene_gn=ANKRD37;refGene_tx=XM_003984591.5
B1 16338946 16339099 refGene_gn=ANKRD37;refGene_tx=XM_003984591.5
B1 16341482 16341574 refGene_gn=ANKRD37;refGene_tx=XM_003984591.5
B1 16342115 16342319 refGene_gn=ANKRD37;refGene_tx=XM_003984591.5
B1 16342566 16342567 refGene_gn=ANKRD37;refGene_tx=XM_003984591.5
B1 16342930 16343017 refGene_gn=UFSP2;refGene_tx=XM_006930570.4
B1 16350242 16350367 refGene_gn=UFSP2;refGene_tx=XM_006930570.4
B1 16351095 16351172 refGene_gn=UFSP2;refGene_tx=XM_006930570.4
B1 16353034 16353159 refGene_gn=UFSP2;refGene_tx=XM_006930570.4
B1 16353372 16353537 refGene_gn=UFSP2;refGene_tx=XM_006930570.4
B1 16356490 16356637 refGene_gn=UFSP2;refGene_tx=XM_006930570.4
B1 16358259 16358452 refGene_gn=UFSP2;refGene_tx=XM_006930570.4
B1 16358833 16358991 refGene_gn=UFSP2;refGene_tx=XM_006930570.4
B1 16361641 16361708 refGene_gn=UFSP2;refGene_tx=XM_006930570.4
B1 16361789 16361973 refGene_gn=UFSP2;refGene_tx=XM_006930570.4
B1 16364132 16364211 refGene_gn=UFSP2;refGene_tx=XM_006930570.4
B1 16367116 16367119 refGene_gn=UFSP2;refGene_tx=XM_006930570.4
B1 16401315 16401365 refGene_gn=CCDC110;refGene_tx=XM_006930569.4
B1 16401673 16401859 refGene_gn=CCDC110;refGene_tx=XM_006930569.4
B1 16401861 16403788 refGene_gn=CCDC110;refGene_tx=XM_006930569.4
B1 16404530 16404641 refGene_gn=CCDC110;refGene_tx=XM_006930569.4
B1 16406122 16406188 refGene_gn=CCDC110;refGene_tx=XM_006930569.4
B1 16406454 16406510 refGene_gn=CCDC110;refGene_tx=XM_006930569.4
B1 16423561 16423666 refGene_gn=CCDC110;refGene_tx=XM_006930569.4
B1 16423994 16424004 refGene_gn=CCDC110;refGene_tx=XM_006930569.4
B1 16445181 16445371
refGene_gn=PDLIM3;refGene_tx=XM_003984592.5;refGene_gn=PDLIM3;refGene_tx=XM_003984593.5;refGene_gn=PDLIM3;refGene_tx=XM_003984594.5
B1 16446630 16446742
refGene_gn=PDLIM3;refGene_tx=XM_003984592.5;refGene_gn=PDLIM3;refGene_tx=XM_003984593.5;refGene_gn=PDLIM3;refGene_tx=XM_003984594.5
B1 16448372 16448503
refGene_gn=PDLIM3;refGene_tx=XM_003984592.5;refGene_gn=PDLIM3;refGene_tx=XM_003984593.5;refGene_gn=PDLIM3;refGene_tx=XM_003984594.5
B1 16450102 16450366 refGene_gn=PDLIM3;refGene_tx=XM_003984592.5

B1 16455397 16455465
refGene_gn=PDLIM3;refGene_tx=XM_003984592.5;refGene_gn=PDLIM3;refGene_tx=XM_003984594.5
B1 16455858 16456046 refGene_gn=PDLIM3;refGene_tx=XM_003984593.5
B1 16464532 16464617
refGene_gn=PDLIM3;refGene_tx=XM_003984592.5;refGene_gn=PDLIM3;refGene_tx=XM_003984593.5;refGene_gn=PDLIM3;refGene_tx=XM_003984594.5
B1 16466199 16466351
refGene_gn=PDLIM3;refGene_tx=XM_003984592.5;refGene_gn=PDLIM3;refGene_tx=XM_003984593.5;refGene_gn=PDLIM3;refGene_tx=XM_003984594.5
B1 16474670 16474763
refGene_gn=PDLIM3;refGene_tx=XM_003984592.5;refGene_gn=PDLIM3;refGene_tx=XM_003984593.5;refGene_gn=PDLIM3;refGene_tx=XM_003984594.5
B1 16530717 16530779
refGene_gn=SORBS2;refGene_tx=XM_011281387.3;refGene_gn=SORBS2;refGene_tx=XM_011281388.3;refGene_gn=SORBS2;refGene_tx=XM_011281390.3;refGene_gn=SORBS2;refGene_tx=XM_019830023.2;refGene_gn=SORBS2;refGene_tx=XM_019830025.2;refGene_gn=SORBS2;refGene_tx=XM_019830037.2;refGene_gn=SORBS2;refGene_tx=XM_019830038.2;refGene_gn=SORBS2;refGene_tx=XM_019830039.2;refGene_gn=SORBS2;refGene_tx=XM_019830043.2;refGene_gn=SORBS2;refGene_tx=XM_019830044.2;refGene_gn=SORBS2;refGene_tx=XM_019830045.2;refGene_gn=SORBS2;refGene_tx=XM_019830046.2;refGene_gn=SORBS2;refGene_tx=XM_023252354.1;refGene_gn=SORBS2;refGene_tx=XM_023252355.1;refGene_gn=SORBS2;refGene_tx=XM_023252356.1;refGene_gn=SORBS2;refGene_tx=XM_023252357.1
B1 16532521 16532628
refGene_gn=SORBS2;refGene_tx=XM_011281387.3;refGene_gn=SORBS2;refGene_tx=XM_011281388.3;refGene_gn=SORBS2;refGene_tx=XM_011281390.3;refGene_gn=SORBS2;refGene_tx=XM_019830023.2;refGene_gn=SORBS2;refGene_tx=XM_019830025.2;refGene_gn=SORBS2;refGene_tx=XM_019830037.2;refGene_gn=SORBS2;refGene_tx=XM_019830038.2;refGene_gn=SORBS2;refGene_tx=XM_019830039.2;refGene_gn=SORBS2;refGene_tx=XM_019830043.2;refGene_gn=SORBS2;refGene_tx=XM_019830044.2;refGene_gn=SORBS2;refGene_tx=XM_019830045.2;refGene_gn=SORBS2;refGene_tx=XM_019830046.2;refGene_gn=SORBS2;refGene_tx=XM_023252354.1;refGene_gn=SORBS2;refGene_tx=XM_023252355.1;refGene_gn=SORBS2;refGene_tx=XM_023252356.1;refGene_gn=SORBS2;refGene_tx=XM_023252357.1
B1 16536290 16536340
refGene_gn=SORBS2;refGene_tx=XM_011281387.3;refGene_gn=SORBS2;refGene_tx=XM_011281388.3;refGene_gn=SORBS2;refGene_tx=XM_011281390.3;refGene_gn=SORBS2;refGene_tx=XM_019830023.2;refGene_gn=SORBS2;refGene_tx=XM_019830025.2;refGene_gn=SORBS2;refGene_tx=XM_019830037.2;refGene_gn=SORBS2;refGene_tx=XM_019830038.2;refGene_gn=SORBS2;refGene_tx=XM_019830039.2;refGene_gn=SORBS2;refGene_tx=XM_019830043.2;refGene_gn=SORBS2;refGene_tx=XM_019830044.2;refGene_gn=SORBS2;refGene_tx=XM_019830045.2;refGene_gn=SORBS2;refGene_tx=XM_019830046.2;refGene_gn=SORBS2;refGene_tx=XM_023252354.1;refGene_gn=SORBS2;refGene_tx=XM_023252355.1;refGene_gn=SORBS2;refGene_tx=XM_023252356.1;refGene_gn=SORBS2;refGene_tx=XM_023252357.1
B1 16546074 16546308

refGene_gn=SORBS2;refGene_tx=XM_023252324.1;refGene_gn=SORBS2;refGene_tx=XM_023252325.1;refGene_gn=SORBS2;refGene_tx=XM_023252326.1;refGene_gn=SORBS2;refGene_tx=XM_023252327.1;refGene_gn=SORBS2;refGene_tx=XM_023252328.1;refGene_gn=SORBS2;refGene_tx=XM_023252329.1;refGene_gn=SORBS2;refGene_tx=XM_023252330.1;refGene_gn=SORBS2;refGene_tx=XM_023252331.1;refGene_gn=SORBS2;refGene_tx=XM_023252332.1;refGene_gn=SORBS2;refGene_tx=XM_023252333.1;refGene_gn=SORBS2;refGene_tx=XM_023252334.1;refGene_gn=SORBS2;refGene_tx=XM_023252335.1;refGene_gn=SORBS2;refGene_tx=XM_023252336.1;refGene_gn=SORBS2;refGene_tx=XM_023252337.1;refGene_gn=SORBS2;refGene_tx=XM_023252338.1;refGene_gn=SORBS2;refGene_tx=XM_023252339.1;refGene_gn=SORBS2;refGene_tx=XM_023252340.1;refGene_gn=SORBS2;refGene_tx=XM_023252341.1;refGene_gn=SORBS2;refGene_tx=XM_023252342.1;refGene_gn=SORBS2;refGene_tx=XM_023252343.1;refGene_gn=SORBS2;refGene_tx=XM_023252344.1;refGene_gn=SORBS2;refGene_tx=XM_023252345.1;refGene_gn=SORBS2;refGene_tx=XM_023252346.1;refGene_gn=SORBS2;refGene_tx=XM_023252347.1;refGene_gn=SORBS2;refGene_tx=XM_023252348.1;refGene_gn=SORBS2;refGene_tx=XM_023252349.1;refGene_gn=SORBS2;refGene_tx=XM_023252350.1;refGene_gn=SORBS2;refGene_tx=XM_023252351.1;refGene_gn=SORBS2;refGene_tx=XM_023252352.1;refGene_gn=SORBS2;refGene_tx=XM_023252353.1
B1 16547792 16548554
refGene_gn=SORBS2;refGene_tx=XM_023252324.1;refGene_gn=SORBS2;refGene_tx=XM_023252325.1;refGene_gn=SORBS2;refGene_tx=XM_023252326.1;refGene_gn=SORBS2;refGene_tx=XM_023252327.1;refGene_gn=SORBS2;refGene_tx=XM_023252328.1;refGene_gn=SORBS2;refGene_tx=XM_023252329.1;refGene_gn=SORBS2;refGene_tx=XM_023252330.1;refGene_gn=SORBS2;refGene_tx=XM_023252331.1;refGene_gn=SORBS2;refGene_tx=XM_023252332.1;refGene_gn=SORBS2;refGene_tx=XM_023252333.1;refGene_gn=SORBS2;refGene_tx=XM_023252334.1;refGene_gn=SORBS2;refGene_tx=XM_023252335.1;refGene_gn=SORBS2;refGene_tx=XM_023252336.1;refGene_gn=SORBS2;refGene_tx=XM_023252337.1;refGene_gn=SORBS2;refGene_tx=XM_023252338.1;refGene_gn=SORBS2;refGene_tx=XM_023252339.1;refGene_gn=SORBS2;refGene_tx=XM_023252340.1;refGene_gn=SORBS2;refGene_tx=XM_023252341.1;refGene_gn=SORBS2;refGene_tx=XM_023252342.1;refGene_gn=SORBS2;refGene_tx=XM_023252343.1;refGene_gn=SORBS2;refGene_tx=XM_023252344.1;refGene_gn=SORBS2;refGene_tx=XM_023252345.1;refGene_gn=SORBS2;refGene_tx=XM_023252346.1;refGene_gn=SORBS2;refGene_tx=XM_023252347.1;refGene_gn=SORBS2;refGene_tx=XM_023252348.1;refGene_gn=SORBS2;refGene_tx=XM_023252349.1;refGene_gn=SORBS2;refGene_tx=XM_023252350.1;refGene_gn=SORBS2;refGene_tx=XM_023252351.1;refGene_gn=SORBS2;refGene_tx=XM_023252352.1;refGene_gn=SORBS2;refGene_tx=XM_023252353.1
B1 16552133 16552337
refGene_gn=SORBS2;refGene_tx=XM_011281387.3;refGene_gn=SORBS2;refGene_tx=XM_011281388.3;refGene_gn=SORBS2;refGene_tx=XM_011281390.3;refGene_gn=SORBS2;refGene_tx=XM_019830023.2;refGene_gn=SORBS2;refGene_tx=XM_019830025.2;refGene_gn=SORBS2;refGene_tx=XM_019830037.2;refGene_gn=SORBS2;refGene_tx=XM_019830038.2;refGene_gn=SORBS2;refGene_tx=XM_019830039.2;refGene_gn=SORBS2;refGene_tx=XM_019830043.2;refGene_gn=SORBS2;refGene_tx=XM_019830044.2;refGene_gn=SORBS2;refGene_tx=XM_019830045.2;refGene_gn=SORBS2;refGene_tx=XM_019830046.2;refGene_gn=SORBS2;refGene_tx=XM_023252324.1;refGene_gn=SORBS2;refGene_tx=XM_023252325.1;refGene_gn=SORBS2;refGene_tx=XM_023252326.1;refGene_gn=SORBS2;refGene_tx=XM_02325232

7.1;refGene_gn=SORBS2;refGene_tx=XM_023252328.1;refGene_gn=SORBS2;refGene_tx=XM_023252329.1;refGene_gn=SORBS2;refGene_tx=XM_023252330.1;refGene_gn=SORBS2;refGene_tx=XM_023252331.1;refGene_gn=SORBS2;refGene_tx=XM_023252332.1;refGene_gn=SORBS2;refGene_tx=XM_023252333.1;refGene_gn=SORBS2;refGene_tx=XM_023252334.1;refGene_gn=SORBS2;refGene_tx=XM_023252335.1;refGene_gn=SORBS2;refGene_tx=XM_023252336.1;refGene_gn=SORBS2;refGene_tx=XM_023252337.1;refGene_gn=SORBS2;refGene_tx=XM_023252338.1;refGene_gn=SORBS2;refGene_tx=XM_023252339.1;refGene_gn=SORBS2;refGene_tx=XM_023252340.1;refGene_gn=SORBS2;refGene_tx=XM_023252341.1;refGene_gn=SORBS2;refGene_tx=XM_023252342.1;refGene_gn=SORBS2;refGene_tx=XM_023252343.1;refGene_gn=SORBS2;refGene_tx=XM_023252344.1;refGene_gn=SORBS2;refGene_tx=XM_023252345.1;refGene_gn=SORBS2;refGene_tx=XM_023252346.1;refGene_gn=SORBS2;refGene_tx=XM_023252347.1;refGene_gn=SORBS2;refGene_tx=XM_023252348.1;refGene_gn=SORBS2;refGene_tx=XM_023252349.1;refGene_gn=SORBS2;refGene_tx=XM_023252350.1;refGene_gn=SORBS2;refGene_tx=XM_023252351.1;refGene_gn=SORBS2;refGene_tx=XM_023252352.1;refGene_gn=SORBS2;refGene_tx=XM_023252353.1;refGene_gn=SORBS2;refGene_tx=XM_023252354.1;refGene_gn=SORBS2;refGene_tx=XM_023252355.1;refGene_gn=SORBS2;refGene_tx=XM_023252356.1;refGene_gn=SORBS2;refGene_tx=XM_023252357.1
B1 16554285 16554414
refGene_gn=SORBS2;refGene_tx=XM_011281387.3;refGene_gn=SORBS2;refGene_tx=XM_011281388.3;refGene_gn=SORBS2;refGene_tx=XM_011281390.3;refGene_gn=SORBS2;refGene_tx=XM_019830023.2;refGene_gn=SORBS2;refGene_tx=XM_019830025.2;refGene_gn=SORBS2;refGene_tx=XM_019830037.2;refGene_gn=SORBS2;refGene_tx=XM_019830038.2;refGene_gn=SORBS2;refGene_tx=XM_019830039.2;refGene_gn=SORBS2;refGene_tx=XM_019830043.2;refGene_gn=SORBS2;refGene_tx=XM_019830044.2;refGene_gn=SORBS2;refGene_tx=XM_019830045.2;refGene_gn=SORBS2;refGene_tx=XM_019830046.2;refGene_gn=SORBS2;refGene_tx=XM_023252324.1;refGene_gn=SORBS2;refGene_tx=XM_023252325.1;refGene_gn=SORBS2;refGene_tx=XM_023252326.1;refGene_gn=SORBS2;refGene_tx=XM_023252327.1;refGene_gn=SORBS2;refGene_tx=XM_023252328.1;refGene_gn=SORBS2;refGene_tx=XM_023252329.1;refGene_gn=SORBS2;refGene_tx=XM_023252330.1;refGene_gn=SORBS2;refGene_tx=XM_023252331.1;refGene_gn=SORBS2;refGene_tx=XM_023252332.1;refGene_gn=SORBS2;refGene_tx=XM_023252333.1;refGene_gn=SORBS2;refGene_tx=XM_023252334.1;refGene_gn=SORBS2;refGene_tx=XM_023252335.1;refGene_gn=SORBS2;refGene_tx=XM_023252336.1;refGene_gn=SORBS2;refGene_tx=XM_023252337.1;refGene_gn=SORBS2;refGene_tx=XM_023252338.1;refGene_gn=SORBS2;refGene_tx=XM_023252339.1;refGene_gn=SORBS2;refGene_tx=XM_023252340.1;refGene_gn=SORBS2;refGene_tx=XM_023252341.1;refGene_gn=SORBS2;refGene_tx=XM_023252342.1;refGene_gn=SORBS2;refGene_tx=XM_023252343.1;refGene_gn=SORBS2;refGene_tx=XM_023252344.1;refGene_gn=SORBS2;refGene_tx=XM_023252345.1;refGene_gn=SORBS2;refGene_tx=XM_023252346.1;refGene_gn=SORBS2;refGene_tx=XM_023252347.1;refGene_gn=SORBS2;refGene_tx=XM_023252348.1;refGene_gn=SORBS2;refGene_tx=XM_023252349.1;refGene_gn=SORBS2;refGene_tx=XM_023252350.1;refGene_gn=SORBS2;refGene_tx=XM_023252351.1;refGene_gn=SORBS2;refGene_tx=XM_023252352.1;refGene_gn=SORBS2;refGene_tx=XM_023252353.1;refGene_gn=SORBS2;refGene_tx=XM_023252354.1;refGene_gn=SORBS2;refGene_tx=XM_023252355.1;refGene_gn=SORBS2;refGene_tx=XM_023252356.1;refGene_gn=SORBS2;refGene_tx=XM_023252357.1
B1 16554499 16554614

refGene_gn=SORBS2;refGene_tx=XM_011281387.3;refGene_gn=SORBS2;refGene_
tx=XM_011281388.3;refGene_gn=SORBS2;refGene_tx=XM_011281390.3;refGene_
gn=SORBS2;refGene_tx=XM_019830023.2;refGene_gn=SORBS2;refGene_tx=XM_01
9830025.2;refGene_gn=SORBS2;refGene_tx=XM_019830037.2;refGene_gn=SORBS
2;refGene_tx=XM_019830038.2;refGene_gn=SORBS2;refGene_tx=XM_019830039.
2;refGene_gn=SORBS2;refGene_tx=XM_019830043.2;refGene_gn=SORBS2;refGen
e_tx=XM_019830044.2;refGene_gn=SORBS2;refGene_tx=XM_019830045.2;refGen
e_gn=SORBS2;refGene_tx=XM_019830046.2;refGene_gn=SORBS2;refGene_tx=XM_
023252324.1;refGene_gn=SORBS2;refGene_tx=XM_023252325.1;refGene_gn=SOR
BS2;refGene_tx=XM_023252326.1;refGene_gn=SORBS2;refGene_tx=XM_02325232
7.1;refGene_gn=SORBS2;refGene_tx=XM_023252328.1;refGene_gn=SORBS2;refG
ene_tx=XM_023252329.1;refGene_gn=SORBS2;refGene_tx=XM_023252330.1;refG
ene_gn=SORBS2;refGene_tx=XM_023252331.1;refGene_gn=SORBS2;refGene_tx=X
M_023252332.1;refGene_gn=SORBS2;refGene_tx=XM_023252333.1;refGene_gn=S
ORBS2;refGene_tx=XM_023252334.1;refGene_gn=SORBS2;refGene_tx=XM_023252
335.1;refGene_gn=SORBS2;refGene_tx=XM_023252336.1;refGene_gn=SORBS2;re
fGene_tx=XM_023252337.1;refGene_gn=SORBS2;refGene_tx=XM_023252338.1;re
fGene_gn=SORBS2;refGene_tx=XM_023252339.1;refGene_gn=SORBS2;refGene_tx
=XM_023252340.1;refGene_gn=SORBS2;refGene_tx=XM_023252341.1;refGene_gn
=SORBS2;refGene_tx=XM_023252342.1;refGene_gn=SORBS2;refGene_tx=XM_0232
52343.1;refGene_gn=SORBS2;refGene_tx=XM_023252344.1;refGene_gn=SORBS2;
refGene_tx=XM_023252345.1;refGene_gn=SORBS2;refGene_tx=XM_023252346.1;
refGene_gn=SORBS2;refGene_tx=XM_023252347.1;refGene_gn=SORBS2;refGene_
tx=XM_023252348.1;refGene_gn=SORBS2;refGene_tx=XM_023252349.1;refGene_
gn=SORBS2;refGene_tx=XM_023252350.1;refGene_gn=SORBS2;refGene_tx=XM_02
3252351.1;refGene_gn=SORBS2;refGene_tx=XM_023252352.1;refGene_gn=SORBS
2;refGene_tx=XM_023252353.1;refGene_gn=SORBS2;refGene_tx=XM_023252354.
1;refGene_gn=SORBS2;refGene_tx=XM_023252355.1;refGene_gn=SORBS2;refGen
e_tx=XM_023252356.1;refGene_gn=SORBS2;refGene_tx=XM_023252357.1
B1 16559412 16559459
refGene_gn=SORBS2;refGene_tx=XM_011281387.3;refGene_gn=SORBS2;refGene_
tx=XM_011281388.3;refGene_gn=SORBS2;refGene_tx=XM_011281390.3;refGene_
gn=SORBS2;refGene_tx=XM_019830023.2;refGene_gn=SORBS2;refGene_tx=XM_01
9830025.2;refGene_gn=SORBS2;refGene_tx=XM_019830037.2;refGene_gn=SORBS
2;refGene_tx=XM_019830038.2;refGene_gn=SORBS2;refGene_tx=XM_019830039.
2;refGene_gn=SORBS2;refGene_tx=XM_019830043.2;refGene_gn=SORBS2;refGen
e_tx=XM_019830044.2;refGene_gn=SORBS2;refGene_tx=XM_019830045.2;refGen
e_gn=SORBS2;refGene_tx=XM_019830046.2;refGene_gn=SORBS2;refGene_tx=XM_
023252324.1;refGene_gn=SORBS2;refGene_tx=XM_023252325.1;refGene_gn=SOR
BS2;refGene_tx=XM_023252326.1;refGene_gn=SORBS2;refGene_tx=XM_02325232
7.1;refGene_gn=SORBS2;refGene_tx=XM_023252328.1;refGene_gn=SORBS2;refG
ene_tx=XM_023252329.1;refGene_gn=SORBS2;refGene_tx=XM_023252330.1;refG
ene_gn=SORBS2;refGene_tx=XM_023252331.1;refGene_gn=SORBS2;refGene_tx=X
M_023252332.1;refGene_gn=SORBS2;refGene_tx=XM_023252333.1;refGene_gn=S
ORBS2;refGene_tx=XM_023252334.1;refGene_gn=SORBS2;refGene_tx=XM_023252
335.1;refGene_gn=SORBS2;refGene_tx=XM_023252336.1;refGene_gn=SORBS2;re
fGene_tx=XM_023252337.1;refGene_gn=SORBS2;refGene_tx=XM_023252338.1;re
fGene_gn=SORBS2;refGene_tx=XM_023252339.1;refGene_gn=SORBS2;refGene_tx
=XM_023252340.1;refGene_gn=SORBS2;refGene_tx=XM_023252341.1;refGene_gn
=SORBS2;refGene_tx=XM_023252342.1;refGene_gn=SORBS2;refGene_tx=XM_0232

52343.1;refGene_gn=SORBS2;refGene_tx=XM_023252344.1;refGene_gn=SORBS2;refGene_tx=XM_023252345.1;refGene_gn=SORBS2;refGene_tx=XM_023252346.1;refGene_gn=SORBS2;refGene_tx=XM_023252347.1;refGene_gn=SORBS2;refGene_tx=XM_023252348.1;refGene_gn=SORBS2;refGene_tx=XM_023252349.1;refGene_gn=SORBS2;refGene_tx=XM_023252350.1;refGene_gn=SORBS2;refGene_tx=XM_023252351.1;refGene_gn=SORBS2;refGene_tx=XM_023252352.1;refGene_gn=SORBS2;refGene_tx=XM_023252353.1;refGene_gn=SORBS2;refGene_tx=XM_023252354.1;refGene_gn=SORBS2;refGene_tx=XM_023252355.1;refGene_gn=SORBS2;refGene_tx=XM_023252356.1;refGene_gn=SORBS2;refGene_tx=XM_023252357.1

B1 16561061 16561150

refGene_gn=SORBS2;refGene_tx=XM_011281387.3;refGene_gn=SORBS2;refGene_tx=XM_011281388.3;refGene_gn=SORBS2;refGene_tx=XM_011281390.3;refGene_gn=SORBS2;refGene_tx=XM_019830023.2;refGene_gn=SORBS2;refGene_tx=XM_019830025.2;refGene_gn=SORBS2;refGene_tx=XM_019830037.2;refGene_gn=SORBS2;refGene_tx=XM_019830038.2;refGene_gn=SORBS2;refGene_tx=XM_019830039.2;refGene_gn=SORBS2;refGene_tx=XM_019830043.2;refGene_gn=SORBS2;refGene_tx=XM_019830044.2;refGene_gn=SORBS2;refGene_tx=XM_019830045.2;refGene_gn=SORBS2;refGene_tx=XM_019830046.2;refGene_gn=SORBS2;refGene_tx=XM_023252324.1;refGene_gn=SORBS2;refGene_tx=XM_023252325.1;refGene_gn=SORBS2;refGene_tx=XM_023252326.1;refGene_gn=SORBS2;refGene_tx=XM_023252327.1;refGene_gn=SORBS2;refGene_tx=XM_023252328.1;refGene_gn=SORBS2;refGene_tx=XM_023252329.1;refGene_gn=SORBS2;refGene_tx=XM_023252330.1;refGene_gn=SORBS2;refGene_tx=XM_023252331.1;refGene_gn=SORBS2;refGene_tx=XM_023252332.1;refGene_gn=SORBS2;refGene_tx=XM_023252333.1;refGene_gn=SORBS2;refGene_tx=XM_023252334.1;refGene_gn=SORBS2;refGene_tx=XM_023252335.1;refGene_gn=SORBS2;refGene_tx=XM_023252336.1;refGene_gn=SORBS2;refGene_tx=XM_023252337.1;refGene_gn=SORBS2;refGene_tx=XM_023252338.1;refGene_gn=SORBS2;refGene_tx=XM_023252339.1;refGene_gn=SORBS2;refGene_tx=XM_023252340.1;refGene_gn=SORBS2;refGene_tx=XM_023252341.1;refGene_gn=SORBS2;refGene_tx=XM_023252342.1;refGene_gn=SORBS2;refGene_tx=XM_023252343.1;refGene_gn=SORBS2;refGene_tx=XM_023252344.1;refGene_gn=SORBS2;refGene_tx=XM_023252345.1;refGene_gn=SORBS2;refGene_tx=XM_023252346.1;refGene_gn=SORBS2;refGene_tx=XM_023252347.1;refGene_gn=SORBS2;refGene_tx=XM_023252348.1;refGene_gn=SORBS2;refGene_tx=XM_023252349.1;refGene_gn=SORBS2;refGene_tx=XM_023252350.1;refGene_gn=SORBS2;refGene_tx=XM_023252351.1;refGene_gn=SORBS2;refGene_tx=XM_023252352.1;refGene_gn=SORBS2;refGene_tx=XM_023252353.1;refGene_gn=SORBS2;refGene_tx=XM_023252354.1;refGene_gn=SORBS2;refGene_tx=XM_023252355.1;refGene_gn=SORBS2;refGene_tx=XM_023252356.1;refGene_gn=SORBS2;refGene_tx=XM_023252357.1

B1 16564379 16565963

refGene_gn=SORBS2;refGene_tx=XM_011281387.3;refGene_gn=SORBS2;refGene_tx=XM_011281388.3;refGene_gn=SORBS2;refGene_tx=XM_019830023.2;refGene_gn=SORBS2;refGene_tx=XM_019830025.2;refGene_gn=SORBS2;refGene_tx=XM_019830037.2;refGene_gn=SORBS2;refGene_tx=XM_023252324.1;refGene_gn=SORBS2;refGene_tx=XM_023252325.1;refGene_gn=SORBS2;refGene_tx=XM_023252326.1;refGene_gn=SORBS2;refGene_tx=XM_023252327.1;refGene_gn=SORBS2;refGene_tx=XM_023252328.1;refGene_gn=SORBS2;refGene_tx=XM_023252329.1;refGene_gn=SORBS2;refGene_tx=XM_023252330.1;refGene_gn=SORBS2;refGene_tx=XM_023252331.1;refGene_gn=SORBS2;refGene_tx=XM_023252332.1;refGene_gn=SORBS2;refGene_tx=XM_023252333.1;refGene_gn=SORBS2;refGene_tx=XM_02325233

4.1;refGene_gn=SORBS2;refGene_tx=XM_023252335.1;refGene_gn=SORBS2;refGene_tx=XM_023252336.1;refGene_gn=SORBS2;refGene_tx=XM_023252337.1;refGene_gn=SORBS2;refGene_tx=XM_023252338.1;refGene_gn=SORBS2;refGene_tx=XM_023252339.1;refGene_gn=SORBS2;refGene_tx=XM_023252340.1;refGene_gn=SORBS2;refGene_tx=XM_023252341.1;refGene_gn=SORBS2;refGene_tx=XM_023252342.1;refGene_gn=SORBS2;refGene_tx=XM_023252343.1;refGene_gn=SORBS2;refGene_tx=XM_023252344.1;refGene_gn=SORBS2;refGene_tx=XM_023252345.1;refGene_gn=SORBS2;refGene_tx=XM_023252346.1;refGene_gn=SORBS2;refGene_tx=XM_023252347.1;refGene_gn=SORBS2;refGene_tx=XM_023252348.1;refGene_gn=SORBS2;refGene_tx=XM_023252349.1;refGene_gn=SORBS2;refGene_tx=XM_023252350.1;refGene_gn=SORBS2;refGene_tx=XM_023252351.1;refGene_gn=SORBS2;refGene_tx=XM_023252352.1;refGene_gn=SORBS2;refGene_tx=XM_023252354.1;refGene_gn=SORBS2;refGene_tx=XM_023252355.1
B1 16568351 16568536
refGene_gn=SORBS2;refGene_tx=XM_011281387.3;refGene_gn=SORBS2;refGene_tx=XM_011281388.3;refGene_gn=SORBS2;refGene_tx=XM_011281390.3;refGene_gn=SORBS2;refGene_tx=XM_019830023.2;refGene_gn=SORBS2;refGene_tx=XM_019830025.2;refGene_gn=SORBS2;refGene_tx=XM_019830037.2;refGene_gn=SORBS2;refGene_tx=XM_019830038.2;refGene_gn=SORBS2;refGene_tx=XM_019830039.2;refGene_gn=SORBS2;refGene_tx=XM_019830043.2;refGene_gn=SORBS2;refGene_tx=XM_019830044.2;refGene_gn=SORBS2;refGene_tx=XM_019830045.2;refGene_gn=SORBS2;refGene_tx=XM_019830046.2;refGene_gn=SORBS2;refGene_tx=XM_023252324.1;refGene_gn=SORBS2;refGene_tx=XM_023252325.1;refGene_gn=SORBS2;refGene_tx=XM_023252326.1;refGene_gn=SORBS2;refGene_tx=XM_023252327.1;refGene_gn=SORBS2;refGene_tx=XM_023252328.1;refGene_gn=SORBS2;refGene_tx=XM_023252329.1;refGene_gn=SORBS2;refGene_tx=XM_023252330.1;refGene_gn=SORBS2;refGene_tx=XM_023252331.1;refGene_gn=SORBS2;refGene_tx=XM_023252332.1;refGene_gn=SORBS2;refGene_tx=XM_023252333.1;refGene_gn=SORBS2;refGene_tx=XM_023252334.1;refGene_gn=SORBS2;refGene_tx=XM_023252335.1;refGene_gn=SORBS2;refGene_tx=XM_023252336.1;refGene_gn=SORBS2;refGene_tx=XM_023252337.1;refGene_gn=SORBS2;refGene_tx=XM_023252338.1;refGene_gn=SORBS2;refGene_tx=XM_023252339.1;refGene_gn=SORBS2;refGene_tx=XM_023252340.1;refGene_gn=SORBS2;refGene_tx=XM_023252341.1;refGene_gn=SORBS2;refGene_tx=XM_023252342.1;refGene_gn=SORBS2;refGene_tx=XM_023252343.1;refGene_gn=SORBS2;refGene_tx=XM_023252344.1;refGene_gn=SORBS2;refGene_tx=XM_023252345.1;refGene_gn=SORBS2;refGene_tx=XM_023252346.1;refGene_gn=SORBS2;refGene_tx=XM_023252347.1;refGene_gn=SORBS2;refGene_tx=XM_023252348.1;refGene_gn=SORBS2;refGene_tx=XM_023252349.1;refGene_gn=SORBS2;refGene_tx=XM_023252350.1;refGene_gn=SORBS2;refGene_tx=XM_023252351.1;refGene_gn=SORBS2;refGene_tx=XM_023252352.1;refGene_gn=SORBS2;refGene_tx=XM_023252353.1;refGene_gn=SORBS2;refGene_tx=XM_023252354.1;refGene_gn=SORBS2;refGene_tx=XM_023252355.1;refGene_gn=SORBS2;refGene_tx=XM_023252356.1;refGene_gn=SORBS2;refGene_tx=XM_023252357.1
B1 16571133 16571183
refGene_gn=SORBS2;refGene_tx=XM_011281387.3;refGene_gn=SORBS2;refGene_tx=XM_011281388.3;refGene_gn=SORBS2;refGene_tx=XM_011281390.3;refGene_gn=SORBS2;refGene_tx=XM_019830023.2;refGene_gn=SORBS2;refGene_tx=XM_019830025.2;refGene_gn=SORBS2;refGene_tx=XM_019830037.2;refGene_gn=SORBS2;refGene_tx=XM_019830038.2;refGene_gn=SORBS2;refGene_tx=XM_019830039.2;refGene_gn=SORBS2;refGene_tx=XM_019830043.2;refGene_gn=SORBS2;refGen

e_tx=XM_019830044.2;refGene_gn=SORBS2;refGene_tx=XM_019830045.2;refGene_gn=SORBS2;refGene_tx=XM_019830046.2;refGene_gn=SORBS2;refGene_tx=XM_023252324.1;refGene_gn=SORBS2;refGene_tx=XM_023252325.1;refGene_gn=SORBS2;refGene_tx=XM_023252326.1;refGene_gn=SORBS2;refGene_tx=XM_023252327.1;refGene_gn=SORBS2;refGene_tx=XM_023252328.1;refGene_gn=SORBS2;refGene_tx=XM_023252329.1;refGene_gn=SORBS2;refGene_tx=XM_023252330.1;refGene_gn=SORBS2;refGene_tx=XM_023252331.1;refGene_gn=SORBS2;refGene_tx=XM_023252332.1;refGene_gn=SORBS2;refGene_tx=XM_023252333.1;refGene_gn=SORBS2;refGene_tx=XM_023252334.1;refGene_gn=SORBS2;refGene_tx=XM_023252335.1;refGene_gn=SORBS2;refGene_tx=XM_023252336.1;refGene_gn=SORBS2;refGene_tx=XM_023252337.1;refGene_gn=SORBS2;refGene_tx=XM_023252338.1;refGene_gn=SORBS2;refGene_tx=XM_023252339.1;refGene_gn=SORBS2;refGene_tx=XM_023252340.1;refGene_gn=SORBS2;refGene_tx=XM_023252341.1;refGene_gn=SORBS2;refGene_tx=XM_023252342.1;refGene_gn=SORBS2;refGene_tx=XM_023252343.1;refGene_gn=SORBS2;refGene_tx=XM_023252344.1;refGene_gn=SORBS2;refGene_tx=XM_023252345.1;refGene_gn=SORBS2;refGene_tx=XM_023252346.1;refGene_gn=SORBS2;refGene_tx=XM_023252347.1;refGene_gn=SORBS2;refGene_tx=XM_023252348.1;refGene_gn=SORBS2;refGene_tx=XM_023252349.1;refGene_gn=SORBS2;refGene_tx=XM_023252350.1;refGene_gn=SORBS2;refGene_tx=XM_023252351.1;refGene_gn=SORBS2;refGene_tx=XM_023252352.1;refGene_gn=SORBS2;refGene_tx=XM_023252353.1;refGene_gn=SORBS2;refGene_tx=XM_023252354.1;refGene_gn=SORBS2;refGene_tx=XM_023252355.1;refGene_gn=SORBS2;refGene_tx=XM_023252356.1;refGene_gn=SORBS2;refGene_tx=XM_023252357.1

B1 16575636 16575693

refGene_gn=SORBS2;refGene_tx=XM_019830023.2;refGene_gn=SORBS2;refGene_tx=XM_019830025.2;refGene_gn=SORBS2;refGene_tx=XM_019830037.2;refGene_gn=SORBS2;refGene_tx=XM_019830038.2;refGene_gn=SORBS2;refGene_tx=XM_019830039.2;refGene_gn=SORBS2;refGene_tx=XM_019830043.2;refGene_gn=SORBS2;refGene_tx=XM_019830044.2;refGene_gn=SORBS2;refGene_tx=XM_019830045.2;refGene_gn=SORBS2;refGene_tx=XM_019830046.2;refGene_gn=SORBS2;refGene_tx=XM_023252324.1;refGene_gn=SORBS2;refGene_tx=XM_023252325.1;refGene_gn=SORBS2;refGene_tx=XM_023252326.1;refGene_gn=SORBS2;refGene_tx=XM_023252327.1;refGene_gn=SORBS2;refGene_tx=XM_023252328.1;refGene_gn=SORBS2;refGene_tx=XM_023252329.1;refGene_gn=SORBS2;refGene_tx=XM_023252330.1;refGene_gn=SORBS2;refGene_tx=XM_023252331.1;refGene_gn=SORBS2;refGene_tx=XM_023252332.1;refGene_gn=SORBS2;refGene_tx=XM_023252333.1;refGene_gn=SORBS2;refGene_tx=XM_023252334.1;refGene_gn=SORBS2;refGene_tx=XM_023252335.1;refGene_gn=SORBS2;refGene_tx=XM_023252336.1;refGene_gn=SORBS2;refGene_tx=XM_023252337.1;refGene_gn=SORBS2;refGene_tx=XM_023252338.1;refGene_gn=SORBS2;refGene_tx=XM_023252339.1;refGene_gn=SORBS2;refGene_tx=XM_023252341.1;refGene_gn=SORBS2;refGene_tx=XM_023252343.1;refGene_gn=SORBS2;refGene_tx=XM_023252344.1;refGene_gn=SORBS2;refGene_tx=XM_023252345.1;refGene_gn=SORBS2;refGene_tx=XM_023252346.1;refGene_gn=SORBS2;refGene_tx=XM_023252347.1;refGene_gn=SORBS2;refGene_tx=XM_023252348.1;refGene_gn=SORBS2;refGene_tx=XM_023252349.1;refGene_gn=SORBS2;refGene_tx=XM_023252350.1;refGene_gn=SORBS2;refGene_tx=XM_023252351.1;refGene_gn=SORBS2;refGene_tx=XM_023252352.1;refGene_gn=SORBS2;refGene_tx=XM_023252353.1;refGene_gn=SORBS2;refGene_tx=XM_023252356.1

B1 16577432 16577516

refGene_gn=SORBS2;refGene_tx=XM_019830023.2;refGene_gn=SORBS2;refGene_

tx=XM_019830025.2;refGene_gn=SORBS2;refGene_tx=XM_019830038.2;refGene_
gn=SORBS2;refGene_tx=XM_019830039.2;refGene_gn=SORBS2;refGene_tx=XM_01
9830043.2;refGene_gn=SORBS2;refGene_tx=XM_019830045.2;refGene_gn=SORBS
2;refGene_tx=XM_019830046.2;refGene_gn=SORBS2;refGene_tx=XM_023252324.
1;refGene_gn=SORBS2;refGene_tx=XM_023252325.1;refGene_gn=SORBS2;refGen
e_tx=XM_023252326.1;refGene_gn=SORBS2;refGene_tx=XM_023252327.1;refGen
e_gn=SORBS2;refGene_tx=XM_023252329.1;refGene_gn=SORBS2;refGene_tx=XM_
023252330.1;refGene_gn=SORBS2;refGene_tx=XM_023252331.1;refGene_gn=SOR
BS2;refGene_tx=XM_023252332.1;refGene_gn=SORBS2;refGene_tx=XM_02325233
3.1;refGene_gn=SORBS2;refGene_tx=XM_023252334.1;refGene_gn=SORBS2;refG
ene_tx=XM_023252335.1;refGene_gn=SORBS2;refGene_tx=XM_023252336.1;refG
ene_gn=SORBS2;refGene_tx=XM_023252337.1;refGene_gn=SORBS2;refGene_tx=X
M_023252338.1;refGene_gn=SORBS2;refGene_tx=XM_023252341.1;refGene_gn=S
ORBS2;refGene_tx=XM_023252343.1;refGene_gn=SORBS2;refGene_tx=XM_023252
344.1;refGene_gn=SORBS2;refGene_tx=XM_023252345.1;refGene_gn=SORBS2;re
fGene_tx=XM_023252346.1;refGene_gn=SORBS2;refGene_tx=XM_023252347.1;re
fGene_gn=SORBS2;refGene_tx=XM_023252348.1;refGene_gn=SORBS2;refGene_tx
=XM_023252349.1;refGene_gn=SORBS2;refGene_tx=XM_023252350.1;refGene_gn
=SORBS2;refGene_tx=XM_023252351.1;refGene_gn=SORBS2;refGene_tx=XM_0232
52352.1;refGene_gn=SORBS2;refGene_tx=XM_023252353.1;refGene_gn=SORBS2;
refGene_tx=XM_023252356.1
B1 16578243 16578402
refGene_gn=SORBS2;refGene_tx=XM_019830023.2;refGene_gn=SORBS2;refGene_
tx=XM_019830025.2;refGene_gn=SORBS2;refGene_tx=XM_019830038.2;refGene_
gn=SORBS2;refGene_tx=XM_019830039.2;refGene_gn=SORBS2;refGene_tx=XM_01
9830045.2;refGene_gn=SORBS2;refGene_tx=XM_019830046.2;refGene_gn=SORBS
2;refGene_tx=XM_023252324.1;refGene_gn=SORBS2;refGene_tx=XM_023252325.
1;refGene_gn=SORBS2;refGene_tx=XM_023252326.1;refGene_gn=SORBS2;refGen
e_tx=XM_023252327.1;refGene_gn=SORBS2;refGene_tx=XM_023252328.1;refGen
e_gn=SORBS2;refGene_tx=XM_023252329.1;refGene_gn=SORBS2;refGene_tx=XM_
023252330.1;refGene_gn=SORBS2;refGene_tx=XM_023252331.1;refGene_gn=SOR
BS2;refGene_tx=XM_023252332.1;refGene_gn=SORBS2;refGene_tx=XM_02325233
3.1;refGene_gn=SORBS2;refGene_tx=XM_023252334.1;refGene_gn=SORBS2;refG
ene_tx=XM_023252335.1;refGene_gn=SORBS2;refGene_tx=XM_023252337.1;refG
ene_gn=SORBS2;refGene_tx=XM_023252338.1;refGene_gn=SORBS2;refGene_tx=X
M_023252341.1;refGene_gn=SORBS2;refGene_tx=XM_023252343.1;refGene_gn=S
ORBS2;refGene_tx=XM_023252344.1;refGene_gn=SORBS2;refGene_tx=XM_023252
345.1;refGene_gn=SORBS2;refGene_tx=XM_023252346.1;refGene_gn=SORBS2;re
fGene_tx=XM_023252347.1;refGene_gn=SORBS2;refGene_tx=XM_023252348.1;re
fGene_gn=SORBS2;refGene_tx=XM_023252349.1;refGene_gn=SORBS2;refGene_tx
=XM_023252350.1;refGene_gn=SORBS2;refGene_tx=XM_023252351.1;refGene_gn
=SORBS2;refGene_tx=XM_023252352.1;refGene_gn=SORBS2;refGene_tx=XM_0232
52353.1
B1 16585657 16585772
refGene_gn=SORBS2;refGene_tx=XM_011281387.3;refGene_gn=SORBS2;refGene_
tx=XM_011281388.3;refGene_gn=SORBS2;refGene_tx=XM_011281390.3;refGene_
gn=SORBS2;refGene_tx=XM_019830023.2;refGene_gn=SORBS2;refGene_tx=XM_01
9830025.2;refGene_gn=SORBS2;refGene_tx=XM_019830037.2;refGene_gn=SORBS
2;refGene_tx=XM_019830038.2;refGene_gn=SORBS2;refGene_tx=XM_019830039.
2;refGene_gn=SORBS2;refGene_tx=XM_019830043.2;refGene_gn=SORBS2;refGen

e_tx=XM_019830044.2;refGene_gn=SORBS2;refGene_tx=XM_019830045.2;refGene_gn=SORBS2;refGene_tx=XM_019830046.2;refGene_gn=SORBS2;refGene_tx=XM_023252324.1;refGene_gn=SORBS2;refGene_tx=XM_023252325.1;refGene_gn=SORBS2;refGene_tx=XM_023252326.1;refGene_gn=SORBS2;refGene_tx=XM_023252327.1;refGene_gn=SORBS2;refGene_tx=XM_023252328.1;refGene_gn=SORBS2;refGene_tx=XM_023252329.1;refGene_gn=SORBS2;refGene_tx=XM_023252330.1;refGene_gn=SORBS2;refGene_tx=XM_023252331.1;refGene_gn=SORBS2;refGene_tx=XM_023252332.1;refGene_gn=SORBS2;refGene_tx=XM_023252333.1;refGene_gn=SORBS2;refGene_tx=XM_023252334.1;refGene_gn=SORBS2;refGene_tx=XM_023252335.1;refGene_gn=SORBS2;refGene_tx=XM_023252336.1;refGene_gn=SORBS2;refGene_tx=XM_023252337.1;refGene_gn=SORBS2;refGene_tx=XM_023252338.1;refGene_gn=SORBS2;refGene_tx=XM_023252339.1;refGene_gn=SORBS2;refGene_tx=XM_023252340.1;refGene_gn=SORBS2;refGene_tx=XM_023252341.1;refGene_gn=SORBS2;refGene_tx=XM_023252342.1;refGene_gn=SORBS2;refGene_tx=XM_023252343.1;refGene_gn=SORBS2;refGene_tx=XM_023252344.1;refGene_gn=SORBS2;refGene_tx=XM_023252345.1;refGene_gn=SORBS2;refGene_tx=XM_023252346.1;refGene_gn=SORBS2;refGene_tx=XM_023252347.1;refGene_gn=SORBS2;refGene_tx=XM_023252348.1;refGene_gn=SORBS2;refGene_tx=XM_023252349.1;refGene_gn=SORBS2;refGene_tx=XM_023252350.1;refGene_gn=SORBS2;refGene_tx=XM_023252351.1;refGene_gn=SORBS2;refGene_tx=XM_023252352.1;refGene_gn=SORBS2;refGene_tx=XM_023252353.1;refGene_gn=SORBS2;refGene_tx=XM_023252354.1;refGene_gn=SORBS2;refGene_tx=XM_023252355.1;refGene_gn=SORBS2;refGene_tx=XM_023252356.1;refGene_gn=SORBS2;refGene_tx=XM_023252357.1

B1 16588596 16588786

refGene_gn=SORBS2;refGene_tx=XM_011281387.3;refGene_gn=SORBS2;refGene_tx=XM_011281388.3;refGene_gn=SORBS2;refGene_tx=XM_011281390.3;refGene_gn=SORBS2;refGene_tx=XM_019830023.2;refGene_gn=SORBS2;refGene_tx=XM_019830025.2;refGene_gn=SORBS2;refGene_tx=XM_019830037.2;refGene_gn=SORBS2;refGene_tx=XM_019830038.2;refGene_gn=SORBS2;refGene_tx=XM_019830039.2;refGene_gn=SORBS2;refGene_tx=XM_019830043.2;refGene_gn=SORBS2;refGene_tx=XM_019830044.2;refGene_gn=SORBS2;refGene_tx=XM_019830045.2;refGene_gn=SORBS2;refGene_tx=XM_019830046.2;refGene_gn=SORBS2;refGene_tx=XM_023252324.1;refGene_gn=SORBS2;refGene_tx=XM_023252325.1;refGene_gn=SORBS2;refGene_tx=XM_023252326.1;refGene_gn=SORBS2;refGene_tx=XM_023252327.1;refGene_gn=SORBS2;refGene_tx=XM_023252328.1;refGene_gn=SORBS2;refGene_tx=XM_023252329.1;refGene_gn=SORBS2;refGene_tx=XM_023252330.1;refGene_gn=SORBS2;refGene_tx=XM_023252331.1;refGene_gn=SORBS2;refGene_tx=XM_023252332.1;refGene_gn=SORBS2;refGene_tx=XM_023252333.1;refGene_gn=SORBS2;refGene_tx=XM_023252334.1;refGene_gn=SORBS2;refGene_tx=XM_023252335.1;refGene_gn=SORBS2;refGene_tx=XM_023252336.1;refGene_gn=SORBS2;refGene_tx=XM_023252337.1;refGene_gn=SORBS2;refGene_tx=XM_023252338.1;refGene_gn=SORBS2;refGene_tx=XM_023252339.1;refGene_gn=SORBS2;refGene_tx=XM_023252340.1;refGene_gn=SORBS2;refGene_tx=XM_023252341.1;refGene_gn=SORBS2;refGene_tx=XM_023252342.1;refGene_gn=SORBS2;refGene_tx=XM_023252343.1;refGene_gn=SORBS2;refGene_tx=XM_023252344.1;refGene_gn=SORBS2;refGene_tx=XM_023252345.1;refGene_gn=SORBS2;refGene_tx=XM_023252346.1;refGene_gn=SORBS2;refGene_tx=XM_023252347.1;refGene_gn=SORBS2;refGene_tx=XM_023252348.1;refGene_gn=SORBS2;refGene_tx=XM_023252349.1;refGene_gn=SORBS2;refGene_tx=XM_023252350.1;refGene_gn=SORBS2;refGene_tx=XM_023252351.1;refGene_gn=SORBS2;refGene_tx=XM_023252352.1;refGene_gn=SORBS

2;refGene_tx=XM_023252353.1;refGene_gn=SORBS2;refGene_tx=XM_023252354.1;refGene_gn=SORBS2;refGene_tx=XM_023252355.1;refGene_gn=SORBS2;refGene_tx=XM_023252356.1;refGene_gn=SORBS2;refGene_tx=XM_023252357.1
B1 16591553 16591598
refGene_gn=SORBS2;refGene_tx=XM_011281387.3;refGene_gn=SORBS2;refGene_tx=XM_019830023.2;refGene_gn=SORBS2;refGene_tx=XM_019830025.2;refGene_gn=SORBS2;refGene_tx=XM_019830037.2;refGene_gn=SORBS2;refGene_tx=XM_019830038.2;refGene_gn=SORBS2;refGene_tx=XM_019830039.2;refGene_gn=SORBS2;refGene_tx=XM_019830043.2;refGene_gn=SORBS2;refGene_tx=XM_019830044.2;refGene_gn=SORBS2;refGene_tx=XM_019830045.2;refGene_gn=SORBS2;refGene_tx=XM_019830046.2;refGene_gn=SORBS2;refGene_tx=XM_023252324.1;refGene_gn=SORBS2;refGene_tx=XM_023252325.1;refGene_gn=SORBS2;refGene_tx=XM_023252326.1;refGene_gn=SORBS2;refGene_tx=XM_023252327.1;refGene_gn=SORBS2;refGene_tx=XM_023252328.1;refGene_gn=SORBS2;refGene_tx=XM_023252329.1;refGene_gn=SORBS2;refGene_tx=XM_023252330.1;refGene_gn=SORBS2;refGene_tx=XM_023252331.1;refGene_gn=SORBS2;refGene_tx=XM_023252332.1;refGene_gn=SORBS2;refGene_tx=XM_023252334.1;refGene_gn=SORBS2;refGene_tx=XM_023252335.1;refGene_gn=SORBS2;refGene_tx=XM_023252336.1;refGene_gn=SORBS2;refGene_tx=XM_023252337.1;refGene_gn=SORBS2;refGene_tx=XM_023252338.1;refGene_gn=SORBS2;refGene_tx=XM_023252339.1;refGene_gn=SORBS2;refGene_tx=XM_023252340.1;refGene_gn=SORBS2;refGene_tx=XM_023252341.1;refGene_gn=SORBS2;refGene_tx=XM_023252342.1;refGene_gn=SORBS2;refGene_tx=XM_023252343.1;refGene_gn=SORBS2;refGene_tx=XM_023252344.1;refGene_gn=SORBS2;refGene_tx=XM_023252345.1;refGene_gn=SORBS2;refGene_tx=XM_023252346.1;refGene_gn=SORBS2;refGene_tx=XM_023252347.1;refGene_gn=SORBS2;refGene_tx=XM_023252348.1;refGene_gn=SORBS2;refGene_tx=XM_023252349.1;refGene_gn=SORBS2;refGene_tx=XM_023252350.1;refGene_gn=SORBS2;refGene_tx=XM_023252351.1;refGene_gn=SORBS2;refGene_tx=XM_023252353.1;refGene_gn=SORBS2;refGene_tx=XM_023252354.1;refGene_gn=SORBS2;refGene_tx=XM_023252356.1;refGene_gn=SORBS2;refGene_tx=XM_023252357.1
B1 16592289 16592356
refGene_gn=SORBS2;refGene_tx=XM_011281387.3;refGene_gn=SORBS2;refGene_tx=XM_011281388.3;refGene_gn=SORBS2;refGene_tx=XM_011281390.3;refGene_gn=SORBS2;refGene_tx=XM_019830023.2;refGene_gn=SORBS2;refGene_tx=XM_019830025.2;refGene_gn=SORBS2;refGene_tx=XM_019830037.2;refGene_gn=SORBS2;refGene_tx=XM_019830038.2;refGene_gn=SORBS2;refGene_tx=XM_019830039.2;refGene_gn=SORBS2;refGene_tx=XM_019830043.2;refGene_gn=SORBS2;refGene_tx=XM_019830044.2;refGene_gn=SORBS2;refGene_tx=XM_019830045.2;refGene_gn=SORBS2;refGene_tx=XM_019830046.2;refGene_gn=SORBS2;refGene_tx=XM_023252324.1;refGene_gn=SORBS2;refGene_tx=XM_023252325.1;refGene_gn=SORBS2;refGene_tx=XM_023252326.1;refGene_gn=SORBS2;refGene_tx=XM_023252327.1;refGene_gn=SORBS2;refGene_tx=XM_023252328.1;refGene_gn=SORBS2;refGene_tx=XM_023252329.1;refGene_gn=SORBS2;refGene_tx=XM_023252330.1;refGene_gn=SORBS2;refGene_tx=XM_023252331.1;refGene_gn=SORBS2;refGene_tx=XM_023252332.1;refGene_gn=SORBS2;refGene_tx=XM_023252333.1;refGene_gn=SORBS2;refGene_tx=XM_023252334.1;refGene_gn=SORBS2;refGene_tx=XM_023252335.1;refGene_gn=SORBS2;refGene_tx=XM_023252336.1;refGene_gn=SORBS2;refGene_tx=XM_023252337.1;refGene_gn=SORBS2;refGene_tx=XM_023252338.1;refGene_gn=SORBS2;refGene_tx=XM_023252339.1;refGene_gn=SORBS2;refGene_tx=XM_023252340.1;refGene_gn=SORBS2;refGene_tx=XM_023252341.1;refGene_gn

=SORBS2;refGene_tx=XM_023252342.1;refGene_gn=SORBS2;refGene_tx=XM_023252343.1;refGene_gn=SORBS2;refGene_tx=XM_023252344.1;refGene_gn=SORBS2;refGene_tx=XM_023252345.1;refGene_gn=SORBS2;refGene_tx=XM_023252346.1;refGene_gn=SORBS2;refGene_tx=XM_023252347.1;refGene_gn=SORBS2;refGene_tx=XM_023252348.1;refGene_gn=SORBS2;refGene_tx=XM_023252349.1;refGene_gn=SORBS2;refGene_tx=XM_023252350.1;refGene_gn=SORBS2;refGene_tx=XM_023252351.1;refGene_gn=SORBS2;refGene_tx=XM_023252352.1;refGene_gn=SORBS2;refGene_tx=XM_023252353.1;refGene_gn=SORBS2;refGene_tx=XM_023252354.1;refGene_gn=SORBS2;refGene_tx=XM_023252355.1;refGene_gn=SORBS2;refGene_tx=XM_023252356.1;refGene_gn=SORBS2;refGene_tx=XM_023252357.1
B1 16596474 16596498
refGene_gn=SORBS2;refGene_tx=XM_023252351.1;refGene_gn=SORBS2;refGene_tx=XM_023252352.1
B1 16597253 16597426
refGene_gn=SORBS2;refGene_tx=XM_011281387.3;refGene_gn=SORBS2;refGene_tx=XM_011281388.3;refGene_gn=SORBS2;refGene_tx=XM_011281390.3;refGene_gn=SORBS2;refGene_tx=XM_019830023.2;refGene_gn=SORBS2;refGene_tx=XM_019830025.2;refGene_gn=SORBS2;refGene_tx=XM_019830037.2;refGene_gn=SORBS2;refGene_tx=XM_019830038.2;refGene_gn=SORBS2;refGene_tx=XM_019830039.2;refGene_gn=SORBS2;refGene_tx=XM_019830043.2;refGene_gn=SORBS2;refGene_tx=XM_019830044.2;refGene_gn=SORBS2;refGene_tx=XM_019830045.2;refGene_gn=SORBS2;refGene_tx=XM_019830046.2;refGene_gn=SORBS2;refGene_tx=XM_023252324.1;refGene_gn=SORBS2;refGene_tx=XM_023252325.1;refGene_gn=SORBS2;refGene_tx=XM_023252326.1;refGene_gn=SORBS2;refGene_tx=XM_023252327.1;refGene_gn=SORBS2;refGene_tx=XM_023252328.1;refGene_gn=SORBS2;refGene_tx=XM_023252329.1;refGene_gn=SORBS2;refGene_tx=XM_023252330.1;refGene_gn=SORBS2;refGene_tx=XM_023252331.1;refGene_gn=SORBS2;refGene_tx=XM_023252332.1;refGene_gn=SORBS2;refGene_tx=XM_023252333.1;refGene_gn=SORBS2;refGene_tx=XM_023252334.1;refGene_gn=SORBS2;refGene_tx=XM_023252335.1;refGene_gn=SORBS2;refGene_tx=XM_023252336.1;refGene_gn=SORBS2;refGene_tx=XM_023252337.1;refGene_gn=SORBS2;refGene_tx=XM_023252338.1;refGene_gn=SORBS2;refGene_tx=XM_023252339.1;refGene_gn=SORBS2;refGene_tx=XM_023252340.1;refGene_gn=SORBS2;refGene_tx=XM_023252341.1;refGene_gn=SORBS2;refGene_tx=XM_023252342.1;refGene_gn=SORBS2;refGene_tx=XM_023252343.1;refGene_gn=SORBS2;refGene_tx=XM_023252344.1;refGene_gn=SORBS2;refGene_tx=XM_023252345.1;refGene_gn=SORBS2;refGene_tx=XM_023252346.1;refGene_gn=SORBS2;refGene_tx=XM_023252347.1;refGene_gn=SORBS2;refGene_tx=XM_023252348.1;refGene_gn=SORBS2;refGene_tx=XM_023252349.1;refGene_gn=SORBS2;refGene_tx=XM_023252350.1;refGene_gn=SORBS2;refGene_tx=XM_023252353.1;refGene_gn=SORBS2;refGene_tx=XM_023252354.1;refGene_gn=SORBS2;refGene_tx=XM_023252355.1;refGene_gn=SORBS2;refGene_tx=XM_023252356.1;refGene_gn=SORBS2;refGene_tx=XM_023252357.1
B1 16602053 16602192
refGene_gn=SORBS2;refGene_tx=XM_011281387.3;refGene_gn=SORBS2;refGene_tx=XM_011281388.3;refGene_gn=SORBS2;refGene_tx=XM_011281390.3;refGene_gn=SORBS2;refGene_tx=XM_019830023.2;refGene_gn=SORBS2;refGene_tx=XM_019830025.2;refGene_gn=SORBS2;refGene_tx=XM_019830037.2;refGene_gn=SORBS2;refGene_tx=XM_019830038.2;refGene_gn=SORBS2;refGene_tx=XM_019830039.2;refGene_gn=SORBS2;refGene_tx=XM_019830043.2;refGene_gn=SORBS2;refGene_tx=XM_019830044.2;refGene_gn=SORBS2;refGene_tx=XM_019830045.2;refGen

e_gn=SORBS2;refGene_tx=XM_019830046.2;refGene_gn=SORBS2;refGene_tx=XM_023252324.1;refGene_gn=SORBS2;refGene_tx=XM_023252325.1;refGene_gn=SORBS2;refGene_tx=XM_023252326.1;refGene_gn=SORBS2;refGene_tx=XM_023252327.1;refGene_gn=SORBS2;refGene_tx=XM_023252328.1;refGene_gn=SORBS2;refGene_tx=XM_023252329.1;refGene_gn=SORBS2;refGene_tx=XM_023252330.1;refGene_gn=SORBS2;refGene_tx=XM_023252331.1;refGene_gn=SORBS2;refGene_tx=XM_023252332.1;refGene_gn=SORBS2;refGene_tx=XM_023252333.1;refGene_gn=SORBS2;refGene_tx=XM_023252334.1;refGene_gn=SORBS2;refGene_tx=XM_023252335.1;refGene_gn=SORBS2;refGene_tx=XM_023252336.1;refGene_gn=SORBS2;refGene_tx=XM_023252337.1;refGene_gn=SORBS2;refGene_tx=XM_023252338.1;refGene_gn=SORBS2;refGene_tx=XM_023252339.1;refGene_gn=SORBS2;refGene_tx=XM_023252340.1;refGene_gn=SORBS2;refGene_tx=XM_023252341.1;refGene_gn=SORBS2;refGene_tx=XM_023252342.1;refGene_gn=SORBS2;refGene_tx=XM_023252343.1;refGene_gn=SORBS2;refGene_tx=XM_023252344.1;refGene_gn=SORBS2;refGene_tx=XM_023252345.1;refGene_gn=SORBS2;refGene_tx=XM_023252346.1;refGene_gn=SORBS2;refGene_tx=XM_023252347.1;refGene_gn=SORBS2;refGene_tx=XM_023252348.1;refGene_gn=SORBS2;refGene_tx=XM_023252353.1;refGene_gn=SORBS2;refGene_tx=XM_023252354.1;refGene_gn=SORBS2;refGene_tx=XM_023252355.1;refGene_gn=SORBS2;refGene_tx=XM_023252356.1;refGene_gn=SORBS2;refGene_tx=XM_023252357.1
B1 16606227 16606263
refGene_gn=SORBS2;refGene_tx=XM_023252331.1;refGene_gn=SORBS2;refGene_tx=XM_023252333.1;refGene_gn=SORBS2;refGene_tx=XM_023252334.1;refGene_gn=SORBS2;refGene_tx=XM_023252336.1;refGene_gn=SORBS2;refGene_tx=XM_023252337.1;refGene_gn=SORBS2;refGene_tx=XM_023252339.1;refGene_gn=SORBS2;refGene_tx=XM_023252340.1;refGene_gn=SORBS2;refGene_tx=XM_023252354.1;refGene_gn=SORBS2;refGene_tx=XM_023252355.1;refGene_gn=SORBS2;refGene_tx=XM_023252356.1;refGene_gn=SORBS2;refGene_tx=XM_023252357.1
B1 16617379 16618021
refGene_gn=SORBS2;refGene_tx=XM_019830023.2;refGene_gn=SORBS2;refGene_tx=XM_019830025.2;refGene_gn=SORBS2;refGene_tx=XM_019830038.2;refGene_gn=SORBS2;refGene_tx=XM_019830039.2;refGene_gn=SORBS2;refGene_tx=XM_023252324.1;refGene_gn=SORBS2;refGene_tx=XM_023252325.1;refGene_gn=SORBS2;refGene_tx=XM_023252326.1;refGene_gn=SORBS2;refGene_tx=XM_023252327.1;refGene_gn=SORBS2;refGene_tx=XM_023252328.1;refGene_gn=SORBS2;refGene_tx=XM_023252329.1;refGene_gn=SORBS2;refGene_tx=XM_023252330.1;refGene_gn=SORBS2;refGene_tx=XM_023252353.1
B1 16618836 16618961
refGene_gn=SORBS2;refGene_tx=XM_011281387.3;refGene_gn=SORBS2;refGene_tx=XM_011281388.3;refGene_gn=SORBS2;refGene_tx=XM_011281390.3;refGene_gn=SORBS2;refGene_tx=XM_019830023.2;refGene_gn=SORBS2;refGene_tx=XM_019830025.2;refGene_gn=SORBS2;refGene_tx=XM_019830037.2;refGene_gn=SORBS2;refGene_tx=XM_019830038.2;refGene_gn=SORBS2;refGene_tx=XM_019830039.2;refGene_gn=SORBS2;refGene_tx=XM_019830043.2;refGene_gn=SORBS2;refGene_tx=XM_019830044.2;refGene_gn=SORBS2;refGene_tx=XM_019830045.2;refGene_gn=SORBS2;refGene_tx=XM_019830046.2;refGene_gn=SORBS2;refGene_tx=XM_023252324.1;refGene_gn=SORBS2;refGene_tx=XM_023252325.1;refGene_gn=SORBS2;refGene_tx=XM_023252326.1;refGene_gn=SORBS2;refGene_tx=XM_023252327.1;refGene_gn=SORBS2;refGene_tx=XM_023252328.1;refGene_gn=SORBS2;refGene_tx=XM_023252329.1;refGene_gn=SORBS2;refGene_tx=XM_023252330.1;refG

ene_gn=SORBS2;refGene_tx=XM_023252331.1;refGene_gn=SORBS2;refGene_tx=XM_023252332.1;refGene_gn=SORBS2;refGene_tx=XM_023252333.1;refGene_gn=SORBS2;refGene_tx=XM_023252334.1;refGene_gn=SORBS2;refGene_tx=XM_023252335.1;refGene_gn=SORBS2;refGene_tx=XM_023252336.1;refGene_gn=SORBS2;refGene_tx=XM_023252337.1;refGene_gn=SORBS2;refGene_tx=XM_023252338.1;refGene_gn=SORBS2;refGene_tx=XM_023252339.1;refGene_gn=SORBS2;refGene_tx=XM_023252340.1;refGene_gn=SORBS2;refGene_tx=XM_023252342.1;refGene_gn=SORBS2;refGene_tx=XM_023252353.1;refGene_gn=SORBS2;refGene_tx=XM_023252354.1;refGene_gn=SORBS2;refGene_tx=XM_023252355.1;refGene_gn=SORBS2;refGene_tx=XM_023252356.1;refGene_gn=SORBS2;refGene_tx=XM_023252357.1
B1 16619205 16619232
refGene_gn=SORBS2;refGene_tx=XM_011281387.3;refGene_gn=SORBS2;refGene_tx=XM_011281388.3;refGene_gn=SORBS2;refGene_tx=XM_011281390.3;refGene_gn=SORBS2;refGene_tx=XM_019830023.2;refGene_gn=SORBS2;refGene_tx=XM_019830025.2;refGene_gn=SORBS2;refGene_tx=XM_019830037.2;refGene_gn=SORBS2;refGene_tx=XM_019830038.2;refGene_gn=SORBS2;refGene_tx=XM_019830039.2;refGene_gn=SORBS2;refGene_tx=XM_019830043.2;refGene_gn=SORBS2;refGene_tx=XM_019830044.2;refGene_gn=SORBS2;refGene_tx=XM_019830045.2;refGene_gn=SORBS2;refGene_tx=XM_019830046.2;refGene_gn=SORBS2;refGene_tx=XM_023252324.1;refGene_gn=SORBS2;refGene_tx=XM_023252325.1;refGene_gn=SORBS2;refGene_tx=XM_023252326.1;refGene_gn=SORBS2;refGene_tx=XM_023252327.1;refGene_gn=SORBS2;refGene_tx=XM_023252328.1;refGene_gn=SORBS2;refGene_tx=XM_023252329.1;refGene_gn=SORBS2;refGene_tx=XM_023252330.1;refGene_gn=SORBS2;refGene_tx=XM_023252331.1;refGene_gn=SORBS2;refGene_tx=XM_023252332.1;refGene_gn=SORBS2;refGene_tx=XM_023252333.1;refGene_gn=SORBS2;refGene_tx=XM_023252334.1;refGene_gn=SORBS2;refGene_tx=XM_023252335.1;refGene_gn=SORBS2;refGene_tx=XM_023252336.1;refGene_gn=SORBS2;refGene_tx=XM_023252337.1;refGene_gn=SORBS2;refGene_tx=XM_023252338.1;refGene_gn=SORBS2;refGene_tx=XM_023252339.1;refGene_gn=SORBS2;refGene_tx=XM_023252340.1;refGene_gn=SORBS2;refGene_tx=XM_023252342.1;refGene_gn=SORBS2;refGene_tx=XM_023252353.1;refGene_gn=SORBS2;refGene_tx=XM_023252354.1;refGene_gn=SORBS2;refGene_tx=XM_023252355.1;refGene_gn=SORBS2;refGene_tx=XM_023252356.1;refGene_gn=SORBS2;refGene_tx=XM_023252357.1
B1 16625380 16625473
refGene_gn=SORBS2;refGene_tx=XM_011281387.3;refGene_gn=SORBS2;refGene_tx=XM_011281388.3;refGene_gn=SORBS2;refGene_tx=XM_011281390.3;refGene_gn=SORBS2;refGene_tx=XM_019830023.2;refGene_gn=SORBS2;refGene_tx=XM_019830025.2;refGene_gn=SORBS2;refGene_tx=XM_019830037.2;refGene_gn=SORBS2;refGene_tx=XM_019830038.2;refGene_gn=SORBS2;refGene_tx=XM_019830039.2;refGene_gn=SORBS2;refGene_tx=XM_019830043.2;refGene_gn=SORBS2;refGene_tx=XM_019830044.2;refGene_gn=SORBS2;refGene_tx=XM_019830045.2;refGene_gn=SORBS2;refGene_tx=XM_019830046.2;refGene_gn=SORBS2;refGene_tx=XM_023252324.1;refGene_gn=SORBS2;refGene_tx=XM_023252325.1;refGene_gn=SORBS2;refGene_tx=XM_023252326.1;refGene_gn=SORBS2;refGene_tx=XM_023252327.1;refGene_gn=SORBS2;refGene_tx=XM_023252328.1;refGene_gn=SORBS2;refGene_tx=XM_023252331.1;refGene_gn=SORBS2;refGene_tx=XM_023252332.1;refGene_gn=SORBS2;refGene_tx=XM_023252333.1;refGene_gn=SORBS2;refGene_tx=XM_023252334.1;refGene_gn=SORBS2;refGene_tx=XM_023252335.1;refGene_gn=SORBS2;refGene_tx=XM_023252336.1;refGene_gn=SORBS2;refGene_tx=XM_023252339.1;refGene_gn=SORBS2;refGene_tx=XM_023252340.1;refGene_gn=SORBS2;re

fGene_tx=XM_023252342.1;refGene_gn=SORBS2;refGene_tx=XM_023252353.1;re
fGene_gn=SORBS2;refGene_tx=XM_023252354.1;refGene_gn=SORBS2;refGene_tx
=XM_023252355.1;refGene_gn=SORBS2;refGene_tx=XM_023252356.1;refGene_gn
=SORBS2;refGene_tx=XM_023252357.1
B1 16629853 16629863
refGene_gn=SORBS2;refGene_tx=XM_011281387.3;refGene_gn=SORBS2;refGene_
tx=XM_011281388.3;refGene_gn=SORBS2;refGene_tx=XM_011281390.3;refGene_
gn=SORBS2;refGene_tx=XM_019830023.2;refGene_gn=SORBS2;refGene_tx=XM_01
9830025.2;refGene_gn=SORBS2;refGene_tx=XM_019830037.2;refGene_gn=SORBS
2;refGene_tx=XM_019830038.2;refGene_gn=SORBS2;refGene_tx=XM_019830039.
2;refGene_gn=SORBS2;refGene_tx=XM_019830043.2;refGene_gn=SORBS2;refGen
e_tx=XM_019830044.2;refGene_gn=SORBS2;refGene_tx=XM_019830045.2;refGen
e_gn=SORBS2;refGene_tx=XM_019830046.2;refGene_gn=SORBS2;refGene_tx=XM_
023252324.1;refGene_gn=SORBS2;refGene_tx=XM_023252325.1;refGene_gn=SOR
BS2;refGene_tx=XM_023252327.1;refGene_gn=SORBS2;refGene_tx=XM_02325232
8.1;refGene_gn=SORBS2;refGene_tx=XM_023252329.1;refGene_gn=SORBS2;refG
ene_tx=XM_023252330.1;refGene_gn=SORBS2;refGene_tx=XM_023252331.1;refG
ene_gn=SORBS2;refGene_tx=XM_023252332.1;refGene_gn=SORBS2;refGene_tx=X
M_023252333.1;refGene_gn=SORBS2;refGene_tx=XM_023252334.1;refGene_gn=S
ORBS2;refGene_tx=XM_023252335.1;refGene_gn=SORBS2;refGene_tx=XM_023252
336.1;refGene_gn=SORBS2;refGene_tx=XM_023252337.1;refGene_gn=SORBS2;re
fGene_tx=XM_023252338.1;refGene_gn=SORBS2;refGene_tx=XM_023252339.1;re
fGene_gn=SORBS2;refGene_tx=XM_023252340.1;refGene_gn=SORBS2;refGene_tx
=XM_023252342.1;refGene_gn=SORBS2;refGene_tx=XM_023252353.1;refGene_gn
=SORBS2;refGene_tx=XM_023252354.1;refGene_gn=SORBS2;refGene_tx=XM_0232
52355.1;refGene_gn=SORBS2;refGene_tx=XM_023252356.1;refGene_gn=SORBS2;
refGene_tx=XM_023252357.1
B1 17007783 17008224
refGene_gn=TLR3;refGene_tx=NM_001079829.1;refGene_gn=TLR3;refGene_tx=X
M_023252208.1
B1 17010375 17010567
refGene_gn=TLR3;refGene_tx=NM_001079829.1;refGene_gn=TLR3;refGene_tx=X
M_023252208.1
B1 17013086 17014939
refGene_gn=TLR3;refGene_tx=NM_001079829.1;refGene_gn=TLR3;refGene_tx=X
M_023252208.1
B1 17015166 17015395
refGene_gn=TLR3;refGene_tx=NM_001079829.1;refGene_gn=TLR3;refGene_tx=X
M_023252208.1
B1 17030834 17030932
refGene_gn=FAM149A;refGene_tx=XM_011281393.3;refGene_gn=FAM149A;refGen
e_tx=XM_011281394.3;refGene_gn=FAM149A;refGene_tx=XM_019830048.2
B1 17064871 17064982
refGene_gn=FAM149A;refGene_tx=XM_011281393.3;refGene_gn=FAM149A;refGen
e_tx=XM_011281394.3;refGene_gn=FAM149A;refGene_tx=XM_019830048.2
B1 17065485 17065624
refGene_gn=FAM149A;refGene_tx=XM_011281393.3;refGene_gn=FAM149A;refGen
e_tx=XM_011281394.3;refGene_gn=FAM149A;refGene_tx=XM_019830048.2
B1 17068329 17068472
refGene_gn=FAM149A;refGene_tx=XM_011281393.3;refGene_gn=FAM149A;refGen

e_tx=XM_011281394.3;refGene_gn=FAM149A;refGene_tx=XM_019830048.2;refGene_gn=FAM149A;refGene_tx=XM_023252368.1
B1 17070022 17070148
refGene_gn=FAM149A;refGene_tx=XM_011281393.3;refGene_gn=FAM149A;refGene_tx=XM_011281394.3;refGene_gn=FAM149A;refGene_tx=XM_019830048.2;refGene_gn=FAM149A;refGene_tx=XM_023252368.1
B1 17071848 17072019
refGene_gn=FAM149A;refGene_tx=XM_011281393.3;refGene_gn=FAM149A;refGene_tx=XM_011281394.3;refGene_gn=FAM149A;refGene_tx=XM_019830048.2;refGene_gn=FAM149A;refGene_tx=XM_023252368.1
B1 17073106 17073297
refGene_gn=FAM149A;refGene_tx=XM_011281393.3;refGene_gn=FAM149A;refGene_tx=XM_011281394.3;refGene_gn=FAM149A;refGene_tx=XM_019830048.2;refGene_gn=FAM149A;refGene_tx=XM_023252368.1
B1 17074095 17074247
refGene_gn=FAM149A;refGene_tx=XM_011281393.3;refGene_gn=FAM149A;refGene_tx=XM_011281394.3;refGene_gn=FAM149A;refGene_tx=XM_019830048.2;refGene_gn=FAM149A;refGene_tx=XM_023252368.1
B1 17078324 17078428
refGene_gn=FAM149A;refGene_tx=XM_011281393.3;refGene_gn=FAM149A;refGene_tx=XM_011281394.3;refGene_gn=FAM149A;refGene_tx=XM_019830048.2;refGene_gn=FAM149A;refGene_tx=XM_023252368.1
B1 17078907 17079117
refGene_gn=FAM149A;refGene_tx=XM_011281393.3;refGene_gn=FAM149A;refGene_tx=XM_011281394.3;refGene_gn=FAM149A;refGene_tx=XM_019830048.2;refGene_gn=FAM149A;refGene_tx=XM_023252368.1
B1 17082440 17082561
refGene_gn=FAM149A;refGene_tx=XM_011281393.3;refGene_gn=FAM149A;refGene_tx=XM_011281394.3;refGene_gn=FAM149A;refGene_tx=XM_019830048.2;refGene_gn=FAM149A;refGene_tx=XM_023252368.1
B1 17083808 17083937
refGene_gn=FAM149A;refGene_tx=XM_011281393.3;refGene_gn=FAM149A;refGene_tx=XM_011281394.3;refGene_gn=FAM149A;refGene_tx=XM_019830048.2;refGene_gn=FAM149A;refGene_tx=XM_023252368.1
B1 17084268 17084347
refGene_gn=FAM149A;refGene_tx=XM_011281393.3;refGene_gn=FAM149A;refGene_tx=XM_011281394.3;refGene_gn=FAM149A;refGene_tx=XM_019830048.2;refGene_gn=FAM149A;refGene_tx=XM_023252368.1
B1 17087722 17087906
refGene_gn=FAM149A;refGene_tx=XM_011281393.3;refGene_gn=FAM149A;refGene_tx=XM_011281394.3;refGene_gn=FAM149A;refGene_tx=XM_023252368.1
B1 17097047 17097212
refGene_gn=LOC101097556;refGene_tx=XM_023252373.1
B1 17097213 17097262
refGene_gn=LOC101097556;refGene_tx=XM_023252373.1
B1 17102463 17102576
refGene_gn=LOC101097556;refGene_tx=XM_023252373.1
B1 17104728 17104814
refGene_gn=LOC101097556;refGene_tx=XM_023252373.1
B1 17105871 17106062

refGene_gn=LOC101097556;refGene_tx=XM_023252373.1
B1 17106456 17106526
refGene_gn=LOC101097556;refGene_tx=XM_023252373.1
B1 17108099 17108226
refGene_gn=LOC101097556;refGene_tx=XM_023252373.1
B1 17109653 17109839
refGene_gn=LOC101097556;refGene_tx=XM_023252373.1
B1 17112184 17112287
refGene_gn=LOC101097556;refGene_tx=XM_023252373.1
B1 17115557 17116103
refGene_gn=LOC102902016;refGene_tx=XM_019828268.2
B1 17116572 17116707
refGene_gn=LOC101097556;refGene_tx=XM_023252373.1
B1 17116801 17116981
refGene_gn=LOC101097556;refGene_tx=XM_023252373.1
B1 17118422 17118595
refGene_gn=LOC101097556;refGene_tx=XM_023252373.1
B1 17128186 17128244
refGene_gn=KLKB1;refGene_tx=XM_019828262.2;refGene_gn=KLKB1;refGene_tx=XM_023252369.1;refGene_gn=KLKB1;refGene_tx=XM_023252370.1;refGene_gn=KLKB1;refGene_tx=XM_023252372.1
B1 17132716 17132879
refGene_gn=KLKB1;refGene_tx=XM_019828262.2;refGene_gn=KLKB1;refGene_tx=XM_019828263.2;refGene_gn=KLKB1;refGene_tx=XM_019828264.2;refGene_gn=KLKB1;refGene_tx=XM_019828265.2;refGene_gn=KLKB1;refGene_tx=XM_023252369.1;refGene_gn=KLKB1;refGene_tx=XM_023252370.1;refGene_gn=KLKB1;refGene_tx=XM_023252372.1
B1 17134106 17134213
refGene_gn=KLKB1;refGene_tx=XM_019828262.2;refGene_gn=KLKB1;refGene_tx=XM_019828263.2;refGene_gn=KLKB1;refGene_tx=XM_019828264.2;refGene_gn=KLKB1;refGene_tx=XM_019828265.2;refGene_gn=KLKB1;refGene_tx=XM_023252369.1;refGene_gn=KLKB1;refGene_tx=XM_023252370.1;refGene_gn=KLKB1;refGene_tx=XM_023252372.1
B1 17137336 17137496
refGene_gn=KLKB1;refGene_tx=XM_019828262.2;refGene_gn=KLKB1;refGene_tx=XM_019828263.2;refGene_gn=KLKB1;refGene_tx=XM_019828264.2;refGene_gn=KLKB1;refGene_tx=XM_019828265.2;refGene_gn=KLKB1;refGene_tx=XM_023252369.1;refGene_gn=KLKB1;refGene_tx=XM_023252370.1;refGene_gn=KLKB1;refGene_tx=XM_023252372.1
B1 17137899 17138009
refGene_gn=KLKB1;refGene_tx=XM_019828262.2;refGene_gn=KLKB1;refGene_tx=XM_019828263.2;refGene_gn=KLKB1;refGene_tx=XM_019828264.2;refGene_gn=KLKB1;refGene_tx=XM_019828265.2;refGene_gn=KLKB1;refGene_tx=XM_023252369.1;refGene_gn=KLKB1;refGene_tx=XM_023252370.1;refGene_gn=KLKB1;refGene_tx=XM_023252372.1
B1 17143336 17143496
refGene_gn=KLKB1;refGene_tx=XM_019828262.2;refGene_gn=KLKB1;refGene_tx=XM_019828263.2;refGene_gn=KLKB1;refGene_tx=XM_019828264.2;refGene_gn=KLKB1;refGene_tx=XM_019828265.2;refGene_gn=KLKB1;refGene_tx=XM_023252369.1;refGene_gn=KLKB1;refGene_tx=XM_023252370.1;refGene_gn=KLKB1;refGe

ne_tx=XM_023252372.1
B1 17144710 17144820
refGene_gn=KLKB1;refGene_tx=XM_019828262.2;refGene_gn=KLKB1;refGene_tx=XM_019828263.2;refGene_gn=KLKB1;refGene_tx=XM_019828264.2;refGene_gn=KLKB1;refGene_tx=XM_019828265.2;refGene_gn=KLKB1;refGene_tx=XM_023252369.1;refGene_gn=KLKB1;refGene_tx=XM_023252370.1
B1 17144958 17145121
refGene_gn=KLKB1;refGene_tx=XM_019828262.2;refGene_gn=KLKB1;refGene_tx=XM_019828263.2;refGene_gn=KLKB1;refGene_tx=XM_019828264.2;refGene_gn=KLKB1;refGene_tx=XM_019828265.2;refGene_gn=KLKB1;refGene_tx=XM_023252369.1;refGene_gn=KLKB1;refGene_tx=XM_023252370.1
B1 17145220 17145333
refGene_gn=KLKB1;refGene_tx=XM_019828262.2;refGene_gn=KLKB1;refGene_tx=XM_019828263.2;refGene_gn=KLKB1;refGene_tx=XM_019828264.2;refGene_gn=KLKB1;refGene_tx=XM_019828265.2;refGene_gn=KLKB1;refGene_tx=XM_023252369.1;refGene_gn=KLKB1;refGene_tx=XM_023252370.1;refGene_gn=KLKB1;refGene_tx=XM_023252372.1
B1 17145497 17145666
refGene_gn=KLKB1;refGene_tx=XM_019828262.2;refGene_gn=KLKB1;refGene_tx=XM_019828263.2;refGene_gn=KLKB1;refGene_tx=XM_019828264.2;refGene_gn=KLKB1;refGene_tx=XM_019828265.2;refGene_gn=KLKB1;refGene_tx=XM_023252369.1;refGene_gn=KLKB1;refGene_tx=XM_023252370.1;refGene_gn=KLKB1;refGene_tx=XM_023252372.1
B1 17149912 17150088
refGene_gn=KLKB1;refGene_tx=XM_019828262.2;refGene_gn=KLKB1;refGene_tx=XM_019828263.2;refGene_gn=KLKB1;refGene_tx=XM_019828264.2;refGene_gn=KLKB1;refGene_tx=XM_019828265.2;refGene_gn=KLKB1;refGene_tx=XM_023252369.1;refGene_gn=KLKB1;refGene_tx=XM_023252370.1;refGene_gn=KLKB1;refGene_tx=XM_023252372.1
B1 17151037 17151133
refGene_gn=KLKB1;refGene_tx=XM_019828262.2;refGene_gn=KLKB1;refGene_tx=XM_019828263.2;refGene_gn=KLKB1;refGene_tx=XM_019828264.2;refGene_gn=KLKB1;refGene_tx=XM_019828265.2;refGene_gn=KLKB1;refGene_tx=XM_023252369.1;refGene_gn=KLKB1;refGene_tx=XM_023252370.1;refGene_gn=KLKB1;refGene_tx=XM_023252372.1
B1 17153191 17153331
refGene_gn=KLKB1;refGene_tx=XM_019828262.2;refGene_gn=KLKB1;refGene_tx=XM_019828263.2;refGene_gn=KLKB1;refGene_tx=XM_019828264.2;refGene_gn=KLKB1;refGene_tx=XM_019828265.2;refGene_gn=KLKB1;refGene_tx=XM_023252369.1;refGene_gn=KLKB1;refGene_tx=XM_023252370.1;refGene_gn=KLKB1;refGene_tx=XM_023252372.1
B1 17153923 17154103
refGene_gn=KLKB1;refGene_tx=XM_019828262.2;refGene_gn=KLKB1;refGene_tx=XM_019828263.2;refGene_gn=KLKB1;refGene_tx=XM_019828264.2;refGene_gn=KLKB1;refGene_tx=XM_019828265.2;refGene_gn=KLKB1;refGene_tx=XM_023252369.1;refGene_gn=KLKB1;refGene_tx=XM_023252372.1
B1 17154178 17154196
refGene_gn=KLKB1;refGene_tx=XM_023252369.1;refGene_gn=KLKB1;refGene_tx=XM_023252370.1
B1 17160619 17160674

refGene_gn=LOC101094340;refGene_tx=XM_003984601.4;refGene_gn=LOC101094340;refGene_tx=XM_019828269.2
B1 17162632 17162795
refGene_gn=LOC101094340;refGene_tx=XM_003984601.4;refGene_gn=LOC101094340;refGene_tx=XM_019828269.2;refGene_gn=LOC101094340;refGene_tx=XM_019828270.2;refGene_gn=LOC101094340;refGene_tx=XM_019828271.2
B1 17164307 17164414
refGene_gn=LOC101094340;refGene_tx=XM_003984601.4;refGene_gn=LOC101094340;refGene_tx=XM_019828269.2;refGene_gn=LOC101094340;refGene_tx=XM_019828270.2;refGene_gn=LOC101094340;refGene_tx=XM_019828271.2;refGene_gn=LOC101094340;refGene_tx=XM_019828272.2
B1 17165288 17165448
refGene_gn=LOC101094340;refGene_tx=XM_003984601.4;refGene_gn=LOC101094340;refGene_tx=XM_019828269.2;refGene_gn=LOC101094340;refGene_tx=XM_019828270.2;refGene_gn=LOC101094340;refGene_tx=XM_019828271.2;refGene_gn=LOC101094340;refGene_tx=XM_019828272.2;refGene_gn=LOC101094340;refGene_tx=XM_019828274.2
B1 17166629 17166739
refGene_gn=LOC101094340;refGene_tx=XM_003984601.4;refGene_gn=LOC101094340;refGene_tx=XM_019828269.2;refGene_gn=LOC101094340;refGene_tx=XM_019828270.2;refGene_gn=LOC101094340;refGene_tx=XM_019828271.2;refGene_gn=LOC101094340;refGene_tx=XM_019828272.2;refGene_gn=LOC101094340;refGene_tx=XM_019828274.2
B1 17167437 17167597
refGene_gn=LOC101094340;refGene_tx=XM_003984601.4;refGene_gn=LOC101094340;refGene_tx=XM_019828269.2;refGene_gn=LOC101094340;refGene_tx=XM_019828270.2;refGene_gn=LOC101094340;refGene_tx=XM_019828271.2;refGene_gn=LOC101094340;refGene_tx=XM_019828272.2;refGene_gn=LOC101094340;refGene_tx=XM_019828274.2
B1 17169936 17170046
refGene_gn=LOC101094340;refGene_tx=XM_003984601.4;refGene_gn=LOC101094340;refGene_tx=XM_019828269.2;refGene_gn=LOC101094340;refGene_tx=XM_019828270.2;refGene_gn=LOC101094340;refGene_tx=XM_019828271.2;refGene_gn=LOC101094340;refGene_tx=XM_019828272.2;refGene_gn=LOC101094340;refGene_tx=XM_019828274.2
B1 17170146 17170309
refGene_gn=LOC101094340;refGene_tx=XM_003984601.4;refGene_gn=LOC101094340;refGene_tx=XM_019828269.2;refGene_gn=LOC101094340;refGene_tx=XM_019828270.2;refGene_gn=LOC101094340;refGene_tx=XM_019828271.2;refGene_gn=LOC101094340;refGene_tx=XM_019828272.2;refGene_gn=LOC101094340;refGene_tx=XM_019828274.2
B1 17170398 17170505
refGene_gn=LOC101094340;refGene_tx=XM_003984601.4;refGene_gn=LOC101094340;refGene_tx=XM_019828269.2;refGene_gn=LOC101094340;refGene_tx=XM_019828270.2;refGene_gn=LOC101094340;refGene_tx=XM_019828271.2;refGene_gn=LOC101094340;refGene_tx=XM_019828272.2;refGene_gn=LOC101094340;refGene_tx=XM_019828274.2
B1 17173967 17174136
refGene_gn=LOC101094340;refGene_tx=XM_003984601.4;refGene_gn=LOC101094340;refGene_tx=XM_019828269.2;refGene_gn=LOC101094340;refGene_tx=XM_01

9828270.2;refGene_gn=LOC101094340;refGene_tx=XM_019828271.2;refGene_gn=LOC101094340;refGene_tx=XM_019828272.2;refGene_gn=LOC101094340;refGene_tx=XM_019828274.2
B1 17175338 17175514
refGene_gn=LOC101094340;refGene_tx=XM_003984601.4;refGene_gn=LOC101094340;refGene_tx=XM_019828269.2;refGene_gn=LOC101094340;refGene_tx=XM_019828270.2;refGene_gn=LOC101094340;refGene_tx=XM_019828271.2;refGene_gn=LOC101094340;refGene_tx=XM_019828272.2;refGene_gn=LOC101094340;refGene_tx=XM_019828274.2
B1 17176088 17176184
refGene_gn=LOC101094340;refGene_tx=XM_003984601.4;refGene_gn=LOC101094340;refGene_tx=XM_019828269.2;refGene_gn=LOC101094340;refGene_tx=XM_019828270.2;refGene_gn=LOC101094340;refGene_tx=XM_019828271.2;refGene_gn=LOC101094340;refGene_tx=XM_019828272.2;refGene_gn=LOC101094340;refGene_tx=XM_019828274.2
B1 17176708 17176848
refGene_gn=LOC101094340;refGene_tx=XM_003984601.4;refGene_gn=LOC101094340;refGene_tx=XM_019828269.2;refGene_gn=LOC101094340;refGene_tx=XM_019828270.2;refGene_gn=LOC101094340;refGene_tx=XM_019828271.2;refGene_gn=LOC101094340;refGene_tx=XM_019828272.2;refGene_gn=LOC101094340;refGene_tx=XM_019828274.2
B1 17177329 17177491
refGene_gn=LOC101094340;refGene_tx=XM_003984601.4;refGene_gn=LOC101094340;refGene_tx=XM_019828269.2;refGene_gn=LOC101094340;refGene_tx=XM_019828270.2;refGene_gn=LOC101094340;refGene_tx=XM_019828271.2;refGene_gn=LOC101094340;refGene_tx=XM_019828272.2;refGene_gn=LOC101094340;refGene_tx=XM_019828274.2
B1 17381939 17382808 refGene_gn=MTNR1A;refGene_tx=XM_023252374.1
B1 17422644 17422870 refGene_gn=MTNR1A;refGene_tx=XM_023252374.1
B1 17444453 17445082 refGene_gn=FAT1;refGene_tx=XM_023252375.1
B1 17450420 17450558 refGene_gn=FAT1;refGene_tx=XM_023252375.1
B1 17451384 17452016 refGene_gn=FAT1;refGene_tx=XM_023252375.1
B1 17453896 17454007 refGene_gn=FAT1;refGene_tx=XM_023252375.1
B1 17454190 17454344 refGene_gn=FAT1;refGene_tx=XM_023252375.1
B1 17455699 17456162 refGene_gn=FAT1;refGene_tx=XM_023252375.1
B1 17456777 17456935 refGene_gn=FAT1;refGene_tx=XM_023252375.1
B1 17458669 17458801 refGene_gn=FAT1;refGene_tx=XM_023252375.1
B1 17458954 17459756 refGene_gn=FAT1;refGene_tx=XM_023252375.1
B1 17460384 17460582 refGene_gn=FAT1;refGene_tx=XM_023252375.1
B1 17461376 17461520 refGene_gn=FAT1;refGene_tx=XM_023252375.1
B1 17463515 17463653 refGene_gn=FAT1;refGene_tx=XM_023252375.1
B1 17463905 17464120 refGene_gn=FAT1;refGene_tx=XM_023252375.1
B1 17466165 17466343 refGene_gn=FAT1;refGene_tx=XM_023252375.1
B1 17466344 17466556 refGene_gn=FAT1;refGene_tx=XM_023252375.1
B1 17467376 17467610 refGene_gn=FAT1;refGene_tx=XM_023252375.1
B1 17468611 17468765 refGene_gn=FAT1;refGene_tx=XM_023252375.1
B1 17471180 17471377 refGene_gn=FAT1;refGene_tx=XM_023252375.1
B1 17472090 17476158 refGene_gn=FAT1;refGene_tx=XM_023252375.1
B1 17481551 17481762 refGene_gn=FAT1;refGene_tx=XM_023252375.1
B1 17481904 17482180 refGene_gn=FAT1;refGene_tx=XM_023252375.1

B1 17487376 17487516 refGene_gn=FAT1;refGene_tx=XM_023252375.1
B1 17490225 17490436 refGene_gn=FAT1;refGene_tx=XM_023252375.1
B1 17491017 17491347 refGene_gn=FAT1;refGene_tx=XM_023252375.1
B1 17494829 17494891 refGene_gn=FAT1;refGene_tx=XM_023252375.1
B1 17521179 17521494 refGene_gn=FAT1;refGene_tx=XM_023252375.1
B1 17558355 17561614 refGene_gn=FAT1;refGene_tx=XM_023252375.1
B1 18655364 18656276
refGene_gn=ZFP42;refGene_tx=XM_003984616.3;refGene_gn=ZFP42;refGene_tx
=XM_019828288.2
B1 18747374 18747964
refGene_gn=TRIML2;refGene_tx=XM_019828286.2;refGene_gn=TRIML2;refGene_
tx=XM_023252377.1
B1 18754392 18754493
refGene_gn=TRIML2;refGene_tx=XM_019828286.2;refGene_gn=TRIML2;refGene_
tx=XM_023252377.1
B1 18754662 18754685
refGene_gn=TRIML2;refGene_tx=XM_019828286.2;refGene_gn=TRIML2;refGene_
tx=XM_023252377.1
B1 18757083 18757224
refGene_gn=TRIML2;refGene_tx=XM_019828286.2;refGene_gn=TRIML2;refGene_
tx=XM_023252377.1
B1 18758558 18758747
refGene_gn=TRIML2;refGene_tx=XM_019828286.2;refGene_gn=TRIML2;refGene_
tx=XM_023252377.1
B1 18761306 18761402
refGene_gn=TRIML2;refGene_tx=XM_019828286.2;refGene_gn=TRIML2;refGene_
tx=XM_023252377.1
B1 18761676 18761865
refGene_gn=TRIML2;refGene_tx=XM_019828286.2;refGene_gn=TRIML2;refGene_
tx=XM_023252377.1
B1 18787446 18787872
refGene_gn=TRIML1;refGene_tx=XM_003984614.4;refGene_gn=TRIML1;refGene_
tx=XM_019828282.2;refGene_gn=TRIML1;refGene_tx=XM_019828284.2
B1 18788115 18788211
refGene_gn=TRIML1;refGene_tx=XM_003984614.4;refGene_gn=TRIML1;refGene_
tx=XM_019828282.2;refGene_gn=TRIML1;refGene_tx=XM_019828284.2
B1 18789391 18789622
refGene_gn=TRIML1;refGene_tx=XM_003984614.4;refGene_gn=TRIML1;refGene_
tx=XM_019828282.2;refGene_gn=TRIML1;refGene_tx=XM_019828284.2
B1 18790869 18790892
refGene_gn=TRIML1;refGene_tx=XM_003984614.4;refGene_gn=TRIML1;refGene_
tx=XM_019828282.2;refGene_gn=TRIML1;refGene_tx=XM_019828284.2
B1 18791070 18791231
refGene_gn=TRIML1;refGene_tx=XM_003984614.4;refGene_gn=TRIML1;refGene_
tx=XM_019828282.2;refGene_gn=TRIML1;refGene_tx=XM_019828284.2
B1 18792957 18793508
refGene_gn=TRIML1;refGene_tx=XM_003984614.4;refGene_gn=TRIML1;refGene_
tx=XM_019828282.2;refGene_gn=TRIML1;refGene_tx=XM_019828284.2
B1 20542231 20542293
refGene_gn=FRG1;refGene_tx=XM_003984618.5;refGene_gn=FRG1;refGene_tx=X

M_006930586.3
B1 20544538 20544609
refGene_gn=FRG1;refGene_tx=XM_003984618.5;refGene_gn=FRG1;refGene_tx=X
M_006930586.3
B1 20551565 20551691
refGene_gn=FRG1;refGene_tx=XM_003984618.5;refGene_gn=FRG1;refGene_tx=X
M_006930586.3
B1 20552943 20553001
refGene_gn=FRG1;refGene_tx=XM_003984618.5;refGene_gn=FRG1;refGene_tx=X
M_006930586.3
B1 20554417 20554532
refGene_gn=FRG1;refGene_tx=XM_003984618.5;refGene_gn=FRG1;refGene_tx=X
M_006930586.3
B1 20558541 20558646
refGene_gn=FRG1;refGene_tx=XM_003984618.5;refGene_gn=FRG1;refGene_tx=X
M_006930586.3
B1 20561239 20561331
refGene_gn=FRG1;refGene_tx=XM_003984618.5;refGene_gn=FRG1;refGene_tx=X
M_006930586.3
B1 20561825 20561936
refGene_gn=FRG1;refGene_tx=XM_003984618.5;refGene_gn=FRG1;refGene_tx=X
M_006930586.3
B1 20563118 20563155
refGene_gn=FRG1;refGene_tx=XM_003984618.5;refGene_gn=FRG1;refGene_tx=X
M_006930586.3
B1 20580834 20580912 refGene_gn=ASAH1;refGene_tx=XM_006930587.3
B1 20604610 20604657 refGene_gn=ASAH1;refGene_tx=XM_006930587.3
B1 20613029 20613120 refGene_gn=ASAH1;refGene_tx=XM_006930587.3
B1 20613514 20613601 refGene_gn=ASAH1;refGene_tx=XM_006930587.3
B1 20616407 20616486 refGene_gn=ASAH1;refGene_tx=XM_006930587.3
B1 20621086 20621161 refGene_gn=ASAH1;refGene_tx=XM_006930587.3
B1 20622025 20622071 refGene_gn=ASAH1;refGene_tx=XM_006930587.3
B1 20622678 20622823 refGene_gn=ASAH1;refGene_tx=XM_006930587.3
B1 20623369 20623424 refGene_gn=ASAH1;refGene_tx=XM_006930587.3
B1 20623550 20623632 refGene_gn=ASAH1;refGene_tx=XM_006930587.3
B1 20627489 20627621 refGene_gn=ASAH1;refGene_tx=XM_006930587.3
B1 20627725 20627849 refGene_gn=ASAH1;refGene_tx=XM_006930587.3
B1 20628275 20628332 refGene_gn=ASAH1;refGene_tx=XM_006930587.3
B1 20629236 20629326 refGene_gn=ASAH1;refGene_tx=XM_006930587.3
B1 20633610 20633708
refGene_gn=PCM1;refGene_tx=XM_023252378.1;refGene_gn=PCM1;refGene_tx=X
M_023252379.1;refGene_gn=PCM1;refGene_tx=XM_023252380.1;refGene_gn=PCM
1;refGene_tx=XM_023252381.1;refGene_gn=PCM1;refGene_tx=XM_023252382.1;
refGene_gn=PCM1;refGene_tx=XM_023252383.1;refGene_gn=PCM1;refGene_tx=X
M_023252384.1;refGene_gn=PCM1;refGene_tx=XM_023252385.1;refGene_gn=PCM
1;refGene_tx=XM_023252387.1;refGene_gn=PCM1;refGene_tx=XM_023252388.1;
refGene_gn=PCM1;refGene_tx=XM_023252389.1;refGene_gn=PCM1;refGene_tx=X
M_023252390.1;refGene_gn=PCM1;refGene_tx=XM_023252391.1;refGene_gn=PCM
1;refGene_tx=XM_023252392.1;refGene_gn=PCM1;refGene_tx=XM_023252393.1;
refGene_gn=PCM1;refGene_tx=XM_023252394.1;refGene_gn=PCM1;refGene_tx=X

M_023252395.1
B1 20636718 20636744
refGene_gn=PCM1;refGene_tx=XM_023252386.1;refGene_gn=PCM1;refGene_tx=X
M_023252397.1;refGene_gn=PCM1;refGene_tx=XM_023252398.1;refGene_gn=PCM
1;refGene_tx=XM_023252399.1;refGene_gn=PCM1;refGene_tx=XM_023252400.1
B1 20637998 20638116
refGene_gn=PCM1;refGene_tx=XM_023252378.1;refGene_gn=PCM1;refGene_tx=X
M_023252379.1;refGene_gn=PCM1;refGene_tx=XM_023252380.1;refGene_gn=PCM
1;refGene_tx=XM_023252381.1;refGene_gn=PCM1;refGene_tx=XM_023252382.1;
refGene_gn=PCM1;refGene_tx=XM_023252383.1;refGene_gn=PCM1;refGene_tx=X
M_023252384.1;refGene_gn=PCM1;refGene_tx=XM_023252385.1;refGene_gn=PCM
1;refGene_tx=XM_023252386.1;refGene_gn=PCM1;refGene_tx=XM_023252387.1;
refGene_gn=PCM1;refGene_tx=XM_023252388.1;refGene_gn=PCM1;refGene_tx=X
M_023252389.1;refGene_gn=PCM1;refGene_tx=XM_023252390.1;refGene_gn=PCM
1;refGene_tx=XM_023252391.1;refGene_gn=PCM1;refGene_tx=XM_023252392.1;
refGene_gn=PCM1;refGene_tx=XM_023252393.1;refGene_gn=PCM1;refGene_tx=X
M_023252394.1;refGene_gn=PCM1;refGene_tx=XM_023252395.1;refGene_gn=PCM
1;refGene_tx=XM_023252397.1;refGene_gn=PCM1;refGene_tx=XM_023252398.1;
refGene_gn=PCM1;refGene_tx=XM_023252399.1;refGene_gn=PCM1;refGene_tx=X
M_023252400.1
B1 20638225 20638318
refGene_gn=PCM1;refGene_tx=XM_023252378.1;refGene_gn=PCM1;refGene_tx=X
M_023252379.1;refGene_gn=PCM1;refGene_tx=XM_023252380.1;refGene_gn=PCM
1;refGene_tx=XM_023252381.1;refGene_gn=PCM1;refGene_tx=XM_023252382.1;
refGene_gn=PCM1;refGene_tx=XM_023252383.1;refGene_gn=PCM1;refGene_tx=X
M_023252384.1;refGene_gn=PCM1;refGene_tx=XM_023252385.1;refGene_gn=PCM
1;refGene_tx=XM_023252386.1;refGene_gn=PCM1;refGene_tx=XM_023252387.1;
refGene_gn=PCM1;refGene_tx=XM_023252388.1;refGene_gn=PCM1;refGene_tx=X
M_023252389.1;refGene_gn=PCM1;refGene_tx=XM_023252390.1;refGene_gn=PCM
1;refGene_tx=XM_023252391.1;refGene_gn=PCM1;refGene_tx=XM_023252392.1;
refGene_gn=PCM1;refGene_tx=XM_023252393.1;refGene_gn=PCM1;refGene_tx=X
M_023252394.1;refGene_gn=PCM1;refGene_tx=XM_023252395.1;refGene_gn=PCM
1;refGene_tx=XM_023252397.1;refGene_gn=PCM1;refGene_tx=XM_023252398.1;
refGene_gn=PCM1;refGene_tx=XM_023252399.1;refGene_gn=PCM1;refGene_tx=X
M_023252400.1
B1 20640005 20640262
refGene_gn=PCM1;refGene_tx=XM_023252378.1;refGene_gn=PCM1;refGene_tx=X
M_023252379.1;refGene_gn=PCM1;refGene_tx=XM_023252380.1;refGene_gn=PCM
1;refGene_tx=XM_023252381.1;refGene_gn=PCM1;refGene_tx=XM_023252382.1;
refGene_gn=PCM1;refGene_tx=XM_023252383.1;refGene_gn=PCM1;refGene_tx=X
M_023252384.1;refGene_gn=PCM1;refGene_tx=XM_023252385.1;refGene_gn=PCM
1;refGene_tx=XM_023252386.1;refGene_gn=PCM1;refGene_tx=XM_023252387.1;
refGene_gn=PCM1;refGene_tx=XM_023252388.1;refGene_gn=PCM1;refGene_tx=X
M_023252389.1;refGene_gn=PCM1;refGene_tx=XM_023252390.1;refGene_gn=PCM
1;refGene_tx=XM_023252391.1;refGene_gn=PCM1;refGene_tx=XM_023252392.1;
refGene_gn=PCM1;refGene_tx=XM_023252393.1;refGene_gn=PCM1;refGene_tx=X
M_023252394.1;refGene_gn=PCM1;refGene_tx=XM_023252395.1;refGene_gn=PCM
1;refGene_tx=XM_023252396.1;refGene_gn=PCM1;refGene_tx=XM_023252397.1;
refGene_gn=PCM1;refGene_tx=XM_023252398.1;refGene_gn=PCM1;refGene_tx=X
M_023252399.1;refGene_gn=PCM1;refGene_tx=XM_023252400.1

B1 20640815 20640891
refGene_gn=PCM1;refGene_tx=XM_023252378.1;refGene_gn=PCM1;refGene_tx=XM_023252379.1;refGene_gn=PCM1;refGene_tx=XM_023252380.1;refGene_gn=PCM1;refGene_tx=XM_023252381.1;refGene_gn=PCM1;refGene_tx=XM_023252382.1;refGene_gn=PCM1;refGene_tx=XM_023252383.1;refGene_gn=PCM1;refGene_tx=XM_023252384.1;refGene_gn=PCM1;refGene_tx=XM_023252385.1;refGene_gn=PCM1;refGene_tx=XM_023252386.1;refGene_gn=PCM1;refGene_tx=XM_023252387.1;refGene_gn=PCM1;refGene_tx=XM_023252388.1;refGene_gn=PCM1;refGene_tx=XM_023252389.1;refGene_gn=PCM1;refGene_tx=XM_023252390.1;refGene_gn=PCM1;refGene_tx=XM_023252391.1;refGene_gn=PCM1;refGene_tx=XM_023252392.1;refGene_gn=PCM1;refGene_tx=XM_023252393.1;refGene_gn=PCM1;refGene_tx=XM_023252394.1;refGene_gn=PCM1;refGene_tx=XM_023252395.1;refGene_gn=PCM1;refGene_tx=XM_023252397.1;refGene_gn=PCM1;refGene_tx=XM_023252398.1;refGene_gn=PCM1;refGene_tx=XM_023252399.1;refGene_gn=PCM1;refGene_tx=XM_023252400.1
B1 20643638 20643799
refGene_gn=PCM1;refGene_tx=XM_023252378.1;refGene_gn=PCM1;refGene_tx=XM_023252379.1;refGene_gn=PCM1;refGene_tx=XM_023252380.1;refGene_gn=PCM1;refGene_tx=XM_023252381.1;refGene_gn=PCM1;refGene_tx=XM_023252382.1;refGene_gn=PCM1;refGene_tx=XM_023252383.1;refGene_gn=PCM1;refGene_tx=XM_023252384.1;refGene_gn=PCM1;refGene_tx=XM_023252385.1;refGene_gn=PCM1;refGene_tx=XM_023252386.1;refGene_gn=PCM1;refGene_tx=XM_023252387.1;refGene_gn=PCM1;refGene_tx=XM_023252388.1;refGene_gn=PCM1;refGene_tx=XM_023252389.1;refGene_gn=PCM1;refGene_tx=XM_023252390.1;refGene_gn=PCM1;refGene_tx=XM_023252391.1;refGene_gn=PCM1;refGene_tx=XM_023252392.1;refGene_gn=PCM1;refGene_tx=XM_023252393.1;refGene_gn=PCM1;refGene_tx=XM_023252394.1;refGene_gn=PCM1;refGene_tx=XM_023252395.1;refGene_gn=PCM1;refGene_tx=XM_023252396.1;refGene_gn=PCM1;refGene_tx=XM_023252397.1;refGene_gn=PCM1;refGene_tx=XM_023252398.1;refGene_gn=PCM1;refGene_tx=XM_023252399.1;refGene_gn=PCM1;refGene_tx=XM_023252400.1
B1 20644094 20644230
refGene_gn=PCM1;refGene_tx=XM_023252378.1;refGene_gn=PCM1;refGene_tx=XM_023252379.1;refGene_gn=PCM1;refGene_tx=XM_023252380.1;refGene_gn=PCM1;refGene_tx=XM_023252381.1;refGene_gn=PCM1;refGene_tx=XM_023252382.1;refGene_gn=PCM1;refGene_tx=XM_023252383.1;refGene_gn=PCM1;refGene_tx=XM_023252384.1;refGene_gn=PCM1;refGene_tx=XM_023252385.1;refGene_gn=PCM1;refGene_tx=XM_023252386.1;refGene_gn=PCM1;refGene_tx=XM_023252387.1;refGene_gn=PCM1;refGene_tx=XM_023252388.1;refGene_gn=PCM1;refGene_tx=XM_023252389.1;refGene_gn=PCM1;refGene_tx=XM_023252390.1;refGene_gn=PCM1;refGene_tx=XM_023252391.1;refGene_gn=PCM1;refGene_tx=XM_023252392.1;refGene_gn=PCM1;refGene_tx=XM_023252393.1;refGene_gn=PCM1;refGene_tx=XM_023252394.1;refGene_gn=PCM1;refGene_tx=XM_023252395.1;refGene_gn=PCM1;refGene_tx=XM_023252396.1;refGene_gn=PCM1;refGene_tx=XM_023252397.1;refGene_gn=PCM1;refGene_tx=XM_023252398.1;refGene_gn=PCM1;refGene_tx=XM_023252399.1;refGene_gn=PCM1;refGene_tx=XM_023252400.1
B1 20644457 20644520
refGene_gn=PCM1;refGene_tx=XM_023252378.1;refGene_gn=PCM1;refGene_tx=XM_023252379.1;refGene_gn=PCM1;refGene_tx=XM_023252380.1;refGene_gn=PCM1;refGene_tx=XM_023252381.1;refGene_gn=PCM1;refGene_tx=XM_023252382.1;refGene_gn=PCM1;refGene_tx=XM_023252383.1;refGene_gn=PCM1;refGene_tx=X

M_023252384.1;refGene_gn=PCM1;refGene_tx=XM_023252385.1;refGene_gn=PCM1;refGene_tx=XM_023252386.1;refGene_gn=PCM1;refGene_tx=XM_023252387.1;refGene_gn=PCM1;refGene_tx=XM_023252388.1;refGene_gn=PCM1;refGene_tx=XM_023252389.1;refGene_gn=PCM1;refGene_tx=XM_023252390.1;refGene_gn=PCM1;refGene_tx=XM_023252391.1;refGene_gn=PCM1;refGene_tx=XM_023252392.1;refGene_gn=PCM1;refGene_tx=XM_023252393.1;refGene_gn=PCM1;refGene_tx=XM_023252394.1;refGene_gn=PCM1;refGene_tx=XM_023252395.1;refGene_gn=PCM1;refGene_tx=XM_023252396.1;refGene_gn=PCM1;refGene_tx=XM_023252397.1;refGene_gn=PCM1;refGene_tx=XM_023252398.1;refGene_gn=PCM1;refGene_tx=XM_023252399.1;refGene_gn=PCM1;refGene_tx=XM_023252400.1
B1 20646119 20646317
refGene_gn=PCM1;refGene_tx=XM_023252378.1;refGene_gn=PCM1;refGene_tx=XM_023252379.1;refGene_gn=PCM1;refGene_tx=XM_023252380.1;refGene_gn=PCM1;refGene_tx=XM_023252381.1;refGene_gn=PCM1;refGene_tx=XM_023252382.1;refGene_gn=PCM1;refGene_tx=XM_023252383.1;refGene_gn=PCM1;refGene_tx=XM_023252384.1;refGene_gn=PCM1;refGene_tx=XM_023252385.1;refGene_gn=PCM1;refGene_tx=XM_023252386.1;refGene_gn=PCM1;refGene_tx=XM_023252387.1;refGene_gn=PCM1;refGene_tx=XM_023252388.1;refGene_gn=PCM1;refGene_tx=XM_023252389.1;refGene_gn=PCM1;refGene_tx=XM_023252390.1;refGene_gn=PCM1;refGene_tx=XM_023252391.1;refGene_gn=PCM1;refGene_tx=XM_023252392.1;refGene_gn=PCM1;refGene_tx=XM_023252393.1;refGene_gn=PCM1;refGene_tx=XM_023252394.1;refGene_gn=PCM1;refGene_tx=XM_023252395.1;refGene_gn=PCM1;refGene_tx=XM_023252396.1;refGene_gn=PCM1;refGene_tx=XM_023252397.1;refGene_gn=PCM1;refGene_tx=XM_023252398.1;refGene_gn=PCM1;refGene_tx=XM_023252399.1;refGene_gn=PCM1;refGene_tx=XM_023252400.1
B1 20647315 20647450
refGene_gn=PCM1;refGene_tx=XM_023252378.1;refGene_gn=PCM1;refGene_tx=XM_023252379.1;refGene_gn=PCM1;refGene_tx=XM_023252380.1;refGene_gn=PCM1;refGene_tx=XM_023252381.1;refGene_gn=PCM1;refGene_tx=XM_023252382.1;refGene_gn=PCM1;refGene_tx=XM_023252383.1;refGene_gn=PCM1;refGene_tx=XM_023252384.1;refGene_gn=PCM1;refGene_tx=XM_023252385.1;refGene_gn=PCM1;refGene_tx=XM_023252386.1;refGene_gn=PCM1;refGene_tx=XM_023252387.1;refGene_gn=PCM1;refGene_tx=XM_023252388.1;refGene_gn=PCM1;refGene_tx=XM_023252389.1;refGene_gn=PCM1;refGene_tx=XM_023252390.1;refGene_gn=PCM1;refGene_tx=XM_023252391.1;refGene_gn=PCM1;refGene_tx=XM_023252392.1;refGene_gn=PCM1;refGene_tx=XM_023252393.1;refGene_gn=PCM1;refGene_tx=XM_023252394.1;refGene_gn=PCM1;refGene_tx=XM_023252395.1;refGene_gn=PCM1;refGene_tx=XM_023252396.1;refGene_gn=PCM1;refGene_tx=XM_023252397.1;refGene_gn=PCM1;refGene_tx=XM_023252398.1;refGene_gn=PCM1;refGene_tx=XM_023252399.1;refGene_gn=PCM1;refGene_tx=XM_023252400.1
B1 20649608 20649742
refGene_gn=PCM1;refGene_tx=XM_023252378.1;refGene_gn=PCM1;refGene_tx=XM_023252379.1;refGene_gn=PCM1;refGene_tx=XM_023252380.1;refGene_gn=PCM1;refGene_tx=XM_023252381.1;refGene_gn=PCM1;refGene_tx=XM_023252382.1;refGene_gn=PCM1;refGene_tx=XM_023252383.1;refGene_gn=PCM1;refGene_tx=XM_023252384.1;refGene_gn=PCM1;refGene_tx=XM_023252385.1;refGene_gn=PCM1;refGene_tx=XM_023252386.1;refGene_gn=PCM1;refGene_tx=XM_023252387.1;refGene_gn=PCM1;refGene_tx=XM_023252388.1;refGene_gn=PCM1;refGene_tx=XM_023252389.1;refGene_gn=PCM1;refGene_tx=XM_023252390.1;refGene_gn=PCM1;refGene_tx=XM_023252391.1;refGene_gn=PCM1;refGene_tx=XM_023252392.1;

refGene_gn=PCM1;refGene_tx=XM_023252393.1;refGene_gn=PCM1;refGene_tx=XM_023252394.1;refGene_gn=PCM1;refGene_tx=XM_023252395.1;refGene_gn=PCM1;refGene_tx=XM_023252396.1;refGene_gn=PCM1;refGene_tx=XM_023252397.1;refGene_gn=PCM1;refGene_tx=XM_023252398.1;refGene_gn=PCM1;refGene_tx=XM_023252399.1;refGene_gn=PCM1;refGene_tx=XM_023252400.1
B1 20649999 20650158
refGene_gn=PCM1;refGene_tx=XM_023252378.1;refGene_gn=PCM1;refGene_tx=XM_023252379.1;refGene_gn=PCM1;refGene_tx=XM_023252380.1;refGene_gn=PCM1;refGene_tx=XM_023252381.1;refGene_gn=PCM1;refGene_tx=XM_023252382.1;refGene_gn=PCM1;refGene_tx=XM_023252383.1;refGene_gn=PCM1;refGene_tx=XM_023252384.1;refGene_gn=PCM1;refGene_tx=XM_023252385.1;refGene_gn=PCM1;refGene_tx=XM_023252386.1;refGene_gn=PCM1;refGene_tx=XM_023252387.1;refGene_gn=PCM1;refGene_tx=XM_023252388.1;refGene_gn=PCM1;refGene_tx=XM_023252389.1;refGene_gn=PCM1;refGene_tx=XM_023252390.1;refGene_gn=PCM1;refGene_tx=XM_023252391.1;refGene_gn=PCM1;refGene_tx=XM_023252392.1;refGene_gn=PCM1;refGene_tx=XM_023252393.1;refGene_gn=PCM1;refGene_tx=XM_023252394.1;refGene_gn=PCM1;refGene_tx=XM_023252395.1;refGene_gn=PCM1;refGene_tx=XM_023252396.1;refGene_gn=PCM1;refGene_tx=XM_023252397.1;refGene_gn=PCM1;refGene_tx=XM_023252398.1;refGene_gn=PCM1;refGene_tx=XM_023252399.1;refGene_gn=PCM1;refGene_tx=XM_023252400.1
B1 20651520 20651641
refGene_gn=PCM1;refGene_tx=XM_023252378.1;refGene_gn=PCM1;refGene_tx=XM_023252379.1;refGene_gn=PCM1;refGene_tx=XM_023252380.1;refGene_gn=PCM1;refGene_tx=XM_023252381.1;refGene_gn=PCM1;refGene_tx=XM_023252382.1;refGene_gn=PCM1;refGene_tx=XM_023252383.1;refGene_gn=PCM1;refGene_tx=XM_023252384.1;refGene_gn=PCM1;refGene_tx=XM_023252385.1;refGene_gn=PCM1;refGene_tx=XM_023252386.1;refGene_gn=PCM1;refGene_tx=XM_023252387.1;refGene_gn=PCM1;refGene_tx=XM_023252388.1;refGene_gn=PCM1;refGene_tx=XM_023252389.1;refGene_gn=PCM1;refGene_tx=XM_023252390.1;refGene_gn=PCM1;refGene_tx=XM_023252391.1;refGene_gn=PCM1;refGene_tx=XM_023252392.1;refGene_gn=PCM1;refGene_tx=XM_023252393.1;refGene_gn=PCM1;refGene_tx=XM_023252394.1;refGene_gn=PCM1;refGene_tx=XM_023252395.1;refGene_gn=PCM1;refGene_tx=XM_023252396.1;refGene_gn=PCM1;refGene_tx=XM_023252397.1;refGene_gn=PCM1;refGene_tx=XM_023252398.1;refGene_gn=PCM1;refGene_tx=XM_023252399.1;refGene_gn=PCM1;refGene_tx=XM_023252400.1
B1 20655630 20655762
refGene_gn=PCM1;refGene_tx=XM_023252378.1;refGene_gn=PCM1;refGene_tx=XM_023252379.1;refGene_gn=PCM1;refGene_tx=XM_023252380.1;refGene_gn=PCM1;refGene_tx=XM_023252381.1;refGene_gn=PCM1;refGene_tx=XM_023252382.1;refGene_gn=PCM1;refGene_tx=XM_023252383.1;refGene_gn=PCM1;refGene_tx=XM_023252384.1;refGene_gn=PCM1;refGene_tx=XM_023252385.1;refGene_gn=PCM1;refGene_tx=XM_023252386.1;refGene_gn=PCM1;refGene_tx=XM_023252387.1;refGene_gn=PCM1;refGene_tx=XM_023252388.1;refGene_gn=PCM1;refGene_tx=XM_023252389.1;refGene_gn=PCM1;refGene_tx=XM_023252390.1;refGene_gn=PCM1;refGene_tx=XM_023252391.1;refGene_gn=PCM1;refGene_tx=XM_023252392.1;refGene_gn=PCM1;refGene_tx=XM_023252393.1;refGene_gn=PCM1;refGene_tx=XM_023252394.1;refGene_gn=PCM1;refGene_tx=XM_023252395.1;refGene_gn=PCM1;refGene_tx=XM_023252396.1;refGene_gn=PCM1;refGene_tx=XM_023252397.1;refGene_gn=PCM1;refGene_tx=XM_023252398.1;refGene_gn=PCM1;refGene_tx=XM_023252399.1;refGene_gn=PCM1;refGene_tx=XM_023252400.1

B1 20656104 20656277
refGene_gn=PCM1;refGene_tx=XM_023252378.1;refGene_gn=PCM1;refGene_tx=XM_023252379.1;refGene_gn=PCM1;refGene_tx=XM_023252380.1;refGene_gn=PCM1;refGene_tx=XM_023252381.1;refGene_gn=PCM1;refGene_tx=XM_023252382.1;refGene_gn=PCM1;refGene_tx=XM_023252383.1;refGene_gn=PCM1;refGene_tx=XM_023252384.1;refGene_gn=PCM1;refGene_tx=XM_023252385.1;refGene_gn=PCM1;refGene_tx=XM_023252386.1;refGene_gn=PCM1;refGene_tx=XM_023252387.1;refGene_gn=PCM1;refGene_tx=XM_023252388.1;refGene_gn=PCM1;refGene_tx=XM_023252389.1;refGene_gn=PCM1;refGene_tx=XM_023252390.1;refGene_gn=PCM1;refGene_tx=XM_023252391.1;refGene_gn=PCM1;refGene_tx=XM_023252392.1;refGene_gn=PCM1;refGene_tx=XM_023252393.1;refGene_gn=PCM1;refGene_tx=XM_023252394.1;refGene_gn=PCM1;refGene_tx=XM_023252395.1;refGene_gn=PCM1;refGene_tx=XM_023252396.1;refGene_gn=PCM1;refGene_tx=XM_023252397.1;refGene_gn=PCM1;refGene_tx=XM_023252398.1;refGene_gn=PCM1;refGene_tx=XM_023252399.1;refGene_gn=PCM1;refGene_tx=XM_023252400.1
B1 20659057 20659114
refGene_gn=PCM1;refGene_tx=XM_023252378.1;refGene_gn=PCM1;refGene_tx=XM_023252379.1;refGene_gn=PCM1;refGene_tx=XM_023252380.1;refGene_gn=PCM1;refGene_tx=XM_023252381.1;refGene_gn=PCM1;refGene_tx=XM_023252382.1;refGene_gn=PCM1;refGene_tx=XM_023252383.1;refGene_gn=PCM1;refGene_tx=XM_023252384.1;refGene_gn=PCM1;refGene_tx=XM_023252386.1;refGene_gn=PCM1;refGene_tx=XM_023252387.1;refGene_gn=PCM1;refGene_tx=XM_023252388.1;refGene_gn=PCM1;refGene_tx=XM_023252389.1;refGene_gn=PCM1;refGene_tx=XM_023252390.1;refGene_gn=PCM1;refGene_tx=XM_023252392.1;refGene_gn=PCM1;refGene_tx=XM_023252393.1;refGene_gn=PCM1;refGene_tx=XM_023252394.1;refGene_gn=PCM1;refGene_tx=XM_023252396.1
B1 20661370 20661535
refGene_gn=PCM1;refGene_tx=XM_023252378.1;refGene_gn=PCM1;refGene_tx=XM_023252379.1;refGene_gn=PCM1;refGene_tx=XM_023252380.1;refGene_gn=PCM1;refGene_tx=XM_023252381.1;refGene_gn=PCM1;refGene_tx=XM_023252382.1;refGene_gn=PCM1;refGene_tx=XM_023252383.1;refGene_gn=PCM1;refGene_tx=XM_023252384.1;refGene_gn=PCM1;refGene_tx=XM_023252385.1;refGene_gn=PCM1;refGene_tx=XM_023252386.1;refGene_gn=PCM1;refGene_tx=XM_023252387.1;refGene_gn=PCM1;refGene_tx=XM_023252388.1;refGene_gn=PCM1;refGene_tx=XM_023252390.1;refGene_gn=PCM1;refGene_tx=XM_023252392.1;refGene_gn=PCM1;refGene_tx=XM_023252393.1;refGene_gn=PCM1;refGene_tx=XM_023252394.1;refGene_gn=PCM1;refGene_tx=XM_023252395.1;refGene_gn=PCM1;refGene_tx=XM_023252396.1;refGene_gn=PCM1;refGene_tx=XM_023252397.1;refGene_gn=PCM1;refGene_tx=XM_023252398.1
B1 20668604 20668963
refGene_gn=PCM1;refGene_tx=XM_023252378.1;refGene_gn=PCM1;refGene_tx=XM_023252379.1;refGene_gn=PCM1;refGene_tx=XM_023252380.1;refGene_gn=PCM1;refGene_tx=XM_023252381.1;refGene_gn=PCM1;refGene_tx=XM_023252382.1;refGene_gn=PCM1;refGene_tx=XM_023252383.1;refGene_gn=PCM1;refGene_tx=XM_023252384.1;refGene_gn=PCM1;refGene_tx=XM_023252385.1;refGene_gn=PCM1;refGene_tx=XM_023252386.1;refGene_gn=PCM1;refGene_tx=XM_023252387.1;refGene_gn=PCM1;refGene_tx=XM_023252388.1;refGene_gn=PCM1;refGene_tx=XM_023252389.1;refGene_gn=PCM1;refGene_tx=XM_023252390.1;refGene_gn=PCM1;refGene_tx=XM_023252391.1;refGene_gn=PCM1;refGene_tx=XM_023252392.1;refGene_gn=PCM1;refGene_tx=XM_023252393.1;refGene_gn=PCM1;refGene_tx=X

M_023252394.1;refGene_gn=PCM1;refGene_tx=XM_023252395.1;refGene_gn=PCM1;refGene_tx=XM_023252396.1;refGene_gn=PCM1;refGene_tx=XM_023252397.1;refGene_gn=PCM1;refGene_tx=XM_023252398.1;refGene_gn=PCM1;refGene_tx=XM_023252399.1;refGene_gn=PCM1;refGene_tx=XM_023252400.1
B1 20673792 20673964
refGene_gn=PCM1;refGene_tx=XM_023252378.1;refGene_gn=PCM1;refGene_tx=XM_023252379.1;refGene_gn=PCM1;refGene_tx=XM_023252380.1;refGene_gn=PCM1;refGene_tx=XM_023252381.1;refGene_gn=PCM1;refGene_tx=XM_023252382.1;refGene_gn=PCM1;refGene_tx=XM_023252383.1;refGene_gn=PCM1;refGene_tx=XM_023252384.1;refGene_gn=PCM1;refGene_tx=XM_023252385.1;refGene_gn=PCM1;refGene_tx=XM_023252386.1;refGene_gn=PCM1;refGene_tx=XM_023252387.1;refGene_gn=PCM1;refGene_tx=XM_023252388.1;refGene_gn=PCM1;refGene_tx=XM_023252389.1;refGene_gn=PCM1;refGene_tx=XM_023252390.1;refGene_gn=PCM1;refGene_tx=XM_023252391.1;refGene_gn=PCM1;refGene_tx=XM_023252392.1;refGene_gn=PCM1;refGene_tx=XM_023252393.1;refGene_gn=PCM1;refGene_tx=XM_023252394.1;refGene_gn=PCM1;refGene_tx=XM_023252395.1;refGene_gn=PCM1;refGene_tx=XM_023252396.1;refGene_gn=PCM1;refGene_tx=XM_023252397.1;refGene_gn=PCM1;refGene_tx=XM_023252398.1;refGene_gn=PCM1;refGene_tx=XM_023252399.1;refGene_gn=PCM1;refGene_tx=XM_023252400.1
B1 20675996 20676187
refGene_gn=PCM1;refGene_tx=XM_023252378.1;refGene_gn=PCM1;refGene_tx=XM_023252379.1;refGene_gn=PCM1;refGene_tx=XM_023252380.1;refGene_gn=PCM1;refGene_tx=XM_023252381.1;refGene_gn=PCM1;refGene_tx=XM_023252382.1;refGene_gn=PCM1;refGene_tx=XM_023252383.1;refGene_gn=PCM1;refGene_tx=XM_023252384.1;refGene_gn=PCM1;refGene_tx=XM_023252385.1;refGene_gn=PCM1;refGene_tx=XM_023252386.1;refGene_gn=PCM1;refGene_tx=XM_023252387.1;refGene_gn=PCM1;refGene_tx=XM_023252388.1;refGene_gn=PCM1;refGene_tx=XM_023252389.1;refGene_gn=PCM1;refGene_tx=XM_023252390.1;refGene_gn=PCM1;refGene_tx=XM_023252391.1;refGene_gn=PCM1;refGene_tx=XM_023252392.1;refGene_gn=PCM1;refGene_tx=XM_023252393.1;refGene_gn=PCM1;refGene_tx=XM_023252394.1;refGene_gn=PCM1;refGene_tx=XM_023252395.1;refGene_gn=PCM1;refGene_tx=XM_023252396.1;refGene_gn=PCM1;refGene_tx=XM_023252397.1;refGene_gn=PCM1;refGene_tx=XM_023252398.1;refGene_gn=PCM1;refGene_tx=XM_023252399.1;refGene_gn=PCM1;refGene_tx=XM_023252400.1
B1 20676704 20676850
refGene_gn=PCM1;refGene_tx=XM_023252378.1;refGene_gn=PCM1;refGene_tx=XM_023252379.1;refGene_gn=PCM1;refGene_tx=XM_023252380.1;refGene_gn=PCM1;refGene_tx=XM_023252381.1;refGene_gn=PCM1;refGene_tx=XM_023252382.1;refGene_gn=PCM1;refGene_tx=XM_023252383.1;refGene_gn=PCM1;refGene_tx=XM_023252384.1;refGene_gn=PCM1;refGene_tx=XM_023252385.1;refGene_gn=PCM1;refGene_tx=XM_023252386.1;refGene_gn=PCM1;refGene_tx=XM_023252387.1;refGene_gn=PCM1;refGene_tx=XM_023252388.1;refGene_gn=PCM1;refGene_tx=XM_023252389.1;refGene_gn=PCM1;refGene_tx=XM_023252390.1;refGene_gn=PCM1;refGene_tx=XM_023252391.1;refGene_gn=PCM1;refGene_tx=XM_023252392.1;refGene_gn=PCM1;refGene_tx=XM_023252393.1;refGene_gn=PCM1;refGene_tx=XM_023252394.1;refGene_gn=PCM1;refGene_tx=XM_023252395.1;refGene_gn=PCM1;refGene_tx=XM_023252396.1;refGene_gn=PCM1;refGene_tx=XM_023252397.1;refGene_gn=PCM1;refGene_tx=XM_023252398.1;refGene_gn=PCM1;refGene_tx=XM_023252399.1;refGene_gn=PCM1;refGene_tx=XM_023252400.1
B1 20676955 20677175

refGene_gn=PCM1;refGene_tx=XM_023252378.1;refGene_gn=PCM1;refGene_tx=XM_023252379.1;refGene_gn=PCM1;refGene_tx=XM_023252380.1;refGene_gn=PCM1;refGene_tx=XM_023252381.1;refGene_gn=PCM1;refGene_tx=XM_023252382.1;refGene_gn=PCM1;refGene_tx=XM_023252383.1;refGene_gn=PCM1;refGene_tx=XM_023252384.1;refGene_gn=PCM1;refGene_tx=XM_023252385.1;refGene_gn=PCM1;refGene_tx=XM_023252386.1;refGene_gn=PCM1;refGene_tx=XM_023252387.1;refGene_gn=PCM1;refGene_tx=XM_023252388.1;refGene_gn=PCM1;refGene_tx=XM_023252389.1;refGene_gn=PCM1;refGene_tx=XM_023252390.1;refGene_gn=PCM1;refGene_tx=XM_023252391.1;refGene_gn=PCM1;refGene_tx=XM_023252392.1;refGene_gn=PCM1;refGene_tx=XM_023252393.1;refGene_gn=PCM1;refGene_tx=XM_023252394.1;refGene_gn=PCM1;refGene_tx=XM_023252395.1;refGene_gn=PCM1;refGene_tx=XM_023252396.1;refGene_gn=PCM1;refGene_tx=XM_023252397.1;refGene_gn=PCM1;refGene_tx=XM_023252398.1;refGene_gn=PCM1;refGene_tx=XM_023252399.1;refGene_gn=PCM1;refGene_tx=XM_023252400.1
B1 20678517 20678718
refGene_gn=PCM1;refGene_tx=XM_023252378.1;refGene_gn=PCM1;refGene_tx=XM_023252379.1;refGene_gn=PCM1;refGene_tx=XM_023252380.1;refGene_gn=PCM1;refGene_tx=XM_023252381.1;refGene_gn=PCM1;refGene_tx=XM_023252382.1;refGene_gn=PCM1;refGene_tx=XM_023252383.1;refGene_gn=PCM1;refGene_tx=XM_023252384.1;refGene_gn=PCM1;refGene_tx=XM_023252385.1;refGene_gn=PCM1;refGene_tx=XM_023252386.1;refGene_gn=PCM1;refGene_tx=XM_023252387.1;refGene_gn=PCM1;refGene_tx=XM_023252388.1;refGene_gn=PCM1;refGene_tx=XM_023252389.1;refGene_gn=PCM1;refGene_tx=XM_023252390.1;refGene_gn=PCM1;refGene_tx=XM_023252391.1;refGene_gn=PCM1;refGene_tx=XM_023252392.1;refGene_gn=PCM1;refGene_tx=XM_023252393.1;refGene_gn=PCM1;refGene_tx=XM_023252394.1;refGene_gn=PCM1;refGene_tx=XM_023252395.1;refGene_gn=PCM1;refGene_tx=XM_023252396.1;refGene_gn=PCM1;refGene_tx=XM_023252397.1;refGene_gn=PCM1;refGene_tx=XM_023252398.1;refGene_gn=PCM1;refGene_tx=XM_023252399.1;refGene_gn=PCM1;refGene_tx=XM_023252400.1
B1 20680733 20680924
refGene_gn=PCM1;refGene_tx=XM_023252378.1;refGene_gn=PCM1;refGene_tx=XM_023252379.1;refGene_gn=PCM1;refGene_tx=XM_023252380.1;refGene_gn=PCM1;refGene_tx=XM_023252381.1;refGene_gn=PCM1;refGene_tx=XM_023252382.1;refGene_gn=PCM1;refGene_tx=XM_023252383.1;refGene_gn=PCM1;refGene_tx=XM_023252384.1;refGene_gn=PCM1;refGene_tx=XM_023252385.1;refGene_gn=PCM1;refGene_tx=XM_023252386.1;refGene_gn=PCM1;refGene_tx=XM_023252387.1;refGene_gn=PCM1;refGene_tx=XM_023252388.1;refGene_gn=PCM1;refGene_tx=XM_023252389.1;refGene_gn=PCM1;refGene_tx=XM_023252390.1;refGene_gn=PCM1;refGene_tx=XM_023252391.1;refGene_gn=PCM1;refGene_tx=XM_023252392.1;refGene_gn=PCM1;refGene_tx=XM_023252393.1;refGene_gn=PCM1;refGene_tx=XM_023252394.1;refGene_gn=PCM1;refGene_tx=XM_023252395.1;refGene_gn=PCM1;refGene_tx=XM_023252396.1;refGene_gn=PCM1;refGene_tx=XM_023252397.1;refGene_gn=PCM1;refGene_tx=XM_023252398.1;refGene_gn=PCM1;refGene_tx=XM_023252399.1;refGene_gn=PCM1;refGene_tx=XM_023252400.1
B1 20681428 20681569
refGene_gn=PCM1;refGene_tx=XM_023252378.1;refGene_gn=PCM1;refGene_tx=XM_023252379.1;refGene_gn=PCM1;refGene_tx=XM_023252380.1;refGene_gn=PCM1;refGene_tx=XM_023252381.1;refGene_gn=PCM1;refGene_tx=XM_023252382.1;refGene_gn=PCM1;refGene_tx=XM_023252383.1;refGene_gn=PCM1;refGene_tx=XM_023252384.1;refGene_gn=PCM1;refGene_tx=XM_023252385.1;refGene_gn=PCM

1;refGene_tx=XM_023252386.1;refGene_gn=PCM1;refGene_tx=XM_023252387.1;
refGene_gn=PCM1;refGene_tx=XM_023252389.1;refGene_gn=PCM1;refGene_tx=X
M_023252390.1;refGene_gn=PCM1;refGene_tx=XM_023252391.1;refGene_gn=PCM
1;refGene_tx=XM_023252392.1;refGene_gn=PCM1;refGene_tx=XM_023252393.1;
refGene_gn=PCM1;refGene_tx=XM_023252394.1;refGene_gn=PCM1;refGene_tx=X
M_023252395.1;refGene_gn=PCM1;refGene_tx=XM_023252396.1;refGene_gn=PCM
1;refGene_tx=XM_023252397.1;refGene_gn=PCM1;refGene_tx=XM_023252398.1;
refGene_gn=PCM1;refGene_tx=XM_023252399.1;refGene_gn=PCM1;refGene_tx=X
M_023252400.1
B1 20682768 20682898
refGene_gn=PCM1;refGene_tx=XM_023252378.1;refGene_gn=PCM1;refGene_tx=X
M_023252379.1;refGene_gn=PCM1;refGene_tx=XM_023252380.1;refGene_gn=PCM
1;refGene_tx=XM_023252381.1;refGene_gn=PCM1;refGene_tx=XM_023252382.1;
refGene_gn=PCM1;refGene_tx=XM_023252383.1;refGene_gn=PCM1;refGene_tx=X
M_023252384.1;refGene_gn=PCM1;refGene_tx=XM_023252385.1;refGene_gn=PCM
1;refGene_tx=XM_023252386.1;refGene_gn=PCM1;refGene_tx=XM_023252387.1;
refGene_gn=PCM1;refGene_tx=XM_023252388.1;refGene_gn=PCM1;refGene_tx=X
M_023252389.1;refGene_gn=PCM1;refGene_tx=XM_023252390.1;refGene_gn=PCM
1;refGene_tx=XM_023252391.1;refGene_gn=PCM1;refGene_tx=XM_023252392.1;
refGene_gn=PCM1;refGene_tx=XM_023252393.1;refGene_gn=PCM1;refGene_tx=X
M_023252394.1;refGene_gn=PCM1;refGene_tx=XM_023252395.1;refGene_gn=PCM
1;refGene_tx=XM_023252396.1;refGene_gn=PCM1;refGene_tx=XM_023252397.1;
refGene_gn=PCM1;refGene_tx=XM_023252398.1;refGene_gn=PCM1;refGene_tx=X
M_023252399.1;refGene_gn=PCM1;refGene_tx=XM_023252400.1
B1 20683037 20683192
refGene_gn=PCM1;refGene_tx=XM_023252378.1;refGene_gn=PCM1;refGene_tx=X
M_023252379.1;refGene_gn=PCM1;refGene_tx=XM_023252380.1;refGene_gn=PCM
1;refGene_tx=XM_023252381.1;refGene_gn=PCM1;refGene_tx=XM_023252382.1;
refGene_gn=PCM1;refGene_tx=XM_023252383.1;refGene_gn=PCM1;refGene_tx=X
M_023252384.1;refGene_gn=PCM1;refGene_tx=XM_023252385.1;refGene_gn=PCM
1;refGene_tx=XM_023252386.1;refGene_gn=PCM1;refGene_tx=XM_023252387.1;
refGene_gn=PCM1;refGene_tx=XM_023252388.1;refGene_gn=PCM1;refGene_tx=X
M_023252389.1;refGene_gn=PCM1;refGene_tx=XM_023252390.1;refGene_gn=PCM
1;refGene_tx=XM_023252391.1;refGene_gn=PCM1;refGene_tx=XM_023252392.1;
refGene_gn=PCM1;refGene_tx=XM_023252393.1;refGene_gn=PCM1;refGene_tx=X
M_023252394.1;refGene_gn=PCM1;refGene_tx=XM_023252395.1;refGene_gn=PCM
1;refGene_tx=XM_023252396.1;refGene_gn=PCM1;refGene_tx=XM_023252397.1;
refGene_gn=PCM1;refGene_tx=XM_023252398.1;refGene_gn=PCM1;refGene_tx=X
M_023252399.1;refGene_gn=PCM1;refGene_tx=XM_023252400.1
B1 20684308 20684544
refGene_gn=PCM1;refGene_tx=XM_023252378.1;refGene_gn=PCM1;refGene_tx=X
M_023252379.1;refGene_gn=PCM1;refGene_tx=XM_023252380.1;refGene_gn=PCM
1;refGene_tx=XM_023252381.1;refGene_gn=PCM1;refGene_tx=XM_023252382.1;
refGene_gn=PCM1;refGene_tx=XM_023252383.1;refGene_gn=PCM1;refGene_tx=X
M_023252384.1;refGene_gn=PCM1;refGene_tx=XM_023252385.1;refGene_gn=PCM
1;refGene_tx=XM_023252386.1;refGene_gn=PCM1;refGene_tx=XM_023252387.1;
refGene_gn=PCM1;refGene_tx=XM_023252388.1;refGene_gn=PCM1;refGene_tx=X
M_023252389.1;refGene_gn=PCM1;refGene_tx=XM_023252390.1;refGene_gn=PCM
1;refGene_tx=XM_023252391.1;refGene_gn=PCM1;refGene_tx=XM_023252392.1;
refGene_gn=PCM1;refGene_tx=XM_023252393.1;refGene_gn=PCM1;refGene_tx=X

M_023252394.1;refGene_gn=PCM1;refGene_tx=XM_023252395.1;refGene_gn=PCM1;refGene_tx=XM_023252396.1;refGene_gn=PCM1;refGene_tx=XM_023252397.1;refGene_gn=PCM1;refGene_tx=XM_023252398.1;refGene_gn=PCM1;refGene_tx=XM_023252399.1;refGene_gn=PCM1;refGene_tx=XM_023252400.1
B1 20684659 20684817
refGene_gn=PCM1;refGene_tx=XM_023252378.1;refGene_gn=PCM1;refGene_tx=XM_023252379.1;refGene_gn=PCM1;refGene_tx=XM_023252380.1;refGene_gn=PCM1;refGene_tx=XM_023252381.1;refGene_gn=PCM1;refGene_tx=XM_023252382.1;refGene_gn=PCM1;refGene_tx=XM_023252383.1;refGene_gn=PCM1;refGene_tx=XM_023252384.1;refGene_gn=PCM1;refGene_tx=XM_023252385.1;refGene_gn=PCM1;refGene_tx=XM_023252386.1;refGene_gn=PCM1;refGene_tx=XM_023252387.1;refGene_gn=PCM1;refGene_tx=XM_023252388.1;refGene_gn=PCM1;refGene_tx=XM_023252389.1;refGene_gn=PCM1;refGene_tx=XM_023252390.1;refGene_gn=PCM1;refGene_tx=XM_023252391.1;refGene_gn=PCM1;refGene_tx=XM_023252392.1;refGene_gn=PCM1;refGene_tx=XM_023252393.1;refGene_gn=PCM1;refGene_tx=XM_023252394.1;refGene_gn=PCM1;refGene_tx=XM_023252395.1;refGene_gn=PCM1;refGene_tx=XM_023252396.1;refGene_gn=PCM1;refGene_tx=XM_023252397.1;refGene_gn=PCM1;refGene_tx=XM_023252398.1;refGene_gn=PCM1;refGene_tx=XM_023252399.1;refGene_gn=PCM1;refGene_tx=XM_023252400.1
B1 20685344 20685518
refGene_gn=PCM1;refGene_tx=XM_023252378.1;refGene_gn=PCM1;refGene_tx=XM_023252379.1;refGene_gn=PCM1;refGene_tx=XM_023252380.1;refGene_gn=PCM1;refGene_tx=XM_023252381.1;refGene_gn=PCM1;refGene_tx=XM_023252382.1;refGene_gn=PCM1;refGene_tx=XM_023252383.1;refGene_gn=PCM1;refGene_tx=XM_023252384.1;refGene_gn=PCM1;refGene_tx=XM_023252385.1;refGene_gn=PCM1;refGene_tx=XM_023252386.1;refGene_gn=PCM1;refGene_tx=XM_023252387.1;refGene_gn=PCM1;refGene_tx=XM_023252388.1;refGene_gn=PCM1;refGene_tx=XM_023252389.1;refGene_gn=PCM1;refGene_tx=XM_023252390.1;refGene_gn=PCM1;refGene_tx=XM_023252391.1;refGene_gn=PCM1;refGene_tx=XM_023252392.1;refGene_gn=PCM1;refGene_tx=XM_023252393.1;refGene_gn=PCM1;refGene_tx=XM_023252394.1;refGene_gn=PCM1;refGene_tx=XM_023252395.1;refGene_gn=PCM1;refGene_tx=XM_023252396.1;refGene_gn=PCM1;refGene_tx=XM_023252397.1;refGene_gn=PCM1;refGene_tx=XM_023252398.1;refGene_gn=PCM1;refGene_tx=XM_023252399.1;refGene_gn=PCM1;refGene_tx=XM_023252400.1
B1 20686671 20686870
refGene_gn=PCM1;refGene_tx=XM_023252378.1;refGene_gn=PCM1;refGene_tx=XM_023252379.1;refGene_gn=PCM1;refGene_tx=XM_023252380.1;refGene_gn=PCM1;refGene_tx=XM_023252381.1;refGene_gn=PCM1;refGene_tx=XM_023252382.1;refGene_gn=PCM1;refGene_tx=XM_023252383.1;refGene_gn=PCM1;refGene_tx=XM_023252384.1;refGene_gn=PCM1;refGene_tx=XM_023252385.1;refGene_gn=PCM1;refGene_tx=XM_023252386.1;refGene_gn=PCM1;refGene_tx=XM_023252387.1;refGene_gn=PCM1;refGene_tx=XM_023252388.1;refGene_gn=PCM1;refGene_tx=XM_023252389.1;refGene_gn=PCM1;refGene_tx=XM_023252390.1;refGene_gn=PCM1;refGene_tx=XM_023252391.1;refGene_gn=PCM1;refGene_tx=XM_023252392.1;refGene_gn=PCM1;refGene_tx=XM_023252393.1;refGene_gn=PCM1;refGene_tx=XM_023252394.1;refGene_gn=PCM1;refGene_tx=XM_023252395.1;refGene_gn=PCM1;refGene_tx=XM_023252396.1;refGene_gn=PCM1;refGene_tx=XM_023252397.1;refGene_gn=PCM1;refGene_tx=XM_023252398.1;refGene_gn=PCM1;refGene_tx=XM_023252399.1;refGene_gn=PCM1;refGene_tx=XM_023252400.1
B1 20688641 20688858

refGene_gn=PCM1;refGene_tx=XM_023252378.1;refGene_gn=PCM1;refGene_tx=XM_023252379.1;refGene_gn=PCM1;refGene_tx=XM_023252380.1;refGene_gn=PCM1;refGene_tx=XM_023252381.1;refGene_gn=PCM1;refGene_tx=XM_023252382.1;refGene_gn=PCM1;refGene_tx=XM_023252383.1;refGene_gn=PCM1;refGene_tx=XM_023252384.1;refGene_gn=PCM1;refGene_tx=XM_023252385.1;refGene_gn=PCM1;refGene_tx=XM_023252386.1;refGene_gn=PCM1;refGene_tx=XM_023252387.1;refGene_gn=PCM1;refGene_tx=XM_023252388.1;refGene_gn=PCM1;refGene_tx=XM_023252389.1;refGene_gn=PCM1;refGene_tx=XM_023252390.1;refGene_gn=PCM1;refGene_tx=XM_023252391.1;refGene_gn=PCM1;refGene_tx=XM_023252392.1;refGene_gn=PCM1;refGene_tx=XM_023252393.1;refGene_gn=PCM1;refGene_tx=XM_023252394.1;refGene_gn=PCM1;refGene_tx=XM_023252395.1;refGene_gn=PCM1;refGene_tx=XM_023252396.1;refGene_gn=PCM1;refGene_tx=XM_023252397.1;refGene_gn=PCM1;refGene_tx=XM_023252398.1;refGene_gn=PCM1;refGene_tx=XM_023252399.1;refGene_gn=PCM1;refGene_tx=XM_023252400.1
B1 20690356 20690466
refGene_gn=PCM1;refGene_tx=XM_023252378.1;refGene_gn=PCM1;refGene_tx=XM_023252379.1;refGene_gn=PCM1;refGene_tx=XM_023252380.1;refGene_gn=PCM1;refGene_tx=XM_023252381.1;refGene_gn=PCM1;refGene_tx=XM_023252382.1;refGene_gn=PCM1;refGene_tx=XM_023252383.1;refGene_gn=PCM1;refGene_tx=XM_023252384.1;refGene_gn=PCM1;refGene_tx=XM_023252385.1;refGene_gn=PCM1;refGene_tx=XM_023252386.1;refGene_gn=PCM1;refGene_tx=XM_023252387.1;refGene_gn=PCM1;refGene_tx=XM_023252388.1;refGene_gn=PCM1;refGene_tx=XM_023252389.1;refGene_gn=PCM1;refGene_tx=XM_023252390.1;refGene_gn=PCM1;refGene_tx=XM_023252391.1;refGene_gn=PCM1;refGene_tx=XM_023252392.1;refGene_gn=PCM1;refGene_tx=XM_023252393.1;refGene_gn=PCM1;refGene_tx=XM_023252394.1;refGene_gn=PCM1;refGene_tx=XM_023252395.1;refGene_gn=PCM1;refGene_tx=XM_023252396.1;refGene_gn=PCM1;refGene_tx=XM_023252397.1;refGene_gn=PCM1;refGene_tx=XM_023252398.1;refGene_gn=PCM1;refGene_tx=XM_023252399.1;refGene_gn=PCM1;refGene_tx=XM_023252400.1
B1 20693026 20693116
refGene_gn=PCM1;refGene_tx=XM_023252378.1;refGene_gn=PCM1;refGene_tx=XM_023252379.1;refGene_gn=PCM1;refGene_tx=XM_023252380.1;refGene_gn=PCM1;refGene_tx=XM_023252381.1;refGene_gn=PCM1;refGene_tx=XM_023252382.1;refGene_gn=PCM1;refGene_tx=XM_023252383.1;refGene_gn=PCM1;refGene_tx=XM_023252384.1;refGene_gn=PCM1;refGene_tx=XM_023252385.1;refGene_gn=PCM1;refGene_tx=XM_023252386.1;refGene_gn=PCM1;refGene_tx=XM_023252388.1;refGene_gn=PCM1;refGene_tx=XM_023252389.1;refGene_gn=PCM1;refGene_tx=XM_023252390.1;refGene_gn=PCM1;refGene_tx=XM_023252391.1;refGene_gn=PCM1;refGene_tx=XM_023252393.1;refGene_gn=PCM1;refGene_tx=XM_023252396.1
B1 20693269 20693447
refGene_gn=PCM1;refGene_tx=XM_023252378.1;refGene_gn=PCM1;refGene_tx=XM_023252379.1;refGene_gn=PCM1;refGene_tx=XM_023252380.1;refGene_gn=PCM1;refGene_tx=XM_023252381.1;refGene_gn=PCM1;refGene_tx=XM_023252382.1;refGene_gn=PCM1;refGene_tx=XM_023252383.1;refGene_gn=PCM1;refGene_tx=XM_023252384.1;refGene_gn=PCM1;refGene_tx=XM_023252385.1;refGene_gn=PCM1;refGene_tx=XM_023252386.1;refGene_gn=PCM1;refGene_tx=XM_023252387.1;refGene_gn=PCM1;refGene_tx=XM_023252388.1;refGene_gn=PCM1;refGene_tx=XM_023252389.1;refGene_gn=PCM1;refGene_tx=XM_023252390.1;refGene_gn=PCM1;refGene_tx=XM_023252391.1;refGene_gn=PCM1;refGene_tx=XM_023252392.1;refGene_gn=PCM1;refGene_tx=XM_023252393.1;refGene_gn=PCM1;refGene_tx=X

M_023252394.1;refGene_gn=PCM1;refGene_tx=XM_023252395.1;refGene_gn=PCM1;refGene_tx=XM_023252396.1;refGene_gn=PCM1;refGene_tx=XM_023252397.1;refGene_gn=PCM1;refGene_tx=XM_023252398.1;refGene_gn=PCM1;refGene_tx=XM_023252399.1;refGene_gn=PCM1;refGene_tx=XM_023252400.1
B1 20698189 20698372
refGene_gn=PCM1;refGene_tx=XM_023252378.1;refGene_gn=PCM1;refGene_tx=XM_023252379.1;refGene_gn=PCM1;refGene_tx=XM_023252380.1;refGene_gn=PCM1;refGene_tx=XM_023252381.1;refGene_gn=PCM1;refGene_tx=XM_023252382.1;refGene_gn=PCM1;refGene_tx=XM_023252383.1;refGene_gn=PCM1;refGene_tx=XM_023252384.1;refGene_gn=PCM1;refGene_tx=XM_023252385.1;refGene_gn=PCM1;refGene_tx=XM_023252386.1;refGene_gn=PCM1;refGene_tx=XM_023252387.1;refGene_gn=PCM1;refGene_tx=XM_023252388.1;refGene_gn=PCM1;refGene_tx=XM_023252389.1;refGene_gn=PCM1;refGene_tx=XM_023252391.1;refGene_gn=PCM1;refGene_tx=XM_023252396.1
B1 20699824 20699941
refGene_gn=PCM1;refGene_tx=XM_023252378.1;refGene_gn=PCM1;refGene_tx=XM_023252379.1;refGene_gn=PCM1;refGene_tx=XM_023252380.1;refGene_gn=PCM1;refGene_tx=XM_023252381.1;refGene_gn=PCM1;refGene_tx=XM_023252382.1;refGene_gn=PCM1;refGene_tx=XM_023252383.1;refGene_gn=PCM1;refGene_tx=XM_023252384.1;refGene_gn=PCM1;refGene_tx=XM_023252385.1;refGene_gn=PCM1;refGene_tx=XM_023252386.1;refGene_gn=PCM1;refGene_tx=XM_023252387.1;refGene_gn=PCM1;refGene_tx=XM_023252388.1;refGene_gn=PCM1;refGene_tx=XM_023252389.1;refGene_gn=PCM1;refGene_tx=XM_023252390.1;refGene_gn=PCM1;refGene_tx=XM_023252391.1;refGene_gn=PCM1;refGene_tx=XM_023252392.1;refGene_gn=PCM1;refGene_tx=XM_023252396.1;refGene_gn=PCM1;refGene_tx=XM_023252397.1;refGene_gn=PCM1;refGene_tx=XM_023252399.1
B1 20700126 20700297
refGene_gn=PCM1;refGene_tx=XM_023252378.1;refGene_gn=PCM1;refGene_tx=XM_023252379.1;refGene_gn=PCM1;refGene_tx=XM_023252380.1;refGene_gn=PCM1;refGene_tx=XM_023252381.1;refGene_gn=PCM1;refGene_tx=XM_023252382.1;refGene_gn=PCM1;refGene_tx=XM_023252383.1;refGene_gn=PCM1;refGene_tx=XM_023252384.1;refGene_gn=PCM1;refGene_tx=XM_023252385.1;refGene_gn=PCM1;refGene_tx=XM_023252386.1;refGene_gn=PCM1;refGene_tx=XM_023252387.1;refGene_gn=PCM1;refGene_tx=XM_023252388.1;refGene_gn=PCM1;refGene_tx=XM_023252389.1;refGene_gn=PCM1;refGene_tx=XM_023252390.1;refGene_gn=PCM1;refGene_tx=XM_023252391.1;refGene_gn=PCM1;refGene_tx=XM_023252392.1;refGene_gn=PCM1;refGene_tx=XM_023252393.1;refGene_gn=PCM1;refGene_tx=XM_023252394.1;refGene_gn=PCM1;refGene_tx=XM_023252395.1;refGene_gn=PCM1;refGene_tx=XM_023252396.1;refGene_gn=PCM1;refGene_tx=XM_023252397.1;refGene_gn=PCM1;refGene_tx=XM_023252398.1;refGene_gn=PCM1;refGene_tx=XM_023252399.1;refGene_gn=PCM1;refGene_tx=XM_023252400.1
B1 20701016 20701286
refGene_gn=PCM1;refGene_tx=XM_023252378.1;refGene_gn=PCM1;refGene_tx=XM_023252379.1;refGene_gn=PCM1;refGene_tx=XM_023252380.1;refGene_gn=PCM1;refGene_tx=XM_023252381.1;refGene_gn=PCM1;refGene_tx=XM_023252382.1;refGene_gn=PCM1;refGene_tx=XM_023252383.1;refGene_gn=PCM1;refGene_tx=XM_023252384.1;refGene_gn=PCM1;refGene_tx=XM_023252385.1;refGene_gn=PCM1;refGene_tx=XM_023252386.1;refGene_gn=PCM1;refGene_tx=XM_023252387.1;refGene_gn=PCM1;refGene_tx=XM_023252388.1;refGene_gn=PCM1;refGene_tx=XM_023252389.1;refGene_gn=PCM1;refGene_tx=XM_023252390.1;refGene_gn=PCM

1;refGene_tx=XM_023252391.1;refGene_gn=PCM1;refGene_tx=XM_023252392.1;
refGene_gn=PCM1;refGene_tx=XM_023252393.1;refGene_gn=PCM1;refGene_tx=X
M_023252394.1;refGene_gn=PCM1;refGene_tx=XM_023252395.1;refGene_gn=PCM
1;refGene_tx=XM_023252396.1;refGene_gn=PCM1;refGene_tx=XM_023252397.1;
refGene_gn=PCM1;refGene_tx=XM_023252398.1;refGene_gn=PCM1;refGene_tx=X
M_023252399.1;refGene_gn=PCM1;refGene_tx=XM_023252400.1
B1 20702294 20702540
refGene_gn=PCM1;refGene_tx=XM_023252378.1;refGene_gn=PCM1;refGene_tx=X
M_023252379.1;refGene_gn=PCM1;refGene_tx=XM_023252380.1;refGene_gn=PCM
1;refGene_tx=XM_023252381.1;refGene_gn=PCM1;refGene_tx=XM_023252382.1;
refGene_gn=PCM1;refGene_tx=XM_023252383.1;refGene_gn=PCM1;refGene_tx=X
M_023252384.1;refGene_gn=PCM1;refGene_tx=XM_023252385.1;refGene_gn=PCM
1;refGene_tx=XM_023252386.1;refGene_gn=PCM1;refGene_tx=XM_023252387.1;
refGene_gn=PCM1;refGene_tx=XM_023252388.1;refGene_gn=PCM1;refGene_tx=X
M_023252389.1;refGene_gn=PCM1;refGene_tx=XM_023252390.1;refGene_gn=PCM
1;refGene_tx=XM_023252391.1;refGene_gn=PCM1;refGene_tx=XM_023252392.1;
refGene_gn=PCM1;refGene_tx=XM_023252393.1;refGene_gn=PCM1;refGene_tx=X
M_023252394.1;refGene_gn=PCM1;refGene_tx=XM_023252395.1;refGene_gn=PCM
1;refGene_tx=XM_023252396.1;refGene_gn=PCM1;refGene_tx=XM_023252397.1;
refGene_gn=PCM1;refGene_tx=XM_023252398.1;refGene_gn=PCM1;refGene_tx=X
M_023252399.1;refGene_gn=PCM1;refGene_tx=XM_023252400.1
B1 20704222 20704318
refGene_gn=PCM1;refGene_tx=XM_023252378.1;refGene_gn=PCM1;refGene_tx=X
M_023252379.1;refGene_gn=PCM1;refGene_tx=XM_023252380.1;refGene_gn=PCM
1;refGene_tx=XM_023252381.1;refGene_gn=PCM1;refGene_tx=XM_023252382.1;
refGene_gn=PCM1;refGene_tx=XM_023252383.1;refGene_gn=PCM1;refGene_tx=X
M_023252384.1;refGene_gn=PCM1;refGene_tx=XM_023252385.1;refGene_gn=PCM
1;refGene_tx=XM_023252386.1;refGene_gn=PCM1;refGene_tx=XM_023252387.1;
refGene_gn=PCM1;refGene_tx=XM_023252388.1;refGene_gn=PCM1;refGene_tx=X
M_023252389.1;refGene_gn=PCM1;refGene_tx=XM_023252390.1;refGene_gn=PCM
1;refGene_tx=XM_023252391.1;refGene_gn=PCM1;refGene_tx=XM_023252392.1;
refGene_gn=PCM1;refGene_tx=XM_023252393.1;refGene_gn=PCM1;refGene_tx=X
M_023252394.1;refGene_gn=PCM1;refGene_tx=XM_023252395.1;refGene_gn=PCM
1;refGene_tx=XM_023252396.1;refGene_gn=PCM1;refGene_tx=XM_023252397.1;
refGene_gn=PCM1;refGene_tx=XM_023252398.1;refGene_gn=PCM1;refGene_tx=X
M_023252399.1;refGene_gn=PCM1;refGene_tx=XM_023252400.1
B1 20752062 20752162 refGene_gn=FGL1;refGene_tx=XM_019828299.2
B1 20753750 20753830
refGene_gn=FGL1;refGene_tx=XM_011281397.3;refGene_gn=FGL1;refGene_tx=X
M_019828299.2;refGene_gn=FGL1;refGene_tx=XM_019828300.2
B1 20756017 20756198
refGene_gn=FGL1;refGene_tx=XM_011281397.3;refGene_gn=FGL1;refGene_tx=X
M_019828299.2;refGene_gn=FGL1;refGene_tx=XM_019828300.2
B1 20763700 20763860
refGene_gn=FGL1;refGene_tx=XM_011281397.3;refGene_gn=FGL1;refGene_tx=X
M_019828299.2;refGene_gn=FGL1;refGene_tx=XM_019828300.2
B1 20764113 20764211
refGene_gn=FGL1;refGene_tx=XM_011281397.3;refGene_gn=FGL1;refGene_tx=X
M_019828299.2;refGene_gn=FGL1;refGene_tx=XM_019828300.2
B1 20767450 20767539

refGene_gn=FGL1;refGene_tx=XM_011281397.3;refGene_gn=FGL1;refGene_tx=XM_019828299.2;refGene_gn=FGL1;refGene_tx=XM_019828300.2
B1 20767731 20767919
refGene_gn=FGL1;refGene_tx=XM_011281397.3;refGene_gn=FGL1;refGene_tx=XM_019828299.2;refGene_gn=FGL1;refGene_tx=XM_019828300.2
B1 20771652 20771812
refGene_gn=FGL1;refGene_tx=XM_011281397.3;refGene_gn=FGL1;refGene_tx=XM_019828299.2;refGene_gn=FGL1;refGene_tx=XM_019828300.2
B1 20873367 20875452 refGene_gn=MTUS1;refGene_tx=XM_003984620.5
B1 20888913 20889109 refGene_gn=MTUS1;refGene_tx=XM_003984620.5
B1 20910105 20910203 refGene_gn=MTUS1;refGene_tx=XM_003984620.5
B1 20910204 20910268 refGene_gn=MTUS1;refGene_tx=XM_003984620.5
B1 20919584 20919719 refGene_gn=MTUS1;refGene_tx=XM_003984620.5
B1 20923992 20924031 refGene_gn=MTUS1;refGene_tx=XM_003984620.5
B1 20954400 20954615 refGene_gn=MTUS1;refGene_tx=XM_003984620.5
B1 20962117 20962184 refGene_gn=MTUS1;refGene_tx=XM_003984620.5
B1 20978409 20978624 refGene_gn=MTUS1;refGene_tx=XM_003984620.5
B1 20979429 20979535 refGene_gn=MTUS1;refGene_tx=XM_003984620.5
B1 20980812 20980886 refGene_gn=MTUS1;refGene_tx=XM_003984620.5
B1 20981031 20981127 refGene_gn=MTUS1;refGene_tx=XM_003984620.5
B1 20984463 20984580 refGene_gn=MTUS1;refGene_tx=XM_003984620.5
B1 20988530 20988628 refGene_gn=MTUS1;refGene_tx=XM_003984620.5
B1 20989392 20989606 refGene_gn=MTUS1;refGene_tx=XM_003984620.5
B1 20992612 20992801
refGene_gn=PDGFRL;refGene_tx=XM_003984634.5;refGene_gn=PDGFRL;refGene_tx=XM_019828306.2
B1 20994424 20994564
refGene_gn=PDGFRL;refGene_tx=XM_003984634.5;refGene_gn=PDGFRL;refGene_tx=XM_019828306.2
B1 20996755 20997049
refGene_gn=PDGFRL;refGene_tx=XM_003984634.5;refGene_gn=PDGFRL;refGene_tx=XM_019828306.2
B1 21001153 21001305
refGene_gn=PDGFRL;refGene_tx=XM_003984634.5;refGene_gn=PDGFRL;refGene_tx=XM_019828306.2
B1 21029424 21029492 refGene_gn=PDGFRL;refGene_tx=XM_019828306.2
B1 21033041 21033104 refGene_gn=PDGFRL;refGene_tx=XM_019828306.2
B1 21042781 21043079 refGene_gn=PDGFRL;refGene_tx=XM_003984634.5
B1 21063422 21063477 refGene_gn=PDGFRL;refGene_tx=XM_003984634.5
B1 21077959 21078156
refGene_gn=SLC7A2;refGene_tx=XM_003984623.5;refGene_gn=SLC7A2;refGene_tx=XM_019828307.2;refGene_gn=SLC7A2;refGene_tx=XM_019828308.2;refGene_gn=SLC7A2;refGene_tx=XM_023252401.1;refGene_gn=SLC7A2;refGene_tx=XM_023252402.1
B1 21081973 21082082
refGene_gn=SLC7A2;refGene_tx=XM_003984623.5;refGene_gn=SLC7A2;refGene_tx=XM_019828307.2;refGene_gn=SLC7A2;refGene_tx=XM_019828308.2;refGene_gn=SLC7A2;refGene_tx=XM_023252401.1;refGene_gn=SLC7A2;refGene_tx=XM_023252402.1
B1 21082824 21082991

refGene_gn=SLC7A2;refGene_tx=XM_003984623.5;refGene_gn=SLC7A2;refGene_tx=XM_019828307.2;refGene_gn=SLC7A2;refGene_tx=XM_019828308.2;refGene_gn=SLC7A2;refGene_tx=XM_023252401.1;refGene_gn=SLC7A2;refGene_tx=XM_023252402.1
B1 21083962 21084168
refGene_gn=SLC7A2;refGene_tx=XM_003984623.5;refGene_gn=SLC7A2;refGene_tx=XM_019828307.2;refGene_gn=SLC7A2;refGene_tx=XM_019828308.2;refGene_gn=SLC7A2;refGene_tx=XM_023252401.1;refGene_gn=SLC7A2;refGene_tx=XM_023252402.1
B1 21085816 21085919
refGene_gn=SLC7A2;refGene_tx=XM_003984623.5;refGene_gn=SLC7A2;refGene_tx=XM_019828307.2;refGene_gn=SLC7A2;refGene_tx=XM_019828308.2;refGene_gn=SLC7A2;refGene_tx=XM_023252401.1;refGene_gn=SLC7A2;refGene_tx=XM_023252402.1
B1 21088367 21088504 refGene_gn=SLC7A2;refGene_tx=XM_023252402.1
B1 21088780 21088920
refGene_gn=SLC7A2;refGene_tx=XM_003984623.5;refGene_gn=SLC7A2;refGene_tx=XM_019828307.2;refGene_gn=SLC7A2;refGene_tx=XM_019828308.2;refGene_gn=SLC7A2;refGene_tx=XM_023252401.1
B1 21090793 21091016
refGene_gn=SLC7A2;refGene_tx=XM_003984623.5;refGene_gn=SLC7A2;refGene_tx=XM_019828307.2;refGene_gn=SLC7A2;refGene_tx=XM_019828308.2;refGene_gn=SLC7A2;refGene_tx=XM_023252401.1;refGene_gn=SLC7A2;refGene_tx=XM_023252402.1
B1 21092506 21092640
refGene_gn=SLC7A2;refGene_tx=XM_003984623.5;refGene_gn=SLC7A2;refGene_tx=XM_019828307.2;refGene_gn=SLC7A2;refGene_tx=XM_019828308.2;refGene_gn=SLC7A2;refGene_tx=XM_023252401.1;refGene_gn=SLC7A2;refGene_tx=XM_023252402.1
B1 21094025 21094191
refGene_gn=SLC7A2;refGene_tx=XM_003984623.5;refGene_gn=SLC7A2;refGene_tx=XM_019828307.2;refGene_gn=SLC7A2;refGene_tx=XM_019828308.2;refGene_gn=SLC7A2;refGene_tx=XM_023252401.1;refGene_gn=SLC7A2;refGene_tx=XM_023252402.1
B1 21098328 21098484
refGene_gn=SLC7A2;refGene_tx=XM_003984623.5;refGene_gn=SLC7A2;refGene_tx=XM_019828307.2;refGene_gn=SLC7A2;refGene_tx=XM_019828308.2;refGene_gn=SLC7A2;refGene_tx=XM_023252401.1;refGene_gn=SLC7A2;refGene_tx=XM_023252402.1
B1 21099299 21099675
refGene_gn=SLC7A2;refGene_tx=XM_003984623.5;refGene_gn=SLC7A2;refGene_tx=XM_019828307.2;refGene_gn=SLC7A2;refGene_tx=XM_019828308.2;refGene_gn=SLC7A2;refGene_tx=XM_023252401.1;refGene_gn=SLC7A2;refGene_tx=XM_023252402.1
B1 21203220 21203245 refGene_gn=MTMR7;refGene_tx=XM_019828309.2
B1 21206412 21206444 refGene_gn=MTMR7;refGene_tx=XM_019828309.2
B1 21223346 21223370 refGene_gn=MTMR7;refGene_tx=XM_003984625.5
B1 21264248 21264371
refGene_gn=MTMR7;refGene_tx=XM_003984625.5;refGene_gn=MTMR7;refGene_tx=XM_019828309.2

B1 21265699 21265862
refGene_gn=MTMR7;refGene_tx=XM_003984625.5;refGene_gn=MTMR7;refGene_tx
=XM_019828309.2
B1 21274253 21274411
refGene_gn=MTMR7;refGene_tx=XM_003984625.5;refGene_gn=MTMR7;refGene_tx
=XM_019828309.2
B1 21284223 21284352
refGene_gn=MTMR7;refGene_tx=XM_003984625.5;refGene_gn=MTMR7;refGene_tx
=XM_019828309.2
B1 21293045 21293180
refGene_gn=MTMR7;refGene_tx=XM_003984625.5;refGene_gn=MTMR7;refGene_tx
=XM_019828309.2
B1 21303146 21303279
refGene_gn=MTMR7;refGene_tx=XM_003984625.5;refGene_gn=MTMR7;refGene_tx
=XM_019828309.2
B1 21319634 21319744
refGene_gn=MTMR7;refGene_tx=XM_003984625.5;refGene_gn=MTMR7;refGene_tx
=XM_019828309.2
B1 21320250 21320376
refGene_gn=MTMR7;refGene_tx=XM_003984625.5;refGene_gn=MTMR7;refGene_tx
=XM_019828309.2
B1 21322166 21322216
refGene_gn=MTMR7;refGene_tx=XM_003984625.5;refGene_gn=MTMR7;refGene_tx
=XM_019828309.2
B1 21324537 21324738
refGene_gn=MTMR7;refGene_tx=XM_003984625.5;refGene_gn=MTMR7;refGene_tx
=XM_019828309.2
B1 21326016 21326157
refGene_gn=MTMR7;refGene_tx=XM_003984625.5;refGene_gn=MTMR7;refGene_tx
=XM_019828309.2
B1 21327652 21327779
refGene_gn=MTMR7;refGene_tx=XM_003984625.5;refGene_gn=MTMR7;refGene_tx
=XM_019828309.2
B1 21330874 21331237
refGene_gn=MTMR7;refGene_tx=XM_003984625.5;refGene_gn=MTMR7;refGene_tx
=XM_019828309.2
B1 21352592 21352673
refGene_gn=VPS37A;refGene_tx=XM_011281400.3;refGene_gn=VPS37A;refGene_
tx=XM_019828311.2
B1 21353621 21353765
refGene_gn=VPS37A;refGene_tx=XM_011281400.3;refGene_gn=VPS37A;refGene_
tx=XM_019828311.2
B1 21357784 21357853
refGene_gn=VPS37A;refGene_tx=XM_011281400.3;refGene_gn=VPS37A;refGene_
tx=XM_019828311.2
B1 21357930 21357989
refGene_gn=VPS37A;refGene_tx=XM_011281400.3;refGene_gn=VPS37A;refGene_
tx=XM_019828311.2
B1 21358070 21358198
refGene_gn=VPS37A;refGene_tx=XM_011281400.3;refGene_gn=VPS37A;refGene_

tx=XM_019828311.2
B1 21361531 21361602
refGene_gn=VPS37A;refGene_tx=XM_011281400.3;refGene_gn=VPS37A;refGene_
tx=XM_019828311.2
B1 21363391 21363617
refGene_gn=VPS37A;refGene_tx=XM_011281400.3;refGene_gn=VPS37A;refGene_
tx=XM_019828311.2
B1 21372920 21373021
refGene_gn=VPS37A;refGene_tx=XM_011281400.3;refGene_gn=VPS37A;refGene_
tx=XM_019828311.2
B1 21373624 21373739
refGene_gn=VPS37A;refGene_tx=XM_011281400.3;refGene_gn=VPS37A;refGene_
tx=XM_019828311.2
B1 21375604 21375679
refGene_gn=VPS37A;refGene_tx=XM_011281400.3;refGene_gn=VPS37A;refGene_
tx=XM_019828311.2
B1 21391024 21391032 refGene_gn=VPS37A;refGene_tx=XM_019828311.2
B1 21395742 21395867 refGene_gn=VPS37A;refGene_tx=XM_011281400.3
B1 21398036 21398153
refGene_gn=CNOT7;refGene_tx=XM_003984626.5;refGene_gn=CNOT7;refGene_tx
=XM_019828312.2;refGene_gn=CNOT7;refGene_tx=XM_019828313.2;refGene_gn=
CNOT7;refGene_tx=XM_019828314.2
B1 21399991 21400185
refGene_gn=CNOT7;refGene_tx=XM_003984626.5;refGene_gn=CNOT7;refGene_tx
=XM_019828312.2;refGene_gn=CNOT7;refGene_tx=XM_019828313.2
B1 21404632 21404661 refGene_gn=CNOT7;refGene_tx=XM_023252404.1
B1 21404983 21405145
refGene_gn=CNOT7;refGene_tx=XM_003984626.5;refGene_gn=CNOT7;refGene_tx
=XM_019828312.2;refGene_gn=CNOT7;refGene_tx=XM_019828314.2;refGene_gn=
CNOT7;refGene_tx=XM_023252404.1
B1 21408067 21408212
refGene_gn=CNOT7;refGene_tx=XM_003984626.5;refGene_gn=CNOT7;refGene_tx
=XM_019828312.2;refGene_gn=CNOT7;refGene_tx=XM_019828313.2;refGene_gn=
CNOT7;refGene_tx=XM_019828314.2;refGene_gn=CNOT7;refGene_tx=XM_0232524
04.1
B1 21410515 21410626
refGene_gn=CNOT7;refGene_tx=XM_003984626.5;refGene_gn=CNOT7;refGene_tx
=XM_019828312.2;refGene_gn=CNOT7;refGene_tx=XM_019828313.2;refGene_gn=
CNOT7;refGene_tx=XM_019828314.2;refGene_gn=CNOT7;refGene_tx=XM_0232524
04.1
B1 21412189 21412318
refGene_gn=CNOT7;refGene_tx=XM_003984626.5;refGene_gn=CNOT7;refGene_tx
=XM_019828312.2;refGene_gn=CNOT7;refGene_tx=XM_019828313.2;refGene_gn=
CNOT7;refGene_tx=XM_019828314.2;refGene_gn=CNOT7;refGene_tx=XM_0232524
04.1
B1 21422412 21422453
refGene_gn=ZDHHC2;refGene_tx=XM_003984627.5;refGene_gn=ZDHHC2;refGene_
tx=XM_023252406.1
B1 21424690 21424803
refGene_gn=ZDHHC2;refGene_tx=XM_003984627.5;refGene_gn=ZDHHC2;refGene_

tx=XM_023252406.1
B1 21428495 21428588
refGene_gn=ZDHHC2;refGene_tx=XM_003984627.5;refGene_gn=ZDHHC2;refGene_tx=XM_023252406.1
B1 21428886 21429013
refGene_gn=ZDHHC2;refGene_tx=XM_003984627.5;refGene_gn=ZDHHC2;refGene_tx=XM_023252406.1
B1 21431457 21431590
refGene_gn=ZDHHC2;refGene_tx=XM_003984627.5;refGene_gn=ZDHHC2;refGene_tx=XM_023252406.1
B1 21433914 21434035
refGene_gn=ZDHHC2;refGene_tx=XM_003984627.5;refGene_gn=ZDHHC2;refGene_tx=XM_023252406.1
B1 21442483 21442516
refGene_gn=ZDHHC2;refGene_tx=XM_003984627.5;refGene_gn=ZDHHC2;refGene_tx=XM_023252406.1
B1 21443261 21443331
refGene_gn=ZDHHC2;refGene_tx=XM_003984627.5;refGene_gn=ZDHHC2;refGene_tx=XM_023252406.1
B1 21450706 21450827
refGene_gn=ZDHHC2;refGene_tx=XM_003984627.5;refGene_gn=ZDHHC2;refGene_tx=XM_023252406.1
B1 21461324 21461419
refGene_gn=ZDHHC2;refGene_tx=XM_003984627.5;refGene_gn=ZDHHC2;refGene_tx=XM_023252406.1
B1 21462509 21462536
refGene_gn=ZDHHC2;refGene_tx=XM_003984627.5;refGene_gn=ZDHHC2;refGene_tx=XM_023252406.1
B1 21471505 21471539 refGene_gn=ZDHHC2;refGene_tx=XM_003984627.5
B1 21486242 21486369 refGene_gn=ZDHHC2;refGene_tx=XM_023252406.1
B1 21526873 21526942
refGene_gn=MICU3;refGene_tx=XM_006930595.4;refGene_gn=MICU3;refGene_tx=XM_023252407.1;refGene_gn=MICU3;refGene_tx=XM_023252408.1
B1 21527994 21528152
refGene_gn=MICU3;refGene_tx=XM_006930595.4;refGene_gn=MICU3;refGene_tx=XM_023252407.1;refGene_gn=MICU3;refGene_tx=XM_023252408.1
B1 21530200 21530309
refGene_gn=MICU3;refGene_tx=XM_006930595.4;refGene_gn=MICU3;refGene_tx=XM_023252407.1;refGene_gn=MICU3;refGene_tx=XM_023252408.1
B1 21535908 21536055 refGene_gn=MICU3;refGene_tx=XM_023252409.1
B1 21540648 21540820
refGene_gn=MICU3;refGene_tx=XM_006930595.4;refGene_gn=MICU3;refGene_tx=XM_023252407.1;refGene_gn=MICU3;refGene_tx=XM_023252408.1;refGene_gn=MICU3;refGene_tx=XM_023252409.1
B1 21541288 21541389
refGene_gn=MICU3;refGene_tx=XM_006930595.4;refGene_gn=MICU3;refGene_tx=XM_023252407.1;refGene_gn=MICU3;refGene_tx=XM_023252408.1;refGene_gn=MICU3;refGene_tx=XM_023252409.1
B1 21547174 21547270
refGene_gn=MICU3;refGene_tx=XM_006930595.4;refGene_gn=MICU3;refGene_tx

=XM_023252407.1;refGene_gn=MICU3;refGene_tx=XM_023252408.1;refGene_gn=MICU3;refGene_tx=XM_023252409.1
B1 21558384 21558423 refGene_gn=MICU3;refGene_tx=XM_006930595.4;refGene_gn=MICU3;refGene_tx=XM_023252408.1;refGene_gn=MICU3;refGene_tx=XM_023252409.1
B1 21561547 21561619 refGene_gn=MICU3;refGene_tx=XM_006930595.4;refGene_gn=MICU3;refGene_tx=XM_023252407.1;refGene_gn=MICU3;refGene_tx=XM_023252408.1;refGene_gn=MICU3;refGene_tx=XM_023252409.1
B1 21564197 21564280 refGene_gn=MICU3;refGene_tx=XM_006930595.4;refGene_gn=MICU3;refGene_tx=XM_023252407.1;refGene_gn=MICU3;refGene_tx=XM_023252408.1;refGene_gn=MICU3;refGene_tx=XM_023252409.1
B1 21567100 21567148 refGene_gn=MICU3;refGene_tx=XM_006930595.4;refGene_gn=MICU3;refGene_tx=XM_023252407.1;refGene_gn=MICU3;refGene_tx=XM_023252409.1
B1 21570926 21571005 refGene_gn=MICU3;refGene_tx=XM_006930595.4;refGene_gn=MICU3;refGene_tx=XM_023252407.1;refGene_gn=MICU3;refGene_tx=XM_023252408.1;refGene_gn=MICU3;refGene_tx=XM_023252409.1
B1 21576727 21576759 refGene_gn=MICU3;refGene_tx=XM_006930595.4;refGene_gn=MICU3;refGene_tx=XM_023252407.1;refGene_gn=MICU3;refGene_tx=XM_023252408.1;refGene_gn=MICU3;refGene_tx=XM_023252409.1
B1 21583796 21583950 refGene_gn=MICU3;refGene_tx=XM_006930595.4;refGene_gn=MICU3;refGene_tx=XM_023252407.1;refGene_gn=MICU3;refGene_tx=XM_023252408.1;refGene_gn=MICU3;refGene_tx=XM_023252409.1
B1 21633971 21634355 refGene_gn=MICU3;refGene_tx=XM_006930595.4;refGene_gn=MICU3;refGene_tx=XM_023252407.1;refGene_gn=MICU3;refGene_tx=XM_023252408.1;refGene_gn=MICU3;refGene_tx=XM_023252409.1
B1 21654730 21655016 refGene_gn=FGF20;refGene_tx=XM_023252220.1
B1 21660446 21660550 refGene_gn=FGF20;refGene_tx=XM_023252220.1
B1 21661689 21661935 refGene_gn=FGF20;refGene_tx=XM_023252220.1
B1 22088963 22089194 refGene_gn=LOC111560024;refGene_tx=XM_023252221.1
B1 22092811 22092844 refGene_gn=LOC111560024;refGene_tx=XM_023252221.1
B1 22484130 22484233 refGene_gn=MSR1;refGene_tx=XM_023252410.1;refGene_gn=MSR1;refGene_tx=XM_023252411.1;refGene_gn=MSR1;refGene_tx=XM_023252412.1
B1 22486471 22486585 refGene_gn=MSR1;refGene_tx=XM_023252410.1;refGene_gn=MSR1;refGene_tx=XM_023252411.1;refGene_gn=MSR1;refGene_tx=XM_023252412.1
B1 22492514 22492927 refGene_gn=MSR1;refGene_tx=XM_023252410.1;refGene_gn=MSR1;refGene_tx=XM_023252411.1;refGene_gn=MSR1;refGene_tx=XM_023252412.1
B1 22497689 22497876 refGene_gn=MSR1;refGene_tx=XM_023252410.1;refGene_gn=MSR1;refGene_tx=X

M_023252411.1;refGene_gn=MSR1;refGene_tx=XM_023252412.1
B1 22507851 22507932
refGene_gn=MSR1;refGene_tx=XM_023252410.1;refGene_gn=MSR1;refGene_tx=XM_023252411.1;refGene_gn=MSR1;refGene_tx=XM_023252412.1
B1 22514235 22514325
refGene_gn=MSR1;refGene_tx=XM_023252410.1;refGene_gn=MSR1;refGene_tx=XM_023252411.1;refGene_gn=MSR1;refGene_tx=XM_023252412.1
B1 22523043 22523097
refGene_gn=MSR1;refGene_tx=XM_023252410.1;refGene_gn=MSR1;refGene_tx=XM_023252411.1;refGene_gn=MSR1;refGene_tx=XM_023252412.1
B1 22525296 22525340 refGene_gn=MSR1;refGene_tx=XM_023252412.1
B1 22541523 22541712
refGene_gn=MSR1;refGene_tx=XM_023252410.1;refGene_gn=MSR1;refGene_tx=XM_023252411.1
B1 22553974 22554108
refGene_gn=MSR1;refGene_tx=XM_023252410.1;refGene_gn=MSR1;refGene_tx=XM_023252411.1
B1 22913607 22913641 refGene_gn=TUSC3;refGene_tx=XM_019828315.2
B1 22932367 22932383 refGene_gn=TUSC3;refGene_tx=XM_019828316.2
B1 22938861 22938874 refGene_gn=TUSC3;refGene_tx=XM_019828317.2
B1 22943203 22943294
refGene_gn=TUSC3;refGene_tx=XM_019828315.2;refGene_gn=TUSC3;refGene_tx=XM_019828316.2;refGene_gn=TUSC3;refGene_tx=XM_019828317.2
B1 22946812 22946887
refGene_gn=TUSC3;refGene_tx=XM_019828315.2;refGene_gn=TUSC3;refGene_tx=XM_019828316.2;refGene_gn=TUSC3;refGene_tx=XM_019828317.2
B1 22964907 22964971
refGene_gn=TUSC3;refGene_tx=XM_019828315.2;refGene_gn=TUSC3;refGene_tx=XM_019828316.2;refGene_gn=TUSC3;refGene_tx=XM_019828317.2
B1 23025622 23025712
refGene_gn=TUSC3;refGene_tx=XM_019828315.2;refGene_gn=TUSC3;refGene_tx=XM_019828316.2;refGene_gn=TUSC3;refGene_tx=XM_019828317.2
B1 23050692 23050833
refGene_gn=TUSC3;refGene_tx=XM_019828315.2;refGene_gn=TUSC3;refGene_tx=XM_019828316.2;refGene_gn=TUSC3;refGene_tx=XM_019828317.2
B1 23055059 23055200
refGene_gn=TUSC3;refGene_tx=XM_019828315.2;refGene_gn=TUSC3;refGene_tx=XM_019828316.2;refGene_gn=TUSC3;refGene_tx=XM_019828317.2
B1 23059785 23059903
refGene_gn=TUSC3;refGene_tx=XM_019828315.2;refGene_gn=TUSC3;refGene_tx=XM_019828316.2;refGene_gn=TUSC3;refGene_tx=XM_019828317.2
B1 23090681 23090851
refGene_gn=TUSC3;refGene_tx=XM_019828315.2;refGene_gn=TUSC3;refGene_tx=XM_019828316.2;refGene_gn=TUSC3;refGene_tx=XM_019828317.2
B1 23200682 23200997
refGene_gn=TUSC3;refGene_tx=XM_019828315.2;refGene_gn=TUSC3;refGene_tx=XM_019828316.2;refGene_gn=TUSC3;refGene_tx=XM_019828317.2
B1 23536498 23536537
refGene_gn=SGCZ;refGene_tx=XM_019828324.2;refGene_gn=SGCZ;refGene_tx=XM_023252415.1

B1 24152102 24152297
refGene_gn=SGCZ;refGene_tx=XM_019828324.2;refGene_gn=SGCZ;refGene_tx=XM_023252413.1;refGene_gn=SGCZ;refGene_tx=XM_023252414.1;refGene_gn=SGCZ;refGene_tx=XM_023252415.1
B1 24426341 24426443
refGene_gn=SGCZ;refGene_tx=XM_019828324.2;refGene_gn=SGCZ;refGene_tx=XM_023252413.1;refGene_gn=SGCZ;refGene_tx=XM_023252414.1;refGene_gn=SGCZ;refGene_tx=XM_023252415.1
B1 24512741 24512829
refGene_gn=SGCZ;refGene_tx=XM_019828324.2;refGene_gn=SGCZ;refGene_tx=XM_023252413.1;refGene_gn=SGCZ;refGene_tx=XM_023252414.1;refGene_gn=SGCZ;refGene_tx=XM_023252415.1
B1 24571641 24571764
refGene_gn=SGCZ;refGene_tx=XM_019828324.2;refGene_gn=SGCZ;refGene_tx=XM_023252413.1;refGene_gn=SGCZ;refGene_tx=XM_023252414.1;refGene_gn=SGCZ;refGene_tx=XM_023252415.1
B1 24607399 24607472
refGene_gn=SGCZ;refGene_tx=XM_019828324.2;refGene_gn=SGCZ;refGene_tx=XM_023252413.1;refGene_gn=SGCZ;refGene_tx=XM_023252414.1;refGene_gn=SGCZ;refGene_tx=XM_023252415.1
B1 24609658 24609800 refGene_gn=SGCZ;refGene_tx=XM_023252415.1
B1 24621033 24621157
refGene_gn=SGCZ;refGene_tx=XM_019828324.2;refGene_gn=SGCZ;refGene_tx=XM_023252413.1;refGene_gn=SGCZ;refGene_tx=XM_023252414.1
B1 24635241 24635436
refGene_gn=SGCZ;refGene_tx=XM_019828324.2;refGene_gn=SGCZ;refGene_tx=XM_023252413.1;refGene_gn=SGCZ;refGene_tx=XM_023252414.1
B1 25095690 25096605
refGene_gn=CB1H8orf48;refGene_tx=XM_011281405.3
B1 25166056 25167073
refGene_gn=DLC1;refGene_tx=XM_019828335.2;refGene_gn=DLC1;refGene_tx=XM_019828336.2;refGene_gn=DLC1;refGene_tx=XM_019828337.2;refGene_gn=DLC1;refGene_tx=XM_019828341.2;refGene_gn=DLC1;refGene_tx=XM_019828342.2;refGene_gn=DLC1;refGene_tx=XM_019828346.2;refGene_gn=DLC1;refGene_tx=XM_023252416.1;refGene_gn=DLC1;refGene_tx=XM_023252417.1;refGene_gn=DLC1;refGene_tx=XM_023252418.1
B1 25258319 25258469
refGene_gn=DLC1;refGene_tx=XM_019828335.2;refGene_gn=DLC1;refGene_tx=XM_019828336.2;refGene_gn=DLC1;refGene_tx=XM_019828337.2;refGene_gn=DLC1;refGene_tx=XM_019828341.2;refGene_gn=DLC1;refGene_tx=XM_019828346.2;refGene_gn=DLC1;refGene_tx=XM_023252416.1;refGene_gn=DLC1;refGene_tx=XM_023252417.1;refGene_gn=DLC1;refGene_tx=XM_023252418.1;refGene_gn=DLC1;refGene_tx=XM_023252419.1
B1 25276950 25277091
refGene_gn=DLC1;refGene_tx=XM_019828335.2;refGene_gn=DLC1;refGene_tx=XM_019828336.2;refGene_gn=DLC1;refGene_tx=XM_019828337.2;refGene_gn=DLC1;refGene_tx=XM_019828341.2;refGene_gn=DLC1;refGene_tx=XM_019828342.2;refGene_gn=DLC1;refGene_tx=XM_019828346.2;refGene_gn=DLC1;refGene_tx=XM_023252416.1;refGene_gn=DLC1;refGene_tx=XM_023252417.1;refGene_gn=DLC1;refGene_tx=XM_023252418.1;refGene_gn=DLC1;refGene_tx=XM_023252419.1

B1 25363109 25363143
refGene_gn=DLC1;refGene_tx=XM_019828335.2;refGene_gn=DLC1;refGene_tx=XM_019828336.2;refGene_gn=DLC1;refGene_tx=XM_019828337.2;refGene_gn=DLC1;refGene_tx=XM_019828341.2;refGene_gn=DLC1;refGene_tx=XM_019828342.2;refGene_gn=DLC1;refGene_tx=XM_019828346.2;refGene_gn=DLC1;refGene_tx=XM_023252416.1;refGene_gn=DLC1;refGene_tx=XM_023252417.1;refGene_gn=DLC1;refGene_tx=XM_023252418.1;refGene_gn=DLC1;refGene_tx=XM_023252419.1
B1 25392614 25392753 refGene_gn=DLC1;refGene_tx=XM_019828343.2
B1 25529305 25529342 refGene_gn=DLC1;refGene_tx=XM_011281421.3
B1 25545788 25545860
refGene_gn=DLC1;refGene_tx=XM_011281421.3;refGene_gn=DLC1;refGene_tx=XM_019828335.2;refGene_gn=DLC1;refGene_tx=XM_019828336.2;refGene_gn=DLC1;refGene_tx=XM_019828337.2;refGene_gn=DLC1;refGene_tx=XM_019828341.2;refGene_gn=DLC1;refGene_tx=XM_019828342.2;refGene_gn=DLC1;refGene_tx=XM_019828343.2;refGene_gn=DLC1;refGene_tx=XM_019828346.2;refGene_gn=DLC1;refGene_tx=XM_023252416.1;refGene_gn=DLC1;refGene_tx=XM_023252417.1;refGene_gn=DLC1;refGene_tx=XM_023252418.1;refGene_gn=DLC1;refGene_tx=XM_023252419.1
B1 25548286 25548332 refGene_gn=DLC1;refGene_tx=XM_019828344.2
B1 25551780 25551862
refGene_gn=DLC1;refGene_tx=XM_011281421.3;refGene_gn=DLC1;refGene_tx=XM_019828335.2;refGene_gn=DLC1;refGene_tx=XM_019828336.2;refGene_gn=DLC1;refGene_tx=XM_019828337.2;refGene_gn=DLC1;refGene_tx=XM_019828341.2;refGene_gn=DLC1;refGene_tx=XM_019828342.2;refGene_gn=DLC1;refGene_tx=XM_019828343.2;refGene_gn=DLC1;refGene_tx=XM_019828344.2;refGene_gn=DLC1;refGene_tx=XM_019828346.2;refGene_gn=DLC1;refGene_tx=XM_023252416.1;refGene_gn=DLC1;refGene_tx=XM_023252417.1;refGene_gn=DLC1;refGene_tx=XM_023252418.1;refGene_gn=DLC1;refGene_tx=XM_023252419.1
B1 25557413 25557432 refGene_gn=DLC1;refGene_tx=XM_019828346.2
B1 25558911 25558975
refGene_gn=DLC1;refGene_tx=XM_011281421.3;refGene_gn=DLC1;refGene_tx=XM_011281422.3;refGene_gn=DLC1;refGene_tx=XM_019828335.2;refGene_gn=DLC1;refGene_tx=XM_019828336.2;refGene_gn=DLC1;refGene_tx=XM_019828337.2;refGene_gn=DLC1;refGene_tx=XM_019828341.2;refGene_gn=DLC1;refGene_tx=XM_019828342.2;refGene_gn=DLC1;refGene_tx=XM_019828343.2;refGene_gn=DLC1;refGene_tx=XM_019828344.2;refGene_gn=DLC1;refGene_tx=XM_019828345.2;refGene_gn=DLC1;refGene_tx=XM_023252416.1;refGene_gn=DLC1;refGene_tx=XM_023252417.1;refGene_gn=DLC1;refGene_tx=XM_023252418.1;refGene_gn=DLC1;refGene_tx=XM_023252419.1
B1 25561286 25562710
refGene_gn=DLC1;refGene_tx=XM_011281421.3;refGene_gn=DLC1;refGene_tx=XM_011281422.3;refGene_gn=DLC1;refGene_tx=XM_019828335.2;refGene_gn=DLC1;refGene_tx=XM_019828336.2;refGene_gn=DLC1;refGene_tx=XM_019828337.2;refGene_gn=DLC1;refGene_tx=XM_019828341.2;refGene_gn=DLC1;refGene_tx=XM_019828342.2;refGene_gn=DLC1;refGene_tx=XM_019828343.2;refGene_gn=DLC1;refGene_tx=XM_019828344.2;refGene_gn=DLC1;refGene_tx=XM_019828345.2;refGene_gn=DLC1;refGene_tx=XM_023252416.1;refGene_gn=DLC1;refGene_tx=XM_023252417.1;refGene_gn=DLC1;refGene_tx=XM_023252418.1;refGene_gn=DLC1;refGene_tx=XM_023252419.1
B1 25563222 25563399

refGene_gn=DLC1;refGene_tx=XM_011281421.3;refGene_gn=DLC1;refGene_tx=XM_011281422.3;refGene_gn=DLC1;refGene_tx=XM_019828335.2;refGene_gn=DLC1;refGene_tx=XM_019828336.2;refGene_gn=DLC1;refGene_tx=XM_019828337.2;refGene_gn=DLC1;refGene_tx=XM_019828341.2;refGene_gn=DLC1;refGene_tx=XM_019828342.2;refGene_gn=DLC1;refGene_tx=XM_019828343.2;refGene_gn=DLC1;refGene_tx=XM_019828344.2;refGene_gn=DLC1;refGene_tx=XM_019828345.2;refGene_gn=DLC1;refGene_tx=XM_023252416.1;refGene_gn=DLC1;refGene_tx=XM_023252417.1;refGene_gn=DLC1;refGene_tx=XM_023252418.1;refGene_gn=DLC1;refGene_tx=XM_023252419.1
B1 25565853 25566013
refGene_gn=DLC1;refGene_tx=XM_011281421.3;refGene_gn=DLC1;refGene_tx=XM_011281422.3;refGene_gn=DLC1;refGene_tx=XM_019828335.2;refGene_gn=DLC1;refGene_tx=XM_019828336.2;refGene_gn=DLC1;refGene_tx=XM_019828337.2;refGene_gn=DLC1;refGene_tx=XM_019828341.2;refGene_gn=DLC1;refGene_tx=XM_019828342.2;refGene_gn=DLC1;refGene_tx=XM_019828343.2;refGene_gn=DLC1;refGene_tx=XM_019828344.2;refGene_gn=DLC1;refGene_tx=XM_019828345.2;refGene_gn=DLC1;refGene_tx=XM_023252416.1;refGene_gn=DLC1;refGene_tx=XM_023252417.1;refGene_gn=DLC1;refGene_tx=XM_023252418.1;refGene_gn=DLC1;refGene_tx=XM_023252419.1
B1 25566134 25566333
refGene_gn=DLC1;refGene_tx=XM_011281421.3;refGene_gn=DLC1;refGene_tx=XM_011281422.3;refGene_gn=DLC1;refGene_tx=XM_019828335.2;refGene_gn=DLC1;refGene_tx=XM_019828336.2;refGene_gn=DLC1;refGene_tx=XM_019828337.2;refGene_gn=DLC1;refGene_tx=XM_019828341.2;refGene_gn=DLC1;refGene_tx=XM_019828342.2;refGene_gn=DLC1;refGene_tx=XM_019828343.2;refGene_gn=DLC1;refGene_tx=XM_019828344.2;refGene_gn=DLC1;refGene_tx=XM_019828345.2;refGene_gn=DLC1;refGene_tx=XM_023252416.1;refGene_gn=DLC1;refGene_tx=XM_023252417.1;refGene_gn=DLC1;refGene_tx=XM_023252418.1;refGene_gn=DLC1;refGene_tx=XM_023252419.1
B1 25567875 25568089
refGene_gn=DLC1;refGene_tx=XM_011281421.3;refGene_gn=DLC1;refGene_tx=XM_011281422.3;refGene_gn=DLC1;refGene_tx=XM_019828335.2;refGene_gn=DLC1;refGene_tx=XM_019828336.2;refGene_gn=DLC1;refGene_tx=XM_019828337.2;refGene_gn=DLC1;refGene_tx=XM_019828341.2;refGene_gn=DLC1;refGene_tx=XM_019828342.2;refGene_gn=DLC1;refGene_tx=XM_019828343.2;refGene_gn=DLC1;refGene_tx=XM_019828344.2;refGene_gn=DLC1;refGene_tx=XM_019828345.2;refGene_gn=DLC1;refGene_tx=XM_023252416.1;refGene_gn=DLC1;refGene_tx=XM_023252417.1;refGene_gn=DLC1;refGene_tx=XM_023252418.1;refGene_gn=DLC1;refGene_tx=XM_023252419.1
B1 25569694 25569809
refGene_gn=DLC1;refGene_tx=XM_011281421.3;refGene_gn=DLC1;refGene_tx=XM_011281422.3;refGene_gn=DLC1;refGene_tx=XM_019828335.2;refGene_gn=DLC1;refGene_tx=XM_019828336.2;refGene_gn=DLC1;refGene_tx=XM_019828337.2;refGene_gn=DLC1;refGene_tx=XM_019828341.2;refGene_gn=DLC1;refGene_tx=XM_019828342.2;refGene_gn=DLC1;refGene_tx=XM_019828343.2;refGene_gn=DLC1;refGene_tx=XM_019828344.2;refGene_gn=DLC1;refGene_tx=XM_019828345.2;refGene_gn=DLC1;refGene_tx=XM_023252416.1;refGene_gn=DLC1;refGene_tx=XM_023252417.1;refGene_gn=DLC1;refGene_tx=XM_023252418.1;refGene_gn=DLC1;refGene_tx=XM_023252419.1
B1 25570371 25570590

refGene_gn=DLC1;refGene_tx=XM_011281421.3;refGene_gn=DLC1;refGene_tx=XM_011281422.3;refGene_gn=DLC1;refGene_tx=XM_019828335.2;refGene_gn=DLC1;refGene_tx=XM_019828336.2;refGene_gn=DLC1;refGene_tx=XM_019828337.2;refGene_gn=DLC1;refGene_tx=XM_019828341.2;refGene_gn=DLC1;refGene_tx=XM_019828342.2;refGene_gn=DLC1;refGene_tx=XM_019828343.2;refGene_gn=DLC1;refGene_tx=XM_019828344.2;refGene_gn=DLC1;refGene_tx=XM_019828345.2;refGene_gn=DLC1;refGene_tx=XM_023252416.1;refGene_gn=DLC1;refGene_tx=XM_023252417.1;refGene_gn=DLC1;refGene_tx=XM_023252418.1;refGene_gn=DLC1;refGene_tx=XM_023252419.1
B1 25571409 25571627
refGene_gn=DLC1;refGene_tx=XM_011281421.3;refGene_gn=DLC1;refGene_tx=XM_011281422.3;refGene_gn=DLC1;refGene_tx=XM_019828335.2;refGene_gn=DLC1;refGene_tx=XM_019828336.2;refGene_gn=DLC1;refGene_tx=XM_019828337.2;refGene_gn=DLC1;refGene_tx=XM_019828341.2;refGene_gn=DLC1;refGene_tx=XM_019828342.2;refGene_gn=DLC1;refGene_tx=XM_019828343.2;refGene_gn=DLC1;refGene_tx=XM_019828344.2;refGene_gn=DLC1;refGene_tx=XM_019828345.2;refGene_gn=DLC1;refGene_tx=XM_023252416.1;refGene_gn=DLC1;refGene_tx=XM_023252417.1;refGene_gn=DLC1;refGene_tx=XM_023252418.1;refGene_gn=DLC1;refGene_tx=XM_023252419.1
B1 25572359 25572533
refGene_gn=DLC1;refGene_tx=XM_011281421.3;refGene_gn=DLC1;refGene_tx=XM_011281422.3;refGene_gn=DLC1;refGene_tx=XM_019828335.2;refGene_gn=DLC1;refGene_tx=XM_019828336.2;refGene_gn=DLC1;refGene_tx=XM_019828337.2;refGene_gn=DLC1;refGene_tx=XM_019828341.2;refGene_gn=DLC1;refGene_tx=XM_019828342.2;refGene_gn=DLC1;refGene_tx=XM_019828343.2;refGene_gn=DLC1;refGene_tx=XM_019828344.2;refGene_gn=DLC1;refGene_tx=XM_019828345.2;refGene_gn=DLC1;refGene_tx=XM_023252416.1;refGene_gn=DLC1;refGene_tx=XM_023252417.1;refGene_gn=DLC1;refGene_tx=XM_023252418.1;refGene_gn=DLC1;refGene_tx=XM_023252419.1
B1 25573286 25573407
refGene_gn=DLC1;refGene_tx=XM_011281421.3;refGene_gn=DLC1;refGene_tx=XM_011281422.3;refGene_gn=DLC1;refGene_tx=XM_019828335.2;refGene_gn=DLC1;refGene_tx=XM_019828336.2;refGene_gn=DLC1;refGene_tx=XM_019828337.2;refGene_gn=DLC1;refGene_tx=XM_019828341.2;refGene_gn=DLC1;refGene_tx=XM_019828342.2;refGene_gn=DLC1;refGene_tx=XM_019828343.2;refGene_gn=DLC1;refGene_tx=XM_019828344.2;refGene_gn=DLC1;refGene_tx=XM_019828345.2;refGene_gn=DLC1;refGene_tx=XM_023252416.1;refGene_gn=DLC1;refGene_tx=XM_023252417.1;refGene_gn=DLC1;refGene_tx=XM_023252418.1;refGene_gn=DLC1;refGene_tx=XM_023252419.1
B1 25632025 25633056
refGene_gn=KIAA1456;refGene_tx=XM_003984641.4;refGene_gn=KIAA1456;refGene_tx=XM_019828348.2;refGene_gn=KIAA1456;refGene_tx=XM_019828349.2;refGene_gn=KIAA1456;refGene_tx=XM_019828350.2;refGene_gn=KIAA1456;refGene_tx=XM_019828351.2;refGene_gn=KIAA1456;refGene_tx=XM_023252420.1
B1 25639511 25639685
refGene_gn=KIAA1456;refGene_tx=XM_003984641.4;refGene_gn=KIAA1456;refGene_tx=XM_019828348.2;refGene_gn=KIAA1456;refGene_tx=XM_019828349.2;refGene_gn=KIAA1456;refGene_tx=XM_019828350.2;refGene_gn=KIAA1456;refGene_tx=XM_023252420.1
B1 25645751 25645905

refGene_gn=KIAA1456;refGene_tx=XM_003984641.4;refGene_gn=KIAA1456;refGene_tx=XM_019828348.2;refGene_gn=KIAA1456;refGene_tx=XM_019828349.2;refGene_gn=KIAA1456;refGene_tx=XM_019828350.2;refGene_gn=KIAA1456;refGene_tx=XM_023252420.1
B1 25885343 25886058 refGene_gn=LONRF1;refGene_tx=XM_023252421.1
B1 25895923 25896042 refGene_gn=LONRF1;refGene_tx=XM_023252421.1
B1 25897022 25897145 refGene_gn=LONRF1;refGene_tx=XM_023252421.1
B1 25907846 25907996 refGene_gn=LONRF1;refGene_tx=XM_023252421.1
B1 25908487 25908728 refGene_gn=LONRF1;refGene_tx=XM_023252421.1
B1 25908820 25908917 refGene_gn=LONRF1;refGene_tx=XM_023252421.1
B1 25910776 25910891 refGene_gn=LONRF1;refGene_tx=XM_023252421.1
B1 25913761 25913883 refGene_gn=LONRF1;refGene_tx=XM_023252421.1
B1 25916069 25916228 refGene_gn=LONRF1;refGene_tx=XM_023252421.1
B1 25916338 25916501 refGene_gn=LONRF1;refGene_tx=XM_023252421.1
B1 25919092 25919245 refGene_gn=LONRF1;refGene_tx=XM_023252421.1
B1 25921549 25921708 refGene_gn=LONRF1;refGene_tx=XM_023252421.1
B1 26023386 26024529
refGene_gn=PRAG1;refGene_tx=XM_003984642.5;refGene_gn=PRAG1;refGene_tx=XM_019828352.2
B1 26031607 26032359
refGene_gn=PRAG1;refGene_tx=XM_003984642.5;refGene_gn=PRAG1;refGene_tx=XM_019828352.2
B1 26044036 26044194
refGene_gn=PRAG1;refGene_tx=XM_003984642.5;refGene_gn=PRAG1;refGene_tx=XM_019828352.2
B1 26065210 26067033
refGene_gn=PRAG1;refGene_tx=XM_003984642.5;refGene_gn=PRAG1;refGene_tx=XM_019828352.2
B1 26069027 26069357 refGene_gn=PRAG1;refGene_tx=XM_003984642.5
B1 26338821 26338842
refGene_gn=LOC111560217;refGene_tx=XM_023253386.1
B1 26340697 26340798
refGene_gn=LOC111560217;refGene_tx=XM_023253386.1
B1 26341026 26341636
refGene_gn=LOC111560217;refGene_tx=XM_023253386.1
B1 26341694 26342570 refGene_gn=CLDN23;refGene_tx=XM_023252422.1
B1 26350960 26350987
refGene_gn=LOC101088073;refGene_tx=XM_023252423.1
B1 26351702 26351811
refGene_gn=LOC101088073;refGene_tx=XM_023252423.1
B1 26353422 26353900
refGene_gn=LOC101088073;refGene_tx=XM_023252423.1
B1 26355756 26355993
refGene_gn=LOC101088073;refGene_tx=XM_023252423.1
B1 26357289 26357400
refGene_gn=LOC101088073;refGene_tx=XM_023252423.1
B1 26358744 26358870
refGene_gn=LOC101088073;refGene_tx=XM_023252423.1
B1 26359292 26359842
refGene_gn=LOC101088073;refGene_tx=XM_023252423.1

B1 26361297 26361498
refGene_gn=LOC101088073;refGene_tx=XM_023252423.1
B1 26361674 26361839
refGene_gn=LOC101088073;refGene_tx=XM_023252423.1
B1 26361925 26362127
refGene_gn=LOC101088073;refGene_tx=XM_023252423.1
B1 26363560 26363744
refGene_gn=LOC101088073;refGene_tx=XM_023252423.1
B1 26364138 26364357
refGene_gn=LOC101088073;refGene_tx=XM_023252423.1
B1 26364933 26365120
refGene_gn=LOC101088073;refGene_tx=XM_023252423.1
B1 26366365 26366550
refGene_gn=LOC101088073;refGene_tx=XM_023252423.1
B1 26366766 26366896
refGene_gn=LOC101088073;refGene_tx=XM_023252423.1
B1 26368438 26368677
refGene_gn=LOC101088073;refGene_tx=XM_023252423.1
B1 26370441 26370657
refGene_gn=LOC101088073;refGene_tx=XM_023252423.1
B1 26371443 26371638
refGene_gn=LOC101088073;refGene_tx=XM_023252423.1
B1 26372866 26373079
refGene_gn=LOC101088073;refGene_tx=XM_023252423.1
B1 26373699 26374011
refGene_gn=LOC101088073;refGene_tx=XM_023252423.1
B1 26379409 26379634
refGene_gn=LOC101088073;refGene_tx=XM_023252423.1
B1 26381498 26381624
refGene_gn=LOC101088073;refGene_tx=XM_023252423.1
B1 26382421 26382647
refGene_gn=LOC101088073;refGene_tx=XM_023252423.1
B1 26383799 26383968
refGene_gn=LOC101088073;refGene_tx=XM_023252423.1
B1 26384266 26384435
refGene_gn=LOC101088073;refGene_tx=XM_023252423.1
B1 26385129 26385333
refGene_gn=LOC101088073;refGene_tx=XM_023252423.1
B1 26385910 26386036
refGene_gn=LOC101088073;refGene_tx=XM_023252423.1
B1 26387167 26387287
refGene_gn=LOC101088073;refGene_tx=XM_023252423.1
B1 26388511 26388625
refGene_gn=LOC101088073;refGene_tx=XM_023252423.1
B1 26389504 26389744
refGene_gn=LOC101088073;refGene_tx=XM_023252423.1
B1 26390188 26390308
refGene_gn=LOC101088073;refGene_tx=XM_023252423.1
B1 26391953 26392067
refGene_gn=LOC101088073;refGene_tx=XM_023252423.1

B1 26394809 26394885
refGene_gn=LOC101088073;refGene_tx=XM_023252423.1
B1 26463183 26463217 refGene_gn=MFHAS1;refGene_tx=XM_023252425.1
B1 26467261 26467289 refGene_gn=MFHAS1;refGene_tx=XM_023252426.1
B1 26470751 26470776
refGene_gn=MFHAS1;refGene_tx=XM_023252427.1;refGene_gn=MFHAS1;refGene_tx=XM_023252428.1
B1 26473402 26473533
refGene_gn=MFHAS1;refGene_tx=XM_019828356.2;refGene_gn=MFHAS1;refGene_tx=XM_023252425.1;refGene_gn=MFHAS1;refGene_tx=XM_023252426.1;refGene_gn=MFHAS1;refGene_tx=XM_023252427.1;refGene_gn=MFHAS1;refGene_tx=XM_023252428.1
B1 26559447 26562445
refGene_gn=MFHAS1;refGene_tx=XM_019828356.2;refGene_gn=MFHAS1;refGene_tx=XM_023252425.1;refGene_gn=MFHAS1;refGene_tx=XM_023252426.1;refGene_gn=MFHAS1;refGene_tx=XM_023252427.1;refGene_gn=MFHAS1;refGene_tx=XM_023252428.1
B1 26648063 26648171 refGene_gn=ERI1;refGene_tx=XM_003984644.4
B1 26650115 26650157 refGene_gn=ERI1;refGene_tx=XM_019828357.2
B1 26652763 26652942
refGene_gn=ERI1;refGene_tx=XM_003984644.4;refGene_gn=ERI1;refGene_tx=XM_019828357.2;refGene_gn=ERI1;refGene_tx=XM_023252429.1
B1 26655268 26655479
refGene_gn=ERI1;refGene_tx=XM_003984644.4;refGene_gn=ERI1;refGene_tx=XM_019828357.2;refGene_gn=ERI1;refGene_tx=XM_023252429.1
B1 26659229 26659313
refGene_gn=ERI1;refGene_tx=XM_003984644.4;refGene_gn=ERI1;refGene_tx=XM_019828357.2;refGene_gn=ERI1;refGene_tx=XM_023252429.1
B1 26660620 26660730
refGene_gn=ERI1;refGene_tx=XM_003984644.4;refGene_gn=ERI1;refGene_tx=XM_019828357.2;refGene_gn=ERI1;refGene_tx=XM_023252429.1
B1 26662664 26662779
refGene_gn=ERI1;refGene_tx=XM_003984644.4;refGene_gn=ERI1;refGene_tx=XM_019828357.2;refGene_gn=ERI1;refGene_tx=XM_023252429.1
B1 26672721 26672964
refGene_gn=ERI1;refGene_tx=XM_003984644.4;refGene_gn=ERI1;refGene_tx=XM_019828357.2;refGene_gn=ERI1;refGene_tx=XM_023252429.1
B1 26778581 26779464
refGene_gn=PPP1R3B;refGene_tx=XM_003984645.5;refGene_gn=PPP1R3B;refGene_tx=XM_006930598.4
B1 26785809 26785841 refGene_gn=PPP1R3B;refGene_tx=XM_006930598.4
B1 27120942 27121615
refGene_gn=TNKS;refGene_tx=XM_011281425.3;refGene_gn=TNKS;refGene_tx=XM_023252430.1
B1 27154647 27154872
refGene_gn=TNKS;refGene_tx=XM_011281425.3;refGene_gn=TNKS;refGene_tx=XM_023252430.1
B1 27185343 27185439
refGene_gn=TNKS;refGene_tx=XM_011281425.3;refGene_gn=TNKS;refGene_tx=XM_023252430.1

B1 27236320 27236357
refGene_gn=TNKS;refGene_tx=XM_011281425.3;refGene_gn=TNKS;refGene_tx=XM_023252430.1
B1 27237117 27237193
refGene_gn=TNKS;refGene_tx=XM_011281425.3;refGene_gn=TNKS;refGene_tx=XM_023252430.1
B1 27260014 27260109
refGene_gn=TNKS;refGene_tx=XM_011281425.3;refGene_gn=TNKS;refGene_tx=XM_023252430.1
B1 27262296 27262363
refGene_gn=TNKS;refGene_tx=XM_011281425.3;refGene_gn=TNKS;refGene_tx=XM_023252430.1
B1 27262677 27262864
refGene_gn=TNKS;refGene_tx=XM_011281425.3;refGene_gn=TNKS;refGene_tx=XM_023252430.1
B1 27268374 27268496
refGene_gn=TNKS;refGene_tx=XM_011281425.3;refGene_gn=TNKS;refGene_tx=XM_023252430.1
B1 27270141 27270233
refGene_gn=TNKS;refGene_tx=XM_011281425.3;refGene_gn=TNKS;refGene_tx=XM_023252430.1
B1 27270325 27270404
refGene_gn=TNKS;refGene_tx=XM_011281425.3;refGene_gn=TNKS;refGene_tx=XM_023252430.1
B1 27283482 27283654
refGene_gn=TNKS;refGene_tx=XM_011281425.3;refGene_gn=TNKS;refGene_tx=XM_023252430.1
B1 27290271 27290351
refGene_gn=TNKS;refGene_tx=XM_011281425.3;refGene_gn=TNKS;refGene_tx=XM_023252430.1
B1 27294023 27294169
refGene_gn=TNKS;refGene_tx=XM_011281425.3;refGene_gn=TNKS;refGene_tx=XM_023252430.1
B1 27296373 27296539
refGene_gn=TNKS;refGene_tx=XM_011281425.3;refGene_gn=TNKS;refGene_tx=XM_023252430.1
B1 27298002 27298222
refGene_gn=TNKS;refGene_tx=XM_011281425.3;refGene_gn=TNKS;refGene_tx=XM_023252430.1
B1 27298510 27298620
refGene_gn=TNKS;refGene_tx=XM_011281425.3;refGene_gn=TNKS;refGene_tx=XM_023252430.1
B1 27306051 27306240
refGene_gn=TNKS;refGene_tx=XM_011281425.3;refGene_gn=TNKS;refGene_tx=XM_023252430.1
B1 27309265 27309503
refGene_gn=TNKS;refGene_tx=XM_011281425.3;refGene_gn=TNKS;refGene_tx=XM_023252430.1
B1 27310092 27310175
refGene_gn=TNKS;refGene_tx=XM_011281425.3;refGene_gn=TNKS;refGene_tx=X

M_023252430.1
B1 27318325 27318446
refGene_gn=TNKS;refGene_tx=XM_011281425.3;refGene_gn=TNKS;refGene_tx=X
M_023252430.1
B1 27319665 27319763
refGene_gn=TNKS;refGene_tx=XM_011281425.3;refGene_gn=TNKS;refGene_tx=X
M_023252430.1
B1 27321340 27321415
refGene_gn=TNKS;refGene_tx=XM_011281425.3;refGene_gn=TNKS;refGene_tx=X
M_023252430.1
B1 27322273 27322379
refGene_gn=TNKS;refGene_tx=XM_011281425.3;refGene_gn=TNKS;refGene_tx=X
M_023252430.1
B1 27322961 27323148
refGene_gn=TNKS;refGene_tx=XM_011281425.3;refGene_gn=TNKS;refGene_tx=X
M_023252430.1
B1 27325807 27325964
refGene_gn=TNKS;refGene_tx=XM_011281425.3;refGene_gn=TNKS;refGene_tx=X
M_023252430.1
B1 27335562 27335649
refGene_gn=TNKS;refGene_tx=XM_011281425.3;refGene_gn=TNKS;refGene_tx=X
M_023252430.1
B1 27355279 27355665 refGene_gn=DUSP4;refGene_tx=XM_003984646.5
B1 27356300 27356520 refGene_gn=DUSP4;refGene_tx=XM_003984646.5
B1 27357527 27357673 refGene_gn=DUSP4;refGene_tx=XM_003984646.5
B1 27366453 27366886 refGene_gn=DUSP4;refGene_tx=XM_003984646.5
B1 27943478 27944070
refGene_gn=LOC101093179;refGene_tx=XM_003984659.2
B1 27966612 27966697
refGene_gn=LOC101093179;refGene_tx=XM_003984659.2
B1 27969703 27970157
refGene_gn=LOC101093179;refGene_tx=XM_003984659.2
B1 28012772 28012798
refGene_gn=SARAF;refGene_tx=XM_003984647.5;refGene_gn=SARAF;refGene_tx
=XM_006930599.4
B1 28015261 28015416
refGene_gn=SARAF;refGene_tx=XM_003984647.5;refGene_gn=SARAF;refGene_tx
=XM_006930599.4
B1 28016051 28016181
refGene_gn=SARAF;refGene_tx=XM_003984647.5;refGene_gn=SARAF;refGene_tx
=XM_006930599.4
B1 28018057 28018475
refGene_gn=SARAF;refGene_tx=XM_003984647.5;refGene_gn=SARAF;refGene_tx
=XM_006930599.4
B1 28020973 28021152
refGene_gn=SARAF;refGene_tx=XM_003984647.5;refGene_gn=SARAF;refGene_tx
=XM_006930599.4
B1 28031114 28031217
refGene_gn=SARAF;refGene_tx=XM_003984647.5;refGene_gn=SARAF;refGene_tx
=XM_006930599.4

B1 28037312 28037328 refGene_gn=LEPROTL1;refGene_tx=XM_023252431.1
B1 28044640 28044716 refGene_gn=LEPROTL1;refGene_tx=XM_023252431.1
B1 28046417 28046604 refGene_gn=LEPROTL1;refGene_tx=XM_023252431.1
B1 28047565 28047682 refGene_gn=LEPROTL1;refGene_tx=XM_023252431.1
B1 28072241 28073205 refGene_gn=MBOAT4;refGene_tx=XM_003984661.3
B1 28076354 28076579 refGene_gn=MBOAT4;refGene_tx=XM_003984661.3
B1 28080110 28080229 refGene_gn=MBOAT4;refGene_tx=XM_003984661.3
B1 28082883 28082906 refGene_gn=DCTN6;refGene_tx=XM_003984648.4
B1 28092044 28092109 refGene_gn=DCTN6;refGene_tx=XM_003984648.4
B1 28094786 28094883 refGene_gn=DCTN6;refGene_tx=XM_023252432.1
B1 28101739 28101845
refGene_gn=DCTN6;refGene_tx=XM_003984648.4;refGene_gn=DCTN6;refGene_tx=XM_023252432.1
B1 28103464 28103553
refGene_gn=DCTN6;refGene_tx=XM_003984648.4;refGene_gn=DCTN6;refGene_tx=XM_023252432.1
B1 28104832 28104880
refGene_gn=DCTN6;refGene_tx=XM_003984648.4;refGene_gn=DCTN6;refGene_tx=XM_023252432.1
B1 28105892 28106035
refGene_gn=DCTN6;refGene_tx=XM_003984648.4;refGene_gn=DCTN6;refGene_tx=XM_023252432.1
B1 28108446 28108545
refGene_gn=DCTN6;refGene_tx=XM_003984648.4;refGene_gn=DCTN6;refGene_tx=XM_023252432.1
B1 28113888 28114097
refGene_gn=LOC101093930;refGene_tx=XM_011281426.2
B1 28114322 28114623
refGene_gn=LOC101093930;refGene_tx=XM_011281426.2
B1 28183709 28184015
refGene_gn=LOC101089601;refGene_tx=XM_011281465.1
B1 28261478 28261544
refGene_gn=RBPMS;refGene_tx=XM_003984649.4;refGene_gn=RBPMS;refGene_tx=XM_003984650.4;refGene_gn=RBPMS;refGene_tx=XM_019828359.2;refGene_gn=RBPMS;refGene_tx=XM_019828364.2;refGene_gn=RBPMS;refGene_tx=XM_023252433.1;refGene_gn=RBPMS;refGene_tx=XM_023252434.1;refGene_gn=RBPMS;refGene_tx=XM_023252435.1;refGene_gn=RBPMS;refGene_tx=XM_023252436.1;refGene_gn=RBPMS;refGene_tx=XM_023252437.1
B1 28265998 28266022 refGene_gn=RBPMS;refGene_tx=XM_019828362.1
B1 28320578 28320632 refGene_gn=RBPMS;refGene_tx=XM_019828361.1
B1 28339656 28339734
refGene_gn=RBPMS;refGene_tx=XM_003984649.4;refGene_gn=RBPMS;refGene_tx=XM_003984650.4;refGene_gn=RBPMS;refGene_tx=XM_019828359.2;refGene_gn=RBPMS;refGene_tx=XM_019828361.1;refGene_gn=RBPMS;refGene_tx=XM_019828362.1;refGene_gn=RBPMS;refGene_tx=XM_023252433.1;refGene_gn=RBPMS;refGene_tx=XM_023252434.1;refGene_gn=RBPMS;refGene_tx=XM_023252435.1;refGene_gn=RBPMS;refGene_tx=XM_023252436.1;refGene_gn=RBPMS;refGene_tx=XM_023252437.1
B1 28342699 28342738
refGene_gn=RBPMS;refGene_tx=XM_003984649.4;refGene_gn=RBPMS;refGene_tx

=XM_003984650.4;refGene_gn=RBPMS;refGene_tx=XM_019828359.2;refGene_gn=RBPMS;refGene_tx=XM_019828361.1;refGene_gn=RBPMS;refGene_tx=XM_019828362.1;refGene_gn=RBPMS;refGene_tx=XM_023252433.1;refGene_gn=RBPMS;refGene_tx=XM_023252434.1;refGene_gn=RBPMS;refGene_tx=XM_023252435.1;refGene_gn=RBPMS;refGene_tx=XM_023252436.1;refGene_gn=RBPMS;refGene_tx=XM_023252437.1
B1 28343986 28344049
refGene_gn=RBPMS;refGene_tx=XM_003984649.4;refGene_gn=RBPMS;refGene_tx=XM_003984650.4;refGene_gn=RBPMS;refGene_tx=XM_019828359.2;refGene_gn=RBPMS;refGene_tx=XM_019828361.1;refGene_gn=RBPMS;refGene_tx=XM_019828362.1;refGene_gn=RBPMS;refGene_tx=XM_023252433.1;refGene_gn=RBPMS;refGene_tx=XM_023252434.1;refGene_gn=RBPMS;refGene_tx=XM_023252435.1;refGene_gn=RBPMS;refGene_tx=XM_023252436.1;refGene_gn=RBPMS;refGene_tx=XM_023252437.1
B1 28368575 28368686 refGene_gn=RBPMS;refGene_tx=XM_019828363.2
B1 28370869 28371020
refGene_gn=RBPMS;refGene_tx=XM_003984649.4;refGene_gn=RBPMS;refGene_tx=XM_003984650.4;refGene_gn=RBPMS;refGene_tx=XM_019828359.2;refGene_gn=RBPMS;refGene_tx=XM_019828361.1;refGene_gn=RBPMS;refGene_tx=XM_019828362.1;refGene_gn=RBPMS;refGene_tx=XM_019828363.2;refGene_gn=RBPMS;refGene_tx=XM_019828364.2;refGene_gn=RBPMS;refGene_tx=XM_023252433.1;refGene_gn=RBPMS;refGene_tx=XM_023252434.1;refGene_gn=RBPMS;refGene_tx=XM_023252435.1;refGene_gn=RBPMS;refGene_tx=XM_023252436.1;refGene_gn=RBPMS;refGene_tx=XM_023252437.1
B1 28411082 28411213
refGene_gn=RBPMS;refGene_tx=XM_003984649.4;refGene_gn=RBPMS;refGene_tx=XM_003984650.4;refGene_gn=RBPMS;refGene_tx=XM_019828359.2;refGene_gn=RBPMS;refGene_tx=XM_019828361.1;refGene_gn=RBPMS;refGene_tx=XM_019828362.1;refGene_gn=RBPMS;refGene_tx=XM_019828363.2;refGene_gn=RBPMS;refGene_tx=XM_019828364.2;refGene_gn=RBPMS;refGene_tx=XM_023252433.1;refGene_gn=RBPMS;refGene_tx=XM_023252434.1;refGene_gn=RBPMS;refGene_tx=XM_023252435.1;refGene_gn=RBPMS;refGene_tx=XM_023252436.1
B1 28413911 28414043 refGene_gn=RBPMS;refGene_tx=XM_003984649.4
B1 28422078 28422212
refGene_gn=RBPMS;refGene_tx=XM_019828359.2;refGene_gn=RBPMS;refGene_tx=XM_019828361.1;refGene_gn=RBPMS;refGene_tx=XM_019828362.1;refGene_gn=RBPMS;refGene_tx=XM_019828363.2;refGene_gn=RBPMS;refGene_tx=XM_019828364.2;refGene_gn=RBPMS;refGene_tx=XM_023252433.1;refGene_gn=RBPMS;refGene_tx=XM_023252434.1;refGene_gn=RBPMS;refGene_tx=XM_023252435.1
B1 28424068 28424138
refGene_gn=RBPMS;refGene_tx=XM_003984650.4;refGene_gn=RBPMS;refGene_tx=XM_019828359.2;refGene_gn=RBPMS;refGene_tx=XM_019828361.1;refGene_gn=RBPMS;refGene_tx=XM_019828362.1;refGene_gn=RBPMS;refGene_tx=XM_019828363.2;refGene_gn=RBPMS;refGene_tx=XM_019828364.2;refGene_gn=RBPMS;refGene_tx=XM_023252433.1;refGene_gn=RBPMS;refGene_tx=XM_023252434.1;refGene_gn=RBPMS;refGene_tx=XM_023252435.1;refGene_gn=RBPMS;refGene_tx=XM_023252436.1;refGene_gn=RBPMS;refGene_tx=XM_023252437.1
B1 28431219 28431234
refGene_gn=RBPMS;refGene_tx=XM_019828359.2;refGene_gn=RBPMS;refGene_tx=XM_019828361.1;refGene_gn=RBPMS;refGene_tx=XM_019828362.1;refGene_gn=

RBPMS;refGene_tx=XM_019828363.2;refGene_gn=RBPMS;refGene_tx=XM_019828364.2;refGene_gn=RBPMS;refGene_tx=XM_023252433.1;refGene_gn=RBPMS;refGene_tx=XM_023252434.1
B1 28437462 28437583 refGene_gn=RBPMS;refGene_tx=XM_023252435.1;refGene_gn=RBPMS;refGene_tx=XM_023252437.1
B1 28452399 28452516 refGene_gn=GTF2E2;refGene_tx=XM_003984651.4;refGene_gn=GTF2E2;refGene_tx=XM_006930602.3
B1 28453773 28453889 refGene_gn=GTF2E2;refGene_tx=XM_003984651.4;refGene_gn=GTF2E2;refGene_tx=XM_006930602.3
B1 28470802 28470870 refGene_gn=GTF2E2;refGene_tx=XM_006930603.4
B1 28477077 28477171 refGene_gn=GTF2E2;refGene_tx=XM_003984651.4;refGene_gn=GTF2E2;refGene_tx=XM_006930602.3;refGene_gn=GTF2E2;refGene_tx=XM_006930603.4
B1 28484925 28485108 refGene_gn=GTF2E2;refGene_tx=XM_003984651.4;refGene_gn=GTF2E2;refGene_tx=XM_006930602.3;refGene_gn=GTF2E2;refGene_tx=XM_006930603.4
B1 28486694 28486802 refGene_gn=GTF2E2;refGene_tx=XM_003984651.4;refGene_gn=GTF2E2;refGene_tx=XM_006930602.3;refGene_gn=GTF2E2;refGene_tx=XM_006930603.4
B1 28500343 28500435 refGene_gn=GTF2E2;refGene_tx=XM_003984651.4;refGene_gn=GTF2E2;refGene_tx=XM_006930602.3;refGene_gn=GTF2E2;refGene_tx=XM_006930603.4
B1 28512778 28513075 refGene_gn=SMIM18;refGene_tx=XM_006930601.4;refGene_gn=SMIM18;refGene_tx=XM_019828365.2;refGene_gn=SMIM18;refGene_tx=XM_019828366.1
B1 28522664 28522830 refGene_gn=GTF2E2;refGene_tx=XM_003984651.4;refGene_gn=GTF2E2;refGene_tx=XM_006930602.3;refGene_gn=GTF2E2;refGene_tx=XM_006930603.4
B1 28547580 28547730 refGene_gn=GSR;refGene_tx=XM_023252439.1
B1 28549820 28549954 refGene_gn=GSR;refGene_tx=XM_023252439.1
B1 28551893 28552025 refGene_gn=GSR;refGene_tx=XM_023252439.1
B1 28554756 28554868 refGene_gn=GSR;refGene_tx=XM_023252439.1
B1 28558581 28558740 refGene_gn=GSR;refGene_tx=XM_023252439.1
B1 28560003 28560090 refGene_gn=GSR;refGene_tx=XM_023252439.1
B1 28563611 28563711 refGene_gn=GSR;refGene_tx=XM_023252439.1
B1 28567448 28567503 refGene_gn=GSR;refGene_tx=XM_023252439.1
B1 28570407 28570555 refGene_gn=GSR;refGene_tx=XM_023252439.1
B1 28572882 28572952 refGene_gn=GSR;refGene_tx=XM_023252439.1
B1 28574116 28574205 refGene_gn=GSR;refGene_tx=XM_023252439.1
B1 28576937 28576964 refGene_gn=GSR;refGene_tx=XM_023252439.1
B1 28592150 28592450 refGene_gn=GSR;refGene_tx=XM_023252439.1
B1 28603451 28603539 refGene_gn=UBXN8;refGene_tx=XM_003984652.5
B1 28611253 28611360 refGene_gn=UBXN8;refGene_tx=XM_003984652.5
B1 28611361 28611377 refGene_gn=UBXN8;refGene_tx=XM_003984652.5
B1 28614661 28614732 refGene_gn=UBXN8;refGene_tx=XM_003984652.5
B1 28620185 28620308 refGene_gn=UBXN8;refGene_tx=XM_003984652.5
B1 28622876 28623017 refGene_gn=UBXN8;refGene_tx=XM_003984652.5

B1 28624816 28624858 refGene_gn=UBXN8;refGene_tx=XM_003984652.5
B1 28628690 28628765 refGene_gn=UBXN8;refGene_tx=XM_003984652.5
B1 28631109 28631277 refGene_gn=UBXN8;refGene_tx=XM_003984652.5
B1 28657442 28657515 refGene_gn=PPP2CB;refGene_tx=XM_006930608.3
B1 28661427 28661546 refGene_gn=PPP2CB;refGene_tx=XM_006930608.3
B1 28663465 28663627 refGene_gn=PPP2CB;refGene_tx=XM_006930608.3
B1 28663737 28663827 refGene_gn=PPP2CB;refGene_tx=XM_006930608.3
B1 28665855 28666029 refGene_gn=PPP2CB;refGene_tx=XM_006930608.3
B1 28667635 28667845 refGene_gn=PPP2CB;refGene_tx=XM_006930608.3
B1 28691702 28691804 refGene_gn=PPP2CB;refGene_tx=XM_006930608.3
B1 28714058 28714096
refGene_gn=TEX15;refGene_tx=XM_019828368.2;refGene_gn=TEX15;refGene_tx
=XM_023252440.1
B1 28717288 28718406
refGene_gn=TEX15;refGene_tx=XM_019828368.2;refGene_gn=TEX15;refGene_tx
=XM_023252440.1
B1 28720488 28720547
refGene_gn=TEX15;refGene_tx=XM_019828368.2;refGene_gn=TEX15;refGene_tx
=XM_023252440.1
B1 28723105 28730442
refGene_gn=TEX15;refGene_tx=XM_019828368.2;refGene_gn=TEX15;refGene_tx
=XM_023252440.1
B1 28738987 28739150 refGene_gn=TEX15;refGene_tx=XM_019828368.2
B1 28741530 28741677 refGene_gn=TEX15;refGene_tx=XM_019828368.2
B1 28749839 28750077 refGene_gn=TEX15;refGene_tx=XM_019828368.2
B1 28765742 28765908 refGene_gn=TEX15;refGene_tx=XM_019828368.2
B1 28778781 28778920 refGene_gn=TEX15;refGene_tx=XM_019828368.2
B1 28898407 28898422 refGene_gn=PURG;refGene_tx=XM_023252442.1
B1 28933019 28934057
refGene_gn=PURG;refGene_tx=XM_023252442.1;refGene_gn=PURG;refGene_tx=X
M_023252443.1
B1 28973620 28973716 refGene_gn=WRN;refGene_tx=XM_023252444.1
B1 28974574 28974687 refGene_gn=WRN;refGene_tx=XM_023252444.1
B1 28978244 28978390 refGene_gn=WRN;refGene_tx=XM_023252444.1
B1 28979610 28979759 refGene_gn=WRN;refGene_tx=XM_023252444.1
B1 28980682 28980832 refGene_gn=WRN;refGene_tx=XM_023252444.1
B1 28982847 28982917 refGene_gn=WRN;refGene_tx=XM_023252444.1
B1 28989481 28989596 refGene_gn=WRN;refGene_tx=XM_023252444.1
B1 28993470 28993894 refGene_gn=WRN;refGene_tx=XM_023252444.1
B1 28996515 28996596 refGene_gn=WRN;refGene_tx=XM_023252444.1
B1 28999457 28999593 refGene_gn=WRN;refGene_tx=XM_023252444.1
B1 29001149 29001225 refGene_gn=WRN;refGene_tx=XM_023252444.1
B1 29002679 29002747 refGene_gn=WRN;refGene_tx=XM_023252444.1
B1 29003035 29003144 refGene_gn=WRN;refGene_tx=XM_023252444.1
B1 29003695 29003764 refGene_gn=WRN;refGene_tx=XM_023252444.1
B1 29008210 29008293 refGene_gn=WRN;refGene_tx=XM_023252444.1
B1 29012008 29012058 refGene_gn=WRN;refGene_tx=XM_023252444.1
B1 29012059 29012116 refGene_gn=WRN;refGene_tx=XM_023252444.1
B1 29023182 29023370 refGene_gn=WRN;refGene_tx=XM_023252444.1
B1 29028908 29029083 refGene_gn=WRN;refGene_tx=XM_023252444.1

B1	29033690	29033872	refGene_gn=WRN;refGene_tx=XM_023252444.1
B1	29039422	29039524	refGene_gn=WRN;refGene_tx=XM_023252444.1
B1	29039808	29039901	refGene_gn=WRN;refGene_tx=XM_023252444.1
B1	29045244	29045386	refGene_gn=WRN;refGene_tx=XM_023252444.1
B1	29052897	29053068	refGene_gn=WRN;refGene_tx=XM_023252444.1
B1	29053145	29053243	refGene_gn=WRN;refGene_tx=XM_023252444.1
B1	29054103	29054176	refGene_gn=WRN;refGene_tx=XM_023252444.1
B1	29055363	29055440	refGene_gn=WRN;refGene_tx=XM_023252444.1
B1	29060383	29060459	refGene_gn=WRN;refGene_tx=XM_023252444.1
B1	29060697	29060810	refGene_gn=WRN;refGene_tx=XM_023252444.1
B1	29063103	29063218	refGene_gn=WRN;refGene_tx=XM_023252444.1
B1	29066590	29066722	refGene_gn=WRN;refGene_tx=XM_023252444.1
B1	29069313	29069476	refGene_gn=WRN;refGene_tx=XM_023252444.1
B1	29077264	29077461	refGene_gn=WRN;refGene_tx=XM_023252444.1
B1	29082653	29082773	refGene_gn=WRN;refGene_tx=XM_023252444.1
B1	29490296	29491029
refGene_gn=NRG1;refGene_tx=XM_019828378.2;refGene_gn=NRG1;refGene_tx=XM_019828379.2;refGene_gn=NRG1;refGene_tx=XM_019828382.2;refGene_gn=NRG1;refGene_tx=XM_019828386.2;refGene_gn=NRG1;refGene_tx=XM_023252445.1;refGene_gn=NRG1;refGene_tx=XM_023252447.1
B1	30384581	30384678
refGene_gn=NRG1;refGene_tx=XM_011281428.3;refGene_gn=NRG1;refGene_tx=XM_019828380.2;refGene_gn=NRG1;refGene_tx=XM_019828381.2;refGene_gn=NRG1;refGene_tx=XM_019828387.1;refGene_gn=NRG1;refGene_tx=XM_023252448.1
B1	30432520	30432698
refGene_gn=NRG1;refGene_tx=XM_011281428.3;refGene_gn=NRG1;refGene_tx=XM_019828378.2;refGene_gn=NRG1;refGene_tx=XM_019828379.2;refGene_gn=NRG1;refGene_tx=XM_019828380.2;refGene_gn=NRG1;refGene_tx=XM_019828381.2;refGene_gn=NRG1;refGene_tx=XM_019828382.2;refGene_gn=NRG1;refGene_tx=XM_019828386.2;refGene_gn=NRG1;refGene_tx=XM_019828387.1;refGene_gn=NRG1;refGene_tx=XM_023252445.1;refGene_gn=NRG1;refGene_tx=XM_023252447.1;refGene_gn=NRG1;refGene_tx=XM_023252448.1
B1	30443157	30443279
refGene_gn=NRG1;refGene_tx=XM_011281428.3;refGene_gn=NRG1;refGene_tx=XM_019828378.2;refGene_gn=NRG1;refGene_tx=XM_019828379.2;refGene_gn=NRG1;refGene_tx=XM_019828380.2;refGene_gn=NRG1;refGene_tx=XM_019828381.2;refGene_gn=NRG1;refGene_tx=XM_019828382.2;refGene_gn=NRG1;refGene_tx=XM_019828386.2;refGene_gn=NRG1;refGene_tx=XM_019828387.1;refGene_gn=NRG1;refGene_tx=XM_023252445.1;refGene_gn=NRG1;refGene_tx=XM_023252447.1;refGene_gn=NRG1;refGene_tx=XM_023252448.1
B1	30451490	30451541
refGene_gn=NRG1;refGene_tx=XM_011281428.3;refGene_gn=NRG1;refGene_tx=XM_019828378.2;refGene_gn=NRG1;refGene_tx=XM_019828379.2;refGene_gn=NRG1;refGene_tx=XM_019828380.2;refGene_gn=NRG1;refGene_tx=XM_019828381.2;refGene_gn=NRG1;refGene_tx=XM_019828386.2;refGene_gn=NRG1;refGene_tx=XM_019828387.1;refGene_gn=NRG1;refGene_tx=XM_023252445.1;refGene_gn=NRG1;refGene_tx=XM_023252447.1
B1	30453859	30453910
refGene_gn=NRG1;refGene_tx=XM_011281428.3;refGene_gn=NRG1;refGene_tx=XM_019828378.2;refGene_gn=NRG1;refGene_tx=XM_019828379.2;refGene_gn=NRG

1;refGene_tx=XM_019828380.2;refGene_gn=NRG1;refGene_tx=XM_019828381.2;
refGene_gn=NRG1;refGene_tx=XM_019828386.2;refGene_gn=NRG1;refGene_tx=X
M_019828387.1;refGene_gn=NRG1;refGene_tx=XM_023252445.1;refGene_gn=NRG
1;refGene_tx=XM_023252447.1
B1 30483767 30484434
refGene_gn=NRG1;refGene_tx=XM_019828376.2;refGene_gn=NRG1;refGene_tx=X
M_019828377.2;refGene_gn=NRG1;refGene_tx=XM_019828383.1;refGene_gn=NRG
1;refGene_tx=XM_019828384.1;refGene_gn=NRG1;refGene_tx=XM_019828385.1
B1 30571527 30571657
refGene_gn=NRG1;refGene_tx=XM_011281428.3;refGene_gn=NRG1;refGene_tx=X
M_019828376.2;refGene_gn=NRG1;refGene_tx=XM_019828377.2;refGene_gn=NRG
1;refGene_tx=XM_019828378.2;refGene_gn=NRG1;refGene_tx=XM_019828379.2;
refGene_gn=NRG1;refGene_tx=XM_019828380.2;refGene_gn=NRG1;refGene_tx=X
M_019828381.2;refGene_gn=NRG1;refGene_tx=XM_019828382.2;refGene_gn=NRG
1;refGene_tx=XM_019828383.1;refGene_gn=NRG1;refGene_tx=XM_019828384.1;
refGene_gn=NRG1;refGene_tx=XM_019828385.1;refGene_gn=NRG1;refGene_tx=X
M_019828386.2;refGene_gn=NRG1;refGene_tx=XM_019828387.1;refGene_gn=NRG
1;refGene_tx=XM_023252445.1;refGene_gn=NRG1;refGene_tx=XM_023252447.1;
refGene_gn=NRG1;refGene_tx=XM_023252448.1;refGene_gn=NRG1;refGene_tx=X
M_023252449.1
B1 30584104 30584172
refGene_gn=NRG1;refGene_tx=XM_019828377.2;refGene_gn=NRG1;refGene_tx=X
M_019828378.2;refGene_gn=NRG1;refGene_tx=XM_019828380.2;refGene_gn=NRG
1;refGene_tx=XM_019828384.1;refGene_gn=NRG1;refGene_tx=XM_019828387.1
B1 30584777 30584836
refGene_gn=NRG1;refGene_tx=XM_011281428.3;refGene_gn=NRG1;refGene_tx=X
M_019828376.2;refGene_gn=NRG1;refGene_tx=XM_019828379.2;refGene_gn=NRG
1;refGene_tx=XM_019828381.2;refGene_gn=NRG1;refGene_tx=XM_019828382.2;
refGene_gn=NRG1;refGene_tx=XM_019828383.1;refGene_gn=NRG1;refGene_tx=X
M_019828385.1;refGene_gn=NRG1;refGene_tx=XM_019828386.2;refGene_gn=NRG
1;refGene_tx=XM_023252445.1;refGene_gn=NRG1;refGene_tx=XM_023252447.1;
refGene_gn=NRG1;refGene_tx=XM_023252448.1;refGene_gn=NRG1;refGene_tx=X
M_023252449.1
B1 30591221 30591245
refGene_gn=NRG1;refGene_tx=XM_011281428.3;refGene_gn=NRG1;refGene_tx=X
M_019828376.2;refGene_gn=NRG1;refGene_tx=XM_019828382.2;refGene_gn=NRG
1;refGene_tx=XM_019828383.1;refGene_gn=NRG1;refGene_tx=XM_019828385.1;
refGene_gn=NRG1;refGene_tx=XM_023252445.1;refGene_gn=NRG1;refGene_tx=X
M_023252447.1;refGene_gn=NRG1;refGene_tx=XM_023252449.1
B1 30595895 30595998
refGene_gn=NRG1;refGene_tx=XM_011281428.3;refGene_gn=NRG1;refGene_tx=X
M_019828376.2;refGene_gn=NRG1;refGene_tx=XM_019828377.2;refGene_gn=NRG
1;refGene_tx=XM_019828378.2;refGene_gn=NRG1;refGene_tx=XM_019828379.2;
refGene_gn=NRG1;refGene_tx=XM_019828380.2;refGene_gn=NRG1;refGene_tx=X
M_019828381.2;refGene_gn=NRG1;refGene_tx=XM_019828382.2;refGene_gn=NRG
1;refGene_tx=XM_019828383.1;refGene_gn=NRG1;refGene_tx=XM_019828384.1;
refGene_gn=NRG1;refGene_tx=XM_019828385.1;refGene_gn=NRG1;refGene_tx=X
M_019828386.2;refGene_gn=NRG1;refGene_tx=XM_019828387.1;refGene_gn=NRG
1;refGene_tx=XM_019828388.2;refGene_gn=NRG1;refGene_tx=XM_023252445.1;
refGene_gn=NRG1;refGene_tx=XM_023252447.1;refGene_gn=NRG1;refGene_tx=X

M_023252448.1;refGene_gn=NRG1;refGene_tx=XM_023252449.1
B1 30597733 30597860
refGene_gn=NRG1;refGene_tx=XM_011281428.3;refGene_gn=NRG1;refGene_tx=XM_019828376.2;refGene_gn=NRG1;refGene_tx=XM_019828377.2;refGene_gn=NRG1;refGene_tx=XM_019828378.2;refGene_gn=NRG1;refGene_tx=XM_019828379.2;refGene_gn=NRG1;refGene_tx=XM_019828380.2;refGene_gn=NRG1;refGene_tx=XM_019828381.2;refGene_gn=NRG1;refGene_tx=XM_019828382.2;refGene_gn=NRG1;refGene_tx=XM_019828383.1;refGene_gn=NRG1;refGene_tx=XM_019828384.1;refGene_gn=NRG1;refGene_tx=XM_019828385.1;refGene_gn=NRG1;refGene_tx=XM_019828386.2;refGene_gn=NRG1;refGene_tx=XM_019828387.1;refGene_gn=NRG1;refGene_tx=XM_019828388.2;refGene_gn=NRG1;refGene_tx=XM_023252445.1;refGene_gn=NRG1;refGene_tx=XM_023252447.1;refGene_gn=NRG1;refGene_tx=XM_023252448.1;refGene_gn=NRG1;refGene_tx=XM_023252449.1
B1 30600245 30600376
refGene_gn=NRG1;refGene_tx=XM_011281428.3;refGene_gn=NRG1;refGene_tx=XM_019828376.2;refGene_gn=NRG1;refGene_tx=XM_019828377.2;refGene_gn=NRG1;refGene_tx=XM_019828378.2;refGene_gn=NRG1;refGene_tx=XM_019828379.2;refGene_gn=NRG1;refGene_tx=XM_019828380.2;refGene_gn=NRG1;refGene_tx=XM_019828381.2;refGene_gn=NRG1;refGene_tx=XM_019828382.2;refGene_gn=NRG1;refGene_tx=XM_019828383.1;refGene_gn=NRG1;refGene_tx=XM_019828384.1;refGene_gn=NRG1;refGene_tx=XM_019828385.1;refGene_gn=NRG1;refGene_tx=XM_019828386.2;refGene_gn=NRG1;refGene_tx=XM_019828387.1;refGene_gn=NRG1;refGene_tx=XM_019828388.2;refGene_gn=NRG1;refGene_tx=XM_023252445.1;refGene_gn=NRG1;refGene_tx=XM_023252447.1;refGene_gn=NRG1;refGene_tx=XM_023252448.1;refGene_gn=NRG1;refGene_tx=XM_023252449.1
B1 30601185 30601396
refGene_gn=NRG1;refGene_tx=XM_011281428.3;refGene_gn=NRG1;refGene_tx=XM_019828376.2;refGene_gn=NRG1;refGene_tx=XM_019828377.2;refGene_gn=NRG1;refGene_tx=XM_019828378.2;refGene_gn=NRG1;refGene_tx=XM_019828379.2;refGene_gn=NRG1;refGene_tx=XM_019828380.2;refGene_gn=NRG1;refGene_tx=XM_019828381.2;refGene_gn=NRG1;refGene_tx=XM_019828382.2;refGene_gn=NRG1;refGene_tx=XM_019828383.1;refGene_gn=NRG1;refGene_tx=XM_019828384.1;refGene_gn=NRG1;refGene_tx=XM_019828385.1;refGene_gn=NRG1;refGene_tx=XM_019828386.2;refGene_gn=NRG1;refGene_tx=XM_019828387.1;refGene_gn=NRG1;refGene_tx=XM_019828388.2;refGene_gn=NRG1;refGene_tx=XM_023252445.1;refGene_gn=NRG1;refGene_tx=XM_023252447.1;refGene_gn=NRG1;refGene_tx=XM_023252448.1;refGene_gn=NRG1;refGene_tx=XM_023252449.1
B1 30603626 30603747
refGene_gn=NRG1;refGene_tx=XM_019828383.1;refGene_gn=NRG1;refGene_tx=XM_019828384.1;refGene_gn=NRG1;refGene_tx=XM_019828386.2;refGene_gn=NRG1;refGene_tx=XM_019828387.1;refGene_gn=NRG1;refGene_tx=XM_023252447.1
B1 30604156 30604808
refGene_gn=NRG1;refGene_tx=XM_011281428.3;refGene_gn=NRG1;refGene_tx=XM_019828376.2;refGene_gn=NRG1;refGene_tx=XM_019828377.2;refGene_gn=NRG1;refGene_tx=XM_019828378.2;refGene_gn=NRG1;refGene_tx=XM_019828379.2;refGene_gn=NRG1;refGene_tx=XM_019828380.2;refGene_gn=NRG1;refGene_tx=XM_019828381.2;refGene_gn=NRG1;refGene_tx=XM_019828382.2;refGene_gn=NRG1;refGene_tx=XM_019828388.2;refGene_gn=NRG1;refGene_tx=XM_023252445.1;refGene_gn=NRG1;refGene_tx=XM_023252448.1;refGene_gn=NRG1;refGene_tx=XM_023252449.1

B1 30966197 30966218
refGene_gn=LOC101100463;refGene_tx=XM_019828394.2
B1 30966220 30966229
refGene_gn=LOC101100463;refGene_tx=XM_019828395.2
B1 30969357 30969416
refGene_gn=LOC101100463;refGene_tx=XM_019828394.2;refGene_gn=LOC101100463;refGene_tx=XM_019828395.2
B1 31040414 31040518
refGene_gn=LOC101100463;refGene_tx=XM_019828390.2;refGene_gn=LOC101100463;refGene_tx=XM_019828393.2
B1 31058944 31059243
refGene_gn=LOC101100463;refGene_tx=XM_019828390.2;refGene_gn=LOC101100463;refGene_tx=XM_019828393.2;refGene_gn=LOC101100463;refGene_tx=XM_019828394.2;refGene_gn=LOC101100463;refGene_tx=XM_019828395.2;refGene_gn=LOC101100463;refGene_tx=XM_019828396.2
B1 31060189 31060416
refGene_gn=LOC101100463;refGene_tx=XM_019828390.2;refGene_gn=LOC101100463;refGene_tx=XM_019828393.2;refGene_gn=LOC101100463;refGene_tx=XM_019828394.2;refGene_gn=LOC101100463;refGene_tx=XM_019828395.2;refGene_gn=LOC101100463;refGene_tx=XM_019828396.2
B1 31072728 31072953 refGene_gn=FUT10;refGene_tx=XM_003984668.5
B1 31083266 31084102 refGene_gn=FUT10;refGene_tx=XM_003984668.5
B1 31137984 31138273 refGene_gn=FUT10;refGene_tx=XM_003984668.5
B1 31143438 31143519 refGene_gn=FUT10;refGene_tx=XM_003984668.5
B1 31155664 31155679 refGene_gn=MAK16;refGene_tx=XM_003984669.4
B1 31158392 31158442 refGene_gn=MAK16;refGene_tx=XM_003984669.4
B1 31158534 31158644 refGene_gn=MAK16;refGene_tx=XM_003984669.4
B1 31158750 31158815 refGene_gn=MAK16;refGene_tx=XM_003984669.4
B1 31159006 31159158 refGene_gn=MAK16;refGene_tx=XM_003984669.4
B1 31160386 31160441 refGene_gn=MAK16;refGene_tx=XM_003984669.4
B1 31163954 31164029 refGene_gn=MAK16;refGene_tx=XM_003984669.4
B1 31164729 31164834 refGene_gn=MAK16;refGene_tx=XM_003984669.4
B1 31165551 31165617 refGene_gn=MAK16;refGene_tx=XM_003984669.4
B1 31165784 31165979 refGene_gn=MAK16;refGene_tx=XM_003984669.4
B1 31166506 31166611 refGene_gn=TTI2;refGene_tx=XM_003984670.4
B1 31166790 31166850 refGene_gn=TTI2;refGene_tx=XM_006930614.3
B1 31167499 31167662
refGene_gn=TTI2;refGene_tx=XM_003984670.4;refGene_gn=TTI2;refGene_tx=XM_006930614.3
B1 31169420 31169564
refGene_gn=TTI2;refGene_tx=XM_003984670.4;refGene_gn=TTI2;refGene_tx=XM_006930614.3
B1 31169641 31169829
refGene_gn=TTI2;refGene_tx=XM_003984670.4;refGene_gn=TTI2;refGene_tx=XM_006930614.3
B1 31171963 31172056
refGene_gn=TTI2;refGene_tx=XM_003984670.4;refGene_gn=TTI2;refGene_tx=XM_006930614.3
B1 31173136 31173323
refGene_gn=TTI2;refGene_tx=XM_003984670.4;refGene_gn=TTI2;refGene_tx=X

M_006930614.3
B1 31174440 31175084
refGene_gn=TTI2;refGene_tx=XM_003984670.4;refGene_gn=TTI2;refGene_tx=XM_006930614.3
B1 31388690 31389573
refGene_gn=ADRA1A;refGene_tx=XM_006930617.4;refGene_gn=ADRA1A;refGene_tx=XM_006930618.4;refGene_gn=ADRA1A;refGene_tx=XM_011281429.3;refGene_gn=ADRA1A;refGene_tx=XM_011281431.3;refGene_gn=ADRA1A;refGene_tx=XM_011281433.3;refGene_gn=ADRA1A;refGene_tx=XM_019828400.1;refGene_gn=ADRA1A;refGene_tx=XM_019828402.2
B1 31412355 31412468 refGene_gn=ADRA1A;refGene_tx=XM_019828402.2
B1 31491627 31492145
refGene_gn=ADRA1A;refGene_tx=XM_006930617.4;refGene_gn=ADRA1A;refGene_tx=XM_006930618.4;refGene_gn=ADRA1A;refGene_tx=XM_011281429.3;refGene_gn=ADRA1A;refGene_tx=XM_011281431.3;refGene_gn=ADRA1A;refGene_tx=XM_011281433.3;refGene_gn=ADRA1A;refGene_tx=XM_019828400.1
B1 31496990 31496999
refGene_gn=ADRA1A;refGene_tx=XM_006930617.4;refGene_gn=ADRA1A;refGene_tx=XM_006930618.4
B1 31606057 31606149 refGene_gn=DPYSL2;refGene_tx=XM_011281434.3
B1 31607743 31607909 refGene_gn=DPYSL2;refGene_tx=XM_011281434.3
B1 31608744 31608924 refGene_gn=DPYSL2;refGene_tx=XM_011281434.3
B1 31613929 31614100 refGene_gn=DPYSL2;refGene_tx=XM_011281434.3
B1 31617057 31617138 refGene_gn=DPYSL2;refGene_tx=XM_011281434.3
B1 31617139 31617200 refGene_gn=DPYSL2;refGene_tx=XM_011281434.3
B1 31617667 31617824 refGene_gn=DPYSL2;refGene_tx=XM_011281434.3
B1 31626582 31626703 refGene_gn=DPYSL2;refGene_tx=XM_011281434.3
B1 31631181 31631250 refGene_gn=DPYSL2;refGene_tx=XM_011281434.3
B1 31631779 31631860 refGene_gn=DPYSL2;refGene_tx=XM_011281434.3
B1 31632372 31632434 refGene_gn=DPYSL2;refGene_tx=XM_011281434.3
B1 31635351 31635516 refGene_gn=DPYSL2;refGene_tx=XM_011281434.3
B1 31671473 31671658 refGene_gn=DPYSL2;refGene_tx=XM_011281434.3
B1 31673412 31673501 refGene_gn=DPYSL2;refGene_tx=XM_011281434.3
B1 31737868 31738222 refGene_gn=DPYSL2;refGene_tx=XM_011281434.3
B1 31742324 31743419
refGene_gn=PNMA2;refGene_tx=XM_011281435.2;refGene_gn=PNMA2;refGene_tx=XM_019828405.1
B1 31790470 31790522
refGene_gn=LOC102900107;refGene_tx=XM_011281437.3;refGene_gn=LOC102900107;refGene_tx=XM_019828413.2;refGene_gn=LOC102900107;refGene_tx=XM_019828414.2;refGene_gn=LOC102900107;refGene_tx=XM_023252454.1;refGene_gn=LOC102900107;refGene_tx=XM_023252455.1;refGene_gn=LOC102900107;refGene_tx=XM_023252456.1
B1 31792250 31792789
refGene_gn=LOC102900107;refGene_tx=XM_006930620.3;refGene_gn=LOC102900107;refGene_tx=XM_006930621.3;refGene_gn=LOC102900107;refGene_tx=XM_011281437.3;refGene_gn=LOC102900107;refGene_tx=XM_019828413.2;refGene_gn=LOC102900107;refGene_tx=XM_019828414.2;refGene_gn=LOC102900107;refGene_tx=XM_019828416.2;refGene_gn=LOC102900107;refGene_tx=XM_023252454.1;
refGene_gn=LOC102900107;refGene_tx=XM_023252455.1;refGene_gn=LOC102900

107;refGene_tx=XM_023252456.1
B1 31792890 31792986
refGene_gn=LOC102900107;refGene_tx=XM_006930621.3;refGene_gn=LOC102900107;refGene_tx=XM_011281437.3;refGene_gn=LOC102900107;refGene_tx=XM_019828413.2;refGene_gn=LOC102900107;refGene_tx=XM_019828414.2
B1 31795182 31795336
refGene_gn=LOC102900107;refGene_tx=XM_006930620.3;refGene_gn=LOC102900107;refGene_tx=XM_006930621.3;refGene_gn=LOC102900107;refGene_tx=XM_011281437.3;refGene_gn=LOC102900107;refGene_tx=XM_019828413.2;refGene_gn=LOC102900107;refGene_tx=XM_019828414.2;refGene_gn=LOC102900107;refGene_tx=XM_019828416.2;refGene_gn=LOC102900107;refGene_tx=XM_023252454.1;refGene_gn=LOC102900107;refGene_tx=XM_023252455.1;refGene_gn=LOC102900107;refGene_tx=XM_023252456.1
B1 31797687 31797832
refGene_gn=LOC102900107;refGene_tx=XM_006930620.3;refGene_gn=LOC102900107;refGene_tx=XM_006930621.3;refGene_gn=LOC102900107;refGene_tx=XM_011281437.3;refGene_gn=LOC102900107;refGene_tx=XM_019828413.2;refGene_gn=LOC102900107;refGene_tx=XM_019828414.2;refGene_gn=LOC102900107;refGene_tx=XM_019828416.2;refGene_gn=LOC102900107;refGene_tx=XM_023252454.1;refGene_gn=LOC102900107;refGene_tx=XM_023252455.1;refGene_gn=LOC102900107;refGene_tx=XM_023252456.1
B1 31835812 31835861
refGene_gn=BNIP3L;refGene_tx=XM_003984675.4;refGene_gn=BNIP3L;refGene_tx=XM_006930622.3
B1 31838817 31838967
refGene_gn=BNIP3L;refGene_tx=XM_003984675.4;refGene_gn=BNIP3L;refGene_tx=XM_006930622.3
B1 31839098 31839202
refGene_gn=BNIP3L;refGene_tx=XM_003984675.4;refGene_gn=BNIP3L;refGene_tx=XM_006930622.3
B1 31844060 31844133
refGene_gn=BNIP3L;refGene_tx=XM_003984675.4;refGene_gn=BNIP3L;refGene_tx=XM_006930622.3
B1 31848694 31848878
refGene_gn=BNIP3L;refGene_tx=XM_003984675.4;refGene_gn=BNIP3L;refGene_tx=XM_006930622.3
B1 31855859 31855959 refGene_gn=BNIP3L;refGene_tx=XM_003984675.4
B1 31889946 31890226
refGene_gn=PPP2R2A;refGene_tx=XM_011281440.3;refGene_gn=PPP2R2A;refGene_tx=XM_011281441.3
B1 31893720 31893812
refGene_gn=PPP2R2A;refGene_tx=XM_011281440.3;refGene_gn=PPP2R2A;refGene_tx=XM_011281441.3
B1 31896431 31896601
refGene_gn=PPP2R2A;refGene_tx=XM_011281440.3;refGene_gn=PPP2R2A;refGene_tx=XM_011281441.3
B1 31897523 31897688
refGene_gn=PPP2R2A;refGene_tx=XM_011281440.3;refGene_gn=PPP2R2A;refGene_tx=XM_011281441.3
B1 31899319 31899497

refGene_gn=PPP2R2A;refGene_tx=XM_011281440.3;refGene_gn=PPP2R2A;refGene_tx=XM_011281441.3
B1 31900169 31900282
refGene_gn=PPP2R2A;refGene_tx=XM_011281440.3;refGene_gn=PPP2R2A;refGene_tx=XM_011281441.3
B1 31907253 31907419
refGene_gn=PPP2R2A;refGene_tx=XM_011281440.3;refGene_gn=PPP2R2A;refGene_tx=XM_011281441.3
B1 31924043 31924141
refGene_gn=PPP2R2A;refGene_tx=XM_011281440.3;refGene_gn=PPP2R2A;refGene_tx=XM_011281441.3
B1 31937587 31937603 refGene_gn=PPP2R2A;refGene_tx=XM_011281441.3
B1 31971847 31971922 refGene_gn=PPP2R2A;refGene_tx=XM_011281440.3
B1 31973787 31973794 refGene_gn=PPP2R2A;refGene_tx=XM_011281440.3
B1 32174411 32174542
refGene_gn=EBF2;refGene_tx=XM_023252458.1;refGene_gn=EBF2;refGene_tx=XM_023252459.1
B1 32176964 32177121
refGene_gn=EBF2;refGene_tx=XM_023252458.1;refGene_gn=EBF2;refGene_tx=XM_023252459.1
B1 32178169 32178233
refGene_gn=EBF2;refGene_tx=XM_023252458.1;refGene_gn=EBF2;refGene_tx=XM_023252459.1
B1 32178537 32178593
refGene_gn=EBF2;refGene_tx=XM_023252458.1;refGene_gn=EBF2;refGene_tx=XM_023252459.1
B1 32179101 32179175
refGene_gn=EBF2;refGene_tx=XM_023252458.1;refGene_gn=EBF2;refGene_tx=XM_023252459.1
B1 32185617 32185686
refGene_gn=EBF2;refGene_tx=XM_023252458.1;refGene_gn=EBF2;refGene_tx=XM_023252459.1
B1 32312960 32313042
refGene_gn=EBF2;refGene_tx=XM_023252458.1;refGene_gn=EBF2;refGene_tx=XM_023252459.1
B1 32330431 32330549
refGene_gn=EBF2;refGene_tx=XM_023252458.1;refGene_gn=EBF2;refGene_tx=XM_023252459.1
B1 32332063 32332194
refGene_gn=EBF2;refGene_tx=XM_023252458.1;refGene_gn=EBF2;refGene_tx=XM_023252459.1
B1 32333086 32333213
refGene_gn=EBF2;refGene_tx=XM_023252458.1;refGene_gn=EBF2;refGene_tx=XM_023252459.1
B1 32354822 32354911
refGene_gn=EBF2;refGene_tx=XM_023252458.1;refGene_gn=EBF2;refGene_tx=XM_023252459.1
B1 32356168 32356234
refGene_gn=EBF2;refGene_tx=XM_023252458.1;refGene_gn=EBF2;refGene_tx=XM_023252459.1

B1 32356317 32356495
refGene_gn=EBF2;refGene_tx=XM_023252458.1;refGene_gn=EBF2;refGene_tx=XM_023252459.1
B1 32358146 32358332
refGene_gn=EBF2;refGene_tx=XM_023252458.1;refGene_gn=EBF2;refGene_tx=XM_023252459.1
B1 32367092 32367260 refGene_gn=EBF2;refGene_tx=XM_023252458.1
B1 32371977 32372009
refGene_gn=EBF2;refGene_tx=XM_023252458.1;refGene_gn=EBF2;refGene_tx=XM_023252459.1
B1 32425617 32425752
refGene_gn=LOC101080279;refGene_tx=XM_023252222.1
B1 32425753 32425820
refGene_gn=LOC101080279;refGene_tx=XM_023252222.1
B1 32425822 32426682
refGene_gn=LOC101080279;refGene_tx=XM_023252222.1
B1 32685350 32686549
refGene_gn=CDCA2;refGene_tx=XM_006930628.4;refGene_gn=CDCA2;refGene_tx=XM_006930631.4;refGene_gn=CDCA2;refGene_tx=XM_011281443.3;refGene_gn=CDCA2;refGene_tx=XM_011281444.3;refGene_gn=CDCA2;refGene_tx=XM_019828419.2;refGene_gn=CDCA2;refGene_tx=XM_023252460.1;refGene_gn=CDCA2;refGene_tx=XM_023252461.1
B1 32688366 32688538
refGene_gn=CDCA2;refGene_tx=XM_006930628.4;refGene_gn=CDCA2;refGene_tx=XM_006930631.4;refGene_gn=CDCA2;refGene_tx=XM_011281443.3;refGene_gn=CDCA2;refGene_tx=XM_011281444.3;refGene_gn=CDCA2;refGene_tx=XM_019828419.2;refGene_gn=CDCA2;refGene_tx=XM_023252460.1;refGene_gn=CDCA2;refGene_tx=XM_023252461.1
B1 32698202 32698340
refGene_gn=CDCA2;refGene_tx=XM_006930628.4;refGene_gn=CDCA2;refGene_tx=XM_006930631.4;refGene_gn=CDCA2;refGene_tx=XM_011281443.3;refGene_gn=CDCA2;refGene_tx=XM_011281444.3;refGene_gn=CDCA2;refGene_tx=XM_019828419.2;refGene_gn=CDCA2;refGene_tx=XM_023252460.1;refGene_gn=CDCA2;refGene_tx=XM_023252461.1
B1 32699249 32699338
refGene_gn=CDCA2;refGene_tx=XM_006930628.4;refGene_gn=CDCA2;refGene_tx=XM_006930631.4;refGene_gn=CDCA2;refGene_tx=XM_011281443.3;refGene_gn=CDCA2;refGene_tx=XM_011281444.3;refGene_gn=CDCA2;refGene_tx=XM_019828419.2;refGene_gn=CDCA2;refGene_tx=XM_023252460.1;refGene_gn=CDCA2;refGene_tx=XM_023252461.1
B1 32700896 32700975
refGene_gn=CDCA2;refGene_tx=XM_006930628.4;refGene_gn=CDCA2;refGene_tx=XM_006930631.4;refGene_gn=CDCA2;refGene_tx=XM_011281443.3;refGene_gn=CDCA2;refGene_tx=XM_011281444.3;refGene_gn=CDCA2;refGene_tx=XM_019828419.2;refGene_gn=CDCA2;refGene_tx=XM_023252460.1;refGene_gn=CDCA2;refGene_tx=XM_023252461.1
B1 32705202 32705447
refGene_gn=CDCA2;refGene_tx=XM_006930628.4;refGene_gn=CDCA2;refGene_tx=XM_006930631.4;refGene_gn=CDCA2;refGene_tx=XM_011281443.3;refGene_gn=CDCA2;refGene_tx=XM_011281444.3;refGene_gn=CDCA2;refGene_tx=XM_0198284

19.2;refGene_gn=CDCA2;refGene_tx=XM_023252460.1;refGene_gn=CDCA2;refGene_tx=XM_023252461.1
B1 32705875 32705960
refGene_gn=CDCA2;refGene_tx=XM_006930628.4;refGene_gn=CDCA2;refGene_tx=XM_006930631.4;refGene_gn=CDCA2;refGene_tx=XM_011281443.3;refGene_gn=CDCA2;refGene_tx=XM_011281444.3;refGene_gn=CDCA2;refGene_tx=XM_019828419.2;refGene_gn=CDCA2;refGene_tx=XM_023252460.1;refGene_gn=CDCA2;refGene_tx=XM_023252461.1
B1 32708206 32708415
refGene_gn=CDCA2;refGene_tx=XM_006930628.4;refGene_gn=CDCA2;refGene_tx=XM_006930631.4;refGene_gn=CDCA2;refGene_tx=XM_011281443.3;refGene_gn=CDCA2;refGene_tx=XM_011281444.3;refGene_gn=CDCA2;refGene_tx=XM_019828419.2;refGene_gn=CDCA2;refGene_tx=XM_023252460.1;refGene_gn=CDCA2;refGene_tx=XM_023252461.1
B1 32709228 32709279
refGene_gn=CDCA2;refGene_tx=XM_006930628.4;refGene_gn=CDCA2;refGene_tx=XM_006930631.4;refGene_gn=CDCA2;refGene_tx=XM_011281443.3;refGene_gn=CDCA2;refGene_tx=XM_019828419.2;refGene_gn=CDCA2;refGene_tx=XM_023252460.1
B1 32716345 32716430
refGene_gn=CDCA2;refGene_tx=XM_006930628.4;refGene_gn=CDCA2;refGene_tx=XM_006930631.4;refGene_gn=CDCA2;refGene_tx=XM_011281443.3;refGene_gn=CDCA2;refGene_tx=XM_011281444.3;refGene_gn=CDCA2;refGene_tx=XM_019828419.2;refGene_gn=CDCA2;refGene_tx=XM_023252460.1;refGene_gn=CDCA2;refGene_tx=XM_023252461.1
B1 32717933 32718133
refGene_gn=CDCA2;refGene_tx=XM_006930628.4;refGene_gn=CDCA2;refGene_tx=XM_006930631.4;refGene_gn=CDCA2;refGene_tx=XM_011281443.3;refGene_gn=CDCA2;refGene_tx=XM_011281444.3;refGene_gn=CDCA2;refGene_tx=XM_019828419.2
B1 32719366 32719517
refGene_gn=CDCA2;refGene_tx=XM_006930628.4;refGene_gn=CDCA2;refGene_tx=XM_006930631.4;refGene_gn=CDCA2;refGene_tx=XM_011281443.3;refGene_gn=CDCA2;refGene_tx=XM_011281444.3;refGene_gn=CDCA2;refGene_tx=XM_019828419.2
B1 32732253 32732405
refGene_gn=CDCA2;refGene_tx=XM_006930628.4;refGene_gn=CDCA2;refGene_tx=XM_011281443.3;refGene_gn=CDCA2;refGene_tx=XM_011281444.3;refGene_gn=CDCA2;refGene_tx=XM_023252460.1;refGene_gn=CDCA2;refGene_tx=XM_023252461.1
B1 32733549 32733723
refGene_gn=CDCA2;refGene_tx=XM_006930628.4;refGene_gn=CDCA2;refGene_tx=XM_011281443.3;refGene_gn=CDCA2;refGene_tx=XM_011281444.3;refGene_gn=CDCA2;refGene_tx=XM_023252460.1;refGene_gn=CDCA2;refGene_tx=XM_023252461.1
B1 32733815 32733879
refGene_gn=CDCA2;refGene_tx=XM_006930628.4;refGene_gn=CDCA2;refGene_tx=XM_011281443.3;refGene_gn=CDCA2;refGene_tx=XM_011281444.3;refGene_gn=CDCA2;refGene_tx=XM_023252460.1;refGene_gn=CDCA2;refGene_tx=XM_023252461.1

B1 32735508 32735556 refGene_gn=KCTD9;refGene_tx=XM_003984680.5
B1 32747764 32747886 refGene_gn=KCTD9;refGene_tx=XM_003984680.5
B1 32750450 32750494 refGene_gn=KCTD9;refGene_tx=XM_003984680.5
B1 32753939 32754036 refGene_gn=KCTD9;refGene_tx=XM_003984680.5
B1 32754684 32754743 refGene_gn=KCTD9;refGene_tx=XM_003984680.5
B1 32754947 32755076 refGene_gn=KCTD9;refGene_tx=XM_003984680.5
B1 32759412 32759480 refGene_gn=KCTD9;refGene_tx=XM_003984680.5
B1 32759567 32759663 refGene_gn=KCTD9;refGene_tx=XM_003984680.5
B1 32760587 32760737 refGene_gn=KCTD9;refGene_tx=XM_003984680.5
B1 32767899 32768005 refGene_gn=KCTD9;refGene_tx=XM_003984680.5
B1 32768716 32768850 refGene_gn=KCTD9;refGene_tx=XM_003984680.5
B1 32771083 32771200 refGene_gn=KCTD9;refGene_tx=XM_003984680.5
B1 32777342 32777344 refGene_gn=GNRH1;refGene_tx=XM_023252462.1
B1 32777572 32777714
refGene_gn=GNRH1;refGene_tx=XM_003984681.3;refGene_gn=GNRH1;refGene_tx
=XM_023252462.1
B1 32779858 32779954
refGene_gn=GNRH1;refGene_tx=XM_003984681.3;refGene_gn=GNRH1;refGene_tx
=XM_023252462.1
B1 32781626 32781668
refGene_gn=GNRH1;refGene_tx=XM_003984681.3;refGene_gn=GNRH1;refGene_tx
=XM_023252462.1
B1 32789642 32789747
refGene_gn=DOCK5;refGene_tx=XM_023252463.1;refGene_gn=DOCK5;refGene_tx
=XM_023252465.1
B1 32790328 32790432
refGene_gn=DOCK5;refGene_tx=XM_023252463.1;refGene_gn=DOCK5;refGene_tx
=XM_023252465.1
B1 32791409 32791548
refGene_gn=DOCK5;refGene_tx=XM_023252463.1;refGene_gn=DOCK5;refGene_tx
=XM_023252465.1
B1 32792244 32792413
refGene_gn=DOCK5;refGene_tx=XM_023252463.1;refGene_gn=DOCK5;refGene_tx
=XM_023252465.1
B1 32799456 32799463 refGene_gn=DOCK5;refGene_tx=XM_023252464.1
B1 32800144 32800311
refGene_gn=DOCK5;refGene_tx=XM_023252463.1;refGene_gn=DOCK5;refGene_tx
=XM_023252464.1;refGene_gn=DOCK5;refGene_tx=XM_023252465.1
B1 32802475 32802613
refGene_gn=DOCK5;refGene_tx=XM_023252463.1;refGene_gn=DOCK5;refGene_tx
=XM_023252464.1;refGene_gn=DOCK5;refGene_tx=XM_023252465.1
B1 32803017 32803101
refGene_gn=DOCK5;refGene_tx=XM_023252463.1;refGene_gn=DOCK5;refGene_tx
=XM_023252464.1;refGene_gn=DOCK5;refGene_tx=XM_023252465.1
B1 32807160 32807337
refGene_gn=DOCK5;refGene_tx=XM_023252463.1;refGene_gn=DOCK5;refGene_tx
=XM_023252464.1;refGene_gn=DOCK5;refGene_tx=XM_023252465.1
B1 32809873 32809960
refGene_gn=DOCK5;refGene_tx=XM_023252463.1;refGene_gn=DOCK5;refGene_tx
=XM_023252464.1;refGene_gn=DOCK5;refGene_tx=XM_023252465.1

B1 32810382 32810467
refGene_gn=DOCK5;refGene_tx=XM_023252463.1;refGene_gn=DOCK5;refGene_tx=XM_023252464.1;refGene_gn=DOCK5;refGene_tx=XM_023252465.1
B1 32811483 32811565
refGene_gn=DOCK5;refGene_tx=XM_023252463.1;refGene_gn=DOCK5;refGene_tx=XM_023252464.1;refGene_gn=DOCK5;refGene_tx=XM_023252465.1
B1 32814586 32814728
refGene_gn=DOCK5;refGene_tx=XM_023252463.1;refGene_gn=DOCK5;refGene_tx=XM_023252464.1;refGene_gn=DOCK5;refGene_tx=XM_023252465.1
B1 32820757 32820862
refGene_gn=DOCK5;refGene_tx=XM_023252463.1;refGene_gn=DOCK5;refGene_tx=XM_023252464.1;refGene_gn=DOCK5;refGene_tx=XM_023252465.1
B1 32822568 32822658
refGene_gn=DOCK5;refGene_tx=XM_023252463.1;refGene_gn=DOCK5;refGene_tx=XM_023252464.1;refGene_gn=DOCK5;refGene_tx=XM_023252465.1
B1 32827912 32828032
refGene_gn=DOCK5;refGene_tx=XM_023252463.1;refGene_gn=DOCK5;refGene_tx=XM_023252464.1;refGene_gn=DOCK5;refGene_tx=XM_023252465.1
B1 32831818 32831909
refGene_gn=DOCK5;refGene_tx=XM_023252463.1;refGene_gn=DOCK5;refGene_tx=XM_023252464.1;refGene_gn=DOCK5;refGene_tx=XM_023252465.1
B1 32832771 32832812
refGene_gn=DOCK5;refGene_tx=XM_023252463.1;refGene_gn=DOCK5;refGene_tx=XM_023252464.1;refGene_gn=DOCK5;refGene_tx=XM_023252465.1
B1 32833591 32833751
refGene_gn=DOCK5;refGene_tx=XM_023252463.1;refGene_gn=DOCK5;refGene_tx=XM_023252464.1;refGene_gn=DOCK5;refGene_tx=XM_023252465.1
B1 32836176 32836262
refGene_gn=DOCK5;refGene_tx=XM_023252463.1;refGene_gn=DOCK5;refGene_tx=XM_023252464.1;refGene_gn=DOCK5;refGene_tx=XM_023252465.1
B1 32837025 32837180
refGene_gn=DOCK5;refGene_tx=XM_023252463.1;refGene_gn=DOCK5;refGene_tx=XM_023252464.1;refGene_gn=DOCK5;refGene_tx=XM_023252465.1
B1 32837746 32837805
refGene_gn=DOCK5;refGene_tx=XM_023252463.1;refGene_gn=DOCK5;refGene_tx=XM_023252464.1;refGene_gn=DOCK5;refGene_tx=XM_023252465.1
B1 32841963 32842064
refGene_gn=DOCK5;refGene_tx=XM_023252463.1;refGene_gn=DOCK5;refGene_tx=XM_023252464.1;refGene_gn=DOCK5;refGene_tx=XM_023252465.1
B1 32844123 32844202
refGene_gn=DOCK5;refGene_tx=XM_023252463.1;refGene_gn=DOCK5;refGene_tx=XM_023252464.1;refGene_gn=DOCK5;refGene_tx=XM_023252465.1
B1 32845472 32845567
refGene_gn=DOCK5;refGene_tx=XM_023252463.1;refGene_gn=DOCK5;refGene_tx=XM_023252464.1;refGene_gn=DOCK5;refGene_tx=XM_023252465.1
B1 32849476 32849575
refGene_gn=DOCK5;refGene_tx=XM_023252463.1;refGene_gn=DOCK5;refGene_tx=XM_023252464.1;refGene_gn=DOCK5;refGene_tx=XM_023252465.1
B1 32853743 32853839
refGene_gn=DOCK5;refGene_tx=XM_023252463.1;refGene_gn=DOCK5;refGene_tx

=XM_023252464.1;refGene_gn=DOCK5;refGene_tx=XM_023252465.1
B1	32858381	32858518
refGene_gn=DOCK5;refGene_tx=XM_023252463.1;refGene_gn=DOCK5;refGene_tx=XM_023252464.1;refGene_gn=DOCK5;refGene_tx=XM_023252465.1
B1	32860856	32860963
refGene_gn=DOCK5;refGene_tx=XM_023252463.1;refGene_gn=DOCK5;refGene_tx=XM_023252464.1;refGene_gn=DOCK5;refGene_tx=XM_023252465.1
B1	32861549	32861620
refGene_gn=DOCK5;refGene_tx=XM_023252463.1;refGene_gn=DOCK5;refGene_tx=XM_023252464.1;refGene_gn=DOCK5;refGene_tx=XM_023252465.1
B1	32863090	32863202
refGene_gn=DOCK5;refGene_tx=XM_023252463.1;refGene_gn=DOCK5;refGene_tx=XM_023252464.1;refGene_gn=DOCK5;refGene_tx=XM_023252465.1
B1	32866212	32866347
refGene_gn=DOCK5;refGene_tx=XM_023252463.1;refGene_gn=DOCK5;refGene_tx=XM_023252464.1;refGene_gn=DOCK5;refGene_tx=XM_023252465.1
B1	32868385	32868486
refGene_gn=DOCK5;refGene_tx=XM_023252463.1;refGene_gn=DOCK5;refGene_tx=XM_023252464.1;refGene_gn=DOCK5;refGene_tx=XM_023252465.1
B1	32870634	32870724
refGene_gn=DOCK5;refGene_tx=XM_023252463.1;refGene_gn=DOCK5;refGene_tx=XM_023252464.1;refGene_gn=DOCK5;refGene_tx=XM_023252465.1
B1	32871289	32871387
refGene_gn=DOCK5;refGene_tx=XM_023252463.1;refGene_gn=DOCK5;refGene_tx=XM_023252464.1;refGene_gn=DOCK5;refGene_tx=XM_023252465.1
B1	32879043	32879227
refGene_gn=DOCK5;refGene_tx=XM_023252463.1;refGene_gn=DOCK5;refGene_tx=XM_023252464.1;refGene_gn=DOCK5;refGene_tx=XM_023252465.1
B1	32880481	32880585
refGene_gn=DOCK5;refGene_tx=XM_023252463.1;refGene_gn=DOCK5;refGene_tx=XM_023252464.1;refGene_gn=DOCK5;refGene_tx=XM_023252465.1
B1	32883678	32883751
refGene_gn=DOCK5;refGene_tx=XM_023252463.1;refGene_gn=DOCK5;refGene_tx=XM_023252464.1;refGene_gn=DOCK5;refGene_tx=XM_023252465.1
B1	32885525	32885624
refGene_gn=DOCK5;refGene_tx=XM_023252463.1;refGene_gn=DOCK5;refGene_tx=XM_023252464.1;refGene_gn=DOCK5;refGene_tx=XM_023252465.1
B1	32887187	32887312
refGene_gn=DOCK5;refGene_tx=XM_023252463.1;refGene_gn=DOCK5;refGene_tx=XM_023252464.1;refGene_gn=DOCK5;refGene_tx=XM_023252465.1
B1	32891976	32892102
refGene_gn=DOCK5;refGene_tx=XM_023252463.1;refGene_gn=DOCK5;refGene_tx=XM_023252464.1;refGene_gn=DOCK5;refGene_tx=XM_023252465.1
B1	32894474	32894617
refGene_gn=DOCK5;refGene_tx=XM_023252463.1;refGene_gn=DOCK5;refGene_tx=XM_023252464.1;refGene_gn=DOCK5;refGene_tx=XM_023252465.1
B1	32898611	32898684
refGene_gn=DOCK5;refGene_tx=XM_023252463.1;refGene_gn=DOCK5;refGene_tx=XM_023252464.1;refGene_gn=DOCK5;refGene_tx=XM_023252465.1
B1	32901483	32901613

refGene_gn=DOCK5;refGene_tx=XM_023252463.1;refGene_gn=DOCK5;refGene_tx=XM_023252464.1;refGene_gn=DOCK5;refGene_tx=XM_023252465.1
B1 32905865 32905947
refGene_gn=DOCK5;refGene_tx=XM_023252463.1;refGene_gn=DOCK5;refGene_tx=XM_023252464.1;refGene_gn=DOCK5;refGene_tx=XM_023252465.1
B1 32907086 32907244
refGene_gn=DOCK5;refGene_tx=XM_023252463.1;refGene_gn=DOCK5;refGene_tx=XM_023252464.1;refGene_gn=DOCK5;refGene_tx=XM_023252465.1
B1 32908581 32908662 refGene_gn=DOCK5;refGene_tx=XM_023252465.1
B1 32909386 32909522
refGene_gn=DOCK5;refGene_tx=XM_023252463.1;refGene_gn=DOCK5;refGene_tx=XM_023252464.1
B1 32912731 32912880
refGene_gn=DOCK5;refGene_tx=XM_023252463.1;refGene_gn=DOCK5;refGene_tx=XM_023252464.1
B1 32923665 32923762
refGene_gn=DOCK5;refGene_tx=XM_023252463.1;refGene_gn=DOCK5;refGene_tx=XM_023252464.1
B1 32925821 32925877
refGene_gn=DOCK5;refGene_tx=XM_023252463.1;refGene_gn=DOCK5;refGene_tx=XM_023252464.1
B1 32931060 32931101
refGene_gn=DOCK5;refGene_tx=XM_023252463.1;refGene_gn=DOCK5;refGene_tx=XM_023252464.1
B1 32952905 32952989
refGene_gn=DOCK5;refGene_tx=XM_023252463.1;refGene_gn=DOCK5;refGene_tx=XM_023252464.1
B1 33014969 33015012
refGene_gn=DOCK5;refGene_tx=XM_023252463.1;refGene_gn=DOCK5;refGene_tx=XM_023252464.1
B1 33235742 33236789 refGene_gn=NEFL;refGene_tx=XM_003984682.5
B1 33237861 33237986 refGene_gn=NEFL;refGene_tx=XM_003984682.5
B1 33238458 33238778 refGene_gn=NEFL;refGene_tx=XM_003984682.5
B1 33239342 33239482 refGene_gn=NEFL;refGene_tx=XM_003984682.5
B1 33272837 33274218 refGene_gn=NEFM;refGene_tx=XM_023252466.1
B1 33275515 33275640 refGene_gn=NEFM;refGene_tx=XM_023252466.1
B1 33276361 33277441 refGene_gn=NEFM;refGene_tx=XM_023252466.1
B1 33767790 33767835
refGene_gn=ADAM7;refGene_tx=XM_023252467.1;refGene_gn=ADAM7;refGene_tx=XM_023252468.1
B1 33769240 33769297
refGene_gn=ADAM7;refGene_tx=XM_023252467.1;refGene_gn=ADAM7;refGene_tx=XM_023252468.1;refGene_gn=ADAM7;refGene_tx=XM_023252469.1
B1 33769957 33770023
refGene_gn=ADAM7;refGene_tx=XM_023252467.1;refGene_gn=ADAM7;refGene_tx=XM_023252469.1
B1 33772813 33772913
refGene_gn=ADAM7;refGene_tx=XM_023252467.1;refGene_gn=ADAM7;refGene_tx=XM_023252468.1;refGene_gn=ADAM7;refGene_tx=XM_023252469.1
B1 33773280 33773386

refGene_gn=ADAM7;refGene_tx=XM_023252467.1;refGene_gn=ADAM7;refGene_tx=XM_023252468.1;refGene_gn=ADAM7;refGene_tx=XM_023252469.1
B1 33773889 33773968
refGene_gn=ADAM7;refGene_tx=XM_023252467.1;refGene_gn=ADAM7;refGene_tx=XM_023252468.1;refGene_gn=ADAM7;refGene_tx=XM_023252469.1
B1 33775221 33775302
refGene_gn=ADAM7;refGene_tx=XM_023252467.1;refGene_gn=ADAM7;refGene_tx=XM_023252468.1;refGene_gn=ADAM7;refGene_tx=XM_023252469.1
B1 33781867 33782063
refGene_gn=ADAM7;refGene_tx=XM_023252467.1;refGene_gn=ADAM7;refGene_tx=XM_023252468.1;refGene_gn=ADAM7;refGene_tx=XM_023252469.1
B1 33783062 33783165
refGene_gn=ADAM7;refGene_tx=XM_023252467.1;refGene_gn=ADAM7;refGene_tx=XM_023252468.1;refGene_gn=ADAM7;refGene_tx=XM_023252469.1
B1 33783581 33783777
refGene_gn=ADAM7;refGene_tx=XM_023252467.1;refGene_gn=ADAM7;refGene_tx=XM_023252468.1;refGene_gn=ADAM7;refGene_tx=XM_023252469.1
B1 33784780 33784870
refGene_gn=ADAM7;refGene_tx=XM_023252467.1;refGene_gn=ADAM7;refGene_tx=XM_023252468.1;refGene_gn=ADAM7;refGene_tx=XM_023252469.1
B1 33786793 33786968
refGene_gn=ADAM7;refGene_tx=XM_023252467.1;refGene_gn=ADAM7;refGene_tx=XM_023252468.1;refGene_gn=ADAM7;refGene_tx=XM_023252469.1
B1 33788925 33789056
refGene_gn=ADAM7;refGene_tx=XM_023252467.1;refGene_gn=ADAM7;refGene_tx=XM_023252468.1;refGene_gn=ADAM7;refGene_tx=XM_023252469.1
B1 33790665 33790750
refGene_gn=ADAM7;refGene_tx=XM_023252467.1;refGene_gn=ADAM7;refGene_tx=XM_023252468.1;refGene_gn=ADAM7;refGene_tx=XM_023252469.1
B1 33793051 33793221
refGene_gn=ADAM7;refGene_tx=XM_023252467.1;refGene_gn=ADAM7;refGene_tx=XM_023252468.1;refGene_gn=ADAM7;refGene_tx=XM_023252469.1
B1 33801124 33801196
refGene_gn=ADAM7;refGene_tx=XM_023252467.1;refGene_gn=ADAM7;refGene_tx=XM_023252468.1;refGene_gn=ADAM7;refGene_tx=XM_023252469.1
B1 33806868 33806922
refGene_gn=ADAM7;refGene_tx=XM_023252467.1;refGene_gn=ADAM7;refGene_tx=XM_023252468.1;refGene_gn=ADAM7;refGene_tx=XM_023252469.1
B1 33808778 33808971
refGene_gn=ADAM7;refGene_tx=XM_023252467.1;refGene_gn=ADAM7;refGene_tx=XM_023252468.1;refGene_gn=ADAM7;refGene_tx=XM_023252469.1
B1 33809931 33810008
refGene_gn=ADAM7;refGene_tx=XM_023252467.1;refGene_gn=ADAM7;refGene_tx=XM_023252468.1;refGene_gn=ADAM7;refGene_tx=XM_023252469.1
B1 33811547 33811626
refGene_gn=ADAM7;refGene_tx=XM_023252467.1;refGene_gn=ADAM7;refGene_tx=XM_023252468.1;refGene_gn=ADAM7;refGene_tx=XM_023252469.1
B1 33826681 33826758
refGene_gn=ADAM7;refGene_tx=XM_023252467.1;refGene_gn=ADAM7;refGene_tx=XM_023252468.1;refGene_gn=ADAM7;refGene_tx=XM_023252469.1

B1	33831251	33831355
refGene_gn=ADAM7;refGene_tx=XM_023252467.1;refGene_gn=ADAM7;refGene_tx
=XM_023252468.1;refGene_gn=ADAM7;refGene_tx=XM_023252469.1
B1	33832069	33832121
refGene_gn=ADAM7;refGene_tx=XM_023252467.1;refGene_gn=ADAM7;refGene_tx
=XM_023252468.1;refGene_gn=ADAM7;refGene_tx=XM_023252469.1
B1	33878338	33878345	refGene_gn=ADAMDEC1;refGene_tx=XM_019829988.2
B1	33879627	33879713	refGene_gn=ADAMDEC1;refGene_tx=XM_019829988.2
B1	33883397	33883575	refGene_gn=ADAMDEC1;refGene_tx=XM_019829988.2
B1	33885028	33885159	refGene_gn=ADAMDEC1;refGene_tx=XM_019829988.2
B1	33885877	33885959	refGene_gn=ADAMDEC1;refGene_tx=XM_019829988.2
B1	33886318	33886488	refGene_gn=ADAMDEC1;refGene_tx=XM_019829988.2
B1	33886782	33886854	refGene_gn=ADAMDEC1;refGene_tx=XM_019829988.2
B1	33887751	33887814	refGene_gn=ADAMDEC1;refGene_tx=XM_019829988.2
B1	33888030	33888217	refGene_gn=ADAMDEC1;refGene_tx=XM_019829988.2
B1	33889274	33889351	refGene_gn=ADAMDEC1;refGene_tx=XM_019829988.2
B1	33890810	33890889	refGene_gn=ADAMDEC1;refGene_tx=XM_019829988.2
B1	33892070	33892147	refGene_gn=ADAMDEC1;refGene_tx=XM_019829988.2
B1	33893592	33893708	refGene_gn=ADAMDEC1;refGene_tx=XM_019829988.2
B1	33895436	33895485	refGene_gn=ADAMDEC1;refGene_tx=XM_019829988.2
B1	33926597	33926618
refGene_gn=ADAM28;refGene_tx=XM_006930633.4;refGene_gn=ADAM28;refGene_
tx=XM_023252470.1;refGene_gn=ADAM28;refGene_tx=XM_023252471.1
B1	33927414	33927474
refGene_gn=ADAM28;refGene_tx=XM_006930633.4;refGene_gn=ADAM28;refGene_
tx=XM_023252470.1;refGene_gn=ADAM28;refGene_tx=XM_023252471.1
B1	33928752	33928818
refGene_gn=ADAM28;refGene_tx=XM_006930633.4;refGene_gn=ADAM28;refGene_
tx=XM_023252470.1;refGene_gn=ADAM28;refGene_tx=XM_023252471.1
B1	33929331	33929397
refGene_gn=ADAM28;refGene_tx=XM_006930633.4;refGene_gn=ADAM28;refGene_
tx=XM_023252470.1;refGene_gn=ADAM28;refGene_tx=XM_023252471.1;refGene_
gn=ADAM28;refGene_tx=XM_023252472.1
B1	33930069	33930148
refGene_gn=ADAM28;refGene_tx=XM_006930633.4;refGene_gn=ADAM28;refGene_
tx=XM_023252470.1;refGene_gn=ADAM28;refGene_tx=XM_023252471.1;refGene_
gn=ADAM28;refGene_tx=XM_023252472.1
B1	33931181	33931290
refGene_gn=ADAM28;refGene_tx=XM_006930633.4;refGene_gn=ADAM28;refGene_
tx=XM_023252470.1;refGene_gn=ADAM28;refGene_tx=XM_023252471.1;refGene_
gn=ADAM28;refGene_tx=XM_023252472.1;refGene_gn=ADAM28;refGene_tx=XM_02
3252473.1
B1	33945029	33945112
refGene_gn=ADAM28;refGene_tx=XM_006930633.4;refGene_gn=ADAM28;refGene_
tx=XM_023252470.1;refGene_gn=ADAM28;refGene_tx=XM_023252471.1;refGene_
gn=ADAM28;refGene_tx=XM_023252472.1
B1	33945420	33945501
refGene_gn=ADAM28;refGene_tx=XM_006930633.4;refGene_gn=ADAM28;refGene_
tx=XM_023252470.1;refGene_gn=ADAM28;refGene_tx=XM_023252471.1;refGene_
gn=ADAM28;refGene_tx=XM_023252472.1;refGene_gn=ADAM28;refGene_tx=XM_02

3252473.1
B1 33946454 33946614
refGene_gn=ADAM28;refGene_tx=XM_006930633.4;refGene_gn=ADAM28;refGene_tx=XM_023252470.1;refGene_gn=ADAM28;refGene_tx=XM_023252471.1;refGene_gn=ADAM28;refGene_tx=XM_023252472.1;refGene_gn=ADAM28;refGene_tx=XM_023252473.1
B1 33949495 33949598
refGene_gn=ADAM28;refGene_tx=XM_006930633.4;refGene_gn=ADAM28;refGene_tx=XM_023252470.1;refGene_gn=ADAM28;refGene_tx=XM_023252471.1;refGene_gn=ADAM28;refGene_tx=XM_023252472.1;refGene_gn=ADAM28;refGene_tx=XM_023252473.1
B1 33954495 33954691
refGene_gn=ADAM28;refGene_tx=XM_006930633.4;refGene_gn=ADAM28;refGene_tx=XM_023252470.1;refGene_gn=ADAM28;refGene_tx=XM_023252471.1;refGene_gn=ADAM28;refGene_tx=XM_023252472.1;refGene_gn=ADAM28;refGene_tx=XM_023252473.1
B1 33956929 33957019
refGene_gn=ADAM28;refGene_tx=XM_006930633.4;refGene_gn=ADAM28;refGene_tx=XM_023252470.1;refGene_gn=ADAM28;refGene_tx=XM_023252471.1;refGene_gn=ADAM28;refGene_tx=XM_023252472.1;refGene_gn=ADAM28;refGene_tx=XM_023252473.1
B1 33959274 33959452
refGene_gn=ADAM28;refGene_tx=XM_006930633.4;refGene_gn=ADAM28;refGene_tx=XM_023252470.1;refGene_gn=ADAM28;refGene_tx=XM_023252471.1;refGene_gn=ADAM28;refGene_tx=XM_023252472.1;refGene_gn=ADAM28;refGene_tx=XM_023252473.1
B1 33960224 33960355
refGene_gn=ADAM28;refGene_tx=XM_006930633.4;refGene_gn=ADAM28;refGene_tx=XM_023252470.1;refGene_gn=ADAM28;refGene_tx=XM_023252471.1;refGene_gn=ADAM28;refGene_tx=XM_023252472.1;refGene_gn=ADAM28;refGene_tx=XM_023252473.1
B1 33968315 33968400
refGene_gn=ADAM28;refGene_tx=XM_006930633.4;refGene_gn=ADAM28;refGene_tx=XM_023252470.1;refGene_gn=ADAM28;refGene_tx=XM_023252471.1;refGene_gn=ADAM28;refGene_tx=XM_023252472.1;refGene_gn=ADAM28;refGene_tx=XM_023252473.1
B1 33971301 33971471
refGene_gn=ADAM28;refGene_tx=XM_006930633.4;refGene_gn=ADAM28;refGene_tx=XM_023252470.1;refGene_gn=ADAM28;refGene_tx=XM_023252471.1;refGene_gn=ADAM28;refGene_tx=XM_023252472.1;refGene_gn=ADAM28;refGene_tx=XM_023252473.1
B1 33976666 33976738
refGene_gn=ADAM28;refGene_tx=XM_006930633.4;refGene_gn=ADAM28;refGene_tx=XM_023252472.1;refGene_gn=ADAM28;refGene_tx=XM_023252473.1
B1 33977646 33977718
refGene_gn=ADAM28;refGene_tx=XM_006930633.4;refGene_gn=ADAM28;refGene_tx=XM_023252470.1;refGene_gn=ADAM28;refGene_tx=XM_023252472.1;refGene_gn=ADAM28;refGene_tx=XM_023252473.1
B1 33980391 33980584
refGene_gn=ADAM28;refGene_tx=XM_006930633.4;refGene_gn=ADAM28;refGene_

tx=XM_023252470.1;refGene_gn=ADAM28;refGene_tx=XM_023252471.1;refGene_gn=ADAM28;refGene_tx=XM_023252472.1;refGene_gn=ADAM28;refGene_tx=XM_023252473.1
B1 33982361 33982438 refGene_gn=ADAM28;refGene_tx=XM_006930633.4;refGene_gn=ADAM28;refGene_tx=XM_023252470.1;refGene_gn=ADAM28;refGene_tx=XM_023252471.1;refGene_gn=ADAM28;refGene_tx=XM_023252472.1;refGene_gn=ADAM28;refGene_tx=XM_023252473.1
B1 33983844 33983923 refGene_gn=ADAM28;refGene_tx=XM_006930633.4;refGene_gn=ADAM28;refGene_tx=XM_023252470.1;refGene_gn=ADAM28;refGene_tx=XM_023252471.1;refGene_gn=ADAM28;refGene_tx=XM_023252472.1;refGene_gn=ADAM28;refGene_tx=XM_023252473.1
B1 33986700 33986777 refGene_gn=ADAM28;refGene_tx=XM_006930633.4;refGene_gn=ADAM28;refGene_tx=XM_023252470.1;refGene_gn=ADAM28;refGene_tx=XM_023252471.1;refGene_gn=ADAM28;refGene_tx=XM_023252472.1;refGene_gn=ADAM28;refGene_tx=XM_023252473.1
B1 33993636 33993740 refGene_gn=ADAM28;refGene_tx=XM_006930633.4;refGene_gn=ADAM28;refGene_tx=XM_023252470.1;refGene_gn=ADAM28;refGene_tx=XM_023252471.1;refGene_gn=ADAM28;refGene_tx=XM_023252472.1;refGene_gn=ADAM28;refGene_tx=XM_023252473.1
B1 33999312 33999370 refGene_gn=ADAM28;refGene_tx=XM_006930633.4;refGene_gn=ADAM28;refGene_tx=XM_023252470.1;refGene_gn=ADAM28;refGene_tx=XM_023252471.1;refGene_gn=ADAM28;refGene_tx=XM_023252472.1;refGene_gn=ADAM28;refGene_tx=XM_023252473.1
B1 34154267 34154291 refGene_gn=LOC102901501;refGene_tx=XM_023253387.1
B1 34154292 34154363 refGene_gn=LOC102901501;refGene_tx=XM_023253387.1
B1 34154365 34154533 refGene_gn=LOC102901501;refGene_tx=XM_023253387.1
B1 34154535 34154827 refGene_gn=LOC102901501;refGene_tx=XM_023253387.1
B1 34154828 34155187 refGene_gn=LOC102901501;refGene_tx=XM_023253387.1
B1 34155189 34155320 refGene_gn=LOC102901501;refGene_tx=XM_023253387.1
B1 34155595 34155776 refGene_gn=LOC102901501;refGene_tx=XM_023253387.1
B1 34155778 34156012 refGene_gn=LOC102901501;refGene_tx=XM_023253387.1
B1 34156014 34156197 refGene_gn=LOC102901501;refGene_tx=XM_023253387.1
B1 34156199 34156285 refGene_gn=LOC102901501;refGene_tx=XM_023253387.1
B1 34156286 34156390 refGene_gn=LOC102901501;refGene_tx=XM_023253387.1

B1 34156391 34156430
refGene_gn=LOC102901501;refGene_tx=XM_023253387.1
B1 34156679 34157206
refGene_gn=LOC102901501;refGene_tx=XM_023253387.1
B1 34157208 34157328
refGene_gn=LOC102901501;refGene_tx=XM_023253387.1
B1 34157330 34157489
refGene_gn=LOC102901501;refGene_tx=XM_023253387.1
B1 34157491 34157547
refGene_gn=LOC102901501;refGene_tx=XM_023253387.1
B1 34283236 34283295
refGene_gn=LOC102901501;refGene_tx=XM_023253387.1
B1 34302613 34302806
refGene_gn=LOC102901501;refGene_tx=XM_023253387.1
B1 34322044 34322247
refGene_gn=LOC102901501;refGene_tx=XM_023253387.1
B1 34515675 34515793
refGene_gn=STC1;refGene_tx=XM_003984690.5;refGene_gn=STC1;refGene_tx=XM_006930637.4
B1 34517795 34517938
refGene_gn=STC1;refGene_tx=XM_003984690.5;refGene_gn=STC1;refGene_tx=XM_006930637.4
B1 34518655 34518867
refGene_gn=STC1;refGene_tx=XM_003984690.5;refGene_gn=STC1;refGene_tx=XM_006930637.4
B1 34524997 34525268
refGene_gn=STC1;refGene_tx=XM_003984690.5;refGene_gn=STC1;refGene_tx=XM_006930637.4
B1 34525403 34525522 refGene_gn=STC1;refGene_tx=XM_006930637.4
B1 34677424 34677704 refGene_gn=NKX2-6;refGene_tx=XM_023252223.1
B1 34680642 34681268 refGene_gn=NKX2-6;refGene_tx=XM_023252223.1
B1 34712176 34712453 refGene_gn=NKX3-1;refGene_tx=XM_023252474.1
B1 34713545 34713964 refGene_gn=NKX3-1;refGene_tx=XM_023252474.1
B1 34822106 34822627
refGene_gn=SLC25A37;refGene_tx=XM_003984691.5;refGene_gn=SLC25A37;refGene_tx=XM_019828422.2;refGene_gn=SLC25A37;refGene_tx=XM_019828423.2;refGene_gn=SLC25A37;refGene_tx=XM_019828424.2;refGene_gn=SLC25A37;refGene_tx=XM_019828425.2;refGene_gn=SLC25A37;refGene_tx=XM_019828426.2;refGene_gn=SLC25A37;refGene_tx=XM_019828427.2;refGene_gn=SLC25A37;refGene_tx=XM_019828428.2;refGene_gn=SLC25A37;refGene_tx=XM_023252475.1
B1 34824699 34824756
refGene_gn=SLC25A37;refGene_tx=XM_003984691.5;refGene_gn=SLC25A37;refGene_tx=XM_019828422.2;refGene_gn=SLC25A37;refGene_tx=XM_019828423.2;refGene_gn=SLC25A37;refGene_tx=XM_019828424.2;refGene_gn=SLC25A37;refGene_tx=XM_019828425.2;refGene_gn=SLC25A37;refGene_tx=XM_019828426.2;refGene_gn=SLC25A37;refGene_tx=XM_019828427.2;refGene_gn=SLC25A37;refGene_tx=XM_019828428.2;refGene_gn=SLC25A37;refGene_tx=XM_023252475.1
B1 34825560 34825597 refGene_gn=SLC25A37;refGene_tx=XM_019828426.2
B1 34826793 34827022
refGene_gn=SLC25A37;refGene_tx=XM_003984691.5;refGene_gn=SLC25A37;refG

ene_tx=XM_019828422.2;refGene_gn=SLC25A37;refGene_tx=XM_019828423.2;refGene_gn=SLC25A37;refGene_tx=XM_019828424.2;refGene_gn=SLC25A37;refGene_tx=XM_019828425.2;refGene_gn=SLC25A37;refGene_tx=XM_023252475.1
B1 34861324 34861546 refGene_gn=SLC25A37;refGene_tx=XM_003984691.5
B1 34938316 34938324
refGene_gn=ENTPD4;refGene_tx=XM_023252477.1;refGene_gn=ENTPD4;refGene_tx=XM_023252478.1;refGene_gn=ENTPD4;refGene_tx=XM_023252479.1;refGene_gn=ENTPD4;refGene_tx=XM_023252480.1;refGene_gn=ENTPD4;refGene_tx=XM_023252481.1;refGene_gn=ENTPD4;refGene_tx=XM_023252482.1;refGene_gn=ENTPD4;refGene_tx=XM_023252483.1
B1 34940029 34940227
refGene_gn=ENTPD4;refGene_tx=XM_023252477.1;refGene_gn=ENTPD4;refGene_tx=XM_023252478.1;refGene_gn=ENTPD4;refGene_tx=XM_023252479.1;refGene_gn=ENTPD4;refGene_tx=XM_023252480.1;refGene_gn=ENTPD4;refGene_tx=XM_023252481.1;refGene_gn=ENTPD4;refGene_tx=XM_023252482.1;refGene_gn=ENTPD4;refGene_tx=XM_023252483.1
B1 34940737 34940943
refGene_gn=ENTPD4;refGene_tx=XM_023252477.1;refGene_gn=ENTPD4;refGene_tx=XM_023252478.1;refGene_gn=ENTPD4;refGene_tx=XM_023252479.1;refGene_gn=ENTPD4;refGene_tx=XM_023252480.1;refGene_gn=ENTPD4;refGene_tx=XM_023252481.1;refGene_gn=ENTPD4;refGene_tx=XM_023252482.1;refGene_gn=ENTPD4;refGene_tx=XM_023252483.1
B1 34943476 34943627
refGene_gn=ENTPD4;refGene_tx=XM_023252477.1;refGene_gn=ENTPD4;refGene_tx=XM_023252478.1;refGene_gn=ENTPD4;refGene_tx=XM_023252479.1;refGene_gn=ENTPD4;refGene_tx=XM_023252480.1;refGene_gn=ENTPD4;refGene_tx=XM_023252481.1;refGene_gn=ENTPD4;refGene_tx=XM_023252482.1;refGene_gn=ENTPD4;refGene_tx=XM_023252483.1
B1 34944399 34944503
refGene_gn=ENTPD4;refGene_tx=XM_023252477.1;refGene_gn=ENTPD4;refGene_tx=XM_023252478.1;refGene_gn=ENTPD4;refGene_tx=XM_023252479.1;refGene_gn=ENTPD4;refGene_tx=XM_023252480.1;refGene_gn=ENTPD4;refGene_tx=XM_023252481.1;refGene_gn=ENTPD4;refGene_tx=XM_023252482.1;refGene_gn=ENTPD4;refGene_tx=XM_023252483.1
B1 34953083 34953143
refGene_gn=ENTPD4;refGene_tx=XM_023252477.1;refGene_gn=ENTPD4;refGene_tx=XM_023252478.1;refGene_gn=ENTPD4;refGene_tx=XM_023252479.1;refGene_gn=ENTPD4;refGene_tx=XM_023252480.1;refGene_gn=ENTPD4;refGene_tx=XM_023252481.1;refGene_gn=ENTPD4;refGene_tx=XM_023252482.1;refGene_gn=ENTPD4;refGene_tx=XM_023252483.1
B1 34953420 34953575
refGene_gn=ENTPD4;refGene_tx=XM_023252477.1;refGene_gn=ENTPD4;refGene_tx=XM_023252478.1;refGene_gn=ENTPD4;refGene_tx=XM_023252479.1;refGene_gn=ENTPD4;refGene_tx=XM_023252480.1;refGene_gn=ENTPD4;refGene_tx=XM_023252481.1;refGene_gn=ENTPD4;refGene_tx=XM_023252482.1;refGene_gn=ENTPD4;refGene_tx=XM_023252483.1
B1 34955238 34955405
refGene_gn=ENTPD4;refGene_tx=XM_023252477.1;refGene_gn=ENTPD4;refGene_tx=XM_023252478.1;refGene_gn=ENTPD4;refGene_tx=XM_023252479.1;refGene_gn=ENTPD4;refGene_tx=XM_023252480.1;refGene_gn=ENTPD4;refGene_tx=XM_02

3252481.1;refGene_gn=ENTPD4;refGene_tx=XM_023252482.1;refGene_gn=ENTPD4;refGene_tx=XM_023252483.1
B1 34958577 34958902
refGene_gn=ENTPD4;refGene_tx=XM_023252477.1;refGene_gn=ENTPD4;refGene_tx=XM_023252478.1;refGene_gn=ENTPD4;refGene_tx=XM_023252479.1;refGene_gn=ENTPD4;refGene_tx=XM_023252480.1;refGene_gn=ENTPD4;refGene_tx=XM_023252481.1;refGene_gn=ENTPD4;refGene_tx=XM_023252482.1;refGene_gn=ENTPD4;refGene_tx=XM_023252483.1
B1 34961139 34961225
refGene_gn=ENTPD4;refGene_tx=XM_023252477.1;refGene_gn=ENTPD4;refGene_tx=XM_023252478.1;refGene_gn=ENTPD4;refGene_tx=XM_023252479.1;refGene_gn=ENTPD4;refGene_tx=XM_023252480.1;refGene_gn=ENTPD4;refGene_tx=XM_023252481.1;refGene_gn=ENTPD4;refGene_tx=XM_023252482.1;refGene_gn=ENTPD4;refGene_tx=XM_023252483.1
B1 34961441 34961495
refGene_gn=ENTPD4;refGene_tx=XM_023252477.1;refGene_gn=ENTPD4;refGene_tx=XM_023252478.1;refGene_gn=ENTPD4;refGene_tx=XM_023252479.1;refGene_gn=ENTPD4;refGene_tx=XM_023252482.1;refGene_gn=ENTPD4;refGene_tx=XM_023252483.1
B1 34962100 34962262
refGene_gn=ENTPD4;refGene_tx=XM_023252477.1;refGene_gn=ENTPD4;refGene_tx=XM_023252478.1;refGene_gn=ENTPD4;refGene_tx=XM_023252479.1;refGene_gn=ENTPD4;refGene_tx=XM_023252480.1;refGene_gn=ENTPD4;refGene_tx=XM_023252481.1;refGene_gn=ENTPD4;refGene_tx=XM_023252482.1;refGene_gn=ENTPD4;refGene_tx=XM_023252483.1
B1 34963456 34963679
refGene_gn=ENTPD4;refGene_tx=XM_023252477.1;refGene_gn=ENTPD4;refGene_tx=XM_023252478.1;refGene_gn=ENTPD4;refGene_tx=XM_023252479.1;refGene_gn=ENTPD4;refGene_tx=XM_023252480.1;refGene_gn=ENTPD4;refGene_tx=XM_023252481.1
B1 34973353 34973384 refGene_gn=ENTPD4;refGene_tx=XM_023252482.1
B1 34973656 34973678 refGene_gn=ENTPD4;refGene_tx=XM_023252483.1
B1 35028499 35028854 refGene_gn=LOXL2;refGene_tx=XM_003984692.4
B1 35035119 35035295 refGene_gn=LOXL2;refGene_tx=XM_003984692.4
B1 35049297 35049509 refGene_gn=LOXL2;refGene_tx=XM_003984692.4
B1 35056041 35056264 refGene_gn=LOXL2;refGene_tx=XM_003984692.4
B1 35059854 35060038 refGene_gn=LOXL2;refGene_tx=XM_003984692.4
B1 35066215 35066367 refGene_gn=LOXL2;refGene_tx=XM_003984692.4
B1 35072907 35073075 refGene_gn=LOXL2;refGene_tx=XM_003984692.4
B1 35075146 35075309 refGene_gn=LOXL2;refGene_tx=XM_003984692.4
B1 35080197 35080441 refGene_gn=LOXL2;refGene_tx=XM_003984692.4
B1 35086317 35086433 refGene_gn=LOXL2;refGene_tx=XM_003984692.4
B1 35087576 35087713 refGene_gn=LOXL2;refGene_tx=XM_003984692.4
B1 35089559 35089671 refGene_gn=LOXL2;refGene_tx=XM_003984692.4
B1 35090420 35090500 refGene_gn=LOXL2;refGene_tx=XM_003984692.4
B1 35092543 35092680
refGene_gn=R3HCC1;refGene_tx=XM_023252484.1;refGene_gn=R3HCC1;refGene_tx=XM_023252485.1;refGene_gn=R3HCC1;refGene_tx=XM_023252486.1
B1 35093682 35093778
refGene_gn=R3HCC1;refGene_tx=XM_023252484.1;refGene_gn=R3HCC1;refGene_

tx=XM_023252485.1
B1 35095105 35095176
refGene_gn=R3HCC1;refGene_tx=XM_023252484.1;refGene_gn=R3HCC1;refGene_tx=XM_023252485.1;refGene_gn=R3HCC1;refGene_tx=XM_023252486.1
B1 35096592 35096765
refGene_gn=R3HCC1;refGene_tx=XM_023252484.1;refGene_gn=R3HCC1;refGene_tx=XM_023252485.1;refGene_gn=R3HCC1;refGene_tx=XM_023252486.1
B1 35097744 35098339
refGene_gn=R3HCC1;refGene_tx=XM_023252484.1;refGene_gn=R3HCC1;refGene_tx=XM_023252485.1;refGene_gn=R3HCC1;refGene_tx=XM_023252486.1
B1 35098989 35099127
refGene_gn=R3HCC1;refGene_tx=XM_023252484.1;refGene_gn=R3HCC1;refGene_tx=XM_023252485.1;refGene_gn=R3HCC1;refGene_tx=XM_023252486.1
B1 35099540 35099668
refGene_gn=R3HCC1;refGene_tx=XM_023252484.1;refGene_gn=R3HCC1;refGene_tx=XM_023252485.1;refGene_gn=R3HCC1;refGene_tx=XM_023252486.1
B1 35109310 35109316
refGene_gn=R3HCC1;refGene_tx=XM_023252484.1;refGene_gn=R3HCC1;refGene_tx=XM_023252486.1
B1 35111500 35111524 refGene_gn=R3HCC1;refGene_tx=XM_023252485.1
B1 35140075 35140128 refGene_gn=CHMP7;refGene_tx=XM_003984693.5
B1 35140360 35140540 refGene_gn=CHMP7;refGene_tx=XM_003984693.5
B1 35141026 35141087 refGene_gn=CHMP7;refGene_tx=XM_003984693.5
B1 35141317 35141416 refGene_gn=CHMP7;refGene_tx=XM_003984693.5
B1 35141702 35141822 refGene_gn=CHMP7;refGene_tx=XM_003984693.5
B1 35142072 35142121 refGene_gn=CHMP7;refGene_tx=XM_003984693.5
B1 35142993 35143127 refGene_gn=CHMP7;refGene_tx=XM_003984693.5
B1 35143263 35143449 refGene_gn=CHMP7;refGene_tx=XM_003984693.5
B1 35149426 35149598 refGene_gn=CHMP7;refGene_tx=XM_003984693.5
B1 35151590 35151889 refGene_gn=CHMP7;refGene_tx=XM_003984693.5
B1 35213284 35213502
refGene_gn=RHOBTB2;refGene_tx=XM_006930639.4;refGene_gn=RHOBTB2;refGene_tx=XM_023252487.1
B1 35214807 35214913
refGene_gn=RHOBTB2;refGene_tx=XM_006930639.4;refGene_gn=RHOBTB2;refGene_tx=XM_023252487.1
B1 35215732 35215821
refGene_gn=RHOBTB2;refGene_tx=XM_006930639.4;refGene_gn=RHOBTB2;refGene_tx=XM_023252487.1
B1 35219546 35219697
refGene_gn=RHOBTB2;refGene_tx=XM_006930639.4;refGene_gn=RHOBTB2;refGene_tx=XM_023252487.1
B1 35221353 35221472
refGene_gn=RHOBTB2;refGene_tx=XM_006930639.4;refGene_gn=RHOBTB2;refGene_tx=XM_023252487.1
B1 35221779 35222801
refGene_gn=RHOBTB2;refGene_tx=XM_006930639.4;refGene_gn=RHOBTB2;refGene_tx=XM_023252487.1
B1 35223429 35223615
refGene_gn=RHOBTB2;refGene_tx=XM_006930639.4;refGene_gn=RHOBTB2;refGen

e_tx=XM_023252487.1
B1 35224114 35224218
refGene_gn=RHOBTB2;refGene_tx=XM_006930639.4;refGene_gn=RHOBTB2;refGene_tx=XM_023252487.1
B1 35224906 35225108
refGene_gn=RHOBTB2;refGene_tx=XM_006930639.4;refGene_gn=RHOBTB2;refGene_tx=XM_023252487.1
B1 35232705 35232761 refGene_gn=RHOBTB2;refGene_tx=XM_023252487.1
B1 35286917 35287048
refGene_gn=PEBP4;refGene_tx=XM_003984698.4;refGene_gn=PEBP4;refGene_tx=XM_006930640.4;refGene_gn=PEBP4;refGene_tx=XM_019828434.2;refGene_gn=PEBP4;refGene_tx=XM_019828435.2
B1 35294113 35294240
refGene_gn=PEBP4;refGene_tx=XM_003984698.4;refGene_gn=PEBP4;refGene_tx=XM_006930640.4;refGene_gn=PEBP4;refGene_tx=XM_019828434.2;refGene_gn=PEBP4;refGene_tx=XM_019828435.2
B1 35393472 35393571
refGene_gn=PEBP4;refGene_tx=XM_003984698.4;refGene_gn=PEBP4;refGene_tx=XM_006930640.4;refGene_gn=PEBP4;refGene_tx=XM_019828434.2;refGene_gn=PEBP4;refGene_tx=XM_019828435.2
B1 35475962 35476008
refGene_gn=PEBP4;refGene_tx=XM_003984698.4;refGene_gn=PEBP4;refGene_tx=XM_006930640.4;refGene_gn=PEBP4;refGene_tx=XM_019828434.2;refGene_gn=PEBP4;refGene_tx=XM_019828435.2
B1 35478265 35478379
refGene_gn=PEBP4;refGene_tx=XM_003984698.4;refGene_gn=PEBP4;refGene_tx=XM_006930640.4;refGene_gn=PEBP4;refGene_tx=XM_019828434.2;refGene_gn=PEBP4;refGene_tx=XM_019828435.2
B1 35489156 35489311
refGene_gn=PEBP4;refGene_tx=XM_003984698.4;refGene_gn=PEBP4;refGene_tx=XM_006930640.4;refGene_gn=PEBP4;refGene_tx=XM_019828434.2;refGene_gn=PEBP4;refGene_tx=XM_019828435.2
B1 35506714 35506868 refGene_gn=EGR3;refGene_tx=XM_003984699.5
B1 35507308 35507348 refGene_gn=EGR3;refGene_tx=XM_023252488.1
B1 35508181 35509191
refGene_gn=EGR3;refGene_tx=XM_003984699.5;refGene_gn=EGR3;refGene_tx=XM_023252488.1
B1 35530264 35530272 refGene_gn=BIN3;refGene_tx=XM_003984702.4
B1 35552717 35552766 refGene_gn=BIN3;refGene_tx=XM_003984702.4
B1 35559473 35559514 refGene_gn=BIN3;refGene_tx=XM_003984702.4
B1 35559943 35560005 refGene_gn=BIN3;refGene_tx=XM_003984702.4
B1 35563147 35563284 refGene_gn=BIN3;refGene_tx=XM_003984702.4
B1 35563841 35563882 refGene_gn=BIN3;refGene_tx=XM_003984702.4
B1 35568282 35568424 refGene_gn=BIN3;refGene_tx=XM_003984702.4
B1 35568639 35568774 refGene_gn=BIN3;refGene_tx=XM_003984702.4
B1 35571489 35571636 refGene_gn=BIN3;refGene_tx=XM_003984702.4
B1 35573420 35573465
refGene_gn=CCAR2;refGene_tx=XM_006930641.4;refGene_gn=CCAR2;refGene_tx=XM_011281449.2;refGene_gn=CCAR2;refGene_tx=XM_011281450.2;refGene_gn=CCAR2;refGene_tx=XM_019828431.2;refGene_gn=CCAR2;refGene_tx=XM_0198284

32.2;refGene_gn=CCAR2;refGene_tx=XM_023252489.1
B1 35573719 35573925
refGene_gn=CCAR2;refGene_tx=XM_006930641.4;refGene_gn=CCAR2;refGene_tx=XM_011281449.2;refGene_gn=CCAR2;refGene_tx=XM_011281450.2;refGene_gn=CCAR2;refGene_tx=XM_019828431.2;refGene_gn=CCAR2;refGene_tx=XM_019828432.2;refGene_gn=CCAR2;refGene_tx=XM_023252489.1
B1 35574078 35574267
refGene_gn=CCAR2;refGene_tx=XM_006930641.4;refGene_gn=CCAR2;refGene_tx=XM_011281449.2;refGene_gn=CCAR2;refGene_tx=XM_011281450.2;refGene_gn=CCAR2;refGene_tx=XM_019828431.2;refGene_gn=CCAR2;refGene_tx=XM_019828432.2;refGene_gn=CCAR2;refGene_tx=XM_023252489.1
B1 35574365 35574477
refGene_gn=CCAR2;refGene_tx=XM_006930641.4;refGene_gn=CCAR2;refGene_tx=XM_011281449.2;refGene_gn=CCAR2;refGene_tx=XM_011281450.2;refGene_gn=CCAR2;refGene_tx=XM_019828431.2;refGene_gn=CCAR2;refGene_tx=XM_019828432.2;refGene_gn=CCAR2;refGene_tx=XM_023252489.1
B1 35574653 35574800
refGene_gn=CCAR2;refGene_tx=XM_006930641.4;refGene_gn=CCAR2;refGene_tx=XM_011281449.2;refGene_gn=CCAR2;refGene_tx=XM_011281450.2;refGene_gn=CCAR2;refGene_tx=XM_019828431.2;refGene_gn=CCAR2;refGene_tx=XM_019828432.2;refGene_gn=CCAR2;refGene_tx=XM_023252489.1
B1 35575160 35575243
refGene_gn=CCAR2;refGene_tx=XM_006930641.4;refGene_gn=CCAR2;refGene_tx=XM_011281449.2;refGene_gn=CCAR2;refGene_tx=XM_011281450.2;refGene_gn=CCAR2;refGene_tx=XM_019828431.2;refGene_gn=CCAR2;refGene_tx=XM_019828432.2;refGene_gn=CCAR2;refGene_tx=XM_023252489.1
B1 35575362 35575507
refGene_gn=CCAR2;refGene_tx=XM_006930641.4;refGene_gn=CCAR2;refGene_tx=XM_011281449.2;refGene_gn=CCAR2;refGene_tx=XM_011281450.2;refGene_gn=CCAR2;refGene_tx=XM_019828431.2;refGene_gn=CCAR2;refGene_tx=XM_019828432.2;refGene_gn=CCAR2;refGene_tx=XM_023252489.1
B1 35576108 35576330
refGene_gn=CCAR2;refGene_tx=XM_006930641.4;refGene_gn=CCAR2;refGene_tx=XM_011281449.2;refGene_gn=CCAR2;refGene_tx=XM_011281450.2;refGene_gn=CCAR2;refGene_tx=XM_019828431.2;refGene_gn=CCAR2;refGene_tx=XM_019828432.2;refGene_gn=CCAR2;refGene_tx=XM_023252489.1
B1 35576438 35576669
refGene_gn=CCAR2;refGene_tx=XM_006930641.4;refGene_gn=CCAR2;refGene_tx=XM_011281449.2;refGene_gn=CCAR2;refGene_tx=XM_011281450.2;refGene_gn=CCAR2;refGene_tx=XM_019828431.2;refGene_gn=CCAR2;refGene_tx=XM_019828432.2;refGene_gn=CCAR2;refGene_tx=XM_023252489.1
B1 35576756 35576928
refGene_gn=CCAR2;refGene_tx=XM_006930641.4;refGene_gn=CCAR2;refGene_tx=XM_011281449.2;refGene_gn=CCAR2;refGene_tx=XM_011281450.2;refGene_gn=CCAR2;refGene_tx=XM_019828431.2;refGene_gn=CCAR2;refGene_tx=XM_019828432.2;refGene_gn=CCAR2;refGene_tx=XM_023252489.1
B1 35577380 35577544
refGene_gn=CCAR2;refGene_tx=XM_006930641.4;refGene_gn=CCAR2;refGene_tx=XM_011281449.2;refGene_gn=CCAR2;refGene_tx=XM_011281450.2;refGene_gn=CCAR2;refGene_tx=XM_019828431.2;refGene_gn=CCAR2;refGene_tx=XM_0198284

32.2;refGene_gn=CCAR2;refGene_tx=XM_023252489.1
B1 35577909 35578023
refGene_gn=CCAR2;refGene_tx=XM_006930641.4;refGene_gn=CCAR2;refGene_tx=XM_011281449.2;refGene_gn=CCAR2;refGene_tx=XM_011281450.2;refGene_gn=CCAR2;refGene_tx=XM_019828431.2;refGene_gn=CCAR2;refGene_tx=XM_019828432.2;refGene_gn=CCAR2;refGene_tx=XM_023252489.1
B1 35578110 35578333
refGene_gn=CCAR2;refGene_tx=XM_006930641.4;refGene_gn=CCAR2;refGene_tx=XM_011281449.2;refGene_gn=CCAR2;refGene_tx=XM_011281450.2;refGene_gn=CCAR2;refGene_tx=XM_019828431.2;refGene_gn=CCAR2;refGene_tx=XM_019828432.2;refGene_gn=CCAR2;refGene_tx=XM_023252489.1
B1 35580331 35580451
refGene_gn=CCAR2;refGene_tx=XM_006930641.4;refGene_gn=CCAR2;refGene_tx=XM_011281449.2;refGene_gn=CCAR2;refGene_tx=XM_011281450.2;refGene_gn=CCAR2;refGene_tx=XM_019828431.2;refGene_gn=CCAR2;refGene_tx=XM_019828432.2;refGene_gn=CCAR2;refGene_tx=XM_023252489.1
B1 35584410 35584507
refGene_gn=CCAR2;refGene_tx=XM_006930641.4;refGene_gn=CCAR2;refGene_tx=XM_011281449.2;refGene_gn=CCAR2;refGene_tx=XM_011281450.2;refGene_gn=CCAR2;refGene_tx=XM_019828431.2;refGene_gn=CCAR2;refGene_tx=XM_019828432.2;refGene_gn=CCAR2;refGene_tx=XM_023252489.1
B1 35584688 35584818
refGene_gn=CCAR2;refGene_tx=XM_006930641.4;refGene_gn=CCAR2;refGene_tx=XM_011281449.2;refGene_gn=CCAR2;refGene_tx=XM_011281450.2;refGene_gn=CCAR2;refGene_tx=XM_019828431.2;refGene_gn=CCAR2;refGene_tx=XM_019828432.2;refGene_gn=CCAR2;refGene_tx=XM_023252489.1
B1 35584988 35585103
refGene_gn=CCAR2;refGene_tx=XM_006930641.4;refGene_gn=CCAR2;refGene_tx=XM_011281449.2;refGene_gn=CCAR2;refGene_tx=XM_011281450.2;refGene_gn=CCAR2;refGene_tx=XM_019828431.2;refGene_gn=CCAR2;refGene_tx=XM_019828432.2;refGene_gn=CCAR2;refGene_tx=XM_023252489.1
B1 35585307 35585399
refGene_gn=CCAR2;refGene_tx=XM_006930641.4;refGene_gn=CCAR2;refGene_tx=XM_011281449.2;refGene_gn=CCAR2;refGene_tx=XM_011281450.2;refGene_gn=CCAR2;refGene_tx=XM_019828431.2;refGene_gn=CCAR2;refGene_tx=XM_019828432.2;refGene_gn=CCAR2;refGene_tx=XM_023252489.1
B1 35585827 35585919
refGene_gn=CCAR2;refGene_tx=XM_006930641.4;refGene_gn=CCAR2;refGene_tx=XM_011281449.2;refGene_gn=CCAR2;refGene_tx=XM_011281450.2;refGene_gn=CCAR2;refGene_tx=XM_019828431.2;refGene_gn=CCAR2;refGene_tx=XM_019828432.2;refGene_gn=CCAR2;refGene_tx=XM_023252489.1
B1 35586197 35586255
refGene_gn=CCAR2;refGene_tx=XM_006930641.4;refGene_gn=CCAR2;refGene_tx=XM_011281449.2;refGene_gn=CCAR2;refGene_tx=XM_011281450.2;refGene_gn=CCAR2;refGene_tx=XM_019828431.2;refGene_gn=CCAR2;refGene_tx=XM_019828432.2;refGene_gn=CCAR2;refGene_tx=XM_023252489.1
B1 35589177 35589289
refGene_gn=CB1H8orf58;refGene_tx=XM_006930643.4;refGene_gn=CB1H8orf58;refGene_tx=XM_006930644.4;refGene_gn=CB1H8orf58;refGene_tx=XM_019828437.2;refGene_gn=CB1H8orf58;refGene_tx=XM_023252490.1;refGene_gn=CB1H8or

f58;refGene_tx=XM_023252491.1;refGene_gn=CB1H8orf58;refGene_tx=XM_0232
52492.1
B1 35589833 35589940
refGene_gn=CB1H8orf58;refGene_tx=XM_006930643.4;refGene_gn=CB1H8orf58;
refGene_tx=XM_006930644.4;refGene_gn=CB1H8orf58;refGene_tx=XM_01982843
7.2;refGene_gn=CB1H8orf58;refGene_tx=XM_023252490.1;refGene_gn=CB1H8or
f58;refGene_tx=XM_023252491.1;refGene_gn=CB1H8orf58;refGene_tx=XM_0232
52492.1
B1 35590157 35590270
refGene_gn=CB1H8orf58;refGene_tx=XM_006930643.4;refGene_gn=CB1H8orf58;
refGene_tx=XM_006930644.4;refGene_gn=CB1H8orf58;refGene_tx=XM_01982843
7.2;refGene_gn=CB1H8orf58;refGene_tx=XM_023252490.1;refGene_gn=CB1H8or
f58;refGene_tx=XM_023252491.1;refGene_gn=CB1H8orf58;refGene_tx=XM_0232
52492.1
B1 35590403 35590539
refGene_gn=CB1H8orf58;refGene_tx=XM_006930643.4;refGene_gn=CB1H8orf58;
refGene_tx=XM_006930644.4;refGene_gn=CB1H8orf58;refGene_tx=XM_01982843
7.2;refGene_gn=CB1H8orf58;refGene_tx=XM_023252490.1;refGene_gn=CB1H8or
f58;refGene_tx=XM_023252491.1;refGene_gn=CB1H8orf58;refGene_tx=XM_0232
52492.1
B1 35590647 35590812
refGene_gn=CB1H8orf58;refGene_tx=XM_006930643.4;refGene_gn=CB1H8orf58;
refGene_tx=XM_006930644.4;refGene_gn=CB1H8orf58;refGene_tx=XM_02325249
0.1;refGene_gn=CB1H8orf58;refGene_tx=XM_023252491.1;refGene_gn=CB1H8or
f58;refGene_tx=XM_023252492.1
B1 35591171 35591840
refGene_gn=CB1H8orf58;refGene_tx=XM_006930643.4;refGene_gn=CB1H8orf58;
refGene_tx=XM_006930644.4;refGene_gn=CB1H8orf58;refGene_tx=XM_01982843
7.2;refGene_gn=CB1H8orf58;refGene_tx=XM_023252490.1;refGene_gn=CB1H8or
f58;refGene_tx=XM_023252491.1;refGene_gn=CB1H8orf58;refGene_tx=XM_0232
52492.1
B1 35592845 35592894
refGene_gn=CB1H8orf58;refGene_tx=XM_023252490.1;refGene_gn=CB1H8orf58;
refGene_tx=XM_023252492.1
B1 35599009 35599187
refGene_gn=PDLIM2;refGene_tx=XM_006930645.4;refGene_gn=PDLIM2;refGene_
tx=XM_006930646.4;refGene_gn=PDLIM2;refGene_tx=XM_019828438.2
B1 35600743 35600861
refGene_gn=PDLIM2;refGene_tx=XM_006930645.4;refGene_gn=PDLIM2;refGene_
tx=XM_006930646.4;refGene_gn=PDLIM2;refGene_tx=XM_019828438.2
B1 35602303 35602449
refGene_gn=PDLIM2;refGene_tx=XM_006930645.4;refGene_gn=PDLIM2;refGene_
tx=XM_006930646.4;refGene_gn=PDLIM2;refGene_tx=XM_019828438.2
B1 35602696 35602785
refGene_gn=PDLIM2;refGene_tx=XM_006930645.4;refGene_gn=PDLIM2;refGene_
tx=XM_006930646.4;refGene_gn=PDLIM2;refGene_tx=XM_019828438.2
B1 35606290 35606369
refGene_gn=PDLIM2;refGene_tx=XM_006930645.4;refGene_gn=PDLIM2;refGene_
tx=XM_006930646.4;refGene_gn=PDLIM2;refGene_tx=XM_019828438.2
B1 35606524 35606676

refGene_gn=PDLIM2;refGene_tx=XM_006930645.4;refGene_gn=PDLIM2;refGene_tx=XM_006930646.4;refGene_gn=PDLIM2;refGene_tx=XM_019828438.2
B1 35606785 35606855
refGene_gn=PDLIM2;refGene_tx=XM_006930645.4;refGene_gn=PDLIM2;refGene_tx=XM_006930646.4;refGene_gn=PDLIM2;refGene_tx=XM_019828438.2
B1 35609586 35609738
refGene_gn=PDLIM2;refGene_tx=XM_006930645.4;refGene_gn=PDLIM2;refGene_tx=XM_006930646.4;refGene_gn=PDLIM2;refGene_tx=XM_019828438.2
B1 35610459 35610552
refGene_gn=PDLIM2;refGene_tx=XM_006930645.4;refGene_gn=PDLIM2;refGene_tx=XM_006930646.4;refGene_gn=PDLIM2;refGene_tx=XM_019828438.2
B1 35616275 35616337
refGene_gn=SORBS3;refGene_tx=XM_023252493.1;refGene_gn=SORBS3;refGene_tx=XM_023252494.1;refGene_gn=SORBS3;refGene_tx=XM_023252495.1;refGene_gn=SORBS3;refGene_tx=XM_023252496.1;refGene_gn=SORBS3;refGene_tx=XM_023252497.1;refGene_gn=SORBS3;refGene_tx=XM_023252498.1;refGene_gn=SORBS3;refGene_tx=XM_023252499.1;refGene_gn=SORBS3;refGene_tx=XM_023252500.1;refGene_gn=SORBS3;refGene_tx=XM_023252501.1;refGene_gn=SORBS3;refGene_tx=XM_023252502.1
B1 35618968 35619075
refGene_gn=SORBS3;refGene_tx=XM_023252493.1;refGene_gn=SORBS3;refGene_tx=XM_023252494.1;refGene_gn=SORBS3;refGene_tx=XM_023252495.1;refGene_gn=SORBS3;refGene_tx=XM_023252496.1;refGene_gn=SORBS3;refGene_tx=XM_023252497.1;refGene_gn=SORBS3;refGene_tx=XM_023252498.1;refGene_gn=SORBS3;refGene_tx=XM_023252499.1;refGene_gn=SORBS3;refGene_tx=XM_023252500.1;refGene_gn=SORBS3;refGene_tx=XM_023252501.1;refGene_gn=SORBS3;refGene_tx=XM_023252502.1
B1 35619650 35619751
refGene_gn=SORBS3;refGene_tx=XM_023252493.1;refGene_gn=SORBS3;refGene_tx=XM_023252494.1;refGene_gn=SORBS3;refGene_tx=XM_023252495.1;refGene_gn=SORBS3;refGene_tx=XM_023252496.1;refGene_gn=SORBS3;refGene_tx=XM_023252497.1;refGene_gn=SORBS3;refGene_tx=XM_023252498.1;refGene_gn=SORBS3;refGene_tx=XM_023252499.1;refGene_gn=SORBS3;refGene_tx=XM_023252500.1;refGene_gn=SORBS3;refGene_tx=XM_023252501.1;refGene_gn=SORBS3;refGene_tx=XM_023252502.1
B1 35620554 35620860
refGene_gn=SORBS3;refGene_tx=XM_023252493.1;refGene_gn=SORBS3;refGene_tx=XM_023252494.1;refGene_gn=SORBS3;refGene_tx=XM_023252495.1;refGene_gn=SORBS3;refGene_tx=XM_023252496.1;refGene_gn=SORBS3;refGene_tx=XM_023252497.1;refGene_gn=SORBS3;refGene_tx=XM_023252498.1;refGene_gn=SORBS3;refGene_tx=XM_023252499.1;refGene_gn=SORBS3;refGene_tx=XM_023252500.1;refGene_gn=SORBS3;refGene_tx=XM_023252501.1;refGene_gn=SORBS3;refGene_tx=XM_023252502.1
B1 35622044 35622170
refGene_gn=SORBS3;refGene_tx=XM_023252493.1;refGene_gn=SORBS3;refGene_tx=XM_023252494.1;refGene_gn=SORBS3;refGene_tx=XM_023252495.1;refGene_gn=SORBS3;refGene_tx=XM_023252496.1;refGene_gn=SORBS3;refGene_tx=XM_023252497.1;refGene_gn=SORBS3;refGene_tx=XM_023252498.1;refGene_gn=SORBS3;refGene_tx=XM_023252499.1;refGene_gn=SORBS3;refGene_tx=XM_023252500.1;refGene_gn=SORBS3;refGene_tx=XM_023252501.1;refGene_gn=SORBS3;refGen

e_tx=XM_023252502.1
B1 35623570 35623685
refGene_gn=SORBS3;refGene_tx=XM_023252493.1;refGene_gn=SORBS3;refGene_tx=XM_023252494.1;refGene_gn=SORBS3;refGene_tx=XM_023252495.1;refGene_gn=SORBS3;refGene_tx=XM_023252496.1;refGene_gn=SORBS3;refGene_tx=XM_023252497.1;refGene_gn=SORBS3;refGene_tx=XM_023252498.1;refGene_gn=SORBS3;refGene_tx=XM_023252499.1;refGene_gn=SORBS3;refGene_tx=XM_023252500.1;refGene_gn=SORBS3;refGene_tx=XM_023252501.1;refGene_gn=SORBS3;refGene_tx=XM_023252502.1
B1 35623870 35623917
refGene_gn=SORBS3;refGene_tx=XM_023252493.1;refGene_gn=SORBS3;refGene_tx=XM_023252494.1;refGene_gn=SORBS3;refGene_tx=XM_023252495.1;refGene_gn=SORBS3;refGene_tx=XM_023252496.1;refGene_gn=SORBS3;refGene_tx=XM_023252497.1;refGene_gn=SORBS3;refGene_tx=XM_023252498.1;refGene_gn=SORBS3;refGene_tx=XM_023252499.1;refGene_gn=SORBS3;refGene_tx=XM_023252500.1;refGene_gn=SORBS3;refGene_tx=XM_023252501.1;refGene_gn=SORBS3;refGene_tx=XM_023252502.1
B1 35624043 35624096
refGene_gn=SORBS3;refGene_tx=XM_023252493.1;refGene_gn=SORBS3;refGene_tx=XM_023252494.1;refGene_gn=SORBS3;refGene_tx=XM_023252495.1;refGene_gn=SORBS3;refGene_tx=XM_023252496.1;refGene_gn=SORBS3;refGene_tx=XM_023252497.1;refGene_gn=SORBS3;refGene_tx=XM_023252498.1;refGene_gn=SORBS3;refGene_tx=XM_023252499.1;refGene_gn=SORBS3;refGene_tx=XM_023252500.1;refGene_gn=SORBS3;refGene_tx=XM_023252501.1;refGene_gn=SORBS3;refGene_tx=XM_023252502.1
B1 35624331 35624471
refGene_gn=SORBS3;refGene_tx=XM_023252493.1;refGene_gn=SORBS3;refGene_tx=XM_023252494.1;refGene_gn=SORBS3;refGene_tx=XM_023252495.1;refGene_gn=SORBS3;refGene_tx=XM_023252496.1;refGene_gn=SORBS3;refGene_tx=XM_023252497.1;refGene_gn=SORBS3;refGene_tx=XM_023252498.1;refGene_gn=SORBS3;refGene_tx=XM_023252499.1;refGene_gn=SORBS3;refGene_tx=XM_023252500.1;refGene_gn=SORBS3;refGene_tx=XM_023252501.1;refGene_gn=SORBS3;refGene_tx=XM_023252502.1
B1 35624874 35624921
refGene_gn=SORBS3;refGene_tx=XM_023252493.1;refGene_gn=SORBS3;refGene_tx=XM_023252494.1;refGene_gn=SORBS3;refGene_tx=XM_023252496.1;refGene_gn=SORBS3;refGene_tx=XM_023252497.1;refGene_gn=SORBS3;refGene_tx=XM_023252498.1;refGene_gn=SORBS3;refGene_tx=XM_023252499.1
B1 35625469 35625556
refGene_gn=SORBS3;refGene_tx=XM_023252493.1;refGene_gn=SORBS3;refGene_tx=XM_023252494.1;refGene_gn=SORBS3;refGene_tx=XM_023252495.1;refGene_gn=SORBS3;refGene_tx=XM_023252496.1;refGene_gn=SORBS3;refGene_tx=XM_023252497.1;refGene_gn=SORBS3;refGene_tx=XM_023252498.1;refGene_gn=SORBS3;refGene_tx=XM_023252499.1
B1 35626357 35626411
refGene_gn=SORBS3;refGene_tx=XM_023252493.1;refGene_gn=SORBS3;refGene_tx=XM_023252494.1;refGene_gn=SORBS3;refGene_tx=XM_023252495.1;refGene_gn=SORBS3;refGene_tx=XM_023252496.1;refGene_gn=SORBS3;refGene_tx=XM_023252497.1;refGene_gn=SORBS3;refGene_tx=XM_023252498.1;refGene_gn=SORBS3;refGene_tx=XM_023252499.1

B1 35626510 35626597
refGene_gn=SORBS3;refGene_tx=XM_023252493.1;refGene_gn=SORBS3;refGene_tx=XM_023252494.1;refGene_gn=SORBS3;refGene_tx=XM_023252495.1;refGene_gn=SORBS3;refGene_tx=XM_023252496.1;refGene_gn=SORBS3;refGene_tx=XM_023252497.1;refGene_gn=SORBS3;refGene_tx=XM_023252498.1;refGene_gn=SORBS3;refGene_tx=XM_023252499.1
B1 35626847 35626935
refGene_gn=SORBS3;refGene_tx=XM_023252493.1;refGene_gn=SORBS3;refGene_tx=XM_023252494.1;refGene_gn=SORBS3;refGene_tx=XM_023252495.1;refGene_gn=SORBS3;refGene_tx=XM_023252496.1;refGene_gn=SORBS3;refGene_tx=XM_023252497.1;refGene_gn=SORBS3;refGene_tx=XM_023252498.1;refGene_gn=SORBS3;refGene_tx=XM_023252499.1
B1 35628619 35628713
refGene_gn=SORBS3;refGene_tx=XM_023252493.1;refGene_gn=SORBS3;refGene_tx=XM_023252494.1;refGene_gn=SORBS3;refGene_tx=XM_023252495.1;refGene_gn=SORBS3;refGene_tx=XM_023252496.1;refGene_gn=SORBS3;refGene_tx=XM_023252497.1;refGene_gn=SORBS3;refGene_tx=XM_023252498.1;refGene_gn=SORBS3;refGene_tx=XM_023252499.1
B1 35629145 35629184
refGene_gn=SORBS3;refGene_tx=XM_023252493.1;refGene_gn=SORBS3;refGene_tx=XM_023252494.1;refGene_gn=SORBS3;refGene_tx=XM_023252495.1;refGene_gn=SORBS3;refGene_tx=XM_023252496.1;refGene_gn=SORBS3;refGene_tx=XM_023252497.1;refGene_gn=SORBS3;refGene_tx=XM_023252498.1;refGene_gn=SORBS3;refGene_tx=XM_023252499.1
B1 35631594 35631658
refGene_gn=SORBS3;refGene_tx=XM_023252493.1;refGene_gn=SORBS3;refGene_tx=XM_023252494.1;refGene_gn=SORBS3;refGene_tx=XM_023252495.1;refGene_gn=SORBS3;refGene_tx=XM_023252496.1;refGene_gn=SORBS3;refGene_tx=XM_023252497.1;refGene_gn=SORBS3;refGene_tx=XM_023252498.1;refGene_gn=SORBS3;refGene_tx=XM_023252499.1
B1 35632527 35632721
refGene_gn=SORBS3;refGene_tx=XM_023252493.1;refGene_gn=SORBS3;refGene_tx=XM_023252494.1;refGene_gn=SORBS3;refGene_tx=XM_023252495.1;refGene_gn=SORBS3;refGene_tx=XM_023252496.1;refGene_gn=SORBS3;refGene_tx=XM_023252497.1;refGene_gn=SORBS3;refGene_tx=XM_023252498.1;refGene_gn=SORBS3;refGene_tx=XM_023252499.1
B1 35634413 35634531
refGene_gn=SORBS3;refGene_tx=XM_023252493.1;refGene_gn=SORBS3;refGene_tx=XM_023252494.1;refGene_gn=SORBS3;refGene_tx=XM_023252495.1;refGene_gn=SORBS3;refGene_tx=XM_023252496.1;refGene_gn=SORBS3;refGene_tx=XM_023252497.1;refGene_gn=SORBS3;refGene_tx=XM_023252498.1
B1 35634778 35634935
refGene_gn=SORBS3;refGene_tx=XM_023252493.1;refGene_gn=SORBS3;refGene_tx=XM_023252494.1;refGene_gn=SORBS3;refGene_tx=XM_023252495.1;refGene_gn=SORBS3;refGene_tx=XM_023252496.1;refGene_gn=SORBS3;refGene_tx=XM_023252497.1;refGene_gn=SORBS3;refGene_tx=XM_023252498.1
B1 35640231 35640338
refGene_gn=SORBS3;refGene_tx=XM_023252493.1;refGene_gn=SORBS3;refGene_tx=XM_023252494.1;refGene_gn=SORBS3;refGene_tx=XM_023252495.1;refGene_gn=SORBS3;refGene_tx=XM_023252496.1

B1 35648832 35648843 refGene_gn=PPP3CC;refGene_tx=XM_019828439.2
B1 35649250 35649471
refGene_gn=PPP3CC;refGene_tx=XM_019828439.2;refGene_gn=PPP3CC;refGene_tx=XM_023252503.1;refGene_gn=PPP3CC;refGene_tx=XM_023252504.1;refGene_gn=PPP3CC;refGene_tx=XM_023252505.1;refGene_gn=PPP3CC;refGene_tx=XM_023252506.1
B1 35650826 35650856
refGene_gn=PPP3CC;refGene_tx=XM_019828439.2;refGene_gn=PPP3CC;refGene_tx=XM_023252503.1;refGene_gn=PPP3CC;refGene_tx=XM_023252504.1
B1 35657660 35657758
refGene_gn=PPP3CC;refGene_tx=XM_019828439.2;refGene_gn=PPP3CC;refGene_tx=XM_023252503.1;refGene_gn=PPP3CC;refGene_tx=XM_023252504.1;refGene_gn=PPP3CC;refGene_tx=XM_023252505.1;refGene_gn=PPP3CC;refGene_tx=XM_023252506.1
B1 35658377 35658459
refGene_gn=PPP3CC;refGene_tx=XM_019828439.2;refGene_gn=PPP3CC;refGene_tx=XM_023252503.1;refGene_gn=PPP3CC;refGene_tx=XM_023252504.1;refGene_gn=PPP3CC;refGene_tx=XM_023252505.1;refGene_gn=PPP3CC;refGene_tx=XM_023252506.1
B1 35659373 35659400
refGene_gn=PPP3CC;refGene_tx=XM_019828439.2;refGene_gn=PPP3CC;refGene_tx=XM_023252503.1;refGene_gn=PPP3CC;refGene_tx=XM_023252505.1
B1 35671694 35671766
refGene_gn=PPP3CC;refGene_tx=XM_019828439.2;refGene_gn=PPP3CC;refGene_tx=XM_023252503.1;refGene_gn=PPP3CC;refGene_tx=XM_023252504.1;refGene_gn=PPP3CC;refGene_tx=XM_023252505.1;refGene_gn=PPP3CC;refGene_tx=XM_023252506.1
B1 35672495 35672621
refGene_gn=PPP3CC;refGene_tx=XM_019828439.2;refGene_gn=PPP3CC;refGene_tx=XM_023252503.1;refGene_gn=PPP3CC;refGene_tx=XM_023252504.1;refGene_gn=PPP3CC;refGene_tx=XM_023252505.1;refGene_gn=PPP3CC;refGene_tx=XM_023252506.1
B1 35676352 35676447
refGene_gn=PPP3CC;refGene_tx=XM_019828439.2;refGene_gn=PPP3CC;refGene_tx=XM_023252503.1;refGene_gn=PPP3CC;refGene_tx=XM_023252504.1;refGene_gn=PPP3CC;refGene_tx=XM_023252505.1;refGene_gn=PPP3CC;refGene_tx=XM_023252506.1
B1 35676533 35676611
refGene_gn=PPP3CC;refGene_tx=XM_019828439.2;refGene_gn=PPP3CC;refGene_tx=XM_023252503.1;refGene_gn=PPP3CC;refGene_tx=XM_023252504.1;refGene_gn=PPP3CC;refGene_tx=XM_023252505.1;refGene_gn=PPP3CC;refGene_tx=XM_023252506.1
B1 35681920 35682060
refGene_gn=PPP3CC;refGene_tx=XM_019828439.2;refGene_gn=PPP3CC;refGene_tx=XM_023252503.1;refGene_gn=PPP3CC;refGene_tx=XM_023252504.1;refGene_gn=PPP3CC;refGene_tx=XM_023252505.1;refGene_gn=PPP3CC;refGene_tx=XM_023252506.1
B1 35683478 35683624
refGene_gn=PPP3CC;refGene_tx=XM_019828439.2;refGene_gn=PPP3CC;refGene_tx=XM_023252503.1;refGene_gn=PPP3CC;refGene_tx=XM_023252504.1;refGene_

gn=PPP3CC;refGene_tx=XM_023252505.1;refGene_gn=PPP3CC;refGene_tx=XM_02
3252506.1
B1 35695860 35695972
refGene_gn=PPP3CC;refGene_tx=XM_019828439.2;refGene_gn=PPP3CC;refGene_
tx=XM_023252503.1;refGene_gn=PPP3CC;refGene_tx=XM_023252504.1;refGene_
gn=PPP3CC;refGene_tx=XM_023252505.1;refGene_gn=PPP3CC;refGene_tx=XM_02
3252506.1
B1 35708712 35708837
refGene_gn=PPP3CC;refGene_tx=XM_019828439.2;refGene_gn=PPP3CC;refGene_
tx=XM_023252503.1;refGene_gn=PPP3CC;refGene_tx=XM_023252504.1;refGene_
gn=PPP3CC;refGene_tx=XM_023252505.1;refGene_gn=PPP3CC;refGene_tx=XM_02
3252506.1
B1 35709161 35709359
refGene_gn=PPP3CC;refGene_tx=XM_019828439.2;refGene_gn=PPP3CC;refGene_
tx=XM_023252503.1;refGene_gn=PPP3CC;refGene_tx=XM_023252504.1;refGene_
gn=PPP3CC;refGene_tx=XM_023252505.1;refGene_gn=PPP3CC;refGene_tx=XM_02
3252506.1
B1 35748670 35748719
refGene_gn=PPP3CC;refGene_tx=XM_019828439.2;refGene_gn=PPP3CC;refGene_
tx=XM_023252503.1;refGene_gn=PPP3CC;refGene_tx=XM_023252504.1;refGene_
gn=PPP3CC;refGene_tx=XM_023252505.1;refGene_gn=PPP3CC;refGene_tx=XM_02
3252506.1
B1 35765439 35765586
refGene_gn=SLC39A14;refGene_tx=XM_023252507.1;refGene_gn=SLC39A14;refG
ene_tx=XM_023252508.1
B1 35767078 35767263
refGene_gn=SLC39A14;refGene_tx=XM_023252507.1;refGene_gn=SLC39A14;refG
ene_tx=XM_023252508.1
B1 35768293 35768501
refGene_gn=SLC39A14;refGene_tx=XM_023252507.1;refGene_gn=SLC39A14;refG
ene_tx=XM_023252508.1
B1 35768619 35768808
refGene_gn=SLC39A14;refGene_tx=XM_023252507.1;refGene_gn=SLC39A14;refG
ene_tx=XM_023252508.1
B1 35769548 35769671
refGene_gn=SLC39A14;refGene_tx=XM_023252507.1;refGene_gn=SLC39A14;refG
ene_tx=XM_023252508.1
B1 35771641 35771811 refGene_gn=SLC39A14;refGene_tx=XM_023252507.1
B1 35773820 35773990 refGene_gn=SLC39A14;refGene_tx=XM_023252508.1
B1 35774797 35774984
refGene_gn=SLC39A14;refGene_tx=XM_023252507.1;refGene_gn=SLC39A14;refG
ene_tx=XM_023252508.1
B1 35776748 35777015
refGene_gn=SLC39A14;refGene_tx=XM_023252507.1;refGene_gn=SLC39A14;refG
ene_tx=XM_023252508.1
B1 35821859 35822016
refGene_gn=PIWIL2;refGene_tx=XM_003984708.5;refGene_gn=PIWIL2;refGene_
tx=XM_019828440.2;refGene_gn=PIWIL2;refGene_tx=XM_019828442.2
B1 35823793 35823901
refGene_gn=PIWIL2;refGene_tx=XM_003984708.5;refGene_gn=PIWIL2;refGene_

tx=XM_019828440.2;refGene_gn=PIWIL2;refGene_tx=XM_019828442.2
B1 35824476 35824730
refGene_gn=PIWIL2;refGene_tx=XM_003984708.5;refGene_gn=PIWIL2;refGene_tx=XM_019828440.2;refGene_gn=PIWIL2;refGene_tx=XM_019828442.2
B1 35863045 35863151
refGene_gn=PIWIL2;refGene_tx=XM_003984708.5;refGene_gn=PIWIL2;refGene_tx=XM_019828440.2;refGene_gn=PIWIL2;refGene_tx=XM_019828442.2
B1 35864027 35864116
refGene_gn=PIWIL2;refGene_tx=XM_003984708.5;refGene_gn=PIWIL2;refGene_tx=XM_019828440.2;refGene_gn=PIWIL2;refGene_tx=XM_019828442.2
B1 35864702 35864819
refGene_gn=PIWIL2;refGene_tx=XM_003984708.5;refGene_gn=PIWIL2;refGene_tx=XM_019828440.2;refGene_gn=PIWIL2;refGene_tx=XM_019828442.2
B1 35865408 35865510
refGene_gn=PIWIL2;refGene_tx=XM_003984708.5;refGene_gn=PIWIL2;refGene_tx=XM_019828440.2;refGene_gn=PIWIL2;refGene_tx=XM_019828442.2
B1 35867606 35867795
refGene_gn=PIWIL2;refGene_tx=XM_003984708.5;refGene_gn=PIWIL2;refGene_tx=XM_019828440.2;refGene_gn=PIWIL2;refGene_tx=XM_019828442.2
B1 35868731 35868845
refGene_gn=PIWIL2;refGene_tx=XM_003984708.5;refGene_gn=PIWIL2;refGene_tx=XM_019828440.2;refGene_gn=PIWIL2;refGene_tx=XM_019828442.2
B1 35873830 35873971
refGene_gn=PIWIL2;refGene_tx=XM_003984708.5;refGene_gn=PIWIL2;refGene_tx=XM_019828440.2;refGene_gn=PIWIL2;refGene_tx=XM_019828442.2
B1 35875421 35875511
refGene_gn=PIWIL2;refGene_tx=XM_003984708.5;refGene_gn=PIWIL2;refGene_tx=XM_019828440.2;refGene_gn=PIWIL2;refGene_tx=XM_019828442.2
B1 35876379 35876464
refGene_gn=PIWIL2;refGene_tx=XM_003984708.5;refGene_gn=PIWIL2;refGene_tx=XM_019828440.2;refGene_gn=PIWIL2;refGene_tx=XM_019828442.2
B1 35877059 35877248
refGene_gn=PIWIL2;refGene_tx=XM_003984708.5;refGene_gn=PIWIL2;refGene_tx=XM_019828440.2;refGene_gn=PIWIL2;refGene_tx=XM_019828442.2
B1 35887479 35887593
refGene_gn=PIWIL2;refGene_tx=XM_003984708.5;refGene_gn=PIWIL2;refGene_tx=XM_019828440.2;refGene_gn=PIWIL2;refGene_tx=XM_019828442.2
B1 35887899 35887980
refGene_gn=PIWIL2;refGene_tx=XM_003984708.5;refGene_gn=PIWIL2;refGene_tx=XM_019828440.2;refGene_gn=PIWIL2;refGene_tx=XM_019828442.2
B1 35888811 35888936
refGene_gn=PIWIL2;refGene_tx=XM_003984708.5;refGene_gn=PIWIL2;refGene_tx=XM_019828440.2;refGene_gn=PIWIL2;refGene_tx=XM_019828442.2
B1 35889429 35889550
refGene_gn=PIWIL2;refGene_tx=XM_003984708.5;refGene_gn=PIWIL2;refGene_tx=XM_019828440.2;refGene_gn=PIWIL2;refGene_tx=XM_019828442.2
B1 35893146 35893257
refGene_gn=PIWIL2;refGene_tx=XM_003984708.5;refGene_gn=PIWIL2;refGene_tx=XM_019828440.2;refGene_gn=PIWIL2;refGene_tx=XM_019828442.2
B1 35893870 35894077

refGene_gn=PIWIL2;refGene_tx=XM_003984708.5;refGene_gn=PIWIL2;refGene_tx=XM_019828440.2;refGene_gn=PIWIL2;refGene_tx=XM_019828442.2
B1	35894566 35894705
refGene_gn=PIWIL2;refGene_tx=XM_003984708.5;refGene_gn=PIWIL2;refGene_tx=XM_019828440.2;refGene_gn=PIWIL2;refGene_tx=XM_019828442.2
B1	35894857 35894945
refGene_gn=PIWIL2;refGene_tx=XM_003984708.5;refGene_gn=PIWIL2;refGene_tx=XM_019828440.2;refGene_gn=PIWIL2;refGene_tx=XM_019828442.2
B1	35896436 35896625
refGene_gn=PIWIL2;refGene_tx=XM_003984708.5;refGene_gn=PIWIL2;refGene_tx=XM_019828440.2;refGene_gn=PIWIL2;refGene_tx=XM_019828442.2
B1	35927038 35927161 refGene_gn=POLR3D;refGene_tx=XM_006930651.3
B1	35927261 35927414 refGene_gn=POLR3D;refGene_tx=XM_006930651.3
B1	35927842 35928108 refGene_gn=POLR3D;refGene_tx=XM_006930651.3
B1	35928480 35928649 refGene_gn=POLR3D;refGene_tx=XM_006930651.3
B1	35928831 35928956 refGene_gn=POLR3D;refGene_tx=XM_006930651.3
B1	35929115 35929267 refGene_gn=POLR3D;refGene_tx=XM_006930651.3
B1	35929894 35929938 refGene_gn=POLR3D;refGene_tx=XM_006930651.3
B1	35930894 35931059 refGene_gn=POLR3D;refGene_tx=XM_006930651.3
B1	35948054 35948219 refGene_gn=PHYHIP;refGene_tx=XM_003984711.4
B1	35949471 35949646 refGene_gn=PHYHIP;refGene_tx=XM_003984711.4
B1	35951501 35951619 refGene_gn=PHYHIP;refGene_tx=XM_003984711.4
B1	35953324 35953859 refGene_gn=PHYHIP;refGene_tx=XM_003984711.4
B1	35961230 35961365
refGene_gn=BMP1;refGene_tx=XM_023252509.1;refGene_gn=BMP1;refGene_tx=XM_023252510.1
B1	35963283 35963534
refGene_gn=BMP1;refGene_tx=XM_023252509.1;refGene_gn=BMP1;refGene_tx=XM_023252510.1
B1	35965141 35965355
refGene_gn=BMP1;refGene_tx=XM_023252509.1;refGene_gn=BMP1;refGene_tx=XM_023252510.1
B1	35965760 35965888
refGene_gn=BMP1;refGene_tx=XM_023252509.1;refGene_gn=BMP1;refGene_tx=XM_023252510.1
B1	35969042 35969168
refGene_gn=BMP1;refGene_tx=XM_023252509.1;refGene_gn=BMP1;refGene_tx=XM_023252510.1
B1	35969745 35969831 refGene_gn=BMP1;refGene_tx=XM_023252511.1
B1	35973157 35973338
refGene_gn=BMP1;refGene_tx=XM_023252509.1;refGene_gn=BMP1;refGene_tx=XM_023252510.1;refGene_gn=BMP1;refGene_tx=XM_023252511.1
B1	35973722 35973883
refGene_gn=BMP1;refGene_tx=XM_023252509.1;refGene_gn=BMP1;refGene_tx=XM_023252510.1;refGene_gn=BMP1;refGene_tx=XM_023252511.1
B1	35974608 35974734
refGene_gn=BMP1;refGene_tx=XM_023252509.1;refGene_gn=BMP1;refGene_tx=XM_023252510.1;refGene_gn=BMP1;refGene_tx=XM_023252511.1
B1	35977795 35977991
refGene_gn=BMP1;refGene_tx=XM_023252509.1;refGene_gn=BMP1;refGene_tx=X

M_023252510.1;refGene_gn=BMP1;refGene_tx=XM_023252511.1
B1 35978089 35978235
refGene_gn=BMP1;refGene_tx=XM_023252509.1;refGene_gn=BMP1;refGene_tx=XM_023252510.1;refGene_gn=BMP1;refGene_tx=XM_023252511.1
B1 35978482 35978599
refGene_gn=BMP1;refGene_tx=XM_023252509.1;refGene_gn=BMP1;refGene_tx=XM_023252510.1;refGene_gn=BMP1;refGene_tx=XM_023252511.1
B1 35982005 35982108
refGene_gn=BMP1;refGene_tx=XM_023252509.1;refGene_gn=BMP1;refGene_tx=XM_023252510.1;refGene_gn=BMP1;refGene_tx=XM_023252511.1
B1 35990325 35990441
refGene_gn=BMP1;refGene_tx=XM_023252509.1;refGene_gn=BMP1;refGene_tx=XM_023252510.1;refGene_gn=BMP1;refGene_tx=XM_023252511.1
B1 35991005 35991130
refGene_gn=BMP1;refGene_tx=XM_023252509.1;refGene_gn=BMP1;refGene_tx=XM_023252510.1;refGene_gn=BMP1;refGene_tx=XM_023252511.1
B1 35992587 35992693
refGene_gn=BMP1;refGene_tx=XM_023252509.1;refGene_gn=BMP1;refGene_tx=XM_023252510.1;refGene_gn=BMP1;refGene_tx=XM_023252511.1
B1 35993488 35993704
refGene_gn=BMP1;refGene_tx=XM_023252509.1;refGene_gn=BMP1;refGene_tx=XM_023252510.1;refGene_gn=BMP1;refGene_tx=XM_023252511.1
B1 35993971 35994089
refGene_gn=BMP1;refGene_tx=XM_023252509.1;refGene_gn=BMP1;refGene_tx=XM_023252510.1;refGene_gn=BMP1;refGene_tx=XM_023252511.1
B1 35994278 35994443
refGene_gn=BMP1;refGene_tx=XM_023252509.1;refGene_gn=BMP1;refGene_tx=XM_023252511.1
B1 35996536 35996650
refGene_gn=BMP1;refGene_tx=XM_023252509.1;refGene_gn=BMP1;refGene_tx=XM_023252511.1
B1 36003934 36004088
refGene_gn=BMP1;refGene_tx=XM_023252509.1;refGene_gn=BMP1;refGene_tx=XM_023252511.1
B1 36005507 36005645
refGene_gn=SFTPC;refGene_tx=XM_003984712.3;refGene_gn=SFTPC;refGene_tx=XM_019828445.1;refGene_gn=SFTPC;refGene_tx=XM_019828446.1
B1 36005769 36005853
refGene_gn=SFTPC;refGene_tx=XM_019828445.1;refGene_gn=SFTPC;refGene_tx=XM_019828446.1
B1 36006021 36006132
refGene_gn=SFTPC;refGene_tx=XM_003984712.3;refGene_gn=SFTPC;refGene_tx=XM_019828445.1;refGene_gn=SFTPC;refGene_tx=XM_019828446.1
B1 36006386 36006509
refGene_gn=SFTPC;refGene_tx=XM_003984712.3;refGene_gn=SFTPC;refGene_tx=XM_019828445.1;refGene_gn=SFTPC;refGene_tx=XM_019828446.1
B1 36006838 36006997
refGene_gn=SFTPC;refGene_tx=XM_003984712.3;refGene_gn=SFTPC;refGene_tx=XM_019828445.1
B1 36007696 36007765

refGene_gn=SFTPC;refGene_tx=XM_003984712.3;refGene_gn=SFTPC;refGene_tx=XM_019828445.1;refGene_gn=SFTPC;refGene_tx=XM_019828446.1
B1 36014194 36014400
refGene_gn=LGI3;refGene_tx=XM_003984713.5;refGene_gn=LGI3;refGene_tx=XM_019828447.2
B1 36015308 36015380
refGene_gn=LGI3;refGene_tx=XM_003984713.5;refGene_gn=LGI3;refGene_tx=XM_019828447.2
B1 36016231 36016303
refGene_gn=LGI3;refGene_tx=XM_003984713.5;refGene_gn=LGI3;refGene_tx=XM_019828447.2
B1 36016647 36016719
refGene_gn=LGI3;refGene_tx=XM_003984713.5;refGene_gn=LGI3;refGene_tx=XM_019828447.2
B1 36016822 36016894
refGene_gn=LGI3;refGene_tx=XM_003984713.5;refGene_gn=LGI3;refGene_tx=XM_019828447.2
B1 36017656 36017826
refGene_gn=LGI3;refGene_tx=XM_003984713.5;refGene_gn=LGI3;refGene_tx=XM_019828447.2
B1 36017974 36018139 refGene_gn=LGI3;refGene_tx=XM_003984713.5
B1 36020297 36021115
refGene_gn=LGI3;refGene_tx=XM_003984713.5;refGene_gn=LGI3;refGene_tx=XM_019828447.2
B1 36026241 36026273 refGene_gn=REEP4;refGene_tx=XM_011281451.3
B1 36027040 36027113 refGene_gn=REEP4;refGene_tx=XM_011281451.3
B1 36027497 36027574 refGene_gn=REEP4;refGene_tx=XM_011281451.3
B1 36027668 36027789 refGene_gn=REEP4;refGene_tx=XM_011281451.3
B1 36028216 36028330 refGene_gn=REEP4;refGene_tx=XM_011281451.3
B1 36028736 36028872 refGene_gn=REEP4;refGene_tx=XM_011281451.3
B1 36029032 36029187 refGene_gn=REEP4;refGene_tx=XM_011281451.3
B1 36029273 36029339 refGene_gn=REEP4;refGene_tx=XM_011281451.3
B1 36038798 36039410
refGene_gn=HR;refGene_tx=XM_011281452.3;refGene_gn=HR;refGene_tx=XM_023252512.1
B1 36040141 36040931
refGene_gn=HR;refGene_tx=XM_011281452.3;refGene_gn=HR;refGene_tx=XM_023252512.1
B1 36042612 36042766
refGene_gn=HR;refGene_tx=XM_011281452.3;refGene_gn=HR;refGene_tx=XM_023252512.1
B1 36042845 36043030
refGene_gn=HR;refGene_tx=XM_011281452.3;refGene_gn=HR;refGene_tx=XM_023252512.1
B1 36044452 36044620
refGene_gn=HR;refGene_tx=XM_011281452.3;refGene_gn=HR;refGene_tx=XM_023252512.1
B1 36045394 36045484
refGene_gn=HR;refGene_tx=XM_011281452.3;refGene_gn=HR;refGene_tx=XM_023252512.1

B1 36045660 36045776
refGene_gn=HR;refGene_tx=XM_011281452.3;refGene_gn=HR;refGene_tx=XM_023252512.1
B1 36046581 36046663
refGene_gn=HR;refGene_tx=XM_011281452.3;refGene_gn=HR;refGene_tx=XM_023252512.1
B1 36047002 36047166
refGene_gn=HR;refGene_tx=XM_011281452.3;refGene_gn=HR;refGene_tx=XM_023252512.1
B1 36047283 36047520
refGene_gn=HR;refGene_tx=XM_011281452.3;refGene_gn=HR;refGene_tx=XM_023252512.1
B1 36047732 36047898
refGene_gn=HR;refGene_tx=XM_011281452.3;refGene_gn=HR;refGene_tx=XM_023252512.1
B1 36048089 36048159
refGene_gn=HR;refGene_tx=XM_011281452.3;refGene_gn=HR;refGene_tx=XM_023252512.1
B1 36048348 36048479
refGene_gn=HR;refGene_tx=XM_011281452.3;refGene_gn=HR;refGene_tx=XM_023252512.1
B1 36048598 36048718
refGene_gn=HR;refGene_tx=XM_011281452.3;refGene_gn=HR;refGene_tx=XM_023252512.1
B1 36048817 36048933
refGene_gn=HR;refGene_tx=XM_011281452.3;refGene_gn=HR;refGene_tx=XM_023252512.1
B1 36050932 36051097
refGene_gn=HR;refGene_tx=XM_011281452.3;refGene_gn=HR;refGene_tx=XM_023252512.1
B1 36051529 36051658
refGene_gn=HR;refGene_tx=XM_011281452.3;refGene_gn=HR;refGene_tx=XM_023252512.1
B1 36051812 36051875
refGene_gn=HR;refGene_tx=XM_011281452.3;refGene_gn=HR;refGene_tx=XM_023252512.1
B1 36057503 36057665 refGene_gn=NUDT18;refGene_tx=XM_003984715.4
B1 36058522 36058736 refGene_gn=NUDT18;refGene_tx=XM_003984715.4
B1 36058999 36059595 refGene_gn=NUDT18;refGene_tx=XM_003984715.4
B1 36063102 36063241
refGene_gn=FAM160B2;refGene_tx=XM_023252513.1;refGene_gn=FAM160B2;refGene_tx=XM_023252514.1;refGene_gn=FAM160B2;refGene_tx=XM_023252515.1;refGene_gn=FAM160B2;refGene_tx=XM_023252516.1;refGene_gn=FAM160B2;refGene_tx=XM_023252517.1;refGene_gn=FAM160B2;refGene_tx=XM_023252518.1;refGene_gn=FAM160B2;refGene_tx=XM_023252519.1;refGene_gn=FAM160B2;refGene_tx=XM_023252520.1;refGene_gn=FAM160B2;refGene_tx=XM_023252521.1;refGene_gn=FAM160B2;refGene_tx=XM_023252522.1
B1 36063469 36063570
refGene_gn=FAM160B2;refGene_tx=XM_023252513.1;refGene_gn=FAM160B2;refGene_tx=XM_023252514.1;refGene_gn=FAM160B2;refGene_tx=XM_023252515.1;re

fGene_gn=FAM160B2;refGene_tx=XM_023252516.1;refGene_gn=FAM160B2;refGen
e_tx=XM_023252517.1;refGene_gn=FAM160B2;refGene_tx=XM_023252518.1;refG
ene_gn=FAM160B2;refGene_tx=XM_023252519.1;refGene_gn=FAM160B2;refGene_
tx=XM_023252520.1;refGene_gn=FAM160B2;refGene_tx=XM_023252521.1;refGen
e_gn=FAM160B2;refGene_tx=XM_023252522.1
B1 36063785 36063926
refGene_gn=FAM160B2;refGene_tx=XM_023252513.1;refGene_gn=FAM160B2;refG
ene_tx=XM_023252514.1;refGene_gn=FAM160B2;refGene_tx=XM_023252515.1;re
fGene_gn=FAM160B2;refGene_tx=XM_023252516.1;refGene_gn=FAM160B2;refGen
e_tx=XM_023252517.1;refGene_gn=FAM160B2;refGene_tx=XM_023252518.1;refG
ene_gn=FAM160B2;refGene_tx=XM_023252519.1;refGene_gn=FAM160B2;refGene_
tx=XM_023252520.1;refGene_gn=FAM160B2;refGene_tx=XM_023252521.1;refGen
e_gn=FAM160B2;refGene_tx=XM_023252522.1
B1 36064266 36064449
refGene_gn=FAM160B2;refGene_tx=XM_023252513.1;refGene_gn=FAM160B2;refG
ene_tx=XM_023252514.1;refGene_gn=FAM160B2;refGene_tx=XM_023252515.1;re
fGene_gn=FAM160B2;refGene_tx=XM_023252516.1;refGene_gn=FAM160B2;refGen
e_tx=XM_023252517.1;refGene_gn=FAM160B2;refGene_tx=XM_023252518.1;refG
ene_gn=FAM160B2;refGene_tx=XM_023252519.1;refGene_gn=FAM160B2;refGene_
tx=XM_023252520.1;refGene_gn=FAM160B2;refGene_tx=XM_023252521.1;refGen
e_gn=FAM160B2;refGene_tx=XM_023252522.1
B1 36064653 36064759
refGene_gn=FAM160B2;refGene_tx=XM_023252513.1;refGene_gn=FAM160B2;refG
ene_tx=XM_023252514.1;refGene_gn=FAM160B2;refGene_tx=XM_023252515.1;re
fGene_gn=FAM160B2;refGene_tx=XM_023252516.1;refGene_gn=FAM160B2;refGen
e_tx=XM_023252517.1;refGene_gn=FAM160B2;refGene_tx=XM_023252518.1;refG
ene_gn=FAM160B2;refGene_tx=XM_023252519.1;refGene_gn=FAM160B2;refGene_
tx=XM_023252520.1;refGene_gn=FAM160B2;refGene_tx=XM_023252521.1;refGen
e_gn=FAM160B2;refGene_tx=XM_023252522.1
B1 36064918 36065047
refGene_gn=FAM160B2;refGene_tx=XM_023252513.1;refGene_gn=FAM160B2;refG
ene_tx=XM_023252514.1;refGene_gn=FAM160B2;refGene_tx=XM_023252515.1;re
fGene_gn=FAM160B2;refGene_tx=XM_023252516.1;refGene_gn=FAM160B2;refGen
e_tx=XM_023252517.1;refGene_gn=FAM160B2;refGene_tx=XM_023252518.1;refG
ene_gn=FAM160B2;refGene_tx=XM_023252519.1;refGene_gn=FAM160B2;refGene_
tx=XM_023252520.1;refGene_gn=FAM160B2;refGene_tx=XM_023252521.1;refGen
e_gn=FAM160B2;refGene_tx=XM_023252522.1
B1 36065164 36065310
refGene_gn=FAM160B2;refGene_tx=XM_023252513.1;refGene_gn=FAM160B2;refG
ene_tx=XM_023252514.1;refGene_gn=FAM160B2;refGene_tx=XM_023252515.1;re
fGene_gn=FAM160B2;refGene_tx=XM_023252516.1;refGene_gn=FAM160B2;refGen
e_tx=XM_023252517.1;refGene_gn=FAM160B2;refGene_tx=XM_023252518.1;refG
ene_gn=FAM160B2;refGene_tx=XM_023252519.1;refGene_gn=FAM160B2;refGene_
tx=XM_023252520.1;refGene_gn=FAM160B2;refGene_tx=XM_023252521.1;refGen
e_gn=FAM160B2;refGene_tx=XM_023252522.1
B1 36065796 36065986
refGene_gn=FAM160B2;refGene_tx=XM_023252513.1;refGene_gn=FAM160B2;refG
ene_tx=XM_023252514.1;refGene_gn=FAM160B2;refGene_tx=XM_023252515.1;re
fGene_gn=FAM160B2;refGene_tx=XM_023252516.1;refGene_gn=FAM160B2;refGen
e_tx=XM_023252517.1;refGene_gn=FAM160B2;refGene_tx=XM_023252518.1;refG

ene_gn=FAM160B2;refGene_tx=XM_023252519.1;refGene_gn=FAM160B2;refGene_tx=XM_023252520.1;refGene_gn=FAM160B2;refGene_tx=XM_023252521.1;refGene_gn=FAM160B2;refGene_tx=XM_023252522.1
B1 36066541 36066618
refGene_gn=FAM160B2;refGene_tx=XM_023252513.1;refGene_gn=FAM160B2;refGene_tx=XM_023252514.1;refGene_gn=FAM160B2;refGene_tx=XM_023252515.1;refGene_gn=FAM160B2;refGene_tx=XM_023252516.1;refGene_gn=FAM160B2;refGene_tx=XM_023252517.1;refGene_gn=FAM160B2;refGene_tx=XM_023252518.1;refGene_gn=FAM160B2;refGene_tx=XM_023252519.1;refGene_gn=FAM160B2;refGene_tx=XM_023252520.1;refGene_gn=FAM160B2;refGene_tx=XM_023252521.1;refGene_gn=FAM160B2;refGene_tx=XM_023252522.1
B1 36066747 36066943
refGene_gn=FAM160B2;refGene_tx=XM_023252513.1;refGene_gn=FAM160B2;refGene_tx=XM_023252514.1;refGene_gn=FAM160B2;refGene_tx=XM_023252515.1;refGene_gn=FAM160B2;refGene_tx=XM_023252516.1;refGene_gn=FAM160B2;refGene_tx=XM_023252517.1;refGene_gn=FAM160B2;refGene_tx=XM_023252518.1;refGene_gn=FAM160B2;refGene_tx=XM_023252519.1;refGene_gn=FAM160B2;refGene_tx=XM_023252520.1;refGene_gn=FAM160B2;refGene_tx=XM_023252521.1;refGene_gn=FAM160B2;refGene_tx=XM_023252522.1
B1 36067232 36067429
refGene_gn=FAM160B2;refGene_tx=XM_023252513.1;refGene_gn=FAM160B2;refGene_tx=XM_023252514.1;refGene_gn=FAM160B2;refGene_tx=XM_023252515.1;refGene_gn=FAM160B2;refGene_tx=XM_023252516.1;refGene_gn=FAM160B2;refGene_tx=XM_023252517.1;refGene_gn=FAM160B2;refGene_tx=XM_023252518.1;refGene_gn=FAM160B2;refGene_tx=XM_023252519.1;refGene_gn=FAM160B2;refGene_tx=XM_023252520.1;refGene_gn=FAM160B2;refGene_tx=XM_023252521.1;refGene_gn=FAM160B2;refGene_tx=XM_023252522.1
B1 36067532 36067760
refGene_gn=FAM160B2;refGene_tx=XM_023252513.1;refGene_gn=FAM160B2;refGene_tx=XM_023252514.1;refGene_gn=FAM160B2;refGene_tx=XM_023252515.1;refGene_gn=FAM160B2;refGene_tx=XM_023252516.1;refGene_gn=FAM160B2;refGene_tx=XM_023252517.1;refGene_gn=FAM160B2;refGene_tx=XM_023252518.1;refGene_gn=FAM160B2;refGene_tx=XM_023252519.1;refGene_gn=FAM160B2;refGene_tx=XM_023252520.1;refGene_gn=FAM160B2;refGene_tx=XM_023252521.1;refGene_gn=FAM160B2;refGene_tx=XM_023252522.1
B1 36067985 36068108
refGene_gn=FAM160B2;refGene_tx=XM_023252513.1;refGene_gn=FAM160B2;refGene_tx=XM_023252514.1;refGene_gn=FAM160B2;refGene_tx=XM_023252515.1;refGene_gn=FAM160B2;refGene_tx=XM_023252516.1;refGene_gn=FAM160B2;refGene_tx=XM_023252517.1;refGene_gn=FAM160B2;refGene_tx=XM_023252518.1;refGene_gn=FAM160B2;refGene_tx=XM_023252519.1;refGene_gn=FAM160B2;refGene_tx=XM_023252520.1;refGene_gn=FAM160B2;refGene_tx=XM_023252521.1;refGene_gn=FAM160B2;refGene_tx=XM_023252522.1
B1 36068203 36068308
refGene_gn=FAM160B2;refGene_tx=XM_023252513.1;refGene_gn=FAM160B2;refGene_tx=XM_023252514.1;refGene_gn=FAM160B2;refGene_tx=XM_023252515.1;refGene_gn=FAM160B2;refGene_tx=XM_023252516.1;refGene_gn=FAM160B2;refGene_tx=XM_023252517.1;refGene_gn=FAM160B2;refGene_tx=XM_023252518.1;refGene_gn=FAM160B2;refGene_tx=XM_023252519.1;refGene_gn=FAM160B2;refGene_tx=XM_023252520.1;refGene_gn=FAM160B2;refGene_tx=XM_023252521.1;refGen

e_gn=FAM160B2;refGene_tx=XM_023252522.1
B1	36068677	36068683	refGene_gn=FAM160B2;refGene_tx=XM_023252521.1
B1	36071060	36071233
refGene_gn=FAM160B2;refGene_tx=XM_023252513.1;refGene_gn=FAM160B2;refGene_tx=XM_023252514.1;refGene_gn=FAM160B2;refGene_tx=XM_023252515.1;refGene_gn=FAM160B2;refGene_tx=XM_023252516.1;refGene_gn=FAM160B2;refGene_tx=XM_023252517.1;refGene_gn=FAM160B2;refGene_tx=XM_023252518.1;refGene_gn=FAM160B2;refGene_tx=XM_023252519.1;refGene_gn=FAM160B2;refGene_tx=XM_023252520.1
B1	36074145	36074224
refGene_gn=FAM160B2;refGene_tx=XM_023252513.1;refGene_gn=FAM160B2;refGene_tx=XM_023252514.1;refGene_gn=FAM160B2;refGene_tx=XM_023252515.1;refGene_gn=FAM160B2;refGene_tx=XM_023252516.1;refGene_gn=FAM160B2;refGene_tx=XM_023252517.1;refGene_gn=FAM160B2;refGene_tx=XM_023252518.1;refGene_gn=FAM160B2;refGene_tx=XM_023252519.1
B1	36078493	36078538
refGene_gn=FAM160B2;refGene_tx=XM_023252513.1;refGene_gn=FAM160B2;refGene_tx=XM_023252514.1;refGene_gn=FAM160B2;refGene_tx=XM_023252515.1;refGene_gn=FAM160B2;refGene_tx=XM_023252516.1;refGene_gn=FAM160B2;refGene_tx=XM_023252517.1;refGene_gn=FAM160B2;refGene_tx=XM_023252518.1;refGene_gn=FAM160B2;refGene_tx=XM_023252519.1
B1	36084185	36084299
refGene_gn=DMTN;refGene_tx=XM_019828448.2;refGene_gn=DMTN;refGene_tx=XM_019828449.2;refGene_gn=DMTN;refGene_tx=XM_019828450.2;refGene_gn=DMTN;refGene_tx=XM_019828451.2;refGene_gn=DMTN;refGene_tx=XM_019828452.2;refGene_gn=DMTN;refGene_tx=XM_019828453.2;refGene_gn=DMTN;refGene_tx=XM_019828454.2;refGene_gn=DMTN;refGene_tx=XM_023252523.1;refGene_gn=DMTN;refGene_tx=XM_023252524.1;refGene_gn=DMTN;refGene_tx=XM_023252525.1
B1	36084463	36084544
refGene_gn=DMTN;refGene_tx=XM_019828448.2;refGene_gn=DMTN;refGene_tx=XM_019828449.2;refGene_gn=DMTN;refGene_tx=XM_019828450.2;refGene_gn=DMTN;refGene_tx=XM_019828451.2;refGene_gn=DMTN;refGene_tx=XM_019828452.2;refGene_gn=DMTN;refGene_tx=XM_019828453.2;refGene_gn=DMTN;refGene_tx=XM_019828454.2;refGene_gn=DMTN;refGene_tx=XM_023252523.1;refGene_gn=DMTN;refGene_tx=XM_023252524.1;refGene_gn=DMTN;refGene_tx=XM_023252525.1
B1	36084817	36084883
refGene_gn=DMTN;refGene_tx=XM_019828448.2;refGene_gn=DMTN;refGene_tx=XM_019828449.2;refGene_gn=DMTN;refGene_tx=XM_019828451.2;refGene_gn=DMTN;refGene_tx=XM_019828452.2;refGene_gn=DMTN;refGene_tx=XM_023252523.1;refGene_gn=DMTN;refGene_tx=XM_023252524.1;refGene_gn=DMTN;refGene_tx=XM_023252525.1
B1	36085061	36085069
refGene_gn=DMTN;refGene_tx=XM_019828448.2;refGene_gn=DMTN;refGene_tx=XM_019828449.2;refGene_gn=DMTN;refGene_tx=XM_019828450.2;refGene_gn=DMTN;refGene_tx=XM_019828451.2;refGene_gn=DMTN;refGene_tx=XM_019828452.2;refGene_gn=DMTN;refGene_tx=XM_019828453.2;refGene_gn=DMTN;refGene_tx=XM_019828454.2;refGene_gn=DMTN;refGene_tx=XM_023252523.1;refGene_gn=DMTN;refGene_tx=XM_023252524.1;refGene_gn=DMTN;refGene_tx=XM_023252525.1
B1	36085255	36085304
refGene_gn=DMTN;refGene_tx=XM_019828448.2;refGene_gn=DMTN;refGene_tx=X

M_019828449.2;refGene_gn=DMTN;refGene_tx=XM_019828450.2;refGene_gn=DMTN;refGene_tx=XM_019828451.2;refGene_gn=DMTN;refGene_tx=XM_019828452.2;refGene_gn=DMTN;refGene_tx=XM_019828453.2;refGene_gn=DMTN;refGene_tx=XM_019828454.2;refGene_gn=DMTN;refGene_tx=XM_023252523.1;refGene_gn=DMTN;refGene_tx=XM_023252524.1;refGene_gn=DMTN;refGene_tx=XM_023252525.1
B1 36085472 36085537
refGene_gn=DMTN;refGene_tx=XM_019828448.2;refGene_gn=DMTN;refGene_tx=XM_019828449.2;refGene_gn=DMTN;refGene_tx=XM_019828450.2;refGene_gn=DMTN;refGene_tx=XM_019828451.2;refGene_gn=DMTN;refGene_tx=XM_019828452.2;refGene_gn=DMTN;refGene_tx=XM_019828453.2;refGene_gn=DMTN;refGene_tx=XM_019828454.2;refGene_gn=DMTN;refGene_tx=XM_023252523.1;refGene_gn=DMTN;refGene_tx=XM_023252524.1;refGene_gn=DMTN;refGene_tx=XM_023252525.1
B1 36089498 36089604
refGene_gn=DMTN;refGene_tx=XM_019828448.2;refGene_gn=DMTN;refGene_tx=XM_019828449.2;refGene_gn=DMTN;refGene_tx=XM_019828450.2;refGene_gn=DMTN;refGene_tx=XM_019828451.2;refGene_gn=DMTN;refGene_tx=XM_019828452.2;refGene_gn=DMTN;refGene_tx=XM_019828453.2;refGene_gn=DMTN;refGene_tx=XM_019828454.2;refGene_gn=DMTN;refGene_tx=XM_023252523.1;refGene_gn=DMTN;refGene_tx=XM_023252524.1;refGene_gn=DMTN;refGene_tx=XM_023252525.1
B1 36090221 36090346
refGene_gn=DMTN;refGene_tx=XM_019828448.2;refGene_gn=DMTN;refGene_tx=XM_019828449.2;refGene_gn=DMTN;refGene_tx=XM_019828450.2;refGene_gn=DMTN;refGene_tx=XM_019828451.2;refGene_gn=DMTN;refGene_tx=XM_019828452.2;refGene_gn=DMTN;refGene_tx=XM_019828453.2;refGene_gn=DMTN;refGene_tx=XM_019828454.2;refGene_gn=DMTN;refGene_tx=XM_023252523.1;refGene_gn=DMTN;refGene_tx=XM_023252524.1;refGene_gn=DMTN;refGene_tx=XM_023252525.1
B1 36090856 36091009
refGene_gn=DMTN;refGene_tx=XM_019828448.2;refGene_gn=DMTN;refGene_tx=XM_019828449.2;refGene_gn=DMTN;refGene_tx=XM_019828450.2;refGene_gn=DMTN;refGene_tx=XM_019828451.2;refGene_gn=DMTN;refGene_tx=XM_019828452.2;refGene_gn=DMTN;refGene_tx=XM_019828453.2;refGene_gn=DMTN;refGene_tx=XM_019828454.2;refGene_gn=DMTN;refGene_tx=XM_023252523.1;refGene_gn=DMTN;refGene_tx=XM_023252524.1;refGene_gn=DMTN;refGene_tx=XM_023252525.1
B1 36091239 36091296
refGene_gn=DMTN;refGene_tx=XM_019828448.2;refGene_gn=DMTN;refGene_tx=XM_019828449.2;refGene_gn=DMTN;refGene_tx=XM_019828450.2;refGene_gn=DMTN;refGene_tx=XM_019828451.2;refGene_gn=DMTN;refGene_tx=XM_019828452.2;refGene_gn=DMTN;refGene_tx=XM_019828453.2;refGene_gn=DMTN;refGene_tx=XM_019828454.2;refGene_gn=DMTN;refGene_tx=XM_023252523.1;refGene_gn=DMTN;refGene_tx=XM_023252524.1;refGene_gn=DMTN;refGene_tx=XM_023252525.1
B1 36091656 36091756
refGene_gn=DMTN;refGene_tx=XM_019828448.2;refGene_gn=DMTN;refGene_tx=XM_019828449.2;refGene_gn=DMTN;refGene_tx=XM_019828450.2;refGene_gn=DMTN;refGene_tx=XM_019828451.2;refGene_gn=DMTN;refGene_tx=XM_019828452.2;refGene_gn=DMTN;refGene_tx=XM_019828453.2;refGene_gn=DMTN;refGene_tx=XM_019828454.2;refGene_gn=DMTN;refGene_tx=XM_023252523.1;refGene_gn=DMTN;refGene_tx=XM_023252524.1;refGene_gn=DMTN;refGene_tx=XM_023252525.1
B1 36092086 36092131
refGene_gn=DMTN;refGene_tx=XM_019828448.2;refGene_gn=DMTN;refGene_tx=XM_019828450.2;refGene_gn=DMTN;refGene_tx=XM_019828451.2;refGene_gn=DMT

N;refGene_tx=XM_019828453.2;refGene_gn=DMTN;refGene_tx=XM_023252523.1;
refGene_gn=DMTN;refGene_tx=XM_023252524.1;refGene_gn=DMTN;refGene_tx=X
M_023252525.1
B1 36093058 36093214
refGene_gn=DMTN;refGene_tx=XM_019828448.2;refGene_gn=DMTN;refGene_tx=X
M_019828449.2;refGene_gn=DMTN;refGene_tx=XM_019828450.2;refGene_gn=DMT
N;refGene_tx=XM_019828451.2;refGene_gn=DMTN;refGene_tx=XM_019828452.2;
refGene_gn=DMTN;refGene_tx=XM_019828453.2;refGene_gn=DMTN;refGene_tx=X
M_019828454.2;refGene_gn=DMTN;refGene_tx=XM_023252523.1;refGene_gn=DMT
N;refGene_tx=XM_023252524.1;refGene_gn=DMTN;refGene_tx=XM_023252525.1
B1 36093583 36093658
refGene_gn=DMTN;refGene_tx=XM_019828448.2;refGene_gn=DMTN;refGene_tx=X
M_019828449.2;refGene_gn=DMTN;refGene_tx=XM_019828450.2;refGene_gn=DMT
N;refGene_tx=XM_023252523.1;refGene_gn=DMTN;refGene_tx=XM_023252524.1;
refGene_gn=DMTN;refGene_tx=XM_023252525.1
B1 36093937 36093955
refGene_gn=DMTN;refGene_tx=XM_019828448.2;refGene_gn=DMTN;refGene_tx=X
M_019828449.2;refGene_gn=DMTN;refGene_tx=XM_019828450.2;refGene_gn=DMT
N;refGene_tx=XM_019828451.2;refGene_gn=DMTN;refGene_tx=XM_019828452.2;
refGene_gn=DMTN;refGene_tx=XM_019828453.2;refGene_gn=DMTN;refGene_tx=X
M_019828454.2;refGene_gn=DMTN;refGene_tx=XM_023252523.1;refGene_gn=DMT
N;refGene_tx=XM_023252524.1;refGene_gn=DMTN;refGene_tx=XM_023252525.1
B1 36111182 36111476
refGene_gn=FGF17;refGene_tx=XM_003984716.5;refGene_gn=FGF17;refGene_tx
=XM_019828455.2;refGene_gn=FGF17;refGene_tx=XM_019828460.2
B1 36112532 36112639
refGene_gn=FGF17;refGene_tx=XM_003984716.5;refGene_gn=FGF17;refGene_tx
=XM_019828455.2;refGene_gn=FGF17;refGene_tx=XM_019828460.2
B1 36112879 36113057
refGene_gn=FGF17;refGene_tx=XM_003984716.5;refGene_gn=FGF17;refGene_tx
=XM_019828455.2;refGene_gn=FGF17;refGene_tx=XM_019828460.2
B1 36116059 36116096
refGene_gn=FGF17;refGene_tx=XM_003984716.5;refGene_gn=FGF17;refGene_tx
=XM_019828455.2
B1 36116269 36116403
refGene_gn=FGF17;refGene_tx=XM_003984716.5;refGene_gn=FGF17;refGene_tx
=XM_019828455.2
B1 36119459 36119495
refGene_gn=FGF17;refGene_tx=XM_003984716.5;refGene_gn=FGF17;refGene_tx
=XM_019828455.2
B1 36121806 36121851
refGene_gn=NPM2;refGene_tx=XM_023252526.1;refGene_gn=NPM2;refGene_tx=X
M_023252527.1
B1 36121932 36121966
refGene_gn=NPM2;refGene_tx=XM_023252526.1;refGene_gn=NPM2;refGene_tx=X
M_023252527.1;refGene_gn=NPM2;refGene_tx=XM_023252528.1
B1 36122814 36122849
refGene_gn=NPM2;refGene_tx=XM_023252526.1;refGene_gn=NPM2;refGene_tx=X
M_023252527.1
B1 36123018 36123203

refGene_gn=NPM2;refGene_tx=XM_023252526.1;refGene_gn=NPM2;refGene_tx=XM_023252527.1;refGene_gn=NPM2;refGene_tx=XM_023252528.1
B1 36123993 36124084
refGene_gn=NPM2;refGene_tx=XM_023252526.1;refGene_gn=NPM2;refGene_tx=XM_023252527.1;refGene_gn=NPM2;refGene_tx=XM_023252528.1
B1 36134999 36135125
refGene_gn=NPM2;refGene_tx=XM_023252526.1;refGene_gn=NPM2;refGene_tx=XM_023252527.1;refGene_gn=NPM2;refGene_tx=XM_023252528.1
B1 36135270 36135356
refGene_gn=NPM2;refGene_tx=XM_023252526.1;refGene_gn=NPM2;refGene_tx=XM_023252527.1;refGene_gn=NPM2;refGene_tx=XM_023252528.1
B1 36135457 36135605
refGene_gn=NPM2;refGene_tx=XM_023252526.1;refGene_gn=NPM2;refGene_tx=XM_023252527.1;refGene_gn=NPM2;refGene_tx=XM_023252528.1
B1 36147440 36147534
refGene_gn=XPO7;refGene_tx=XM_019828465.2;refGene_gn=XPO7;refGene_tx=XM_019828466.2
B1 36148530 36148658
refGene_gn=XPO7;refGene_tx=XM_019828465.2;refGene_gn=XPO7;refGene_tx=XM_019828466.2
B1 36149326 36149425
refGene_gn=XPO7;refGene_tx=XM_019828465.2;refGene_gn=XPO7;refGene_tx=XM_019828466.2
B1 36150538 36150699
refGene_gn=XPO7;refGene_tx=XM_019828465.2;refGene_gn=XPO7;refGene_tx=XM_019828466.2
B1 36152716 36152855
refGene_gn=XPO7;refGene_tx=XM_019828465.2;refGene_gn=XPO7;refGene_tx=XM_019828466.2
B1 36153097 36153312
refGene_gn=XPO7;refGene_tx=XM_019828465.2;refGene_gn=XPO7;refGene_tx=XM_019828466.2
B1 36153537 36153620
refGene_gn=XPO7;refGene_tx=XM_019828465.2;refGene_gn=XPO7;refGene_tx=XM_019828466.2
B1 36155997 36156105
refGene_gn=XPO7;refGene_tx=XM_019828465.2;refGene_gn=XPO7;refGene_tx=XM_019828466.2
B1 36156814 36156903
refGene_gn=XPO7;refGene_tx=XM_019828465.2;refGene_gn=XPO7;refGene_tx=XM_019828466.2
B1 36159427 36159534
refGene_gn=XPO7;refGene_tx=XM_019828465.2;refGene_gn=XPO7;refGene_tx=XM_019828466.2
B1 36160506 36160615
refGene_gn=XPO7;refGene_tx=XM_019828465.2;refGene_gn=XPO7;refGene_tx=XM_019828466.2
B1 36161022 36161086
refGene_gn=XPO7;refGene_tx=XM_019828465.2;refGene_gn=XPO7;refGene_tx=XM_019828466.2

B1 36161999 36162080
refGene_gn=XPO7;refGene_tx=XM_019828465.2;refGene_gn=XPO7;refGene_tx=XM_019828466.2
B1 36163219 36163293
refGene_gn=XPO7;refGene_tx=XM_019828465.2;refGene_gn=XPO7;refGene_tx=XM_019828466.2
B1 36163806 36163942
refGene_gn=XPO7;refGene_tx=XM_019828465.2;refGene_gn=XPO7;refGene_tx=XM_019828466.2
B1 36166806 36166912
refGene_gn=XPO7;refGene_tx=XM_019828465.2;refGene_gn=XPO7;refGene_tx=XM_019828466.2
B1 36167600 36167794
refGene_gn=XPO7;refGene_tx=XM_019828465.2;refGene_gn=XPO7;refGene_tx=XM_019828466.2
B1 36169768 36169941
refGene_gn=XPO7;refGene_tx=XM_019828465.2;refGene_gn=XPO7;refGene_tx=XM_019828466.2
B1 36171118 36171265
refGene_gn=XPO7;refGene_tx=XM_019828465.2;refGene_gn=XPO7;refGene_tx=XM_019828466.2
B1 36174451 36174571
refGene_gn=XPO7;refGene_tx=XM_019828465.2;refGene_gn=XPO7;refGene_tx=XM_019828466.2
B1 36176235 36176309
refGene_gn=XPO7;refGene_tx=XM_019828465.2;refGene_gn=XPO7;refGene_tx=XM_019828466.2
B1 36176888 36177054
refGene_gn=XPO7;refGene_tx=XM_019828465.2;refGene_gn=XPO7;refGene_tx=XM_019828466.2
B1 36178434 36178539
refGene_gn=XPO7;refGene_tx=XM_019828465.2;refGene_gn=XPO7;refGene_tx=XM_019828466.2
B1 36181219 36181312
refGene_gn=XPO7;refGene_tx=XM_019828465.2;refGene_gn=XPO7;refGene_tx=XM_019828466.2
B1 36182608 36182775
refGene_gn=XPO7;refGene_tx=XM_019828465.2;refGene_gn=XPO7;refGene_tx=XM_019828466.2
B1 36183362 36183456
refGene_gn=XPO7;refGene_tx=XM_019828465.2;refGene_gn=XPO7;refGene_tx=XM_019828466.2
B1 36185500 36185647
refGene_gn=XPO7;refGene_tx=XM_019828465.2;refGene_gn=XPO7;refGene_tx=XM_019828466.2
B1 36186173 36186194
refGene_gn=XPO7;refGene_tx=XM_019828465.2;refGene_gn=XPO7;refGene_tx=XM_019828466.2
B1 36243798 36243861 refGene_gn=DOK2;refGene_tx=XM_003984731.5
B1 36244837 36245119 refGene_gn=DOK2;refGene_tx=XM_003984731.5

B1 36245370 36245458 refGene_gn=DOK2;refGene_tx=XM_003984731.5
B1 36246366 36246551
refGene_gn=DOK2;refGene_tx=XM_003984731.5;refGene_gn=DOK2;refGene_tx=XM_019828468.2
B1 36247050 36247671
refGene_gn=DOK2;refGene_tx=XM_003984731.5;refGene_gn=DOK2;refGene_tx=XM_019828468.2
B1 36343172 36343212 refGene_gn=GFRA2;refGene_tx=XM_011281463.3
B1 36348448 36348763 refGene_gn=GFRA2;refGene_tx=XM_011281463.3
B1 36354337 36354421 refGene_gn=GFRA2;refGene_tx=XM_011281463.3
B1 36373701 36373797 refGene_gn=GFRA2;refGene_tx=XM_011281463.3
B1 36373799 36374055 refGene_gn=GFRA2;refGene_tx=XM_011281463.3
B1 36412154 36412264 refGene_gn=GFRA2;refGene_tx=XM_011281463.3
B1 36413014 36413155 refGene_gn=GFRA2;refGene_tx=XM_011281463.3
B1 36415547 36415720 refGene_gn=GFRA2;refGene_tx=XM_011281463.3
B1 36425246 36425300 refGene_gn=GFRA2;refGene_tx=XM_011281463.3
B1 36426399 36426522 refGene_gn=GFRA2;refGene_tx=XM_011281463.3
B1 36510985 36511141
refGene_gn=LOC109498816;refGene_tx=XM_019828472.2
B1 37783324 37783669
refGene_gn=LZTS1;refGene_tx=XM_003984723.4;refGene_gn=LZTS1;refGene_tx=XM_006930658.3
B1 37785187 37785991
refGene_gn=LZTS1;refGene_tx=XM_003984723.4;refGene_gn=LZTS1;refGene_tx=XM_006930658.3
B1 37788563 37789235
refGene_gn=LZTS1;refGene_tx=XM_003984723.4;refGene_gn=LZTS1;refGene_tx=XM_006930658.3
B1 37810691 37810831 refGene_gn=ATP6V1B2;refGene_tx=XM_003984724.5
B1 37812871 37813001 refGene_gn=ATP6V1B2;refGene_tx=XM_003984724.5
B1 37814067 37814172 refGene_gn=ATP6V1B2;refGene_tx=XM_003984724.5
B1 37815395 37815478 refGene_gn=ATP6V1B2;refGene_tx=XM_003984724.5
B1 37817342 37817493 refGene_gn=ATP6V1B2;refGene_tx=XM_003984724.5
B1 37819081 37819205 refGene_gn=ATP6V1B2;refGene_tx=XM_003984724.5
B1 37819810 37819908 refGene_gn=ATP6V1B2;refGene_tx=XM_003984724.5
B1 37820259 37820361 refGene_gn=ATP6V1B2;refGene_tx=XM_003984724.5
B1 37820891 37821031 refGene_gn=ATP6V1B2;refGene_tx=XM_003984724.5
B1 37821484 37821562 refGene_gn=ATP6V1B2;refGene_tx=XM_003984724.5
B1 37821691 37821785 refGene_gn=ATP6V1B2;refGene_tx=XM_003984724.5
B1 37822530 37822629 refGene_gn=ATP6V1B2;refGene_tx=XM_003984724.5
B1 37827690 37827746 refGene_gn=ATP6V1B2;refGene_tx=XM_003984724.5
B1 37834292 37834428 refGene_gn=ATP6V1B2;refGene_tx=XM_003984724.5
B1 37857906 37858030
refGene_gn=SLC18A1;refGene_tx=XM_006930662.3;refGene_gn=SLC18A1;refGene_tx=XM_019829989.2;refGene_gn=SLC18A1;refGene_tx=XM_019829990.2
B1 37858663 37859027
refGene_gn=SLC18A1;refGene_tx=XM_006930662.3;refGene_gn=SLC18A1;refGene_tx=XM_019829989.2;refGene_gn=SLC18A1;refGene_tx=XM_019829990.2
B1 37859623 37859682
refGene_gn=SLC18A1;refGene_tx=XM_006930662.3;refGene_gn=SLC18A1;refGen

e_tx=XM_019829989.2;refGene_gn=SLC18A1;refGene_tx=XM_019829990.2
B1 37862280 37862364
refGene_gn=SLC18A1;refGene_tx=XM_006930662.3;refGene_gn=SLC18A1;refGene_tx=XM_019829989.2;refGene_gn=SLC18A1;refGene_tx=XM_019829990.2
B1 37863497 37863590
refGene_gn=SLC18A1;refGene_tx=XM_006930662.3;refGene_gn=SLC18A1;refGene_tx=XM_019829989.2;refGene_gn=SLC18A1;refGene_tx=XM_019829990.2
B1 37865412 37865502
refGene_gn=SLC18A1;refGene_tx=XM_006930662.3;refGene_gn=SLC18A1;refGene_tx=XM_019829989.2;refGene_gn=SLC18A1;refGene_tx=XM_019829990.2
B1 37865750 37865794
refGene_gn=SLC18A1;refGene_tx=XM_006930662.3;refGene_gn=SLC18A1;refGene_tx=XM_019829989.2;refGene_gn=SLC18A1;refGene_tx=XM_019829990.2
B1 37868070 37868131
refGene_gn=SLC18A1;refGene_tx=XM_006930662.3;refGene_gn=SLC18A1;refGene_tx=XM_019829989.2;refGene_gn=SLC18A1;refGene_tx=XM_019829990.2
B1 37868214 37868310
refGene_gn=SLC18A1;refGene_tx=XM_006930662.3;refGene_gn=SLC18A1;refGene_tx=XM_019829990.2
B1 37874918 37874997
refGene_gn=SLC18A1;refGene_tx=XM_006930662.3;refGene_gn=SLC18A1;refGene_tx=XM_019829989.2;refGene_gn=SLC18A1;refGene_tx=XM_019829990.2
B1 37875896 37875948
refGene_gn=SLC18A1;refGene_tx=XM_006930662.3;refGene_gn=SLC18A1;refGene_tx=XM_019829989.2;refGene_gn=SLC18A1;refGene_tx=XM_019829990.2
B1 37877668 37877732
refGene_gn=SLC18A1;refGene_tx=XM_006930662.3;refGene_gn=SLC18A1;refGene_tx=XM_019829989.2;refGene_gn=SLC18A1;refGene_tx=XM_019829990.2
B1 37877991 37878111
refGene_gn=SLC18A1;refGene_tx=XM_006930662.3;refGene_gn=SLC18A1;refGene_tx=XM_019829989.2;refGene_gn=SLC18A1;refGene_tx=XM_019829990.2
B1 37878330 37878464
refGene_gn=SLC18A1;refGene_tx=XM_006930662.3;refGene_gn=SLC18A1;refGene_tx=XM_019829989.2
B1 37879355 37879514
refGene_gn=SLC18A1;refGene_tx=XM_006930662.3;refGene_gn=SLC18A1;refGene_tx=XM_019829989.2;refGene_gn=SLC18A1;refGene_tx=XM_019829990.2
B1 38074369 38074370 refGene_gn=LPL;refGene_tx=NM_001042567.1
B1 38076636 38076741 refGene_gn=LPL;refGene_tx=NM_001042567.1
B1 38078534 38078717 refGene_gn=LPL;refGene_tx=NM_001042567.1
B1 38079916 38080037 refGene_gn=LPL;refGene_tx=NM_001042567.1
B1 38082234 38082477 refGene_gn=LPL;refGene_tx=NM_001042567.1
B1 38083702 38083936 refGene_gn=LPL;refGene_tx=NM_001042567.1
B1 38084693 38084805 refGene_gn=LPL;refGene_tx=NM_001042567.1
B1 38086308 38086488 refGene_gn=LPL;refGene_tx=NM_001042567.1
B1 38090951 38091121 refGene_gn=LPL;refGene_tx=NM_001042567.1
B1 38098342 38098430 refGene_gn=LPL;refGene_tx=NM_001042567.1
B1 40446257 40447130
refGene_gn=LOC101095575;refGene_tx=XM_004001593.5
B1 40502962 40503178

refGene_gn=PSD3;refGene_tx=XM_023252529.1;refGene_gn=PSD3;refGene_tx=XM_023252530.1;refGene_gn=PSD3;refGene_tx=XM_023252531.1;refGene_gn=PSD3;refGene_tx=XM_023252532.1;refGene_gn=PSD3;refGene_tx=XM_023252533.1;refGene_gn=PSD3;refGene_tx=XM_023252537.1;refGene_gn=PSD3;refGene_tx=XM_023252538.1;refGene_gn=PSD3;refGene_tx=XM_023252539.1;refGene_gn=PSD3;refGene_tx=XM_023252540.1;refGene_gn=PSD3;refGene_tx=XM_023252541.1
B1 40533108 40533211 refGene_gn=PSD3;refGene_tx=XM_023252533.1
B1 40542164 40542185 refGene_gn=PSD3;refGene_tx=XM_023252535.1
B1 40543384 40543396 refGene_gn=PSD3;refGene_tx=XM_023252536.1
B1 40545438 40545489 refGene_gn=PSD3;refGene_tx=XM_023252534.1
B1 40549107 40549251
refGene_gn=PSD3;refGene_tx=XM_023252529.1;refGene_gn=PSD3;refGene_tx=XM_023252530.1;refGene_gn=PSD3;refGene_tx=XM_023252531.1;refGene_gn=PSD3;refGene_tx=XM_023252532.1;refGene_gn=PSD3;refGene_tx=XM_023252533.1;refGene_gn=PSD3;refGene_tx=XM_023252534.1;refGene_gn=PSD3;refGene_tx=XM_023252535.1;refGene_gn=PSD3;refGene_tx=XM_023252536.1;refGene_gn=PSD3;refGene_tx=XM_023252537.1;refGene_gn=PSD3;refGene_tx=XM_023252538.1;refGene_gn=PSD3;refGene_tx=XM_023252539.1;refGene_gn=PSD3;refGene_tx=XM_023252540.1;refGene_gn=PSD3;refGene_tx=XM_023252541.1
B1 40563845 40563990
refGene_gn=PSD3;refGene_tx=XM_023252529.1;refGene_gn=PSD3;refGene_tx=XM_023252530.1;refGene_gn=PSD3;refGene_tx=XM_023252531.1;refGene_gn=PSD3;refGene_tx=XM_023252532.1;refGene_gn=PSD3;refGene_tx=XM_023252533.1;refGene_gn=PSD3;refGene_tx=XM_023252534.1;refGene_gn=PSD3;refGene_tx=XM_023252535.1;refGene_gn=PSD3;refGene_tx=XM_023252536.1;refGene_gn=PSD3;refGene_tx=XM_023252537.1;refGene_gn=PSD3;refGene_tx=XM_023252538.1;refGene_gn=PSD3;refGene_tx=XM_023252539.1;refGene_gn=PSD3;refGene_tx=XM_023252540.1;refGene_gn=PSD3;refGene_tx=XM_023252541.1
B1 40567062 40567220
refGene_gn=PSD3;refGene_tx=XM_023252529.1;refGene_gn=PSD3;refGene_tx=XM_023252530.1;refGene_gn=PSD3;refGene_tx=XM_023252531.1;refGene_gn=PSD3;refGene_tx=XM_023252532.1;refGene_gn=PSD3;refGene_tx=XM_023252533.1;refGene_gn=PSD3;refGene_tx=XM_023252534.1;refGene_gn=PSD3;refGene_tx=XM_023252535.1;refGene_gn=PSD3;refGene_tx=XM_023252536.1;refGene_gn=PSD3;refGene_tx=XM_023252537.1;refGene_gn=PSD3;refGene_tx=XM_023252538.1;refGene_gn=PSD3;refGene_tx=XM_023252539.1;refGene_gn=PSD3;refGene_tx=XM_023252540.1;refGene_gn=PSD3;refGene_tx=XM_023252541.1
B1 40607224 40607295
refGene_gn=PSD3;refGene_tx=XM_023252529.1;refGene_gn=PSD3;refGene_tx=XM_023252530.1;refGene_gn=PSD3;refGene_tx=XM_023252531.1;refGene_gn=PSD3;refGene_tx=XM_023252532.1;refGene_gn=PSD3;refGene_tx=XM_023252533.1;refGene_gn=PSD3;refGene_tx=XM_023252534.1;refGene_gn=PSD3;refGene_tx=XM_023252535.1;refGene_gn=PSD3;refGene_tx=XM_023252536.1;refGene_gn=PSD3;refGene_tx=XM_023252537.1;refGene_gn=PSD3;refGene_tx=XM_023252538.1;refGene_gn=PSD3;refGene_tx=XM_023252539.1;refGene_gn=PSD3;refGene_tx=XM_023252540.1;refGene_gn=PSD3;refGene_tx=XM_023252541.1
B1 40676465 40676659
refGene_gn=PSD3;refGene_tx=XM_023252529.1;refGene_gn=PSD3;refGene_tx=XM_023252530.1;refGene_gn=PSD3;refGene_tx=XM_023252531.1;refGene_gn=PSD3;refGene_tx=XM_023252532.1;refGene_gn=PSD3;refGene_tx=XM_023252533.1;

refGene_gn=PSD3;refGene_tx=XM_023252534.1;refGene_gn=PSD3;refGene_tx=XM_023252535.1;refGene_gn=PSD3;refGene_tx=XM_023252536.1;refGene_gn=PSD3;refGene_tx=XM_023252537.1;refGene_gn=PSD3;refGene_tx=XM_023252538.1;refGene_gn=PSD3;refGene_tx=XM_023252539.1;refGene_gn=PSD3;refGene_tx=XM_023252540.1;refGene_gn=PSD3;refGene_tx=XM_023252541.1
B1 40720881 40720925
refGene_gn=PSD3;refGene_tx=XM_023252529.1;refGene_gn=PSD3;refGene_tx=XM_023252530.1;refGene_gn=PSD3;refGene_tx=XM_023252531.1;refGene_gn=PSD3;refGene_tx=XM_023252532.1;refGene_gn=PSD3;refGene_tx=XM_023252533.1;refGene_gn=PSD3;refGene_tx=XM_023252534.1;refGene_gn=PSD3;refGene_tx=XM_023252535.1;refGene_gn=PSD3;refGene_tx=XM_023252536.1;refGene_gn=PSD3;refGene_tx=XM_023252537.1;refGene_gn=PSD3;refGene_tx=XM_023252538.1;refGene_gn=PSD3;refGene_tx=XM_023252539.1;refGene_gn=PSD3;refGene_tx=XM_023252540.1;refGene_gn=PSD3;refGene_tx=XM_023252541.1
B1 40827181 40827271
refGene_gn=PSD3;refGene_tx=XM_023252529.1;refGene_gn=PSD3;refGene_tx=XM_023252530.1;refGene_gn=PSD3;refGene_tx=XM_023252531.1;refGene_gn=PSD3;refGene_tx=XM_023252532.1;refGene_gn=PSD3;refGene_tx=XM_023252533.1;refGene_gn=PSD3;refGene_tx=XM_023252534.1;refGene_gn=PSD3;refGene_tx=XM_023252535.1;refGene_gn=PSD3;refGene_tx=XM_023252536.1;refGene_gn=PSD3;refGene_tx=XM_023252537.1;refGene_gn=PSD3;refGene_tx=XM_023252538.1;refGene_gn=PSD3;refGene_tx=XM_023252539.1;refGene_gn=PSD3;refGene_tx=XM_023252540.1;refGene_gn=PSD3;refGene_tx=XM_023252541.1
B1 40853470 40853529
refGene_gn=PSD3;refGene_tx=XM_023252529.1;refGene_gn=PSD3;refGene_tx=XM_023252530.1;refGene_gn=PSD3;refGene_tx=XM_023252531.1;refGene_gn=PSD3;refGene_tx=XM_023252532.1;refGene_gn=PSD3;refGene_tx=XM_023252533.1;refGene_gn=PSD3;refGene_tx=XM_023252534.1;refGene_gn=PSD3;refGene_tx=XM_023252535.1;refGene_gn=PSD3;refGene_tx=XM_023252536.1;refGene_gn=PSD3;refGene_tx=XM_023252537.1;refGene_gn=PSD3;refGene_tx=XM_023252538.1;refGene_gn=PSD3;refGene_tx=XM_023252539.1;refGene_gn=PSD3;refGene_tx=XM_023252540.1;refGene_gn=PSD3;refGene_tx=XM_023252541.1
B1 40856496 40856609
refGene_gn=PSD3;refGene_tx=XM_023252529.1;refGene_gn=PSD3;refGene_tx=XM_023252530.1;refGene_gn=PSD3;refGene_tx=XM_023252531.1;refGene_gn=PSD3;refGene_tx=XM_023252532.1;refGene_gn=PSD3;refGene_tx=XM_023252533.1;refGene_gn=PSD3;refGene_tx=XM_023252534.1;refGene_gn=PSD3;refGene_tx=XM_023252535.1;refGene_gn=PSD3;refGene_tx=XM_023252536.1;refGene_gn=PSD3;refGene_tx=XM_023252537.1;refGene_gn=PSD3;refGene_tx=XM_023252538.1;refGene_gn=PSD3;refGene_tx=XM_023252539.1;refGene_gn=PSD3;refGene_tx=XM_023252540.1;refGene_gn=PSD3;refGene_tx=XM_023252541.1
B1 40858584 40858665
refGene_gn=PSD3;refGene_tx=XM_023252529.1;refGene_gn=PSD3;refGene_tx=XM_023252530.1;refGene_gn=PSD3;refGene_tx=XM_023252531.1;refGene_gn=PSD3;refGene_tx=XM_023252532.1;refGene_gn=PSD3;refGene_tx=XM_023252533.1;refGene_gn=PSD3;refGene_tx=XM_023252534.1;refGene_gn=PSD3;refGene_tx=XM_023252535.1;refGene_gn=PSD3;refGene_tx=XM_023252536.1;refGene_gn=PSD3;refGene_tx=XM_023252537.1;refGene_gn=PSD3;refGene_tx=XM_023252538.1;refGene_gn=PSD3;refGene_tx=XM_023252539.1;refGene_gn=PSD3;refGene_tx=XM_023252540.1;refGene_gn=PSD3;refGene_tx=XM_023252541.1

B1 40858765 40858960
refGene_gn=PSD3;refGene_tx=XM_023252529.1;refGene_gn=PSD3;refGene_tx=XM_023252530.1;refGene_gn=PSD3;refGene_tx=XM_023252531.1;refGene_gn=PSD3;refGene_tx=XM_023252532.1;refGene_gn=PSD3;refGene_tx=XM_023252533.1;refGene_gn=PSD3;refGene_tx=XM_023252534.1;refGene_gn=PSD3;refGene_tx=XM_023252535.1;refGene_gn=PSD3;refGene_tx=XM_023252536.1;refGene_gn=PSD3;refGene_tx=XM_023252537.1;refGene_gn=PSD3;refGene_tx=XM_023252538.1;refGene_gn=PSD3;refGene_tx=XM_023252539.1;refGene_gn=PSD3;refGene_tx=XM_023252540.1;refGene_gn=PSD3;refGene_tx=XM_023252541.1
B1 40864850 40864897 refGene_gn=PSD3;refGene_tx=XM_023252541.1
B1 40925888 40925924
refGene_gn=PSD3;refGene_tx=XM_023252529.1;refGene_gn=PSD3;refGene_tx=XM_023252530.1;refGene_gn=PSD3;refGene_tx=XM_023252531.1;refGene_gn=PSD3;refGene_tx=XM_023252533.1;refGene_gn=PSD3;refGene_tx=XM_023252534.1;refGene_gn=PSD3;refGene_tx=XM_023252535.1;refGene_gn=PSD3;refGene_tx=XM_023252536.1;refGene_gn=PSD3;refGene_tx=XM_023252537.1;refGene_gn=PSD3;refGene_tx=XM_023252538.1;refGene_gn=PSD3;refGene_tx=XM_023252539.1;refGene_gn=PSD3;refGene_tx=XM_023252540.1
B1 40930821 40931217
refGene_gn=PSD3;refGene_tx=XM_023252529.1;refGene_gn=PSD3;refGene_tx=XM_023252530.1;refGene_gn=PSD3;refGene_tx=XM_023252531.1;refGene_gn=PSD3;refGene_tx=XM_023252532.1;refGene_gn=PSD3;refGene_tx=XM_023252533.1;refGene_gn=PSD3;refGene_tx=XM_023252534.1;refGene_gn=PSD3;refGene_tx=XM_023252535.1;refGene_gn=PSD3;refGene_tx=XM_023252536.1;refGene_gn=PSD3;refGene_tx=XM_023252537.1;refGene_gn=PSD3;refGene_tx=XM_023252538.1;refGene_gn=PSD3;refGene_tx=XM_023252539.1;refGene_gn=PSD3;refGene_tx=XM_023252540.1
B1 40933842 40934923
refGene_gn=PSD3;refGene_tx=XM_023252529.1;refGene_gn=PSD3;refGene_tx=XM_023252530.1;refGene_gn=PSD3;refGene_tx=XM_023252531.1;refGene_gn=PSD3;refGene_tx=XM_023252532.1;refGene_gn=PSD3;refGene_tx=XM_023252533.1;refGene_gn=PSD3;refGene_tx=XM_023252534.1;refGene_gn=PSD3;refGene_tx=XM_023252535.1;refGene_gn=PSD3;refGene_tx=XM_023252536.1;refGene_gn=PSD3;refGene_tx=XM_023252537.1;refGene_gn=PSD3;refGene_tx=XM_023252538.1;refGene_gn=PSD3;refGene_tx=XM_023252539.1;refGene_gn=PSD3;refGene_tx=XM_023252540.1
B1 40949286 40949317 refGene_gn=PSD3;refGene_tx=XM_023252538.1
B1 40999765 40999874
refGene_gn=PSD3;refGene_tx=XM_023252529.1;refGene_gn=PSD3;refGene_tx=XM_023252530.1;refGene_gn=PSD3;refGene_tx=XM_023252531.1;refGene_gn=PSD3;refGene_tx=XM_023252532.1;refGene_gn=PSD3;refGene_tx=XM_023252533.1;refGene_gn=PSD3;refGene_tx=XM_023252534.1;refGene_gn=PSD3;refGene_tx=XM_023252535.1;refGene_gn=PSD3;refGene_tx=XM_023252536.1;refGene_gn=PSD3;refGene_tx=XM_023252537.1
B1 41075580 41075601 refGene_gn=PSD3;refGene_tx=XM_023252537.1
B1 41150489 41151254
refGene_gn=PSD3;refGene_tx=XM_023252529.1;refGene_gn=PSD3;refGene_tx=XM_023252530.1;refGene_gn=PSD3;refGene_tx=XM_023252531.1;refGene_gn=PSD3;refGene_tx=XM_023252532.1;refGene_gn=PSD3;refGene_tx=XM_023252533.1;refGene_gn=PSD3;refGene_tx=XM_023252534.1;refGene_gn=PSD3;refGene_tx=X

M_023252535.1;refGene_gn=PSD3;refGene_tx=XM_023252536.1
B1 41402180 41402361
refGene_gn=SH2D4A;refGene_tx=XM_003984734.5;refGene_gn=SH2D4A;refGene_tx=XM_006930668.3;refGene_gn=SH2D4A;refGene_tx=XM_011281481.3;refGene_gn=SH2D4A;refGene_tx=XM_011281482.3
B1 41413129 41413289
refGene_gn=SH2D4A;refGene_tx=XM_003984734.5;refGene_gn=SH2D4A;refGene_tx=XM_006930668.3;refGene_gn=SH2D4A;refGene_tx=XM_011281481.3;refGene_gn=SH2D4A;refGene_tx=XM_011281482.3
B1 41414674 41414846
refGene_gn=SH2D4A;refGene_tx=XM_003984734.5;refGene_gn=SH2D4A;refGene_tx=XM_006930668.3;refGene_gn=SH2D4A;refGene_tx=XM_011281481.3;refGene_gn=SH2D4A;refGene_tx=XM_011281482.3
B1 41430910 41430991
refGene_gn=SH2D4A;refGene_tx=XM_003984734.5;refGene_gn=SH2D4A;refGene_tx=XM_006930668.3;refGene_gn=SH2D4A;refGene_tx=XM_011281481.3
B1 41434974 41435086
refGene_gn=SH2D4A;refGene_tx=XM_003984734.5;refGene_gn=SH2D4A;refGene_tx=XM_006930668.3;refGene_gn=SH2D4A;refGene_tx=XM_011281481.3;refGene_gn=SH2D4A;refGene_tx=XM_011281482.3
B1 41436915 41437126
refGene_gn=SH2D4A;refGene_tx=XM_003984734.5;refGene_gn=SH2D4A;refGene_tx=XM_006930668.3;refGene_gn=SH2D4A;refGene_tx=XM_011281481.3;refGene_gn=SH2D4A;refGene_tx=XM_011281482.3
B1 41439865 41439996
refGene_gn=SH2D4A;refGene_tx=XM_003984734.5;refGene_gn=SH2D4A;refGene_tx=XM_006930668.3;refGene_gn=SH2D4A;refGene_tx=XM_011281481.3;refGene_gn=SH2D4A;refGene_tx=XM_011281482.3
B1 41456693 41456914
refGene_gn=SH2D4A;refGene_tx=XM_003984734.5;refGene_gn=SH2D4A;refGene_tx=XM_006930668.3;refGene_gn=SH2D4A;refGene_tx=XM_011281481.3;refGene_gn=SH2D4A;refGene_tx=XM_011281482.3
B1 41457404 41457497
refGene_gn=SH2D4A;refGene_tx=XM_003984734.5;refGene_gn=SH2D4A;refGene_tx=XM_006930668.3;refGene_gn=SH2D4A;refGene_tx=XM_011281481.3;refGene_gn=SH2D4A;refGene_tx=XM_011281482.3
B1 41468814 41469104
refGene_gn=CSGALNACT1;refGene_tx=XM_006930671.3;refGene_gn=CSGALNACT1;refGene_tx=XM_006930676.4;refGene_gn=CSGALNACT1;refGene_tx=XM_006930677.3;refGene_gn=CSGALNACT1;refGene_tx=XM_019828473.2;refGene_gn=CSGALNACT1;refGene_tx=XM_019828474.2;refGene_gn=CSGALNACT1;refGene_tx=XM_023252542.1;refGene_gn=CSGALNACT1;refGene_tx=XM_023252543.1;refGene_gn=CSGALNACT1;refGene_tx=XM_023252544.1
B1 41471739 41471821
refGene_gn=CSGALNACT1;refGene_tx=XM_006930671.3;refGene_gn=CSGALNACT1;refGene_tx=XM_006930676.4;refGene_gn=CSGALNACT1;refGene_tx=XM_006930677.3;refGene_gn=CSGALNACT1;refGene_tx=XM_019828473.2;refGene_gn=CSGALNACT1;refGene_tx=XM_019828474.2;refGene_gn=CSGALNACT1;refGene_tx=XM_023252542.1;refGene_gn=CSGALNACT1;refGene_tx=XM_023252543.1;refGene_gn=CSGALNACT1;refGene_tx=XM_023252544.1

B1 41480700 41480795
refGene_gn=CSGALNACT1;refGene_tx=XM_006930671.3;refGene_gn=CSGALNACT1;
refGene_tx=XM_006930676.4;refGene_gn=CSGALNACT1;refGene_tx=XM_00693067
7.3;refGene_gn=CSGALNACT1;refGene_tx=XM_019828473.2;refGene_gn=CSGALNA
CT1;refGene_tx=XM_019828474.2;refGene_gn=CSGALNACT1;refGene_tx=XM_0232
52542.1;refGene_gn=CSGALNACT1;refGene_tx=XM_023252543.1;refGene_gn=CSG
ALNACT1;refGene_tx=XM_023252544.1
B1 41481950 41482129
refGene_gn=CSGALNACT1;refGene_tx=XM_006930671.3;refGene_gn=CSGALNACT1;
refGene_tx=XM_006930676.4;refGene_gn=CSGALNACT1;refGene_tx=XM_00693067
7.3;refGene_gn=CSGALNACT1;refGene_tx=XM_019828473.2;refGene_gn=CSGALNA
CT1;refGene_tx=XM_019828474.2;refGene_gn=CSGALNACT1;refGene_tx=XM_0232
52542.1;refGene_gn=CSGALNACT1;refGene_tx=XM_023252543.1;refGene_gn=CSG
ALNACT1;refGene_tx=XM_023252544.1
B1 41497554 41497656
refGene_gn=CSGALNACT1;refGene_tx=XM_006930671.3;refGene_gn=CSGALNACT1;
refGene_tx=XM_006930676.4;refGene_gn=CSGALNACT1;refGene_tx=XM_00693067
7.3;refGene_gn=CSGALNACT1;refGene_tx=XM_019828473.2;refGene_gn=CSGALNA
CT1;refGene_tx=XM_019828474.2;refGene_gn=CSGALNACT1;refGene_tx=XM_0232
52542.1;refGene_gn=CSGALNACT1;refGene_tx=XM_023252543.1;refGene_gn=CSG
ALNACT1;refGene_tx=XM_023252544.1
B1 41513101 41513318
refGene_gn=CSGALNACT1;refGene_tx=XM_006930671.3;refGene_gn=CSGALNACT1;
refGene_tx=XM_006930676.4;refGene_gn=CSGALNACT1;refGene_tx=XM_00693067
7.3;refGene_gn=CSGALNACT1;refGene_tx=XM_019828473.2;refGene_gn=CSGALNA
CT1;refGene_tx=XM_019828474.2;refGene_gn=CSGALNACT1;refGene_tx=XM_0232
52542.1;refGene_gn=CSGALNACT1;refGene_tx=XM_023252543.1;refGene_gn=CSG
ALNACT1;refGene_tx=XM_023252544.1
B1 41560677 41561308
refGene_gn=CSGALNACT1;refGene_tx=XM_006930671.3;refGene_gn=CSGALNACT1;
refGene_tx=XM_006930676.4;refGene_gn=CSGALNACT1;refGene_tx=XM_00693067
7.3;refGene_gn=CSGALNACT1;refGene_tx=XM_019828473.2;refGene_gn=CSGALNA
CT1;refGene_tx=XM_019828474.2;refGene_gn=CSGALNACT1;refGene_tx=XM_0232
52542.1;refGene_gn=CSGALNACT1;refGene_tx=XM_023252543.1;refGene_gn=CSG
ALNACT1;refGene_tx=XM_023252544.1
B1 41876261 41876390
refGene_gn=INTS10;refGene_tx=XM_003984736.5;refGene_gn=INTS10;refGene_
tx=XM_019828475.2
B1 41876948 41877016
refGene_gn=INTS10;refGene_tx=XM_003984736.5;refGene_gn=INTS10;refGene_
tx=XM_019828475.2;refGene_gn=INTS10;refGene_tx=XM_019828476.1;refGene_
gn=INTS10;refGene_tx=XM_019828477.1
B1 41878243 41878347
refGene_gn=INTS10;refGene_tx=XM_003984736.5;refGene_gn=INTS10;refGene_
tx=XM_019828475.2;refGene_gn=INTS10;refGene_tx=XM_019828476.1;refGene_
gn=INTS10;refGene_tx=XM_019828477.1
B1 41879048 41879188
refGene_gn=INTS10;refGene_tx=XM_003984736.5;refGene_gn=INTS10;refGene_
tx=XM_019828475.2;refGene_gn=INTS10;refGene_tx=XM_019828476.1;refGene_
gn=INTS10;refGene_tx=XM_019828477.1

B1 41881144 41881226
refGene_gn=INTS10;refGene_tx=XM_003984736.5;refGene_gn=INTS10;refGene_tx=XM_019828475.2;refGene_gn=INTS10;refGene_tx=XM_019828476.1;refGene_gn=INTS10;refGene_tx=XM_019828477.1
B1 41881723 41881864
refGene_gn=INTS10;refGene_tx=XM_003984736.5;refGene_gn=INTS10;refGene_tx=XM_019828475.2;refGene_gn=INTS10;refGene_tx=XM_019828476.1;refGene_gn=INTS10;refGene_tx=XM_019828477.1
B1 41882289 41882461
refGene_gn=INTS10;refGene_tx=XM_003984736.5;refGene_gn=INTS10;refGene_tx=XM_019828475.2;refGene_gn=INTS10;refGene_tx=XM_019828476.1;refGene_gn=INTS10;refGene_tx=XM_019828477.1
B1 41882890 41883060
refGene_gn=INTS10;refGene_tx=XM_003984736.5;refGene_gn=INTS10;refGene_tx=XM_019828475.2;refGene_gn=INTS10;refGene_tx=XM_019828476.1;refGene_gn=INTS10;refGene_tx=XM_019828477.1
B1 41883847 41883981
refGene_gn=INTS10;refGene_tx=XM_003984736.5;refGene_gn=INTS10;refGene_tx=XM_019828475.2;refGene_gn=INTS10;refGene_tx=XM_019828476.1;refGene_gn=INTS10;refGene_tx=XM_019828477.1
B1 41888650 41888804
refGene_gn=INTS10;refGene_tx=XM_003984736.5;refGene_gn=INTS10;refGene_tx=XM_019828475.2;refGene_gn=INTS10;refGene_tx=XM_019828476.1;refGene_gn=INTS10;refGene_tx=XM_019828477.1
B1 41890604 41890687
refGene_gn=INTS10;refGene_tx=XM_003984736.5;refGene_gn=INTS10;refGene_tx=XM_019828475.2;refGene_gn=INTS10;refGene_tx=XM_019828476.1;refGene_gn=INTS10;refGene_tx=XM_019828477.1
B1 41891760 41891913
refGene_gn=INTS10;refGene_tx=XM_003984736.5;refGene_gn=INTS10;refGene_tx=XM_019828475.2;refGene_gn=INTS10;refGene_tx=XM_019828476.1;refGene_gn=INTS10;refGene_tx=XM_019828477.1
B1 41894902 41895011
refGene_gn=INTS10;refGene_tx=XM_003984736.5;refGene_gn=INTS10;refGene_tx=XM_019828475.2;refGene_gn=INTS10;refGene_tx=XM_019828476.1;refGene_gn=INTS10;refGene_tx=XM_019828477.1
B1 41900060 41900140
refGene_gn=INTS10;refGene_tx=XM_003984736.5;refGene_gn=INTS10;refGene_tx=XM_019828475.2;refGene_gn=INTS10;refGene_tx=XM_019828476.1;refGene_gn=INTS10;refGene_tx=XM_019828477.1
B1 41901476 41901639
refGene_gn=INTS10;refGene_tx=XM_003984736.5;refGene_gn=INTS10;refGene_tx=XM_019828475.2;refGene_gn=INTS10;refGene_tx=XM_019828476.1;refGene_gn=INTS10;refGene_tx=XM_019828477.1
B1 41902689 41902783
refGene_gn=INTS10;refGene_tx=XM_003984736.5;refGene_gn=INTS10;refGene_tx=XM_019828475.2;refGene_gn=INTS10;refGene_tx=XM_019828476.1;refGene_gn=INTS10;refGene_tx=XM_019828477.1
B1 41905733 41905811
refGene_gn=INTS10;refGene_tx=XM_019828475.2;refGene_gn=INTS10;refGene_

tx=XM_019828476.1;refGene_gn=INTS10;refGene_tx=XM_019828477.1
B1 41908012 41908169
refGene_gn=INTS10;refGene_tx=XM_003984736.5;refGene_gn=INTS10;refGene_tx=XM_019828475.2;refGene_gn=INTS10;refGene_tx=XM_019828476.1;refGene_gn=INTS10;refGene_tx=XM_019828477.1
B1 41914467 41914649
refGene_gn=HGSNAT;refGene_tx=XM_023252545.1;refGene_gn=HGSNAT;refGene_tx=XM_023252546.1;refGene_gn=HGSNAT;refGene_tx=XM_023252547.1
B1 41915375 41915488
refGene_gn=HGSNAT;refGene_tx=XM_023252545.1;refGene_gn=HGSNAT;refGene_tx=XM_023252546.1;refGene_gn=HGSNAT;refGene_tx=XM_023252547.1
B1 41915583 41915654
refGene_gn=HGSNAT;refGene_tx=XM_023252545.1;refGene_gn=HGSNAT;refGene_tx=XM_023252546.1;refGene_gn=HGSNAT;refGene_tx=XM_023252547.1
B1 41916270 41916348
refGene_gn=HGSNAT;refGene_tx=XM_023252545.1;refGene_gn=HGSNAT;refGene_tx=XM_023252546.1;refGene_gn=HGSNAT;refGene_tx=XM_023252547.1
B1 41918538 41918625
refGene_gn=HGSNAT;refGene_tx=XM_023252545.1;refGene_gn=HGSNAT;refGene_tx=XM_023252546.1;refGene_gn=HGSNAT;refGene_tx=XM_023252547.1
B1 41920022 41920149
refGene_gn=HGSNAT;refGene_tx=XM_023252545.1;refGene_gn=HGSNAT;refGene_tx=XM_023252546.1;refGene_gn=HGSNAT;refGene_tx=XM_023252547.1
B1 41920530 41920652
refGene_gn=HGSNAT;refGene_tx=XM_023252545.1;refGene_gn=HGSNAT;refGene_tx=XM_023252546.1;refGene_gn=HGSNAT;refGene_tx=XM_023252547.1
B1 41923145 41923261
refGene_gn=HGSNAT;refGene_tx=XM_023252545.1;refGene_gn=HGSNAT;refGene_tx=XM_023252546.1;refGene_gn=HGSNAT;refGene_tx=XM_023252547.1
B1 41924412 41924573
refGene_gn=HGSNAT;refGene_tx=XM_023252545.1;refGene_gn=HGSNAT;refGene_tx=XM_023252546.1;refGene_gn=HGSNAT;refGene_tx=XM_023252547.1
B1 41925948 41925979
refGene_gn=HGSNAT;refGene_tx=XM_023252545.1;refGene_gn=HGSNAT;refGene_tx=XM_023252546.1;refGene_gn=HGSNAT;refGene_tx=XM_023252547.1
B1 41926963 41927040
refGene_gn=HGSNAT;refGene_tx=XM_023252545.1;refGene_gn=HGSNAT;refGene_tx=XM_023252546.1
B1 41929162 41929272
refGene_gn=HGSNAT;refGene_tx=XM_023252545.1;refGene_gn=HGSNAT;refGene_tx=XM_023252546.1
B1 41929957 41930027
refGene_gn=HGSNAT;refGene_tx=XM_023252545.1;refGene_gn=HGSNAT;refGene_tx=XM_023252546.1
B1 41932206 41932276
refGene_gn=HGSNAT;refGene_tx=XM_023252545.1;refGene_gn=HGSNAT;refGene_tx=XM_023252546.1
B1 41935357 41935479
refGene_gn=HGSNAT;refGene_tx=XM_023252545.1;refGene_gn=HGSNAT;refGene_tx=XM_023252546.1;refGene_gn=HGSNAT;refGene_tx=XM_023252547.1

B1 41935681 41935818
refGene_gn=HGSNAT;refGene_tx=XM_023252545.1;refGene_gn=HGSNAT;refGene_tx=XM_023252546.1;refGene_gn=HGSNAT;refGene_tx=XM_023252547.1
B1 41945243 41945362
refGene_gn=HGSNAT;refGene_tx=XM_023252545.1;refGene_gn=HGSNAT;refGene_tx=XM_023252546.1;refGene_gn=HGSNAT;refGene_tx=XM_023252547.1
B1 41965650 41965870
refGene_gn=HGSNAT;refGene_tx=XM_023252545.1;refGene_gn=HGSNAT;refGene_tx=XM_023252546.1;refGene_gn=HGSNAT;refGene_tx=XM_023252547.1
B1 41978742 41979513
refGene_gn=POMK;refGene_tx=XM_011281485.3;refGene_gn=POMK;refGene_tx=XM_011281488.3;refGene_gn=POMK;refGene_tx=XM_019828480.2;refGene_gn=POMK;refGene_tx=XM_019828481.2;refGene_gn=POMK;refGene_tx=XM_019828482.2;refGene_gn=POMK;refGene_tx=XM_019828483.2;refGene_gn=POMK;refGene_tx=XM_019828484.2;refGene_gn=POMK;refGene_tx=XM_019828485.2;refGene_gn=POMK;refGene_tx=XM_019828486.2;refGene_gn=POMK;refGene_tx=XM_019828488.2;refGene_gn=POMK;refGene_tx=XM_019828489.2;refGene_gn=POMK;refGene_tx=XM_019828490.2;refGene_gn=POMK;refGene_tx=XM_023252548.1
B1 41992161 41992443
refGene_gn=POMK;refGene_tx=XM_011281485.3;refGene_gn=POMK;refGene_tx=XM_011281488.3;refGene_gn=POMK;refGene_tx=XM_019828480.2;refGene_gn=POMK;refGene_tx=XM_019828481.2;refGene_gn=POMK;refGene_tx=XM_019828482.2;refGene_gn=POMK;refGene_tx=XM_019828483.2;refGene_gn=POMK;refGene_tx=XM_019828484.2;refGene_gn=POMK;refGene_tx=XM_019828485.2;refGene_gn=POMK;refGene_tx=XM_019828486.2;refGene_gn=POMK;refGene_tx=XM_019828488.2;refGene_gn=POMK;refGene_tx=XM_019828489.2;refGene_gn=POMK;refGene_tx=XM_019828490.2;refGene_gn=POMK;refGene_tx=XM_023252548.1
B1 42009893 42010016 refGene_gn=FNTA;refGene_tx=XM_003984738.5
B1 42010291 42010463 refGene_gn=FNTA;refGene_tx=XM_003984738.5
B1 42011880 42011943 refGene_gn=FNTA;refGene_tx=XM_003984738.5
B1 42017272 42017421 refGene_gn=FNTA;refGene_tx=XM_003984738.5
B1 42018975 42019102 refGene_gn=FNTA;refGene_tx=XM_003984738.5
B1 42022137 42022242 refGene_gn=FNTA;refGene_tx=XM_003984738.5
B1 42025878 42025993 refGene_gn=FNTA;refGene_tx=XM_003984738.5
B1 42037414 42037500 refGene_gn=FNTA;refGene_tx=XM_003984738.5
B1 42039530 42039736 refGene_gn=FNTA;refGene_tx=XM_003984738.5
B1 42062510 42062651
refGene_gn=HOOK3;refGene_tx=XM_019828491.2;refGene_gn=HOOK3;refGene_tx=XM_019828492.2;refGene_gn=HOOK3;refGene_tx=XM_019828493.2;refGene_gn=HOOK3;refGene_tx=XM_019828495.2;refGene_gn=HOOK3;refGene_tx=XM_019828496.2;refGene_gn=HOOK3;refGene_tx=XM_023252550.1;refGene_gn=HOOK3;refGene_tx=XM_023252551.1
B1 42065135 42065207
refGene_gn=HOOK3;refGene_tx=XM_019828491.2;refGene_gn=HOOK3;refGene_tx=XM_019828492.2;refGene_gn=HOOK3;refGene_tx=XM_019828493.2;refGene_gn=HOOK3;refGene_tx=XM_019828495.2;refGene_gn=HOOK3;refGene_tx=XM_019828496.2;refGene_gn=HOOK3;refGene_tx=XM_023252550.1;refGene_gn=HOOK3;refGene_tx=XM_023252551.1
B1 42065374 42065479
refGene_gn=HOOK3;refGene_tx=XM_019828491.2;refGene_gn=HOOK3;refGene_tx

=XM_019828492.2;refGene_gn=HOOK3;refGene_tx=XM_019828493.2;refGene_gn=HOOK3;refGene_tx=XM_019828495.2;refGene_gn=HOOK3;refGene_tx=XM_019828496.2;refGene_gn=HOOK3;refGene_tx=XM_023252550.1;refGene_gn=HOOK3;refGene_tx=XM_023252551.1
B1 42068360 42068461 refGene_gn=HOOK3;refGene_tx=XM_019828491.2;refGene_gn=HOOK3;refGene_tx=XM_019828492.2;refGene_gn=HOOK3;refGene_tx=XM_019828493.2;refGene_gn=HOOK3;refGene_tx=XM_019828495.2;refGene_gn=HOOK3;refGene_tx=XM_019828496.2;refGene_gn=HOOK3;refGene_tx=XM_023252550.1;refGene_gn=HOOK3;refGene_tx=XM_023252551.1
B1 42069027 42069110 refGene_gn=HOOK3;refGene_tx=XM_019828491.2;refGene_gn=HOOK3;refGene_tx=XM_019828492.2;refGene_gn=HOOK3;refGene_tx=XM_019828493.2;refGene_gn=HOOK3;refGene_tx=XM_019828495.2;refGene_gn=HOOK3;refGene_tx=XM_019828496.2;refGene_gn=HOOK3;refGene_tx=XM_023252550.1;refGene_gn=HOOK3;refGene_tx=XM_023252551.1
B1 42072494 42072529 refGene_gn=HOOK3;refGene_tx=XM_019828491.2;refGene_gn=HOOK3;refGene_tx=XM_019828492.2;refGene_gn=HOOK3;refGene_tx=XM_019828493.2;refGene_gn=HOOK3;refGene_tx=XM_019828495.2;refGene_gn=HOOK3;refGene_tx=XM_019828496.2;refGene_gn=HOOK3;refGene_tx=XM_023252550.1;refGene_gn=HOOK3;refGene_tx=XM_023252551.1
B1 42074442 42074448 refGene_gn=HOOK3;refGene_tx=XM_019828491.2;refGene_gn=HOOK3;refGene_tx=XM_019828493.2;refGene_gn=HOOK3;refGene_tx=XM_019828495.2;refGene_gn=HOOK3;refGene_tx=XM_019828496.2;refGene_gn=HOOK3;refGene_tx=XM_023252550.1
B1 42074650 42074698 refGene_gn=HOOK3;refGene_tx=XM_019828498.2
B1 42077174 42077262 refGene_gn=HOOK3;refGene_tx=XM_019828491.2;refGene_gn=HOOK3;refGene_tx=XM_019828492.2;refGene_gn=HOOK3;refGene_tx=XM_019828493.2;refGene_gn=HOOK3;refGene_tx=XM_019828495.2;refGene_gn=HOOK3;refGene_tx=XM_019828496.2;refGene_gn=HOOK3;refGene_tx=XM_019828498.2;refGene_gn=HOOK3;refGene_tx=XM_023252550.1;refGene_gn=HOOK3;refGene_tx=XM_023252551.1
B1 42090091 42090232 refGene_gn=HOOK3;refGene_tx=XM_019828491.2;refGene_gn=HOOK3;refGene_tx=XM_019828492.2;refGene_gn=HOOK3;refGene_tx=XM_019828493.2;refGene_gn=HOOK3;refGene_tx=XM_019828495.2;refGene_gn=HOOK3;refGene_tx=XM_019828496.2;refGene_gn=HOOK3;refGene_tx=XM_019828498.2;refGene_gn=HOOK3;refGene_tx=XM_023252550.1;refGene_gn=HOOK3;refGene_tx=XM_023252551.1
B1 42092802 42092872 refGene_gn=HOOK3;refGene_tx=XM_019828491.2;refGene_gn=HOOK3;refGene_tx=XM_019828492.2;refGene_gn=HOOK3;refGene_tx=XM_019828493.2;refGene_gn=HOOK3;refGene_tx=XM_019828495.2;refGene_gn=HOOK3;refGene_tx=XM_019828496.2;refGene_gn=HOOK3;refGene_tx=XM_019828498.2;refGene_gn=HOOK3;refGene_tx=XM_023252550.1;refGene_gn=HOOK3;refGene_tx=XM_023252551.1
B1 42100229 42100317 refGene_gn=HOOK3;refGene_tx=XM_019828491.2;refGene_gn=HOOK3;refGene_tx=XM_019828492.2;refGene_gn=HOOK3;refGene_tx=XM_019828493.2;refGene_gn=HOOK3;refGene_tx=XM_019828495.2;refGene_gn=HOOK3;refGene_tx=XM_0198284

96.2;refGene_gn=HOOK3;refGene_tx=XM_019828498.2;refGene_gn=HOOK3;refGene_tx=XM_023252550.1;refGene_gn=HOOK3;refGene_tx=XM_023252551.1
B1 42101042 42101153
refGene_gn=HOOK3;refGene_tx=XM_019828491.2;refGene_gn=HOOK3;refGene_tx=XM_019828492.2;refGene_gn=HOOK3;refGene_tx=XM_019828493.2;refGene_gn=HOOK3;refGene_tx=XM_019828495.2;refGene_gn=HOOK3;refGene_tx=XM_019828496.2;refGene_gn=HOOK3;refGene_tx=XM_019828498.2;refGene_gn=HOOK3;refGene_tx=XM_023252550.1;refGene_gn=HOOK3;refGene_tx=XM_023252551.1
B1 42105392 42105594
refGene_gn=HOOK3;refGene_tx=XM_019828491.2;refGene_gn=HOOK3;refGene_tx=XM_019828492.2;refGene_gn=HOOK3;refGene_tx=XM_019828493.2;refGene_gn=HOOK3;refGene_tx=XM_019828495.2;refGene_gn=HOOK3;refGene_tx=XM_019828496.2;refGene_gn=HOOK3;refGene_tx=XM_019828498.2;refGene_gn=HOOK3;refGene_tx=XM_023252550.1;refGene_gn=HOOK3;refGene_tx=XM_023252551.1
B1 42107168 42107309
refGene_gn=HOOK3;refGene_tx=XM_019828491.2;refGene_gn=HOOK3;refGene_tx=XM_019828492.2;refGene_gn=HOOK3;refGene_tx=XM_019828493.2;refGene_gn=HOOK3;refGene_tx=XM_019828495.2;refGene_gn=HOOK3;refGene_tx=XM_019828496.2;refGene_gn=HOOK3;refGene_tx=XM_019828498.2;refGene_gn=HOOK3;refGene_tx=XM_023252550.1;refGene_gn=HOOK3;refGene_tx=XM_023252551.1
B1 42113821 42113985
refGene_gn=HOOK3;refGene_tx=XM_019828491.2;refGene_gn=HOOK3;refGene_tx=XM_019828492.2;refGene_gn=HOOK3;refGene_tx=XM_019828493.2;refGene_gn=HOOK3;refGene_tx=XM_019828495.2;refGene_gn=HOOK3;refGene_tx=XM_019828496.2;refGene_gn=HOOK3;refGene_tx=XM_019828498.2;refGene_gn=HOOK3;refGene_tx=XM_023252550.1;refGene_gn=HOOK3;refGene_tx=XM_023252551.1
B1 42121422 42121506
refGene_gn=HOOK3;refGene_tx=XM_019828491.2;refGene_gn=HOOK3;refGene_tx=XM_019828492.2;refGene_gn=HOOK3;refGene_tx=XM_019828493.2;refGene_gn=HOOK3;refGene_tx=XM_019828495.2;refGene_gn=HOOK3;refGene_tx=XM_019828496.2;refGene_gn=HOOK3;refGene_tx=XM_019828498.2;refGene_gn=HOOK3;refGene_tx=XM_023252550.1;refGene_gn=HOOK3;refGene_tx=XM_023252551.1
B1 42123132 42123195
refGene_gn=HOOK3;refGene_tx=XM_019828491.2;refGene_gn=HOOK3;refGene_tx=XM_019828492.2;refGene_gn=HOOK3;refGene_tx=XM_019828493.2;refGene_gn=HOOK3;refGene_tx=XM_019828495.2;refGene_gn=HOOK3;refGene_tx=XM_019828496.2;refGene_gn=HOOK3;refGene_tx=XM_019828498.2;refGene_gn=HOOK3;refGene_tx=XM_023252550.1;refGene_gn=HOOK3;refGene_tx=XM_023252551.1
B1 42132088 42132156
refGene_gn=HOOK3;refGene_tx=XM_019828491.2;refGene_gn=HOOK3;refGene_tx=XM_019828492.2;refGene_gn=HOOK3;refGene_tx=XM_019828493.2;refGene_gn=HOOK3;refGene_tx=XM_019828495.2;refGene_gn=HOOK3;refGene_tx=XM_019828496.2;refGene_gn=HOOK3;refGene_tx=XM_019828498.2;refGene_gn=HOOK3;refGene_tx=XM_023252550.1;refGene_gn=HOOK3;refGene_tx=XM_023252551.1
B1 42136177 42136310
refGene_gn=HOOK3;refGene_tx=XM_019828491.2;refGene_gn=HOOK3;refGene_tx=XM_019828492.2;refGene_gn=HOOK3;refGene_tx=XM_019828493.2;refGene_gn=HOOK3;refGene_tx=XM_019828495.2;refGene_gn=HOOK3;refGene_tx=XM_019828496.2;refGene_gn=HOOK3;refGene_tx=XM_019828498.2;refGene_gn=HOOK3;refGene_tx=XM_023252550.1;refGene_gn=HOOK3;refGene_tx=XM_023252551.1

B1 42137790 42137841
refGene_gn=HOOK3;refGene_tx=XM_019828491.2;refGene_gn=HOOK3;refGene_tx
=XM_019828492.2;refGene_gn=HOOK3;refGene_tx=XM_019828493.2;refGene_gn=
HOOK3;refGene_tx=XM_019828495.2;refGene_gn=HOOK3;refGene_tx=XM_0198284
96.2;refGene_gn=HOOK3;refGene_tx=XM_019828498.2;refGene_gn=HOOK3;refGe
ne_tx=XM_023252550.1;refGene_gn=HOOK3;refGene_tx=XM_023252551.1
B1 42141008 42141081
refGene_gn=HOOK3;refGene_tx=XM_019828491.2;refGene_gn=HOOK3;refGene_tx
=XM_019828492.2;refGene_gn=HOOK3;refGene_tx=XM_019828493.2;refGene_gn=
HOOK3;refGene_tx=XM_019828495.2;refGene_gn=HOOK3;refGene_tx=XM_0198284
96.2;refGene_gn=HOOK3;refGene_tx=XM_019828498.2;refGene_gn=HOOK3;refGe
ne_tx=XM_023252550.1;refGene_gn=HOOK3;refGene_tx=XM_023252551.1
B1 42154011 42154110
refGene_gn=HOOK3;refGene_tx=XM_019828491.2;refGene_gn=HOOK3;refGene_tx
=XM_019828492.2;refGene_gn=HOOK3;refGene_tx=XM_019828495.2;refGene_gn=
HOOK3;refGene_tx=XM_019828496.2;refGene_gn=HOOK3;refGene_tx=XM_0198284
98.2
B1 42161108 42161194
refGene_gn=HOOK3;refGene_tx=XM_019828491.2;refGene_gn=HOOK3;refGene_tx
=XM_019828492.2;refGene_gn=HOOK3;refGene_tx=XM_019828493.2;refGene_gn=
HOOK3;refGene_tx=XM_019828495.2;refGene_gn=HOOK3;refGene_tx=XM_0198284
96.2;refGene_gn=HOOK3;refGene_tx=XM_019828498.2;refGene_gn=HOOK3;refGe
ne_tx=XM_023252550.1;refGene_gn=HOOK3;refGene_tx=XM_023252551.1
B1 42161300 42161441
refGene_gn=HOOK3;refGene_tx=XM_019828491.2;refGene_gn=HOOK3;refGene_tx
=XM_019828492.2;refGene_gn=HOOK3;refGene_tx=XM_019828493.2;refGene_gn=
HOOK3;refGene_tx=XM_019828498.2
B1 42169160 42169217
refGene_gn=HOOK3;refGene_tx=XM_019828496.2;refGene_gn=HOOK3;refGene_tx
=XM_023252550.1;refGene_gn=HOOK3;refGene_tx=XM_023252551.1
B1 42176576 42176713
refGene_gn=RNF170;refGene_tx=XM_003984739.5;refGene_gn=RNF170;refGene_
tx=XM_006930684.4;refGene_gn=RNF170;refGene_tx=XM_023252552.1;refGene_
gn=RNF170;refGene_tx=XM_023252553.1
B1 42196475 42196551
refGene_gn=RNF170;refGene_tx=XM_003984739.5;refGene_gn=RNF170;refGene_
tx=XM_006930684.4;refGene_gn=RNF170;refGene_tx=XM_023252552.1;refGene_
gn=RNF170;refGene_tx=XM_023252553.1
B1 42199718 42199827
refGene_gn=RNF170;refGene_tx=XM_003984739.5;refGene_gn=RNF170;refGene_
tx=XM_003984740.4;refGene_gn=RNF170;refGene_tx=XM_006930684.4;refGene_
gn=RNF170;refGene_tx=XM_023252552.1;refGene_gn=RNF170;refGene_tx=XM_02
3252553.1;refGene_gn=RNF170;refGene_tx=XM_023252554.1
B1 42203793 42203867
refGene_gn=RNF170;refGene_tx=XM_003984739.5;refGene_gn=RNF170;refGene_
tx=XM_003984740.4;refGene_gn=RNF170;refGene_tx=XM_006930684.4;refGene_
gn=RNF170;refGene_tx=XM_023252552.1;refGene_gn=RNF170;refGene_tx=XM_02
3252553.1;refGene_gn=RNF170;refGene_tx=XM_023252554.1
B1 42208387 42208498
refGene_gn=RNF170;refGene_tx=XM_003984739.5;refGene_gn=RNF170;refGene_

tx=XM_003984740.4;refGene_gn=RNF170;refGene_tx=XM_006930684.4;refGene_gn=RNF170;refGene_tx=XM_023252552.1;refGene_gn=RNF170;refGene_tx=XM_023252553.1;refGene_gn=RNF170;refGene_tx=XM_023252554.1
B1 42212559 42212829 refGene_gn=RNF170;refGene_tx=XM_003984739.5;refGene_gn=RNF170;refGene_tx=XM_003984740.4;refGene_gn=RNF170;refGene_tx=XM_006930684.4;refGene_gn=RNF170;refGene_tx=XM_023252552.1;refGene_gn=RNF170;refGene_tx=XM_023252553.1;refGene_gn=RNF170;refGene_tx=XM_023252554.1
B1 42225158 42225229 refGene_gn=THAP1;refGene_tx=XM_003984741.5
B1 42225559 42225627 refGene_gn=THAP1;refGene_tx=XM_006930685.4
B1 42228863 42229059 refGene_gn=THAP1;refGene_tx=XM_003984741.5;refGene_gn=THAP1;refGene_tx=XM_006930685.4
B1 42233680 42234055 refGene_gn=THAP1;refGene_tx=XM_003984741.5;refGene_gn=THAP1;refGene_tx=XM_006930685.4
B1 42283595 42283781 refGene_gn=LOC101097968;refGene_tx=XM_003984742.3
B1 42284078 42284288 refGene_gn=LOC101097968;refGene_tx=XM_003984742.3
B1 42285639 42285718 refGene_gn=CHRNA6;refGene_tx=XM_003984743.4
B1 42287389 42287529 refGene_gn=CHRNA6;refGene_tx=XM_003984743.4
B1 42296354 42296390 refGene_gn=CHRNA6;refGene_tx=XM_023252556.1
B1 42296641 42296686 refGene_gn=CHRNA6;refGene_tx=XM_003984743.4;refGene_gn=CHRNA6;refGene_tx=XM_019828502.2;refGene_gn=CHRNA6;refGene_tx=XM_019828503.2;refGene_gn=CHRNA6;refGene_tx=XM_023252556.1
B1 42297607 42297717 refGene_gn=CHRNA6;refGene_tx=XM_003984743.4;refGene_gn=CHRNA6;refGene_tx=XM_019828502.2;refGene_gn=CHRNA6;refGene_tx=XM_019828503.2;refGene_gn=CHRNA6;refGene_tx=XM_023252556.1
B1 42297818 42298797 refGene_gn=CHRNA6;refGene_tx=XM_003984743.4;refGene_gn=CHRNA6;refGene_tx=XM_019828502.2;refGene_gn=CHRNA6;refGene_tx=XM_019828503.2;refGene_gn=CHRNA6;refGene_tx=XM_023252556.1
B1 42300141 42300273 refGene_gn=CHRNA6;refGene_tx=XM_003984743.4;refGene_gn=CHRNA6;refGene_tx=XM_019828502.2;refGene_gn=CHRNA6;refGene_tx=XM_019828503.2;refGene_gn=CHRNA6;refGene_tx=XM_023252556.1
B1 42321952 42322087 refGene_gn=CHRNB3;refGene_tx=XM_023252558.1;refGene_gn=CHRNB3;refGene_tx=XM_023252560.1
B1 42324108 42325129 refGene_gn=CHRNB3;refGene_tx=XM_019828504.1;refGene_gn=CHRNB3;refGene_tx=XM_023252557.1;refGene_gn=CHRNB3;refGene_tx=XM_023252558.1;refGene_gn=CHRNB3;refGene_tx=XM_023252559.1;refGene_gn=CHRNB3;refGene_tx=XM_023252560.1
B1 42325349 42325459 refGene_gn=CHRNB3;refGene_tx=XM_019828504.1;refGene_gn=CHRNB3;refGene_tx=XM_023252557.1;refGene_gn=CHRNB3;refGene_tx=XM_023252558.1;refGene_

gn=CHRNB3;refGene_tx=XM_023252559.1;refGene_gn=CHRNB3;refGene_tx=XM_02
3252560.1
B1	42343133 42343178
refGene_gn=CHRNB3;refGene_tx=XM_023252557.1;refGene_gn=CHRNB3;refGene_
tx=XM_023252558.1;refGene_gn=CHRNB3;refGene_tx=XM_023252559.1;refGene_
gn=CHRNB3;refGene_tx=XM_023252560.1
B1	42344350 42344523
refGene_gn=CHRNB3;refGene_tx=XM_019828504.1;refGene_gn=CHRNB3;refGene_
tx=XM_023252557.1;refGene_gn=CHRNB3;refGene_tx=XM_023252558.1;refGene_
gn=CHRNB3;refGene_tx=XM_023252559.1;refGene_gn=CHRNB3;refGene_tx=XM_02
3252560.1
B1	42353467 42353522
refGene_gn=CHRNB3;refGene_tx=XM_019828504.1;refGene_gn=CHRNB3;refGene_
tx=XM_023252557.1;refGene_gn=CHRNB3;refGene_tx=XM_023252558.1;refGene_
gn=CHRNB3;refGene_tx=XM_023252559.1;refGene_gn=CHRNB3;refGene_tx=XM_02
3252560.1
B1	42448159 42448200 refGene_gn=SMIM19;refGene_tx=XM_023252564.1
B1	42461676 42461741
refGene_gn=SMIM19;refGene_tx=XM_019828509.2;refGene_gn=SMIM19;refGene_
tx=XM_023252561.1;refGene_gn=SMIM19;refGene_tx=XM_023252562.1;refGene_
gn=SMIM19;refGene_tx=XM_023252563.1
B1	42465733 42465858
refGene_gn=SMIM19;refGene_tx=XM_019828509.2;refGene_gn=SMIM19;refGene_
tx=XM_023252561.1;refGene_gn=SMIM19;refGene_tx=XM_023252562.1;refGene_
gn=SMIM19;refGene_tx=XM_023252563.1;refGene_gn=SMIM19;refGene_tx=XM_02
3252564.1
B1	42466658 42466747 refGene_gn=SMIM19;refGene_tx=XM_019828509.2
B1	42470876 42471010
refGene_gn=SMIM19;refGene_tx=XM_023252561.1;refGene_gn=SMIM19;refGene_
tx=XM_023252562.1;refGene_gn=SMIM19;refGene_tx=XM_023252563.1;refGene_
gn=SMIM19;refGene_tx=XM_023252564.1
B1	42532795 42533084
refGene_gn=SLC20A2;refGene_tx=NM_001009839.1;refGene_gn=SLC20A2;refGen
e_tx=XM_006930663.4;refGene_gn=SLC20A2;refGene_tx=XM_006930665.4;refGe
ne_gn=SLC20A2;refGene_tx=XM_006930666.4;refGene_gn=SLC20A2;refGene_tx=
XM_019828149.2;refGene_gn=SLC20A2;refGene_tx=XM_019828150.2;refGene_gn
=SLC20A2;refGene_tx=XM_023252209.1;refGene_gn=SLC20A2;refGene_tx=XM_02
3252210.1;refGene_gn=SLC20A2;refGene_tx=XM_023252211.1;refGene_gn=SLC2
0A2;refGene_tx=XM_023252212.1;refGene_gn=SLC20A2;refGene_tx=XM_0232522
13.1
B1	42538027 42538168
refGene_gn=SLC20A2;refGene_tx=NM_001009839.1;refGene_gn=SLC20A2;refGen
e_tx=XM_006930663.4;refGene_gn=SLC20A2;refGene_tx=XM_006930665.4;refGe
ne_gn=SLC20A2;refGene_tx=XM_006930666.4;refGene_gn=SLC20A2;refGene_tx=
XM_019828149.2;refGene_gn=SLC20A2;refGene_tx=XM_019828150.2;refGene_gn
=SLC20A2;refGene_tx=XM_023252209.1;refGene_gn=SLC20A2;refGene_tx=XM_02
3252210.1;refGene_gn=SLC20A2;refGene_tx=XM_023252211.1;refGene_gn=SLC2
0A2;refGene_tx=XM_023252212.1;refGene_gn=SLC20A2;refGene_tx=XM_0232522
13.1
B1	42539754 42539840

refGene_gn=SLC20A2;refGene_tx=NM_001009839.1;refGene_gn=SLC20A2;refGen
e_tx=XM_006930663.4;refGene_gn=SLC20A2;refGene_tx=XM_006930665.4;refGe
ne_gn=SLC20A2;refGene_tx=XM_006930666.4;refGene_gn=SLC20A2;refGene_tx=
XM_019828149.2;refGene_gn=SLC20A2;refGene_tx=XM_019828150.2;refGene_gn
=SLC20A2;refGene_tx=XM_023252209.1;refGene_gn=SLC20A2;refGene_tx=XM_02
3252210.1;refGene_gn=SLC20A2;refGene_tx=XM_023252211.1;refGene_gn=SLC2
0A2;refGene_tx=XM_023252212.1;refGene_gn=SLC20A2;refGene_tx=XM_0232522
13.1
B1 42542840 42542937
refGene_gn=SLC20A2;refGene_tx=NM_001009839.1;refGene_gn=SLC20A2;refGen
e_tx=XM_006930663.4;refGene_gn=SLC20A2;refGene_tx=XM_006930665.4;refGe
ne_gn=SLC20A2;refGene_tx=XM_006930666.4;refGene_gn=SLC20A2;refGene_tx=
XM_019828149.2;refGene_gn=SLC20A2;refGene_tx=XM_019828150.2;refGene_gn
=SLC20A2;refGene_tx=XM_023252209.1;refGene_gn=SLC20A2;refGene_tx=XM_02
3252210.1;refGene_gn=SLC20A2;refGene_tx=XM_023252211.1;refGene_gn=SLC2
0A2;refGene_tx=XM_023252212.1;refGene_gn=SLC20A2;refGene_tx=XM_0232522
13.1
B1 42553517 42553634
refGene_gn=SLC20A2;refGene_tx=NM_001009839.1;refGene_gn=SLC20A2;refGen
e_tx=XM_006930663.4;refGene_gn=SLC20A2;refGene_tx=XM_006930665.4;refGe
ne_gn=SLC20A2;refGene_tx=XM_006930666.4;refGene_gn=SLC20A2;refGene_tx=
XM_019828149.2;refGene_gn=SLC20A2;refGene_tx=XM_019828150.2;refGene_gn
=SLC20A2;refGene_tx=XM_023252209.1;refGene_gn=SLC20A2;refGene_tx=XM_02
3252210.1;refGene_gn=SLC20A2;refGene_tx=XM_023252211.1;refGene_gn=SLC2
0A2;refGene_tx=XM_023252212.1;refGene_gn=SLC20A2;refGene_tx=XM_0232522
13.1
B1 42558640 42558844
refGene_gn=SLC20A2;refGene_tx=NM_001009839.1;refGene_gn=SLC20A2;refGen
e_tx=XM_006930663.4;refGene_gn=SLC20A2;refGene_tx=XM_006930665.4;refGe
ne_gn=SLC20A2;refGene_tx=XM_006930666.4;refGene_gn=SLC20A2;refGene_tx=
XM_019828149.2;refGene_gn=SLC20A2;refGene_tx=XM_019828150.2;refGene_gn
=SLC20A2;refGene_tx=XM_023252209.1;refGene_gn=SLC20A2;refGene_tx=XM_02
3252210.1;refGene_gn=SLC20A2;refGene_tx=XM_023252211.1;refGene_gn=SLC2
0A2;refGene_tx=XM_023252212.1;refGene_gn=SLC20A2;refGene_tx=XM_0232522
13.1
B1 42560390 42560982
refGene_gn=SLC20A2;refGene_tx=NM_001009839.1;refGene_gn=SLC20A2;refGen
e_tx=XM_006930663.4;refGene_gn=SLC20A2;refGene_tx=XM_006930665.4;refGe
ne_gn=SLC20A2;refGene_tx=XM_006930666.4;refGene_gn=SLC20A2;refGene_tx=
XM_019828149.2;refGene_gn=SLC20A2;refGene_tx=XM_019828150.2;refGene_gn
=SLC20A2;refGene_tx=XM_023252209.1;refGene_gn=SLC20A2;refGene_tx=XM_02
3252210.1;refGene_gn=SLC20A2;refGene_tx=XM_023252211.1;refGene_gn=SLC2
0A2;refGene_tx=XM_023252212.1;refGene_gn=SLC20A2;refGene_tx=XM_0232522
13.1
B1 42569617 42569803
refGene_gn=SLC20A2;refGene_tx=NM_001009839.1;refGene_gn=SLC20A2;refGen
e_tx=XM_006930663.4;refGene_gn=SLC20A2;refGene_tx=XM_006930665.4;refGe
ne_gn=SLC20A2;refGene_tx=XM_006930666.4;refGene_gn=SLC20A2;refGene_tx=
XM_019828149.2;refGene_gn=SLC20A2;refGene_tx=XM_019828150.2;refGene_gn
=SLC20A2;refGene_tx=XM_023252209.1;refGene_gn=SLC20A2;refGene_tx=XM_02

3252210.1;refGene_gn=SLC20A2;refGene_tx=XM_023252211.1;refGene_gn=SLC20A2;refGene_tx=XM_023252212.1;refGene_gn=SLC20A2;refGene_tx=XM_023252213.1
B1 42570840 42570925 refGene_gn=SLC20A2;refGene_tx=NM_001009839.1;refGene_gn=SLC20A2;refGene_tx=XM_006930663.4;refGene_gn=SLC20A2;refGene_tx=XM_006930665.4;refGene_gn=SLC20A2;refGene_tx=XM_006930666.4;refGene_gn=SLC20A2;refGene_tx=XM_019828149.2;refGene_gn=SLC20A2;refGene_tx=XM_019828150.2;refGene_gn=SLC20A2;refGene_tx=XM_023252209.1;refGene_gn=SLC20A2;refGene_tx=XM_023252210.1;refGene_gn=SLC20A2;refGene_tx=XM_023252211.1;refGene_gn=SLC20A2;refGene_tx=XM_023252212.1;refGene_gn=SLC20A2;refGene_tx=XM_023252213.1
B1 42582943 42583108 refGene_gn=SLC20A2;refGene_tx=NM_001009839.1;refGene_gn=SLC20A2;refGene_tx=XM_006930663.4;refGene_gn=SLC20A2;refGene_tx=XM_006930665.4;refGene_gn=SLC20A2;refGene_tx=XM_006930666.4;refGene_gn=SLC20A2;refGene_tx=XM_019828149.2;refGene_gn=SLC20A2;refGene_tx=XM_019828150.2;refGene_gn=SLC20A2;refGene_tx=XM_023252209.1;refGene_gn=SLC20A2;refGene_tx=XM_023252210.1;refGene_gn=SLC20A2;refGene_tx=XM_023252211.1;refGene_gn=SLC20A2;refGene_tx=XM_023252212.1;refGene_gn=SLC20A2;refGene_tx=XM_023252213.1
B1 42594111 42594203 refGene_gn=VDAC3;refGene_tx=XM_023252565.1;refGene_gn=VDAC3;refGene_tx=XM_023252566.1
B1 42594651 42594709 refGene_gn=VDAC3;refGene_tx=XM_023252565.1;refGene_gn=VDAC3;refGene_tx=XM_023252566.1
B1 42596103 42596254 refGene_gn=VDAC3;refGene_tx=XM_023252565.1;refGene_gn=VDAC3;refGene_tx=XM_023252566.1
B1 42597528 42597756 refGene_gn=VDAC3;refGene_tx=XM_023252565.1;refGene_gn=VDAC3;refGene_tx=XM_023252566.1
B1 42599320 42599373 refGene_gn=VDAC3;refGene_tx=XM_023252565.1;refGene_gn=VDAC3;refGene_tx=XM_023252566.1
B1 42600206 42600359 refGene_gn=VDAC3;refGene_tx=XM_023252565.1;refGene_gn=VDAC3;refGene_tx=XM_023252566.1
B1 42602796 42602799 refGene_gn=VDAC3;refGene_tx=XM_023252565.1
B1 42604001 42604051 refGene_gn=VDAC3;refGene_tx=XM_023252565.1;refGene_gn=VDAC3;refGene_tx=XM_023252566.1
B1 42604753 42604820 refGene_gn=VDAC3;refGene_tx=XM_023252565.1;refGene_gn=VDAC3;refGene_tx=XM_023252566.1
B1 42620782 42620893 refGene_gn=DKK4;refGene_tx=XM_023252567.1
B1 42621404 42621555 refGene_gn=DKK4;refGene_tx=XM_023252567.1
B1 42622029 42622182 refGene_gn=DKK4;refGene_tx=XM_023252567.1
B1 42623237 42623488 refGene_gn=DKK4;refGene_tx=XM_023252567.1

B1 42625322 42625417 refGene_gn=POLB;refGene_tx=XM_003984750.5
B1 42626682 42626822 refGene_gn=POLB;refGene_tx=XM_003984750.5
B1 42627322 42627387 refGene_gn=POLB;refGene_tx=XM_003984750.5
B1 42633715 42633802 refGene_gn=POLB;refGene_tx=XM_003984750.5
B1 42634919 42634990 refGene_gn=POLB;refGene_tx=XM_003984750.5
B1 42639890 42639963 refGene_gn=POLB;refGene_tx=XM_003984750.5
B1 42640060 42640115 refGene_gn=POLB;refGene_tx=XM_003984750.5
B1 42641717 42641769 refGene_gn=POLB;refGene_tx=XM_003984750.5
B1 42643931 42643981 refGene_gn=POLB;refGene_tx=XM_003984750.5
B1 42646445 42646504 refGene_gn=POLB;refGene_tx=XM_003984750.5
B1 42647849 42647924 refGene_gn=POLB;refGene_tx=XM_003984750.5
B1 42653016 42653083 refGene_gn=POLB;refGene_tx=XM_003984750.5
B1 42655398 42655456 refGene_gn=POLB;refGene_tx=XM_003984750.5
B1 42655795 42655856 refGene_gn=POLB;refGene_tx=XM_003984750.5
B1 42678757 42678823
refGene_gn=IKBKB;refGene_tx=XM_003984751.5;refGene_gn=IKBKB;refGene_tx=XM_011281493.3;refGene_gn=IKBKB;refGene_tx=XM_011281494.3;refGene_gn=IKBKB;refGene_tx=XM_019828511.2
B1 42679869 42679960
refGene_gn=IKBKB;refGene_tx=XM_003984751.5;refGene_gn=IKBKB;refGene_tx=XM_011281493.3;refGene_gn=IKBKB;refGene_tx=XM_011281494.3;refGene_gn=IKBKB;refGene_tx=XM_019828511.2
B1 42682344 42682472
refGene_gn=IKBKB;refGene_tx=XM_003984751.5;refGene_gn=IKBKB;refGene_tx=XM_011281493.3;refGene_gn=IKBKB;refGene_tx=XM_011281494.3;refGene_gn=IKBKB;refGene_tx=XM_019828511.2
B1 42685487 42685635
refGene_gn=IKBKB;refGene_tx=XM_003984751.5;refGene_gn=IKBKB;refGene_tx=XM_011281493.3;refGene_gn=IKBKB;refGene_tx=XM_011281494.3;refGene_gn=IKBKB;refGene_tx=XM_019828511.2
B1 42685843 42685943
refGene_gn=IKBKB;refGene_tx=XM_003984751.5;refGene_gn=IKBKB;refGene_tx=XM_011281493.3;refGene_gn=IKBKB;refGene_tx=XM_011281494.3;refGene_gn=IKBKB;refGene_tx=XM_019828511.2
B1 42686050 42686100
refGene_gn=IKBKB;refGene_tx=XM_003984751.5;refGene_gn=IKBKB;refGene_tx=XM_011281493.3;refGene_gn=IKBKB;refGene_tx=XM_011281494.3;refGene_gn=IKBKB;refGene_tx=XM_019828511.2
B1 42687523 42687645
refGene_gn=IKBKB;refGene_tx=XM_003984751.5;refGene_gn=IKBKB;refGene_tx=XM_011281493.3;refGene_gn=IKBKB;refGene_tx=XM_011281494.3;refGene_gn=IKBKB;refGene_tx=XM_019828511.2
B1 42688682 42688744
refGene_gn=IKBKB;refGene_tx=XM_003984751.5;refGene_gn=IKBKB;refGene_tx=XM_011281493.3;refGene_gn=IKBKB;refGene_tx=XM_011281494.3;refGene_gn=IKBKB;refGene_tx=XM_019828511.2
B1 42688919 42689071
refGene_gn=IKBKB;refGene_tx=XM_003984751.5;refGene_gn=IKBKB;refGene_tx=XM_011281493.3;refGene_gn=IKBKB;refGene_tx=XM_011281494.3;refGene_gn=IKBKB;refGene_tx=XM_019828511.2

B1 42689625 42689749
refGene_gn=IKBKB;refGene_tx=XM_003984751.5;refGene_gn=IKBKB;refGene_tx
=XM_011281493.3;refGene_gn=IKBKB;refGene_tx=XM_011281494.3;refGene_gn=
IKBKB;refGene_tx=XM_019828511.2
B1 42691093 42691208
refGene_gn=IKBKB;refGene_tx=XM_003984751.5;refGene_gn=IKBKB;refGene_tx
=XM_011281493.3;refGene_gn=IKBKB;refGene_tx=XM_011281494.3;refGene_gn=
IKBKB;refGene_tx=XM_019828511.2
B1 42692068 42692263
refGene_gn=IKBKB;refGene_tx=XM_003984751.5;refGene_gn=IKBKB;refGene_tx
=XM_011281493.3;refGene_gn=IKBKB;refGene_tx=XM_011281494.3;refGene_gn=
IKBKB;refGene_tx=XM_019828511.2
B1 42692628 42692758
refGene_gn=IKBKB;refGene_tx=XM_003984751.5;refGene_gn=IKBKB;refGene_tx
=XM_011281493.3;refGene_gn=IKBKB;refGene_tx=XM_011281494.3;refGene_gn=
IKBKB;refGene_tx=XM_019828511.2
B1 42693922 42694030
refGene_gn=IKBKB;refGene_tx=XM_003984751.5;refGene_gn=IKBKB;refGene_tx
=XM_011281493.3;refGene_gn=IKBKB;refGene_tx=XM_011281494.3;refGene_gn=
IKBKB;refGene_tx=XM_019828511.2
B1 42697844 42697969
refGene_gn=IKBKB;refGene_tx=XM_003984751.5;refGene_gn=IKBKB;refGene_tx
=XM_011281493.3;refGene_gn=IKBKB;refGene_tx=XM_011281494.3;refGene_gn=
IKBKB;refGene_tx=XM_019828511.2
B1 42700040 42700130
refGene_gn=IKBKB;refGene_tx=XM_003984751.5;refGene_gn=IKBKB;refGene_tx
=XM_011281493.3;refGene_gn=IKBKB;refGene_tx=XM_011281494.3;refGene_gn=
IKBKB;refGene_tx=XM_019828511.2
B1 42701188 42701277
refGene_gn=IKBKB;refGene_tx=XM_003984751.5;refGene_gn=IKBKB;refGene_tx
=XM_011281493.3;refGene_gn=IKBKB;refGene_tx=XM_011281494.3;refGene_gn=
IKBKB;refGene_tx=XM_019828511.2
B1 42726409 42726440 refGene_gn=IKBKB;refGene_tx=XM_019828511.2
B1 42726467 42726537
refGene_gn=IKBKB;refGene_tx=XM_003984751.5;refGene_gn=IKBKB;refGene_tx
=XM_011281493.3;refGene_gn=IKBKB;refGene_tx=XM_011281494.3
B1 42729128 42729246
refGene_gn=IKBKB;refGene_tx=XM_003984751.5;refGene_gn=IKBKB;refGene_tx
=XM_011281493.3;refGene_gn=IKBKB;refGene_tx=XM_011281494.3
B1 42730403 42730498
refGene_gn=IKBKB;refGene_tx=XM_003984751.5;refGene_gn=IKBKB;refGene_tx
=XM_011281493.3;refGene_gn=IKBKB;refGene_tx=XM_011281494.3
B1 42731646 42731682 refGene_gn=IKBKB;refGene_tx=XM_011281494.3
B1 42749960 42750065
refGene_gn=IKBKB;refGene_tx=XM_003984751.5;refGene_gn=IKBKB;refGene_tx
=XM_011281493.3
B1 42810086 42810158 refGene_gn=PLAT;refGene_tx=XM_011281496.3
B1 42811032 42811075 refGene_gn=PLAT;refGene_tx=XM_011281496.3
B1 42812848 42812986 refGene_gn=PLAT;refGene_tx=XM_011281496.3
B1 42813561 42813672 refGene_gn=PLAT;refGene_tx=XM_011281496.3

B1 42814032 42814207 refGene_gn=PLAT;refGene_tx=XM_011281496.3
B1 42814872 42814964 refGene_gn=PLAT;refGene_tx=XM_011281496.3
B1 42816415 42816590 refGene_gn=PLAT;refGene_tx=XM_011281496.3
B1 42817127 42817213 refGene_gn=PLAT;refGene_tx=XM_011281496.3
B1 42818812 42819008 refGene_gn=PLAT;refGene_tx=XM_011281496.3
B1 42819363 42819500 refGene_gn=PLAT;refGene_tx=XM_011281496.3
B1 42819663 42819804 refGene_gn=PLAT;refGene_tx=XM_011281496.3
B1 42820685 42820852 refGene_gn=PLAT;refGene_tx=XM_011281496.3
B1 42822422 42822581 refGene_gn=PLAT;refGene_tx=XM_011281496.3
B1 42829280 42829381
refGene_gn=AP3M2;refGene_tx=XM_003984753.5;refGene_gn=AP3M2;refGene_tx
=XM_023252570.1
B1 42830530 42830675
refGene_gn=AP3M2;refGene_tx=XM_003984753.5;refGene_gn=AP3M2;refGene_tx
=XM_023252570.1
B1 42830949 42831157
refGene_gn=AP3M2;refGene_tx=XM_003984753.5;refGene_gn=AP3M2;refGene_tx
=XM_023252570.1
B1 42833016 42833150
refGene_gn=AP3M2;refGene_tx=XM_003984753.5;refGene_gn=AP3M2;refGene_tx
=XM_023252570.1
B1 42833405 42833491
refGene_gn=AP3M2;refGene_tx=XM_003984753.5;refGene_gn=AP3M2;refGene_tx
=XM_023252570.1
B1 42840629 42840767
refGene_gn=AP3M2;refGene_tx=XM_003984753.5;refGene_gn=AP3M2;refGene_tx
=XM_023252570.1
B1 42845101 42845273
refGene_gn=AP3M2;refGene_tx=XM_003984753.5;refGene_gn=AP3M2;refGene_tx
=XM_023252570.1
B1 42848813 42849086
refGene_gn=AP3M2;refGene_tx=XM_003984753.5;refGene_gn=AP3M2;refGene_tx
=XM_023252570.1
B1 42940741 42941341
refGene_gn=KAT6A;refGene_tx=XM_011281497.3;refGene_gn=KAT6A;refGene_tx
=XM_011281498.3;refGene_gn=KAT6A;refGene_tx=XM_023252568.1;refGene_gn=
KAT6A;refGene_tx=XM_023252569.1
B1 42996693 42996802
refGene_gn=KAT6A;refGene_tx=XM_011281497.3;refGene_gn=KAT6A;refGene_tx
=XM_011281498.3;refGene_gn=KAT6A;refGene_tx=XM_023252568.1;refGene_gn=
KAT6A;refGene_tx=XM_023252569.1
B1 43000920 43001036
refGene_gn=KAT6A;refGene_tx=XM_011281497.3;refGene_gn=KAT6A;refGene_tx
=XM_011281498.3;refGene_gn=KAT6A;refGene_tx=XM_023252568.1;refGene_gn=
KAT6A;refGene_tx=XM_023252569.1
B1 43001629 43001711
refGene_gn=KAT6A;refGene_tx=XM_011281497.3;refGene_gn=KAT6A;refGene_tx
=XM_011281498.3;refGene_gn=KAT6A;refGene_tx=XM_023252568.1;refGene_gn=
KAT6A;refGene_tx=XM_023252569.1
B1 43004201 43004337

refGene_gn=KAT6A;refGene_tx=XM_011281497.3;refGene_gn=KAT6A;refGene_tx=XM_011281498.3;refGene_gn=KAT6A;refGene_tx=XM_023252568.1;refGene_gn=KAT6A;refGene_tx=XM_023252569.1
B1 43005700 43006017
refGene_gn=KAT6A;refGene_tx=XM_011281497.3;refGene_gn=KAT6A;refGene_tx=XM_011281498.3;refGene_gn=KAT6A;refGene_tx=XM_023252568.1;refGene_gn=KAT6A;refGene_tx=XM_023252569.1
B1 43011313 43011432
refGene_gn=KAT6A;refGene_tx=XM_011281497.3;refGene_gn=KAT6A;refGene_tx=XM_011281498.3;refGene_gn=KAT6A;refGene_tx=XM_023252568.1;refGene_gn=KAT6A;refGene_tx=XM_023252569.1
B1 43025719 43025835
refGene_gn=KAT6A;refGene_tx=XM_011281497.3;refGene_gn=KAT6A;refGene_tx=XM_011281498.3;refGene_gn=KAT6A;refGene_tx=XM_023252568.1;refGene_gn=KAT6A;refGene_tx=XM_023252569.1
B1 43033494 43033636
refGene_gn=KAT6A;refGene_tx=XM_011281497.3;refGene_gn=KAT6A;refGene_tx=XM_011281498.3;refGene_gn=KAT6A;refGene_tx=XM_023252568.1;refGene_gn=KAT6A;refGene_tx=XM_023252569.1
B1 43035125 43035287
refGene_gn=KAT6A;refGene_tx=XM_011281497.3;refGene_gn=KAT6A;refGene_tx=XM_011281498.3;refGene_gn=KAT6A;refGene_tx=XM_023252568.1;refGene_gn=KAT6A;refGene_tx=XM_023252569.1
B1 43036367 43036461
refGene_gn=KAT6A;refGene_tx=XM_011281497.3;refGene_gn=KAT6A;refGene_tx=XM_011281498.3;refGene_gn=KAT6A;refGene_tx=XM_023252568.1;refGene_gn=KAT6A;refGene_tx=XM_023252569.1
B1 43038428 43038660
refGene_gn=KAT6A;refGene_tx=XM_011281497.3;refGene_gn=KAT6A;refGene_tx=XM_011281498.3;refGene_gn=KAT6A;refGene_tx=XM_023252568.1;refGene_gn=KAT6A;refGene_tx=XM_023252569.1
B1 43039378 43039586
refGene_gn=KAT6A;refGene_tx=XM_011281497.3;refGene_gn=KAT6A;refGene_tx=XM_011281498.3;refGene_gn=KAT6A;refGene_tx=XM_023252568.1;refGene_gn=KAT6A;refGene_tx=XM_023252569.1
B1 43040769 43041372
refGene_gn=KAT6A;refGene_tx=XM_011281497.3;refGene_gn=KAT6A;refGene_tx=XM_011281498.3;refGene_gn=KAT6A;refGene_tx=XM_023252568.1;refGene_gn=KAT6A;refGene_tx=XM_023252569.1
B1 43044134 43044447
refGene_gn=KAT6A;refGene_tx=XM_011281497.3;refGene_gn=KAT6A;refGene_tx=XM_011281498.3;refGene_gn=KAT6A;refGene_tx=XM_023252568.1;refGene_gn=KAT6A;refGene_tx=XM_023252569.1
B1 43046515 43049199
refGene_gn=KAT6A;refGene_tx=XM_011281497.3;refGene_gn=KAT6A;refGene_tx=XM_011281498.3;refGene_gn=KAT6A;refGene_tx=XM_023252568.1;refGene_gn=KAT6A;refGene_tx=XM_023252569.1
B1 43083597 43083723 refGene_gn=ANK1;refGene_tx=XM_023252571.1
B1 43202126 43202228 refGene_gn=ANK1;refGene_tx=XM_023252571.1
B1 43223108 43223207 refGene_gn=ANK1;refGene_tx=XM_023252571.1

B1 43227651 43227750 refGene_gn=ANK1;refGene_tx=XM_023252571.1
B1 43228837 43228936 refGene_gn=ANK1;refGene_tx=XM_023252571.1
B1 43229882 43230068 refGene_gn=ANK1;refGene_tx=XM_023252571.1
B1 43231152 43231251 refGene_gn=ANK1;refGene_tx=XM_023252571.1
B1 43231946 43232045 refGene_gn=ANK1;refGene_tx=XM_023252571.1
B1 43232339 43232438 refGene_gn=ANK1;refGene_tx=XM_023252571.1
B1 43235913 43236111 refGene_gn=ANK1;refGene_tx=XM_023252571.1
B1 43237508 43237607 refGene_gn=ANK1;refGene_tx=XM_023252571.1
B1 43238009 43238108 refGene_gn=ANK1;refGene_tx=XM_023252571.1
B1 43238616 43238715 refGene_gn=ANK1;refGene_tx=XM_023252571.1
B1 43240279 43240477 refGene_gn=ANK1;refGene_tx=XM_023252571.1
B1 43241037 43241136 refGene_gn=ANK1;refGene_tx=XM_023252571.1
B1 43241622 43241721 refGene_gn=ANK1;refGene_tx=XM_023252571.1
B1 43245390 43245588 refGene_gn=ANK1;refGene_tx=XM_023252571.1
B1 43247858 43247957 refGene_gn=ANK1;refGene_tx=XM_023252571.1
B1 43248080 43248179 refGene_gn=ANK1;refGene_tx=XM_023252571.1
B1 43248422 43248521 refGene_gn=ANK1;refGene_tx=XM_023252571.1
B1 43249660 43249753 refGene_gn=ANK1;refGene_tx=XM_023252571.1
B1 43250189 43250262 refGene_gn=ANK1;refGene_tx=XM_023252571.1
B1 43252198 43252292 refGene_gn=ANK1;refGene_tx=XM_023252571.1
B1 43253727 43253806 refGene_gn=ANK1;refGene_tx=XM_023252571.1
B1 43255801 43255899 refGene_gn=ANK1;refGene_tx=XM_023252571.1
B1 43255999 43256171 refGene_gn=ANK1;refGene_tx=XM_023252571.1
B1 43256172 43256225 refGene_gn=ANK1;refGene_tx=XM_023252571.1
B1 43257501 43257656 refGene_gn=ANK1;refGene_tx=XM_023252571.1
B1 43257958 43258170 refGene_gn=ANK1;refGene_tx=XM_023252571.1
B1 43258755 43258960 refGene_gn=ANK1;refGene_tx=XM_023252571.1
B1 43259282 43259379 refGene_gn=ANK1;refGene_tx=XM_023252571.1
B1 43259633 43259862 refGene_gn=ANK1;refGene_tx=XM_023252571.1
B1 43261043 43261169 refGene_gn=ANK1;refGene_tx=XM_023252571.1
B1 43261288 43261408 refGene_gn=ANK1;refGene_tx=XM_023252571.1
B1 43262672 43262751 refGene_gn=ANK1;refGene_tx=XM_023252571.1
B1 43263030 43263105 refGene_gn=ANK1;refGene_tx=XM_023252571.1
B1 43264880 43265012 refGene_gn=ANK1;refGene_tx=XM_023252571.1
B1 43265826 43265973 refGene_gn=ANK1;refGene_tx=XM_023252571.1
B1 43284390 43284949 refGene_gn=ANK1;refGene_tx=XM_023252571.1
B1 43287936 43288234 refGene_gn=ANK1;refGene_tx=XM_023252571.1
B1 43292278 43292362 refGene_gn=ANK1;refGene_tx=XM_023252571.1
B1 43293375 43293516 refGene_gn=ANK1;refGene_tx=XM_023252571.1
B1 43299983 43300010 refGene_gn=ANK1;refGene_tx=XM_023252571.1
B1 43305230 43305612 refGene_gn=NKX6-3;refGene_tx=XM_003984760.4
B1 43307554 43307724 refGene_gn=NKX6-3;refGene_tx=XM_003984760.4
B1 43309090 43309330 refGene_gn=NKX6-3;refGene_tx=XM_003984760.4
B1 43327117 43327226
refGene_gn=GPAT4;refGene_tx=XM_003984761.5;refGene_gn=GPAT4;refGene_tx
=XM_019828518.2
B1 43329204 43329284 refGene_gn=GPAT4;refGene_tx=XM_003984761.5
B1 43329371 43329500
refGene_gn=GPAT4;refGene_tx=XM_003984761.5;refGene_gn=GPAT4;refGene_tx
=XM_019828518.2

B1 43333797 43333883
refGene_gn=GPAT4;refGene_tx=XM_003984761.5;refGene_gn=GPAT4;refGene_tx=XM_019828518.2
B1 43334431 43334487
refGene_gn=GPAT4;refGene_tx=XM_003984761.5;refGene_gn=GPAT4;refGene_tx=XM_019828518.2
B1 43335815 43335931
refGene_gn=GPAT4;refGene_tx=XM_003984761.5;refGene_gn=GPAT4;refGene_tx=XM_019828518.2
B1 43336539 43336633
refGene_gn=GPAT4;refGene_tx=XM_003984761.5;refGene_gn=GPAT4;refGene_tx=XM_019828518.2
B1 43336816 43336906
refGene_gn=GPAT4;refGene_tx=XM_003984761.5;refGene_gn=GPAT4;refGene_tx=XM_019828518.2
B1 43337662 43337737
refGene_gn=GPAT4;refGene_tx=XM_003984761.5;refGene_gn=GPAT4;refGene_tx=XM_019828518.2
B1 43338540 43338841
refGene_gn=GPAT4;refGene_tx=XM_003984761.5;refGene_gn=GPAT4;refGene_tx=XM_019828518.2
B1 43339002 43339072
refGene_gn=GPAT4;refGene_tx=XM_003984761.5;refGene_gn=GPAT4;refGene_tx=XM_019828518.2
B1 43346736 43346901
refGene_gn=GPAT4;refGene_tx=XM_003984761.5;refGene_gn=GPAT4;refGene_tx=XM_019828518.2
B1 43393544 43393641
refGene_gn=GINS4;refGene_tx=XM_003984762.4;refGene_gn=GINS4;refGene_tx=XM_023252572.1
B1 43393723 43393814
refGene_gn=GINS4;refGene_tx=XM_003984762.4;refGene_gn=GINS4;refGene_tx=XM_023252572.1
B1 43396453 43396542
refGene_gn=GINS4;refGene_tx=XM_003984762.4;refGene_gn=GINS4;refGene_tx=XM_023252572.1
B1 43396708 43396806
refGene_gn=GINS4;refGene_tx=XM_003984762.4;refGene_gn=GINS4;refGene_tx=XM_023252572.1
B1 43398241 43398355
refGene_gn=GINS4;refGene_tx=XM_003984762.4;refGene_gn=GINS4;refGene_tx=XM_023252572.1
B1 43399181 43399268 refGene_gn=GINS4;refGene_tx=XM_003984762.4
B1 43403204 43403300
refGene_gn=GINS4;refGene_tx=XM_003984762.4;refGene_gn=GINS4;refGene_tx=XM_023252572.1
B1 43418678 43418906
refGene_gn=GOLGA7;refGene_tx=XM_019828519.2;refGene_gn=GOLGA7;refGene_tx=XM_019828520.1
B1 43421574 43421627

refGene_gn=GOLGA7;refGene_tx=XM_003984763.5;refGene_gn=GOLGA7;refGene_tx=XM_006930700.4
B1 43422692 43422794
refGene_gn=GOLGA7;refGene_tx=XM_003984763.5;refGene_gn=GOLGA7;refGene_tx=XM_006930700.4
B1 43429544 43429697
refGene_gn=GOLGA7;refGene_tx=XM_003984763.5;refGene_gn=GOLGA7;refGene_tx=XM_006930700.4;refGene_gn=GOLGA7;refGene_tx=XM_019828519.2;refGene_gn=GOLGA7;refGene_tx=XM_019828520.1
B1 43434678 43434789
refGene_gn=GOLGA7;refGene_tx=XM_003984763.5;refGene_gn=GOLGA7;refGene_tx=XM_006930700.4;refGene_gn=GOLGA7;refGene_tx=XM_019828519.2;refGene_gn=GOLGA7;refGene_tx=XM_019828520.1
B1 43620166 43620710
refGene_gn=SFRP1;refGene_tx=XM_011281505.3;refGene_gn=SFRP1;refGene_tx=XM_019828521.1
B1 43625766 43625844
refGene_gn=SFRP1;refGene_tx=XM_011281505.3;refGene_gn=SFRP1;refGene_tx=XM_019828521.1
B1 43629016 43629101 refGene_gn=SFRP1;refGene_tx=XM_019828521.1
B1 43660149 43660472
refGene_gn=SFRP1;refGene_tx=XM_011281505.3;refGene_gn=SFRP1;refGene_tx=XM_019828521.1
B1 43979246 43979348
refGene_gn=ZMAT4;refGene_tx=XM_011281506.2;refGene_gn=ZMAT4;refGene_tx=XM_019828523.1
B1 44030770 44030860
refGene_gn=ZMAT4;refGene_tx=XM_011281506.2;refGene_gn=ZMAT4;refGene_tx=XM_019828523.1
B1 44099553 44099710
refGene_gn=ZMAT4;refGene_tx=XM_011281506.2;refGene_gn=ZMAT4;refGene_tx=XM_019828523.1
B1 44126866 44127094
refGene_gn=ZMAT4;refGene_tx=XM_011281506.2;refGene_gn=ZMAT4;refGene_tx=XM_019828523.1
B1 44210500 44210597
refGene_gn=ZMAT4;refGene_tx=XM_011281506.2;refGene_gn=ZMAT4;refGene_tx=XM_019828523.1
B1 44240387 44240539 refGene_gn=ZMAT4;refGene_tx=XM_019828523.1
B1 44258569 44258585
refGene_gn=ZMAT4;refGene_tx=XM_011281506.2;refGene_gn=ZMAT4;refGene_tx=XM_019828523.1
B1 44574706 44575105
refGene_gn=LOC109499346;refGene_tx=XM_023252224.1
B1 44579996 44580320 refGene_gn=TCIM;refGene_tx=XM_003984767.5
B1 44679456 44679812 refGene_gn=IDO2;refGene_tx=XM_023252573.1
B1 44681234 44681383 refGene_gn=IDO2;refGene_tx=XM_023252573.1
B1 44687589 44687641 refGene_gn=IDO2;refGene_tx=XM_023252573.1
B1 44702105 44702223 refGene_gn=IDO2;refGene_tx=XM_023252573.1
B1 44704302 44704402 refGene_gn=IDO2;refGene_tx=XM_023252573.1

B1 44708553 44708568 refGene_gn=IDO2;refGene_tx=XM_023252573.1
B1 44710860 44710979 refGene_gn=IDO2;refGene_tx=XM_023252573.1
B1 44713399 44713519 refGene_gn=IDO2;refGene_tx=XM_023252573.1
B1 44723368 44723464 refGene_gn=IDO2;refGene_tx=XM_023252573.1
B1 44751927 44752026 refGene_gn=IDO2;refGene_tx=XM_023252573.1
B1 44768716 44769072 refGene_gn=IDO1;refGene_tx=XM_003984814.3
B1 44770032 44770181 refGene_gn=IDO1;refGene_tx=XM_003984814.3
B1 44770840 44770892 refGene_gn=IDO1;refGene_tx=XM_003984814.3
B1 44771965 44772083 refGene_gn=IDO1;refGene_tx=XM_003984814.3
B1 44772626 44772726 refGene_gn=IDO1;refGene_tx=XM_003984814.3
B1 44774250 44774265 refGene_gn=IDO1;refGene_tx=XM_003984814.3
B1 44775413 44775532 refGene_gn=IDO1;refGene_tx=XM_003984814.3
B1 44778494 44778614 refGene_gn=IDO1;refGene_tx=XM_003984814.3
B1 44778730 44778826 refGene_gn=IDO1;refGene_tx=XM_003984814.3
B1 44782202 44782301 refGene_gn=IDO1;refGene_tx=XM_003984814.3
B1 44821471 44821535
refGene_gn=LOC102899536;refGene_tx=XM_023252574.1;refGene_gn=LOC102899536;refGene_tx=XM_023252575.1;refGene_gn=LOC102899536;refGene_tx=XM_023252576.1
B1 44822064 44822144
refGene_gn=LOC102899536;refGene_tx=XM_023252574.1;refGene_gn=LOC102899536;refGene_tx=XM_023252575.1;refGene_gn=LOC102899536;refGene_tx=XM_023252576.1
B1 44831398 44831454
refGene_gn=LOC102899536;refGene_tx=XM_019828527.2;refGene_gn=LOC102899536;refGene_tx=XM_023252574.1;refGene_gn=LOC102899536;refGene_tx=XM_023252575.1;refGene_gn=LOC102899536;refGene_tx=XM_023252576.1
B1 44835886 44835965
refGene_gn=LOC102899536;refGene_tx=XM_019828527.2;refGene_gn=LOC102899536;refGene_tx=XM_023252574.1;refGene_gn=LOC102899536;refGene_tx=XM_023252575.1;refGene_gn=LOC102899536;refGene_tx=XM_023252576.1
B1 44854531 44854608
refGene_gn=LOC102899536;refGene_tx=XM_019828527.2;refGene_gn=LOC102899536;refGene_tx=XM_023252574.1;refGene_gn=LOC102899536;refGene_tx=XM_023252575.1;refGene_gn=LOC102899536;refGene_tx=XM_023252576.1
B1 44855122 44855300
refGene_gn=LOC102899536;refGene_tx=XM_019828527.2;refGene_gn=LOC102899536;refGene_tx=XM_019828528.2;refGene_gn=LOC102899536;refGene_tx=XM_023252574.1;refGene_gn=LOC102899536;refGene_tx=XM_023252575.1;refGene_gn=LOC102899536;refGene_tx=XM_023252576.1
B1 44861483 44861546
refGene_gn=LOC102899536;refGene_tx=XM_019828527.2;refGene_gn=LOC102899536;refGene_tx=XM_019828528.2;refGene_gn=LOC102899536;refGene_tx=XM_023252574.1;refGene_gn=LOC102899536;refGene_tx=XM_023252575.1;refGene_gn=LOC102899536;refGene_tx=XM_023252576.1
B1 44865276 44865348
refGene_gn=LOC102899536;refGene_tx=XM_019828527.2;refGene_gn=LOC102899536;refGene_tx=XM_019828528.2;refGene_gn=LOC102899536;refGene_tx=XM_023252574.1;refGene_gn=LOC102899536;refGene_tx=XM_023252575.1;refGene_gn=LOC102899536;refGene_tx=XM_023252576.1

B1 44865576 44865743
refGene_gn=LOC102899536;refGene_tx=XM_019828527.2;refGene_gn=LOC102899536;refGene_tx=XM_019828528.2;refGene_gn=LOC102899536;refGene_tx=XM_023252574.1;refGene_gn=LOC102899536;refGene_tx=XM_023252575.1;refGene_gn=LOC102899536;refGene_tx=XM_023252576.1
B1 44866796 44866878
refGene_gn=LOC102899536;refGene_tx=XM_019828527.2;refGene_gn=LOC102899536;refGene_tx=XM_019828528.2;refGene_gn=LOC102899536;refGene_tx=XM_023252574.1;refGene_gn=LOC102899536;refGene_tx=XM_023252575.1;refGene_gn=LOC102899536;refGene_tx=XM_023252576.1
B1 44873366 44873506
refGene_gn=LOC102899536;refGene_tx=XM_019828527.2;refGene_gn=LOC102899536;refGene_tx=XM_019828528.2;refGene_gn=LOC102899536;refGene_tx=XM_023252574.1;refGene_gn=LOC102899536;refGene_tx=XM_023252575.1;refGene_gn=LOC102899536;refGene_tx=XM_023252576.1
B1 44880168 44880355
refGene_gn=LOC102899536;refGene_tx=XM_019828527.2;refGene_gn=LOC102899536;refGene_tx=XM_019828528.2;refGene_gn=LOC102899536;refGene_tx=XM_023252574.1;refGene_gn=LOC102899536;refGene_tx=XM_023252575.1;refGene_gn=LOC102899536;refGene_tx=XM_023252576.1
B1 44899159 44899258
refGene_gn=LOC102899536;refGene_tx=XM_019828527.2;refGene_gn=LOC102899536;refGene_tx=XM_019828528.2;refGene_gn=LOC102899536;refGene_tx=XM_023252574.1;refGene_gn=LOC102899536;refGene_tx=XM_023252575.1;refGene_gn=LOC102899536;refGene_tx=XM_023252576.1
B1 44900866 44901062
refGene_gn=LOC102899536;refGene_tx=XM_019828527.2;refGene_gn=LOC102899536;refGene_tx=XM_019828528.2;refGene_gn=LOC102899536;refGene_tx=XM_023252574.1;refGene_gn=LOC102899536;refGene_tx=XM_023252575.1;refGene_gn=LOC102899536;refGene_tx=XM_023252576.1
B1 44907799 44907890
refGene_gn=LOC102899536;refGene_tx=XM_019828527.2;refGene_gn=LOC102899536;refGene_tx=XM_019828528.2;refGene_gn=LOC102899536;refGene_tx=XM_023252574.1;refGene_gn=LOC102899536;refGene_tx=XM_023252575.1;refGene_gn=LOC102899536;refGene_tx=XM_023252576.1
B1 44908152 44908342
refGene_gn=LOC102899536;refGene_tx=XM_019828527.2;refGene_gn=LOC102899536;refGene_tx=XM_019828528.2;refGene_gn=LOC102899536;refGene_tx=XM_023252574.1;refGene_gn=LOC102899536;refGene_tx=XM_023252575.1;refGene_gn=LOC102899536;refGene_tx=XM_023252576.1
B1 44914777 44914864
refGene_gn=LOC102899536;refGene_tx=XM_019828527.2;refGene_gn=LOC102899536;refGene_tx=XM_019828528.2;refGene_gn=LOC102899536;refGene_tx=XM_023252574.1;refGene_gn=LOC102899536;refGene_tx=XM_023252575.1;refGene_gn=LOC102899536;refGene_tx=XM_023252576.1
B1 44917387 44917507
refGene_gn=LOC102899536;refGene_tx=XM_019828527.2;refGene_gn=LOC102899536;refGene_tx=XM_019828528.2;refGene_gn=LOC102899536;refGene_tx=XM_023252574.1;refGene_gn=LOC102899536;refGene_tx=XM_023252575.1;refGene_gn=LOC102899536;refGene_tx=XM_023252576.1

B1 44925876 44926039
refGene_gn=LOC102899536;refGene_tx=XM_019828527.2;refGene_gn=LOC102899536;refGene_tx=XM_019828528.2;refGene_gn=LOC102899536;refGene_tx=XM_023252574.1;refGene_gn=LOC102899536;refGene_tx=XM_023252576.1
B1 44928408 44928451
refGene_gn=LOC102899536;refGene_tx=XM_019828527.2;refGene_gn=LOC102899536;refGene_tx=XM_019828528.2;refGene_gn=LOC102899536;refGene_tx=XM_023252574.1;refGene_gn=LOC102899536;refGene_tx=XM_023252576.1
B1 44933624 44933679
refGene_gn=LOC101097234;refGene_tx=XM_006930702.3;refGene_gn=LOC101097234;refGene_tx=XM_023252577.1
B1 44934293 44934370
refGene_gn=LOC101097234;refGene_tx=XM_006930702.3;refGene_gn=LOC101097234;refGene_tx=XM_023252577.1
B1 44946071 44946127
refGene_gn=LOC101097234;refGene_tx=XM_006930702.3;refGene_gn=LOC101097234;refGene_tx=XM_023252577.1
B1 44949123 44949202
refGene_gn=LOC101097234;refGene_tx=XM_006930702.3;refGene_gn=LOC101097234;refGene_tx=XM_023252577.1
B1 44955522 44955599
refGene_gn=LOC101097234;refGene_tx=XM_006930702.3;refGene_gn=LOC101097234;refGene_tx=XM_023252577.1
B1 44956711 44956889
refGene_gn=LOC101097234;refGene_tx=XM_006930702.3;refGene_gn=LOC101097234;refGene_tx=XM_023252577.1
B1 44968755 44968821
refGene_gn=LOC101097234;refGene_tx=XM_006930702.3
B1 44975116 44975188
refGene_gn=LOC101097234;refGene_tx=XM_006930702.3;refGene_gn=LOC101097234;refGene_tx=XM_023252577.1
B1 44975378 44975545
refGene_gn=LOC101097234;refGene_tx=XM_006930702.3;refGene_gn=LOC101097234;refGene_tx=XM_023252577.1
B1 44976572 44976654
refGene_gn=LOC101097234;refGene_tx=XM_006930702.3;refGene_gn=LOC101097234;refGene_tx=XM_023252577.1
B1 44978935 44979072
refGene_gn=LOC101097234;refGene_tx=XM_006930702.3;refGene_gn=LOC101097234;refGene_tx=XM_023252577.1
B1 44982734 44982918
refGene_gn=LOC101097234;refGene_tx=XM_006930702.3;refGene_gn=LOC101097234;refGene_tx=XM_023252577.1
B1 44992776 44992872
refGene_gn=LOC101097234;refGene_tx=XM_006930702.3;refGene_gn=LOC101097234;refGene_tx=XM_023252577.1
B1 44995697 44995896
refGene_gn=LOC101097234;refGene_tx=XM_006930702.3;refGene_gn=LOC101097234;refGene_tx=XM_023252577.1
B1 44999642 44999748

refGene_gn=LOC101097234;refGene_tx=XM_006930702.3;refGene_gn=LOC101097234;refGene_tx=XM_023252577.1
B1 45012809 45012999
refGene_gn=LOC101097234;refGene_tx=XM_006930702.3;refGene_gn=LOC101097234;refGene_tx=XM_023252577.1
B1 45031987 45032068
refGene_gn=LOC101097234;refGene_tx=XM_006930702.3;refGene_gn=LOC101097234;refGene_tx=XM_023252577.1
B1 45052907 45053022
refGene_gn=LOC101097234;refGene_tx=XM_006930702.3;refGene_gn=LOC101097234;refGene_tx=XM_023252577.1
B1 45061437 45061597
refGene_gn=LOC101097234;refGene_tx=XM_006930702.3;refGene_gn=LOC101097234;refGene_tx=XM_023252577.1
B1 45072415 45072461
refGene_gn=LOC101097234;refGene_tx=XM_006930702.3;refGene_gn=LOC101097234;refGene_tx=XM_023252577.1
B1 45098923 45098957
refGene_gn=ADAM2;refGene_tx=XM_011281508.3;refGene_gn=ADAM2;refGene_tx=XM_023252578.1
B1 45100168 45100328
refGene_gn=ADAM2;refGene_tx=XM_011281508.3;refGene_gn=ADAM2;refGene_tx=XM_023252578.1
B1 45108909 45109051
refGene_gn=ADAM2;refGene_tx=XM_011281508.3;refGene_gn=ADAM2;refGene_tx=XM_023252578.1
B1 45109274 45109352
refGene_gn=ADAM2;refGene_tx=XM_011281508.3;refGene_gn=ADAM2;refGene_tx=XM_023252578.1
B1 45118960 45119144
refGene_gn=ADAM2;refGene_tx=XM_011281508.3;refGene_gn=ADAM2;refGene_tx=XM_023252578.1
B1 45133974 45134080
refGene_gn=ADAM2;refGene_tx=XM_011281508.3;refGene_gn=ADAM2;refGene_tx=XM_023252578.1
B1 45145833 45146029
refGene_gn=ADAM2;refGene_tx=XM_011281508.3;refGene_gn=ADAM2;refGene_tx=XM_023252578.1
B1 45146129 45146228
refGene_gn=ADAM2;refGene_tx=XM_011281508.3;refGene_gn=ADAM2;refGene_tx=XM_023252578.1
B1 45150149 45150330
refGene_gn=ADAM2;refGene_tx=XM_011281508.3;refGene_gn=ADAM2;refGene_tx=XM_023252578.1
B1 45165391 45165528
refGene_gn=ADAM2;refGene_tx=XM_011281508.3;refGene_gn=ADAM2;refGene_tx=XM_023252578.1
B1 45185736 45185818
refGene_gn=ADAM2;refGene_tx=XM_011281508.3;refGene_gn=ADAM2;refGene_tx=XM_023252578.1

B1 45186843 45187010
refGene_gn=ADAM2;refGene_tx=XM_011281508.3;refGene_gn=ADAM2;refGene_tx
=XM_023252578.1
B1 45187663 45187735
refGene_gn=ADAM2;refGene_tx=XM_011281508.3;refGene_gn=ADAM2;refGene_tx
=XM_023252578.1
B1 45202707 45202764
refGene_gn=ADAM2;refGene_tx=XM_011281508.3;refGene_gn=ADAM2;refGene_tx
=XM_023252578.1
B1 45207263 45207432
refGene_gn=ADAM2;refGene_tx=XM_011281508.3;refGene_gn=ADAM2;refGene_tx
=XM_023252578.1
B1 45207850 45207927
refGene_gn=ADAM2;refGene_tx=XM_011281508.3;refGene_gn=ADAM2;refGene_tx
=XM_023252578.1
B1 45213643 45213688 refGene_gn=ADAM2;refGene_tx=XM_023252578.1
B1 45218408 45218487 refGene_gn=ADAM2;refGene_tx=XM_011281508.3
B1 45229448 45229504 refGene_gn=ADAM2;refGene_tx=XM_011281508.3
B1 45233688 45233765 refGene_gn=ADAM2;refGene_tx=XM_011281508.3
B1 45234254 45234309 refGene_gn=ADAM2;refGene_tx=XM_011281508.3
B1 45257301 45257350
refGene_gn=LOC101097721;refGene_tx=XM_011281509.3;refGene_gn=LOC101097
721;refGene_tx=XM_011281510.3;refGene_gn=LOC101097721;refGene_tx=XM_01
9828530.1;refGene_gn=LOC101097721;refGene_tx=XM_019828531.1
B1 45260627 45260790
refGene_gn=LOC101097721;refGene_tx=XM_011281509.3;refGene_gn=LOC101097
721;refGene_tx=XM_011281510.3;refGene_gn=LOC101097721;refGene_tx=XM_01
1281511.3;refGene_gn=LOC101097721;refGene_tx=XM_019828530.1;refGene_gn
=LOC101097721;refGene_tx=XM_019828531.1
B1 45299711 45299826
refGene_gn=LOC101097721;refGene_tx=XM_011281509.3;refGene_gn=LOC101097
721;refGene_tx=XM_011281510.3;refGene_gn=LOC101097721;refGene_tx=XM_01
9828530.1;refGene_gn=LOC101097721;refGene_tx=XM_019828531.1
B1 45303896 45303983
refGene_gn=LOC101097721;refGene_tx=XM_011281509.3;refGene_gn=LOC101097
721;refGene_tx=XM_011281510.3;refGene_gn=LOC101097721;refGene_tx=XM_01
1281511.3;refGene_gn=LOC101097721;refGene_tx=XM_019828530.1;refGene_gn
=LOC101097721;refGene_tx=XM_019828531.1
B1 45313344 45313534
refGene_gn=LOC101097721;refGene_tx=XM_011281509.3;refGene_gn=LOC101097
721;refGene_tx=XM_011281510.3;refGene_gn=LOC101097721;refGene_tx=XM_01
1281511.3;refGene_gn=LOC101097721;refGene_tx=XM_019828530.1;refGene_gn
=LOC101097721;refGene_tx=XM_019828531.1
B1 45314048 45314139
refGene_gn=LOC101097721;refGene_tx=XM_011281509.3;refGene_gn=LOC101097
721;refGene_tx=XM_011281510.3;refGene_gn=LOC101097721;refGene_tx=XM_01
1281511.3;refGene_gn=LOC101097721;refGene_tx=XM_019828530.1;refGene_gn
=LOC101097721;refGene_tx=XM_019828531.1
B1 45314617 45314813
refGene_gn=LOC101097721;refGene_tx=XM_011281509.3;refGene_gn=LOC101097

721;refGene_tx=XM_011281510.3;refGene_gn=LOC101097721;refGene_tx=XM_011281511.3;refGene_gn=LOC101097721;refGene_tx=XM_019828530.1;refGene_gn=LOC101097721;refGene_tx=XM_019828531.1
B1 45316385 45316484
refGene_gn=LOC101097721;refGene_tx=XM_011281509.3;refGene_gn=LOC101097721;refGene_tx=XM_011281510.3;refGene_gn=LOC101097721;refGene_tx=XM_011281511.3;refGene_gn=LOC101097721;refGene_tx=XM_019828530.1;refGene_gn=LOC101097721;refGene_tx=XM_019828531.1
B1 45330885 45331072
refGene_gn=LOC101097721;refGene_tx=XM_011281509.3;refGene_gn=LOC101097721;refGene_tx=XM_011281510.3;refGene_gn=LOC101097721;refGene_tx=XM_011281511.3;refGene_gn=LOC101097721;refGene_tx=XM_019828530.1;refGene_gn=LOC101097721;refGene_tx=XM_019828531.1
B1 45343008 45343145
refGene_gn=LOC101097721;refGene_tx=XM_011281509.3;refGene_gn=LOC101097721;refGene_tx=XM_011281510.3;refGene_gn=LOC101097721;refGene_tx=XM_011281511.3;refGene_gn=LOC101097721;refGene_tx=XM_019828530.1;refGene_gn=LOC101097721;refGene_tx=XM_019828531.1
B1 45345069 45345151
refGene_gn=LOC101097721;refGene_tx=XM_011281509.3;refGene_gn=LOC101097721;refGene_tx=XM_011281510.3;refGene_gn=LOC101097721;refGene_tx=XM_011281511.3;refGene_gn=LOC101097721;refGene_tx=XM_019828530.1;refGene_gn=LOC101097721;refGene_tx=XM_019828531.1
B1 45346741 45346908
refGene_gn=LOC101097721;refGene_tx=XM_011281509.3;refGene_gn=LOC101097721;refGene_tx=XM_011281510.3;refGene_gn=LOC101097721;refGene_tx=XM_011281511.3;refGene_gn=LOC101097721;refGene_tx=XM_019828530.1;refGene_gn=LOC101097721;refGene_tx=XM_019828531.1
B1 45347144 45347216
refGene_gn=LOC101097721;refGene_tx=XM_011281509.3;refGene_gn=LOC101097721;refGene_tx=XM_011281510.3;refGene_gn=LOC101097721;refGene_tx=XM_011281511.3;refGene_gn=LOC101097721;refGene_tx=XM_019828530.1;refGene_gn=LOC101097721;refGene_tx=XM_019828531.1
B1 45352667 45352733
refGene_gn=LOC101097721;refGene_tx=XM_011281509.3;refGene_gn=LOC101097721;refGene_tx=XM_011281510.3;refGene_gn=LOC101097721;refGene_tx=XM_011281511.3;refGene_gn=LOC101097721;refGene_tx=XM_019828530.1;refGene_gn=LOC101097721;refGene_tx=XM_019828531.1
B1 45363245 45363423
refGene_gn=LOC101097721;refGene_tx=XM_011281509.3;refGene_gn=LOC101097721;refGene_tx=XM_011281510.3;refGene_gn=LOC101097721;refGene_tx=XM_011281511.3;refGene_gn=LOC101097721;refGene_tx=XM_019828530.1;refGene_gn=LOC101097721;refGene_tx=XM_019828531.1
B1 45364126 45364203
refGene_gn=LOC101097721;refGene_tx=XM_011281509.3;refGene_gn=LOC101097721;refGene_tx=XM_011281510.3;refGene_gn=LOC101097721;refGene_tx=XM_011281511.3
B1 45390270 45390349
refGene_gn=LOC101097721;refGene_tx=XM_011281509.3;refGene_gn=LOC101097721;refGene_tx=XM_011281510.3;refGene_gn=LOC101097721;refGene_tx=XM_01

1281511.3
B1 45397333 45397389
refGene_gn=LOC101097721;refGene_tx=XM_011281509.3;refGene_gn=LOC101097721;refGene_tx=XM_011281510.3;refGene_gn=LOC101097721;refGene_tx=XM_011281511.3
B1 45402009 45402089
refGene_gn=LOC101097721;refGene_tx=XM_011281509.3;refGene_gn=LOC101097721;refGene_tx=XM_011281510.3;refGene_gn=LOC101097721;refGene_tx=XM_011281511.3
B1 45402645 45402709
refGene_gn=LOC101097721;refGene_tx=XM_011281509.3;refGene_gn=LOC101097721;refGene_tx=XM_011281510.3;refGene_gn=LOC101097721;refGene_tx=XM_011281511.3
B1 45428082 45428095
refGene_gn=LOC101097969;refGene_tx=XM_006930748.4;refGene_gn=LOC101097969;refGene_tx=XM_019828533.2
B1 45438089 45438140
refGene_gn=LOC101097969;refGene_tx=XM_006930748.4;refGene_gn=LOC101097969;refGene_tx=XM_019828533.2
B1 45440093 45440253
refGene_gn=LOC101097969;refGene_tx=XM_006930748.4;refGene_gn=LOC101097969;refGene_tx=XM_019828533.2
B1 45449518 45449630
refGene_gn=LOC101097969;refGene_tx=XM_006930748.4;refGene_gn=LOC101097969;refGene_tx=XM_019828533.2
B1 45450328 45450415
refGene_gn=LOC101097969;refGene_tx=XM_006930748.4;refGene_gn=LOC101097969;refGene_tx=XM_019828533.2
B1 45462559 45462779
refGene_gn=LOC101097969;refGene_tx=XM_006930748.4;refGene_gn=LOC101097969;refGene_tx=XM_019828533.2
B1 45463344 45463435
refGene_gn=LOC101097969;refGene_tx=XM_006930748.4;refGene_gn=LOC101097969;refGene_tx=XM_019828533.2
B1 45472213 45472409
refGene_gn=LOC101097969;refGene_tx=XM_006930748.4;refGene_gn=LOC101097969;refGene_tx=XM_019828533.2
B1 45488407 45488503
refGene_gn=LOC101097969;refGene_tx=XM_006930748.4;refGene_gn=LOC101097969;refGene_tx=XM_019828533.2
B1 45493531 45493721
refGene_gn=LOC101097969;refGene_tx=XM_006930748.4;refGene_gn=LOC101097969;refGene_tx=XM_019828533.2
B1 45496478 45496615
refGene_gn=LOC101097969;refGene_tx=XM_006930748.4;refGene_gn=LOC101097969;refGene_tx=XM_019828533.2
B1 45515632 45515714
refGene_gn=LOC101097969;refGene_tx=XM_006930748.4;refGene_gn=LOC101097969;refGene_tx=XM_019828533.2
B1 45516458 45516622

refGene_gn=LOC101097969;refGene_tx=XM_006930748.4;refGene_gn=LOC101097969;refGene_tx=XM_019828533.2
B1 45516803 45516875
refGene_gn=LOC101097969;refGene_tx=XM_006930748.4;refGene_gn=LOC101097969;refGene_tx=XM_019828533.2
B1 45536094 45536160
refGene_gn=LOC101097969;refGene_tx=XM_006930748.4;refGene_gn=LOC101097969;refGene_tx=XM_019828533.2
B1 45537038 45537213
refGene_gn=LOC101097969;refGene_tx=XM_006930748.4;refGene_gn=LOC101097969;refGene_tx=XM_019828533.2
B1 45538032 45538109
refGene_gn=LOC101097969;refGene_tx=XM_006930748.4
B1 45538885 45538964
refGene_gn=LOC101097969;refGene_tx=XM_006930748.4
B1 45541372 45541428
refGene_gn=LOC101097969;refGene_tx=XM_006930748.4
B1 45556050 45556130
refGene_gn=LOC101097969;refGene_tx=XM_006930748.4
B1 45556620 45556675
refGene_gn=LOC101097969;refGene_tx=XM_006930748.4
B1 45585806 45585813
refGene_gn=ADAM32;refGene_tx=XM_011281548.3;refGene_gn=ADAM32;refGene_tx=XM_019828534.2;refGene_gn=ADAM32;refGene_tx=XM_019828535.2;refGene_gn=ADAM32;refGene_tx=XM_019828537.1;refGene_gn=ADAM32;refGene_tx=XM_019828539.1;refGene_gn=ADAM32;refGene_tx=XM_023252579.1;refGene_gn=ADAM32;refGene_tx=XM_023252580.1
B1 45586676 45586715
refGene_gn=ADAM32;refGene_tx=XM_011281548.3;refGene_gn=ADAM32;refGene_tx=XM_019828534.2;refGene_gn=ADAM32;refGene_tx=XM_019828535.2;refGene_gn=ADAM32;refGene_tx=XM_019828537.1;refGene_gn=ADAM32;refGene_tx=XM_019828539.1;refGene_gn=ADAM32;refGene_tx=XM_023252579.1;refGene_gn=ADAM32;refGene_tx=XM_023252580.1
B1 45588994 45589033
refGene_gn=ADAM32;refGene_tx=XM_011281548.3;refGene_gn=ADAM32;refGene_tx=XM_019828534.2;refGene_gn=ADAM32;refGene_tx=XM_019828535.2;refGene_gn=ADAM32;refGene_tx=XM_019828537.1;refGene_gn=ADAM32;refGene_tx=XM_019828539.1;refGene_gn=ADAM32;refGene_tx=XM_023252579.1;refGene_gn=ADAM32;refGene_tx=XM_023252580.1
B1 45593441 45593480
refGene_gn=ADAM32;refGene_tx=XM_011281548.3;refGene_gn=ADAM32;refGene_tx=XM_019828534.2;refGene_gn=ADAM32;refGene_tx=XM_019828535.2;refGene_gn=ADAM32;refGene_tx=XM_019828537.1;refGene_gn=ADAM32;refGene_tx=XM_019828539.1;refGene_gn=ADAM32;refGene_tx=XM_023252579.1;refGene_gn=ADAM32;refGene_tx=XM_023252580.1
B1 45605825 45605858
refGene_gn=ADAM32;refGene_tx=XM_011281548.3;refGene_gn=ADAM32;refGene_tx=XM_019828534.2;refGene_gn=ADAM32;refGene_tx=XM_019828537.1;refGene_gn=ADAM32;refGene_tx=XM_019828539.1;refGene_gn=ADAM32;refGene_tx=XM_023252579.1;refGene_gn=ADAM32;refGene_tx=XM_023252580.1

B1 45612837 45613000
refGene_gn=ADAM32;refGene_tx=XM_011281548.3;refGene_gn=ADAM32;refGene_tx=XM_019828534.2;refGene_gn=ADAM32;refGene_tx=XM_019828535.2;refGene_gn=ADAM32;refGene_tx=XM_019828537.1;refGene_gn=ADAM32;refGene_tx=XM_019828539.1;refGene_gn=ADAM32;refGene_tx=XM_023252579.1;refGene_gn=ADAM32;refGene_tx=XM_023252580.1
B1 45613576 45613676
refGene_gn=ADAM32;refGene_tx=XM_011281548.3;refGene_gn=ADAM32;refGene_tx=XM_019828534.2;refGene_gn=ADAM32;refGene_tx=XM_019828535.2;refGene_gn=ADAM32;refGene_tx=XM_019828537.1;refGene_gn=ADAM32;refGene_tx=XM_019828539.1;refGene_gn=ADAM32;refGene_tx=XM_023252579.1;refGene_gn=ADAM32;refGene_tx=XM_023252580.1
B1 45633685 45633769
refGene_gn=ADAM32;refGene_tx=XM_011281548.3;refGene_gn=ADAM32;refGene_tx=XM_019828534.2;refGene_gn=ADAM32;refGene_tx=XM_019828535.2;refGene_gn=ADAM32;refGene_tx=XM_019828537.1;refGene_gn=ADAM32;refGene_tx=XM_019828539.1;refGene_gn=ADAM32;refGene_tx=XM_023252579.1;refGene_gn=ADAM32;refGene_tx=XM_023252580.1
B1 45636200 45636384
refGene_gn=ADAM32;refGene_tx=XM_011281548.3;refGene_gn=ADAM32;refGene_tx=XM_019828534.2;refGene_gn=ADAM32;refGene_tx=XM_019828535.2;refGene_gn=ADAM32;refGene_tx=XM_019828537.1;refGene_gn=ADAM32;refGene_tx=XM_019828539.1;refGene_gn=ADAM32;refGene_tx=XM_023252579.1;refGene_gn=ADAM32;refGene_tx=XM_023252580.1
B1 45638220 45638326
refGene_gn=ADAM32;refGene_tx=XM_011281548.3;refGene_gn=ADAM32;refGene_tx=XM_019828534.2;refGene_gn=ADAM32;refGene_tx=XM_019828535.2;refGene_gn=ADAM32;refGene_tx=XM_019828537.1;refGene_gn=ADAM32;refGene_tx=XM_019828539.1;refGene_gn=ADAM32;refGene_tx=XM_023252579.1;refGene_gn=ADAM32;refGene_tx=XM_023252580.1
B1 45688157 45688353
refGene_gn=ADAM32;refGene_tx=XM_011281548.3;refGene_gn=ADAM32;refGene_tx=XM_019828534.2;refGene_gn=ADAM32;refGene_tx=XM_019828535.2;refGene_gn=ADAM32;refGene_tx=XM_019828537.1;refGene_gn=ADAM32;refGene_tx=XM_019828539.1;refGene_gn=ADAM32;refGene_tx=XM_023252579.1;refGene_gn=ADAM32;refGene_tx=XM_023252580.1
B1 45689581 45689674
refGene_gn=ADAM32;refGene_tx=XM_011281548.3;refGene_gn=ADAM32;refGene_tx=XM_019828534.2;refGene_gn=ADAM32;refGene_tx=XM_019828535.2;refGene_gn=ADAM32;refGene_tx=XM_019828537.1;refGene_gn=ADAM32;refGene_tx=XM_019828539.1;refGene_gn=ADAM32;refGene_tx=XM_023252579.1;refGene_gn=ADAM32;refGene_tx=XM_023252580.1
B1 45696558 45696736
refGene_gn=ADAM32;refGene_tx=XM_011281548.3;refGene_gn=ADAM32;refGene_tx=XM_019828534.2;refGene_gn=ADAM32;refGene_tx=XM_019828535.2;refGene_gn=ADAM32;refGene_tx=XM_019828537.1;refGene_gn=ADAM32;refGene_tx=XM_019828539.1;refGene_gn=ADAM32;refGene_tx=XM_023252579.1;refGene_gn=ADAM32;refGene_tx=XM_023252580.1
B1 45705124 45705261
refGene_gn=ADAM32;refGene_tx=XM_011281548.3;refGene_gn=ADAM32;refGene_

tx=XM_019828534.2;refGene_gn=ADAM32;refGene_tx=XM_019828535.2;refGene_gn=ADAM32;refGene_tx=XM_019828537.1;refGene_gn=ADAM32;refGene_tx=XM_019828539.1;refGene_gn=ADAM32;refGene_tx=XM_023252579.1;refGene_gn=ADAM32;refGene_tx=XM_023252580.1
B1 45712652 45712734
refGene_gn=ADAM32;refGene_tx=XM_011281548.3;refGene_gn=ADAM32;refGene_tx=XM_019828534.2;refGene_gn=ADAM32;refGene_tx=XM_019828535.2;refGene_gn=ADAM32;refGene_tx=XM_019828537.1;refGene_gn=ADAM32;refGene_tx=XM_019828539.1;refGene_gn=ADAM32;refGene_tx=XM_023252579.1;refGene_gn=ADAM32;refGene_tx=XM_023252580.1
B1 45728951 45729118
refGene_gn=ADAM32;refGene_tx=XM_011281548.3;refGene_gn=ADAM32;refGene_tx=XM_019828534.2;refGene_gn=ADAM32;refGene_tx=XM_019828535.2;refGene_gn=ADAM32;refGene_tx=XM_019828537.1;refGene_gn=ADAM32;refGene_tx=XM_019828539.1;refGene_gn=ADAM32;refGene_tx=XM_023252579.1;refGene_gn=ADAM32;refGene_tx=XM_023252580.1
B1 45729323 45729395
refGene_gn=ADAM32;refGene_tx=XM_011281548.3;refGene_gn=ADAM32;refGene_tx=XM_019828534.2;refGene_gn=ADAM32;refGene_tx=XM_019828535.2;refGene_gn=ADAM32;refGene_tx=XM_019828537.1;refGene_gn=ADAM32;refGene_tx=XM_019828539.1;refGene_gn=ADAM32;refGene_tx=XM_023252579.1;refGene_gn=ADAM32;refGene_tx=XM_023252580.1
B1 45732225 45732294
refGene_gn=ADAM32;refGene_tx=XM_011281548.3;refGene_gn=ADAM32;refGene_tx=XM_019828534.2;refGene_gn=ADAM32;refGene_tx=XM_019828535.2;refGene_gn=ADAM32;refGene_tx=XM_019828537.1;refGene_gn=ADAM32;refGene_tx=XM_019828539.1;refGene_gn=ADAM32;refGene_tx=XM_023252579.1;refGene_gn=ADAM32;refGene_tx=XM_023252580.1
B1 45737002 45737180
refGene_gn=ADAM32;refGene_tx=XM_011281548.3;refGene_gn=ADAM32;refGene_tx=XM_019828534.2;refGene_gn=ADAM32;refGene_tx=XM_019828535.2;refGene_gn=ADAM32;refGene_tx=XM_019828537.1;refGene_gn=ADAM32;refGene_tx=XM_019828539.1;refGene_gn=ADAM32;refGene_tx=XM_023252579.1;refGene_gn=ADAM32;refGene_tx=XM_023252580.1
B1 45738421 45738498
refGene_gn=ADAM32;refGene_tx=XM_011281548.3;refGene_gn=ADAM32;refGene_tx=XM_019828534.2;refGene_gn=ADAM32;refGene_tx=XM_019828535.2;refGene_gn=ADAM32;refGene_tx=XM_019828537.1;refGene_gn=ADAM32;refGene_tx=XM_023252579.1
B1 45741173 45741249
refGene_gn=ADAM32;refGene_tx=XM_011281548.3;refGene_gn=ADAM32;refGene_tx=XM_019828534.2;refGene_gn=ADAM32;refGene_tx=XM_019828535.2;refGene_gn=ADAM32;refGene_tx=XM_019828537.1;refGene_gn=ADAM32;refGene_tx=XM_019828539.1;refGene_gn=ADAM32;refGene_tx=XM_023252579.1;refGene_gn=ADAM32;refGene_tx=XM_023252580.1
B1 45743653 45743715
refGene_gn=ADAM32;refGene_tx=XM_019828534.2;refGene_gn=ADAM32;refGene_tx=XM_019828535.2
B1 45754476 45754556
refGene_gn=ADAM32;refGene_tx=XM_019828534.2;refGene_gn=ADAM32;refGene_

tx=XM_019828535.2;refGene_gn=ADAM32;refGene_tx=XM_019828537.1
B1	45767366	45767424
refGene_gn=ADAM32;refGene_tx=XM_019828534.2;refGene_gn=ADAM32;refGene_
tx=XM_019828535.2
B1	45773100	45773194	refGene_gn=ADAM9;refGene_tx=XM_003984768.5
B1	45774839	45774907	refGene_gn=ADAM9;refGene_tx=XM_003984768.5
B1	45779848	45779936	refGene_gn=ADAM9;refGene_tx=XM_003984768.5
B1	45781660	45781802	refGene_gn=ADAM9;refGene_tx=XM_003984768.5
B1	45789677	45789783	refGene_gn=ADAM9;refGene_tx=XM_003984768.5
B1	45790072	45790153	refGene_gn=ADAM9;refGene_tx=XM_003984768.5
B1	45796805	45796928	refGene_gn=ADAM9;refGene_tx=XM_019828541.2
B1	45821829	45822013
refGene_gn=ADAM9;refGene_tx=XM_003984768.5;refGene_gn=ADAM9;refGene_tx
=XM_019828541.2
B1	45830588	45830694
refGene_gn=ADAM9;refGene_tx=XM_003984768.5;refGene_gn=ADAM9;refGene_tx
=XM_019828541.2
B1	45838161	45838357
refGene_gn=ADAM9;refGene_tx=XM_003984768.5;refGene_gn=ADAM9;refGene_tx
=XM_019828541.2
B1	45839335	45839428
refGene_gn=ADAM9;refGene_tx=XM_003984768.5;refGene_gn=ADAM9;refGene_tx
=XM_019828541.2
B1	45847517	45847689
refGene_gn=ADAM9;refGene_tx=XM_003984768.5;refGene_gn=ADAM9;refGene_tx
=XM_019828541.2
B1	45868162	45868296
refGene_gn=ADAM9;refGene_tx=XM_003984768.5;refGene_gn=ADAM9;refGene_tx
=XM_019828541.2
B1	45868800	45868882
refGene_gn=ADAM9;refGene_tx=XM_003984768.5;refGene_gn=ADAM9;refGene_tx
=XM_019828541.2
B1	45873956	45874126
refGene_gn=ADAM9;refGene_tx=XM_003984768.5;refGene_gn=ADAM9;refGene_tx
=XM_019828541.2
B1	45875251	45875323
refGene_gn=ADAM9;refGene_tx=XM_003984768.5;refGene_gn=ADAM9;refGene_tx
=XM_019828541.2
B1	45877654	45877720
refGene_gn=ADAM9;refGene_tx=XM_003984768.5;refGene_gn=ADAM9;refGene_tx
=XM_019828541.2
B1	45878807	45879003
refGene_gn=ADAM9;refGene_tx=XM_003984768.5;refGene_gn=ADAM9;refGene_tx
=XM_019828541.2
B1	45880361	45880438
refGene_gn=ADAM9;refGene_tx=XM_003984768.5;refGene_gn=ADAM9;refGene_tx
=XM_019828541.2
B1	45887407	45887486
refGene_gn=ADAM9;refGene_tx=XM_003984768.5;refGene_gn=ADAM9;refGene_tx
=XM_019828541.2

B1 45900725 45900784
refGene_gn=ADAM9;refGene_tx=XM_003984768.5;refGene_gn=ADAM9;refGene_tx
=XM_019828541.2
B1 45904094 45904192
refGene_gn=ADAM9;refGene_tx=XM_003984768.5;refGene_gn=ADAM9;refGene_tx
=XM_019828541.2
B1 45915226 45915323
refGene_gn=ADAM9;refGene_tx=XM_003984768.5;refGene_gn=ADAM9;refGene_tx
=XM_019828541.2
B1 45915930 45916157 refGene_gn=TM2D2;refGene_tx=XM_003984769.5
B1 45916945 45917033 refGene_gn=TM2D2;refGene_tx=XM_003984769.5
B1 45918663 45918779 refGene_gn=TM2D2;refGene_tx=XM_003984769.5
B1 45920207 45920421 refGene_gn=TM2D2;refGene_tx=XM_003984769.5
B1 45923368 45923534 refGene_gn=HTRA4;refGene_tx=XM_023252584.1
B1 45927250 45927346 refGene_gn=HTRA4;refGene_tx=XM_023252584.1
B1 45927762 45927820 refGene_gn=HTRA4;refGene_tx=XM_023252584.1
B1 45928442 45928557 refGene_gn=HTRA4;refGene_tx=XM_023252584.1
B1 45930336 45930369 refGene_gn=HTRA4;refGene_tx=XM_023252584.1
B1 45930791 45930986 refGene_gn=HTRA4;refGene_tx=XM_023252584.1
B1 45931862 45932067 refGene_gn=HTRA4;refGene_tx=XM_023252584.1
B1 45932850 45932950 refGene_gn=HTRA4;refGene_tx=XM_023252584.1
B1 45933404 45933882 refGene_gn=HTRA4;refGene_tx=XM_023252584.1
B1 45937591 45937954
refGene_gn=PLEKHA2;refGene_tx=XM_019828543.2;refGene_gn=PLEKHA2;refGen
e_tx=XM_023252581.1;refGene_gn=PLEKHA2;refGene_tx=XM_023252582.1;refGe
ne_gn=PLEKHA2;refGene_tx=XM_023252583.1
B1 45938395 45938473
refGene_gn=PLEKHA2;refGene_tx=XM_019828543.2;refGene_gn=PLEKHA2;refGen
e_tx=XM_023252581.1;refGene_gn=PLEKHA2;refGene_tx=XM_023252582.1;refGe
ne_gn=PLEKHA2;refGene_tx=XM_023252583.1
B1 45946494 45946558
refGene_gn=PLEKHA2;refGene_tx=XM_019828543.2;refGene_gn=PLEKHA2;refGen
e_tx=XM_023252581.1;refGene_gn=PLEKHA2;refGene_tx=XM_023252582.1;refGe
ne_gn=PLEKHA2;refGene_tx=XM_023252583.1
B1 45950030 45950101
refGene_gn=PLEKHA2;refGene_tx=XM_019828543.2;refGene_gn=PLEKHA2;refGen
e_tx=XM_023252581.1;refGene_gn=PLEKHA2;refGene_tx=XM_023252582.1;refGe
ne_gn=PLEKHA2;refGene_tx=XM_023252583.1
B1 45950480 45950549
refGene_gn=PLEKHA2;refGene_tx=XM_019828543.2;refGene_gn=PLEKHA2;refGen
e_tx=XM_023252581.1;refGene_gn=PLEKHA2;refGene_tx=XM_023252582.1;refGe
ne_gn=PLEKHA2;refGene_tx=XM_023252583.1
B1 45950872 45951019
refGene_gn=PLEKHA2;refGene_tx=XM_019828543.2;refGene_gn=PLEKHA2;refGen
e_tx=XM_023252581.1;refGene_gn=PLEKHA2;refGene_tx=XM_023252582.1;refGe
ne_gn=PLEKHA2;refGene_tx=XM_023252583.1
B1 45951588 45951729
refGene_gn=PLEKHA2;refGene_tx=XM_019828543.2;refGene_gn=PLEKHA2;refGen
e_tx=XM_023252581.1;refGene_gn=PLEKHA2;refGene_tx=XM_023252582.1;refGe
ne_gn=PLEKHA2;refGene_tx=XM_023252583.1

B1 45959005 45959103
refGene_gn=PLEKHA2;refGene_tx=XM_019828543.2;refGene_gn=PLEKHA2;refGen
e_tx=XM_023252581.1;refGene_gn=PLEKHA2;refGene_tx=XM_023252582.1;refGe
ne_gn=PLEKHA2;refGene_tx=XM_023252583.1
B1 45960518 45960567
refGene_gn=PLEKHA2;refGene_tx=XM_019828543.2;refGene_gn=PLEKHA2;refGen
e_tx=XM_023252581.1;refGene_gn=PLEKHA2;refGene_tx=XM_023252582.1;refGe
ne_gn=PLEKHA2;refGene_tx=XM_023252583.1
B1 45966000 45966057
refGene_gn=PLEKHA2;refGene_tx=XM_019828543.2;refGene_gn=PLEKHA2;refGen
e_tx=XM_023252581.1;refGene_gn=PLEKHA2;refGene_tx=XM_023252582.1;refGe
ne_gn=PLEKHA2;refGene_tx=XM_023252583.1
B1 45966269 45966365
refGene_gn=PLEKHA2;refGene_tx=XM_019828543.2;refGene_gn=PLEKHA2;refGen
e_tx=XM_023252581.1;refGene_gn=PLEKHA2;refGene_tx=XM_023252582.1
B1 45979665 45979806
refGene_gn=PLEKHA2;refGene_tx=XM_019828543.2;refGene_gn=PLEKHA2;refGen
e_tx=XM_023252581.1;refGene_gn=PLEKHA2;refGene_tx=XM_023252582.1;refGe
ne_gn=PLEKHA2;refGene_tx=XM_023252583.1
B1 46044056 46044125
refGene_gn=TACC1;refGene_tx=XM_003984771.5;refGene_gn=TACC1;refGene_tx
=XM_011281518.3;refGene_gn=TACC1;refGene_tx=XM_011281519.3;refGene_gn=
TACC1;refGene_tx=XM_019828544.2;refGene_gn=TACC1;refGene_tx=XM_0198285
47.2;refGene_gn=TACC1;refGene_tx=XM_023252585.1;refGene_gn=TACC1;refGe
ne_tx=XM_023252586.1;refGene_gn=TACC1;refGene_tx=XM_023252587.1;refGen
e_gn=TACC1;refGene_tx=XM_023252588.1;refGene_gn=TACC1;refGene_tx=XM_02
3252589.1;refGene_gn=TACC1;refGene_tx=XM_023252590.1;refGene_gn=TACC1;
refGene_tx=XM_023252591.1;refGene_gn=TACC1;refGene_tx=XM_023252592.1;r
efGene_gn=TACC1;refGene_tx=XM_023252593.1
B1 46044834 46044879 refGene_gn=TACC1;refGene_tx=XM_019828545.2
B1 46045210 46045331
refGene_gn=TACC1;refGene_tx=XM_003984771.5;refGene_gn=TACC1;refGene_tx
=XM_011281518.3;refGene_gn=TACC1;refGene_tx=XM_011281519.3;refGene_gn=
TACC1;refGene_tx=XM_019828544.2;refGene_gn=TACC1;refGene_tx=XM_0198285
45.2;refGene_gn=TACC1;refGene_tx=XM_019828547.2;refGene_gn=TACC1;refGe
ne_tx=XM_023252585.1;refGene_gn=TACC1;refGene_tx=XM_023252586.1;refGen
e_gn=TACC1;refGene_tx=XM_023252587.1;refGene_gn=TACC1;refGene_tx=XM_02
3252588.1;refGene_gn=TACC1;refGene_tx=XM_023252589.1;refGene_gn=TACC1;
refGene_tx=XM_023252590.1;refGene_gn=TACC1;refGene_tx=XM_023252591.1;r
efGene_gn=TACC1;refGene_tx=XM_023252592.1;refGene_gn=TACC1;refGene_tx=
XM_023252593.1
B1 46048431 46048538
refGene_gn=TACC1;refGene_tx=XM_003984771.5;refGene_gn=TACC1;refGene_tx
=XM_011281518.3;refGene_gn=TACC1;refGene_tx=XM_011281519.3;refGene_gn=
TACC1;refGene_tx=XM_019828544.2;refGene_gn=TACC1;refGene_tx=XM_0198285
45.2;refGene_gn=TACC1;refGene_tx=XM_019828547.2;refGene_gn=TACC1;refGe
ne_tx=XM_023252585.1;refGene_gn=TACC1;refGene_tx=XM_023252586.1;refGen
e_gn=TACC1;refGene_tx=XM_023252587.1;refGene_gn=TACC1;refGene_tx=XM_02
3252588.1;refGene_gn=TACC1;refGene_tx=XM_023252589.1;refGene_gn=TACC1;
refGene_tx=XM_023252590.1;refGene_gn=TACC1;refGene_tx=XM_023252591.1;r

efGene_gn=TACC1;refGene_tx=XM_023252592.1;refGene_gn=TACC1;refGene_tx=XM_023252593.1
B1 46049349 46049510
refGene_gn=TACC1;refGene_tx=XM_003984771.5;refGene_gn=TACC1;refGene_tx=XM_011281518.3;refGene_gn=TACC1;refGene_tx=XM_011281519.3;refGene_gn=TACC1;refGene_tx=XM_019828544.2;refGene_gn=TACC1;refGene_tx=XM_019828545.2;refGene_gn=TACC1;refGene_tx=XM_019828547.2;refGene_gn=TACC1;refGene_tx=XM_023252585.1;refGene_gn=TACC1;refGene_tx=XM_023252586.1;refGene_gn=TACC1;refGene_tx=XM_023252587.1;refGene_gn=TACC1;refGene_tx=XM_023252588.1;refGene_gn=TACC1;refGene_tx=XM_023252589.1;refGene_gn=TACC1;refGene_tx=XM_023252590.1;refGene_gn=TACC1;refGene_tx=XM_023252591.1;refGene_gn=TACC1;refGene_tx=XM_023252592.1;refGene_gn=TACC1;refGene_tx=XM_023252593.1
B1 46050910 46050954
refGene_gn=TACC1;refGene_tx=XM_003984771.5;refGene_gn=TACC1;refGene_tx=XM_011281518.3;refGene_gn=TACC1;refGene_tx=XM_011281519.3;refGene_gn=TACC1;refGene_tx=XM_019828544.2;refGene_gn=TACC1;refGene_tx=XM_019828545.2;refGene_gn=TACC1;refGene_tx=XM_019828547.2;refGene_gn=TACC1;refGene_tx=XM_023252585.1;refGene_gn=TACC1;refGene_tx=XM_023252586.1;refGene_gn=TACC1;refGene_tx=XM_023252587.1;refGene_gn=TACC1;refGene_tx=XM_023252588.1;refGene_gn=TACC1;refGene_tx=XM_023252589.1;refGene_gn=TACC1;refGene_tx=XM_023252590.1;refGene_gn=TACC1;refGene_tx=XM_023252591.1;refGene_gn=TACC1;refGene_tx=XM_023252592.1;refGene_gn=TACC1;refGene_tx=XM_023252593.1
B1 46052067 46052144
refGene_gn=TACC1;refGene_tx=XM_003984771.5;refGene_gn=TACC1;refGene_tx=XM_011281518.3;refGene_gn=TACC1;refGene_tx=XM_011281519.3;refGene_gn=TACC1;refGene_tx=XM_019828544.2;refGene_gn=TACC1;refGene_tx=XM_019828545.2;refGene_gn=TACC1;refGene_tx=XM_019828547.2;refGene_gn=TACC1;refGene_tx=XM_023252585.1;refGene_gn=TACC1;refGene_tx=XM_023252586.1;refGene_gn=TACC1;refGene_tx=XM_023252587.1;refGene_gn=TACC1;refGene_tx=XM_023252588.1;refGene_gn=TACC1;refGene_tx=XM_023252589.1;refGene_gn=TACC1;refGene_tx=XM_023252590.1;refGene_gn=TACC1;refGene_tx=XM_023252591.1;refGene_gn=TACC1;refGene_tx=XM_023252592.1;refGene_gn=TACC1;refGene_tx=XM_023252593.1
B1 46053762 46053888
refGene_gn=TACC1;refGene_tx=XM_003984771.5;refGene_gn=TACC1;refGene_tx=XM_011281518.3;refGene_gn=TACC1;refGene_tx=XM_011281519.3;refGene_gn=TACC1;refGene_tx=XM_019828544.2;refGene_gn=TACC1;refGene_tx=XM_019828545.2;refGene_gn=TACC1;refGene_tx=XM_019828547.2;refGene_gn=TACC1;refGene_tx=XM_023252585.1;refGene_gn=TACC1;refGene_tx=XM_023252586.1;refGene_gn=TACC1;refGene_tx=XM_023252587.1;refGene_gn=TACC1;refGene_tx=XM_023252588.1;refGene_gn=TACC1;refGene_tx=XM_023252589.1;refGene_gn=TACC1;refGene_tx=XM_023252590.1;refGene_gn=TACC1;refGene_tx=XM_023252591.1;refGene_gn=TACC1;refGene_tx=XM_023252592.1;refGene_gn=TACC1;refGene_tx=XM_023252593.1
B1 46062301 46062354
refGene_gn=TACC1;refGene_tx=XM_003984771.5;refGene_gn=TACC1;refGene_tx=XM_011281518.3;refGene_gn=TACC1;refGene_tx=XM_011281519.3;refGene_gn=TACC1;refGene_tx=XM_019828544.2;refGene_gn=TACC1;refGene_tx=XM_0198285

45.2;refGene_gn=TACC1;refGene_tx=XM_019828547.2;refGene_gn=TACC1;refGene_tx=XM_023252585.1;refGene_gn=TACC1;refGene_tx=XM_023252586.1;refGene_gn=TACC1;refGene_tx=XM_023252587.1;refGene_gn=TACC1;refGene_tx=XM_023252588.1;refGene_gn=TACC1;refGene_tx=XM_023252589.1;refGene_gn=TACC1;refGene_tx=XM_023252590.1;refGene_gn=TACC1;refGene_tx=XM_023252591.1;refGene_gn=TACC1;refGene_tx=XM_023252592.1;refGene_gn=TACC1;refGene_tx=XM_023252593.1
B1 46064919 46065127
refGene_gn=TACC1;refGene_tx=XM_003984771.5;refGene_gn=TACC1;refGene_tx=XM_011281518.3;refGene_gn=TACC1;refGene_tx=XM_011281519.3;refGene_gn=TACC1;refGene_tx=XM_019828544.2;refGene_gn=TACC1;refGene_tx=XM_019828545.2;refGene_gn=TACC1;refGene_tx=XM_019828547.2;refGene_gn=TACC1;refGene_tx=XM_023252585.1;refGene_gn=TACC1;refGene_tx=XM_023252586.1;refGene_gn=TACC1;refGene_tx=XM_023252587.1;refGene_gn=TACC1;refGene_tx=XM_023252588.1;refGene_gn=TACC1;refGene_tx=XM_023252589.1;refGene_gn=TACC1;refGene_tx=XM_023252590.1;refGene_gn=TACC1;refGene_tx=XM_023252591.1;refGene_gn=TACC1;refGene_tx=XM_023252592.1;refGene_gn=TACC1;refGene_tx=XM_023252593.1
B1 46067003 46067064
refGene_gn=TACC1;refGene_tx=XM_003984771.5;refGene_gn=TACC1;refGene_tx=XM_011281518.3;refGene_gn=TACC1;refGene_tx=XM_011281519.3;refGene_gn=TACC1;refGene_tx=XM_019828544.2;refGene_gn=TACC1;refGene_tx=XM_019828545.2;refGene_gn=TACC1;refGene_tx=XM_019828547.2;refGene_gn=TACC1;refGene_tx=XM_023252585.1;refGene_gn=TACC1;refGene_tx=XM_023252586.1;refGene_gn=TACC1;refGene_tx=XM_023252587.1;refGene_gn=TACC1;refGene_tx=XM_023252588.1;refGene_gn=TACC1;refGene_tx=XM_023252589.1;refGene_gn=TACC1;refGene_tx=XM_023252590.1;refGene_gn=TACC1;refGene_tx=XM_023252591.1;refGene_gn=TACC1;refGene_tx=XM_023252592.1;refGene_gn=TACC1;refGene_tx=XM_023252593.1
B1 46068364 46068400
refGene_gn=TACC1;refGene_tx=XM_019828544.2;refGene_gn=TACC1;refGene_tx=XM_019828545.2;refGene_gn=TACC1;refGene_tx=XM_019828547.2;refGene_gn=TACC1;refGene_tx=XM_023252585.1;refGene_gn=TACC1;refGene_tx=XM_023252588.1;refGene_gn=TACC1;refGene_tx=XM_023252591.1
B1 46070689 46071791
refGene_gn=TACC1;refGene_tx=XM_003984771.5;refGene_gn=TACC1;refGene_tx=XM_011281518.3;refGene_gn=TACC1;refGene_tx=XM_019828544.2;refGene_gn=TACC1;refGene_tx=XM_019828545.2;refGene_gn=TACC1;refGene_tx=XM_019828547.2;refGene_gn=TACC1;refGene_tx=XM_023252585.1;refGene_gn=TACC1;refGene_tx=XM_023252586.1;refGene_gn=TACC1;refGene_tx=XM_023252587.1;refGene_gn=TACC1;refGene_tx=XM_023252588.1;refGene_gn=TACC1;refGene_tx=XM_023252589.1;refGene_gn=TACC1;refGene_tx=XM_023252590.1;refGene_gn=TACC1;refGene_tx=XM_023252591.1
B1 46096749 46096865
refGene_gn=TACC1;refGene_tx=XM_003984771.5;refGene_gn=TACC1;refGene_tx=XM_011281518.3;refGene_gn=TACC1;refGene_tx=XM_019828544.2;refGene_gn=TACC1;refGene_tx=XM_019828545.2;refGene_gn=TACC1;refGene_tx=XM_019828547.2;refGene_gn=TACC1;refGene_tx=XM_023252585.1;refGene_gn=TACC1;refGene_tx=XM_023252586.1;refGene_gn=TACC1;refGene_tx=XM_023252587.1;refGene_gn=TACC1;refGene_tx=XM_023252588.1;refGene_gn=TACC1;refGene_tx=XM_02

3252589.1;refGene_gn=TACC1;refGene_tx=XM_023252590.1
B1 46097817 46097978
refGene_gn=TACC1;refGene_tx=XM_003984771.5;refGene_gn=TACC1;refGene_tx=XM_011281518.3;refGene_gn=TACC1;refGene_tx=XM_011281519.3;refGene_gn=TACC1;refGene_tx=XM_019828544.2;refGene_gn=TACC1;refGene_tx=XM_019828545.2;refGene_gn=TACC1;refGene_tx=XM_019828547.2;refGene_gn=TACC1;refGene_tx=XM_023252588.1;refGene_gn=TACC1;refGene_tx=XM_023252589.1;refGene_gn=TACC1;refGene_tx=XM_023252590.1;refGene_gn=TACC1;refGene_tx=XM_023252592.1
B1 46130961 46131038
refGene_gn=TACC1;refGene_tx=XM_023252585.1;refGene_gn=TACC1;refGene_tx=XM_023252586.1;refGene_gn=TACC1;refGene_tx=XM_023252587.1;refGene_gn=TACC1;refGene_tx=XM_023252593.1
B1 46398379 46398467
refGene_gn=FGFR1;refGene_tx=XM_011281520.3;refGene_gn=FGFR1;refGene_tx=XM_011281521.3;refGene_gn=FGFR1;refGene_tx=XM_011281522.3;refGene_gn=FGFR1;refGene_tx=XM_011281523.3;refGene_gn=FGFR1;refGene_tx=XM_011281524.3;refGene_gn=FGFR1;refGene_tx=XM_011281525.3;refGene_gn=FGFR1;refGene_tx=XM_023252594.1
B1 46425224 46425491
refGene_gn=FGFR1;refGene_tx=XM_011281520.3;refGene_gn=FGFR1;refGene_tx=XM_011281521.3;refGene_gn=FGFR1;refGene_tx=XM_011281522.3;refGene_gn=FGFR1;refGene_tx=XM_023252594.1
B1 46426825 46426915
refGene_gn=FGFR1;refGene_tx=XM_011281520.3;refGene_gn=FGFR1;refGene_tx=XM_011281521.3;refGene_gn=FGFR1;refGene_tx=XM_011281522.3;refGene_gn=FGFR1;refGene_tx=XM_011281523.3;refGene_gn=FGFR1;refGene_tx=XM_011281524.3;refGene_gn=FGFR1;refGene_tx=XM_011281525.3;refGene_gn=FGFR1;refGene_tx=XM_023252594.1
B1 46427159 46427332
refGene_gn=FGFR1;refGene_tx=XM_011281520.3;refGene_gn=FGFR1;refGene_tx=XM_011281521.3;refGene_gn=FGFR1;refGene_tx=XM_011281522.3;refGene_gn=FGFR1;refGene_tx=XM_011281523.3;refGene_gn=FGFR1;refGene_tx=XM_011281524.3;refGene_gn=FGFR1;refGene_tx=XM_011281525.3;refGene_gn=FGFR1;refGene_tx=XM_023252594.1
B1 46429820 46429944
refGene_gn=FGFR1;refGene_tx=XM_011281520.3;refGene_gn=FGFR1;refGene_tx=XM_011281521.3;refGene_gn=FGFR1;refGene_tx=XM_011281522.3;refGene_gn=FGFR1;refGene_tx=XM_011281523.3;refGene_gn=FGFR1;refGene_tx=XM_011281524.3;refGene_gn=FGFR1;refGene_tx=XM_011281525.3;refGene_gn=FGFR1;refGene_tx=XM_023252594.1
B1 46430923 46431114
refGene_gn=FGFR1;refGene_tx=XM_011281520.3;refGene_gn=FGFR1;refGene_tx=XM_011281521.3;refGene_gn=FGFR1;refGene_tx=XM_011281522.3;refGene_gn=FGFR1;refGene_tx=XM_011281523.3;refGene_gn=FGFR1;refGene_tx=XM_011281524.3;refGene_gn=FGFR1;refGene_tx=XM_011281525.3;refGene_gn=FGFR1;refGene_tx=XM_023252594.1
B1 46433073 46433218
refGene_gn=FGFR1;refGene_tx=XM_011281520.3;refGene_gn=FGFR1;refGene_tx=XM_011281521.3;refGene_gn=FGFR1;refGene_tx=XM_011281522.3;refGene_gn=

FGFR1;refGene_tx=XM_011281523.3;refGene_gn=FGFR1;refGene_tx=XM_0112815
24.3;refGene_gn=FGFR1;refGene_tx=XM_011281525.3;refGene_gn=FGFR1;refGe
ne_tx=XM_023252594.1
B1 46435229 46435432
refGene_gn=FGFR1;refGene_tx=XM_011281520.3;refGene_gn=FGFR1;refGene_tx
=XM_011281521.3;refGene_gn=FGFR1;refGene_tx=XM_011281522.3;refGene_gn=
FGFR1;refGene_tx=XM_011281523.3;refGene_gn=FGFR1;refGene_tx=XM_0112815
24.3;refGene_gn=FGFR1;refGene_tx=XM_011281525.3;refGene_gn=FGFR1;refGe
ne_tx=XM_023252594.1
B1 46436297 46436443
refGene_gn=FGFR1;refGene_tx=XM_011281520.3;refGene_gn=FGFR1;refGene_tx
=XM_011281521.3;refGene_gn=FGFR1;refGene_tx=XM_011281522.3;refGene_gn=
FGFR1;refGene_tx=XM_011281523.3;refGene_gn=FGFR1;refGene_tx=XM_0112815
24.3;refGene_gn=FGFR1;refGene_tx=XM_011281525.3;refGene_gn=FGFR1;refGe
ne_tx=XM_023252594.1
B1 46436632 46436754
refGene_gn=FGFR1;refGene_tx=XM_011281520.3;refGene_gn=FGFR1;refGene_tx
=XM_011281521.3;refGene_gn=FGFR1;refGene_tx=XM_011281522.3;refGene_gn=
FGFR1;refGene_tx=XM_011281523.3;refGene_gn=FGFR1;refGene_tx=XM_0112815
24.3;refGene_gn=FGFR1;refGene_tx=XM_011281525.3;refGene_gn=FGFR1;refGe
ne_tx=XM_023252594.1
B1 46437225 46437336
refGene_gn=FGFR1;refGene_tx=XM_011281520.3;refGene_gn=FGFR1;refGene_tx
=XM_011281521.3;refGene_gn=FGFR1;refGene_tx=XM_011281522.3;refGene_gn=
FGFR1;refGene_tx=XM_011281523.3;refGene_gn=FGFR1;refGene_tx=XM_0112815
24.3;refGene_gn=FGFR1;refGene_tx=XM_011281525.3;refGene_gn=FGFR1;refGe
ne_tx=XM_023252594.1
B1 46437829 46438020
refGene_gn=FGFR1;refGene_tx=XM_011281520.3;refGene_gn=FGFR1;refGene_tx
=XM_011281521.3;refGene_gn=FGFR1;refGene_tx=XM_011281522.3;refGene_gn=
FGFR1;refGene_tx=XM_011281523.3;refGene_gn=FGFR1;refGene_tx=XM_0112815
24.3;refGene_gn=FGFR1;refGene_tx=XM_011281525.3;refGene_gn=FGFR1;refGe
ne_tx=XM_023252594.1
B1 46438679 46438802
refGene_gn=FGFR1;refGene_tx=XM_011281520.3;refGene_gn=FGFR1;refGene_tx
=XM_011281521.3;refGene_gn=FGFR1;refGene_tx=XM_011281522.3;refGene_gn=
FGFR1;refGene_tx=XM_011281523.3;refGene_gn=FGFR1;refGene_tx=XM_0112815
24.3;refGene_gn=FGFR1;refGene_tx=XM_011281525.3;refGene_gn=FGFR1;refGe
ne_tx=XM_023252594.1
B1 46439026 46439097
refGene_gn=FGFR1;refGene_tx=XM_011281520.3;refGene_gn=FGFR1;refGene_tx
=XM_011281521.3;refGene_gn=FGFR1;refGene_tx=XM_011281522.3;refGene_gn=
FGFR1;refGene_tx=XM_011281523.3;refGene_gn=FGFR1;refGene_tx=XM_0112815
24.3;refGene_gn=FGFR1;refGene_tx=XM_011281525.3;refGene_gn=FGFR1;refGe
ne_tx=XM_023252594.1
B1 46439406 46439544
refGene_gn=FGFR1;refGene_tx=XM_011281520.3;refGene_gn=FGFR1;refGene_tx
=XM_011281521.3;refGene_gn=FGFR1;refGene_tx=XM_011281522.3;refGene_gn=
FGFR1;refGene_tx=XM_011281523.3;refGene_gn=FGFR1;refGene_tx=XM_0112815
24.3;refGene_gn=FGFR1;refGene_tx=XM_011281525.3;refGene_gn=FGFR1;refGe

ne_tx=XM_023252594.1
B1 46439722 46439828
refGene_gn=FGFR1;refGene_tx=XM_011281520.3;refGene_gn=FGFR1;refGene_tx=XM_011281521.3;refGene_gn=FGFR1;refGene_tx=XM_011281522.3;refGene_gn=FGFR1;refGene_tx=XM_011281523.3;refGene_gn=FGFR1;refGene_tx=XM_011281524.3;refGene_gn=FGFR1;refGene_tx=XM_011281525.3;refGene_gn=FGFR1;refGene_tx=XM_023252594.1
B1 46439938 46440115
refGene_gn=FGFR1;refGene_tx=XM_011281520.3;refGene_gn=FGFR1;refGene_tx=XM_011281521.3;refGene_gn=FGFR1;refGene_tx=XM_011281522.3;refGene_gn=FGFR1;refGene_tx=XM_011281523.3;refGene_gn=FGFR1;refGene_tx=XM_011281524.3;refGene_gn=FGFR1;refGene_tx=XM_011281525.3;refGene_gn=FGFR1;refGene_tx=XM_023252594.1
B1 46445122 46445185
refGene_gn=LETM2;refGene_tx=XM_006930710.3;refGene_gn=LETM2;refGene_tx=XM_006930711.3;refGene_gn=LETM2;refGene_tx=XM_006930712.4;refGene_gn=LETM2;refGene_tx=XM_006930713.3;refGene_gn=LETM2;refGene_tx=XM_006930714.3;refGene_gn=LETM2;refGene_tx=XM_019828555.2;refGene_gn=LETM2;refGene_tx=XM_023252596.1
B1 46446183 46446297
refGene_gn=LETM2;refGene_tx=XM_006930710.3;refGene_gn=LETM2;refGene_tx=XM_006930711.3;refGene_gn=LETM2;refGene_tx=XM_006930712.4;refGene_gn=LETM2;refGene_tx=XM_006930714.3;refGene_gn=LETM2;refGene_tx=XM_019828555.2;refGene_gn=LETM2;refGene_tx=XM_023252595.1;refGene_gn=LETM2;refGene_tx=XM_023252596.1
B1 46446616 46446709
refGene_gn=LETM2;refGene_tx=XM_006930710.3;refGene_gn=LETM2;refGene_tx=XM_006930711.3;refGene_gn=LETM2;refGene_tx=XM_006930712.4;refGene_gn=LETM2;refGene_tx=XM_006930713.3;refGene_gn=LETM2;refGene_tx=XM_006930714.3;refGene_gn=LETM2;refGene_tx=XM_019828555.2;refGene_gn=LETM2;refGene_tx=XM_023252595.1;refGene_gn=LETM2;refGene_tx=XM_023252596.1
B1 46449001 46449115
refGene_gn=LETM2;refGene_tx=XM_006930710.3;refGene_gn=LETM2;refGene_tx=XM_006930711.3;refGene_gn=LETM2;refGene_tx=XM_006930712.4;refGene_gn=LETM2;refGene_tx=XM_006930713.3;refGene_gn=LETM2;refGene_tx=XM_006930714.3;refGene_gn=LETM2;refGene_tx=XM_019828555.2;refGene_gn=LETM2;refGene_tx=XM_023252595.1;refGene_gn=LETM2;refGene_tx=XM_023252596.1
B1 46451966 46452086
refGene_gn=LETM2;refGene_tx=XM_006930710.3;refGene_gn=LETM2;refGene_tx=XM_006930711.3;refGene_gn=LETM2;refGene_tx=XM_006930712.4;refGene_gn=LETM2;refGene_tx=XM_006930713.3;refGene_gn=LETM2;refGene_tx=XM_006930714.3;refGene_gn=LETM2;refGene_tx=XM_019828555.2;refGene_gn=LETM2;refGene_tx=XM_023252595.1;refGene_gn=LETM2;refGene_tx=XM_023252596.1
B1 46453199 46453400
refGene_gn=LETM2;refGene_tx=XM_006930710.3;refGene_gn=LETM2;refGene_tx=XM_006930711.3;refGene_gn=LETM2;refGene_tx=XM_006930712.4;refGene_gn=LETM2;refGene_tx=XM_006930713.3;refGene_gn=LETM2;refGene_tx=XM_006930714.3;refGene_gn=LETM2;refGene_tx=XM_019828555.2;refGene_gn=LETM2;refGene_tx=XM_023252595.1;refGene_gn=LETM2;refGene_tx=XM_023252596.1
B1 46454059 46454197

refGene_gn=LETM2;refGene_tx=XM_006930710.3;refGene_gn=LETM2;refGene_tx=XM_006930711.3;refGene_gn=LETM2;refGene_tx=XM_006930712.4;refGene_gn=LETM2;refGene_tx=XM_006930713.3;refGene_gn=LETM2;refGene_tx=XM_006930714.3;refGene_gn=LETM2;refGene_tx=XM_019828555.2;refGene_gn=LETM2;refGene_tx=XM_023252595.1;refGene_gn=LETM2;refGene_tx=XM_023252596.1
B1 46458398 46458542
refGene_gn=LETM2;refGene_tx=XM_006930710.3;refGene_gn=LETM2;refGene_tx=XM_006930711.3;refGene_gn=LETM2;refGene_tx=XM_006930712.4;refGene_gn=LETM2;refGene_tx=XM_006930713.3;refGene_gn=LETM2;refGene_tx=XM_006930714.3;refGene_gn=LETM2;refGene_tx=XM_019828555.2;refGene_gn=LETM2;refGene_tx=XM_023252595.1;refGene_gn=LETM2;refGene_tx=XM_023252596.1
B1 46459153 46459607
refGene_gn=LETM2;refGene_tx=XM_006930710.3;refGene_gn=LETM2;refGene_tx=XM_006930711.3;refGene_gn=LETM2;refGene_tx=XM_006930712.4;refGene_gn=LETM2;refGene_tx=XM_006930713.3;refGene_gn=LETM2;refGene_tx=XM_006930714.3;refGene_gn=LETM2;refGene_tx=XM_019828555.2;refGene_gn=LETM2;refGene_tx=XM_023252595.1;refGene_gn=LETM2;refGene_tx=XM_023252596.1
B1 46461405 46461452
refGene_gn=LETM2;refGene_tx=XM_006930710.3;refGene_gn=LETM2;refGene_tx=XM_006930711.3;refGene_gn=LETM2;refGene_tx=XM_006930712.4;refGene_gn=LETM2;refGene_tx=XM_006930713.3;refGene_gn=LETM2;refGene_tx=XM_023252595.1
B1 46512700 46513375
refGene_gn=NSD3;refGene_tx=XM_003984777.4;refGene_gn=NSD3;refGene_tx=XM_023252597.1;refGene_gn=NSD3;refGene_tx=XM_023252598.1;refGene_gn=NSD3;refGene_tx=XM_023252599.1;refGene_gn=NSD3;refGene_tx=XM_023252600.1
B1 46521950 46522022
refGene_gn=NSD3;refGene_tx=XM_003984777.4;refGene_gn=NSD3;refGene_tx=XM_023252597.1;refGene_gn=NSD3;refGene_tx=XM_023252598.1;refGene_gn=NSD3;refGene_tx=XM_023252599.1;refGene_gn=NSD3;refGene_tx=XM_023252600.1
B1 46522610 46522773
refGene_gn=NSD3;refGene_tx=XM_003984777.4;refGene_gn=NSD3;refGene_tx=XM_023252597.1;refGene_gn=NSD3;refGene_tx=XM_023252598.1;refGene_gn=NSD3;refGene_tx=XM_023252599.1;refGene_gn=NSD3;refGene_tx=XM_023252600.1
B1 46528180 46528335
refGene_gn=NSD3;refGene_tx=XM_003984777.4;refGene_gn=NSD3;refGene_tx=XM_023252597.1;refGene_gn=NSD3;refGene_tx=XM_023252598.1;refGene_gn=NSD3;refGene_tx=XM_023252599.1;refGene_gn=NSD3;refGene_tx=XM_023252600.1
B1 46530675 46531191
refGene_gn=NSD3;refGene_tx=XM_003984777.4;refGene_gn=NSD3;refGene_tx=XM_023252597.1;refGene_gn=NSD3;refGene_tx=XM_023252598.1;refGene_gn=NSD3;refGene_tx=XM_023252599.1;refGene_gn=NSD3;refGene_tx=XM_023252600.1
B1 46533474 46533601
refGene_gn=NSD3;refGene_tx=XM_003984777.4;refGene_gn=NSD3;refGene_tx=XM_023252597.1;refGene_gn=NSD3;refGene_tx=XM_023252598.1;refGene_gn=NSD3;refGene_tx=XM_023252599.1;refGene_gn=NSD3;refGene_tx=XM_023252600.1
B1 46537651 46537752
refGene_gn=NSD3;refGene_tx=XM_003984777.4;refGene_gn=NSD3;refGene_tx=XM_023252597.1;refGene_gn=NSD3;refGene_tx=XM_023252598.1;refGene_gn=NSD3;refGene_tx=XM_023252599.1;refGene_gn=NSD3;refGene_tx=XM_023252600.1

B1	46539884 46539930
refGene_gn=NSD3;refGene_tx=XM_003984777.4;refGene_gn=NSD3;refGene_tx=XM_023252597.1;refGene_gn=NSD3;refGene_tx=XM_023252598.1;refGene_gn=NSD3;refGene_tx=XM_023252599.1;refGene_gn=NSD3;refGene_tx=XM_023252600.1
B1	46540839 46540922 refGene_gn=NSD3;refGene_tx=XM_003984777.4
B1	46542861 46542992
refGene_gn=NSD3;refGene_tx=XM_023252597.1;refGene_gn=NSD3;refGene_tx=XM_023252598.1;refGene_gn=NSD3;refGene_tx=XM_023252599.1;refGene_gn=NSD3;refGene_tx=XM_023252600.1
B1	46543371 46543500
refGene_gn=NSD3;refGene_tx=XM_023252597.1;refGene_gn=NSD3;refGene_tx=XM_023252598.1;refGene_gn=NSD3;refGene_tx=XM_023252599.1;refGene_gn=NSD3;refGene_tx=XM_023252600.1
B1	46544686 46544813
refGene_gn=NSD3;refGene_tx=XM_023252597.1;refGene_gn=NSD3;refGene_tx=XM_023252598.1;refGene_gn=NSD3;refGene_tx=XM_023252599.1;refGene_gn=NSD3;refGene_tx=XM_023252600.1
B1	46553238 46553436
refGene_gn=NSD3;refGene_tx=XM_023252597.1;refGene_gn=NSD3;refGene_tx=XM_023252598.1;refGene_gn=NSD3;refGene_tx=XM_023252599.1
B1	46554190 46554364
refGene_gn=NSD3;refGene_tx=XM_023252597.1;refGene_gn=NSD3;refGene_tx=XM_023252598.1;refGene_gn=NSD3;refGene_tx=XM_023252599.1;refGene_gn=NSD3;refGene_tx=XM_023252600.1
B1	46558552 46558699
refGene_gn=NSD3;refGene_tx=XM_023252597.1;refGene_gn=NSD3;refGene_tx=XM_023252598.1;refGene_gn=NSD3;refGene_tx=XM_023252600.1
B1	46561987 46562171
refGene_gn=NSD3;refGene_tx=XM_023252597.1;refGene_gn=NSD3;refGene_tx=XM_023252598.1;refGene_gn=NSD3;refGene_tx=XM_023252599.1;refGene_gn=NSD3;refGene_tx=XM_023252600.1
B1	46566563 46566766
refGene_gn=NSD3;refGene_tx=XM_023252597.1;refGene_gn=NSD3;refGene_tx=XM_023252598.1;refGene_gn=NSD3;refGene_tx=XM_023252599.1;refGene_gn=NSD3;refGene_tx=XM_023252600.1
B1	46567508 46567621
refGene_gn=NSD3;refGene_tx=XM_023252597.1;refGene_gn=NSD3;refGene_tx=XM_023252598.1;refGene_gn=NSD3;refGene_tx=XM_023252599.1;refGene_gn=NSD3;refGene_tx=XM_023252600.1
B1	46568214 46568484
refGene_gn=NSD3;refGene_tx=XM_023252597.1;refGene_gn=NSD3;refGene_tx=XM_023252598.1;refGene_gn=NSD3;refGene_tx=XM_023252599.1;refGene_gn=NSD3;refGene_tx=XM_023252600.1
B1	46573792 46573909
refGene_gn=NSD3;refGene_tx=XM_023252597.1;refGene_gn=NSD3;refGene_tx=XM_023252598.1;refGene_gn=NSD3;refGene_tx=XM_023252599.1;refGene_gn=NSD3;refGene_tx=XM_023252600.1
B1	46577315 46577457
refGene_gn=NSD3;refGene_tx=XM_023252597.1;refGene_gn=NSD3;refGene_tx=XM_023252598.1;refGene_gn=NSD3;refGene_tx=XM_023252599.1;refGene_gn=NSD

3;refGene_tx=XM_023252600.1
B1 46578496 46578603
refGene_gn=NSD3;refGene_tx=XM_023252597.1;refGene_gn=NSD3;refGene_tx=XM_023252598.1;refGene_gn=NSD3;refGene_tx=XM_023252599.1;refGene_gn=NSD3;refGene_tx=XM_023252600.1
B1 46579364 46579569
refGene_gn=NSD3;refGene_tx=XM_023252597.1;refGene_gn=NSD3;refGene_tx=XM_023252598.1;refGene_gn=NSD3;refGene_tx=XM_023252599.1;refGene_gn=NSD3;refGene_tx=XM_023252600.1
B1 46579987 46580229
refGene_gn=NSD3;refGene_tx=XM_023252597.1;refGene_gn=NSD3;refGene_tx=XM_023252598.1;refGene_gn=NSD3;refGene_tx=XM_023252599.1;refGene_gn=NSD3;refGene_tx=XM_023252600.1
B1 46586101 46586242
refGene_gn=PLPP5;refGene_tx=XM_023252601.1;refGene_gn=PLPP5;refGene_tx=XM_023252602.1;refGene_gn=PLPP5;refGene_tx=XM_023252604.1;refGene_gn=PLPP5;refGene_tx=XM_023252606.1
B1 46586694 46586768
refGene_gn=PLPP5;refGene_tx=XM_023252603.1;refGene_gn=PLPP5;refGene_tx=XM_023252605.1;refGene_gn=PLPP5;refGene_tx=XM_023252609.1
B1 46586961 46587070
refGene_gn=PLPP5;refGene_tx=XM_023252601.1;refGene_gn=PLPP5;refGene_tx=XM_023252602.1;refGene_gn=PLPP5;refGene_tx=XM_023252603.1;refGene_gn=PLPP5;refGene_tx=XM_023252604.1;refGene_gn=PLPP5;refGene_tx=XM_023252605.1;refGene_gn=PLPP5;refGene_tx=XM_023252606.1;refGene_gn=PLPP5;refGene_tx=XM_023252607.1;refGene_gn=PLPP5;refGene_tx=XM_023252608.1;refGene_gn=PLPP5;refGene_tx=XM_023252609.1
B1 46587509 46587600
refGene_gn=PLPP5;refGene_tx=XM_023252601.1;refGene_gn=PLPP5;refGene_tx=XM_023252602.1;refGene_gn=PLPP5;refGene_tx=XM_023252603.1;refGene_gn=PLPP5;refGene_tx=XM_023252604.1;refGene_gn=PLPP5;refGene_tx=XM_023252605.1;refGene_gn=PLPP5;refGene_tx=XM_023252606.1;refGene_gn=PLPP5;refGene_tx=XM_023252607.1;refGene_gn=PLPP5;refGene_tx=XM_023252608.1;refGene_gn=PLPP5;refGene_tx=XM_023252609.1
B1 46588022 46588086
refGene_gn=PLPP5;refGene_tx=XM_023252601.1;refGene_gn=PLPP5;refGene_tx=XM_023252602.1;refGene_gn=PLPP5;refGene_tx=XM_023252603.1;refGene_gn=PLPP5;refGene_tx=XM_023252604.1;refGene_gn=PLPP5;refGene_tx=XM_023252605.1;refGene_gn=PLPP5;refGene_tx=XM_023252606.1;refGene_gn=PLPP5;refGene_tx=XM_023252607.1;refGene_gn=PLPP5;refGene_tx=XM_023252608.1;refGene_gn=PLPP5;refGene_tx=XM_023252609.1
B1 46588615 46588740
refGene_gn=PLPP5;refGene_tx=XM_023252601.1;refGene_gn=PLPP5;refGene_tx=XM_023252602.1;refGene_gn=PLPP5;refGene_tx=XM_023252603.1;refGene_gn=PLPP5;refGene_tx=XM_023252604.1;refGene_gn=PLPP5;refGene_tx=XM_023252605.1;refGene_gn=PLPP5;refGene_tx=XM_023252606.1;refGene_gn=PLPP5;refGene_tx=XM_023252607.1;refGene_gn=PLPP5;refGene_tx=XM_023252608.1;refGene_gn=PLPP5;refGene_tx=XM_023252609.1
B1 46589393 46589580
refGene_gn=PLPP5;refGene_tx=XM_023252601.1;refGene_gn=PLPP5;refGene_tx

=XM_023252602.1;refGene_gn=PLPP5;refGene_tx=XM_023252603.1;refGene_gn=PLPP5;refGene_tx=XM_023252604.1;refGene_gn=PLPP5;refGene_tx=XM_023252605.1;refGene_gn=PLPP5;refGene_tx=XM_023252606.1;refGene_gn=PLPP5;refGene_tx=XM_023252607.1;refGene_gn=PLPP5;refGene_tx=XM_023252608.1;refGene_gn=PLPP5;refGene_tx=XM_023252609.1
B1 46590454 46590501 refGene_gn=PLPP5;refGene_tx=XM_023252606.1
B1 46590945 46591106
refGene_gn=PLPP5;refGene_tx=XM_023252601.1;refGene_gn=PLPP5;refGene_tx=XM_023252602.1;refGene_gn=PLPP5;refGene_tx=XM_023252603.1;refGene_gn=PLPP5;refGene_tx=XM_023252605.1;refGene_gn=PLPP5;refGene_tx=XM_023252607.1;refGene_gn=PLPP5;refGene_tx=XM_023252608.1
B1 46591979 46592050 refGene_gn=PLPP5;refGene_tx=XM_023252604.1
B1 46595220 46595302 refGene_gn=DDHD2;refGene_tx=XM_023253388.1
B1 46603682 46603842 refGene_gn=DDHD2;refGene_tx=XM_023253388.1
B1 46604536 46604707 refGene_gn=DDHD2;refGene_tx=XM_023253388.1
B1 46604845 46604948 refGene_gn=DDHD2;refGene_tx=XM_023253388.1
B1 46605426 46605582 refGene_gn=DDHD2;refGene_tx=XM_023253388.1
B1 46605693 46605822 refGene_gn=DDHD2;refGene_tx=XM_023253388.1
B1 46606864 46606960 refGene_gn=DDHD2;refGene_tx=XM_023253388.1
B1 46608050 46608173 refGene_gn=DDHD2;refGene_tx=XM_023253388.1
B1 46609014 46609082 refGene_gn=DDHD2;refGene_tx=XM_023253388.1
B1 46609328 46609537 refGene_gn=DDHD2;refGene_tx=XM_023253388.1
B1 46612047 46612183 refGene_gn=DDHD2;refGene_tx=XM_023253388.1
B1 46613126 46613216 refGene_gn=DDHD2;refGene_tx=XM_023253388.1
B1 46615349 46615470 refGene_gn=DDHD2;refGene_tx=XM_023253388.1
B1 46616018 46616108 refGene_gn=DDHD2;refGene_tx=XM_023253388.1
B1 46616660 46616725 refGene_gn=DDHD2;refGene_tx=XM_023253388.1
B1 46618546 46618787 refGene_gn=DDHD2;refGene_tx=XM_023253388.1
B1 46633166 46633652
refGene_gn=BAG4;refGene_tx=XM_003984780.5;refGene_gn=BAG4;refGene_tx=XM_019828561.2
B1 46634379 46634634
refGene_gn=BAG4;refGene_tx=XM_003984780.5;refGene_gn=BAG4;refGene_tx=XM_019828561.2
B1 46635860 46636115 refGene_gn=BAG4;refGene_tx=XM_003984780.5
B1 46646660 46646765
refGene_gn=BAG4;refGene_tx=XM_003984780.5;refGene_gn=BAG4;refGene_tx=XM_019828561.2
B1 46663893 46664163
refGene_gn=BAG4;refGene_tx=XM_003984780.5;refGene_gn=BAG4;refGene_tx=XM_019828561.2
B1 46664706 46664752 refGene_gn=LSM1;refGene_tx=XM_003984782.5
B1 46668959 46669028 refGene_gn=LSM1;refGene_tx=XM_003984782.5
B1 46670383 46670499 refGene_gn=LSM1;refGene_tx=XM_003984782.5
B1 46674434 46674605 refGene_gn=LSM1;refGene_tx=XM_003984782.5
B1 46683589 46683653 refGene_gn=STAR;refGene_tx=NM_001246196.1
B1 46684993 46685107 refGene_gn=STAR;refGene_tx=NM_001246196.1
B1 46685434 46685562 refGene_gn=STAR;refGene_tx=NM_001246196.1
B1 46686199 46686358 refGene_gn=STAR;refGene_tx=NM_001246196.1
B1 46686498 46686683 refGene_gn=STAR;refGene_tx=NM_001246196.1

B1	46687251	46687345	refGene_gn=STAR;refGene_tx=NM_001246196.1
B1	46687839	46687953	refGene_gn=STAR;refGene_tx=NM_001246196.1
B1	46691692	46691800
refGene_gn=ASH2L;refGene_tx=XM_003984783.5;refGene_gn=ASH2L;refGene_tx=XM_023252610.1;refGene_gn=ASH2L;refGene_tx=XM_023252611.1
B1	46691894	46691954
refGene_gn=ASH2L;refGene_tx=XM_003984783.5;refGene_gn=ASH2L;refGene_tx=XM_023252610.1;refGene_gn=ASH2L;refGene_tx=XM_023252611.1
B1	46692689	46692788
refGene_gn=ASH2L;refGene_tx=XM_003984783.5;refGene_gn=ASH2L;refGene_tx=XM_023252610.1;refGene_gn=ASH2L;refGene_tx=XM_023252611.1
B1	46694080	46694173
refGene_gn=ASH2L;refGene_tx=XM_003984783.5;refGene_gn=ASH2L;refGene_tx=XM_023252610.1;refGene_gn=ASH2L;refGene_tx=XM_023252611.1
B1	46698913	46699107
refGene_gn=ASH2L;refGene_tx=XM_003984783.5;refGene_gn=ASH2L;refGene_tx=XM_023252610.1;refGene_gn=ASH2L;refGene_tx=XM_023252611.1
B1	46699404	46699572
refGene_gn=ASH2L;refGene_tx=XM_003984783.5;refGene_gn=ASH2L;refGene_tx=XM_023252610.1;refGene_gn=ASH2L;refGene_tx=XM_023252611.1
B1	46707520	46707738
refGene_gn=ASH2L;refGene_tx=XM_003984783.5;refGene_gn=ASH2L;refGene_tx=XM_023252610.1;refGene_gn=ASH2L;refGene_tx=XM_023252611.1
B1	46708483	46708577
refGene_gn=ASH2L;refGene_tx=XM_003984783.5;refGene_gn=ASH2L;refGene_tx=XM_023252610.1;refGene_gn=ASH2L;refGene_tx=XM_023252611.1
B1	46710030	46710048
refGene_gn=ASH2L;refGene_tx=XM_023252610.1;refGene_gn=ASH2L;refGene_tx=XM_023252611.1
B1	46710781	46710857
refGene_gn=ASH2L;refGene_tx=XM_003984783.5;refGene_gn=ASH2L;refGene_tx=XM_023252610.1;refGene_gn=ASH2L;refGene_tx=XM_023252611.1
B1	46712217	46712313
refGene_gn=ASH2L;refGene_tx=XM_003984783.5;refGene_gn=ASH2L;refGene_tx=XM_023252610.1;refGene_gn=ASH2L;refGene_tx=XM_023252611.1
B1	46712946	46713042
refGene_gn=ASH2L;refGene_tx=XM_003984783.5;refGene_gn=ASH2L;refGene_tx=XM_023252610.1;refGene_gn=ASH2L;refGene_tx=XM_023252611.1
B1	46717596	46717691
refGene_gn=ASH2L;refGene_tx=XM_003984783.5;refGene_gn=ASH2L;refGene_tx=XM_023252610.1;refGene_gn=ASH2L;refGene_tx=XM_023252611.1
B1	46717977	46718066
refGene_gn=ASH2L;refGene_tx=XM_003984783.5;refGene_gn=ASH2L;refGene_tx=XM_023252610.1;refGene_gn=ASH2L;refGene_tx=XM_023252611.1
B1	46721984	46722130
refGene_gn=ASH2L;refGene_tx=XM_003984783.5;refGene_gn=ASH2L;refGene_tx=XM_023252610.1;refGene_gn=ASH2L;refGene_tx=XM_023252611.1
B1	46722261	46722328
refGene_gn=ASH2L;refGene_tx=XM_003984783.5;refGene_gn=ASH2L;refGene_tx=XM_023252610.1

B1 46723004 46723192
refGene_gn=ASH2L;refGene_tx=XM_003984783.5;refGene_gn=ASH2L;refGene_tx=XM_023252610.1
B1 46740235 46740267 refGene_gn=EIF4EBP1;refGene_tx=XM_003984784.3
B1 46743165 46743345 refGene_gn=EIF4EBP1;refGene_tx=XM_003984784.3
B1 46763192 46763337 refGene_gn=EIF4EBP1;refGene_tx=XM_003984784.3
B1 46802625 46803799
refGene_gn=ADRB3;refGene_tx=NM_001009336.1;refGene_gn=ADRB3;refGene_tx=XM_019828152.2
B1 46804500 46804523 refGene_gn=ADRB3;refGene_tx=NM_001009336.1
B1 46823282 46823305 refGene_gn=ADRB3;refGene_tx=XM_019828152.2
B1 46825181 46825296
refGene_gn=GOT1L1;refGene_tx=XM_003984822.4;refGene_gn=GOT1L1;refGene_tx=XM_019828562.2
B1 46826058 46826240
refGene_gn=GOT1L1;refGene_tx=XM_003984822.4;refGene_gn=GOT1L1;refGene_tx=XM_019828562.2
B1 46826979 46827091
refGene_gn=GOT1L1;refGene_tx=XM_003984822.4;refGene_gn=GOT1L1;refGene_tx=XM_019828562.2
B1 46827338 46827448
refGene_gn=GOT1L1;refGene_tx=XM_003984822.4;refGene_gn=GOT1L1;refGene_tx=XM_019828562.2
B1 46827653 46827746
refGene_gn=GOT1L1;refGene_tx=XM_003984822.4;refGene_gn=GOT1L1;refGene_tx=XM_019828562.2
B1 46827839 46827990
refGene_gn=GOT1L1;refGene_tx=XM_003984822.4;refGene_gn=GOT1L1;refGene_tx=XM_019828562.2
B1 46828496 46828662
refGene_gn=GOT1L1;refGene_tx=XM_003984822.4;refGene_gn=GOT1L1;refGene_tx=XM_019828562.2
B1 46829164 46829307
refGene_gn=GOT1L1;refGene_tx=XM_003984822.4;refGene_gn=GOT1L1;refGene_tx=XM_019828562.2
B1 46829554 46829703
refGene_gn=GOT1L1;refGene_tx=XM_003984822.4;refGene_gn=GOT1L1;refGene_tx=XM_019828562.2
B1 46865864 46866241
refGene_gn=RAB11FIP1;refGene_tx=XM_003984823.5;refGene_gn=RAB11FIP1;refGene_tx=XM_023252616.1
B1 46880831 46881274
refGene_gn=RAB11FIP1;refGene_tx=XM_003984823.5;refGene_gn=RAB11FIP1;refGene_tx=XM_023252615.1;refGene_gn=RAB11FIP1;refGene_tx=XM_023252616.1
B1 46882572 46883377
refGene_gn=RAB11FIP1;refGene_tx=XM_003984823.5;refGene_gn=RAB11FIP1;refGene_tx=XM_023252615.1;refGene_gn=RAB11FIP1;refGene_tx=XM_023252616.1
B1 46884899 46886783
refGene_gn=RAB11FIP1;refGene_tx=XM_003984823.5;refGene_gn=RAB11FIP1;refGene_tx=XM_023252615.1

B1 46887552 46887661
refGene_gn=RAB11FIP1;refGene_tx=XM_003984823.5;refGene_gn=RAB11FIP1;refGene_tx=XM_023252615.1;refGene_gn=RAB11FIP1;refGene_tx=XM_023252616.1
B1 46896824 46897043
refGene_gn=RAB11FIP1;refGene_tx=XM_003984823.5;refGene_gn=RAB11FIP1;refGene_tx=XM_023252615.1;refGene_gn=RAB11FIP1;refGene_tx=XM_023252616.1
B1 46903323 46903329 refGene_gn=ADGRA2;refGene_tx=XM_023252613.1
B1 46908638 46908792 refGene_gn=BRF2;refGene_tx=XM_003984785.4
B1 46909705 46909765 refGene_gn=BRF2;refGene_tx=XM_003984785.4
B1 46910223 46910545 refGene_gn=BRF2;refGene_tx=XM_003984785.4
B1 46911503 46912233 refGene_gn=BRF2;refGene_tx=XM_003984785.4
B1 46914441 46915708
refGene_gn=ADGRA2;refGene_tx=XM_023252612.1;refGene_gn=ADGRA2;refGene_tx=XM_023252613.1;refGene_gn=ADGRA2;refGene_tx=XM_023252614.1
B1 46915938 46916028
refGene_gn=ADGRA2;refGene_tx=XM_023252612.1;refGene_gn=ADGRA2;refGene_tx=XM_023252613.1;refGene_gn=ADGRA2;refGene_tx=XM_023252614.1
B1 46916541 46916687
refGene_gn=ADGRA2;refGene_tx=XM_023252612.1;refGene_gn=ADGRA2;refGene_tx=XM_023252613.1;refGene_gn=ADGRA2;refGene_tx=XM_023252614.1
B1 46917147 46917271
refGene_gn=ADGRA2;refGene_tx=XM_023252612.1;refGene_gn=ADGRA2;refGene_tx=XM_023252613.1;refGene_gn=ADGRA2;refGene_tx=XM_023252614.1
B1 46917583 46917711
refGene_gn=ADGRA2;refGene_tx=XM_023252612.1;refGene_gn=ADGRA2;refGene_tx=XM_023252613.1;refGene_gn=ADGRA2;refGene_tx=XM_023252614.1
B1 46918379 46918588
refGene_gn=ADGRA2;refGene_tx=XM_023252612.1;refGene_gn=ADGRA2;refGene_tx=XM_023252613.1;refGene_gn=ADGRA2;refGene_tx=XM_023252614.1
B1 46919513 46919730
refGene_gn=ADGRA2;refGene_tx=XM_023252612.1;refGene_gn=ADGRA2;refGene_tx=XM_023252613.1;refGene_gn=ADGRA2;refGene_tx=XM_023252614.1
B1 46919926 46920151
refGene_gn=ADGRA2;refGene_tx=XM_023252612.1;refGene_gn=ADGRA2;refGene_tx=XM_023252613.1
B1 46920934 46921096
refGene_gn=ADGRA2;refGene_tx=XM_023252612.1;refGene_gn=ADGRA2;refGene_tx=XM_023252613.1;refGene_gn=ADGRA2;refGene_tx=XM_023252614.1
B1 46921221 46921371
refGene_gn=ADGRA2;refGene_tx=XM_023252612.1;refGene_gn=ADGRA2;refGene_tx=XM_023252613.1;refGene_gn=ADGRA2;refGene_tx=XM_023252614.1
B1 46921878 46922077
refGene_gn=ADGRA2;refGene_tx=XM_023252612.1;refGene_gn=ADGRA2;refGene_tx=XM_023252613.1;refGene_gn=ADGRA2;refGene_tx=XM_023252614.1
B1 46922669 46922834
refGene_gn=ADGRA2;refGene_tx=XM_023252612.1;refGene_gn=ADGRA2;refGene_tx=XM_023252613.1;refGene_gn=ADGRA2;refGene_tx=XM_023252614.1
B1 46923346 46923560
refGene_gn=ADGRA2;refGene_tx=XM_023252612.1;refGene_gn=ADGRA2;refGene_tx=XM_023252613.1;refGene_gn=ADGRA2;refGene_tx=XM_023252614.1

B1	46923976	46924140	refGene_gn=ADGRA2;refGene_tx=XM_023252612.1;refGene_gn=ADGRA2;refGene_tx=XM_023252613.1;refGene_gn=ADGRA2;refGene_tx=XM_023252614.1
B1	46924426	46924498	refGene_gn=ADGRA2;refGene_tx=XM_023252612.1;refGene_gn=ADGRA2;refGene_tx=XM_023252613.1;refGene_gn=ADGRA2;refGene_tx=XM_023252614.1
B1	46924653	46924725	refGene_gn=ADGRA2;refGene_tx=XM_023252612.1;refGene_gn=ADGRA2;refGene_tx=XM_023252613.1;refGene_gn=ADGRA2;refGene_tx=XM_023252614.1
B1	46924982	46925054	refGene_gn=ADGRA2;refGene_tx=XM_023252612.1;refGene_gn=ADGRA2;refGene_tx=XM_023252613.1;refGene_gn=ADGRA2;refGene_tx=XM_023252614.1
B1	46935027	46935099	refGene_gn=ADGRA2;refGene_tx=XM_023252612.1;refGene_gn=ADGRA2;refGene_tx=XM_023252613.1;refGene_gn=ADGRA2;refGene_tx=XM_023252614.1
B1	46947325	46947588	refGene_gn=ADGRA2;refGene_tx=XM_023252612.1;refGene_gn=ADGRA2;refGene_tx=XM_023252613.1;refGene_gn=ADGRA2;refGene_tx=XM_023252614.1
B1	46965543	46965675	refGene_gn=PLPBP;refGene_tx=XM_006930724.4;refGene_gn=PLPBP;refGene_tx=XM_006930725.4
B1	46967290	46967389	refGene_gn=PLPBP;refGene_tx=XM_006930724.4;refGene_gn=PLPBP;refGene_tx=XM_006930725.4
B1	46967787	46967930	refGene_gn=PLPBP;refGene_tx=XM_006930724.4;refGene_gn=PLPBP;refGene_tx=XM_006930725.4
B1	46969832	46969967	refGene_gn=PLPBP;refGene_tx=XM_006930724.4;refGene_gn=PLPBP;refGene_tx=XM_006930725.4
B1	46979930	46980006	refGene_gn=PLPBP;refGene_tx=XM_006930724.4;refGene_gn=PLPBP;refGene_tx=XM_006930725.4
B1	46980635	46980671	refGene_gn=PLPBP;refGene_tx=XM_006930724.4;refGene_gn=PLPBP;refGene_tx=XM_006930725.4
B1	46980749	46980857	refGene_gn=PLPBP;refGene_tx=XM_006930724.4;refGene_gn=PLPBP;refGene_tx=XM_006930725.4
B1	46983036	46983135	refGene_gn=PLPBP;refGene_tx=XM_006930724.4
B1	46991462	46991657	refGene_gn=ERLIN2;refGene_tx=XM_003984787.5;refGene_gn=ERLIN2;refGene_tx=XM_006930726.4
B1	46992049	46992129	refGene_gn=ERLIN2;refGene_tx=XM_003984787.5;refGene_gn=ERLIN2;refGene_tx=XM_006930726.4
B1	46993804	46993894	refGene_gn=ERLIN2;refGene_tx=XM_003984787.5;refGene_gn=ERLIN2;refGene_tx=XM_006930726.4
B1	46994698	46994790

refGene_gn=ERLIN2;refGene_tx=XM_003984787.5;refGene_gn=ERLIN2;refGene_tx=XM_006930726.4
B1 46995325 46995384
refGene_gn=ERLIN2;refGene_tx=XM_003984787.5;refGene_gn=ERLIN2;refGene_tx=XM_006930726.4
B1 46995543 46995617
refGene_gn=ERLIN2;refGene_tx=XM_003984787.5;refGene_gn=ERLIN2;refGene_tx=XM_006930726.4
B1 46999447 46999573
refGene_gn=ERLIN2;refGene_tx=XM_003984787.5;refGene_gn=ERLIN2;refGene_tx=XM_006930726.4
B1 46999727 46999789
refGene_gn=ERLIN2;refGene_tx=XM_003984787.5;refGene_gn=ERLIN2;refGene_tx=XM_006930726.4
B1 47001661 47001708
refGene_gn=ERLIN2;refGene_tx=XM_003984787.5;refGene_gn=ERLIN2;refGene_tx=XM_006930726.4
B1 47002912 47002994
refGene_gn=ERLIN2;refGene_tx=XM_003984787.5;refGene_gn=ERLIN2;refGene_tx=XM_006930726.4
B1 47004508 47004615
refGene_gn=ERLIN2;refGene_tx=XM_003984787.5;refGene_gn=ERLIN2;refGene_tx=XM_006930726.4
B1 47036538 47038074 refGene_gn=ZNF703;refGene_tx=XM_023252617.1
B1 47039092 47039335 refGene_gn=ZNF703;refGene_tx=XM_023252617.1
B1 47716738 47717150
refGene_gn=KCNU1;refGene_tx=XM_019828564.2;refGene_gn=KCNU1;refGene_tx=XM_019828566.2
B1 47719394 47719504
refGene_gn=KCNU1;refGene_tx=XM_019828564.2;refGene_gn=KCNU1;refGene_tx=XM_019828566.2
B1 47721278 47721470
refGene_gn=KCNU1;refGene_tx=XM_019828564.2;refGene_gn=KCNU1;refGene_tx=XM_019828566.2
B1 47730203 47730343
refGene_gn=KCNU1;refGene_tx=XM_019828564.2;refGene_gn=KCNU1;refGene_tx=XM_019828566.2
B1 47733192 47733264
refGene_gn=KCNU1;refGene_tx=XM_019828564.2;refGene_gn=KCNU1;refGene_tx=XM_019828566.2
B1 47737843 47738030
refGene_gn=KCNU1;refGene_tx=XM_019828564.2;refGene_gn=KCNU1;refGene_tx=XM_019828566.2
B1 47739402 47739627
refGene_gn=KCNU1;refGene_tx=XM_019828564.2;refGene_gn=KCNU1;refGene_tx=XM_019828566.2
B1 47742842 47742939
refGene_gn=KCNU1;refGene_tx=XM_019828564.2;refGene_gn=KCNU1;refGene_tx=XM_019828566.2
B1 47785237 47785352

refGene_gn=KCNU1;refGene_tx=XM_019828564.2;refGene_gn=KCNU1;refGene_tx=XM_019828566.2
B1 47809718 47809816
refGene_gn=KCNU1;refGene_tx=XM_019828564.2;refGene_gn=KCNU1;refGene_tx=XM_019828566.2
B1 47809951 47810041
refGene_gn=KCNU1;refGene_tx=XM_019828564.2;refGene_gn=KCNU1;refGene_tx=XM_019828566.2
B1 47813829 47813898
refGene_gn=KCNU1;refGene_tx=XM_019828564.2;refGene_gn=KCNU1;refGene_tx=XM_019828566.2
B1 47814251 47814364
refGene_gn=KCNU1;refGene_tx=XM_019828564.2;refGene_gn=KCNU1;refGene_tx=XM_019828566.2
B1 47818906 47819059
refGene_gn=KCNU1;refGene_tx=XM_019828564.2;refGene_gn=KCNU1;refGene_tx=XM_019828566.2
B1 47822827 47822897
refGene_gn=KCNU1;refGene_tx=XM_019828564.2;refGene_gn=KCNU1;refGene_tx=XM_019828566.2
B1 47824204 47824287
refGene_gn=KCNU1;refGene_tx=XM_019828564.2;refGene_gn=KCNU1;refGene_tx=XM_019828566.2
B1 47825370 47825476
refGene_gn=KCNU1;refGene_tx=XM_019828564.2;refGene_gn=KCNU1;refGene_tx=XM_019828566.2
B1 47827841 47827952
refGene_gn=KCNU1;refGene_tx=XM_019828564.2;refGene_gn=KCNU1;refGene_tx=XM_019828566.2
B1 47829578 47829670
refGene_gn=KCNU1;refGene_tx=XM_019828564.2;refGene_gn=KCNU1;refGene_tx=XM_019828566.2
B1 47830785 47830956 refGene_gn=KCNU1;refGene_tx=XM_019828564.2
B1 47837199 47837275
refGene_gn=KCNU1;refGene_tx=XM_019828564.2;refGene_gn=KCNU1;refGene_tx=XM_019828566.2
B1 47838277 47838353
refGene_gn=KCNU1;refGene_tx=XM_019828564.2;refGene_gn=KCNU1;refGene_tx=XM_019828566.2
B1 47839339 47839451
refGene_gn=KCNU1;refGene_tx=XM_019828564.2;refGene_gn=KCNU1;refGene_tx=XM_019828566.2
B1 47841781 47841872
refGene_gn=KCNU1;refGene_tx=XM_019828564.2;refGene_gn=KCNU1;refGene_tx=XM_019828566.2
B1 47842953 47843015
refGene_gn=KCNU1;refGene_tx=XM_019828564.2;refGene_gn=KCNU1;refGene_tx=XM_019828566.2
B1 47860943 47861063
refGene_gn=KCNU1;refGene_tx=XM_019828564.2;refGene_gn=KCNU1;refGene_tx

=XM_019828566.2
B1	47863899	47864046
refGene_gn=KCNU1;refGene_tx=XM_019828564.2;refGene_gn=KCNU1;refGene_tx
=XM_019828566.2
B1	48613647	48613774	refGene_gn=UNC5D;refGene_tx=XM_023252623.1
B1	48633818	48634023
refGene_gn=UNC5D;refGene_tx=XM_019828578.2;refGene_gn=UNC5D;refGene_tx
=XM_023252618.1;refGene_gn=UNC5D;refGene_tx=XM_023252619.1;refGene_gn=
UNC5D;refGene_tx=XM_023252620.1;refGene_gn=UNC5D;refGene_tx=XM_0232526
21.1;refGene_gn=UNC5D;refGene_tx=XM_023252622.1;refGene_gn=UNC5D;refGe
ne_tx=XM_023252624.1;refGene_gn=UNC5D;refGene_tx=XM_023252625.1;refGen
e_gn=UNC5D;refGene_tx=XM_023252627.1;refGene_gn=UNC5D;refGene_tx=XM_02
3252628.1
B1	48640074	48640090	refGene_gn=UNC5D;refGene_tx=XM_023252626.1
B1	48647388	48647567
refGene_gn=UNC5D;refGene_tx=XM_019828578.2;refGene_gn=UNC5D;refGene_tx
=XM_023252618.1;refGene_gn=UNC5D;refGene_tx=XM_023252619.1;refGene_gn=
UNC5D;refGene_tx=XM_023252620.1;refGene_gn=UNC5D;refGene_tx=XM_0232526
21.1;refGene_gn=UNC5D;refGene_tx=XM_023252622.1;refGene_gn=UNC5D;refGe
ne_tx=XM_023252623.1;refGene_gn=UNC5D;refGene_tx=XM_023252624.1;refGen
e_gn=UNC5D;refGene_tx=XM_023252625.1;refGene_gn=UNC5D;refGene_tx=XM_02
3252626.1;refGene_gn=UNC5D;refGene_tx=XM_023252627.1;refGene_gn=UNC5D;
refGene_tx=XM_023252628.1
B1	48653505	48653670
refGene_gn=UNC5D;refGene_tx=XM_019828578.2;refGene_gn=UNC5D;refGene_tx
=XM_023252618.1;refGene_gn=UNC5D;refGene_tx=XM_023252619.1;refGene_gn=
UNC5D;refGene_tx=XM_023252620.1;refGene_gn=UNC5D;refGene_tx=XM_0232526
21.1;refGene_gn=UNC5D;refGene_tx=XM_023252622.1;refGene_gn=UNC5D;refGe
ne_tx=XM_023252623.1;refGene_gn=UNC5D;refGene_tx=XM_023252624.1;refGen
e_gn=UNC5D;refGene_tx=XM_023252625.1;refGene_gn=UNC5D;refGene_tx=XM_02
3252626.1;refGene_gn=UNC5D;refGene_tx=XM_023252627.1;refGene_gn=UNC5D;
refGene_tx=XM_023252628.1
B1	48660004	48660154
refGene_gn=UNC5D;refGene_tx=XM_019828578.2;refGene_gn=UNC5D;refGene_tx
=XM_023252618.1;refGene_gn=UNC5D;refGene_tx=XM_023252619.1;refGene_gn=
UNC5D;refGene_tx=XM_023252620.1;refGene_gn=UNC5D;refGene_tx=XM_0232526
21.1;refGene_gn=UNC5D;refGene_tx=XM_023252622.1;refGene_gn=UNC5D;refGe
ne_tx=XM_023252623.1;refGene_gn=UNC5D;refGene_tx=XM_023252624.1;refGen
e_gn=UNC5D;refGene_tx=XM_023252625.1;refGene_gn=UNC5D;refGene_tx=XM_02
3252626.1;refGene_gn=UNC5D;refGene_tx=XM_023252627.1;refGene_gn=UNC5D;
refGene_tx=XM_023252628.1
B1	48669051	48669279
refGene_gn=UNC5D;refGene_tx=XM_019828578.2;refGene_gn=UNC5D;refGene_tx
=XM_023252618.1;refGene_gn=UNC5D;refGene_tx=XM_023252619.1;refGene_gn=
UNC5D;refGene_tx=XM_023252620.1;refGene_gn=UNC5D;refGene_tx=XM_0232526
21.1;refGene_gn=UNC5D;refGene_tx=XM_023252622.1;refGene_gn=UNC5D;refGe
ne_tx=XM_023252623.1;refGene_gn=UNC5D;refGene_tx=XM_023252624.1;refGen
e_gn=UNC5D;refGene_tx=XM_023252625.1;refGene_gn=UNC5D;refGene_tx=XM_02
3252626.1;refGene_gn=UNC5D;refGene_tx=XM_023252627.1;refGene_gn=UNC5D;
refGene_tx=XM_023252628.1

B1 48670935 48671104
refGene_gn=UNC5D;refGene_tx=XM_019828578.2;refGene_gn=UNC5D;refGene_tx=XM_023252618.1;refGene_gn=UNC5D;refGene_tx=XM_023252619.1;refGene_gn=UNC5D;refGene_tx=XM_023252620.1;refGene_gn=UNC5D;refGene_tx=XM_023252621.1;refGene_gn=UNC5D;refGene_tx=XM_023252622.1;refGene_gn=UNC5D;refGene_tx=XM_023252623.1;refGene_gn=UNC5D;refGene_tx=XM_023252624.1;refGene_gn=UNC5D;refGene_tx=XM_023252625.1;refGene_gn=UNC5D;refGene_tx=XM_023252626.1;refGene_gn=UNC5D;refGene_tx=XM_023252627.1;refGene_gn=UNC5D;refGene_tx=XM_023252628.1
B1 48691084 48691169
refGene_gn=UNC5D;refGene_tx=XM_019828578.2;refGene_gn=UNC5D;refGene_tx=XM_023252618.1;refGene_gn=UNC5D;refGene_tx=XM_023252619.1;refGene_gn=UNC5D;refGene_tx=XM_023252620.1;refGene_gn=UNC5D;refGene_tx=XM_023252621.1;refGene_gn=UNC5D;refGene_tx=XM_023252622.1;refGene_gn=UNC5D;refGene_tx=XM_023252623.1;refGene_gn=UNC5D;refGene_tx=XM_023252624.1;refGene_gn=UNC5D;refGene_tx=XM_023252625.1;refGene_gn=UNC5D;refGene_tx=XM_023252626.1;refGene_gn=UNC5D;refGene_tx=XM_023252627.1;refGene_gn=UNC5D;refGene_tx=XM_023252628.1
B1 48696056 48696434
refGene_gn=UNC5D;refGene_tx=XM_019828578.2;refGene_gn=UNC5D;refGene_tx=XM_023252618.1;refGene_gn=UNC5D;refGene_tx=XM_023252619.1;refGene_gn=UNC5D;refGene_tx=XM_023252620.1;refGene_gn=UNC5D;refGene_tx=XM_023252621.1;refGene_gn=UNC5D;refGene_tx=XM_023252622.1;refGene_gn=UNC5D;refGene_tx=XM_023252623.1;refGene_gn=UNC5D;refGene_tx=XM_023252624.1;refGene_gn=UNC5D;refGene_tx=XM_023252625.1;refGene_gn=UNC5D;refGene_tx=XM_023252626.1;refGene_gn=UNC5D;refGene_tx=XM_023252627.1;refGene_gn=UNC5D;refGene_tx=XM_023252628.1
B1 48700062 48700248
refGene_gn=UNC5D;refGene_tx=XM_019828578.2;refGene_gn=UNC5D;refGene_tx=XM_023252618.1;refGene_gn=UNC5D;refGene_tx=XM_023252619.1;refGene_gn=UNC5D;refGene_tx=XM_023252620.1;refGene_gn=UNC5D;refGene_tx=XM_023252621.1;refGene_gn=UNC5D;refGene_tx=XM_023252622.1;refGene_gn=UNC5D;refGene_tx=XM_023252623.1;refGene_gn=UNC5D;refGene_tx=XM_023252624.1;refGene_gn=UNC5D;refGene_tx=XM_023252625.1;refGene_gn=UNC5D;refGene_tx=XM_023252626.1;refGene_gn=UNC5D;refGene_tx=XM_023252627.1;refGene_gn=UNC5D;refGene_tx=XM_023252628.1
B1 48701009 48701024
refGene_gn=UNC5D;refGene_tx=XM_019828578.2;refGene_gn=UNC5D;refGene_tx=XM_023252618.1;refGene_gn=UNC5D;refGene_tx=XM_023252620.1;refGene_gn=UNC5D;refGene_tx=XM_023252623.1;refGene_gn=UNC5D;refGene_tx=XM_023252624.1;refGene_gn=UNC5D;refGene_tx=XM_023252626.1
B1 48715886 48715919
refGene_gn=UNC5D;refGene_tx=XM_019828578.2;refGene_gn=UNC5D;refGene_tx=XM_023252618.1;refGene_gn=UNC5D;refGene_tx=XM_023252619.1;refGene_gn=UNC5D;refGene_tx=XM_023252620.1;refGene_gn=UNC5D;refGene_tx=XM_023252622.1;refGene_gn=UNC5D;refGene_tx=XM_023252623.1;refGene_gn=UNC5D;refGene_tx=XM_023252624.1;refGene_gn=UNC5D;refGene_tx=XM_023252625.1;refGene_gn=UNC5D;refGene_tx=XM_023252626.1;refGene_gn=UNC5D;refGene_tx=XM_023252628.1
B1 48733136 48733301

refGene_gn=UNC5D;refGene_tx=XM_019828578.2;refGene_gn=UNC5D;refGene_tx=XM_023252618.1;refGene_gn=UNC5D;refGene_tx=XM_023252619.1;refGene_gn=UNC5D;refGene_tx=XM_023252620.1;refGene_gn=UNC5D;refGene_tx=XM_023252621.1;refGene_gn=UNC5D;refGene_tx=XM_023252622.1;refGene_gn=UNC5D;refGene_tx=XM_023252623.1;refGene_gn=UNC5D;refGene_tx=XM_023252624.1;refGene_gn=UNC5D;refGene_tx=XM_023252625.1;refGene_gn=UNC5D;refGene_tx=XM_023252626.1;refGene_gn=UNC5D;refGene_tx=XM_023252627.1;refGene_gn=UNC5D;refGene_tx=XM_023252628.1
B1 48735045 48735213
refGene_gn=UNC5D;refGene_tx=XM_019828578.2;refGene_gn=UNC5D;refGene_tx=XM_023252618.1;refGene_gn=UNC5D;refGene_tx=XM_023252619.1;refGene_gn=UNC5D;refGene_tx=XM_023252620.1;refGene_gn=UNC5D;refGene_tx=XM_023252621.1;refGene_gn=UNC5D;refGene_tx=XM_023252622.1;refGene_gn=UNC5D;refGene_tx=XM_023252623.1;refGene_gn=UNC5D;refGene_tx=XM_023252626.1
B1 48736089 48736270
refGene_gn=UNC5D;refGene_tx=XM_019828578.2;refGene_gn=UNC5D;refGene_tx=XM_023252618.1;refGene_gn=UNC5D;refGene_tx=XM_023252619.1;refGene_gn=UNC5D;refGene_tx=XM_023252620.1;refGene_gn=UNC5D;refGene_tx=XM_023252621.1;refGene_gn=UNC5D;refGene_tx=XM_023252622.1;refGene_gn=UNC5D;refGene_tx=XM_023252623.1;refGene_gn=UNC5D;refGene_tx=XM_023252624.1;refGene_gn=UNC5D;refGene_tx=XM_023252625.1;refGene_gn=UNC5D;refGene_tx=XM_023252626.1;refGene_gn=UNC5D;refGene_tx=XM_023252627.1;refGene_gn=UNC5D;refGene_tx=XM_023252628.1
B1 48822661 48822765
refGene_gn=UNC5D;refGene_tx=XM_019828578.2;refGene_gn=UNC5D;refGene_tx=XM_023252618.1;refGene_gn=UNC5D;refGene_tx=XM_023252619.1;refGene_gn=UNC5D;refGene_tx=XM_023252620.1;refGene_gn=UNC5D;refGene_tx=XM_023252621.1;refGene_gn=UNC5D;refGene_tx=XM_023252622.1;refGene_gn=UNC5D;refGene_tx=XM_023252623.1;refGene_gn=UNC5D;refGene_tx=XM_023252624.1;refGene_gn=UNC5D;refGene_tx=XM_023252625.1;refGene_gn=UNC5D;refGene_tx=XM_023252626.1;refGene_gn=UNC5D;refGene_tx=XM_023252627.1;refGene_gn=UNC5D;refGene_tx=XM_023252628.1
B1 48848676 48848820
refGene_gn=UNC5D;refGene_tx=XM_019828578.2;refGene_gn=UNC5D;refGene_tx=XM_023252618.1;refGene_gn=UNC5D;refGene_tx=XM_023252619.1;refGene_gn=UNC5D;refGene_tx=XM_023252620.1;refGene_gn=UNC5D;refGene_tx=XM_023252621.1;refGene_gn=UNC5D;refGene_tx=XM_023252622.1;refGene_gn=UNC5D;refGene_tx=XM_023252623.1;refGene_gn=UNC5D;refGene_tx=XM_023252624.1;refGene_gn=UNC5D;refGene_tx=XM_023252625.1;refGene_gn=UNC5D;refGene_tx=XM_023252626.1;refGene_gn=UNC5D;refGene_tx=XM_023252627.1;refGene_gn=UNC5D;refGene_tx=XM_023252628.1
B1 48862325 48862544
refGene_gn=UNC5D;refGene_tx=XM_019828578.2;refGene_gn=UNC5D;refGene_tx=XM_023252618.1;refGene_gn=UNC5D;refGene_tx=XM_023252619.1;refGene_gn=UNC5D;refGene_tx=XM_023252620.1;refGene_gn=UNC5D;refGene_tx=XM_023252621.1;refGene_gn=UNC5D;refGene_tx=XM_023252622.1;refGene_gn=UNC5D;refGene_tx=XM_023252623.1;refGene_gn=UNC5D;refGene_tx=XM_023252624.1;refGene_gn=UNC5D;refGene_tx=XM_023252625.1;refGene_gn=UNC5D;refGene_tx=XM_023252626.1;refGene_gn=UNC5D;refGene_tx=XM_023252627.1;refGene_gn=UNC5D;refGene_tx=XM_023252628.1

B1 49179625 49179728
refGene_gn=UNC5D;refGene_tx=XM_023252618.1;refGene_gn=UNC5D;refGene_tx=XM_023252619.1;refGene_gn=UNC5D;refGene_tx=XM_023252620.1;refGene_gn=UNC5D;refGene_tx=XM_023252621.1;refGene_gn=UNC5D;refGene_tx=XM_023252622.1;refGene_gn=UNC5D;refGene_tx=XM_023252623.1;refGene_gn=UNC5D;refGene_tx=XM_023252624.1;refGene_gn=UNC5D;refGene_tx=XM_023252625.1;refGene_gn=UNC5D;refGene_tx=XM_023252626.1;refGene_gn=UNC5D;refGene_tx=XM_023252627.1;refGene_gn=UNC5D;refGene_tx=XM_023252628.1
B1 50623237 50623458
refGene_gn=DUSP26;refGene_tx=XM_003984789.5;refGene_gn=DUSP26;refGene_tx=XM_006930727.4
B1 50624920 50625135
refGene_gn=DUSP26;refGene_tx=XM_003984789.5;refGene_gn=DUSP26;refGene_tx=XM_006930727.4
B1 50625636 50625836
refGene_gn=DUSP26;refGene_tx=XM_003984789.5;refGene_gn=DUSP26;refGene_tx=XM_006930727.4
B1 50639316 50639341
refGene_gn=RNF122;refGene_tx=XM_003984790.5;refGene_gn=RNF122;refGene_tx=XM_011281534.3
B1 50640167 50640176 refGene_gn=RNF122;refGene_tx=XM_011281534.3
B1 50643164 50643288 refGene_gn=RNF122;refGene_tx=XM_019828583.2
B1 50644055 50644212
refGene_gn=RNF122;refGene_tx=XM_003984790.5;refGene_gn=RNF122;refGene_tx=XM_011281534.3;refGene_gn=RNF122;refGene_tx=XM_019828583.2;refGene_gn=RNF122;refGene_tx=XM_019828585.2
B1 50649637 50649683
refGene_gn=RNF122;refGene_tx=XM_003984790.5;refGene_gn=RNF122;refGene_tx=XM_011281534.3;refGene_gn=RNF122;refGene_tx=XM_019828583.2;refGene_gn=RNF122;refGene_tx=XM_019828585.2
B1 50650200 50650242
refGene_gn=RNF122;refGene_tx=XM_003984790.5;refGene_gn=RNF122;refGene_tx=XM_011281534.3;refGene_gn=RNF122;refGene_tx=XM_019828583.2;refGene_gn=RNF122;refGene_tx=XM_019828585.2
B1 50651450 50651533
refGene_gn=RNF122;refGene_tx=XM_003984790.5;refGene_gn=RNF122;refGene_tx=XM_011281534.3;refGene_gn=RNF122;refGene_tx=XM_019828583.2;refGene_gn=RNF122;refGene_tx=XM_019828585.2
B1 50651808 50651923
refGene_gn=RNF122;refGene_tx=XM_003984790.5;refGene_gn=RNF122;refGene_tx=XM_011281534.3;refGene_gn=RNF122;refGene_tx=XM_019828583.2;refGene_gn=RNF122;refGene_tx=XM_019828585.2
B1 50695377 50695437
refGene_gn=STMN4;refGene_tx=XM_003984791.4;refGene_gn=STMN4;refGene_tx=XM_011281535.2
B1 50697725 50697746
refGene_gn=STMN4;refGene_tx=XM_019828586.1;refGene_gn=STMN4;refGene_tx=XM_019828587.1
B1 50698432 50698624
refGene_gn=STMN4;refGene_tx=XM_003984791.4;refGene_gn=STMN4;refGene_tx

=XM_011281535.2;refGene_gn=STMN4;refGene_tx=XM_019828586.1;refGene_gn=STMN4;refGene_tx=XM_019828587.1
B1 50699732 50699941
refGene_gn=STMN4;refGene_tx=XM_003984791.4;refGene_gn=STMN4;refGene_tx=XM_011281535.2;refGene_gn=STMN4;refGene_tx=XM_019828586.1;refGene_gn=STMN4;refGene_tx=XM_019828587.1
B1 50700353 50700434
refGene_gn=STMN4;refGene_tx=XM_011281535.2;refGene_gn=STMN4;refGene_tx=XM_019828586.1
B1 50701100 50701196
refGene_gn=STMN4;refGene_tx=XM_003984791.4;refGene_gn=STMN4;refGene_tx=XM_011281535.2;refGene_gn=STMN4;refGene_tx=XM_019828586.1;refGene_gn=STMN4;refGene_tx=XM_019828587.1
B1 50702495 50702508
refGene_gn=STMN4;refGene_tx=XM_003984791.4;refGene_gn=STMN4;refGene_tx=XM_011281535.2;refGene_gn=STMN4;refGene_tx=XM_019828586.1;refGene_gn=STMN4;refGene_tx=XM_019828587.1
B1 50747431 50748009
refGene_gn=TRIM35;refGene_tx=XM_003984794.5;refGene_gn=TRIM35;refGene_tx=XM_019828588.2
B1 50749424 50749543
refGene_gn=TRIM35;refGene_tx=XM_003984794.5;refGene_gn=TRIM35;refGene_tx=XM_019828588.2
B1 50750357 50750380
refGene_gn=TRIM35;refGene_tx=XM_003984794.5;refGene_gn=TRIM35;refGene_tx=XM_019828588.2
B1 50754031 50754262 refGene_gn=TRIM35;refGene_tx=XM_003984794.5
B1 50758814 50758910
refGene_gn=TRIM35;refGene_tx=XM_003984794.5;refGene_gn=TRIM35;refGene_tx=XM_019828588.2
B1 50771972 50772407
refGene_gn=TRIM35;refGene_tx=XM_003984794.5;refGene_gn=TRIM35;refGene_tx=XM_019828588.2
B1 50846574 50846778
refGene_gn=PTK2B;refGene_tx=XM_011281537.2;refGene_gn=PTK2B;refGene_tx=XM_011281538.2;refGene_gn=PTK2B;refGene_tx=XM_011281539.2;refGene_gn=PTK2B;refGene_tx=XM_019828589.1;refGene_gn=PTK2B;refGene_tx=XM_019828591.1
B1 50874696 50874741 refGene_gn=PTK2B;refGene_tx=XM_019828590.2
B1 50875998 50876177
refGene_gn=PTK2B;refGene_tx=XM_011281537.2;refGene_gn=PTK2B;refGene_tx=XM_011281538.2;refGene_gn=PTK2B;refGene_tx=XM_011281539.2;refGene_gn=PTK2B;refGene_tx=XM_019828589.1;refGene_gn=PTK2B;refGene_tx=XM_019828590.2;refGene_gn=PTK2B;refGene_tx=XM_019828591.1
B1 50876720 50876808
refGene_gn=PTK2B;refGene_tx=XM_011281537.2;refGene_gn=PTK2B;refGene_tx=XM_011281538.2;refGene_gn=PTK2B;refGene_tx=XM_011281539.2;refGene_gn=PTK2B;refGene_tx=XM_019828589.1;refGene_gn=PTK2B;refGene_tx=XM_019828590.2;refGene_gn=PTK2B;refGene_tx=XM_019828591.1
B1 50878011 50878091

refGene_gn=PTK2B;refGene_tx=XM_011281537.2;refGene_gn=PTK2B;refGene_tx
=XM_011281538.2;refGene_gn=PTK2B;refGene_tx=XM_011281539.2;refGene_gn=
PTK2B;refGene_tx=XM_019828589.1;refGene_gn=PTK2B;refGene_tx=XM_0198285
90.2;refGene_gn=PTK2B;refGene_tx=XM_019828591.1
B1 50880213 50880276
refGene_gn=PTK2B;refGene_tx=XM_011281537.2;refGene_gn=PTK2B;refGene_tx
=XM_011281538.2;refGene_gn=PTK2B;refGene_tx=XM_011281539.2;refGene_gn=
PTK2B;refGene_tx=XM_019828589.1;refGene_gn=PTK2B;refGene_tx=XM_0198285
90.2;refGene_gn=PTK2B;refGene_tx=XM_019828591.1
B1 50880492 50880547
refGene_gn=PTK2B;refGene_tx=XM_011281537.2;refGene_gn=PTK2B;refGene_tx
=XM_011281538.2;refGene_gn=PTK2B;refGene_tx=XM_011281539.2;refGene_gn=
PTK2B;refGene_tx=XM_019828589.1;refGene_gn=PTK2B;refGene_tx=XM_0198285
90.2;refGene_gn=PTK2B;refGene_tx=XM_019828591.1
B1 50880957 50881098
refGene_gn=PTK2B;refGene_tx=XM_011281537.2;refGene_gn=PTK2B;refGene_tx
=XM_011281538.2;refGene_gn=PTK2B;refGene_tx=XM_011281539.2;refGene_gn=
PTK2B;refGene_tx=XM_019828589.1;refGene_gn=PTK2B;refGene_tx=XM_0198285
90.2;refGene_gn=PTK2B;refGene_tx=XM_019828591.1
B1 50881512 50881587
refGene_gn=PTK2B;refGene_tx=XM_011281537.2;refGene_gn=PTK2B;refGene_tx
=XM_011281538.2;refGene_gn=PTK2B;refGene_tx=XM_011281539.2;refGene_gn=
PTK2B;refGene_tx=XM_019828589.1;refGene_gn=PTK2B;refGene_tx=XM_0198285
90.2;refGene_gn=PTK2B;refGene_tx=XM_019828591.1
B1 50882434 50882536
refGene_gn=PTK2B;refGene_tx=XM_011281537.2;refGene_gn=PTK2B;refGene_tx
=XM_011281538.2;refGene_gn=PTK2B;refGene_tx=XM_011281539.2;refGene_gn=
PTK2B;refGene_tx=XM_019828589.1;refGene_gn=PTK2B;refGene_tx=XM_0198285
90.2;refGene_gn=PTK2B;refGene_tx=XM_019828591.1
B1 50883258 50883376
refGene_gn=PTK2B;refGene_tx=XM_011281537.2;refGene_gn=PTK2B;refGene_tx
=XM_011281538.2;refGene_gn=PTK2B;refGene_tx=XM_011281539.2;refGene_gn=
PTK2B;refGene_tx=XM_019828589.1;refGene_gn=PTK2B;refGene_tx=XM_0198285
90.2;refGene_gn=PTK2B;refGene_tx=XM_019828591.1
B1 50883928 50883968
refGene_gn=PTK2B;refGene_tx=XM_011281537.2;refGene_gn=PTK2B;refGene_tx
=XM_011281538.2;refGene_gn=PTK2B;refGene_tx=XM_011281539.2;refGene_gn=
PTK2B;refGene_tx=XM_019828589.1;refGene_gn=PTK2B;refGene_tx=XM_0198285
90.2;refGene_gn=PTK2B;refGene_tx=XM_019828591.1
B1 50884074 50884143
refGene_gn=PTK2B;refGene_tx=XM_011281538.2;refGene_gn=PTK2B;refGene_tx
=XM_019828589.1;refGene_gn=PTK2B;refGene_tx=XM_019828590.2;refGene_gn=
PTK2B;refGene_tx=XM_019828591.1
B1 50884397 50884444
refGene_gn=PTK2B;refGene_tx=XM_011281537.2;refGene_gn=PTK2B;refGene_tx
=XM_011281538.2;refGene_gn=PTK2B;refGene_tx=XM_011281539.2;refGene_gn=
PTK2B;refGene_tx=XM_019828589.1;refGene_gn=PTK2B;refGene_tx=XM_0198285
90.2;refGene_gn=PTK2B;refGene_tx=XM_019828591.1
B1 50885334 50885385
refGene_gn=PTK2B;refGene_tx=XM_011281537.2;refGene_gn=PTK2B;refGene_tx

=XM_011281538.2;refGene_gn=PTK2B;refGene_tx=XM_011281539.2;refGene_gn=PTK2B;refGene_tx=XM_019828589.1;refGene_gn=PTK2B;refGene_tx=XM_019828590.2;refGene_gn=PTK2B;refGene_tx=XM_019828591.1
B1 50885862 50885960
refGene_gn=PTK2B;refGene_tx=XM_011281537.2;refGene_gn=PTK2B;refGene_tx=XM_011281538.2;refGene_gn=PTK2B;refGene_tx=XM_011281539.2;refGene_gn=PTK2B;refGene_tx=XM_019828589.1;refGene_gn=PTK2B;refGene_tx=XM_019828590.2;refGene_gn=PTK2B;refGene_tx=XM_019828591.1
B1 50886747 50886832
refGene_gn=PTK2B;refGene_tx=XM_011281537.2;refGene_gn=PTK2B;refGene_tx=XM_011281538.2;refGene_gn=PTK2B;refGene_tx=XM_011281539.2;refGene_gn=PTK2B;refGene_tx=XM_019828589.1;refGene_gn=PTK2B;refGene_tx=XM_019828590.2;refGene_gn=PTK2B;refGene_tx=XM_019828591.1
B1 50887072 50887173
refGene_gn=PTK2B;refGene_tx=XM_011281537.2;refGene_gn=PTK2B;refGene_tx=XM_011281538.2;refGene_gn=PTK2B;refGene_tx=XM_011281539.2;refGene_gn=PTK2B;refGene_tx=XM_019828589.1;refGene_gn=PTK2B;refGene_tx=XM_019828590.2;refGene_gn=PTK2B;refGene_tx=XM_019828591.1
B1 50887422 50887538
refGene_gn=PTK2B;refGene_tx=XM_011281537.2;refGene_gn=PTK2B;refGene_tx=XM_011281538.2;refGene_gn=PTK2B;refGene_tx=XM_011281539.2;refGene_gn=PTK2B;refGene_tx=XM_019828589.1;refGene_gn=PTK2B;refGene_tx=XM_019828590.2;refGene_gn=PTK2B;refGene_tx=XM_019828591.1
B1 50889505 50889606
refGene_gn=PTK2B;refGene_tx=XM_011281537.2;refGene_gn=PTK2B;refGene_tx=XM_011281538.2;refGene_gn=PTK2B;refGene_tx=XM_011281539.2;refGene_gn=PTK2B;refGene_tx=XM_019828589.1;refGene_gn=PTK2B;refGene_tx=XM_019828590.2;refGene_gn=PTK2B;refGene_tx=XM_019828591.1
B1 50890008 50890098
refGene_gn=PTK2B;refGene_tx=XM_011281537.2;refGene_gn=PTK2B;refGene_tx=XM_011281538.2;refGene_gn=PTK2B;refGene_tx=XM_011281539.2;refGene_gn=PTK2B;refGene_tx=XM_019828589.1;refGene_gn=PTK2B;refGene_tx=XM_019828590.2;refGene_gn=PTK2B;refGene_tx=XM_019828591.1
B1 50890981 50891186
refGene_gn=PTK2B;refGene_tx=XM_011281537.2;refGene_gn=PTK2B;refGene_tx=XM_011281538.2;refGene_gn=PTK2B;refGene_tx=XM_011281539.2;refGene_gn=PTK2B;refGene_tx=XM_019828589.1;refGene_gn=PTK2B;refGene_tx=XM_019828590.2;refGene_gn=PTK2B;refGene_tx=XM_019828591.1
B1 50893785 50893894
refGene_gn=PTK2B;refGene_tx=XM_011281537.2;refGene_gn=PTK2B;refGene_tx=XM_011281538.2;refGene_gn=PTK2B;refGene_tx=XM_011281539.2;refGene_gn=PTK2B;refGene_tx=XM_019828589.1;refGene_gn=PTK2B;refGene_tx=XM_019828590.2;refGene_gn=PTK2B;refGene_tx=XM_019828591.1
B1 50895122 50895188
refGene_gn=PTK2B;refGene_tx=XM_011281537.2;refGene_gn=PTK2B;refGene_tx=XM_011281538.2;refGene_gn=PTK2B;refGene_tx=XM_011281539.2;refGene_gn=PTK2B;refGene_tx=XM_019828589.1;refGene_gn=PTK2B;refGene_tx=XM_019828590.2;refGene_gn=PTK2B;refGene_tx=XM_019828591.1
B1 50896444 50896609
refGene_gn=PTK2B;refGene_tx=XM_011281537.2;refGene_gn=PTK2B;refGene_tx

=XM_019828589.1;refGene_gn=PTK2B;refGene_tx=XM_019828590.2;refGene_gn=PTK2B;refGene_tx=XM_019828591.1
B1 50901617 50901764
refGene_gn=PTK2B;refGene_tx=XM_011281537.2;refGene_gn=PTK2B;refGene_tx=XM_011281538.2;refGene_gn=PTK2B;refGene_tx=XM_011281539.2;refGene_gn=PTK2B;refGene_tx=XM_019828589.1;refGene_gn=PTK2B;refGene_tx=XM_019828590.2
B1 50902051 50902087
refGene_gn=PTK2B;refGene_tx=XM_011281537.2;refGene_gn=PTK2B;refGene_tx=XM_011281538.2;refGene_gn=PTK2B;refGene_tx=XM_011281539.2;refGene_gn=PTK2B;refGene_tx=XM_019828589.1;refGene_gn=PTK2B;refGene_tx=XM_019828590.2
B1 50902457 50902482
refGene_gn=PTK2B;refGene_tx=XM_011281537.2;refGene_gn=PTK2B;refGene_tx=XM_011281538.2;refGene_gn=PTK2B;refGene_tx=XM_011281539.2;refGene_gn=PTK2B;refGene_tx=XM_019828589.1;refGene_gn=PTK2B;refGene_tx=XM_019828590.2
B1 50904172 50904219
refGene_gn=PTK2B;refGene_tx=XM_011281537.2;refGene_gn=PTK2B;refGene_tx=XM_011281538.2;refGene_gn=PTK2B;refGene_tx=XM_011281539.2;refGene_gn=PTK2B;refGene_tx=XM_019828589.1;refGene_gn=PTK2B;refGene_tx=XM_019828590.2
B1 50904895 50905033
refGene_gn=PTK2B;refGene_tx=XM_011281537.2;refGene_gn=PTK2B;refGene_tx=XM_011281538.2;refGene_gn=PTK2B;refGene_tx=XM_011281539.2;refGene_gn=PTK2B;refGene_tx=XM_019828589.1;refGene_gn=PTK2B;refGene_tx=XM_019828590.2
B1 50905287 50905368
refGene_gn=PTK2B;refGene_tx=XM_011281537.2;refGene_gn=PTK2B;refGene_tx=XM_011281538.2;refGene_gn=PTK2B;refGene_tx=XM_011281539.2;refGene_gn=PTK2B;refGene_tx=XM_019828589.1;refGene_gn=PTK2B;refGene_tx=XM_019828590.2
B1 50907285 50907501
refGene_gn=PTK2B;refGene_tx=XM_011281537.2;refGene_gn=PTK2B;refGene_tx=XM_011281538.2;refGene_gn=PTK2B;refGene_tx=XM_011281539.2;refGene_gn=PTK2B;refGene_tx=XM_019828589.1;refGene_gn=PTK2B;refGene_tx=XM_019828590.2
B1 50910387 50910513 refGene_gn=CHRNA2;refGene_tx=XM_023252629.1
B1 50911713 50912728 refGene_gn=CHRNA2;refGene_tx=XM_023252629.1
B1 50916457 50916567 refGene_gn=CHRNA2;refGene_tx=XM_023252629.1
B1 50920055 50920100 refGene_gn=CHRNA2;refGene_tx=XM_023252629.1
B1 50920476 50920628 refGene_gn=CHRNA2;refGene_tx=XM_023252629.1
B1 50921825 50921934 refGene_gn=CHRNA2;refGene_tx=XM_023252629.1
B1 50944857 50944958 refGene_gn=EPHX2;refGene_tx=XM_003984798.3
B1 50960209 50960294 refGene_gn=EPHX2;refGene_tx=XM_003984798.3
B1 50966633 50966790 refGene_gn=EPHX2;refGene_tx=XM_003984798.3
B1 50967993 50968184 refGene_gn=EPHX2;refGene_tx=XM_003984798.3
B1 50969886 50970009 refGene_gn=EPHX2;refGene_tx=XM_003984798.3
B1 50973065 50973140 refGene_gn=EPHX2;refGene_tx=XM_003984798.3
B1 50977147 50977243 refGene_gn=EPHX2;refGene_tx=XM_003984798.3

B1 50977787 50977866 refGene_gn=EPHX2;refGene_tx=XM_003984798.3
B1 50980746 50980781 refGene_gn=EPHX2;refGene_tx=XM_003984798.3
B1 50987033 50987060 refGene_gn=EPHX2;refGene_tx=XM_003984798.3
B1 50989502 50989588 refGene_gn=EPHX2;refGene_tx=XM_003984798.3
B1 50993389 50993501 refGene_gn=EPHX2;refGene_tx=XM_003984798.3
B1 51008990 51009062 refGene_gn=EPHX2;refGene_tx=XM_003984798.3
B1 51012280 51012314 refGene_gn=EPHX2;refGene_tx=XM_003984798.3
B1 51014185 51014288 refGene_gn=EPHX2;refGene_tx=XM_003984798.3
B1 51015160 51015230 refGene_gn=EPHX2;refGene_tx=XM_003984798.3
B1 51019720 51019801 refGene_gn=EPHX2;refGene_tx=XM_003984798.3
B1 51021414 51021473 refGene_gn=EPHX2;refGene_tx=XM_003984798.3
B1 51021646 51021725 refGene_gn=EPHX2;refGene_tx=XM_003984798.3
B1 51029966 51029969
refGene_gn=LOC101092602;refGene_tx=XM_019828593.1
B1 51030890 51030992
refGene_gn=LOC101092602;refGene_tx=XM_019828593.1
B1 51033772 51033898
refGene_gn=LOC101092602;refGene_tx=XM_019828593.1
B1 51035797 51035892
refGene_gn=LOC101092602;refGene_tx=XM_019828593.1
B1 51040318 51040409
refGene_gn=LOC101092602;refGene_tx=XM_019828593.1
B1 51043957 51044143
refGene_gn=LOC101092602;refGene_tx=XM_019828593.1
B1 51053523 51053631
refGene_gn=LOC101092602;refGene_tx=XM_019828593.1
B1 51054470 51054547
refGene_gn=LOC101092602;refGene_tx=XM_019828593.1
B1 51060051 51060201
refGene_gn=LOC101092602;refGene_tx=XM_019828593.1
B1 51061059 51061224
refGene_gn=LOC101092602;refGene_tx=XM_019828593.1
B1 51061719 51061810
refGene_gn=LOC101092602;refGene_tx=XM_019828593.1
B1 51062583 51062712
refGene_gn=LOC101092602;refGene_tx=XM_019828593.1
B1 51072320 51072330 refGene_gn=CLU;refGene_tx=XM_003984800.5
B1 51072585 51072761 refGene_gn=CLU;refGene_tx=XM_003984800.5
B1 51073729 51073959 refGene_gn=CLU;refGene_tx=XM_003984800.5
B1 51077385 51077490 refGene_gn=CLU;refGene_tx=XM_003984800.5
B1 51078537 51078946 refGene_gn=CLU;refGene_tx=XM_003984800.5
B1 51080460 51080631 refGene_gn=CLU;refGene_tx=XM_003984800.5
B1 51084087 51084236 refGene_gn=CLU;refGene_tx=XM_003984800.5
B1 51085573 51085667 refGene_gn=CLU;refGene_tx=XM_003984800.5
B1 51102799 51102806 refGene_gn=SCARA3;refGene_tx=XM_011281541.2
B1 51128356 51128455 refGene_gn=SCARA3;refGene_tx=XM_011281541.2
B1 51129858 51129978 refGene_gn=SCARA3;refGene_tx=XM_011281541.2
B1 51135198 51135297 refGene_gn=SCARA3;refGene_tx=XM_011281541.2
B1 51136283 51137327 refGene_gn=SCARA3;refGene_tx=XM_011281541.2
B1 51153488 51153940 refGene_gn=SCARA3;refGene_tx=XM_011281541.2

B1 51222533 51222563
refGene_gn=CCDC25;refGene_tx=XM_003984829.5;refGene_gn=CCDC25;refGene_tx=XM_011281542.3;refGene_gn=CCDC25;refGene_tx=XM_019828595.2
B1 51227222 51227268
refGene_gn=CCDC25;refGene_tx=XM_003984829.5;refGene_gn=CCDC25;refGene_tx=XM_011281542.3;refGene_gn=CCDC25;refGene_tx=XM_019828595.2
B1 51235019 51235222
refGene_gn=CCDC25;refGene_tx=XM_003984829.5;refGene_gn=CCDC25;refGene_tx=XM_011281542.3;refGene_gn=CCDC25;refGene_tx=XM_019828595.2
B1 51235664 51235768
refGene_gn=CCDC25;refGene_tx=XM_003984829.5;refGene_gn=CCDC25;refGene_tx=XM_011281542.3;refGene_gn=CCDC25;refGene_tx=XM_019828595.2
B1 51239318 51239394
refGene_gn=CCDC25;refGene_tx=XM_003984829.5;refGene_gn=CCDC25;refGene_tx=XM_011281542.3;refGene_gn=CCDC25;refGene_tx=XM_019828595.2
B1 51244513 51244565
refGene_gn=CCDC25;refGene_tx=XM_003984829.5;refGene_gn=CCDC25;refGene_tx=XM_011281542.3;refGene_gn=CCDC25;refGene_tx=XM_019828595.2
B1 51249766 51249806
refGene_gn=CCDC25;refGene_tx=XM_003984829.5;refGene_gn=CCDC25;refGene_tx=XM_011281542.3;refGene_gn=CCDC25;refGene_tx=XM_019828595.2
B1 51254993 51255081
refGene_gn=CCDC25;refGene_tx=XM_003984829.5;refGene_gn=CCDC25;refGene_tx=XM_011281542.3
B1 51260080 51260108
refGene_gn=CCDC25;refGene_tx=XM_003984829.5;refGene_gn=CCDC25;refGene_tx=XM_019828595.2
B1 51262881 51262934 refGene_gn=ESCO2;refGene_tx=XM_006930741.3
B1 51264609 51265423 refGene_gn=ESCO2;refGene_tx=XM_006930741.3
B1 51273087 51273181 refGene_gn=ESCO2;refGene_tx=XM_006930741.3
B1 51277110 51277168 refGene_gn=ESCO2;refGene_tx=XM_006930741.3
B1 51280553 51280677 refGene_gn=ESCO2;refGene_tx=XM_006930741.3
B1 51281457 51281589 refGene_gn=ESCO2;refGene_tx=XM_006930741.3
B1 51282720 51282810 refGene_gn=ESCO2;refGene_tx=XM_006930741.3
B1 51283441 51283585 refGene_gn=ESCO2;refGene_tx=XM_006930741.3
B1 51286749 51286916 refGene_gn=ESCO2;refGene_tx=XM_006930741.3
B1 51289739 51289872 refGene_gn=ESCO2;refGene_tx=XM_006930741.3
B1 51297869 51298069
refGene_gn=PBK;refGene_tx=XM_006930742.4;refGene_gn=PBK;refGene_tx=XM_019828597.2;refGene_gn=PBK;refGene_tx=XM_019828598.2;refGene_gn=PBK;refGene_tx=XM_019828599.1;refGene_gn=PBK;refGene_tx=XM_019828600.2;refGene_gn=PBK;refGene_tx=XM_019828601.2
B1 51298824 51299004
refGene_gn=PBK;refGene_tx=XM_006930742.4;refGene_gn=PBK;refGene_tx=XM_019828597.2;refGene_gn=PBK;refGene_tx=XM_019828598.2;refGene_gn=PBK;refGene_tx=XM_019828599.1;refGene_gn=PBK;refGene_tx=XM_019828600.2
B1 51306984 51307114
refGene_gn=PBK;refGene_tx=XM_006930742.4;refGene_gn=PBK;refGene_tx=XM_019828597.2;refGene_gn=PBK;refGene_tx=XM_019828598.2;refGene_gn=PBK;refGene_tx=XM_019828599.1;refGene_gn=PBK;refGene_tx=XM_019828600.2;refGe

ne_gn=PBK;refGene_tx=XM_019828601.2
B1 51309045 51309215
refGene_gn=PBK;refGene_tx=XM_006930742.4;refGene_gn=PBK;refGene_tx=XM_019828597.2;refGene_gn=PBK;refGene_tx=XM_019828598.2;refGene_gn=PBK;refGene_tx=XM_019828599.1;refGene_gn=PBK;refGene_tx=XM_019828600.2;refGene_gn=PBK;refGene_tx=XM_019828601.2
B1 51309746 51309889
refGene_gn=PBK;refGene_tx=XM_006930742.4;refGene_gn=PBK;refGene_tx=XM_019828597.2;refGene_gn=PBK;refGene_tx=XM_019828598.2;refGene_gn=PBK;refGene_tx=XM_019828599.1;refGene_gn=PBK;refGene_tx=XM_019828600.2;refGene_gn=PBK;refGene_tx=XM_019828601.2
B1 51315328 51315422
refGene_gn=PBK;refGene_tx=XM_006930742.4;refGene_gn=PBK;refGene_tx=XM_019828597.2;refGene_gn=PBK;refGene_tx=XM_019828598.2;refGene_gn=PBK;refGene_tx=XM_019828599.1;refGene_gn=PBK;refGene_tx=XM_019828600.2;refGene_gn=PBK;refGene_tx=XM_019828601.2
B1 51321080 51321138
refGene_gn=PBK;refGene_tx=XM_006930742.4;refGene_gn=PBK;refGene_tx=XM_019828597.2;refGene_gn=PBK;refGene_tx=XM_019828598.2;refGene_gn=PBK;refGene_tx=XM_019828599.1;refGene_gn=PBK;refGene_tx=XM_019828600.2;refGene_gn=PBK;refGene_tx=XM_019828601.2
B1 51368684 51368821
refGene_gn=SCARA5;refGene_tx=XM_003984802.5;refGene_gn=SCARA5;refGene_tx=XM_019828609.1
B1 51378774 51378972
refGene_gn=SCARA5;refGene_tx=XM_003984802.5;refGene_gn=SCARA5;refGene_tx=XM_019828609.1
B1 51402979 51403036
refGene_gn=SCARA5;refGene_tx=XM_003984802.5;refGene_gn=SCARA5;refGene_tx=XM_019828609.1
B1 51405857 51405956
refGene_gn=SCARA5;refGene_tx=XM_003984802.5;refGene_gn=SCARA5;refGene_tx=XM_019828609.1
B1 51408028 51408109
refGene_gn=SCARA5;refGene_tx=XM_003984802.5;refGene_gn=SCARA5;refGene_tx=XM_019828609.1
B1 51418180 51418855
refGene_gn=SCARA5;refGene_tx=XM_003984802.5;refGene_gn=SCARA5;refGene_tx=XM_019828609.1
B1 51462997 51463126
refGene_gn=SCARA5;refGene_tx=XM_003984802.5;refGene_gn=SCARA5;refGene_tx=XM_019828609.1
B1 51488264 51488376
refGene_gn=SCARA5;refGene_tx=XM_003984802.5;refGene_gn=SCARA5;refGene_tx=XM_019828609.1
B1 51528748 51528888
refGene_gn=NUGGC;refGene_tx=XM_003984831.4;refGene_gn=NUGGC;refGene_tx=XM_019828602.2;refGene_gn=NUGGC;refGene_tx=XM_019828604.2;refGene_gn=NUGGC;refGene_tx=XM_019828605.2;refGene_gn=NUGGC;refGene_tx=XM_019828606.1;refGene_gn=NUGGC;refGene_tx=XM_019828607.2;refGene_gn=NUGGC;refGe

ne_tx=XM_019828608.2;refGene_gn=NUGGC;refGene_tx=XM_023252630.1
B1 51531203 51531294
refGene_gn=NUGGC;refGene_tx=XM_003984831.4;refGene_gn=NUGGC;refGene_tx=XM_019828602.2;refGene_gn=NUGGC;refGene_tx=XM_019828604.2;refGene_gn=NUGGC;refGene_tx=XM_019828605.2;refGene_gn=NUGGC;refGene_tx=XM_019828606.1;refGene_gn=NUGGC;refGene_tx=XM_019828607.2;refGene_gn=NUGGC;refGene_tx=XM_019828608.2;refGene_gn=NUGGC;refGene_tx=XM_023252630.1
B1 51533541 51533678
refGene_gn=NUGGC;refGene_tx=XM_003984831.4;refGene_gn=NUGGC;refGene_tx=XM_019828602.2;refGene_gn=NUGGC;refGene_tx=XM_019828604.2;refGene_gn=NUGGC;refGene_tx=XM_019828605.2;refGene_gn=NUGGC;refGene_tx=XM_019828606.1;refGene_gn=NUGGC;refGene_tx=XM_019828607.2
B1 51534384 51534493
refGene_gn=NUGGC;refGene_tx=XM_003984831.4;refGene_gn=NUGGC;refGene_tx=XM_019828602.2;refGene_gn=NUGGC;refGene_tx=XM_019828604.2;refGene_gn=NUGGC;refGene_tx=XM_019828605.2;refGene_gn=NUGGC;refGene_tx=XM_019828606.1;refGene_gn=NUGGC;refGene_tx=XM_019828607.2;refGene_gn=NUGGC;refGene_tx=XM_019828608.2
B1 51535903 51536042
refGene_gn=NUGGC;refGene_tx=XM_003984831.4;refGene_gn=NUGGC;refGene_tx=XM_019828602.2;refGene_gn=NUGGC;refGene_tx=XM_019828604.2;refGene_gn=NUGGC;refGene_tx=XM_019828605.2;refGene_gn=NUGGC;refGene_tx=XM_019828606.1;refGene_gn=NUGGC;refGene_tx=XM_019828607.2;refGene_gn=NUGGC;refGene_tx=XM_019828608.2;refGene_gn=NUGGC;refGene_tx=XM_023252630.1
B1 51537759 51537917
refGene_gn=NUGGC;refGene_tx=XM_003984831.4;refGene_gn=NUGGC;refGene_tx=XM_019828602.2;refGene_gn=NUGGC;refGene_tx=XM_019828604.2;refGene_gn=NUGGC;refGene_tx=XM_019828605.2;refGene_gn=NUGGC;refGene_tx=XM_019828606.1;refGene_gn=NUGGC;refGene_tx=XM_019828607.2;refGene_gn=NUGGC;refGene_tx=XM_019828608.2;refGene_gn=NUGGC;refGene_tx=XM_023252630.1
B1 51541622 51541787
refGene_gn=NUGGC;refGene_tx=XM_003984831.4;refGene_gn=NUGGC;refGene_tx=XM_019828602.2;refGene_gn=NUGGC;refGene_tx=XM_019828604.2;refGene_gn=NUGGC;refGene_tx=XM_019828605.2;refGene_gn=NUGGC;refGene_tx=XM_019828606.1;refGene_gn=NUGGC;refGene_tx=XM_019828607.2;refGene_gn=NUGGC;refGene_tx=XM_019828608.2;refGene_gn=NUGGC;refGene_tx=XM_023252630.1
B1 51546167 51546301
refGene_gn=NUGGC;refGene_tx=XM_003984831.4;refGene_gn=NUGGC;refGene_tx=XM_019828602.2;refGene_gn=NUGGC;refGene_tx=XM_019828604.2;refGene_gn=NUGGC;refGene_tx=XM_019828605.2;refGene_gn=NUGGC;refGene_tx=XM_019828606.1;refGene_gn=NUGGC;refGene_tx=XM_019828607.2;refGene_gn=NUGGC;refGene_tx=XM_019828608.2;refGene_gn=NUGGC;refGene_tx=XM_023252630.1
B1 51547508 51547614
refGene_gn=NUGGC;refGene_tx=XM_003984831.4;refGene_gn=NUGGC;refGene_tx=XM_019828602.2;refGene_gn=NUGGC;refGene_tx=XM_019828604.2;refGene_gn=NUGGC;refGene_tx=XM_019828605.2;refGene_gn=NUGGC;refGene_tx=XM_019828606.1;refGene_gn=NUGGC;refGene_tx=XM_019828607.2;refGene_gn=NUGGC;refGene_tx=XM_019828608.2;refGene_gn=NUGGC;refGene_tx=XM_023252630.1
B1 51556162 51556252
refGene_gn=NUGGC;refGene_tx=XM_003984831.4;refGene_gn=NUGGC;refGene_tx

=XM_019828602.2;refGene_gn=NUGGC;refGene_tx=XM_019828604.2;refGene_gn=NUGGC;refGene_tx=XM_019828605.2;refGene_gn=NUGGC;refGene_tx=XM_019828606.1;refGene_gn=NUGGC;refGene_tx=XM_019828607.2;refGene_gn=NUGGC;refGene_tx=XM_019828608.2;refGene_gn=NUGGC;refGene_tx=XM_023252630.1
B1 51556540 51556559
refGene_gn=NUGGC;refGene_tx=XM_003984831.4;refGene_gn=NUGGC;refGene_tx=XM_019828602.2;refGene_gn=NUGGC;refGene_tx=XM_019828604.2;refGene_gn=NUGGC;refGene_tx=XM_019828605.2;refGene_gn=NUGGC;refGene_tx=XM_019828606.1;refGene_gn=NUGGC;refGene_tx=XM_019828607.2;refGene_gn=NUGGC;refGene_tx=XM_019828608.2;refGene_gn=NUGGC;refGene_tx=XM_023252630.1
B1 51558128 51558304
refGene_gn=NUGGC;refGene_tx=XM_003984831.4;refGene_gn=NUGGC;refGene_tx=XM_019828602.2;refGene_gn=NUGGC;refGene_tx=XM_019828604.2;refGene_gn=NUGGC;refGene_tx=XM_019828605.2;refGene_gn=NUGGC;refGene_tx=XM_019828606.1;refGene_gn=NUGGC;refGene_tx=XM_019828607.2;refGene_gn=NUGGC;refGene_tx=XM_019828608.2;refGene_gn=NUGGC;refGene_tx=XM_023252630.1
B1 51563983 51564193
refGene_gn=NUGGC;refGene_tx=XM_003984831.4;refGene_gn=NUGGC;refGene_tx=XM_019828602.2;refGene_gn=NUGGC;refGene_tx=XM_019828604.2;refGene_gn=NUGGC;refGene_tx=XM_019828605.2;refGene_gn=NUGGC;refGene_tx=XM_019828606.1;refGene_gn=NUGGC;refGene_tx=XM_019828607.2;refGene_gn=NUGGC;refGene_tx=XM_019828608.2;refGene_gn=NUGGC;refGene_tx=XM_023252630.1
B1 51568316 51568547
refGene_gn=NUGGC;refGene_tx=XM_003984831.4;refGene_gn=NUGGC;refGene_tx=XM_019828602.2;refGene_gn=NUGGC;refGene_tx=XM_019828604.2;refGene_gn=NUGGC;refGene_tx=XM_019828605.2;refGene_gn=NUGGC;refGene_tx=XM_019828606.1;refGene_gn=NUGGC;refGene_tx=XM_019828607.2;refGene_gn=NUGGC;refGene_tx=XM_019828608.2;refGene_gn=NUGGC;refGene_tx=XM_023252630.1
B1 51569028 51569251
refGene_gn=NUGGC;refGene_tx=XM_003984831.4;refGene_gn=NUGGC;refGene_tx=XM_019828602.2;refGene_gn=NUGGC;refGene_tx=XM_019828604.2;refGene_gn=NUGGC;refGene_tx=XM_019828605.2;refGene_gn=NUGGC;refGene_tx=XM_019828606.1;refGene_gn=NUGGC;refGene_tx=XM_019828607.2;refGene_gn=NUGGC;refGene_tx=XM_019828608.2;refGene_gn=NUGGC;refGene_tx=XM_023252630.1
B1 51570327 51570436
refGene_gn=NUGGC;refGene_tx=XM_003984831.4;refGene_gn=NUGGC;refGene_tx=XM_019828602.2;refGene_gn=NUGGC;refGene_tx=XM_019828604.2;refGene_gn=NUGGC;refGene_tx=XM_019828605.2;refGene_gn=NUGGC;refGene_tx=XM_019828606.1;refGene_gn=NUGGC;refGene_tx=XM_019828607.2;refGene_gn=NUGGC;refGene_tx=XM_019828608.2;refGene_gn=NUGGC;refGene_tx=XM_023252630.1
B1 51571048 51571153
refGene_gn=NUGGC;refGene_tx=XM_003984831.4;refGene_gn=NUGGC;refGene_tx=XM_019828602.2;refGene_gn=NUGGC;refGene_tx=XM_019828604.2;refGene_gn=NUGGC;refGene_tx=XM_019828605.2;refGene_gn=NUGGC;refGene_tx=XM_019828606.1;refGene_gn=NUGGC;refGene_tx=XM_019828607.2;refGene_gn=NUGGC;refGene_tx=XM_019828608.2;refGene_gn=NUGGC;refGene_tx=XM_023252630.1
B1 51573845 51573888
refGene_gn=NUGGC;refGene_tx=XM_003984831.4;refGene_gn=NUGGC;refGene_tx=XM_019828602.2;refGene_gn=NUGGC;refGene_tx=XM_019828604.2;refGene_gn=NUGGC;refGene_tx=XM_019828605.2;refGene_gn=NUGGC;refGene_tx=XM_0198286

06.1;refGene_gn=NUGGC;refGene_tx=XM_019828607.2;refGene_gn=NUGGC;refGene_tx=XM_019828608.2;refGene_gn=NUGGC;refGene_tx=XM_023252630.1
B1 51585055 51585074 refGene_gn=ELP3;refGene_tx=XM_019828610.1
B1 51588095 51588195 refGene_gn=ELP3;refGene_tx=XM_019828610.1
B1 51591159 51591298 refGene_gn=ELP3;refGene_tx=XM_019828610.1
B1 51607931 51608002 refGene_gn=ELP3;refGene_tx=XM_019828610.1
B1 51608799 51608863 refGene_gn=ELP3;refGene_tx=XM_019828610.1
B1 51610250 51610319 refGene_gn=ELP3;refGene_tx=XM_019828610.1
B1 51613203 51613358 refGene_gn=ELP3;refGene_tx=XM_019828610.1
B1 51629761 51629923 refGene_gn=ELP3;refGene_tx=XM_019828610.1
B1 51632317 51632444 refGene_gn=ELP3;refGene_tx=XM_019828610.1
B1 51641034 51641228 refGene_gn=ELP3;refGene_tx=XM_019828610.1
B1 51654549 51654640 refGene_gn=ELP3;refGene_tx=XM_019828610.1
B1 51657383 51657449 refGene_gn=ELP3;refGene_tx=XM_019828610.1
B1 51659075 51659303 refGene_gn=ELP3;refGene_tx=XM_019828610.1
B1 51660228 51660310 refGene_gn=ELP3;refGene_tx=XM_019828610.1
B1 51679371 51679448 refGene_gn=ELP3;refGene_tx=XM_019828610.1
B1 51860676 51860802
refGene_gn=PNOC;refGene_tx=XM_003984805.3;refGene_gn=PNOC;refGene_tx=XM_019828611.2;refGene_gn=PNOC;refGene_tx=XM_019828613.1
B1 51869599 51870004
refGene_gn=PNOC;refGene_tx=XM_003984805.3;refGene_gn=PNOC;refGene_tx=XM_019828611.2;refGene_gn=PNOC;refGene_tx=XM_019828613.1
B1 51873009 51873095
refGene_gn=PNOC;refGene_tx=XM_003984805.3;refGene_gn=PNOC;refGene_tx=XM_019828611.2
B1 51878850 51878962 refGene_gn=ZNF395;refGene_tx=XM_023252631.1
B1 51879262 51879366 refGene_gn=ZNF395;refGene_tx=XM_023252631.1
B1 51880246 51880339 refGene_gn=ZNF395;refGene_tx=XM_023252631.1
B1 51881922 51882199 refGene_gn=ZNF395;refGene_tx=XM_023252631.1
B1 51882937 51883038 refGene_gn=ZNF395;refGene_tx=XM_023252631.1
B1 51883502 51883738 refGene_gn=ZNF395;refGene_tx=XM_023252631.1
B1 51887367 51887477 refGene_gn=ZNF395;refGene_tx=XM_023252631.1
B1 51889365 51889571 refGene_gn=ZNF395;refGene_tx=XM_023252631.1
B1 51891182 51891416 refGene_gn=ZNF395;refGene_tx=XM_023252631.1
B1 51959867 51960050
refGene_gn=FBXO16;refGene_tx=XM_011281554.3;refGene_gn=FBXO16;refGene_tx=XM_019828615.2;refGene_gn=FBXO16;refGene_tx=XM_019828617.2;refGene_gn=FBXO16;refGene_tx=XM_019828618.2
B1 51960551 51960577 refGene_gn=FBXO16;refGene_tx=XM_011281554.3
B1 51969186 51969189 refGene_gn=FBXO16;refGene_tx=XM_019828619.2
B1 51972855 51972958
refGene_gn=FBXO16;refGene_tx=XM_011281554.3;refGene_gn=FBXO16;refGene_tx=XM_019828615.2;refGene_gn=FBXO16;refGene_tx=XM_019828617.2;refGene_gn=FBXO16;refGene_tx=XM_019828618.2;refGene_gn=FBXO16;refGene_tx=XM_019828619.2
B1 51980265 51980495
refGene_gn=FBXO16;refGene_tx=XM_011281554.3;refGene_gn=FBXO16;refGene_tx=XM_019828615.2;refGene_gn=FBXO16;refGene_tx=XM_019828617.2;refGene_gn=FBXO16;refGene_tx=XM_019828618.2;refGene_gn=FBXO16;refGene_tx=XM_01

9828619.2
B1	51986090	51986255
refGene_gn=FBXO16;refGene_tx=XM_011281554.3;refGene_gn=FBXO16;refGene_tx=XM_019828615.2;refGene_gn=FBXO16;refGene_tx=XM_019828617.2;refGene_gn=FBXO16;refGene_tx=XM_019828619.2
B1	51989329	51989536
refGene_gn=FBXO16;refGene_tx=XM_011281554.3;refGene_gn=FBXO16;refGene_tx=XM_019828615.2;refGene_gn=FBXO16;refGene_tx=XM_019828617.2;refGene_gn=FBXO16;refGene_tx=XM_019828618.2;refGene_gn=FBXO16;refGene_tx=XM_019828619.2
B1	52002031	52002067
refGene_gn=FBXO16;refGene_tx=XM_011281554.3;refGene_gn=FBXO16;refGene_tx=XM_019828615.2;refGene_gn=FBXO16;refGene_tx=XM_019828618.2;refGene_gn=FBXO16;refGene_tx=XM_019828619.2
B1	52010100	52010199
refGene_gn=FBXO16;refGene_tx=XM_011281554.3;refGene_gn=FBXO16;refGene_tx=XM_019828615.2;refGene_gn=FBXO16;refGene_tx=XM_019828617.2;refGene_gn=FBXO16;refGene_tx=XM_019828618.2;refGene_gn=FBXO16;refGene_tx=XM_019828619.2
B1	52029447	52029636	refGene_gn=FZD3;refGene_tx=XM_003984807.5
B1	52055111	52055308
refGene_gn=FZD3;refGene_tx=XM_003984807.5;refGene_gn=FZD3;refGene_tx=XM_019828620.2;refGene_gn=FZD3;refGene_tx=XM_019828621.2
B1	52059420	52060438
refGene_gn=FZD3;refGene_tx=XM_003984807.5;refGene_gn=FZD3;refGene_tx=XM_019828620.2;refGene_gn=FZD3;refGene_tx=XM_019828621.2
B1	52097098	52097247
refGene_gn=FZD3;refGene_tx=XM_003984807.5;refGene_gn=FZD3;refGene_tx=XM_019828620.2;refGene_gn=FZD3;refGene_tx=XM_019828621.2
B1	52101703	52101937
refGene_gn=FZD3;refGene_tx=XM_003984807.5;refGene_gn=FZD3;refGene_tx=XM_019828620.2;refGene_gn=FZD3;refGene_tx=XM_019828621.2
B1	52113923	52114137
refGene_gn=FZD3;refGene_tx=XM_003984807.5;refGene_gn=FZD3;refGene_tx=XM_019828620.2;refGene_gn=FZD3;refGene_tx=XM_019828621.2
B1	52261784	52263932
refGene_gn=EXTL3;refGene_tx=XM_003984808.4;refGene_gn=EXTL3;refGene_tx=XM_019828622.2;refGene_gn=EXTL3;refGene_tx=XM_019828623.2
B1	52276066	52276194
refGene_gn=EXTL3;refGene_tx=XM_003984808.4;refGene_gn=EXTL3;refGene_tx=XM_019828622.2;refGene_gn=EXTL3;refGene_tx=XM_019828623.2
B1	52280760	52280905
refGene_gn=EXTL3;refGene_tx=XM_003984808.4;refGene_gn=EXTL3;refGene_tx=XM_019828622.2;refGene_gn=EXTL3;refGene_tx=XM_019828623.2
B1	52288806	52288935
refGene_gn=EXTL3;refGene_tx=XM_003984808.4;refGene_gn=EXTL3;refGene_tx=XM_019828622.2;refGene_gn=EXTL3;refGene_tx=XM_019828623.2
B1	52294996	52295206
refGene_gn=EXTL3;refGene_tx=XM_003984808.4;refGene_gn=EXTL3;refGene_tx=XM_019828622.2;refGene_gn=EXTL3;refGene_tx=XM_019828623.2

B1 52310305 52310482
refGene_gn=INTS9;refGene_tx=XM_006930745.4;refGene_gn=INTS9;refGene_tx
=XM_019828624.2
B1 52314283 52314421
refGene_gn=INTS9;refGene_tx=XM_006930745.4;refGene_gn=INTS9;refGene_tx
=XM_019828624.2
B1 52315148 52315226 refGene_gn=INTS9;refGene_tx=XM_023252633.1
B1 52315704 52315803
refGene_gn=INTS9;refGene_tx=XM_006930745.4;refGene_gn=INTS9;refGene_tx
=XM_019828624.2;refGene_gn=INTS9;refGene_tx=XM_023252633.1
B1 52319428 52319596
refGene_gn=INTS9;refGene_tx=XM_006930745.4;refGene_gn=INTS9;refGene_tx
=XM_019828624.2;refGene_gn=INTS9;refGene_tx=XM_023252633.1
B1 52321629 52321754
refGene_gn=INTS9;refGene_tx=XM_006930745.4;refGene_gn=INTS9;refGene_tx
=XM_019828624.2;refGene_gn=INTS9;refGene_tx=XM_023252633.1
B1 52324752 52324924
refGene_gn=INTS9;refGene_tx=XM_006930745.4;refGene_gn=INTS9;refGene_tx
=XM_019828624.2;refGene_gn=INTS9;refGene_tx=XM_023252633.1
B1 52328382 52328443
refGene_gn=INTS9;refGene_tx=XM_006930745.4;refGene_gn=INTS9;refGene_tx
=XM_019828624.2;refGene_gn=INTS9;refGene_tx=XM_023252633.1
B1 52332995 52333176
refGene_gn=INTS9;refGene_tx=XM_006930745.4;refGene_gn=INTS9;refGene_tx
=XM_019828624.2;refGene_gn=INTS9;refGene_tx=XM_023252633.1
B1 52335432 52335544
refGene_gn=INTS9;refGene_tx=XM_006930745.4;refGene_gn=INTS9;refGene_tx
=XM_019828624.2;refGene_gn=INTS9;refGene_tx=XM_023252633.1
B1 52346138 52346273
refGene_gn=INTS9;refGene_tx=XM_006930745.4;refGene_gn=INTS9;refGene_tx
=XM_019828624.2;refGene_gn=INTS9;refGene_tx=XM_023252633.1
B1 52347108 52347229
refGene_gn=INTS9;refGene_tx=XM_006930745.4;refGene_gn=INTS9;refGene_tx
=XM_019828624.2;refGene_gn=INTS9;refGene_tx=XM_023252633.1
B1 52362475 52362562
refGene_gn=INTS9;refGene_tx=XM_006930745.4;refGene_gn=INTS9;refGene_tx
=XM_019828624.2;refGene_gn=INTS9;refGene_tx=XM_023252633.1
B1 52364555 52364695
refGene_gn=INTS9;refGene_tx=XM_006930745.4;refGene_gn=INTS9;refGene_tx
=XM_019828624.2;refGene_gn=INTS9;refGene_tx=XM_023252633.1
B1 52369171 52369234
refGene_gn=INTS9;refGene_tx=XM_006930745.4;refGene_gn=INTS9;refGene_tx
=XM_019828624.2;refGene_gn=INTS9;refGene_tx=XM_023252633.1
B1 52372804 52372865
refGene_gn=INTS9;refGene_tx=XM_006930745.4;refGene_gn=INTS9;refGene_tx
=XM_023252633.1
B1 52381388 52381516
refGene_gn=INTS9;refGene_tx=XM_006930745.4;refGene_gn=INTS9;refGene_tx
=XM_019828624.2;refGene_gn=INTS9;refGene_tx=XM_023252633.1
B1 52412996 52413005

refGene_gn=INTS9;refGene_tx=XM_006930745.4;refGene_gn=INTS9;refGene_tx=XM_023252633.1
B1 52492191 52492214
refGene_gn=HMBOX1;refGene_tx=XM_011281556.3;refGene_gn=HMBOX1;refGene_tx=XM_011281557.3;refGene_gn=HMBOX1;refGene_tx=XM_019828625.2;refGene_gn=HMBOX1;refGene_tx=XM_019828626.2;refGene_gn=HMBOX1;refGene_tx=XM_019828627.2;refGene_gn=HMBOX1;refGene_tx=XM_019828628.2;refGene_gn=HMBOX1;refGene_tx=XM_023252634.1;refGene_gn=HMBOX1;refGene_tx=XM_023252635.1;refGene_gn=HMBOX1;refGene_tx=XM_023252636.1;refGene_gn=HMBOX1;refGene_tx=XM_023252637.1;refGene_gn=HMBOX1;refGene_tx=XM_023252638.1;refGene_gn=HMBOX1;refGene_tx=XM_023252639.1;refGene_gn=HMBOX1;refGene_tx=XM_023252640.1
B1 52507866 52508343
refGene_gn=HMBOX1;refGene_tx=XM_011281556.3;refGene_gn=HMBOX1;refGene_tx=XM_011281557.3;refGene_gn=HMBOX1;refGene_tx=XM_019828625.2;refGene_gn=HMBOX1;refGene_tx=XM_019828626.2;refGene_gn=HMBOX1;refGene_tx=XM_019828627.2;refGene_gn=HMBOX1;refGene_tx=XM_019828628.2;refGene_gn=HMBOX1;refGene_tx=XM_023252634.1;refGene_gn=HMBOX1;refGene_tx=XM_023252635.1;refGene_gn=HMBOX1;refGene_tx=XM_023252636.1;refGene_gn=HMBOX1;refGene_tx=XM_023252637.1;refGene_gn=HMBOX1;refGene_tx=XM_023252638.1;refGene_gn=HMBOX1;refGene_tx=XM_023252639.1;refGene_gn=HMBOX1;refGene_tx=XM_023252640.1
B1 52516243 52516329
refGene_gn=HMBOX1;refGene_tx=XM_011281556.3;refGene_gn=HMBOX1;refGene_tx=XM_011281557.3;refGene_gn=HMBOX1;refGene_tx=XM_019828625.2;refGene_gn=HMBOX1;refGene_tx=XM_019828626.2;refGene_gn=HMBOX1;refGene_tx=XM_019828627.2;refGene_gn=HMBOX1;refGene_tx=XM_019828628.2;refGene_gn=HMBOX1;refGene_tx=XM_023252634.1;refGene_gn=HMBOX1;refGene_tx=XM_023252635.1;refGene_gn=HMBOX1;refGene_tx=XM_023252636.1;refGene_gn=HMBOX1;refGene_tx=XM_023252637.1;refGene_gn=HMBOX1;refGene_tx=XM_023252638.1;refGene_gn=HMBOX1;refGene_tx=XM_023252639.1;refGene_gn=HMBOX1;refGene_tx=XM_023252640.1
B1 52549168 52549279
refGene_gn=HMBOX1;refGene_tx=XM_011281556.3;refGene_gn=HMBOX1;refGene_tx=XM_011281557.3;refGene_gn=HMBOX1;refGene_tx=XM_019828625.2;refGene_gn=HMBOX1;refGene_tx=XM_019828626.2;refGene_gn=HMBOX1;refGene_tx=XM_019828627.2;refGene_gn=HMBOX1;refGene_tx=XM_019828628.2;refGene_gn=HMBOX1;refGene_tx=XM_023252634.1;refGene_gn=HMBOX1;refGene_tx=XM_023252635.1;refGene_gn=HMBOX1;refGene_tx=XM_023252636.1;refGene_gn=HMBOX1;refGene_tx=XM_023252637.1;refGene_gn=HMBOX1;refGene_tx=XM_023252638.1;refGene_gn=HMBOX1;refGene_tx=XM_023252639.1;refGene_gn=HMBOX1;refGene_tx=XM_023252640.1
B1 52560790 52560944
refGene_gn=HMBOX1;refGene_tx=XM_011281556.3;refGene_gn=HMBOX1;refGene_tx=XM_011281557.3;refGene_gn=HMBOX1;refGene_tx=XM_019828625.2;refGene_gn=HMBOX1;refGene_tx=XM_019828626.2;refGene_gn=HMBOX1;refGene_tx=XM_019828627.2;refGene_gn=HMBOX1;refGene_tx=XM_019828628.2;refGene_gn=HMBOX1;refGene_tx=XM_023252634.1;refGene_gn=HMBOX1;refGene_tx=XM_023252635.1;refGene_gn=HMBOX1;refGene_tx=XM_023252636.1;refGene_gn=HMBOX1;refGene_tx=XM_023252637.1;refGene_gn=HMBOX1;refGene_tx=XM_023252638.1;refGen

e_gn=HMBOX1;refGene_tx=XM_023252639.1;refGene_gn=HMBOX1;refGene_tx=XM_023252640.1
B1 52594608 52594691 refGene_gn=HMBOX1;refGene_tx=XM_011281556.3;refGene_gn=HMBOX1;refGene_tx=XM_011281557.3;refGene_gn=HMBOX1;refGene_tx=XM_019828625.2;refGene_gn=HMBOX1;refGene_tx=XM_019828626.2;refGene_gn=HMBOX1;refGene_tx=XM_019828627.2;refGene_gn=HMBOX1;refGene_tx=XM_019828628.2;refGene_gn=HMBOX1;refGene_tx=XM_023252634.1;refGene_gn=HMBOX1;refGene_tx=XM_023252635.1;refGene_gn=HMBOX1;refGene_tx=XM_023252636.1;refGene_gn=HMBOX1;refGene_tx=XM_023252637.1;refGene_gn=HMBOX1;refGene_tx=XM_023252638.1;refGene_gn=HMBOX1;refGene_tx=XM_023252639.1;refGene_gn=HMBOX1;refGene_tx=XM_023252640.1
B1 52596272 52596368 refGene_gn=HMBOX1;refGene_tx=XM_011281556.3;refGene_gn=HMBOX1;refGene_tx=XM_011281557.3;refGene_gn=HMBOX1;refGene_tx=XM_019828625.2;refGene_gn=HMBOX1;refGene_tx=XM_019828626.2;refGene_gn=HMBOX1;refGene_tx=XM_019828627.2;refGene_gn=HMBOX1;refGene_tx=XM_019828628.2;refGene_gn=HMBOX1;refGene_tx=XM_023252634.1;refGene_gn=HMBOX1;refGene_tx=XM_023252635.1;refGene_gn=HMBOX1;refGene_tx=XM_023252636.1;refGene_gn=HMBOX1;refGene_tx=XM_023252637.1;refGene_gn=HMBOX1;refGene_tx=XM_023252638.1;refGene_gn=HMBOX1;refGene_tx=XM_023252639.1;refGene_gn=HMBOX1;refGene_tx=XM_023252640.1
B1 52597485 52597580 refGene_gn=HMBOX1;refGene_tx=XM_011281556.3;refGene_gn=HMBOX1;refGene_tx=XM_011281557.3;refGene_gn=HMBOX1;refGene_tx=XM_019828625.2;refGene_gn=HMBOX1;refGene_tx=XM_019828626.2;refGene_gn=HMBOX1;refGene_tx=XM_019828627.2;refGene_gn=HMBOX1;refGene_tx=XM_019828628.2;refGene_gn=HMBOX1;refGene_tx=XM_023252634.1;refGene_gn=HMBOX1;refGene_tx=XM_023252635.1;refGene_gn=HMBOX1;refGene_tx=XM_023252636.1;refGene_gn=HMBOX1;refGene_tx=XM_023252637.1;refGene_gn=HMBOX1;refGene_tx=XM_023252638.1;refGene_gn=HMBOX1;refGene_tx=XM_023252639.1;refGene_gn=HMBOX1;refGene_tx=XM_023252640.1
B1 52597820 52597916 refGene_gn=HMBOX1;refGene_tx=XM_019828625.2;refGene_gn=HMBOX1;refGene_tx=XM_019828626.2;refGene_gn=HMBOX1;refGene_tx=XM_019828627.2;refGene_gn=HMBOX1;refGene_tx=XM_023252634.1;refGene_gn=HMBOX1;refGene_tx=XM_023252635.1;refGene_gn=HMBOX1;refGene_tx=XM_023252636.1;refGene_gn=HMBOX1;refGene_tx=XM_023252637.1;refGene_gn=HMBOX1;refGene_tx=XM_023252638.1;refGene_gn=HMBOX1;refGene_tx=XM_023252639.1;refGene_gn=HMBOX1;refGene_tx=XM_023252640.1
B1 52599585 52599723 refGene_gn=HMBOX1;refGene_tx=XM_011281556.3;refGene_gn=HMBOX1;refGene_tx=XM_011281557.3;refGene_gn=HMBOX1;refGene_tx=XM_019828625.2;refGene_gn=HMBOX1;refGene_tx=XM_019828626.2;refGene_gn=HMBOX1;refGene_tx=XM_019828627.2;refGene_gn=HMBOX1;refGene_tx=XM_019828628.2;refGene_gn=HMBOX1;refGene_tx=XM_023252634.1;refGene_gn=HMBOX1;refGene_tx=XM_023252635.1;refGene_gn=HMBOX1;refGene_tx=XM_023252636.1;refGene_gn=HMBOX1;refGene_tx=XM_023252637.1;refGene_gn=HMBOX1;refGene_tx=XM_023252638.1
B1 52620572 52620862 refGene_gn=KIF13B;refGene_tx=XM_023252641.1
B1 52622988 52623991 refGene_gn=KIF13B;refGene_tx=XM_023252641.1

B1	52627558	52627621	refGene_gn=KIF13B;refGene_tx=XM_023252641.1
B1	52648283	52648417	refGene_gn=KIF13B;refGene_tx=XM_023252641.1
B1	52656058	52656170	refGene_gn=KIF13B;refGene_tx=XM_023252641.1
B1	52667708	52667762	refGene_gn=KIF13B;refGene_tx=XM_023252641.1
B1	52669070	52669148	refGene_gn=KIF13B;refGene_tx=XM_023252641.1
B1	52669934	52670087	refGene_gn=KIF13B;refGene_tx=XM_023252641.1
B1	52673650	52673743	refGene_gn=KIF13B;refGene_tx=XM_023252641.1
B1	52676807	52676984	refGene_gn=KIF13B;refGene_tx=XM_023252641.1
B1	52678559	52678684	refGene_gn=KIF13B;refGene_tx=XM_023252641.1
B1	52680310	52680366	refGene_gn=KIF13B;refGene_tx=XM_023252641.1
B1	52681120	52681247	refGene_gn=KIF13B;refGene_tx=XM_023252641.1
B1	52681826	52681926	refGene_gn=KIF13B;refGene_tx=XM_023252641.1
B1	52683463	52683493	refGene_gn=KIF13B;refGene_tx=XM_023252641.1
B1	52684121	52684268	refGene_gn=KIF13B;refGene_tx=XM_023252641.1
B1	52689590	52689723	refGene_gn=KIF13B;refGene_tx=XM_023252641.1
B1	52691154	52691312	refGene_gn=KIF13B;refGene_tx=XM_023252641.1
B1	52693191	52693302	refGene_gn=KIF13B;refGene_tx=XM_023252641.1
B1	52693303	52693366	refGene_gn=KIF13B;refGene_tx=XM_023252641.1
B1	52699222	52699351	refGene_gn=KIF13B;refGene_tx=XM_023252641.1
B1	52699633	52699783	refGene_gn=KIF13B;refGene_tx=XM_023252641.1
B1	52701174	52701321	refGene_gn=KIF13B;refGene_tx=XM_023252641.1
B1	52704397	52704560	refGene_gn=KIF13B;refGene_tx=XM_023252641.1
B1	52705447	52705658	refGene_gn=KIF13B;refGene_tx=XM_023252641.1
B1	52706169	52706360	refGene_gn=KIF13B;refGene_tx=XM_023252641.1
B1	52707900	52707987	refGene_gn=KIF13B;refGene_tx=XM_023252641.1
B1	52711141	52711272	refGene_gn=KIF13B;refGene_tx=XM_023252641.1
B1	52721871	52722006	refGene_gn=KIF13B;refGene_tx=XM_023252641.1
B1	52724855	52724966	refGene_gn=KIF13B;refGene_tx=XM_023252641.1
B1	52726329	52726542	refGene_gn=KIF13B;refGene_tx=XM_023252641.1
B1	52734459	52734571	refGene_gn=KIF13B;refGene_tx=XM_023252641.1
B1	52735785	52735898	refGene_gn=KIF13B;refGene_tx=XM_023252641.1
B1	52738069	52738204	refGene_gn=KIF13B;refGene_tx=XM_023252641.1
B1	52741180	52741268	refGene_gn=KIF13B;refGene_tx=XM_023252641.1
B1	52744772	52744953	refGene_gn=KIF13B;refGene_tx=XM_023252641.1
B1	52746742	52746835	refGene_gn=KIF13B;refGene_tx=XM_023252641.1
B1	52748279	52748340	refGene_gn=KIF13B;refGene_tx=XM_023252641.1
B1	52752931	52752944	refGene_gn=KIF13B;refGene_tx=XM_023252641.1
B1	52807310	52807404	refGene_gn=KIF13B;refGene_tx=XM_023252641.1
B1	52823399	52823454	refGene_gn=KIF13B;refGene_tx=XM_023252641.1
B1	53049830	53049966
refGene_gn=MSRA;refGene_tx=XM_011281559.3;refGene_gn=MSRA;refGene_tx=XM_011281560.3
B1	53095748	53095761	refGene_gn=MSRA;refGene_tx=XM_019828634.1
B1	53096120	53096178	refGene_gn=MSRA;refGene_tx=XM_019828632.1
B1	53114628	53114665	refGene_gn=MSRA;refGene_tx=XM_019828633.1
B1	53228802	53228871
refGene_gn=MSRA;refGene_tx=XM_011281559.3;refGene_gn=MSRA;refGene_tx=XM_019828632.1;refGene_gn=MSRA;refGene_tx=XM_019828633.1;refGene_gn=MSRA;refGene_tx=XM_019828634.1
B1	53275188	53275308

refGene_gn=MSRA;refGene_tx=XM_011281559.3;refGene_gn=MSRA;refGene_tx=XM_011281560.3;refGene_gn=MSRA;refGene_tx=XM_019828632.1;refGene_gn=MSRA;refGene_tx=XM_019828633.1;refGene_gn=MSRA;refGene_tx=XM_019828634.1
B1 53352991 53353096
refGene_gn=MSRA;refGene_tx=XM_011281559.3;refGene_gn=MSRA;refGene_tx=XM_011281560.3;refGene_gn=MSRA;refGene_tx=XM_019828632.1;refGene_gn=MSRA;refGene_tx=XM_019828633.1;refGene_gn=MSRA;refGene_tx=XM_019828634.1
B1 53396589 53396696
refGene_gn=MSRA;refGene_tx=XM_011281559.3;refGene_gn=MSRA;refGene_tx=XM_011281560.3;refGene_gn=MSRA;refGene_tx=XM_019828632.1;refGene_gn=MSRA;refGene_tx=XM_019828633.1;refGene_gn=MSRA;refGene_tx=XM_019828634.1
B1 53504217 53504382
refGene_gn=MSRA;refGene_tx=XM_011281559.3;refGene_gn=MSRA;refGene_tx=XM_011281560.3;refGene_gn=MSRA;refGene_tx=XM_019828632.1;refGene_gn=MSRA;refGene_tx=XM_019828633.1;refGene_gn=MSRA;refGene_tx=XM_019828634.1
B1 53571491 53571617
refGene_gn=LOC105260490;refGene_tx=XM_023252647.1
B1 53572260 53572533
refGene_gn=LOC105260490;refGene_tx=XM_023252646.1;refGene_gn=LOC105260490;refGene_tx=XM_023252648.1;refGene_gn=LOC105260490;refGene_tx=XM_023252649.1
B1 53575964 53576107
refGene_gn=LOC105260490;refGene_tx=XM_023252646.1;refGene_gn=LOC105260490;refGene_tx=XM_023252647.1;refGene_gn=LOC105260490;refGene_tx=XM_023252648.1;refGene_gn=LOC105260490;refGene_tx=XM_023252649.1
B1 53577429 53577856
refGene_gn=LOC105260490;refGene_tx=XM_023252646.1;refGene_gn=LOC105260490;refGene_tx=XM_023252647.1;refGene_gn=LOC105260490;refGene_tx=XM_023252648.1;refGene_gn=LOC105260490;refGene_tx=XM_023252649.1
B1 53589436 53589463
refGene_gn=LOC102900749;refGene_tx=XM_019828647.2
B1 53615738 53616035
refGene_gn=LOC102900749;refGene_tx=XM_019828641.2;refGene_gn=LOC102900749;refGene_tx=XM_019828642.2;refGene_gn=LOC102900749;refGene_tx=XM_023252643.1;refGene_gn=LOC102900749;refGene_tx=XM_023252644.1
B1 53616214 53616357
refGene_gn=LOC102900749;refGene_tx=XM_019828641.2;refGene_gn=LOC102900749;refGene_tx=XM_019828642.2;refGene_gn=LOC102900749;refGene_tx=XM_019828647.2;refGene_gn=LOC102900749;refGene_tx=XM_023252643.1;refGene_gn=LOC102900749;refGene_tx=XM_023252644.1
B1 53617344 53617604
refGene_gn=LOC102900749;refGene_tx=XM_019828641.2;refGene_gn=LOC102900749;refGene_tx=XM_019828642.2;refGene_gn=LOC102900749;refGene_tx=XM_019828647.2;refGene_gn=LOC102900749;refGene_tx=XM_023252643.1;refGene_gn=LOC102900749;refGene_tx=XM_023252644.1
B1 53618237 53618430
refGene_gn=LOC102900749;refGene_tx=XM_019828641.2;refGene_gn=LOC102900749;refGene_tx=XM_019828642.2;refGene_gn=LOC102900749;refGene_tx=XM_019828647.2;refGene_gn=LOC102900749;refGene_tx=XM_023252643.1;refGene_gn=LOC102900749;refGene_tx=XM_023252644.1

B1 53626883 53626944
refGene_gn=LOC102900749;refGene_tx=XM_023252644.1
B1 53627582 53627697
refGene_gn=LOC102900749;refGene_tx=XM_019828641.2;refGene_gn=LOC102900749;refGene_tx=XM_019828642.2;refGene_gn=LOC102900749;refGene_tx=XM_019828647.2;refGene_gn=LOC102900749;refGene_tx=XM_023252643.1
B1 53645959 53646038
refGene_gn=LOC101085119;refGene_tx=XM_019828651.1;refGene_gn=LOC101085119;refGene_tx=XM_019828653.2;refGene_gn=LOC101085119;refGene_tx=XM_019828654.2
B1 53648309 53648358
refGene_gn=LOC101085119;refGene_tx=XM_023252645.1
B1 53650128 53650321
refGene_gn=LOC101085119;refGene_tx=XM_019828651.1;refGene_gn=LOC101085119;refGene_tx=XM_019828652.2;refGene_gn=LOC101085119;refGene_tx=XM_019828653.2;refGene_gn=LOC101085119;refGene_tx=XM_019828654.2;refGene_gn=LOC101085119;refGene_tx=XM_023252645.1
B1 53651630 53651881
refGene_gn=LOC101085119;refGene_tx=XM_019828651.1;refGene_gn=LOC101085119;refGene_tx=XM_019828652.2;refGene_gn=LOC101085119;refGene_tx=XM_019828653.2;refGene_gn=LOC101085119;refGene_tx=XM_019828654.2;refGene_gn=LOC101085119;refGene_tx=XM_019828655.2;refGene_gn=LOC101085119;refGene_tx=XM_023252645.1
B1 53653280 53653423
refGene_gn=LOC101085119;refGene_tx=XM_019828651.1;refGene_gn=LOC101085119;refGene_tx=XM_019828652.2;refGene_gn=LOC101085119;refGene_tx=XM_019828653.2;refGene_gn=LOC101085119;refGene_tx=XM_019828654.2;refGene_gn=LOC101085119;refGene_tx=XM_019828655.2;refGene_gn=LOC101085119;refGene_tx=XM_023252645.1
B1 53662760 53663063
refGene_gn=LOC101085119;refGene_tx=XM_019828651.1;refGene_gn=LOC101085119;refGene_tx=XM_019828652.2;refGene_gn=LOC101085119;refGene_tx=XM_019828653.2;refGene_gn=LOC101085119;refGene_tx=XM_019828654.2;refGene_gn=LOC101085119;refGene_tx=XM_019828655.2;refGene_gn=LOC101085119;refGene_tx=XM_023252645.1
B1 53666728 53666743
refGene_gn=LOC101085119;refGene_tx=XM_019828653.2
B1 53668142 53668148
refGene_gn=LOC101085119;refGene_tx=XM_019828654.2
B1 53749846 53755002
refGene_gn=RP1L1;refGene_tx=XM_003984847.5;refGene_gn=RP1L1;refGene_tx=XM_019828657.1;refGene_gn=RP1L1;refGene_tx=XM_019828659.2;refGene_gn=RP1L1;refGene_tx=XM_019828660.1;refGene_gn=RP1L1;refGene_tx=XM_019828662.1;refGene_gn=RP1L1;refGene_tx=XM_019828663.2;refGene_gn=RP1L1;refGene_tx=XM_019828665.1
B1 53756241 53756383
refGene_gn=RP1L1;refGene_tx=XM_003984847.5;refGene_gn=RP1L1;refGene_tx=XM_019828657.1;refGene_gn=RP1L1;refGene_tx=XM_019828659.2;refGene_gn=RP1L1;refGene_tx=XM_019828660.1;refGene_gn=RP1L1;refGene_tx=XM_019828662.1;refGene_gn=RP1L1;refGene_tx=XM_019828663.2

B1 53762056 53762665
refGene_gn=RP1L1;refGene_tx=XM_003984847.5;refGene_gn=RP1L1;refGene_tx=XM_019828657.1;refGene_gn=RP1L1;refGene_tx=XM_019828659.2;refGene_gn=RP1L1;refGene_tx=XM_019828660.1;refGene_gn=RP1L1;refGene_tx=XM_019828662.1;refGene_gn=RP1L1;refGene_tx=XM_019828663.2
B1 53815559 53815607
refGene_gn=CB1H8orf74;refGene_tx=XM_003984838.4
B1 53817155 53817348
refGene_gn=CB1H8orf74;refGene_tx=XM_003984838.4;refGene_gn=CB1H8orf74;refGene_tx=XM_023252650.1
B1 53839787 53840208
refGene_gn=CB1H8orf74;refGene_tx=XM_003984838.4;refGene_gn=CB1H8orf74;refGene_tx=XM_023252650.1
B1 53842728 53842965
refGene_gn=CB1H8orf74;refGene_tx=XM_003984838.4;refGene_gn=CB1H8orf74;refGene_tx=XM_023252650.1
B1 53870799 53871722 refGene_gn=SOX7;refGene_tx=XM_011281561.3
B1 53875651 53875889 refGene_gn=SOX7;refGene_tx=XM_011281561.3
B1 53913680 53914010 refGene_gn=PINX1;refGene_tx=XM_006930758.4
B1 53914011 53914149 refGene_gn=PINX1;refGene_tx=XM_006930758.4
B1 53970274 53970351 refGene_gn=PINX1;refGene_tx=XM_006930758.4
B1 53976890 53976983 refGene_gn=PINX1;refGene_tx=XM_006930758.4
B1 53991346 53991425 refGene_gn=PINX1;refGene_tx=XM_006930758.4
B1 53993235 53993328 refGene_gn=PINX1;refGene_tx=XM_006930758.4
B1 53994995 53995105 refGene_gn=PINX1;refGene_tx=XM_006930758.4
B1 54001568 54001587 refGene_gn=PINX1;refGene_tx=XM_006930758.4
B1 54067453 54068418 refGene_gn=XKR6;refGene_tx=XM_023252651.1
B1 54093141 54093338 refGene_gn=XKR6;refGene_tx=XM_023252651.1
B1 54387127 54387864 refGene_gn=XKR6;refGene_tx=XM_023252651.1
B1 54468392 54468574
refGene_gn=MTMR9;refGene_tx=XM_011281563.3;refGene_gn=MTMR9;refGene_tx=XM_019828675.2
B1 54484255 54484364
refGene_gn=MTMR9;refGene_tx=XM_011281563.3;refGene_gn=MTMR9;refGene_tx=XM_019828675.2
B1 54488709 54488835
refGene_gn=MTMR9;refGene_tx=XM_011281563.3;refGene_gn=MTMR9;refGene_tx=XM_019828675.2
B1 54499110 54499284
refGene_gn=MTMR9;refGene_tx=XM_011281563.3;refGene_gn=MTMR9;refGene_tx=XM_019828675.2
B1 54501843 54502061
refGene_gn=MTMR9;refGene_tx=XM_011281563.3;refGene_gn=MTMR9;refGene_tx=XM_019828675.2
B1 54503024 54503186
refGene_gn=MTMR9;refGene_tx=XM_011281563.3;refGene_gn=MTMR9;refGene_tx=XM_019828675.2
B1 54505735 54505877
refGene_gn=MTMR9;refGene_tx=XM_011281563.3;refGene_gn=MTMR9;refGene_tx=XM_019828675.2

B1 54507671 54507892
refGene_gn=MTMR9;refGene_tx=XM_011281563.3;refGene_gn=MTMR9;refGene_tx=XM_019828675.2
B1 54518297 54518449 refGene_gn=MTMR9;refGene_tx=XM_011281563.3
B1 54524610 54524774
refGene_gn=MTMR9;refGene_tx=XM_011281563.3;refGene_gn=MTMR9;refGene_tx=XM_019828675.2
B1 54550315 54550499
refGene_gn=LOC101083586;refGene_tx=XM_006930761.3;refGene_gn=LOC101083586;refGene_tx=XM_019828676.1;refGene_gn=LOC101083586;refGene_tx=XM_019828677.1
B1 54552545 54552588
refGene_gn=LOC101083586;refGene_tx=XM_006930761.3;refGene_gn=LOC101083586;refGene_tx=XM_019828676.1
B1 54552887 54552958
refGene_gn=LOC101083586;refGene_tx=XM_006930761.3;refGene_gn=LOC101083586;refGene_tx=XM_019828676.1
B1 54554709 54554859
refGene_gn=LOC101083586;refGene_tx=XM_006930761.3;refGene_gn=LOC101083586;refGene_tx=XM_019828676.1;refGene_gn=LOC101083586;refGene_tx=XM_019828677.1
B1 54554951 54555124
refGene_gn=LOC101083586;refGene_tx=XM_006930761.3;refGene_gn=LOC101083586;refGene_tx=XM_019828676.1;refGene_gn=LOC101083586;refGene_tx=XM_019828677.1
B1 54555965 54556056
refGene_gn=LOC101083586;refGene_tx=XM_006930761.3;refGene_gn=LOC101083586;refGene_tx=XM_019828676.1;refGene_gn=LOC101083586;refGene_tx=XM_019828677.1
B1 54558144 54558507
refGene_gn=LOC101083586;refGene_tx=XM_006930761.3;refGene_gn=LOC101083586;refGene_tx=XM_019828676.1;refGene_gn=LOC101083586;refGene_tx=XM_019828677.1
B1 54559082 54559222
refGene_gn=LOC101083586;refGene_tx=XM_006930761.3;refGene_gn=LOC101083586;refGene_tx=XM_019828676.1;refGene_gn=LOC101083586;refGene_tx=XM_019828677.1
B1 54624324 54624588
refGene_gn=FAM167A;refGene_tx=XM_003984841.5;refGene_gn=FAM167A;refGene_tx=XM_019828678.2;refGene_gn=FAM167A;refGene_tx=XM_023252653.1;refGene_gn=FAM167A;refGene_tx=XM_023252654.1
B1 54644695 54645061
refGene_gn=FAM167A;refGene_tx=XM_003984841.5;refGene_gn=FAM167A;refGene_tx=XM_019828678.2;refGene_gn=FAM167A;refGene_tx=XM_023252653.1;refGene_gn=FAM167A;refGene_tx=XM_023252654.1
B1 54755220 54755328
refGene_gn=BLK;refGene_tx=XM_003984842.4;refGene_gn=BLK;refGene_tx=XM_019828679.1;refGene_gn=BLK;refGene_tx=XM_019828680.1
B1 54757054 54757106
refGene_gn=BLK;refGene_tx=XM_003984842.4;refGene_gn=BLK;refGene_tx=XM_

019828679.1;refGene_gn=BLK;refGene_tx=XM_019828680.1
B1 54759203 54759297
refGene_gn=BLK;refGene_tx=XM_003984842.4;refGene_gn=BLK;refGene_tx=XM_019828679.1;refGene_gn=BLK;refGene_tx=XM_019828680.1
B1 54760252 54760351
refGene_gn=BLK;refGene_tx=XM_003984842.4;refGene_gn=BLK;refGene_tx=XM_019828679.1;refGene_gn=BLK;refGene_tx=XM_019828680.1
B1 54761651 54761755
refGene_gn=BLK;refGene_tx=XM_003984842.4;refGene_gn=BLK;refGene_tx=XM_019828679.1;refGene_gn=BLK;refGene_tx=XM_019828680.1
B1 54766676 54766823
refGene_gn=BLK;refGene_tx=XM_003984842.4;refGene_gn=BLK;refGene_tx=XM_019828679.1;refGene_gn=BLK;refGene_tx=XM_019828680.1
B1 54767259 54767412
refGene_gn=BLK;refGene_tx=XM_003984842.4;refGene_gn=BLK;refGene_tx=XM_019828679.1;refGene_gn=BLK;refGene_tx=XM_019828680.1
B1 54768254 54768434
refGene_gn=BLK;refGene_tx=XM_003984842.4;refGene_gn=BLK;refGene_tx=XM_019828679.1;refGene_gn=BLK;refGene_tx=XM_019828680.1
B1 54769604 54769681
refGene_gn=BLK;refGene_tx=XM_003984842.4;refGene_gn=BLK;refGene_tx=XM_019828679.1;refGene_gn=BLK;refGene_tx=XM_019828680.1
B1 54772515 54772666
refGene_gn=BLK;refGene_tx=XM_003984842.4;refGene_gn=BLK;refGene_tx=XM_019828679.1;refGene_gn=BLK;refGene_tx=XM_019828680.1
B1 54773880 54774012
refGene_gn=BLK;refGene_tx=XM_003984842.4;refGene_gn=BLK;refGene_tx=XM_019828679.1;refGene_gn=BLK;refGene_tx=XM_019828680.1
B1 54774739 54774945
refGene_gn=BLK;refGene_tx=XM_003984842.4;refGene_gn=BLK;refGene_tx=XM_019828679.1;refGene_gn=BLK;refGene_tx=XM_019828680.1
B1 54936361 54936977
refGene_gn=GATA4;refGene_tx=XM_011281564.3;refGene_gn=GATA4;refGene_tx=XM_019828684.2;refGene_gn=GATA4;refGene_tx=XM_023252656.1
B1 54975019 54975189
refGene_gn=GATA4;refGene_tx=XM_011281564.3;refGene_gn=GATA4;refGene_tx=XM_019828684.2;refGene_gn=GATA4;refGene_tx=XM_023252656.1
B1 54976910 54977036
refGene_gn=GATA4;refGene_tx=XM_011281564.3;refGene_gn=GATA4;refGene_tx=XM_019828684.2;refGene_gn=GATA4;refGene_tx=XM_023252656.1
B1 54980191 54980279
refGene_gn=GATA4;refGene_tx=XM_011281564.3;refGene_gn=GATA4;refGene_tx=XM_019828684.2;refGene_gn=GATA4;refGene_tx=XM_023252656.1
B1 54982151 54982300
refGene_gn=GATA4;refGene_tx=XM_011281564.3;refGene_gn=GATA4;refGene_tx=XM_019828684.2;refGene_gn=GATA4;refGene_tx=XM_023252656.1
B1 54983531 54983714
refGene_gn=GATA4;refGene_tx=XM_011281564.3;refGene_gn=GATA4;refGene_tx=XM_019828684.2;refGene_gn=GATA4;refGene_tx=XM_023252656.1
B1 54996543 54996681

refGene_gn=NEIL2;refGene_tx=XM_003984843.4;refGene_gn=NEIL2;refGene_tx
=XM_006930763.4
B1 54998156 54998288 refGene_gn=NEIL2;refGene_tx=XM_006930764.1
B1 55001701 55002054
refGene_gn=NEIL2;refGene_tx=XM_003984843.4;refGene_gn=NEIL2;refGene_tx
=XM_006930763.4;refGene_gn=NEIL2;refGene_tx=XM_006930764.1
B1 55006835 55007032
refGene_gn=NEIL2;refGene_tx=XM_003984843.4;refGene_gn=NEIL2;refGene_tx
=XM_006930763.4;refGene_gn=NEIL2;refGene_tx=XM_006930764.1
B1 55008957 55009271
refGene_gn=NEIL2;refGene_tx=XM_003984843.4;refGene_gn=NEIL2;refGene_tx
=XM_006930763.4;refGene_gn=NEIL2;refGene_tx=XM_006930764.1
B1 55019004 55019103 refGene_gn=FDFT1;refGene_tx=XM_003984844.5
B1 55028169 55028570
refGene_gn=FDFT1;refGene_tx=XM_003984844.5;refGene_gn=FDFT1;refGene_tx
=XM_023252658.1
B1 55029406 55029590
refGene_gn=FDFT1;refGene_tx=XM_003984844.5;refGene_gn=FDFT1;refGene_tx
=XM_023252658.1
B1 55035971 55036100
refGene_gn=FDFT1;refGene_tx=XM_003984844.5;refGene_gn=FDFT1;refGene_tx
=XM_023252658.1
B1 55039246 55039438
refGene_gn=FDFT1;refGene_tx=XM_003984844.5;refGene_gn=FDFT1;refGene_tx
=XM_023252658.1
B1 55046949 55047126
refGene_gn=FDFT1;refGene_tx=XM_003984844.5;refGene_gn=FDFT1;refGene_tx
=XM_023252658.1
B1 55048202 55048355
refGene_gn=FDFT1;refGene_tx=XM_003984844.5;refGene_gn=FDFT1;refGene_tx
=XM_023252658.1
B1 55056628 55056850
refGene_gn=FDFT1;refGene_tx=XM_003984844.5;refGene_gn=FDFT1;refGene_tx
=XM_023252658.1
B1 55062309 55062407
refGene_gn=CTSB;refGene_tx=XM_003984845.4;refGene_gn=CTSB;refGene_tx=X
M_006930765.3;refGene_gn=CTSB;refGene_tx=XM_019828685.1
B1 55062895 55063024
refGene_gn=CTSB;refGene_tx=XM_003984845.4;refGene_gn=CTSB;refGene_tx=X
M_006930765.3;refGene_gn=CTSB;refGene_tx=XM_019828685.1
B1 55064011 55064128
refGene_gn=CTSB;refGene_tx=XM_003984845.4;refGene_gn=CTSB;refGene_tx=X
M_006930765.3;refGene_gn=CTSB;refGene_tx=XM_019828685.1
B1 55064652 55064796
refGene_gn=CTSB;refGene_tx=XM_003984845.4;refGene_gn=CTSB;refGene_tx=X
M_006930765.3;refGene_gn=CTSB;refGene_tx=XM_019828685.1
B1 55065051 55065137
refGene_gn=CTSB;refGene_tx=XM_003984845.4;refGene_gn=CTSB;refGene_tx=X
M_006930765.3;refGene_gn=CTSB;refGene_tx=XM_019828685.1
B1 55065950 55066069

refGene_gn=CTSB;refGene_tx=XM_003984845.4;refGene_gn=CTSB;refGene_tx=XM_006930765.3
B1 55067287 55067402
refGene_gn=CTSB;refGene_tx=XM_003984845.4;refGene_gn=CTSB;refGene_tx=XM_006930765.3;refGene_gn=CTSB;refGene_tx=XM_019828685.1
B1 55068179 55068265
refGene_gn=CTSB;refGene_tx=XM_003984845.4;refGene_gn=CTSB;refGene_tx=XM_006930765.3
B1 55069246 55069372
refGene_gn=CTSB;refGene_tx=XM_003984845.4;refGene_gn=CTSB;refGene_tx=XM_006930765.3
B1 55169225 55169398 refGene_gn=DEFB136;refGene_tx=XM_019829911.2
B1 55169850 55169905 refGene_gn=DEFB136;refGene_tx=XM_019829911.2
B1 55209862 55210005 refGene_gn=DEFB134;refGene_tx=XM_011281571.2
B1 55212216 55212274 refGene_gn=DEFB134;refGene_tx=XM_011281571.2
B1 55224038 55224182
refGene_gn=LOC109499398;refGene_tx=XM_019829991.1
B1 55552107 55554429
refGene_gn=LOC101087152;refGene_tx=XM_003984851.4
B1 55627263 55627757
refGene_gn=LOC111560220;refGene_tx=XM_023253389.1
B1 55627848 55628134
refGene_gn=LOC111560220;refGene_tx=XM_023253389.1
B1 55697285 55700168
refGene_gn=ADAM29;refGene_tx=XM_006930770.4;refGene_gn=ADAM29;refGene_tx=XM_011281565.3;refGene_gn=ADAM29;refGene_tx=XM_019828686.2
B1 55850913 55850984
refGene_gn=GLRA3;refGene_tx=XM_023252660.1;refGene_gn=GLRA3;refGene_tx=XM_023252661.1
B1 55892539 55892667
refGene_gn=GLRA3;refGene_tx=XM_023252660.1;refGene_gn=GLRA3;refGene_tx=XM_023252661.1
B1 55918386 55918454
refGene_gn=GLRA3;refGene_tx=XM_023252660.1;refGene_gn=GLRA3;refGene_tx=XM_023252661.1
B1 55968956 55969180
refGene_gn=GLRA3;refGene_tx=XM_023252660.1;refGene_gn=GLRA3;refGene_tx=XM_023252661.1
B1 55981562 55981645
refGene_gn=GLRA3;refGene_tx=XM_023252660.1;refGene_gn=GLRA3;refGene_tx=XM_023252661.1
B1 56004949 56005087
refGene_gn=GLRA3;refGene_tx=XM_023252660.1;refGene_gn=GLRA3;refGene_tx=XM_023252661.1
B1 56013111 56013326
refGene_gn=GLRA3;refGene_tx=XM_023252660.1;refGene_gn=GLRA3;refGene_tx=XM_023252661.1
B1 56031886 56032030
refGene_gn=GLRA3;refGene_tx=XM_023252660.1;refGene_gn=GLRA3;refGene_tx=XM_023252661.1

B1 56034267 56034312 refGene_gn=GLRA3;refGene_tx=XM_023252660.1
B1 56048122 56048401
refGene_gn=GLRA3;refGene_tx=XM_023252660.1;refGene_gn=GLRA3;refGene_tx
=XM_023252661.1
B1 56190822 56190915 refGene_gn=HPGD;refGene_tx=XM_011281573.3
B1 56191236 56191360 refGene_gn=HPGD;refGene_tx=XM_011281573.3
B1 56194998 56195105 refGene_gn=HPGD;refGene_tx=XM_011281573.3
B1 56203196 56203293 refGene_gn=HPGD;refGene_tx=XM_011281573.3
B1 56214491 56214568 refGene_gn=HPGD;refGene_tx=XM_011281573.3
B1 56216795 56216959 refGene_gn=HPGD;refGene_tx=XM_011281573.3
B1 56217978 56218019 refGene_gn=HPGD;refGene_tx=XM_011281573.3
B1 56218021 56218116 refGene_gn=HPGD;refGene_tx=XM_011281573.3
B1 56395463 56395506
refGene_gn=CEP44;refGene_tx=XM_011281574.3;refGene_gn=CEP44;refGene_tx
=XM_011281575.3;refGene_gn=CEP44;refGene_tx=XM_011281576.3;refGene_gn=
CEP44;refGene_tx=XM_011281577.3;refGene_gn=CEP44;refGene_tx=XM_0112815
78.3;refGene_gn=CEP44;refGene_tx=XM_019828693.2;refGene_gn=CEP44;refGe
ne_tx=XM_019828694.2;refGene_gn=CEP44;refGene_tx=XM_019828695.2;refGen
e_gn=CEP44;refGene_tx=XM_019828696.2
B1 56396266 56396304
refGene_gn=CEP44;refGene_tx=XM_011281574.3;refGene_gn=CEP44;refGene_tx
=XM_011281575.3;refGene_gn=CEP44;refGene_tx=XM_011281576.3;refGene_gn=
CEP44;refGene_tx=XM_011281577.3;refGene_gn=CEP44;refGene_tx=XM_0198286
93.2;refGene_gn=CEP44;refGene_tx=XM_019828694.2;refGene_gn=CEP44;refGe
ne_tx=XM_019828695.2;refGene_gn=CEP44;refGene_tx=XM_019828696.2
B1 56396556 56396681
refGene_gn=CEP44;refGene_tx=XM_011281574.3;refGene_gn=CEP44;refGene_tx
=XM_011281575.3;refGene_gn=CEP44;refGene_tx=XM_011281576.3;refGene_gn=
CEP44;refGene_tx=XM_011281577.3;refGene_gn=CEP44;refGene_tx=XM_0112815
78.3;refGene_gn=CEP44;refGene_tx=XM_019828693.2;refGene_gn=CEP44;refGe
ne_tx=XM_019828694.2;refGene_gn=CEP44;refGene_tx=XM_019828695.2;refGen
e_gn=CEP44;refGene_tx=XM_019828696.2
B1 56401920 56401996
refGene_gn=CEP44;refGene_tx=XM_011281574.3;refGene_gn=CEP44;refGene_tx
=XM_011281575.3;refGene_gn=CEP44;refGene_tx=XM_011281576.3;refGene_gn=
CEP44;refGene_tx=XM_011281577.3;refGene_gn=CEP44;refGene_tx=XM_0112815
78.3;refGene_gn=CEP44;refGene_tx=XM_019828693.2;refGene_gn=CEP44;refGe
ne_tx=XM_019828694.2;refGene_gn=CEP44;refGene_tx=XM_019828695.2;refGen
e_gn=CEP44;refGene_tx=XM_019828696.2
B1 56402737 56402944
refGene_gn=CEP44;refGene_tx=XM_011281574.3;refGene_gn=CEP44;refGene_tx
=XM_011281575.3;refGene_gn=CEP44;refGene_tx=XM_011281576.3;refGene_gn=
CEP44;refGene_tx=XM_011281577.3;refGene_gn=CEP44;refGene_tx=XM_0112815
78.3;refGene_gn=CEP44;refGene_tx=XM_019828693.2;refGene_gn=CEP44;refGe
ne_tx=XM_019828694.2;refGene_gn=CEP44;refGene_tx=XM_019828695.2
B1 56404217 56404388
refGene_gn=CEP44;refGene_tx=XM_011281574.3;refGene_gn=CEP44;refGene_tx
=XM_011281575.3;refGene_gn=CEP44;refGene_tx=XM_011281576.3;refGene_gn=
CEP44;refGene_tx=XM_011281577.3;refGene_gn=CEP44;refGene_tx=XM_0112815
78.3;refGene_gn=CEP44;refGene_tx=XM_019828693.2;refGene_gn=CEP44;refGe

ne_tx=XM_019828694.2;refGene_gn=CEP44;refGene_tx=XM_019828695.2;refGene_gn=CEP44;refGene_tx=XM_019828696.2
B1 56406824 56406947
refGene_gn=CEP44;refGene_tx=XM_011281574.3;refGene_gn=CEP44;refGene_tx=XM_011281575.3;refGene_gn=CEP44;refGene_tx=XM_011281576.3;refGene_gn=CEP44;refGene_tx=XM_011281577.3;refGene_gn=CEP44;refGene_tx=XM_011281578.3;refGene_gn=CEP44;refGene_tx=XM_019828693.2;refGene_gn=CEP44;refGene_tx=XM_019828694.2;refGene_gn=CEP44;refGene_tx=XM_019828695.2;refGene_gn=CEP44;refGene_tx=XM_019828696.2
B1 56407769 56407916
refGene_gn=CEP44;refGene_tx=XM_011281574.3;refGene_gn=CEP44;refGene_tx=XM_011281575.3;refGene_gn=CEP44;refGene_tx=XM_011281576.3;refGene_gn=CEP44;refGene_tx=XM_011281577.3;refGene_gn=CEP44;refGene_tx=XM_011281578.3;refGene_gn=CEP44;refGene_tx=XM_019828693.2;refGene_gn=CEP44;refGene_tx=XM_019828694.2;refGene_gn=CEP44;refGene_tx=XM_019828695.2;refGene_gn=CEP44;refGene_tx=XM_019828696.2
B1 56409071 56409219
refGene_gn=CEP44;refGene_tx=XM_011281574.3;refGene_gn=CEP44;refGene_tx=XM_011281575.3;refGene_gn=CEP44;refGene_tx=XM_011281576.3;refGene_gn=CEP44;refGene_tx=XM_011281577.3;refGene_gn=CEP44;refGene_tx=XM_011281578.3;refGene_gn=CEP44;refGene_tx=XM_019828693.2;refGene_gn=CEP44;refGene_tx=XM_019828694.2;refGene_gn=CEP44;refGene_tx=XM_019828695.2;refGene_gn=CEP44;refGene_tx=XM_019828696.2
B1 56412296 56412385
refGene_gn=CEP44;refGene_tx=XM_011281574.3;refGene_gn=CEP44;refGene_tx=XM_011281575.3;refGene_gn=CEP44;refGene_tx=XM_011281576.3;refGene_gn=CEP44;refGene_tx=XM_011281577.3;refGene_gn=CEP44;refGene_tx=XM_011281578.3;refGene_gn=CEP44;refGene_tx=XM_019828693.2;refGene_gn=CEP44;refGene_tx=XM_019828694.2;refGene_gn=CEP44;refGene_tx=XM_019828695.2;refGene_gn=CEP44;refGene_tx=XM_019828696.2
B1 56439722 56440051 refGene_gn=FBXO8;refGene_tx=XM_003984855.4
B1 56443109 56443236 refGene_gn=FBXO8;refGene_tx=XM_003984855.4
B1 56461632 56461751 refGene_gn=FBXO8;refGene_tx=XM_003984855.4
B1 56462926 56463123 refGene_gn=FBXO8;refGene_tx=XM_003984855.4
B1 56464268 56464456 refGene_gn=FBXO8;refGene_tx=XM_003984855.4
B1 57116092 57116241
refGene_gn=LOC109499352;refGene_tx=XM_019829912.1
B1 57116882 57116922
refGene_gn=LOC109499352;refGene_tx=XM_019829912.1
B1 57117330 57117449
refGene_gn=LOC109499352;refGene_tx=XM_019829912.1
B1 57123296 57123408
refGene_gn=LOC109499352;refGene_tx=XM_019829912.1
B1 57123519 57123725
refGene_gn=LOC109499352;refGene_tx=XM_019829912.1
B1 57124188 57124320
refGene_gn=LOC109499352;refGene_tx=XM_019829912.1
B1 57124411 57124466
refGene_gn=LOC109499352;refGene_tx=XM_019829912.1
B1 57125084 57125639

refGene_gn=HAND2;refGene_tx=XM_019828697.2;refGene_gn=HAND2;refGene_tx=XM_023252662.1
B1 57126324 57126381 refGene_gn=HAND2;refGene_tx=XM_019828697.2
B1 57127001 57127100
refGene_gn=HAND2;refGene_tx=XM_019828697.2;refGene_gn=HAND2;refGene_tx=XM_023252662.1
B1 57285357 57285599
refGene_gn=SCRG1;refGene_tx=XM_011281579.3;refGene_gn=SCRG1;refGene_tx=XM_019828703.1
B1 57287939 57287994
refGene_gn=SCRG1;refGene_tx=XM_011281579.3;refGene_gn=SCRG1;refGene_tx=XM_019828703.1
B1 57297209 57297332
refGene_gn=SAP30;refGene_tx=XM_003984857.5;refGene_gn=SAP30;refGene_tx=XM_006930772.4
B1 57299599 57299698
refGene_gn=SAP30;refGene_tx=XM_003984857.5;refGene_gn=SAP30;refGene_tx=XM_006930772.4
B1 57300135 57300261 refGene_gn=SAP30;refGene_tx=XM_003984857.5
B1 57301951 57302269
refGene_gn=SAP30;refGene_tx=XM_003984857.5;refGene_gn=SAP30;refGene_tx=XM_006930772.4
B1 57334697 57334847
refGene_gn=HMGB2;refGene_tx=XM_003984858.5;refGene_gn=HMGB2;refGene_tx=XM_006930773.4
B1 57335151 57335297
refGene_gn=HMGB2;refGene_tx=XM_003984858.5;refGene_gn=HMGB2;refGene_tx=XM_006930773.4
B1 57335372 57335547
refGene_gn=HMGB2;refGene_tx=XM_003984858.5;refGene_gn=HMGB2;refGene_tx=XM_006930773.4
B1 57336099 57336261
refGene_gn=HMGB2;refGene_tx=XM_003984858.5;refGene_gn=HMGB2;refGene_tx=XM_006930773.4
B1 57347092 57347230
refGene_gn=GALNT7;refGene_tx=XM_023252664.1;refGene_gn=GALNT7;refGene_tx=XM_023252665.1
B1 57350559 57350688
refGene_gn=GALNT7;refGene_tx=XM_023252664.1;refGene_gn=GALNT7;refGene_tx=XM_023252665.1
B1 57351704 57351803
refGene_gn=GALNT7;refGene_tx=XM_023252664.1;refGene_gn=GALNT7;refGene_tx=XM_023252665.1
B1 57354958 57355177
refGene_gn=GALNT7;refGene_tx=XM_023252664.1;refGene_gn=GALNT7;refGene_tx=XM_023252665.1
B1 57361108 57361231
refGene_gn=GALNT7;refGene_tx=XM_023252664.1;refGene_gn=GALNT7;refGene_tx=XM_023252665.1
B1 57362963 57363081

refGene_gn=GALNT7;refGene_tx=XM_023252664.1;refGene_gn=GALNT7;refGene_tx=XM_023252665.1
B1 57365856 57366039 refGene_gn=GALNT7;refGene_tx=XM_023252665.1
B1 57366180 57366363 refGene_gn=GALNT7;refGene_tx=XM_023252664.1
B1 57368638 57368718
refGene_gn=GALNT7;refGene_tx=XM_023252664.1;refGene_gn=GALNT7;refGene_tx=XM_023252665.1
B1 57368968 57369099
refGene_gn=GALNT7;refGene_tx=XM_023252664.1;refGene_gn=GALNT7;refGene_tx=XM_023252665.1
B1 57374078 57374245
refGene_gn=GALNT7;refGene_tx=XM_023252664.1;refGene_gn=GALNT7;refGene_tx=XM_023252665.1
B1 57411858 57412319
refGene_gn=GALNT7;refGene_tx=XM_023252664.1;refGene_gn=GALNT7;refGene_tx=XM_023252665.1
B1 57493817 57493943
refGene_gn=GALNT7;refGene_tx=XM_023252664.1;refGene_gn=GALNT7;refGene_tx=XM_023252665.1
B1 57596881 57597049 refGene_gn=GALNTL6;refGene_tx=XM_003984859.5
B1 57606158 57606188 refGene_gn=GALNTL6;refGene_tx=XM_019828706.2
B1 57609384 57609480 refGene_gn=GALNTL6;refGene_tx=XM_023252666.1
B1 57615749 57615899
refGene_gn=GALNTL6;refGene_tx=XM_003984859.5;refGene_gn=GALNTL6;refGene_tx=XM_019828706.2;refGene_gn=GALNTL6;refGene_tx=XM_023252666.1
B1 57624188 57624305
refGene_gn=GALNTL6;refGene_tx=XM_003984859.5;refGene_gn=GALNTL6;refGene_tx=XM_019828706.2;refGene_gn=GALNTL6;refGene_tx=XM_023252666.1
B1 57673691 57673913
refGene_gn=GALNTL6;refGene_tx=XM_003984859.5;refGene_gn=GALNTL6;refGene_tx=XM_019828706.2;refGene_gn=GALNTL6;refGene_tx=XM_023252666.1
B1 57707355 57707463
refGene_gn=GALNTL6;refGene_tx=XM_003984859.5;refGene_gn=GALNTL6;refGene_tx=XM_019828706.2;refGene_gn=GALNTL6;refGene_tx=XM_023252666.1
B1 57732330 57732448
refGene_gn=GALNTL6;refGene_tx=XM_003984859.5;refGene_gn=GALNTL6;refGene_tx=XM_019828706.2;refGene_gn=GALNTL6;refGene_tx=XM_023252666.1
B1 57821716 57821900
refGene_gn=GALNTL6;refGene_tx=XM_003984859.5;refGene_gn=GALNTL6;refGene_tx=XM_019828706.2;refGene_gn=GALNTL6;refGene_tx=XM_023252666.1
B1 57826021 57826207
refGene_gn=GALNTL6;refGene_tx=XM_003984859.5;refGene_gn=GALNTL6;refGene_tx=XM_019828706.2;refGene_gn=GALNTL6;refGene_tx=XM_023252666.1
B1 57906702 57906745
refGene_gn=LOC111556190;refGene_tx=XM_023252667.1
B1 57923907 57924152
refGene_gn=LOC111556190;refGene_tx=XM_023252667.1
B1 58262279 58262446
refGene_gn=GALNTL6;refGene_tx=XM_003984859.5;refGene_gn=GALNTL6;refGene_tx=XM_019828706.2;refGene_gn=GALNTL6;refGene_tx=XM_023252666.1

B1 58284644 58284783
refGene_gn=GALNTL6;refGene_tx=XM_003984859.5;refGene_gn=GALNTL6;refGene_tx=XM_019828706.2;refGene_gn=GALNTL6;refGene_tx=XM_023252666.1
B1 58367432 58367541
refGene_gn=GALNTL6;refGene_tx=XM_003984859.5;refGene_gn=GALNTL6;refGene_tx=XM_019828706.2;refGene_gn=GALNTL6;refGene_tx=XM_023252666.1
B1 58807712 58807850
refGene_gn=GALNTL6;refGene_tx=XM_003984859.5;refGene_gn=GALNTL6;refGene_tx=XM_019828706.2;refGene_gn=GALNTL6;refGene_tx=XM_023252666.1
B1 60574286 60574353
refGene_gn=AADAT;refGene_tx=XM_003984863.4;refGene_gn=AADAT;refGene_tx=XM_006930777.4;refGene_gn=AADAT;refGene_tx=XM_019828714.2;refGene_gn=AADAT;refGene_tx=XM_023252669.1;refGene_gn=AADAT;refGene_tx=XM_023252670.1;refGene_gn=AADAT;refGene_tx=XM_023252671.1
B1 60575041 60575210
refGene_gn=AADAT;refGene_tx=XM_003984863.4;refGene_gn=AADAT;refGene_tx=XM_006930777.4;refGene_gn=AADAT;refGene_tx=XM_019828714.2;refGene_gn=AADAT;refGene_tx=XM_023252669.1;refGene_gn=AADAT;refGene_tx=XM_023252670.1;refGene_gn=AADAT;refGene_tx=XM_023252671.1
B1 60576214 60576347
refGene_gn=AADAT;refGene_tx=XM_003984863.4;refGene_gn=AADAT;refGene_tx=XM_006930777.4;refGene_gn=AADAT;refGene_tx=XM_019828714.2;refGene_gn=AADAT;refGene_tx=XM_023252669.1;refGene_gn=AADAT;refGene_tx=XM_023252670.1;refGene_gn=AADAT;refGene_tx=XM_023252671.1
B1 60580079 60580154
refGene_gn=AADAT;refGene_tx=XM_003984863.4;refGene_gn=AADAT;refGene_tx=XM_006930777.4;refGene_gn=AADAT;refGene_tx=XM_019828714.2;refGene_gn=AADAT;refGene_tx=XM_023252669.1;refGene_gn=AADAT;refGene_tx=XM_023252670.1;refGene_gn=AADAT;refGene_tx=XM_023252671.1
B1 60584493 60584703
refGene_gn=AADAT;refGene_tx=XM_003984863.4;refGene_gn=AADAT;refGene_tx=XM_006930777.4;refGene_gn=AADAT;refGene_tx=XM_019828714.2;refGene_gn=AADAT;refGene_tx=XM_023252669.1;refGene_gn=AADAT;refGene_tx=XM_023252670.1;refGene_gn=AADAT;refGene_tx=XM_023252671.1
B1 60587676 60587742
refGene_gn=AADAT;refGene_tx=XM_003984863.4;refGene_gn=AADAT;refGene_tx=XM_006930777.4;refGene_gn=AADAT;refGene_tx=XM_019828714.2;refGene_gn=AADAT;refGene_tx=XM_023252669.1;refGene_gn=AADAT;refGene_tx=XM_023252670.1;refGene_gn=AADAT;refGene_tx=XM_023252671.1
B1 60589068 60589151
refGene_gn=AADAT;refGene_tx=XM_003984863.4;refGene_gn=AADAT;refGene_tx=XM_006930777.4;refGene_gn=AADAT;refGene_tx=XM_019828714.2;refGene_gn=AADAT;refGene_tx=XM_023252669.1;refGene_gn=AADAT;refGene_tx=XM_023252670.1;refGene_gn=AADAT;refGene_tx=XM_023252671.1
B1 60589605 60589702
refGene_gn=AADAT;refGene_tx=XM_003984863.4;refGene_gn=AADAT;refGene_tx=XM_006930777.4;refGene_gn=AADAT;refGene_tx=XM_019828714.2;refGene_gn=AADAT;refGene_tx=XM_023252669.1;refGene_gn=AADAT;refGene_tx=XM_023252670.1;refGene_gn=AADAT;refGene_tx=XM_023252671.1
B1 60591390 60591452

refGene_gn=AADAT;refGene_tx=XM_003984863.4;refGene_gn=AADAT;refGene_tx=XM_006930777.4;refGene_gn=AADAT;refGene_tx=XM_019828714.2;refGene_gn=AADAT;refGene_tx=XM_023252669.1;refGene_gn=AADAT;refGene_tx=XM_023252670.1;refGene_gn=AADAT;refGene_tx=XM_023252671.1
B1 60591723 60591788
refGene_gn=AADAT;refGene_tx=XM_003984863.4;refGene_gn=AADAT;refGene_tx=XM_006930777.4;refGene_gn=AADAT;refGene_tx=XM_019828714.2;refGene_gn=AADAT;refGene_tx=XM_023252669.1;refGene_gn=AADAT;refGene_tx=XM_023252670.1;refGene_gn=AADAT;refGene_tx=XM_023252671.1
B1 60593086 60593193
refGene_gn=AADAT;refGene_tx=XM_003984863.4;refGene_gn=AADAT;refGene_tx=XM_006930777.4;refGene_gn=AADAT;refGene_tx=XM_019828714.2;refGene_gn=AADAT;refGene_tx=XM_023252669.1;refGene_gn=AADAT;refGene_tx=XM_023252670.1;refGene_gn=AADAT;refGene_tx=XM_023252671.1
B1 60595669 60595771
refGene_gn=AADAT;refGene_tx=XM_003984863.4;refGene_gn=AADAT;refGene_tx=XM_006930777.4;refGene_gn=AADAT;refGene_tx=XM_019828714.2;refGene_gn=AADAT;refGene_tx=XM_023252669.1;refGene_gn=AADAT;refGene_tx=XM_023252670.1;refGene_gn=AADAT;refGene_tx=XM_023252671.1
B1 60596726 60596768 refGene_gn=AADAT;refGene_tx=XM_003984863.4
B1 60601834 60601882 refGene_gn=AADAT;refGene_tx=XM_019828714.2
B1 60605676 60605760
refGene_gn=AADAT;refGene_tx=XM_006930777.4;refGene_gn=AADAT;refGene_tx=XM_023252669.1;refGene_gn=AADAT;refGene_tx=XM_023252670.1;refGene_gn=AADAT;refGene_tx=XM_023252671.1
B1 60648575 60648864
refGene_gn=MFAP3L;refGene_tx=XM_019828718.1;refGene_gn=MFAP3L;refGene_tx=XM_023252672.1;refGene_gn=MFAP3L;refGene_tx=XM_023252673.1;refGene_gn=MFAP3L;refGene_tx=XM_023252674.1;refGene_gn=MFAP3L;refGene_tx=XM_023252675.1;refGene_gn=MFAP3L;refGene_tx=XM_023252676.1
B1 60661679 60662512
refGene_gn=MFAP3L;refGene_tx=XM_019828718.1;refGene_gn=MFAP3L;refGene_tx=XM_023252672.1;refGene_gn=MFAP3L;refGene_tx=XM_023252673.1;refGene_gn=MFAP3L;refGene_tx=XM_023252674.1;refGene_gn=MFAP3L;refGene_tx=XM_023252675.1;refGene_gn=MFAP3L;refGene_tx=XM_023252676.1
B1 60669088 60669121 refGene_gn=MFAP3L;refGene_tx=XM_023252676.1
B1 60681791 60681896
refGene_gn=MFAP3L;refGene_tx=XM_019828718.1;refGene_gn=MFAP3L;refGene_tx=XM_023252672.1;refGene_gn=MFAP3L;refGene_tx=XM_023252673.1;refGene_gn=MFAP3L;refGene_tx=XM_023252674.1
B1 60715577 60715641 refGene_gn=MFAP3L;refGene_tx=XM_023252675.1
B1 60758778 60758783 refGene_gn=MFAP3L;refGene_tx=XM_023252675.1
B1 60888754 60888802
refGene_gn=LOC101092264;refGene_tx=XM_003984866.5
B1 60893490 60893650
refGene_gn=LOC101092264;refGene_tx=XM_003984866.5
B1 60895897 60896087
refGene_gn=LOC101092264;refGene_tx=XM_003984866.5
B1 60897249 60897348
refGene_gn=LOC101092264;refGene_tx=XM_003984866.5

B1 60905792 60905943
refGene_gn=LOC101092264;refGene_tx=XM_003984866.5
B1 60911096 60911184
refGene_gn=LOC101092264;refGene_tx=XM_003984866.5
B1 60915866 60916039
refGene_gn=LOC101092264;refGene_tx=XM_003984866.5
B1 60918242 60918374
refGene_gn=LOC101092264;refGene_tx=XM_003984866.5
B1 60927143 60927302
refGene_gn=CLCN3;refGene_tx=XM_003984867.5;refGene_gn=CLCN3;refGene_tx=XM_011281581.3;refGene_gn=CLCN3;refGene_tx=XM_011281582.3;refGene_gn=CLCN3;refGene_tx=XM_011281583.3
B1 60929034 60929110
refGene_gn=CLCN3;refGene_tx=XM_003984867.5;refGene_gn=CLCN3;refGene_tx=XM_011281581.3
B1 60934103 60934320
refGene_gn=CLCN3;refGene_tx=XM_003984867.5;refGene_gn=CLCN3;refGene_tx=XM_011281581.3;refGene_gn=CLCN3;refGene_tx=XM_011281582.3;refGene_gn=CLCN3;refGene_tx=XM_011281583.3
B1 60941063 60941462
refGene_gn=CLCN3;refGene_tx=XM_003984867.5;refGene_gn=CLCN3;refGene_tx=XM_011281581.3;refGene_gn=CLCN3;refGene_tx=XM_011281582.3;refGene_gn=CLCN3;refGene_tx=XM_011281583.3
B1 60944292 60944479
refGene_gn=CLCN3;refGene_tx=XM_003984867.5;refGene_gn=CLCN3;refGene_tx=XM_011281581.3;refGene_gn=CLCN3;refGene_tx=XM_011281582.3;refGene_gn=CLCN3;refGene_tx=XM_011281583.3
B1 60947420 60947966
refGene_gn=CLCN3;refGene_tx=XM_003984867.5;refGene_gn=CLCN3;refGene_tx=XM_011281581.3;refGene_gn=CLCN3;refGene_tx=XM_011281582.3;refGene_gn=CLCN3;refGene_tx=XM_011281583.3
B1 60948865 60948946
refGene_gn=CLCN3;refGene_tx=XM_003984867.5;refGene_gn=CLCN3;refGene_tx=XM_011281581.3;refGene_gn=CLCN3;refGene_tx=XM_011281582.3;refGene_gn=CLCN3;refGene_tx=XM_011281583.3
B1 60951360 60951567
refGene_gn=CLCN3;refGene_tx=XM_003984867.5;refGene_gn=CLCN3;refGene_tx=XM_011281581.3;refGene_gn=CLCN3;refGene_tx=XM_011281582.3;refGene_gn=CLCN3;refGene_tx=XM_011281583.3
B1 60953397 60953520
refGene_gn=CLCN3;refGene_tx=XM_003984867.5;refGene_gn=CLCN3;refGene_tx=XM_011281581.3;refGene_gn=CLCN3;refGene_tx=XM_011281582.3;refGene_gn=CLCN3;refGene_tx=XM_011281583.3
B1 60954500 60954688
refGene_gn=CLCN3;refGene_tx=XM_003984867.5;refGene_gn=CLCN3;refGene_tx=XM_011281581.3;refGene_gn=CLCN3;refGene_tx=XM_011281582.3;refGene_gn=CLCN3;refGene_tx=XM_011281583.3
B1 60956014 60956114
refGene_gn=CLCN3;refGene_tx=XM_003984867.5;refGene_gn=CLCN3;refGene_tx=XM_011281581.3;refGene_gn=CLCN3;refGene_tx=XM_011281582.3;refGene_gn=

CLCN3;refGene_tx=XM_011281583.3
B1 60964136 60964294
refGene_gn=CLCN3;refGene_tx=XM_003984867.5;refGene_gn=CLCN3;refGene_tx=XM_011281581.3;refGene_gn=CLCN3;refGene_tx=XM_011281582.3;refGene_gn=CLCN3;refGene_tx=XM_011281583.3
B1 60981107 60981186
refGene_gn=CLCN3;refGene_tx=XM_011281581.3;refGene_gn=CLCN3;refGene_tx=XM_011281583.3
B1 61009344 61009504
refGene_gn=CLCN3;refGene_tx=XM_003984867.5;refGene_gn=CLCN3;refGene_tx=XM_011281582.3
B1 61041757 61041874
refGene_gn=NEK1;refGene_tx=XM_019828725.2;refGene_gn=NEK1;refGene_tx=XM_019828726.2;refGene_gn=NEK1;refGene_tx=XM_023252678.1;refGene_gn=NEK1;refGene_tx=XM_023252679.1;refGene_gn=NEK1;refGene_tx=XM_023252680.1;refGene_gn=NEK1;refGene_tx=XM_023252681.1
B1 61042304 61042401
refGene_gn=NEK1;refGene_tx=XM_019828725.2;refGene_gn=NEK1;refGene_tx=XM_019828726.2;refGene_gn=NEK1;refGene_tx=XM_023252678.1;refGene_gn=NEK1;refGene_tx=XM_023252679.1;refGene_gn=NEK1;refGene_tx=XM_023252680.1;refGene_gn=NEK1;refGene_tx=XM_023252681.1
B1 61058475 61058573
refGene_gn=NEK1;refGene_tx=XM_019828725.2;refGene_gn=NEK1;refGene_tx=XM_019828726.2;refGene_gn=NEK1;refGene_tx=XM_023252678.1;refGene_gn=NEK1;refGene_tx=XM_023252679.1;refGene_gn=NEK1;refGene_tx=XM_023252680.1;refGene_gn=NEK1;refGene_tx=XM_023252681.1
B1 61066909 61066993
refGene_gn=NEK1;refGene_tx=XM_019828725.2;refGene_gn=NEK1;refGene_tx=XM_019828726.2;refGene_gn=NEK1;refGene_tx=XM_023252678.1;refGene_gn=NEK1;refGene_tx=XM_023252679.1;refGene_gn=NEK1;refGene_tx=XM_023252680.1;refGene_gn=NEK1;refGene_tx=XM_023252681.1
B1 61069949 61070017
refGene_gn=NEK1;refGene_tx=XM_019828725.2;refGene_gn=NEK1;refGene_tx=XM_019828726.2;refGene_gn=NEK1;refGene_tx=XM_023252678.1;refGene_gn=NEK1;refGene_tx=XM_023252679.1;refGene_gn=NEK1;refGene_tx=XM_023252680.1;refGene_gn=NEK1;refGene_tx=XM_023252681.1
B1 61070133 61070220
refGene_gn=NEK1;refGene_tx=XM_019828725.2;refGene_gn=NEK1;refGene_tx=XM_019828726.2;refGene_gn=NEK1;refGene_tx=XM_023252678.1;refGene_gn=NEK1;refGene_tx=XM_023252679.1;refGene_gn=NEK1;refGene_tx=XM_023252680.1;refGene_gn=NEK1;refGene_tx=XM_023252681.1
B1 61073553 61073608
refGene_gn=NEK1;refGene_tx=XM_019828725.2;refGene_gn=NEK1;refGene_tx=XM_019828726.2;refGene_gn=NEK1;refGene_tx=XM_023252678.1;refGene_gn=NEK1;refGene_tx=XM_023252679.1;refGene_gn=NEK1;refGene_tx=XM_023252680.1;refGene_gn=NEK1;refGene_tx=XM_023252681.1
B1 61076158 61076359
refGene_gn=NEK1;refGene_tx=XM_019828725.2;refGene_gn=NEK1;refGene_tx=XM_019828726.2;refGene_gn=NEK1;refGene_tx=XM_023252678.1;refGene_gn=NEK1;refGene_tx=XM_023252679.1;refGene_gn=NEK1;refGene_tx=XM_023252680.1;

refGene_gn=NEK1;refGene_tx=XM_023252681.1
B1 61082642 61082703
refGene_gn=NEK1;refGene_tx=XM_019828725.2;refGene_gn=NEK1;refGene_tx=XM_019828726.2;refGene_gn=NEK1;refGene_tx=XM_023252678.1;refGene_gn=NEK1;refGene_tx=XM_023252679.1;refGene_gn=NEK1;refGene_tx=XM_023252680.1;refGene_gn=NEK1;refGene_tx=XM_023252681.1
B1 61084952 61085104
refGene_gn=NEK1;refGene_tx=XM_019828725.2;refGene_gn=NEK1;refGene_tx=XM_019828726.2;refGene_gn=NEK1;refGene_tx=XM_023252678.1;refGene_gn=NEK1;refGene_tx=XM_023252679.1;refGene_gn=NEK1;refGene_tx=XM_023252680.1;refGene_gn=NEK1;refGene_tx=XM_023252681.1
B1 61100422 61100482
refGene_gn=NEK1;refGene_tx=XM_019828725.2;refGene_gn=NEK1;refGene_tx=XM_019828726.2;refGene_gn=NEK1;refGene_tx=XM_023252678.1;refGene_gn=NEK1;refGene_tx=XM_023252679.1;refGene_gn=NEK1;refGene_tx=XM_023252680.1;refGene_gn=NEK1;refGene_tx=XM_023252681.1
B1 61100734 61100794
refGene_gn=NEK1;refGene_tx=XM_019828725.2;refGene_gn=NEK1;refGene_tx=XM_019828726.2;refGene_gn=NEK1;refGene_tx=XM_023252678.1;refGene_gn=NEK1;refGene_tx=XM_023252679.1;refGene_gn=NEK1;refGene_tx=XM_023252680.1;refGene_gn=NEK1;refGene_tx=XM_023252681.1
B1 61100888 61100936
refGene_gn=NEK1;refGene_tx=XM_019828725.2;refGene_gn=NEK1;refGene_tx=XM_019828726.2;refGene_gn=NEK1;refGene_tx=XM_023252678.1;refGene_gn=NEK1;refGene_tx=XM_023252679.1;refGene_gn=NEK1;refGene_tx=XM_023252680.1;refGene_gn=NEK1;refGene_tx=XM_023252681.1
B1 61101098 61101173
refGene_gn=NEK1;refGene_tx=XM_019828725.2;refGene_gn=NEK1;refGene_tx=XM_019828726.2;refGene_gn=NEK1;refGene_tx=XM_023252678.1;refGene_gn=NEK1;refGene_tx=XM_023252679.1;refGene_gn=NEK1;refGene_tx=XM_023252680.1;refGene_gn=NEK1;refGene_tx=XM_023252681.1
B1 61106767 61106931
refGene_gn=NEK1;refGene_tx=XM_019828725.2;refGene_gn=NEK1;refGene_tx=XM_019828726.2;refGene_gn=NEK1;refGene_tx=XM_023252678.1;refGene_gn=NEK1;refGene_tx=XM_023252679.1;refGene_gn=NEK1;refGene_tx=XM_023252680.1;refGene_gn=NEK1;refGene_tx=XM_023252681.1
B1 61127058 61127161
refGene_gn=NEK1;refGene_tx=XM_019828725.2;refGene_gn=NEK1;refGene_tx=XM_019828726.2;refGene_gn=NEK1;refGene_tx=XM_023252678.1;refGene_gn=NEK1;refGene_tx=XM_023252679.1;refGene_gn=NEK1;refGene_tx=XM_023252680.1;refGene_gn=NEK1;refGene_tx=XM_023252681.1
B1 61155703 61155787
refGene_gn=NEK1;refGene_tx=XM_023252678.1;refGene_gn=NEK1;refGene_tx=XM_023252679.1;refGene_gn=NEK1;refGene_tx=XM_023252680.1;refGene_gn=NEK1;refGene_tx=XM_023252681.1
B1 61156215 61156299
refGene_gn=NEK1;refGene_tx=XM_019828725.2;refGene_gn=NEK1;refGene_tx=XM_019828726.2;refGene_gn=NEK1;refGene_tx=XM_023252678.1;refGene_gn=NEK1;refGene_tx=XM_023252679.1;refGene_gn=NEK1;refGene_tx=XM_023252680.1;refGene_gn=NEK1;refGene_tx=XM_023252681.1

B1 61156765 61156843
refGene_gn=NEK1;refGene_tx=XM_019828725.2;refGene_gn=NEK1;refGene_tx=XM_019828726.2;refGene_gn=NEK1;refGene_tx=XM_023252678.1;refGene_gn=NEK1;refGene_tx=XM_023252679.1;refGene_gn=NEK1;refGene_tx=XM_023252680.1;refGene_gn=NEK1;refGene_tx=XM_023252681.1
B1 61157415 61157511
refGene_gn=NEK1;refGene_tx=XM_019828725.2;refGene_gn=NEK1;refGene_tx=XM_019828726.2;refGene_gn=NEK1;refGene_tx=XM_023252678.1;refGene_gn=NEK1;refGene_tx=XM_023252679.1;refGene_gn=NEK1;refGene_tx=XM_023252680.1;refGene_gn=NEK1;refGene_tx=XM_023252681.1
B1 61173364 61173496
refGene_gn=NEK1;refGene_tx=XM_019828725.2;refGene_gn=NEK1;refGene_tx=XM_019828726.2;refGene_gn=NEK1;refGene_tx=XM_023252678.1;refGene_gn=NEK1;refGene_tx=XM_023252679.1;refGene_gn=NEK1;refGene_tx=XM_023252680.1;refGene_gn=NEK1;refGene_tx=XM_023252681.1
B1 61175385 61175451
refGene_gn=NEK1;refGene_tx=XM_019828725.2;refGene_gn=NEK1;refGene_tx=XM_019828726.2;refGene_gn=NEK1;refGene_tx=XM_023252678.1;refGene_gn=NEK1;refGene_tx=XM_023252679.1;refGene_gn=NEK1;refGene_tx=XM_023252680.1;refGene_gn=NEK1;refGene_tx=XM_023252681.1
B1 61175530 61175759
refGene_gn=NEK1;refGene_tx=XM_019828725.2;refGene_gn=NEK1;refGene_tx=XM_019828726.2;refGene_gn=NEK1;refGene_tx=XM_023252678.1;refGene_gn=NEK1;refGene_tx=XM_023252679.1;refGene_gn=NEK1;refGene_tx=XM_023252680.1;refGene_gn=NEK1;refGene_tx=XM_023252681.1
B1 61191810 61191963
refGene_gn=NEK1;refGene_tx=XM_019828725.2;refGene_gn=NEK1;refGene_tx=XM_019828726.2;refGene_gn=NEK1;refGene_tx=XM_023252678.1;refGene_gn=NEK1;refGene_tx=XM_023252679.1;refGene_gn=NEK1;refGene_tx=XM_023252680.1
B1 61208325 61208514
refGene_gn=NEK1;refGene_tx=XM_019828725.2;refGene_gn=NEK1;refGene_tx=XM_019828726.2;refGene_gn=NEK1;refGene_tx=XM_023252678.1;refGene_gn=NEK1;refGene_tx=XM_023252679.1;refGene_gn=NEK1;refGene_tx=XM_023252680.1;refGene_gn=NEK1;refGene_tx=XM_023252681.1
B1 61211841 61211962
refGene_gn=NEK1;refGene_tx=XM_019828725.2;refGene_gn=NEK1;refGene_tx=XM_019828726.2;refGene_gn=NEK1;refGene_tx=XM_023252678.1;refGene_gn=NEK1;refGene_tx=XM_023252679.1;refGene_gn=NEK1;refGene_tx=XM_023252680.1;refGene_gn=NEK1;refGene_tx=XM_023252681.1
B1 61219038 61219127
refGene_gn=NEK1;refGene_tx=XM_019828725.2;refGene_gn=NEK1;refGene_tx=XM_019828726.2;refGene_gn=NEK1;refGene_tx=XM_023252678.1;refGene_gn=NEK1;refGene_tx=XM_023252679.1;refGene_gn=NEK1;refGene_tx=XM_023252680.1;refGene_gn=NEK1;refGene_tx=XM_023252681.1
B1 61220032 61220280
refGene_gn=NEK1;refGene_tx=XM_019828725.2;refGene_gn=NEK1;refGene_tx=XM_019828726.2;refGene_gn=NEK1;refGene_tx=XM_023252678.1;refGene_gn=NEK1;refGene_tx=XM_023252679.1;refGene_gn=NEK1;refGene_tx=XM_023252680.1;refGene_gn=NEK1;refGene_tx=XM_023252681.1
B1 61228389 61228541

refGene_gn=NEK1;refGene_tx=XM_019828725.2;refGene_gn=NEK1;refGene_tx=XM_019828726.2;refGene_gn=NEK1;refGene_tx=XM_023252678.1;refGene_gn=NEK1;refGene_tx=XM_023252679.1;refGene_gn=NEK1;refGene_tx=XM_023252680.1;refGene_gn=NEK1;refGene_tx=XM_023252681.1
B1 61231762 61231971
refGene_gn=NEK1;refGene_tx=XM_019828725.2;refGene_gn=NEK1;refGene_tx=XM_019828726.2;refGene_gn=NEK1;refGene_tx=XM_023252678.1;refGene_gn=NEK1;refGene_tx=XM_023252679.1;refGene_gn=NEK1;refGene_tx=XM_023252680.1;refGene_gn=NEK1;refGene_tx=XM_023252681.1
B1 61233691 61233822
refGene_gn=NEK1;refGene_tx=XM_019828725.2;refGene_gn=NEK1;refGene_tx=XM_019828726.2;refGene_gn=NEK1;refGene_tx=XM_023252678.1;refGene_gn=NEK1;refGene_tx=XM_023252679.1;refGene_gn=NEK1;refGene_tx=XM_023252680.1;refGene_gn=NEK1;refGene_tx=XM_023252681.1
B1 61234099 61234232
refGene_gn=NEK1;refGene_tx=XM_019828725.2;refGene_gn=NEK1;refGene_tx=XM_019828726.2;refGene_gn=NEK1;refGene_tx=XM_023252678.1;refGene_gn=NEK1;refGene_tx=XM_023252679.1;refGene_gn=NEK1;refGene_tx=XM_023252680.1;refGene_gn=NEK1;refGene_tx=XM_023252681.1
B1 61249680 61249735
refGene_gn=NEK1;refGene_tx=XM_019828725.2;refGene_gn=NEK1;refGene_tx=XM_019828726.2;refGene_gn=NEK1;refGene_tx=XM_023252678.1;refGene_gn=NEK1;refGene_tx=XM_023252679.1;refGene_gn=NEK1;refGene_tx=XM_023252681.1
B1 61258887 61258901
refGene_gn=NEK1;refGene_tx=XM_019828725.2;refGene_gn=NEK1;refGene_tx=XM_019828726.2;refGene_gn=NEK1;refGene_tx=XM_023252678.1;refGene_gn=NEK1;refGene_tx=XM_023252679.1;refGene_gn=NEK1;refGene_tx=XM_023252680.1;refGene_gn=NEK1;refGene_tx=XM_023252681.1
B1 61359870 61360263 refGene_gn=SH3RF1;refGene_tx=XM_023252683.1
B1 61472744 61473020 refGene_gn=SH3RF1;refGene_tx=XM_023252683.1
B1 61474115 61474211 refGene_gn=SH3RF1;refGene_tx=XM_023252683.1
B1 61499930 61500233 refGene_gn=SH3RF1;refGene_tx=XM_023252683.1
B1 61505116 61505227 refGene_gn=SH3RF1;refGene_tx=XM_023252683.1
B1 61513306 61513476 refGene_gn=SH3RF1;refGene_tx=XM_023252683.1
B1 61514652 61514823 refGene_gn=SH3RF1;refGene_tx=XM_023252683.1
B1 61517254 61517514 refGene_gn=SH3RF1;refGene_tx=XM_023252683.1
B1 61518447 61518809 refGene_gn=SH3RF1;refGene_tx=XM_023252683.1
B1 61528343 61528702 refGene_gn=SH3RF1;refGene_tx=XM_023252683.1
B1 61537375 61537544 refGene_gn=SH3RF1;refGene_tx=XM_023252683.1
B1 61634031 61634173
refGene_gn=CBR4;refGene_tx=XM_003984869.5;refGene_gn=CBR4;refGene_tx=XM_019828728.2;refGene_gn=CBR4;refGene_tx=XM_023252688.1
B1 61635351 61635364 refGene_gn=CBR4;refGene_tx=XM_019828729.2
B1 61636465 61636586
refGene_gn=CBR4;refGene_tx=XM_003984869.5;refGene_gn=CBR4;refGene_tx=XM_019828728.2;refGene_gn=CBR4;refGene_tx=XM_019828729.2;refGene_gn=CBR4;refGene_tx=XM_023252688.1
B1 61637227 61637364
refGene_gn=CBR4;refGene_tx=XM_003984869.5;refGene_gn=CBR4;refGene_tx=XM_019828728.2;refGene_gn=CBR4;refGene_tx=XM_019828729.2;refGene_gn=CBR

4;refGene_tx=XM_023252688.1
B1 61640168 61640303
refGene_gn=CBR4;refGene_tx=XM_003984869.5;refGene_gn=CBR4;refGene_tx=XM_019828728.2;refGene_gn=CBR4;refGene_tx=XM_019828729.2
B1 61655623 61655802
refGene_gn=CBR4;refGene_tx=XM_003984869.5;refGene_gn=CBR4;refGene_tx=XM_019828729.2;refGene_gn=CBR4;refGene_tx=XM_023252688.1
B1 61713928 61714101
refGene_gn=PALLD;refGene_tx=XM_023252684.1;refGene_gn=PALLD;refGene_tx=XM_023252685.1
B1 61715079 61715093 refGene_gn=PALLD;refGene_tx=XM_023252686.1
B1 61715253 61715387
refGene_gn=PALLD;refGene_tx=XM_023252684.1;refGene_gn=PALLD;refGene_tx=XM_023252685.1;refGene_gn=PALLD;refGene_tx=XM_023252686.1
B1 61715915 61716081
refGene_gn=PALLD;refGene_tx=XM_023252684.1;refGene_gn=PALLD;refGene_tx=XM_023252685.1;refGene_gn=PALLD;refGene_tx=XM_023252686.1
B1 61718886 61719094
refGene_gn=PALLD;refGene_tx=XM_023252684.1;refGene_gn=PALLD;refGene_tx=XM_023252685.1;refGene_gn=PALLD;refGene_tx=XM_023252686.1
B1 61721825 61721958
refGene_gn=PALLD;refGene_tx=XM_023252684.1;refGene_gn=PALLD;refGene_tx=XM_023252685.1;refGene_gn=PALLD;refGene_tx=XM_023252686.1
B1 61723854 61723949
refGene_gn=PALLD;refGene_tx=XM_023252684.1;refGene_gn=PALLD;refGene_tx=XM_023252685.1;refGene_gn=PALLD;refGene_tx=XM_023252686.1
B1 61732829 61732951 refGene_gn=CBR4;refGene_tx=XM_019828728.2
B1 61733137 61733287
refGene_gn=PALLD;refGene_tx=XM_023252684.1;refGene_gn=PALLD;refGene_tx=XM_023252685.1;refGene_gn=PALLD;refGene_tx=XM_023252686.1
B1 61737198 61737420
refGene_gn=PALLD;refGene_tx=XM_023252684.1;refGene_gn=PALLD;refGene_tx=XM_023252685.1;refGene_gn=PALLD;refGene_tx=XM_023252686.1
B1 61738575 61738626
refGene_gn=PALLD;refGene_tx=XM_023252684.1;refGene_gn=PALLD;refGene_tx=XM_023252686.1
B1 61739600 61739699
refGene_gn=PALLD;refGene_tx=XM_023252684.1;refGene_gn=PALLD;refGene_tx=XM_023252685.1;refGene_gn=PALLD;refGene_tx=XM_023252686.1
B1 61743369 61743505
refGene_gn=PALLD;refGene_tx=XM_023252684.1;refGene_gn=PALLD;refGene_tx=XM_023252685.1;refGene_gn=PALLD;refGene_tx=XM_023252686.1
B1 61753999 61754680
refGene_gn=PALLD;refGene_tx=XM_023252684.1;refGene_gn=PALLD;refGene_tx=XM_023252685.1;refGene_gn=PALLD;refGene_tx=XM_023252686.1
B1 61910990 61911333
refGene_gn=PALLD;refGene_tx=XM_023252684.1;refGene_gn=PALLD;refGene_tx=XM_023252685.1;refGene_gn=PALLD;refGene_tx=XM_023252686.1
B1 61913548 61913668
refGene_gn=PALLD;refGene_tx=XM_023252684.1;refGene_gn=PALLD;refGene_tx

=XM_023252685.1;refGene_gn=PALLD;refGene_tx=XM_023252686.1
B1	61929573	61929597
refGene_gn=PALLD;refGene_tx=XM_023252684.1;refGene_gn=PALLD;refGene_tx=XM_023252685.1;refGene_gn=PALLD;refGene_tx=XM_023252686.1
B1	61930122	61930264
refGene_gn=PALLD;refGene_tx=XM_023252684.1;refGene_gn=PALLD;refGene_tx=XM_023252685.1;refGene_gn=PALLD;refGene_tx=XM_023252686.1
B1	61934584	61934659
refGene_gn=PALLD;refGene_tx=XM_023252684.1;refGene_gn=PALLD;refGene_tx=XM_023252685.1;refGene_gn=PALLD;refGene_tx=XM_023252686.1
B1	61937547	61937653
refGene_gn=PALLD;refGene_tx=XM_023252684.1;refGene_gn=PALLD;refGene_tx=XM_023252685.1;refGene_gn=PALLD;refGene_tx=XM_023252686.1
B1	61939123	61939190
refGene_gn=PALLD;refGene_tx=XM_023252684.1;refGene_gn=PALLD;refGene_tx=XM_023252685.1;refGene_gn=PALLD;refGene_tx=XM_023252686.1
B1	61949067	61949246
refGene_gn=PALLD;refGene_tx=XM_023252684.1;refGene_gn=PALLD;refGene_tx=XM_023252685.1;refGene_gn=PALLD;refGene_tx=XM_023252686.1
B1	62099489	62100400
refGene_gn=PALLD;refGene_tx=XM_023252684.1;refGene_gn=PALLD;refGene_tx=XM_023252685.1;refGene_gn=PALLD;refGene_tx=XM_023252686.1
B1	62206351	62206355
refGene_gn=DDX60;refGene_tx=XM_006930786.4;refGene_gn=DDX60;refGene_tx=XM_019828730.2;refGene_gn=DDX60;refGene_tx=XM_019828731.2;refGene_gn=DDX60;refGene_tx=XM_019828733.2
B1	62206487	62206557
refGene_gn=DDX60;refGene_tx=XM_006930786.4;refGene_gn=DDX60;refGene_tx=XM_019828730.2;refGene_gn=DDX60;refGene_tx=XM_019828731.2;refGene_gn=DDX60;refGene_tx=XM_019828733.2
B1	62207471	62207661
refGene_gn=DDX60;refGene_tx=XM_006930786.4;refGene_gn=DDX60;refGene_tx=XM_019828730.2;refGene_gn=DDX60;refGene_tx=XM_019828731.2;refGene_gn=DDX60;refGene_tx=XM_019828733.2
B1	62208726	62209068
refGene_gn=DDX60;refGene_tx=XM_006930786.4;refGene_gn=DDX60;refGene_tx=XM_019828730.2;refGene_gn=DDX60;refGene_tx=XM_019828731.2;refGene_gn=DDX60;refGene_tx=XM_019828733.2
B1	62211059	62211176
refGene_gn=DDX60;refGene_tx=XM_006930786.4;refGene_gn=DDX60;refGene_tx=XM_019828730.2;refGene_gn=DDX60;refGene_tx=XM_019828731.2
B1	62213714	62213876
refGene_gn=DDX60;refGene_tx=XM_006930786.4;refGene_gn=DDX60;refGene_tx=XM_019828730.2;refGene_gn=DDX60;refGene_tx=XM_019828731.2;refGene_gn=DDX60;refGene_tx=XM_019828733.2
B1	62216158	62216320
refGene_gn=DDX60;refGene_tx=XM_006930786.4;refGene_gn=DDX60;refGene_tx=XM_019828730.2;refGene_gn=DDX60;refGene_tx=XM_019828731.2;refGene_gn=DDX60;refGene_tx=XM_019828733.2
B1	62223744	62223886

refGene_gn=DDX60;refGene_tx=XM_006930786.4;refGene_gn=DDX60;refGene_tx=XM_019828730.2;refGene_gn=DDX60;refGene_tx=XM_019828731.2;refGene_gn=DDX60;refGene_tx=XM_019828733.2
B1 62224845 62225001
refGene_gn=DDX60;refGene_tx=XM_006930786.4;refGene_gn=DDX60;refGene_tx=XM_019828730.2;refGene_gn=DDX60;refGene_tx=XM_019828731.2;refGene_gn=DDX60;refGene_tx=XM_019828733.2
B1 62228943 62229049
refGene_gn=DDX60;refGene_tx=XM_006930786.4;refGene_gn=DDX60;refGene_tx=XM_019828730.2;refGene_gn=DDX60;refGene_tx=XM_019828731.2;refGene_gn=DDX60;refGene_tx=XM_019828733.2
B1 62229997 62230113
refGene_gn=DDX60;refGene_tx=XM_006930786.4;refGene_gn=DDX60;refGene_tx=XM_019828730.2;refGene_gn=DDX60;refGene_tx=XM_019828731.2;refGene_gn=DDX60;refGene_tx=XM_019828733.2
B1 62231598 62231759
refGene_gn=DDX60;refGene_tx=XM_006930786.4;refGene_gn=DDX60;refGene_tx=XM_019828730.2;refGene_gn=DDX60;refGene_tx=XM_019828731.2;refGene_gn=DDX60;refGene_tx=XM_019828733.2
B1 62235204 62235457
refGene_gn=DDX60;refGene_tx=XM_006930786.4;refGene_gn=DDX60;refGene_tx=XM_019828730.2;refGene_gn=DDX60;refGene_tx=XM_019828731.2;refGene_gn=DDX60;refGene_tx=XM_019828733.2
B1 62237021 62237188
refGene_gn=DDX60;refGene_tx=XM_006930786.4;refGene_gn=DDX60;refGene_tx=XM_019828730.2;refGene_gn=DDX60;refGene_tx=XM_019828731.2;refGene_gn=DDX60;refGene_tx=XM_019828733.2
B1 62237734 62237893
refGene_gn=DDX60;refGene_tx=XM_006930786.4;refGene_gn=DDX60;refGene_tx=XM_019828730.2;refGene_gn=DDX60;refGene_tx=XM_019828731.2;refGene_gn=DDX60;refGene_tx=XM_019828733.2
B1 62239669 62239819
refGene_gn=DDX60;refGene_tx=XM_006930786.4;refGene_gn=DDX60;refGene_tx=XM_019828730.2;refGene_gn=DDX60;refGene_tx=XM_019828731.2;refGene_gn=DDX60;refGene_tx=XM_019828733.2
B1 62242827 62242947
refGene_gn=DDX60;refGene_tx=XM_006930786.4;refGene_gn=DDX60;refGene_tx=XM_019828730.2;refGene_gn=DDX60;refGene_tx=XM_019828731.2;refGene_gn=DDX60;refGene_tx=XM_019828733.2
B1 62244099 62244195
refGene_gn=DDX60;refGene_tx=XM_006930786.4;refGene_gn=DDX60;refGene_tx=XM_019828730.2;refGene_gn=DDX60;refGene_tx=XM_019828731.2;refGene_gn=DDX60;refGene_tx=XM_019828733.2
B1 62245792 62245908
refGene_gn=DDX60;refGene_tx=XM_006930786.4;refGene_gn=DDX60;refGene_tx=XM_019828730.2;refGene_gn=DDX60;refGene_tx=XM_019828731.2;refGene_gn=DDX60;refGene_tx=XM_019828733.2
B1 62246972 62247103
refGene_gn=DDX60;refGene_tx=XM_006930786.4;refGene_gn=DDX60;refGene_tx=XM_019828730.2;refGene_gn=DDX60;refGene_tx=XM_019828731.2;refGene_gn=

DDX60;refGene_tx=XM_019828733.2
B1 62247189 62247299
refGene_gn=DDX60;refGene_tx=XM_006930786.4;refGene_gn=DDX60;refGene_tx=XM_019828730.2;refGene_gn=DDX60;refGene_tx=XM_019828731.2;refGene_gn=DDX60;refGene_tx=XM_019828733.2
B1 62253907 62254012
refGene_gn=DDX60;refGene_tx=XM_006930786.4;refGene_gn=DDX60;refGene_tx=XM_019828730.2;refGene_gn=DDX60;refGene_tx=XM_019828731.2;refGene_gn=DDX60;refGene_tx=XM_019828733.2
B1 62254575 62254704
refGene_gn=DDX60;refGene_tx=XM_006930786.4;refGene_gn=DDX60;refGene_tx=XM_019828730.2;refGene_gn=DDX60;refGene_tx=XM_019828731.2;refGene_gn=DDX60;refGene_tx=XM_019828733.2
B1 62256000 62256125
refGene_gn=DDX60;refGene_tx=XM_006930786.4;refGene_gn=DDX60;refGene_tx=XM_019828730.2;refGene_gn=DDX60;refGene_tx=XM_019828731.2;refGene_gn=DDX60;refGene_tx=XM_019828733.2
B1 62260429 62260588
refGene_gn=DDX60;refGene_tx=XM_006930786.4;refGene_gn=DDX60;refGene_tx=XM_019828730.2;refGene_gn=DDX60;refGene_tx=XM_019828731.2;refGene_gn=DDX60;refGene_tx=XM_019828733.2
B1 62263683 62263831
refGene_gn=DDX60;refGene_tx=XM_006930786.4;refGene_gn=DDX60;refGene_tx=XM_019828730.2;refGene_gn=DDX60;refGene_tx=XM_019828731.2;refGene_gn=DDX60;refGene_tx=XM_019828733.2
B1 62266012 62266165
refGene_gn=DDX60;refGene_tx=XM_006930786.4;refGene_gn=DDX60;refGene_tx=XM_019828730.2;refGene_gn=DDX60;refGene_tx=XM_019828731.2;refGene_gn=DDX60;refGene_tx=XM_019828733.2
B1 62268332 62268437
refGene_gn=DDX60;refGene_tx=XM_006930786.4;refGene_gn=DDX60;refGene_tx=XM_019828730.2;refGene_gn=DDX60;refGene_tx=XM_019828731.2;refGene_gn=DDX60;refGene_tx=XM_019828733.2
B1 62273469 62273670
refGene_gn=DDX60;refGene_tx=XM_006930786.4;refGene_gn=DDX60;refGene_tx=XM_019828730.2;refGene_gn=DDX60;refGene_tx=XM_019828731.2;refGene_gn=DDX60;refGene_tx=XM_019828733.2
B1 62278954 62279059
refGene_gn=DDX60;refGene_tx=XM_006930786.4;refGene_gn=DDX60;refGene_tx=XM_019828730.2;refGene_gn=DDX60;refGene_tx=XM_019828731.2;refGene_gn=DDX60;refGene_tx=XM_019828733.2
B1 62279356 62279498
refGene_gn=DDX60;refGene_tx=XM_006930786.4;refGene_gn=DDX60;refGene_tx=XM_019828730.2;refGene_gn=DDX60;refGene_tx=XM_019828731.2;refGene_gn=DDX60;refGene_tx=XM_019828733.2
B1 62281350 62281472
refGene_gn=DDX60;refGene_tx=XM_006930786.4;refGene_gn=DDX60;refGene_tx=XM_019828730.2;refGene_gn=DDX60;refGene_tx=XM_019828731.2;refGene_gn=DDX60;refGene_tx=XM_019828733.2
B1 62288758 62288906

refGene_gn=DDX60;refGene_tx=XM_006930786.4;refGene_gn=DDX60;refGene_tx=XM_019828730.2;refGene_gn=DDX60;refGene_tx=XM_019828731.2;refGene_gn=DDX60;refGene_tx=XM_019828733.2
B1 62290225 62290368
refGene_gn=DDX60;refGene_tx=XM_006930786.4;refGene_gn=DDX60;refGene_tx=XM_019828730.2;refGene_gn=DDX60;refGene_tx=XM_019828731.2;refGene_gn=DDX60;refGene_tx=XM_019828733.2
B1 62294773 62294925
refGene_gn=DDX60;refGene_tx=XM_006930786.4;refGene_gn=DDX60;refGene_tx=XM_019828730.2;refGene_gn=DDX60;refGene_tx=XM_019828731.2;refGene_gn=DDX60;refGene_tx=XM_019828733.2
B1 62302448 62302511
refGene_gn=DDX60;refGene_tx=XM_006930786.4;refGene_gn=DDX60;refGene_tx=XM_019828730.2;refGene_gn=DDX60;refGene_tx=XM_019828731.2;refGene_gn=DDX60;refGene_tx=XM_019828733.2
B1 62305737 62305837
refGene_gn=DDX60;refGene_tx=XM_006930786.4;refGene_gn=DDX60;refGene_tx=XM_019828730.2;refGene_gn=DDX60;refGene_tx=XM_019828731.2;refGene_gn=DDX60;refGene_tx=XM_019828733.2
B1 62370575 62370644 refGene_gn=ANXA10;refGene_tx=XM_011281585.3
B1 62372858 62372981 refGene_gn=ANXA10;refGene_tx=XM_011281585.3
B1 62376415 62376474 refGene_gn=ANXA10;refGene_tx=XM_011281585.3
B1 62378808 62378904 refGene_gn=ANXA10;refGene_tx=XM_011281585.3
B1 62379553 62379647 refGene_gn=ANXA10;refGene_tx=XM_011281585.3
B1 62379742 62379796 refGene_gn=ANXA10;refGene_tx=XM_011281585.3
B1 62395029 62395109 refGene_gn=ANXA10;refGene_tx=XM_011281585.3
B1 62396974 62397065 refGene_gn=ANXA10;refGene_tx=XM_011281585.3
B1 62399039 62399153 refGene_gn=ANXA10;refGene_tx=XM_011281585.3
B1 62422065 62422160 refGene_gn=ANXA10;refGene_tx=XM_011281585.3
B1 62431896 62431978 refGene_gn=ANXA10;refGene_tx=XM_011281585.3
B1 62468645 62468663 refGene_gn=ANXA10;refGene_tx=XM_011281585.3
B1 62929181 62929247
refGene_gn=LOC101080450;refGene_tx=XM_023253390.1
B1 62952341 62952593
refGene_gn=LOC101080450;refGene_tx=XM_023253390.1
B1 62952595 62953147
refGene_gn=LOC101080450;refGene_tx=XM_023253390.1
B1 63299953 63300142
refGene_gn=SPOCK3;refGene_tx=XM_023252690.1;refGene_gn=SPOCK3;refGene_tx=XM_023252691.1;refGene_gn=SPOCK3;refGene_tx=XM_023252692.1;refGene_gn=SPOCK3;refGene_tx=XM_023252693.1
B1 63464136 63464145
refGene_gn=SPOCK3;refGene_tx=XM_023252690.1;refGene_gn=SPOCK3;refGene_tx=XM_023252691.1
B1 63464236 63464282
refGene_gn=SPOCK3;refGene_tx=XM_023252690.1;refGene_gn=SPOCK3;refGene_tx=XM_023252691.1;refGene_gn=SPOCK3;refGene_tx=XM_023252692.1;refGene_gn=SPOCK3;refGene_tx=XM_023252693.1
B1 63517449 63517564
refGene_gn=SPOCK3;refGene_tx=XM_023252690.1;refGene_gn=SPOCK3;refGene_

tx=XM_023252691.1;refGene_gn=SPOCK3;refGene_tx=XM_023252692.1;refGene_gn=SPOCK3;refGene_tx=XM_023252693.1
B1 63614112 63614236
refGene_gn=SPOCK3;refGene_tx=XM_023252690.1;refGene_gn=SPOCK3;refGene_tx=XM_023252691.1;refGene_gn=SPOCK3;refGene_tx=XM_023252692.1;refGene_gn=SPOCK3;refGene_tx=XM_023252693.1
B1 63634753 63634868
refGene_gn=SPOCK3;refGene_tx=XM_023252690.1;refGene_gn=SPOCK3;refGene_tx=XM_023252691.1;refGene_gn=SPOCK3;refGene_tx=XM_023252692.1;refGene_gn=SPOCK3;refGene_tx=XM_023252693.1
B1 63739949 63740069
refGene_gn=SPOCK3;refGene_tx=XM_023252690.1;refGene_gn=SPOCK3;refGene_tx=XM_023252691.1;refGene_gn=SPOCK3;refGene_tx=XM_023252692.1;refGene_gn=SPOCK3;refGene_tx=XM_023252693.1
B1 63773876 63774109
refGene_gn=SPOCK3;refGene_tx=XM_023252690.1;refGene_gn=SPOCK3;refGene_tx=XM_023252691.1;refGene_gn=SPOCK3;refGene_tx=XM_023252692.1;refGene_gn=SPOCK3;refGene_tx=XM_023252693.1
B1 63778725 63778788
refGene_gn=SPOCK3;refGene_tx=XM_023252690.1;refGene_gn=SPOCK3;refGene_tx=XM_023252691.1;refGene_gn=SPOCK3;refGene_tx=XM_023252692.1
B1 63781790 63781928
refGene_gn=SPOCK3;refGene_tx=XM_023252690.1;refGene_gn=SPOCK3;refGene_tx=XM_023252691.1;refGene_gn=SPOCK3;refGene_tx=XM_023252692.1
B1 63785153 63785323
refGene_gn=SPOCK3;refGene_tx=XM_023252690.1;refGene_gn=SPOCK3;refGene_tx=XM_023252691.1;refGene_gn=SPOCK3;refGene_tx=XM_023252692.1
B1 64316475 64316610
refGene_gn=TLL1;refGene_tx=XM_023252694.1;refGene_gn=TLL1;refGene_tx=XM_023252695.1;refGene_gn=TLL1;refGene_tx=XM_023252696.1;refGene_gn=TLL1;refGene_tx=XM_023252697.1
B1 64317817 64318068
refGene_gn=TLL1;refGene_tx=XM_023252694.1;refGene_gn=TLL1;refGene_tx=XM_023252695.1;refGene_gn=TLL1;refGene_tx=XM_023252696.1;refGene_gn=TLL1;refGene_tx=XM_023252697.1
B1 64326741 64326955
refGene_gn=TLL1;refGene_tx=XM_023252694.1;refGene_gn=TLL1;refGene_tx=XM_023252695.1;refGene_gn=TLL1;refGene_tx=XM_023252696.1;refGene_gn=TLL1;refGene_tx=XM_023252697.1
B1 64338390 64338518
refGene_gn=TLL1;refGene_tx=XM_023252694.1;refGene_gn=TLL1;refGene_tx=XM_023252695.1;refGene_gn=TLL1;refGene_tx=XM_023252696.1;refGene_gn=TLL1;refGene_tx=XM_023252697.1
B1 64340650 64340776
refGene_gn=TLL1;refGene_tx=XM_023252694.1;refGene_gn=TLL1;refGene_tx=XM_023252695.1;refGene_gn=TLL1;refGene_tx=XM_023252696.1;refGene_gn=TLL1;refGene_tx=XM_023252697.1
B1 64350687 64350868
refGene_gn=TLL1;refGene_tx=XM_023252694.1;refGene_gn=TLL1;refGene_tx=XM_023252695.1;refGene_gn=TLL1;refGene_tx=XM_023252696.1;refGene_gn=TLL

1;refGene_tx=XM_023252697.1
B1 64360356 64360517
refGene_gn=TLL1;refGene_tx=XM_023252694.1;refGene_gn=TLL1;refGene_tx=XM_023252695.1;refGene_gn=TLL1;refGene_tx=XM_023252696.1;refGene_gn=TLL1;refGene_tx=XM_023252697.1
B1 64362547 64362673
refGene_gn=TLL1;refGene_tx=XM_023252694.1;refGene_gn=TLL1;refGene_tx=XM_023252695.1;refGene_gn=TLL1;refGene_tx=XM_023252696.1;refGene_gn=TLL1;refGene_tx=XM_023252697.1
B1 64364740 64364936
refGene_gn=TLL1;refGene_tx=XM_023252694.1;refGene_gn=TLL1;refGene_tx=XM_023252695.1;refGene_gn=TLL1;refGene_tx=XM_023252696.1;refGene_gn=TLL1;refGene_tx=XM_023252697.1
B1 64377959 64378105
refGene_gn=TLL1;refGene_tx=XM_023252694.1;refGene_gn=TLL1;refGene_tx=XM_023252695.1;refGene_gn=TLL1;refGene_tx=XM_023252696.1;refGene_gn=TLL1;refGene_tx=XM_023252697.1
B1 64379275 64379392
refGene_gn=TLL1;refGene_tx=XM_023252694.1;refGene_gn=TLL1;refGene_tx=XM_023252695.1;refGene_gn=TLL1;refGene_tx=XM_023252696.1;refGene_gn=TLL1;refGene_tx=XM_023252697.1
B1 64381900 64382003
refGene_gn=TLL1;refGene_tx=XM_023252694.1;refGene_gn=TLL1;refGene_tx=XM_023252695.1;refGene_gn=TLL1;refGene_tx=XM_023252696.1;refGene_gn=TLL1;refGene_tx=XM_023252697.1
B1 64395770 64395886
refGene_gn=TLL1;refGene_tx=XM_023252694.1;refGene_gn=TLL1;refGene_tx=XM_023252695.1;refGene_gn=TLL1;refGene_tx=XM_023252696.1;refGene_gn=TLL1;refGene_tx=XM_023252697.1
B1 64406505 64406630
refGene_gn=TLL1;refGene_tx=XM_023252694.1;refGene_gn=TLL1;refGene_tx=XM_023252695.1;refGene_gn=TLL1;refGene_tx=XM_023252696.1;refGene_gn=TLL1;refGene_tx=XM_023252697.1
B1 64414858 64414964
refGene_gn=TLL1;refGene_tx=XM_023252694.1;refGene_gn=TLL1;refGene_tx=XM_023252695.1;refGene_gn=TLL1;refGene_tx=XM_023252696.1;refGene_gn=TLL1;refGene_tx=XM_023252697.1
B1 64426850 64427029
refGene_gn=TLL1;refGene_tx=XM_023252694.1;refGene_gn=TLL1;refGene_tx=XM_023252695.1;refGene_gn=TLL1;refGene_tx=XM_023252696.1;refGene_gn=TLL1;refGene_tx=XM_023252697.1
B1 64442500 64442618
refGene_gn=TLL1;refGene_tx=XM_023252694.1;refGene_gn=TLL1;refGene_tx=XM_023252695.1;refGene_gn=TLL1;refGene_tx=XM_023252696.1;refGene_gn=TLL1;refGene_tx=XM_023252697.1
B1 64443147 64443300
refGene_gn=TLL1;refGene_tx=XM_023252694.1;refGene_gn=TLL1;refGene_tx=XM_023252695.1;refGene_gn=TLL1;refGene_tx=XM_023252696.1
B1 64445035 64445116
refGene_gn=TLL1;refGene_tx=XM_023252694.1;refGene_gn=TLL1;refGene_tx=X

M_023252695.1;refGene_gn=TLL1;refGene_tx=XM_023252696.1
B1 64447217 64447325
refGene_gn=TLL1;refGene_tx=XM_023252694.1;refGene_gn=TLL1;refGene_tx=XM_023252695.1
B1 64500958 64500980 refGene_gn=TLL1;refGene_tx=XM_023252695.1
B1 64532940 64533109 refGene_gn=TLL1;refGene_tx=XM_023252694.1
B1 64906585 64906684 refGene_gn=CPE;refGene_tx=XM_011281641.3
B1 64908534 64908653 refGene_gn=CPE;refGene_tx=XM_011281641.3
B1 64911719 64911819 refGene_gn=CPE;refGene_tx=XM_011281641.3
B1 64916551 64916691 refGene_gn=CPE;refGene_tx=XM_011281641.3
B1 64927403 64927586 refGene_gn=CPE;refGene_tx=XM_011281641.3
B1 64929071 64929189 refGene_gn=CPE;refGene_tx=XM_011281641.3
B1 64938176 64938344 refGene_gn=CPE;refGene_tx=XM_011281641.3
B1 64940272 64940469 refGene_gn=CPE;refGene_tx=XM_011281641.3
B1 65031392 65031699 refGene_gn=CPE;refGene_tx=XM_011281641.3
B1 65055667 65055863
refGene_gn=MSMO1;refGene_tx=XM_003984882.5;refGene_gn=MSMO1;refGene_tx=XM_019828736.2
B1 65057234 65057389
refGene_gn=MSMO1;refGene_tx=XM_003984882.5;refGene_gn=MSMO1;refGene_tx=XM_019828736.2
B1 65058780 65058907
refGene_gn=MSMO1;refGene_tx=XM_003984882.5;refGene_gn=MSMO1;refGene_tx=XM_019828736.2
B1 65059984 65060133
refGene_gn=MSMO1;refGene_tx=XM_003984882.5;refGene_gn=MSMO1;refGene_tx=XM_019828736.2
B1 65063794 65064049
refGene_gn=MSMO1;refGene_tx=XM_003984882.5;refGene_gn=MSMO1;refGene_tx=XM_019828736.2
B1 65074827 65074856
refGene_gn=KLHL2;refGene_tx=XM_023252700.1;refGene_gn=KLHL2;refGene_tx=XM_023252701.1;refGene_gn=KLHL2;refGene_tx=XM_023252702.1
B1 65076009 65076153
refGene_gn=KLHL2;refGene_tx=XM_023252700.1;refGene_gn=KLHL2;refGene_tx=XM_023252702.1
B1 65078960 65079101
refGene_gn=KLHL2;refGene_tx=XM_023252700.1;refGene_gn=KLHL2;refGene_tx=XM_023252701.1;refGene_gn=KLHL2;refGene_tx=XM_023252702.1
B1 65079716 65079845
refGene_gn=KLHL2;refGene_tx=XM_023252700.1;refGene_gn=KLHL2;refGene_tx=XM_023252701.1;refGene_gn=KLHL2;refGene_tx=XM_023252702.1
B1 65081138 65081240
refGene_gn=KLHL2;refGene_tx=XM_023252700.1;refGene_gn=KLHL2;refGene_tx=XM_023252701.1;refGene_gn=KLHL2;refGene_tx=XM_023252702.1
B1 65081870 65082068
refGene_gn=KLHL2;refGene_tx=XM_023252700.1;refGene_gn=KLHL2;refGene_tx=XM_023252701.1;refGene_gn=KLHL2;refGene_tx=XM_023252702.1
B1 65087259 65087377
refGene_gn=KLHL2;refGene_tx=XM_023252700.1;refGene_gn=KLHL2;refGene_tx

=XM_023252701.1;refGene_gn=KLHL2;refGene_tx=XM_023252702.1
B1	65093699	65093849
refGene_gn=KLHL2;refGene_tx=XM_023252700.1;refGene_gn=KLHL2;refGene_tx=XM_023252701.1;refGene_gn=KLHL2;refGene_tx=XM_023252702.1
B1	65095061	65095178
refGene_gn=KLHL2;refGene_tx=XM_023252700.1;refGene_gn=KLHL2;refGene_tx=XM_023252701.1;refGene_gn=KLHL2;refGene_tx=XM_023252702.1
B1	65101290	65101400
refGene_gn=KLHL2;refGene_tx=XM_023252700.1;refGene_gn=KLHL2;refGene_tx=XM_023252701.1;refGene_gn=KLHL2;refGene_tx=XM_023252702.1
B1	65120925	65121088
refGene_gn=KLHL2;refGene_tx=XM_023252700.1;refGene_gn=KLHL2;refGene_tx=XM_023252701.1;refGene_gn=KLHL2;refGene_tx=XM_023252702.1
B1	65143323	65143445
refGene_gn=KLHL2;refGene_tx=XM_023252700.1;refGene_gn=KLHL2;refGene_tx=XM_023252701.1
B1	65167495	65167602
refGene_gn=KLHL2;refGene_tx=XM_023252700.1;refGene_gn=KLHL2;refGene_tx=XM_023252701.1;refGene_gn=KLHL2;refGene_tx=XM_023252702.1
B1	65178491	65178617
refGene_gn=KLHL2;refGene_tx=XM_023252700.1;refGene_gn=KLHL2;refGene_tx=XM_023252701.1
B1	65191474	65191500
refGene_gn=KLHL2;refGene_tx=XM_023252700.1;refGene_gn=KLHL2;refGene_tx=XM_023252701.1
B1	65282197	65282224
refGene_gn=TMEM192;refGene_tx=XM_003984901.5;refGene_gn=TMEM192;refGene_tx=XM_019828739.2;refGene_gn=TMEM192;refGene_tx=XM_019828740.2
B1	65291239	65291386
refGene_gn=TMEM192;refGene_tx=XM_003984901.5;refGene_gn=TMEM192;refGene_tx=XM_019828739.2;refGene_gn=TMEM192;refGene_tx=XM_019828740.2
B1	65297495	65297760
refGene_gn=TMEM192;refGene_tx=XM_003984901.5;refGene_gn=TMEM192;refGene_tx=XM_019828739.2;refGene_gn=TMEM192;refGene_tx=XM_019828740.2
B1	65303321	65303456
refGene_gn=TMEM192;refGene_tx=XM_003984901.5;refGene_gn=TMEM192;refGene_tx=XM_019828739.2;refGene_gn=TMEM192;refGene_tx=XM_019828740.2
B1	65309932	65310035
refGene_gn=TMEM192;refGene_tx=XM_003984901.5;refGene_gn=TMEM192;refGene_tx=XM_019828739.2;refGene_gn=TMEM192;refGene_tx=XM_019828740.2
B1	65310587	65310726
refGene_gn=TMEM192;refGene_tx=XM_003984901.5;refGene_gn=TMEM192;refGene_tx=XM_019828739.2
B1	65312590	65312621 refGene_gn=TMEM192;refGene_tx=XM_019828740.2
B1	65332370	65333780
refGene_gn=LOC101097889;refGene_tx=XM_019828738.2
B1	65359045	65360461
refGene_gn=LOC101098136;refGene_tx=XM_003984887.5;refGene_gn=LOC101098136;refGene_tx=XM_023252704.1
B1	65396037	65397423

refGene_gn=LOC101082488;refGene_tx=XM_003984902.1
B1 65458644 65460057
refGene_gn=LOC101082747;refGene_tx=XM_003984903.1
B1 65476327 65477749
refGene_gn=LOC101098376;refGene_tx=XM_023252225.1
B1 65513474 65513563 refGene_gn=APELA;refGene_tx=XM_011281590.3
B1 65514931 65515007 refGene_gn=APELA;refGene_tx=XM_011281590.3
B1 66500832 66500845
refGene_gn=MARCH1;refGene_tx=XM_023252706.1;refGene_gn=MARCH1;refGene_tx=XM_023252712.1
B1 66578028 66578177
refGene_gn=MARCH1;refGene_tx=XM_023252706.1;refGene_gn=MARCH1;refGene_tx=XM_023252707.1;refGene_gn=MARCH1;refGene_tx=XM_023252708.1;refGene_gn=MARCH1;refGene_tx=XM_023252712.1
B1 66725577 66725628
refGene_gn=MARCH1;refGene_tx=XM_023252706.1;refGene_gn=MARCH1;refGene_tx=XM_023252707.1;refGene_gn=MARCH1;refGene_tx=XM_023252708.1;refGene_gn=MARCH1;refGene_tx=XM_023252712.1
B1 66802538 66802843
refGene_gn=MARCH1;refGene_tx=XM_023252705.1;refGene_gn=MARCH1;refGene_tx=XM_023252706.1;refGene_gn=MARCH1;refGene_tx=XM_023252707.1;refGene_gn=MARCH1;refGene_tx=XM_023252708.1;refGene_gn=MARCH1;refGene_tx=XM_023252709.1;refGene_gn=MARCH1;refGene_tx=XM_023252711.1;refGene_gn=MARCH1;refGene_tx=XM_023252712.1
B1 66803122 66803884
refGene_gn=MARCH1;refGene_tx=XM_023252705.1;refGene_gn=MARCH1;refGene_tx=XM_023252706.1;refGene_gn=MARCH1;refGene_tx=XM_023252707.1;refGene_gn=MARCH1;refGene_tx=XM_023252708.1;refGene_gn=MARCH1;refGene_tx=XM_023252709.1;refGene_gn=MARCH1;refGene_tx=XM_023252710.1
B1 66835041 66835222
refGene_gn=MARCH1;refGene_tx=XM_023252705.1;refGene_gn=MARCH1;refGene_tx=XM_023252706.1;refGene_gn=MARCH1;refGene_tx=XM_023252707.1;refGene_gn=MARCH1;refGene_tx=XM_023252708.1;refGene_gn=MARCH1;refGene_tx=XM_023252709.1;refGene_gn=MARCH1;refGene_tx=XM_023252710.1;refGene_gn=MARCH1;refGene_tx=XM_023252711.1;refGene_gn=MARCH1;refGene_tx=XM_023252712.1
B1 66864898 66864968 refGene_gn=TMA16;refGene_tx=XM_019828756.2
B1 66875829 66875977
refGene_gn=MARCH1;refGene_tx=XM_023252705.1;refGene_gn=MARCH1;refGene_tx=XM_023252706.1;refGene_gn=MARCH1;refGene_tx=XM_023252707.1;refGene_gn=MARCH1;refGene_tx=XM_023252708.1;refGene_gn=MARCH1;refGene_tx=XM_023252709.1;refGene_gn=MARCH1;refGene_tx=XM_023252710.1;refGene_gn=MARCH1;refGene_tx=XM_023252711.1;refGene_gn=MARCH1;refGene_tx=XM_023252712.1
B1 66885467 66885523 refGene_gn=MARCH1;refGene_tx=XM_023252709.1
B1 66887926 66888225
refGene_gn=MARCH1;refGene_tx=XM_023252705.1;refGene_gn=MARCH1;refGene_tx=XM_023252706.1;refGene_gn=MARCH1;refGene_tx=XM_023252707.1;refGene_gn=MARCH1;refGene_tx=XM_023252708.1;refGene_gn=MARCH1;refGene_tx=XM_023252710.1;refGene_gn=MARCH1;refGene_tx=XM_023252711.1;refGene_gn=MARCH

1;refGene_tx=XM_023252712.1
B1 66898421 66898602 refGene_gn=TMA16;refGene_tx=XM_003984890.5
B1 66900046 66900089
refGene_gn=TMA16;refGene_tx=XM_003984890.5;refGene_gn=TMA16;refGene_tx
=XM_019828756.2
B1 66903042 66903191
refGene_gn=TMA16;refGene_tx=XM_003984890.5;refGene_gn=TMA16;refGene_tx
=XM_019828756.2
B1 66904350 66904435
refGene_gn=TMA16;refGene_tx=XM_003984890.5;refGene_gn=TMA16;refGene_tx
=XM_019828756.2
B1 66905592 66905630
refGene_gn=TMA16;refGene_tx=XM_003984890.5;refGene_gn=TMA16;refGene_tx
=XM_019828756.2
B1 66911725 66911838
refGene_gn=TMA16;refGene_tx=XM_003984890.5;refGene_gn=TMA16;refGene_tx
=XM_019828756.2
B1 66925575 66925578
refGene_gn=TMA16;refGene_tx=XM_003984890.5;refGene_gn=TMA16;refGene_tx
=XM_019828756.2
B1 66934073 66935951 refGene_gn=TKTL2;refGene_tx=XM_003984891.4
B1 66936319 66936508
refGene_gn=LOC111560221;refGene_tx=XM_023253391.1
B1 66971755 66973544
refGene_gn=LOC111560221;refGene_tx=XM_023253391.1
B1 67049408 67050749 refGene_gn=NPY5R;refGene_tx=XM_011281591.3
B1 67073899 67074595
refGene_gn=NPY1R;refGene_tx=XM_003984893.5;refGene_gn=NPY1R;refGene_tx
=XM_006930791.3;refGene_gn=NPY1R;refGene_tx=XM_006930792.4;refGene_gn=
NPY1R;refGene_tx=XM_019828758.1;refGene_gn=NPY1R;refGene_tx=XM_0198287
59.1
B1 67074699 67075152
refGene_gn=NPY1R;refGene_tx=XM_003984893.5;refGene_gn=NPY1R;refGene_tx
=XM_006930791.3;refGene_gn=NPY1R;refGene_tx=XM_006930792.4;refGene_gn=
NPY1R;refGene_tx=XM_019828758.1;refGene_gn=NPY1R;refGene_tx=XM_0198287
59.1
B1 67185641 67185970 refGene_gn=NAF1;refGene_tx=XM_003984894.5
B1 67190777 67190946 refGene_gn=NAF1;refGene_tx=XM_003984894.5
B1 67206001 67206095 refGene_gn=NAF1;refGene_tx=XM_003984894.5
B1 67209617 67209700 refGene_gn=NAF1;refGene_tx=XM_003984894.5
B1 67216488 67216649 refGene_gn=NAF1;refGene_tx=XM_003984894.5
B1 67220057 67220109 refGene_gn=NAF1;refGene_tx=XM_003984894.5
B1 67224508 67224611 refGene_gn=NAF1;refGene_tx=XM_003984894.5
B1 67228063 67228527 refGene_gn=NAF1;refGene_tx=XM_003984894.5
B1 68177945 68178071 refGene_gn=FSTL5;refGene_tx=XM_023252713.1
B1 68257318 68257352 refGene_gn=FSTL5;refGene_tx=XM_023252713.1
B1 68393858 68394107 refGene_gn=FSTL5;refGene_tx=XM_023252713.1
B1 68537650 68537847 refGene_gn=FSTL5;refGene_tx=XM_023252713.1
B1 68553209 68553330 refGene_gn=FSTL5;refGene_tx=XM_023252713.1
B1 68646236 68646403 refGene_gn=FSTL5;refGene_tx=XM_023252713.1

B1 68708519 68708640 refGene_gn=FSTL5;refGene_tx=XM_023252713.1
B1 68747463 68747625 refGene_gn=FSTL5;refGene_tx=XM_023252713.1
B1 68751989 68752124 refGene_gn=FSTL5;refGene_tx=XM_023252713.1
B1 68788306 68788333 refGene_gn=FSTL5;refGene_tx=XM_023252713.1
B1 68797353 68797472 refGene_gn=FSTL5;refGene_tx=XM_023252713.1
B1 68818709 68818859 refGene_gn=FSTL5;refGene_tx=XM_023252713.1
B1 68835708 68835795 refGene_gn=FSTL5;refGene_tx=XM_023252713.1
B1 68835796 68835817 refGene_gn=FSTL5;refGene_tx=XM_023252713.1
B1 68842036 68842161 refGene_gn=FSTL5;refGene_tx=XM_023252713.1
B1 68894043 68894746 refGene_gn=FSTL5;refGene_tx=XM_023252713.1
B1 70779813 70779839
refGene_gn=RAPGEF2;refGene_tx=XM_011281593.3;refGene_gn=RAPGEF2;refGene_tx=XM_019828768.2;refGene_gn=RAPGEF2;refGene_tx=XM_019828769.2;refGene_gn=RAPGEF2;refGene_tx=XM_019828770.2;refGene_gn=RAPGEF2;refGene_tx=XM_019828771.2;refGene_gn=RAPGEF2;refGene_tx=XM_019828772.2;refGene_gn=RAPGEF2;refGene_tx=XM_023252714.1;refGene_gn=RAPGEF2;refGene_tx=XM_023252715.1;refGene_gn=RAPGEF2;refGene_tx=XM_023252716.1;refGene_gn=RAPGEF2;refGene_tx=XM_023252717.1;refGene_gn=RAPGEF2;refGene_tx=XM_023252718.1;refGene_gn=RAPGEF2;refGene_tx=XM_023252719.1
B1 70780635 70780941
refGene_gn=RAPGEF2;refGene_tx=XM_011281593.3;refGene_gn=RAPGEF2;refGene_tx=XM_019828768.2;refGene_gn=RAPGEF2;refGene_tx=XM_019828769.2;refGene_gn=RAPGEF2;refGene_tx=XM_019828770.2;refGene_gn=RAPGEF2;refGene_tx=XM_019828771.2;refGene_gn=RAPGEF2;refGene_tx=XM_019828772.2;refGene_gn=RAPGEF2;refGene_tx=XM_023252714.1;refGene_gn=RAPGEF2;refGene_tx=XM_023252715.1;refGene_gn=RAPGEF2;refGene_tx=XM_023252716.1;refGene_gn=RAPGEF2;refGene_tx=XM_023252717.1;refGene_gn=RAPGEF2;refGene_tx=XM_023252718.1;refGene_gn=RAPGEF2;refGene_tx=XM_023252719.1
B1 70782774 70783334
refGene_gn=RAPGEF2;refGene_tx=XM_011281593.3;refGene_gn=RAPGEF2;refGene_tx=XM_019828768.2;refGene_gn=RAPGEF2;refGene_tx=XM_019828769.2;refGene_gn=RAPGEF2;refGene_tx=XM_019828770.2;refGene_gn=RAPGEF2;refGene_tx=XM_019828771.2;refGene_gn=RAPGEF2;refGene_tx=XM_019828772.2;refGene_gn=RAPGEF2;refGene_tx=XM_023252714.1;refGene_gn=RAPGEF2;refGene_tx=XM_023252715.1;refGene_gn=RAPGEF2;refGene_tx=XM_023252716.1;refGene_gn=RAPGEF2;refGene_tx=XM_023252717.1;refGene_gn=RAPGEF2;refGene_tx=XM_023252718.1;refGene_gn=RAPGEF2;refGene_tx=XM_023252719.1
B1 70783895 70784121
refGene_gn=RAPGEF2;refGene_tx=XM_011281593.3;refGene_gn=RAPGEF2;refGene_tx=XM_019828768.2;refGene_gn=RAPGEF2;refGene_tx=XM_019828769.2;refGene_gn=RAPGEF2;refGene_tx=XM_019828770.2;refGene_gn=RAPGEF2;refGene_tx=XM_019828771.2;refGene_gn=RAPGEF2;refGene_tx=XM_019828772.2;refGene_gn=RAPGEF2;refGene_tx=XM_023252714.1;refGene_gn=RAPGEF2;refGene_tx=XM_023252715.1;refGene_gn=RAPGEF2;refGene_tx=XM_023252716.1;refGene_gn=RAPGEF2;refGene_tx=XM_023252717.1;refGene_gn=RAPGEF2;refGene_tx=XM_023252718.1;refGene_gn=RAPGEF2;refGene_tx=XM_023252719.1
B1 70785329 70785452
refGene_gn=RAPGEF2;refGene_tx=XM_011281593.3;refGene_gn=RAPGEF2;refGene_tx=XM_019828768.2;refGene_gn=RAPGEF2;refGene_tx=XM_019828769.2;refGene_gn=RAPGEF2;refGene_tx=XM_019828770.2;refGene_gn=RAPGEF2;refGene_tx=

XM_019828771.2;refGene_gn=RAPGEF2;refGene_tx=XM_023252714.1;refGene_gn=RAPGEF2;refGene_tx=XM_023252715.1;refGene_gn=RAPGEF2;refGene_tx=XM_023252716.1;refGene_gn=RAPGEF2;refGene_tx=XM_023252719.1
B1 70786215 70786368
refGene_gn=RAPGEF2;refGene_tx=XM_011281593.3;refGene_gn=RAPGEF2;refGene_tx=XM_019828768.2;refGene_gn=RAPGEF2;refGene_tx=XM_019828769.2;refGene_gn=RAPGEF2;refGene_tx=XM_019828770.2;refGene_gn=RAPGEF2;refGene_tx=XM_019828771.2;refGene_gn=RAPGEF2;refGene_tx=XM_019828772.2;refGene_gn=RAPGEF2;refGene_tx=XM_023252714.1;refGene_gn=RAPGEF2;refGene_tx=XM_023252715.1;refGene_gn=RAPGEF2;refGene_tx=XM_023252716.1;refGene_gn=RAPGEF2;refGene_tx=XM_023252717.1;refGene_gn=RAPGEF2;refGene_tx=XM_023252718.1;refGene_gn=RAPGEF2;refGene_tx=XM_023252719.1
B1 70788305 70788512
refGene_gn=RAPGEF2;refGene_tx=XM_011281593.3;refGene_gn=RAPGEF2;refGene_tx=XM_019828768.2;refGene_gn=RAPGEF2;refGene_tx=XM_019828769.2;refGene_gn=RAPGEF2;refGene_tx=XM_019828770.2;refGene_gn=RAPGEF2;refGene_tx=XM_019828771.2;refGene_gn=RAPGEF2;refGene_tx=XM_019828772.2;refGene_gn=RAPGEF2;refGene_tx=XM_023252714.1;refGene_gn=RAPGEF2;refGene_tx=XM_023252715.1;refGene_gn=RAPGEF2;refGene_tx=XM_023252716.1;refGene_gn=RAPGEF2;refGene_tx=XM_023252717.1;refGene_gn=RAPGEF2;refGene_tx=XM_023252718.1;refGene_gn=RAPGEF2;refGene_tx=XM_023252719.1
B1 70789925 70790149
refGene_gn=RAPGEF2;refGene_tx=XM_011281593.3;refGene_gn=RAPGEF2;refGene_tx=XM_019828768.2;refGene_gn=RAPGEF2;refGene_tx=XM_019828769.2;refGene_gn=RAPGEF2;refGene_tx=XM_019828770.2;refGene_gn=RAPGEF2;refGene_tx=XM_019828771.2;refGene_gn=RAPGEF2;refGene_tx=XM_019828772.2;refGene_gn=RAPGEF2;refGene_tx=XM_023252714.1;refGene_gn=RAPGEF2;refGene_tx=XM_023252715.1;refGene_gn=RAPGEF2;refGene_tx=XM_023252716.1;refGene_gn=RAPGEF2;refGene_tx=XM_023252717.1;refGene_gn=RAPGEF2;refGene_tx=XM_023252718.1;refGene_gn=RAPGEF2;refGene_tx=XM_023252719.1
B1 70791140 70791164
refGene_gn=RAPGEF2;refGene_tx=XM_011281593.3;refGene_gn=RAPGEF2;refGene_tx=XM_019828768.2;refGene_gn=RAPGEF2;refGene_tx=XM_019828769.2;refGene_gn=RAPGEF2;refGene_tx=XM_019828770.2;refGene_gn=RAPGEF2;refGene_tx=XM_019828771.2;refGene_gn=RAPGEF2;refGene_tx=XM_019828772.2;refGene_gn=RAPGEF2;refGene_tx=XM_023252714.1;refGene_gn=RAPGEF2;refGene_tx=XM_023252716.1;refGene_gn=RAPGEF2;refGene_tx=XM_023252717.1;refGene_gn=RAPGEF2;refGene_tx=XM_023252718.1;refGene_gn=RAPGEF2;refGene_tx=XM_023252719.1
B1 70791779 70791903
refGene_gn=RAPGEF2;refGene_tx=XM_011281593.3;refGene_gn=RAPGEF2;refGene_tx=XM_019828768.2;refGene_gn=RAPGEF2;refGene_tx=XM_019828769.2;refGene_gn=RAPGEF2;refGene_tx=XM_019828770.2;refGene_gn=RAPGEF2;refGene_tx=XM_019828771.2;refGene_gn=RAPGEF2;refGene_tx=XM_019828772.2;refGene_gn=RAPGEF2;refGene_tx=XM_023252714.1;refGene_gn=RAPGEF2;refGene_tx=XM_023252715.1;refGene_gn=RAPGEF2;refGene_tx=XM_023252716.1;refGene_gn=RAPGEF2;refGene_tx=XM_023252717.1;refGene_gn=RAPGEF2;refGene_tx=XM_023252718.1;refGene_gn=RAPGEF2;refGene_tx=XM_023252719.1
B1 70791995 70792207
refGene_gn=RAPGEF2;refGene_tx=XM_011281593.3;refGene_gn=RAPGEF2;refGen

e_tx=XM_019828768.2;refGene_gn=RAPGEF2;refGene_tx=XM_019828769.2;refGene_gn=RAPGEF2;refGene_tx=XM_019828770.2;refGene_gn=RAPGEF2;refGene_tx=XM_019828771.2;refGene_gn=RAPGEF2;refGene_tx=XM_019828772.2;refGene_gn=RAPGEF2;refGene_tx=XM_023252714.1;refGene_gn=RAPGEF2;refGene_tx=XM_023252715.1;refGene_gn=RAPGEF2;refGene_tx=XM_023252716.1;refGene_gn=RAPGEF2;refGene_tx=XM_023252717.1;refGene_gn=RAPGEF2;refGene_tx=XM_023252718.1;refGene_gn=RAPGEF2;refGene_tx=XM_023252719.1
B1 70793088 70793472
refGene_gn=RAPGEF2;refGene_tx=XM_011281593.3;refGene_gn=RAPGEF2;refGene_tx=XM_019828768.2;refGene_gn=RAPGEF2;refGene_tx=XM_019828769.2;refGene_gn=RAPGEF2;refGene_tx=XM_019828770.2;refGene_gn=RAPGEF2;refGene_tx=XM_019828771.2;refGene_gn=RAPGEF2;refGene_tx=XM_019828772.2;refGene_gn=RAPGEF2;refGene_tx=XM_023252714.1;refGene_gn=RAPGEF2;refGene_tx=XM_023252715.1;refGene_gn=RAPGEF2;refGene_tx=XM_023252716.1;refGene_gn=RAPGEF2;refGene_tx=XM_023252717.1;refGene_gn=RAPGEF2;refGene_tx=XM_023252718.1;refGene_gn=RAPGEF2;refGene_tx=XM_023252719.1
B1 70795631 70795872
refGene_gn=RAPGEF2;refGene_tx=XM_011281593.3;refGene_gn=RAPGEF2;refGene_tx=XM_019828768.2;refGene_gn=RAPGEF2;refGene_tx=XM_019828769.2;refGene_gn=RAPGEF2;refGene_tx=XM_019828770.2;refGene_gn=RAPGEF2;refGene_tx=XM_019828771.2;refGene_gn=RAPGEF2;refGene_tx=XM_019828772.2;refGene_gn=RAPGEF2;refGene_tx=XM_023252714.1;refGene_gn=RAPGEF2;refGene_tx=XM_023252715.1;refGene_gn=RAPGEF2;refGene_tx=XM_023252716.1;refGene_gn=RAPGEF2;refGene_tx=XM_023252717.1;refGene_gn=RAPGEF2;refGene_tx=XM_023252718.1;refGene_gn=RAPGEF2;refGene_tx=XM_023252719.1
B1 70796486 70796644
refGene_gn=RAPGEF2;refGene_tx=XM_011281593.3;refGene_gn=RAPGEF2;refGene_tx=XM_019828768.2;refGene_gn=RAPGEF2;refGene_tx=XM_019828769.2;refGene_gn=RAPGEF2;refGene_tx=XM_019828770.2;refGene_gn=RAPGEF2;refGene_tx=XM_019828771.2;refGene_gn=RAPGEF2;refGene_tx=XM_019828772.2;refGene_gn=RAPGEF2;refGene_tx=XM_023252714.1;refGene_gn=RAPGEF2;refGene_tx=XM_023252715.1;refGene_gn=RAPGEF2;refGene_tx=XM_023252716.1;refGene_gn=RAPGEF2;refGene_tx=XM_023252717.1;refGene_gn=RAPGEF2;refGene_tx=XM_023252718.1;refGene_gn=RAPGEF2;refGene_tx=XM_023252719.1
B1 70800661 70800908
refGene_gn=RAPGEF2;refGene_tx=XM_011281593.3;refGene_gn=RAPGEF2;refGene_tx=XM_019828768.2;refGene_gn=RAPGEF2;refGene_tx=XM_019828769.2;refGene_gn=RAPGEF2;refGene_tx=XM_019828770.2;refGene_gn=RAPGEF2;refGene_tx=XM_019828771.2;refGene_gn=RAPGEF2;refGene_tx=XM_019828772.2;refGene_gn=RAPGEF2;refGene_tx=XM_023252714.1;refGene_gn=RAPGEF2;refGene_tx=XM_023252715.1;refGene_gn=RAPGEF2;refGene_tx=XM_023252716.1;refGene_gn=RAPGEF2;refGene_tx=XM_023252717.1;refGene_gn=RAPGEF2;refGene_tx=XM_023252718.1;refGene_gn=RAPGEF2;refGene_tx=XM_023252719.1
B1 70801377 70801486
refGene_gn=RAPGEF2;refGene_tx=XM_011281593.3;refGene_gn=RAPGEF2;refGene_tx=XM_019828768.2;refGene_gn=RAPGEF2;refGene_tx=XM_019828769.2;refGene_gn=RAPGEF2;refGene_tx=XM_019828770.2;refGene_gn=RAPGEF2;refGene_tx=XM_019828771.2;refGene_gn=RAPGEF2;refGene_tx=XM_019828772.2;refGene_gn=RAPGEF2;refGene_tx=XM_023252714.1;refGene_gn=RAPGEF2;refGene_tx=XM_023252715.1;refGene_gn=RAPGEF2;refGene_tx=XM_023252716.1;refGene_gn=RAPG

EF2;refGene_tx=XM_023252717.1;refGene_gn=RAPGEF2;refGene_tx=XM_023252718.1;refGene_gn=RAPGEF2;refGene_tx=XM_023252719.1
B1 70801582 70801786
refGene_gn=RAPGEF2;refGene_tx=XM_011281593.3;refGene_gn=RAPGEF2;refGene_tx=XM_019828768.2;refGene_gn=RAPGEF2;refGene_tx=XM_019828769.2;refGene_gn=RAPGEF2;refGene_tx=XM_019828770.2;refGene_gn=RAPGEF2;refGene_tx=XM_019828771.2;refGene_gn=RAPGEF2;refGene_tx=XM_019828772.2;refGene_gn=RAPGEF2;refGene_tx=XM_023252714.1;refGene_gn=RAPGEF2;refGene_tx=XM_023252715.1;refGene_gn=RAPGEF2;refGene_tx=XM_023252716.1;refGene_gn=RAPGEF2;refGene_tx=XM_023252717.1;refGene_gn=RAPGEF2;refGene_tx=XM_023252718.1;refGene_gn=RAPGEF2;refGene_tx=XM_023252719.1
B1 70801874 70801982
refGene_gn=RAPGEF2;refGene_tx=XM_011281593.3;refGene_gn=RAPGEF2;refGene_tx=XM_019828768.2;refGene_gn=RAPGEF2;refGene_tx=XM_019828769.2;refGene_gn=RAPGEF2;refGene_tx=XM_019828770.2;refGene_gn=RAPGEF2;refGene_tx=XM_019828771.2;refGene_gn=RAPGEF2;refGene_tx=XM_019828772.2;refGene_gn=RAPGEF2;refGene_tx=XM_023252714.1;refGene_gn=RAPGEF2;refGene_tx=XM_023252715.1;refGene_gn=RAPGEF2;refGene_tx=XM_023252716.1;refGene_gn=RAPGEF2;refGene_tx=XM_023252717.1;refGene_gn=RAPGEF2;refGene_tx=XM_023252718.1;refGene_gn=RAPGEF2;refGene_tx=XM_023252719.1
B1 70804546 70804711
refGene_gn=RAPGEF2;refGene_tx=XM_011281593.3;refGene_gn=RAPGEF2;refGene_tx=XM_019828768.2;refGene_gn=RAPGEF2;refGene_tx=XM_019828769.2;refGene_gn=RAPGEF2;refGene_tx=XM_019828770.2;refGene_gn=RAPGEF2;refGene_tx=XM_019828771.2;refGene_gn=RAPGEF2;refGene_tx=XM_019828772.2;refGene_gn=RAPGEF2;refGene_tx=XM_023252714.1;refGene_gn=RAPGEF2;refGene_tx=XM_023252715.1;refGene_gn=RAPGEF2;refGene_tx=XM_023252716.1;refGene_gn=RAPGEF2;refGene_tx=XM_023252717.1;refGene_gn=RAPGEF2;refGene_tx=XM_023252718.1;refGene_gn=RAPGEF2;refGene_tx=XM_023252719.1
B1 70804989 70805142
refGene_gn=RAPGEF2;refGene_tx=XM_011281593.3;refGene_gn=RAPGEF2;refGene_tx=XM_019828768.2;refGene_gn=RAPGEF2;refGene_tx=XM_019828769.2;refGene_gn=RAPGEF2;refGene_tx=XM_019828770.2;refGene_gn=RAPGEF2;refGene_tx=XM_019828771.2;refGene_gn=RAPGEF2;refGene_tx=XM_019828772.2;refGene_gn=RAPGEF2;refGene_tx=XM_023252714.1;refGene_gn=RAPGEF2;refGene_tx=XM_023252715.1;refGene_gn=RAPGEF2;refGene_tx=XM_023252716.1;refGene_gn=RAPGEF2;refGene_tx=XM_023252717.1;refGene_gn=RAPGEF2;refGene_tx=XM_023252718.1;refGene_gn=RAPGEF2;refGene_tx=XM_023252719.1
B1 70811565 70811724
refGene_gn=RAPGEF2;refGene_tx=XM_011281593.3;refGene_gn=RAPGEF2;refGene_tx=XM_019828768.2;refGene_gn=RAPGEF2;refGene_tx=XM_019828769.2;refGene_gn=RAPGEF2;refGene_tx=XM_019828770.2;refGene_gn=RAPGEF2;refGene_tx=XM_019828771.2;refGene_gn=RAPGEF2;refGene_tx=XM_019828772.2;refGene_gn=RAPGEF2;refGene_tx=XM_023252714.1;refGene_gn=RAPGEF2;refGene_tx=XM_023252715.1;refGene_gn=RAPGEF2;refGene_tx=XM_023252716.1;refGene_gn=RAPGEF2;refGene_tx=XM_023252717.1;refGene_gn=RAPGEF2;refGene_tx=XM_023252718.1;refGene_gn=RAPGEF2;refGene_tx=XM_023252719.1
B1 70812350 70812487
refGene_gn=RAPGEF2;refGene_tx=XM_011281593.3;refGene_gn=RAPGEF2;refGene_tx=XM_019828768.2;refGene_gn=RAPGEF2;refGene_tx=XM_019828769.2;refGe

ne_gn=RAPGEF2;refGene_tx=XM_019828770.2;refGene_gn=RAPGEF2;refGene_tx=XM_019828771.2;refGene_gn=RAPGEF2;refGene_tx=XM_019828772.2;refGene_gn=RAPGEF2;refGene_tx=XM_023252714.1;refGene_gn=RAPGEF2;refGene_tx=XM_023252715.1;refGene_gn=RAPGEF2;refGene_tx=XM_023252716.1;refGene_gn=RAPGEF2;refGene_tx=XM_023252717.1;refGene_gn=RAPGEF2;refGene_tx=XM_023252718.1;refGene_gn=RAPGEF2;refGene_tx=XM_023252719.1
B1 70818919 70819097
refGene_gn=RAPGEF2;refGene_tx=XM_011281593.3;refGene_gn=RAPGEF2;refGene_tx=XM_019828768.2;refGene_gn=RAPGEF2;refGene_tx=XM_019828769.2;refGene_gn=RAPGEF2;refGene_tx=XM_019828770.2;refGene_gn=RAPGEF2;refGene_tx=XM_019828771.2;refGene_gn=RAPGEF2;refGene_tx=XM_019828772.2;refGene_gn=RAPGEF2;refGene_tx=XM_023252714.1;refGene_gn=RAPGEF2;refGene_tx=XM_023252715.1;refGene_gn=RAPGEF2;refGene_tx=XM_023252716.1;refGene_gn=RAPGEF2;refGene_tx=XM_023252717.1;refGene_gn=RAPGEF2;refGene_tx=XM_023252718.1;refGene_gn=RAPGEF2;refGene_tx=XM_023252719.1
B1 70829682 70829814
refGene_gn=RAPGEF2;refGene_tx=XM_011281593.3;refGene_gn=RAPGEF2;refGene_tx=XM_019828768.2;refGene_gn=RAPGEF2;refGene_tx=XM_019828769.2;refGene_gn=RAPGEF2;refGene_tx=XM_019828770.2;refGene_gn=RAPGEF2;refGene_tx=XM_019828771.2;refGene_gn=RAPGEF2;refGene_tx=XM_019828772.2;refGene_gn=RAPGEF2;refGene_tx=XM_023252714.1;refGene_gn=RAPGEF2;refGene_tx=XM_023252715.1;refGene_gn=RAPGEF2;refGene_tx=XM_023252716.1;refGene_gn=RAPGEF2;refGene_tx=XM_023252717.1;refGene_gn=RAPGEF2;refGene_tx=XM_023252718.1;refGene_gn=RAPGEF2;refGene_tx=XM_023252719.1
B1 70868210 70868270
refGene_gn=RAPGEF2;refGene_tx=XM_019828770.2;refGene_gn=RAPGEF2;refGene_tx=XM_019828772.2;refGene_gn=RAPGEF2;refGene_tx=XM_023252719.1
B1 70869113 70869167 refGene_gn=RAPGEF2;refGene_tx=XM_019828771.2
B1 70892649 70892667
refGene_gn=RAPGEF2;refGene_tx=XM_011281593.3;refGene_gn=RAPGEF2;refGene_tx=XM_019828768.2;refGene_gn=RAPGEF2;refGene_tx=XM_019828769.2;refGene_gn=RAPGEF2;refGene_tx=XM_023252715.1;refGene_gn=RAPGEF2;refGene_tx=XM_023252716.1;refGene_gn=RAPGEF2;refGene_tx=XM_023252717.1
B1 70895057 70895225
refGene_gn=RAPGEF2;refGene_tx=XM_011281593.3;refGene_gn=RAPGEF2;refGene_tx=XM_019828768.2;refGene_gn=RAPGEF2;refGene_tx=XM_019828769.2;refGene_gn=RAPGEF2;refGene_tx=XM_023252714.1;refGene_gn=RAPGEF2;refGene_tx=XM_023252715.1;refGene_gn=RAPGEF2;refGene_tx=XM_023252716.1;refGene_gn=RAPGEF2;refGene_tx=XM_023252717.1;refGene_gn=RAPGEF2;refGene_tx=XM_023252718.1
B1 70897026 70897102
refGene_gn=RAPGEF2;refGene_tx=XM_011281593.3;refGene_gn=RAPGEF2;refGene_tx=XM_019828768.2;refGene_gn=RAPGEF2;refGene_tx=XM_019828769.2;refGene_gn=RAPGEF2;refGene_tx=XM_023252714.1;refGene_gn=RAPGEF2;refGene_tx=XM_023252715.1;refGene_gn=RAPGEF2;refGene_tx=XM_023252716.1;refGene_gn=RAPGEF2;refGene_tx=XM_023252717.1;refGene_gn=RAPGEF2;refGene_tx=XM_023252718.1
B1 70905749 70905760 refGene_gn=RAPGEF2;refGene_tx=XM_019828768.2
B1 70924618 70924702
refGene_gn=RAPGEF2;refGene_tx=XM_011281593.3;refGene_gn=RAPGEF2;refGen

e_tx=XM_023252714.1;refGene_gn=RAPGEF2;refGene_tx=XM_023252715.1;refGene_gn=RAPGEF2;refGene_tx=XM_023252716.1;refGene_gn=RAPGEF2;refGene_tx=XM_023252717.1;refGene_gn=RAPGEF2;refGene_tx=XM_023252718.1
B1 70939815 70939872
refGene_gn=RAPGEF2;refGene_tx=XM_011281593.3;refGene_gn=RAPGEF2;refGene_tx=XM_023252714.1;refGene_gn=RAPGEF2;refGene_tx=XM_023252715.1;refGene_gn=RAPGEF2;refGene_tx=XM_023252716.1;refGene_gn=RAPGEF2;refGene_tx=XM_023252717.1;refGene_gn=RAPGEF2;refGene_tx=XM_023252718.1
B1 70951720 70951791
refGene_gn=RAPGEF2;refGene_tx=XM_011281593.3;refGene_gn=RAPGEF2;refGene_tx=XM_023252714.1;refGene_gn=RAPGEF2;refGene_tx=XM_023252715.1;refGene_gn=RAPGEF2;refGene_tx=XM_023252716.1;refGene_gn=RAPGEF2;refGene_tx=XM_023252717.1;refGene_gn=RAPGEF2;refGene_tx=XM_023252718.1
B1 71025100 71025169
refGene_gn=RAPGEF2;refGene_tx=XM_011281593.3;refGene_gn=RAPGEF2;refGene_tx=XM_023252714.1;refGene_gn=RAPGEF2;refGene_tx=XM_023252715.1;refGene_gn=RAPGEF2;refGene_tx=XM_023252716.1;refGene_gn=RAPGEF2;refGene_tx=XM_023252717.1;refGene_gn=RAPGEF2;refGene_tx=XM_023252718.1
B1 71076352 71076479
refGene_gn=CB1H4orf45;refGene_tx=XM_019828785.2
B1 71128904 71129137
refGene_gn=CB1H4orf45;refGene_tx=XM_019828785.2
B1 71143238 71143297
refGene_gn=CB1H4orf45;refGene_tx=XM_019828785.2
B1 71172795 71172982
refGene_gn=CB1H4orf45;refGene_tx=XM_019828785.2
B1 71183858 71183937
refGene_gn=FNIP2;refGene_tx=XM_019828773.2;refGene_gn=FNIP2;refGene_tx=XM_023252720.1;refGene_gn=FNIP2;refGene_tx=XM_023252721.1;refGene_gn=FNIP2;refGene_tx=XM_023252722.1;refGene_gn=FNIP2;refGene_tx=XM_023252723.1
B1 71185882 71185998
refGene_gn=FNIP2;refGene_tx=XM_019828773.2;refGene_gn=FNIP2;refGene_tx=XM_023252720.1;refGene_gn=FNIP2;refGene_tx=XM_023252721.1;refGene_gn=FNIP2;refGene_tx=XM_023252722.1
B1 71187778 71187826
refGene_gn=CB1H4orf45;refGene_tx=XM_019828785.2
B1 71189472 71189673
refGene_gn=FNIP2;refGene_tx=XM_019828773.2;refGene_gn=FNIP2;refGene_tx=XM_023252720.1;refGene_gn=FNIP2;refGene_tx=XM_023252721.1;refGene_gn=FNIP2;refGene_tx=XM_023252722.1;refGene_gn=FNIP2;refGene_tx=XM_023252723.1
B1 71204563 71204720
refGene_gn=FNIP2;refGene_tx=XM_019828773.2;refGene_gn=FNIP2;refGene_tx=XM_023252720.1;refGene_gn=FNIP2;refGene_tx=XM_023252721.1;refGene_gn=FNIP2;refGene_tx=XM_023252722.1;refGene_gn=FNIP2;refGene_tx=XM_023252723.1
B1 71205615 71206969
refGene_gn=FNIP2;refGene_tx=XM_019828773.2;refGene_gn=FNIP2;refGene_tx=XM_023252720.1;refGene_gn=FNIP2;refGene_tx=XM_023252721.1;refGene_gn=

FNIP2;refGene_tx=XM_023252722.1;refGene_gn=FNIP2;refGene_tx=XM_0232527
23.1
B1 71214589 71214759
refGene_gn=FNIP2;refGene_tx=XM_019828773.2;refGene_gn=FNIP2;refGene_tx
=XM_023252720.1;refGene_gn=FNIP2;refGene_tx=XM_023252721.1;refGene_gn=
FNIP2;refGene_tx=XM_023252722.1;refGene_gn=FNIP2;refGene_tx=XM_0232527
23.1
B1 71214876 71215023
refGene_gn=FNIP2;refGene_tx=XM_019828773.2;refGene_gn=FNIP2;refGene_tx
=XM_023252720.1;refGene_gn=FNIP2;refGene_tx=XM_023252721.1;refGene_gn=
FNIP2;refGene_tx=XM_023252722.1;refGene_gn=FNIP2;refGene_tx=XM_0232527
23.1
B1 71218035 71218124
refGene_gn=FNIP2;refGene_tx=XM_019828773.2;refGene_gn=FNIP2;refGene_tx
=XM_023252720.1;refGene_gn=FNIP2;refGene_tx=XM_023252721.1;refGene_gn=
FNIP2;refGene_tx=XM_023252722.1;refGene_gn=FNIP2;refGene_tx=XM_0232527
23.1
B1 71218568 71218770
refGene_gn=FNIP2;refGene_tx=XM_019828773.2;refGene_gn=FNIP2;refGene_tx
=XM_023252720.1;refGene_gn=FNIP2;refGene_tx=XM_023252721.1;refGene_gn=
FNIP2;refGene_tx=XM_023252722.1;refGene_gn=FNIP2;refGene_tx=XM_0232527
23.1
B1 71226823 71226953
refGene_gn=FNIP2;refGene_tx=XM_019828773.2;refGene_gn=FNIP2;refGene_tx
=XM_023252720.1;refGene_gn=FNIP2;refGene_tx=XM_023252721.1;refGene_gn=
FNIP2;refGene_tx=XM_023252722.1;refGene_gn=FNIP2;refGene_tx=XM_0232527
23.1
B1 71236291 71236363
refGene_gn=FNIP2;refGene_tx=XM_019828773.2;refGene_gn=FNIP2;refGene_tx
=XM_023252720.1;refGene_gn=FNIP2;refGene_tx=XM_023252721.1;refGene_gn=
FNIP2;refGene_tx=XM_023252722.1;refGene_gn=FNIP2;refGene_tx=XM_0232527
23.1
B1 71237942 71238032
refGene_gn=FNIP2;refGene_tx=XM_019828773.2;refGene_gn=FNIP2;refGene_tx
=XM_023252720.1;refGene_gn=FNIP2;refGene_tx=XM_023252723.1
B1 71238208 71238309
refGene_gn=FNIP2;refGene_tx=XM_019828773.2;refGene_gn=FNIP2;refGene_tx
=XM_023252720.1;refGene_gn=FNIP2;refGene_tx=XM_023252721.1;refGene_gn=
FNIP2;refGene_tx=XM_023252722.1;refGene_gn=FNIP2;refGene_tx=XM_0232527
23.1
B1 71239839 71239911
refGene_gn=FNIP2;refGene_tx=XM_019828773.2;refGene_gn=FNIP2;refGene_tx
=XM_023252720.1;refGene_gn=FNIP2;refGene_tx=XM_023252721.1;refGene_gn=
FNIP2;refGene_tx=XM_023252722.1;refGene_gn=FNIP2;refGene_tx=XM_0232527
23.1
B1 71240012 71240113
refGene_gn=FNIP2;refGene_tx=XM_019828773.2;refGene_gn=FNIP2;refGene_tx
=XM_023252720.1;refGene_gn=FNIP2;refGene_tx=XM_023252721.1;refGene_gn=
FNIP2;refGene_tx=XM_023252722.1;refGene_gn=FNIP2;refGene_tx=XM_0232527
23.1

B1 71243117 71243258
refGene_gn=FNIP2;refGene_tx=XM_019828773.2;refGene_gn=FNIP2;refGene_tx=XM_023252720.1;refGene_gn=FNIP2;refGene_tx=XM_023252721.1;refGene_gn=FNIP2;refGene_tx=XM_023252722.1;refGene_gn=FNIP2;refGene_tx=XM_023252723.1
B1 71246154 71246281
refGene_gn=FNIP2;refGene_tx=XM_019828773.2;refGene_gn=FNIP2;refGene_tx=XM_023252720.1;refGene_gn=FNIP2;refGene_tx=XM_023252721.1;refGene_gn=FNIP2;refGene_tx=XM_023252722.1;refGene_gn=FNIP2;refGene_tx=XM_023252723.1
B1 71275898 71276074
refGene_gn=FNIP2;refGene_tx=XM_019828773.2;refGene_gn=FNIP2;refGene_tx=XM_023252721.1;refGene_gn=FNIP2;refGene_tx=XM_023252723.1
B1 71314264 71314377
refGene_gn=FNIP2;refGene_tx=XM_023252720.1;refGene_gn=FNIP2;refGene_tx=XM_023252722.1
B1 71347615 71347700 refGene_gn=PPID;refGene_tx=XM_003984906.4
B1 71348942 71349083 refGene_gn=PPID;refGene_tx=XM_003984906.4
B1 71351296 71351403 refGene_gn=PPID;refGene_tx=XM_003984906.4
B1 71353368 71353557 refGene_gn=PPID;refGene_tx=XM_003984906.4
B1 71354460 71354583 refGene_gn=PPID;refGene_tx=XM_003984906.4
B1 71354725 71354832 refGene_gn=PPID;refGene_tx=XM_003984906.4
B1 71356584 71356726 refGene_gn=PPID;refGene_tx=XM_003984906.4
B1 71359560 71359647 refGene_gn=PPID;refGene_tx=XM_003984906.4
B1 71359743 71359786 refGene_gn=PPID;refGene_tx=XM_003984906.4
B1 71360535 71360624 refGene_gn=PPID;refGene_tx=XM_003984906.4
B1 71361559 71361723 refGene_gn=ETFDH;refGene_tx=NM_001290236.1
B1 71362966 71363188 refGene_gn=ETFDH;refGene_tx=NM_001290236.1
B1 71363582 71363765 refGene_gn=ETFDH;refGene_tx=NM_001290236.1
B1 71366091 71366260 refGene_gn=ETFDH;refGene_tx=NM_001290236.1
B1 71369933 71370077 refGene_gn=ETFDH;refGene_tx=NM_001290236.1
B1 71372769 71372910 refGene_gn=ETFDH;refGene_tx=NM_001290236.1
B1 71374491 71374638 refGene_gn=ETFDH;refGene_tx=NM_001290236.1
B1 71377997 71378075 refGene_gn=ETFDH;refGene_tx=NM_001290236.1
B1 71380909 71381028 refGene_gn=ETFDH;refGene_tx=NM_001290236.1
B1 71381666 71381748 refGene_gn=ETFDH;refGene_tx=NM_001290236.1
B1 71383100 71383330 refGene_gn=ETFDH;refGene_tx=NM_001290236.1
B1 71383746 71383887 refGene_gn=ETFDH;refGene_tx=NM_001290236.1
B1 71392539 71392573 refGene_gn=ETFDH;refGene_tx=NM_001290236.1
B1 71392956 71393080
refGene_gn=CB1H4orf46;refGene_tx=XM_023252730.1;refGene_gn=CB1H4orf46;refGene_tx=XM_023252731.1
B1 71393250 71393465
refGene_gn=CB1H4orf46;refGene_tx=XM_023252730.1;refGene_gn=CB1H4orf46;refGene_tx=XM_023252731.1
B1 71395194 71395350
refGene_gn=CB1H4orf46;refGene_tx=XM_023252730.1;refGene_gn=CB1H4orf46;refGene_tx=XM_023252731.1
B1 71405304 71405606
refGene_gn=RXFP1;refGene_tx=XM_019828775.2;refGene_gn=RXFP1;refGene_tx

=XM_019828782.2;refGene_gn=RXFP1;refGene_tx=XM_019828784.2;refGene_gn=RXFP1;refGene_tx=XM_023252724.1;refGene_gn=RXFP1;refGene_tx=XM_023252725.1;refGene_gn=RXFP1;refGene_tx=XM_023252726.1;refGene_gn=RXFP1;refGene_tx=XM_023252727.1;refGene_gn=RXFP1;refGene_tx=XM_023252728.1;refGene_gn=RXFP1;refGene_tx=XM_023252729.1
B1 71407476 71407695
refGene_gn=RXFP1;refGene_tx=XM_019828775.2;refGene_gn=RXFP1;refGene_tx=XM_019828782.2;refGene_gn=RXFP1;refGene_tx=XM_019828784.2;refGene_gn=RXFP1;refGene_tx=XM_023252724.1;refGene_gn=RXFP1;refGene_tx=XM_023252725.1;refGene_gn=RXFP1;refGene_tx=XM_023252726.1;refGene_gn=RXFP1;refGene_tx=XM_023252727.1;refGene_gn=RXFP1;refGene_tx=XM_023252728.1;refGene_gn=RXFP1;refGene_tx=XM_023252729.1
B1 71409593 71410004
refGene_gn=RXFP1;refGene_tx=XM_019828775.2;refGene_gn=RXFP1;refGene_tx=XM_019828782.2;refGene_gn=RXFP1;refGene_tx=XM_019828784.2;refGene_gn=RXFP1;refGene_tx=XM_023252724.1;refGene_gn=RXFP1;refGene_tx=XM_023252725.1;refGene_gn=RXFP1;refGene_tx=XM_023252726.1;refGene_gn=RXFP1;refGene_tx=XM_023252727.1;refGene_gn=RXFP1;refGene_tx=XM_023252728.1;refGene_gn=RXFP1;refGene_tx=XM_023252729.1
B1 71412063 71412293
refGene_gn=RXFP1;refGene_tx=XM_019828775.2;refGene_gn=RXFP1;refGene_tx=XM_019828782.2;refGene_gn=RXFP1;refGene_tx=XM_019828784.2;refGene_gn=RXFP1;refGene_tx=XM_023252724.1;refGene_gn=RXFP1;refGene_tx=XM_023252725.1;refGene_gn=RXFP1;refGene_tx=XM_023252726.1;refGene_gn=RXFP1;refGene_tx=XM_023252727.1;refGene_gn=RXFP1;refGene_tx=XM_023252728.1;refGene_gn=RXFP1;refGene_tx=XM_023252729.1
B1 71414041 71414113
refGene_gn=RXFP1;refGene_tx=XM_019828775.2;refGene_gn=RXFP1;refGene_tx=XM_019828782.2;refGene_gn=RXFP1;refGene_tx=XM_019828784.2;refGene_gn=RXFP1;refGene_tx=XM_023252725.1;refGene_gn=RXFP1;refGene_tx=XM_023252726.1;refGene_gn=RXFP1;refGene_tx=XM_023252727.1;refGene_gn=RXFP1;refGene_tx=XM_023252728.1;refGene_gn=RXFP1;refGene_tx=XM_023252729.1
B1 71414956 71415028
refGene_gn=RXFP1;refGene_tx=XM_019828775.2;refGene_gn=RXFP1;refGene_tx=XM_019828782.2;refGene_gn=RXFP1;refGene_tx=XM_019828784.2;refGene_gn=RXFP1;refGene_tx=XM_023252724.1;refGene_gn=RXFP1;refGene_tx=XM_023252725.1;refGene_gn=RXFP1;refGene_tx=XM_023252726.1;refGene_gn=RXFP1;refGene_tx=XM_023252727.1;refGene_gn=RXFP1;refGene_tx=XM_023252728.1;refGene_gn=RXFP1;refGene_tx=XM_023252729.1
B1 71418178 71418250
refGene_gn=RXFP1;refGene_tx=XM_019828775.2;refGene_gn=RXFP1;refGene_tx=XM_019828782.2;refGene_gn=RXFP1;refGene_tx=XM_019828784.2;refGene_gn=RXFP1;refGene_tx=XM_023252724.1;refGene_gn=RXFP1;refGene_tx=XM_023252725.1;refGene_gn=RXFP1;refGene_tx=XM_023252726.1;refGene_gn=RXFP1;refGene_tx=XM_023252727.1;refGene_gn=RXFP1;refGene_tx=XM_023252728.1;refGene_gn=RXFP1;refGene_tx=XM_023252729.1
B1 71421819 71421891
refGene_gn=RXFP1;refGene_tx=XM_019828775.2;refGene_gn=RXFP1;refGene_tx=XM_019828782.2;refGene_gn=RXFP1;refGene_tx=XM_019828784.2;refGene_gn=RXFP1;refGene_tx=XM_023252724.1;refGene_gn=RXFP1;refGene_tx=XM_0232527

25.1;refGene_gn=RXFP1;refGene_tx=XM_023252727.1;refGene_gn=RXFP1;refGene_tx=XM_023252728.1;refGene_gn=RXFP1;refGene_tx=XM_023252729.1
B1 71424425 71424497
refGene_gn=RXFP1;refGene_tx=XM_019828775.2;refGene_gn=RXFP1;refGene_tx=XM_019828782.2;refGene_gn=RXFP1;refGene_tx=XM_019828784.2;refGene_gn=RXFP1;refGene_tx=XM_023252724.1;refGene_gn=RXFP1;refGene_tx=XM_023252726.1;refGene_gn=RXFP1;refGene_tx=XM_023252727.1;refGene_gn=RXFP1;refGene_tx=XM_023252728.1;refGene_gn=RXFP1;refGene_tx=XM_023252729.1
B1 71434559 71434634
refGene_gn=RXFP1;refGene_tx=XM_019828775.2;refGene_gn=RXFP1;refGene_tx=XM_019828782.2;refGene_gn=RXFP1;refGene_tx=XM_019828784.2;refGene_gn=RXFP1;refGene_tx=XM_023252724.1;refGene_gn=RXFP1;refGene_tx=XM_023252725.1;refGene_gn=RXFP1;refGene_tx=XM_023252726.1;refGene_gn=RXFP1;refGene_tx=XM_023252727.1;refGene_gn=RXFP1;refGene_tx=XM_023252728.1;refGene_gn=RXFP1;refGene_tx=XM_023252729.1
B1 71435249 71435321
refGene_gn=RXFP1;refGene_tx=XM_019828775.2;refGene_gn=RXFP1;refGene_tx=XM_019828782.2;refGene_gn=RXFP1;refGene_tx=XM_019828784.2;refGene_gn=RXFP1;refGene_tx=XM_023252724.1;refGene_gn=RXFP1;refGene_tx=XM_023252725.1;refGene_gn=RXFP1;refGene_tx=XM_023252726.1;refGene_gn=RXFP1;refGene_tx=XM_023252728.1
B1 71435407 71435479
refGene_gn=RXFP1;refGene_tx=XM_019828775.2;refGene_gn=RXFP1;refGene_tx=XM_019828782.2;refGene_gn=RXFP1;refGene_tx=XM_019828784.2;refGene_gn=RXFP1;refGene_tx=XM_023252724.1;refGene_gn=RXFP1;refGene_tx=XM_023252725.1;refGene_gn=RXFP1;refGene_tx=XM_023252726.1;refGene_gn=RXFP1;refGene_tx=XM_023252727.1;refGene_gn=RXFP1;refGene_tx=XM_023252728.1;refGene_gn=RXFP1;refGene_tx=XM_023252729.1
B1 71439751 71439823
refGene_gn=RXFP1;refGene_tx=XM_019828775.2;refGene_gn=RXFP1;refGene_tx=XM_019828782.2;refGene_gn=RXFP1;refGene_tx=XM_019828784.2;refGene_gn=RXFP1;refGene_tx=XM_023252724.1;refGene_gn=RXFP1;refGene_tx=XM_023252725.1;refGene_gn=RXFP1;refGene_tx=XM_023252726.1;refGene_gn=RXFP1;refGene_tx=XM_023252727.1
B1 71450500 71450572
refGene_gn=RXFP1;refGene_tx=XM_019828775.2;refGene_gn=RXFP1;refGene_tx=XM_019828782.2;refGene_gn=RXFP1;refGene_tx=XM_019828784.2;refGene_gn=RXFP1;refGene_tx=XM_023252724.1;refGene_gn=RXFP1;refGene_tx=XM_023252725.1;refGene_gn=RXFP1;refGene_tx=XM_023252726.1;refGene_gn=RXFP1;refGene_tx=XM_023252727.1;refGene_gn=RXFP1;refGene_tx=XM_023252728.1;refGene_gn=RXFP1;refGene_tx=XM_023252729.1
B1 71455878 71456065
refGene_gn=RXFP1;refGene_tx=XM_019828775.2;refGene_gn=RXFP1;refGene_tx=XM_019828782.2;refGene_gn=RXFP1;refGene_tx=XM_023252724.1;refGene_gn=RXFP1;refGene_tx=XM_023252725.1;refGene_gn=RXFP1;refGene_tx=XM_023252726.1;refGene_gn=RXFP1;refGene_tx=XM_023252727.1;refGene_gn=RXFP1;refGene_tx=XM_023252728.1;refGene_gn=RXFP1;refGene_tx=XM_023252729.1
B1 71459618 71459717
refGene_gn=RXFP1;refGene_tx=XM_019828775.2;refGene_gn=RXFP1;refGene_tx=XM_019828782.2;refGene_gn=RXFP1;refGene_tx=XM_019828784.2;refGene_gn=

RXFP1;refGene_tx=XM_023252724.1;refGene_gn=RXFP1;refGene_tx=XM_0232527
25.1;refGene_gn=RXFP1;refGene_tx=XM_023252726.1;refGene_gn=RXFP1;refGe
ne_tx=XM_023252727.1;refGene_gn=RXFP1;refGene_tx=XM_023252728.1;refGen
e_gn=RXFP1;refGene_tx=XM_023252729.1
B1 71465070 71465192
refGene_gn=CB1H4orf46;refGene_tx=XM_023252730.1
B1 71471081 71471171
refGene_gn=CB1H4orf46;refGene_tx=XM_023252730.1
B1 71481390 71481528
refGene_gn=RXFP1;refGene_tx=XM_019828775.2;refGene_gn=RXFP1;refGene_tx
=XM_019828782.2;refGene_gn=RXFP1;refGene_tx=XM_023252724.1;refGene_gn=
RXFP1;refGene_tx=XM_023252725.1;refGene_gn=RXFP1;refGene_tx=XM_0232527
26.1;refGene_gn=RXFP1;refGene_tx=XM_023252727.1;refGene_gn=RXFP1;refGe
ne_tx=XM_023252728.1;refGene_gn=RXFP1;refGene_tx=XM_023252729.1
B1 71518494 71518543
refGene_gn=RXFP1;refGene_tx=XM_019828775.2;refGene_gn=RXFP1;refGene_tx
=XM_019828782.2;refGene_gn=RXFP1;refGene_tx=XM_023252724.1;refGene_gn=
RXFP1;refGene_tx=XM_023252725.1;refGene_gn=RXFP1;refGene_tx=XM_0232527
26.1;refGene_gn=RXFP1;refGene_tx=XM_023252727.1;refGene_gn=RXFP1;refGe
ne_tx=XM_023252728.1;refGene_gn=RXFP1;refGene_tx=XM_023252729.1
B1 71738089 71738173
refGene_gn=TMEM144;refGene_tx=XM_011281596.3;refGene_gn=TMEM144;refGen
e_tx=XM_019828789.2;refGene_gn=TMEM144;refGene_tx=XM_019828791.2;refGe
ne_gn=TMEM144;refGene_tx=XM_019828794.2;refGene_gn=TMEM144;refGene_tx=
XM_019828798.2;refGene_gn=TMEM144;refGene_tx=XM_019828799.2;refGene_gn
=TMEM144;refGene_tx=XM_019828800.2;refGene_gn=TMEM144;refGene_tx=XM_02
3252732.1;refGene_gn=TMEM144;refGene_tx=XM_023252733.1;refGene_gn=TMEM
144;refGene_tx=XM_023252734.1;refGene_gn=TMEM144;refGene_tx=XM_0232527
35.1;refGene_gn=TMEM144;refGene_tx=XM_023252736.1;refGene_gn=TMEM144;r
efGene_tx=XM_023252737.1;refGene_gn=TMEM144;refGene_tx=XM_023252738.1;
refGene_gn=TMEM144;refGene_tx=XM_023252739.1;refGene_gn=TMEM144;refGen
e_tx=XM_023252740.1;refGene_gn=TMEM144;refGene_tx=XM_023252741.1
B1 71744777 71744831
refGene_gn=TMEM144;refGene_tx=XM_011281596.3;refGene_gn=TMEM144;refGen
e_tx=XM_019828789.2;refGene_gn=TMEM144;refGene_tx=XM_019828791.2;refGe
ne_gn=TMEM144;refGene_tx=XM_019828794.2;refGene_gn=TMEM144;refGene_tx=
XM_019828798.2;refGene_gn=TMEM144;refGene_tx=XM_019828799.2;refGene_gn
=TMEM144;refGene_tx=XM_019828800.2;refGene_gn=TMEM144;refGene_tx=XM_02
3252732.1;refGene_gn=TMEM144;refGene_tx=XM_023252733.1;refGene_gn=TMEM
144;refGene_tx=XM_023252734.1;refGene_gn=TMEM144;refGene_tx=XM_0232527
35.1;refGene_gn=TMEM144;refGene_tx=XM_023252736.1;refGene_gn=TMEM144;r
efGene_tx=XM_023252737.1;refGene_gn=TMEM144;refGene_tx=XM_023252738.1;
refGene_gn=TMEM144;refGene_tx=XM_023252739.1;refGene_gn=TMEM144;refGen
e_tx=XM_023252740.1;refGene_gn=TMEM144;refGene_tx=XM_023252741.1
B1 71748081 71748179
refGene_gn=TMEM144;refGene_tx=XM_011281596.3;refGene_gn=TMEM144;refGen
e_tx=XM_019828789.2;refGene_gn=TMEM144;refGene_tx=XM_019828791.2;refGe
ne_gn=TMEM144;refGene_tx=XM_019828794.2;refGene_gn=TMEM144;refGene_tx=
XM_019828798.2;refGene_gn=TMEM144;refGene_tx=XM_019828799.2;refGene_gn
=TMEM144;refGene_tx=XM_019828800.2;refGene_gn=TMEM144;refGene_tx=XM_02

3252732.1;refGene_gn=TMEM144;refGene_tx=XM_023252733.1;refGene_gn=TMEM144;refGene_tx=XM_023252734.1;refGene_gn=TMEM144;refGene_tx=XM_023252735.1;refGene_gn=TMEM144;refGene_tx=XM_023252736.1;refGene_gn=TMEM144;refGene_tx=XM_023252737.1;refGene_gn=TMEM144;refGene_tx=XM_023252738.1;refGene_gn=TMEM144;refGene_tx=XM_023252739.1;refGene_gn=TMEM144;refGene_tx=XM_023252740.1;refGene_gn=TMEM144;refGene_tx=XM_023252741.1
B1 71749291 71749411
refGene_gn=TMEM144;refGene_tx=XM_011281596.3;refGene_gn=TMEM144;refGene_tx=XM_019828789.2;refGene_gn=TMEM144;refGene_tx=XM_019828791.2;refGene_gn=TMEM144;refGene_tx=XM_019828794.2;refGene_gn=TMEM144;refGene_tx=XM_019828798.2;refGene_gn=TMEM144;refGene_tx=XM_019828799.2;refGene_gn=TMEM144;refGene_tx=XM_019828800.2;refGene_gn=TMEM144;refGene_tx=XM_023252732.1;refGene_gn=TMEM144;refGene_tx=XM_023252733.1;refGene_gn=TMEM144;refGene_tx=XM_023252734.1;refGene_gn=TMEM144;refGene_tx=XM_023252735.1;refGene_gn=TMEM144;refGene_tx=XM_023252736.1;refGene_gn=TMEM144;refGene_tx=XM_023252737.1;refGene_gn=TMEM144;refGene_tx=XM_023252738.1;refGene_gn=TMEM144;refGene_tx=XM_023252739.1;refGene_gn=TMEM144;refGene_tx=XM_023252740.1;refGene_gn=TMEM144;refGene_tx=XM_023252741.1
B1 71751976 71752095
refGene_gn=TMEM144;refGene_tx=XM_011281596.3;refGene_gn=TMEM144;refGene_tx=XM_019828789.2;refGene_gn=TMEM144;refGene_tx=XM_019828791.2;refGene_gn=TMEM144;refGene_tx=XM_019828794.2;refGene_gn=TMEM144;refGene_tx=XM_019828798.2;refGene_gn=TMEM144;refGene_tx=XM_019828799.2;refGene_gn=TMEM144;refGene_tx=XM_019828800.2;refGene_gn=TMEM144;refGene_tx=XM_023252732.1;refGene_gn=TMEM144;refGene_tx=XM_023252733.1;refGene_gn=TMEM144;refGene_tx=XM_023252734.1;refGene_gn=TMEM144;refGene_tx=XM_023252735.1;refGene_gn=TMEM144;refGene_tx=XM_023252736.1;refGene_gn=TMEM144;refGene_tx=XM_023252737.1;refGene_gn=TMEM144;refGene_tx=XM_023252738.1;refGene_gn=TMEM144;refGene_tx=XM_023252739.1;refGene_gn=TMEM144;refGene_tx=XM_023252740.1;refGene_gn=TMEM144;refGene_tx=XM_023252741.1
B1 71754295 71754372
refGene_gn=TMEM144;refGene_tx=XM_011281596.3;refGene_gn=TMEM144;refGene_tx=XM_019828789.2;refGene_gn=TMEM144;refGene_tx=XM_019828791.2;refGene_gn=TMEM144;refGene_tx=XM_019828794.2;refGene_gn=TMEM144;refGene_tx=XM_019828798.2;refGene_gn=TMEM144;refGene_tx=XM_019828799.2;refGene_gn=TMEM144;refGene_tx=XM_019828800.2;refGene_gn=TMEM144;refGene_tx=XM_023252732.1;refGene_gn=TMEM144;refGene_tx=XM_023252733.1;refGene_gn=TMEM144;refGene_tx=XM_023252734.1;refGene_gn=TMEM144;refGene_tx=XM_023252735.1;refGene_gn=TMEM144;refGene_tx=XM_023252736.1;refGene_gn=TMEM144;refGene_tx=XM_023252737.1;refGene_gn=TMEM144;refGene_tx=XM_023252738.1;refGene_gn=TMEM144;refGene_tx=XM_023252739.1;refGene_gn=TMEM144;refGene_tx=XM_023252740.1;refGene_gn=TMEM144;refGene_tx=XM_023252741.1
B1 71756277 71756359
refGene_gn=TMEM144;refGene_tx=XM_011281596.3;refGene_gn=TMEM144;refGene_tx=XM_019828789.2;refGene_gn=TMEM144;refGene_tx=XM_019828791.2;refGene_gn=TMEM144;refGene_tx=XM_019828794.2;refGene_gn=TMEM144;refGene_tx=XM_019828798.2;refGene_gn=TMEM144;refGene_tx=XM_019828799.2;refGene_gn=TMEM144;refGene_tx=XM_019828800.2;refGene_gn=TMEM144;refGene_tx=XM_023252732.1;refGene_gn=TMEM144;refGene_tx=XM_023252733.1;refGene_gn=TMEM144;refGene_tx=XM_023252734.1;refGene_gn=TMEM144;refGene_tx=XM_0232527

35.1;refGene_gn=TMEM144;refGene_tx=XM_023252736.1;refGene_gn=TMEM144;refGene_tx=XM_023252737.1;refGene_gn=TMEM144;refGene_tx=XM_023252738.1;refGene_gn=TMEM144;refGene_tx=XM_023252739.1;refGene_gn=TMEM144;refGene_tx=XM_023252740.1;refGene_gn=TMEM144;refGene_tx=XM_023252741.1
B1 71762452 71762533
refGene_gn=TMEM144;refGene_tx=XM_011281596.3;refGene_gn=TMEM144;refGene_tx=XM_019828789.2;refGene_gn=TMEM144;refGene_tx=XM_019828791.2;refGene_gn=TMEM144;refGene_tx=XM_019828794.2;refGene_gn=TMEM144;refGene_tx=XM_019828798.2;refGene_gn=TMEM144;refGene_tx=XM_019828799.2;refGene_gn=TMEM144;refGene_tx=XM_019828800.2;refGene_gn=TMEM144;refGene_tx=XM_023252732.1;refGene_gn=TMEM144;refGene_tx=XM_023252733.1;refGene_gn=TMEM144;refGene_tx=XM_023252734.1;refGene_gn=TMEM144;refGene_tx=XM_023252735.1;refGene_gn=TMEM144;refGene_tx=XM_023252736.1;refGene_gn=TMEM144;refGene_tx=XM_023252737.1;refGene_gn=TMEM144;refGene_tx=XM_023252738.1;refGene_gn=TMEM144;refGene_tx=XM_023252739.1;refGene_gn=TMEM144;refGene_tx=XM_023252740.1;refGene_gn=TMEM144;refGene_tx=XM_023252741.1
B1 71764443 71764543
refGene_gn=TMEM144;refGene_tx=XM_011281596.3;refGene_gn=TMEM144;refGene_tx=XM_019828789.2;refGene_gn=TMEM144;refGene_tx=XM_019828791.2;refGene_gn=TMEM144;refGene_tx=XM_019828794.2;refGene_gn=TMEM144;refGene_tx=XM_019828798.2;refGene_gn=TMEM144;refGene_tx=XM_019828799.2;refGene_gn=TMEM144;refGene_tx=XM_019828800.2;refGene_gn=TMEM144;refGene_tx=XM_023252732.1;refGene_gn=TMEM144;refGene_tx=XM_023252733.1;refGene_gn=TMEM144;refGene_tx=XM_023252734.1;refGene_gn=TMEM144;refGene_tx=XM_023252735.1;refGene_gn=TMEM144;refGene_tx=XM_023252736.1;refGene_gn=TMEM144;refGene_tx=XM_023252737.1;refGene_gn=TMEM144;refGene_tx=XM_023252738.1;refGene_gn=TMEM144;refGene_tx=XM_023252739.1;refGene_gn=TMEM144;refGene_tx=XM_023252740.1;refGene_gn=TMEM144;refGene_tx=XM_023252741.1
B1 71766537 71766660
refGene_gn=TMEM144;refGene_tx=XM_011281596.3;refGene_gn=TMEM144;refGene_tx=XM_019828789.2;refGene_gn=TMEM144;refGene_tx=XM_019828791.2;refGene_gn=TMEM144;refGene_tx=XM_019828794.2;refGene_gn=TMEM144;refGene_tx=XM_019828798.2;refGene_gn=TMEM144;refGene_tx=XM_019828799.2;refGene_gn=TMEM144;refGene_tx=XM_019828800.2;refGene_gn=TMEM144;refGene_tx=XM_023252732.1;refGene_gn=TMEM144;refGene_tx=XM_023252733.1;refGene_gn=TMEM144;refGene_tx=XM_023252734.1;refGene_gn=TMEM144;refGene_tx=XM_023252735.1;refGene_gn=TMEM144;refGene_tx=XM_023252736.1;refGene_gn=TMEM144;refGene_tx=XM_023252737.1;refGene_gn=TMEM144;refGene_tx=XM_023252738.1;refGene_gn=TMEM144;refGene_tx=XM_023252739.1;refGene_gn=TMEM144;refGene_tx=XM_023252740.1;refGene_gn=TMEM144;refGene_tx=XM_023252741.1
B1 71767522 71767568 refGene_gn=TMEM144;refGene_tx=XM_019828800.2
B1 71769034 71769143
refGene_gn=TMEM144;refGene_tx=XM_011281596.3;refGene_gn=TMEM144;refGene_tx=XM_019828789.2;refGene_gn=TMEM144;refGene_tx=XM_019828791.2;refGene_gn=TMEM144;refGene_tx=XM_019828794.2;refGene_gn=TMEM144;refGene_tx=XM_019828798.2;refGene_gn=TMEM144;refGene_tx=XM_019828799.2;refGene_gn=TMEM144;refGene_tx=XM_023252732.1;refGene_gn=TMEM144;refGene_tx=XM_023252733.1;refGene_gn=TMEM144;refGene_tx=XM_023252734.1;refGene_gn=TMEM144;refGene_tx=XM_023252735.1;refGene_gn=TMEM144;refGene_tx=XM_023252736.1;refGene_gn=TMEM144;refGene_tx=XM_023252737.1;refGene_gn=TMEM144;r

efGene_tx=XM_023252738.1;refGene_gn=TMEM144;refGene_tx=XM_023252739.1;refGene_gn=TMEM144;refGene_tx=XM_023252740.1;refGene_gn=TMEM144;refGene_tx=XM_023252741.1
B1 71814138 71815036 refGene_gn=FAM198B;refGene_tx=XM_003984914.5;refGene_gn=FAM198B;refGene_tx=XM_011281597.3
B1 71831013 71831228 refGene_gn=FAM198B;refGene_tx=XM_003984914.5;refGene_gn=FAM198B;refGene_tx=XM_011281597.3
B1 71855154 71855381 refGene_gn=FAM198B;refGene_tx=XM_003984914.5;refGene_gn=FAM198B;refGene_tx=XM_011281597.3
B1 71858242 71858450 refGene_gn=FAM198B;refGene_tx=XM_003984914.5;refGene_gn=FAM198B;refGene_tx=XM_011281597.3
B1 72559114 72559279 refGene_gn=GRIA2;refGene_tx=XM_011281602.3
B1 72559673 72559919 refGene_gn=GRIA2;refGene_tx=XM_011281598.3;refGene_gn=GRIA2;refGene_tx=XM_011281599.3;refGene_gn=GRIA2;refGene_tx=XM_011281602.3
B1 72561066 72561181 refGene_gn=GRIA2;refGene_tx=XM_011281599.3;refGene_gn=GRIA2;refGene_tx=XM_011281600.2;refGene_gn=GRIA2;refGene_tx=XM_011281602.3
B1 72561597 72561722 refGene_gn=GRIA2;refGene_tx=XM_011281598.3;refGene_gn=GRIA2;refGene_tx=XM_011281600.2;refGene_gn=GRIA2;refGene_tx=XM_011281602.3
B1 72562606 72562854 refGene_gn=GRIA2;refGene_tx=XM_011281598.3;refGene_gn=GRIA2;refGene_tx=XM_011281599.3;refGene_gn=GRIA2;refGene_tx=XM_011281600.2;refGene_gn=GRIA2;refGene_tx=XM_011281602.3
B1 72586880 72587079 refGene_gn=GRIA2;refGene_tx=XM_011281598.3;refGene_gn=GRIA2;refGene_tx=XM_011281599.3;refGene_gn=GRIA2;refGene_tx=XM_011281600.2;refGene_gn=GRIA2;refGene_tx=XM_011281602.3
B1 72591651 72592022 refGene_gn=GRIA2;refGene_tx=XM_011281598.3;refGene_gn=GRIA2;refGene_tx=XM_011281599.3;refGene_gn=GRIA2;refGene_tx=XM_011281600.2;refGene_gn=GRIA2;refGene_tx=XM_011281602.3
B1 72592528 72592735 refGene_gn=GRIA2;refGene_tx=XM_011281598.3;refGene_gn=GRIA2;refGene_tx=XM_011281599.3;refGene_gn=GRIA2;refGene_tx=XM_011281600.2;refGene_gn=GRIA2;refGene_tx=XM_011281602.3
B1 72594241 72594352 refGene_gn=GRIA2;refGene_tx=XM_011281598.3;refGene_gn=GRIA2;refGene_tx=XM_011281599.3;refGene_gn=GRIA2;refGene_tx=XM_011281600.2;refGene_gn=GRIA2;refGene_tx=XM_011281602.3
B1 72595029 72595134 refGene_gn=GRIA2;refGene_tx=XM_011281598.3;refGene_gn=GRIA2;refGene_tx=XM_011281599.3;refGene_gn=GRIA2;refGene_tx=XM_011281600.2;refGene_gn=GRIA2;refGene_tx=XM_011281602.3
B1 72595383 72595551

refGene_gn=GRIA2;refGene_tx=XM_011281598.3;refGene_gn=GRIA2;refGene_tx=XM_011281599.3;refGene_gn=GRIA2;refGene_tx=XM_011281600.2;refGene_gn=GRIA2;refGene_tx=XM_011281602.3
B1 72612344 72612506
refGene_gn=GRIA2;refGene_tx=XM_011281598.3;refGene_gn=GRIA2;refGene_tx=XM_011281599.3;refGene_gn=GRIA2;refGene_tx=XM_011281600.2;refGene_gn=GRIA2;refGene_tx=XM_011281602.3
B1 72617365 72617419
refGene_gn=GRIA2;refGene_tx=XM_011281598.3;refGene_gn=GRIA2;refGene_tx=XM_011281599.3;refGene_gn=GRIA2;refGene_tx=XM_011281600.2;refGene_gn=GRIA2;refGene_tx=XM_011281602.3
B1 72621784 72621981
refGene_gn=GRIA2;refGene_tx=XM_011281598.3;refGene_gn=GRIA2;refGene_tx=XM_011281599.3;refGene_gn=GRIA2;refGene_tx=XM_011281600.2;refGene_gn=GRIA2;refGene_tx=XM_011281602.3
B1 72634967 72635207
refGene_gn=GRIA2;refGene_tx=XM_011281598.3;refGene_gn=GRIA2;refGene_tx=XM_011281599.3;refGene_gn=GRIA2;refGene_tx=XM_011281600.2;refGene_gn=GRIA2;refGene_tx=XM_011281602.3
B1 72715603 72715744
refGene_gn=GRIA2;refGene_tx=XM_011281598.3;refGene_gn=GRIA2;refGene_tx=XM_011281599.3;refGene_gn=GRIA2;refGene_tx=XM_011281600.2;refGene_gn=GRIA2;refGene_tx=XM_011281602.3
B1 72716267 72716355
refGene_gn=GRIA2;refGene_tx=XM_011281598.3;refGene_gn=GRIA2;refGene_tx=XM_011281599.3;refGene_gn=GRIA2;refGene_tx=XM_011281600.2;refGene_gn=GRIA2;refGene_tx=XM_011281602.3
B1 72762080 72762377
refGene_gn=GLRB;refGene_tx=XM_006930801.4;refGene_gn=GLRB;refGene_tx=XM_019828802.1;refGene_gn=GLRB;refGene_tx=XM_019828803.2;refGene_gn=GLRB;refGene_tx=XM_023252742.1
B1 72775514 72775813
refGene_gn=GLRB;refGene_tx=XM_006930801.4;refGene_gn=GLRB;refGene_tx=XM_019828802.1;refGene_gn=GLRB;refGene_tx=XM_019828803.2;refGene_gn=GLRB;refGene_tx=XM_023252742.1
B1 72789213 72789366
refGene_gn=GLRB;refGene_tx=XM_006930801.4;refGene_gn=GLRB;refGene_tx=XM_019828802.1;refGene_gn=GLRB;refGene_tx=XM_019828803.2;refGene_gn=GLRB;refGene_tx=XM_023252742.1
B1 72795020 72795161
refGene_gn=GLRB;refGene_tx=XM_006930801.4;refGene_gn=GLRB;refGene_tx=XM_019828802.1;refGene_gn=GLRB;refGene_tx=XM_019828803.2;refGene_gn=GLRB;refGene_tx=XM_023252742.1
B1 72797001 72797084
refGene_gn=GLRB;refGene_tx=XM_006930801.4;refGene_gn=GLRB;refGene_tx=XM_019828802.1;refGene_gn=GLRB;refGene_tx=XM_019828803.2;refGene_gn=GLRB;refGene_tx=XM_023252742.1
B1 72797191 72797421
refGene_gn=GLRB;refGene_tx=XM_006930801.4;refGene_gn=GLRB;refGene_tx=XM_019828802.1;refGene_gn=GLRB;refGene_tx=XM_019828803.2;refGene_gn=GLR

B;refGene_tx=XM_023252742.1
B1 72817900 72817968
refGene_gn=GLRB;refGene_tx=XM_006930801.4;refGene_gn=GLRB;refGene_tx=XM_019828802.1;refGene_gn=GLRB;refGene_tx=XM_019828803.2;refGene_gn=GLRB;refGene_tx=XM_023252742.1
B1 72819763 72819870
refGene_gn=GLRB;refGene_tx=XM_019828802.1;refGene_gn=GLRB;refGene_tx=XM_023252742.1
B1 72822469 72822543 refGene_gn=GLRB;refGene_tx=XM_019828802.1
B1 72854887 72855009 refGene_gn=GLRB;refGene_tx=XM_023252742.1
B1 72947099 72947217 refGene_gn=PDGFC;refGene_tx=XM_003984917.5
B1 73059662 73059718 refGene_gn=PDGFC;refGene_tx=XM_003984917.5
B1 73059719 73059859 refGene_gn=PDGFC;refGene_tx=XM_003984917.5
B1 73095159 73095340 refGene_gn=PDGFC;refGene_tx=XM_003984917.5
B1 73137559 73137767 refGene_gn=PDGFC;refGene_tx=XM_003984917.5
B1 73142811 73142954 refGene_gn=PDGFC;refGene_tx=XM_003984917.5
B1 73142955 73143030 refGene_gn=PDGFC;refGene_tx=XM_003984917.5
B1 73149620 73149737 refGene_gn=PDGFC;refGene_tx=XM_003984917.5
B1 73922461 73922590 refGene_gn=CTSO;refGene_tx=XM_023252745.1
B1 73925241 73925350 refGene_gn=CTSO;refGene_tx=XM_023252745.1
B1 73925989 73926129 refGene_gn=CTSO;refGene_tx=XM_023252745.1
B1 73928339 73928507 refGene_gn=CTSO;refGene_tx=XM_023252745.1
B1 73929745 73929867 refGene_gn=CTSO;refGene_tx=XM_023252745.1
B1 73935883 73936047 refGene_gn=CTSO;refGene_tx=XM_023252745.1
B1 73936874 73936967 refGene_gn=CTSO;refGene_tx=XM_023252745.1
B1 73939339 73939374 refGene_gn=CTSO;refGene_tx=XM_023252745.1
B1 73944106 73944260
refGene_gn=TDO2;refGene_tx=XM_023252743.1;refGene_gn=TDO2;refGene_tx=XM_023252744.1
B1 73945957 73946048
refGene_gn=TDO2;refGene_tx=XM_023252743.1;refGene_gn=TDO2;refGene_tx=XM_023252744.1
B1 73946949 73947029
refGene_gn=TDO2;refGene_tx=XM_023252743.1;refGene_gn=TDO2;refGene_tx=XM_023252744.1
B1 73948525 73948583
refGene_gn=TDO2;refGene_tx=XM_023252743.1;refGene_gn=TDO2;refGene_tx=XM_023252744.1
B1 73950047 73950159
refGene_gn=TDO2;refGene_tx=XM_023252743.1;refGene_gn=TDO2;refGene_tx=XM_023252744.1
B1 73953335 73953443
refGene_gn=TDO2;refGene_tx=XM_023252743.1;refGene_gn=TDO2;refGene_tx=XM_023252744.1
B1 73954692 73954879
refGene_gn=TDO2;refGene_tx=XM_023252743.1;refGene_gn=TDO2;refGene_tx=XM_023252744.1
B1 73956129 73956257
refGene_gn=TDO2;refGene_tx=XM_023252743.1;refGene_gn=TDO2;refGene_tx=XM_023252744.1

B1 73958042 73958132 refGene_gn=TDO2;refGene_tx=XM_023252744.1
B1 73958834 73958905 refGene_gn=TDO2;refGene_tx=XM_023252743.1
B1 73961132 73961223 refGene_gn=TDO2;refGene_tx=XM_023252743.1
B1 73963085 73963191 refGene_gn=TDO2;refGene_tx=XM_023252743.1
B1 73963410 73963445 refGene_gn=TDO2;refGene_tx=XM_023252743.1
B1 73988995 73989035 refGene_gn=ASIC5;refGene_tx=XM_003984922.4
B1 73991755 73991821 refGene_gn=ASIC5;refGene_tx=XM_003984922.4
B1 73991823 73992061 refGene_gn=ASIC5;refGene_tx=XM_003984922.4
B1 73997612 73997784 refGene_gn=ASIC5;refGene_tx=XM_003984922.4
B1 73997785 73997851 refGene_gn=ASIC5;refGene_tx=XM_003984922.4
B1 73999297 73999423 refGene_gn=ASIC5;refGene_tx=XM_003984922.4
B1 74002001 74002151 refGene_gn=ASIC5;refGene_tx=XM_003984922.4
B1 74003087 74003235 refGene_gn=ASIC5;refGene_tx=XM_003984922.4
B1 74005740 74005797 refGene_gn=ASIC5;refGene_tx=XM_003984922.4
B1 74008210 74008311 refGene_gn=ASIC5;refGene_tx=XM_003984922.4
B1 74008312 74008380 refGene_gn=ASIC5;refGene_tx=XM_003984922.4
B1 74015901 74015993 refGene_gn=ASIC5;refGene_tx=XM_003984922.4
B1 74018374 74018562 refGene_gn=ASIC5;refGene_tx=XM_003984922.4
B1 74054060 74054084 refGene_gn=GUCY1B1;refGene_tx=XM_023252746.1
B1 74055201 74055328 refGene_gn=GUCY1B1;refGene_tx=XM_023252746.1
B1 74055670 74055825 refGene_gn=GUCY1B1;refGene_tx=XM_023252746.1
B1 74056624 74056765 refGene_gn=GUCY1B1;refGene_tx=XM_023252746.1
B1 74057796 74058034 refGene_gn=GUCY1B1;refGene_tx=XM_023252746.1
B1 74059787 74059985 refGene_gn=GUCY1B1;refGene_tx=XM_023252746.1
B1 74063264 74063398 refGene_gn=GUCY1B1;refGene_tx=XM_023252746.1
B1 74064370 74064477 refGene_gn=GUCY1B1;refGene_tx=XM_023252746.1
B1 74064479 74064486 refGene_gn=GUCY1B1;refGene_tx=XM_023252746.1
B1 74064896 74065127 refGene_gn=GUCY1B1;refGene_tx=XM_023252746.1
B1 74068642 74068840 refGene_gn=GUCY1B1;refGene_tx=XM_023252746.1
B1 74081361 74081480 refGene_gn=GUCY1B1;refGene_tx=XM_023252746.1
B1 74082958 74083059 refGene_gn=GUCY1B1;refGene_tx=XM_023252746.1
B1 74099648 74099722 refGene_gn=GUCY1B1;refGene_tx=XM_023252746.1
B1 74100401 74100404 refGene_gn=GUCY1B1;refGene_tx=XM_023252746.1
B1 74140316 74140518
refGene_gn=GUCY1A1;refGene_tx=XM_011281605.3;refGene_gn=GUCY1A1;refGene_tx=XM_011281606.3;refGene_gn=GUCY1A1;refGene_tx=XM_011281607.3;refGene_gn=GUCY1A1;refGene_tx=XM_019828807.2
B1 74151071 74151226
refGene_gn=GUCY1A1;refGene_tx=XM_011281605.3;refGene_gn=GUCY1A1;refGene_tx=XM_011281606.3;refGene_gn=GUCY1A1;refGene_tx=XM_011281607.3;refGene_gn=GUCY1A1;refGene_tx=XM_019828807.2
B1 74155729 74155873
refGene_gn=GUCY1A1;refGene_tx=XM_011281605.3;refGene_gn=GUCY1A1;refGene_tx=XM_011281606.3;refGene_gn=GUCY1A1;refGene_tx=XM_011281607.3;refGene_gn=GUCY1A1;refGene_tx=XM_019828807.2
B1 74159997 74160483
refGene_gn=GUCY1A1;refGene_tx=XM_011281605.3;refGene_gn=GUCY1A1;refGene_tx=XM_011281606.3;refGene_gn=GUCY1A1;refGene_tx=XM_011281607.3;refGene_gn=GUCY1A1;refGene_tx=XM_019828807.2
B1 74162119 74162829

refGene_gn=GUCY1A1;refGene_tx=XM_011281605.3;refGene_gn=GUCY1A1;refGene_tx=XM_011281606.3;refGene_gn=GUCY1A1;refGene_tx=XM_011281607.3;refGene_gn=GUCY1A1;refGene_tx=XM_019828807.2
B1 74164140 74164199
refGene_gn=GUCY1A1;refGene_tx=XM_011281605.3;refGene_gn=GUCY1A1;refGene_tx=XM_011281606.3;refGene_gn=GUCY1A1;refGene_tx=XM_011281607.3;refGene_gn=GUCY1A1;refGene_tx=XM_019828807.2
B1 74167390 74167452
refGene_gn=GUCY1A1;refGene_tx=XM_011281605.3;refGene_gn=GUCY1A1;refGene_tx=XM_011281606.3;refGene_gn=GUCY1A1;refGene_tx=XM_011281607.3;refGene_gn=GUCY1A1;refGene_tx=XM_019828807.2
B1 74171034 74171289
refGene_gn=GUCY1A1;refGene_tx=XM_011281605.3;refGene_gn=GUCY1A1;refGene_tx=XM_011281606.3;refGene_gn=GUCY1A1;refGene_tx=XM_011281607.3;refGene_gn=GUCY1A1;refGene_tx=XM_019828807.2
B1 74468460 74468535
refGene_gn=MAP9;refGene_tx=XM_019828808.2;refGene_gn=MAP9;refGene_tx=XM_023252747.1;refGene_gn=MAP9;refGene_tx=XM_023252748.1;refGene_gn=MAP9;refGene_tx=XM_023252749.1
B1 74469283 74469368
refGene_gn=MAP9;refGene_tx=XM_019828808.2;refGene_gn=MAP9;refGene_tx=XM_023252747.1;refGene_gn=MAP9;refGene_tx=XM_023252748.1;refGene_gn=MAP9;refGene_tx=XM_023252749.1
B1 74470509 74470830
refGene_gn=MAP9;refGene_tx=XM_019828808.2;refGene_gn=MAP9;refGene_tx=XM_023252747.1;refGene_gn=MAP9;refGene_tx=XM_023252748.1;refGene_gn=MAP9;refGene_tx=XM_023252749.1
B1 74476980 74477210
refGene_gn=MAP9;refGene_tx=XM_019828808.2;refGene_gn=MAP9;refGene_tx=XM_023252747.1;refGene_gn=MAP9;refGene_tx=XM_023252748.1;refGene_gn=MAP9;refGene_tx=XM_023252749.1
B1 74485407 74485501
refGene_gn=MAP9;refGene_tx=XM_019828808.2;refGene_gn=MAP9;refGene_tx=XM_023252747.1;refGene_gn=MAP9;refGene_tx=XM_023252748.1;refGene_gn=MAP9;refGene_tx=XM_023252749.1
B1 74487271 74487516
refGene_gn=MAP9;refGene_tx=XM_019828808.2;refGene_gn=MAP9;refGene_tx=XM_023252747.1;refGene_gn=MAP9;refGene_tx=XM_023252748.1;refGene_gn=MAP9;refGene_tx=XM_023252749.1
B1 74489369 74489440
refGene_gn=MAP9;refGene_tx=XM_019828808.2;refGene_gn=MAP9;refGene_tx=XM_023252747.1;refGene_gn=MAP9;refGene_tx=XM_023252748.1;refGene_gn=MAP9;refGene_tx=XM_023252749.1
B1 74490462 74490631
refGene_gn=MAP9;refGene_tx=XM_019828808.2;refGene_gn=MAP9;refGene_tx=XM_023252747.1;refGene_gn=MAP9;refGene_tx=XM_023252748.1;refGene_gn=MAP9;refGene_tx=XM_023252749.1
B1 74491681 74491771
refGene_gn=MAP9;refGene_tx=XM_019828808.2;refGene_gn=MAP9;refGene_tx=XM_023252747.1;refGene_gn=MAP9;refGene_tx=XM_023252748.1;refGene_gn=MAP

9;refGene_tx=XM_023252749.1
B1	74493659	74493821
refGene_gn=MAP9;refGene_tx=XM_019828808.2;refGene_gn=MAP9;refGene_tx=XM_023252747.1;refGene_gn=MAP9;refGene_tx=XM_023252748.1;refGene_gn=MAP9;refGene_tx=XM_023252749.1
B1	74493951	74494097
refGene_gn=MAP9;refGene_tx=XM_019828808.2;refGene_gn=MAP9;refGene_tx=XM_023252747.1;refGene_gn=MAP9;refGene_tx=XM_023252748.1;refGene_gn=MAP9;refGene_tx=XM_023252749.1
B1	74494276	74494409
refGene_gn=MAP9;refGene_tx=XM_019828808.2;refGene_gn=MAP9;refGene_tx=XM_023252747.1;refGene_gn=MAP9;refGene_tx=XM_023252748.1;refGene_gn=MAP9;refGene_tx=XM_023252749.1
B1	74501436	74501559
refGene_gn=MAP9;refGene_tx=XM_019828808.2;refGene_gn=MAP9;refGene_tx=XM_023252747.1;refGene_gn=MAP9;refGene_tx=XM_023252748.1;refGene_gn=MAP9;refGene_tx=XM_023252749.1
B1	74616982	74618131	refGene_gn=NPY2R;refGene_tx=XM_006930805.3
B1	74965360	74965560
refGene_gn=RBM46;refGene_tx=XM_006930807.3;refGene_gn=RBM46;refGene_tx=XM_006930808.3;refGene_gn=RBM46;refGene_tx=XM_019828809.2
B1	74974021	74974077
refGene_gn=RBM46;refGene_tx=XM_023252750.1;refGene_gn=RBM46;refGene_tx=XM_023252751.1
B1	74977652	74977672	refGene_gn=RBM46;refGene_tx=XM_023252752.1
B1	74994658	74995441
refGene_gn=RBM46;refGene_tx=XM_006930807.3;refGene_gn=RBM46;refGene_tx=XM_006930808.3;refGene_gn=RBM46;refGene_tx=XM_019828809.2;refGene_gn=RBM46;refGene_tx=XM_023252750.1;refGene_gn=RBM46;refGene_tx=XM_023252751.1;refGene_gn=RBM46;refGene_tx=XM_023252752.1
B1	74995983	74996451
refGene_gn=RBM46;refGene_tx=XM_006930807.3;refGene_gn=RBM46;refGene_tx=XM_006930808.3;refGene_gn=RBM46;refGene_tx=XM_019828809.2;refGene_gn=RBM46;refGene_tx=XM_023252750.1;refGene_gn=RBM46;refGene_tx=XM_023252751.1;refGene_gn=RBM46;refGene_tx=XM_023252752.1
B1	74999580	74999731
refGene_gn=RBM46;refGene_tx=XM_006930807.3;refGene_gn=RBM46;refGene_tx=XM_006930808.3;refGene_gn=RBM46;refGene_tx=XM_019828809.2;refGene_gn=RBM46;refGene_tx=XM_023252750.1;refGene_gn=RBM46;refGene_tx=XM_023252751.1;refGene_gn=RBM46;refGene_tx=XM_023252752.1
B1	75044783	75044936
refGene_gn=LRAT;refGene_tx=XM_006930809.4;refGene_gn=LRAT;refGene_tx=XM_006930810.3;refGene_gn=LRAT;refGene_tx=XM_019828817.2
B1	75045645	75045735	refGene_gn=LRAT;refGene_tx=XM_019828818.1
B1	75049174	75049714
refGene_gn=LRAT;refGene_tx=XM_006930809.4;refGene_gn=LRAT;refGene_tx=XM_006930810.3;refGene_gn=LRAT;refGene_tx=XM_019828817.2;refGene_gn=LRAT;refGene_tx=XM_019828818.1
B1	75180513	75180591	refGene_gn=FGG;refGene_tx=XM_003984925.4
B1	75180696	75180741	refGene_gn=FGG;refGene_tx=XM_003984925.4

B1 75181164 75181348 refGene_gn=FGG;refGene_tx=XM_003984925.4
B1 75181455 75181549 refGene_gn=FGG;refGene_tx=XM_003984925.4
B1 75183074 75183205 refGene_gn=FGG;refGene_tx=XM_003984925.4
B1 75183556 75183690 refGene_gn=FGG;refGene_tx=XM_003984925.4
B1 75184563 75184748 refGene_gn=FGG;refGene_tx=XM_003984925.4
B1 75186221 75186499 refGene_gn=FGG;refGene_tx=XM_003984925.4
B1 75187839 75188009 refGene_gn=FGG;refGene_tx=XM_003984925.4
B1 75188507 75188522 refGene_gn=FGG;refGene_tx=XM_003984925.4
B1 75200623 75200677 refGene_gn=FGA;refGene_tx=XM_019828819.2
B1 75202955 75203081 refGene_gn=FGA;refGene_tx=XM_019828819.2
B1 75203546 75203730 refGene_gn=FGA;refGene_tx=XM_019828819.2
B1 75204827 75204973 refGene_gn=FGA;refGene_tx=XM_019828819.2
B1 75205562 75207036 refGene_gn=FGA;refGene_tx=XM_019828819.2
B1 75207801 75208523 refGene_gn=FGA;refGene_tx=XM_019828819.2
B1 75221386 75221618 refGene_gn=FGB;refGene_tx=XM_019828820.1
B1 75222232 75222518 refGene_gn=FGB;refGene_tx=XM_019828820.1
B1 75223019 75223145 refGene_gn=FGB;refGene_tx=XM_019828820.1
B1 75223501 75223615 refGene_gn=FGB;refGene_tx=XM_019828820.1
B1 75224146 75224374 refGene_gn=FGB;refGene_tx=XM_019828820.1
B1 75225330 75225514 refGene_gn=FGB;refGene_tx=XM_019828820.1
B1 75225993 75226185 refGene_gn=FGB;refGene_tx=XM_019828820.1
B1 75228538 75228559 refGene_gn=FGB;refGene_tx=XM_019828820.1
B1 75228774 75228849 refGene_gn=FGB;refGene_tx=XM_019828820.1
B1 75237488 75237497 refGene_gn=PLRG1;refGene_tx=XM_011281610.3
B1 75239047 75239154 refGene_gn=PLRG1;refGene_tx=XM_011281610.3
B1 75240190 75240333
refGene_gn=PLRG1;refGene_tx=XM_011281610.3;refGene_gn=PLRG1;refGene_tx=XM_011281611.3
B1 75241033 75241087
refGene_gn=PLRG1;refGene_tx=XM_011281610.3;refGene_gn=PLRG1;refGene_tx=XM_011281611.3
B1 75242196 75242290
refGene_gn=PLRG1;refGene_tx=XM_011281610.3;refGene_gn=PLRG1;refGene_tx=XM_011281611.3
B1 75242489 75242577
refGene_gn=PLRG1;refGene_tx=XM_011281610.3;refGene_gn=PLRG1;refGene_tx=XM_011281611.3
B1 75244182 75244284
refGene_gn=PLRG1;refGene_tx=XM_011281610.3;refGene_gn=PLRG1;refGene_tx=XM_011281611.3
B1 75245269 75245362
refGene_gn=PLRG1;refGene_tx=XM_011281610.3;refGene_gn=PLRG1;refGene_tx=XM_011281611.3
B1 75247893 75248043
refGene_gn=PLRG1;refGene_tx=XM_011281610.3;refGene_gn=PLRG1;refGene_tx=XM_011281611.3
B1 75248132 75248234
refGene_gn=PLRG1;refGene_tx=XM_011281610.3;refGene_gn=PLRG1;refGene_tx=XM_011281611.3
B1 75248769 75248872

refGene_gn=PLRG1;refGene_tx=XM_011281610.3;refGene_gn=PLRG1;refGene_tx=XM_011281611.3
B1 75250246 75250355
refGene_gn=PLRG1;refGene_tx=XM_011281610.3;refGene_gn=PLRG1;refGene_tx=XM_011281611.3
B1 75251306 75251446
refGene_gn=PLRG1;refGene_tx=XM_011281610.3;refGene_gn=PLRG1;refGene_tx=XM_011281611.3
B1 75251947 75252141
refGene_gn=PLRG1;refGene_tx=XM_011281610.3;refGene_gn=PLRG1;refGene_tx=XM_011281611.3
B1 75253096 75253156
refGene_gn=PLRG1;refGene_tx=XM_011281610.3;refGene_gn=PLRG1;refGene_tx=XM_011281611.3
B1 75316243 75318262 refGene_gn=DCHS2;refGene_tx=XM_023252753.1
B1 75436848 75437040 refGene_gn=DCHS2;refGene_tx=XM_023252753.1
B1 75452141 75452373 refGene_gn=DCHS2;refGene_tx=XM_023252753.1
B1 75468568 75468805 refGene_gn=DCHS2;refGene_tx=XM_023252753.1
B1 75470115 75471132 refGene_gn=DCHS2;refGene_tx=XM_023252753.1
B1 75473796 75473984 refGene_gn=DCHS2;refGene_tx=XM_023252753.1
B1 75475278 75475378 refGene_gn=DCHS2;refGene_tx=XM_023252753.1
B1 75479758 75479916 refGene_gn=DCHS2;refGene_tx=XM_023252753.1
B1 75481258 75482102 refGene_gn=DCHS2;refGene_tx=XM_023252753.1
B1 75484898 75485138 refGene_gn=DCHS2;refGene_tx=XM_023252753.1
B1 75499292 75499427 refGene_gn=DCHS2;refGene_tx=XM_023252753.1
B1 75499636 75499846 refGene_gn=DCHS2;refGene_tx=XM_023252753.1
B1 75505134 75505992 refGene_gn=DCHS2;refGene_tx=XM_023252753.1
B1 75525483 75525597 refGene_gn=DCHS2;refGene_tx=XM_023252753.1
B1 75535938 75536150 refGene_gn=DCHS2;refGene_tx=XM_023252753.1
B1 75540540 75540692 refGene_gn=DCHS2;refGene_tx=XM_023252753.1
B1 75550009 75550140 refGene_gn=DCHS2;refGene_tx=XM_023252753.1
B1 75552140 75552427 refGene_gn=DCHS2;refGene_tx=XM_023252753.1
B1 75553329 75553462 refGene_gn=DCHS2;refGene_tx=XM_023252753.1
B1 75555577 75558195 refGene_gn=DCHS2;refGene_tx=XM_023252753.1
B1 75944311 75944576 refGene_gn=SFRP2;refGene_tx=XM_003984929.4
B1 75944578 75944809 refGene_gn=SFRP2;refGene_tx=XM_003984929.4
B1 75947222 75947303 refGene_gn=SFRP2;refGene_tx=XM_003984929.4
B1 75951349 75951654 refGene_gn=SFRP2;refGene_tx=XM_003984929.4
B1 75980921 75980927 refGene_gn=RNF175;refGene_tx=XM_011281613.3
B1 75984200 75984365
refGene_gn=RNF175;refGene_tx=XM_019828822.1;refGene_gn=RNF175;refGene_tx=XM_019828823.2
B1 75998982 75999137
refGene_gn=RNF175;refGene_tx=XM_011281613.3;refGene_gn=RNF175;refGene_tx=XM_019828822.1;refGene_gn=RNF175;refGene_tx=XM_019828823.2
B1 76004225 76004333
refGene_gn=RNF175;refGene_tx=XM_011281613.3;refGene_gn=RNF175;refGene_tx=XM_019828822.1;refGene_gn=RNF175;refGene_tx=XM_019828823.2
B1 76007205 76007326
refGene_gn=RNF175;refGene_tx=XM_011281613.3;refGene_gn=RNF175;refGene_

tx=XM_019828822.1;refGene_gn=RNF175;refGene_tx=XM_019828823.2
B1	76012740	76012874
refGene_gn=RNF175;refGene_tx=XM_011281613.3;refGene_gn=RNF175;refGene_tx=XM_019828822.1;refGene_gn=RNF175;refGene_tx=XM_019828823.2
B1	76016119	76016221
refGene_gn=RNF175;refGene_tx=XM_011281613.3;refGene_gn=RNF175;refGene_tx=XM_019828822.1
B1	76018375	76018496
refGene_gn=RNF175;refGene_tx=XM_011281613.3;refGene_gn=RNF175;refGene_tx=XM_019828822.1;refGene_gn=RNF175;refGene_tx=XM_019828823.2
B1	76023403	76025758	refGene_gn=TLR2;refGene_tx=XM_003984930.5
B1	76071412	76071688	refGene_gn=TMEM131L;refGene_tx=XM_023252754.1
B1	76073137	76073277	refGene_gn=TMEM131L;refGene_tx=XM_023252754.1
B1	76075186	76075275	refGene_gn=TMEM131L;refGene_tx=XM_023252754.1
B1	76076566	76076687	refGene_gn=TMEM131L;refGene_tx=XM_023252754.1
B1	76084611	76084694	refGene_gn=TMEM131L;refGene_tx=XM_023252754.1
B1	76085572	76085651	refGene_gn=TMEM131L;refGene_tx=XM_023252754.1
B1	76090844	76091033	refGene_gn=TMEM131L;refGene_tx=XM_023252754.1
B1	76092221	76092388	refGene_gn=TMEM131L;refGene_tx=XM_023252754.1
B1	76095598	76095720	refGene_gn=TMEM131L;refGene_tx=XM_023252754.1
B1	76104426	76104575	refGene_gn=TMEM131L;refGene_tx=XM_023252754.1
B1	76110767	76111393	refGene_gn=TMEM131L;refGene_tx=XM_023252754.1
B1	76111774	76111924	refGene_gn=TMEM131L;refGene_tx=XM_023252754.1
B1	76112736	76112922	refGene_gn=TMEM131L;refGene_tx=XM_023252754.1
B1	76113132	76113319	refGene_gn=TMEM131L;refGene_tx=XM_023252754.1
B1	76115573	76115716	refGene_gn=TMEM131L;refGene_tx=XM_023252754.1
B1	76117778	76117906	refGene_gn=TMEM131L;refGene_tx=XM_023252754.1
B1	76121136	76121209	refGene_gn=TMEM131L;refGene_tx=XM_023252754.1
B1	76122213	76122323	refGene_gn=TMEM131L;refGene_tx=XM_023252754.1
B1	76124722	76124864	refGene_gn=TMEM131L;refGene_tx=XM_023252754.1
B1	76126202	76126320	refGene_gn=TMEM131L;refGene_tx=XM_023252754.1
B1	76127386	76127441	refGene_gn=TMEM131L;refGene_tx=XM_023252754.1
B1	76127442	76127457	refGene_gn=TMEM131L;refGene_tx=XM_023252754.1
B1	76128781	76128952	refGene_gn=TMEM131L;refGene_tx=XM_023252754.1
B1	76129670	76129824	refGene_gn=TMEM131L;refGene_tx=XM_023252754.1
B1	76130323	76130417	refGene_gn=TMEM131L;refGene_tx=XM_023252754.1
B1	76132723	76132832	refGene_gn=TMEM131L;refGene_tx=XM_023252754.1
B1	76133144	76133203	refGene_gn=TMEM131L;refGene_tx=XM_023252754.1
B1	76134810	76134964	refGene_gn=TMEM131L;refGene_tx=XM_023252754.1
B1	76135171	76135249	refGene_gn=TMEM131L;refGene_tx=XM_023252754.1
B1	76155557	76155668	refGene_gn=TMEM131L;refGene_tx=XM_023252754.1
B1	76157011	76157128	refGene_gn=TMEM131L;refGene_tx=XM_023252754.1
B1	76158175	76158299	refGene_gn=TMEM131L;refGene_tx=XM_023252754.1
B1	76163081	76163150	refGene_gn=TMEM131L;refGene_tx=XM_023252754.1
B1	76233831	76233875	refGene_gn=TMEM131L;refGene_tx=XM_023252754.1
B1	76243467	76243538	refGene_gn=TMEM131L;refGene_tx=XM_023252754.1
B1	76244238	76244362	refGene_gn=TMEM131L;refGene_tx=XM_023252754.1
B1	76299251	76299358
refGene_gn=MND1;refGene_tx=XM_011281614.3;refGene_gn=MND1;refGene_tx=XM_023252755.1;refGene_gn=MND1;refGene_tx=XM_023252756.1

B1 76307790 76307835
refGene_gn=MND1;refGene_tx=XM_011281614.3;refGene_gn=MND1;refGene_tx=XM_023252755.1;refGene_gn=MND1;refGene_tx=XM_023252756.1
B1 76318199 76318314
refGene_gn=MND1;refGene_tx=XM_011281614.3;refGene_gn=MND1;refGene_tx=XM_023252755.1;refGene_gn=MND1;refGene_tx=XM_023252756.1
B1 76321364 76321439
refGene_gn=MND1;refGene_tx=XM_011281614.3;refGene_gn=MND1;refGene_tx=XM_023252755.1;refGene_gn=MND1;refGene_tx=XM_023252756.1
B1 76351992 76352141
refGene_gn=MND1;refGene_tx=XM_011281614.3;refGene_gn=MND1;refGene_tx=XM_023252755.1;refGene_gn=MND1;refGene_tx=XM_023252756.1
B1 76355357 76355415
refGene_gn=MND1;refGene_tx=XM_011281614.3;refGene_gn=MND1;refGene_tx=XM_023252755.1;refGene_gn=MND1;refGene_tx=XM_023252756.1
B1 76357308 76357374
refGene_gn=MND1;refGene_tx=XM_011281614.3;refGene_gn=MND1;refGene_tx=XM_023252755.1;refGene_gn=MND1;refGene_tx=XM_023252756.1
B1 76363395 76363401 refGene_gn=MND1;refGene_tx=XM_011281614.3
B1 76363908 76363911 refGene_gn=MND1;refGene_tx=XM_023252755.1
B1 76376883 76377036
refGene_gn=TRIM2;refGene_tx=XM_011281615.3;refGene_gn=TRIM2;refGene_tx=XM_019828831.2;refGene_gn=TRIM2;refGene_tx=XM_019828832.2;refGene_gn=TRIM2;refGene_tx=XM_019828833.2;refGene_gn=TRIM2;refGene_tx=XM_019828834.2;refGene_gn=TRIM2;refGene_tx=XM_023252757.1;refGene_gn=TRIM2;refGene_tx=XM_023252758.1;refGene_gn=TRIM2;refGene_tx=XM_023252759.1
B1 76381306 76381447
refGene_gn=TRIM2;refGene_tx=XM_011281615.3;refGene_gn=TRIM2;refGene_tx=XM_019828831.2;refGene_gn=TRIM2;refGene_tx=XM_019828832.2;refGene_gn=TRIM2;refGene_tx=XM_019828833.2;refGene_gn=TRIM2;refGene_tx=XM_019828834.2;refGene_gn=TRIM2;refGene_tx=XM_023252757.1;refGene_gn=TRIM2;refGene_tx=XM_023252758.1;refGene_gn=TRIM2;refGene_tx=XM_023252759.1
B1 76385721 76385792
refGene_gn=TRIM2;refGene_tx=XM_011281615.3;refGene_gn=TRIM2;refGene_tx=XM_019828831.2;refGene_gn=TRIM2;refGene_tx=XM_019828832.2;refGene_gn=TRIM2;refGene_tx=XM_019828833.2;refGene_gn=TRIM2;refGene_tx=XM_019828834.2;refGene_gn=TRIM2;refGene_tx=XM_023252757.1;refGene_gn=TRIM2;refGene_tx=XM_023252758.1;refGene_gn=TRIM2;refGene_tx=XM_023252759.1
B1 76386836 76387017
refGene_gn=TRIM2;refGene_tx=XM_011281615.3;refGene_gn=TRIM2;refGene_tx=XM_019828831.2;refGene_gn=TRIM2;refGene_tx=XM_019828832.2;refGene_gn=TRIM2;refGene_tx=XM_019828833.2;refGene_gn=TRIM2;refGene_tx=XM_019828834.2;refGene_gn=TRIM2;refGene_tx=XM_023252757.1;refGene_gn=TRIM2;refGene_tx=XM_023252758.1;refGene_gn=TRIM2;refGene_tx=XM_023252759.1
B1 76392178 76392181 refGene_gn=TRIM2;refGene_tx=XM_011281618.3
B1 76393271 76393439
refGene_gn=TRIM2;refGene_tx=XM_011281615.3;refGene_gn=TRIM2;refGene_tx=XM_011281618.3;refGene_gn=TRIM2;refGene_tx=XM_019828831.2;refGene_gn=TRIM2;refGene_tx=XM_019828832.2;refGene_gn=TRIM2;refGene_tx=XM_019828833.2;refGene_gn=TRIM2;refGene_tx=XM_019828834.2;refGene_gn=TRIM2;refGe

ne_tx=XM_023252757.1;refGene_gn=TRIM2;refGene_tx=XM_023252758.1;refGene_gn=TRIM2;refGene_tx=XM_023252759.1
B1	76393675	76393779
refGene_gn=TRIM2;refGene_tx=XM_011281615.3;refGene_gn=TRIM2;refGene_tx=XM_011281618.3;refGene_gn=TRIM2;refGene_tx=XM_019828831.2;refGene_gn=TRIM2;refGene_tx=XM_019828832.2;refGene_gn=TRIM2;refGene_tx=XM_019828833.2;refGene_gn=TRIM2;refGene_tx=XM_019828834.2;refGene_gn=TRIM2;refGene_tx=XM_023252757.1;refGene_gn=TRIM2;refGene_tx=XM_023252758.1;refGene_gn=TRIM2;refGene_tx=XM_023252759.1
B1	76409619	76410343
refGene_gn=TRIM2;refGene_tx=XM_011281615.3;refGene_gn=TRIM2;refGene_tx=XM_011281618.3;refGene_gn=TRIM2;refGene_tx=XM_019828831.2;refGene_gn=TRIM2;refGene_tx=XM_019828832.2;refGene_gn=TRIM2;refGene_tx=XM_019828833.2;refGene_gn=TRIM2;refGene_tx=XM_019828834.2;refGene_gn=TRIM2;refGene_tx=XM_023252757.1;refGene_gn=TRIM2;refGene_tx=XM_023252758.1;refGene_gn=TRIM2;refGene_tx=XM_023252759.1
B1	76411204	76411385
refGene_gn=TRIM2;refGene_tx=XM_011281615.3;refGene_gn=TRIM2;refGene_tx=XM_011281618.3;refGene_gn=TRIM2;refGene_tx=XM_019828831.2;refGene_gn=TRIM2;refGene_tx=XM_019828832.2;refGene_gn=TRIM2;refGene_tx=XM_019828833.2;refGene_gn=TRIM2;refGene_tx=XM_019828834.2;refGene_gn=TRIM2;refGene_tx=XM_023252757.1;refGene_gn=TRIM2;refGene_tx=XM_023252758.1;refGene_gn=TRIM2;refGene_tx=XM_023252759.1
B1	76412560	76412712
refGene_gn=TRIM2;refGene_tx=XM_011281615.3;refGene_gn=TRIM2;refGene_tx=XM_011281618.3;refGene_gn=TRIM2;refGene_tx=XM_019828831.2;refGene_gn=TRIM2;refGene_tx=XM_019828832.2;refGene_gn=TRIM2;refGene_tx=XM_019828833.2;refGene_gn=TRIM2;refGene_tx=XM_019828834.2;refGene_gn=TRIM2;refGene_tx=XM_023252757.1;refGene_gn=TRIM2;refGene_tx=XM_023252758.1;refGene_gn=TRIM2;refGene_tx=XM_023252759.1
B1	76423925	76424163
refGene_gn=TRIM2;refGene_tx=XM_011281615.3;refGene_gn=TRIM2;refGene_tx=XM_011281618.3;refGene_gn=TRIM2;refGene_tx=XM_019828831.2;refGene_gn=TRIM2;refGene_tx=XM_019828832.2;refGene_gn=TRIM2;refGene_tx=XM_019828833.2;refGene_gn=TRIM2;refGene_tx=XM_019828834.2;refGene_gn=TRIM2;refGene_tx=XM_023252757.1;refGene_gn=TRIM2;refGene_tx=XM_023252759.1
B1	76430072	76430257
refGene_gn=TRIM2;refGene_tx=XM_011281615.3;refGene_gn=TRIM2;refGene_tx=XM_011281618.3;refGene_gn=TRIM2;refGene_tx=XM_019828831.2;refGene_gn=TRIM2;refGene_tx=XM_019828832.2;refGene_gn=TRIM2;refGene_tx=XM_019828833.2;refGene_gn=TRIM2;refGene_tx=XM_019828834.2;refGene_gn=TRIM2;refGene_tx=XM_023252757.1;refGene_gn=TRIM2;refGene_tx=XM_023252758.1;refGene_gn=TRIM2;refGene_tx=XM_023252759.1
B1	76440239	76440242
refGene_gn=TRIM2;refGene_tx=XM_019828833.2;refGene_gn=TRIM2;refGene_tx=XM_019828834.2
B1	76494238	76494268
refGene_gn=TRIM2;refGene_tx=XM_011281615.3;refGene_gn=TRIM2;refGene_tx=XM_011281618.3;refGene_gn=TRIM2;refGene_tx=XM_019828831.2;refGene_gn=TRIM2;refGene_tx=XM_019828832.2

B1 76702476 76704498
refGene_gn=FHDC1;refGene_tx=XM_003984933.5;refGene_gn=FHDC1;refGene_tx=XM_019828837.2;refGene_gn=FHDC1;refGene_tx=XM_019828838.2;refGene_gn=FHDC1;refGene_tx=XM_023252760.1
B1 76706742 76706907
refGene_gn=FHDC1;refGene_tx=XM_003984933.5;refGene_gn=FHDC1;refGene_tx=XM_019828837.2;refGene_gn=FHDC1;refGene_tx=XM_019828838.2;refGene_gn=FHDC1;refGene_tx=XM_023252760.1
B1 76710677 76710795
refGene_gn=FHDC1;refGene_tx=XM_003984933.5;refGene_gn=FHDC1;refGene_tx=XM_019828837.2;refGene_gn=FHDC1;refGene_tx=XM_019828838.2;refGene_gn=FHDC1;refGene_tx=XM_023252760.1
B1 76712883 76712954
refGene_gn=FHDC1;refGene_tx=XM_003984933.5;refGene_gn=FHDC1;refGene_tx=XM_019828837.2;refGene_gn=FHDC1;refGene_tx=XM_019828838.2;refGene_gn=FHDC1;refGene_tx=XM_023252760.1
B1 76714962 76715070
refGene_gn=FHDC1;refGene_tx=XM_003984933.5;refGene_gn=FHDC1;refGene_tx=XM_019828837.2;refGene_gn=FHDC1;refGene_tx=XM_019828838.2;refGene_gn=FHDC1;refGene_tx=XM_023252760.1
B1 76715170 76715241
refGene_gn=FHDC1;refGene_tx=XM_003984933.5;refGene_gn=FHDC1;refGene_tx=XM_019828837.2;refGene_gn=FHDC1;refGene_tx=XM_019828838.2;refGene_gn=FHDC1;refGene_tx=XM_023252760.1
B1 76719169 76719270
refGene_gn=FHDC1;refGene_tx=XM_003984933.5;refGene_gn=FHDC1;refGene_tx=XM_019828837.2;refGene_gn=FHDC1;refGene_tx=XM_019828838.2;refGene_gn=FHDC1;refGene_tx=XM_023252760.1
B1 76719357 76719443
refGene_gn=FHDC1;refGene_tx=XM_003984933.5;refGene_gn=FHDC1;refGene_tx=XM_019828837.2;refGene_gn=FHDC1;refGene_tx=XM_019828838.2;refGene_gn=FHDC1;refGene_tx=XM_023252760.1
B1 76724952 76725055
refGene_gn=FHDC1;refGene_tx=XM_003984933.5;refGene_gn=FHDC1;refGene_tx=XM_019828837.2;refGene_gn=FHDC1;refGene_tx=XM_019828838.2;refGene_gn=FHDC1;refGene_tx=XM_023252760.1
B1 76725709 76725771
refGene_gn=FHDC1;refGene_tx=XM_003984933.5;refGene_gn=FHDC1;refGene_tx=XM_019828837.2;refGene_gn=FHDC1;refGene_tx=XM_019828838.2;refGene_gn=FHDC1;refGene_tx=XM_023252760.1
B1 76734866 76735352
refGene_gn=FHDC1;refGene_tx=XM_003984933.5;refGene_gn=FHDC1;refGene_tx=XM_019828837.2;refGene_gn=FHDC1;refGene_tx=XM_019828838.2;refGene_gn=FHDC1;refGene_tx=XM_023252760.1
B1 76760213 76760369
refGene_gn=ARFIP1;refGene_tx=XM_006930817.3;refGene_gn=ARFIP1;refGene_tx=XM_019828849.2
B1 76775811 76775871 refGene_gn=ARFIP1;refGene_tx=XM_006930819.4
B1 76780303 76780462
refGene_gn=ARFIP1;refGene_tx=XM_019828844.2;refGene_gn=ARFIP1;refGene_

tx=XM_019828845.2;refGene_gn=ARFIP1;refGene_tx=XM_019828846.2;refGene_gn=ARFIP1;refGene_tx=XM_019828847.2;refGene_gn=ARFIP1;refGene_tx=XM_019828848.2;refGene_gn=ARFIP1;refGene_tx=XM_019828851.2
B1 76788349 76788524
refGene_gn=ARFIP1;refGene_tx=XM_006930817.3;refGene_gn=ARFIP1;refGene_tx=XM_006930819.4;refGene_gn=ARFIP1;refGene_tx=XM_019828844.2;refGene_gn=ARFIP1;refGene_tx=XM_019828845.2;refGene_gn=ARFIP1;refGene_tx=XM_019828846.2;refGene_gn=ARFIP1;refGene_tx=XM_019828847.2;refGene_gn=ARFIP1;refGene_tx=XM_019828848.2;refGene_gn=ARFIP1;refGene_tx=XM_019828849.2;refGene_gn=ARFIP1;refGene_tx=XM_019828851.2
B1 76794964 76795122
refGene_gn=ARFIP1;refGene_tx=XM_006930817.3;refGene_gn=ARFIP1;refGene_tx=XM_006930819.4;refGene_gn=ARFIP1;refGene_tx=XM_019828844.2;refGene_gn=ARFIP1;refGene_tx=XM_019828845.2;refGene_gn=ARFIP1;refGene_tx=XM_019828846.2;refGene_gn=ARFIP1;refGene_tx=XM_019828847.2;refGene_gn=ARFIP1;refGene_tx=XM_019828848.2;refGene_gn=ARFIP1;refGene_tx=XM_019828849.2;refGene_gn=ARFIP1;refGene_tx=XM_019828851.2
B1 76803084 76803306
refGene_gn=ARFIP1;refGene_tx=XM_006930817.3;refGene_gn=ARFIP1;refGene_tx=XM_006930819.4;refGene_gn=ARFIP1;refGene_tx=XM_019828844.2;refGene_gn=ARFIP1;refGene_tx=XM_019828845.2;refGene_gn=ARFIP1;refGene_tx=XM_019828846.2;refGene_gn=ARFIP1;refGene_tx=XM_019828847.2;refGene_gn=ARFIP1;refGene_tx=XM_019828848.2;refGene_gn=ARFIP1;refGene_tx=XM_019828849.2
B1 76812456 76812569
refGene_gn=ARFIP1;refGene_tx=XM_006930817.3;refGene_gn=ARFIP1;refGene_tx=XM_006930819.4;refGene_gn=ARFIP1;refGene_tx=XM_019828844.2;refGene_gn=ARFIP1;refGene_tx=XM_019828845.2;refGene_gn=ARFIP1;refGene_tx=XM_019828846.2;refGene_gn=ARFIP1;refGene_tx=XM_019828847.2;refGene_gn=ARFIP1;refGene_tx=XM_019828848.2;refGene_gn=ARFIP1;refGene_tx=XM_019828849.2;refGene_gn=ARFIP1;refGene_tx=XM_019828851.2
B1 76814161 76814257
refGene_gn=ARFIP1;refGene_tx=XM_006930817.3;refGene_gn=ARFIP1;refGene_tx=XM_006930819.4;refGene_gn=ARFIP1;refGene_tx=XM_019828844.2;refGene_gn=ARFIP1;refGene_tx=XM_019828845.2;refGene_gn=ARFIP1;refGene_tx=XM_019828846.2;refGene_gn=ARFIP1;refGene_tx=XM_019828847.2;refGene_gn=ARFIP1;refGene_tx=XM_019828851.2
B1 76820519 76820628
refGene_gn=ARFIP1;refGene_tx=XM_006930817.3;refGene_gn=ARFIP1;refGene_tx=XM_006930819.4;refGene_gn=ARFIP1;refGene_tx=XM_019828844.2;refGene_gn=ARFIP1;refGene_tx=XM_019828845.2;refGene_gn=ARFIP1;refGene_tx=XM_019828846.2;refGene_gn=ARFIP1;refGene_tx=XM_019828847.2;refGene_gn=ARFIP1;refGene_tx=XM_019828848.2;refGene_gn=ARFIP1;refGene_tx=XM_019828849.2;refGene_gn=ARFIP1;refGene_tx=XM_019828851.2
B1 76838238 76838340
refGene_gn=ARFIP1;refGene_tx=XM_006930817.3;refGene_gn=ARFIP1;refGene_tx=XM_006930819.4;refGene_gn=ARFIP1;refGene_tx=XM_019828844.2;refGene_gn=ARFIP1;refGene_tx=XM_019828845.2;refGene_gn=ARFIP1;refGene_tx=XM_019828846.2;refGene_gn=ARFIP1;refGene_tx=XM_019828847.2;refGene_gn=ARFIP1;refGene_tx=XM_019828848.2;refGene_gn=ARFIP1;refGene_tx=XM_019828849.

2;refGene_gn=ARFIP1;refGene_tx=XM_019828851.2
B1 76872870 76872948
refGene_gn=ARFIP1;refGene_tx=XM_006930817.3;refGene_gn=ARFIP1;refGene_tx=XM_006930819.4;refGene_gn=ARFIP1;refGene_tx=XM_019828844.2;refGene_gn=ARFIP1;refGene_tx=XM_019828845.2;refGene_gn=ARFIP1;refGene_tx=XM_019828848.2;refGene_gn=ARFIP1;refGene_tx=XM_019828849.2;refGene_gn=ARFIP1;refGene_tx=XM_019828851.2
B1 76895605 76897141
refGene_gn=TIGD4;refGene_tx=XM_003984937.4;refGene_gn=TIGD4;refGene_tx=XM_019828839.2;refGene_gn=TIGD4;refGene_tx=XM_019828840.2
B1 76973937 76974025
refGene_gn=TMEM154;refGene_tx=XM_019828852.2;refGene_gn=TMEM154;refGene_tx=XM_019828853.2;refGene_gn=TMEM154;refGene_tx=XM_019828854.2;refGene_gn=TMEM154;refGene_tx=XM_019828855.2
B1 76997156 76997417
refGene_gn=TMEM154;refGene_tx=XM_019828852.2;refGene_gn=TMEM154;refGene_tx=XM_019828853.2;refGene_gn=TMEM154;refGene_tx=XM_019828854.2;refGene_gn=TMEM154;refGene_tx=XM_019828855.2
B1 76997508 76997547
refGene_gn=TMEM154;refGene_tx=XM_019828852.2;refGene_gn=TMEM154;refGene_tx=XM_019828853.2;refGene_gn=TMEM154;refGene_tx=XM_019828854.2;refGene_gn=TMEM154;refGene_tx=XM_019828855.2
B1 77003754 77003782
refGene_gn=TMEM154;refGene_tx=XM_019828852.2;refGene_gn=TMEM154;refGene_tx=XM_019828853.2;refGene_gn=TMEM154;refGene_tx=XM_019828854.2;refGene_gn=TMEM154;refGene_tx=XM_019828855.2
B1 77005031 77005117
refGene_gn=TMEM154;refGene_tx=XM_019828852.2;refGene_gn=TMEM154;refGene_tx=XM_019828853.2;refGene_gn=TMEM154;refGene_tx=XM_019828854.2;refGene_gn=TMEM154;refGene_tx=XM_019828855.2
B1 77007667 77007725
refGene_gn=TMEM154;refGene_tx=XM_019828852.2;refGene_gn=TMEM154;refGene_tx=XM_019828853.2;refGene_gn=TMEM154;refGene_tx=XM_019828854.2;refGene_gn=TMEM154;refGene_tx=XM_019828855.2
B1 77017157 77017173 refGene_gn=TMEM154;refGene_tx=XM_019828853.2
B1 77017200 77017282 refGene_gn=TMEM154;refGene_tx=XM_019828852.2
B1 77020927 77020943
refGene_gn=TMEM154;refGene_tx=XM_019828854.2;refGene_gn=TMEM154;refGene_tx=XM_019828855.2
B1 77237461 77237974
refGene_gn=FBXW7;refGene_tx=XM_003984939.5;refGene_gn=FBXW7;refGene_tx=XM_006930821.3;refGene_gn=FBXW7;refGene_tx=XM_019828857.2
B1 77269780 77269927 refGene_gn=FBXW7;refGene_tx=XM_003984940.5
B1 77299007 77299268 refGene_gn=FBXW7;refGene_tx=XM_003984938.5
B1 77301681 77301764
refGene_gn=FBXW7;refGene_tx=XM_003984938.5;refGene_gn=FBXW7;refGene_tx=XM_003984939.5;refGene_gn=FBXW7;refGene_tx=XM_003984940.5;refGene_gn=FBXW7;refGene_tx=XM_006930821.3;refGene_gn=FBXW7;refGene_tx=XM_019828857.2
B1 77305007 77305149

refGene_gn=FBXW7;refGene_tx=XM_003984938.5;refGene_gn=FBXW7;refGene_tx=XM_003984939.5;refGene_gn=FBXW7;refGene_tx=XM_003984940.5;refGene_gn=FBXW7;refGene_tx=XM_006930821.3;refGene_gn=FBXW7;refGene_tx=XM_019828857.2
B1 77313695 77313830
refGene_gn=FBXW7;refGene_tx=XM_003984938.5;refGene_gn=FBXW7;refGene_tx=XM_003984939.5;refGene_gn=FBXW7;refGene_tx=XM_003984940.5;refGene_gn=FBXW7;refGene_tx=XM_006930821.3;refGene_gn=FBXW7;refGene_tx=XM_019828857.2
B1 77319681 77319805
refGene_gn=FBXW7;refGene_tx=XM_003984938.5;refGene_gn=FBXW7;refGene_tx=XM_003984939.5;refGene_gn=FBXW7;refGene_tx=XM_003984940.5;refGene_gn=FBXW7;refGene_tx=XM_006930821.3;refGene_gn=FBXW7;refGene_tx=XM_019828857.2
B1 77322143 77322280
refGene_gn=FBXW7;refGene_tx=XM_003984938.5;refGene_gn=FBXW7;refGene_tx=XM_003984939.5;refGene_gn=FBXW7;refGene_tx=XM_003984940.5;refGene_gn=FBXW7;refGene_tx=XM_006930821.3;refGene_gn=FBXW7;refGene_tx=XM_019828857.2
B1 77323234 77323348
refGene_gn=FBXW7;refGene_tx=XM_003984938.5;refGene_gn=FBXW7;refGene_tx=XM_003984939.5;refGene_gn=FBXW7;refGene_tx=XM_003984940.5;refGene_gn=FBXW7;refGene_tx=XM_006930821.3;refGene_gn=FBXW7;refGene_tx=XM_019828857.2
B1 77324571 77324753
refGene_gn=FBXW7;refGene_tx=XM_003984938.5;refGene_gn=FBXW7;refGene_tx=XM_003984939.5;refGene_gn=FBXW7;refGene_tx=XM_003984940.5;refGene_gn=FBXW7;refGene_tx=XM_006930821.3;refGene_gn=FBXW7;refGene_tx=XM_019828857.2
B1 77326781 77327007
refGene_gn=FBXW7;refGene_tx=XM_003984938.5;refGene_gn=FBXW7;refGene_tx=XM_003984939.5;refGene_gn=FBXW7;refGene_tx=XM_003984940.5;refGene_gn=FBXW7;refGene_tx=XM_006930821.3;refGene_gn=FBXW7;refGene_tx=XM_019828857.2
B1 77328906 77329117
refGene_gn=FBXW7;refGene_tx=XM_003984938.5;refGene_gn=FBXW7;refGene_tx=XM_003984939.5;refGene_gn=FBXW7;refGene_tx=XM_003984940.5;refGene_gn=FBXW7;refGene_tx=XM_006930821.3;refGene_gn=FBXW7;refGene_tx=XM_019828857.2
B1 77330201 77330470
refGene_gn=FBXW7;refGene_tx=XM_003984938.5;refGene_gn=FBXW7;refGene_tx=XM_003984939.5;refGene_gn=FBXW7;refGene_tx=XM_003984940.5;refGene_gn=FBXW7;refGene_tx=XM_006930821.3;refGene_gn=FBXW7;refGene_tx=XM_019828857.2
B1 77858265 77858441
refGene_gn=GATB;refGene_tx=XM_003984942.5;refGene_gn=GATB;refGene_tx=XM_006930822.4;refGene_gn=GATB;refGene_tx=XM_019828860.2;refGene_gn=GATB;refGene_tx=XM_019828861.2
B1 77860335 77860486
refGene_gn=GATB;refGene_tx=XM_003984942.5;refGene_gn=GATB;refGene_tx=X

M_006930822.4;refGene_gn=GATB;refGene_tx=XM_019828860.2;refGene_gn=GATB;refGene_tx=XM_019828861.2
B1 77890536 77890650
refGene_gn=GATB;refGene_tx=XM_003984942.5;refGene_gn=GATB;refGene_tx=XM_006930822.4;refGene_gn=GATB;refGene_tx=XM_019828860.2;refGene_gn=GATB;refGene_tx=XM_019828861.2
B1 77893228 77893427
refGene_gn=GATB;refGene_tx=XM_003984942.5;refGene_gn=GATB;refGene_tx=XM_006930822.4;refGene_gn=GATB;refGene_tx=XM_019828860.2;refGene_gn=GATB;refGene_tx=XM_019828861.2
B1 77893883 77894006
refGene_gn=GATB;refGene_tx=XM_003984942.5;refGene_gn=GATB;refGene_tx=XM_006930822.4;refGene_gn=GATB;refGene_tx=XM_019828860.2;refGene_gn=GATB;refGene_tx=XM_019828861.2
B1 77902500 77902614
refGene_gn=GATB;refGene_tx=XM_003984942.5;refGene_gn=GATB;refGene_tx=XM_006930822.4;refGene_gn=GATB;refGene_tx=XM_019828860.2;refGene_gn=GATB;refGene_tx=XM_019828861.2
B1 77904944 77905029
refGene_gn=GATB;refGene_tx=XM_003984942.5;refGene_gn=GATB;refGene_tx=XM_006930822.4;refGene_gn=GATB;refGene_tx=XM_019828860.2;refGene_gn=GATB;refGene_tx=XM_019828861.2
B1 77905744 77905804
refGene_gn=GATB;refGene_tx=XM_003984942.5;refGene_gn=GATB;refGene_tx=XM_019828860.2;refGene_gn=GATB;refGene_tx=XM_019828861.2
B1 77907759 77907949
refGene_gn=GATB;refGene_tx=XM_003984942.5;refGene_gn=GATB;refGene_tx=XM_006930822.4;refGene_gn=GATB;refGene_tx=XM_019828860.2;refGene_gn=GATB;refGene_tx=XM_019828861.2
B1 77919734 77919868
refGene_gn=GATB;refGene_tx=XM_003984942.5;refGene_gn=GATB;refGene_tx=XM_006930822.4;refGene_gn=GATB;refGene_tx=XM_019828860.2;refGene_gn=GATB;refGene_tx=XM_019828861.2
B1 77927240 77927319
refGene_gn=GATB;refGene_tx=XM_003984942.5;refGene_gn=GATB;refGene_tx=XM_006930822.4;refGene_gn=GATB;refGene_tx=XM_019828860.2;refGene_gn=GATB;refGene_tx=XM_019828861.2
B1 77940569 77940704
refGene_gn=GATB;refGene_tx=XM_003984942.5;refGene_gn=GATB;refGene_tx=XM_006930822.4;refGene_gn=GATB;refGene_tx=XM_019828860.2;refGene_gn=GATB;refGene_tx=XM_019828861.2
B1 77942215 77942344
refGene_gn=GATB;refGene_tx=XM_003984942.5;refGene_gn=GATB;refGene_tx=XM_006930822.4;refGene_gn=GATB;refGene_tx=XM_019828860.2;refGene_gn=GATB;refGene_tx=XM_019828861.2
B1 77950130 77950384 refGene_gn=FAM160A1;refGene_tx=XM_006930824.4
B1 77956571 77956710 refGene_gn=FAM160A1;refGene_tx=XM_006930824.4
B1 77957069 77957248 refGene_gn=FAM160A1;refGene_tx=XM_006930824.4
B1 77963931 77964226 refGene_gn=FAM160A1;refGene_tx=XM_006930824.4
B1 77964227 77965078 refGene_gn=FAM160A1;refGene_tx=XM_006930824.4

B1 77968237 77968428 refGene_gn=FAM160A1;refGene_tx=XM_006930824.4
B1 77977506 77977586 refGene_gn=FAM160A1;refGene_tx=XM_006930824.4
B1 77978736 77978904 refGene_gn=FAM160A1;refGene_tx=XM_006930824.4
B1 78013981 78014068 refGene_gn=FAM160A1;refGene_tx=XM_006930824.4
B1 78016092 78016251 refGene_gn=FAM160A1;refGene_tx=XM_006930824.4
B1 78027056 78027683 refGene_gn=FAM160A1;refGene_tx=XM_006930824.4
B1 78039539 78039644 refGene_gn=FAM160A1;refGene_tx=XM_006930824.4
B1 78313138 78313250 refGene_gn=SH3D19;refGene_tx=XM_023252763.1
B1 78405149 78405189 refGene_gn=SH3D19;refGene_tx=XM_023252763.1
B1 78443532 78443573 refGene_gn=SH3D19;refGene_tx=XM_023252763.1
B1 78448762 78448778 refGene_gn=SH3D19;refGene_tx=XM_011281621.3
B1 78450101 78450144
refGene_gn=SH3D19;refGene_tx=XM_011281621.3;refGene_gn=SH3D19;refGene_tx=XM_023252763.1
B1 78451975 78452114
refGene_gn=SH3D19;refGene_tx=XM_011281621.3;refGene_gn=SH3D19;refGene_tx=XM_023252763.1
B1 78452522 78452685
refGene_gn=SH3D19;refGene_tx=XM_011281621.3;refGene_gn=SH3D19;refGene_tx=XM_023252763.1
B1 78453407 78454409
refGene_gn=SH3D19;refGene_tx=XM_011281619.3;refGene_gn=SH3D19;refGene_tx=XM_011281620.3;refGene_gn=SH3D19;refGene_tx=XM_011281621.3;refGene_gn=SH3D19;refGene_tx=XM_011281622.3;refGene_gn=SH3D19;refGene_tx=XM_023252763.1
B1 78463433 78463541
refGene_gn=SH3D19;refGene_tx=XM_011281619.3;refGene_gn=SH3D19;refGene_tx=XM_011281620.3;refGene_gn=SH3D19;refGene_tx=XM_011281621.3;refGene_gn=SH3D19;refGene_tx=XM_011281622.3;refGene_gn=SH3D19;refGene_tx=XM_023252763.1
B1 78468027 78468140
refGene_gn=SH3D19;refGene_tx=XM_011281619.3;refGene_gn=SH3D19;refGene_tx=XM_011281620.3;refGene_gn=SH3D19;refGene_tx=XM_011281621.3;refGene_gn=SH3D19;refGene_tx=XM_011281622.3;refGene_gn=SH3D19;refGene_tx=XM_023252763.1
B1 78473713 78473775
refGene_gn=SH3D19;refGene_tx=XM_011281619.3;refGene_gn=SH3D19;refGene_tx=XM_011281620.3;refGene_gn=SH3D19;refGene_tx=XM_011281621.3;refGene_gn=SH3D19;refGene_tx=XM_011281622.3;refGene_gn=SH3D19;refGene_tx=XM_023252763.1
B1 78474823 78475088
refGene_gn=SH3D19;refGene_tx=XM_011281619.3;refGene_gn=SH3D19;refGene_tx=XM_011281620.3;refGene_gn=SH3D19;refGene_tx=XM_011281621.3;refGene_gn=SH3D19;refGene_tx=XM_011281622.3;refGene_gn=SH3D19;refGene_tx=XM_023252763.1
B1 78477493 78477562
refGene_gn=SH3D19;refGene_tx=XM_011281619.3;refGene_gn=SH3D19;refGene_tx=XM_011281620.3;refGene_gn=SH3D19;refGene_tx=XM_011281621.3;refGene_gn=SH3D19;refGene_tx=XM_011281622.3;refGene_gn=SH3D19;refGene_tx=XM_023252763.1

B1 78477728 78477869
refGene_gn=SH3D19;refGene_tx=XM_011281619.3;refGene_gn=SH3D19;refGene_tx=XM_011281620.3;refGene_gn=SH3D19;refGene_tx=XM_011281621.3;refGene_gn=SH3D19;refGene_tx=XM_011281622.3;refGene_gn=SH3D19;refGene_tx=XM_023252763.1
B1 78483426 78483508
refGene_gn=SH3D19;refGene_tx=XM_011281619.3;refGene_gn=SH3D19;refGene_tx=XM_011281620.3;refGene_gn=SH3D19;refGene_tx=XM_011281621.3;refGene_gn=SH3D19;refGene_tx=XM_011281622.3;refGene_gn=SH3D19;refGene_tx=XM_023252763.1
B1 78484912 78485043
refGene_gn=SH3D19;refGene_tx=XM_011281619.3;refGene_gn=SH3D19;refGene_tx=XM_011281620.3;refGene_gn=SH3D19;refGene_tx=XM_011281621.3;refGene_gn=SH3D19;refGene_tx=XM_011281622.3;refGene_gn=SH3D19;refGene_tx=XM_023252763.1
B1 78486886 78486945
refGene_gn=SH3D19;refGene_tx=XM_011281619.3;refGene_gn=SH3D19;refGene_tx=XM_011281620.3;refGene_gn=SH3D19;refGene_tx=XM_011281621.3;refGene_gn=SH3D19;refGene_tx=XM_011281622.3;refGene_gn=SH3D19;refGene_tx=XM_023252763.1
B1 78489083 78489286
refGene_gn=SH3D19;refGene_tx=XM_011281619.3;refGene_gn=SH3D19;refGene_tx=XM_011281620.3;refGene_gn=SH3D19;refGene_tx=XM_011281621.3;refGene_gn=SH3D19;refGene_tx=XM_011281622.3;refGene_gn=SH3D19;refGene_tx=XM_023252763.1
B1 78490063 78490119
refGene_gn=SH3D19;refGene_tx=XM_011281619.3;refGene_gn=SH3D19;refGene_tx=XM_011281620.3;refGene_gn=SH3D19;refGene_tx=XM_011281621.3;refGene_gn=SH3D19;refGene_tx=XM_011281622.3;refGene_gn=SH3D19;refGene_tx=XM_023252763.1
B1 78494217 78494404
refGene_gn=SH3D19;refGene_tx=XM_011281619.3;refGene_gn=SH3D19;refGene_tx=XM_011281620.3;refGene_gn=SH3D19;refGene_tx=XM_011281621.3;refGene_gn=SH3D19;refGene_tx=XM_011281622.3;refGene_gn=SH3D19;refGene_tx=XM_023252763.1
B1 78494839 78494865
refGene_gn=SH3D19;refGene_tx=XM_011281619.3;refGene_gn=SH3D19;refGene_tx=XM_011281620.3;refGene_gn=SH3D19;refGene_tx=XM_011281621.3;refGene_gn=SH3D19;refGene_tx=XM_011281622.3;refGene_gn=SH3D19;refGene_tx=XM_023252763.1
B1 78514996 78515118 refGene_gn=RPS3A;refGene_tx=XM_003984944.3
B1 78515287 78515397 refGene_gn=RPS3A;refGene_tx=XM_003984944.3
B1 78515973 78516182 refGene_gn=RPS3A;refGene_tx=XM_003984944.3
B1 78517650 78517838 refGene_gn=RPS3A;refGene_tx=XM_003984944.3
B1 78518214 78518318 refGene_gn=RPS3A;refGene_tx=XM_003984944.3
B1 78519134 78519196 refGene_gn=RPS3A;refGene_tx=XM_003984944.3
B1 78526186 78526484
refGene_gn=LOC101080451;refGene_tx=XM_019829917.2
B1 78526758 78526928
refGene_gn=LOC101080451;refGene_tx=XM_019829917.2

B1 78589285 78589498
refGene_gn=LRBA;refGene_tx=XM_011281623.3;refGene_gn=LRBA;refGene_tx=XM_023252764.1;refGene_gn=LRBA;refGene_tx=XM_023252765.1;refGene_gn=LRBA;refGene_tx=XM_023252766.1;refGene_gn=LRBA;refGene_tx=XM_023252767.1
B1 78639352 78639527
refGene_gn=LOC109499331;refGene_tx=XM_019829887.1
B1 78656892 78656921
refGene_gn=LOC109499331;refGene_tx=XM_019829887.1
B1 78677751 78677983
refGene_gn=LRBA;refGene_tx=XM_011281623.3;refGene_gn=LRBA;refGene_tx=XM_023252764.1;refGene_gn=LRBA;refGene_tx=XM_023252765.1;refGene_gn=LRBA;refGene_tx=XM_023252766.1;refGene_gn=LRBA;refGene_tx=XM_023252767.1
B1 78678196 78678297
refGene_gn=LRBA;refGene_tx=XM_011281623.3;refGene_gn=LRBA;refGene_tx=XM_023252764.1;refGene_gn=LRBA;refGene_tx=XM_023252765.1;refGene_gn=LRBA;refGene_tx=XM_023252766.1;refGene_gn=LRBA;refGene_tx=XM_023252767.1
B1 78685442 78685538
refGene_gn=LRBA;refGene_tx=XM_011281623.3;refGene_gn=LRBA;refGene_tx=XM_023252764.1;refGene_gn=LRBA;refGene_tx=XM_023252765.1;refGene_gn=LRBA;refGene_tx=XM_023252766.1;refGene_gn=LRBA;refGene_tx=XM_023252767.1
B1 78696979 78697101
refGene_gn=LRBA;refGene_tx=XM_011281623.3;refGene_gn=LRBA;refGene_tx=XM_023252764.1;refGene_gn=LRBA;refGene_tx=XM_023252765.1;refGene_gn=LRBA;refGene_tx=XM_023252766.1;refGene_gn=LRBA;refGene_tx=XM_023252767.1
B1 78697186 78697313
refGene_gn=LRBA;refGene_tx=XM_011281623.3;refGene_gn=LRBA;refGene_tx=XM_023252764.1;refGene_gn=LRBA;refGene_tx=XM_023252765.1;refGene_gn=LRBA;refGene_tx=XM_023252766.1;refGene_gn=LRBA;refGene_tx=XM_023252767.1
B1 78697998 78698118
refGene_gn=LRBA;refGene_tx=XM_011281623.3;refGene_gn=LRBA;refGene_tx=XM_023252764.1;refGene_gn=LRBA;refGene_tx=XM_023252765.1;refGene_gn=LRBA;refGene_tx=XM_023252766.1;refGene_gn=LRBA;refGene_tx=XM_023252767.1
B1 78699897 78700044
refGene_gn=LRBA;refGene_tx=XM_011281623.3;refGene_gn=LRBA;refGene_tx=XM_023252764.1;refGene_gn=LRBA;refGene_tx=XM_023252765.1;refGene_gn=LRBA;refGene_tx=XM_023252766.1;refGene_gn=LRBA;refGene_tx=XM_023252767.1
B1 78712811 78713009
refGene_gn=LRBA;refGene_tx=XM_011281623.3;refGene_gn=LRBA;refGene_tx=XM_023252764.1;refGene_gn=LRBA;refGene_tx=XM_023252765.1;refGene_gn=LRBA;refGene_tx=XM_023252766.1;refGene_gn=LRBA;refGene_tx=XM_023252767.1
B1 78713206 78713340
refGene_gn=LRBA;refGene_tx=XM_011281623.3;refGene_gn=LRBA;refGene_tx=XM_023252764.1;refGene_gn=LRBA;refGene_tx=XM_023252765.1;refGene_gn=LRBA;refGene_tx=XM_023252766.1;refGene_gn=LRBA;refGene_tx=XM_023252767.1
B1 78715702 78715811
refGene_gn=LRBA;refGene_tx=XM_011281623.3;refGene_gn=LRBA;refGene_tx=XM_023252764.1;refGene_gn=LRBA;refGene_tx=XM_023252765.1;refGene_gn=LRBA;refGene_tx=XM_023252766.1;refGene_gn=LRBA;refGene_tx=XM_023252767.1
B1 78716118 78716271
refGene_gn=LRBA;refGene_tx=XM_011281623.3;refGene_gn=LRBA;refGene_tx=X

M_023252764.1;refGene_gn=LRBA;refGene_tx=XM_023252765.1;refGene_gn=LRBA;refGene_tx=XM_023252766.1;refGene_gn=LRBA;refGene_tx=XM_023252767.1
B1 78723369 78723538
refGene_gn=LRBA;refGene_tx=XM_011281623.3;refGene_gn=LRBA;refGene_tx=XM_023252764.1;refGene_gn=LRBA;refGene_tx=XM_023252765.1;refGene_gn=LRBA;refGene_tx=XM_023252766.1;refGene_gn=LRBA;refGene_tx=XM_023252767.1
B1 78729087 78729167
refGene_gn=LRBA;refGene_tx=XM_011281623.3;refGene_gn=LRBA;refGene_tx=XM_023252764.1;refGene_gn=LRBA;refGene_tx=XM_023252765.1;refGene_gn=LRBA;refGene_tx=XM_023252766.1;refGene_gn=LRBA;refGene_tx=XM_023252767.1
B1 78731013 78731076
refGene_gn=LRBA;refGene_tx=XM_011281623.3;refGene_gn=LRBA;refGene_tx=XM_023252764.1;refGene_gn=LRBA;refGene_tx=XM_023252765.1;refGene_gn=LRBA;refGene_tx=XM_023252766.1;refGene_gn=LRBA;refGene_tx=XM_023252767.1
B1 78733084 78733182
refGene_gn=LRBA;refGene_tx=XM_011281623.3;refGene_gn=LRBA;refGene_tx=XM_023252764.1;refGene_gn=LRBA;refGene_tx=XM_023252765.1;refGene_gn=LRBA;refGene_tx=XM_023252766.1;refGene_gn=LRBA;refGene_tx=XM_023252767.1
B1 78750584 78750677
refGene_gn=LRBA;refGene_tx=XM_011281623.3;refGene_gn=LRBA;refGene_tx=XM_023252764.1;refGene_gn=LRBA;refGene_tx=XM_023252765.1;refGene_gn=LRBA;refGene_tx=XM_023252766.1;refGene_gn=LRBA;refGene_tx=XM_023252767.1
B1 78752085 78752194
refGene_gn=LRBA;refGene_tx=XM_011281623.3;refGene_gn=LRBA;refGene_tx=XM_023252764.1;refGene_gn=LRBA;refGene_tx=XM_023252765.1;refGene_gn=LRBA;refGene_tx=XM_023252766.1;refGene_gn=LRBA;refGene_tx=XM_023252767.1
B1 78752599 78752681
refGene_gn=LRBA;refGene_tx=XM_011281623.3;refGene_gn=LRBA;refGene_tx=XM_023252764.1;refGene_gn=LRBA;refGene_tx=XM_023252765.1;refGene_gn=LRBA;refGene_tx=XM_023252766.1;refGene_gn=LRBA;refGene_tx=XM_023252767.1
B1 78755368 78755492
refGene_gn=LRBA;refGene_tx=XM_011281623.3;refGene_gn=LRBA;refGene_tx=XM_023252764.1;refGene_gn=LRBA;refGene_tx=XM_023252765.1;refGene_gn=LRBA;refGene_tx=XM_023252766.1;refGene_gn=LRBA;refGene_tx=XM_023252767.1
B1 78755783 78755976
refGene_gn=LRBA;refGene_tx=XM_011281623.3;refGene_gn=LRBA;refGene_tx=XM_023252764.1;refGene_gn=LRBA;refGene_tx=XM_023252765.1;refGene_gn=LRBA;refGene_tx=XM_023252766.1;refGene_gn=LRBA;refGene_tx=XM_023252767.1
B1 78788348 78789404
refGene_gn=LRBA;refGene_tx=XM_011281623.3;refGene_gn=LRBA;refGene_tx=XM_023252764.1;refGene_gn=LRBA;refGene_tx=XM_023252765.1;refGene_gn=LRBA;refGene_tx=XM_023252766.1;refGene_gn=LRBA;refGene_tx=XM_023252767.1
B1 78790607 78790786
refGene_gn=LRBA;refGene_tx=XM_011281623.3;refGene_gn=LRBA;refGene_tx=XM_023252764.1;refGene_gn=LRBA;refGene_tx=XM_023252765.1;refGene_gn=LRBA;refGene_tx=XM_023252766.1;refGene_gn=LRBA;refGene_tx=XM_023252767.1
B1 78791603 78791757
refGene_gn=LRBA;refGene_tx=XM_011281623.3;refGene_gn=LRBA;refGene_tx=XM_023252764.1;refGene_gn=LRBA;refGene_tx=XM_023252765.1;refGene_gn=LRBA;refGene_tx=XM_023252766.1;refGene_gn=LRBA;refGene_tx=XM_023252767.1

B1 78792401 78792582
refGene_gn=LRBA;refGene_tx=XM_011281623.3;refGene_gn=LRBA;refGene_tx=XM_023252764.1;refGene_gn=LRBA;refGene_tx=XM_023252765.1;refGene_gn=LRBA;refGene_tx=XM_023252766.1;refGene_gn=LRBA;refGene_tx=XM_023252767.1
B1 78796465 78796593
refGene_gn=LRBA;refGene_tx=XM_011281623.3;refGene_gn=LRBA;refGene_tx=XM_023252764.1;refGene_gn=LRBA;refGene_tx=XM_023252765.1;refGene_gn=LRBA;refGene_tx=XM_023252766.1;refGene_gn=LRBA;refGene_tx=XM_023252767.1
B1 78797093 78797201
refGene_gn=LRBA;refGene_tx=XM_011281623.3;refGene_gn=LRBA;refGene_tx=XM_023252764.1;refGene_gn=LRBA;refGene_tx=XM_023252765.1;refGene_gn=LRBA;refGene_tx=XM_023252766.1;refGene_gn=LRBA;refGene_tx=XM_023252767.1
B1 78801437 78801597
refGene_gn=LRBA;refGene_tx=XM_011281623.3;refGene_gn=LRBA;refGene_tx=XM_023252764.1;refGene_gn=LRBA;refGene_tx=XM_023252765.1;refGene_gn=LRBA;refGene_tx=XM_023252766.1;refGene_gn=LRBA;refGene_tx=XM_023252767.1
B1 78804736 78805184
refGene_gn=LRBA;refGene_tx=XM_011281623.3;refGene_gn=LRBA;refGene_tx=XM_023252764.1;refGene_gn=LRBA;refGene_tx=XM_023252765.1;refGene_gn=LRBA;refGene_tx=XM_023252766.1;refGene_gn=LRBA;refGene_tx=XM_023252767.1
B1 78812296 78812430
refGene_gn=LRBA;refGene_tx=XM_011281623.3;refGene_gn=LRBA;refGene_tx=XM_023252764.1;refGene_gn=LRBA;refGene_tx=XM_023252765.1;refGene_gn=LRBA;refGene_tx=XM_023252766.1;refGene_gn=LRBA;refGene_tx=XM_023252767.1
B1 78825135 78825214
refGene_gn=LRBA;refGene_tx=XM_011281623.3;refGene_gn=LRBA;refGene_tx=XM_023252764.1;refGene_gn=LRBA;refGene_tx=XM_023252765.1;refGene_gn=LRBA;refGene_tx=XM_023252766.1;refGene_gn=LRBA;refGene_tx=XM_023252767.1
B1 78826036 78826170
refGene_gn=LRBA;refGene_tx=XM_011281623.3;refGene_gn=LRBA;refGene_tx=XM_023252764.1;refGene_gn=LRBA;refGene_tx=XM_023252765.1;refGene_gn=LRBA;refGene_tx=XM_023252766.1;refGene_gn=LRBA;refGene_tx=XM_023252767.1
B1 78833346 78833408
refGene_gn=LRBA;refGene_tx=XM_011281623.3;refGene_gn=LRBA;refGene_tx=XM_023252764.1;refGene_gn=LRBA;refGene_tx=XM_023252765.1;refGene_gn=LRBA;refGene_tx=XM_023252766.1;refGene_gn=LRBA;refGene_tx=XM_023252767.1
B1 78865496 78865561
refGene_gn=LRBA;refGene_tx=XM_011281623.3;refGene_gn=LRBA;refGene_tx=XM_023252764.1;refGene_gn=LRBA;refGene_tx=XM_023252765.1;refGene_gn=LRBA;refGene_tx=XM_023252766.1;refGene_gn=LRBA;refGene_tx=XM_023252767.1
B1 78912142 78912251
refGene_gn=LRBA;refGene_tx=XM_011281623.3;refGene_gn=LRBA;refGene_tx=XM_023252764.1;refGene_gn=LRBA;refGene_tx=XM_023252765.1;refGene_gn=LRBA;refGene_tx=XM_023252766.1;refGene_gn=LRBA;refGene_tx=XM_023252767.1
B1 78978382 78978549
refGene_gn=LRBA;refGene_tx=XM_011281623.3;refGene_gn=LRBA;refGene_tx=XM_023252764.1;refGene_gn=LRBA;refGene_tx=XM_023252765.1;refGene_gn=LRBA;refGene_tx=XM_023252766.1;refGene_gn=LRBA;refGene_tx=XM_023252767.1
B1 79065354 79065515
refGene_gn=LRBA;refGene_tx=XM_011281623.3;refGene_gn=LRBA;refGene_tx=X

M_023252764.1;refGene_gn=LRBA;refGene_tx=XM_023252765.1;refGene_gn=LRBA;refGene_tx=XM_023252766.1;refGene_gn=LRBA;refGene_tx=XM_023252767.1
B1 79073343 79073490
refGene_gn=LRBA;refGene_tx=XM_011281623.3;refGene_gn=LRBA;refGene_tx=XM_023252764.1;refGene_gn=LRBA;refGene_tx=XM_023252765.1;refGene_gn=LRBA;refGene_tx=XM_023252766.1;refGene_gn=LRBA;refGene_tx=XM_023252767.1
B1 79076028 79076165
refGene_gn=LRBA;refGene_tx=XM_011281623.3;refGene_gn=LRBA;refGene_tx=XM_023252764.1;refGene_gn=LRBA;refGene_tx=XM_023252765.1;refGene_gn=LRBA;refGene_tx=XM_023252766.1;refGene_gn=LRBA;refGene_tx=XM_023252767.1
B1 79080221 79081301 refGene_gn=MAB21L2;refGene_tx=XM_006930833.2
B1 79160911 79161029
refGene_gn=LRBA;refGene_tx=XM_011281623.3;refGene_gn=LRBA;refGene_tx=XM_023252764.1;refGene_gn=LRBA;refGene_tx=XM_023252765.1;refGene_gn=LRBA;refGene_tx=XM_023252766.1;refGene_gn=LRBA;refGene_tx=XM_023252767.1
B1 79163147 79163250
refGene_gn=LRBA;refGene_tx=XM_011281623.3;refGene_gn=LRBA;refGene_tx=XM_023252764.1;refGene_gn=LRBA;refGene_tx=XM_023252765.1;refGene_gn=LRBA;refGene_tx=XM_023252766.1;refGene_gn=LRBA;refGene_tx=XM_023252767.1
B1 79168083 79168199
refGene_gn=LRBA;refGene_tx=XM_011281623.3;refGene_gn=LRBA;refGene_tx=XM_023252764.1;refGene_gn=LRBA;refGene_tx=XM_023252765.1;refGene_gn=LRBA;refGene_tx=XM_023252766.1;refGene_gn=LRBA;refGene_tx=XM_023252767.1
B1 79173141 79173254
refGene_gn=LRBA;refGene_tx=XM_011281623.3;refGene_gn=LRBA;refGene_tx=XM_023252764.1;refGene_gn=LRBA;refGene_tx=XM_023252765.1;refGene_gn=LRBA;refGene_tx=XM_023252766.1;refGene_gn=LRBA;refGene_tx=XM_023252767.1
B1 79201309 79201450
refGene_gn=LRBA;refGene_tx=XM_011281623.3;refGene_gn=LRBA;refGene_tx=XM_023252764.1;refGene_gn=LRBA;refGene_tx=XM_023252765.1;refGene_gn=LRBA;refGene_tx=XM_023252766.1;refGene_gn=LRBA;refGene_tx=XM_023252767.1
B1 79202487 79202607
refGene_gn=LRBA;refGene_tx=XM_011281623.3;refGene_gn=LRBA;refGene_tx=XM_023252764.1;refGene_gn=LRBA;refGene_tx=XM_023252765.1;refGene_gn=LRBA;refGene_tx=XM_023252766.1;refGene_gn=LRBA;refGene_tx=XM_023252767.1
B1 79221196 79221349
refGene_gn=LRBA;refGene_tx=XM_011281623.3;refGene_gn=LRBA;refGene_tx=XM_023252764.1;refGene_gn=LRBA;refGene_tx=XM_023252765.1;refGene_gn=LRBA;refGene_tx=XM_023252766.1;refGene_gn=LRBA;refGene_tx=XM_023252767.1
B1 79286193 79286361
refGene_gn=LRBA;refGene_tx=XM_011281623.3;refGene_gn=LRBA;refGene_tx=XM_023252764.1;refGene_gn=LRBA;refGene_tx=XM_023252765.1;refGene_gn=LRBA;refGene_tx=XM_023252766.1;refGene_gn=LRBA;refGene_tx=XM_023252767.1
B1 79317038 79317128
refGene_gn=LRBA;refGene_tx=XM_011281623.3;refGene_gn=LRBA;refGene_tx=XM_023252764.1;refGene_gn=LRBA;refGene_tx=XM_023252765.1;refGene_gn=LRBA;refGene_tx=XM_023252766.1;refGene_gn=LRBA;refGene_tx=XM_023252767.1
B1 79321501 79321694
refGene_gn=LRBA;refGene_tx=XM_011281623.3;refGene_gn=LRBA;refGene_tx=XM_023252764.1;refGene_gn=LRBA;refGene_tx=XM_023252765.1;refGene_gn=LRB

A;refGene_tx=XM_023252766.1;refGene_gn=LRBA;refGene_tx=XM_023252767.1
B1 79326104 79326167
refGene_gn=LRBA;refGene_tx=XM_011281623.3;refGene_gn=LRBA;refGene_tx=XM_023252764.1;refGene_gn=LRBA;refGene_tx=XM_023252765.1;refGene_gn=LRBA;refGene_tx=XM_023252766.1;refGene_gn=LRBA;refGene_tx=XM_023252767.1
B1 79333224 79333380
refGene_gn=LRBA;refGene_tx=XM_011281623.3;refGene_gn=LRBA;refGene_tx=XM_023252764.1;refGene_gn=LRBA;refGene_tx=XM_023252765.1;refGene_gn=LRBA;refGene_tx=XM_023252766.1;refGene_gn=LRBA;refGene_tx=XM_023252767.1
B1 79339732 79339900
refGene_gn=LRBA;refGene_tx=XM_011281623.3;refGene_gn=LRBA;refGene_tx=XM_023252764.1;refGene_gn=LRBA;refGene_tx=XM_023252765.1;refGene_gn=LRBA;refGene_tx=XM_023252766.1;refGene_gn=LRBA;refGene_tx=XM_023252767.1
B1 79354605 79354707
refGene_gn=LRBA;refGene_tx=XM_011281623.3;refGene_gn=LRBA;refGene_tx=XM_023252764.1;refGene_gn=LRBA;refGene_tx=XM_023252765.1;refGene_gn=LRBA;refGene_tx=XM_023252766.1;refGene_gn=LRBA;refGene_tx=XM_023252767.1
B1 79358061 79358258
refGene_gn=LRBA;refGene_tx=XM_011281623.3;refGene_gn=LRBA;refGene_tx=XM_023252764.1;refGene_gn=LRBA;refGene_tx=XM_023252765.1;refGene_gn=LRBA;refGene_tx=XM_023252766.1;refGene_gn=LRBA;refGene_tx=XM_023252767.1
B1 79362276 79362428
refGene_gn=LRBA;refGene_tx=XM_011281623.3;refGene_gn=LRBA;refGene_tx=XM_023252764.1;refGene_gn=LRBA;refGene_tx=XM_023252765.1;refGene_gn=LRBA;refGene_tx=XM_023252766.1;refGene_gn=LRBA;refGene_tx=XM_023252767.1
B1 79368254 79368345
refGene_gn=LRBA;refGene_tx=XM_011281623.3;refGene_gn=LRBA;refGene_tx=XM_023252764.1;refGene_gn=LRBA;refGene_tx=XM_023252765.1;refGene_gn=LRBA;refGene_tx=XM_023252766.1;refGene_gn=LRBA;refGene_tx=XM_023252767.1
B1 79377137 79377365
refGene_gn=DCLK2;refGene_tx=XM_011281624.3;refGene_gn=DCLK2;refGene_tx=XM_011281625.3;refGene_gn=DCLK2;refGene_tx=XM_011281626.3;refGene_gn=DCLK2;refGene_tx=XM_011281627.3;refGene_gn=DCLK2;refGene_tx=XM_019828866.2;refGene_gn=DCLK2;refGene_tx=XM_019828868.2
B1 79378800 79378813 refGene_gn=DCLK2;refGene_tx=XM_019828868.2
B1 79380321 79380387 refGene_gn=DCLK2;refGene_tx=XM_019828867.2
B1 79382684 79382801
refGene_gn=DCLK2;refGene_tx=XM_011281624.3;refGene_gn=DCLK2;refGene_tx=XM_011281625.3;refGene_gn=DCLK2;refGene_tx=XM_011281626.3;refGene_gn=DCLK2;refGene_tx=XM_011281627.3;refGene_gn=DCLK2;refGene_tx=XM_019828866.2;refGene_gn=DCLK2;refGene_tx=XM_019828867.2;refGene_gn=DCLK2;refGene_tx=XM_019828868.2
B1 79383964 79384045
refGene_gn=DCLK2;refGene_tx=XM_011281624.3;refGene_gn=DCLK2;refGene_tx=XM_011281625.3;refGene_gn=DCLK2;refGene_tx=XM_011281626.3;refGene_gn=DCLK2;refGene_tx=XM_011281627.3;refGene_gn=DCLK2;refGene_tx=XM_019828866.2;refGene_gn=DCLK2;refGene_tx=XM_019828867.2;refGene_gn=DCLK2;refGene_tx=XM_019828868.2
B1 79384667 79384764
refGene_gn=DCLK2;refGene_tx=XM_011281624.3;refGene_gn=DCLK2;refGene_tx

=XM_011281625.3;refGene_gn=DCLK2;refGene_tx=XM_011281626.3;refGene_gn=DCLK2;refGene_tx=XM_011281627.3;refGene_gn=DCLK2;refGene_tx=XM_019828866.2;refGene_gn=DCLK2;refGene_tx=XM_019828867.2;refGene_gn=DCLK2;refGene_tx=XM_019828868.2
B1 79390813 79390891
refGene_gn=DCLK2;refGene_tx=XM_011281624.3;refGene_gn=DCLK2;refGene_tx=XM_011281625.3;refGene_gn=DCLK2;refGene_tx=XM_011281626.3;refGene_gn=DCLK2;refGene_tx=XM_011281627.3;refGene_gn=DCLK2;refGene_tx=XM_019828866.2;refGene_gn=DCLK2;refGene_tx=XM_019828867.2;refGene_gn=DCLK2;refGene_tx=XM_019828868.2
B1 79391315 79391449
refGene_gn=DCLK2;refGene_tx=XM_011281624.3;refGene_gn=DCLK2;refGene_tx=XM_011281625.3;refGene_gn=DCLK2;refGene_tx=XM_011281626.3;refGene_gn=DCLK2;refGene_tx=XM_011281627.3;refGene_gn=DCLK2;refGene_tx=XM_019828866.2;refGene_gn=DCLK2;refGene_tx=XM_019828867.2;refGene_gn=DCLK2;refGene_tx=XM_019828868.2
B1 79397931 79398078
refGene_gn=DCLK2;refGene_tx=XM_011281624.3;refGene_gn=DCLK2;refGene_tx=XM_011281625.3;refGene_gn=DCLK2;refGene_tx=XM_011281626.3;refGene_gn=DCLK2;refGene_tx=XM_011281627.3;refGene_gn=DCLK2;refGene_tx=XM_019828866.2;refGene_gn=DCLK2;refGene_tx=XM_019828867.2;refGene_gn=DCLK2;refGene_tx=XM_019828868.2
B1 79398320 79398440
refGene_gn=DCLK2;refGene_tx=XM_011281624.3;refGene_gn=DCLK2;refGene_tx=XM_011281625.3;refGene_gn=DCLK2;refGene_tx=XM_011281626.3;refGene_gn=DCLK2;refGene_tx=XM_011281627.3;refGene_gn=DCLK2;refGene_tx=XM_019828867.2;refGene_gn=DCLK2;refGene_tx=XM_019828868.2
B1 79406164 79406222
refGene_gn=DCLK2;refGene_tx=XM_011281624.3;refGene_gn=DCLK2;refGene_tx=XM_011281625.3;refGene_gn=DCLK2;refGene_tx=XM_011281626.3;refGene_gn=DCLK2;refGene_tx=XM_011281627.3;refGene_gn=DCLK2;refGene_tx=XM_019828866.2;refGene_gn=DCLK2;refGene_tx=XM_019828867.2;refGene_gn=DCLK2;refGene_tx=XM_019828868.2
B1 79408237 79408346
refGene_gn=DCLK2;refGene_tx=XM_011281624.3;refGene_gn=DCLK2;refGene_tx=XM_011281625.3;refGene_gn=DCLK2;refGene_tx=XM_011281626.3;refGene_gn=DCLK2;refGene_tx=XM_011281627.3;refGene_gn=DCLK2;refGene_tx=XM_019828866.2;refGene_gn=DCLK2;refGene_tx=XM_019828867.2;refGene_gn=DCLK2;refGene_tx=XM_019828868.2
B1 79409182 79409258
refGene_gn=DCLK2;refGene_tx=XM_011281624.3;refGene_gn=DCLK2;refGene_tx=XM_011281625.3;refGene_gn=DCLK2;refGene_tx=XM_011281626.3;refGene_gn=DCLK2;refGene_tx=XM_011281627.3;refGene_gn=DCLK2;refGene_tx=XM_019828866.2;refGene_gn=DCLK2;refGene_tx=XM_019828867.2;refGene_gn=DCLK2;refGene_tx=XM_019828868.2
B1 79427644 79427739
refGene_gn=DCLK2;refGene_tx=XM_011281624.3;refGene_gn=DCLK2;refGene_tx=XM_011281625.3;refGene_gn=DCLK2;refGene_tx=XM_011281626.3;refGene_gn=DCLK2;refGene_tx=XM_011281627.3;refGene_gn=DCLK2;refGene_tx=XM_019828866.2;refGene_gn=DCLK2;refGene_tx=XM_019828867.2;refGene_gn=DCLK2;refGe

ne_tx=XM_019828868.2
B1	79432515 79432566
refGene_gn=DCLK2;refGene_tx=XM_011281624.3;refGene_gn=DCLK2;refGene_tx=XM_011281625.3;refGene_gn=DCLK2;refGene_tx=XM_019828866.2;refGene_gn=DCLK2;refGene_tx=XM_019828867.2;refGene_gn=DCLK2;refGene_tx=XM_019828868.2
B1	79433450 79433552
refGene_gn=DCLK2;refGene_tx=XM_011281624.3;refGene_gn=DCLK2;refGene_tx=XM_011281625.3;refGene_gn=DCLK2;refGene_tx=XM_011281626.3;refGene_gn=DCLK2;refGene_tx=XM_011281627.3;refGene_gn=DCLK2;refGene_tx=XM_019828866.2;refGene_gn=DCLK2;refGene_tx=XM_019828867.2;refGene_gn=DCLK2;refGene_tx=XM_019828868.2
B1	79436515 79436618
refGene_gn=DCLK2;refGene_tx=XM_011281624.3;refGene_gn=DCLK2;refGene_tx=XM_011281625.3;refGene_gn=DCLK2;refGene_tx=XM_011281626.3;refGene_gn=DCLK2;refGene_tx=XM_011281627.3;refGene_gn=DCLK2;refGene_tx=XM_019828866.2;refGene_gn=DCLK2;refGene_tx=XM_019828867.2;refGene_gn=DCLK2;refGene_tx=XM_019828868.2
B1	79516436 79516771
refGene_gn=DCLK2;refGene_tx=XM_011281624.3;refGene_gn=DCLK2;refGene_tx=XM_011281625.3;refGene_gn=DCLK2;refGene_tx=XM_011281626.3;refGene_gn=DCLK2;refGene_tx=XM_011281627.3;refGene_gn=DCLK2;refGene_tx=XM_019828866.2;refGene_gn=DCLK2;refGene_tx=XM_019828867.2;refGene_gn=DCLK2;refGene_tx=XM_019828868.2
B1	79536739 79537160
refGene_gn=DCLK2;refGene_tx=XM_011281624.3;refGene_gn=DCLK2;refGene_tx=XM_011281625.3;refGene_gn=DCLK2;refGene_tx=XM_011281626.3;refGene_gn=DCLK2;refGene_tx=XM_011281627.3;refGene_gn=DCLK2;refGene_tx=XM_019828866.2;refGene_gn=DCLK2;refGene_tx=XM_019828867.2;refGene_gn=DCLK2;refGene_tx=XM_019828868.2
B1	79836244 79836263
refGene_gn=IQCM;refGene_tx=XM_019828871.2;refGene_gn=IQCM;refGene_tx=XM_023252769.1;refGene_gn=IQCM;refGene_tx=XM_023252770.1;refGene_gn=IQCM;refGene_tx=XM_023252771.1
B1	79845087 79845170
refGene_gn=IQCM;refGene_tx=XM_019828871.2;refGene_gn=IQCM;refGene_tx=XM_023252769.1;refGene_gn=IQCM;refGene_tx=XM_023252770.1;refGene_gn=IQCM;refGene_tx=XM_023252771.1
B1	79847451 79847719
refGene_gn=IQCM;refGene_tx=XM_019828871.2;refGene_gn=IQCM;refGene_tx=XM_023252769.1;refGene_gn=IQCM;refGene_tx=XM_023252770.1;refGene_gn=IQCM;refGene_tx=XM_023252771.1
B1	79889039 79889130
refGene_gn=IQCM;refGene_tx=XM_019828871.2;refGene_gn=IQCM;refGene_tx=XM_023252769.1;refGene_gn=IQCM;refGene_tx=XM_023252770.1;refGene_gn=IQCM;refGene_tx=XM_023252771.1
B1	79893956 79894048
refGene_gn=IQCM;refGene_tx=XM_019828871.2;refGene_gn=IQCM;refGene_tx=XM_023252769.1;refGene_gn=IQCM;refGene_tx=XM_023252770.1;refGene_gn=IQCM;refGene_tx=XM_023252771.1

B1 79947134 79947250
refGene_gn=IQCM;refGene_tx=XM_019828871.2;refGene_gn=IQCM;refGene_tx=XM_023252769.1;refGene_gn=IQCM;refGene_tx=XM_023252770.1;refGene_gn=IQCM;refGene_tx=XM_023252771.1
B1 79983259 79983315
refGene_gn=IQCM;refGene_tx=XM_019828871.2;refGene_gn=IQCM;refGene_tx=XM_023252769.1;refGene_gn=IQCM;refGene_tx=XM_023252770.1;refGene_gn=IQCM;refGene_tx=XM_023252771.1
B1 80013940 80014139
refGene_gn=IQCM;refGene_tx=XM_019828871.2;refGene_gn=IQCM;refGene_tx=XM_023252769.1;refGene_gn=IQCM;refGene_tx=XM_023252770.1;refGene_gn=IQCM;refGene_tx=XM_023252771.1
B1 80031985 80032130
refGene_gn=IQCM;refGene_tx=XM_019828871.2;refGene_gn=IQCM;refGene_tx=XM_023252769.1;refGene_gn=IQCM;refGene_tx=XM_023252770.1;refGene_gn=IQCM;refGene_tx=XM_023252771.1
B1 80048349 80048484
refGene_gn=IQCM;refGene_tx=XM_019828871.2;refGene_gn=IQCM;refGene_tx=XM_023252769.1;refGene_gn=IQCM;refGene_tx=XM_023252770.1;refGene_gn=IQCM;refGene_tx=XM_023252771.1
B1 80144361 80144523
refGene_gn=IQCM;refGene_tx=XM_019828871.2;refGene_gn=IQCM;refGene_tx=XM_023252769.1;refGene_gn=IQCM;refGene_tx=XM_023252770.1;refGene_gn=IQCM;refGene_tx=XM_023252771.1
B1 80220813 80220923
refGene_gn=IQCM;refGene_tx=XM_019828871.2;refGene_gn=IQCM;refGene_tx=XM_023252769.1;refGene_gn=IQCM;refGene_tx=XM_023252770.1;refGene_gn=IQCM;refGene_tx=XM_023252771.1
B1 81046624 81048381
refGene_gn=NR3C2;refGene_tx=XM_003984956.5;refGene_gn=NR3C2;refGene_tx=XM_019828874.2;refGene_gn=NR3C2;refGene_tx=XM_019828875.2;refGene_gn=NR3C2;refGene_tx=XM_019828876.2
B1 81211015 81211167
refGene_gn=NR3C2;refGene_tx=XM_003984956.5;refGene_gn=NR3C2;refGene_tx=XM_019828874.2;refGene_gn=NR3C2;refGene_tx=XM_019828875.2;refGene_gn=NR3C2;refGene_tx=XM_019828876.2
B1 81287023 81287140
refGene_gn=NR3C2;refGene_tx=XM_003984956.5;refGene_gn=NR3C2;refGene_tx=XM_019828874.2;refGene_gn=NR3C2;refGene_tx=XM_019828875.2;refGene_gn=NR3C2;refGene_tx=XM_019828876.2
B1 81318632 81318980
refGene_gn=NR3C2;refGene_tx=XM_003984956.5;refGene_gn=NR3C2;refGene_tx=XM_019828874.2;refGene_gn=NR3C2;refGene_tx=XM_019828875.2;refGene_gn=NR3C2;refGene_tx=XM_019828876.2
B1 81320752 81320897
refGene_gn=NR3C2;refGene_tx=XM_003984956.5;refGene_gn=NR3C2;refGene_tx=XM_019828874.2;refGene_gn=NR3C2;refGene_tx=XM_019828875.2;refGene_gn=NR3C2;refGene_tx=XM_019828876.2
B1 81346596 81346727
refGene_gn=NR3C2;refGene_tx=XM_003984956.5;refGene_gn=NR3C2;refGene_tx

=XM_019828874.2;refGene_gn=NR3C2;refGene_tx=XM_019828875.2;refGene_gn=NR3C2;refGene_tx=XM_019828876.2
B1 81352346 81352504
refGene_gn=NR3C2;refGene_tx=XM_003984956.5;refGene_gn=NR3C2;refGene_tx=XM_019828874.2;refGene_gn=NR3C2;refGene_tx=XM_019828875.2;refGene_gn=NR3C2;refGene_tx=XM_019828876.2
B1 81387075 81387231
refGene_gn=NR3C2;refGene_tx=XM_003984956.5;refGene_gn=NR3C2;refGene_tx=XM_019828874.2;refGene_gn=NR3C2;refGene_tx=XM_019828875.2;refGene_gn=NR3C2;refGene_tx=XM_019828876.2
B1 81397803 81397892
refGene_gn=ARHGAP10;refGene_tx=XM_003984958.4;refGene_gn=ARHGAP10;refGene_tx=XM_006930835.3
B1 81407156 81407248
refGene_gn=ARHGAP10;refGene_tx=XM_003984958.4;refGene_gn=ARHGAP10;refGene_tx=XM_006930835.3
B1 81408414 81408567
refGene_gn=ARHGAP10;refGene_tx=XM_003984958.4;refGene_gn=ARHGAP10;refGene_tx=XM_006930835.3
B1 81425873 81426033
refGene_gn=ARHGAP10;refGene_tx=XM_003984958.4;refGene_gn=ARHGAP10;refGene_tx=XM_006930835.3
B1 81451950 81452101
refGene_gn=ARHGAP10;refGene_tx=XM_003984958.4;refGene_gn=ARHGAP10;refGene_tx=XM_006930835.3
B1 81499995 81500155
refGene_gn=ARHGAP10;refGene_tx=XM_003984958.4;refGene_gn=ARHGAP10;refGene_tx=XM_006930835.3
B1 81501813 81501919
refGene_gn=ARHGAP10;refGene_tx=XM_003984958.4;refGene_gn=ARHGAP10;refGene_tx=XM_006930835.3
B1 81513126 81513185
refGene_gn=ARHGAP10;refGene_tx=XM_003984958.4;refGene_gn=ARHGAP10;refGene_tx=XM_006930835.3
B1 81522425 81522513
refGene_gn=ARHGAP10;refGene_tx=XM_003984958.4;refGene_gn=ARHGAP10;refGene_tx=XM_006930835.3
B1 81529274 81529349
refGene_gn=ARHGAP10;refGene_tx=XM_003984958.4;refGene_gn=ARHGAP10;refGene_tx=XM_006930835.3
B1 81551903 81551969
refGene_gn=ARHGAP10;refGene_tx=XM_003984958.4;refGene_gn=ARHGAP10;refGene_tx=XM_006930835.3
B1 81554756 81554802
refGene_gn=ARHGAP10;refGene_tx=XM_003984958.4;refGene_gn=ARHGAP10;refGene_tx=XM_006930835.3
B1 81556782 81556864
refGene_gn=ARHGAP10;refGene_tx=XM_003984958.4;refGene_gn=ARHGAP10;refGene_tx=XM_006930835.3
B1 81585013 81585108

refGene_gn=ARHGAP10;refGene_tx=XM_003984958.4;refGene_gn=ARHGAP10;refGene_tx=XM_006930835.3
B1	81587288	81587395
refGene_gn=ARHGAP10;refGene_tx=XM_003984958.4;refGene_gn=ARHGAP10;refGene_tx=XM_006930835.3
B1	81590620	81590750
refGene_gn=ARHGAP10;refGene_tx=XM_003984958.4;refGene_gn=ARHGAP10;refGene_tx=XM_006930835.3
B1	81601494	81601599	refGene_gn=ARHGAP10;refGene_tx=XM_003984958.4
B1	81603390	81603501	refGene_gn=ARHGAP10;refGene_tx=XM_003984958.4
B1	81610298	81610400
refGene_gn=ARHGAP10;refGene_tx=XM_003984958.4;refGene_gn=ARHGAP10;refGene_tx=XM_006930835.3
B1	81618635	81618707
refGene_gn=ARHGAP10;refGene_tx=XM_003984958.4;refGene_gn=ARHGAP10;refGene_tx=XM_006930835.3
B1	81641143	81641205
refGene_gn=ARHGAP10;refGene_tx=XM_003984958.4;refGene_gn=ARHGAP10;refGene_tx=XM_006930835.3
B1	81641278	81641374
refGene_gn=ARHGAP10;refGene_tx=XM_003984958.4;refGene_gn=ARHGAP10;refGene_tx=XM_006930835.3
B1	81721584	81721680	refGene_gn=PRMT9;refGene_tx=XM_019828877.2
B1	81722114	81722268
refGene_gn=ARHGAP10;refGene_tx=XM_003984958.4;refGene_gn=ARHGAP10;refGene_tx=XM_006930835.3
B1	81777547	81777736	refGene_gn=PRMT9;refGene_tx=XM_003984959.5
B1	81780086	81780235
refGene_gn=PRMT9;refGene_tx=XM_003984959.5;refGene_gn=PRMT9;refGene_tx=XM_019828877.2;refGene_gn=PRMT9;refGene_tx=XM_019828878.2
B1	81784831	81785068
refGene_gn=PRMT9;refGene_tx=XM_003984959.5;refGene_gn=PRMT9;refGene_tx=XM_019828877.2;refGene_gn=PRMT9;refGene_tx=XM_019828878.2
B1	81785933	81786101
refGene_gn=PRMT9;refGene_tx=XM_003984959.5;refGene_gn=PRMT9;refGene_tx=XM_019828877.2;refGene_gn=PRMT9;refGene_tx=XM_019828878.2
B1	81788323	81788426
refGene_gn=PRMT9;refGene_tx=XM_003984959.5;refGene_gn=PRMT9;refGene_tx=XM_019828877.2;refGene_gn=PRMT9;refGene_tx=XM_019828878.2
B1	81791064	81791171
refGene_gn=PRMT9;refGene_tx=XM_003984959.5;refGene_gn=PRMT9;refGene_tx=XM_019828877.2;refGene_gn=PRMT9;refGene_tx=XM_019828878.2;refGene_gn=PRMT9;refGene_tx=XM_023252772.1;refGene_gn=PRMT9;refGene_tx=XM_023252774.1
B1	81794730	81794923
refGene_gn=PRMT9;refGene_tx=XM_003984959.5;refGene_gn=PRMT9;refGene_tx=XM_019828877.2;refGene_gn=PRMT9;refGene_tx=XM_019828878.2;refGene_gn=PRMT9;refGene_tx=XM_023252772.1;refGene_gn=PRMT9;refGene_tx=XM_023252774.1
B1	81797036	81797220

refGene_gn=PRMT9;refGene_tx=XM_003984959.5;refGene_gn=PRMT9;refGene_tx=XM_019828877.2;refGene_gn=PRMT9;refGene_tx=XM_019828878.2;refGene_gn=PRMT9;refGene_tx=XM_023252772.1;refGene_gn=PRMT9;refGene_tx=XM_023252774.1
B1 81800328 81801046
refGene_gn=PRMT9;refGene_tx=XM_003984959.5;refGene_gn=PRMT9;refGene_tx=XM_019828877.2;refGene_gn=PRMT9;refGene_tx=XM_019828878.2;refGene_gn=PRMT9;refGene_tx=XM_023252772.1;refGene_gn=PRMT9;refGene_tx=XM_023252774.1
B1 81810745 81810899
refGene_gn=PRMT9;refGene_tx=XM_003984959.5;refGene_gn=PRMT9;refGene_tx=XM_019828877.2;refGene_gn=PRMT9;refGene_tx=XM_019828878.2;refGene_gn=PRMT9;refGene_tx=XM_023252772.1;refGene_gn=PRMT9;refGene_tx=XM_023252774.1
B1 81813989 81814112
refGene_gn=PRMT9;refGene_tx=XM_003984959.5;refGene_gn=PRMT9;refGene_tx=XM_019828877.2;refGene_gn=PRMT9;refGene_tx=XM_019828878.2;refGene_gn=PRMT9;refGene_tx=XM_023252772.1;refGene_gn=PRMT9;refGene_tx=XM_023252774.1
B1 81814410 81814626
refGene_gn=PRMT9;refGene_tx=XM_003984959.5;refGene_gn=PRMT9;refGene_tx=XM_019828877.2;refGene_gn=PRMT9;refGene_tx=XM_019828878.2;refGene_gn=PRMT9;refGene_tx=XM_023252772.1;refGene_gn=PRMT9;refGene_tx=XM_023252774.1
B1 81816919 81817185 refGene_gn=TMEM184C;refGene_tx=XM_003984960.5
B1 81817393 81817565 refGene_gn=TMEM184C;refGene_tx=XM_003984960.5
B1 81818039 81818139 refGene_gn=TMEM184C;refGene_tx=XM_003984960.5
B1 81819498 81819611 refGene_gn=TMEM184C;refGene_tx=XM_003984960.5
B1 81822234 81822328 refGene_gn=TMEM184C;refGene_tx=XM_003984960.5
B1 81823429 81823504 refGene_gn=TMEM184C;refGene_tx=XM_003984960.5
B1 81833006 81833212 refGene_gn=TMEM184C;refGene_tx=XM_003984960.5
B1 81833951 81833988 refGene_gn=TMEM184C;refGene_tx=XM_003984960.5
B1 81834086 81834217 refGene_gn=TMEM184C;refGene_tx=XM_003984960.5
B1 81841014 81841233 refGene_gn=TMEM184C;refGene_tx=XM_003984960.5
B1 81921985 81922126 refGene_gn=EDNRA;refGene_tx=XM_003984961.5
B1 81925395 81925504 refGene_gn=EDNRA;refGene_tx=XM_003984961.5
B1 81925930 81926064 refGene_gn=EDNRA;refGene_tx=XM_003984961.5
B1 81930282 81930435 refGene_gn=EDNRA;refGene_tx=XM_003984961.5
B1 81933531 81933730 refGene_gn=EDNRA;refGene_tx=XM_003984961.5
B1 81948423 81948551 refGene_gn=EDNRA;refGene_tx=XM_003984961.5
B1 81976053 81976473 refGene_gn=EDNRA;refGene_tx=XM_003984961.5
B1 82511025 82511117
refGene_gn=TTC29;refGene_tx=XM_006930836.3;refGene_gn=TTC29;refGene_tx=XM_006930837.3;refGene_gn=TTC29;refGene_tx=XM_019828882.2;refGene_gn=TTC29;refGene_tx=XM_019828883.2;refGene_gn=TTC29;refGene_tx=XM_023252775.1
B1 82512896 82512980
refGene_gn=TTC29;refGene_tx=XM_006930836.3;refGene_gn=TTC29;refGene_tx=XM_006930837.3;refGene_gn=TTC29;refGene_tx=XM_019828882.2;refGene_gn=TTC29;refGene_tx=XM_019828883.2;refGene_gn=TTC29;refGene_tx=XM_0232527

75.1
B1 82549205 82549429
refGene_gn=TTC29;refGene_tx=XM_006930836.3;refGene_gn=TTC29;refGene_tx=XM_006930837.3;refGene_gn=TTC29;refGene_tx=XM_019828882.2;refGene_gn=TTC29;refGene_tx=XM_019828883.2;refGene_gn=TTC29;refGene_tx=XM_023252775.1
B1 82554866 82555052
refGene_gn=TTC29;refGene_tx=XM_006930836.3;refGene_gn=TTC29;refGene_tx=XM_006930837.3;refGene_gn=TTC29;refGene_tx=XM_019828882.2;refGene_gn=TTC29;refGene_tx=XM_019828883.2;refGene_gn=TTC29;refGene_tx=XM_023252775.1
B1 82595059 82595272
refGene_gn=TTC29;refGene_tx=XM_006930836.3;refGene_gn=TTC29;refGene_tx=XM_006930837.3;refGene_gn=TTC29;refGene_tx=XM_019828882.2;refGene_gn=TTC29;refGene_tx=XM_019828883.2;refGene_gn=TTC29;refGene_tx=XM_023252775.1
B1 82601510 82601596
refGene_gn=TTC29;refGene_tx=XM_006930836.3;refGene_gn=TTC29;refGene_tx=XM_006930837.3;refGene_gn=TTC29;refGene_tx=XM_019828882.2;refGene_gn=TTC29;refGene_tx=XM_019828883.2;refGene_gn=TTC29;refGene_tx=XM_023252775.1
B1 82634175 82634267
refGene_gn=TTC29;refGene_tx=XM_006930836.3;refGene_gn=TTC29;refGene_tx=XM_006930837.3;refGene_gn=TTC29;refGene_tx=XM_019828882.2;refGene_gn=TTC29;refGene_tx=XM_019828883.2;refGene_gn=TTC29;refGene_tx=XM_023252775.1
B1 82652390 82652514
refGene_gn=TTC29;refGene_tx=XM_006930836.3;refGene_gn=TTC29;refGene_tx=XM_006930837.3;refGene_gn=TTC29;refGene_tx=XM_019828882.2;refGene_gn=TTC29;refGene_tx=XM_019828883.2;refGene_gn=TTC29;refGene_tx=XM_023252775.1
B1 82661786 82662015
refGene_gn=TTC29;refGene_tx=XM_006930836.3;refGene_gn=TTC29;refGene_tx=XM_006930837.3;refGene_gn=TTC29;refGene_tx=XM_019828882.2;refGene_gn=TTC29;refGene_tx=XM_019828883.2;refGene_gn=TTC29;refGene_tx=XM_023252775.1
B1 82745530 82745597
refGene_gn=TTC29;refGene_tx=XM_006930836.3;refGene_gn=TTC29;refGene_tx=XM_006930837.3;refGene_gn=TTC29;refGene_tx=XM_019828882.2;refGene_gn=TTC29;refGene_tx=XM_019828883.2;refGene_gn=TTC29;refGene_tx=XM_023252775.1
B1 82745881 82745912
refGene_gn=TTC29;refGene_tx=XM_006930836.3;refGene_gn=TTC29;refGene_tx=XM_006930837.3;refGene_gn=TTC29;refGene_tx=XM_019828882.2;refGene_gn=TTC29;refGene_tx=XM_019828883.2;refGene_gn=TTC29;refGene_tx=XM_023252775.1
B1 82807084 82808026 refGene_gn=POU4F2;refGene_tx=XM_011281629.3
B1 82808489 82808774 refGene_gn=POU4F2;refGene_tx=XM_011281629.3
B1 82878107 82878207
refGene_gn=SLC10A7;refGene_tx=XM_003984966.4;refGene_gn=SLC10A7;refGen

e_tx=XM_006930838.3;refGene_gn=SLC10A7;refGene_tx=XM_019828885.2;refGene_gn=SLC10A7;refGene_tx=XM_019828886.2;refGene_gn=SLC10A7;refGene_tx=XM_019828891.2
B1 82886952 82887035
refGene_gn=SLC10A7;refGene_tx=XM_003984966.4;refGene_gn=SLC10A7;refGene_tx=XM_006930838.3;refGene_gn=SLC10A7;refGene_tx=XM_019828885.2;refGene_gn=SLC10A7;refGene_tx=XM_019828886.2;refGene_gn=SLC10A7;refGene_tx=XM_019828891.2
B1 82893066 82893203
refGene_gn=SLC10A7;refGene_tx=XM_003984966.4;refGene_gn=SLC10A7;refGene_tx=XM_006930838.3;refGene_gn=SLC10A7;refGene_tx=XM_019828885.2;refGene_gn=SLC10A7;refGene_tx=XM_019828886.2
B1 82898127 82898203
refGene_gn=SLC10A7;refGene_tx=XM_003984966.4;refGene_gn=SLC10A7;refGene_tx=XM_006930838.3;refGene_gn=SLC10A7;refGene_tx=XM_019828885.2;refGene_gn=SLC10A7;refGene_tx=XM_019828886.2;refGene_gn=SLC10A7;refGene_tx=XM_019828891.2
B1 82955888 82955927
refGene_gn=SLC10A7;refGene_tx=XM_003984966.4;refGene_gn=SLC10A7;refGene_tx=XM_019828885.2;refGene_gn=SLC10A7;refGene_tx=XM_019828886.2
B1 83064167 83064203
refGene_gn=SLC10A7;refGene_tx=XM_003984966.4;refGene_gn=SLC10A7;refGene_tx=XM_006930838.3;refGene_gn=SLC10A7;refGene_tx=XM_019828885.2;refGene_gn=SLC10A7;refGene_tx=XM_019828886.2
B1 83075492 83075576 refGene_gn=SLC10A7;refGene_tx=XM_019828886.2
B1 83083506 83083590
refGene_gn=SLC10A7;refGene_tx=XM_003984966.4;refGene_gn=SLC10A7;refGene_tx=XM_006930838.3;refGene_gn=SLC10A7;refGene_tx=XM_019828885.2
B1 83093123 83093289
refGene_gn=SLC10A7;refGene_tx=XM_003984966.4;refGene_gn=SLC10A7;refGene_tx=XM_006930838.3;refGene_gn=SLC10A7;refGene_tx=XM_019828885.2
B1 83094237 83094289
refGene_gn=SLC10A7;refGene_tx=XM_003984966.4;refGene_gn=SLC10A7;refGene_tx=XM_006930838.3;refGene_gn=SLC10A7;refGene_tx=XM_019828885.2
B1 83104997 83105071
refGene_gn=SLC10A7;refGene_tx=XM_003984966.4;refGene_gn=SLC10A7;refGene_tx=XM_006930838.3;refGene_gn=SLC10A7;refGene_tx=XM_019828885.2
B1 83111411 83111416 refGene_gn=SLC10A7;refGene_tx=XM_019828885.2
B1 83130732 83130878
refGene_gn=SLC10A7;refGene_tx=XM_003984966.4;refGene_gn=SLC10A7;refGene_tx=XM_006930838.3
B1 83133173 83133203
refGene_gn=SLC10A7;refGene_tx=XM_003984966.4;refGene_gn=SLC10A7;refGene_tx=XM_006930838.3
B1 83154425 83155034
refGene_gn=LOC109498950;refGene_tx=XM_019828894.2;refGene_gn=LOC109498950;refGene_tx=XM_019828895.2;refGene_gn=LOC109498950;refGene_tx=XM_023252776.1
B1 83155847 83155911
refGene_gn=LOC109498950;refGene_tx=XM_019828894.2;refGene_gn=LOC109498

950;refGene_tx=XM_019828895.2;refGene_gn=LOC109498950;refGene_tx=XM_023252776.1
B1 83156434 83156747 refGene_gn=LOC109498950;refGene_tx=XM_019828894.2;refGene_gn=LOC109498950;refGene_tx=XM_019828895.2;refGene_gn=LOC109498950;refGene_tx=XM_023252776.1
B1 83159716 83159880 refGene_gn=LOC109498950;refGene_tx=XM_019828894.2;refGene_gn=LOC109498950;refGene_tx=XM_019828895.2;refGene_gn=LOC109498950;refGene_tx=XM_023252776.1
B1 83162031 83162254 refGene_gn=LOC109498950;refGene_tx=XM_019828894.2;refGene_gn=LOC109498950;refGene_tx=XM_019828895.2;refGene_gn=LOC109498950;refGene_tx=XM_023252776.1
B1 83165506 83165714 refGene_gn=LOC109498950;refGene_tx=XM_019828894.2;refGene_gn=LOC109498950;refGene_tx=XM_023252776.1
B1 83187473 83187508 refGene_gn=LSM6;refGene_tx=XM_003984968.3;refGene_gn=LSM6;refGene_tx=XM_006930839.2
B1 83189693 83189807 refGene_gn=LSM6;refGene_tx=XM_003984968.3;refGene_gn=LSM6;refGene_tx=XM_006930839.2
B1 83193402 83193496 refGene_gn=LSM6;refGene_tx=XM_003984968.3;refGene_gn=LSM6;refGene_tx=XM_006930839.2
B1 83391238 83391281 refGene_gn=ZNF827;refGene_tx=XM_019828897.2
B1 83422364 83423308 refGene_gn=ZNF827;refGene_tx=XM_019828897.2
B1 83423309 83423418 refGene_gn=ZNF827;refGene_tx=XM_019828897.2
B1 83433210 83433383 refGene_gn=ZNF827;refGene_tx=XM_019828897.2
B1 83438972 83439453 refGene_gn=ZNF827;refGene_tx=XM_019828897.2
B1 83456219 83456453 refGene_gn=ZNF827;refGene_tx=XM_019828897.2
B1 83482797 83483037 refGene_gn=ZNF827;refGene_tx=XM_019828897.2
B1 83485605 83485663 refGene_gn=ZNF827;refGene_tx=XM_019828897.2
B1 83504074 83504178 refGene_gn=ZNF827;refGene_tx=XM_019828897.2
B1 83546355 83546493 refGene_gn=ZNF827;refGene_tx=XM_019828897.2
B1 83547966 83548138 refGene_gn=ZNF827;refGene_tx=XM_019828897.2
B1 83549456 83549623 refGene_gn=ZNF827;refGene_tx=XM_019828897.2
B1 83558071 83558263 refGene_gn=ZNF827;refGene_tx=XM_019828897.2
B1 83558611 83558789 refGene_gn=ZNF827;refGene_tx=XM_019828897.2
B1 83560805 83560817 refGene_gn=ZNF827;refGene_tx=XM_019828897.2
B1 83562336 83563156 refGene_gn=ZNF827;refGene_tx=XM_019828897.2
B1 83629764 83630055 refGene_gn=MMAA;refGene_tx=XM_006930842.3;refGene_gn=MMAA;refGene_tx=XM_019828905.2;refGene_gn=MMAA;refGene_tx=XM_019828906.2;refGene_gn=MMAA;refGene_tx=XM_023252777.1;refGene_gn=MMAA;refGene_tx=XM_023252778.1
B1 83631092 83631242 refGene_gn=MMAA;refGene_tx=XM_006930842.3;refGene_gn=MMAA;refGene_tx=XM_019828905.2;refGene_gn=MMAA;refGene_tx=XM_019828906.2;refGene_gn=MMAA;refGene_tx=XM_023252777.1;refGene_gn=MMAA;refGene_tx=XM_023252778.1

B1 83632063 83632149
refGene_gn=MMAA;refGene_tx=XM_006930842.3;refGene_gn=MMAA;refGene_tx=X
M_019828905.2;refGene_gn=MMAA;refGene_tx=XM_019828906.2;refGene_gn=MMA
A;refGene_tx=XM_023252777.1;refGene_gn=MMAA;refGene_tx=XM_023252778.1
B1 83635035 83635206
refGene_gn=MMAA;refGene_tx=XM_006930842.3;refGene_gn=MMAA;refGene_tx=X
M_019828905.2;refGene_gn=MMAA;refGene_tx=XM_019828906.2;refGene_gn=MMA
A;refGene_tx=XM_023252777.1;refGene_gn=MMAA;refGene_tx=XM_023252778.1
B1 83638228 83638351
refGene_gn=MMAA;refGene_tx=XM_006930842.3;refGene_gn=MMAA;refGene_tx=X
M_019828905.2;refGene_gn=MMAA;refGene_tx=XM_019828906.2;refGene_gn=MMA
A;refGene_tx=XM_023252777.1;refGene_gn=MMAA;refGene_tx=XM_023252778.1
B1 83640540 83640985
refGene_gn=MMAA;refGene_tx=XM_006930842.3;refGene_gn=MMAA;refGene_tx=X
M_019828905.2;refGene_gn=MMAA;refGene_tx=XM_019828906.2;refGene_gn=MMA
A;refGene_tx=XM_023252777.1;refGene_gn=MMAA;refGene_tx=XM_023252778.1
B1 83700244 83700388 refGene_gn=SMAD1;refGene_tx=XM_023252779.1
B1 83704210 83704467 refGene_gn=SMAD1;refGene_tx=XM_023252779.1
B1 83709906 83710128 refGene_gn=SMAD1;refGene_tx=XM_023252779.1
B1 83713284 83713401 refGene_gn=SMAD1;refGene_tx=XM_023252779.1
B1 83715966 83716224 refGene_gn=SMAD1;refGene_tx=XM_023252779.1
B1 83745742 83746142 refGene_gn=SMAD1;refGene_tx=XM_023252779.1
B1 84033633 84033792 refGene_gn=OTUD4;refGene_tx=XM_023252780.1
B1 84039647 84039731 refGene_gn=OTUD4;refGene_tx=XM_023252780.1
B1 84042178 84042229 refGene_gn=OTUD4;refGene_tx=XM_023252780.1
B1 84048556 84048603 refGene_gn=OTUD4;refGene_tx=XM_023252780.1
B1 84049293 84049366 refGene_gn=OTUD4;refGene_tx=XM_023252780.1
B1 84050668 84050750 refGene_gn=OTUD4;refGene_tx=XM_023252780.1
B1 84052349 84052482 refGene_gn=OTUD4;refGene_tx=XM_023252780.1
B1 84056166 84056224 refGene_gn=OTUD4;refGene_tx=XM_023252780.1
B1 84056525 84056643 refGene_gn=OTUD4;refGene_tx=XM_023252780.1
B1 84056732 84056797 refGene_gn=OTUD4;refGene_tx=XM_023252780.1
B1 84059467 84059562 refGene_gn=OTUD4;refGene_tx=XM_023252780.1
B1 84060817 84060918 refGene_gn=OTUD4;refGene_tx=XM_023252780.1
B1 84061032 84061219 refGene_gn=OTUD4;refGene_tx=XM_023252780.1
B1 84065547 84065710 refGene_gn=OTUD4;refGene_tx=XM_023252780.1
B1 84067584 84067708 refGene_gn=OTUD4;refGene_tx=XM_023252780.1
B1 84068006 84068062 refGene_gn=OTUD4;refGene_tx=XM_023252780.1
B1 84068581 84068659 refGene_gn=OTUD4;refGene_tx=XM_023252780.1
B1 84070400 84070539 refGene_gn=OTUD4;refGene_tx=XM_023252780.1
B1 84071313 84071574 refGene_gn=OTUD4;refGene_tx=XM_023252780.1
B1 84072540 84072581 refGene_gn=OTUD4;refGene_tx=XM_023252780.1
B1 84073987 84075196 refGene_gn=OTUD4;refGene_tx=XM_023252780.1
B1 84084747 84084795 refGene_gn=ABCE1;refGene_tx=XM_003984975.4
B1 84086118 84086230 refGene_gn=ABCE1;refGene_tx=XM_003984975.4
B1 84087634 84087757 refGene_gn=ABCE1;refGene_tx=XM_003984975.4
B1 84088114 84088257 refGene_gn=ABCE1;refGene_tx=XM_003984975.4
B1 84088343 84088454 refGene_gn=ABCE1;refGene_tx=XM_003984975.4
B1 84091050 84091109 refGene_gn=ABCE1;refGene_tx=XM_003984975.4
B1 84091206 84091266 refGene_gn=ABCE1;refGene_tx=XM_003984975.4

B1 84092247 84092469 refGene_gn=ABCE1;refGene_tx=XM_003984975.4
B1 84095403 84095525 refGene_gn=ABCE1;refGene_tx=XM_003984975.4
B1 84100920 84101010 refGene_gn=ABCE1;refGene_tx=XM_003984975.4
B1 84101900 84101997 refGene_gn=ABCE1;refGene_tx=XM_003984975.4
B1 84102246 84102316 refGene_gn=ABCE1;refGene_tx=XM_003984975.4
B1 84102458 84102596 refGene_gn=ABCE1;refGene_tx=XM_003984975.4
B1 84103202 84103320 refGene_gn=ABCE1;refGene_tx=XM_003984975.4
B1 84104316 84104414 refGene_gn=ABCE1;refGene_tx=XM_003984975.4
B1 84107591 84107677 refGene_gn=ABCE1;refGene_tx=XM_003984975.4
B1 84108773 84108876 refGene_gn=ABCE1;refGene_tx=XM_003984975.4
B1 84118903 84119018
refGene_gn=ANAPC10;refGene_tx=XM_003984976.5;refGene_gn=ANAPC10;refGen
e_tx=XM_011281634.3;refGene_gn=ANAPC10;refGene_tx=XM_019828911.1;refGe
ne_gn=ANAPC10;refGene_tx=XM_023252781.1;refGene_gn=ANAPC10;refGene_tx=
XM_023252782.1
B1 84142617 84142708
refGene_gn=ANAPC10;refGene_tx=XM_003984976.5;refGene_gn=ANAPC10;refGen
e_tx=XM_011281634.3;refGene_gn=ANAPC10;refGene_tx=XM_019828911.1;refGe
ne_gn=ANAPC10;refGene_tx=XM_023252781.1;refGene_gn=ANAPC10;refGene_tx=
XM_023252782.1
B1 84167265 84167386
refGene_gn=ANAPC10;refGene_tx=XM_003984976.5;refGene_gn=ANAPC10;refGen
e_tx=XM_011281634.3;refGene_gn=ANAPC10;refGene_tx=XM_019828911.1;refGe
ne_gn=ANAPC10;refGene_tx=XM_023252781.1;refGene_gn=ANAPC10;refGene_tx=
XM_023252782.1
B1 84167988 84168018 refGene_gn=ANAPC10;refGene_tx=XM_019828911.1
B1 84191715 84191748 refGene_gn=ANAPC10;refGene_tx=XM_023252782.1
B1 84218637 84218706
refGene_gn=ANAPC10;refGene_tx=XM_011281634.3;refGene_gn=ANAPC10;refGen
e_tx=XM_023252781.1
B1 84224750 84224981
refGene_gn=ANAPC10;refGene_tx=XM_003984976.5;refGene_gn=ANAPC10;refGen
e_tx=XM_011281634.3;refGene_gn=ANAPC10;refGene_tx=XM_019828911.1;refGe
ne_gn=ANAPC10;refGene_tx=XM_023252781.1
B1 84409696 84409890 refGene_gn=HHIP;refGene_tx=XM_003984977.5
B1 84412605 84412754 refGene_gn=HHIP;refGene_tx=XM_003984977.5
B1 84429772 84429854 refGene_gn=HHIP;refGene_tx=XM_003984977.5
B1 84432974 84433105 refGene_gn=HHIP;refGene_tx=XM_003984977.5
B1 84434071 84434195 refGene_gn=HHIP;refGene_tx=XM_003984977.5
B1 84436462 84436584 refGene_gn=HHIP;refGene_tx=XM_003984977.5
B1 84439867 84440011 refGene_gn=HHIP;refGene_tx=XM_003984977.5
B1 84440619 84440793 refGene_gn=HHIP;refGene_tx=XM_003984977.5
B1 84441179 84441331 refGene_gn=HHIP;refGene_tx=XM_003984977.5
B1 84486066 84486268 refGene_gn=HHIP;refGene_tx=XM_003984977.5
B1 84487197 84487354 refGene_gn=HHIP;refGene_tx=XM_003984977.5
B1 84493308 84493501 refGene_gn=HHIP;refGene_tx=XM_003984977.5
B1 84499266 84499545 refGene_gn=HHIP;refGene_tx=XM_003984977.5
B1 84500015 84500069
refGene_gn=LOC109498955;refGene_tx=XM_023253392.1
B1 84500175 84500482

refGene_gn=LOC109498955;refGene_tx=XM_023253392.1
B1 84502416 84502442
refGene_gn=LOC109498955;refGene_tx=XM_023253392.1
B1 84907444 84907487
refGene_gn=GYPA;refGene_tx=XM_019828917.2;refGene_gn=GYPA;refGene_tx=XM_019828918.2
B1 84910784 84910868 refGene_gn=GYPA;refGene_tx=XM_019828917.2
B1 84931405 84931444
refGene_gn=GYPA;refGene_tx=XM_019828917.2;refGene_gn=GYPA;refGene_tx=XM_019828918.2
B1 84934432 84934468
refGene_gn=GYPA;refGene_tx=XM_019828917.2;refGene_gn=GYPA;refGene_tx=XM_019828918.2
B1 84936073 84936159
refGene_gn=GYPA;refGene_tx=XM_019828917.2;refGene_gn=GYPA;refGene_tx=XM_019828918.2
B1 84938934 84939010
refGene_gn=GYPA;refGene_tx=XM_019828917.2;refGene_gn=GYPA;refGene_tx=XM_019828918.2
B1 84942954 84942962
refGene_gn=GYPA;refGene_tx=XM_019828917.2;refGene_gn=GYPA;refGene_tx=XM_019828918.2
B1 85093540 85098722 refGene_gn=FREM3;refGene_tx=XM_003984978.3
B1 85100794 85100884 refGene_gn=FREM3;refGene_tx=XM_003984978.3
B1 85145618 85145765 refGene_gn=FREM3;refGene_tx=XM_003984978.3
B1 85149177 85149408 refGene_gn=FREM3;refGene_tx=XM_003984978.3
B1 85152363 85152489 refGene_gn=FREM3;refGene_tx=XM_003984978.3
B1 85164138 85164390 refGene_gn=FREM3;refGene_tx=XM_003984978.3
B1 85175669 85175819 refGene_gn=FREM3;refGene_tx=XM_003984978.3
B1 85178991 85179233 refGene_gn=FREM3;refGene_tx=XM_003984978.3
B1 85213406 85213472 refGene_gn=SMARCA5;refGene_tx=XM_003984979.5
B1 85214151 85214259 refGene_gn=SMARCA5;refGene_tx=XM_003984979.5
B1 85216207 85216420 refGene_gn=SMARCA5;refGene_tx=XM_003984979.5
B1 85216844 85216963 refGene_gn=SMARCA5;refGene_tx=XM_003984979.5
B1 85217560 85217693 refGene_gn=SMARCA5;refGene_tx=XM_003984979.5
B1 85218669 85218792 refGene_gn=SMARCA5;refGene_tx=XM_003984979.5
B1 85218891 85219005 refGene_gn=SMARCA5;refGene_tx=XM_003984979.5
B1 85219903 85220014 refGene_gn=SMARCA5;refGene_tx=XM_003984979.5
B1 85220762 85220882 refGene_gn=SMARCA5;refGene_tx=XM_003984979.5
B1 85221279 85221428 refGene_gn=SMARCA5;refGene_tx=XM_003984979.5
B1 85224017 85224150 refGene_gn=SMARCA5;refGene_tx=XM_003984979.5
B1 85225334 85225487 refGene_gn=SMARCA5;refGene_tx=XM_003984979.5
B1 85225568 85225690 refGene_gn=SMARCA5;refGene_tx=XM_003984979.5
B1 85227414 85227641 refGene_gn=SMARCA5;refGene_tx=XM_003984979.5
B1 85229465 85229575 refGene_gn=SMARCA5;refGene_tx=XM_003984979.5
B1 85234847 85234916 refGene_gn=SMARCA5;refGene_tx=XM_003984979.5
B1 85236648 85236780 refGene_gn=SMARCA5;refGene_tx=XM_003984979.5
B1 85236874 85237030 refGene_gn=SMARCA5;refGene_tx=XM_003984979.5
B1 85238162 85238342 refGene_gn=SMARCA5;refGene_tx=XM_003984979.5
B1 85239080 85239181 refGene_gn=SMARCA5;refGene_tx=XM_003984979.5

B1 85240680 85240781 refGene_gn=SMARCA5;refGene_tx=XM_003984979.5
B1 85244242 85244409 refGene_gn=SMARCA5;refGene_tx=XM_003984979.5
B1 85248396 85248471 refGene_gn=SMARCA5;refGene_tx=XM_003984979.5
B1 85251901 85252078 refGene_gn=SMARCA5;refGene_tx=XM_003984979.5
B1 85311790 85311949
refGene_gn=GAB1;refGene_tx=XM_003984980.5;refGene_gn=GAB1;refGene_tx=XM_003984981.5;refGene_gn=GAB1;refGene_tx=XM_006930848.4;refGene_gn=GAB1;refGene_tx=XM_019828924.2
B1 85314827 85314950
refGene_gn=GAB1;refGene_tx=XM_003984980.5;refGene_gn=GAB1;refGene_tx=XM_003984981.5;refGene_gn=GAB1;refGene_tx=XM_006930848.4;refGene_gn=GAB1;refGene_tx=XM_019828924.2
B1 85320117 85320241
refGene_gn=GAB1;refGene_tx=XM_003984980.5;refGene_gn=GAB1;refGene_tx=XM_003984981.5;refGene_gn=GAB1;refGene_tx=XM_006930848.4;refGene_gn=GAB1;refGene_tx=XM_019828924.2
B1 85321255 85321349
refGene_gn=GAB1;refGene_tx=XM_003984980.5;refGene_gn=GAB1;refGene_tx=XM_003984981.5;refGene_gn=GAB1;refGene_tx=XM_006930848.4;refGene_gn=GAB1;refGene_tx=XM_019828924.2
B1 85323046 85323136
refGene_gn=GAB1;refGene_tx=XM_003984981.5;refGene_gn=GAB1;refGene_tx=XM_019828924.2
B1 85331862 85332166
refGene_gn=GAB1;refGene_tx=XM_003984980.5;refGene_gn=GAB1;refGene_tx=XM_003984981.5;refGene_gn=GAB1;refGene_tx=XM_006930848.4;refGene_gn=GAB1;refGene_tx=XM_019828924.2
B1 85332345 85332431
refGene_gn=GAB1;refGene_tx=XM_003984980.5;refGene_gn=GAB1;refGene_tx=XM_003984981.5;refGene_gn=GAB1;refGene_tx=XM_006930848.4;refGene_gn=GAB1;refGene_tx=XM_019828924.2
B1 85333650 85334252
refGene_gn=GAB1;refGene_tx=XM_003984980.5;refGene_gn=GAB1;refGene_tx=XM_003984981.5;refGene_gn=GAB1;refGene_tx=XM_006930848.4;refGene_gn=GAB1;refGene_tx=XM_019828924.2
B1 85337853 85337934
refGene_gn=GAB1;refGene_tx=XM_006930848.4;refGene_gn=GAB1;refGene_tx=XM_019828924.2
B1 85338281 85338507
refGene_gn=GAB1;refGene_tx=XM_003984980.5;refGene_gn=GAB1;refGene_tx=XM_003984981.5;refGene_gn=GAB1;refGene_tx=XM_006930848.4;refGene_gn=GAB1;refGene_tx=XM_019828924.2
B1 85349989 85350284
refGene_gn=GAB1;refGene_tx=XM_003984980.5;refGene_gn=GAB1;refGene_tx=XM_003984981.5;refGene_gn=GAB1;refGene_tx=XM_006930848.4;refGene_gn=GAB1;refGene_tx=XM_019828924.2
B1 85430781 85430853
refGene_gn=GAB1;refGene_tx=XM_003984980.5;refGene_gn=GAB1;refGene_tx=XM_003984981.5;refGene_gn=GAB1;refGene_tx=XM_006930848.4;refGene_gn=GAB1;refGene_tx=XM_019828924.2

B1 85509219 85509381 refGene_gn=USP38;refGene_tx=XM_003984982.5
B1 85514274 85515628 refGene_gn=USP38;refGene_tx=XM_003984982.5
B1 85516748 85516855 refGene_gn=USP38;refGene_tx=XM_003984982.5
B1 85519196 85519290 refGene_gn=USP38;refGene_tx=XM_003984982.5
B1 85522111 85522305 refGene_gn=USP38;refGene_tx=XM_003984982.5
B1 85525884 85526043 refGene_gn=USP38;refGene_tx=XM_003984982.5
B1 85529660 85529762 refGene_gn=USP38;refGene_tx=XM_003984982.5
B1 85531705 85531835 refGene_gn=USP38;refGene_tx=XM_003984982.5
B1 85538572 85538708 refGene_gn=USP38;refGene_tx=XM_003984982.5
B1 85540308 85540990 refGene_gn=USP38;refGene_tx=XM_003984982.5
B1 86240515 86240606
refGene_gn=INPP4B;refGene_tx=XM_019828925.2;refGene_gn=INPP4B;refGene_tx=XM_019828926.2;refGene_gn=INPP4B;refGene_tx=XM_019828927.1;refGene_gn=INPP4B;refGene_tx=XM_019828929.1;refGene_gn=INPP4B;refGene_tx=XM_019828930.1;refGene_gn=INPP4B;refGene_tx=XM_019828931.2
B1 86242823 86242868
refGene_gn=INPP4B;refGene_tx=XM_019828925.2;refGene_gn=INPP4B;refGene_tx=XM_019828926.2;refGene_gn=INPP4B;refGene_tx=XM_019828927.1;refGene_gn=INPP4B;refGene_tx=XM_019828929.1;refGene_gn=INPP4B;refGene_tx=XM_019828930.1;refGene_gn=INPP4B;refGene_tx=XM_019828931.2
B1 86267718 86267837
refGene_gn=INPP4B;refGene_tx=XM_019828925.2;refGene_gn=INPP4B;refGene_tx=XM_019828926.2;refGene_gn=INPP4B;refGene_tx=XM_019828927.1;refGene_gn=INPP4B;refGene_tx=XM_019828929.1;refGene_gn=INPP4B;refGene_tx=XM_019828930.1;refGene_gn=INPP4B;refGene_tx=XM_019828931.2
B1 86269868 86269985
refGene_gn=INPP4B;refGene_tx=XM_019828925.2;refGene_gn=INPP4B;refGene_tx=XM_019828926.2;refGene_gn=INPP4B;refGene_tx=XM_019828927.1;refGene_gn=INPP4B;refGene_tx=XM_019828929.1;refGene_gn=INPP4B;refGene_tx=XM_019828930.1;refGene_gn=INPP4B;refGene_tx=XM_019828931.2
B1 86346978 86347029
refGene_gn=INPP4B;refGene_tx=XM_019828925.2;refGene_gn=INPP4B;refGene_tx=XM_019828926.2;refGene_gn=INPP4B;refGene_tx=XM_019828927.1;refGene_gn=INPP4B;refGene_tx=XM_019828929.1;refGene_gn=INPP4B;refGene_tx=XM_019828930.1;refGene_gn=INPP4B;refGene_tx=XM_019828931.2
B1 86357632 86357712
refGene_gn=INPP4B;refGene_tx=XM_019828925.2;refGene_gn=INPP4B;refGene_tx=XM_019828926.2;refGene_gn=INPP4B;refGene_tx=XM_019828927.1;refGene_gn=INPP4B;refGene_tx=XM_019828929.1;refGene_gn=INPP4B;refGene_tx=XM_019828930.1;refGene_gn=INPP4B;refGene_tx=XM_019828931.2
B1 86393083 86393195
refGene_gn=INPP4B;refGene_tx=XM_019828925.2;refGene_gn=INPP4B;refGene_tx=XM_019828926.2;refGene_gn=INPP4B;refGene_tx=XM_019828927.1;refGene_gn=INPP4B;refGene_tx=XM_019828929.1;refGene_gn=INPP4B;refGene_tx=XM_019828930.1;refGene_gn=INPP4B;refGene_tx=XM_019828931.2
B1 86402227 86402300
refGene_gn=INPP4B;refGene_tx=XM_019828925.2;refGene_gn=INPP4B;refGene_tx=XM_019828926.2;refGene_gn=INPP4B;refGene_tx=XM_019828927.1;refGene_gn=INPP4B;refGene_tx=XM_019828929.1;refGene_gn=INPP4B;refGene_tx=XM_019828930.1;refGene_gn=INPP4B;refGene_tx=XM_019828931.2

B1 86414693 86414841
refGene_gn=INPP4B;refGene_tx=XM_019828925.2;refGene_gn=INPP4B;refGene_tx=XM_019828926.2;refGene_gn=INPP4B;refGene_tx=XM_019828927.1;refGene_gn=INPP4B;refGene_tx=XM_019828929.1;refGene_gn=INPP4B;refGene_tx=XM_019828930.1;refGene_gn=INPP4B;refGene_tx=XM_019828931.2
B1 86438244 86438375
refGene_gn=INPP4B;refGene_tx=XM_019828925.2;refGene_gn=INPP4B;refGene_tx=XM_019828926.2;refGene_gn=INPP4B;refGene_tx=XM_019828927.1;refGene_gn=INPP4B;refGene_tx=XM_019828929.1;refGene_gn=INPP4B;refGene_tx=XM_019828930.1;refGene_gn=INPP4B;refGene_tx=XM_019828931.2
B1 86438759 86438864
refGene_gn=INPP4B;refGene_tx=XM_019828925.2;refGene_gn=INPP4B;refGene_tx=XM_019828926.2;refGene_gn=INPP4B;refGene_tx=XM_019828927.1;refGene_gn=INPP4B;refGene_tx=XM_019828929.1;refGene_gn=INPP4B;refGene_tx=XM_019828930.1;refGene_gn=INPP4B;refGene_tx=XM_019828931.2
B1 86453656 86453765
refGene_gn=INPP4B;refGene_tx=XM_019828925.2;refGene_gn=INPP4B;refGene_tx=XM_019828926.2;refGene_gn=INPP4B;refGene_tx=XM_019828927.1;refGene_gn=INPP4B;refGene_tx=XM_019828929.1;refGene_gn=INPP4B;refGene_tx=XM_019828930.1;refGene_gn=INPP4B;refGene_tx=XM_019828931.2
B1 86477127 86477305
refGene_gn=INPP4B;refGene_tx=XM_019828925.2;refGene_gn=INPP4B;refGene_tx=XM_019828926.2;refGene_gn=INPP4B;refGene_tx=XM_019828927.1;refGene_gn=INPP4B;refGene_tx=XM_019828929.1;refGene_gn=INPP4B;refGene_tx=XM_019828930.1;refGene_gn=INPP4B;refGene_tx=XM_019828931.2
B1 86488767 86488971
refGene_gn=INPP4B;refGene_tx=XM_019828925.2;refGene_gn=INPP4B;refGene_tx=XM_019828926.2;refGene_gn=INPP4B;refGene_tx=XM_019828927.1;refGene_gn=INPP4B;refGene_tx=XM_019828929.1;refGene_gn=INPP4B;refGene_tx=XM_019828930.1;refGene_gn=INPP4B;refGene_tx=XM_019828931.2
B1 86504299 86504459
refGene_gn=INPP4B;refGene_tx=XM_019828925.2;refGene_gn=INPP4B;refGene_tx=XM_019828926.2;refGene_gn=INPP4B;refGene_tx=XM_019828927.1;refGene_gn=INPP4B;refGene_tx=XM_019828929.1;refGene_gn=INPP4B;refGene_tx=XM_019828930.1;refGene_gn=INPP4B;refGene_tx=XM_019828931.2
B1 86524363 86524536
refGene_gn=INPP4B;refGene_tx=XM_019828925.2;refGene_gn=INPP4B;refGene_tx=XM_019828926.2;refGene_gn=INPP4B;refGene_tx=XM_019828927.1;refGene_gn=INPP4B;refGene_tx=XM_019828929.1;refGene_gn=INPP4B;refGene_tx=XM_019828930.1;refGene_gn=INPP4B;refGene_tx=XM_019828931.2
B1 86525381 86525505
refGene_gn=INPP4B;refGene_tx=XM_019828925.2;refGene_gn=INPP4B;refGene_tx=XM_019828926.2;refGene_gn=INPP4B;refGene_tx=XM_019828927.1;refGene_gn=INPP4B;refGene_tx=XM_019828929.1;refGene_gn=INPP4B;refGene_tx=XM_019828930.1;refGene_gn=INPP4B;refGene_tx=XM_019828931.2
B1 86526623 86526741
refGene_gn=INPP4B;refGene_tx=XM_019828925.2;refGene_gn=INPP4B;refGene_tx=XM_019828926.2;refGene_gn=INPP4B;refGene_tx=XM_019828927.1;refGene_gn=INPP4B;refGene_tx=XM_019828929.1;refGene_gn=INPP4B;refGene_tx=XM_019828930.1;refGene_gn=INPP4B;refGene_tx=XM_019828931.2

B1 86531653 86531794
refGene_gn=INPP4B;refGene_tx=XM_019828925.2;refGene_gn=INPP4B;refGene_tx=XM_019828926.2;refGene_gn=INPP4B;refGene_tx=XM_019828927.1;refGene_gn=INPP4B;refGene_tx=XM_019828929.1;refGene_gn=INPP4B;refGene_tx=XM_019828930.1;refGene_gn=INPP4B;refGene_tx=XM_019828931.2
B1 86536427 86536525
refGene_gn=INPP4B;refGene_tx=XM_019828925.2;refGene_gn=INPP4B;refGene_tx=XM_019828926.2;refGene_gn=INPP4B;refGene_tx=XM_019828927.1;refGene_gn=INPP4B;refGene_tx=XM_019828929.1;refGene_gn=INPP4B;refGene_tx=XM_019828930.1;refGene_gn=INPP4B;refGene_tx=XM_019828931.2
B1 86556682 86556795
refGene_gn=INPP4B;refGene_tx=XM_019828925.2;refGene_gn=INPP4B;refGene_tx=XM_019828926.2;refGene_gn=INPP4B;refGene_tx=XM_019828927.1;refGene_gn=INPP4B;refGene_tx=XM_019828929.1;refGene_gn=INPP4B;refGene_tx=XM_019828930.1;refGene_gn=INPP4B;refGene_tx=XM_019828931.2
B1 86560696 86560851
refGene_gn=INPP4B;refGene_tx=XM_019828925.2;refGene_gn=INPP4B;refGene_tx=XM_019828926.2;refGene_gn=INPP4B;refGene_tx=XM_019828927.1;refGene_gn=INPP4B;refGene_tx=XM_019828929.1;refGene_gn=INPP4B;refGene_tx=XM_019828930.1;refGene_gn=INPP4B;refGene_tx=XM_019828931.2
B1 86586509 86586531 refGene_gn=INPP4B;refGene_tx=XM_019828931.2
B1 86607987 86608162
refGene_gn=INPP4B;refGene_tx=XM_019828925.2;refGene_gn=INPP4B;refGene_tx=XM_019828926.2;refGene_gn=INPP4B;refGene_tx=XM_019828927.1
B1 86609395 86609528
refGene_gn=INPP4B;refGene_tx=XM_019828929.1;refGene_gn=INPP4B;refGene_tx=XM_019828930.1
B1 86777374 86777485
refGene_gn=IL15;refGene_tx=NM_001009207.1;refGene_gn=IL15;refGene_tx=XM_006930830.3;refGene_gn=IL15;refGene_tx=XM_006930831.3
B1 86780426 86780564
refGene_gn=IL15;refGene_tx=NM_001009207.1;refGene_gn=IL15;refGene_tx=XM_006930830.3;refGene_gn=IL15;refGene_tx=XM_006930831.3
B1 86782521 86782566
refGene_gn=IL15;refGene_tx=NM_001009207.1;refGene_gn=IL15;refGene_tx=XM_006930830.3;refGene_gn=IL15;refGene_tx=XM_006930831.3
B1 86787829 86787914
refGene_gn=IL15;refGene_tx=NM_001009207.1;refGene_gn=IL15;refGene_tx=XM_006930830.3;refGene_gn=IL15;refGene_tx=XM_006930831.3
B1 86789731 86789829
refGene_gn=IL15;refGene_tx=NM_001009207.1;refGene_gn=IL15;refGene_tx=XM_006930830.3;refGene_gn=IL15;refGene_tx=XM_006930831.3
B1 86790780 86790792
refGene_gn=IL15;refGene_tx=NM_001009207.1;refGene_gn=IL15;refGene_tx=XM_006930830.3;refGene_gn=IL15;refGene_tx=XM_006930831.3
B1 87191546 87191821
refGene_gn=ZNF330;refGene_tx=XM_011281654.3;refGene_gn=ZNF330;refGene_tx=XM_011281655.3;refGene_gn=ZNF330;refGene_tx=XM_019828932.2
B1 87193004 87193122
refGene_gn=ZNF330;refGene_tx=XM_011281654.3;refGene_gn=ZNF330;refGene_

tx=XM_011281655.3;refGene_gn=ZNF330;refGene_tx=XM_019828932.2
B1 87194177 87194224
refGene_gn=ZNF330;refGene_tx=XM_011281654.3;refGene_gn=ZNF330;refGene_tx=XM_011281655.3;refGene_gn=ZNF330;refGene_tx=XM_019828932.2
B1 87198637 87198742
refGene_gn=ZNF330;refGene_tx=XM_011281654.3;refGene_gn=ZNF330;refGene_tx=XM_011281655.3;refGene_gn=ZNF330;refGene_tx=XM_019828932.2
B1 87199181 87199308
refGene_gn=ZNF330;refGene_tx=XM_011281654.3;refGene_gn=ZNF330;refGene_tx=XM_011281655.3;refGene_gn=ZNF330;refGene_tx=XM_019828932.2
B1 87204943 87205023
refGene_gn=ZNF330;refGene_tx=XM_011281654.3;refGene_gn=ZNF330;refGene_tx=XM_011281655.3;refGene_gn=ZNF330;refGene_tx=XM_019828932.2
B1 87206944 87207015
refGene_gn=ZNF330;refGene_tx=XM_011281654.3;refGene_gn=ZNF330;refGene_tx=XM_011281655.3;refGene_gn=ZNF330;refGene_tx=XM_019828932.2
B1 87207109 87207129
refGene_gn=ZNF330;refGene_tx=XM_011281654.3;refGene_gn=ZNF330;refGene_tx=XM_011281655.3;refGene_gn=ZNF330;refGene_tx=XM_019828932.2
B1 87209007 87209127
refGene_gn=ZNF330;refGene_tx=XM_011281654.3;refGene_gn=ZNF330;refGene_tx=XM_011281655.3;refGene_gn=ZNF330;refGene_tx=XM_019828932.2
B1 87299578 87300062
refGene_gn=RNF150;refGene_tx=XM_019828933.2;refGene_gn=RNF150;refGene_tx=XM_019828934.2;refGene_gn=RNF150;refGene_tx=XM_019828935.2
B1 87455634 87455885
refGene_gn=RNF150;refGene_tx=XM_019828933.2;refGene_gn=RNF150;refGene_tx=XM_019828934.2;refGene_gn=RNF150;refGene_tx=XM_019828935.2
B1 87477102 87477174
refGene_gn=RNF150;refGene_tx=XM_019828933.2;refGene_gn=RNF150;refGene_tx=XM_019828934.2;refGene_gn=RNF150;refGene_tx=XM_019828935.2
B1 87479373 87479456
refGene_gn=RNF150;refGene_tx=XM_019828933.2;refGene_gn=RNF150;refGene_tx=XM_019828934.2;refGene_gn=RNF150;refGene_tx=XM_019828935.2
B1 87505059 87505156
refGene_gn=RNF150;refGene_tx=XM_019828933.2;refGene_gn=RNF150;refGene_tx=XM_019828934.2;refGene_gn=RNF150;refGene_tx=XM_019828935.2
B1 87512066 87512277
refGene_gn=RNF150;refGene_tx=XM_019828933.2;refGene_gn=RNF150;refGene_tx=XM_019828934.2;refGene_gn=RNF150;refGene_tx=XM_019828935.2
B1 87524970 87525032 refGene_gn=RNF150;refGene_tx=XM_019828935.2
B1 87554806 87554925 refGene_gn=RNF150;refGene_tx=XM_019828933.2
B1 87595004 87595093 refGene_gn=RNF150;refGene_tx=XM_019828934.2
B1 87655350 87655480
refGene_gn=TBC1D9;refGene_tx=XM_023252784.1;refGene_gn=TBC1D9;refGene_tx=XM_023252785.1
B1 87704408 87704519
refGene_gn=TBC1D9;refGene_tx=XM_023252784.1;refGene_gn=TBC1D9;refGene_tx=XM_023252785.1
B1 87717717 87717836

refGene_gn=TBC1D9;refGene_tx=XM_023252784.1;refGene_gn=TBC1D9;refGene_tx=XM_023252785.1
B1 87724154 87724383
refGene_gn=TBC1D9;refGene_tx=XM_023252784.1;refGene_gn=TBC1D9;refGene_tx=XM_023252785.1
B1 87724814 87725076
refGene_gn=TBC1D9;refGene_tx=XM_023252784.1;refGene_gn=TBC1D9;refGene_tx=XM_023252785.1
B1 87726857 87727065
refGene_gn=TBC1D9;refGene_tx=XM_023252784.1;refGene_gn=TBC1D9;refGene_tx=XM_023252785.1
B1 87733053 87733260
refGene_gn=TBC1D9;refGene_tx=XM_023252784.1;refGene_gn=TBC1D9;refGene_tx=XM_023252785.1
B1 87734114 87734288
refGene_gn=TBC1D9;refGene_tx=XM_023252784.1;refGene_gn=TBC1D9;refGene_tx=XM_023252785.1
B1 87734875 87735026
refGene_gn=TBC1D9;refGene_tx=XM_023252784.1;refGene_gn=TBC1D9;refGene_tx=XM_023252785.1
B1 87737877 87738092
refGene_gn=TBC1D9;refGene_tx=XM_023252784.1;refGene_gn=TBC1D9;refGene_tx=XM_023252785.1
B1 87740456 87740574
refGene_gn=TBC1D9;refGene_tx=XM_023252784.1;refGene_gn=TBC1D9;refGene_tx=XM_023252785.1
B1 87742344 87742630
refGene_gn=TBC1D9;refGene_tx=XM_023252784.1;refGene_gn=TBC1D9;refGene_tx=XM_023252785.1
B1 87742947 87743077
refGene_gn=TBC1D9;refGene_tx=XM_023252784.1;refGene_gn=TBC1D9;refGene_tx=XM_023252785.1
B1 87753122 87753221
refGene_gn=TBC1D9;refGene_tx=XM_023252784.1;refGene_gn=TBC1D9;refGene_tx=XM_023252785.1
B1 87753368 87753437
refGene_gn=TBC1D9;refGene_tx=XM_023252784.1;refGene_gn=TBC1D9;refGene_tx=XM_023252785.1
B1 87758010 87758251
refGene_gn=TBC1D9;refGene_tx=XM_023252784.1;refGene_gn=TBC1D9;refGene_tx=XM_023252785.1
B1 87763278 87763344
refGene_gn=TBC1D9;refGene_tx=XM_023252784.1;refGene_gn=TBC1D9;refGene_tx=XM_023252785.1
B1 87764345 87764432
refGene_gn=TBC1D9;refGene_tx=XM_023252784.1;refGene_gn=TBC1D9;refGene_tx=XM_023252785.1
B1 87768551 87768626
refGene_gn=TBC1D9;refGene_tx=XM_023252784.1;refGene_gn=TBC1D9;refGene_tx=XM_023252785.1

B1 87768719 87768823
refGene_gn=TBC1D9;refGene_tx=XM_023252784.1;refGene_gn=TBC1D9;refGene_tx=XM_023252785.1
B1 87770194 87770917
refGene_gn=TBC1D9;refGene_tx=XM_023252784.1;refGene_gn=TBC1D9;refGene_tx=XM_023252785.1
B1 87801613 87801739 refGene_gn=UCP1;refGene_tx=XM_003984985.4
B1 87802382 87802581 refGene_gn=UCP1;refGene_tx=XM_003984985.4
B1 87804084 87804285 refGene_gn=UCP1;refGene_tx=XM_003984985.4
B1 87804384 87804486 refGene_gn=UCP1;refGene_tx=XM_003984985.4
B1 87805735 87805916 refGene_gn=UCP1;refGene_tx=XM_003984985.4
B1 87808508 87808623 refGene_gn=UCP1;refGene_tx=XM_003984985.4
B1 87816425 87816571
refGene_gn=ELMOD2;refGene_tx=XM_003984986.5;refGene_gn=ELMOD2;refGene_tx=XM_019828939.2;refGene_gn=ELMOD2;refGene_tx=XM_019828940.2;refGene_gn=ELMOD2;refGene_tx=XM_019828941.2;refGene_gn=ELMOD2;refGene_tx=XM_019828942.2;refGene_gn=ELMOD2;refGene_tx=XM_019828944.2;refGene_gn=ELMOD2;refGene_tx=XM_019828946.1
B1 87823330 87823464
refGene_gn=ELMOD2;refGene_tx=XM_003984986.5;refGene_gn=ELMOD2;refGene_tx=XM_019828939.2;refGene_gn=ELMOD2;refGene_tx=XM_019828940.2;refGene_gn=ELMOD2;refGene_tx=XM_019828941.2;refGene_gn=ELMOD2;refGene_tx=XM_019828942.2;refGene_gn=ELMOD2;refGene_tx=XM_019828944.2;refGene_gn=ELMOD2;refGene_tx=XM_019828946.1
B1 87824959 87825028
refGene_gn=ELMOD2;refGene_tx=XM_003984986.5;refGene_gn=ELMOD2;refGene_tx=XM_019828939.2;refGene_gn=ELMOD2;refGene_tx=XM_019828940.2;refGene_gn=ELMOD2;refGene_tx=XM_019828941.2;refGene_gn=ELMOD2;refGene_tx=XM_019828942.2;refGene_gn=ELMOD2;refGene_tx=XM_019828944.2;refGene_gn=ELMOD2;refGene_tx=XM_019828946.1
B1 87828261 87828395
refGene_gn=ELMOD2;refGene_tx=XM_003984986.5;refGene_gn=ELMOD2;refGene_tx=XM_019828939.2;refGene_gn=ELMOD2;refGene_tx=XM_019828940.2;refGene_gn=ELMOD2;refGene_tx=XM_019828941.2;refGene_gn=ELMOD2;refGene_tx=XM_019828942.2;refGene_gn=ELMOD2;refGene_tx=XM_019828944.2;refGene_gn=ELMOD2;refGene_tx=XM_019828946.1
B1 87830916 87831046
refGene_gn=ELMOD2;refGene_tx=XM_003984986.5;refGene_gn=ELMOD2;refGene_tx=XM_019828939.2;refGene_gn=ELMOD2;refGene_tx=XM_019828940.2;refGene_gn=ELMOD2;refGene_tx=XM_019828941.2;refGene_gn=ELMOD2;refGene_tx=XM_019828942.2;refGene_gn=ELMOD2;refGene_tx=XM_019828944.2;refGene_gn=ELMOD2;refGene_tx=XM_019828946.1
B1 87831292 87831390
refGene_gn=ELMOD2;refGene_tx=XM_003984986.5;refGene_gn=ELMOD2;refGene_tx=XM_019828939.2;refGene_gn=ELMOD2;refGene_tx=XM_019828940.2;refGene_gn=ELMOD2;refGene_tx=XM_019828941.2;refGene_gn=ELMOD2;refGene_tx=XM_019828942.2;refGene_gn=ELMOD2;refGene_tx=XM_019828944.2;refGene_gn=ELMOD2;refGene_tx=XM_019828946.1
B1 87837045 87837057 refGene_gn=ELMOD2;refGene_tx=XM_019828946.1
B1 87840966 87840995

refGene_gn=ELMOD2;refGene_tx=XM_003984986.5;refGene_gn=ELMOD2;refGene_tx=XM_019828939.2;refGene_gn=ELMOD2;refGene_tx=XM_019828940.2;refGene_gn=ELMOD2;refGene_tx=XM_019828941.2;refGene_gn=ELMOD2;refGene_tx=XM_019828942.2;refGene_gn=ELMOD2;refGene_tx=XM_019828944.2
B1 87842905 87843047
refGene_gn=ELMOD2;refGene_tx=XM_003984986.5;refGene_gn=ELMOD2;refGene_tx=XM_019828939.2;refGene_gn=ELMOD2;refGene_tx=XM_019828940.2;refGene_gn=ELMOD2;refGene_tx=XM_019828941.2;refGene_gn=ELMOD2;refGene_tx=XM_019828942.2;refGene_gn=ELMOD2;refGene_tx=XM_019828944.2
B1 87857602 87857696
refGene_gn=MGAT4D;refGene_tx=XM_023252787.1;refGene_gn=MGAT4D;refGene_tx=XM_023252789.1
B1 87878171 87878330
refGene_gn=MGAT4D;refGene_tx=XM_023252787.1;refGene_gn=MGAT4D;refGene_tx=XM_023252789.1
B1 87880751 87880892
refGene_gn=MGAT4D;refGene_tx=XM_023252787.1;refGene_gn=MGAT4D;refGene_tx=XM_023252789.1
B1 87885069 87885203
refGene_gn=MGAT4D;refGene_tx=XM_023252787.1;refGene_gn=MGAT4D;refGene_tx=XM_023252789.1
B1 87894083 87894191
refGene_gn=MGAT4D;refGene_tx=XM_023252787.1;refGene_gn=MGAT4D;refGene_tx=XM_023252789.1
B1 87896353 87896386 refGene_gn=MGAT4D;refGene_tx=XM_023252787.1
B1 87904439 87904448 refGene_gn=MGAT4D;refGene_tx=XM_023252789.1
B1 87930904 87931048 refGene_gn=CLGN;refGene_tx=XM_003984987.5
B1 87933592 87933666 refGene_gn=CLGN;refGene_tx=XM_003984987.5
B1 87934331 87934390 refGene_gn=CLGN;refGene_tx=XM_003984987.5
B1 87938847 87938989 refGene_gn=CLGN;refGene_tx=XM_003984987.5
B1 87942740 87942822 refGene_gn=CLGN;refGene_tx=XM_003984987.5
B1 87944230 87944423 refGene_gn=CLGN;refGene_tx=XM_003984987.5
B1 87948483 87948673 refGene_gn=CLGN;refGene_tx=XM_003984987.5
B1 87954601 87954715 refGene_gn=CLGN;refGene_tx=XM_003984987.5
B1 87954837 87954988 refGene_gn=CLGN;refGene_tx=XM_003984987.5
B1 87957272 87957488 refGene_gn=CLGN;refGene_tx=XM_003984987.5
B1 87958809 87958935 refGene_gn=CLGN;refGene_tx=XM_003984987.5
B1 87959152 87959312 refGene_gn=CLGN;refGene_tx=XM_003984987.5
B1 87960391 87960492 refGene_gn=CLGN;refGene_tx=XM_003984987.5
B1 87962921 87963002 refGene_gn=CLGN;refGene_tx=XM_003984987.5
B1 87971544 87971687 refGene_gn=SCOC;refGene_tx=XM_023252790.1
B1 87973585 87973669 refGene_gn=SCOC;refGene_tx=XM_023252790.1
B1 87973991 87974063 refGene_gn=SCOC;refGene_tx=XM_023252790.1
B1 87979898 87980007 refGene_gn=SCOC;refGene_tx=XM_023252790.1
B1 87980008 87980089 refGene_gn=SCOC;refGene_tx=XM_023252790.1
B1 88175105 88175576
refGene_gn=MAML3;refGene_tx=XM_003984990.4;refGene_gn=MAML3;refGene_tx=XM_006930850.3
B1 88418601 88420218
refGene_gn=MAML3;refGene_tx=XM_003984990.4;refGene_gn=MAML3;refGene_tx

=XM_006930850.3
B1 88580920 88581166
refGene_gn=MAML3;refGene_tx=XM_003984990.4;refGene_gn=MAML3;refGene_tx
=XM_006930850.3
B1 88585270 88585355
refGene_gn=MAML3;refGene_tx=XM_003984990.4;refGene_gn=MAML3;refGene_tx
=XM_006930850.3
B1 88590154 88591152
refGene_gn=MAML3;refGene_tx=XM_003984990.4;refGene_gn=MAML3;refGene_tx
=XM_006930850.3
B1 88599296 88599426 refGene_gn=MGST2;refGene_tx=XM_003984991.5
B1 88599913 88599995 refGene_gn=MGST2;refGene_tx=XM_003984991.5
B1 88603330 88603401 refGene_gn=MGST2;refGene_tx=XM_003984991.5
B1 88619650 88619750 refGene_gn=MGST2;refGene_tx=XM_003984991.5
B1 88625204 88625262 refGene_gn=MGST2;refGene_tx=XM_003984991.5
B1 88724015 88724084
refGene_gn=SETD7;refGene_tx=XM_003984992.5;refGene_gn=SETD7;refGene_tx
=XM_006930851.4
B1 88724758 88724891
refGene_gn=SETD7;refGene_tx=XM_003984992.5;refGene_gn=SETD7;refGene_tx
=XM_006930851.4
B1 88732456 88732586
refGene_gn=SETD7;refGene_tx=XM_003984992.5;refGene_gn=SETD7;refGene_tx
=XM_006930851.4
B1 88746747 88746949
refGene_gn=SETD7;refGene_tx=XM_003984992.5;refGene_gn=SETD7;refGene_tx
=XM_006930851.4
B1 88751852 88752042
refGene_gn=SETD7;refGene_tx=XM_003984992.5;refGene_gn=SETD7;refGene_tx
=XM_006930851.4
B1 88757804 88757886
refGene_gn=SETD7;refGene_tx=XM_003984992.5;refGene_gn=SETD7;refGene_tx
=XM_006930851.4
B1 88761496 88761614
refGene_gn=SETD7;refGene_tx=XM_003984992.5;refGene_gn=SETD7;refGene_tx
=XM_006930851.4
B1 88764659 88764817
refGene_gn=SETD7;refGene_tx=XM_003984992.5;refGene_gn=SETD7;refGene_tx
=XM_006930851.4
B1 88769273 88769454 refGene_gn=SETD7;refGene_tx=XM_003984992.5
B1 88775784 88775959 refGene_gn=SETD7;refGene_tx=XM_006930851.4
B1 88810896 88811337
refGene_gn=RAB33B;refGene_tx=XM_023252792.1;refGene_gn=RAB33B;refGene_
tx=XM_023252793.1
B1 88814086 88814092 refGene_gn=RAB33B;refGene_tx=XM_023252797.1
B1 88820265 88820286 refGene_gn=RAB33B;refGene_tx=XM_023252796.1
B1 88822602 88822635 refGene_gn=RAB33B;refGene_tx=XM_023252794.1
B1 88823324 88823351 refGene_gn=RAB33B;refGene_tx=XM_023252795.1
B1 88823593 88823861
refGene_gn=RAB33B;refGene_tx=XM_023252792.1;refGene_gn=RAB33B;refGene_

tx=XM_023252793.1;refGene_gn=RAB33B;refGene_tx=XM_023252794.1;refGene_gn=RAB33B;refGene_tx=XM_023252795.1;refGene_gn=RAB33B;refGene_tx=XM_023252796.1;refGene_gn=RAB33B;refGene_tx=XM_023252797.1
B1 88824056 88824406
refGene_gn=RAB33B;refGene_tx=XM_023252792.1;refGene_gn=RAB33B;refGene_tx=XM_023252794.1;refGene_gn=RAB33B;refGene_tx=XM_023252795.1;refGene_gn=RAB33B;refGene_tx=XM_023252796.1;refGene_gn=RAB33B;refGene_tx=XM_023252797.1
B1 88865660 88865861
refGene_gn=NAA15;refGene_tx=XM_003984993.5;refGene_gn=NAA15;refGene_tx=XM_019828951.2;refGene_gn=NAA15;refGene_tx=XM_023252791.1
B1 88867212 88867310
refGene_gn=NAA15;refGene_tx=XM_003984993.5;refGene_gn=NAA15;refGene_tx=XM_019828951.2;refGene_gn=NAA15;refGene_tx=XM_023252791.1
B1 88868929 88869076
refGene_gn=NAA15;refGene_tx=XM_003984993.5;refGene_gn=NAA15;refGene_tx=XM_019828951.2;refGene_gn=NAA15;refGene_tx=XM_023252791.1
B1 88874611 88874710
refGene_gn=NAA15;refGene_tx=XM_003984993.5;refGene_gn=NAA15;refGene_tx=XM_019828951.2;refGene_gn=NAA15;refGene_tx=XM_023252791.1
B1 88876636 88876745
refGene_gn=NAA15;refGene_tx=XM_003984993.5;refGene_gn=NAA15;refGene_tx=XM_019828951.2;refGene_gn=NAA15;refGene_tx=XM_023252791.1
B1 88881370 88881564
refGene_gn=NAA15;refGene_tx=XM_003984993.5;refGene_gn=NAA15;refGene_tx=XM_019828951.2;refGene_gn=NAA15;refGene_tx=XM_023252791.1
B1 88890953 88891167
refGene_gn=NAA15;refGene_tx=XM_003984993.5;refGene_gn=NAA15;refGene_tx=XM_019828951.2;refGene_gn=NAA15;refGene_tx=XM_023252791.1
B1 88892250 88892379
refGene_gn=NAA15;refGene_tx=XM_003984993.5;refGene_gn=NAA15;refGene_tx=XM_019828951.2;refGene_gn=NAA15;refGene_tx=XM_023252791.1
B1 88893188 88893341
refGene_gn=NAA15;refGene_tx=XM_003984993.5;refGene_gn=NAA15;refGene_tx=XM_019828951.2;refGene_gn=NAA15;refGene_tx=XM_023252791.1
B1 88894551 88894721
refGene_gn=NAA15;refGene_tx=XM_003984993.5;refGene_gn=NAA15;refGene_tx=XM_019828951.2;refGene_gn=NAA15;refGene_tx=XM_023252791.1
B1 88897694 88897767
refGene_gn=NAA15;refGene_tx=XM_003984993.5;refGene_gn=NAA15;refGene_tx=XM_019828951.2;refGene_gn=NAA15;refGene_tx=XM_023252791.1
B1 88899604 88899711
refGene_gn=NAA15;refGene_tx=XM_003984993.5;refGene_gn=NAA15;refGene_tx=XM_019828951.2;refGene_gn=NAA15;refGene_tx=XM_023252791.1
B1 88899857 88899953
refGene_gn=NAA15;refGene_tx=XM_003984993.5;refGene_gn=NAA15;refGene_tx=XM_019828951.2;refGene_gn=NAA15;refGene_tx=XM_023252791.1
B1 88901554 88901674
refGene_gn=NAA15;refGene_tx=XM_003984993.5;refGene_gn=NAA15;refGene_tx=XM_019828951.2;refGene_gn=NAA15;refGene_tx=XM_023252791.1

B1 88904991 88905145
refGene_gn=NAA15;refGene_tx=XM_003984993.5;refGene_gn=NAA15;refGene_tx=XM_019828951.2;refGene_gn=NAA15;refGene_tx=XM_023252791.1
B1 88906584 88906719
refGene_gn=NAA15;refGene_tx=XM_003984993.5;refGene_gn=NAA15;refGene_tx=XM_019828951.2;refGene_gn=NAA15;refGene_tx=XM_023252791.1
B1 88907531 88907689
refGene_gn=NAA15;refGene_tx=XM_003984993.5;refGene_gn=NAA15;refGene_tx=XM_019828951.2;refGene_gn=NAA15;refGene_tx=XM_023252791.1
B1 88910797 88910902
refGene_gn=NAA15;refGene_tx=XM_003984993.5;refGene_gn=NAA15;refGene_tx=XM_019828951.2;refGene_gn=NAA15;refGene_tx=XM_023252791.1
B1 88912861 88912946
refGene_gn=NAA15;refGene_tx=XM_003984993.5;refGene_gn=NAA15;refGene_tx=XM_023252791.1
B1 88947531 88947585
refGene_gn=NAA15;refGene_tx=XM_003984993.5;refGene_gn=NAA15;refGene_tx=XM_023252791.1
B1 88951597 88951664 refGene_gn=NDUFC1;refGene_tx=XM_003984994.3
B1 88952012 88952116 refGene_gn=NDUFC1;refGene_tx=XM_003984994.3
B1 88954086 88954146 refGene_gn=NDUFC1;refGene_tx=XM_003984994.3
B1 88962161 88962243 refGene_gn=MGARP;refGene_tx=XM_011281693.3
B1 88966290 88966394
refGene_gn=MGARP;refGene_tx=XM_011281693.3;refGene_gn=MGARP;refGene_tx=XM_023252798.1;refGene_gn=MGARP;refGene_tx=XM_023252799.1
B1 88972295 88972389
refGene_gn=MGARP;refGene_tx=XM_011281693.3;refGene_gn=MGARP;refGene_tx=XM_023252798.1;refGene_gn=MGARP;refGene_tx=XM_023252799.1
B1 88973756 88974202
refGene_gn=MGARP;refGene_tx=XM_011281693.3;refGene_gn=MGARP;refGene_tx=XM_023252798.1;refGene_gn=MGARP;refGene_tx=XM_023252799.1
B1 89060418 89060490
refGene_gn=ELF2;refGene_tx=XM_023252800.1;refGene_gn=ELF2;refGene_tx=XM_023252801.1;refGene_gn=ELF2;refGene_tx=XM_023252802.1;refGene_gn=ELF2;refGene_tx=XM_023252803.1;refGene_gn=ELF2;refGene_tx=XM_023252804.1
B1 89071420 89071586
refGene_gn=ELF2;refGene_tx=XM_023252800.1;refGene_gn=ELF2;refGene_tx=XM_023252801.1;refGene_gn=ELF2;refGene_tx=XM_023252802.1;refGene_gn=ELF2;refGene_tx=XM_023252803.1;refGene_gn=ELF2;refGene_tx=XM_023252804.1
B1 89101577 89101635
refGene_gn=ELF2;refGene_tx=XM_003984995.5;refGene_gn=ELF2;refGene_tx=XM_003984998.5;refGene_gn=ELF2;refGene_tx=XM_006930854.4;refGene_gn=ELF2;refGene_tx=XM_006930855.4;refGene_gn=ELF2;refGene_tx=XM_019828955.2
B1 89110935 89111049
refGene_gn=ELF2;refGene_tx=XM_003984995.5;refGene_gn=ELF2;refGene_tx=XM_003984998.5;refGene_gn=ELF2;refGene_tx=XM_006930854.4;refGene_gn=ELF2;refGene_tx=XM_006930855.4;refGene_gn=ELF2;refGene_tx=XM_019828955.2;refGene_gn=ELF2;refGene_tx=XM_019828956.2;refGene_gn=ELF2;refGene_tx=XM_023252800.1;refGene_gn=ELF2;refGene_tx=XM_023252801.1;refGene_gn=ELF2;refGene_tx=XM_023252802.1;refGene_gn=ELF2;refGene_tx=XM_023252803.1;

refGene_gn=ELF2;refGene_tx=XM_023252804.1
B1 89112907 89113081
refGene_gn=ELF2;refGene_tx=XM_003984995.5;refGene_gn=ELF2;refGene_tx=XM_003984998.5;refGene_gn=ELF2;refGene_tx=XM_006930854.4;refGene_gn=ELF2;refGene_tx=XM_006930855.4;refGene_gn=ELF2;refGene_tx=XM_019828955.2;refGene_gn=ELF2;refGene_tx=XM_019828956.2;refGene_gn=ELF2;refGene_tx=XM_023252800.1;refGene_gn=ELF2;refGene_tx=XM_023252801.1;refGene_gn=ELF2;refGene_tx=XM_023252802.1;refGene_gn=ELF2;refGene_tx=XM_023252803.1;refGene_gn=ELF2;refGene_tx=XM_023252804.1
B1 89115621 89115660
refGene_gn=ELF2;refGene_tx=XM_019828955.2;refGene_gn=ELF2;refGene_tx=XM_019828956.2;refGene_gn=ELF2;refGene_tx=XM_023252800.1;refGene_gn=ELF2;refGene_tx=XM_023252801.1
B1 89116014 89116101
refGene_gn=ELF2;refGene_tx=XM_003984995.5;refGene_gn=ELF2;refGene_tx=XM_003984998.5;refGene_gn=ELF2;refGene_tx=XM_019828955.2;refGene_gn=ELF2;refGene_tx=XM_019828956.2;refGene_gn=ELF2;refGene_tx=XM_023252800.1;refGene_gn=ELF2;refGene_tx=XM_023252801.1;refGene_gn=ELF2;refGene_tx=XM_023252802.1;refGene_gn=ELF2;refGene_tx=XM_023252803.1
B1 89120342 89120535
refGene_gn=ELF2;refGene_tx=XM_003984995.5;refGene_gn=ELF2;refGene_tx=XM_003984998.5;refGene_gn=ELF2;refGene_tx=XM_006930854.4;refGene_gn=ELF2;refGene_tx=XM_006930855.4;refGene_gn=ELF2;refGene_tx=XM_019828955.2;refGene_gn=ELF2;refGene_tx=XM_019828956.2;refGene_gn=ELF2;refGene_tx=XM_023252800.1;refGene_gn=ELF2;refGene_tx=XM_023252801.1;refGene_gn=ELF2;refGene_tx=XM_023252802.1;refGene_gn=ELF2;refGene_tx=XM_023252803.1;refGene_gn=ELF2;refGene_tx=XM_023252804.1
B1 89121150 89121498
refGene_gn=ELF2;refGene_tx=XM_003984995.5;refGene_gn=ELF2;refGene_tx=XM_003984998.5;refGene_gn=ELF2;refGene_tx=XM_006930854.4;refGene_gn=ELF2;refGene_tx=XM_006930855.4;refGene_gn=ELF2;refGene_tx=XM_019828955.2;refGene_gn=ELF2;refGene_tx=XM_019828956.2;refGene_gn=ELF2;refGene_tx=XM_023252800.1;refGene_gn=ELF2;refGene_tx=XM_023252801.1;refGene_gn=ELF2;refGene_tx=XM_023252802.1;refGene_gn=ELF2;refGene_tx=XM_023252803.1;refGene_gn=ELF2;refGene_tx=XM_023252804.1
B1 89121874 89122502
refGene_gn=ELF2;refGene_tx=XM_003984995.5;refGene_gn=ELF2;refGene_tx=XM_003984998.5;refGene_gn=ELF2;refGene_tx=XM_006930854.4;refGene_gn=ELF2;refGene_tx=XM_006930855.4;refGene_gn=ELF2;refGene_tx=XM_019828955.2;refGene_gn=ELF2;refGene_tx=XM_019828956.2;refGene_gn=ELF2;refGene_tx=XM_023252800.1;refGene_gn=ELF2;refGene_tx=XM_023252801.1;refGene_gn=ELF2;refGene_tx=XM_023252802.1;refGene_gn=ELF2;refGene_tx=XM_023252803.1;refGene_gn=ELF2;refGene_tx=XM_023252804.1
B1 89133697 89134533
refGene_gn=NOCT;refGene_tx=XM_011281691.3;refGene_gn=NOCT;refGene_tx=XM_019828960.2;refGene_gn=NOCT;refGene_tx=XM_023252806.1;refGene_gn=NOCT;refGene_tx=XM_023252807.1;refGene_gn=NOCT;refGene_tx=XM_023252808.1
B1 89136229 89136499
refGene_gn=NOCT;refGene_tx=XM_011281691.3;refGene_gn=NOCT;refGene_tx=XM_019828960.2;refGene_gn=NOCT;refGene_tx=XM_023252806.1;refGene_gn=NOC

T;refGene_tx=XM_023252807.1;refGene_gn=NOCT;refGene_tx=XM_023252808.1
B1	89148830	89148966	refGene_gn=NOCT;refGene_tx=XM_011281691.3
B1	89160305	89160495	refGene_gn=NOCT;refGene_tx=XM_023252806.1
B1	89869852	89870129	refGene_gn=SLC7A11;refGene_tx=XM_011281688.3
B1	89873969	89874096	refGene_gn=SLC7A11;refGene_tx=XM_011281688.3
B1	89878880	89878996	refGene_gn=SLC7A11;refGene_tx=XM_011281688.3
B1	89887235	89887361	refGene_gn=SLC7A11;refGene_tx=XM_011281688.3
B1	89890860	89890960	refGene_gn=SLC7A11;refGene_tx=XM_011281688.3
B1	89898540	89898585	refGene_gn=SLC7A11;refGene_tx=XM_011281688.3
B1	89925844	89925968	refGene_gn=SLC7A11;refGene_tx=XM_011281688.3
B1	89927867	89927971	refGene_gn=SLC7A11;refGene_tx=XM_011281688.3
B1	89928853	89928950	refGene_gn=SLC7A11;refGene_tx=XM_011281688.3
B1	89930459	89930609	refGene_gn=SLC7A11;refGene_tx=XM_011281688.3
B1	89931863	89932041	refGene_gn=SLC7A11;refGene_tx=XM_011281688.3
B1	89943989	89944057	refGene_gn=SLC7A11;refGene_tx=XM_011281688.3
B1	90585282	90587760
refGene_gn=PCDH18;refGene_tx=XM_011281686.3;refGene_gn=PCDH18;refGene_tx=XM_011281687.3
B1	90588598	90588687
refGene_gn=PCDH18;refGene_tx=XM_011281686.3;refGene_gn=PCDH18;refGene_tx=XM_011281687.3
B1	90588777	90588941
refGene_gn=PCDH18;refGene_tx=XM_011281686.3;refGene_gn=PCDH18;refGene_tx=XM_011281687.3
B1	90596633	90597301
refGene_gn=PCDH18;refGene_tx=XM_011281686.3;refGene_gn=PCDH18;refGene_tx=XM_011281687.3
B1	93566245	93567358	refGene_gn=PABPC4L;refGene_tx=XM_023252811.1
B1	94408483	94408551	refGene_gn=PCDH10;refGene_tx=XM_011281684.3
B1	94430506	94430526	refGene_gn=PCDH10;refGene_tx=XM_011281685.3
B1	94450564	94450605	refGene_gn=PCDH10;refGene_tx=XM_006930866.4
B1	94456216	94456522
refGene_gn=PCDH10;refGene_tx=XM_006930866.4;refGene_gn=PCDH10;refGene_tx=XM_011281684.3;refGene_gn=PCDH10;refGene_tx=XM_011281685.3
B1	94464498	94464605
refGene_gn=PCDH10;refGene_tx=XM_006930866.4;refGene_gn=PCDH10;refGene_tx=XM_011281684.3;refGene_gn=PCDH10;refGene_tx=XM_011281685.3
B1	94465155	94465214
refGene_gn=PCDH10;refGene_tx=XM_006930866.4;refGene_gn=PCDH10;refGene_tx=XM_011281684.3;refGene_gn=PCDH10;refGene_tx=XM_011281685.3
B1	94466789	94469420
refGene_gn=PCDH10;refGene_tx=XM_006930866.4;refGene_gn=PCDH10;refGene_tx=XM_011281684.3;refGene_gn=PCDH10;refGene_tx=XM_011281685.3
B1	97837787	97837893
refGene_gn=CB1H4orf33;refGene_tx=XM_003985011.5;refGene_gn=CB1H4orf33;refGene_tx=XM_019828964.2
B1	97840072	97840272
refGene_gn=CB1H4orf33;refGene_tx=XM_003985011.5;refGene_gn=CB1H4orf33;refGene_tx=XM_019828964.2
B1	97840631	97840683

refGene_gn=CB1H4orf33;refGene_tx=XM_003985011.5;refGene_gn=CB1H4orf33;refGene_tx=XM_019828964.2
B1 97843195 97843256 refGene_gn=CB1H4orf33;refGene_tx=XM_003985011.5;refGene_gn=CB1H4orf33;refGene_tx=XM_019828964.2
B1 97848160 97848341 refGene_gn=CB1H4orf33;refGene_tx=XM_003985011.5;refGene_gn=CB1H4orf33;refGene_tx=XM_019828964.2
B1 97859579 97859613 refGene_gn=SCLT1;refGene_tx=XM_006930868.3
B1 97875538 97875606 refGene_gn=SCLT1;refGene_tx=XM_006930868.3
B1 97889385 97889444 refGene_gn=SCLT1;refGene_tx=XM_006930868.3
B1 97890303 97890376 refGene_gn=SCLT1;refGene_tx=XM_006930868.3
B1 97899520 97899576 refGene_gn=SCLT1;refGene_tx=XM_006930868.3
B1 97911433 97911569 refGene_gn=SCLT1;refGene_tx=XM_006930868.3
B1 97919952 97920075 refGene_gn=SCLT1;refGene_tx=XM_006930868.3
B1 97922227 97922293 refGene_gn=SCLT1;refGene_tx=XM_006930868.3
B1 97927003 97927074 refGene_gn=SCLT1;refGene_tx=XM_006930868.3
B1 97972390 97972481 refGene_gn=SCLT1;refGene_tx=XM_006930868.3
B1 97983199 97983291 refGene_gn=SCLT1;refGene_tx=XM_006930868.3
B1 97992794 97992972 refGene_gn=SCLT1;refGene_tx=XM_006930868.3
B1 97998236 97998335 refGene_gn=SCLT1;refGene_tx=XM_006930868.3
B1 98015227 98015299 refGene_gn=SCLT1;refGene_tx=XM_006930868.3
B1 98022745 98022820 refGene_gn=SCLT1;refGene_tx=XM_006930868.3
B1 98025096 98025242 refGene_gn=SCLT1;refGene_tx=XM_006930868.3
B1 98031133 98031326 refGene_gn=SCLT1;refGene_tx=XM_006930868.3
B1 98037613 98037810 refGene_gn=SCLT1;refGene_tx=XM_006930868.3
B1 98080554 98080633 refGene_gn=SCLT1;refGene_tx=XM_006930868.3
B1 98088192 98088288 refGene_gn=SCLT1;refGene_tx=XM_006930868.3
B1 98092999 98093062 refGene_gn=SCLT1;refGene_tx=XM_006930868.3
B1 98104963 98105874 refGene_gn=JADE1;refGene_tx=XM_003985012.5;refGene_gn=JADE1;refGene_tx=XM_003985013.5;refGene_gn=JADE1;refGene_tx=XM_006930869.4;refGene_gn=JADE1;refGene_tx=XM_006930870.4;refGene_gn=JADE1;refGene_tx=XM_019828965.2;refGene_gn=JADE1;refGene_tx=XM_019828966.2;refGene_gn=JADE1;refGene_tx=XM_019828967.2;refGene_gn=JADE1;refGene_tx=XM_019828968.2;refGene_gn=JADE1;refGene_tx=XM_019828969.2;refGene_gn=JADE1;refGene_tx=XM_019828971.2;refGene_gn=JADE1;refGene_tx=XM_023252812.1;refGene_gn=JADE1;refGene_tx=XM_023252813.1
B1 98109446 98109564 refGene_gn=JADE1;refGene_tx=XM_003985012.5;refGene_gn=JADE1;refGene_tx=XM_003985013.5;refGene_gn=JADE1;refGene_tx=XM_006930869.4;refGene_gn=JADE1;refGene_tx=XM_006930870.4;refGene_gn=JADE1;refGene_tx=XM_019828965.2;refGene_gn=JADE1;refGene_tx=XM_019828966.2;refGene_gn=JADE1;refGene_tx=XM_019828967.2;refGene_gn=JADE1;refGene_tx=XM_019828968.2;refGene_gn=JADE1;refGene_tx=XM_019828969.2;refGene_gn=JADE1;refGene_tx=XM_019828971.2;refGene_gn=JADE1;refGene_tx=XM_023252812.1;refGene_gn=JADE1;refGene_tx=XM_023252813.1
B1 98114955 98115477 refGene_gn=JADE1;refGene_tx=XM_003985012.5;refGene_gn=JADE1;refGene_tx=XM_003985013.5;refGene_gn=JADE1;refGene_tx=XM_006930869.4;refGene_gn=

JADE1;refGene_tx=XM_006930870.4;refGene_gn=JADE1;refGene_tx=XM_0198289
65.2;refGene_gn=JADE1;refGene_tx=XM_019828966.2;refGene_gn=JADE1;refGe
ne_tx=XM_019828967.2;refGene_gn=JADE1;refGene_tx=XM_019828968.2;refGen
e_gn=JADE1;refGene_tx=XM_019828969.2;refGene_gn=JADE1;refGene_tx=XM_01
9828971.2;refGene_gn=JADE1;refGene_tx=XM_023252812.1;refGene_gn=JADE1;
refGene_tx=XM_023252813.1
B1 98119024 98119141
refGene_gn=JADE1;refGene_tx=XM_003985012.5;refGene_gn=JADE1;refGene_tx
=XM_003985013.5;refGene_gn=JADE1;refGene_tx=XM_006930869.4;refGene_gn=
JADE1;refGene_tx=XM_006930870.4;refGene_gn=JADE1;refGene_tx=XM_0198289
65.2;refGene_gn=JADE1;refGene_tx=XM_019828966.2;refGene_gn=JADE1;refGe
ne_tx=XM_019828967.2;refGene_gn=JADE1;refGene_tx=XM_019828968.2;refGen
e_gn=JADE1;refGene_tx=XM_019828969.2;refGene_gn=JADE1;refGene_tx=XM_01
9828971.2;refGene_gn=JADE1;refGene_tx=XM_023252812.1;refGene_gn=JADE1;
refGene_tx=XM_023252813.1
B1 98120463 98120631
refGene_gn=JADE1;refGene_tx=XM_003985012.5;refGene_gn=JADE1;refGene_tx
=XM_003985013.5;refGene_gn=JADE1;refGene_tx=XM_006930869.4;refGene_gn=
JADE1;refGene_tx=XM_006930870.4;refGene_gn=JADE1;refGene_tx=XM_0198289
65.2;refGene_gn=JADE1;refGene_tx=XM_019828966.2;refGene_gn=JADE1;refGe
ne_tx=XM_019828967.2;refGene_gn=JADE1;refGene_tx=XM_019828968.2;refGen
e_gn=JADE1;refGene_tx=XM_019828969.2;refGene_gn=JADE1;refGene_tx=XM_01
9828971.2;refGene_gn=JADE1;refGene_tx=XM_023252812.1;refGene_gn=JADE1;
refGene_tx=XM_023252813.1
B1 98124357 98124569
refGene_gn=JADE1;refGene_tx=XM_003985012.5;refGene_gn=JADE1;refGene_tx
=XM_003985013.5;refGene_gn=JADE1;refGene_tx=XM_006930869.4;refGene_gn=
JADE1;refGene_tx=XM_006930870.4;refGene_gn=JADE1;refGene_tx=XM_0198289
65.2;refGene_gn=JADE1;refGene_tx=XM_019828966.2;refGene_gn=JADE1;refGe
ne_tx=XM_019828967.2;refGene_gn=JADE1;refGene_tx=XM_019828968.2;refGen
e_gn=JADE1;refGene_tx=XM_019828969.2;refGene_gn=JADE1;refGene_tx=XM_01
9828971.2;refGene_gn=JADE1;refGene_tx=XM_023252812.1;refGene_gn=JADE1;
refGene_tx=XM_023252813.1
B1 98127136 98127324
refGene_gn=JADE1;refGene_tx=XM_003985012.5;refGene_gn=JADE1;refGene_tx
=XM_003985013.5;refGene_gn=JADE1;refGene_tx=XM_006930869.4;refGene_gn=
JADE1;refGene_tx=XM_006930870.4;refGene_gn=JADE1;refGene_tx=XM_0198289
65.2;refGene_gn=JADE1;refGene_tx=XM_019828966.2;refGene_gn=JADE1;refGe
ne_tx=XM_019828967.2;refGene_gn=JADE1;refGene_tx=XM_019828968.2;refGen
e_gn=JADE1;refGene_tx=XM_019828969.2;refGene_gn=JADE1;refGene_tx=XM_01
9828971.2;refGene_gn=JADE1;refGene_tx=XM_023252812.1;refGene_gn=JADE1;
refGene_tx=XM_023252813.1
B1 98129236 98129394
refGene_gn=JADE1;refGene_tx=XM_003985012.5;refGene_gn=JADE1;refGene_tx
=XM_003985013.5;refGene_gn=JADE1;refGene_tx=XM_006930869.4;refGene_gn=
JADE1;refGene_tx=XM_006930870.4;refGene_gn=JADE1;refGene_tx=XM_0198289
65.2;refGene_gn=JADE1;refGene_tx=XM_019828966.2;refGene_gn=JADE1;refGe
ne_tx=XM_019828967.2;refGene_gn=JADE1;refGene_tx=XM_019828968.2;refGen
e_gn=JADE1;refGene_tx=XM_019828969.2;refGene_gn=JADE1;refGene_tx=XM_01
9828971.2;refGene_gn=JADE1;refGene_tx=XM_023252812.1;refGene_gn=JADE1;

refGene_tx=XM_023252813.1
B1 98132038 98132081
refGene_gn=JADE1;refGene_tx=XM_019828967.2;refGene_gn=JADE1;refGene_tx=XM_019828968.2;refGene_gn=JADE1;refGene_tx=XM_019828969.2;refGene_gn=JADE1;refGene_tx=XM_019828971.2
B1 98132422 98132508
refGene_gn=JADE1;refGene_tx=XM_003985012.5;refGene_gn=JADE1;refGene_tx=XM_003985013.5;refGene_gn=JADE1;refGene_tx=XM_006930869.4;refGene_gn=JADE1;refGene_tx=XM_006930870.4;refGene_gn=JADE1;refGene_tx=XM_019828965.2;refGene_gn=JADE1;refGene_tx=XM_019828966.2;refGene_gn=JADE1;refGene_tx=XM_019828967.2;refGene_gn=JADE1;refGene_tx=XM_019828968.2;refGene_gn=JADE1;refGene_tx=XM_019828969.2;refGene_gn=JADE1;refGene_tx=XM_019828971.2;refGene_gn=JADE1;refGene_tx=XM_023252812.1;refGene_gn=JADE1;refGene_tx=XM_023252813.1
B1 98139590 98139660 refGene_gn=JADE1;refGene_tx=XM_019828965.2
B1 98143596 98143674
refGene_gn=JADE1;refGene_tx=XM_003985012.5;refGene_gn=JADE1;refGene_tx=XM_003985013.5;refGene_gn=JADE1;refGene_tx=XM_006930869.4;refGene_gn=JADE1;refGene_tx=XM_006930870.4;refGene_gn=JADE1;refGene_tx=XM_019828965.2;refGene_gn=JADE1;refGene_tx=XM_019828966.2;refGene_gn=JADE1;refGene_tx=XM_019828967.2;refGene_gn=JADE1;refGene_tx=XM_019828968.2;refGene_gn=JADE1;refGene_tx=XM_019828969.2;refGene_gn=JADE1;refGene_tx=XM_019828971.2;refGene_gn=JADE1;refGene_tx=XM_023252812.1;refGene_gn=JADE1;refGene_tx=XM_023252813.1
B1 98159258 98159277 refGene_gn=JADE1;refGene_tx=XM_023252812.1
B1 98160693 98160706 refGene_gn=JADE1;refGene_tx=XM_006930869.4
B1 98650152 98650666
refGene_gn=PGRMC2;refGene_tx=XM_003985015.4;refGene_gn=PGRMC2;refGene_tx=XM_019828973.2
B1 98666011 98666167
refGene_gn=PGRMC2;refGene_tx=XM_003985015.4;refGene_gn=PGRMC2;refGene_tx=XM_019828973.2
B1 98667138 98667166 refGene_gn=PGRMC2;refGene_tx=XM_019828973.2
B1 98667292 98667390
refGene_gn=PGRMC2;refGene_tx=XM_003985015.4;refGene_gn=PGRMC2;refGene_tx=XM_019828973.2
B1 98706592 98706751
refGene_gn=LARP1B;refGene_tx=XM_019828979.2;refGene_gn=LARP1B;refGene_tx=XM_023252816.1;refGene_gn=LARP1B;refGene_tx=XM_023252817.1;refGene_gn=LARP1B;refGene_tx=XM_023252818.1;refGene_gn=LARP1B;refGene_tx=XM_023252821.1;refGene_gn=LARP1B;refGene_tx=XM_023252822.1
B1 98707941 98707974 refGene_gn=LARP1B;refGene_tx=XM_023252820.1
B1 98712829 98712967
refGene_gn=LARP1B;refGene_tx=XM_023252815.1;refGene_gn=LARP1B;refGene_tx=XM_023252819.1
B1 98714763 98714891
refGene_gn=LARP1B;refGene_tx=XM_019828979.2;refGene_gn=LARP1B;refGene_tx=XM_023252815.1;refGene_gn=LARP1B;refGene_tx=XM_023252816.1;refGene_gn=LARP1B;refGene_tx=XM_023252817.1;refGene_gn=LARP1B;refGene_tx=XM_023252818.1;refGene_gn=LARP1B;refGene_tx=XM_023252819.1;refGene_gn=LARP1

B;refGene_tx=XM_023252820.1;refGene_gn=LARP1B;refGene_tx=XM_023252821.1;refGene_gn=LARP1B;refGene_tx=XM_023252822.1
B1	98715608 98715718 refGene_gn=LARP1B;refGene_tx=XM_019828979.2;refGene_gn=LARP1B;refGene_tx=XM_023252815.1;refGene_gn=LARP1B;refGene_tx=XM_023252816.1;refGene_gn=LARP1B;refGene_tx=XM_023252817.1;refGene_gn=LARP1B;refGene_tx=XM_023252818.1;refGene_gn=LARP1B;refGene_tx=XM_023252819.1;refGene_gn=LARP1B;refGene_tx=XM_023252820.1;refGene_gn=LARP1B;refGene_tx=XM_023252821.1;refGene_gn=LARP1B;refGene_tx=XM_023252822.1
B1	98720331 98720476 refGene_gn=LARP1B;refGene_tx=XM_019828979.2;refGene_gn=LARP1B;refGene_tx=XM_023252815.1;refGene_gn=LARP1B;refGene_tx=XM_023252816.1;refGene_gn=LARP1B;refGene_tx=XM_023252817.1;refGene_gn=LARP1B;refGene_tx=XM_023252818.1;refGene_gn=LARP1B;refGene_tx=XM_023252819.1;refGene_gn=LARP1B;refGene_tx=XM_023252820.1;refGene_gn=LARP1B;refGene_tx=XM_023252821.1;refGene_gn=LARP1B;refGene_tx=XM_023252822.1
B1	98720864 98721025 refGene_gn=LARP1B;refGene_tx=XM_019828979.2;refGene_gn=LARP1B;refGene_tx=XM_023252815.1;refGene_gn=LARP1B;refGene_tx=XM_023252816.1;refGene_gn=LARP1B;refGene_tx=XM_023252817.1;refGene_gn=LARP1B;refGene_tx=XM_023252818.1;refGene_gn=LARP1B;refGene_tx=XM_023252819.1;refGene_gn=LARP1B;refGene_tx=XM_023252820.1;refGene_gn=LARP1B;refGene_tx=XM_023252821.1;refGene_gn=LARP1B;refGene_tx=XM_023252822.1
B1	98733094 98733201 refGene_gn=LARP1B;refGene_tx=XM_019828979.2;refGene_gn=LARP1B;refGene_tx=XM_023252815.1;refGene_gn=LARP1B;refGene_tx=XM_023252816.1;refGene_gn=LARP1B;refGene_tx=XM_023252817.1;refGene_gn=LARP1B;refGene_tx=XM_023252818.1;refGene_gn=LARP1B;refGene_tx=XM_023252819.1;refGene_gn=LARP1B;refGene_tx=XM_023252820.1;refGene_gn=LARP1B;refGene_tx=XM_023252821.1;refGene_gn=LARP1B;refGene_tx=XM_023252822.1
B1	98733656 98733868 refGene_gn=LARP1B;refGene_tx=XM_019828979.2;refGene_gn=LARP1B;refGene_tx=XM_023252815.1;refGene_gn=LARP1B;refGene_tx=XM_023252816.1;refGene_gn=LARP1B;refGene_tx=XM_023252817.1;refGene_gn=LARP1B;refGene_tx=XM_023252818.1;refGene_gn=LARP1B;refGene_tx=XM_023252819.1;refGene_gn=LARP1B;refGene_tx=XM_023252820.1;refGene_gn=LARP1B;refGene_tx=XM_023252821.1;refGene_gn=LARP1B;refGene_tx=XM_023252822.1;refGene_gn=LARP1B;refGene_tx=XM_023252824.1
B1	98748719 98748843 refGene_gn=LARP1B;refGene_tx=XM_019828979.2;refGene_gn=LARP1B;refGene_tx=XM_023252815.1;refGene_gn=LARP1B;refGene_tx=XM_023252816.1;refGene_gn=LARP1B;refGene_tx=XM_023252817.1;refGene_gn=LARP1B;refGene_tx=XM_023252818.1;refGene_gn=LARP1B;refGene_tx=XM_023252819.1;refGene_gn=LARP1B;refGene_tx=XM_023252820.1;refGene_gn=LARP1B;refGene_tx=XM_023252821.1;refGene_gn=LARP1B;refGene_tx=XM_023252822.1
B1	98785590 98785653 refGene_gn=LARP1B;refGene_tx=XM_023252823.1
B1	98790000 98790366 refGene_gn=LARP1B;refGene_tx=XM_019828979.2;refGene_gn=LARP1B;refGene_tx=XM_023252815.1;refGene_gn=LARP1B;refGene_tx=XM_023252816.1;refGene_gn=LARP1B;refGene_tx=XM_023252817.1;refGene_gn=LARP1B;refGene_tx=XM_02

3252818.1;refGene_gn=LARP1B;refGene_tx=XM_023252819.1;refGene_gn=LARP1B;refGene_tx=XM_023252820.1;refGene_gn=LARP1B;refGene_tx=XM_023252821.1;refGene_gn=LARP1B;refGene_tx=XM_023252822.1;refGene_gn=LARP1B;refGene_tx=XM_023252823.1;refGene_gn=LARP1B;refGene_tx=XM_023252824.1
B1 98793840 98794013
refGene_gn=LARP1B;refGene_tx=XM_019828979.2;refGene_gn=LARP1B;refGene_tx=XM_023252815.1;refGene_gn=LARP1B;refGene_tx=XM_023252816.1;refGene_gn=LARP1B;refGene_tx=XM_023252817.1;refGene_gn=LARP1B;refGene_tx=XM_023252818.1;refGene_gn=LARP1B;refGene_tx=XM_023252819.1;refGene_gn=LARP1B;refGene_tx=XM_023252820.1;refGene_gn=LARP1B;refGene_tx=XM_023252821.1;refGene_gn=LARP1B;refGene_tx=XM_023252822.1;refGene_gn=LARP1B;refGene_tx=XM_023252823.1;refGene_gn=LARP1B;refGene_tx=XM_023252824.1
B1 98798339 98798359 refGene_gn=LARP1B;refGene_tx=XM_023252825.1
B1 98798944 98799119
refGene_gn=LARP1B;refGene_tx=XM_019828979.2;refGene_gn=LARP1B;refGene_tx=XM_023252815.1;refGene_gn=LARP1B;refGene_tx=XM_023252816.1;refGene_gn=LARP1B;refGene_tx=XM_023252817.1;refGene_gn=LARP1B;refGene_tx=XM_023252818.1;refGene_gn=LARP1B;refGene_tx=XM_023252819.1;refGene_gn=LARP1B;refGene_tx=XM_023252820.1;refGene_gn=LARP1B;refGene_tx=XM_023252821.1;refGene_gn=LARP1B;refGene_tx=XM_023252822.1;refGene_gn=LARP1B;refGene_tx=XM_023252823.1;refGene_gn=LARP1B;refGene_tx=XM_023252824.1;refGene_gn=LARP1B;refGene_tx=XM_023252825.1
B1 98807786 98807931
refGene_gn=LARP1B;refGene_tx=XM_019828979.2;refGene_gn=LARP1B;refGene_tx=XM_023252815.1;refGene_gn=LARP1B;refGene_tx=XM_023252816.1;refGene_gn=LARP1B;refGene_tx=XM_023252817.1;refGene_gn=LARP1B;refGene_tx=XM_023252818.1;refGene_gn=LARP1B;refGene_tx=XM_023252819.1;refGene_gn=LARP1B;refGene_tx=XM_023252820.1;refGene_gn=LARP1B;refGene_tx=XM_023252821.1;refGene_gn=LARP1B;refGene_tx=XM_023252822.1;refGene_gn=LARP1B;refGene_tx=XM_023252823.1;refGene_gn=LARP1B;refGene_tx=XM_023252824.1;refGene_gn=LARP1B;refGene_tx=XM_023252825.1
B1 98811605 98811771
refGene_gn=LARP1B;refGene_tx=XM_019828979.2;refGene_gn=LARP1B;refGene_tx=XM_023252815.1;refGene_gn=LARP1B;refGene_tx=XM_023252816.1;refGene_gn=LARP1B;refGene_tx=XM_023252817.1;refGene_gn=LARP1B;refGene_tx=XM_023252818.1;refGene_gn=LARP1B;refGene_tx=XM_023252819.1;refGene_gn=LARP1B;refGene_tx=XM_023252820.1;refGene_gn=LARP1B;refGene_tx=XM_023252821.1;refGene_gn=LARP1B;refGene_tx=XM_023252822.1;refGene_gn=LARP1B;refGene_tx=XM_023252823.1;refGene_gn=LARP1B;refGene_tx=XM_023252824.1;refGene_gn=LARP1B;refGene_tx=XM_023252825.1
B1 98812008 98812152
refGene_gn=LARP1B;refGene_tx=XM_019828979.2;refGene_gn=LARP1B;refGene_tx=XM_023252815.1;refGene_gn=LARP1B;refGene_tx=XM_023252816.1;refGene_gn=LARP1B;refGene_tx=XM_023252817.1;refGene_gn=LARP1B;refGene_tx=XM_023252818.1;refGene_gn=LARP1B;refGene_tx=XM_023252819.1;refGene_gn=LARP1B;refGene_tx=XM_023252820.1;refGene_gn=LARP1B;refGene_tx=XM_023252821.1;refGene_gn=LARP1B;refGene_tx=XM_023252822.1;refGene_gn=LARP1B;refGene_tx=XM_023252823.1;refGene_gn=LARP1B;refGene_tx=XM_023252824.1;refGene_gn=LARP1B;refGene_tx=XM_023252825.1
B1 98817067 98817208

refGene_gn=LARP1B;refGene_tx=XM_019828979.2;refGene_gn=LARP1B;refGene_tx=XM_023252815.1;refGene_gn=LARP1B;refGene_tx=XM_023252816.1;refGene_gn=LARP1B;refGene_tx=XM_023252817.1;refGene_gn=LARP1B;refGene_tx=XM_023252818.1;refGene_gn=LARP1B;refGene_tx=XM_023252819.1;refGene_gn=LARP1B;refGene_tx=XM_023252820.1;refGene_gn=LARP1B;refGene_tx=XM_023252821.1;refGene_gn=LARP1B;refGene_tx=XM_023252822.1;refGene_gn=LARP1B;refGene_tx=XM_023252823.1;refGene_gn=LARP1B;refGene_tx=XM_023252824.1;refGene_gn=LARP1B;refGene_tx=XM_023252825.1
B1 98820221 98820396
refGene_gn=LARP1B;refGene_tx=XM_019828979.2;refGene_gn=LARP1B;refGene_tx=XM_023252815.1;refGene_gn=LARP1B;refGene_tx=XM_023252816.1;refGene_gn=LARP1B;refGene_tx=XM_023252817.1;refGene_gn=LARP1B;refGene_tx=XM_023252818.1;refGene_gn=LARP1B;refGene_tx=XM_023252819.1;refGene_gn=LARP1B;refGene_tx=XM_023252820.1;refGene_gn=LARP1B;refGene_tx=XM_023252821.1;refGene_gn=LARP1B;refGene_tx=XM_023252822.1;refGene_gn=LARP1B;refGene_tx=XM_023252823.1;refGene_gn=LARP1B;refGene_tx=XM_023252824.1;refGene_gn=LARP1B;refGene_tx=XM_023252825.1
B1 98822547 98822607
refGene_gn=LARP1B;refGene_tx=XM_019828979.2;refGene_gn=LARP1B;refGene_tx=XM_023252815.1;refGene_gn=LARP1B;refGene_tx=XM_023252817.1;refGene_gn=LARP1B;refGene_tx=XM_023252818.1;refGene_gn=LARP1B;refGene_tx=XM_023252819.1;refGene_gn=LARP1B;refGene_tx=XM_023252820.1;refGene_gn=LARP1B;refGene_tx=XM_023252821.1;refGene_gn=LARP1B;refGene_tx=XM_023252822.1;refGene_gn=LARP1B;refGene_tx=XM_023252823.1;refGene_gn=LARP1B;refGene_tx=XM_023252824.1;refGene_gn=LARP1B;refGene_tx=XM_023252825.1
B1 98823017 98823076
refGene_gn=LARP1B;refGene_tx=XM_019828979.2;refGene_gn=LARP1B;refGene_tx=XM_023252815.1;refGene_gn=LARP1B;refGene_tx=XM_023252816.1;refGene_gn=LARP1B;refGene_tx=XM_023252817.1;refGene_gn=LARP1B;refGene_tx=XM_023252818.1;refGene_gn=LARP1B;refGene_tx=XM_023252819.1;refGene_gn=LARP1B;refGene_tx=XM_023252820.1;refGene_gn=LARP1B;refGene_tx=XM_023252823.1;refGene_gn=LARP1B;refGene_tx=XM_023252824.1;refGene_gn=LARP1B;refGene_tx=XM_023252825.1
B1 98833886 98834385
refGene_gn=LARP1B;refGene_tx=XM_019828979.2;refGene_gn=LARP1B;refGene_tx=XM_023252815.1;refGene_gn=LARP1B;refGene_tx=XM_023252816.1;refGene_gn=LARP1B;refGene_tx=XM_023252817.1;refGene_gn=LARP1B;refGene_tx=XM_023252818.1;refGene_gn=LARP1B;refGene_tx=XM_023252819.1;refGene_gn=LARP1B;refGene_tx=XM_023252820.1;refGene_gn=LARP1B;refGene_tx=XM_023252823.1;refGene_gn=LARP1B;refGene_tx=XM_023252824.1;refGene_gn=LARP1B;refGene_tx=XM_023252825.1
B1 98862827 98862879
refGene_gn=ABHD18;refGene_tx=XM_006930871.4;refGene_gn=ABHD18;refGene_tx=XM_023252827.1
B1 98864382 98864545
refGene_gn=ABHD18;refGene_tx=XM_006930871.4;refGene_gn=ABHD18;refGene_tx=XM_023252827.1
B1 98865247 98865626
refGene_gn=ABHD18;refGene_tx=XM_006930871.4;refGene_gn=ABHD18;refGene_tx=XM_023252827.1

B1 98869120 98869222
refGene_gn=ABHD18;refGene_tx=XM_006930871.4;refGene_gn=ABHD18;refGene_tx=XM_023252827.1
B1 98869880 98869970
refGene_gn=ABHD18;refGene_tx=XM_006930871.4;refGene_gn=ABHD18;refGene_tx=XM_023252827.1
B1 98872033 98872172
refGene_gn=ABHD18;refGene_tx=XM_006930871.4;refGene_gn=ABHD18;refGene_tx=XM_023252827.1
B1 98877171 98877199
refGene_gn=ABHD18;refGene_tx=XM_006930871.4;refGene_gn=ABHD18;refGene_tx=XM_023252827.1
B1 98878999 98879084
refGene_gn=ABHD18;refGene_tx=XM_006930871.4;refGene_gn=ABHD18;refGene_tx=XM_023252827.1
B1 98879188 98879267 refGene_gn=ABHD18;refGene_tx=XM_006930871.4
B1 98883544 98883645 refGene_gn=ABHD18;refGene_tx=XM_006930871.4
B1 98887328 98887413 refGene_gn=ABHD18;refGene_tx=XM_006930871.4
B1 98888048 98888140 refGene_gn=ABHD18;refGene_tx=XM_006930871.4
B1 98899283 98899348
refGene_gn=MFSD8;refGene_tx=XM_006930873.4;refGene_gn=MFSD8;refGene_tx=XM_011281678.3;refGene_gn=MFSD8;refGene_tx=XM_019828986.2;refGene_gn=MFSD8;refGene_tx=XM_019828987.2;refGene_gn=MFSD8;refGene_tx=XM_019828988.1;refGene_gn=MFSD8;refGene_tx=XM_019828989.1
B1 98913910 98914002
refGene_gn=MFSD8;refGene_tx=XM_006930873.4;refGene_gn=MFSD8;refGene_tx=XM_011281678.3;refGene_gn=MFSD8;refGene_tx=XM_019828986.2;refGene_gn=MFSD8;refGene_tx=XM_019828987.2;refGene_gn=MFSD8;refGene_tx=XM_019828988.1;refGene_gn=MFSD8;refGene_tx=XM_019828989.1
B1 98920024 98920068
refGene_gn=MFSD8;refGene_tx=XM_006930873.4;refGene_gn=MFSD8;refGene_tx=XM_011281678.3;refGene_gn=MFSD8;refGene_tx=XM_019828986.2;refGene_gn=MFSD8;refGene_tx=XM_019828987.2;refGene_gn=MFSD8;refGene_tx=XM_019828988.1;refGene_gn=MFSD8;refGene_tx=XM_019828989.1
B1 98925911 98926152
refGene_gn=MFSD8;refGene_tx=XM_006930873.4;refGene_gn=MFSD8;refGene_tx=XM_011281678.3;refGene_gn=MFSD8;refGene_tx=XM_019828986.2;refGene_gn=MFSD8;refGene_tx=XM_019828987.2;refGene_gn=MFSD8;refGene_tx=XM_019828988.1;refGene_gn=MFSD8;refGene_tx=XM_019828989.1
B1 98927018 98927132
refGene_gn=MFSD8;refGene_tx=XM_006930873.4;refGene_gn=MFSD8;refGene_tx=XM_011281678.3;refGene_gn=MFSD8;refGene_tx=XM_019828986.2;refGene_gn=MFSD8;refGene_tx=XM_019828987.2;refGene_gn=MFSD8;refGene_tx=XM_019828988.1;refGene_gn=MFSD8;refGene_tx=XM_019828989.1
B1 98930059 98930204
refGene_gn=MFSD8;refGene_tx=XM_006930873.4;refGene_gn=MFSD8;refGene_tx=XM_011281678.3;refGene_gn=MFSD8;refGene_tx=XM_019828986.2;refGene_gn=MFSD8;refGene_tx=XM_019828987.2;refGene_gn=MFSD8;refGene_tx=XM_019828988.1
B1 98930935 98930991

refGene_gn=MFSD8;refGene_tx=XM_006930873.4;refGene_gn=MFSD8;refGene_tx=XM_011281678.3;refGene_gn=MFSD8;refGene_tx=XM_019828986.2;refGene_gn=MFSD8;refGene_tx=XM_019828987.2;refGene_gn=MFSD8;refGene_tx=XM_019828988.1
B1 98935053 98935162 refGene_gn=MFSD8;refGene_tx=XM_006930873.4;refGene_gn=MFSD8;refGene_tx=XM_011281678.3;refGene_gn=MFSD8;refGene_tx=XM_019828986.2;refGene_gn=MFSD8;refGene_tx=XM_019828987.2;refGene_gn=MFSD8;refGene_tx=XM_019828988.1;refGene_gn=MFSD8;refGene_tx=XM_019828989.1
B1 98937244 98937379 refGene_gn=MFSD8;refGene_tx=XM_006930873.4;refGene_gn=MFSD8;refGene_tx=XM_011281678.3;refGene_gn=MFSD8;refGene_tx=XM_019828986.2;refGene_gn=MFSD8;refGene_tx=XM_019828988.1;refGene_gn=MFSD8;refGene_tx=XM_019828989.1
B1 98940476 98940580 refGene_gn=MFSD8;refGene_tx=XM_006930873.4;refGene_gn=MFSD8;refGene_tx=XM_011281678.3;refGene_gn=MFSD8;refGene_tx=XM_019828986.2;refGene_gn=MFSD8;refGene_tx=XM_019828987.2;refGene_gn=MFSD8;refGene_tx=XM_019828988.1;refGene_gn=MFSD8;refGene_tx=XM_019828989.1
B1 98940667 98940915 refGene_gn=MFSD8;refGene_tx=XM_006930873.4;refGene_gn=MFSD8;refGene_tx=XM_011281678.3;refGene_gn=MFSD8;refGene_tx=XM_019828986.2;refGene_gn=MFSD8;refGene_tx=XM_019828987.2;refGene_gn=MFSD8;refGene_tx=XM_019828988.1;refGene_gn=MFSD8;refGene_tx=XM_019828989.1
B1 98945893 98946100 refGene_gn=MFSD8;refGene_tx=XM_006930873.4;refGene_gn=MFSD8;refGene_tx=XM_019828986.2;refGene_gn=MFSD8;refGene_tx=XM_019828987.2;refGene_gn=MFSD8;refGene_tx=XM_019828989.1
B1 98957454 98957523 refGene_gn=MFSD8;refGene_tx=XM_011281678.3;refGene_gn=MFSD8;refGene_tx=XM_019828988.1
B1 98972857 98972960 refGene_gn=PLK4;refGene_tx=XM_004001097.4
B1 98973369 98973476 refGene_gn=PLK4;refGene_tx=XM_004001097.4
B1 98975267 98975408 refGene_gn=PLK4;refGene_tx=XM_004001097.4
B1 98976683 98976831 refGene_gn=PLK4;refGene_tx=XM_004001097.4
B1 98976978 98977070 refGene_gn=PLK4;refGene_tx=XM_004001097.4
B1 98977163 98977297 refGene_gn=PLK4;refGene_tx=XM_004001097.4
B1 98978024 98978174 refGene_gn=PLK4;refGene_tx=XM_004001097.4
B1 98979160 98979263 refGene_gn=PLK4;refGene_tx=XM_004001097.4
B1 98979824 98979929 refGene_gn=PLK4;refGene_tx=XM_004001097.4
B1 98980754 98981122 refGene_gn=PLK4;refGene_tx=XM_004001097.4
B1 98983427 98983528 refGene_gn=PLK4;refGene_tx=XM_004001097.4
B1 98984560 98985581 refGene_gn=PLK4;refGene_tx=XM_004001097.4
B1 98988624 98988739 refGene_gn=PLK4;refGene_tx=XM_004001097.4
B1 98988821 98988917 refGene_gn=PLK4;refGene_tx=XM_004001097.4
B1 98989723 98989819 refGene_gn=PLK4;refGene_tx=XM_004001097.4
B1 98990476 98990506 refGene_gn=PLK4;refGene_tx=XM_004001097.4
B1 99042111 99042303 refGene_gn=HSPA4L;refGene_tx=XM_004001096.5;refGene_gn=HSPA4L;refGene_tx=XM_019828991.2

B1 99044614 99044776
refGene_gn=HSPA4L;refGene_tx=XM_004001096.5;refGene_gn=HSPA4L;refGene_tx=XM_019828991.2
B1 99048428 99048548
refGene_gn=HSPA4L;refGene_tx=XM_004001096.5;refGene_gn=HSPA4L;refGene_tx=XM_019828991.2
B1 99050702 99050810
refGene_gn=HSPA4L;refGene_tx=XM_004001096.5;refGene_gn=HSPA4L;refGene_tx=XM_019828991.2
B1 99051101 99051227
refGene_gn=HSPA4L;refGene_tx=XM_004001096.5;refGene_gn=HSPA4L;refGene_tx=XM_019828991.2
B1 99054817 99054955
refGene_gn=HSPA4L;refGene_tx=XM_004001096.5;refGene_gn=HSPA4L;refGene_tx=XM_019828991.2
B1 99056475 99056574
refGene_gn=HSPA4L;refGene_tx=XM_004001096.5;refGene_gn=HSPA4L;refGene_tx=XM_019828991.2
B1 99061687 99061887
refGene_gn=HSPA4L;refGene_tx=XM_004001096.5;refGene_gn=HSPA4L;refGene_tx=XM_019828991.2
B1 99063484 99063618
refGene_gn=HSPA4L;refGene_tx=XM_004001096.5;refGene_gn=HSPA4L;refGene_tx=XM_019828991.2
B1 99066489 99066596
refGene_gn=HSPA4L;refGene_tx=XM_004001096.5;refGene_gn=HSPA4L;refGene_tx=XM_019828991.2
B1 99067080 99067232
refGene_gn=HSPA4L;refGene_tx=XM_004001096.5;refGene_gn=HSPA4L;refGene_tx=XM_019828991.2
B1 99067843 99067920
refGene_gn=HSPA4L;refGene_tx=XM_004001096.5;refGene_gn=HSPA4L;refGene_tx=XM_019828991.2
B1 99068064 99068309
refGene_gn=HSPA4L;refGene_tx=XM_004001096.5;refGene_gn=HSPA4L;refGene_tx=XM_019828991.2
B1 99070370 99070504
refGene_gn=HSPA4L;refGene_tx=XM_004001096.5;refGene_gn=HSPA4L;refGene_tx=XM_019828991.2
B1 99070858 99070958
refGene_gn=HSPA4L;refGene_tx=XM_004001096.5;refGene_gn=HSPA4L;refGene_tx=XM_019828991.2
B1 99074576 99074699
refGene_gn=HSPA4L;refGene_tx=XM_004001096.5;refGene_gn=HSPA4L;refGene_tx=XM_019828991.2
B1 99076648 99076789
refGene_gn=HSPA4L;refGene_tx=XM_004001096.5;refGene_gn=HSPA4L;refGene_tx=XM_019828991.2
B1 99078510 99078568
refGene_gn=HSPA4L;refGene_tx=XM_004001096.5;refGene_gn=HSPA4L;refGene_

tx=XM_019828991.2
B1 99090857 99090964
refGene_gn=HSPA4L;refGene_tx=XM_004001096.5;refGene_gn=HSPA4L;refGene_
tx=XM_019828991.2
B1 99104168 99104369 refGene_gn=SLC25A31;refGene_tx=XM_004001095.5
B1 99108555 99108681 refGene_gn=SLC25A31;refGene_tx=XM_004001095.5
B1 99110171 99110326 refGene_gn=SLC25A31;refGene_tx=XM_004001095.5
B1 99112913 99113031 refGene_gn=SLC25A31;refGene_tx=XM_004001095.5
B1 99121850 99121978 refGene_gn=SLC25A31;refGene_tx=XM_004001095.5
B1 99131939 99132183 refGene_gn=SLC25A31;refGene_tx=XM_004001095.5
B1 99138509 99138641 refGene_gn=INTU;refGene_tx=XM_023253393.1
B1 99162648 99162892 refGene_gn=INTU;refGene_tx=XM_023253393.1
B1 99166937 99167035 refGene_gn=INTU;refGene_tx=XM_023253393.1
B1 99178235 99178721 refGene_gn=INTU;refGene_tx=XM_023253393.1
B1 99179320 99179542 refGene_gn=INTU;refGene_tx=XM_023253393.1
B1 99180350 99180413 refGene_gn=INTU;refGene_tx=XM_023253393.1
B1 99181087 99181141 refGene_gn=INTU;refGene_tx=XM_023253393.1
B1 99182853 99183043 refGene_gn=INTU;refGene_tx=XM_023253393.1
B1 99190091 99190169 refGene_gn=INTU;refGene_tx=XM_023253393.1
B1 99197945 99198035 refGene_gn=INTU;refGene_tx=XM_023253393.1
B1 99203951 99204070 refGene_gn=INTU;refGene_tx=XM_023253393.1
B1 99208102 99208306 refGene_gn=INTU;refGene_tx=XM_023253393.1
B1 99218553 99218639 refGene_gn=INTU;refGene_tx=XM_023253393.1
B1 99228538 99229077 refGene_gn=INTU;refGene_tx=XM_023253393.1
B1 99242411 99242557 refGene_gn=INTU;refGene_tx=XM_023253393.1
B1 101295475 101297340
refGene_gn=FAT4;refGene_tx=XM_011281676.2
B1 101300126 101300388
refGene_gn=FAT4;refGene_tx=XM_011281676.2
B1 101304707 101304925
refGene_gn=FAT4;refGene_tx=XM_011281676.2
B1 101306801 101306926
refGene_gn=FAT4;refGene_tx=XM_011281676.2
B1 101310019 101310199
refGene_gn=FAT4;refGene_tx=XM_011281676.2
B1 101311076 101311162
refGene_gn=FAT4;refGene_tx=XM_011281676.2
B1 101315654 101315962
refGene_gn=FAT4;refGene_tx=XM_011281676.2
B1 101319897 101320002
refGene_gn=FAT4;refGene_tx=XM_011281676.2
B1 101331048 101335398
refGene_gn=FAT4;refGene_tx=XM_011281676.2
B1 101337375 101337626
refGene_gn=FAT4;refGene_tx=XM_011281676.2
B1 101348001 101348182
refGene_gn=FAT4;refGene_tx=XM_011281676.2
B1 101366988 101367163
refGene_gn=FAT4;refGene_tx=XM_011281676.2
B1 101368233 101369156

refGene_gn=FAT4;refGene_tx=XM_011281676.2
B1 101375916 101376267
refGene_gn=FAT4;refGene_tx=XM_011281676.2
B1 101377790 101378052
refGene_gn=FAT4;refGene_tx=XM_011281676.2
B1 101392791 101392923
refGene_gn=FAT4;refGene_tx=XM_011281676.2
B1 101465159 101470334
refGene_gn=FAT4;refGene_tx=XM_011281676.2
B1 102026490 102027002
refGene_gn=ANKRD50;refGene_tx=XM_006930874.4
B1 102046474 102046704
refGene_gn=ANKRD50;refGene_tx=XM_006930874.4;refGene_gn=ANKRD50;refGene_tx=XM_019828994.2
B1 102048608 102052156
refGene_gn=ANKRD50;refGene_tx=XM_006930874.4;refGene_gn=ANKRD50;refGene_tx=XM_019828994.2
B1 103165720 103166680
refGene_gn=SPRY1;refGene_tx=XM_006930876.4;refGene_gn=SPRY1;refGene_tx=XM_023252828.1
B1 103240651 103240828
refGene_gn=SPATA5;refGene_tx=XM_003985028.4;refGene_gn=SPATA5;refGene_tx=XM_011281675.3;refGene_gn=SPATA5;refGene_tx=XM_019828995.2
B1 103265845 103265980
refGene_gn=SPATA5;refGene_tx=XM_011281675.3;refGene_gn=SPATA5;refGene_tx=XM_019828995.2
B1 103314737 103314902
refGene_gn=SPATA5;refGene_tx=XM_003985028.4;refGene_gn=SPATA5;refGene_tx=XM_011281675.3;refGene_gn=SPATA5;refGene_tx=XM_019828995.2
B1 103475634 103475761
refGene_gn=SPATA5;refGene_tx=XM_003985028.4;refGene_gn=SPATA5;refGene_tx=XM_011281675.3;refGene_gn=SPATA5;refGene_tx=XM_019828995.2
B1 103511294 103511374
refGene_gn=SPATA5;refGene_tx=XM_003985028.4;refGene_gn=SPATA5;refGene_tx=XM_011281675.3;refGene_gn=SPATA5;refGene_tx=XM_019828995.2
B1 103513475 103513529
refGene_gn=SPATA5;refGene_tx=XM_003985028.4;refGene_gn=SPATA5;refGene_tx=XM_011281675.3;refGene_gn=SPATA5;refGene_tx=XM_019828995.2
B1 103532180 103532390
refGene_gn=SPATA5;refGene_tx=XM_003985028.4;refGene_gn=SPATA5;refGene_tx=XM_011281675.3;refGene_gn=SPATA5;refGene_tx=XM_019828995.2
B1 103559715 103559870
refGene_gn=SPATA5;refGene_tx=XM_003985028.4;refGene_gn=SPATA5;refGene_tx=XM_011281675.3;refGene_gn=SPATA5;refGene_tx=XM_019828995.2
B1 103590985 103591241
refGene_gn=SPATA5;refGene_tx=XM_003985028.4;refGene_gn=SPATA5;refGene_tx=XM_011281675.3;refGene_gn=SPATA5;refGene_tx=XM_019828995.2
B1 103593982 103594106
refGene_gn=SPATA5;refGene_tx=XM_003985028.4;refGene_gn=SPATA5;refGene_tx=XM_011281675.3;refGene_gn=SPATA5;refGene_tx=XM_019828995.2

B1	103596784	103596841
refGene_gn=SPATA5;refGene_tx=XM_003985028.4;refGene_gn=SPATA5;refGene_tx=XM_011281675.3;refGene_gn=SPATA5;refGene_tx=XM_019828995.2
B1	103597111	103597241
refGene_gn=SPATA5;refGene_tx=XM_003985028.4;refGene_gn=SPATA5;refGene_tx=XM_011281675.3;refGene_gn=SPATA5;refGene_tx=XM_019828995.2
B1	103598189	103598828
refGene_gn=SPATA5;refGene_tx=XM_003985028.4;refGene_gn=SPATA5;refGene_tx=XM_011281675.3;refGene_gn=SPATA5;refGene_tx=XM_019828995.2
B1	103599418	103599477
refGene_gn=SPATA5;refGene_tx=XM_003985028.4;refGene_gn=SPATA5;refGene_tx=XM_011281675.3;refGene_gn=SPATA5;refGene_tx=XM_019828995.2
B1	103603028	103603186
refGene_gn=SPATA5;refGene_tx=XM_003985028.4;refGene_gn=SPATA5;refGene_tx=XM_011281675.3;refGene_gn=SPATA5;refGene_tx=XM_019828995.2
B1	103604486	103604608
refGene_gn=SPATA5;refGene_tx=XM_003985028.4;refGene_gn=SPATA5;refGene_tx=XM_011281675.3;refGene_gn=SPATA5;refGene_tx=XM_019828995.2
B1	103609355	103609515
refGene_gn=SPATA5;refGene_tx=XM_003985028.4;refGene_gn=SPATA5;refGene_tx=XM_011281675.3
B1	103610062	103610300
refGene_gn=NUDT6;refGene_tx=XM_023252832.1
B1	103613747	103613951
refGene_gn=NUDT6;refGene_tx=XM_023252832.1
B1	103618566	103618622
refGene_gn=NUDT6;refGene_tx=XM_023252832.1
B1	103619835	103619853
refGene_gn=NUDT6;refGene_tx=XM_023252833.1
B1	103645676	103645731
refGene_gn=NUDT6;refGene_tx=XM_023252832.1;refGene_gn=NUDT6;refGene_tx=XM_023252833.1;refGene_gn=NUDT6;refGene_tx=XM_023252834.1
B1	103649921	103650319
refGene_gn=NUDT6;refGene_tx=XM_023252832.1;refGene_gn=NUDT6;refGene_tx=XM_023252833.1;refGene_gn=NUDT6;refGene_tx=XM_023252834.1
B1	103650755	103650941
refGene_gn=FGF2;refGene_tx=XM_023253395.1
B1	103658973	103659077
refGene_gn=FGF2;refGene_tx=XM_023253395.1
B1	103711941	103712011
refGene_gn=FGF2;refGene_tx=XM_023253395.1
B1	103712012	103712272
refGene_gn=FGF2;refGene_tx=XM_023253395.1
B1	103712274	103712383
refGene_gn=FGF2;refGene_tx=XM_023253395.1
B1	103712385	103712572
refGene_gn=FGF2;refGene_tx=XM_023253395.1
B1	103712676	103712771
refGene_gn=FGF2;refGene_tx=XM_023253395.1
B1	103782465	103784607

refGene_gn=BBS12;refGene_tx=XM_006930879.4;refGene_gn=BBS12;refGene_tx=XM_019828999.2;refGene_gn=BBS12;refGene_tx=XM_023252830.1
B1 103795598 103795601
refGene_gn=LOC101090350;refGene_tx=XM_019829000.2
B1 103796782 103796920
refGene_gn=LOC101090350;refGene_tx=XM_019829000.2
B1 103797294 103797423
refGene_gn=LOC101090350;refGene_tx=XM_019829000.2
B1 103798028 103798157
refGene_gn=LOC101090350;refGene_tx=XM_019829000.2
B1 103798378 103798468
refGene_gn=LOC101090350;refGene_tx=XM_019829000.2
B1 103887616 103887784
refGene_gn=IL21;refGene_tx=XM_003985030.4
B1 103887892 103887928
refGene_gn=IL21;refGene_tx=XM_003985030.4
B1 103892256 103892412
refGene_gn=IL21;refGene_tx=XM_003985030.4
B1 103894205 103894283
refGene_gn=IL21;refGene_tx=XM_003985030.4
B1 103894379 103894403
refGene_gn=IL21;refGene_tx=XM_003985030.4
B1 103987465 103987566
refGene_gn=ADAD1;refGene_tx=XM_019829001.2;refGene_gn=ADAD1;refGene_tx=XM_019829002.2;refGene_gn=ADAD1;refGene_tx=XM_019829003.2
B1 104012959 104013106
refGene_gn=IL2;refGene_tx=NM_001043337.1;refGene_gn=IL2;refGene_tx=XM_023252215.1
B1 104013196 104013256
refGene_gn=IL2;refGene_tx=NM_001043337.1;refGene_gn=IL2;refGene_tx=XM_023252215.1
B1 104015398 104015545
refGene_gn=IL2;refGene_tx=NM_001043337.1;refGene_gn=IL2;refGene_tx=XM_023252215.1
B1 104017387 104017498
refGene_gn=IL2;refGene_tx=XM_023252215.1
B1 104018555 104018666
refGene_gn=IL2;refGene_tx=NM_001043337.1
B1 104040441 104040555
refGene_gn=ADAD1;refGene_tx=XM_011281674.3;refGene_gn=ADAD1;refGene_tx=XM_019829001.2;refGene_gn=ADAD1;refGene_tx=XM_019829002.2
B1 104049830 104049963
refGene_gn=ADAD1;refGene_tx=XM_011281674.3;refGene_gn=ADAD1;refGene_tx=XM_019829001.2;refGene_gn=ADAD1;refGene_tx=XM_019829002.2;refGene_gn=ADAD1;refGene_tx=XM_019829003.2
B1 104054695 104054933
refGene_gn=ADAD1;refGene_tx=XM_011281674.3;refGene_gn=ADAD1;refGene_tx=XM_019829001.2;refGene_gn=ADAD1;refGene_tx=XM_019829002.2;refGene_gn=ADAD1;refGene_tx=XM_019829003.2
B1 104067569 104067799

refGene_gn=ADAD1;refGene_tx=XM_011281674.3;refGene_gn=ADAD1;refGene_tx
=XM_019829001.2;refGene_gn=ADAD1;refGene_tx=XM_019829002.2;refGene_gn=
ADAD1;refGene_tx=XM_019829003.2
B1 104069844 104070015
refGene_gn=ADAD1;refGene_tx=XM_011281674.3;refGene_gn=ADAD1;refGene_tx
=XM_019829001.2;refGene_gn=ADAD1;refGene_tx=XM_019829002.2;refGene_gn=
ADAD1;refGene_tx=XM_019829003.2
B1 104072369 104072493
refGene_gn=ADAD1;refGene_tx=XM_011281674.3;refGene_gn=ADAD1;refGene_tx
=XM_019829001.2;refGene_gn=ADAD1;refGene_tx=XM_019829002.2;refGene_gn=
ADAD1;refGene_tx=XM_019829003.2
B1 104079786 104079912
refGene_gn=ADAD1;refGene_tx=XM_011281674.3;refGene_gn=ADAD1;refGene_tx
=XM_019829001.2;refGene_gn=ADAD1;refGene_tx=XM_019829002.2;refGene_gn=
ADAD1;refGene_tx=XM_019829003.2
B1 104084133 104084202
refGene_gn=ADAD1;refGene_tx=XM_011281674.3;refGene_gn=ADAD1;refGene_tx
=XM_019829001.2;refGene_gn=ADAD1;refGene_tx=XM_019829002.2;refGene_gn=
ADAD1;refGene_tx=XM_019829003.2
B1 104090603 104090771
refGene_gn=ADAD1;refGene_tx=XM_011281674.3;refGene_gn=ADAD1;refGene_tx
=XM_019829001.2;refGene_gn=ADAD1;refGene_tx=XM_019829002.2;refGene_gn=
ADAD1;refGene_tx=XM_019829003.2
B1 104093843 104094032
refGene_gn=ADAD1;refGene_tx=XM_011281674.3;refGene_gn=ADAD1;refGene_tx
=XM_019829001.2;refGene_gn=ADAD1;refGene_tx=XM_019829002.2;refGene_gn=
ADAD1;refGene_tx=XM_019829003.2
B1 104094761 104094941
refGene_gn=ADAD1;refGene_tx=XM_011281674.3;refGene_gn=ADAD1;refGene_tx
=XM_019829001.2;refGene_gn=ADAD1;refGene_tx=XM_019829002.2;refGene_gn=
ADAD1;refGene_tx=XM_019829003.2
B1 104095506 104095558
refGene_gn=ADAD1;refGene_tx=XM_011281674.3;refGene_gn=ADAD1;refGene_tx
=XM_019829001.2;refGene_gn=ADAD1;refGene_tx=XM_019829003.2
B1 104128955 104129172
refGene_gn=KIAA1109;refGene_tx=XM_023252835.1
B1 104131499 104131665
refGene_gn=KIAA1109;refGene_tx=XM_023252835.1
B1 104134787 104134922
refGene_gn=KIAA1109;refGene_tx=XM_023252835.1
B1 104135341 104135536
refGene_gn=KIAA1109;refGene_tx=XM_023252835.1
B1 104137286 104137490
refGene_gn=KIAA1109;refGene_tx=XM_023252835.1
B1 104138177 104138429
refGene_gn=KIAA1109;refGene_tx=XM_023252835.1
B1 104141475 104141693
refGene_gn=KIAA1109;refGene_tx=XM_023252835.1
B1 104141924 104142117
refGene_gn=KIAA1109;refGene_tx=XM_023252835.1

B1 104142214 104142372
refGene_gn=KIAA1109;refGene_tx=XM_023252835.1
B1 104142752 104142903
refGene_gn=KIAA1109;refGene_tx=XM_023252835.1
B1 104144025 104144283
refGene_gn=KIAA1109;refGene_tx=XM_023252835.1
B1 104145043 104145181
refGene_gn=KIAA1109;refGene_tx=XM_023252835.1
B1 104147377 104147557
refGene_gn=KIAA1109;refGene_tx=XM_023252835.1
B1 104149060 104149261
refGene_gn=KIAA1109;refGene_tx=XM_023252835.1
B1 104152847 104153049
refGene_gn=KIAA1109;refGene_tx=XM_023252835.1
B1 104154375 104154515
refGene_gn=KIAA1109;refGene_tx=XM_023252835.1
B1 104155019 104155181
refGene_gn=KIAA1109;refGene_tx=XM_023252835.1
B1 104156897 104157099
refGene_gn=KIAA1109;refGene_tx=XM_023252835.1
B1 104157714 104157895
refGene_gn=KIAA1109;refGene_tx=XM_023252835.1
B1 104159345 104159561
refGene_gn=KIAA1109;refGene_tx=XM_023252835.1
B1 104160126 104160380
refGene_gn=KIAA1109;refGene_tx=XM_023252835.1
B1 104164348 104164439
refGene_gn=KIAA1109;refGene_tx=XM_023252835.1
B1 104165103 104165213
refGene_gn=KIAA1109;refGene_tx=XM_023252835.1
B1 104167568 104167692
refGene_gn=KIAA1109;refGene_tx=XM_023252835.1
B1 104169011 104169142
refGene_gn=KIAA1109;refGene_tx=XM_023252835.1
B1 104170272 104170499
refGene_gn=KIAA1109;refGene_tx=XM_023252835.1
B1 104172329 104172389
refGene_gn=KIAA1109;refGene_tx=XM_023252835.1
B1 104175890 104176089
refGene_gn=KIAA1109;refGene_tx=XM_023252835.1
B1 104177175 104177387
refGene_gn=KIAA1109;refGene_tx=XM_023252835.1
B1 104178930 104179099
refGene_gn=KIAA1109;refGene_tx=XM_023252835.1
B1 104179699 104179885
refGene_gn=KIAA1109;refGene_tx=XM_023252835.1
B1 104180639 104180737
refGene_gn=KIAA1109;refGene_tx=XM_023252835.1
B1 104188661 104188767
refGene_gn=KIAA1109;refGene_tx=XM_023252835.1

B1 104191152 104191397
refGene_gn=KIAA1109;refGene_tx=XM_023252835.1
B1 104196250 104196489
refGene_gn=KIAA1109;refGene_tx=XM_023252835.1
B1 104198181 104198388
refGene_gn=KIAA1109;refGene_tx=XM_023252835.1
B1 104201712 104201763
refGene_gn=KIAA1109;refGene_tx=XM_023252835.1
B1 104203716 104203810
refGene_gn=KIAA1109;refGene_tx=XM_023252835.1
B1 104203965 104203971
refGene_gn=KIAA1109;refGene_tx=XM_023252835.1
B1 104204663 104204903
refGene_gn=KIAA1109;refGene_tx=XM_023252835.1
B1 104205312 104206019
refGene_gn=KIAA1109;refGene_tx=XM_023252835.1
B1 104207922 104208098
refGene_gn=KIAA1109;refGene_tx=XM_023252835.1
B1 104211378 104211554
refGene_gn=KIAA1109;refGene_tx=XM_023252835.1
B1 104213344 104213503
refGene_gn=KIAA1109;refGene_tx=XM_023252835.1
B1 104213973 104214193
refGene_gn=KIAA1109;refGene_tx=XM_023252835.1
B1 104216398 104216541
refGene_gn=KIAA1109;refGene_tx=XM_023252835.1
B1 104220294 104220535
refGene_gn=KIAA1109;refGene_tx=XM_023252835.1
B1 104222181 104222334
refGene_gn=KIAA1109;refGene_tx=XM_023252835.1
B1 104222509 104222660
refGene_gn=KIAA1109;refGene_tx=XM_023252835.1
B1 104223111 104223263
refGene_gn=KIAA1109;refGene_tx=XM_023252835.1
B1 104225806 104226018
refGene_gn=KIAA1109;refGene_tx=XM_023252835.1
B1 104226690 104226888
refGene_gn=KIAA1109;refGene_tx=XM_023252835.1
B1 104229628 104229828
refGene_gn=KIAA1109;refGene_tx=XM_023252835.1
B1 104230158 104230286
refGene_gn=KIAA1109;refGene_tx=XM_023252835.1
B1 104230630 104230777
refGene_gn=KIAA1109;refGene_tx=XM_023252835.1
B1 104232587 104232708
refGene_gn=KIAA1109;refGene_tx=XM_023252835.1
B1 104233177 104233351
refGene_gn=KIAA1109;refGene_tx=XM_023252835.1
B1 104235193 104235264
refGene_gn=KIAA1109;refGene_tx=XM_023252835.1

B1 104237458 104238296
refGene_gn=KIAA1109;refGene_tx=XM_023252835.1
B1 104239331 104239590
refGene_gn=KIAA1109;refGene_tx=XM_023252835.1
B1 104242650 104242901
refGene_gn=KIAA1109;refGene_tx=XM_023252835.1
B1 104249091 104249372
refGene_gn=KIAA1109;refGene_tx=XM_023252835.1
B1 104250063 104250183
refGene_gn=KIAA1109;refGene_tx=XM_023252835.1
B1 104257222 104257351
refGene_gn=KIAA1109;refGene_tx=XM_023252835.1
B1 104259528 104259708
refGene_gn=KIAA1109;refGene_tx=XM_023252835.1
B1 104267621 104267759
refGene_gn=KIAA1109;refGene_tx=XM_023252835.1
B1 104268556 104268784
refGene_gn=KIAA1109;refGene_tx=XM_023252835.1
B1 104275672 104275881
refGene_gn=KIAA1109;refGene_tx=XM_023252835.1
B1 104277860 104277967
refGene_gn=KIAA1109;refGene_tx=XM_023252835.1
B1 104278113 104278292
refGene_gn=KIAA1109;refGene_tx=XM_023252835.1
B1 104280988 104281086
refGene_gn=KIAA1109;refGene_tx=XM_023252835.1
B1 104281333 104281489
refGene_gn=KIAA1109;refGene_tx=XM_023252835.1
B1 104288760 104288878
refGene_gn=KIAA1109;refGene_tx=XM_023252835.1
B1 104290182 104290274
refGene_gn=KIAA1109;refGene_tx=XM_023252835.1
B1 104292308 104292387
refGene_gn=KIAA1109;refGene_tx=XM_023252835.1
B1 104292792 104292949
refGene_gn=KIAA1109;refGene_tx=XM_023252835.1
B1 104296978 104297129
refGene_gn=KIAA1109;refGene_tx=XM_023252835.1
B1 104298985 104299065
refGene_gn=KIAA1109;refGene_tx=XM_023252835.1
B1 104301235 104301429
refGene_gn=KIAA1109;refGene_tx=XM_023252835.1
B1 104301803 104301885
refGene_gn=KIAA1109;refGene_tx=XM_023252835.1
B1 104304921 104305108
refGene_gn=KIAA1109;refGene_tx=XM_023252835.1
B1 104319387 104319452
refGene_gn=KIAA1109;refGene_tx=XM_023252835.1
B1 104321045 104321124
refGene_gn=KIAA1109;refGene_tx=XM_023252835.1

B1 104322144 104322261
refGene_gn=KIAA1109;refGene_tx=XM_023252835.1
B1 104325157 104325254
refGene_gn=KIAA1109;refGene_tx=XM_023252835.1
B1 104522686 104522913
refGene_gn=TRPC3;refGene_tx=XM_023252836.1;refGene_gn=TRPC3;refGene_tx
=XM_023252837.1;refGene_gn=TRPC3;refGene_tx=XM_023252838.1
B1 104540200 104540972
refGene_gn=TRPC3;refGene_tx=XM_023252836.1;refGene_gn=TRPC3;refGene_tx
=XM_023252837.1;refGene_gn=TRPC3;refGene_tx=XM_023252838.1
B1 104547640 104547829
refGene_gn=TRPC3;refGene_tx=XM_023252836.1;refGene_gn=TRPC3;refGene_tx
=XM_023252837.1;refGene_gn=TRPC3;refGene_tx=XM_023252838.1
B1 104556343 104556508
refGene_gn=TRPC3;refGene_tx=XM_023252836.1;refGene_gn=TRPC3;refGene_tx
=XM_023252837.1;refGene_gn=TRPC3;refGene_tx=XM_023252838.1
B1 104559223 104559440
refGene_gn=TRPC3;refGene_tx=XM_023252836.1;refGene_gn=TRPC3;refGene_tx
=XM_023252837.1;refGene_gn=TRPC3;refGene_tx=XM_023252838.1
B1 104560821 104561055
refGene_gn=TRPC3;refGene_tx=XM_023252836.1;refGene_gn=TRPC3;refGene_tx
=XM_023252837.1;refGene_gn=TRPC3;refGene_tx=XM_023252838.1
B1 104563352 104563617
refGene_gn=TRPC3;refGene_tx=XM_023252836.1;refGene_gn=TRPC3;refGene_tx
=XM_023252837.1;refGene_gn=TRPC3;refGene_tx=XM_023252838.1
B1 104566360 104566556
refGene_gn=TRPC3;refGene_tx=XM_023252836.1;refGene_gn=TRPC3;refGene_tx
=XM_023252837.1;refGene_gn=TRPC3;refGene_tx=XM_023252838.1
B1 104567867 104568077
refGene_gn=TRPC3;refGene_tx=XM_023252836.1;refGene_gn=TRPC3;refGene_tx
=XM_023252837.1;refGene_gn=TRPC3;refGene_tx=XM_023252838.1
B1 104569402 104569450
refGene_gn=TRPC3;refGene_tx=XM_023252836.1
B1 104571502 104571586
refGene_gn=TRPC3;refGene_tx=XM_023252836.1;refGene_gn=TRPC3;refGene_tx
=XM_023252837.1
B1 104591236 104591312
refGene_gn=TRPC3;refGene_tx=XM_023252836.1;refGene_gn=TRPC3;refGene_tx
=XM_023252837.1;refGene_gn=TRPC3;refGene_tx=XM_023252838.1
B1 104593979 104594122
refGene_gn=TRPC3;refGene_tx=XM_023252836.1;refGene_gn=TRPC3;refGene_tx
=XM_023252837.1;refGene_gn=TRPC3;refGene_tx=XM_023252838.1
B1 104602982 104603018
refGene_gn=BBS7;refGene_tx=XM_003985033.5;refGene_gn=BBS7;refGene_tx=X
M_006930881.4
B1 104604947 104605013
refGene_gn=BBS7;refGene_tx=XM_003985033.5;refGene_gn=BBS7;refGene_tx=X
M_006930881.4
B1 104608993 104609056
refGene_gn=BBS7;refGene_tx=XM_003985033.5;refGene_gn=BBS7;refGene_tx=X

M_006930881.4
B1 104610802 104610978
refGene_gn=BBS7;refGene_tx=XM_003985033.5;refGene_gn=BBS7;refGene_tx=X
M_006930881.4
B1 104613339 104613526
refGene_gn=BBS7;refGene_tx=XM_003985033.5;refGene_gn=BBS7;refGene_tx=X
M_006930881.4
B1 104615937 104616010
refGene_gn=BBS7;refGene_tx=XM_003985033.5;refGene_gn=BBS7;refGene_tx=X
M_006930881.4
B1 104616675 104616792
refGene_gn=BBS7;refGene_tx=XM_003985033.5;refGene_gn=BBS7;refGene_tx=X
M_006930881.4
B1 104618432 104618563
refGene_gn=BBS7;refGene_tx=XM_003985033.5;refGene_gn=BBS7;refGene_tx=X
M_006930881.4
B1 104619817 104619902
refGene_gn=BBS7;refGene_tx=XM_003985033.5;refGene_gn=BBS7;refGene_tx=X
M_006930881.4
B1 104620886 104620989
refGene_gn=BBS7;refGene_tx=XM_003985033.5;refGene_gn=BBS7;refGene_tx=X
M_006930881.4
B1 104622604 104622797
refGene_gn=BBS7;refGene_tx=XM_003985033.5;refGene_gn=BBS7;refGene_tx=X
M_006930881.4
B1 104624021 104624096
refGene_gn=BBS7;refGene_tx=XM_003985033.5;refGene_gn=BBS7;refGene_tx=X
M_006930881.4
B1 104627106 104627172
refGene_gn=BBS7;refGene_tx=XM_003985033.5
B1 104631412 104631552
refGene_gn=BBS7;refGene_tx=XM_003985033.5;refGene_gn=BBS7;refGene_tx=X
M_006930881.4
B1 104632438 104632603
refGene_gn=BBS7;refGene_tx=XM_003985033.5;refGene_gn=BBS7;refGene_tx=X
M_006930881.4
B1 104635647 104635757
refGene_gn=BBS7;refGene_tx=XM_003985033.5;refGene_gn=BBS7;refGene_tx=X
M_006930881.4
B1 104635874 104635978
refGene_gn=BBS7;refGene_tx=XM_003985033.5;refGene_gn=BBS7;refGene_tx=X
M_006930881.4
B1 104636109 104636233
refGene_gn=BBS7;refGene_tx=XM_003985033.5;refGene_gn=BBS7;refGene_tx=X
M_006930881.4
B1 104638336 104638470
refGene_gn=BBS7;refGene_tx=XM_003985033.5;refGene_gn=BBS7;refGene_tx=X
M_006930881.4
B1 104640732 104640945
refGene_gn=CCNA2;refGene_tx=XM_003985035.5

B1 104641721 104641965
refGene_gn=CCNA2;refGene_tx=XM_003985035.5
B1 104643691 104643804
refGene_gn=CCNA2;refGene_tx=XM_003985035.5
B1 104643999 104644223
refGene_gn=CCNA2;refGene_tx=XM_003985035.5
B1 104644870 104645078
refGene_gn=CCNA2;refGene_tx=XM_003985035.5
B1 104645498 104645612
refGene_gn=CCNA2;refGene_tx=XM_003985035.5
B1 104646052 104646186
refGene_gn=CCNA2;refGene_tx=XM_003985035.5
B1 104646531 104646580
refGene_gn=CCNA2;refGene_tx=XM_003985035.5
B1 104647330 104647415
refGene_gn=EXOSC9;refGene_tx=XM_023253396.1
B1 104647810 104647889
refGene_gn=EXOSC9;refGene_tx=XM_023253396.1
B1 104649886 104650086
refGene_gn=EXOSC9;refGene_tx=XM_023253396.1
B1 104650826 104650973
refGene_gn=EXOSC9;refGene_tx=XM_023253396.1
B1 104651671 104651760
refGene_gn=EXOSC9;refGene_tx=XM_023253396.1
B1 104653022 104653234
refGene_gn=EXOSC9;refGene_tx=XM_023253396.1
B1 104657551 104657708
refGene_gn=EXOSC9;refGene_tx=XM_023253396.1
B1 104658603 104658706
refGene_gn=EXOSC9;refGene_tx=XM_023253396.1
B1 104658819 104658939
refGene_gn=EXOSC9;refGene_tx=XM_023253396.1
B1 104659838 104659933
refGene_gn=EXOSC9;refGene_tx=XM_023253396.1
B1 104660278 104660344
refGene_gn=EXOSC9;refGene_tx=XM_023253396.1
B1 104703224 104703358
refGene_gn=TMEM155;refGene_tx=XM_019829006.1
B1 104704893 104705062
refGene_gn=TMEM155;refGene_tx=XM_019829006.1
B1 104753692 104753701
refGene_gn=ANXA5;refGene_tx=XM_006930883.4
B1 104765626 104765708
refGene_gn=ANXA5;refGene_tx=XM_006930883.4
B1 104766983 104767078
refGene_gn=ANXA5;refGene_tx=XM_006930883.4
B1 104768209 104768323
refGene_gn=ANXA5;refGene_tx=XM_006930883.4
B1 104770192 104770283
refGene_gn=ANXA5;refGene_tx=XM_006930883.4

B1 104772125 104772205
refGene_gn=ANXA5;refGene_tx=XM_006930883.4
B1 104772594 104772651
refGene_gn=ANXA5;refGene_tx=XM_006930883.4
B1 104778564 104778658
refGene_gn=ANXA5;refGene_tx=XM_006930883.4
B1 104779784 104779880
refGene_gn=ANXA5;refGene_tx=XM_006930883.4
B1 104781398 104781457
refGene_gn=ANXA5;refGene_tx=XM_006930883.4
B1 104781669 104781792
refGene_gn=ANXA5;refGene_tx=XM_006930883.4
B1 104782987 104783050
refGene_gn=ANXA5;refGene_tx=XM_006930883.4
B1 105184592 105184932
refGene_gn=QRFPR;refGene_tx=XM_011281672.3;refGene_gn=QRFPR;refGene_tx=XM_019829007.2
B1 105219645 105219804
refGene_gn=QRFPR;refGene_tx=XM_011281672.3;refGene_gn=QRFPR;refGene_tx=XM_019829007.2
B1 105222478 105222543
refGene_gn=QRFPR;refGene_tx=XM_011281672.3;refGene_gn=QRFPR;refGene_tx=XM_019829007.2
B1 105226245 105226481
refGene_gn=QRFPR;refGene_tx=XM_011281672.3;refGene_gn=QRFPR;refGene_tx=XM_019829007.2
B1 105227671 105227769
refGene_gn=QRFPR;refGene_tx=XM_011281672.3;refGene_gn=QRFPR;refGene_tx=XM_019829007.2
B1 105228531 105228872
refGene_gn=QRFPR;refGene_tx=XM_011281672.3;refGene_gn=QRFPR;refGene_tx=XM_019829007.2
B1 105394647 105394713
refGene_gn=TNIP3;refGene_tx=XM_006930884.3;refGene_gn=TNIP3;refGene_tx=XM_023252839.1
B1 105398429 105398510
refGene_gn=TNIP3;refGene_tx=XM_006930884.3;refGene_gn=TNIP3;refGene_tx=XM_023252839.1
B1 105400200 105400266
refGene_gn=TNIP3;refGene_tx=XM_006930884.3;refGene_gn=TNIP3;refGene_tx=XM_023252839.1
B1 105401710 105401842
refGene_gn=TNIP3;refGene_tx=XM_006930884.3;refGene_gn=TNIP3;refGene_tx=XM_023252839.1
B1 105405039 105405168
refGene_gn=TNIP3;refGene_tx=XM_006930884.3;refGene_gn=TNIP3;refGene_tx=XM_023252839.1
B1 105409867 105409987
refGene_gn=TNIP3;refGene_tx=XM_006930884.3;refGene_gn=TNIP3;refGene_tx=XM_023252839.1

B1 105412876 105413002
refGene_gn=TNIP3;refGene_tx=XM_006930884.3;refGene_gn=TNIP3;refGene_tx=XM_023252839.1
B1 105417072 105417123
refGene_gn=TNIP3;refGene_tx=XM_006930884.3;refGene_gn=TNIP3;refGene_tx=XM_023252839.1
B1 105418025 105418124
refGene_gn=TNIP3;refGene_tx=XM_006930884.3;refGene_gn=TNIP3;refGene_tx=XM_023252839.1
B1 105421275 105421336
refGene_gn=TNIP3;refGene_tx=XM_006930884.3;refGene_gn=TNIP3;refGene_tx=XM_023252839.1
B1 105431139 105431171
refGene_gn=TNIP3;refGene_tx=XM_006930884.3;refGene_gn=TNIP3;refGene_tx=XM_023252839.1
B1 105507867 105508055
refGene_gn=NDNF;refGene_tx=XM_003985039.4;refGene_gn=NDNF;refGene_tx=XM_006930885.3
B1 105516652 105516777
refGene_gn=NDNF;refGene_tx=XM_003985039.4;refGene_gn=NDNF;refGene_tx=XM_006930885.3
B1 105518689 105520083
refGene_gn=NDNF;refGene_tx=XM_003985039.4;refGene_gn=NDNF;refGene_tx=XM_006930885.3
B1 105599365 105599458
refGene_gn=PRDM5;refGene_tx=XM_023252840.1;refGene_gn=PRDM5;refGene_tx=XM_023252841.1;refGene_gn=PRDM5;refGene_tx=XM_023252842.1;refGene_gn=PRDM5;refGene_tx=XM_023252843.1
B1 105620992 105621076
refGene_gn=PRDM5;refGene_tx=XM_023252840.1;refGene_gn=PRDM5;refGene_tx=XM_023252841.1;refGene_gn=PRDM5;refGene_tx=XM_023252842.1;refGene_gn=PRDM5;refGene_tx=XM_023252843.1
B1 105669860 105669983
refGene_gn=PRDM5;refGene_tx=XM_023252840.1;refGene_gn=PRDM5;refGene_tx=XM_023252841.1;refGene_gn=PRDM5;refGene_tx=XM_023252842.1;refGene_gn=PRDM5;refGene_tx=XM_023252843.1
B1 105693346 105693521
refGene_gn=PRDM5;refGene_tx=XM_023252840.1;refGene_gn=PRDM5;refGene_tx=XM_023252841.1;refGene_gn=PRDM5;refGene_tx=XM_023252842.1;refGene_gn=PRDM5;refGene_tx=XM_023252843.1
B1 105695805 105695980
refGene_gn=PRDM5;refGene_tx=XM_023252840.1;refGene_gn=PRDM5;refGene_tx=XM_023252841.1;refGene_gn=PRDM5;refGene_tx=XM_023252842.1;refGene_gn=PRDM5;refGene_tx=XM_023252843.1
B1 105697109 105697202
refGene_gn=PRDM5;refGene_tx=XM_023252840.1;refGene_gn=PRDM5;refGene_tx=XM_023252841.1;refGene_gn=PRDM5;refGene_tx=XM_023252842.1;refGene_gn=PRDM5;refGene_tx=XM_023252843.1
B1 105697463 105697585
refGene_gn=PRDM5;refGene_tx=XM_023252840.1;refGene_gn=PRDM5;refGene_tx

=XM_023252841.1;refGene_gn=PRDM5;refGene_tx=XM_023252842.1;refGene_gn=PRDM5;refGene_tx=XM_023252843.1
B1	105701513	105701593
refGene_gn=PRDM5;refGene_tx=XM_023252840.1;refGene_gn=PRDM5;refGene_tx=XM_023252841.1;refGene_gn=PRDM5;refGene_tx=XM_023252842.1;refGene_gn=PRDM5;refGene_tx=XM_023252843.1
B1	105705559	105705644
refGene_gn=PRDM5;refGene_tx=XM_023252840.1;refGene_gn=PRDM5;refGene_tx=XM_023252841.1;refGene_gn=PRDM5;refGene_tx=XM_023252843.1
B1	105706897	105707055
refGene_gn=PRDM5;refGene_tx=XM_023252840.1;refGene_gn=PRDM5;refGene_tx=XM_023252841.1;refGene_gn=PRDM5;refGene_tx=XM_023252843.1
B1	105725799	105725893
refGene_gn=PRDM5;refGene_tx=XM_023252840.1;refGene_gn=PRDM5;refGene_tx=XM_023252841.1;refGene_gn=PRDM5;refGene_tx=XM_023252842.1;refGene_gn=PRDM5;refGene_tx=XM_023252843.1
B1	105730847	105731008
refGene_gn=PRDM5;refGene_tx=XM_023252840.1;refGene_gn=PRDM5;refGene_tx=XM_023252841.1;refGene_gn=PRDM5;refGene_tx=XM_023252842.1;refGene_gn=PRDM5;refGene_tx=XM_023252843.1
B1	105735048	105735142
refGene_gn=PRDM5;refGene_tx=XM_023252840.1;refGene_gn=PRDM5;refGene_tx=XM_023252841.1;refGene_gn=PRDM5;refGene_tx=XM_023252842.1;refGene_gn=PRDM5;refGene_tx=XM_023252843.1
B1	105757766	105757852
refGene_gn=PRDM5;refGene_tx=XM_023252840.1;refGene_gn=PRDM5;refGene_tx=XM_023252841.1;refGene_gn=PRDM5;refGene_tx=XM_023252842.1
B1	105795552	105795657
refGene_gn=PRDM5;refGene_tx=XM_023252840.1;refGene_gn=PRDM5;refGene_tx=XM_023252842.1;refGene_gn=PRDM5;refGene_tx=XM_023252843.1
B1	105805311	105805476
refGene_gn=PRDM5;refGene_tx=XM_023252840.1;refGene_gn=PRDM5;refGene_tx=XM_023252841.1;refGene_gn=PRDM5;refGene_tx=XM_023252842.1
B1	106024917	106025916
refGene_gn=LOC101092105;refGene_tx=XM_011281671.3
B1	106349708	106349781
refGene_gn=MAD2L1;refGene_tx=XM_003985050.5
B1	106350696	106350843
refGene_gn=MAD2L1;refGene_tx=XM_003985050.5
B1	106354016	106354137
refGene_gn=MAD2L1;refGene_tx=XM_003985050.5
B1	106355406	106355510
refGene_gn=MAD2L1;refGene_tx=XM_003985050.5
B1	106356178	106356351
refGene_gn=MAD2L1;refGene_tx=XM_003985050.5
B1	106366016	106366081
refGene_gn=LOC101093602;refGene_tx=XM_023252844.1;refGene_gn=LOC101093602;refGene_tx=XM_023252845.1
B1	106366995	106367163
refGene_gn=LOC101093602;refGene_tx=XM_023252844.1;refGene_gn=LOC101093

602;refGene_tx=XM_023252845.1
B1 106368775 106368925
refGene_gn=LOC101093602;refGene_tx=XM_023252844.1;refGene_gn=LOC101093602;refGene_tx=XM_023252845.1
B1 106371365 106371407
refGene_gn=LOC101093602;refGene_tx=XM_023252844.1;refGene_gn=LOC101093602;refGene_tx=XM_023252845.1
B1 106372266 106372351
refGene_gn=LOC101093602;refGene_tx=XM_023252844.1;refGene_gn=LOC101093602;refGene_tx=XM_023252845.1
B1 106376508 106376586
refGene_gn=LOC101093602;refGene_tx=XM_023252844.1;refGene_gn=LOC101093602;refGene_tx=XM_023252845.1
B1 106379067 106379134
refGene_gn=LOC101093602;refGene_tx=XM_023252844.1;refGene_gn=LOC101093602;refGene_tx=XM_023252845.1
B1 106381702 106381891
refGene_gn=LOC101093602;refGene_tx=XM_023252844.1;refGene_gn=LOC101093602;refGene_tx=XM_023252845.1
B1 106385056 106385226
refGene_gn=LOC101093602;refGene_tx=XM_023252844.1;refGene_gn=LOC101093602;refGene_tx=XM_023252845.1
B1 106386583 106386745
refGene_gn=LOC101093602;refGene_tx=XM_023252844.1;refGene_gn=LOC101093602;refGene_tx=XM_023252845.1
B1 106388663 106388773
refGene_gn=LOC101093602;refGene_tx=XM_023252844.1;refGene_gn=LOC101093602;refGene_tx=XM_023252845.1
B1 106389297 106389487
refGene_gn=LOC101093602;refGene_tx=XM_023252844.1;refGene_gn=LOC101093602;refGene_tx=XM_023252845.1
B1 106390945 106391057
refGene_gn=LOC101093602;refGene_tx=XM_023252844.1;refGene_gn=LOC101093602;refGene_tx=XM_023252845.1
B1 106392311 106392457
refGene_gn=LOC101093602;refGene_tx=XM_023252844.1;refGene_gn=LOC101093602;refGene_tx=XM_023252845.1
B1 106654181 106654303
refGene_gn=PDE5A;refGene_tx=XM_011281658.3;refGene_gn=PDE5A;refGene_tx=XM_019828144.2
B1 106655693 106655728
refGene_gn=PDE5A;refGene_tx=XM_019828141.2
B1 106674206 106674795
refGene_gn=PDE5A;refGene_tx=NM_001204776.2;refGene_gn=PDE5A;refGene_tx=XM_011281658.3;refGene_gn=PDE5A;refGene_tx=XM_019828141.2;refGene_gn=PDE5A;refGene_tx=XM_019828142.2;refGene_gn=PDE5A;refGene_tx=XM_019828144.2
B1 106686133 106686223
refGene_gn=PDE5A;refGene_tx=NM_001204776.2;refGene_gn=PDE5A;refGene_tx=XM_011281658.3;refGene_gn=PDE5A;refGene_tx=XM_019828141.2;refGene_gn=

PDE5A;refGene_tx=XM_019828142.2;refGene_gn=PDE5A;refGene_tx=XM_0198281
44.2
B1 106712109 106712181
refGene_gn=PDE5A;refGene_tx=NM_001204776.2;refGene_gn=PDE5A;refGene_tx
=XM_011281658.3;refGene_gn=PDE5A;refGene_tx=XM_019828141.2;refGene_gn=
PDE5A;refGene_tx=XM_019828142.2;refGene_gn=PDE5A;refGene_tx=XM_0198281
44.2
B1 106713823 106713913
refGene_gn=PDE5A;refGene_tx=NM_001204776.2;refGene_gn=PDE5A;refGene_tx
=XM_011281658.3;refGene_gn=PDE5A;refGene_tx=XM_019828141.2;refGene_gn=
PDE5A;refGene_tx=XM_019828142.2;refGene_gn=PDE5A;refGene_tx=XM_0198281
44.2
B1 106716253 106716391
refGene_gn=PDE5A;refGene_tx=NM_001204776.2;refGene_gn=PDE5A;refGene_tx
=XM_011281658.3;refGene_gn=PDE5A;refGene_tx=XM_019828141.2;refGene_gn=
PDE5A;refGene_tx=XM_019828142.2;refGene_gn=PDE5A;refGene_tx=XM_0198281
44.2
B1 106718412 106718480
refGene_gn=PDE5A;refGene_tx=NM_001204776.2;refGene_gn=PDE5A;refGene_tx
=XM_011281658.3;refGene_gn=PDE5A;refGene_tx=XM_019828141.2;refGene_gn=
PDE5A;refGene_tx=XM_019828142.2;refGene_gn=PDE5A;refGene_tx=XM_0198281
44.2
B1 106725802 106725911
refGene_gn=PDE5A;refGene_tx=NM_001204776.2;refGene_gn=PDE5A;refGene_tx
=XM_011281658.3;refGene_gn=PDE5A;refGene_tx=XM_019828141.2;refGene_gn=
PDE5A;refGene_tx=XM_019828142.2;refGene_gn=PDE5A;refGene_tx=XM_0198281
44.2
B1 106726871 106726959
refGene_gn=PDE5A;refGene_tx=NM_001204776.2;refGene_gn=PDE5A;refGene_tx
=XM_011281658.3;refGene_gn=PDE5A;refGene_tx=XM_019828141.2;refGene_gn=
PDE5A;refGene_tx=XM_019828142.2;refGene_gn=PDE5A;refGene_tx=XM_0198281
44.2
B1 106737092 106737268
refGene_gn=PDE5A;refGene_tx=NM_001204776.2;refGene_gn=PDE5A;refGene_tx
=XM_011281658.3;refGene_gn=PDE5A;refGene_tx=XM_019828141.2;refGene_gn=
PDE5A;refGene_tx=XM_019828142.2;refGene_gn=PDE5A;refGene_tx=XM_0198281
44.2
B1 106740602 106740662
refGene_gn=PDE5A;refGene_tx=NM_001204776.2;refGene_gn=PDE5A;refGene_tx
=XM_011281658.3;refGene_gn=PDE5A;refGene_tx=XM_019828141.2;refGene_gn=
PDE5A;refGene_tx=XM_019828142.2;refGene_gn=PDE5A;refGene_tx=XM_0198281
44.2
B1 106759148 106759295
refGene_gn=PDE5A;refGene_tx=NM_001204776.2;refGene_gn=PDE5A;refGene_tx
=XM_011281658.3;refGene_gn=PDE5A;refGene_tx=XM_019828141.2;refGene_gn=
PDE5A;refGene_tx=XM_019828142.2
B1 106764369 106764495
refGene_gn=PDE5A;refGene_tx=NM_001204776.2;refGene_gn=PDE5A;refGene_tx
=XM_011281658.3;refGene_gn=PDE5A;refGene_tx=XM_019828141.2;refGene_gn=
PDE5A;refGene_tx=XM_019828142.2;refGene_gn=PDE5A;refGene_tx=XM_0198281

44.2
B1	106766510	106766605
refGene_gn=PDE5A;refGene_tx=NM_001204776.2;refGene_gn=PDE5A;refGene_tx=XM_011281658.3;refGene_gn=PDE5A;refGene_tx=XM_019828141.2;refGene_gn=PDE5A;refGene_tx=XM_019828142.2;refGene_gn=PDE5A;refGene_tx=XM_019828144.2
B1	106775644	106775732
refGene_gn=PDE5A;refGene_tx=NM_001204776.2;refGene_gn=PDE5A;refGene_tx=XM_011281658.3;refGene_gn=PDE5A;refGene_tx=XM_019828141.2;refGene_gn=PDE5A;refGene_tx=XM_019828142.2;refGene_gn=PDE5A;refGene_tx=XM_019828144.2
B1	106780227	106780328
refGene_gn=PDE5A;refGene_tx=NM_001204776.2;refGene_gn=PDE5A;refGene_tx=XM_011281658.3;refGene_gn=PDE5A;refGene_tx=XM_019828141.2;refGene_gn=PDE5A;refGene_tx=XM_019828142.2;refGene_gn=PDE5A;refGene_tx=XM_019828144.2
B1	106781994	106782072
refGene_gn=PDE5A;refGene_tx=NM_001204776.2;refGene_gn=PDE5A;refGene_tx=XM_011281658.3;refGene_gn=PDE5A;refGene_tx=XM_019828141.2;refGene_gn=PDE5A;refGene_tx=XM_019828142.2;refGene_gn=PDE5A;refGene_tx=XM_019828144.2
B1	106784153	106784217
refGene_gn=PDE5A;refGene_tx=NM_001204776.2;refGene_gn=PDE5A;refGene_tx=XM_011281658.3;refGene_gn=PDE5A;refGene_tx=XM_019828141.2;refGene_gn=PDE5A;refGene_tx=XM_019828142.2;refGene_gn=PDE5A;refGene_tx=XM_019828144.2
B1	106789302	106789377
refGene_gn=PDE5A;refGene_tx=NM_001204776.2;refGene_gn=PDE5A;refGene_tx=XM_011281658.3;refGene_gn=PDE5A;refGene_tx=XM_019828141.2;refGene_gn=PDE5A;refGene_tx=XM_019828142.2;refGene_gn=PDE5A;refGene_tx=XM_019828144.2
B1	106790789	106790873
refGene_gn=PDE5A;refGene_tx=NM_001204776.2;refGene_gn=PDE5A;refGene_tx=XM_011281658.3;refGene_gn=PDE5A;refGene_tx=XM_019828141.2;refGene_gn=PDE5A;refGene_tx=XM_019828142.2;refGene_gn=PDE5A;refGene_tx=XM_019828144.2
B1	106792496	106792634
refGene_gn=PDE5A;refGene_tx=NM_001204776.2;refGene_gn=PDE5A;refGene_tx=XM_011281658.3;refGene_gn=PDE5A;refGene_tx=XM_019828141.2;refGene_gn=PDE5A;refGene_tx=XM_019828142.2;refGene_gn=PDE5A;refGene_tx=XM_019828144.2
B1	106828071	106828138
refGene_gn=FABP2;refGene_tx=XM_003985051.4
B1	106829350	106829523
refGene_gn=FABP2;refGene_tx=XM_003985051.4
B1	106831160	106831268
refGene_gn=FABP2;refGene_tx=XM_003985051.4
B1	106831647	106831698
refGene_gn=FABP2;refGene_tx=XM_003985051.4
B1	106858462	106858668

refGene_gn=CB1H4orf3;refGene_tx=XM_003985052.4;refGene_gn=CB1H4orf3;refGene_tx=XM_006930892.2
B1 106860093 106860094
refGene_gn=CB1H4orf3;refGene_tx=XM_003985052.4
B1 106860113 106860114
refGene_gn=CB1H4orf3;refGene_tx=XM_006930892.2
B1 106864870 106865858
refGene_gn=USP53;refGene_tx=XM_011281669.3;refGene_gn=USP53;refGene_tx=XM_011281670.3
B1 106867821 106867918
refGene_gn=USP53;refGene_tx=XM_011281669.3;refGene_gn=USP53;refGene_tx=XM_011281670.3
B1 106876259 106876336
refGene_gn=USP53;refGene_tx=XM_011281669.3;refGene_gn=USP53;refGene_tx=XM_011281670.3
B1 106877641 106878380
refGene_gn=USP53;refGene_tx=XM_011281669.3;refGene_gn=USP53;refGene_tx=XM_011281670.3
B1 106879730 106879877
refGene_gn=USP53;refGene_tx=XM_011281669.3;refGene_gn=USP53;refGene_tx=XM_011281670.3
B1 106881352 106881505
refGene_gn=USP53;refGene_tx=XM_011281669.3;refGene_gn=USP53;refGene_tx=XM_011281670.3
B1 106882583 106882746
refGene_gn=USP53;refGene_tx=XM_011281669.3;refGene_gn=USP53;refGene_tx=XM_011281670.3
B1 106890813 106890963
refGene_gn=USP53;refGene_tx=XM_011281669.3;refGene_gn=USP53;refGene_tx=XM_011281670.3
B1 106891749 106891896
refGene_gn=USP53;refGene_tx=XM_011281669.3
B1 106892893 106892999
refGene_gn=USP53;refGene_tx=XM_011281669.3;refGene_gn=USP53;refGene_tx=XM_011281670.3
B1 106895821 106895904
refGene_gn=USP53;refGene_tx=XM_011281669.3;refGene_gn=USP53;refGene_tx=XM_011281670.3
B1 106896059 106896173
refGene_gn=USP53;refGene_tx=XM_011281669.3;refGene_gn=USP53;refGene_tx=XM_011281670.3
B1 106897511 106897646
refGene_gn=USP53;refGene_tx=XM_011281669.3;refGene_gn=USP53;refGene_tx=XM_011281670.3
B1 106901402 106901495
refGene_gn=USP53;refGene_tx=XM_011281669.3;refGene_gn=USP53;refGene_tx=XM_011281670.3
B1 106912537 106912681
refGene_gn=USP53;refGene_tx=XM_011281669.3;refGene_gn=USP53;refGene_tx=XM_011281670.3

B1 106945259 106945494
refGene_gn=MYOZ2;refGene_tx=XM_003985054.3
B1 106958730 106958914
refGene_gn=MYOZ2;refGene_tx=XM_003985054.3
B1 106964119 106964249
refGene_gn=MYOZ2;refGene_tx=XM_003985054.3
B1 106969685 106969832
refGene_gn=MYOZ2;refGene_tx=XM_003985054.3
B1 106969833 106969856
refGene_gn=MYOZ2;refGene_tx=XM_003985054.3
B1 106985106 106985182
refGene_gn=MYOZ2;refGene_tx=XM_003985054.3
B1 107046911 107047448
refGene_gn=SYNPO2;refGene_tx=XM_006930894.4;refGene_gn=SYNPO2;refGene_tx=XM_023252846.1
B1 107073316 107075499
refGene_gn=SYNPO2;refGene_tx=XM_006930894.4
B1 107077944 107078756
refGene_gn=SYNPO2;refGene_tx=XM_006930894.4
B1 107082177 107082329
refGene_gn=SYNPO2;refGene_tx=XM_006930894.4;refGene_gn=SYNPO2;refGene_tx=XM_023252846.1
B1 107216995 107217100
refGene_gn=SYNPO2;refGene_tx=XM_006930894.4;refGene_gn=SYNPO2;refGene_tx=XM_023252846.1
B1 107268968 107269086
refGene_gn=SEC24D;refGene_tx=XM_011281667.3;refGene_gn=SEC24D;refGene_tx=XM_019829009.2;refGene_gn=SEC24D;refGene_tx=XM_019829010.2
B1 107279842 107279972
refGene_gn=SEC24D;refGene_tx=XM_011281667.3;refGene_gn=SEC24D;refGene_tx=XM_019829009.2;refGene_gn=SEC24D;refGene_tx=XM_019829010.2
B1 107286496 107286645
refGene_gn=SEC24D;refGene_tx=XM_011281667.3;refGene_gn=SEC24D;refGene_tx=XM_019829009.2;refGene_gn=SEC24D;refGene_tx=XM_019829010.2
B1 107288138 107288414
refGene_gn=SEC24D;refGene_tx=XM_011281667.3;refGene_gn=SEC24D;refGene_tx=XM_019829009.2;refGene_gn=SEC24D;refGene_tx=XM_019829010.2
B1 107288684 107288815
refGene_gn=SEC24D;refGene_tx=XM_011281667.3;refGene_gn=SEC24D;refGene_tx=XM_019829009.2;refGene_gn=SEC24D;refGene_tx=XM_019829010.2
B1 107297595 107297707
refGene_gn=SEC24D;refGene_tx=XM_011281667.3;refGene_gn=SEC24D;refGene_tx=XM_019829009.2;refGene_gn=SEC24D;refGene_tx=XM_019829010.2
B1 107304026 107304154
refGene_gn=SEC24D;refGene_tx=XM_011281667.3;refGene_gn=SEC24D;refGene_tx=XM_019829009.2;refGene_gn=SEC24D;refGene_tx=XM_019829010.2
B1 107327646 107327785
refGene_gn=SEC24D;refGene_tx=XM_011281667.3;refGene_gn=SEC24D;refGene_tx=XM_019829009.2;refGene_gn=SEC24D;refGene_tx=XM_019829010.2
B1 107331115 107331231

refGene_gn=SEC24D;refGene_tx=XM_011281667.3;refGene_gn=SEC24D;refGene_tx=XM_019829009.2;refGene_gn=SEC24D;refGene_tx=XM_019829010.2
B1 107338398 107338523
refGene_gn=SEC24D;refGene_tx=XM_011281667.3;refGene_gn=SEC24D;refGene_tx=XM_019829009.2;refGene_gn=SEC24D;refGene_tx=XM_019829010.2
B1 107343163 107343355
refGene_gn=SEC24D;refGene_tx=XM_011281667.3;refGene_gn=SEC24D;refGene_tx=XM_019829009.2;refGene_gn=SEC24D;refGene_tx=XM_019829010.2
B1 107343946 107344040
refGene_gn=SEC24D;refGene_tx=XM_011281667.3;refGene_gn=SEC24D;refGene_tx=XM_019829009.2;refGene_gn=SEC24D;refGene_tx=XM_019829010.2
B1 107351675 107351792
refGene_gn=SEC24D;refGene_tx=XM_011281667.3;refGene_gn=SEC24D;refGene_tx=XM_019829009.2;refGene_gn=SEC24D;refGene_tx=XM_019829010.2
B1 107352631 107352802
refGene_gn=SEC24D;refGene_tx=XM_011281667.3;refGene_gn=SEC24D;refGene_tx=XM_019829009.2;refGene_gn=SEC24D;refGene_tx=XM_019829010.2
B1 107354050 107354147
refGene_gn=SEC24D;refGene_tx=XM_011281667.3;refGene_gn=SEC24D;refGene_tx=XM_019829009.2;refGene_gn=SEC24D;refGene_tx=XM_019829010.2
B1 107354276 107354422
refGene_gn=SEC24D;refGene_tx=XM_011281667.3;refGene_gn=SEC24D;refGene_tx=XM_019829009.2;refGene_gn=SEC24D;refGene_tx=XM_019829010.2
B1 107355414 107355553
refGene_gn=SEC24D;refGene_tx=XM_011281667.3;refGene_gn=SEC24D;refGene_tx=XM_019829009.2;refGene_gn=SEC24D;refGene_tx=XM_019829010.2
B1 107356362 107356481
refGene_gn=SEC24D;refGene_tx=XM_011281667.3;refGene_gn=SEC24D;refGene_tx=XM_019829009.2;refGene_gn=SEC24D;refGene_tx=XM_019829010.2
B1 107360980 107361160
refGene_gn=SEC24D;refGene_tx=XM_011281667.3;refGene_gn=SEC24D;refGene_tx=XM_019829009.2;refGene_gn=SEC24D;refGene_tx=XM_019829010.2
B1 107363474 107363666
refGene_gn=SEC24D;refGene_tx=XM_011281667.3;refGene_gn=SEC24D;refGene_tx=XM_019829009.2;refGene_gn=SEC24D;refGene_tx=XM_019829010.2
B1 107366403 107366493
refGene_gn=SEC24D;refGene_tx=XM_011281667.3;refGene_gn=SEC24D;refGene_tx=XM_019829009.2;refGene_gn=SEC24D;refGene_tx=XM_019829010.2
B1 107372614 107372755
refGene_gn=SEC24D;refGene_tx=XM_011281667.3;refGene_gn=SEC24D;refGene_tx=XM_019829009.2;refGene_gn=SEC24D;refGene_tx=XM_019829010.2
B1 107384569 107384874
refGene_gn=METTL14;refGene_tx=XM_003985057.3;refGene_gn=METTL14;refGene_tx=XM_019829012.2
B1 107386255 107386466
refGene_gn=METTL14;refGene_tx=XM_003985057.3;refGene_gn=METTL14;refGene_tx=XM_019829012.2
B1 107387646 107387763
refGene_gn=METTL14;refGene_tx=XM_003985057.3;refGene_gn=METTL14;refGene_tx=XM_019829012.2

B1 107389813 107389906
refGene_gn=METTL14;refGene_tx=XM_003985057.3;refGene_gn=METTL14;refGene_tx=XM_019829012.2
B1 107395108 107395250
refGene_gn=METTL14;refGene_tx=XM_003985057.3;refGene_gn=METTL14;refGene_tx=XM_019829012.2
B1 107396444 107396535
refGene_gn=METTL14;refGene_tx=XM_003985057.3;refGene_gn=METTL14;refGene_tx=XM_019829012.2
B1 107400682 107400770
refGene_gn=METTL14;refGene_tx=XM_003985057.3;refGene_gn=METTL14;refGene_tx=XM_019829012.2
B1 107401122 107401203
refGene_gn=METTL14;refGene_tx=XM_003985057.3;refGene_gn=METTL14;refGene_tx=XM_019829012.2
B1 107403988 107404076
refGene_gn=METTL14;refGene_tx=XM_003985057.3;refGene_gn=METTL14;refGene_tx=XM_019829012.2
B1 107404990 107405079
refGene_gn=METTL14;refGene_tx=XM_003985057.3
B1 107409101 107409167
refGene_gn=METTL14;refGene_tx=XM_003985057.3
B1 107481569 107482080
refGene_gn=PRSS12;refGene_tx=XM_023252847.1;refGene_gn=PRSS12;refGene_tx=XM_023252848.1
B1 107494627 107494766
refGene_gn=PRSS12;refGene_tx=XM_023252847.1;refGene_gn=PRSS12;refGene_tx=XM_023252848.1
B1 107496992 107497171
refGene_gn=PRSS12;refGene_tx=XM_023252847.1;refGene_gn=PRSS12;refGene_tx=XM_023252848.1
B1 107501890 107502041
refGene_gn=PRSS12;refGene_tx=XM_023252847.1;refGene_gn=PRSS12;refGene_tx=XM_023252848.1
B1 107513147 107513326
refGene_gn=PRSS12;refGene_tx=XM_023252847.1;refGene_gn=PRSS12;refGene_tx=XM_023252848.1
B1 107515050 107515192
refGene_gn=PRSS12;refGene_tx=XM_023252847.1;refGene_gn=PRSS12;refGene_tx=XM_023252848.1
B1 107517940 107518137
refGene_gn=PRSS12;refGene_tx=XM_023252847.1;refGene_gn=PRSS12;refGene_tx=XM_023252848.1
B1 107525079 107525221
refGene_gn=PRSS12;refGene_tx=XM_023252847.1;refGene_gn=PRSS12;refGene_tx=XM_023252848.1
B1 107538676 107538882
refGene_gn=PRSS12;refGene_tx=XM_023252847.1;refGene_gn=PRSS12;refGene_tx=XM_023252848.1
B1 107541886 107541965

refGene_gn=PRSS12;refGene_tx=XM_023252847.1;refGene_gn=PRSS12;refGene_tx=XM_023252848.1
B1 107542670 107542793
refGene_gn=PRSS12;refGene_tx=XM_023252847.1;refGene_gn=PRSS12;refGene_tx=XM_023252848.1
B1 107556430 107556711
refGene_gn=PRSS12;refGene_tx=XM_023252847.1
B1 107558014 107558322
refGene_gn=PRSS12;refGene_tx=XM_023252847.1;refGene_gn=PRSS12;refGene_tx=XM_023252848.1
B1 107591557 107591677
refGene_gn=NDST3;refGene_tx=XM_006930896.4;refGene_gn=NDST3;refGene_tx=XM_019829011.2;refGene_gn=NDST3;refGene_tx=XM_023252849.1
B1 107596807 107596910
refGene_gn=NDST3;refGene_tx=XM_006930896.4;refGene_gn=NDST3;refGene_tx=XM_019829011.2;refGene_gn=NDST3;refGene_tx=XM_023252849.1
B1 107606612 107606722
refGene_gn=NDST3;refGene_tx=XM_006930896.4;refGene_gn=NDST3;refGene_tx=XM_019829011.2;refGene_gn=NDST3;refGene_tx=XM_023252849.1
B1 107608243 107608414
refGene_gn=NDST3;refGene_tx=XM_006930896.4;refGene_gn=NDST3;refGene_tx=XM_019829011.2;refGene_gn=NDST3;refGene_tx=XM_023252849.1
B1 107611411 107611586
refGene_gn=NDST3;refGene_tx=XM_006930896.4;refGene_gn=NDST3;refGene_tx=XM_019829011.2;refGene_gn=NDST3;refGene_tx=XM_023252849.1
B1 107615879 107616003
refGene_gn=NDST3;refGene_tx=XM_006930896.4;refGene_gn=NDST3;refGene_tx=XM_019829011.2;refGene_gn=NDST3;refGene_tx=XM_023252849.1
B1 107620044 107620141
refGene_gn=NDST3;refGene_tx=XM_006930896.4;refGene_gn=NDST3;refGene_tx=XM_019829011.2;refGene_gn=NDST3;refGene_tx=XM_023252849.1
B1 107622377 107622560
refGene_gn=NDST3;refGene_tx=XM_006930896.4;refGene_gn=NDST3;refGene_tx=XM_019829011.2;refGene_gn=NDST3;refGene_tx=XM_023252849.1
B1 107677520 107677649
refGene_gn=NDST3;refGene_tx=XM_006930896.4;refGene_gn=NDST3;refGene_tx=XM_019829011.2;refGene_gn=NDST3;refGene_tx=XM_023252849.1
B1 107684970 107685156
refGene_gn=NDST3;refGene_tx=XM_006930896.4;refGene_gn=NDST3;refGene_tx=XM_019829011.2;refGene_gn=NDST3;refGene_tx=XM_023252849.1
B1 107705276 107705431
refGene_gn=NDST3;refGene_tx=XM_006930896.4;refGene_gn=NDST3;refGene_tx=XM_019829011.2;refGene_gn=NDST3;refGene_tx=XM_023252849.1
B1 107711499 107711587
refGene_gn=NDST3;refGene_tx=XM_006930896.4;refGene_gn=NDST3;refGene_tx=XM_019829011.2
B1 107759827 107760850
refGene_gn=NDST3;refGene_tx=XM_006930896.4;refGene_gn=NDST3;refGene_tx=XM_019829011.2
B1 108672279 108673392

refGene_gn=TRAM1L1;refGene_tx=XM_003985060.5
B1 110760912 110761890
refGene_gn=LOC101097478;refGene_tx=XM_023252850.1;refGene_gn=LOC101097
478;refGene_tx=XM_023252851.1
B1 110862099 110862187
refGene_gn=LOC101097478;refGene_tx=XM_023252850.1;refGene_gn=LOC101097
478;refGene_tx=XM_023252851.1
B1 110872760 110872915
refGene_gn=LOC101097478;refGene_tx=XM_023252850.1;refGene_gn=LOC101097
478;refGene_tx=XM_023252851.1
B1 110901450 110901636
refGene_gn=LOC101097478;refGene_tx=XM_023252850.1;refGene_gn=LOC101097
478;refGene_tx=XM_023252851.1
B1 110903538 110903667
refGene_gn=LOC101097478;refGene_tx=XM_023252850.1;refGene_gn=LOC101097
478;refGene_tx=XM_023252851.1
B1 110963467 110963650
refGene_gn=LOC101097478;refGene_tx=XM_023252850.1;refGene_gn=LOC101097
478;refGene_tx=XM_023252851.1
B1 110981935 110982032
refGene_gn=LOC101097478;refGene_tx=XM_023252850.1;refGene_gn=LOC101097
478;refGene_tx=XM_023252851.1
B1 110986561 110986685
refGene_gn=LOC101097478;refGene_tx=XM_023252850.1;refGene_gn=LOC101097
478;refGene_tx=XM_023252851.1
B1 110988736 110988911
refGene_gn=LOC101097478;refGene_tx=XM_023252850.1;refGene_gn=LOC101097
478;refGene_tx=XM_023252851.1
B1 110994401 110994572
refGene_gn=LOC101097478;refGene_tx=XM_023252850.1;refGene_gn=LOC101097
478;refGene_tx=XM_023252851.1
B1 110995720 110995830
refGene_gn=LOC101097478;refGene_tx=XM_023252850.1;refGene_gn=LOC101097
478;refGene_tx=XM_023252851.1
B1 110999548 110999651
refGene_gn=LOC101097478;refGene_tx=XM_023252850.1;refGene_gn=LOC101097
478;refGene_tx=XM_023252851.1
B1 111000797 111000838
refGene_gn=LOC101097478;refGene_tx=XM_023252851.1
B1 111002583 111002757
refGene_gn=LOC101097478;refGene_tx=XM_023252850.1;refGene_gn=LOC101097
478;refGene_tx=XM_023252851.1
B1 111127519 111127883
refGene_gn=UGT8;refGene_tx=XM_019829015.2
B1 111130119 111130339
refGene_gn=UGT8;refGene_tx=XM_019829015.2
B1 111132902 111132979
refGene_gn=UGT8;refGene_tx=XM_019829015.2
B1 111134692 111134835
refGene_gn=UGT8;refGene_tx=XM_019829015.2

B1 111180952 111181285
refGene_gn=UGT8;refGene_tx=XM_019829015.2
B1 111181286 111181775
refGene_gn=UGT8;refGene_tx=XM_019829015.2
B1 111677446 111677844
refGene_gn=ARSJ;refGene_tx=XM_011281663.3
B1 111757060 111758465
refGene_gn=ARSJ;refGene_tx=XM_011281663.3
B1 111873788 111873853
refGene_gn=CAMK2D;refGene_tx=XM_006930898.4;refGene_gn=CAMK2D;refGene_tx=XM_006930899.4;refGene_gn=CAMK2D;refGene_tx=XM_006930900.4;refGene_gn=CAMK2D;refGene_tx=XM_006930901.4;refGene_gn=CAMK2D;refGene_tx=XM_011281662.3;refGene_gn=CAMK2D;refGene_tx=XM_019829018.2;refGene_gn=CAMK2D;refGene_tx=XM_019829019.2;refGene_gn=CAMK2D;refGene_tx=XM_019829020.2;refGene_gn=CAMK2D;refGene_tx=XM_019829021.2;refGene_gn=CAMK2D;refGene_tx=XM_019829022.2;refGene_gn=CAMK2D;refGene_tx=XM_019829023.2;refGene_gn=CAMK2D;refGene_tx=XM_019829024.2;refGene_gn=CAMK2D;refGene_tx=XM_019829025.2;refGene_gn=CAMK2D;refGene_tx=XM_019829026.2;refGene_gn=CAMK2D;refGene_tx=XM_019829027.2;refGene_gn=CAMK2D;refGene_tx=XM_019829028.2;refGene_gn=CAMK2D;refGene_tx=XM_019829029.2;refGene_gn=CAMK2D;refGene_tx=XM_019829030.2;refGene_gn=CAMK2D;refGene_tx=XM_023252852.1
B1 111875479 111875574
refGene_gn=CAMK2D;refGene_tx=XM_006930898.4;refGene_gn=CAMK2D;refGene_tx=XM_006930899.4;refGene_gn=CAMK2D;refGene_tx=XM_006930900.4;refGene_gn=CAMK2D;refGene_tx=XM_006930901.4;refGene_gn=CAMK2D;refGene_tx=XM_011281662.3;refGene_gn=CAMK2D;refGene_tx=XM_019829018.2;refGene_gn=CAMK2D;refGene_tx=XM_019829019.2;refGene_gn=CAMK2D;refGene_tx=XM_019829020.2;refGene_gn=CAMK2D;refGene_tx=XM_019829021.2;refGene_gn=CAMK2D;refGene_tx=XM_019829022.2;refGene_gn=CAMK2D;refGene_tx=XM_019829023.2;refGene_gn=CAMK2D;refGene_tx=XM_019829024.2;refGene_gn=CAMK2D;refGene_tx=XM_019829025.2;refGene_gn=CAMK2D;refGene_tx=XM_019829026.2;refGene_gn=CAMK2D;refGene_tx=XM_019829027.2;refGene_gn=CAMK2D;refGene_tx=XM_019829028.2;refGene_gn=CAMK2D;refGene_tx=XM_019829029.2;refGene_gn=CAMK2D;refGene_tx=XM_019829030.2;refGene_gn=CAMK2D;refGene_tx=XM_023252852.1
B1 111975654 111975714
refGene_gn=CAMK2D;refGene_tx=XM_006930898.4;refGene_gn=CAMK2D;refGene_tx=XM_006930899.4;refGene_gn=CAMK2D;refGene_tx=XM_006930900.4;refGene_gn=CAMK2D;refGene_tx=XM_006930901.4;refGene_gn=CAMK2D;refGene_tx=XM_011281662.3;refGene_gn=CAMK2D;refGene_tx=XM_019829018.2;refGene_gn=CAMK2D;refGene_tx=XM_019829019.2;refGene_gn=CAMK2D;refGene_tx=XM_019829020.2;refGene_gn=CAMK2D;refGene_tx=XM_019829021.2;refGene_gn=CAMK2D;refGene_tx=XM_019829022.2;refGene_gn=CAMK2D;refGene_tx=XM_019829023.2;refGene_gn=CAMK2D;refGene_tx=XM_019829024.2;refGene_gn=CAMK2D;refGene_tx=XM_019829025.2;refGene_gn=CAMK2D;refGene_tx=XM_019829026.2;refGene_gn=CAMK2D;refGene_tx=XM_019829027.2;refGene_gn=CAMK2D;refGene_tx=XM_019829028.2;refGene_gn=CAMK2D;refGene_tx=XM_019829029.2;refGene_gn=CAMK2D;refGene_tx=XM_019829030.2;refGene_gn=CAMK2D;refGene_tx=XM_023252852.1
B1 112032482 112032537
refGene_gn=CAMK2D;refGene_tx=XM_006930898.4;refGene_gn=CAMK2D;refGene_tx=XM_006930899.4;refGene_gn=CAMK2D;refGene_tx=XM_006930900.4;refGene_

gn=CAMK2D;refGene_tx=XM_006930901.4;refGene_gn=CAMK2D;refGene_tx=XM_01
1281662.3;refGene_gn=CAMK2D;refGene_tx=XM_019829018.2;refGene_gn=CAMK2
D;refGene_tx=XM_019829019.2;refGene_gn=CAMK2D;refGene_tx=XM_019829020.
2;refGene_gn=CAMK2D;refGene_tx=XM_019829021.2;refGene_gn=CAMK2D;refGen
e_tx=XM_019829022.2;refGene_gn=CAMK2D;refGene_tx=XM_019829023.2;refGen
e_gn=CAMK2D;refGene_tx=XM_019829024.2;refGene_gn=CAMK2D;refGene_tx=XM_
019829025.2;refGene_gn=CAMK2D;refGene_tx=XM_019829026.2;refGene_gn=CAM
K2D;refGene_tx=XM_019829027.2;refGene_gn=CAMK2D;refGene_tx=XM_01982902
8.2;refGene_gn=CAMK2D;refGene_tx=XM_019829029.2;refGene_gn=CAMK2D;refG
ene_tx=XM_019829030.2;refGene_gn=CAMK2D;refGene_tx=XM_023252852.1
B1	112096952	112097018
refGene_gn=CAMK2D;refGene_tx=XM_006930898.4;refGene_gn=CAMK2D;refGene_
tx=XM_006930899.4;refGene_gn=CAMK2D;refGene_tx=XM_006930900.4;refGene_
gn=CAMK2D;refGene_tx=XM_006930901.4;refGene_gn=CAMK2D;refGene_tx=XM_01
1281662.3;refGene_gn=CAMK2D;refGene_tx=XM_019829018.2;refGene_gn=CAMK2
D;refGene_tx=XM_019829019.2;refGene_gn=CAMK2D;refGene_tx=XM_019829020.
2;refGene_gn=CAMK2D;refGene_tx=XM_019829021.2;refGene_gn=CAMK2D;refGen
e_tx=XM_019829022.2;refGene_gn=CAMK2D;refGene_tx=XM_019829023.2;refGen
e_gn=CAMK2D;refGene_tx=XM_019829024.2;refGene_gn=CAMK2D;refGene_tx=XM_
019829025.2;refGene_gn=CAMK2D;refGene_tx=XM_019829026.2;refGene_gn=CAM
K2D;refGene_tx=XM_019829027.2;refGene_gn=CAMK2D;refGene_tx=XM_01982902
8.2;refGene_gn=CAMK2D;refGene_tx=XM_019829029.2;refGene_gn=CAMK2D;refG
ene_tx=XM_019829030.2;refGene_gn=CAMK2D;refGene_tx=XM_023252852.1
B1	112100125	112100198
refGene_gn=CAMK2D;refGene_tx=XM_006930898.4;refGene_gn=CAMK2D;refGene_
tx=XM_006930899.4;refGene_gn=CAMK2D;refGene_tx=XM_006930900.4;refGene_
gn=CAMK2D;refGene_tx=XM_006930901.4;refGene_gn=CAMK2D;refGene_tx=XM_01
1281662.3;refGene_gn=CAMK2D;refGene_tx=XM_019829018.2;refGene_gn=CAMK2
D;refGene_tx=XM_019829020.2;refGene_gn=CAMK2D;refGene_tx=XM_019829023.
2;refGene_gn=CAMK2D;refGene_tx=XM_019829026.2;refGene_gn=CAMK2D;refGen
e_tx=XM_019829027.2;refGene_gn=CAMK2D;refGene_tx=XM_019829028.2;refGen
e_gn=CAMK2D;refGene_tx=XM_023252852.1
B1	112101143	112101216
refGene_gn=CAMK2D;refGene_tx=XM_019829019.2;refGene_gn=CAMK2D;refGene_
tx=XM_019829021.2;refGene_gn=CAMK2D;refGene_tx=XM_019829022.2;refGene_
gn=CAMK2D;refGene_tx=XM_019829024.2;refGene_gn=CAMK2D;refGene_tx=XM_01
9829025.2;refGene_gn=CAMK2D;refGene_tx=XM_019829029.2;refGene_gn=CAMK2
D;refGene_tx=XM_019829030.2
B1	112110654	112110757
refGene_gn=CAMK2D;refGene_tx=XM_006930898.4;refGene_gn=CAMK2D;refGene_
tx=XM_006930899.4;refGene_gn=CAMK2D;refGene_tx=XM_006930900.4;refGene_
gn=CAMK2D;refGene_tx=XM_006930901.4;refGene_gn=CAMK2D;refGene_tx=XM_01
1281662.3;refGene_gn=CAMK2D;refGene_tx=XM_019829018.2;refGene_gn=CAMK2
D;refGene_tx=XM_019829019.2;refGene_gn=CAMK2D;refGene_tx=XM_019829020.
2;refGene_gn=CAMK2D;refGene_tx=XM_019829021.2;refGene_gn=CAMK2D;refGen
e_tx=XM_019829022.2;refGene_gn=CAMK2D;refGene_tx=XM_019829023.2;refGen
e_gn=CAMK2D;refGene_tx=XM_019829024.2;refGene_gn=CAMK2D;refGene_tx=XM_
019829025.2;refGene_gn=CAMK2D;refGene_tx=XM_019829026.2;refGene_gn=CAM
K2D;refGene_tx=XM_019829027.2;refGene_gn=CAMK2D;refGene_tx=XM_01982902
8.2;refGene_gn=CAMK2D;refGene_tx=XM_019829029.2;refGene_gn=CAMK2D;refG

ene_tx=XM_019829030.2;refGene_gn=CAMK2D;refGene_tx=XM_023252852.1
B1 112118255 112118339
refGene_gn=CAMK2D;refGene_tx=XM_006930898.4;refGene_gn=CAMK2D;refGene_tx=XM_006930899.4;refGene_gn=CAMK2D;refGene_tx=XM_006930900.4;refGene_gn=CAMK2D;refGene_tx=XM_006930901.4;refGene_gn=CAMK2D;refGene_tx=XM_011281662.3;refGene_gn=CAMK2D;refGene_tx=XM_019829018.2;refGene_gn=CAMK2D;refGene_tx=XM_019829019.2;refGene_gn=CAMK2D;refGene_tx=XM_019829020.2;refGene_gn=CAMK2D;refGene_tx=XM_019829021.2;refGene_gn=CAMK2D;refGene_tx=XM_019829022.2;refGene_gn=CAMK2D;refGene_tx=XM_019829023.2;refGene_gn=CAMK2D;refGene_tx=XM_019829024.2;refGene_gn=CAMK2D;refGene_tx=XM_019829025.2;refGene_gn=CAMK2D;refGene_tx=XM_019829026.2;refGene_gn=CAMK2D;refGene_tx=XM_019829027.2;refGene_gn=CAMK2D;refGene_tx=XM_019829028.2;refGene_gn=CAMK2D;refGene_tx=XM_019829029.2;refGene_gn=CAMK2D;refGene_tx=XM_019829030.2;refGene_gn=CAMK2D;refGene_tx=XM_023252852.1
B1 112133500 112133595
refGene_gn=CAMK2D;refGene_tx=XM_006930898.4;refGene_gn=CAMK2D;refGene_tx=XM_006930899.4;refGene_gn=CAMK2D;refGene_tx=XM_006930900.4;refGene_gn=CAMK2D;refGene_tx=XM_006930901.4;refGene_gn=CAMK2D;refGene_tx=XM_011281662.3;refGene_gn=CAMK2D;refGene_tx=XM_019829018.2;refGene_gn=CAMK2D;refGene_tx=XM_019829019.2;refGene_gn=CAMK2D;refGene_tx=XM_019829020.2;refGene_gn=CAMK2D;refGene_tx=XM_019829021.2;refGene_gn=CAMK2D;refGene_tx=XM_019829022.2;refGene_gn=CAMK2D;refGene_tx=XM_019829023.2;refGene_gn=CAMK2D;refGene_tx=XM_019829024.2;refGene_gn=CAMK2D;refGene_tx=XM_019829025.2;refGene_gn=CAMK2D;refGene_tx=XM_019829026.2;refGene_gn=CAMK2D;refGene_tx=XM_019829027.2;refGene_gn=CAMK2D;refGene_tx=XM_019829028.2;refGene_gn=CAMK2D;refGene_tx=XM_019829029.2;refGene_gn=CAMK2D;refGene_tx=XM_019829030.2;refGene_gn=CAMK2D;refGene_tx=XM_023252852.1
B1 112135953 112136076
refGene_gn=CAMK2D;refGene_tx=XM_006930898.4;refGene_gn=CAMK2D;refGene_tx=XM_006930899.4;refGene_gn=CAMK2D;refGene_tx=XM_006930900.4;refGene_gn=CAMK2D;refGene_tx=XM_006930901.4;refGene_gn=CAMK2D;refGene_tx=XM_011281662.3;refGene_gn=CAMK2D;refGene_tx=XM_019829018.2;refGene_gn=CAMK2D;refGene_tx=XM_019829019.2;refGene_gn=CAMK2D;refGene_tx=XM_019829020.2;refGene_gn=CAMK2D;refGene_tx=XM_019829021.2;refGene_gn=CAMK2D;refGene_tx=XM_019829022.2;refGene_gn=CAMK2D;refGene_tx=XM_019829023.2;refGene_gn=CAMK2D;refGene_tx=XM_019829024.2;refGene_gn=CAMK2D;refGene_tx=XM_019829025.2;refGene_gn=CAMK2D;refGene_tx=XM_019829026.2;refGene_gn=CAMK2D;refGene_tx=XM_019829027.2;refGene_gn=CAMK2D;refGene_tx=XM_019829028.2;refGene_gn=CAMK2D;refGene_tx=XM_019829029.2;refGene_gn=CAMK2D;refGene_tx=XM_019829030.2;refGene_gn=CAMK2D;refGene_tx=XM_023252852.1
B1 112136942 112137026
refGene_gn=CAMK2D;refGene_tx=XM_006930898.4;refGene_gn=CAMK2D;refGene_tx=XM_006930899.4;refGene_gn=CAMK2D;refGene_tx=XM_006930900.4;refGene_gn=CAMK2D;refGene_tx=XM_006930901.4;refGene_gn=CAMK2D;refGene_tx=XM_011281662.3;refGene_gn=CAMK2D;refGene_tx=XM_019829018.2;refGene_gn=CAMK2D;refGene_tx=XM_019829019.2;refGene_gn=CAMK2D;refGene_tx=XM_019829020.2;refGene_gn=CAMK2D;refGene_tx=XM_019829021.2;refGene_gn=CAMK2D;refGene_tx=XM_019829022.2;refGene_gn=CAMK2D;refGene_tx=XM_019829023.2;refGene_gn=CAMK2D;refGene_tx=XM_019829024.2;refGene_gn=CAMK2D;refGene_tx=XM_019829025.2;refGene_gn=CAMK2D;refGene_tx=XM_019829026.2;refGene_gn=CAM

K2D;refGene_tx=XM_019829027.2;refGene_gn=CAMK2D;refGene_tx=XM_01982902
8.2;refGene_gn=CAMK2D;refGene_tx=XM_019829029.2;refGene_gn=CAMK2D;refG
ene_tx=XM_019829030.2;refGene_gn=CAMK2D;refGene_tx=XM_023252852.1
B1 112137474 112137517
refGene_gn=CAMK2D;refGene_tx=XM_006930898.4;refGene_gn=CAMK2D;refGene_
tx=XM_006930899.4;refGene_gn=CAMK2D;refGene_tx=XM_006930900.4;refGene_
gn=CAMK2D;refGene_tx=XM_006930901.4;refGene_gn=CAMK2D;refGene_tx=XM_01
1281662.3;refGene_gn=CAMK2D;refGene_tx=XM_019829018.2;refGene_gn=CAMK2
D;refGene_tx=XM_019829019.2;refGene_gn=CAMK2D;refGene_tx=XM_019829020.
2;refGene_gn=CAMK2D;refGene_tx=XM_019829021.2;refGene_gn=CAMK2D;refGen
e_tx=XM_019829022.2;refGene_gn=CAMK2D;refGene_tx=XM_019829023.2;refGen
e_gn=CAMK2D;refGene_tx=XM_019829024.2;refGene_gn=CAMK2D;refGene_tx=XM_
019829025.2;refGene_gn=CAMK2D;refGene_tx=XM_019829026.2;refGene_gn=CAM
K2D;refGene_tx=XM_019829027.2;refGene_gn=CAMK2D;refGene_tx=XM_01982902
8.2;refGene_gn=CAMK2D;refGene_tx=XM_019829029.2;refGene_gn=CAMK2D;refG
ene_tx=XM_019829030.2;refGene_gn=CAMK2D;refGene_tx=XM_023252852.1
B1 112140914 112140952
refGene_gn=CAMK2D;refGene_tx=XM_006930898.4;refGene_gn=CAMK2D;refGene_
tx=XM_006930899.4;refGene_gn=CAMK2D;refGene_tx=XM_006930900.4;refGene_
gn=CAMK2D;refGene_tx=XM_006930901.4;refGene_gn=CAMK2D;refGene_tx=XM_01
1281662.3;refGene_gn=CAMK2D;refGene_tx=XM_019829018.2;refGene_gn=CAMK2
D;refGene_tx=XM_019829019.2;refGene_gn=CAMK2D;refGene_tx=XM_019829020.
2;refGene_gn=CAMK2D;refGene_tx=XM_019829021.2;refGene_gn=CAMK2D;refGen
e_tx=XM_019829022.2;refGene_gn=CAMK2D;refGene_tx=XM_019829023.2;refGen
e_gn=CAMK2D;refGene_tx=XM_019829024.2;refGene_gn=CAMK2D;refGene_tx=XM_
019829025.2;refGene_gn=CAMK2D;refGene_tx=XM_019829026.2;refGene_gn=CAM
K2D;refGene_tx=XM_019829027.2;refGene_gn=CAMK2D;refGene_tx=XM_01982902
8.2;refGene_gn=CAMK2D;refGene_tx=XM_019829029.2;refGene_gn=CAMK2D;refG
ene_tx=XM_019829030.2;refGene_gn=CAMK2D;refGene_tx=XM_023252852.1
B1 112142569 112142602
refGene_gn=CAMK2D;refGene_tx=XM_019829018.2;refGene_gn=CAMK2D;refGene_
tx=XM_019829019.2;refGene_gn=CAMK2D;refGene_tx=XM_019829020.2;refGene_
gn=CAMK2D;refGene_tx=XM_019829021.2;refGene_gn=CAMK2D;refGene_tx=XM_01
9829023.2;refGene_gn=CAMK2D;refGene_tx=XM_019829026.2;refGene_gn=CAMK2
D;refGene_tx=XM_019829027.2;refGene_gn=CAMK2D;refGene_tx=XM_019829030.
2
B1 112145508 112145568
refGene_gn=CAMK2D;refGene_tx=XM_006930898.4;refGene_gn=CAMK2D;refGene_
tx=XM_006930899.4;refGene_gn=CAMK2D;refGene_tx=XM_019829018.2;refGene_
gn=CAMK2D;refGene_tx=XM_019829019.2;refGene_gn=CAMK2D;refGene_tx=XM_01
9829020.2;refGene_gn=CAMK2D;refGene_tx=XM_019829021.2;refGene_gn=CAMK2
D;refGene_tx=XM_019829022.2;refGene_gn=CAMK2D;refGene_tx=XM_019829023.
2
B1 112147584 112147626
refGene_gn=CAMK2D;refGene_tx=XM_006930898.4;refGene_gn=CAMK2D;refGene_
tx=XM_006930899.4;refGene_gn=CAMK2D;refGene_tx=XM_006930900.4;refGene_
gn=CAMK2D;refGene_tx=XM_006930901.4;refGene_gn=CAMK2D;refGene_tx=XM_01
9829018.2;refGene_gn=CAMK2D;refGene_tx=XM_019829019.2;refGene_gn=CAMK2
D;refGene_tx=XM_019829020.2;refGene_gn=CAMK2D;refGene_tx=XM_019829021.
2;refGene_gn=CAMK2D;refGene_tx=XM_019829022.2;refGene_gn=CAMK2D;refGen

e_tx=XM_019829023.2;refGene_gn=CAMK2D;refGene_tx=XM_019829024.2;refGen
e_gn=CAMK2D;refGene_tx=XM_019829025.2
B1 112149993 112150042
refGene_gn=CAMK2D;refGene_tx=XM_006930898.4;refGene_gn=CAMK2D;refGene_
tx=XM_006930899.4;refGene_gn=CAMK2D;refGene_tx=XM_006930900.4;refGene_
gn=CAMK2D;refGene_tx=XM_006930901.4;refGene_gn=CAMK2D;refGene_tx=XM_01
1281662.3;refGene_gn=CAMK2D;refGene_tx=XM_019829018.2;refGene_gn=CAMK2
D;refGene_tx=XM_019829019.2;refGene_gn=CAMK2D;refGene_tx=XM_019829020.
2;refGene_gn=CAMK2D;refGene_tx=XM_019829021.2;refGene_gn=CAMK2D;refGen
e_tx=XM_019829022.2;refGene_gn=CAMK2D;refGene_tx=XM_019829023.2;refGen
e_gn=CAMK2D;refGene_tx=XM_019829024.2;refGene_gn=CAMK2D;refGene_tx=XM_
019829025.2;refGene_gn=CAMK2D;refGene_tx=XM_019829026.2;refGene_gn=CAM
K2D;refGene_tx=XM_019829027.2;refGene_gn=CAMK2D;refGene_tx=XM_01982902
8.2;refGene_gn=CAMK2D;refGene_tx=XM_019829029.2;refGene_gn=CAMK2D;refG
ene_tx=XM_019829030.2;refGene_gn=CAMK2D;refGene_tx=XM_023252852.1
B1 112177425 112177501
refGene_gn=CAMK2D;refGene_tx=XM_006930898.4;refGene_gn=CAMK2D;refGene_
tx=XM_006930899.4;refGene_gn=CAMK2D;refGene_tx=XM_006930900.4;refGene_
gn=CAMK2D;refGene_tx=XM_006930901.4;refGene_gn=CAMK2D;refGene_tx=XM_01
1281662.3;refGene_gn=CAMK2D;refGene_tx=XM_019829018.2;refGene_gn=CAMK2
D;refGene_tx=XM_019829019.2;refGene_gn=CAMK2D;refGene_tx=XM_019829020.
2;refGene_gn=CAMK2D;refGene_tx=XM_019829021.2;refGene_gn=CAMK2D;refGen
e_tx=XM_019829022.2;refGene_gn=CAMK2D;refGene_tx=XM_019829023.2;refGen
e_gn=CAMK2D;refGene_tx=XM_019829024.2;refGene_gn=CAMK2D;refGene_tx=XM_
019829025.2;refGene_gn=CAMK2D;refGene_tx=XM_019829026.2;refGene_gn=CAM
K2D;refGene_tx=XM_019829027.2;refGene_gn=CAMK2D;refGene_tx=XM_01982902
8.2;refGene_gn=CAMK2D;refGene_tx=XM_019829029.2;refGene_gn=CAMK2D;refG
ene_tx=XM_019829030.2;refGene_gn=CAMK2D;refGene_tx=XM_023252852.1
B1 112182324 112182419
refGene_gn=CAMK2D;refGene_tx=XM_006930898.4;refGene_gn=CAMK2D;refGene_
tx=XM_006930899.4;refGene_gn=CAMK2D;refGene_tx=XM_006930900.4;refGene_
gn=CAMK2D;refGene_tx=XM_006930901.4;refGene_gn=CAMK2D;refGene_tx=XM_01
1281662.3;refGene_gn=CAMK2D;refGene_tx=XM_019829018.2;refGene_gn=CAMK2
D;refGene_tx=XM_019829019.2;refGene_gn=CAMK2D;refGene_tx=XM_019829020.
2;refGene_gn=CAMK2D;refGene_tx=XM_019829021.2;refGene_gn=CAMK2D;refGen
e_tx=XM_019829022.2;refGene_gn=CAMK2D;refGene_tx=XM_019829023.2;refGen
e_gn=CAMK2D;refGene_tx=XM_019829024.2;refGene_gn=CAMK2D;refGene_tx=XM_
019829025.2;refGene_gn=CAMK2D;refGene_tx=XM_019829026.2;refGene_gn=CAM
K2D;refGene_tx=XM_019829027.2;refGene_gn=CAMK2D;refGene_tx=XM_01982902
8.2;refGene_gn=CAMK2D;refGene_tx=XM_019829029.2;refGene_gn=CAMK2D;refG
ene_tx=XM_019829030.2;refGene_gn=CAMK2D;refGene_tx=XM_023252852.1
B1 112184416 112184649
refGene_gn=CAMK2D;refGene_tx=XM_006930898.4;refGene_gn=CAMK2D;refGene_
tx=XM_006930899.4;refGene_gn=CAMK2D;refGene_tx=XM_006930900.4;refGene_
gn=CAMK2D;refGene_tx=XM_006930901.4;refGene_gn=CAMK2D;refGene_tx=XM_01
1281662.3;refGene_gn=CAMK2D;refGene_tx=XM_019829018.2;refGene_gn=CAMK2
D;refGene_tx=XM_019829019.2;refGene_gn=CAMK2D;refGene_tx=XM_019829020.
2;refGene_gn=CAMK2D;refGene_tx=XM_019829021.2;refGene_gn=CAMK2D;refGen
e_tx=XM_019829022.2;refGene_gn=CAMK2D;refGene_tx=XM_019829023.2;refGen
e_gn=CAMK2D;refGene_tx=XM_019829024.2;refGene_gn=CAMK2D;refGene_tx=XM_

019829025.2;refGene_gn=CAMK2D;refGene_tx=XM_019829026.2;refGene_gn=CAMK2D;refGene_tx=XM_019829027.2;refGene_gn=CAMK2D;refGene_tx=XM_019829028.2;refGene_gn=CAMK2D;refGene_tx=XM_019829029.2;refGene_gn=CAMK2D;refGene_tx=XM_019829030.2;refGene_gn=CAMK2D;refGene_tx=XM_023252852.1
B1 112186088 112186155
refGene_gn=CAMK2D;refGene_tx=XM_006930898.4;refGene_gn=CAMK2D;refGene_tx=XM_006930899.4;refGene_gn=CAMK2D;refGene_tx=XM_006930900.4;refGene_gn=CAMK2D;refGene_tx=XM_006930901.4;refGene_gn=CAMK2D;refGene_tx=XM_011281662.3;refGene_gn=CAMK2D;refGene_tx=XM_019829018.2;refGene_gn=CAMK2D;refGene_tx=XM_019829019.2;refGene_gn=CAMK2D;refGene_tx=XM_019829020.2;refGene_gn=CAMK2D;refGene_tx=XM_019829021.2;refGene_gn=CAMK2D;refGene_tx=XM_019829022.2;refGene_gn=CAMK2D;refGene_tx=XM_019829024.2;refGene_gn=CAMK2D;refGene_tx=XM_019829025.2;refGene_gn=CAMK2D;refGene_tx=XM_019829026.2;refGene_gn=CAMK2D;refGene_tx=XM_019829027.2;refGene_gn=CAMK2D;refGene_tx=XM_019829028.2;refGene_gn=CAMK2D;refGene_tx=XM_019829030.2
B1 112187346 112187350
refGene_gn=CAMK2D;refGene_tx=XM_019829023.2
B1 112286487 112286671
refGene_gn=ANK2;refGene_tx=XM_019829050.2
B1 112289644 112289736
refGene_gn=ANK2;refGene_tx=XM_019829050.2
B1 112291225 112291408
refGene_gn=ANK2;refGene_tx=XM_019829050.2
B1 112292682 112292847
refGene_gn=ANK2;refGene_tx=XM_019829050.2
B1 112292950 112293034
refGene_gn=ANK2;refGene_tx=XM_019829050.2
B1 112295980 112296269
refGene_gn=ANK2;refGene_tx=XM_019829050.2
B1 112297890 112298176
refGene_gn=ANK2;refGene_tx=XM_019829050.2
B1 112301260 112301404
refGene_gn=ANK2;refGene_tx=XM_019829050.2
B1 112302346 112302478
refGene_gn=ANK2;refGene_tx=XM_019829050.2
B1 112305659 112305734
refGene_gn=ANK2;refGene_tx=XM_019829050.2
B1 112307290 112312384
refGene_gn=ANK2;refGene_tx=XM_019829050.2
B1 112312386 112313538
refGene_gn=ANK2;refGene_tx=XM_019829050.2
B1 112316561 112316583
refGene_gn=ANK2;refGene_tx=XM_019829050.2
B1 112318775 112318808
refGene_gn=ANK2;refGene_tx=XM_019829050.2
B1 112320459 112320582
refGene_gn=ANK2;refGene_tx=XM_019829050.2
B1 112323065 112323191
refGene_gn=ANK2;refGene_tx=XM_019829050.2

B1 112323992 112324221
refGene_gn=ANK2;refGene_tx=XM_019829050.2
B1 112326442 112326539
refGene_gn=ANK2;refGene_tx=XM_019829050.2
B1 112329571 112329776
refGene_gn=ANK2;refGene_tx=XM_019829050.2
B1 112330352 112330564
refGene_gn=ANK2;refGene_tx=XM_019829050.2
B1 112332806 112332961
refGene_gn=ANK2;refGene_tx=XM_019829050.2
B1 112333954 112334053
refGene_gn=ANK2;refGene_tx=XM_019829050.2
B1 112335580 112335805
refGene_gn=ANK2;refGene_tx=XM_019829050.2
B1 112347276 112347380
refGene_gn=ANK2;refGene_tx=XM_019829050.2
B1 112348010 112348113
refGene_gn=ANK2;refGene_tx=XM_019829050.2
B1 112354979 112355124
refGene_gn=ANK2;refGene_tx=XM_019829050.2
B1 112362598 112362671
refGene_gn=ANK2;refGene_tx=XM_019829050.2
B1 112372015 112372114
refGene_gn=ANK2;refGene_tx=XM_019829050.2
B1 112373050 112373149
refGene_gn=ANK2;refGene_tx=XM_019829050.2
B1 112378541 112378640
refGene_gn=ANK2;refGene_tx=XM_019829050.2
B1 112379350 112379449
refGene_gn=ANK2;refGene_tx=XM_019829050.2
B1 112384687 112384885
refGene_gn=ANK2;refGene_tx=XM_019829050.2
B1 112389908 112390007
refGene_gn=ANK2;refGene_tx=XM_019829050.2
B1 112390620 112390719
refGene_gn=ANK2;refGene_tx=XM_019829050.2
B1 112393618 112393816
refGene_gn=ANK2;refGene_tx=XM_019829050.2
B1 112400038 112400137
refGene_gn=ANK2;refGene_tx=XM_019829050.2
B1 112405594 112405693
refGene_gn=ANK2;refGene_tx=XM_019829050.2
B1 112405859 112405958
refGene_gn=ANK2;refGene_tx=XM_019829050.2
B1 112408137 112408335
refGene_gn=ANK2;refGene_tx=XM_019829050.2
B1 112412826 112412925
refGene_gn=ANK2;refGene_tx=XM_019829050.2
B1 112419774 112419873
refGene_gn=ANK2;refGene_tx=XM_019829050.2

B1	112421113	112421212
refGene_gn=ANK2;refGene_tx=XM_019829050.2
B1	112424366	112424390
refGene_gn=ANK2;refGene_tx=XM_019829050.2
B1	112424835	112425021
refGene_gn=ANK2;refGene_tx=XM_019829050.2
B1	112429880	112429979
refGene_gn=ANK2;refGene_tx=XM_019829050.2
B1	112458564	112458663
refGene_gn=ANK2;refGene_tx=XM_019829050.2
B1	112460052	112460151
refGene_gn=ANK2;refGene_tx=XM_019829050.2
B1	112496472	112496574
refGene_gn=ANK2;refGene_tx=XM_019829050.2
B1	112627233	112627317
refGene_gn=ANK2;refGene_tx=XM_019829050.2
B1	113013849	113013930
refGene_gn=LARP7;refGene_tx=XM_023252853.1;refGene_gn=LARP7;refGene_tx=XM_023252854.1;refGene_gn=LARP7;refGene_tx=XM_023252855.1
B1	113017664	113017756
refGene_gn=LARP7;refGene_tx=XM_023252853.1;refGene_gn=LARP7;refGene_tx=XM_023252854.1;refGene_gn=LARP7;refGene_tx=XM_023252855.1
B1	113017967	113018127
refGene_gn=LARP7;refGene_tx=XM_023252853.1;refGene_gn=LARP7;refGene_tx=XM_023252854.1;refGene_gn=LARP7;refGene_tx=XM_023252855.1
B1	113021226	113021345
refGene_gn=LARP7;refGene_tx=XM_023252853.1;refGene_gn=LARP7;refGene_tx=XM_023252854.1;refGene_gn=LARP7;refGene_tx=XM_023252855.1
B1	113022133	113022282
refGene_gn=LARP7;refGene_tx=XM_023252853.1;refGene_gn=LARP7;refGene_tx=XM_023252854.1;refGene_gn=LARP7;refGene_tx=XM_023252855.1
B1	113023959	113024104
refGene_gn=LARP7;refGene_tx=XM_023252853.1;refGene_gn=LARP7;refGene_tx=XM_023252854.1;refGene_gn=LARP7;refGene_tx=XM_023252855.1
B1	113024199	113024553
refGene_gn=LARP7;refGene_tx=XM_023252853.1;refGene_gn=LARP7;refGene_tx=XM_023252854.1;refGene_gn=LARP7;refGene_tx=XM_023252855.1
B1	113024623	113024717
refGene_gn=LARP7;refGene_tx=XM_023252853.1;refGene_gn=LARP7;refGene_tx=XM_023252854.1;refGene_gn=LARP7;refGene_tx=XM_023252855.1
B1	113024797	113024962
refGene_gn=LARP7;refGene_tx=XM_023252853.1;refGene_gn=LARP7;refGene_tx=XM_023252854.1;refGene_gn=LARP7;refGene_tx=XM_023252855.1
B1	113025102	113025186
refGene_gn=LARP7;refGene_tx=XM_023252853.1;refGene_gn=LARP7;refGene_tx=XM_023252854.1;refGene_gn=LARP7;refGene_tx=XM_023252855.1
B1	113025255	113025356
refGene_gn=LARP7;refGene_tx=XM_023252853.1;refGene_gn=LARP7;refGene_tx=XM_023252854.1;refGene_gn=LARP7;refGene_tx=XM_023252855.1
B1	113025923	113026125

refGene_gn=LARP7;refGene_tx=XM_023252853.1;refGene_gn=LARP7;refGene_tx=XM_023252854.1;refGene_gn=LARP7;refGene_tx=XM_023252855.1
B1 113039203 113039224
refGene_gn=ZGRF1;refGene_tx=XM_023252856.1
B1 113040873 113040954
refGene_gn=ZGRF1;refGene_tx=XM_023252856.1
B1 113049817 113049877
refGene_gn=ZGRF1;refGene_tx=XM_023252856.1
B1 113052101 113052287
refGene_gn=ZGRF1;refGene_tx=XM_023252856.1
B1 113052604 113054879
refGene_gn=ZGRF1;refGene_tx=XM_023252856.1
B1 113060785 113060859
refGene_gn=ZGRF1;refGene_tx=XM_023252856.1
B1 113064625 113064676
refGene_gn=ZGRF1;refGene_tx=XM_023252856.1
B1 113067755 113067839
refGene_gn=ZGRF1;refGene_tx=XM_023252856.1
B1 113069863 113070049
refGene_gn=ZGRF1;refGene_tx=XM_023252856.1
B1 113079769 113079923
refGene_gn=ZGRF1;refGene_tx=XM_023252856.1
B1 113083369 113084013
refGene_gn=ZGRF1;refGene_tx=XM_023252856.1
B1 113085151 113085290
refGene_gn=ZGRF1;refGene_tx=XM_023252856.1
B1 113086076 113086261
refGene_gn=ZGRF1;refGene_tx=XM_023252856.1
B1 113087423 113087620
refGene_gn=ZGRF1;refGene_tx=XM_023252856.1
B1 113089339 113089479
refGene_gn=ZGRF1;refGene_tx=XM_023252856.1
B1 113097337 113097481
refGene_gn=ZGRF1;refGene_tx=XM_023252856.1
B1 113099012 113099127
refGene_gn=ZGRF1;refGene_tx=XM_023252856.1
B1 113101516 113101776
refGene_gn=ZGRF1;refGene_tx=XM_023252856.1
B1 113103569 113103729
refGene_gn=ZGRF1;refGene_tx=XM_023252856.1
B1 113104982 113105060
refGene_gn=ZGRF1;refGene_tx=XM_023252856.1
B1 113105506 113105654
refGene_gn=ZGRF1;refGene_tx=XM_023252856.1
B1 113109243 113109371
refGene_gn=ZGRF1;refGene_tx=XM_023252856.1
B1 113111571 113111695
refGene_gn=ZGRF1;refGene_tx=XM_023252856.1
B1 113122368 113122545
refGene_gn=ZGRF1;refGene_tx=XM_023252856.1

B1	113122677	113122812
refGene_gn=ZGRF1;refGene_tx=XM_023252856.1
B1	113123506	113123768
refGene_gn=ZGRF1;refGene_tx=XM_023252856.1
B1	113123920	113124066
refGene_gn=ZGRF1;refGene_tx=XM_023252856.1
B1	113151792	113152611
refGene_gn=NEUROG2;refGene_tx=XM_023252857.1
B1	113196200	113196400
refGene_gn=ALPK1;refGene_tx=XM_019829070.1;refGene_gn=ALPK1;refGene_tx=XM_019829071.1;refGene_gn=ALPK1;refGene_tx=XM_019829072.1;refGene_gn=ALPK1;refGene_tx=XM_019829073.2
B1	113199160	113199347
refGene_gn=ALPK1;refGene_tx=XM_019829070.1;refGene_gn=ALPK1;refGene_tx=XM_019829071.1;refGene_gn=ALPK1;refGene_tx=XM_019829072.1;refGene_gn=ALPK1;refGene_tx=XM_019829073.2
B1	113200565	113200728
refGene_gn=ALPK1;refGene_tx=XM_019829070.1;refGene_gn=ALPK1;refGene_tx=XM_019829071.1;refGene_gn=ALPK1;refGene_tx=XM_019829072.1;refGene_gn=ALPK1;refGene_tx=XM_019829073.2
B1	113205101	113205255
refGene_gn=ALPK1;refGene_tx=XM_019829070.1;refGene_gn=ALPK1;refGene_tx=XM_019829071.1;refGene_gn=ALPK1;refGene_tx=XM_019829072.1;refGene_gn=ALPK1;refGene_tx=XM_019829073.2
B1	113207410	113209526
refGene_gn=ALPK1;refGene_tx=XM_019829070.1;refGene_gn=ALPK1;refGene_tx=XM_019829071.1;refGene_gn=ALPK1;refGene_tx=XM_019829072.1;refGene_gn=ALPK1;refGene_tx=XM_019829073.2
B1	113210466	113210571
refGene_gn=ALPK1;refGene_tx=XM_019829070.1;refGene_gn=ALPK1;refGene_tx=XM_019829071.1;refGene_gn=ALPK1;refGene_tx=XM_019829072.1;refGene_gn=ALPK1;refGene_tx=XM_019829073.2
B1	113212868	113212964
refGene_gn=ALPK1;refGene_tx=XM_019829070.1;refGene_gn=ALPK1;refGene_tx=XM_019829071.1;refGene_gn=ALPK1;refGene_tx=XM_019829072.1;refGene_gn=ALPK1;refGene_tx=XM_019829073.2
B1	113214072	113214149
refGene_gn=ALPK1;refGene_tx=XM_019829070.1;refGene_gn=ALPK1;refGene_tx=XM_019829071.1;refGene_gn=ALPK1;refGene_tx=XM_019829072.1;refGene_gn=ALPK1;refGene_tx=XM_019829073.2
B1	113215453	113215540
refGene_gn=ALPK1;refGene_tx=XM_019829070.1;refGene_gn=ALPK1;refGene_tx=XM_019829071.1;refGene_gn=ALPK1;refGene_tx=XM_019829072.1;refGene_gn=ALPK1;refGene_tx=XM_019829073.2
B1	113217226	113217286
refGene_gn=ALPK1;refGene_tx=XM_019829070.1;refGene_gn=ALPK1;refGene_tx=XM_019829071.1;refGene_gn=ALPK1;refGene_tx=XM_019829072.1;refGene_gn=ALPK1;refGene_tx=XM_019829073.2
B1	113226505	113226704
refGene_gn=ALPK1;refGene_tx=XM_019829070.1;refGene_gn=ALPK1;refGene_tx

=XM_019829071.1;refGene_gn=ALPK1;refGene_tx=XM_019829072.1;refGene_gn=ALPK1;refGene_tx=XM_019829073.2
B1 113257034 113257189
refGene_gn=ALPK1;refGene_tx=XM_019829070.1;refGene_gn=ALPK1;refGene_tx=XM_019829071.1;refGene_gn=ALPK1;refGene_tx=XM_019829072.1;refGene_gn=ALPK1;refGene_tx=XM_019829073.2
B1 113263115 113263236
refGene_gn=ALPK1;refGene_tx=XM_019829070.1;refGene_gn=ALPK1;refGene_tx=XM_019829071.1;refGene_gn=ALPK1;refGene_tx=XM_019829072.1;refGene_gn=ALPK1;refGene_tx=XM_019829073.2
B1 113339126 113339684
refGene_gn=TIFA;refGene_tx=XM_003985074.5;refGene_gn=TIFA;refGene_tx=XM_019829075.2;refGene_gn=TIFA;refGene_tx=XM_019829077.2
B1 113344576 113344842
refGene_gn=AP1AR;refGene_tx=XM_011281713.3;refGene_gn=AP1AR;refGene_tx=XM_019829078.2
B1 113346257 113346386
refGene_gn=AP1AR;refGene_tx=XM_011281713.3;refGene_gn=AP1AR;refGene_tx=XM_019829078.2
B1 113347632 113347706
refGene_gn=AP1AR;refGene_tx=XM_011281713.3;refGene_gn=AP1AR;refGene_tx=XM_019829078.2
B1 113348365 113348424
refGene_gn=AP1AR;refGene_tx=XM_011281713.3;refGene_gn=AP1AR;refGene_tx=XM_019829078.2
B1 113350682 113350781
refGene_gn=AP1AR;refGene_tx=XM_011281713.3
B1 113351975 113352072
refGene_gn=AP1AR;refGene_tx=XM_011281713.3;refGene_gn=AP1AR;refGene_tx=XM_019829078.2
B1 113354709 113354735
refGene_gn=AP1AR;refGene_tx=XM_011281713.3;refGene_gn=AP1AR;refGene_tx=XM_019829078.2
B1 113357495 113357522
refGene_gn=AP1AR;refGene_tx=XM_011281713.3;refGene_gn=AP1AR;refGene_tx=XM_019829078.2
B1 113359325 113359374
refGene_gn=AP1AR;refGene_tx=XM_011281713.3;refGene_gn=AP1AR;refGene_tx=XM_019829078.2
B1 113382771 113382854
refGene_gn=AP1AR;refGene_tx=XM_011281713.3;refGene_gn=AP1AR;refGene_tx=XM_019829078.2
B1 113437678 113437924
refGene_gn=FAM241A;refGene_tx=XM_023252859.1
B1 113476726 113476882
refGene_gn=FAM241A;refGene_tx=XM_023252859.1
B1 114664243 114664315
refGene_gn=LOC101086141;refGene_tx=XM_023252232.1
B1 114667165 114667720
refGene_gn=LOC101086141;refGene_tx=XM_023252232.1

B1 114850450 114850458
refGene_gn=PITX2;refGene_tx=XM_019829080.2
B1 114851103 114851159
refGene_gn=PITX2;refGene_tx=XM_003985077.3;refGene_gn=PITX2;refGene_tx=XM_003985078.4;refGene_gn=PITX2;refGene_tx=XM_019829080.2;refGene_gn=PITX2;refGene_tx=XM_019829081.1;refGene_gn=PITX2;refGene_tx=XM_019829082.1
B1 114851640 114851778
refGene_gn=PITX2;refGene_tx=XM_003985077.3;refGene_gn=PITX2;refGene_tx=XM_019829080.2;refGene_gn=PITX2;refGene_tx=XM_019829081.1
B1 114861883 114862091
refGene_gn=PITX2;refGene_tx=XM_003985075.3
B1 114862952 114863158
refGene_gn=PITX2;refGene_tx=XM_003985075.3;refGene_gn=PITX2;refGene_tx=XM_003985077.3;refGene_gn=PITX2;refGene_tx=XM_003985078.4;refGene_gn=PITX2;refGene_tx=XM_019829080.2;refGene_gn=PITX2;refGene_tx=XM_019829081.1;refGene_gn=PITX2;refGene_tx=XM_019829082.1
B1 114865635 114866199
refGene_gn=PITX2;refGene_tx=XM_003985075.3;refGene_gn=PITX2;refGene_tx=XM_003985077.3;refGene_gn=PITX2;refGene_tx=XM_003985078.4;refGene_gn=PITX2;refGene_tx=XM_019829080.2;refGene_gn=PITX2;refGene_tx=XM_019829081.1;refGene_gn=PITX2;refGene_tx=XM_019829082.1
B1 114919582 114919735
refGene_gn=ENPEP;refGene_tx=XM_003985081.5
B1 114921133 114921212
refGene_gn=ENPEP;refGene_tx=XM_003985081.5
B1 114928440 114928581
refGene_gn=ENPEP;refGene_tx=XM_003985081.5
B1 114930237 114930408
refGene_gn=ENPEP;refGene_tx=XM_003985081.5
B1 114930496 114930601
refGene_gn=ENPEP;refGene_tx=XM_003985081.5
B1 114930707 114930781
refGene_gn=ENPEP;refGene_tx=XM_003985081.5
B1 114931782 114931933
refGene_gn=ENPEP;refGene_tx=XM_003985081.5
B1 114939024 114939080
refGene_gn=ENPEP;refGene_tx=XM_003985081.5
B1 114939210 114939347
refGene_gn=ENPEP;refGene_tx=XM_003985081.5
B1 114949462 114949542
refGene_gn=ENPEP;refGene_tx=XM_003985081.5
B1 114960128 114960280
refGene_gn=ENPEP;refGene_tx=XM_003985081.5
B1 114960410 114960476
refGene_gn=ENPEP;refGene_tx=XM_003985081.5
B1 114966541 114966607
refGene_gn=ENPEP;refGene_tx=XM_003985081.5
B1 114968639 114968774
refGene_gn=ENPEP;refGene_tx=XM_003985081.5

B1	114972102	114972216
refGene_gn=ENPEP;refGene_tx=XM_003985081.5
B1	114972881	114973036
refGene_gn=ENPEP;refGene_tx=XM_003985081.5
B1	114975214	114975335
refGene_gn=ENPEP;refGene_tx=XM_003985081.5
B1	114990121	114990253
refGene_gn=ENPEP;refGene_tx=XM_003985081.5
B1	114992297	114992439
refGene_gn=ENPEP;refGene_tx=XM_003985081.5
B1	115005148	115005777
refGene_gn=ENPEP;refGene_tx=XM_003985081.5
B1	115199530	115199619
refGene_gn=ELOVL6;refGene_tx=XM_003985082.5;refGene_gn=ELOVL6;refGene_tx=XM_019829086.2;refGene_gn=ELOVL6;refGene_tx=XM_019829087.2;refGene_gn=ELOVL6;refGene_tx=XM_023252860.1
B1	115289415	115289547
refGene_gn=ELOVL6;refGene_tx=XM_003985082.5;refGene_gn=ELOVL6;refGene_tx=XM_019829086.2;refGene_gn=ELOVL6;refGene_tx=XM_019829087.2;refGene_gn=ELOVL6;refGene_tx=XM_019829088.2;refGene_gn=ELOVL6;refGene_tx=XM_023252860.1
B1	115334443	115334595
refGene_gn=ELOVL6;refGene_tx=XM_003985082.5;refGene_gn=ELOVL6;refGene_tx=XM_019829086.2;refGene_gn=ELOVL6;refGene_tx=XM_019829087.2;refGene_gn=ELOVL6;refGene_tx=XM_019829088.2;refGene_gn=ELOVL6;refGene_tx=XM_023252860.1
B1	115341779	115342204
refGene_gn=ELOVL6;refGene_tx=XM_003985082.5;refGene_gn=ELOVL6;refGene_tx=XM_019829086.2;refGene_gn=ELOVL6;refGene_tx=XM_019829087.2;refGene_gn=ELOVL6;refGene_tx=XM_019829088.2;refGene_gn=ELOVL6;refGene_tx=XM_023252860.1
B1	115393359	115393610
refGene_gn=EGF;refGene_tx=NM_001009381.1;refGene_gn=EGF;refGene_tx=XM_006930906.3;refGene_gn=EGF;refGene_tx=XM_006930907.3;refGene_gn=EGF;refGene_tx=XM_006930908.3;refGene_gn=EGF;refGene_tx=XM_023252214.1
B1	115395539	115395645
refGene_gn=EGF;refGene_tx=NM_001009381.1;refGene_gn=EGF;refGene_tx=XM_006930906.3;refGene_gn=EGF;refGene_tx=XM_006930907.3;refGene_gn=EGF;refGene_tx=XM_006930908.3;refGene_gn=EGF;refGene_tx=XM_023252214.1
B1	115398946	115399058
refGene_gn=EGF;refGene_tx=NM_001009381.1;refGene_gn=EGF;refGene_tx=XM_006930906.3;refGene_gn=EGF;refGene_tx=XM_006930907.3;refGene_gn=EGF;refGene_tx=XM_006930908.3;refGene_gn=EGF;refGene_tx=XM_023252214.1
B1	115403305	115403470
refGene_gn=EGF;refGene_tx=NM_001009381.1;refGene_gn=EGF;refGene_tx=XM_006930906.3;refGene_gn=EGF;refGene_tx=XM_006930907.3;refGene_gn=EGF;refGene_tx=XM_023252214.1
B1	115409467	115409612
refGene_gn=EGF;refGene_tx=NM_001009381.1;refGene_gn=EGF;refGene_tx=XM_006930906.3;refGene_gn=EGF;refGene_tx=XM_006930907.3;refGene_gn=EGF;re

fGene_tx=XM_006930908.3;refGene_gn=EGF;refGene_tx=XM_023252214.1
B1	115410966	115411089
refGene_gn=EGF;refGene_tx=NM_001009381.1;refGene_gn=EGF;refGene_tx=XM_006930906.3;refGene_gn=EGF;refGene_tx=XM_006930907.3;refGene_gn=EGF;refGene_tx=XM_006930908.3;refGene_gn=EGF;refGene_tx=XM_023252214.1
B1	115415834	115415960
refGene_gn=EGF;refGene_tx=NM_001009381.1;refGene_gn=EGF;refGene_tx=XM_006930906.3;refGene_gn=EGF;refGene_tx=XM_006930907.3;refGene_gn=EGF;refGene_tx=XM_006930908.3;refGene_gn=EGF;refGene_tx=XM_023252214.1
B1	115416604	115416721
refGene_gn=EGF;refGene_tx=NM_001009381.1;refGene_gn=EGF;refGene_tx=XM_006930906.3;refGene_gn=EGF;refGene_tx=XM_006930907.3;refGene_gn=EGF;refGene_tx=XM_006930908.3;refGene_gn=EGF;refGene_tx=XM_023252214.1
B1	115421603	115421726
refGene_gn=EGF;refGene_tx=NM_001009381.1;refGene_gn=EGF;refGene_tx=XM_006930906.3;refGene_gn=EGF;refGene_tx=XM_006930908.3;refGene_gn=EGF;refGene_tx=XM_023252214.1
B1	115424961	115425114
refGene_gn=EGF;refGene_tx=NM_001009381.1;refGene_gn=EGF;refGene_tx=XM_006930906.3;refGene_gn=EGF;refGene_tx=XM_006930907.3;refGene_gn=EGF;refGene_tx=XM_006930908.3;refGene_gn=EGF;refGene_tx=XM_023252214.1
B1	115426013	115426181
refGene_gn=EGF;refGene_tx=NM_001009381.1;refGene_gn=EGF;refGene_tx=XM_006930906.3;refGene_gn=EGF;refGene_tx=XM_006930907.3;refGene_gn=EGF;refGene_tx=XM_006930908.3;refGene_gn=EGF;refGene_tx=XM_023252214.1
B1	115429843	115430067
refGene_gn=EGF;refGene_tx=NM_001009381.1;refGene_gn=EGF;refGene_tx=XM_006930906.3;refGene_gn=EGF;refGene_tx=XM_006930907.3;refGene_gn=EGF;refGene_tx=XM_006930908.3;refGene_gn=EGF;refGene_tx=XM_023252214.1
B1	115431558	115431663
refGene_gn=EGF;refGene_tx=NM_001009381.1;refGene_gn=EGF;refGene_tx=XM_006930906.3;refGene_gn=EGF;refGene_tx=XM_006930907.3;refGene_gn=EGF;refGene_tx=XM_006930908.3;refGene_gn=EGF;refGene_tx=XM_023252214.1
B1	115436068	115436217
refGene_gn=EGF;refGene_tx=NM_001009381.1;refGene_gn=EGF;refGene_tx=XM_006930906.3;refGene_gn=EGF;refGene_tx=XM_006930907.3;refGene_gn=EGF;refGene_tx=XM_006930908.3;refGene_gn=EGF;refGene_tx=XM_023252214.1
B1	115438748	115438885
refGene_gn=EGF;refGene_tx=NM_001009381.1;refGene_gn=EGF;refGene_tx=XM_006930906.3;refGene_gn=EGF;refGene_tx=XM_006930907.3;refGene_gn=EGF;refGene_tx=XM_006930908.3;refGene_gn=EGF;refGene_tx=XM_023252214.1
B1	115439946	115440072
refGene_gn=EGF;refGene_tx=NM_001009381.1;refGene_gn=EGF;refGene_tx=XM_006930906.3;refGene_gn=EGF;refGene_tx=XM_006930907.3;refGene_gn=EGF;refGene_tx=XM_006930908.3;refGene_gn=EGF;refGene_tx=XM_023252214.1
B1	115440968	115441091
refGene_gn=EGF;refGene_tx=NM_001009381.1;refGene_gn=EGF;refGene_tx=XM_006930906.3;refGene_gn=EGF;refGene_tx=XM_006930907.3;refGene_gn=EGF;refGene_tx=XM_006930908.3;refGene_gn=EGF;refGene_tx=XM_023252214.1
B1	115442123	115442246

refGene_gn=EGF;refGene_tx=NM_001009381.1;refGene_gn=EGF;refGene_tx=XM_006930906.3;refGene_gn=EGF;refGene_tx=XM_006930907.3;refGene_gn=EGF;refGene_tx=XM_006930908.3;refGene_gn=EGF;refGene_tx=XM_023252214.1
B1 115443555 115443681
refGene_gn=EGF;refGene_tx=NM_001009381.1;refGene_gn=EGF;refGene_tx=XM_006930906.3;refGene_gn=EGF;refGene_tx=XM_006930907.3;refGene_gn=EGF;refGene_tx=XM_006930908.3;refGene_gn=EGF;refGene_tx=XM_023252214.1
B1 115459829 115460032
refGene_gn=EGF;refGene_tx=NM_001009381.1;refGene_gn=EGF;refGene_tx=XM_006930906.3;refGene_gn=EGF;refGene_tx=XM_006930907.3;refGene_gn=EGF;refGene_tx=XM_006930908.3;refGene_gn=EGF;refGene_tx=XM_023252214.1
B1 115460876 115461104
refGene_gn=EGF;refGene_tx=NM_001009381.1;refGene_gn=EGF;refGene_tx=XM_006930906.3;refGene_gn=EGF;refGene_tx=XM_006930907.3;refGene_gn=EGF;refGene_tx=XM_006930908.3;refGene_gn=EGF;refGene_tx=XM_023252214.1
B1 115461466 115461648
refGene_gn=EGF;refGene_tx=NM_001009381.1;refGene_gn=EGF;refGene_tx=XM_006930906.3;refGene_gn=EGF;refGene_tx=XM_006930907.3;refGene_gn=EGF;refGene_tx=XM_006930908.3;refGene_gn=EGF;refGene_tx=XM_023252214.1
B1 115464384 115464584
refGene_gn=EGF;refGene_tx=NM_001009381.1;refGene_gn=EGF;refGene_tx=XM_006930906.3;refGene_gn=EGF;refGene_tx=XM_006930907.3;refGene_gn=EGF;refGene_tx=XM_006930908.3;refGene_gn=EGF;refGene_tx=XM_023252214.1
B1 115489395 115489522
refGene_gn=EGF;refGene_tx=NM_001009381.1;refGene_gn=EGF;refGene_tx=XM_006930906.3;refGene_gn=EGF;refGene_tx=XM_006930907.3;refGene_gn=EGF;refGene_tx=XM_006930908.3
B1 115521560 115522717
refGene_gn=LRIT3;refGene_tx=XM_003985083.3
B1 115524080 115524386
refGene_gn=LRIT3;refGene_tx=XM_003985083.3
B1 115533413 115533886
refGene_gn=LRIT3;refGene_tx=XM_003985083.3
B1 115536812 115536931
refGene_gn=LRIT3;refGene_tx=XM_003985083.3
B1 115539044 115539159
refGene_gn=RRH;refGene_tx=XM_003985084.3
B1 115540819 115540998
refGene_gn=RRH;refGene_tx=XM_003985084.3
B1 115543948 115544044
refGene_gn=RRH;refGene_tx=XM_019829091.2
B1 115545325 115545494
refGene_gn=RRH;refGene_tx=XM_003985084.3;refGene_gn=RRH;refGene_tx=XM_019829091.2
B1 115546378 115546532
refGene_gn=RRH;refGene_tx=XM_003985084.3;refGene_gn=RRH;refGene_tx=XM_019829091.2
B1 115547466 115547566
refGene_gn=RRH;refGene_tx=XM_003985084.3;refGene_gn=RRH;refGene_tx=XM_019829091.2

B1 115549233 115549424
refGene_gn=RRH;refGene_tx=XM_003985084.3;refGene_gn=RRH;refGene_tx=XM_019829091.2
B1 115555175 115555281
refGene_gn=RRH;refGene_tx=XM_003985084.3;refGene_gn=RRH;refGene_tx=XM_019829091.2
B1 115558065 115558079
refGene_gn=GAR1;refGene_tx=XM_003985085.5;refGene_gn=GAR1;refGene_tx=XM_006930912.3;refGene_gn=GAR1;refGene_tx=XM_006930913.4;refGene_gn=GAR1;refGene_tx=XM_006930914.3;refGene_gn=GAR1;refGene_tx=XM_011281712.3;refGene_gn=GAR1;refGene_tx=XM_019829092.2;refGene_gn=GAR1;refGene_tx=XM_019829093.2
B1 115558412 115558478
refGene_gn=GAR1;refGene_tx=XM_003985085.5;refGene_gn=GAR1;refGene_tx=XM_006930912.3;refGene_gn=GAR1;refGene_tx=XM_006930913.4;refGene_gn=GAR1;refGene_tx=XM_006930914.3;refGene_gn=GAR1;refGene_tx=XM_011281712.3;refGene_gn=GAR1;refGene_tx=XM_019829092.2;refGene_gn=GAR1;refGene_tx=XM_019829093.2
B1 115559930 115560069
refGene_gn=GAR1;refGene_tx=XM_003985085.5;refGene_gn=GAR1;refGene_tx=XM_006930912.3;refGene_gn=GAR1;refGene_tx=XM_006930913.4;refGene_gn=GAR1;refGene_tx=XM_006930914.3;refGene_gn=GAR1;refGene_tx=XM_011281712.3;refGene_gn=GAR1;refGene_tx=XM_019829092.2;refGene_gn=GAR1;refGene_tx=XM_019829093.2
B1 115563325 115563385
refGene_gn=GAR1;refGene_tx=XM_003985085.5;refGene_gn=GAR1;refGene_tx=XM_006930912.3;refGene_gn=GAR1;refGene_tx=XM_006930913.4;refGene_gn=GAR1;refGene_tx=XM_006930914.3;refGene_gn=GAR1;refGene_tx=XM_011281712.3;refGene_gn=GAR1;refGene_tx=XM_019829092.2;refGene_gn=GAR1;refGene_tx=XM_019829093.2
B1 115563815 115563970
refGene_gn=GAR1;refGene_tx=XM_003985085.5;refGene_gn=GAR1;refGene_tx=XM_006930912.3;refGene_gn=GAR1;refGene_tx=XM_006930913.4;refGene_gn=GAR1;refGene_tx=XM_006930914.3;refGene_gn=GAR1;refGene_tx=XM_011281712.3;refGene_gn=GAR1;refGene_tx=XM_019829092.2;refGene_gn=GAR1;refGene_tx=XM_019829093.2
B1 115565414 115565621
refGene_gn=GAR1;refGene_tx=XM_003985085.5;refGene_gn=GAR1;refGene_tx=XM_006930912.3;refGene_gn=GAR1;refGene_tx=XM_006930913.4;refGene_gn=GAR1;refGene_tx=XM_006930914.3;refGene_gn=GAR1;refGene_tx=XM_011281712.3;refGene_gn=GAR1;refGene_tx=XM_019829092.2;refGene_gn=GAR1;refGene_tx=XM_019829093.2
B1 115566299 115566717
refGene_gn=GAR1;refGene_tx=XM_019829093.2
B1 115585877 115585889
refGene_gn=CFI;refGene_tx=XM_019829089.1
B1 115587271 115587328
refGene_gn=CFI;refGene_tx=XM_006930915.4;refGene_gn=CFI;refGene_tx=XM_011281710.3
B1 115602235 115602277

refGene_gn=CFI;refGene_tx=XM_006930915.4;refGene_gn=CFI;refGene_tx=XM_011281710.3;refGene_gn=CFI;refGene_tx=XM_019829089.1
B1 115611628 115611908
refGene_gn=CFI;refGene_tx=XM_006930915.4;refGene_gn=CFI;refGene_tx=XM_011281710.3;refGene_gn=CFI;refGene_tx=XM_019829089.1
B1 115613701 115613855
refGene_gn=CFI;refGene_tx=XM_006930915.4;refGene_gn=CFI;refGene_tx=XM_011281710.3;refGene_gn=CFI;refGene_tx=XM_019829089.1
B1 115617120 115617302
refGene_gn=CFI;refGene_tx=XM_006930915.4;refGene_gn=CFI;refGene_tx=XM_011281710.3;refGene_gn=CFI;refGene_tx=XM_019829089.1
B1 115618164 115618278
refGene_gn=CFI;refGene_tx=XM_006930915.4;refGene_gn=CFI;refGene_tx=XM_011281710.3;refGene_gn=CFI;refGene_tx=XM_019829089.1
B1 115618378 115618489
refGene_gn=CFI;refGene_tx=XM_006930915.4;refGene_gn=CFI;refGene_tx=XM_011281710.3;refGene_gn=CFI;refGene_tx=XM_019829089.1
B1 115625334 115625382
refGene_gn=CFI;refGene_tx=XM_006930915.4;refGene_gn=CFI;refGene_tx=XM_019829089.1
B1 115628654 115628678
refGene_gn=CFI;refGene_tx=XM_006930915.4;refGene_gn=CFI;refGene_tx=XM_011281710.3;refGene_gn=CFI;refGene_tx=XM_019829089.1
B1 115631491 115631527
refGene_gn=CFI;refGene_tx=XM_006930915.4;refGene_gn=CFI;refGene_tx=XM_011281710.3;refGene_gn=CFI;refGene_tx=XM_019829089.1
B1 115634038 115634142
refGene_gn=CFI;refGene_tx=XM_006930915.4;refGene_gn=CFI;refGene_tx=XM_011281710.3;refGene_gn=CFI;refGene_tx=XM_019829089.1
B1 115634310 115634414
refGene_gn=CFI;refGene_tx=XM_006930915.4;refGene_gn=CFI;refGene_tx=XM_011281710.3;refGene_gn=CFI;refGene_tx=XM_019829089.1
B1 115637152 115637430
refGene_gn=CFI;refGene_tx=XM_006930915.4;refGene_gn=CFI;refGene_tx=XM_011281710.3;refGene_gn=CFI;refGene_tx=XM_019829089.1
B1 115640871 115640976
refGene_gn=CFI;refGene_tx=XM_006930915.4;refGene_gn=CFI;refGene_tx=XM_011281710.3;refGene_gn=CFI;refGene_tx=XM_019829089.1
B1 115642806 115643024
refGene_gn=CFI;refGene_tx=XM_006930915.4;refGene_gn=CFI;refGene_tx=XM_011281710.3;refGene_gn=CFI;refGene_tx=XM_019829089.1
B1 115660707 115660918
refGene_gn=PLA2G12A;refGene_tx=XM_006930916.3;refGene_gn=PLA2G12A;refGene_tx=XM_006930918.4
B1 115677077 115677154
refGene_gn=PLA2G12A;refGene_tx=XM_006930916.3;refGene_gn=PLA2G12A;refGene_tx=XM_006930918.4
B1 115678550 115678716
refGene_gn=PLA2G12A;refGene_tx=XM_006930916.3;refGene_gn=PLA2G12A;refGene_tx=XM_006930918.4

B1	115680800	115680919
refGene_gn=PLA2G12A;refGene_tx=XM_006930918.4
B1	115688473	115688564
refGene_gn=PLA2G12A;refGene_tx=XM_023252861.1
B1	115688762	115688954
refGene_gn=MCUB;refGene_tx=XM_023252865.1;refGene_gn=MCUB;refGene_tx=XM_023252866.1;refGene_gn=MCUB;refGene_tx=XM_023252867.1;refGene_gn=MCUB;refGene_tx=XM_023252868.1;refGene_gn=MCUB;refGene_tx=XM_023252869.1
B1	115694239	115694282
refGene_gn=PLA2G12A;refGene_tx=XM_003985087.5;refGene_gn=PLA2G12A;refGene_tx=XM_006930916.3;refGene_gn=PLA2G12A;refGene_tx=XM_019829095.1;refGene_gn=PLA2G12A;refGene_tx=XM_023252861.1;refGene_gn=PLA2G12A;refGene_tx=XM_023252862.1;refGene_gn=PLA2G12A;refGene_tx=XM_023252863.1
B1	115694784	115694931
refGene_gn=PLA2G12A;refGene_tx=XM_003985087.5;refGene_gn=PLA2G12A;refGene_tx=XM_006930916.3;refGene_gn=PLA2G12A;refGene_tx=XM_019829095.1;refGene_gn=PLA2G12A;refGene_tx=XM_023252861.1;refGene_gn=PLA2G12A;refGene_tx=XM_023252862.1;refGene_gn=PLA2G12A;refGene_tx=XM_023252863.1;refGene_gn=PLA2G12A;refGene_tx=XM_023252864.1
B1	115695554	115695631
refGene_gn=PLA2G12A;refGene_tx=XM_003985087.5;refGene_gn=PLA2G12A;refGene_tx=XM_006930916.3;refGene_gn=PLA2G12A;refGene_tx=XM_019829095.1;refGene_gn=PLA2G12A;refGene_tx=XM_023252861.1;refGene_gn=PLA2G12A;refGene_tx=XM_023252862.1;refGene_gn=PLA2G12A;refGene_tx=XM_023252863.1;refGene_gn=PLA2G12A;refGene_tx=XM_023252864.1
B1	115697613	115697789
refGene_gn=PLA2G12A;refGene_tx=XM_003985087.5;refGene_gn=PLA2G12A;refGene_tx=XM_006930916.3;refGene_gn=PLA2G12A;refGene_tx=XM_019829095.1;refGene_gn=PLA2G12A;refGene_tx=XM_023252861.1;refGene_gn=PLA2G12A;refGene_tx=XM_023252862.1;refGene_gn=PLA2G12A;refGene_tx=XM_023252863.1;refGene_gn=PLA2G12A;refGene_tx=XM_023252864.1
B1	115701089	115701249
refGene_gn=PLA2G12A;refGene_tx=XM_003985087.5;refGene_gn=PLA2G12A;refGene_tx=XM_006930916.3;refGene_gn=PLA2G12A;refGene_tx=XM_019829095.1;refGene_gn=PLA2G12A;refGene_tx=XM_023252861.1;refGene_gn=PLA2G12A;refGene_tx=XM_023252862.1;refGene_gn=PLA2G12A;refGene_tx=XM_023252863.1;refGene_gn=PLA2G12A;refGene_tx=XM_023252864.1
B1	115701880	115702119
refGene_gn=PLA2G12A;refGene_tx=XM_003985087.5;refGene_gn=PLA2G12A;refGene_tx=XM_006930916.3;refGene_gn=PLA2G12A;refGene_tx=XM_019829095.1;refGene_gn=PLA2G12A;refGene_tx=XM_023252861.1;refGene_gn=PLA2G12A;refGene_tx=XM_023252862.1;refGene_gn=PLA2G12A;refGene_tx=XM_023252863.1;refGene_gn=PLA2G12A;refGene_tx=XM_023252864.1
B1	115705711	115705828
refGene_gn=MCUB;refGene_tx=XM_023252865.1;refGene_gn=MCUB;refGene_tx=XM_023252866.1;refGene_gn=MCUB;refGene_tx=XM_023252867.1;refGene_gn=MCUB;refGene_tx=XM_023252868.1;refGene_gn=MCUB;refGene_tx=XM_023252869.1
B1	115706650	115706854
refGene_gn=MCUB;refGene_tx=XM_023252865.1;refGene_gn=MCUB;refGene_tx=XM_023252866.1;refGene_gn=MCUB;refGene_tx=XM_023252867.1;refGene_gn=MCU

B;refGene_tx=XM_023252868.1
B1 115708238 115708399
refGene_gn=MCUB;refGene_tx=XM_023252865.1;refGene_gn=MCUB;refGene_tx=XM_023252866.1;refGene_gn=MCUB;refGene_tx=XM_023252867.1;refGene_gn=MCUB;refGene_tx=XM_023252868.1;refGene_gn=MCUB;refGene_tx=XM_023252869.1
B1 115720893 115720998
refGene_gn=MCUB;refGene_tx=XM_023252865.1;refGene_gn=MCUB;refGene_tx=XM_023252866.1;refGene_gn=MCUB;refGene_tx=XM_023252867.1;refGene_gn=MCUB;refGene_tx=XM_023252868.1;refGene_gn=MCUB;refGene_tx=XM_023252869.1
B1 115724136 115724307
refGene_gn=MCUB;refGene_tx=XM_023252865.1;refGene_gn=MCUB;refGene_tx=XM_023252866.1;refGene_gn=MCUB;refGene_tx=XM_023252867.1;refGene_gn=MCUB;refGene_tx=XM_023252868.1;refGene_gn=MCUB;refGene_tx=XM_023252869.1
B1 115724787 115724863
refGene_gn=MCUB;refGene_tx=XM_023252865.1;refGene_gn=MCUB;refGene_tx=XM_023252866.1;refGene_gn=MCUB;refGene_tx=XM_023252867.1;refGene_gn=MCUB;refGene_tx=XM_023252868.1;refGene_gn=MCUB;refGene_tx=XM_023252869.1
B1 115736063 115736145
refGene_gn=MCUB;refGene_tx=XM_023252865.1;refGene_gn=MCUB;refGene_tx=XM_023252869.1
B1 115739613 115739646
refGene_gn=MCUB;refGene_tx=XM_023252868.1
B1 115742029 115742139
refGene_gn=MCUB;refGene_tx=XM_023252865.1;refGene_gn=MCUB;refGene_tx=XM_023252867.1;refGene_gn=MCUB;refGene_tx=XM_023252869.1
B1 115798051 115798153
refGene_gn=MCUB;refGene_tx=XM_023252866.1
B1 115818110 115818225
refGene_gn=SEC24B;refGene_tx=XM_023252870.1
B1 115818471 115818575
refGene_gn=SEC24B;refGene_tx=XM_023252870.1
B1 115821187 115821280
refGene_gn=SEC24B;refGene_tx=XM_023252870.1
B1 115822087 115822192
refGene_gn=SEC24B;refGene_tx=XM_023252870.1
B1 115825127 115825265
refGene_gn=SEC24B;refGene_tx=XM_023252870.1
B1 115826142 115826238
refGene_gn=SEC24B;refGene_tx=XM_023252870.1
B1 115826239 115826319
refGene_gn=SEC24B;refGene_tx=XM_023252870.1
B1 115827769 115827880
refGene_gn=SEC24B;refGene_tx=XM_023252870.1
B1 115828948 115829122
refGene_gn=SEC24B;refGene_tx=XM_023252870.1
B1 115830188 115830347
refGene_gn=SEC24B;refGene_tx=XM_023252870.1
B1 115830771 115830895
refGene_gn=SEC24B;refGene_tx=XM_023252870.1
B1 115833829 115834036

refGene_gn=SEC24B;refGene_tx=XM_023252870.1
B1 115834278 115834334
refGene_gn=SEC24B;refGene_tx=XM_023252870.1
B1 115834931 115835050
refGene_gn=SEC24B;refGene_tx=XM_023252870.1
B1 115837671 115837784
refGene_gn=SEC24B;refGene_tx=XM_023252870.1
B1 115839507 115839617
refGene_gn=SEC24B;refGene_tx=XM_023252870.1
B1 115840319 115840446
refGene_gn=SEC24B;refGene_tx=XM_023252870.1
B1 115843361 115843464
refGene_gn=SEC24B;refGene_tx=XM_023252870.1
B1 115846232 115846417
refGene_gn=SEC24B;refGene_tx=XM_023252870.1
B1 115854885 115855109
refGene_gn=SEC24B;refGene_tx=XM_023252870.1
B1 115858921 115859002
refGene_gn=SEC24B;refGene_tx=XM_023252870.1
B1 115873483 115873588
refGene_gn=SEC24B;refGene_tx=XM_023252870.1
B1 115883504 115883678
refGene_gn=SEC24B;refGene_tx=XM_023252870.1
B1 115894485 115895235
refGene_gn=SEC24B;refGene_tx=XM_023252870.1
B1 115918493 115918620
refGene_gn=SEC24B;refGene_tx=XM_023252870.1
B1 116014227 116014515
refGene_gn=COL25A1;refGene_tx=XM_019829100.2;refGene_gn=COL25A1;refGene_tx=XM_019829102.2;refGene_gn=COL25A1;refGene_tx=XM_019829103.2;refGene_gn=COL25A1;refGene_tx=XM_019829104.2;refGene_gn=COL25A1;refGene_tx=XM_019829105.2;refGene_gn=COL25A1;refGene_tx=XM_019829106.2;refGene_gn=COL25A1;refGene_tx=XM_019829107.2;refGene_gn=COL25A1;refGene_tx=XM_019829108.2;refGene_gn=COL25A1;refGene_tx=XM_019829109.2;refGene_gn=COL25A1;refGene_tx=XM_019829111.2;refGene_gn=COL25A1;refGene_tx=XM_019829113.2;refGene_gn=COL25A1;refGene_tx=XM_019829114.2;refGene_gn=COL25A1;refGene_tx=XM_019829115.2;refGene_gn=COL25A1;refGene_tx=XM_019829116.2;refGene_gn=COL25A1;refGene_tx=XM_019829117.2;refGene_gn=COL25A1;refGene_tx=XM_019829118.2;refGene_gn=COL25A1;refGene_tx=XM_019829119.2
B1 116015606 116015676
refGene_gn=COL25A1;refGene_tx=XM_019829100.2;refGene_gn=COL25A1;refGene_tx=XM_019829102.2;refGene_gn=COL25A1;refGene_tx=XM_019829103.2;refGene_gn=COL25A1;refGene_tx=XM_019829104.2;refGene_gn=COL25A1;refGene_tx=XM_019829105.2;refGene_gn=COL25A1;refGene_tx=XM_019829106.2;refGene_gn=COL25A1;refGene_tx=XM_019829107.2;refGene_gn=COL25A1;refGene_tx=XM_019829108.2;refGene_gn=COL25A1;refGene_tx=XM_019829109.2;refGene_gn=COL25A1;refGene_tx=XM_019829111.2;refGene_gn=COL25A1;refGene_tx=XM_019829113.2;refGene_gn=COL25A1;refGene_tx=XM_019829114.2;refGene_gn=COL25A1;refGene_tx=XM_019829115.2;refGene_gn=COL25A1;refGene_tx=XM_019829116.2;refGene_gn=COL25A1;refGene_tx=XM_019829117.2;refGene_gn=COL25A1;refGen

e_tx=XM_019829118.2;refGene_gn=COL25A1;refGene_tx=XM_019829119.2
B1 116263354 116263399
refGene_gn=COL25A1;refGene_tx=XM_019829100.2;refGene_gn=COL25A1;refGene_tx=XM_019829102.2;refGene_gn=COL25A1;refGene_tx=XM_019829103.2;refGene_gn=COL25A1;refGene_tx=XM_019829104.2;refGene_gn=COL25A1;refGene_tx=XM_019829105.2;refGene_gn=COL25A1;refGene_tx=XM_019829106.2;refGene_gn=COL25A1;refGene_tx=XM_019829107.2;refGene_gn=COL25A1;refGene_tx=XM_019829108.2;refGene_gn=COL25A1;refGene_tx=XM_019829109.2;refGene_gn=COL25A1;refGene_tx=XM_019829111.2;refGene_gn=COL25A1;refGene_tx=XM_019829113.2;refGene_gn=COL25A1;refGene_tx=XM_019829114.2;refGene_gn=COL25A1;refGene_tx=XM_019829115.2;refGene_gn=COL25A1;refGene_tx=XM_019829116.2;refGene_gn=COL25A1;refGene_tx=XM_019829117.2;refGene_gn=COL25A1;refGene_tx=XM_019829118.2;refGene_gn=COL25A1;refGene_tx=XM_019829119.2
B1 116265450 116265458
refGene_gn=COL25A1;refGene_tx=XM_019829100.2;refGene_gn=COL25A1;refGene_tx=XM_019829102.2;refGene_gn=COL25A1;refGene_tx=XM_019829103.2;refGene_gn=COL25A1;refGene_tx=XM_019829104.2;refGene_gn=COL25A1;refGene_tx=XM_019829105.2;refGene_gn=COL25A1;refGene_tx=XM_019829106.2;refGene_gn=COL25A1;refGene_tx=XM_019829107.2;refGene_gn=COL25A1;refGene_tx=XM_019829108.2;refGene_gn=COL25A1;refGene_tx=XM_019829109.2;refGene_gn=COL25A1;refGene_tx=XM_019829111.2;refGene_gn=COL25A1;refGene_tx=XM_019829113.2;refGene_gn=COL25A1;refGene_tx=XM_019829114.2;refGene_gn=COL25A1;refGene_tx=XM_019829115.2;refGene_gn=COL25A1;refGene_tx=XM_019829116.2;refGene_gn=COL25A1;refGene_tx=XM_019829117.2;refGene_gn=COL25A1;refGene_tx=XM_019829118.2;refGene_gn=COL25A1;refGene_tx=XM_019829119.2
B1 116301937 116301955
refGene_gn=COL25A1;refGene_tx=XM_019829100.2;refGene_gn=COL25A1;refGene_tx=XM_019829102.2;refGene_gn=COL25A1;refGene_tx=XM_019829103.2;refGene_gn=COL25A1;refGene_tx=XM_019829105.2;refGene_gn=COL25A1;refGene_tx=XM_019829106.2;refGene_gn=COL25A1;refGene_tx=XM_019829108.2;refGene_gn=COL25A1;refGene_tx=XM_019829109.2;refGene_gn=COL25A1;refGene_tx=XM_019829111.2;refGene_gn=COL25A1;refGene_tx=XM_019829113.2;refGene_gn=COL25A1;refGene_tx=XM_019829114.2;refGene_gn=COL25A1;refGene_tx=XM_019829115.2;refGene_gn=COL25A1;refGene_tx=XM_019829116.2;refGene_gn=COL25A1;refGene_tx=XM_019829117.2;refGene_gn=COL25A1;refGene_tx=XM_019829118.2;refGene_gn=COL25A1;refGene_tx=XM_019829119.2
B1 116326475 116326502
refGene_gn=COL25A1;refGene_tx=XM_019829100.2;refGene_gn=COL25A1;refGene_tx=XM_019829102.2;refGene_gn=COL25A1;refGene_tx=XM_019829103.2;refGene_gn=COL25A1;refGene_tx=XM_019829104.2;refGene_gn=COL25A1;refGene_tx=XM_019829105.2;refGene_gn=COL25A1;refGene_tx=XM_019829106.2;refGene_gn=COL25A1;refGene_tx=XM_019829107.2;refGene_gn=COL25A1;refGene_tx=XM_019829108.2;refGene_gn=COL25A1;refGene_tx=XM_019829109.2;refGene_gn=COL25A1;refGene_tx=XM_019829111.2;refGene_gn=COL25A1;refGene_tx=XM_019829113.2;refGene_gn=COL25A1;refGene_tx=XM_019829114.2;refGene_gn=COL25A1;refGene_tx=XM_019829115.2;refGene_gn=COL25A1;refGene_tx=XM_019829116.2;refGene_gn=COL25A1;refGene_tx=XM_019829117.2;refGene_gn=COL25A1;refGene_tx=XM_019829118.2;refGene_gn=COL25A1;refGene_tx=XM_019829119.2
B1 116326648 116326675
refGene_gn=COL25A1;refGene_tx=XM_019829100.2;refGene_gn=COL25A1;refGen

e_tx=XM_019829102.2;refGene_gn=COL25A1;refGene_tx=XM_019829103.2;refGene_gn=COL25A1;refGene_tx=XM_019829104.2;refGene_gn=COL25A1;refGene_tx=XM_019829105.2;refGene_gn=COL25A1;refGene_tx=XM_019829106.2;refGene_gn=COL25A1;refGene_tx=XM_019829107.2;refGene_gn=COL25A1;refGene_tx=XM_019829108.2;refGene_gn=COL25A1;refGene_tx=XM_019829109.2;refGene_gn=COL25A1;refGene_tx=XM_019829111.2;refGene_gn=COL25A1;refGene_tx=XM_019829113.2;refGene_gn=COL25A1;refGene_tx=XM_019829114.2;refGene_gn=COL25A1;refGene_tx=XM_019829115.2;refGene_gn=COL25A1;refGene_tx=XM_019829116.2;refGene_gn=COL25A1;refGene_tx=XM_019829117.2;refGene_gn=COL25A1;refGene_tx=XM_019829118.2;refGene_gn=COL25A1;refGene_tx=XM_019829119.2
B1 116359889 116359949
refGene_gn=COL25A1;refGene_tx=XM_019829103.2;refGene_gn=COL25A1;refGene_tx=XM_019829107.2;refGene_gn=COL25A1;refGene_tx=XM_019829115.2
B1 116360710 116360782
refGene_gn=COL25A1;refGene_tx=XM_019829100.2;refGene_gn=COL25A1;refGene_tx=XM_019829102.2;refGene_gn=COL25A1;refGene_tx=XM_019829104.2;refGene_gn=COL25A1;refGene_tx=XM_019829105.2;refGene_gn=COL25A1;refGene_tx=XM_019829106.2;refGene_gn=COL25A1;refGene_tx=XM_019829108.2;refGene_gn=COL25A1;refGene_tx=XM_019829109.2;refGene_gn=COL25A1;refGene_tx=XM_019829111.2;refGene_gn=COL25A1;refGene_tx=XM_019829113.2;refGene_gn=COL25A1;refGene_tx=XM_019829114.2;refGene_gn=COL25A1;refGene_tx=XM_019829116.2;refGene_gn=COL25A1;refGene_tx=XM_019829117.2;refGene_gn=COL25A1;refGene_tx=XM_019829118.2;refGene_gn=COL25A1;refGene_tx=XM_019829119.2
B1 116361516 116361624
refGene_gn=COL25A1;refGene_tx=XM_019829100.2;refGene_gn=COL25A1;refGene_tx=XM_019829102.2;refGene_gn=COL25A1;refGene_tx=XM_019829103.2;refGene_gn=COL25A1;refGene_tx=XM_019829104.2;refGene_gn=COL25A1;refGene_tx=XM_019829105.2;refGene_gn=COL25A1;refGene_tx=XM_019829106.2;refGene_gn=COL25A1;refGene_tx=XM_019829107.2;refGene_gn=COL25A1;refGene_tx=XM_019829108.2;refGene_gn=COL25A1;refGene_tx=XM_019829109.2;refGene_gn=COL25A1;refGene_tx=XM_019829111.2;refGene_gn=COL25A1;refGene_tx=XM_019829113.2;refGene_gn=COL25A1;refGene_tx=XM_019829114.2;refGene_gn=COL25A1;refGene_tx=XM_019829115.2;refGene_gn=COL25A1;refGene_tx=XM_019829116.2;refGene_gn=COL25A1;refGene_tx=XM_019829117.2;refGene_gn=COL25A1;refGene_tx=XM_019829118.2;refGene_gn=COL25A1;refGene_tx=XM_019829119.2
B1 116363968 116364004
refGene_gn=COL25A1;refGene_tx=XM_019829100.2;refGene_gn=COL25A1;refGene_tx=XM_019829102.2;refGene_gn=COL25A1;refGene_tx=XM_019829103.2;refGene_gn=COL25A1;refGene_tx=XM_019829104.2;refGene_gn=COL25A1;refGene_tx=XM_019829105.2;refGene_gn=COL25A1;refGene_tx=XM_019829106.2;refGene_gn=COL25A1;refGene_tx=XM_019829107.2;refGene_gn=COL25A1;refGene_tx=XM_019829108.2;refGene_gn=COL25A1;refGene_tx=XM_019829109.2;refGene_gn=COL25A1;refGene_tx=XM_019829111.2;refGene_gn=COL25A1;refGene_tx=XM_019829113.2;refGene_gn=COL25A1;refGene_tx=XM_019829114.2;refGene_gn=COL25A1;refGene_tx=XM_019829115.2;refGene_gn=COL25A1;refGene_tx=XM_019829116.2;refGene_gn=COL25A1;refGene_tx=XM_019829117.2;refGene_gn=COL25A1;refGene_tx=XM_019829118.2;refGene_gn=COL25A1;refGene_tx=XM_019829119.2
B1 116378893 116378920
refGene_gn=COL25A1;refGene_tx=XM_019829100.2;refGene_gn=COL25A1;refGene_tx=XM_019829102.2;refGene_gn=COL25A1;refGene_tx=XM_019829103.2;refGe

ne_gn=COL25A1;refGene_tx=XM_019829104.2;refGene_gn=COL25A1;refGene_tx=XM_019829105.2;refGene_gn=COL25A1;refGene_tx=XM_019829106.2;refGene_gn=COL25A1;refGene_tx=XM_019829107.2;refGene_gn=COL25A1;refGene_tx=XM_019829108.2;refGene_gn=COL25A1;refGene_tx=XM_019829109.2;refGene_gn=COL25A1;refGene_tx=XM_019829111.2;refGene_gn=COL25A1;refGene_tx=XM_019829113.2;refGene_gn=COL25A1;refGene_tx=XM_019829114.2;refGene_gn=COL25A1;refGene_tx=XM_019829115.2;refGene_gn=COL25A1;refGene_tx=XM_019829116.2;refGene_gn=COL25A1;refGene_tx=XM_019829117.2;refGene_gn=COL25A1;refGene_tx=XM_019829118.2;refGene_gn=COL25A1;refGene_tx=XM_019829119.2
B1 116381166 116381211
refGene_gn=COL25A1;refGene_tx=XM_019829100.2;refGene_gn=COL25A1;refGene_tx=XM_019829102.2;refGene_gn=COL25A1;refGene_tx=XM_019829103.2;refGene_gn=COL25A1;refGene_tx=XM_019829104.2;refGene_gn=COL25A1;refGene_tx=XM_019829105.2;refGene_gn=COL25A1;refGene_tx=XM_019829106.2;refGene_gn=COL25A1;refGene_tx=XM_019829107.2;refGene_gn=COL25A1;refGene_tx=XM_019829108.2;refGene_gn=COL25A1;refGene_tx=XM_019829109.2;refGene_gn=COL25A1;refGene_tx=XM_019829111.2;refGene_gn=COL25A1;refGene_tx=XM_019829113.2;refGene_gn=COL25A1;refGene_tx=XM_019829114.2;refGene_gn=COL25A1;refGene_tx=XM_019829115.2;refGene_gn=COL25A1;refGene_tx=XM_019829116.2;refGene_gn=COL25A1;refGene_tx=XM_019829117.2;refGene_gn=COL25A1;refGene_tx=XM_019829118.2;refGene_gn=COL25A1;refGene_tx=XM_019829119.2
B1 116397527 116397581
refGene_gn=COL25A1;refGene_tx=XM_019829100.2;refGene_gn=COL25A1;refGene_tx=XM_019829102.2;refGene_gn=COL25A1;refGene_tx=XM_019829103.2;refGene_gn=COL25A1;refGene_tx=XM_019829104.2;refGene_gn=COL25A1;refGene_tx=XM_019829105.2;refGene_gn=COL25A1;refGene_tx=XM_019829106.2;refGene_gn=COL25A1;refGene_tx=XM_019829107.2;refGene_gn=COL25A1;refGene_tx=XM_019829108.2;refGene_gn=COL25A1;refGene_tx=XM_019829109.2;refGene_gn=COL25A1;refGene_tx=XM_019829111.2;refGene_gn=COL25A1;refGene_tx=XM_019829113.2;refGene_gn=COL25A1;refGene_tx=XM_019829114.2;refGene_gn=COL25A1;refGene_tx=XM_019829115.2;refGene_gn=COL25A1;refGene_tx=XM_019829116.2;refGene_gn=COL25A1;refGene_tx=XM_019829117.2;refGene_gn=COL25A1;refGene_tx=XM_019829118.2;refGene_gn=COL25A1;refGene_tx=XM_019829119.2
B1 116399896 116399923
refGene_gn=COL25A1;refGene_tx=XM_019829100.2;refGene_gn=COL25A1;refGene_tx=XM_019829102.2;refGene_gn=COL25A1;refGene_tx=XM_019829103.2;refGene_gn=COL25A1;refGene_tx=XM_019829104.2;refGene_gn=COL25A1;refGene_tx=XM_019829105.2;refGene_gn=COL25A1;refGene_tx=XM_019829106.2;refGene_gn=COL25A1;refGene_tx=XM_019829107.2;refGene_gn=COL25A1;refGene_tx=XM_019829108.2;refGene_gn=COL25A1;refGene_tx=XM_019829109.2;refGene_gn=COL25A1;refGene_tx=XM_019829111.2;refGene_gn=COL25A1;refGene_tx=XM_019829113.2;refGene_gn=COL25A1;refGene_tx=XM_019829114.2;refGene_gn=COL25A1;refGene_tx=XM_019829115.2;refGene_gn=COL25A1;refGene_tx=XM_019829116.2;refGene_gn=COL25A1;refGene_tx=XM_019829117.2;refGene_gn=COL25A1;refGene_tx=XM_019829118.2;refGene_gn=COL25A1;refGene_tx=XM_019829119.2
B1 116402319 116402364
refGene_gn=COL25A1;refGene_tx=XM_019829100.2;refGene_gn=COL25A1;refGene_tx=XM_019829102.2;refGene_gn=COL25A1;refGene_tx=XM_019829103.2;refGene_gn=COL25A1;refGene_tx=XM_019829104.2;refGene_gn=COL25A1;refGene_tx=XM_019829105.2;refGene_gn=COL25A1;refGene_tx=XM_019829106.2;refGene_gn

=COL25A1;refGene_tx=XM_019829107.2;refGene_gn=COL25A1;refGene_tx=XM_01
9829108.2;refGene_gn=COL25A1;refGene_tx=XM_019829109.2;refGene_gn=COL2
5A1;refGene_tx=XM_019829111.2;refGene_gn=COL25A1;refGene_tx=XM_0198291
13.2;refGene_gn=COL25A1;refGene_tx=XM_019829114.2;refGene_gn=COL25A1;r
efGene_tx=XM_019829115.2;refGene_gn=COL25A1;refGene_tx=XM_019829116.2;
refGene_gn=COL25A1;refGene_tx=XM_019829117.2;refGene_gn=COL25A1;refGen
e_tx=XM_019829118.2;refGene_gn=COL25A1;refGene_tx=XM_019829119.2
B1 116409721 116409754
refGene_gn=COL25A1;refGene_tx=XM_019829100.2;refGene_gn=COL25A1;refGen
e_tx=XM_019829102.2;refGene_gn=COL25A1;refGene_tx=XM_019829103.2;refGe
ne_gn=COL25A1;refGene_tx=XM_019829104.2;refGene_gn=COL25A1;refGene_tx=
XM_019829105.2;refGene_gn=COL25A1;refGene_tx=XM_019829106.2;refGene_gn
=COL25A1;refGene_tx=XM_019829107.2;refGene_gn=COL25A1;refGene_tx=XM_01
9829108.2;refGene_gn=COL25A1;refGene_tx=XM_019829109.2;refGene_gn=COL2
5A1;refGene_tx=XM_019829111.2;refGene_gn=COL25A1;refGene_tx=XM_0198291
13.2;refGene_gn=COL25A1;refGene_tx=XM_019829114.2;refGene_gn=COL25A1;r
efGene_tx=XM_019829115.2;refGene_gn=COL25A1;refGene_tx=XM_019829116.2;
refGene_gn=COL25A1;refGene_tx=XM_019829117.2;refGene_gn=COL25A1;refGen
e_tx=XM_019829118.2;refGene_gn=COL25A1;refGene_tx=XM_019829119.2
B1 116410199 116410235
refGene_gn=COL25A1;refGene_tx=XM_019829100.2;refGene_gn=COL25A1;refGen
e_tx=XM_019829102.2;refGene_gn=COL25A1;refGene_tx=XM_019829103.2;refGe
ne_gn=COL25A1;refGene_tx=XM_019829104.2;refGene_gn=COL25A1;refGene_tx=
XM_019829105.2;refGene_gn=COL25A1;refGene_tx=XM_019829106.2;refGene_gn
=COL25A1;refGene_tx=XM_019829107.2;refGene_gn=COL25A1;refGene_tx=XM_01
9829108.2;refGene_gn=COL25A1;refGene_tx=XM_019829109.2;refGene_gn=COL2
5A1;refGene_tx=XM_019829111.2;refGene_gn=COL25A1;refGene_tx=XM_0198291
13.2;refGene_gn=COL25A1;refGene_tx=XM_019829114.2;refGene_gn=COL25A1;r
efGene_tx=XM_019829115.2;refGene_gn=COL25A1;refGene_tx=XM_019829116.2;
refGene_gn=COL25A1;refGene_tx=XM_019829117.2;refGene_gn=COL25A1;refGen
e_tx=XM_019829118.2;refGene_gn=COL25A1;refGene_tx=XM_019829119.2
B1 116414972 116415017
refGene_gn=COL25A1;refGene_tx=XM_019829100.2;refGene_gn=COL25A1;refGen
e_tx=XM_019829102.2;refGene_gn=COL25A1;refGene_tx=XM_019829103.2;refGe
ne_gn=COL25A1;refGene_tx=XM_019829104.2;refGene_gn=COL25A1;refGene_tx=
XM_019829105.2;refGene_gn=COL25A1;refGene_tx=XM_019829106.2;refGene_gn
=COL25A1;refGene_tx=XM_019829107.2;refGene_gn=COL25A1;refGene_tx=XM_01
9829108.2;refGene_gn=COL25A1;refGene_tx=XM_019829109.2;refGene_gn=COL2
5A1;refGene_tx=XM_019829113.2;refGene_gn=COL25A1;refGene_tx=XM_0198291
14.2;refGene_gn=COL25A1;refGene_tx=XM_019829115.2;refGene_gn=COL25A1;r
efGene_tx=XM_019829116.2;refGene_gn=COL25A1;refGene_tx=XM_019829117.2;
refGene_gn=COL25A1;refGene_tx=XM_019829118.2
B1 116428481 116428544
refGene_gn=COL25A1;refGene_tx=XM_019829100.2;refGene_gn=COL25A1;refGen
e_tx=XM_019829102.2;refGene_gn=COL25A1;refGene_tx=XM_019829103.2;refGe
ne_gn=COL25A1;refGene_tx=XM_019829104.2;refGene_gn=COL25A1;refGene_tx=
XM_019829105.2;refGene_gn=COL25A1;refGene_tx=XM_019829106.2;refGene_gn
=COL25A1;refGene_tx=XM_019829107.2;refGene_gn=COL25A1;refGene_tx=XM_01
9829108.2;refGene_gn=COL25A1;refGene_tx=XM_019829109.2;refGene_gn=COL2
5A1;refGene_tx=XM_019829111.2;refGene_gn=COL25A1;refGene_tx=XM_0198291

13.2;refGene_gn=COL25A1;refGene_tx=XM_019829114.2;refGene_gn=COL25A1;refGene_tx=XM_019829115.2;refGene_gn=COL25A1;refGene_tx=XM_019829116.2;refGene_gn=COL25A1;refGene_tx=XM_019829117.2;refGene_gn=COL25A1;refGene_tx=XM_019829118.2;refGene_gn=COL25A1;refGene_tx=XM_019829119.2
B1 116432677 116432746
refGene_gn=COL25A1;refGene_tx=XM_019829100.2;refGene_gn=COL25A1;refGene_tx=XM_019829102.2;refGene_gn=COL25A1;refGene_tx=XM_019829103.2;refGene_gn=COL25A1;refGene_tx=XM_019829104.2;refGene_gn=COL25A1;refGene_tx=XM_019829105.2;refGene_gn=COL25A1;refGene_tx=XM_019829106.2;refGene_gn=COL25A1;refGene_tx=XM_019829107.2;refGene_gn=COL25A1;refGene_tx=XM_019829108.2;refGene_gn=COL25A1;refGene_tx=XM_019829109.2;refGene_gn=COL25A1;refGene_tx=XM_019829111.2;refGene_gn=COL25A1;refGene_tx=XM_019829113.2;refGene_gn=COL25A1;refGene_tx=XM_019829114.2;refGene_gn=COL25A1;refGene_tx=XM_019829115.2;refGene_gn=COL25A1;refGene_tx=XM_019829116.2;refGene_gn=COL25A1;refGene_tx=XM_019829117.2;refGene_gn=COL25A1;refGene_tx=XM_019829118.2;refGene_gn=COL25A1;refGene_tx=XM_019829119.2
B1 116433737 116433782
refGene_gn=COL25A1;refGene_tx=XM_019829100.2;refGene_gn=COL25A1;refGene_tx=XM_019829102.2;refGene_gn=COL25A1;refGene_tx=XM_019829103.2;refGene_gn=COL25A1;refGene_tx=XM_019829104.2;refGene_gn=COL25A1;refGene_tx=XM_019829105.2;refGene_gn=COL25A1;refGene_tx=XM_019829106.2;refGene_gn=COL25A1;refGene_tx=XM_019829107.2;refGene_gn=COL25A1;refGene_tx=XM_019829108.2;refGene_gn=COL25A1;refGene_tx=XM_019829111.2;refGene_gn=COL25A1;refGene_tx=XM_019829113.2;refGene_gn=COL25A1;refGene_tx=XM_019829114.2;refGene_gn=COL25A1;refGene_tx=XM_019829116.2;refGene_gn=COL25A1;refGene_tx=XM_019829117.2;refGene_gn=COL25A1;refGene_tx=XM_019829118.2;refGene_gn=COL25A1;refGene_tx=XM_019829119.2
B1 116435304 116435349
refGene_gn=COL25A1;refGene_tx=XM_019829100.2;refGene_gn=COL25A1;refGene_tx=XM_019829102.2;refGene_gn=COL25A1;refGene_tx=XM_019829103.2;refGene_gn=COL25A1;refGene_tx=XM_019829104.2;refGene_gn=COL25A1;refGene_tx=XM_019829105.2;refGene_gn=COL25A1;refGene_tx=XM_019829106.2;refGene_gn=COL25A1;refGene_tx=XM_019829107.2;refGene_gn=COL25A1;refGene_tx=XM_019829108.2;refGene_gn=COL25A1;refGene_tx=XM_019829109.2;refGene_gn=COL25A1;refGene_tx=XM_019829111.2;refGene_gn=COL25A1;refGene_tx=XM_019829113.2;refGene_gn=COL25A1;refGene_tx=XM_019829114.2;refGene_gn=COL25A1;refGene_tx=XM_019829115.2;refGene_gn=COL25A1;refGene_tx=XM_019829116.2;refGene_gn=COL25A1;refGene_tx=XM_019829117.2;refGene_gn=COL25A1;refGene_tx=XM_019829118.2;refGene_gn=COL25A1;refGene_tx=XM_019829119.2
B1 116436263 116436341
refGene_gn=COL25A1;refGene_tx=XM_019829100.2;refGene_gn=COL25A1;refGene_tx=XM_019829102.2;refGene_gn=COL25A1;refGene_tx=XM_019829103.2;refGene_gn=COL25A1;refGene_tx=XM_019829104.2;refGene_gn=COL25A1;refGene_tx=XM_019829105.2;refGene_gn=COL25A1;refGene_tx=XM_019829106.2;refGene_gn=COL25A1;refGene_tx=XM_019829107.2;refGene_gn=COL25A1;refGene_tx=XM_019829108.2;refGene_gn=COL25A1;refGene_tx=XM_019829109.2;refGene_gn=COL25A1;refGene_tx=XM_019829111.2;refGene_gn=COL25A1;refGene_tx=XM_019829113.2;refGene_gn=COL25A1;refGene_tx=XM_019829114.2;refGene_gn=COL25A1;refGene_tx=XM_019829115.2;refGene_gn=COL25A1;refGene_tx=XM_019829116.2;refGene_gn=COL25A1;refGene_tx=XM_019829117.2;refGene_gn=COL25A1;refGen

e_tx=XM_019829118.2;refGene_gn=COL25A1;refGene_tx=XM_019829119.2
B1 116442312 116442336
refGene_gn=COL25A1;refGene_tx=XM_019829100.2;refGene_gn=COL25A1;refGene_tx=XM_019829102.2;refGene_gn=COL25A1;refGene_tx=XM_019829103.2;refGene_gn=COL25A1;refGene_tx=XM_019829104.2;refGene_gn=COL25A1;refGene_tx=XM_019829106.2;refGene_gn=COL25A1;refGene_tx=XM_019829107.2;refGene_gn=COL25A1;refGene_tx=XM_019829108.2;refGene_gn=COL25A1;refGene_tx=XM_019829109.2;refGene_gn=COL25A1;refGene_tx=XM_019829111.2;refGene_gn=COL25A1;refGene_tx=XM_019829113.2;refGene_gn=COL25A1;refGene_tx=XM_019829114.2;refGene_gn=COL25A1;refGene_tx=XM_019829116.2;refGene_gn=COL25A1;refGene_tx=XM_019829117.2;refGene_gn=COL25A1;refGene_tx=XM_019829118.2;refGene_gn=COL25A1;refGene_tx=XM_019829119.2
B1 116442948 116442993
refGene_gn=COL25A1;refGene_tx=XM_019829100.2;refGene_gn=COL25A1;refGene_tx=XM_019829102.2;refGene_gn=COL25A1;refGene_tx=XM_019829103.2;refGene_gn=COL25A1;refGene_tx=XM_019829104.2;refGene_gn=COL25A1;refGene_tx=XM_019829105.2;refGene_gn=COL25A1;refGene_tx=XM_019829106.2;refGene_gn=COL25A1;refGene_tx=XM_019829107.2;refGene_gn=COL25A1;refGene_tx=XM_019829108.2;refGene_gn=COL25A1;refGene_tx=XM_019829109.2;refGene_gn=COL25A1;refGene_tx=XM_019829111.2;refGene_gn=COL25A1;refGene_tx=XM_019829113.2;refGene_gn=COL25A1;refGene_tx=XM_019829114.2;refGene_gn=COL25A1;refGene_tx=XM_019829115.2;refGene_gn=COL25A1;refGene_tx=XM_019829116.2;refGene_gn=COL25A1;refGene_tx=XM_019829117.2;refGene_gn=COL25A1;refGene_tx=XM_019829118.2;refGene_gn=COL25A1;refGene_tx=XM_019829119.2
B1 116446420 116446465
refGene_gn=COL25A1;refGene_tx=XM_019829100.2;refGene_gn=COL25A1;refGene_tx=XM_019829102.2;refGene_gn=COL25A1;refGene_tx=XM_019829103.2;refGene_gn=COL25A1;refGene_tx=XM_019829104.2;refGene_gn=COL25A1;refGene_tx=XM_019829105.2;refGene_gn=COL25A1;refGene_tx=XM_019829106.2;refGene_gn=COL25A1;refGene_tx=XM_019829107.2;refGene_gn=COL25A1;refGene_tx=XM_019829108.2;refGene_gn=COL25A1;refGene_tx=XM_019829109.2;refGene_gn=COL25A1;refGene_tx=XM_019829111.2;refGene_gn=COL25A1;refGene_tx=XM_019829113.2;refGene_gn=COL25A1;refGene_tx=XM_019829114.2;refGene_gn=COL25A1;refGene_tx=XM_019829115.2;refGene_gn=COL25A1;refGene_tx=XM_019829116.2;refGene_gn=COL25A1;refGene_tx=XM_019829117.2;refGene_gn=COL25A1;refGene_tx=XM_019829118.2;refGene_gn=COL25A1;refGene_tx=XM_019829119.2
B1 116449195 116449276
refGene_gn=COL25A1;refGene_tx=XM_019829100.2;refGene_gn=COL25A1;refGene_tx=XM_019829102.2;refGene_gn=COL25A1;refGene_tx=XM_019829103.2;refGene_gn=COL25A1;refGene_tx=XM_019829104.2;refGene_gn=COL25A1;refGene_tx=XM_019829105.2;refGene_gn=COL25A1;refGene_tx=XM_019829106.2;refGene_gn=COL25A1;refGene_tx=XM_019829107.2;refGene_gn=COL25A1;refGene_tx=XM_019829108.2;refGene_gn=COL25A1;refGene_tx=XM_019829109.2;refGene_gn=COL25A1;refGene_tx=XM_019829111.2;refGene_gn=COL25A1;refGene_tx=XM_019829113.2;refGene_gn=COL25A1;refGene_tx=XM_019829114.2;refGene_gn=COL25A1;refGene_tx=XM_019829115.2;refGene_gn=COL25A1;refGene_tx=XM_019829116.2;refGene_gn=COL25A1;refGene_tx=XM_019829117.2;refGene_gn=COL25A1;refGene_tx=XM_019829118.2;refGene_gn=COL25A1;refGene_tx=XM_019829119.2
B1 116450700 116450763
refGene_gn=COL25A1;refGene_tx=XM_019829100.2;refGene_gn=COL25A1;refGen

e_tx=XM_019829102.2;refGene_gn=COL25A1;refGene_tx=XM_019829103.2;refGene_gn=COL25A1;refGene_tx=XM_019829104.2;refGene_gn=COL25A1;refGene_tx=XM_019829105.2;refGene_gn=COL25A1;refGene_tx=XM_019829106.2;refGene_gn=COL25A1;refGene_tx=XM_019829107.2;refGene_gn=COL25A1;refGene_tx=XM_019829108.2;refGene_gn=COL25A1;refGene_tx=XM_019829109.2;refGene_gn=COL25A1;refGene_tx=XM_019829111.2;refGene_gn=COL25A1;refGene_tx=XM_019829113.2;refGene_gn=COL25A1;refGene_tx=XM_019829114.2;refGene_gn=COL25A1;refGene_tx=XM_019829115.2;refGene_gn=COL25A1;refGene_tx=XM_019829116.2;refGene_gn=COL25A1;refGene_tx=XM_019829117.2;refGene_gn=COL25A1;refGene_tx=XM_019829118.2;refGene_gn=COL25A1;refGene_tx=XM_019829119.2
B1 116451405 116451456
refGene_gn=COL25A1;refGene_tx=XM_019829100.2;refGene_gn=COL25A1;refGene_tx=XM_019829102.2;refGene_gn=COL25A1;refGene_tx=XM_019829103.2;refGene_gn=COL25A1;refGene_tx=XM_019829104.2;refGene_gn=COL25A1;refGene_tx=XM_019829105.2;refGene_gn=COL25A1;refGene_tx=XM_019829106.2;refGene_gn=COL25A1;refGene_tx=XM_019829107.2;refGene_gn=COL25A1;refGene_tx=XM_019829108.2;refGene_gn=COL25A1;refGene_tx=XM_019829109.2;refGene_gn=COL25A1;refGene_tx=XM_019829111.2;refGene_gn=COL25A1;refGene_tx=XM_019829113.2;refGene_gn=COL25A1;refGene_tx=XM_019829114.2;refGene_gn=COL25A1;refGene_tx=XM_019829115.2;refGene_gn=COL25A1;refGene_tx=XM_019829116.2;refGene_gn=COL25A1;refGene_tx=XM_019829117.2;refGene_gn=COL25A1;refGene_tx=XM_019829118.2;refGene_gn=COL25A1;refGene_tx=XM_019829119.2
B1 116453445 116453472
refGene_gn=COL25A1;refGene_tx=XM_019829100.2;refGene_gn=COL25A1;refGene_tx=XM_019829102.2;refGene_gn=COL25A1;refGene_tx=XM_019829103.2;refGene_gn=COL25A1;refGene_tx=XM_019829104.2;refGene_gn=COL25A1;refGene_tx=XM_019829105.2;refGene_gn=COL25A1;refGene_tx=XM_019829106.2;refGene_gn=COL25A1;refGene_tx=XM_019829107.2;refGene_gn=COL25A1;refGene_tx=XM_019829108.2;refGene_gn=COL25A1;refGene_tx=XM_019829109.2;refGene_gn=COL25A1;refGene_tx=XM_019829111.2;refGene_gn=COL25A1;refGene_tx=XM_019829113.2;refGene_gn=COL25A1;refGene_tx=XM_019829114.2;refGene_gn=COL25A1;refGene_tx=XM_019829115.2;refGene_gn=COL25A1;refGene_tx=XM_019829116.2;refGene_gn=COL25A1;refGene_tx=XM_019829117.2;refGene_gn=COL25A1;refGene_tx=XM_019829118.2;refGene_gn=COL25A1;refGene_tx=XM_019829119.2
B1 116456490 116456571
refGene_gn=COL25A1;refGene_tx=XM_019829100.2;refGene_gn=COL25A1;refGene_tx=XM_019829102.2;refGene_gn=COL25A1;refGene_tx=XM_019829103.2;refGene_gn=COL25A1;refGene_tx=XM_019829104.2;refGene_gn=COL25A1;refGene_tx=XM_019829105.2;refGene_gn=COL25A1;refGene_tx=XM_019829106.2;refGene_gn=COL25A1;refGene_tx=XM_019829107.2;refGene_gn=COL25A1;refGene_tx=XM_019829108.2;refGene_gn=COL25A1;refGene_tx=XM_019829109.2;refGene_gn=COL25A1;refGene_tx=XM_019829111.2;refGene_gn=COL25A1;refGene_tx=XM_019829115.2
B1 116459075 116459111
refGene_gn=COL25A1;refGene_tx=XM_019829100.2;refGene_gn=COL25A1;refGene_tx=XM_019829102.2;refGene_gn=COL25A1;refGene_tx=XM_019829103.2;refGene_gn=COL25A1;refGene_tx=XM_019829104.2;refGene_gn=COL25A1;refGene_tx=XM_019829105.2;refGene_gn=COL25A1;refGene_tx=XM_019829106.2;refGene_gn=COL25A1;refGene_tx=XM_019829107.2;refGene_gn=COL25A1;refGene_tx=XM_019829108.2;refGene_gn=COL25A1;refGene_tx=XM_019829109.2;refGene_gn=COL2

5A1;refGene_tx=XM_019829111.2;refGene_gn=COL25A1;refGene_tx=XM_0198291
13.2;refGene_gn=COL25A1;refGene_tx=XM_019829114.2;refGene_gn=COL25A1;r
efGene_tx=XM_019829115.2;refGene_gn=COL25A1;refGene_tx=XM_019829119.2
B1 116460697 116460751
refGene_gn=COL25A1;refGene_tx=XM_019829100.2;refGene_gn=COL25A1;refGen
e_tx=XM_019829102.2;refGene_gn=COL25A1;refGene_tx=XM_019829103.2;refGe
ne_gn=COL25A1;refGene_tx=XM_019829104.2;refGene_gn=COL25A1;refGene_tx=
XM_019829105.2;refGene_gn=COL25A1;refGene_tx=XM_019829106.2;refGene_gn
=COL25A1;refGene_tx=XM_019829107.2;refGene_gn=COL25A1;refGene_tx=XM_01
9829108.2;refGene_gn=COL25A1;refGene_tx=XM_019829109.2;refGene_gn=COL2
5A1;refGene_tx=XM_019829111.2;refGene_gn=COL25A1;refGene_tx=XM_0198291
13.2;refGene_gn=COL25A1;refGene_tx=XM_019829114.2;refGene_gn=COL25A1;r
efGene_tx=XM_019829115.2;refGene_gn=COL25A1;refGene_tx=XM_019829116.2;
refGene_gn=COL25A1;refGene_tx=XM_019829117.2;refGene_gn=COL25A1;refGen
e_tx=XM_019829118.2;refGene_gn=COL25A1;refGene_tx=XM_019829119.2
B1 116465755 116465809
refGene_gn=COL25A1;refGene_tx=XM_019829100.2;refGene_gn=COL25A1;refGen
e_tx=XM_019829102.2;refGene_gn=COL25A1;refGene_tx=XM_019829103.2;refGe
ne_gn=COL25A1;refGene_tx=XM_019829104.2;refGene_gn=COL25A1;refGene_tx=
XM_019829105.2;refGene_gn=COL25A1;refGene_tx=XM_019829106.2;refGene_gn
=COL25A1;refGene_tx=XM_019829107.2;refGene_gn=COL25A1;refGene_tx=XM_01
9829108.2;refGene_gn=COL25A1;refGene_tx=XM_019829109.2;refGene_gn=COL2
5A1;refGene_tx=XM_019829111.2;refGene_gn=COL25A1;refGene_tx=XM_0198291
13.2;refGene_gn=COL25A1;refGene_tx=XM_019829114.2;refGene_gn=COL25A1;r
efGene_tx=XM_019829115.2;refGene_gn=COL25A1;refGene_tx=XM_019829116.2;
refGene_gn=COL25A1;refGene_tx=XM_019829117.2;refGene_gn=COL25A1;refGen
e_tx=XM_019829118.2;refGene_gn=COL25A1;refGene_tx=XM_019829119.2
B1 116467262 116467289
refGene_gn=COL25A1;refGene_tx=XM_019829100.2;refGene_gn=COL25A1;refGen
e_tx=XM_019829102.2;refGene_gn=COL25A1;refGene_tx=XM_019829103.2;refGe
ne_gn=COL25A1;refGene_tx=XM_019829104.2;refGene_gn=COL25A1;refGene_tx=
XM_019829105.2;refGene_gn=COL25A1;refGene_tx=XM_019829107.2;refGene_gn
=COL25A1;refGene_tx=XM_019829109.2;refGene_gn=COL25A1;refGene_tx=XM_01
9829111.2;refGene_gn=COL25A1;refGene_tx=XM_019829113.2;refGene_gn=COL2
5A1;refGene_tx=XM_019829114.2;refGene_gn=COL25A1;refGene_tx=XM_0198291
16.2;refGene_gn=COL25A1;refGene_tx=XM_019829117.2
B1 116468175 116468229
refGene_gn=COL25A1;refGene_tx=XM_019829100.2;refGene_gn=COL25A1;refGen
e_tx=XM_019829102.2;refGene_gn=COL25A1;refGene_tx=XM_019829103.2;refGe
ne_gn=COL25A1;refGene_tx=XM_019829104.2;refGene_gn=COL25A1;refGene_tx=
XM_019829105.2;refGene_gn=COL25A1;refGene_tx=XM_019829106.2;refGene_gn
=COL25A1;refGene_tx=XM_019829107.2;refGene_gn=COL25A1;refGene_tx=XM_01
9829108.2;refGene_gn=COL25A1;refGene_tx=XM_019829109.2;refGene_gn=COL2
5A1;refGene_tx=XM_019829111.2;refGene_gn=COL25A1;refGene_tx=XM_0198291
13.2;refGene_gn=COL25A1;refGene_tx=XM_019829114.2;refGene_gn=COL25A1;r
efGene_tx=XM_019829115.2;refGene_gn=COL25A1;refGene_tx=XM_019829116.2;
refGene_gn=COL25A1;refGene_tx=XM_019829117.2;refGene_gn=COL25A1;refGen
e_tx=XM_019829118.2;refGene_gn=COL25A1;refGene_tx=XM_019829119.2
B1 116472027 116472114
refGene_gn=COL25A1;refGene_tx=XM_019829100.2;refGene_gn=COL25A1;refGen

e_tx=XM_019829103.2;refGene_gn=COL25A1;refGene_tx=XM_019829104.2;refGene_gn=COL25A1;refGene_tx=XM_019829105.2;refGene_gn=COL25A1;refGene_tx=XM_019829106.2;refGene_gn=COL25A1;refGene_tx=XM_019829107.2;refGene_gn=COL25A1;refGene_tx=XM_019829109.2;refGene_gn=COL25A1;refGene_tx=XM_019829111.2;refGene_gn=COL25A1;refGene_tx=XM_019829113.2;refGene_gn=COL25A1;refGene_tx=XM_019829115.2;refGene_gn=COL25A1;refGene_tx=XM_019829116.2
B1 116472597 116472675
refGene_gn=COL25A1;refGene_tx=XM_019829102.2;refGene_gn=COL25A1;refGene_tx=XM_019829108.2;refGene_gn=COL25A1;refGene_tx=XM_019829114.2;refGene_gn=COL25A1;refGene_tx=XM_019829117.2;refGene_gn=COL25A1;refGene_tx=XM_019829118.2;refGene_gn=COL25A1;refGene_tx=XM_019829119.2
B1 116474514 116474553
refGene_gn=COL25A1;refGene_tx=XM_019829100.2;refGene_gn=COL25A1;refGene_tx=XM_019829102.2;refGene_gn=COL25A1;refGene_tx=XM_019829103.2;refGene_gn=COL25A1;refGene_tx=XM_019829104.2;refGene_gn=COL25A1;refGene_tx=XM_019829105.2;refGene_gn=COL25A1;refGene_tx=XM_019829106.2;refGene_gn=COL25A1;refGene_tx=XM_019829107.2;refGene_gn=COL25A1;refGene_tx=XM_019829108.2;refGene_gn=COL25A1;refGene_tx=XM_019829109.2;refGene_gn=COL25A1;refGene_tx=XM_019829111.2;refGene_gn=COL25A1;refGene_tx=XM_019829113.2;refGene_gn=COL25A1;refGene_tx=XM_019829114.2;refGene_gn=COL25A1;refGene_tx=XM_019829115.2;refGene_gn=COL25A1;refGene_tx=XM_019829116.2;refGene_gn=COL25A1;refGene_tx=XM_019829117.2;refGene_gn=COL25A1;refGene_tx=XM_019829118.2;refGene_gn=COL25A1;refGene_tx=XM_019829119.2
B1 116478300 116478303
refGene_gn=COL25A1;refGene_tx=XM_019829100.2;refGene_gn=COL25A1;refGene_tx=XM_019829102.2;refGene_gn=COL25A1;refGene_tx=XM_019829103.2;refGene_gn=COL25A1;refGene_tx=XM_019829104.2;refGene_gn=COL25A1;refGene_tx=XM_019829105.2;refGene_gn=COL25A1;refGene_tx=XM_019829106.2;refGene_gn=COL25A1;refGene_tx=XM_019829107.2;refGene_gn=COL25A1;refGene_tx=XM_019829108.2;refGene_gn=COL25A1;refGene_tx=XM_019829109.2;refGene_gn=COL25A1;refGene_tx=XM_019829111.2;refGene_gn=COL25A1;refGene_tx=XM_019829113.2;refGene_gn=COL25A1;refGene_tx=XM_019829114.2;refGene_gn=COL25A1;refGene_tx=XM_019829115.2;refGene_gn=COL25A1;refGene_tx=XM_019829116.2;refGene_gn=COL25A1;refGene_tx=XM_019829117.2;refGene_gn=COL25A1;refGene_tx=XM_019829118.2;refGene_gn=COL25A1;refGene_tx=XM_019829119.2
B1 116504178 116504303
refGene_gn=ETNPPL;refGene_tx=XM_019829121.1
B1 116511894 116512094
refGene_gn=ETNPPL;refGene_tx=XM_003985096.5;refGene_gn=ETNPPL;refGene_tx=XM_023252872.1;refGene_gn=ETNPPL;refGene_tx=XM_023252873.1;refGene_gn=ETNPPL;refGene_tx=XM_023252874.1
B1 116514930 116515049
refGene_gn=ETNPPL;refGene_tx=XM_003985096.5;refGene_gn=ETNPPL;refGene_tx=XM_019829121.1;refGene_gn=ETNPPL;refGene_tx=XM_023252872.1;refGene_gn=ETNPPL;refGene_tx=XM_023252873.1;refGene_gn=ETNPPL;refGene_tx=XM_023252874.1
B1 116515327 116515487
refGene_gn=ETNPPL;refGene_tx=XM_003985096.5;refGene_gn=ETNPPL;refGene_tx=XM_019829121.1;refGene_gn=ETNPPL;refGene_tx=XM_023252872.1;refGene_

gn=ETNPPL;refGene_tx=XM_023252873.1;refGene_gn=ETNPPL;refGene_tx=XM_023252874.1
B1	116517947	116518022	refGene_gn=ETNPPL;refGene_tx=XM_003985096.5;refGene_gn=ETNPPL;refGene_tx=XM_019829121.1;refGene_gn=ETNPPL;refGene_tx=XM_023252872.1;refGene_gn=ETNPPL;refGene_tx=XM_023252873.1;refGene_gn=ETNPPL;refGene_tx=XM_023252874.1
B1	116522001	116522092	refGene_gn=ETNPPL;refGene_tx=XM_003985096.5;refGene_gn=ETNPPL;refGene_tx=XM_019829121.1;refGene_gn=ETNPPL;refGene_tx=XM_023252872.1;refGene_gn=ETNPPL;refGene_tx=XM_023252873.1;refGene_gn=ETNPPL;refGene_tx=XM_023252874.1
B1	116522751	116522868	refGene_gn=ETNPPL;refGene_tx=XM_003985096.5;refGene_gn=ETNPPL;refGene_tx=XM_019829121.1;refGene_gn=ETNPPL;refGene_tx=XM_023252872.1;refGene_gn=ETNPPL;refGene_tx=XM_023252873.1;refGene_gn=ETNPPL;refGene_tx=XM_023252874.1
B1	116524960	116525043	refGene_gn=ETNPPL;refGene_tx=XM_003985096.5;refGene_gn=ETNPPL;refGene_tx=XM_019829121.1;refGene_gn=ETNPPL;refGene_tx=XM_023252872.1;refGene_gn=ETNPPL;refGene_tx=XM_023252873.1;refGene_gn=ETNPPL;refGene_tx=XM_023252874.1
B1	116525636	116525862	refGene_gn=ETNPPL;refGene_tx=XM_003985096.5;refGene_gn=ETNPPL;refGene_tx=XM_019829121.1;refGene_gn=ETNPPL;refGene_tx=XM_023252872.1;refGene_gn=ETNPPL;refGene_tx=XM_023252873.1;refGene_gn=ETNPPL;refGene_tx=XM_023252874.1
B1	116526756	116526911	refGene_gn=ETNPPL;refGene_tx=XM_003985096.5;refGene_gn=ETNPPL;refGene_tx=XM_019829121.1;refGene_gn=ETNPPL;refGene_tx=XM_023252872.1;refGene_gn=ETNPPL;refGene_tx=XM_023252873.1;refGene_gn=ETNPPL;refGene_tx=XM_023252874.1
B1	116527811	116527901	refGene_gn=ETNPPL;refGene_tx=XM_003985096.5;refGene_gn=ETNPPL;refGene_tx=XM_019829121.1;refGene_gn=ETNPPL;refGene_tx=XM_023252872.1;refGene_gn=ETNPPL;refGene_tx=XM_023252873.1;refGene_gn=ETNPPL;refGene_tx=XM_023252874.1
B1	116528334	116528465	refGene_gn=ETNPPL;refGene_tx=XM_003985096.5;refGene_gn=ETNPPL;refGene_tx=XM_019829121.1;refGene_gn=ETNPPL;refGene_tx=XM_023252872.1;refGene_gn=ETNPPL;refGene_tx=XM_023252873.1;refGene_gn=ETNPPL;refGene_tx=XM_023252874.1
B1	116530526	116530594	refGene_gn=ETNPPL;refGene_tx=XM_003985096.5;refGene_gn=ETNPPL;refGene_tx=XM_019829121.1;refGene_gn=ETNPPL;refGene_tx=XM_023252872.1;refGene_gn=ETNPPL;refGene_tx=XM_023252873.1;refGene_gn=ETNPPL;refGene_tx=XM_023252874.1
B1	116532559	116532688	refGene_gn=ETNPPL;refGene_tx=XM_003985096.5;refGene_gn=ETNPPL;refGene_tx=XM_019829121.1

B1 116536552 116536642
refGene_gn=ETNPPL;refGene_tx=XM_023252872.1
B1 116546897 116546915
refGene_gn=ETNPPL;refGene_tx=XM_023252874.1
B1 116552076 116552106
refGene_gn=ETNPPL;refGene_tx=XM_023252873.1
B1 116552872 116552879
refGene_gn=LOC109499033;refGene_tx=XM_023253397.1
B1 116553469 116553556
refGene_gn=LOC109499033;refGene_tx=XM_023253397.1
B1 116554748 116554960
refGene_gn=LOC109499033;refGene_tx=XM_023253397.1
B1 116559386 116559590
refGene_gn=LOC109499033;refGene_tx=XM_023253397.1
B1 116570881 116571163
refGene_gn=LOC109499033;refGene_tx=XM_023253397.1
B1 116586694 116586713
refGene_gn=OSTC;refGene_tx=XM_011281705.3
B1 116589882 116590080
refGene_gn=OSTC;refGene_tx=XM_011281705.3
B1 116592344 116592438
refGene_gn=OSTC;refGene_tx=XM_011281705.3
B1 116596318 116596457
refGene_gn=OSTC;refGene_tx=XM_011281705.3
B1 116619628 116619713
refGene_gn=RPL34;refGene_tx=XM_019829126.1
B1 116622018 116622122
refGene_gn=RPL34;refGene_tx=XM_019829126.1
B1 116622434 116622534
refGene_gn=RPL34;refGene_tx=XM_019829126.1
B1 116622617 116622682
refGene_gn=RPL34;refGene_tx=XM_019829126.1
B1 117003028 117003244
refGene_gn=LEF1;refGene_tx=XM_006930926.4;refGene_gn=LEF1;refGene_tx=XM_011281702.3;refGene_gn=LEF1;refGene_tx=XM_011281703.3;refGene_gn=LEF1;refGene_tx=XM_011281704.3
B1 117004533 117004542
refGene_gn=LEF1;refGene_tx=XM_019829129.2;refGene_gn=LEF1;refGene_tx=XM_019829130.2
B1 117005708 117005775
refGene_gn=LEF1;refGene_tx=XM_006930926.4;refGene_gn=LEF1;refGene_tx=XM_011281702.3;refGene_gn=LEF1;refGene_tx=XM_011281703.3;refGene_gn=LEF1;refGene_tx=XM_011281704.3;refGene_gn=LEF1;refGene_tx=XM_019829129.2;refGene_gn=LEF1;refGene_tx=XM_019829130.2
B1 117007203 117007337
refGene_gn=LEF1;refGene_tx=XM_006930926.4;refGene_gn=LEF1;refGene_tx=XM_011281702.3;refGene_gn=LEF1;refGene_tx=XM_011281703.3;refGene_gn=LEF1;refGene_tx=XM_011281704.3;refGene_gn=LEF1;refGene_tx=XM_019829129.2;refGene_gn=LEF1;refGene_tx=XM_019829130.2
B1 117086039 117086172

refGene_gn=LEF1;refGene_tx=XM_006930926.4;refGene_gn=LEF1;refGene_tx=XM_011281702.3;refGene_gn=LEF1;refGene_tx=XM_011281703.3;refGene_gn=LEF1;refGene_tx=XM_011281704.3;refGene_gn=LEF1;refGene_tx=XM_019829129.2;refGene_gn=LEF1;refGene_tx=XM_019829130.2
B1 117092442 117092533
refGene_gn=LEF1;refGene_tx=XM_006930926.4;refGene_gn=LEF1;refGene_tx=XM_011281702.3;refGene_gn=LEF1;refGene_tx=XM_011281703.3;refGene_gn=LEF1;refGene_tx=XM_011281704.3;refGene_gn=LEF1;refGene_tx=XM_019829129.2;refGene_gn=LEF1;refGene_tx=XM_019829130.2
B1 117094118 117094202
refGene_gn=LEF1;refGene_tx=XM_006930926.4;refGene_gn=LEF1;refGene_tx=XM_011281702.3;refGene_gn=LEF1;refGene_tx=XM_019829129.2;refGene_gn=LEF1;refGene_tx=XM_019829130.2
B1 117096281 117096404
refGene_gn=LEF1;refGene_tx=XM_006930926.4;refGene_gn=LEF1;refGene_tx=XM_011281702.3;refGene_gn=LEF1;refGene_tx=XM_011281703.3;refGene_gn=LEF1;refGene_tx=XM_011281704.3;refGene_gn=LEF1;refGene_tx=XM_019829129.2;refGene_gn=LEF1;refGene_tx=XM_019829130.2
B1 117097443 117097606
refGene_gn=LEF1;refGene_tx=XM_006930926.4;refGene_gn=LEF1;refGene_tx=XM_011281702.3;refGene_gn=LEF1;refGene_tx=XM_011281703.3;refGene_gn=LEF1;refGene_tx=XM_011281704.3;refGene_gn=LEF1;refGene_tx=XM_019829129.2;refGene_gn=LEF1;refGene_tx=XM_019829130.2
B1 117102499 117102607
refGene_gn=LEF1;refGene_tx=XM_006930926.4;refGene_gn=LEF1;refGene_tx=XM_011281702.3;refGene_gn=LEF1;refGene_tx=XM_011281703.3;refGene_gn=LEF1;refGene_tx=XM_011281704.3;refGene_gn=LEF1;refGene_tx=XM_019829129.2;refGene_gn=LEF1;refGene_tx=XM_019829130.2
B1 117108062 117108111
refGene_gn=LEF1;refGene_tx=XM_006930926.4;refGene_gn=LEF1;refGene_tx=XM_011281702.3;refGene_gn=LEF1;refGene_tx=XM_011281703.3;refGene_gn=LEF1;refGene_tx=XM_011281704.3;refGene_gn=LEF1;refGene_tx=XM_019829129.2;refGene_gn=LEF1;refGene_tx=XM_019829130.2
B1 117108746 117108781
refGene_gn=LEF1;refGene_tx=XM_006930926.4;refGene_gn=LEF1;refGene_tx=XM_011281704.3;refGene_gn=LEF1;refGene_tx=XM_019829130.2
B1 117121773 117121853
refGene_gn=LEF1;refGene_tx=XM_011281702.3;refGene_gn=LEF1;refGene_tx=XM_011281703.3;refGene_gn=LEF1;refGene_tx=XM_019829129.2
B1 117134097 117134216
refGene_gn=HADH;refGene_tx=XM_003985100.5
B1 117135179 117135296
refGene_gn=HADH;refGene_tx=XM_003985100.5
B1 117141008 117141081
refGene_gn=HADH;refGene_tx=XM_003985100.5
B1 117145118 117145208
refGene_gn=HADH;refGene_tx=XM_003985100.5
B1 117148511 117148638
refGene_gn=HADH;refGene_tx=XM_003985100.5
B1 117152594 117152752

refGene_gn=HADH;refGene_tx=XM_003985100.5
B1	117158576	117158705
refGene_gn=HADH;refGene_tx=XM_003985100.5
B1	117183587	117183719
refGene_gn=HADH;refGene_tx=XM_003985100.5
B1	117198574	117198753
refGene_gn=LOC101089856;refGene_tx=XM_019829131.2
B1	117199045	117199068
refGene_gn=LOC101089856;refGene_tx=XM_023252875.1
B1	117199513	117199681
refGene_gn=LOC101089856;refGene_tx=XM_019829131.2;refGene_gn=LOC101089
856;refGene_tx=XM_023252875.1
B1	117201475	117201637
refGene_gn=LOC101089856;refGene_tx=XM_019829131.2;refGene_gn=LOC101089
856;refGene_tx=XM_023252875.1
B1	117203598	117204234
refGene_gn=LOC101089856;refGene_tx=XM_019829131.2;refGene_gn=LOC101089
856;refGene_tx=XM_023252875.1
B1	117222206	117222696
refGene_gn=LOC101089856;refGene_tx=XM_019829131.2;refGene_gn=LOC101089
856;refGene_tx=XM_023252875.1
B1	117247842	117248046
refGene_gn=SGMS2;refGene_tx=XM_019829135.2;refGene_gn=SGMS2;refGene_tx
=XM_019829136.2;refGene_gn=SGMS2;refGene_tx=XM_019829137.2;refGene_gn=
SGMS2;refGene_tx=XM_019829138.2;refGene_gn=SGMS2;refGene_tx=XM_0198291
39.2;refGene_gn=SGMS2;refGene_tx=XM_023252878.1
B1	117249595	117249762
refGene_gn=SGMS2;refGene_tx=XM_019829135.2;refGene_gn=SGMS2;refGene_tx
=XM_019829136.2;refGene_gn=SGMS2;refGene_tx=XM_019829137.2;refGene_gn=
SGMS2;refGene_tx=XM_019829138.2;refGene_gn=SGMS2;refGene_tx=XM_0198291
39.2;refGene_gn=SGMS2;refGene_tx=XM_023252878.1
B1	117254656	117254810
refGene_gn=SGMS2;refGene_tx=XM_019829135.2;refGene_gn=SGMS2;refGene_tx
=XM_019829136.2;refGene_gn=SGMS2;refGene_tx=XM_019829137.2;refGene_gn=
SGMS2;refGene_tx=XM_019829138.2;refGene_gn=SGMS2;refGene_tx=XM_0198291
39.2;refGene_gn=SGMS2;refGene_tx=XM_023252878.1
B1	117260389	117260507
refGene_gn=SGMS2;refGene_tx=XM_019829135.2;refGene_gn=SGMS2;refGene_tx
=XM_019829136.2;refGene_gn=SGMS2;refGene_tx=XM_019829137.2;refGene_gn=
SGMS2;refGene_tx=XM_019829138.2;refGene_gn=SGMS2;refGene_tx=XM_0198291
39.2;refGene_gn=SGMS2;refGene_tx=XM_023252878.1
B1	117264186	117264641
refGene_gn=SGMS2;refGene_tx=XM_019829135.2;refGene_gn=SGMS2;refGene_tx
=XM_019829136.2;refGene_gn=SGMS2;refGene_tx=XM_019829137.2;refGene_gn=
SGMS2;refGene_tx=XM_019829138.2;refGene_gn=SGMS2;refGene_tx=XM_0198291
39.2;refGene_gn=SGMS2;refGene_tx=XM_023252878.1
B1	117422183	117422243
refGene_gn=PAPSS1;refGene_tx=XM_003985102.5
B1	117422570	117422681
refGene_gn=PAPSS1;refGene_tx=XM_023252876.1

B1	117436818	117436933
refGene_gn=PAPSS1;refGene_tx=XM_003985102.5;refGene_gn=PAPSS1;refGene_tx=XM_023252876.1;refGene_gn=PAPSS1;refGene_tx=XM_023252877.1
B1	117443779	117444015
refGene_gn=PAPSS1;refGene_tx=XM_003985102.5;refGene_gn=PAPSS1;refGene_tx=XM_023252876.1;refGene_gn=PAPSS1;refGene_tx=XM_023252877.1
B1	117449412	117449551
refGene_gn=PAPSS1;refGene_tx=XM_003985102.5;refGene_gn=PAPSS1;refGene_tx=XM_023252876.1;refGene_gn=PAPSS1;refGene_tx=XM_023252877.1
B1	117453936	117454055
refGene_gn=PAPSS1;refGene_tx=XM_003985102.5;refGene_gn=PAPSS1;refGene_tx=XM_023252876.1;refGene_gn=PAPSS1;refGene_tx=XM_023252877.1
B1	117471608	117471722
refGene_gn=PAPSS1;refGene_tx=XM_003985102.5;refGene_gn=PAPSS1;refGene_tx=XM_023252876.1;refGene_gn=PAPSS1;refGene_tx=XM_023252877.1
B1	117473916	117474028
refGene_gn=PAPSS1;refGene_tx=XM_003985102.5;refGene_gn=PAPSS1;refGene_tx=XM_023252876.1;refGene_gn=PAPSS1;refGene_tx=XM_023252877.1
B1	117476333	117476539
refGene_gn=PAPSS1;refGene_tx=XM_003985102.5;refGene_gn=PAPSS1;refGene_tx=XM_023252876.1;refGene_gn=PAPSS1;refGene_tx=XM_023252877.1
B1	117480375	117480511
refGene_gn=PAPSS1;refGene_tx=XM_003985102.5;refGene_gn=PAPSS1;refGene_tx=XM_023252876.1;refGene_gn=PAPSS1;refGene_tx=XM_023252877.1
B1	117493817	117494086
refGene_gn=PAPSS1;refGene_tx=XM_003985102.5;refGene_gn=PAPSS1;refGene_tx=XM_023252876.1;refGene_gn=PAPSS1;refGene_tx=XM_023252877.1
B1	117512023	117512253
refGene_gn=PAPSS1;refGene_tx=XM_003985102.5;refGene_gn=PAPSS1;refGene_tx=XM_023252876.1;refGene_gn=PAPSS1;refGene_tx=XM_023252877.1
B1	117526412	117526551
refGene_gn=PAPSS1;refGene_tx=XM_003985102.5;refGene_gn=PAPSS1;refGene_tx=XM_023252876.1;refGene_gn=PAPSS1;refGene_tx=XM_023252877.1
B1	118003515	118003737
refGene_gn=DKK2;refGene_tx=XM_003985106.5
B1	118098414	118098565
refGene_gn=DKK2;refGene_tx=XM_003985106.5
B1	118099747	118099903
refGene_gn=DKK2;refGene_tx=XM_003985106.5
B1	118102461	118102712
refGene_gn=DKK2;refGene_tx=XM_003985106.5
B1	118587314	118587734
refGene_gn=GIMD1;refGene_tx=XM_011281812.3
B1	118596872	118597133
refGene_gn=GIMD1;refGene_tx=XM_011281812.3
B1	118607231	118607398
refGene_gn=AIMP1;refGene_tx=XM_006930928.4;refGene_gn=AIMP1;refGene_tx=XM_019829140.2;refGene_gn=AIMP1;refGene_tx=XM_019829141.2
B1	118620653	118620822
refGene_gn=AIMP1;refGene_tx=XM_006930928.4;refGene_gn=AIMP1;refGene_tx

=XM_019829140.2;refGene_gn=AIMP1;refGene_tx=XM_019829141.2
B1 118626804 118627010
refGene_gn=AIMP1;refGene_tx=XM_006930928.4;refGene_gn=AIMP1;refGene_tx
=XM_019829140.2;refGene_gn=AIMP1;refGene_tx=XM_019829141.2
B1 118629632 118629797
refGene_gn=AIMP1;refGene_tx=XM_006930928.4;refGene_gn=AIMP1;refGene_tx
=XM_019829140.2;refGene_gn=AIMP1;refGene_tx=XM_019829141.2
B1 118632963 118633077
refGene_gn=AIMP1;refGene_tx=XM_006930928.4;refGene_gn=AIMP1;refGene_tx
=XM_019829140.2;refGene_gn=AIMP1;refGene_tx=XM_019829141.2
B1 118635561 118635695
refGene_gn=AIMP1;refGene_tx=XM_006930928.4;refGene_gn=AIMP1;refGene_tx
=XM_019829140.2;refGene_gn=AIMP1;refGene_tx=XM_019829141.2
B1 118645794 118645796
refGene_gn=AIMP1;refGene_tx=XM_006930928.4
B1 118652370 118652563
refGene_gn=TBCK;refGene_tx=XM_023252879.1;refGene_gn=TBCK;refGene_tx=X
M_023252880.1;refGene_gn=TBCK;refGene_tx=XM_023252881.1;refGene_gn=TBC
K;refGene_tx=XM_023252882.1
B1 118653264 118653342
refGene_gn=TBCK;refGene_tx=XM_023252879.1;refGene_gn=TBCK;refGene_tx=X
M_023252880.1;refGene_gn=TBCK;refGene_tx=XM_023252882.1
B1 118668615 118668688
refGene_gn=TBCK;refGene_tx=XM_023252879.1;refGene_gn=TBCK;refGene_tx=X
M_023252880.1;refGene_gn=TBCK;refGene_tx=XM_023252881.1;refGene_gn=TBC
K;refGene_tx=XM_023252882.1
B1 118712615 118712730
refGene_gn=TBCK;refGene_tx=XM_023252879.1;refGene_gn=TBCK;refGene_tx=X
M_023252880.1;refGene_gn=TBCK;refGene_tx=XM_023252881.1;refGene_gn=TBC
K;refGene_tx=XM_023252882.1;refGene_gn=TBCK;refGene_tx=XM_023252883.1
B1 118714718 118714792
refGene_gn=TBCK;refGene_tx=XM_023252879.1;refGene_gn=TBCK;refGene_tx=X
M_023252880.1;refGene_gn=TBCK;refGene_tx=XM_023252881.1;refGene_gn=TBC
K;refGene_tx=XM_023252882.1;refGene_gn=TBCK;refGene_tx=XM_023252883.1
B1 118722119 118722261
refGene_gn=TBCK;refGene_tx=XM_023252879.1;refGene_gn=TBCK;refGene_tx=X
M_023252880.1;refGene_gn=TBCK;refGene_tx=XM_023252881.1;refGene_gn=TBC
K;refGene_tx=XM_023252882.1;refGene_gn=TBCK;refGene_tx=XM_023252883.1;
refGene_gn=TBCK;refGene_tx=XM_023252884.1
B1 118727831 118727892
refGene_gn=TBCK;refGene_tx=XM_023252879.1;refGene_gn=TBCK;refGene_tx=X
M_023252880.1;refGene_gn=TBCK;refGene_tx=XM_023252881.1;refGene_gn=TBC
K;refGene_tx=XM_023252882.1;refGene_gn=TBCK;refGene_tx=XM_023252883.1;
refGene_gn=TBCK;refGene_tx=XM_023252884.1
B1 118731097 118731159
refGene_gn=TBCK;refGene_tx=XM_023252879.1;refGene_gn=TBCK;refGene_tx=X
M_023252880.1;refGene_gn=TBCK;refGene_tx=XM_023252881.1;refGene_gn=TBC
K;refGene_tx=XM_023252882.1;refGene_gn=TBCK;refGene_tx=XM_023252883.1;
refGene_gn=TBCK;refGene_tx=XM_023252884.1
B1 118731416 118731478

refGene_gn=TBCK;refGene_tx=XM_023252879.1;refGene_gn=TBCK;refGene_tx=XM_023252880.1;refGene_gn=TBCK;refGene_tx=XM_023252881.1;refGene_gn=TBCK;refGene_tx=XM_023252882.1;refGene_gn=TBCK;refGene_tx=XM_023252883.1;refGene_gn=TBCK;refGene_tx=XM_023252884.1
B1 118732293 118732442
refGene_gn=TBCK;refGene_tx=XM_023252879.1;refGene_gn=TBCK;refGene_tx=XM_023252880.1;refGene_gn=TBCK;refGene_tx=XM_023252881.1;refGene_gn=TBCK;refGene_tx=XM_023252882.1;refGene_gn=TBCK;refGene_tx=XM_023252883.1;refGene_gn=TBCK;refGene_tx=XM_023252884.1
B1 118733276 118733415
refGene_gn=TBCK;refGene_tx=XM_023252879.1;refGene_gn=TBCK;refGene_tx=XM_023252880.1;refGene_gn=TBCK;refGene_tx=XM_023252881.1;refGene_gn=TBCK;refGene_tx=XM_023252882.1;refGene_gn=TBCK;refGene_tx=XM_023252883.1;refGene_gn=TBCK;refGene_tx=XM_023252884.1
B1 118740543 118740643
refGene_gn=TBCK;refGene_tx=XM_023252879.1;refGene_gn=TBCK;refGene_tx=XM_023252880.1;refGene_gn=TBCK;refGene_tx=XM_023252881.1;refGene_gn=TBCK;refGene_tx=XM_023252882.1;refGene_gn=TBCK;refGene_tx=XM_023252883.1;refGene_gn=TBCK;refGene_tx=XM_023252884.1
B1 118746553 118746603
refGene_gn=TBCK;refGene_tx=XM_023252879.1;refGene_gn=TBCK;refGene_tx=XM_023252880.1;refGene_gn=TBCK;refGene_tx=XM_023252881.1;refGene_gn=TBCK;refGene_tx=XM_023252882.1;refGene_gn=TBCK;refGene_tx=XM_023252883.1;refGene_gn=TBCK;refGene_tx=XM_023252884.1
B1 118746784 118746914
refGene_gn=TBCK;refGene_tx=XM_023252879.1;refGene_gn=TBCK;refGene_tx=XM_023252880.1;refGene_gn=TBCK;refGene_tx=XM_023252881.1;refGene_gn=TBCK;refGene_tx=XM_023252882.1;refGene_gn=TBCK;refGene_tx=XM_023252883.1;refGene_gn=TBCK;refGene_tx=XM_023252884.1
B1 118747889 118747988
refGene_gn=TBCK;refGene_tx=XM_023252879.1;refGene_gn=TBCK;refGene_tx=XM_023252880.1;refGene_gn=TBCK;refGene_tx=XM_023252881.1;refGene_gn=TBCK;refGene_tx=XM_023252882.1;refGene_gn=TBCK;refGene_tx=XM_023252883.1;refGene_gn=TBCK;refGene_tx=XM_023252884.1
B1 118749614 118749677
refGene_gn=TBCK;refGene_tx=XM_023252879.1;refGene_gn=TBCK;refGene_tx=XM_023252880.1;refGene_gn=TBCK;refGene_tx=XM_023252881.1;refGene_gn=TBCK;refGene_tx=XM_023252882.1;refGene_gn=TBCK;refGene_tx=XM_023252883.1;refGene_gn=TBCK;refGene_tx=XM_023252884.1
B1 118750435 118750562
refGene_gn=TBCK;refGene_tx=XM_023252879.1;refGene_gn=TBCK;refGene_tx=XM_023252880.1;refGene_gn=TBCK;refGene_tx=XM_023252881.1;refGene_gn=TBCK;refGene_tx=XM_023252882.1;refGene_gn=TBCK;refGene_tx=XM_023252883.1;refGene_gn=TBCK;refGene_tx=XM_023252884.1
B1 118752083 118752134
refGene_gn=TBCK;refGene_tx=XM_023252879.1;refGene_gn=TBCK;refGene_tx=XM_023252880.1;refGene_gn=TBCK;refGene_tx=XM_023252881.1;refGene_gn=TBCK;refGene_tx=XM_023252882.1;refGene_gn=TBCK;refGene_tx=XM_023252883.1;refGene_gn=TBCK;refGene_tx=XM_023252884.1
B1 118753448 118753532

refGene_gn=TBCK;refGene_tx=XM_023252879.1;refGene_gn=TBCK;refGene_tx=XM_023252880.1;refGene_gn=TBCK;refGene_tx=XM_023252881.1;refGene_gn=TBCK;refGene_tx=XM_023252883.1;refGene_gn=TBCK;refGene_tx=XM_023252884.1
B1 118763038 118763124
refGene_gn=TBCK;refGene_tx=XM_023252879.1;refGene_gn=TBCK;refGene_tx=XM_023252880.1;refGene_gn=TBCK;refGene_tx=XM_023252881.1;refGene_gn=TBCK;refGene_tx=XM_023252882.1;refGene_gn=TBCK;refGene_tx=XM_023252883.1;refGene_gn=TBCK;refGene_tx=XM_023252884.1
B1 118788739 118788776
refGene_gn=TBCK;refGene_tx=XM_023252879.1;refGene_gn=TBCK;refGene_tx=XM_023252880.1;refGene_gn=TBCK;refGene_tx=XM_023252881.1;refGene_gn=TBCK;refGene_tx=XM_023252882.1;refGene_gn=TBCK;refGene_tx=XM_023252883.1;refGene_gn=TBCK;refGene_tx=XM_023252884.1
B1 118790074 118790236
refGene_gn=TBCK;refGene_tx=XM_023252879.1;refGene_gn=TBCK;refGene_tx=XM_023252880.1;refGene_gn=TBCK;refGene_tx=XM_023252881.1;refGene_gn=TBCK;refGene_tx=XM_023252882.1;refGene_gn=TBCK;refGene_tx=XM_023252883.1;refGene_gn=TBCK;refGene_tx=XM_023252884.1
B1 118802620 118802796
refGene_gn=TBCK;refGene_tx=XM_023252879.1;refGene_gn=TBCK;refGene_tx=XM_023252880.1;refGene_gn=TBCK;refGene_tx=XM_023252881.1;refGene_gn=TBCK;refGene_tx=XM_023252882.1;refGene_gn=TBCK;refGene_tx=XM_023252883.1;refGene_gn=TBCK;refGene_tx=XM_023252884.1
B1 118833272 118833448
refGene_gn=TBCK;refGene_tx=XM_023252879.1;refGene_gn=TBCK;refGene_tx=XM_023252880.1;refGene_gn=TBCK;refGene_tx=XM_023252881.1;refGene_gn=TBCK;refGene_tx=XM_023252882.1;refGene_gn=TBCK;refGene_tx=XM_023252883.1;refGene_gn=TBCK;refGene_tx=XM_023252884.1
B1 118857221 118857381
refGene_gn=TBCK;refGene_tx=XM_023252879.1;refGene_gn=TBCK;refGene_tx=XM_023252880.1;refGene_gn=TBCK;refGene_tx=XM_023252881.1;refGene_gn=TBCK;refGene_tx=XM_023252882.1;refGene_gn=TBCK;refGene_tx=XM_023252883.1;refGene_gn=TBCK;refGene_tx=XM_023252884.1
B1 118880012 118880123
refGene_gn=TBCK;refGene_tx=XM_023252879.1;refGene_gn=TBCK;refGene_tx=XM_023252880.1;refGene_gn=TBCK;refGene_tx=XM_023252881.1;refGene_gn=TBCK;refGene_tx=XM_023252882.1;refGene_gn=TBCK;refGene_tx=XM_023252883.1;refGene_gn=TBCK;refGene_tx=XM_023252884.1
B1 118945070 118945273
refGene_gn=NPNT;refGene_tx=XM_019829143.2
B1 118946628 118946739
refGene_gn=NPNT;refGene_tx=XM_011281807.3;refGene_gn=NPNT;refGene_tx=XM_011281808.3;refGene_gn=NPNT;refGene_tx=XM_011281809.3;refGene_gn=NPNT;refGene_tx=XM_011281810.3
B1 118948103 118948360
refGene_gn=NPNT;refGene_tx=XM_011281807.3;refGene_gn=NPNT;refGene_tx=XM_011281808.3;refGene_gn=NPNT;refGene_tx=XM_011281809.3;refGene_gn=NPNT;refGene_tx=XM_011281810.3
B1 118956769 118956868
refGene_gn=NPNT;refGene_tx=XM_011281807.3;refGene_gn=NPNT;refGene_tx=X

M_011281808.3;refGene_gn=NPNT;refGene_tx=XM_011281809.3;refGene_gn=NPNT;refGene_tx=XM_011281810.3;refGene_gn=NPNT;refGene_tx=XM_019829143.2
B1 118957341 118957428
refGene_gn=NPNT;refGene_tx=XM_011281807.3;refGene_gn=NPNT;refGene_tx=XM_011281808.3;refGene_gn=NPNT;refGene_tx=XM_011281809.3;refGene_gn=NPNT;refGene_tx=XM_011281810.3;refGene_gn=NPNT;refGene_tx=XM_019829143.2
B1 118976630 118977008
refGene_gn=NPNT;refGene_tx=XM_011281807.3;refGene_gn=NPNT;refGene_tx=XM_011281808.3;refGene_gn=NPNT;refGene_tx=XM_011281809.3;refGene_gn=NPNT;refGene_tx=XM_011281810.3;refGene_gn=NPNT;refGene_tx=XM_019829143.2
B1 118979168 118979291
refGene_gn=NPNT;refGene_tx=XM_011281807.3;refGene_gn=NPNT;refGene_tx=XM_011281808.3;refGene_gn=NPNT;refGene_tx=XM_011281809.3;refGene_gn=NPNT;refGene_tx=XM_011281810.3;refGene_gn=NPNT;refGene_tx=XM_019829143.2
B1 118979584 118979719
refGene_gn=NPNT;refGene_tx=XM_011281807.3;refGene_gn=NPNT;refGene_tx=XM_011281808.3;refGene_gn=NPNT;refGene_tx=XM_011281809.3;refGene_gn=NPNT;refGene_tx=XM_011281810.3;refGene_gn=NPNT;refGene_tx=XM_019829143.2
B1 118981322 118981442
refGene_gn=NPNT;refGene_tx=XM_011281807.3;refGene_gn=NPNT;refGene_tx=XM_011281808.3;refGene_gn=NPNT;refGene_tx=XM_011281809.3;refGene_gn=NPNT;refGene_tx=XM_011281810.3;refGene_gn=NPNT;refGene_tx=XM_019829143.2
B1 118983621 118983741
refGene_gn=NPNT;refGene_tx=XM_011281807.3;refGene_gn=NPNT;refGene_tx=XM_011281808.3;refGene_gn=NPNT;refGene_tx=XM_011281809.3;refGene_gn=NPNT;refGene_tx=XM_011281810.3;refGene_gn=NPNT;refGene_tx=XM_019829143.2
B1 118988778 118988871
refGene_gn=NPNT;refGene_tx=XM_011281807.3;refGene_gn=NPNT;refGene_tx=XM_011281808.3;refGene_gn=NPNT;refGene_tx=XM_019829143.2
B1 118993140 118993233
refGene_gn=NPNT;refGene_tx=XM_011281807.3;refGene_gn=NPNT;refGene_tx=XM_011281808.3;refGene_gn=NPNT;refGene_tx=XM_011281809.3;refGene_gn=NPNT;refGene_tx=XM_011281810.3;refGene_gn=NPNT;refGene_tx=XM_019829143.2
B1 119008739 119008790
refGene_gn=NPNT;refGene_tx=XM_011281807.3;refGene_gn=NPNT;refGene_tx=XM_011281809.3;refGene_gn=NPNT;refGene_tx=XM_019829143.2
B1 119023931 119024032
refGene_gn=NPNT;refGene_tx=XM_011281807.3;refGene_gn=NPNT;refGene_tx=XM_011281808.3;refGene_gn=NPNT;refGene_tx=XM_011281809.3;refGene_gn=NPNT;refGene_tx=XM_011281810.3;refGene_gn=NPNT;refGene_tx=XM_019829143.2
B1 119026223 119026294
refGene_gn=NPNT;refGene_tx=XM_011281807.3;refGene_gn=NPNT;refGene_tx=XM_011281808.3;refGene_gn=NPNT;refGene_tx=XM_011281809.3;refGene_gn=NPNT;refGene_tx=XM_011281810.3;refGene_gn=NPNT;refGene_tx=XM_019829143.2
B1 119063134 119063271
refGene_gn=GSTCD;refGene_tx=XM_006930933.3
B1 119066096 119066166
refGene_gn=GSTCD;refGene_tx=XM_006930933.3
B1 119069987 119070018
refGene_gn=GSTCD;refGene_tx=XM_006930933.3

B1 119072631 119072765
refGene_gn=GSTCD;refGene_tx=XM_006930933.3
B1 119090146 119090275
refGene_gn=GSTCD;refGene_tx=XM_006930933.3
B1 119092587 119092632
refGene_gn=GSTCD;refGene_tx=XM_006930933.3
B1 119092794 119092910
refGene_gn=GSTCD;refGene_tx=XM_006930933.3
B1 119169059 119169153
refGene_gn=GSTCD;refGene_tx=XM_006930933.3
B1 119174192 119174444
refGene_gn=GSTCD;refGene_tx=XM_006930933.3
B1 119182907 119183363
refGene_gn=GSTCD;refGene_tx=XM_006930933.3
B1 119184457 119184874
refGene_gn=GSTCD;refGene_tx=XM_006930933.3
B1 119203042 119203198
refGene_gn=INTS12;refGene_tx=XM_003985117.3;refGene_gn=INTS12;refGene_tx=XM_006930934.4;refGene_gn=INTS12;refGene_tx=XM_011281806.2
B1 119207293 119207446
refGene_gn=INTS12;refGene_tx=XM_003985117.3;refGene_gn=INTS12;refGene_tx=XM_006930934.4;refGene_gn=INTS12;refGene_tx=XM_011281806.2
B1 119209193 119209381
refGene_gn=INTS12;refGene_tx=XM_003985117.3;refGene_gn=INTS12;refGene_tx=XM_006930934.4;refGene_gn=INTS12;refGene_tx=XM_011281806.2
B1 119211575 119211735
refGene_gn=INTS12;refGene_tx=XM_003985117.3;refGene_gn=INTS12;refGene_tx=XM_006930934.4;refGene_gn=INTS12;refGene_tx=XM_011281806.2
B1 119216422 119216569
refGene_gn=INTS12;refGene_tx=XM_003985117.3;refGene_gn=INTS12;refGene_tx=XM_006930934.4;refGene_gn=INTS12;refGene_tx=XM_011281806.2
B1 119219135 119219720
refGene_gn=INTS12;refGene_tx=XM_003985117.3;refGene_gn=INTS12;refGene_tx=XM_006930934.4;refGene_gn=INTS12;refGene_tx=XM_011281806.2
B1 119226925 119227111
refGene_gn=ARHGEF38;refGene_tx=XM_003985118.4
B1 119236938 119237198
refGene_gn=ARHGEF38;refGene_tx=XM_003985118.4
B1 119237362 119237561
refGene_gn=ARHGEF38;refGene_tx=XM_003985118.4
B1 119238367 119238511
refGene_gn=ARHGEF38;refGene_tx=XM_003985118.4
B1 119241917 119242229
refGene_gn=ARHGEF38;refGene_tx=XM_003985118.4
B1 119245922 119246045
refGene_gn=ARHGEF38;refGene_tx=XM_003985118.4
B1 119247790 119247895
refGene_gn=ARHGEF38;refGene_tx=XM_003985118.4
B1 119253341 119253475
refGene_gn=ARHGEF38;refGene_tx=XM_003985118.4

B1	119256145	119256345
refGene_gn=ARHGEF38;refGene_tx=XM_003985118.4
B1	119267084	119267102
refGene_gn=ARHGEF38;refGene_tx=XM_003985118.4
B1	119273940	119274016
refGene_gn=ARHGEF38;refGene_tx=XM_003985118.4
B1	119274018	119274087
refGene_gn=ARHGEF38;refGene_tx=XM_003985118.4
B1	119293353	119293477
refGene_gn=ARHGEF38;refGene_tx=XM_003985118.4
B1	119317955	119318143
refGene_gn=ARHGEF38;refGene_tx=XM_003985118.4
B1	119354838	119355034
refGene_gn=ARHGEF38;refGene_tx=XM_003985118.4
B1	119429967	119430142
refGene_gn=PPA2;refGene_tx=XM_006930935.4;refGene_gn=PPA2;refGene_tx=XM_006930936.4
B1	119430332	119430468
refGene_gn=PPA2;refGene_tx=XM_023252885.1
B1	119446115	119446180
refGene_gn=PPA2;refGene_tx=XM_006930935.4;refGene_gn=PPA2;refGene_tx=XM_006930936.4;refGene_gn=PPA2;refGene_tx=XM_023252885.1
B1	119448079	119448124
refGene_gn=PPA2;refGene_tx=XM_006930935.4
B1	119452716	119452770
refGene_gn=PPA2;refGene_tx=XM_006930935.4;refGene_gn=PPA2;refGene_tx=XM_006930936.4;refGene_gn=PPA2;refGene_tx=XM_023252885.1
B1	119454201	119454321
refGene_gn=PPA2;refGene_tx=XM_006930935.4;refGene_gn=PPA2;refGene_tx=XM_006930936.4;refGene_gn=PPA2;refGene_tx=XM_023252885.1
B1	119462564	119462651
refGene_gn=PPA2;refGene_tx=XM_006930935.4;refGene_gn=PPA2;refGene_tx=XM_006930936.4;refGene_gn=PPA2;refGene_tx=XM_023252885.1
B1	119474320	119474447
refGene_gn=PPA2;refGene_tx=XM_006930935.4;refGene_gn=PPA2;refGene_tx=XM_006930936.4;refGene_gn=PPA2;refGene_tx=XM_023252885.1
B1	119492493	119492621
refGene_gn=PPA2;refGene_tx=XM_006930935.4;refGene_gn=PPA2;refGene_tx=XM_006930936.4;refGene_gn=PPA2;refGene_tx=XM_023252885.1
B1	119496352	119496438
refGene_gn=PPA2;refGene_tx=XM_006930935.4;refGene_gn=PPA2;refGene_tx=XM_006930936.4;refGene_gn=PPA2;refGene_tx=XM_023252885.1
B1	119503952	119504022
refGene_gn=PPA2;refGene_tx=XM_006930935.4;refGene_gn=PPA2;refGene_tx=XM_006930936.4;refGene_gn=PPA2;refGene_tx=XM_023252885.1
B1	119520376	119520413
refGene_gn=PPA2;refGene_tx=XM_006930935.4;refGene_gn=PPA2;refGene_tx=XM_006930936.4;refGene_gn=PPA2;refGene_tx=XM_023252885.1
B1	119521482	119521517
refGene_gn=PPA2;refGene_tx=XM_006930935.4;refGene_gn=PPA2;refGene_tx=X

M_006930936.4;refGene_gn=PPA2;refGene_tx=XM_023252885.1
B1 119602197 119603189
refGene_gn=TET2;refGene_tx=XM_006930937.4
B1 119603191 119603716
refGene_gn=TET2;refGene_tx=XM_006930937.4
B1 119605637 119605992
refGene_gn=TET2;refGene_tx=XM_006930937.4
B1 119608450 119608588
refGene_gn=TET2;refGene_tx=XM_006930937.4
B1 119616279 119616369
refGene_gn=TET2;refGene_tx=XM_006930937.4
B1 119619455 119619606
refGene_gn=TET2;refGene_tx=XM_006930937.4
B1 119641244 119641453
refGene_gn=TET2;refGene_tx=XM_006930937.4
B1 119642089 119642183
refGene_gn=TET2;refGene_tx=XM_006930937.4
B1 119643580 119643671
refGene_gn=TET2;refGene_tx=XM_006930937.4
B1 119645312 119648721
refGene_gn=TET2;refGene_tx=XM_006930937.4
B1 120279348 120280425
refGene_gn=CXXC4;refGene_tx=XM_019829149.2;refGene_gn=CXXC4;refGene_tx
=XM_019829151.2
B1 120298614 120298659
refGene_gn=CXXC4;refGene_tx=XM_019829149.2;refGene_gn=CXXC4;refGene_tx
=XM_019829151.2
B1 120887619 120888167
refGene_gn=TACR3;refGene_tx=XM_003985120.4
B1 120926887 120927076
refGene_gn=TACR3;refGene_tx=XM_003985120.4
B1 120929677 120929828
refGene_gn=TACR3;refGene_tx=XM_003985120.4
B1 120974874 120975071
refGene_gn=TACR3;refGene_tx=XM_003985120.4
B1 120977463 120977776
refGene_gn=TACR3;refGene_tx=XM_003985120.4
B1 121297896 121297952
refGene_gn=CENPE;refGene_tx=XM_019829159.2;refGene_gn=CENPE;refGene_tx
=XM_023252886.1;refGene_gn=CENPE;refGene_tx=XM_023252887.1;refGene_gn=
CENPE;refGene_tx=XM_023252888.1;refGene_gn=CENPE;refGene_tx=XM_0232528
89.1;refGene_gn=CENPE;refGene_tx=XM_023252890.1;refGene_gn=CENPE;refGe
ne_tx=XM_023252891.1
B1 121299711 121299803
refGene_gn=CENPE;refGene_tx=XM_019829159.2;refGene_gn=CENPE;refGene_tx
=XM_023252886.1;refGene_gn=CENPE;refGene_tx=XM_023252887.1;refGene_gn=
CENPE;refGene_tx=XM_023252888.1;refGene_gn=CENPE;refGene_tx=XM_0232528
89.1;refGene_gn=CENPE;refGene_tx=XM_023252890.1;refGene_gn=CENPE;refGe
ne_tx=XM_023252891.1
B1 121302431 121302521

refGene_gn=CENPE;refGene_tx=XM_019829159.2;refGene_gn=CENPE;refGene_tx=XM_023252886.1;refGene_gn=CENPE;refGene_tx=XM_023252887.1;refGene_gn=CENPE;refGene_tx=XM_023252888.1;refGene_gn=CENPE;refGene_tx=XM_023252889.1;refGene_gn=CENPE;refGene_tx=XM_023252890.1;refGene_gn=CENPE;refGene_tx=XM_023252891.1
B1 121302638 121302757
refGene_gn=CENPE;refGene_tx=XM_019829159.2;refGene_gn=CENPE;refGene_tx=XM_023252886.1;refGene_gn=CENPE;refGene_tx=XM_023252887.1;refGene_gn=CENPE;refGene_tx=XM_023252888.1;refGene_gn=CENPE;refGene_tx=XM_023252889.1;refGene_gn=CENPE;refGene_tx=XM_023252890.1;refGene_gn=CENPE;refGene_tx=XM_023252891.1
B1 121303925 121304045
refGene_gn=CENPE;refGene_tx=XM_019829159.2;refGene_gn=CENPE;refGene_tx=XM_023252886.1;refGene_gn=CENPE;refGene_tx=XM_023252887.1;refGene_gn=CENPE;refGene_tx=XM_023252888.1;refGene_gn=CENPE;refGene_tx=XM_023252889.1;refGene_gn=CENPE;refGene_tx=XM_023252890.1;refGene_gn=CENPE;refGene_tx=XM_023252891.1
B1 121304441 121304523
refGene_gn=CENPE;refGene_tx=XM_019829159.2;refGene_gn=CENPE;refGene_tx=XM_023252886.1;refGene_gn=CENPE;refGene_tx=XM_023252887.1;refGene_gn=CENPE;refGene_tx=XM_023252888.1;refGene_gn=CENPE;refGene_tx=XM_023252889.1;refGene_gn=CENPE;refGene_tx=XM_023252890.1;refGene_gn=CENPE;refGene_tx=XM_023252891.1
B1 121304675 121304743
refGene_gn=CENPE;refGene_tx=XM_019829159.2;refGene_gn=CENPE;refGene_tx=XM_023252886.1;refGene_gn=CENPE;refGene_tx=XM_023252887.1;refGene_gn=CENPE;refGene_tx=XM_023252888.1;refGene_gn=CENPE;refGene_tx=XM_023252889.1;refGene_gn=CENPE;refGene_tx=XM_023252890.1;refGene_gn=CENPE;refGene_tx=XM_023252891.1
B1 121304816 121304882
refGene_gn=CENPE;refGene_tx=XM_019829159.2;refGene_gn=CENPE;refGene_tx=XM_023252886.1;refGene_gn=CENPE;refGene_tx=XM_023252887.1;refGene_gn=CENPE;refGene_tx=XM_023252888.1;refGene_gn=CENPE;refGene_tx=XM_023252889.1;refGene_gn=CENPE;refGene_tx=XM_023252890.1;refGene_gn=CENPE;refGene_tx=XM_023252891.1
B1 121309061 121309113
refGene_gn=CENPE;refGene_tx=XM_019829159.2;refGene_gn=CENPE;refGene_tx=XM_023252886.1;refGene_gn=CENPE;refGene_tx=XM_023252887.1;refGene_gn=CENPE;refGene_tx=XM_023252888.1;refGene_gn=CENPE;refGene_tx=XM_023252889.1;refGene_gn=CENPE;refGene_tx=XM_023252890.1;refGene_gn=CENPE;refGene_tx=XM_023252891.1
B1 121310854 121310942
refGene_gn=CENPE;refGene_tx=XM_019829159.2;refGene_gn=CENPE;refGene_tx=XM_023252886.1;refGene_gn=CENPE;refGene_tx=XM_023252887.1;refGene_gn=CENPE;refGene_tx=XM_023252888.1;refGene_gn=CENPE;refGene_tx=XM_023252889.1;refGene_gn=CENPE;refGene_tx=XM_023252890.1;refGene_gn=CENPE;refGene_tx=XM_023252891.1
B1 121311290 121311420
refGene_gn=CENPE;refGene_tx=XM_019829159.2;refGene_gn=CENPE;refGene_tx=XM_023252886.1;refGene_gn=CENPE;refGene_tx=XM_023252887.1;refGene_gn=

CENPE;refGene_tx=XM_023252888.1;refGene_gn=CENPE;refGene_tx=XM_023252889.1;refGene_gn=CENPE;refGene_tx=XM_023252890.1;refGene_gn=CENPE;refGene_tx=XM_023252891.1
B1 121313173 121313293
refGene_gn=CENPE;refGene_tx=XM_019829159.2;refGene_gn=CENPE;refGene_tx=XM_023252886.1;refGene_gn=CENPE;refGene_tx=XM_023252887.1;refGene_gn=CENPE;refGene_tx=XM_023252888.1;refGene_gn=CENPE;refGene_tx=XM_023252889.1;refGene_gn=CENPE;refGene_tx=XM_023252890.1;refGene_gn=CENPE;refGene_tx=XM_023252891.1
B1 121316677 121316836
refGene_gn=CENPE;refGene_tx=XM_019829159.2;refGene_gn=CENPE;refGene_tx=XM_023252886.1;refGene_gn=CENPE;refGene_tx=XM_023252887.1;refGene_gn=CENPE;refGene_tx=XM_023252888.1;refGene_gn=CENPE;refGene_tx=XM_023252889.1;refGene_gn=CENPE;refGene_tx=XM_023252890.1;refGene_gn=CENPE;refGene_tx=XM_023252891.1
B1 121320184 121320329
refGene_gn=CENPE;refGene_tx=XM_019829159.2;refGene_gn=CENPE;refGene_tx=XM_023252886.1;refGene_gn=CENPE;refGene_tx=XM_023252887.1;refGene_gn=CENPE;refGene_tx=XM_023252888.1;refGene_gn=CENPE;refGene_tx=XM_023252889.1;refGene_gn=CENPE;refGene_tx=XM_023252890.1;refGene_gn=CENPE;refGene_tx=XM_023252891.1
B1 121321245 121321334
refGene_gn=CENPE;refGene_tx=XM_019829159.2;refGene_gn=CENPE;refGene_tx=XM_023252886.1;refGene_gn=CENPE;refGene_tx=XM_023252887.1;refGene_gn=CENPE;refGene_tx=XM_023252888.1;refGene_gn=CENPE;refGene_tx=XM_023252889.1;refGene_gn=CENPE;refGene_tx=XM_023252890.1;refGene_gn=CENPE;refGene_tx=XM_023252891.1
B1 121323213 121323381
refGene_gn=CENPE;refGene_tx=XM_019829159.2;refGene_gn=CENPE;refGene_tx=XM_023252886.1;refGene_gn=CENPE;refGene_tx=XM_023252887.1;refGene_gn=CENPE;refGene_tx=XM_023252888.1;refGene_gn=CENPE;refGene_tx=XM_023252889.1;refGene_gn=CENPE;refGene_tx=XM_023252890.1;refGene_gn=CENPE;refGene_tx=XM_023252891.1
B1 121327166 121327241
refGene_gn=CENPE;refGene_tx=XM_019829159.2;refGene_gn=CENPE;refGene_tx=XM_023252886.1;refGene_gn=CENPE;refGene_tx=XM_023252887.1;refGene_gn=CENPE;refGene_tx=XM_023252889.1;refGene_gn=CENPE;refGene_tx=XM_023252890.1;refGene_gn=CENPE;refGene_tx=XM_023252891.1
B1 121327466 121327586
refGene_gn=CENPE;refGene_tx=XM_019829159.2;refGene_gn=CENPE;refGene_tx=XM_023252886.1;refGene_gn=CENPE;refGene_tx=XM_023252887.1;refGene_gn=CENPE;refGene_tx=XM_023252888.1;refGene_gn=CENPE;refGene_tx=XM_023252889.1;refGene_gn=CENPE;refGene_tx=XM_023252890.1;refGene_gn=CENPE;refGene_tx=XM_023252891.1
B1 121329310 121329433
refGene_gn=CENPE;refGene_tx=XM_019829159.2;refGene_gn=CENPE;refGene_tx=XM_023252886.1;refGene_gn=CENPE;refGene_tx=XM_023252887.1;refGene_gn=CENPE;refGene_tx=XM_023252888.1;refGene_gn=CENPE;refGene_tx=XM_023252889.1;refGene_gn=CENPE;refGene_tx=XM_023252890.1;refGene_gn=CENPE;refGene_tx=XM_023252891.1

B1 121329514 121329680
refGene_gn=CENPE;refGene_tx=XM_019829159.2;refGene_gn=CENPE;refGene_tx=XM_023252886.1;refGene_gn=CENPE;refGene_tx=XM_023252887.1;refGene_gn=CENPE;refGene_tx=XM_023252888.1;refGene_gn=CENPE;refGene_tx=XM_023252889.1;refGene_gn=CENPE;refGene_tx=XM_023252890.1;refGene_gn=CENPE;refGene_tx=XM_023252891.1
B1 121330084 121330239
refGene_gn=CENPE;refGene_tx=XM_019829159.2;refGene_gn=CENPE;refGene_tx=XM_023252886.1;refGene_gn=CENPE;refGene_tx=XM_023252887.1;refGene_gn=CENPE;refGene_tx=XM_023252888.1;refGene_gn=CENPE;refGene_tx=XM_023252889.1;refGene_gn=CENPE;refGene_tx=XM_023252890.1;refGene_gn=CENPE;refGene_tx=XM_023252891.1
B1 121331822 121332134
refGene_gn=CENPE;refGene_tx=XM_019829159.2;refGene_gn=CENPE;refGene_tx=XM_023252886.1;refGene_gn=CENPE;refGene_tx=XM_023252887.1;refGene_gn=CENPE;refGene_tx=XM_023252888.1;refGene_gn=CENPE;refGene_tx=XM_023252889.1;refGene_gn=CENPE;refGene_tx=XM_023252890.1;refGene_gn=CENPE;refGene_tx=XM_023252891.1
B1 121332247 121332520
refGene_gn=CENPE;refGene_tx=XM_019829159.2;refGene_gn=CENPE;refGene_tx=XM_023252886.1;refGene_gn=CENPE;refGene_tx=XM_023252887.1;refGene_gn=CENPE;refGene_tx=XM_023252888.1;refGene_gn=CENPE;refGene_tx=XM_023252889.1;refGene_gn=CENPE;refGene_tx=XM_023252890.1;refGene_gn=CENPE;refGene_tx=XM_023252891.1
B1 121332674 121332833
refGene_gn=CENPE;refGene_tx=XM_019829159.2;refGene_gn=CENPE;refGene_tx=XM_023252886.1;refGene_gn=CENPE;refGene_tx=XM_023252887.1;refGene_gn=CENPE;refGene_tx=XM_023252888.1;refGene_gn=CENPE;refGene_tx=XM_023252889.1;refGene_gn=CENPE;refGene_tx=XM_023252890.1;refGene_gn=CENPE;refGene_tx=XM_023252891.1
B1 121336593 121336797
refGene_gn=CENPE;refGene_tx=XM_019829159.2;refGene_gn=CENPE;refGene_tx=XM_023252886.1;refGene_gn=CENPE;refGene_tx=XM_023252887.1;refGene_gn=CENPE;refGene_tx=XM_023252888.1;refGene_gn=CENPE;refGene_tx=XM_023252889.1;refGene_gn=CENPE;refGene_tx=XM_023252890.1;refGene_gn=CENPE;refGene_tx=XM_023252891.1
B1 121338661 121338820
refGene_gn=CENPE;refGene_tx=XM_019829159.2;refGene_gn=CENPE;refGene_tx=XM_023252886.1;refGene_gn=CENPE;refGene_tx=XM_023252887.1;refGene_gn=CENPE;refGene_tx=XM_023252888.1;refGene_gn=CENPE;refGene_tx=XM_023252889.1;refGene_gn=CENPE;refGene_tx=XM_023252890.1;refGene_gn=CENPE;refGene_tx=XM_023252891.1
B1 121340520 121340811
refGene_gn=CENPE;refGene_tx=XM_019829159.2;refGene_gn=CENPE;refGene_tx=XM_023252886.1;refGene_gn=CENPE;refGene_tx=XM_023252887.1;refGene_gn=CENPE;refGene_tx=XM_023252888.1;refGene_gn=CENPE;refGene_tx=XM_023252889.1;refGene_gn=CENPE;refGene_tx=XM_023252890.1;refGene_gn=CENPE;refGene_tx=XM_023252891.1
B1 121340930 121341089
refGene_gn=CENPE;refGene_tx=XM_019829159.2;refGene_gn=CENPE;refGene_tx

=XM_023252886.1;refGene_gn=CENPE;refGene_tx=XM_023252887.1;refGene_gn=CENPE;refGene_tx=XM_023252888.1;refGene_gn=CENPE;refGene_tx=XM_023252889.1;refGene_gn=CENPE;refGene_tx=XM_023252890.1;refGene_gn=CENPE;refGene_tx=XM_023252891.1
B1 121342066 121342357
refGene_gn=CENPE;refGene_tx=XM_019829159.2;refGene_gn=CENPE;refGene_tx=XM_023252886.1;refGene_gn=CENPE;refGene_tx=XM_023252887.1;refGene_gn=CENPE;refGene_tx=XM_023252888.1;refGene_gn=CENPE;refGene_tx=XM_023252889.1;refGene_gn=CENPE;refGene_tx=XM_023252890.1;refGene_gn=CENPE;refGene_tx=XM_023252891.1
B1 121343889 121344180
refGene_gn=CENPE;refGene_tx=XM_019829159.2;refGene_gn=CENPE;refGene_tx=XM_023252886.1;refGene_gn=CENPE;refGene_tx=XM_023252887.1;refGene_gn=CENPE;refGene_tx=XM_023252888.1;refGene_gn=CENPE;refGene_tx=XM_023252889.1;refGene_gn=CENPE;refGene_tx=XM_023252890.1;refGene_gn=CENPE;refGene_tx=XM_023252891.1
B1 121344334 121344493
refGene_gn=CENPE;refGene_tx=XM_019829159.2;refGene_gn=CENPE;refGene_tx=XM_023252886.1;refGene_gn=CENPE;refGene_tx=XM_023252887.1;refGene_gn=CENPE;refGene_tx=XM_023252888.1;refGene_gn=CENPE;refGene_tx=XM_023252889.1;refGene_gn=CENPE;refGene_tx=XM_023252890.1;refGene_gn=CENPE;refGene_tx=XM_023252891.1
B1 121344708 121344996
refGene_gn=CENPE;refGene_tx=XM_019829159.2;refGene_gn=CENPE;refGene_tx=XM_023252886.1;refGene_gn=CENPE;refGene_tx=XM_023252887.1;refGene_gn=CENPE;refGene_tx=XM_023252888.1;refGene_gn=CENPE;refGene_tx=XM_023252889.1;refGene_gn=CENPE;refGene_tx=XM_023252890.1;refGene_gn=CENPE;refGene_tx=XM_023252891.1
B1 121345147 121345252
refGene_gn=CENPE;refGene_tx=XM_019829159.2;refGene_gn=CENPE;refGene_tx=XM_023252886.1;refGene_gn=CENPE;refGene_tx=XM_023252887.1;refGene_gn=CENPE;refGene_tx=XM_023252888.1;refGene_gn=CENPE;refGene_tx=XM_023252889.1;refGene_gn=CENPE;refGene_tx=XM_023252890.1;refGene_gn=CENPE;refGene_tx=XM_023252891.1
B1 121345428 121345716
refGene_gn=CENPE;refGene_tx=XM_019829159.2;refGene_gn=CENPE;refGene_tx=XM_023252886.1;refGene_gn=CENPE;refGene_tx=XM_023252887.1;refGene_gn=CENPE;refGene_tx=XM_023252888.1;refGene_gn=CENPE;refGene_tx=XM_023252889.1;refGene_gn=CENPE;refGene_tx=XM_023252890.1;refGene_gn=CENPE;refGene_tx=XM_023252891.1
B1 121346303 121346462
refGene_gn=CENPE;refGene_tx=XM_019829159.2;refGene_gn=CENPE;refGene_tx=XM_023252886.1;refGene_gn=CENPE;refGene_tx=XM_023252887.1;refGene_gn=CENPE;refGene_tx=XM_023252888.1;refGene_gn=CENPE;refGene_tx=XM_023252889.1;refGene_gn=CENPE;refGene_tx=XM_023252890.1;refGene_gn=CENPE;refGene_tx=XM_023252891.1
B1 121346855 121347137
refGene_gn=CENPE;refGene_tx=XM_019829159.2;refGene_gn=CENPE;refGene_tx=XM_023252886.1;refGene_gn=CENPE;refGene_tx=XM_023252887.1;refGene_gn=CENPE;refGene_tx=XM_023252888.1;refGene_gn=CENPE;refGene_tx=XM_0232528

89.1;refGene_gn=CENPE;refGene_tx=XM_023252890.1;refGene_gn=CENPE;refGene_tx=XM_023252891.1
B1	121347782	121347941
refGene_gn=CENPE;refGene_tx=XM_019829159.2;refGene_gn=CENPE;refGene_tx=XM_023252886.1;refGene_gn=CENPE;refGene_tx=XM_023252887.1;refGene_gn=CENPE;refGene_tx=XM_023252888.1;refGene_gn=CENPE;refGene_tx=XM_023252889.1;refGene_gn=CENPE;refGene_tx=XM_023252890.1;refGene_gn=CENPE;refGene_tx=XM_023252891.1
B1	121349067	121349337
refGene_gn=CENPE;refGene_tx=XM_019829159.2;refGene_gn=CENPE;refGene_tx=XM_023252886.1;refGene_gn=CENPE;refGene_tx=XM_023252887.1;refGene_gn=CENPE;refGene_tx=XM_023252888.1;refGene_gn=CENPE;refGene_tx=XM_023252889.1;refGene_gn=CENPE;refGene_tx=XM_023252890.1;refGene_gn=CENPE;refGene_tx=XM_023252891.1
B1	121349967	121350126
refGene_gn=CENPE;refGene_tx=XM_019829159.2;refGene_gn=CENPE;refGene_tx=XM_023252886.1;refGene_gn=CENPE;refGene_tx=XM_023252887.1;refGene_gn=CENPE;refGene_tx=XM_023252888.1;refGene_gn=CENPE;refGene_tx=XM_023252889.1
B1	121350290	121350581
refGene_gn=CENPE;refGene_tx=XM_019829159.2;refGene_gn=CENPE;refGene_tx=XM_023252886.1;refGene_gn=CENPE;refGene_tx=XM_023252887.1;refGene_gn=CENPE;refGene_tx=XM_023252888.1;refGene_gn=CENPE;refGene_tx=XM_023252889.1;refGene_gn=CENPE;refGene_tx=XM_023252890.1
B1	121351808	121351913
refGene_gn=CENPE;refGene_tx=XM_019829159.2;refGene_gn=CENPE;refGene_tx=XM_023252886.1;refGene_gn=CENPE;refGene_tx=XM_023252887.1;refGene_gn=CENPE;refGene_tx=XM_023252888.1;refGene_gn=CENPE;refGene_tx=XM_023252889.1;refGene_gn=CENPE;refGene_tx=XM_023252890.1;refGene_gn=CENPE;refGene_tx=XM_023252891.1
B1	121353856	121354075
refGene_gn=CENPE;refGene_tx=XM_019829159.2;refGene_gn=CENPE;refGene_tx=XM_023252886.1;refGene_gn=CENPE;refGene_tx=XM_023252887.1;refGene_gn=CENPE;refGene_tx=XM_023252888.1;refGene_gn=CENPE;refGene_tx=XM_023252889.1;refGene_gn=CENPE;refGene_tx=XM_023252890.1;refGene_gn=CENPE;refGene_tx=XM_023252891.1
B1	121356166	121356364
refGene_gn=CENPE;refGene_tx=XM_019829159.2;refGene_gn=CENPE;refGene_tx=XM_023252886.1;refGene_gn=CENPE;refGene_tx=XM_023252887.1;refGene_gn=CENPE;refGene_tx=XM_023252888.1;refGene_gn=CENPE;refGene_tx=XM_023252889.1;refGene_gn=CENPE;refGene_tx=XM_023252890.1;refGene_gn=CENPE;refGene_tx=XM_023252891.1
B1	121357054	121357258
refGene_gn=CENPE;refGene_tx=XM_019829159.2;refGene_gn=CENPE;refGene_tx=XM_023252886.1;refGene_gn=CENPE;refGene_tx=XM_023252887.1;refGene_gn=CENPE;refGene_tx=XM_023252888.1;refGene_gn=CENPE;refGene_tx=XM_023252889.1;refGene_gn=CENPE;refGene_tx=XM_023252890.1;refGene_gn=CENPE;refGene_tx=XM_023252891.1
B1	121359808	121360027
refGene_gn=CENPE;refGene_tx=XM_019829159.2;refGene_gn=CENPE;refGene_tx

=XM_023252886.1;refGene_gn=CENPE;refGene_tx=XM_023252887.1;refGene_gn=CENPE;refGene_tx=XM_023252888.1;refGene_gn=CENPE;refGene_tx=XM_023252889.1;refGene_gn=CENPE;refGene_tx=XM_023252890.1;refGene_gn=CENPE;refGene_tx=XM_023252891.1
B1 121363496 121363682
refGene_gn=CENPE;refGene_tx=XM_019829159.2;refGene_gn=CENPE;refGene_tx=XM_023252886.1;refGene_gn=CENPE;refGene_tx=XM_023252887.1;refGene_gn=CENPE;refGene_tx=XM_023252888.1;refGene_gn=CENPE;refGene_tx=XM_023252889.1;refGene_gn=CENPE;refGene_tx=XM_023252890.1;refGene_gn=CENPE;refGene_tx=XM_023252891.1
B1 121369834 121369947
refGene_gn=CENPE;refGene_tx=XM_019829159.2;refGene_gn=CENPE;refGene_tx=XM_023252886.1;refGene_gn=CENPE;refGene_tx=XM_023252887.1;refGene_gn=CENPE;refGene_tx=XM_023252888.1;refGene_gn=CENPE;refGene_tx=XM_023252889.1;refGene_gn=CENPE;refGene_tx=XM_023252890.1;refGene_gn=CENPE;refGene_tx=XM_023252891.1
B1 121371298 121371396
refGene_gn=CENPE;refGene_tx=XM_019829159.2;refGene_gn=CENPE;refGene_tx=XM_023252886.1;refGene_gn=CENPE;refGene_tx=XM_023252887.1;refGene_gn=CENPE;refGene_tx=XM_023252888.1;refGene_gn=CENPE;refGene_tx=XM_023252889.1;refGene_gn=CENPE;refGene_tx=XM_023252890.1;refGene_gn=CENPE;refGene_tx=XM_023252891.1
B1 121373025 121373209
refGene_gn=CENPE;refGene_tx=XM_019829159.2;refGene_gn=CENPE;refGene_tx=XM_023252886.1;refGene_gn=CENPE;refGene_tx=XM_023252887.1;refGene_gn=CENPE;refGene_tx=XM_023252888.1;refGene_gn=CENPE;refGene_tx=XM_023252889.1;refGene_gn=CENPE;refGene_tx=XM_023252890.1;refGene_gn=CENPE;refGene_tx=XM_023252891.1
B1 121376048 121376335
refGene_gn=CENPE;refGene_tx=XM_019829159.2;refGene_gn=CENPE;refGene_tx=XM_023252886.1;refGene_gn=CENPE;refGene_tx=XM_023252887.1;refGene_gn=CENPE;refGene_tx=XM_023252888.1;refGene_gn=CENPE;refGene_tx=XM_023252889.1;refGene_gn=CENPE;refGene_tx=XM_023252890.1;refGene_gn=CENPE;refGene_tx=XM_023252891.1
B1 121379109 121379204
refGene_gn=CENPE;refGene_tx=XM_019829159.2;refGene_gn=CENPE;refGene_tx=XM_023252886.1;refGene_gn=CENPE;refGene_tx=XM_023252888.1;refGene_gn=CENPE;refGene_tx=XM_023252889.1;refGene_gn=CENPE;refGene_tx=XM_023252890.1;refGene_gn=CENPE;refGene_tx=XM_023252891.1
B1 121383017 121383097
refGene_gn=CENPE;refGene_tx=XM_023252887.1
B1 121387031 121387106
refGene_gn=BDH2;refGene_tx=XM_003985122.4
B1 121388014 121388093
refGene_gn=BDH2;refGene_tx=XM_003985122.4
B1 121390191 121390288
refGene_gn=BDH2;refGene_tx=XM_003985122.4
B1 121392771 121392880
refGene_gn=BDH2;refGene_tx=XM_003985122.4
B1 121399615 121399676

refGene_gn=BDH2;refGene_tx=XM_003985122.4
B1	121401132	121401246
refGene_gn=BDH2;refGene_tx=XM_003985122.4
B1	121404517	121404576
refGene_gn=BDH2;refGene_tx=XM_003985122.4
B1	121405161	121405254
refGene_gn=BDH2;refGene_tx=XM_003985122.4
B1	121407256	121407310
refGene_gn=BDH2;refGene_tx=XM_003985122.4
B1	121425165	121425255
refGene_gn=SLC9B2;refGene_tx=XM_003985123.5;refGene_gn=SLC9B2;refGene_tx=XM_019829162.2
B1	121425938	121426119
refGene_gn=SLC9B2;refGene_tx=XM_003985123.5;refGene_gn=SLC9B2;refGene_tx=XM_019829162.2
B1	121432113	121432284
refGene_gn=SLC9B2;refGene_tx=XM_003985123.5;refGene_gn=SLC9B2;refGene_tx=XM_019829162.2;refGene_gn=SLC9B2;refGene_tx=XM_019829163.2
B1	121436206	121436349
refGene_gn=SLC9B2;refGene_tx=XM_003985123.5;refGene_gn=SLC9B2;refGene_tx=XM_019829162.2;refGene_gn=SLC9B2;refGene_tx=XM_019829163.2
B1	121437430	121437558
refGene_gn=SLC9B2;refGene_tx=XM_003985123.5;refGene_gn=SLC9B2;refGene_tx=XM_019829162.2;refGene_gn=SLC9B2;refGene_tx=XM_019829163.2
B1	121439297	121439473
refGene_gn=SLC9B2;refGene_tx=XM_003985123.5;refGene_gn=SLC9B2;refGene_tx=XM_019829162.2;refGene_gn=SLC9B2;refGene_tx=XM_019829163.2
B1	121443115	121443222
refGene_gn=SLC9B2;refGene_tx=XM_003985123.5;refGene_gn=SLC9B2;refGene_tx=XM_019829162.2;refGene_gn=SLC9B2;refGene_tx=XM_019829163.2
B1	121446348	121446498
refGene_gn=SLC9B2;refGene_tx=XM_003985123.5;refGene_gn=SLC9B2;refGene_tx=XM_019829162.2;refGene_gn=SLC9B2;refGene_tx=XM_019829163.2
B1	121454255	121454364
refGene_gn=SLC9B2;refGene_tx=XM_003985123.5;refGene_gn=SLC9B2;refGene_tx=XM_019829162.2;refGene_gn=SLC9B2;refGene_tx=XM_019829163.2
B1	121458258	121458395
refGene_gn=SLC9B2;refGene_tx=XM_003985123.5;refGene_gn=SLC9B2;refGene_tx=XM_019829163.2
B1	121461111	121461363
refGene_gn=SLC9B2;refGene_tx=XM_003985123.5;refGene_gn=SLC9B2;refGene_tx=XM_019829162.2;refGene_gn=SLC9B2;refGene_tx=XM_019829163.2
B1	121494544	121494613
refGene_gn=SLC9B1;refGene_tx=XM_023252892.1;refGene_gn=SLC9B1;refGene_tx=XM_023252893.1;refGene_gn=SLC9B1;refGene_tx=XM_023252894.1;refGene_gn=SLC9B1;refGene_tx=XM_023252895.1;refGene_gn=SLC9B1;refGene_tx=XM_023252896.1;refGene_gn=SLC9B1;refGene_tx=XM_023252897.1;refGene_gn=SLC9B1;refGene_tx=XM_023252898.1;refGene_gn=SLC9B1;refGene_tx=XM_023252899.1
B1	121496658	121496797

refGene_gn=SLC9B1;refGene_tx=XM_023252892.1;refGene_gn=SLC9B1;refGene_tx=XM_023252893.1;refGene_gn=SLC9B1;refGene_tx=XM_023252894.1;refGene_gn=SLC9B1;refGene_tx=XM_023252895.1;refGene_gn=SLC9B1;refGene_tx=XM_023252896.1;refGene_gn=SLC9B1;refGene_tx=XM_023252897.1;refGene_gn=SLC9B1;refGene_tx=XM_023252898.1;refGene_gn=SLC9B1;refGene_tx=XM_023252899.1
B1 121511206 121511377
refGene_gn=SLC9B1;refGene_tx=XM_023252892.1;refGene_gn=SLC9B1;refGene_tx=XM_023252893.1;refGene_gn=SLC9B1;refGene_tx=XM_023252894.1;refGene_gn=SLC9B1;refGene_tx=XM_023252895.1;refGene_gn=SLC9B1;refGene_tx=XM_023252896.1;refGene_gn=SLC9B1;refGene_tx=XM_023252897.1;refGene_gn=SLC9B1;refGene_tx=XM_023252898.1;refGene_gn=SLC9B1;refGene_tx=XM_023252899.1;refGene_gn=SLC9B1;refGene_tx=XM_023252900.1
B1 121512589 121512732
refGene_gn=SLC9B1;refGene_tx=XM_023252892.1;refGene_gn=SLC9B1;refGene_tx=XM_023252893.1;refGene_gn=SLC9B1;refGene_tx=XM_023252894.1;refGene_gn=SLC9B1;refGene_tx=XM_023252895.1;refGene_gn=SLC9B1;refGene_tx=XM_023252896.1;refGene_gn=SLC9B1;refGene_tx=XM_023252897.1;refGene_gn=SLC9B1;refGene_tx=XM_023252898.1;refGene_gn=SLC9B1;refGene_tx=XM_023252899.1;refGene_gn=SLC9B1;refGene_tx=XM_023252900.1
B1 121516539 121516667
refGene_gn=SLC9B1;refGene_tx=XM_023252892.1;refGene_gn=SLC9B1;refGene_tx=XM_023252893.1;refGene_gn=SLC9B1;refGene_tx=XM_023252894.1;refGene_gn=SLC9B1;refGene_tx=XM_023252895.1;refGene_gn=SLC9B1;refGene_tx=XM_023252896.1;refGene_gn=SLC9B1;refGene_tx=XM_023252897.1;refGene_gn=SLC9B1;refGene_tx=XM_023252898.1;refGene_gn=SLC9B1;refGene_tx=XM_023252899.1;refGene_gn=SLC9B1;refGene_tx=XM_023252900.1
B1 121536173 121536361
refGene_gn=SLC9B1;refGene_tx=XM_023252892.1;refGene_gn=SLC9B1;refGene_tx=XM_023252893.1;refGene_gn=SLC9B1;refGene_tx=XM_023252894.1;refGene_gn=SLC9B1;refGene_tx=XM_023252895.1;refGene_gn=SLC9B1;refGene_tx=XM_023252896.1;refGene_gn=SLC9B1;refGene_tx=XM_023252897.1;refGene_gn=SLC9B1;refGene_tx=XM_023252898.1;refGene_gn=SLC9B1;refGene_tx=XM_023252899.1;refGene_gn=SLC9B1;refGene_tx=XM_023252900.1
B1 121540397 121540504
refGene_gn=SLC9B1;refGene_tx=XM_023252892.1;refGene_gn=SLC9B1;refGene_tx=XM_023252893.1;refGene_gn=SLC9B1;refGene_tx=XM_023252894.1;refGene_gn=SLC9B1;refGene_tx=XM_023252895.1;refGene_gn=SLC9B1;refGene_tx=XM_023252896.1;refGene_gn=SLC9B1;refGene_tx=XM_023252897.1;refGene_gn=SLC9B1;refGene_tx=XM_023252898.1;refGene_gn=SLC9B1;refGene_tx=XM_023252899.1;refGene_gn=SLC9B1;refGene_tx=XM_023252900.1
B1 121541337 121541487
refGene_gn=SLC9B1;refGene_tx=XM_023252892.1;refGene_gn=SLC9B1;refGene_tx=XM_023252893.1;refGene_gn=SLC9B1;refGene_tx=XM_023252894.1;refGene_gn=SLC9B1;refGene_tx=XM_023252895.1;refGene_gn=SLC9B1;refGene_tx=XM_023252896.1;refGene_gn=SLC9B1;refGene_tx=XM_023252897.1;refGene_gn=SLC9B1;refGene_tx=XM_023252898.1;refGene_gn=SLC9B1;refGene_tx=XM_023252899.1;refGene_gn=SLC9B1;refGene_tx=XM_023252900.1
B1 121547009 121547118
refGene_gn=SLC9B1;refGene_tx=XM_023252892.1;refGene_gn=SLC9B1;refGene_

tx=XM_023252893.1;refGene_gn=SLC9B1;refGene_tx=XM_023252894.1;refGene_gn=SLC9B1;refGene_tx=XM_023252895.1;refGene_gn=SLC9B1;refGene_tx=XM_023252896.1;refGene_gn=SLC9B1;refGene_tx=XM_023252897.1;refGene_gn=SLC9B1;refGene_tx=XM_023252898.1;refGene_gn=SLC9B1;refGene_tx=XM_023252899.1;refGene_gn=SLC9B1;refGene_tx=XM_023252900.1
B1	121547690	121547827
refGene_gn=SLC9B1;refGene_tx=XM_023252892.1;refGene_gn=SLC9B1;refGene_tx=XM_023252893.1;refGene_gn=SLC9B1;refGene_tx=XM_023252894.1;refGene_gn=SLC9B1;refGene_tx=XM_023252895.1;refGene_gn=SLC9B1;refGene_tx=XM_023252896.1;refGene_gn=SLC9B1;refGene_tx=XM_023252897.1;refGene_gn=SLC9B1;refGene_tx=XM_023252898.1;refGene_gn=SLC9B1;refGene_tx=XM_023252899.1;refGene_gn=SLC9B1;refGene_tx=XM_023252900.1
B1	121549261	121549474
refGene_gn=SLC9B1;refGene_tx=XM_023252892.1;refGene_gn=SLC9B1;refGene_tx=XM_023252893.1;refGene_gn=SLC9B1;refGene_tx=XM_023252894.1;refGene_gn=SLC9B1;refGene_tx=XM_023252895.1;refGene_gn=SLC9B1;refGene_tx=XM_023252896.1;refGene_gn=SLC9B1;refGene_tx=XM_023252897.1;refGene_gn=SLC9B1;refGene_tx=XM_023252898.1;refGene_gn=SLC9B1;refGene_tx=XM_023252899.1;refGene_gn=SLC9B1;refGene_tx=XM_023252900.1
B1	121570826	121570916
refGene_gn=CISD2;refGene_tx=XM_003985124.4
B1	121572503	121572718
refGene_gn=CISD2;refGene_tx=XM_003985124.4
B1	121583071	121583174
refGene_gn=CISD2;refGene_tx=XM_003985124.4
B1	121631757	121631781
refGene_gn=UBE2D3;refGene_tx=XM_003985128.4;refGene_gn=UBE2D3;refGene_tx=XM_006930938.3;refGene_gn=UBE2D3;refGene_tx=XM_006930939.3;refGene_gn=UBE2D3;refGene_tx=XM_006930940.3
B1	121646428	121646492
refGene_gn=UBE2D3;refGene_tx=XM_003985128.4;refGene_gn=UBE2D3;refGene_tx=XM_006930938.3;refGene_gn=UBE2D3;refGene_tx=XM_006930939.3;refGene_gn=UBE2D3;refGene_tx=XM_006930940.3
B1	121646579	121646611
refGene_gn=UBE2D3;refGene_tx=XM_003985128.4;refGene_gn=UBE2D3;refGene_tx=XM_006930938.3;refGene_gn=UBE2D3;refGene_tx=XM_006930939.3;refGene_gn=UBE2D3;refGene_tx=XM_006930940.3
B1	121653065	121653143
refGene_gn=UBE2D3;refGene_tx=XM_003985128.4;refGene_gn=UBE2D3;refGene_tx=XM_006930938.3;refGene_gn=UBE2D3;refGene_tx=XM_006930939.3;refGene_gn=UBE2D3;refGene_tx=XM_006930940.3
B1	121654348	121654454
refGene_gn=UBE2D3;refGene_tx=XM_003985128.4;refGene_gn=UBE2D3;refGene_tx=XM_006930938.3;refGene_gn=UBE2D3;refGene_tx=XM_006930939.3;refGene_gn=UBE2D3;refGene_tx=XM_006930940.3
B1	121656227	121656321
refGene_gn=UBE2D3;refGene_tx=XM_003985128.4;refGene_gn=UBE2D3;refGene_tx=XM_006930938.3;refGene_gn=UBE2D3;refGene_tx=XM_006930939.3;refGene_gn=UBE2D3;refGene_tx=XM_006930940.3
B1	121656741	121656790

refGene_gn=UBE2D3;refGene_tx=XM_006930938.3;refGene_gn=UBE2D3;refGene_tx=XM_006930939.3;refGene_gn=UBE2D3;refGene_tx=XM_006930940.3
B1 121658326 121658372
refGene_gn=UBE2D3;refGene_tx=XM_003985128.4
B1 121686724 121686901
refGene_gn=MANBA;refGene_tx=XM_023252901.1
B1 121706062 121706157
refGene_gn=MANBA;refGene_tx=XM_023252901.1
B1 121710011 121710117
refGene_gn=MANBA;refGene_tx=XM_023252901.1
B1 121710947 121711118
refGene_gn=MANBA;refGene_tx=XM_023252901.1
B1 121718027 121718151
refGene_gn=MANBA;refGene_tx=XM_023252901.1
B1 121734119 121734295
refGene_gn=MANBA;refGene_tx=XM_023252901.1
B1 121735396 121735507
refGene_gn=MANBA;refGene_tx=XM_023252901.1
B1 121753350 121753502
refGene_gn=MANBA;refGene_tx=XM_023252901.1
B1 121758019 121758137
refGene_gn=MANBA;refGene_tx=XM_023252901.1
B1 121759359 121759446
refGene_gn=MANBA;refGene_tx=XM_023252901.1
B1 121763048 121763216
refGene_gn=MANBA;refGene_tx=XM_023252901.1
B1 121772256 121772475
refGene_gn=MANBA;refGene_tx=XM_023252901.1
B1 121779784 121779949
refGene_gn=MANBA;refGene_tx=XM_023252901.1
B1 121792493 121792560
refGene_gn=MANBA;refGene_tx=XM_023252901.1
B1 121792562 121792637
refGene_gn=MANBA;refGene_tx=XM_023252901.1
B1 121796019 121796162
refGene_gn=MANBA;refGene_tx=XM_023252901.1
B1 121796969 121797227
refGene_gn=MANBA;refGene_tx=XM_023252901.1
B1 121800248 121800473
refGene_gn=MANBA;refGene_tx=XM_023252901.1
B1 121815942 121816100
refGene_gn=NFKB1;refGene_tx=XM_023252903.1;refGene_gn=NFKB1;refGene_tx=XM_023252904.1;refGene_gn=NFKB1;refGene_tx=XM_023252905.1;refGene_gn=NFKB1;refGene_tx=XM_023252906.1;refGene_gn=NFKB1;refGene_tx=XM_023252907.1
B1 121819771 121819940
refGene_gn=NFKB1;refGene_tx=XM_023252903.1;refGene_gn=NFKB1;refGene_tx=XM_023252904.1;refGene_gn=NFKB1;refGene_tx=XM_023252905.1;refGene_gn=NFKB1;refGene_tx=XM_023252906.1;refGene_gn=NFKB1;refGene_tx=XM_023252907.1

B1 121820684 121820857
refGene_gn=NFKB1;refGene_tx=XM_023252903.1;refGene_gn=NFKB1;refGene_tx=XM_023252904.1;refGene_gn=NFKB1;refGene_tx=XM_023252905.1;refGene_gn=NFKB1;refGene_tx=XM_023252906.1;refGene_gn=NFKB1;refGene_tx=XM_023252907.1
B1 121821175 121821242
refGene_gn=NFKB1;refGene_tx=XM_023252903.1;refGene_gn=NFKB1;refGene_tx=XM_023252904.1;refGene_gn=NFKB1;refGene_tx=XM_023252905.1;refGene_gn=NFKB1;refGene_tx=XM_023252906.1;refGene_gn=NFKB1;refGene_tx=XM_023252907.1
B1 121822632 121822757
refGene_gn=NFKB1;refGene_tx=XM_023252903.1;refGene_gn=NFKB1;refGene_tx=XM_023252904.1;refGene_gn=NFKB1;refGene_tx=XM_023252905.1;refGene_gn=NFKB1;refGene_tx=XM_023252906.1;refGene_gn=NFKB1;refGene_tx=XM_023252907.1
B1 121825531 121825634
refGene_gn=NFKB1;refGene_tx=XM_023252903.1;refGene_gn=NFKB1;refGene_tx=XM_023252904.1;refGene_gn=NFKB1;refGene_tx=XM_023252905.1;refGene_gn=NFKB1;refGene_tx=XM_023252906.1;refGene_gn=NFKB1;refGene_tx=XM_023252907.1
B1 121825970 121826140
refGene_gn=NFKB1;refGene_tx=XM_023252903.1;refGene_gn=NFKB1;refGene_tx=XM_023252904.1;refGene_gn=NFKB1;refGene_tx=XM_023252905.1;refGene_gn=NFKB1;refGene_tx=XM_023252906.1;refGene_gn=NFKB1;refGene_tx=XM_023252907.1
B1 121826594 121826796
refGene_gn=NFKB1;refGene_tx=XM_023252903.1;refGene_gn=NFKB1;refGene_tx=XM_023252904.1;refGene_gn=NFKB1;refGene_tx=XM_023252905.1;refGene_gn=NFKB1;refGene_tx=XM_023252906.1;refGene_gn=NFKB1;refGene_tx=XM_023252907.1
B1 121831895 121832010
refGene_gn=NFKB1;refGene_tx=XM_023252903.1;refGene_gn=NFKB1;refGene_tx=XM_023252904.1;refGene_gn=NFKB1;refGene_tx=XM_023252905.1;refGene_gn=NFKB1;refGene_tx=XM_023252906.1;refGene_gn=NFKB1;refGene_tx=XM_023252907.1
B1 121835453 121835595
refGene_gn=NFKB1;refGene_tx=XM_023252903.1;refGene_gn=NFKB1;refGene_tx=XM_023252904.1;refGene_gn=NFKB1;refGene_tx=XM_023252905.1;refGene_gn=NFKB1;refGene_tx=XM_023252906.1;refGene_gn=NFKB1;refGene_tx=XM_023252907.1
B1 121836735 121836921
refGene_gn=NFKB1;refGene_tx=XM_023252903.1;refGene_gn=NFKB1;refGene_tx=XM_023252904.1;refGene_gn=NFKB1;refGene_tx=XM_023252905.1;refGene_gn=NFKB1;refGene_tx=XM_023252906.1;refGene_gn=NFKB1;refGene_tx=XM_023252907.1
B1 121838930 121839020
refGene_gn=NFKB1;refGene_tx=XM_023252903.1;refGene_gn=NFKB1;refGene_tx=XM_023252904.1;refGene_gn=NFKB1;refGene_tx=XM_023252905.1;refGene_gn=NFKB1;refGene_tx=XM_023252906.1;refGene_gn=NFKB1;refGene_tx=XM_023252907.1

B1 121840349 121840493
refGene_gn=NFKB1;refGene_tx=XM_023252903.1;refGene_gn=NFKB1;refGene_tx=XM_023252904.1;refGene_gn=NFKB1;refGene_tx=XM_023252905.1;refGene_gn=NFKB1;refGene_tx=XM_023252906.1;refGene_gn=NFKB1;refGene_tx=XM_023252907.1
B1 121847744 121847883
refGene_gn=NFKB1;refGene_tx=XM_023252903.1;refGene_gn=NFKB1;refGene_tx=XM_023252904.1;refGene_gn=NFKB1;refGene_tx=XM_023252905.1;refGene_gn=NFKB1;refGene_tx=XM_023252906.1;refGene_gn=NFKB1;refGene_tx=XM_023252907.1
B1 121850015 121850107
refGene_gn=NFKB1;refGene_tx=XM_023252903.1;refGene_gn=NFKB1;refGene_tx=XM_023252904.1;refGene_gn=NFKB1;refGene_tx=XM_023252905.1;refGene_gn=NFKB1;refGene_tx=XM_023252906.1;refGene_gn=NFKB1;refGene_tx=XM_023252907.1
B1 121852588 121852693
refGene_gn=NFKB1;refGene_tx=XM_023252903.1;refGene_gn=NFKB1;refGene_tx=XM_023252904.1;refGene_gn=NFKB1;refGene_tx=XM_023252905.1;refGene_gn=NFKB1;refGene_tx=XM_023252906.1;refGene_gn=NFKB1;refGene_tx=XM_023252907.1
B1 121854645 121854804
refGene_gn=NFKB1;refGene_tx=XM_023252903.1;refGene_gn=NFKB1;refGene_tx=XM_023252904.1;refGene_gn=NFKB1;refGene_tx=XM_023252905.1;refGene_gn=NFKB1;refGene_tx=XM_023252906.1;refGene_gn=NFKB1;refGene_tx=XM_023252907.1
B1 121856838 121857002
refGene_gn=NFKB1;refGene_tx=XM_023252903.1;refGene_gn=NFKB1;refGene_tx=XM_023252904.1;refGene_gn=NFKB1;refGene_tx=XM_023252905.1;refGene_gn=NFKB1;refGene_tx=XM_023252906.1;refGene_gn=NFKB1;refGene_tx=XM_023252907.1
B1 121867440 121867589
refGene_gn=NFKB1;refGene_tx=XM_023252903.1;refGene_gn=NFKB1;refGene_tx=XM_023252904.1;refGene_gn=NFKB1;refGene_tx=XM_023252905.1;refGene_gn=NFKB1;refGene_tx=XM_023252906.1;refGene_gn=NFKB1;refGene_tx=XM_023252907.1
B1 121879876 121879885
refGene_gn=NFKB1;refGene_tx=XM_023252907.1
B1 121896582 121896681
refGene_gn=NFKB1;refGene_tx=XM_023252903.1;refGene_gn=NFKB1;refGene_tx=XM_023252904.1;refGene_gn=NFKB1;refGene_tx=XM_023252905.1;refGene_gn=NFKB1;refGene_tx=XM_023252906.1
B1 121900574 121900615
refGene_gn=NFKB1;refGene_tx=XM_023252903.1;refGene_gn=NFKB1;refGene_tx=XM_023252904.1;refGene_gn=NFKB1;refGene_tx=XM_023252905.1;refGene_gn=NFKB1;refGene_tx=XM_023252906.1
B1 121904950 121905026
refGene_gn=NFKB1;refGene_tx=XM_023252903.1;refGene_gn=NFKB1;refGene_tx=XM_023252904.1;refGene_gn=NFKB1;refGene_tx=XM_023252905.1;refGene_gn=NFKB1;refGene_tx=XM_023252906.1
B1 121909432 121909471

refGene_gn=NFKB1;refGene_tx=XM_023252903.1;refGene_gn=NFKB1;refGene_tx=XM_023252904.1
B1 122067481 122067700
refGene_gn=SLC39A8;refGene_tx=XM_006930941.4;refGene_gn=SLC39A8;refGene_tx=XM_019829164.2;refGene_gn=SLC39A8;refGene_tx=XM_023252908.1
B1 122093868 122094031
refGene_gn=SLC39A8;refGene_tx=XM_006930941.4;refGene_gn=SLC39A8;refGene_tx=XM_019829164.2;refGene_gn=SLC39A8;refGene_tx=XM_023252908.1
B1 122102069 122102239
refGene_gn=SLC39A8;refGene_tx=XM_006930941.4;refGene_gn=SLC39A8;refGene_tx=XM_019829164.2;refGene_gn=SLC39A8;refGene_tx=XM_023252908.1
B1 122103332 122103455
refGene_gn=SLC39A8;refGene_tx=XM_006930941.4;refGene_gn=SLC39A8;refGene_tx=XM_019829164.2;refGene_gn=SLC39A8;refGene_tx=XM_023252908.1
B1 122103935 122104103
refGene_gn=SLC39A8;refGene_tx=XM_006930941.4;refGene_gn=SLC39A8;refGene_tx=XM_019829164.2;refGene_gn=SLC39A8;refGene_tx=XM_023252908.1
B1 122141836 122142044
refGene_gn=SLC39A8;refGene_tx=XM_006930941.4;refGene_gn=SLC39A8;refGene_tx=XM_019829164.2;refGene_gn=SLC39A8;refGene_tx=XM_023252908.1
B1 122142464 122142649
refGene_gn=SLC39A8;refGene_tx=XM_006930941.4;refGene_gn=SLC39A8;refGene_tx=XM_019829164.2;refGene_gn=SLC39A8;refGene_tx=XM_023252908.1
B1 122145081 122145231
refGene_gn=SLC39A8;refGene_tx=XM_006930941.4;refGene_gn=SLC39A8;refGene_tx=XM_019829164.2;refGene_gn=SLC39A8;refGene_tx=XM_023252908.1
B1 122280844 122281021
refGene_gn=BANK1;refGene_tx=XM_023252909.1
B1 122281293 122281311
refGene_gn=BANK1;refGene_tx=XM_023252909.1
B1 122282589 122282645
refGene_gn=BANK1;refGene_tx=XM_023252909.1
B1 122283662 122283692
refGene_gn=BANK1;refGene_tx=XM_023252909.1
B1 122288982 122289046
refGene_gn=BANK1;refGene_tx=XM_023252909.1
B1 122293703 122293879
refGene_gn=BANK1;refGene_tx=XM_023252909.1
B1 122306177 122306243
refGene_gn=BANK1;refGene_tx=XM_023252909.1
B1 122320233 122320530
refGene_gn=BANK1;refGene_tx=XM_023252909.1
B1 122325442 122325754
refGene_gn=BANK1;refGene_tx=XM_023252909.1
B1 122329249 122329325
refGene_gn=BANK1;refGene_tx=XM_023252909.1
B1 122440876 122441073
refGene_gn=BANK1;refGene_tx=XM_023252909.1
B1 122469978 122470084
refGene_gn=BANK1;refGene_tx=XM_023252909.1

B1	122492549	122492689
refGene_gn=BANK1;refGene_tx=XM_023252909.1
B1	122501922	122502061
refGene_gn=BANK1;refGene_tx=XM_023252909.1
B1	122510323	122510478
refGene_gn=BANK1;refGene_tx=XM_023252909.1
B1	122557505	122557904
refGene_gn=BANK1;refGene_tx=XM_023252909.1
B1	122592248	122592318
refGene_gn=BANK1;refGene_tx=XM_023252909.1
B1	122927671	122927729
refGene_gn=PPP3CA;refGene_tx=XM_003985136.4;refGene_gn=PPP3CA;refGene_tx=XM_003985137.4
B1	123072472	123072494
refGene_gn=PPP3CA;refGene_tx=XM_019829166.2
B1	123092391	123092592
refGene_gn=PPP3CA;refGene_tx=XM_003985136.4;refGene_gn=PPP3CA;refGene_tx=XM_003985137.4;refGene_gn=PPP3CA;refGene_tx=XM_019829166.2
B1	123174630	123174755
refGene_gn=PPP3CA;refGene_tx=XM_003985136.4;refGene_gn=PPP3CA;refGene_tx=XM_003985137.4;refGene_gn=PPP3CA;refGene_tx=XM_019829166.2
B1	123183626	123183738
refGene_gn=PPP3CA;refGene_tx=XM_003985136.4;refGene_gn=PPP3CA;refGene_tx=XM_003985137.4;refGene_gn=PPP3CA;refGene_tx=XM_019829166.2
B1	123185283	123185429
refGene_gn=PPP3CA;refGene_tx=XM_003985136.4;refGene_gn=PPP3CA;refGene_tx=XM_003985137.4;refGene_gn=PPP3CA;refGene_tx=XM_019829166.2
B1	123187932	123188072
refGene_gn=PPP3CA;refGene_tx=XM_003985136.4;refGene_gn=PPP3CA;refGene_tx=XM_003985137.4;refGene_gn=PPP3CA;refGene_tx=XM_019829166.2
B1	123198342	123198420
refGene_gn=PPP3CA;refGene_tx=XM_003985136.4;refGene_gn=PPP3CA;refGene_tx=XM_003985137.4;refGene_gn=PPP3CA;refGene_tx=XM_019829166.2
B1	123200880	123200975
refGene_gn=PPP3CA;refGene_tx=XM_003985136.4;refGene_gn=PPP3CA;refGene_tx=XM_003985137.4;refGene_gn=PPP3CA;refGene_tx=XM_019829166.2
B1	123217959	123218085
refGene_gn=PPP3CA;refGene_tx=XM_003985136.4;refGene_gn=PPP3CA;refGene_tx=XM_003985137.4;refGene_gn=PPP3CA;refGene_tx=XM_019829166.2
B1	123220266	123220341
refGene_gn=PPP3CA;refGene_tx=XM_003985136.4;refGene_gn=PPP3CA;refGene_tx=XM_003985137.4;refGene_gn=PPP3CA;refGene_tx=XM_019829166.2
B1	123240202	123240287
refGene_gn=PPP3CA;refGene_tx=XM_003985136.4;refGene_gn=PPP3CA;refGene_tx=XM_003985137.4;refGene_gn=PPP3CA;refGene_tx=XM_019829166.2
B1	123245545	123245643
refGene_gn=PPP3CA;refGene_tx=XM_003985136.4;refGene_gn=PPP3CA;refGene_tx=XM_003985137.4;refGene_gn=PPP3CA;refGene_tx=XM_019829166.2
B1	123248715	123248745
refGene_gn=PPP3CA;refGene_tx=XM_003985136.4;refGene_gn=PPP3CA;refGene_

tx=XM_019829166.2
B1 123251932 123252129
refGene_gn=PPP3CA;refGene_tx=XM_003985136.4;refGene_gn=PPP3CA;refGene_tx=XM_003985137.4;refGene_gn=PPP3CA;refGene_tx=XM_019829166.2
B1 123648558 123648625
refGene_gn=EMCN;refGene_tx=XM_003985140.4;refGene_gn=EMCN;refGene_tx=XM_006930944.3
B1 123686278 123686398
refGene_gn=EMCN;refGene_tx=XM_003985140.4;refGene_gn=EMCN;refGene_tx=XM_006930944.3
B1 123692550 123692628
refGene_gn=EMCN;refGene_tx=XM_003985140.4;refGene_gn=EMCN;refGene_tx=XM_006930944.3
B1 123703796 123703907
refGene_gn=EMCN;refGene_tx=XM_003985140.4;refGene_gn=EMCN;refGene_tx=XM_006930944.3
B1 123716050 123716089
refGene_gn=EMCN;refGene_tx=XM_006930944.3
B1 123727890 123727983
refGene_gn=EMCN;refGene_tx=XM_003985140.4;refGene_gn=EMCN;refGene_tx=XM_006930944.3
B1 123728459 123728513
refGene_gn=EMCN;refGene_tx=XM_003985140.4;refGene_gn=EMCN;refGene_tx=XM_006930944.3
B1 123730448 123730544
refGene_gn=EMCN;refGene_tx=XM_003985140.4;refGene_gn=EMCN;refGene_tx=XM_006930944.3
B1 123734983 123735008
refGene_gn=EMCN;refGene_tx=XM_003985140.4;refGene_gn=EMCN;refGene_tx=XM_006930944.3
B1 123736217 123736279
refGene_gn=EMCN;refGene_tx=XM_003985140.4;refGene_gn=EMCN;refGene_tx=XM_006930944.3
B1 123743349 123743384
refGene_gn=EMCN;refGene_tx=XM_003985140.4;refGene_gn=EMCN;refGene_tx=XM_006930944.3
B1 123930106 123930197
refGene_gn=DDIT4L;refGene_tx=XM_003985141.5
B1 123932326 123932817
refGene_gn=DDIT4L;refGene_tx=XM_003985141.5
B1 124141214 124141217
refGene_gn=LOC101087401;refGene_tx=XM_023252911.1
B1 124141651 124141729
refGene_gn=LOC101087401;refGene_tx=XM_023252911.1
B1 124142013 124142127
refGene_gn=LOC101087401;refGene_tx=XM_023252911.1
B1 124142478 124142608
refGene_gn=LOC101087401;refGene_tx=XM_023252911.1
B1 124142881 124142943
refGene_gn=LOC101087401;refGene_tx=XM_023252911.1

B1 124144919 124145052
refGene_gn=DNAJB14;refGene_tx=XM_023252912.1;refGene_gn=DNAJB14;refGene_tx=XM_023252913.1
B1 124160745 124160917
refGene_gn=DNAJB14;refGene_tx=XM_023252912.1
B1 124169091 124169237
refGene_gn=DNAJB14;refGene_tx=XM_023252912.1;refGene_gn=DNAJB14;refGene_tx=XM_023252913.1
B1 124177792 124177978
refGene_gn=DNAJB14;refGene_tx=XM_023252912.1;refGene_gn=DNAJB14;refGene_tx=XM_023252913.1
B1 124179792 124179887
refGene_gn=DNAJB14;refGene_tx=XM_023252912.1;refGene_gn=DNAJB14;refGene_tx=XM_023252913.1
B1 124180673 124180783
refGene_gn=DNAJB14;refGene_tx=XM_023252912.1;refGene_gn=DNAJB14;refGene_tx=XM_023252913.1
B1 124181624 124181797
refGene_gn=DNAJB14;refGene_tx=XM_023252912.1;refGene_gn=DNAJB14;refGene_tx=XM_023252913.1
B1 124186176 124186301
refGene_gn=DNAJB14;refGene_tx=XM_023252912.1;refGene_gn=DNAJB14;refGene_tx=XM_023252913.1
B1 124197563 124197572
refGene_gn=LAMTOR3;refGene_tx=XM_003985143.5;refGene_gn=LAMTOR3;refGene_tx=XM_019829168.2
B1 124200691 124200726
refGene_gn=LAMTOR3;refGene_tx=XM_003985143.5;refGene_gn=LAMTOR3;refGene_tx=XM_019829168.2
B1 124206941 124207000
refGene_gn=LAMTOR3;refGene_tx=XM_003985143.5;refGene_gn=LAMTOR3;refGene_tx=XM_019829168.2
B1 124208590 124208724
refGene_gn=LAMTOR3;refGene_tx=XM_003985143.5;refGene_gn=LAMTOR3;refGene_tx=XM_019829168.2
B1 124211058 124211122
refGene_gn=LAMTOR3;refGene_tx=XM_003985143.5;refGene_gn=LAMTOR3;refGene_tx=XM_019829168.2
B1 124213925 124213999
refGene_gn=LAMTOR3;refGene_tx=XM_003985143.5
B1 124216616 124216678
refGene_gn=LAMTOR3;refGene_tx=XM_019829168.2
B1 124224721 124224790
refGene_gn=DAPP1;refGene_tx=XM_003985144.5
B1 124227028 124227116
refGene_gn=DAPP1;refGene_tx=XM_003985144.5
B1 124229736 124229822
refGene_gn=DAPP1;refGene_tx=XM_003985144.5
B1 124231172 124231235
refGene_gn=DAPP1;refGene_tx=XM_003985144.5

B1	124232488	124232536
refGene_gn=DAPP1;refGene_tx=XM_003985144.5
B1	124238409	124238540
refGene_gn=DAPP1;refGene_tx=XM_003985144.5
B1	124251134	124251268
refGene_gn=DAPP1;refGene_tx=XM_003985144.5
B1	124255204	124255327
refGene_gn=DAPP1;refGene_tx=XM_003985144.5
B1	124276697	124276798
refGene_gn=DAPP1;refGene_tx=XM_003985144.5
B1	124393765	124399171
refGene_gn=LOC102902241;refGene_tx=XM_006930945.4
B1	124428830	124429002
refGene_gn=MTTP;refGene_tx=XM_019829170.2;refGene_gn=MTTP;refGene_tx=XM_019829171.2
B1	124430499	124430670
refGene_gn=MTTP;refGene_tx=XM_019829170.2;refGene_gn=MTTP;refGene_tx=XM_019829171.2
B1	124432850	124432975
refGene_gn=MTTP;refGene_tx=XM_019829170.2;refGene_gn=MTTP;refGene_tx=XM_019829171.2
B1	124436835	124437063
refGene_gn=MTTP;refGene_tx=XM_019829170.2;refGene_gn=MTTP;refGene_tx=XM_019829171.2
B1	124438687	124438809
refGene_gn=MTTP;refGene_tx=XM_019829170.2;refGene_gn=MTTP;refGene_tx=XM_019829171.2
B1	124438914	124439012
refGene_gn=MTTP;refGene_tx=XM_019829170.2;refGene_gn=MTTP;refGene_tx=XM_019829171.2
B1	124441247	124441459
refGene_gn=MTTP;refGene_tx=XM_019829170.2;refGene_gn=MTTP;refGene_tx=XM_019829171.2
B1	124442953	124443166
refGene_gn=MTTP;refGene_tx=XM_019829170.2;refGene_gn=MTTP;refGene_tx=XM_019829171.2
B1	124448837	124448945
refGene_gn=MTTP;refGene_tx=XM_019829170.2;refGene_gn=MTTP;refGene_tx=XM_019829171.2
B1	124450293	124450462
refGene_gn=MTTP;refGene_tx=XM_019829170.2;refGene_gn=MTTP;refGene_tx=XM_019829171.2
B1	124455064	124455222
refGene_gn=MTTP;refGene_tx=XM_019829170.2;refGene_gn=MTTP;refGene_tx=XM_019829171.2
B1	124457367	124457518
refGene_gn=MTTP;refGene_tx=XM_019829170.2;refGene_gn=MTTP;refGene_tx=XM_019829171.2
B1	124459260	124459400
refGene_gn=MTTP;refGene_tx=XM_019829170.2;refGene_gn=MTTP;refGene_tx=X

M_019829171.2
B1 124459724 124459841
refGene_gn=MTTP;refGene_tx=XM_019829170.2;refGene_gn=MTTP;refGene_tx=X
M_019829171.2
B1 124461040 124461148
refGene_gn=MTTP;refGene_tx=XM_019829170.2;refGene_gn=MTTP;refGene_tx=X
M_019829171.2
B1 124467156 124467300
refGene_gn=MTTP;refGene_tx=XM_019829170.2;refGene_gn=MTTP;refGene_tx=X
M_019829171.2
B1 124468606 124468794
refGene_gn=MTTP;refGene_tx=XM_019829170.2
B1 124474736 124474797
refGene_gn=MTTP;refGene_tx=XM_019829170.2
B1 124493473 124493658
refGene_gn=TRMT10A;refGene_tx=XM_006930947.4;refGene_gn=TRMT10A;refGen
e_tx=XM_006930949.4;refGene_gn=TRMT10A;refGene_tx=XM_019829173.2;refGe
ne_gn=TRMT10A;refGene_tx=XM_019829174.2;refGene_gn=TRMT10A;refGene_tx=
XM_019829175.2
B1 124494492 124494655
refGene_gn=TRMT10A;refGene_tx=XM_006930947.4;refGene_gn=TRMT10A;refGen
e_tx=XM_006930949.4;refGene_gn=TRMT10A;refGene_tx=XM_019829173.2;refGe
ne_gn=TRMT10A;refGene_tx=XM_019829174.2;refGene_gn=TRMT10A;refGene_tx=
XM_019829175.2
B1 124495612 124495684
refGene_gn=TRMT10A;refGene_tx=XM_006930947.4;refGene_gn=TRMT10A;refGen
e_tx=XM_006930949.4;refGene_gn=TRMT10A;refGene_tx=XM_019829173.2;refGe
ne_gn=TRMT10A;refGene_tx=XM_019829174.2;refGene_gn=TRMT10A;refGene_tx=
XM_019829175.2
B1 124502981 124503056
refGene_gn=TRMT10A;refGene_tx=XM_006930947.4;refGene_gn=TRMT10A;refGen
e_tx=XM_006930949.4;refGene_gn=TRMT10A;refGene_tx=XM_019829173.2;refGe
ne_gn=TRMT10A;refGene_tx=XM_019829174.2;refGene_gn=TRMT10A;refGene_tx=
XM_019829175.2
B1 124504984 124505134
refGene_gn=TRMT10A;refGene_tx=XM_006930947.4;refGene_gn=TRMT10A;refGen
e_tx=XM_006930949.4;refGene_gn=TRMT10A;refGene_tx=XM_019829173.2;refGe
ne_gn=TRMT10A;refGene_tx=XM_019829174.2;refGene_gn=TRMT10A;refGene_tx=
XM_019829175.2
B1 124507042 124507148
refGene_gn=TRMT10A;refGene_tx=XM_006930947.4;refGene_gn=TRMT10A;refGen
e_tx=XM_006930949.4;refGene_gn=TRMT10A;refGene_tx=XM_019829173.2;refGe
ne_gn=TRMT10A;refGene_tx=XM_019829174.2;refGene_gn=TRMT10A;refGene_tx=
XM_019829175.2
B1 124508376 124508645
refGene_gn=TRMT10A;refGene_tx=XM_006930947.4;refGene_gn=TRMT10A;refGen
e_tx=XM_006930949.4;refGene_gn=TRMT10A;refGene_tx=XM_019829173.2;refGe
ne_gn=TRMT10A;refGene_tx=XM_019829174.2;refGene_gn=TRMT10A;refGene_tx=
XM_019829175.2
B1 124523147 124523347

refGene_gn=CB1H4orf17;refGene_tx=XM_003985146.4;refGene_gn=CB1H4orf17;
refGene_tx=XM_006930952.3;refGene_gn=CB1H4orf17;refGene_tx=XM_02325291
4.1
B1 124524820 124524864
refGene_gn=CB1H4orf17;refGene_tx=XM_003985146.4;refGene_gn=CB1H4orf17;
refGene_tx=XM_006930952.3;refGene_gn=CB1H4orf17;refGene_tx=XM_02325291
4.1
B1 124525801 124525988
refGene_gn=CB1H4orf17;refGene_tx=XM_003985146.4;refGene_gn=CB1H4orf17;
refGene_tx=XM_006930952.3;refGene_gn=CB1H4orf17;refGene_tx=XM_02325291
4.1
B1 124527388 124527470
refGene_gn=CB1H4orf17;refGene_tx=XM_003985146.4;refGene_gn=CB1H4orf17;
refGene_tx=XM_006930952.3;refGene_gn=CB1H4orf17;refGene_tx=XM_02325291
4.1
B1 124536462 124536606
refGene_gn=CB1H4orf17;refGene_tx=XM_003985146.4;refGene_gn=CB1H4orf17;
refGene_tx=XM_006930952.3
B1 124541865 124541930
refGene_gn=CB1H4orf17;refGene_tx=XM_003985146.4;refGene_gn=CB1H4orf17;
refGene_tx=XM_006930952.3;refGene_gn=CB1H4orf17;refGene_tx=XM_02325291
4.1
B1 124544502 124544712
refGene_gn=CB1H4orf17;refGene_tx=XM_003985146.4;refGene_gn=CB1H4orf17;
refGene_tx=XM_006930952.3;refGene_gn=CB1H4orf17;refGene_tx=XM_02325291
4.1
B1 124551330 124551457
refGene_gn=CB1H4orf17;refGene_tx=XM_003985146.4;refGene_gn=CB1H4orf17;
refGene_tx=XM_006930952.3;refGene_gn=CB1H4orf17;refGene_tx=XM_02325291
4.1
B1 124661832 124661850
refGene_gn=LOC101100387;refGene_tx=XM_003985148.5
B1 124664798 124664900
refGene_gn=LOC101100387;refGene_tx=XM_003985148.5
B1 124665360 124665499
refGene_gn=LOC101100387;refGene_tx=XM_003985148.5;refGene_gn=LOC101100
387;refGene_tx=XM_006930953.4
B1 124667364 124667452
refGene_gn=LOC101100387;refGene_tx=XM_003985148.5;refGene_gn=LOC101100
387;refGene_tx=XM_006930953.4
B1 124667549 124667769
refGene_gn=LOC101100387;refGene_tx=XM_003985148.5;refGene_gn=LOC101100
387;refGene_tx=XM_006930953.4
B1 124670011 124670272
refGene_gn=LOC101100387;refGene_tx=XM_003985148.5;refGene_gn=LOC101100
387;refGene_tx=XM_006930953.4
B1 124671960 124672096
refGene_gn=LOC101100387;refGene_tx=XM_003985148.5;refGene_gn=LOC101100
387;refGene_tx=XM_006930953.4
B1 124672934 124673073

refGene_gn=LOC101100387;refGene_tx=XM_003985148.5;refGene_gn=LOC101100387;refGene_tx=XM_006930953.4
B1 124675109 124675134
refGene_gn=LOC101100387;refGene_tx=XM_003985148.5;refGene_gn=LOC101100387;refGene_tx=XM_006930953.4
B1 124731991 124732027
refGene_gn=LOC101100805;refGene_tx=XM_003985149.3
B1 124734616 124734718
refGene_gn=LOC101100805;refGene_tx=XM_003985149.3
B1 124737091 124737230
refGene_gn=LOC101100805;refGene_tx=XM_003985149.3
B1 124738674 124738762
refGene_gn=LOC101100805;refGene_tx=XM_003985149.3
B1 124738904 124739124
refGene_gn=LOC101100805;refGene_tx=XM_003985149.3
B1 124739303 124739564
refGene_gn=LOC101100805;refGene_tx=XM_003985149.3
B1 124740667 124740803
refGene_gn=LOC101100805;refGene_tx=XM_003985149.3
B1 124741602 124741741
refGene_gn=LOC101100805;refGene_tx=XM_003985149.3
B1 124743274 124743299
refGene_gn=LOC101100805;refGene_tx=XM_003985149.3
B1 124796354 124796372
refGene_gn=LOC101101050;refGene_tx=XM_003985150.4
B1 124797653 124797755
refGene_gn=LOC101101050;refGene_tx=XM_003985150.4
B1 124798808 124798950
refGene_gn=LOC101101050;refGene_tx=XM_003985150.4
B1 124800403 124800491
refGene_gn=LOC101101050;refGene_tx=XM_003985150.4
B1 124802612 124802844
refGene_gn=LOC101101050;refGene_tx=XM_003985150.4
B1 124806745 124807006
refGene_gn=LOC101101050;refGene_tx=XM_003985150.4
B1 124810728 124810864
refGene_gn=LOC101101050;refGene_tx=XM_003985150.4
B1 124811347 124811486
refGene_gn=LOC101101050;refGene_tx=XM_003985150.4
B1 124813519 124813544
refGene_gn=LOC101101050;refGene_tx=XM_003985150.4
B1 124834999 124835011
refGene_gn=ADH5;refGene_tx=XM_003985151.3
B1 124839377 124839479
refGene_gn=ADH5;refGene_tx=XM_003985151.3
B1 124841078 124841220
refGene_gn=ADH5;refGene_tx=XM_003985151.3
B1 124841735 124841823
refGene_gn=ADH5;refGene_tx=XM_003985151.3
B1 124845703 124845923

refGene_gn=ADH5;refGene_tx=XM_003985151.3
B1 124846074 124846335
refGene_gn=ADH5;refGene_tx=XM_003985151.3
B1 124846994 124847130
refGene_gn=ADH5;refGene_tx=XM_003985151.3
B1 124849956 124850095
refGene_gn=ADH5;refGene_tx=XM_003985151.3
B1 124850437 124850462
refGene_gn=ADH5;refGene_tx=XM_003985151.3
B1 124860672 124860836
refGene_gn=METAP1;refGene_tx=XM_023252915.1;refGene_gn=METAP1;refGene_tx=XM_023252916.1
B1 124865160 124865226
refGene_gn=METAP1;refGene_tx=XM_023252915.1;refGene_gn=METAP1;refGene_tx=XM_023252916.1
B1 124869738 124869882
refGene_gn=METAP1;refGene_tx=XM_023252915.1;refGene_gn=METAP1;refGene_tx=XM_023252916.1
B1 124872957 124873089
refGene_gn=METAP1;refGene_tx=XM_023252915.1;refGene_gn=METAP1;refGene_tx=XM_023252916.1
B1 124874601 124874740
refGene_gn=METAP1;refGene_tx=XM_023252915.1;refGene_gn=METAP1;refGene_tx=XM_023252916.1
B1 124876794 124876878
refGene_gn=METAP1;refGene_tx=XM_023252915.1;refGene_gn=METAP1;refGene_tx=XM_023252916.1
B1 124878019 124878111
refGene_gn=METAP1;refGene_tx=XM_023252915.1;refGene_gn=METAP1;refGene_tx=XM_023252916.1
B1 124879282 124879343
refGene_gn=METAP1;refGene_tx=XM_023252915.1;refGene_gn=METAP1;refGene_tx=XM_023252916.1
B1 124880287 124880400
refGene_gn=METAP1;refGene_tx=XM_023252915.1;refGene_gn=METAP1;refGene_tx=XM_023252916.1
B1 124887414 124887466
refGene_gn=METAP1;refGene_tx=XM_023252915.1;refGene_gn=METAP1;refGene_tx=XM_023252916.1
B1 124917355 124917469
refGene_gn=METAP1;refGene_tx=XM_023252915.1;refGene_gn=METAP1;refGene_tx=XM_023252916.1
B1 124973648 124973666
refGene_gn=EIF4E;refGene_tx=XM_003985152.5
B1 124974297 124974354
refGene_gn=EIF4E;refGene_tx=XM_019829177.2
B1 124997729 124997836
refGene_gn=EIF4E;refGene_tx=XM_003985152.5;refGene_gn=EIF4E;refGene_tx=XM_019829177.2
B1 125008747 125008843

refGene_gn=EIF4E;refGene_tx=XM_003985152.5;refGene_gn=EIF4E;refGene_tx=XM_019829177.2
B1 125012020 125012084
refGene_gn=EIF4E;refGene_tx=XM_003985152.5;refGene_gn=EIF4E;refGene_tx=XM_019829177.2
B1 125012793 125012907
refGene_gn=EIF4E;refGene_tx=XM_003985152.5;refGene_gn=EIF4E;refGene_tx=XM_019829177.2
B1 125014240 125014380
refGene_gn=EIF4E;refGene_tx=XM_003985152.5;refGene_gn=EIF4E;refGene_tx=XM_019829177.2
B1 125016973 125017088
refGene_gn=EIF4E;refGene_tx=XM_003985152.5;refGene_gn=EIF4E;refGene_tx=XM_019829177.2
B1 125215616 125215697
refGene_gn=TSPAN5;refGene_tx=XM_011281794.2;refGene_gn=TSPAN5;refGene_tx=XM_019829180.2
B1 125355940 125355991
refGene_gn=TSPAN5;refGene_tx=XM_011281794.2;refGene_gn=TSPAN5;refGene_tx=XM_019829180.2
B1 125368268 125368310
refGene_gn=TSPAN5;refGene_tx=XM_023252918.1
B1 125375092 125375239
refGene_gn=TSPAN5;refGene_tx=XM_011281794.2;refGene_gn=TSPAN5;refGene_tx=XM_023252918.1
B1 125379209 125379380
refGene_gn=TSPAN5;refGene_tx=XM_011281794.2;refGene_gn=TSPAN5;refGene_tx=XM_019829180.2;refGene_gn=TSPAN5;refGene_tx=XM_023252918.1
B1 125382315 125382441
refGene_gn=TSPAN5;refGene_tx=XM_011281794.2;refGene_gn=TSPAN5;refGene_tx=XM_019829180.2;refGene_gn=TSPAN5;refGene_tx=XM_023252918.1
B1 125384335 125384383
refGene_gn=TSPAN5;refGene_tx=XM_011281794.2;refGene_gn=TSPAN5;refGene_tx=XM_019829180.2;refGene_gn=TSPAN5;refGene_tx=XM_023252918.1
B1 125384481 125384598
refGene_gn=TSPAN5;refGene_tx=XM_011281794.2;refGene_gn=TSPAN5;refGene_tx=XM_019829180.2;refGene_gn=TSPAN5;refGene_tx=XM_023252918.1
B1 125386941 125387007
refGene_gn=TSPAN5;refGene_tx=XM_011281794.2;refGene_gn=TSPAN5;refGene_tx=XM_019829180.2;refGene_gn=TSPAN5;refGene_tx=XM_023252918.1
B1 125409103 125409231
refGene_gn=RAP1GDS1;refGene_tx=XM_019829185.2;refGene_gn=RAP1GDS1;refGene_tx=XM_019829186.2;refGene_gn=RAP1GDS1;refGene_tx=XM_023252919.1;refGene_gn=RAP1GDS1;refGene_tx=XM_023252920.1;refGene_gn=RAP1GDS1;refGene_tx=XM_023252921.1
B1 125413255 125413384
refGene_gn=RAP1GDS1;refGene_tx=XM_019829185.2;refGene_gn=RAP1GDS1;refGene_tx=XM_019829186.2;refGene_gn=RAP1GDS1;refGene_tx=XM_023252919.1;refGene_gn=RAP1GDS1;refGene_tx=XM_023252920.1;refGene_gn=RAP1GDS1;refGene_tx=XM_023252921.1

B1 125422651 125422778
refGene_gn=RAP1GDS1;refGene_tx=XM_019829185.2;refGene_gn=RAP1GDS1;refGene_tx=XM_019829186.2;refGene_gn=RAP1GDS1;refGene_tx=XM_023252919.1;refGene_gn=RAP1GDS1;refGene_tx=XM_023252920.1;refGene_gn=RAP1GDS1;refGene_tx=XM_023252921.1
B1 125429052 125429192
refGene_gn=RAP1GDS1;refGene_tx=XM_019829185.2;refGene_gn=RAP1GDS1;refGene_tx=XM_019829186.2;refGene_gn=RAP1GDS1;refGene_tx=XM_023252919.1;refGene_gn=RAP1GDS1;refGene_tx=XM_023252920.1;refGene_gn=RAP1GDS1;refGene_tx=XM_023252921.1
B1 125429951 125430077
refGene_gn=RAP1GDS1;refGene_tx=XM_019829185.2;refGene_gn=RAP1GDS1;refGene_tx=XM_019829186.2;refGene_gn=RAP1GDS1;refGene_tx=XM_023252919.1;refGene_gn=RAP1GDS1;refGene_tx=XM_023252920.1;refGene_gn=RAP1GDS1;refGene_tx=XM_023252921.1
B1 125431053 125431188
refGene_gn=RAP1GDS1;refGene_tx=XM_019829185.2;refGene_gn=RAP1GDS1;refGene_tx=XM_019829186.2;refGene_gn=RAP1GDS1;refGene_tx=XM_023252919.1;refGene_gn=RAP1GDS1;refGene_tx=XM_023252920.1;refGene_gn=RAP1GDS1;refGene_tx=XM_023252921.1
B1 125432653 125432785
refGene_gn=RAP1GDS1;refGene_tx=XM_019829185.2;refGene_gn=RAP1GDS1;refGene_tx=XM_019829186.2;refGene_gn=RAP1GDS1;refGene_tx=XM_023252919.1;refGene_gn=RAP1GDS1;refGene_tx=XM_023252920.1;refGene_gn=RAP1GDS1;refGene_tx=XM_023252921.1
B1 125433216 125433360
refGene_gn=RAP1GDS1;refGene_tx=XM_019829185.2;refGene_gn=RAP1GDS1;refGene_tx=XM_019829186.2;refGene_gn=RAP1GDS1;refGene_tx=XM_023252919.1;refGene_gn=RAP1GDS1;refGene_tx=XM_023252920.1;refGene_gn=RAP1GDS1;refGene_tx=XM_023252921.1
B1 125447573 125447699
refGene_gn=RAP1GDS1;refGene_tx=XM_019829185.2;refGene_gn=RAP1GDS1;refGene_tx=XM_019829186.2;refGene_gn=RAP1GDS1;refGene_tx=XM_023252919.1;refGene_gn=RAP1GDS1;refGene_tx=XM_023252920.1;refGene_gn=RAP1GDS1;refGene_tx=XM_023252921.1
B1 125461941 125462070
refGene_gn=RAP1GDS1;refGene_tx=XM_019829185.2;refGene_gn=RAP1GDS1;refGene_tx=XM_019829186.2;refGene_gn=RAP1GDS1;refGene_tx=XM_023252919.1;refGene_gn=RAP1GDS1;refGene_tx=XM_023252920.1;refGene_gn=RAP1GDS1;refGene_tx=XM_023252921.1
B1 125475314 125475461
refGene_gn=RAP1GDS1;refGene_tx=XM_019829185.2;refGene_gn=RAP1GDS1;refGene_tx=XM_019829186.2;refGene_gn=RAP1GDS1;refGene_tx=XM_023252919.1;refGene_gn=RAP1GDS1;refGene_tx=XM_023252920.1
B1 125510822 125510948
refGene_gn=RAP1GDS1;refGene_tx=XM_019829185.2;refGene_gn=RAP1GDS1;refGene_tx=XM_019829186.2;refGene_gn=RAP1GDS1;refGene_tx=XM_023252919.1;refGene_gn=RAP1GDS1;refGene_tx=XM_023252920.1;refGene_gn=RAP1GDS1;refGene_tx=XM_023252921.1
B1 125520434 125520557

refGene_gn=RAP1GDS1;refGene_tx=XM_023252919.1;refGene_gn=RAP1GDS1;refGene_tx=XM_023252920.1;refGene_gn=RAP1GDS1;refGene_tx=XM_023252921.1
B1 125558364 125558472
refGene_gn=RAP1GDS1;refGene_tx=XM_023252919.1;refGene_gn=RAP1GDS1;refGene_tx=XM_023252920.1;refGene_gn=RAP1GDS1;refGene_tx=XM_023252921.1
B1 125584344 125584348
refGene_gn=RAP1GDS1;refGene_tx=XM_023252919.1;refGene_gn=RAP1GDS1;refGene_tx=XM_023252920.1;refGene_gn=RAP1GDS1;refGene_tx=XM_023252921.1
B1 125689376 125689485
refGene_gn=STPG2;refGene_tx=XM_023252922.1
B1 125698665 125698778
refGene_gn=STPG2;refGene_tx=XM_023252922.1
B1 125704838 125704997
refGene_gn=STPG2;refGene_tx=XM_023252922.1
B1 125707791 125707956
refGene_gn=STPG2;refGene_tx=XM_023252922.1
B1 125719872 125719985
refGene_gn=STPG2;refGene_tx=XM_023252922.1
B1 125723691 125723803
refGene_gn=STPG2;refGene_tx=XM_023252922.1
B1 125819758 125819866
refGene_gn=STPG2;refGene_tx=XM_023252922.1
B1 125838005 125838165
refGene_gn=STPG2;refGene_tx=XM_023252922.1
B1 125846441 125846602
refGene_gn=STPG2;refGene_tx=XM_023252922.1
B1 125860932 125861025
refGene_gn=STPG2;refGene_tx=XM_023252922.1
B1 125930722 125930882
refGene_gn=STPG2;refGene_tx=XM_023252922.1
B1 126014275 126014391
refGene_gn=STPG2;refGene_tx=XM_023252922.1
B1 126249878 126250027
refGene_gn=STPG2;refGene_tx=XM_023252922.1
B1 126253839 126253840
refGene_gn=STPG2;refGene_tx=XM_023252922.1
B1 127706900 127708076
refGene_gn=PDHA2;refGene_tx=XM_011281793.3
B1 127934438 127934562
refGene_gn=UNC5C;refGene_tx=XM_011281792.3;refGene_gn=UNC5C;refGene_tx=XM_019829187.2
B1 128135596 128135818
refGene_gn=UNC5C;refGene_tx=XM_011281792.3;refGene_gn=UNC5C;refGene_tx=XM_019829187.2
B1 128165962 128166106
refGene_gn=UNC5C;refGene_tx=XM_011281792.3;refGene_gn=UNC5C;refGene_tx=XM_019829187.2
B1 128188898 128189002
refGene_gn=UNC5C;refGene_tx=XM_011281792.3;refGene_gn=UNC5C;refGene_tx=XM_019829187.2

B1 128215465 128215646
refGene_gn=UNC5C;refGene_tx=XM_011281792.3;refGene_gn=UNC5C;refGene_tx=XM_019829187.2
B1 128219221 128219389
refGene_gn=UNC5C;refGene_tx=XM_011281792.3;refGene_gn=UNC5C;refGene_tx=XM_019829187.2
B1 128221971 128222136
refGene_gn=UNC5C;refGene_tx=XM_011281792.3;refGene_gn=UNC5C;refGene_tx=XM_019829187.2
B1 128241352 128241409
refGene_gn=UNC5C;refGene_tx=XM_019829187.2
B1 128243426 128243618
refGene_gn=UNC5C;refGene_tx=XM_011281792.3;refGene_gn=UNC5C;refGene_tx=XM_019829187.2
B1 128244349 128244694
refGene_gn=UNC5C;refGene_tx=XM_011281792.3;refGene_gn=UNC5C;refGene_tx=XM_019829187.2
B1 128247580 128247668
refGene_gn=UNC5C;refGene_tx=XM_011281792.3;refGene_gn=UNC5C;refGene_tx=XM_019829187.2
B1 128257786 128257955
refGene_gn=UNC5C;refGene_tx=XM_011281792.3;refGene_gn=UNC5C;refGene_tx=XM_019829187.2
B1 128261896 128262130
refGene_gn=UNC5C;refGene_tx=XM_011281792.3;refGene_gn=UNC5C;refGene_tx=XM_019829187.2
B1 128276665 128276815
refGene_gn=UNC5C;refGene_tx=XM_011281792.3;refGene_gn=UNC5C;refGene_tx=XM_019829187.2
B1 128279074 128279239
refGene_gn=UNC5C;refGene_tx=XM_011281792.3;refGene_gn=UNC5C;refGene_tx=XM_019829187.2
B1 128285683 128285862
refGene_gn=UNC5C;refGene_tx=XM_011281792.3;refGene_gn=UNC5C;refGene_tx=XM_019829187.2
B1 128286633 128286799
refGene_gn=UNC5C;refGene_tx=XM_011281792.3;refGene_gn=UNC5C;refGene_tx=XM_019829187.2
B1 128300843 128300969
refGene_gn=BMPR1B;refGene_tx=XM_006930956.4;refGene_gn=BMPR1B;refGene_tx=XM_019829189.2;refGene_gn=BMPR1B;refGene_tx=XM_019829199.2;refGene_gn=BMPR1B;refGene_tx=XM_019829200.2;refGene_gn=BMPR1B;refGene_tx=XM_019829204.2;refGene_gn=BMPR1B;refGene_tx=XM_019829207.2;refGene_gn=BMPR1B;refGene_tx=XM_019829209.2;refGene_gn=BMPR1B;refGene_tx=XM_019829210.1;refGene_gn=BMPR1B;refGene_tx=XM_019829211.2;refGene_gn=BMPR1B;refGene_tx=XM_023252924.1;refGene_gn=BMPR1B;refGene_tx=XM_023252925.1;refGene_gn=BMPR1B;refGene_tx=XM_023252926.1;refGene_gn=BMPR1B;refGene_tx=XM_023252927.1;refGene_gn=BMPR1B;refGene_tx=XM_023252928.1;refGene_gn=BMPR1B;refGene_tx=XM_023252929.1;refGene_gn=BMPR1B;refGene_tx=XM_023252930.1

B1	128302767	128302898
refGene_gn=BMPR1B;refGene_tx=XM_006930956.4;refGene_gn=BMPR1B;refGene_tx=XM_019829189.2;refGene_gn=BMPR1B;refGene_tx=XM_019829199.2;refGene_gn=BMPR1B;refGene_tx=XM_019829200.2;refGene_gn=BMPR1B;refGene_tx=XM_019829204.2;refGene_gn=BMPR1B;refGene_tx=XM_019829207.2;refGene_gn=BMPR1B;refGene_tx=XM_019829209.2;refGene_gn=BMPR1B;refGene_tx=XM_019829210.1;refGene_gn=BMPR1B;refGene_tx=XM_019829211.2;refGene_gn=BMPR1B;refGene_tx=XM_023252924.1;refGene_gn=BMPR1B;refGene_tx=XM_023252925.1;refGene_gn=BMPR1B;refGene_tx=XM_023252926.1;refGene_gn=BMPR1B;refGene_tx=XM_023252927.1;refGene_gn=BMPR1B;refGene_tx=XM_023252928.1;refGene_gn=BMPR1B;refGene_tx=XM_023252929.1;refGene_gn=BMPR1B;refGene_tx=XM_023252930.1
B1	128306323	128306499
refGene_gn=BMPR1B;refGene_tx=XM_006930956.4;refGene_gn=BMPR1B;refGene_tx=XM_019829189.2;refGene_gn=BMPR1B;refGene_tx=XM_019829199.2;refGene_gn=BMPR1B;refGene_tx=XM_019829200.2;refGene_gn=BMPR1B;refGene_tx=XM_019829204.2;refGene_gn=BMPR1B;refGene_tx=XM_019829207.2;refGene_gn=BMPR1B;refGene_tx=XM_019829209.2;refGene_gn=BMPR1B;refGene_tx=XM_019829210.1;refGene_gn=BMPR1B;refGene_tx=XM_019829211.2;refGene_gn=BMPR1B;refGene_tx=XM_023252924.1;refGene_gn=BMPR1B;refGene_tx=XM_023252925.1;refGene_gn=BMPR1B;refGene_tx=XM_023252926.1;refGene_gn=BMPR1B;refGene_tx=XM_023252927.1;refGene_gn=BMPR1B;refGene_tx=XM_023252928.1;refGene_gn=BMPR1B;refGene_tx=XM_023252929.1;refGene_gn=BMPR1B;refGene_tx=XM_023252930.1
B1	128317879	128318177
refGene_gn=BMPR1B;refGene_tx=XM_006930956.4;refGene_gn=BMPR1B;refGene_tx=XM_019829189.2;refGene_gn=BMPR1B;refGene_tx=XM_019829199.2;refGene_gn=BMPR1B;refGene_tx=XM_019829200.2;refGene_gn=BMPR1B;refGene_tx=XM_019829204.2;refGene_gn=BMPR1B;refGene_tx=XM_019829207.2;refGene_gn=BMPR1B;refGene_tx=XM_019829209.2;refGene_gn=BMPR1B;refGene_tx=XM_019829210.1;refGene_gn=BMPR1B;refGene_tx=XM_019829211.2;refGene_gn=BMPR1B;refGene_tx=XM_023252924.1;refGene_gn=BMPR1B;refGene_tx=XM_023252925.1;refGene_gn=BMPR1B;refGene_tx=XM_023252926.1;refGene_gn=BMPR1B;refGene_tx=XM_023252927.1;refGene_gn=BMPR1B;refGene_tx=XM_023252928.1;refGene_gn=BMPR1B;refGene_tx=XM_023252929.1;refGene_gn=BMPR1B;refGene_tx=XM_023252930.1
B1	128318979	128319172
refGene_gn=BMPR1B;refGene_tx=XM_006930956.4;refGene_gn=BMPR1B;refGene_tx=XM_019829189.2;refGene_gn=BMPR1B;refGene_tx=XM_019829199.2;refGene_gn=BMPR1B;refGene_tx=XM_019829200.2;refGene_gn=BMPR1B;refGene_tx=XM_019829204.2;refGene_gn=BMPR1B;refGene_tx=XM_019829207.2;refGene_gn=BMPR1B;refGene_tx=XM_019829209.2;refGene_gn=BMPR1B;refGene_tx=XM_019829210.1;refGene_gn=BMPR1B;refGene_tx=XM_019829211.2;refGene_gn=BMPR1B;refGene_tx=XM_023252924.1;refGene_gn=BMPR1B;refGene_tx=XM_023252925.1;refGene_gn=BMPR1B;refGene_tx=XM_023252926.1;refGene_gn=BMPR1B;refGene_tx=XM_023252927.1;refGene_gn=BMPR1B;refGene_tx=XM_023252928.1;refGene_gn=BMPR1B;refGene_tx=XM_023252929.1;refGene_gn=BMPR1B;refGene_tx=XM_023252930.1
B1	128323413	128323552
refGene_gn=BMPR1B;refGene_tx=XM_006930956.4;refGene_gn=BMPR1B;refGene_

tx=XM_019829189.2;refGene_gn=BMPR1B;refGene_tx=XM_019829199.2;refGene_
gn=BMPR1B;refGene_tx=XM_019829200.2;refGene_gn=BMPR1B;refGene_tx=XM_01
9829204.2;refGene_gn=BMPR1B;refGene_tx=XM_019829207.2;refGene_gn=BMPR1
B;refGene_tx=XM_019829209.2;refGene_gn=BMPR1B;refGene_tx=XM_019829210.
1;refGene_gn=BMPR1B;refGene_tx=XM_019829211.2;refGene_gn=BMPR1B;refGen
e_tx=XM_023252924.1;refGene_gn=BMPR1B;refGene_tx=XM_023252925.1;refGen
e_gn=BMPR1B;refGene_tx=XM_023252926.1;refGene_gn=BMPR1B;refGene_tx=XM_
023252927.1;refGene_gn=BMPR1B;refGene_tx=XM_023252928.1;refGene_gn=BMP
R1B;refGene_tx=XM_023252929.1;refGene_gn=BMPR1B;refGene_tx=XM_02325293
0.1
B1 128324618 128324715
refGene_gn=BMPR1B;refGene_tx=XM_006930956.4;refGene_gn=BMPR1B;refGene_
tx=XM_019829189.2;refGene_gn=BMPR1B;refGene_tx=XM_019829199.2;refGene_
gn=BMPR1B;refGene_tx=XM_019829200.2;refGene_gn=BMPR1B;refGene_tx=XM_01
9829204.2;refGene_gn=BMPR1B;refGene_tx=XM_019829207.2;refGene_gn=BMPR1
B;refGene_tx=XM_019829209.2;refGene_gn=BMPR1B;refGene_tx=XM_019829210.
1;refGene_gn=BMPR1B;refGene_tx=XM_019829211.2;refGene_gn=BMPR1B;refGen
e_tx=XM_023252924.1;refGene_gn=BMPR1B;refGene_tx=XM_023252925.1;refGen
e_gn=BMPR1B;refGene_tx=XM_023252926.1;refGene_gn=BMPR1B;refGene_tx=XM_
023252927.1;refGene_gn=BMPR1B;refGene_tx=XM_023252928.1;refGene_gn=BMP
R1B;refGene_tx=XM_023252929.1;refGene_gn=BMPR1B;refGene_tx=XM_02325293
0.1
B1 128334550 128334653
refGene_gn=BMPR1B;refGene_tx=XM_006930956.4;refGene_gn=BMPR1B;refGene_
tx=XM_019829189.2;refGene_gn=BMPR1B;refGene_tx=XM_019829199.2;refGene_
gn=BMPR1B;refGene_tx=XM_019829200.2;refGene_gn=BMPR1B;refGene_tx=XM_01
9829204.2;refGene_gn=BMPR1B;refGene_tx=XM_019829207.2;refGene_gn=BMPR1
B;refGene_tx=XM_019829209.2;refGene_gn=BMPR1B;refGene_tx=XM_019829210.
1;refGene_gn=BMPR1B;refGene_tx=XM_019829211.2;refGene_gn=BMPR1B;refGen
e_tx=XM_023252924.1;refGene_gn=BMPR1B;refGene_tx=XM_023252925.1;refGen
e_gn=BMPR1B;refGene_tx=XM_023252926.1;refGene_gn=BMPR1B;refGene_tx=XM_
023252927.1;refGene_gn=BMPR1B;refGene_tx=XM_023252928.1;refGene_gn=BMP
R1B;refGene_tx=XM_023252929.1;refGene_gn=BMPR1B;refGene_tx=XM_02325293
0.1
B1 128335428 128335531
refGene_gn=BMPR1B;refGene_tx=XM_006930956.4;refGene_gn=BMPR1B;refGene_
tx=XM_019829189.2;refGene_gn=BMPR1B;refGene_tx=XM_019829199.2;refGene_
gn=BMPR1B;refGene_tx=XM_019829200.2;refGene_gn=BMPR1B;refGene_tx=XM_01
9829204.2;refGene_gn=BMPR1B;refGene_tx=XM_019829207.2;refGene_gn=BMPR1
B;refGene_tx=XM_019829209.2;refGene_gn=BMPR1B;refGene_tx=XM_019829210.
1;refGene_gn=BMPR1B;refGene_tx=XM_019829211.2;refGene_gn=BMPR1B;refGen
e_tx=XM_023252924.1;refGene_gn=BMPR1B;refGene_tx=XM_023252925.1;refGen
e_gn=BMPR1B;refGene_tx=XM_023252926.1;refGene_gn=BMPR1B;refGene_tx=XM_
023252927.1;refGene_gn=BMPR1B;refGene_tx=XM_023252928.1;refGene_gn=BMP
R1B;refGene_tx=XM_023252929.1;refGene_gn=BMPR1B;refGene_tx=XM_02325293
0.1
B1 128345768 128345928
refGene_gn=BMPR1B;refGene_tx=XM_006930956.4;refGene_gn=BMPR1B;refGene_
tx=XM_019829189.2;refGene_gn=BMPR1B;refGene_tx=XM_019829199.2;refGene_
gn=BMPR1B;refGene_tx=XM_019829200.2;refGene_gn=BMPR1B;refGene_tx=XM_01

9829204.2;refGene_gn=BMPR1B;refGene_tx=XM_019829207.2;refGene_gn=BMPR1B;refGene_tx=XM_019829209.2;refGene_gn=BMPR1B;refGene_tx=XM_019829210.1;refGene_gn=BMPR1B;refGene_tx=XM_019829211.2;refGene_gn=BMPR1B;refGene_tx=XM_023252924.1;refGene_gn=BMPR1B;refGene_tx=XM_023252925.1;refGene_gn=BMPR1B;refGene_tx=XM_023252926.1;refGene_gn=BMPR1B;refGene_tx=XM_023252927.1;refGene_gn=BMPR1B;refGene_tx=XM_023252928.1;refGene_gn=BMPR1B;refGene_tx=XM_023252929.1;refGene_gn=BMPR1B;refGene_tx=XM_023252930.1
B1 128409154 128409350
refGene_gn=BMPR1B;refGene_tx=XM_023252924.1
B1 128410886 128410941
refGene_gn=BMPR1B;refGene_tx=XM_019829189.2;refGene_gn=BMPR1B;refGene_tx=XM_023252924.1
B1 128812023 128812113
refGene_gn=PDLIM5;refGene_tx=XM_019829215.2;refGene_gn=PDLIM5;refGene_tx=XM_019829217.1;refGene_gn=PDLIM5;refGene_tx=XM_019829218.2;refGene_gn=PDLIM5;refGene_tx=XM_019829219.2;refGene_gn=PDLIM5;refGene_tx=XM_019829220.2;refGene_gn=PDLIM5;refGene_tx=XM_019829221.1;refGene_gn=PDLIM5;refGene_tx=XM_019829222.2;refGene_gn=PDLIM5;refGene_tx=XM_019829223.2;refGene_gn=PDLIM5;refGene_tx=XM_019829224.2;refGene_gn=PDLIM5;refGene_tx=XM_019829225.2;refGene_gn=PDLIM5;refGene_tx=XM_019829226.2;refGene_gn=PDLIM5;refGene_tx=XM_019829228.2;refGene_gn=PDLIM5;refGene_tx=XM_019829229.2;refGene_gn=PDLIM5;refGene_tx=XM_019829231.2;refGene_gn=PDLIM5;refGene_tx=XM_023252931.1
B1 128813499 128813615
refGene_gn=PDLIM5;refGene_tx=XM_019829215.2;refGene_gn=PDLIM5;refGene_tx=XM_019829217.1;refGene_gn=PDLIM5;refGene_tx=XM_019829218.2;refGene_gn=PDLIM5;refGene_tx=XM_019829219.2;refGene_gn=PDLIM5;refGene_tx=XM_019829220.2;refGene_gn=PDLIM5;refGene_tx=XM_019829221.1;refGene_gn=PDLIM5;refGene_tx=XM_019829222.2;refGene_gn=PDLIM5;refGene_tx=XM_019829223.2;refGene_gn=PDLIM5;refGene_tx=XM_019829224.2;refGene_gn=PDLIM5;refGene_tx=XM_019829225.2;refGene_gn=PDLIM5;refGene_tx=XM_019829226.2;refGene_gn=PDLIM5;refGene_tx=XM_019829228.2;refGene_gn=PDLIM5;refGene_tx=XM_019829229.2;refGene_gn=PDLIM5;refGene_tx=XM_019829231.2;refGene_gn=PDLIM5;refGene_tx=XM_023252931.1
B1 128817023 128817144
refGene_gn=PDLIM5;refGene_tx=XM_019829215.2;refGene_gn=PDLIM5;refGene_tx=XM_019829217.1;refGene_gn=PDLIM5;refGene_tx=XM_019829218.2;refGene_gn=PDLIM5;refGene_tx=XM_019829219.2;refGene_gn=PDLIM5;refGene_tx=XM_019829220.2;refGene_gn=PDLIM5;refGene_tx=XM_019829221.1;refGene_gn=PDLIM5;refGene_tx=XM_019829222.2;refGene_gn=PDLIM5;refGene_tx=XM_019829223.2;refGene_gn=PDLIM5;refGene_tx=XM_019829224.2;refGene_gn=PDLIM5;refGene_tx=XM_019829225.2;refGene_gn=PDLIM5;refGene_tx=XM_019829226.2;refGene_gn=PDLIM5;refGene_tx=XM_019829228.2;refGene_gn=PDLIM5;refGene_tx=XM_019829229.2;refGene_gn=PDLIM5;refGene_tx=XM_019829231.2;refGene_gn=PDLIM5;refGene_tx=XM_023252931.1
B1 128819622 128819803
refGene_gn=PDLIM5;refGene_tx=XM_019829215.2;refGene_gn=PDLIM5;refGene_tx=XM_019829217.1;refGene_gn=PDLIM5;refGene_tx=XM_019829218.2;refGene_gn=PDLIM5;refGene_tx=XM_019829219.2;refGene_gn=PDLIM5;refGene_tx=XM_01

9829220.2;refGene_gn=PDLIM5;refGene_tx=XM_019829221.1;refGene_gn=PDLIM5;refGene_tx=XM_019829222.2;refGene_gn=PDLIM5;refGene_tx=XM_019829223.2;refGene_gn=PDLIM5;refGene_tx=XM_019829224.2;refGene_gn=PDLIM5;refGene_tx=XM_019829225.2;refGene_gn=PDLIM5;refGene_tx=XM_019829226.2;refGene_gn=PDLIM5;refGene_tx=XM_019829228.2;refGene_gn=PDLIM5;refGene_tx=XM_019829229.2;refGene_gn=PDLIM5;refGene_tx=XM_019829231.2;refGene_gn=PDLIM5;refGene_tx=XM_023252931.1
B1 128834010 128834185
refGene_gn=PDLIM5;refGene_tx=XM_019829215.2;refGene_gn=PDLIM5;refGene_tx=XM_019829217.1;refGene_gn=PDLIM5;refGene_tx=XM_019829218.2;refGene_gn=PDLIM5;refGene_tx=XM_019829219.2;refGene_gn=PDLIM5;refGene_tx=XM_019829220.2;refGene_gn=PDLIM5;refGene_tx=XM_019829221.1;refGene_gn=PDLIM5;refGene_tx=XM_019829222.2;refGene_gn=PDLIM5;refGene_tx=XM_019829223.2;refGene_gn=PDLIM5;refGene_tx=XM_019829224.2;refGene_gn=PDLIM5;refGene_tx=XM_019829225.2;refGene_gn=PDLIM5;refGene_tx=XM_019829226.2;refGene_gn=PDLIM5;refGene_tx=XM_019829228.2;refGene_gn=PDLIM5;refGene_tx=XM_019829229.2;refGene_gn=PDLIM5;refGene_tx=XM_019829231.2;refGene_gn=PDLIM5;refGene_tx=XM_023252931.1
B1 128863189 128863377
refGene_gn=PDLIM5;refGene_tx=XM_019829215.2;refGene_gn=PDLIM5;refGene_tx=XM_019829217.1;refGene_gn=PDLIM5;refGene_tx=XM_019829218.2;refGene_gn=PDLIM5;refGene_tx=XM_019829219.2;refGene_gn=PDLIM5;refGene_tx=XM_019829220.2;refGene_gn=PDLIM5;refGene_tx=XM_019829221.1;refGene_gn=PDLIM5;refGene_tx=XM_019829222.2;refGene_gn=PDLIM5;refGene_tx=XM_019829223.2;refGene_gn=PDLIM5;refGene_tx=XM_019829224.2;refGene_gn=PDLIM5;refGene_tx=XM_019829225.2;refGene_gn=PDLIM5;refGene_tx=XM_019829226.2;refGene_gn=PDLIM5;refGene_tx=XM_019829228.2;refGene_gn=PDLIM5;refGene_tx=XM_019829229.2;refGene_gn=PDLIM5;refGene_tx=XM_019829231.2;refGene_gn=PDLIM5;refGene_tx=XM_023252931.1
B1 128870835 128870967
refGene_gn=PDLIM5;refGene_tx=XM_019829231.2
B1 128871921 128872059
refGene_gn=PDLIM5;refGene_tx=XM_019829231.2
B1 128874023 128874140
refGene_gn=PDLIM5;refGene_tx=XM_019829231.2
B1 128899200 128899294
refGene_gn=PDLIM5;refGene_tx=XM_011281790.3;refGene_gn=PDLIM5;refGene_tx=XM_019829230.2
B1 128899853 128899890
refGene_gn=PDLIM5;refGene_tx=XM_011281790.3;refGene_gn=PDLIM5;refGene_tx=XM_019829215.2;refGene_gn=PDLIM5;refGene_tx=XM_019829217.1;refGene_gn=PDLIM5;refGene_tx=XM_019829218.2;refGene_gn=PDLIM5;refGene_tx=XM_019829219.2;refGene_gn=PDLIM5;refGene_tx=XM_019829220.2;refGene_gn=PDLIM5;refGene_tx=XM_019829221.1;refGene_gn=PDLIM5;refGene_tx=XM_019829222.2;refGene_gn=PDLIM5;refGene_tx=XM_019829223.2;refGene_gn=PDLIM5;refGene_tx=XM_019829224.2;refGene_gn=PDLIM5;refGene_tx=XM_019829225.2;refGene_gn=PDLIM5;refGene_tx=XM_019829226.2;refGene_gn=PDLIM5;refGene_tx=XM_019829228.2;refGene_gn=PDLIM5;refGene_tx=XM_019829229.2;refGene_gn=PDLIM5;refGene_tx=XM_019829230.2;refGene_gn=PDLIM5;refGene_tx=XM_019829231.2;refGene_gn=PDLIM5;refGene_tx=XM_023252931.1

B1 128900247 128900420
refGene_gn=PDLIM5;refGene_tx=XM_011281790.3;refGene_gn=PDLIM5;refGene_tx=XM_019829215.2;refGene_gn=PDLIM5;refGene_tx=XM_019829217.1;refGene_gn=PDLIM5;refGene_tx=XM_019829218.2;refGene_gn=PDLIM5;refGene_tx=XM_019829219.2;refGene_gn=PDLIM5;refGene_tx=XM_019829220.2;refGene_gn=PDLIM5;refGene_tx=XM_019829221.1;refGene_gn=PDLIM5;refGene_tx=XM_019829222.2;refGene_gn=PDLIM5;refGene_tx=XM_019829223.2;refGene_gn=PDLIM5;refGene_tx=XM_019829224.2;refGene_gn=PDLIM5;refGene_tx=XM_019829225.2;refGene_gn=PDLIM5;refGene_tx=XM_019829226.2;refGene_gn=PDLIM5;refGene_tx=XM_019829228.2;refGene_gn=PDLIM5;refGene_tx=XM_019829229.2;refGene_gn=PDLIM5;refGene_tx=XM_019829230.2;refGene_gn=PDLIM5;refGene_tx=XM_019829231.2;refGene_gn=PDLIM5;refGene_tx=XM_023252931.1
B1 128900677 128900695
refGene_gn=PDLIM5;refGene_tx=XM_011281790.3;refGene_gn=PDLIM5;refGene_tx=XM_019829215.2;refGene_gn=PDLIM5;refGene_tx=XM_019829217.1;refGene_gn=PDLIM5;refGene_tx=XM_019829218.2;refGene_gn=PDLIM5;refGene_tx=XM_019829220.2;refGene_gn=PDLIM5;refGene_tx=XM_019829221.1;refGene_gn=PDLIM5;refGene_tx=XM_019829222.2;refGene_gn=PDLIM5;refGene_tx=XM_019829223.2;refGene_gn=PDLIM5;refGene_tx=XM_019829225.2;refGene_gn=PDLIM5;refGene_tx=XM_019829226.2;refGene_gn=PDLIM5;refGene_tx=XM_019829228.2;refGene_gn=PDLIM5;refGene_tx=XM_019829230.2;refGene_gn=PDLIM5;refGene_tx=XM_019829231.2;refGene_gn=PDLIM5;refGene_tx=XM_023252931.1
B1 128903055 128903070
refGene_gn=PDLIM5;refGene_tx=XM_011281790.3;refGene_gn=PDLIM5;refGene_tx=XM_019829215.2;refGene_gn=PDLIM5;refGene_tx=XM_019829217.1;refGene_gn=PDLIM5;refGene_tx=XM_019829220.2;refGene_gn=PDLIM5;refGene_tx=XM_019829221.1;refGene_gn=PDLIM5;refGene_tx=XM_019829222.2;refGene_gn=PDLIM5;refGene_tx=XM_019829225.2;refGene_gn=PDLIM5;refGene_tx=XM_019829226.2;refGene_gn=PDLIM5;refGene_tx=XM_019829228.2;refGene_gn=PDLIM5;refGene_tx=XM_023252931.1
B1 128906514 128906531
refGene_gn=PDLIM5;refGene_tx=XM_011281790.3;refGene_gn=PDLIM5;refGene_tx=XM_019829225.2;refGene_gn=PDLIM5;refGene_tx=XM_019829228.2
B1 128909477 128909896
refGene_gn=PDLIM5;refGene_tx=XM_019829215.2;refGene_gn=PDLIM5;refGene_tx=XM_019829217.1;refGene_gn=PDLIM5;refGene_tx=XM_019829218.2;refGene_gn=PDLIM5;refGene_tx=XM_019829219.2;refGene_gn=PDLIM5;refGene_tx=XM_019829220.2;refGene_gn=PDLIM5;refGene_tx=XM_019829221.1;refGene_gn=PDLIM5;refGene_tx=XM_019829222.2;refGene_gn=PDLIM5;refGene_tx=XM_019829223.2;refGene_gn=PDLIM5;refGene_tx=XM_019829224.2;refGene_gn=PDLIM5;refGene_tx=XM_019829226.2;refGene_gn=PDLIM5;refGene_tx=XM_019829230.2;refGene_gn=PDLIM5;refGene_tx=XM_019829231.2;refGene_gn=PDLIM5;refGene_tx=XM_023252931.1
B1 128911472 128911515
refGene_gn=PDLIM5;refGene_tx=XM_011281790.3;refGene_gn=PDLIM5;refGene_tx=XM_019829215.2;refGene_gn=PDLIM5;refGene_tx=XM_019829217.1;refGene_gn=PDLIM5;refGene_tx=XM_019829218.2;refGene_gn=PDLIM5;refGene_tx=XM_019829219.2;refGene_gn=PDLIM5;refGene_tx=XM_019829220.2;refGene_gn=PDLIM5;refGene_tx=XM_019829221.1;refGene_gn=PDLIM5;refGene_tx=XM_019829222.2;refGene_gn=PDLIM5;refGene_tx=XM_019829223.2;refGene_gn=PDLIM5;refGen

e_tx=XM_019829224.2;refGene_gn=PDLIM5;refGene_tx=XM_019829225.2;refGene_gn=PDLIM5;refGene_tx=XM_019829226.2;refGene_gn=PDLIM5;refGene_tx=XM_019829228.2;refGene_gn=PDLIM5;refGene_tx=XM_019829230.2;refGene_gn=PDLIM5;refGene_tx=XM_019829231.2;refGene_gn=PDLIM5;refGene_tx=XM_023252931.1
B1 128915653 128915790
refGene_gn=PDLIM5;refGene_tx=XM_019829220.2;refGene_gn=PDLIM5;refGene_tx=XM_023252931.1
B1 128966520 128966672
refGene_gn=PDLIM5;refGene_tx=XM_011281790.3;refGene_gn=PDLIM5;refGene_tx=XM_019829215.2;refGene_gn=PDLIM5;refGene_tx=XM_019829217.1;refGene_gn=PDLIM5;refGene_tx=XM_019829218.2;refGene_gn=PDLIM5;refGene_tx=XM_019829219.2;refGene_gn=PDLIM5;refGene_tx=XM_019829222.2;refGene_gn=PDLIM5;refGene_tx=XM_019829223.2;refGene_gn=PDLIM5;refGene_tx=XM_019829224.2;refGene_gn=PDLIM5;refGene_tx=XM_019829225.2;refGene_gn=PDLIM5;refGene_tx=XM_019829230.2;refGene_gn=PDLIM5;refGene_tx=XM_019829231.2
B1 129034178 129034274
refGene_gn=PDLIM5;refGene_tx=XM_011281790.3;refGene_gn=PDLIM5;refGene_tx=XM_019829215.2;refGene_gn=PDLIM5;refGene_tx=XM_019829217.1;refGene_gn=PDLIM5;refGene_tx=XM_019829218.2;refGene_gn=PDLIM5;refGene_tx=XM_019829219.2;refGene_gn=PDLIM5;refGene_tx=XM_019829221.1;refGene_gn=PDLIM5;refGene_tx=XM_019829222.2;refGene_gn=PDLIM5;refGene_tx=XM_019829223.2;refGene_gn=PDLIM5;refGene_tx=XM_019829224.2;refGene_gn=PDLIM5;refGene_tx=XM_019829225.2;refGene_gn=PDLIM5;refGene_tx=XM_019829226.2;refGene_gn=PDLIM5;refGene_tx=XM_019829228.2;refGene_gn=PDLIM5;refGene_tx=XM_019829230.2;refGene_gn=PDLIM5;refGene_tx=XM_019829231.2
B1 129156450 129156583
refGene_gn=HPGDS;refGene_tx=XM_003985161.5
B1 129164739 129164832
refGene_gn=HPGDS;refGene_tx=XM_003985161.5
B1 129165667 129165777
refGene_gn=HPGDS;refGene_tx=XM_003985161.5
B1 129171525 129171624
refGene_gn=HPGDS;refGene_tx=XM_003985161.5
B1 129176222 129176387
refGene_gn=HPGDS;refGene_tx=XM_003985161.5
B1 129181787 129181849
refGene_gn=SMARCAD1;refGene_tx=XM_006930958.3;refGene_gn=SMARCAD1;refGene_tx=XM_019829232.2;refGene_gn=SMARCAD1;refGene_tx=XM_019829235.1;refGene_gn=SMARCAD1;refGene_tx=XM_019829237.1;refGene_gn=SMARCAD1;refGene_tx=XM_019829238.2
B1 129186050 129186160
refGene_gn=SMARCAD1;refGene_tx=XM_006930958.3;refGene_gn=SMARCAD1;refGene_tx=XM_019829232.2;refGene_gn=SMARCAD1;refGene_tx=XM_019829235.1;refGene_gn=SMARCAD1;refGene_tx=XM_019829237.1;refGene_gn=SMARCAD1;refGene_tx=XM_019829238.2
B1 129187101 129187284
refGene_gn=SMARCAD1;refGene_tx=XM_006930958.3;refGene_gn=SMARCAD1;refGene_tx=XM_019829232.2;refGene_gn=SMARCAD1;refGene_tx=XM_019829235.1;refGene_gn=SMARCAD1;refGene_tx=XM_019829237.1;refGene_gn=SMARCAD1;refGen

e_tx=XM_019829238.2
B1	129188186	129188305
refGene_gn=SMARCAD1;refGene_tx=XM_006930958.3;refGene_gn=SMARCAD1;refGene_tx=XM_019829232.2;refGene_gn=SMARCAD1;refGene_tx=XM_019829235.1;refGene_gn=SMARCAD1;refGene_tx=XM_019829237.1;refGene_gn=SMARCAD1;refGene_tx=XM_019829238.2
B1	129188924	129189113
refGene_gn=SMARCAD1;refGene_tx=XM_006930958.3;refGene_gn=SMARCAD1;refGene_tx=XM_019829232.2;refGene_gn=SMARCAD1;refGene_tx=XM_019829235.1;refGene_gn=SMARCAD1;refGene_tx=XM_019829237.1;refGene_gn=SMARCAD1;refGene_tx=XM_019829238.2
B1	129191891	129192013
refGene_gn=SMARCAD1;refGene_tx=XM_006930958.3;refGene_gn=SMARCAD1;refGene_tx=XM_019829232.2;refGene_gn=SMARCAD1;refGene_tx=XM_019829235.1;refGene_gn=SMARCAD1;refGene_tx=XM_019829237.1;refGene_gn=SMARCAD1;refGene_tx=XM_019829238.2
B1	129192331	129192449
refGene_gn=SMARCAD1;refGene_tx=XM_006930958.3;refGene_gn=SMARCAD1;refGene_tx=XM_019829232.2;refGene_gn=SMARCAD1;refGene_tx=XM_019829235.1;refGene_gn=SMARCAD1;refGene_tx=XM_019829237.1;refGene_gn=SMARCAD1;refGene_tx=XM_019829238.2
B1	129192547	129192643
refGene_gn=SMARCAD1;refGene_tx=XM_006930958.3;refGene_gn=SMARCAD1;refGene_tx=XM_019829232.2;refGene_gn=SMARCAD1;refGene_tx=XM_019829235.1;refGene_gn=SMARCAD1;refGene_tx=XM_019829237.1;refGene_gn=SMARCAD1;refGene_tx=XM_019829238.2
B1	129194105	129194243
refGene_gn=SMARCAD1;refGene_tx=XM_006930958.3;refGene_gn=SMARCAD1;refGene_tx=XM_019829232.2;refGene_gn=SMARCAD1;refGene_tx=XM_019829235.1;refGene_gn=SMARCAD1;refGene_tx=XM_019829237.1;refGene_gn=SMARCAD1;refGene_tx=XM_019829238.2
B1	129194755	129194891
refGene_gn=SMARCAD1;refGene_tx=XM_006930958.3;refGene_gn=SMARCAD1;refGene_tx=XM_019829232.2;refGene_gn=SMARCAD1;refGene_tx=XM_019829235.1;refGene_gn=SMARCAD1;refGene_tx=XM_019829237.1;refGene_gn=SMARCAD1;refGene_tx=XM_019829238.2
B1	129196361	129196437
refGene_gn=SMARCAD1;refGene_tx=XM_006930958.3;refGene_gn=SMARCAD1;refGene_tx=XM_019829232.2;refGene_gn=SMARCAD1;refGene_tx=XM_019829235.1;refGene_gn=SMARCAD1;refGene_tx=XM_019829237.1;refGene_gn=SMARCAD1;refGene_tx=XM_019829238.2
B1	129196538	129196598
refGene_gn=SMARCAD1;refGene_tx=XM_006930958.3;refGene_gn=SMARCAD1;refGene_tx=XM_019829232.2;refGene_gn=SMARCAD1;refGene_tx=XM_019829235.1;refGene_gn=SMARCAD1;refGene_tx=XM_019829237.1;refGene_gn=SMARCAD1;refGene_tx=XM_019829238.2
B1	129197297	129197397
refGene_gn=SMARCAD1;refGene_tx=XM_006930958.3;refGene_gn=SMARCAD1;refGene_tx=XM_019829232.2;refGene_gn=SMARCAD1;refGene_tx=XM_019829235.1;refGene_gn=SMARCAD1;refGene_tx=XM_019829237.1;refGene_gn=SMARCAD1;refGen

e_tx=XM_019829238.2
B1 129199793 129199884
refGene_gn=SMARCAD1;refGene_tx=XM_006930958.3;refGene_gn=SMARCAD1;refGene_tx=XM_019829232.2;refGene_gn=SMARCAD1;refGene_tx=XM_019829235.1;refGene_gn=SMARCAD1;refGene_tx=XM_019829237.1;refGene_gn=SMARCAD1;refGene_tx=XM_019829238.2
B1 129205561 129205761
refGene_gn=SMARCAD1;refGene_tx=XM_006930958.3;refGene_gn=SMARCAD1;refGene_tx=XM_019829232.2;refGene_gn=SMARCAD1;refGene_tx=XM_019829235.1;refGene_gn=SMARCAD1;refGene_tx=XM_019829237.1;refGene_gn=SMARCAD1;refGene_tx=XM_019829238.2
B1 129215060 129215452
refGene_gn=SMARCAD1;refGene_tx=XM_006930958.3;refGene_gn=SMARCAD1;refGene_tx=XM_019829232.2;refGene_gn=SMARCAD1;refGene_tx=XM_019829235.1;refGene_gn=SMARCAD1;refGene_tx=XM_019829237.1
B1 129216877 129216959
refGene_gn=SMARCAD1;refGene_tx=XM_006930958.3;refGene_gn=SMARCAD1;refGene_tx=XM_019829232.2;refGene_gn=SMARCAD1;refGene_tx=XM_019829235.1;refGene_gn=SMARCAD1;refGene_tx=XM_019829237.1
B1 129218691 129218793
refGene_gn=SMARCAD1;refGene_tx=XM_006930958.3;refGene_gn=SMARCAD1;refGene_tx=XM_019829232.2;refGene_gn=SMARCAD1;refGene_tx=XM_019829235.1;refGene_gn=SMARCAD1;refGene_tx=XM_019829237.1
B1 129226504 129226608
refGene_gn=SMARCAD1;refGene_tx=XM_006930958.3;refGene_gn=SMARCAD1;refGene_tx=XM_019829232.2;refGene_gn=SMARCAD1;refGene_tx=XM_019829235.1;refGene_gn=SMARCAD1;refGene_tx=XM_019829237.1
B1 129231888 129231955
refGene_gn=SMARCAD1;refGene_tx=XM_006930958.3;refGene_gn=SMARCAD1;refGene_tx=XM_019829232.2;refGene_gn=SMARCAD1;refGene_tx=XM_019829235.1;refGene_gn=SMARCAD1;refGene_tx=XM_019829237.1
B1 129234800 129234972
refGene_gn=SMARCAD1;refGene_tx=XM_006930958.3;refGene_gn=SMARCAD1;refGene_tx=XM_019829232.2;refGene_gn=SMARCAD1;refGene_tx=XM_019829235.1;refGene_gn=SMARCAD1;refGene_tx=XM_019829237.1
B1 129241049 129241227
refGene_gn=SMARCAD1;refGene_tx=XM_006930958.3;refGene_gn=SMARCAD1;refGene_tx=XM_019829232.2;refGene_gn=SMARCAD1;refGene_tx=XM_019829235.1;refGene_gn=SMARCAD1;refGene_tx=XM_019829237.1
B1 129258319 129258509
refGene_gn=SMARCAD1;refGene_tx=XM_006930958.3;refGene_gn=SMARCAD1;refGene_tx=XM_019829232.2;refGene_gn=SMARCAD1;refGene_tx=XM_019829235.1;refGene_gn=SMARCAD1;refGene_tx=XM_019829237.1
B1 129573980 129575051
refGene_gn=ATOH1;refGene_tx=XM_004001418.3
B1 129629371 129629794
refGene_gn=GRID2;refGene_tx=XM_019829242.1
B1 129632429 129632670
refGene_gn=GRID2;refGene_tx=XM_019829242.1
B1 129679914 129680203

refGene_gn=GRID2;refGene_tx=XM_019829243.1
B1 129743078 129743245
refGene_gn=GRID2;refGene_tx=XM_019829242.1;refGene_gn=GRID2;refGene_tx=XM_019829243.1
B1 129854394 129854590
refGene_gn=GRID2;refGene_tx=XM_019829242.1;refGene_gn=GRID2;refGene_tx=XM_019829243.1
B1 129878181 129878320
refGene_gn=GRID2;refGene_tx=XM_019829242.1;refGene_gn=GRID2;refGene_tx=XM_019829243.1
B1 129929112 129929425
refGene_gn=GRID2;refGene_tx=XM_019829242.1;refGene_gn=GRID2;refGene_tx=XM_019829243.1
B1 129971075 129971273
refGene_gn=GRID2;refGene_tx=XM_019829242.1;refGene_gn=GRID2;refGene_tx=XM_019829243.1
B1 130004396 130004498
refGene_gn=GRID2;refGene_tx=XM_019829242.1;refGene_gn=GRID2;refGene_tx=XM_019829243.1
B1 130170562 130170682
refGene_gn=GRID2;refGene_tx=XM_019829242.1;refGene_gn=GRID2;refGene_tx=XM_019829243.1
B1 130188071 130188233
refGene_gn=GRID2;refGene_tx=XM_019829242.1;refGene_gn=GRID2;refGene_tx=XM_019829243.1
B1 130196433 130196607
refGene_gn=GRID2;refGene_tx=XM_019829242.1;refGene_gn=GRID2;refGene_tx=XM_019829243.1
B1 130209995 130210049
refGene_gn=GRID2;refGene_tx=XM_019829242.1;refGene_gn=GRID2;refGene_tx=XM_019829243.1
B1 130301144 130301350
refGene_gn=GRID2;refGene_tx=XM_019829242.1;refGene_gn=GRID2;refGene_tx=XM_019829243.1
B1 130318307 130318592
refGene_gn=GRID2;refGene_tx=XM_019829242.1;refGene_gn=GRID2;refGene_tx=XM_019829243.1
B1 130782549 130782705
refGene_gn=GRID2;refGene_tx=XM_019829242.1;refGene_gn=GRID2;refGene_tx=XM_019829243.1
B1 131077970 131078058
refGene_gn=GRID2;refGene_tx=XM_019829242.1;refGene_gn=GRID2;refGene_tx=XM_019829243.1
B1 131709095 131709581
refGene_gn=CCSER1;refGene_tx=XM_011281786.3;refGene_gn=CCSER1;refGene_tx=XM_019829245.1;refGene_gn=CCSER1;refGene_tx=XM_023252933.1
B1 131991556 131991602
refGene_gn=CCSER1;refGene_tx=XM_019829245.1
B1 132142290 132142335
refGene_gn=CCSER1;refGene_tx=XM_011281786.3;refGene_gn=CCSER1;refGene_

tx=XM_019829245.1;refGene_gn=CCSER1;refGene_tx=XM_023252933.1
B1	132313797	132313857
refGene_gn=CCSER1;refGene_tx=XM_011281787.2
B1	132333453	132333531
refGene_gn=CCSER1;refGene_tx=XM_011281786.3;refGene_gn=CCSER1;refGene_tx=XM_011281787.2;refGene_gn=CCSER1;refGene_tx=XM_019829245.1;refGene_gn=CCSER1;refGene_tx=XM_023252933.1
B1	132423296	132423380
refGene_gn=CCSER1;refGene_tx=XM_011281786.3;refGene_gn=CCSER1;refGene_tx=XM_011281787.2;refGene_gn=CCSER1;refGene_tx=XM_019829245.1;refGene_gn=CCSER1;refGene_tx=XM_023252933.1
B1	132497392	132497470
refGene_gn=CCSER1;refGene_tx=XM_011281786.3;refGene_gn=CCSER1;refGene_tx=XM_011281787.2;refGene_gn=CCSER1;refGene_tx=XM_019829245.1;refGene_gn=CCSER1;refGene_tx=XM_023252933.1
B1	132578347	132578555
refGene_gn=CCSER1;refGene_tx=XM_011281786.3;refGene_gn=CCSER1;refGene_tx=XM_011281787.2;refGene_gn=CCSER1;refGene_tx=XM_019829245.1;refGene_gn=CCSER1;refGene_tx=XM_023252933.1
B1	132720216	132720337
refGene_gn=CCSER1;refGene_tx=XM_011281786.3;refGene_gn=CCSER1;refGene_tx=XM_011281787.2;refGene_gn=CCSER1;refGene_tx=XM_019829245.1;refGene_gn=CCSER1;refGene_tx=XM_023252933.1
B1	132776317	132776411
refGene_gn=CCSER1;refGene_tx=XM_011281786.3;refGene_gn=CCSER1;refGene_tx=XM_011281787.2;refGene_gn=CCSER1;refGene_tx=XM_019829245.1;refGene_gn=CCSER1;refGene_tx=XM_023252933.1
B1	132856571	132856756
refGene_gn=CCSER1;refGene_tx=XM_011281786.3;refGene_gn=CCSER1;refGene_tx=XM_011281787.2;refGene_gn=CCSER1;refGene_tx=XM_019829245.1;refGene_gn=CCSER1;refGene_tx=XM_023252933.1
B1	132862131	132863455
refGene_gn=CCSER1;refGene_tx=XM_011281786.3;refGene_gn=CCSER1;refGene_tx=XM_011281787.2;refGene_gn=CCSER1;refGene_tx=XM_019829245.1;refGene_gn=CCSER1;refGene_tx=XM_023252933.1
B1	133184960	133185382
refGene_gn=MMRN1;refGene_tx=XM_006930965.4
B1	133186562	133186715
refGene_gn=MMRN1;refGene_tx=XM_006930965.4
B1	133204607	133206596
refGene_gn=MMRN1;refGene_tx=XM_006930965.4
B1	133214234	133214408
refGene_gn=MMRN1;refGene_tx=XM_006930965.4
B1	133216260	133216365
refGene_gn=MMRN1;refGene_tx=XM_006930965.4
B1	133226777	133226884
refGene_gn=MMRN1;refGene_tx=XM_006930965.4
B1	133229847	133229967
refGene_gn=MMRN1;refGene_tx=XM_006930965.4
B1	133248988	133249563

refGene_gn=MMRN1;refGene_tx=XM_006930965.4
B1 133308817 133308938
refGene_gn=SNCA;refGene_tx=XM_003985175.5;refGene_gn=SNCA;refGene_tx=XM_006930966.4;refGene_gn=SNCA;refGene_tx=XM_006930967.4;refGene_gn=SNCA;refGene_tx=XM_006930969.4
B1 133319449 133319491
refGene_gn=SNCA;refGene_tx=XM_003985175.5;refGene_gn=SNCA;refGene_tx=XM_006930966.4;refGene_gn=SNCA;refGene_tx=XM_006930967.4;refGene_gn=SNCA;refGene_tx=XM_006930969.4
B1 133327280 133327423
refGene_gn=SNCA;refGene_tx=XM_003985175.5;refGene_gn=SNCA;refGene_tx=XM_006930966.4;refGene_gn=SNCA;refGene_tx=XM_006930967.4;refGene_gn=SNCA;refGene_tx=XM_006930969.4
B1 133451386 133451470
refGene_gn=SNCA;refGene_tx=XM_003985175.5;refGene_gn=SNCA;refGene_tx=XM_006930966.4;refGene_gn=SNCA;refGene_tx=XM_006930967.4;refGene_gn=SNCA;refGene_tx=XM_006930969.4
B1 133460671 133460704
refGene_gn=SNCA;refGene_tx=XM_003985175.5;refGene_gn=SNCA;refGene_tx=XM_006930966.4;refGene_gn=SNCA;refGene_tx=XM_006930967.4;refGene_gn=SNCA;refGene_tx=XM_006930969.4
B1 133928670 133930989
refGene_gn=GPRIN3;refGene_tx=XM_003985177.3;refGene_gn=GPRIN3;refGene_tx=XM_019829247.1
B1 134074218 134075796
refGene_gn=TIGD2;refGene_tx=XM_006930970.4
B1 134105917 134106022
refGene_gn=FAM13A;refGene_tx=XM_003985178.5;refGene_gn=FAM13A;refGene_tx=XM_019829252.2;refGene_gn=FAM13A;refGene_tx=XM_023252934.1;refGene_gn=FAM13A;refGene_tx=XM_023252935.1;refGene_gn=FAM13A;refGene_tx=XM_023252936.1;refGene_gn=FAM13A;refGene_tx=XM_023252937.1;refGene_gn=FAM13A;refGene_tx=XM_023252938.1
B1 134138724 134138914
refGene_gn=FAM13A;refGene_tx=XM_003985178.5;refGene_gn=FAM13A;refGene_tx=XM_019829252.2;refGene_gn=FAM13A;refGene_tx=XM_023252934.1;refGene_gn=FAM13A;refGene_tx=XM_023252935.1;refGene_gn=FAM13A;refGene_tx=XM_023252936.1;refGene_gn=FAM13A;refGene_tx=XM_023252937.1;refGene_gn=FAM13A;refGene_tx=XM_023252938.1;refGene_gn=FAM13A;refGene_tx=XM_023252939.1
B1 134146441 134146651
refGene_gn=FAM13A;refGene_tx=XM_003985178.5;refGene_gn=FAM13A;refGene_tx=XM_019829252.2;refGene_gn=FAM13A;refGene_tx=XM_023252934.1;refGene_gn=FAM13A;refGene_tx=XM_023252935.1;refGene_gn=FAM13A;refGene_tx=XM_023252936.1;refGene_gn=FAM13A;refGene_tx=XM_023252937.1;refGene_gn=FAM13A;refGene_tx=XM_023252938.1;refGene_gn=FAM13A;refGene_tx=XM_023252939.1
B1 134168347 134168525
refGene_gn=FAM13A;refGene_tx=XM_003985178.5;refGene_gn=FAM13A;refGene_tx=XM_019829252.2;refGene_gn=FAM13A;refGene_tx=XM_023252934.1;refGene_gn=FAM13A;refGene_tx=XM_023252935.1;refGene_gn=FAM13A;refGene_tx=XM_02

3252936.1;refGene_gn=FAM13A;refGene_tx=XM_023252937.1;refGene_gn=FAM13A;refGene_tx=XM_023252938.1;refGene_gn=FAM13A;refGene_tx=XM_023252939.1
B1 134229722 134229876
refGene_gn=FAM13A;refGene_tx=XM_003985178.5;refGene_gn=FAM13A;refGene_tx=XM_019829252.2;refGene_gn=FAM13A;refGene_tx=XM_023252934.1;refGene_gn=FAM13A;refGene_tx=XM_023252935.1;refGene_gn=FAM13A;refGene_tx=XM_023252936.1;refGene_gn=FAM13A;refGene_tx=XM_023252937.1;refGene_gn=FAM13A;refGene_tx=XM_023252938.1;refGene_gn=FAM13A;refGene_tx=XM_023252939.1
B1 134259676 134259754
refGene_gn=FAM13A;refGene_tx=XM_003985178.5;refGene_gn=FAM13A;refGene_tx=XM_023252934.1;refGene_gn=FAM13A;refGene_tx=XM_023252935.1;refGene_gn=FAM13A;refGene_tx=XM_023252936.1;refGene_gn=FAM13A;refGene_tx=XM_023252937.1;refGene_gn=FAM13A;refGene_tx=XM_023252938.1;refGene_gn=FAM13A;refGene_tx=XM_023252939.1
B1 134307524 134307688
refGene_gn=FAM13A;refGene_tx=XM_003985178.5;refGene_gn=FAM13A;refGene_tx=XM_019829252.2;refGene_gn=FAM13A;refGene_tx=XM_023252934.1;refGene_gn=FAM13A;refGene_tx=XM_023252935.1;refGene_gn=FAM13A;refGene_tx=XM_023252936.1;refGene_gn=FAM13A;refGene_tx=XM_023252937.1;refGene_gn=FAM13A;refGene_tx=XM_023252938.1;refGene_gn=FAM13A;refGene_tx=XM_023252939.1
B1 134333588 134333617
refGene_gn=FAM13A;refGene_tx=XM_006930971.4;refGene_gn=FAM13A;refGene_tx=XM_006930972.3
B1 134349600 134349642
refGene_gn=FAM13A;refGene_tx=XM_003985178.5;refGene_gn=FAM13A;refGene_tx=XM_006930971.4;refGene_gn=FAM13A;refGene_tx=XM_006930972.3;refGene_gn=FAM13A;refGene_tx=XM_019829252.2;refGene_gn=FAM13A;refGene_tx=XM_023252934.1;refGene_gn=FAM13A;refGene_tx=XM_023252936.1;refGene_gn=FAM13A;refGene_tx=XM_023252937.1;refGene_gn=FAM13A;refGene_tx=XM_023252938.1;refGene_gn=FAM13A;refGene_tx=XM_023252939.1
B1 134366849 134366891
refGene_gn=FAM13A;refGene_tx=XM_003985178.5;refGene_gn=FAM13A;refGene_tx=XM_006930971.4;refGene_gn=FAM13A;refGene_tx=XM_019829252.2;refGene_gn=FAM13A;refGene_tx=XM_023252934.1;refGene_gn=FAM13A;refGene_tx=XM_023252935.1;refGene_gn=FAM13A;refGene_tx=XM_023252937.1;refGene_gn=FAM13A;refGene_tx=XM_023252938.1;refGene_gn=FAM13A;refGene_tx=XM_023252939.1
B1 134369531 134369711
refGene_gn=FAM13A;refGene_tx=XM_003985178.5;refGene_gn=FAM13A;refGene_tx=XM_006930971.4;refGene_gn=FAM13A;refGene_tx=XM_006930972.3;refGene_gn=FAM13A;refGene_tx=XM_019829252.2;refGene_gn=FAM13A;refGene_tx=XM_023252934.1;refGene_gn=FAM13A;refGene_tx=XM_023252935.1;refGene_gn=FAM13A;refGene_tx=XM_023252936.1;refGene_gn=FAM13A;refGene_tx=XM_023252937.1;refGene_gn=FAM13A;refGene_tx=XM_023252938.1;refGene_gn=FAM13A;refGene_tx=XM_023252939.1
B1 134375881 134376068
refGene_gn=FAM13A;refGene_tx=XM_003985178.5;refGene_gn=FAM13A;refGene_

tx=XM_006930971.4;refGene_gn=FAM13A;refGene_tx=XM_006930972.3;refGene_gn=FAM13A;refGene_tx=XM_019829252.2;refGene_gn=FAM13A;refGene_tx=XM_023252934.1;refGene_gn=FAM13A;refGene_tx=XM_023252935.1;refGene_gn=FAM13A;refGene_tx=XM_023252936.1;refGene_gn=FAM13A;refGene_tx=XM_023252937.1;refGene_gn=FAM13A;refGene_tx=XM_023252938.1;refGene_gn=FAM13A;refGene_tx=XM_023252939.1
B1 134388687 134388764
refGene_gn=FAM13A;refGene_tx=XM_003985178.5;refGene_gn=FAM13A;refGene_tx=XM_006930971.4;refGene_gn=FAM13A;refGene_tx=XM_006930972.3;refGene_gn=FAM13A;refGene_tx=XM_019829252.2;refGene_gn=FAM13A;refGene_tx=XM_023252934.1;refGene_gn=FAM13A;refGene_tx=XM_023252935.1;refGene_gn=FAM13A;refGene_tx=XM_023252936.1;refGene_gn=FAM13A;refGene_tx=XM_023252937.1;refGene_gn=FAM13A;refGene_tx=XM_023252938.1;refGene_gn=FAM13A;refGene_tx=XM_023252939.1
B1 134389141 134389184
refGene_gn=FAM13A;refGene_tx=XM_003985178.5;refGene_gn=FAM13A;refGene_tx=XM_006930971.4;refGene_gn=FAM13A;refGene_tx=XM_006930972.3;refGene_gn=FAM13A;refGene_tx=XM_019829252.2;refGene_gn=FAM13A;refGene_tx=XM_023252934.1;refGene_gn=FAM13A;refGene_tx=XM_023252935.1;refGene_gn=FAM13A;refGene_tx=XM_023252936.1;refGene_gn=FAM13A;refGene_tx=XM_023252937.1;refGene_gn=FAM13A;refGene_tx=XM_023252938.1;refGene_gn=FAM13A;refGene_tx=XM_023252939.1
B1 134398609 134398757
refGene_gn=FAM13A;refGene_tx=XM_003985178.5;refGene_gn=FAM13A;refGene_tx=XM_006930971.4;refGene_gn=FAM13A;refGene_tx=XM_006930972.3;refGene_gn=FAM13A;refGene_tx=XM_019829252.2;refGene_gn=FAM13A;refGene_tx=XM_023252934.1;refGene_gn=FAM13A;refGene_tx=XM_023252935.1;refGene_gn=FAM13A;refGene_tx=XM_023252936.1;refGene_gn=FAM13A;refGene_tx=XM_023252937.1;refGene_gn=FAM13A;refGene_tx=XM_023252938.1;refGene_gn=FAM13A;refGene_tx=XM_023252939.1
B1 134407561 134407772
refGene_gn=FAM13A;refGene_tx=XM_003985178.5;refGene_gn=FAM13A;refGene_tx=XM_006930971.4;refGene_gn=FAM13A;refGene_tx=XM_006930972.3;refGene_gn=FAM13A;refGene_tx=XM_019829252.2;refGene_gn=FAM13A;refGene_tx=XM_023252934.1;refGene_gn=FAM13A;refGene_tx=XM_023252935.1;refGene_gn=FAM13A;refGene_tx=XM_023252936.1;refGene_gn=FAM13A;refGene_tx=XM_023252937.1;refGene_gn=FAM13A;refGene_tx=XM_023252938.1;refGene_gn=FAM13A;refGene_tx=XM_023252939.1
B1 134408244 134408383
refGene_gn=FAM13A;refGene_tx=XM_003985178.5;refGene_gn=FAM13A;refGene_tx=XM_006930971.4;refGene_gn=FAM13A;refGene_tx=XM_006930972.3;refGene_gn=FAM13A;refGene_tx=XM_019829252.2;refGene_gn=FAM13A;refGene_tx=XM_023252934.1;refGene_gn=FAM13A;refGene_tx=XM_023252935.1;refGene_gn=FAM13A;refGene_tx=XM_023252936.1;refGene_gn=FAM13A;refGene_tx=XM_023252937.1;refGene_gn=FAM13A;refGene_tx=XM_023252938.1;refGene_gn=FAM13A;refGene_tx=XM_023252939.1
B1 134409115 134409197
refGene_gn=FAM13A;refGene_tx=XM_003985178.5;refGene_gn=FAM13A;refGene_tx=XM_006930971.4;refGene_gn=FAM13A;refGene_tx=XM_006930972.3;refGene_gn=FAM13A;refGene_tx=XM_019829252.2;refGene_gn=FAM13A;refGene_tx=XM_02

3252934.1;refGene_gn=FAM13A;refGene_tx=XM_023252935.1;refGene_gn=FAM13A;refGene_tx=XM_023252936.1;refGene_gn=FAM13A;refGene_tx=XM_023252937.1;refGene_gn=FAM13A;refGene_tx=XM_023252938.1;refGene_gn=FAM13A;refGene_tx=XM_023252939.1
B1 134410015 134410236 refGene_gn=FAM13A;refGene_tx=XM_003985178.5;refGene_gn=FAM13A;refGene_tx=XM_006930971.4;refGene_gn=FAM13A;refGene_tx=XM_006930972.3;refGene_gn=FAM13A;refGene_tx=XM_019829252.2;refGene_gn=FAM13A;refGene_tx=XM_023252934.1;refGene_gn=FAM13A;refGene_tx=XM_023252935.1;refGene_gn=FAM13A;refGene_tx=XM_023252936.1;refGene_gn=FAM13A;refGene_tx=XM_023252937.1;refGene_gn=FAM13A;refGene_tx=XM_023252938.1;refGene_gn=FAM13A;refGene_tx=XM_023252939.1
B1 134410893 134410977 refGene_gn=FAM13A;refGene_tx=XM_003985178.5;refGene_gn=FAM13A;refGene_tx=XM_006930971.4;refGene_gn=FAM13A;refGene_tx=XM_006930972.3;refGene_gn=FAM13A;refGene_tx=XM_019829252.2;refGene_gn=FAM13A;refGene_tx=XM_023252934.1;refGene_gn=FAM13A;refGene_tx=XM_023252935.1;refGene_gn=FAM13A;refGene_tx=XM_023252936.1;refGene_gn=FAM13A;refGene_tx=XM_023252937.1;refGene_gn=FAM13A;refGene_tx=XM_023252938.1;refGene_gn=FAM13A;refGene_tx=XM_023252939.1
B1 134424238 134424334 refGene_gn=FAM13A;refGene_tx=XM_003985178.5;refGene_gn=FAM13A;refGene_tx=XM_006930971.4;refGene_gn=FAM13A;refGene_tx=XM_006930972.3;refGene_gn=FAM13A;refGene_tx=XM_019829252.2;refGene_gn=FAM13A;refGene_tx=XM_023252934.1;refGene_gn=FAM13A;refGene_tx=XM_023252935.1;refGene_gn=FAM13A;refGene_tx=XM_023252936.1;refGene_gn=FAM13A;refGene_tx=XM_023252937.1;refGene_gn=FAM13A;refGene_tx=XM_023252938.1;refGene_gn=FAM13A;refGene_tx=XM_023252939.1
B1 134426536 134426620 refGene_gn=FAM13A;refGene_tx=XM_003985178.5;refGene_gn=FAM13A;refGene_tx=XM_006930971.4;refGene_gn=FAM13A;refGene_tx=XM_006930972.3;refGene_gn=FAM13A;refGene_tx=XM_019829252.2;refGene_gn=FAM13A;refGene_tx=XM_023252934.1;refGene_gn=FAM13A;refGene_tx=XM_023252935.1;refGene_gn=FAM13A;refGene_tx=XM_023252936.1;refGene_gn=FAM13A;refGene_tx=XM_023252938.1;refGene_gn=FAM13A;refGene_tx=XM_023252939.1
B1 134432154 134432351 refGene_gn=FAM13A;refGene_tx=XM_003985178.5;refGene_gn=FAM13A;refGene_tx=XM_006930971.4;refGene_gn=FAM13A;refGene_tx=XM_006930972.3;refGene_gn=FAM13A;refGene_tx=XM_019829252.2;refGene_gn=FAM13A;refGene_tx=XM_023252934.1;refGene_gn=FAM13A;refGene_tx=XM_023252935.1;refGene_gn=FAM13A;refGene_tx=XM_023252936.1;refGene_gn=FAM13A;refGene_tx=XM_023252937.1;refGene_gn=FAM13A;refGene_tx=XM_023252938.1;refGene_gn=FAM13A;refGene_tx=XM_023252939.1
B1 134432919 134433021 refGene_gn=FAM13A;refGene_tx=XM_003985178.5;refGene_gn=FAM13A;refGene_tx=XM_006930971.4;refGene_gn=FAM13A;refGene_tx=XM_006930972.3;refGene_gn=FAM13A;refGene_tx=XM_019829252.2;refGene_gn=FAM13A;refGene_tx=XM_023252934.1;refGene_gn=FAM13A;refGene_tx=XM_023252935.1;refGene_gn=FAM13A;refGene_tx=XM_023252936.1;refGene_gn=FAM13A;refGene_tx=XM_023252937.1;refGene_gn=FAM13A;refGene_tx=XM_023252938.1;refGene_gn=FAM13A;refGen

e_tx=XM_023252939.1
B1	134435477	134435604
refGene_gn=FAM13A;refGene_tx=XM_003985178.5;refGene_gn=FAM13A;refGene_tx=XM_006930971.4;refGene_gn=FAM13A;refGene_tx=XM_006930972.3;refGene_gn=FAM13A;refGene_tx=XM_019829252.2;refGene_gn=FAM13A;refGene_tx=XM_023252934.1;refGene_gn=FAM13A;refGene_tx=XM_023252935.1;refGene_gn=FAM13A;refGene_tx=XM_023252936.1;refGene_gn=FAM13A;refGene_tx=XM_023252937.1;refGene_gn=FAM13A;refGene_tx=XM_023252939.1
B1	134439269	134439297
refGene_gn=FAM13A;refGene_tx=XM_023252938.1
B1	134460696	134460905
refGene_gn=HERC3;refGene_tx=XM_003985179.4;refGene_gn=HERC3;refGene_tx=XM_023252940.1;refGene_gn=HERC3;refGene_tx=XM_023252941.1
B1	134463076	134463179
refGene_gn=HERC3;refGene_tx=XM_003985179.4;refGene_gn=HERC3;refGene_tx=XM_023252940.1;refGene_gn=HERC3;refGene_tx=XM_023252941.1
B1	134463414	134463598
refGene_gn=HERC3;refGene_tx=XM_003985179.4;refGene_gn=HERC3;refGene_tx=XM_023252940.1;refGene_gn=HERC3;refGene_tx=XM_023252941.1
B1	134474393	134474984
refGene_gn=NAP1L5;refGene_tx=XM_011281785.3
B1	134483100	134483183
refGene_gn=HERC3;refGene_tx=XM_003985179.4;refGene_gn=HERC3;refGene_tx=XM_023252940.1;refGene_gn=HERC3;refGene_tx=XM_023252941.1
B1	134483677	134483744
refGene_gn=HERC3;refGene_tx=XM_003985179.4;refGene_gn=HERC3;refGene_tx=XM_023252940.1;refGene_gn=HERC3;refGene_tx=XM_023252941.1
B1	134486348	134486515
refGene_gn=HERC3;refGene_tx=XM_003985179.4;refGene_gn=HERC3;refGene_tx=XM_023252940.1;refGene_gn=HERC3;refGene_tx=XM_023252941.1
B1	134487879	134488023
refGene_gn=HERC3;refGene_tx=XM_003985179.4;refGene_gn=HERC3;refGene_tx=XM_023252940.1;refGene_gn=HERC3;refGene_tx=XM_023252941.1
B1	134489692	134489863
refGene_gn=HERC3;refGene_tx=XM_003985179.4;refGene_gn=HERC3;refGene_tx=XM_023252940.1;refGene_gn=HERC3;refGene_tx=XM_023252941.1
B1	134491084	134491174
refGene_gn=HERC3;refGene_tx=XM_003985179.4;refGene_gn=HERC3;refGene_tx=XM_023252940.1;refGene_gn=HERC3;refGene_tx=XM_023252941.1
B1	134491265	134491289
refGene_gn=HERC3;refGene_tx=XM_003985179.4;refGene_gn=HERC3;refGene_tx=XM_023252940.1;refGene_gn=HERC3;refGene_tx=XM_023252941.1
B1	134497096	134497210
refGene_gn=HERC3;refGene_tx=XM_003985179.4;refGene_gn=HERC3;refGene_tx=XM_023252940.1;refGene_gn=HERC3;refGene_tx=XM_023252941.1
B1	134497342	134497506
refGene_gn=HERC3;refGene_tx=XM_003985179.4;refGene_gn=HERC3;refGene_tx=XM_023252940.1;refGene_gn=HERC3;refGene_tx=XM_023252941.1
B1	134499285	134499475
refGene_gn=HERC3;refGene_tx=XM_003985179.4;refGene_gn=HERC3;refGene_tx

=XM_023252940.1;refGene_gn=HERC3;refGene_tx=XM_023252941.1
B1 134500350 134500462
refGene_gn=HERC3;refGene_tx=XM_003985179.4;refGene_gn=HERC3;refGene_tx
=XM_023252940.1;refGene_gn=HERC3;refGene_tx=XM_023252941.1
B1 134502791 134502851
refGene_gn=HERC3;refGene_tx=XM_003985179.4;refGene_gn=HERC3;refGene_tx
=XM_023252940.1;refGene_gn=HERC3;refGene_tx=XM_023252941.1
B1 134504414 134504539
refGene_gn=HERC3;refGene_tx=XM_003985179.4;refGene_gn=HERC3;refGene_tx
=XM_023252940.1;refGene_gn=HERC3;refGene_tx=XM_023252941.1
B1 134508116 134508193
refGene_gn=HERC3;refGene_tx=XM_003985179.4;refGene_gn=HERC3;refGene_tx
=XM_023252940.1;refGene_gn=HERC3;refGene_tx=XM_023252941.1
B1 134510404 134510565
refGene_gn=HERC3;refGene_tx=XM_003985179.4;refGene_gn=HERC3;refGene_tx
=XM_023252940.1;refGene_gn=HERC3;refGene_tx=XM_023252941.1
B1 134511148 134511279
refGene_gn=HERC3;refGene_tx=XM_003985179.4;refGene_gn=HERC3;refGene_tx
=XM_023252940.1;refGene_gn=HERC3;refGene_tx=XM_023252941.1
B1 134512031 134512123
refGene_gn=HERC3;refGene_tx=XM_003985179.4;refGene_gn=HERC3;refGene_tx
=XM_023252940.1;refGene_gn=HERC3;refGene_tx=XM_023252941.1
B1 134513296 134513518
refGene_gn=HERC3;refGene_tx=XM_003985179.4;refGene_gn=HERC3;refGene_tx
=XM_023252940.1;refGene_gn=HERC3;refGene_tx=XM_023252941.1
B1 134514346 134514423
refGene_gn=HERC3;refGene_tx=XM_003985179.4;refGene_gn=HERC3;refGene_tx
=XM_023252940.1;refGene_gn=HERC3;refGene_tx=XM_023252941.1
B1 134518212 134518372
refGene_gn=HERC3;refGene_tx=XM_003985179.4;refGene_gn=HERC3;refGene_tx
=XM_023252940.1;refGene_gn=HERC3;refGene_tx=XM_023252941.1
B1 134558811 134559037
refGene_gn=HERC3;refGene_tx=XM_003985179.4;refGene_gn=HERC3;refGene_tx
=XM_023252940.1;refGene_gn=HERC3;refGene_tx=XM_023252941.1
B1 134570519 134570540
refGene_gn=LOC109499364;refGene_tx=XM_019829933.1
B1 134570913 134571539
refGene_gn=LOC109499364;refGene_tx=XM_019829933.1
B1 134575582 134575740
refGene_gn=LOC109499364;refGene_tx=XM_019829933.1
B1 134576635 134576739
refGene_gn=LOC109499364;refGene_tx=XM_019829933.1
B1 134625924 134626130
refGene_gn=PYURF;refGene_tx=XM_003985180.5
B1 134627523 134627665
refGene_gn=PYURF;refGene_tx=XM_003985180.5
B1 134627758 134627974
refGene_gn=PIGY;refGene_tx=XM_011281783.3
B1 134635630 134635836
refGene_gn=HERC5;refGene_tx=XM_023252942.1;refGene_gn=HERC5;refGene_tx

=XM_023252943.1
B1	134636906	134637009
refGene_gn=HERC5;refGene_tx=XM_023252942.1;refGene_gn=HERC5;refGene_tx=XM_023252943.1
B1	134637094	134637278
refGene_gn=HERC5;refGene_tx=XM_023252942.1;refGene_gn=HERC5;refGene_tx=XM_023252943.1
B1	134638851	134638922
refGene_gn=HERC5;refGene_tx=XM_023252942.1;refGene_gn=HERC5;refGene_tx=XM_023252943.1
B1	134639792	134639859
refGene_gn=HERC5;refGene_tx=XM_023252942.1;refGene_gn=HERC5;refGene_tx=XM_023252943.1
B1	134643923	134644090
refGene_gn=HERC5;refGene_tx=XM_023252942.1;refGene_gn=HERC5;refGene_tx=XM_023252943.1
B1	134644670	134644814
refGene_gn=HERC5;refGene_tx=XM_023252942.1;refGene_gn=HERC5;refGene_tx=XM_023252943.1
B1	134647652	134647823
refGene_gn=HERC5;refGene_tx=XM_023252942.1;refGene_gn=HERC5;refGene_tx=XM_023252943.1
B1	134650290	134650401
refGene_gn=HERC5;refGene_tx=XM_023252942.1;refGene_gn=HERC5;refGene_tx=XM_023252943.1
B1	134651472	134651589
refGene_gn=HERC5;refGene_tx=XM_023252942.1;refGene_gn=HERC5;refGene_tx=XM_023252943.1
B1	134658458	134658613
refGene_gn=HERC5;refGene_tx=XM_023252942.1;refGene_gn=HERC5;refGene_tx=XM_023252943.1
B1	134660175	134660365
refGene_gn=HERC5;refGene_tx=XM_023252942.1;refGene_gn=HERC5;refGene_tx=XM_023252943.1
B1	134663571	134663665
refGene_gn=HERC5;refGene_tx=XM_023252942.1;refGene_gn=HERC5;refGene_tx=XM_023252943.1
B1	134666238	134666298
refGene_gn=HERC5;refGene_tx=XM_023252942.1;refGene_gn=HERC5;refGene_tx=XM_023252943.1
B1	134668134	134668238
refGene_gn=HERC5;refGene_tx=XM_023252942.1;refGene_gn=HERC5;refGene_tx=XM_023252943.1
B1	134669208	134669285
refGene_gn=HERC5;refGene_tx=XM_023252942.1;refGene_gn=HERC5;refGene_tx=XM_023252943.1
B1	134670907	134671053
refGene_gn=HERC5;refGene_tx=XM_023252942.1;refGene_gn=HERC5;refGene_tx=XM_023252943.1
B1	134673362	134673493

refGene_gn=HERC5;refGene_tx=XM_023252942.1;refGene_gn=HERC5;refGene_tx=XM_023252943.1
B1	134673743	134673835
refGene_gn=HERC5;refGene_tx=XM_023252942.1;refGene_gn=HERC5;refGene_tx=XM_023252943.1
B1	134675045	134675267
refGene_gn=HERC5;refGene_tx=XM_023252942.1;refGene_gn=HERC5;refGene_tx=XM_023252943.1
B1	134678805	134678882
refGene_gn=HERC5;refGene_tx=XM_023252942.1;refGene_gn=HERC5;refGene_tx=XM_023252943.1
B1	134679546	134679667
refGene_gn=HERC5;refGene_tx=XM_023252942.1;refGene_gn=HERC5;refGene_tx=XM_023252943.1
B1	134681656	134681939
refGene_gn=HERC5;refGene_tx=XM_023252942.1
B1	134687106	134687330
refGene_gn=HERC6;refGene_tx=XM_023252944.1;refGene_gn=HERC6;refGene_tx=XM_023252945.1
B1	134690927	134691030
refGene_gn=HERC6;refGene_tx=XM_023252944.1;refGene_gn=HERC6;refGene_tx=XM_023252945.1
B1	134691120	134691304
refGene_gn=HERC6;refGene_tx=XM_023252944.1;refGene_gn=HERC6;refGene_tx=XM_023252945.1
B1	134692976	134693047
refGene_gn=HERC6;refGene_tx=XM_023252944.1;refGene_gn=HERC6;refGene_tx=XM_023252945.1;refGene_gn=HERC6;refGene_tx=XM_023252946.1
B1	134693898	134693965
refGene_gn=HERC6;refGene_tx=XM_023252944.1;refGene_gn=HERC6;refGene_tx=XM_023252945.1
B1	134695244	134695411
refGene_gn=HERC6;refGene_tx=XM_023252944.1;refGene_gn=HERC6;refGene_tx=XM_023252945.1;refGene_gn=HERC6;refGene_tx=XM_023252946.1
B1	134696712	134696856
refGene_gn=HERC6;refGene_tx=XM_023252944.1;refGene_gn=HERC6;refGene_tx=XM_023252945.1;refGene_gn=HERC6;refGene_tx=XM_023252946.1
B1	134700309	134700471
refGene_gn=HERC6;refGene_tx=XM_023252944.1;refGene_gn=HERC6;refGene_tx=XM_023252945.1;refGene_gn=HERC6;refGene_tx=XM_023252946.1
B1	134701751	134701859
refGene_gn=HERC6;refGene_tx=XM_023252944.1;refGene_gn=HERC6;refGene_tx=XM_023252945.1;refGene_gn=HERC6;refGene_tx=XM_023252946.1
B1	134702162	134702276
refGene_gn=HERC6;refGene_tx=XM_023252944.1;refGene_gn=HERC6;refGene_tx=XM_023252945.1;refGene_gn=HERC6;refGene_tx=XM_023252946.1
B1	134708479	134708640
refGene_gn=HERC6;refGene_tx=XM_023252944.1;refGene_gn=HERC6;refGene_tx=XM_023252945.1;refGene_gn=HERC6;refGene_tx=XM_023252946.1
B1	134711035	134711225

refGene_gn=HERC6;refGene_tx=XM_023252944.1;refGene_gn=HERC6;refGene_tx=XM_023252945.1;refGene_gn=HERC6;refGene_tx=XM_023252946.1
B1 134715487 134715581
refGene_gn=HERC6;refGene_tx=XM_023252944.1;refGene_gn=HERC6;refGene_tx=XM_023252945.1;refGene_gn=HERC6;refGene_tx=XM_023252946.1
B1 134718732 134718792
refGene_gn=HERC6;refGene_tx=XM_023252944.1;refGene_gn=HERC6;refGene_tx=XM_023252945.1;refGene_gn=HERC6;refGene_tx=XM_023252946.1
B1 134719706 134719828
refGene_gn=HERC6;refGene_tx=XM_023252944.1;refGene_gn=HERC6;refGene_tx=XM_023252945.1;refGene_gn=HERC6;refGene_tx=XM_023252946.1
B1 134728170 134728238
refGene_gn=HERC6;refGene_tx=XM_023252944.1;refGene_gn=HERC6;refGene_tx=XM_023252945.1;refGene_gn=HERC6;refGene_tx=XM_023252946.1
B1 134728895 134729038
refGene_gn=HERC6;refGene_tx=XM_023252944.1;refGene_gn=HERC6;refGene_tx=XM_023252945.1;refGene_gn=HERC6;refGene_tx=XM_023252946.1
B1 134736403 134736531
refGene_gn=HERC6;refGene_tx=XM_023252944.1;refGene_gn=HERC6;refGene_tx=XM_023252945.1;refGene_gn=HERC6;refGene_tx=XM_023252946.1
B1 134738478 134738573
refGene_gn=HERC6;refGene_tx=XM_023252944.1;refGene_gn=HERC6;refGene_tx=XM_023252945.1;refGene_gn=HERC6;refGene_tx=XM_023252946.1
B1 134739913 134740141
refGene_gn=HERC6;refGene_tx=XM_023252944.1;refGene_gn=HERC6;refGene_tx=XM_023252945.1;refGene_gn=HERC6;refGene_tx=XM_023252946.1
B1 134743480 134743557
refGene_gn=HERC6;refGene_tx=XM_023252944.1;refGene_gn=HERC6;refGene_tx=XM_023252945.1;refGene_gn=HERC6;refGene_tx=XM_023252946.1
B1 134745404 134745564
refGene_gn=HERC6;refGene_tx=XM_023252944.1;refGene_gn=HERC6;refGene_tx=XM_023252945.1;refGene_gn=HERC6;refGene_tx=XM_023252946.1
B1 134747392 134747591
refGene_gn=HERC6;refGene_tx=XM_023252944.1;refGene_gn=HERC6;refGene_tx=XM_023252945.1;refGene_gn=HERC6;refGene_tx=XM_023252946.1
B1 134819035 134819475
refGene_gn=PPM1K;refGene_tx=XM_006930974.4
B1 134820587 134820688
refGene_gn=PPM1K;refGene_tx=XM_006930974.4
B1 134828390 134828556
refGene_gn=PPM1K;refGene_tx=XM_006930974.4
B1 134828708 134828853
refGene_gn=PPM1K;refGene_tx=XM_006930974.4
B1 134830798 134830933
refGene_gn=PPM1K;refGene_tx=XM_006930974.4
B1 134833685 134833817
refGene_gn=PPM1K;refGene_tx=XM_006930974.4
B1 134932836 134933039
refGene_gn=ABCG2;refGene_tx=XM_023252947.1
B1 134939537 134939597

refGene_gn=ABCG2;refGene_tx=XM_023252947.1
B1 134940212 134940327
refGene_gn=ABCG2;refGene_tx=XM_023252947.1
B1 134940911 134941064
refGene_gn=ABCG2;refGene_tx=XM_023252947.1
B1 134943725 134943883
refGene_gn=ABCG2;refGene_tx=XM_023252947.1
B1 134949932 134950084
refGene_gn=ABCG2;refGene_tx=XM_023252947.1
B1 134951730 134951835
refGene_gn=ABCG2;refGene_tx=XM_023252947.1
B1 134953079 134953327
refGene_gn=ABCG2;refGene_tx=XM_023252947.1
B1 134958868 134958889
refGene_gn=ABCG2;refGene_tx=XM_023252947.1
B1 134958890 134958952
refGene_gn=ABCG2;refGene_tx=XM_023252947.1
B1 134964303 134964393
refGene_gn=ABCG2;refGene_tx=XM_023252947.1
B1 134965787 134965912
refGene_gn=ABCG2;refGene_tx=XM_023252947.1
B1 134967147 134967302
refGene_gn=ABCG2;refGene_tx=XM_023252947.1
B1 134972915 134973005
refGene_gn=ABCG2;refGene_tx=XM_023252947.1
B1 134973894 134973977
refGene_gn=ABCG2;refGene_tx=XM_023252947.1
B1 134975581 134975729
refGene_gn=ABCG2;refGene_tx=XM_023252947.1
B1 134986177 134986414
refGene_gn=PKD2;refGene_tx=XM_011281830.3
B1 134986783 134986931
refGene_gn=PKD2;refGene_tx=XM_011281830.3
B1 134991016 134991180
refGene_gn=PKD2;refGene_tx=XM_011281830.3
B1 134992140 134992258
refGene_gn=PKD2;refGene_tx=XM_011281830.3
B1 134992517 134992639
refGene_gn=PKD2;refGene_tx=XM_011281830.3
B1 134994875 134994974
refGene_gn=PKD2;refGene_tx=XM_011281830.3
B1 134998817 134998938
refGene_gn=PKD2;refGene_tx=XM_011281830.3
B1 135000074 135000256
refGene_gn=PKD2;refGene_tx=XM_011281830.3
B1 135003151 135003319
refGene_gn=PKD2;refGene_tx=XM_011281830.3
B1 135007605 135007834
refGene_gn=PKD2;refGene_tx=XM_011281830.3
B1 135010446 135010671

refGene_gn=PKD2;refGene_tx=XM_011281830.3
B1 135015296 135015547
refGene_gn=PKD2;refGene_tx=XM_011281830.3
B1 135017638 135017772
refGene_gn=PKD2;refGene_tx=XM_011281830.3
B1 135028099 135028213
refGene_gn=PKD2;refGene_tx=XM_011281830.3
B1 135035887 135036488
refGene_gn=PKD2;refGene_tx=XM_011281830.3
B1 135059207 135059612
refGene_gn=SPP1;refGene_tx=XM_003985184.5;refGene_gn=SPP1;refGene_tx=XM_006930977.3
B1 135060309 135060588
refGene_gn=SPP1;refGene_tx=XM_003985184.5;refGene_gn=SPP1;refGene_tx=XM_006930977.3
B1 135061639 135061681
refGene_gn=SPP1;refGene_tx=XM_003985184.5;refGene_gn=SPP1;refGene_tx=XM_006930977.3
B1 135061975 135062056
refGene_gn=SPP1;refGene_tx=XM_003985184.5
B1 135065156 135065195
refGene_gn=SPP1;refGene_tx=XM_003985184.5;refGene_gn=SPP1;refGene_tx=XM_006930977.3
B1 135065306 135065360
refGene_gn=SPP1;refGene_tx=XM_003985184.5;refGene_gn=SPP1;refGene_tx=XM_006930977.3
B1 135204076 135205537
refGene_gn=MEPE;refGene_tx=XM_023252235.1
B1 135210729 135210822
refGene_gn=MEPE;refGene_tx=XM_023252235.1
B1 135211766 135211820
refGene_gn=MEPE;refGene_tx=XM_023252235.1
B1 135216248 135216302
refGene_gn=MEPE;refGene_tx=XM_023252235.1
B1 135233846 135234389
refGene_gn=IBSP;refGene_tx=XM_003985189.3
B1 135234986 135235157
refGene_gn=IBSP;refGene_tx=XM_003985189.3
B1 135240453 135240519
refGene_gn=IBSP;refGene_tx=XM_003985189.3
B1 135244428 135244506
refGene_gn=IBSP;refGene_tx=XM_003985189.3
B1 135244601 135244652
refGene_gn=IBSP;refGene_tx=XM_003985189.3
B1 135244733 135244787
refGene_gn=IBSP;refGene_tx=XM_003985189.3
B1 135380307 135381678
refGene_gn=DMP1;refGene_tx=XM_019829256.1
B1 135383531 135383579
refGene_gn=DMP1;refGene_tx=XM_019829256.1

B1	135383768	135383801
refGene_gn=DMP1;refGene_tx=XM_019829256.1
B1	135386034	135386082
refGene_gn=DMP1;refGene_tx=XM_019829256.1
B1	135386530	135386584
refGene_gn=DMP1;refGene_tx=XM_019829256.1
B1	135411350	135412362
refGene_gn=DSPP;refGene_tx=XM_023252948.1
B1	135414730	135415235
refGene_gn=DSPP;refGene_tx=XM_023252948.1
B1	135415717	135416713
refGene_gn=DSPP;refGene_tx=XM_023252948.1
B1	135416848	135416935
refGene_gn=DSPP;refGene_tx=XM_023252948.1
B1	135418562	135418613
refGene_gn=DSPP;refGene_tx=XM_023252948.1
B1	135501534	135501563
refGene_gn=NUDT9;refGene_tx=XM_019829262.2
B1	135513570	135513604
refGene_gn=SPARCL1;refGene_tx=XM_003985193.4
B1	135528595	135528660
refGene_gn=SPARCL1;refGene_tx=XM_003985193.4;refGene_gn=SPARCL1;refGene_tx=XM_011281774.2;refGene_gn=SPARCL1;refGene_tx=XM_011281775.2;refGene_gn=SPARCL1;refGene_tx=XM_011281776.2;refGene_gn=SPARCL1;refGene_tx=XM_019829259.1;refGene_gn=SPARCL1;refGene_tx=XM_019829260.2
B1	135532445	135532622
refGene_gn=SPARCL1;refGene_tx=XM_003985193.4;refGene_gn=SPARCL1;refGene_tx=XM_011281774.2;refGene_gn=SPARCL1;refGene_tx=XM_011281775.2;refGene_gn=SPARCL1;refGene_tx=XM_011281776.2;refGene_gn=SPARCL1;refGene_tx=XM_019829259.1;refGene_gn=SPARCL1;refGene_tx=XM_019829260.2
B1	135533005	135534010
refGene_gn=SPARCL1;refGene_tx=XM_003985193.4;refGene_gn=SPARCL1;refGene_tx=XM_011281774.2;refGene_gn=SPARCL1;refGene_tx=XM_011281775.2;refGene_gn=SPARCL1;refGene_tx=XM_011281776.2;refGene_gn=SPARCL1;refGene_tx=XM_019829259.1;refGene_gn=SPARCL1;refGene_tx=XM_019829260.2
B1	135535311	135535384
refGene_gn=SPARCL1;refGene_tx=XM_003985193.4;refGene_gn=SPARCL1;refGene_tx=XM_011281774.2;refGene_gn=SPARCL1;refGene_tx=XM_011281775.2;refGene_gn=SPARCL1;refGene_tx=XM_011281776.2;refGene_gn=SPARCL1;refGene_tx=XM_019829259.1;refGene_gn=SPARCL1;refGene_tx=XM_019829260.2
B1	135536284	135536403
refGene_gn=SPARCL1;refGene_tx=XM_003985193.4;refGene_gn=SPARCL1;refGene_tx=XM_011281774.2;refGene_gn=SPARCL1;refGene_tx=XM_011281775.2;refGene_gn=SPARCL1;refGene_tx=XM_011281776.2;refGene_gn=SPARCL1;refGene_tx=XM_019829259.1;refGene_gn=SPARCL1;refGene_tx=XM_019829260.2
B1	135537114	135537235
refGene_gn=SPARCL1;refGene_tx=XM_003985193.4;refGene_gn=SPARCL1;refGene_tx=XM_011281774.2;refGene_gn=SPARCL1;refGene_tx=XM_011281775.2;refGene_gn=SPARCL1;refGene_tx=XM_011281776.2;refGene_gn=SPARCL1;refGene_tx=XM_019829259.1;refGene_gn=SPARCL1;refGene_tx=XM_019829260.2

B1	135538952	135539089
refGene_gn=SPARCL1;refGene_tx=XM_003985193.4;refGene_gn=SPARCL1;refGene_tx=XM_011281774.2;refGene_gn=SPARCL1;refGene_tx=XM_011281775.2;refGene_gn=SPARCL1;refGene_tx=XM_011281776.2;refGene_gn=SPARCL1;refGene_tx=XM_019829259.1;refGene_gn=SPARCL1;refGene_tx=XM_019829260.2
B1	135540990	135541139
refGene_gn=SPARCL1;refGene_tx=XM_003985193.4;refGene_gn=SPARCL1;refGene_tx=XM_011281774.2;refGene_gn=SPARCL1;refGene_tx=XM_011281775.2;refGene_gn=SPARCL1;refGene_tx=XM_011281776.2;refGene_gn=SPARCL1;refGene_tx=XM_019829259.1;refGene_gn=SPARCL1;refGene_tx=XM_019829260.2
B1	135541756	135541905
refGene_gn=SPARCL1;refGene_tx=XM_003985193.4;refGene_gn=SPARCL1;refGene_tx=XM_011281774.2;refGene_gn=SPARCL1;refGene_tx=XM_011281775.2;refGene_gn=SPARCL1;refGene_tx=XM_011281776.2;refGene_gn=SPARCL1;refGene_tx=XM_019829259.1;refGene_gn=SPARCL1;refGene_tx=XM_019829260.2
B1	135543455	135543469
refGene_gn=NUDT9;refGene_tx=XM_011281779.2
B1	135547547	135547576
refGene_gn=SPARCL1;refGene_tx=XM_003985193.4;refGene_gn=SPARCL1;refGene_tx=XM_011281774.2;refGene_gn=SPARCL1;refGene_tx=XM_011281775.2;refGene_gn=SPARCL1;refGene_tx=XM_011281776.2;refGene_gn=SPARCL1;refGene_tx=XM_019829259.1;refGene_gn=SPARCL1;refGene_tx=XM_019829260.2
B1	135556323	135556355
refGene_gn=NUDT9;refGene_tx=XM_011281778.2
B1	135561225	135561404
refGene_gn=NUDT9;refGene_tx=XM_011281777.3;refGene_gn=NUDT9;refGene_tx=XM_019829261.2;refGene_gn=NUDT9;refGene_tx=XM_023252949.1
B1	135562730	135562815
refGene_gn=NUDT9;refGene_tx=XM_011281777.3;refGene_gn=NUDT9;refGene_tx=XM_011281778.2;refGene_gn=NUDT9;refGene_tx=XM_011281779.2;refGene_gn=NUDT9;refGene_tx=XM_019829261.2;refGene_gn=NUDT9;refGene_tx=XM_019829262.2;refGene_gn=NUDT9;refGene_tx=XM_023252949.1
B1	135564326	135564473
refGene_gn=NUDT9;refGene_tx=XM_011281777.3;refGene_gn=NUDT9;refGene_tx=XM_011281778.2;refGene_gn=NUDT9;refGene_tx=XM_011281779.2;refGene_gn=NUDT9;refGene_tx=XM_019829261.2;refGene_gn=NUDT9;refGene_tx=XM_019829262.2;refGene_gn=NUDT9;refGene_tx=XM_023252949.1
B1	135566460	135566572
refGene_gn=NUDT9;refGene_tx=XM_011281777.3;refGene_gn=NUDT9;refGene_tx=XM_011281778.2;refGene_gn=NUDT9;refGene_tx=XM_011281779.2;refGene_gn=NUDT9;refGene_tx=XM_019829261.2;refGene_gn=NUDT9;refGene_tx=XM_019829262.2;refGene_gn=NUDT9;refGene_tx=XM_023252949.1
B1	135579518	135579605
refGene_gn=NUDT9;refGene_tx=XM_011281777.3;refGene_gn=NUDT9;refGene_tx=XM_011281778.2;refGene_gn=NUDT9;refGene_tx=XM_011281779.2;refGene_gn=NUDT9;refGene_tx=XM_019829261.2;refGene_gn=NUDT9;refGene_tx=XM_019829262.2;refGene_gn=NUDT9;refGene_tx=XM_023252949.1
B1	135581772	135581868
refGene_gn=NUDT9;refGene_tx=XM_011281777.3;refGene_gn=NUDT9;refGene_tx=XM_011281778.2;refGene_gn=NUDT9;refGene_tx=XM_011281779.2;refGene_gn=

NUDT9;refGene_tx=XM_019829261.2;refGene_gn=NUDT9;refGene_tx=XM_0198292
62.2;refGene_gn=NUDT9;refGene_tx=XM_023252949.1
B1 135584370 135584610
refGene_gn=NUDT9;refGene_tx=XM_011281777.3;refGene_gn=NUDT9;refGene_tx
=XM_011281778.2;refGene_gn=NUDT9;refGene_tx=XM_011281779.2;refGene_gn=
NUDT9;refGene_tx=XM_019829261.2;refGene_gn=NUDT9;refGene_tx=XM_0198292
62.2;refGene_gn=NUDT9;refGene_tx=XM_023252949.1
B1 135596642 135596749
refGene_gn=NUDT9;refGene_tx=XM_011281777.3;refGene_gn=NUDT9;refGene_tx
=XM_011281778.2;refGene_gn=NUDT9;refGene_tx=XM_011281779.2;refGene_gn=
NUDT9;refGene_tx=XM_019829262.2
B1 135608919 135609129
refGene_gn=HSD17B11;refGene_tx=XM_003985194.5
B1 135616265 135616373
refGene_gn=HSD17B11;refGene_tx=XM_003985194.5
B1 135618460 135618592
refGene_gn=HSD17B11;refGene_tx=XM_003985194.5
B1 135620680 135620787
refGene_gn=HSD17B11;refGene_tx=XM_003985194.5
B1 135626821 135626959
refGene_gn=HSD17B11;refGene_tx=XM_003985194.5
B1 135636996 135637113
refGene_gn=HSD17B11;refGene_tx=XM_003985194.5
B1 135642121 135642212
refGene_gn=HSD17B11;refGene_tx=XM_003985194.5
B1 135650782 135650992
refGene_gn=HSD17B13;refGene_tx=XM_003985204.4
B1 135656714 135656822
refGene_gn=HSD17B13;refGene_tx=XM_003985204.4
B1 135657730 135657862
refGene_gn=HSD17B13;refGene_tx=XM_003985204.4
B1 135666097 135666204
refGene_gn=HSD17B13;refGene_tx=XM_003985204.4
B1 135668249 135668387
refGene_gn=HSD17B13;refGene_tx=XM_003985204.4
B1 135671712 135671829
refGene_gn=HSD17B13;refGene_tx=XM_003985204.4
B1 135677183 135677274
refGene_gn=HSD17B13;refGene_tx=XM_003985204.4
B1 135797452 135797665
refGene_gn=KLHL8;refGene_tx=XM_003985196.5;refGene_gn=KLHL8;refGene_tx
=XM_006930979.4
B1 135806564 135807113
refGene_gn=KLHL8;refGene_tx=XM_003985196.5;refGene_gn=KLHL8;refGene_tx
=XM_006930979.4
B1 135808837 135809024
refGene_gn=KLHL8;refGene_tx=XM_003985196.5;refGene_gn=KLHL8;refGene_tx
=XM_006930979.4
B1 135811386 135811530
refGene_gn=KLHL8;refGene_tx=XM_003985196.5;refGene_gn=KLHL8;refGene_tx

=XM_006930979.4
B1	135812483	135812595
refGene_gn=KLHL8;refGene_tx=XM_003985196.5;refGene_gn=KLHL8;refGene_tx
=XM_006930979.4
B1	135818537	135818706
refGene_gn=KLHL8;refGene_tx=XM_003985196.5;refGene_gn=KLHL8;refGene_tx
=XM_006930979.4
B1	135818927	135819087
refGene_gn=KLHL8;refGene_tx=XM_003985196.5;refGene_gn=KLHL8;refGene_tx
=XM_006930979.4
B1	135821155	135821357
refGene_gn=KLHL8;refGene_tx=XM_003985196.5;refGene_gn=KLHL8;refGene_tx
=XM_006930979.4
B1	135821597	135821721
refGene_gn=KLHL8;refGene_tx=XM_003985196.5;refGene_gn=KLHL8;refGene_tx
=XM_006930979.4
B1	135855863	135855985
refGene_gn=AFF1;refGene_tx=XM_011281773.2;refGene_gn=AFF1;refGene_tx=X
M_019829263.2;refGene_gn=AFF1;refGene_tx=XM_023252950.1
B1	135857008	135857214
refGene_gn=AFF1;refGene_tx=XM_011281773.2;refGene_gn=AFF1;refGene_tx=X
M_019829263.2;refGene_gn=AFF1;refGene_tx=XM_023252950.1
B1	135859078	135859216
refGene_gn=AFF1;refGene_tx=XM_011281773.2;refGene_gn=AFF1;refGene_tx=X
M_019829263.2;refGene_gn=AFF1;refGene_tx=XM_023252950.1
B1	135859667	135859736
refGene_gn=AFF1;refGene_tx=XM_011281773.2;refGene_gn=AFF1;refGene_tx=X
M_019829263.2;refGene_gn=AFF1;refGene_tx=XM_023252950.1
B1	135860324	135860461
refGene_gn=AFF1;refGene_tx=XM_011281773.2;refGene_gn=AFF1;refGene_tx=X
M_019829263.2;refGene_gn=AFF1;refGene_tx=XM_023252950.1
B1	135863739	135863800
refGene_gn=AFF1;refGene_tx=XM_011281773.2;refGene_gn=AFF1;refGene_tx=X
M_019829263.2;refGene_gn=AFF1;refGene_tx=XM_023252950.1
B1	135863990	135864082
refGene_gn=AFF1;refGene_tx=XM_011281773.2;refGene_gn=AFF1;refGene_tx=X
M_019829263.2;refGene_gn=AFF1;refGene_tx=XM_023252950.1
B1	135864784	135865028
refGene_gn=AFF1;refGene_tx=XM_011281773.2;refGene_gn=AFF1;refGene_tx=X
M_019829263.2;refGene_gn=AFF1;refGene_tx=XM_023252950.1
B1	135866058	135866162
refGene_gn=AFF1;refGene_tx=XM_011281773.2;refGene_gn=AFF1;refGene_tx=X
M_019829263.2;refGene_gn=AFF1;refGene_tx=XM_023252950.1
B1	135876087	135877062
refGene_gn=AFF1;refGene_tx=XM_011281773.2;refGene_gn=AFF1;refGene_tx=X
M_019829263.2;refGene_gn=AFF1;refGene_tx=XM_023252950.1
B1	135882673	135882830
refGene_gn=AFF1;refGene_tx=XM_011281773.2;refGene_gn=AFF1;refGene_tx=X
M_019829263.2;refGene_gn=AFF1;refGene_tx=XM_023252950.1
B1	135884856	135884894

refGene_gn=AFF1;refGene_tx=XM_011281773.2;refGene_gn=AFF1;refGene_tx=XM_019829263.2;refGene_gn=AFF1;refGene_tx=XM_023252950.1
B1 135885016 135885071
refGene_gn=AFF1;refGene_tx=XM_011281773.2;refGene_gn=AFF1;refGene_tx=XM_019829263.2;refGene_gn=AFF1;refGene_tx=XM_023252950.1
B1 135897407 135897459
refGene_gn=AFF1;refGene_tx=XM_011281773.2;refGene_gn=AFF1;refGene_tx=XM_019829263.2;refGene_gn=AFF1;refGene_tx=XM_023252950.1
B1 135900902 135900939
refGene_gn=AFF1;refGene_tx=XM_011281773.2;refGene_gn=AFF1;refGene_tx=XM_019829263.2;refGene_gn=AFF1;refGene_tx=XM_023252950.1
B1 135901926 135902013
refGene_gn=AFF1;refGene_tx=XM_011281773.2;refGene_gn=AFF1;refGene_tx=XM_019829263.2;refGene_gn=AFF1;refGene_tx=XM_023252950.1
B1 135907459 135907504
refGene_gn=AFF1;refGene_tx=XM_011281773.2;refGene_gn=AFF1;refGene_tx=XM_019829263.2;refGene_gn=AFF1;refGene_tx=XM_023252950.1
B1 135947002 135947923
refGene_gn=AFF1;refGene_tx=XM_011281773.2;refGene_gn=AFF1;refGene_tx=XM_019829263.2;refGene_gn=AFF1;refGene_tx=XM_023252950.1
B1 135948328 135948449
refGene_gn=AFF1;refGene_tx=XM_011281773.2;refGene_gn=AFF1;refGene_tx=XM_019829263.2;refGene_gn=AFF1;refGene_tx=XM_023252950.1
B1 135985224 135985241
refGene_gn=AFF1;refGene_tx=XM_011281773.2
B1 136044402 136044464
refGene_gn=AFF1;refGene_tx=XM_019829263.2;refGene_gn=AFF1;refGene_tx=XM_023252950.1
B1 136106102 136106167
refGene_gn=CB1H4orf36;refGene_tx=XM_023252952.1;refGene_gn=CB1H4orf36;refGene_tx=XM_023252954.1;refGene_gn=CB1H4orf36;refGene_tx=XM_023252955.1;refGene_gn=CB1H4orf36;refGene_tx=XM_023252956.1;refGene_gn=CB1H4orf36;refGene_tx=XM_023252957.1;refGene_gn=CB1H4orf36;refGene_tx=XM_023252958.1
B1 136109343 136109498
refGene_gn=CB1H4orf36;refGene_tx=XM_023252952.1;refGene_gn=CB1H4orf36;refGene_tx=XM_023252954.1;refGene_gn=CB1H4orf36;refGene_tx=XM_023252955.1;refGene_gn=CB1H4orf36;refGene_tx=XM_023252956.1;refGene_gn=CB1H4orf36;refGene_tx=XM_023252957.1;refGene_gn=CB1H4orf36;refGene_tx=XM_023252958.1
B1 136109732 136109866
refGene_gn=CB1H4orf36;refGene_tx=XM_023252952.1;refGene_gn=CB1H4orf36;refGene_tx=XM_023252954.1;refGene_gn=CB1H4orf36;refGene_tx=XM_023252955.1;refGene_gn=CB1H4orf36;refGene_tx=XM_023252956.1;refGene_gn=CB1H4orf36;refGene_tx=XM_023252957.1;refGene_gn=CB1H4orf36;refGene_tx=XM_023252958.1
B1 136128358 136128735
refGene_gn=SLC10A6;refGene_tx=XM_003985207.4;refGene_gn=SLC10A6;refGene_tx=XM_006930984.2;refGene_gn=SLC10A6;refGene_tx=XM_023252951.1
B1 136142594 136142713

refGene_gn=SLC10A6;refGene_tx=XM_003985207.4;refGene_gn=SLC10A6;refGene_tx=XM_006930984.2;refGene_gn=SLC10A6;refGene_tx=XM_023252951.1
B1 136147032 136147121
refGene_gn=SLC10A6;refGene_tx=XM_003985207.4;refGene_gn=SLC10A6;refGene_tx=XM_006930984.2;refGene_gn=SLC10A6;refGene_tx=XM_023252951.1
B1 136150189 136150365
refGene_gn=SLC10A6;refGene_tx=XM_003985207.4;refGene_gn=SLC10A6;refGene_tx=XM_006930984.2;refGene_gn=SLC10A6;refGene_tx=XM_023252951.1
B1 136153181 136153339
refGene_gn=SLC10A6;refGene_tx=XM_003985207.4;refGene_gn=SLC10A6;refGene_tx=XM_006930984.2;refGene_gn=SLC10A6;refGene_tx=XM_023252951.1
B1 136156904 136156909
refGene_gn=SLC10A6;refGene_tx=XM_023252951.1
B1 136157566 136157804
refGene_gn=SLC10A6;refGene_tx=XM_003985207.4;refGene_gn=SLC10A6;refGene_tx=XM_006930984.2
B1 136167421 136167517
refGene_gn=PTPN13;refGene_tx=XM_011281764.3
B1 136170297 136170360
refGene_gn=PTPN13;refGene_tx=XM_011281764.3
B1 136173823 136174039
refGene_gn=PTPN13;refGene_tx=XM_011281764.3
B1 136175020 136175358
refGene_gn=PTPN13;refGene_tx=XM_011281764.3
B1 136178175 136178266
refGene_gn=PTPN13;refGene_tx=XM_011281764.3
B1 136179746 136179895
refGene_gn=PTPN13;refGene_tx=XM_011281764.3
B1 136181421 136181528
refGene_gn=PTPN13;refGene_tx=XM_011281764.3
B1 136184150 136184206
refGene_gn=PTPN13;refGene_tx=XM_011281764.3
B1 136190934 136191023
refGene_gn=PTPN13;refGene_tx=XM_011281764.3
B1 136191124 136191262
refGene_gn=PTPN13;refGene_tx=XM_011281764.3
B1 136191934 136192028
refGene_gn=PTPN13;refGene_tx=XM_011281764.3
B1 136194212 136194274
refGene_gn=PTPN13;refGene_tx=XM_011281764.3
B1 136196943 136197014
refGene_gn=PTPN13;refGene_tx=XM_011281764.3
B1 136200459 136200670
refGene_gn=PTPN13;refGene_tx=XM_011281764.3
B1 136200770 136200942
refGene_gn=PTPN13;refGene_tx=XM_011281764.3
B1 136201455 136201614
refGene_gn=PTPN13;refGene_tx=XM_011281764.3
B1 136203459 136203643
refGene_gn=PTPN13;refGene_tx=XM_011281764.3

B1 136204825 136205184
refGene_gn=PTPN13;refGene_tx=XM_011281764.3
B1 136207185 136207284
refGene_gn=PTPN13;refGene_tx=XM_011281764.3
B1 136207403 136207618
refGene_gn=PTPN13;refGene_tx=XM_011281764.3
B1 136209444 136209604
refGene_gn=PTPN13;refGene_tx=XM_011281764.3
B1 136210308 136210394
refGene_gn=PTPN13;refGene_tx=XM_011281764.3
B1 136211251 136211345
refGene_gn=PTPN13;refGene_tx=XM_011281764.3
B1 136212587 136212719
refGene_gn=PTPN13;refGene_tx=XM_011281764.3
B1 136213777 136214241
refGene_gn=PTPN13;refGene_tx=XM_011281764.3
B1 136219213 136219345
refGene_gn=PTPN13;refGene_tx=XM_011281764.3
B1 136219519 136219627
refGene_gn=PTPN13;refGene_tx=XM_011281764.3
B1 136219942 136220032
refGene_gn=PTPN13;refGene_tx=XM_011281764.3
B1 136227976 136228033
refGene_gn=PTPN13;refGene_tx=XM_011281764.3
B1 136229543 136229641
refGene_gn=PTPN13;refGene_tx=XM_011281764.3
B1 136229778 136230196
refGene_gn=PTPN13;refGene_tx=XM_011281764.3
B1 136234897 136235060
refGene_gn=PTPN13;refGene_tx=XM_011281764.3
B1 136237775 136237958
refGene_gn=PTPN13;refGene_tx=XM_011281764.3
B1 136240848 136240911
refGene_gn=PTPN13;refGene_tx=XM_011281764.3
B1 136240912 136241002
refGene_gn=PTPN13;refGene_tx=XM_011281764.3
B1 136242380 136242519
refGene_gn=PTPN13;refGene_tx=XM_011281764.3
B1 136243024 136243178
refGene_gn=PTPN13;refGene_tx=XM_011281764.3
B1 136245178 136245353
refGene_gn=PTPN13;refGene_tx=XM_011281764.3
B1 136245483 136245558
refGene_gn=PTPN13;refGene_tx=XM_011281764.3
B1 136248785 136249008
refGene_gn=PTPN13;refGene_tx=XM_011281764.3
B1 136255744 136255838
refGene_gn=PTPN13;refGene_tx=XM_011281764.3
B1 136256457 136256553
refGene_gn=PTPN13;refGene_tx=XM_011281764.3

B1 136276812 136277376
refGene_gn=PTPN13;refGene_tx=XM_011281764.3
B1 136288420 136288508
refGene_gn=PTPN13;refGene_tx=XM_011281764.3
B1 136292780 136292966
refGene_gn=PTPN13;refGene_tx=XM_011281764.3
B1 136295440 136295506
refGene_gn=PTPN13;refGene_tx=XM_011281764.3
B1 136307841 136308020
refGene_gn=PTPN13;refGene_tx=XM_011281764.3
B1 136340905 136341020
refGene_gn=PTPN13;refGene_tx=XM_011281764.3
B1 136798858 136798924
refGene_gn=MAPK10;refGene_tx=XM_011281754.3;refGene_gn=MAPK10;refGene_tx=XM_011281756.3;refGene_gn=MAPK10;refGene_tx=XM_011281760.3;refGene_gn=MAPK10;refGene_tx=XM_011281762.3;refGene_gn=MAPK10;refGene_tx=XM_019829271.2;refGene_gn=MAPK10;refGene_tx=XM_019829272.2;refGene_gn=MAPK10;refGene_tx=XM_019829273.2
B1 136822842 136823012
refGene_gn=MAPK10;refGene_tx=XM_011281754.3;refGene_gn=MAPK10;refGene_tx=XM_011281756.3;refGene_gn=MAPK10;refGene_tx=XM_011281758.3;refGene_gn=MAPK10;refGene_tx=XM_011281760.3;refGene_gn=MAPK10;refGene_tx=XM_011281762.3;refGene_gn=MAPK10;refGene_tx=XM_019829271.2;refGene_gn=MAPK10;refGene_tx=XM_019829272.2;refGene_gn=MAPK10;refGene_tx=XM_019829273.2;refGene_gn=MAPK10;refGene_tx=XM_019829274.2
B1 136872465 136872595
refGene_gn=MAPK10;refGene_tx=XM_011281754.3;refGene_gn=MAPK10;refGene_tx=XM_011281756.3;refGene_gn=MAPK10;refGene_tx=XM_011281758.3;refGene_gn=MAPK10;refGene_tx=XM_011281760.3;refGene_gn=MAPK10;refGene_tx=XM_011281762.3;refGene_gn=MAPK10;refGene_tx=XM_019829271.2;refGene_gn=MAPK10;refGene_tx=XM_019829272.2;refGene_gn=MAPK10;refGene_tx=XM_019829273.2;refGene_gn=MAPK10;refGene_tx=XM_019829274.2
B1 136876889 136876948
refGene_gn=MAPK10;refGene_tx=XM_011281754.3;refGene_gn=MAPK10;refGene_tx=XM_011281756.3;refGene_gn=MAPK10;refGene_tx=XM_011281758.3;refGene_gn=MAPK10;refGene_tx=XM_011281760.3;refGene_gn=MAPK10;refGene_tx=XM_011281762.3;refGene_gn=MAPK10;refGene_tx=XM_019829271.2;refGene_gn=MAPK10;refGene_tx=XM_019829272.2;refGene_gn=MAPK10;refGene_tx=XM_019829273.2;refGene_gn=MAPK10;refGene_tx=XM_019829274.2
B1 136877974 136878113
refGene_gn=MAPK10;refGene_tx=XM_011281754.3;refGene_gn=MAPK10;refGene_tx=XM_011281756.3;refGene_gn=MAPK10;refGene_tx=XM_011281758.3;refGene_gn=MAPK10;refGene_tx=XM_011281760.3;refGene_gn=MAPK10;refGene_tx=XM_011281762.3;refGene_gn=MAPK10;refGene_tx=XM_019829271.2;refGene_gn=MAPK10;refGene_tx=XM_019829272.2;refGene_gn=MAPK10;refGene_tx=XM_019829273.2;refGene_gn=MAPK10;refGene_tx=XM_019829274.2
B1 136878804 136878970
refGene_gn=MAPK10;refGene_tx=XM_011281754.3;refGene_gn=MAPK10;refGene_tx=XM_011281756.3;refGene_gn=MAPK10;refGene_tx=XM_011281758.3;refGene_gn=MAPK10;refGene_tx=XM_011281760.3;refGene_gn=MAPK10;refGene_tx=XM_01

1281762.3;refGene_gn=MAPK10;refGene_tx=XM_019829271.2;refGene_gn=MAPK1
0;refGene_tx=XM_019829272.2;refGene_gn=MAPK10;refGene_tx=XM_019829273.
2;refGene_gn=MAPK10;refGene_tx=XM_019829274.2
B1	136881412	136881484
refGene_gn=MAPK10;refGene_tx=XM_011281754.3;refGene_gn=MAPK10;refGene_
tx=XM_011281756.3;refGene_gn=MAPK10;refGene_tx=XM_011281758.3;refGene_
gn=MAPK10;refGene_tx=XM_011281760.3;refGene_gn=MAPK10;refGene_tx=XM_01
9829271.2;refGene_gn=MAPK10;refGene_tx=XM_019829272.2;refGene_gn=MAPK1
0;refGene_tx=XM_019829273.2;refGene_gn=MAPK10;refGene_tx=XM_019829274.
2
B1	136891431	136891503
refGene_gn=MAPK10;refGene_tx=XM_011281762.3
B1	136906567	136906750
refGene_gn=MAPK10;refGene_tx=XM_011281754.3;refGene_gn=MAPK10;refGene_
tx=XM_011281756.3;refGene_gn=MAPK10;refGene_tx=XM_011281758.3;refGene_
gn=MAPK10;refGene_tx=XM_011281760.3;refGene_gn=MAPK10;refGene_tx=XM_01
1281762.3;refGene_gn=MAPK10;refGene_tx=XM_019829271.2;refGene_gn=MAPK1
0;refGene_tx=XM_019829272.2;refGene_gn=MAPK10;refGene_tx=XM_019829273.
2;refGene_gn=MAPK10;refGene_tx=XM_019829274.2
B1	136910942	136911067
refGene_gn=MAPK10;refGene_tx=XM_011281754.3;refGene_gn=MAPK10;refGene_
tx=XM_011281756.3;refGene_gn=MAPK10;refGene_tx=XM_011281758.3;refGene_
gn=MAPK10;refGene_tx=XM_011281760.3;refGene_gn=MAPK10;refGene_tx=XM_01
1281762.3;refGene_gn=MAPK10;refGene_tx=XM_019829271.2;refGene_gn=MAPK1
0;refGene_tx=XM_019829272.2;refGene_gn=MAPK10;refGene_tx=XM_019829273.
2;refGene_gn=MAPK10;refGene_tx=XM_019829274.2
B1	136940743	136940807
refGene_gn=MAPK10;refGene_tx=XM_011281754.3;refGene_gn=MAPK10;refGene_
tx=XM_011281756.3;refGene_gn=MAPK10;refGene_tx=XM_011281758.3;refGene_
gn=MAPK10;refGene_tx=XM_011281760.3;refGene_gn=MAPK10;refGene_tx=XM_01
1281762.3;refGene_gn=MAPK10;refGene_tx=XM_019829271.2;refGene_gn=MAPK1
0;refGene_tx=XM_019829272.2;refGene_gn=MAPK10;refGene_tx=XM_019829273.
2;refGene_gn=MAPK10;refGene_tx=XM_019829274.2
B1	136942509	136942587
refGene_gn=MAPK10;refGene_tx=XM_011281754.3;refGene_gn=MAPK10;refGene_
tx=XM_011281756.3;refGene_gn=MAPK10;refGene_tx=XM_011281758.3;refGene_
gn=MAPK10;refGene_tx=XM_011281760.3;refGene_gn=MAPK10;refGene_tx=XM_01
1281762.3;refGene_gn=MAPK10;refGene_tx=XM_019829271.2;refGene_gn=MAPK1
0;refGene_tx=XM_019829272.2;refGene_gn=MAPK10;refGene_tx=XM_019829273.
2;refGene_gn=MAPK10;refGene_tx=XM_019829274.2
B1	136957140	136957288
refGene_gn=MAPK10;refGene_tx=XM_011281754.3;refGene_gn=MAPK10;refGene_
tx=XM_011281756.3;refGene_gn=MAPK10;refGene_tx=XM_011281758.3;refGene_
gn=MAPK10;refGene_tx=XM_011281760.3;refGene_gn=MAPK10;refGene_tx=XM_01
1281762.3;refGene_gn=MAPK10;refGene_tx=XM_019829271.2;refGene_gn=MAPK1
0;refGene_tx=XM_019829272.2;refGene_gn=MAPK10;refGene_tx=XM_019829273.
2;refGene_gn=MAPK10;refGene_tx=XM_019829274.2
B1	136973773	136974017
refGene_gn=ARHGAP24;refGene_tx=XM_003985210.3;refGene_gn=ARHGAP24;refG
ene_tx=XM_003985212.3;refGene_gn=ARHGAP24;refGene_tx=XM_003985213.4;re

fGene_gn=ARHGAP24;refGene_tx=XM_006930991.4;refGene_gn=ARHGAP24;refGene_tx=XM_019829275.2;refGene_gn=ARHGAP24;refGene_tx=XM_019829276.2;refGene_gn=ARHGAP24;refGene_tx=XM_019829277.2;refGene_gn=ARHGAP24;refGene_tx=XM_019829278.2;refGene_gn=ARHGAP24;refGene_tx=XM_023252961.1
B1 136978088 136979166
refGene_gn=ARHGAP24;refGene_tx=XM_003985210.3;refGene_gn=ARHGAP24;refGene_tx=XM_003985212.3;refGene_gn=ARHGAP24;refGene_tx=XM_003985213.4;refGene_gn=ARHGAP24;refGene_tx=XM_006930991.4;refGene_gn=ARHGAP24;refGene_tx=XM_019829275.2;refGene_gn=ARHGAP24;refGene_tx=XM_019829276.2;refGene_gn=ARHGAP24;refGene_tx=XM_019829277.2;refGene_gn=ARHGAP24;refGene_tx=XM_019829278.2;refGene_gn=ARHGAP24;refGene_tx=XM_023252961.1
B1 136991093 136991215
refGene_gn=ARHGAP24;refGene_tx=XM_003985210.3;refGene_gn=ARHGAP24;refGene_tx=XM_003985212.3;refGene_gn=ARHGAP24;refGene_tx=XM_003985213.4;refGene_gn=ARHGAP24;refGene_tx=XM_006930991.4;refGene_gn=ARHGAP24;refGene_tx=XM_019829275.2;refGene_gn=ARHGAP24;refGene_tx=XM_019829276.2;refGene_gn=ARHGAP24;refGene_tx=XM_019829277.2;refGene_gn=ARHGAP24;refGene_tx=XM_019829278.2;refGene_gn=ARHGAP24;refGene_tx=XM_023252961.1
B1 136993745 136993819
refGene_gn=ARHGAP24;refGene_tx=XM_003985210.3;refGene_gn=ARHGAP24;refGene_tx=XM_003985212.3;refGene_gn=ARHGAP24;refGene_tx=XM_003985213.4;refGene_gn=ARHGAP24;refGene_tx=XM_006930991.4;refGene_gn=ARHGAP24;refGene_tx=XM_019829275.2;refGene_gn=ARHGAP24;refGene_tx=XM_019829276.2;refGene_gn=ARHGAP24;refGene_tx=XM_019829277.2;refGene_gn=ARHGAP24;refGene_tx=XM_019829278.2;refGene_gn=ARHGAP24;refGene_tx=XM_023252961.1
B1 136996682 136996815
refGene_gn=ARHGAP24;refGene_tx=XM_003985210.3;refGene_gn=ARHGAP24;refGene_tx=XM_003985212.3;refGene_gn=ARHGAP24;refGene_tx=XM_003985213.4;refGene_gn=ARHGAP24;refGene_tx=XM_006930991.4;refGene_gn=ARHGAP24;refGene_tx=XM_019829275.2;refGene_gn=ARHGAP24;refGene_tx=XM_019829276.2;refGene_gn=ARHGAP24;refGene_tx=XM_019829277.2;refGene_gn=ARHGAP24;refGene_tx=XM_019829278.2;refGene_gn=ARHGAP24;refGene_tx=XM_023252961.1
B1 137025155 137025363
refGene_gn=ARHGAP24;refGene_tx=XM_003985210.3;refGene_gn=ARHGAP24;refGene_tx=XM_003985212.3;refGene_gn=ARHGAP24;refGene_tx=XM_003985213.4;refGene_gn=ARHGAP24;refGene_tx=XM_006930991.4;refGene_gn=ARHGAP24;refGene_tx=XM_019829275.2;refGene_gn=ARHGAP24;refGene_tx=XM_019829276.2;refGene_gn=ARHGAP24;refGene_tx=XM_019829277.2;refGene_gn=ARHGAP24;refGene_tx=XM_019829278.2;refGene_gn=ARHGAP24;refGene_tx=XM_023252961.1
B1 137038023 137038138
refGene_gn=ARHGAP24;refGene_tx=XM_019829277.2
B1 137045366 137045489
refGene_gn=ARHGAP24;refGene_tx=XM_003985210.3;refGene_gn=ARHGAP24;refGene_tx=XM_003985212.3;refGene_gn=ARHGAP24;refGene_tx=XM_003985213.4;refGene_gn=ARHGAP24;refGene_tx=XM_006930991.4;refGene_gn=ARHGAP24;refGene_tx=XM_019829275.2;refGene_gn=ARHGAP24;refGene_tx=XM_019829276.2;refGene_gn=ARHGAP24;refGene_tx=XM_019829278.2;refGene_gn=ARHGAP24;refGene_tx=XM_023252961.1
B1 137257546 137257634
refGene_gn=ARHGAP24;refGene_tx=XM_003985210.3;refGene_gn=ARHGAP24;refG

ene_tx=XM_019829275.2;refGene_gn=ARHGAP24;refGene_tx=XM_019829276.2;refGene_gn=ARHGAP24;refGene_tx=XM_023252961.1
B1 137399494 137399674
refGene_gn=ARHGAP24;refGene_tx=XM_003985210.3;refGene_gn=ARHGAP24;refGene_tx=XM_019829275.2;refGene_gn=ARHGAP24;refGene_tx=XM_019829276.2;refGene_gn=ARHGAP24;refGene_tx=XM_023252961.1
B1 137920452 137920520
refGene_gn=LOC109499444;refGene_tx=XM_023253399.1
B1 137921556 137922106
refGene_gn=LOC109499444;refGene_tx=XM_023253399.1
B1 138014051 138014231
refGene_gn=WDFY3;refGene_tx=XM_023252963.1;refGene_gn=WDFY3;refGene_tx=XM_023252964.1;refGene_gn=WDFY3;refGene_tx=XM_023252965.1;refGene_gn=WDFY3;refGene_tx=XM_023252966.1;refGene_gn=WDFY3;refGene_tx=XM_023252967.1;refGene_gn=WDFY3;refGene_tx=XM_023252968.1;refGene_gn=WDFY3;refGene_tx=XM_023252969.1;refGene_gn=WDFY3;refGene_tx=XM_023252970.1;refGene_gn=WDFY3;refGene_tx=XM_023252971.1;refGene_gn=WDFY3;refGene_tx=XM_023252972.1
B1 138025298 138025422
refGene_gn=WDFY3;refGene_tx=XM_023252963.1;refGene_gn=WDFY3;refGene_tx=XM_023252964.1;refGene_gn=WDFY3;refGene_tx=XM_023252965.1;refGene_gn=WDFY3;refGene_tx=XM_023252966.1;refGene_gn=WDFY3;refGene_tx=XM_023252967.1;refGene_gn=WDFY3;refGene_tx=XM_023252968.1;refGene_gn=WDFY3;refGene_tx=XM_023252969.1;refGene_gn=WDFY3;refGene_tx=XM_023252970.1;refGene_gn=WDFY3;refGene_tx=XM_023252971.1;refGene_gn=WDFY3;refGene_tx=XM_023252972.1
B1 138033483 138033537
refGene_gn=WDFY3;refGene_tx=XM_023252963.1;refGene_gn=WDFY3;refGene_tx=XM_023252964.1;refGene_gn=WDFY3;refGene_tx=XM_023252965.1;refGene_gn=WDFY3;refGene_tx=XM_023252966.1;refGene_gn=WDFY3;refGene_tx=XM_023252967.1;refGene_gn=WDFY3;refGene_tx=XM_023252969.1;refGene_gn=WDFY3;refGene_tx=XM_023252971.1
B1 138037151 138037261
refGene_gn=WDFY3;refGene_tx=XM_023252963.1;refGene_gn=WDFY3;refGene_tx=XM_023252964.1;refGene_gn=WDFY3;refGene_tx=XM_023252965.1;refGene_gn=WDFY3;refGene_tx=XM_023252966.1;refGene_gn=WDFY3;refGene_tx=XM_023252967.1;refGene_gn=WDFY3;refGene_tx=XM_023252968.1;refGene_gn=WDFY3;refGene_tx=XM_023252969.1;refGene_gn=WDFY3;refGene_tx=XM_023252970.1;refGene_gn=WDFY3;refGene_tx=XM_023252971.1;refGene_gn=WDFY3;refGene_tx=XM_023252972.1;refGene_gn=WDFY3;refGene_tx=XM_023252973.1
B1 138041637 138041799
refGene_gn=WDFY3;refGene_tx=XM_023252963.1;refGene_gn=WDFY3;refGene_tx=XM_023252964.1;refGene_gn=WDFY3;refGene_tx=XM_023252965.1;refGene_gn=WDFY3;refGene_tx=XM_023252966.1;refGene_gn=WDFY3;refGene_tx=XM_023252967.1;refGene_gn=WDFY3;refGene_tx=XM_023252968.1;refGene_gn=WDFY3;refGene_tx=XM_023252969.1;refGene_gn=WDFY3;refGene_tx=XM_023252970.1;refGene_gn=WDFY3;refGene_tx=XM_023252971.1;refGene_gn=WDFY3;refGene_tx=XM_023252972.1;refGene_gn=WDFY3;refGene_tx=XM_023252973.1
B1 138051595 138051788
refGene_gn=WDFY3;refGene_tx=XM_023252963.1;refGene_gn=WDFY3;refGene_tx

=XM_023252964.1;refGene_gn=WDFY3;refGene_tx=XM_023252965.1;refGene_gn=
WDFY3;refGene_tx=XM_023252966.1;refGene_gn=WDFY3;refGene_tx=XM_0232529
67.1;refGene_gn=WDFY3;refGene_tx=XM_023252968.1;refGene_gn=WDFY3;refGe
ne_tx=XM_023252969.1;refGene_gn=WDFY3;refGene_tx=XM_023252970.1;refGen
e_gn=WDFY3;refGene_tx=XM_023252971.1;refGene_gn=WDFY3;refGene_tx=XM_02
3252972.1;refGene_gn=WDFY3;refGene_tx=XM_023252973.1
B1 138053024 138053211
refGene_gn=WDFY3;refGene_tx=XM_023252963.1;refGene_gn=WDFY3;refGene_tx
=XM_023252964.1;refGene_gn=WDFY3;refGene_tx=XM_023252965.1;refGene_gn=
WDFY3;refGene_tx=XM_023252966.1;refGene_gn=WDFY3;refGene_tx=XM_0232529
67.1;refGene_gn=WDFY3;refGene_tx=XM_023252968.1;refGene_gn=WDFY3;refGe
ne_tx=XM_023252969.1;refGene_gn=WDFY3;refGene_tx=XM_023252970.1;refGen
e_gn=WDFY3;refGene_tx=XM_023252971.1;refGene_gn=WDFY3;refGene_tx=XM_02
3252972.1;refGene_gn=WDFY3;refGene_tx=XM_023252973.1
B1 138055524 138055691
refGene_gn=WDFY3;refGene_tx=XM_023252963.1;refGene_gn=WDFY3;refGene_tx
=XM_023252964.1;refGene_gn=WDFY3;refGene_tx=XM_023252965.1;refGene_gn=
WDFY3;refGene_tx=XM_023252966.1;refGene_gn=WDFY3;refGene_tx=XM_0232529
67.1;refGene_gn=WDFY3;refGene_tx=XM_023252968.1;refGene_gn=WDFY3;refGe
ne_tx=XM_023252969.1;refGene_gn=WDFY3;refGene_tx=XM_023252970.1;refGen
e_gn=WDFY3;refGene_tx=XM_023252971.1;refGene_gn=WDFY3;refGene_tx=XM_02
3252972.1;refGene_gn=WDFY3;refGene_tx=XM_023252973.1
B1 138061397 138061865
refGene_gn=WDFY3;refGene_tx=XM_023252963.1;refGene_gn=WDFY3;refGene_tx
=XM_023252964.1;refGene_gn=WDFY3;refGene_tx=XM_023252965.1;refGene_gn=
WDFY3;refGene_tx=XM_023252966.1;refGene_gn=WDFY3;refGene_tx=XM_0232529
67.1;refGene_gn=WDFY3;refGene_tx=XM_023252968.1;refGene_gn=WDFY3;refGe
ne_tx=XM_023252969.1;refGene_gn=WDFY3;refGene_tx=XM_023252970.1;refGen
e_gn=WDFY3;refGene_tx=XM_023252971.1;refGene_gn=WDFY3;refGene_tx=XM_02
3252972.1;refGene_gn=WDFY3;refGene_tx=XM_023252973.1
B1 138062844 138062946
refGene_gn=WDFY3;refGene_tx=XM_023252963.1;refGene_gn=WDFY3;refGene_tx
=XM_023252964.1;refGene_gn=WDFY3;refGene_tx=XM_023252965.1;refGene_gn=
WDFY3;refGene_tx=XM_023252966.1;refGene_gn=WDFY3;refGene_tx=XM_0232529
67.1;refGene_gn=WDFY3;refGene_tx=XM_023252968.1;refGene_gn=WDFY3;refGe
ne_tx=XM_023252969.1;refGene_gn=WDFY3;refGene_tx=XM_023252970.1;refGen
e_gn=WDFY3;refGene_tx=XM_023252971.1;refGene_gn=WDFY3;refGene_tx=XM_02
3252972.1;refGene_gn=WDFY3;refGene_tx=XM_023252973.1
B1 138065029 138065223
refGene_gn=WDFY3;refGene_tx=XM_023252963.1;refGene_gn=WDFY3;refGene_tx
=XM_023252964.1;refGene_gn=WDFY3;refGene_tx=XM_023252965.1;refGene_gn=
WDFY3;refGene_tx=XM_023252966.1;refGene_gn=WDFY3;refGene_tx=XM_0232529
67.1;refGene_gn=WDFY3;refGene_tx=XM_023252968.1;refGene_gn=WDFY3;refGe
ne_tx=XM_023252969.1;refGene_gn=WDFY3;refGene_tx=XM_023252970.1;refGen
e_gn=WDFY3;refGene_tx=XM_023252971.1;refGene_gn=WDFY3;refGene_tx=XM_02
3252972.1;refGene_gn=WDFY3;refGene_tx=XM_023252973.1
B1 138077542 138078000
refGene_gn=WDFY3;refGene_tx=XM_023252963.1;refGene_gn=WDFY3;refGene_tx
=XM_023252964.1;refGene_gn=WDFY3;refGene_tx=XM_023252965.1;refGene_gn=
WDFY3;refGene_tx=XM_023252966.1;refGene_gn=WDFY3;refGene_tx=XM_0232529

67.1;refGene_gn=WDFY3;refGene_tx=XM_023252968.1;refGene_gn=WDFY3;refGene_tx=XM_023252969.1;refGene_gn=WDFY3;refGene_tx=XM_023252970.1;refGene_gn=WDFY3;refGene_tx=XM_023252971.1;refGene_gn=WDFY3;refGene_tx=XM_023252972.1;refGene_gn=WDFY3;refGene_tx=XM_023252973.1
B1 138079415 138079499
refGene_gn=WDFY3;refGene_tx=XM_023252963.1;refGene_gn=WDFY3;refGene_tx=XM_023252964.1;refGene_gn=WDFY3;refGene_tx=XM_023252965.1;refGene_gn=WDFY3;refGene_tx=XM_023252966.1;refGene_gn=WDFY3;refGene_tx=XM_023252967.1;refGene_gn=WDFY3;refGene_tx=XM_023252968.1;refGene_gn=WDFY3;refGene_tx=XM_023252969.1;refGene_gn=WDFY3;refGene_tx=XM_023252970.1;refGene_gn=WDFY3;refGene_tx=XM_023252972.1;refGene_gn=WDFY3;refGene_tx=XM_023252973.1
B1 138084777 138084955
refGene_gn=WDFY3;refGene_tx=XM_023252963.1;refGene_gn=WDFY3;refGene_tx=XM_023252964.1;refGene_gn=WDFY3;refGene_tx=XM_023252965.1;refGene_gn=WDFY3;refGene_tx=XM_023252966.1;refGene_gn=WDFY3;refGene_tx=XM_023252967.1;refGene_gn=WDFY3;refGene_tx=XM_023252968.1;refGene_gn=WDFY3;refGene_tx=XM_023252969.1;refGene_gn=WDFY3;refGene_tx=XM_023252970.1;refGene_gn=WDFY3;refGene_tx=XM_023252971.1;refGene_gn=WDFY3;refGene_tx=XM_023252972.1;refGene_gn=WDFY3;refGene_tx=XM_023252973.1
B1 138085742 138085957
refGene_gn=WDFY3;refGene_tx=XM_023252963.1;refGene_gn=WDFY3;refGene_tx=XM_023252964.1;refGene_gn=WDFY3;refGene_tx=XM_023252965.1;refGene_gn=WDFY3;refGene_tx=XM_023252966.1;refGene_gn=WDFY3;refGene_tx=XM_023252967.1;refGene_gn=WDFY3;refGene_tx=XM_023252968.1;refGene_gn=WDFY3;refGene_tx=XM_023252969.1;refGene_gn=WDFY3;refGene_tx=XM_023252970.1;refGene_gn=WDFY3;refGene_tx=XM_023252971.1;refGene_gn=WDFY3;refGene_tx=XM_023252972.1;refGene_gn=WDFY3;refGene_tx=XM_023252973.1
B1 138088259 138088372
refGene_gn=WDFY3;refGene_tx=XM_023252963.1;refGene_gn=WDFY3;refGene_tx=XM_023252964.1;refGene_gn=WDFY3;refGene_tx=XM_023252965.1;refGene_gn=WDFY3;refGene_tx=XM_023252966.1;refGene_gn=WDFY3;refGene_tx=XM_023252967.1;refGene_gn=WDFY3;refGene_tx=XM_023252968.1;refGene_gn=WDFY3;refGene_tx=XM_023252969.1;refGene_gn=WDFY3;refGene_tx=XM_023252970.1;refGene_gn=WDFY3;refGene_tx=XM_023252971.1;refGene_gn=WDFY3;refGene_tx=XM_023252972.1;refGene_gn=WDFY3;refGene_tx=XM_023252973.1
B1 138090658 138090890
refGene_gn=WDFY3;refGene_tx=XM_023252963.1;refGene_gn=WDFY3;refGene_tx=XM_023252964.1;refGene_gn=WDFY3;refGene_tx=XM_023252965.1;refGene_gn=WDFY3;refGene_tx=XM_023252966.1;refGene_gn=WDFY3;refGene_tx=XM_023252967.1;refGene_gn=WDFY3;refGene_tx=XM_023252968.1;refGene_gn=WDFY3;refGene_tx=XM_023252969.1;refGene_gn=WDFY3;refGene_tx=XM_023252970.1;refGene_gn=WDFY3;refGene_tx=XM_023252971.1;refGene_gn=WDFY3;refGene_tx=XM_023252972.1;refGene_gn=WDFY3;refGene_tx=XM_023252973.1
B1 138092358 138092459
refGene_gn=WDFY3;refGene_tx=XM_023252963.1;refGene_gn=WDFY3;refGene_tx=XM_023252964.1;refGene_gn=WDFY3;refGene_tx=XM_023252965.1;refGene_gn=WDFY3;refGene_tx=XM_023252966.1;refGene_gn=WDFY3;refGene_tx=XM_023252967.1;refGene_gn=WDFY3;refGene_tx=XM_023252968.1;refGene_gn=WDFY3;refGene_tx=XM_023252969.1;refGene_gn=WDFY3;refGene_tx=XM_023252970.1;refGen

e_gn=WDFY3;refGene_tx=XM_023252971.1;refGene_gn=WDFY3;refGene_tx=XM_023252972.1;refGene_gn=WDFY3;refGene_tx=XM_023252973.1
B1 138092598 138092817 refGene_gn=WDFY3;refGene_tx=XM_023252963.1;refGene_gn=WDFY3;refGene_tx=XM_023252964.1;refGene_gn=WDFY3;refGene_tx=XM_023252965.1;refGene_gn=WDFY3;refGene_tx=XM_023252966.1;refGene_gn=WDFY3;refGene_tx=XM_023252967.1;refGene_gn=WDFY3;refGene_tx=XM_023252968.1;refGene_gn=WDFY3;refGene_tx=XM_023252969.1;refGene_gn=WDFY3;refGene_tx=XM_023252970.1;refGene_gn=WDFY3;refGene_tx=XM_023252971.1;refGene_gn=WDFY3;refGene_tx=XM_023252972.1;refGene_gn=WDFY3;refGene_tx=XM_023252973.1
B1 138097755 138097937 refGene_gn=WDFY3;refGene_tx=XM_023252963.1;refGene_gn=WDFY3;refGene_tx=XM_023252964.1;refGene_gn=WDFY3;refGene_tx=XM_023252965.1;refGene_gn=WDFY3;refGene_tx=XM_023252966.1;refGene_gn=WDFY3;refGene_tx=XM_023252967.1;refGene_gn=WDFY3;refGene_tx=XM_023252968.1;refGene_gn=WDFY3;refGene_tx=XM_023252969.1;refGene_gn=WDFY3;refGene_tx=XM_023252970.1;refGene_gn=WDFY3;refGene_tx=XM_023252971.1;refGene_gn=WDFY3;refGene_tx=XM_023252972.1;refGene_gn=WDFY3;refGene_tx=XM_023252973.1
B1 138099762 138099994 refGene_gn=WDFY3;refGene_tx=XM_023252963.1;refGene_gn=WDFY3;refGene_tx=XM_023252964.1;refGene_gn=WDFY3;refGene_tx=XM_023252965.1;refGene_gn=WDFY3;refGene_tx=XM_023252966.1;refGene_gn=WDFY3;refGene_tx=XM_023252967.1;refGene_gn=WDFY3;refGene_tx=XM_023252968.1;refGene_gn=WDFY3;refGene_tx=XM_023252969.1;refGene_gn=WDFY3;refGene_tx=XM_023252970.1;refGene_gn=WDFY3;refGene_tx=XM_023252971.1;refGene_gn=WDFY3;refGene_tx=XM_023252972.1;refGene_gn=WDFY3;refGene_tx=XM_023252973.1
B1 138101865 138102026 refGene_gn=WDFY3;refGene_tx=XM_023252963.1;refGene_gn=WDFY3;refGene_tx=XM_023252964.1;refGene_gn=WDFY3;refGene_tx=XM_023252965.1;refGene_gn=WDFY3;refGene_tx=XM_023252966.1;refGene_gn=WDFY3;refGene_tx=XM_023252967.1;refGene_gn=WDFY3;refGene_tx=XM_023252968.1;refGene_gn=WDFY3;refGene_tx=XM_023252969.1;refGene_gn=WDFY3;refGene_tx=XM_023252970.1;refGene_gn=WDFY3;refGene_tx=XM_023252971.1;refGene_gn=WDFY3;refGene_tx=XM_023252972.1;refGene_gn=WDFY3;refGene_tx=XM_023252973.1
B1 138104273 138104385 refGene_gn=WDFY3;refGene_tx=XM_023252963.1;refGene_gn=WDFY3;refGene_tx=XM_023252964.1;refGene_gn=WDFY3;refGene_tx=XM_023252965.1;refGene_gn=WDFY3;refGene_tx=XM_023252966.1;refGene_gn=WDFY3;refGene_tx=XM_023252967.1;refGene_gn=WDFY3;refGene_tx=XM_023252968.1;refGene_gn=WDFY3;refGene_tx=XM_023252969.1;refGene_gn=WDFY3;refGene_tx=XM_023252970.1;refGene_gn=WDFY3;refGene_tx=XM_023252971.1;refGene_gn=WDFY3;refGene_tx=XM_023252972.1;refGene_gn=WDFY3;refGene_tx=XM_023252973.1
B1 138106122 138106313 refGene_gn=WDFY3;refGene_tx=XM_023252963.1;refGene_gn=WDFY3;refGene_tx=XM_023252964.1;refGene_gn=WDFY3;refGene_tx=XM_023252965.1;refGene_gn=WDFY3;refGene_tx=XM_023252966.1;refGene_gn=WDFY3;refGene_tx=XM_023252967.1;refGene_gn=WDFY3;refGene_tx=XM_023252968.1;refGene_gn=WDFY3;refGene_tx=XM_023252969.1;refGene_gn=WDFY3;refGene_tx=XM_023252970.1;refGene_gn=WDFY3;refGene_tx=XM_023252971.1;refGene_gn=WDFY3;refGene_tx=XM_023252972.1;refGene_gn=WDFY3;refGene_tx=XM_023252973.1

B1 138107964 138108117
refGene_gn=WDFY3;refGene_tx=XM_023252963.1;refGene_gn=WDFY3;refGene_tx=XM_023252964.1;refGene_gn=WDFY3;refGene_tx=XM_023252965.1;refGene_gn=WDFY3;refGene_tx=XM_023252966.1;refGene_gn=WDFY3;refGene_tx=XM_023252967.1;refGene_gn=WDFY3;refGene_tx=XM_023252968.1;refGene_gn=WDFY3;refGene_tx=XM_023252969.1;refGene_gn=WDFY3;refGene_tx=XM_023252970.1;refGene_gn=WDFY3;refGene_tx=XM_023252971.1;refGene_gn=WDFY3;refGene_tx=XM_023252972.1;refGene_gn=WDFY3;refGene_tx=XM_023252973.1
B1 138111745 138111819
refGene_gn=WDFY3;refGene_tx=XM_023252963.1;refGene_gn=WDFY3;refGene_tx=XM_023252964.1;refGene_gn=WDFY3;refGene_tx=XM_023252965.1;refGene_gn=WDFY3;refGene_tx=XM_023252966.1;refGene_gn=WDFY3;refGene_tx=XM_023252967.1;refGene_gn=WDFY3;refGene_tx=XM_023252968.1;refGene_gn=WDFY3;refGene_tx=XM_023252969.1;refGene_gn=WDFY3;refGene_tx=XM_023252970.1;refGene_gn=WDFY3;refGene_tx=XM_023252971.1;refGene_gn=WDFY3;refGene_tx=XM_023252972.1;refGene_gn=WDFY3;refGene_tx=XM_023252973.1
B1 138111905 138112067
refGene_gn=WDFY3;refGene_tx=XM_023252963.1;refGene_gn=WDFY3;refGene_tx=XM_023252964.1;refGene_gn=WDFY3;refGene_tx=XM_023252965.1;refGene_gn=WDFY3;refGene_tx=XM_023252966.1;refGene_gn=WDFY3;refGene_tx=XM_023252967.1;refGene_gn=WDFY3;refGene_tx=XM_023252968.1;refGene_gn=WDFY3;refGene_tx=XM_023252969.1;refGene_gn=WDFY3;refGene_tx=XM_023252970.1;refGene_gn=WDFY3;refGene_tx=XM_023252971.1;refGene_gn=WDFY3;refGene_tx=XM_023252972.1;refGene_gn=WDFY3;refGene_tx=XM_023252973.1
B1 138113463 138113558
refGene_gn=WDFY3;refGene_tx=XM_023252963.1;refGene_gn=WDFY3;refGene_tx=XM_023252964.1;refGene_gn=WDFY3;refGene_tx=XM_023252965.1;refGene_gn=WDFY3;refGene_tx=XM_023252966.1;refGene_gn=WDFY3;refGene_tx=XM_023252967.1;refGene_gn=WDFY3;refGene_tx=XM_023252968.1;refGene_gn=WDFY3;refGene_tx=XM_023252969.1;refGene_gn=WDFY3;refGene_tx=XM_023252970.1;refGene_gn=WDFY3;refGene_tx=XM_023252971.1;refGene_gn=WDFY3;refGene_tx=XM_023252972.1;refGene_gn=WDFY3;refGene_tx=XM_023252973.1
B1 138117273 138117394
refGene_gn=WDFY3;refGene_tx=XM_023252963.1;refGene_gn=WDFY3;refGene_tx=XM_023252964.1;refGene_gn=WDFY3;refGene_tx=XM_023252965.1;refGene_gn=WDFY3;refGene_tx=XM_023252966.1;refGene_gn=WDFY3;refGene_tx=XM_023252967.1;refGene_gn=WDFY3;refGene_tx=XM_023252968.1;refGene_gn=WDFY3;refGene_tx=XM_023252969.1;refGene_gn=WDFY3;refGene_tx=XM_023252970.1;refGene_gn=WDFY3;refGene_tx=XM_023252971.1;refGene_gn=WDFY3;refGene_tx=XM_023252972.1;refGene_gn=WDFY3;refGene_tx=XM_023252973.1
B1 138117623 138117841
refGene_gn=WDFY3;refGene_tx=XM_023252963.1;refGene_gn=WDFY3;refGene_tx=XM_023252964.1;refGene_gn=WDFY3;refGene_tx=XM_023252965.1;refGene_gn=WDFY3;refGene_tx=XM_023252966.1;refGene_gn=WDFY3;refGene_tx=XM_023252967.1;refGene_gn=WDFY3;refGene_tx=XM_023252968.1;refGene_gn=WDFY3;refGene_tx=XM_023252969.1;refGene_gn=WDFY3;refGene_tx=XM_023252970.1;refGene_gn=WDFY3;refGene_tx=XM_023252971.1;refGene_gn=WDFY3;refGene_tx=XM_023252972.1;refGene_gn=WDFY3;refGene_tx=XM_023252973.1
B1 138121325 138121561
refGene_gn=WDFY3;refGene_tx=XM_023252963.1;refGene_gn=WDFY3;refGene_tx

=XM_023252964.1;refGene_gn=WDFY3;refGene_tx=XM_023252965.1;refGene_gn=WDFY3;refGene_tx=XM_023252966.1;refGene_gn=WDFY3;refGene_tx=XM_023252967.1;refGene_gn=WDFY3;refGene_tx=XM_023252968.1;refGene_gn=WDFY3;refGene_tx=XM_023252969.1;refGene_gn=WDFY3;refGene_tx=XM_023252970.1;refGene_gn=WDFY3;refGene_tx=XM_023252971.1;refGene_gn=WDFY3;refGene_tx=XM_023252972.1;refGene_gn=WDFY3;refGene_tx=XM_023252973.1
B1 138122768 138122903
refGene_gn=WDFY3;refGene_tx=XM_023252963.1;refGene_gn=WDFY3;refGene_tx=XM_023252964.1;refGene_gn=WDFY3;refGene_tx=XM_023252965.1;refGene_gn=WDFY3;refGene_tx=XM_023252966.1;refGene_gn=WDFY3;refGene_tx=XM_023252967.1;refGene_gn=WDFY3;refGene_tx=XM_023252968.1;refGene_gn=WDFY3;refGene_tx=XM_023252969.1;refGene_gn=WDFY3;refGene_tx=XM_023252970.1;refGene_gn=WDFY3;refGene_tx=XM_023252971.1;refGene_gn=WDFY3;refGene_tx=XM_023252972.1;refGene_gn=WDFY3;refGene_tx=XM_023252973.1
B1 138125352 138125541
refGene_gn=WDFY3;refGene_tx=XM_023252963.1;refGene_gn=WDFY3;refGene_tx=XM_023252964.1;refGene_gn=WDFY3;refGene_tx=XM_023252965.1;refGene_gn=WDFY3;refGene_tx=XM_023252966.1;refGene_gn=WDFY3;refGene_tx=XM_023252967.1;refGene_gn=WDFY3;refGene_tx=XM_023252968.1;refGene_gn=WDFY3;refGene_tx=XM_023252969.1;refGene_gn=WDFY3;refGene_tx=XM_023252970.1;refGene_gn=WDFY3;refGene_tx=XM_023252971.1;refGene_gn=WDFY3;refGene_tx=XM_023252972.1;refGene_gn=WDFY3;refGene_tx=XM_023252973.1
B1 138126904 138127138
refGene_gn=WDFY3;refGene_tx=XM_023252963.1;refGene_gn=WDFY3;refGene_tx=XM_023252964.1;refGene_gn=WDFY3;refGene_tx=XM_023252965.1;refGene_gn=WDFY3;refGene_tx=XM_023252966.1;refGene_gn=WDFY3;refGene_tx=XM_023252967.1;refGene_gn=WDFY3;refGene_tx=XM_023252968.1;refGene_gn=WDFY3;refGene_tx=XM_023252969.1;refGene_gn=WDFY3;refGene_tx=XM_023252970.1;refGene_gn=WDFY3;refGene_tx=XM_023252971.1;refGene_gn=WDFY3;refGene_tx=XM_023252972.1;refGene_gn=WDFY3;refGene_tx=XM_023252973.1
B1 138133231 138133331
refGene_gn=WDFY3;refGene_tx=XM_023252963.1;refGene_gn=WDFY3;refGene_tx=XM_023252964.1;refGene_gn=WDFY3;refGene_tx=XM_023252965.1;refGene_gn=WDFY3;refGene_tx=XM_023252966.1;refGene_gn=WDFY3;refGene_tx=XM_023252967.1;refGene_gn=WDFY3;refGene_tx=XM_023252968.1;refGene_gn=WDFY3;refGene_tx=XM_023252969.1;refGene_gn=WDFY3;refGene_tx=XM_023252970.1;refGene_gn=WDFY3;refGene_tx=XM_023252971.1;refGene_gn=WDFY3;refGene_tx=XM_023252972.1;refGene_gn=WDFY3;refGene_tx=XM_023252973.1
B1 138135528 138135689
refGene_gn=WDFY3;refGene_tx=XM_023252963.1;refGene_gn=WDFY3;refGene_tx=XM_023252964.1;refGene_gn=WDFY3;refGene_tx=XM_023252965.1;refGene_gn=WDFY3;refGene_tx=XM_023252966.1;refGene_gn=WDFY3;refGene_tx=XM_023252967.1;refGene_gn=WDFY3;refGene_tx=XM_023252968.1;refGene_gn=WDFY3;refGene_tx=XM_023252969.1;refGene_gn=WDFY3;refGene_tx=XM_023252970.1;refGene_gn=WDFY3;refGene_tx=XM_023252971.1;refGene_gn=WDFY3;refGene_tx=XM_023252972.1;refGene_gn=WDFY3;refGene_tx=XM_023252973.1
B1 138136784 138137014
refGene_gn=WDFY3;refGene_tx=XM_023252963.1;refGene_gn=WDFY3;refGene_tx=XM_023252964.1;refGene_gn=WDFY3;refGene_tx=XM_023252965.1;refGene_gn=WDFY3;refGene_tx=XM_023252966.1;refGene_gn=WDFY3;refGene_tx=XM_0232529

67.1;refGene_gn=WDFY3;refGene_tx=XM_023252968.1;refGene_gn=WDFY3;refGene_tx=XM_023252969.1;refGene_gn=WDFY3;refGene_tx=XM_023252970.1;refGene_gn=WDFY3;refGene_tx=XM_023252971.1;refGene_gn=WDFY3;refGene_tx=XM_023252972.1;refGene_gn=WDFY3;refGene_tx=XM_023252973.1
B1 138138580 138138690
refGene_gn=WDFY3;refGene_tx=XM_023252963.1;refGene_gn=WDFY3;refGene_tx=XM_023252964.1;refGene_gn=WDFY3;refGene_tx=XM_023252965.1;refGene_gn=WDFY3;refGene_tx=XM_023252966.1;refGene_gn=WDFY3;refGene_tx=XM_023252967.1;refGene_gn=WDFY3;refGene_tx=XM_023252968.1;refGene_gn=WDFY3;refGene_tx=XM_023252969.1;refGene_gn=WDFY3;refGene_tx=XM_023252970.1;refGene_gn=WDFY3;refGene_tx=XM_023252971.1;refGene_gn=WDFY3;refGene_tx=XM_023252972.1;refGene_gn=WDFY3;refGene_tx=XM_023252973.1
B1 138140695 138140878
refGene_gn=WDFY3;refGene_tx=XM_023252963.1;refGene_gn=WDFY3;refGene_tx=XM_023252964.1;refGene_gn=WDFY3;refGene_tx=XM_023252965.1;refGene_gn=WDFY3;refGene_tx=XM_023252966.1;refGene_gn=WDFY3;refGene_tx=XM_023252967.1;refGene_gn=WDFY3;refGene_tx=XM_023252968.1;refGene_gn=WDFY3;refGene_tx=XM_023252969.1;refGene_gn=WDFY3;refGene_tx=XM_023252970.1;refGene_gn=WDFY3;refGene_tx=XM_023252971.1;refGene_gn=WDFY3;refGene_tx=XM_023252972.1;refGene_gn=WDFY3;refGene_tx=XM_023252973.1
B1 138141773 138141931
refGene_gn=WDFY3;refGene_tx=XM_023252963.1;refGene_gn=WDFY3;refGene_tx=XM_023252964.1;refGene_gn=WDFY3;refGene_tx=XM_023252965.1;refGene_gn=WDFY3;refGene_tx=XM_023252966.1;refGene_gn=WDFY3;refGene_tx=XM_023252967.1;refGene_gn=WDFY3;refGene_tx=XM_023252968.1;refGene_gn=WDFY3;refGene_tx=XM_023252969.1;refGene_gn=WDFY3;refGene_tx=XM_023252970.1;refGene_gn=WDFY3;refGene_tx=XM_023252971.1;refGene_gn=WDFY3;refGene_tx=XM_023252972.1;refGene_gn=WDFY3;refGene_tx=XM_023252973.1
B1 138143025 138143103
refGene_gn=WDFY3;refGene_tx=XM_023252963.1;refGene_gn=WDFY3;refGene_tx=XM_023252964.1;refGene_gn=WDFY3;refGene_tx=XM_023252965.1;refGene_gn=WDFY3;refGene_tx=XM_023252966.1;refGene_gn=WDFY3;refGene_tx=XM_023252967.1;refGene_gn=WDFY3;refGene_tx=XM_023252968.1;refGene_gn=WDFY3;refGene_tx=XM_023252969.1;refGene_gn=WDFY3;refGene_tx=XM_023252970.1;refGene_gn=WDFY3;refGene_tx=XM_023252971.1;refGene_gn=WDFY3;refGene_tx=XM_023252972.1;refGene_gn=WDFY3;refGene_tx=XM_023252973.1
B1 138144736 138144964
refGene_gn=WDFY3;refGene_tx=XM_023252963.1;refGene_gn=WDFY3;refGene_tx=XM_023252964.1;refGene_gn=WDFY3;refGene_tx=XM_023252965.1;refGene_gn=WDFY3;refGene_tx=XM_023252966.1;refGene_gn=WDFY3;refGene_tx=XM_023252967.1;refGene_gn=WDFY3;refGene_tx=XM_023252968.1;refGene_gn=WDFY3;refGene_tx=XM_023252969.1;refGene_gn=WDFY3;refGene_tx=XM_023252970.1;refGene_gn=WDFY3;refGene_tx=XM_023252971.1;refGene_gn=WDFY3;refGene_tx=XM_023252972.1;refGene_gn=WDFY3;refGene_tx=XM_023252973.1
B1 138151229 138151283
refGene_gn=WDFY3;refGene_tx=XM_023252963.1;refGene_gn=WDFY3;refGene_tx=XM_023252964.1;refGene_gn=WDFY3;refGene_tx=XM_023252965.1;refGene_gn=WDFY3;refGene_tx=XM_023252966.1;refGene_gn=WDFY3;refGene_tx=XM_023252967.1;refGene_gn=WDFY3;refGene_tx=XM_023252968.1;refGene_gn=WDFY3;refGene_tx=XM_023252970.1;refGene_gn=WDFY3;refGene_tx=XM_023252971.1;refGen

e_gn=WDFY3;refGene_tx=XM_023252973.1
B1	138153544	138153713
refGene_gn=WDFY3;refGene_tx=XM_023252963.1;refGene_gn=WDFY3;refGene_tx=XM_023252964.1;refGene_gn=WDFY3;refGene_tx=XM_023252965.1;refGene_gn=WDFY3;refGene_tx=XM_023252966.1;refGene_gn=WDFY3;refGene_tx=XM_023252967.1;refGene_gn=WDFY3;refGene_tx=XM_023252968.1;refGene_gn=WDFY3;refGene_tx=XM_023252969.1;refGene_gn=WDFY3;refGene_tx=XM_023252970.1;refGene_gn=WDFY3;refGene_tx=XM_023252971.1;refGene_gn=WDFY3;refGene_tx=XM_023252972.1;refGene_gn=WDFY3;refGene_tx=XM_023252973.1
B1	138156746	138156910
refGene_gn=WDFY3;refGene_tx=XM_023252963.1;refGene_gn=WDFY3;refGene_tx=XM_023252964.1;refGene_gn=WDFY3;refGene_tx=XM_023252965.1;refGene_gn=WDFY3;refGene_tx=XM_023252966.1;refGene_gn=WDFY3;refGene_tx=XM_023252967.1;refGene_gn=WDFY3;refGene_tx=XM_023252968.1;refGene_gn=WDFY3;refGene_tx=XM_023252969.1;refGene_gn=WDFY3;refGene_tx=XM_023252970.1;refGene_gn=WDFY3;refGene_tx=XM_023252971.1;refGene_gn=WDFY3;refGene_tx=XM_023252972.1;refGene_gn=WDFY3;refGene_tx=XM_023252973.1
B1	138159472	138159621
refGene_gn=WDFY3;refGene_tx=XM_023252963.1;refGene_gn=WDFY3;refGene_tx=XM_023252964.1;refGene_gn=WDFY3;refGene_tx=XM_023252965.1;refGene_gn=WDFY3;refGene_tx=XM_023252966.1;refGene_gn=WDFY3;refGene_tx=XM_023252967.1;refGene_gn=WDFY3;refGene_tx=XM_023252968.1;refGene_gn=WDFY3;refGene_tx=XM_023252969.1;refGene_gn=WDFY3;refGene_tx=XM_023252970.1;refGene_gn=WDFY3;refGene_tx=XM_023252971.1;refGene_gn=WDFY3;refGene_tx=XM_023252972.1;refGene_gn=WDFY3;refGene_tx=XM_023252973.1
B1	138160739	138160860
refGene_gn=WDFY3;refGene_tx=XM_023252963.1;refGene_gn=WDFY3;refGene_tx=XM_023252964.1;refGene_gn=WDFY3;refGene_tx=XM_023252965.1;refGene_gn=WDFY3;refGene_tx=XM_023252966.1;refGene_gn=WDFY3;refGene_tx=XM_023252967.1;refGene_gn=WDFY3;refGene_tx=XM_023252968.1;refGene_gn=WDFY3;refGene_tx=XM_023252969.1;refGene_gn=WDFY3;refGene_tx=XM_023252970.1;refGene_gn=WDFY3;refGene_tx=XM_023252971.1;refGene_gn=WDFY3;refGene_tx=XM_023252972.1;refGene_gn=WDFY3;refGene_tx=XM_023252973.1
B1	138161672	138161758
refGene_gn=WDFY3;refGene_tx=XM_023252963.1;refGene_gn=WDFY3;refGene_tx=XM_023252964.1;refGene_gn=WDFY3;refGene_tx=XM_023252965.1;refGene_gn=WDFY3;refGene_tx=XM_023252966.1;refGene_gn=WDFY3;refGene_tx=XM_023252967.1;refGene_gn=WDFY3;refGene_tx=XM_023252968.1;refGene_gn=WDFY3;refGene_tx=XM_023252969.1;refGene_gn=WDFY3;refGene_tx=XM_023252970.1;refGene_gn=WDFY3;refGene_tx=XM_023252971.1;refGene_gn=WDFY3;refGene_tx=XM_023252972.1;refGene_gn=WDFY3;refGene_tx=XM_023252973.1
B1	138163466	138163547
refGene_gn=WDFY3;refGene_tx=XM_023252963.1;refGene_gn=WDFY3;refGene_tx=XM_023252964.1;refGene_gn=WDFY3;refGene_tx=XM_023252965.1;refGene_gn=WDFY3;refGene_tx=XM_023252966.1;refGene_gn=WDFY3;refGene_tx=XM_023252967.1;refGene_gn=WDFY3;refGene_tx=XM_023252968.1;refGene_gn=WDFY3;refGene_tx=XM_023252969.1;refGene_gn=WDFY3;refGene_tx=XM_023252970.1;refGene_gn=WDFY3;refGene_tx=XM_023252971.1;refGene_gn=WDFY3;refGene_tx=XM_023252972.1;refGene_gn=WDFY3;refGene_tx=XM_023252973.1
B1	138166775	138166830

refGene_gn=WDFY3;refGene_tx=XM_023252963.1;refGene_gn=WDFY3;refGene_tx
=XM_023252964.1;refGene_gn=WDFY3;refGene_tx=XM_023252965.1;refGene_gn=
WDFY3;refGene_tx=XM_023252966.1;refGene_gn=WDFY3;refGene_tx=XM_0232529
67.1;refGene_gn=WDFY3;refGene_tx=XM_023252968.1;refGene_gn=WDFY3;refGe
ne_tx=XM_023252969.1;refGene_gn=WDFY3;refGene_tx=XM_023252970.1;refGen
e_gn=WDFY3;refGene_tx=XM_023252971.1;refGene_gn=WDFY3;refGene_tx=XM_02
3252972.1;refGene_gn=WDFY3;refGene_tx=XM_023252973.1
B1 138167115 138167235
refGene_gn=WDFY3;refGene_tx=XM_023252963.1;refGene_gn=WDFY3;refGene_tx
=XM_023252964.1;refGene_gn=WDFY3;refGene_tx=XM_023252965.1;refGene_gn=
WDFY3;refGene_tx=XM_023252966.1;refGene_gn=WDFY3;refGene_tx=XM_0232529
67.1;refGene_gn=WDFY3;refGene_tx=XM_023252968.1;refGene_gn=WDFY3;refGe
ne_tx=XM_023252969.1;refGene_gn=WDFY3;refGene_tx=XM_023252970.1;refGen
e_gn=WDFY3;refGene_tx=XM_023252971.1;refGene_gn=WDFY3;refGene_tx=XM_02
3252972.1;refGene_gn=WDFY3;refGene_tx=XM_023252973.1
B1 138169054 138169172
refGene_gn=WDFY3;refGene_tx=XM_023252963.1;refGene_gn=WDFY3;refGene_tx
=XM_023252964.1;refGene_gn=WDFY3;refGene_tx=XM_023252965.1;refGene_gn=
WDFY3;refGene_tx=XM_023252966.1;refGene_gn=WDFY3;refGene_tx=XM_0232529
67.1;refGene_gn=WDFY3;refGene_tx=XM_023252968.1;refGene_gn=WDFY3;refGe
ne_tx=XM_023252969.1;refGene_gn=WDFY3;refGene_tx=XM_023252970.1;refGen
e_gn=WDFY3;refGene_tx=XM_023252971.1;refGene_gn=WDFY3;refGene_tx=XM_02
3252972.1;refGene_gn=WDFY3;refGene_tx=XM_023252973.1
B1 138170130 138170237
refGene_gn=WDFY3;refGene_tx=XM_023252963.1;refGene_gn=WDFY3;refGene_tx
=XM_023252964.1;refGene_gn=WDFY3;refGene_tx=XM_023252965.1;refGene_gn=
WDFY3;refGene_tx=XM_023252966.1;refGene_gn=WDFY3;refGene_tx=XM_0232529
67.1;refGene_gn=WDFY3;refGene_tx=XM_023252968.1;refGene_gn=WDFY3;refGe
ne_tx=XM_023252969.1;refGene_gn=WDFY3;refGene_tx=XM_023252970.1;refGen
e_gn=WDFY3;refGene_tx=XM_023252971.1;refGene_gn=WDFY3;refGene_tx=XM_02
3252972.1;refGene_gn=WDFY3;refGene_tx=XM_023252973.1
B1 138171790 138171944
refGene_gn=WDFY3;refGene_tx=XM_023252963.1;refGene_gn=WDFY3;refGene_tx
=XM_023252964.1;refGene_gn=WDFY3;refGene_tx=XM_023252965.1;refGene_gn=
WDFY3;refGene_tx=XM_023252966.1;refGene_gn=WDFY3;refGene_tx=XM_0232529
67.1;refGene_gn=WDFY3;refGene_tx=XM_023252968.1;refGene_gn=WDFY3;refGe
ne_tx=XM_023252969.1;refGene_gn=WDFY3;refGene_tx=XM_023252970.1;refGen
e_gn=WDFY3;refGene_tx=XM_023252971.1;refGene_gn=WDFY3;refGene_tx=XM_02
3252972.1;refGene_gn=WDFY3;refGene_tx=XM_023252973.1
B1 138176743 138176835
refGene_gn=WDFY3;refGene_tx=XM_023252963.1;refGene_gn=WDFY3;refGene_tx
=XM_023252964.1;refGene_gn=WDFY3;refGene_tx=XM_023252965.1;refGene_gn=
WDFY3;refGene_tx=XM_023252966.1;refGene_gn=WDFY3;refGene_tx=XM_0232529
67.1;refGene_gn=WDFY3;refGene_tx=XM_023252968.1;refGene_gn=WDFY3;refGe
ne_tx=XM_023252969.1;refGene_gn=WDFY3;refGene_tx=XM_023252970.1;refGen
e_gn=WDFY3;refGene_tx=XM_023252971.1;refGene_gn=WDFY3;refGene_tx=XM_02
3252972.1;refGene_gn=WDFY3;refGene_tx=XM_023252973.1
B1 138177060 138177273
refGene_gn=WDFY3;refGene_tx=XM_023252963.1;refGene_gn=WDFY3;refGene_tx
=XM_023252964.1;refGene_gn=WDFY3;refGene_tx=XM_023252965.1;refGene_gn=

WDFY3;refGene_tx=XM_023252966.1;refGene_gn=WDFY3;refGene_tx=XM_0232529
67.1;refGene_gn=WDFY3;refGene_tx=XM_023252968.1;refGene_gn=WDFY3;refGe
ne_tx=XM_023252969.1;refGene_gn=WDFY3;refGene_tx=XM_023252970.1;refGen
e_gn=WDFY3;refGene_tx=XM_023252971.1;refGene_gn=WDFY3;refGene_tx=XM_02
3252972.1;refGene_gn=WDFY3;refGene_tx=XM_023252973.1
B1 138179797 138179945
refGene_gn=WDFY3;refGene_tx=XM_023252963.1;refGene_gn=WDFY3;refGene_tx
=XM_023252964.1;refGene_gn=WDFY3;refGene_tx=XM_023252965.1;refGene_gn=
WDFY3;refGene_tx=XM_023252966.1;refGene_gn=WDFY3;refGene_tx=XM_0232529
67.1;refGene_gn=WDFY3;refGene_tx=XM_023252968.1;refGene_gn=WDFY3;refGe
ne_tx=XM_023252969.1;refGene_gn=WDFY3;refGene_tx=XM_023252970.1;refGen
e_gn=WDFY3;refGene_tx=XM_023252971.1;refGene_gn=WDFY3;refGene_tx=XM_02
3252972.1;refGene_gn=WDFY3;refGene_tx=XM_023252973.1
B1 138181182 138181337
refGene_gn=WDFY3;refGene_tx=XM_023252963.1;refGene_gn=WDFY3;refGene_tx
=XM_023252964.1;refGene_gn=WDFY3;refGene_tx=XM_023252965.1;refGene_gn=
WDFY3;refGene_tx=XM_023252966.1;refGene_gn=WDFY3;refGene_tx=XM_0232529
67.1;refGene_gn=WDFY3;refGene_tx=XM_023252968.1;refGene_gn=WDFY3;refGe
ne_tx=XM_023252969.1;refGene_gn=WDFY3;refGene_tx=XM_023252970.1;refGen
e_gn=WDFY3;refGene_tx=XM_023252971.1;refGene_gn=WDFY3;refGene_tx=XM_02
3252972.1;refGene_gn=WDFY3;refGene_tx=XM_023252973.1
B1 138182201 138182360
refGene_gn=WDFY3;refGene_tx=XM_023252963.1;refGene_gn=WDFY3;refGene_tx
=XM_023252964.1;refGene_gn=WDFY3;refGene_tx=XM_023252965.1;refGene_gn=
WDFY3;refGene_tx=XM_023252966.1;refGene_gn=WDFY3;refGene_tx=XM_0232529
67.1;refGene_gn=WDFY3;refGene_tx=XM_023252968.1;refGene_gn=WDFY3;refGe
ne_tx=XM_023252969.1;refGene_gn=WDFY3;refGene_tx=XM_023252970.1;refGen
e_gn=WDFY3;refGene_tx=XM_023252971.1;refGene_gn=WDFY3;refGene_tx=XM_02
3252972.1;refGene_gn=WDFY3;refGene_tx=XM_023252973.1
B1 138184702 138184882
refGene_gn=WDFY3;refGene_tx=XM_023252963.1;refGene_gn=WDFY3;refGene_tx
=XM_023252964.1;refGene_gn=WDFY3;refGene_tx=XM_023252965.1;refGene_gn=
WDFY3;refGene_tx=XM_023252966.1;refGene_gn=WDFY3;refGene_tx=XM_0232529
67.1;refGene_gn=WDFY3;refGene_tx=XM_023252968.1;refGene_gn=WDFY3;refGe
ne_tx=XM_023252969.1;refGene_gn=WDFY3;refGene_tx=XM_023252970.1;refGen
e_gn=WDFY3;refGene_tx=XM_023252971.1;refGene_gn=WDFY3;refGene_tx=XM_02
3252972.1;refGene_gn=WDFY3;refGene_tx=XM_023252973.1
B1 138188271 138188454
refGene_gn=WDFY3;refGene_tx=XM_023252963.1;refGene_gn=WDFY3;refGene_tx
=XM_023252964.1;refGene_gn=WDFY3;refGene_tx=XM_023252965.1;refGene_gn=
WDFY3;refGene_tx=XM_023252966.1;refGene_gn=WDFY3;refGene_tx=XM_0232529
67.1;refGene_gn=WDFY3;refGene_tx=XM_023252968.1;refGene_gn=WDFY3;refGe
ne_tx=XM_023252969.1;refGene_gn=WDFY3;refGene_tx=XM_023252970.1;refGen
e_gn=WDFY3;refGene_tx=XM_023252971.1;refGene_gn=WDFY3;refGene_tx=XM_02
3252972.1;refGene_gn=WDFY3;refGene_tx=XM_023252973.1
B1 138189604 138189701
refGene_gn=WDFY3;refGene_tx=XM_023252963.1;refGene_gn=WDFY3;refGene_tx
=XM_023252964.1;refGene_gn=WDFY3;refGene_tx=XM_023252965.1;refGene_gn=
WDFY3;refGene_tx=XM_023252966.1;refGene_gn=WDFY3;refGene_tx=XM_0232529
67.1;refGene_gn=WDFY3;refGene_tx=XM_023252968.1;refGene_gn=WDFY3;refGe

ne_tx=XM_023252969.1;refGene_gn=WDFY3;refGene_tx=XM_023252970.1;refGen
e_gn=WDFY3;refGene_tx=XM_023252971.1;refGene_gn=WDFY3;refGene_tx=XM_02
3252972.1;refGene_gn=WDFY3;refGene_tx=XM_023252973.1
B1 138192211 138192535
refGene_gn=WDFY3;refGene_tx=XM_023252963.1;refGene_gn=WDFY3;refGene_tx
=XM_023252964.1;refGene_gn=WDFY3;refGene_tx=XM_023252965.1;refGene_gn=
WDFY3;refGene_tx=XM_023252966.1;refGene_gn=WDFY3;refGene_tx=XM_0232529
67.1;refGene_gn=WDFY3;refGene_tx=XM_023252968.1;refGene_gn=WDFY3;refGe
ne_tx=XM_023252969.1;refGene_gn=WDFY3;refGene_tx=XM_023252970.1;refGen
e_gn=WDFY3;refGene_tx=XM_023252971.1;refGene_gn=WDFY3;refGene_tx=XM_02
3252972.1;refGene_gn=WDFY3;refGene_tx=XM_023252973.1
B1 138193076 138193188
refGene_gn=WDFY3;refGene_tx=XM_023252963.1;refGene_gn=WDFY3;refGene_tx
=XM_023252964.1;refGene_gn=WDFY3;refGene_tx=XM_023252965.1;refGene_gn=
WDFY3;refGene_tx=XM_023252966.1;refGene_gn=WDFY3;refGene_tx=XM_0232529
67.1;refGene_gn=WDFY3;refGene_tx=XM_023252968.1;refGene_gn=WDFY3;refGe
ne_tx=XM_023252969.1;refGene_gn=WDFY3;refGene_tx=XM_023252970.1;refGen
e_gn=WDFY3;refGene_tx=XM_023252971.1;refGene_gn=WDFY3;refGene_tx=XM_02
3252972.1;refGene_gn=WDFY3;refGene_tx=XM_023252973.1
B1 138193581 138193779
refGene_gn=WDFY3;refGene_tx=XM_023252963.1;refGene_gn=WDFY3;refGene_tx
=XM_023252964.1;refGene_gn=WDFY3;refGene_tx=XM_023252965.1;refGene_gn=
WDFY3;refGene_tx=XM_023252966.1;refGene_gn=WDFY3;refGene_tx=XM_0232529
67.1;refGene_gn=WDFY3;refGene_tx=XM_023252968.1;refGene_gn=WDFY3;refGe
ne_tx=XM_023252969.1;refGene_gn=WDFY3;refGene_tx=XM_023252970.1;refGen
e_gn=WDFY3;refGene_tx=XM_023252971.1;refGene_gn=WDFY3;refGene_tx=XM_02
3252972.1;refGene_gn=WDFY3;refGene_tx=XM_023252973.1
B1 138197121 138197245
refGene_gn=WDFY3;refGene_tx=XM_023252963.1;refGene_gn=WDFY3;refGene_tx
=XM_023252964.1;refGene_gn=WDFY3;refGene_tx=XM_023252965.1;refGene_gn=
WDFY3;refGene_tx=XM_023252966.1;refGene_gn=WDFY3;refGene_tx=XM_0232529
67.1;refGene_gn=WDFY3;refGene_tx=XM_023252968.1;refGene_gn=WDFY3;refGe
ne_tx=XM_023252969.1;refGene_gn=WDFY3;refGene_tx=XM_023252970.1;refGen
e_gn=WDFY3;refGene_tx=XM_023252971.1;refGene_gn=WDFY3;refGene_tx=XM_02
3252972.1;refGene_gn=WDFY3;refGene_tx=XM_023252973.1
B1 138220622 138220752
refGene_gn=CDS1;refGene_tx=XM_003985217.5
B1 138224642 138224746
refGene_gn=CDS1;refGene_tx=XM_003985217.5
B1 138227238 138227358
refGene_gn=CDS1;refGene_tx=XM_003985217.5
B1 138228825 138228978
refGene_gn=CDS1;refGene_tx=XM_003985217.5
B1 138231922 138231991
refGene_gn=CDS1;refGene_tx=XM_003985217.5
B1 138234640 138234728
refGene_gn=CDS1;refGene_tx=XM_003985217.5
B1 138236026 138236109
refGene_gn=CDS1;refGene_tx=XM_003985217.5
B1 138237763 138237822

refGene_gn=CDS1;refGene_tx=XM_003985217.5
B1 138254723 138254863
refGene_gn=CDS1;refGene_tx=XM_003985217.5
B1 138258141 138258239
refGene_gn=CDS1;refGene_tx=XM_003985217.5
B1 138264853 138264950
refGene_gn=CDS1;refGene_tx=XM_003985217.5
B1 138272029 138272157
refGene_gn=CDS1;refGene_tx=XM_003985217.5
B1 138289720 138289837
refGene_gn=CDS1;refGene_tx=XM_003985217.5
B1 138374809 138375479
refGene_gn=NKX6-1;refGene_tx=XM_019829291.2
B1 138377218 138377391
refGene_gn=NKX6-1;refGene_tx=XM_019829291.2
B1 138379623 138379884
refGene_gn=NKX6-1;refGene_tx=XM_019829291.2
B1 139235590 139235687
refGene_gn=GPAT3;refGene_tx=XM_003985219.5;refGene_gn=GPAT3;refGene_tx=XM_006930996.4;refGene_gn=GPAT3;refGene_tx=XM_006930997.3;refGene_gn=GPAT3;refGene_tx=XM_019829293.2
B1 139240485 139240565
refGene_gn=GPAT3;refGene_tx=XM_003985219.5;refGene_gn=GPAT3;refGene_tx=XM_006930996.4;refGene_gn=GPAT3;refGene_tx=XM_006930997.3;refGene_gn=GPAT3;refGene_tx=XM_019829293.2
B1 139241045 139241174
refGene_gn=GPAT3;refGene_tx=XM_003985219.5;refGene_gn=GPAT3;refGene_tx=XM_006930996.4;refGene_gn=GPAT3;refGene_tx=XM_006930997.3;refGene_gn=GPAT3;refGene_tx=XM_019829293.2
B1 139242669 139242755
refGene_gn=GPAT3;refGene_tx=XM_003985219.5;refGene_gn=GPAT3;refGene_tx=XM_006930996.4;refGene_gn=GPAT3;refGene_tx=XM_006930997.3;refGene_gn=GPAT3;refGene_tx=XM_019829293.2
B1 139243278 139243334
refGene_gn=GPAT3;refGene_tx=XM_003985219.5;refGene_gn=GPAT3;refGene_tx=XM_006930996.4;refGene_gn=GPAT3;refGene_tx=XM_006930997.3;refGene_gn=GPAT3;refGene_tx=XM_019829293.2
B1 139244516 139244632
refGene_gn=GPAT3;refGene_tx=XM_003985219.5;refGene_gn=GPAT3;refGene_tx=XM_006930996.4;refGene_gn=GPAT3;refGene_tx=XM_006930997.3;refGene_gn=GPAT3;refGene_tx=XM_019829293.2
B1 139248636 139248730
refGene_gn=GPAT3;refGene_tx=XM_003985219.5;refGene_gn=GPAT3;refGene_tx=XM_006930996.4;refGene_gn=GPAT3;refGene_tx=XM_006930997.3;refGene_gn=GPAT3;refGene_tx=XM_019829293.2
B1 139250853 139250943
refGene_gn=GPAT3;refGene_tx=XM_003985219.5;refGene_gn=GPAT3;refGene_tx=XM_006930996.4;refGene_gn=GPAT3;refGene_tx=XM_006930997.3;refGene_gn=GPAT3;refGene_tx=XM_019829293.2
B1 139251465 139251540

refGene_gn=GPAT3;refGene_tx=XM_003985219.5;refGene_gn=GPAT3;refGene_tx
=XM_006930996.4;refGene_gn=GPAT3;refGene_tx=XM_006930997.3
B1 139255265 139255536
refGene_gn=GPAT3;refGene_tx=XM_003985219.5;refGene_gn=GPAT3;refGene_tx
=XM_006930996.4;refGene_gn=GPAT3;refGene_tx=XM_006930997.3;refGene_gn=
GPAT3;refGene_tx=XM_019829293.2
B1 139289402 139289469
refGene_gn=GPAT3;refGene_tx=XM_003985219.5;refGene_gn=GPAT3;refGene_tx
=XM_006930996.4;refGene_gn=GPAT3;refGene_tx=XM_006930997.3;refGene_gn=
GPAT3;refGene_tx=XM_019829293.2
B1 139297502 139297643
refGene_gn=GPAT3;refGene_tx=XM_003985219.5;refGene_gn=GPAT3;refGene_tx
=XM_006930996.4;refGene_gn=GPAT3;refGene_tx=XM_006930997.3;refGene_gn=
GPAT3;refGene_tx=XM_019829293.2
B1 139336488 139336575
refGene_gn=ABRAXAS1;refGene_tx=XM_006930998.4
B1 139338762 139338853
refGene_gn=ABRAXAS1;refGene_tx=XM_006930998.4;refGene_gn=ABRAXAS1;refG
ene_tx=XM_023252974.1
B1 139342821 139342858
refGene_gn=ABRAXAS1;refGene_tx=XM_006930998.4;refGene_gn=ABRAXAS1;refG
ene_tx=XM_023252974.1
B1 139343929 139343996
refGene_gn=ABRAXAS1;refGene_tx=XM_006930998.4;refGene_gn=ABRAXAS1;refG
ene_tx=XM_023252974.1
B1 139345519 139345713
refGene_gn=ABRAXAS1;refGene_tx=XM_006930998.4;refGene_gn=ABRAXAS1;refG
ene_tx=XM_019829294.2;refGene_gn=ABRAXAS1;refGene_tx=XM_019829295.2;re
fGene_gn=ABRAXAS1;refGene_tx=XM_023252974.1
B1 139347081 139347201
refGene_gn=ABRAXAS1;refGene_tx=XM_006930998.4;refGene_gn=ABRAXAS1;refG
ene_tx=XM_019829294.2;refGene_gn=ABRAXAS1;refGene_tx=XM_019829295.2;re
fGene_gn=ABRAXAS1;refGene_tx=XM_023252974.1
B1 139348139 139348224
refGene_gn=ABRAXAS1;refGene_tx=XM_006930998.4;refGene_gn=ABRAXAS1;refG
ene_tx=XM_019829294.2;refGene_gn=ABRAXAS1;refGene_tx=XM_019829295.2;re
fGene_gn=ABRAXAS1;refGene_tx=XM_023252974.1
B1 139350848 139350963
refGene_gn=ABRAXAS1;refGene_tx=XM_006930998.4;refGene_gn=ABRAXAS1;refG
ene_tx=XM_019829294.2;refGene_gn=ABRAXAS1;refGene_tx=XM_019829295.2;re
fGene_gn=ABRAXAS1;refGene_tx=XM_023252974.1
B1 139351350 139351784
refGene_gn=ABRAXAS1;refGene_tx=XM_006930998.4;refGene_gn=ABRAXAS1;refG
ene_tx=XM_019829294.2;refGene_gn=ABRAXAS1;refGene_tx=XM_019829295.2;re
fGene_gn=ABRAXAS1;refGene_tx=XM_023252974.1
B1 139355790 139355867
refGene_gn=MRPS18C;refGene_tx=XM_011281748.3;refGene_gn=MRPS18C;refGen
e_tx=XM_023252976.1
B1 139355952 139356012
refGene_gn=MRPS18C;refGene_tx=XM_011281748.3;refGene_gn=MRPS18C;refGen

e_tx=XM_023252976.1
B1 139356537 139356595
refGene_gn=MRPS18C;refGene_tx=XM_011281748.3;refGene_gn=MRPS18C;refGene_tx=XM_023252976.1
B1 139357594 139357678
refGene_gn=MRPS18C;refGene_tx=XM_011281748.3
B1 139358985 139359038
refGene_gn=MRPS18C;refGene_tx=XM_011281748.3;refGene_gn=MRPS18C;refGene_tx=XM_023252976.1
B1 139360248 139360348
refGene_gn=MRPS18C;refGene_tx=XM_011281748.3;refGene_gn=MRPS18C;refGene_tx=XM_023252976.1
B1 139360780 139360966
refGene_gn=HELQ;refGene_tx=XM_006930999.4;refGene_gn=HELQ;refGene_tx=XM_019829296.2;refGene_gn=HELQ;refGene_tx=XM_019829298.2;refGene_gn=HELQ;refGene_tx=XM_019829299.2;refGene_gn=HELQ;refGene_tx=XM_019829300.2
B1 139361140 139361221
refGene_gn=HELQ;refGene_tx=XM_019829297.2
B1 139362445 139363160
refGene_gn=HELQ;refGene_tx=XM_006930999.4;refGene_gn=HELQ;refGene_tx=XM_019829296.2;refGene_gn=HELQ;refGene_tx=XM_019829297.2;refGene_gn=HELQ;refGene_tx=XM_019829298.2;refGene_gn=HELQ;refGene_tx=XM_019829299.2;refGene_gn=HELQ;refGene_tx=XM_019829300.2
B1 139366899 139367078
refGene_gn=HELQ;refGene_tx=XM_006930999.4;refGene_gn=HELQ;refGene_tx=XM_019829296.2;refGene_gn=HELQ;refGene_tx=XM_019829297.2;refGene_gn=HELQ;refGene_tx=XM_019829298.2;refGene_gn=HELQ;refGene_tx=XM_019829299.2;refGene_gn=HELQ;refGene_tx=XM_019829300.2;refGene_gn=HELQ;refGene_tx=XM_023252975.1
B1 139368217 139368418
refGene_gn=HELQ;refGene_tx=XM_006930999.4;refGene_gn=HELQ;refGene_tx=XM_019829296.2;refGene_gn=HELQ;refGene_tx=XM_019829297.2;refGene_gn=HELQ;refGene_tx=XM_019829298.2;refGene_gn=HELQ;refGene_tx=XM_019829299.2;refGene_gn=HELQ;refGene_tx=XM_019829300.2;refGene_gn=HELQ;refGene_tx=XM_023252975.1
B1 139368884 139368957
refGene_gn=HELQ;refGene_tx=XM_006930999.4;refGene_gn=HELQ;refGene_tx=XM_019829296.2;refGene_gn=HELQ;refGene_tx=XM_019829297.2;refGene_gn=HELQ;refGene_tx=XM_019829298.2;refGene_gn=HELQ;refGene_tx=XM_019829299.2;refGene_gn=HELQ;refGene_tx=XM_019829300.2;refGene_gn=HELQ;refGene_tx=XM_023252975.1
B1 139371435 139371546
refGene_gn=HELQ;refGene_tx=XM_006930999.4;refGene_gn=HELQ;refGene_tx=XM_011281749.2;refGene_gn=HELQ;refGene_tx=XM_019829296.2;refGene_gn=HELQ;refGene_tx=XM_019829297.2;refGene_gn=HELQ;refGene_tx=XM_019829298.2;refGene_gn=HELQ;refGene_tx=XM_019829299.2;refGene_gn=HELQ;refGene_tx=XM_019829300.2;refGene_gn=HELQ;refGene_tx=XM_023252975.1
B1 139372999 139373098
refGene_gn=HELQ;refGene_tx=XM_006930999.4;refGene_gn=HELQ;refGene_tx=XM_011281749.2;refGene_gn=HELQ;refGene_tx=XM_019829296.2;refGene_gn=HEL

Q;refGene_tx=XM_019829297.2;refGene_gn=HELQ;refGene_tx=XM_019829298.2;
refGene_gn=HELQ;refGene_tx=XM_019829299.2;refGene_gn=HELQ;refGene_tx=X
M_019829300.2;refGene_gn=HELQ;refGene_tx=XM_023252975.1
B1 139374609 139374755
refGene_gn=HELQ;refGene_tx=XM_006930999.4;refGene_gn=HELQ;refGene_tx=X
M_011281749.2;refGene_gn=HELQ;refGene_tx=XM_019829296.2;refGene_gn=HEL
Q;refGene_tx=XM_019829297.2;refGene_gn=HELQ;refGene_tx=XM_019829298.2;
refGene_gn=HELQ;refGene_tx=XM_019829299.2;refGene_gn=HELQ;refGene_tx=X
M_019829300.2;refGene_gn=HELQ;refGene_tx=XM_023252975.1
B1 139377101 139377341
refGene_gn=HELQ;refGene_tx=XM_006930999.4;refGene_gn=HELQ;refGene_tx=X
M_011281749.2;refGene_gn=HELQ;refGene_tx=XM_019829296.2;refGene_gn=HEL
Q;refGene_tx=XM_019829297.2;refGene_gn=HELQ;refGene_tx=XM_019829298.2;
refGene_gn=HELQ;refGene_tx=XM_019829299.2;refGene_gn=HELQ;refGene_tx=X
M_019829300.2;refGene_gn=HELQ;refGene_tx=XM_023252975.1
B1 139381456 139381598
refGene_gn=HELQ;refGene_tx=XM_006930999.4;refGene_gn=HELQ;refGene_tx=X
M_011281749.2;refGene_gn=HELQ;refGene_tx=XM_019829296.2;refGene_gn=HEL
Q;refGene_tx=XM_019829297.2;refGene_gn=HELQ;refGene_tx=XM_019829298.2;
refGene_gn=HELQ;refGene_tx=XM_019829299.2;refGene_gn=HELQ;refGene_tx=X
M_019829300.2;refGene_gn=HELQ;refGene_tx=XM_023252975.1
B1 139382044 139382149
refGene_gn=HELQ;refGene_tx=XM_006930999.4;refGene_gn=HELQ;refGene_tx=X
M_011281749.2;refGene_gn=HELQ;refGene_tx=XM_019829296.2;refGene_gn=HEL
Q;refGene_tx=XM_019829297.2;refGene_gn=HELQ;refGene_tx=XM_019829298.2;
refGene_gn=HELQ;refGene_tx=XM_019829299.2;refGene_gn=HELQ;refGene_tx=X
M_019829300.2;refGene_gn=HELQ;refGene_tx=XM_023252975.1
B1 139388043 139388266
refGene_gn=HELQ;refGene_tx=XM_006930999.4;refGene_gn=HELQ;refGene_tx=X
M_011281749.2;refGene_gn=HELQ;refGene_tx=XM_019829296.2;refGene_gn=HEL
Q;refGene_tx=XM_019829297.2;refGene_gn=HELQ;refGene_tx=XM_019829298.2;
refGene_gn=HELQ;refGene_tx=XM_019829299.2;refGene_gn=HELQ;refGene_tx=X
M_019829300.2;refGene_gn=HELQ;refGene_tx=XM_023252975.1
B1 139389977 139390135
refGene_gn=HELQ;refGene_tx=XM_006930999.4;refGene_gn=HELQ;refGene_tx=X
M_011281749.2;refGene_gn=HELQ;refGene_tx=XM_019829296.2;refGene_gn=HEL
Q;refGene_tx=XM_019829297.2;refGene_gn=HELQ;refGene_tx=XM_019829298.2;
refGene_gn=HELQ;refGene_tx=XM_019829299.2;refGene_gn=HELQ;refGene_tx=X
M_019829300.2;refGene_gn=HELQ;refGene_tx=XM_023252975.1
B1 139391338 139391437
refGene_gn=HELQ;refGene_tx=XM_006930999.4;refGene_gn=HELQ;refGene_tx=X
M_011281749.2;refGene_gn=HELQ;refGene_tx=XM_019829296.2;refGene_gn=HEL
Q;refGene_tx=XM_019829297.2;refGene_gn=HELQ;refGene_tx=XM_019829298.2;
refGene_gn=HELQ;refGene_tx=XM_019829299.2;refGene_gn=HELQ;refGene_tx=X
M_019829300.2;refGene_gn=HELQ;refGene_tx=XM_023252975.1
B1 139393801 139393951
refGene_gn=HELQ;refGene_tx=XM_019829300.2
B1 139394389 139394563
refGene_gn=HELQ;refGene_tx=XM_006930999.4;refGene_gn=HELQ;refGene_tx=X
M_011281749.2;refGene_gn=HELQ;refGene_tx=XM_019829296.2;refGene_gn=HEL

Q;refGene_tx=XM_019829297.2;refGene_gn=HELQ;refGene_tx=XM_019829298.2;
refGene_gn=HELQ;refGene_tx=XM_019829299.2;refGene_gn=HELQ;refGene_tx=X
M_023252975.1
B1 139397598 139397712
refGene_gn=HELQ;refGene_tx=XM_006930999.4;refGene_gn=HELQ;refGene_tx=X
M_011281749.2;refGene_gn=HELQ;refGene_tx=XM_019829296.2;refGene_gn=HEL
Q;refGene_tx=XM_019829297.2;refGene_gn=HELQ;refGene_tx=XM_023252975.1
B1 139398603 139398738
refGene_gn=HELQ;refGene_tx=XM_006930999.4;refGene_gn=HELQ;refGene_tx=X
M_011281749.2;refGene_gn=HELQ;refGene_tx=XM_019829296.2;refGene_gn=HEL
Q;refGene_tx=XM_019829297.2;refGene_gn=HELQ;refGene_tx=XM_019829298.2;
refGene_gn=HELQ;refGene_tx=XM_023252975.1
B1 139399003 139399048
refGene_gn=HELQ;refGene_tx=XM_019829296.2
B1 139401914 139402019
refGene_gn=HELQ;refGene_tx=XM_006930999.4;refGene_gn=HELQ;refGene_tx=X
M_011281749.2;refGene_gn=HELQ;refGene_tx=XM_019829297.2;refGene_gn=HEL
Q;refGene_tx=XM_019829298.2;refGene_gn=HELQ;refGene_tx=XM_019829299.2;
refGene_gn=HELQ;refGene_tx=XM_023252975.1
B1 139441282 139441509
refGene_gn=HPSE;refGene_tx=XM_023252977.1;refGene_gn=HPSE;refGene_tx=X
M_023252978.1
B1 139455239 139455385
refGene_gn=HPSE;refGene_tx=XM_023252977.1;refGene_gn=HPSE;refGene_tx=X
M_023252978.1
B1 139457363 139457489
refGene_gn=HPSE;refGene_tx=XM_023252977.1;refGene_gn=HPSE;refGene_tx=X
M_023252978.1
B1 139463062 139463236
refGene_gn=HPSE;refGene_tx=XM_023252977.1;refGene_gn=HPSE;refGene_tx=X
M_023252978.1
B1 139465469 139465638
refGene_gn=HPSE;refGene_tx=XM_023252977.1;refGene_gn=HPSE;refGene_tx=X
M_023252978.1
B1 139465965 139466013
refGene_gn=HPSE;refGene_tx=XM_023252977.1;refGene_gn=HPSE;refGene_tx=X
M_023252978.1
B1 139466607 139466701
refGene_gn=HPSE;refGene_tx=XM_023252977.1
B1 139467567 139467674
refGene_gn=HPSE;refGene_tx=XM_023252977.1
B1 139469372 139469487
refGene_gn=HPSE;refGene_tx=XM_023252977.1;refGene_gn=HPSE;refGene_tx=X
M_023252978.1
B1 139473262 139473381
refGene_gn=HPSE;refGene_tx=XM_023252977.1;refGene_gn=HPSE;refGene_tx=X
M_023252978.1
B1 139474243 139474390
refGene_gn=HPSE;refGene_tx=XM_023252977.1;refGene_gn=HPSE;refGene_tx=X
M_023252978.1

B1 139479793 139479953
refGene_gn=HPSE;refGene_tx=XM_023252977.1;refGene_gn=HPSE;refGene_tx=XM_023252978.1
B1 139492748 139492995
refGene_gn=COQ2;refGene_tx=XM_003985223.3;refGene_gn=COQ2;refGene_tx=XM_006931000.2;refGene_gn=COQ2;refGene_tx=XM_006931001.3
B1 139497791 139497958
refGene_gn=COQ2;refGene_tx=XM_003985223.3;refGene_gn=COQ2;refGene_tx=XM_006931000.2;refGene_gn=COQ2;refGene_tx=XM_006931001.3
B1 139506254 139506376
refGene_gn=COQ2;refGene_tx=XM_003985223.3;refGene_gn=COQ2;refGene_tx=XM_006931000.2;refGene_gn=COQ2;refGene_tx=XM_006931001.3
B1 139507637 139507723
refGene_gn=COQ2;refGene_tx=XM_003985223.3;refGene_gn=COQ2;refGene_tx=XM_006931000.2;refGene_gn=COQ2;refGene_tx=XM_006931001.3
B1 139509123 139509257
refGene_gn=COQ2;refGene_tx=XM_003985223.3;refGene_gn=COQ2;refGene_tx=XM_006931000.2;refGene_gn=COQ2;refGene_tx=XM_006931001.3
B1 139511202 139511391
refGene_gn=COQ2;refGene_tx=XM_003985223.3;refGene_gn=COQ2;refGene_tx=XM_006931000.2;refGene_gn=COQ2;refGene_tx=XM_006931001.3
B1 139513757 139513922
refGene_gn=COQ2;refGene_tx=XM_003985223.3;refGene_gn=COQ2;refGene_tx=XM_006931000.2
B1 139514647 139514680
refGene_gn=COQ2;refGene_tx=XM_006931001.3
B1 139584846 139584946
refGene_gn=LOC101090352;refGene_tx=XM_019829303.2;refGene_gn=LOC101090352;refGene_tx=XM_019829304.1
B1 139586731 139586856
refGene_gn=LOC101090352;refGene_tx=XM_019829303.2;refGene_gn=LOC101090352;refGene_tx=XM_019829304.1
B1 139591444 139591549
refGene_gn=LOC101090352;refGene_tx=XM_019829303.2;refGene_gn=LOC101090352;refGene_tx=XM_019829304.1
B1 139642820 139642932
refGene_gn=LOC101084956;refGene_tx=XM_003985224.4
B1 139645319 139645444
refGene_gn=LOC101084956;refGene_tx=XM_003985224.4
B1 139652117 139652222
refGene_gn=LOC101084956;refGene_tx=XM_003985224.4
B1 139662502 139662636
refGene_gn=COPS4;refGene_tx=XM_003985225.5;refGene_gn=COPS4;refGene_tx=XM_003985226.5
B1 139668623 139668708
refGene_gn=COPS4;refGene_tx=XM_003985225.5
B1 139670808 139670924
refGene_gn=COPS4;refGene_tx=XM_003985225.5;refGene_gn=COPS4;refGene_tx=XM_003985226.5
B1 139677029 139677200

refGene_gn=COPS4;refGene_tx=XM_003985225.5;refGene_gn=COPS4;refGene_tx=XM_003985226.5
B1	139681108	139681259
refGene_gn=COPS4;refGene_tx=XM_003985225.5;refGene_gn=COPS4;refGene_tx=XM_003985226.5
B1	139681424	139681578
refGene_gn=COPS4;refGene_tx=XM_003985225.5;refGene_gn=COPS4;refGene_tx=XM_003985226.5
B1	139686182	139686286
refGene_gn=COPS4;refGene_tx=XM_003985225.5;refGene_gn=COPS4;refGene_tx=XM_003985226.5
B1	139687241	139687393
refGene_gn=COPS4;refGene_tx=XM_003985225.5;refGene_gn=COPS4;refGene_tx=XM_003985226.5
B1	139689232	139689312
refGene_gn=COPS4;refGene_tx=XM_003985225.5;refGene_gn=COPS4;refGene_tx=XM_003985226.5
B1	139704565	139704639
refGene_gn=COPS4;refGene_tx=XM_003985225.5;refGene_gn=COPS4;refGene_tx=XM_003985226.5
B1	139733756	139734440
refGene_gn=LIN54;refGene_tx=XM_003985227.4;refGene_gn=LIN54;refGene_tx=XM_006931004.3;refGene_gn=LIN54;refGene_tx=XM_006931005.4;refGene_gn=LIN54;refGene_tx=XM_011281747.2;refGene_gn=LIN54;refGene_tx=XM_023252980.1;refGene_gn=LIN54;refGene_tx=XM_023252981.1
B1	139738950	139739074
refGene_gn=LIN54;refGene_tx=XM_003985227.4;refGene_gn=LIN54;refGene_tx=XM_006931004.3;refGene_gn=LIN54;refGene_tx=XM_006931005.4;refGene_gn=LIN54;refGene_tx=XM_023252980.1;refGene_gn=LIN54;refGene_tx=XM_023252981.1
B1	139746023	139746166
refGene_gn=LIN54;refGene_tx=XM_003985227.4;refGene_gn=LIN54;refGene_tx=XM_006931005.4;refGene_gn=LIN54;refGene_tx=XM_023252980.1;refGene_gn=LIN54;refGene_tx=XM_023252981.1
B1	139774105	139774322
refGene_gn=LIN54;refGene_tx=XM_003985227.4;refGene_gn=LIN54;refGene_tx=XM_006931005.4;refGene_gn=LIN54;refGene_tx=XM_011281747.2;refGene_gn=LIN54;refGene_tx=XM_023252980.1;refGene_gn=LIN54;refGene_tx=XM_023252981.1
B1	139779180	139779254
refGene_gn=LIN54;refGene_tx=XM_003985227.4;refGene_gn=LIN54;refGene_tx=XM_006931004.3;refGene_gn=LIN54;refGene_tx=XM_006931005.4;refGene_gn=LIN54;refGene_tx=XM_011281747.2;refGene_gn=LIN54;refGene_tx=XM_023252980.1;refGene_gn=LIN54;refGene_tx=XM_023252981.1
B1	139779398	139779596
refGene_gn=LIN54;refGene_tx=XM_003985227.4;refGene_gn=LIN54;refGene_tx=XM_006931004.3;refGene_gn=LIN54;refGene_tx=XM_006931005.4;refGene_gn=LIN54;refGene_tx=XM_011281747.2;refGene_gn=LIN54;refGene_tx=XM_023252980.1;refGene_gn=LIN54;refGene_tx=XM_023252981.1
B1	139780732	139780824

refGene_gn=LIN54;refGene_tx=XM_003985227.4;refGene_gn=LIN54;refGene_tx=XM_006931004.3;refGene_gn=LIN54;refGene_tx=XM_006931005.4;refGene_gn=LIN54;refGene_tx=XM_011281747.2;refGene_gn=LIN54;refGene_tx=XM_023252980.1;refGene_gn=LIN54;refGene_tx=XM_023252981.1
B1 139781547 139781619
refGene_gn=LIN54;refGene_tx=XM_003985227.4;refGene_gn=LIN54;refGene_tx=XM_006931004.3;refGene_gn=LIN54;refGene_tx=XM_006931005.4;refGene_gn=LIN54;refGene_tx=XM_011281747.2;refGene_gn=LIN54;refGene_tx=XM_023252980.1;refGene_gn=LIN54;refGene_tx=XM_023252981.1
B1 139782835 139782938
refGene_gn=LIN54;refGene_tx=XM_003985227.4;refGene_gn=LIN54;refGene_tx=XM_006931004.3;refGene_gn=LIN54;refGene_tx=XM_006931005.4;refGene_gn=LIN54;refGene_tx=XM_011281747.2;refGene_gn=LIN54;refGene_tx=XM_023252980.1;refGene_gn=LIN54;refGene_tx=XM_023252981.1
B1 139783090 139783228
refGene_gn=LIN54;refGene_tx=XM_003985227.4;refGene_gn=LIN54;refGene_tx=XM_006931004.3;refGene_gn=LIN54;refGene_tx=XM_006931005.4;refGene_gn=LIN54;refGene_tx=XM_011281747.2;refGene_gn=LIN54;refGene_tx=XM_023252980.1;refGene_gn=LIN54;refGene_tx=XM_023252981.1
B1 139787406 139787609
refGene_gn=LIN54;refGene_tx=XM_003985227.4;refGene_gn=LIN54;refGene_tx=XM_006931004.3;refGene_gn=LIN54;refGene_tx=XM_006931005.4;refGene_gn=LIN54;refGene_tx=XM_011281747.2;refGene_gn=LIN54;refGene_tx=XM_023252980.1;refGene_gn=LIN54;refGene_tx=XM_023252981.1
B1 139789315 139789517
refGene_gn=LIN54;refGene_tx=XM_003985227.4;refGene_gn=LIN54;refGene_tx=XM_006931004.3;refGene_gn=LIN54;refGene_tx=XM_006931005.4;refGene_gn=LIN54;refGene_tx=XM_011281747.2;refGene_gn=LIN54;refGene_tx=XM_023252980.1;refGene_gn=LIN54;refGene_tx=XM_023252981.1
B1 139797522 139799500
refGene_gn=THAP9;refGene_tx=XM_003985230.4;refGene_gn=THAP9;refGene_tx=XM_019829306.1
B1 139807851 139808002
refGene_gn=THAP9;refGene_tx=XM_003985230.4;refGene_gn=THAP9;refGene_tx=XM_019829306.1
B1 139809418 139809722
refGene_gn=THAP9;refGene_tx=XM_003985230.4;refGene_gn=THAP9;refGene_tx=XM_019829306.1
B1 139811131 139811327
refGene_gn=THAP9;refGene_tx=XM_003985230.4
B1 139816845 139816925
refGene_gn=THAP9;refGene_tx=XM_003985230.4
B1 139839244 139839323
refGene_gn=SEC31A;refGene_tx=XM_003985231.5;refGene_gn=SEC31A;refGene_tx=XM_003985232.5;refGene_gn=SEC31A;refGene_tx=XM_003985233.5;refGene_gn=SEC31A;refGene_tx=XM_003985234.5;refGene_gn=SEC31A;refGene_tx=XM_003985236.5;refGene_gn=SEC31A;refGene_tx=XM_006931006.4;refGene_gn=SEC31A;refGene_tx=XM_006931013.4;refGene_gn=SEC31A;refGene_tx=XM_006931016.4;refGene_gn=SEC31A;refGene_tx=XM_006931017.4;refGene_gn=SEC31A;refGene_tx=XM_019829307.2;refGene_gn=SEC31A;refGene_tx=XM_019829308.2;refGen

e_gn=SEC31A;refGene_tx=XM_019829309.2;refGene_gn=SEC31A;refGene_tx=XM_019829310.1;refGene_gn=SEC31A;refGene_tx=XM_019829311.2;refGene_gn=SEC31A;refGene_tx=XM_019829312.2;refGene_gn=SEC31A;refGene_tx=XM_019829313.1;refGene_gn=SEC31A;refGene_tx=XM_019829314.2;refGene_gn=SEC31A;refGene_tx=XM_019829315.1;refGene_gn=SEC31A;refGene_tx=XM_019829316.2;refGene_gn=SEC31A;refGene_tx=XM_019829317.1;refGene_gn=SEC31A;refGene_tx=XM_019829318.2

B1 139839988 139840112

refGene_gn=SEC31A;refGene_tx=XM_003985231.5;refGene_gn=SEC31A;refGene_tx=XM_003985232.5;refGene_gn=SEC31A;refGene_tx=XM_003985233.5;refGene_gn=SEC31A;refGene_tx=XM_003985234.5;refGene_gn=SEC31A;refGene_tx=XM_003985236.5;refGene_gn=SEC31A;refGene_tx=XM_006931006.4;refGene_gn=SEC31A;refGene_tx=XM_006931013.4;refGene_gn=SEC31A;refGene_tx=XM_006931016.4;refGene_gn=SEC31A;refGene_tx=XM_006931017.4;refGene_gn=SEC31A;refGene_tx=XM_019829307.2;refGene_gn=SEC31A;refGene_tx=XM_019829308.2;refGene_gn=SEC31A;refGene_tx=XM_019829309.2;refGene_gn=SEC31A;refGene_tx=XM_019829310.1;refGene_gn=SEC31A;refGene_tx=XM_019829311.2;refGene_gn=SEC31A;refGene_tx=XM_019829312.2;refGene_gn=SEC31A;refGene_tx=XM_019829313.1;refGene_gn=SEC31A;refGene_tx=XM_019829314.2;refGene_gn=SEC31A;refGene_tx=XM_019829315.1;refGene_gn=SEC31A;refGene_tx=XM_019829316.2;refGene_gn=SEC31A;refGene_tx=XM_019829317.1;refGene_gn=SEC31A;refGene_tx=XM_019829318.2

B1 139841982 139842181

refGene_gn=SEC31A;refGene_tx=XM_003985231.5;refGene_gn=SEC31A;refGene_tx=XM_003985232.5;refGene_gn=SEC31A;refGene_tx=XM_003985233.5;refGene_gn=SEC31A;refGene_tx=XM_003985234.5;refGene_gn=SEC31A;refGene_tx=XM_003985236.5;refGene_gn=SEC31A;refGene_tx=XM_006931006.4;refGene_gn=SEC31A;refGene_tx=XM_006931013.4;refGene_gn=SEC31A;refGene_tx=XM_006931016.4;refGene_gn=SEC31A;refGene_tx=XM_006931017.4;refGene_gn=SEC31A;refGene_tx=XM_019829307.2;refGene_gn=SEC31A;refGene_tx=XM_019829308.2;refGene_gn=SEC31A;refGene_tx=XM_019829309.2;refGene_gn=SEC31A;refGene_tx=XM_019829310.1;refGene_gn=SEC31A;refGene_tx=XM_019829311.2;refGene_gn=SEC31A;refGene_tx=XM_019829312.2;refGene_gn=SEC31A;refGene_tx=XM_019829313.1;refGene_gn=SEC31A;refGene_tx=XM_019829314.2;refGene_gn=SEC31A;refGene_tx=XM_019829315.1;refGene_gn=SEC31A;refGene_tx=XM_019829316.2;refGene_gn=SEC31A;refGene_tx=XM_019829317.1;refGene_gn=SEC31A;refGene_tx=XM_019829318.2

B1 139844351 139844447

refGene_gn=SEC31A;refGene_tx=XM_003985231.5;refGene_gn=SEC31A;refGene_tx=XM_003985232.5;refGene_gn=SEC31A;refGene_tx=XM_003985233.5;refGene_gn=SEC31A;refGene_tx=XM_003985234.5;refGene_gn=SEC31A;refGene_tx=XM_003985236.5;refGene_gn=SEC31A;refGene_tx=XM_006931006.4;refGene_gn=SEC31A;refGene_tx=XM_006931013.4;refGene_gn=SEC31A;refGene_tx=XM_006931016.4;refGene_gn=SEC31A;refGene_tx=XM_006931017.4;refGene_gn=SEC31A;refGene_tx=XM_019829307.2;refGene_gn=SEC31A;refGene_tx=XM_019829308.2;refGene_gn=SEC31A;refGene_tx=XM_019829309.2;refGene_gn=SEC31A;refGene_tx=XM_019829310.1;refGene_gn=SEC31A;refGene_tx=XM_019829311.2;refGene_gn=SEC31A;refGene_tx=XM_019829312.2;refGene_gn=SEC31A;refGene_tx=XM_019829313.1;refGene_gn=SEC31A;refGene_tx=XM_019829314.2;refGene_gn=SEC31A;refGene_tx=XM_019829315.1;refGene_gn=SEC31A;refGene_tx=XM_019829316.2;refG

ene_gn=SEC31A;refGene_tx=XM_019829317.1;refGene_gn=SEC31A;refGene_tx=XM_019829318.2
B1 139845349 139845490
refGene_gn=SEC31A;refGene_tx=XM_003985231.5;refGene_gn=SEC31A;refGene_tx=XM_003985232.5;refGene_gn=SEC31A;refGene_tx=XM_003985233.5;refGene_gn=SEC31A;refGene_tx=XM_003985234.5;refGene_gn=SEC31A;refGene_tx=XM_003985236.5;refGene_gn=SEC31A;refGene_tx=XM_006931006.4;refGene_gn=SEC31A;refGene_tx=XM_006931013.4;refGene_gn=SEC31A;refGene_tx=XM_006931016.4;refGene_gn=SEC31A;refGene_tx=XM_006931017.4;refGene_gn=SEC31A;refGene_tx=XM_019829307.2;refGene_gn=SEC31A;refGene_tx=XM_019829308.2;refGene_gn=SEC31A;refGene_tx=XM_019829309.2;refGene_gn=SEC31A;refGene_tx=XM_019829310.1;refGene_gn=SEC31A;refGene_tx=XM_019829311.2;refGene_gn=SEC31A;refGene_tx=XM_019829312.2;refGene_gn=SEC31A;refGene_tx=XM_019829313.1;refGene_gn=SEC31A;refGene_tx=XM_019829314.2;refGene_gn=SEC31A;refGene_tx=XM_019829315.1;refGene_gn=SEC31A;refGene_tx=XM_019829316.2;refGene_gn=SEC31A;refGene_tx=XM_019829317.1;refGene_gn=SEC31A;refGene_tx=XM_019829318.2
B1 139846837 139846980
refGene_gn=SEC31A;refGene_tx=XM_003985231.5;refGene_gn=SEC31A;refGene_tx=XM_003985232.5;refGene_gn=SEC31A;refGene_tx=XM_003985233.5;refGene_gn=SEC31A;refGene_tx=XM_003985234.5;refGene_gn=SEC31A;refGene_tx=XM_003985236.5;refGene_gn=SEC31A;refGene_tx=XM_006931006.4;refGene_gn=SEC31A;refGene_tx=XM_006931013.4;refGene_gn=SEC31A;refGene_tx=XM_006931016.4;refGene_gn=SEC31A;refGene_tx=XM_006931017.4;refGene_gn=SEC31A;refGene_tx=XM_019829307.2;refGene_gn=SEC31A;refGene_tx=XM_019829308.2;refGene_gn=SEC31A;refGene_tx=XM_019829309.2;refGene_gn=SEC31A;refGene_tx=XM_019829310.1;refGene_gn=SEC31A;refGene_tx=XM_019829311.2;refGene_gn=SEC31A;refGene_tx=XM_019829312.2;refGene_gn=SEC31A;refGene_tx=XM_019829313.1;refGene_gn=SEC31A;refGene_tx=XM_019829314.2;refGene_gn=SEC31A;refGene_tx=XM_019829315.1;refGene_gn=SEC31A;refGene_tx=XM_019829316.2;refGene_gn=SEC31A;refGene_tx=XM_019829317.1;refGene_gn=SEC31A;refGene_tx=XM_019829318.2
B1 139850837 139850937
refGene_gn=SEC31A;refGene_tx=XM_003985231.5;refGene_gn=SEC31A;refGene_tx=XM_003985232.5;refGene_gn=SEC31A;refGene_tx=XM_003985233.5;refGene_gn=SEC31A;refGene_tx=XM_003985234.5;refGene_gn=SEC31A;refGene_tx=XM_003985236.5;refGene_gn=SEC31A;refGene_tx=XM_006931006.4;refGene_gn=SEC31A;refGene_tx=XM_006931013.4;refGene_gn=SEC31A;refGene_tx=XM_006931016.4;refGene_gn=SEC31A;refGene_tx=XM_006931017.4;refGene_gn=SEC31A;refGene_tx=XM_019829307.2;refGene_gn=SEC31A;refGene_tx=XM_019829308.2;refGene_gn=SEC31A;refGene_tx=XM_019829309.2;refGene_gn=SEC31A;refGene_tx=XM_019829310.1;refGene_gn=SEC31A;refGene_tx=XM_019829311.2;refGene_gn=SEC31A;refGene_tx=XM_019829312.2;refGene_gn=SEC31A;refGene_tx=XM_019829313.1;refGene_gn=SEC31A;refGene_tx=XM_019829314.2;refGene_gn=SEC31A;refGene_tx=XM_019829315.1;refGene_gn=SEC31A;refGene_tx=XM_019829316.2;refGene_gn=SEC31A;refGene_tx=XM_019829317.1;refGene_gn=SEC31A;refGene_tx=XM_019829318.2
B1 139853970 139854132
refGene_gn=SEC31A;refGene_tx=XM_003985231.5;refGene_gn=SEC31A;refGene_tx=XM_003985232.5;refGene_gn=SEC31A;refGene_tx=XM_003985233.5;refGene_

gn=SEC31A;refGene_tx=XM_003985234.5;refGene_gn=SEC31A;refGene_tx=XM_00
3985236.5;refGene_gn=SEC31A;refGene_tx=XM_006931006.4;refGene_gn=SEC31
A;refGene_tx=XM_006931013.4;refGene_gn=SEC31A;refGene_tx=XM_006931016.
4;refGene_gn=SEC31A;refGene_tx=XM_006931017.4;refGene_gn=SEC31A;refGen
e_tx=XM_019829307.2;refGene_gn=SEC31A;refGene_tx=XM_019829308.2;refGen
e_gn=SEC31A;refGene_tx=XM_019829309.2;refGene_gn=SEC31A;refGene_tx=XM_
019829310.1;refGene_gn=SEC31A;refGene_tx=XM_019829311.2;refGene_gn=SEC
31A;refGene_tx=XM_019829312.2;refGene_gn=SEC31A;refGene_tx=XM_01982931
3.1;refGene_gn=SEC31A;refGene_tx=XM_019829314.2;refGene_gn=SEC31A;refG
ene_tx=XM_019829315.1;refGene_gn=SEC31A;refGene_tx=XM_019829316.2;refG
ene_gn=SEC31A;refGene_tx=XM_019829317.1;refGene_gn=SEC31A;refGene_tx=X
M_019829318.2
B1 139854324 139854477
refGene_gn=SEC31A;refGene_tx=XM_003985231.5;refGene_gn=SEC31A;refGene_
tx=XM_003985232.5;refGene_gn=SEC31A;refGene_tx=XM_003985233.5;refGene_
gn=SEC31A;refGene_tx=XM_003985234.5;refGene_gn=SEC31A;refGene_tx=XM_00
3985236.5;refGene_gn=SEC31A;refGene_tx=XM_006931006.4;refGene_gn=SEC31
A;refGene_tx=XM_006931013.4;refGene_gn=SEC31A;refGene_tx=XM_006931016.
4;refGene_gn=SEC31A;refGene_tx=XM_006931017.4;refGene_gn=SEC31A;refGen
e_tx=XM_019829307.2;refGene_gn=SEC31A;refGene_tx=XM_019829308.2;refGen
e_gn=SEC31A;refGene_tx=XM_019829309.2;refGene_gn=SEC31A;refGene_tx=XM_
019829310.1;refGene_gn=SEC31A;refGene_tx=XM_019829311.2;refGene_gn=SEC
31A;refGene_tx=XM_019829312.2;refGene_gn=SEC31A;refGene_tx=XM_01982931
3.1;refGene_gn=SEC31A;refGene_tx=XM_019829314.2;refGene_gn=SEC31A;refG
ene_tx=XM_019829315.1;refGene_gn=SEC31A;refGene_tx=XM_019829316.2;refG
ene_gn=SEC31A;refGene_tx=XM_019829317.1;refGene_gn=SEC31A;refGene_tx=X
M_019829318.2
B1 139860226 139860463
refGene_gn=SEC31A;refGene_tx=XM_003985231.5;refGene_gn=SEC31A;refGene_
tx=XM_003985232.5;refGene_gn=SEC31A;refGene_tx=XM_003985233.5;refGene_
gn=SEC31A;refGene_tx=XM_003985234.5;refGene_gn=SEC31A;refGene_tx=XM_00
3985236.5;refGene_gn=SEC31A;refGene_tx=XM_006931006.4;refGene_gn=SEC31
A;refGene_tx=XM_006931013.4;refGene_gn=SEC31A;refGene_tx=XM_006931016.
4;refGene_gn=SEC31A;refGene_tx=XM_006931017.4;refGene_gn=SEC31A;refGen
e_tx=XM_019829307.2;refGene_gn=SEC31A;refGene_tx=XM_019829308.2;refGen
e_gn=SEC31A;refGene_tx=XM_019829309.2;refGene_gn=SEC31A;refGene_tx=XM_
019829310.1;refGene_gn=SEC31A;refGene_tx=XM_019829311.2;refGene_gn=SEC
31A;refGene_tx=XM_019829312.2;refGene_gn=SEC31A;refGene_tx=XM_01982931
3.1;refGene_gn=SEC31A;refGene_tx=XM_019829314.2;refGene_gn=SEC31A;refG
ene_tx=XM_019829315.1;refGene_gn=SEC31A;refGene_tx=XM_019829316.2;refG
ene_gn=SEC31A;refGene_tx=XM_019829317.1;refGene_gn=SEC31A;refGene_tx=X
M_019829318.2
B1 139861468 139861543
refGene_gn=SEC31A;refGene_tx=XM_003985231.5;refGene_gn=SEC31A;refGene_
tx=XM_003985232.5;refGene_gn=SEC31A;refGene_tx=XM_003985233.5;refGene_
gn=SEC31A;refGene_tx=XM_003985234.5;refGene_gn=SEC31A;refGene_tx=XM_00
3985236.5;refGene_gn=SEC31A;refGene_tx=XM_006931006.4;refGene_gn=SEC31
A;refGene_tx=XM_006931013.4;refGene_gn=SEC31A;refGene_tx=XM_006931016.
4;refGene_gn=SEC31A;refGene_tx=XM_006931017.4;refGene_gn=SEC31A;refGen
e_tx=XM_019829307.2;refGene_gn=SEC31A;refGene_tx=XM_019829308.2;refGen

e_gn=SEC31A;refGene_tx=XM_019829309.2;refGene_gn=SEC31A;refGene_tx=XM_
019829310.1;refGene_gn=SEC31A;refGene_tx=XM_019829311.2;refGene_gn=SEC
31A;refGene_tx=XM_019829312.2;refGene_gn=SEC31A;refGene_tx=XM_01982931
3.1;refGene_gn=SEC31A;refGene_tx=XM_019829314.2;refGene_gn=SEC31A;refG
ene_tx=XM_019829315.1;refGene_gn=SEC31A;refGene_tx=XM_019829316.2;refG
ene_gn=SEC31A;refGene_tx=XM_019829317.1;refGene_gn=SEC31A;refGene_tx=X
M_019829318.2
B1 139863618 139863657
refGene_gn=SEC31A;refGene_tx=XM_003985231.5;refGene_gn=SEC31A;refGene_
tx=XM_003985232.5;refGene_gn=SEC31A;refGene_tx=XM_003985233.5;refGene_
gn=SEC31A;refGene_tx=XM_006931006.4;refGene_gn=SEC31A;refGene_tx=XM_00
6931013.4;refGene_gn=SEC31A;refGene_tx=XM_006931016.4;refGene_gn=SEC31
A;refGene_tx=XM_019829307.2;refGene_gn=SEC31A;refGene_tx=XM_019829308.
2;refGene_gn=SEC31A;refGene_tx=XM_019829309.2;refGene_gn=SEC31A;refGen
e_tx=XM_019829310.1;refGene_gn=SEC31A;refGene_tx=XM_019829311.2;refGen
e_gn=SEC31A;refGene_tx=XM_019829312.2;refGene_gn=SEC31A;refGene_tx=XM_
019829314.2;refGene_gn=SEC31A;refGene_tx=XM_019829315.1;refGene_gn=SEC
31A;refGene_tx=XM_019829316.2;refGene_gn=SEC31A;refGene_tx=XM_01982931
8.2
B1 139864163 139864241
refGene_gn=SEC31A;refGene_tx=XM_003985231.5;refGene_gn=SEC31A;refGene_
tx=XM_003985232.5;refGene_gn=SEC31A;refGene_tx=XM_003985233.5;refGene_
gn=SEC31A;refGene_tx=XM_006931006.4;refGene_gn=SEC31A;refGene_tx=XM_00
6931013.4;refGene_gn=SEC31A;refGene_tx=XM_006931016.4;refGene_gn=SEC31
A;refGene_tx=XM_019829307.2;refGene_gn=SEC31A;refGene_tx=XM_019829308.
2;refGene_gn=SEC31A;refGene_tx=XM_019829309.2;refGene_gn=SEC31A;refGen
e_tx=XM_019829310.1;refGene_gn=SEC31A;refGene_tx=XM_019829311.2;refGen
e_gn=SEC31A;refGene_tx=XM_019829312.2;refGene_gn=SEC31A;refGene_tx=XM_
019829313.1;refGene_gn=SEC31A;refGene_tx=XM_019829314.2;refGene_gn=SEC
31A;refGene_tx=XM_019829316.2;refGene_gn=SEC31A;refGene_tx=XM_01982931
8.2
B1 139866617 139866693
refGene_gn=SEC31A;refGene_tx=XM_003985231.5;refGene_gn=SEC31A;refGene_
tx=XM_003985232.5;refGene_gn=SEC31A;refGene_tx=XM_003985233.5;refGene_
gn=SEC31A;refGene_tx=XM_003985234.5;refGene_gn=SEC31A;refGene_tx=XM_00
3985236.5;refGene_gn=SEC31A;refGene_tx=XM_006931006.4;refGene_gn=SEC31
A;refGene_tx=XM_006931013.4;refGene_gn=SEC31A;refGene_tx=XM_006931016.
4;refGene_gn=SEC31A;refGene_tx=XM_006931017.4;refGene_gn=SEC31A;refGen
e_tx=XM_019829307.2;refGene_gn=SEC31A;refGene_tx=XM_019829308.2;refGen
e_gn=SEC31A;refGene_tx=XM_019829309.2;refGene_gn=SEC31A;refGene_tx=XM_
019829310.1;refGene_gn=SEC31A;refGene_tx=XM_019829311.2;refGene_gn=SEC
31A;refGene_tx=XM_019829312.2;refGene_gn=SEC31A;refGene_tx=XM_01982931
3.1;refGene_gn=SEC31A;refGene_tx=XM_019829314.2;refGene_gn=SEC31A;refG
ene_tx=XM_019829315.1;refGene_gn=SEC31A;refGene_tx=XM_019829316.2;refG
ene_gn=SEC31A;refGene_tx=XM_019829317.1;refGene_gn=SEC31A;refGene_tx=X
M_019829318.2
B1 139867318 139867497
refGene_gn=SEC31A;refGene_tx=XM_003985231.5;refGene_gn=SEC31A;refGene_
tx=XM_003985232.5;refGene_gn=SEC31A;refGene_tx=XM_003985233.5;refGene_
gn=SEC31A;refGene_tx=XM_003985234.5;refGene_gn=SEC31A;refGene_tx=XM_00

3985236.5;refGene_gn=SEC31A;refGene_tx=XM_006931006.4;refGene_gn=SEC31
A;refGene_tx=XM_006931013.4;refGene_gn=SEC31A;refGene_tx=XM_006931016.
4;refGene_gn=SEC31A;refGene_tx=XM_006931017.4;refGene_gn=SEC31A;refGen
e_tx=XM_019829307.2;refGene_gn=SEC31A;refGene_tx=XM_019829308.2;refGen
e_gn=SEC31A;refGene_tx=XM_019829309.2;refGene_gn=SEC31A;refGene_tx=XM_
019829310.1;refGene_gn=SEC31A;refGene_tx=XM_019829311.2;refGene_gn=SEC
31A;refGene_tx=XM_019829312.2;refGene_gn=SEC31A;refGene_tx=XM_01982931
3.1;refGene_gn=SEC31A;refGene_tx=XM_019829314.2;refGene_gn=SEC31A;refG
ene_tx=XM_019829315.1;refGene_gn=SEC31A;refGene_tx=XM_019829316.2;refG
ene_gn=SEC31A;refGene_tx=XM_019829317.1;refGene_gn=SEC31A;refGene_tx=X
M_019829318.2
B1 139868384 139868511
refGene_gn=SEC31A;refGene_tx=XM_003985231.5;refGene_gn=SEC31A;refGene_
tx=XM_003985232.5;refGene_gn=SEC31A;refGene_tx=XM_003985233.5;refGene_
gn=SEC31A;refGene_tx=XM_003985234.5;refGene_gn=SEC31A;refGene_tx=XM_00
3985236.5;refGene_gn=SEC31A;refGene_tx=XM_006931006.4;refGene_gn=SEC31
A;refGene_tx=XM_006931013.4;refGene_gn=SEC31A;refGene_tx=XM_006931016.
4;refGene_gn=SEC31A;refGene_tx=XM_006931017.4;refGene_gn=SEC31A;refGen
e_tx=XM_019829307.2;refGene_gn=SEC31A;refGene_tx=XM_019829308.2;refGen
e_gn=SEC31A;refGene_tx=XM_019829309.2;refGene_gn=SEC31A;refGene_tx=XM_
019829310.1;refGene_gn=SEC31A;refGene_tx=XM_019829311.2;refGene_gn=SEC
31A;refGene_tx=XM_019829312.2;refGene_gn=SEC31A;refGene_tx=XM_01982931
3.1;refGene_gn=SEC31A;refGene_tx=XM_019829314.2;refGene_gn=SEC31A;refG
ene_tx=XM_019829315.1;refGene_gn=SEC31A;refGene_tx=XM_019829316.2;refG
ene_gn=SEC31A;refGene_tx=XM_019829317.1;refGene_gn=SEC31A;refGene_tx=X
M_019829318.2
B1 139869470 139869616
refGene_gn=SEC31A;refGene_tx=XM_003985231.5;refGene_gn=SEC31A;refGene_
tx=XM_003985232.5;refGene_gn=SEC31A;refGene_tx=XM_003985233.5;refGene_
gn=SEC31A;refGene_tx=XM_003985234.5;refGene_gn=SEC31A;refGene_tx=XM_00
3985236.5;refGene_gn=SEC31A;refGene_tx=XM_006931006.4;refGene_gn=SEC31
A;refGene_tx=XM_006931013.4;refGene_gn=SEC31A;refGene_tx=XM_006931016.
4;refGene_gn=SEC31A;refGene_tx=XM_006931017.4;refGene_gn=SEC31A;refGen
e_tx=XM_019829307.2;refGene_gn=SEC31A;refGene_tx=XM_019829308.2;refGen
e_gn=SEC31A;refGene_tx=XM_019829309.2;refGene_gn=SEC31A;refGene_tx=XM_
019829310.1;refGene_gn=SEC31A;refGene_tx=XM_019829311.2;refGene_gn=SEC
31A;refGene_tx=XM_019829312.2;refGene_gn=SEC31A;refGene_tx=XM_01982931
3.1;refGene_gn=SEC31A;refGene_tx=XM_019829314.2;refGene_gn=SEC31A;refG
ene_tx=XM_019829315.1;refGene_gn=SEC31A;refGene_tx=XM_019829316.2;refG
ene_gn=SEC31A;refGene_tx=XM_019829317.1;refGene_gn=SEC31A;refGene_tx=X
M_019829318.2
B1 139870890 139871064
refGene_gn=SEC31A;refGene_tx=XM_003985231.5;refGene_gn=SEC31A;refGene_
tx=XM_003985232.5;refGene_gn=SEC31A;refGene_tx=XM_003985233.5;refGene_
gn=SEC31A;refGene_tx=XM_003985234.5;refGene_gn=SEC31A;refGene_tx=XM_00
3985236.5;refGene_gn=SEC31A;refGene_tx=XM_006931006.4;refGene_gn=SEC31
A;refGene_tx=XM_006931013.4;refGene_gn=SEC31A;refGene_tx=XM_006931016.
4;refGene_gn=SEC31A;refGene_tx=XM_006931017.4;refGene_gn=SEC31A;refGen
e_tx=XM_019829307.2;refGene_gn=SEC31A;refGene_tx=XM_019829308.2;refGen
e_gn=SEC31A;refGene_tx=XM_019829309.2;refGene_gn=SEC31A;refGene_tx=XM_

019829310.1;refGene_gn=SEC31A;refGene_tx=XM_019829311.2;refGene_gn=SEC31A;refGene_tx=XM_019829312.2;refGene_gn=SEC31A;refGene_tx=XM_019829313.1;refGene_gn=SEC31A;refGene_tx=XM_019829314.2;refGene_gn=SEC31A;refGene_tx=XM_019829315.1;refGene_gn=SEC31A;refGene_tx=XM_019829316.2;refGene_gn=SEC31A;refGene_tx=XM_019829317.1;refGene_gn=SEC31A;refGene_tx=XM_019829318.2
B1 139873406 139873580
refGene_gn=SEC31A;refGene_tx=XM_003985231.5;refGene_gn=SEC31A;refGene_tx=XM_003985232.5;refGene_gn=SEC31A;refGene_tx=XM_003985233.5;refGene_gn=SEC31A;refGene_tx=XM_003985234.5;refGene_gn=SEC31A;refGene_tx=XM_003985236.5;refGene_gn=SEC31A;refGene_tx=XM_006931006.4;refGene_gn=SEC31A;refGene_tx=XM_006931013.4;refGene_gn=SEC31A;refGene_tx=XM_006931016.4;refGene_gn=SEC31A;refGene_tx=XM_006931017.4;refGene_gn=SEC31A;refGene_tx=XM_019829307.2;refGene_gn=SEC31A;refGene_tx=XM_019829308.2;refGene_gn=SEC31A;refGene_tx=XM_019829309.2;refGene_gn=SEC31A;refGene_tx=XM_019829310.1;refGene_gn=SEC31A;refGene_tx=XM_019829311.2;refGene_gn=SEC31A;refGene_tx=XM_019829312.2;refGene_gn=SEC31A;refGene_tx=XM_019829313.1;refGene_gn=SEC31A;refGene_tx=XM_019829314.2;refGene_gn=SEC31A;refGene_tx=XM_019829315.1;refGene_gn=SEC31A;refGene_tx=XM_019829316.2;refGene_gn=SEC31A;refGene_tx=XM_019829317.1;refGene_gn=SEC31A;refGene_tx=XM_019829318.2
B1 139876557 139876650
refGene_gn=SEC31A;refGene_tx=XM_006931006.4;refGene_gn=SEC31A;refGene_tx=XM_006931013.4;refGene_gn=SEC31A;refGene_tx=XM_006931016.4;refGene_gn=SEC31A;refGene_tx=XM_019829307.2;refGene_gn=SEC31A;refGene_tx=XM_019829308.2;refGene_gn=SEC31A;refGene_tx=XM_019829309.2;refGene_gn=SEC31A;refGene_tx=XM_019829310.1;refGene_gn=SEC31A;refGene_tx=XM_019829311.2;refGene_gn=SEC31A;refGene_tx=XM_019829312.2;refGene_gn=SEC31A;refGene_tx=XM_019829313.1;refGene_gn=SEC31A;refGene_tx=XM_019829314.2;refGene_gn=SEC31A;refGene_tx=XM_019829315.1;refGene_gn=SEC31A;refGene_tx=XM_019829317.1;refGene_gn=SEC31A;refGene_tx=XM_019829318.2
B1 139877325 139877452
refGene_gn=SEC31A;refGene_tx=XM_003985231.5;refGene_gn=SEC31A;refGene_tx=XM_003985232.5;refGene_gn=SEC31A;refGene_tx=XM_003985233.5;refGene_gn=SEC31A;refGene_tx=XM_003985234.5;refGene_gn=SEC31A;refGene_tx=XM_003985236.5;refGene_gn=SEC31A;refGene_tx=XM_006931006.4;refGene_gn=SEC31A;refGene_tx=XM_006931013.4;refGene_gn=SEC31A;refGene_tx=XM_006931016.4;refGene_gn=SEC31A;refGene_tx=XM_006931017.4;refGene_gn=SEC31A;refGene_tx=XM_019829307.2;refGene_gn=SEC31A;refGene_tx=XM_019829308.2;refGene_gn=SEC31A;refGene_tx=XM_019829309.2;refGene_gn=SEC31A;refGene_tx=XM_019829310.1;refGene_gn=SEC31A;refGene_tx=XM_019829311.2;refGene_gn=SEC31A;refGene_tx=XM_019829312.2;refGene_gn=SEC31A;refGene_tx=XM_019829313.1;refGene_gn=SEC31A;refGene_tx=XM_019829314.2;refGene_gn=SEC31A;refGene_tx=XM_019829315.1;refGene_gn=SEC31A;refGene_tx=XM_019829316.2;refGene_gn=SEC31A;refGene_tx=XM_019829317.1;refGene_gn=SEC31A;refGene_tx=XM_019829318.2
B1 139879017 139879359
refGene_gn=SEC31A;refGene_tx=XM_003985231.5;refGene_gn=SEC31A;refGene_tx=XM_003985232.5;refGene_gn=SEC31A;refGene_tx=XM_003985234.5;refGene_gn=SEC31A;refGene_tx=XM_003985236.5;refGene_gn=SEC31A;refGene_tx=XM_00

6931006.4;refGene_gn=SEC31A;refGene_tx=XM_006931013.4;refGene_gn=SEC31A;refGene_tx=XM_019829307.2;refGene_gn=SEC31A;refGene_tx=XM_019829308.2;refGene_gn=SEC31A;refGene_tx=XM_019829309.2;refGene_gn=SEC31A;refGene_tx=XM_019829310.1;refGene_gn=SEC31A;refGene_tx=XM_019829311.2;refGene_gn=SEC31A;refGene_tx=XM_019829312.2;refGene_gn=SEC31A;refGene_tx=XM_019829313.1;refGene_gn=SEC31A;refGene_tx=XM_019829314.2;refGene_gn=SEC31A;refGene_tx=XM_019829315.1;refGene_gn=SEC31A;refGene_tx=XM_019829316.2;refGene_gn=SEC31A;refGene_tx=XM_019829317.1;refGene_gn=SEC31A;refGene_tx=XM_019829318.2
B1 139885915 139885954
refGene_gn=SEC31A;refGene_tx=XM_019829307.2;refGene_gn=SEC31A;refGene_tx=XM_019829308.2;refGene_gn=SEC31A;refGene_tx=XM_019829309.2;refGene_gn=SEC31A;refGene_tx=XM_019829310.1;refGene_gn=SEC31A;refGene_tx=XM_019829311.2;refGene_gn=SEC31A;refGene_tx=XM_019829312.2;refGene_gn=SEC31A;refGene_tx=XM_019829313.1;refGene_gn=SEC31A;refGene_tx=XM_019829314.2;refGene_gn=SEC31A;refGene_tx=XM_019829315.1;refGene_gn=SEC31A;refGene_tx=XM_019829316.2;refGene_gn=SEC31A;refGene_tx=XM_019829317.1;refGene_gn=SEC31A;refGene_tx=XM_019829318.2
B1 139887594 139887653
refGene_gn=SEC31A;refGene_tx=XM_003985231.5;refGene_gn=SEC31A;refGene_tx=XM_003985232.5;refGene_gn=SEC31A;refGene_tx=XM_003985233.5;refGene_gn=SEC31A;refGene_tx=XM_003985234.5;refGene_gn=SEC31A;refGene_tx=XM_003985236.5;refGene_gn=SEC31A;refGene_tx=XM_006931006.4;refGene_gn=SEC31A;refGene_tx=XM_006931013.4;refGene_gn=SEC31A;refGene_tx=XM_006931016.4;refGene_gn=SEC31A;refGene_tx=XM_006931017.4;refGene_gn=SEC31A;refGene_tx=XM_019829307.2;refGene_gn=SEC31A;refGene_tx=XM_019829308.2;refGene_gn=SEC31A;refGene_tx=XM_019829309.2;refGene_gn=SEC31A;refGene_tx=XM_019829310.1;refGene_gn=SEC31A;refGene_tx=XM_019829311.2;refGene_gn=SEC31A;refGene_tx=XM_019829312.2;refGene_gn=SEC31A;refGene_tx=XM_019829313.1;refGene_gn=SEC31A;refGene_tx=XM_019829314.2;refGene_gn=SEC31A;refGene_tx=XM_019829315.1;refGene_gn=SEC31A;refGene_tx=XM_019829316.2;refGene_gn=SEC31A;refGene_tx=XM_019829317.1;refGene_gn=SEC31A;refGene_tx=XM_019829318.2
B1 139888882 139889146
refGene_gn=SEC31A;refGene_tx=XM_003985231.5;refGene_gn=SEC31A;refGene_tx=XM_003985232.5;refGene_gn=SEC31A;refGene_tx=XM_003985233.5;refGene_gn=SEC31A;refGene_tx=XM_003985234.5;refGene_gn=SEC31A;refGene_tx=XM_003985236.5;refGene_gn=SEC31A;refGene_tx=XM_006931006.4;refGene_gn=SEC31A;refGene_tx=XM_006931013.4;refGene_gn=SEC31A;refGene_tx=XM_006931016.4;refGene_gn=SEC31A;refGene_tx=XM_006931017.4;refGene_gn=SEC31A;refGene_tx=XM_019829307.2;refGene_gn=SEC31A;refGene_tx=XM_019829308.2;refGene_gn=SEC31A;refGene_tx=XM_019829309.2;refGene_gn=SEC31A;refGene_tx=XM_019829310.1;refGene_gn=SEC31A;refGene_tx=XM_019829311.2;refGene_gn=SEC31A;refGene_tx=XM_019829312.2;refGene_gn=SEC31A;refGene_tx=XM_019829313.1;refGene_gn=SEC31A;refGene_tx=XM_019829314.2;refGene_gn=SEC31A;refGene_tx=XM_019829315.1;refGene_gn=SEC31A;refGene_tx=XM_019829316.2;refGene_gn=SEC31A;refGene_tx=XM_019829317.1;refGene_gn=SEC31A;refGene_tx=XM_019829318.2
B1 139891656 139891776
refGene_gn=SEC31A;refGene_tx=XM_003985231.5;refGene_gn=SEC31A;refGene_

tx=XM_003985232.5;refGene_gn=SEC31A;refGene_tx=XM_003985233.5;refGene_
gn=SEC31A;refGene_tx=XM_003985234.5;refGene_gn=SEC31A;refGene_tx=XM_00
3985236.5;refGene_gn=SEC31A;refGene_tx=XM_006931006.4;refGene_gn=SEC31
A;refGene_tx=XM_006931013.4;refGene_gn=SEC31A;refGene_tx=XM_006931016.
4;refGene_gn=SEC31A;refGene_tx=XM_006931017.4;refGene_gn=SEC31A;refGen
e_tx=XM_019829307.2;refGene_gn=SEC31A;refGene_tx=XM_019829308.2;refGen
e_gn=SEC31A;refGene_tx=XM_019829309.2;refGene_gn=SEC31A;refGene_tx=XM_
019829310.1;refGene_gn=SEC31A;refGene_tx=XM_019829311.2;refGene_gn=SEC
31A;refGene_tx=XM_019829312.2;refGene_gn=SEC31A;refGene_tx=XM_01982931
3.1;refGene_gn=SEC31A;refGene_tx=XM_019829314.2;refGene_gn=SEC31A;refG
ene_tx=XM_019829315.1;refGene_gn=SEC31A;refGene_tx=XM_019829316.2;refG
ene_gn=SEC31A;refGene_tx=XM_019829317.1
B1 139896945 139897017
refGene_gn=SEC31A;refGene_tx=XM_003985231.5;refGene_gn=SEC31A;refGene_
tx=XM_003985232.5;refGene_gn=SEC31A;refGene_tx=XM_003985233.5;refGene_
gn=SEC31A;refGene_tx=XM_003985234.5;refGene_gn=SEC31A;refGene_tx=XM_00
3985236.5;refGene_gn=SEC31A;refGene_tx=XM_006931006.4;refGene_gn=SEC31
A;refGene_tx=XM_006931013.4;refGene_gn=SEC31A;refGene_tx=XM_006931016.
4;refGene_gn=SEC31A;refGene_tx=XM_006931017.4;refGene_gn=SEC31A;refGen
e_tx=XM_019829307.2;refGene_gn=SEC31A;refGene_tx=XM_019829308.2;refGen
e_gn=SEC31A;refGene_tx=XM_019829309.2;refGene_gn=SEC31A;refGene_tx=XM_
019829310.1;refGene_gn=SEC31A;refGene_tx=XM_019829311.2;refGene_gn=SEC
31A;refGene_tx=XM_019829312.2;refGene_gn=SEC31A;refGene_tx=XM_01982931
3.1;refGene_gn=SEC31A;refGene_tx=XM_019829314.2;refGene_gn=SEC31A;refG
ene_tx=XM_019829315.1;refGene_gn=SEC31A;refGene_tx=XM_019829316.2;refG
ene_gn=SEC31A;refGene_tx=XM_019829317.1;refGene_gn=SEC31A;refGene_tx=X
M_019829318.2
B1 139898078 139898258
refGene_gn=SEC31A;refGene_tx=XM_003985231.5;refGene_gn=SEC31A;refGene_
tx=XM_003985232.5;refGene_gn=SEC31A;refGene_tx=XM_003985233.5;refGene_
gn=SEC31A;refGene_tx=XM_003985234.5;refGene_gn=SEC31A;refGene_tx=XM_00
3985236.5;refGene_gn=SEC31A;refGene_tx=XM_006931006.4;refGene_gn=SEC31
A;refGene_tx=XM_006931013.4;refGene_gn=SEC31A;refGene_tx=XM_006931016.
4;refGene_gn=SEC31A;refGene_tx=XM_006931017.4;refGene_gn=SEC31A;refGen
e_tx=XM_019829307.2;refGene_gn=SEC31A;refGene_tx=XM_019829308.2;refGen
e_gn=SEC31A;refGene_tx=XM_019829309.2;refGene_gn=SEC31A;refGene_tx=XM_
019829310.1;refGene_gn=SEC31A;refGene_tx=XM_019829311.2;refGene_gn=SEC
31A;refGene_tx=XM_019829312.2;refGene_gn=SEC31A;refGene_tx=XM_01982931
3.1;refGene_gn=SEC31A;refGene_tx=XM_019829314.2;refGene_gn=SEC31A;refG
ene_tx=XM_019829315.1;refGene_gn=SEC31A;refGene_tx=XM_019829316.2;refG
ene_gn=SEC31A;refGene_tx=XM_019829317.1;refGene_gn=SEC31A;refGene_tx=X
M_019829318.2
B1 139915220 139915452
refGene_gn=SCD5;refGene_tx=XM_023252983.1
B1 140013100 140013231
refGene_gn=SCD5;refGene_tx=XM_023252983.1
B1 140032943 140033149
refGene_gn=SCD5;refGene_tx=XM_023252983.1
B1 140066626 140066859
refGene_gn=SCD5;refGene_tx=XM_023252983.1

B1	140070951	140071142
refGene_gn=SCD5;refGene_tx=XM_023252983.1
B1	140190352	140190432
refGene_gn=TMEM150C;refGene_tx=XM_003985237.5;refGene_gn=TMEM150C;refGene_tx=XM_023252984.1
B1	140191275	140191329
refGene_gn=TMEM150C;refGene_tx=XM_003985237.5;refGene_gn=TMEM150C;refGene_tx=XM_023252984.1
B1	140191445	140191478
refGene_gn=TMEM150C;refGene_tx=XM_003985237.5;refGene_gn=TMEM150C;refGene_tx=XM_023252984.1
B1	140192173	140192241
refGene_gn=TMEM150C;refGene_tx=XM_003985237.5;refGene_gn=TMEM150C;refGene_tx=XM_023252984.1
B1	140199191	140199319
refGene_gn=TMEM150C;refGene_tx=XM_003985237.5;refGene_gn=TMEM150C;refGene_tx=XM_023252984.1
B1	140202758	140202936
refGene_gn=TMEM150C;refGene_tx=XM_003985237.5;refGene_gn=TMEM150C;refGene_tx=XM_023252984.1
B1	140205683	140205892
refGene_gn=TMEM150C;refGene_tx=XM_003985237.5;refGene_gn=TMEM150C;refGene_tx=XM_023252984.1
B1	140222731	140222871
refGene_gn=ENOPH1;refGene_tx=XM_004001353.5;refGene_gn=ENOPH1;refGene_tx=XM_023252987.1;refGene_gn=ENOPH1;refGene_tx=XM_023252988.1;refGene_gn=ENOPH1;refGene_tx=XM_023252989.1
B1	140228617	140228741
refGene_gn=ENOPH1;refGene_tx=XM_004001353.5;refGene_gn=ENOPH1;refGene_tx=XM_023252987.1;refGene_gn=ENOPH1;refGene_tx=XM_023252988.1;refGene_gn=ENOPH1;refGene_tx=XM_023252989.1
B1	140230594	140230727
refGene_gn=ENOPH1;refGene_tx=XM_004001353.5;refGene_gn=ENOPH1;refGene_tx=XM_023252987.1;refGene_gn=ENOPH1;refGene_tx=XM_023252988.1;refGene_gn=ENOPH1;refGene_tx=XM_023252989.1
B1	140233048	140233251
refGene_gn=ENOPH1;refGene_tx=XM_004001353.5;refGene_gn=ENOPH1;refGene_tx=XM_023252987.1;refGene_gn=ENOPH1;refGene_tx=XM_023252988.1;refGene_gn=ENOPH1;refGene_tx=XM_023252989.1
B1	140234422	140234428
refGene_gn=LOC101088078;refGene_tx=XM_023252985.1
B1	140234494	140235316
refGene_gn=LOC101088078;refGene_tx=XM_023252985.1
B1	140235317	140235500
refGene_gn=LOC101088078;refGene_tx=XM_023252985.1
B1	140236550	140236652
refGene_gn=ENOPH1;refGene_tx=XM_004001353.5
B1	140258672	140258756
refGene_gn=ENOPH1;refGene_tx=XM_004001353.5;refGene_gn=ENOPH1;refGene_tx=XM_023252987.1

B1	140259930	140260373
refGene_gn=HNRNPDL;refGene_tx=XM_003985238.5
B1	140261157	140261326
refGene_gn=HNRNPDL;refGene_tx=XM_003985238.5
B1	140261424	140261586
refGene_gn=HNRNPDL;refGene_tx=XM_003985238.5
B1	140262032	140262164
refGene_gn=HNRNPDL;refGene_tx=XM_003985238.5
B1	140262309	140262424
refGene_gn=HNRNPDL;refGene_tx=XM_003985238.5
B1	140263138	140263197
refGene_gn=HNRNPDL;refGene_tx=XM_003985238.5
B1	140263198	140263310
refGene_gn=HNRNPDL;refGene_tx=XM_003985238.5
B1	140263647	140263718
refGene_gn=HNRNPDL;refGene_tx=XM_003985238.5
B1	140315357	140315590
refGene_gn=HNRNPD;refGene_tx=XM_003985246.5;refGene_gn=HNRNPD;refGene_tx=XM_023252990.1;refGene_gn=HNRNPD;refGene_tx=XM_023252991.1;refGene_gn=HNRNPD;refGene_tx=XM_023252992.1
B1	140317436	140317493
refGene_gn=HNRNPD;refGene_tx=XM_003985246.5;refGene_gn=HNRNPD;refGene_tx=XM_023252991.1
B1	140327989	140328158
refGene_gn=HNRNPD;refGene_tx=XM_003985246.5;refGene_gn=HNRNPD;refGene_tx=XM_023252990.1;refGene_gn=HNRNPD;refGene_tx=XM_023252991.1;refGene_gn=HNRNPD;refGene_tx=XM_023252992.1
B1	140328727	140328889
refGene_gn=HNRNPD;refGene_tx=XM_003985246.5;refGene_gn=HNRNPD;refGene_tx=XM_023252990.1;refGene_gn=HNRNPD;refGene_tx=XM_023252991.1;refGene_gn=HNRNPD;refGene_tx=XM_023252992.1
B1	140329933	140330065
refGene_gn=HNRNPD;refGene_tx=XM_003985246.5;refGene_gn=HNRNPD;refGene_tx=XM_023252990.1;refGene_gn=HNRNPD;refGene_tx=XM_023252991.1;refGene_gn=HNRNPD;refGene_tx=XM_023252992.1
B1	140330524	140330624
refGene_gn=HNRNPD;refGene_tx=XM_003985246.5;refGene_gn=HNRNPD;refGene_tx=XM_023252990.1;refGene_gn=HNRNPD;refGene_tx=XM_023252991.1;refGene_gn=HNRNPD;refGene_tx=XM_023252992.1
B1	140330732	140330879
refGene_gn=HNRNPD;refGene_tx=XM_003985246.5;refGene_gn=HNRNPD;refGene_tx=XM_023252990.1
B1	140331984	140332052
refGene_gn=HNRNPD;refGene_tx=XM_003985246.5;refGene_gn=HNRNPD;refGene_tx=XM_023252990.1;refGene_gn=HNRNPD;refGene_tx=XM_023252991.1;refGene_gn=HNRNPD;refGene_tx=XM_023252992.1
B1	140343330	140343342
refGene_gn=LOC102901761;refGene_tx=XM_006931023.4
B1	140344930	140344957
refGene_gn=LOC102901761;refGene_tx=XM_006931022.3

B1 140344985 140345123
refGene_gn=LOC102901761;refGene_tx=XM_006931021.3;refGene_gn=LOC102901761;refGene_tx=XM_006931022.3;refGene_gn=LOC102901761;refGene_tx=XM_006931023.4;refGene_gn=LOC102901761;refGene_tx=XM_023252994.1;refGene_gn=LOC102901761;refGene_tx=XM_023252995.1
B1 140355493 140355652
refGene_gn=LOC102901761;refGene_tx=XM_006931021.3;refGene_gn=LOC102901761;refGene_tx=XM_006931022.3;refGene_gn=LOC102901761;refGene_tx=XM_006931023.4;refGene_gn=LOC102901761;refGene_tx=XM_023252994.1;refGene_gn=LOC102901761;refGene_tx=XM_023252995.1
B1 140366006 140366108
refGene_gn=LOC102901761;refGene_tx=XM_006931021.3;refGene_gn=LOC102901761;refGene_tx=XM_006931022.3;refGene_gn=LOC102901761;refGene_tx=XM_006931023.4;refGene_gn=LOC102901761;refGene_tx=XM_023252994.1
B1 140368326 140368423
refGene_gn=LOC102901761;refGene_tx=XM_006931021.3;refGene_gn=LOC102901761;refGene_tx=XM_006931022.3;refGene_gn=LOC102901761;refGene_tx=XM_006931023.4;refGene_gn=LOC102901761;refGene_tx=XM_023252994.1;refGene_gn=LOC102901761;refGene_tx=XM_023252995.1
B1 140372747 140372818
refGene_gn=LOC102901761;refGene_tx=XM_006931021.3;refGene_gn=LOC102901761;refGene_tx=XM_006931022.3;refGene_gn=LOC102901761;refGene_tx=XM_006931023.4;refGene_gn=LOC102901761;refGene_tx=XM_023252994.1;refGene_gn=LOC102901761;refGene_tx=XM_023252995.1
B1 140377896 140377974
refGene_gn=LOC102901761;refGene_tx=XM_006931021.3;refGene_gn=LOC102901761;refGene_tx=XM_006931022.3;refGene_gn=LOC102901761;refGene_tx=XM_006931023.4;refGene_gn=LOC102901761;refGene_tx=XM_023252995.1
B1 140581826 140581895
refGene_gn=RASGEF1B;refGene_tx=XM_023252996.1
B1 140684986 140685034
refGene_gn=RASGEF1B;refGene_tx=XM_023252996.1
B1 140685205 140685310
refGene_gn=RASGEF1B;refGene_tx=XM_019829321.2
B1 140782177 140782231
refGene_gn=RASGEF1B;refGene_tx=XM_019829321.2;refGene_gn=RASGEF1B;refGene_tx=XM_023252996.1
B1 140981840 140981927
refGene_gn=RASGEF1B;refGene_tx=XM_019829321.2;refGene_gn=RASGEF1B;refGene_tx=XM_023252996.1
B1 141117964 141118147
refGene_gn=RASGEF1B;refGene_tx=XM_003985241.4;refGene_gn=RASGEF1B;refGene_tx=XM_006931026.3;refGene_gn=RASGEF1B;refGene_tx=XM_006931027.4;refGene_gn=RASGEF1B;refGene_tx=XM_006931028.4;refGene_gn=RASGEF1B;refGene_tx=XM_019829321.2;refGene_gn=RASGEF1B;refGene_tx=XM_023252996.1
B1 141119842 141119965
refGene_gn=RASGEF1B;refGene_tx=XM_003985241.4;refGene_gn=RASGEF1B;refGene_tx=XM_006931026.3;refGene_gn=RASGEF1B;refGene_tx=XM_006931027.4;refGene_gn=RASGEF1B;refGene_tx=XM_006931028.4;refGene_gn=RASGEF1B;refGene_tx=XM_019829321.2;refGene_gn=RASGEF1B;refGene_tx=XM_023252996.1

B1 141120585 141120723
refGene_gn=RASGEF1B;refGene_tx=XM_003985241.4;refGene_gn=RASGEF1B;refGene_tx=XM_006931026.3;refGene_gn=RASGEF1B;refGene_tx=XM_006931027.4;refGene_gn=RASGEF1B;refGene_tx=XM_006931028.4;refGene_gn=RASGEF1B;refGene_tx=XM_019829321.2;refGene_gn=RASGEF1B;refGene_tx=XM_023252996.1
B1 141126652 141126865
refGene_gn=RASGEF1B;refGene_tx=XM_003985241.4;refGene_gn=RASGEF1B;refGene_tx=XM_006931026.3;refGene_gn=RASGEF1B;refGene_tx=XM_006931027.4;refGene_gn=RASGEF1B;refGene_tx=XM_006931028.4;refGene_gn=RASGEF1B;refGene_tx=XM_019829321.2;refGene_gn=RASGEF1B;refGene_tx=XM_023252996.1
B1 141127340 141127415
refGene_gn=RASGEF1B;refGene_tx=XM_003985241.4;refGene_gn=RASGEF1B;refGene_tx=XM_006931026.3;refGene_gn=RASGEF1B;refGene_tx=XM_006931027.4;refGene_gn=RASGEF1B;refGene_tx=XM_006931028.4;refGene_gn=RASGEF1B;refGene_tx=XM_019829321.2;refGene_gn=RASGEF1B;refGene_tx=XM_023252996.1
B1 141129001 141129097
refGene_gn=RASGEF1B;refGene_tx=XM_003985241.4;refGene_gn=RASGEF1B;refGene_tx=XM_006931026.3;refGene_gn=RASGEF1B;refGene_tx=XM_006931027.4;refGene_gn=RASGEF1B;refGene_tx=XM_006931028.4;refGene_gn=RASGEF1B;refGene_tx=XM_019829321.2;refGene_gn=RASGEF1B;refGene_tx=XM_023252996.1
B1 141129220 141129323
refGene_gn=RASGEF1B;refGene_tx=XM_003985241.4;refGene_gn=RASGEF1B;refGene_tx=XM_006931026.3;refGene_gn=RASGEF1B;refGene_tx=XM_006931027.4;refGene_gn=RASGEF1B;refGene_tx=XM_006931028.4;refGene_gn=RASGEF1B;refGene_tx=XM_019829321.2;refGene_gn=RASGEF1B;refGene_tx=XM_023252996.1
B1 141130771 141130851
refGene_gn=RASGEF1B;refGene_tx=XM_003985241.4;refGene_gn=RASGEF1B;refGene_tx=XM_006931026.3;refGene_gn=RASGEF1B;refGene_tx=XM_006931027.4;refGene_gn=RASGEF1B;refGene_tx=XM_006931028.4;refGene_gn=RASGEF1B;refGene_tx=XM_019829321.2;refGene_gn=RASGEF1B;refGene_tx=XM_023252996.1
B1 141131945 141132041
refGene_gn=RASGEF1B;refGene_tx=XM_003985241.4;refGene_gn=RASGEF1B;refGene_tx=XM_006931026.3;refGene_gn=RASGEF1B;refGene_tx=XM_006931027.4;refGene_gn=RASGEF1B;refGene_tx=XM_006931028.4;refGene_gn=RASGEF1B;refGene_tx=XM_019829321.2;refGene_gn=RASGEF1B;refGene_tx=XM_023252996.1
B1 141137076 141137172
refGene_gn=RASGEF1B;refGene_tx=XM_003985241.4;refGene_gn=RASGEF1B;refGene_tx=XM_006931026.3;refGene_gn=RASGEF1B;refGene_tx=XM_006931027.4;refGene_gn=RASGEF1B;refGene_tx=XM_006931028.4;refGene_gn=RASGEF1B;refGene_tx=XM_019829321.2;refGene_gn=RASGEF1B;refGene_tx=XM_023252996.1
B1 141137759 141137883
refGene_gn=RASGEF1B;refGene_tx=XM_003985241.4;refGene_gn=RASGEF1B;refGene_tx=XM_006931026.3;refGene_gn=RASGEF1B;refGene_tx=XM_006931027.4;refGene_gn=RASGEF1B;refGene_tx=XM_006931028.4;refGene_gn=RASGEF1B;refGene_tx=XM_019829321.2;refGene_gn=RASGEF1B;refGene_tx=XM_023252996.1
B1 141139275 141139348
refGene_gn=RASGEF1B;refGene_tx=XM_003985241.4;refGene_gn=RASGEF1B;refGene_tx=XM_006931026.3;refGene_gn=RASGEF1B;refGene_tx=XM_006931027.4;refGene_gn=RASGEF1B;refGene_tx=XM_006931028.4;refGene_gn=RASGEF1B;refGene_tx=XM_019829321.2;refGene_gn=RASGEF1B;refGene_tx=XM_023252996.1

B1 141142686 141142711
refGene_gn=RASGEF1B;refGene_tx=XM_003985241.4;refGene_gn=RASGEF1B;refGene_tx=XM_006931026.3;refGene_gn=RASGEF1B;refGene_tx=XM_006931027.4;refGene_gn=RASGEF1B;refGene_tx=XM_006931028.4;refGene_gn=RASGEF1B;refGene_tx=XM_019829321.2;refGene_gn=RASGEF1B;refGene_tx=XM_023252996.1
B1 141322348 141322809
refGene_gn=PRKG2;refGene_tx=XM_023252998.1;refGene_gn=PRKG2;refGene_tx=XM_023252999.1;refGene_gn=PRKG2;refGene_tx=XM_023253000.1
B1 141345347 141345514
refGene_gn=PRKG2;refGene_tx=XM_023252998.1;refGene_gn=PRKG2;refGene_tx=XM_023252999.1;refGene_gn=PRKG2;refGene_tx=XM_023253000.1
B1 141350084 141350198
refGene_gn=PRKG2;refGene_tx=XM_023252998.1;refGene_gn=PRKG2;refGene_tx=XM_023252999.1;refGene_gn=PRKG2;refGene_tx=XM_023253000.1
B1 141351827 141351933
refGene_gn=PRKG2;refGene_tx=XM_023252998.1;refGene_gn=PRKG2;refGene_tx=XM_023252999.1;refGene_gn=PRKG2;refGene_tx=XM_023253000.1
B1 141355202 141355266
refGene_gn=PRKG2;refGene_tx=XM_023252998.1;refGene_gn=PRKG2;refGene_tx=XM_023252999.1;refGene_gn=PRKG2;refGene_tx=XM_023253000.1
B1 141363006 141363084
refGene_gn=PRKG2;refGene_tx=XM_023252998.1;refGene_gn=PRKG2;refGene_tx=XM_023252999.1;refGene_gn=PRKG2;refGene_tx=XM_023253000.1
B1 141364589 141364684
refGene_gn=PRKG2;refGene_tx=XM_023252998.1;refGene_gn=PRKG2;refGene_tx=XM_023252999.1;refGene_gn=PRKG2;refGene_tx=XM_023253000.1
B1 141368214 141368283
refGene_gn=PRKG2;refGene_tx=XM_023252998.1;refGene_gn=PRKG2;refGene_tx=XM_023252999.1;refGene_gn=PRKG2;refGene_tx=XM_023253000.1
B1 141372949 141373048
refGene_gn=PRKG2;refGene_tx=XM_023252998.1;refGene_gn=PRKG2;refGene_tx=XM_023252999.1;refGene_gn=PRKG2;refGene_tx=XM_023253000.1
B1 141374450 141374604
refGene_gn=PRKG2;refGene_tx=XM_023252998.1;refGene_gn=PRKG2;refGene_tx=XM_023252999.1;refGene_gn=PRKG2;refGene_tx=XM_023253000.1
B1 141376583 141376720
refGene_gn=PRKG2;refGene_tx=XM_023252998.1;refGene_gn=PRKG2;refGene_tx=XM_023252999.1;refGene_gn=PRKG2;refGene_tx=XM_023253000.1;refGene_gn=PRKG2;refGene_tx=XM_023253001.1
B1 141378792 141378882
refGene_gn=PRKG2;refGene_tx=XM_023252998.1;refGene_gn=PRKG2;refGene_tx=XM_023252999.1;refGene_gn=PRKG2;refGene_tx=XM_023253000.1;refGene_gn=PRKG2;refGene_tx=XM_023253001.1
B1 141381181 141381323
refGene_gn=PRKG2;refGene_tx=XM_023252998.1;refGene_gn=PRKG2;refGene_tx=XM_023252999.1;refGene_gn=PRKG2;refGene_tx=XM_023253000.1;refGene_gn=PRKG2;refGene_tx=XM_023253001.1
B1 141398178 141398342
refGene_gn=PRKG2;refGene_tx=XM_023252998.1;refGene_gn=PRKG2;refGene_tx=XM_023252999.1;refGene_gn=PRKG2;refGene_tx=XM_023253000.1;refGene_gn=

PRKG2;refGene_tx=XM_023253001.1
B1 141402658 141402781
refGene_gn=PRKG2;refGene_tx=XM_023252998.1;refGene_gn=PRKG2;refGene_tx=XM_023252999.1;refGene_gn=PRKG2;refGene_tx=XM_023253000.1;refGene_gn=PRKG2;refGene_tx=XM_023253001.1
B1 141403826 141403889
refGene_gn=PRKG2;refGene_tx=XM_023252998.1;refGene_gn=PRKG2;refGene_tx=XM_023252999.1;refGene_gn=PRKG2;refGene_tx=XM_023253000.1;refGene_gn=PRKG2;refGene_tx=XM_023253001.1
B1 141411537 141411604
refGene_gn=PRKG2;refGene_tx=XM_023252998.1;refGene_gn=PRKG2;refGene_tx=XM_023252999.1;refGene_gn=PRKG2;refGene_tx=XM_023253000.1;refGene_gn=PRKG2;refGene_tx=XM_023253001.1
B1 141414644 141414740
refGene_gn=PRKG2;refGene_tx=XM_023252998.1;refGene_gn=PRKG2;refGene_tx=XM_023252999.1;refGene_gn=PRKG2;refGene_tx=XM_023253000.1;refGene_gn=PRKG2;refGene_tx=XM_023253001.1
B1 141450343 141450535
refGene_gn=BMP3;refGene_tx=XM_006931036.4
B1 141457742 141458653
refGene_gn=BMP3;refGene_tx=XM_006931036.4
B1 141473920 141474245
refGene_gn=BMP3;refGene_tx=XM_006931036.4
B1 141533403 141533499
refGene_gn=CB1H4orf22;refGene_tx=XM_003985248.4;refGene_gn=CB1H4orf22;refGene_tx=XM_019829323.2
B1 141550242 141550372
refGene_gn=CB1H4orf22;refGene_tx=XM_003985248.4;refGene_gn=CB1H4orf22;refGene_tx=XM_019829323.2
B1 141613181 141613324
refGene_gn=CB1H4orf22;refGene_tx=XM_003985248.4;refGene_gn=CB1H4orf22;refGene_tx=XM_019829323.2
B1 141895353 141895444
refGene_gn=CB1H4orf22;refGene_tx=XM_003985248.4;refGene_gn=CB1H4orf22;refGene_tx=XM_019829323.2
B1 142108797 142108928
refGene_gn=CB1H4orf22;refGene_tx=XM_003985248.4;refGene_gn=CB1H4orf22;refGene_tx=XM_019829323.2
B1 142121775 142121886
refGene_gn=CB1H4orf22;refGene_tx=XM_003985248.4;refGene_gn=CB1H4orf22;refGene_tx=XM_019829323.2
B1 142165613 142165961
refGene_gn=FGF5;refGene_tx=NM_001114546.1;refGene_gn=FGF5;refGene_tx=XM_006931032.3;refGene_gn=FGF5;refGene_tx=XM_019828138.2
B1 142173991 142174044
refGene_gn=FGF5;refGene_tx=XM_019828138.2
B1 142177859 142177963
refGene_gn=FGF5;refGene_tx=NM_001114546.1;refGene_gn=FGF5;refGene_tx=XM_019828138.2
B1 142185640 142186001

refGene_gn=FGF5;refGene_tx=NM_001114546.1;refGene_gn=FGF5;refGene_tx=XM_006931032.3;refGene_gn=FGF5;refGene_tx=XM_019828138.2
B1 142244818 142246431
refGene_gn=PRDM8;refGene_tx=XM_023253003.1;refGene_gn=PRDM8;refGene_tx=XM_023253004.1;refGene_gn=PRDM8;refGene_tx=XM_023253005.1;refGene_gn=PRDM8;refGene_tx=XM_023253006.1;refGene_gn=PRDM8;refGene_tx=XM_023253008.1;refGene_gn=PRDM8;refGene_tx=XM_023253009.1
B1 142246816 142247048
refGene_gn=PRDM8;refGene_tx=XM_023253003.1;refGene_gn=PRDM8;refGene_tx=XM_023253004.1;refGene_gn=PRDM8;refGene_tx=XM_023253005.1;refGene_gn=PRDM8;refGene_tx=XM_023253006.1;refGene_gn=PRDM8;refGene_tx=XM_023253008.1;refGene_gn=PRDM8;refGene_tx=XM_023253009.1
B1 142248043 142248264
refGene_gn=PRDM8;refGene_tx=XM_023253003.1;refGene_gn=PRDM8;refGene_tx=XM_023253004.1;refGene_gn=PRDM8;refGene_tx=XM_023253005.1;refGene_gn=PRDM8;refGene_tx=XM_023253006.1;refGene_gn=PRDM8;refGene_tx=XM_023253008.1;refGene_gn=PRDM8;refGene_tx=XM_023253009.1
B1 142249795 142249796
refGene_gn=PRDM8;refGene_tx=XM_023253003.1;refGene_gn=PRDM8;refGene_tx=XM_023253009.1
B1 142371033 142371185
refGene_gn=ANTXR2;refGene_tx=XM_011281741.2
B1 142371981 142372053
refGene_gn=ANTXR2;refGene_tx=XM_011281741.2
B1 142374118 142374190
refGene_gn=ANTXR2;refGene_tx=XM_011281741.2
B1 142390451 142390533
refGene_gn=ANTXR2;refGene_tx=XM_011281741.2
B1 142390988 142391096
refGene_gn=ANTXR2;refGene_tx=XM_011281741.2
B1 142391232 142391301
refGene_gn=ANTXR2;refGene_tx=XM_011281741.2
B1 142392051 142392132
refGene_gn=ANTXR2;refGene_tx=XM_011281741.2
B1 142414372 142414433
refGene_gn=ANTXR2;refGene_tx=XM_011281741.2
B1 142416490 142416589
refGene_gn=ANTXR2;refGene_tx=XM_011281741.2
B1 142418845 142418915
refGene_gn=ANTXR2;refGene_tx=XM_011281741.2
B1 142431889 142431968
refGene_gn=ANTXR2;refGene_tx=XM_011281741.2
B1 142445234 142445330
refGene_gn=ANTXR2;refGene_tx=XM_011281741.2
B1 142461036 142461081
refGene_gn=ANTXR2;refGene_tx=XM_011281741.2
B1 142461928 142462021
refGene_gn=ANTXR2;refGene_tx=XM_011281741.2
B1 142468219 142468387
refGene_gn=ANTXR2;refGene_tx=XM_011281741.2

B1 142468701 142468782
refGene_gn=ANTXR2;refGene_tx=XM_011281741.2
B1 142521612 142521651
refGene_gn=ANTXR2;refGene_tx=XM_011281741.2
B1 142942690 142944352
refGene_gn=GK2;refGene_tx=XM_004001309.5
B1 143009690 143010371
refGene_gn=NAA11;refGene_tx=XM_006931033.3
B1 143400518 143400703
refGene_gn=PAQR3;refGene_tx=XM_003985250.5;refGene_gn=PAQR3;refGene_tx=XM_023253015.1
B1 143404704 143404867
refGene_gn=PAQR3;refGene_tx=XM_003985250.5;refGene_gn=PAQR3;refGene_tx=XM_023253015.1
B1 143409293 143409449
refGene_gn=PAQR3;refGene_tx=XM_003985250.5;refGene_gn=PAQR3;refGene_tx=XM_023253015.1;refGene_gn=PAQR3;refGene_tx=XM_023253016.1
B1 143412811 143413009
refGene_gn=PAQR3;refGene_tx=XM_003985250.5;refGene_gn=PAQR3;refGene_tx=XM_023253015.1;refGene_gn=PAQR3;refGene_tx=XM_023253016.1
B1 143415577 143415668
refGene_gn=PAQR3;refGene_tx=XM_003985250.5;refGene_gn=PAQR3;refGene_tx=XM_023253015.1;refGene_gn=PAQR3;refGene_tx=XM_023253016.1
B1 143419085 143419228
refGene_gn=PAQR3;refGene_tx=XM_003985250.5;refGene_gn=PAQR3;refGene_tx=XM_023253015.1;refGene_gn=PAQR3;refGene_tx=XM_023253016.1
B1 143427590 143429026
refGene_gn=BMP2K;refGene_tx=XM_023253010.1;refGene_gn=BMP2K;refGene_tx=XM_023253012.1;refGene_gn=BMP2K;refGene_tx=XM_023253013.1;refGene_gn=BMP2K;refGene_tx=XM_023253014.1
B1 143444215 143444326
refGene_gn=BMP2K;refGene_tx=XM_023253010.1;refGene_gn=BMP2K;refGene_tx=XM_023253012.1;refGene_gn=BMP2K;refGene_tx=XM_023253013.1
B1 143451519 143451677
refGene_gn=BMP2K;refGene_tx=XM_023253010.1;refGene_gn=BMP2K;refGene_tx=XM_023253012.1;refGene_gn=BMP2K;refGene_tx=XM_023253013.1;refGene_gn=BMP2K;refGene_tx=XM_023253014.1
B1 143456145 143456330
refGene_gn=BMP2K;refGene_tx=XM_023253010.1;refGene_gn=BMP2K;refGene_tx=XM_023253012.1;refGene_gn=BMP2K;refGene_tx=XM_023253013.1;refGene_gn=BMP2K;refGene_tx=XM_023253014.1
B1 143457322 143457421
refGene_gn=BMP2K;refGene_tx=XM_023253010.1;refGene_gn=BMP2K;refGene_tx=XM_023253012.1;refGene_gn=BMP2K;refGene_tx=XM_023253013.1;refGene_gn=BMP2K;refGene_tx=XM_023253014.1
B1 143458966 143459178
refGene_gn=BMP2K;refGene_tx=XM_023253010.1;refGene_gn=BMP2K;refGene_tx=XM_023253012.1;refGene_gn=BMP2K;refGene_tx=XM_023253013.1;refGene_gn=BMP2K;refGene_tx=XM_023253014.1
B1 143463176 143463337

refGene_gn=BMP2K;refGene_tx=XM_023253010.1;refGene_gn=BMP2K;refGene_tx=XM_023253012.1;refGene_gn=BMP2K;refGene_tx=XM_023253013.1;refGene_gn=BMP2K;refGene_tx=XM_023253014.1
B1 143467437 143467517
refGene_gn=BMP2K;refGene_tx=XM_023253010.1;refGene_gn=BMP2K;refGene_tx=XM_023253012.1;refGene_gn=BMP2K;refGene_tx=XM_023253013.1;refGene_gn=BMP2K;refGene_tx=XM_023253014.1
B1 143468966 143469070
refGene_gn=BMP2K;refGene_tx=XM_023253010.1;refGene_gn=BMP2K;refGene_tx=XM_023253012.1;refGene_gn=BMP2K;refGene_tx=XM_023253013.1;refGene_gn=BMP2K;refGene_tx=XM_023253014.1
B1 143481808 143481941
refGene_gn=BMP2K;refGene_tx=XM_023253010.1;refGene_gn=BMP2K;refGene_tx=XM_023253012.1;refGene_gn=BMP2K;refGene_tx=XM_023253013.1;refGene_gn=BMP2K;refGene_tx=XM_023253014.1
B1 143486393 143486475
refGene_gn=BMP2K;refGene_tx=XM_023253010.1;refGene_gn=BMP2K;refGene_tx=XM_023253012.1;refGene_gn=BMP2K;refGene_tx=XM_023253013.1;refGene_gn=BMP2K;refGene_tx=XM_023253014.1
B1 143488667 143488789
refGene_gn=BMP2K;refGene_tx=XM_023253010.1;refGene_gn=BMP2K;refGene_tx=XM_023253012.1;refGene_gn=BMP2K;refGene_tx=XM_023253013.1;refGene_gn=BMP2K;refGene_tx=XM_023253014.1
B1 143491260 143491403
refGene_gn=BMP2K;refGene_tx=XM_023253010.1;refGene_gn=BMP2K;refGene_tx=XM_023253012.1;refGene_gn=BMP2K;refGene_tx=XM_023253013.1;refGene_gn=BMP2K;refGene_tx=XM_023253014.1
B1 143498882 143498988
refGene_gn=BMP2K;refGene_tx=XM_023253010.1;refGene_gn=BMP2K;refGene_tx=XM_023253012.1;refGene_gn=BMP2K;refGene_tx=XM_023253013.1;refGene_gn=BMP2K;refGene_tx=XM_023253014.1
B1 143503388 143503507
refGene_gn=BMP2K;refGene_tx=XM_023253010.1;refGene_gn=BMP2K;refGene_tx=XM_023253012.1;refGene_gn=BMP2K;refGene_tx=XM_023253013.1;refGene_gn=BMP2K;refGene_tx=XM_023253014.1
B1 143555237 143555412
refGene_gn=BMP2K;refGene_tx=XM_023253010.1;refGene_gn=BMP2K;refGene_tx=XM_023253012.1;refGene_gn=BMP2K;refGene_tx=XM_023253013.1;refGene_gn=BMP2K;refGene_tx=XM_023253014.1
B1 143684038 143684098
refGene_gn=ANXA3;refGene_tx=XM_019829326.2;refGene_gn=ANXA3;refGene_tx=XM_023253018.1
B1 143692075 143692198
refGene_gn=ANXA3;refGene_tx=XM_019829326.2;refGene_gn=ANXA3;refGene_tx=XM_023253018.1
B1 143694244 143694303
refGene_gn=ANXA3;refGene_tx=XM_019829326.2;refGene_gn=ANXA3;refGene_tx=XM_023253018.1
B1 143695230 143695326
refGene_gn=ANXA3;refGene_tx=XM_019829326.2;refGene_gn=ANXA3;refGene_tx

=XM_023253018.1
B1	143696992	143697086
refGene_gn=ANXA3;refGene_tx=XM_019829326.2;refGene_gn=ANXA3;refGene_tx
=XM_023253018.1
B1	143697520	143697577
refGene_gn=ANXA3;refGene_tx=XM_019829326.2;refGene_gn=ANXA3;refGene_tx
=XM_023253018.1
B1	143700669	143700749
refGene_gn=ANXA3;refGene_tx=XM_019829326.2;refGene_gn=ANXA3;refGene_tx
=XM_023253018.1
B1	143704611	143704702
refGene_gn=ANXA3;refGene_tx=XM_019829326.2;refGene_gn=ANXA3;refGene_tx
=XM_023253018.1
B1	143708362	143708476
refGene_gn=ANXA3;refGene_tx=XM_019829326.2;refGene_gn=ANXA3;refGene_tx
=XM_023253018.1
B1	143711610	143711705
refGene_gn=ANXA3;refGene_tx=XM_019829326.2;refGene_gn=ANXA3;refGene_tx
=XM_023253018.1
B1	143716426	143716514
refGene_gn=ANXA3;refGene_tx=XM_019829326.2;refGene_gn=ANXA3;refGene_tx
=XM_023253018.1
B1	143733619	143733634
refGene_gn=ANXA3;refGene_tx=XM_019829326.2;refGene_gn=ANXA3;refGene_tx
=XM_023253018.1
B1	143748138	143748732
refGene_gn=FRAS1;refGene_tx=XM_023253019.1
B1	143750088	143750235
refGene_gn=FRAS1;refGene_tx=XM_023253019.1
B1	143751604	143751810
refGene_gn=FRAS1;refGene_tx=XM_023253019.1
B1	143754659	143754826
refGene_gn=FRAS1;refGene_tx=XM_023253019.1
B1	143762690	143762807
refGene_gn=FRAS1;refGene_tx=XM_023253019.1
B1	143766598	143766758
refGene_gn=FRAS1;refGene_tx=XM_023253019.1
B1	143767740	143767848
refGene_gn=FRAS1;refGene_tx=XM_023253019.1
B1	143769454	143769605
refGene_gn=FRAS1;refGene_tx=XM_023253019.1
B1	143772337	143772552
refGene_gn=FRAS1;refGene_tx=XM_023253019.1
B1	143774477	143774638
refGene_gn=FRAS1;refGene_tx=XM_023253019.1
B1	143777228	143777461
refGene_gn=FRAS1;refGene_tx=XM_023253019.1
B1	143780256	143780532
refGene_gn=FRAS1;refGene_tx=XM_023253019.1
B1	143781712	143781900

refGene_gn=FRAS1;refGene_tx=XM_023253019.1
B1 143783465 143783666
refGene_gn=FRAS1;refGene_tx=XM_023253019.1
B1 143785602 143785759
refGene_gn=FRAS1;refGene_tx=XM_023253019.1
B1 143789801 143790007
refGene_gn=FRAS1;refGene_tx=XM_023253019.1
B1 143796984 143797132
refGene_gn=FRAS1;refGene_tx=XM_023253019.1
B1 143797701 143797862
refGene_gn=FRAS1;refGene_tx=XM_023253019.1
B1 143799428 143799773
refGene_gn=FRAS1;refGene_tx=XM_023253019.1
B1 143801011 143801258
refGene_gn=FRAS1;refGene_tx=XM_023253019.1
B1 143803512 143803681
refGene_gn=FRAS1;refGene_tx=XM_023253019.1
B1 143805941 143806101
refGene_gn=FRAS1;refGene_tx=XM_023253019.1
B1 143807512 143807663
refGene_gn=FRAS1;refGene_tx=XM_023253019.1
B1 143809540 143809654
refGene_gn=FRAS1;refGene_tx=XM_023253019.1
B1 143811655 143811883
refGene_gn=FRAS1;refGene_tx=XM_023253019.1
B1 143813738 143813879
refGene_gn=FRAS1;refGene_tx=XM_023253019.1
B1 143814155 143814280
refGene_gn=FRAS1;refGene_tx=XM_023253019.1
B1 143822767 143822947
refGene_gn=FRAS1;refGene_tx=XM_023253019.1
B1 143823220 143823340
refGene_gn=FRAS1;refGene_tx=XM_023253019.1
B1 143824770 143824959
refGene_gn=FRAS1;refGene_tx=XM_023253019.1
B1 143827173 143827437
refGene_gn=FRAS1;refGene_tx=XM_023253019.1
B1 143828641 143828795
refGene_gn=FRAS1;refGene_tx=XM_023253019.1
B1 143829786 143829977
refGene_gn=FRAS1;refGene_tx=XM_023253019.1
B1 143833536 143833672
refGene_gn=FRAS1;refGene_tx=XM_023253019.1
B1 143835243 143835406
refGene_gn=FRAS1;refGene_tx=XM_023253019.1
B1 143835815 143835964
refGene_gn=FRAS1;refGene_tx=XM_023253019.1
B1 143841648 143841896
refGene_gn=FRAS1;refGene_tx=XM_023253019.1
B1 143843866 143843992

refGene_gn=FRAS1;refGene_tx=XM_023253019.1
B1	143845324	143845456
refGene_gn=FRAS1;refGene_tx=XM_023253019.1
B1	143853956	143853989
refGene_gn=FRAS1;refGene_tx=XM_023253019.1
B1	143856331	143856469
refGene_gn=FRAS1;refGene_tx=XM_023253019.1
B1	143858996	143859111
refGene_gn=FRAS1;refGene_tx=XM_023253019.1
B1	143867552	143867669
refGene_gn=FRAS1;refGene_tx=XM_023253019.1
B1	143872559	143872738
refGene_gn=FRAS1;refGene_tx=XM_023253019.1
B1	143879411	143879565
refGene_gn=FRAS1;refGene_tx=XM_023253019.1
B1	143889452	143889779
refGene_gn=FRAS1;refGene_tx=XM_023253019.1
B1	143892899	143892984
refGene_gn=FRAS1;refGene_tx=XM_023253019.1
B1	143896713	143896984
refGene_gn=FRAS1;refGene_tx=XM_023253019.1
B1	143900192	143900330
refGene_gn=FRAS1;refGene_tx=XM_023253019.1
B1	143901861	143902002
refGene_gn=FRAS1;refGene_tx=XM_023253019.1
B1	143902917	143903058
refGene_gn=FRAS1;refGene_tx=XM_023253019.1
B1	143907906	143908053
refGene_gn=FRAS1;refGene_tx=XM_023253019.1
B1	143913609	143913756
refGene_gn=FRAS1;refGene_tx=XM_023253019.1
B1	143914048	143914201
refGene_gn=FRAS1;refGene_tx=XM_023253019.1
B1	143933642	143933786
refGene_gn=FRAS1;refGene_tx=XM_023253019.1
B1	143936340	143936481
refGene_gn=FRAS1;refGene_tx=XM_023253019.1
B1	143950861	143951038
refGene_gn=FRAS1;refGene_tx=XM_023253019.1
B1	143952395	143952536
refGene_gn=FRAS1;refGene_tx=XM_023253019.1
B1	143954129	143954270
refGene_gn=FRAS1;refGene_tx=XM_023253019.1
B1	143958528	143958672
refGene_gn=FRAS1;refGene_tx=XM_023253019.1
B1	143974162	143974297
refGene_gn=FRAS1;refGene_tx=XM_023253019.1
B1	143975609	143975753
refGene_gn=FRAS1;refGene_tx=XM_023253019.1
B1	143977268	143977416

refGene_gn=FRAS1;refGene_tx=XM_023253019.1
B1 143979029 143979065
refGene_gn=FRAS1;refGene_tx=XM_023253019.1
B1 143980917 143981007
refGene_gn=FRAS1;refGene_tx=XM_023253019.1
B1 143985703 143985895
refGene_gn=FRAS1;refGene_tx=XM_023253019.1
B1 143986135 143986237
refGene_gn=FRAS1;refGene_tx=XM_023253019.1
B1 143988351 143988435
refGene_gn=FRAS1;refGene_tx=XM_023253019.1
B1 143998436 143998570
refGene_gn=FRAS1;refGene_tx=XM_023253019.1
B1 144001143 144001303
refGene_gn=FRAS1;refGene_tx=XM_023253019.1
B1 144010484 144010577
refGene_gn=FRAS1;refGene_tx=XM_023253019.1
B1 144018476 144018584
refGene_gn=FRAS1;refGene_tx=XM_023253019.1
B1 144165437 144165469
refGene_gn=FRAS1;refGene_tx=XM_023253019.1
B1 144173877 144173953
refGene_gn=FRAS1;refGene_tx=XM_023253019.1
B1 144295061 144295180
refGene_gn=MRPL1;refGene_tx=XM_003985258.5;refGene_gn=MRPL1;refGene_tx
=XM_006931041.4;refGene_gn=MRPL1;refGene_tx=XM_019829331.1
B1 144305571 144305653
refGene_gn=MRPL1;refGene_tx=XM_003985258.5;refGene_gn=MRPL1;refGene_tx
=XM_006931041.4;refGene_gn=MRPL1;refGene_tx=XM_019829331.1
B1 144338313 144338420
refGene_gn=MRPL1;refGene_tx=XM_003985258.5;refGene_gn=MRPL1;refGene_tx
=XM_006931041.4;refGene_gn=MRPL1;refGene_tx=XM_019829331.1
B1 144360016 144360128
refGene_gn=MRPL1;refGene_tx=XM_003985258.5;refGene_gn=MRPL1;refGene_tx
=XM_006931041.4;refGene_gn=MRPL1;refGene_tx=XM_019829331.1
B1 144362811 144362883
refGene_gn=MRPL1;refGene_tx=XM_003985258.5;refGene_gn=MRPL1;refGene_tx
=XM_006931041.4;refGene_gn=MRPL1;refGene_tx=XM_019829331.1
B1 144364247 144364331
refGene_gn=MRPL1;refGene_tx=XM_003985258.5;refGene_gn=MRPL1;refGene_tx
=XM_006931041.4;refGene_gn=MRPL1;refGene_tx=XM_019829331.1
B1 144367576 144367835
refGene_gn=MRPL1;refGene_tx=XM_003985258.5;refGene_gn=MRPL1;refGene_tx
=XM_006931041.4;refGene_gn=MRPL1;refGene_tx=XM_019829331.1
B1 144378919 144379031
refGene_gn=MRPL1;refGene_tx=XM_003985258.5;refGene_gn=MRPL1;refGene_tx
=XM_006931041.4;refGene_gn=MRPL1;refGene_tx=XM_019829331.1
B1 144392405 144392442
refGene_gn=MRPL1;refGene_tx=XM_006931041.4
B1 144392760 144392791

refGene_gn=MRPL1;refGene_tx=XM_003985258.5
B1 144393219 144393229
refGene_gn=MRPL1;refGene_tx=XM_019829331.1
B1 144442171 144442176
refGene_gn=CNOT6L;refGene_tx=XM_006931042.4;refGene_gn=CNOT6L;refGene_tx=XM_023253021.1
B1 144442242 144442247
refGene_gn=CNOT6L;refGene_tx=XM_019829328.2
B1 144487030 144487152
refGene_gn=CNOT6L;refGene_tx=XM_006931042.4;refGene_gn=CNOT6L;refGene_tx=XM_019829328.2;refGene_gn=CNOT6L;refGene_tx=XM_023253020.1;refGene_gn=CNOT6L;refGene_tx=XM_023253021.1
B1 144490177 144490364
refGene_gn=CNOT6L;refGene_tx=XM_006931042.4;refGene_gn=CNOT6L;refGene_tx=XM_019829328.2;refGene_gn=CNOT6L;refGene_tx=XM_023253020.1;refGene_gn=CNOT6L;refGene_tx=XM_023253021.1
B1 144491860 144491946
refGene_gn=CNOT6L;refGene_tx=XM_006931042.4;refGene_gn=CNOT6L;refGene_tx=XM_019829328.2;refGene_gn=CNOT6L;refGene_tx=XM_023253020.1;refGene_gn=CNOT6L;refGene_tx=XM_023253021.1
B1 144515138 144515228
refGene_gn=CNOT6L;refGene_tx=XM_006931042.4;refGene_gn=CNOT6L;refGene_tx=XM_019829328.2;refGene_gn=CNOT6L;refGene_tx=XM_023253020.1;refGene_gn=CNOT6L;refGene_tx=XM_023253021.1
B1 144526381 144526450
refGene_gn=CNOT6L;refGene_tx=XM_006931042.4;refGene_gn=CNOT6L;refGene_tx=XM_019829328.2;refGene_gn=CNOT6L;refGene_tx=XM_023253020.1;refGene_gn=CNOT6L;refGene_tx=XM_023253021.1
B1 144528981 144529139
refGene_gn=CNOT6L;refGene_tx=XM_006931042.4;refGene_gn=CNOT6L;refGene_tx=XM_019829328.2;refGene_gn=CNOT6L;refGene_tx=XM_023253020.1;refGene_gn=CNOT6L;refGene_tx=XM_023253021.1
B1 144531189 144531344
refGene_gn=CNOT6L;refGene_tx=XM_006931042.4;refGene_gn=CNOT6L;refGene_tx=XM_019829328.2;refGene_gn=CNOT6L;refGene_tx=XM_023253020.1;refGene_gn=CNOT6L;refGene_tx=XM_023253021.1
B1 144544009 144544161
refGene_gn=CNOT6L;refGene_tx=XM_006931042.4;refGene_gn=CNOT6L;refGene_tx=XM_019829328.2;refGene_gn=CNOT6L;refGene_tx=XM_023253020.1;refGene_gn=CNOT6L;refGene_tx=XM_023253021.1
B1 144548161 144548389
refGene_gn=CNOT6L;refGene_tx=XM_006931042.4;refGene_gn=CNOT6L;refGene_tx=XM_019829328.2;refGene_gn=CNOT6L;refGene_tx=XM_023253020.1;refGene_gn=CNOT6L;refGene_tx=XM_023253021.1
B1 144550543 144550746
refGene_gn=CNOT6L;refGene_tx=XM_006931042.4;refGene_gn=CNOT6L;refGene_tx=XM_019829328.2;refGene_gn=CNOT6L;refGene_tx=XM_023253020.1;refGene_gn=CNOT6L;refGene_tx=XM_023253021.1
B1 144553765 144553871
refGene_gn=CNOT6L;refGene_tx=XM_023253021.1

B1 144555743 144555956
refGene_gn=CNOT6L;refGene_tx=XM_006931042.4;refGene_gn=CNOT6L;refGene_tx=XM_019829328.2;refGene_gn=CNOT6L;refGene_tx=XM_023253020.1;refGene_gn=CNOT6L;refGene_tx=XM_023253021.1
B1 144698879 144698904
refGene_gn=CXCL13;refGene_tx=XM_003985260.4
B1 144699420 144699501
refGene_gn=CXCL13;refGene_tx=XM_003985260.4
B1 144702331 144702464
refGene_gn=CXCL13;refGene_tx=XM_003985260.4
B1 144704112 144704176
refGene_gn=CXCL13;refGene_tx=XM_003985260.4
B1 145030314 145030435
refGene_gn=CCNG2;refGene_tx=XM_003985261.3;refGene_gn=CCNG2;refGene_tx=XM_006931043.2;refGene_gn=CCNG2;refGene_tx=XM_011281738.2
B1 145031591 145031797
refGene_gn=CCNG2;refGene_tx=XM_003985261.3;refGene_gn=CCNG2;refGene_tx=XM_006931043.2;refGene_gn=CCNG2;refGene_tx=XM_011281738.2
B1 145033735 145033834
refGene_gn=CCNG2;refGene_tx=XM_003985261.3;refGene_gn=CCNG2;refGene_tx=XM_006931043.2;refGene_gn=CCNG2;refGene_tx=XM_011281738.2
B1 145033920 145033999
refGene_gn=CCNG2;refGene_tx=XM_003985261.3;refGene_gn=CCNG2;refGene_tx=XM_006931043.2;refGene_gn=CCNG2;refGene_tx=XM_011281738.2
B1 145034188 145034439
refGene_gn=CCNG2;refGene_tx=XM_003985261.3;refGene_gn=CCNG2;refGene_tx=XM_006931043.2;refGene_gn=CCNG2;refGene_tx=XM_011281738.2
B1 145035745 145035883
refGene_gn=CCNG2;refGene_tx=XM_003985261.3;refGene_gn=CCNG2;refGene_tx=XM_006931043.2;refGene_gn=CCNG2;refGene_tx=XM_011281738.2
B1 145036481 145036619
refGene_gn=CCNG2;refGene_tx=XM_003985261.3;refGene_gn=CCNG2;refGene_tx=XM_006931043.2;refGene_gn=CCNG2;refGene_tx=XM_011281738.2
B1 145111122 145111236
refGene_gn=CCNI;refGene_tx=XM_003985264.5
B1 145124840 145124969
refGene_gn=CCNI;refGene_tx=XM_003985264.5
B1 145127943 145128018
refGene_gn=CCNI;refGene_tx=XM_003985264.5
B1 145128171 145128312
refGene_gn=CCNI;refGene_tx=XM_003985264.5
B1 145128602 145128833
refGene_gn=CCNI;refGene_tx=XM_003985264.5
B1 145135637 145136087
refGene_gn=CCNI;refGene_tx=XM_003985264.5
B1 145143432 145143436
refGene_gn=SEPT11;refGene_tx=XM_006931045.3
B1 145144106 145144122
refGene_gn=SEPT11;refGene_tx=XM_006931044.3
B1 145146145 145146170

refGene_gn=SEPT11;refGene_tx=XM_011281732.2;refGene_gn=SEPT11;refGene_tx=XM_011281733.2;refGene_gn=SEPT11;refGene_tx=XM_011281734.2;refGene_gn=SEPT11;refGene_tx=XM_019829332.2
B1 145148680 145148696
refGene_gn=SEPT11;refGene_tx=XM_003985263.4
B1 145152265 145152453
refGene_gn=SEPT11;refGene_tx=XM_003985263.4;refGene_gn=SEPT11;refGene_tx=XM_006931044.3;refGene_gn=SEPT11;refGene_tx=XM_006931045.3;refGene_gn=SEPT11;refGene_tx=XM_011281732.2;refGene_gn=SEPT11;refGene_tx=XM_011281733.2;refGene_gn=SEPT11;refGene_tx=XM_011281734.2;refGene_gn=SEPT11;refGene_tx=XM_019829332.2
B1 145153898 145154031
refGene_gn=SEPT11;refGene_tx=XM_003985263.4;refGene_gn=SEPT11;refGene_tx=XM_006931044.3;refGene_gn=SEPT11;refGene_tx=XM_006931045.3;refGene_gn=SEPT11;refGene_tx=XM_011281732.2;refGene_gn=SEPT11;refGene_tx=XM_011281733.2;refGene_gn=SEPT11;refGene_tx=XM_011281734.2;refGene_gn=SEPT11;refGene_tx=XM_019829332.2
B1 145159854 145160023
refGene_gn=SEPT11;refGene_tx=XM_003985263.4;refGene_gn=SEPT11;refGene_tx=XM_006931044.3;refGene_gn=SEPT11;refGene_tx=XM_006931045.3;refGene_gn=SEPT11;refGene_tx=XM_011281732.2;refGene_gn=SEPT11;refGene_tx=XM_011281733.2;refGene_gn=SEPT11;refGene_tx=XM_011281734.2;refGene_gn=SEPT11;refGene_tx=XM_019829332.2
B1 145161269 145161366
refGene_gn=SEPT11;refGene_tx=XM_003985263.4;refGene_gn=SEPT11;refGene_tx=XM_006931044.3;refGene_gn=SEPT11;refGene_tx=XM_006931045.3;refGene_gn=SEPT11;refGene_tx=XM_011281732.2;refGene_gn=SEPT11;refGene_tx=XM_011281733.2;refGene_gn=SEPT11;refGene_tx=XM_011281734.2;refGene_gn=SEPT11;refGene_tx=XM_019829332.2
B1 145166193 145166355
refGene_gn=SEPT11;refGene_tx=XM_003985263.4;refGene_gn=SEPT11;refGene_tx=XM_006931044.3;refGene_gn=SEPT11;refGene_tx=XM_006931045.3;refGene_gn=SEPT11;refGene_tx=XM_011281732.2;refGene_gn=SEPT11;refGene_tx=XM_011281733.2;refGene_gn=SEPT11;refGene_tx=XM_011281734.2;refGene_gn=SEPT11;refGene_tx=XM_019829332.2
B1 145169428 145169615
refGene_gn=SEPT11;refGene_tx=XM_003985263.4;refGene_gn=SEPT11;refGene_tx=XM_006931044.3;refGene_gn=SEPT11;refGene_tx=XM_006931045.3;refGene_gn=SEPT11;refGene_tx=XM_011281732.2;refGene_gn=SEPT11;refGene_tx=XM_011281733.2;refGene_gn=SEPT11;refGene_tx=XM_011281734.2;refGene_gn=SEPT11;refGene_tx=XM_019829332.2
B1 145176378 145176574
refGene_gn=SEPT11;refGene_tx=XM_003985263.4;refGene_gn=SEPT11;refGene_tx=XM_006931044.3;refGene_gn=SEPT11;refGene_tx=XM_006931045.3;refGene_gn=SEPT11;refGene_tx=XM_011281732.2;refGene_gn=SEPT11;refGene_tx=XM_011281733.2;refGene_gn=SEPT11;refGene_tx=XM_011281734.2
B1 145187182 145187297
refGene_gn=SEPT11;refGene_tx=XM_003985263.4;refGene_gn=SEPT11;refGene_tx=XM_006931044.3;refGene_gn=SEPT11;refGene_tx=XM_006931045.3;refGene_gn=SEPT11;refGene_tx=XM_011281732.2;refGene_gn=SEPT11;refGene_tx=XM_01

1281733.2;refGene_gn=SEPT11;refGene_tx=XM_011281734.2
B1 145231506 145231533
refGene_gn=SEPT11;refGene_tx=XM_003985263.4;refGene_gn=SEPT11;refGene_tx=XM_006931044.3;refGene_gn=SEPT11;refGene_tx=XM_006931045.3;refGene_gn=SEPT11;refGene_tx=XM_011281732.2;refGene_gn=SEPT11;refGene_tx=XM_011281733.2;refGene_gn=SEPT11;refGene_tx=XM_011281734.2
B1 145268314 145270663
refGene_gn=SOWAHB;refGene_tx=XM_011281731.3
B1 145363844 145364213
refGene_gn=SHROOM3;refGene_tx=XM_023253022.1;refGene_gn=SHROOM3;refGene_tx=XM_023253023.1;refGene_gn=SHROOM3;refGene_tx=XM_023253024.1;refGene_gn=SHROOM3;refGene_tx=XM_023253025.1;refGene_gn=SHROOM3;refGene_tx=XM_023253027.1;refGene_gn=SHROOM3;refGene_tx=XM_023253028.1
B1 145369423 145369696
refGene_gn=SHROOM3;refGene_tx=XM_023253022.1;refGene_gn=SHROOM3;refGene_tx=XM_023253023.1;refGene_gn=SHROOM3;refGene_tx=XM_023253024.1;refGene_gn=SHROOM3;refGene_tx=XM_023253025.1;refGene_gn=SHROOM3;refGene_tx=XM_023253027.1;refGene_gn=SHROOM3;refGene_tx=XM_023253028.1
B1 145380621 145380772
refGene_gn=SHROOM3;refGene_tx=XM_023253022.1;refGene_gn=SHROOM3;refGene_tx=XM_023253023.1;refGene_gn=SHROOM3;refGene_tx=XM_023253024.1;refGene_gn=SHROOM3;refGene_tx=XM_023253025.1;refGene_gn=SHROOM3;refGene_tx=XM_023253027.1;refGene_gn=SHROOM3;refGene_tx=XM_023253028.1
B1 145384097 145384589
refGene_gn=SHROOM3;refGene_tx=XM_023253022.1;refGene_gn=SHROOM3;refGene_tx=XM_023253023.1;refGene_gn=SHROOM3;refGene_tx=XM_023253024.1;refGene_gn=SHROOM3;refGene_tx=XM_023253025.1;refGene_gn=SHROOM3;refGene_tx=XM_023253027.1;refGene_gn=SHROOM3;refGene_tx=XM_023253028.1
B1 145385747 145386644
refGene_gn=SHROOM3;refGene_tx=XM_023253022.1;refGene_gn=SHROOM3;refGene_tx=XM_023253023.1;refGene_gn=SHROOM3;refGene_tx=XM_023253024.1;refGene_gn=SHROOM3;refGene_tx=XM_023253025.1;refGene_gn=SHROOM3;refGene_tx=XM_023253027.1;refGene_gn=SHROOM3;refGene_tx=XM_023253028.1
B1 145392067 145392141
refGene_gn=SHROOM3;refGene_tx=XM_023253022.1;refGene_gn=SHROOM3;refGene_tx=XM_023253023.1;refGene_gn=SHROOM3;refGene_tx=XM_023253024.1;refGene_gn=SHROOM3;refGene_tx=XM_023253025.1;refGene_gn=SHROOM3;refGene_tx=XM_023253027.1;refGene_gn=SHROOM3;refGene_tx=XM_023253028.1
B1 145399327 145402526
refGene_gn=SHROOM3;refGene_tx=XM_023253022.1;refGene_gn=SHROOM3;refGene_tx=XM_023253023.1;refGene_gn=SHROOM3;refGene_tx=XM_023253024.1;refGene_gn=SHROOM3;refGene_tx=XM_023253025.1;refGene_gn=SHROOM3;refGene_tx=XM_023253027.1;refGene_gn=SHROOM3;refGene_tx=XM_023253028.1
B1 145409622 145409754
refGene_gn=SHROOM3;refGene_tx=XM_023253022.1;refGene_gn=SHROOM3;refGene_tx=XM_023253023.1;refGene_gn=SHROOM3;refGene_tx=XM_023253024.1;refGene_gn=SHROOM3;refGene_tx=XM_023253025.1;refGene_gn=SHROOM3;refGene_tx=XM_023253027.1;refGene_gn=SHROOM3;refGene_tx=XM_023253028.1
B1 145423243 145423323
refGene_gn=SHROOM3;refGene_tx=XM_023253024.1

B1 145428584 145428716
refGene_gn=SHROOM3;refGene_tx=XM_023253022.1;refGene_gn=SHROOM3;refGene_tx=XM_023253023.1
B1 145586734 145586889
refGene_gn=SHROOM3;refGene_tx=XM_023253022.1;refGene_gn=SHROOM3;refGene_tx=XM_023253023.1
B1 145699625 145699793
refGene_gn=SHROOM3;refGene_tx=XM_023253022.1;refGene_gn=SHROOM3;refGene_tx=XM_023253023.1
B1 145732816 145732890
refGene_gn=CCDC158;refGene_tx=XM_023253029.1
B1 145738959 145738964
refGene_gn=CCDC158;refGene_tx=XM_023253029.1
B1 145739195 145739404
refGene_gn=CCDC158;refGene_tx=XM_023253029.1
B1 145743404 145743514
refGene_gn=CCDC158;refGene_tx=XM_023253029.1
B1 145743626 145743697
refGene_gn=CCDC158;refGene_tx=XM_023253029.1
B1 145743699 145743819
refGene_gn=CCDC158;refGene_tx=XM_023253029.1
B1 145743821 145743952
refGene_gn=CCDC158;refGene_tx=XM_023253029.1
B1 145744306 145744400
refGene_gn=CCDC158;refGene_tx=XM_023253029.1
B1 145745261 145745303
refGene_gn=CCDC158;refGene_tx=XM_023253029.1
B1 145745305 145745360
refGene_gn=CCDC158;refGene_tx=XM_023253029.1
B1 145746497 145746526
refGene_gn=CCDC158;refGene_tx=XM_023253029.1
B1 145746528 145746611
refGene_gn=CCDC158;refGene_tx=XM_023253029.1
B1 145753308 145753428
refGene_gn=CCDC158;refGene_tx=XM_023253029.1
B1 145755410 145755420
refGene_gn=CCDC158;refGene_tx=XM_023253029.1
B1 145755422 145755607
refGene_gn=CCDC158;refGene_tx=XM_023253029.1
B1 145756749 145757232
refGene_gn=CCDC158;refGene_tx=XM_023253029.1
B1 145764788 145764978
refGene_gn=CCDC158;refGene_tx=XM_023253029.1
B1 145767021 145767174
refGene_gn=CCDC158;refGene_tx=XM_023253029.1
B1 145769693 145769806
refGene_gn=CCDC158;refGene_tx=XM_023253029.1
B1 145771623 145771782
refGene_gn=CCDC158;refGene_tx=XM_023253029.1
B1 145772938 145773031

refGene_gn=CCDC158;refGene_tx=XM_023253029.1
B1 145773820 145773946
refGene_gn=CCDC158;refGene_tx=XM_023253029.1
B1 145810382 145810543
refGene_gn=CCDC158;refGene_tx=XM_023253029.1
B1 145811878 145811938
refGene_gn=CCDC158;refGene_tx=XM_023253029.1
B1 145812831 145812891
refGene_gn=CCDC158;refGene_tx=XM_023253029.1
B1 145816641 145816709
refGene_gn=CCDC158;refGene_tx=XM_023253029.1
B1 145819351 145819510
refGene_gn=CCDC158;refGene_tx=XM_023253029.1
B1 145821294 145821402
refGene_gn=CCDC158;refGene_tx=XM_023253029.1
B1 145823801 145823878
refGene_gn=CCDC158;refGene_tx=XM_023253029.1
B1 145827397 145828257
refGene_gn=STBD1;refGene_tx=XM_003985320.4
B1 145830377 145830579
refGene_gn=STBD1;refGene_tx=XM_003985320.4
B1 145914288 145914405
refGene_gn=SCARB2;refGene_tx=XM_003985268.4
B1 145936690 145936848
refGene_gn=SCARB2;refGene_tx=XM_003985268.4
B1 145950853 145951001
refGene_gn=SCARB2;refGene_tx=XM_003985268.4
B1 145952085 145952274
refGene_gn=SCARB2;refGene_tx=XM_003985268.4
B1 145960921 145961013
refGene_gn=SCARB2;refGene_tx=XM_003985268.4
B1 145969727 145969847
refGene_gn=SCARB2;refGene_tx=XM_003985268.4
B1 145972526 145972696
refGene_gn=SCARB2;refGene_tx=XM_003985268.4
B1 145977717 145977836
refGene_gn=SCARB2;refGene_tx=XM_003985268.4
B1 145979121 145979195
refGene_gn=SCARB2;refGene_tx=XM_003985268.4
B1 145981674 145981726
refGene_gn=SCARB2;refGene_tx=XM_003985268.4
B1 145985116 145985275
refGene_gn=SCARB2;refGene_tx=XM_003985268.4
B1 145986771 145986810
refGene_gn=SCARB2;refGene_tx=XM_003985268.4
B1 145996267 145996334
refGene_gn=NUP54;refGene_tx=XM_003985270.4;refGene_gn=NUP54;refGene_tx
=XM_019829347.1
B1 145996990 145997152
refGene_gn=NUP54;refGene_tx=XM_019829347.1

B1	145999955	146000039
refGene_gn=NUP54;refGene_tx=XM_003985270.4;refGene_gn=NUP54;refGene_tx
=XM_019829347.1
B1	146000131	146000278
refGene_gn=NUP54;refGene_tx=XM_003985270.4;refGene_gn=NUP54;refGene_tx
=XM_019829347.1
B1	146009011	146009238
refGene_gn=NUP54;refGene_tx=XM_003985270.4;refGene_gn=NUP54;refGene_tx
=XM_019829347.1
B1	146011002	146011190
refGene_gn=NUP54;refGene_tx=XM_003985270.4;refGene_gn=NUP54;refGene_tx
=XM_019829347.1
B1	146011928	146012125
refGene_gn=NUP54;refGene_tx=XM_003985270.4;refGene_gn=NUP54;refGene_tx
=XM_019829347.1
B1	146013753	146013808
refGene_gn=NUP54;refGene_tx=XM_003985270.4;refGene_gn=NUP54;refGene_tx
=XM_019829347.1
B1	146014249	146014343
refGene_gn=NUP54;refGene_tx=XM_003985270.4;refGene_gn=NUP54;refGene_tx
=XM_019829347.1
B1	146019523	146019631
refGene_gn=NUP54;refGene_tx=XM_003985270.4;refGene_gn=NUP54;refGene_tx
=XM_019829347.1
B1	146023618	146023738
refGene_gn=NUP54;refGene_tx=XM_003985270.4;refGene_gn=NUP54;refGene_tx
=XM_019829347.1
B1	146024022	146024133
refGene_gn=NUP54;refGene_tx=XM_003985270.4;refGene_gn=NUP54;refGene_tx
=XM_019829347.1
B1	146026157	146026286
refGene_gn=NUP54;refGene_tx=XM_003985270.4;refGene_gn=NUP54;refGene_tx
=XM_019829347.1
B1	146044672	146044812
refGene_gn=ART3;refGene_tx=XM_006931050.2;refGene_gn=ART3;refGene_tx=X
M_019829348.2;refGene_gn=ART3;refGene_tx=XM_023253031.1;refGene_gn=ART
3;refGene_tx=XM_023253032.1;refGene_gn=ART3;refGene_tx=XM_023253033.1;
refGene_gn=ART3;refGene_tx=XM_023253034.1;refGene_gn=ART3;refGene_tx=X
M_023253035.1
B1	146046439	146046472
refGene_gn=ART3;refGene_tx=XM_006931050.2;refGene_gn=ART3;refGene_tx=X
M_019829348.2;refGene_gn=ART3;refGene_tx=XM_023253031.1;refGene_gn=ART
3;refGene_tx=XM_023253032.1;refGene_gn=ART3;refGene_tx=XM_023253033.1;
refGene_gn=ART3;refGene_tx=XM_023253034.1;refGene_gn=ART3;refGene_tx=X
M_023253035.1
B1	146047780	146047813
refGene_gn=ART3;refGene_tx=XM_006931050.2;refGene_gn=ART3;refGene_tx=X
M_019829348.2;refGene_gn=ART3;refGene_tx=XM_023253031.1;refGene_gn=ART
3;refGene_tx=XM_023253032.1;refGene_gn=ART3;refGene_tx=XM_023253033.1
B1	146048701	146048734

refGene_gn=ART3;refGene_tx=XM_006931050.2;refGene_gn=ART3;refGene_tx=XM_019829348.2;refGene_gn=ART3;refGene_tx=XM_023253031.1
B1 146049349 146049382
refGene_gn=ART3;refGene_tx=XM_006931050.2;refGene_gn=ART3;refGene_tx=XM_019829348.2;refGene_gn=ART3;refGene_tx=XM_023253031.1;refGene_gn=ART3;refGene_tx=XM_023253032.1;refGene_gn=ART3;refGene_tx=XM_023253033.1;refGene_gn=ART3;refGene_tx=XM_023253034.1;refGene_gn=ART3;refGene_tx=XM_023253035.1
B1 146050443 146050476
refGene_gn=ART3;refGene_tx=XM_006931050.2;refGene_gn=ART3;refGene_tx=XM_019829348.2;refGene_gn=ART3;refGene_tx=XM_023253031.1;refGene_gn=ART3;refGene_tx=XM_023253032.1;refGene_gn=ART3;refGene_tx=XM_023253033.1;refGene_gn=ART3;refGene_tx=XM_023253034.1;refGene_gn=ART3;refGene_tx=XM_023253035.1
B1 146052173 146052203
refGene_gn=ART3;refGene_tx=XM_006931050.2;refGene_gn=ART3;refGene_tx=XM_019829348.2;refGene_gn=ART3;refGene_tx=XM_023253032.1;refGene_gn=ART3;refGene_tx=XM_023253034.1
B1 146053178 146053211
refGene_gn=ART3;refGene_tx=XM_006931050.2;refGene_gn=ART3;refGene_tx=XM_019829348.2;refGene_gn=ART3;refGene_tx=XM_023253031.1;refGene_gn=ART3;refGene_tx=XM_023253032.1;refGene_gn=ART3;refGene_tx=XM_023253033.1;refGene_gn=ART3;refGene_tx=XM_023253034.1;refGene_gn=ART3;refGene_tx=XM_023253035.1
B1 146054144 146054177
refGene_gn=ART3;refGene_tx=XM_006931050.2;refGene_gn=ART3;refGene_tx=XM_019829348.2;refGene_gn=ART3;refGene_tx=XM_023253031.1;refGene_gn=ART3;refGene_tx=XM_023253032.1;refGene_gn=ART3;refGene_tx=XM_023253033.1;refGene_gn=ART3;refGene_tx=XM_023253034.1;refGene_gn=ART3;refGene_tx=XM_023253035.1
B1 146072026 146072738
refGene_gn=ART3;refGene_tx=XM_006931050.2;refGene_gn=ART3;refGene_tx=XM_019829348.2;refGene_gn=ART3;refGene_tx=XM_023253031.1;refGene_gn=ART3;refGene_tx=XM_023253032.1;refGene_gn=ART3;refGene_tx=XM_023253033.1;refGene_gn=ART3;refGene_tx=XM_023253034.1;refGene_gn=ART3;refGene_tx=XM_023253035.1
B1 146078771 146078840
refGene_gn=ART3;refGene_tx=XM_006931050.2;refGene_gn=ART3;refGene_tx=XM_019829348.2;refGene_gn=ART3;refGene_tx=XM_023253031.1;refGene_gn=ART3;refGene_tx=XM_023253032.1;refGene_gn=ART3;refGene_tx=XM_023253033.1;refGene_gn=ART3;refGene_tx=XM_023253034.1;refGene_gn=ART3;refGene_tx=XM_023253035.1
B1 146137331 146137392
refGene_gn=CXCL11;refGene_tx=XM_023253037.1
B1 146137969 146138096
refGene_gn=CXCL11;refGene_tx=XM_023253037.1
B1 146138211 146138284
refGene_gn=CXCL11;refGene_tx=XM_023253037.1
B1 146138515 146138557
refGene_gn=CXCL11;refGene_tx=XM_023253037.1

B1 146169769 146169830
refGene_gn=CXCL10;refGene_tx=XM_003985274.3
B1 146170337 146170464
refGene_gn=CXCL10;refGene_tx=XM_003985274.3
B1 146170682 146170772
refGene_gn=CXCL10;refGene_tx=XM_003985274.3
B1 146171228 146171247
refGene_gn=CXCL10;refGene_tx=XM_003985274.3
B1 146196272 146196336
refGene_gn=CXCL9;refGene_tx=XM_003985275.4
B1 146198478 146198605
refGene_gn=CXCL9;refGene_tx=XM_003985275.4
B1 146199935 146200020
refGene_gn=CXCL9;refGene_tx=XM_003985275.4
B1 146201075 146201180
refGene_gn=CXCL9;refGene_tx=XM_003985275.4
B1 146218500 146218590
refGene_gn=SDAD1;refGene_tx=XM_011281730.2
B1 146220266 146220371
refGene_gn=SDAD1;refGene_tx=XM_011281730.2
B1 146221095 146221194
refGene_gn=SDAD1;refGene_tx=XM_011281730.2
B1 146223853 146223964
refGene_gn=SDAD1;refGene_tx=XM_011281730.2
B1 146225562 146225634
refGene_gn=SDAD1;refGene_tx=XM_011281730.2
B1 146225713 146225814
refGene_gn=SDAD1;refGene_tx=XM_011281730.2
B1 146227473 146227531
refGene_gn=SDAD1;refGene_tx=XM_011281730.2
B1 146228127 146228202
refGene_gn=SDAD1;refGene_tx=XM_011281730.2
B1 146229576 146229678
refGene_gn=SDAD1;refGene_tx=XM_011281730.2
B1 146230234 146230304
refGene_gn=SDAD1;refGene_tx=XM_011281730.2
B1 146230925 146231029
refGene_gn=SDAD1;refGene_tx=XM_011281730.2
B1 146233034 146233092
refGene_gn=SDAD1;refGene_tx=XM_011281730.2
B1 146234173 146234232
refGene_gn=SDAD1;refGene_tx=XM_011281730.2
B1 146234829 146234906
refGene_gn=SDAD1;refGene_tx=XM_011281730.2
B1 146235076 146235174
refGene_gn=SDAD1;refGene_tx=XM_011281730.2
B1 146235482 146235559
refGene_gn=SDAD1;refGene_tx=XM_011281730.2
B1 146236025 146236152
refGene_gn=SDAD1;refGene_tx=XM_011281730.2

B1 146237426 146237521
refGene_gn=SDAD1;refGene_tx=XM_011281730.2
B1 146237656 146237847
refGene_gn=SDAD1;refGene_tx=XM_011281730.2
B1 146237956 146238041
refGene_gn=SDAD1;refGene_tx=XM_011281730.2
B1 146238873 146239035
refGene_gn=SDAD1;refGene_tx=XM_011281730.2
B1 146243113 146243161
refGene_gn=SDAD1;refGene_tx=XM_011281730.2
B1 146251574 146251777
refGene_gn=NAAA;refGene_tx=XM_019829942.2
B1 146252452 146252617
refGene_gn=NAAA;refGene_tx=XM_019829942.2
B1 146255053 146255180
refGene_gn=NAAA;refGene_tx=XM_019829942.2
B1 146259607 146259698
refGene_gn=NAAA;refGene_tx=XM_019829942.2
B1 146262991 146263068
refGene_gn=NAAA;refGene_tx=XM_019829942.2
B1 146267664 146267837
refGene_gn=NAAA;refGene_tx=XM_019829942.2
B1 146267979 146268042
refGene_gn=NAAA;refGene_tx=XM_019829942.2
B1 146269246 146269313
refGene_gn=NAAA;refGene_tx=XM_019829942.2
B1 146269966 146269995
refGene_gn=NAAA;refGene_tx=XM_019829942.2
B1 146273742 146273818
refGene_gn=NAAA;refGene_tx=XM_019829942.2
B1 146294044 146294099
refGene_gn=PPEF2;refGene_tx=XM_003985322.5;refGene_gn=PPEF2;refGene_tx
=XM_023253038.1
B1 146298794 146298922
refGene_gn=PPEF2;refGene_tx=XM_003985322.5;refGene_gn=PPEF2;refGene_tx
=XM_023253038.1;refGene_gn=PPEF2;refGene_tx=XM_023253039.1
B1 146299072 146299130
refGene_gn=PPEF2;refGene_tx=XM_003985322.5;refGene_gn=PPEF2;refGene_tx
=XM_023253038.1;refGene_gn=PPEF2;refGene_tx=XM_023253039.1
B1 146300333 146300509
refGene_gn=PPEF2;refGene_tx=XM_003985322.5;refGene_gn=PPEF2;refGene_tx
=XM_023253038.1;refGene_gn=PPEF2;refGene_tx=XM_023253039.1
B1 146301810 146301925
refGene_gn=PPEF2;refGene_tx=XM_003985322.5;refGene_gn=PPEF2;refGene_tx
=XM_023253038.1;refGene_gn=PPEF2;refGene_tx=XM_023253039.1
B1 146304480 146304527
refGene_gn=PPEF2;refGene_tx=XM_003985322.5;refGene_gn=PPEF2;refGene_tx
=XM_023253038.1;refGene_gn=PPEF2;refGene_tx=XM_023253039.1
B1 146305530 146305697
refGene_gn=PPEF2;refGene_tx=XM_003985322.5;refGene_gn=PPEF2;refGene_tx

=XM_023253038.1;refGene_gn=PPEF2;refGene_tx=XM_023253039.1
B1 146306177 146306214
refGene_gn=PPEF2;refGene_tx=XM_003985322.5;refGene_gn=PPEF2;refGene_tx
=XM_023253038.1;refGene_gn=PPEF2;refGene_tx=XM_023253039.1
B1 146306306 146306456
refGene_gn=PPEF2;refGene_tx=XM_003985322.5;refGene_gn=PPEF2;refGene_tx
=XM_023253038.1;refGene_gn=PPEF2;refGene_tx=XM_023253039.1
B1 146309998 146310382
refGene_gn=PPEF2;refGene_tx=XM_003985322.5;refGene_gn=PPEF2;refGene_tx
=XM_023253038.1;refGene_gn=PPEF2;refGene_tx=XM_023253039.1
B1 146312208 146312394
refGene_gn=PPEF2;refGene_tx=XM_003985322.5;refGene_gn=PPEF2;refGene_tx
=XM_023253038.1;refGene_gn=PPEF2;refGene_tx=XM_023253039.1
B1 146313499 146313642
refGene_gn=PPEF2;refGene_tx=XM_003985322.5;refGene_gn=PPEF2;refGene_tx
=XM_023253038.1;refGene_gn=PPEF2;refGene_tx=XM_023253039.1
B1 146316927 146317034
refGene_gn=PPEF2;refGene_tx=XM_003985322.5;refGene_gn=PPEF2;refGene_tx
=XM_023253038.1;refGene_gn=PPEF2;refGene_tx=XM_023253039.1
B1 146318572 146318736
refGene_gn=PPEF2;refGene_tx=XM_003985322.5;refGene_gn=PPEF2;refGene_tx
=XM_023253038.1;refGene_gn=PPEF2;refGene_tx=XM_023253039.1
B1 146319460 146319548
refGene_gn=PPEF2;refGene_tx=XM_003985322.5;refGene_gn=PPEF2;refGene_tx
=XM_023253038.1;refGene_gn=PPEF2;refGene_tx=XM_023253039.1
B1 146322216 146322470
refGene_gn=PPEF2;refGene_tx=XM_003985322.5;refGene_gn=PPEF2;refGene_tx
=XM_023253038.1;refGene_gn=PPEF2;refGene_tx=XM_023253039.1
B1 146391401 146391491
refGene_gn=USO1;refGene_tx=XM_023253040.1;refGene_gn=USO1;refGene_tx=X
M_023253041.1
B1 146392223 146392439
refGene_gn=USO1;refGene_tx=XM_023253040.1;refGene_gn=USO1;refGene_tx=X
M_023253041.1
B1 146393406 146393514
refGene_gn=USO1;refGene_tx=XM_023253040.1;refGene_gn=USO1;refGene_tx=X
M_023253041.1
B1 146395552 146395651
refGene_gn=USO1;refGene_tx=XM_023253040.1;refGene_gn=USO1;refGene_tx=X
M_023253041.1
B1 146398886 146398976
refGene_gn=USO1;refGene_tx=XM_023253040.1;refGene_gn=USO1;refGene_tx=X
M_023253041.1
B1 146399653 146399817
refGene_gn=USO1;refGene_tx=XM_023253040.1;refGene_gn=USO1;refGene_tx=X
M_023253041.1
B1 146400791 146400930
refGene_gn=USO1;refGene_tx=XM_023253040.1;refGene_gn=USO1;refGene_tx=X
M_023253041.1
B1 146403543 146403665

refGene_gn=USO1;refGene_tx=XM_023253040.1;refGene_gn=USO1;refGene_tx=XM_023253041.1
B1 146403944 146404126
refGene_gn=USO1;refGene_tx=XM_023253040.1;refGene_gn=USO1;refGene_tx=XM_023253041.1
B1 146404475 146404594
refGene_gn=USO1;refGene_tx=XM_023253040.1;refGene_gn=USO1;refGene_tx=XM_023253041.1
B1 146404908 146405019
refGene_gn=USO1;refGene_tx=XM_023253040.1;refGene_gn=USO1;refGene_tx=XM_023253041.1
B1 146410983 146411004
refGene_gn=USO1;refGene_tx=XM_023253040.1
B1 146412772 146412984
refGene_gn=USO1;refGene_tx=XM_023253040.1;refGene_gn=USO1;refGene_tx=XM_023253041.1
B1 146413387 146413542
refGene_gn=USO1;refGene_tx=XM_023253040.1;refGene_gn=USO1;refGene_tx=XM_023253041.1
B1 146413940 146414029
refGene_gn=USO1;refGene_tx=XM_023253040.1;refGene_gn=USO1;refGene_tx=XM_023253041.1
B1 146415919 146416060
refGene_gn=USO1;refGene_tx=XM_023253040.1;refGene_gn=USO1;refGene_tx=XM_023253041.1
B1 146419696 146419875
refGene_gn=USO1;refGene_tx=XM_023253040.1;refGene_gn=USO1;refGene_tx=XM_023253041.1
B1 146432623 146432744
refGene_gn=USO1;refGene_tx=XM_023253040.1;refGene_gn=USO1;refGene_tx=XM_023253041.1
B1 146435663 146435719
refGene_gn=USO1;refGene_tx=XM_023253040.1;refGene_gn=USO1;refGene_tx=XM_023253041.1
B1 146435875 146435978
refGene_gn=USO1;refGene_tx=XM_023253040.1;refGene_gn=USO1;refGene_tx=XM_023253041.1
B1 146436386 146436487
refGene_gn=USO1;refGene_tx=XM_023253040.1;refGene_gn=USO1;refGene_tx=XM_023253041.1
B1 146450801 146450878
refGene_gn=USO1;refGene_tx=XM_023253040.1;refGene_gn=USO1;refGene_tx=XM_023253041.1
B1 146460102 146460167
refGene_gn=USO1;refGene_tx=XM_023253040.1;refGene_gn=USO1;refGene_tx=XM_023253041.1
B1 146460244 146460331
refGene_gn=USO1;refGene_tx=XM_023253040.1;refGene_gn=USO1;refGene_tx=XM_023253041.1
B1 146485865 146485931

refGene_gn=USO1;refGene_tx=XM_023253040.1;refGene_gn=USO1;refGene_tx=XM_023253041.1
B1 146547612 146547707
refGene_gn=G3BP2;refGene_tx=XM_003985278.5;refGene_gn=G3BP2;refGene_tx=XM_003985279.5;refGene_gn=G3BP2;refGene_tx=XM_006931055.4;refGene_gn=G3BP2;refGene_tx=XM_006931056.4;refGene_gn=G3BP2;refGene_tx=XM_023253042.1
B1 146549882 146549964
refGene_gn=G3BP2;refGene_tx=XM_003985278.5;refGene_gn=G3BP2;refGene_tx=XM_003985279.5;refGene_gn=G3BP2;refGene_tx=XM_006931055.4;refGene_gn=G3BP2;refGene_tx=XM_006931056.4;refGene_gn=G3BP2;refGene_tx=XM_023253042.1
B1 146550787 146550961
refGene_gn=G3BP2;refGene_tx=XM_003985278.5;refGene_gn=G3BP2;refGene_tx=XM_003985279.5;refGene_gn=G3BP2;refGene_tx=XM_006931055.4;refGene_gn=G3BP2;refGene_tx=XM_006931056.4;refGene_gn=G3BP2;refGene_tx=XM_023253042.1
B1 146552085 146552176
refGene_gn=G3BP2;refGene_tx=XM_003985278.5;refGene_gn=G3BP2;refGene_tx=XM_003985279.5;refGene_gn=G3BP2;refGene_tx=XM_006931055.4;refGene_gn=G3BP2;refGene_tx=XM_006931056.4;refGene_gn=G3BP2;refGene_tx=XM_023253042.1
B1 146553321 146553424
refGene_gn=G3BP2;refGene_tx=XM_003985278.5;refGene_gn=G3BP2;refGene_tx=XM_003985279.5;refGene_gn=G3BP2;refGene_tx=XM_006931055.4;refGene_gn=G3BP2;refGene_tx=XM_006931056.4;refGene_gn=G3BP2;refGene_tx=XM_023253042.1
B1 146553977 146554158
refGene_gn=G3BP2;refGene_tx=XM_003985278.5;refGene_gn=G3BP2;refGene_tx=XM_003985279.5;refGene_gn=G3BP2;refGene_tx=XM_006931055.4;refGene_gn=G3BP2;refGene_tx=XM_006931056.4;refGene_gn=G3BP2;refGene_tx=XM_023253042.1
B1 146555132 146555231
refGene_gn=G3BP2;refGene_tx=XM_003985278.5;refGene_gn=G3BP2;refGene_tx=XM_006931055.4;refGene_gn=G3BP2;refGene_tx=XM_006931056.4;refGene_gn=G3BP2;refGene_tx=XM_023253042.1
B1 146559803 146559906
refGene_gn=G3BP2;refGene_tx=XM_003985278.5;refGene_gn=G3BP2;refGene_tx=XM_003985279.5;refGene_gn=G3BP2;refGene_tx=XM_006931055.4;refGene_gn=G3BP2;refGene_tx=XM_006931056.4;refGene_gn=G3BP2;refGene_tx=XM_023253042.1
B1 146561115 146561244
refGene_gn=G3BP2;refGene_tx=XM_003985278.5;refGene_gn=G3BP2;refGene_tx=XM_003985279.5;refGene_gn=G3BP2;refGene_tx=XM_006931055.4;refGene_gn=G3BP2;refGene_tx=XM_006931056.4;refGene_gn=G3BP2;refGene_tx=XM_023253042.1
B1 146561796 146561915
refGene_gn=G3BP2;refGene_tx=XM_003985278.5;refGene_gn=G3BP2;refGene_tx=XM_003985279.5;refGene_gn=G3BP2;refGene_tx=XM_006931055.4;refGene_gn=G3BP2;refGene_tx=XM_006931056.4;refGene_gn=G3BP2;refGene_tx=XM_0232530

42.1
B1	146562401	146562674
refGene_gn=G3BP2;refGene_tx=XM_003985278.5;refGene_gn=G3BP2;refGene_tx=XM_003985279.5;refGene_gn=G3BP2;refGene_tx=XM_006931055.4;refGene_gn=G3BP2;refGene_tx=XM_006931056.4;refGene_gn=G3BP2;refGene_tx=XM_023253042.1
B1	146580669	146580837
refGene_gn=CDKL2;refGene_tx=XM_006931059.4;refGene_gn=CDKL2;refGene_tx=XM_019829355.2
B1	146597018	146597213
refGene_gn=CDKL2;refGene_tx=XM_006931059.4;refGene_gn=CDKL2;refGene_tx=XM_019829355.2
B1	146599150	146599329
refGene_gn=CDKL2;refGene_tx=XM_006931059.4;refGene_gn=CDKL2;refGene_tx=XM_019829355.2
B1	146601523	146601636
refGene_gn=CDKL2;refGene_tx=XM_006931059.4;refGene_gn=CDKL2;refGene_tx=XM_019829355.2
B1	146603130	146603270
refGene_gn=CDKL2;refGene_tx=XM_006931059.4;refGene_gn=CDKL2;refGene_tx=XM_019829355.2
B1	146612441	146612530
refGene_gn=CDKL2;refGene_tx=XM_006931059.4;refGene_gn=CDKL2;refGene_tx=XM_019829355.2
B1	146615023	146615159
refGene_gn=CDKL2;refGene_tx=XM_006931059.4;refGene_gn=CDKL2;refGene_tx=XM_019829355.2
B1	146616031	146616321
refGene_gn=CDKL2;refGene_tx=XM_006931059.4;refGene_gn=CDKL2;refGene_tx=XM_019829355.2
B1	146616927	146617021
refGene_gn=CDKL2;refGene_tx=XM_006931059.4;refGene_gn=CDKL2;refGene_tx=XM_019829355.2
B1	146620123	146620247
refGene_gn=CDKL2;refGene_tx=XM_006931059.4;refGene_gn=CDKL2;refGene_tx=XM_019829355.2
B1	146620621	146620728
refGene_gn=CDKL2;refGene_tx=XM_006931059.4;refGene_gn=CDKL2;refGene_tx=XM_019829355.2
B1	146626594	146626660
refGene_gn=CDKL2;refGene_tx=XM_006931059.4;refGene_gn=CDKL2;refGene_tx=XM_019829355.2
B1	146644960	146645355
refGene_gn=ODAPH;refGene_tx=XM_019829356.2
B1	146653620	146653687
refGene_gn=ODAPH;refGene_tx=XM_019829356.2
B1	146671469	146671724
refGene_gn=THAP6;refGene_tx=XM_003985281.4;refGene_gn=THAP6;refGene_tx=XM_003985282.4;refGene_gn=THAP6;refGene_tx=XM_006931060.3;refGene_gn=THAP6;refGene_tx=XM_019829360.1

B1 146677958 146678084
refGene_gn=THAP6;refGene_tx=XM_003985281.4;refGene_gn=THAP6;refGene_tx=XM_003985282.4;refGene_gn=THAP6;refGene_tx=XM_006931060.3;refGene_gn=THAP6;refGene_tx=XM_019829360.1
B1 146683145 146683353
refGene_gn=THAP6;refGene_tx=XM_003985281.4;refGene_gn=THAP6;refGene_tx=XM_003985282.4;refGene_gn=THAP6;refGene_tx=XM_006931060.3;refGene_gn=THAP6;refGene_tx=XM_019829360.1
B1 146684495 146684575
refGene_gn=THAP6;refGene_tx=XM_003985281.4;refGene_gn=THAP6;refGene_tx=XM_003985282.4;refGene_gn=THAP6;refGene_tx=XM_006931060.3
B1 146685695 146685794
refGene_gn=RCHY1;refGene_tx=XM_003985283.5
B1 146691526 146691646
refGene_gn=RCHY1;refGene_tx=XM_003985283.5
B1 146691870 146691986
refGene_gn=RCHY1;refGene_tx=XM_003985283.5
B1 146702485 146702564
refGene_gn=RCHY1;refGene_tx=XM_003985283.5;refGene_gn=RCHY1;refGene_tx=XM_019829357.1;refGene_gn=RCHY1;refGene_tx=XM_019829358.1
B1 146704137 146704182
refGene_gn=RCHY1;refGene_tx=XM_003985283.5;refGene_gn=RCHY1;refGene_tx=XM_019829357.1;refGene_gn=RCHY1;refGene_tx=XM_019829358.1
B1 146704288 146704347
refGene_gn=RCHY1;refGene_tx=XM_003985283.5;refGene_gn=RCHY1;refGene_tx=XM_019829357.1;refGene_gn=RCHY1;refGene_tx=XM_019829358.1
B1 146704436 146704463
refGene_gn=RCHY1;refGene_tx=XM_003985283.5;refGene_gn=RCHY1;refGene_tx=XM_019829357.1;refGene_gn=RCHY1;refGene_tx=XM_019829358.1
B1 146705319 146705440
refGene_gn=RCHY1;refGene_tx=XM_003985283.5;refGene_gn=RCHY1;refGene_tx=XM_019829357.1;refGene_gn=RCHY1;refGene_tx=XM_019829358.1
B1 146707601 146707730
refGene_gn=RCHY1;refGene_tx=XM_003985283.5;refGene_gn=RCHY1;refGene_tx=XM_019829357.1;refGene_gn=RCHY1;refGene_tx=XM_019829358.1
B1 147030288 147030373
refGene_gn=PARM1;refGene_tx=XM_023253044.1;refGene_gn=PARM1;refGene_tx=XM_023253045.1
B1 147043720 147043799
refGene_gn=PARM1;refGene_tx=XM_023253044.1;refGene_gn=PARM1;refGene_tx=XM_023253045.1
B1 147062166 147062320
refGene_gn=PARM1;refGene_tx=XM_023253043.1
B1 147064279 147064331
refGene_gn=PARM1;refGene_tx=XM_023253043.1;refGene_gn=PARM1;refGene_tx=XM_023253045.1
B1 147065736 147066468
refGene_gn=PARM1;refGene_tx=XM_023253043.1;refGene_gn=PARM1;refGene_tx=XM_023253044.1;refGene_gn=PARM1;refGene_tx=XM_023253045.1
B1 147139911 147139954

refGene_gn=PARM1;refGene_tx=XM_023253043.1;refGene_gn=PARM1;refGene_tx=XM_023253044.1;refGene_gn=PARM1;refGene_tx=XM_023253045.1
B1 147256916 147256980
refGene_gn=BTC;refGene_tx=XM_019829362.2
B1 147273244 147273298
refGene_gn=BTC;refGene_tx=XM_019829362.2
B1 147282413 147282512
refGene_gn=BTC;refGene_tx=XM_019829362.2
B1 147293929 147294047
refGene_gn=BTC;refGene_tx=XM_019829362.2
B1 147299258 147299405
refGene_gn=BTC;refGene_tx=XM_019829362.2
B1 147301744 147301853
refGene_gn=BTC;refGene_tx=XM_019829362.2
B1 147572654 147572748
refGene_gn=AREG;refGene_tx=XM_006931061.4;refGene_gn=AREG;refGene_tx=XM_019829363.2
B1 147575295 147575451
refGene_gn=AREG;refGene_tx=XM_006931061.4;refGene_gn=AREG;refGene_tx=XM_019829363.2
B1 147576640 147576842
refGene_gn=AREG;refGene_tx=XM_006931061.4;refGene_gn=AREG;refGene_tx=XM_019829363.2
B1 147579445 147579691
refGene_gn=AREG;refGene_tx=XM_006931061.4;refGene_gn=AREG;refGene_tx=XM_019829363.2
B1 147580698 147580972
refGene_gn=AREG;refGene_tx=XM_006931061.4;refGene_gn=AREG;refGene_tx=XM_019829363.2
B1 147638709 147638791
refGene_gn=EREG;refGene_tx=XM_023253401.1
B1 147641045 147641195
refGene_gn=EREG;refGene_tx=XM_023253401.1
B1 147642060 147642184
refGene_gn=EREG;refGene_tx=XM_023253401.1
B1 147644664 147644751
refGene_gn=EREG;refGene_tx=XM_023253401.1
B1 147660489 147660686
refGene_gn=EREG;refGene_tx=XM_023253401.1
B1 147660803 147660844
refGene_gn=EREG;refGene_tx=XM_023253401.1
B1 147696738 147696793
refGene_gn=EPGN;refGene_tx=XM_019829365.2
B1 147698443 147698596
refGene_gn=EPGN;refGene_tx=XM_019829365.2
B1 147699334 147699455
refGene_gn=EPGN;refGene_tx=XM_019829365.2
B1 147702172 147702262
refGene_gn=EPGN;refGene_tx=XM_019829365.2
B1 147702834 147702877

refGene_gn=EPGN;refGene_tx=XM_019829365.2
B1 147709885 147709998
refGene_gn=MTHFD2L;refGene_tx=XM_023253046.1;refGene_gn=MTHFD2L;refGene_tx=XM_023253047.1
B1 147737520 147737646
refGene_gn=MTHFD2L;refGene_tx=XM_023253046.1;refGene_gn=MTHFD2L;refGene_tx=XM_023253047.1
B1 147782288 147782381
refGene_gn=MTHFD2L;refGene_tx=XM_023253046.1;refGene_gn=MTHFD2L;refGene_tx=XM_023253047.1
B1 147803881 147803989
refGene_gn=MTHFD2L;refGene_tx=XM_023253046.1
B1 147805187 147805340
refGene_gn=MTHFD2L;refGene_tx=XM_023253046.1;refGene_gn=MTHFD2L;refGene_tx=XM_023253047.1
B1 147835229 147835352
refGene_gn=MTHFD2L;refGene_tx=XM_023253046.1;refGene_gn=MTHFD2L;refGene_tx=XM_023253047.1
B1 147836374 147836559
refGene_gn=MTHFD2L;refGene_tx=XM_023253046.1;refGene_gn=MTHFD2L;refGene_tx=XM_023253047.1
B1 147852415 147852539
refGene_gn=MTHFD2L;refGene_tx=XM_023253048.1
B1 147854601 147854744
refGene_gn=MTHFD2L;refGene_tx=XM_023253046.1;refGene_gn=MTHFD2L;refGene_tx=XM_023253047.1;refGene_gn=MTHFD2L;refGene_tx=XM_023253048.1
B1 147887867 147887883
refGene_gn=LOC102901432;refGene_tx=XM_023253049.1
B1 147888669 147888753
refGene_gn=LOC102901432;refGene_tx=XM_023253049.1
B1 147888861 147888985
refGene_gn=LOC102901432;refGene_tx=XM_023253049.1
B1 147889091 147889179
refGene_gn=LOC102901432;refGene_tx=XM_023253049.1
B1 147900094 147900110
refGene_gn=LOC111560177;refGene_tx=XM_023253050.1
B1 147900883 147900967
refGene_gn=LOC111560177;refGene_tx=XM_023253050.1
B1 147901075 147901199
refGene_gn=LOC111560177;refGene_tx=XM_023253050.1
B1 147901304 147901392
refGene_gn=LOC111560177;refGene_tx=XM_023253050.1
B1 147988552 147988565
refGene_gn=LOC101093854;refGene_tx=XM_023253051.1
B1 147989918 147990002
refGene_gn=LOC101093854;refGene_tx=XM_023253051.1
B1 147994976 147995106
refGene_gn=LOC101093854;refGene_tx=XM_023253051.1
B1 147995762 147995826
refGene_gn=LOC101093854;refGene_tx=XM_023253051.1

B1 148006788 148006830
refGene_gn=LOC111560178;refGene_tx=XM_023253053.1
B1 148006831 148006877
refGene_gn=LOC111560178;refGene_tx=XM_023253053.1
B1 148006983 148007107
refGene_gn=LOC111560178;refGene_tx=XM_023253053.1
B1 148007211 148007299
refGene_gn=LOC111560178;refGene_tx=XM_023253053.1
B1 148024305 148024393
refGene_gn=LOC101094096;refGene_tx=XM_011281728.3
B1 148024511 148024635
refGene_gn=LOC101094096;refGene_tx=XM_011281728.3
B1 148024939 148025042
refGene_gn=LOC101094096;refGene_tx=XM_011281728.3
B1 148030534 148030637
refGene_gn=LOC101094344;refGene_tx=XM_003985329.4
B1 148030789 148030925
refGene_gn=LOC101094344;refGene_tx=XM_003985329.4
B1 148031262 148031371
refGene_gn=LOC101094344;refGene_tx=XM_003985329.4
B1 148041270 148041289
refGene_gn=CXCL6;refGene_tx=XM_003985330.5
B1 148041657 148041741
refGene_gn=CXCL6;refGene_tx=XM_003985330.5
B1 148041864 148041997
refGene_gn=CXCL6;refGene_tx=XM_003985330.5
B1 148042160 148042371
refGene_gn=CXCL6;refGene_tx=XM_003985330.5
B1 148127850 148127872
refGene_gn=CXCL8;refGene_tx=NM_001009281.1
B1 148128350 148128434
refGene_gn=CXCL8;refGene_tx=NM_001009281.1
B1 148128719 148128855
refGene_gn=CXCL8;refGene_tx=NM_001009281.1
B1 148130253 148130317
refGene_gn=CXCL8;refGene_tx=NM_001009281.1
B1 148226095 148226289
refGene_gn=RASSF6;refGene_tx=XM_019829371.2;refGene_gn=RASSF6;refGene_tx=XM_019829372.2
B1 148237661 148237760
refGene_gn=RASSF6;refGene_tx=XM_019829371.2
B1 148249663 148249713
refGene_gn=RASSF6;refGene_tx=XM_019829375.1
B1 148252896 148252975
refGene_gn=RASSF6;refGene_tx=XM_019829371.2;refGene_gn=RASSF6;refGene_tx=XM_019829372.2;refGene_gn=RASSF6;refGene_tx=XM_019829375.1
B1 148257349 148257489
refGene_gn=RASSF6;refGene_tx=XM_019829371.2;refGene_gn=RASSF6;refGene_tx=XM_019829372.2;refGene_gn=RASSF6;refGene_tx=XM_019829375.1
B1 148263147 148263245

refGene_gn=RASSF6;refGene_tx=XM_019829371.2;refGene_gn=RASSF6;refGene_tx=XM_019829372.2;refGene_gn=RASSF6;refGene_tx=XM_019829375.1
B1 148264652 148264837
refGene_gn=RASSF6;refGene_tx=XM_019829371.2;refGene_gn=RASSF6;refGene_tx=XM_019829372.2;refGene_gn=RASSF6;refGene_tx=XM_019829375.1
B1 148268658 148268760
refGene_gn=RASSF6;refGene_tx=XM_019829371.2;refGene_gn=RASSF6;refGene_tx=XM_019829372.2;refGene_gn=RASSF6;refGene_tx=XM_019829375.1
B1 148269106 148269158
refGene_gn=RASSF6;refGene_tx=XM_019829371.2;refGene_gn=RASSF6;refGene_tx=XM_019829372.2;refGene_gn=RASSF6;refGene_tx=XM_019829375.1
B1 148274045 148274164
refGene_gn=RASSF6;refGene_tx=XM_019829371.2;refGene_gn=RASSF6;refGene_tx=XM_019829372.2;refGene_gn=RASSF6;refGene_tx=XM_019829375.1
B1 148274270 148274368
refGene_gn=RASSF6;refGene_tx=XM_019829371.2;refGene_gn=RASSF6;refGene_tx=XM_019829372.2;refGene_gn=RASSF6;refGene_tx=XM_019829375.1
B1 148274469 148274545
refGene_gn=RASSF6;refGene_tx=XM_019829371.2;refGene_gn=RASSF6;refGene_tx=XM_019829372.2;refGene_gn=RASSF6;refGene_tx=XM_019829375.1
B1 148282331 148282379
refGene_gn=LOC101095098;refGene_tx=XM_003985331.3;refGene_gn=LOC101095098;refGene_tx=XM_019829376.2;refGene_gn=LOC101095098;refGene_tx=XM_019829377.2;refGene_gn=LOC101095098;refGene_tx=XM_019829378.2;refGene_gn=LOC101095098;refGene_tx=XM_019829379.2;refGene_gn=LOC101095098;refGene_tx=XM_019829380.2
B1 148288982 148289115
refGene_gn=LOC101095098;refGene_tx=XM_003985331.3;refGene_gn=LOC101095098;refGene_tx=XM_019829376.2;refGene_gn=LOC101095098;refGene_tx=XM_019829377.2;refGene_gn=LOC101095098;refGene_tx=XM_019829378.2;refGene_gn=LOC101095098;refGene_tx=XM_019829380.2
B1 148297178 148297405
refGene_gn=LOC101095098;refGene_tx=XM_003985331.3;refGene_gn=LOC101095098;refGene_tx=XM_019829376.2;refGene_gn=LOC101095098;refGene_tx=XM_019829377.2;refGene_gn=LOC101095098;refGene_tx=XM_019829378.2
B1 148299904 148300043
refGene_gn=LOC101095098;refGene_tx=XM_003985331.3;refGene_gn=LOC101095098;refGene_tx=XM_019829376.2;refGene_gn=LOC101095098;refGene_tx=XM_019829377.2;refGene_gn=LOC101095098;refGene_tx=XM_019829378.2;refGene_gn=LOC101095098;refGene_tx=XM_019829379.2
B1 148302236 148302334
refGene_gn=LOC101095098;refGene_tx=XM_003985331.3;refGene_gn=LOC101095098;refGene_tx=XM_019829376.2;refGene_gn=LOC101095098;refGene_tx=XM_019829377.2;refGene_gn=LOC101095098;refGene_tx=XM_019829378.2;refGene_gn=LOC101095098;refGene_tx=XM_019829379.2;refGene_gn=LOC101095098;refGene_tx=XM_019829380.2
B1 148304206 148304339
refGene_gn=LOC101095098;refGene_tx=XM_003985331.3;refGene_gn=LOC101095098;refGene_tx=XM_019829376.2;refGene_gn=LOC101095098;refGene_tx=XM_019829377.2;refGene_gn=LOC101095098;refGene_tx=XM_019829379.2;refGene_gn

=LOC101095098;refGene_tx=XM_019829380.2
B1 148305842 148306057
refGene_gn=LOC101095098;refGene_tx=XM_003985331.3;refGene_gn=LOC101095098;refGene_tx=XM_019829376.2;refGene_gn=LOC101095098;refGene_tx=XM_019829377.2;refGene_gn=LOC101095098;refGene_tx=XM_019829379.2;refGene_gn=LOC101095098;refGene_tx=XM_019829380.2
B1 148312759 148312889
refGene_gn=LOC101095098;refGene_tx=XM_003985331.3;refGene_gn=LOC101095098;refGene_tx=XM_019829376.2;refGene_gn=LOC101095098;refGene_tx=XM_019829377.2;refGene_gn=LOC101095098;refGene_tx=XM_019829378.2;refGene_gn=LOC101095098;refGene_tx=XM_019829379.2;refGene_gn=LOC101095098;refGene_tx=XM_019829380.2
B1 148314368 148314466
refGene_gn=LOC101095098;refGene_tx=XM_003985331.3;refGene_gn=LOC101095098;refGene_tx=XM_019829376.2;refGene_gn=LOC101095098;refGene_tx=XM_019829377.2;refGene_gn=LOC101095098;refGene_tx=XM_019829378.2;refGene_gn=LOC101095098;refGene_tx=XM_019829379.2;refGene_gn=LOC101095098;refGene_tx=XM_019829380.2
B1 148317462 148317595
refGene_gn=LOC101095098;refGene_tx=XM_003985331.3;refGene_gn=LOC101095098;refGene_tx=XM_019829376.2;refGene_gn=LOC101095098;refGene_tx=XM_019829377.2;refGene_gn=LOC101095098;refGene_tx=XM_019829378.2;refGene_gn=LOC101095098;refGene_tx=XM_019829379.2;refGene_gn=LOC101095098;refGene_tx=XM_019829380.2
B1 148327868 148328080
refGene_gn=LOC101095098;refGene_tx=XM_003985331.3;refGene_gn=LOC101095098;refGene_tx=XM_019829376.2;refGene_gn=LOC101095098;refGene_tx=XM_019829377.2;refGene_gn=LOC101095098;refGene_tx=XM_019829378.2;refGene_gn=LOC101095098;refGene_tx=XM_019829379.2;refGene_gn=LOC101095098;refGene_tx=XM_019829380.2
B1 148329408 148329550
refGene_gn=LOC101095098;refGene_tx=XM_003985331.3;refGene_gn=LOC101095098;refGene_tx=XM_019829376.2;refGene_gn=LOC101095098;refGene_tx=XM_019829378.2;refGene_gn=LOC101095098;refGene_tx=XM_019829379.2;refGene_gn=LOC101095098;refGene_tx=XM_019829380.2
B1 148329689 148329741
refGene_gn=LOC101095098;refGene_tx=XM_003985331.3;refGene_gn=LOC101095098;refGene_tx=XM_019829376.2;refGene_gn=LOC101095098;refGene_tx=XM_019829378.2;refGene_gn=LOC101095098;refGene_tx=XM_019829379.2;refGene_gn=LOC101095098;refGene_tx=XM_019829380.2
B1 148330890 148330975
refGene_gn=LOC101095098;refGene_tx=XM_003985331.3;refGene_gn=LOC101095098;refGene_tx=XM_019829376.2;refGene_gn=LOC101095098;refGene_tx=XM_019829378.2;refGene_gn=LOC101095098;refGene_tx=XM_019829379.2;refGene_gn=LOC101095098;refGene_tx=XM_019829380.2
B1 148340535 148340562
refGene_gn=AFM;refGene_tx=XM_019829381.2
B1 148341728 148341861
refGene_gn=AFM;refGene_tx=XM_019829381.2
B1 148342707 148342931

refGene_gn=AFM;refGene_tx=XM_019829381.2
B1 148343708 148343841
refGene_gn=AFM;refGene_tx=XM_019829381.2
B1 148345085 148345183
refGene_gn=AFM;refGene_tx=XM_019829381.2
B1 148347881 148348014
refGene_gn=AFM;refGene_tx=XM_019829381.2
B1 148351467 148351682
refGene_gn=AFM;refGene_tx=XM_019829381.2
B1 148354235 148354365
refGene_gn=AFM;refGene_tx=XM_019829381.2
B1 148355124 148355222
refGene_gn=AFM;refGene_tx=XM_019829381.2
B1 148355826 148355959
refGene_gn=AFM;refGene_tx=XM_019829381.2
B1 148356758 148356970
refGene_gn=AFM;refGene_tx=XM_019829381.2
B1 148357672 148357805
refGene_gn=AFM;refGene_tx=XM_019829381.2
B1 148358335 148358384
refGene_gn=AFM;refGene_tx=XM_019829381.2
B1 148361650 148361738
refGene_gn=AFM;refGene_tx=XM_019829381.2
B1 148371925 148371970
refGene_gn=AFP;refGene_tx=XM_003985288.2
B1 148373263 148373396
refGene_gn=AFP;refGene_tx=XM_003985288.2
B1 148374757 148374981
refGene_gn=AFP;refGene_tx=XM_003985288.2
B1 148377250 148377389
refGene_gn=AFP;refGene_tx=XM_003985288.2
B1 148378205 148378303
refGene_gn=AFP;refGene_tx=XM_003985288.2
B1 148379070 148379203
refGene_gn=AFP;refGene_tx=XM_003985288.2
B1 148380805 148381020
refGene_gn=AFP;refGene_tx=XM_003985288.2
B1 148382730 148382860
refGene_gn=AFP;refGene_tx=XM_003985288.2
B1 148385714 148385812
refGene_gn=AFP;refGene_tx=XM_003985288.2
B1 148386748 148386881
refGene_gn=AFP;refGene_tx=XM_003985288.2
B1 148387728 148387940
refGene_gn=AFP;refGene_tx=XM_003985288.2
B1 148389949 148390082
refGene_gn=AFP;refGene_tx=XM_003985288.2
B1 148391014 148391066
refGene_gn=AFP;refGene_tx=XM_003985288.2
B1 148391839 148391924

refGene_gn=AFP;refGene_tx=XM_003985288.2
B1 148407732 148407774
refGene_gn=ALB;refGene_tx=NM_001009961.2
B1 148408393 148408526
refGene_gn=ALB;refGene_tx=NM_001009961.2
B1 148409979 148410203
refGene_gn=ALB;refGene_tx=NM_001009961.2
B1 148410792 148410931
refGene_gn=ALB;refGene_tx=NM_001009961.2
B1 148411978 148412076
refGene_gn=ALB;refGene_tx=NM_001009961.2
B1 148413369 148413502
refGene_gn=ALB;refGene_tx=NM_001009961.2
B1 148414927 148415142
refGene_gn=ALB;refGene_tx=NM_001009961.2
B1 148416069 148416199
refGene_gn=ALB;refGene_tx=NM_001009961.2
B1 148417793 148417891
refGene_gn=ALB;refGene_tx=NM_001009961.2
B1 148418512 148418645
refGene_gn=ALB;refGene_tx=NM_001009961.2
B1 148419217 148419429
refGene_gn=ALB;refGene_tx=NM_001009961.2
B1 148421181 148421314
refGene_gn=ALB;refGene_tx=NM_001009961.2
B1 148422176 148422234
refGene_gn=ALB;refGene_tx=NM_001009961.2
B1 148422982 148423061
refGene_gn=ALB;refGene_tx=NM_001009961.2
B1 148532098 148532512
refGene_gn=ANKRD17;refGene_tx=XM_023253055.1;refGene_gn=ANKRD17;refGen
e_tx=XM_023253056.1;refGene_gn=ANKRD17;refGene_tx=XM_023253057.1
B1 148538304 148538313
refGene_gn=ANKRD17;refGene_tx=XM_023253055.1;refGene_gn=ANKRD17;refGen
e_tx=XM_023253056.1;refGene_gn=ANKRD17;refGene_tx=XM_023253057.1
B1 148567666 148567720
refGene_gn=ANKRD17;refGene_tx=XM_003985289.5;refGene_gn=ANKRD17;refGen
e_tx=XM_006931066.4;refGene_gn=ANKRD17;refGene_tx=XM_019829382.2;refGe
ne_gn=ANKRD17;refGene_tx=XM_019829383.2
B1 148613533 148613687
refGene_gn=ANKRD17;refGene_tx=XM_003985289.5;refGene_gn=ANKRD17;refGen
e_tx=XM_006931066.4;refGene_gn=ANKRD17;refGene_tx=XM_019829382.2;refGe
ne_gn=ANKRD17;refGene_tx=XM_019829383.2;refGene_gn=ANKRD17;refGene_tx=
XM_023253055.1;refGene_gn=ANKRD17;refGene_tx=XM_023253056.1;refGene_gn
=ANKRD17;refGene_tx=XM_023253057.1
B1 148620050 148620207
refGene_gn=ANKRD17;refGene_tx=XM_003985289.5;refGene_gn=ANKRD17;refGen
e_tx=XM_006931066.4;refGene_gn=ANKRD17;refGene_tx=XM_019829382.2;refGe
ne_gn=ANKRD17;refGene_tx=XM_019829383.2;refGene_gn=ANKRD17;refGene_tx=
XM_023253055.1;refGene_gn=ANKRD17;refGene_tx=XM_023253056.1;refGene_gn

=ANKRD17;refGene_tx=XM_023253057.1
B1	148623871	148624019
refGene_gn=ANKRD17;refGene_tx=XM_003985289.5;refGene_gn=ANKRD17;refGene_tx=XM_006931066.4;refGene_gn=ANKRD17;refGene_tx=XM_019829382.2;refGene_gn=ANKRD17;refGene_tx=XM_019829383.2;refGene_gn=ANKRD17;refGene_tx=XM_023253055.1;refGene_gn=ANKRD17;refGene_tx=XM_023253056.1;refGene_gn=ANKRD17;refGene_tx=XM_023253057.1
B1	148624264	148624412
refGene_gn=ANKRD17;refGene_tx=XM_003985289.5;refGene_gn=ANKRD17;refGene_tx=XM_006931066.4;refGene_gn=ANKRD17;refGene_tx=XM_019829382.2;refGene_gn=ANKRD17;refGene_tx=XM_019829383.2;refGene_gn=ANKRD17;refGene_tx=XM_023253055.1;refGene_gn=ANKRD17;refGene_tx=XM_023253056.1;refGene_gn=ANKRD17;refGene_tx=XM_023253057.1
B1	148625572	148625806
refGene_gn=ANKRD17;refGene_tx=XM_003985289.5;refGene_gn=ANKRD17;refGene_tx=XM_006931066.4;refGene_gn=ANKRD17;refGene_tx=XM_019829382.2;refGene_gn=ANKRD17;refGene_tx=XM_019829383.2;refGene_gn=ANKRD17;refGene_tx=XM_023253055.1;refGene_gn=ANKRD17;refGene_tx=XM_023253056.1;refGene_gn=ANKRD17;refGene_tx=XM_023253057.1
B1	148628459	148628554
refGene_gn=ANKRD17;refGene_tx=XM_003985289.5;refGene_gn=ANKRD17;refGene_tx=XM_006931066.4;refGene_gn=ANKRD17;refGene_tx=XM_019829382.2;refGene_gn=ANKRD17;refGene_tx=XM_019829383.2;refGene_gn=ANKRD17;refGene_tx=XM_023253055.1;refGene_gn=ANKRD17;refGene_tx=XM_023253056.1;refGene_gn=ANKRD17;refGene_tx=XM_023253057.1
B1	148630495	148630733
refGene_gn=ANKRD17;refGene_tx=XM_003985289.5;refGene_gn=ANKRD17;refGene_tx=XM_006931066.4;refGene_gn=ANKRD17;refGene_tx=XM_019829382.2;refGene_gn=ANKRD17;refGene_tx=XM_019829383.2;refGene_gn=ANKRD17;refGene_tx=XM_023253055.1;refGene_gn=ANKRD17;refGene_tx=XM_023253056.1;refGene_gn=ANKRD17;refGene_tx=XM_023253057.1
B1	148631763	148631955
refGene_gn=ANKRD17;refGene_tx=XM_003985289.5;refGene_gn=ANKRD17;refGene_tx=XM_006931066.4;refGene_gn=ANKRD17;refGene_tx=XM_019829382.2;refGene_gn=ANKRD17;refGene_tx=XM_019829383.2;refGene_gn=ANKRD17;refGene_tx=XM_023253055.1;refGene_gn=ANKRD17;refGene_tx=XM_023253056.1;refGene_gn=ANKRD17;refGene_tx=XM_023253057.1
B1	148632328	148632438
refGene_gn=ANKRD17;refGene_tx=XM_003985289.5;refGene_gn=ANKRD17;refGene_tx=XM_006931066.4;refGene_gn=ANKRD17;refGene_tx=XM_019829382.2;refGene_gn=ANKRD17;refGene_tx=XM_019829383.2;refGene_gn=ANKRD17;refGene_tx=XM_023253055.1;refGene_gn=ANKRD17;refGene_tx=XM_023253056.1;refGene_gn=ANKRD17;refGene_tx=XM_023253057.1
B1	148634123	148634211
refGene_gn=ANKRD17;refGene_tx=XM_003985289.5;refGene_gn=ANKRD17;refGene_tx=XM_006931066.4;refGene_gn=ANKRD17;refGene_tx=XM_019829382.2;refGene_gn=ANKRD17;refGene_tx=XM_019829383.2;refGene_gn=ANKRD17;refGene_tx=XM_023253055.1;refGene_gn=ANKRD17;refGene_tx=XM_023253056.1;refGene_gn=ANKRD17;refGene_tx=XM_023253057.1
B1	148635936	148636064

refGene_gn=ANKRD17;refGene_tx=XM_003985289.5;refGene_gn=ANKRD17;refGen
e_tx=XM_006931066.4;refGene_gn=ANKRD17;refGene_tx=XM_019829382.2;refGe
ne_gn=ANKRD17;refGene_tx=XM_019829383.2;refGene_gn=ANKRD17;refGene_tx=
XM_023253055.1;refGene_gn=ANKRD17;refGene_tx=XM_023253056.1;refGene_gn
=ANKRD17;refGene_tx=XM_023253057.1
B1 148636317 148636461
refGene_gn=ANKRD17;refGene_tx=XM_003985289.5;refGene_gn=ANKRD17;refGen
e_tx=XM_006931066.4;refGene_gn=ANKRD17;refGene_tx=XM_019829382.2;refGe
ne_gn=ANKRD17;refGene_tx=XM_019829383.2;refGene_gn=ANKRD17;refGene_tx=
XM_023253055.1;refGene_gn=ANKRD17;refGene_tx=XM_023253056.1;refGene_gn
=ANKRD17;refGene_tx=XM_023253057.1
B1 148636873 148636976
refGene_gn=ANKRD17;refGene_tx=XM_003985289.5;refGene_gn=ANKRD17;refGen
e_tx=XM_006931066.4;refGene_gn=ANKRD17;refGene_tx=XM_019829382.2;refGe
ne_gn=ANKRD17;refGene_tx=XM_019829383.2;refGene_gn=ANKRD17;refGene_tx=
XM_023253055.1;refGene_gn=ANKRD17;refGene_tx=XM_023253056.1;refGene_gn
=ANKRD17;refGene_tx=XM_023253057.1
B1 148638450 148639203
refGene_gn=ANKRD17;refGene_tx=XM_003985289.5;refGene_gn=ANKRD17;refGen
e_tx=XM_019829382.2;refGene_gn=ANKRD17;refGene_tx=XM_023253055.1;refGe
ne_gn=ANKRD17;refGene_tx=XM_023253056.1
B1 148643652 148643801
refGene_gn=ANKRD17;refGene_tx=XM_003985289.5;refGene_gn=ANKRD17;refGen
e_tx=XM_006931066.4;refGene_gn=ANKRD17;refGene_tx=XM_019829382.2;refGe
ne_gn=ANKRD17;refGene_tx=XM_019829383.2;refGene_gn=ANKRD17;refGene_tx=
XM_023253055.1;refGene_gn=ANKRD17;refGene_tx=XM_023253056.1;refGene_gn
=ANKRD17;refGene_tx=XM_023253057.1
B1 148650193 148650305
refGene_gn=ANKRD17;refGene_tx=XM_003985289.5;refGene_gn=ANKRD17;refGen
e_tx=XM_006931066.4;refGene_gn=ANKRD17;refGene_tx=XM_019829382.2;refGe
ne_gn=ANKRD17;refGene_tx=XM_019829383.2;refGene_gn=ANKRD17;refGene_tx=
XM_023253055.1;refGene_gn=ANKRD17;refGene_tx=XM_023253056.1;refGene_gn
=ANKRD17;refGene_tx=XM_023253057.1
B1 148650455 148650601
refGene_gn=ANKRD17;refGene_tx=XM_003985289.5;refGene_gn=ANKRD17;refGen
e_tx=XM_006931066.4;refGene_gn=ANKRD17;refGene_tx=XM_019829382.2;refGe
ne_gn=ANKRD17;refGene_tx=XM_019829383.2;refGene_gn=ANKRD17;refGene_tx=
XM_023253055.1;refGene_gn=ANKRD17;refGene_tx=XM_023253056.1;refGene_gn
=ANKRD17;refGene_tx=XM_023253057.1
B1 148655251 148655394
refGene_gn=ANKRD17;refGene_tx=XM_003985289.5;refGene_gn=ANKRD17;refGen
e_tx=XM_006931066.4;refGene_gn=ANKRD17;refGene_tx=XM_019829382.2;refGe
ne_gn=ANKRD17;refGene_tx=XM_019829383.2;refGene_gn=ANKRD17;refGene_tx=
XM_023253055.1;refGene_gn=ANKRD17;refGene_tx=XM_023253056.1;refGene_gn
=ANKRD17;refGene_tx=XM_023253057.1
B1 148655936 148656150
refGene_gn=ANKRD17;refGene_tx=XM_003985289.5;refGene_gn=ANKRD17;refGen
e_tx=XM_006931066.4;refGene_gn=ANKRD17;refGene_tx=XM_019829382.2;refGe
ne_gn=ANKRD17;refGene_tx=XM_019829383.2;refGene_gn=ANKRD17;refGene_tx=
XM_023253055.1;refGene_gn=ANKRD17;refGene_tx=XM_023253056.1;refGene_gn

=ANKRD17;refGene_tx=XM_023253057.1
B1 148656648 148656824
refGene_gn=ANKRD17;refGene_tx=XM_003985289.5;refGene_gn=ANKRD17;refGene_tx=XM_006931066.4;refGene_gn=ANKRD17;refGene_tx=XM_019829382.2;refGene_gn=ANKRD17;refGene_tx=XM_019829383.2;refGene_gn=ANKRD17;refGene_tx=XM_023253055.1;refGene_gn=ANKRD17;refGene_tx=XM_023253056.1;refGene_gn=ANKRD17;refGene_tx=XM_023253057.1
B1 148657976 148658139
refGene_gn=ANKRD17;refGene_tx=XM_003985289.5;refGene_gn=ANKRD17;refGene_tx=XM_006931066.4;refGene_gn=ANKRD17;refGene_tx=XM_019829382.2;refGene_gn=ANKRD17;refGene_tx=XM_019829383.2;refGene_gn=ANKRD17;refGene_tx=XM_023253055.1;refGene_gn=ANKRD17;refGene_tx=XM_023253056.1;refGene_gn=ANKRD17;refGene_tx=XM_023253057.1
B1 148660430 148660526
refGene_gn=ANKRD17;refGene_tx=XM_003985289.5;refGene_gn=ANKRD17;refGene_tx=XM_006931066.4;refGene_gn=ANKRD17;refGene_tx=XM_019829382.2;refGene_gn=ANKRD17;refGene_tx=XM_019829383.2;refGene_gn=ANKRD17;refGene_tx=XM_023253055.1;refGene_gn=ANKRD17;refGene_tx=XM_023253056.1;refGene_gn=ANKRD17;refGene_tx=XM_023253057.1
B1 148661769 148661886
refGene_gn=ANKRD17;refGene_tx=XM_003985289.5;refGene_gn=ANKRD17;refGene_tx=XM_006931066.4;refGene_gn=ANKRD17;refGene_tx=XM_019829382.2;refGene_gn=ANKRD17;refGene_tx=XM_019829383.2;refGene_gn=ANKRD17;refGene_tx=XM_023253055.1;refGene_gn=ANKRD17;refGene_tx=XM_023253056.1;refGene_gn=ANKRD17;refGene_tx=XM_023253057.1
B1 148674172 148674344
refGene_gn=ANKRD17;refGene_tx=XM_003985289.5;refGene_gn=ANKRD17;refGene_tx=XM_006931066.4;refGene_gn=ANKRD17;refGene_tx=XM_019829382.2;refGene_gn=ANKRD17;refGene_tx=XM_019829383.2;refGene_gn=ANKRD17;refGene_tx=XM_023253055.1;refGene_gn=ANKRD17;refGene_tx=XM_023253056.1;refGene_gn=ANKRD17;refGene_tx=XM_023253057.1
B1 148675778 148676226
refGene_gn=ANKRD17;refGene_tx=XM_003985289.5;refGene_gn=ANKRD17;refGene_tx=XM_006931066.4;refGene_gn=ANKRD17;refGene_tx=XM_019829382.2;refGene_gn=ANKRD17;refGene_tx=XM_019829383.2;refGene_gn=ANKRD17;refGene_tx=XM_023253055.1;refGene_gn=ANKRD17;refGene_tx=XM_023253056.1;refGene_gn=ANKRD17;refGene_tx=XM_023253057.1
B1 148676709 148676865
refGene_gn=ANKRD17;refGene_tx=XM_003985289.5;refGene_gn=ANKRD17;refGene_tx=XM_006931066.4;refGene_gn=ANKRD17;refGene_tx=XM_019829382.2;refGene_gn=ANKRD17;refGene_tx=XM_019829383.2;refGene_gn=ANKRD17;refGene_tx=XM_023253055.1;refGene_gn=ANKRD17;refGene_tx=XM_023253056.1;refGene_gn=ANKRD17;refGene_tx=XM_023253057.1
B1 148678562 148678712
refGene_gn=ANKRD17;refGene_tx=XM_003985289.5;refGene_gn=ANKRD17;refGene_tx=XM_006931066.4;refGene_gn=ANKRD17;refGene_tx=XM_019829382.2;refGene_gn=ANKRD17;refGene_tx=XM_019829383.2;refGene_gn=ANKRD17;refGene_tx=XM_023253055.1;refGene_gn=ANKRD17;refGene_tx=XM_023253056.1;refGene_gn=ANKRD17;refGene_tx=XM_023253057.1
B1 148680046 148681683

refGene_gn=ANKRD17;refGene_tx=XM_003985289.5;refGene_gn=ANKRD17;refGene_tx=XM_006931066.4;refGene_gn=ANKRD17;refGene_tx=XM_019829382.2;refGene_gn=ANKRD17;refGene_tx=XM_019829383.2;refGene_gn=ANKRD17;refGene_tx=XM_023253055.1;refGene_gn=ANKRD17;refGene_tx=XM_023253056.1;refGene_gn=ANKRD17;refGene_tx=XM_023253057.1
B1 148686531 148686729
refGene_gn=ANKRD17;refGene_tx=XM_003985289.5;refGene_gn=ANKRD17;refGene_tx=XM_006931066.4;refGene_gn=ANKRD17;refGene_tx=XM_019829382.2;refGene_gn=ANKRD17;refGene_tx=XM_019829383.2;refGene_gn=ANKRD17;refGene_tx=XM_023253055.1;refGene_gn=ANKRD17;refGene_tx=XM_023253056.1;refGene_gn=ANKRD17;refGene_tx=XM_023253057.1
B1 148695217 148695466
refGene_gn=ANKRD17;refGene_tx=XM_003985289.5;refGene_gn=ANKRD17;refGene_tx=XM_006931066.4;refGene_gn=ANKRD17;refGene_tx=XM_019829382.2;refGene_gn=ANKRD17;refGene_tx=XM_019829383.2;refGene_gn=ANKRD17;refGene_tx=XM_023253055.1;refGene_gn=ANKRD17;refGene_tx=XM_023253056.1;refGene_gn=ANKRD17;refGene_tx=XM_023253057.1
B1 148695987 148696166
refGene_gn=ANKRD17;refGene_tx=XM_003985289.5;refGene_gn=ANKRD17;refGene_tx=XM_006931066.4;refGene_gn=ANKRD17;refGene_tx=XM_019829382.2;refGene_gn=ANKRD17;refGene_tx=XM_019829383.2;refGene_gn=ANKRD17;refGene_tx=XM_023253055.1;refGene_gn=ANKRD17;refGene_tx=XM_023253056.1;refGene_gn=ANKRD17;refGene_tx=XM_023253057.1
B1 148696574 148696739
refGene_gn=ANKRD17;refGene_tx=XM_003985289.5;refGene_gn=ANKRD17;refGene_tx=XM_006931066.4;refGene_gn=ANKRD17;refGene_tx=XM_019829382.2;refGene_gn=ANKRD17;refGene_tx=XM_019829383.2;refGene_gn=ANKRD17;refGene_tx=XM_023253055.1;refGene_gn=ANKRD17;refGene_tx=XM_023253056.1;refGene_gn=ANKRD17;refGene_tx=XM_023253057.1
B1 148697379 148697439
refGene_gn=ANKRD17;refGene_tx=XM_003985289.5;refGene_gn=ANKRD17;refGene_tx=XM_006931066.4;refGene_gn=ANKRD17;refGene_tx=XM_019829382.2;refGene_gn=ANKRD17;refGene_tx=XM_019829383.2;refGene_gn=ANKRD17;refGene_tx=XM_023253055.1;refGene_gn=ANKRD17;refGene_tx=XM_023253056.1;refGene_gn=ANKRD17;refGene_tx=XM_023253057.1
B1 148703831 148704173
refGene_gn=LOC101095576;refGene_tx=XM_003985332.4;refGene_gn=LOC101095576;refGene_tx=XM_006931068.3;refGene_gn=LOC101095576;refGene_tx=XM_019829384.2;refGene_gn=LOC101095576;refGene_tx=XM_023253058.1
B1 148705318 148705419
refGene_gn=LOC101095576;refGene_tx=XM_003985332.4;refGene_gn=LOC101095576;refGene_tx=XM_006931068.3;refGene_gn=LOC101095576;refGene_tx=XM_019829384.2;refGene_gn=LOC101095576;refGene_tx=XM_023253058.1
B1 148707885 148708049
refGene_gn=LOC101095576;refGene_tx=XM_003985332.4;refGene_gn=LOC101095576;refGene_tx=XM_006931068.3;refGene_gn=LOC101095576;refGene_tx=XM_019829384.2;refGene_gn=LOC101095576;refGene_tx=XM_023253058.1
B1 148709022 148709147
refGene_gn=LOC101095576;refGene_tx=XM_003985332.4;refGene_gn=LOC101095576;refGene_tx=XM_006931068.3;refGene_gn=LOC101095576;refGene_tx=XM_01

9829384.2;refGene_gn=LOC101095576;refGene_tx=XM_023253058.1
B1 148710916 148711024
refGene_gn=LOC101095576;refGene_tx=XM_003985332.4;refGene_gn=LOC101095576;refGene_tx=XM_006931068.3;refGene_gn=LOC101095576;refGene_tx=XM_019829384.2;refGene_gn=LOC101095576;refGene_tx=XM_023253058.1
B1 148713851 148714025
refGene_gn=LOC101095576;refGene_tx=XM_003985332.4;refGene_gn=LOC101095576;refGene_tx=XM_006931068.3;refGene_gn=LOC101095576;refGene_tx=XM_019829384.2
B1 148719194 148719275
refGene_gn=LOC101095576;refGene_tx=XM_023253058.1
B1 149075420 149075489
refGene_gn=ADAMTS3;refGene_tx=XM_011281725.3
B1 149076774 149076802
refGene_gn=ADAMTS3;refGene_tx=XM_011281725.3
B1 149096215 149096625
refGene_gn=ADAMTS3;refGene_tx=XM_011281725.3
B1 149191072 149191229
refGene_gn=ADAMTS3;refGene_tx=XM_011281725.3
B1 149274986 149275186
refGene_gn=ADAMTS3;refGene_tx=XM_011281725.3
B1 149298673 149298757
refGene_gn=ADAMTS3;refGene_tx=XM_011281725.3
B1 149300611 149300768
refGene_gn=ADAMTS3;refGene_tx=XM_011281725.3
B1 149301518 149301624
refGene_gn=ADAMTS3;refGene_tx=XM_011281725.3
B1 149302018 149302162
refGene_gn=ADAMTS3;refGene_tx=XM_011281725.3
B1 149302804 149302937
refGene_gn=ADAMTS3;refGene_tx=XM_011281725.3
B1 149305218 149305332
refGene_gn=ADAMTS3;refGene_tx=XM_011281725.3
B1 149307598 149307744
refGene_gn=ADAMTS3;refGene_tx=XM_011281725.3
B1 149308884 149309060
refGene_gn=ADAMTS3;refGene_tx=XM_011281725.3
B1 149310666 149310800
refGene_gn=ADAMTS3;refGene_tx=XM_011281725.3
B1 149312626 149312750
refGene_gn=ADAMTS3;refGene_tx=XM_011281725.3
B1 149317061 149317142
refGene_gn=ADAMTS3;refGene_tx=XM_011281725.3
B1 149319091 149319255
refGene_gn=ADAMTS3;refGene_tx=XM_011281725.3
B1 149324877 149325043
refGene_gn=ADAMTS3;refGene_tx=XM_011281725.3
B1 149327599 149327732
refGene_gn=ADAMTS3;refGene_tx=XM_011281725.3
B1 149331565 149331719

refGene_gn=ADAMTS3;refGene_tx=XM_011281725.3
B1	149331720	149331774
refGene_gn=ADAMTS3;refGene_tx=XM_011281725.3
B1	149333774	149333892
refGene_gn=ADAMTS3;refGene_tx=XM_011281725.3
B1	149339345	149339902
refGene_gn=ADAMTS3;refGene_tx=XM_011281725.3
B1	149457662	149458512
refGene_gn=NPFFR2;refGene_tx=XM_003985291.5;refGene_gn=NPFFR2;refGene_tx=XM_023253059.1
B1	149464897	149464997
refGene_gn=NPFFR2;refGene_tx=XM_003985291.5;refGene_gn=NPFFR2;refGene_tx=XM_023253059.1
B1	149476486	149476814
refGene_gn=NPFFR2;refGene_tx=XM_003985291.5;refGene_gn=NPFFR2;refGene_tx=XM_023253059.1
B1	149611162	149611295
refGene_gn=LOC101096061;refGene_tx=XM_023253402.1
B1	149614733	149614831
refGene_gn=LOC101096061;refGene_tx=XM_023253402.1
B1	149628594	149628684
refGene_gn=LOC101096061;refGene_tx=XM_023253402.1
B1	149638839	149638942
refGene_gn=LOC101096061;refGene_tx=XM_023253402.1
B1	149639334	149639666
refGene_gn=LOC101096061;refGene_tx=XM_023253402.1
B1	149639667	149639908
refGene_gn=LOC101096061;refGene_tx=XM_023253402.1
B1	149639910	149640089
refGene_gn=LOC101096061;refGene_tx=XM_023253402.1
B1	149797308	149797366
refGene_gn=GC;refGene_tx=XM_006931069.4;refGene_gn=GC;refGene_tx=XM_019829395.2
B1	149810388	149810458
refGene_gn=GC;refGene_tx=XM_006931069.4;refGene_gn=GC;refGene_tx=XM_019829395.2
B1	149811021	149811154
refGene_gn=GC;refGene_tx=XM_006931069.4;refGene_gn=GC;refGene_tx=XM_019829395.2
B1	149813406	149813618
refGene_gn=GC;refGene_tx=XM_006931069.4;refGene_gn=GC;refGene_tx=XM_019829395.2
B1	149814737	149814870
refGene_gn=GC;refGene_tx=XM_006931069.4;refGene_gn=GC;refGene_tx=XM_019829395.2
B1	149815161	149815256
refGene_gn=GC;refGene_tx=XM_006931069.4;refGene_gn=GC;refGene_tx=XM_019829395.2
B1	149818577	149818707
refGene_gn=GC;refGene_tx=XM_006931069.4;refGene_gn=GC;refGene_tx=XM_01

9829395.2
B1	149819787	149819990
refGene_gn=GC;refGene_tx=XM_006931069.4;refGene_gn=GC;refGene_tx=XM_01
9829395.2
B1	149820715	149820845
refGene_gn=GC;refGene_tx=XM_006931069.4;refGene_gn=GC;refGene_tx=XM_01
9829395.2
B1	149821292	149821390
refGene_gn=GC;refGene_tx=XM_006931069.4;refGene_gn=GC;refGene_tx=XM_01
9829395.2
B1	149823179	149823312
refGene_gn=GC;refGene_tx=XM_006931069.4;refGene_gn=GC;refGene_tx=XM_01
9829395.2
B1	149827101	149827131
refGene_gn=GC;refGene_tx=XM_006931069.4;refGene_gn=GC;refGene_tx=XM_01
9829395.2
B1	149967853	149967897
refGene_gn=SLC4A4;refGene_tx=XM_011281724.3;refGene_gn=SLC4A4;refGene_
tx=XM_019829396.2;refGene_gn=SLC4A4;refGene_tx=XM_019829397.2;refGene_
gn=SLC4A4;refGene_tx=XM_019829398.2;refGene_gn=SLC4A4;refGene_tx=XM_01
9829399.2;refGene_gn=SLC4A4;refGene_tx=XM_023253060.1
B1	149970522	149970619
refGene_gn=SLC4A4;refGene_tx=XM_011281724.3;refGene_gn=SLC4A4;refGene_
tx=XM_019829396.2;refGene_gn=SLC4A4;refGene_tx=XM_019829397.2;refGene_
gn=SLC4A4;refGene_tx=XM_019829398.2;refGene_gn=SLC4A4;refGene_tx=XM_01
9829399.2;refGene_gn=SLC4A4;refGene_tx=XM_023253060.1
B1	149974156	149974318
refGene_gn=SLC4A4;refGene_tx=XM_011281724.3;refGene_gn=SLC4A4;refGene_
tx=XM_019829396.2;refGene_gn=SLC4A4;refGene_tx=XM_019829397.2;refGene_
gn=SLC4A4;refGene_tx=XM_019829398.2;refGene_gn=SLC4A4;refGene_tx=XM_01
9829399.2;refGene_gn=SLC4A4;refGene_tx=XM_023253060.1
B1	149976947	149977121
refGene_gn=SLC4A4;refGene_tx=XM_011281724.3;refGene_gn=SLC4A4;refGene_
tx=XM_019829396.2;refGene_gn=SLC4A4;refGene_tx=XM_019829397.2;refGene_
gn=SLC4A4;refGene_tx=XM_019829398.2;refGene_gn=SLC4A4;refGene_tx=XM_01
9829399.2;refGene_gn=SLC4A4;refGene_tx=XM_023253060.1
B1	149979839	149979908
refGene_gn=SLC4A4;refGene_tx=XM_011281724.3;refGene_gn=SLC4A4;refGene_
tx=XM_019829396.2;refGene_gn=SLC4A4;refGene_tx=XM_019829397.2;refGene_
gn=SLC4A4;refGene_tx=XM_019829398.2;refGene_gn=SLC4A4;refGene_tx=XM_01
9829399.2;refGene_gn=SLC4A4;refGene_tx=XM_023253060.1
B1	149988842	149988915
refGene_gn=SLC4A4;refGene_tx=XM_011281724.3;refGene_gn=SLC4A4;refGene_
tx=XM_019829396.2;refGene_gn=SLC4A4;refGene_tx=XM_019829397.2;refGene_
gn=SLC4A4;refGene_tx=XM_019829398.2;refGene_gn=SLC4A4;refGene_tx=XM_01
9829399.2;refGene_gn=SLC4A4;refGene_tx=XM_023253060.1
B1	149990070	149990249
refGene_gn=SLC4A4;refGene_tx=XM_011281724.3;refGene_gn=SLC4A4;refGene_
tx=XM_019829396.2;refGene_gn=SLC4A4;refGene_tx=XM_019829397.2;refGene_
gn=SLC4A4;refGene_tx=XM_019829398.2;refGene_gn=SLC4A4;refGene_tx=XM_01

9829399.2;refGene_gn=SLC4A4;refGene_tx=XM_023253060.1
B1	150004504	150004666
refGene_gn=SLC4A4;refGene_tx=XM_011281724.3;refGene_gn=SLC4A4;refGene_tx=XM_019829396.2;refGene_gn=SLC4A4;refGene_tx=XM_019829397.2;refGene_gn=SLC4A4;refGene_tx=XM_019829398.2;refGene_gn=SLC4A4;refGene_tx=XM_019829399.2;refGene_gn=SLC4A4;refGene_tx=XM_023253060.1
B1	150006523	150006637
refGene_gn=SLC4A4;refGene_tx=XM_011281724.3;refGene_gn=SLC4A4;refGene_tx=XM_019829396.2;refGene_gn=SLC4A4;refGene_tx=XM_019829397.2;refGene_gn=SLC4A4;refGene_tx=XM_019829398.2;refGene_gn=SLC4A4;refGene_tx=XM_019829399.2;refGene_gn=SLC4A4;refGene_tx=XM_023253060.1
B1	150019870	150020062
refGene_gn=SLC4A4;refGene_tx=XM_011281724.3;refGene_gn=SLC4A4;refGene_tx=XM_019829396.2;refGene_gn=SLC4A4;refGene_tx=XM_019829397.2;refGene_gn=SLC4A4;refGene_tx=XM_019829398.2;refGene_gn=SLC4A4;refGene_tx=XM_019829399.2;refGene_gn=SLC4A4;refGene_tx=XM_023253060.1
B1	150035070	150035141
refGene_gn=SLC4A4;refGene_tx=XM_011281724.3;refGene_gn=SLC4A4;refGene_tx=XM_019829396.2;refGene_gn=SLC4A4;refGene_tx=XM_019829397.2;refGene_gn=SLC4A4;refGene_tx=XM_019829398.2;refGene_gn=SLC4A4;refGene_tx=XM_019829399.2;refGene_gn=SLC4A4;refGene_tx=XM_023253060.1
B1	150047187	150047459
refGene_gn=SLC4A4;refGene_tx=XM_011281724.3;refGene_gn=SLC4A4;refGene_tx=XM_019829396.2;refGene_gn=SLC4A4;refGene_tx=XM_019829397.2;refGene_gn=SLC4A4;refGene_tx=XM_019829398.2;refGene_gn=SLC4A4;refGene_tx=XM_019829399.2;refGene_gn=SLC4A4;refGene_tx=XM_023253060.1
B1	150054478	150054612
refGene_gn=SLC4A4;refGene_tx=XM_011281724.3;refGene_gn=SLC4A4;refGene_tx=XM_019829396.2;refGene_gn=SLC4A4;refGene_tx=XM_019829397.2;refGene_gn=SLC4A4;refGene_tx=XM_019829398.2;refGene_gn=SLC4A4;refGene_tx=XM_019829399.2;refGene_gn=SLC4A4;refGene_tx=XM_023253060.1
B1	150068408	150068583
refGene_gn=SLC4A4;refGene_tx=XM_011281724.3;refGene_gn=SLC4A4;refGene_tx=XM_019829396.2;refGene_gn=SLC4A4;refGene_tx=XM_019829397.2;refGene_gn=SLC4A4;refGene_tx=XM_019829398.2;refGene_gn=SLC4A4;refGene_tx=XM_019829399.2;refGene_gn=SLC4A4;refGene_tx=XM_023253060.1
B1	150071370	150071490
refGene_gn=SLC4A4;refGene_tx=XM_011281724.3;refGene_gn=SLC4A4;refGene_tx=XM_019829396.2;refGene_gn=SLC4A4;refGene_tx=XM_019829397.2;refGene_gn=SLC4A4;refGene_tx=XM_019829398.2;refGene_gn=SLC4A4;refGene_tx=XM_019829399.2;refGene_gn=SLC4A4;refGene_tx=XM_023253060.1
B1	150072124	150072279
refGene_gn=SLC4A4;refGene_tx=XM_011281724.3;refGene_gn=SLC4A4;refGene_tx=XM_019829396.2;refGene_gn=SLC4A4;refGene_tx=XM_019829397.2;refGene_gn=SLC4A4;refGene_tx=XM_019829398.2;refGene_gn=SLC4A4;refGene_tx=XM_019829399.2;refGene_gn=SLC4A4;refGene_tx=XM_023253060.1
B1	150074536	150074624
refGene_gn=SLC4A4;refGene_tx=XM_011281724.3;refGene_gn=SLC4A4;refGene_tx=XM_019829396.2;refGene_gn=SLC4A4;refGene_tx=XM_019829397.2;refGene_gn=SLC4A4;refGene_tx=XM_019829398.2;refGene_gn=SLC4A4;refGene_tx=XM_01

9829399.2;refGene_gn=SLC4A4;refGene_tx=XM_023253060.1
B1 150081092 150081250
refGene_gn=SLC4A4;refGene_tx=XM_011281724.3;refGene_gn=SLC4A4;refGene_tx=XM_019829396.2;refGene_gn=SLC4A4;refGene_tx=XM_019829397.2;refGene_gn=SLC4A4;refGene_tx=XM_019829398.2;refGene_gn=SLC4A4;refGene_tx=XM_019829399.2;refGene_gn=SLC4A4;refGene_tx=XM_023253060.1
B1 150113395 150113472
refGene_gn=SLC4A4;refGene_tx=XM_011281724.3;refGene_gn=SLC4A4;refGene_tx=XM_019829396.2;refGene_gn=SLC4A4;refGene_tx=XM_019829397.2;refGene_gn=SLC4A4;refGene_tx=XM_019829398.2;refGene_gn=SLC4A4;refGene_tx=XM_019829399.2;refGene_gn=SLC4A4;refGene_tx=XM_023253060.1
B1 150143661 150143757
refGene_gn=SLC4A4;refGene_tx=XM_019829396.2;refGene_gn=SLC4A4;refGene_tx=XM_019829397.2;refGene_gn=SLC4A4;refGene_tx=XM_019829398.2;refGene_gn=SLC4A4;refGene_tx=XM_023253060.1
B1 150155553 150155733
refGene_gn=SLC4A4;refGene_tx=XM_011281724.3;refGene_gn=SLC4A4;refGene_tx=XM_019829396.2;refGene_gn=SLC4A4;refGene_tx=XM_019829397.2;refGene_gn=SLC4A4;refGene_tx=XM_019829398.2;refGene_gn=SLC4A4;refGene_tx=XM_019829399.2;refGene_gn=SLC4A4;refGene_tx=XM_023253060.1
B1 150162675 150162836
refGene_gn=SLC4A4;refGene_tx=XM_011281724.3;refGene_gn=SLC4A4;refGene_tx=XM_019829396.2;refGene_gn=SLC4A4;refGene_tx=XM_019829397.2;refGene_gn=SLC4A4;refGene_tx=XM_019829398.2;refGene_gn=SLC4A4;refGene_tx=XM_019829399.2;refGene_gn=SLC4A4;refGene_tx=XM_023253060.1
B1 150175253 150175389
refGene_gn=SLC4A4;refGene_tx=XM_011281724.3;refGene_gn=SLC4A4;refGene_tx=XM_019829396.2;refGene_gn=SLC4A4;refGene_tx=XM_019829397.2;refGene_gn=SLC4A4;refGene_tx=XM_019829398.2;refGene_gn=SLC4A4;refGene_tx=XM_019829399.2;refGene_gn=SLC4A4;refGene_tx=XM_023253060.1
B1 150251413 150251593
refGene_gn=SLC4A4;refGene_tx=XM_011281724.3;refGene_gn=SLC4A4;refGene_tx=XM_019829396.2;refGene_gn=SLC4A4;refGene_tx=XM_019829397.2;refGene_gn=SLC4A4;refGene_tx=XM_019829398.2;refGene_gn=SLC4A4;refGene_tx=XM_019829399.2;refGene_gn=SLC4A4;refGene_tx=XM_023253060.1
B1 150269290 150269364
refGene_gn=SLC4A4;refGene_tx=XM_011281724.3;refGene_gn=SLC4A4;refGene_tx=XM_019829396.2;refGene_gn=SLC4A4;refGene_tx=XM_019829397.2;refGene_gn=SLC4A4;refGene_tx=XM_019829398.2;refGene_gn=SLC4A4;refGene_tx=XM_019829399.2;refGene_gn=SLC4A4;refGene_tx=XM_023253060.1
B1 150269714 150269806
refGene_gn=SLC4A4;refGene_tx=XM_011281724.3;refGene_gn=SLC4A4;refGene_tx=XM_019829396.2;refGene_gn=SLC4A4;refGene_tx=XM_019829397.2;refGene_gn=SLC4A4;refGene_tx=XM_019829398.2;refGene_gn=SLC4A4;refGene_tx=XM_019829399.2
B1 150413198 150413225
refGene_gn=DCK;refGene_tx=XM_003985295.5
B1 150415467 150415558
refGene_gn=DCK;refGene_tx=XM_003985295.5
B1 150416339 150416455

refGene_gn=DCK;refGene_tx=XM_003985295.5
B1 150417870 150418018
refGene_gn=DCK;refGene_tx=XM_003985295.5
B1 150419405 150419599
refGene_gn=DCK;refGene_tx=XM_003985295.5
B1 150438102 150438218
refGene_gn=DCK;refGene_tx=XM_003985295.5
B1 150443872 150443990
refGene_gn=DCK;refGene_tx=XM_003985295.5
B1 150457055 150457133
refGene_gn=MOB1B;refGene_tx=XM_006931070.4;refGene_gn=MOB1B;refGene_tx
=XM_023253061.1
B1 150460398 150460562
refGene_gn=MOB1B;refGene_tx=XM_006931070.4
B1 150464046 150464180
refGene_gn=MOB1B;refGene_tx=XM_006931070.4;refGene_gn=MOB1B;refGene_tx
=XM_023253061.1
B1 150469779 150469873
refGene_gn=MOB1B;refGene_tx=XM_006931070.4;refGene_gn=MOB1B;refGene_tx
=XM_023253061.1
B1 150474303 150474470
refGene_gn=MOB1B;refGene_tx=XM_006931070.4;refGene_gn=MOB1B;refGene_tx
=XM_023253061.1
B1 150505257 150505271
refGene_gn=MOB1B;refGene_tx=XM_006931070.4;refGene_gn=MOB1B;refGene_tx
=XM_023253061.1
B1 150543590 150543944
refGene_gn=GRSF1;refGene_tx=XM_023253062.1
B1 150544234 150544237
refGene_gn=GRSF1;refGene_tx=XM_003985297.5
B1 150546412 150546569
refGene_gn=GRSF1;refGene_tx=XM_003985297.5;refGene_gn=GRSF1;refGene_tx
=XM_023253062.1
B1 150547613 150547769
refGene_gn=GRSF1;refGene_tx=XM_003985297.5;refGene_gn=GRSF1;refGene_tx
=XM_023253062.1
B1 150548123 150548267
refGene_gn=GRSF1;refGene_tx=XM_003985297.5;refGene_gn=GRSF1;refGene_tx
=XM_023253062.1
B1 150548954 150549090
refGene_gn=GRSF1;refGene_tx=XM_003985297.5;refGene_gn=GRSF1;refGene_tx
=XM_023253062.1
B1 150551241 150551423
refGene_gn=GRSF1;refGene_tx=XM_003985297.5;refGene_gn=GRSF1;refGene_tx
=XM_023253062.1
B1 150552607 150552729
refGene_gn=GRSF1;refGene_tx=XM_003985297.5;refGene_gn=GRSF1;refGene_tx
=XM_023253062.1
B1 150553900 150554036
refGene_gn=GRSF1;refGene_tx=XM_003985297.5;refGene_gn=GRSF1;refGene_tx

=XM_023253062.1
B1 150554647 150554697
refGene_gn=GRSF1;refGene_tx=XM_003985297.5;refGene_gn=GRSF1;refGene_tx=XM_023253062.1
B1 150567970 150568114
refGene_gn=RUFY3;refGene_tx=XM_003985298.5;refGene_gn=RUFY3;refGene_tx=XM_011281722.2;refGene_gn=RUFY3;refGene_tx=XM_019829403.2;refGene_gn=RUFY3;refGene_tx=XM_023253063.1
B1 150569899 150569968
refGene_gn=RUFY3;refGene_tx=XM_003985298.5;refGene_gn=RUFY3;refGene_tx=XM_011281722.2;refGene_gn=RUFY3;refGene_tx=XM_019829403.2;refGene_gn=RUFY3;refGene_tx=XM_023253063.1
B1 150570754 150570782
refGene_gn=RUFY3;refGene_tx=XM_003985298.5;refGene_gn=RUFY3;refGene_tx=XM_011281722.2;refGene_gn=RUFY3;refGene_tx=XM_019829403.2
B1 150571461 150571474
refGene_gn=RUFY3;refGene_tx=XM_023253064.1
B1 150573637 150573702
refGene_gn=RUFY3;refGene_tx=XM_003985298.5;refGene_gn=RUFY3;refGene_tx=XM_011281722.2;refGene_gn=RUFY3;refGene_tx=XM_019829403.2;refGene_gn=RUFY3;refGene_tx=XM_023253063.1;refGene_gn=RUFY3;refGene_tx=XM_023253064.1
B1 150575382 150575482
refGene_gn=RUFY3;refGene_tx=XM_003985298.5;refGene_gn=RUFY3;refGene_tx=XM_011281722.2;refGene_gn=RUFY3;refGene_tx=XM_019829403.2;refGene_gn=RUFY3;refGene_tx=XM_023253063.1;refGene_gn=RUFY3;refGene_tx=XM_023253064.1
B1 150576823 150576943
refGene_gn=RUFY3;refGene_tx=XM_003985298.5;refGene_gn=RUFY3;refGene_tx=XM_011281722.2;refGene_gn=RUFY3;refGene_tx=XM_019829403.2;refGene_gn=RUFY3;refGene_tx=XM_023253063.1;refGene_gn=RUFY3;refGene_tx=XM_023253064.1
B1 150579450 150579523
refGene_gn=RUFY3;refGene_tx=XM_003985299.5;refGene_gn=RUFY3;refGene_tx=XM_011281723.3;refGene_gn=RUFY3;refGene_tx=XM_023253065.1
B1 150581096 150581194
refGene_gn=RUFY3;refGene_tx=XM_003985298.5;refGene_gn=RUFY3;refGene_tx=XM_003985299.5;refGene_gn=RUFY3;refGene_tx=XM_011281722.2;refGene_gn=RUFY3;refGene_tx=XM_011281723.3;refGene_gn=RUFY3;refGene_tx=XM_019829403.2;refGene_gn=RUFY3;refGene_tx=XM_023253063.1;refGene_gn=RUFY3;refGene_tx=XM_023253064.1;refGene_gn=RUFY3;refGene_tx=XM_023253065.1
B1 150581692 150581860
refGene_gn=RUFY3;refGene_tx=XM_003985298.5;refGene_gn=RUFY3;refGene_tx=XM_003985299.5;refGene_gn=RUFY3;refGene_tx=XM_011281722.2;refGene_gn=RUFY3;refGene_tx=XM_011281723.3;refGene_gn=RUFY3;refGene_tx=XM_019829403.2;refGene_gn=RUFY3;refGene_tx=XM_023253063.1;refGene_gn=RUFY3;refGene_tx=XM_023253064.1;refGene_gn=RUFY3;refGene_tx=XM_023253065.1
B1 150584281 150584365
refGene_gn=RUFY3;refGene_tx=XM_003985298.5;refGene_gn=RUFY3;refGene_tx=XM_003985299.5;refGene_gn=RUFY3;refGene_tx=XM_011281722.2;refGene_gn=

RUFY3;refGene_tx=XM_011281723.3;refGene_gn=RUFY3;refGene_tx=XM_019829403.2;refGene_gn=RUFY3;refGene_tx=XM_023253063.1;refGene_gn=RUFY3;refGene_tx=XM_023253064.1;refGene_gn=RUFY3;refGene_tx=XM_023253065.1
B1 150585476 150585569
refGene_gn=RUFY3;refGene_tx=XM_003985298.5;refGene_gn=RUFY3;refGene_tx=XM_003985299.5;refGene_gn=RUFY3;refGene_tx=XM_011281722.2;refGene_gn=RUFY3;refGene_tx=XM_011281723.3;refGene_gn=RUFY3;refGene_tx=XM_019829403.2;refGene_gn=RUFY3;refGene_tx=XM_023253063.1;refGene_gn=RUFY3;refGene_tx=XM_023253064.1;refGene_gn=RUFY3;refGene_tx=XM_023253065.1
B1 150588622 150588692
refGene_gn=RUFY3;refGene_tx=XM_003985298.5;refGene_gn=RUFY3;refGene_tx=XM_003985299.5;refGene_gn=RUFY3;refGene_tx=XM_011281722.2;refGene_gn=RUFY3;refGene_tx=XM_011281723.3;refGene_gn=RUFY3;refGene_tx=XM_019829403.2;refGene_gn=RUFY3;refGene_tx=XM_023253063.1;refGene_gn=RUFY3;refGene_tx=XM_023253064.1;refGene_gn=RUFY3;refGene_tx=XM_023253065.1
B1 150589953 150590019
refGene_gn=RUFY3;refGene_tx=XM_003985298.5;refGene_gn=RUFY3;refGene_tx=XM_003985299.5;refGene_gn=RUFY3;refGene_tx=XM_011281722.2;refGene_gn=RUFY3;refGene_tx=XM_011281723.3;refGene_gn=RUFY3;refGene_tx=XM_019829403.2;refGene_gn=RUFY3;refGene_tx=XM_023253063.1;refGene_gn=RUFY3;refGene_tx=XM_023253064.1;refGene_gn=RUFY3;refGene_tx=XM_023253065.1
B1 150590920 150590982
refGene_gn=RUFY3;refGene_tx=XM_003985298.5;refGene_gn=RUFY3;refGene_tx=XM_003985299.5;refGene_gn=RUFY3;refGene_tx=XM_011281722.2;refGene_gn=RUFY3;refGene_tx=XM_011281723.3;refGene_gn=RUFY3;refGene_tx=XM_019829403.2;refGene_gn=RUFY3;refGene_tx=XM_023253063.1;refGene_gn=RUFY3;refGene_tx=XM_023253064.1;refGene_gn=RUFY3;refGene_tx=XM_023253065.1
B1 150592327 150592451
refGene_gn=RUFY3;refGene_tx=XM_003985298.5;refGene_gn=RUFY3;refGene_tx=XM_003985299.5;refGene_gn=RUFY3;refGene_tx=XM_011281722.2;refGene_gn=RUFY3;refGene_tx=XM_011281723.3;refGene_gn=RUFY3;refGene_tx=XM_019829403.2;refGene_gn=RUFY3;refGene_tx=XM_023253063.1;refGene_gn=RUFY3;refGene_tx=XM_023253064.1;refGene_gn=RUFY3;refGene_tx=XM_023253065.1
B1 150594143 150594245
refGene_gn=RUFY3;refGene_tx=XM_003985298.5;refGene_gn=RUFY3;refGene_tx=XM_003985299.5;refGene_gn=RUFY3;refGene_tx=XM_011281722.2;refGene_gn=RUFY3;refGene_tx=XM_011281723.3;refGene_gn=RUFY3;refGene_tx=XM_019829403.2;refGene_gn=RUFY3;refGene_tx=XM_023253063.1;refGene_gn=RUFY3;refGene_tx=XM_023253064.1;refGene_gn=RUFY3;refGene_tx=XM_023253065.1
B1 150595077 150595195
refGene_gn=RUFY3;refGene_tx=XM_003985298.5;refGene_gn=RUFY3;refGene_tx=XM_003985299.5;refGene_gn=RUFY3;refGene_tx=XM_011281722.2;refGene_gn=RUFY3;refGene_tx=XM_011281723.3;refGene_gn=RUFY3;refGene_tx=XM_019829403.2;refGene_gn=RUFY3;refGene_tx=XM_023253063.1;refGene_gn=RUFY3;refGene_tx=XM_023253064.1;refGene_gn=RUFY3;refGene_tx=XM_023253065.1
B1 150595998 150596172
refGene_gn=RUFY3;refGene_tx=XM_003985298.5;refGene_gn=RUFY3;refGene_tx=XM_003985299.5;refGene_gn=RUFY3;refGene_tx=XM_011281722.2;refGene_gn=RUFY3;refGene_tx=XM_011281723.3;refGene_gn=RUFY3;refGene_tx=XM_019829403.2;refGene_gn=RUFY3;refGene_tx=XM_023253063.1;refGene_gn=RUFY3;refGe

ne_tx=XM_023253064.1;refGene_gn=RUFY3;refGene_tx=XM_023253065.1
B1	150614127	150614181
refGene_gn=RUFY3;refGene_tx=XM_011281723.3;refGene_gn=RUFY3;refGene_tx=XM_019829403.2
B1	150630940	150631118
refGene_gn=RUFY3;refGene_tx=XM_003985298.5;refGene_gn=RUFY3;refGene_tx=XM_003985299.5;refGene_gn=RUFY3;refGene_tx=XM_011281723.3;refGene_gn=RUFY3;refGene_tx=XM_019829403.2
B1	150647518	150647873
refGene_gn=RUFY3;refGene_tx=XM_011281722.2;refGene_gn=RUFY3;refGene_tx=XM_023253063.1;refGene_gn=RUFY3;refGene_tx=XM_023253064.1;refGene_gn=RUFY3;refGene_tx=XM_023253065.1
B1	150659775	150661221
refGene_gn=UTP3;refGene_tx=XM_003985300.5
B1	150687551	150687615
refGene_gn=JCHAIN;refGene_tx=XM_003985301.5
B1	150691264	150691391
refGene_gn=JCHAIN;refGene_tx=XM_003985301.5
B1	150693473	150693554
refGene_gn=JCHAIN;refGene_tx=XM_003985301.5
B1	150694955	150695160
refGene_gn=JCHAIN;refGene_tx=XM_003985301.5
B1	150706346	150709205
refGene_gn=ENAM;refGene_tx=XM_003985302.2
B1	150711185	150711236
refGene_gn=ENAM;refGene_tx=XM_003985302.2
B1	150712695	150712758
refGene_gn=ENAM;refGene_tx=XM_003985302.2
B1	150713867	150714140
refGene_gn=ENAM;refGene_tx=XM_003985302.2
B1	150715533	150715575
refGene_gn=ENAM;refGene_tx=XM_003985302.2
B1	150716288	150716333
refGene_gn=ENAM;refGene_tx=XM_003985302.2
B1	150716455	150716524
refGene_gn=ENAM;refGene_tx=XM_003985302.2
B1	150718317	150718371
refGene_gn=ENAM;refGene_tx=XM_003985302.2
B1	150747815	150748352
refGene_gn=AMBN;refGene_tx=XM_023253403.1
B1	150750615	150750660
refGene_gn=AMBN;refGene_tx=XM_023253403.1
B1	150751120	150751165
refGene_gn=AMBN;refGene_tx=XM_023253403.1
B1	150751260	150751320
refGene_gn=AMBN;refGene_tx=XM_023253403.1
B1	150751821	150751860
refGene_gn=AMBN;refGene_tx=XM_023253403.1
B1	150752748	150752985
refGene_gn=AMBN;refGene_tx=XM_023253403.1

B1 150754389 150754500
refGene_gn=AMBN;refGene_tx=XM_023253403.1
B1 150755005 150755053
refGene_gn=AMBN;refGene_tx=XM_023253403.1
B1 150756370 150756421
refGene_gn=AMBN;refGene_tx=XM_023253403.1
B1 150759166 150759235
refGene_gn=AMBN;refGene_tx=XM_023253403.1
B1 150768546 150768712
refGene_gn=AMBN;refGene_tx=XM_023253403.1
B1 150768776 150768955
refGene_gn=AMBN;refGene_tx=XM_023253403.1
B1 150769438 150769641
refGene_gn=AMBN;refGene_tx=XM_023253403.1
B1 150776188 150776345
refGene_gn=AMBN;refGene_tx=XM_023253403.1
B1 150804585 150804596
refGene_gn=AMTN;refGene_tx=XM_003985336.4;refGene_gn=AMTN;refGene_tx=XM_006931073.3
B1 150805298 150805563
refGene_gn=AMTN;refGene_tx=XM_003985336.4;refGene_gn=AMTN;refGene_tx=XM_006931073.3
B1 150808257 150808284
refGene_gn=AMTN;refGene_tx=XM_003985336.4
B1 150809412 150809448
refGene_gn=AMTN;refGene_tx=XM_003985336.4;refGene_gn=AMTN;refGene_tx=XM_006931073.3
B1 150813978 150814068
refGene_gn=AMTN;refGene_tx=XM_003985336.4;refGene_gn=AMTN;refGene_tx=XM_006931073.3
B1 150814991 150815057
refGene_gn=AMTN;refGene_tx=XM_003985336.4;refGene_gn=AMTN;refGene_tx=XM_006931073.3
B1 150816181 150816265
refGene_gn=AMTN;refGene_tx=XM_003985336.4;refGene_gn=AMTN;refGene_tx=XM_006931073.3
B1 150819352 150819406
refGene_gn=AMTN;refGene_tx=XM_003985336.4;refGene_gn=AMTN;refGene_tx=XM_006931073.3
B1 150936759 150938382
refGene_gn=LOC102900847;refGene_tx=XM_023253404.1
B1 150942773 150942827
refGene_gn=LOC102900847;refGene_tx=XM_023253404.1
B1 150978302 150979511
refGene_gn=CABS1;refGene_tx=XM_019829406.2
B1 151039081 151039543
refGene_gn=CSN3;refGene_tx=XM_003985304.1
B1 151040710 151040743
refGene_gn=CSN3;refGene_tx=XM_003985304.1
B1 151044232 151044286

refGene_gn=CSN3;refGene_tx=XM_003985304.1
B1	151064154	151064325
refGene_gn=FDCSP;refGene_tx=XM_019829408.2
B1	151065368	151065401
refGene_gn=FDCSP;refGene_tx=XM_019829408.2
B1	151065838	151065908
refGene_gn=FDCSP;refGene_tx=XM_019829408.2
B1	151070211	151070234
refGene_gn=FDCSP;refGene_tx=XM_019829408.2
B1	151090766	151090778
refGene_gn=ODAM;refGene_tx=XM_023253405.1
B1	151090912	151091059
refGene_gn=ODAM;refGene_tx=XM_023253405.1
B1	151091489	151091561
refGene_gn=ODAM;refGene_tx=XM_023253405.1
B1	151092564	151092612
refGene_gn=ODAM;refGene_tx=XM_023253405.1
B1	151093423	151093528
refGene_gn=ODAM;refGene_tx=XM_023253405.1
B1	151095576	151095624
refGene_gn=ODAM;refGene_tx=XM_023253405.1
B1	151096953	151097187
refGene_gn=ODAM;refGene_tx=XM_023253405.1
B1	151097734	151097782
refGene_gn=ODAM;refGene_tx=XM_023253405.1
B1	151098312	151098354
refGene_gn=ODAM;refGene_tx=XM_023253405.1
B1	151098442	151098493
refGene_gn=ODAM;refGene_tx=XM_023253405.1
B1	151154363	151154831
refGene_gn=PRR27;refGene_tx=XM_023252237.1
B1	151155257	151155314
refGene_gn=PRR27;refGene_tx=XM_023252237.1
B1	151225201	151225303
refGene_gn=LOC102901003;refGene_tx=XM_023253406.1
B1	151227201	151227330
refGene_gn=LOC102901003;refGene_tx=XM_023253406.1
B1	151228719	151228764
refGene_gn=LOC102901003;refGene_tx=XM_023253406.1
B1	151228857	151228881
refGene_gn=LOC102901003;refGene_tx=XM_023253406.1
B1	151229994	151230021
refGene_gn=LOC102901003;refGene_tx=XM_023253406.1
B1	151231113	151231155
refGene_gn=LOC102901003;refGene_tx=XM_023253406.1
B1	151231245	151231269
refGene_gn=LOC102901003;refGene_tx=XM_023253406.1
B1	151232171	151232198
refGene_gn=LOC102901003;refGene_tx=XM_023253406.1
B1	151233514	151233577

refGene_gn=LOC102901003;refGene_tx=XM_023253406.1
B1 151237167 151237251
refGene_gn=LOC102901003;refGene_tx=XM_023253406.1
B1 151300165 151300216
refGene_gn=CSN2;refGene_tx=XM_003985340.1
B1 151300931 151300958
refGene_gn=CSN2;refGene_tx=XM_003985340.1
B1 151301055 151301082
refGene_gn=CSN2;refGene_tx=XM_003985340.1
B1 151302126 151302150
refGene_gn=CSN2;refGene_tx=XM_003985340.1
B1 151302246 151302291
refGene_gn=CSN2;refGene_tx=XM_003985340.1
B1 151303594 151304155
refGene_gn=CSN2;refGene_tx=XM_003985340.1
B1 151316746 151316905
refGene_gn=CSN1S1;refGene_tx=XM_006931074.3
B1 151317902 151317944
refGene_gn=CSN1S1;refGene_tx=XM_006931074.3
B1 151318380 151318398
refGene_gn=CSN1S1;refGene_tx=XM_006931074.3
B1 151319763 151319805
refGene_gn=CSN1S1;refGene_tx=XM_006931074.3
B1 151320807 151320852
refGene_gn=CSN1S1;refGene_tx=XM_006931074.3
B1 151320984 151321008
refGene_gn=CSN1S1;refGene_tx=XM_006931074.3
B1 151321816 151321849
refGene_gn=CSN1S1;refGene_tx=XM_006931074.3
B1 151323109 151323133
refGene_gn=CSN1S1;refGene_tx=XM_006931074.3
B1 151324327 151324369
refGene_gn=CSN1S1;refGene_tx=XM_006931074.3
B1 151324455 151324479
refGene_gn=CSN1S1;refGene_tx=XM_006931074.3
B1 151325106 151325130
refGene_gn=CSN1S1;refGene_tx=XM_006931074.3
B1 151325638 151325677
refGene_gn=CSN1S1;refGene_tx=XM_006931074.3
B1 151326341 151326362
refGene_gn=CSN1S1;refGene_tx=XM_006931074.3
B1 151326831 151326864
refGene_gn=CSN1S1;refGene_tx=XM_006931074.3
B1 151328670 151328721
refGene_gn=CSN1S1;refGene_tx=XM_006931074.3
B1 151398740 151398894
refGene_gn=SULT1E1;refGene_tx=XM_003985305.3;refGene_gn=SULT1E1;refGen
e_tx=XM_006931075.2
B1 151401624 151401750
refGene_gn=SULT1E1;refGene_tx=XM_003985305.3;refGene_gn=SULT1E1;refGen

e_tx=XM_006931075.2
B1	151402890	151402988
refGene_gn=SULT1E1;refGene_tx=XM_003985305.3;refGene_gn=SULT1E1;refGene_tx=XM_006931075.2
B1	151406781	151406908
refGene_gn=SULT1E1;refGene_tx=XM_003985305.3;refGene_gn=SULT1E1;refGene_tx=XM_006931075.2
B1	151408803	151408898
refGene_gn=SULT1E1;refGene_tx=XM_003985305.3;refGene_gn=SULT1E1;refGene_tx=XM_006931075.2
B1	151411872	151412053
refGene_gn=SULT1E1;refGene_tx=XM_003985305.3;refGene_gn=SULT1E1;refGene_tx=XM_006931075.2
B1	151413946	151414059
refGene_gn=SULT1E1;refGene_tx=XM_003985305.3;refGene_gn=SULT1E1;refGene_tx=XM_006931075.2
B1	151453127	151453275
refGene_gn=LOC101086557;refGene_tx=XM_023253066.1
B1	151462733	151462859
refGene_gn=LOC101086557;refGene_tx=XM_023253066.1
B1	151463665	151463763
refGene_gn=LOC101086557;refGene_tx=XM_023253066.1
B1	151469516	151469643
refGene_gn=LOC101086557;refGene_tx=XM_023253066.1
B1	151470997	151471092
refGene_gn=LOC101086557;refGene_tx=XM_023253066.1
B1	151472640	151472821
refGene_gn=LOC101086557;refGene_tx=XM_023253066.1
B1	151476268	151476318
refGene_gn=LOC101086557;refGene_tx=XM_023253066.1
B1	151476319	151476382
refGene_gn=LOC101086557;refGene_tx=XM_023253066.1
B1	151504097	151504245
refGene_gn=SULT1B1;refGene_tx=XM_023253068.1;refGene_gn=SULT1B1;refGene_tx=XM_023253069.1
B1	151505036	151505184
refGene_gn=SULT1B1;refGene_tx=XM_011281852.2
B1	151505953	151506082
refGene_gn=SULT1B1;refGene_tx=XM_011281852.2;refGene_gn=SULT1B1;refGene_tx=XM_023253068.1;refGene_gn=SULT1B1;refGene_tx=XM_023253069.1
B1	151509263	151509361
refGene_gn=SULT1B1;refGene_tx=XM_011281852.2;refGene_gn=SULT1B1;refGene_tx=XM_023253068.1;refGene_gn=SULT1B1;refGene_tx=XM_023253069.1
B1	151511297	151511424
refGene_gn=SULT1B1;refGene_tx=XM_011281852.2;refGene_gn=SULT1B1;refGene_tx=XM_023253068.1;refGene_gn=SULT1B1;refGene_tx=XM_023253069.1
B1	151512045	151512140
refGene_gn=SULT1B1;refGene_tx=XM_011281852.2;refGene_gn=SULT1B1;refGene_tx=XM_023253068.1;refGene_gn=SULT1B1;refGene_tx=XM_023253069.1
B1	151515779	151515960

refGene_gn=SULT1B1;refGene_tx=XM_011281852.2;refGene_gn=SULT1B1;refGene_tx=XM_023253068.1;refGene_gn=SULT1B1;refGene_tx=XM_023253069.1
B1	151520013	151520126
refGene_gn=SULT1B1;refGene_tx=XM_011281852.2;refGene_gn=SULT1B1;refGene_tx=XM_023253068.1;refGene_gn=SULT1B1;refGene_tx=XM_023253069.1
B1	151563009	151563727
refGene_gn=LOC101087059;refGene_tx=XM_003985308.3;refGene_gn=LOC101087059;refGene_tx=XM_003985310.3;refGene_gn=LOC101087059;refGene_tx=XM_011281850.2
B1	151575482	151576224
refGene_gn=LOC101087059;refGene_tx=XM_003985309.3;refGene_gn=LOC101087059;refGene_tx=XM_003985310.3
B1	151594345	151595069
refGene_gn=LOC101087059;refGene_tx=XM_011281849.2
B1	151603632	151603781
refGene_gn=LOC101087059;refGene_tx=XM_003985308.3;refGene_gn=LOC101087059;refGene_tx=XM_003985309.3;refGene_gn=LOC101087059;refGene_tx=XM_003985310.3;refGene_gn=LOC101087059;refGene_tx=XM_011281849.2;refGene_gn=LOC101087059;refGene_tx=XM_011281850.2
B1	151606207	151606339
refGene_gn=LOC101087059;refGene_tx=XM_003985308.3;refGene_gn=LOC101087059;refGene_tx=XM_003985309.3;refGene_gn=LOC101087059;refGene_tx=XM_011281849.2
B1	151608485	151608573
refGene_gn=LOC101087059;refGene_tx=XM_003985308.3;refGene_gn=LOC101087059;refGene_tx=XM_003985309.3;refGene_gn=LOC101087059;refGene_tx=XM_003985310.3;refGene_gn=LOC101087059;refGene_tx=XM_011281849.2;refGene_gn=LOC101087059;refGene_tx=XM_011281850.2
B1	151609824	151610044
refGene_gn=LOC101087059;refGene_tx=XM_003985308.3;refGene_gn=LOC101087059;refGene_tx=XM_003985309.3;refGene_gn=LOC101087059;refGene_tx=XM_003985310.3;refGene_gn=LOC101087059;refGene_tx=XM_011281849.2;refGene_gn=LOC101087059;refGene_tx=XM_011281850.2
B1	151614367	151614647
refGene_gn=LOC101087059;refGene_tx=XM_003985308.3;refGene_gn=LOC101087059;refGene_tx=XM_003985309.3;refGene_gn=LOC101087059;refGene_tx=XM_003985310.3;refGene_gn=LOC101087059;refGene_tx=XM_011281849.2;refGene_gn=LOC101087059;refGene_tx=XM_011281850.2
B1	151710326	151710606
refGene_gn=LOC101087662;refGene_tx=XM_003985311.4;refGene_gn=LOC101087662;refGene_tx=XM_019829415.1;refGene_gn=LOC101087662;refGene_tx=XM_019829416.2;refGene_gn=LOC101087662;refGene_tx=XM_023253070.1
B1	151712756	151712976
refGene_gn=LOC101087662;refGene_tx=XM_003985311.4;refGene_gn=LOC101087662;refGene_tx=XM_019829415.1;refGene_gn=LOC101087662;refGene_tx=XM_019829416.2;refGene_gn=LOC101087662;refGene_tx=XM_023253070.1
B1	151714758	151714846
refGene_gn=LOC101087662;refGene_tx=XM_003985311.4;refGene_gn=LOC101087662;refGene_tx=XM_019829415.1;refGene_gn=LOC101087662;refGene_tx=XM_019829416.2;refGene_gn=LOC101087662;refGene_tx=XM_023253070.1

B1 151723456 151723588
refGene_gn=LOC101087662;refGene_tx=XM_003985311.4;refGene_gn=LOC101087662;refGene_tx=XM_019829415.1;refGene_gn=LOC101087662;refGene_tx=XM_019829416.2;refGene_gn=LOC101087662;refGene_tx=XM_023253070.1
B1 151728813 151728962
refGene_gn=LOC101087662;refGene_tx=XM_003985311.4;refGene_gn=LOC101087662;refGene_tx=XM_019829415.1;refGene_gn=LOC101087662;refGene_tx=XM_019829416.2;refGene_gn=LOC101087662;refGene_tx=XM_023253070.1
B1 151735083 151735804
refGene_gn=LOC101087662;refGene_tx=XM_003985311.4;refGene_gn=LOC101087662;refGene_tx=XM_023253070.1
B1 151765227 151765452
refGene_gn=YTHDC1;refGene_tx=XM_003985312.4;refGene_gn=YTHDC1;refGene_tx=XM_003985313.4;refGene_gn=YTHDC1;refGene_tx=XM_006931083.3;refGene_gn=YTHDC1;refGene_tx=XM_011281848.3;refGene_gn=YTHDC1;refGene_tx=XM_019829412.2;refGene_gn=YTHDC1;refGene_tx=XM_019829413.2;refGene_gn=YTHDC1;refGene_tx=XM_019829414.2
B1 151767772 151767907
refGene_gn=YTHDC1;refGene_tx=XM_003985312.4;refGene_gn=YTHDC1;refGene_tx=XM_003985313.4;refGene_gn=YTHDC1;refGene_tx=XM_006931083.3;refGene_gn=YTHDC1;refGene_tx=XM_011281848.3;refGene_gn=YTHDC1;refGene_tx=XM_019829412.2;refGene_gn=YTHDC1;refGene_tx=XM_019829413.2;refGene_gn=YTHDC1;refGene_tx=XM_019829414.2
B1 151769521 151769584
refGene_gn=YTHDC1;refGene_tx=XM_003985312.4;refGene_gn=YTHDC1;refGene_tx=XM_003985313.4;refGene_gn=YTHDC1;refGene_tx=XM_006931083.3;refGene_gn=YTHDC1;refGene_tx=XM_011281848.3;refGene_gn=YTHDC1;refGene_tx=XM_019829412.2;refGene_gn=YTHDC1;refGene_tx=XM_019829413.2;refGene_gn=YTHDC1;refGene_tx=XM_019829414.2
B1 151769693 151769757
refGene_gn=YTHDC1;refGene_tx=XM_003985312.4;refGene_gn=YTHDC1;refGene_tx=XM_003985313.4;refGene_gn=YTHDC1;refGene_tx=XM_006931083.3;refGene_gn=YTHDC1;refGene_tx=XM_011281848.3;refGene_gn=YTHDC1;refGene_tx=XM_019829412.2;refGene_gn=YTHDC1;refGene_tx=XM_019829413.2;refGene_gn=YTHDC1;refGene_tx=XM_019829414.2
B1 151769829 151769866
refGene_gn=YTHDC1;refGene_tx=XM_003985312.4;refGene_gn=YTHDC1;refGene_tx=XM_003985313.4;refGene_gn=YTHDC1;refGene_tx=XM_006931083.3;refGene_gn=YTHDC1;refGene_tx=XM_011281848.3;refGene_gn=YTHDC1;refGene_tx=XM_019829412.2;refGene_gn=YTHDC1;refGene_tx=XM_019829413.2;refGene_gn=YTHDC1;refGene_tx=XM_019829414.2
B1 151771093 151771176
refGene_gn=YTHDC1;refGene_tx=XM_003985312.4;refGene_gn=YTHDC1;refGene_tx=XM_003985313.4;refGene_gn=YTHDC1;refGene_tx=XM_006931083.3;refGene_gn=YTHDC1;refGene_tx=XM_011281848.3;refGene_gn=YTHDC1;refGene_tx=XM_019829412.2;refGene_gn=YTHDC1;refGene_tx=XM_019829413.2;refGene_gn=YTHDC1;refGene_tx=XM_019829414.2
B1 151773691 151773858
refGene_gn=YTHDC1;refGene_tx=XM_003985312.4;refGene_gn=YTHDC1;refGene_tx=XM_003985313.4;refGene_gn=YTHDC1;refGene_tx=XM_006931083.3;refGene_

gn=YTHDC1;refGene_tx=XM_011281848.3;refGene_gn=YTHDC1;refGene_tx=XM_019829412.2;refGene_gn=YTHDC1;refGene_tx=XM_019829413.2;refGene_gn=YTHDC1;refGene_tx=XM_019829414.2
B1 151774803 151774888
refGene_gn=YTHDC1;refGene_tx=XM_003985312.4;refGene_gn=YTHDC1;refGene_tx=XM_003985313.4;refGene_gn=YTHDC1;refGene_tx=XM_006931083.3;refGene_gn=YTHDC1;refGene_tx=XM_011281848.3;refGene_gn=YTHDC1;refGene_tx=XM_019829412.2;refGene_gn=YTHDC1;refGene_tx=XM_019829413.2;refGene_gn=YTHDC1;refGene_tx=XM_019829414.2
B1 151781331 151781449
refGene_gn=YTHDC1;refGene_tx=XM_003985312.4;refGene_gn=YTHDC1;refGene_tx=XM_003985313.4;refGene_gn=YTHDC1;refGene_tx=XM_006931083.3;refGene_gn=YTHDC1;refGene_tx=XM_011281848.3;refGene_gn=YTHDC1;refGene_tx=XM_019829412.2;refGene_gn=YTHDC1;refGene_tx=XM_019829413.2;refGene_gn=YTHDC1;refGene_tx=XM_019829414.2
B1 151781527 151781636
refGene_gn=YTHDC1;refGene_tx=XM_003985312.4;refGene_gn=YTHDC1;refGene_tx=XM_003985313.4;refGene_gn=YTHDC1;refGene_tx=XM_006931083.3;refGene_gn=YTHDC1;refGene_tx=XM_011281848.3;refGene_gn=YTHDC1;refGene_tx=XM_019829412.2;refGene_gn=YTHDC1;refGene_tx=XM_019829413.2;refGene_gn=YTHDC1;refGene_tx=XM_019829414.2
B1 151783120 151783215
refGene_gn=YTHDC1;refGene_tx=XM_003985312.4;refGene_gn=YTHDC1;refGene_tx=XM_003985313.4;refGene_gn=YTHDC1;refGene_tx=XM_006931083.3;refGene_gn=YTHDC1;refGene_tx=XM_011281848.3;refGene_gn=YTHDC1;refGene_tx=XM_019829412.2;refGene_gn=YTHDC1;refGene_tx=XM_019829413.2;refGene_gn=YTHDC1;refGene_tx=XM_019829414.2
B1 151783775 151783829
refGene_gn=YTHDC1;refGene_tx=XM_003985312.4;refGene_gn=YTHDC1;refGene_tx=XM_006931083.3;refGene_gn=YTHDC1;refGene_tx=XM_011281848.3;refGene_gn=YTHDC1;refGene_tx=XM_019829412.2;refGene_gn=YTHDC1;refGene_tx=XM_019829413.2
B1 151784298 151784388
refGene_gn=YTHDC1;refGene_tx=XM_003985312.4;refGene_gn=YTHDC1;refGene_tx=XM_003985313.4;refGene_gn=YTHDC1;refGene_tx=XM_006931083.3;refGene_gn=YTHDC1;refGene_tx=XM_011281848.3;refGene_gn=YTHDC1;refGene_tx=XM_019829412.2;refGene_gn=YTHDC1;refGene_tx=XM_019829413.2;refGene_gn=YTHDC1;refGene_tx=XM_019829414.2
B1 151788586 151789016
refGene_gn=YTHDC1;refGene_tx=XM_003985312.4;refGene_gn=YTHDC1;refGene_tx=XM_003985313.4;refGene_gn=YTHDC1;refGene_tx=XM_006931083.3;refGene_gn=YTHDC1;refGene_tx=XM_011281848.3;refGene_gn=YTHDC1;refGene_tx=XM_019829412.2;refGene_gn=YTHDC1;refGene_tx=XM_019829413.2;refGene_gn=YTHDC1;refGene_tx=XM_019829414.2
B1 151789137 151789466
refGene_gn=YTHDC1;refGene_tx=XM_003985312.4;refGene_gn=YTHDC1;refGene_tx=XM_003985313.4;refGene_gn=YTHDC1;refGene_tx=XM_006931083.3;refGene_gn=YTHDC1;refGene_tx=XM_011281848.3;refGene_gn=YTHDC1;refGene_tx=XM_019829412.2;refGene_gn=YTHDC1;refGene_tx=XM_019829413.2;refGene_gn=YTHDC1;refGene_tx=XM_019829414.2

B1 151790086 151790188
refGene_gn=YTHDC1;refGene_tx=XM_003985312.4;refGene_gn=YTHDC1;refGene_tx=XM_003985313.4;refGene_gn=YTHDC1;refGene_tx=XM_006931083.3;refGene_gn=YTHDC1;refGene_tx=XM_019829412.2;refGene_gn=YTHDC1;refGene_tx=XM_019829413.2;refGene_gn=YTHDC1;refGene_tx=XM_019829414.2
B1 151794676 151794712
refGene_gn=YTHDC1;refGene_tx=XM_019829412.2;refGene_gn=YTHDC1;refGene_tx=XM_019829413.2;refGene_gn=YTHDC1;refGene_tx=XM_019829414.2
B1 151799366 151799394
refGene_gn=YTHDC1;refGene_tx=XM_003985312.4;refGene_gn=YTHDC1;refGene_tx=XM_003985313.4;refGene_gn=YTHDC1;refGene_tx=XM_006931083.3;refGene_gn=YTHDC1;refGene_tx=XM_019829412.2;refGene_gn=YTHDC1;refGene_tx=XM_019829413.2;refGene_gn=YTHDC1;refGene_tx=XM_019829414.2
B1 151874134 151874142
refGene_gn=LOC101088329;refGene_tx=XM_019829425.1;refGene_gn=LOC101088329;refGene_tx=XM_019829426.1
B1 151889269 151889394
refGene_gn=LOC101088329;refGene_tx=XM_019829425.1;refGene_gn=LOC101088329;refGene_tx=XM_019829426.1
B1 151893910 151894032
refGene_gn=LOC101088329;refGene_tx=XM_019829425.1;refGene_gn=LOC101088329;refGene_tx=XM_019829426.1
B1 151896614 151896682
refGene_gn=LOC101088329;refGene_tx=XM_019829425.1;refGene_gn=LOC101088329;refGene_tx=XM_019829426.1
B1 151899162 151899326
refGene_gn=LOC101088329;refGene_tx=XM_019829425.1;refGene_gn=LOC101088329;refGene_tx=XM_019829426.1
B1 151901152 151901197
refGene_gn=LOC101088329;refGene_tx=XM_019829425.1
B1 151902739 151902778
refGene_gn=LOC101088329;refGene_tx=XM_019829425.1;refGene_gn=LOC101088329;refGene_tx=XM_019829426.1
B1 151904888 151905066
refGene_gn=LOC101088329;refGene_tx=XM_019829425.1;refGene_gn=LOC101088329;refGene_tx=XM_019829426.1
B1 151906466 151906726
refGene_gn=LOC101088329;refGene_tx=XM_019829425.1;refGene_gn=LOC101088329;refGene_tx=XM_019829426.1
B1 151907315 151907458
refGene_gn=LOC101088329;refGene_tx=XM_019829425.1;refGene_gn=LOC101088329;refGene_tx=XM_019829426.1
B1 151915888 151916050
refGene_gn=LOC101088329;refGene_tx=XM_019829425.1;refGene_gn=LOC101088329;refGene_tx=XM_019829426.1
B1 151923531 151923623
refGene_gn=LOC101099042;refGene_tx=XM_019829945.1
B1 151944864 151944983
refGene_gn=LOC101099042;refGene_tx=XM_019829945.1
B1 151949391 151949507

refGene_gn=LOC101099042;refGene_tx=XM_019829945.1
B1 151949592 151949664
refGene_gn=LOC101099042;refGene_tx=XM_019829945.1
B1 151951021 151951202
refGene_gn=LOC101099042;refGene_tx=XM_019829945.1
B1 151952596 151952635
refGene_gn=LOC101099042;refGene_tx=XM_019829945.1
B1 151953450 151953625
refGene_gn=LOC101099042;refGene_tx=XM_019829945.1
B1 151953705 151953788
refGene_gn=LOC101099042;refGene_tx=XM_019829945.1
B1 151961257 151961400
refGene_gn=LOC101099042;refGene_tx=XM_019829945.1
B1 151962044 151962206
refGene_gn=LOC101099042;refGene_tx=XM_019829945.1
B1 151976059 151976108
refGene_gn=LOC101099294;refGene_tx=XM_023253408.1
B1 151978404 151978575
refGene_gn=LOC101099294;refGene_tx=XM_023253408.1
B1 151981630 151981749
refGene_gn=LOC101099294;refGene_tx=XM_023253408.1
B1 151981845 151981913
refGene_gn=LOC101099294;refGene_tx=XM_023253408.1
B1 151984205 151984366
refGene_gn=LOC101099294;refGene_tx=XM_023253408.1
B1 151985070 151985109
refGene_gn=LOC101099294;refGene_tx=XM_023253408.1
B1 151985636 151985811
refGene_gn=LOC101099294;refGene_tx=XM_023253408.1
B1 151986922 151987182
refGene_gn=LOC101099294;refGene_tx=XM_023253408.1
B1 151987866 151988009
refGene_gn=LOC101099294;refGene_tx=XM_023253408.1
B1 151988866 151989028
refGene_gn=LOC101099294;refGene_tx=XM_023253408.1
B1 152033017 152033028
refGene_gn=LOC101099548;refGene_tx=XM_003985344.3;refGene_gn=LOC101099
548;refGene_tx=XM_019829424.2
B1 152065084 152065236
refGene_gn=LOC101099548;refGene_tx=XM_003985344.3;refGene_gn=LOC101099
548;refGene_tx=XM_019829424.2
B1 152072825 152072944
refGene_gn=LOC101099548;refGene_tx=XM_003985344.3;refGene_gn=LOC101099
548;refGene_tx=XM_019829424.2
B1 152086211 152086279
refGene_gn=LOC101099548;refGene_tx=XM_003985344.3;refGene_gn=LOC101099
548;refGene_tx=XM_019829424.2
B1 152087878 152088042
refGene_gn=LOC101099548;refGene_tx=XM_003985344.3;refGene_gn=LOC101099
548;refGene_tx=XM_019829424.2

B1 152090588 152090627
refGene_gn=LOC101099548;refGene_tx=XM_003985344.3;refGene_gn=LOC101099548;refGene_tx=XM_019829424.2
B1 152095435 152095637
refGene_gn=LOC101099548;refGene_tx=XM_003985344.3;refGene_gn=LOC101099548;refGene_tx=XM_019829424.2
B1 152097550 152097810
refGene_gn=LOC101099548;refGene_tx=XM_003985344.3;refGene_gn=LOC101099548;refGene_tx=XM_019829424.2
B1 152099989 152100132
refGene_gn=LOC101099548;refGene_tx=XM_003985344.3;refGene_gn=LOC101099548;refGene_tx=XM_019829424.2
B1 152100438 152100474
refGene_gn=LOC101099548;refGene_tx=XM_019829424.2
B1 152104010 152104169
refGene_gn=LOC101099548;refGene_tx=XM_003985344.3
B1 152131656 152131709
refGene_gn=LOC101099798;refGene_tx=XM_023253409.1
B1 152133701 152133823
refGene_gn=LOC101099798;refGene_tx=XM_023253409.1
B1 152135699 152135818
refGene_gn=LOC101099798;refGene_tx=XM_023253409.1
B1 152138848 152138916
refGene_gn=LOC101099798;refGene_tx=XM_023253409.1
B1 152139550 152139711
refGene_gn=LOC101099798;refGene_tx=XM_023253409.1
B1 152152497 152152536
refGene_gn=LOC101099798;refGene_tx=XM_023253409.1
B1 152153414 152153586
refGene_gn=LOC101099798;refGene_tx=XM_023253409.1
B1 152154704 152154964
refGene_gn=LOC101099798;refGene_tx=XM_023253409.1
B1 152158597 152158740
refGene_gn=LOC101099798;refGene_tx=XM_023253409.1
B1 152161244 152161403
refGene_gn=LOC101099798;refGene_tx=XM_023253409.1
B1 152177163 152177174
refGene_gn=TMPRSS11A;refGene_tx=XM_019829427.2
B1 152190926 152191051
refGene_gn=TMPRSS11A;refGene_tx=XM_019829427.2
B1 152193550 152193669
refGene_gn=TMPRSS11A;refGene_tx=XM_019829427.2
B1 152208258 152208326
refGene_gn=TMPRSS11A;refGene_tx=XM_019829427.2
B1 152210259 152210420
refGene_gn=TMPRSS11A;refGene_tx=XM_019829427.2
B1 152213415 152213454
refGene_gn=TMPRSS11A;refGene_tx=XM_019829427.2
B1 152215375 152215547
refGene_gn=TMPRSS11A;refGene_tx=XM_019829427.2

B1 152222872 152223132
refGene_gn=TMPRSS11A;refGene_tx=XM_019829427.2
B1 152227674 152227817
refGene_gn=TMPRSS11A;refGene_tx=XM_019829427.2
B1 152229563 152229725
refGene_gn=TMPRSS11A;refGene_tx=XM_019829427.2
B1 152277768 152277895
refGene_gn=TMPRSS11D;refGene_tx=XM_023253410.1
B1 152280026 152280145
refGene_gn=TMPRSS11D;refGene_tx=XM_023253410.1
B1 152287856 152287924
refGene_gn=TMPRSS11D;refGene_tx=XM_023253410.1
B1 152294258 152294416
refGene_gn=TMPRSS11D;refGene_tx=XM_023253410.1
B1 152298194 152298233
refGene_gn=TMPRSS11D;refGene_tx=XM_023253410.1
B1 152301045 152301223
refGene_gn=TMPRSS11D;refGene_tx=XM_023253410.1
B1 152306695 152306955
refGene_gn=TMPRSS11D;refGene_tx=XM_023253410.1
B1 152308078 152308221
refGene_gn=TMPRSS11D;refGene_tx=XM_023253410.1
B1 152311250 152311412
refGene_gn=TMPRSS11D;refGene_tx=XM_023253410.1
B1 152328218 152328274
refGene_gn=LOC101088575;refGene_tx=XM_011281844.2
B1 152336947 152337066
refGene_gn=LOC101088575;refGene_tx=XM_011281844.2
B1 152343805 152343924
refGene_gn=LOC101088575;refGene_tx=XM_011281844.2
B1 152348726 152348794
refGene_gn=LOC101088575;refGene_tx=XM_011281844.2
B1 152352330 152352494
refGene_gn=LOC101088575;refGene_tx=XM_011281844.2
B1 152368618 152368657
refGene_gn=LOC101088575;refGene_tx=XM_011281844.2
B1 152372167 152372339
refGene_gn=LOC101088575;refGene_tx=XM_011281844.2
B1 152373540 152373800
refGene_gn=LOC101088575;refGene_tx=XM_011281844.2
B1 152375284 152375427
refGene_gn=LOC101088575;refGene_tx=XM_011281844.2
B1 152380469 152380631
refGene_gn=LOC101088575;refGene_tx=XM_011281844.2
B1 152391618 152392137
refGene_gn=GNRHR;refGene_tx=NM_001287228.1;refGene_gn=GNRHR;refGene_tx
=XM_019828145.1
B1 152409729 152409949
refGene_gn=GNRHR;refGene_tx=NM_001287228.1;refGene_gn=GNRHR;refGene_tx
=XM_019828145.1

B1	152412387	152412632
refGene_gn=GNRHR;refGene_tx=NM_001287228.1;refGene_gn=GNRHR;refGene_tx=XM_019828145.1
B1	152429215	152429286
refGene_gn=UBA6;refGene_tx=XM_023253071.1
B1	152432687	152432750
refGene_gn=UBA6;refGene_tx=XM_023253071.1
B1	152456709	152456804
refGene_gn=UBA6;refGene_tx=XM_023253071.1;refGene_gn=UBA6;refGene_tx=XM_023253072.1
B1	152457335	152457364
refGene_gn=UBA6;refGene_tx=XM_023253071.1;refGene_gn=UBA6;refGene_tx=XM_023253072.1
B1	152461857	152461952
refGene_gn=UBA6;refGene_tx=XM_023253071.1;refGene_gn=UBA6;refGene_tx=XM_023253072.1
B1	152462474	152462586
refGene_gn=UBA6;refGene_tx=XM_023253071.1;refGene_gn=UBA6;refGene_tx=XM_023253072.1
B1	152470088	152470169
refGene_gn=UBA6;refGene_tx=XM_023253071.1;refGene_gn=UBA6;refGene_tx=XM_023253072.1
B1	152475456	152475579
refGene_gn=UBA6;refGene_tx=XM_023253071.1;refGene_gn=UBA6;refGene_tx=XM_023253072.1
B1	152478130	152478254
refGene_gn=UBA6;refGene_tx=XM_023253071.1;refGene_gn=UBA6;refGene_tx=XM_023253072.1
B1	152480215	152480319
refGene_gn=UBA6;refGene_tx=XM_023253071.1;refGene_gn=UBA6;refGene_tx=XM_023253072.1
B1	152480933	152480996
refGene_gn=UBA6;refGene_tx=XM_023253071.1;refGene_gn=UBA6;refGene_tx=XM_023253072.1
B1	152481739	152481816
refGene_gn=UBA6;refGene_tx=XM_023253071.1;refGene_gn=UBA6;refGene_tx=XM_023253072.1
B1	152482434	152482501
refGene_gn=UBA6;refGene_tx=XM_023253071.1;refGene_gn=UBA6;refGene_tx=XM_023253072.1
B1	152488751	152488895
refGene_gn=UBA6;refGene_tx=XM_023253071.1;refGene_gn=UBA6;refGene_tx=XM_023253072.1
B1	152490074	152490142
refGene_gn=UBA6;refGene_tx=XM_023253071.1;refGene_gn=UBA6;refGene_tx=XM_023253072.1
B1	152491252	152491331
refGene_gn=UBA6;refGene_tx=XM_023253071.1;refGene_gn=UBA6;refGene_tx=XM_023253072.1
B1	152492481	152492562

refGene_gn=UBA6;refGene_tx=XM_023253071.1;refGene_gn=UBA6;refGene_tx=XM_023253072.1
B1 152496904 152496982
refGene_gn=UBA6;refGene_tx=XM_023253071.1;refGene_gn=UBA6;refGene_tx=XM_023253072.1
B1 152498327 152498509
refGene_gn=UBA6;refGene_tx=XM_023253071.1;refGene_gn=UBA6;refGene_tx=XM_023253072.1
B1 152500001 152500107
refGene_gn=UBA6;refGene_tx=XM_023253071.1;refGene_gn=UBA6;refGene_tx=XM_023253072.1
B1 152502015 152502105
refGene_gn=UBA6;refGene_tx=XM_023253071.1;refGene_gn=UBA6;refGene_tx=XM_023253072.1
B1 152502201 152502282
refGene_gn=UBA6;refGene_tx=XM_023253071.1;refGene_gn=UBA6;refGene_tx=XM_023253072.1
B1 152503060 152503189
refGene_gn=UBA6;refGene_tx=XM_023253071.1;refGene_gn=UBA6;refGene_tx=XM_023253072.1
B1 152504762 152504814
refGene_gn=UBA6;refGene_tx=XM_023253071.1;refGene_gn=UBA6;refGene_tx=XM_023253072.1
B1 152504902 152504966
refGene_gn=UBA6;refGene_tx=XM_023253071.1;refGene_gn=UBA6;refGene_tx=XM_023253072.1
B1 152505787 152505857
refGene_gn=UBA6;refGene_tx=XM_023253071.1;refGene_gn=UBA6;refGene_tx=XM_023253072.1
B1 152506392 152506464
refGene_gn=UBA6;refGene_tx=XM_023253071.1;refGene_gn=UBA6;refGene_tx=XM_023253072.1
B1 152509313 152509431
refGene_gn=UBA6;refGene_tx=XM_023253071.1;refGene_gn=UBA6;refGene_tx=XM_023253072.1
B1 152510863 152511057
refGene_gn=UBA6;refGene_tx=XM_023253071.1;refGene_gn=UBA6;refGene_tx=XM_023253072.1
B1 152512096 152512224
refGene_gn=UBA6;refGene_tx=XM_023253071.1;refGene_gn=UBA6;refGene_tx=XM_023253072.1
B1 152515300 152515388
refGene_gn=UBA6;refGene_tx=XM_023253071.1;refGene_gn=UBA6;refGene_tx=XM_023253072.1
B1 152515552 152515647
refGene_gn=UBA6;refGene_tx=XM_023253071.1;refGene_gn=UBA6;refGene_tx=XM_023253072.1
B1 152518692 152518828
refGene_gn=UBA6;refGene_tx=XM_023253071.1;refGene_gn=UBA6;refGene_tx=XM_023253072.1

B1 152533733 152533763
refGene_gn=STAP1;refGene_tx=XM_011281843.3;refGene_gn=STAP1;refGene_tx=XM_019829431.2
B1 152535240 152535438
refGene_gn=STAP1;refGene_tx=XM_011281843.3;refGene_gn=STAP1;refGene_tx=XM_019829431.2;refGene_gn=STAP1;refGene_tx=XM_019829433.2
B1 152539225 152539295
refGene_gn=STAP1;refGene_tx=XM_011281843.3;refGene_gn=STAP1;refGene_tx=XM_019829431.2;refGene_gn=STAP1;refGene_tx=XM_019829433.2
B1 152547101 152547230
refGene_gn=STAP1;refGene_tx=XM_011281843.3;refGene_gn=STAP1;refGene_tx=XM_019829431.2;refGene_gn=STAP1;refGene_tx=XM_019829433.2
B1 152548719 152548886
refGene_gn=STAP1;refGene_tx=XM_011281843.3;refGene_gn=STAP1;refGene_tx=XM_019829431.2;refGene_gn=STAP1;refGene_tx=XM_019829433.2
B1 152551798 152551855
refGene_gn=STAP1;refGene_tx=XM_011281843.3;refGene_gn=STAP1;refGene_tx=XM_019829431.2;refGene_gn=STAP1;refGene_tx=XM_019829433.2
B1 152554332 152554446
refGene_gn=STAP1;refGene_tx=XM_011281843.3;refGene_gn=STAP1;refGene_tx=XM_019829431.2;refGene_gn=STAP1;refGene_tx=XM_019829433.2
B1 152559515 152559587
refGene_gn=STAP1;refGene_tx=XM_011281843.3;refGene_gn=STAP1;refGene_tx=XM_019829431.2;refGene_gn=STAP1;refGene_tx=XM_019829433.2
B1 152568760 152568880
refGene_gn=STAP1;refGene_tx=XM_011281843.3;refGene_gn=STAP1;refGene_tx=XM_019829431.2;refGene_gn=STAP1;refGene_tx=XM_019829433.2
B1 152580928 152580946
refGene_gn=CENPC;refGene_tx=XM_006931085.4;refGene_gn=CENPC;refGene_tx=XM_019829418.2;refGene_gn=CENPC;refGene_tx=XM_019829419.2
B1 152585165 152585212
refGene_gn=CENPC;refGene_tx=XM_006931085.4;refGene_gn=CENPC;refGene_tx=XM_019829418.2;refGene_gn=CENPC;refGene_tx=XM_019829419.2
B1 152589893 152589964
refGene_gn=CENPC;refGene_tx=XM_006931085.4;refGene_gn=CENPC;refGene_tx=XM_019829418.2;refGene_gn=CENPC;refGene_tx=XM_019829419.2
B1 152590922 152591017
refGene_gn=CENPC;refGene_tx=XM_006931085.4;refGene_gn=CENPC;refGene_tx=XM_019829418.2;refGene_gn=CENPC;refGene_tx=XM_019829419.2
B1 152595390 152595496
refGene_gn=CENPC;refGene_tx=XM_006931085.4;refGene_gn=CENPC;refGene_tx=XM_019829418.2;refGene_gn=CENPC;refGene_tx=XM_019829419.2
B1 152600419 152600705
refGene_gn=CENPC;refGene_tx=XM_006931085.4;refGene_gn=CENPC;refGene_tx=XM_019829418.2;refGene_gn=CENPC;refGene_tx=XM_019829419.2
B1 152601631 152601847
refGene_gn=CENPC;refGene_tx=XM_006931085.4;refGene_gn=CENPC;refGene_tx=XM_019829418.2;refGene_gn=CENPC;refGene_tx=XM_019829419.2
B1 152604822 152605436
refGene_gn=CENPC;refGene_tx=XM_006931085.4;refGene_gn=CENPC;refGene_tx

=XM_019829418.2;refGene_gn=CENPC;refGene_tx=XM_019829419.2
B1 152610859 152611051
refGene_gn=CENPC;refGene_tx=XM_006931085.4;refGene_gn=CENPC;refGene_tx
=XM_019829418.2;refGene_gn=CENPC;refGene_tx=XM_019829419.2
B1 152617045 152617337
refGene_gn=CENPC;refGene_tx=XM_006931085.4;refGene_gn=CENPC;refGene_tx
=XM_019829418.2;refGene_gn=CENPC;refGene_tx=XM_019829419.2
B1 152628362 152628506
refGene_gn=CENPC;refGene_tx=XM_006931085.4;refGene_gn=CENPC;refGene_tx
=XM_019829418.2;refGene_gn=CENPC;refGene_tx=XM_019829419.2
B1 152630880 152630963
refGene_gn=CENPC;refGene_tx=XM_006931085.4;refGene_gn=CENPC;refGene_tx
=XM_019829418.2;refGene_gn=CENPC;refGene_tx=XM_019829419.2
B1 152636209 152636269
refGene_gn=CENPC;refGene_tx=XM_006931085.4;refGene_gn=CENPC;refGene_tx
=XM_019829419.2
B1 152641980 152642085
refGene_gn=CENPC;refGene_tx=XM_006931085.4;refGene_gn=CENPC;refGene_tx
=XM_019829418.2;refGene_gn=CENPC;refGene_tx=XM_019829419.2
B1 152649827 152649956
refGene_gn=CENPC;refGene_tx=XM_006931085.4;refGene_gn=CENPC;refGene_tx
=XM_019829418.2
B1 152651904 152652000
refGene_gn=CENPC;refGene_tx=XM_006931085.4;refGene_gn=CENPC;refGene_tx
=XM_019829418.2;refGene_gn=CENPC;refGene_tx=XM_019829419.2
B1 152653670 152653825
refGene_gn=CENPC;refGene_tx=XM_006931085.4;refGene_gn=CENPC;refGene_tx
=XM_019829418.2;refGene_gn=CENPC;refGene_tx=XM_019829419.2
B1 152660137 152660228
refGene_gn=CENPC;refGene_tx=XM_006931085.4;refGene_gn=CENPC;refGene_tx
=XM_019829418.2;refGene_gn=CENPC;refGene_tx=XM_019829419.2
B1 152661378 152661449
refGene_gn=CENPC;refGene_tx=XM_006931085.4;refGene_gn=CENPC;refGene_tx
=XM_019829418.2;refGene_gn=CENPC;refGene_tx=XM_019829419.2
B1 153326334 153328670
refGene_gn=LOC111560225;refGene_tx=XM_023253411.1
B1 153329603 153329792
refGene_gn=LOC111560225;refGene_tx=XM_023253411.1
B1 154392879 154393075
refGene_gn=EPHA5;refGene_tx=XM_023253073.1;refGene_gn=EPHA5;refGene_tx
=XM_023253074.1;refGene_gn=EPHA5;refGene_tx=XM_023253075.1;refGene_gn=
EPHA5;refGene_tx=XM_023253076.1;refGene_gn=EPHA5;refGene_tx=XM_0232530
77.1;refGene_gn=EPHA5;refGene_tx=XM_023253078.1;refGene_gn=EPHA5;refGe
ne_tx=XM_023253079.1
B1 154426272 154426337
refGene_gn=EPHA5;refGene_tx=XM_023253073.1;refGene_gn=EPHA5;refGene_tx
=XM_023253074.1;refGene_gn=EPHA5;refGene_tx=XM_023253075.1;refGene_gn=
EPHA5;refGene_tx=XM_023253076.1;refGene_gn=EPHA5;refGene_tx=XM_0232530
77.1;refGene_gn=EPHA5;refGene_tx=XM_023253078.1;refGene_gn=EPHA5;refGe
ne_tx=XM_023253079.1

B1 154473383 154474047
refGene_gn=EPHA5;refGene_tx=XM_023253073.1;refGene_gn=EPHA5;refGene_tx=XM_023253074.1;refGene_gn=EPHA5;refGene_tx=XM_023253075.1;refGene_gn=EPHA5;refGene_tx=XM_023253076.1;refGene_gn=EPHA5;refGene_tx=XM_023253077.1;refGene_gn=EPHA5;refGene_tx=XM_023253078.1;refGene_gn=EPHA5;refGene_tx=XM_023253079.1
B1 154577583 154577739
refGene_gn=EPHA5;refGene_tx=XM_023253073.1;refGene_gn=EPHA5;refGene_tx=XM_023253074.1;refGene_gn=EPHA5;refGene_tx=XM_023253075.1;refGene_gn=EPHA5;refGene_tx=XM_023253076.1;refGene_gn=EPHA5;refGene_tx=XM_023253077.1;refGene_gn=EPHA5;refGene_tx=XM_023253078.1
B1 154581739 154582075
refGene_gn=EPHA5;refGene_tx=XM_023253073.1;refGene_gn=EPHA5;refGene_tx=XM_023253074.1;refGene_gn=EPHA5;refGene_tx=XM_023253075.1;refGene_gn=EPHA5;refGene_tx=XM_023253076.1
B1 154650268 154650393
refGene_gn=EPHA5;refGene_tx=XM_023253073.1;refGene_gn=EPHA5;refGene_tx=XM_023253074.1;refGene_gn=EPHA5;refGene_tx=XM_023253075.1;refGene_gn=EPHA5;refGene_tx=XM_023253076.1;refGene_gn=EPHA5;refGene_tx=XM_023253077.1;refGene_gn=EPHA5;refGene_tx=XM_023253078.1;refGene_gn=EPHA5;refGene_tx=XM_023253079.1
B1 154657954 154658114
refGene_gn=EPHA5;refGene_tx=XM_023253073.1;refGene_gn=EPHA5;refGene_tx=XM_023253074.1;refGene_gn=EPHA5;refGene_tx=XM_023253075.1;refGene_gn=EPHA5;refGene_tx=XM_023253076.1;refGene_gn=EPHA5;refGene_tx=XM_023253077.1;refGene_gn=EPHA5;refGene_tx=XM_023253078.1;refGene_gn=EPHA5;refGene_tx=XM_023253079.1
B1 154666329 154666435
refGene_gn=EPHA5;refGene_tx=XM_023253073.1;refGene_gn=EPHA5;refGene_tx=XM_023253074.1;refGene_gn=EPHA5;refGene_tx=XM_023253075.1;refGene_gn=EPHA5;refGene_tx=XM_023253076.1;refGene_gn=EPHA5;refGene_tx=XM_023253077.1;refGene_gn=EPHA5;refGene_tx=XM_023253078.1;refGene_gn=EPHA5;refGene_tx=XM_023253079.1
B1 154692004 154692070
refGene_gn=EPHA5;refGene_tx=XM_023253073.1;refGene_gn=EPHA5;refGene_tx=XM_023253074.1;refGene_gn=EPHA5;refGene_tx=XM_023253077.1;refGene_gn=EPHA5;refGene_tx=XM_023253078.1;refGene_gn=EPHA5;refGene_tx=XM_023253079.1
B1 154701353 154701421
refGene_gn=EPHA5;refGene_tx=XM_023253073.1;refGene_gn=EPHA5;refGene_tx=XM_023253074.1;refGene_gn=EPHA5;refGene_tx=XM_023253075.1;refGene_gn=EPHA5;refGene_tx=XM_023253076.1;refGene_gn=EPHA5;refGene_tx=XM_023253077.1;refGene_gn=EPHA5;refGene_tx=XM_023253078.1;refGene_gn=EPHA5;refGene_tx=XM_023253079.1
B1 154702962 154703088
refGene_gn=EPHA5;refGene_tx=XM_023253073.1;refGene_gn=EPHA5;refGene_tx=XM_023253074.1;refGene_gn=EPHA5;refGene_tx=XM_023253075.1;refGene_gn=EPHA5;refGene_tx=XM_023253076.1;refGene_gn=EPHA5;refGene_tx=XM_023253077.1;refGene_gn=EPHA5;refGene_tx=XM_023253078.1;refGene_gn=EPHA5;refGene_tx=XM_023253079.1

B1	154703823	154704009
refGene_gn=EPHA5;refGene_tx=XM_023253073.1;refGene_gn=EPHA5;refGene_tx=XM_023253074.1;refGene_gn=EPHA5;refGene_tx=XM_023253075.1;refGene_gn=EPHA5;refGene_tx=XM_023253076.1;refGene_gn=EPHA5;refGene_tx=XM_023253077.1;refGene_gn=EPHA5;refGene_tx=XM_023253078.1;refGene_gn=EPHA5;refGene_tx=XM_023253079.1
B1	154718815	154718877
refGene_gn=EPHA5;refGene_tx=XM_023253073.1;refGene_gn=EPHA5;refGene_tx=XM_023253074.1;refGene_gn=EPHA5;refGene_tx=XM_023253075.1;refGene_gn=EPHA5;refGene_tx=XM_023253076.1;refGene_gn=EPHA5;refGene_tx=XM_023253077.1;refGene_gn=EPHA5;refGene_tx=XM_023253078.1;refGene_gn=EPHA5;refGene_tx=XM_023253079.1
B1	154720293	154720503
refGene_gn=EPHA5;refGene_tx=XM_023253073.1;refGene_gn=EPHA5;refGene_tx=XM_023253074.1;refGene_gn=EPHA5;refGene_tx=XM_023253075.1;refGene_gn=EPHA5;refGene_tx=XM_023253076.1;refGene_gn=EPHA5;refGene_tx=XM_023253077.1;refGene_gn=EPHA5;refGene_tx=XM_023253078.1;refGene_gn=EPHA5;refGene_tx=XM_023253079.1
B1	154723530	154723680
refGene_gn=EPHA5;refGene_tx=XM_023253073.1;refGene_gn=EPHA5;refGene_tx=XM_023253074.1;refGene_gn=EPHA5;refGene_tx=XM_023253075.1;refGene_gn=EPHA5;refGene_tx=XM_023253076.1;refGene_gn=EPHA5;refGene_tx=XM_023253077.1;refGene_gn=EPHA5;refGene_tx=XM_023253078.1;refGene_gn=EPHA5;refGene_tx=XM_023253079.1
B1	154732815	154733009
refGene_gn=EPHA5;refGene_tx=XM_023253073.1;refGene_gn=EPHA5;refGene_tx=XM_023253074.1;refGene_gn=EPHA5;refGene_tx=XM_023253075.1;refGene_gn=EPHA5;refGene_tx=XM_023253076.1;refGene_gn=EPHA5;refGene_tx=XM_023253077.1;refGene_gn=EPHA5;refGene_tx=XM_023253078.1;refGene_gn=EPHA5;refGene_tx=XM_023253079.1
B1	154736549	154736709
refGene_gn=EPHA5;refGene_tx=XM_023253073.1;refGene_gn=EPHA5;refGene_tx=XM_023253074.1;refGene_gn=EPHA5;refGene_tx=XM_023253075.1;refGene_gn=EPHA5;refGene_tx=XM_023253076.1;refGene_gn=EPHA5;refGene_tx=XM_023253077.1;refGene_gn=EPHA5;refGene_tx=XM_023253078.1;refGene_gn=EPHA5;refGene_tx=XM_023253079.1
B1	154744808	154744914
refGene_gn=EPHA5;refGene_tx=XM_023253073.1;refGene_gn=EPHA5;refGene_tx=XM_023253074.1;refGene_gn=EPHA5;refGene_tx=XM_023253075.1;refGene_gn=EPHA5;refGene_tx=XM_023253076.1;refGene_gn=EPHA5;refGene_tx=XM_023253077.1;refGene_gn=EPHA5;refGene_tx=XM_023253079.1
B1	155435215	155435449
refGene_gn=TECRL;refGene_tx=XM_003985351.3
B1	155461552	155461604
refGene_gn=TECRL;refGene_tx=XM_003985351.3
B1	155490716	155490761
refGene_gn=TECRL;refGene_tx=XM_003985351.3
B1	155496004	155496108
refGene_gn=TECRL;refGene_tx=XM_003985351.3
B1	155505653	155505769

refGene_gn=TECRL;refGene_tx=XM_003985351.3
B1	155509719	155509825
refGene_gn=TECRL;refGene_tx=XM_003985351.3
B1	155511136	155511209
refGene_gn=TECRL;refGene_tx=XM_003985351.3
B1	155515872	155515916
refGene_gn=TECRL;refGene_tx=XM_003985351.3
B1	155527269	155527327
refGene_gn=TECRL;refGene_tx=XM_003985351.3
B1	155530758	155530844
refGene_gn=TECRL;refGene_tx=XM_003985351.3
B1	155531387	155531433
refGene_gn=TECRL;refGene_tx=XM_003985351.3
B1	155532289	155532417
refGene_gn=TECRL;refGene_tx=XM_003985351.3
B1	157353763	157354563
refGene_gn=ADGRL3;refGene_tx=XM_011281860.3
B1	157380512	157380609
refGene_gn=ADGRL3;refGene_tx=XM_011281860.3
B1	157388235	157388361
refGene_gn=ADGRL3;refGene_tx=XM_011281860.3
B1	157399138	157399307
refGene_gn=ADGRL3;refGene_tx=XM_011281860.3
B1	157401871	157401898
refGene_gn=ADGRL3;refGene_tx=XM_011281860.3
B1	157427248	157427340
refGene_gn=ADGRL3;refGene_tx=XM_011281860.3
B1	157429408	157429475
refGene_gn=ADGRL3;refGene_tx=XM_011281860.3
B1	157441311	157441532
refGene_gn=ADGRL3;refGene_tx=XM_011281860.3
B1	157444617	157444827
refGene_gn=ADGRL3;refGene_tx=XM_011281860.3
B1	157466530	157466707
refGene_gn=ADGRL3;refGene_tx=XM_011281860.3
B1	157467704	157467752
refGene_gn=ADGRL3;refGene_tx=XM_011281860.3
B1	157467753	157467914
refGene_gn=ADGRL3;refGene_tx=XM_011281860.3
B1	157474579	157474702
refGene_gn=ADGRL3;refGene_tx=XM_011281860.3
B1	157475615	157475799
refGene_gn=ADGRL3;refGene_tx=XM_011281860.3
B1	157492866	157492905
refGene_gn=ADGRL3;refGene_tx=XM_011281860.3
B1	157495913	157496099
refGene_gn=ADGRL3;refGene_tx=XM_011281860.3
B1	157505309	157505413
refGene_gn=ADGRL3;refGene_tx=XM_011281860.3
B1	157507490	157507793

refGene_gn=ADGRL3;refGene_tx=XM_011281860.3
B1 157586253 157586334
refGene_gn=ADGRL3;refGene_tx=XM_011281860.3
B1 157633833 157634634
refGene_gn=ADGRL3;refGene_tx=XM_011281860.3
B1 157636757 157636772
refGene_gn=ADGRL3;refGene_tx=XM_011281860.3
B1 157684758 157684868
refGene_gn=ADGRL3;refGene_tx=XM_011281860.3
B1 157767629 157767843
refGene_gn=ADGRL3;refGene_tx=XM_011281860.3
B1 157849040 157849244
refGene_gn=ADGRL3;refGene_tx=XM_011281860.3
B1 157868605 157868660
refGene_gn=ADGRL3;refGene_tx=XM_011281860.3
B1 161873242 161873717
refGene_gn=IGFBP7;refGene_tx=XM_003985355.4
B1 161935837 161935947
refGene_gn=IGFBP7;refGene_tx=XM_003985355.4
B1 161944408 161944525
refGene_gn=IGFBP7;refGene_tx=XM_003985355.4
B1 161945254 161945381
refGene_gn=IGFBP7;refGene_tx=XM_003985355.4
B1 161946384 161946404
refGene_gn=IGFBP7;refGene_tx=XM_003985355.4
B1 161946748 161946838
refGene_gn=POLR2B;refGene_tx=XM_003985353.5;refGene_gn=POLR2B;refGene_
tx=XM_006931102.4;refGene_gn=POLR2B;refGene_tx=XM_019829457.2
B1 161947289 161947485
refGene_gn=POLR2B;refGene_tx=XM_003985353.5;refGene_gn=POLR2B;refGene_
tx=XM_006931102.4;refGene_gn=POLR2B;refGene_tx=XM_019829457.2
B1 161951160 161951321
refGene_gn=POLR2B;refGene_tx=XM_003985353.5;refGene_gn=POLR2B;refGene_
tx=XM_006931102.4;refGene_gn=POLR2B;refGene_tx=XM_019829457.2
B1 161951704 161951818
refGene_gn=POLR2B;refGene_tx=XM_003985353.5;refGene_gn=POLR2B;refGene_
tx=XM_006931102.4;refGene_gn=POLR2B;refGene_tx=XM_019829457.2
B1 161952792 161952900
refGene_gn=POLR2B;refGene_tx=XM_003985353.5;refGene_gn=POLR2B;refGene_
tx=XM_006931102.4;refGene_gn=POLR2B;refGene_tx=XM_019829457.2
B1 161953137 161953227
refGene_gn=POLR2B;refGene_tx=XM_003985353.5;refGene_gn=POLR2B;refGene_
tx=XM_006931102.4;refGene_gn=POLR2B;refGene_tx=XM_019829457.2
B1 161953319 161953570
refGene_gn=POLR2B;refGene_tx=XM_003985353.5;refGene_gn=POLR2B;refGene_
tx=XM_006931102.4;refGene_gn=POLR2B;refGene_tx=XM_019829457.2
B1 161954811 161954906
refGene_gn=POLR2B;refGene_tx=XM_003985353.5;refGene_gn=POLR2B;refGene_
tx=XM_006931102.4;refGene_gn=POLR2B;refGene_tx=XM_019829457.2
B1 161955897 161955994

refGene_gn=POLR2B;refGene_tx=XM_003985353.5;refGene_gn=POLR2B;refGene_tx=XM_006931102.4;refGene_gn=POLR2B;refGene_tx=XM_019829457.2
B1 161964341 161964510
refGene_gn=POLR2B;refGene_tx=XM_003985353.5;refGene_gn=POLR2B;refGene_tx=XM_006931102.4;refGene_gn=POLR2B;refGene_tx=XM_019829457.2
B1 161964828 161965027
refGene_gn=POLR2B;refGene_tx=XM_003985353.5;refGene_gn=POLR2B;refGene_tx=XM_006931102.4;refGene_gn=POLR2B;refGene_tx=XM_019829457.2
B1 161966131 161966286
refGene_gn=POLR2B;refGene_tx=XM_003985353.5;refGene_gn=POLR2B;refGene_tx=XM_006931102.4;refGene_gn=POLR2B;refGene_tx=XM_019829457.2
B1 161968391 161968503
refGene_gn=POLR2B;refGene_tx=XM_003985353.5;refGene_gn=POLR2B;refGene_tx=XM_006931102.4;refGene_gn=POLR2B;refGene_tx=XM_019829457.2
B1 161968605 161968745
refGene_gn=POLR2B;refGene_tx=XM_003985353.5;refGene_gn=POLR2B;refGene_tx=XM_006931102.4;refGene_gn=POLR2B;refGene_tx=XM_019829457.2
B1 161968974 161969118
refGene_gn=POLR2B;refGene_tx=XM_003985353.5;refGene_gn=POLR2B;refGene_tx=XM_006931102.4;refGene_gn=POLR2B;refGene_tx=XM_019829457.2
B1 161969766 161969953
refGene_gn=POLR2B;refGene_tx=XM_003985353.5;refGene_gn=POLR2B;refGene_tx=XM_006931102.4;refGene_gn=POLR2B;refGene_tx=XM_019829457.2
B1 161970728 161970848
refGene_gn=POLR2B;refGene_tx=XM_003985353.5;refGene_gn=POLR2B;refGene_tx=XM_006931102.4;refGene_gn=POLR2B;refGene_tx=XM_019829457.2
B1 161971022 161971219
refGene_gn=POLR2B;refGene_tx=XM_003985353.5;refGene_gn=POLR2B;refGene_tx=XM_006931102.4;refGene_gn=POLR2B;refGene_tx=XM_019829457.2
B1 161973898 161974063
refGene_gn=POLR2B;refGene_tx=XM_003985353.5;refGene_gn=POLR2B;refGene_tx=XM_006931102.4;refGene_gn=POLR2B;refGene_tx=XM_019829457.2
B1 161975546 161975705
refGene_gn=POLR2B;refGene_tx=XM_003985353.5;refGene_gn=POLR2B;refGene_tx=XM_006931102.4;refGene_gn=POLR2B;refGene_tx=XM_019829457.2
B1 161975887 161976107
refGene_gn=POLR2B;refGene_tx=XM_003985353.5;refGene_gn=POLR2B;refGene_tx=XM_006931102.4
B1 161976223 161976336
refGene_gn=POLR2B;refGene_tx=XM_003985353.5;refGene_gn=POLR2B;refGene_tx=XM_006931102.4
B1 161979667 161979818
refGene_gn=POLR2B;refGene_tx=XM_003985353.5;refGene_gn=POLR2B;refGene_tx=XM_006931102.4
B1 161981922 161981995
refGene_gn=POLR2B;refGene_tx=XM_003985353.5;refGene_gn=POLR2B;refGene_tx=XM_006931102.4
B1 161988415 161988434
refGene_gn=POLR2B;refGene_tx=XM_003985353.5;refGene_gn=POLR2B;refGene_tx=XM_006931102.4

B1 161989189 161990336
refGene_gn=NOA1;refGene_tx=XM_003985356.2
B1 161991650 161991815
refGene_gn=NOA1;refGene_tx=XM_003985356.2
B1 161993117 161993323
refGene_gn=NOA1;refGene_tx=XM_003985356.2
B1 161996152 161996284
refGene_gn=NOA1;refGene_tx=XM_003985356.2
B1 161997443 161997560
refGene_gn=NOA1;refGene_tx=XM_003985356.2
B1 162001179 162001300
refGene_gn=NOA1;refGene_tx=XM_003985356.2
B1 162002082 162002285
refGene_gn=NOA1;refGene_tx=XM_003985356.2
B1 162020707 162022866
refGene_gn=REST;refGene_tx=XM_019829458.2;refGene_gn=REST;refGene_tx=XM_019829460.2;refGene_gn=REST;refGene_tx=XM_023253080.1
B1 162030284 162030368
refGene_gn=REST;refGene_tx=XM_019829458.2;refGene_gn=REST;refGene_tx=XM_019829460.2;refGene_gn=REST;refGene_tx=XM_023253080.1
B1 162036185 162037080
refGene_gn=REST;refGene_tx=XM_019829458.2;refGene_gn=REST;refGene_tx=XM_019829460.2;refGene_gn=REST;refGene_tx=XM_023253080.1
B1 162122736 162122791
refGene_gn=SPINK2;refGene_tx=XM_006931104.4
B1 162123863 162123916
refGene_gn=SPINK2;refGene_tx=XM_006931104.4
B1 162132820 162132930
refGene_gn=SPINK2;refGene_tx=XM_006931104.4
B1 162134212 162134258
refGene_gn=SPINK2;refGene_tx=XM_006931104.4
B1 162238834 162238896
refGene_gn=HOPX;refGene_tx=XM_019829461.1
B1 162262333 162262490
refGene_gn=HOPX;refGene_tx=XM_011281867.3;refGene_gn=HOPX;refGene_tx=XM_011281868.2;refGene_gn=HOPX;refGene_tx=XM_011281869.3;refGene_gn=HOPX;refGene_tx=XM_019829461.1
B1 162269047 162269125
refGene_gn=HOPX;refGene_tx=XM_011281867.3;refGene_gn=HOPX;refGene_tx=XM_011281868.2;refGene_gn=HOPX;refGene_tx=XM_011281869.3;refGene_gn=HOPX;refGene_tx=XM_019829461.1
B1 162299609 162299724
refGene_gn=THEGL;refGene_tx=XM_023253081.1
B1 162300953 162301129
refGene_gn=THEGL;refGene_tx=XM_023253081.1
B1 162303539 162303671
refGene_gn=THEGL;refGene_tx=XM_023253081.1
B1 162310308 162310423
refGene_gn=THEGL;refGene_tx=XM_023253081.1
B1 162324616 162324754

refGene_gn=THEGL;refGene_tx=XM_023253081.1
B1 162327012 162327141
refGene_gn=THEGL;refGene_tx=XM_023253081.1
B1 162335132 162335212
refGene_gn=THEGL;refGene_tx=XM_023253081.1
B1 162348405 162348425
refGene_gn=THEGL;refGene_tx=XM_023253081.1
B1 162355494 162356065
refGene_gn=THEGL;refGene_tx=XM_023253081.1
B1 162367649 162367829
refGene_gn=ARL9;refGene_tx=XM_019829948.1
B1 162378912 162379088
refGene_gn=ARL9;refGene_tx=XM_019829948.1
B1 162381730 162381893
refGene_gn=ARL9;refGene_tx=XM_019829948.1
B1 162385223 162385365
refGene_gn=ARL9;refGene_tx=XM_019829948.1
B1 162385640 162385687
refGene_gn=ARL9;refGene_tx=XM_019829948.1
B1 162388993 162389171
refGene_gn=SRP72;refGene_tx=XM_011281872.3;refGene_gn=SRP72;refGene_tx=XM_019829467.2
B1 162389819 162389979
refGene_gn=SRP72;refGene_tx=XM_011281872.3;refGene_gn=SRP72;refGene_tx=XM_019829467.2
B1 162392597 162392635
refGene_gn=SRP72;refGene_tx=XM_011281872.3;refGene_gn=SRP72;refGene_tx=XM_019829467.2
B1 162396133 162396271
refGene_gn=SRP72;refGene_tx=XM_011281872.3;refGene_gn=SRP72;refGene_tx=XM_019829467.2
B1 162397200 162397278
refGene_gn=SRP72;refGene_tx=XM_011281872.3;refGene_gn=SRP72;refGene_tx=XM_019829467.2
B1 162397404 162397508
refGene_gn=SRP72;refGene_tx=XM_011281872.3;refGene_gn=SRP72;refGene_tx=XM_019829467.2
B1 162398551 162398647
refGene_gn=SRP72;refGene_tx=XM_011281872.3;refGene_gn=SRP72;refGene_tx=XM_019829467.2
B1 162400246 162400311
refGene_gn=SRP72;refGene_tx=XM_011281872.3;refGene_gn=SRP72;refGene_tx=XM_019829467.2
B1 162402151 162402224
refGene_gn=SRP72;refGene_tx=XM_011281872.3;refGene_gn=SRP72;refGene_tx=XM_019829467.2
B1 162403043 162403172
refGene_gn=SRP72;refGene_tx=XM_011281872.3;refGene_gn=SRP72;refGene_tx=XM_019829467.2
B1 162404098 162404230

refGene_gn=SRP72;refGene_tx=XM_011281872.3;refGene_gn=SRP72;refGene_tx
=XM_019829467.2
B1 162406593 162406651
refGene_gn=SRP72;refGene_tx=XM_011281872.3;refGene_gn=SRP72;refGene_tx
=XM_019829467.2
B1 162406744 162406869
refGene_gn=SRP72;refGene_tx=XM_011281872.3;refGene_gn=SRP72;refGene_tx
=XM_019829467.2
B1 162407515 162407547
refGene_gn=SRP72;refGene_tx=XM_011281872.3;refGene_gn=SRP72;refGene_tx
=XM_019829467.2
B1 162409545 162409657
refGene_gn=SRP72;refGene_tx=XM_011281872.3;refGene_gn=SRP72;refGene_tx
=XM_019829467.2
B1 162409736 162409880
refGene_gn=SRP72;refGene_tx=XM_011281872.3;refGene_gn=SRP72;refGene_tx
=XM_019829467.2
B1 162411135 162411259
refGene_gn=SRP72;refGene_tx=XM_011281872.3;refGene_gn=SRP72;refGene_tx
=XM_019829467.2
B1 162414067 162414188
refGene_gn=SRP72;refGene_tx=XM_011281872.3;refGene_gn=SRP72;refGene_tx
=XM_019829467.2
B1 162415614 162415627
refGene_gn=SRP72;refGene_tx=XM_019829467.2
B1 162416304 162416416
refGene_gn=SRP72;refGene_tx=XM_011281872.3
B1 162424057 162424224
refGene_gn=LOC101084705;refGene_tx=XM_003985360.4;refGene_gn=LOC101084
705;refGene_tx=XM_019829466.1;refGene_gn=LOC101084705;refGene_tx=XM_02
3253082.1
B1 162430513 162430672
refGene_gn=LOC101084705;refGene_tx=XM_003985360.4;refGene_gn=LOC101084
705;refGene_tx=XM_019829466.1;refGene_gn=LOC101084705;refGene_tx=XM_02
3253082.1
B1 162432217 162432398
refGene_gn=LOC101084705;refGene_tx=XM_003985360.4;refGene_gn=LOC101084
705;refGene_tx=XM_019829466.1;refGene_gn=LOC101084705;refGene_tx=XM_02
3253082.1
B1 162433786 162433870
refGene_gn=LOC101084705;refGene_tx=XM_003985360.4;refGene_gn=LOC101084
705;refGene_tx=XM_019829466.1;refGene_gn=LOC101084705;refGene_tx=XM_02
3253082.1
B1 162434553 162434667
refGene_gn=LOC101084705;refGene_tx=XM_003985360.4;refGene_gn=LOC101084
705;refGene_tx=XM_019829466.1;refGene_gn=LOC101084705;refGene_tx=XM_02
3253082.1
B1 162434765 162434945
refGene_gn=LOC101084705;refGene_tx=XM_003985360.4;refGene_gn=LOC101084
705;refGene_tx=XM_019829466.1;refGene_gn=LOC101084705;refGene_tx=XM_02

3253082.1
B1	162437939	162438118
refGene_gn=LOC101084705;refGene_tx=XM_003985360.4;refGene_gn=LOC101084705;refGene_tx=XM_019829466.1;refGene_gn=LOC101084705;refGene_tx=XM_023253082.1
B1	162445855	162446053
refGene_gn=LOC101084705;refGene_tx=XM_003985360.4;refGene_gn=LOC101084705;refGene_tx=XM_019829466.1;refGene_gn=LOC101084705;refGene_tx=XM_023253082.1
B1	162449928	162449995
refGene_gn=LOC101084705;refGene_tx=XM_003985360.4;refGene_gn=LOC101084705;refGene_tx=XM_019829466.1;refGene_gn=LOC101084705;refGene_tx=XM_023253082.1
B1	162450810	162450938
refGene_gn=PPAT;refGene_tx=XM_003985361.4
B1	162454063	162454200
refGene_gn=LOC101084705;refGene_tx=XM_003985360.4;refGene_gn=LOC101084705;refGene_tx=XM_019829466.1
B1	162456563	162456629
refGene_gn=LOC101084705;refGene_tx=XM_003985360.4;refGene_gn=LOC101084705;refGene_tx=XM_019829466.1
B1	162456927	162457299
refGene_gn=LOC101084705;refGene_tx=XM_003985360.4;refGene_gn=LOC101084705;refGene_tx=XM_019829466.1
B1	162457648	162457720
refGene_gn=LOC101084705;refGene_tx=XM_003985360.4;refGene_gn=LOC101084705;refGene_tx=XM_019829466.1
B1	162464471	162464561
refGene_gn=LOC101084705;refGene_tx=XM_003985360.4;refGene_gn=LOC101084705;refGene_tx=XM_019829466.1
B1	162465680	162466115
refGene_gn=LOC101084705;refGene_tx=XM_019829466.1
B1	162467605	162467692
refGene_gn=LOC101084705;refGene_tx=XM_003985360.4;refGene_gn=LOC101084705;refGene_tx=XM_019829466.1
B1	162472157	162472284
refGene_gn=LOC101084705;refGene_tx=XM_003985360.4;refGene_gn=LOC101084705;refGene_tx=XM_019829466.1
B1	162474395	162474462
refGene_gn=PPAT;refGene_tx=XM_003985361.4
B1	162475261	162475468
refGene_gn=PPAT;refGene_tx=XM_003985361.4
B1	162477221	162477334
refGene_gn=PPAT;refGene_tx=XM_003985361.4
B1	162477444	162477590
refGene_gn=PPAT;refGene_tx=XM_003985361.4
B1	162478038	162478111
refGene_gn=PPAT;refGene_tx=XM_003985361.4
B1	162478899	162479051
refGene_gn=PPAT;refGene_tx=XM_003985361.4

B1 162479443 162479571
refGene_gn=PPAT;refGene_tx=XM_003985361.4
B1 162481189 162481411
refGene_gn=PPAT;refGene_tx=XM_003985361.4
B1 162482620 162482741
refGene_gn=PPAT;refGene_tx=XM_003985361.4
B1 162484012 162484350
refGene_gn=PPAT;refGene_tx=XM_003985361.4
B1 162497866 162498096
refGene_gn=AASDH;refGene_tx=XM_006931107.3;refGene_gn=AASDH;refGene_tx=XM_019829475.2;refGene_gn=AASDH;refGene_tx=XM_019829476.2;refGene_gn=AASDH;refGene_tx=XM_019829477.1
B1 162499154 162499275
refGene_gn=AASDH;refGene_tx=XM_006931107.3;refGene_gn=AASDH;refGene_tx=XM_019829475.2;refGene_gn=AASDH;refGene_tx=XM_019829476.2;refGene_gn=AASDH;refGene_tx=XM_019829477.1;refGene_gn=AASDH;refGene_tx=XM_019829479.1
B1 162512212 162512526
refGene_gn=AASDH;refGene_tx=XM_006931107.3;refGene_gn=AASDH;refGene_tx=XM_006931109.4;refGene_gn=AASDH;refGene_tx=XM_019829475.2;refGene_gn=AASDH;refGene_tx=XM_019829476.2;refGene_gn=AASDH;refGene_tx=XM_019829477.1;refGene_gn=AASDH;refGene_tx=XM_019829478.1
B1 162516162 162516355
refGene_gn=AASDH;refGene_tx=XM_006931107.3;refGene_gn=AASDH;refGene_tx=XM_019829475.2;refGene_gn=AASDH;refGene_tx=XM_019829476.2;refGene_gn=AASDH;refGene_tx=XM_019829477.1;refGene_gn=AASDH;refGene_tx=XM_019829478.1;refGene_gn=AASDH;refGene_tx=XM_019829479.1
B1 162523321 162523563
refGene_gn=AASDH;refGene_tx=XM_006931107.3;refGene_gn=AASDH;refGene_tx=XM_006931109.4;refGene_gn=AASDH;refGene_tx=XM_019829475.2;refGene_gn=AASDH;refGene_tx=XM_019829476.2;refGene_gn=AASDH;refGene_tx=XM_019829477.1;refGene_gn=AASDH;refGene_tx=XM_019829478.1;refGene_gn=AASDH;refGene_tx=XM_019829479.1
B1 162523935 162524042
refGene_gn=AASDH;refGene_tx=XM_006931107.3;refGene_gn=AASDH;refGene_tx=XM_006931109.4;refGene_gn=AASDH;refGene_tx=XM_019829475.2;refGene_gn=AASDH;refGene_tx=XM_019829476.2;refGene_gn=AASDH;refGene_tx=XM_019829477.1;refGene_gn=AASDH;refGene_tx=XM_019829478.1;refGene_gn=AASDH;refGene_tx=XM_019829479.1
B1 162524769 162524942
refGene_gn=AASDH;refGene_tx=XM_006931107.3;refGene_gn=AASDH;refGene_tx=XM_006931109.4;refGene_gn=AASDH;refGene_tx=XM_019829475.2;refGene_gn=AASDH;refGene_tx=XM_019829476.2;refGene_gn=AASDH;refGene_tx=XM_019829477.1;refGene_gn=AASDH;refGene_tx=XM_019829478.1;refGene_gn=AASDH;refGene_tx=XM_019829479.1
B1 162525369 162525562
refGene_gn=AASDH;refGene_tx=XM_006931107.3;refGene_gn=AASDH;refGene_tx=XM_006931109.4;refGene_gn=AASDH;refGene_tx=XM_019829475.2;refGene_gn=AASDH;refGene_tx=XM_019829476.2;refGene_gn=AASDH;refGene_tx=XM_019829477.1;refGene_gn=AASDH;refGene_tx=XM_019829478.1;refGene_gn=AASDH;refGe

ne_tx=XM_019829479.1
B1	162526179	162526295
refGene_gn=AASDH;refGene_tx=XM_006931107.3;refGene_gn=AASDH;refGene_tx=XM_006931109.4;refGene_gn=AASDH;refGene_tx=XM_019829475.2;refGene_gn=AASDH;refGene_tx=XM_019829476.2;refGene_gn=AASDH;refGene_tx=XM_019829477.1;refGene_gn=AASDH;refGene_tx=XM_019829478.1;refGene_gn=AASDH;refGene_tx=XM_019829479.1
B1	162528370	162529163
refGene_gn=AASDH;refGene_tx=XM_006931107.3;refGene_gn=AASDH;refGene_tx=XM_006931109.4;refGene_gn=AASDH;refGene_tx=XM_019829475.2;refGene_gn=AASDH;refGene_tx=XM_019829476.2;refGene_gn=AASDH;refGene_tx=XM_019829477.1;refGene_gn=AASDH;refGene_tx=XM_019829478.1;refGene_gn=AASDH;refGene_tx=XM_019829479.1
B1	162533315	162533479
refGene_gn=AASDH;refGene_tx=XM_006931107.3;refGene_gn=AASDH;refGene_tx=XM_006931109.4;refGene_gn=AASDH;refGene_tx=XM_019829475.2;refGene_gn=AASDH;refGene_tx=XM_019829476.2;refGene_gn=AASDH;refGene_tx=XM_019829477.1;refGene_gn=AASDH;refGene_tx=XM_019829478.1;refGene_gn=AASDH;refGene_tx=XM_019829479.1
B1	162534574	162534697
refGene_gn=AASDH;refGene_tx=XM_006931107.3;refGene_gn=AASDH;refGene_tx=XM_006931109.4;refGene_gn=AASDH;refGene_tx=XM_019829475.2;refGene_gn=AASDH;refGene_tx=XM_019829476.2;refGene_gn=AASDH;refGene_tx=XM_019829477.1;refGene_gn=AASDH;refGene_tx=XM_019829478.1;refGene_gn=AASDH;refGene_tx=XM_019829479.1
B1	162535521	162535653
refGene_gn=AASDH;refGene_tx=XM_006931107.3;refGene_gn=AASDH;refGene_tx=XM_006931109.4;refGene_gn=AASDH;refGene_tx=XM_019829475.2;refGene_gn=AASDH;refGene_tx=XM_019829476.2;refGene_gn=AASDH;refGene_tx=XM_019829477.1;refGene_gn=AASDH;refGene_tx=XM_019829478.1;refGene_gn=AASDH;refGene_tx=XM_019829479.1
B1	162537705	162538095
refGene_gn=AASDH;refGene_tx=XM_006931107.3;refGene_gn=AASDH;refGene_tx=XM_006931109.4;refGene_gn=AASDH;refGene_tx=XM_019829475.2;refGene_gn=AASDH;refGene_tx=XM_019829476.2;refGene_gn=AASDH;refGene_tx=XM_019829477.1;refGene_gn=AASDH;refGene_tx=XM_019829478.1;refGene_gn=AASDH;refGene_tx=XM_019829479.1
B1	162538365	162538427
refGene_gn=AASDH;refGene_tx=XM_019829477.1
B1	162543860	162544021
refGene_gn=KIAA1211;refGene_tx=XM_019829471.1;refGene_gn=KIAA1211;refGene_tx=XM_023253083.1;refGene_gn=KIAA1211;refGene_tx=XM_023253084.1;refGene_gn=KIAA1211;refGene_tx=XM_023253085.1;refGene_gn=KIAA1211;refGene_tx=XM_023253086.1;refGene_gn=KIAA1211;refGene_tx=XM_023253087.1;refGene_gn=KIAA1211;refGene_tx=XM_023253088.1;refGene_gn=KIAA1211;refGene_tx=XM_023253089.1;refGene_gn=KIAA1211;refGene_tx=XM_023253090.1
B1	162552610	162552773
refGene_gn=KIAA1211;refGene_tx=XM_019829471.1;refGene_gn=KIAA1211;refGene_tx=XM_023253083.1;refGene_gn=KIAA1211;refGene_tx=XM_023253084.1;refGene_gn=KIAA1211;refGene_tx=XM_023253085.1;refGene_gn=KIAA1211;refGen

e_tx=XM_023253086.1;refGene_gn=KIAA1211;refGene_tx=XM_023253087.1;refGene_gn=KIAA1211;refGene_tx=XM_023253088.1;refGene_gn=KIAA1211;refGene_tx=XM_023253089.1;refGene_gn=KIAA1211;refGene_tx=XM_023253090.1
B1 162553345 162553536
refGene_gn=KIAA1211;refGene_tx=XM_019829471.1;refGene_gn=KIAA1211;refGene_tx=XM_023253083.1;refGene_gn=KIAA1211;refGene_tx=XM_023253084.1;refGene_gn=KIAA1211;refGene_tx=XM_023253085.1;refGene_gn=KIAA1211;refGene_tx=XM_023253086.1;refGene_gn=KIAA1211;refGene_tx=XM_023253087.1;refGene_gn=KIAA1211;refGene_tx=XM_023253088.1;refGene_gn=KIAA1211;refGene_tx=XM_023253089.1;refGene_gn=KIAA1211;refGene_tx=XM_023253090.1
B1 162560558 162563379
refGene_gn=KIAA1211;refGene_tx=XM_019829471.1;refGene_gn=KIAA1211;refGene_tx=XM_023253083.1;refGene_gn=KIAA1211;refGene_tx=XM_023253084.1;refGene_gn=KIAA1211;refGene_tx=XM_023253085.1;refGene_gn=KIAA1211;refGene_tx=XM_023253086.1;refGene_gn=KIAA1211;refGene_tx=XM_023253087.1;refGene_gn=KIAA1211;refGene_tx=XM_023253088.1;refGene_gn=KIAA1211;refGene_tx=XM_023253089.1;refGene_gn=KIAA1211;refGene_tx=XM_023253090.1
B1 162564056 162564239
refGene_gn=KIAA1211;refGene_tx=XM_019829471.1;refGene_gn=KIAA1211;refGene_tx=XM_023253083.1;refGene_gn=KIAA1211;refGene_tx=XM_023253084.1;refGene_gn=KIAA1211;refGene_tx=XM_023253085.1;refGene_gn=KIAA1211;refGene_tx=XM_023253086.1;refGene_gn=KIAA1211;refGene_tx=XM_023253087.1;refGene_gn=KIAA1211;refGene_tx=XM_023253088.1;refGene_gn=KIAA1211;refGene_tx=XM_023253089.1;refGene_gn=KIAA1211;refGene_tx=XM_023253090.1
B1 162566637 162566706
refGene_gn=KIAA1211;refGene_tx=XM_019829471.1;refGene_gn=KIAA1211;refGene_tx=XM_023253083.1;refGene_gn=KIAA1211;refGene_tx=XM_023253084.1;refGene_gn=KIAA1211;refGene_tx=XM_023253085.1;refGene_gn=KIAA1211;refGene_tx=XM_023253086.1;refGene_gn=KIAA1211;refGene_tx=XM_023253087.1;refGene_gn=KIAA1211;refGene_tx=XM_023253088.1;refGene_gn=KIAA1211;refGene_tx=XM_023253089.1;refGene_gn=KIAA1211;refGene_tx=XM_023253090.1
B1 162569373 162569538
refGene_gn=KIAA1211;refGene_tx=XM_019829471.1;refGene_gn=KIAA1211;refGene_tx=XM_023253083.1;refGene_gn=KIAA1211;refGene_tx=XM_023253084.1;refGene_gn=KIAA1211;refGene_tx=XM_023253085.1;refGene_gn=KIAA1211;refGene_tx=XM_023253086.1;refGene_gn=KIAA1211;refGene_tx=XM_023253087.1;refGene_gn=KIAA1211;refGene_tx=XM_023253088.1;refGene_gn=KIAA1211;refGene_tx=XM_023253089.1;refGene_gn=KIAA1211;refGene_tx=XM_023253090.1
B1 162575001 162575023
refGene_gn=KIAA1211;refGene_tx=XM_023253088.1;refGene_gn=KIAA1211;refGene_tx=XM_023253089.1
B1 162577001 162577121
refGene_gn=KIAA1211;refGene_tx=XM_019829471.1;refGene_gn=KIAA1211;refGene_tx=XM_023253083.1;refGene_gn=KIAA1211;refGene_tx=XM_023253084.1;refGene_gn=KIAA1211;refGene_tx=XM_023253085.1;refGene_gn=KIAA1211;refGene_tx=XM_023253086.1;refGene_gn=KIAA1211;refGene_tx=XM_023253087.1;refGene_gn=KIAA1211;refGene_tx=XM_023253088.1
B1 162853068 162853171
refGene_gn=CEP135;refGene_tx=XM_019829474.2;refGene_gn=CEP135;refGene_tx=XM_023253091.1;refGene_gn=CEP135;refGene_tx=XM_023253092.1;refGene_

gn=CEP135;refGene_tx=XM_023253093.1;refGene_gn=CEP135;refGene_tx=XM_023253094.1;refGene_gn=CEP135;refGene_tx=XM_023253095.1
B1 162855931 162856036
refGene_gn=CEP135;refGene_tx=XM_019829474.2;refGene_gn=CEP135;refGene_tx=XM_023253091.1;refGene_gn=CEP135;refGene_tx=XM_023253092.1;refGene_gn=CEP135;refGene_tx=XM_023253093.1;refGene_gn=CEP135;refGene_tx=XM_023253094.1;refGene_gn=CEP135;refGene_tx=XM_023253095.1
B1 162856926 162857129
refGene_gn=CEP135;refGene_tx=XM_019829474.2;refGene_gn=CEP135;refGene_tx=XM_023253091.1;refGene_gn=CEP135;refGene_tx=XM_023253092.1;refGene_gn=CEP135;refGene_tx=XM_023253093.1;refGene_gn=CEP135;refGene_tx=XM_023253094.1;refGene_gn=CEP135;refGene_tx=XM_023253095.1
B1 162857966 162858176
refGene_gn=CEP135;refGene_tx=XM_019829474.2;refGene_gn=CEP135;refGene_tx=XM_023253091.1;refGene_gn=CEP135;refGene_tx=XM_023253092.1;refGene_gn=CEP135;refGene_tx=XM_023253093.1;refGene_gn=CEP135;refGene_tx=XM_023253094.1
B1 162864406 162864592
refGene_gn=CEP135;refGene_tx=XM_019829474.2;refGene_gn=CEP135;refGene_tx=XM_023253091.1;refGene_gn=CEP135;refGene_tx=XM_023253092.1;refGene_gn=CEP135;refGene_tx=XM_023253093.1;refGene_gn=CEP135;refGene_tx=XM_023253094.1;refGene_gn=CEP135;refGene_tx=XM_023253095.1
B1 162864940 162865051
refGene_gn=CEP135;refGene_tx=XM_019829474.2;refGene_gn=CEP135;refGene_tx=XM_023253091.1;refGene_gn=CEP135;refGene_tx=XM_023253092.1;refGene_gn=CEP135;refGene_tx=XM_023253093.1;refGene_gn=CEP135;refGene_tx=XM_023253094.1;refGene_gn=CEP135;refGene_tx=XM_023253095.1
B1 162865852 162866021
refGene_gn=CEP135;refGene_tx=XM_019829474.2;refGene_gn=CEP135;refGene_tx=XM_023253091.1;refGene_gn=CEP135;refGene_tx=XM_023253092.1;refGene_gn=CEP135;refGene_tx=XM_023253093.1;refGene_gn=CEP135;refGene_tx=XM_023253094.1;refGene_gn=CEP135;refGene_tx=XM_023253095.1
B1 162866746 162866802
refGene_gn=CEP135;refGene_tx=XM_019829474.2;refGene_gn=CEP135;refGene_tx=XM_023253091.1;refGene_gn=CEP135;refGene_tx=XM_023253092.1;refGene_gn=CEP135;refGene_tx=XM_023253093.1;refGene_gn=CEP135;refGene_tx=XM_023253094.1;refGene_gn=CEP135;refGene_tx=XM_023253095.1
B1 162867429 162867584
refGene_gn=CEP135;refGene_tx=XM_019829474.2;refGene_gn=CEP135;refGene_tx=XM_023253091.1;refGene_gn=CEP135;refGene_tx=XM_023253092.1;refGene_gn=CEP135;refGene_tx=XM_023253093.1;refGene_gn=CEP135;refGene_tx=XM_023253094.1;refGene_gn=CEP135;refGene_tx=XM_023253095.1
B1 162867657 162867773
refGene_gn=CEP135;refGene_tx=XM_019829474.2;refGene_gn=CEP135;refGene_tx=XM_023253091.1;refGene_gn=CEP135;refGene_tx=XM_023253092.1;refGene_gn=CEP135;refGene_tx=XM_023253093.1;refGene_gn=CEP135;refGene_tx=XM_023253094.1;refGene_gn=CEP135;refGene_tx=XM_023253095.1
B1 162871133 162871285
refGene_gn=CEP135;refGene_tx=XM_019829474.2;refGene_gn=CEP135;refGene_tx=XM_023253091.1;refGene_gn=CEP135;refGene_tx=XM_023253092.1;refGene_

gn=CEP135;refGene_tx=XM_023253093.1;refGene_gn=CEP135;refGene_tx=XM_023253094.1;refGene_gn=CEP135;refGene_tx=XM_023253095.1
B1 162879594 162879672
refGene_gn=CEP135;refGene_tx=XM_019829474.2;refGene_gn=CEP135;refGene_tx=XM_023253091.1;refGene_gn=CEP135;refGene_tx=XM_023253092.1;refGene_gn=CEP135;refGene_tx=XM_023253093.1;refGene_gn=CEP135;refGene_tx=XM_023253094.1;refGene_gn=CEP135;refGene_tx=XM_023253095.1
B1 162884180 162884333
refGene_gn=CEP135;refGene_tx=XM_019829474.2;refGene_gn=CEP135;refGene_tx=XM_023253091.1;refGene_gn=CEP135;refGene_tx=XM_023253092.1;refGene_gn=CEP135;refGene_tx=XM_023253093.1;refGene_gn=CEP135;refGene_tx=XM_023253094.1;refGene_gn=CEP135;refGene_tx=XM_023253095.1
B1 162885091 162885244
refGene_gn=CEP135;refGene_tx=XM_019829474.2;refGene_gn=CEP135;refGene_tx=XM_023253091.1;refGene_gn=CEP135;refGene_tx=XM_023253092.1;refGene_gn=CEP135;refGene_tx=XM_023253093.1;refGene_gn=CEP135;refGene_tx=XM_023253094.1;refGene_gn=CEP135;refGene_tx=XM_023253095.1
B1 162890027 162890251
refGene_gn=CEP135;refGene_tx=XM_019829474.2;refGene_gn=CEP135;refGene_tx=XM_023253091.1;refGene_gn=CEP135;refGene_tx=XM_023253092.1;refGene_gn=CEP135;refGene_tx=XM_023253093.1;refGene_gn=CEP135;refGene_tx=XM_023253094.1;refGene_gn=CEP135;refGene_tx=XM_023253095.1
B1 162893332 162893471
refGene_gn=CEP135;refGene_tx=XM_019829474.2;refGene_gn=CEP135;refGene_tx=XM_023253091.1;refGene_gn=CEP135;refGene_tx=XM_023253092.1;refGene_gn=CEP135;refGene_tx=XM_023253093.1;refGene_gn=CEP135;refGene_tx=XM_023253094.1;refGene_gn=CEP135;refGene_tx=XM_023253095.1
B1 162895623 162895689
refGene_gn=CEP135;refGene_tx=XM_019829474.2;refGene_gn=CEP135;refGene_tx=XM_023253091.1;refGene_gn=CEP135;refGene_tx=XM_023253092.1;refGene_gn=CEP135;refGene_tx=XM_023253093.1;refGene_gn=CEP135;refGene_tx=XM_023253094.1;refGene_gn=CEP135;refGene_tx=XM_023253095.1
B1 162899032 162899248
refGene_gn=CEP135;refGene_tx=XM_019829474.2;refGene_gn=CEP135;refGene_tx=XM_023253091.1;refGene_gn=CEP135;refGene_tx=XM_023253092.1;refGene_gn=CEP135;refGene_tx=XM_023253093.1;refGene_gn=CEP135;refGene_tx=XM_023253094.1;refGene_gn=CEP135;refGene_tx=XM_023253095.1
B1 162900884 162901013
refGene_gn=CEP135;refGene_tx=XM_019829474.2;refGene_gn=CEP135;refGene_tx=XM_023253091.1;refGene_gn=CEP135;refGene_tx=XM_023253092.1;refGene_gn=CEP135;refGene_tx=XM_023253093.1;refGene_gn=CEP135;refGene_tx=XM_023253094.1;refGene_gn=CEP135;refGene_tx=XM_023253095.1
B1 162904823 162904908
refGene_gn=CEP135;refGene_tx=XM_019829474.2;refGene_gn=CEP135;refGene_tx=XM_023253091.1;refGene_gn=CEP135;refGene_tx=XM_023253092.1;refGene_gn=CEP135;refGene_tx=XM_023253093.1;refGene_gn=CEP135;refGene_tx=XM_023253094.1;refGene_gn=CEP135;refGene_tx=XM_023253095.1
B1 162906799 162906851
refGene_gn=CEP135;refGene_tx=XM_019829474.2
B1 162907283 162907425

refGene_gn=CEP135;refGene_tx=XM_019829474.2;refGene_gn=CEP135;refGene_tx=XM_023253091.1;refGene_gn=CEP135;refGene_tx=XM_023253092.1;refGene_gn=CEP135;refGene_tx=XM_023253093.1;refGene_gn=CEP135;refGene_tx=XM_023253094.1;refGene_gn=CEP135;refGene_tx=XM_023253095.1
B1 162910258 162910426
refGene_gn=CEP135;refGene_tx=XM_023253091.1;refGene_gn=CEP135;refGene_tx=XM_023253092.1;refGene_gn=CEP135;refGene_tx=XM_023253093.1;refGene_gn=CEP135;refGene_tx=XM_023253094.1;refGene_gn=CEP135;refGene_tx=XM_023253095.1
B1 162911803 162911994
refGene_gn=CEP135;refGene_tx=XM_023253091.1;refGene_gn=CEP135;refGene_tx=XM_023253092.1;refGene_gn=CEP135;refGene_tx=XM_023253093.1;refGene_gn=CEP135;refGene_tx=XM_023253094.1;refGene_gn=CEP135;refGene_tx=XM_023253095.1
B1 162913382 162913495
refGene_gn=CEP135;refGene_tx=XM_023253091.1;refGene_gn=CEP135;refGene_tx=XM_023253092.1;refGene_gn=CEP135;refGene_tx=XM_023253093.1;refGene_gn=CEP135;refGene_tx=XM_023253094.1;refGene_gn=CEP135;refGene_tx=XM_023253095.1
B1 162951044 162951197
refGene_gn=EXOC1;refGene_tx=XM_006931112.4;refGene_gn=EXOC1;refGene_tx=XM_011281877.3;refGene_gn=EXOC1;refGene_tx=XM_019829480.2;refGene_gn=EXOC1;refGene_tx=XM_019829481.2;refGene_gn=EXOC1;refGene_tx=XM_019829482.2;refGene_gn=EXOC1;refGene_tx=XM_019829483.2;refGene_gn=EXOC1;refGene_tx=XM_019829484.2;refGene_gn=EXOC1;refGene_tx=XM_019829485.2
B1 162953043 162953238
refGene_gn=EXOC1;refGene_tx=XM_006931112.4;refGene_gn=EXOC1;refGene_tx=XM_011281877.3;refGene_gn=EXOC1;refGene_tx=XM_019829480.2;refGene_gn=EXOC1;refGene_tx=XM_019829481.2;refGene_gn=EXOC1;refGene_tx=XM_019829482.2;refGene_gn=EXOC1;refGene_tx=XM_019829483.2;refGene_gn=EXOC1;refGene_tx=XM_019829484.2;refGene_gn=EXOC1;refGene_tx=XM_019829485.2
B1 162955598 162955798
refGene_gn=EXOC1;refGene_tx=XM_006931112.4;refGene_gn=EXOC1;refGene_tx=XM_011281877.3;refGene_gn=EXOC1;refGene_tx=XM_019829480.2;refGene_gn=EXOC1;refGene_tx=XM_019829481.2;refGene_gn=EXOC1;refGene_tx=XM_019829482.2;refGene_gn=EXOC1;refGene_tx=XM_019829483.2;refGene_gn=EXOC1;refGene_tx=XM_019829484.2;refGene_gn=EXOC1;refGene_tx=XM_019829485.2
B1 162957625 162957809
refGene_gn=EXOC1;refGene_tx=XM_006931112.4;refGene_gn=EXOC1;refGene_tx=XM_011281877.3;refGene_gn=EXOC1;refGene_tx=XM_019829480.2;refGene_gn=EXOC1;refGene_tx=XM_019829481.2;refGene_gn=EXOC1;refGene_tx=XM_019829482.2;refGene_gn=EXOC1;refGene_tx=XM_019829483.2;refGene_gn=EXOC1;refGene_tx=XM_019829484.2;refGene_gn=EXOC1;refGene_tx=XM_019829485.2
B1 162965824 162966053
refGene_gn=EXOC1;refGene_tx=XM_006931112.4;refGene_gn=EXOC1;refGene_tx=XM_011281877.3;refGene_gn=EXOC1;refGene_tx=XM_019829480.2;refGene_gn=EXOC1;refGene_tx=XM_019829481.2;refGene_gn=EXOC1;refGene_tx=XM_019829482.2;refGene_gn=EXOC1;refGene_tx=XM_019829483.2;refGene_gn=EXOC1;refGene_tx=XM_019829484.2;refGene_gn=EXOC1;refGene_tx=XM_019829485.2
B1 162966602 162966679

refGene_gn=EXOC1;refGene_tx=XM_006931112.4;refGene_gn=EXOC1;refGene_tx=XM_011281877.3;refGene_gn=EXOC1;refGene_tx=XM_019829480.2;refGene_gn=EXOC1;refGene_tx=XM_019829481.2;refGene_gn=EXOC1;refGene_tx=XM_019829482.2;refGene_gn=EXOC1;refGene_tx=XM_019829483.2;refGene_gn=EXOC1;refGene_tx=XM_019829484.2;refGene_gn=EXOC1;refGene_tx=XM_019829485.2
B1 162968697 162968805
refGene_gn=EXOC1;refGene_tx=XM_006931112.4;refGene_gn=EXOC1;refGene_tx=XM_011281877.3;refGene_gn=EXOC1;refGene_tx=XM_019829480.2;refGene_gn=EXOC1;refGene_tx=XM_019829481.2;refGene_gn=EXOC1;refGene_tx=XM_019829482.2;refGene_gn=EXOC1;refGene_tx=XM_019829483.2;refGene_gn=EXOC1;refGene_tx=XM_019829484.2;refGene_gn=EXOC1;refGene_tx=XM_019829485.2
B1 162969348 162969512
refGene_gn=EXOC1;refGene_tx=XM_006931112.4;refGene_gn=EXOC1;refGene_tx=XM_011281877.3;refGene_gn=EXOC1;refGene_tx=XM_019829480.2;refGene_gn=EXOC1;refGene_tx=XM_019829481.2;refGene_gn=EXOC1;refGene_tx=XM_019829482.2;refGene_gn=EXOC1;refGene_tx=XM_019829483.2;refGene_gn=EXOC1;refGene_tx=XM_019829484.2;refGene_gn=EXOC1;refGene_tx=XM_019829485.2
B1 162971026 162971071
refGene_gn=EXOC1;refGene_tx=XM_006931112.4;refGene_gn=EXOC1;refGene_tx=XM_019829480.2;refGene_gn=EXOC1;refGene_tx=XM_019829481.2;refGene_gn=EXOC1;refGene_tx=XM_019829482.2;refGene_gn=EXOC1;refGene_tx=XM_019829484.2
B1 162975411 162975517
refGene_gn=EXOC1;refGene_tx=XM_006931112.4;refGene_gn=EXOC1;refGene_tx=XM_011281877.3;refGene_gn=EXOC1;refGene_tx=XM_019829480.2;refGene_gn=EXOC1;refGene_tx=XM_019829481.2;refGene_gn=EXOC1;refGene_tx=XM_019829482.2;refGene_gn=EXOC1;refGene_tx=XM_019829483.2;refGene_gn=EXOC1;refGene_tx=XM_019829484.2;refGene_gn=EXOC1;refGene_tx=XM_019829485.2
B1 162980958 162981108
refGene_gn=EXOC1;refGene_tx=XM_006931112.4;refGene_gn=EXOC1;refGene_tx=XM_011281877.3;refGene_gn=EXOC1;refGene_tx=XM_019829480.2;refGene_gn=EXOC1;refGene_tx=XM_019829481.2;refGene_gn=EXOC1;refGene_tx=XM_019829482.2;refGene_gn=EXOC1;refGene_tx=XM_019829483.2;refGene_gn=EXOC1;refGene_tx=XM_019829484.2;refGene_gn=EXOC1;refGene_tx=XM_019829485.2
B1 162988129 162988183
refGene_gn=EXOC1;refGene_tx=XM_019829480.2;refGene_gn=EXOC1;refGene_tx=XM_019829481.2;refGene_gn=EXOC1;refGene_tx=XM_019829482.2;refGene_gn=EXOC1;refGene_tx=XM_019829483.2
B1 162988936 162989046
refGene_gn=EXOC1;refGene_tx=XM_006931112.4;refGene_gn=EXOC1;refGene_tx=XM_011281877.3;refGene_gn=EXOC1;refGene_tx=XM_019829480.2;refGene_gn=EXOC1;refGene_tx=XM_019829481.2;refGene_gn=EXOC1;refGene_tx=XM_019829482.2;refGene_gn=EXOC1;refGene_tx=XM_019829483.2;refGene_gn=EXOC1;refGene_tx=XM_019829484.2;refGene_gn=EXOC1;refGene_tx=XM_019829485.2
B1 162989907 162990040
refGene_gn=EXOC1;refGene_tx=XM_006931112.4;refGene_gn=EXOC1;refGene_tx=XM_011281877.3;refGene_gn=EXOC1;refGene_tx=XM_019829480.2;refGene_gn=EXOC1;refGene_tx=XM_019829481.2;refGene_gn=EXOC1;refGene_tx=XM_019829482.2;refGene_gn=EXOC1;refGene_tx=XM_019829483.2;refGene_gn=EXOC1;refGene_tx=XM_019829484.2;refGene_gn=EXOC1;refGene_tx=XM_019829485.2

B1 162990244 162990472
refGene_gn=EXOC1;refGene_tx=XM_006931112.4;refGene_gn=EXOC1;refGene_tx=XM_011281877.3;refGene_gn=EXOC1;refGene_tx=XM_019829480.2;refGene_gn=EXOC1;refGene_tx=XM_019829481.2;refGene_gn=EXOC1;refGene_tx=XM_019829482.2;refGene_gn=EXOC1;refGene_tx=XM_019829483.2;refGene_gn=EXOC1;refGene_tx=XM_019829484.2;refGene_gn=EXOC1;refGene_tx=XM_019829485.2
B1 162993012 162993200
refGene_gn=EXOC1;refGene_tx=XM_006931112.4;refGene_gn=EXOC1;refGene_tx=XM_011281877.3;refGene_gn=EXOC1;refGene_tx=XM_019829480.2;refGene_gn=EXOC1;refGene_tx=XM_019829481.2;refGene_gn=EXOC1;refGene_tx=XM_019829482.2;refGene_gn=EXOC1;refGene_tx=XM_019829483.2;refGene_gn=EXOC1;refGene_tx=XM_019829484.2;refGene_gn=EXOC1;refGene_tx=XM_019829485.2
B1 162994725 162994746
refGene_gn=EXOC1;refGene_tx=XM_019829480.2;refGene_gn=EXOC1;refGene_tx=XM_019829481.2;refGene_gn=EXOC1;refGene_tx=XM_019829483.2;refGene_gn=EXOC1;refGene_tx=XM_019829484.2;refGene_gn=EXOC1;refGene_tx=XM_019829485.2
B1 162997625 162997785
refGene_gn=EXOC1;refGene_tx=XM_006931112.4;refGene_gn=EXOC1;refGene_tx=XM_011281877.3;refGene_gn=EXOC1;refGene_tx=XM_019829480.2;refGene_gn=EXOC1;refGene_tx=XM_019829481.2;refGene_gn=EXOC1;refGene_tx=XM_019829482.2;refGene_gn=EXOC1;refGene_tx=XM_019829483.2;refGene_gn=EXOC1;refGene_tx=XM_019829484.2;refGene_gn=EXOC1;refGene_tx=XM_019829485.2
B1 163000859 163000990
refGene_gn=EXOC1;refGene_tx=XM_006931112.4;refGene_gn=EXOC1;refGene_tx=XM_011281877.3;refGene_gn=EXOC1;refGene_tx=XM_019829480.2;refGene_gn=EXOC1;refGene_tx=XM_019829481.2;refGene_gn=EXOC1;refGene_tx=XM_019829482.2;refGene_gn=EXOC1;refGene_tx=XM_019829483.2;refGene_gn=EXOC1;refGene_tx=XM_019829484.2;refGene_gn=EXOC1;refGene_tx=XM_019829485.2
B1 163007312 163007436
refGene_gn=EXOC1;refGene_tx=XM_006931112.4;refGene_gn=EXOC1;refGene_tx=XM_011281877.3;refGene_gn=EXOC1;refGene_tx=XM_019829480.2;refGene_gn=EXOC1;refGene_tx=XM_019829481.2;refGene_gn=EXOC1;refGene_tx=XM_019829482.2;refGene_gn=EXOC1;refGene_tx=XM_019829483.2;refGene_gn=EXOC1;refGene_tx=XM_019829484.2;refGene_gn=EXOC1;refGene_tx=XM_019829485.2
B1 163035194 163035461
refGene_gn=LOC109499159;refGene_tx=XM_019829488.1
B1 163041313 163041444
refGene_gn=LOC109499159;refGene_tx=XM_019829488.1
B1 163051409 163051530
refGene_gn=LOC109499159;refGene_tx=XM_019829488.1
B1 163222844 163222953
refGene_gn=NMU;refGene_tx=XM_023253099.1
B1 163229915 163229974
refGene_gn=NMU;refGene_tx=XM_023253099.1
B1 163244044 163244092
refGene_gn=NMU;refGene_tx=XM_023253099.1
B1 163248031 163248091
refGene_gn=NMU;refGene_tx=XM_023253099.1
B1 163249578 163249608

refGene_gn=NMU;refGene_tx=XM_023253099.1
B1 163249690 163249741
refGene_gn=NMU;refGene_tx=XM_023253099.1
B1 163251573 163251648
refGene_gn=NMU;refGene_tx=XM_023253099.1
B1 163254584 163254638
refGene_gn=NMU;refGene_tx=XM_023253099.1
B1 163255933 163255969
refGene_gn=NMU;refGene_tx=XM_023253099.1
B1 163262482 163262488
refGene_gn=PDCL2;refGene_tx=XM_006931113.4
B1 163273933 163274054
refGene_gn=PDCL2;refGene_tx=XM_006931113.4
B1 163274617 163274708
refGene_gn=PDCL2;refGene_tx=XM_006931113.4
B1 163287586 163287730
refGene_gn=PDCL2;refGene_tx=XM_006931113.4
B1 163294222 163294431
refGene_gn=PDCL2;refGene_tx=XM_006931113.4
B1 163297865 163298023
refGene_gn=PDCL2;refGene_tx=XM_006931113.4
B1 163362649 163362696
refGene_gn=CLOCK;refGene_tx=XM_006931114.4;refGene_gn=CLOCK;refGene_tx
=XM_019829486.2;refGene_gn=CLOCK;refGene_tx=XM_023253096.1;refGene_gn=
CLOCK;refGene_tx=XM_023253097.1
B1 163365093 163365153
refGene_gn=CLOCK;refGene_tx=XM_006931114.4;refGene_gn=CLOCK;refGene_tx
=XM_019829486.2;refGene_gn=CLOCK;refGene_tx=XM_023253096.1;refGene_gn=
CLOCK;refGene_tx=XM_023253097.1
B1 163368963 163369112
refGene_gn=CLOCK;refGene_tx=XM_006931114.4;refGene_gn=CLOCK;refGene_tx
=XM_019829486.2;refGene_gn=CLOCK;refGene_tx=XM_023253096.1;refGene_gn=
CLOCK;refGene_tx=XM_023253097.1
B1 163372194 163372286
refGene_gn=CLOCK;refGene_tx=XM_006931114.4;refGene_gn=CLOCK;refGene_tx
=XM_019829486.2;refGene_gn=CLOCK;refGene_tx=XM_023253096.1;refGene_gn=
CLOCK;refGene_tx=XM_023253097.1
B1 163383952 163384042
refGene_gn=CLOCK;refGene_tx=XM_006931114.4;refGene_gn=CLOCK;refGene_tx
=XM_019829486.2;refGene_gn=CLOCK;refGene_tx=XM_019829487.2;refGene_gn=
CLOCK;refGene_tx=XM_023253096.1;refGene_gn=CLOCK;refGene_tx=XM_0232530
97.1
B1 163389856 163389977
refGene_gn=CLOCK;refGene_tx=XM_006931114.4;refGene_gn=CLOCK;refGene_tx
=XM_019829486.2;refGene_gn=CLOCK;refGene_tx=XM_019829487.2;refGene_gn=
CLOCK;refGene_tx=XM_023253096.1;refGene_gn=CLOCK;refGene_tx=XM_0232530
97.1
B1 163393344 163393458
refGene_gn=CLOCK;refGene_tx=XM_006931114.4;refGene_gn=CLOCK;refGene_tx
=XM_019829486.2;refGene_gn=CLOCK;refGene_tx=XM_019829487.2;refGene_gn=

CLOCK;refGene_tx=XM_023253096.1;refGene_gn=CLOCK;refGene_tx=XM_0232530
97.1
B1 163393596 163393715
refGene_gn=CLOCK;refGene_tx=XM_006931114.4;refGene_gn=CLOCK;refGene_tx
=XM_019829486.2;refGene_gn=CLOCK;refGene_tx=XM_019829487.2;refGene_gn=
CLOCK;refGene_tx=XM_023253096.1;refGene_gn=CLOCK;refGene_tx=XM_0232530
97.1
B1 163396186 163396269
refGene_gn=CLOCK;refGene_tx=XM_006931114.4;refGene_gn=CLOCK;refGene_tx
=XM_019829486.2;refGene_gn=CLOCK;refGene_tx=XM_019829487.2;refGene_gn=
CLOCK;refGene_tx=XM_023253096.1;refGene_gn=CLOCK;refGene_tx=XM_0232530
97.1
B1 163396473 163396580
refGene_gn=CLOCK;refGene_tx=XM_006931114.4;refGene_gn=CLOCK;refGene_tx
=XM_019829486.2;refGene_gn=CLOCK;refGene_tx=XM_019829487.2;refGene_gn=
CLOCK;refGene_tx=XM_023253096.1;refGene_gn=CLOCK;refGene_tx=XM_0232530
97.1
B1 163398583 163398731
refGene_gn=CLOCK;refGene_tx=XM_006931114.4;refGene_gn=CLOCK;refGene_tx
=XM_019829486.2;refGene_gn=CLOCK;refGene_tx=XM_019829487.2;refGene_gn=
CLOCK;refGene_tx=XM_023253096.1;refGene_gn=CLOCK;refGene_tx=XM_0232530
97.1
B1 163399295 163399371
refGene_gn=CLOCK;refGene_tx=XM_006931114.4;refGene_gn=CLOCK;refGene_tx
=XM_019829486.2;refGene_gn=CLOCK;refGene_tx=XM_019829487.2;refGene_gn=
CLOCK;refGene_tx=XM_023253096.1;refGene_gn=CLOCK;refGene_tx=XM_0232530
97.1
B1 163405617 163405759
refGene_gn=CLOCK;refGene_tx=XM_006931114.4;refGene_gn=CLOCK;refGene_tx
=XM_019829486.2;refGene_gn=CLOCK;refGene_tx=XM_019829487.2;refGene_gn=
CLOCK;refGene_tx=XM_023253096.1;refGene_gn=CLOCK;refGene_tx=XM_0232530
97.1
B1 163407921 163408022
refGene_gn=CLOCK;refGene_tx=XM_006931114.4;refGene_gn=CLOCK;refGene_tx
=XM_019829486.2;refGene_gn=CLOCK;refGene_tx=XM_019829487.2;refGene_gn=
CLOCK;refGene_tx=XM_023253096.1;refGene_gn=CLOCK;refGene_tx=XM_0232530
97.1
B1 163408716 163408806
refGene_gn=CLOCK;refGene_tx=XM_006931114.4;refGene_gn=CLOCK;refGene_tx
=XM_019829486.2;refGene_gn=CLOCK;refGene_tx=XM_019829487.2;refGene_gn=
CLOCK;refGene_tx=XM_023253096.1;refGene_gn=CLOCK;refGene_tx=XM_0232530
97.1
B1 163412335 163412488
refGene_gn=CLOCK;refGene_tx=XM_006931114.4;refGene_gn=CLOCK;refGene_tx
=XM_019829486.2;refGene_gn=CLOCK;refGene_tx=XM_019829487.2;refGene_gn=
CLOCK;refGene_tx=XM_023253096.1;refGene_gn=CLOCK;refGene_tx=XM_0232530
97.1
B1 163418732 163418942
refGene_gn=CLOCK;refGene_tx=XM_006931114.4;refGene_gn=CLOCK;refGene_tx
=XM_019829486.2;refGene_gn=CLOCK;refGene_tx=XM_019829487.2;refGene_gn=

CLOCK;refGene_tx=XM_023253096.1;refGene_gn=CLOCK;refGene_tx=XM_0232530
97.1
B1 163419716 163419922
refGene_gn=CLOCK;refGene_tx=XM_006931114.4;refGene_gn=CLOCK;refGene_tx
=XM_019829486.2;refGene_gn=CLOCK;refGene_tx=XM_019829487.2;refGene_gn=
CLOCK;refGene_tx=XM_023253096.1;refGene_gn=CLOCK;refGene_tx=XM_0232530
97.1
B1 163424024 163424271
refGene_gn=CLOCK;refGene_tx=XM_006931114.4;refGene_gn=CLOCK;refGene_tx
=XM_019829486.2;refGene_gn=CLOCK;refGene_tx=XM_019829487.2;refGene_gn=
CLOCK;refGene_tx=XM_023253096.1;refGene_gn=CLOCK;refGene_tx=XM_0232530
97.1
B1 163428935 163429115
refGene_gn=CLOCK;refGene_tx=XM_006931114.4;refGene_gn=CLOCK;refGene_tx
=XM_019829486.2;refGene_gn=CLOCK;refGene_tx=XM_019829487.2;refGene_gn=
CLOCK;refGene_tx=XM_023253096.1;refGene_gn=CLOCK;refGene_tx=XM_0232530
97.1
B1 163438564 163438641
refGene_gn=TMEM165;refGene_tx=XM_003985371.5
B1 163439318 163439424
refGene_gn=TMEM165;refGene_tx=XM_003985371.5
B1 163443514 163443697
refGene_gn=TMEM165;refGene_tx=XM_003985371.5
B1 163444458 163444634
refGene_gn=TMEM165;refGene_tx=XM_003985371.5
B1 163452289 163452515
refGene_gn=TMEM165;refGene_tx=XM_003985371.5
B1 163465336 163465543
refGene_gn=TMEM165;refGene_tx=XM_003985371.5
B1 163486396 163486656
refGene_gn=SRD5A3;refGene_tx=XM_006931118.2;refGene_gn=SRD5A3;refGene_
tx=XM_023253098.1
B1 163488199 163488334
refGene_gn=SRD5A3;refGene_tx=XM_006931118.2
B1 163492059 163492257
refGene_gn=SRD5A3;refGene_tx=XM_006931118.2;refGene_gn=SRD5A3;refGene_
tx=XM_023253098.1
B1 163493780 163493923
refGene_gn=SRD5A3;refGene_tx=XM_006931118.2;refGene_gn=SRD5A3;refGene_
tx=XM_023253098.1
B1 163500895 163501116
refGene_gn=SRD5A3;refGene_tx=XM_006931118.2;refGene_gn=SRD5A3;refGene_
tx=XM_023253098.1
B1 163649141 163649208
refGene_gn=KDR;refGene_tx=XM_011281879.3
B1 163653411 163653505
refGene_gn=KDR;refGene_tx=XM_011281879.3
B1 163655489 163655686
refGene_gn=KDR;refGene_tx=XM_011281879.3
B1 163658929 163659060

refGene_gn=KDR;refGene_tx=XM_011281879.3
B1 163659290 163659459
refGene_gn=KDR;refGene_tx=XM_011281879.3
B1 163660049 163660189
refGene_gn=KDR;refGene_tx=XM_011281879.3
B1 163660894 163661072
refGene_gn=KDR;refGene_tx=XM_011281879.3
B1 163662499 163662614
refGene_gn=KDR;refGene_tx=XM_011281879.3
B1 163662706 163662870
refGene_gn=KDR;refGene_tx=XM_011281879.3
B1 163664963 163665120
refGene_gn=KDR;refGene_tx=XM_011281879.3
B1 163665986 163666110
refGene_gn=KDR;refGene_tx=XM_011281879.3
B1 163666655 163666764
refGene_gn=KDR;refGene_tx=XM_011281879.3
B1 163668104 163668446
refGene_gn=KDR;refGene_tx=XM_011281879.3
B1 163670522 163670669
refGene_gn=KDR;refGene_tx=XM_011281879.3
B1 163670996 163671128
refGene_gn=KDR;refGene_tx=XM_011281879.3
B1 163674805 163674912
refGene_gn=KDR;refGene_tx=XM_011281879.3
B1 163675345 163675481
refGene_gn=KDR;refGene_tx=XM_011281879.3
B1 163675852 163675957
refGene_gn=KDR;refGene_tx=XM_011281879.3
B1 163677279 163677393
refGene_gn=KDR;refGene_tx=XM_011281879.3
B1 163677969 163678058
refGene_gn=KDR;refGene_tx=XM_011281879.3
B1 163679108 163679262
refGene_gn=KDR;refGene_tx=XM_011281879.3
B1 163681163 163681261
refGene_gn=KDR;refGene_tx=XM_011281879.3
B1 163683669 163683792
refGene_gn=KDR;refGene_tx=XM_011281879.3
B1 163683943 163684055
refGene_gn=KDR;refGene_tx=XM_011281879.3
B1 163684243 163684343
refGene_gn=KDR;refGene_tx=XM_011281879.3
B1 163684743 163684849
refGene_gn=KDR;refGene_tx=XM_011281879.3
B1 163685949 163686101
refGene_gn=KDR;refGene_tx=XM_011281879.3
B1 163691124 163691224
refGene_gn=KDR;refGene_tx=XM_011281879.3
B1 163691957 163692043

refGene_gn=KDR;refGene_tx=XM_011281879.3
B1	163693862	163694085
refGene_gn=KDR;refGene_tx=XM_011281879.3
B1	163953935	163954067
refGene_gn=KIT;refGene_tx=NM_001009837.3
B1	163955171	163955277
refGene_gn=KIT;refGene_tx=NM_001009837.3
B1	163955641	163955741
refGene_gn=KIT;refGene_tx=NM_001009837.3
B1	163955852	163955964
refGene_gn=KIT;refGene_tx=NM_001009837.3
B1	163959981	163960104
refGene_gn=KIT;refGene_tx=NM_001009837.3
B1	163961160	163961288
refGene_gn=KIT;refGene_tx=NM_001009837.3
B1	163961778	163961870
refGene_gn=KIT;refGene_tx=NM_001009837.3
B1	163963706	163963857
refGene_gn=KIT;refGene_tx=NM_001009837.3
B1	163965022	163965133
refGene_gn=KIT;refGene_tx=NM_001009837.3
B1	163965222	163965327
refGene_gn=KIT;refGene_tx=NM_001009837.3
B1	163965609	163965736
refGene_gn=KIT;refGene_tx=NM_001009837.3
B1	163965833	163965940
refGene_gn=KIT;refGene_tx=NM_001009837.3
B1	163967162	163967356
refGene_gn=KIT;refGene_tx=NM_001009837.3
B1	163968897	163969012
refGene_gn=KIT;refGene_tx=NM_001009837.3
B1	163984089	163984205
refGene_gn=KIT;refGene_tx=NM_001009837.3
B1	163986391	163986581
refGene_gn=KIT;refGene_tx=NM_001009837.3
B1	163991692	163991864
refGene_gn=KIT;refGene_tx=NM_001009837.3
B1	163996874	163996878
refGene_gn=KIT;refGene_tx=NM_001009837.3
B1	163996879	163997012
refGene_gn=KIT;refGene_tx=NM_001009837.3
B1	163998111	163998393
refGene_gn=KIT;refGene_tx=NM_001009837.3
B1	164001217	164001487
refGene_gn=KIT;refGene_tx=NM_001009837.3
B1	164039201	164039268
refGene_gn=KIT;refGene_tx=NM_001009837.3
B1	164397129	164397277
refGene_gn=PDGFRA;refGene_tx=XM_003985377.4;refGene_gn=PDGFRA;refGene_
tx=XM_006931120.3;refGene_gn=PDGFRA;refGene_tx=XM_019829489.2;refGene_

gn=PDGFRA;refGene_tx=XM_019829490.2;refGene_gn=PDGFRA;refGene_tx=XM_023253101.1;refGene_gn=PDGFRA;refGene_tx=XM_023253102.1
B1 164403171 164403413
refGene_gn=PDGFRA;refGene_tx=XM_003985377.4;refGene_gn=PDGFRA;refGene_tx=XM_006931120.3;refGene_gn=PDGFRA;refGene_tx=XM_019829489.2;refGene_gn=PDGFRA;refGene_tx=XM_019829490.2;refGene_gn=PDGFRA;refGene_tx=XM_023253101.1;refGene_gn=PDGFRA;refGene_tx=XM_023253102.1
B1 164404960 164405066
refGene_gn=PDGFRA;refGene_tx=XM_003985377.4;refGene_gn=PDGFRA;refGene_tx=XM_006931120.3;refGene_gn=PDGFRA;refGene_tx=XM_019829489.2;refGene_gn=PDGFRA;refGene_tx=XM_019829490.2;refGene_gn=PDGFRA;refGene_tx=XM_023253101.1;refGene_gn=PDGFRA;refGene_tx=XM_023253102.1
B1 164405182 164405282
refGene_gn=PDGFRA;refGene_tx=XM_003985377.4;refGene_gn=PDGFRA;refGene_tx=XM_006931120.3;refGene_gn=PDGFRA;refGene_tx=XM_019829489.2;refGene_gn=PDGFRA;refGene_tx=XM_019829490.2;refGene_gn=PDGFRA;refGene_tx=XM_023253101.1;refGene_gn=PDGFRA;refGene_tx=XM_023253102.1
B1 164406553 164406665
refGene_gn=PDGFRA;refGene_tx=XM_003985377.4;refGene_gn=PDGFRA;refGene_tx=XM_006931120.3;refGene_gn=PDGFRA;refGene_tx=XM_019829489.2;refGene_gn=PDGFRA;refGene_tx=XM_019829490.2;refGene_gn=PDGFRA;refGene_tx=XM_023253101.1;refGene_gn=PDGFRA;refGene_tx=XM_023253102.1
B1 164408708 164408831
refGene_gn=PDGFRA;refGene_tx=XM_003985377.4;refGene_gn=PDGFRA;refGene_tx=XM_006931120.3;refGene_gn=PDGFRA;refGene_tx=XM_019829489.2;refGene_gn=PDGFRA;refGene_tx=XM_019829490.2;refGene_gn=PDGFRA;refGene_tx=XM_023253101.1;refGene_gn=PDGFRA;refGene_tx=XM_023253102.1
B1 164409171 164409287
refGene_gn=PDGFRA;refGene_tx=XM_003985377.4;refGene_gn=PDGFRA;refGene_tx=XM_006931120.3;refGene_gn=PDGFRA;refGene_tx=XM_019829489.2;refGene_gn=PDGFRA;refGene_tx=XM_019829490.2;refGene_gn=PDGFRA;refGene_tx=XM_023253101.1;refGene_gn=PDGFRA;refGene_tx=XM_023253102.1
B1 164411624 164411791
refGene_gn=PDGFRA;refGene_tx=XM_003985377.4;refGene_gn=PDGFRA;refGene_tx=XM_006931120.3;refGene_gn=PDGFRA;refGene_tx=XM_019829489.2;refGene_gn=PDGFRA;refGene_tx=XM_019829490.2;refGene_gn=PDGFRA;refGene_tx=XM_023253101.1;refGene_gn=PDGFRA;refGene_tx=XM_023253102.1
B1 164412567 164412721
refGene_gn=PDGFRA;refGene_tx=XM_003985377.4;refGene_gn=PDGFRA;refGene_tx=XM_006931120.3;refGene_gn=PDGFRA;refGene_tx=XM_019829489.2;refGene_gn=PDGFRA;refGene_tx=XM_019829490.2;refGene_gn=PDGFRA;refGene_tx=XM_023253101.1;refGene_gn=PDGFRA;refGene_tx=XM_023253102.1
B1 164413937 164414048
refGene_gn=PDGFRA;refGene_tx=XM_003985377.4;refGene_gn=PDGFRA;refGene_tx=XM_006931120.3;refGene_gn=PDGFRA;refGene_tx=XM_019829489.2;refGene_gn=PDGFRA;refGene_tx=XM_019829490.2;refGene_gn=PDGFRA;refGene_tx=XM_023253101.1;refGene_gn=PDGFRA;refGene_tx=XM_023253102.1
B1 164414437 164414542
refGene_gn=PDGFRA;refGene_tx=XM_003985377.4;refGene_gn=PDGFRA;refGene_tx=XM_006931120.3;refGene_gn=PDGFRA;refGene_tx=XM_019829489.2;refGene_

gn=PDGFRA;refGene_tx=XM_019829490.2;refGene_gn=PDGFRA;refGene_tx=XM_023253101.1;refGene_gn=PDGFRA;refGene_tx=XM_023253102.1
B1 164416957 164417090
refGene_gn=PDGFRA;refGene_tx=XM_003985377.4;refGene_gn=PDGFRA;refGene_tx=XM_006931120.3;refGene_gn=PDGFRA;refGene_tx=XM_019829489.2;refGene_gn=PDGFRA;refGene_tx=XM_019829490.2;refGene_gn=PDGFRA;refGene_tx=XM_023253101.1;refGene_gn=PDGFRA;refGene_tx=XM_023253102.1
B1 164417295 164417390
refGene_gn=PDGFRA;refGene_tx=XM_003985377.4;refGene_gn=PDGFRA;refGene_tx=XM_006931120.3;refGene_gn=PDGFRA;refGene_tx=XM_019829489.2;refGene_gn=PDGFRA;refGene_tx=XM_019829490.2;refGene_gn=PDGFRA;refGene_tx=XM_023253101.1;refGene_gn=PDGFRA;refGene_tx=XM_023253102.1
B1 164418154 164418348
refGene_gn=PDGFRA;refGene_tx=XM_003985377.4;refGene_gn=PDGFRA;refGene_tx=XM_006931120.3;refGene_gn=PDGFRA;refGene_tx=XM_019829489.2;refGene_gn=PDGFRA;refGene_tx=XM_019829490.2;refGene_gn=PDGFRA;refGene_tx=XM_023253101.1;refGene_gn=PDGFRA;refGene_tx=XM_023253102.1
B1 164420263 164420390
refGene_gn=PDGFRA;refGene_tx=XM_003985377.4;refGene_gn=PDGFRA;refGene_tx=XM_006931120.3;refGene_gn=PDGFRA;refGene_tx=XM_019829489.2;refGene_gn=PDGFRA;refGene_tx=XM_019829490.2;refGene_gn=PDGFRA;refGene_tx=XM_023253101.1;refGene_gn=PDGFRA;refGene_tx=XM_023253102.1
B1 164421730 164421846
refGene_gn=PDGFRA;refGene_tx=XM_003985377.4;refGene_gn=PDGFRA;refGene_tx=XM_006931120.3;refGene_gn=PDGFRA;refGene_tx=XM_019829489.2;refGene_gn=PDGFRA;refGene_tx=XM_019829490.2;refGene_gn=PDGFRA;refGene_tx=XM_023253101.1;refGene_gn=PDGFRA;refGene_tx=XM_023253102.1
B1 164425648 164425838
refGene_gn=PDGFRA;refGene_tx=XM_003985377.4;refGene_gn=PDGFRA;refGene_tx=XM_006931120.3;refGene_gn=PDGFRA;refGene_tx=XM_019829489.2;refGene_gn=PDGFRA;refGene_tx=XM_019829490.2;refGene_gn=PDGFRA;refGene_tx=XM_023253101.1;refGene_gn=PDGFRA;refGene_tx=XM_023253102.1
B1 164425930 164426102
refGene_gn=PDGFRA;refGene_tx=XM_003985377.4;refGene_gn=PDGFRA;refGene_tx=XM_006931120.3;refGene_gn=PDGFRA;refGene_tx=XM_019829489.2;refGene_gn=PDGFRA;refGene_tx=XM_019829490.2;refGene_gn=PDGFRA;refGene_tx=XM_023253101.1;refGene_gn=PDGFRA;refGene_tx=XM_023253102.1
B1 164428191 164428322
refGene_gn=PDGFRA;refGene_tx=XM_003985377.4;refGene_gn=PDGFRA;refGene_tx=XM_006931120.3;refGene_gn=PDGFRA;refGene_tx=XM_019829489.2;refGene_gn=PDGFRA;refGene_tx=XM_019829490.2;refGene_gn=PDGFRA;refGene_tx=XM_023253101.1;refGene_gn=PDGFRA;refGene_tx=XM_023253102.1
B1 164429618 164429879
refGene_gn=PDGFRA;refGene_tx=XM_003985377.4;refGene_gn=PDGFRA;refGene_tx=XM_006931120.3;refGene_gn=PDGFRA;refGene_tx=XM_019829489.2;refGene_gn=PDGFRA;refGene_tx=XM_019829490.2;refGene_gn=PDGFRA;refGene_tx=XM_023253101.1;refGene_gn=PDGFRA;refGene_tx=XM_023253102.1
B1 164433617 164433935
refGene_gn=PDGFRA;refGene_tx=XM_003985377.4;refGene_gn=PDGFRA;refGene_tx=XM_006931120.3;refGene_gn=PDGFRA;refGene_tx=XM_019829489.2;refGene_

gn=PDGFRA;refGene_tx=XM_019829490.2;refGene_gn=PDGFRA;refGene_tx=XM_023253101.1;refGene_gn=PDGFRA;refGene_tx=XM_023253102.1
B1 164435670 164435719
refGene_gn=PDGFRA;refGene_tx=XM_003985377.4;refGene_gn=PDGFRA;refGene_tx=XM_006931120.3;refGene_gn=PDGFRA;refGene_tx=XM_019829489.2;refGene_gn=PDGFRA;refGene_tx=XM_019829490.2;refGene_gn=PDGFRA;refGene_tx=XM_023253101.1;refGene_gn=PDGFRA;refGene_tx=XM_023253102.1
B1 164538658 164538999
refGene_gn=GSX2;refGene_tx=XM_019829492.2
B1 164539682 164540259
refGene_gn=GSX2;refGene_tx=XM_019829492.2
B1 164576578 164576697
refGene_gn=CHIC2;refGene_tx=XM_003985378.5
B1 164581032 164581073
refGene_gn=CHIC2;refGene_tx=XM_006931121.4
B1 164599611 164599666
refGene_gn=CHIC2;refGene_tx=XM_003985378.5;refGene_gn=CHIC2;refGene_tx=XM_006931121.4
B1 164599814 164599970
refGene_gn=CHIC2;refGene_tx=XM_003985378.5;refGene_gn=CHIC2;refGene_tx=XM_006931121.4
B1 164624617 164624674
refGene_gn=CHIC2;refGene_tx=XM_003985378.5;refGene_gn=CHIC2;refGene_tx=XM_006931121.4
B1 164624840 164624900
refGene_gn=CHIC2;refGene_tx=XM_003985378.5;refGene_gn=CHIC2;refGene_tx=XM_006931121.4
B1 164628369 164628420
refGene_gn=CHIC2;refGene_tx=XM_003985378.5;refGene_gn=CHIC2;refGene_tx=XM_006931121.4
B1 165014028 165014420
refGene_gn=LNX1;refGene_tx=XM_011281882.3;refGene_gn=LNX1;refGene_tx=XM_019829496.2
B1 165023569 165023661
refGene_gn=LNX1;refGene_tx=XM_011281883.3
B1 165073423 165073473
refGene_gn=LNX1;refGene_tx=XM_019829497.2
B1 165075166 165075339
refGene_gn=LNX1;refGene_tx=XM_023253103.1
B1 165083170 165083412
refGene_gn=LNX1;refGene_tx=XM_011281882.3;refGene_gn=LNX1;refGene_tx=XM_011281883.3;refGene_gn=LNX1;refGene_tx=XM_019829496.2;refGene_gn=LNX1;refGene_tx=XM_019829497.2;refGene_gn=LNX1;refGene_tx=XM_023253103.1
B1 165084159 165084312
refGene_gn=LNX1;refGene_tx=XM_011281882.3;refGene_gn=LNX1;refGene_tx=XM_011281883.3;refGene_gn=LNX1;refGene_tx=XM_019829496.2;refGene_gn=LNX1;refGene_tx=XM_019829497.2;refGene_gn=LNX1;refGene_tx=XM_023253103.1
B1 165089372 165089575
refGene_gn=LNX1;refGene_tx=XM_011281882.3;refGene_gn=LNX1;refGene_tx=XM_011281883.3;refGene_gn=LNX1;refGene_tx=XM_019829496.2;refGene_gn=LNX

1;refGene_tx=XM_019829497.2;refGene_gn=LNX1;refGene_tx=XM_023253103.1
B1	165091815	165092187
refGene_gn=LNX1;refGene_tx=XM_011281882.3;refGene_gn=LNX1;refGene_tx=XM_011281883.3;refGene_gn=LNX1;refGene_tx=XM_019829496.2;refGene_gn=LNX1;refGene_tx=XM_019829497.2;refGene_gn=LNX1;refGene_tx=XM_023253103.1
B1	165104918	165105053
refGene_gn=LNX1;refGene_tx=XM_011281882.3;refGene_gn=LNX1;refGene_tx=XM_011281883.3;refGene_gn=LNX1;refGene_tx=XM_019829496.2;refGene_gn=LNX1;refGene_tx=XM_019829497.2;refGene_gn=LNX1;refGene_tx=XM_023253103.1
B1	165112372	165112550
refGene_gn=LNX1;refGene_tx=XM_011281882.3;refGene_gn=LNX1;refGene_tx=XM_011281883.3;refGene_gn=LNX1;refGene_tx=XM_019829496.2;refGene_gn=LNX1;refGene_tx=XM_019829497.2;refGene_gn=LNX1;refGene_tx=XM_023253103.1
B1	165114658	165114887
refGene_gn=LNX1;refGene_tx=XM_011281882.3;refGene_gn=LNX1;refGene_tx=XM_011281883.3;refGene_gn=LNX1;refGene_tx=XM_019829496.2;refGene_gn=LNX1;refGene_tx=XM_019829497.2;refGene_gn=LNX1;refGene_tx=XM_023253103.1
B1	165119335	165119494
refGene_gn=LNX1;refGene_tx=XM_011281882.3;refGene_gn=LNX1;refGene_tx=XM_011281883.3;refGene_gn=LNX1;refGene_tx=XM_019829496.2;refGene_gn=LNX1;refGene_tx=XM_019829497.2;refGene_gn=LNX1;refGene_tx=XM_023253103.1
B1	165119857	165119993
refGene_gn=LNX1;refGene_tx=XM_011281882.3;refGene_gn=LNX1;refGene_tx=XM_011281883.3;refGene_gn=LNX1;refGene_tx=XM_019829496.2;refGene_gn=LNX1;refGene_tx=XM_019829497.2;refGene_gn=LNX1;refGene_tx=XM_023253103.1
B1	165121371	165121519
refGene_gn=FIP1L1;refGene_tx=XM_003985382.4;refGene_gn=FIP1L1;refGene_tx=XM_003985383.5;refGene_gn=FIP1L1;refGene_tx=XM_003985384.5;refGene_gn=FIP1L1;refGene_tx=XM_006931123.4;refGene_gn=FIP1L1;refGene_tx=XM_006931124.4;refGene_gn=FIP1L1;refGene_tx=XM_006931125.4;refGene_gn=FIP1L1;refGene_tx=XM_006931126.4;refGene_gn=FIP1L1;refGene_tx=XM_011281880.3;refGene_gn=FIP1L1;refGene_tx=XM_011281881.3;refGene_gn=FIP1L1;refGene_tx=XM_019829498.2;refGene_gn=FIP1L1;refGene_tx=XM_019829499.2
B1	165122089	165122227
refGene_gn=FIP1L1;refGene_tx=XM_003985382.4;refGene_gn=FIP1L1;refGene_tx=XM_003985383.5;refGene_gn=FIP1L1;refGene_tx=XM_003985384.5;refGene_gn=FIP1L1;refGene_tx=XM_006931123.4;refGene_gn=FIP1L1;refGene_tx=XM_006931124.4;refGene_gn=FIP1L1;refGene_tx=XM_006931125.4;refGene_gn=FIP1L1;refGene_tx=XM_006931126.4;refGene_gn=FIP1L1;refGene_tx=XM_011281880.3;refGene_gn=FIP1L1;refGene_tx=XM_011281881.3;refGene_gn=FIP1L1;refGene_tx=XM_019829498.2;refGene_gn=FIP1L1;refGene_tx=XM_019829499.2
B1	165129647	165129861
refGene_gn=FIP1L1;refGene_tx=XM_003985382.4;refGene_gn=FIP1L1;refGene_tx=XM_003985383.5;refGene_gn=FIP1L1;refGene_tx=XM_003985384.5;refGene_gn=FIP1L1;refGene_tx=XM_006931123.4;refGene_gn=FIP1L1;refGene_tx=XM_006931124.4;refGene_gn=FIP1L1;refGene_tx=XM_006931125.4;refGene_gn=FIP1L1;refGene_tx=XM_006931126.4;refGene_gn=FIP1L1;refGene_tx=XM_011281880.3;refGene_gn=FIP1L1;refGene_tx=XM_011281881.3;refGene_gn=FIP1L1;refGene_tx=XM_019829498.2;refGene_gn=FIP1L1;refGene_tx=XM_019829499.2
B1	165137406	165137462

refGene_gn=FIP1L1;refGene_tx=XM_003985382.4;refGene_gn=FIP1L1;refGene_tx=XM_003985383.5;refGene_gn=FIP1L1;refGene_tx=XM_003985384.5;refGene_gn=FIP1L1;refGene_tx=XM_006931123.4;refGene_gn=FIP1L1;refGene_tx=XM_006931124.4;refGene_gn=FIP1L1;refGene_tx=XM_006931125.4;refGene_gn=FIP1L1;refGene_tx=XM_006931126.4;refGene_gn=FIP1L1;refGene_tx=XM_011281880.3;refGene_gn=FIP1L1;refGene_tx=XM_011281881.3;refGene_gn=FIP1L1;refGene_tx=XM_019829498.2;refGene_gn=FIP1L1;refGene_tx=XM_019829499.2
B1 165139280 165139335
refGene_gn=FIP1L1;refGene_tx=XM_003985382.4;refGene_gn=FIP1L1;refGene_tx=XM_003985383.5;refGene_gn=FIP1L1;refGene_tx=XM_003985384.5;refGene_gn=FIP1L1;refGene_tx=XM_006931123.4;refGene_gn=FIP1L1;refGene_tx=XM_006931124.4;refGene_gn=FIP1L1;refGene_tx=XM_006931125.4;refGene_gn=FIP1L1;refGene_tx=XM_006931126.4;refGene_gn=FIP1L1;refGene_tx=XM_011281880.3;refGene_gn=FIP1L1;refGene_tx=XM_011281881.3;refGene_gn=FIP1L1;refGene_tx=XM_019829498.2;refGene_gn=FIP1L1;refGene_tx=XM_019829499.2
B1 165141360 165141387
refGene_gn=FIP1L1;refGene_tx=XM_003985383.5;refGene_gn=FIP1L1;refGene_tx=XM_006931123.4;refGene_gn=FIP1L1;refGene_tx=XM_006931124.4;refGene_gn=FIP1L1;refGene_tx=XM_006931125.4;refGene_gn=FIP1L1;refGene_tx=XM_006931126.4;refGene_gn=FIP1L1;refGene_tx=XM_011281881.3;refGene_gn=FIP1L1;refGene_tx=XM_019829498.2;refGene_gn=FIP1L1;refGene_tx=XM_019829499.2
B1 165155572 165155729
refGene_gn=FIP1L1;refGene_tx=XM_003985382.4;refGene_gn=FIP1L1;refGene_tx=XM_003985383.5;refGene_gn=FIP1L1;refGene_tx=XM_003985384.5;refGene_gn=FIP1L1;refGene_tx=XM_006931123.4;refGene_gn=FIP1L1;refGene_tx=XM_006931124.4;refGene_gn=FIP1L1;refGene_tx=XM_006931125.4;refGene_gn=FIP1L1;refGene_tx=XM_006931126.4;refGene_gn=FIP1L1;refGene_tx=XM_011281880.3;refGene_gn=FIP1L1;refGene_tx=XM_011281881.3;refGene_gn=FIP1L1;refGene_tx=XM_019829498.2;refGene_gn=FIP1L1;refGene_tx=XM_019829499.2
B1 165157634 165157728
refGene_gn=FIP1L1;refGene_tx=XM_003985382.4;refGene_gn=FIP1L1;refGene_tx=XM_003985383.5;refGene_gn=FIP1L1;refGene_tx=XM_003985384.5;refGene_gn=FIP1L1;refGene_tx=XM_006931123.4;refGene_gn=FIP1L1;refGene_tx=XM_006931124.4;refGene_gn=FIP1L1;refGene_tx=XM_006931125.4;refGene_gn=FIP1L1;refGene_tx=XM_006931126.4;refGene_gn=FIP1L1;refGene_tx=XM_011281880.3;refGene_gn=FIP1L1;refGene_tx=XM_011281881.3;refGene_gn=FIP1L1;refGene_tx=XM_019829498.2;refGene_gn=FIP1L1;refGene_tx=XM_019829499.2
B1 165168101 165168209
refGene_gn=FIP1L1;refGene_tx=XM_003985382.4;refGene_gn=FIP1L1;refGene_tx=XM_003985383.5;refGene_gn=FIP1L1;refGene_tx=XM_006931123.4;refGene_gn=FIP1L1;refGene_tx=XM_006931124.4;refGene_gn=FIP1L1;refGene_tx=XM_019829498.2;refGene_gn=FIP1L1;refGene_tx=XM_019829499.2
B1 165174680 165174790
refGene_gn=FIP1L1;refGene_tx=XM_003985382.4;refGene_gn=FIP1L1;refGene_tx=XM_003985383.5;refGene_gn=FIP1L1;refGene_tx=XM_003985384.5;refGene_gn=FIP1L1;refGene_tx=XM_006931123.4;refGene_gn=FIP1L1;refGene_tx=XM_006931124.4;refGene_gn=FIP1L1;refGene_tx=XM_006931125.4;refGene_gn=FIP1L1;refGene_tx=XM_006931126.4;refGene_gn=FIP1L1;refGene_tx=XM_011281880.3;refGene_gn=FIP1L1;refGene_tx=XM_011281881.3;refGene_gn=FIP1L1;refGen

e_tx=XM_019829498.2;refGene_gn=FIP1L1;refGene_tx=XM_019829499.2
B1 165183153 165183222
refGene_gn=FIP1L1;refGene_tx=XM_003985382.4;refGene_gn=FIP1L1;refGene_tx=XM_003985383.5;refGene_gn=FIP1L1;refGene_tx=XM_006931123.4;refGene_gn=FIP1L1;refGene_tx=XM_006931125.4;refGene_gn=FIP1L1;refGene_tx=XM_019829498.2
B1 165183519 165183650
refGene_gn=FIP1L1;refGene_tx=XM_003985382.4;refGene_gn=FIP1L1;refGene_tx=XM_003985383.5;refGene_gn=FIP1L1;refGene_tx=XM_003985384.5;refGene_gn=FIP1L1;refGene_tx=XM_006931123.4;refGene_gn=FIP1L1;refGene_tx=XM_006931124.4;refGene_gn=FIP1L1;refGene_tx=XM_006931125.4;refGene_gn=FIP1L1;refGene_tx=XM_006931126.4;refGene_gn=FIP1L1;refGene_tx=XM_011281880.3;refGene_gn=FIP1L1;refGene_tx=XM_011281881.3;refGene_gn=FIP1L1;refGene_tx=XM_019829498.2;refGene_gn=FIP1L1;refGene_tx=XM_019829499.2
B1 165184110 165184218
refGene_gn=FIP1L1;refGene_tx=XM_003985382.4;refGene_gn=FIP1L1;refGene_tx=XM_003985383.5;refGene_gn=FIP1L1;refGene_tx=XM_003985384.5;refGene_gn=FIP1L1;refGene_tx=XM_006931123.4;refGene_gn=FIP1L1;refGene_tx=XM_006931124.4;refGene_gn=FIP1L1;refGene_tx=XM_006931125.4;refGene_gn=FIP1L1;refGene_tx=XM_006931126.4;refGene_gn=FIP1L1;refGene_tx=XM_011281880.3;refGene_gn=FIP1L1;refGene_tx=XM_011281881.3;refGene_gn=FIP1L1;refGene_tx=XM_019829498.2;refGene_gn=FIP1L1;refGene_tx=XM_019829499.2
B1 165184762 165184827
refGene_gn=FIP1L1;refGene_tx=XM_003985382.4;refGene_gn=FIP1L1;refGene_tx=XM_003985383.5;refGene_gn=FIP1L1;refGene_tx=XM_003985384.5;refGene_gn=FIP1L1;refGene_tx=XM_006931123.4;refGene_gn=FIP1L1;refGene_tx=XM_006931124.4;refGene_gn=FIP1L1;refGene_tx=XM_006931125.4;refGene_gn=FIP1L1;refGene_tx=XM_006931126.4;refGene_gn=FIP1L1;refGene_tx=XM_011281880.3;refGene_gn=FIP1L1;refGene_tx=XM_011281881.3;refGene_gn=FIP1L1;refGene_tx=XM_019829498.2;refGene_gn=FIP1L1;refGene_tx=XM_019829499.2
B1 165189695 165189799
refGene_gn=FIP1L1;refGene_tx=XM_003985382.4;refGene_gn=FIP1L1;refGene_tx=XM_003985383.5;refGene_gn=FIP1L1;refGene_tx=XM_003985384.5;refGene_gn=FIP1L1;refGene_tx=XM_006931123.4;refGene_gn=FIP1L1;refGene_tx=XM_006931124.4;refGene_gn=FIP1L1;refGene_tx=XM_006931125.4;refGene_gn=FIP1L1;refGene_tx=XM_006931126.4;refGene_gn=FIP1L1;refGene_tx=XM_011281880.3;refGene_gn=FIP1L1;refGene_tx=XM_011281881.3;refGene_gn=FIP1L1;refGene_tx=XM_019829498.2;refGene_gn=FIP1L1;refGene_tx=XM_019829499.2
B1 165190904 165190962
refGene_gn=FIP1L1;refGene_tx=XM_003985382.4;refGene_gn=FIP1L1;refGene_tx=XM_003985383.5;refGene_gn=FIP1L1;refGene_tx=XM_003985384.5;refGene_gn=FIP1L1;refGene_tx=XM_006931123.4;refGene_gn=FIP1L1;refGene_tx=XM_006931124.4;refGene_gn=FIP1L1;refGene_tx=XM_006931125.4;refGene_gn=FIP1L1;refGene_tx=XM_006931126.4;refGene_gn=FIP1L1;refGene_tx=XM_011281880.3;refGene_gn=FIP1L1;refGene_tx=XM_011281881.3;refGene_gn=FIP1L1;refGene_tx=XM_019829498.2;refGene_gn=FIP1L1;refGene_tx=XM_019829499.2
B1 165199203 165199243
refGene_gn=FIP1L1;refGene_tx=XM_003985382.4;refGene_gn=FIP1L1;refGene_tx=XM_003985383.5;refGene_gn=FIP1L1;refGene_tx=XM_003985384.5;refGene_gn=FIP1L1;refGene_tx=XM_006931123.4;refGene_gn=FIP1L1;refGene_tx=XM_00

6931124.4;refGene_gn=FIP1L1;refGene_tx=XM_006931125.4;refGene_gn=FIP1L1;refGene_tx=XM_006931126.4;refGene_gn=FIP1L1;refGene_tx=XM_011281880.3;refGene_gn=FIP1L1;refGene_tx=XM_011281881.3;refGene_gn=FIP1L1;refGene_tx=XM_019829498.2;refGene_gn=FIP1L1;refGene_tx=XM_019829499.2
B1 165199346 165199391
refGene_gn=FIP1L1;refGene_tx=XM_003985382.4;refGene_gn=FIP1L1;refGene_tx=XM_006931123.4;refGene_gn=FIP1L1;refGene_tx=XM_006931124.4;refGene_gn=FIP1L1;refGene_tx=XM_006931125.4;refGene_gn=FIP1L1;refGene_tx=XM_006931126.4;refGene_gn=FIP1L1;refGene_tx=XM_011281880.3;refGene_gn=FIP1L1;refGene_tx=XM_019829498.2;refGene_gn=FIP1L1;refGene_tx=XM_019829499.2
B1 165200555 165200640
refGene_gn=FIP1L1;refGene_tx=XM_003985382.4;refGene_gn=FIP1L1;refGene_tx=XM_003985383.5;refGene_gn=FIP1L1;refGene_tx=XM_003985384.5;refGene_gn=FIP1L1;refGene_tx=XM_006931123.4;refGene_gn=FIP1L1;refGene_tx=XM_006931124.4;refGene_gn=FIP1L1;refGene_tx=XM_006931125.4;refGene_gn=FIP1L1;refGene_tx=XM_006931126.4;refGene_gn=FIP1L1;refGene_tx=XM_011281880.3;refGene_gn=FIP1L1;refGene_tx=XM_011281881.3;refGene_gn=FIP1L1;refGene_tx=XM_019829498.2;refGene_gn=FIP1L1;refGene_tx=XM_019829499.2
B1 165208642 165209480
refGene_gn=SCFD2;refGene_tx=XM_023253104.1;refGene_gn=SCFD2;refGene_tx=XM_023253105.1;refGene_gn=SCFD2;refGene_tx=XM_023253106.1;refGene_gn=SCFD2;refGene_tx=XM_023253107.1
B1 165223324 165223493
refGene_gn=SCFD2;refGene_tx=XM_023253104.1;refGene_gn=SCFD2;refGene_tx=XM_023253105.1;refGene_gn=SCFD2;refGene_tx=XM_023253106.1;refGene_gn=SCFD2;refGene_tx=XM_023253107.1
B1 165266983 165267111
refGene_gn=SCFD2;refGene_tx=XM_023253104.1;refGene_gn=SCFD2;refGene_tx=XM_023253105.1;refGene_gn=SCFD2;refGene_tx=XM_023253106.1;refGene_gn=SCFD2;refGene_tx=XM_023253107.1
B1 165284605 165284781
refGene_gn=SCFD2;refGene_tx=XM_023253104.1;refGene_gn=SCFD2;refGene_tx=XM_023253105.1;refGene_gn=SCFD2;refGene_tx=XM_023253106.1;refGene_gn=SCFD2;refGene_tx=XM_023253107.1
B1 165377078 165377328
refGene_gn=SCFD2;refGene_tx=XM_023253104.1;refGene_gn=SCFD2;refGene_tx=XM_023253105.1;refGene_gn=SCFD2;refGene_tx=XM_023253106.1;refGene_gn=SCFD2;refGene_tx=XM_023253107.1
B1 165486038 165486160
refGene_gn=SCFD2;refGene_tx=XM_023253107.1
B1 165495539 165495697
refGene_gn=SCFD2;refGene_tx=XM_023253106.1
B1 165578108 165578254
refGene_gn=SCFD2;refGene_tx=XM_023253104.1;refGene_gn=SCFD2;refGene_tx=XM_023253105.1
B1 165593058 165593193
refGene_gn=SCFD2;refGene_tx=XM_023253104.1;refGene_gn=SCFD2;refGene_tx=XM_023253105.1
B1 165604284 165604404

refGene_gn=SCFD2;refGene_tx=XM_023253104.1
B1	165613258	165613351
refGene_gn=SCFD2;refGene_tx=XM_023253104.1;refGene_gn=SCFD2;refGene_tx=XM_023253105.1
B1	165621470	165621941
refGene_gn=RASL11B;refGene_tx=XM_003985387.5
B1	165622715	165622792
refGene_gn=RASL11B;refGene_tx=XM_003985387.5
B1	165623929	165623986
refGene_gn=RASL11B;refGene_tx=XM_003985387.5
B1	165624611	165624753
refGene_gn=RASL11B;refGene_tx=XM_003985387.5
B1	165695547	165697284
refGene_gn=ERVMER34-1;refGene_tx=XM_019829500.2;refGene_gn=ERVMER34-1;refGene_tx=XM_023253108.1
B1	165761725	165761761
refGene_gn=USP46;refGene_tx=XM_023253109.1;refGene_gn=USP46;refGene_tx=XM_023253111.1
B1	165764294	165764309
refGene_gn=USP46;refGene_tx=XM_023253110.1
B1	165784343	165784424
refGene_gn=USP46;refGene_tx=XM_023253109.1;refGene_gn=USP46;refGene_tx=XM_023253110.1;refGene_gn=USP46;refGene_tx=XM_023253111.1
B1	165787442	165787656
refGene_gn=USP46;refGene_tx=XM_023253109.1;refGene_gn=USP46;refGene_tx=XM_023253110.1;refGene_gn=USP46;refGene_tx=XM_023253111.1
B1	165789091	165789321
refGene_gn=USP46;refGene_tx=XM_023253109.1;refGene_gn=USP46;refGene_tx=XM_023253110.1;refGene_gn=USP46;refGene_tx=XM_023253111.1
B1	165802867	165802944
refGene_gn=USP46;refGene_tx=XM_023253109.1;refGene_gn=USP46;refGene_tx=XM_023253110.1;refGene_gn=USP46;refGene_tx=XM_023253111.1
B1	165811428	165811512
refGene_gn=USP46;refGene_tx=XM_023253109.1;refGene_gn=USP46;refGene_tx=XM_023253110.1;refGene_gn=USP46;refGene_tx=XM_023253111.1
B1	165813572	165813770
refGene_gn=USP46;refGene_tx=XM_023253109.1;refGene_gn=USP46;refGene_tx=XM_023253110.1
B1	165816926	165817005
refGene_gn=USP46;refGene_tx=XM_023253109.1;refGene_gn=USP46;refGene_tx=XM_023253110.1;refGene_gn=USP46;refGene_tx=XM_023253111.1
B1	165818044	165818146
refGene_gn=USP46;refGene_tx=XM_023253109.1;refGene_gn=USP46;refGene_tx=XM_023253110.1;refGene_gn=USP46;refGene_tx=XM_023253111.1
B1	166204738	166204746
refGene_gn=SPATA18;refGene_tx=XM_019829507.2;refGene_gn=SPATA18;refGene_tx=XM_023253112.1;refGene_gn=SPATA18;refGene_tx=XM_023253113.1;refGene_gn=SPATA18;refGene_tx=XM_023253114.1;refGene_gn=SPATA18;refGene_tx=XM_023253115.1
B1	166205059	166205105

refGene_gn=SPATA18;refGene_tx=XM_019829507.2;refGene_gn=SPATA18;refGene_tx=XM_023253112.1;refGene_gn=SPATA18;refGene_tx=XM_023253113.1;refGene_gn=SPATA18;refGene_tx=XM_023253114.1;refGene_gn=SPATA18;refGene_tx=XM_023253115.1
B1 166208269 166208353
refGene_gn=SPATA18;refGene_tx=XM_019829507.2;refGene_gn=SPATA18;refGene_tx=XM_023253112.1;refGene_gn=SPATA18;refGene_tx=XM_023253113.1;refGene_gn=SPATA18;refGene_tx=XM_023253114.1;refGene_gn=SPATA18;refGene_tx=XM_023253115.1
B1 166210080 166210204
refGene_gn=SPATA18;refGene_tx=XM_019829507.2;refGene_gn=SPATA18;refGene_tx=XM_023253112.1;refGene_gn=SPATA18;refGene_tx=XM_023253113.1;refGene_gn=SPATA18;refGene_tx=XM_023253114.1;refGene_gn=SPATA18;refGene_tx=XM_023253115.1
B1 166213613 166213789
refGene_gn=SPATA18;refGene_tx=XM_019829507.2;refGene_gn=SPATA18;refGene_tx=XM_023253112.1;refGene_gn=SPATA18;refGene_tx=XM_023253113.1;refGene_gn=SPATA18;refGene_tx=XM_023253114.1;refGene_gn=SPATA18;refGene_tx=XM_023253115.1
B1 166214604 166214763
refGene_gn=SPATA18;refGene_tx=XM_019829507.2;refGene_gn=SPATA18;refGene_tx=XM_023253112.1;refGene_gn=SPATA18;refGene_tx=XM_023253113.1;refGene_gn=SPATA18;refGene_tx=XM_023253114.1;refGene_gn=SPATA18;refGene_tx=XM_023253115.1
B1 166216208 166216473
refGene_gn=SPATA18;refGene_tx=XM_019829507.2;refGene_gn=SPATA18;refGene_tx=XM_023253112.1;refGene_gn=SPATA18;refGene_tx=XM_023253113.1;refGene_gn=SPATA18;refGene_tx=XM_023253114.1;refGene_gn=SPATA18;refGene_tx=XM_023253115.1
B1 166218275 166218527
refGene_gn=SPATA18;refGene_tx=XM_019829507.2;refGene_gn=SPATA18;refGene_tx=XM_023253112.1;refGene_gn=SPATA18;refGene_tx=XM_023253113.1;refGene_gn=SPATA18;refGene_tx=XM_023253114.1;refGene_gn=SPATA18;refGene_tx=XM_023253115.1
B1 166221794 166221890
refGene_gn=SPATA18;refGene_tx=XM_023253112.1;refGene_gn=SPATA18;refGene_tx=XM_023253114.1;refGene_gn=SPATA18;refGene_tx=XM_023253115.1
B1 166224236 166224238
refGene_gn=SPATA18;refGene_tx=XM_019829507.2
B1 166224247 166224351
refGene_gn=SPATA18;refGene_tx=XM_023253112.1;refGene_gn=SPATA18;refGene_tx=XM_023253113.1;refGene_gn=SPATA18;refGene_tx=XM_023253114.1;refGene_gn=SPATA18;refGene_tx=XM_023253115.1
B1 166228239 166228355
refGene_gn=SPATA18;refGene_tx=XM_023253112.1;refGene_gn=SPATA18;refGene_tx=XM_023253113.1
B1 166228461 166228567
refGene_gn=SPATA18;refGene_tx=XM_023253112.1;refGene_gn=SPATA18;refGene_tx=XM_023253113.1
B1 166243391 166243478

refGene_gn=SPATA18;refGene_tx=XM_023253112.1;refGene_gn=SPATA18;refGene_tx=XM_023253113.1
B1 166257157 166257196
refGene_gn=SGCB;refGene_tx=XM_023253126.1
B1 166262666 166262876
refGene_gn=SGCB;refGene_tx=XM_023253126.1
B1 166266845 166267031
refGene_gn=SGCB;refGene_tx=XM_023253126.1
B1 166267681 166267873
refGene_gn=SGCB;refGene_tx=XM_023253126.1
B1 166268184 166268316
refGene_gn=SGCB;refGene_tx=XM_023253126.1
B1 166271874 166272078
refGene_gn=SGCB;refGene_tx=XM_023253126.1
B1 166277424 166277920
refGene_gn=LRRC66;refGene_tx=XM_023253116.1
B1 166289890 166290060
refGene_gn=LRRC66;refGene_tx=XM_023253116.1
B1 166295800 166295990
refGene_gn=LRRC66;refGene_tx=XM_023253116.1
B1 166297243 166297384
refGene_gn=LRRC66;refGene_tx=XM_023253116.1
B1 166297386 166299479
refGene_gn=LRRC66;refGene_tx=XM_023253116.1
B1 166354032 166354088
refGene_gn=DCUN1D4;refGene_tx=XM_019829511.2;refGene_gn=DCUN1D4;refGene_tx=XM_023253117.1;refGene_gn=DCUN1D4;refGene_tx=XM_023253119.1;refGene_gn=DCUN1D4;refGene_tx=XM_023253121.1;refGene_gn=DCUN1D4;refGene_tx=XM_023253122.1;refGene_gn=DCUN1D4;refGene_tx=XM_023253123.1;refGene_gn=DCUN1D4;refGene_tx=XM_023253124.1;refGene_gn=DCUN1D4;refGene_tx=XM_023253125.1
B1 166354197 166354300
refGene_gn=DCUN1D4;refGene_tx=XM_019829511.2;refGene_gn=DCUN1D4;refGene_tx=XM_023253117.1;refGene_gn=DCUN1D4;refGene_tx=XM_023253119.1;refGene_gn=DCUN1D4;refGene_tx=XM_023253121.1;refGene_gn=DCUN1D4;refGene_tx=XM_023253122.1;refGene_gn=DCUN1D4;refGene_tx=XM_023253123.1;refGene_gn=DCUN1D4;refGene_tx=XM_023253124.1;refGene_gn=DCUN1D4;refGene_tx=XM_023253125.1
B1 166356428 166356533
refGene_gn=DCUN1D4;refGene_tx=XM_019829511.2;refGene_gn=DCUN1D4;refGene_tx=XM_023253117.1;refGene_gn=DCUN1D4;refGene_tx=XM_023253121.1;refGene_gn=DCUN1D4;refGene_tx=XM_023253122.1;refGene_gn=DCUN1D4;refGene_tx=XM_023253123.1;refGene_gn=DCUN1D4;refGene_tx=XM_023253124.1;refGene_gn=DCUN1D4;refGene_tx=XM_023253125.1
B1 166358724 166358862
refGene_gn=DCUN1D4;refGene_tx=XM_023253120.1
B1 166368901 166369010
refGene_gn=DCUN1D4;refGene_tx=XM_019829511.2;refGene_gn=DCUN1D4;refGene_tx=XM_023253117.1;refGene_gn=DCUN1D4;refGene_tx=XM_023253119.1;refGene_gn=DCUN1D4;refGene_tx=XM_023253120.1;refGene_gn=DCUN1D4;refGene_tx=

XM_023253121.1;refGene_gn=DCUN1D4;refGene_tx=XM_023253122.1;refGene_gn=DCUN1D4;refGene_tx=XM_023253123.1;refGene_gn=DCUN1D4;refGene_tx=XM_023253124.1;refGene_gn=DCUN1D4;refGene_tx=XM_023253125.1
B1 166375962 166376054
refGene_gn=DCUN1D4;refGene_tx=XM_019829511.2;refGene_gn=DCUN1D4;refGene_tx=XM_023253117.1;refGene_gn=DCUN1D4;refGene_tx=XM_023253119.1;refGene_gn=DCUN1D4;refGene_tx=XM_023253120.1;refGene_gn=DCUN1D4;refGene_tx=XM_023253121.1;refGene_gn=DCUN1D4;refGene_tx=XM_023253122.1;refGene_gn=DCUN1D4;refGene_tx=XM_023253123.1;refGene_gn=DCUN1D4;refGene_tx=XM_023253124.1;refGene_gn=DCUN1D4;refGene_tx=XM_023253125.1
B1 166381738 166381809
refGene_gn=DCUN1D4;refGene_tx=XM_019829511.2;refGene_gn=DCUN1D4;refGene_tx=XM_023253117.1;refGene_gn=DCUN1D4;refGene_tx=XM_023253119.1;refGene_gn=DCUN1D4;refGene_tx=XM_023253120.1;refGene_gn=DCUN1D4;refGene_tx=XM_023253121.1;refGene_gn=DCUN1D4;refGene_tx=XM_023253122.1;refGene_gn=DCUN1D4;refGene_tx=XM_023253123.1;refGene_gn=DCUN1D4;refGene_tx=XM_023253124.1;refGene_gn=DCUN1D4;refGene_tx=XM_023253125.1
B1 166392123 166392215
refGene_gn=DCUN1D4;refGene_tx=XM_019829511.2;refGene_gn=DCUN1D4;refGene_tx=XM_023253117.1;refGene_gn=DCUN1D4;refGene_tx=XM_023253119.1;refGene_gn=DCUN1D4;refGene_tx=XM_023253120.1;refGene_gn=DCUN1D4;refGene_tx=XM_023253121.1;refGene_gn=DCUN1D4;refGene_tx=XM_023253122.1;refGene_gn=DCUN1D4;refGene_tx=XM_023253123.1;refGene_gn=DCUN1D4;refGene_tx=XM_023253124.1;refGene_gn=DCUN1D4;refGene_tx=XM_023253125.1
B1 166395343 166395458
refGene_gn=DCUN1D4;refGene_tx=XM_019829511.2;refGene_gn=DCUN1D4;refGene_tx=XM_023253117.1;refGene_gn=DCUN1D4;refGene_tx=XM_023253119.1;refGene_gn=DCUN1D4;refGene_tx=XM_023253120.1;refGene_gn=DCUN1D4;refGene_tx=XM_023253121.1;refGene_gn=DCUN1D4;refGene_tx=XM_023253122.1;refGene_gn=DCUN1D4;refGene_tx=XM_023253123.1;refGene_gn=DCUN1D4;refGene_tx=XM_023253124.1;refGene_gn=DCUN1D4;refGene_tx=XM_023253125.1
B1 166406280 166406320
refGene_gn=DCUN1D4;refGene_tx=XM_019829511.2;refGene_gn=DCUN1D4;refGene_tx=XM_023253117.1;refGene_gn=DCUN1D4;refGene_tx=XM_023253119.1;refGene_gn=DCUN1D4;refGene_tx=XM_023253120.1
B1 166406491 166406562
refGene_gn=DCUN1D4;refGene_tx=XM_019829511.2;refGene_gn=DCUN1D4;refGene_tx=XM_023253117.1;refGene_gn=DCUN1D4;refGene_tx=XM_023253119.1;refGene_gn=DCUN1D4;refGene_tx=XM_023253120.1
B1 166427864 166427889
refGene_gn=DCUN1D4;refGene_tx=XM_023253117.1;refGene_gn=DCUN1D4;refGene_tx=XM_023253119.1;refGene_gn=DCUN1D4;refGene_tx=XM_023253120.1
B1 166437652 166437713
refGene_gn=DCUN1D4;refGene_tx=XM_019829511.2
B1 166437909 166437915
refGene_gn=DCUN1D4;refGene_tx=XM_019829511.2
B1 166473129 166473208
refGene_gn=CWH43;refGene_tx=XM_023253127.1
B1 166476488 166476644
refGene_gn=CWH43;refGene_tx=XM_023253127.1

B1	166482774	166482836
refGene_gn=CWH43;refGene_tx=XM_023253127.1
B1	166487832	166487977
refGene_gn=CWH43;refGene_tx=XM_023253127.1
B1	166493292	166493442
refGene_gn=CWH43;refGene_tx=XM_023253127.1
B1	166495095	166495231
refGene_gn=CWH43;refGene_tx=XM_023253127.1
B1	166496295	166496401
refGene_gn=CWH43;refGene_tx=XM_023253127.1
B1	166503395	166503475
refGene_gn=CWH43;refGene_tx=XM_023253127.1
B1	166507692	166507815
refGene_gn=CWH43;refGene_tx=XM_023253127.1
B1	166510379	166510637
refGene_gn=CWH43;refGene_tx=XM_023253127.1
B1	166515745	166515784
refGene_gn=CWH43;refGene_tx=XM_023253127.1
B1	166515786	166515833
refGene_gn=CWH43;refGene_tx=XM_023253127.1
B1	166520930	166521132
refGene_gn=CWH43;refGene_tx=XM_023253127.1
B1	166523635	166523790
refGene_gn=CWH43;refGene_tx=XM_023253127.1
B1	166524172	166524293
refGene_gn=CWH43;refGene_tx=XM_023253127.1
B1	166528494	166528686
refGene_gn=CWH43;refGene_tx=XM_023253127.1
B1	166530538	166530581
refGene_gn=CWH43;refGene_tx=XM_023253127.1
B1	166609453	166609519
refGene_gn=OCIAD2;refGene_tx=XM_011281889.2
B1	166613823	166613920
refGene_gn=OCIAD2;refGene_tx=XM_011281889.2
B1	166615566	166615620
refGene_gn=OCIAD2;refGene_tx=XM_011281889.2
B1	166618931	166618979
refGene_gn=OCIAD2;refGene_tx=XM_011281889.2
B1	166620170	166620220
refGene_gn=OCIAD2;refGene_tx=XM_011281889.2
B1	166620222	166620276
refGene_gn=OCIAD2;refGene_tx=XM_011281889.2
B1	166620278	166620292
refGene_gn=OCIAD2;refGene_tx=XM_011281889.2
B1	166622308	166622390
refGene_gn=OCIAD2;refGene_tx=XM_011281889.2
B1	166631074	166631112
refGene_gn=OCIAD1;refGene_tx=XM_003985391.5;refGene_gn=OCIAD1;refGene_tx=XM_019829529.2;refGene_gn=OCIAD1;refGene_tx=XM_019829530.2;refGene_gn=OCIAD1;refGene_tx=XM_019829531.2

B1 166633809 166633962
refGene_gn=OCIAD1;refGene_tx=XM_003985391.5;refGene_gn=OCIAD1;refGene_tx=XM_019829529.2;refGene_gn=OCIAD1;refGene_tx=XM_019829531.2
B1 166638081 166638274
refGene_gn=OCIAD1;refGene_tx=XM_003985391.5;refGene_gn=OCIAD1;refGene_tx=XM_019829529.2;refGene_gn=OCIAD1;refGene_tx=XM_019829530.2;refGene_gn=OCIAD1;refGene_tx=XM_019829531.2;refGene_gn=OCIAD1;refGene_tx=XM_019829532.2;refGene_gn=OCIAD1;refGene_tx=XM_019829533.2
B1 166639294 166639430
refGene_gn=OCIAD1;refGene_tx=XM_003985391.5;refGene_gn=OCIAD1;refGene_tx=XM_019829529.2;refGene_gn=OCIAD1;refGene_tx=XM_019829530.2;refGene_gn=OCIAD1;refGene_tx=XM_019829531.2;refGene_gn=OCIAD1;refGene_tx=XM_019829532.2;refGene_gn=OCIAD1;refGene_tx=XM_019829533.2
B1 166640411 166640459
refGene_gn=OCIAD1;refGene_tx=XM_003985391.5;refGene_gn=OCIAD1;refGene_tx=XM_019829529.2;refGene_gn=OCIAD1;refGene_tx=XM_019829530.2;refGene_gn=OCIAD1;refGene_tx=XM_019829531.2;refGene_gn=OCIAD1;refGene_tx=XM_019829532.2;refGene_gn=OCIAD1;refGene_tx=XM_019829533.2
B1 166646695 166646749
refGene_gn=OCIAD1;refGene_tx=XM_003985391.5;refGene_gn=OCIAD1;refGene_tx=XM_019829529.2;refGene_gn=OCIAD1;refGene_tx=XM_019829530.2;refGene_gn=OCIAD1;refGene_tx=XM_019829531.2;refGene_gn=OCIAD1;refGene_tx=XM_019829532.2;refGene_gn=OCIAD1;refGene_tx=XM_019829533.2
B1 166655513 166655594
refGene_gn=OCIAD1;refGene_tx=XM_003985391.5;refGene_gn=OCIAD1;refGene_tx=XM_019829529.2;refGene_gn=OCIAD1;refGene_tx=XM_019829530.2;refGene_gn=OCIAD1;refGene_tx=XM_019829531.2;refGene_gn=OCIAD1;refGene_tx=XM_019829532.2;refGene_gn=OCIAD1;refGene_tx=XM_019829533.2
B1 166655997 166656055
refGene_gn=OCIAD1;refGene_tx=XM_003985391.5;refGene_gn=OCIAD1;refGene_tx=XM_019829529.2;refGene_gn=OCIAD1;refGene_tx=XM_019829530.2;refGene_gn=OCIAD1;refGene_tx=XM_019829531.2;refGene_gn=OCIAD1;refGene_tx=XM_019829532.2;refGene_gn=OCIAD1;refGene_tx=XM_019829533.2
B1 166818084 166818204
refGene_gn=FRYL;refGene_tx=XM_019829524.2;refGene_gn=FRYL;refGene_tx=XM_019829525.2;refGene_gn=FRYL;refGene_tx=XM_019829526.2;refGene_gn=FRYL;refGene_tx=XM_019829527.2;refGene_gn=FRYL;refGene_tx=XM_023253128.1;refGene_gn=FRYL;refGene_tx=XM_023253129.1;refGene_gn=FRYL;refGene_tx=XM_023253130.1;refGene_gn=FRYL;refGene_tx=XM_023253131.1;refGene_gn=FRYL;refGene_tx=XM_023253132.1
B1 166831664 166831718
refGene_gn=FRYL;refGene_tx=XM_019829524.2;refGene_gn=FRYL;refGene_tx=XM_019829525.2;refGene_gn=FRYL;refGene_tx=XM_019829526.2;refGene_gn=FRYL;refGene_tx=XM_019829527.2;refGene_gn=FRYL;refGene_tx=XM_023253128.1;refGene_gn=FRYL;refGene_tx=XM_023253129.1;refGene_gn=FRYL;refGene_tx=XM_023253130.1;refGene_gn=FRYL;refGene_tx=XM_023253131.1;refGene_gn=FRYL;refGene_tx=XM_023253132.1
B1 166834731 166834871
refGene_gn=FRYL;refGene_tx=XM_019829524.2;refGene_gn=FRYL;refGene_tx=XM_019829525.2;refGene_gn=FRYL;refGene_tx=XM_019829526.2;refGene_gn=FRY

L;refGene_tx=XM_019829527.2;refGene_gn=FRYL;refGene_tx=XM_023253128.1;
refGene_gn=FRYL;refGene_tx=XM_023253129.1;refGene_gn=FRYL;refGene_tx=X
M_023253130.1;refGene_gn=FRYL;refGene_tx=XM_023253131.1;refGene_gn=FRY
L;refGene_tx=XM_023253132.1
B1 166835780 166835877
refGene_gn=FRYL;refGene_tx=XM_019829524.2;refGene_gn=FRYL;refGene_tx=X
M_019829525.2;refGene_gn=FRYL;refGene_tx=XM_019829526.2;refGene_gn=FRY
L;refGene_tx=XM_019829527.2;refGene_gn=FRYL;refGene_tx=XM_023253128.1;
refGene_gn=FRYL;refGene_tx=XM_023253129.1;refGene_gn=FRYL;refGene_tx=X
M_023253130.1;refGene_gn=FRYL;refGene_tx=XM_023253131.1;refGene_gn=FRY
L;refGene_tx=XM_023253132.1
B1 166847514 166847594
refGene_gn=FRYL;refGene_tx=XM_019829524.2;refGene_gn=FRYL;refGene_tx=X
M_019829525.2;refGene_gn=FRYL;refGene_tx=XM_019829526.2;refGene_gn=FRY
L;refGene_tx=XM_019829527.2;refGene_gn=FRYL;refGene_tx=XM_023253128.1;
refGene_gn=FRYL;refGene_tx=XM_023253129.1;refGene_gn=FRYL;refGene_tx=X
M_023253130.1;refGene_gn=FRYL;refGene_tx=XM_023253131.1;refGene_gn=FRY
L;refGene_tx=XM_023253132.1
B1 166848193 166848274
refGene_gn=FRYL;refGene_tx=XM_019829524.2;refGene_gn=FRYL;refGene_tx=X
M_019829525.2;refGene_gn=FRYL;refGene_tx=XM_019829526.2;refGene_gn=FRY
L;refGene_tx=XM_019829527.2;refGene_gn=FRYL;refGene_tx=XM_023253128.1;
refGene_gn=FRYL;refGene_tx=XM_023253129.1;refGene_gn=FRYL;refGene_tx=X
M_023253130.1;refGene_gn=FRYL;refGene_tx=XM_023253131.1;refGene_gn=FRY
L;refGene_tx=XM_023253132.1
B1 166850061 166850230
refGene_gn=FRYL;refGene_tx=XM_019829524.2;refGene_gn=FRYL;refGene_tx=X
M_019829525.2;refGene_gn=FRYL;refGene_tx=XM_019829526.2;refGene_gn=FRY
L;refGene_tx=XM_019829527.2;refGene_gn=FRYL;refGene_tx=XM_023253128.1;
refGene_gn=FRYL;refGene_tx=XM_023253129.1;refGene_gn=FRYL;refGene_tx=X
M_023253130.1;refGene_gn=FRYL;refGene_tx=XM_023253131.1;refGene_gn=FRY
L;refGene_tx=XM_023253132.1
B1 166850571 166850664
refGene_gn=FRYL;refGene_tx=XM_019829524.2;refGene_gn=FRYL;refGene_tx=X
M_019829525.2;refGene_gn=FRYL;refGene_tx=XM_019829526.2;refGene_gn=FRY
L;refGene_tx=XM_019829527.2;refGene_gn=FRYL;refGene_tx=XM_023253128.1;
refGene_gn=FRYL;refGene_tx=XM_023253129.1;refGene_gn=FRYL;refGene_tx=X
M_023253130.1;refGene_gn=FRYL;refGene_tx=XM_023253131.1;refGene_gn=FRY
L;refGene_tx=XM_023253132.1
B1 166855068 166855167
refGene_gn=FRYL;refGene_tx=XM_019829524.2;refGene_gn=FRYL;refGene_tx=X
M_019829525.2;refGene_gn=FRYL;refGene_tx=XM_019829526.2;refGene_gn=FRY
L;refGene_tx=XM_019829527.2;refGene_gn=FRYL;refGene_tx=XM_023253128.1;
refGene_gn=FRYL;refGene_tx=XM_023253129.1;refGene_gn=FRYL;refGene_tx=X
M_023253130.1;refGene_gn=FRYL;refGene_tx=XM_023253131.1;refGene_gn=FRY
L;refGene_tx=XM_023253132.1
B1 166856132 166856234
refGene_gn=FRYL;refGene_tx=XM_019829524.2;refGene_gn=FRYL;refGene_tx=X
M_019829525.2;refGene_gn=FRYL;refGene_tx=XM_019829526.2;refGene_gn=FRY
L;refGene_tx=XM_019829527.2;refGene_gn=FRYL;refGene_tx=XM_023253128.1;

refGene_gn=FRYL;refGene_tx=XM_023253129.1;refGene_gn=FRYL;refGene_tx=XM_023253130.1;refGene_gn=FRYL;refGene_tx=XM_023253131.1;refGene_gn=FRYL;refGene_tx=XM_023253132.1
B1 166859311 166859356
refGene_gn=FRYL;refGene_tx=XM_019829524.2;refGene_gn=FRYL;refGene_tx=XM_019829525.2;refGene_gn=FRYL;refGene_tx=XM_019829526.2;refGene_gn=FRYL;refGene_tx=XM_019829527.2;refGene_gn=FRYL;refGene_tx=XM_023253128.1;refGene_gn=FRYL;refGene_tx=XM_023253129.1
B1 166861741 166861845
refGene_gn=FRYL;refGene_tx=XM_019829524.2;refGene_gn=FRYL;refGene_tx=XM_019829525.2;refGene_gn=FRYL;refGene_tx=XM_019829526.2;refGene_gn=FRYL;refGene_tx=XM_019829527.2;refGene_gn=FRYL;refGene_tx=XM_023253128.1;refGene_gn=FRYL;refGene_tx=XM_023253129.1;refGene_gn=FRYL;refGene_tx=XM_023253130.1;refGene_gn=FRYL;refGene_tx=XM_023253131.1;refGene_gn=FRYL;refGene_tx=XM_023253132.1
B1 166862039 166862148
refGene_gn=FRYL;refGene_tx=XM_019829524.2;refGene_gn=FRYL;refGene_tx=XM_019829525.2;refGene_gn=FRYL;refGene_tx=XM_019829526.2;refGene_gn=FRYL;refGene_tx=XM_019829527.2;refGene_gn=FRYL;refGene_tx=XM_023253128.1;refGene_gn=FRYL;refGene_tx=XM_023253129.1;refGene_gn=FRYL;refGene_tx=XM_023253130.1;refGene_gn=FRYL;refGene_tx=XM_023253131.1;refGene_gn=FRYL;refGene_tx=XM_023253132.1
B1 166863910 166863997
refGene_gn=FRYL;refGene_tx=XM_019829524.2;refGene_gn=FRYL;refGene_tx=XM_019829525.2;refGene_gn=FRYL;refGene_tx=XM_019829526.2;refGene_gn=FRYL;refGene_tx=XM_019829527.2;refGene_gn=FRYL;refGene_tx=XM_023253128.1;refGene_gn=FRYL;refGene_tx=XM_023253129.1;refGene_gn=FRYL;refGene_tx=XM_023253130.1;refGene_gn=FRYL;refGene_tx=XM_023253131.1;refGene_gn=FRYL;refGene_tx=XM_023253132.1
B1 166866829 166867001
refGene_gn=FRYL;refGene_tx=XM_019829524.2;refGene_gn=FRYL;refGene_tx=XM_019829525.2;refGene_gn=FRYL;refGene_tx=XM_019829526.2;refGene_gn=FRYL;refGene_tx=XM_019829527.2;refGene_gn=FRYL;refGene_tx=XM_023253128.1;refGene_gn=FRYL;refGene_tx=XM_023253129.1;refGene_gn=FRYL;refGene_tx=XM_023253130.1;refGene_gn=FRYL;refGene_tx=XM_023253131.1;refGene_gn=FRYL;refGene_tx=XM_023253132.1
B1 166867455 166867588
refGene_gn=FRYL;refGene_tx=XM_019829524.2;refGene_gn=FRYL;refGene_tx=XM_019829525.2;refGene_gn=FRYL;refGene_tx=XM_019829526.2;refGene_gn=FRYL;refGene_tx=XM_019829527.2;refGene_gn=FRYL;refGene_tx=XM_023253128.1;refGene_gn=FRYL;refGene_tx=XM_023253129.1;refGene_gn=FRYL;refGene_tx=XM_023253130.1;refGene_gn=FRYL;refGene_tx=XM_023253131.1;refGene_gn=FRYL;refGene_tx=XM_023253132.1
B1 166870483 166870591
refGene_gn=FRYL;refGene_tx=XM_019829524.2;refGene_gn=FRYL;refGene_tx=XM_019829525.2;refGene_gn=FRYL;refGene_tx=XM_019829526.2;refGene_gn=FRYL;refGene_tx=XM_019829527.2;refGene_gn=FRYL;refGene_tx=XM_023253128.1;refGene_gn=FRYL;refGene_tx=XM_023253129.1;refGene_gn=FRYL;refGene_tx=XM_023253130.1;refGene_gn=FRYL;refGene_tx=XM_023253131.1;refGene_gn=FRYL;refGene_tx=XM_023253132.1

B1	166873561	166873799
refGene_gn=FRYL;refGene_tx=XM_019829524.2;refGene_gn=FRYL;refGene_tx=XM_019829525.2;refGene_gn=FRYL;refGene_tx=XM_019829526.2;refGene_gn=FRYL;refGene_tx=XM_019829527.2;refGene_gn=FRYL;refGene_tx=XM_023253128.1;refGene_gn=FRYL;refGene_tx=XM_023253129.1;refGene_gn=FRYL;refGene_tx=XM_023253130.1;refGene_gn=FRYL;refGene_tx=XM_023253131.1;refGene_gn=FRYL;refGene_tx=XM_023253132.1
B1	166874910	166875096
refGene_gn=FRYL;refGene_tx=XM_019829524.2;refGene_gn=FRYL;refGene_tx=XM_019829525.2;refGene_gn=FRYL;refGene_tx=XM_019829526.2;refGene_gn=FRYL;refGene_tx=XM_019829527.2;refGene_gn=FRYL;refGene_tx=XM_023253128.1;refGene_gn=FRYL;refGene_tx=XM_023253129.1;refGene_gn=FRYL;refGene_tx=XM_023253130.1;refGene_gn=FRYL;refGene_tx=XM_023253131.1;refGene_gn=FRYL;refGene_tx=XM_023253132.1
B1	166875357	166875444
refGene_gn=FRYL;refGene_tx=XM_019829524.2;refGene_gn=FRYL;refGene_tx=XM_019829525.2;refGene_gn=FRYL;refGene_tx=XM_019829526.2;refGene_gn=FRYL;refGene_tx=XM_019829527.2;refGene_gn=FRYL;refGene_tx=XM_023253128.1;refGene_gn=FRYL;refGene_tx=XM_023253129.1;refGene_gn=FRYL;refGene_tx=XM_023253130.1;refGene_gn=FRYL;refGene_tx=XM_023253131.1;refGene_gn=FRYL;refGene_tx=XM_023253132.1
B1	166876574	166876843
refGene_gn=FRYL;refGene_tx=XM_019829524.2;refGene_gn=FRYL;refGene_tx=XM_019829525.2;refGene_gn=FRYL;refGene_tx=XM_019829526.2;refGene_gn=FRYL;refGene_tx=XM_019829527.2;refGene_gn=FRYL;refGene_tx=XM_023253128.1;refGene_gn=FRYL;refGene_tx=XM_023253129.1;refGene_gn=FRYL;refGene_tx=XM_023253130.1;refGene_gn=FRYL;refGene_tx=XM_023253131.1;refGene_gn=FRYL;refGene_tx=XM_023253132.1
B1	166879557	166879753
refGene_gn=FRYL;refGene_tx=XM_019829524.2;refGene_gn=FRYL;refGene_tx=XM_019829525.2;refGene_gn=FRYL;refGene_tx=XM_019829526.2;refGene_gn=FRYL;refGene_tx=XM_019829527.2;refGene_gn=FRYL;refGene_tx=XM_023253128.1;refGene_gn=FRYL;refGene_tx=XM_023253129.1;refGene_gn=FRYL;refGene_tx=XM_023253130.1;refGene_gn=FRYL;refGene_tx=XM_023253131.1;refGene_gn=FRYL;refGene_tx=XM_023253132.1
B1	166880579	166880704
refGene_gn=FRYL;refGene_tx=XM_019829524.2;refGene_gn=FRYL;refGene_tx=XM_019829525.2;refGene_gn=FRYL;refGene_tx=XM_019829526.2;refGene_gn=FRYL;refGene_tx=XM_019829527.2;refGene_gn=FRYL;refGene_tx=XM_023253128.1;refGene_gn=FRYL;refGene_tx=XM_023253129.1;refGene_gn=FRYL;refGene_tx=XM_023253130.1;refGene_gn=FRYL;refGene_tx=XM_023253131.1;refGene_gn=FRYL;refGene_tx=XM_023253132.1
B1	166883198	166883256
refGene_gn=FRYL;refGene_tx=XM_019829524.2;refGene_gn=FRYL;refGene_tx=XM_019829525.2;refGene_gn=FRYL;refGene_tx=XM_019829526.2;refGene_gn=FRYL;refGene_tx=XM_019829527.2;refGene_gn=FRYL;refGene_tx=XM_023253128.1;refGene_gn=FRYL;refGene_tx=XM_023253129.1;refGene_gn=FRYL;refGene_tx=XM_023253130.1;refGene_gn=FRYL;refGene_tx=XM_023253131.1;refGene_gn=FRYL;refGene_tx=XM_023253132.1
B1	166885522	166885614

refGene_gn=FRYL;refGene_tx=XM_019829524.2;refGene_gn=FRYL;refGene_tx=XM_019829525.2;refGene_gn=FRYL;refGene_tx=XM_019829526.2;refGene_gn=FRYL;refGene_tx=XM_019829527.2;refGene_gn=FRYL;refGene_tx=XM_023253128.1;refGene_gn=FRYL;refGene_tx=XM_023253129.1;refGene_gn=FRYL;refGene_tx=XM_023253130.1;refGene_gn=FRYL;refGene_tx=XM_023253131.1;refGene_gn=FRYL;refGene_tx=XM_023253132.1
B1 166889909 166890082
refGene_gn=FRYL;refGene_tx=XM_019829524.2;refGene_gn=FRYL;refGene_tx=XM_019829525.2;refGene_gn=FRYL;refGene_tx=XM_019829526.2;refGene_gn=FRYL;refGene_tx=XM_019829527.2;refGene_gn=FRYL;refGene_tx=XM_023253128.1;refGene_gn=FRYL;refGene_tx=XM_023253129.1;refGene_gn=FRYL;refGene_tx=XM_023253130.1;refGene_gn=FRYL;refGene_tx=XM_023253131.1;refGene_gn=FRYL;refGene_tx=XM_023253132.1
B1 166892120 166892281
refGene_gn=FRYL;refGene_tx=XM_019829524.2;refGene_gn=FRYL;refGene_tx=XM_019829525.2;refGene_gn=FRYL;refGene_tx=XM_019829526.2;refGene_gn=FRYL;refGene_tx=XM_019829527.2;refGene_gn=FRYL;refGene_tx=XM_023253128.1;refGene_gn=FRYL;refGene_tx=XM_023253129.1;refGene_gn=FRYL;refGene_tx=XM_023253130.1;refGene_gn=FRYL;refGene_tx=XM_023253131.1;refGene_gn=FRYL;refGene_tx=XM_023253132.1
B1 166892804 166892915
refGene_gn=FRYL;refGene_tx=XM_019829524.2;refGene_gn=FRYL;refGene_tx=XM_019829525.2;refGene_gn=FRYL;refGene_tx=XM_019829526.2;refGene_gn=FRYL;refGene_tx=XM_019829527.2;refGene_gn=FRYL;refGene_tx=XM_023253128.1;refGene_gn=FRYL;refGene_tx=XM_023253129.1;refGene_gn=FRYL;refGene_tx=XM_023253130.1;refGene_gn=FRYL;refGene_tx=XM_023253131.1;refGene_gn=FRYL;refGene_tx=XM_023253132.1
B1 166893712 166893867
refGene_gn=FRYL;refGene_tx=XM_019829524.2;refGene_gn=FRYL;refGene_tx=XM_019829525.2;refGene_gn=FRYL;refGene_tx=XM_019829526.2;refGene_gn=FRYL;refGene_tx=XM_019829527.2;refGene_gn=FRYL;refGene_tx=XM_023253128.1;refGene_gn=FRYL;refGene_tx=XM_023253129.1;refGene_gn=FRYL;refGene_tx=XM_023253130.1;refGene_gn=FRYL;refGene_tx=XM_023253131.1;refGene_gn=FRYL;refGene_tx=XM_023253132.1
B1 166894153 166894253
refGene_gn=FRYL;refGene_tx=XM_019829524.2;refGene_gn=FRYL;refGene_tx=XM_019829525.2;refGene_gn=FRYL;refGene_tx=XM_019829526.2;refGene_gn=FRYL;refGene_tx=XM_019829527.2;refGene_gn=FRYL;refGene_tx=XM_023253128.1;refGene_gn=FRYL;refGene_tx=XM_023253129.1;refGene_gn=FRYL;refGene_tx=XM_023253130.1;refGene_gn=FRYL;refGene_tx=XM_023253131.1;refGene_gn=FRYL;refGene_tx=XM_023253132.1
B1 166895730 166895899
refGene_gn=FRYL;refGene_tx=XM_019829524.2;refGene_gn=FRYL;refGene_tx=XM_019829525.2;refGene_gn=FRYL;refGene_tx=XM_019829526.2;refGene_gn=FRYL;refGene_tx=XM_019829527.2;refGene_gn=FRYL;refGene_tx=XM_023253128.1;refGene_gn=FRYL;refGene_tx=XM_023253129.1;refGene_gn=FRYL;refGene_tx=XM_023253130.1;refGene_gn=FRYL;refGene_tx=XM_023253131.1;refGene_gn=FRYL;refGene_tx=XM_023253132.1
B1 166901621 166901881
refGene_gn=FRYL;refGene_tx=XM_019829524.2;refGene_gn=FRYL;refGene_tx=X

M_019829525.2;refGene_gn=FRYL;refGene_tx=XM_019829526.2;refGene_gn=FRY
L;refGene_tx=XM_019829527.2;refGene_gn=FRYL;refGene_tx=XM_023253128.1;
refGene_gn=FRYL;refGene_tx=XM_023253129.1;refGene_gn=FRYL;refGene_tx=X
M_023253130.1;refGene_gn=FRYL;refGene_tx=XM_023253131.1;refGene_gn=FRY
L;refGene_tx=XM_023253132.1
B1 166902242 166902383
refGene_gn=FRYL;refGene_tx=XM_019829524.2;refGene_gn=FRYL;refGene_tx=X
M_019829525.2;refGene_gn=FRYL;refGene_tx=XM_019829526.2;refGene_gn=FRY
L;refGene_tx=XM_019829527.2;refGene_gn=FRYL;refGene_tx=XM_023253128.1;
refGene_gn=FRYL;refGene_tx=XM_023253129.1;refGene_gn=FRYL;refGene_tx=X
M_023253130.1;refGene_gn=FRYL;refGene_tx=XM_023253131.1;refGene_gn=FRY
L;refGene_tx=XM_023253132.1
B1 166905175 166905344
refGene_gn=FRYL;refGene_tx=XM_019829524.2;refGene_gn=FRYL;refGene_tx=X
M_019829525.2;refGene_gn=FRYL;refGene_tx=XM_019829526.2;refGene_gn=FRY
L;refGene_tx=XM_019829527.2;refGene_gn=FRYL;refGene_tx=XM_023253128.1;
refGene_gn=FRYL;refGene_tx=XM_023253129.1;refGene_gn=FRYL;refGene_tx=X
M_023253130.1;refGene_gn=FRYL;refGene_tx=XM_023253131.1;refGene_gn=FRY
L;refGene_tx=XM_023253132.1
B1 166908013 166908098
refGene_gn=FRYL;refGene_tx=XM_019829524.2;refGene_gn=FRYL;refGene_tx=X
M_019829525.2;refGene_gn=FRYL;refGene_tx=XM_019829526.2;refGene_gn=FRY
L;refGene_tx=XM_019829527.2;refGene_gn=FRYL;refGene_tx=XM_023253128.1;
refGene_gn=FRYL;refGene_tx=XM_023253129.1;refGene_gn=FRYL;refGene_tx=X
M_023253130.1;refGene_gn=FRYL;refGene_tx=XM_023253131.1;refGene_gn=FRY
L;refGene_tx=XM_023253132.1
B1 166908658 166908771
refGene_gn=FRYL;refGene_tx=XM_019829524.2;refGene_gn=FRYL;refGene_tx=X
M_019829525.2;refGene_gn=FRYL;refGene_tx=XM_019829526.2;refGene_gn=FRY
L;refGene_tx=XM_019829527.2;refGene_gn=FRYL;refGene_tx=XM_023253128.1;
refGene_gn=FRYL;refGene_tx=XM_023253129.1;refGene_gn=FRYL;refGene_tx=X
M_023253130.1;refGene_gn=FRYL;refGene_tx=XM_023253131.1;refGene_gn=FRY
L;refGene_tx=XM_023253132.1
B1 166909752 166909903
refGene_gn=FRYL;refGene_tx=XM_019829524.2;refGene_gn=FRYL;refGene_tx=X
M_019829525.2;refGene_gn=FRYL;refGene_tx=XM_019829526.2;refGene_gn=FRY
L;refGene_tx=XM_019829527.2;refGene_gn=FRYL;refGene_tx=XM_023253128.1;
refGene_gn=FRYL;refGene_tx=XM_023253129.1;refGene_gn=FRYL;refGene_tx=X
M_023253130.1;refGene_gn=FRYL;refGene_tx=XM_023253131.1;refGene_gn=FRY
L;refGene_tx=XM_023253132.1
B1 166910566 166910670
refGene_gn=FRYL;refGene_tx=XM_019829524.2;refGene_gn=FRYL;refGene_tx=X
M_019829525.2;refGene_gn=FRYL;refGene_tx=XM_019829526.2;refGene_gn=FRY
L;refGene_tx=XM_019829527.2;refGene_gn=FRYL;refGene_tx=XM_023253128.1;
refGene_gn=FRYL;refGene_tx=XM_023253129.1;refGene_gn=FRYL;refGene_tx=X
M_023253130.1;refGene_gn=FRYL;refGene_tx=XM_023253131.1;refGene_gn=FRY
L;refGene_tx=XM_023253132.1
B1 166911854 166912040
refGene_gn=FRYL;refGene_tx=XM_019829524.2;refGene_gn=FRYL;refGene_tx=X
M_019829525.2;refGene_gn=FRYL;refGene_tx=XM_019829526.2;refGene_gn=FRY

L;refGene_tx=XM_019829527.2;refGene_gn=FRYL;refGene_tx=XM_023253128.1;
refGene_gn=FRYL;refGene_tx=XM_023253129.1;refGene_gn=FRYL;refGene_tx=X
M_023253130.1;refGene_gn=FRYL;refGene_tx=XM_023253131.1;refGene_gn=FRY
L;refGene_tx=XM_023253132.1
B1 166913098 166913303
refGene_gn=FRYL;refGene_tx=XM_019829524.2;refGene_gn=FRYL;refGene_tx=X
M_019829525.2;refGene_gn=FRYL;refGene_tx=XM_019829526.2;refGene_gn=FRY
L;refGene_tx=XM_019829527.2;refGene_gn=FRYL;refGene_tx=XM_023253128.1;
refGene_gn=FRYL;refGene_tx=XM_023253129.1;refGene_gn=FRYL;refGene_tx=X
M_023253130.1;refGene_gn=FRYL;refGene_tx=XM_023253131.1;refGene_gn=FRY
L;refGene_tx=XM_023253132.1
B1 166914527 166914649
refGene_gn=FRYL;refGene_tx=XM_019829524.2;refGene_gn=FRYL;refGene_tx=X
M_019829525.2;refGene_gn=FRYL;refGene_tx=XM_019829526.2;refGene_gn=FRY
L;refGene_tx=XM_019829527.2;refGene_gn=FRYL;refGene_tx=XM_023253128.1;
refGene_gn=FRYL;refGene_tx=XM_023253129.1;refGene_gn=FRYL;refGene_tx=X
M_023253130.1;refGene_gn=FRYL;refGene_tx=XM_023253131.1;refGene_gn=FRY
L;refGene_tx=XM_023253132.1
B1 166915393 166915584
refGene_gn=FRYL;refGene_tx=XM_019829524.2;refGene_gn=FRYL;refGene_tx=X
M_019829525.2;refGene_gn=FRYL;refGene_tx=XM_019829526.2;refGene_gn=FRY
L;refGene_tx=XM_019829527.2;refGene_gn=FRYL;refGene_tx=XM_023253128.1;
refGene_gn=FRYL;refGene_tx=XM_023253129.1;refGene_gn=FRYL;refGene_tx=X
M_023253130.1;refGene_gn=FRYL;refGene_tx=XM_023253131.1;refGene_gn=FRY
L;refGene_tx=XM_023253132.1
B1 166916902 166916997
refGene_gn=FRYL;refGene_tx=XM_019829524.2;refGene_gn=FRYL;refGene_tx=X
M_019829525.2;refGene_gn=FRYL;refGene_tx=XM_019829526.2;refGene_gn=FRY
L;refGene_tx=XM_019829527.2;refGene_gn=FRYL;refGene_tx=XM_023253128.1;
refGene_gn=FRYL;refGene_tx=XM_023253129.1;refGene_gn=FRYL;refGene_tx=X
M_023253130.1;refGene_gn=FRYL;refGene_tx=XM_023253131.1;refGene_gn=FRY
L;refGene_tx=XM_023253132.1
B1 166918117 166918725
refGene_gn=FRYL;refGene_tx=XM_019829524.2;refGene_gn=FRYL;refGene_tx=X
M_019829525.2;refGene_gn=FRYL;refGene_tx=XM_019829526.2;refGene_gn=FRY
L;refGene_tx=XM_019829527.2;refGene_gn=FRYL;refGene_tx=XM_023253128.1;
refGene_gn=FRYL;refGene_tx=XM_023253129.1;refGene_gn=FRYL;refGene_tx=X
M_023253130.1;refGene_gn=FRYL;refGene_tx=XM_023253131.1;refGene_gn=FRY
L;refGene_tx=XM_023253132.1
B1 166919062 166919160
refGene_gn=FRYL;refGene_tx=XM_019829524.2;refGene_gn=FRYL;refGene_tx=X
M_019829525.2;refGene_gn=FRYL;refGene_tx=XM_019829526.2;refGene_gn=FRY
L;refGene_tx=XM_019829527.2;refGene_gn=FRYL;refGene_tx=XM_023253128.1;
refGene_gn=FRYL;refGene_tx=XM_023253129.1;refGene_gn=FRYL;refGene_tx=X
M_023253130.1;refGene_gn=FRYL;refGene_tx=XM_023253131.1;refGene_gn=FRY
L;refGene_tx=XM_023253132.1
B1 166924041 166924212
refGene_gn=FRYL;refGene_tx=XM_019829524.2;refGene_gn=FRYL;refGene_tx=X
M_019829525.2;refGene_gn=FRYL;refGene_tx=XM_019829526.2;refGene_gn=FRY
L;refGene_tx=XM_019829527.2;refGene_gn=FRYL;refGene_tx=XM_023253128.1;

refGene_gn=FRYL;refGene_tx=XM_023253129.1;refGene_gn=FRYL;refGene_tx=XM_023253130.1;refGene_gn=FRYL;refGene_tx=XM_023253131.1;refGene_gn=FRYL;refGene_tx=XM_023253132.1
B1 166926406 166926547
refGene_gn=FRYL;refGene_tx=XM_019829524.2;refGene_gn=FRYL;refGene_tx=XM_019829525.2;refGene_gn=FRYL;refGene_tx=XM_019829526.2;refGene_gn=FRYL;refGene_tx=XM_019829527.2;refGene_gn=FRYL;refGene_tx=XM_023253128.1;refGene_gn=FRYL;refGene_tx=XM_023253129.1;refGene_gn=FRYL;refGene_tx=XM_023253130.1;refGene_gn=FRYL;refGene_tx=XM_023253131.1;refGene_gn=FRYL;refGene_tx=XM_023253132.1
B1 166930136 166930334
refGene_gn=FRYL;refGene_tx=XM_019829524.2;refGene_gn=FRYL;refGene_tx=XM_019829525.2;refGene_gn=FRYL;refGene_tx=XM_019829526.2;refGene_gn=FRYL;refGene_tx=XM_019829527.2;refGene_gn=FRYL;refGene_tx=XM_023253128.1;refGene_gn=FRYL;refGene_tx=XM_023253129.1;refGene_gn=FRYL;refGene_tx=XM_023253130.1;refGene_gn=FRYL;refGene_tx=XM_023253131.1;refGene_gn=FRYL;refGene_tx=XM_023253132.1
B1 166934734 166934896
refGene_gn=FRYL;refGene_tx=XM_019829524.2;refGene_gn=FRYL;refGene_tx=XM_019829525.2;refGene_gn=FRYL;refGene_tx=XM_019829526.2;refGene_gn=FRYL;refGene_tx=XM_019829527.2;refGene_gn=FRYL;refGene_tx=XM_023253128.1;refGene_gn=FRYL;refGene_tx=XM_023253129.1;refGene_gn=FRYL;refGene_tx=XM_023253130.1;refGene_gn=FRYL;refGene_tx=XM_023253131.1;refGene_gn=FRYL;refGene_tx=XM_023253132.1
B1 166935024 166935099
refGene_gn=FRYL;refGene_tx=XM_019829524.2;refGene_gn=FRYL;refGene_tx=XM_019829525.2;refGene_gn=FRYL;refGene_tx=XM_019829526.2;refGene_gn=FRYL;refGene_tx=XM_019829527.2;refGene_gn=FRYL;refGene_tx=XM_023253128.1;refGene_gn=FRYL;refGene_tx=XM_023253129.1;refGene_gn=FRYL;refGene_tx=XM_023253130.1;refGene_gn=FRYL;refGene_tx=XM_023253131.1;refGene_gn=FRYL;refGene_tx=XM_023253132.1
B1 166935444 166935621
refGene_gn=FRYL;refGene_tx=XM_019829524.2;refGene_gn=FRYL;refGene_tx=XM_019829525.2;refGene_gn=FRYL;refGene_tx=XM_019829526.2;refGene_gn=FRYL;refGene_tx=XM_019829527.2;refGene_gn=FRYL;refGene_tx=XM_023253128.1;refGene_gn=FRYL;refGene_tx=XM_023253129.1;refGene_gn=FRYL;refGene_tx=XM_023253130.1;refGene_gn=FRYL;refGene_tx=XM_023253131.1;refGene_gn=FRYL;refGene_tx=XM_023253132.1
B1 166939118 166939127
refGene_gn=FRYL;refGene_tx=XM_019829524.2;refGene_gn=FRYL;refGene_tx=XM_019829525.2;refGene_gn=FRYL;refGene_tx=XM_019829526.2;refGene_gn=FRYL;refGene_tx=XM_019829527.2;refGene_gn=FRYL;refGene_tx=XM_023253129.1;refGene_gn=FRYL;refGene_tx=XM_023253130.1
B1 166939507 166939711
refGene_gn=FRYL;refGene_tx=XM_019829524.2;refGene_gn=FRYL;refGene_tx=XM_019829525.2;refGene_gn=FRYL;refGene_tx=XM_019829526.2;refGene_gn=FRYL;refGene_tx=XM_019829527.2;refGene_gn=FRYL;refGene_tx=XM_023253128.1;refGene_gn=FRYL;refGene_tx=XM_023253129.1;refGene_gn=FRYL;refGene_tx=XM_023253130.1;refGene_gn=FRYL;refGene_tx=XM_023253131.1;refGene_gn=FRYL;refGene_tx=XM_023253132.1

B1	166943011	166943179
refGene_gn=FRYL;refGene_tx=XM_019829524.2;refGene_gn=FRYL;refGene_tx=XM_019829525.2;refGene_gn=FRYL;refGene_tx=XM_019829526.2;refGene_gn=FRYL;refGene_tx=XM_019829527.2;refGene_gn=FRYL;refGene_tx=XM_023253128.1;refGene_gn=FRYL;refGene_tx=XM_023253129.1;refGene_gn=FRYL;refGene_tx=XM_023253130.1;refGene_gn=FRYL;refGene_tx=XM_023253131.1;refGene_gn=FRYL;refGene_tx=XM_023253132.1
B1	166949613	166949864
refGene_gn=FRYL;refGene_tx=XM_019829524.2;refGene_gn=FRYL;refGene_tx=XM_019829525.2;refGene_gn=FRYL;refGene_tx=XM_019829526.2;refGene_gn=FRYL;refGene_tx=XM_019829527.2;refGene_gn=FRYL;refGene_tx=XM_023253128.1;refGene_gn=FRYL;refGene_tx=XM_023253129.1;refGene_gn=FRYL;refGene_tx=XM_023253130.1;refGene_gn=FRYL;refGene_tx=XM_023253131.1;refGene_gn=FRYL;refGene_tx=XM_023253132.1
B1	166953172	166953380
refGene_gn=FRYL;refGene_tx=XM_019829524.2;refGene_gn=FRYL;refGene_tx=XM_019829525.2;refGene_gn=FRYL;refGene_tx=XM_019829526.2;refGene_gn=FRYL;refGene_tx=XM_019829527.2;refGene_gn=FRYL;refGene_tx=XM_023253128.1;refGene_gn=FRYL;refGene_tx=XM_023253129.1;refGene_gn=FRYL;refGene_tx=XM_023253130.1;refGene_gn=FRYL;refGene_tx=XM_023253131.1;refGene_gn=FRYL;refGene_tx=XM_023253132.1
B1	166954535	166954547
refGene_gn=FRYL;refGene_tx=XM_019829527.2
B1	166955116	166955266
refGene_gn=FRYL;refGene_tx=XM_019829524.2;refGene_gn=FRYL;refGene_tx=XM_019829525.2;refGene_gn=FRYL;refGene_tx=XM_019829526.2;refGene_gn=FRYL;refGene_tx=XM_023253128.1;refGene_gn=FRYL;refGene_tx=XM_023253129.1;refGene_gn=FRYL;refGene_tx=XM_023253130.1;refGene_gn=FRYL;refGene_tx=XM_023253131.1;refGene_gn=FRYL;refGene_tx=XM_023253132.1
B1	166956041	166956140
refGene_gn=FRYL;refGene_tx=XM_019829524.2;refGene_gn=FRYL;refGene_tx=XM_019829525.2;refGene_gn=FRYL;refGene_tx=XM_019829526.2;refGene_gn=FRYL;refGene_tx=XM_023253128.1;refGene_gn=FRYL;refGene_tx=XM_023253129.1;refGene_gn=FRYL;refGene_tx=XM_023253130.1;refGene_gn=FRYL;refGene_tx=XM_023253131.1;refGene_gn=FRYL;refGene_tx=XM_023253132.1
B1	166962651	166962720
refGene_gn=FRYL;refGene_tx=XM_019829524.2;refGene_gn=FRYL;refGene_tx=XM_019829525.2;refGene_gn=FRYL;refGene_tx=XM_019829526.2;refGene_gn=FRYL;refGene_tx=XM_023253128.1;refGene_gn=FRYL;refGene_tx=XM_023253129.1;refGene_gn=FRYL;refGene_tx=XM_023253130.1;refGene_gn=FRYL;refGene_tx=XM_023253131.1;refGene_gn=FRYL;refGene_tx=XM_023253132.1
B1	166965320	166965338
refGene_gn=FRYL;refGene_tx=XM_019829524.2;refGene_gn=FRYL;refGene_tx=XM_019829525.2;refGene_gn=FRYL;refGene_tx=XM_019829526.2;refGene_gn=FRYL;refGene_tx=XM_023253128.1;refGene_gn=FRYL;refGene_tx=XM_023253130.1;refGene_gn=FRYL;refGene_tx=XM_023253131.1
B1	166966070	166966181
refGene_gn=FRYL;refGene_tx=XM_019829524.2;refGene_gn=FRYL;refGene_tx=XM_019829525.2;refGene_gn=FRYL;refGene_tx=XM_019829526.2;refGene_gn=FRYL;refGene_tx=XM_023253128.1;refGene_gn=FRYL;refGene_tx=XM_023253129.1;

refGene_gn=FRYL;refGene_tx=XM_023253130.1;refGene_gn=FRYL;refGene_tx=XM_023253131.1;refGene_gn=FRYL;refGene_tx=XM_023253132.1
B1 166967267 166967458
refGene_gn=FRYL;refGene_tx=XM_019829524.2;refGene_gn=FRYL;refGene_tx=XM_019829525.2;refGene_gn=FRYL;refGene_tx=XM_019829526.2;refGene_gn=FRYL;refGene_tx=XM_023253128.1;refGene_gn=FRYL;refGene_tx=XM_023253129.1;refGene_gn=FRYL;refGene_tx=XM_023253130.1;refGene_gn=FRYL;refGene_tx=XM_023253131.1;refGene_gn=FRYL;refGene_tx=XM_023253132.1
B1 166967783 166968042
refGene_gn=FRYL;refGene_tx=XM_019829524.2;refGene_gn=FRYL;refGene_tx=XM_019829525.2;refGene_gn=FRYL;refGene_tx=XM_019829526.2;refGene_gn=FRYL;refGene_tx=XM_023253128.1;refGene_gn=FRYL;refGene_tx=XM_023253129.1;refGene_gn=FRYL;refGene_tx=XM_023253130.1;refGene_gn=FRYL;refGene_tx=XM_023253131.1;refGene_gn=FRYL;refGene_tx=XM_023253132.1
B1 166973286 166973430
refGene_gn=ZAR1;refGene_tx=XM_023252238.1
B1 166974779 166974854
refGene_gn=ZAR1;refGene_tx=XM_023252238.1
B1 166975145 166975238
refGene_gn=ZAR1;refGene_tx=XM_023252238.1
B1 166977102 166978056
refGene_gn=ZAR1;refGene_tx=XM_023252238.1
B1 166979461 166979974
refGene_gn=SLC10A4;refGene_tx=XM_023253133.1
B1 166983416 166983627
refGene_gn=SLC10A4;refGene_tx=XM_023253133.1
B1 166984390 166984986
refGene_gn=SLC10A4;refGene_tx=XM_023253133.1
B1 167051728 167051795
refGene_gn=SLAIN2;refGene_tx=XM_023253134.1;refGene_gn=SLAIN2;refGene_tx=XM_023253135.1
B1 167052893 167053212
refGene_gn=SLAIN2;refGene_tx=XM_023253134.1;refGene_gn=SLAIN2;refGene_tx=XM_023253135.1
B1 167067269 167067347
refGene_gn=SLAIN2;refGene_tx=XM_023253134.1
B1 167076964 167077102
refGene_gn=SLAIN2;refGene_tx=XM_023253134.1;refGene_gn=SLAIN2;refGene_tx=XM_023253135.1
B1 167077534 167077894
refGene_gn=SLAIN2;refGene_tx=XM_023253134.1;refGene_gn=SLAIN2;refGene_tx=XM_023253135.1
B1 167081389 167081551
refGene_gn=SLAIN2;refGene_tx=XM_023253134.1;refGene_gn=SLAIN2;refGene_tx=XM_023253135.1
B1 167083248 167083413
refGene_gn=SLAIN2;refGene_tx=XM_023253134.1;refGene_gn=SLAIN2;refGene_tx=XM_023253135.1
B1 167089829 167089978
refGene_gn=SLAIN2;refGene_tx=XM_023253134.1;refGene_gn=SLAIN2;refGene_

tx=XM_023253135.1
B1	167121275	167121664
refGene_gn=SLAIN2;refGene_tx=XM_023253134.1;refGene_gn=SLAIN2;refGene_tx=XM_023253135.1
B1	167203069	167203105
refGene_gn=TEC;refGene_tx=XM_011281891.3;refGene_gn=TEC;refGene_tx=XM_019829536.2
B1	167224938	167225124
refGene_gn=TEC;refGene_tx=XM_011281891.3;refGene_gn=TEC;refGene_tx=XM_019829536.2;refGene_gn=TEC;refGene_tx=XM_023253137.1
B1	167280066	167280171
refGene_gn=TEC;refGene_tx=XM_011281891.3;refGene_gn=TEC;refGene_tx=XM_019829536.2;refGene_gn=TEC;refGene_tx=XM_023253137.1
B1	167281773	167281855
refGene_gn=TEC;refGene_tx=XM_011281891.3;refGene_gn=TEC;refGene_tx=XM_019829536.2;refGene_gn=TEC;refGene_tx=XM_023253137.1
B1	167282505	167282634
refGene_gn=TEC;refGene_tx=XM_011281891.3;refGene_gn=TEC;refGene_tx=XM_019829536.2;refGene_gn=TEC;refGene_tx=XM_023253137.1
B1	167284772	167284813
refGene_gn=TEC;refGene_tx=XM_011281891.3;refGene_gn=TEC;refGene_tx=XM_019829536.2;refGene_gn=TEC;refGene_tx=XM_023253137.1
B1	167285843	167286019
refGene_gn=TEC;refGene_tx=XM_011281891.3;refGene_gn=TEC;refGene_tx=XM_019829536.2;refGene_gn=TEC;refGene_tx=XM_023253137.1
B1	167289940	167290006
refGene_gn=TEC;refGene_tx=XM_011281891.3;refGene_gn=TEC;refGene_tx=XM_019829536.2;refGene_gn=TEC;refGene_tx=XM_023253137.1
B1	167295385	167295440
refGene_gn=TEC;refGene_tx=XM_011281891.3;refGene_gn=TEC;refGene_tx=XM_019829536.2;refGene_gn=TEC;refGene_tx=XM_023253137.1
B1	167298595	167298675
refGene_gn=TEC;refGene_tx=XM_011281891.3;refGene_gn=TEC;refGene_tx=XM_019829536.2;refGene_gn=TEC;refGene_tx=XM_023253137.1
B1	167299874	167300008
refGene_gn=TEC;refGene_tx=XM_011281891.3;refGene_gn=TEC;refGene_tx=XM_019829536.2;refGene_gn=TEC;refGene_tx=XM_023253137.1
B1	167303285	167303360
refGene_gn=TEC;refGene_tx=XM_011281891.3;refGene_gn=TEC;refGene_tx=XM_019829536.2;refGene_gn=TEC;refGene_tx=XM_023253137.1
B1	167304121	167304293
refGene_gn=TEC;refGene_tx=XM_011281891.3;refGene_gn=TEC;refGene_tx=XM_019829536.2;refGene_gn=TEC;refGene_tx=XM_023253137.1
B1	167304401	167304618
refGene_gn=TEC;refGene_tx=XM_011281891.3;refGene_gn=TEC;refGene_tx=XM_019829536.2;refGene_gn=TEC;refGene_tx=XM_023253137.1
B1	167310660	167310725
refGene_gn=TEC;refGene_tx=XM_011281891.3;refGene_gn=TEC;refGene_tx=XM_019829536.2;refGene_gn=TEC;refGene_tx=XM_023253137.1
B1	167314270	167314389

refGene_gn=TEC;refGene_tx=XM_011281891.3;refGene_gn=TEC;refGene_tx=XM_019829536.2;refGene_gn=TEC;refGene_tx=XM_023253137.1
B1 167314475 167314633
refGene_gn=TEC;refGene_tx=XM_011281891.3;refGene_gn=TEC;refGene_tx=XM_019829536.2;refGene_gn=TEC;refGene_tx=XM_023253137.1
B1 167315787 167315871
refGene_gn=TEC;refGene_tx=XM_011281891.3;refGene_gn=TEC;refGene_tx=XM_019829536.2;refGene_gn=TEC;refGene_tx=XM_023253137.1
B1 167318595 167318683
refGene_gn=TXK;refGene_tx=XM_023253138.1
B1 167338165 167338236
refGene_gn=TXK;refGene_tx=XM_003985397.5;refGene_gn=TXK;refGene_tx=XM_023253138.1
B1 167339246 167339349
refGene_gn=TXK;refGene_tx=XM_003985397.5;refGene_gn=TXK;refGene_tx=XM_023253138.1
B1 167340378 167340584
refGene_gn=TXK;refGene_tx=XM_003985397.5;refGene_gn=TXK;refGene_tx=XM_023253138.1
B1 167343230 167343296
refGene_gn=TXK;refGene_tx=XM_003985397.5;refGene_gn=TXK;refGene_tx=XM_023253138.1
B1 167349824 167349879
refGene_gn=TXK;refGene_tx=XM_003985397.5;refGene_gn=TXK;refGene_tx=XM_023253138.1
B1 167359694 167359774
refGene_gn=TXK;refGene_tx=XM_003985397.5;refGene_gn=TXK;refGene_tx=XM_023253138.1
B1 167360947 167361075
refGene_gn=TXK;refGene_tx=XM_003985397.5;refGene_gn=TXK;refGene_tx=XM_023253138.1
B1 167365806 167365881
refGene_gn=TXK;refGene_tx=XM_003985397.5;refGene_gn=TXK;refGene_tx=XM_023253138.1
B1 167368810 167368982
refGene_gn=TXK;refGene_tx=XM_003985397.5;refGene_gn=TXK;refGene_tx=XM_023253138.1
B1 167371818 167372035
refGene_gn=TXK;refGene_tx=XM_003985397.5;refGene_gn=TXK;refGene_tx=XM_023253138.1
B1 167374612 167374677
refGene_gn=TXK;refGene_tx=XM_003985397.5;refGene_gn=TXK;refGene_tx=XM_023253138.1
B1 167377404 167377523
refGene_gn=TXK;refGene_tx=XM_003985397.5;refGene_gn=TXK;refGene_tx=XM_023253138.1
B1 167379034 167379192
refGene_gn=TXK;refGene_tx=XM_003985397.5;refGene_gn=TXK;refGene_tx=XM_023253138.1
B1 167381951 167382020

refGene_gn=TXK;refGene_tx=XM_003985397.5;refGene_gn=TXK;refGene_tx=XM_023253138.1
B1 167401156 167401776
refGene_gn=NIPAL1;refGene_tx=XM_003985451.3
B1 167402298 167402459
refGene_gn=NIPAL1;refGene_tx=XM_003985451.3
B1 167403965 167404056
refGene_gn=NIPAL1;refGene_tx=XM_003985451.3
B1 167412461 167412518
refGene_gn=NIPAL1;refGene_tx=XM_003985451.3
B1 167415611 167415894
refGene_gn=NIPAL1;refGene_tx=XM_003985451.3
B1 167461903 167462010
refGene_gn=CNGA1;refGene_tx=XM_006931133.3
B1 167462865 167462988
refGene_gn=CNGA1;refGene_tx=XM_006931133.3
B1 167464434 167464497
refGene_gn=CNGA1;refGene_tx=XM_006931133.3
B1 167469056 167469098
refGene_gn=CNGA1;refGene_tx=XM_006931133.3
B1 167469187 167469298
refGene_gn=CNGA1;refGene_tx=XM_006931133.3
B1 167471243 167471351
refGene_gn=CNGA1;refGene_tx=XM_006931133.3
B1 167472104 167472211
refGene_gn=CNGA1;refGene_tx=XM_006931133.3
B1 167481512 167482927
refGene_gn=CNGA1;refGene_tx=XM_006931133.3
B1 167499176 167499420
refGene_gn=NFXL1;refGene_tx=XM_003985399.4;refGene_gn=NFXL1;refGene_tx=XM_019829539.2;refGene_gn=NFXL1;refGene_tx=XM_019829540.2;refGene_gn=NFXL1;refGene_tx=XM_019829542.1
B1 167502691 167502865
refGene_gn=NFXL1;refGene_tx=XM_003985399.4;refGene_gn=NFXL1;refGene_tx=XM_019829539.2;refGene_gn=NFXL1;refGene_tx=XM_019829540.2;refGene_gn=NFXL1;refGene_tx=XM_019829542.1
B1 167513973 167514083
refGene_gn=NFXL1;refGene_tx=XM_003985399.4;refGene_gn=NFXL1;refGene_tx=XM_019829539.2;refGene_gn=NFXL1;refGene_tx=XM_019829540.2;refGene_gn=NFXL1;refGene_tx=XM_019829542.1
B1 167516135 167516269
refGene_gn=NFXL1;refGene_tx=XM_003985399.4;refGene_gn=NFXL1;refGene_tx=XM_019829539.2;refGene_gn=NFXL1;refGene_tx=XM_019829540.2;refGene_gn=NFXL1;refGene_tx=XM_019829542.1
B1 167519384 167519563
refGene_gn=NFXL1;refGene_tx=XM_003985399.4;refGene_gn=NFXL1;refGene_tx=XM_019829539.2;refGene_gn=NFXL1;refGene_tx=XM_019829540.2;refGene_gn=NFXL1;refGene_tx=XM_019829542.1
B1 167519797 167519959
refGene_gn=NFXL1;refGene_tx=XM_003985399.4;refGene_gn=NFXL1;refGene_tx

=XM_019829539.2;refGene_gn=NFXL1;refGene_tx=XM_019829540.2;refGene_gn=NFXL1;refGene_tx=XM_019829542.1
B1 167520297 167520398
refGene_gn=NFXL1;refGene_tx=XM_003985399.4;refGene_gn=NFXL1;refGene_tx=XM_019829539.2;refGene_gn=NFXL1;refGene_tx=XM_019829540.2;refGene_gn=NFXL1;refGene_tx=XM_019829542.1
B1 167521045 167521160
refGene_gn=NFXL1;refGene_tx=XM_003985399.4;refGene_gn=NFXL1;refGene_tx=XM_019829539.2;refGene_gn=NFXL1;refGene_tx=XM_019829540.2;refGene_gn=NFXL1;refGene_tx=XM_019829542.1
B1 167522294 167522419
refGene_gn=NFXL1;refGene_tx=XM_003985399.4;refGene_gn=NFXL1;refGene_tx=XM_019829539.2;refGene_gn=NFXL1;refGene_tx=XM_019829540.2;refGene_gn=NFXL1;refGene_tx=XM_019829542.1
B1 167523103 167523226
refGene_gn=NFXL1;refGene_tx=XM_003985399.4;refGene_gn=NFXL1;refGene_tx=XM_019829539.2;refGene_gn=NFXL1;refGene_tx=XM_019829540.2;refGene_gn=NFXL1;refGene_tx=XM_019829542.1
B1 167527916 167528007
refGene_gn=NFXL1;refGene_tx=XM_003985399.4;refGene_gn=NFXL1;refGene_tx=XM_019829539.2;refGene_gn=NFXL1;refGene_tx=XM_019829540.2;refGene_gn=NFXL1;refGene_tx=XM_019829542.1
B1 167533679 167533800
refGene_gn=NFXL1;refGene_tx=XM_003985399.4;refGene_gn=NFXL1;refGene_tx=XM_019829539.2;refGene_gn=NFXL1;refGene_tx=XM_019829540.2;refGene_gn=NFXL1;refGene_tx=XM_019829542.1
B1 167534026 167534186
refGene_gn=NFXL1;refGene_tx=XM_003985399.4;refGene_gn=NFXL1;refGene_tx=XM_019829539.2;refGene_gn=NFXL1;refGene_tx=XM_019829540.2;refGene_gn=NFXL1;refGene_tx=XM_019829542.1
B1 167534923 167535015
refGene_gn=NFXL1;refGene_tx=XM_003985399.4;refGene_gn=NFXL1;refGene_tx=XM_019829539.2;refGene_gn=NFXL1;refGene_tx=XM_019829540.2;refGene_gn=NFXL1;refGene_tx=XM_019829542.1
B1 167543009 167543031
refGene_gn=NFXL1;refGene_tx=XM_003985399.4;refGene_gn=NFXL1;refGene_tx=XM_019829539.2;refGene_gn=NFXL1;refGene_tx=XM_019829540.2;refGene_gn=NFXL1;refGene_tx=XM_019829542.1
B1 167543464 167543605
refGene_gn=NFXL1;refGene_tx=XM_003985399.4;refGene_gn=NFXL1;refGene_tx=XM_019829539.2;refGene_gn=NFXL1;refGene_tx=XM_019829540.2;refGene_gn=NFXL1;refGene_tx=XM_019829542.1
B1 167549505 167549672
refGene_gn=NFXL1;refGene_tx=XM_003985399.4;refGene_gn=NFXL1;refGene_tx=XM_019829539.2;refGene_gn=NFXL1;refGene_tx=XM_019829540.2;refGene_gn=NFXL1;refGene_tx=XM_019829542.1
B1 167555608 167555678
refGene_gn=NFXL1;refGene_tx=XM_003985399.4;refGene_gn=NFXL1;refGene_tx=XM_019829539.2;refGene_gn=NFXL1;refGene_tx=XM_019829540.2;refGene_gn=NFXL1;refGene_tx=XM_019829542.1

B1 167561983 167562088
refGene_gn=NFXL1;refGene_tx=XM_003985399.4;refGene_gn=NFXL1;refGene_tx=XM_019829539.2;refGene_gn=NFXL1;refGene_tx=XM_019829540.2;refGene_gn=NFXL1;refGene_tx=XM_019829542.1
B1 167566178 167566265
refGene_gn=NFXL1;refGene_tx=XM_003985399.4;refGene_gn=NFXL1;refGene_tx=XM_019829539.2;refGene_gn=NFXL1;refGene_tx=XM_019829540.2;refGene_gn=NFXL1;refGene_tx=XM_019829542.1
B1 167567062 167567116
refGene_gn=NFXL1;refGene_tx=XM_003985399.4;refGene_gn=NFXL1;refGene_tx=XM_019829539.2;refGene_gn=NFXL1;refGene_tx=XM_019829540.2;refGene_gn=NFXL1;refGene_tx=XM_019829542.1
B1 167569924 167570098
refGene_gn=NFXL1;refGene_tx=XM_003985399.4;refGene_gn=NFXL1;refGene_tx=XM_019829539.2;refGene_gn=NFXL1;refGene_tx=XM_019829540.2;refGene_gn=NFXL1;refGene_tx=XM_019829542.1
B1 167578181 167578253
refGene_gn=CORIN;refGene_tx=XM_019829549.2;refGene_gn=CORIN;refGene_tx=XM_019829551.2
B1 167578482 167578761
refGene_gn=CORIN;refGene_tx=XM_003985401.5;refGene_gn=CORIN;refGene_tx=XM_019829546.2;refGene_gn=CORIN;refGene_tx=XM_019829548.2;refGene_gn=CORIN;refGene_tx=XM_023253140.1
B1 167611371 167611516
refGene_gn=CORIN;refGene_tx=XM_003985401.5;refGene_gn=CORIN;refGene_tx=XM_019829546.2;refGene_gn=CORIN;refGene_tx=XM_019829548.2;refGene_gn=CORIN;refGene_tx=XM_019829549.2;refGene_gn=CORIN;refGene_tx=XM_019829551.2;refGene_gn=CORIN;refGene_tx=XM_023253140.1
B1 167629051 167629252
refGene_gn=CORIN;refGene_tx=XM_003985401.5;refGene_gn=CORIN;refGene_tx=XM_019829546.2;refGene_gn=CORIN;refGene_tx=XM_019829548.2;refGene_gn=CORIN;refGene_tx=XM_019829549.2;refGene_gn=CORIN;refGene_tx=XM_019829551.2;refGene_gn=CORIN;refGene_tx=XM_023253140.1
B1 167655273 167655481
refGene_gn=CORIN;refGene_tx=XM_003985401.5;refGene_gn=CORIN;refGene_tx=XM_019829546.2;refGene_gn=CORIN;refGene_tx=XM_019829548.2;refGene_gn=CORIN;refGene_tx=XM_019829549.2;refGene_gn=CORIN;refGene_tx=XM_019829551.2;refGene_gn=CORIN;refGene_tx=XM_023253140.1
B1 167676714 167676896
refGene_gn=CORIN;refGene_tx=XM_003985401.5;refGene_gn=CORIN;refGene_tx=XM_019829546.2;refGene_gn=CORIN;refGene_tx=XM_019829548.2;refGene_gn=CORIN;refGene_tx=XM_019829549.2;refGene_gn=CORIN;refGene_tx=XM_019829551.2;refGene_gn=CORIN;refGene_tx=XM_023253140.1
B1 167737375 167737489
refGene_gn=CORIN;refGene_tx=XM_003985401.5;refGene_gn=CORIN;refGene_tx=XM_019829546.2;refGene_gn=CORIN;refGene_tx=XM_019829548.2;refGene_gn=CORIN;refGene_tx=XM_019829549.2;refGene_gn=CORIN;refGene_tx=XM_019829551.2;refGene_gn=CORIN;refGene_tx=XM_023253140.1
B1 167748814 167748922
refGene_gn=CORIN;refGene_tx=XM_003985401.5;refGene_gn=CORIN;refGene_tx

=XM_019829546.2;refGene_gn=CORIN;refGene_tx=XM_019829548.2;refGene_gn=
CORIN;refGene_tx=XM_019829549.2;refGene_gn=CORIN;refGene_tx=XM_0198295
51.2
B1 167750587 167750698
refGene_gn=CORIN;refGene_tx=XM_003985401.5;refGene_gn=CORIN;refGene_tx
=XM_019829546.2;refGene_gn=CORIN;refGene_tx=XM_019829548.2;refGene_gn=
CORIN;refGene_tx=XM_019829549.2;refGene_gn=CORIN;refGene_tx=XM_0198295
51.2;refGene_gn=CORIN;refGene_tx=XM_023253140.1
B1 167752390 167752507
refGene_gn=CORIN;refGene_tx=XM_003985401.5;refGene_gn=CORIN;refGene_tx
=XM_019829546.2;refGene_gn=CORIN;refGene_tx=XM_019829548.2;refGene_gn=
CORIN;refGene_tx=XM_019829549.2;refGene_gn=CORIN;refGene_tx=XM_0198295
51.2;refGene_gn=CORIN;refGene_tx=XM_023253140.1
B1 167756977 167757085
refGene_gn=CORIN;refGene_tx=XM_003985401.5;refGene_gn=CORIN;refGene_tx
=XM_019829546.2;refGene_gn=CORIN;refGene_tx=XM_019829548.2;refGene_gn=
CORIN;refGene_tx=XM_019829549.2;refGene_gn=CORIN;refGene_tx=XM_0198295
51.2;refGene_gn=CORIN;refGene_tx=XM_023253140.1
B1 167765464 167765696
refGene_gn=CORIN;refGene_tx=XM_003985401.5;refGene_gn=CORIN;refGene_tx
=XM_019829546.2;refGene_gn=CORIN;refGene_tx=XM_019829548.2;refGene_gn=
CORIN;refGene_tx=XM_019829549.2;refGene_gn=CORIN;refGene_tx=XM_0198295
51.2;refGene_gn=CORIN;refGene_tx=XM_023253140.1
B1 167772004 167772150
refGene_gn=CORIN;refGene_tx=XM_003985401.5;refGene_gn=CORIN;refGene_tx
=XM_019829546.2;refGene_gn=CORIN;refGene_tx=XM_019829548.2;refGene_gn=
CORIN;refGene_tx=XM_019829549.2;refGene_gn=CORIN;refGene_tx=XM_0198295
51.2;refGene_gn=CORIN;refGene_tx=XM_023253140.1
B1 167772918 167773026
refGene_gn=CORIN;refGene_tx=XM_003985401.5;refGene_gn=CORIN;refGene_tx
=XM_019829546.2;refGene_gn=CORIN;refGene_tx=XM_019829549.2;refGene_gn=
CORIN;refGene_tx=XM_019829551.2;refGene_gn=CORIN;refGene_tx=XM_0232531
40.1
B1 167773489 167773561
refGene_gn=CORIN;refGene_tx=XM_019829546.2;refGene_gn=CORIN;refGene_tx
=XM_019829549.2;refGene_gn=CORIN;refGene_tx=XM_023253140.1
B1 167782689 167782803
refGene_gn=CORIN;refGene_tx=XM_003985401.5;refGene_gn=CORIN;refGene_tx
=XM_019829546.2;refGene_gn=CORIN;refGene_tx=XM_019829548.2;refGene_gn=
CORIN;refGene_tx=XM_019829549.2;refGene_gn=CORIN;refGene_tx=XM_0198295
51.2;refGene_gn=CORIN;refGene_tx=XM_023253140.1
B1 167784658 167784769
refGene_gn=CORIN;refGene_tx=XM_003985401.5;refGene_gn=CORIN;refGene_tx
=XM_019829546.2;refGene_gn=CORIN;refGene_tx=XM_019829548.2;refGene_gn=
CORIN;refGene_tx=XM_019829549.2;refGene_gn=CORIN;refGene_tx=XM_0198295
51.2;refGene_gn=CORIN;refGene_tx=XM_023253140.1
B1 167786863 167786993
refGene_gn=CORIN;refGene_tx=XM_003985401.5;refGene_gn=CORIN;refGene_tx
=XM_019829546.2;refGene_gn=CORIN;refGene_tx=XM_019829548.2;refGene_gn=
CORIN;refGene_tx=XM_019829549.2;refGene_gn=CORIN;refGene_tx=XM_0198295

51.2;refGene_gn=CORIN;refGene_tx=XM_023253140.1
B1 167807080 167807197
refGene_gn=CORIN;refGene_tx=XM_003985401.5;refGene_gn=CORIN;refGene_tx=XM_019829546.2;refGene_gn=CORIN;refGene_tx=XM_019829548.2;refGene_gn=CORIN;refGene_tx=XM_019829549.2;refGene_gn=CORIN;refGene_tx=XM_019829551.2;refGene_gn=CORIN;refGene_tx=XM_023253140.1
B1 167809513 167809563
refGene_gn=CORIN;refGene_tx=XM_003985401.5;refGene_gn=CORIN;refGene_tx=XM_019829546.2;refGene_gn=CORIN;refGene_tx=XM_019829548.2;refGene_gn=CORIN;refGene_tx=XM_019829549.2;refGene_gn=CORIN;refGene_tx=XM_019829551.2;refGene_gn=CORIN;refGene_tx=XM_023253140.1
B1 167809711 167809886
refGene_gn=CORIN;refGene_tx=XM_003985401.5;refGene_gn=CORIN;refGene_tx=XM_019829546.2;refGene_gn=CORIN;refGene_tx=XM_019829548.2;refGene_gn=CORIN;refGene_tx=XM_019829549.2;refGene_gn=CORIN;refGene_tx=XM_019829551.2;refGene_gn=CORIN;refGene_tx=XM_023253140.1
B1 167824093 167824365
refGene_gn=CORIN;refGene_tx=XM_003985401.5;refGene_gn=CORIN;refGene_tx=XM_019829546.2;refGene_gn=CORIN;refGene_tx=XM_019829548.2;refGene_gn=CORIN;refGene_tx=XM_019829549.2;refGene_gn=CORIN;refGene_tx=XM_019829551.2;refGene_gn=CORIN;refGene_tx=XM_023253140.1
B1 167826493 167826627
refGene_gn=CORIN;refGene_tx=XM_003985401.5;refGene_gn=CORIN;refGene_tx=XM_019829546.2;refGene_gn=CORIN;refGene_tx=XM_019829548.2;refGene_gn=CORIN;refGene_tx=XM_019829549.2;refGene_gn=CORIN;refGene_tx=XM_019829551.2;refGene_gn=CORIN;refGene_tx=XM_023253140.1
B1 167835585 167835768
refGene_gn=CORIN;refGene_tx=XM_003985401.5;refGene_gn=CORIN;refGene_tx=XM_019829546.2;refGene_gn=CORIN;refGene_tx=XM_019829548.2;refGene_gn=CORIN;refGene_tx=XM_019829549.2;refGene_gn=CORIN;refGene_tx=XM_019829551.2;refGene_gn=CORIN;refGene_tx=XM_023253140.1
B1 167839860 167840191
refGene_gn=ATP10D;refGene_tx=XM_003985402.5;refGene_gn=ATP10D;refGene_tx=XM_019829544.2;refGene_gn=ATP10D;refGene_tx=XM_023253139.1
B1 167844273 167844461
refGene_gn=ATP10D;refGene_tx=XM_003985402.5;refGene_gn=ATP10D;refGene_tx=XM_019829544.2;refGene_gn=ATP10D;refGene_tx=XM_023253139.1
B1 167848824 167848929
refGene_gn=ATP10D;refGene_tx=XM_003985402.5;refGene_gn=ATP10D;refGene_tx=XM_019829544.2;refGene_gn=ATP10D;refGene_tx=XM_023253139.1
B1 167850140 167850221
refGene_gn=ATP10D;refGene_tx=XM_003985402.5;refGene_gn=ATP10D;refGene_tx=XM_019829544.2;refGene_gn=ATP10D;refGene_tx=XM_023253139.1
B1 167855010 167855211
refGene_gn=ATP10D;refGene_tx=XM_003985402.5;refGene_gn=ATP10D;refGene_tx=XM_019829544.2;refGene_gn=ATP10D;refGene_tx=XM_023253139.1
B1 167857901 167858027
refGene_gn=ATP10D;refGene_tx=XM_003985402.5;refGene_gn=ATP10D;refGene_tx=XM_019829544.2;refGene_gn=ATP10D;refGene_tx=XM_023253139.1
B1 167859750 167859827

refGene_gn=ATP10D;refGene_tx=XM_003985402.5;refGene_gn=ATP10D;refGene_tx=XM_019829544.2;refGene_gn=ATP10D;refGene_tx=XM_023253139.1
B1 167862704 167863014
refGene_gn=ATP10D;refGene_tx=XM_003985402.5;refGene_gn=ATP10D;refGene_tx=XM_019829544.2;refGene_gn=ATP10D;refGene_tx=XM_023253139.1
B1 167867262 167867447
refGene_gn=ATP10D;refGene_tx=XM_003985402.5;refGene_gn=ATP10D;refGene_tx=XM_019829544.2;refGene_gn=ATP10D;refGene_tx=XM_023253139.1
B1 167869515 167869642
refGene_gn=ATP10D;refGene_tx=XM_003985402.5;refGene_gn=ATP10D;refGene_tx=XM_019829544.2;refGene_gn=ATP10D;refGene_tx=XM_023253139.1
B1 167871015 167871122
refGene_gn=ATP10D;refGene_tx=XM_003985402.5;refGene_gn=ATP10D;refGene_tx=XM_019829544.2;refGene_gn=ATP10D;refGene_tx=XM_023253139.1
B1 167871742 167872352
refGene_gn=ATP10D;refGene_tx=XM_003985402.5;refGene_gn=ATP10D;refGene_tx=XM_019829544.2;refGene_gn=ATP10D;refGene_tx=XM_023253139.1
B1 167874802 167874991
refGene_gn=ATP10D;refGene_tx=XM_003985402.5;refGene_gn=ATP10D;refGene_tx=XM_019829544.2;refGene_gn=ATP10D;refGene_tx=XM_023253139.1
B1 167882126 167882365
refGene_gn=ATP10D;refGene_tx=XM_003985402.5;refGene_gn=ATP10D;refGene_tx=XM_019829544.2;refGene_gn=ATP10D;refGene_tx=XM_023253139.1
B1 167889893 167890146
refGene_gn=ATP10D;refGene_tx=XM_003985402.5;refGene_gn=ATP10D;refGene_tx=XM_019829544.2;refGene_gn=ATP10D;refGene_tx=XM_023253139.1
B1 167890267 167890395
refGene_gn=ATP10D;refGene_tx=XM_003985402.5;refGene_gn=ATP10D;refGene_tx=XM_019829544.2;refGene_gn=ATP10D;refGene_tx=XM_023253139.1
B1 167890798 167890930
refGene_gn=ATP10D;refGene_tx=XM_003985402.5;refGene_gn=ATP10D;refGene_tx=XM_019829544.2
B1 167891210 167891317
refGene_gn=ATP10D;refGene_tx=XM_003985402.5;refGene_gn=ATP10D;refGene_tx=XM_019829544.2;refGene_gn=ATP10D;refGene_tx=XM_023253139.1
B1 167898836 167898922
refGene_gn=ATP10D;refGene_tx=XM_003985402.5;refGene_gn=ATP10D;refGene_tx=XM_019829544.2;refGene_gn=ATP10D;refGene_tx=XM_023253139.1
B1 167900913 167901118
refGene_gn=ATP10D;refGene_tx=XM_003985402.5;refGene_gn=ATP10D;refGene_tx=XM_019829544.2;refGene_gn=ATP10D;refGene_tx=XM_023253139.1
B1 167918049 167918244
refGene_gn=ATP10D;refGene_tx=XM_003985402.5;refGene_gn=ATP10D;refGene_tx=XM_019829544.2;refGene_gn=ATP10D;refGene_tx=XM_023253139.1
B1 167920381 167920671
refGene_gn=ATP10D;refGene_tx=XM_003985402.5;refGene_gn=ATP10D;refGene_tx=XM_023253139.1
B1 167968904 167968970
refGene_gn=COMMD8;refGene_tx=XM_023253141.1
B1 167971735 167971765

refGene_gn=COMMD8;refGene_tx=XM_023253142.1
B1 167973666 167973822
refGene_gn=COMMD8;refGene_tx=XM_023253141.1;refGene_gn=COMMD8;refGene_tx=XM_023253142.1
B1 167977926 167978079
refGene_gn=COMMD8;refGene_tx=XM_023253141.1;refGene_gn=COMMD8;refGene_tx=XM_023253142.1
B1 167981105 167981261
refGene_gn=COMMD8;refGene_tx=XM_023253141.1;refGene_gn=COMMD8;refGene_tx=XM_023253142.1
B1 167982324 167982345
refGene_gn=COMMD8;refGene_tx=XM_023253141.1;refGene_gn=COMMD8;refGene_tx=XM_023253142.1
B1 168001960 168002311
refGene_gn=GABRB1;refGene_tx=XM_011281892.3
B1 168019327 168019572
refGene_gn=GABRB1;refGene_tx=XM_011281892.3
B1 168023127 168023280
refGene_gn=GABRB1;refGene_tx=XM_011281892.3
B1 168023388 168023526
refGene_gn=GABRB1;refGene_tx=XM_011281892.3
B1 168103317 168103400
refGene_gn=GABRB1;refGene_tx=XM_011281892.3
B1 168243584 168243805
refGene_gn=GABRB1;refGene_tx=XM_011281892.3
B1 168381963 168382031
refGene_gn=GABRB1;refGene_tx=XM_011281892.3
B1 168382454 168382546
refGene_gn=GABRB1;refGene_tx=XM_011281892.3
B1 168382712 168382792
refGene_gn=GABRB1;refGene_tx=XM_011281892.3
B1 168415146 168415232
refGene_gn=GABRA4;refGene_tx=XM_003985405.5
B1 168415655 168415774
refGene_gn=GABRA4;refGene_tx=XM_003985405.5
B1 168432134 168432202
refGene_gn=GABRA4;refGene_tx=XM_003985405.5
B1 168433214 168433435
refGene_gn=GABRA4;refGene_tx=XM_003985405.5
B1 168433609 168433692
refGene_gn=GABRA4;refGene_tx=XM_003985405.5
B1 168436237 168436381
refGene_gn=GABRA4;refGene_tx=XM_003985405.5
B1 168437128 168437281
refGene_gn=GABRA4;refGene_tx=XM_003985405.5
B1 168443604 168443864
refGene_gn=GABRA4;refGene_tx=XM_003985405.5
B1 168481060 168481591
refGene_gn=GABRA4;refGene_tx=XM_003985405.5
B1 168654390 168654630

refGene_gn=LOC101098629;refGene_tx=XM_006931136.3;refGene_gn=LOC101098629;refGene_tx=XM_019829553.2;refGene_gn=LOC101098629;refGene_tx=XM_019829554.2;refGene_gn=LOC101098629;refGene_tx=XM_023253143.1;refGene_gn=LOC101098629;refGene_tx=XM_023253144.1;refGene_gn=LOC101098629;refGene_tx=XM_023253145.1
B1 168921535 168921606
refGene_gn=GABRA2;refGene_tx=XM_011281893.3;refGene_gn=GABRA2;refGene_tx=XM_019829558.2
B1 168924234 168924350
refGene_gn=GABRA2;refGene_tx=XM_011281893.3;refGene_gn=GABRA2;refGene_tx=XM_019829558.2
B1 168969875 168969943
refGene_gn=GABRA2;refGene_tx=XM_011281893.3;refGene_gn=GABRA2;refGene_tx=XM_019829558.2;refGene_gn=GABRA2;refGene_tx=XM_023253146.1
B1 168990949 168991170
refGene_gn=GABRA2;refGene_tx=XM_011281893.3;refGene_gn=GABRA2;refGene_tx=XM_019829558.2;refGene_gn=GABRA2;refGene_tx=XM_023253146.1
B1 168993812 168993895
refGene_gn=GABRA2;refGene_tx=XM_011281893.3;refGene_gn=GABRA2;refGene_tx=XM_019829558.2;refGene_gn=GABRA2;refGene_tx=XM_023253146.1
B1 168998408 168998552
refGene_gn=GABRA2;refGene_tx=XM_011281893.3;refGene_gn=GABRA2;refGene_tx=XM_019829558.2;refGene_gn=GABRA2;refGene_tx=XM_023253146.1
B1 169000077 169000230
refGene_gn=GABRA2;refGene_tx=XM_011281893.3;refGene_gn=GABRA2;refGene_tx=XM_019829558.2;refGene_gn=GABRA2;refGene_tx=XM_023253146.1
B1 169029455 169029658
refGene_gn=GABRA2;refGene_tx=XM_011281893.3;refGene_gn=GABRA2;refGene_tx=XM_019829558.2;refGene_gn=GABRA2;refGene_tx=XM_023253146.1
B1 169035729 169035909
refGene_gn=GABRA2;refGene_tx=XM_011281893.3;refGene_gn=GABRA2;refGene_tx=XM_023253146.1
B1 169041139 169041436
refGene_gn=GABRA2;refGene_tx=XM_011281893.3;refGene_gn=GABRA2;refGene_tx=XM_019829558.2;refGene_gn=GABRA2;refGene_tx=XM_023253146.1
B1 169144518 169144622
refGene_gn=GABRG1;refGene_tx=XM_003985410.5
B1 169157653 169157802
refGene_gn=GABRG1;refGene_tx=XM_003985410.5
B1 169169086 169169154
refGene_gn=GABRG1;refGene_tx=XM_003985410.5
B1 169190368 169190589
refGene_gn=GABRG1;refGene_tx=XM_003985410.5
B1 169191468 169191551
refGene_gn=GABRG1;refGene_tx=XM_003985410.5
B1 169194698 169194836
refGene_gn=GABRG1;refGene_tx=XM_003985410.5
B1 169194958 169195111
refGene_gn=GABRG1;refGene_tx=XM_003985410.5
B1 169199883 169200095

refGene_gn=GABRG1;refGene_tx=XM_003985410.5
B1 169210622 169210892
refGene_gn=GABRG1;refGene_tx=XM_003985410.5
B1 170297046 170297170
refGene_gn=GNPDA2;refGene_tx=XM_003985411.4
B1 170302107 170302209
refGene_gn=GNPDA2;refGene_tx=XM_003985411.4
B1 170303689 170303872
refGene_gn=GNPDA2;refGene_tx=XM_003985411.4
B1 170309781 170309966
refGene_gn=GNPDA2;refGene_tx=XM_003985411.4
B1 170315839 170316014
refGene_gn=GNPDA2;refGene_tx=XM_003985411.4
B1 170321493 170321555
refGene_gn=GNPDA2;refGene_tx=XM_003985411.4
B1 170325405 170325543
refGene_gn=GUF1;refGene_tx=XM_011281894.3;refGene_gn=GUF1;refGene_tx=XM_011281895.3;refGene_gn=GUF1;refGene_tx=XM_019829559.2
B1 170326548 170326585
refGene_gn=GUF1;refGene_tx=XM_011281894.3;refGene_gn=GUF1;refGene_tx=XM_011281895.3;refGene_gn=GUF1;refGene_tx=XM_019829559.2
B1 170328172 170328292
refGene_gn=GUF1;refGene_tx=XM_011281894.3;refGene_gn=GUF1;refGene_tx=XM_011281895.3;refGene_gn=GUF1;refGene_tx=XM_019829559.2
B1 170329707 170329809
refGene_gn=GUF1;refGene_tx=XM_011281894.3;refGene_gn=GUF1;refGene_tx=XM_011281895.3;refGene_gn=GUF1;refGene_tx=XM_019829559.2
B1 170332827 170332961
refGene_gn=GUF1;refGene_tx=XM_011281894.3;refGene_gn=GUF1;refGene_tx=XM_011281895.3;refGene_gn=GUF1;refGene_tx=XM_019829559.2
B1 170333858 170334002
refGene_gn=GUF1;refGene_tx=XM_011281894.3;refGene_gn=GUF1;refGene_tx=XM_011281895.3;refGene_gn=GUF1;refGene_tx=XM_019829559.2
B1 170334823 170334956
refGene_gn=GUF1;refGene_tx=XM_011281894.3;refGene_gn=GUF1;refGene_tx=XM_011281895.3;refGene_gn=GUF1;refGene_tx=XM_019829559.2
B1 170335310 170335434
refGene_gn=GUF1;refGene_tx=XM_011281894.3;refGene_gn=GUF1;refGene_tx=XM_011281895.3;refGene_gn=GUF1;refGene_tx=XM_019829559.2
B1 170336939 170337079
refGene_gn=GUF1;refGene_tx=XM_011281894.3;refGene_gn=GUF1;refGene_tx=XM_011281895.3;refGene_gn=GUF1;refGene_tx=XM_019829559.2
B1 170338933 170339137
refGene_gn=GUF1;refGene_tx=XM_011281894.3;refGene_gn=GUF1;refGene_tx=XM_011281895.3;refGene_gn=GUF1;refGene_tx=XM_019829559.2
B1 170340143 170340208
refGene_gn=GUF1;refGene_tx=XM_011281894.3;refGene_gn=GUF1;refGene_tx=XM_011281895.3;refGene_gn=GUF1;refGene_tx=XM_019829559.2
B1 170342031 170342115
refGene_gn=GUF1;refGene_tx=XM_011281894.3;refGene_gn=GUF1;refGene_tx=X

M_011281895.3;refGene_gn=GUF1;refGene_tx=XM_019829559.2
B1 170343159 170343237
refGene_gn=GUF1;refGene_tx=XM_011281894.3;refGene_gn=GUF1;refGene_tx=XM_011281895.3;refGene_gn=GUF1;refGene_tx=XM_019829559.2
B1 170344108 170344189
refGene_gn=GUF1;refGene_tx=XM_011281894.3;refGene_gn=GUF1;refGene_tx=XM_011281895.3;refGene_gn=GUF1;refGene_tx=XM_019829559.2
B1 170344459 170344608
refGene_gn=GUF1;refGene_tx=XM_011281894.3;refGene_gn=GUF1;refGene_tx=XM_011281895.3;refGene_gn=GUF1;refGene_tx=XM_019829559.2
B1 170344951 170345063
refGene_gn=GUF1;refGene_tx=XM_011281894.3;refGene_gn=GUF1;refGene_tx=XM_011281895.3;refGene_gn=GUF1;refGene_tx=XM_019829559.2
B1 170346520 170346685
refGene_gn=GUF1;refGene_tx=XM_011281894.3
B1 170368487 170368603
refGene_gn=YIPF7;refGene_tx=XM_023253147.1
B1 170377551 170377712
refGene_gn=YIPF7;refGene_tx=XM_023253147.1
B1 170385233 170385379
refGene_gn=YIPF7;refGene_tx=XM_023253147.1
B1 170390005 170390187
refGene_gn=YIPF7;refGene_tx=XM_023253147.1
B1 170393621 170393784
refGene_gn=YIPF7;refGene_tx=XM_023253147.1
B1 170589686 170590656
refGene_gn=KCTD8;refGene_tx=XM_023253149.1;refGene_gn=KCTD8;refGene_tx=XM_023253150.1
B1 170845390 170845851
refGene_gn=KCTD8;refGene_tx=XM_023253149.1
B1 171752055 171752235
refGene_gn=GRXCR1;refGene_tx=XM_003985413.3;refGene_gn=GRXCR1;refGene_tx=XM_019829568.2
B1 171761382 171761448
refGene_gn=GRXCR1;refGene_tx=XM_003985413.3;refGene_gn=GRXCR1;refGene_tx=XM_019829568.2
B1 171762959 171763036
refGene_gn=TRIM61;refGene_tx=XM_019829569.1
B1 171763037 171763236
refGene_gn=TRIM61;refGene_tx=XM_019829569.1
B1 171763237 171763290
refGene_gn=TRIM61;refGene_tx=XM_019829569.1
B1 171763291 171763475
refGene_gn=TRIM61;refGene_tx=XM_019829569.1
B1 171810775 171811018
refGene_gn=GRXCR1;refGene_tx=XM_003985413.3;refGene_gn=GRXCR1;refGene_tx=XM_019829568.2
B1 171878098 171878608
refGene_gn=GRXCR1;refGene_tx=XM_003985413.3
B1 172060248 172060318

refGene_gn=GRXCR1;refGene_tx=XM_019829568.2
B1 172061123 172061395
refGene_gn=GRXCR1;refGene_tx=XM_019829568.2
B1 172062097 172062146
refGene_gn=ATP8A1;refGene_tx=XM_023253151.1
B1 172084336 172084400
refGene_gn=ATP8A1;refGene_tx=XM_019829562.2;refGene_gn=ATP8A1;refGene_tx=XM_019829563.2;refGene_gn=ATP8A1;refGene_tx=XM_019829564.2
B1 172096671 172096786
refGene_gn=ATP8A1;refGene_tx=XM_019829562.2;refGene_gn=ATP8A1;refGene_tx=XM_019829563.2;refGene_gn=ATP8A1;refGene_tx=XM_019829564.2;refGene_gn=ATP8A1;refGene_tx=XM_023253151.1
B1 172098023 172098123
refGene_gn=ATP8A1;refGene_tx=XM_019829562.2;refGene_gn=ATP8A1;refGene_tx=XM_019829563.2;refGene_gn=ATP8A1;refGene_tx=XM_019829564.2;refGene_gn=ATP8A1;refGene_tx=XM_023253151.1
B1 172099286 172099385
refGene_gn=ATP8A1;refGene_tx=XM_019829562.2;refGene_gn=ATP8A1;refGene_tx=XM_019829563.2;refGene_gn=ATP8A1;refGene_tx=XM_019829564.2;refGene_gn=ATP8A1;refGene_tx=XM_023253151.1
B1 172105880 172105926
refGene_gn=ATP8A1;refGene_tx=XM_019829562.2;refGene_gn=ATP8A1;refGene_tx=XM_019829563.2;refGene_gn=ATP8A1;refGene_tx=XM_019829564.2;refGene_gn=ATP8A1;refGene_tx=XM_023253151.1
B1 172121537 172121578
refGene_gn=ATP8A1;refGene_tx=XM_019829562.2;refGene_gn=ATP8A1;refGene_tx=XM_019829563.2;refGene_gn=ATP8A1;refGene_tx=XM_019829564.2;refGene_gn=ATP8A1;refGene_tx=XM_019829566.2;refGene_gn=ATP8A1;refGene_tx=XM_023253151.1
B1 172127620 172127694
refGene_gn=ATP8A1;refGene_tx=XM_019829562.2;refGene_gn=ATP8A1;refGene_tx=XM_019829564.2;refGene_gn=ATP8A1;refGene_tx=XM_023253151.1
B1 172131011 172131085
refGene_gn=ATP8A1;refGene_tx=XM_019829563.2
B1 172133602 172133672
refGene_gn=ATP8A1;refGene_tx=XM_019829562.2;refGene_gn=ATP8A1;refGene_tx=XM_019829563.2;refGene_gn=ATP8A1;refGene_tx=XM_019829564.2;refGene_gn=ATP8A1;refGene_tx=XM_019829566.2;refGene_gn=ATP8A1;refGene_tx=XM_019829567.2;refGene_gn=ATP8A1;refGene_tx=XM_023253151.1
B1 172134727 172134855
refGene_gn=ATP8A1;refGene_tx=XM_019829562.2;refGene_gn=ATP8A1;refGene_tx=XM_019829563.2;refGene_gn=ATP8A1;refGene_tx=XM_019829564.2;refGene_gn=ATP8A1;refGene_tx=XM_019829566.2;refGene_gn=ATP8A1;refGene_tx=XM_019829567.2;refGene_gn=ATP8A1;refGene_tx=XM_023253151.1
B1 172139249 172139361
refGene_gn=ATP8A1;refGene_tx=XM_019829562.2;refGene_gn=ATP8A1;refGene_tx=XM_019829563.2;refGene_gn=ATP8A1;refGene_tx=XM_019829564.2;refGene_gn=ATP8A1;refGene_tx=XM_019829566.2;refGene_gn=ATP8A1;refGene_tx=XM_019829567.2;refGene_gn=ATP8A1;refGene_tx=XM_023253151.1
B1 172140639 172140805

refGene_gn=ATP8A1;refGene_tx=XM_019829562.2;refGene_gn=ATP8A1;refGene_tx=XM_019829563.2;refGene_gn=ATP8A1;refGene_tx=XM_019829564.2;refGene_gn=ATP8A1;refGene_tx=XM_019829566.2;refGene_gn=ATP8A1;refGene_tx=XM_019829567.2;refGene_gn=ATP8A1;refGene_tx=XM_023253151.1
B1 172142064 172142192
refGene_gn=ATP8A1;refGene_tx=XM_019829562.2;refGene_gn=ATP8A1;refGene_tx=XM_019829563.2;refGene_gn=ATP8A1;refGene_tx=XM_019829564.2;refGene_gn=ATP8A1;refGene_tx=XM_019829566.2;refGene_gn=ATP8A1;refGene_tx=XM_019829567.2;refGene_gn=ATP8A1;refGene_tx=XM_023253151.1
B1 172144796 172144874
refGene_gn=ATP8A1;refGene_tx=XM_019829562.2;refGene_gn=ATP8A1;refGene_tx=XM_019829563.2;refGene_gn=ATP8A1;refGene_tx=XM_019829564.2;refGene_gn=ATP8A1;refGene_tx=XM_019829566.2;refGene_gn=ATP8A1;refGene_tx=XM_019829567.2;refGene_gn=ATP8A1;refGene_tx=XM_023253151.1
B1 172146079 172146168
refGene_gn=ATP8A1;refGene_tx=XM_019829562.2;refGene_gn=ATP8A1;refGene_tx=XM_019829563.2;refGene_gn=ATP8A1;refGene_tx=XM_019829564.2;refGene_gn=ATP8A1;refGene_tx=XM_019829566.2;refGene_gn=ATP8A1;refGene_tx=XM_019829567.2;refGene_gn=ATP8A1;refGene_tx=XM_023253151.1
B1 172152371 172152416
refGene_gn=ATP8A1;refGene_tx=XM_019829562.2;refGene_gn=ATP8A1;refGene_tx=XM_019829563.2;refGene_gn=ATP8A1;refGene_tx=XM_019829566.2;refGene_gn=ATP8A1;refGene_tx=XM_019829567.2;refGene_gn=ATP8A1;refGene_tx=XM_023253151.1
B1 172161703 172161776
refGene_gn=ATP8A1;refGene_tx=XM_019829562.2;refGene_gn=ATP8A1;refGene_tx=XM_019829563.2;refGene_gn=ATP8A1;refGene_tx=XM_019829564.2;refGene_gn=ATP8A1;refGene_tx=XM_019829566.2;refGene_gn=ATP8A1;refGene_tx=XM_019829567.2;refGene_gn=ATP8A1;refGene_tx=XM_023253151.1
B1 172163869 172163975
refGene_gn=ATP8A1;refGene_tx=XM_019829562.2;refGene_gn=ATP8A1;refGene_tx=XM_019829563.2;refGene_gn=ATP8A1;refGene_tx=XM_019829564.2;refGene_gn=ATP8A1;refGene_tx=XM_019829566.2;refGene_gn=ATP8A1;refGene_tx=XM_019829567.2;refGene_gn=ATP8A1;refGene_tx=XM_023253151.1
B1 172165397 172165480
refGene_gn=ATP8A1;refGene_tx=XM_019829562.2;refGene_gn=ATP8A1;refGene_tx=XM_019829563.2;refGene_gn=ATP8A1;refGene_tx=XM_019829564.2;refGene_gn=ATP8A1;refGene_tx=XM_019829566.2;refGene_gn=ATP8A1;refGene_tx=XM_019829567.2;refGene_gn=ATP8A1;refGene_tx=XM_023253151.1
B1 172170139 172170189
refGene_gn=ATP8A1;refGene_tx=XM_019829562.2;refGene_gn=ATP8A1;refGene_tx=XM_019829563.2;refGene_gn=ATP8A1;refGene_tx=XM_019829564.2;refGene_gn=ATP8A1;refGene_tx=XM_019829566.2;refGene_gn=ATP8A1;refGene_tx=XM_019829567.2;refGene_gn=ATP8A1;refGene_tx=XM_023253151.1
B1 172174562 172174632
refGene_gn=ATP8A1;refGene_tx=XM_019829562.2;refGene_gn=ATP8A1;refGene_tx=XM_019829563.2;refGene_gn=ATP8A1;refGene_tx=XM_019829564.2;refGene_gn=ATP8A1;refGene_tx=XM_019829566.2;refGene_gn=ATP8A1;refGene_tx=XM_019829567.2;refGene_gn=ATP8A1;refGene_tx=XM_023253151.1
B1 172179914 172179999

refGene_gn=ATP8A1;refGene_tx=XM_019829562.2;refGene_gn=ATP8A1;refGene_tx=XM_019829563.2;refGene_gn=ATP8A1;refGene_tx=XM_019829564.2;refGene_gn=ATP8A1;refGene_tx=XM_019829566.2;refGene_gn=ATP8A1;refGene_tx=XM_019829567.2;refGene_gn=ATP8A1;refGene_tx=XM_023253151.1
B1 172182423 172182563
refGene_gn=ATP8A1;refGene_tx=XM_019829562.2;refGene_gn=ATP8A1;refGene_tx=XM_019829563.2;refGene_gn=ATP8A1;refGene_tx=XM_019829564.2;refGene_gn=ATP8A1;refGene_tx=XM_019829566.2;refGene_gn=ATP8A1;refGene_tx=XM_019829567.2;refGene_gn=ATP8A1;refGene_tx=XM_023253151.1
B1 172196894 172197033
refGene_gn=ATP8A1;refGene_tx=XM_019829562.2;refGene_gn=ATP8A1;refGene_tx=XM_019829563.2;refGene_gn=ATP8A1;refGene_tx=XM_019829564.2;refGene_gn=ATP8A1;refGene_tx=XM_019829566.2;refGene_gn=ATP8A1;refGene_tx=XM_019829567.2;refGene_gn=ATP8A1;refGene_tx=XM_023253151.1
B1 172200575 172200640
refGene_gn=ATP8A1;refGene_tx=XM_019829562.2;refGene_gn=ATP8A1;refGene_tx=XM_019829563.2;refGene_gn=ATP8A1;refGene_tx=XM_019829564.2;refGene_gn=ATP8A1;refGene_tx=XM_019829566.2;refGene_gn=ATP8A1;refGene_tx=XM_019829567.2;refGene_gn=ATP8A1;refGene_tx=XM_023253151.1
B1 172219245 172219418
refGene_gn=ATP8A1;refGene_tx=XM_019829562.2;refGene_gn=ATP8A1;refGene_tx=XM_019829563.2;refGene_gn=ATP8A1;refGene_tx=XM_019829564.2;refGene_gn=ATP8A1;refGene_tx=XM_019829566.2;refGene_gn=ATP8A1;refGene_tx=XM_019829567.2;refGene_gn=ATP8A1;refGene_tx=XM_023253151.1
B1 172238312 172238496
refGene_gn=ATP8A1;refGene_tx=XM_019829562.2;refGene_gn=ATP8A1;refGene_tx=XM_019829563.2;refGene_gn=ATP8A1;refGene_tx=XM_019829564.2;refGene_gn=ATP8A1;refGene_tx=XM_019829566.2;refGene_gn=ATP8A1;refGene_tx=XM_019829567.2;refGene_gn=ATP8A1;refGene_tx=XM_023253151.1
B1 172238589 172238700
refGene_gn=ATP8A1;refGene_tx=XM_019829562.2;refGene_gn=ATP8A1;refGene_tx=XM_019829563.2;refGene_gn=ATP8A1;refGene_tx=XM_019829564.2;refGene_gn=ATP8A1;refGene_tx=XM_019829566.2;refGene_gn=ATP8A1;refGene_tx=XM_019829567.2;refGene_gn=ATP8A1;refGene_tx=XM_023253151.1
B1 172245281 172245356
refGene_gn=ATP8A1;refGene_tx=XM_019829562.2;refGene_gn=ATP8A1;refGene_tx=XM_019829563.2;refGene_gn=ATP8A1;refGene_tx=XM_019829564.2;refGene_gn=ATP8A1;refGene_tx=XM_019829566.2;refGene_gn=ATP8A1;refGene_tx=XM_019829567.2;refGene_gn=ATP8A1;refGene_tx=XM_023253151.1
B1 172245460 172245583
refGene_gn=ATP8A1;refGene_tx=XM_019829562.2;refGene_gn=ATP8A1;refGene_tx=XM_019829563.2;refGene_gn=ATP8A1;refGene_tx=XM_019829564.2;refGene_gn=ATP8A1;refGene_tx=XM_019829566.2;refGene_gn=ATP8A1;refGene_tx=XM_019829567.2;refGene_gn=ATP8A1;refGene_tx=XM_023253151.1
B1 172249068 172249147
refGene_gn=ATP8A1;refGene_tx=XM_019829562.2;refGene_gn=ATP8A1;refGene_tx=XM_019829563.2;refGene_gn=ATP8A1;refGene_tx=XM_019829564.2;refGene_gn=ATP8A1;refGene_tx=XM_019829566.2;refGene_gn=ATP8A1;refGene_tx=XM_019829567.2;refGene_gn=ATP8A1;refGene_tx=XM_023253151.1
B1 172254667 172254729

refGene_gn=ATP8A1;refGene_tx=XM_019829562.2;refGene_gn=ATP8A1;refGene_tx=XM_019829563.2;refGene_gn=ATP8A1;refGene_tx=XM_019829564.2;refGene_gn=ATP8A1;refGene_tx=XM_019829566.2;refGene_gn=ATP8A1;refGene_tx=XM_019829567.2;refGene_gn=ATP8A1;refGene_tx=XM_023253151.1
B1 172257864 172257921
refGene_gn=ATP8A1;refGene_tx=XM_019829562.2;refGene_gn=ATP8A1;refGene_tx=XM_019829563.2;refGene_gn=ATP8A1;refGene_tx=XM_019829564.2;refGene_gn=ATP8A1;refGene_tx=XM_019829566.2;refGene_gn=ATP8A1;refGene_tx=XM_019829567.2;refGene_gn=ATP8A1;refGene_tx=XM_023253151.1
B1 172258848 172258956
refGene_gn=ATP8A1;refGene_tx=XM_019829562.2;refGene_gn=ATP8A1;refGene_tx=XM_019829563.2;refGene_gn=ATP8A1;refGene_tx=XM_019829564.2;refGene_gn=ATP8A1;refGene_tx=XM_019829566.2;refGene_gn=ATP8A1;refGene_tx=XM_019829567.2;refGene_gn=ATP8A1;refGene_tx=XM_023253151.1
B1 172279402 172279491
refGene_gn=ATP8A1;refGene_tx=XM_019829562.2;refGene_gn=ATP8A1;refGene_tx=XM_019829563.2;refGene_gn=ATP8A1;refGene_tx=XM_019829564.2;refGene_gn=ATP8A1;refGene_tx=XM_019829566.2;refGene_gn=ATP8A1;refGene_tx=XM_019829567.2;refGene_gn=ATP8A1;refGene_tx=XM_023253151.1
B1 172280212 172280305
refGene_gn=ATP8A1;refGene_tx=XM_019829562.2;refGene_gn=ATP8A1;refGene_tx=XM_019829563.2;refGene_gn=ATP8A1;refGene_tx=XM_019829564.2;refGene_gn=ATP8A1;refGene_tx=XM_019829566.2;refGene_gn=ATP8A1;refGene_tx=XM_019829567.2;refGene_gn=ATP8A1;refGene_tx=XM_023253151.1
B1 172287864 172287956
refGene_gn=ATP8A1;refGene_tx=XM_019829562.2;refGene_gn=ATP8A1;refGene_tx=XM_019829563.2;refGene_gn=ATP8A1;refGene_tx=XM_019829564.2;refGene_gn=ATP8A1;refGene_tx=XM_019829566.2;refGene_gn=ATP8A1;refGene_tx=XM_019829567.2;refGene_gn=ATP8A1;refGene_tx=XM_023253151.1
B1 172289141 172289239
refGene_gn=ATP8A1;refGene_tx=XM_019829562.2;refGene_gn=ATP8A1;refGene_tx=XM_019829563.2;refGene_gn=ATP8A1;refGene_tx=XM_019829564.2;refGene_gn=ATP8A1;refGene_tx=XM_019829566.2;refGene_gn=ATP8A1;refGene_tx=XM_019829567.2;refGene_gn=ATP8A1;refGene_tx=XM_023253151.1
B1 172301130 172301570
refGene_gn=SHISA3;refGene_tx=XM_023253152.1
B1 172304893 172305458
refGene_gn=SHISA3;refGene_tx=XM_023253152.1
B1 172498565 172499055
refGene_gn=BEND4;refGene_tx=XM_011281896.3;refGene_gn=BEND4;refGene_tx=XM_023253153.1;refGene_gn=BEND4;refGene_tx=XM_023253154.1;refGene_gn=BEND4;refGene_tx=XM_023253155.1
B1 172504822 172504960
refGene_gn=BEND4;refGene_tx=XM_011281896.3;refGene_gn=BEND4;refGene_tx=XM_023253154.1;refGene_gn=BEND4;refGene_tx=XM_023253155.1
B1 172506450 172507017
refGene_gn=BEND4;refGene_tx=XM_011281896.3;refGene_gn=BEND4;refGene_tx=XM_023253153.1;refGene_gn=BEND4;refGene_tx=XM_023253154.1;refGene_gn=BEND4;refGene_tx=XM_023253155.1
B1 172519956 172520048

refGene_gn=BEND4;refGene_tx=XM_011281896.3;refGene_gn=BEND4;refGene_tx=XM_023253153.1;refGene_gn=BEND4;refGene_tx=XM_023253154.1;refGene_gn=BEND4;refGene_tx=XM_023253155.1
B1 172526686 172526927
refGene_gn=BEND4;refGene_tx=XM_011281896.3;refGene_gn=BEND4;refGene_tx=XM_023253153.1;refGene_gn=BEND4;refGene_tx=XM_023253154.1
B1 172530378 172530596
refGene_gn=BEND4;refGene_tx=XM_011281896.3;refGene_gn=BEND4;refGene_tx=XM_023253153.1;refGene_gn=BEND4;refGene_tx=XM_023253154.1;refGene_gn=BEND4;refGene_tx=XM_023253155.1
B1 172566225 172566270
refGene_gn=SLC30A9;refGene_tx=XM_023253156.1;refGene_gn=SLC30A9;refGene_tx=XM_023253157.1
B1 172573522 172573636
refGene_gn=SLC30A9;refGene_tx=XM_023253156.1;refGene_gn=SLC30A9;refGene_tx=XM_023253157.1
B1 172575477 172575607
refGene_gn=SLC30A9;refGene_tx=XM_023253156.1;refGene_gn=SLC30A9;refGene_tx=XM_023253157.1
B1 172577979 172578145
refGene_gn=SLC30A9;refGene_tx=XM_023253156.1;refGene_gn=SLC30A9;refGene_tx=XM_023253157.1
B1 172580743 172580851
refGene_gn=SLC30A9;refGene_tx=XM_023253156.1;refGene_gn=SLC30A9;refGene_tx=XM_023253157.1
B1 172581317 172581389
refGene_gn=SLC30A9;refGene_tx=XM_023253156.1;refGene_gn=SLC30A9;refGene_tx=XM_023253157.1
B1 172582208 172582248
refGene_gn=SLC30A9;refGene_tx=XM_023253156.1;refGene_gn=SLC30A9;refGene_tx=XM_023253157.1
B1 172584820 172584956
refGene_gn=SLC30A9;refGene_tx=XM_023253156.1;refGene_gn=SLC30A9;refGene_tx=XM_023253157.1
B1 172587190 172587246
refGene_gn=SLC30A9;refGene_tx=XM_023253156.1;refGene_gn=SLC30A9;refGene_tx=XM_023253157.1
B1 172596610 172596713
refGene_gn=SLC30A9;refGene_tx=XM_023253156.1;refGene_gn=SLC30A9;refGene_tx=XM_023253157.1
B1 172599647 172599715
refGene_gn=SLC30A9;refGene_tx=XM_023253156.1;refGene_gn=SLC30A9;refGene_tx=XM_023253157.1
B1 172606050 172606109
refGene_gn=SLC30A9;refGene_tx=XM_023253156.1;refGene_gn=SLC30A9;refGene_tx=XM_023253157.1
B1 172611608 172611691
refGene_gn=SLC30A9;refGene_tx=XM_023253156.1;refGene_gn=SLC30A9;refGene_tx=XM_023253157.1
B1 172612073 172612166

refGene_gn=SLC30A9;refGene_tx=XM_023253156.1;refGene_gn=SLC30A9;refGene_tx=XM_023253157.1
B1 172614167 172614267
refGene_gn=SLC30A9;refGene_tx=XM_023253156.1;refGene_gn=SLC30A9;refGene_tx=XM_023253157.1
B1 172616529 172616592
refGene_gn=SLC30A9;refGene_tx=XM_023253156.1;refGene_gn=SLC30A9;refGene_tx=XM_023253157.1
B1 172629005 172629173
refGene_gn=SLC30A9;refGene_tx=XM_023253156.1;refGene_gn=SLC30A9;refGene_tx=XM_023253157.1
B1 172650069 172650181
refGene_gn=SLC30A9;refGene_tx=XM_023253156.1;refGene_gn=SLC30A9;refGene_tx=XM_023253157.1
B1 172673292 172673422
refGene_gn=TMEM33;refGene_tx=XM_023253159.1;refGene_gn=TMEM33;refGene_tx=XM_023253160.1;refGene_gn=TMEM33;refGene_tx=XM_023253161.1
B1 172679214 172679298
refGene_gn=TMEM33;refGene_tx=XM_023253159.1;refGene_gn=TMEM33;refGene_tx=XM_023253160.1;refGene_gn=TMEM33;refGene_tx=XM_023253161.1
B1 172683422 172683556
refGene_gn=TMEM33;refGene_tx=XM_023253159.1;refGene_gn=TMEM33;refGene_tx=XM_023253160.1;refGene_gn=TMEM33;refGene_tx=XM_023253161.1
B1 172684974 172685042
refGene_gn=TMEM33;refGene_tx=XM_023253159.1;refGene_gn=TMEM33;refGene_tx=XM_023253160.1;refGene_gn=TMEM33;refGene_tx=XM_023253161.1
B1 172688833 172689021
refGene_gn=TMEM33;refGene_tx=XM_023253159.1;refGene_gn=TMEM33;refGene_tx=XM_023253160.1;refGene_gn=TMEM33;refGene_tx=XM_023253161.1
B1 172689495 172689590
refGene_gn=TMEM33;refGene_tx=XM_023253159.1;refGene_gn=TMEM33;refGene_tx=XM_023253160.1;refGene_gn=TMEM33;refGene_tx=XM_023253161.1
B1 172693152 172693197
refGene_gn=TMEM33;refGene_tx=XM_023253159.1;refGene_gn=TMEM33;refGene_tx=XM_023253160.1
B1 172854507 172854748
refGene_gn=PHOX2B;refGene_tx=XM_023252239.1
B1 172855595 172855783
refGene_gn=PHOX2B;refGene_tx=XM_023252239.1
B1 172856777 172857296
refGene_gn=PHOX2B;refGene_tx=XM_023252239.1
B1 172919277 172919303
refGene_gn=LIMCH1;refGene_tx=XM_019829582.2;refGene_gn=LIMCH1;refGene_tx=XM_019829583.2;refGene_gn=LIMCH1;refGene_tx=XM_019829584.2;refGene_gn=LIMCH1;refGene_tx=XM_019829585.2;refGene_gn=LIMCH1;refGene_tx=XM_019829586.2;refGene_gn=LIMCH1;refGene_tx=XM_023253162.1;refGene_gn=LIMCH1;refGene_tx=XM_023253163.1;refGene_gn=LIMCH1;refGene_tx=XM_023253165.1;refGene_gn=LIMCH1;refGene_tx=XM_023253166.1;refGene_gn=LIMCH1;refGene_tx=XM_023253167.1;refGene_gn=LIMCH1;refGene_tx=XM_023253168.1;refGene_gn=LIMCH1;refGene_tx=XM_023253169.1;refGene_gn=LIMCH1;refGene_tx=XM_

023253170.1;refGene_gn=LIMCH1;refGene_tx=XM_023253171.1
B1 172924917 172925020
refGene_gn=LIMCH1;refGene_tx=XM_019829582.2;refGene_gn=LIMCH1;refGene_tx=XM_019829583.2;refGene_gn=LIMCH1;refGene_tx=XM_019829584.2;refGene_gn=LIMCH1;refGene_tx=XM_019829585.2;refGene_gn=LIMCH1;refGene_tx=XM_019829586.2;refGene_gn=LIMCH1;refGene_tx=XM_023253162.1;refGene_gn=LIMCH1;refGene_tx=XM_023253163.1;refGene_gn=LIMCH1;refGene_tx=XM_023253165.1;refGene_gn=LIMCH1;refGene_tx=XM_023253166.1;refGene_gn=LIMCH1;refGene_tx=XM_023253167.1;refGene_gn=LIMCH1;refGene_tx=XM_023253168.1;refGene_gn=LIMCH1;refGene_tx=XM_023253169.1;refGene_gn=LIMCH1;refGene_tx=XM_023253170.1;refGene_gn=LIMCH1;refGene_tx=XM_023253171.1
B1 172927465 172927574
refGene_gn=LIMCH1;refGene_tx=XM_019829582.2;refGene_gn=LIMCH1;refGene_tx=XM_019829583.2;refGene_gn=LIMCH1;refGene_tx=XM_019829584.2;refGene_gn=LIMCH1;refGene_tx=XM_019829585.2;refGene_gn=LIMCH1;refGene_tx=XM_019829586.2;refGene_gn=LIMCH1;refGene_tx=XM_023253162.1;refGene_gn=LIMCH1;refGene_tx=XM_023253163.1;refGene_gn=LIMCH1;refGene_tx=XM_023253165.1;refGene_gn=LIMCH1;refGene_tx=XM_023253166.1;refGene_gn=LIMCH1;refGene_tx=XM_023253167.1;refGene_gn=LIMCH1;refGene_tx=XM_023253168.1;refGene_gn=LIMCH1;refGene_tx=XM_023253169.1;refGene_gn=LIMCH1;refGene_tx=XM_023253170.1;refGene_gn=LIMCH1;refGene_tx=XM_023253171.1
B1 172929420 172929498
refGene_gn=LIMCH1;refGene_tx=XM_019829582.2;refGene_gn=LIMCH1;refGene_tx=XM_019829583.2;refGene_gn=LIMCH1;refGene_tx=XM_023253162.1;refGene_gn=LIMCH1;refGene_tx=XM_023253163.1;refGene_gn=LIMCH1;refGene_tx=XM_023253165.1;refGene_gn=LIMCH1;refGene_tx=XM_023253166.1;refGene_gn=LIMCH1;refGene_tx=XM_023253167.1
B1 172931858 172931982
refGene_gn=LIMCH1;refGene_tx=XM_019829582.2;refGene_gn=LIMCH1;refGene_tx=XM_019829583.2;refGene_gn=LIMCH1;refGene_tx=XM_019829584.2;refGene_gn=LIMCH1;refGene_tx=XM_019829585.2;refGene_gn=LIMCH1;refGene_tx=XM_019829586.2;refGene_gn=LIMCH1;refGene_tx=XM_023253162.1;refGene_gn=LIMCH1;refGene_tx=XM_023253163.1;refGene_gn=LIMCH1;refGene_tx=XM_023253165.1;refGene_gn=LIMCH1;refGene_tx=XM_023253166.1;refGene_gn=LIMCH1;refGene_tx=XM_023253167.1;refGene_gn=LIMCH1;refGene_tx=XM_023253168.1;refGene_gn=LIMCH1;refGene_tx=XM_023253169.1;refGene_gn=LIMCH1;refGene_tx=XM_023253170.1;refGene_gn=LIMCH1;refGene_tx=XM_023253171.1
B1 172932252 172932374
refGene_gn=LIMCH1;refGene_tx=XM_019829582.2;refGene_gn=LIMCH1;refGene_tx=XM_019829583.2;refGene_gn=LIMCH1;refGene_tx=XM_019829584.2;refGene_gn=LIMCH1;refGene_tx=XM_019829585.2;refGene_gn=LIMCH1;refGene_tx=XM_019829586.2;refGene_gn=LIMCH1;refGene_tx=XM_023253162.1;refGene_gn=LIMCH1;refGene_tx=XM_023253163.1;refGene_gn=LIMCH1;refGene_tx=XM_023253165.1;refGene_gn=LIMCH1;refGene_tx=XM_023253166.1;refGene_gn=LIMCH1;refGene_tx=XM_023253167.1;refGene_gn=LIMCH1;refGene_tx=XM_023253168.1;refGene_gn=LIMCH1;refGene_tx=XM_023253169.1;refGene_gn=LIMCH1;refGene_tx=XM_023253170.1;refGene_gn=LIMCH1;refGene_tx=XM_023253171.1
B1 172934120 172934251
refGene_gn=LIMCH1;refGene_tx=XM_019829582.2;refGene_gn=LIMCH1;refGene_tx=XM_019829583.2;refGene_gn=LIMCH1;refGene_tx=XM_019829584.2;refGene_

gn=LIMCH1;refGene_tx=XM_019829585.2;refGene_gn=LIMCH1;refGene_tx=XM_01
9829586.2;refGene_gn=LIMCH1;refGene_tx=XM_023253162.1;refGene_gn=LIMCH
1;refGene_tx=XM_023253163.1;refGene_gn=LIMCH1;refGene_tx=XM_023253165.
1;refGene_gn=LIMCH1;refGene_tx=XM_023253166.1;refGene_gn=LIMCH1;refGen
e_tx=XM_023253167.1;refGene_gn=LIMCH1;refGene_tx=XM_023253168.1;refGen
e_gn=LIMCH1;refGene_tx=XM_023253169.1;refGene_gn=LIMCH1;refGene_tx=XM_
023253170.1;refGene_gn=LIMCH1;refGene_tx=XM_023253171.1
B1 172935619 172935724
refGene_gn=LIMCH1;refGene_tx=XM_019829582.2;refGene_gn=LIMCH1;refGene_
tx=XM_019829583.2;refGene_gn=LIMCH1;refGene_tx=XM_019829584.2;refGene_
gn=LIMCH1;refGene_tx=XM_019829585.2;refGene_gn=LIMCH1;refGene_tx=XM_01
9829586.2;refGene_gn=LIMCH1;refGene_tx=XM_023253162.1;refGene_gn=LIMCH
1;refGene_tx=XM_023253163.1;refGene_gn=LIMCH1;refGene_tx=XM_023253165.
1;refGene_gn=LIMCH1;refGene_tx=XM_023253166.1;refGene_gn=LIMCH1;refGen
e_tx=XM_023253167.1;refGene_gn=LIMCH1;refGene_tx=XM_023253168.1;refGen
e_gn=LIMCH1;refGene_tx=XM_023253169.1;refGene_gn=LIMCH1;refGene_tx=XM_
023253170.1;refGene_gn=LIMCH1;refGene_tx=XM_023253171.1
B1 172936589 172936682
refGene_gn=LIMCH1;refGene_tx=XM_019829582.2;refGene_gn=LIMCH1;refGene_
tx=XM_019829583.2;refGene_gn=LIMCH1;refGene_tx=XM_019829584.2;refGene_
gn=LIMCH1;refGene_tx=XM_019829585.2;refGene_gn=LIMCH1;refGene_tx=XM_01
9829586.2;refGene_gn=LIMCH1;refGene_tx=XM_023253162.1;refGene_gn=LIMCH
1;refGene_tx=XM_023253163.1;refGene_gn=LIMCH1;refGene_tx=XM_023253165.
1;refGene_gn=LIMCH1;refGene_tx=XM_023253166.1;refGene_gn=LIMCH1;refGen
e_tx=XM_023253167.1;refGene_gn=LIMCH1;refGene_tx=XM_023253168.1;refGen
e_gn=LIMCH1;refGene_tx=XM_023253169.1;refGene_gn=LIMCH1;refGene_tx=XM_
023253170.1;refGene_gn=LIMCH1;refGene_tx=XM_023253171.1
B1 172939849 172939933
refGene_gn=LIMCH1;refGene_tx=XM_019829582.2;refGene_gn=LIMCH1;refGene_
tx=XM_019829583.2;refGene_gn=LIMCH1;refGene_tx=XM_019829584.2;refGene_
gn=LIMCH1;refGene_tx=XM_019829585.2;refGene_gn=LIMCH1;refGene_tx=XM_01
9829586.2;refGene_gn=LIMCH1;refGene_tx=XM_023253162.1;refGene_gn=LIMCH
1;refGene_tx=XM_023253163.1;refGene_gn=LIMCH1;refGene_tx=XM_023253165.
1;refGene_gn=LIMCH1;refGene_tx=XM_023253166.1;refGene_gn=LIMCH1;refGen
e_tx=XM_023253167.1;refGene_gn=LIMCH1;refGene_tx=XM_023253168.1;refGen
e_gn=LIMCH1;refGene_tx=XM_023253169.1;refGene_gn=LIMCH1;refGene_tx=XM_
023253170.1;refGene_gn=LIMCH1;refGene_tx=XM_023253171.1
B1 172944398 172944439
refGene_gn=LIMCH1;refGene_tx=XM_019829582.2;refGene_gn=LIMCH1;refGene_
tx=XM_019829583.2;refGene_gn=LIMCH1;refGene_tx=XM_019829584.2;refGene_
gn=LIMCH1;refGene_tx=XM_019829585.2;refGene_gn=LIMCH1;refGene_tx=XM_01
9829586.2;refGene_gn=LIMCH1;refGene_tx=XM_023253162.1;refGene_gn=LIMCH
1;refGene_tx=XM_023253163.1;refGene_gn=LIMCH1;refGene_tx=XM_023253165.
1;refGene_gn=LIMCH1;refGene_tx=XM_023253166.1;refGene_gn=LIMCH1;refGen
e_tx=XM_023253167.1;refGene_gn=LIMCH1;refGene_tx=XM_023253168.1;refGen
e_gn=LIMCH1;refGene_tx=XM_023253169.1;refGene_gn=LIMCH1;refGene_tx=XM_
023253170.1;refGene_gn=LIMCH1;refGene_tx=XM_023253171.1
B1 172945192 172945267
refGene_gn=LIMCH1;refGene_tx=XM_023253163.1;refGene_gn=LIMCH1;refGene_
tx=XM_023253168.1

B1 172949403 172949509
refGene_gn=LIMCH1;refGene_tx=XM_019829582.2;refGene_gn=LIMCH1;refGene_tx=XM_019829583.2;refGene_gn=LIMCH1;refGene_tx=XM_019829584.2;refGene_gn=LIMCH1;refGene_tx=XM_019829585.2;refGene_gn=LIMCH1;refGene_tx=XM_019829586.2;refGene_gn=LIMCH1;refGene_tx=XM_023253162.1;refGene_gn=LIMCH1;refGene_tx=XM_023253163.1;refGene_gn=LIMCH1;refGene_tx=XM_023253165.1;refGene_gn=LIMCH1;refGene_tx=XM_023253166.1;refGene_gn=LIMCH1;refGene_tx=XM_023253167.1;refGene_gn=LIMCH1;refGene_tx=XM_023253168.1;refGene_gn=LIMCH1;refGene_tx=XM_023253169.1;refGene_gn=LIMCH1;refGene_tx=XM_023253170.1;refGene_gn=LIMCH1;refGene_tx=XM_023253171.1
B1 172954155 172954319
refGene_gn=LIMCH1;refGene_tx=XM_019829582.2;refGene_gn=LIMCH1;refGene_tx=XM_019829583.2;refGene_gn=LIMCH1;refGene_tx=XM_019829584.2;refGene_gn=LIMCH1;refGene_tx=XM_019829585.2;refGene_gn=LIMCH1;refGene_tx=XM_019829586.2;refGene_gn=LIMCH1;refGene_tx=XM_023253162.1;refGene_gn=LIMCH1;refGene_tx=XM_023253163.1;refGene_gn=LIMCH1;refGene_tx=XM_023253165.1;refGene_gn=LIMCH1;refGene_tx=XM_023253166.1;refGene_gn=LIMCH1;refGene_tx=XM_023253167.1;refGene_gn=LIMCH1;refGene_tx=XM_023253168.1;refGene_gn=LIMCH1;refGene_tx=XM_023253169.1;refGene_gn=LIMCH1;refGene_tx=XM_023253170.1;refGene_gn=LIMCH1;refGene_tx=XM_023253171.1
B1 172955945 172956036
refGene_gn=LIMCH1;refGene_tx=XM_019829582.2;refGene_gn=LIMCH1;refGene_tx=XM_019829583.2;refGene_gn=LIMCH1;refGene_tx=XM_019829584.2;refGene_gn=LIMCH1;refGene_tx=XM_019829585.2;refGene_gn=LIMCH1;refGene_tx=XM_019829586.2;refGene_gn=LIMCH1;refGene_tx=XM_023253162.1;refGene_gn=LIMCH1;refGene_tx=XM_023253163.1;refGene_gn=LIMCH1;refGene_tx=XM_023253165.1;refGene_gn=LIMCH1;refGene_tx=XM_023253166.1;refGene_gn=LIMCH1;refGene_tx=XM_023253167.1;refGene_gn=LIMCH1;refGene_tx=XM_023253168.1;refGene_gn=LIMCH1;refGene_tx=XM_023253169.1;refGene_gn=LIMCH1;refGene_tx=XM_023253170.1;refGene_gn=LIMCH1;refGene_tx=XM_023253171.1
B1 172971428 172971644
refGene_gn=LIMCH1;refGene_tx=XM_019829582.2;refGene_gn=LIMCH1;refGene_tx=XM_019829583.2;refGene_gn=LIMCH1;refGene_tx=XM_019829584.2;refGene_gn=LIMCH1;refGene_tx=XM_019829585.2;refGene_gn=LIMCH1;refGene_tx=XM_019829586.2;refGene_gn=LIMCH1;refGene_tx=XM_023253162.1;refGene_gn=LIMCH1;refGene_tx=XM_023253163.1;refGene_gn=LIMCH1;refGene_tx=XM_023253165.1;refGene_gn=LIMCH1;refGene_tx=XM_023253166.1;refGene_gn=LIMCH1;refGene_tx=XM_023253167.1;refGene_gn=LIMCH1;refGene_tx=XM_023253168.1;refGene_gn=LIMCH1;refGene_tx=XM_023253169.1;refGene_gn=LIMCH1;refGene_tx=XM_023253170.1;refGene_gn=LIMCH1;refGene_tx=XM_023253171.1
B1 172975512 172975921
refGene_gn=LIMCH1;refGene_tx=XM_019829582.2;refGene_gn=LIMCH1;refGene_tx=XM_019829583.2;refGene_gn=LIMCH1;refGene_tx=XM_019829584.2;refGene_gn=LIMCH1;refGene_tx=XM_019829585.2;refGene_gn=LIMCH1;refGene_tx=XM_019829586.2;refGene_gn=LIMCH1;refGene_tx=XM_023253162.1;refGene_gn=LIMCH1;refGene_tx=XM_023253163.1;refGene_gn=LIMCH1;refGene_tx=XM_023253165.1;refGene_gn=LIMCH1;refGene_tx=XM_023253166.1;refGene_gn=LIMCH1;refGene_tx=XM_023253167.1;refGene_gn=LIMCH1;refGene_tx=XM_023253168.1;refGene_gn=LIMCH1;refGene_tx=XM_023253169.1;refGene_gn=LIMCH1;refGene_tx=XM_023253170.1;refGene_gn=LIMCH1;refGene_tx=XM_023253171.1

B1 172976131 172976289
refGene_gn=LIMCH1;refGene_tx=XM_019829582.2;refGene_gn=LIMCH1;refGene_tx=XM_019829583.2;refGene_gn=LIMCH1;refGene_tx=XM_019829584.2;refGene_gn=LIMCH1;refGene_tx=XM_019829585.2;refGene_gn=LIMCH1;refGene_tx=XM_019829586.2;refGene_gn=LIMCH1;refGene_tx=XM_023253162.1;refGene_gn=LIMCH1;refGene_tx=XM_023253163.1;refGene_gn=LIMCH1;refGene_tx=XM_023253165.1;refGene_gn=LIMCH1;refGene_tx=XM_023253166.1;refGene_gn=LIMCH1;refGene_tx=XM_023253167.1;refGene_gn=LIMCH1;refGene_tx=XM_023253168.1;refGene_gn=LIMCH1;refGene_tx=XM_023253169.1;refGene_gn=LIMCH1;refGene_tx=XM_023253170.1;refGene_gn=LIMCH1;refGene_tx=XM_023253171.1
B1 172977709 172977836
refGene_gn=LIMCH1;refGene_tx=XM_019829582.2;refGene_gn=LIMCH1;refGene_tx=XM_019829583.2;refGene_gn=LIMCH1;refGene_tx=XM_019829584.2;refGene_gn=LIMCH1;refGene_tx=XM_019829585.2;refGene_gn=LIMCH1;refGene_tx=XM_019829586.2;refGene_gn=LIMCH1;refGene_tx=XM_023253162.1;refGene_gn=LIMCH1;refGene_tx=XM_023253163.1;refGene_gn=LIMCH1;refGene_tx=XM_023253165.1;refGene_gn=LIMCH1;refGene_tx=XM_023253166.1;refGene_gn=LIMCH1;refGene_tx=XM_023253167.1;refGene_gn=LIMCH1;refGene_tx=XM_023253168.1;refGene_gn=LIMCH1;refGene_tx=XM_023253169.1;refGene_gn=LIMCH1;refGene_tx=XM_023253170.1;refGene_gn=LIMCH1;refGene_tx=XM_023253171.1
B1 172982900 172982936
refGene_gn=LIMCH1;refGene_tx=XM_019829582.2;refGene_gn=LIMCH1;refGene_tx=XM_019829584.2;refGene_gn=LIMCH1;refGene_tx=XM_019829585.2;refGene_gn=LIMCH1;refGene_tx=XM_023253162.1;refGene_gn=LIMCH1;refGene_tx=XM_023253163.1;refGene_gn=LIMCH1;refGene_tx=XM_023253165.1;refGene_gn=LIMCH1;refGene_tx=XM_023253166.1;refGene_gn=LIMCH1;refGene_tx=XM_023253169.1;refGene_gn=LIMCH1;refGene_tx=XM_023253170.1
B1 172988169 172988430
refGene_gn=LIMCH1;refGene_tx=XM_023253162.1;refGene_gn=LIMCH1;refGene_tx=XM_023253163.1
B1 172988872 172988984
refGene_gn=LIMCH1;refGene_tx=XM_023253162.1;refGene_gn=LIMCH1;refGene_tx=XM_023253163.1
B1 172989093 172989209
refGene_gn=LIMCH1;refGene_tx=XM_023253162.1;refGene_gn=LIMCH1;refGene_tx=XM_023253163.1
B1 172990577 172990907
refGene_gn=LIMCH1;refGene_tx=XM_023253162.1;refGene_gn=LIMCH1;refGene_tx=XM_023253163.1
B1 172991682 172991925
refGene_gn=LIMCH1;refGene_tx=XM_023253162.1;refGene_gn=LIMCH1;refGene_tx=XM_023253163.1
B1 172994408 172994711
refGene_gn=LIMCH1;refGene_tx=XM_023253162.1;refGene_gn=LIMCH1;refGene_tx=XM_023253163.1
B1 173002767 173003034
refGene_gn=LIMCH1;refGene_tx=XM_023253162.1;refGene_gn=LIMCH1;refGene_tx=XM_023253163.1
B1 173004121 173004374
refGene_gn=LIMCH1;refGene_tx=XM_019829582.2;refGene_gn=LIMCH1;refGene_

tx=XM_019829583.2;refGene_gn=LIMCH1;refGene_tx=XM_019829584.2;refGene_gn=LIMCH1;refGene_tx=XM_019829585.2;refGene_gn=LIMCH1;refGene_tx=XM_019829586.2;refGene_gn=LIMCH1;refGene_tx=XM_023253162.1;refGene_gn=LIMCH1;refGene_tx=XM_023253163.1;refGene_gn=LIMCH1;refGene_tx=XM_023253165.1;refGene_gn=LIMCH1;refGene_tx=XM_023253166.1;refGene_gn=LIMCH1;refGene_tx=XM_023253167.1;refGene_gn=LIMCH1;refGene_tx=XM_023253168.1;refGene_gn=LIMCH1;refGene_tx=XM_023253169.1;refGene_gn=LIMCH1;refGene_tx=XM_023253170.1;refGene_gn=LIMCH1;refGene_tx=XM_023253171.1
B1 173010475 173010671
refGene_gn=LIMCH1;refGene_tx=XM_019829582.2;refGene_gn=LIMCH1;refGene_tx=XM_019829583.2;refGene_gn=LIMCH1;refGene_tx=XM_019829584.2;refGene_gn=LIMCH1;refGene_tx=XM_019829585.2;refGene_gn=LIMCH1;refGene_tx=XM_019829586.2;refGene_gn=LIMCH1;refGene_tx=XM_023253162.1;refGene_gn=LIMCH1;refGene_tx=XM_023253163.1;refGene_gn=LIMCH1;refGene_tx=XM_023253165.1;refGene_gn=LIMCH1;refGene_tx=XM_023253166.1;refGene_gn=LIMCH1;refGene_tx=XM_023253167.1;refGene_gn=LIMCH1;refGene_tx=XM_023253168.1;refGene_gn=LIMCH1;refGene_tx=XM_023253169.1;refGene_gn=LIMCH1;refGene_tx=XM_023253170.1;refGene_gn=LIMCH1;refGene_tx=XM_023253171.1
B1 173011047 173011071
refGene_gn=LIMCH1;refGene_tx=XM_019829582.2;refGene_gn=LIMCH1;refGene_tx=XM_019829583.2;refGene_gn=LIMCH1;refGene_tx=XM_019829584.2;refGene_gn=LIMCH1;refGene_tx=XM_019829585.2;refGene_gn=LIMCH1;refGene_tx=XM_019829586.2;refGene_gn=LIMCH1;refGene_tx=XM_023253163.1
B1 173017876 173017987
refGene_gn=LIMCH1;refGene_tx=XM_023253162.1;refGene_gn=LIMCH1;refGene_tx=XM_023253165.1;refGene_gn=LIMCH1;refGene_tx=XM_023253166.1;refGene_gn=LIMCH1;refGene_tx=XM_023253167.1;refGene_gn=LIMCH1;refGene_tx=XM_023253168.1;refGene_gn=LIMCH1;refGene_tx=XM_023253169.1;refGene_gn=LIMCH1;refGene_tx=XM_023253170.1;refGene_gn=LIMCH1;refGene_tx=XM_023253171.1
B1 173020206 173020237
refGene_gn=LIMCH1;refGene_tx=XM_023253162.1;refGene_gn=LIMCH1;refGene_tx=XM_023253165.1;refGene_gn=LIMCH1;refGene_tx=XM_023253166.1;refGene_gn=LIMCH1;refGene_tx=XM_023253167.1;refGene_gn=LIMCH1;refGene_tx=XM_023253168.1;refGene_gn=LIMCH1;refGene_tx=XM_023253169.1;refGene_gn=LIMCH1;refGene_tx=XM_023253170.1;refGene_gn=LIMCH1;refGene_tx=XM_023253171.1
B1 173025790 173025897
refGene_gn=LIMCH1;refGene_tx=XM_023253162.1;refGene_gn=LIMCH1;refGene_tx=XM_023253165.1;refGene_gn=LIMCH1;refGene_tx=XM_023253166.1;refGene_gn=LIMCH1;refGene_tx=XM_023253167.1;refGene_gn=LIMCH1;refGene_tx=XM_023253168.1;refGene_gn=LIMCH1;refGene_tx=XM_023253169.1;refGene_gn=LIMCH1;refGene_tx=XM_023253170.1;refGene_gn=LIMCH1;refGene_tx=XM_023253171.1
B1 173097361 173097431
refGene_gn=LIMCH1;refGene_tx=XM_023253162.1;refGene_gn=LIMCH1;refGene_tx=XM_023253165.1;refGene_gn=LIMCH1;refGene_tx=XM_023253166.1;refGene_gn=LIMCH1;refGene_tx=XM_023253167.1;refGene_gn=LIMCH1;refGene_tx=XM_023253168.1;refGene_gn=LIMCH1;refGene_tx=XM_023253169.1;refGene_gn=LIMCH1;refGene_tx=XM_023253170.1;refGene_gn=LIMCH1;refGene_tx=XM_023253171.

1
B1	173120632	173120703
refGene_gn=LIMCH1;refGene_tx=XM_023253162.1;refGene_gn=LIMCH1;refGene_tx=XM_023253165.1;refGene_gn=LIMCH1;refGene_tx=XM_023253166.1;refGene_gn=LIMCH1;refGene_tx=XM_023253167.1;refGene_gn=LIMCH1;refGene_tx=XM_023253168.1;refGene_gn=LIMCH1;refGene_tx=XM_023253169.1;refGene_gn=LIMCH1;refGene_tx=XM_023253170.1;refGene_gn=LIMCH1;refGene_tx=XM_023253171.1
B1	173249173	173249269
refGene_gn=LIMCH1;refGene_tx=XM_023253162.1;refGene_gn=LIMCH1;refGene_tx=XM_023253165.1;refGene_gn=LIMCH1;refGene_tx=XM_023253166.1;refGene_gn=LIMCH1;refGene_tx=XM_023253167.1;refGene_gn=LIMCH1;refGene_tx=XM_023253168.1;refGene_gn=LIMCH1;refGene_tx=XM_023253169.1;refGene_gn=LIMCH1;refGene_tx=XM_023253170.1;refGene_gn=LIMCH1;refGene_tx=XM_023253171.1
B1	173313553	173313640
refGene_gn=UCHL1;refGene_tx=XM_011281913.2
B1	173318845	173318904
refGene_gn=UCHL1;refGene_tx=XM_011281913.2
B1	173320052	173320119
refGene_gn=UCHL1;refGene_tx=XM_011281913.2
B1	173320993	173321041
refGene_gn=UCHL1;refGene_tx=XM_011281913.2
B1	173321119	173321205
refGene_gn=UCHL1;refGene_tx=XM_011281913.2
B1	173323285	173323436
refGene_gn=UCHL1;refGene_tx=XM_011281913.2
B1	173326193	173326322
refGene_gn=UCHL1;refGene_tx=XM_011281913.2
B1	173326797	173326809
refGene_gn=UCHL1;refGene_tx=XM_011281913.2
B1	173326926	173326959
refGene_gn=UCHL1;refGene_tx=XM_011281913.2
B1	173510584	173510630
refGene_gn=APBB2;refGene_tx=XM_023253173.1;refGene_gn=APBB2;refGene_tx=XM_023253174.1;refGene_gn=APBB2;refGene_tx=XM_023253175.1;refGene_gn=APBB2;refGene_tx=XM_023253176.1;refGene_gn=APBB2;refGene_tx=XM_023253177.1;refGene_gn=APBB2;refGene_tx=XM_023253178.1;refGene_gn=APBB2;refGene_tx=XM_023253179.1;refGene_gn=APBB2;refGene_tx=XM_023253181.1;refGene_gn=APBB2;refGene_tx=XM_023253182.1;refGene_gn=APBB2;refGene_tx=XM_023253183.1;refGene_gn=APBB2;refGene_tx=XM_023253184.1
B1	173547237	173547256
refGene_gn=APBB2;refGene_tx=XM_019829594.2;refGene_gn=APBB2;refGene_tx=XM_023253180.1
B1	173558062	173558878
refGene_gn=APBB2;refGene_tx=XM_019829594.2;refGene_gn=APBB2;refGene_tx=XM_023253173.1;refGene_gn=APBB2;refGene_tx=XM_023253174.1;refGene_gn=APBB2;refGene_tx=XM_023253175.1;refGene_gn=APBB2;refGene_tx=XM_023253176.1;refGene_gn=APBB2;refGene_tx=XM_023253177.1;refGene_gn=APBB2;refGene_tx=XM_023253178.1;refGene_gn=APBB2;refGene_tx=XM_023253179.1;refGen

e_gn=APBB2;refGene_tx=XM_023253180.1;refGene_gn=APBB2;refGene_tx=XM_02
3253181.1;refGene_gn=APBB2;refGene_tx=XM_023253182.1;refGene_gn=APBB2;
refGene_tx=XM_023253183.1;refGene_gn=APBB2;refGene_tx=XM_023253184.1
B1 173624808 173625017
refGene_gn=APBB2;refGene_tx=XM_019829594.2;refGene_gn=APBB2;refGene_tx
=XM_023253173.1;refGene_gn=APBB2;refGene_tx=XM_023253174.1;refGene_gn=
APBB2;refGene_tx=XM_023253175.1;refGene_gn=APBB2;refGene_tx=XM_0232531
76.1;refGene_gn=APBB2;refGene_tx=XM_023253177.1;refGene_gn=APBB2;refGe
ne_tx=XM_023253178.1;refGene_gn=APBB2;refGene_tx=XM_023253179.1;refGen
e_gn=APBB2;refGene_tx=XM_023253180.1;refGene_gn=APBB2;refGene_tx=XM_02
3253181.1;refGene_gn=APBB2;refGene_tx=XM_023253182.1;refGene_gn=APBB2;
refGene_tx=XM_023253183.1;refGene_gn=APBB2;refGene_tx=XM_023253184.1
B1 173634566 173634629
refGene_gn=APBB2;refGene_tx=XM_019829594.2;refGene_gn=APBB2;refGene_tx
=XM_023253173.1;refGene_gn=APBB2;refGene_tx=XM_023253174.1;refGene_gn=
APBB2;refGene_tx=XM_023253175.1;refGene_gn=APBB2;refGene_tx=XM_0232531
76.1;refGene_gn=APBB2;refGene_tx=XM_023253177.1;refGene_gn=APBB2;refGe
ne_tx=XM_023253178.1;refGene_gn=APBB2;refGene_tx=XM_023253179.1;refGen
e_gn=APBB2;refGene_tx=XM_023253180.1
B1 173635022 173635108
refGene_gn=APBB2;refGene_tx=XM_019829594.2;refGene_gn=APBB2;refGene_tx
=XM_023253173.1;refGene_gn=APBB2;refGene_tx=XM_023253174.1;refGene_gn=
APBB2;refGene_tx=XM_023253175.1;refGene_gn=APBB2;refGene_tx=XM_0232531
76.1;refGene_gn=APBB2;refGene_tx=XM_023253177.1;refGene_gn=APBB2;refGe
ne_tx=XM_023253178.1;refGene_gn=APBB2;refGene_tx=XM_023253179.1;refGen
e_gn=APBB2;refGene_tx=XM_023253180.1;refGene_gn=APBB2;refGene_tx=XM_02
3253181.1;refGene_gn=APBB2;refGene_tx=XM_023253182.1;refGene_gn=APBB2;
refGene_tx=XM_023253183.1;refGene_gn=APBB2;refGene_tx=XM_023253184.1
B1 173635202 173635263
refGene_gn=APBB2;refGene_tx=XM_019829594.2;refGene_gn=APBB2;refGene_tx
=XM_023253173.1;refGene_gn=APBB2;refGene_tx=XM_023253174.1;refGene_gn=
APBB2;refGene_tx=XM_023253175.1;refGene_gn=APBB2;refGene_tx=XM_0232531
76.1;refGene_gn=APBB2;refGene_tx=XM_023253177.1;refGene_gn=APBB2;refGe
ne_tx=XM_023253178.1;refGene_gn=APBB2;refGene_tx=XM_023253179.1;refGen
e_gn=APBB2;refGene_tx=XM_023253180.1;refGene_gn=APBB2;refGene_tx=XM_02
3253181.1;refGene_gn=APBB2;refGene_tx=XM_023253182.1;refGene_gn=APBB2;
refGene_tx=XM_023253183.1;refGene_gn=APBB2;refGene_tx=XM_023253184.1
B1 173675492 173675639
refGene_gn=APBB2;refGene_tx=XM_019829594.2;refGene_gn=APBB2;refGene_tx
=XM_023253173.1;refGene_gn=APBB2;refGene_tx=XM_023253174.1;refGene_gn=
APBB2;refGene_tx=XM_023253175.1;refGene_gn=APBB2;refGene_tx=XM_0232531
76.1;refGene_gn=APBB2;refGene_tx=XM_023253177.1;refGene_gn=APBB2;refGe
ne_tx=XM_023253178.1;refGene_gn=APBB2;refGene_tx=XM_023253179.1;refGen
e_gn=APBB2;refGene_tx=XM_023253180.1;refGene_gn=APBB2;refGene_tx=XM_02
3253181.1;refGene_gn=APBB2;refGene_tx=XM_023253182.1;refGene_gn=APBB2;
refGene_tx=XM_023253183.1;refGene_gn=APBB2;refGene_tx=XM_023253184.1
B1 173679056 173679184
refGene_gn=APBB2;refGene_tx=XM_019829594.2;refGene_gn=APBB2;refGene_tx
=XM_023253173.1;refGene_gn=APBB2;refGene_tx=XM_023253174.1;refGene_gn=
APBB2;refGene_tx=XM_023253175.1;refGene_gn=APBB2;refGene_tx=XM_0232531

76.1;refGene_gn=APBB2;refGene_tx=XM_023253177.1;refGene_gn=APBB2;refGene_tx=XM_023253178.1;refGene_gn=APBB2;refGene_tx=XM_023253179.1;refGene_gn=APBB2;refGene_tx=XM_023253180.1;refGene_gn=APBB2;refGene_tx=XM_023253181.1;refGene_gn=APBB2;refGene_tx=XM_023253182.1;refGene_gn=APBB2;refGene_tx=XM_023253183.1;refGene_gn=APBB2;refGene_tx=XM_023253184.1
B1 173725905 173725911
refGene_gn=APBB2;refGene_tx=XM_019829594.2;refGene_gn=APBB2;refGene_tx=XM_023253173.1;refGene_gn=APBB2;refGene_tx=XM_023253174.1;refGene_gn=APBB2;refGene_tx=XM_023253175.1;refGene_gn=APBB2;refGene_tx=XM_023253176.1;refGene_gn=APBB2;refGene_tx=XM_023253177.1;refGene_gn=APBB2;refGene_tx=XM_023253180.1;refGene_gn=APBB2;refGene_tx=XM_023253181.1
B1 173736574 173736689
refGene_gn=APBB2;refGene_tx=XM_019829594.2;refGene_gn=APBB2;refGene_tx=XM_023253173.1;refGene_gn=APBB2;refGene_tx=XM_023253174.1;refGene_gn=APBB2;refGene_tx=XM_023253175.1;refGene_gn=APBB2;refGene_tx=XM_023253176.1;refGene_gn=APBB2;refGene_tx=XM_023253177.1;refGene_gn=APBB2;refGene_tx=XM_023253178.1;refGene_gn=APBB2;refGene_tx=XM_023253179.1;refGene_gn=APBB2;refGene_tx=XM_023253180.1;refGene_gn=APBB2;refGene_tx=XM_023253181.1;refGene_gn=APBB2;refGene_tx=XM_023253182.1;refGene_gn=APBB2;refGene_tx=XM_023253183.1;refGene_gn=APBB2;refGene_tx=XM_023253184.1
B1 173739272 173739360
refGene_gn=APBB2;refGene_tx=XM_006931155.3;refGene_gn=APBB2;refGene_tx=XM_019829594.2;refGene_gn=APBB2;refGene_tx=XM_019829600.1;refGene_gn=APBB2;refGene_tx=XM_019829601.1;refGene_gn=APBB2;refGene_tx=XM_023253173.1;refGene_gn=APBB2;refGene_tx=XM_023253174.1;refGene_gn=APBB2;refGene_tx=XM_023253175.1;refGene_gn=APBB2;refGene_tx=XM_023253176.1;refGene_gn=APBB2;refGene_tx=XM_023253177.1;refGene_gn=APBB2;refGene_tx=XM_023253178.1;refGene_gn=APBB2;refGene_tx=XM_023253179.1;refGene_gn=APBB2;refGene_tx=XM_023253180.1;refGene_gn=APBB2;refGene_tx=XM_023253181.1;refGene_gn=APBB2;refGene_tx=XM_023253182.1;refGene_gn=APBB2;refGene_tx=XM_023253183.1;refGene_gn=APBB2;refGene_tx=XM_023253184.1
B1 173740446 173740530
refGene_gn=APBB2;refGene_tx=XM_006931155.3;refGene_gn=APBB2;refGene_tx=XM_019829594.2;refGene_gn=APBB2;refGene_tx=XM_019829600.1;refGene_gn=APBB2;refGene_tx=XM_019829601.1;refGene_gn=APBB2;refGene_tx=XM_023253173.1;refGene_gn=APBB2;refGene_tx=XM_023253174.1;refGene_gn=APBB2;refGene_tx=XM_023253175.1;refGene_gn=APBB2;refGene_tx=XM_023253176.1;refGene_gn=APBB2;refGene_tx=XM_023253177.1;refGene_gn=APBB2;refGene_tx=XM_023253178.1;refGene_gn=APBB2;refGene_tx=XM_023253179.1;refGene_gn=APBB2;refGene_tx=XM_023253180.1;refGene_gn=APBB2;refGene_tx=XM_023253181.1;refGene_gn=APBB2;refGene_tx=XM_023253182.1;refGene_gn=APBB2;refGene_tx=XM_023253183.1;refGene_gn=APBB2;refGene_tx=XM_023253184.1
B1 173742558 173742674
refGene_gn=APBB2;refGene_tx=XM_006931155.3;refGene_gn=APBB2;refGene_tx=XM_019829594.2;refGene_gn=APBB2;refGene_tx=XM_019829600.1;refGene_gn=APBB2;refGene_tx=XM_019829601.1;refGene_gn=APBB2;refGene_tx=XM_023253173.1;refGene_gn=APBB2;refGene_tx=XM_023253174.1;refGene_gn=APBB2;refGene_tx=XM_023253175.1;refGene_gn=APBB2;refGene_tx=XM_023253176.1;refGene_gn=APBB2;refGene_tx=XM_023253177.1;refGene_gn=APBB2;refGene_tx=XM_023253178.1;refGene_gn=APBB2;refGene_tx=XM_023253179.1;refGene_gn=APBB2;

refGene_tx=XM_023253180.1;refGene_gn=APBB2;refGene_tx=XM_023253181.1;r
efGene_gn=APBB2;refGene_tx=XM_023253182.1;refGene_gn=APBB2;refGene_tx=
XM_023253183.1;refGene_gn=APBB2;refGene_tx=XM_023253184.1
B1 173744352 173744532
refGene_gn=APBB2;refGene_tx=XM_006931155.3;refGene_gn=APBB2;refGene_tx
=XM_019829594.2;refGene_gn=APBB2;refGene_tx=XM_019829600.1;refGene_gn=
APBB2;refGene_tx=XM_019829601.1;refGene_gn=APBB2;refGene_tx=XM_0232531
73.1;refGene_gn=APBB2;refGene_tx=XM_023253174.1;refGene_gn=APBB2;refGe
ne_tx=XM_023253175.1;refGene_gn=APBB2;refGene_tx=XM_023253176.1;refGen
e_gn=APBB2;refGene_tx=XM_023253177.1;refGene_gn=APBB2;refGene_tx=XM_02
3253178.1;refGene_gn=APBB2;refGene_tx=XM_023253179.1;refGene_gn=APBB2;
refGene_tx=XM_023253180.1;refGene_gn=APBB2;refGene_tx=XM_023253181.1;r
efGene_gn=APBB2;refGene_tx=XM_023253182.1;refGene_gn=APBB2;refGene_tx=
XM_023253183.1;refGene_gn=APBB2;refGene_tx=XM_023253184.1
B1 173750292 173750460
refGene_gn=APBB2;refGene_tx=XM_006931155.3;refGene_gn=APBB2;refGene_tx
=XM_019829594.2;refGene_gn=APBB2;refGene_tx=XM_019829600.1;refGene_gn=
APBB2;refGene_tx=XM_019829601.1;refGene_gn=APBB2;refGene_tx=XM_0232531
73.1;refGene_gn=APBB2;refGene_tx=XM_023253174.1;refGene_gn=APBB2;refGe
ne_tx=XM_023253175.1;refGene_gn=APBB2;refGene_tx=XM_023253176.1;refGen
e_gn=APBB2;refGene_tx=XM_023253177.1;refGene_gn=APBB2;refGene_tx=XM_02
3253178.1;refGene_gn=APBB2;refGene_tx=XM_023253179.1;refGene_gn=APBB2;
refGene_tx=XM_023253180.1;refGene_gn=APBB2;refGene_tx=XM_023253181.1;r
efGene_gn=APBB2;refGene_tx=XM_023253182.1;refGene_gn=APBB2;refGene_tx=
XM_023253183.1;refGene_gn=APBB2;refGene_tx=XM_023253184.1
B1 173756833 173757472
refGene_gn=NSUN7;refGene_tx=XM_003985419.4;refGene_gn=NSUN7;refGene_tx
=XM_006931156.4;refGene_gn=NSUN7;refGene_tx=XM_006931157.3;refGene_gn=
NSUN7;refGene_tx=XM_019829603.2
B1 173758308 173758432
refGene_gn=NSUN7;refGene_tx=XM_003985419.4;refGene_gn=NSUN7;refGene_tx
=XM_006931156.4;refGene_gn=NSUN7;refGene_tx=XM_006931157.3
B1 173765982 173766100
refGene_gn=NSUN7;refGene_tx=XM_003985419.4;refGene_gn=NSUN7;refGene_tx
=XM_006931156.4;refGene_gn=NSUN7;refGene_tx=XM_006931157.3;refGene_gn=
NSUN7;refGene_tx=XM_019829603.2
B1 173771310 173771412
refGene_gn=NSUN7;refGene_tx=XM_003985419.4;refGene_gn=NSUN7;refGene_tx
=XM_006931156.4;refGene_gn=NSUN7;refGene_tx=XM_006931157.3;refGene_gn=
NSUN7;refGene_tx=XM_019829603.2
B1 173776812 173776956
refGene_gn=NSUN7;refGene_tx=XM_003985419.4;refGene_gn=NSUN7;refGene_tx
=XM_006931156.4;refGene_gn=NSUN7;refGene_tx=XM_006931157.3;refGene_gn=
NSUN7;refGene_tx=XM_019829603.2
B1 173782783 173782994
refGene_gn=NSUN7;refGene_tx=XM_003985419.4;refGene_gn=NSUN7;refGene_tx
=XM_006931156.4;refGene_gn=NSUN7;refGene_tx=XM_006931157.3;refGene_gn=
NSUN7;refGene_tx=XM_019829603.2
B1 173784537 173784721
refGene_gn=NSUN7;refGene_tx=XM_003985419.4;refGene_gn=NSUN7;refGene_tx

=XM_006931156.4;refGene_gn=NSUN7;refGene_tx=XM_006931157.3;refGene_gn=NSUN7;refGene_tx=XM_019829603.2
B1 173785710 173785863
refGene_gn=NSUN7;refGene_tx=XM_003985419.4;refGene_gn=NSUN7;refGene_tx=XM_006931156.4;refGene_gn=NSUN7;refGene_tx=XM_006931157.3;refGene_gn=NSUN7;refGene_tx=XM_019829603.2
B1 173797671 173797802
refGene_gn=NSUN7;refGene_tx=XM_003985419.4;refGene_gn=NSUN7;refGene_tx=XM_006931156.4;refGene_gn=NSUN7;refGene_tx=XM_006931157.3;refGene_gn=NSUN7;refGene_tx=XM_019829603.2
B1 173798407 173798466
refGene_gn=NSUN7;refGene_tx=XM_003985419.4;refGene_gn=NSUN7;refGene_tx=XM_006931156.4;refGene_gn=NSUN7;refGene_tx=XM_006931157.3;refGene_gn=NSUN7;refGene_tx=XM_019829603.2
B1 173808141 173808454
refGene_gn=NSUN7;refGene_tx=XM_003985419.4;refGene_gn=NSUN7;refGene_tx=XM_006931156.4;refGene_gn=NSUN7;refGene_tx=XM_006931157.3;refGene_gn=NSUN7;refGene_tx=XM_019829603.2
B1 174054831 174055957
refGene_gn=RBM47;refGene_tx=XM_019829604.2;refGene_gn=RBM47;refGene_tx=XM_019829605.1;refGene_gn=RBM47;refGene_tx=XM_019829606.2;refGene_gn=RBM47;refGene_tx=XM_019829607.2
B1 174056823 174057030
refGene_gn=RBM47;refGene_tx=XM_019829604.2;refGene_gn=RBM47;refGene_tx=XM_019829605.1;refGene_gn=RBM47;refGene_tx=XM_019829606.2;refGene_gn=RBM47;refGene_tx=XM_019829607.2
B1 174059901 174060098
refGene_gn=RBM47;refGene_tx=XM_019829604.2;refGene_gn=RBM47;refGene_tx=XM_019829605.1;refGene_gn=RBM47;refGene_tx=XM_019829606.2;refGene_gn=RBM47;refGene_tx=XM_019829607.2
B1 174065049 174065289
refGene_gn=RBM47;refGene_tx=XM_019829604.2;refGene_gn=RBM47;refGene_tx=XM_019829605.1;refGene_gn=RBM47;refGene_tx=XM_019829606.2;refGene_gn=RBM47;refGene_tx=XM_019829607.2
B1 174122869 174123411
refGene_gn=CHRNA9;refGene_tx=XM_023253185.1
B1 174126519 174127052
refGene_gn=CHRNA9;refGene_tx=XM_023253185.1
B1 174134056 174134211
refGene_gn=CHRNA9;refGene_tx=XM_023253185.1
B1 174135140 174135286
refGene_gn=CHRNA9;refGene_tx=XM_023253185.1
B1 174135543 174135607
refGene_gn=CHRNA9;refGene_tx=XM_023253185.1
B1 174208321 174208897
refGene_gn=RHOH;refGene_tx=XM_003985423.5;refGene_gn=RHOH;refGene_tx=XM_006931159.4;refGene_gn=RHOH;refGene_tx=XM_019829611.2;refGene_gn=RHOH;refGene_tx=XM_019829613.2;refGene_gn=RHOH;refGene_tx=XM_019829614.1;refGene_gn=RHOH;refGene_tx=XM_019829615.1;refGene_gn=RHOH;refGene_tx=XM_019829616.1

B1 174279552 174279688
refGene_gn=N4BP2;refGene_tx=XM_019829617.2;refGene_gn=N4BP2;refGene_tx=XM_019829619.2;refGene_gn=N4BP2;refGene_tx=XM_019829621.2;refGene_gn=N4BP2;refGene_tx=XM_019829622.2
B1 174286622 174286668
refGene_gn=N4BP2;refGene_tx=XM_011281916.3;refGene_gn=N4BP2;refGene_tx=XM_019829618.2;refGene_gn=N4BP2;refGene_tx=XM_023253186.1;refGene_gn=N4BP2;refGene_tx=XM_023253187.1
B1 174287901 174288025
refGene_gn=N4BP2;refGene_tx=XM_011281916.3;refGene_gn=N4BP2;refGene_tx=XM_019829617.2;refGene_gn=N4BP2;refGene_tx=XM_019829618.2;refGene_gn=N4BP2;refGene_tx=XM_019829619.2;refGene_gn=N4BP2;refGene_tx=XM_019829621.2;refGene_gn=N4BP2;refGene_tx=XM_019829622.2;refGene_gn=N4BP2;refGene_tx=XM_023253186.1;refGene_gn=N4BP2;refGene_tx=XM_023253187.1
B1 174295866 174296035
refGene_gn=N4BP2;refGene_tx=XM_011281916.3;refGene_gn=N4BP2;refGene_tx=XM_019829617.2;refGene_gn=N4BP2;refGene_tx=XM_019829618.2;refGene_gn=N4BP2;refGene_tx=XM_019829619.2;refGene_gn=N4BP2;refGene_tx=XM_019829621.2;refGene_gn=N4BP2;refGene_tx=XM_019829622.2;refGene_gn=N4BP2;refGene_tx=XM_023253186.1;refGene_gn=N4BP2;refGene_tx=XM_023253187.1
B1 174297235 174297424
refGene_gn=N4BP2;refGene_tx=XM_011281916.3;refGene_gn=N4BP2;refGene_tx=XM_019829617.2;refGene_gn=N4BP2;refGene_tx=XM_019829618.2;refGene_gn=N4BP2;refGene_tx=XM_019829619.2;refGene_gn=N4BP2;refGene_tx=XM_019829621.2;refGene_gn=N4BP2;refGene_tx=XM_019829622.2;refGene_gn=N4BP2;refGene_tx=XM_023253186.1;refGene_gn=N4BP2;refGene_tx=XM_023253187.1
B1 174302875 174303014
refGene_gn=N4BP2;refGene_tx=XM_011281916.3;refGene_gn=N4BP2;refGene_tx=XM_019829617.2;refGene_gn=N4BP2;refGene_tx=XM_019829618.2;refGene_gn=N4BP2;refGene_tx=XM_019829619.2;refGene_gn=N4BP2;refGene_tx=XM_019829621.2;refGene_gn=N4BP2;refGene_tx=XM_019829622.2;refGene_gn=N4BP2;refGene_tx=XM_023253186.1;refGene_gn=N4BP2;refGene_tx=XM_023253187.1
B1 174306769 174306888
refGene_gn=N4BP2;refGene_tx=XM_011281916.3;refGene_gn=N4BP2;refGene_tx=XM_019829617.2;refGene_gn=N4BP2;refGene_tx=XM_019829618.2;refGene_gn=N4BP2;refGene_tx=XM_019829619.2;refGene_gn=N4BP2;refGene_tx=XM_019829621.2;refGene_gn=N4BP2;refGene_tx=XM_019829622.2;refGene_gn=N4BP2;refGene_tx=XM_019829623.2;refGene_gn=N4BP2;refGene_tx=XM_023253186.1;refGene_gn=N4BP2;refGene_tx=XM_023253187.1
B1 174307443 174307580
refGene_gn=N4BP2;refGene_tx=XM_019829623.2
B1 174309662 174309859
refGene_gn=N4BP2;refGene_tx=XM_011281916.3;refGene_gn=N4BP2;refGene_tx=XM_019829617.2;refGene_gn=N4BP2;refGene_tx=XM_019829618.2;refGene_gn=N4BP2;refGene_tx=XM_019829619.2;refGene_gn=N4BP2;refGene_tx=XM_019829621.2;refGene_gn=N4BP2;refGene_tx=XM_019829622.2;refGene_gn=N4BP2;refGene_tx=XM_019829623.2;refGene_gn=N4BP2;refGene_tx=XM_023253186.1;refGene_gn=N4BP2;refGene_tx=XM_023253187.1
B1 174311371 174311417
refGene_gn=N4BP2;refGene_tx=XM_011281916.3;refGene_gn=N4BP2;refGene_tx

=XM_019829617.2;refGene_gn=N4BP2;refGene_tx=XM_019829618.2;refGene_gn=N4BP2;refGene_tx=XM_019829619.2;refGene_gn=N4BP2;refGene_tx=XM_019829621.2;refGene_gn=N4BP2;refGene_tx=XM_019829622.2;refGene_gn=N4BP2;refGene_tx=XM_019829623.2;refGene_gn=N4BP2;refGene_tx=XM_023253186.1;refGene_gn=N4BP2;refGene_tx=XM_023253187.1
B1 174311936 174312022
refGene_gn=N4BP2;refGene_tx=XM_011281916.3;refGene_gn=N4BP2;refGene_tx=XM_019829617.2;refGene_gn=N4BP2;refGene_tx=XM_019829618.2;refGene_gn=N4BP2;refGene_tx=XM_019829619.2;refGene_gn=N4BP2;refGene_tx=XM_019829621.2;refGene_gn=N4BP2;refGene_tx=XM_019829622.2;refGene_gn=N4BP2;refGene_tx=XM_019829623.2;refGene_gn=N4BP2;refGene_tx=XM_023253186.1;refGene_gn=N4BP2;refGene_tx=XM_023253187.1
B1 174312628 174315021
refGene_gn=N4BP2;refGene_tx=XM_011281916.3;refGene_gn=N4BP2;refGene_tx=XM_019829617.2;refGene_gn=N4BP2;refGene_tx=XM_019829618.2;refGene_gn=N4BP2;refGene_tx=XM_019829619.2;refGene_gn=N4BP2;refGene_tx=XM_019829621.2;refGene_gn=N4BP2;refGene_tx=XM_019829622.2;refGene_gn=N4BP2;refGene_tx=XM_019829623.2;refGene_gn=N4BP2;refGene_tx=XM_023253186.1;refGene_gn=N4BP2;refGene_tx=XM_023253187.1
B1 174316569 174316725
refGene_gn=N4BP2;refGene_tx=XM_011281916.3;refGene_gn=N4BP2;refGene_tx=XM_019829617.2;refGene_gn=N4BP2;refGene_tx=XM_019829618.2;refGene_gn=N4BP2;refGene_tx=XM_019829619.2;refGene_gn=N4BP2;refGene_tx=XM_019829621.2;refGene_gn=N4BP2;refGene_tx=XM_019829622.2;refGene_gn=N4BP2;refGene_tx=XM_019829623.2;refGene_gn=N4BP2;refGene_tx=XM_023253186.1;refGene_gn=N4BP2;refGene_tx=XM_023253187.1
B1 174321281 174321358
refGene_gn=N4BP2;refGene_tx=XM_011281916.3;refGene_gn=N4BP2;refGene_tx=XM_019829617.2;refGene_gn=N4BP2;refGene_tx=XM_019829618.2;refGene_gn=N4BP2;refGene_tx=XM_019829619.2;refGene_gn=N4BP2;refGene_tx=XM_019829621.2;refGene_gn=N4BP2;refGene_tx=XM_019829622.2;refGene_gn=N4BP2;refGene_tx=XM_019829623.2;refGene_gn=N4BP2;refGene_tx=XM_023253186.1;refGene_gn=N4BP2;refGene_tx=XM_023253187.1
B1 174326469 174326558
refGene_gn=N4BP2;refGene_tx=XM_011281916.3;refGene_gn=N4BP2;refGene_tx=XM_019829617.2;refGene_gn=N4BP2;refGene_tx=XM_019829618.2;refGene_gn=N4BP2;refGene_tx=XM_019829619.2;refGene_gn=N4BP2;refGene_tx=XM_019829621.2;refGene_gn=N4BP2;refGene_tx=XM_019829622.2;refGene_gn=N4BP2;refGene_tx=XM_019829623.2;refGene_gn=N4BP2;refGene_tx=XM_023253186.1;refGene_gn=N4BP2;refGene_tx=XM_023253187.1
B1 174333640 174333765
refGene_gn=N4BP2;refGene_tx=XM_011281916.3;refGene_gn=N4BP2;refGene_tx=XM_019829617.2;refGene_gn=N4BP2;refGene_tx=XM_019829618.2;refGene_gn=N4BP2;refGene_tx=XM_019829619.2;refGene_gn=N4BP2;refGene_tx=XM_019829621.2;refGene_gn=N4BP2;refGene_tx=XM_019829622.2;refGene_gn=N4BP2;refGene_tx=XM_019829623.2;refGene_gn=N4BP2;refGene_tx=XM_023253186.1;refGene_gn=N4BP2;refGene_tx=XM_023253187.1
B1 174334348 174335489
refGene_gn=N4BP2;refGene_tx=XM_011281916.3;refGene_gn=N4BP2;refGene_tx=XM_019829617.2;refGene_gn=N4BP2;refGene_tx=XM_019829618.2;refGene_gn=

N4BP2;refGene_tx=XM_019829619.2;refGene_gn=N4BP2;refGene_tx=XM_0198296
21.2;refGene_gn=N4BP2;refGene_tx=XM_019829622.2;refGene_gn=N4BP2;refGe
ne_tx=XM_019829623.2;refGene_gn=N4BP2;refGene_tx=XM_023253186.1;refGen
e_gn=N4BP2;refGene_tx=XM_023253187.1
B1 174339339 174339568
refGene_gn=N4BP2;refGene_tx=XM_011281916.3;refGene_gn=N4BP2;refGene_tx
=XM_019829617.2;refGene_gn=N4BP2;refGene_tx=XM_019829618.2;refGene_gn=
N4BP2;refGene_tx=XM_019829623.2;refGene_gn=N4BP2;refGene_tx=XM_0232531
86.1;refGene_gn=N4BP2;refGene_tx=XM_023253187.1
B1 174442924 174443062
refGene_gn=PDS5A;refGene_tx=XM_023253188.1
B1 174486543 174486747
refGene_gn=PDS5A;refGene_tx=XM_023253188.1
B1 174487837 174487924
refGene_gn=PDS5A;refGene_tx=XM_023253188.1
B1 174488422 174488520
refGene_gn=PDS5A;refGene_tx=XM_023253188.1
B1 174493041 174493168
refGene_gn=PDS5A;refGene_tx=XM_023253188.1
B1 174495052 174495133
refGene_gn=PDS5A;refGene_tx=XM_023253188.1
B1 174497053 174497194
refGene_gn=PDS5A;refGene_tx=XM_023253188.1
B1 174500004 174500120
refGene_gn=PDS5A;refGene_tx=XM_023253188.1
B1 174502471 174502566
refGene_gn=PDS5A;refGene_tx=XM_023253188.1
B1 174503300 174503446
refGene_gn=PDS5A;refGene_tx=XM_023253188.1
B1 174505593 174505745
refGene_gn=PDS5A;refGene_tx=XM_023253188.1
B1 174507222 174507336
refGene_gn=PDS5A;refGene_tx=XM_023253188.1
B1 174509594 174509676
refGene_gn=PDS5A;refGene_tx=XM_023253188.1
B1 174510995 174511044
refGene_gn=PDS5A;refGene_tx=XM_023253188.1
B1 174511228 174511368
refGene_gn=PDS5A;refGene_tx=XM_023253188.1
B1 174517971 174518087
refGene_gn=PDS5A;refGene_tx=XM_023253188.1
B1 174528292 174528398
refGene_gn=PDS5A;refGene_tx=XM_023253188.1
B1 174530099 174530260
refGene_gn=PDS5A;refGene_tx=XM_023253188.1
B1 174534509 174534633
refGene_gn=PDS5A;refGene_tx=XM_023253188.1
B1 174535460 174535619
refGene_gn=PDS5A;refGene_tx=XM_023253188.1
B1 174538253 174538322

refGene_gn=PDS5A;refGene_tx=XM_023253188.1
B1	174539427	174539564
refGene_gn=PDS5A;refGene_tx=XM_023253188.1
B1	174542669	174542793
refGene_gn=PDS5A;refGene_tx=XM_023253188.1
B1	174543109	174543314
refGene_gn=PDS5A;refGene_tx=XM_023253188.1
B1	174543829	174543944
refGene_gn=PDS5A;refGene_tx=XM_023253188.1
B1	174558000	174558133
refGene_gn=PDS5A;refGene_tx=XM_023253188.1
B1	174558757	174558877
refGene_gn=PDS5A;refGene_tx=XM_023253188.1
B1	174563164	174563227
refGene_gn=PDS5A;refGene_tx=XM_023253188.1
B1	174563971	174564117
refGene_gn=PDS5A;refGene_tx=XM_023253188.1
B1	174565662	174565771
refGene_gn=PDS5A;refGene_tx=XM_023253188.1
B1	174571298	174571651
refGene_gn=PDS5A;refGene_tx=XM_023253188.1
B1	174577114	174577118
refGene_gn=PDS5A;refGene_tx=XM_023253188.1
B1	174619658	174619733
refGene_gn=UBE2K;refGene_tx=XM_003985425.4
B1	174620069	174620198
refGene_gn=UBE2K;refGene_tx=XM_003985425.4
B1	174623441	174623541
refGene_gn=UBE2K;refGene_tx=XM_003985425.4
B1	174634876	174634959
refGene_gn=UBE2K;refGene_tx=XM_003985425.4
B1	174644056	174644115
refGene_gn=UBE2K;refGene_tx=XM_003985425.4
B1	174648283	174648377
refGene_gn=UBE2K;refGene_tx=XM_003985425.4
B1	174689739	174689802
refGene_gn=UBE2K;refGene_tx=XM_003985425.4
B1	174797481	174797556
refGene_gn=SMIM14;refGene_tx=XM_003985429.5;refGene_gn=SMIM14;refGene_tx=XM_023253189.1
B1	174820620	174820669
refGene_gn=SMIM14;refGene_tx=XM_003985429.5;refGene_gn=SMIM14;refGene_tx=XM_023253189.1
B1	174835746	174835889
refGene_gn=SMIM14;refGene_tx=XM_003985429.5;refGene_gn=SMIM14;refGene_tx=XM_023253189.1
B1	174838001	174838034
refGene_gn=SMIM14;refGene_tx=XM_003985429.5;refGene_gn=SMIM14;refGene_tx=XM_023253189.1
B1	174862225	174862387

refGene_gn=UGDH;refGene_tx=XM_003985430.5
B1 174867460 174867562
refGene_gn=UGDH;refGene_tx=XM_003985430.5
B1 174872735 174872936
refGene_gn=UGDH;refGene_tx=XM_003985430.5
B1 174873043 174873241
refGene_gn=UGDH;refGene_tx=XM_003985430.5
B1 174873709 174873857
refGene_gn=UGDH;refGene_tx=XM_003985430.5
B1 174877522 174877617
refGene_gn=UGDH;refGene_tx=XM_003985430.5
B1 174879346 174879477
refGene_gn=UGDH;refGene_tx=XM_003985430.5
B1 174879724 174879858
refGene_gn=UGDH;refGene_tx=XM_003985430.5
B1 174881296 174881388
refGene_gn=UGDH;refGene_tx=XM_003985430.5
B1 174881497 174881608
refGene_gn=UGDH;refGene_tx=XM_003985430.5
B1 174884086 174884197
refGene_gn=UGDH;refGene_tx=XM_003985430.5
B1 174892811 174892867
refGene_gn=LIAS;refGene_tx=XM_003985433.4
B1 174894957 174895069
refGene_gn=LIAS;refGene_tx=XM_003985433.4
B1 174895833 174895904
refGene_gn=LIAS;refGene_tx=XM_003985433.4
B1 174897497 174897643
refGene_gn=LIAS;refGene_tx=XM_003985433.4
B1 174899513 174899642
refGene_gn=LIAS;refGene_tx=XM_003985433.4
B1 174901834 174901892
refGene_gn=LIAS;refGene_tx=XM_003985433.4
B1 174901978 174902135
refGene_gn=LIAS;refGene_tx=XM_003985433.4
B1 174904108 174904193
refGene_gn=LIAS;refGene_tx=XM_003985433.4;refGene_gn=LIAS;refGene_tx=XM_023253190.1
B1 174905533 174905627
refGene_gn=LIAS;refGene_tx=XM_003985433.4
B1 174906569 174906742
refGene_gn=LIAS;refGene_tx=XM_003985433.4;refGene_gn=LIAS;refGene_tx=XM_023253190.1
B1 174907821 174907866
refGene_gn=LIAS;refGene_tx=XM_003985433.4;refGene_gn=LIAS;refGene_tx=XM_023253190.1
B1 174908567 174908613
refGene_gn=RPL9;refGene_tx=XM_003985435.4
B1 174908708 174908824
refGene_gn=RPL9;refGene_tx=XM_003985435.4

B1 174909353 174909449
refGene_gn=RPL9;refGene_tx=XM_003985435.4
B1 174910815 174910948
refGene_gn=RPL9;refGene_tx=XM_003985435.4
B1 174912213 174912294
refGene_gn=RPL9;refGene_tx=XM_003985435.4
B1 174912520 174912627
refGene_gn=RPL9;refGene_tx=XM_003985435.4
B1 174919471 174919836
refGene_gn=KLB;refGene_tx=XM_003985434.5
B1 174922364 174923508
refGene_gn=KLB;refGene_tx=XM_003985434.5
B1 174929549 174929818
refGene_gn=KLB;refGene_tx=XM_003985434.5
B1 174932279 174932790
refGene_gn=KLB;refGene_tx=XM_003985434.5
B1 174952752 174953532
refGene_gn=KLB;refGene_tx=XM_003985434.5
B1 174953533 174953578
refGene_gn=KLB;refGene_tx=XM_003985434.5
B1 174978995 174978998
refGene_gn=RFC1;refGene_tx=XM_011281918.3
B1 174993478 174993607
refGene_gn=RFC1;refGene_tx=XM_011281918.3
B1 175013310 175013386
refGene_gn=RFC1;refGene_tx=XM_011281918.3
B1 175015201 175015312
refGene_gn=RFC1;refGene_tx=XM_011281918.3
B1 175030860 175031093
refGene_gn=RFC1;refGene_tx=XM_011281918.3
B1 175032230 175032308
refGene_gn=RFC1;refGene_tx=XM_011281918.3
B1 175036453 175036531
refGene_gn=RFC1;refGene_tx=XM_011281918.3
B1 175037701 175037762
refGene_gn=RFC1;refGene_tx=XM_011281918.3
B1 175037764 175037788
refGene_gn=RFC1;refGene_tx=XM_011281918.3
B1 175038575 175038859
refGene_gn=RFC1;refGene_tx=XM_011281918.3
B1 175041265 175041373
refGene_gn=RFC1;refGene_tx=XM_011281918.3
B1 175043211 175043391
refGene_gn=RFC1;refGene_tx=XM_011281918.3
B1 175044559 175044664
refGene_gn=RFC1;refGene_tx=XM_011281918.3
B1 175047072 175047472
refGene_gn=RFC1;refGene_tx=XM_011281918.3
B1 175049038 175049148
refGene_gn=RFC1;refGene_tx=XM_011281918.3

B1 175050771 175050886
refGene_gn=RFC1;refGene_tx=XM_011281918.3
B1 175052433 175052525
refGene_gn=RFC1;refGene_tx=XM_011281918.3
B1 175052726 175052864
refGene_gn=RFC1;refGene_tx=XM_011281918.3
B1 175053004 175053100
refGene_gn=RFC1;refGene_tx=XM_011281918.3
B1 175053224 175053323
refGene_gn=RFC1;refGene_tx=XM_011281918.3
B1 175054668 175054823
refGene_gn=RFC1;refGene_tx=XM_011281918.3
B1 175054932 175055050
refGene_gn=RFC1;refGene_tx=XM_011281918.3
B1 175056914 175057060
refGene_gn=RFC1;refGene_tx=XM_011281918.3
B1 175060459 175060673
refGene_gn=RFC1;refGene_tx=XM_011281918.3
B1 175061801 175061993
refGene_gn=RFC1;refGene_tx=XM_011281918.3
B1 175062607 175062620
refGene_gn=WDR19;refGene_tx=XM_019829626.1
B1 175062963 175063047
refGene_gn=RFC1;refGene_tx=XM_011281918.3
B1 175082581 175082693
refGene_gn=WDR19;refGene_tx=XM_003985437.4;refGene_gn=WDR19;refGene_tx=XM_006931161.3;refGene_gn=WDR19;refGene_tx=XM_019829627.2
B1 175083000 175083077
refGene_gn=WDR19;refGene_tx=XM_003985437.4;refGene_gn=WDR19;refGene_tx=XM_006931161.3;refGene_gn=WDR19;refGene_tx=XM_019829626.1;refGene_gn=WDR19;refGene_tx=XM_019829627.2
B1 175083660 175083784
refGene_gn=WDR19;refGene_tx=XM_003985437.4;refGene_gn=WDR19;refGene_tx=XM_006931161.3;refGene_gn=WDR19;refGene_tx=XM_019829626.1;refGene_gn=WDR19;refGene_tx=XM_019829627.2
B1 175084670 175084821
refGene_gn=WDR19;refGene_tx=XM_003985437.4;refGene_gn=WDR19;refGene_tx=XM_006931161.3;refGene_gn=WDR19;refGene_tx=XM_019829626.1;refGene_gn=WDR19;refGene_tx=XM_019829627.2
B1 175086508 175086590
refGene_gn=WDR19;refGene_tx=XM_003985437.4;refGene_gn=WDR19;refGene_tx=XM_006931161.3;refGene_gn=WDR19;refGene_tx=XM_019829626.1;refGene_gn=WDR19;refGene_tx=XM_019829627.2
B1 175089999 175090124
refGene_gn=WDR19;refGene_tx=XM_003985437.4;refGene_gn=WDR19;refGene_tx=XM_006931161.3;refGene_gn=WDR19;refGene_tx=XM_019829626.1;refGene_gn=WDR19;refGene_tx=XM_019829627.2
B1 175091763 175091860
refGene_gn=WDR19;refGene_tx=XM_003985437.4;refGene_gn=WDR19;refGene_tx=XM_006931161.3;refGene_gn=WDR19;refGene_tx=XM_019829626.1;refGene_gn=

WDR19;refGene_tx=XM_019829627.2
B1 175093123 175093201
refGene_gn=WDR19;refGene_tx=XM_003985437.4;refGene_gn=WDR19;refGene_tx=XM_006931161.3;refGene_gn=WDR19;refGene_tx=XM_019829626.1;refGene_gn=WDR19;refGene_tx=XM_019829627.2
B1 175102787 175102856
refGene_gn=WDR19;refGene_tx=XM_003985437.4;refGene_gn=WDR19;refGene_tx=XM_006931161.3;refGene_gn=WDR19;refGene_tx=XM_019829626.1;refGene_gn=WDR19;refGene_tx=XM_019829627.2
B1 175105128 175105241
refGene_gn=WDR19;refGene_tx=XM_003985437.4;refGene_gn=WDR19;refGene_tx=XM_006931161.3;refGene_gn=WDR19;refGene_tx=XM_019829626.1;refGene_gn=WDR19;refGene_tx=XM_019829627.2
B1 175107423 175107548
refGene_gn=WDR19;refGene_tx=XM_003985437.4;refGene_gn=WDR19;refGene_tx=XM_006931161.3;refGene_gn=WDR19;refGene_tx=XM_019829626.1;refGene_gn=WDR19;refGene_tx=XM_019829627.2
B1 175107863 175108010
refGene_gn=WDR19;refGene_tx=XM_003985437.4;refGene_gn=WDR19;refGene_tx=XM_006931161.3;refGene_gn=WDR19;refGene_tx=XM_019829626.1;refGene_gn=WDR19;refGene_tx=XM_019829627.2
B1 175109200 175109284
refGene_gn=WDR19;refGene_tx=XM_003985437.4;refGene_gn=WDR19;refGene_tx=XM_006931161.3;refGene_gn=WDR19;refGene_tx=XM_019829626.1;refGene_gn=WDR19;refGene_tx=XM_019829627.2
B1 175109735 175109818
refGene_gn=WDR19;refGene_tx=XM_003985437.4;refGene_gn=WDR19;refGene_tx=XM_006931161.3;refGene_gn=WDR19;refGene_tx=XM_019829626.1;refGene_gn=WDR19;refGene_tx=XM_019829627.2
B1 175109898 175110039
refGene_gn=WDR19;refGene_tx=XM_003985437.4;refGene_gn=WDR19;refGene_tx=XM_006931161.3;refGene_gn=WDR19;refGene_tx=XM_019829626.1;refGene_gn=WDR19;refGene_tx=XM_019829627.2
B1 175112601 175112659
refGene_gn=WDR19;refGene_tx=XM_003985437.4;refGene_gn=WDR19;refGene_tx=XM_006931161.3;refGene_gn=WDR19;refGene_tx=XM_019829626.1;refGene_gn=WDR19;refGene_tx=XM_019829627.2
B1 175117370 175117480
refGene_gn=WDR19;refGene_tx=XM_003985437.4;refGene_gn=WDR19;refGene_tx=XM_006931161.3;refGene_gn=WDR19;refGene_tx=XM_019829626.1;refGene_gn=WDR19;refGene_tx=XM_019829627.2
B1 175119967 175120078
refGene_gn=WDR19;refGene_tx=XM_003985437.4;refGene_gn=WDR19;refGene_tx=XM_006931161.3;refGene_gn=WDR19;refGene_tx=XM_019829626.1;refGene_gn=WDR19;refGene_tx=XM_019829627.2
B1 175120273 175120433
refGene_gn=WDR19;refGene_tx=XM_003985437.4;refGene_gn=WDR19;refGene_tx=XM_006931161.3;refGene_gn=WDR19;refGene_tx=XM_019829626.1;refGene_gn=WDR19;refGene_tx=XM_019829627.2
B1 175122679 175122884

refGene_gn=WDR19;refGene_tx=XM_003985437.4;refGene_gn=WDR19;refGene_tx=XM_006931161.3;refGene_gn=WDR19;refGene_tx=XM_019829626.1;refGene_gn=WDR19;refGene_tx=XM_019829627.2
B1 175123009 175123157
refGene_gn=WDR19;refGene_tx=XM_003985437.4;refGene_gn=WDR19;refGene_tx=XM_006931161.3;refGene_gn=WDR19;refGene_tx=XM_019829626.1;refGene_gn=WDR19;refGene_tx=XM_019829627.2
B1 175127959 175128109
refGene_gn=WDR19;refGene_tx=XM_003985437.4;refGene_gn=WDR19;refGene_tx=XM_006931161.3;refGene_gn=WDR19;refGene_tx=XM_019829626.1;refGene_gn=WDR19;refGene_tx=XM_019829627.2
B1 175135709 175135832
refGene_gn=WDR19;refGene_tx=XM_003985437.4;refGene_gn=WDR19;refGene_tx=XM_006931161.3;refGene_gn=WDR19;refGene_tx=XM_019829626.1;refGene_gn=WDR19;refGene_tx=XM_019829627.2
B1 175136531 175136638
refGene_gn=WDR19;refGene_tx=XM_003985437.4;refGene_gn=WDR19;refGene_tx=XM_006931161.3;refGene_gn=WDR19;refGene_tx=XM_019829626.1;refGene_gn=WDR19;refGene_tx=XM_019829627.2
B1 175137486 175137601
refGene_gn=WDR19;refGene_tx=XM_003985437.4;refGene_gn=WDR19;refGene_tx=XM_006931161.3;refGene_gn=WDR19;refGene_tx=XM_019829626.1;refGene_gn=WDR19;refGene_tx=XM_019829627.2
B1 175137693 175137866
refGene_gn=WDR19;refGene_tx=XM_003985437.4;refGene_gn=WDR19;refGene_tx=XM_006931161.3;refGene_gn=WDR19;refGene_tx=XM_019829626.1;refGene_gn=WDR19;refGene_tx=XM_019829627.2
B1 175138328 175138399
refGene_gn=WDR19;refGene_tx=XM_003985437.4;refGene_gn=WDR19;refGene_tx=XM_006931161.3;refGene_gn=WDR19;refGene_tx=XM_019829626.1;refGene_gn=WDR19;refGene_tx=XM_019829627.2
B1 175147530 175147704
refGene_gn=WDR19;refGene_tx=XM_003985437.4;refGene_gn=WDR19;refGene_tx=XM_006931161.3;refGene_gn=WDR19;refGene_tx=XM_019829626.1;refGene_gn=WDR19;refGene_tx=XM_019829627.2
B1 175148227 175148340
refGene_gn=WDR19;refGene_tx=XM_003985437.4;refGene_gn=WDR19;refGene_tx=XM_006931161.3;refGene_gn=WDR19;refGene_tx=XM_019829626.1;refGene_gn=WDR19;refGene_tx=XM_019829627.2
B1 175150189 175150270
refGene_gn=WDR19;refGene_tx=XM_003985437.4;refGene_gn=WDR19;refGene_tx=XM_006931161.3;refGene_gn=WDR19;refGene_tx=XM_019829626.1;refGene_gn=WDR19;refGene_tx=XM_019829627.2
B1 175155314 175155430
refGene_gn=WDR19;refGene_tx=XM_003985437.4;refGene_gn=WDR19;refGene_tx=XM_006931161.3;refGene_gn=WDR19;refGene_tx=XM_019829626.1;refGene_gn=WDR19;refGene_tx=XM_019829627.2
B1 175158900 175159016
refGene_gn=WDR19;refGene_tx=XM_003985437.4;refGene_gn=WDR19;refGene_tx=XM_006931161.3;refGene_gn=WDR19;refGene_tx=XM_019829626.1;refGene_gn=

WDR19;refGene_tx=XM_019829627.2
B1 175171631 175171757
refGene_gn=WDR19;refGene_tx=XM_003985437.4;refGene_gn=WDR19;refGene_tx=XM_006931161.3;refGene_gn=WDR19;refGene_tx=XM_019829626.1;refGene_gn=WDR19;refGene_tx=XM_019829627.2
B1 175173593 175173659
refGene_gn=WDR19;refGene_tx=XM_003985437.4;refGene_gn=WDR19;refGene_tx=XM_006931161.3;refGene_gn=WDR19;refGene_tx=XM_019829626.1
B1 175174186 175174278
refGene_gn=WDR19;refGene_tx=XM_003985437.4;refGene_gn=WDR19;refGene_tx=XM_006931161.3;refGene_gn=WDR19;refGene_tx=XM_019829626.1
B1 175176586 175176592
refGene_gn=WDR19;refGene_tx=XM_003985437.4;refGene_gn=WDR19;refGene_tx=XM_006931161.3;refGene_gn=WDR19;refGene_tx=XM_019829626.1
B1 175209493 175209550
refGene_gn=KLHL5;refGene_tx=XM_006931162.4;refGene_gn=KLHL5;refGene_tx=XM_011281919.3;refGene_gn=KLHL5;refGene_tx=XM_019829630.2;refGene_gn=KLHL5;refGene_tx=XM_019829631.2;refGene_gn=KLHL5;refGene_tx=XM_019829632.2;refGene_gn=KLHL5;refGene_tx=XM_023253191.1;refGene_gn=KLHL5;refGene_tx=XM_023253192.1;refGene_gn=KLHL5;refGene_tx=XM_023253193.1;refGene_gn=KLHL5;refGene_tx=XM_023253194.1
B1 175211099 175211141
refGene_gn=KLHL5;refGene_tx=XM_019829628.1
B1 175213612 175213784
refGene_gn=KLHL5;refGene_tx=XM_006931162.4;refGene_gn=KLHL5;refGene_tx=XM_011281919.3;refGene_gn=KLHL5;refGene_tx=XM_019829628.1;refGene_gn=KLHL5;refGene_tx=XM_019829630.2;refGene_gn=KLHL5;refGene_tx=XM_019829631.2;refGene_gn=KLHL5;refGene_tx=XM_019829632.2;refGene_gn=KLHL5;refGene_tx=XM_023253191.1;refGene_gn=KLHL5;refGene_tx=XM_023253192.1;refGene_gn=KLHL5;refGene_tx=XM_023253193.1;refGene_gn=KLHL5;refGene_tx=XM_023253194.1
B1 175217773 175217986
refGene_gn=KLHL5;refGene_tx=XM_006931162.4;refGene_gn=KLHL5;refGene_tx=XM_011281919.3;refGene_gn=KLHL5;refGene_tx=XM_019829628.1;refGene_gn=KLHL5;refGene_tx=XM_019829630.2;refGene_gn=KLHL5;refGene_tx=XM_019829631.2;refGene_gn=KLHL5;refGene_tx=XM_019829632.2;refGene_gn=KLHL5;refGene_tx=XM_023253191.1;refGene_gn=KLHL5;refGene_tx=XM_023253192.1;refGene_gn=KLHL5;refGene_tx=XM_023253193.1;refGene_gn=KLHL5;refGene_tx=XM_023253194.1
B1 175227229 175227392
refGene_gn=KLHL5;refGene_tx=XM_006931162.4;refGene_gn=KLHL5;refGene_tx=XM_011281919.3;refGene_gn=KLHL5;refGene_tx=XM_019829628.1;refGene_gn=KLHL5;refGene_tx=XM_019829630.2;refGene_gn=KLHL5;refGene_tx=XM_019829631.2;refGene_gn=KLHL5;refGene_tx=XM_019829632.2;refGene_gn=KLHL5;refGene_tx=XM_023253191.1;refGene_gn=KLHL5;refGene_tx=XM_023253192.1;refGene_gn=KLHL5;refGene_tx=XM_023253193.1;refGene_gn=KLHL5;refGene_tx=XM_023253194.1
B1 175232560 175232785
refGene_gn=KLHL5;refGene_tx=XM_006931162.4;refGene_gn=KLHL5;refGene_tx=XM_011281919.3;refGene_gn=KLHL5;refGene_tx=XM_019829628.1;refGene_gn=

KLHL5;refGene_tx=XM_019829630.2;refGene_gn=KLHL5;refGene_tx=XM_0198296
31.2;refGene_gn=KLHL5;refGene_tx=XM_019829632.2;refGene_gn=KLHL5;refGe
ne_tx=XM_023253191.1;refGene_gn=KLHL5;refGene_tx=XM_023253192.1;refGen
e_gn=KLHL5;refGene_tx=XM_023253193.1;refGene_gn=KLHL5;refGene_tx=XM_02
3253194.1
B1 175239408 175239595
refGene_gn=KLHL5;refGene_tx=XM_006931162.4;refGene_gn=KLHL5;refGene_tx
=XM_011281919.3;refGene_gn=KLHL5;refGene_tx=XM_019829628.1;refGene_gn=
KLHL5;refGene_tx=XM_019829630.2;refGene_gn=KLHL5;refGene_tx=XM_0198296
31.2;refGene_gn=KLHL5;refGene_tx=XM_019829632.2;refGene_gn=KLHL5;refGe
ne_tx=XM_023253191.1;refGene_gn=KLHL5;refGene_tx=XM_023253192.1;refGen
e_gn=KLHL5;refGene_tx=XM_023253193.1;refGene_gn=KLHL5;refGene_tx=XM_02
3253194.1
B1 175255828 175256041
refGene_gn=KLHL5;refGene_tx=XM_006931162.4;refGene_gn=KLHL5;refGene_tx
=XM_011281919.3;refGene_gn=KLHL5;refGene_tx=XM_019829628.1;refGene_gn=
KLHL5;refGene_tx=XM_019829630.2;refGene_gn=KLHL5;refGene_tx=XM_0198296
31.2;refGene_gn=KLHL5;refGene_tx=XM_019829632.2;refGene_gn=KLHL5;refGe
ne_tx=XM_023253191.1;refGene_gn=KLHL5;refGene_tx=XM_023253192.1;refGen
e_gn=KLHL5;refGene_tx=XM_023253193.1;refGene_gn=KLHL5;refGene_tx=XM_02
3253194.1
B1 175265307 175265504
refGene_gn=KLHL5;refGene_tx=XM_006931162.4;refGene_gn=KLHL5;refGene_tx
=XM_011281919.3;refGene_gn=KLHL5;refGene_tx=XM_019829628.1;refGene_gn=
KLHL5;refGene_tx=XM_019829630.2;refGene_gn=KLHL5;refGene_tx=XM_0198296
31.2;refGene_gn=KLHL5;refGene_tx=XM_019829632.2;refGene_gn=KLHL5;refGe
ne_tx=XM_023253191.1;refGene_gn=KLHL5;refGene_tx=XM_023253192.1;refGen
e_gn=KLHL5;refGene_tx=XM_023253193.1;refGene_gn=KLHL5;refGene_tx=XM_02
3253194.1
B1 175266036 175266173
refGene_gn=KLHL5;refGene_tx=XM_006931162.4;refGene_gn=KLHL5;refGene_tx
=XM_011281919.3;refGene_gn=KLHL5;refGene_tx=XM_019829628.1;refGene_gn=
KLHL5;refGene_tx=XM_019829630.2;refGene_gn=KLHL5;refGene_tx=XM_0198296
31.2;refGene_gn=KLHL5;refGene_tx=XM_019829632.2;refGene_gn=KLHL5;refGe
ne_tx=XM_023253191.1;refGene_gn=KLHL5;refGene_tx=XM_023253192.1;refGen
e_gn=KLHL5;refGene_tx=XM_023253193.1;refGene_gn=KLHL5;refGene_tx=XM_02
3253194.1
B1 175269204 175269387
refGene_gn=KLHL5;refGene_tx=XM_006931162.4;refGene_gn=KLHL5;refGene_tx
=XM_011281919.3;refGene_gn=KLHL5;refGene_tx=XM_019829628.1;refGene_gn=
KLHL5;refGene_tx=XM_019829630.2;refGene_gn=KLHL5;refGene_tx=XM_0198296
31.2;refGene_gn=KLHL5;refGene_tx=XM_019829632.2;refGene_gn=KLHL5;refGe
ne_tx=XM_023253191.1;refGene_gn=KLHL5;refGene_tx=XM_023253192.1;refGen
e_gn=KLHL5;refGene_tx=XM_023253193.1;refGene_gn=KLHL5;refGene_tx=XM_02
3253194.1
B1 175281935 175282034
refGene_gn=KLHL5;refGene_tx=XM_011281919.3;refGene_gn=KLHL5;refGene_tx
=XM_019829628.1;refGene_gn=KLHL5;refGene_tx=XM_023253192.1
B1 175283134 175283514
refGene_gn=KLHL5;refGene_tx=XM_006931162.4;refGene_gn=KLHL5;refGene_tx

=XM_011281919.3;refGene_gn=KLHL5;refGene_tx=XM_019829628.1;refGene_gn=KLHL5;refGene_tx=XM_023253191.1
B1	175296615	175296665
refGene_gn=KLHL5;refGene_tx=XM_023253192.1;refGene_gn=KLHL5;refGene_tx=XM_023253193.1
B1	175328398	175328486
refGene_gn=TMEM156;refGene_tx=XM_003985466.4;refGene_gn=TMEM156;refGene_tx=XM_006931166.3
B1	175354441	175354711
refGene_gn=TMEM156;refGene_tx=XM_003985466.4;refGene_gn=TMEM156;refGene_tx=XM_006931166.3
B1	175356243	175356504
refGene_gn=TMEM156;refGene_tx=XM_003985466.4;refGene_gn=TMEM156;refGene_tx=XM_006931166.3
B1	175359406	175359526
refGene_gn=TMEM156;refGene_tx=XM_003985466.4;refGene_gn=TMEM156;refGene_tx=XM_006931166.3
B1	175362849	175362933
refGene_gn=TMEM156;refGene_tx=XM_003985466.4;refGene_gn=TMEM156;refGene_tx=XM_006931166.3
B1	175377367	175377470
refGene_gn=TMEM156;refGene_tx=XM_003985466.4;refGene_gn=TMEM156;refGene_tx=XM_006931166.3
B1	175380554	175380567
refGene_gn=TMEM156;refGene_tx=XM_003985466.4;refGene_gn=TMEM156;refGene_tx=XM_006931166.3
B1	175399867	175399969
refGene_gn=FAM114A1;refGene_tx=XM_003985439.3;refGene_gn=FAM114A1;refGene_tx=XM_019829634.1;refGene_gn=FAM114A1;refGene_tx=XM_023253196.1;refGene_gn=FAM114A1;refGene_tx=XM_023253197.1;refGene_gn=FAM114A1;refGene_tx=XM_023253198.1
B1	175401796	175401850
refGene_gn=FAM114A1;refGene_tx=XM_003985439.3;refGene_gn=FAM114A1;refGene_tx=XM_019829634.1;refGene_gn=FAM114A1;refGene_tx=XM_023253196.1;refGene_gn=FAM114A1;refGene_tx=XM_023253197.1;refGene_gn=FAM114A1;refGene_tx=XM_023253198.1
B1	175406185	175406258
refGene_gn=FAM114A1;refGene_tx=XM_003985439.3;refGene_gn=FAM114A1;refGene_tx=XM_019829634.1;refGene_gn=FAM114A1;refGene_tx=XM_023253196.1;refGene_gn=FAM114A1;refGene_tx=XM_023253197.1;refGene_gn=FAM114A1;refGene_tx=XM_023253198.1
B1	175409123	175409263
refGene_gn=FAM114A1;refGene_tx=XM_003985439.3;refGene_gn=FAM114A1;refGene_tx=XM_019829634.1;refGene_gn=FAM114A1;refGene_tx=XM_023253196.1;refGene_gn=FAM114A1;refGene_tx=XM_023253197.1;refGene_gn=FAM114A1;refGene_tx=XM_023253198.1
B1	175410566	175410728
refGene_gn=FAM114A1;refGene_tx=XM_003985439.3;refGene_gn=FAM114A1;refGene_tx=XM_019829634.1;refGene_gn=FAM114A1;refGene_tx=XM_023253196.1;refGene_gn=FAM114A1;refGene_tx=XM_023253197.1;refGene_gn=FAM114A1;refGen

e_tx=XM_023253198.1
B1 175412422 175412514
refGene_gn=FAM114A1;refGene_tx=XM_003985439.3;refGene_gn=FAM114A1;refGene_tx=XM_019829634.1;refGene_gn=FAM114A1;refGene_tx=XM_023253196.1;refGene_gn=FAM114A1;refGene_tx=XM_023253197.1;refGene_gn=FAM114A1;refGene_tx=XM_023253198.1
B1 175417909 175418033
refGene_gn=FAM114A1;refGene_tx=XM_003985439.3;refGene_gn=FAM114A1;refGene_tx=XM_019829634.1;refGene_gn=FAM114A1;refGene_tx=XM_023253196.1;refGene_gn=FAM114A1;refGene_tx=XM_023253197.1;refGene_gn=FAM114A1;refGene_tx=XM_023253198.1
B1 175424217 175424370
refGene_gn=FAM114A1;refGene_tx=XM_003985439.3;refGene_gn=FAM114A1;refGene_tx=XM_019829634.1;refGene_gn=FAM114A1;refGene_tx=XM_023253196.1;refGene_gn=FAM114A1;refGene_tx=XM_023253197.1;refGene_gn=FAM114A1;refGene_tx=XM_023253198.1
B1 175428256 175428391
refGene_gn=FAM114A1;refGene_tx=XM_003985439.3;refGene_gn=FAM114A1;refGene_tx=XM_019829634.1;refGene_gn=FAM114A1;refGene_tx=XM_023253196.1;refGene_gn=FAM114A1;refGene_tx=XM_023253197.1;refGene_gn=FAM114A1;refGene_tx=XM_023253198.1
B1 175431241 175431345
refGene_gn=FAM114A1;refGene_tx=XM_003985439.3;refGene_gn=FAM114A1;refGene_tx=XM_019829634.1;refGene_gn=FAM114A1;refGene_tx=XM_023253196.1;refGene_gn=FAM114A1;refGene_tx=XM_023253197.1;refGene_gn=FAM114A1;refGene_tx=XM_023253198.1
B1 175431470 175431584
refGene_gn=FAM114A1;refGene_tx=XM_003985439.3;refGene_gn=FAM114A1;refGene_tx=XM_019829634.1;refGene_gn=FAM114A1;refGene_tx=XM_023253196.1;refGene_gn=FAM114A1;refGene_tx=XM_023253197.1
B1 175444647 175444735
refGene_gn=FAM114A1;refGene_tx=XM_003985439.3;refGene_gn=FAM114A1;refGene_tx=XM_019829634.1;refGene_gn=FAM114A1;refGene_tx=XM_023253196.1;refGene_gn=FAM114A1;refGene_tx=XM_023253197.1
B1 175456723 175457068
refGene_gn=FAM114A1;refGene_tx=XM_003985439.3;refGene_gn=FAM114A1;refGene_tx=XM_019829634.1;refGene_gn=FAM114A1;refGene_tx=XM_023253196.1;refGene_gn=FAM114A1;refGene_tx=XM_023253197.1
B1 175493241 175495656
refGene_gn=TLR6;refGene_tx=XM_003985467.4
B1 175521623 175523993
refGene_gn=TLR1;refGene_tx=XM_003985440.3;refGene_gn=TLR1;refGene_tx=XM_011281921.2;refGene_gn=TLR1;refGene_tx=XM_019829647.2
B1 175541602 175541754
refGene_gn=TLR10;refGene_tx=XM_019829643.1;refGene_gn=TLR10;refGene_tx=XM_019829644.1;refGene_gn=TLR10;refGene_tx=XM_019829645.1
B1 175541963 175542050
refGene_gn=TLR10;refGene_tx=XM_019829643.1;refGene_gn=TLR10;refGene_tx=XM_019829645.1
B1 175548041 175548139

refGene_gn=TLR10;refGene_tx=XM_019829643.1;refGene_gn=TLR10;refGene_tx=XM_019829644.1;refGene_gn=TLR10;refGene_tx=XM_019829645.1
B1 175548364 175548479
refGene_gn=TLR10;refGene_tx=XM_019829643.1;refGene_gn=TLR10;refGene_tx=XM_019829644.1
B1 175548953 175549008
refGene_gn=TLR10;refGene_tx=XM_019829643.1;refGene_gn=TLR10;refGene_tx=XM_019829644.1;refGene_gn=TLR10;refGene_tx=XM_019829645.1
B1 175549355 175551785
refGene_gn=TLR10;refGene_tx=XM_006931168.2;refGene_gn=TLR10;refGene_tx=XM_019829635.1;refGene_gn=TLR10;refGene_tx=XM_019829636.1;refGene_gn=TLR10;refGene_tx=XM_019829638.1;refGene_gn=TLR10;refGene_tx=XM_019829639.1;refGene_gn=TLR10;refGene_tx=XM_019829640.2
B1 175617555 175617737
refGene_gn=KLF3;refGene_tx=XM_006931169.4;refGene_gn=KLF3;refGene_tx=XM_023253200.1;refGene_gn=KLF3;refGene_tx=XM_023253201.1
B1 175619779 175619940
refGene_gn=KLF3;refGene_tx=XM_006931169.4;refGene_gn=KLF3;refGene_tx=XM_023253200.1;refGene_gn=KLF3;refGene_tx=XM_023253201.1
B1 175624922 175625073
refGene_gn=KLF3;refGene_tx=XM_006931169.4;refGene_gn=KLF3;refGene_tx=XM_023253200.1;refGene_gn=KLF3;refGene_tx=XM_023253201.1
B1 175625749 175626236
refGene_gn=KLF3;refGene_tx=XM_006931169.4;refGene_gn=KLF3;refGene_tx=XM_023253200.1;refGene_gn=KLF3;refGene_tx=XM_023253201.1
B1 175633646 175633703
refGene_gn=KLF3;refGene_tx=XM_006931169.4;refGene_gn=KLF3;refGene_tx=XM_023253200.1;refGene_gn=KLF3;refGene_tx=XM_023253201.1
B1 175647729 175648035
refGene_gn=LOC111560191;refGene_tx=XM_023253202.1
B1 175648297 175648360
refGene_gn=LOC111560191;refGene_tx=XM_023253202.1
B1 176085125 176085350
refGene_gn=TBC1D1;refGene_tx=XM_023253204.1;refGene_gn=TBC1D1;refGene_tx=XM_023253205.1;refGene_gn=TBC1D1;refGene_tx=XM_023253206.1;refGene_gn=TBC1D1;refGene_tx=XM_023253207.1;refGene_gn=TBC1D1;refGene_tx=XM_023253208.1
B1 176089000 176089174
refGene_gn=TBC1D1;refGene_tx=XM_023253204.1;refGene_gn=TBC1D1;refGene_tx=XM_023253205.1;refGene_gn=TBC1D1;refGene_tx=XM_023253206.1;refGene_gn=TBC1D1;refGene_tx=XM_023253207.1;refGene_gn=TBC1D1;refGene_tx=XM_023253208.1;refGene_gn=TBC1D1;refGene_tx=XM_023253209.1
B1 176095696 176095866
refGene_gn=TBC1D1;refGene_tx=XM_023253204.1;refGene_gn=TBC1D1;refGene_tx=XM_023253205.1;refGene_gn=TBC1D1;refGene_tx=XM_023253206.1;refGene_gn=TBC1D1;refGene_tx=XM_023253207.1;refGene_gn=TBC1D1;refGene_tx=XM_023253208.1
B1 176102538 176102698
refGene_gn=TBC1D1;refGene_tx=XM_023253204.1;refGene_gn=TBC1D1;refGene_tx=XM_023253205.1;refGene_gn=TBC1D1;refGene_tx=XM_023253206.1;refGene_

gn=TBC1D1;refGene_tx=XM_023253207.1;refGene_gn=TBC1D1;refGene_tx=XM_02
3253208.1;refGene_gn=TBC1D1;refGene_tx=XM_023253209.1
B1 176104695 176104940
refGene_gn=TBC1D1;refGene_tx=XM_023253204.1;refGene_gn=TBC1D1;refGene_
tx=XM_023253205.1;refGene_gn=TBC1D1;refGene_tx=XM_023253206.1;refGene_
gn=TBC1D1;refGene_tx=XM_023253207.1;refGene_gn=TBC1D1;refGene_tx=XM_02
3253208.1;refGene_gn=TBC1D1;refGene_tx=XM_023253209.1
B1 176119090 176119249
refGene_gn=TBC1D1;refGene_tx=XM_023253204.1;refGene_gn=TBC1D1;refGene_
tx=XM_023253205.1;refGene_gn=TBC1D1;refGene_tx=XM_023253206.1;refGene_
gn=TBC1D1;refGene_tx=XM_023253207.1;refGene_gn=TBC1D1;refGene_tx=XM_02
3253208.1;refGene_gn=TBC1D1;refGene_tx=XM_023253209.1
B1 176124913 176125075
refGene_gn=TBC1D1;refGene_tx=XM_023253204.1;refGene_gn=TBC1D1;refGene_
tx=XM_023253205.1;refGene_gn=TBC1D1;refGene_tx=XM_023253206.1;refGene_
gn=TBC1D1;refGene_tx=XM_023253207.1;refGene_gn=TBC1D1;refGene_tx=XM_02
3253208.1;refGene_gn=TBC1D1;refGene_tx=XM_023253209.1
B1 176131198 176131384
refGene_gn=TBC1D1;refGene_tx=XM_023253204.1;refGene_gn=TBC1D1;refGene_
tx=XM_023253205.1;refGene_gn=TBC1D1;refGene_tx=XM_023253206.1;refGene_
gn=TBC1D1;refGene_tx=XM_023253207.1;refGene_gn=TBC1D1;refGene_tx=XM_02
3253208.1;refGene_gn=TBC1D1;refGene_tx=XM_023253209.1
B1 176168016 176168156
refGene_gn=TBC1D1;refGene_tx=XM_023253204.1;refGene_gn=TBC1D1;refGene_
tx=XM_023253205.1;refGene_gn=TBC1D1;refGene_tx=XM_023253206.1;refGene_
gn=TBC1D1;refGene_tx=XM_023253207.1;refGene_gn=TBC1D1;refGene_tx=XM_02
3253208.1;refGene_gn=TBC1D1;refGene_tx=XM_023253209.1
B1 176169097 176169217
refGene_gn=TBC1D1;refGene_tx=XM_023253204.1;refGene_gn=TBC1D1;refGene_
tx=XM_023253205.1;refGene_gn=TBC1D1;refGene_tx=XM_023253208.1;refGene_
gn=TBC1D1;refGene_tx=XM_023253209.1
B1 176170048 176170210
refGene_gn=TBC1D1;refGene_tx=XM_023253204.1;refGene_gn=TBC1D1;refGene_
tx=XM_023253205.1;refGene_gn=TBC1D1;refGene_tx=XM_023253206.1;refGene_
gn=TBC1D1;refGene_tx=XM_023253208.1;refGene_gn=TBC1D1;refGene_tx=XM_02
3253209.1
B1 176172158 176172439
refGene_gn=TBC1D1;refGene_tx=XM_023253204.1;refGene_gn=TBC1D1;refGene_
tx=XM_023253205.1;refGene_gn=TBC1D1;refGene_tx=XM_023253206.1;refGene_
gn=TBC1D1;refGene_tx=XM_023253207.1;refGene_gn=TBC1D1;refGene_tx=XM_02
3253208.1;refGene_gn=TBC1D1;refGene_tx=XM_023253209.1
B1 176176083 176176170
refGene_gn=TBC1D1;refGene_tx=XM_023253204.1;refGene_gn=TBC1D1;refGene_
tx=XM_023253205.1;refGene_gn=TBC1D1;refGene_tx=XM_023253206.1;refGene_
gn=TBC1D1;refGene_tx=XM_023253207.1;refGene_gn=TBC1D1;refGene_tx=XM_02
3253208.1;refGene_gn=TBC1D1;refGene_tx=XM_023253209.1
B1 176177611 176177740
refGene_gn=TBC1D1;refGene_tx=XM_023253204.1;refGene_gn=TBC1D1;refGene_
tx=XM_023253205.1;refGene_gn=TBC1D1;refGene_tx=XM_023253206.1;refGene_
gn=TBC1D1;refGene_tx=XM_023253207.1;refGene_gn=TBC1D1;refGene_tx=XM_02

3253208.1;refGene_gn=TBC1D1;refGene_tx=XM_023253209.1
B1	176183600	176183711
refGene_gn=TBC1D1;refGene_tx=XM_023253204.1;refGene_gn=TBC1D1;refGene_tx=XM_023253205.1;refGene_gn=TBC1D1;refGene_tx=XM_023253206.1;refGene_gn=TBC1D1;refGene_tx=XM_023253207.1;refGene_gn=TBC1D1;refGene_tx=XM_023253208.1;refGene_gn=TBC1D1;refGene_tx=XM_023253209.1
B1	176190252	176190344
refGene_gn=TBC1D1;refGene_tx=XM_023253204.1;refGene_gn=TBC1D1;refGene_tx=XM_023253205.1;refGene_gn=TBC1D1;refGene_tx=XM_023253206.1;refGene_gn=TBC1D1;refGene_tx=XM_023253207.1;refGene_gn=TBC1D1;refGene_tx=XM_023253208.1;refGene_gn=TBC1D1;refGene_tx=XM_023253209.1
B1	176196430	176196563
refGene_gn=TBC1D1;refGene_tx=XM_023253204.1;refGene_gn=TBC1D1;refGene_tx=XM_023253205.1;refGene_gn=TBC1D1;refGene_tx=XM_023253206.1;refGene_gn=TBC1D1;refGene_tx=XM_023253207.1;refGene_gn=TBC1D1;refGene_tx=XM_023253208.1;refGene_gn=TBC1D1;refGene_tx=XM_023253209.1
B1	176197546	176197651
refGene_gn=TBC1D1;refGene_tx=XM_023253204.1;refGene_gn=TBC1D1;refGene_tx=XM_023253205.1;refGene_gn=TBC1D1;refGene_tx=XM_023253206.1;refGene_gn=TBC1D1;refGene_tx=XM_023253207.1;refGene_gn=TBC1D1;refGene_tx=XM_023253208.1;refGene_gn=TBC1D1;refGene_tx=XM_023253209.1
B1	176200985	176201075
refGene_gn=TBC1D1;refGene_tx=XM_023253204.1;refGene_gn=TBC1D1;refGene_tx=XM_023253205.1;refGene_gn=TBC1D1;refGene_tx=XM_023253206.1;refGene_gn=TBC1D1;refGene_tx=XM_023253207.1;refGene_gn=TBC1D1;refGene_tx=XM_023253208.1;refGene_gn=TBC1D1;refGene_tx=XM_023253209.1
B1	176203899	176204418
refGene_gn=TBC1D1;refGene_tx=XM_023253204.1;refGene_gn=TBC1D1;refGene_tx=XM_023253205.1;refGene_gn=TBC1D1;refGene_tx=XM_023253206.1;refGene_gn=TBC1D1;refGene_tx=XM_023253207.1;refGene_gn=TBC1D1;refGene_tx=XM_023253208.1;refGene_gn=TBC1D1;refGene_tx=XM_023253209.1
B1	176236093	176236123
refGene_gn=TBC1D1;refGene_tx=XM_023253208.1
B1	176301987	176302404
refGene_gn=TBC1D1;refGene_tx=XM_023253204.1;refGene_gn=TBC1D1;refGene_tx=XM_023253205.1;refGene_gn=TBC1D1;refGene_tx=XM_023253206.1;refGene_gn=TBC1D1;refGene_tx=XM_023253207.1;refGene_gn=TBC1D1;refGene_tx=XM_023253209.1
B1	176348707	176348810
refGene_gn=PGM2;refGene_tx=XM_019829651.1
B1	176354850	176354984
refGene_gn=PGM2;refGene_tx=XM_019829651.1
B1	176360787	176360977
refGene_gn=PGM2;refGene_tx=XM_019829651.1
B1	176362064	176362194
refGene_gn=PGM2;refGene_tx=XM_019829651.1
B1	176363639	176363733
refGene_gn=PGM2;refGene_tx=XM_019829651.1
B1	176363821	176364002
refGene_gn=PGM2;refGene_tx=XM_019829651.1

B1 176365102 176365200
refGene_gn=PGM2;refGene_tx=XM_019829651.1
B1 176366392 176366582
refGene_gn=PGM2;refGene_tx=XM_019829651.1
B1 176370520 176370714
refGene_gn=PGM2;refGene_tx=XM_019829651.1
B1 176370849 176370933
refGene_gn=PGM2;refGene_tx=XM_019829651.1
B1 176372872 176372957
refGene_gn=PGM2;refGene_tx=XM_019829651.1
B1 176376670 176376777
refGene_gn=PGM2;refGene_tx=XM_019829651.1
B1 176381211 176381379
refGene_gn=PGM2;refGene_tx=XM_019829651.1
B1 176386414 176386465
refGene_gn=PGM2;refGene_tx=XM_019829651.1
B1 176501802 176501890
refGene_gn=RELL1;refGene_tx=XM_023253210.1;refGene_gn=RELL1;refGene_tx
=XM_023253211.1
B1 176534465 176534690
refGene_gn=RELL1;refGene_tx=XM_023253210.1;refGene_gn=RELL1;refGene_tx
=XM_023253211.1
B1 176537031 176537103
refGene_gn=RELL1;refGene_tx=XM_023253210.1;refGene_gn=RELL1;refGene_tx
=XM_023253211.1
B1 176544329 176544387
refGene_gn=RELL1;refGene_tx=XM_023253210.1;refGene_gn=RELL1;refGene_tx
=XM_023253211.1
B1 176546764 176547001
refGene_gn=RELL1;refGene_tx=XM_023253210.1;refGene_gn=RELL1;refGene_tx
=XM_023253211.1
B1 176552762 176552898
refGene_gn=RELL1;refGene_tx=XM_023253210.1;refGene_gn=RELL1;refGene_tx
=XM_023253211.1
B1 176585603 176585906
refGene_gn=CB1H4orf19;refGene_tx=XM_023253213.1
B1 176585907 176586532
refGene_gn=CB1H4orf19;refGene_tx=XM_023253213.1
B1 176587568 176587600
refGene_gn=CB1H4orf19;refGene_tx=XM_023253213.1
B1 176641442 176641718
refGene_gn=LOC111560193;refGene_tx=XM_023253214.1
B1 176679987 176683920
refGene_gn=NWD2;refGene_tx=XM_003985445.5;refGene_gn=NWD2;refGene_tx=X
M_023253215.1
B1 176688591 176689181
refGene_gn=NWD2;refGene_tx=XM_003985445.5;refGene_gn=NWD2;refGene_tx=X
M_023253215.1
B1 176694055 176694200
refGene_gn=NWD2;refGene_tx=XM_003985445.5;refGene_gn=NWD2;refGene_tx=X

M_023253215.1
B1 176702366 176702570
refGene_gn=NWD2;refGene_tx=XM_003985445.5;refGene_gn=NWD2;refGene_tx=X
M_023253215.1
B1 176768527 176768644
refGene_gn=NWD2;refGene_tx=XM_003985445.5
B1 176785046 176785135
refGene_gn=NWD2;refGene_tx=XM_003985445.5
B1 176859479 176859630
refGene_gn=NWD2;refGene_tx=XM_003985445.5
B1 177596564 177596911
refGene_gn=DTHD1;refGene_tx=XM_003985470.3;refGene_gn=DTHD1;refGene_tx
=XM_019829658.1
B1 177600289 177600347
refGene_gn=DTHD1;refGene_tx=XM_003985470.3;refGene_gn=DTHD1;refGene_tx
=XM_019829658.1
B1 177627324 177627569
refGene_gn=DTHD1;refGene_tx=XM_003985470.3;refGene_gn=DTHD1;refGene_tx
=XM_019829658.1
B1 177634532 177634822
refGene_gn=DTHD1;refGene_tx=XM_003985470.3;refGene_gn=DTHD1;refGene_tx
=XM_019829658.1
B1 177636787 177636952
refGene_gn=DTHD1;refGene_tx=XM_003985470.3;refGene_gn=DTHD1;refGene_tx
=XM_019829658.1
B1 177648150 177648395
refGene_gn=DTHD1;refGene_tx=XM_003985470.3;refGene_gn=DTHD1;refGene_tx
=XM_019829658.1
B1 177650929 177651109
refGene_gn=DTHD1;refGene_tx=XM_003985470.3;refGene_gn=DTHD1;refGene_tx
=XM_019829658.1
B1 177653872 177654200
refGene_gn=DTHD1;refGene_tx=XM_003985470.3;refGene_gn=DTHD1;refGene_tx
=XM_019829658.1
B1 177657914 177658530
refGene_gn=DTHD1;refGene_tx=XM_003985470.3;refGene_gn=DTHD1;refGene_tx
=XM_019829658.1
B1 177661004 177661275
refGene_gn=DTHD1;refGene_tx=XM_003985470.3;refGene_gn=DTHD1;refGene_tx
=XM_019829658.1
B1 177718591 177719499
refGene_gn=ARAP2;refGene_tx=XM_003985471.5;refGene_gn=ARAP2;refGene_tx
=XM_019829654.2;refGene_gn=ARAP2;refGene_tx=XM_019829655.2;refGene_gn=
ARAP2;refGene_tx=XM_019829656.2;refGene_gn=ARAP2;refGene_tx=XM_0198296
57.2
B1 177728599 177728658
refGene_gn=ARAP2;refGene_tx=XM_003985471.5;refGene_gn=ARAP2;refGene_tx
=XM_019829654.2;refGene_gn=ARAP2;refGene_tx=XM_019829655.2;refGene_gn=
ARAP2;refGene_tx=XM_019829656.2;refGene_gn=ARAP2;refGene_tx=XM_0198296
57.2

B1 177729766 177729843
refGene_gn=ARAP2;refGene_tx=XM_003985471.5;refGene_gn=ARAP2;refGene_tx=XM_019829654.2;refGene_gn=ARAP2;refGene_tx=XM_019829655.2;refGene_gn=ARAP2;refGene_tx=XM_019829656.2;refGene_gn=ARAP2;refGene_tx=XM_019829657.2
B1 177730570 177730662
refGene_gn=ARAP2;refGene_tx=XM_003985471.5;refGene_gn=ARAP2;refGene_tx=XM_019829654.2;refGene_gn=ARAP2;refGene_tx=XM_019829655.2;refGene_gn=ARAP2;refGene_tx=XM_019829656.2;refGene_gn=ARAP2;refGene_tx=XM_019829657.2
B1 177732324 177732678
refGene_gn=ARAP2;refGene_tx=XM_003985471.5;refGene_gn=ARAP2;refGene_tx=XM_019829654.2;refGene_gn=ARAP2;refGene_tx=XM_019829655.2;refGene_gn=ARAP2;refGene_tx=XM_019829656.2;refGene_gn=ARAP2;refGene_tx=XM_019829657.2
B1 177748805 177748875
refGene_gn=ARAP2;refGene_tx=XM_003985471.5;refGene_gn=ARAP2;refGene_tx=XM_019829654.2;refGene_gn=ARAP2;refGene_tx=XM_019829655.2;refGene_gn=ARAP2;refGene_tx=XM_019829656.2;refGene_gn=ARAP2;refGene_tx=XM_019829657.2
B1 177756122 177756243
refGene_gn=ARAP2;refGene_tx=XM_003985471.5;refGene_gn=ARAP2;refGene_tx=XM_019829654.2;refGene_gn=ARAP2;refGene_tx=XM_019829655.2;refGene_gn=ARAP2;refGene_tx=XM_019829656.2;refGene_gn=ARAP2;refGene_tx=XM_019829657.2
B1 177765131 177765310
refGene_gn=ARAP2;refGene_tx=XM_003985471.5;refGene_gn=ARAP2;refGene_tx=XM_019829654.2;refGene_gn=ARAP2;refGene_tx=XM_019829655.2;refGene_gn=ARAP2;refGene_tx=XM_019829656.2;refGene_gn=ARAP2;refGene_tx=XM_019829657.2
B1 177777107 177777223
refGene_gn=ARAP2;refGene_tx=XM_003985471.5;refGene_gn=ARAP2;refGene_tx=XM_019829654.2;refGene_gn=ARAP2;refGene_tx=XM_019829655.2;refGene_gn=ARAP2;refGene_tx=XM_019829656.2;refGene_gn=ARAP2;refGene_tx=XM_019829657.2
B1 177780053 177780253
refGene_gn=ARAP2;refGene_tx=XM_003985471.5;refGene_gn=ARAP2;refGene_tx=XM_019829654.2;refGene_gn=ARAP2;refGene_tx=XM_019829655.2;refGene_gn=ARAP2;refGene_tx=XM_019829656.2;refGene_gn=ARAP2;refGene_tx=XM_019829657.2
B1 177783532 177783618
refGene_gn=ARAP2;refGene_tx=XM_003985471.5;refGene_gn=ARAP2;refGene_tx=XM_019829654.2;refGene_gn=ARAP2;refGene_tx=XM_019829655.2;refGene_gn=ARAP2;refGene_tx=XM_019829656.2;refGene_gn=ARAP2;refGene_tx=XM_019829657.2
B1 177784351 177784534
refGene_gn=ARAP2;refGene_tx=XM_003985471.5;refGene_gn=ARAP2;refGene_tx=XM_019829654.2;refGene_gn=ARAP2;refGene_tx=XM_019829655.2;refGene_gn=ARAP2;refGene_tx=XM_019829656.2;refGene_gn=ARAP2;refGene_tx=XM_019829657.2

B1 177789668 177789843
refGene_gn=ARAP2;refGene_tx=XM_003985471.5;refGene_gn=ARAP2;refGene_tx=XM_019829654.2;refGene_gn=ARAP2;refGene_tx=XM_019829655.2;refGene_gn=ARAP2;refGene_tx=XM_019829656.2;refGene_gn=ARAP2;refGene_tx=XM_019829657.2
B1 177790301 177790436
refGene_gn=ARAP2;refGene_tx=XM_003985471.5;refGene_gn=ARAP2;refGene_tx=XM_019829654.2;refGene_gn=ARAP2;refGene_tx=XM_019829655.2;refGene_gn=ARAP2;refGene_tx=XM_019829656.2;refGene_gn=ARAP2;refGene_tx=XM_019829657.2
B1 177803387 177803532
refGene_gn=ARAP2;refGene_tx=XM_003985471.5;refGene_gn=ARAP2;refGene_tx=XM_019829654.2;refGene_gn=ARAP2;refGene_tx=XM_019829655.2;refGene_gn=ARAP2;refGene_tx=XM_019829656.2;refGene_gn=ARAP2;refGene_tx=XM_019829657.2
B1 177805572 177805675
refGene_gn=ARAP2;refGene_tx=XM_003985471.5;refGene_gn=ARAP2;refGene_tx=XM_019829654.2;refGene_gn=ARAP2;refGene_tx=XM_019829655.2;refGene_gn=ARAP2;refGene_tx=XM_019829656.2;refGene_gn=ARAP2;refGene_tx=XM_019829657.2
B1 177806335 177806534
refGene_gn=ARAP2;refGene_tx=XM_003985471.5;refGene_gn=ARAP2;refGene_tx=XM_019829654.2;refGene_gn=ARAP2;refGene_tx=XM_019829655.2;refGene_gn=ARAP2;refGene_tx=XM_019829656.2;refGene_gn=ARAP2;refGene_tx=XM_019829657.2
B1 177806720 177806784
refGene_gn=ARAP2;refGene_tx=XM_003985471.5;refGene_gn=ARAP2;refGene_tx=XM_019829654.2;refGene_gn=ARAP2;refGene_tx=XM_019829655.2;refGene_gn=ARAP2;refGene_tx=XM_019829656.2;refGene_gn=ARAP2;refGene_tx=XM_019829657.2
B1 177827580 177827744
refGene_gn=ARAP2;refGene_tx=XM_003985471.5;refGene_gn=ARAP2;refGene_tx=XM_019829654.2;refGene_gn=ARAP2;refGene_tx=XM_019829655.2;refGene_gn=ARAP2;refGene_tx=XM_019829656.2;refGene_gn=ARAP2;refGene_tx=XM_019829657.2
B1 177831889 177832102
refGene_gn=ARAP2;refGene_tx=XM_003985471.5;refGene_gn=ARAP2;refGene_tx=XM_019829654.2;refGene_gn=ARAP2;refGene_tx=XM_019829655.2;refGene_gn=ARAP2;refGene_tx=XM_019829656.2;refGene_gn=ARAP2;refGene_tx=XM_019829657.2
B1 177835362 177835468
refGene_gn=ARAP2;refGene_tx=XM_003985471.5;refGene_gn=ARAP2;refGene_tx=XM_019829654.2;refGene_gn=ARAP2;refGene_tx=XM_019829655.2;refGene_gn=ARAP2;refGene_tx=XM_019829656.2;refGene_gn=ARAP2;refGene_tx=XM_019829657.2
B1 177841099 177841247
refGene_gn=ARAP2;refGene_tx=XM_003985471.5;refGene_gn=ARAP2;refGene_tx=XM_019829654.2;refGene_gn=ARAP2;refGene_tx=XM_019829655.2;refGene_gn=ARAP2;refGene_tx=XM_019829656.2;refGene_gn=ARAP2;refGene_tx=XM_019829657.2

B1 177842739 177842808
refGene_gn=ARAP2;refGene_tx=XM_003985471.5;refGene_gn=ARAP2;refGene_tx
=XM_019829654.2;refGene_gn=ARAP2;refGene_tx=XM_019829656.2;refGene_gn=
ARAP2;refGene_tx=XM_019829657.2
B1 177845340 177845415
refGene_gn=ARAP2;refGene_tx=XM_003985471.5;refGene_gn=ARAP2;refGene_tx
=XM_019829654.2;refGene_gn=ARAP2;refGene_tx=XM_019829655.2;refGene_gn=
ARAP2;refGene_tx=XM_019829656.2;refGene_gn=ARAP2;refGene_tx=XM_0198296
57.2
B1 177848183 177848301
refGene_gn=ARAP2;refGene_tx=XM_003985471.5;refGene_gn=ARAP2;refGene_tx
=XM_019829654.2;refGene_gn=ARAP2;refGene_tx=XM_019829655.2;refGene_gn=
ARAP2;refGene_tx=XM_019829656.2;refGene_gn=ARAP2;refGene_tx=XM_0198296
57.2
B1 177857305 177857434
refGene_gn=ARAP2;refGene_tx=XM_003985471.5;refGene_gn=ARAP2;refGene_tx
=XM_019829654.2;refGene_gn=ARAP2;refGene_tx=XM_019829655.2;refGene_gn=
ARAP2;refGene_tx=XM_019829656.2;refGene_gn=ARAP2;refGene_tx=XM_0198296
57.2
B1 177858690 177858734
refGene_gn=ARAP2;refGene_tx=XM_019829657.2
B1 177865713 177865790
refGene_gn=ARAP2;refGene_tx=XM_019829656.2
B1 177871336 177871476
refGene_gn=ARAP2;refGene_tx=XM_003985471.5;refGene_gn=ARAP2;refGene_tx
=XM_019829654.2;refGene_gn=ARAP2;refGene_tx=XM_019829655.2
B1 177879155 177879238
refGene_gn=ARAP2;refGene_tx=XM_003985471.5;refGene_gn=ARAP2;refGene_tx
=XM_019829654.2;refGene_gn=ARAP2;refGene_tx=XM_019829655.2
B1 177880311 177880347
refGene_gn=ARAP2;refGene_tx=XM_003985471.5;refGene_gn=ARAP2;refGene_tx
=XM_019829654.2;refGene_gn=ARAP2;refGene_tx=XM_019829655.2
B1 177882279 177882343
refGene_gn=ARAP2;refGene_tx=XM_003985471.5;refGene_gn=ARAP2;refGene_tx
=XM_019829654.2;refGene_gn=ARAP2;refGene_tx=XM_019829655.2
B1 177887665 177887800
refGene_gn=ARAP2;refGene_tx=XM_003985471.5;refGene_gn=ARAP2;refGene_tx
=XM_019829654.2;refGene_gn=ARAP2;refGene_tx=XM_019829655.2
B1 177894155 177894539
refGene_gn=ARAP2;refGene_tx=XM_003985471.5;refGene_gn=ARAP2;refGene_tx
=XM_019829654.2;refGene_gn=ARAP2;refGene_tx=XM_019829655.2
B1 182282882 182283259
refGene_gn=PCDH7;refGene_tx=XM_019829661.2;refGene_gn=PCDH7;refGene_tx
=XM_023253216.1;refGene_gn=PCDH7;refGene_tx=XM_023253217.1;refGene_gn=
PCDH7;refGene_tx=XM_023253218.1;refGene_gn=PCDH7;refGene_tx=XM_0232532
19.1;refGene_gn=PCDH7;refGene_tx=XM_023253220.1;refGene_gn=PCDH7;refGe
ne_tx=XM_023253224.1
B1 182453162 182453189
refGene_gn=PCDH7;refGene_tx=XM_019829661.2
B1 182494214 182494303

refGene_gn=PCDH7;refGene_tx=XM_023253221.1;refGene_gn=PCDH7;refGene_tx=XM_023253222.1
B1 182522425 182522642
refGene_gn=PCDH7;refGene_tx=XM_019829661.2;refGene_gn=PCDH7;refGene_tx=XM_023253216.1;refGene_gn=PCDH7;refGene_tx=XM_023253217.1;refGene_gn=PCDH7;refGene_tx=XM_023253218.1;refGene_gn=PCDH7;refGene_tx=XM_023253219.1;refGene_gn=PCDH7;refGene_tx=XM_023253220.1;refGene_gn=PCDH7;refGene_tx=XM_023253221.1;refGene_gn=PCDH7;refGene_tx=XM_023253222.1
B1 182648112 182648172
refGene_gn=PCDH7;refGene_tx=XM_019829661.2;refGene_gn=PCDH7;refGene_tx=XM_023253216.1;refGene_gn=PCDH7;refGene_tx=XM_023253217.1;refGene_gn=PCDH7;refGene_tx=XM_023253220.1;refGene_gn=PCDH7;refGene_tx=XM_023253221.1;refGene_gn=PCDH7;refGene_tx=XM_023253222.1
B1 182701918 182701954
refGene_gn=PCDH7;refGene_tx=XM_023253223.1
B1 182708182 182711374
refGene_gn=PCDH7;refGene_tx=XM_019829661.2;refGene_gn=PCDH7;refGene_tx=XM_023253216.1;refGene_gn=PCDH7;refGene_tx=XM_023253217.1;refGene_gn=PCDH7;refGene_tx=XM_023253218.1;refGene_gn=PCDH7;refGene_tx=XM_023253219.1;refGene_gn=PCDH7;refGene_tx=XM_023253220.1;refGene_gn=PCDH7;refGene_tx=XM_023253221.1;refGene_gn=PCDH7;refGene_tx=XM_023253222.1;refGene_gn=PCDH7;refGene_tx=XM_023253223.1;refGene_gn=PCDH7;refGene_tx=XM_023253224.1
B1 186080089 186080567
refGene_gn=STIM2;refGene_tx=XM_023253225.1;refGene_gn=STIM2;refGene_tx=XM_023253226.1
B1 186085455 186085729
refGene_gn=STIM2;refGene_tx=XM_023253225.1;refGene_gn=STIM2;refGene_tx=XM_023253226.1
B1 186095474 186095713
refGene_gn=STIM2;refGene_tx=XM_023253225.1;refGene_gn=STIM2;refGene_tx=XM_023253226.1
B1 186095945 186096046
refGene_gn=STIM2;refGene_tx=XM_023253225.1;refGene_gn=STIM2;refGene_tx=XM_023253226.1
B1 186096465 186096489
refGene_gn=STIM2;refGene_tx=XM_023253225.1
B1 186096751 186096919
refGene_gn=STIM2;refGene_tx=XM_023253225.1;refGene_gn=STIM2;refGene_tx=XM_023253226.1
B1 186101799 186101977
refGene_gn=STIM2;refGene_tx=XM_023253225.1;refGene_gn=STIM2;refGene_tx=XM_023253226.1
B1 186102750 186102928
refGene_gn=STIM2;refGene_tx=XM_023253225.1;refGene_gn=STIM2;refGene_tx=XM_023253226.1
B1 186105991 186106107
refGene_gn=STIM2;refGene_tx=XM_023253225.1;refGene_gn=STIM2;refGene_tx=XM_023253226.1
B1 186110047 186110159

refGene_gn=STIM2;refGene_tx=XM_023253225.1;refGene_gn=STIM2;refGene_tx=XM_023253226.1
B1 186141028 186141143
refGene_gn=STIM2;refGene_tx=XM_023253225.1;refGene_gn=STIM2;refGene_tx=XM_023253226.1
B1 186168584 186168715
refGene_gn=STIM2;refGene_tx=XM_023253225.1;refGene_gn=STIM2;refGene_tx=XM_023253226.1
B1 186230880 186231289
refGene_gn=STIM2;refGene_tx=XM_023253225.1;refGene_gn=STIM2;refGene_tx=XM_023253226.1
B1 186328843 186328918
refGene_gn=TBC1D19;refGene_tx=XM_003985476.5;refGene_gn=TBC1D19;refGene_tx=XM_006931184.4
B1 186329945 186330016
refGene_gn=TBC1D19;refGene_tx=XM_003985476.5;refGene_gn=TBC1D19;refGene_tx=XM_006931184.4
B1 186334209 186334235
refGene_gn=TBC1D19;refGene_tx=XM_019829664.2
B1 186334731 186334847
refGene_gn=TBC1D19;refGene_tx=XM_003985476.5;refGene_gn=TBC1D19;refGene_tx=XM_006931184.4;refGene_gn=TBC1D19;refGene_tx=XM_019829664.2
B1 186341913 186342005
refGene_gn=TBC1D19;refGene_tx=XM_003985476.5;refGene_gn=TBC1D19;refGene_tx=XM_006931184.4;refGene_gn=TBC1D19;refGene_tx=XM_019829664.2
B1 186344209 186344319
refGene_gn=TBC1D19;refGene_tx=XM_003985476.5;refGene_gn=TBC1D19;refGene_tx=XM_006931184.4;refGene_gn=TBC1D19;refGene_tx=XM_019829664.2
B1 186347726 186347759
refGene_gn=TBC1D19;refGene_tx=XM_003985476.5;refGene_gn=TBC1D19;refGene_tx=XM_006931184.4;refGene_gn=TBC1D19;refGene_tx=XM_019829664.2
B1 186358352 186358397
refGene_gn=TBC1D19;refGene_tx=XM_003985476.5;refGene_gn=TBC1D19;refGene_tx=XM_006931184.4;refGene_gn=TBC1D19;refGene_tx=XM_019829664.2
B1 186361256 186361341
refGene_gn=TBC1D19;refGene_tx=XM_003985476.5;refGene_gn=TBC1D19;refGene_tx=XM_006931184.4;refGene_gn=TBC1D19;refGene_tx=XM_019829664.2
B1 186386617 186388039
refGene_gn=CAR1;refGene_tx=NM_001305591.1;refGene_gn=CAR1;refGene_tx=XM_023252207.1
B1 186392387 186392450
refGene_gn=TBC1D19;refGene_tx=XM_003985476.5;refGene_gn=TBC1D19;refGene_tx=XM_006931184.4;refGene_gn=TBC1D19;refGene_tx=XM_019829664.2
B1 186395831 186395906
refGene_gn=TBC1D19;refGene_tx=XM_003985476.5;refGene_gn=TBC1D19;refGene_tx=XM_006931184.4;refGene_gn=TBC1D19;refGene_tx=XM_019829664.2
B1 186408073 186408186
refGene_gn=TBC1D19;refGene_tx=XM_003985476.5;refGene_gn=TBC1D19;refGene_tx=XM_006931184.4;refGene_gn=TBC1D19;refGene_tx=XM_019829664.2
B1 186409877 186409916

refGene_gn=TBC1D19;refGene_tx=XM_003985476.5;refGene_gn=TBC1D19;refGene_tx=XM_006931184.4;refGene_gn=TBC1D19;refGene_tx=XM_019829664.2
B1 186416171 186416244
refGene_gn=TBC1D19;refGene_tx=XM_003985476.5;refGene_gn=TBC1D19;refGene_tx=XM_006931184.4;refGene_gn=TBC1D19;refGene_tx=XM_019829664.2
B1 186423973 186424084
refGene_gn=TBC1D19;refGene_tx=XM_003985476.5;refGene_gn=TBC1D19;refGene_tx=XM_006931184.4;refGene_gn=TBC1D19;refGene_tx=XM_019829664.2
B1 186428551 186428598
refGene_gn=TBC1D19;refGene_tx=XM_003985476.5;refGene_gn=TBC1D19;refGene_tx=XM_006931184.4;refGene_gn=TBC1D19;refGene_tx=XM_019829664.2
B1 186429780 186429844
refGene_gn=TBC1D19;refGene_tx=XM_003985476.5;refGene_gn=TBC1D19;refGene_tx=XM_006931184.4;refGene_gn=TBC1D19;refGene_tx=XM_019829664.2
B1 186431083 186431158
refGene_gn=TBC1D19;refGene_tx=XM_003985476.5;refGene_gn=TBC1D19;refGene_tx=XM_006931184.4;refGene_gn=TBC1D19;refGene_tx=XM_019829664.2
B1 186442842 186442918
refGene_gn=TBC1D19;refGene_tx=XM_003985476.5;refGene_gn=TBC1D19;refGene_tx=XM_006931184.4;refGene_gn=TBC1D19;refGene_tx=XM_019829664.2
B1 186448895 186448941
refGene_gn=TBC1D19;refGene_tx=XM_003985476.5;refGene_gn=TBC1D19;refGene_tx=XM_019829664.2
B1 186449905 186449978
refGene_gn=TBC1D19;refGene_tx=XM_003985476.5;refGene_gn=TBC1D19;refGene_tx=XM_019829664.2
B1 186474905 186475004
refGene_gn=TBC1D19;refGene_tx=XM_003985476.5;refGene_gn=TBC1D19;refGene_tx=XM_019829664.2
B1 186560356 186560468
refGene_gn=CCKAR;refGene_tx=XM_003985477.4;refGene_gn=CCKAR;refGene_tx=XM_019829667.2
B1 186561178 186561430
refGene_gn=CCKAR;refGene_tx=XM_003985477.4;refGene_gn=CCKAR;refGene_tx=XM_019829667.2;refGene_gn=CCKAR;refGene_tx=XM_019829668.2
B1 186564392 186564654
refGene_gn=CCKAR;refGene_tx=XM_003985477.4;refGene_gn=CCKAR;refGene_tx=XM_019829667.2;refGene_gn=CCKAR;refGene_tx=XM_019829668.2
B1 186568038 186568166
refGene_gn=CCKAR;refGene_tx=XM_003985477.4;refGene_gn=CCKAR;refGene_tx=XM_019829667.2;refGene_gn=CCKAR;refGene_tx=XM_019829668.2
B1 186569558 186570094
refGene_gn=CCKAR;refGene_tx=XM_003985477.4;refGene_gn=CCKAR;refGene_tx=XM_019829667.2;refGene_gn=CCKAR;refGene_tx=XM_019829668.2
B1 186570212 186570315
refGene_gn=CCKAR;refGene_tx=XM_019829667.2;refGene_gn=CCKAR;refGene_tx=XM_019829668.2
B1 186617577 186617893
refGene_gn=RBPJ;refGene_tx=XM_011281983.3;refGene_gn=RBPJ;refGene_tx=XM_023253227.1;refGene_gn=RBPJ;refGene_tx=XM_023253228.1;refGene_gn=RBP

J;refGene_tx=XM_023253229.1;refGene_gn=RBPJ;refGene_tx=XM_023253230.1;
refGene_gn=RBPJ;refGene_tx=XM_023253231.1;refGene_gn=RBPJ;refGene_tx=X
M_023253232.1;refGene_gn=RBPJ;refGene_tx=XM_023253233.1;refGene_gn=RBP
J;refGene_tx=XM_023253234.1;refGene_gn=RBPJ;refGene_tx=XM_023253235.1
B1 186618050 186618154
refGene_gn=RBPJ;refGene_tx=XM_011281983.3;refGene_gn=RBPJ;refGene_tx=X
M_023253227.1;refGene_gn=RBPJ;refGene_tx=XM_023253228.1;refGene_gn=RBP
J;refGene_tx=XM_023253229.1;refGene_gn=RBPJ;refGene_tx=XM_023253230.1;
refGene_gn=RBPJ;refGene_tx=XM_023253231.1;refGene_gn=RBPJ;refGene_tx=X
M_023253232.1;refGene_gn=RBPJ;refGene_tx=XM_023253233.1;refGene_gn=RBP
J;refGene_tx=XM_023253234.1;refGene_gn=RBPJ;refGene_tx=XM_023253235.1
B1 186618502 186618658
refGene_gn=RBPJ;refGene_tx=XM_011281983.3;refGene_gn=RBPJ;refGene_tx=X
M_023253227.1;refGene_gn=RBPJ;refGene_tx=XM_023253228.1;refGene_gn=RBP
J;refGene_tx=XM_023253229.1;refGene_gn=RBPJ;refGene_tx=XM_023253230.1;
refGene_gn=RBPJ;refGene_tx=XM_023253231.1;refGene_gn=RBPJ;refGene_tx=X
M_023253232.1;refGene_gn=RBPJ;refGene_tx=XM_023253233.1;refGene_gn=RBP
J;refGene_tx=XM_023253234.1;refGene_gn=RBPJ;refGene_tx=XM_023253235.1
B1 186619710 186619851
refGene_gn=RBPJ;refGene_tx=XM_011281983.3;refGene_gn=RBPJ;refGene_tx=X
M_023253227.1;refGene_gn=RBPJ;refGene_tx=XM_023253228.1;refGene_gn=RBP
J;refGene_tx=XM_023253229.1;refGene_gn=RBPJ;refGene_tx=XM_023253230.1;
refGene_gn=RBPJ;refGene_tx=XM_023253231.1;refGene_gn=RBPJ;refGene_tx=X
M_023253232.1;refGene_gn=RBPJ;refGene_tx=XM_023253233.1;refGene_gn=RBP
J;refGene_tx=XM_023253234.1;refGene_gn=RBPJ;refGene_tx=XM_023253235.1
B1 186623031 186623144
refGene_gn=RBPJ;refGene_tx=XM_011281983.3;refGene_gn=RBPJ;refGene_tx=X
M_023253227.1;refGene_gn=RBPJ;refGene_tx=XM_023253228.1;refGene_gn=RBP
J;refGene_tx=XM_023253229.1;refGene_gn=RBPJ;refGene_tx=XM_023253230.1;
refGene_gn=RBPJ;refGene_tx=XM_023253231.1;refGene_gn=RBPJ;refGene_tx=X
M_023253232.1;refGene_gn=RBPJ;refGene_tx=XM_023253233.1;refGene_gn=RBP
J;refGene_tx=XM_023253234.1;refGene_gn=RBPJ;refGene_tx=XM_023253235.1
B1 186623296 186623434
refGene_gn=RBPJ;refGene_tx=XM_011281983.3;refGene_gn=RBPJ;refGene_tx=X
M_023253227.1;refGene_gn=RBPJ;refGene_tx=XM_023253228.1;refGene_gn=RBP
J;refGene_tx=XM_023253229.1;refGene_gn=RBPJ;refGene_tx=XM_023253230.1;
refGene_gn=RBPJ;refGene_tx=XM_023253231.1;refGene_gn=RBPJ;refGene_tx=X
M_023253232.1;refGene_gn=RBPJ;refGene_tx=XM_023253233.1;refGene_gn=RBP
J;refGene_tx=XM_023253234.1;refGene_gn=RBPJ;refGene_tx=XM_023253235.1
B1 186627410 186627585
refGene_gn=RBPJ;refGene_tx=XM_011281983.3;refGene_gn=RBPJ;refGene_tx=X
M_023253227.1;refGene_gn=RBPJ;refGene_tx=XM_023253228.1;refGene_gn=RBP
J;refGene_tx=XM_023253229.1;refGene_gn=RBPJ;refGene_tx=XM_023253230.1;
refGene_gn=RBPJ;refGene_tx=XM_023253231.1;refGene_gn=RBPJ;refGene_tx=X
M_023253232.1;refGene_gn=RBPJ;refGene_tx=XM_023253233.1;refGene_gn=RBP
J;refGene_tx=XM_023253234.1;refGene_gn=RBPJ;refGene_tx=XM_023253235.1
B1 186632643 186632809
refGene_gn=RBPJ;refGene_tx=XM_011281983.3;refGene_gn=RBPJ;refGene_tx=X
M_023253227.1;refGene_gn=RBPJ;refGene_tx=XM_023253228.1;refGene_gn=RBP
J;refGene_tx=XM_023253229.1;refGene_gn=RBPJ;refGene_tx=XM_023253230.1;

refGene_gn=RBPJ;refGene_tx=XM_023253231.1;refGene_gn=RBPJ;refGene_tx=XM_023253232.1;refGene_gn=RBPJ;refGene_tx=XM_023253233.1;refGene_gn=RBPJ;refGene_tx=XM_023253234.1;refGene_gn=RBPJ;refGene_tx=XM_023253235.1
B1 186640101 186640197
refGene_gn=RBPJ;refGene_tx=XM_011281983.3;refGene_gn=RBPJ;refGene_tx=XM_023253227.1;refGene_gn=RBPJ;refGene_tx=XM_023253228.1;refGene_gn=RBPJ;refGene_tx=XM_023253229.1;refGene_gn=RBPJ;refGene_tx=XM_023253230.1;refGene_gn=RBPJ;refGene_tx=XM_023253231.1;refGene_gn=RBPJ;refGene_tx=XM_023253232.1;refGene_gn=RBPJ;refGene_tx=XM_023253233.1;refGene_gn=RBPJ;refGene_tx=XM_023253234.1;refGene_gn=RBPJ;refGene_tx=XM_023253235.1
B1 186654357 186654396
refGene_gn=RBPJ;refGene_tx=XM_011281983.3;refGene_gn=RBPJ;refGene_tx=XM_023253227.1;refGene_gn=RBPJ;refGene_tx=XM_023253228.1;refGene_gn=RBPJ;refGene_tx=XM_023253229.1;refGene_gn=RBPJ;refGene_tx=XM_023253230.1;refGene_gn=RBPJ;refGene_tx=XM_023253235.1
B1 186678849 186678866
refGene_gn=RBPJ;refGene_tx=XM_023253229.1;refGene_gn=RBPJ;refGene_tx=XM_023253230.1
B1 186714007 186714027
refGene_gn=RBPJ;refGene_tx=XM_023253228.1
B1 186714240 186714299
refGene_gn=RBPJ;refGene_tx=XM_023253227.1;refGene_gn=RBPJ;refGene_tx=XM_023253235.1
B1 186832622 186832654
refGene_gn=RBPJ;refGene_tx=XM_011281983.3
B1 187031531 187031569
refGene_gn=SMIM20;refGene_tx=XM_023253237.1
B1 187032309 187032366
refGene_gn=SMIM20;refGene_tx=XM_023253237.1
B1 187037092 187037096
refGene_gn=SMIM20;refGene_tx=XM_023253236.1
B1 187037981 187038145
refGene_gn=SMIM20;refGene_tx=XM_023253236.1
B1 187042441 187042488
refGene_gn=SMIM20;refGene_tx=XM_023253236.1
B1 187043227 187043336
refGene_gn=SMIM20;refGene_tx=XM_023253236.1;refGene_gn=SMIM20;refGene_tx=XM_023253237.1
B1 187092927 187093020
refGene_gn=SEL1L3;refGene_tx=XM_023253238.1
B1 187093021 187093093
refGene_gn=SEL1L3;refGene_tx=XM_023253238.1
B1 187106676 187107247
refGene_gn=SEL1L3;refGene_tx=XM_023253238.1
B1 187117608 187117735
refGene_gn=SEL1L3;refGene_tx=XM_023253238.1
B1 187119384 187119506
refGene_gn=SEL1L3;refGene_tx=XM_023253238.1
B1 187119861 187119977
refGene_gn=SEL1L3;refGene_tx=XM_023253238.1

B1 187122747 187122806
refGene_gn=SEL1L3;refGene_tx=XM_023253238.1
B1 187131098 187131231
refGene_gn=SEL1L3;refGene_tx=XM_023253238.1
B1 187133053 187133186
refGene_gn=SEL1L3;refGene_tx=XM_023253238.1
B1 187134717 187134858
refGene_gn=SEL1L3;refGene_tx=XM_023253238.1
B1 187147385 187147597
refGene_gn=SEL1L3;refGene_tx=XM_023253238.1
B1 187148751 187148931
refGene_gn=SEL1L3;refGene_tx=XM_023253238.1
B1 187157825 187157945
refGene_gn=SEL1L3;refGene_tx=XM_023253238.1
B1 187159690 187159831
refGene_gn=SEL1L3;refGene_tx=XM_023253238.1
B1 187163918 187163981
refGene_gn=SEL1L3;refGene_tx=XM_023253238.1
B1 187167038 187167215
refGene_gn=SEL1L3;refGene_tx=XM_023253238.1
B1 187170413 187170541
refGene_gn=SEL1L3;refGene_tx=XM_023253238.1
B1 187172817 187172901
refGene_gn=SEL1L3;refGene_tx=XM_023253238.1
B1 187180962 187181053
refGene_gn=SEL1L3;refGene_tx=XM_023253238.1
B1 187181185 187181270
refGene_gn=SEL1L3;refGene_tx=XM_023253238.1
B1 187182502 187182612
refGene_gn=SEL1L3;refGene_tx=XM_023253238.1
B1 187188025 187188153
refGene_gn=SEL1L3;refGene_tx=XM_023253238.1
B1 187188991 187189094
refGene_gn=SEL1L3;refGene_tx=XM_023253238.1
B1 187189175 187189248
refGene_gn=SEL1L3;refGene_tx=XM_023253238.1
B1 187197827 187197961
refGene_gn=SEL1L3;refGene_tx=XM_023253238.1
B1 187245466 187246093
refGene_gn=SLC34A2;refGene_tx=XM_003985480.4;refGene_gn=SLC34A2;refGene_tx=XM_019829677.2
B1 187247769 187247894
refGene_gn=SLC34A2;refGene_tx=XM_003985480.4;refGene_gn=SLC34A2;refGene_tx=XM_019829677.2
B1 187247980 187248097
refGene_gn=SLC34A2;refGene_tx=XM_003985480.4;refGene_gn=SLC34A2;refGene_tx=XM_019829677.2
B1 187249521 187249689
refGene_gn=SLC34A2;refGene_tx=XM_003985480.4;refGene_gn=SLC34A2;refGene_tx=XM_019829677.2

B1	187250830	187250951
refGene_gn=SLC34A2;refGene_tx=XM_003985480.4;refGene_gn=SLC34A2;refGene_tx=XM_019829677.2
B1	187251378	187251474
refGene_gn=SLC34A2;refGene_tx=XM_003985480.4;refGene_gn=SLC34A2;refGene_tx=XM_019829677.2
B1	187252258	187252454
refGene_gn=SLC34A2;refGene_tx=XM_003985480.4;refGene_gn=SLC34A2;refGene_tx=XM_019829677.2
B1	187254856	187254968
refGene_gn=SLC34A2;refGene_tx=XM_003985480.4;refGene_gn=SLC34A2;refGene_tx=XM_019829677.2
B1	187256088	187256232
refGene_gn=SLC34A2;refGene_tx=XM_003985480.4;refGene_gn=SLC34A2;refGene_tx=XM_019829677.2
B1	187257706	187257835
refGene_gn=SLC34A2;refGene_tx=XM_003985480.4;refGene_gn=SLC34A2;refGene_tx=XM_019829677.2
B1	187259128	187259263
refGene_gn=SLC34A2;refGene_tx=XM_003985480.4;refGene_gn=SLC34A2;refGene_tx=XM_019829677.2
B1	187259352	187259464
refGene_gn=SLC34A2;refGene_tx=XM_003985480.4;refGene_gn=SLC34A2;refGene_tx=XM_019829677.2
B1	187449341	187449614
refGene_gn=LOC101101466;refGene_tx=XM_011281981.3
B1	187472026	187472251
refGene_gn=ANAPC4;refGene_tx=XM_023253239.1;refGene_gn=ANAPC4;refGene_tx=XM_023253240.1
B1	187472747	187472871
refGene_gn=ANAPC4;refGene_tx=XM_023253239.1;refGene_gn=ANAPC4;refGene_tx=XM_023253240.1
B1	187473563	187473737
refGene_gn=ANAPC4;refGene_tx=XM_023253239.1;refGene_gn=ANAPC4;refGene_tx=XM_023253240.1
B1	187474646	187474721
refGene_gn=ANAPC4;refGene_tx=XM_023253239.1;refGene_gn=ANAPC4;refGene_tx=XM_023253240.1
B1	187475965	187476067
refGene_gn=ANAPC4;refGene_tx=XM_023253239.1;refGene_gn=ANAPC4;refGene_tx=XM_023253240.1
B1	187476164	187476203
refGene_gn=ANAPC4;refGene_tx=XM_023253239.1;refGene_gn=ANAPC4;refGene_tx=XM_023253240.1
B1	187476281	187476343
refGene_gn=ANAPC4;refGene_tx=XM_023253239.1;refGene_gn=ANAPC4;refGene_tx=XM_023253240.1
B1	187477209	187477307
refGene_gn=ANAPC4;refGene_tx=XM_023253239.1;refGene_gn=ANAPC4;refGene_tx=XM_023253240.1

B1	187480684	187480778
refGene_gn=ANAPC4;refGene_tx=XM_023253239.1;refGene_gn=ANAPC4;refGene_tx=XM_023253240.1
B1	187482879	187482936
refGene_gn=ANAPC4;refGene_tx=XM_023253239.1;refGene_gn=ANAPC4;refGene_tx=XM_023253240.1
B1	187483488	187483545
refGene_gn=ANAPC4;refGene_tx=XM_023253239.1;refGene_gn=ANAPC4;refGene_tx=XM_023253240.1
B1	187484832	187484882
refGene_gn=ANAPC4;refGene_tx=XM_023253239.1;refGene_gn=ANAPC4;refGene_tx=XM_023253240.1
B1	187487408	187487464
refGene_gn=ANAPC4;refGene_tx=XM_023253239.1;refGene_gn=ANAPC4;refGene_tx=XM_023253240.1
B1	187490718	187490770
refGene_gn=ANAPC4;refGene_tx=XM_023253239.1;refGene_gn=ANAPC4;refGene_tx=XM_023253240.1
B1	187490851	187490952
refGene_gn=ANAPC4;refGene_tx=XM_023253239.1;refGene_gn=ANAPC4;refGene_tx=XM_023253240.1
B1	187492411	187492488
refGene_gn=ANAPC4;refGene_tx=XM_023253239.1;refGene_gn=ANAPC4;refGene_tx=XM_023253240.1
B1	187492597	187492640
refGene_gn=ANAPC4;refGene_tx=XM_023253239.1;refGene_gn=ANAPC4;refGene_tx=XM_023253240.1
B1	187493363	187493428
refGene_gn=ANAPC4;refGene_tx=XM_023253239.1;refGene_gn=ANAPC4;refGene_tx=XM_023253240.1
B1	187493836	187493923
refGene_gn=ANAPC4;refGene_tx=XM_023253239.1;refGene_gn=ANAPC4;refGene_tx=XM_023253240.1
B1	187495015	187495099
refGene_gn=ANAPC4;refGene_tx=XM_023253239.1;refGene_gn=ANAPC4;refGene_tx=XM_023253240.1
B1	187496693	187496798
refGene_gn=ANAPC4;refGene_tx=XM_023253239.1;refGene_gn=ANAPC4;refGene_tx=XM_023253240.1
B1	187497469	187497554
refGene_gn=ANAPC4;refGene_tx=XM_023253239.1;refGene_gn=ANAPC4;refGene_tx=XM_023253240.1
B1	187498184	187498229
refGene_gn=ANAPC4;refGene_tx=XM_023253239.1;refGene_gn=ANAPC4;refGene_tx=XM_023253240.1
B1	187498321	187498348
refGene_gn=ANAPC4;refGene_tx=XM_023253239.1;refGene_gn=ANAPC4;refGene_tx=XM_023253240.1
B1	187498470	187498545
refGene_gn=ANAPC4;refGene_tx=XM_023253239.1;refGene_gn=ANAPC4;refGene_

tx=XM_023253240.1
B1 187500309 187500442
refGene_gn=ANAPC4;refGene_tx=XM_023253239.1;refGene_gn=ANAPC4;refGene_
tx=XM_023253240.1
B1 187502168 187502274
refGene_gn=ANAPC4;refGene_tx=XM_023253239.1;refGene_gn=ANAPC4;refGene_
tx=XM_023253240.1
B1 187506326 187506455
refGene_gn=ANAPC4;refGene_tx=XM_023253239.1;refGene_gn=ANAPC4;refGene_
tx=XM_023253240.1
B1 187514829 187514965
refGene_gn=ZCCHC4;refGene_tx=XM_003985482.4
B1 187520214 187520359
refGene_gn=ZCCHC4;refGene_tx=XM_003985482.4
B1 187521130 187521182
refGene_gn=ZCCHC4;refGene_tx=XM_003985482.4
B1 187523127 187523203
refGene_gn=ZCCHC4;refGene_tx=XM_003985482.4
B1 187523444 187523566
refGene_gn=ZCCHC4;refGene_tx=XM_003985482.4
B1 187531639 187531740
refGene_gn=ZCCHC4;refGene_tx=XM_003985482.4
B1 187533253 187533404
refGene_gn=ZCCHC4;refGene_tx=XM_003985482.4
B1 187537137 187537210
refGene_gn=ZCCHC4;refGene_tx=XM_003985482.4
B1 187549710 187549791
refGene_gn=ZCCHC4;refGene_tx=XM_003985482.4
B1 187550635 187550911
refGene_gn=ZCCHC4;refGene_tx=XM_003985482.4
B1 187564708 187564791
refGene_gn=ZCCHC4;refGene_tx=XM_003985482.4
B1 187565530 187565649
refGene_gn=ZCCHC4;refGene_tx=XM_003985482.4
B1 187566641 187566768
refGene_gn=ZCCHC4;refGene_tx=XM_003985482.4
B1 187589613 187589787
refGene_gn=PI4K2B;refGene_tx=XM_023253242.1;refGene_gn=PI4K2B;refGene_
tx=XM_023253243.1
B1 187597294 187597354
refGene_gn=PI4K2B;refGene_tx=XM_023253242.1;refGene_gn=PI4K2B;refGene_
tx=XM_023253243.1
B1 187597918 187598052
refGene_gn=PI4K2B;refGene_tx=XM_023253242.1;refGene_gn=PI4K2B;refGene_
tx=XM_023253243.1
B1 187601346 187601446
refGene_gn=PI4K2B;refGene_tx=XM_023253242.1;refGene_gn=PI4K2B;refGene_
tx=XM_023253243.1
B1 187603312 187603371
refGene_gn=PI4K2B;refGene_tx=XM_023253242.1;refGene_gn=PI4K2B;refGene_

tx=XM_023253243.1
B1 187604518 187604672
refGene_gn=PI4K2B;refGene_tx=XM_023253242.1;refGene_gn=PI4K2B;refGene_
tx=XM_023253243.1
B1 187606510 187606642
refGene_gn=PI4K2B;refGene_tx=XM_023253242.1;refGene_gn=PI4K2B;refGene_
tx=XM_023253243.1
B1 187607220 187607421
refGene_gn=PI4K2B;refGene_tx=XM_023253242.1;refGene_gn=PI4K2B;refGene_
tx=XM_023253243.1
B1 187609065 187609202
refGene_gn=PI4K2B;refGene_tx=XM_023253242.1;refGene_gn=PI4K2B;refGene_
tx=XM_023253243.1
B1 187617757 187618064
refGene_gn=PI4K2B;refGene_tx=XM_023253242.1;refGene_gn=PI4K2B;refGene_
tx=XM_023253243.1
B1 187687813 187687927
refGene_gn=SEPSECS;refGene_tx=XM_003985484.5;refGene_gn=SEPSECS;refGen
e_tx=XM_019829679.2
B1 187688736 187688891
refGene_gn=SEPSECS;refGene_tx=XM_003985484.5;refGene_gn=SEPSECS;refGen
e_tx=XM_019829678.2;refGene_gn=SEPSECS;refGene_tx=XM_019829679.2
B1 187691513 187691632
refGene_gn=SEPSECS;refGene_tx=XM_003985484.5;refGene_gn=SEPSECS;refGen
e_tx=XM_019829678.2;refGene_gn=SEPSECS;refGene_tx=XM_019829679.2
B1 187691962 187692121
refGene_gn=SEPSECS;refGene_tx=XM_003985484.5;refGene_gn=SEPSECS;refGen
e_tx=XM_019829678.2;refGene_gn=SEPSECS;refGene_tx=XM_019829679.2
B1 187695044 187695198
refGene_gn=SEPSECS;refGene_tx=XM_003985484.5;refGene_gn=SEPSECS;refGen
e_tx=XM_019829678.2;refGene_gn=SEPSECS;refGene_tx=XM_019829679.2
B1 187698770 187698873
refGene_gn=SEPSECS;refGene_tx=XM_003985484.5;refGene_gn=SEPSECS;refGen
e_tx=XM_019829678.2;refGene_gn=SEPSECS;refGene_tx=XM_019829679.2
B1 187706078 187706208
refGene_gn=SEPSECS;refGene_tx=XM_003985484.5;refGene_gn=SEPSECS;refGen
e_tx=XM_019829678.2;refGene_gn=SEPSECS;refGene_tx=XM_019829679.2
B1 187706330 187706422
refGene_gn=SEPSECS;refGene_tx=XM_003985484.5;refGene_gn=SEPSECS;refGen
e_tx=XM_019829678.2;refGene_gn=SEPSECS;refGene_tx=XM_019829679.2
B1 187717689 187717783
refGene_gn=SEPSECS;refGene_tx=XM_003985484.5;refGene_gn=SEPSECS;refGen
e_tx=XM_019829678.2;refGene_gn=SEPSECS;refGene_tx=XM_019829679.2
B1 187719445 187719536
refGene_gn=SEPSECS;refGene_tx=XM_003985484.5;refGene_gn=SEPSECS;refGen
e_tx=XM_019829678.2;refGene_gn=SEPSECS;refGene_tx=XM_019829679.2
B1 187719614 187719633
refGene_gn=SEPSECS;refGene_tx=XM_019829679.2
B1 187720936 187721240
refGene_gn=SEPSECS;refGene_tx=XM_003985484.5;refGene_gn=SEPSECS;refGen

e_tx=XM_019829678.2
B1	187801258	187801467
refGene_gn=LGI2;refGene_tx=XM_023253245.1
B1	187803359	187803431
refGene_gn=LGI2;refGene_tx=XM_023253245.1
B1	187805014	187805086
refGene_gn=LGI2;refGene_tx=XM_023253245.1
B1	187807444	187807516
refGene_gn=LGI2;refGene_tx=XM_023253245.1
B1	187812410	187812482
refGene_gn=LGI2;refGene_tx=XM_023253245.1
B1	187813419	187813589
refGene_gn=LGI2;refGene_tx=XM_023253245.1
B1	187820232	187820397
refGene_gn=LGI2;refGene_tx=XM_023253245.1
B1	187827777	187828595
refGene_gn=LGI2;refGene_tx=XM_023253245.1
B1	187897041	187897089
refGene_gn=CCDC149;refGene_tx=XM_023253246.1;refGene_gn=CCDC149;refGen
e_tx=XM_023253247.1
B1	187915936	187915942
refGene_gn=CCDC149;refGene_tx=XM_019829682.2
B1	187932436	187932598
refGene_gn=CCDC149;refGene_tx=XM_019829682.2;refGene_gn=CCDC149;refGen
e_tx=XM_019829683.2;refGene_gn=CCDC149;refGene_tx=XM_023253246.1;refGe
ne_gn=CCDC149;refGene_tx=XM_023253247.1;refGene_gn=CCDC149;refGene_tx=
XM_023253249.1
B1	187934566	187934605
refGene_gn=CCDC149;refGene_tx=XM_019829682.2;refGene_gn=CCDC149;refGen
e_tx=XM_019829683.2;refGene_gn=CCDC149;refGene_tx=XM_023253246.1;refGe
ne_gn=CCDC149;refGene_tx=XM_023253247.1;refGene_gn=CCDC149;refGene_tx=
XM_023253249.1
B1	187954097	187954205
refGene_gn=CCDC149;refGene_tx=XM_019829682.2;refGene_gn=CCDC149;refGen
e_tx=XM_019829683.2;refGene_gn=CCDC149;refGene_tx=XM_023253246.1;refGe
ne_gn=CCDC149;refGene_tx=XM_023253247.1;refGene_gn=CCDC149;refGene_tx=
XM_023253249.1
B1	187968305	187968422
refGene_gn=CCDC149;refGene_tx=XM_019829682.2;refGene_gn=CCDC149;refGen
e_tx=XM_019829683.2;refGene_gn=CCDC149;refGene_tx=XM_023253246.1;refGe
ne_gn=CCDC149;refGene_tx=XM_023253247.1;refGene_gn=CCDC149;refGene_tx=
XM_023253249.1
B1	187969092	187969265
refGene_gn=CCDC149;refGene_tx=XM_019829682.2;refGene_gn=CCDC149;refGen
e_tx=XM_019829683.2;refGene_gn=CCDC149;refGene_tx=XM_023253246.1;refGe
ne_gn=CCDC149;refGene_tx=XM_023253247.1;refGene_gn=CCDC149;refGene_tx=
XM_023253249.1
B1	187970009	187970082
refGene_gn=CCDC149;refGene_tx=XM_019829682.2;refGene_gn=CCDC149;refGen
e_tx=XM_019829683.2;refGene_gn=CCDC149;refGene_tx=XM_023253246.1;refGe

ne_gn=CCDC149;refGene_tx=XM_023253247.1;refGene_gn=CCDC149;refGene_tx=XM_023253249.1
B1 187971508 187971593
refGene_gn=CCDC149;refGene_tx=XM_019829682.2;refGene_gn=CCDC149;refGene_tx=XM_019829683.2;refGene_gn=CCDC149;refGene_tx=XM_023253246.1;refGene_gn=CCDC149;refGene_tx=XM_023253247.1;refGene_gn=CCDC149;refGene_tx=XM_023253249.1
B1 187973887 187974032
refGene_gn=CCDC149;refGene_tx=XM_019829682.2;refGene_gn=CCDC149;refGene_tx=XM_019829683.2;refGene_gn=CCDC149;refGene_tx=XM_023253246.1;refGene_gn=CCDC149;refGene_tx=XM_023253247.1;refGene_gn=CCDC149;refGene_tx=XM_023253249.1
B1 187984103 187984180
refGene_gn=CCDC149;refGene_tx=XM_019829682.2;refGene_gn=CCDC149;refGene_tx=XM_019829683.2;refGene_gn=CCDC149;refGene_tx=XM_023253246.1;refGene_gn=CCDC149;refGene_tx=XM_023253247.1;refGene_gn=CCDC149;refGene_tx=XM_023253249.1
B1 187986040 187986073
refGene_gn=CCDC149;refGene_tx=XM_019829682.2;refGene_gn=CCDC149;refGene_tx=XM_019829683.2;refGene_gn=CCDC149;refGene_tx=XM_023253246.1;refGene_gn=CCDC149;refGene_tx=XM_023253249.1
B1 187991845 187991962
refGene_gn=CCDC149;refGene_tx=XM_019829682.2;refGene_gn=CCDC149;refGene_tx=XM_019829683.2;refGene_gn=CCDC149;refGene_tx=XM_023253246.1;refGene_gn=CCDC149;refGene_tx=XM_023253247.1;refGene_gn=CCDC149;refGene_tx=XM_023253249.1
B1 187998052 187998486
refGene_gn=CCDC149;refGene_tx=XM_019829682.2;refGene_gn=CCDC149;refGene_tx=XM_019829683.2;refGene_gn=CCDC149;refGene_tx=XM_023253246.1;refGene_gn=CCDC149;refGene_tx=XM_023253247.1;refGene_gn=CCDC149;refGene_tx=XM_023253249.1
B1 188007021 188007762
refGene_gn=SOD3;refGene_tx=XM_023253250.1
B1 188190655 188190726
refGene_gn=DHX15;refGene_tx=XM_003985486.4
B1 188197498 188197934
refGene_gn=DHX15;refGene_tx=XM_003985486.4
B1 188203304 188203498
refGene_gn=DHX15;refGene_tx=XM_003985486.4
B1 188221148 188221308
refGene_gn=DHX15;refGene_tx=XM_003985486.4
B1 188223602 188223821
refGene_gn=DHX15;refGene_tx=XM_003985486.4
B1 188228805 188228973
refGene_gn=DHX15;refGene_tx=XM_003985486.4
B1 188236233 188236320
refGene_gn=DHX15;refGene_tx=XM_003985486.4
B1 188237559 188237709
refGene_gn=DHX15;refGene_tx=XM_003985486.4
B1 188238110 188238219

refGene_gn=DHX15;refGene_tx=XM_003985486.4
B1	188238958	188239150
refGene_gn=DHX15;refGene_tx=XM_003985486.4
B1	188242037	188242160
refGene_gn=DHX15;refGene_tx=XM_003985486.4
B1	188247474	188247665
refGene_gn=DHX15;refGene_tx=XM_003985486.4
B1	188250070	188250208
refGene_gn=DHX15;refGene_tx=XM_003985486.4
B1	188250209	188250241
refGene_gn=DHX15;refGene_tx=XM_003985486.4
B1	188251706	188251824
refGene_gn=DHX15;refGene_tx=XM_003985486.4
B1	188483785	188483803
refGene_gn=PPARGC1A;refGene_tx=XM_019829686.2
B1	188645693	188645762
refGene_gn=PPARGC1A;refGene_tx=XM_019829685.2
B1	188854081	188854129
refGene_gn=PPARGC1A;refGene_tx=XM_003985487.5
B1	188855678	188855777
refGene_gn=PPARGC1A;refGene_tx=XM_019829684.2;refGene_gn=PPARGC1A;refGene_tx=XM_023253252.1
B1	188859142	188859322
refGene_gn=PPARGC1A;refGene_tx=XM_003985487.5;refGene_gn=PPARGC1A;refGene_tx=XM_019829684.2;refGene_gn=PPARGC1A;refGene_tx=XM_019829685.2;refGene_gn=PPARGC1A;refGene_tx=XM_019829686.2;refGene_gn=PPARGC1A;refGene_tx=XM_023253252.1
B1	188910530	188910725
refGene_gn=PPARGC1A;refGene_tx=XM_003985487.5;refGene_gn=PPARGC1A;refGene_tx=XM_019829684.2;refGene_gn=PPARGC1A;refGene_tx=XM_019829685.2;refGene_gn=PPARGC1A;refGene_tx=XM_019829686.2;refGene_gn=PPARGC1A;refGene_tx=XM_023253252.1
B1	188912765	188912888
refGene_gn=PPARGC1A;refGene_tx=XM_003985487.5;refGene_gn=PPARGC1A;refGene_tx=XM_019829684.2;refGene_gn=PPARGC1A;refGene_tx=XM_019829685.2;refGene_gn=PPARGC1A;refGene_tx=XM_019829686.2;refGene_gn=PPARGC1A;refGene_tx=XM_023253252.1
B1	188913748	188913953
refGene_gn=PPARGC1A;refGene_tx=XM_003985487.5;refGene_gn=PPARGC1A;refGene_tx=XM_019829684.2;refGene_gn=PPARGC1A;refGene_tx=XM_019829685.2;refGene_gn=PPARGC1A;refGene_tx=XM_019829686.2;refGene_gn=PPARGC1A;refGene_tx=XM_023253252.1
B1	188918228	188918274
refGene_gn=PPARGC1A;refGene_tx=XM_003985487.5;refGene_gn=PPARGC1A;refGene_tx=XM_019829684.2;refGene_gn=PPARGC1A;refGene_tx=XM_019829685.2;refGene_gn=PPARGC1A;refGene_tx=XM_019829686.2;refGene_gn=PPARGC1A;refGene_tx=XM_023253252.1
B1	188918383	188918457
refGene_gn=PPARGC1A;refGene_tx=XM_003985487.5;refGene_gn=PPARGC1A;refGene_tx=XM_019829684.2;refGene_gn=PPARGC1A;refGene_tx=XM_019829685.2;re

fGene_gn=PPARGC1A;refGene_tx=XM_019829686.2;refGene_gn=PPARGC1A;refGene_tx=XM_023253252.1
B1 188931721 188932637
refGene_gn=PPARGC1A;refGene_tx=XM_003985487.5;refGene_gn=PPARGC1A;refGene_tx=XM_019829684.2;refGene_gn=PPARGC1A;refGene_tx=XM_019829685.2;refGene_gn=PPARGC1A;refGene_tx=XM_019829686.2;refGene_gn=PPARGC1A;refGene_tx=XM_023253252.1
B1 188933237 188933342
refGene_gn=PPARGC1A;refGene_tx=XM_003985487.5;refGene_gn=PPARGC1A;refGene_tx=XM_019829684.2;refGene_gn=PPARGC1A;refGene_tx=XM_019829685.2;refGene_gn=PPARGC1A;refGene_tx=XM_019829686.2;refGene_gn=PPARGC1A;refGene_tx=XM_023253252.1
B1 188933501 188933622
refGene_gn=PPARGC1A;refGene_tx=XM_003985487.5;refGene_gn=PPARGC1A;refGene_tx=XM_019829684.2;refGene_gn=PPARGC1A;refGene_tx=XM_019829685.2;refGene_gn=PPARGC1A;refGene_tx=XM_019829686.2;refGene_gn=PPARGC1A;refGene_tx=XM_023253252.1
B1 188942820 188942942
refGene_gn=PPARGC1A;refGene_tx=XM_003985487.5;refGene_gn=PPARGC1A;refGene_tx=XM_019829684.2;refGene_gn=PPARGC1A;refGene_tx=XM_019829685.2;refGene_gn=PPARGC1A;refGene_tx=XM_019829686.2;refGene_gn=PPARGC1A;refGene_tx=XM_023253252.1
B1 188943275 188943427
refGene_gn=PPARGC1A;refGene_tx=XM_003985487.5;refGene_gn=PPARGC1A;refGene_tx=XM_019829684.2;refGene_gn=PPARGC1A;refGene_tx=XM_019829685.2;refGene_gn=PPARGC1A;refGene_tx=XM_019829686.2;refGene_gn=PPARGC1A;refGene_tx=XM_023253252.1
B1 188949973 188950077
refGene_gn=PPARGC1A;refGene_tx=XM_003985487.5;refGene_gn=PPARGC1A;refGene_tx=XM_019829684.2;refGene_gn=PPARGC1A;refGene_tx=XM_019829685.2;refGene_gn=PPARGC1A;refGene_tx=XM_019829686.2;refGene_gn=PPARGC1A;refGene_tx=XM_023253252.1
B1 189742838 189742879
refGene_gn=LOC101082399;refGene_tx=XM_019829688.2
B1 189768473 189768676
refGene_gn=LOC101082399;refGene_tx=XM_019829687.1
B1 189821337 189821453
refGene_gn=LOC101082399;refGene_tx=XM_023253253.1
B1 189828622 189828746
refGene_gn=LOC101082399;refGene_tx=XM_019829687.1;refGene_gn=LOC101082399;refGene_tx=XM_019829688.2;refGene_gn=LOC101082399;refGene_tx=XM_023253253.1
B1 189829466 189830263
refGene_gn=LOC101082399;refGene_tx=XM_019829687.1;refGene_gn=LOC101082399;refGene_tx=XM_019829688.2;refGene_gn=LOC101082399;refGene_tx=XM_023253253.1
B1 189844718 189844946
refGene_gn=LOC101082399;refGene_tx=XM_019829687.1;refGene_gn=LOC101082399;refGene_tx=XM_019829688.2;refGene_gn=LOC101082399;refGene_tx=XM_023253253.1

B1 189874328 189874386
refGene_gn=LOC101082399;refGene_tx=XM_019829687.1;refGene_gn=LOC101082
399;refGene_tx=XM_019829688.2;refGene_gn=LOC101082399;refGene_tx=XM_02
3253253.1
B1 190002916 190003161
refGene_gn=ADGRA3;refGene_tx=XM_023253254.1
B1 190039418 190039490
refGene_gn=ADGRA3;refGene_tx=XM_023253254.1
B1 190050662 190050734
refGene_gn=ADGRA3;refGene_tx=XM_023253254.1
B1 190056782 190056854
refGene_gn=ADGRA3;refGene_tx=XM_023253254.1
B1 190064186 190064258
refGene_gn=ADGRA3;refGene_tx=XM_023253254.1
B1 190066097 190066258
refGene_gn=ADGRA3;refGene_tx=XM_023253254.1
B1 190069357 190069571
refGene_gn=ADGRA3;refGene_tx=XM_023253254.1
B1 190074781 190074946
refGene_gn=ADGRA3;refGene_tx=XM_023253254.1
B1 190077388 190077590
refGene_gn=ADGRA3;refGene_tx=XM_023253254.1
B1 190078514 190078670
refGene_gn=ADGRA3;refGene_tx=XM_023253254.1
B1 190099366 190099528
refGene_gn=ADGRA3;refGene_tx=XM_023253254.1
B1 190103183 190103387
refGene_gn=ADGRA3;refGene_tx=XM_023253254.1
B1 190112891 190113105
refGene_gn=ADGRA3;refGene_tx=XM_023253254.1
B1 190113316 190113525
refGene_gn=ADGRA3;refGene_tx=XM_023253254.1
B1 190123806 190123931
refGene_gn=ADGRA3;refGene_tx=XM_023253254.1
B1 190125008 190125132
refGene_gn=ADGRA3;refGene_tx=XM_023253254.1
B1 190134426 190134572
refGene_gn=ADGRA3;refGene_tx=XM_023253254.1
B1 190138074 190138170
refGene_gn=ADGRA3;refGene_tx=XM_023253254.1
B1 190138309 190139552
refGene_gn=ADGRA3;refGene_tx=XM_023253254.1
B1 190503025 190503086
refGene_gn=KCNIP4;refGene_tx=XM_019829694.1;refGene_gn=KCNIP4;refGene_
tx=XM_019829697.1
B1 190750402 190750490
refGene_gn=KCNIP4;refGene_tx=XM_019829696.2
B1 191119306 191119406
refGene_gn=KCNIP4;refGene_tx=XM_019829695.2
B1 191433470 191433582

refGene_gn=KCNIP4;refGene_tx=XM_011281974.2
B1 191527589 191527691
refGene_gn=KCNIP4;refGene_tx=XM_019829694.1
B1 191550381 191550506
refGene_gn=KCNIP4;refGene_tx=XM_011281974.2;refGene_gn=KCNIP4;refGene_tx=XM_019829694.1;refGene_gn=KCNIP4;refGene_tx=XM_019829695.2;refGene_gn=KCNIP4;refGene_tx=XM_019829696.2;refGene_gn=KCNIP4;refGene_tx=XM_019829697.1
B1 191627062 191627132
refGene_gn=KCNIP4;refGene_tx=XM_011281974.2;refGene_gn=KCNIP4;refGene_tx=XM_019829694.1;refGene_gn=KCNIP4;refGene_tx=XM_019829695.2;refGene_gn=KCNIP4;refGene_tx=XM_019829696.2;refGene_gn=KCNIP4;refGene_tx=XM_019829697.1
B1 191635163 191635234
refGene_gn=KCNIP4;refGene_tx=XM_011281974.2;refGene_gn=KCNIP4;refGene_tx=XM_019829694.1;refGene_gn=KCNIP4;refGene_tx=XM_019829695.2;refGene_gn=KCNIP4;refGene_tx=XM_019829696.2;refGene_gn=KCNIP4;refGene_tx=XM_019829697.1
B1 191655596 191655704
refGene_gn=KCNIP4;refGene_tx=XM_011281974.2;refGene_gn=KCNIP4;refGene_tx=XM_019829694.1;refGene_gn=KCNIP4;refGene_tx=XM_019829695.2;refGene_gn=KCNIP4;refGene_tx=XM_019829696.2;refGene_gn=KCNIP4;refGene_tx=XM_019829697.1
B1 191657747 191657852
refGene_gn=KCNIP4;refGene_tx=XM_011281974.2;refGene_gn=KCNIP4;refGene_tx=XM_019829694.1;refGene_gn=KCNIP4;refGene_tx=XM_019829695.2;refGene_gn=KCNIP4;refGene_tx=XM_019829696.2;refGene_gn=KCNIP4;refGene_tx=XM_019829697.1
B1 191658454 191658517
refGene_gn=KCNIP4;refGene_tx=XM_011281974.2;refGene_gn=KCNIP4;refGene_tx=XM_019829694.1;refGene_gn=KCNIP4;refGene_tx=XM_019829695.2;refGene_gn=KCNIP4;refGene_tx=XM_019829696.2;refGene_gn=KCNIP4;refGene_tx=XM_019829697.1
B1 191660299 191660347
refGene_gn=KCNIP4;refGene_tx=XM_011281974.2;refGene_gn=KCNIP4;refGene_tx=XM_019829694.1;refGene_gn=KCNIP4;refGene_tx=XM_019829695.2;refGene_gn=KCNIP4;refGene_tx=XM_019829696.2;refGene_gn=KCNIP4;refGene_tx=XM_019829697.1
B1 191663220 191663277
refGene_gn=PACRGL;refGene_tx=XM_011281977.2;refGene_gn=PACRGL;refGene_tx=XM_019829689.1;refGene_gn=PACRGL;refGene_tx=XM_019829690.1;refGene_gn=PACRGL;refGene_tx=XM_019829693.2;refGene_gn=PACRGL;refGene_tx=XM_023253255.1;refGene_gn=PACRGL;refGene_tx=XM_023253256.1;refGene_gn=PACRGL;refGene_tx=XM_023253257.1
B1 191666101 191666182
refGene_gn=PACRGL;refGene_tx=XM_011281977.2;refGene_gn=PACRGL;refGene_tx=XM_019829689.1;refGene_gn=PACRGL;refGene_tx=XM_019829690.1;refGene_gn=PACRGL;refGene_tx=XM_019829693.2;refGene_gn=PACRGL;refGene_tx=XM_023253255.1;refGene_gn=PACRGL;refGene_tx=XM_023253256.1;refGene_gn=PACRGL;refGene_tx=XM_023253257.1

B1	191667912	191668020
refGene_gn=PACRGL;refGene_tx=XM_011281977.2;refGene_gn=PACRGL;refGene_tx=XM_019829689.1;refGene_gn=PACRGL;refGene_tx=XM_019829690.1;refGene_gn=PACRGL;refGene_tx=XM_019829693.2;refGene_gn=PACRGL;refGene_tx=XM_023253255.1;refGene_gn=PACRGL;refGene_tx=XM_023253256.1;refGene_gn=PACRGL;refGene_tx=XM_023253257.1
B1	191668489	191668624
refGene_gn=PACRGL;refGene_tx=XM_011281977.2;refGene_gn=PACRGL;refGene_tx=XM_019829689.1;refGene_gn=PACRGL;refGene_tx=XM_019829690.1;refGene_gn=PACRGL;refGene_tx=XM_019829693.2;refGene_gn=PACRGL;refGene_tx=XM_023253255.1;refGene_gn=PACRGL;refGene_tx=XM_023253256.1;refGene_gn=PACRGL;refGene_tx=XM_023253257.1
B1	191671323	191671414
refGene_gn=PACRGL;refGene_tx=XM_011281977.2;refGene_gn=PACRGL;refGene_tx=XM_019829689.1;refGene_gn=PACRGL;refGene_tx=XM_019829690.1;refGene_gn=PACRGL;refGene_tx=XM_019829693.2;refGene_gn=PACRGL;refGene_tx=XM_023253255.1;refGene_gn=PACRGL;refGene_tx=XM_023253256.1;refGene_gn=PACRGL;refGene_tx=XM_023253257.1
B1	191673407	191673475
refGene_gn=PACRGL;refGene_tx=XM_011281977.2;refGene_gn=PACRGL;refGene_tx=XM_019829689.1;refGene_gn=PACRGL;refGene_tx=XM_019829690.1;refGene_gn=PACRGL;refGene_tx=XM_019829693.2;refGene_gn=PACRGL;refGene_tx=XM_023253255.1;refGene_gn=PACRGL;refGene_tx=XM_023253256.1;refGene_gn=PACRGL;refGene_tx=XM_023253257.1
B1	191676524	191676679
refGene_gn=PACRGL;refGene_tx=XM_011281977.2;refGene_gn=PACRGL;refGene_tx=XM_019829689.1;refGene_gn=PACRGL;refGene_tx=XM_019829690.1;refGene_gn=PACRGL;refGene_tx=XM_019829693.2;refGene_gn=PACRGL;refGene_tx=XM_023253255.1;refGene_gn=PACRGL;refGene_tx=XM_023253256.1;refGene_gn=PACRGL;refGene_tx=XM_023253257.1
B1	191676807	191676875
refGene_gn=PACRGL;refGene_tx=XM_011281977.2;refGene_gn=PACRGL;refGene_tx=XM_019829689.1;refGene_gn=PACRGL;refGene_tx=XM_019829690.1;refGene_gn=PACRGL;refGene_tx=XM_019829693.2;refGene_gn=PACRGL;refGene_tx=XM_023253255.1;refGene_gn=PACRGL;refGene_tx=XM_023253256.1;refGene_gn=PACRGL;refGene_tx=XM_023253257.1
B1	191678600	191678725
refGene_gn=PACRGL;refGene_tx=XM_023253255.1;refGene_gn=PACRGL;refGene_tx=XM_023253256.1
B1	191734346	191734588
refGene_gn=SLIT2;refGene_tx=XM_023253260.1;refGene_gn=SLIT2;refGene_tx=XM_023253261.1;refGene_gn=SLIT2;refGene_tx=XM_023253262.1;refGene_gn=SLIT2;refGene_tx=XM_023253263.1;refGene_gn=SLIT2;refGene_tx=XM_023253264.1;refGene_gn=SLIT2;refGene_tx=XM_023253265.1;refGene_gn=SLIT2;refGene_tx=XM_023253266.1;refGene_gn=SLIT2;refGene_tx=XM_023253267.1;refGene_gn=SLIT2;refGene_tx=XM_023253268.1;refGene_gn=SLIT2;refGene_tx=XM_023253269.1;refGene_gn=SLIT2;refGene_tx=XM_023253270.1
B1	191735686	191735898
refGene_gn=SLIT2;refGene_tx=XM_023253260.1;refGene_gn=SLIT2;refGene_tx=XM_023253261.1;refGene_gn=SLIT2;refGene_tx=XM_023253262.1;refGene_gn=

SLIT2;refGene_tx=XM_023253263.1;refGene_gn=SLIT2;refGene_tx=XM_0232532
64.1;refGene_gn=SLIT2;refGene_tx=XM_023253265.1;refGene_gn=SLIT2;refGe
ne_tx=XM_023253266.1;refGene_gn=SLIT2;refGene_tx=XM_023253267.1;refGen
e_gn=SLIT2;refGene_tx=XM_023253268.1;refGene_gn=SLIT2;refGene_tx=XM_02
3253269.1;refGene_gn=SLIT2;refGene_tx=XM_023253270.1
B1 191736110 191736402
refGene_gn=SLIT2;refGene_tx=XM_023253260.1;refGene_gn=SLIT2;refGene_tx
=XM_023253261.1;refGene_gn=SLIT2;refGene_tx=XM_023253262.1;refGene_gn=
SLIT2;refGene_tx=XM_023253263.1;refGene_gn=SLIT2;refGene_tx=XM_0232532
64.1;refGene_gn=SLIT2;refGene_tx=XM_023253265.1;refGene_gn=SLIT2;refGe
ne_tx=XM_023253266.1;refGene_gn=SLIT2;refGene_tx=XM_023253267.1;refGen
e_gn=SLIT2;refGene_tx=XM_023253268.1;refGene_gn=SLIT2;refGene_tx=XM_02
3253269.1;refGene_gn=SLIT2;refGene_tx=XM_023253270.1
B1 191742009 191742164
refGene_gn=SLIT2;refGene_tx=XM_023253260.1;refGene_gn=SLIT2;refGene_tx
=XM_023253261.1;refGene_gn=SLIT2;refGene_tx=XM_023253262.1;refGene_gn=
SLIT2;refGene_tx=XM_023253263.1;refGene_gn=SLIT2;refGene_tx=XM_0232532
64.1;refGene_gn=SLIT2;refGene_tx=XM_023253265.1;refGene_gn=SLIT2;refGe
ne_tx=XM_023253266.1;refGene_gn=SLIT2;refGene_tx=XM_023253267.1;refGen
e_gn=SLIT2;refGene_tx=XM_023253268.1;refGene_gn=SLIT2;refGene_tx=XM_02
3253269.1;refGene_gn=SLIT2;refGene_tx=XM_023253270.1
B1 191754606 191754737
refGene_gn=SLIT2;refGene_tx=XM_023253260.1;refGene_gn=SLIT2;refGene_tx
=XM_023253261.1;refGene_gn=SLIT2;refGene_tx=XM_023253262.1;refGene_gn=
SLIT2;refGene_tx=XM_023253263.1;refGene_gn=SLIT2;refGene_tx=XM_0232532
64.1;refGene_gn=SLIT2;refGene_tx=XM_023253265.1;refGene_gn=SLIT2;refGe
ne_tx=XM_023253266.1;refGene_gn=SLIT2;refGene_tx=XM_023253267.1;refGen
e_gn=SLIT2;refGene_tx=XM_023253268.1;refGene_gn=SLIT2;refGene_tx=XM_02
3253269.1;refGene_gn=SLIT2;refGene_tx=XM_023253270.1
B1 191756089 191756330
refGene_gn=SLIT2;refGene_tx=XM_023253260.1;refGene_gn=SLIT2;refGene_tx
=XM_023253261.1;refGene_gn=SLIT2;refGene_tx=XM_023253262.1;refGene_gn=
SLIT2;refGene_tx=XM_023253263.1;refGene_gn=SLIT2;refGene_tx=XM_0232532
64.1;refGene_gn=SLIT2;refGene_tx=XM_023253265.1;refGene_gn=SLIT2;refGe
ne_tx=XM_023253266.1;refGene_gn=SLIT2;refGene_tx=XM_023253267.1;refGen
e_gn=SLIT2;refGene_tx=XM_023253268.1;refGene_gn=SLIT2;refGene_tx=XM_02
3253269.1;refGene_gn=SLIT2;refGene_tx=XM_023253270.1
B1 191756440 191756578
refGene_gn=SLIT2;refGene_tx=XM_023253260.1;refGene_gn=SLIT2;refGene_tx
=XM_023253261.1;refGene_gn=SLIT2;refGene_tx=XM_023253262.1;refGene_gn=
SLIT2;refGene_tx=XM_023253263.1;refGene_gn=SLIT2;refGene_tx=XM_0232532
64.1;refGene_gn=SLIT2;refGene_tx=XM_023253265.1;refGene_gn=SLIT2;refGe
ne_tx=XM_023253266.1;refGene_gn=SLIT2;refGene_tx=XM_023253267.1;refGen
e_gn=SLIT2;refGene_tx=XM_023253268.1;refGene_gn=SLIT2;refGene_tx=XM_02
3253269.1;refGene_gn=SLIT2;refGene_tx=XM_023253270.1
B1 191762822 191762916
refGene_gn=SLIT2;refGene_tx=XM_023253260.1;refGene_gn=SLIT2;refGene_tx
=XM_023253261.1;refGene_gn=SLIT2;refGene_tx=XM_023253262.1;refGene_gn=
SLIT2;refGene_tx=XM_023253263.1;refGene_gn=SLIT2;refGene_tx=XM_0232532
64.1;refGene_gn=SLIT2;refGene_tx=XM_023253265.1;refGene_gn=SLIT2;refGe

ne_tx=XM_023253266.1;refGene_gn=SLIT2;refGene_tx=XM_023253267.1;refGen
e_gn=SLIT2;refGene_tx=XM_023253268.1;refGene_gn=SLIT2;refGene_tx=XM_02
3253269.1;refGene_gn=SLIT2;refGene_tx=XM_023253270.1
B1 191778580 191778607
refGene_gn=SLIT2;refGene_tx=XM_023253260.1;refGene_gn=SLIT2;refGene_tx
=XM_023253261.1;refGene_gn=SLIT2;refGene_tx=XM_023253262.1;refGene_gn=
SLIT2;refGene_tx=XM_023253264.1;refGene_gn=SLIT2;refGene_tx=XM_0232532
65.1;refGene_gn=SLIT2;refGene_tx=XM_023253266.1
B1 191782735 191782875
refGene_gn=SLIT2;refGene_tx=XM_023253260.1;refGene_gn=SLIT2;refGene_tx
=XM_023253261.1;refGene_gn=SLIT2;refGene_tx=XM_023253262.1;refGene_gn=
SLIT2;refGene_tx=XM_023253263.1;refGene_gn=SLIT2;refGene_tx=XM_0232532
64.1;refGene_gn=SLIT2;refGene_tx=XM_023253265.1;refGene_gn=SLIT2;refGe
ne_tx=XM_023253266.1;refGene_gn=SLIT2;refGene_tx=XM_023253267.1;refGen
e_gn=SLIT2;refGene_tx=XM_023253268.1;refGene_gn=SLIT2;refGene_tx=XM_02
3253269.1;refGene_gn=SLIT2;refGene_tx=XM_023253270.1
B1 191784641 191784739
refGene_gn=SLIT2;refGene_tx=XM_023253260.1;refGene_gn=SLIT2;refGene_tx
=XM_023253261.1;refGene_gn=SLIT2;refGene_tx=XM_023253262.1;refGene_gn=
SLIT2;refGene_tx=XM_023253263.1;refGene_gn=SLIT2;refGene_tx=XM_0232532
64.1;refGene_gn=SLIT2;refGene_tx=XM_023253265.1;refGene_gn=SLIT2;refGe
ne_tx=XM_023253266.1;refGene_gn=SLIT2;refGene_tx=XM_023253267.1;refGen
e_gn=SLIT2;refGene_tx=XM_023253268.1;refGene_gn=SLIT2;refGene_tx=XM_02
3253269.1;refGene_gn=SLIT2;refGene_tx=XM_023253270.1
B1 191784878 191785003
refGene_gn=SLIT2;refGene_tx=XM_023253260.1;refGene_gn=SLIT2;refGene_tx
=XM_023253261.1;refGene_gn=SLIT2;refGene_tx=XM_023253262.1;refGene_gn=
SLIT2;refGene_tx=XM_023253263.1;refGene_gn=SLIT2;refGene_tx=XM_0232532
64.1;refGene_gn=SLIT2;refGene_tx=XM_023253265.1;refGene_gn=SLIT2;refGe
ne_tx=XM_023253266.1;refGene_gn=SLIT2;refGene_tx=XM_023253267.1;refGen
e_gn=SLIT2;refGene_tx=XM_023253268.1;refGene_gn=SLIT2;refGene_tx=XM_02
3253269.1;refGene_gn=SLIT2;refGene_tx=XM_023253270.1
B1 191797225 191797258
refGene_gn=SLIT2;refGene_tx=XM_023253260.1;refGene_gn=SLIT2;refGene_tx
=XM_023253261.1;refGene_gn=SLIT2;refGene_tx=XM_023253262.1;refGene_gn=
SLIT2;refGene_tx=XM_023253263.1;refGene_gn=SLIT2;refGene_tx=XM_0232532
65.1
B1 191797601 191797765
refGene_gn=SLIT2;refGene_tx=XM_023253260.1;refGene_gn=SLIT2;refGene_tx
=XM_023253261.1;refGene_gn=SLIT2;refGene_tx=XM_023253262.1;refGene_gn=
SLIT2;refGene_tx=XM_023253263.1;refGene_gn=SLIT2;refGene_tx=XM_0232532
64.1;refGene_gn=SLIT2;refGene_tx=XM_023253265.1;refGene_gn=SLIT2;refGe
ne_tx=XM_023253266.1;refGene_gn=SLIT2;refGene_tx=XM_023253267.1;refGen
e_gn=SLIT2;refGene_tx=XM_023253268.1;refGene_gn=SLIT2;refGene_tx=XM_02
3253269.1;refGene_gn=SLIT2;refGene_tx=XM_023253270.1
B1 191800807 191800879
refGene_gn=SLIT2;refGene_tx=XM_023253260.1;refGene_gn=SLIT2;refGene_tx
=XM_023253261.1;refGene_gn=SLIT2;refGene_tx=XM_023253262.1;refGene_gn=
SLIT2;refGene_tx=XM_023253263.1;refGene_gn=SLIT2;refGene_tx=XM_0232532
64.1;refGene_gn=SLIT2;refGene_tx=XM_023253265.1;refGene_gn=SLIT2;refGe

ne_tx=XM_023253266.1;refGene_gn=SLIT2;refGene_tx=XM_023253267.1;refGene_gn=SLIT2;refGene_tx=XM_023253268.1;refGene_gn=SLIT2;refGene_tx=XM_023253269.1;refGene_gn=SLIT2;refGene_tx=XM_023253270.1
B1 191802511 191802583
refGene_gn=SLIT2;refGene_tx=XM_023253260.1;refGene_gn=SLIT2;refGene_tx=XM_023253261.1;refGene_gn=SLIT2;refGene_tx=XM_023253262.1;refGene_gn=SLIT2;refGene_tx=XM_023253263.1;refGene_gn=SLIT2;refGene_tx=XM_023253264.1;refGene_gn=SLIT2;refGene_tx=XM_023253265.1;refGene_gn=SLIT2;refGene_tx=XM_023253266.1;refGene_gn=SLIT2;refGene_tx=XM_023253267.1;refGene_gn=SLIT2;refGene_tx=XM_023253268.1;refGene_gn=SLIT2;refGene_tx=XM_023253269.1;refGene_gn=SLIT2;refGene_tx=XM_023253270.1
B1 191803061 191803133
refGene_gn=SLIT2;refGene_tx=XM_023253260.1;refGene_gn=SLIT2;refGene_tx=XM_023253261.1;refGene_gn=SLIT2;refGene_tx=XM_023253262.1;refGene_gn=SLIT2;refGene_tx=XM_023253263.1;refGene_gn=SLIT2;refGene_tx=XM_023253264.1;refGene_gn=SLIT2;refGene_tx=XM_023253265.1;refGene_gn=SLIT2;refGene_tx=XM_023253266.1;refGene_gn=SLIT2;refGene_tx=XM_023253267.1;refGene_gn=SLIT2;refGene_tx=XM_023253268.1;refGene_gn=SLIT2;refGene_tx=XM_023253269.1;refGene_gn=SLIT2;refGene_tx=XM_023253270.1
B1 191806447 191806516
refGene_gn=SLIT2;refGene_tx=XM_023253260.1;refGene_gn=SLIT2;refGene_tx=XM_023253261.1;refGene_gn=SLIT2;refGene_tx=XM_023253262.1;refGene_gn=SLIT2;refGene_tx=XM_023253263.1;refGene_gn=SLIT2;refGene_tx=XM_023253264.1;refGene_gn=SLIT2;refGene_tx=XM_023253265.1;refGene_gn=SLIT2;refGene_tx=XM_023253266.1;refGene_gn=SLIT2;refGene_tx=XM_023253267.1;refGene_gn=SLIT2;refGene_tx=XM_023253268.1;refGene_gn=SLIT2;refGene_tx=XM_023253269.1;refGene_gn=SLIT2;refGene_tx=XM_023253270.1
B1 191810140 191810273
refGene_gn=SLIT2;refGene_tx=XM_023253260.1;refGene_gn=SLIT2;refGene_tx=XM_023253261.1;refGene_gn=SLIT2;refGene_tx=XM_023253262.1;refGene_gn=SLIT2;refGene_tx=XM_023253263.1;refGene_gn=SLIT2;refGene_tx=XM_023253264.1;refGene_gn=SLIT2;refGene_tx=XM_023253265.1;refGene_gn=SLIT2;refGene_tx=XM_023253266.1;refGene_gn=SLIT2;refGene_tx=XM_023253267.1;refGene_gn=SLIT2;refGene_tx=XM_023253268.1;refGene_gn=SLIT2;refGene_tx=XM_023253269.1;refGene_gn=SLIT2;refGene_tx=XM_023253270.1
B1 191811615 191811782
refGene_gn=SLIT2;refGene_tx=XM_023253260.1;refGene_gn=SLIT2;refGene_tx=XM_023253261.1;refGene_gn=SLIT2;refGene_tx=XM_023253262.1;refGene_gn=SLIT2;refGene_tx=XM_023253263.1;refGene_gn=SLIT2;refGene_tx=XM_023253264.1;refGene_gn=SLIT2;refGene_tx=XM_023253265.1;refGene_gn=SLIT2;refGene_tx=XM_023253266.1;refGene_gn=SLIT2;refGene_tx=XM_023253267.1;refGene_gn=SLIT2;refGene_tx=XM_023253268.1;refGene_gn=SLIT2;refGene_tx=XM_023253269.1;refGene_gn=SLIT2;refGene_tx=XM_023253270.1
B1 191813337 191813481
refGene_gn=SLIT2;refGene_tx=XM_023253260.1;refGene_gn=SLIT2;refGene_tx=XM_023253261.1;refGene_gn=SLIT2;refGene_tx=XM_023253262.1;refGene_gn=SLIT2;refGene_tx=XM_023253263.1;refGene_gn=SLIT2;refGene_tx=XM_023253264.1;refGene_gn=SLIT2;refGene_tx=XM_023253265.1;refGene_gn=SLIT2;refGene_tx=XM_023253266.1;refGene_gn=SLIT2;refGene_tx=XM_023253267.1;refGene_gn=SLIT2;refGene_tx=XM_023253268.1;refGene_gn=SLIT2;refGene_tx=XM_02

3253269.1;refGene_gn=SLIT2;refGene_tx=XM_023253270.1
B1 191817189 191817333
refGene_gn=SLIT2;refGene_tx=XM_023253260.1;refGene_gn=SLIT2;refGene_tx=XM_023253261.1;refGene_gn=SLIT2;refGene_tx=XM_023253262.1;refGene_gn=SLIT2;refGene_tx=XM_023253263.1;refGene_gn=SLIT2;refGene_tx=XM_023253264.1;refGene_gn=SLIT2;refGene_tx=XM_023253265.1;refGene_gn=SLIT2;refGene_tx=XM_023253266.1;refGene_gn=SLIT2;refGene_tx=XM_023253267.1;refGene_gn=SLIT2;refGene_tx=XM_023253268.1;refGene_gn=SLIT2;refGene_tx=XM_023253269.1;refGene_gn=SLIT2;refGene_tx=XM_023253270.1
B1 191819125 191819200
refGene_gn=SLIT2;refGene_tx=XM_023253260.1;refGene_gn=SLIT2;refGene_tx=XM_023253261.1;refGene_gn=SLIT2;refGene_tx=XM_023253262.1;refGene_gn=SLIT2;refGene_tx=XM_023253263.1;refGene_gn=SLIT2;refGene_tx=XM_023253264.1;refGene_gn=SLIT2;refGene_tx=XM_023253265.1;refGene_gn=SLIT2;refGene_tx=XM_023253266.1;refGene_gn=SLIT2;refGene_tx=XM_023253267.1;refGene_gn=SLIT2;refGene_tx=XM_023253268.1;refGene_gn=SLIT2;refGene_tx=XM_023253269.1;refGene_gn=SLIT2;refGene_tx=XM_023253270.1
B1 191821792 191821943
refGene_gn=SLIT2;refGene_tx=XM_023253260.1;refGene_gn=SLIT2;refGene_tx=XM_023253261.1;refGene_gn=SLIT2;refGene_tx=XM_023253262.1;refGene_gn=SLIT2;refGene_tx=XM_023253263.1;refGene_gn=SLIT2;refGene_tx=XM_023253264.1;refGene_gn=SLIT2;refGene_tx=XM_023253265.1;refGene_gn=SLIT2;refGene_tx=XM_023253266.1;refGene_gn=SLIT2;refGene_tx=XM_023253267.1;refGene_gn=SLIT2;refGene_tx=XM_023253268.1;refGene_gn=SLIT2;refGene_tx=XM_023253269.1;refGene_gn=SLIT2;refGene_tx=XM_023253270.1
B1 191823042 191823060
refGene_gn=SLIT2;refGene_tx=XM_023253260.1;refGene_gn=SLIT2;refGene_tx=XM_023253261.1;refGene_gn=SLIT2;refGene_tx=XM_023253263.1;refGene_gn=SLIT2;refGene_tx=XM_023253264.1
B1 191825496 191825520
refGene_gn=SLIT2;refGene_tx=XM_023253260.1;refGene_gn=SLIT2;refGene_tx=XM_023253261.1;refGene_gn=SLIT2;refGene_tx=XM_023253262.1;refGene_gn=SLIT2;refGene_tx=XM_023253263.1;refGene_gn=SLIT2;refGene_tx=XM_023253264.1;refGene_gn=SLIT2;refGene_tx=XM_023253266.1;refGene_gn=SLIT2;refGene_tx=XM_023253267.1;refGene_gn=SLIT2;refGene_tx=XM_023253268.1
B1 191826492 191826656
refGene_gn=SLIT2;refGene_tx=XM_023253260.1;refGene_gn=SLIT2;refGene_tx=XM_023253261.1;refGene_gn=SLIT2;refGene_tx=XM_023253262.1;refGene_gn=SLIT2;refGene_tx=XM_023253263.1;refGene_gn=SLIT2;refGene_tx=XM_023253264.1;refGene_gn=SLIT2;refGene_tx=XM_023253265.1;refGene_gn=SLIT2;refGene_tx=XM_023253266.1;refGene_gn=SLIT2;refGene_tx=XM_023253267.1;refGene_gn=SLIT2;refGene_tx=XM_023253268.1;refGene_gn=SLIT2;refGene_tx=XM_023253269.1;refGene_gn=SLIT2;refGene_tx=XM_023253270.1
B1 191826768 191826912
refGene_gn=SLIT2;refGene_tx=XM_023253260.1;refGene_gn=SLIT2;refGene_tx=XM_023253261.1;refGene_gn=SLIT2;refGene_tx=XM_023253262.1;refGene_gn=SLIT2;refGene_tx=XM_023253263.1;refGene_gn=SLIT2;refGene_tx=XM_023253264.1;refGene_gn=SLIT2;refGene_tx=XM_023253265.1;refGene_gn=SLIT2;refGene_tx=XM_023253266.1;refGene_gn=SLIT2;refGene_tx=XM_023253267.1;refGene_gn=SLIT2;refGene_tx=XM_023253268.1;refGene_gn=SLIT2;refGene_tx=XM_02

3253269.1;refGene_gn=SLIT2;refGene_tx=XM_023253270.1
B1	191831019	191831091
refGene_gn=SLIT2;refGene_tx=XM_023253260.1;refGene_gn=SLIT2;refGene_tx=XM_023253261.1;refGene_gn=SLIT2;refGene_tx=XM_023253262.1;refGene_gn=SLIT2;refGene_tx=XM_023253263.1;refGene_gn=SLIT2;refGene_tx=XM_023253264.1;refGene_gn=SLIT2;refGene_tx=XM_023253265.1;refGene_gn=SLIT2;refGene_tx=XM_023253266.1;refGene_gn=SLIT2;refGene_tx=XM_023253267.1;refGene_gn=SLIT2;refGene_tx=XM_023253268.1;refGene_gn=SLIT2;refGene_tx=XM_023253269.1;refGene_gn=SLIT2;refGene_tx=XM_023253270.1
B1	191843074	191843146
refGene_gn=SLIT2;refGene_tx=XM_023253260.1;refGene_gn=SLIT2;refGene_tx=XM_023253261.1;refGene_gn=SLIT2;refGene_tx=XM_023253262.1;refGene_gn=SLIT2;refGene_tx=XM_023253263.1;refGene_gn=SLIT2;refGene_tx=XM_023253264.1;refGene_gn=SLIT2;refGene_tx=XM_023253265.1;refGene_gn=SLIT2;refGene_tx=XM_023253266.1;refGene_gn=SLIT2;refGene_tx=XM_023253267.1;refGene_gn=SLIT2;refGene_tx=XM_023253268.1;refGene_gn=SLIT2;refGene_tx=XM_023253269.1;refGene_gn=SLIT2;refGene_tx=XM_023253270.1
B1	191843647	191843719
refGene_gn=SLIT2;refGene_tx=XM_023253260.1;refGene_gn=SLIT2;refGene_tx=XM_023253261.1;refGene_gn=SLIT2;refGene_tx=XM_023253262.1;refGene_gn=SLIT2;refGene_tx=XM_023253263.1;refGene_gn=SLIT2;refGene_tx=XM_023253264.1;refGene_gn=SLIT2;refGene_tx=XM_023253265.1;refGene_gn=SLIT2;refGene_tx=XM_023253266.1;refGene_gn=SLIT2;refGene_tx=XM_023253267.1;refGene_gn=SLIT2;refGene_tx=XM_023253268.1;refGene_gn=SLIT2;refGene_tx=XM_023253269.1;refGene_gn=SLIT2;refGene_tx=XM_023253270.1
B1	191862225	191862364
refGene_gn=SLIT2;refGene_tx=XM_023253260.1;refGene_gn=SLIT2;refGene_tx=XM_023253261.1;refGene_gn=SLIT2;refGene_tx=XM_023253262.1;refGene_gn=SLIT2;refGene_tx=XM_023253263.1;refGene_gn=SLIT2;refGene_tx=XM_023253264.1;refGene_gn=SLIT2;refGene_tx=XM_023253265.1;refGene_gn=SLIT2;refGene_tx=XM_023253266.1;refGene_gn=SLIT2;refGene_tx=XM_023253267.1;refGene_gn=SLIT2;refGene_tx=XM_023253268.1;refGene_gn=SLIT2;refGene_tx=XM_023253269.1;refGene_gn=SLIT2;refGene_tx=XM_023253270.1
B1	191863299	191863311
refGene_gn=SLIT2;refGene_tx=XM_023253260.1;refGene_gn=SLIT2;refGene_tx=XM_023253262.1;refGene_gn=SLIT2;refGene_tx=XM_023253263.1;refGene_gn=SLIT2;refGene_tx=XM_023253264.1;refGene_gn=SLIT2;refGene_tx=XM_023253265.1;refGene_gn=SLIT2;refGene_tx=XM_023253266.1;refGene_gn=SLIT2;refGene_tx=XM_023253267.1;refGene_gn=SLIT2;refGene_tx=XM_023253269.1
B1	191864491	191864655
refGene_gn=SLIT2;refGene_tx=XM_023253260.1;refGene_gn=SLIT2;refGene_tx=XM_023253261.1;refGene_gn=SLIT2;refGene_tx=XM_023253262.1;refGene_gn=SLIT2;refGene_tx=XM_023253263.1;refGene_gn=SLIT2;refGene_tx=XM_023253264.1;refGene_gn=SLIT2;refGene_tx=XM_023253265.1;refGene_gn=SLIT2;refGene_tx=XM_023253266.1;refGene_gn=SLIT2;refGene_tx=XM_023253267.1;refGene_gn=SLIT2;refGene_tx=XM_023253268.1;refGene_gn=SLIT2;refGene_tx=XM_023253269.1;refGene_gn=SLIT2;refGene_tx=XM_023253270.1
B1	191867468	191867540
refGene_gn=SLIT2;refGene_tx=XM_023253260.1;refGene_gn=SLIT2;refGene_tx=XM_023253261.1;refGene_gn=SLIT2;refGene_tx=XM_023253262.1;refGene_gn=

SLIT2;refGene_tx=XM_023253263.1;refGene_gn=SLIT2;refGene_tx=XM_0232532
64.1;refGene_gn=SLIT2;refGene_tx=XM_023253265.1;refGene_gn=SLIT2;refGe
ne_tx=XM_023253266.1;refGene_gn=SLIT2;refGene_tx=XM_023253267.1;refGen
e_gn=SLIT2;refGene_tx=XM_023253268.1;refGene_gn=SLIT2;refGene_tx=XM_02
3253269.1;refGene_gn=SLIT2;refGene_tx=XM_023253270.1
B1 191873282 191873354
refGene_gn=SLIT2;refGene_tx=XM_023253260.1;refGene_gn=SLIT2;refGene_tx
=XM_023253261.1;refGene_gn=SLIT2;refGene_tx=XM_023253262.1;refGene_gn=
SLIT2;refGene_tx=XM_023253263.1;refGene_gn=SLIT2;refGene_tx=XM_0232532
64.1;refGene_gn=SLIT2;refGene_tx=XM_023253265.1;refGene_gn=SLIT2;refGe
ne_tx=XM_023253266.1;refGene_gn=SLIT2;refGene_tx=XM_023253267.1;refGen
e_gn=SLIT2;refGene_tx=XM_023253268.1;refGene_gn=SLIT2;refGene_tx=XM_02
3253269.1;refGene_gn=SLIT2;refGene_tx=XM_023253270.1
B1 191885652 191885724
refGene_gn=SLIT2;refGene_tx=XM_023253260.1;refGene_gn=SLIT2;refGene_tx
=XM_023253261.1;refGene_gn=SLIT2;refGene_tx=XM_023253262.1;refGene_gn=
SLIT2;refGene_tx=XM_023253263.1;refGene_gn=SLIT2;refGene_tx=XM_0232532
64.1;refGene_gn=SLIT2;refGene_tx=XM_023253265.1;refGene_gn=SLIT2;refGe
ne_tx=XM_023253266.1;refGene_gn=SLIT2;refGene_tx=XM_023253267.1;refGen
e_gn=SLIT2;refGene_tx=XM_023253268.1;refGene_gn=SLIT2;refGene_tx=XM_02
3253269.1;refGene_gn=SLIT2;refGene_tx=XM_023253270.1
B1 192090921 192090993
refGene_gn=SLIT2;refGene_tx=XM_023253260.1;refGene_gn=SLIT2;refGene_tx
=XM_023253261.1;refGene_gn=SLIT2;refGene_tx=XM_023253262.1;refGene_gn=
SLIT2;refGene_tx=XM_023253263.1;refGene_gn=SLIT2;refGene_tx=XM_0232532
64.1;refGene_gn=SLIT2;refGene_tx=XM_023253265.1;refGene_gn=SLIT2;refGe
ne_tx=XM_023253266.1;refGene_gn=SLIT2;refGene_tx=XM_023253267.1;refGen
e_gn=SLIT2;refGene_tx=XM_023253268.1;refGene_gn=SLIT2;refGene_tx=XM_02
3253269.1;refGene_gn=SLIT2;refGene_tx=XM_023253270.1
B1 192103021 192103093
refGene_gn=SLIT2;refGene_tx=XM_023253260.1;refGene_gn=SLIT2;refGene_tx
=XM_023253261.1;refGene_gn=SLIT2;refGene_tx=XM_023253262.1;refGene_gn=
SLIT2;refGene_tx=XM_023253263.1;refGene_gn=SLIT2;refGene_tx=XM_0232532
64.1;refGene_gn=SLIT2;refGene_tx=XM_023253265.1;refGene_gn=SLIT2;refGe
ne_tx=XM_023253266.1;refGene_gn=SLIT2;refGene_tx=XM_023253267.1;refGen
e_gn=SLIT2;refGene_tx=XM_023253268.1;refGene_gn=SLIT2;refGene_tx=XM_02
3253269.1;refGene_gn=SLIT2;refGene_tx=XM_023253270.1
B1 192104262 192104334
refGene_gn=SLIT2;refGene_tx=XM_023253260.1;refGene_gn=SLIT2;refGene_tx
=XM_023253261.1;refGene_gn=SLIT2;refGene_tx=XM_023253262.1;refGene_gn=
SLIT2;refGene_tx=XM_023253263.1;refGene_gn=SLIT2;refGene_tx=XM_0232532
64.1;refGene_gn=SLIT2;refGene_tx=XM_023253265.1;refGene_gn=SLIT2;refGe
ne_tx=XM_023253266.1;refGene_gn=SLIT2;refGene_tx=XM_023253267.1;refGen
e_gn=SLIT2;refGene_tx=XM_023253268.1;refGene_gn=SLIT2;refGene_tx=XM_02
3253269.1;refGene_gn=SLIT2;refGene_tx=XM_023253270.1
B1 192107090 192107269
refGene_gn=SLIT2;refGene_tx=XM_023253260.1;refGene_gn=SLIT2;refGene_tx
=XM_023253261.1;refGene_gn=SLIT2;refGene_tx=XM_023253262.1;refGene_gn=
SLIT2;refGene_tx=XM_023253263.1;refGene_gn=SLIT2;refGene_tx=XM_0232532
64.1;refGene_gn=SLIT2;refGene_tx=XM_023253265.1;refGene_gn=SLIT2;refGe

ne_tx=XM_023253266.1;refGene_gn=SLIT2;refGene_tx=XM_023253267.1;refGene_gn=SLIT2;refGene_tx=XM_023253268.1;refGene_gn=SLIT2;refGene_tx=XM_023253269.1;refGene_gn=SLIT2;refGene_tx=XM_023253270.1
B1 194052092 194052243
refGene_gn=LCORL;refGene_tx=XM_003985500.4;refGene_gn=LCORL;refGene_tx=XM_023253275.1;refGene_gn=LCORL;refGene_tx=XM_023253280.1
B1 194052972 194053306
refGene_gn=LCORL;refGene_tx=XM_023253272.1;refGene_gn=LCORL;refGene_tx=XM_023253273.1;refGene_gn=LCORL;refGene_tx=XM_023253274.1;refGene_gn=LCORL;refGene_tx=XM_023253279.1
B1 194102828 194102894
refGene_gn=LCORL;refGene_tx=XM_003985500.4;refGene_gn=LCORL;refGene_tx=XM_023253272.1;refGene_gn=LCORL;refGene_tx=XM_023253273.1;refGene_gn=LCORL;refGene_tx=XM_023253274.1;refGene_gn=LCORL;refGene_tx=XM_023253275.1;refGene_gn=LCORL;refGene_tx=XM_023253277.1;refGene_gn=LCORL;refGene_tx=XM_023253278.1;refGene_gn=LCORL;refGene_tx=XM_023253279.1;refGene_gn=LCORL;refGene_tx=XM_023253280.1
B1 194113414 194113494
refGene_gn=LCORL;refGene_tx=XM_003985500.4;refGene_gn=LCORL;refGene_tx=XM_023253272.1;refGene_gn=LCORL;refGene_tx=XM_023253273.1;refGene_gn=LCORL;refGene_tx=XM_023253274.1;refGene_gn=LCORL;refGene_tx=XM_023253275.1;refGene_gn=LCORL;refGene_tx=XM_023253279.1;refGene_gn=LCORL;refGene_tx=XM_023253280.1
B1 194114370 194114500
refGene_gn=LCORL;refGene_tx=XM_003985500.4;refGene_gn=LCORL;refGene_tx=XM_023253272.1;refGene_gn=LCORL;refGene_tx=XM_023253273.1;refGene_gn=LCORL;refGene_tx=XM_023253274.1;refGene_gn=LCORL;refGene_tx=XM_023253275.1;refGene_gn=LCORL;refGene_tx=XM_023253277.1;refGene_gn=LCORL;refGene_tx=XM_023253278.1;refGene_gn=LCORL;refGene_tx=XM_023253279.1;refGene_gn=LCORL;refGene_tx=XM_023253280.1
B1 194160777 194161029
refGene_gn=LCORL;refGene_tx=XM_003985500.4;refGene_gn=LCORL;refGene_tx=XM_023253272.1;refGene_gn=LCORL;refGene_tx=XM_023253273.1;refGene_gn=LCORL;refGene_tx=XM_023253274.1;refGene_gn=LCORL;refGene_tx=XM_023253275.1;refGene_gn=LCORL;refGene_tx=XM_023253277.1;refGene_gn=LCORL;refGene_tx=XM_023253278.1;refGene_gn=LCORL;refGene_tx=XM_023253279.1;refGene_gn=LCORL;refGene_tx=XM_023253280.1
B1 194185200 194185294
refGene_gn=LCORL;refGene_tx=XM_003985500.4;refGene_gn=LCORL;refGene_tx=XM_023253272.1;refGene_gn=LCORL;refGene_tx=XM_023253273.1;refGene_gn=LCORL;refGene_tx=XM_023253274.1;refGene_gn=LCORL;refGene_tx=XM_023253275.1;refGene_gn=LCORL;refGene_tx=XM_023253277.1;refGene_gn=LCORL;refGene_tx=XM_023253278.1;refGene_gn=LCORL;refGene_tx=XM_023253279.1;refGene_gn=LCORL;refGene_tx=XM_023253280.1
B1 194186651 194187684
refGene_gn=LCORL;refGene_tx=XM_003985500.4
B1 194193357 194198171
refGene_gn=LCORL;refGene_tx=XM_006931197.4;refGene_gn=LCORL;refGene_tx=XM_023253272.1;refGene_gn=LCORL;refGene_tx=XM_023253273.1;refGene_gn=LCORL;refGene_tx=XM_023253274.1;refGene_gn=LCORL;refGene_tx=XM_0232532

75.1;refGene_gn=LCORL;refGene_tx=XM_023253277.1;refGene_gn=LCORL;refGene_tx=XM_023253278.1;refGene_gn=LCORL;refGene_tx=XM_023253279.1
B1 194204664 194204690
refGene_gn=LCORL;refGene_tx=XM_006931197.4;refGene_gn=LCORL;refGene_tx=XM_023253272.1;refGene_gn=LCORL;refGene_tx=XM_023253273.1;refGene_gn=LCORL;refGene_tx=XM_023253275.1;refGene_gn=LCORL;refGene_tx=XM_023253277.1;refGene_gn=LCORL;refGene_tx=XM_023253278.1
B1 194217336 194217517
refGene_gn=LCORL;refGene_tx=XM_023253274.1;refGene_gn=LCORL;refGene_tx=XM_023253279.1;refGene_gn=LCORL;refGene_tx=XM_023253280.1
B1 194220003 194220127
refGene_gn=NCAPG;refGene_tx=XM_003985501.5;refGene_gn=NCAPG;refGene_tx=XM_019829714.2;refGene_gn=NCAPG;refGene_tx=XM_019829715.2
B1 194221062 194221132
refGene_gn=NCAPG;refGene_tx=XM_003985501.5;refGene_gn=NCAPG;refGene_tx=XM_019829714.2;refGene_gn=NCAPG;refGene_tx=XM_019829715.2
B1 194222561 194222648
refGene_gn=NCAPG;refGene_tx=XM_003985501.5;refGene_gn=NCAPG;refGene_tx=XM_019829714.2;refGene_gn=NCAPG;refGene_tx=XM_019829715.2
B1 194223045 194223184
refGene_gn=NCAPG;refGene_tx=XM_003985501.5;refGene_gn=NCAPG;refGene_tx=XM_019829714.2;refGene_gn=NCAPG;refGene_tx=XM_019829715.2
B1 194223643 194223805
refGene_gn=NCAPG;refGene_tx=XM_003985501.5;refGene_gn=NCAPG;refGene_tx=XM_019829714.2;refGene_gn=NCAPG;refGene_tx=XM_019829715.2
B1 194226475 194226650
refGene_gn=NCAPG;refGene_tx=XM_003985501.5;refGene_gn=NCAPG;refGene_tx=XM_019829714.2
B1 194226916 194227098
refGene_gn=NCAPG;refGene_tx=XM_003985501.5;refGene_gn=NCAPG;refGene_tx=XM_019829714.2
B1 194236859 194237087
refGene_gn=NCAPG;refGene_tx=XM_003985501.5;refGene_gn=NCAPG;refGene_tx=XM_019829714.2;refGene_gn=NCAPG;refGene_tx=XM_019829715.2
B1 194243909 194244029
refGene_gn=NCAPG;refGene_tx=XM_003985501.5;refGene_gn=NCAPG;refGene_tx=XM_019829714.2;refGene_gn=NCAPG;refGene_tx=XM_019829715.2
B1 194247441 194247552
refGene_gn=NCAPG;refGene_tx=XM_003985501.5;refGene_gn=NCAPG;refGene_tx=XM_019829714.2;refGene_gn=NCAPG;refGene_tx=XM_019829715.2
B1 194249754 194249934
refGene_gn=NCAPG;refGene_tx=XM_003985501.5;refGene_gn=NCAPG;refGene_tx=XM_019829714.2;refGene_gn=NCAPG;refGene_tx=XM_019829715.2
B1 194250254 194250341
refGene_gn=NCAPG;refGene_tx=XM_003985501.5;refGene_gn=NCAPG;refGene_tx=XM_019829714.2;refGene_gn=NCAPG;refGene_tx=XM_019829715.2
B1 194252103 194252227
refGene_gn=NCAPG;refGene_tx=XM_003985501.5;refGene_gn=NCAPG;refGene_tx=XM_019829714.2;refGene_gn=NCAPG;refGene_tx=XM_019829715.2
B1 194253202 194253343

refGene_gn=NCAPG;refGene_tx=XM_003985501.5;refGene_gn=NCAPG;refGene_tx=XM_019829715.2
B1 194259884 194260034
refGene_gn=NCAPG;refGene_tx=XM_003985501.5;refGene_gn=NCAPG;refGene_tx=XM_019829714.2;refGene_gn=NCAPG;refGene_tx=XM_019829715.2
B1 194260722 194260915
refGene_gn=NCAPG;refGene_tx=XM_003985501.5;refGene_gn=NCAPG;refGene_tx=XM_019829714.2;refGene_gn=NCAPG;refGene_tx=XM_019829715.2
B1 194261939 194262024
refGene_gn=NCAPG;refGene_tx=XM_003985501.5;refGene_gn=NCAPG;refGene_tx=XM_019829714.2;refGene_gn=NCAPG;refGene_tx=XM_019829715.2
B1 194262272 194262418
refGene_gn=NCAPG;refGene_tx=XM_003985501.5;refGene_gn=NCAPG;refGene_tx=XM_019829714.2;refGene_gn=NCAPG;refGene_tx=XM_019829715.2
B1 194265057 194265286
refGene_gn=NCAPG;refGene_tx=XM_003985501.5;refGene_gn=NCAPG;refGene_tx=XM_019829714.2;refGene_gn=NCAPG;refGene_tx=XM_019829715.2
B1 194265762 194265966
refGene_gn=NCAPG;refGene_tx=XM_003985501.5;refGene_gn=NCAPG;refGene_tx=XM_019829714.2;refGene_gn=NCAPG;refGene_tx=XM_019829715.2
B1 194267155 194267266
refGene_gn=NCAPG;refGene_tx=XM_003985501.5;refGene_gn=NCAPG;refGene_tx=XM_019829714.2;refGene_gn=NCAPG;refGene_tx=XM_019829715.2
B1 194276614 194277265
refGene_gn=DCAF16;refGene_tx=XM_006931199.4;refGene_gn=DCAF16;refGene_tx=XM_019829716.2
B1 194302470 194303039
refGene_gn=FAM184B;refGene_tx=XM_019829717.2;refGene_gn=FAM184B;refGene_tx=XM_023253281.1
B1 194303253 194303400
refGene_gn=FAM184B;refGene_tx=XM_003985503.5;refGene_gn=FAM184B;refGene_tx=XM_019829717.2;refGene_gn=FAM184B;refGene_tx=XM_023253281.1
B1 194379933 194380686
refGene_gn=FAM184B;refGene_tx=XM_003985503.5;refGene_gn=FAM184B;refGene_tx=XM_019829717.2;refGene_gn=FAM184B;refGene_tx=XM_023253281.1
B1 194381441 194381577
refGene_gn=FAM184B;refGene_tx=XM_003985503.5;refGene_gn=FAM184B;refGene_tx=XM_019829717.2;refGene_gn=FAM184B;refGene_tx=XM_023253281.1
B1 194382903 194383043
refGene_gn=FAM184B;refGene_tx=XM_003985503.5;refGene_gn=FAM184B;refGene_tx=XM_019829717.2;refGene_gn=FAM184B;refGene_tx=XM_023253281.1
B1 194383571 194383778
refGene_gn=FAM184B;refGene_tx=XM_003985503.5;refGene_gn=FAM184B;refGene_tx=XM_019829717.2;refGene_gn=FAM184B;refGene_tx=XM_023253281.1
B1 194404701 194404812
refGene_gn=FAM184B;refGene_tx=XM_003985503.5;refGene_gn=FAM184B;refGene_tx=XM_019829717.2;refGene_gn=FAM184B;refGene_tx=XM_023253281.1
B1 194408854 194408962
refGene_gn=FAM184B;refGene_tx=XM_003985503.5;refGene_gn=FAM184B;refGene_tx=XM_019829717.2;refGene_gn=FAM184B;refGene_tx=XM_023253281.1

B1 194422018 194422122
refGene_gn=FAM184B;refGene_tx=XM_003985503.5;refGene_gn=FAM184B;refGene_tx=XM_019829717.2;refGene_gn=FAM184B;refGene_tx=XM_023253281.1
B1 194432711 194432841
refGene_gn=FAM184B;refGene_tx=XM_003985503.5;refGene_gn=FAM184B;refGene_tx=XM_019829717.2;refGene_gn=FAM184B;refGene_tx=XM_023253281.1
B1 194433580 194433793
refGene_gn=FAM184B;refGene_tx=XM_003985503.5;refGene_gn=FAM184B;refGene_tx=XM_019829717.2;refGene_gn=FAM184B;refGene_tx=XM_023253281.1
B1 194443047 194443201
refGene_gn=FAM184B;refGene_tx=XM_003985503.5;refGene_gn=FAM184B;refGene_tx=XM_019829717.2;refGene_gn=FAM184B;refGene_tx=XM_023253281.1
B1 194446463 194446627
refGene_gn=FAM184B;refGene_tx=XM_003985503.5;refGene_gn=FAM184B;refGene_tx=XM_019829717.2;refGene_gn=FAM184B;refGene_tx=XM_023253281.1
B1 194449186 194449359
refGene_gn=FAM184B;refGene_tx=XM_003985503.5;refGene_gn=FAM184B;refGene_tx=XM_019829717.2;refGene_gn=FAM184B;refGene_tx=XM_023253281.1
B1 194450437 194450584
refGene_gn=FAM184B;refGene_tx=XM_003985503.5;refGene_gn=FAM184B;refGene_tx=XM_019829717.2;refGene_gn=FAM184B;refGene_tx=XM_023253281.1
B1 194452499 194452617
refGene_gn=FAM184B;refGene_tx=XM_003985503.5;refGene_gn=FAM184B;refGene_tx=XM_019829717.2;refGene_gn=FAM184B;refGene_tx=XM_023253281.1
B1 194454151 194454309
refGene_gn=FAM184B;refGene_tx=XM_003985503.5;refGene_gn=FAM184B;refGene_tx=XM_019829717.2;refGene_gn=FAM184B;refGene_tx=XM_023253281.1
B1 194455329 194455529
refGene_gn=FAM184B;refGene_tx=XM_003985503.5;refGene_gn=FAM184B;refGene_tx=XM_019829717.2
B1 194457273 194457367
refGene_gn=FAM184B;refGene_tx=XM_003985503.5;refGene_gn=FAM184B;refGene_tx=XM_019829717.2
B1 194464125 194464323
refGene_gn=MED28;refGene_tx=XM_003985504.4;refGene_gn=MED28;refGene_tx=XM_006931201.2;refGene_gn=MED28;refGene_tx=XM_006931202.2
B1 194465807 194465920
refGene_gn=MED28;refGene_tx=XM_003985504.4;refGene_gn=MED28;refGene_tx=XM_006931201.2;refGene_gn=MED28;refGene_tx=XM_006931202.2
B1 194466592 194466659
refGene_gn=MED28;refGene_tx=XM_003985504.4;refGene_gn=MED28;refGene_tx=XM_006931201.2;refGene_gn=MED28;refGene_tx=XM_006931202.2
B1 194470997 194471147
refGene_gn=MED28;refGene_tx=XM_003985504.4;refGene_gn=MED28;refGene_tx=XM_006931201.2;refGene_gn=MED28;refGene_tx=XM_006931202.2
B1 194474677 194474867
refGene_gn=LAP3;refGene_tx=XM_023253283.1
B1 194475570 194475680
refGene_gn=LAP3;refGene_tx=XM_023253283.1
B1 194477758 194477838

refGene_gn=LAP3;refGene_tx=XM_023253283.1
B1 194484611 194484714
refGene_gn=LAP3;refGene_tx=XM_023253283.1
B1 194485800 194485889
refGene_gn=LAP3;refGene_tx=XM_023253283.1
B1 194487321 194487446
refGene_gn=LAP3;refGene_tx=XM_023253283.1
B1 194491664 194491823
refGene_gn=LAP3;refGene_tx=XM_023253283.1
B1 194494462 194494627
refGene_gn=LAP3;refGene_tx=XM_023253283.1
B1 194496501 194496661
refGene_gn=LAP3;refGene_tx=XM_023253283.1
B1 194497731 194497837
refGene_gn=LAP3;refGene_tx=XM_023253283.1
B1 194498539 194498594
refGene_gn=LAP3;refGene_tx=XM_023253283.1
B1 194499921 194500037
refGene_gn=LAP3;refGene_tx=XM_023253283.1
B1 194502391 194502493
refGene_gn=LAP3;refGene_tx=XM_023253283.1
B1 194539911 194540177
refGene_gn=CLRN2;refGene_tx=XM_003985506.4
B1 194540822 194541002
refGene_gn=CLRN2;refGene_tx=XM_003985506.4
B1 194546618 194546871
refGene_gn=CLRN2;refGene_tx=XM_003985506.4
B1 194557346 194557445
refGene_gn=QDPR;refGene_tx=XM_023253284.1
B1 194564068 194564161
refGene_gn=QDPR;refGene_tx=XM_023253284.1
B1 194567772 194567869
refGene_gn=QDPR;refGene_tx=XM_023253284.1
B1 194570334 194570475
refGene_gn=QDPR;refGene_tx=XM_023253284.1
B1 194574785 194574894
refGene_gn=QDPR;refGene_tx=XM_023253284.1
B1 194576071 194576155
refGene_gn=QDPR;refGene_tx=XM_023253284.1
B1 194579133 194579239
refGene_gn=QDPR;refGene_tx=XM_023253284.1
B1 194978292 194978389
refGene_gn=LOC111560232;refGene_tx=XM_023253412.1
B1 194988026 194988152
refGene_gn=LOC111560232;refGene_tx=XM_023253412.1
B1 194991156 194991267
refGene_gn=LOC111560232;refGene_tx=XM_023253412.1
B1 195003579 195003594
refGene_gn=LOC111560232;refGene_tx=XM_023253412.1
B1 195050834 195051004

refGene_gn=LOC111560232;refGene_tx=XM_023253412.1
B1 195117626 195117758
refGene_gn=LDB2;refGene_tx=XM_003985531.4;refGene_gn=LDB2;refGene_tx=XM_003985532.4;refGene_gn=LDB2;refGene_tx=XM_006931220.3;refGene_gn=LDB2;refGene_tx=XM_006931221.3;refGene_gn=LDB2;refGene_tx=XM_006931222.3;refGene_gn=LDB2;refGene_tx=XM_006931223.4;refGene_gn=LDB2;refGene_tx=XM_006931224.3;refGene_gn=LDB2;refGene_tx=XM_006931225.3;refGene_gn=LDB2;refGene_tx=XM_006931226.3;refGene_gn=LDB2;refGene_tx=XM_019829721.2
B1 195225926 195225929
refGene_gn=LDB2;refGene_tx=XM_019829720.2
B1 195257990 195258093
refGene_gn=LDB2;refGene_tx=XM_003985531.4;refGene_gn=LDB2;refGene_tx=XM_003985532.4;refGene_gn=LDB2;refGene_tx=XM_006931220.3;refGene_gn=LDB2;refGene_tx=XM_006931221.3;refGene_gn=LDB2;refGene_tx=XM_006931222.3;refGene_gn=LDB2;refGene_tx=XM_006931223.4;refGene_gn=LDB2;refGene_tx=XM_006931224.3;refGene_gn=LDB2;refGene_tx=XM_006931225.3;refGene_gn=LDB2;refGene_tx=XM_006931226.3;refGene_gn=LDB2;refGene_tx=XM_019829720.2;refGene_gn=LDB2;refGene_tx=XM_019829721.2
B1 195414888 195415061
refGene_gn=LDB2;refGene_tx=XM_003985531.4;refGene_gn=LDB2;refGene_tx=XM_003985532.4;refGene_gn=LDB2;refGene_tx=XM_006931220.3;refGene_gn=LDB2;refGene_tx=XM_006931221.3;refGene_gn=LDB2;refGene_tx=XM_006931222.3;refGene_gn=LDB2;refGene_tx=XM_006931223.4;refGene_gn=LDB2;refGene_tx=XM_006931224.3;refGene_gn=LDB2;refGene_tx=XM_006931225.3;refGene_gn=LDB2;refGene_tx=XM_006931226.3;refGene_gn=LDB2;refGene_tx=XM_019829720.2;refGene_gn=LDB2;refGene_tx=XM_019829721.2
B1 195422188 195422311
refGene_gn=LDB2;refGene_tx=XM_003985531.4;refGene_gn=LDB2;refGene_tx=XM_003985532.4;refGene_gn=LDB2;refGene_tx=XM_006931220.3;refGene_gn=LDB2;refGene_tx=XM_006931221.3;refGene_gn=LDB2;refGene_tx=XM_006931222.3;refGene_gn=LDB2;refGene_tx=XM_006931223.4;refGene_gn=LDB2;refGene_tx=XM_006931224.3;refGene_gn=LDB2;refGene_tx=XM_006931225.3;refGene_gn=LDB2;refGene_tx=XM_006931226.3;refGene_gn=LDB2;refGene_tx=XM_019829720.2;refGene_gn=LDB2;refGene_tx=XM_019829721.2
B1 195425114 195425198
refGene_gn=LDB2;refGene_tx=XM_003985531.4;refGene_gn=LDB2;refGene_tx=XM_003985532.4;refGene_gn=LDB2;refGene_tx=XM_006931220.3;refGene_gn=LDB2;refGene_tx=XM_006931221.3;refGene_gn=LDB2;refGene_tx=XM_006931222.3;refGene_gn=LDB2;refGene_tx=XM_006931223.4;refGene_gn=LDB2;refGene_tx=XM_006931224.3;refGene_gn=LDB2;refGene_tx=XM_006931225.3;refGene_gn=LDB2;refGene_tx=XM_006931226.3;refGene_gn=LDB2;refGene_tx=XM_019829720.2;refGene_gn=LDB2;refGene_tx=XM_019829721.2
B1 195488757 195488881
refGene_gn=LDB2;refGene_tx=XM_003985531.4;refGene_gn=LDB2;refGene_tx=XM_003985532.4;refGene_gn=LDB2;refGene_tx=XM_006931220.3;refGene_gn=LDB2;refGene_tx=XM_006931221.3;refGene_gn=LDB2;refGene_tx=XM_006931222.3;refGene_gn=LDB2;refGene_tx=XM_006931223.4;refGene_gn=LDB2;refGene_tx=XM_006931224.3;refGene_gn=LDB2;refGene_tx=XM_006931225.3;refGene_gn=LDB2;refGene_tx=XM_006931226.3;refGene_gn=LDB2;refGene_tx=XM_019829720.2;refGene_gn=LDB2;refGene_tx=XM_019829721.2

B1 195492499 195492651
refGene_gn=LDB2;refGene_tx=XM_003985531.4;refGene_gn=LDB2;refGene_tx=XM_003985532.4;refGene_gn=LDB2;refGene_tx=XM_006931220.3;refGene_gn=LDB2;refGene_tx=XM_006931221.3;refGene_gn=LDB2;refGene_tx=XM_006931222.3;refGene_gn=LDB2;refGene_tx=XM_006931223.4;refGene_gn=LDB2;refGene_tx=XM_006931224.3;refGene_gn=LDB2;refGene_tx=XM_006931225.3;refGene_gn=LDB2;refGene_tx=XM_006931226.3;refGene_gn=LDB2;refGene_tx=XM_019829720.2;refGene_gn=LDB2;refGene_tx=XM_019829721.2
B1 195495189 195495294
refGene_gn=LDB2;refGene_tx=XM_003985532.4;refGene_gn=LDB2;refGene_tx=XM_006931223.4;refGene_gn=LDB2;refGene_tx=XM_006931224.3;refGene_gn=LDB2;refGene_tx=XM_006931225.3
B1 195498023 195498074
refGene_gn=LDB2;refGene_tx=XM_006931226.3;refGene_gn=LDB2;refGene_tx=XM_019829721.2
B1 195498288 195498519
refGene_gn=LDB2;refGene_tx=XM_003985531.4;refGene_gn=LDB2;refGene_tx=XM_006931220.3;refGene_gn=LDB2;refGene_tx=XM_006931221.3;refGene_gn=LDB2;refGene_tx=XM_006931222.3;refGene_gn=LDB2;refGene_tx=XM_019829720.2
B1 195755489 195755691
refGene_gn=TAPT1;refGene_tx=XM_023253285.1
B1 195767182 195767313
refGene_gn=TAPT1;refGene_tx=XM_023253285.1
B1 195783513 195783632
refGene_gn=TAPT1;refGene_tx=XM_023253285.1
B1 195794682 195794845
refGene_gn=TAPT1;refGene_tx=XM_023253285.1
B1 195797282 195797418
refGene_gn=TAPT1;refGene_tx=XM_023253285.1
B1 195798720 195798818
refGene_gn=TAPT1;refGene_tx=XM_023253285.1
B1 195798997 195799067
refGene_gn=TAPT1;refGene_tx=XM_023253285.1
B1 195800318 195800399
refGene_gn=TAPT1;refGene_tx=XM_023253285.1
B1 195808729 195808839
refGene_gn=TAPT1;refGene_tx=XM_023253285.1
B1 195810219 195810279
refGene_gn=TAPT1;refGene_tx=XM_023253285.1
B1 195810653 195810722
refGene_gn=TAPT1;refGene_tx=XM_023253285.1
B1 195815846 195815923
refGene_gn=TAPT1;refGene_tx=XM_023253285.1
B1 195819412 195819573
refGene_gn=TAPT1;refGene_tx=XM_023253285.1
B1 195822543 195822761
refGene_gn=TAPT1;refGene_tx=XM_023253285.1
B1 195895259 195895482
refGene_gn=PROM1;refGene_tx=XM_019829725.2;refGene_gn=PROM1;refGene_tx=XM_019829726.2;refGene_gn=PROM1;refGene_tx=XM_019829727.2;refGene_gn=

PROM1;refGene_tx=XM_019829728.2;refGene_gn=PROM1;refGene_tx=XM_0198297
33.2;refGene_gn=PROM1;refGene_tx=XM_023253286.1
B1 195933458 195933514
refGene_gn=PROM1;refGene_tx=XM_019829725.2;refGene_gn=PROM1;refGene_tx
=XM_019829726.2;refGene_gn=PROM1;refGene_tx=XM_019829727.2;refGene_gn=
PROM1;refGene_tx=XM_019829728.2;refGene_gn=PROM1;refGene_tx=XM_0198297
33.2;refGene_gn=PROM1;refGene_tx=XM_023253286.1
B1 195941914 195941941
refGene_gn=PROM1;refGene_tx=XM_019829725.2;refGene_gn=PROM1;refGene_tx
=XM_019829726.2;refGene_gn=PROM1;refGene_tx=XM_019829727.2;refGene_gn=
PROM1;refGene_tx=XM_019829728.2;refGene_gn=PROM1;refGene_tx=XM_0232532
86.1
B1 195944449 195944655
refGene_gn=PROM1;refGene_tx=XM_019829725.2;refGene_gn=PROM1;refGene_tx
=XM_019829726.2;refGene_gn=PROM1;refGene_tx=XM_019829727.2;refGene_gn=
PROM1;refGene_tx=XM_019829728.2;refGene_gn=PROM1;refGene_tx=XM_0198297
33.2;refGene_gn=PROM1;refGene_tx=XM_023253286.1
B1 195953507 195953628
refGene_gn=PROM1;refGene_tx=XM_019829725.2;refGene_gn=PROM1;refGene_tx
=XM_019829726.2;refGene_gn=PROM1;refGene_tx=XM_019829727.2;refGene_gn=
PROM1;refGene_tx=XM_019829728.2;refGene_gn=PROM1;refGene_tx=XM_0198297
33.2;refGene_gn=PROM1;refGene_tx=XM_023253286.1
B1 195955696 195955760
refGene_gn=PROM1;refGene_tx=XM_019829725.2;refGene_gn=PROM1;refGene_tx
=XM_019829726.2;refGene_gn=PROM1;refGene_tx=XM_019829727.2;refGene_gn=
PROM1;refGene_tx=XM_019829728.2;refGene_gn=PROM1;refGene_tx=XM_0198297
33.2;refGene_gn=PROM1;refGene_tx=XM_023253286.1
B1 195957416 195957506
refGene_gn=PROM1;refGene_tx=XM_019829725.2;refGene_gn=PROM1;refGene_tx
=XM_019829726.2;refGene_gn=PROM1;refGene_tx=XM_019829727.2;refGene_gn=
PROM1;refGene_tx=XM_019829728.2;refGene_gn=PROM1;refGene_tx=XM_0198297
33.2;refGene_gn=PROM1;refGene_tx=XM_023253286.1
B1 195962618 195962836
refGene_gn=PROM1;refGene_tx=XM_019829725.2;refGene_gn=PROM1;refGene_tx
=XM_019829726.2;refGene_gn=PROM1;refGene_tx=XM_019829727.2;refGene_gn=
PROM1;refGene_tx=XM_019829728.2;refGene_gn=PROM1;refGene_tx=XM_0198297
33.2;refGene_gn=PROM1;refGene_tx=XM_023253286.1
B1 195964894 195964969
refGene_gn=PROM1;refGene_tx=XM_019829725.2;refGene_gn=PROM1;refGene_tx
=XM_019829726.2;refGene_gn=PROM1;refGene_tx=XM_019829727.2;refGene_gn=
PROM1;refGene_tx=XM_019829728.2;refGene_gn=PROM1;refGene_tx=XM_0198297
33.2;refGene_gn=PROM1;refGene_tx=XM_023253286.1
B1 195968002 195968066
refGene_gn=PROM1;refGene_tx=XM_019829725.2;refGene_gn=PROM1;refGene_tx
=XM_019829726.2;refGene_gn=PROM1;refGene_tx=XM_019829727.2;refGene_gn=
PROM1;refGene_tx=XM_019829728.2;refGene_gn=PROM1;refGene_tx=XM_0198297
33.2;refGene_gn=PROM1;refGene_tx=XM_023253286.1
B1 195971233 195971393
refGene_gn=PROM1;refGene_tx=XM_019829725.2;refGene_gn=PROM1;refGene_tx
=XM_019829726.2;refGene_gn=PROM1;refGene_tx=XM_019829727.2;refGene_gn=

PROM1;refGene_tx=XM_019829728.2;refGene_gn=PROM1;refGene_tx=XM_0198297
33.2;refGene_gn=PROM1;refGene_tx=XM_023253286.1
B1 195974131 195974284
refGene_gn=PROM1;refGene_tx=XM_019829725.2;refGene_gn=PROM1;refGene_tx
=XM_019829726.2;refGene_gn=PROM1;refGene_tx=XM_019829727.2;refGene_gn=
PROM1;refGene_tx=XM_019829728.2;refGene_gn=PROM1;refGene_tx=XM_0198297
33.2;refGene_gn=PROM1;refGene_tx=XM_023253286.1
B1 195979708 195979832
refGene_gn=PROM1;refGene_tx=XM_019829725.2;refGene_gn=PROM1;refGene_tx
=XM_019829726.2;refGene_gn=PROM1;refGene_tx=XM_019829727.2;refGene_gn=
PROM1;refGene_tx=XM_019829728.2;refGene_gn=PROM1;refGene_tx=XM_0198297
33.2;refGene_gn=PROM1;refGene_tx=XM_023253286.1
B1 195981610 195981714
refGene_gn=PROM1;refGene_tx=XM_019829725.2;refGene_gn=PROM1;refGene_tx
=XM_019829726.2;refGene_gn=PROM1;refGene_tx=XM_019829727.2;refGene_gn=
PROM1;refGene_tx=XM_019829728.2;refGene_gn=PROM1;refGene_tx=XM_0198297
33.2;refGene_gn=PROM1;refGene_tx=XM_023253286.1
B1 195983554 195983639
refGene_gn=PROM1;refGene_tx=XM_019829725.2;refGene_gn=PROM1;refGene_tx
=XM_019829726.2;refGene_gn=PROM1;refGene_tx=XM_019829727.2;refGene_gn=
PROM1;refGene_tx=XM_019829728.2;refGene_gn=PROM1;refGene_tx=XM_0198297
33.2;refGene_gn=PROM1;refGene_tx=XM_023253286.1
B1 195986214 195986361
refGene_gn=PROM1;refGene_tx=XM_019829725.2;refGene_gn=PROM1;refGene_tx
=XM_019829726.2;refGene_gn=PROM1;refGene_tx=XM_019829727.2;refGene_gn=
PROM1;refGene_tx=XM_019829728.2;refGene_gn=PROM1;refGene_tx=XM_0198297
33.2;refGene_gn=PROM1;refGene_tx=XM_023253286.1
B1 195987019 195987091
refGene_gn=PROM1;refGene_tx=XM_019829725.2;refGene_gn=PROM1;refGene_tx
=XM_019829726.2;refGene_gn=PROM1;refGene_tx=XM_019829727.2;refGene_gn=
PROM1;refGene_tx=XM_019829728.2;refGene_gn=PROM1;refGene_tx=XM_0198297
33.2;refGene_gn=PROM1;refGene_tx=XM_023253286.1
B1 195988168 195988261
refGene_gn=PROM1;refGene_tx=XM_019829725.2;refGene_gn=PROM1;refGene_tx
=XM_019829726.2;refGene_gn=PROM1;refGene_tx=XM_019829727.2;refGene_gn=
PROM1;refGene_tx=XM_019829728.2;refGene_gn=PROM1;refGene_tx=XM_0198297
33.2;refGene_gn=PROM1;refGene_tx=XM_023253286.1
B1 195988898 195988916
refGene_gn=PROM1;refGene_tx=XM_019829725.2;refGene_gn=PROM1;refGene_tx
=XM_019829726.2;refGene_gn=PROM1;refGene_tx=XM_019829727.2;refGene_gn=
PROM1;refGene_tx=XM_019829728.2
B1 195990172 195990226
refGene_gn=PROM1;refGene_tx=XM_019829725.2;refGene_gn=PROM1;refGene_tx
=XM_019829726.2;refGene_gn=PROM1;refGene_tx=XM_019829727.2;refGene_gn=
PROM1;refGene_tx=XM_019829728.2;refGene_gn=PROM1;refGene_tx=XM_0198297
33.2;refGene_gn=PROM1;refGene_tx=XM_023253286.1
B1 195992764 195992845
refGene_gn=PROM1;refGene_tx=XM_019829725.2;refGene_gn=PROM1;refGene_tx
=XM_019829726.2;refGene_gn=PROM1;refGene_tx=XM_019829727.2;refGene_gn=
PROM1;refGene_tx=XM_019829728.2;refGene_gn=PROM1;refGene_tx=XM_0198297

33.2;refGene_gn=PROM1;refGene_tx=XM_023253286.1
B1 195993183 195993252
refGene_gn=PROM1;refGene_tx=XM_019829725.2;refGene_gn=PROM1;refGene_tx=XM_019829726.2;refGene_gn=PROM1;refGene_tx=XM_019829727.2;refGene_gn=PROM1;refGene_tx=XM_019829728.2;refGene_gn=PROM1;refGene_tx=XM_019829733.2;refGene_gn=PROM1;refGene_tx=XM_023253286.1
B1 195996337 195996430
refGene_gn=PROM1;refGene_tx=XM_019829725.2;refGene_gn=PROM1;refGene_tx=XM_019829726.2;refGene_gn=PROM1;refGene_tx=XM_019829727.2;refGene_gn=PROM1;refGene_tx=XM_019829728.2;refGene_gn=PROM1;refGene_tx=XM_019829733.2;refGene_gn=PROM1;refGene_tx=XM_023253286.1
B1 195998871 195998987
refGene_gn=PROM1;refGene_tx=XM_019829725.2;refGene_gn=PROM1;refGene_tx=XM_019829726.2;refGene_gn=PROM1;refGene_tx=XM_019829727.2;refGene_gn=PROM1;refGene_tx=XM_019829728.2;refGene_gn=PROM1;refGene_tx=XM_019829733.2;refGene_gn=PROM1;refGene_tx=XM_023253286.1
B1 195999212 195999249
refGene_gn=PROM1;refGene_tx=XM_019829728.2
B1 195999912 195999967
refGene_gn=PROM1;refGene_tx=XM_019829725.2;refGene_gn=PROM1;refGene_tx=XM_019829726.2;refGene_gn=PROM1;refGene_tx=XM_019829727.2;refGene_gn=PROM1;refGene_tx=XM_019829733.2;refGene_gn=PROM1;refGene_tx=XM_023253286.1
B1 196086711 196087476
refGene_gn=FGFBP1;refGene_tx=XM_006931219.4
B1 196183153 196183217
refGene_gn=CD38;refGene_tx=XM_023253287.1
B1 196186459 196186546
refGene_gn=CD38;refGene_tx=XM_023253287.1
B1 196186884 196186977
refGene_gn=CD38;refGene_tx=XM_023253287.1
B1 196188693 196188767
refGene_gn=CD38;refGene_tx=XM_023253287.1
B1 196188924 196189080
refGene_gn=CD38;refGene_tx=XM_023253288.1
B1 196192308 196192394
refGene_gn=CD38;refGene_tx=XM_023253287.1;refGene_gn=CD38;refGene_tx=XM_023253288.1
B1 196197894 196198030
refGene_gn=CD38;refGene_tx=XM_023253287.1;refGene_gn=CD38;refGene_tx=XM_023253288.1
B1 196201992 196202122
refGene_gn=CD38;refGene_tx=XM_023253287.1;refGene_gn=CD38;refGene_tx=XM_023253288.1
B1 196241343 196241597
refGene_gn=CD38;refGene_tx=XM_023253287.1;refGene_gn=CD38;refGene_tx=XM_023253288.1
B1 196244714 196245225
refGene_gn=LOC111560057;refGene_tx=XM_023252240.1
B1 196247631 196247798

refGene_gn=LOC111560057;refGene_tx=XM_023252240.1
B1 196287949 196288055
refGene_gn=BST1;refGene_tx=XM_023253289.1
B1 196298598 196298662
refGene_gn=BST1;refGene_tx=XM_023253290.1;refGene_gn=BST1;refGene_tx=XM_023253291.1
B1 196298740 196298800
refGene_gn=BST1;refGene_tx=XM_023253289.1;refGene_gn=BST1;refGene_tx=XM_023253290.1;refGene_gn=BST1;refGene_tx=XM_023253291.1
B1 196302392 196302479
refGene_gn=BST1;refGene_tx=XM_023253289.1;refGene_gn=BST1;refGene_tx=XM_023253290.1;refGene_gn=BST1;refGene_tx=XM_023253291.1
B1 196305133 196305226
refGene_gn=BST1;refGene_tx=XM_023253289.1;refGene_gn=BST1;refGene_tx=XM_023253290.1;refGene_gn=BST1;refGene_tx=XM_023253291.1
B1 196305578 196305655
refGene_gn=BST1;refGene_tx=XM_023253289.1;refGene_gn=BST1;refGene_tx=XM_023253290.1;refGene_gn=BST1;refGene_tx=XM_023253291.1
B1 196309580 196309663
refGene_gn=BST1;refGene_tx=XM_023253289.1;refGene_gn=BST1;refGene_tx=XM_023253290.1;refGene_gn=BST1;refGene_tx=XM_023253291.1
B1 196314143 196314279
refGene_gn=BST1;refGene_tx=XM_023253289.1;refGene_gn=BST1;refGene_tx=XM_023253290.1;refGene_gn=BST1;refGene_tx=XM_023253291.1
B1 196316013 196316140
refGene_gn=BST1;refGene_tx=XM_023253289.1;refGene_gn=BST1;refGene_tx=XM_023253290.1;refGene_gn=BST1;refGene_tx=XM_023253291.1
B1 196318104 196318277
refGene_gn=BST1;refGene_tx=XM_023253289.1;refGene_gn=BST1;refGene_tx=XM_023253290.1;refGene_gn=BST1;refGene_tx=XM_023253291.1
B1 196371254 196371338
refGene_gn=FBXL5;refGene_tx=XM_023253294.1
B1 196380963 196381179
refGene_gn=FBXL5;refGene_tx=XM_023253293.1;refGene_gn=FBXL5;refGene_tx=XM_023253294.1
B1 196385166 196385262
refGene_gn=FBXL5;refGene_tx=XM_023253293.1;refGene_gn=FBXL5;refGene_tx=XM_023253294.1
B1 196387417 196387604
refGene_gn=FBXL5;refGene_tx=XM_023253293.1;refGene_gn=FBXL5;refGene_tx=XM_023253294.1
B1 196389966 196390149
refGene_gn=FBXL5;refGene_tx=XM_023253293.1;refGene_gn=FBXL5;refGene_tx=XM_023253294.1
B1 196392120 196392246
refGene_gn=FBXL5;refGene_tx=XM_023253293.1;refGene_gn=FBXL5;refGene_tx=XM_023253294.1
B1 196395983 196396132
refGene_gn=FBXL5;refGene_tx=XM_023253293.1;refGene_gn=FBXL5;refGene_tx=XM_023253294.1

B1	196396754	196396837
refGene_gn=FBXL5;refGene_tx=XM_023253293.1;refGene_gn=FBXL5;refGene_tx
=XM_023253294.1
B1	196397741	196398467
refGene_gn=FBXL5;refGene_tx=XM_023253293.1;refGene_gn=FBXL5;refGene_tx
=XM_023253294.1
B1	196410456	196410605
refGene_gn=FBXL5;refGene_tx=XM_023253293.1;refGene_gn=FBXL5;refGene_tx
=XM_023253294.1
B1	196417649	196417726
refGene_gn=FBXL5;refGene_tx=XM_023253293.1;refGene_gn=FBXL5;refGene_tx
=XM_023253294.1
B1	196423707	196423896
refGene_gn=CC2D2A;refGene_tx=XM_023253295.1;refGene_gn=CC2D2A;refGene_
tx=XM_023253296.1;refGene_gn=CC2D2A;refGene_tx=XM_023253297.1;refGene_
gn=CC2D2A;refGene_tx=XM_023253298.1;refGene_gn=CC2D2A;refGene_tx=XM_02
3253299.1;refGene_gn=CC2D2A;refGene_tx=XM_023253300.1;refGene_gn=CC2D2
A;refGene_tx=XM_023253301.1;refGene_gn=CC2D2A;refGene_tx=XM_023253302.
1;refGene_gn=CC2D2A;refGene_tx=XM_023253303.1;refGene_gn=CC2D2A;refGen
e_tx=XM_023253304.1
B1	196425274	196425452
refGene_gn=CC2D2A;refGene_tx=XM_023253295.1;refGene_gn=CC2D2A;refGene_
tx=XM_023253296.1;refGene_gn=CC2D2A;refGene_tx=XM_023253297.1;refGene_
gn=CC2D2A;refGene_tx=XM_023253298.1;refGene_gn=CC2D2A;refGene_tx=XM_02
3253299.1;refGene_gn=CC2D2A;refGene_tx=XM_023253300.1;refGene_gn=CC2D2
A;refGene_tx=XM_023253301.1;refGene_gn=CC2D2A;refGene_tx=XM_023253302.
1;refGene_gn=CC2D2A;refGene_tx=XM_023253303.1;refGene_gn=CC2D2A;refGen
e_tx=XM_023253304.1
B1	196427152	196427211
refGene_gn=CC2D2A;refGene_tx=XM_023253295.1;refGene_gn=CC2D2A;refGene_
tx=XM_023253296.1;refGene_gn=CC2D2A;refGene_tx=XM_023253297.1;refGene_
gn=CC2D2A;refGene_tx=XM_023253298.1;refGene_gn=CC2D2A;refGene_tx=XM_02
3253299.1;refGene_gn=CC2D2A;refGene_tx=XM_023253300.1;refGene_gn=CC2D2
A;refGene_tx=XM_023253301.1;refGene_gn=CC2D2A;refGene_tx=XM_023253302.
1;refGene_gn=CC2D2A;refGene_tx=XM_023253303.1;refGene_gn=CC2D2A;refGen
e_tx=XM_023253304.1
B1	196428790	196428913
refGene_gn=CC2D2A;refGene_tx=XM_023253295.1;refGene_gn=CC2D2A;refGene_
tx=XM_023253296.1;refGene_gn=CC2D2A;refGene_tx=XM_023253297.1;refGene_
gn=CC2D2A;refGene_tx=XM_023253298.1;refGene_gn=CC2D2A;refGene_tx=XM_02
3253299.1;refGene_gn=CC2D2A;refGene_tx=XM_023253300.1;refGene_gn=CC2D2
A;refGene_tx=XM_023253301.1;refGene_gn=CC2D2A;refGene_tx=XM_023253302.
1;refGene_gn=CC2D2A;refGene_tx=XM_023253303.1;refGene_gn=CC2D2A;refGen
e_tx=XM_023253304.1
B1	196438499	196438634
refGene_gn=CC2D2A;refGene_tx=XM_023253295.1;refGene_gn=CC2D2A;refGene_
tx=XM_023253296.1;refGene_gn=CC2D2A;refGene_tx=XM_023253297.1;refGene_
gn=CC2D2A;refGene_tx=XM_023253298.1;refGene_gn=CC2D2A;refGene_tx=XM_02
3253299.1;refGene_gn=CC2D2A;refGene_tx=XM_023253300.1;refGene_gn=CC2D2
A;refGene_tx=XM_023253301.1;refGene_gn=CC2D2A;refGene_tx=XM_023253302.

1
B1 196440073 196440187
refGene_gn=CC2D2A;refGene_tx=XM_023253295.1;refGene_gn=CC2D2A;refGene_tx=XM_023253296.1;refGene_gn=CC2D2A;refGene_tx=XM_023253297.1;refGene_gn=CC2D2A;refGene_tx=XM_023253298.1;refGene_gn=CC2D2A;refGene_tx=XM_023253299.1;refGene_gn=CC2D2A;refGene_tx=XM_023253300.1;refGene_gn=CC2D2A;refGene_tx=XM_023253301.1;refGene_gn=CC2D2A;refGene_tx=XM_023253302.1
B1 196442012 196442102
refGene_gn=CC2D2A;refGene_tx=XM_023253295.1;refGene_gn=CC2D2A;refGene_tx=XM_023253296.1;refGene_gn=CC2D2A;refGene_tx=XM_023253297.1;refGene_gn=CC2D2A;refGene_tx=XM_023253298.1;refGene_gn=CC2D2A;refGene_tx=XM_023253299.1;refGene_gn=CC2D2A;refGene_tx=XM_023253300.1;refGene_gn=CC2D2A;refGene_tx=XM_023253301.1;refGene_gn=CC2D2A;refGene_tx=XM_023253302.1;refGene_gn=CC2D2A;refGene_tx=XM_023253303.1;refGene_gn=CC2D2A;refGene_tx=XM_023253304.1
B1 196449517 196449721
refGene_gn=CC2D2A;refGene_tx=XM_023253295.1;refGene_gn=CC2D2A;refGene_tx=XM_023253296.1;refGene_gn=CC2D2A;refGene_tx=XM_023253297.1;refGene_gn=CC2D2A;refGene_tx=XM_023253298.1;refGene_gn=CC2D2A;refGene_tx=XM_023253299.1;refGene_gn=CC2D2A;refGene_tx=XM_023253300.1;refGene_gn=CC2D2A;refGene_tx=XM_023253301.1;refGene_gn=CC2D2A;refGene_tx=XM_023253303.1
B1 196452967 196452988
refGene_gn=CC2D2A;refGene_tx=XM_023253295.1;refGene_gn=CC2D2A;refGene_tx=XM_023253296.1;refGene_gn=CC2D2A;refGene_tx=XM_023253298.1;refGene_gn=CC2D2A;refGene_tx=XM_023253299.1;refGene_gn=CC2D2A;refGene_tx=XM_023253301.1;refGene_gn=CC2D2A;refGene_tx=XM_023253303.1
B1 196455767 196455944
refGene_gn=CC2D2A;refGene_tx=XM_023253295.1;refGene_gn=CC2D2A;refGene_tx=XM_023253296.1;refGene_gn=CC2D2A;refGene_tx=XM_023253297.1;refGene_gn=CC2D2A;refGene_tx=XM_023253298.1;refGene_gn=CC2D2A;refGene_tx=XM_023253299.1;refGene_gn=CC2D2A;refGene_tx=XM_023253300.1;refGene_gn=CC2D2A;refGene_tx=XM_023253301.1;refGene_gn=CC2D2A;refGene_tx=XM_023253302.1;refGene_gn=CC2D2A;refGene_tx=XM_023253303.1;refGene_gn=CC2D2A;refGene_tx=XM_023253304.1
B1 196458824 196458923
refGene_gn=CC2D2A;refGene_tx=XM_023253295.1;refGene_gn=CC2D2A;refGene_tx=XM_023253296.1;refGene_gn=CC2D2A;refGene_tx=XM_023253297.1;refGene_gn=CC2D2A;refGene_tx=XM_023253298.1;refGene_gn=CC2D2A;refGene_tx=XM_023253299.1;refGene_gn=CC2D2A;refGene_tx=XM_023253300.1;refGene_gn=CC2D2A;refGene_tx=XM_023253301.1;refGene_gn=CC2D2A;refGene_tx=XM_023253302.1;refGene_gn=CC2D2A;refGene_tx=XM_023253303.1;refGene_gn=CC2D2A;refGene_tx=XM_023253304.1
B1 196459514 196459611
refGene_gn=CC2D2A;refGene_tx=XM_023253295.1;refGene_gn=CC2D2A;refGene_tx=XM_023253296.1;refGene_gn=CC2D2A;refGene_tx=XM_023253297.1;refGene_gn=CC2D2A;refGene_tx=XM_023253298.1;refGene_gn=CC2D2A;refGene_tx=XM_023253299.1;refGene_gn=CC2D2A;refGene_tx=XM_023253300.1;refGene_gn=CC2D2A;refGene_tx=XM_023253301.1;refGene_gn=CC2D2A;refGene_tx=XM_023253302.

1;refGene_gn=CC2D2A;refGene_tx=XM_023253303.1;refGene_gn=CC2D2A;refGene_tx=XM_023253304.1
B1 196460869 196460979
refGene_gn=CC2D2A;refGene_tx=XM_023253295.1;refGene_gn=CC2D2A;refGene_tx=XM_023253296.1;refGene_gn=CC2D2A;refGene_tx=XM_023253297.1;refGene_gn=CC2D2A;refGene_tx=XM_023253298.1;refGene_gn=CC2D2A;refGene_tx=XM_023253299.1;refGene_gn=CC2D2A;refGene_tx=XM_023253300.1;refGene_gn=CC2D2A;refGene_tx=XM_023253301.1;refGene_gn=CC2D2A;refGene_tx=XM_023253302.1;refGene_gn=CC2D2A;refGene_tx=XM_023253303.1;refGene_gn=CC2D2A;refGene_tx=XM_023253304.1
B1 196461169 196461275
refGene_gn=CC2D2A;refGene_tx=XM_023253295.1;refGene_gn=CC2D2A;refGene_tx=XM_023253296.1;refGene_gn=CC2D2A;refGene_tx=XM_023253297.1;refGene_gn=CC2D2A;refGene_tx=XM_023253298.1;refGene_gn=CC2D2A;refGene_tx=XM_023253299.1;refGene_gn=CC2D2A;refGene_tx=XM_023253300.1;refGene_gn=CC2D2A;refGene_tx=XM_023253301.1;refGene_gn=CC2D2A;refGene_tx=XM_023253302.1;refGene_gn=CC2D2A;refGene_tx=XM_023253303.1;refGene_gn=CC2D2A;refGene_tx=XM_023253304.1
B1 196464371 196464539
refGene_gn=CC2D2A;refGene_tx=XM_023253295.1;refGene_gn=CC2D2A;refGene_tx=XM_023253296.1;refGene_gn=CC2D2A;refGene_tx=XM_023253297.1;refGene_gn=CC2D2A;refGene_tx=XM_023253298.1;refGene_gn=CC2D2A;refGene_tx=XM_023253299.1;refGene_gn=CC2D2A;refGene_tx=XM_023253300.1;refGene_gn=CC2D2A;refGene_tx=XM_023253301.1;refGene_gn=CC2D2A;refGene_tx=XM_023253302.1;refGene_gn=CC2D2A;refGene_tx=XM_023253303.1;refGene_gn=CC2D2A;refGene_tx=XM_023253304.1
B1 196466655 196466670
refGene_gn=CC2D2A;refGene_tx=XM_023253295.1;refGene_gn=CC2D2A;refGene_tx=XM_023253297.1;refGene_gn=CC2D2A;refGene_tx=XM_023253298.1;refGene_gn=CC2D2A;refGene_tx=XM_023253299.1;refGene_gn=CC2D2A;refGene_tx=XM_023253301.1;refGene_gn=CC2D2A;refGene_tx=XM_023253302.1;refGene_gn=CC2D2A;refGene_tx=XM_023253303.1;refGene_gn=CC2D2A;refGene_tx=XM_023253304.1
B1 196467621 196467713
refGene_gn=CC2D2A;refGene_tx=XM_023253295.1;refGene_gn=CC2D2A;refGene_tx=XM_023253296.1;refGene_gn=CC2D2A;refGene_tx=XM_023253297.1;refGene_gn=CC2D2A;refGene_tx=XM_023253298.1;refGene_gn=CC2D2A;refGene_tx=XM_023253299.1;refGene_gn=CC2D2A;refGene_tx=XM_023253300.1;refGene_gn=CC2D2A;refGene_tx=XM_023253301.1;refGene_gn=CC2D2A;refGene_tx=XM_023253302.1;refGene_gn=CC2D2A;refGene_tx=XM_023253303.1;refGene_gn=CC2D2A;refGene_tx=XM_023253304.1
B1 196469021 196469114
refGene_gn=CC2D2A;refGene_tx=XM_023253295.1;refGene_gn=CC2D2A;refGene_tx=XM_023253296.1;refGene_gn=CC2D2A;refGene_tx=XM_023253297.1;refGene_gn=CC2D2A;refGene_tx=XM_023253298.1;refGene_gn=CC2D2A;refGene_tx=XM_023253299.1;refGene_gn=CC2D2A;refGene_tx=XM_023253300.1;refGene_gn=CC2D2A;refGene_tx=XM_023253301.1;refGene_gn=CC2D2A;refGene_tx=XM_023253302.1;refGene_gn=CC2D2A;refGene_tx=XM_023253303.1;refGene_gn=CC2D2A;refGene_tx=XM_023253304.1
B1 196471148 196471352

refGene_gn=CC2D2A;refGene_tx=XM_023253295.1;refGene_gn=CC2D2A;refGene_tx=XM_023253296.1;refGene_gn=CC2D2A;refGene_tx=XM_023253297.1;refGene_gn=CC2D2A;refGene_tx=XM_023253298.1;refGene_gn=CC2D2A;refGene_tx=XM_023253299.1;refGene_gn=CC2D2A;refGene_tx=XM_023253300.1;refGene_gn=CC2D2A;refGene_tx=XM_023253301.1;refGene_gn=CC2D2A;refGene_tx=XM_023253302.1;refGene_gn=CC2D2A;refGene_tx=XM_023253303.1;refGene_gn=CC2D2A;refGene_tx=XM_023253304.1
B1 196475754 196475893
refGene_gn=CC2D2A;refGene_tx=XM_023253295.1;refGene_gn=CC2D2A;refGene_tx=XM_023253296.1;refGene_gn=CC2D2A;refGene_tx=XM_023253297.1;refGene_gn=CC2D2A;refGene_tx=XM_023253298.1;refGene_gn=CC2D2A;refGene_tx=XM_023253299.1;refGene_gn=CC2D2A;refGene_tx=XM_023253300.1;refGene_gn=CC2D2A;refGene_tx=XM_023253301.1;refGene_gn=CC2D2A;refGene_tx=XM_023253302.1;refGene_gn=CC2D2A;refGene_tx=XM_023253303.1;refGene_gn=CC2D2A;refGene_tx=XM_023253304.1
B1 196476865 196476925
refGene_gn=CC2D2A;refGene_tx=XM_023253295.1;refGene_gn=CC2D2A;refGene_tx=XM_023253296.1;refGene_gn=CC2D2A;refGene_tx=XM_023253297.1;refGene_gn=CC2D2A;refGene_tx=XM_023253299.1;refGene_gn=CC2D2A;refGene_tx=XM_023253301.1;refGene_gn=CC2D2A;refGene_tx=XM_023253302.1;refGene_gn=CC2D2A;refGene_tx=XM_023253303.1;refGene_gn=CC2D2A;refGene_tx=XM_023253304.1
B1 196478765 196478913
refGene_gn=CC2D2A;refGene_tx=XM_023253295.1;refGene_gn=CC2D2A;refGene_tx=XM_023253296.1;refGene_gn=CC2D2A;refGene_tx=XM_023253297.1;refGene_gn=CC2D2A;refGene_tx=XM_023253298.1;refGene_gn=CC2D2A;refGene_tx=XM_023253299.1;refGene_gn=CC2D2A;refGene_tx=XM_023253300.1;refGene_gn=CC2D2A;refGene_tx=XM_023253301.1;refGene_gn=CC2D2A;refGene_tx=XM_023253302.1;refGene_gn=CC2D2A;refGene_tx=XM_023253303.1;refGene_gn=CC2D2A;refGene_tx=XM_023253304.1
B1 196481617 196481774
refGene_gn=CC2D2A;refGene_tx=XM_023253295.1;refGene_gn=CC2D2A;refGene_tx=XM_023253296.1;refGene_gn=CC2D2A;refGene_tx=XM_023253297.1;refGene_gn=CC2D2A;refGene_tx=XM_023253298.1;refGene_gn=CC2D2A;refGene_tx=XM_023253299.1;refGene_gn=CC2D2A;refGene_tx=XM_023253300.1;refGene_gn=CC2D2A;refGene_tx=XM_023253301.1;refGene_gn=CC2D2A;refGene_tx=XM_023253302.1;refGene_gn=CC2D2A;refGene_tx=XM_023253303.1;refGene_gn=CC2D2A;refGene_tx=XM_023253304.1
B1 196495133 196495311
refGene_gn=CC2D2A;refGene_tx=XM_023253295.1;refGene_gn=CC2D2A;refGene_tx=XM_023253296.1;refGene_gn=CC2D2A;refGene_tx=XM_023253297.1;refGene_gn=CC2D2A;refGene_tx=XM_023253298.1;refGene_gn=CC2D2A;refGene_tx=XM_023253299.1;refGene_gn=CC2D2A;refGene_tx=XM_023253300.1;refGene_gn=CC2D2A;refGene_tx=XM_023253301.1;refGene_gn=CC2D2A;refGene_tx=XM_023253302.1;refGene_gn=CC2D2A;refGene_tx=XM_023253303.1;refGene_gn=CC2D2A;refGene_tx=XM_023253304.1
B1 196499095 196499334
refGene_gn=CC2D2A;refGene_tx=XM_023253295.1;refGene_gn=CC2D2A;refGene_tx=XM_023253296.1;refGene_gn=CC2D2A;refGene_tx=XM_023253297.1;refGene_gn=CC2D2A;refGene_tx=XM_023253298.1;refGene_gn=CC2D2A;refGene_tx=XM_02

3253299.1;refGene_gn=CC2D2A;refGene_tx=XM_023253300.1;refGene_gn=CC2D2A;refGene_tx=XM_023253301.1;refGene_gn=CC2D2A;refGene_tx=XM_023253302.1;refGene_gn=CC2D2A;refGene_tx=XM_023253303.1;refGene_gn=CC2D2A;refGene_tx=XM_023253304.1
B1 196500189 196500346
refGene_gn=CC2D2A;refGene_tx=XM_023253295.1;refGene_gn=CC2D2A;refGene_tx=XM_023253296.1;refGene_gn=CC2D2A;refGene_tx=XM_023253297.1;refGene_gn=CC2D2A;refGene_tx=XM_023253298.1;refGene_gn=CC2D2A;refGene_tx=XM_023253299.1;refGene_gn=CC2D2A;refGene_tx=XM_023253300.1;refGene_gn=CC2D2A;refGene_tx=XM_023253301.1;refGene_gn=CC2D2A;refGene_tx=XM_023253302.1;refGene_gn=CC2D2A;refGene_tx=XM_023253303.1;refGene_gn=CC2D2A;refGene_tx=XM_023253304.1
B1 196503565 196503706
refGene_gn=CC2D2A;refGene_tx=XM_023253295.1;refGene_gn=CC2D2A;refGene_tx=XM_023253296.1;refGene_gn=CC2D2A;refGene_tx=XM_023253297.1;refGene_gn=CC2D2A;refGene_tx=XM_023253298.1;refGene_gn=CC2D2A;refGene_tx=XM_023253299.1;refGene_gn=CC2D2A;refGene_tx=XM_023253300.1;refGene_gn=CC2D2A;refGene_tx=XM_023253301.1;refGene_gn=CC2D2A;refGene_tx=XM_023253302.1;refGene_gn=CC2D2A;refGene_tx=XM_023253303.1;refGene_gn=CC2D2A;refGene_tx=XM_023253304.1
B1 196508082 196508189
refGene_gn=CC2D2A;refGene_tx=XM_023253295.1;refGene_gn=CC2D2A;refGene_tx=XM_023253296.1;refGene_gn=CC2D2A;refGene_tx=XM_023253297.1;refGene_gn=CC2D2A;refGene_tx=XM_023253298.1;refGene_gn=CC2D2A;refGene_tx=XM_023253299.1;refGene_gn=CC2D2A;refGene_tx=XM_023253300.1;refGene_gn=CC2D2A;refGene_tx=XM_023253301.1;refGene_gn=CC2D2A;refGene_tx=XM_023253302.1;refGene_gn=CC2D2A;refGene_tx=XM_023253303.1;refGene_gn=CC2D2A;refGene_tx=XM_023253304.1
B1 196509356 196509566
refGene_gn=CC2D2A;refGene_tx=XM_023253295.1;refGene_gn=CC2D2A;refGene_tx=XM_023253296.1;refGene_gn=CC2D2A;refGene_tx=XM_023253297.1;refGene_gn=CC2D2A;refGene_tx=XM_023253298.1;refGene_gn=CC2D2A;refGene_tx=XM_023253299.1;refGene_gn=CC2D2A;refGene_tx=XM_023253300.1;refGene_gn=CC2D2A;refGene_tx=XM_023253301.1;refGene_gn=CC2D2A;refGene_tx=XM_023253302.1;refGene_gn=CC2D2A;refGene_tx=XM_023253303.1;refGene_gn=CC2D2A;refGene_tx=XM_023253304.1
B1 196516253 196516385
refGene_gn=CC2D2A;refGene_tx=XM_023253295.1;refGene_gn=CC2D2A;refGene_tx=XM_023253296.1;refGene_gn=CC2D2A;refGene_tx=XM_023253297.1;refGene_gn=CC2D2A;refGene_tx=XM_023253298.1;refGene_gn=CC2D2A;refGene_tx=XM_023253299.1;refGene_gn=CC2D2A;refGene_tx=XM_023253300.1;refGene_gn=CC2D2A;refGene_tx=XM_023253301.1;refGene_gn=CC2D2A;refGene_tx=XM_023253302.1;refGene_gn=CC2D2A;refGene_tx=XM_023253303.1;refGene_gn=CC2D2A;refGene_tx=XM_023253304.1
B1 196517486 196517623
refGene_gn=CC2D2A;refGene_tx=XM_023253295.1;refGene_gn=CC2D2A;refGene_tx=XM_023253296.1;refGene_gn=CC2D2A;refGene_tx=XM_023253297.1;refGene_gn=CC2D2A;refGene_tx=XM_023253298.1;refGene_gn=CC2D2A;refGene_tx=XM_023253299.1;refGene_gn=CC2D2A;refGene_tx=XM_023253300.1;refGene_gn=CC2D2A;refGene_tx=XM_023253301.1;refGene_gn=CC2D2A;refGene_tx=XM_023253302.

1;refGene_gn=CC2D2A;refGene_tx=XM_023253303.1;refGene_gn=CC2D2A;refGene_tx=XM_023253304.1
B1	196523707	196523870
refGene_gn=CC2D2A;refGene_tx=XM_023253295.1;refGene_gn=CC2D2A;refGene_tx=XM_023253296.1;refGene_gn=CC2D2A;refGene_tx=XM_023253297.1;refGene_gn=CC2D2A;refGene_tx=XM_023253298.1;refGene_gn=CC2D2A;refGene_tx=XM_023253299.1;refGene_gn=CC2D2A;refGene_tx=XM_023253300.1;refGene_gn=CC2D2A;refGene_tx=XM_023253301.1;refGene_gn=CC2D2A;refGene_tx=XM_023253302.1;refGene_gn=CC2D2A;refGene_tx=XM_023253303.1;refGene_gn=CC2D2A;refGene_tx=XM_023253304.1
B1	196533316	196533487
refGene_gn=CC2D2A;refGene_tx=XM_023253295.1;refGene_gn=CC2D2A;refGene_tx=XM_023253296.1;refGene_gn=CC2D2A;refGene_tx=XM_023253297.1;refGene_gn=CC2D2A;refGene_tx=XM_023253298.1;refGene_gn=CC2D2A;refGene_tx=XM_023253299.1;refGene_gn=CC2D2A;refGene_tx=XM_023253300.1;refGene_gn=CC2D2A;refGene_tx=XM_023253301.1;refGene_gn=CC2D2A;refGene_tx=XM_023253302.1;refGene_gn=CC2D2A;refGene_tx=XM_023253303.1;refGene_gn=CC2D2A;refGene_tx=XM_023253304.1
B1	196534531	196534633
refGene_gn=CC2D2A;refGene_tx=XM_023253295.1;refGene_gn=CC2D2A;refGene_tx=XM_023253296.1;refGene_gn=CC2D2A;refGene_tx=XM_023253297.1;refGene_gn=CC2D2A;refGene_tx=XM_023253298.1;refGene_gn=CC2D2A;refGene_tx=XM_023253299.1;refGene_gn=CC2D2A;refGene_tx=XM_023253300.1;refGene_gn=CC2D2A;refGene_tx=XM_023253301.1;refGene_gn=CC2D2A;refGene_tx=XM_023253302.1;refGene_gn=CC2D2A;refGene_tx=XM_023253303.1;refGene_gn=CC2D2A;refGene_tx=XM_023253304.1
B1	196542526	196542628
refGene_gn=CC2D2A;refGene_tx=XM_023253295.1;refGene_gn=CC2D2A;refGene_tx=XM_023253296.1;refGene_gn=CC2D2A;refGene_tx=XM_023253297.1;refGene_gn=CC2D2A;refGene_tx=XM_023253298.1;refGene_gn=CC2D2A;refGene_tx=XM_023253299.1;refGene_gn=CC2D2A;refGene_tx=XM_023253300.1;refGene_gn=CC2D2A;refGene_tx=XM_023253301.1;refGene_gn=CC2D2A;refGene_tx=XM_023253302.1;refGene_gn=CC2D2A;refGene_tx=XM_023253303.1;refGene_gn=CC2D2A;refGene_tx=XM_023253304.1
B1	196542945	196543034
refGene_gn=CC2D2A;refGene_tx=XM_023253295.1;refGene_gn=CC2D2A;refGene_tx=XM_023253296.1;refGene_gn=CC2D2A;refGene_tx=XM_023253297.1;refGene_gn=CC2D2A;refGene_tx=XM_023253298.1;refGene_gn=CC2D2A;refGene_tx=XM_023253299.1;refGene_gn=CC2D2A;refGene_tx=XM_023253300.1;refGene_gn=CC2D2A;refGene_tx=XM_023253301.1;refGene_gn=CC2D2A;refGene_tx=XM_023253302.1;refGene_gn=CC2D2A;refGene_tx=XM_023253303.1;refGene_gn=CC2D2A;refGene_tx=XM_023253304.1
B1	196563951	196564075
refGene_gn=CC2D2A;refGene_tx=XM_023253295.1;refGene_gn=CC2D2A;refGene_tx=XM_023253296.1;refGene_gn=CC2D2A;refGene_tx=XM_023253297.1;refGene_gn=CC2D2A;refGene_tx=XM_023253298.1;refGene_gn=CC2D2A;refGene_tx=XM_023253299.1;refGene_gn=CC2D2A;refGene_tx=XM_023253300.1;refGene_gn=CC2D2A;refGene_tx=XM_023253301.1;refGene_gn=CC2D2A;refGene_tx=XM_023253302.1;refGene_gn=CC2D2A;refGene_tx=XM_023253303.1;refGene_gn=CC2D2A;refGene_tx=XM_023253304.1

B1 196565213 196565276
refGene_gn=CC2D2A;refGene_tx=XM_023253299.1
B1 196565901 196565985
refGene_gn=CC2D2A;refGene_tx=XM_023253295.1;refGene_gn=CC2D2A;refGene_tx=XM_023253296.1;refGene_gn=CC2D2A;refGene_tx=XM_023253297.1;refGene_gn=CC2D2A;refGene_tx=XM_023253298.1;refGene_gn=CC2D2A;refGene_tx=XM_023253300.1;refGene_gn=CC2D2A;refGene_tx=XM_023253302.1;refGene_gn=CC2D2A;refGene_tx=XM_023253303.1;refGene_gn=CC2D2A;refGene_tx=XM_023253304.1
B1 196568057 196568096
refGene_gn=CC2D2A;refGene_tx=XM_023253295.1;refGene_gn=CC2D2A;refGene_tx=XM_023253296.1;refGene_gn=CC2D2A;refGene_tx=XM_023253297.1;refGene_gn=CC2D2A;refGene_tx=XM_023253298.1;refGene_gn=CC2D2A;refGene_tx=XM_023253300.1;refGene_gn=CC2D2A;refGene_tx=XM_023253302.1;refGene_gn=CC2D2A;refGene_tx=XM_023253303.1;refGene_gn=CC2D2A;refGene_tx=XM_023253304.1
B1 196597688 196598320
refGene_gn=C1QTNF7;refGene_tx=XM_003985524.5;refGene_gn=C1QTNF7;refGene_tx=XM_003985525.4;refGene_gn=C1QTNF7;refGene_tx=XM_006931216.3;refGene_gn=C1QTNF7;refGene_tx=XM_006931217.3
B1 196604759 196605005
refGene_gn=C1QTNF7;refGene_tx=XM_003985524.5;refGene_gn=C1QTNF7;refGene_tx=XM_003985525.4;refGene_gn=C1QTNF7;refGene_tx=XM_006931216.3;refGene_gn=C1QTNF7;refGene_tx=XM_006931217.3
B1 196704036 196704064
refGene_gn=C1QTNF7;refGene_tx=XM_003985524.5
B1 196904924 196905152
refGene_gn=CPEB2;refGene_tx=XM_023253305.1
B1 196911258 196911440
refGene_gn=CPEB2;refGene_tx=XM_023253305.1
B1 196913909 196914024
refGene_gn=CPEB2;refGene_tx=XM_023253305.1
B1 196914333 196914452
refGene_gn=CPEB2;refGene_tx=XM_023253305.1
B1 196918955 196919045
refGene_gn=CPEB2;refGene_tx=XM_023253305.1
B1 196920256 196920427
refGene_gn=CPEB2;refGene_tx=XM_023253305.1
B1 196935071 196935095
refGene_gn=CPEB2;refGene_tx=XM_023253305.1
B1 196942967 196943018
refGene_gn=CPEB2;refGene_tx=XM_023253305.1
B1 196960093 196960184
refGene_gn=CPEB2;refGene_tx=XM_023253305.1
B1 196969200 196969290
refGene_gn=CPEB2;refGene_tx=XM_023253305.1
B1 196970041 196970323
refGene_gn=CPEB2;refGene_tx=XM_023253305.1
B1 196973232 196974600
refGene_gn=CPEB2;refGene_tx=XM_023253305.1

B1 196974601 196974928
refGene_gn=CPEB2;refGene_tx=XM_023253305.1
B1 198207170 198207401
refGene_gn=BOD1L1;refGene_tx=XM_019829965.2;refGene_gn=BOD1L1;refGene_tx=XM_023253306.1
B1 198213877 198214002
refGene_gn=BOD1L1;refGene_tx=XM_019829965.2;refGene_gn=BOD1L1;refGene_tx=XM_023253306.1
B1 198218718 198218909
refGene_gn=BOD1L1;refGene_tx=XM_019829965.2;refGene_gn=BOD1L1;refGene_tx=XM_023253306.1
B1 198219620 198220235
refGene_gn=BOD1L1;refGene_tx=XM_019829965.2;refGene_gn=BOD1L1;refGene_tx=XM_023253306.1
B1 198220416 198220566
refGene_gn=BOD1L1;refGene_tx=XM_019829965.2;refGene_gn=BOD1L1;refGene_tx=XM_023253306.1
B1 198222676 198222843
refGene_gn=BOD1L1;refGene_tx=XM_019829965.2;refGene_gn=BOD1L1;refGene_tx=XM_023253306.1
B1 198225141 198225253
refGene_gn=BOD1L1;refGene_tx=XM_019829965.2;refGene_gn=BOD1L1;refGene_tx=XM_023253306.1
B1 198225908 198226047
refGene_gn=BOD1L1;refGene_tx=XM_019829965.2;refGene_gn=BOD1L1;refGene_tx=XM_023253306.1
B1 198226733 198226806
refGene_gn=BOD1L1;refGene_tx=XM_019829965.2;refGene_gn=BOD1L1;refGene_tx=XM_023253306.1
B1 198228799 198234782
refGene_gn=BOD1L1;refGene_tx=XM_019829965.2;refGene_gn=BOD1L1;refGene_tx=XM_023253306.1
B1 198237062 198237127
refGene_gn=BOD1L1;refGene_tx=XM_019829965.2;refGene_gn=BOD1L1;refGene_tx=XM_023253306.1
B1 198237995 198238080
refGene_gn=BOD1L1;refGene_tx=XM_019829965.2;refGene_gn=BOD1L1;refGene_tx=XM_023253306.1
B1 198242233 198242277
refGene_gn=BOD1L1;refGene_tx=XM_019829965.2;refGene_gn=BOD1L1;refGene_tx=XM_023253306.1
B1 198243777 198243838
refGene_gn=BOD1L1;refGene_tx=XM_019829965.2;refGene_gn=BOD1L1;refGene_tx=XM_023253306.1
B1 198245401 198245472
refGene_gn=BOD1L1;refGene_tx=XM_019829965.2;refGene_gn=BOD1L1;refGene_tx=XM_023253306.1
B1 198246093 198246166
refGene_gn=BOD1L1;refGene_tx=XM_019829965.2;refGene_gn=BOD1L1;refGene_tx=XM_023253306.1

B1 198247393 198247467
refGene_gn=BOD1L1;refGene_tx=XM_019829965.2;refGene_gn=BOD1L1;refGene_tx=XM_023253306.1
B1 198249091 198249176
refGene_gn=BOD1L1;refGene_tx=XM_019829965.2;refGene_gn=BOD1L1;refGene_tx=XM_023253306.1
B1 198249542 198249616
refGene_gn=BOD1L1;refGene_tx=XM_019829965.2;refGene_gn=BOD1L1;refGene_tx=XM_023253306.1
B1 198250579 198250655
refGene_gn=BOD1L1;refGene_tx=XM_019829965.2;refGene_gn=BOD1L1;refGene_tx=XM_023253306.1
B1 198250855 198250890
refGene_gn=BOD1L1;refGene_tx=XM_019829965.2;refGene_gn=BOD1L1;refGene_tx=XM_023253306.1
B1 198251951 198251997
refGene_gn=BOD1L1;refGene_tx=XM_019829965.2;refGene_gn=BOD1L1;refGene_tx=XM_023253306.1
B1 198253787 198253837
refGene_gn=BOD1L1;refGene_tx=XM_019829965.2;refGene_gn=BOD1L1;refGene_tx=XM_023253306.1
B1 198253934 198254019
refGene_gn=BOD1L1;refGene_tx=XM_019829965.2;refGene_gn=BOD1L1;refGene_tx=XM_023253306.1
B1 198254432 198254586
refGene_gn=BOD1L1;refGene_tx=XM_019829965.2;refGene_gn=BOD1L1;refGene_tx=XM_023253306.1
B1 198257185 198257326
refGene_gn=BOD1L1;refGene_tx=XM_019829965.2
B1 198260134 198260252
refGene_gn=BOD1L1;refGene_tx=XM_019829965.2;refGene_gn=BOD1L1;refGene_tx=XM_023253306.1
B1 198281829 198282295
refGene_gn=NKX3-2;refGene_tx=XM_023253307.1
B1 198283823 198284359
refGene_gn=NKX3-2;refGene_tx=XM_023253307.1
B1 198339287 198339362
refGene_gn=RAB28;refGene_tx=XM_003985516.5;refGene_gn=RAB28;refGene_tx=XM_003985517.4;refGene_gn=RAB28;refGene_tx=XM_006931210.2;refGene_gn=RAB28;refGene_tx=XM_006931211.3;refGene_gn=RAB28;refGene_tx=XM_006931212.4;refGene_gn=RAB28;refGene_tx=XM_019829743.2
B1 198343980 198344077
refGene_gn=RAB28;refGene_tx=XM_003985516.5;refGene_gn=RAB28;refGene_tx=XM_003985517.4;refGene_gn=RAB28;refGene_tx=XM_006931210.2;refGene_gn=RAB28;refGene_tx=XM_006931211.3;refGene_gn=RAB28;refGene_tx=XM_006931212.4;refGene_gn=RAB28;refGene_tx=XM_019829743.2
B1 198349278 198349367
refGene_gn=RAB28;refGene_tx=XM_003985516.5;refGene_gn=RAB28;refGene_tx=XM_003985517.4;refGene_gn=RAB28;refGene_tx=XM_006931210.2;refGene_gn=RAB28;refGene_tx=XM_006931211.3;refGene_gn=RAB28;refGene_tx=XM_0069312

12.4;refGene_gn=RAB28;refGene_tx=XM_019829743.2
B1 198361249 198361379
refGene_gn=RAB28;refGene_tx=XM_003985516.5;refGene_gn=RAB28;refGene_tx=XM_003985517.4;refGene_gn=RAB28;refGene_tx=XM_006931210.2;refGene_gn=RAB28;refGene_tx=XM_006931211.3;refGene_gn=RAB28;refGene_tx=XM_006931212.4;refGene_gn=RAB28;refGene_tx=XM_019829743.2
B1 198410753 198410857
refGene_gn=RAB28;refGene_tx=XM_003985516.5;refGene_gn=RAB28;refGene_tx=XM_003985517.4;refGene_gn=RAB28;refGene_tx=XM_006931210.2;refGene_gn=RAB28;refGene_tx=XM_006931211.3;refGene_gn=RAB28;refGene_tx=XM_006931212.4;refGene_gn=RAB28;refGene_tx=XM_019829743.2
B1 198415246 198415324
refGene_gn=RAB28;refGene_tx=XM_003985516.5;refGene_gn=RAB28;refGene_tx=XM_003985517.4;refGene_gn=RAB28;refGene_tx=XM_006931210.2;refGene_gn=RAB28;refGene_tx=XM_006931211.3;refGene_gn=RAB28;refGene_tx=XM_006931212.4;refGene_gn=RAB28;refGene_tx=XM_019829743.2
B1 198423926 198424016
refGene_gn=RAB28;refGene_tx=XM_003985517.4;refGene_gn=RAB28;refGene_tx=XM_006931211.3
B1 198425486 198425579
refGene_gn=RAB28;refGene_tx=XM_003985516.5
B1 198425700 198425796
refGene_gn=RAB28;refGene_tx=XM_006931210.2
B1 198426238 198426448
refGene_gn=RAB28;refGene_tx=XM_019829743.2
B1 198427481 198427556
refGene_gn=RAB28;refGene_tx=XM_006931212.4
B1 200056621 200057587
refGene_gn=HS3ST1;refGene_tx=XM_003985515.5;refGene_gn=HS3ST1;refGene_tx=XM_011281965.3
B1 200766488 200766545
refGene_gn=CLNK;refGene_tx=XM_023253310.1
B1 200771957 200772102
refGene_gn=CLNK;refGene_tx=XM_023253310.1
B1 200773292 200773399
refGene_gn=CLNK;refGene_tx=XM_023253310.1
B1 200779062 200779108
refGene_gn=CLNK;refGene_tx=XM_023253310.1
B1 200793327 200793353
refGene_gn=CLNK;refGene_tx=XM_023253310.1
B1 200793566 200793586
refGene_gn=CLNK;refGene_tx=XM_023253310.1
B1 200794607 200794718
refGene_gn=CLNK;refGene_tx=XM_023253310.1
B1 200802062 200802090
refGene_gn=CLNK;refGene_tx=XM_023253310.1
B1 200804996 200805015
refGene_gn=CLNK;refGene_tx=XM_023253310.1
B1 200807619 200807701
refGene_gn=CLNK;refGene_tx=XM_023253310.1

B1 200809261 200809302
refGene_gn=CLNK;refGene_tx=XM_023253310.1
B1 200815523 200815657
refGene_gn=CLNK;refGene_tx=XM_023253310.1
B1 200819677 200819755
refGene_gn=CLNK;refGene_tx=XM_023253310.1
B1 200823580 200823736
refGene_gn=CLNK;refGene_tx=XM_023253310.1
B1 200832368 200832515
refGene_gn=CLNK;refGene_tx=XM_023253310.1
B1 200868190 200871436
refGene_gn=ZNF518B;refGene_tx=XM_003985514.5;refGene_gn=ZNF518B;refGene_tx=XM_019829746.2;refGene_gn=ZNF518B;refGene_tx=XM_023253308.1;refGene_gn=ZNF518B;refGene_tx=XM_023253309.1
B1 200873458 200873542
refGene_gn=ZNF518B;refGene_tx=XM_023253309.1
B1 201074235 201074251
refGene_gn=WDR1;refGene_tx=XM_023253311.1
B1 201074698 201074820
refGene_gn=WDR1;refGene_tx=XM_023253311.1
B1 201087538 201087629
refGene_gn=WDR1;refGene_tx=XM_023253311.1
B1 201092235 201092383
refGene_gn=WDR1;refGene_tx=XM_023253311.1
B1 201093424 201093605
refGene_gn=WDR1;refGene_tx=XM_023253311.1
B1 201101134 201101212
refGene_gn=WDR1;refGene_tx=XM_023253311.1
B1 201101747 201101828
refGene_gn=WDR1;refGene_tx=XM_023253311.1
B1 201102051 201102285
refGene_gn=WDR1;refGene_tx=XM_023253311.1
B1 201104876 201104964
refGene_gn=WDR1;refGene_tx=XM_023253311.1
B1 201106693 201106850
refGene_gn=WDR1;refGene_tx=XM_023253311.1
B1 201108136 201108224
refGene_gn=WDR1;refGene_tx=XM_023253311.1
B1 201111758 201111869
refGene_gn=WDR1;refGene_tx=XM_023253311.1
B1 201114018 201114192
refGene_gn=WDR1;refGene_tx=XM_023253311.1
B1 201115919 201116064
refGene_gn=WDR1;refGene_tx=XM_023253311.1
B1 201117632 201117739
refGene_gn=WDR1;refGene_tx=XM_023253311.1
B1 201166837 201166900
refGene_gn=SLC2A9;refGene_tx=XM_003985513.5;refGene_gn=SLC2A9;refGene_tx=XM_019829751.2;refGene_gn=SLC2A9;refGene_tx=XM_019829752.2
B1 201171619 201171730

refGene_gn=SLC2A9;refGene_tx=XM_019829750.2
B1	201174212	201174311
refGene_gn=SLC2A9;refGene_tx=XM_003985513.5;refGene_gn=SLC2A9;refGene_tx=XM_019829750.2;refGene_gn=SLC2A9;refGene_tx=XM_019829751.2;refGene_gn=SLC2A9;refGene_tx=XM_019829752.2
B1	201199537	201199698
refGene_gn=SLC2A9;refGene_tx=XM_003985513.5;refGene_gn=SLC2A9;refGene_tx=XM_019829750.2;refGene_gn=SLC2A9;refGene_tx=XM_019829751.2;refGene_gn=SLC2A9;refGene_tx=XM_019829752.2
B1	201216535	201216660
refGene_gn=SLC2A9;refGene_tx=XM_003985513.5;refGene_gn=SLC2A9;refGene_tx=XM_019829750.2;refGene_gn=SLC2A9;refGene_tx=XM_019829751.2;refGene_gn=SLC2A9;refGene_tx=XM_019829752.2
B1	201231169	201231315
refGene_gn=SLC2A9;refGene_tx=XM_003985513.5;refGene_gn=SLC2A9;refGene_tx=XM_019829750.2;refGene_gn=SLC2A9;refGene_tx=XM_019829751.2;refGene_gn=SLC2A9;refGene_tx=XM_019829752.2
B1	201271611	201271744
refGene_gn=SLC2A9;refGene_tx=XM_003985513.5;refGene_gn=SLC2A9;refGene_tx=XM_019829750.2;refGene_gn=SLC2A9;refGene_tx=XM_019829751.2;refGene_gn=SLC2A9;refGene_tx=XM_019829752.2
B1	201305046	201305234
refGene_gn=SLC2A9;refGene_tx=XM_003985513.5;refGene_gn=SLC2A9;refGene_tx=XM_019829750.2;refGene_gn=SLC2A9;refGene_tx=XM_019829751.2;refGene_gn=SLC2A9;refGene_tx=XM_019829752.2
B1	201316938	201317049
refGene_gn=SLC2A9;refGene_tx=XM_003985513.5;refGene_gn=SLC2A9;refGene_tx=XM_019829750.2;refGene_gn=SLC2A9;refGene_tx=XM_019829751.2;refGene_gn=SLC2A9;refGene_tx=XM_019829752.2
B1	201330014	201330116
refGene_gn=SLC2A9;refGene_tx=XM_003985513.5;refGene_gn=SLC2A9;refGene_tx=XM_019829750.2;refGene_gn=SLC2A9;refGene_tx=XM_019829751.2;refGene_gn=SLC2A9;refGene_tx=XM_019829752.2
B1	201333589	201333665
refGene_gn=SLC2A9;refGene_tx=XM_003985513.5;refGene_gn=SLC2A9;refGene_tx=XM_019829750.2;refGene_gn=SLC2A9;refGene_tx=XM_019829751.2;refGene_gn=SLC2A9;refGene_tx=XM_019829752.2
B1	201366908	201367036
refGene_gn=SLC2A9;refGene_tx=XM_003985513.5;refGene_gn=SLC2A9;refGene_tx=XM_019829750.2;refGene_gn=SLC2A9;refGene_tx=XM_019829751.2;refGene_gn=SLC2A9;refGene_tx=XM_019829752.2
B1	201384350	201384605
refGene_gn=SLC2A9;refGene_tx=XM_003985513.5;refGene_gn=SLC2A9;refGene_tx=XM_019829750.2;refGene_gn=SLC2A9;refGene_tx=XM_019829751.2;refGene_gn=SLC2A9;refGene_tx=XM_019829752.2
B1	201422334	201423759
refGene_gn=DRD5;refGene_tx=XM_003985519.5
B1	201438064	201438235
refGene_gn=OTOP1;refGene_tx=XM_023253312.1
B1	201445979	201446908

refGene_gn=OTOP1;refGene_tx=XM_023253312.1
B1	201450842	201450973
refGene_gn=OTOP1;refGene_tx=XM_023253312.1
B1	201457569	201457628
refGene_gn=OTOP1;refGene_tx=XM_023253312.1
B1	201461578	201461715
refGene_gn=OTOP1;refGene_tx=XM_023253312.1
B1	201472470	201472867
refGene_gn=OTOP1;refGene_tx=XM_023253312.1
B1	201487748	201487848
refGene_gn=TMEM128;refGene_tx=XM_003985511.3
B1	201490669	201490828
refGene_gn=TMEM128;refGene_tx=XM_003985511.3
B1	201494514	201494656
refGene_gn=TMEM128;refGene_tx=XM_003985511.3
B1	201496178	201496275
refGene_gn=TMEM128;refGene_tx=XM_003985511.3
B1	201508923	201509058
refGene_gn=LYAR;refGene_tx=XM_006931205.4;refGene_gn=LYAR;refGene_tx=XM_006931206.4;refGene_gn=LYAR;refGene_tx=XM_006931207.4;refGene_gn=LYAR;refGene_tx=XM_006931208.4;refGene_gn=LYAR;refGene_tx=XM_023253314.1
B1	201509784	201509870
refGene_gn=LYAR;refGene_tx=XM_006931205.4;refGene_gn=LYAR;refGene_tx=XM_006931206.4;refGene_gn=LYAR;refGene_tx=XM_006931207.4;refGene_gn=LYAR;refGene_tx=XM_006931208.4;refGene_gn=LYAR;refGene_tx=XM_023253314.1
B1	201514031	201514118
refGene_gn=LYAR;refGene_tx=XM_006931205.4;refGene_gn=LYAR;refGene_tx=XM_006931206.4;refGene_gn=LYAR;refGene_tx=XM_006931207.4;refGene_gn=LYAR;refGene_tx=XM_006931208.4;refGene_gn=LYAR;refGene_tx=XM_023253314.1
B1	201514421	201514839
refGene_gn=LYAR;refGene_tx=XM_006931205.4;refGene_gn=LYAR;refGene_tx=XM_006931206.4;refGene_gn=LYAR;refGene_tx=XM_006931207.4;refGene_gn=LYAR;refGene_tx=XM_006931208.4;refGene_gn=LYAR;refGene_tx=XM_023253314.1
B1	201518196	201518280
refGene_gn=LYAR;refGene_tx=XM_006931205.4;refGene_gn=LYAR;refGene_tx=XM_006931206.4;refGene_gn=LYAR;refGene_tx=XM_006931207.4;refGene_gn=LYAR;refGene_tx=XM_006931208.4;refGene_gn=LYAR;refGene_tx=XM_023253314.1
B1	201518396	201518504
refGene_gn=LYAR;refGene_tx=XM_006931205.4;refGene_gn=LYAR;refGene_tx=XM_006931206.4;refGene_gn=LYAR;refGene_tx=XM_006931207.4;refGene_gn=LYAR;refGene_tx=XM_006931208.4;refGene_gn=LYAR;refGene_tx=XM_023253314.1
B1	201522787	201522902
refGene_gn=LYAR;refGene_tx=XM_006931205.4;refGene_gn=LYAR;refGene_tx=XM_006931206.4;refGene_gn=LYAR;refGene_tx=XM_006931207.4;refGene_gn=LYAR;refGene_tx=XM_006931208.4;refGene_gn=LYAR;refGene_tx=XM_023253314.1
B1	201524276	201524398
refGene_gn=LYAR;refGene_tx=XM_006931205.4;refGene_gn=LYAR;refGene_tx=XM_006931206.4;refGene_gn=LYAR;refGene_tx=XM_006931207.4;refGene_gn=LYAR;refGene_tx=XM_006931208.4;refGene_gn=LYAR;refGene_tx=XM_023253314.1
B1	201538866	201539018

refGene_gn=ZBTB49;refGene_tx=XM_023253313.1
B1 201541010 201542116
refGene_gn=ZBTB49;refGene_tx=XM_023253313.1
B1 201544610 201544657
refGene_gn=ZBTB49;refGene_tx=XM_023253313.1
B1 201551834 201551908
refGene_gn=ZBTB49;refGene_tx=XM_023253313.1
B1 201553461 201553544
refGene_gn=ZBTB49;refGene_tx=XM_023253313.1
B1 201553635 201553797
refGene_gn=ZBTB49;refGene_tx=XM_023253313.1
B1 201557369 201557599
refGene_gn=ZBTB49;refGene_tx=XM_023253313.1
B1 201557600 201558047
refGene_gn=ZBTB49;refGene_tx=XM_023253313.1
B1 201605645 201605774
refGene_gn=NSG1;refGene_tx=XM_023253315.1
B1 201609306 201609423
refGene_gn=NSG1;refGene_tx=XM_023253315.1
B1 201627642 201627753
refGene_gn=NSG1;refGene_tx=XM_023253315.1
B1 201634520 201634724
refGene_gn=NSG1;refGene_tx=XM_023253315.1
B1 201637147 201637243
refGene_gn=STX18;refGene_tx=XM_003985508.5
B1 201637946 201638027
refGene_gn=STX18;refGene_tx=XM_003985508.5
B1 201639673 201639743
refGene_gn=STX18;refGene_tx=XM_003985508.5
B1 201641061 201641120
refGene_gn=STX18;refGene_tx=XM_003985508.5
B1 201650304 201650393
refGene_gn=STX18;refGene_tx=XM_003985508.5
B1 201659318 201659434
refGene_gn=STX18;refGene_tx=XM_003985508.5
B1 201673555 201673622
refGene_gn=STX18;refGene_tx=XM_003985508.5
B1 201674025 201674103
refGene_gn=STX18;refGene_tx=XM_003985508.5
B1 201675855 201675971
refGene_gn=STX18;refGene_tx=XM_003985508.5
B1 201686983 201687051
refGene_gn=STX18;refGene_tx=XM_003985508.5
B1 201760158 201760326
refGene_gn=STX18;refGene_tx=XM_003985508.5
B1 202048751 202049220
refGene_gn=MSX1;refGene_tx=XM_019829968.2
B1 202051560 202052003
refGene_gn=MSX1;refGene_tx=XM_019829968.2
B1 202176941 202177025

refGene_gn=CYTL1;refGene_tx=XM_003985603.3
B1	202178164	202178293
refGene_gn=CYTL1;refGene_tx=XM_003985603.3
B1	202178425	202178470
refGene_gn=CYTL1;refGene_tx=XM_003985603.3
B1	202180051	202180201
refGene_gn=CYTL1;refGene_tx=XM_003985603.3
B1	202206240	202206292
refGene_gn=STK32B;refGene_tx=XM_019829756.2;refGene_gn=STK32B;refGene_tx=XM_019829757.2;refGene_gn=STK32B;refGene_tx=XM_019829758.2;refGene_gn=STK32B;refGene_tx=XM_019829759.2;refGene_gn=STK32B;refGene_tx=XM_023253316.1
B1	202299319	202299375
refGene_gn=STK32B;refGene_tx=XM_019829756.2;refGene_gn=STK32B;refGene_tx=XM_019829757.2;refGene_gn=STK32B;refGene_tx=XM_019829758.2;refGene_gn=STK32B;refGene_tx=XM_023253316.1
B1	202329811	202329963
refGene_gn=STK32B;refGene_tx=XM_019829756.2;refGene_gn=STK32B;refGene_tx=XM_019829757.2;refGene_gn=STK32B;refGene_tx=XM_019829758.2;refGene_gn=STK32B;refGene_tx=XM_019829759.2;refGene_gn=STK32B;refGene_tx=XM_019829761.2;refGene_gn=STK32B;refGene_tx=XM_019829763.2;refGene_gn=STK32B;refGene_tx=XM_019829764.2;refGene_gn=STK32B;refGene_tx=XM_019829765.2;refGene_gn=STK32B;refGene_tx=XM_023253316.1;refGene_gn=STK32B;refGene_tx=XM_023253317.1
B1	202470700	202470874
refGene_gn=STK32B;refGene_tx=XM_019829756.2;refGene_gn=STK32B;refGene_tx=XM_019829757.2;refGene_gn=STK32B;refGene_tx=XM_019829758.2;refGene_gn=STK32B;refGene_tx=XM_019829759.2;refGene_gn=STK32B;refGene_tx=XM_019829761.2;refGene_gn=STK32B;refGene_tx=XM_019829763.2;refGene_gn=STK32B;refGene_tx=XM_019829764.2;refGene_gn=STK32B;refGene_tx=XM_019829765.2;refGene_gn=STK32B;refGene_tx=XM_023253316.1;refGene_gn=STK32B;refGene_tx=XM_023253317.1
B1	202526637	202526675
refGene_gn=STK32B;refGene_tx=XM_019829756.2;refGene_gn=STK32B;refGene_tx=XM_019829757.2;refGene_gn=STK32B;refGene_tx=XM_019829758.2;refGene_gn=STK32B;refGene_tx=XM_019829759.2;refGene_gn=STK32B;refGene_tx=XM_019829761.2;refGene_gn=STK32B;refGene_tx=XM_019829763.2;refGene_gn=STK32B;refGene_tx=XM_019829764.2;refGene_gn=STK32B;refGene_tx=XM_019829765.2;refGene_gn=STK32B;refGene_tx=XM_023253316.1;refGene_gn=STK32B;refGene_tx=XM_023253317.1
B1	202543020	202543110
refGene_gn=STK32B;refGene_tx=XM_019829756.2;refGene_gn=STK32B;refGene_tx=XM_019829757.2;refGene_gn=STK32B;refGene_tx=XM_019829758.2;refGene_gn=STK32B;refGene_tx=XM_019829759.2;refGene_gn=STK32B;refGene_tx=XM_019829761.2;refGene_gn=STK32B;refGene_tx=XM_019829763.2;refGene_gn=STK32B;refGene_tx=XM_019829764.2;refGene_gn=STK32B;refGene_tx=XM_019829765.2;refGene_gn=STK32B;refGene_tx=XM_023253317.1
B1	202568600	202568704
refGene_gn=STK32B;refGene_tx=XM_019829756.2;refGene_gn=STK32B;refGene_tx=XM_019829757.2;refGene_gn=STK32B;refGene_tx=XM_019829758.2;refGene_

gn=STK32B;refGene_tx=XM_019829759.2;refGene_gn=STK32B;refGene_tx=XM_01
9829761.2;refGene_gn=STK32B;refGene_tx=XM_019829763.2;refGene_gn=STK32
B;refGene_tx=XM_019829764.2;refGene_gn=STK32B;refGene_tx=XM_019829765.
2;refGene_gn=STK32B;refGene_tx=XM_023253316.1;refGene_gn=STK32B;refGen
e_tx=XM_023253317.1
B1 202577500 202577617
refGene_gn=STK32B;refGene_tx=XM_019829756.2;refGene_gn=STK32B;refGene_
tx=XM_019829757.2;refGene_gn=STK32B;refGene_tx=XM_019829758.2;refGene_
gn=STK32B;refGene_tx=XM_019829759.2;refGene_gn=STK32B;refGene_tx=XM_01
9829761.2;refGene_gn=STK32B;refGene_tx=XM_019829763.2;refGene_gn=STK32
B;refGene_tx=XM_019829764.2;refGene_gn=STK32B;refGene_tx=XM_019829765.
2;refGene_gn=STK32B;refGene_tx=XM_023253316.1;refGene_gn=STK32B;refGen
e_tx=XM_023253317.1
B1 202579519 202579645
refGene_gn=STK32B;refGene_tx=XM_019829756.2;refGene_gn=STK32B;refGene_
tx=XM_019829757.2;refGene_gn=STK32B;refGene_tx=XM_019829758.2;refGene_
gn=STK32B;refGene_tx=XM_019829759.2;refGene_gn=STK32B;refGene_tx=XM_01
9829761.2;refGene_gn=STK32B;refGene_tx=XM_019829763.2;refGene_gn=STK32
B;refGene_tx=XM_019829764.2;refGene_gn=STK32B;refGene_tx=XM_019829765.
2;refGene_gn=STK32B;refGene_tx=XM_023253316.1;refGene_gn=STK32B;refGen
e_tx=XM_023253317.1
B1 202582400 202582532
refGene_gn=STK32B;refGene_tx=XM_019829756.2;refGene_gn=STK32B;refGene_
tx=XM_019829757.2;refGene_gn=STK32B;refGene_tx=XM_019829758.2;refGene_
gn=STK32B;refGene_tx=XM_019829759.2;refGene_gn=STK32B;refGene_tx=XM_01
9829761.2;refGene_gn=STK32B;refGene_tx=XM_019829763.2;refGene_gn=STK32
B;refGene_tx=XM_019829764.2;refGene_gn=STK32B;refGene_tx=XM_019829765.
2;refGene_gn=STK32B;refGene_tx=XM_023253316.1;refGene_gn=STK32B;refGen
e_tx=XM_023253317.1
B1 202583245 202583310
refGene_gn=STK32B;refGene_tx=XM_019829756.2;refGene_gn=STK32B;refGene_
tx=XM_019829757.2;refGene_gn=STK32B;refGene_tx=XM_019829758.2;refGene_
gn=STK32B;refGene_tx=XM_019829759.2;refGene_gn=STK32B;refGene_tx=XM_01
9829761.2;refGene_gn=STK32B;refGene_tx=XM_019829763.2;refGene_gn=STK32
B;refGene_tx=XM_019829764.2;refGene_gn=STK32B;refGene_tx=XM_019829765.
2;refGene_gn=STK32B;refGene_tx=XM_023253316.1;refGene_gn=STK32B;refGen
e_tx=XM_023253317.1
B1 202585136 202585176
refGene_gn=STK32B;refGene_tx=XM_019829758.2
B1 202585476 202585615
refGene_gn=STK32B;refGene_tx=XM_019829757.2
B1 202600589 202600728
refGene_gn=STK32B;refGene_tx=XM_019829756.2;refGene_gn=STK32B;refGene_
tx=XM_019829759.2;refGene_gn=STK32B;refGene_tx=XM_019829761.2;refGene_
gn=STK32B;refGene_tx=XM_019829763.2;refGene_gn=STK32B;refGene_tx=XM_01
9829764.2;refGene_gn=STK32B;refGene_tx=XM_019829765.2;refGene_gn=STK32
B;refGene_tx=XM_023253316.1;refGene_gn=STK32B;refGene_tx=XM_023253317.
1
B1 202637265 202637536
refGene_gn=EVC2;refGene_tx=XM_023253318.1

B1 202640176 202640278
refGene_gn=EVC2;refGene_tx=XM_023253318.1
B1 202642950 202643147
refGene_gn=EVC2;refGene_tx=XM_023253318.1
B1 202645598 202645686
refGene_gn=EVC2;refGene_tx=XM_023253318.1
B1 202647122 202647337
refGene_gn=EVC2;refGene_tx=XM_023253318.1
B1 202657351 202657579
refGene_gn=EVC2;refGene_tx=XM_023253318.1
B1 202672059 202672182
refGene_gn=EVC2;refGene_tx=XM_023253318.1
B1 202674298 202674503
refGene_gn=EVC2;refGene_tx=XM_023253318.1
B1 202679939 202680394
refGene_gn=EVC2;refGene_tx=XM_023253318.1
B1 202684447 202684607
refGene_gn=EVC2;refGene_tx=XM_023253318.1
B1 202687397 202687576
refGene_gn=EVC2;refGene_tx=XM_023253318.1
B1 202689756 202689993
refGene_gn=EVC2;refGene_tx=XM_023253318.1
B1 202697778 202697927
refGene_gn=EVC2;refGene_tx=XM_023253318.1
B1 202697929 202698102
refGene_gn=EVC2;refGene_tx=XM_023253318.1
B1 202724604 202724744
refGene_gn=EVC2;refGene_tx=XM_023253318.1
B1 202728093 202728228
refGene_gn=EVC2;refGene_tx=XM_023253318.1
B1 202744625 202744679
refGene_gn=EVC2;refGene_tx=XM_023253318.1
B1 202750083 202750193
refGene_gn=EVC2;refGene_tx=XM_023253318.1
B1 202757001 202757188
refGene_gn=EVC2;refGene_tx=XM_023253318.1
B1 202759281 202759350
refGene_gn=EVC2;refGene_tx=XM_023253318.1
B1 202763047 202763214
refGene_gn=EVC2;refGene_tx=XM_023253318.1
B1 202766471 202766526
refGene_gn=EVC2;refGene_tx=XM_023253318.1
B1 202772676 202772877
refGene_gn=EVC2;refGene_tx=XM_023253318.1
B1 202775345 202775519
refGene_gn=EVC;refGene_tx=XM_023253319.1;refGene_gn=EVC;refGene_tx=XM_
023253320.1
B1 202778288 202778293
refGene_gn=EVC;refGene_tx=XM_023253321.1
B1 202781839 202781962

refGene_gn=EVC;refGene_tx=XM_023253319.1;refGene_gn=EVC;refGene_tx=XM_023253320.1;refGene_gn=EVC;refGene_tx=XM_023253321.1
B1 202785588 202785600
refGene_gn=EVC;refGene_tx=XM_023253322.1
B1 202787997 202788115
refGene_gn=EVC;refGene_tx=XM_023253319.1;refGene_gn=EVC;refGene_tx=XM_023253320.1;refGene_gn=EVC;refGene_tx=XM_023253321.1;refGene_gn=EVC;refGene_tx=XM_023253322.1
B1 202790145 202790378
refGene_gn=EVC;refGene_tx=XM_023253319.1;refGene_gn=EVC;refGene_tx=XM_023253320.1;refGene_gn=EVC;refGene_tx=XM_023253321.1;refGene_gn=EVC;refGene_tx=XM_023253322.1
B1 202791649 202791734
refGene_gn=EVC;refGene_tx=XM_023253319.1;refGene_gn=EVC;refGene_tx=XM_023253320.1;refGene_gn=EVC;refGene_tx=XM_023253321.1;refGene_gn=EVC;refGene_tx=XM_023253322.1
B1 202797578 202797677
refGene_gn=EVC;refGene_tx=XM_023253319.1;refGene_gn=EVC;refGene_tx=XM_023253320.1;refGene_gn=EVC;refGene_tx=XM_023253321.1;refGene_gn=EVC;refGene_tx=XM_023253322.1
B1 202800795 202800933
refGene_gn=EVC;refGene_tx=XM_023253319.1;refGene_gn=EVC;refGene_tx=XM_023253320.1;refGene_gn=EVC;refGene_tx=XM_023253321.1;refGene_gn=EVC;refGene_tx=XM_023253322.1
B1 202803698 202803857
refGene_gn=EVC;refGene_tx=XM_023253319.1;refGene_gn=EVC;refGene_tx=XM_023253320.1;refGene_gn=EVC;refGene_tx=XM_023253321.1;refGene_gn=EVC;refGene_tx=XM_023253322.1
B1 202808888 202809105
refGene_gn=EVC;refGene_tx=XM_023253319.1;refGene_gn=EVC;refGene_tx=XM_023253320.1;refGene_gn=EVC;refGene_tx=XM_023253321.1;refGene_gn=EVC;refGene_tx=XM_023253322.1
B1 202811628 202811777
refGene_gn=EVC;refGene_tx=XM_023253319.1;refGene_gn=EVC;refGene_tx=XM_023253320.1;refGene_gn=EVC;refGene_tx=XM_023253321.1;refGene_gn=EVC;refGene_tx=XM_023253322.1
B1 202814060 202814159
refGene_gn=EVC;refGene_tx=XM_023253319.1;refGene_gn=EVC;refGene_tx=XM_023253320.1;refGene_gn=EVC;refGene_tx=XM_023253321.1;refGene_gn=EVC;refGene_tx=XM_023253322.1
B1 202822666 202822879
refGene_gn=EVC;refGene_tx=XM_023253319.1;refGene_gn=EVC;refGene_tx=XM_023253320.1;refGene_gn=EVC;refGene_tx=XM_023253321.1;refGene_gn=EVC;refGene_tx=XM_023253322.1
B1 202831119 202831229
refGene_gn=EVC;refGene_tx=XM_023253319.1;refGene_gn=EVC;refGene_tx=XM_023253320.1;refGene_gn=EVC;refGene_tx=XM_023253321.1;refGene_gn=EVC;refGene_tx=XM_023253322.1
B1 202834640 202834851
refGene_gn=EVC;refGene_tx=XM_023253319.1;refGene_gn=EVC;refGene_tx=XM_

023253320.1;refGene_gn=EVC;refGene_tx=XM_023253321.1;refGene_gn=EVC;refGene_tx=XM_023253322.1
B1 202835775 202835982
refGene_gn=EVC;refGene_tx=XM_023253319.1;refGene_gn=EVC;refGene_tx=XM_023253320.1;refGene_gn=EVC;refGene_tx=XM_023253321.1;refGene_gn=EVC;refGene_tx=XM_023253322.1
B1 202838622 202838767
refGene_gn=EVC;refGene_tx=XM_023253319.1;refGene_gn=EVC;refGene_tx=XM_023253320.1;refGene_gn=EVC;refGene_tx=XM_023253321.1;refGene_gn=EVC;refGene_tx=XM_023253322.1
B1 202840793 202840905
refGene_gn=EVC;refGene_tx=XM_023253319.1;refGene_gn=EVC;refGene_tx=XM_023253320.1;refGene_gn=EVC;refGene_tx=XM_023253321.1;refGene_gn=EVC;refGene_tx=XM_023253322.1
B1 202845633 202845760
refGene_gn=EVC;refGene_tx=XM_023253319.1;refGene_gn=EVC;refGene_tx=XM_023253320.1;refGene_gn=EVC;refGene_tx=XM_023253321.1;refGene_gn=EVC;refGene_tx=XM_023253322.1
B1 202847285 202847379
refGene_gn=EVC;refGene_tx=XM_023253319.1;refGene_gn=EVC;refGene_tx=XM_023253320.1;refGene_gn=EVC;refGene_tx=XM_023253321.1;refGene_gn=EVC;refGene_tx=XM_023253322.1
B1 202848040 202848152
refGene_gn=EVC;refGene_tx=XM_023253319.1;refGene_gn=EVC;refGene_tx=XM_023253320.1;refGene_gn=EVC;refGene_tx=XM_023253321.1;refGene_gn=EVC;refGene_tx=XM_023253322.1
B1 202848688 202848773
refGene_gn=EVC;refGene_tx=XM_023253319.1;refGene_gn=EVC;refGene_tx=XM_023253320.1;refGene_gn=EVC;refGene_tx=XM_023253321.1;refGene_gn=EVC;refGene_tx=XM_023253322.1
B1 202855378 202855470
refGene_gn=CRMP1;refGene_tx=XM_023253323.1;refGene_gn=CRMP1;refGene_tx=XM_023253324.1;refGene_gn=CRMP1;refGene_tx=XM_023253325.1
B1 202858090 202858256
refGene_gn=CRMP1;refGene_tx=XM_023253323.1;refGene_gn=CRMP1;refGene_tx=XM_023253324.1;refGene_gn=CRMP1;refGene_tx=XM_023253325.1
B1 202860318 202860498
refGene_gn=CRMP1;refGene_tx=XM_023253323.1;refGene_gn=CRMP1;refGene_tx=XM_023253324.1;refGene_gn=CRMP1;refGene_tx=XM_023253325.1
B1 202869500 202869671
refGene_gn=CRMP1;refGene_tx=XM_023253323.1;refGene_gn=CRMP1;refGene_tx=XM_023253324.1;refGene_gn=CRMP1;refGene_tx=XM_023253325.1
B1 202870344 202870486
refGene_gn=CRMP1;refGene_tx=XM_023253323.1;refGene_gn=CRMP1;refGene_tx=XM_023253324.1;refGene_gn=CRMP1;refGene_tx=XM_023253325.1
B1 202871752 202871909
refGene_gn=CRMP1;refGene_tx=XM_023253323.1;refGene_gn=CRMP1;refGene_tx=XM_023253324.1;refGene_gn=CRMP1;refGene_tx=XM_023253325.1
B1 202874415 202874536
refGene_gn=CRMP1;refGene_tx=XM_023253323.1;refGene_gn=CRMP1;refGene_tx

=XM_023253324.1;refGene_gn=CRMP1;refGene_tx=XM_023253325.1
B1 202875667 202875736
refGene_gn=CRMP1;refGene_tx=XM_023253323.1;refGene_gn=CRMP1;refGene_tx=XM_023253324.1;refGene_gn=CRMP1;refGene_tx=XM_023253325.1
B1 202880809 202880890
refGene_gn=CRMP1;refGene_tx=XM_023253323.1;refGene_gn=CRMP1;refGene_tx=XM_023253324.1;refGene_gn=CRMP1;refGene_tx=XM_023253325.1
B1 202882808 202882870
refGene_gn=CRMP1;refGene_tx=XM_023253323.1;refGene_gn=CRMP1;refGene_tx=XM_023253324.1;refGene_gn=CRMP1;refGene_tx=XM_023253325.1
B1 202896575 202896740
refGene_gn=CRMP1;refGene_tx=XM_023253323.1;refGene_gn=CRMP1;refGene_tx=XM_023253324.1;refGene_gn=CRMP1;refGene_tx=XM_023253325.1
B1 202900653 202900838
refGene_gn=CRMP1;refGene_tx=XM_023253323.1;refGene_gn=CRMP1;refGene_tx=XM_023253324.1;refGene_gn=CRMP1;refGene_tx=XM_023253325.1
B1 202905447 202905536
refGene_gn=CRMP1;refGene_tx=XM_023253323.1;refGene_gn=CRMP1;refGene_tx=XM_023253324.1;refGene_gn=CRMP1;refGene_tx=XM_023253325.1
B1 202923654 202923789
refGene_gn=CRMP1;refGene_tx=XM_023253325.1
B1 202927343 202927382
refGene_gn=CRMP1;refGene_tx=XM_023253324.1
B1 202931663 202932044
refGene_gn=CRMP1;refGene_tx=XM_023253323.1
B1 202944538 202944601
refGene_gn=CB1H4orf50;refGene_tx=XM_019829788.2
B1 203000966 203001218
refGene_gn=CB1H4orf50;refGene_tx=XM_023253326.1;refGene_gn=CB1H4orf50;refGene_tx=XM_023253327.1;refGene_gn=CB1H4orf50;refGene_tx=XM_023253328.1
B1 203006002 203006130
refGene_gn=CB1H4orf50;refGene_tx=XM_019829788.2;refGene_gn=CB1H4orf50;refGene_tx=XM_023253326.1;refGene_gn=CB1H4orf50;refGene_tx=XM_023253327.1;refGene_gn=CB1H4orf50;refGene_tx=XM_023253328.1
B1 203008017 203008066
refGene_gn=CB1H4orf50;refGene_tx=XM_019829788.2;refGene_gn=CB1H4orf50;refGene_tx=XM_023253326.1;refGene_gn=CB1H4orf50;refGene_tx=XM_023253327.1;refGene_gn=CB1H4orf50;refGene_tx=XM_023253328.1
B1 203014663 203014846
refGene_gn=CB1H4orf50;refGene_tx=XM_019829788.2;refGene_gn=CB1H4orf50;refGene_tx=XM_023253326.1;refGene_gn=CB1H4orf50;refGene_tx=XM_023253327.1;refGene_gn=CB1H4orf50;refGene_tx=XM_023253328.1
B1 203016007 203016064
refGene_gn=CB1H4orf50;refGene_tx=XM_019829788.2;refGene_gn=CB1H4orf50;refGene_tx=XM_023253326.1;refGene_gn=CB1H4orf50;refGene_tx=XM_023253327.1;refGene_gn=CB1H4orf50;refGene_tx=XM_023253328.1
B1 203019889 203020057
refGene_gn=CB1H4orf50;refGene_tx=XM_019829788.2;refGene_gn=CB1H4orf50;refGene_tx=XM_023253326.1;refGene_gn=CB1H4orf50;refGene_tx=XM_02325332

7.1;refGene_gn=CB1H4orf50;refGene_tx=XM_023253328.1
B1 203027601 203027934
refGene_gn=CB1H4orf50;refGene_tx=XM_019829788.2;refGene_gn=CB1H4orf50;
refGene_tx=XM_023253326.1;refGene_gn=CB1H4orf50;refGene_tx=XM_02325332
7.1;refGene_gn=CB1H4orf50;refGene_tx=XM_023253328.1
B1 203055927 203055957
refGene_gn=JAKMIP1;refGene_tx=XM_023253413.1
B1 203057162 203057207
refGene_gn=JAKMIP1;refGene_tx=XM_023253413.1
B1 203061006 203061072
refGene_gn=JAKMIP1;refGene_tx=XM_023253413.1
B1 203069078 203069282
refGene_gn=JAKMIP1;refGene_tx=XM_023253413.1
B1 203072234 203072312
refGene_gn=JAKMIP1;refGene_tx=XM_023253413.1
B1 203073576 203073645
refGene_gn=JAKMIP1;refGene_tx=XM_023253413.1
B1 203079257 203079323
refGene_gn=JAKMIP1;refGene_tx=XM_023253413.1
B1 203079905 203079959
refGene_gn=JAKMIP1;refGene_tx=XM_023253413.1
B1 203080642 203080744
refGene_gn=JAKMIP1;refGene_tx=XM_023253413.1
B1 203083424 203083523
refGene_gn=JAKMIP1;refGene_tx=XM_023253413.1
B1 203085161 203085224
refGene_gn=JAKMIP1;refGene_tx=XM_023253413.1
B1 203088328 203088412
refGene_gn=JAKMIP1;refGene_tx=XM_023253413.1
B1 203089667 203089796
refGene_gn=JAKMIP1;refGene_tx=XM_023253413.1
B1 203091036 203091165
refGene_gn=JAKMIP1;refGene_tx=XM_023253413.1
B1 203104422 203104482
refGene_gn=JAKMIP1;refGene_tx=XM_023253413.1
B1 203108233 203108388
refGene_gn=JAKMIP1;refGene_tx=XM_023253413.1
B1 203111964 203112084
refGene_gn=JAKMIP1;refGene_tx=XM_023253413.1
B1 203112914 203113124
refGene_gn=JAKMIP1;refGene_tx=XM_023253413.1
B1 203124175 203124670
refGene_gn=JAKMIP1;refGene_tx=XM_023253413.1
B1 203129757 203129886
refGene_gn=JAKMIP1;refGene_tx=XM_023253413.1
B1 203257428 203257660
refGene_gn=WFS1;refGene_tx=XM_023253329.1;refGene_gn=WFS1;refGene_tx=X
M_023253330.1
B1 203266202 203266285
refGene_gn=WFS1;refGene_tx=XM_023253329.1;refGene_gn=WFS1;refGene_tx=X

M_023253330.1
B1 203266926 203267071
refGene_gn=WFS1;refGene_tx=XM_023253329.1;refGene_gn=WFS1;refGene_tx=X
M_023253330.1
B1 203269083 203269254
refGene_gn=WFS1;refGene_tx=XM_023253329.1;refGene_gn=WFS1;refGene_tx=X
M_023253330.1
B1 203270760 203270841
refGene_gn=WFS1;refGene_tx=XM_023253329.1;refGene_gn=WFS1;refGene_tx=X
M_023253330.1
B1 203273437 203273583
refGene_gn=WFS1;refGene_tx=XM_023253329.1;refGene_gn=WFS1;refGene_tx=X
M_023253330.1
B1 203277625 203279434
refGene_gn=WFS1;refGene_tx=XM_023253329.1;refGene_gn=WFS1;refGene_tx=X
M_023253330.1
B1 203292668 203292960
refGene_gn=PPP2R2C;refGene_tx=XM_019829791.2;refGene_gn=PPP2R2C;refGen
e_tx=XM_023253331.1;refGene_gn=PPP2R2C;refGene_tx=XM_023253332.1
B1 203301682 203301774
refGene_gn=PPP2R2C;refGene_tx=XM_019829791.2;refGene_gn=PPP2R2C;refGen
e_tx=XM_023253331.1;refGene_gn=PPP2R2C;refGene_tx=XM_023253332.1
B1 203305484 203305654
refGene_gn=PPP2R2C;refGene_tx=XM_019829791.2;refGene_gn=PPP2R2C;refGen
e_tx=XM_023253331.1;refGene_gn=PPP2R2C;refGene_tx=XM_023253332.1
B1 203317161 203317326
refGene_gn=PPP2R2C;refGene_tx=XM_019829791.2;refGene_gn=PPP2R2C;refGen
e_tx=XM_023253331.1;refGene_gn=PPP2R2C;refGene_tx=XM_023253332.1
B1 203333538 203333716
refGene_gn=PPP2R2C;refGene_tx=XM_019829791.2;refGene_gn=PPP2R2C;refGen
e_tx=XM_023253331.1;refGene_gn=PPP2R2C;refGene_tx=XM_023253332.1
B1 203339964 203340077
refGene_gn=PPP2R2C;refGene_tx=XM_019829791.2;refGene_gn=PPP2R2C;refGen
e_tx=XM_023253331.1;refGene_gn=PPP2R2C;refGene_tx=XM_023253332.1
B1 203342331 203342497
refGene_gn=PPP2R2C;refGene_tx=XM_019829791.2;refGene_gn=PPP2R2C;refGen
e_tx=XM_023253331.1;refGene_gn=PPP2R2C;refGene_tx=XM_023253332.1
B1 203344522 203344620
refGene_gn=PPP2R2C;refGene_tx=XM_019829791.2;refGene_gn=PPP2R2C;refGen
e_tx=XM_023253331.1;refGene_gn=PPP2R2C;refGene_tx=XM_023253332.1
B1 203393150 203393196
refGene_gn=PPP2R2C;refGene_tx=XM_019829791.2
B1 203429606 203429634
refGene_gn=PPP2R2C;refGene_tx=XM_023253332.1
B1 203429683 203429753
refGene_gn=PPP2R2C;refGene_tx=XM_023253331.1
B1 203491827 203491840
refGene_gn=MAN2B2;refGene_tx=XM_023253414.1
B1 203493983 203494156
refGene_gn=MAN2B2;refGene_tx=XM_023253414.1

B1 203494924 203495013
refGene_gn=MAN2B2;refGene_tx=XM_023253414.1
B1 203495415 203495423
refGene_gn=MAN2B2;refGene_tx=XM_023253414.1
B1 203495904 203496098
refGene_gn=MAN2B2;refGene_tx=XM_023253414.1
B1 203500472 203500550
refGene_gn=MAN2B2;refGene_tx=XM_023253414.1
B1 203500552 203500649
refGene_gn=MAN2B2;refGene_tx=XM_023253414.1
B1 203500651 203500675
refGene_gn=MAN2B2;refGene_tx=XM_023253414.1
B1 203502502 203502692
refGene_gn=MAN2B2;refGene_tx=XM_023253414.1
B1 203502816 203502907
refGene_gn=MAN2B2;refGene_tx=XM_023253414.1
B1 203503359 203503907
refGene_gn=MAN2B2;refGene_tx=XM_023253414.1
B1 203504734 203504825
refGene_gn=MAN2B2;refGene_tx=XM_023253414.1
B1 203505118 203505514
refGene_gn=MAN2B2;refGene_tx=XM_023253414.1
B1 203506380 203506435
refGene_gn=MAN2B2;refGene_tx=XM_023253414.1
B1 203506437 203506601
refGene_gn=MAN2B2;refGene_tx=XM_023253414.1
B1 203506603 203506656
refGene_gn=MAN2B2;refGene_tx=XM_023253414.1
B1 203509424 203509577
refGene_gn=MAN2B2;refGene_tx=XM_023253414.1
B1 203510298 203510398
refGene_gn=MAN2B2;refGene_tx=XM_023253414.1
B1 203510399 203510564
refGene_gn=MAN2B2;refGene_tx=XM_023253414.1
B1 203511652 203511763
refGene_gn=MAN2B2;refGene_tx=XM_023253414.1
B1 203511838 203512010
refGene_gn=MAN2B2;refGene_tx=XM_023253414.1
B1 203515951 203516131
refGene_gn=MAN2B2;refGene_tx=XM_023253414.1
B1 203516977 203517165
refGene_gn=MAN2B2;refGene_tx=XM_023253414.1
B1 203534048 203534432
refGene_gn=MRFAP1;refGene_tx=XM_003985553.5;refGene_gn=MRFAP1;refGene_
tx=XM_019829795.2
B1 203553159 203553495
refGene_gn=LOC111560210;refGene_tx=XM_023253334.1
B1 203565447 203565783
refGene_gn=LOC111560209;refGene_tx=XM_023253333.1
B1 203580856 203580994

refGene_gn=S100P;refGene_tx=XM_023253335.1
B1	203583308	203583458
refGene_gn=S100P;refGene_tx=XM_023253335.1
B1	203595202	203595586
refGene_gn=LOC111560059;refGene_tx=XM_023252241.1
B1	203598515	203599172
refGene_gn=BLOC1S4;refGene_tx=XM_011282003.3
B1	203780785	203781507
refGene_gn=TBC1D14;refGene_tx=XM_011281956.3;refGene_gn=TBC1D14;refGene_tx=XM_023253336.1;refGene_gn=TBC1D14;refGene_tx=XM_023253337.1;refGene_gn=TBC1D14;refGene_tx=XM_023253338.1
B1	203788988	203789074
refGene_gn=TBC1D14;refGene_tx=XM_019829799.2
B1	203808246	203808281
refGene_gn=TBC1D14;refGene_tx=XM_006931262.3
B1	203819311	203819432
refGene_gn=TBC1D14;refGene_tx=XM_006931262.3;refGene_gn=TBC1D14;refGene_tx=XM_011281956.3;refGene_gn=TBC1D14;refGene_tx=XM_019829799.2;refGene_gn=TBC1D14;refGene_tx=XM_023253336.1;refGene_gn=TBC1D14;refGene_tx=XM_023253337.1;refGene_gn=TBC1D14;refGene_tx=XM_023253338.1
B1	203839002	203839005
refGene_gn=TBC1D14;refGene_tx=XM_006931263.3
B1	203846984	203847103
refGene_gn=TBC1D14;refGene_tx=XM_006931262.3;refGene_gn=TBC1D14;refGene_tx=XM_006931263.3;refGene_gn=TBC1D14;refGene_tx=XM_011281956.3;refGene_gn=TBC1D14;refGene_tx=XM_019829799.2;refGene_gn=TBC1D14;refGene_tx=XM_023253336.1;refGene_gn=TBC1D14;refGene_tx=XM_023253337.1;refGene_gn=TBC1D14;refGene_tx=XM_023253338.1
B1	203848866	203848949
refGene_gn=TBC1D14;refGene_tx=XM_006931262.3;refGene_gn=TBC1D14;refGene_tx=XM_006931263.3;refGene_gn=TBC1D14;refGene_tx=XM_006931264.3;refGene_gn=TBC1D14;refGene_tx=XM_011281956.3;refGene_gn=TBC1D14;refGene_tx=XM_019829799.2;refGene_gn=TBC1D14;refGene_tx=XM_023253336.1;refGene_gn=TBC1D14;refGene_tx=XM_023253337.1;refGene_gn=TBC1D14;refGene_tx=XM_023253338.1
B1	203850120	203850238
refGene_gn=TBC1D14;refGene_tx=XM_006931262.3;refGene_gn=TBC1D14;refGene_tx=XM_006931263.3;refGene_gn=TBC1D14;refGene_tx=XM_006931264.3;refGene_gn=TBC1D14;refGene_tx=XM_011281956.3;refGene_gn=TBC1D14;refGene_tx=XM_019829799.2;refGene_gn=TBC1D14;refGene_tx=XM_023253336.1;refGene_gn=TBC1D14;refGene_tx=XM_023253337.1;refGene_gn=TBC1D14;refGene_tx=XM_023253338.1
B1	203852218	203852325
refGene_gn=TBC1D14;refGene_tx=XM_006931262.3;refGene_gn=TBC1D14;refGene_tx=XM_006931263.3;refGene_gn=TBC1D14;refGene_tx=XM_006931264.3;refGene_gn=TBC1D14;refGene_tx=XM_011281956.3;refGene_gn=TBC1D14;refGene_tx=XM_019829799.2;refGene_gn=TBC1D14;refGene_tx=XM_023253336.1;refGene_gn=TBC1D14;refGene_tx=XM_023253337.1;refGene_gn=TBC1D14;refGene_tx=XM_023253338.1
B1	203855383	203855464

refGene_gn=TBC1D14;refGene_tx=XM_006931262.3;refGene_gn=TBC1D14;refGene_tx=XM_006931263.3;refGene_gn=TBC1D14;refGene_tx=XM_006931264.3;refGene_gn=TBC1D14;refGene_tx=XM_011281956.3;refGene_gn=TBC1D14;refGene_tx=XM_019829799.2;refGene_gn=TBC1D14;refGene_tx=XM_023253336.1;refGene_gn=TBC1D14;refGene_tx=XM_023253337.1;refGene_gn=TBC1D14;refGene_tx=XM_023253338.1
B1 203856391 203856486
refGene_gn=TBC1D14;refGene_tx=XM_006931262.3;refGene_gn=TBC1D14;refGene_tx=XM_006931263.3;refGene_gn=TBC1D14;refGene_tx=XM_006931264.3;refGene_gn=TBC1D14;refGene_tx=XM_011281956.3;refGene_gn=TBC1D14;refGene_tx=XM_019829799.2;refGene_gn=TBC1D14;refGene_tx=XM_023253336.1;refGene_gn=TBC1D14;refGene_tx=XM_023253337.1;refGene_gn=TBC1D14;refGene_tx=XM_023253338.1
B1 203859496 203859568
refGene_gn=TBC1D14;refGene_tx=XM_006931262.3;refGene_gn=TBC1D14;refGene_tx=XM_006931263.3;refGene_gn=TBC1D14;refGene_tx=XM_006931264.3;refGene_gn=TBC1D14;refGene_tx=XM_011281956.3;refGene_gn=TBC1D14;refGene_tx=XM_019829799.2;refGene_gn=TBC1D14;refGene_tx=XM_023253338.1
B1 203860279 203860408
refGene_gn=TBC1D14;refGene_tx=XM_006931262.3;refGene_gn=TBC1D14;refGene_tx=XM_006931263.3;refGene_gn=TBC1D14;refGene_tx=XM_006931264.3;refGene_gn=TBC1D14;refGene_tx=XM_011281956.3;refGene_gn=TBC1D14;refGene_tx=XM_019829799.2;refGene_gn=TBC1D14;refGene_tx=XM_023253336.1;refGene_gn=TBC1D14;refGene_tx=XM_023253338.1
B1 203864188 203864298
refGene_gn=TBC1D14;refGene_tx=XM_006931262.3;refGene_gn=TBC1D14;refGene_tx=XM_006931263.3;refGene_gn=TBC1D14;refGene_tx=XM_006931264.3;refGene_gn=TBC1D14;refGene_tx=XM_011281956.3;refGene_gn=TBC1D14;refGene_tx=XM_019829799.2;refGene_gn=TBC1D14;refGene_tx=XM_023253336.1;refGene_gn=TBC1D14;refGene_tx=XM_023253337.1;refGene_gn=TBC1D14;refGene_tx=XM_023253338.1
B1 203875462 203875721
refGene_gn=TBC1D14;refGene_tx=XM_006931262.3;refGene_gn=TBC1D14;refGene_tx=XM_006931263.3;refGene_gn=TBC1D14;refGene_tx=XM_006931264.3;refGene_gn=TBC1D14;refGene_tx=XM_011281956.3;refGene_gn=TBC1D14;refGene_tx=XM_019829799.2;refGene_gn=TBC1D14;refGene_tx=XM_023253336.1;refGene_gn=TBC1D14;refGene_tx=XM_023253337.1;refGene_gn=TBC1D14;refGene_tx=XM_023253338.1
B1 203879689 203879755
refGene_gn=TBC1D14;refGene_tx=XM_006931262.3;refGene_gn=TBC1D14;refGene_tx=XM_006931263.3;refGene_gn=TBC1D14;refGene_tx=XM_006931264.3;refGene_gn=TBC1D14;refGene_tx=XM_011281956.3;refGene_gn=TBC1D14;refGene_tx=XM_019829799.2;refGene_gn=TBC1D14;refGene_tx=XM_023253336.1;refGene_gn=TBC1D14;refGene_tx=XM_023253337.1;refGene_gn=TBC1D14;refGene_tx=XM_023253338.1
B1 203887820 203888066
refGene_gn=CCDC96;refGene_tx=XM_023253339.1
B1 203888067 203889561
refGene_gn=CCDC96;refGene_tx=XM_023253339.1
B1 203890140 203890410

refGene_gn=TADA2B;refGene_tx=XM_023253340.1
B1	203903109	203904102
refGene_gn=TADA2B;refGene_tx=XM_023253340.1
B1	203908514	203908861
refGene_gn=GRPEL1;refGene_tx=XM_011281955.3
B1	203911355	203911437
refGene_gn=GRPEL1;refGene_tx=XM_011281955.3
B1	203912674	203912837
refGene_gn=GRPEL1;refGene_tx=XM_011281955.3
B1	203915331	203915393
refGene_gn=GRPEL1;refGene_tx=XM_011281955.3
B1	204009643	204010135
refGene_gn=SORCS2;refGene_tx=XM_023253426.1
B1	204197405	204197473
refGene_gn=SORCS2;refGene_tx=XM_023253426.1
B1	204293027	204293127
refGene_gn=SORCS2;refGene_tx=XM_023253426.1
B1	204388365	204388530
refGene_gn=SORCS2;refGene_tx=XM_023253426.1
B1	204398901	204398975
refGene_gn=SORCS2;refGene_tx=XM_023253426.1
B1	204403079	204403144
refGene_gn=SORCS2;refGene_tx=XM_023253426.1
B1	204406501	204406620
refGene_gn=SORCS2;refGene_tx=XM_023253426.1
B1	204409913	204410003
refGene_gn=SORCS2;refGene_tx=XM_023253426.1
B1	204417389	204417569
refGene_gn=SORCS2;refGene_tx=XM_023253426.1
B1	204422193	204422340
refGene_gn=SORCS2;refGene_tx=XM_023253426.1
B1	204429183	204429286
refGene_gn=SORCS2;refGene_tx=XM_023253426.1
B1	204433146	204433223
refGene_gn=SORCS2;refGene_tx=XM_023253426.1
B1	204438933	204439025
refGene_gn=SORCS2;refGene_tx=XM_023253426.1
B1	204439912	204440020
refGene_gn=SORCS2;refGene_tx=XM_023253426.1
B1	204445112	204445233
refGene_gn=SORCS2;refGene_tx=XM_023253426.1
B1	204446179	204446313
refGene_gn=SORCS2;refGene_tx=XM_023253426.1
B1	204447034	204447163
refGene_gn=SORCS2;refGene_tx=XM_023253426.1
B1	204448897	204449069
refGene_gn=SORCS2;refGene_tx=XM_023253426.1
B1	204454537	204454724
refGene_gn=SORCS2;refGene_tx=XM_023253426.1
B1	204455029	204455163

refGene_gn=SORCS2;refGene_tx=XM_023253426.1
B1	204456077	204456201
refGene_gn=SORCS2;refGene_tx=XM_023253426.1
B1	204457235	204457348
refGene_gn=SORCS2;refGene_tx=XM_023253426.1
B1	204458319	204458445
refGene_gn=SORCS2;refGene_tx=XM_023253426.1
B1	204461045	204461145
refGene_gn=SORCS2;refGene_tx=XM_023253426.1
B1	204461763	204461866
refGene_gn=SORCS2;refGene_tx=XM_023253426.1
B1	204466869	204466973
refGene_gn=SORCS2;refGene_tx=XM_023253426.1
B1	204470555	204470620
refGene_gn=SORCS2;refGene_tx=XM_023253426.1
B1	204486348	204486375
refGene_gn=AFAP1;refGene_tx=XM_019829803.2;refGene_gn=AFAP1;refGene_tx=XM_019829804.2;refGene_gn=AFAP1;refGene_tx=XM_019829806.2;refGene_gn=AFAP1;refGene_tx=XM_023253427.1;refGene_gn=AFAP1;refGene_tx=XM_023253428.1;refGene_gn=AFAP1;refGene_tx=XM_023253429.1;refGene_gn=AFAP1;refGene_tx=XM_023253430.1;refGene_gn=AFAP1;refGene_tx=XM_023253431.1
B1	204488682	204488844
refGene_gn=AFAP1;refGene_tx=XM_019829803.2;refGene_gn=AFAP1;refGene_tx=XM_019829804.2;refGene_gn=AFAP1;refGene_tx=XM_019829806.2;refGene_gn=AFAP1;refGene_tx=XM_023253427.1;refGene_gn=AFAP1;refGene_tx=XM_023253428.1;refGene_gn=AFAP1;refGene_tx=XM_023253429.1;refGene_gn=AFAP1;refGene_tx=XM_023253430.1;refGene_gn=AFAP1;refGene_tx=XM_023253431.1
B1	204491008	204491199
refGene_gn=AFAP1;refGene_tx=XM_019829803.2;refGene_gn=AFAP1;refGene_tx=XM_019829804.2;refGene_gn=AFAP1;refGene_tx=XM_019829806.2;refGene_gn=AFAP1;refGene_tx=XM_023253427.1;refGene_gn=AFAP1;refGene_tx=XM_023253428.1;refGene_gn=AFAP1;refGene_tx=XM_023253429.1;refGene_gn=AFAP1;refGene_tx=XM_023253430.1;refGene_gn=AFAP1;refGene_tx=XM_023253431.1
B1	204492817	204492982
refGene_gn=AFAP1;refGene_tx=XM_019829803.2;refGene_gn=AFAP1;refGene_tx=XM_019829804.2;refGene_gn=AFAP1;refGene_tx=XM_019829806.2;refGene_gn=AFAP1;refGene_tx=XM_023253427.1;refGene_gn=AFAP1;refGene_tx=XM_023253428.1;refGene_gn=AFAP1;refGene_tx=XM_023253429.1;refGene_gn=AFAP1;refGene_tx=XM_023253430.1;refGene_gn=AFAP1;refGene_tx=XM_023253431.1
B1	204496729	204496844
refGene_gn=AFAP1;refGene_tx=XM_019829803.2;refGene_gn=AFAP1;refGene_tx=XM_019829804.2;refGene_gn=AFAP1;refGene_tx=XM_019829806.2;refGene_gn=AFAP1;refGene_tx=XM_023253427.1;refGene_gn=AFAP1;refGene_tx=XM_023253428.1;refGene_gn=AFAP1;refGene_tx=XM_023253429.1;refGene_gn=AFAP1;refGene_tx=XM_023253430.1;refGene_gn=AFAP1;refGene_tx=XM_023253431.1
B1	204498968	204499226
refGene_gn=AFAP1;refGene_tx=XM_019829804.2;refGene_gn=AFAP1;refGene_tx=XM_019829806.2;refGene_gn=AFAP1;refGene_tx=XM_023253427.1;refGene_gn=AFAP1;refGene_tx=XM_023253428.1;refGene_gn=AFAP1;refGene_tx=XM_023253429.1;refGene_gn=AFAP1;refGene_tx=XM_023253430.1;refGene_gn=AFAP1;refGe

ne_tx=XM_023253431.1
B1 204502609 204502727
refGene_gn=AFAP1;refGene_tx=XM_019829803.2;refGene_gn=AFAP1;refGene_tx=XM_019829804.2;refGene_gn=AFAP1;refGene_tx=XM_019829806.2;refGene_gn=AFAP1;refGene_tx=XM_023253427.1;refGene_gn=AFAP1;refGene_tx=XM_023253428.1;refGene_gn=AFAP1;refGene_tx=XM_023253429.1;refGene_gn=AFAP1;refGene_tx=XM_023253430.1;refGene_gn=AFAP1;refGene_tx=XM_023253431.1
B1 204509296 204509442
refGene_gn=AFAP1;refGene_tx=XM_019829803.2;refGene_gn=AFAP1;refGene_tx=XM_019829804.2;refGene_gn=AFAP1;refGene_tx=XM_019829806.2;refGene_gn=AFAP1;refGene_tx=XM_023253427.1;refGene_gn=AFAP1;refGene_tx=XM_023253428.1;refGene_gn=AFAP1;refGene_tx=XM_023253429.1;refGene_gn=AFAP1;refGene_tx=XM_023253430.1;refGene_gn=AFAP1;refGene_tx=XM_023253431.1
B1 204513795 204514007
refGene_gn=AFAP1;refGene_tx=XM_019829803.2;refGene_gn=AFAP1;refGene_tx=XM_019829804.2;refGene_gn=AFAP1;refGene_tx=XM_019829806.2;refGene_gn=AFAP1;refGene_tx=XM_023253427.1;refGene_gn=AFAP1;refGene_tx=XM_023253428.1;refGene_gn=AFAP1;refGene_tx=XM_023253429.1;refGene_gn=AFAP1;refGene_tx=XM_023253430.1;refGene_gn=AFAP1;refGene_tx=XM_023253431.1
B1 204520524 204520674
refGene_gn=AFAP1;refGene_tx=XM_019829803.2;refGene_gn=AFAP1;refGene_tx=XM_019829804.2;refGene_gn=AFAP1;refGene_tx=XM_019829806.2;refGene_gn=AFAP1;refGene_tx=XM_023253427.1;refGene_gn=AFAP1;refGene_tx=XM_023253428.1;refGene_gn=AFAP1;refGene_tx=XM_023253429.1;refGene_gn=AFAP1;refGene_tx=XM_023253430.1;refGene_gn=AFAP1;refGene_tx=XM_023253431.1
B1 204525827 204525909
refGene_gn=AFAP1;refGene_tx=XM_019829803.2;refGene_gn=AFAP1;refGene_tx=XM_019829804.2;refGene_gn=AFAP1;refGene_tx=XM_019829806.2;refGene_gn=AFAP1;refGene_tx=XM_023253427.1;refGene_gn=AFAP1;refGene_tx=XM_023253428.1;refGene_gn=AFAP1;refGene_tx=XM_023253429.1;refGene_gn=AFAP1;refGene_tx=XM_023253430.1;refGene_gn=AFAP1;refGene_tx=XM_023253431.1
B1 204528541 204528637
refGene_gn=AFAP1;refGene_tx=XM_019829803.2;refGene_gn=AFAP1;refGene_tx=XM_019829804.2;refGene_gn=AFAP1;refGene_tx=XM_019829806.2;refGene_gn=AFAP1;refGene_tx=XM_023253427.1;refGene_gn=AFAP1;refGene_tx=XM_023253428.1;refGene_gn=AFAP1;refGene_tx=XM_023253429.1;refGene_gn=AFAP1;refGene_tx=XM_023253430.1;refGene_gn=AFAP1;refGene_tx=XM_023253431.1
B1 204541165 204541345
refGene_gn=AFAP1;refGene_tx=XM_019829803.2;refGene_gn=AFAP1;refGene_tx=XM_019829804.2;refGene_gn=AFAP1;refGene_tx=XM_019829806.2;refGene_gn=AFAP1;refGene_tx=XM_023253427.1;refGene_gn=AFAP1;refGene_tx=XM_023253428.1;refGene_gn=AFAP1;refGene_tx=XM_023253429.1;refGene_gn=AFAP1;refGene_tx=XM_023253430.1;refGene_gn=AFAP1;refGene_tx=XM_023253431.1
B1 204544719 204544931
refGene_gn=AFAP1;refGene_tx=XM_019829803.2;refGene_gn=AFAP1;refGene_tx=XM_019829804.2;refGene_gn=AFAP1;refGene_tx=XM_019829806.2;refGene_gn=AFAP1;refGene_tx=XM_023253427.1;refGene_gn=AFAP1;refGene_tx=XM_023253428.1;refGene_gn=AFAP1;refGene_tx=XM_023253429.1;refGene_gn=AFAP1;refGene_tx=XM_023253430.1;refGene_gn=AFAP1;refGene_tx=XM_023253431.1
B1 204553999 204554108

refGene_gn=AFAP1;refGene_tx=XM_019829803.2;refGene_gn=AFAP1;refGene_tx=XM_019829804.2;refGene_gn=AFAP1;refGene_tx=XM_019829806.2;refGene_gn=AFAP1;refGene_tx=XM_023253427.1;refGene_gn=AFAP1;refGene_tx=XM_023253428.1;refGene_gn=AFAP1;refGene_tx=XM_023253429.1;refGene_gn=AFAP1;refGene_tx=XM_023253430.1;refGene_gn=AFAP1;refGene_tx=XM_023253431.1
B1 204565486 204565584
refGene_gn=AFAP1;refGene_tx=XM_019829803.2;refGene_gn=AFAP1;refGene_tx=XM_019829804.2;refGene_gn=AFAP1;refGene_tx=XM_019829806.2;refGene_gn=AFAP1;refGene_tx=XM_023253427.1;refGene_gn=AFAP1;refGene_tx=XM_023253428.1;refGene_gn=AFAP1;refGene_tx=XM_023253429.1;refGene_gn=AFAP1;refGene_tx=XM_023253430.1;refGene_gn=AFAP1;refGene_tx=XM_023253431.1
B1 204567437 204567566
refGene_gn=AFAP1;refGene_tx=XM_019829803.2;refGene_gn=AFAP1;refGene_tx=XM_019829804.2;refGene_gn=AFAP1;refGene_tx=XM_019829806.2;refGene_gn=AFAP1;refGene_tx=XM_023253427.1;refGene_gn=AFAP1;refGene_tx=XM_023253428.1;refGene_gn=AFAP1;refGene_tx=XM_023253429.1;refGene_gn=AFAP1;refGene_tx=XM_023253430.1;refGene_gn=AFAP1;refGene_tx=XM_023253431.1
B1 204573739 204573770
refGene_gn=AFAP1;refGene_tx=XM_019829803.2;refGene_gn=AFAP1;refGene_tx=XM_019829804.2;refGene_gn=AFAP1;refGene_tx=XM_023253427.1;refGene_gn=AFAP1;refGene_tx=XM_023253429.1;refGene_gn=AFAP1;refGene_tx=XM_023253430.1
B1 204574142 204574330
refGene_gn=AFAP1;refGene_tx=XM_023253428.1
B1 204617995 204618072
refGene_gn=AFAP1;refGene_tx=XM_023253428.1
B1 204626976 204627063
refGene_gn=LOC111560233;refGene_tx=XM_023253415.1
B1 204628550 204628652
refGene_gn=LOC111560233;refGene_tx=XM_023253415.1
B1 204628654 204629327
refGene_gn=LOC111560233;refGene_tx=XM_023253415.1
B1 204629388 204629531
refGene_gn=LOC111560233;refGene_tx=XM_023253415.1
B1 204629533 204629890
refGene_gn=LOC111560233;refGene_tx=XM_023253415.1
B1 204639568 204639682
refGene_gn=ABLIM2;refGene_tx=XM_023253441.1;refGene_gn=ABLIM2;refGene_tx=XM_023253442.1;refGene_gn=ABLIM2;refGene_tx=XM_023253443.1;refGene_gn=ABLIM2;refGene_tx=XM_023253444.1;refGene_gn=ABLIM2;refGene_tx=XM_023253445.1;refGene_gn=ABLIM2;refGene_tx=XM_023253446.1;refGene_gn=ABLIM2;refGene_tx=XM_023253447.1;refGene_gn=ABLIM2;refGene_tx=XM_023253448.1;refGene_gn=ABLIM2;refGene_tx=XM_023253449.1;refGene_gn=ABLIM2;refGene_tx=XM_023253450.1;refGene_gn=ABLIM2;refGene_tx=XM_023253451.1;refGene_gn=ABLIM2;refGene_tx=XM_023253452.1;refGene_gn=ABLIM2;refGene_tx=XM_023253453.1;refGene_gn=ABLIM2;refGene_tx=XM_023253454.1;refGene_gn=ABLIM2;refGene_tx=XM_023253455.1;refGene_gn=ABLIM2;refGene_tx=XM_023253456.1;refGene_gn=ABLIM2;refGene_tx=XM_023253457.1;refGene_gn=ABLIM2;refGene_tx=XM_023253458.1;refGene_gn=ABLIM2;refGene_tx=XM_023253459.1;refGene_gn=ABLIM2;refGene_tx=XM_023253460.1;refGene_gn=ABLIM2;refGene_tx=X

M_023253461.1
B1	204652372	204652453
refGene_gn=ABLIM2;refGene_tx=XM_023253441.1;refGene_gn=ABLIM2;refGene_tx=XM_023253442.1;refGene_gn=ABLIM2;refGene_tx=XM_023253443.1;refGene_gn=ABLIM2;refGene_tx=XM_023253444.1;refGene_gn=ABLIM2;refGene_tx=XM_023253445.1;refGene_gn=ABLIM2;refGene_tx=XM_023253446.1;refGene_gn=ABLIM2;refGene_tx=XM_023253447.1;refGene_gn=ABLIM2;refGene_tx=XM_023253448.1;refGene_gn=ABLIM2;refGene_tx=XM_023253449.1;refGene_gn=ABLIM2;refGene_tx=XM_023253450.1;refGene_gn=ABLIM2;refGene_tx=XM_023253451.1;refGene_gn=ABLIM2;refGene_tx=XM_023253452.1;refGene_gn=ABLIM2;refGene_tx=XM_023253453.1;refGene_gn=ABLIM2;refGene_tx=XM_023253454.1;refGene_gn=ABLIM2;refGene_tx=XM_023253455.1;refGene_gn=ABLIM2;refGene_tx=XM_023253456.1;refGene_gn=ABLIM2;refGene_tx=XM_023253457.1;refGene_gn=ABLIM2;refGene_tx=XM_023253458.1;refGene_gn=ABLIM2;refGene_tx=XM_023253459.1;refGene_gn=ABLIM2;refGene_tx=XM_023253460.1;refGene_gn=ABLIM2;refGene_tx=XM_023253461.1
B1	204652662	204652670
refGene_gn=ABLIM2;refGene_tx=XM_023253441.1;refGene_gn=ABLIM2;refGene_tx=XM_023253442.1;refGene_gn=ABLIM2;refGene_tx=XM_023253443.1;refGene_gn=ABLIM2;refGene_tx=XM_023253444.1;refGene_gn=ABLIM2;refGene_tx=XM_023253445.1;refGene_gn=ABLIM2;refGene_tx=XM_023253446.1;refGene_gn=ABLIM2;refGene_tx=XM_023253447.1;refGene_gn=ABLIM2;refGene_tx=XM_023253448.1;refGene_gn=ABLIM2;refGene_tx=XM_023253449.1;refGene_gn=ABLIM2;refGene_tx=XM_023253450.1;refGene_gn=ABLIM2;refGene_tx=XM_023253451.1;refGene_gn=ABLIM2;refGene_tx=XM_023253452.1;refGene_gn=ABLIM2;refGene_tx=XM_023253453.1;refGene_gn=ABLIM2;refGene_tx=XM_023253454.1;refGene_gn=ABLIM2;refGene_tx=XM_023253455.1;refGene_gn=ABLIM2;refGene_tx=XM_023253456.1;refGene_gn=ABLIM2;refGene_tx=XM_023253457.1;refGene_gn=ABLIM2;refGene_tx=XM_023253458.1;refGene_gn=ABLIM2;refGene_tx=XM_023253459.1;refGene_gn=ABLIM2;refGene_tx=XM_023253460.1;refGene_gn=ABLIM2;refGene_tx=XM_023253461.1
B1	204654101	204654156
refGene_gn=ABLIM2;refGene_tx=XM_023253441.1;refGene_gn=ABLIM2;refGene_tx=XM_023253442.1;refGene_gn=ABLIM2;refGene_tx=XM_023253443.1;refGene_gn=ABLIM2;refGene_tx=XM_023253444.1;refGene_gn=ABLIM2;refGene_tx=XM_023253445.1;refGene_gn=ABLIM2;refGene_tx=XM_023253447.1;refGene_gn=ABLIM2;refGene_tx=XM_023253448.1;refGene_gn=ABLIM2;refGene_tx=XM_023253449.1;refGene_gn=ABLIM2;refGene_tx=XM_023253450.1;refGene_gn=ABLIM2;refGene_tx=XM_023253451.1;refGene_gn=ABLIM2;refGene_tx=XM_023253452.1;refGene_gn=ABLIM2;refGene_tx=XM_023253454.1;refGene_gn=ABLIM2;refGene_tx=XM_023253457.1;refGene_gn=ABLIM2;refGene_tx=XM_023253458.1;refGene_gn=ABLIM2;refGene_tx=XM_023253459.1
B1	204659065	204659127
refGene_gn=ABLIM2;refGene_tx=XM_023253441.1;refGene_gn=ABLIM2;refGene_tx=XM_023253442.1;refGene_gn=ABLIM2;refGene_tx=XM_023253443.1;refGene_gn=ABLIM2;refGene_tx=XM_023253444.1;refGene_gn=ABLIM2;refGene_tx=XM_023253445.1;refGene_gn=ABLIM2;refGene_tx=XM_023253447.1;refGene_gn=ABLIM2;refGene_tx=XM_023253448.1;refGene_gn=ABLIM2;refGene_tx=XM_023253449.1;refGene_gn=ABLIM2;refGene_tx=XM_023253450.1;refGene_gn=ABLIM2;refGene_tx=XM_023253451.1;refGene_gn=ABLIM2;refGene_tx=XM_023253452.1;refGen

e_gn=ABLIM2;refGene_tx=XM_023253454.1;refGene_gn=ABLIM2;refGene_tx=XM_023253457.1;refGene_gn=ABLIM2;refGene_tx=XM_023253458.1;refGene_gn=ABLIM2;refGene_tx=XM_023253459.1
B1 204670478 204670637
refGene_gn=ABLIM2;refGene_tx=XM_023253441.1;refGene_gn=ABLIM2;refGene_tx=XM_023253442.1;refGene_gn=ABLIM2;refGene_tx=XM_023253443.1;refGene_gn=ABLIM2;refGene_tx=XM_023253444.1;refGene_gn=ABLIM2;refGene_tx=XM_023253445.1;refGene_gn=ABLIM2;refGene_tx=XM_023253446.1;refGene_gn=ABLIM2;refGene_tx=XM_023253447.1;refGene_gn=ABLIM2;refGene_tx=XM_023253448.1;refGene_gn=ABLIM2;refGene_tx=XM_023253449.1;refGene_gn=ABLIM2;refGene_tx=XM_023253450.1;refGene_gn=ABLIM2;refGene_tx=XM_023253451.1;refGene_gn=ABLIM2;refGene_tx=XM_023253452.1;refGene_gn=ABLIM2;refGene_tx=XM_023253453.1;refGene_gn=ABLIM2;refGene_tx=XM_023253454.1;refGene_gn=ABLIM2;refGene_tx=XM_023253455.1;refGene_gn=ABLIM2;refGene_tx=XM_023253456.1;refGene_gn=ABLIM2;refGene_tx=XM_023253457.1;refGene_gn=ABLIM2;refGene_tx=XM_023253458.1;refGene_gn=ABLIM2;refGene_tx=XM_023253459.1;refGene_gn=ABLIM2;refGene_tx=XM_023253460.1;refGene_gn=ABLIM2;refGene_tx=XM_023253461.1;refGene_gn=ABLIM2;refGene_tx=XM_023253462.1
B1 204671362 204671415
refGene_gn=ABLIM2;refGene_tx=XM_023253441.1;refGene_gn=ABLIM2;refGene_tx=XM_023253442.1;refGene_gn=ABLIM2;refGene_tx=XM_023253443.1;refGene_gn=ABLIM2;refGene_tx=XM_023253444.1;refGene_gn=ABLIM2;refGene_tx=XM_023253445.1;refGene_gn=ABLIM2;refGene_tx=XM_023253446.1;refGene_gn=ABLIM2;refGene_tx=XM_023253447.1;refGene_gn=ABLIM2;refGene_tx=XM_023253448.1;refGene_gn=ABLIM2;refGene_tx=XM_023253449.1;refGene_gn=ABLIM2;refGene_tx=XM_023253450.1;refGene_gn=ABLIM2;refGene_tx=XM_023253451.1;refGene_gn=ABLIM2;refGene_tx=XM_023253452.1;refGene_gn=ABLIM2;refGene_tx=XM_023253453.1;refGene_gn=ABLIM2;refGene_tx=XM_023253454.1;refGene_gn=ABLIM2;refGene_tx=XM_023253455.1;refGene_gn=ABLIM2;refGene_tx=XM_023253456.1;refGene_gn=ABLIM2;refGene_tx=XM_023253457.1;refGene_gn=ABLIM2;refGene_tx=XM_023253458.1;refGene_gn=ABLIM2;refGene_tx=XM_023253459.1;refGene_gn=ABLIM2;refGene_tx=XM_023253460.1;refGene_gn=ABLIM2;refGene_tx=XM_023253461.1;refGene_gn=ABLIM2;refGene_tx=XM_023253462.1
B1 204672448 204672568
refGene_gn=ABLIM2;refGene_tx=XM_023253441.1;refGene_gn=ABLIM2;refGene_tx=XM_023253442.1;refGene_gn=ABLIM2;refGene_tx=XM_023253443.1;refGene_gn=ABLIM2;refGene_tx=XM_023253444.1;refGene_gn=ABLIM2;refGene_tx=XM_023253445.1;refGene_gn=ABLIM2;refGene_tx=XM_023253446.1
B1 204677617 204677671
refGene_gn=ABLIM2;refGene_tx=XM_023253441.1;refGene_gn=ABLIM2;refGene_tx=XM_023253442.1;refGene_gn=ABLIM2;refGene_tx=XM_023253443.1;refGene_gn=ABLIM2;refGene_tx=XM_023253444.1;refGene_gn=ABLIM2;refGene_tx=XM_023253445.1;refGene_gn=ABLIM2;refGene_tx=XM_023253446.1;refGene_gn=ABLIM2;refGene_tx=XM_023253447.1;refGene_gn=ABLIM2;refGene_tx=XM_023253448.1;refGene_gn=ABLIM2;refGene_tx=XM_023253449.1;refGene_gn=ABLIM2;refGene_tx=XM_023253450.1;refGene_gn=ABLIM2;refGene_tx=XM_023253453.1;refGene_gn=ABLIM2;refGene_tx=XM_023253454.1;refGene_gn=ABLIM2;refGene_tx=XM_023253455.1;refGene_gn=ABLIM2;refGene_tx=XM_023253456.1
B1 204678111 204678213
refGene_gn=ABLIM2;refGene_tx=XM_023253441.1;refGene_gn=ABLIM2;refGene_

tx=XM_023253442.1;refGene_gn=ABLIM2;refGene_tx=XM_023253443.1;refGene_gn=ABLIM2;refGene_tx=XM_023253444.1;refGene_gn=ABLIM2;refGene_tx=XM_023253445.1;refGene_gn=ABLIM2;refGene_tx=XM_023253446.1;refGene_gn=ABLIM2;refGene_tx=XM_023253447.1;refGene_gn=ABLIM2;refGene_tx=XM_023253448.1;refGene_gn=ABLIM2;refGene_tx=XM_023253449.1;refGene_gn=ABLIM2;refGene_tx=XM_023253450.1;refGene_gn=ABLIM2;refGene_tx=XM_023253451.1;refGene_gn=ABLIM2;refGene_tx=XM_023253452.1;refGene_gn=ABLIM2;refGene_tx=XM_023253453.1;refGene_gn=ABLIM2;refGene_tx=XM_023253454.1;refGene_gn=ABLIM2;refGene_tx=XM_023253455.1;refGene_gn=ABLIM2;refGene_tx=XM_023253456.1
B1 204684606 204684705
refGene_gn=ABLIM2;refGene_tx=XM_023253441.1;refGene_gn=ABLIM2;refGene_tx=XM_023253442.1;refGene_gn=ABLIM2;refGene_tx=XM_023253443.1;refGene_gn=ABLIM2;refGene_tx=XM_023253444.1;refGene_gn=ABLIM2;refGene_tx=XM_023253446.1;refGene_gn=ABLIM2;refGene_tx=XM_023253447.1;refGene_gn=ABLIM2;refGene_tx=XM_023253448.1;refGene_gn=ABLIM2;refGene_tx=XM_023253449.1;refGene_gn=ABLIM2;refGene_tx=XM_023253450.1;refGene_gn=ABLIM2;refGene_tx=XM_023253451.1;refGene_gn=ABLIM2;refGene_tx=XM_023253452.1;refGene_gn=ABLIM2;refGene_tx=XM_023253453.1;refGene_gn=ABLIM2;refGene_tx=XM_023253455.1
B1 204689466 204689587
refGene_gn=ABLIM2;refGene_tx=XM_023253441.1;refGene_gn=ABLIM2;refGene_tx=XM_023253442.1;refGene_gn=ABLIM2;refGene_tx=XM_023253443.1;refGene_gn=ABLIM2;refGene_tx=XM_023253444.1;refGene_gn=ABLIM2;refGene_tx=XM_023253445.1;refGene_gn=ABLIM2;refGene_tx=XM_023253446.1;refGene_gn=ABLIM2;refGene_tx=XM_023253447.1;refGene_gn=ABLIM2;refGene_tx=XM_023253448.1;refGene_gn=ABLIM2;refGene_tx=XM_023253449.1;refGene_gn=ABLIM2;refGene_tx=XM_023253450.1;refGene_gn=ABLIM2;refGene_tx=XM_023253451.1;refGene_gn=ABLIM2;refGene_tx=XM_023253452.1;refGene_gn=ABLIM2;refGene_tx=XM_023253453.1;refGene_gn=ABLIM2;refGene_tx=XM_023253454.1;refGene_gn=ABLIM2;refGene_tx=XM_023253455.1;refGene_gn=ABLIM2;refGene_tx=XM_023253456.1;refGene_gn=ABLIM2;refGene_tx=XM_023253457.1;refGene_gn=ABLIM2;refGene_tx=XM_023253458.1;refGene_gn=ABLIM2;refGene_tx=XM_023253459.1;refGene_gn=ABLIM2;refGene_tx=XM_023253460.1;refGene_gn=ABLIM2;refGene_tx=XM_023253461.1;refGene_gn=ABLIM2;refGene_tx=XM_023253462.1
B1 204691431 204691464
refGene_gn=ABLIM2;refGene_tx=XM_023253441.1;refGene_gn=ABLIM2;refGene_tx=XM_023253442.1;refGene_gn=ABLIM2;refGene_tx=XM_023253443.1;refGene_gn=ABLIM2;refGene_tx=XM_023253445.1;refGene_gn=ABLIM2;refGene_tx=XM_023253446.1;refGene_gn=ABLIM2;refGene_tx=XM_023253447.1;refGene_gn=ABLIM2;refGene_tx=XM_023253448.1;refGene_gn=ABLIM2;refGene_tx=XM_023253451.1;refGene_gn=ABLIM2;refGene_tx=XM_023253452.1;refGene_gn=ABLIM2;refGene_tx=XM_023253453.1;refGene_gn=ABLIM2;refGene_tx=XM_023253457.1;refGene_gn=ABLIM2;refGene_tx=XM_023253458.1;refGene_gn=ABLIM2;refGene_tx=XM_023253460.1;refGene_gn=ABLIM2;refGene_tx=XM_023253462.1
B1 204693925 204694072
refGene_gn=ABLIM2;refGene_tx=XM_023253441.1;refGene_gn=ABLIM2;refGene_tx=XM_023253442.1;refGene_gn=ABLIM2;refGene_tx=XM_023253443.1;refGene_gn=ABLIM2;refGene_tx=XM_023253444.1;refGene_gn=ABLIM2;refGene_tx=XM_023253445.1;refGene_gn=ABLIM2;refGene_tx=XM_023253446.1;refGene_gn=ABLIM

2;refGene_tx=XM_023253447.1;refGene_gn=ABLIM2;refGene_tx=XM_023253448.1;refGene_gn=ABLIM2;refGene_tx=XM_023253449.1;refGene_gn=ABLIM2;refGene_tx=XM_023253450.1;refGene_gn=ABLIM2;refGene_tx=XM_023253451.1;refGene_gn=ABLIM2;refGene_tx=XM_023253452.1;refGene_gn=ABLIM2;refGene_tx=XM_023253453.1;refGene_gn=ABLIM2;refGene_tx=XM_023253454.1;refGene_gn=ABLIM2;refGene_tx=XM_023253455.1;refGene_gn=ABLIM2;refGene_tx=XM_023253456.1;refGene_gn=ABLIM2;refGene_tx=XM_023253457.1;refGene_gn=ABLIM2;refGene_tx=XM_023253458.1;refGene_gn=ABLIM2;refGene_tx=XM_023253459.1;refGene_gn=ABLIM2;refGene_tx=XM_023253460.1;refGene_gn=ABLIM2;refGene_tx=XM_023253461.1;refGene_gn=ABLIM2;refGene_tx=XM_023253462.1
B1 204701575 204701653
refGene_gn=ABLIM2;refGene_tx=XM_023253441.1;refGene_gn=ABLIM2;refGene_tx=XM_023253442.1;refGene_gn=ABLIM2;refGene_tx=XM_023253443.1;refGene_gn=ABLIM2;refGene_tx=XM_023253444.1;refGene_gn=ABLIM2;refGene_tx=XM_023253445.1;refGene_gn=ABLIM2;refGene_tx=XM_023253446.1;refGene_gn=ABLIM2;refGene_tx=XM_023253447.1;refGene_gn=ABLIM2;refGene_tx=XM_023253448.1;refGene_gn=ABLIM2;refGene_tx=XM_023253449.1;refGene_gn=ABLIM2;refGene_tx=XM_023253450.1;refGene_gn=ABLIM2;refGene_tx=XM_023253451.1;refGene_gn=ABLIM2;refGene_tx=XM_023253452.1;refGene_gn=ABLIM2;refGene_tx=XM_023253453.1;refGene_gn=ABLIM2;refGene_tx=XM_023253454.1;refGene_gn=ABLIM2;refGene_tx=XM_023253455.1;refGene_gn=ABLIM2;refGene_tx=XM_023253456.1;refGene_gn=ABLIM2;refGene_tx=XM_023253457.1;refGene_gn=ABLIM2;refGene_tx=XM_023253458.1;refGene_gn=ABLIM2;refGene_tx=XM_023253459.1;refGene_gn=ABLIM2;refGene_tx=XM_023253460.1;refGene_gn=ABLIM2;refGene_tx=XM_023253461.1;refGene_gn=ABLIM2;refGene_tx=XM_023253462.1
B1 204707588 204707647
refGene_gn=ABLIM2;refGene_tx=XM_023253441.1;refGene_gn=ABLIM2;refGene_tx=XM_023253442.1;refGene_gn=ABLIM2;refGene_tx=XM_023253443.1;refGene_gn=ABLIM2;refGene_tx=XM_023253444.1;refGene_gn=ABLIM2;refGene_tx=XM_023253445.1;refGene_gn=ABLIM2;refGene_tx=XM_023253446.1;refGene_gn=ABLIM2;refGene_tx=XM_023253447.1;refGene_gn=ABLIM2;refGene_tx=XM_023253448.1;refGene_gn=ABLIM2;refGene_tx=XM_023253449.1;refGene_gn=ABLIM2;refGene_tx=XM_023253450.1;refGene_gn=ABLIM2;refGene_tx=XM_023253451.1;refGene_gn=ABLIM2;refGene_tx=XM_023253452.1;refGene_gn=ABLIM2;refGene_tx=XM_023253453.1;refGene_gn=ABLIM2;refGene_tx=XM_023253454.1;refGene_gn=ABLIM2;refGene_tx=XM_023253455.1;refGene_gn=ABLIM2;refGene_tx=XM_023253456.1;refGene_gn=ABLIM2;refGene_tx=XM_023253457.1;refGene_gn=ABLIM2;refGene_tx=XM_023253458.1;refGene_gn=ABLIM2;refGene_tx=XM_023253459.1;refGene_gn=ABLIM2;refGene_tx=XM_023253460.1;refGene_gn=ABLIM2;refGene_tx=XM_023253461.1;refGene_gn=ABLIM2;refGene_tx=XM_023253462.1
B1 204710090 204710178
refGene_gn=ABLIM2;refGene_tx=XM_023253441.1;refGene_gn=ABLIM2;refGene_tx=XM_023253442.1;refGene_gn=ABLIM2;refGene_tx=XM_023253443.1;refGene_gn=ABLIM2;refGene_tx=XM_023253444.1;refGene_gn=ABLIM2;refGene_tx=XM_023253445.1;refGene_gn=ABLIM2;refGene_tx=XM_023253446.1;refGene_gn=ABLIM2;refGene_tx=XM_023253447.1;refGene_gn=ABLIM2;refGene_tx=XM_023253448.1;refGene_gn=ABLIM2;refGene_tx=XM_023253449.1;refGene_gn=ABLIM2;refGene_tx=XM_023253450.1;refGene_gn=ABLIM2;refGene_tx=XM_023253451.1;refGene_gn=ABLIM2;refGene_tx=XM_023253452.1;refGene_gn=ABLIM2;refGene_tx=XM_023253453.1;refGene_gn=ABLIM2;refGene_tx=XM_023253454.1;refGene_gn=ABL

IM2;refGene_tx=XM_023253455.1;refGene_gn=ABLIM2;refGene_tx=XM_02325345
6.1;refGene_gn=ABLIM2;refGene_tx=XM_023253457.1;refGene_gn=ABLIM2;refG
ene_tx=XM_023253458.1;refGene_gn=ABLIM2;refGene_tx=XM_023253459.1;refG
ene_gn=ABLIM2;refGene_tx=XM_023253460.1;refGene_gn=ABLIM2;refGene_tx=X
M_023253461.1;refGene_gn=ABLIM2;refGene_tx=XM_023253462.1
B1 204720223 204720317
refGene_gn=ABLIM2;refGene_tx=XM_023253441.1;refGene_gn=ABLIM2;refGene_
tx=XM_023253442.1;refGene_gn=ABLIM2;refGene_tx=XM_023253443.1;refGene_
gn=ABLIM2;refGene_tx=XM_023253444.1;refGene_gn=ABLIM2;refGene_tx=XM_02
3253445.1;refGene_gn=ABLIM2;refGene_tx=XM_023253446.1;refGene_gn=ABLIM
2;refGene_tx=XM_023253447.1;refGene_gn=ABLIM2;refGene_tx=XM_023253448.
1;refGene_gn=ABLIM2;refGene_tx=XM_023253449.1;refGene_gn=ABLIM2;refGen
e_tx=XM_023253450.1;refGene_gn=ABLIM2;refGene_tx=XM_023253451.1;refGen
e_gn=ABLIM2;refGene_tx=XM_023253452.1;refGene_gn=ABLIM2;refGene_tx=XM_
023253453.1;refGene_gn=ABLIM2;refGene_tx=XM_023253454.1;refGene_gn=ABL
IM2;refGene_tx=XM_023253455.1;refGene_gn=ABLIM2;refGene_tx=XM_02325345
6.1;refGene_gn=ABLIM2;refGene_tx=XM_023253457.1;refGene_gn=ABLIM2;refG
ene_tx=XM_023253458.1;refGene_gn=ABLIM2;refGene_tx=XM_023253459.1;refG
ene_gn=ABLIM2;refGene_tx=XM_023253460.1;refGene_gn=ABLIM2;refGene_tx=X
M_023253461.1;refGene_gn=ABLIM2;refGene_tx=XM_023253462.1
B1 204722565 204722692
refGene_gn=ABLIM2;refGene_tx=XM_023253441.1;refGene_gn=ABLIM2;refGene_
tx=XM_023253442.1;refGene_gn=ABLIM2;refGene_tx=XM_023253443.1;refGene_
gn=ABLIM2;refGene_tx=XM_023253444.1;refGene_gn=ABLIM2;refGene_tx=XM_02
3253445.1;refGene_gn=ABLIM2;refGene_tx=XM_023253446.1;refGene_gn=ABLIM
2;refGene_tx=XM_023253447.1;refGene_gn=ABLIM2;refGene_tx=XM_023253448.
1;refGene_gn=ABLIM2;refGene_tx=XM_023253449.1;refGene_gn=ABLIM2;refGen
e_tx=XM_023253450.1;refGene_gn=ABLIM2;refGene_tx=XM_023253451.1;refGen
e_gn=ABLIM2;refGene_tx=XM_023253452.1;refGene_gn=ABLIM2;refGene_tx=XM_
023253453.1;refGene_gn=ABLIM2;refGene_tx=XM_023253454.1;refGene_gn=ABL
IM2;refGene_tx=XM_023253455.1;refGene_gn=ABLIM2;refGene_tx=XM_02325345
6.1;refGene_gn=ABLIM2;refGene_tx=XM_023253457.1;refGene_gn=ABLIM2;refG
ene_tx=XM_023253458.1;refGene_gn=ABLIM2;refGene_tx=XM_023253459.1;refG
ene_gn=ABLIM2;refGene_tx=XM_023253460.1;refGene_gn=ABLIM2;refGene_tx=X
M_023253461.1;refGene_gn=ABLIM2;refGene_tx=XM_023253462.1
B1 204728554 204728670
refGene_gn=ABLIM2;refGene_tx=XM_023253441.1;refGene_gn=ABLIM2;refGene_
tx=XM_023253442.1;refGene_gn=ABLIM2;refGene_tx=XM_023253443.1;refGene_
gn=ABLIM2;refGene_tx=XM_023253444.1;refGene_gn=ABLIM2;refGene_tx=XM_02
3253445.1;refGene_gn=ABLIM2;refGene_tx=XM_023253446.1;refGene_gn=ABLIM
2;refGene_tx=XM_023253447.1;refGene_gn=ABLIM2;refGene_tx=XM_023253448.
1;refGene_gn=ABLIM2;refGene_tx=XM_023253449.1;refGene_gn=ABLIM2;refGen
e_tx=XM_023253450.1;refGene_gn=ABLIM2;refGene_tx=XM_023253451.1;refGen
e_gn=ABLIM2;refGene_tx=XM_023253452.1;refGene_gn=ABLIM2;refGene_tx=XM_
023253453.1;refGene_gn=ABLIM2;refGene_tx=XM_023253454.1;refGene_gn=ABL
IM2;refGene_tx=XM_023253455.1;refGene_gn=ABLIM2;refGene_tx=XM_02325345
6.1;refGene_gn=ABLIM2;refGene_tx=XM_023253457.1;refGene_gn=ABLIM2;refG
ene_tx=XM_023253458.1;refGene_gn=ABLIM2;refGene_tx=XM_023253459.1;refG
ene_gn=ABLIM2;refGene_tx=XM_023253460.1;refGene_gn=ABLIM2;refGene_tx=X
M_023253461.1;refGene_gn=ABLIM2;refGene_tx=XM_023253462.1

B1	204735355	204735539
refGene_gn=ABLIM2;refGene_tx=XM_023253441.1;refGene_gn=ABLIM2;refGene_tx=XM_023253442.1;refGene_gn=ABLIM2;refGene_tx=XM_023253443.1;refGene_gn=ABLIM2;refGene_tx=XM_023253444.1;refGene_gn=ABLIM2;refGene_tx=XM_023253445.1;refGene_gn=ABLIM2;refGene_tx=XM_023253446.1;refGene_gn=ABLIM2;refGene_tx=XM_023253447.1;refGene_gn=ABLIM2;refGene_tx=XM_023253448.1;refGene_gn=ABLIM2;refGene_tx=XM_023253449.1;refGene_gn=ABLIM2;refGene_tx=XM_023253450.1;refGene_gn=ABLIM2;refGene_tx=XM_023253451.1;refGene_gn=ABLIM2;refGene_tx=XM_023253452.1;refGene_gn=ABLIM2;refGene_tx=XM_023253453.1;refGene_gn=ABLIM2;refGene_tx=XM_023253454.1;refGene_gn=ABLIM2;refGene_tx=XM_023253455.1;refGene_gn=ABLIM2;refGene_tx=XM_023253456.1;refGene_gn=ABLIM2;refGene_tx=XM_023253457.1;refGene_gn=ABLIM2;refGene_tx=XM_023253458.1;refGene_gn=ABLIM2;refGene_tx=XM_023253459.1;refGene_gn=ABLIM2;refGene_tx=XM_023253460.1;refGene_gn=ABLIM2;refGene_tx=XM_023253461.1;refGene_gn=ABLIM2;refGene_tx=XM_023253462.1
B1	204742285	204742429
refGene_gn=ABLIM2;refGene_tx=XM_023253441.1;refGene_gn=ABLIM2;refGene_tx=XM_023253442.1;refGene_gn=ABLIM2;refGene_tx=XM_023253443.1;refGene_gn=ABLIM2;refGene_tx=XM_023253444.1;refGene_gn=ABLIM2;refGene_tx=XM_023253445.1;refGene_gn=ABLIM2;refGene_tx=XM_023253446.1;refGene_gn=ABLIM2;refGene_tx=XM_023253447.1;refGene_gn=ABLIM2;refGene_tx=XM_023253448.1;refGene_gn=ABLIM2;refGene_tx=XM_023253449.1;refGene_gn=ABLIM2;refGene_tx=XM_023253450.1;refGene_gn=ABLIM2;refGene_tx=XM_023253451.1;refGene_gn=ABLIM2;refGene_tx=XM_023253452.1;refGene_gn=ABLIM2;refGene_tx=XM_023253453.1;refGene_gn=ABLIM2;refGene_tx=XM_023253454.1;refGene_gn=ABLIM2;refGene_tx=XM_023253455.1;refGene_gn=ABLIM2;refGene_tx=XM_023253456.1;refGene_gn=ABLIM2;refGene_tx=XM_023253457.1;refGene_gn=ABLIM2;refGene_tx=XM_023253458.1;refGene_gn=ABLIM2;refGene_tx=XM_023253459.1;refGene_gn=ABLIM2;refGene_tx=XM_023253460.1;refGene_gn=ABLIM2;refGene_tx=XM_023253461.1;refGene_gn=ABLIM2;refGene_tx=XM_023253462.1
B1	204757781	204757800
refGene_gn=ABLIM2;refGene_tx=XM_023253441.1;refGene_gn=ABLIM2;refGene_tx=XM_023253442.1;refGene_gn=ABLIM2;refGene_tx=XM_023253444.1;refGene_gn=ABLIM2;refGene_tx=XM_023253445.1;refGene_gn=ABLIM2;refGene_tx=XM_023253446.1;refGene_gn=ABLIM2;refGene_tx=XM_023253447.1;refGene_gn=ABLIM2;refGene_tx=XM_023253448.1;refGene_gn=ABLIM2;refGene_tx=XM_023253449.1;refGene_gn=ABLIM2;refGene_tx=XM_023253450.1;refGene_gn=ABLIM2;refGene_tx=XM_023253451.1;refGene_gn=ABLIM2;refGene_tx=XM_023253452.1;refGene_gn=ABLIM2;refGene_tx=XM_023253453.1;refGene_gn=ABLIM2;refGene_tx=XM_023253454.1;refGene_gn=ABLIM2;refGene_tx=XM_023253455.1;refGene_gn=ABLIM2;refGene_tx=XM_023253456.1;refGene_gn=ABLIM2;refGene_tx=XM_023253457.1;refGene_gn=ABLIM2;refGene_tx=XM_023253458.1;refGene_gn=ABLIM2;refGene_tx=XM_023253459.1;refGene_gn=ABLIM2;refGene_tx=XM_023253460.1;refGene_gn=ABLIM2;refGene_tx=XM_023253461.1;refGene_gn=ABLIM2;refGene_tx=XM_023253462.1
B1	204782206	204782216
refGene_gn=ABLIM2;refGene_tx=XM_023253443.1
B1	204821618	204821808
refGene_gn=SH3TC1;refGene_tx=XM_023253417.1;refGene_gn=SH3TC1;refGene_tx=XM_023253418.1;refGene_gn=SH3TC1;refGene_tx=XM_023253419.1;refGene_

gn=SH3TC1;refGene_tx=XM_023253420.1;refGene_gn=SH3TC1;refGene_tx=XM_02
3253421.1;refGene_gn=SH3TC1;refGene_tx=XM_023253422.1;refGene_gn=SH3TC
1;refGene_tx=XM_023253424.1
B1 204823433 204823508
refGene_gn=SH3TC1;refGene_tx=XM_023253417.1;refGene_gn=SH3TC1;refGene_
tx=XM_023253418.1;refGene_gn=SH3TC1;refGene_tx=XM_023253419.1;refGene_
gn=SH3TC1;refGene_tx=XM_023253420.1;refGene_gn=SH3TC1;refGene_tx=XM_02
3253421.1;refGene_gn=SH3TC1;refGene_tx=XM_023253422.1;refGene_gn=SH3TC
1;refGene_tx=XM_023253423.1;refGene_gn=SH3TC1;refGene_tx=XM_023253424.
1
B1 204825585 204825713
refGene_gn=SH3TC1;refGene_tx=XM_023253417.1;refGene_gn=SH3TC1;refGene_
tx=XM_023253418.1;refGene_gn=SH3TC1;refGene_tx=XM_023253419.1;refGene_
gn=SH3TC1;refGene_tx=XM_023253420.1;refGene_gn=SH3TC1;refGene_tx=XM_02
3253421.1;refGene_gn=SH3TC1;refGene_tx=XM_023253422.1;refGene_gn=SH3TC
1;refGene_tx=XM_023253423.1;refGene_gn=SH3TC1;refGene_tx=XM_023253424.
1;refGene_gn=SH3TC1;refGene_tx=XM_023253425.1
B1 204826437 204826543
refGene_gn=SH3TC1;refGene_tx=XM_023253417.1;refGene_gn=SH3TC1;refGene_
tx=XM_023253418.1;refGene_gn=SH3TC1;refGene_tx=XM_023253419.1;refGene_
gn=SH3TC1;refGene_tx=XM_023253420.1;refGene_gn=SH3TC1;refGene_tx=XM_02
3253421.1;refGene_gn=SH3TC1;refGene_tx=XM_023253422.1;refGene_gn=SH3TC
1;refGene_tx=XM_023253423.1;refGene_gn=SH3TC1;refGene_tx=XM_023253424.
1;refGene_gn=SH3TC1;refGene_tx=XM_023253425.1
B1 204827295 204827442
refGene_gn=SH3TC1;refGene_tx=XM_023253417.1;refGene_gn=SH3TC1;refGene_
tx=XM_023253418.1;refGene_gn=SH3TC1;refGene_tx=XM_023253419.1;refGene_
gn=SH3TC1;refGene_tx=XM_023253420.1;refGene_gn=SH3TC1;refGene_tx=XM_02
3253421.1;refGene_gn=SH3TC1;refGene_tx=XM_023253422.1;refGene_gn=SH3TC
1;refGene_tx=XM_023253423.1;refGene_gn=SH3TC1;refGene_tx=XM_023253424.
1;refGene_gn=SH3TC1;refGene_tx=XM_023253425.1
B1 204828092 204828297
refGene_gn=SH3TC1;refGene_tx=XM_023253417.1;refGene_gn=SH3TC1;refGene_
tx=XM_023253418.1;refGene_gn=SH3TC1;refGene_tx=XM_023253419.1;refGene_
gn=SH3TC1;refGene_tx=XM_023253420.1;refGene_gn=SH3TC1;refGene_tx=XM_02
3253421.1;refGene_gn=SH3TC1;refGene_tx=XM_023253422.1;refGene_gn=SH3TC
1;refGene_tx=XM_023253423.1;refGene_gn=SH3TC1;refGene_tx=XM_023253424.
1;refGene_gn=SH3TC1;refGene_tx=XM_023253425.1
B1 204829088 204829165
refGene_gn=SH3TC1;refGene_tx=XM_023253417.1;refGene_gn=SH3TC1;refGene_
tx=XM_023253418.1;refGene_gn=SH3TC1;refGene_tx=XM_023253419.1;refGene_
gn=SH3TC1;refGene_tx=XM_023253420.1;refGene_gn=SH3TC1;refGene_tx=XM_02
3253421.1;refGene_gn=SH3TC1;refGene_tx=XM_023253422.1;refGene_gn=SH3TC
1;refGene_tx=XM_023253423.1;refGene_gn=SH3TC1;refGene_tx=XM_023253424.
1;refGene_gn=SH3TC1;refGene_tx=XM_023253425.1
B1 204829902 204830098
refGene_gn=SH3TC1;refGene_tx=XM_023253417.1;refGene_gn=SH3TC1;refGene_
tx=XM_023253418.1;refGene_gn=SH3TC1;refGene_tx=XM_023253419.1;refGene_
gn=SH3TC1;refGene_tx=XM_023253420.1;refGene_gn=SH3TC1;refGene_tx=XM_02
3253421.1;refGene_gn=SH3TC1;refGene_tx=XM_023253422.1;refGene_gn=SH3TC

1;refGene_tx=XM_023253423.1;refGene_gn=SH3TC1;refGene_tx=XM_023253424.
1;refGene_gn=SH3TC1;refGene_tx=XM_023253425.1
B1 204832135 204832269
refGene_gn=SH3TC1;refGene_tx=XM_023253417.1;refGene_gn=SH3TC1;refGene_
tx=XM_023253418.1;refGene_gn=SH3TC1;refGene_tx=XM_023253419.1;refGene_
gn=SH3TC1;refGene_tx=XM_023253420.1;refGene_gn=SH3TC1;refGene_tx=XM_02
3253421.1;refGene_gn=SH3TC1;refGene_tx=XM_023253422.1;refGene_gn=SH3TC
1;refGene_tx=XM_023253423.1;refGene_gn=SH3TC1;refGene_tx=XM_023253424.
1;refGene_gn=SH3TC1;refGene_tx=XM_023253425.1
B1 204834112 204834154
refGene_gn=SH3TC1;refGene_tx=XM_023253417.1;refGene_gn=SH3TC1;refGene_
tx=XM_023253418.1;refGene_gn=SH3TC1;refGene_tx=XM_023253419.1;refGene_
gn=SH3TC1;refGene_tx=XM_023253420.1;refGene_gn=SH3TC1;refGene_tx=XM_02
3253421.1;refGene_gn=SH3TC1;refGene_tx=XM_023253422.1;refGene_gn=SH3TC
1;refGene_tx=XM_023253423.1;refGene_gn=SH3TC1;refGene_tx=XM_023253424.
1;refGene_gn=SH3TC1;refGene_tx=XM_023253425.1
B1 204835104 204835188
refGene_gn=SH3TC1;refGene_tx=XM_023253417.1;refGene_gn=SH3TC1;refGene_
tx=XM_023253418.1;refGene_gn=SH3TC1;refGene_tx=XM_023253419.1;refGene_
gn=SH3TC1;refGene_tx=XM_023253420.1;refGene_gn=SH3TC1;refGene_tx=XM_02
3253421.1;refGene_gn=SH3TC1;refGene_tx=XM_023253423.1;refGene_gn=SH3TC
1;refGene_tx=XM_023253424.1;refGene_gn=SH3TC1;refGene_tx=XM_023253425.
1
B1 204835835 204837494
refGene_gn=SH3TC1;refGene_tx=XM_023253417.1;refGene_gn=SH3TC1;refGene_
tx=XM_023253418.1;refGene_gn=SH3TC1;refGene_tx=XM_023253419.1;refGene_
gn=SH3TC1;refGene_tx=XM_023253420.1;refGene_gn=SH3TC1;refGene_tx=XM_02
3253421.1;refGene_gn=SH3TC1;refGene_tx=XM_023253422.1;refGene_gn=SH3TC
1;refGene_tx=XM_023253423.1;refGene_gn=SH3TC1;refGene_tx=XM_023253424.
1;refGene_gn=SH3TC1;refGene_tx=XM_023253425.1
B1 204839480 204839661
refGene_gn=SH3TC1;refGene_tx=XM_023253417.1;refGene_gn=SH3TC1;refGene_
tx=XM_023253418.1;refGene_gn=SH3TC1;refGene_tx=XM_023253419.1;refGene_
gn=SH3TC1;refGene_tx=XM_023253420.1;refGene_gn=SH3TC1;refGene_tx=XM_02
3253421.1;refGene_gn=SH3TC1;refGene_tx=XM_023253422.1;refGene_gn=SH3TC
1;refGene_tx=XM_023253423.1;refGene_gn=SH3TC1;refGene_tx=XM_023253424.
1;refGene_gn=SH3TC1;refGene_tx=XM_023253425.1
B1 204840668 204840819
refGene_gn=SH3TC1;refGene_tx=XM_023253417.1;refGene_gn=SH3TC1;refGene_
tx=XM_023253418.1;refGene_gn=SH3TC1;refGene_tx=XM_023253419.1;refGene_
gn=SH3TC1;refGene_tx=XM_023253420.1;refGene_gn=SH3TC1;refGene_tx=XM_02
3253421.1;refGene_gn=SH3TC1;refGene_tx=XM_023253422.1;refGene_gn=SH3TC
1;refGene_tx=XM_023253423.1;refGene_gn=SH3TC1;refGene_tx=XM_023253424.
1;refGene_gn=SH3TC1;refGene_tx=XM_023253425.1
B1 204842663 204842786
refGene_gn=SH3TC1;refGene_tx=XM_023253417.1;refGene_gn=SH3TC1;refGene_
tx=XM_023253418.1;refGene_gn=SH3TC1;refGene_tx=XM_023253419.1;refGene_
gn=SH3TC1;refGene_tx=XM_023253420.1;refGene_gn=SH3TC1;refGene_tx=XM_02
3253421.1;refGene_gn=SH3TC1;refGene_tx=XM_023253422.1;refGene_gn=SH3TC
1;refGene_tx=XM_023253423.1;refGene_gn=SH3TC1;refGene_tx=XM_023253424.

1;refGene_gn=SH3TC1;refGene_tx=XM_023253425.1
B1	204843431	204843582
refGene_gn=SH3TC1;refGene_tx=XM_023253417.1;refGene_gn=SH3TC1;refGene_tx=XM_023253418.1;refGene_gn=SH3TC1;refGene_tx=XM_023253419.1;refGene_gn=SH3TC1;refGene_tx=XM_023253420.1;refGene_gn=SH3TC1;refGene_tx=XM_023253421.1;refGene_gn=SH3TC1;refGene_tx=XM_023253422.1;refGene_gn=SH3TC1;refGene_tx=XM_023253423.1;refGene_gn=SH3TC1;refGene_tx=XM_023253424.1;refGene_gn=SH3TC1;refGene_tx=XM_023253425.1
B1	204844170	204844343
refGene_gn=SH3TC1;refGene_tx=XM_023253424.1
B1	204844375	204844572
refGene_gn=SH3TC1;refGene_tx=XM_023253417.1;refGene_gn=SH3TC1;refGene_tx=XM_023253418.1;refGene_gn=SH3TC1;refGene_tx=XM_023253419.1;refGene_gn=SH3TC1;refGene_tx=XM_023253420.1;refGene_gn=SH3TC1;refGene_tx=XM_023253421.1;refGene_gn=SH3TC1;refGene_tx=XM_023253422.1;refGene_gn=SH3TC1;refGene_tx=XM_023253423.1;refGene_gn=SH3TC1;refGene_tx=XM_023253425.1
B1	204846938	204847196
refGene_gn=SH3TC1;refGene_tx=XM_023253417.1;refGene_gn=SH3TC1;refGene_tx=XM_023253418.1;refGene_gn=SH3TC1;refGene_tx=XM_023253419.1;refGene_gn=SH3TC1;refGene_tx=XM_023253420.1;refGene_gn=SH3TC1;refGene_tx=XM_023253421.1;refGene_gn=SH3TC1;refGene_tx=XM_023253422.1;refGene_gn=SH3TC1;refGene_tx=XM_023253423.1;refGene_gn=SH3TC1;refGene_tx=XM_023253425.1
B1	204871528	204872553
refGene_gn=HTRA3;refGene_tx=XM_023253435.1;refGene_gn=HTRA3;refGene_tx=XM_023253436.1;refGene_gn=HTRA3;refGene_tx=XM_023253437.1;refGene_gn=HTRA3;refGene_tx=XM_023253438.1;refGene_gn=HTRA3;refGene_tx=XM_023253439.1;refGene_gn=HTRA3;refGene_tx=XM_023253440.1
B1	204883252	204883357
refGene_gn=HTRA3;refGene_tx=XM_023253435.1;refGene_gn=HTRA3;refGene_tx=XM_023253436.1;refGene_gn=HTRA3;refGene_tx=XM_023253437.1;refGene_gn=HTRA3;refGene_tx=XM_023253438.1;refGene_gn=HTRA3;refGene_tx=XM_023253439.1;refGene_gn=HTRA3;refGene_tx=XM_023253440.1
B1	204886119	204886342
refGene_gn=HTRA3;refGene_tx=XM_023253435.1;refGene_gn=HTRA3;refGene_tx=XM_023253436.1;refGene_gn=HTRA3;refGene_tx=XM_023253437.1;refGene_gn=HTRA3;refGene_tx=XM_023253438.1;refGene_gn=HTRA3;refGene_tx=XM_023253439.1;refGene_gn=HTRA3;refGene_tx=XM_023253440.1
B1	204888832	204889027
refGene_gn=HTRA3;refGene_tx=XM_023253435.1;refGene_gn=HTRA3;refGene_tx=XM_023253436.1;refGene_gn=HTRA3;refGene_tx=XM_023253437.1;refGene_gn=HTRA3;refGene_tx=XM_023253438.1;refGene_gn=HTRA3;refGene_tx=XM_023253439.1;refGene_gn=HTRA3;refGene_tx=XM_023253440.1
B1	204889646	204889679
refGene_gn=HTRA3;refGene_tx=XM_023253435.1;refGene_gn=HTRA3;refGene_tx=XM_023253436.1;refGene_gn=HTRA3;refGene_tx=XM_023253437.1;refGene_gn=HTRA3;refGene_tx=XM_023253438.1;refGene_gn=HTRA3;refGene_tx=XM_023253439.1;refGene_gn=HTRA3;refGene_tx=XM_023253440.1
B1	204890933	204891048

refGene_gn=HTRA3;refGene_tx=XM_023253435.1;refGene_gn=HTRA3;refGene_tx=XM_023253436.1;refGene_gn=HTRA3;refGene_tx=XM_023253437.1;refGene_gn=HTRA3;refGene_tx=XM_023253438.1;refGene_gn=HTRA3;refGene_tx=XM_023253439.1;refGene_gn=HTRA3;refGene_tx=XM_023253440.1
B1 204891890 204892101
refGene_gn=HTRA3;refGene_tx=XM_023253435.1;refGene_gn=HTRA3;refGene_tx=XM_023253438.1
B1 204892207 204892426
refGene_gn=HTRA3;refGene_tx=XM_023253435.1;refGene_gn=HTRA3;refGene_tx=XM_023253437.1;refGene_gn=HTRA3;refGene_tx=XM_023253439.1
B1 204898869 204898918
refGene_gn=HTRA3;refGene_tx=XM_023253436.1;refGene_gn=HTRA3;refGene_tx=XM_023253440.1
B1 204899369 204899465
refGene_gn=HTRA3;refGene_tx=XM_023253436.1;refGene_gn=HTRA3;refGene_tx=XM_023253440.1
B1 204901024 204901190
refGene_gn=HTRA3;refGene_tx=XM_023253436.1;refGene_gn=HTRA3;refGene_tx=XM_023253440.1
B1 204945504 204945624
refGene_gn=ACOX3;refGene_tx=XM_003985589.5;refGene_gn=ACOX3;refGene_tx=XM_019829807.2;refGene_gn=ACOX3;refGene_tx=XM_023253434.1
B1 204948184 204948328
refGene_gn=ACOX3;refGene_tx=XM_003985589.5;refGene_gn=ACOX3;refGene_tx=XM_019829807.2;refGene_gn=ACOX3;refGene_tx=XM_023253434.1
B1 204950423 204950491
refGene_gn=ACOX3;refGene_tx=XM_003985589.5;refGene_gn=ACOX3;refGene_tx=XM_023253434.1
B1 204951524 204951699
refGene_gn=ACOX3;refGene_tx=XM_003985589.5;refGene_gn=ACOX3;refGene_tx=XM_023253434.1
B1 204955943 204956059
refGene_gn=ACOX3;refGene_tx=XM_003985589.5;refGene_gn=ACOX3;refGene_tx=XM_019829807.2;refGene_gn=ACOX3;refGene_tx=XM_023253434.1
B1 204961010 204961124
refGene_gn=ACOX3;refGene_tx=XM_003985589.5;refGene_gn=ACOX3;refGene_tx=XM_019829807.2;refGene_gn=ACOX3;refGene_tx=XM_023253434.1
B1 204961287 204961410
refGene_gn=ACOX3;refGene_tx=XM_003985589.5;refGene_gn=ACOX3;refGene_tx=XM_019829807.2;refGene_gn=ACOX3;refGene_tx=XM_023253434.1
B1 204963651 204963772
refGene_gn=ACOX3;refGene_tx=XM_003985589.5;refGene_gn=ACOX3;refGene_tx=XM_019829807.2;refGene_gn=ACOX3;refGene_tx=XM_023253434.1
B1 204965208 204965331
refGene_gn=ACOX3;refGene_tx=XM_003985589.5;refGene_gn=ACOX3;refGene_tx=XM_019829807.2;refGene_gn=ACOX3;refGene_tx=XM_023253434.1
B1 204966992 204967175
refGene_gn=ACOX3;refGene_tx=XM_003985589.5;refGene_gn=ACOX3;refGene_tx=XM_019829807.2;refGene_gn=ACOX3;refGene_tx=XM_023253434.1
B1 204968810 204968907

refGene_gn=ACOX3;refGene_tx=XM_003985589.5;refGene_gn=ACOX3;refGene_tx=XM_019829807.2;refGene_gn=ACOX3;refGene_tx=XM_023253434.1
B1 204971565 204971654
refGene_gn=ACOX3;refGene_tx=XM_003985589.5;refGene_gn=ACOX3;refGene_tx=XM_019829807.2;refGene_gn=ACOX3;refGene_tx=XM_023253434.1
B1 204975986 204976130
refGene_gn=ACOX3;refGene_tx=XM_003985589.5;refGene_gn=ACOX3;refGene_tx=XM_019829807.2;refGene_gn=ACOX3;refGene_tx=XM_023253434.1
B1 204978491 204978581
refGene_gn=ACOX3;refGene_tx=XM_003985589.5;refGene_gn=ACOX3;refGene_tx=XM_019829807.2;refGene_gn=ACOX3;refGene_tx=XM_023253434.1
B1 204978945 204979020
refGene_gn=ACOX3;refGene_tx=XM_003985589.5;refGene_gn=ACOX3;refGene_tx=XM_019829807.2;refGene_gn=ACOX3;refGene_tx=XM_023253434.1
B1 204980130 204980364
refGene_gn=ACOX3;refGene_tx=XM_003985589.5;refGene_gn=ACOX3;refGene_tx=XM_019829807.2;refGene_gn=ACOX3;refGene_tx=XM_023253434.1
B1 204980697 204980841
refGene_gn=ACOX3;refGene_tx=XM_003985589.5;refGene_gn=ACOX3;refGene_tx=XM_019829807.2;refGene_gn=ACOX3;refGene_tx=XM_023253434.1
B1 204997199 204997788
refGene_gn=TRMT44;refGene_tx=XM_011281996.3;refGene_gn=TRMT44;refGene_tx=XM_023253432.1
B1 204998565 204998680
refGene_gn=TRMT44;refGene_tx=XM_011281996.3;refGene_gn=TRMT44;refGene_tx=XM_023253432.1;refGene_gn=TRMT44;refGene_tx=XM_023253433.1
B1 205005762 205005982
refGene_gn=TRMT44;refGene_tx=XM_011281996.3;refGene_gn=TRMT44;refGene_tx=XM_023253432.1;refGene_gn=TRMT44;refGene_tx=XM_023253433.1
B1 205007439 205007508
refGene_gn=TRMT44;refGene_tx=XM_011281996.3;refGene_gn=TRMT44;refGene_tx=XM_023253432.1;refGene_gn=TRMT44;refGene_tx=XM_023253433.1
B1 205008288 205008396
refGene_gn=TRMT44;refGene_tx=XM_011281996.3;refGene_gn=TRMT44;refGene_tx=XM_023253432.1;refGene_gn=TRMT44;refGene_tx=XM_023253433.1
B1 205010110 205010182
refGene_gn=TRMT44;refGene_tx=XM_011281996.3;refGene_gn=TRMT44;refGene_tx=XM_023253432.1;refGene_gn=TRMT44;refGene_tx=XM_023253433.1
B1 205014358 205014465
refGene_gn=TRMT44;refGene_tx=XM_011281996.3;refGene_gn=TRMT44;refGene_tx=XM_023253432.1;refGene_gn=TRMT44;refGene_tx=XM_023253433.1
B1 205015518 205015702
refGene_gn=TRMT44;refGene_tx=XM_011281996.3;refGene_gn=TRMT44;refGene_tx=XM_023253432.1;refGene_gn=TRMT44;refGene_tx=XM_023253433.1
B1 205017598 205018007
refGene_gn=TRMT44;refGene_tx=XM_011281996.3;refGene_gn=TRMT44;refGene_tx=XM_023253432.1;refGene_gn=TRMT44;refGene_tx=XM_023253433.1
B1 205020179 205020296
refGene_gn=TRMT44;refGene_tx=XM_011281996.3;refGene_gn=TRMT44;refGene_tx=XM_023253432.1;refGene_gn=TRMT44;refGene_tx=XM_023253433.1

B1	205026245	205026475
refGene_gn=TRMT44;refGene_tx=XM_011281996.3;refGene_gn=TRMT44;refGene_tx=XM_023253432.1;refGene_gn=TRMT44;refGene_tx=XM_023253433.1
B1	205046884	205047799
refGene_gn=LOC109499301;refGene_tx=XM_023253463.1;refGene_gn=LOC109499301;refGene_tx=XM_023253464.1
B1	205095634	205095719
refGene_gn=CPZ;refGene_tx=XM_003985548.5
B1	205099533	205099746
refGene_gn=CPZ;refGene_tx=XM_003985548.5
B1	205104937	205105134
refGene_gn=CPZ;refGene_tx=XM_003985548.5
B1	205105695	205105857
refGene_gn=CPZ;refGene_tx=XM_003985548.5
B1	205106319	205106475
refGene_gn=CPZ;refGene_tx=XM_003985548.5
B1	205110875	205111011
refGene_gn=CPZ;refGene_tx=XM_003985548.5
B1	205112937	205113077
refGene_gn=CPZ;refGene_tx=XM_003985548.5
B1	205115887	205115987
refGene_gn=CPZ;refGene_tx=XM_003985548.5
B1	205116193	205116561
refGene_gn=CPZ;refGene_tx=XM_003985548.5
B1	205300634	205301293
refGene_gn=HMX1;refGene_tx=XM_023253341.1
B1	205304012	205304406
refGene_gn=HMX1;refGene_tx=XM_023253341.1
B1	205442136	205443567
refGene_gn=ADRA2C;refGene_tx=XM_023253342.1
B1	205638925	205639138
refGene_gn=LRPAP1;refGene_tx=XM_023253347.1
B1	205642351	205642496
refGene_gn=LRPAP1;refGene_tx=XM_023253347.1
B1	205646651	205646776
refGene_gn=LRPAP1;refGene_tx=XM_023253347.1
B1	205647572	205647693
refGene_gn=LRPAP1;refGene_tx=XM_023253347.1
B1	205647883	205648045
refGene_gn=LRPAP1;refGene_tx=XM_023253347.1
B1	205648989	205649072
refGene_gn=LRPAP1;refGene_tx=XM_023253347.1
B1	205650496	205650673
refGene_gn=LRPAP1;refGene_tx=XM_023253347.1
B1	205651947	205652010
refGene_gn=LRPAP1;refGene_tx=XM_023253347.1
B1	205665927	205666064
refGene_gn=DOK7;refGene_tx=XM_023253343.1;refGene_gn=DOK7;refGene_tx=XM_023253344.1
B1	205666295	205666424

refGene_gn=DOK7;refGene_tx=XM_023253343.1;refGene_gn=DOK7;refGene_tx=XM_023253344.1
B1	205670540	205670583
refGene_gn=DOK7;refGene_tx=XM_023253343.1;refGene_gn=DOK7;refGene_tx=XM_023253344.1
B1	205674490	205675197
refGene_gn=DOK7;refGene_tx=XM_023253343.1;refGene_gn=DOK7;refGene_tx=XM_023253344.1;refGene_gn=DOK7;refGene_tx=XM_023253345.1
B1	205675963	205676083
refGene_gn=DOK7;refGene_tx=XM_023253343.1;refGene_gn=DOK7;refGene_tx=XM_023253344.1;refGene_gn=DOK7;refGene_tx=XM_023253345.1
B1	205678159	205678279
refGene_gn=DOK7;refGene_tx=XM_023253343.1;refGene_gn=DOK7;refGene_tx=XM_023253344.1;refGene_gn=DOK7;refGene_tx=XM_023253345.1
B1	205686500	205686701
refGene_gn=DOK7;refGene_tx=XM_023253343.1;refGene_gn=DOK7;refGene_tx=XM_023253344.1;refGene_gn=DOK7;refGene_tx=XM_023253345.1
B1	205687655	205687886
refGene_gn=DOK7;refGene_tx=XM_023253343.1;refGene_gn=DOK7;refGene_tx=XM_023253344.1;refGene_gn=DOK7;refGene_tx=XM_023253345.1
B1	205695170	205695216
refGene_gn=DOK7;refGene_tx=XM_023253343.1;refGene_gn=DOK7;refGene_tx=XM_023253344.1;refGene_gn=DOK7;refGene_tx=XM_023253345.1
B1	205695287	205695341
refGene_gn=DOK7;refGene_tx=XM_023253343.1;refGene_gn=DOK7;refGene_tx=XM_023253344.1;refGene_gn=DOK7;refGene_tx=XM_023253345.1
B1	205703518	205703701
refGene_gn=HGFAC;refGene_tx=XM_011281954.3
B1	205704756	205704905
refGene_gn=HGFAC;refGene_tx=XM_011281954.3
B1	205704991	205705132
refGene_gn=HGFAC;refGene_tx=XM_011281954.3
B1	205705430	205705570
refGene_gn=HGFAC;refGene_tx=XM_011281954.3
B1	205707607	205707863
refGene_gn=HGFAC;refGene_tx=XM_011281954.3
B1	205708460	205708546
refGene_gn=HGFAC;refGene_tx=XM_011281954.3
B1	205708750	205708772
refGene_gn=HGFAC;refGene_tx=XM_011281954.3
B1	205708774	205708924
refGene_gn=HGFAC;refGene_tx=XM_011281954.3
B1	205709018	205709129
refGene_gn=HGFAC;refGene_tx=XM_011281954.3
B1	205709218	205709350
refGene_gn=HGFAC;refGene_tx=XM_011281954.3
B1	205709433	205709556
refGene_gn=HGFAC;refGene_tx=XM_011281954.3
B1	205710234	205710314
refGene_gn=HGFAC;refGene_tx=XM_011281954.3

B1	205710549	205710646
refGene_gn=HGFAC;refGene_tx=XM_011281954.3
B1	205710798	205710976
refGene_gn=HGFAC;refGene_tx=XM_011281954.3
B1	205711523	205711706
refGene_gn=HGFAC;refGene_tx=XM_011281954.3
B1	205714054	205714281
refGene_gn=RGS12;refGene_tx=XM_023253348.1;refGene_gn=RGS12;refGene_tx=XM_023253349.1;refGene_gn=RGS12;refGene_tx=XM_023253351.1
B1	205722218	205722680
refGene_gn=RGS12;refGene_tx=XM_023253348.1;refGene_gn=RGS12;refGene_tx=XM_023253349.1;refGene_gn=RGS12;refGene_tx=XM_023253350.1;refGene_gn=RGS12;refGene_tx=XM_023253351.1
B1	205724052	205724206
refGene_gn=RGS12;refGene_tx=XM_023253348.1;refGene_gn=RGS12;refGene_tx=XM_023253349.1;refGene_gn=RGS12;refGene_tx=XM_023253351.1
B1	205724549	205724620
refGene_gn=RGS12;refGene_tx=XM_023253348.1;refGene_gn=RGS12;refGene_tx=XM_023253349.1;refGene_gn=RGS12;refGene_tx=XM_023253350.1;refGene_gn=RGS12;refGene_tx=XM_023253351.1
B1	205725971	205726068
refGene_gn=RGS12;refGene_tx=XM_023253348.1;refGene_gn=RGS12;refGene_tx=XM_023253349.1;refGene_gn=RGS12;refGene_tx=XM_023253350.1;refGene_gn=RGS12;refGene_tx=XM_023253351.1
B1	205728081	205728208
refGene_gn=RGS12;refGene_tx=XM_023253348.1;refGene_gn=RGS12;refGene_tx=XM_023253349.1;refGene_gn=RGS12;refGene_tx=XM_023253350.1;refGene_gn=RGS12;refGene_tx=XM_023253351.1
B1	205728694	205728768
refGene_gn=RGS12;refGene_tx=XM_023253348.1;refGene_gn=RGS12;refGene_tx=XM_023253349.1;refGene_gn=RGS12;refGene_tx=XM_023253350.1;refGene_gn=RGS12;refGene_tx=XM_023253351.1
B1	205729034	205729229
refGene_gn=RGS12;refGene_tx=XM_023253348.1;refGene_gn=RGS12;refGene_tx=XM_023253349.1;refGene_gn=RGS12;refGene_tx=XM_023253350.1;refGene_gn=RGS12;refGene_tx=XM_023253351.1
B1	205731177	205731254
refGene_gn=RGS12;refGene_tx=XM_023253348.1;refGene_gn=RGS12;refGene_tx=XM_023253349.1;refGene_gn=RGS12;refGene_tx=XM_023253350.1;refGene_gn=RGS12;refGene_tx=XM_023253351.1
B1	205732870	205733024
refGene_gn=RGS12;refGene_tx=XM_023253348.1;refGene_gn=RGS12;refGene_tx=XM_023253349.1;refGene_gn=RGS12;refGene_tx=XM_023253350.1;refGene_gn=RGS12;refGene_tx=XM_023253351.1
B1	205733551	205733731
refGene_gn=RGS12;refGene_tx=XM_023253348.1;refGene_gn=RGS12;refGene_tx=XM_023253349.1;refGene_gn=RGS12;refGene_tx=XM_023253350.1;refGene_gn=RGS12;refGene_tx=XM_023253351.1
B1	205734465	205734609
refGene_gn=RGS12;refGene_tx=XM_023253348.1;refGene_gn=RGS12;refGene_tx

=XM_023253349.1;refGene_gn=RGS12;refGene_tx=XM_023253350.1;refGene_gn=RGS12;refGene_tx=XM_023253351.1
B1 205735312 205735405
refGene_gn=RGS12;refGene_tx=XM_023253348.1;refGene_gn=RGS12;refGene_tx=XM_023253349.1;refGene_gn=RGS12;refGene_tx=XM_023253350.1;refGene_gn=RGS12;refGene_tx=XM_023253351.1
B1 205735935 205736105
refGene_gn=RGS12;refGene_tx=XM_023253348.1;refGene_gn=RGS12;refGene_tx=XM_023253349.1;refGene_gn=RGS12;refGene_tx=XM_023253350.1;refGene_gn=RGS12;refGene_tx=XM_023253351.1
B1 205779219 205779273
refGene_gn=RGS12;refGene_tx=XM_023253351.1
B1 205792393 205792510
refGene_gn=RGS12;refGene_tx=XM_023253348.1;refGene_gn=RGS12;refGene_tx=XM_023253350.1
B1 205806901 205808773
refGene_gn=RGS12;refGene_tx=XM_023253348.1;refGene_gn=RGS12;refGene_tx=XM_023253349.1;refGene_gn=RGS12;refGene_tx=XM_023253350.1
B1 205861835 205862076
refGene_gn=MSANTD1;refGene_tx=XM_019829819.2;refGene_gn=MSANTD1;refGene_tx=XM_019829820.2
B1 205864729 205865005
refGene_gn=MSANTD1;refGene_tx=XM_019829819.2;refGene_gn=MSANTD1;refGene_tx=XM_019829820.2
B1 205868543 205868808
refGene_gn=MSANTD1;refGene_tx=XM_019829819.2;refGene_gn=MSANTD1;refGene_tx=XM_019829820.2
B1 205870469 205870515
refGene_gn=MSANTD1;refGene_tx=XM_019829819.2;refGene_gn=MSANTD1;refGene_tx=XM_019829820.2
B1 205870524 205870864
refGene_gn=MSANTD1;refGene_tx=XM_019829824.2
B1 205872140 205872226
refGene_gn=MSANTD1;refGene_tx=XM_019829824.2
B1 205873121 205873370
refGene_gn=MSANTD1;refGene_tx=XM_019829824.2
B1 205877890 205878104
refGene_gn=HTT;refGene_tx=XM_023253466.1;refGene_gn=HTT;refGene_tx=XM_023253467.1
B1 205878984 205879145
refGene_gn=HTT;refGene_tx=XM_023253466.1;refGene_gn=HTT;refGene_tx=XM_023253467.1
B1 205879329 205879492
refGene_gn=HTT;refGene_tx=XM_023253466.1;refGene_gn=HTT;refGene_tx=XM_023253467.1
B1 205881229 205881335
refGene_gn=HTT;refGene_tx=XM_023253466.1;refGene_gn=HTT;refGene_tx=XM_023253467.1
B1 205881748 205881962
refGene_gn=HTT;refGene_tx=XM_023253466.1;refGene_gn=HTT;refGene_tx=XM_

023253467.1
B1 205882089 205882204
refGene_gn=HTT;refGene_tx=XM_023253466.1;refGene_gn=HTT;refGene_tx=XM_
023253467.1
B1 205882368 205882559
refGene_gn=HTT;refGene_tx=XM_023253466.1;refGene_gn=HTT;refGene_tx=XM_
023253467.1
B1 205884718 205884802
refGene_gn=HTT;refGene_tx=XM_023253468.1
B1 205884927 205885083
refGene_gn=HTT;refGene_tx=XM_023253466.1;refGene_gn=HTT;refGene_tx=XM_
023253467.1;refGene_gn=HTT;refGene_tx=XM_023253468.1
B1 205886816 205886946
refGene_gn=HTT;refGene_tx=XM_023253466.1;refGene_gn=HTT;refGene_tx=XM_
023253467.1;refGene_gn=HTT;refGene_tx=XM_023253468.1
B1 205887051 205887182
refGene_gn=HTT;refGene_tx=XM_023253466.1;refGene_gn=HTT;refGene_tx=XM_
023253467.1;refGene_gn=HTT;refGene_tx=XM_023253468.1
B1 205888783 205888866
refGene_gn=HTT;refGene_tx=XM_023253466.1;refGene_gn=HTT;refGene_tx=XM_
023253467.1;refGene_gn=HTT;refGene_tx=XM_023253468.1
B1 205890203 205890343
refGene_gn=HTT;refGene_tx=XM_023253466.1;refGene_gn=HTT;refGene_tx=XM_
023253467.1;refGene_gn=HTT;refGene_tx=XM_023253468.1
B1 205890691 205890846
refGene_gn=HTT;refGene_tx=XM_023253466.1;refGene_gn=HTT;refGene_tx=XM_
023253467.1;refGene_gn=HTT;refGene_tx=XM_023253468.1
B1 205892012 205892113
refGene_gn=HTT;refGene_tx=XM_023253466.1;refGene_gn=HTT;refGene_tx=XM_
023253467.1;refGene_gn=HTT;refGene_tx=XM_023253468.1
B1 205894082 205894209
refGene_gn=HTT;refGene_tx=XM_023253466.1;refGene_gn=HTT;refGene_tx=XM_
023253467.1;refGene_gn=HTT;refGene_tx=XM_023253468.1
B1 205895937 205896125
refGene_gn=HTT;refGene_tx=XM_023253466.1;refGene_gn=HTT;refGene_tx=XM_
023253467.1;refGene_gn=HTT;refGene_tx=XM_023253468.1
B1 205898788 205898884
refGene_gn=HTT;refGene_tx=XM_023253466.1;refGene_gn=HTT;refGene_tx=XM_
023253467.1;refGene_gn=HTT;refGene_tx=XM_023253468.1
B1 205899775 205899953
refGene_gn=HTT;refGene_tx=XM_023253466.1;refGene_gn=HTT;refGene_tx=XM_
023253467.1;refGene_gn=HTT;refGene_tx=XM_023253468.1
B1 205901793 205901939
refGene_gn=HTT;refGene_tx=XM_023253466.1;refGene_gn=HTT;refGene_tx=XM_
023253467.1;refGene_gn=HTT;refGene_tx=XM_023253468.1
B1 205902308 205902522
refGene_gn=HTT;refGene_tx=XM_023253466.1;refGene_gn=HTT;refGene_tx=XM_
023253467.1;refGene_gn=HTT;refGene_tx=XM_023253468.1
B1 205904337 205904460
refGene_gn=HTT;refGene_tx=XM_023253466.1;refGene_gn=HTT;refGene_tx=XM_

023253467.1;refGene_gn=HTT;refGene_tx=XM_023253468.1
B1 205905509 205905648
refGene_gn=HTT;refGene_tx=XM_023253466.1;refGene_gn=HTT;refGene_tx=XM_023253467.1;refGene_gn=HTT;refGene_tx=XM_023253468.1
B1 205906941 205907018
refGene_gn=HTT;refGene_tx=XM_023253466.1;refGene_gn=HTT;refGene_tx=XM_023253467.1;refGene_gn=HTT;refGene_tx=XM_023253468.1
B1 205907280 205907457
refGene_gn=HTT;refGene_tx=XM_023253466.1;refGene_gn=HTT;refGene_tx=XM_023253467.1;refGene_gn=HTT;refGene_tx=XM_023253468.1
B1 205907594 205907774
refGene_gn=HTT;refGene_tx=XM_023253466.1;refGene_gn=HTT;refGene_tx=XM_023253467.1;refGene_gn=HTT;refGene_tx=XM_023253468.1
B1 205910269 205910411
refGene_gn=HTT;refGene_tx=XM_023253466.1;refGene_gn=HTT;refGene_tx=XM_023253467.1;refGene_gn=HTT;refGene_tx=XM_023253468.1
B1 205914960 205915168
refGene_gn=HTT;refGene_tx=XM_023253466.1;refGene_gn=HTT;refGene_tx=XM_023253467.1;refGene_gn=HTT;refGene_tx=XM_023253468.1
B1 205932433 205932576
refGene_gn=HTT;refGene_tx=XM_023253466.1;refGene_gn=HTT;refGene_tx=XM_023253467.1;refGene_gn=HTT;refGene_tx=XM_023253468.1
B1 205933674 205933910
refGene_gn=HTT;refGene_tx=XM_023253466.1;refGene_gn=HTT;refGene_tx=XM_023253467.1;refGene_gn=HTT;refGene_tx=XM_023253468.1
B1 205934199 205934322
refGene_gn=HTT;refGene_tx=XM_023253466.1;refGene_gn=HTT;refGene_tx=XM_023253467.1;refGene_gn=HTT;refGene_tx=XM_023253468.1
B1 205937862 205937979
refGene_gn=HTT;refGene_tx=XM_023253466.1;refGene_gn=HTT;refGene_tx=XM_023253467.1;refGene_gn=HTT;refGene_tx=XM_023253468.1
B1 205939719 205939856
refGene_gn=HTT;refGene_tx=XM_023253466.1;refGene_gn=HTT;refGene_tx=XM_023253467.1;refGene_gn=HTT;refGene_tx=XM_023253468.1
B1 205942464 205942613
refGene_gn=HTT;refGene_tx=XM_023253466.1;refGene_gn=HTT;refGene_tx=XM_023253467.1;refGene_gn=HTT;refGene_tx=XM_023253468.1
B1 205944461 205944517
refGene_gn=HTT;refGene_tx=XM_023253466.1;refGene_gn=HTT;refGene_tx=XM_023253467.1;refGene_gn=HTT;refGene_tx=XM_023253468.1
B1 205945406 205945568
refGene_gn=HTT;refGene_tx=XM_023253466.1;refGene_gn=HTT;refGene_tx=XM_023253467.1;refGene_gn=HTT;refGene_tx=XM_023253468.1
B1 205945685 205945764
refGene_gn=HTT;refGene_tx=XM_023253466.1;refGene_gn=HTT;refGene_tx=XM_023253467.1;refGene_gn=HTT;refGene_tx=XM_023253468.1
B1 205946823 205947047
refGene_gn=HTT;refGene_tx=XM_023253466.1;refGene_gn=HTT;refGene_tx=XM_023253467.1;refGene_gn=HTT;refGene_tx=XM_023253468.1
B1 205947534 205947612

refGene_gn=HTT;refGene_tx=XM_023253466.1;refGene_gn=HTT;refGene_tx=XM_023253467.1;refGene_gn=HTT;refGene_tx=XM_023253468.1
B1 205952235 205952346
refGene_gn=HTT;refGene_tx=XM_023253466.1;refGene_gn=HTT;refGene_tx=XM_023253467.1;refGene_gn=HTT;refGene_tx=XM_023253468.1
B1 205955098 205955226
refGene_gn=HTT;refGene_tx=XM_023253466.1;refGene_gn=HTT;refGene_tx=XM_023253467.1;refGene_gn=HTT;refGene_tx=XM_023253468.1
B1 205956608 205956735
refGene_gn=HTT;refGene_tx=XM_023253466.1;refGene_gn=HTT;refGene_tx=XM_023253467.1;refGene_gn=HTT;refGene_tx=XM_023253468.1
B1 205959079 205959282
refGene_gn=HTT;refGene_tx=XM_023253466.1;refGene_gn=HTT;refGene_tx=XM_023253467.1;refGene_gn=HTT;refGene_tx=XM_023253468.1
B1 205960290 205960442
refGene_gn=HTT;refGene_tx=XM_023253466.1;refGene_gn=HTT;refGene_tx=XM_023253467.1;refGene_gn=HTT;refGene_tx=XM_023253468.1
B1 205961927 205962004
refGene_gn=HTT;refGene_tx=XM_023253466.1;refGene_gn=HTT;refGene_tx=XM_023253467.1;refGene_gn=HTT;refGene_tx=XM_023253468.1
B1 205963413 205963534
refGene_gn=HTT;refGene_tx=XM_023253466.1;refGene_gn=HTT;refGene_tx=XM_023253467.1;refGene_gn=HTT;refGene_tx=XM_023253468.1
B1 205965280 205965427
refGene_gn=HTT;refGene_tx=XM_023253466.1;refGene_gn=HTT;refGene_tx=XM_023253467.1;refGene_gn=HTT;refGene_tx=XM_023253468.1
B1 205968258 205968359
refGene_gn=HTT;refGene_tx=XM_023253466.1;refGene_gn=HTT;refGene_tx=XM_023253467.1;refGene_gn=HTT;refGene_tx=XM_023253468.1
B1 205968607 205968671
refGene_gn=HTT;refGene_tx=XM_023253466.1;refGene_gn=HTT;refGene_tx=XM_023253467.1;refGene_gn=HTT;refGene_tx=XM_023253468.1
B1 205969621 205969761
refGene_gn=HTT;refGene_tx=XM_023253466.1;refGene_gn=HTT;refGene_tx=XM_023253467.1;refGene_gn=HTT;refGene_tx=XM_023253468.1
B1 205970815 205970913
refGene_gn=HTT;refGene_tx=XM_023253466.1;refGene_gn=HTT;refGene_tx=XM_023253467.1;refGene_gn=HTT;refGene_tx=XM_023253468.1
B1 205971003 205971162
refGene_gn=HTT;refGene_tx=XM_023253466.1;refGene_gn=HTT;refGene_tx=XM_023253467.1;refGene_gn=HTT;refGene_tx=XM_023253468.1
B1 205972247 205972385
refGene_gn=HTT;refGene_tx=XM_023253466.1;refGene_gn=HTT;refGene_tx=XM_023253467.1;refGene_gn=HTT;refGene_tx=XM_023253468.1
B1 205972638 205972750
refGene_gn=HTT;refGene_tx=XM_023253466.1;refGene_gn=HTT;refGene_tx=XM_023253467.1;refGene_gn=HTT;refGene_tx=XM_023253468.1
B1 205974551 205974670
refGene_gn=HTT;refGene_tx=XM_023253466.1;refGene_gn=HTT;refGene_tx=XM_023253467.1;refGene_gn=HTT;refGene_tx=XM_023253468.1

B1 205974920 205975044
refGene_gn=HTT;refGene_tx=XM_023253466.1;refGene_gn=HTT;refGene_tx=XM_023253467.1;refGene_gn=HTT;refGene_tx=XM_023253468.1
B1 205977689 205978051
refGene_gn=HTT;refGene_tx=XM_023253466.1;refGene_gn=HTT;refGene_tx=XM_023253467.1;refGene_gn=HTT;refGene_tx=XM_023253468.1
B1 205979411 205979492
refGene_gn=HTT;refGene_tx=XM_023253466.1;refGene_gn=HTT;refGene_tx=XM_023253467.1;refGene_gn=HTT;refGene_tx=XM_023253468.1
B1 205981912 205981960
refGene_gn=HTT;refGene_tx=XM_023253466.1;refGene_gn=HTT;refGene_tx=XM_023253468.1
B1 205983123 205983325
refGene_gn=HTT;refGene_tx=XM_023253466.1;refGene_gn=HTT;refGene_tx=XM_023253467.1;refGene_gn=HTT;refGene_tx=XM_023253468.1
B1 205988641 205988820
refGene_gn=HTT;refGene_tx=XM_023253466.1;refGene_gn=HTT;refGene_tx=XM_023253467.1;refGene_gn=HTT;refGene_tx=XM_023253468.1
B1 205989736 205989878
refGene_gn=HTT;refGene_tx=XM_023253466.1;refGene_gn=HTT;refGene_tx=XM_023253467.1;refGene_gn=HTT;refGene_tx=XM_023253468.1
B1 205992348 205992487
refGene_gn=HTT;refGene_tx=XM_023253466.1;refGene_gn=HTT;refGene_tx=XM_023253467.1;refGene_gn=HTT;refGene_tx=XM_023253468.1
B1 205994058 205994138
refGene_gn=HTT;refGene_tx=XM_023253466.1;refGene_gn=HTT;refGene_tx=XM_023253467.1;refGene_gn=HTT;refGene_tx=XM_023253468.1
B1 205994533 205994593
refGene_gn=HTT;refGene_tx=XM_023253466.1;refGene_gn=HTT;refGene_tx=XM_023253467.1;refGene_gn=HTT;refGene_tx=XM_023253468.1
B1 205999447 205999568
refGene_gn=HTT;refGene_tx=XM_023253466.1;refGene_gn=HTT;refGene_tx=XM_023253467.1;refGene_gn=HTT;refGene_tx=XM_023253468.1
B1 206009095 206009179
refGene_gn=HTT;refGene_tx=XM_023253466.1;refGene_gn=HTT;refGene_tx=XM_023253467.1;refGene_gn=HTT;refGene_tx=XM_023253468.1
B1 206022849 206023094
refGene_gn=HTT;refGene_tx=XM_023253466.1;refGene_gn=HTT;refGene_tx=XM_023253467.1;refGene_gn=HTT;refGene_tx=XM_023253468.1
B1 206035934 206036042
refGene_gn=GRK4;refGene_tx=XM_019829825.2;refGene_gn=GRK4;refGene_tx=XM_023253497.1;refGene_gn=GRK4;refGene_tx=XM_023253498.1;refGene_gn=GRK4;refGene_tx=XM_023253499.1;refGene_gn=GRK4;refGene_tx=XM_023253500.1;refGene_gn=GRK4;refGene_tx=XM_023253501.1
B1 206037093 206037225
refGene_gn=GRK4;refGene_tx=XM_019829825.2;refGene_gn=GRK4;refGene_tx=XM_023253497.1;refGene_gn=GRK4;refGene_tx=XM_023253498.1;refGene_gn=GRK4;refGene_tx=XM_023253499.1;refGene_gn=GRK4;refGene_tx=XM_023253500.1;refGene_gn=GRK4;refGene_tx=XM_023253501.1
B1 206037576 206037714

refGene_gn=GRK4;refGene_tx=XM_019829825.2;refGene_gn=GRK4;refGene_tx=XM_023253497.1;refGene_gn=GRK4;refGene_tx=XM_023253498.1;refGene_gn=GRK4;refGene_tx=XM_023253499.1;refGene_gn=GRK4;refGene_tx=XM_023253500.1;refGene_gn=GRK4;refGene_tx=XM_023253501.1
B1 206039523 206039658
refGene_gn=GRK4;refGene_tx=XM_019829825.2;refGene_gn=GRK4;refGene_tx=XM_023253497.1;refGene_gn=GRK4;refGene_tx=XM_023253498.1;refGene_gn=GRK4;refGene_tx=XM_023253499.1;refGene_gn=GRK4;refGene_tx=XM_023253500.1;refGene_gn=GRK4;refGene_tx=XM_023253501.1
B1 206042787 206042973
refGene_gn=GRK4;refGene_tx=XM_019829825.2;refGene_gn=GRK4;refGene_tx=XM_023253497.1;refGene_gn=GRK4;refGene_tx=XM_023253498.1;refGene_gn=GRK4;refGene_tx=XM_023253499.1;refGene_gn=GRK4;refGene_tx=XM_023253501.1
B1 206043059 206043268
refGene_gn=GRK4;refGene_tx=XM_019829825.2;refGene_gn=GRK4;refGene_tx=XM_023253497.1;refGene_gn=GRK4;refGene_tx=XM_023253498.1;refGene_gn=GRK4;refGene_tx=XM_023253499.1;refGene_gn=GRK4;refGene_tx=XM_023253500.1;refGene_gn=GRK4;refGene_tx=XM_023253501.1
B1 206044237 206044327
refGene_gn=GRK4;refGene_tx=XM_019829825.2;refGene_gn=GRK4;refGene_tx=XM_023253497.1;refGene_gn=GRK4;refGene_tx=XM_023253499.1;refGene_gn=GRK4;refGene_tx=XM_023253500.1;refGene_gn=GRK4;refGene_tx=XM_023253501.1
B1 206058342 206058380
refGene_gn=GRK4;refGene_tx=XM_019829825.2;refGene_gn=GRK4;refGene_tx=XM_023253497.1;refGene_gn=GRK4;refGene_tx=XM_023253498.1;refGene_gn=GRK4;refGene_tx=XM_023253499.1;refGene_gn=GRK4;refGene_tx=XM_023253500.1;refGene_gn=GRK4;refGene_tx=XM_023253501.1
B1 206068297 206068488
refGene_gn=GRK4;refGene_tx=XM_019829825.2;refGene_gn=GRK4;refGene_tx=XM_023253497.1;refGene_gn=GRK4;refGene_tx=XM_023253498.1;refGene_gn=GRK4;refGene_tx=XM_023253499.1;refGene_gn=GRK4;refGene_tx=XM_023253500.1;refGene_gn=GRK4;refGene_tx=XM_023253501.1
B1 206075174 206075315
refGene_gn=GRK4;refGene_tx=XM_019829825.2;refGene_gn=GRK4;refGene_tx=XM_023253497.1;refGene_gn=GRK4;refGene_tx=XM_023253498.1;refGene_gn=GRK4;refGene_tx=XM_023253499.1;refGene_gn=GRK4;refGene_tx=XM_023253500.1;refGene_gn=GRK4;refGene_tx=XM_023253501.1
B1 206079225 206079289
refGene_gn=GRK4;refGene_tx=XM_019829825.2;refGene_gn=GRK4;refGene_tx=XM_023253497.1;refGene_gn=GRK4;refGene_tx=XM_023253498.1;refGene_gn=GRK4;refGene_tx=XM_023253499.1;refGene_gn=GRK4;refGene_tx=XM_023253500.1
B1 206080497 206080590
refGene_gn=GRK4;refGene_tx=XM_019829825.2;refGene_gn=GRK4;refGene_tx=XM_023253497.1;refGene_gn=GRK4;refGene_tx=XM_023253498.1;refGene_gn=GRK4;refGene_tx=XM_023253499.1;refGene_gn=GRK4;refGene_tx=XM_023253500.1
B1 206084776 206084880
refGene_gn=GRK4;refGene_tx=XM_019829825.2;refGene_gn=GRK4;refGene_tx=XM_023253497.1;refGene_gn=GRK4;refGene_tx=XM_023253498.1;refGene_gn=GRK4;refGene_tx=XM_023253499.1;refGene_gn=GRK4;refGene_tx=XM_023253500.1
B1 206085050 206085080

refGene_gn=GRK4;refGene_tx=XM_019829825.2;refGene_gn=GRK4;refGene_tx=XM_023253498.1;refGene_gn=GRK4;refGene_tx=XM_023253499.1;refGene_gn=GRK4;refGene_tx=XM_023253500.1
B1 206091208 206091289
refGene_gn=GRK4;refGene_tx=XM_019829825.2;refGene_gn=GRK4;refGene_tx=XM_023253497.1;refGene_gn=GRK4;refGene_tx=XM_023253498.1;refGene_gn=GRK4;refGene_tx=XM_023253499.1;refGene_gn=GRK4;refGene_tx=XM_023253500.1;refGene_gn=GRK4;refGene_tx=XM_023253501.1
B1 206095091 206095204
refGene_gn=GRK4;refGene_tx=XM_019829825.2;refGene_gn=GRK4;refGene_tx=XM_023253497.1;refGene_gn=GRK4;refGene_tx=XM_023253498.1;refGene_gn=GRK4;refGene_tx=XM_023253499.1;refGene_gn=GRK4;refGene_tx=XM_023253500.1;refGene_gn=GRK4;refGene_tx=XM_023253501.1
B1 206102667 206102763
refGene_gn=GRK4;refGene_tx=XM_019829825.2;refGene_gn=GRK4;refGene_tx=XM_023253497.1;refGene_gn=GRK4;refGene_tx=XM_023253498.1;refGene_gn=GRK4;refGene_tx=XM_023253500.1;refGene_gn=GRK4;refGene_tx=XM_023253501.1
B1 206132412 206132464
refGene_gn=GRK4;refGene_tx=XM_019829825.2;refGene_gn=GRK4;refGene_tx=XM_023253497.1;refGene_gn=GRK4;refGene_tx=XM_023253498.1;refGene_gn=GRK4;refGene_tx=XM_023253499.1;refGene_gn=GRK4;refGene_tx=XM_023253500.1;refGene_gn=GRK4;refGene_tx=XM_023253501.1
B1 206133368 206133563
refGene_gn=NOP14;refGene_tx=XM_003985547.5
B1 206136603 206136738
refGene_gn=NOP14;refGene_tx=XM_003985547.5;refGene_gn=NOP14;refGene_tx=XM_019829832.2
B1 206137257 206137399
refGene_gn=NOP14;refGene_tx=XM_003985547.5;refGene_gn=NOP14;refGene_tx=XM_019829832.2
B1 206141839 206141982
refGene_gn=NOP14;refGene_tx=XM_003985547.5;refGene_gn=NOP14;refGene_tx=XM_019829832.2
B1 206142462 206142597
refGene_gn=NOP14;refGene_tx=XM_003985547.5;refGene_gn=NOP14;refGene_tx=XM_019829832.2
B1 206143584 206143707
refGene_gn=NOP14;refGene_tx=XM_003985547.5;refGene_gn=NOP14;refGene_tx=XM_019829832.2
B1 206144909 206145041
refGene_gn=NOP14;refGene_tx=XM_003985547.5;refGene_gn=NOP14;refGene_tx=XM_019829832.2
B1 206145882 206146132
refGene_gn=NOP14;refGene_tx=XM_003985547.5;refGene_gn=NOP14;refGene_tx=XM_019829832.2
B1 206147061 206147192
refGene_gn=NOP14;refGene_tx=XM_003985547.5;refGene_gn=NOP14;refGene_tx=XM_019829832.2
B1 206147781 206147867
refGene_gn=NOP14;refGene_tx=XM_003985547.5;refGene_gn=NOP14;refGene_tx

=XM_019829832.2
B1 206148878 206149014
refGene_gn=NOP14;refGene_tx=XM_003985547.5;refGene_gn=NOP14;refGene_tx
=XM_019829832.2
B1 206150957 206151059
refGene_gn=NOP14;refGene_tx=XM_003985547.5;refGene_gn=NOP14;refGene_tx
=XM_019829832.2
B1 206151530 206151684
refGene_gn=NOP14;refGene_tx=XM_003985547.5;refGene_gn=NOP14;refGene_tx
=XM_019829832.2
B1 206153301 206153461
refGene_gn=NOP14;refGene_tx=XM_003985547.5;refGene_gn=NOP14;refGene_tx
=XM_019829832.2
B1 206155624 206155772
refGene_gn=NOP14;refGene_tx=XM_003985547.5;refGene_gn=NOP14;refGene_tx
=XM_019829832.2
B1 206157455 206157574
refGene_gn=NOP14;refGene_tx=XM_003985547.5;refGene_gn=NOP14;refGene_tx
=XM_019829832.2
B1 206157723 206157879
refGene_gn=NOP14;refGene_tx=XM_003985547.5;refGene_gn=NOP14;refGene_tx
=XM_019829832.2
B1 206158046 206158146
refGene_gn=NOP14;refGene_tx=XM_003985547.5;refGene_gn=NOP14;refGene_tx
=XM_019829832.2
B1 206159564 206159993
refGene_gn=MFSD10;refGene_tx=XM_023253505.1;refGene_gn=MFSD10;refGene_
tx=XM_023253506.1;refGene_gn=MFSD10;refGene_tx=XM_023253511.1;refGene_
gn=MFSD10;refGene_tx=XM_023253514.1
B1 206160489 206160751
refGene_gn=MFSD10;refGene_tx=XM_023253505.1;refGene_gn=MFSD10;refGene_
tx=XM_023253506.1;refGene_gn=MFSD10;refGene_tx=XM_023253507.1;refGene_
gn=MFSD10;refGene_tx=XM_023253508.1;refGene_gn=MFSD10;refGene_tx=XM_02
3253509.1;refGene_gn=MFSD10;refGene_tx=XM_023253510.1;refGene_gn=MFSD1
0;refGene_tx=XM_023253511.1;refGene_gn=MFSD10;refGene_tx=XM_023253512.
1;refGene_gn=MFSD10;refGene_tx=XM_023253513.1;refGene_gn=MFSD10;refGen
e_tx=XM_023253514.1
B1 206160861 206160958
refGene_gn=MFSD10;refGene_tx=XM_023253505.1;refGene_gn=MFSD10;refGene_
tx=XM_023253506.1;refGene_gn=MFSD10;refGene_tx=XM_023253507.1;refGene_
gn=MFSD10;refGene_tx=XM_023253508.1;refGene_gn=MFSD10;refGene_tx=XM_02
3253509.1;refGene_gn=MFSD10;refGene_tx=XM_023253510.1;refGene_gn=MFSD1
0;refGene_tx=XM_023253511.1;refGene_gn=MFSD10;refGene_tx=XM_023253512.
1;refGene_gn=MFSD10;refGene_tx=XM_023253513.1;refGene_gn=MFSD10;refGen
e_tx=XM_023253514.1
B1 206161125 206161229
refGene_gn=MFSD10;refGene_tx=XM_023253505.1;refGene_gn=MFSD10;refGene_
tx=XM_023253506.1;refGene_gn=MFSD10;refGene_tx=XM_023253507.1;refGene_
gn=MFSD10;refGene_tx=XM_023253508.1;refGene_gn=MFSD10;refGene_tx=XM_02
3253509.1;refGene_gn=MFSD10;refGene_tx=XM_023253510.1;refGene_gn=MFSD1

0;refGene_tx=XM_023253511.1;refGene_gn=MFSD10;refGene_tx=XM_023253512.
1;refGene_gn=MFSD10;refGene_tx=XM_023253513.1;refGene_gn=MFSD10;refGen
e_tx=XM_023253514.1
B1 206161485 206161644
refGene_gn=MFSD10;refGene_tx=XM_023253505.1;refGene_gn=MFSD10;refGene_
tx=XM_023253506.1;refGene_gn=MFSD10;refGene_tx=XM_023253507.1;refGene_
gn=MFSD10;refGene_tx=XM_023253508.1;refGene_gn=MFSD10;refGene_tx=XM_02
3253509.1;refGene_gn=MFSD10;refGene_tx=XM_023253510.1;refGene_gn=MFSD1
0;refGene_tx=XM_023253511.1;refGene_gn=MFSD10;refGene_tx=XM_023253512.
1;refGene_gn=MFSD10;refGene_tx=XM_023253513.1;refGene_gn=MFSD10;refGen
e_tx=XM_023253514.1
B1 206161725 206161892
refGene_gn=MFSD10;refGene_tx=XM_023253505.1;refGene_gn=MFSD10;refGene_
tx=XM_023253506.1;refGene_gn=MFSD10;refGene_tx=XM_023253507.1;refGene_
gn=MFSD10;refGene_tx=XM_023253508.1;refGene_gn=MFSD10;refGene_tx=XM_02
3253509.1;refGene_gn=MFSD10;refGene_tx=XM_023253510.1;refGene_gn=MFSD1
0;refGene_tx=XM_023253511.1;refGene_gn=MFSD10;refGene_tx=XM_023253512.
1;refGene_gn=MFSD10;refGene_tx=XM_023253513.1;refGene_gn=MFSD10;refGen
e_tx=XM_023253514.1
B1 206162128 206162237
refGene_gn=MFSD10;refGene_tx=XM_023253505.1;refGene_gn=MFSD10;refGene_
tx=XM_023253506.1;refGene_gn=MFSD10;refGene_tx=XM_023253507.1;refGene_
gn=MFSD10;refGene_tx=XM_023253508.1;refGene_gn=MFSD10;refGene_tx=XM_02
3253509.1;refGene_gn=MFSD10;refGene_tx=XM_023253510.1;refGene_gn=MFSD1
0;refGene_tx=XM_023253511.1;refGene_gn=MFSD10;refGene_tx=XM_023253512.
1;refGene_gn=MFSD10;refGene_tx=XM_023253513.1;refGene_gn=MFSD10;refGen
e_tx=XM_023253514.1
B1 206162320 206162432
refGene_gn=MFSD10;refGene_tx=XM_023253505.1;refGene_gn=MFSD10;refGene_
tx=XM_023253506.1;refGene_gn=MFSD10;refGene_tx=XM_023253507.1;refGene_
gn=MFSD10;refGene_tx=XM_023253508.1;refGene_gn=MFSD10;refGene_tx=XM_02
3253509.1;refGene_gn=MFSD10;refGene_tx=XM_023253510.1;refGene_gn=MFSD1
0;refGene_tx=XM_023253511.1;refGene_gn=MFSD10;refGene_tx=XM_023253512.
1;refGene_gn=MFSD10;refGene_tx=XM_023253513.1;refGene_gn=MFSD10;refGen
e_tx=XM_023253514.1
B1 206162598 206162707
refGene_gn=MFSD10;refGene_tx=XM_023253505.1;refGene_gn=MFSD10;refGene_
tx=XM_023253506.1;refGene_gn=MFSD10;refGene_tx=XM_023253507.1;refGene_
gn=MFSD10;refGene_tx=XM_023253508.1;refGene_gn=MFSD10;refGene_tx=XM_02
3253509.1;refGene_gn=MFSD10;refGene_tx=XM_023253510.1;refGene_gn=MFSD1
0;refGene_tx=XM_023253511.1;refGene_gn=MFSD10;refGene_tx=XM_023253512.
1;refGene_gn=MFSD10;refGene_tx=XM_023253513.1
B1 206162815 206162906
refGene_gn=MFSD10;refGene_tx=XM_023253505.1;refGene_gn=MFSD10;refGene_
tx=XM_023253506.1;refGene_gn=MFSD10;refGene_tx=XM_023253507.1;refGene_
gn=MFSD10;refGene_tx=XM_023253508.1;refGene_gn=MFSD10;refGene_tx=XM_02
3253509.1;refGene_gn=MFSD10;refGene_tx=XM_023253510.1;refGene_gn=MFSD1
0;refGene_tx=XM_023253511.1;refGene_gn=MFSD10;refGene_tx=XM_023253512.
1;refGene_gn=MFSD10;refGene_tx=XM_023253513.1
B1 206163029 206163253

refGene_gn=MFSD10;refGene_tx=XM_023253505.1;refGene_gn=MFSD10;refGene_tx=XM_023253506.1;refGene_gn=MFSD10;refGene_tx=XM_023253507.1;refGene_gn=MFSD10;refGene_tx=XM_023253508.1;refGene_gn=MFSD10;refGene_tx=XM_023253509.1;refGene_gn=MFSD10;refGene_tx=XM_023253510.1;refGene_gn=MFSD10;refGene_tx=XM_023253511.1;refGene_gn=MFSD10;refGene_tx=XM_023253512.1;refGene_gn=MFSD10;refGene_tx=XM_023253513.1
B1 206163444 206163563
refGene_gn=MFSD10;refGene_tx=XM_023253505.1;refGene_gn=MFSD10;refGene_tx=XM_023253506.1;refGene_gn=MFSD10;refGene_tx=XM_023253507.1;refGene_gn=MFSD10;refGene_tx=XM_023253508.1;refGene_gn=MFSD10;refGene_tx=XM_023253509.1;refGene_gn=MFSD10;refGene_tx=XM_023253510.1;refGene_gn=MFSD10;refGene_tx=XM_023253512.1;refGene_gn=MFSD10;refGene_tx=XM_023253513.1
B1 206165771 206166130
refGene_gn=ADD1;refGene_tx=XM_023253494.1
B1 206168385 206168484
refGene_gn=ADD1;refGene_tx=XM_023253494.1
B1 206171096 206171253
refGene_gn=ADD1;refGene_tx=XM_023253494.1
B1 206175736 206175829
refGene_gn=ADD1;refGene_tx=XM_023253494.1
B1 206176565 206176655
refGene_gn=ADD1;refGene_tx=XM_023253494.1
B1 206177313 206177415
refGene_gn=ADD1;refGene_tx=XM_023253494.1
B1 206179948 206180200
refGene_gn=ADD1;refGene_tx=XM_023253494.1
B1 206185029 206185206
refGene_gn=ADD1;refGene_tx=XM_023253494.1
B1 206185927 206186026
refGene_gn=ADD1;refGene_tx=XM_023253494.1
B1 206186132 206186276
refGene_gn=ADD1;refGene_tx=XM_023253494.1
B1 206188249 206188342
refGene_gn=ADD1;refGene_tx=XM_023253494.1
B1 206188343 206188400
refGene_gn=ADD1;refGene_tx=XM_023253494.1
B1 206188642 206188723
refGene_gn=ADD1;refGene_tx=XM_023253494.1
B1 206197329 206197481
refGene_gn=ADD1;refGene_tx=XM_023253494.1
B1 206205379 206205542
refGene_gn=ADD1;refGene_tx=XM_023253494.1
B1 206210390 206210585
refGene_gn=ADD1;refGene_tx=XM_023253494.1
B1 206265823 206265961
refGene_gn=SH3BP2;refGene_tx=XM_003985541.5;refGene_gn=SH3BP2;refGene_tx=XM_023253519.1;refGene_gn=SH3BP2;refGene_tx=XM_023253520.1;refGene_gn=SH3BP2;refGene_tx=XM_023253521.1;refGene_gn=SH3BP2;refGene_tx=XM_023253522.1

B1	206266182	206266242
refGene_gn=SH3BP2;refGene_tx=XM_003985541.5;refGene_gn=SH3BP2;refGene_tx=XM_023253519.1;refGene_gn=SH3BP2;refGene_tx=XM_023253520.1;refGene_gn=SH3BP2;refGene_tx=XM_023253521.1;refGene_gn=SH3BP2;refGene_tx=XM_023253522.1
B1	206267407	206267489
refGene_gn=SH3BP2;refGene_tx=XM_003985541.5;refGene_gn=SH3BP2;refGene_tx=XM_023253519.1;refGene_gn=SH3BP2;refGene_tx=XM_023253520.1;refGene_gn=SH3BP2;refGene_tx=XM_023253521.1;refGene_gn=SH3BP2;refGene_tx=XM_023253522.1
B1	206267848	206267904
refGene_gn=SH3BP2;refGene_tx=XM_003985541.5;refGene_gn=SH3BP2;refGene_tx=XM_023253519.1;refGene_gn=SH3BP2;refGene_tx=XM_023253520.1;refGene_gn=SH3BP2;refGene_tx=XM_023253521.1;refGene_gn=SH3BP2;refGene_tx=XM_023253522.1
B1	206268143	206268255
refGene_gn=SH3BP2;refGene_tx=XM_003985541.5;refGene_gn=SH3BP2;refGene_tx=XM_023253519.1;refGene_gn=SH3BP2;refGene_tx=XM_023253520.1;refGene_gn=SH3BP2;refGene_tx=XM_023253521.1;refGene_gn=SH3BP2;refGene_tx=XM_023253522.1
B1	206269617	206270269
refGene_gn=SH3BP2;refGene_tx=XM_003985541.5;refGene_gn=SH3BP2;refGene_tx=XM_023253519.1;refGene_gn=SH3BP2;refGene_tx=XM_023253520.1;refGene_gn=SH3BP2;refGene_tx=XM_023253521.1;refGene_gn=SH3BP2;refGene_tx=XM_023253522.1
B1	206271791	206271860
refGene_gn=SH3BP2;refGene_tx=XM_003985541.5;refGene_gn=SH3BP2;refGene_tx=XM_023253519.1;refGene_gn=SH3BP2;refGene_tx=XM_023253520.1;refGene_gn=SH3BP2;refGene_tx=XM_023253521.1;refGene_gn=SH3BP2;refGene_tx=XM_023253522.1
B1	206272163	206272252
refGene_gn=SH3BP2;refGene_tx=XM_003985541.5;refGene_gn=SH3BP2;refGene_tx=XM_023253519.1;refGene_gn=SH3BP2;refGene_tx=XM_023253520.1;refGene_gn=SH3BP2;refGene_tx=XM_023253521.1;refGene_gn=SH3BP2;refGene_tx=XM_023253522.1
B1	206273366	206273437
refGene_gn=SH3BP2;refGene_tx=XM_003985541.5;refGene_gn=SH3BP2;refGene_tx=XM_023253519.1;refGene_gn=SH3BP2;refGene_tx=XM_023253520.1;refGene_gn=SH3BP2;refGene_tx=XM_023253521.1;refGene_gn=SH3BP2;refGene_tx=XM_023253522.1
B1	206273726	206273844
refGene_gn=SH3BP2;refGene_tx=XM_003985541.5;refGene_gn=SH3BP2;refGene_tx=XM_023253519.1;refGene_gn=SH3BP2;refGene_tx=XM_023253520.1;refGene_gn=SH3BP2;refGene_tx=XM_023253521.1;refGene_gn=SH3BP2;refGene_tx=XM_023253522.1
B1	206274561	206274664
refGene_gn=SH3BP2;refGene_tx=XM_003985541.5;refGene_gn=SH3BP2;refGene_tx=XM_023253519.1;refGene_gn=SH3BP2;refGene_tx=XM_023253520.1;refGene_gn=SH3BP2;refGene_tx=XM_023253521.1;refGene_gn=SH3BP2;refGene_tx=XM_023253522.1

B1	206275845	206275985
refGene_gn=SH3BP2;refGene_tx=XM_003985541.5;refGene_gn=SH3BP2;refGene_tx=XM_023253519.1;refGene_gn=SH3BP2;refGene_tx=XM_023253520.1;refGene_gn=SH3BP2;refGene_tx=XM_023253521.1;refGene_gn=SH3BP2;refGene_tx=XM_023253522.1
B1	206277784	206277900
refGene_gn=SH3BP2;refGene_tx=XM_023253519.1
B1	206283289	206283372
refGene_gn=SH3BP2;refGene_tx=XM_003985541.5
B1	206319954	206320233
refGene_gn=TNIP2;refGene_tx=XM_023253523.1;refGene_gn=TNIP2;refGene_tx=XM_023253525.1
B1	206336778	206336832
refGene_gn=TNIP2;refGene_tx=XM_019829839.2
B1	206348714	206349002
refGene_gn=TNIP2;refGene_tx=XM_019829839.2;refGene_gn=TNIP2;refGene_tx=XM_023253523.1;refGene_gn=TNIP2;refGene_tx=XM_023253525.1
B1	206350458	206350548
refGene_gn=TNIP2;refGene_tx=XM_019829839.2;refGene_gn=TNIP2;refGene_tx=XM_023253523.1;refGene_gn=TNIP2;refGene_tx=XM_023253525.1
B1	206351276	206351543
refGene_gn=TNIP2;refGene_tx=XM_019829839.2;refGene_gn=TNIP2;refGene_tx=XM_023253523.1;refGene_gn=TNIP2;refGene_tx=XM_023253525.1
B1	206351659	206351779
refGene_gn=TNIP2;refGene_tx=XM_019829839.2;refGene_gn=TNIP2;refGene_tx=XM_023253523.1;refGene_gn=TNIP2;refGene_tx=XM_023253525.1
B1	206353451	206353718
refGene_gn=TNIP2;refGene_tx=XM_019829839.2;refGene_gn=TNIP2;refGene_tx=XM_023253523.1;refGene_gn=TNIP2;refGene_tx=XM_023253525.1
B1	206363763	206363857
refGene_gn=FAM193A;refGene_tx=XM_023253469.1;refGene_gn=FAM193A;refGene_tx=XM_023253470.1;refGene_gn=FAM193A;refGene_tx=XM_023253471.1;refGene_gn=FAM193A;refGene_tx=XM_023253472.1;refGene_gn=FAM193A;refGene_tx=XM_023253473.1;refGene_gn=FAM193A;refGene_tx=XM_023253474.1;refGene_gn=FAM193A;refGene_tx=XM_023253475.1;refGene_gn=FAM193A;refGene_tx=XM_023253476.1
B1	206373998	206374080
refGene_gn=FAM193A;refGene_tx=XM_023253469.1;refGene_gn=FAM193A;refGene_tx=XM_023253470.1;refGene_gn=FAM193A;refGene_tx=XM_023253471.1;refGene_gn=FAM193A;refGene_tx=XM_023253472.1;refGene_gn=FAM193A;refGene_tx=XM_023253473.1;refGene_gn=FAM193A;refGene_tx=XM_023253474.1;refGene_gn=FAM193A;refGene_tx=XM_023253475.1;refGene_gn=FAM193A;refGene_tx=XM_023253476.1
B1	206381446	206382278
refGene_gn=FAM193A;refGene_tx=XM_023253469.1;refGene_gn=FAM193A;refGene_tx=XM_023253470.1;refGene_gn=FAM193A;refGene_tx=XM_023253471.1;refGene_gn=FAM193A;refGene_tx=XM_023253472.1;refGene_gn=FAM193A;refGene_tx=XM_023253473.1;refGene_gn=FAM193A;refGene_tx=XM_023253474.1;refGene_gn=FAM193A;refGene_tx=XM_023253475.1;refGene_gn=FAM193A;refGene_tx=XM_023253476.1

B1	206384644	206384875
refGene_gn=FAM193A;refGene_tx=XM_023253469.1;refGene_gn=FAM193A;refGen
e_tx=XM_023253470.1;refGene_gn=FAM193A;refGene_tx=XM_023253471.1;refGe
ne_gn=FAM193A;refGene_tx=XM_023253472.1;refGene_gn=FAM193A;refGene_tx=
XM_023253473.1;refGene_gn=FAM193A;refGene_tx=XM_023253474.1;refGene_gn
=FAM193A;refGene_tx=XM_023253475.1;refGene_gn=FAM193A;refGene_tx=XM_02
3253476.1
B1	206385870	206386054
refGene_gn=FAM193A;refGene_tx=XM_023253469.1;refGene_gn=FAM193A;refGen
e_tx=XM_023253470.1;refGene_gn=FAM193A;refGene_tx=XM_023253471.1;refGe
ne_gn=FAM193A;refGene_tx=XM_023253472.1;refGene_gn=FAM193A;refGene_tx=
XM_023253473.1;refGene_gn=FAM193A;refGene_tx=XM_023253474.1;refGene_gn
=FAM193A;refGene_tx=XM_023253475.1;refGene_gn=FAM193A;refGene_tx=XM_02
3253476.1
B1	206386933	206387216
refGene_gn=FAM193A;refGene_tx=XM_023253469.1;refGene_gn=FAM193A;refGen
e_tx=XM_023253470.1;refGene_gn=FAM193A;refGene_tx=XM_023253471.1;refGe
ne_gn=FAM193A;refGene_tx=XM_023253472.1;refGene_gn=FAM193A;refGene_tx=
XM_023253473.1;refGene_gn=FAM193A;refGene_tx=XM_023253474.1;refGene_gn
=FAM193A;refGene_tx=XM_023253475.1;refGene_gn=FAM193A;refGene_tx=XM_02
3253476.1
B1	206388482	206388755
refGene_gn=FAM193A;refGene_tx=XM_023253469.1;refGene_gn=FAM193A;refGen
e_tx=XM_023253470.1;refGene_gn=FAM193A;refGene_tx=XM_023253471.1;refGe
ne_gn=FAM193A;refGene_tx=XM_023253472.1;refGene_gn=FAM193A;refGene_tx=
XM_023253473.1;refGene_gn=FAM193A;refGene_tx=XM_023253474.1;refGene_gn
=FAM193A;refGene_tx=XM_023253475.1;refGene_gn=FAM193A;refGene_tx=XM_02
3253476.1
B1	206389679	206389878
refGene_gn=FAM193A;refGene_tx=XM_023253469.1;refGene_gn=FAM193A;refGen
e_tx=XM_023253470.1;refGene_gn=FAM193A;refGene_tx=XM_023253471.1;refGe
ne_gn=FAM193A;refGene_tx=XM_023253472.1;refGene_gn=FAM193A;refGene_tx=
XM_023253473.1;refGene_gn=FAM193A;refGene_tx=XM_023253474.1;refGene_gn
=FAM193A;refGene_tx=XM_023253475.1;refGene_gn=FAM193A;refGene_tx=XM_02
3253476.1
B1	206399487	206399739
refGene_gn=FAM193A;refGene_tx=XM_023253469.1;refGene_gn=FAM193A;refGen
e_tx=XM_023253470.1;refGene_gn=FAM193A;refGene_tx=XM_023253471.1;refGe
ne_gn=FAM193A;refGene_tx=XM_023253472.1;refGene_gn=FAM193A;refGene_tx=
XM_023253473.1;refGene_gn=FAM193A;refGene_tx=XM_023253474.1;refGene_gn
=FAM193A;refGene_tx=XM_023253475.1;refGene_gn=FAM193A;refGene_tx=XM_02
3253476.1
B1	206404762	206404942
refGene_gn=FAM193A;refGene_tx=XM_023253469.1;refGene_gn=FAM193A;refGen
e_tx=XM_023253470.1;refGene_gn=FAM193A;refGene_tx=XM_023253471.1;refGe
ne_gn=FAM193A;refGene_tx=XM_023253472.1;refGene_gn=FAM193A;refGene_tx=
XM_023253473.1;refGene_gn=FAM193A;refGene_tx=XM_023253474.1;refGene_gn
=FAM193A;refGene_tx=XM_023253475.1;refGene_gn=FAM193A;refGene_tx=XM_02
3253476.1
B1	206405058	206405212

refGene_gn=FAM193A;refGene_tx=XM_023253469.1;refGene_gn=FAM193A;refGene_tx=XM_023253470.1;refGene_gn=FAM193A;refGene_tx=XM_023253471.1;refGene_gn=FAM193A;refGene_tx=XM_023253472.1;refGene_gn=FAM193A;refGene_tx=XM_023253473.1;refGene_gn=FAM193A;refGene_tx=XM_023253474.1;refGene_gn=FAM193A;refGene_tx=XM_023253475.1;refGene_gn=FAM193A;refGene_tx=XM_023253476.1
B1 206407615 206407858
refGene_gn=FAM193A;refGene_tx=XM_023253469.1;refGene_gn=FAM193A;refGene_tx=XM_023253470.1;refGene_gn=FAM193A;refGene_tx=XM_023253471.1;refGene_gn=FAM193A;refGene_tx=XM_023253472.1;refGene_gn=FAM193A;refGene_tx=XM_023253473.1;refGene_gn=FAM193A;refGene_tx=XM_023253474.1;refGene_gn=FAM193A;refGene_tx=XM_023253475.1;refGene_gn=FAM193A;refGene_tx=XM_023253476.1
B1 206407991 206408104
refGene_gn=FAM193A;refGene_tx=XM_023253469.1;refGene_gn=FAM193A;refGene_tx=XM_023253470.1;refGene_gn=FAM193A;refGene_tx=XM_023253471.1;refGene_gn=FAM193A;refGene_tx=XM_023253472.1;refGene_gn=FAM193A;refGene_tx=XM_023253473.1;refGene_gn=FAM193A;refGene_tx=XM_023253474.1;refGene_gn=FAM193A;refGene_tx=XM_023253475.1;refGene_gn=FAM193A;refGene_tx=XM_023253476.1
B1 206409713 206409791
refGene_gn=FAM193A;refGene_tx=XM_023253469.1;refGene_gn=FAM193A;refGene_tx=XM_023253470.1;refGene_gn=FAM193A;refGene_tx=XM_023253471.1;refGene_gn=FAM193A;refGene_tx=XM_023253472.1;refGene_gn=FAM193A;refGene_tx=XM_023253473.1;refGene_gn=FAM193A;refGene_tx=XM_023253474.1;refGene_gn=FAM193A;refGene_tx=XM_023253475.1;refGene_gn=FAM193A;refGene_tx=XM_023253476.1
B1 206416685 206416833
refGene_gn=FAM193A;refGene_tx=XM_023253469.1;refGene_gn=FAM193A;refGene_tx=XM_023253470.1;refGene_gn=FAM193A;refGene_tx=XM_023253471.1;refGene_gn=FAM193A;refGene_tx=XM_023253472.1;refGene_gn=FAM193A;refGene_tx=XM_023253473.1;refGene_gn=FAM193A;refGene_tx=XM_023253474.1;refGene_gn=FAM193A;refGene_tx=XM_023253475.1;refGene_gn=FAM193A;refGene_tx=XM_023253476.1
B1 206421116 206421241
refGene_gn=FAM193A;refGene_tx=XM_023253469.1;refGene_gn=FAM193A;refGene_tx=XM_023253470.1;refGene_gn=FAM193A;refGene_tx=XM_023253471.1;refGene_gn=FAM193A;refGene_tx=XM_023253472.1;refGene_gn=FAM193A;refGene_tx=XM_023253473.1;refGene_gn=FAM193A;refGene_tx=XM_023253474.1;refGene_gn=FAM193A;refGene_tx=XM_023253475.1;refGene_gn=FAM193A;refGene_tx=XM_023253476.1
B1 206429314 206429549
refGene_gn=FAM193A;refGene_tx=XM_023253469.1;refGene_gn=FAM193A;refGene_tx=XM_023253470.1;refGene_gn=FAM193A;refGene_tx=XM_023253471.1;refGene_gn=FAM193A;refGene_tx=XM_023253472.1;refGene_gn=FAM193A;refGene_tx=XM_023253473.1;refGene_gn=FAM193A;refGene_tx=XM_023253474.1;refGene_gn=FAM193A;refGene_tx=XM_023253475.1;refGene_gn=FAM193A;refGene_tx=XM_023253476.1
B1 206432781 206432949
refGene_gn=FAM193A;refGene_tx=XM_023253469.1;refGene_gn=FAM193A;refGen

e_tx=XM_023253470.1;refGene_gn=FAM193A;refGene_tx=XM_023253471.1;refGene_gn=FAM193A;refGene_tx=XM_023253472.1;refGene_gn=FAM193A;refGene_tx=XM_023253473.1;refGene_gn=FAM193A;refGene_tx=XM_023253474.1;refGene_gn=FAM193A;refGene_tx=XM_023253475.1;refGene_gn=FAM193A;refGene_tx=XM_023253476.1
B1 206433435 206433593
refGene_gn=FAM193A;refGene_tx=XM_023253469.1;refGene_gn=FAM193A;refGene_tx=XM_023253470.1;refGene_gn=FAM193A;refGene_tx=XM_023253471.1;refGene_gn=FAM193A;refGene_tx=XM_023253472.1;refGene_gn=FAM193A;refGene_tx=XM_023253473.1;refGene_gn=FAM193A;refGene_tx=XM_023253474.1;refGene_gn=FAM193A;refGene_tx=XM_023253476.1
B1 206458507 206458735
refGene_gn=FAM193A;refGene_tx=XM_023253469.1;refGene_gn=FAM193A;refGene_tx=XM_023253470.1;refGene_gn=FAM193A;refGene_tx=XM_023253471.1;refGene_gn=FAM193A;refGene_tx=XM_023253472.1;refGene_gn=FAM193A;refGene_tx=XM_023253473.1;refGene_gn=FAM193A;refGene_tx=XM_023253474.1;refGene_gn=FAM193A;refGene_tx=XM_023253476.1
B1 206549540 206549690
refGene_gn=RNF4;refGene_tx=XM_003985539.5;refGene_gn=RNF4;refGene_tx=XM_006931241.3;refGene_gn=RNF4;refGene_tx=XM_006931242.4;refGene_gn=RNF4;refGene_tx=XM_019829842.2;refGene_gn=RNF4;refGene_tx=XM_023253532.1
B1 206550271 206550320
refGene_gn=RNF4;refGene_tx=XM_003985539.5;refGene_gn=RNF4;refGene_tx=XM_006931241.3;refGene_gn=RNF4;refGene_tx=XM_006931242.4;refGene_gn=RNF4;refGene_tx=XM_019829842.2;refGene_gn=RNF4;refGene_tx=XM_023253532.1
B1 206550915 206551075
refGene_gn=RNF4;refGene_tx=XM_003985539.5;refGene_gn=RNF4;refGene_tx=XM_006931241.3;refGene_gn=RNF4;refGene_tx=XM_006931242.4;refGene_gn=RNF4;refGene_tx=XM_019829842.2;refGene_gn=RNF4;refGene_tx=XM_023253532.1
B1 206551486 206551496
refGene_gn=RNF4;refGene_tx=XM_003985539.5;refGene_gn=RNF4;refGene_tx=XM_006931241.3;refGene_gn=RNF4;refGene_tx=XM_006931242.4;refGene_gn=RNF4;refGene_tx=XM_019829842.2;refGene_gn=RNF4;refGene_tx=XM_023253532.1
B1 206555524 206555593
refGene_gn=RNF4;refGene_tx=XM_006931241.3;refGene_gn=RNF4;refGene_tx=XM_006931242.4;refGene_gn=RNF4;refGene_tx=XM_019829842.2;refGene_gn=RNF4;refGene_tx=XM_023253532.1
B1 206560078 206560158
refGene_gn=RNF4;refGene_tx=XM_003985539.5;refGene_gn=RNF4;refGene_tx=XM_006931241.3;refGene_gn=RNF4;refGene_tx=XM_006931242.4;refGene_gn=RNF4;refGene_tx=XM_019829842.2;refGene_gn=RNF4;refGene_tx=XM_023253532.1
B1 206562413 206562528
refGene_gn=RNF4;refGene_tx=XM_003985539.5;refGene_gn=RNF4;refGene_tx=XM_006931241.3;refGene_gn=RNF4;refGene_tx=XM_006931242.4;refGene_gn=RNF4;refGene_tx=XM_019829842.2;refGene_gn=RNF4;refGene_tx=XM_023253532.1
B1 206565619 206565628
refGene_gn=RNF4;refGene_tx=XM_003985539.5;refGene_gn=RNF4;refGene_tx=XM_006931241.3;refGene_gn=RNF4;refGene_tx=XM_019829842.2;refGene_gn=RNF4;refGene_tx=XM_023253532.1
B1 206584202 206584319

refGene_gn=CFAP99;refGene_tx=XM_023253502.1;refGene_gn=CFAP99;refGene_tx=XM_023253504.1
B1 206584469 206584632
refGene_gn=CFAP99;refGene_tx=XM_023253502.1;refGene_gn=CFAP99;refGene_tx=XM_023253504.1
B1 206585900 206585967
refGene_gn=CFAP99;refGene_tx=XM_023253503.1
B1 206586257 206586463
refGene_gn=CFAP99;refGene_tx=XM_023253502.1;refGene_gn=CFAP99;refGene_tx=XM_023253503.1;refGene_gn=CFAP99;refGene_tx=XM_023253504.1
B1 206587308 206587460
refGene_gn=CFAP99;refGene_tx=XM_023253502.1;refGene_gn=CFAP99;refGene_tx=XM_023253503.1;refGene_gn=CFAP99;refGene_tx=XM_023253504.1
B1 206587657 206587799
refGene_gn=CFAP99;refGene_tx=XM_023253502.1;refGene_gn=CFAP99;refGene_tx=XM_023253503.1;refGene_gn=CFAP99;refGene_tx=XM_023253504.1
B1 206595217 206595422
refGene_gn=CFAP99;refGene_tx=XM_023253502.1;refGene_gn=CFAP99;refGene_tx=XM_023253503.1;refGene_gn=CFAP99;refGene_tx=XM_023253504.1
B1 206595943 206596032
refGene_gn=CFAP99;refGene_tx=XM_023253502.1;refGene_gn=CFAP99;refGene_tx=XM_023253503.1;refGene_gn=CFAP99;refGene_tx=XM_023253504.1
B1 206596273 206596345
refGene_gn=CFAP99;refGene_tx=XM_023253502.1;refGene_gn=CFAP99;refGene_tx=XM_023253503.1;refGene_gn=CFAP99;refGene_tx=XM_023253504.1
B1 206597237 206597312
refGene_gn=CFAP99;refGene_tx=XM_023253502.1;refGene_gn=CFAP99;refGene_tx=XM_023253503.1;refGene_gn=CFAP99;refGene_tx=XM_023253504.1
B1 206597495 206597576
refGene_gn=CFAP99;refGene_tx=XM_023253502.1;refGene_gn=CFAP99;refGene_tx=XM_023253503.1;refGene_gn=CFAP99;refGene_tx=XM_023253504.1
B1 206599206 206599384
refGene_gn=CFAP99;refGene_tx=XM_023253502.1;refGene_gn=CFAP99;refGene_tx=XM_023253503.1
B1 206600659 206600772
refGene_gn=CFAP99;refGene_tx=XM_023253502.1;refGene_gn=CFAP99;refGene_tx=XM_023253503.1
B1 206604200 206604295
refGene_gn=CFAP99;refGene_tx=XM_023253502.1;refGene_gn=CFAP99;refGene_tx=XM_023253503.1;refGene_gn=CFAP99;refGene_tx=XM_023253504.1
B1 206605488 206605633
refGene_gn=CFAP99;refGene_tx=XM_023253502.1;refGene_gn=CFAP99;refGene_tx=XM_023253503.1;refGene_gn=CFAP99;refGene_tx=XM_023253504.1
B1 206615588 206615699
refGene_gn=CFAP99;refGene_tx=XM_023253502.1;refGene_gn=CFAP99;refGene_tx=XM_023253503.1;refGene_gn=CFAP99;refGene_tx=XM_023253504.1
B1 206624876 206624915
refGene_gn=ZFYVE28;refGene_tx=XM_023253481.1;refGene_gn=ZFYVE28;refGene_tx=XM_023253482.1;refGene_gn=ZFYVE28;refGene_tx=XM_023253483.1
B1 206679692 206679833

refGene_gn=ZFYVE28;refGene_tx=XM_023253481.1;refGene_gn=ZFYVE28;refGene_tx=XM_023253482.1;refGene_gn=ZFYVE28;refGene_tx=XM_023253483.1
B1 206689780 206689918
refGene_gn=ZFYVE28;refGene_tx=XM_019829843.2;refGene_gn=ZFYVE28;refGene_tx=XM_023253481.1;refGene_gn=ZFYVE28;refGene_tx=XM_023253482.1;refGene_gn=ZFYVE28;refGene_tx=XM_023253483.1
B1 206691758 206691961
refGene_gn=ZFYVE28;refGene_tx=XM_019829843.2;refGene_gn=ZFYVE28;refGene_tx=XM_023253481.1;refGene_gn=ZFYVE28;refGene_tx=XM_023253482.1;refGene_gn=ZFYVE28;refGene_tx=XM_023253483.1
B1 206693482 206693572
refGene_gn=ZFYVE28;refGene_tx=XM_019829843.2;refGene_gn=ZFYVE28;refGene_tx=XM_023253481.1;refGene_gn=ZFYVE28;refGene_tx=XM_023253482.1;refGene_gn=ZFYVE28;refGene_tx=XM_023253483.1
B1 206696146 206696236
refGene_gn=ZFYVE28;refGene_tx=XM_019829843.2;refGene_gn=ZFYVE28;refGene_tx=XM_019829845.2;refGene_gn=ZFYVE28;refGene_tx=XM_023253481.1;refGene_gn=ZFYVE28;refGene_tx=XM_023253482.1;refGene_gn=ZFYVE28;refGene_tx=XM_023253483.1
B1 206706360 206706462
refGene_gn=ZFYVE28;refGene_tx=XM_019829843.2;refGene_gn=ZFYVE28;refGene_tx=XM_019829845.2;refGene_gn=ZFYVE28;refGene_tx=XM_023253481.1;refGene_gn=ZFYVE28;refGene_tx=XM_023253482.1;refGene_gn=ZFYVE28;refGene_tx=XM_023253483.1
B1 206712816 206714115
refGene_gn=ZFYVE28;refGene_tx=XM_019829843.2;refGene_gn=ZFYVE28;refGene_tx=XM_019829845.2;refGene_gn=ZFYVE28;refGene_tx=XM_023253481.1;refGene_gn=ZFYVE28;refGene_tx=XM_023253482.1;refGene_gn=ZFYVE28;refGene_tx=XM_023253483.1
B1 206733976 206734110
refGene_gn=ZFYVE28;refGene_tx=XM_019829843.2;refGene_gn=ZFYVE28;refGene_tx=XM_019829845.2;refGene_gn=ZFYVE28;refGene_tx=XM_023253481.1;refGene_gn=ZFYVE28;refGene_tx=XM_023253482.1;refGene_gn=ZFYVE28;refGene_tx=XM_023253483.1
B1 206736305 206736955
refGene_gn=ZFYVE28;refGene_tx=XM_019829843.2;refGene_gn=ZFYVE28;refGene_tx=XM_019829845.2;refGene_gn=ZFYVE28;refGene_tx=XM_023253481.1;refGene_gn=ZFYVE28;refGene_tx=XM_023253482.1;refGene_gn=ZFYVE28;refGene_tx=XM_023253483.1
B1 206737610 206737715
refGene_gn=ZFYVE28;refGene_tx=XM_023253483.1
B1 206739877 206739981
refGene_gn=ZFYVE28;refGene_tx=XM_023253483.1
B1 206740517 206740649
refGene_gn=ZFYVE28;refGene_tx=XM_023253483.1
B1 206747329 206747393
refGene_gn=MXD4;refGene_tx=XM_023253533.1;refGene_gn=MXD4;refGene_tx=XM_023253534.1
B1 206747496 206747596
refGene_gn=MXD4;refGene_tx=XM_023253533.1;refGene_gn=MXD4;refGene_tx=X

M_023253534.1
B1	206751185	206751215
refGene_gn=MXD4;refGene_tx=XM_023253533.1
B1	206756913	206757028
refGene_gn=MXD4;refGene_tx=XM_023253533.1;refGene_gn=MXD4;refGene_tx=X
M_023253534.1
B1	206757997	206758160
refGene_gn=MXD4;refGene_tx=XM_023253533.1;refGene_gn=MXD4;refGene_tx=X
M_023253534.1
B1	206758418	206758576
refGene_gn=MXD4;refGene_tx=XM_023253533.1;refGene_gn=MXD4;refGene_tx=X
M_023253534.1
B1	206768021	206768048
refGene_gn=HAUS3;refGene_tx=XM_019829850.2;refGene_gn=HAUS3;refGene_tx
=XM_023253518.1
B1	206768579	206769488
refGene_gn=HAUS3;refGene_tx=XM_011281940.3;refGene_gn=HAUS3;refGene_tx
=XM_011281941.3;refGene_gn=HAUS3;refGene_tx=XM_019829847.2;refGene_gn=
HAUS3;refGene_tx=XM_019829848.2
B1	206770907	206771347
refGene_gn=HAUS3;refGene_tx=XM_011281940.3;refGene_gn=HAUS3;refGene_tx
=XM_011281941.3;refGene_gn=HAUS3;refGene_tx=XM_019829847.2;refGene_gn=
HAUS3;refGene_tx=XM_019829848.2;refGene_gn=HAUS3;refGene_tx=XM_0198298
50.2;refGene_gn=HAUS3;refGene_tx=XM_023253518.1
B1	206772908	206773137
refGene_gn=HAUS3;refGene_tx=XM_011281940.3;refGene_gn=HAUS3;refGene_tx
=XM_011281941.3;refGene_gn=HAUS3;refGene_tx=XM_019829847.2;refGene_gn=
HAUS3;refGene_tx=XM_019829848.2;refGene_gn=HAUS3;refGene_tx=XM_0198298
50.2;refGene_gn=HAUS3;refGene_tx=XM_023253518.1
B1	206773822	206773928
refGene_gn=HAUS3;refGene_tx=XM_019829848.2
B1	206784654	206784888
refGene_gn=HAUS3;refGene_tx=XM_011281940.3;refGene_gn=HAUS3;refGene_tx
=XM_011281941.3;refGene_gn=HAUS3;refGene_tx=XM_019829847.2;refGene_gn=
HAUS3;refGene_tx=XM_019829848.2;refGene_gn=HAUS3;refGene_tx=XM_0198298
50.2;refGene_gn=HAUS3;refGene_tx=XM_023253518.1
B1	206786716	206786926
refGene_gn=POLN;refGene_tx=XM_023253416.1
B1	206801047	206801141
refGene_gn=POLN;refGene_tx=XM_023253416.1
B1	206801143	206801445
refGene_gn=POLN;refGene_tx=XM_023253416.1
B1	206801447	206801615
refGene_gn=POLN;refGene_tx=XM_023253416.1
B1	206804357	206804413
refGene_gn=POLN;refGene_tx=XM_023253416.1
B1	206810788	206810868
refGene_gn=POLN;refGene_tx=XM_023253416.1
B1	206810870	206810900
refGene_gn=POLN;refGene_tx=XM_023253416.1

B1 206815264 206815422
refGene_gn=POLN;refGene_tx=XM_023253416.1
B1 206817992 206818061
refGene_gn=POLN;refGene_tx=XM_023253416.1
B1 206819576 206819637
refGene_gn=POLN;refGene_tx=XM_023253416.1
B1 206820339 206820404
refGene_gn=POLN;refGene_tx=XM_023253416.1
B1 206833359 206833440
refGene_gn=POLN;refGene_tx=XM_023253416.1
B1 206833592 206833781
refGene_gn=POLN;refGene_tx=XM_023253416.1
B1 206840151 206840208
refGene_gn=POLN;refGene_tx=XM_023253416.1
B1 206840968 206841022
refGene_gn=POLN;refGene_tx=XM_023253416.1
B1 206841700 206841766
refGene_gn=POLN;refGene_tx=XM_023253416.1
B1 206873404 206873462
refGene_gn=POLN;refGene_tx=XM_023253416.1
B1 206874629 206874707
refGene_gn=POLN;refGene_tx=XM_023253416.1
B1 206875584 206875699
refGene_gn=POLN;refGene_tx=XM_023253416.1
B1 206930401 206930550
refGene_gn=POLN;refGene_tx=XM_023253416.1
B1 206934402 206934513
refGene_gn=POLN;refGene_tx=XM_023253416.1
B1 206937272 206937311
refGene_gn=POLN;refGene_tx=XM_023253416.1
B1 206937313 206937350
refGene_gn=POLN;refGene_tx=XM_023253416.1
B1 206941571 206941639
refGene_gn=POLN;refGene_tx=XM_023253416.1
B1 206942930 206942992
refGene_gn=POLN;refGene_tx=XM_023253416.1
B1 206943683 206944100
refGene_gn=POLN;refGene_tx=XM_023253416.1
B1 206950539 206950907
refGene_gn=NAT8L;refGene_tx=XM_019829851.2
B1 206953447 206953612
refGene_gn=NAT8L;refGene_tx=XM_019829851.2
B1 206954952 206955328
refGene_gn=NAT8L;refGene_tx=XM_019829851.2
B1 206961459 206961525
refGene_gn=CB1H4orf48;refGene_tx=XM_011281939.2
B1 206962609 206962698
refGene_gn=CB1H4orf48;refGene_tx=XM_011281939.2
B1 206962884 206963017
refGene_gn=CB1H4orf48;refGene_tx=XM_011281939.2

B1 206982696 206982906
refGene_gn=NELFA;refGene_tx=XM_011281938.3
B1 206984007 206984424
refGene_gn=NELFA;refGene_tx=XM_023253515.1;refGene_gn=NELFA;refGene_tx=XM_023253516.1
B1 207001513 207001685
refGene_gn=NELFA;refGene_tx=XM_011281938.3;refGene_gn=NELFA;refGene_tx=XM_023253515.1;refGene_gn=NELFA;refGene_tx=XM_023253516.1
B1 207002866 207003028
refGene_gn=NELFA;refGene_tx=XM_011281938.3;refGene_gn=NELFA;refGene_tx=XM_023253515.1;refGene_gn=NELFA;refGene_tx=XM_023253516.1
B1 207004270 207004360
refGene_gn=NELFA;refGene_tx=XM_011281938.3;refGene_gn=NELFA;refGene_tx=XM_023253515.1;refGene_gn=NELFA;refGene_tx=XM_023253516.1
B1 207005850 207005981
refGene_gn=NELFA;refGene_tx=XM_011281938.3;refGene_gn=NELFA;refGene_tx=XM_023253515.1;refGene_gn=NELFA;refGene_tx=XM_023253516.1
B1 207006076 207006146
refGene_gn=NELFA;refGene_tx=XM_011281938.3;refGene_gn=NELFA;refGene_tx=XM_023253515.1;refGene_gn=NELFA;refGene_tx=XM_023253516.1
B1 207006474 207006563
refGene_gn=NELFA;refGene_tx=XM_011281938.3;refGene_gn=NELFA;refGene_tx=XM_023253515.1;refGene_gn=NELFA;refGene_tx=XM_023253516.1
B1 207007272 207007384
refGene_gn=NELFA;refGene_tx=XM_011281938.3;refGene_gn=NELFA;refGene_tx=XM_023253515.1;refGene_gn=NELFA;refGene_tx=XM_023253516.1
B1 207008168 207008449
refGene_gn=NELFA;refGene_tx=XM_011281938.3;refGene_gn=NELFA;refGene_tx=XM_023253515.1;refGene_gn=NELFA;refGene_tx=XM_023253516.1
B1 207008547 207008651
refGene_gn=NELFA;refGene_tx=XM_011281938.3;refGene_gn=NELFA;refGene_tx=XM_023253515.1;refGene_gn=NELFA;refGene_tx=XM_023253516.1
B1 207008795 207009066
refGene_gn=NELFA;refGene_tx=XM_011281938.3;refGene_gn=NELFA;refGene_tx=XM_023253515.1;refGene_gn=NELFA;refGene_tx=XM_023253516.1
B1 207013529 207013801
refGene_gn=NSD2;refGene_tx=XM_019829852.2
B1 207014641 207014846
refGene_gn=NSD2;refGene_tx=XM_019829852.2
B1 207015797 207015904
refGene_gn=NSD2;refGene_tx=XM_019829852.2
B1 207016027 207016169
refGene_gn=NSD2;refGene_tx=XM_019829852.2
B1 207024003 207024048
refGene_gn=NSD2;refGene_tx=XM_023253477.1
B1 207024164 207024281
refGene_gn=NSD2;refGene_tx=XM_019829852.2;refGene_gn=NSD2;refGene_tx=XM_023253477.1
B1 207025146 207025416
refGene_gn=NSD2;refGene_tx=XM_019829852.2;refGene_gn=NSD2;refGene_tx=X

M_023253477.1
B1 207026626 207026730
refGene_gn=NSD2;refGene_tx=XM_019829852.2;refGene_gn=NSD2;refGene_tx=XM_023253477.1;refGene_gn=NSD2;refGene_tx=XM_023253478.1
B1 207028104 207028310
refGene_gn=NSD2;refGene_tx=XM_019829852.2;refGene_gn=NSD2;refGene_tx=XM_023253477.1;refGene_gn=NSD2;refGene_tx=XM_023253478.1
B1 207028439 207028596
refGene_gn=NSD2;refGene_tx=XM_019829852.2;refGene_gn=NSD2;refGene_tx=XM_023253477.1;refGene_gn=NSD2;refGene_tx=XM_023253478.1
B1 207028940 207029120
refGene_gn=NSD2;refGene_tx=XM_019829852.2;refGene_gn=NSD2;refGene_tx=XM_023253477.1;refGene_gn=NSD2;refGene_tx=XM_023253478.1
B1 207030521 207030722
refGene_gn=NSD2;refGene_tx=XM_019829852.2;refGene_gn=NSD2;refGene_tx=XM_023253477.1;refGene_gn=NSD2;refGene_tx=XM_023253478.1
B1 207031735 207031859
refGene_gn=NSD2;refGene_tx=XM_019829852.2;refGene_gn=NSD2;refGene_tx=XM_023253477.1;refGene_gn=NSD2;refGene_tx=XM_023253478.1
B1 207032170 207032302
refGene_gn=NSD2;refGene_tx=XM_019829852.2;refGene_gn=NSD2;refGene_tx=XM_023253477.1;refGene_gn=NSD2;refGene_tx=XM_023253478.1
B1 207039969 207040032
refGene_gn=NSD2;refGene_tx=XM_023253480.1
B1 207041945 207042044
refGene_gn=NSD2;refGene_tx=XM_023253479.1
B1 207042647 207042769
refGene_gn=NSD2;refGene_tx=XM_019829852.2;refGene_gn=NSD2;refGene_tx=XM_023253477.1;refGene_gn=NSD2;refGene_tx=XM_023253478.1;refGene_gn=NSD2;refGene_tx=XM_023253479.1;refGene_gn=NSD2;refGene_tx=XM_023253480.1
B1 207043907 207043989
refGene_gn=NSD2;refGene_tx=XM_019829852.2;refGene_gn=NSD2;refGene_tx=XM_023253477.1;refGene_gn=NSD2;refGene_tx=XM_023253478.1;refGene_gn=NSD2;refGene_tx=XM_023253479.1;refGene_gn=NSD2;refGene_tx=XM_023253480.1
B1 207046140 207046259
refGene_gn=NSD2;refGene_tx=XM_019829852.2;refGene_gn=NSD2;refGene_tx=XM_023253477.1;refGene_gn=NSD2;refGene_tx=XM_023253478.1;refGene_gn=NSD2;refGene_tx=XM_023253479.1;refGene_gn=NSD2;refGene_tx=XM_023253480.1
B1 207049078 207049223
refGene_gn=NSD2;refGene_tx=XM_019829852.2;refGene_gn=NSD2;refGene_tx=XM_023253477.1;refGene_gn=NSD2;refGene_tx=XM_023253478.1;refGene_gn=NSD2;refGene_tx=XM_023253479.1;refGene_gn=NSD2;refGene_tx=XM_023253480.1
B1 207057425 207057914
refGene_gn=NSD2;refGene_tx=XM_019829852.2;refGene_gn=NSD2;refGene_tx=XM_023253477.1;refGene_gn=NSD2;refGene_tx=XM_023253478.1;refGene_gn=NSD2;refGene_tx=XM_023253479.1;refGene_gn=NSD2;refGene_tx=XM_023253480.1
B1 207058412 207058579
refGene_gn=NSD2;refGene_tx=XM_019829852.2;refGene_gn=NSD2;refGene_tx=XM_023253477.1;refGene_gn=NSD2;refGene_tx=XM_023253478.1;refGene_gn=NSD2;refGene_tx=XM_023253479.1;refGene_gn=NSD2;refGene_tx=XM_023253480.1

B1 207069048 207069211
refGene_gn=NSD2;refGene_tx=XM_019829852.2;refGene_gn=NSD2;refGene_tx=XM_023253477.1;refGene_gn=NSD2;refGene_tx=XM_023253478.1;refGene_gn=NSD2;refGene_tx=XM_023253479.1;refGene_gn=NSD2;refGene_tx=XM_023253480.1
B1 207071320 207071857
refGene_gn=NSD2;refGene_tx=XM_019829852.2;refGene_gn=NSD2;refGene_tx=XM_023253477.1;refGene_gn=NSD2;refGene_tx=XM_023253478.1;refGene_gn=NSD2;refGene_tx=XM_023253479.1;refGene_gn=NSD2;refGene_tx=XM_023253480.1
B1 207118159 207118238
refGene_gn=LETM1;refGene_tx=XM_023253495.1;refGene_gn=LETM1;refGene_tx=XM_023253496.1
B1 207123376 207123437
refGene_gn=LETM1;refGene_tx=XM_023253495.1;refGene_gn=LETM1;refGene_tx=XM_023253496.1
B1 207124943 207124954
refGene_gn=LETM1;refGene_tx=XM_019829855.2
B1 207129682 207130133
refGene_gn=LETM1;refGene_tx=XM_019829855.2;refGene_gn=LETM1;refGene_tx=XM_023253495.1;refGene_gn=LETM1;refGene_tx=XM_023253496.1
B1 207138137 207138281
refGene_gn=LETM1;refGene_tx=XM_019829855.2;refGene_gn=LETM1;refGene_tx=XM_023253495.1;refGene_gn=LETM1;refGene_tx=XM_023253496.1
B1 207138860 207138998
refGene_gn=LETM1;refGene_tx=XM_019829855.2;refGene_gn=LETM1;refGene_tx=XM_023253495.1;refGene_gn=LETM1;refGene_tx=XM_023253496.1
B1 207139668 207139872
refGene_gn=LETM1;refGene_tx=XM_019829855.2;refGene_gn=LETM1;refGene_tx=XM_023253495.1;refGene_gn=LETM1;refGene_tx=XM_023253496.1
B1 207146602 207146722
refGene_gn=LETM1;refGene_tx=XM_019829855.2;refGene_gn=LETM1;refGene_tx=XM_023253495.1;refGene_gn=LETM1;refGene_tx=XM_023253496.1
B1 207147992 207148124
refGene_gn=LETM1;refGene_tx=XM_019829855.2;refGene_gn=LETM1;refGene_tx=XM_023253495.1;refGene_gn=LETM1;refGene_tx=XM_023253496.1
B1 207148468 207148612
refGene_gn=LETM1;refGene_tx=XM_019829855.2;refGene_gn=LETM1;refGene_tx=XM_023253495.1;refGene_gn=LETM1;refGene_tx=XM_023253496.1
B1 207149186 207149321
refGene_gn=LETM1;refGene_tx=XM_019829855.2;refGene_gn=LETM1;refGene_tx=XM_023253495.1;refGene_gn=LETM1;refGene_tx=XM_023253496.1
B1 207152151 207152286
refGene_gn=LETM1;refGene_tx=XM_019829855.2;refGene_gn=LETM1;refGene_tx=XM_023253495.1;refGene_gn=LETM1;refGene_tx=XM_023253496.1
B1 207153593 207153781
refGene_gn=LETM1;refGene_tx=XM_019829855.2;refGene_gn=LETM1;refGene_tx=XM_023253495.1;refGene_gn=LETM1;refGene_tx=XM_023253496.1
B1 207154769 207154908
refGene_gn=LETM1;refGene_tx=XM_019829855.2;refGene_gn=LETM1;refGene_tx=XM_023253495.1;refGene_gn=LETM1;refGene_tx=XM_023253496.1
B1 207155929 207156079

refGene_gn=LETM1;refGene_tx=XM_019829855.2;refGene_gn=LETM1;refGene_tx=XM_023253495.1;refGene_gn=LETM1;refGene_tx=XM_023253496.1
B1 207164389 207164536
refGene_gn=FGFR3;refGene_tx=XM_023253489.1;refGene_gn=FGFR3;refGene_tx=XM_023253490.1;refGene_gn=FGFR3;refGene_tx=XM_023253491.1;refGene_gn=FGFR3;refGene_tx=XM_023253492.1;refGene_gn=FGFR3;refGene_tx=XM_023253493.1
B1 207164624 207164730
refGene_gn=FGFR3;refGene_tx=XM_023253489.1;refGene_gn=FGFR3;refGene_tx=XM_023253490.1;refGene_gn=FGFR3;refGene_tx=XM_023253491.1;refGene_gn=FGFR3;refGene_tx=XM_023253492.1;refGene_gn=FGFR3;refGene_tx=XM_023253493.1
B1 207164876 207165014
refGene_gn=FGFR3;refGene_tx=XM_023253489.1;refGene_gn=FGFR3;refGene_tx=XM_023253490.1;refGene_gn=FGFR3;refGene_tx=XM_023253491.1;refGene_gn=FGFR3;refGene_tx=XM_023253492.1;refGene_gn=FGFR3;refGene_tx=XM_023253493.1
B1 207165206 207165284
refGene_gn=FGFR3;refGene_tx=XM_023253489.1;refGene_gn=FGFR3;refGene_tx=XM_023253490.1;refGene_gn=FGFR3;refGene_tx=XM_023253491.1;refGene_gn=FGFR3;refGene_tx=XM_023253492.1;refGene_gn=FGFR3;refGene_tx=XM_023253493.1
B1 207165429 207165552
refGene_gn=FGFR3;refGene_tx=XM_023253489.1;refGene_gn=FGFR3;refGene_tx=XM_023253490.1;refGene_gn=FGFR3;refGene_tx=XM_023253491.1;refGene_gn=FGFR3;refGene_tx=XM_023253492.1;refGene_gn=FGFR3;refGene_tx=XM_023253493.1
B1 207165680 207165871
refGene_gn=FGFR3;refGene_tx=XM_023253489.1;refGene_gn=FGFR3;refGene_tx=XM_023253490.1;refGene_gn=FGFR3;refGene_tx=XM_023253491.1;refGene_gn=FGFR3;refGene_tx=XM_023253492.1;refGene_gn=FGFR3;refGene_tx=XM_023253493.1
B1 207165949 207166060
refGene_gn=FGFR3;refGene_tx=XM_023253489.1;refGene_gn=FGFR3;refGene_tx=XM_023253490.1;refGene_gn=FGFR3;refGene_tx=XM_023253491.1;refGene_gn=FGFR3;refGene_tx=XM_023253492.1;refGene_gn=FGFR3;refGene_tx=XM_023253493.1
B1 207166172 207166294
refGene_gn=FGFR3;refGene_tx=XM_023253489.1;refGene_gn=FGFR3;refGene_tx=XM_023253490.1;refGene_gn=FGFR3;refGene_tx=XM_023253491.1;refGene_gn=FGFR3;refGene_tx=XM_023253492.1;refGene_gn=FGFR3;refGene_tx=XM_023253493.1
B1 207166686 207166835
refGene_gn=FGFR3;refGene_tx=XM_023253489.1;refGene_gn=FGFR3;refGene_tx=XM_023253490.1;refGene_gn=FGFR3;refGene_tx=XM_023253491.1;refGene_gn=FGFR3;refGene_tx=XM_023253492.1;refGene_gn=FGFR3;refGene_tx=XM_023253493.1
B1 207167208 207167405
refGene_gn=FGFR3;refGene_tx=XM_023253489.1;refGene_gn=FGFR3;refGene_tx=XM_023253490.1;refGene_gn=FGFR3;refGene_tx=XM_023253491.1;refGene_gn=

FGFR3;refGene_tx=XM_023253492.1;refGene_gn=FGFR3;refGene_tx=XM_0232534
93.1
B1 207167885 207168030
refGene_gn=FGFR3;refGene_tx=XM_023253491.1;refGene_gn=FGFR3;refGene_tx
=XM_023253492.1
B1 207168750 207168901
refGene_gn=FGFR3;refGene_tx=XM_023253489.1;refGene_gn=FGFR3;refGene_tx
=XM_023253490.1;refGene_gn=FGFR3;refGene_tx=XM_023253493.1
B1 207169968 207170159
refGene_gn=FGFR3;refGene_tx=XM_023253489.1;refGene_gn=FGFR3;refGene_tx
=XM_023253490.1;refGene_gn=FGFR3;refGene_tx=XM_023253491.1;refGene_gn=
FGFR3;refGene_tx=XM_023253492.1;refGene_gn=FGFR3;refGene_tx=XM_0232534
93.1
B1 207170240 207170364
refGene_gn=FGFR3;refGene_tx=XM_023253489.1;refGene_gn=FGFR3;refGene_tx
=XM_023253490.1;refGene_gn=FGFR3;refGene_tx=XM_023253491.1;refGene_gn=
FGFR3;refGene_tx=XM_023253492.1;refGene_gn=FGFR3;refGene_tx=XM_0232534
93.1
B1 207170440 207170613
refGene_gn=FGFR3;refGene_tx=XM_023253489.1;refGene_gn=FGFR3;refGene_tx
=XM_023253490.1;refGene_gn=FGFR3;refGene_tx=XM_023253491.1;refGene_gn=
FGFR3;refGene_tx=XM_023253492.1;refGene_gn=FGFR3;refGene_tx=XM_0232534
93.1
B1 207172469 207172523
refGene_gn=FGFR3;refGene_tx=XM_023253489.1;refGene_gn=FGFR3;refGene_tx
=XM_023253490.1;refGene_gn=FGFR3;refGene_tx=XM_023253491.1;refGene_gn=
FGFR3;refGene_tx=XM_023253492.1;refGene_gn=FGFR3;refGene_tx=XM_0232534
93.1
B1 207172713 207172983
refGene_gn=FGFR3;refGene_tx=XM_023253489.1;refGene_gn=FGFR3;refGene_tx
=XM_023253490.1;refGene_gn=FGFR3;refGene_tx=XM_023253491.1;refGene_gn=
FGFR3;refGene_tx=XM_023253492.1;refGene_gn=FGFR3;refGene_tx=XM_0232534
93.1
B1 207178858 207178967
refGene_gn=FGFR3;refGene_tx=XM_023253489.1;refGene_gn=FGFR3;refGene_tx
=XM_023253490.1;refGene_gn=FGFR3;refGene_tx=XM_023253491.1;refGene_gn=
FGFR3;refGene_tx=XM_023253492.1;refGene_gn=FGFR3;refGene_tx=XM_0232534
93.1
B1 207230063 207230129
refGene_gn=TACC3;refGene_tx=XM_023253484.1;refGene_gn=TACC3;refGene_tx
=XM_023253485.1;refGene_gn=TACC3;refGene_tx=XM_023253486.1;refGene_gn=
TACC3;refGene_tx=XM_023253487.1;refGene_gn=TACC3;refGene_tx=XM_0232534
88.1
B1 207230442 207230563
refGene_gn=TACC3;refGene_tx=XM_023253484.1;refGene_gn=TACC3;refGene_tx
=XM_023253485.1;refGene_gn=TACC3;refGene_tx=XM_023253486.1;refGene_gn=
TACC3;refGene_tx=XM_023253487.1;refGene_gn=TACC3;refGene_tx=XM_0232534
88.1
B1 207230712 207230819
refGene_gn=TACC3;refGene_tx=XM_023253484.1;refGene_gn=TACC3;refGene_tx

=XM_023253485.1;refGene_gn=TACC3;refGene_tx=XM_023253486.1;refGene_gn=TACC3;refGene_tx=XM_023253487.1;refGene_gn=TACC3;refGene_tx=XM_023253488.1
B1 207232492 207232653
refGene_gn=TACC3;refGene_tx=XM_023253484.1;refGene_gn=TACC3;refGene_tx=XM_023253485.1;refGene_gn=TACC3;refGene_tx=XM_023253486.1;refGene_gn=TACC3;refGene_tx=XM_023253487.1;refGene_gn=TACC3;refGene_tx=XM_023253488.1
B1 207233196 207233240
refGene_gn=TACC3;refGene_tx=XM_023253484.1;refGene_gn=TACC3;refGene_tx=XM_023253485.1;refGene_gn=TACC3;refGene_tx=XM_023253486.1;refGene_gn=TACC3;refGene_tx=XM_023253487.1;refGene_gn=TACC3;refGene_tx=XM_023253488.1
B1 207233317 207233394
refGene_gn=TACC3;refGene_tx=XM_023253484.1;refGene_gn=TACC3;refGene_tx=XM_023253485.1;refGene_gn=TACC3;refGene_tx=XM_023253486.1;refGene_gn=TACC3;refGene_tx=XM_023253487.1;refGene_gn=TACC3;refGene_tx=XM_023253488.1
B1 207234030 207234147
refGene_gn=TACC3;refGene_tx=XM_023253484.1;refGene_gn=TACC3;refGene_tx=XM_023253485.1;refGene_gn=TACC3;refGene_tx=XM_023253486.1;refGene_gn=TACC3;refGene_tx=XM_023253487.1;refGene_gn=TACC3;refGene_tx=XM_023253488.1
B1 207234351 207234469
refGene_gn=TACC3;refGene_tx=XM_023253484.1;refGene_gn=TACC3;refGene_tx=XM_023253485.1;refGene_gn=TACC3;refGene_tx=XM_023253486.1;refGene_gn=TACC3;refGene_tx=XM_023253487.1;refGene_gn=TACC3;refGene_tx=XM_023253488.1
B1 207234753 207234857
refGene_gn=TACC3;refGene_tx=XM_023253484.1;refGene_gn=TACC3;refGene_tx=XM_023253485.1;refGene_gn=TACC3;refGene_tx=XM_023253486.1;refGene_gn=TACC3;refGene_tx=XM_023253487.1;refGene_gn=TACC3;refGene_tx=XM_023253488.1
B1 207235160 207235213
refGene_gn=TACC3;refGene_tx=XM_023253484.1;refGene_gn=TACC3;refGene_tx=XM_023253485.1;refGene_gn=TACC3;refGene_tx=XM_023253486.1;refGene_gn=TACC3;refGene_tx=XM_023253487.1;refGene_gn=TACC3;refGene_tx=XM_023253488.1
B1 207236878 207237032
refGene_gn=TACC3;refGene_tx=XM_023253484.1;refGene_gn=TACC3;refGene_tx=XM_023253485.1;refGene_gn=TACC3;refGene_tx=XM_023253486.1;refGene_gn=TACC3;refGene_tx=XM_023253487.1;refGene_gn=TACC3;refGene_tx=XM_023253488.1
B1 207237132 207237208
refGene_gn=TACC3;refGene_tx=XM_023253484.1;refGene_gn=TACC3;refGene_tx=XM_023253485.1;refGene_gn=TACC3;refGene_tx=XM_023253486.1;refGene_gn=TACC3;refGene_tx=XM_023253487.1;refGene_gn=TACC3;refGene_tx=XM_023253488.1
B1 207238410 207239343
refGene_gn=TACC3;refGene_tx=XM_023253484.1;refGene_gn=TACC3;refGene_tx

=XM_023253485.1;refGene_gn=TACC3;refGene_tx=XM_023253486.1
B1 207240056 207240199
refGene_gn=TACC3;refGene_tx=XM_023253484.1;refGene_gn=TACC3;refGene_tx=XM_023253485.1;refGene_gn=TACC3;refGene_tx=XM_023253486.1;refGene_gn=TACC3;refGene_tx=XM_023253487.1;refGene_gn=TACC3;refGene_tx=XM_023253488.1
B1 207240354 207240517
refGene_gn=TACC3;refGene_tx=XM_023253484.1;refGene_gn=TACC3;refGene_tx=XM_023253485.1;refGene_gn=TACC3;refGene_tx=XM_023253486.1;refGene_gn=TACC3;refGene_tx=XM_023253487.1;refGene_gn=TACC3;refGene_tx=XM_023253488.1
B1 207240602 207240621
refGene_gn=TMEM129;refGene_tx=XM_019829859.2
B1 207242409 207242423
refGene_gn=TACC3;refGene_tx=XM_023253484.1;refGene_gn=TACC3;refGene_tx=XM_023253485.1;refGene_gn=TACC3;refGene_tx=XM_023253486.1;refGene_gn=TACC3;refGene_tx=XM_023253487.1;refGene_gn=TACC3;refGene_tx=XM_023253488.1
B1 207243168 207243373
refGene_gn=TMEM129;refGene_tx=XM_003985536.5;refGene_gn=TMEM129;refGene_tx=XM_019829860.2;refGene_gn=TMEM129;refGene_tx=XM_019829861.2
B1 207245354 207245829
refGene_gn=TMEM129;refGene_tx=XM_003985536.5;refGene_gn=TMEM129;refGene_tx=XM_019829859.2;refGene_gn=TMEM129;refGene_tx=XM_019829860.2;refGene_gn=TMEM129;refGene_tx=XM_019829861.2
B1 207246355 207246374
refGene_gn=TMEM129;refGene_tx=XM_019829861.2
B1 207246441 207246671
refGene_gn=TMEM129;refGene_tx=XM_003985536.5;refGene_gn=TMEM129;refGene_tx=XM_019829859.2;refGene_gn=TMEM129;refGene_tx=XM_019829860.2
B1 207246800 207247049
refGene_gn=TMEM129;refGene_tx=XM_003985536.5;refGene_gn=TMEM129;refGene_tx=XM_019829859.2
B1 207249451 207249510
refGene_gn=SLBP;refGene_tx=XM_023253526.1;refGene_gn=SLBP;refGene_tx=XM_023253528.1;refGene_gn=SLBP;refGene_tx=XM_023253529.1;refGene_gn=SLBP;refGene_tx=XM_023253530.1
B1 207249646 207249763
refGene_gn=SLBP;refGene_tx=XM_023253526.1;refGene_gn=SLBP;refGene_tx=XM_023253528.1;refGene_gn=SLBP;refGene_tx=XM_023253529.1;refGene_gn=SLBP;refGene_tx=XM_023253530.1
B1 207253894 207253999
refGene_gn=SLBP;refGene_tx=XM_023253526.1;refGene_gn=SLBP;refGene_tx=XM_023253528.1;refGene_gn=SLBP;refGene_tx=XM_023253529.1;refGene_gn=SLBP;refGene_tx=XM_023253530.1
B1 207256667 207256727
refGene_gn=SLBP;refGene_tx=XM_023253526.1;refGene_gn=SLBP;refGene_tx=XM_023253528.1;refGene_gn=SLBP;refGene_tx=XM_023253529.1;refGene_gn=SLBP;refGene_tx=XM_023253530.1
B1 207257017 207257155

refGene_gn=SLBP;refGene_tx=XM_023253526.1;refGene_gn=SLBP;refGene_tx=XM_023253528.1;refGene_gn=SLBP;refGene_tx=XM_023253529.1;refGene_gn=SLBP;refGene_tx=XM_023253530.1
B1 207257791 207257941
refGene_gn=SLBP;refGene_tx=XM_023253526.1;refGene_gn=SLBP;refGene_tx=XM_023253528.1;refGene_gn=SLBP;refGene_tx=XM_023253529.1;refGene_gn=SLBP;refGene_tx=XM_023253530.1
B1 207259514 207259596
refGene_gn=SLBP;refGene_tx=XM_023253526.1;refGene_gn=SLBP;refGene_tx=XM_023253528.1;refGene_gn=SLBP;refGene_tx=XM_023253529.1;refGene_gn=SLBP;refGene_tx=XM_023253530.1
B1 207260995 207261001
refGene_gn=SLBP;refGene_tx=XM_023253530.1
B1 207262611 207262728
refGene_gn=SLBP;refGene_tx=XM_023253528.1
B1 207293108 207293378
refGene_gn=SLBP;refGene_tx=XM_023253526.1
B1 207327417 207327483
refGene_gn=SLBP;refGene_tx=XM_023253529.1
B1 207486997 207487463
refGene_gn=NKX1-1;refGene_tx=XM_023252242.1
B1 207489556 207489768
refGene_gn=NKX1-1;refGene_tx=XM_023252242.1
B1 207489769 207490441
refGene_gn=NKX1-1;refGene_tx=XM_023252242.1
B1 207503179 207503273
refGene_gn=UVSSA;refGene_tx=XM_023253352.1;refGene_gn=UVSSA;refGene_tx=XM_023253354.1
B1 207504884 207505059
refGene_gn=UVSSA;refGene_tx=XM_023253352.1;refGene_gn=UVSSA;refGene_tx=XM_023253354.1
B1 207507842 207507969
refGene_gn=UVSSA;refGene_tx=XM_023253352.1;refGene_gn=UVSSA;refGene_tx=XM_023253354.1
B1 207508392 207508576
refGene_gn=UVSSA;refGene_tx=XM_023253352.1;refGene_gn=UVSSA;refGene_tx=XM_023253354.1
B1 207509235 207509361
refGene_gn=UVSSA;refGene_tx=XM_023253352.1;refGene_gn=UVSSA;refGene_tx=XM_023253354.1
B1 207509737 207509876
refGene_gn=UVSSA;refGene_tx=XM_023253352.1;refGene_gn=UVSSA;refGene_tx=XM_023253354.1
B1 207515719 207515837
refGene_gn=UVSSA;refGene_tx=XM_023253352.1;refGene_gn=UVSSA;refGene_tx=XM_023253354.1
B1 207523851 207523980
refGene_gn=UVSSA;refGene_tx=XM_023253352.1;refGene_gn=UVSSA;refGene_tx=XM_023253354.1
B1 207524110 207524223

refGene_gn=UVSSA;refGene_tx=XM_023253352.1;refGene_gn=UVSSA;refGene_tx=XM_023253354.1
B1 207525101 207525476
refGene_gn=UVSSA;refGene_tx=XM_023253352.1;refGene_gn=UVSSA;refGene_tx=XM_023253354.1
B1 207526154 207526275
refGene_gn=UVSSA;refGene_tx=XM_023253352.1;refGene_gn=UVSSA;refGene_tx=XM_023253354.1
B1 207526800 207527131
refGene_gn=UVSSA;refGene_tx=XM_023253352.1;refGene_gn=UVSSA;refGene_tx=XM_023253354.1
B1 207528112 207528210
refGene_gn=UVSSA;refGene_tx=XM_023253352.1;refGene_gn=UVSSA;refGene_tx=XM_023253354.1
B1 207539201 207539297
refGene_gn=MAEA;refGene_tx=XM_023253355.1;refGene_gn=MAEA;refGene_tx=XM_023253356.1;refGene_gn=MAEA;refGene_tx=XM_023253357.1
B1 207539909 207540105
refGene_gn=MAEA;refGene_tx=XM_023253355.1;refGene_gn=MAEA;refGene_tx=XM_023253356.1;refGene_gn=MAEA;refGene_tx=XM_023253357.1
B1 207540776 207540910
refGene_gn=MAEA;refGene_tx=XM_023253355.1;refGene_gn=MAEA;refGene_tx=XM_023253356.1;refGene_gn=MAEA;refGene_tx=XM_023253357.1
B1 207543641 207543750
refGene_gn=MAEA;refGene_tx=XM_023253355.1;refGene_gn=MAEA;refGene_tx=XM_023253356.1;refGene_gn=MAEA;refGene_tx=XM_023253357.1
B1 207544644 207544721
refGene_gn=MAEA;refGene_tx=XM_023253355.1;refGene_gn=MAEA;refGene_tx=XM_023253356.1
B1 207547321 207547435
refGene_gn=MAEA;refGene_tx=XM_023253355.1
B1 207548065 207548188
refGene_gn=MAEA;refGene_tx=XM_023253355.1;refGene_gn=MAEA;refGene_tx=XM_023253356.1
B1 207552428 207552632
refGene_gn=MAEA;refGene_tx=XM_023253355.1;refGene_gn=MAEA;refGene_tx=XM_023253356.1;refGene_gn=MAEA;refGene_tx=XM_023253357.1
B1 207554546 207554729
refGene_gn=MAEA;refGene_tx=XM_023253355.1;refGene_gn=MAEA;refGene_tx=XM_023253356.1;refGene_gn=MAEA;refGene_tx=XM_023253357.1
B1 207576397 207576466
refGene_gn=MAEA;refGene_tx=XM_023253355.1;refGene_gn=MAEA;refGene_tx=XM_023253356.1;refGene_gn=MAEA;refGene_tx=XM_023253357.1
B1 207596368 207596719
refGene_gn=LOC109499320;refGene_tx=XM_023253358.1
B1 207597946 207598215
refGene_gn=LOC109499320;refGene_tx=XM_023253358.1
B1 207600738 207600790
refGene_gn=LOC109499320;refGene_tx=XM_023253358.1
B1 207611717 207611757

refGene_gn=CTBP1;refGene_tx=XM_023253359.1;refGene_gn=CTBP1;refGene_tx=XM_023253360.1
B1 207621090 207621097
refGene_gn=CTBP1;refGene_tx=XM_011281929.3;refGene_gn=CTBP1;refGene_tx=XM_023253361.1;refGene_gn=CTBP1;refGene_tx=XM_023253362.1
B1 207622520 207622675
refGene_gn=CTBP1;refGene_tx=XM_011281929.3;refGene_gn=CTBP1;refGene_tx=XM_023253359.1;refGene_gn=CTBP1;refGene_tx=XM_023253360.1;refGene_gn=CTBP1;refGene_tx=XM_023253361.1;refGene_gn=CTBP1;refGene_tx=XM_023253362.1
B1 207630284 207630429
refGene_gn=CTBP1;refGene_tx=XM_011281929.3;refGene_gn=CTBP1;refGene_tx=XM_023253359.1;refGene_gn=CTBP1;refGene_tx=XM_023253360.1;refGene_gn=CTBP1;refGene_tx=XM_023253361.1;refGene_gn=CTBP1;refGene_tx=XM_023253362.1
B1 207631559 207631766
refGene_gn=CTBP1;refGene_tx=XM_011281929.3;refGene_gn=CTBP1;refGene_tx=XM_023253359.1;refGene_gn=CTBP1;refGene_tx=XM_023253360.1;refGene_gn=CTBP1;refGene_tx=XM_023253361.1;refGene_gn=CTBP1;refGene_tx=XM_023253362.1
B1 207633457 207633672
refGene_gn=CTBP1;refGene_tx=XM_011281929.3;refGene_gn=CTBP1;refGene_tx=XM_023253359.1;refGene_gn=CTBP1;refGene_tx=XM_023253360.1;refGene_gn=CTBP1;refGene_tx=XM_023253361.1;refGene_gn=CTBP1;refGene_tx=XM_023253362.1
B1 207635562 207635693
refGene_gn=CTBP1;refGene_tx=XM_011281929.3;refGene_gn=CTBP1;refGene_tx=XM_023253359.1;refGene_gn=CTBP1;refGene_tx=XM_023253360.1;refGene_gn=CTBP1;refGene_tx=XM_023253361.1;refGene_gn=CTBP1;refGene_tx=XM_023253362.1
B1 207636326 207636454
refGene_gn=CTBP1;refGene_tx=XM_011281929.3;refGene_gn=CTBP1;refGene_tx=XM_023253359.1;refGene_gn=CTBP1;refGene_tx=XM_023253360.1;refGene_gn=CTBP1;refGene_tx=XM_023253361.1;refGene_gn=CTBP1;refGene_tx=XM_023253362.1
B1 207636980 207637098
refGene_gn=CTBP1;refGene_tx=XM_011281929.3;refGene_gn=CTBP1;refGene_tx=XM_023253359.1;refGene_gn=CTBP1;refGene_tx=XM_023253360.1;refGene_gn=CTBP1;refGene_tx=XM_023253361.1;refGene_gn=CTBP1;refGene_tx=XM_023253362.1
B1 207637462 207637649
refGene_gn=CTBP1;refGene_tx=XM_011281929.3;refGene_gn=CTBP1;refGene_tx=XM_023253359.1;refGene_gn=CTBP1;refGene_tx=XM_023253360.1;refGene_gn=CTBP1;refGene_tx=XM_023253361.1;refGene_gn=CTBP1;refGene_tx=XM_023253362.1
B1 207656610 207657570
refGene_gn=LOC105260525;refGene_tx=XM_023253363.1
B1 207669232 207669455
refGene_gn=SPON2;refGene_tx=XM_023253364.1
B1 207669707 207669931

refGene_gn=SPON2;refGene_tx=XM_023253364.1
B1 207670136 207670328
refGene_gn=SPON2;refGene_tx=XM_023253364.1
B1 207670756 207670931
refGene_gn=SPON2;refGene_tx=XM_023253364.1
B1 207672324 207672509
refGene_gn=SPON2;refGene_tx=XM_023253364.1
B1 207683923 207684116
refGene_gn=LOC101095254;refGene_tx=XM_003985534.3
B1 207697224 207697325
refGene_gn=LOC101095254;refGene_tx=XM_003985534.3
B1 207705028 207705154
refGene_gn=LOC101095254;refGene_tx=XM_003985534.3
B1 207705877 207706024
refGene_gn=LOC101095254;refGene_tx=XM_003985534.3
B1 207706818 207706968
refGene_gn=LOC101095254;refGene_tx=XM_003985534.3
B1 207709029 207709138
refGene_gn=RNF212;refGene_tx=XM_011281927.3
B1 207713769 207713831
refGene_gn=RNF212;refGene_tx=XM_011281927.3
B1 207722942 207723017
refGene_gn=RNF212;refGene_tx=XM_011281927.3
B1 207726544 207726601
refGene_gn=RNF212;refGene_tx=XM_011281927.3
B1 207727460 207727519
refGene_gn=RNF212;refGene_tx=XM_011281927.3
B1 207729303 207729356
refGene_gn=RNF212;refGene_tx=XM_011281927.3
B1 207729443 207729501
refGene_gn=RNF212;refGene_tx=XM_011281927.3
B1 207730468 207730514
refGene_gn=RNF212;refGene_tx=XM_011281927.3
B1 207734367 207734431
refGene_gn=RNF212;refGene_tx=XM_011281927.3
B1 207747749 207747822
refGene_gn=RNF212;refGene_tx=XM_011281927.3
B1 207748092 207748256
refGene_gn=RNF212;refGene_tx=XM_011281927.3
B1 207748437 207748475
refGene_gn=RNF212;refGene_tx=XM_011281927.3
B1 207767454 207767897
refGene_gn=FGFRL1;refGene_tx=XM_023253365.1;refGene_gn=FGFRL1;refGene_tx=XM_023253366.1;refGene_gn=FGFRL1;refGene_tx=XM_023253367.1
B1 207768011 207768365
refGene_gn=FGFRL1;refGene_tx=XM_023253365.1;refGene_gn=FGFRL1;refGene_tx=XM_023253366.1;refGene_gn=FGFRL1;refGene_tx=XM_023253367.1
B1 207768447 207768729
refGene_gn=FGFRL1;refGene_tx=XM_023253365.1;refGene_gn=FGFRL1;refGene_tx=XM_023253366.1;refGene_gn=FGFRL1;refGene_tx=XM_023253367.1

B1 207768824 207768905
refGene_gn=FGFRL1;refGene_tx=XM_023253365.1;refGene_gn=FGFRL1;refGene_tx=XM_023253366.1;refGene_gn=FGFRL1;refGene_tx=XM_023253367.1
B1 207769998 207770271
refGene_gn=FGFRL1;refGene_tx=XM_023253365.1;refGene_gn=FGFRL1;refGene_tx=XM_023253366.1;refGene_gn=FGFRL1;refGene_tx=XM_023253367.1
B1 207780377 207780472
refGene_gn=FGFRL1;refGene_tx=XM_023253365.1;refGene_gn=FGFRL1;refGene_tx=XM_023253366.1;refGene_gn=FGFRL1;refGene_tx=XM_023253367.1
B1 207781987 207781998
refGene_gn=FGFRL1;refGene_tx=XM_023253365.1
B1 207798581 207798688
refGene_gn=IDUA;refGene_tx=NM_001305032.1;refGene_gn=IDUA;refGene_tx=XM_023252206.1
B1 207798836 207798937
refGene_gn=IDUA;refGene_tx=NM_001305032.1;refGene_gn=IDUA;refGene_tx=XM_023252206.1
B1 207799344 207799421
refGene_gn=IDUA;refGene_tx=NM_001305032.1;refGene_gn=IDUA;refGene_tx=XM_023252206.1
B1 207799506 207799632
refGene_gn=IDUA;refGene_tx=NM_001305032.1;refGene_gn=IDUA;refGene_tx=XM_023252206.1
B1 207799708 207799830
refGene_gn=IDUA;refGene_tx=NM_001305032.1;refGene_gn=IDUA;refGene_tx=XM_023252206.1
B1 207799910 207800123
refGene_gn=IDUA;refGene_tx=NM_001305032.1;refGene_gn=IDUA;refGene_tx=XM_023252206.1
B1 207800442 207800659
refGene_gn=IDUA;refGene_tx=NM_001305032.1;refGene_gn=IDUA;refGene_tx=XM_023252206.1
B1 207800761 207800941
refGene_gn=IDUA;refGene_tx=NM_001305032.1;refGene_gn=IDUA;refGene_tx=XM_023252206.1
B1 207801064 207801267
refGene_gn=IDUA;refGene_tx=NM_001305032.1;refGene_gn=IDUA;refGene_tx=XM_023252206.1
B1 207801364 207801460
refGene_gn=IDUA;refGene_tx=NM_001305032.1;refGene_gn=IDUA;refGene_tx=XM_023252206.1
B1 207801731 207801839
refGene_gn=IDUA;refGene_tx=NM_001305032.1;refGene_gn=IDUA;refGene_tx=XM_023252206.1
B1 207802027 207802113
refGene_gn=IDUA;refGene_tx=NM_001305032.1;refGene_gn=IDUA;refGene_tx=XM_023252206.1
B1 207808740 207809358
refGene_gn=LOC102901021;refGene_tx=XM_023253369.1
B1 207809973 207811500

refGene_gn=LOC111560215;refGene_tx=XM_023253368.1
B1 207812371 207812512
refGene_gn=IDUA;refGene_tx=NM_001305032.1;refGene_gn=IDUA;refGene_tx=XM_023252206.1
B1 207813263 207813418
refGene_gn=IDUA;refGene_tx=NM_001305032.1;refGene_gn=IDUA;refGene_tx=XM_023252206.1
B1 207818090 207818121
refGene_gn=LOC102901021;refGene_tx=XM_023253369.1
B1 207818123 207818176
refGene_gn=LOC102901021;refGene_tx=XM_023253369.1
B1 207823469 207823749
refGene_gn=DGKQ;refGene_tx=XM_023253374.1
B1 207826222 207826302
refGene_gn=DGKQ;refGene_tx=XM_023253374.1
B1 207827017 207827117
refGene_gn=DGKQ;refGene_tx=XM_023253374.1
B1 207827505 207827591
refGene_gn=DGKQ;refGene_tx=XM_023253374.1
B1 207827682 207827808
refGene_gn=DGKQ;refGene_tx=XM_023253374.1
B1 207828026 207828252
refGene_gn=DGKQ;refGene_tx=XM_023253374.1
B1 207828397 207828498
refGene_gn=DGKQ;refGene_tx=XM_023253374.1
B1 207828613 207828852
refGene_gn=DGKQ;refGene_tx=XM_023253374.1
B1 207828944 207829035
refGene_gn=DGKQ;refGene_tx=XM_023253374.1
B1 207829129 207829184
refGene_gn=DGKQ;refGene_tx=XM_023253374.1
B1 207829303 207829365
refGene_gn=DGKQ;refGene_tx=XM_023253374.1
B1 207829761 207829912
refGene_gn=DGKQ;refGene_tx=XM_023253374.1
B1 207830323 207830362
refGene_gn=DGKQ;refGene_tx=XM_023253374.1
B1 207830982 207831098
refGene_gn=DGKQ;refGene_tx=XM_023253374.1
B1 207832708 207832860
refGene_gn=DGKQ;refGene_tx=XM_023253374.1
B1 207833103 207833252
refGene_gn=DGKQ;refGene_tx=XM_023253374.1
B1 207833350 207833529
refGene_gn=DGKQ;refGene_tx=XM_023253374.1
B1 207833900 207834001
refGene_gn=DGKQ;refGene_tx=XM_023253374.1
B1 207834164 207834311
refGene_gn=DGKQ;refGene_tx=XM_023253374.1
B1 207834402 207834514

refGene_gn=DGKQ;refGene_tx=XM_023253374.1
B1 207834591 207834744
refGene_gn=DGKQ;refGene_tx=XM_023253374.1
B1 207834973 207835075
refGene_gn=DGKQ;refGene_tx=XM_023253374.1
B1 207836722 207837377
refGene_gn=TMEM175;refGene_tx=XM_019829869.2;refGene_gn=TMEM175;refGene_tx=XM_023253375.1;refGene_gn=TMEM175;refGene_tx=XM_023253376.1;refGene_gn=TMEM175;refGene_tx=XM_023253377.1
B1 207838883 207839019
refGene_gn=TMEM175;refGene_tx=XM_019829869.2;refGene_gn=TMEM175;refGene_tx=XM_023253375.1;refGene_gn=TMEM175;refGene_tx=XM_023253376.1;refGene_gn=TMEM175;refGene_tx=XM_023253377.1
B1 207839168 207839247
refGene_gn=TMEM175;refGene_tx=XM_019829869.2;refGene_gn=TMEM175;refGene_tx=XM_023253375.1;refGene_gn=TMEM175;refGene_tx=XM_023253376.1;refGene_gn=TMEM175;refGene_tx=XM_023253377.1
B1 207843512 207843677
refGene_gn=TMEM175;refGene_tx=XM_019829869.2;refGene_gn=TMEM175;refGene_tx=XM_023253375.1;refGene_gn=TMEM175;refGene_tx=XM_023253376.1;refGene_gn=TMEM175;refGene_tx=XM_023253377.1
B1 207844132 207844216
refGene_gn=TMEM175;refGene_tx=XM_019829869.2;refGene_gn=TMEM175;refGene_tx=XM_023253375.1;refGene_gn=TMEM175;refGene_tx=XM_023253376.1
B1 207844887 207844923
refGene_gn=TMEM175;refGene_tx=XM_019829869.2;refGene_gn=TMEM175;refGene_tx=XM_023253375.1;refGene_gn=TMEM175;refGene_tx=XM_023253376.1;refGene_gn=TMEM175;refGene_tx=XM_023253377.1
B1 207845291 207845343
refGene_gn=TMEM175;refGene_tx=XM_019829869.2;refGene_gn=TMEM175;refGene_tx=XM_023253375.1;refGene_gn=TMEM175;refGene_tx=XM_023253376.1;refGene_gn=TMEM175;refGene_tx=XM_023253377.1
B1 207847349 207847444
refGene_gn=TMEM175;refGene_tx=XM_019829869.2;refGene_gn=TMEM175;refGene_tx=XM_023253375.1;refGene_gn=TMEM175;refGene_tx=XM_023253376.1;refGene_gn=TMEM175;refGene_tx=XM_023253377.1
B1 207849248 207849287
refGene_gn=TMEM175;refGene_tx=XM_023253375.1;refGene_gn=TMEM175;refGene_tx=XM_023253376.1;refGene_gn=TMEM175;refGene_tx=XM_023253377.1
B1 207849562 207849715
refGene_gn=TMEM175;refGene_tx=XM_023253375.1;refGene_gn=TMEM175;refGene_tx=XM_023253376.1;refGene_gn=TMEM175;refGene_tx=XM_023253377.1
B1 207864908 207865053
refGene_gn=GAK;refGene_tx=XM_023253370.1;refGene_gn=GAK;refGene_tx=XM_023253371.1
B1 207874574 207874636
refGene_gn=GAK;refGene_tx=XM_023253370.1;refGene_gn=GAK;refGene_tx=XM_023253371.1
B1 207875278 207875338
refGene_gn=GAK;refGene_tx=XM_023253370.1;refGene_gn=GAK;refGene_tx=XM_

023253371.1;refGene_gn=GAK;refGene_tx=XM_023253372.1;refGene_gn=GAK;refGene_tx=XM_023253373.1
B1 207876256 207876371 refGene_gn=GAK;refGene_tx=XM_023253370.1;refGene_gn=GAK;refGene_tx=XM_023253371.1;refGene_gn=GAK;refGene_tx=XM_023253372.1;refGene_gn=GAK;refGene_tx=XM_023253373.1
B1 207879850 207879993 refGene_gn=GAK;refGene_tx=XM_019829871.2;refGene_gn=GAK;refGene_tx=XM_023253370.1;refGene_gn=GAK;refGene_tx=XM_023253371.1;refGene_gn=GAK;refGene_tx=XM_023253372.1;refGene_gn=GAK;refGene_tx=XM_023253373.1
B1 207884036 207884162 refGene_gn=GAK;refGene_tx=XM_019829871.2;refGene_gn=GAK;refGene_tx=XM_023253370.1;refGene_gn=GAK;refGene_tx=XM_023253371.1;refGene_gn=GAK;refGene_tx=XM_023253372.1;refGene_gn=GAK;refGene_tx=XM_023253373.1
B1 207887410 207887500 refGene_gn=GAK;refGene_tx=XM_019829871.2;refGene_gn=GAK;refGene_tx=XM_023253370.1;refGene_gn=GAK;refGene_tx=XM_023253371.1;refGene_gn=GAK;refGene_tx=XM_023253372.1;refGene_gn=GAK;refGene_tx=XM_023253373.1
B1 207888878 207889014 refGene_gn=GAK;refGene_tx=XM_019829871.2;refGene_gn=GAK;refGene_tx=XM_023253370.1;refGene_gn=GAK;refGene_tx=XM_023253371.1;refGene_gn=GAK;refGene_tx=XM_023253372.1;refGene_gn=GAK;refGene_tx=XM_023253373.1
B1 207889329 207889442 refGene_gn=GAK;refGene_tx=XM_019829871.2;refGene_gn=GAK;refGene_tx=XM_023253370.1;refGene_gn=GAK;refGene_tx=XM_023253371.1;refGene_gn=GAK;refGene_tx=XM_023253372.1;refGene_gn=GAK;refGene_tx=XM_023253373.1
B1 207891304 207891395 refGene_gn=GAK;refGene_tx=XM_019829871.2;refGene_gn=GAK;refGene_tx=XM_023253370.1;refGene_gn=GAK;refGene_tx=XM_023253371.1;refGene_gn=GAK;refGene_tx=XM_023253372.1;refGene_gn=GAK;refGene_tx=XM_023253373.1
B1 207893333 207893457 refGene_gn=GAK;refGene_tx=XM_019829871.2;refGene_gn=GAK;refGene_tx=XM_023253370.1;refGene_gn=GAK;refGene_tx=XM_023253371.1;refGene_gn=GAK;refGene_tx=XM_023253372.1;refGene_gn=GAK;refGene_tx=XM_023253373.1
B1 207900313 207900363 refGene_gn=GAK;refGene_tx=XM_019829871.2;refGene_gn=GAK;refGene_tx=XM_023253370.1;refGene_gn=GAK;refGene_tx=XM_023253371.1;refGene_gn=GAK;refGene_tx=XM_023253372.1;refGene_gn=GAK;refGene_tx=XM_023253373.1
B1 207900924 207901073 refGene_gn=GAK;refGene_tx=XM_019829871.2;refGene_gn=GAK;refGene_tx=XM_023253370.1;refGene_gn=GAK;refGene_tx=XM_023253371.1;refGene_gn=GAK;refGene_tx=XM_023253372.1;refGene_gn=GAK;refGene_tx=XM_023253373.1
B1 207901602 207901725 refGene_gn=GAK;refGene_tx=XM_019829871.2;refGene_gn=GAK;refGene_tx=XM_023253370.1;refGene_gn=GAK;refGene_tx=XM_023253371.1;refGene_gn=GAK;refGene_tx=XM_023253372.1;refGene_gn=GAK;refGene_tx=XM_023253373.1
B1 207902838 207902972 refGene_gn=GAK;refGene_tx=XM_019829871.2;refGene_gn=GAK;refGene_tx=XM_023253370.1;refGene_gn=GAK;refGene_tx=XM_023253371.1;refGene_gn=GAK;refGene_tx=XM_023253372.1;refGene_gn=GAK;refGene_tx=XM_023253373.1

B1	207905174	207905369
refGene_gn=GAK;refGene_tx=XM_019829871.2;refGene_gn=GAK;refGene_tx=XM_023253370.1;refGene_gn=GAK;refGene_tx=XM_023253371.1;refGene_gn=GAK;refGene_tx=XM_023253372.1;refGene_gn=GAK;refGene_tx=XM_023253373.1
B1	207905658	207905776
refGene_gn=GAK;refGene_tx=XM_019829871.2;refGene_gn=GAK;refGene_tx=XM_023253370.1;refGene_gn=GAK;refGene_tx=XM_023253371.1;refGene_gn=GAK;refGene_tx=XM_023253372.1;refGene_gn=GAK;refGene_tx=XM_023253373.1
B1	207906134	207906214
refGene_gn=GAK;refGene_tx=XM_019829871.2;refGene_gn=GAK;refGene_tx=XM_023253370.1;refGene_gn=GAK;refGene_tx=XM_023253371.1;refGene_gn=GAK;refGene_tx=XM_023253372.1;refGene_gn=GAK;refGene_tx=XM_023253373.1
B1	207908785	207908979
refGene_gn=GAK;refGene_tx=XM_019829871.2;refGene_gn=GAK;refGene_tx=XM_023253370.1;refGene_gn=GAK;refGene_tx=XM_023253371.1;refGene_gn=GAK;refGene_tx=XM_023253372.1;refGene_gn=GAK;refGene_tx=XM_023253373.1
B1	207909844	207909991
refGene_gn=GAK;refGene_tx=XM_019829871.2;refGene_gn=GAK;refGene_tx=XM_023253370.1;refGene_gn=GAK;refGene_tx=XM_023253371.1;refGene_gn=GAK;refGene_tx=XM_023253372.1;refGene_gn=GAK;refGene_tx=XM_023253373.1
B1	207910153	207910624
refGene_gn=GAK;refGene_tx=XM_019829871.2;refGene_gn=GAK;refGene_tx=XM_023253370.1;refGene_gn=GAK;refGene_tx=XM_023253371.1;refGene_gn=GAK;refGene_tx=XM_023253372.1;refGene_gn=GAK;refGene_tx=XM_023253373.1
B1	207911057	207911228
refGene_gn=GAK;refGene_tx=XM_019829871.2;refGene_gn=GAK;refGene_tx=XM_023253370.1;refGene_gn=GAK;refGene_tx=XM_023253371.1;refGene_gn=GAK;refGene_tx=XM_023253372.1;refGene_gn=GAK;refGene_tx=XM_023253373.1
B1	207911744	207911855
refGene_gn=GAK;refGene_tx=XM_019829871.2;refGene_gn=GAK;refGene_tx=XM_023253370.1;refGene_gn=GAK;refGene_tx=XM_023253371.1;refGene_gn=GAK;refGene_tx=XM_023253372.1;refGene_gn=GAK;refGene_tx=XM_023253373.1
B1	207912723	207912840
refGene_gn=GAK;refGene_tx=XM_019829871.2;refGene_gn=GAK;refGene_tx=XM_023253370.1;refGene_gn=GAK;refGene_tx=XM_023253371.1;refGene_gn=GAK;refGene_tx=XM_023253372.1;refGene_gn=GAK;refGene_tx=XM_023253373.1
B1	207916552	207916780
refGene_gn=GAK;refGene_tx=XM_019829871.2;refGene_gn=GAK;refGene_tx=XM_023253370.1;refGene_gn=GAK;refGene_tx=XM_023253371.1;refGene_gn=GAK;refGene_tx=XM_023253372.1;refGene_gn=GAK;refGene_tx=XM_023253373.1
B1	207917346	207917495
refGene_gn=GAK;refGene_tx=XM_019829871.2;refGene_gn=GAK;refGene_tx=XM_023253370.1;refGene_gn=GAK;refGene_tx=XM_023253371.1;refGene_gn=GAK;refGene_tx=XM_023253372.1;refGene_gn=GAK;refGene_tx=XM_023253373.1
B1	207918270	207918447
refGene_gn=GAK;refGene_tx=XM_019829871.2;refGene_gn=GAK;refGene_tx=XM_023253370.1;refGene_gn=GAK;refGene_tx=XM_023253371.1;refGene_gn=GAK;refGene_tx=XM_023253372.1;refGene_gn=GAK;refGene_tx=XM_023253373.1
B1	207918542	207918644
refGene_gn=GAK;refGene_tx=XM_019829871.2;refGene_gn=GAK;refGene_tx=XM_

023253370.1;refGene_gn=GAK;refGene_tx=XM_023253371.1;refGene_gn=GAK;refGene_tx=XM_023253372.1;refGene_gn=GAK;refGene_tx=XM_023253373.1
B1 207933405 207933436
refGene_gn=CPLX1;refGene_tx=XM_019829875.2;refGene_gn=CPLX1;refGene_tx=XM_023253378.1;refGene_gn=CPLX1;refGene_tx=XM_023253379.1
B1 207946384 207946547
refGene_gn=CPLX1;refGene_tx=XM_019829873.2
B1 207958904 207959080
refGene_gn=CPLX1;refGene_tx=XM_019829873.2;refGene_gn=CPLX1;refGene_tx=XM_019829875.2;refGene_gn=CPLX1;refGene_tx=XM_023253378.1;refGene_gn=CPLX1;refGene_tx=XM_023253379.1
B1 207962336 207962534
refGene_gn=CPLX1;refGene_tx=XM_019829873.2;refGene_gn=CPLX1;refGene_tx=XM_019829875.2;refGene_gn=CPLX1;refGene_tx=XM_023253378.1;refGene_gn=CPLX1;refGene_tx=XM_023253379.1
B1 207988394 207988442
refGene_gn=PCGF3;refGene_tx=XM_006931299.4;refGene_gn=PCGF3;refGene_tx=XM_006931300.4
B1 207988895 207988976
refGene_gn=PCGF3;refGene_tx=XM_006931299.4;refGene_gn=PCGF3;refGene_tx=XM_006931300.4
B1 207989483 207989600
refGene_gn=PCGF3;refGene_tx=XM_006931301.3
B1 207993835 207993841
refGene_gn=PCGF3;refGene_tx=XM_019829876.2
B1 207994940 207995078
refGene_gn=PCGF3;refGene_tx=XM_006931299.4;refGene_gn=PCGF3;refGene_tx=XM_006931300.4;refGene_gn=PCGF3;refGene_tx=XM_006931301.3;refGene_gn=PCGF3;refGene_tx=XM_019829876.2
B1 208005320 208005409
refGene_gn=PCGF3;refGene_tx=XM_006931299.4;refGene_gn=PCGF3;refGene_tx=XM_006931300.4;refGene_gn=PCGF3;refGene_tx=XM_006931301.3;refGene_gn=PCGF3;refGene_tx=XM_019829876.2
B1 208006347 208006458
refGene_gn=PCGF3;refGene_tx=XM_006931299.4;refGene_gn=PCGF3;refGene_tx=XM_006931300.4;refGene_gn=PCGF3;refGene_tx=XM_006931301.3;refGene_gn=PCGF3;refGene_tx=XM_019829876.2
B1 208015577 208015633
refGene_gn=PCGF3;refGene_tx=XM_006931299.4;refGene_gn=PCGF3;refGene_tx=XM_006931300.4;refGene_gn=PCGF3;refGene_tx=XM_006931301.3;refGene_gn=PCGF3;refGene_tx=XM_019829876.2
B1 208019019 208019116
refGene_gn=PCGF3;refGene_tx=XM_006931299.4;refGene_gn=PCGF3;refGene_tx=XM_006931300.4;refGene_gn=PCGF3;refGene_tx=XM_006931301.3;refGene_gn=PCGF3;refGene_tx=XM_019829876.2
B1 208020460 208020569
refGene_gn=PCGF3;refGene_tx=XM_006931299.4;refGene_gn=PCGF3;refGene_tx=XM_006931300.4;refGene_gn=PCGF3;refGene_tx=XM_006931301.3;refGene_gn=PCGF3;refGene_tx=XM_019829876.2
B1 208054377 208054512

refGene_gn=SLC49A3;refGene_tx=XM_023253380.1;refGene_gn=SLC49A3;refGene_tx=XM_023253381.1
B1 208057471 208057630
refGene_gn=SLC49A3;refGene_tx=XM_023253380.1;refGene_gn=SLC49A3;refGene_tx=XM_023253381.1;refGene_gn=SLC49A3;refGene_tx=XM_023253382.1;refGene_gn=SLC49A3;refGene_tx=XM_023253383.1;refGene_gn=SLC49A3;refGene_tx=XM_023253384.1
B1 208057814 208058028
refGene_gn=SLC49A3;refGene_tx=XM_023253380.1;refGene_gn=SLC49A3;refGene_tx=XM_023253381.1;refGene_gn=SLC49A3;refGene_tx=XM_023253382.1;refGene_gn=SLC49A3;refGene_tx=XM_023253383.1;refGene_gn=SLC49A3;refGene_tx=XM_023253384.1
B1 208058189 208058266
refGene_gn=SLC49A3;refGene_tx=XM_023253380.1;refGene_gn=SLC49A3;refGene_tx=XM_023253381.1;refGene_gn=SLC49A3;refGene_tx=XM_023253382.1;refGene_gn=SLC49A3;refGene_tx=XM_023253383.1;refGene_gn=SLC49A3;refGene_tx=XM_023253384.1
B1 208059265 208059403
refGene_gn=SLC49A3;refGene_tx=XM_023253380.1;refGene_gn=SLC49A3;refGene_tx=XM_023253381.1;refGene_gn=SLC49A3;refGene_tx=XM_023253382.1;refGene_gn=SLC49A3;refGene_tx=XM_023253383.1;refGene_gn=SLC49A3;refGene_tx=XM_023253384.1
B1 208059540 208059657
refGene_gn=SLC49A3;refGene_tx=XM_023253380.1;refGene_gn=SLC49A3;refGene_tx=XM_023253381.1;refGene_gn=SLC49A3;refGene_tx=XM_023253382.1;refGene_gn=SLC49A3;refGene_tx=XM_023253383.1;refGene_gn=SLC49A3;refGene_tx=XM_023253384.1
B1 208060400 208060553
refGene_gn=SLC49A3;refGene_tx=XM_023253380.1;refGene_gn=SLC49A3;refGene_tx=XM_023253381.1;refGene_gn=SLC49A3;refGene_tx=XM_023253382.1;refGene_gn=SLC49A3;refGene_tx=XM_023253383.1;refGene_gn=SLC49A3;refGene_tx=XM_023253384.1
B1 208060794 208060952
refGene_gn=SLC49A3;refGene_tx=XM_023253380.1;refGene_gn=SLC49A3;refGene_tx=XM_023253381.1;refGene_gn=SLC49A3;refGene_tx=XM_023253382.1;refGene_gn=SLC49A3;refGene_tx=XM_023253383.1;refGene_gn=SLC49A3;refGene_tx=XM_023253384.1
B1 208061178 208061288
refGene_gn=SLC49A3;refGene_tx=XM_023253380.1;refGene_gn=SLC49A3;refGene_tx=XM_023253381.1;refGene_gn=SLC49A3;refGene_tx=XM_023253382.1;refGene_gn=SLC49A3;refGene_tx=XM_023253383.1;refGene_gn=SLC49A3;refGene_tx=XM_023253384.1
B1 208061813 208062020
refGene_gn=SLC49A3;refGene_tx=XM_023253380.1;refGene_gn=SLC49A3;refGene_tx=XM_023253381.1;refGene_gn=SLC49A3;refGene_tx=XM_023253382.1;refGene_gn=SLC49A3;refGene_tx=XM_023253383.1;refGene_gn=SLC49A3;refGene_tx=XM_023253384.1
B1 208062900 208063002
refGene_gn=MYL5;refGene_tx=NM_001009874.3
B1 208063289 208063338

refGene_gn=MYL5;refGene_tx=NM_001009874.3
B1 208063957 208064036
refGene_gn=MYL5;refGene_tx=NM_001009874.3
B1 208064506 208064611
refGene_gn=MYL5;refGene_tx=NM_001009874.3
B1 208065381 208065457
refGene_gn=MYL5;refGene_tx=NM_001009874.3
B1 208065645 208065753
refGene_gn=MYL5;refGene_tx=NM_001009874.3
B1 208066303 208066306
refGene_gn=MYL5;refGene_tx=NM_001009874.3
B1 208069877 208069913
refGene_gn=ATP5ME;refGene_tx=XM_006931302.3
B1 208070132 208070187
refGene_gn=ATP5ME;refGene_tx=XM_006931302.3
B1 208070752 208070851
refGene_gn=ATP5ME;refGene_tx=XM_006931302.3
B1 208071333 208071359
refGene_gn=ATP5ME;refGene_tx=XM_006931302.3
B1 208073113 208073175
refGene_gn=PDE6B;refGene_tx=XM_003985610.3;refGene_gn=PDE6B;refGene_tx
=XM_019829878.1
B1 208074058 208074215
refGene_gn=PDE6B;refGene_tx=XM_003985610.3;refGene_gn=PDE6B;refGene_tx
=XM_019829878.1
B1 208075474 208075558
refGene_gn=PDE6B;refGene_tx=XM_003985610.3;refGene_gn=PDE6B;refGene_tx
=XM_019829878.1
B1 208076696 208076771
refGene_gn=PDE6B;refGene_tx=XM_003985610.3;refGene_gn=PDE6B;refGene_tx
=XM_019829878.1
B1 208077094 208077158
refGene_gn=PDE6B;refGene_tx=XM_003985610.3;refGene_gn=PDE6B;refGene_tx
=XM_019829878.1
B1 208077740 208077848
refGene_gn=PDE6B;refGene_tx=XM_003985610.3;refGene_gn=PDE6B;refGene_tx
=XM_019829878.1
B1 208077988 208078089
refGene_gn=PDE6B;refGene_tx=XM_003985610.3;refGene_gn=PDE6B;refGene_tx
=XM_019829878.1
B1 208078422 208078510
refGene_gn=PDE6B;refGene_tx=XM_003985610.3;refGene_gn=PDE6B;refGene_tx
=XM_019829878.1
B1 208078694 208078804
refGene_gn=PDE6B;refGene_tx=XM_003985610.3;refGene_gn=PDE6B;refGene_tx
=XM_019829878.1
B1 208079153 208079261
refGene_gn=PDE6B;refGene_tx=XM_003985610.3;refGene_gn=PDE6B;refGene_tx
=XM_019829878.1
B1 208080474 208080621

refGene_gn=PDE6B;refGene_tx=XM_003985610.3;refGene_gn=PDE6B;refGene_tx
=XM_019829878.1
B1	208081020	208081086
refGene_gn=PDE6B;refGene_tx=XM_003985610.3;refGene_gn=PDE6B;refGene_tx
=XM_019829878.1
B1	208081540	208081684
refGene_gn=PDE6B;refGene_tx=XM_003985610.3;refGene_gn=PDE6B;refGene_tx
=XM_019829878.1
B1	208081976	208082126
refGene_gn=PDE6B;refGene_tx=XM_003985610.3;refGene_gn=PDE6B;refGene_tx
=XM_019829878.1
B1	208082827	208082875
refGene_gn=PDE6B;refGene_tx=XM_003985610.3;refGene_gn=PDE6B;refGene_tx
=XM_019829878.1
B1	208083053	208083120
refGene_gn=PDE6B;refGene_tx=XM_003985610.3;refGene_gn=PDE6B;refGene_tx
=XM_019829878.1
B1	208083995	208084060
refGene_gn=PDE6B;refGene_tx=XM_003985610.3;refGene_gn=PDE6B;refGene_tx
=XM_019829878.1
B1	208085002	208085077
refGene_gn=PDE6B;refGene_tx=XM_003985610.3;refGene_gn=PDE6B;refGene_tx
=XM_019829878.1
B1	208085167	208085308
refGene_gn=PDE6B;refGene_tx=XM_003985610.3;refGene_gn=PDE6B;refGene_tx
=XM_019829878.1
B1	208102247	208102337
refGene_gn=PDE6B;refGene_tx=XM_003985610.3;refGene_gn=PDE6B;refGene_tx
=XM_019829878.1
B1	208102641	208102794
refGene_gn=PDE6B;refGene_tx=XM_003985610.3;refGene_gn=PDE6B;refGene_tx
=XM_019829878.1
B1	208103809	208103983
refGene_gn=PDE6B;refGene_tx=XM_019829878.1
B1	208107012	208107552
refGene_gn=PDE6B;refGene_tx=XM_003985610.3
B1	208152033	208152247
refGene_gn=PIGG;refGene_tx=XM_003985611.3;refGene_gn=PIGG;refGene_tx=X
M_006931303.3;refGene_gn=PIGG;refGene_tx=XM_006931304.2;refGene_gn=PIG
G;refGene_tx=XM_006931305.2;refGene_gn=PIGG;refGene_tx=XM_019829879.1;
refGene_gn=PIGG;refGene_tx=XM_019829880.1;refGene_gn=PIGG;refGene_tx=X
M_019829881.1;refGene_gn=PIGG;refGene_tx=XM_019829883.1;refGene_gn=PIG
G;refGene_tx=XM_019829885.1;refGene_gn=PIGG;refGene_tx=XM_019829886.1
B1	208156593	208156757
refGene_gn=PIGG;refGene_tx=XM_003985611.3;refGene_gn=PIGG;refGene_tx=X
M_006931303.3;refGene_gn=PIGG;refGene_tx=XM_006931304.2;refGene_gn=PIG
G;refGene_tx=XM_006931305.2;refGene_gn=PIGG;refGene_tx=XM_006931306.2;
refGene_gn=PIGG;refGene_tx=XM_019829879.1;refGene_gn=PIGG;refGene_tx=X
M_019829880.1;refGene_gn=PIGG;refGene_tx=XM_019829881.1;refGene_gn=PIG
G;refGene_tx=XM_019829883.1;refGene_gn=PIGG;refGene_tx=XM_019829885.1;

refGene_gn=PIGG;refGene_tx=XM_019829886.1
B1 208160553 208160863
refGene_gn=PIGG;refGene_tx=XM_003985611.3;refGene_gn=PIGG;refGene_tx=XM_006931303.3;refGene_gn=PIGG;refGene_tx=XM_006931304.2;refGene_gn=PIGG;refGene_tx=XM_006931305.2;refGene_gn=PIGG;refGene_tx=XM_019829879.1;refGene_gn=PIGG;refGene_tx=XM_019829880.1;refGene_gn=PIGG;refGene_tx=XM_019829881.1;refGene_gn=PIGG;refGene_tx=XM_019829883.1;refGene_gn=PIGG;refGene_tx=XM_019829885.1;refGene_gn=PIGG;refGene_tx=XM_019829886.1
B1 208163836 208164028
refGene_gn=PIGG;refGene_tx=XM_003985611.3;refGene_gn=PIGG;refGene_tx=XM_006931303.3;refGene_gn=PIGG;refGene_tx=XM_006931304.2;refGene_gn=PIGG;refGene_tx=XM_006931305.2;refGene_gn=PIGG;refGene_tx=XM_006931306.2;refGene_gn=PIGG;refGene_tx=XM_019829879.1;refGene_gn=PIGG;refGene_tx=XM_019829880.1;refGene_gn=PIGG;refGene_tx=XM_019829881.1;refGene_gn=PIGG;refGene_tx=XM_019829883.1;refGene_gn=PIGG;refGene_tx=XM_019829885.1;refGene_gn=PIGG;refGene_tx=XM_019829886.1
B1 208164887 208164903
refGene_gn=PIGG;refGene_tx=XM_019829884.2
B1 208166276 208166734
refGene_gn=PIGG;refGene_tx=XM_003985611.3;refGene_gn=PIGG;refGene_tx=XM_006931303.3;refGene_gn=PIGG;refGene_tx=XM_006931304.2;refGene_gn=PIGG;refGene_tx=XM_006931305.2;refGene_gn=PIGG;refGene_tx=XM_006931306.2;refGene_gn=PIGG;refGene_tx=XM_019829879.1;refGene_gn=PIGG;refGene_tx=XM_019829880.1;refGene_gn=PIGG;refGene_tx=XM_019829881.1;refGene_gn=PIGG;refGene_tx=XM_019829883.1;refGene_gn=PIGG;refGene_tx=XM_019829884.2;refGene_gn=PIGG;refGene_tx=XM_019829885.1;refGene_gn=PIGG;refGene_tx=XM_019829886.1
B1 208168097 208168379
refGene_gn=PIGG;refGene_tx=XM_003985611.3;refGene_gn=PIGG;refGene_tx=XM_006931303.3;refGene_gn=PIGG;refGene_tx=XM_006931305.2;refGene_gn=PIGG;refGene_tx=XM_006931306.2;refGene_gn=PIGG;refGene_tx=XM_019829879.1;refGene_gn=PIGG;refGene_tx=XM_019829880.1;refGene_gn=PIGG;refGene_tx=XM_019829881.1;refGene_gn=PIGG;refGene_tx=XM_019829883.1;refGene_gn=PIGG;refGene_tx=XM_019829884.2;refGene_gn=PIGG;refGene_tx=XM_019829885.1
B1 208168725 208168943
refGene_gn=PIGG;refGene_tx=XM_003985611.3;refGene_gn=PIGG;refGene_tx=XM_006931303.3;refGene_gn=PIGG;refGene_tx=XM_006931304.2;refGene_gn=PIGG;refGene_tx=XM_006931305.2;refGene_gn=PIGG;refGene_tx=XM_006931306.2;refGene_gn=PIGG;refGene_tx=XM_019829879.1;refGene_gn=PIGG;refGene_tx=XM_019829880.1;refGene_gn=PIGG;refGene_tx=XM_019829881.1;refGene_gn=PIGG;refGene_tx=XM_019829883.1;refGene_gn=PIGG;refGene_tx=XM_019829884.2;refGene_gn=PIGG;refGene_tx=XM_019829885.1
B1 208172381 208172594
refGene_gn=PIGG;refGene_tx=XM_003985611.3;refGene_gn=PIGG;refGene_tx=XM_006931303.3;refGene_gn=PIGG;refGene_tx=XM_006931304.2;refGene_gn=PIGG;refGene_tx=XM_006931305.2;refGene_gn=PIGG;refGene_tx=XM_006931306.2;refGene_gn=PIGG;refGene_tx=XM_019829879.1;refGene_gn=PIGG;refGene_tx=XM_019829880.1;refGene_gn=PIGG;refGene_tx=XM_019829881.1;refGene_gn=PIGG;refGene_tx=XM_019829883.1;refGene_gn=PIGG;refGene_tx=XM_019829884.2;refGene_gn=PIGG;refGene_tx=XM_019829886.1

B1	208181594	208181736
refGene_gn=PIGG;refGene_tx=XM_003985611.3;refGene_gn=PIGG;refGene_tx=XM_006931303.3;refGene_gn=PIGG;refGene_tx=XM_006931304.2;refGene_gn=PIGG;refGene_tx=XM_006931305.2;refGene_gn=PIGG;refGene_tx=XM_006931306.2;refGene_gn=PIGG;refGene_tx=XM_019829879.1;refGene_gn=PIGG;refGene_tx=XM_019829880.1;refGene_gn=PIGG;refGene_tx=XM_019829881.1;refGene_gn=PIGG;refGene_tx=XM_019829883.1;refGene_gn=PIGG;refGene_tx=XM_019829884.2
B1	208182652	208182841
refGene_gn=PIGG;refGene_tx=XM_003985611.3;refGene_gn=PIGG;refGene_tx=XM_006931303.3;refGene_gn=PIGG;refGene_tx=XM_006931304.2;refGene_gn=PIGG;refGene_tx=XM_006931305.2;refGene_gn=PIGG;refGene_tx=XM_006931306.2;refGene_gn=PIGG;refGene_tx=XM_019829879.1;refGene_gn=PIGG;refGene_tx=XM_019829880.1;refGene_gn=PIGG;refGene_tx=XM_019829881.1;refGene_gn=PIGG;refGene_tx=XM_019829883.1;refGene_gn=PIGG;refGene_tx=XM_019829884.2
B1	208184610	208184820
refGene_gn=PIGG;refGene_tx=XM_003985611.3;refGene_gn=PIGG;refGene_tx=XM_006931303.3;refGene_gn=PIGG;refGene_tx=XM_006931304.2;refGene_gn=PIGG;refGene_tx=XM_006931305.2;refGene_gn=PIGG;refGene_tx=XM_006931306.2;refGene_gn=PIGG;refGene_tx=XM_019829879.1;refGene_gn=PIGG;refGene_tx=XM_019829880.1;refGene_gn=PIGG;refGene_tx=XM_019829884.2
B1	208193612	208193818
refGene_gn=PIGG;refGene_tx=XM_003985611.3;refGene_gn=PIGG;refGene_tx=XM_006931303.3;refGene_gn=PIGG;refGene_tx=XM_006931304.2;refGene_gn=PIGG;refGene_tx=XM_006931306.2;refGene_gn=PIGG;refGene_tx=XM_019829879.1;refGene_gn=PIGG;refGene_tx=XM_019829880.1;refGene_gn=PIGG;refGene_tx=XM_019829881.1;refGene_gn=PIGG;refGene_tx=XM_019829884.2
B1	208194662	208194816
refGene_gn=PIGG;refGene_tx=XM_003985611.3;refGene_gn=PIGG;refGene_tx=XM_006931304.2;refGene_gn=PIGG;refGene_tx=XM_006931306.2;refGene_gn=PIGG;refGene_tx=XM_019829881.1;refGene_gn=PIGG;refGene_tx=XM_019829884.2
B2	144358	144418
refGene_gn=LOC101084270;refGene_tx=XM_023253563.1;refGene_gn=LOC101084270;refGene_tx=XM_023253564.1;refGene_gn=LOC101084270;refGene_tx=XM_023253565.1
B2	145130	145154
refGene_gn=LOC101084270;refGene_tx=XM_023253563.1;refGene_gn=LOC101084270;refGene_tx=XM_023253564.1
B2	147349	147486
refGene_gn=LOC101084270;refGene_tx=XM_023253563.1;refGene_gn=LOC101084270;refGene_tx=XM_023253564.1;refGene_gn=LOC101084270;refGene_tx=XM_023253565.1
B2	148351	148384
refGene_gn=LOC101084270;refGene_tx=XM_023253563.1;refGene_gn=LOC101084270;refGene_tx=XM_023253564.1;refGene_gn=LOC101084270;refGene_tx=XM_023253565.1
B2	149807	149866
refGene_gn=LOC101084270;refGene_tx=XM_023253563.1;refGene_gn=LOC101084270;refGene_tx=XM_023253564.1;refGene_gn=LOC101084270;refGene_tx=XM_023253565.1
B2	152999	153022

refGene_gn=LOC101084270;refGene_tx=XM_023253563.1;refGene_gn=LOC101084270;refGene_tx=XM_023253564.1;refGene_gn=LOC101084270;refGene_tx=XM_023253565.1
B2 156046 156079 refGene_gn=LOC101084270;refGene_tx=XM_023253563.1;refGene_gn=LOC101084270;refGene_tx=XM_023253564.1;refGene_gn=LOC101084270;refGene_tx=XM_023253565.1
B2 193733 194171 refGene_gn=LOC101096649;refGene_tx=XM_011282019.3
B2 194268 194364 refGene_gn=LOC101096649;refGene_tx=XM_011282019.3
B2 196609 196840 refGene_gn=LOC101096649;refGene_tx=XM_011282019.3
B2 199860 199883 refGene_gn=LOC101096649;refGene_tx=XM_011282019.3
B2 200283 200399 refGene_gn=LOC101096649;refGene_tx=XM_011282019.3
B2 203012 203045 refGene_gn=LOC101096649;refGene_tx=XM_011282019.3
B2 203406 204089 refGene_gn=LOC101096649;refGene_tx=XM_011282019.3
B2 218148 218191 refGene_gn=TRIM15;refGene_tx=XM_023253562.1
B2 218764 218966 refGene_gn=TRIM15;refGene_tx=XM_023253562.1
B2 220011 220044 refGene_gn=TRIM15;refGene_tx=XM_023253562.1
B2 220400 220513 refGene_gn=TRIM15;refGene_tx=XM_023253562.1
B2 221542 221565 refGene_gn=TRIM15;refGene_tx=XM_023253562.1
B2 222315 222543 refGene_gn=TRIM15;refGene_tx=XM_023253562.1
B2 223507 223603 refGene_gn=TRIM15;refGene_tx=XM_023253562.1
B2 224771 225017 refGene_gn=TRIM15;refGene_tx=XM_023253562.1
B2 228026 228455 refGene_gn=LOC101098631;refGene_tx=XM_011282020.3
B2 229411 229507 refGene_gn=LOC101098631;refGene_tx=XM_011282020.3
B2 230034 230265 refGene_gn=LOC101098631;refGene_tx=XM_011282020.3
B2 230797 230820 refGene_gn=LOC101098631;refGene_tx=XM_011282020.3
B2 231091 231207 refGene_gn=LOC101098631;refGene_tx=XM_011282020.3
B2 232837 232870 refGene_gn=LOC101098631;refGene_tx=XM_011282020.3
B2 233206 233724 refGene_gn=LOC101098631;refGene_tx=XM_011282020.3
B2 283122 283485 refGene_gn=RNF39;refGene_tx=XM_023253566.1
B2 284896 284919 refGene_gn=RNF39;refGene_tx=XM_023253566.1
B2 285114 285206 refGene_gn=RNF39;refGene_tx=XM_023253566.1
B2 286740 287315 refGene_gn=RNF39;refGene_tx=XM_023253566.1
B2 288927 289130 refGene_gn=PPP1R11;refGene_tx=XM_003985617.5
B2 291558 291667 refGene_gn=PPP1R11;refGene_tx=XM_003985617.5
B2 292592 292661 refGene_gn=PPP1R11;refGene_tx=XM_003985617.5

B2	295263	295288	refGene_gn=ZNRD1;refGene_tx=XM_003985618.4
B2	296950	297060	refGene_gn=ZNRD1;refGene_tx=XM_003985618.4
B2	297257	297358	refGene_gn=ZNRD1;refGene_tx=XM_003985618.4
B2	297456	297592	refGene_gn=ZNRD1;refGene_tx=XM_003985618.4
B2	300247	300474	refGene_gn=ZFP57;refGene_tx=XM_006931314.4
B2	300721	300797
refGene_gn=ZFP57;refGene_tx=XM_006931314.4;refGene_gn=ZFP57;refGene_tx=XM_006931315.4;refGene_gn=ZFP57;refGene_tx=XM_006931316.4;refGene_gn=ZFP57;refGene_tx=XM_019830076.2
B2	315155	315599
refGene_gn=ZFP57;refGene_tx=XM_006931314.4;refGene_gn=ZFP57;refGene_tx=XM_006931315.4;refGene_gn=ZFP57;refGene_tx=XM_006931316.4;refGene_gn=ZFP57;refGene_tx=XM_019830076.2
B2	316456	316583
refGene_gn=ZFP57;refGene_tx=XM_006931314.4;refGene_gn=ZFP57;refGene_tx=XM_006931315.4;refGene_gn=ZFP57;refGene_tx=XM_006931316.4;refGene_gn=ZFP57;refGene_tx=XM_019830076.2
B2	317004	317088
refGene_gn=ZFP57;refGene_tx=XM_006931314.4;refGene_gn=ZFP57;refGene_tx=XM_006931315.4;refGene_gn=ZFP57;refGene_tx=XM_006931316.4;refGene_gn=ZFP57;refGene_tx=XM_019830076.2
B2	317741	318967
refGene_gn=ZFP57;refGene_tx=XM_006931314.4;refGene_gn=ZFP57;refGene_tx=XM_006931315.4;refGene_gn=ZFP57;refGene_tx=XM_006931316.4;refGene_gn=ZFP57;refGene_tx=XM_019830076.2
B2	319690	319704	refGene_gn=MOG;refGene_tx=XM_019830077.2
B2	319981	320002
refGene_gn=MOG;refGene_tx=XM_019830077.2;refGene_gn=MOG;refGene_tx=XM_019830078.2
B2	320208	320325
refGene_gn=MOG;refGene_tx=XM_019830077.2;refGene_gn=MOG;refGene_tx=XM_019830078.2
B2	321596	321617
refGene_gn=MOG;refGene_tx=XM_019830077.2;refGene_gn=MOG;refGene_tx=XM_019830078.2
B2	321832	321853
refGene_gn=MOG;refGene_tx=XM_019830077.2;refGene_gn=MOG;refGene_tx=XM_019830078.2
B2	322752	322866
refGene_gn=MOG;refGene_tx=XM_019830077.2;refGene_gn=MOG;refGene_tx=XM_019830078.2
B2	326875	327223
refGene_gn=MOG;refGene_tx=XM_019830077.2;refGene_gn=MOG;refGene_tx=XM_019830078.2
B2	328275	328357
refGene_gn=MOG;refGene_tx=XM_019830077.2;refGene_gn=MOG;refGene_tx=XM_019830078.2
B2	348380	348474	refGene_gn=GABBR1;refGene_tx=XM_023253567.1
B2	349062	349266	refGene_gn=GABBR1;refGene_tx=XM_023253567.1
B2	349953	350139	refGene_gn=GABBR1;refGene_tx=XM_023253567.1

B2 351403 351424 refGene_gn=GABBR1;refGene_tx=XM_023253567.1
B2 352542 352703 refGene_gn=GABBR1;refGene_tx=XM_023253567.1
B2 355802 355937 refGene_gn=GABBR1;refGene_tx=XM_023253567.1
B2 356211 356382 refGene_gn=GABBR1;refGene_tx=XM_023253567.1
B2 357316 357418 refGene_gn=GABBR1;refGene_tx=XM_023253567.1
B2 358043 358109 refGene_gn=GABBR1;refGene_tx=XM_023253567.1
B2 358529 358721 refGene_gn=GABBR1;refGene_tx=XM_023253567.1
B2 366044 366287 refGene_gn=GABBR1;refGene_tx=XM_023253567.1
B2 366876 366940 refGene_gn=GABBR1;refGene_tx=XM_023253567.1
B2 368297 368375 refGene_gn=GABBR1;refGene_tx=XM_023253567.1
B2 369881 370032 refGene_gn=GABBR1;refGene_tx=XM_023253567.1
B2 370462 370595 refGene_gn=GABBR1;refGene_tx=XM_023253567.1
B2 371669 371786 refGene_gn=GABBR1;refGene_tx=XM_023253567.1
B2 371876 371984 refGene_gn=GABBR1;refGene_tx=XM_023253567.1
B2 372315 372409 refGene_gn=GABBR1;refGene_tx=XM_023253567.1
B2 372989 373117 refGene_gn=GABBR1;refGene_tx=XM_023253567.1
B2 373547 373676 refGene_gn=GABBR1;refGene_tx=XM_023253567.1
B2 373824 373851 refGene_gn=GABBR1;refGene_tx=XM_023253567.1
B2 373852 373969 refGene_gn=GABBR1;refGene_tx=XM_023253567.1
B2 374169 374343 refGene_gn=GABBR1;refGene_tx=XM_023253567.1
B2 385329 385525
refGene_gn=LOC109500078;refGene_tx=XM_023254636.1
B2 385526 385705
refGene_gn=LOC109500078;refGene_tx=XM_023254636.1
B2 385706 385825
refGene_gn=LOC109500078;refGene_tx=XM_023254636.1
B2 385827 385987
refGene_gn=LOC109500078;refGene_tx=XM_023254636.1
B2 386162 386338
refGene_gn=LOC109500078;refGene_tx=XM_023254636.1
B2 389320 389456
refGene_gn=LOC109500078;refGene_tx=XM_023254636.1
B2 401998 402070 refGene_gn=UBD;refGene_tx=XM_023254637.1
B2 402736 402776 refGene_gn=UBD;refGene_tx=XM_023254637.1
B2 403980 404339 refGene_gn=UBD;refGene_tx=XM_023254637.1
B2 404340 404425 refGene_gn=UBD;refGene_tx=XM_023254637.1
B2 405488 405883 refGene_gn=UBD;refGene_tx=XM_023254637.1
B2 416346 416625
refGene_gn=LOC101100132;refGene_tx=XM_019831549.2
B2 419731 419848
refGene_gn=LOC101100132;refGene_tx=XM_019831549.2
B2 419849 420234
refGene_gn=LOC101100132;refGene_tx=XM_019831549.2
B2 420236 420270
refGene_gn=LOC101100132;refGene_tx=XM_019831549.2
B2 456505 457453
refGene_gn=LOC101100389;refGene_tx=XM_003985627.1
B2 471782 472736
refGene_gn=LOC105260533;refGene_tx=XM_011282042.3
B2 487407 488361

```
refGene_gn=LOC101100889;refGene_tx=XM_011282043.3
B2      501248  502199
refGene_gn=LOC101101134;refGene_tx=XM_023253568.1
B2      526160  527096
refGene_gn=LOC101101385;refGene_tx=XM_003985630.1
B2      550905  551835
refGene_gn=LOC101080455;refGene_tx=XM_003985631.1
B2      580079  581033
refGene_gn=LOC101080711;refGene_tx=XM_003985632.2
B2      617659  618625
refGene_gn=LOC101080967;refGene_tx=XM_003985633.3
B2      635858  636980
refGene_gn=LOC101081467;refGene_tx=XM_003985635.3
B2      660793  661738
refGene_gn=LOC101081729;refGene_tx=XM_003985636.3
B2      670931  671891
refGene_gn=LOC101081980;refGene_tx=XM_003985637.1
B2      687334  688294
refGene_gn=LOC101082234;refGene_tx=XM_003985638.3
B2      716634  717582
refGene_gn=LOC111560402;refGene_tx=XM_023253569.1
B2      727135  728083
refGene_gn=LOC111560403;refGene_tx=XM_023253570.1
B2      778048  778993
refGene_gn=LOC101083004;refGene_tx=XM_006931322.1
B2      807499  808453
refGene_gn=LOC111560562;refGene_tx=XM_023254638.1
B2      932592  933531
refGene_gn=LOC101083769;refGene_tx=XM_023254703.1
B2      1008324 1009269
refGene_gn=LOC101084028;refGene_tx=XM_023254639.1
B2      1024180 1025125
refGene_gn=LOC101084283;refGene_tx=XM_003985644.3
B2      1046079 1047021
refGene_gn=LOC101084538;refGene_tx=XM_011282023.1
B2      1074324 1075278
refGene_gn=LOC101084792;refGene_tx=XM_003985646.2
B2      1106701 1107044
refGene_gn=LOC101097393;refGene_tx=XM_003985619.4
B2      1107046 1107630
refGene_gn=LOC101097393;refGene_tx=XM_003985619.4
B2      1251673 1252343
refGene_gn=LOC101085545;refGene_tx=XM_023254640.1
B2      1252375 1252601
refGene_gn=LOC101085545;refGene_tx=XM_023254640.1
B2      1256818 1257047
refGene_gn=LOC101085545;refGene_tx=XM_023254640.1
B2      1275894 1276839
refGene_gn=LOC101085800;refGene_tx=XM_003985649.2
B2      1297510 1298446
```

refGene_gn=LOC101097641;refGene_tx=XM_003985620.4
B2 1321782 1322712
refGene_gn=LOC101097892;refGene_tx=XM_003985621.3
B2 1333146 1334059
refGene_gn=LOC101086059;refGene_tx=XM_006931324.3
B2 1334060 1334077
refGene_gn=LOC101086059;refGene_tx=XM_006931324.3
B2 1410957 1411871
refGene_gn=LOC101086806;refGene_tx=XM_003985652.3
B2 1419840 1419864
refGene_gn=LOC101086806;refGene_tx=XM_003985652.3
B2 1425138 1425406
refGene_gn=LOC101086806;refGene_tx=XM_003985652.3
B2 1460212 1461145
refGene_gn=LOC101086560;refGene_tx=XM_003985651.4
B2 1573614 1574577
refGene_gn=LOC105260530;refGene_tx=XM_011282026.3
B2 1607898 1608861
refGene_gn=LOC105260531;refGene_tx=XM_011282027.3
B2 1669859 1669868
refGene_gn=ZNF311;refGene_tx=XM_023254665.1;refGene_gn=ZNF311;refGene_tx=XM_023254666.1;refGene_gn=ZNF311;refGene_tx=XM_023254667.1
B2 1670155 1670225
refGene_gn=ZNF311;refGene_tx=XM_023254665.1;refGene_gn=ZNF311;refGene_tx=XM_023254666.1;refGene_gn=ZNF311;refGene_tx=XM_023254667.1;refGene_gn=ZNF311;refGene_tx=XM_023254668.1
B2 1673736 1673813
refGene_gn=ZNF311;refGene_tx=XM_023254665.1;refGene_gn=ZNF311;refGene_tx=XM_023254666.1;refGene_gn=ZNF311;refGene_tx=XM_023254667.1;refGene_gn=ZNF311;refGene_tx=XM_023254668.1
B2 1674154 1674281
refGene_gn=ZNF311;refGene_tx=XM_023254665.1;refGene_gn=ZNF311;refGene_tx=XM_023254666.1;refGene_gn=ZNF311;refGene_tx=XM_023254667.1;refGene_gn=ZNF311;refGene_tx=XM_023254668.1
B2 1674607 1674743
refGene_gn=ZNF311;refGene_tx=XM_023254665.1;refGene_gn=ZNF311;refGene_tx=XM_023254666.1;refGene_gn=ZNF311;refGene_tx=XM_023254667.1;refGene_gn=ZNF311;refGene_tx=XM_023254668.1;refGene_gn=ZNF311;refGene_tx=XM_023254669.1
B2 1676257 1677642
refGene_gn=ZNF311;refGene_tx=XM_023254665.1;refGene_gn=ZNF311;refGene_tx=XM_023254666.1;refGene_gn=ZNF311;refGene_tx=XM_023254667.1;refGene_gn=ZNF311;refGene_tx=XM_023254668.1;refGene_gn=ZNF311;refGene_tx=XM_023254669.1
B2 1721024 1721444 refGene_gn=TRIM27;refGene_tx=XM_023254671.1
B2 1722488 1722584 refGene_gn=TRIM27;refGene_tx=XM_023254671.1
B2 1724981 1725212 refGene_gn=TRIM27;refGene_tx=XM_023254671.1
B2 1731052 1731075 refGene_gn=TRIM27;refGene_tx=XM_023254671.1
B2 1732849 1732965 refGene_gn=TRIM27;refGene_tx=XM_023254671.1
B2 1733097 1733130 refGene_gn=TRIM27;refGene_tx=XM_023254671.1

B2 1734915 1734942 refGene_gn=TRIM27;refGene_tx=XM_023254671.1
B2 1736391 1736987 refGene_gn=TRIM27;refGene_tx=XM_023254671.1
B2 1738993 1739059
refGene_gn=LOC102899754;refGene_tx=XM_023254642.1
B2 1743218 1743399
refGene_gn=LOC102899754;refGene_tx=XM_023254642.1
B2 1743481 1743679
refGene_gn=LOC102899754;refGene_tx=XM_023254642.1
B2 1743960 1744070
refGene_gn=LOC102899754;refGene_tx=XM_023254642.1
B2 1749427 1749433
refGene_gn=LOC102899754;refGene_tx=XM_023254642.1
B2 1750123 1751038
refGene_gn=LOC102899754;refGene_tx=XM_023254642.1
B2 1886831 1888444
refGene_gn=LOC111560563;refGene_tx=XM_023254643.1
B2 1888713 1888913
refGene_gn=LOC111560563;refGene_tx=XM_023254643.1
B2 1888968 1889138
refGene_gn=LOC111560563;refGene_tx=XM_023254643.1
B2 1889315 1889553
refGene_gn=LOC111560563;refGene_tx=XM_023254643.1
B2 1897088 1897146
refGene_gn=LOC111560563;refGene_tx=XM_023254643.1
B2 1901029 1901092
refGene_gn=LOC111560563;refGene_tx=XM_023254643.1
B2 1901343 1901543
refGene_gn=LOC111560563;refGene_tx=XM_023254643.1
B2 1907400 1908243
refGene_gn=LOC111560563;refGene_tx=XM_023254643.1
B2 1908382 1908829
refGene_gn=LOC111560563;refGene_tx=XM_023254643.1;refGene_gn=ZBED9;refGene_tx=XM_011282028.3;refGene_gn=ZBED9;refGene_tx=XM_011282029.3;refGene_gn=ZBED9;refGene_tx=XM_019830082.2;refGene_gn=ZBED9;refGene_tx=XM_019830084.2;refGene_gn=ZBED9;refGene_tx=XM_019830085.2
B2 1909919 1910045 refGene_gn=ZBED9;refGene_tx=XM_019830085.2
B2 1918257 1918405
refGene_gn=ZBED9;refGene_tx=XM_011282028.3;refGene_gn=ZBED9;refGene_tx=XM_019830082.2;refGene_gn=ZBED9;refGene_tx=XM_019830084.2
B2 1919709 1921205
refGene_gn=ZBED9;refGene_tx=XM_011282028.3;refGene_gn=ZBED9;refGene_tx=XM_019830082.2;refGene_gn=ZBED9;refGene_tx=XM_019830084.2
B2 1922267 1924184
refGene_gn=ZBED9;refGene_tx=XM_011282028.3;refGene_gn=ZBED9;refGene_tx=XM_011282029.3;refGene_gn=ZBED9;refGene_tx=XM_019830082.2;refGene_gn=ZBED9;refGene_tx=XM_019830084.2
B2 1963760 1963967
refGene_gn=GPX5;refGene_tx=XM_019830094.1;refGene_gn=GPX5;refGene_tx=XM_019830095.2;refGene_gn=GPX5;refGene_tx=XM_023254695.1
B2 1965188 1965288

refGene_gn=GPX5;refGene_tx=XM_019830094.1;refGene_gn=GPX5;refGene_tx=XM_019830095.2;refGene_gn=GPX5;refGene_tx=XM_023254695.1
B2 1966325 1966443
refGene_gn=GPX5;refGene_tx=XM_019830094.1;refGene_gn=GPX5;refGene_tx=XM_019830095.2;refGene_gn=GPX5;refGene_tx=XM_023254695.1
B2 1969227 1969381
refGene_gn=GPX5;refGene_tx=XM_019830094.1;refGene_gn=GPX5;refGene_tx=XM_019830095.2;refGene_gn=GPX5;refGene_tx=XM_023254695.1
B2 1972087 1972177 refGene_gn=GPX5;refGene_tx=XM_023254695.1
B2 1984870 1984906
refGene_gn=GPX6;refGene_tx=XM_019830099.2;refGene_gn=GPX6;refGene_tx=XM_023254696.1
B2 1988942 1989029 refGene_gn=GPX6;refGene_tx=XM_003985655.5
B2 1992359 1992513
refGene_gn=GPX6;refGene_tx=XM_003985655.5;refGene_gn=GPX6;refGene_tx=XM_019830099.2;refGene_gn=GPX6;refGene_tx=XM_019830102.2;refGene_gn=GPX6;refGene_tx=XM_023254696.1
B2 1997903 1998021
refGene_gn=GPX6;refGene_tx=XM_003985655.5;refGene_gn=GPX6;refGene_tx=XM_019830099.2;refGene_gn=GPX6;refGene_tx=XM_019830102.2;refGene_gn=GPX6;refGene_tx=XM_023254696.1
B2 1998509 1998609
refGene_gn=GPX6;refGene_tx=XM_003985655.5;refGene_gn=GPX6;refGene_tx=XM_019830099.2;refGene_gn=GPX6;refGene_tx=XM_019830102.2;refGene_gn=GPX6;refGene_tx=XM_023254696.1
B2 1999749 1999956
refGene_gn=GPX6;refGene_tx=XM_003985655.5;refGene_gn=GPX6;refGene_tx=XM_019830099.2;refGene_gn=GPX6;refGene_tx=XM_019830102.2;refGene_gn=GPX6;refGene_tx=XM_023254696.1
B2 2025555 2026500
refGene_gn=LOC101090104;refGene_tx=XM_003985665.2
B2 2030740 2030927
refGene_gn=LOC109499510;refGene_tx=XM_023254697.1
B2 2030929 2031129
refGene_gn=LOC109499510;refGene_tx=XM_023254697.1
B2 2031392 2031534
refGene_gn=LOC109499510;refGene_tx=XM_023254697.1
B2 2042176 2042241
refGene_gn=LOC109499510;refGene_tx=XM_023254697.1
B2 2042252 2042660
refGene_gn=ZSCAN23;refGene_tx=XM_023254674.1;refGene_gn=ZSCAN23;refGene_tx=XM_023254675.1;refGene_gn=ZSCAN23;refGene_tx=XM_023254676.1;refGene_gn=ZSCAN23;refGene_tx=XM_023254677.1;refGene_gn=ZSCAN23;refGene_tx=XM_023254678.1;refGene_gn=ZSCAN23;refGene_tx=XM_023254679.1;refGene_gn=ZSCAN23;refGene_tx=XM_023254680.1;refGene_gn=ZSCAN23;refGene_tx=XM_023254681.1
B2 2043134 2043285
refGene_gn=ZSCAN23;refGene_tx=XM_023254674.1;refGene_gn=ZSCAN23;refGene_tx=XM_023254675.1;refGene_gn=ZSCAN23;refGene_tx=XM_023254676.1;refGene_gn=ZSCAN23;refGene_tx=XM_023254677.1;refGene_gn=ZSCAN23;refGene_tx=

XM_023254678.1;refGene_gn=ZSCAN23;refGene_tx=XM_023254679.1;refGene_gn=ZSCAN23;refGene_tx=XM_023254680.1;refGene_gn=ZSCAN23;refGene_tx=XM_023254681.1
B2 2043616 2044323
refGene_gn=ZSCAN23;refGene_tx=XM_023254674.1;refGene_gn=ZSCAN23;refGene_tx=XM_023254675.1;refGene_gn=ZSCAN23;refGene_tx=XM_023254676.1;refGene_gn=ZSCAN23;refGene_tx=XM_023254677.1;refGene_gn=ZSCAN23;refGene_tx=XM_023254678.1;refGene_gn=ZSCAN23;refGene_tx=XM_023254679.1;refGene_gn=ZSCAN23;refGene_tx=XM_023254680.1;refGene_gn=ZSCAN23;refGene_tx=XM_023254681.1
B2 2048938 2048989
refGene_gn=ZSCAN23;refGene_tx=XM_023254676.1;refGene_gn=ZSCAN23;refGene_tx=XM_023254677.1;refGene_gn=ZSCAN23;refGene_tx=XM_023254678.1;refGene_gn=ZSCAN23;refGene_tx=XM_023254679.1;refGene_gn=ZSCAN23;refGene_tx=XM_023254680.1;refGene_gn=ZSCAN23;refGene_tx=XM_023254681.1
B2 2060191 2060602
refGene_gn=ZSCAN12;refGene_tx=XM_006931332.4;refGene_gn=ZSCAN12;refGene_tx=XM_019830086.2
B2 2065149 2065315
refGene_gn=ZSCAN12;refGene_tx=XM_006931332.4;refGene_gn=ZSCAN12;refGene_tx=XM_019830086.2
B2 2067309 2068628
refGene_gn=ZSCAN12;refGene_tx=XM_006931332.4;refGene_gn=ZSCAN12;refGene_tx=XM_019830086.2
B2 2094342 2095205
refGene_gn=LOC101087838;refGene_tx=XM_023254659.1;refGene_gn=LOC101087838;refGene_tx=XM_023254660.1;refGene_gn=LOC101087838;refGene_tx=XM_023254661.1;refGene_gn=LOC101087838;refGene_tx=XM_023254662.1;refGene_gn=LOC101087838;refGene_tx=XM_023254663.1;refGene_gn=LOC101087838;refGene_tx=XM_023254664.1
B2 2096114 2096238
refGene_gn=LOC101087838;refGene_tx=XM_023254659.1;refGene_gn=LOC101087838;refGene_tx=XM_023254660.1;refGene_gn=LOC101087838;refGene_tx=XM_023254661.1;refGene_gn=LOC101087838;refGene_tx=XM_023254662.1;refGene_gn=LOC101087838;refGene_tx=XM_023254663.1;refGene_gn=LOC101087838;refGene_tx=XM_023254664.1
B2 2097212 2097295
refGene_gn=LOC101087838;refGene_tx=XM_023254659.1;refGene_gn=LOC101087838;refGene_tx=XM_023254660.1;refGene_gn=LOC101087838;refGene_tx=XM_023254661.1;refGene_gn=LOC101087838;refGene_tx=XM_023254662.1;refGene_gn=LOC101087838;refGene_tx=XM_023254663.1;refGene_gn=LOC101087838;refGene_tx=XM_023254664.1
B2 2098354 2098502
refGene_gn=LOC101087838;refGene_tx=XM_023254659.1;refGene_gn=LOC101087838;refGene_tx=XM_023254660.1;refGene_gn=LOC101087838;refGene_tx=XM_023254661.1;refGene_gn=LOC101087838;refGene_tx=XM_023254662.1;refGene_gn=LOC101087838;refGene_tx=XM_023254663.1;refGene_gn=LOC101087838;refGene_tx=XM_023254664.1
B2 2100017 2100473
refGene_gn=LOC101087838;refGene_tx=XM_023254659.1;refGene_gn=LOC101087

838;refGene_tx=XM_023254660.1;refGene_gn=LOC101087838;refGene_tx=XM_02
3254661.1;refGene_gn=LOC101087838;refGene_tx=XM_023254662.1;refGene_gn
=LOC101087838;refGene_tx=XM_023254663.1;refGene_gn=LOC101087838;refGen
e_tx=XM_023254664.1
B2	2104879	2104933
refGene_gn=LOC101087838;refGene_tx=XM_023254659.1;refGene_gn=LOC101087
838;refGene_tx=XM_023254660.1;refGene_gn=LOC101087838;refGene_tx=XM_02
3254661.1
B2	2106277	2106355
refGene_gn=LOC101087838;refGene_tx=XM_023254659.1;refGene_gn=LOC101087
838;refGene_tx=XM_023254660.1;refGene_gn=LOC101087838;refGene_tx=XM_02
3254661.1;refGene_gn=LOC101087838;refGene_tx=XM_023254662.1
B2	2106698	2106731
refGene_gn=LOC101087838;refGene_tx=XM_023254663.1
B2	2107487	2107990	refGene_gn=ZSCAN31;refGene_tx=XM_006931335.3
B2	2113422	2113544	refGene_gn=ZSCAN31;refGene_tx=XM_006931338.4
B2	2113668	2114131
refGene_gn=ZSCAN31;refGene_tx=XM_006931335.3;refGene_gn=ZSCAN31;refGen
e_tx=XM_006931338.4;refGene_gn=ZSCAN31;refGene_tx=XM_019831664.1;refGe
ne_gn=ZSCAN31;refGene_tx=XM_019831665.1;refGene_gn=ZSCAN31;refGene_tx=
XM_023254657.1;refGene_gn=ZSCAN31;refGene_tx=XM_023254658.1
B2	2115029	2115180
refGene_gn=ZSCAN31;refGene_tx=XM_006931335.3;refGene_gn=ZSCAN31;refGen
e_tx=XM_006931338.4;refGene_gn=ZSCAN31;refGene_tx=XM_019831664.1;refGe
ne_gn=ZSCAN31;refGene_tx=XM_019831665.1;refGene_gn=ZSCAN31;refGene_tx=
XM_023254657.1;refGene_gn=ZSCAN31;refGene_tx=XM_023254658.1
B2	2115676	2116452
refGene_gn=ZSCAN31;refGene_tx=XM_006931335.3;refGene_gn=ZSCAN31;refGen
e_tx=XM_006931338.4;refGene_gn=ZSCAN31;refGene_tx=XM_019831664.1;refGe
ne_gn=ZSCAN31;refGene_tx=XM_019831665.1;refGene_gn=ZSCAN31;refGene_tx=
XM_023254657.1;refGene_gn=ZSCAN31;refGene_tx=XM_023254658.1
B2	2129572	2129624	refGene_gn=PGBD1;refGene_tx=XM_019831660.1
B2	2129901	2131453
refGene_gn=PGBD1;refGene_tx=XM_006931341.3;refGene_gn=PGBD1;refGene_tx
=XM_006931342.3
B2	2134865	2134962
refGene_gn=PGBD1;refGene_tx=XM_006931341.3;refGene_gn=PGBD1;refGene_tx
=XM_006931342.3;refGene_gn=PGBD1;refGene_tx=XM_019831660.1
B2	2152921	2153078
refGene_gn=PGBD1;refGene_tx=XM_006931341.3;refGene_gn=PGBD1;refGene_tx
=XM_006931342.3;refGene_gn=PGBD1;refGene_tx=XM_019831660.1
B2	2154420	2154807
refGene_gn=PGBD1;refGene_tx=XM_006931341.3;refGene_gn=PGBD1;refGene_tx
=XM_006931342.3;refGene_gn=PGBD1;refGene_tx=XM_019831660.1
B2	2156900	2156995
refGene_gn=LOC109500248;refGene_tx=XM_019831688.1
B2	2157765	2158342
refGene_gn=LOC109500248;refGene_tx=XM_019831688.1
B2	2164955	2165857
refGene_gn=ZSCAN26;refGene_tx=XM_003985657.4;refGene_gn=ZSCAN26;refGen

e_tx=XM_006931345.4;refGene_gn=ZSCAN26;refGene_tx=XM_019831671.1
B2 2169134 2169252
refGene_gn=ZSCAN26;refGene_tx=XM_003985657.4;refGene_gn=ZSCAN26;refGene_tx=XM_006931345.4;refGene_gn=ZSCAN26;refGene_tx=XM_019831671.1
B2 2169574 2170054
refGene_gn=ZSCAN26;refGene_tx=XM_003985657.4;refGene_gn=ZSCAN26;refGene_tx=XM_006931345.4
B2 2171985 2172060 refGene_gn=ZSCAN26;refGene_tx=XM_003985657.4
B2 2179598 2180810 refGene_gn=NKAPL;refGene_tx=XM_003985658.5
B2 2186135 2186561
refGene_gn=LOC102901578;refGene_tx=XM_023254670.1
B2 2188141 2188289
refGene_gn=LOC102901578;refGene_tx=XM_023254670.1
B2 2189348 2189431
refGene_gn=LOC102901578;refGene_tx=XM_023254670.1
B2 2190406 2190530
refGene_gn=LOC102901578;refGene_tx=XM_023254670.1
B2 2191437 2192300
refGene_gn=LOC102901578;refGene_tx=XM_023254670.1
B2 2211274 2211876
refGene_gn=ZSCAN9;refGene_tx=XM_019831673.2;refGene_gn=ZSCAN9;refGene_tx=XM_023254672.1;refGene_gn=ZSCAN9;refGene_tx=XM_023254673.1
B2 2213483 2213582
refGene_gn=ZSCAN9;refGene_tx=XM_019831673.2;refGene_gn=ZSCAN9;refGene_tx=XM_023254672.1
B2 2216707 2216855
refGene_gn=ZSCAN9;refGene_tx=XM_019831673.2;refGene_gn=ZSCAN9;refGene_tx=XM_023254672.1;refGene_gn=ZSCAN9;refGene_tx=XM_023254673.1
B2 2217033 2217450
refGene_gn=ZSCAN9;refGene_tx=XM_019831673.2;refGene_gn=ZSCAN9;refGene_tx=XM_023254672.1;refGene_gn=ZSCAN9;refGene_tx=XM_023254673.1
B2 2241839 2243428
refGene_gn=LOC101091367;refGene_tx=XM_006931347.4
B2 2249167 2249315
refGene_gn=LOC101091367;refGene_tx=XM_006931347.4
B2 2249875 2250277
refGene_gn=LOC101091367;refGene_tx=XM_006931347.4
B2 2280203 2280908 refGene_gn=ZKSCAN8;refGene_tx=XM_023254701.1
B2 2297694 2298713 refGene_gn=ZKSCAN8;refGene_tx=XM_023254701.1
B2 2299302 2299426 refGene_gn=ZKSCAN8;refGene_tx=XM_023254701.1
B2 2299715 2299807 refGene_gn=ZKSCAN8;refGene_tx=XM_023254701.1
B2 2302656 2302795 refGene_gn=ZKSCAN8;refGene_tx=XM_023254701.1
B2 2303396 2303813 refGene_gn=ZKSCAN8;refGene_tx=XM_023254701.1
B2 2312746 2312847
refGene_gn=LOC111560564;refGene_tx=XM_023254644.1
B2 2313417 2314207
refGene_gn=LOC111560564;refGene_tx=XM_023254644.1
B2 2327854 2328375
refGene_gn=ZSCAN16;refGene_tx=XM_006931349.2;refGene_gn=ZSCAN16;refGene_tx=XM_006931350.4;refGene_gn=ZSCAN16;refGene_tx=XM_023254691.1

B2	2332330	2332389	refGene_gn=ZSCAN16;refGene_tx=XM_023254693.1
B2	2335440	2335579
refGene_gn=ZSCAN16;refGene_tx=XM_006931349.2;refGene_gn=ZSCAN16;refGene_tx=XM_023254691.1;refGene_gn=ZSCAN16;refGene_tx=XM_023254693.1
B2	2337831	2338218
refGene_gn=ZSCAN16;refGene_tx=XM_006931349.2;refGene_gn=ZSCAN16;refGene_tx=XM_023254691.1;refGene_gn=ZSCAN16;refGene_tx=XM_023254693.1
B2	2345601	2346538	refGene_gn=ZSCAN16;refGene_tx=XM_006931350.4
B2	2353933	2354081	refGene_gn=ZSCAN16;refGene_tx=XM_006931350.4
B2	2354209	2354608	refGene_gn=ZSCAN16;refGene_tx=XM_006931350.4
B2	2370795	2371700
refGene_gn=ZNF165;refGene_tx=XM_003985660.4;refGene_gn=ZNF165;refGene_tx=XM_006931354.3;refGene_gn=ZNF165;refGene_tx=XM_006931355.4
B2	2373557	2373693
refGene_gn=ZNF165;refGene_tx=XM_003985660.4;refGene_gn=ZNF165;refGene_tx=XM_006931354.3;refGene_gn=ZNF165;refGene_tx=XM_006931355.4
B2	2373978	2374389
refGene_gn=ZNF165;refGene_tx=XM_003985660.4;refGene_gn=ZNF165;refGene_tx=XM_006931354.3;refGene_gn=ZNF165;refGene_tx=XM_006931355.4
B2	2394539	2395481
refGene_gn=LOC101092110;refGene_tx=XM_003985673.3
B2	2864964	2865873
refGene_gn=LOC101101053;refGene_tx=XM_023254646.1
B2	2873738	2874155
refGene_gn=LOC101101053;refGene_tx=XM_023254646.1
B2	2874287	2874539
refGene_gn=LOC101101053;refGene_tx=XM_023254646.1
B2	2884904	2885601
refGene_gn=LOC111560565;refGene_tx=XM_023254647.1
B2	2885603	2885812
refGene_gn=LOC111560565;refGene_tx=XM_023254647.1
B2	2890206	2890393
refGene_gn=LOC111560565;refGene_tx=XM_023254647.1
B2	2895380	2895430
refGene_gn=LOC111560565;refGene_tx=XM_023254647.1
B2	2895825	2895918
refGene_gn=LOC111560565;refGene_tx=XM_023254647.1
B2	2895977	2896403
refGene_gn=LOC111560566;refGene_tx=XM_023254648.1
B2	2907487	2908007
refGene_gn=LOC111560566;refGene_tx=XM_023254648.1
B2	2911237	2911320
refGene_gn=LOC111560566;refGene_tx=XM_023254648.1
B2	2949982	2950912
refGene_gn=LOC101080380;refGene_tx=XM_003985755.1
B2	2973433	2973473
refGene_gn=LOC101081730;refGene_tx=XM_011282061.2
B2	2973691	2974633
refGene_gn=LOC101081730;refGene_tx=XM_003985680.3;refGene_gn=LOC101081730;refGene_tx=XM_011282061.2

B2 3004189 3005131
refGene_gn=LOC101081981;refGene_tx=XM_003985681.1;refGene_gn=LOC101081981;refGene_tx=XM_019831687.2
B2 3011710 3011740
refGene_gn=LOC101081981;refGene_tx=XM_019831687.2
B2 3020379 3020760
refGene_gn=LOC101082235;refGene_tx=XM_003985682.5
B2 3021086 3021479
refGene_gn=LOC105260536;refGene_tx=XM_011282063.2
B2 3022555 3022966
refGene_gn=LOC101080632;refGene_tx=XM_011282064.3
B2 3036694 3037006
refGene_gn=LOC101082491;refGene_tx=XM_003985683.2
B2 3037754 3038165
refGene_gn=LOC101080886;refGene_tx=XM_006931365.3
B2 3041426 3042107
refGene_gn=LOC101082749;refGene_tx=XM_003985684.2
B2 3042659 3043052
refGene_gn=LOC101083005;refGene_tx=XM_011282066.3
B2 3059368 3059749
refGene_gn=LOC101083514;refGene_tx=XM_023254699.1
B2 3060070 3060463
refGene_gn=LOC101083265;refGene_tx=XM_023254698.1
B2 3067183 3067495
refGene_gn=LOC101083933;refGene_tx=XM_003985688.4
B2 3072571 3072883
refGene_gn=LOC101084193;refGene_tx=XM_023254700.1
B2 3077179 3077560
refGene_gn=LOC101084446;refGene_tx=XM_003985690.5
B2 3077882 3078275
refGene_gn=LOC101084707;refGene_tx=XM_003985691.4
B2 3079416 3079827
refGene_gn=LOC101084959;refGene_tx=XM_003985692.4
B2 3082864 3083263
refGene_gn=LOC101085206;refGene_tx=XM_003985693.2
B2 3083584 3083965
refGene_gn=LOC101085463;refGene_tx=XM_003985694.2
B2 3137943 3138048
refGene_gn=LOC109500083;refGene_tx=XM_019831555.1
B2 3139386 3139535
refGene_gn=LOC109500083;refGene_tx=XM_019831555.1
B2 3140065 3140443
refGene_gn=LOC109500083;refGene_tx=XM_019831555.1
B2 3142243 3142259
refGene_gn=LOC109500083;refGene_tx=XM_019831555.1
B2 3303810 3303817 refGene_gn=ZNF184;refGene_tx=XM_006931367.4
B2 3305105 3305121 refGene_gn=ZNF184;refGene_tx=XM_019831658.2
B2 3316202 3316307
refGene_gn=ZNF184;refGene_tx=XM_006931367.4;refGene_gn=ZNF184;refGene_tx=XM_019831657.2;refGene_gn=ZNF184;refGene_tx=XM_019831658.2

B2 3316916 3316946 refGene_gn=ZNF184;refGene_tx=XM_019831659.2
B2 3324040 3324167
refGene_gn=ZNF184;refGene_tx=XM_006931367.4;refGene_gn=ZNF184;refGene_tx=XM_019831657.2;refGene_gn=ZNF184;refGene_tx=XM_019831658.2;refGene_gn=ZNF184;refGene_tx=XM_019831659.2
B2 3324465 3324561
refGene_gn=ZNF184;refGene_tx=XM_006931367.4;refGene_gn=ZNF184;refGene_tx=XM_019831657.2;refGene_gn=ZNF184;refGene_tx=XM_019831658.2;refGene_gn=ZNF184;refGene_tx=XM_019831659.2
B2 3328835 3330793
refGene_gn=ZNF184;refGene_tx=XM_006931367.4;refGene_gn=ZNF184;refGene_tx=XM_019831657.2;refGene_gn=ZNF184;refGene_tx=XM_019831658.2;refGene_gn=ZNF184;refGene_tx=XM_019831659.2
B2 3375266 3376343
refGene_gn=ZNF391;refGene_tx=XM_006931374.3;refGene_gn=ZNF391;refGene_tx=XM_011282067.2;refGene_gn=ZNF391;refGene_tx=XM_023254683.1;refGene_gn=ZNF391;refGene_tx=XM_023254684.1;refGene_gn=ZNF391;refGene_tx=XM_023254685.1;refGene_gn=ZNF391;refGene_tx=XM_023254686.1;refGene_gn=ZNF391;refGene_tx=XM_023254687.1;refGene_gn=ZNF391;refGene_tx=XM_023254688.1;refGene_gn=ZNF391;refGene_tx=XM_023254689.1;refGene_gn=ZNF391;refGene_tx=XM_023254690.1
B2 3406267 3408190
refGene_gn=LOC101081890;refGene_tx=XM_006931381.3;refGene_gn=LOC101081890;refGene_tx=XM_023254654.1;refGene_gn=LOC101081890;refGene_tx=XM_023254655.1;refGene_gn=LOC101081890;refGene_tx=XM_023254656.1
B2 3439089 3442107 refGene_gn=POM121L2;refGene_tx=XM_023253571.1
B2 3482027 3483566
refGene_gn=LOC111560404;refGene_tx=XM_023253572.1
B2 3503060 3503129 refGene_gn=PRSS16;refGene_tx=XM_023253573.1
B2 3503237 3503383 refGene_gn=PRSS16;refGene_tx=XM_023253573.1
B2 3503460 3503640 refGene_gn=PRSS16;refGene_tx=XM_023253573.1
B2 3505485 3505627 refGene_gn=PRSS16;refGene_tx=XM_023253573.1
B2 3506212 3506503 refGene_gn=PRSS16;refGene_tx=XM_023253573.1
B2 3507113 3507161 refGene_gn=PRSS16;refGene_tx=XM_023253573.1
B2 3507259 3507337 refGene_gn=PRSS16;refGene_tx=XM_023253573.1
B2 3507588 3507712 refGene_gn=PRSS16;refGene_tx=XM_023253573.1
B2 3508977 3509107 refGene_gn=PRSS16;refGene_tx=XM_023253573.1
B2 3509728 3509828 refGene_gn=PRSS16;refGene_tx=XM_023253573.1
B2 3510563 3510724 refGene_gn=PRSS16;refGene_tx=XM_023253573.1
B2 3510799 3510872 refGene_gn=PRSS16;refGene_tx=XM_023253573.1
B2 3550241 3550245
refGene_gn=LOC102901244;refGene_tx=XM_023254651.1
B2 3550561 3551027
refGene_gn=LOC102901244;refGene_tx=XM_023254651.1
B2 3551925 3552090
refGene_gn=LOC102901244;refGene_tx=XM_023254651.1
B2 3553340 3553362
refGene_gn=LOC102901244;refGene_tx=XM_023254651.1
B2 3577008 3577395
refGene_gn=LOC101085717;refGene_tx=XM_003985695.3

B2	3577719	3578100
refGene_gn=LOC101086477;refGene_tx=XM_023253574.1
B2	3580563	3580875
refGene_gn=LOC105260538;refGene_tx=XM_011282072.3
B2	3640982	3641341
refGene_gn=FELCATV1R15;refGene_tx=NM_001167487.1
B2	3641342	3641898
refGene_gn=FELCATV1R15;refGene_tx=NM_001167487.1
B2	3650290	3651262
refGene_gn=FELCATV1R14;refGene_tx=NM_001167486.1
B2	3665714	3666599
refGene_gn=LOC105260539;refGene_tx=XM_011282074.3
B2	3666600	3666651
refGene_gn=LOC105260539;refGene_tx=XM_011282074.3
B2	3682694	3683666
refGene_gn=FELCATV1R16;refGene_tx=NM_001167488.1
B2	3770652	3771861
refGene_gn=ZNF322;refGene_tx=XM_003985703.5;refGene_gn=ZNF322;refGene_tx=XM_006931383.3;refGene_gn=ZNF322;refGene_tx=XM_006931384.3;refGene_gn=ZNF322;refGene_tx=XM_006931385.3;refGene_gn=ZNF322;refGene_tx=XM_019831679.2
B2	3789131	3789512	refGene_gn=ABT1;refGene_tx=XM_003985697.5
B2	3789664	3789870	refGene_gn=ABT1;refGene_tx=XM_003985697.5
B2	3790510	3790751	refGene_gn=ABT1;refGene_tx=XM_003985697.5
B2	3828733	3829091
refGene_gn=HMGN4;refGene_tx=XM_006931386.4;refGene_gn=HMGN4;refGene_tx=XM_019831701.2;refGene_gn=HMGN4;refGene_tx=XM_019831702.2;refGene_gn=HMGN4;refGene_tx=XM_023253580.1;refGene_gn=HMGN4;refGene_tx=XM_023253581.1
B2	3833066	3833077	refGene_gn=HMGN4;refGene_tx=XM_019831701.2
B2	3849511	3850185
refGene_gn=BTN1A1;refGene_tx=XM_003985696.3;refGene_gn=BTN1A1;refGene_tx=XM_011282075.2;refGene_gn=BTN1A1;refGene_tx=XM_019831670.1
B2	3850859	3850886
refGene_gn=BTN1A1;refGene_tx=XM_003985696.3;refGene_gn=BTN1A1;refGene_tx=XM_011282075.2;refGene_gn=BTN1A1;refGene_tx=XM_019831670.1
B2	3850980	3851001
refGene_gn=BTN1A1;refGene_tx=XM_003985696.3;refGene_gn=BTN1A1;refGene_tx=XM_011282075.2
B2	3851428	3851578
refGene_gn=BTN1A1;refGene_tx=XM_003985696.3;refGene_gn=BTN1A1;refGene_tx=XM_011282075.2;refGene_gn=BTN1A1;refGene_tx=XM_019831670.1
B2	3852322	3852604
refGene_gn=BTN1A1;refGene_tx=XM_003985696.3;refGene_gn=BTN1A1;refGene_tx=XM_011282075.2;refGene_gn=BTN1A1;refGene_tx=XM_019831670.1
B2	3854033	3854381
refGene_gn=BTN1A1;refGene_tx=XM_003985696.3;refGene_gn=BTN1A1;refGene_tx=XM_011282075.2;refGene_gn=BTN1A1;refGene_tx=XM_019831670.1
B2	3854525	3854604
refGene_gn=BTN1A1;refGene_tx=XM_003985696.3;refGene_gn=BTN1A1;refGene_

tx=XM_011282075.2;refGene_gn=BTN1A1;refGene_tx=XM_019831670.1
B2 3862270 3863806
refGene_gn=LOC102901749;refGene_tx=XM_019831672.2
B2 3872987 3873610
refGene_gn=LOC101082905;refGene_tx=XM_006931388.2;refGene_gn=LOC101082905;refGene_tx=XM_019831666.1;refGene_gn=LOC101082905;refGene_tx=XM_019831667.2;refGene_gn=LOC101082905;refGene_tx=XM_023253575.1
B2 3874724 3874751
refGene_gn=LOC101082905;refGene_tx=XM_006931388.2;refGene_gn=LOC101082905;refGene_tx=XM_019831666.1;refGene_gn=LOC101082905;refGene_tx=XM_019831667.2;refGene_gn=LOC101082905;refGene_tx=XM_023253575.1
B2 3874851 3874872
refGene_gn=LOC101082905;refGene_tx=XM_006931388.2;refGene_gn=LOC101082905;refGene_tx=XM_019831666.1;refGene_gn=LOC101082905;refGene_tx=XM_019831667.2;refGene_gn=LOC101082905;refGene_tx=XM_023253575.1
B2 3875270 3875471
refGene_gn=LOC101082905;refGene_tx=XM_006931388.2;refGene_gn=LOC101082905;refGene_tx=XM_019831666.1;refGene_gn=LOC101082905;refGene_tx=XM_019831667.2;refGene_gn=LOC101082905;refGene_tx=XM_023253575.1
B2 3876802 3877084
refGene_gn=LOC101082905;refGene_tx=XM_006931388.2;refGene_gn=LOC101082905;refGene_tx=XM_019831666.1;refGene_gn=LOC101082905;refGene_tx=XM_019831667.2;refGene_gn=LOC101082905;refGene_tx=XM_023253575.1
B2 3882806 3883154
refGene_gn=LOC101082905;refGene_tx=XM_006931388.2;refGene_gn=LOC101082905;refGene_tx=XM_019831666.1;refGene_gn=LOC101082905;refGene_tx=XM_019831667.2;refGene_gn=LOC101082905;refGene_tx=XM_023253575.1
B2 3883885 3883970
refGene_gn=LOC101082905;refGene_tx=XM_006931388.2;refGene_gn=LOC101082905;refGene_tx=XM_019831666.1;refGene_gn=LOC101082905;refGene_tx=XM_019831667.2;refGene_gn=LOC101082905;refGene_tx=XM_023253575.1
B2 3937891 3938203
refGene_gn=LOC101087478;refGene_tx=XM_023253583.1
B2 3946426 3946807
refGene_gn=LOC101088485;refGene_tx=XM_003985706.5
B2 3947203 3947596
refGene_gn=LOC101088237;refGene_tx=XM_003985705.4
B2 3948612 3949019
refGene_gn=LOC101089008;refGene_tx=XM_023254652.1
B2 3956208 3956589
refGene_gn=LOC101088746;refGene_tx=XM_003985707.4;refGene_gn=LOC101089008;refGene_tx=XM_023254652.1
B2 3957543 3958044
refGene_gn=LOC101089008;refGene_tx=XM_023254652.1
B2 3969244 3969556
refGene_gn=LOC101083921;refGene_tx=XM_011282077.3
B2 3974457 3975126
refGene_gn=LOC101089263;refGene_tx=XM_003985709.5
B2 3975486 3975507
refGene_gn=LOC105260540;refGene_tx=XM_011282078.2

B2	3983000	3983411
refGene_gn=LOC101089514;refGene_tx=XM_006931390.3
B2	3984518	3985055
refGene_gn=LOC102899134;refGene_tx=XM_011282080.3
B2	3987825	3988137
refGene_gn=LOC101089767;refGene_tx=XM_006931391.2
B2	3993513	3993924
refGene_gn=LOC105260541;refGene_tx=XM_019831695.1
B2	3994807	3995179
refGene_gn=LOC105260540;refGene_tx=XM_011282078.2
B2	3995512	3995893
refGene_gn=LOC101090262;refGene_tx=XM_006931392.4
B2	3999573	3999962
refGene_gn=LOC102899210;refGene_tx=XM_019831689.2
B2	4000087	4000099
refGene_gn=LOC101084194;refGene_tx=XM_003985765.4
B2	4001428	4001839
refGene_gn=LOC101084194;refGene_tx=XM_003985765.4
B2	4002023	4002172
refGene_gn=LOC101084447;refGene_tx=XM_011282081.3
B2	4002174	4002640
refGene_gn=LOC101084447;refGene_tx=XM_011282081.3
B2	4003778	4003938
refGene_gn=LOC102899210;refGene_tx=XM_019831689.2
B2	4008403	4008715
refGene_gn=LOC101084708;refGene_tx=XM_023253582.1
B2	4014600	4014981
refGene_gn=LOC101091280;refGene_tx=XM_003985716.5
B2	4035955	4036336
refGene_gn=LOC101091533;refGene_tx=XM_003985717.5;refGene_gn=LOC101091533;refGene_tx=XM_006931393.4
B2	4037250	4037907
refGene_gn=LOC101091785;refGene_tx=XM_003985718.4
B2	4062467	4062860
refGene_gn=LOC101092025;refGene_tx=XM_003985719.4
B2	4063198	4063579
refGene_gn=LOC101092267;refGene_tx=XM_003985720.4
B2	4085682	4086318
refGene_gn=LOC101092516;refGene_tx=XM_003985721.2
B2	4088640	4088952
refGene_gn=LOC101092762;refGene_tx=XM_006931394.4
B2	4096706	4096747
refGene_gn=HFE;refGene_tx=XM_011282083.3;refGene_gn=HFE;refGene_tx=XM_023253576.1;refGene_gn=HFE;refGene_tx=XM_023253577.1
B2	4097513	4097627
refGene_gn=HFE;refGene_tx=XM_011282083.3;refGene_gn=HFE;refGene_tx=XM_023253576.1;refGene_gn=HFE;refGene_tx=XM_023253577.1
B2	4097744	4098020
refGene_gn=HFE;refGene_tx=XM_011282083.3;refGene_gn=HFE;refGene_tx=XM_023253576.1;refGene_gn=HFE;refGene_tx=XM_023253577.1

B2 4098819 4099119
refGene_gn=HFE;refGene_tx=XM_011282083.3;refGene_gn=HFE;refGene_tx=XM_023253576.1;refGene_gn=HFE;refGene_tx=XM_023253577.1
B2 4099291 4099555
refGene_gn=HFE;refGene_tx=XM_011282083.3;refGene_gn=HFE;refGene_tx=XM_023253576.1;refGene_gn=HFE;refGene_tx=XM_023253577.1
B2 4102167 4102261
refGene_gn=HFE;refGene_tx=XM_023253576.1;refGene_gn=HFE;refGene_tx=XM_023253577.1
B2 4119036 4119678
refGene_gn=LOC101093016;refGene_tx=XM_023253578.1
B2 4127389 4127800
refGene_gn=LOC101093269;refGene_tx=XM_003985724.3
B2 4128728 4129121
refGene_gn=LOC101093519;refGene_tx=XM_011282084.3
B2 4129507 4129888
refGene_gn=LOC101093772;refGene_tx=XM_003985725.5
B2 4139122 4139515
refGene_gn=LOC101094019;refGene_tx=XM_003985726.5
B2 4140489 4140831
refGene_gn=LOC101094506;refGene_tx=XM_011282085.3
B2 4145484 4145802
refGene_gn=LOC101094506;refGene_tx=XM_011282085.3
B2 4152014 4152344
refGene_gn=LOC101094758;refGene_tx=XM_003985729.4
B2 4153155 4153482
refGene_gn=LOC101094758;refGene_tx=XM_003985729.4
B2 4155926 4156571
refGene_gn=LOC101085207;refGene_tx=XM_023253584.1
B2 4179164 4179688
refGene_gn=TRIM38;refGene_tx=NM_001290237.1;refGene_gn=TRIM38;refGene_tx=XM_023253539.1;refGene_gn=TRIM38;refGene_tx=XM_023253540.1
B2 4182687 4182800
refGene_gn=TRIM38;refGene_tx=NM_001290237.1;refGene_gn=TRIM38;refGene_tx=XM_023253539.1;refGene_gn=TRIM38;refGene_tx=XM_023253540.1
B2 4182895 4182918
refGene_gn=TRIM38;refGene_tx=NM_001290237.1;refGene_gn=TRIM38;refGene_tx=XM_023253539.1;refGene_gn=TRIM38;refGene_tx=XM_023253540.1
B2 4184165 4184396
refGene_gn=TRIM38;refGene_tx=NM_001290237.1;refGene_gn=TRIM38;refGene_tx=XM_023253539.1
B2 4186332 4186428
refGene_gn=TRIM38;refGene_tx=NM_001290237.1;refGene_gn=TRIM38;refGene_tx=XM_023253539.1;refGene_gn=TRIM38;refGene_tx=XM_023253540.1
B2 4188222 4188633
refGene_gn=TRIM38;refGene_tx=NM_001290237.1;refGene_gn=TRIM38;refGene_tx=XM_023253539.1;refGene_gn=TRIM38;refGene_tx=XM_023253540.1
B2 4207037 4207065
refGene_gn=SLC17A2;refGene_tx=XM_023253585.1;refGene_gn=SLC17A2;refGene_tx=XM_023253586.1

B2	4208661	4208873
refGene_gn=SLC17A2;refGene_tx=XM_023253585.1;refGene_gn=SLC17A2;refGen
e_tx=XM_023253586.1
B2	4210693	4210927	refGene_gn=SLC17A2;refGene_tx=XM_023253585.1
B2	4211011	4211099
refGene_gn=SLC17A2;refGene_tx=XM_023253585.1;refGene_gn=SLC17A2;refGen
e_tx=XM_023253586.1
B2	4211975	4212062
refGene_gn=SLC17A2;refGene_tx=XM_023253585.1;refGene_gn=SLC17A2;refGen
e_tx=XM_023253586.1
B2	4213470	4213589
refGene_gn=SLC17A2;refGene_tx=XM_023253585.1;refGene_gn=SLC17A2;refGen
e_tx=XM_023253586.1
B2	4213701	4213863
refGene_gn=SLC17A2;refGene_tx=XM_023253585.1;refGene_gn=SLC17A2;refGen
e_tx=XM_023253586.1
B2	4214331	4214464
refGene_gn=SLC17A2;refGene_tx=XM_023253585.1;refGene_gn=SLC17A2;refGen
e_tx=XM_023253586.1
B2	4214553	4214701
refGene_gn=SLC17A2;refGene_tx=XM_023253585.1;refGene_gn=SLC17A2;refGen
e_tx=XM_023253586.1
B2	4215986	4216077
refGene_gn=SLC17A2;refGene_tx=XM_023253585.1;refGene_gn=SLC17A2;refGen
e_tx=XM_023253586.1
B2	4217683	4217818
refGene_gn=SLC17A2;refGene_tx=XM_023253585.1;refGene_gn=SLC17A2;refGen
e_tx=XM_023253586.1
B2	4234100	4234191
refGene_gn=SLC17A3;refGene_tx=XM_019830129.2;refGene_gn=SLC17A3;refGen
e_tx=XM_019830130.2;refGene_gn=SLC17A3;refGene_tx=XM_019830132.2;refGe
ne_gn=SLC17A3;refGene_tx=XM_019830133.2
B2	4238865	4239077
refGene_gn=SLC17A3;refGene_tx=XM_019830129.2;refGene_gn=SLC17A3;refGen
e_tx=XM_019830130.2;refGene_gn=SLC17A3;refGene_tx=XM_019830132.2;refGe
ne_gn=SLC17A3;refGene_tx=XM_019830133.2
B2	4239483	4239717
refGene_gn=SLC17A3;refGene_tx=XM_019830129.2;refGene_gn=SLC17A3;refGen
e_tx=XM_019830130.2;refGene_gn=SLC17A3;refGene_tx=XM_019830132.2
B2	4239802	4239890
refGene_gn=SLC17A3;refGene_tx=XM_019830129.2;refGene_gn=SLC17A3;refGen
e_tx=XM_019830130.2;refGene_gn=SLC17A3;refGene_tx=XM_019830132.2;refGe
ne_gn=SLC17A3;refGene_tx=XM_019830133.2
B2	4244483	4244570
refGene_gn=SLC17A3;refGene_tx=XM_019830129.2;refGene_gn=SLC17A3;refGen
e_tx=XM_019830130.2;refGene_gn=SLC17A3;refGene_tx=XM_019830132.2;refGe
ne_gn=SLC17A3;refGene_tx=XM_019830133.2
B2	4247806	4247925
refGene_gn=SLC17A3;refGene_tx=XM_019830129.2;refGene_gn=SLC17A3;refGen
e_tx=XM_019830130.2;refGene_gn=SLC17A3;refGene_tx=XM_019830132.2;refGe

ne_gn=SLC17A3;refGene_tx=XM_019830133.2
B2	4248051	4248213
refGene_gn=SLC17A3;refGene_tx=XM_019830129.2;refGene_gn=SLC17A3;refGene_tx=XM_019830130.2;refGene_gn=SLC17A3;refGene_tx=XM_019830133.2
B2	4248938	4249068
refGene_gn=SLC17A3;refGene_tx=XM_019830129.2;refGene_gn=SLC17A3;refGene_tx=XM_019830130.2;refGene_gn=SLC17A3;refGene_tx=XM_019830132.2;refGene_gn=SLC17A3;refGene_tx=XM_019830133.2
B2	4249168	4249316
refGene_gn=SLC17A3;refGene_tx=XM_019830129.2;refGene_gn=SLC17A3;refGene_tx=XM_019830130.2;refGene_gn=SLC17A3;refGene_tx=XM_019830132.2;refGene_gn=SLC17A3;refGene_tx=XM_019830133.2
B2	4249606	4249697
refGene_gn=SLC17A3;refGene_tx=XM_019830129.2;refGene_gn=SLC17A3;refGene_tx=XM_019830130.2;refGene_gn=SLC17A3;refGene_tx=XM_019830132.2;refGene_gn=SLC17A3;refGene_tx=XM_019830133.2
B2	4250616	4250751
refGene_gn=SLC17A3;refGene_tx=XM_019830129.2;refGene_gn=SLC17A3;refGene_tx=XM_019830130.2;refGene_gn=SLC17A3;refGene_tx=XM_019830132.2;refGene_gn=SLC17A3;refGene_tx=XM_019830133.2
B2	4256387	4256471	refGene_gn=SLC17A1;refGene_tx=XM_003985735.3
B2	4257706	4257788
refGene_gn=SLC17A1;refGene_tx=XM_003985735.3;refGene_gn=SLC17A1;refGene_tx=XM_019830128.2
B2	4261927	4262100
refGene_gn=SLC17A1;refGene_tx=XM_003985735.3;refGene_gn=SLC17A1;refGene_tx=XM_019830128.2
B2	4267248	4267482
refGene_gn=SLC17A1;refGene_tx=XM_003985735.3;refGene_gn=SLC17A1;refGene_tx=XM_019830128.2
B2	4267566	4267654
refGene_gn=SLC17A1;refGene_tx=XM_003985735.3;refGene_gn=SLC17A1;refGene_tx=XM_019830128.2
B2	4268009	4268096
refGene_gn=SLC17A1;refGene_tx=XM_003985735.3;refGene_gn=SLC17A1;refGene_tx=XM_019830128.2
B2	4270225	4270347
refGene_gn=SLC17A1;refGene_tx=XM_003985735.3;refGene_gn=SLC17A1;refGene_tx=XM_019830128.2
B2	4270463	4270622
refGene_gn=SLC17A1;refGene_tx=XM_003985735.3;refGene_gn=SLC17A1;refGene_tx=XM_019830128.2
B2	4271832	4271965
refGene_gn=SLC17A1;refGene_tx=XM_003985735.3;refGene_gn=SLC17A1;refGene_tx=XM_019830128.2
B2	4272055	4272203
refGene_gn=SLC17A1;refGene_tx=XM_003985735.3;refGene_gn=SLC17A1;refGene_tx=XM_019830128.2
B2	4283258	4283349
refGene_gn=SLC17A1;refGene_tx=XM_003985735.3;refGene_gn=SLC17A1;refGen

e_tx=XM_019830128.2
B2	4292902	4293040
refGene_gn=SLC17A1;refGene_tx=XM_003985735.3;refGene_gn=SLC17A1;refGene_tx=XM_019830128.2
B2	4313059	4313283
refGene_gn=SLC17A4;refGene_tx=XM_023253587.1;refGene_gn=SLC17A4;refGene_tx=XM_023253588.1;refGene_gn=SLC17A4;refGene_tx=XM_023253589.1
B2	4314217	4314308
refGene_gn=SLC17A4;refGene_tx=XM_023253587.1;refGene_gn=SLC17A4;refGene_tx=XM_023253588.1;refGene_gn=SLC17A4;refGene_tx=XM_023253589.1
B2	4315135	4315283
refGene_gn=SLC17A4;refGene_tx=XM_023253587.1;refGene_gn=SLC17A4;refGene_tx=XM_023253588.1;refGene_gn=SLC17A4;refGene_tx=XM_023253589.1
B2	4315363	4315496
refGene_gn=SLC17A4;refGene_tx=XM_023253587.1;refGene_gn=SLC17A4;refGene_tx=XM_023253588.1;refGene_gn=SLC17A4;refGene_tx=XM_023253589.1
B2	4317338	4317500
refGene_gn=SLC17A4;refGene_tx=XM_023253587.1;refGene_gn=SLC17A4;refGene_tx=XM_023253588.1;refGene_gn=SLC17A4;refGene_tx=XM_023253589.1
B2	4317623	4317742
refGene_gn=SLC17A4;refGene_tx=XM_023253587.1;refGene_gn=SLC17A4;refGene_tx=XM_023253588.1;refGene_gn=SLC17A4;refGene_tx=XM_023253589.1
B2	4319187	4319274
refGene_gn=SLC17A4;refGene_tx=XM_023253587.1;refGene_gn=SLC17A4;refGene_tx=XM_023253588.1;refGene_gn=SLC17A4;refGene_tx=XM_023253589.1
B2	4319766	4319854
refGene_gn=SLC17A4;refGene_tx=XM_023253587.1;refGene_gn=SLC17A4;refGene_tx=XM_023253588.1;refGene_gn=SLC17A4;refGene_tx=XM_023253589.1
B2	4319951	4320185
refGene_gn=SLC17A4;refGene_tx=XM_023253587.1;refGene_gn=SLC17A4;refGene_tx=XM_023253588.1;refGene_gn=SLC17A4;refGene_tx=XM_023253589.1
B2	4320679	4320885
refGene_gn=SLC17A4;refGene_tx=XM_023253587.1;refGene_gn=SLC17A4;refGene_tx=XM_023253588.1
B2	4327933	4328021
refGene_gn=SLC17A4;refGene_tx=XM_023253587.1;refGene_gn=SLC17A4;refGene_tx=XM_023253588.1;refGene_gn=SLC17A4;refGene_tx=XM_023253589.1
B2	4363425	4363839
refGene_gn=LOC101085975;refGene_tx=XM_003985771.4
B2	4363972	4363984
refGene_gn=LOC101085975;refGene_tx=XM_003985771.4
B2	4364091	4364472
refGene_gn=LOC101096314;refGene_tx=XM_003985736.4
B2	4374269	4374653
refGene_gn=LOC101096566;refGene_tx=XM_003985737.4
B2	4374967	4375363
refGene_gn=LOC101096808;refGene_tx=XM_003985738.5
B2	4414635	4414764
refGene_gn=SCGN;refGene_tx=XM_003985739.3;refGene_gn=SCGN;refGene_tx=XM_023253591.1

B2 4421484 4421553
refGene_gn=SCGN;refGene_tx=XM_003985739.3;refGene_gn=SCGN;refGene_tx=XM_023253591.1
B2 4423500 4423560
refGene_gn=SCGN;refGene_tx=XM_003985739.3;refGene_gn=SCGN;refGene_tx=XM_023253591.1
B2 4423798 4423844
refGene_gn=SCGN;refGene_tx=XM_003985739.3;refGene_gn=SCGN;refGene_tx=XM_023253591.1
B2 4428387 4428443
refGene_gn=SCGN;refGene_tx=XM_003985739.3;refGene_gn=SCGN;refGene_tx=XM_023253591.1
B2 4436625 4436703
refGene_gn=SCGN;refGene_tx=XM_003985739.3;refGene_gn=SCGN;refGene_tx=XM_023253591.1
B2 4437356 4437413
refGene_gn=SCGN;refGene_tx=XM_003985739.3;refGene_gn=SCGN;refGene_tx=XM_023253591.1
B2 4440399 4440489
refGene_gn=SCGN;refGene_tx=XM_003985739.3;refGene_gn=SCGN;refGene_tx=XM_023253591.1
B2 4443750 4443843
refGene_gn=SCGN;refGene_tx=XM_003985739.3;refGene_gn=SCGN;refGene_tx=XM_023253591.1
B2 4448630 4448701
refGene_gn=SCGN;refGene_tx=XM_003985739.3;refGene_gn=SCGN;refGene_tx=XM_023253591.1
B2 4449717 4449799
refGene_gn=SCGN;refGene_tx=XM_003985739.3;refGene_gn=SCGN;refGene_tx=XM_023253591.1
B2 4469396 4469576
refGene_gn=CARMIL1;refGene_tx=XM_011282087.3;refGene_gn=CARMIL1;refGene_tx=XM_019830107.2;refGene_gn=CARMIL1;refGene_tx=XM_019830114.2;refGene_gn=CARMIL1;refGene_tx=XM_019830116.2;refGene_gn=CARMIL1;refGene_tx=XM_023253592.1;refGene_gn=CARMIL1;refGene_tx=XM_023253593.1;refGene_gn=CARMIL1;refGene_tx=XM_023253594.1;refGene_gn=CARMIL1;refGene_tx=XM_023253595.1;refGene_gn=CARMIL1;refGene_tx=XM_023253596.1;refGene_gn=CARMIL1;refGene_tx=XM_023253598.1;refGene_gn=CARMIL1;refGene_tx=XM_023253599.1
B2 4476017 4476167
refGene_gn=CARMIL1;refGene_tx=XM_023253593.1;refGene_gn=CARMIL1;refGene_tx=XM_023253595.1;refGene_gn=CARMIL1;refGene_tx=XM_023253597.1;refGene_gn=CARMIL1;refGene_tx=XM_023253600.1
B2 4476493 4476609
refGene_gn=CARMIL1;refGene_tx=XM_019830107.2;refGene_gn=CARMIL1;refGene_tx=XM_019830114.2;refGene_gn=CARMIL1;refGene_tx=XM_023253592.1;refGene_gn=CARMIL1;refGene_tx=XM_023253594.1;refGene_gn=CARMIL1;refGene_tx=XM_023253596.1;refGene_gn=CARMIL1;refGene_tx=XM_023253597.1;refGene_gn=CARMIL1;refGene_tx=XM_023253598.1;refGene_gn=CARMIL1;refGene_tx=XM_023253600.1;refGene_gn=CARMIL1;refGene_tx=XM_023253601.1;refGene_gn=CARM

IL1;refGene_tx=XM_023253602.1;refGene_gn=CARMIL1;refGene_tx=XM_0232536
03.1
B2 4478447 4478588
refGene_gn=CARMIL1;refGene_tx=XM_011282087.3;refGene_gn=CARMIL1;refGen
e_tx=XM_019830107.2;refGene_gn=CARMIL1;refGene_tx=XM_019830114.2;refGe
ne_gn=CARMIL1;refGene_tx=XM_019830116.2;refGene_gn=CARMIL1;refGene_tx=
XM_023253592.1;refGene_gn=CARMIL1;refGene_tx=XM_023253593.1;refGene_gn
=CARMIL1;refGene_tx=XM_023253594.1;refGene_gn=CARMIL1;refGene_tx=XM_02
3253595.1;refGene_gn=CARMIL1;refGene_tx=XM_023253596.1;refGene_gn=CARM
IL1;refGene_tx=XM_023253597.1;refGene_gn=CARMIL1;refGene_tx=XM_0232535
98.1;refGene_gn=CARMIL1;refGene_tx=XM_023253599.1;refGene_gn=CARMIL1;r
efGene_tx=XM_023253600.1;refGene_gn=CARMIL1;refGene_tx=XM_023253601.1;
refGene_gn=CARMIL1;refGene_tx=XM_023253602.1;refGene_gn=CARMIL1;refGen
e_tx=XM_023253603.1
B2 4481536 4481749
refGene_gn=CARMIL1;refGene_tx=XM_011282087.3;refGene_gn=CARMIL1;refGen
e_tx=XM_019830107.2;refGene_gn=CARMIL1;refGene_tx=XM_019830114.2;refGe
ne_gn=CARMIL1;refGene_tx=XM_019830116.2;refGene_gn=CARMIL1;refGene_tx=
XM_023253592.1;refGene_gn=CARMIL1;refGene_tx=XM_023253593.1;refGene_gn
=CARMIL1;refGene_tx=XM_023253594.1;refGene_gn=CARMIL1;refGene_tx=XM_02
3253595.1;refGene_gn=CARMIL1;refGene_tx=XM_023253596.1;refGene_gn=CARM
IL1;refGene_tx=XM_023253597.1;refGene_gn=CARMIL1;refGene_tx=XM_0232535
98.1;refGene_gn=CARMIL1;refGene_tx=XM_023253599.1;refGene_gn=CARMIL1;r
efGene_tx=XM_023253600.1;refGene_gn=CARMIL1;refGene_tx=XM_023253601.1;
refGene_gn=CARMIL1;refGene_tx=XM_023253602.1;refGene_gn=CARMIL1;refGen
e_tx=XM_023253603.1
B2 4482856 4482932
refGene_gn=CARMIL1;refGene_tx=XM_011282087.3;refGene_gn=CARMIL1;refGen
e_tx=XM_019830107.2;refGene_gn=CARMIL1;refGene_tx=XM_019830114.2;refGe
ne_gn=CARMIL1;refGene_tx=XM_019830116.2;refGene_gn=CARMIL1;refGene_tx=
XM_023253592.1;refGene_gn=CARMIL1;refGene_tx=XM_023253593.1;refGene_gn
=CARMIL1;refGene_tx=XM_023253594.1;refGene_gn=CARMIL1;refGene_tx=XM_02
3253595.1;refGene_gn=CARMIL1;refGene_tx=XM_023253596.1;refGene_gn=CARM
IL1;refGene_tx=XM_023253597.1;refGene_gn=CARMIL1;refGene_tx=XM_0232535
98.1;refGene_gn=CARMIL1;refGene_tx=XM_023253599.1;refGene_gn=CARMIL1;r
efGene_tx=XM_023253600.1;refGene_gn=CARMIL1;refGene_tx=XM_023253601.1;
refGene_gn=CARMIL1;refGene_tx=XM_023253602.1;refGene_gn=CARMIL1;refGen
e_tx=XM_023253603.1
B2 4489320 4489747
refGene_gn=CARMIL1;refGene_tx=XM_011282087.3;refGene_gn=CARMIL1;refGen
e_tx=XM_019830107.2;refGene_gn=CARMIL1;refGene_tx=XM_019830114.2;refGe
ne_gn=CARMIL1;refGene_tx=XM_019830116.2;refGene_gn=CARMIL1;refGene_tx=
XM_023253592.1;refGene_gn=CARMIL1;refGene_tx=XM_023253593.1;refGene_gn
=CARMIL1;refGene_tx=XM_023253594.1;refGene_gn=CARMIL1;refGene_tx=XM_02
3253595.1;refGene_gn=CARMIL1;refGene_tx=XM_023253596.1;refGene_gn=CARM
IL1;refGene_tx=XM_023253597.1;refGene_gn=CARMIL1;refGene_tx=XM_0232535
98.1;refGene_gn=CARMIL1;refGene_tx=XM_023253599.1;refGene_gn=CARMIL1;r
efGene_tx=XM_023253600.1;refGene_gn=CARMIL1;refGene_tx=XM_023253601.1;
refGene_gn=CARMIL1;refGene_tx=XM_023253602.1;refGene_gn=CARMIL1;refGen
e_tx=XM_023253603.1

B2	4496683	4496796
refGene_gn=CARMIL1;refGene_tx=XM_011282087.3;refGene_gn=CARMIL1;refGene_tx=XM_019830107.2;refGene_gn=CARMIL1;refGene_tx=XM_019830114.2;refGene_gn=CARMIL1;refGene_tx=XM_019830116.2;refGene_gn=CARMIL1;refGene_tx=XM_023253592.1;refGene_gn=CARMIL1;refGene_tx=XM_023253593.1;refGene_gn=CARMIL1;refGene_tx=XM_023253594.1;refGene_gn=CARMIL1;refGene_tx=XM_023253595.1;refGene_gn=CARMIL1;refGene_tx=XM_023253596.1;refGene_gn=CARMIL1;refGene_tx=XM_023253597.1;refGene_gn=CARMIL1;refGene_tx=XM_023253598.1;refGene_gn=CARMIL1;refGene_tx=XM_023253599.1;refGene_gn=CARMIL1;refGene_tx=XM_023253600.1;refGene_gn=CARMIL1;refGene_tx=XM_023253601.1;refGene_gn=CARMIL1;refGene_tx=XM_023253602.1;refGene_gn=CARMIL1;refGene_tx=XM_023253603.1
B2	4509963	4510160
refGene_gn=CARMIL1;refGene_tx=XM_011282087.3;refGene_gn=CARMIL1;refGene_tx=XM_019830107.2;refGene_gn=CARMIL1;refGene_tx=XM_019830114.2;refGene_gn=CARMIL1;refGene_tx=XM_019830116.2;refGene_gn=CARMIL1;refGene_tx=XM_023253592.1;refGene_gn=CARMIL1;refGene_tx=XM_023253593.1;refGene_gn=CARMIL1;refGene_tx=XM_023253594.1;refGene_gn=CARMIL1;refGene_tx=XM_023253595.1;refGene_gn=CARMIL1;refGene_tx=XM_023253596.1;refGene_gn=CARMIL1;refGene_tx=XM_023253597.1;refGene_gn=CARMIL1;refGene_tx=XM_023253598.1;refGene_gn=CARMIL1;refGene_tx=XM_023253599.1;refGene_gn=CARMIL1;refGene_tx=XM_023253600.1;refGene_gn=CARMIL1;refGene_tx=XM_023253601.1;refGene_gn=CARMIL1;refGene_tx=XM_023253602.1;refGene_gn=CARMIL1;refGene_tx=XM_023253603.1
B2	4510381	4510448
refGene_gn=CARMIL1;refGene_tx=XM_011282087.3;refGene_gn=CARMIL1;refGene_tx=XM_019830107.2;refGene_gn=CARMIL1;refGene_tx=XM_019830114.2;refGene_gn=CARMIL1;refGene_tx=XM_019830116.2;refGene_gn=CARMIL1;refGene_tx=XM_023253592.1;refGene_gn=CARMIL1;refGene_tx=XM_023253593.1;refGene_gn=CARMIL1;refGene_tx=XM_023253594.1;refGene_gn=CARMIL1;refGene_tx=XM_023253595.1;refGene_gn=CARMIL1;refGene_tx=XM_023253596.1;refGene_gn=CARMIL1;refGene_tx=XM_023253597.1;refGene_gn=CARMIL1;refGene_tx=XM_023253598.1;refGene_gn=CARMIL1;refGene_tx=XM_023253599.1;refGene_gn=CARMIL1;refGene_tx=XM_023253600.1;refGene_gn=CARMIL1;refGene_tx=XM_023253601.1;refGene_gn=CARMIL1;refGene_tx=XM_023253602.1;refGene_gn=CARMIL1;refGene_tx=XM_023253603.1
B2	4514817	4514835
refGene_gn=CARMIL1;refGene_tx=XM_019830107.2;refGene_gn=CARMIL1;refGene_tx=XM_019830114.2;refGene_gn=CARMIL1;refGene_tx=XM_023253592.1;refGene_gn=CARMIL1;refGene_tx=XM_023253593.1;refGene_gn=CARMIL1;refGene_tx=XM_023253594.1;refGene_gn=CARMIL1;refGene_tx=XM_023253595.1;refGene_gn=CARMIL1;refGene_tx=XM_023253597.1;refGene_gn=CARMIL1;refGene_tx=XM_023253598.1;refGene_gn=CARMIL1;refGene_tx=XM_023253599.1;refGene_gn=CARMIL1;refGene_tx=XM_023253600.1;refGene_gn=CARMIL1;refGene_tx=XM_023253601.1;refGene_gn=CARMIL1;refGene_tx=XM_023253602.1;refGene_gn=CARMIL1;refGene_tx=XM_023253603.1
B2	4536800	4536950
refGene_gn=CARMIL1;refGene_tx=XM_011282087.3;refGene_gn=CARMIL1;refGene_tx=XM_019830107.2;refGene_gn=CARMIL1;refGene_tx=XM_019830114.2;refGene_gn=CARMIL1;refGene_tx=XM_019830116.2;refGene_gn=CARMIL1;refGene_tx=

XM_023253592.1;refGene_gn=CARMIL1;refGene_tx=XM_023253593.1;refGene_gn
=CARMIL1;refGene_tx=XM_023253594.1;refGene_gn=CARMIL1;refGene_tx=XM_02
3253595.1;refGene_gn=CARMIL1;refGene_tx=XM_023253596.1;refGene_gn=CARM
IL1;refGene_tx=XM_023253597.1;refGene_gn=CARMIL1;refGene_tx=XM_0232535
98.1;refGene_gn=CARMIL1;refGene_tx=XM_023253599.1;refGene_gn=CARMIL1;r
efGene_tx=XM_023253600.1;refGene_gn=CARMIL1;refGene_tx=XM_023253601.1;
refGene_gn=CARMIL1;refGene_tx=XM_023253602.1;refGene_gn=CARMIL1;refGen
e_tx=XM_023253603.1
B2 4539874 4539962
refGene_gn=CARMIL1;refGene_tx=XM_011282087.3;refGene_gn=CARMIL1;refGen
e_tx=XM_019830107.2;refGene_gn=CARMIL1;refGene_tx=XM_019830114.2;refGe
ne_gn=CARMIL1;refGene_tx=XM_019830116.2;refGene_gn=CARMIL1;refGene_tx=
XM_023253592.1;refGene_gn=CARMIL1;refGene_tx=XM_023253593.1;refGene_gn
=CARMIL1;refGene_tx=XM_023253594.1;refGene_gn=CARMIL1;refGene_tx=XM_02
3253595.1;refGene_gn=CARMIL1;refGene_tx=XM_023253596.1;refGene_gn=CARM
IL1;refGene_tx=XM_023253597.1;refGene_gn=CARMIL1;refGene_tx=XM_0232535
98.1;refGene_gn=CARMIL1;refGene_tx=XM_023253599.1;refGene_gn=CARMIL1;r
efGene_tx=XM_023253600.1;refGene_gn=CARMIL1;refGene_tx=XM_023253601.1;
refGene_gn=CARMIL1;refGene_tx=XM_023253602.1;refGene_gn=CARMIL1;refGen
e_tx=XM_023253603.1
B2 4543036 4543212
refGene_gn=CARMIL1;refGene_tx=XM_011282087.3;refGene_gn=CARMIL1;refGen
e_tx=XM_019830107.2;refGene_gn=CARMIL1;refGene_tx=XM_019830114.2;refGe
ne_gn=CARMIL1;refGene_tx=XM_019830116.2;refGene_gn=CARMIL1;refGene_tx=
XM_023253592.1;refGene_gn=CARMIL1;refGene_tx=XM_023253593.1;refGene_gn
=CARMIL1;refGene_tx=XM_023253594.1;refGene_gn=CARMIL1;refGene_tx=XM_02
3253595.1;refGene_gn=CARMIL1;refGene_tx=XM_023253596.1;refGene_gn=CARM
IL1;refGene_tx=XM_023253597.1;refGene_gn=CARMIL1;refGene_tx=XM_0232535
98.1;refGene_gn=CARMIL1;refGene_tx=XM_023253599.1;refGene_gn=CARMIL1;r
efGene_tx=XM_023253600.1;refGene_gn=CARMIL1;refGene_tx=XM_023253601.1;
refGene_gn=CARMIL1;refGene_tx=XM_023253602.1;refGene_gn=CARMIL1;refGen
e_tx=XM_023253603.1
B2 4553543 4553675
refGene_gn=CARMIL1;refGene_tx=XM_011282087.3;refGene_gn=CARMIL1;refGen
e_tx=XM_019830107.2;refGene_gn=CARMIL1;refGene_tx=XM_019830114.2;refGe
ne_gn=CARMIL1;refGene_tx=XM_019830116.2;refGene_gn=CARMIL1;refGene_tx=
XM_023253592.1;refGene_gn=CARMIL1;refGene_tx=XM_023253593.1;refGene_gn
=CARMIL1;refGene_tx=XM_023253594.1;refGene_gn=CARMIL1;refGene_tx=XM_02
3253595.1;refGene_gn=CARMIL1;refGene_tx=XM_023253596.1;refGene_gn=CARM
IL1;refGene_tx=XM_023253597.1;refGene_gn=CARMIL1;refGene_tx=XM_0232535
98.1;refGene_gn=CARMIL1;refGene_tx=XM_023253599.1;refGene_gn=CARMIL1;r
efGene_tx=XM_023253600.1;refGene_gn=CARMIL1;refGene_tx=XM_023253601.1;
refGene_gn=CARMIL1;refGene_tx=XM_023253602.1;refGene_gn=CARMIL1;refGen
e_tx=XM_023253603.1
B2 4554358 4554487
refGene_gn=CARMIL1;refGene_tx=XM_011282087.3;refGene_gn=CARMIL1;refGen
e_tx=XM_019830107.2;refGene_gn=CARMIL1;refGene_tx=XM_019830114.2;refGe
ne_gn=CARMIL1;refGene_tx=XM_019830116.2;refGene_gn=CARMIL1;refGene_tx=
XM_023253592.1;refGene_gn=CARMIL1;refGene_tx=XM_023253593.1;refGene_gn
=CARMIL1;refGene_tx=XM_023253594.1;refGene_gn=CARMIL1;refGene_tx=XM_02

3253595.1;refGene_gn=CARMIL1;refGene_tx=XM_023253596.1;refGene_gn=CARMIL1;refGene_tx=XM_023253597.1;refGene_gn=CARMIL1;refGene_tx=XM_023253598.1;refGene_gn=CARMIL1;refGene_tx=XM_023253599.1;refGene_gn=CARMIL1;refGene_tx=XM_023253600.1;refGene_gn=CARMIL1;refGene_tx=XM_023253601.1;refGene_gn=CARMIL1;refGene_tx=XM_023253602.1;refGene_gn=CARMIL1;refGene_tx=XM_023253603.1
B2 4562519 4562618
refGene_gn=CARMIL1;refGene_tx=XM_011282087.3;refGene_gn=CARMIL1;refGene_tx=XM_019830107.2;refGene_gn=CARMIL1;refGene_tx=XM_019830114.2;refGene_gn=CARMIL1;refGene_tx=XM_019830116.2;refGene_gn=CARMIL1;refGene_tx=XM_023253592.1;refGene_gn=CARMIL1;refGene_tx=XM_023253593.1;refGene_gn=CARMIL1;refGene_tx=XM_023253594.1;refGene_gn=CARMIL1;refGene_tx=XM_023253595.1;refGene_gn=CARMIL1;refGene_tx=XM_023253596.1;refGene_gn=CARMIL1;refGene_tx=XM_023253597.1;refGene_gn=CARMIL1;refGene_tx=XM_023253599.1;refGene_gn=CARMIL1;refGene_tx=XM_023253600.1;refGene_gn=CARMIL1;refGene_tx=XM_023253601.1;refGene_gn=CARMIL1;refGene_tx=XM_023253602.1;refGene_gn=CARMIL1;refGene_tx=XM_023253603.1
B2 4568735 4568829
refGene_gn=CARMIL1;refGene_tx=XM_011282087.3;refGene_gn=CARMIL1;refGene_tx=XM_019830107.2;refGene_gn=CARMIL1;refGene_tx=XM_019830114.2;refGene_gn=CARMIL1;refGene_tx=XM_019830116.2;refGene_gn=CARMIL1;refGene_tx=XM_023253592.1;refGene_gn=CARMIL1;refGene_tx=XM_023253593.1;refGene_gn=CARMIL1;refGene_tx=XM_023253594.1;refGene_gn=CARMIL1;refGene_tx=XM_023253595.1;refGene_gn=CARMIL1;refGene_tx=XM_023253596.1;refGene_gn=CARMIL1;refGene_tx=XM_023253597.1;refGene_gn=CARMIL1;refGene_tx=XM_023253598.1;refGene_gn=CARMIL1;refGene_tx=XM_023253599.1;refGene_gn=CARMIL1;refGene_tx=XM_023253600.1;refGene_gn=CARMIL1;refGene_tx=XM_023253601.1;refGene_gn=CARMIL1;refGene_tx=XM_023253602.1;refGene_gn=CARMIL1;refGene_tx=XM_023253603.1
B2 4572025 4572094
refGene_gn=CARMIL1;refGene_tx=XM_011282087.3;refGene_gn=CARMIL1;refGene_tx=XM_019830107.2;refGene_gn=CARMIL1;refGene_tx=XM_019830114.2;refGene_gn=CARMIL1;refGene_tx=XM_019830116.2;refGene_gn=CARMIL1;refGene_tx=XM_023253592.1;refGene_gn=CARMIL1;refGene_tx=XM_023253593.1;refGene_gn=CARMIL1;refGene_tx=XM_023253594.1;refGene_gn=CARMIL1;refGene_tx=XM_023253595.1;refGene_gn=CARMIL1;refGene_tx=XM_023253596.1;refGene_gn=CARMIL1;refGene_tx=XM_023253597.1;refGene_gn=CARMIL1;refGene_tx=XM_023253598.1;refGene_gn=CARMIL1;refGene_tx=XM_023253599.1;refGene_gn=CARMIL1;refGene_tx=XM_023253600.1;refGene_gn=CARMIL1;refGene_tx=XM_023253601.1;refGene_gn=CARMIL1;refGene_tx=XM_023253602.1;refGene_gn=CARMIL1;refGene_tx=XM_023253603.1
B2 4573661 4573834
refGene_gn=CARMIL1;refGene_tx=XM_011282087.3;refGene_gn=CARMIL1;refGene_tx=XM_019830107.2;refGene_gn=CARMIL1;refGene_tx=XM_019830114.2;refGene_gn=CARMIL1;refGene_tx=XM_019830116.2;refGene_gn=CARMIL1;refGene_tx=XM_023253592.1;refGene_gn=CARMIL1;refGene_tx=XM_023253593.1;refGene_gn=CARMIL1;refGene_tx=XM_023253594.1;refGene_gn=CARMIL1;refGene_tx=XM_023253595.1;refGene_gn=CARMIL1;refGene_tx=XM_023253596.1;refGene_gn=CARMIL1;refGene_tx=XM_023253597.1;refGene_gn=CARMIL1;refGene_tx=XM_023253598.1;refGene_gn=CARMIL1;refGene_tx=XM_023253599.1;refGene_gn=CARMIL1;r

efGene_tx=XM_023253600.1;refGene_gn=CARMIL1;refGene_tx=XM_023253601.1;
refGene_gn=CARMIL1;refGene_tx=XM_023253602.1;refGene_gn=CARMIL1;refGen
e_tx=XM_023253603.1
B2 4578402 4578457
refGene_gn=CARMIL1;refGene_tx=XM_011282087.3;refGene_gn=CARMIL1;refGen
e_tx=XM_019830107.2;refGene_gn=CARMIL1;refGene_tx=XM_019830114.2;refGe
ne_gn=CARMIL1;refGene_tx=XM_019830116.2;refGene_gn=CARMIL1;refGene_tx=
XM_023253592.1;refGene_gn=CARMIL1;refGene_tx=XM_023253593.1;refGene_gn
=CARMIL1;refGene_tx=XM_023253594.1;refGene_gn=CARMIL1;refGene_tx=XM_02
3253595.1;refGene_gn=CARMIL1;refGene_tx=XM_023253596.1;refGene_gn=CARM
IL1;refGene_tx=XM_023253597.1;refGene_gn=CARMIL1;refGene_tx=XM_0232535
98.1;refGene_gn=CARMIL1;refGene_tx=XM_023253599.1;refGene_gn=CARMIL1;r
efGene_tx=XM_023253600.1;refGene_gn=CARMIL1;refGene_tx=XM_023253601.1;
refGene_gn=CARMIL1;refGene_tx=XM_023253602.1;refGene_gn=CARMIL1;refGen
e_tx=XM_023253603.1
B2 4578558 4578658
refGene_gn=CARMIL1;refGene_tx=XM_011282087.3;refGene_gn=CARMIL1;refGen
e_tx=XM_019830107.2;refGene_gn=CARMIL1;refGene_tx=XM_019830114.2;refGe
ne_gn=CARMIL1;refGene_tx=XM_019830116.2;refGene_gn=CARMIL1;refGene_tx=
XM_023253592.1;refGene_gn=CARMIL1;refGene_tx=XM_023253593.1;refGene_gn
=CARMIL1;refGene_tx=XM_023253594.1;refGene_gn=CARMIL1;refGene_tx=XM_02
3253595.1;refGene_gn=CARMIL1;refGene_tx=XM_023253596.1;refGene_gn=CARM
IL1;refGene_tx=XM_023253597.1;refGene_gn=CARMIL1;refGene_tx=XM_0232535
98.1;refGene_gn=CARMIL1;refGene_tx=XM_023253599.1;refGene_gn=CARMIL1;r
efGene_tx=XM_023253600.1;refGene_gn=CARMIL1;refGene_tx=XM_023253601.1;
refGene_gn=CARMIL1;refGene_tx=XM_023253602.1;refGene_gn=CARMIL1;refGen
e_tx=XM_023253603.1
B2 4579396 4579478
refGene_gn=CARMIL1;refGene_tx=XM_011282087.3;refGene_gn=CARMIL1;refGen
e_tx=XM_019830107.2;refGene_gn=CARMIL1;refGene_tx=XM_019830114.2;refGe
ne_gn=CARMIL1;refGene_tx=XM_019830116.2;refGene_gn=CARMIL1;refGene_tx=
XM_023253592.1;refGene_gn=CARMIL1;refGene_tx=XM_023253593.1;refGene_gn
=CARMIL1;refGene_tx=XM_023253594.1;refGene_gn=CARMIL1;refGene_tx=XM_02
3253595.1;refGene_gn=CARMIL1;refGene_tx=XM_023253596.1;refGene_gn=CARM
IL1;refGene_tx=XM_023253597.1;refGene_gn=CARMIL1;refGene_tx=XM_0232535
98.1;refGene_gn=CARMIL1;refGene_tx=XM_023253599.1;refGene_gn=CARMIL1;r
efGene_tx=XM_023253600.1;refGene_gn=CARMIL1;refGene_tx=XM_023253601.1;
refGene_gn=CARMIL1;refGene_tx=XM_023253602.1;refGene_gn=CARMIL1;refGen
e_tx=XM_023253603.1
B2 4581683 4581695
refGene_gn=CARMIL1;refGene_tx=XM_019830107.2;refGene_gn=CARMIL1;refGen
e_tx=XM_019830114.2;refGene_gn=CARMIL1;refGene_tx=XM_019830116.2;refGe
ne_gn=CARMIL1;refGene_tx=XM_023253592.1;refGene_gn=CARMIL1;refGene_tx=
XM_023253593.1;refGene_gn=CARMIL1;refGene_tx=XM_023253595.1;refGene_gn
=CARMIL1;refGene_tx=XM_023253596.1;refGene_gn=CARMIL1;refGene_tx=XM_02
3253597.1;refGene_gn=CARMIL1;refGene_tx=XM_023253598.1;refGene_gn=CARM
IL1;refGene_tx=XM_023253599.1;refGene_gn=CARMIL1;refGene_tx=XM_0232536
00.1;refGene_gn=CARMIL1;refGene_tx=XM_023253601.1;refGene_gn=CARMIL1;r
efGene_tx=XM_023253602.1;refGene_gn=CARMIL1;refGene_tx=XM_023253603.1
B2 4588155 4588225

refGene_gn=CARMIL1;refGene_tx=XM_011282087.3;refGene_gn=CARMIL1;refGen
e_tx=XM_019830107.2;refGene_gn=CARMIL1;refGene_tx=XM_019830114.2;refGe
ne_gn=CARMIL1;refGene_tx=XM_019830116.2;refGene_gn=CARMIL1;refGene_tx=
XM_023253592.1;refGene_gn=CARMIL1;refGene_tx=XM_023253593.1;refGene_gn
=CARMIL1;refGene_tx=XM_023253594.1;refGene_gn=CARMIL1;refGene_tx=XM_02
3253595.1;refGene_gn=CARMIL1;refGene_tx=XM_023253596.1;refGene_gn=CARM
IL1;refGene_tx=XM_023253597.1;refGene_gn=CARMIL1;refGene_tx=XM_0232535
98.1;refGene_gn=CARMIL1;refGene_tx=XM_023253599.1;refGene_gn=CARMIL1;r
efGene_tx=XM_023253600.1;refGene_gn=CARMIL1;refGene_tx=XM_023253601.1;
refGene_gn=CARMIL1;refGene_tx=XM_023253602.1;refGene_gn=CARMIL1;refGen
e_tx=XM_023253603.1
B2 4593304 4593409
refGene_gn=CARMIL1;refGene_tx=XM_011282087.3;refGene_gn=CARMIL1;refGen
e_tx=XM_019830107.2;refGene_gn=CARMIL1;refGene_tx=XM_019830114.2;refGe
ne_gn=CARMIL1;refGene_tx=XM_019830116.2;refGene_gn=CARMIL1;refGene_tx=
XM_023253592.1;refGene_gn=CARMIL1;refGene_tx=XM_023253593.1;refGene_gn
=CARMIL1;refGene_tx=XM_023253594.1;refGene_gn=CARMIL1;refGene_tx=XM_02
3253595.1;refGene_gn=CARMIL1;refGene_tx=XM_023253596.1;refGene_gn=CARM
IL1;refGene_tx=XM_023253597.1;refGene_gn=CARMIL1;refGene_tx=XM_0232535
98.1;refGene_gn=CARMIL1;refGene_tx=XM_023253599.1;refGene_gn=CARMIL1;r
efGene_tx=XM_023253600.1;refGene_gn=CARMIL1;refGene_tx=XM_023253601.1;
refGene_gn=CARMIL1;refGene_tx=XM_023253602.1;refGene_gn=CARMIL1;refGen
e_tx=XM_023253603.1
B2 4594305 4594382
refGene_gn=CARMIL1;refGene_tx=XM_011282087.3;refGene_gn=CARMIL1;refGen
e_tx=XM_019830107.2;refGene_gn=CARMIL1;refGene_tx=XM_019830114.2;refGe
ne_gn=CARMIL1;refGene_tx=XM_019830116.2;refGene_gn=CARMIL1;refGene_tx=
XM_023253592.1;refGene_gn=CARMIL1;refGene_tx=XM_023253593.1;refGene_gn
=CARMIL1;refGene_tx=XM_023253594.1;refGene_gn=CARMIL1;refGene_tx=XM_02
3253595.1;refGene_gn=CARMIL1;refGene_tx=XM_023253596.1;refGene_gn=CARM
IL1;refGene_tx=XM_023253597.1;refGene_gn=CARMIL1;refGene_tx=XM_0232535
98.1;refGene_gn=CARMIL1;refGene_tx=XM_023253599.1;refGene_gn=CARMIL1;r
efGene_tx=XM_023253600.1;refGene_gn=CARMIL1;refGene_tx=XM_023253601.1;
refGene_gn=CARMIL1;refGene_tx=XM_023253602.1;refGene_gn=CARMIL1;refGen
e_tx=XM_023253603.1
B2 4594516 4594594
refGene_gn=CARMIL1;refGene_tx=XM_011282087.3;refGene_gn=CARMIL1;refGen
e_tx=XM_019830107.2;refGene_gn=CARMIL1;refGene_tx=XM_019830114.2;refGe
ne_gn=CARMIL1;refGene_tx=XM_019830116.2;refGene_gn=CARMIL1;refGene_tx=
XM_023253592.1;refGene_gn=CARMIL1;refGene_tx=XM_023253593.1;refGene_gn
=CARMIL1;refGene_tx=XM_023253594.1;refGene_gn=CARMIL1;refGene_tx=XM_02
3253595.1;refGene_gn=CARMIL1;refGene_tx=XM_023253596.1;refGene_gn=CARM
IL1;refGene_tx=XM_023253597.1;refGene_gn=CARMIL1;refGene_tx=XM_0232535
98.1;refGene_gn=CARMIL1;refGene_tx=XM_023253599.1;refGene_gn=CARMIL1;r
efGene_tx=XM_023253600.1;refGene_gn=CARMIL1;refGene_tx=XM_023253601.1;
refGene_gn=CARMIL1;refGene_tx=XM_023253602.1;refGene_gn=CARMIL1;refGen
e_tx=XM_023253603.1
B2 4597222 4597326
refGene_gn=CARMIL1;refGene_tx=XM_011282087.3;refGene_gn=CARMIL1;refGen
e_tx=XM_019830107.2;refGene_gn=CARMIL1;refGene_tx=XM_019830114.2;refGe

ne_gn=CARMIL1;refGene_tx=XM_019830116.2;refGene_gn=CARMIL1;refGene_tx=XM_023253592.1;refGene_gn=CARMIL1;refGene_tx=XM_023253593.1;refGene_gn=CARMIL1;refGene_tx=XM_023253594.1;refGene_gn=CARMIL1;refGene_tx=XM_023253595.1;refGene_gn=CARMIL1;refGene_tx=XM_023253596.1;refGene_gn=CARMIL1;refGene_tx=XM_023253597.1;refGene_gn=CARMIL1;refGene_tx=XM_023253598.1;refGene_gn=CARMIL1;refGene_tx=XM_023253599.1;refGene_gn=CARMIL1;refGene_tx=XM_023253600.1;refGene_gn=CARMIL1;refGene_tx=XM_023253601.1;refGene_gn=CARMIL1;refGene_tx=XM_023253602.1;refGene_gn=CARMIL1;refGene_tx=XM_023253603.1
B2 4604938 4605025
refGene_gn=CARMIL1;refGene_tx=XM_011282087.3;refGene_gn=CARMIL1;refGene_tx=XM_019830107.2;refGene_gn=CARMIL1;refGene_tx=XM_019830114.2;refGene_gn=CARMIL1;refGene_tx=XM_019830116.2;refGene_gn=CARMIL1;refGene_tx=XM_023253592.1;refGene_gn=CARMIL1;refGene_tx=XM_023253593.1;refGene_gn=CARMIL1;refGene_tx=XM_023253594.1;refGene_gn=CARMIL1;refGene_tx=XM_023253595.1;refGene_gn=CARMIL1;refGene_tx=XM_023253596.1;refGene_gn=CARMIL1;refGene_tx=XM_023253597.1;refGene_gn=CARMIL1;refGene_tx=XM_023253598.1;refGene_gn=CARMIL1;refGene_tx=XM_023253599.1;refGene_gn=CARMIL1;refGene_tx=XM_023253600.1;refGene_gn=CARMIL1;refGene_tx=XM_023253601.1;refGene_gn=CARMIL1;refGene_tx=XM_023253602.1;refGene_gn=CARMIL1;refGene_tx=XM_023253603.1
B2 4616942 4617037
refGene_gn=CARMIL1;refGene_tx=XM_011282087.3;refGene_gn=CARMIL1;refGene_tx=XM_019830107.2;refGene_gn=CARMIL1;refGene_tx=XM_019830114.2;refGene_gn=CARMIL1;refGene_tx=XM_019830116.2;refGene_gn=CARMIL1;refGene_tx=XM_023253592.1;refGene_gn=CARMIL1;refGene_tx=XM_023253593.1;refGene_gn=CARMIL1;refGene_tx=XM_023253594.1;refGene_gn=CARMIL1;refGene_tx=XM_023253595.1;refGene_gn=CARMIL1;refGene_tx=XM_023253596.1;refGene_gn=CARMIL1;refGene_tx=XM_023253597.1;refGene_gn=CARMIL1;refGene_tx=XM_023253598.1;refGene_gn=CARMIL1;refGene_tx=XM_023253599.1;refGene_gn=CARMIL1;refGene_tx=XM_023253600.1;refGene_gn=CARMIL1;refGene_tx=XM_023253601.1;refGene_gn=CARMIL1;refGene_tx=XM_023253602.1;refGene_gn=CARMIL1;refGene_tx=XM_023253603.1
B2 4618228 4618317
refGene_gn=CARMIL1;refGene_tx=XM_011282087.3;refGene_gn=CARMIL1;refGene_tx=XM_019830107.2;refGene_gn=CARMIL1;refGene_tx=XM_019830114.2;refGene_gn=CARMIL1;refGene_tx=XM_019830116.2;refGene_gn=CARMIL1;refGene_tx=XM_023253592.1;refGene_gn=CARMIL1;refGene_tx=XM_023253593.1;refGene_gn=CARMIL1;refGene_tx=XM_023253594.1;refGene_gn=CARMIL1;refGene_tx=XM_023253595.1;refGene_gn=CARMIL1;refGene_tx=XM_023253596.1;refGene_gn=CARMIL1;refGene_tx=XM_023253597.1;refGene_gn=CARMIL1;refGene_tx=XM_023253598.1;refGene_gn=CARMIL1;refGene_tx=XM_023253599.1;refGene_gn=CARMIL1;refGene_tx=XM_023253600.1;refGene_gn=CARMIL1;refGene_tx=XM_023253601.1;refGene_gn=CARMIL1;refGene_tx=XM_023253602.1;refGene_gn=CARMIL1;refGene_tx=XM_023253603.1
B2 4624966 4625042
refGene_gn=CARMIL1;refGene_tx=XM_011282087.3;refGene_gn=CARMIL1;refGene_tx=XM_019830107.2;refGene_gn=CARMIL1;refGene_tx=XM_019830114.2;refGene_gn=CARMIL1;refGene_tx=XM_019830116.2;refGene_gn=CARMIL1;refGene_tx=XM_023253592.1;refGene_gn=CARMIL1;refGene_tx=XM_023253593.1;refGene_gn

=CARMIL1;refGene_tx=XM_023253594.1;refGene_gn=CARMIL1;refGene_tx=XM_023253595.1;refGene_gn=CARMIL1;refGene_tx=XM_023253596.1;refGene_gn=CARMIL1;refGene_tx=XM_023253597.1;refGene_gn=CARMIL1;refGene_tx=XM_023253598.1;refGene_gn=CARMIL1;refGene_tx=XM_023253599.1;refGene_gn=CARMIL1;refGene_tx=XM_023253600.1;refGene_gn=CARMIL1;refGene_tx=XM_023253601.1;refGene_gn=CARMIL1;refGene_tx=XM_023253602.1;refGene_gn=CARMIL1;refGene_tx=XM_023253603.1

B2 4640598 4640672
refGene_gn=CARMIL1;refGene_tx=XM_011282087.3;refGene_gn=CARMIL1;refGene_tx=XM_019830107.2;refGene_gn=CARMIL1;refGene_tx=XM_019830114.2;refGene_gn=CARMIL1;refGene_tx=XM_019830116.2;refGene_gn=CARMIL1;refGene_tx=XM_023253592.1;refGene_gn=CARMIL1;refGene_tx=XM_023253593.1;refGene_gn=CARMIL1;refGene_tx=XM_023253594.1;refGene_gn=CARMIL1;refGene_tx=XM_023253595.1;refGene_gn=CARMIL1;refGene_tx=XM_023253596.1;refGene_gn=CARMIL1;refGene_tx=XM_023253597.1;refGene_gn=CARMIL1;refGene_tx=XM_023253598.1;refGene_gn=CARMIL1;refGene_tx=XM_023253599.1;refGene_gn=CARMIL1;refGene_tx=XM_023253600.1;refGene_gn=CARMIL1;refGene_tx=XM_023253601.1;refGene_gn=CARMIL1;refGene_tx=XM_023253602.1;refGene_gn=CARMIL1;refGene_tx=XM_023253603.1

B2 4640921 4640992
refGene_gn=CARMIL1;refGene_tx=XM_011282087.3;refGene_gn=CARMIL1;refGene_tx=XM_019830107.2;refGene_gn=CARMIL1;refGene_tx=XM_019830114.2;refGene_gn=CARMIL1;refGene_tx=XM_019830116.2;refGene_gn=CARMIL1;refGene_tx=XM_023253592.1;refGene_gn=CARMIL1;refGene_tx=XM_023253593.1;refGene_gn=CARMIL1;refGene_tx=XM_023253594.1;refGene_gn=CARMIL1;refGene_tx=XM_023253595.1;refGene_gn=CARMIL1;refGene_tx=XM_023253596.1;refGene_gn=CARMIL1;refGene_tx=XM_023253597.1;refGene_gn=CARMIL1;refGene_tx=XM_023253598.1;refGene_gn=CARMIL1;refGene_tx=XM_023253599.1;refGene_gn=CARMIL1;refGene_tx=XM_023253600.1;refGene_gn=CARMIL1;refGene_tx=XM_023253601.1;refGene_gn=CARMIL1;refGene_tx=XM_023253602.1;refGene_gn=CARMIL1;refGene_tx=XM_023253603.1

B2 4641329 4641427
refGene_gn=CARMIL1;refGene_tx=XM_011282087.3;refGene_gn=CARMIL1;refGene_tx=XM_019830107.2;refGene_gn=CARMIL1;refGene_tx=XM_019830114.2;refGene_gn=CARMIL1;refGene_tx=XM_019830116.2;refGene_gn=CARMIL1;refGene_tx=XM_023253592.1;refGene_gn=CARMIL1;refGene_tx=XM_023253593.1;refGene_gn=CARMIL1;refGene_tx=XM_023253594.1;refGene_gn=CARMIL1;refGene_tx=XM_023253595.1;refGene_gn=CARMIL1;refGene_tx=XM_023253596.1;refGene_gn=CARMIL1;refGene_tx=XM_023253597.1;refGene_gn=CARMIL1;refGene_tx=XM_023253598.1;refGene_gn=CARMIL1;refGene_tx=XM_023253599.1;refGene_gn=CARMIL1;refGene_tx=XM_023253600.1;refGene_gn=CARMIL1;refGene_tx=XM_023253601.1;refGene_gn=CARMIL1;refGene_tx=XM_023253602.1;refGene_gn=CARMIL1;refGene_tx=XM_023253603.1

B2 4652406 4652528
refGene_gn=CARMIL1;refGene_tx=XM_011282087.3;refGene_gn=CARMIL1;refGene_tx=XM_019830107.2;refGene_gn=CARMIL1;refGene_tx=XM_019830114.2;refGene_gn=CARMIL1;refGene_tx=XM_019830116.2;refGene_gn=CARMIL1;refGene_tx=XM_023253592.1;refGene_gn=CARMIL1;refGene_tx=XM_023253593.1;refGene_gn=CARMIL1;refGene_tx=XM_023253594.1;refGene_gn=CARMIL1;refGene_tx=XM_023253595.1;refGene_gn=CARMIL1;refGene_tx=XM_023253596.1;refGene_gn=CARM

IL1;refGene_tx=XM_023253597.1;refGene_gn=CARMIL1;refGene_tx=XM_0232535
98.1;refGene_gn=CARMIL1;refGene_tx=XM_023253599.1;refGene_gn=CARMIL1;r
efGene_tx=XM_023253600.1;refGene_gn=CARMIL1;refGene_tx=XM_023253601.1;
refGene_gn=CARMIL1;refGene_tx=XM_023253602.1;refGene_gn=CARMIL1;refGen
e_tx=XM_023253603.1
B2 4662322 4662382
refGene_gn=CARMIL1;refGene_tx=XM_011282087.3;refGene_gn=CARMIL1;refGen
e_tx=XM_019830107.2;refGene_gn=CARMIL1;refGene_tx=XM_019830114.2;refGe
ne_gn=CARMIL1;refGene_tx=XM_019830116.2;refGene_gn=CARMIL1;refGene_tx=
XM_023253592.1;refGene_gn=CARMIL1;refGene_tx=XM_023253593.1;refGene_gn
=CARMIL1;refGene_tx=XM_023253594.1;refGene_gn=CARMIL1;refGene_tx=XM_02
3253595.1;refGene_gn=CARMIL1;refGene_tx=XM_023253596.1;refGene_gn=CARM
IL1;refGene_tx=XM_023253597.1;refGene_gn=CARMIL1;refGene_tx=XM_0232535
98.1;refGene_gn=CARMIL1;refGene_tx=XM_023253599.1;refGene_gn=CARMIL1;r
efGene_tx=XM_023253600.1;refGene_gn=CARMIL1;refGene_tx=XM_023253601.1;
refGene_gn=CARMIL1;refGene_tx=XM_023253602.1;refGene_gn=CARMIL1;refGen
e_tx=XM_023253603.1
B2 4667872 4667923
refGene_gn=CARMIL1;refGene_tx=XM_011282087.3;refGene_gn=CARMIL1;refGen
e_tx=XM_019830107.2;refGene_gn=CARMIL1;refGene_tx=XM_019830114.2;refGe
ne_gn=CARMIL1;refGene_tx=XM_019830116.2;refGene_gn=CARMIL1;refGene_tx=
XM_023253592.1;refGene_gn=CARMIL1;refGene_tx=XM_023253593.1;refGene_gn
=CARMIL1;refGene_tx=XM_023253594.1;refGene_gn=CARMIL1;refGene_tx=XM_02
3253595.1;refGene_gn=CARMIL1;refGene_tx=XM_023253596.1;refGene_gn=CARM
IL1;refGene_tx=XM_023253597.1;refGene_gn=CARMIL1;refGene_tx=XM_0232535
98.1;refGene_gn=CARMIL1;refGene_tx=XM_023253599.1;refGene_gn=CARMIL1;r
efGene_tx=XM_023253600.1;refGene_gn=CARMIL1;refGene_tx=XM_023253601.1;
refGene_gn=CARMIL1;refGene_tx=XM_023253602.1;refGene_gn=CARMIL1;refGen
e_tx=XM_023253603.1
B2 4776802 4776900
refGene_gn=CARMIL1;refGene_tx=XM_011282087.3;refGene_gn=CARMIL1;refGen
e_tx=XM_019830107.2;refGene_gn=CARMIL1;refGene_tx=XM_019830114.2;refGe
ne_gn=CARMIL1;refGene_tx=XM_019830116.2;refGene_gn=CARMIL1;refGene_tx=
XM_023253592.1;refGene_gn=CARMIL1;refGene_tx=XM_023253593.1;refGene_gn
=CARMIL1;refGene_tx=XM_023253594.1;refGene_gn=CARMIL1;refGene_tx=XM_02
3253595.1;refGene_gn=CARMIL1;refGene_tx=XM_023253596.1;refGene_gn=CARM
IL1;refGene_tx=XM_023253597.1;refGene_gn=CARMIL1;refGene_tx=XM_0232535
98.1;refGene_gn=CARMIL1;refGene_tx=XM_023253599.1;refGene_gn=CARMIL1;r
efGene_tx=XM_023253600.1;refGene_gn=CARMIL1;refGene_tx=XM_023253601.1;
refGene_gn=CARMIL1;refGene_tx=XM_023253602.1;refGene_gn=CARMIL1;refGen
e_tx=XM_023253603.1
B2 4780936 4780976
refGene_gn=CARMIL1;refGene_tx=XM_011282087.3;refGene_gn=CARMIL1;refGen
e_tx=XM_019830107.2;refGene_gn=CARMIL1;refGene_tx=XM_019830116.2
B2 4781680 4781921
refGene_gn=CARMIL1;refGene_tx=XM_023253592.1;refGene_gn=CARMIL1;refGen
e_tx=XM_023253593.1;refGene_gn=CARMIL1;refGene_tx=XM_023253594.1;refGe
ne_gn=CARMIL1;refGene_tx=XM_023253595.1;refGene_gn=CARMIL1;refGene_tx=
XM_023253596.1;refGene_gn=CARMIL1;refGene_tx=XM_023253597.1;refGene_gn
=CARMIL1;refGene_tx=XM_023253598.1;refGene_gn=CARMIL1;refGene_tx=XM_02

3253599.1;refGene_gn=CARMIL1;refGene_tx=XM_023253600.1;refGene_gn=CARM
IL1;refGene_tx=XM_023253601.1;refGene_gn=CARMIL1;refGene_tx=XM_0232536
02.1;refGene_gn=CARMIL1;refGene_tx=XM_023253603.1
B2 4889051 4889104 refGene_gn=CMAH;refGene_tx=XM_011282052.3
B2 4894248 4894296
refGene_gn=CMAH;refGene_tx=NM_001244985.1;refGene_gn=CMAH;refGene_tx=X
M_011282051.3;refGene_gn=CMAH;refGene_tx=XM_011282052.3
B2 4934909 4935001 refGene_gn=CMAH;refGene_tx=XM_019830061.2
B2 4935217 4935421
refGene_gn=CMAH;refGene_tx=NM_001244985.1;refGene_gn=CMAH;refGene_tx=X
M_011282051.3;refGene_gn=CMAH;refGene_tx=XM_011282052.3;refGene_gn=CMA
H;refGene_tx=XM_019830061.2
B2 4938675 4938767
refGene_gn=CMAH;refGene_tx=NM_001244985.1;refGene_gn=CMAH;refGene_tx=X
M_011282051.3;refGene_gn=CMAH;refGene_tx=XM_011282052.3;refGene_gn=CMA
H;refGene_tx=XM_019830061.2
B2 4947462 4947572
refGene_gn=CMAH;refGene_tx=NM_001244985.1;refGene_gn=CMAH;refGene_tx=X
M_011282051.3;refGene_gn=CMAH;refGene_tx=XM_011282052.3;refGene_gn=CMA
H;refGene_tx=XM_019830061.2
B2 4948876 4949067
refGene_gn=CMAH;refGene_tx=NM_001244985.1;refGene_gn=CMAH;refGene_tx=X
M_011282051.3;refGene_gn=CMAH;refGene_tx=XM_011282052.3;refGene_gn=CMA
H;refGene_tx=XM_019830061.2
B2 4953091 4953239
refGene_gn=CMAH;refGene_tx=NM_001244985.1;refGene_gn=CMAH;refGene_tx=X
M_011282051.3;refGene_gn=CMAH;refGene_tx=XM_011282052.3;refGene_gn=CMA
H;refGene_tx=XM_019830061.2
B2 4953551 4953633
refGene_gn=CMAH;refGene_tx=NM_001244985.1;refGene_gn=CMAH;refGene_tx=X
M_011282051.3;refGene_gn=CMAH;refGene_tx=XM_011282052.3;refGene_gn=CMA
H;refGene_tx=XM_019830061.2
B2 4955264 4955402
refGene_gn=CMAH;refGene_tx=NM_001244985.1;refGene_gn=CMAH;refGene_tx=X
M_011282051.3;refGene_gn=CMAH;refGene_tx=XM_011282052.3;refGene_gn=CMA
H;refGene_tx=XM_019830061.2
B2 4957748 4957884
refGene_gn=CMAH;refGene_tx=NM_001244985.1;refGene_gn=CMAH;refGene_tx=X
M_011282051.3;refGene_gn=CMAH;refGene_tx=XM_011282052.3;refGene_gn=CMA
H;refGene_tx=XM_019830061.2
B2 4974624 4974765
refGene_gn=CMAH;refGene_tx=NM_001244985.1;refGene_gn=CMAH;refGene_tx=X
M_011282051.3;refGene_gn=CMAH;refGene_tx=XM_011282052.3;refGene_gn=CMA
H;refGene_tx=XM_019830061.2
B2 4978865 4979040
refGene_gn=CMAH;refGene_tx=NM_001244985.1;refGene_gn=CMAH;refGene_tx=X
M_011282051.3;refGene_gn=CMAH;refGene_tx=XM_011282052.3;refGene_gn=CMA
H;refGene_tx=XM_019830061.2
B2 4981076 4981193
refGene_gn=CMAH;refGene_tx=NM_001244985.1;refGene_gn=CMAH;refGene_tx=X

M_011282051.3;refGene_gn=CMAH;refGene_tx=XM_011282052.3;refGene_gn=CMAH;refGene_tx=XM_019830061.2
B2 4985704 4985820
refGene_gn=CMAH;refGene_tx=NM_001244985.1;refGene_gn=CMAH;refGene_tx=XM_011282051.3;refGene_gn=CMAH;refGene_tx=XM_011282052.3;refGene_gn=CMAH;refGene_tx=XM_019830061.2
B2 4988265 4988341
refGene_gn=CMAH;refGene_tx=NM_001244985.1;refGene_gn=CMAH;refGene_tx=XM_011282051.3;refGene_gn=CMAH;refGene_tx=XM_011282052.3;refGene_gn=CMAH;refGene_tx=XM_019830061.2
B2 5022402 5022478
refGene_gn=RIPOR2;refGene_tx=XM_023253604.1;refGene_gn=RIPOR2;refGene_tx=XM_023253607.1;refGene_gn=RIPOR2;refGene_tx=XM_023253608.1;refGene_gn=RIPOR2;refGene_tx=XM_023253609.1;refGene_gn=RIPOR2;refGene_tx=XM_023253611.1;refGene_gn=RIPOR2;refGene_tx=XM_023253612.1;refGene_gn=RIPOR2;refGene_tx=XM_023253613.1
B2 5135780 5135841 refGene_gn=RIPOR2;refGene_tx=XM_023253605.1
B2 5191949 5192076
refGene_gn=RIPOR2;refGene_tx=XM_023253604.1;refGene_gn=RIPOR2;refGene_tx=XM_023253605.1;refGene_gn=RIPOR2;refGene_tx=XM_023253607.1;refGene_gn=RIPOR2;refGene_tx=XM_023253608.1;refGene_gn=RIPOR2;refGene_tx=XM_023253609.1;refGene_gn=RIPOR2;refGene_tx=XM_023253610.1;refGene_gn=RIPOR2;refGene_tx=XM_023253611.1;refGene_gn=RIPOR2;refGene_tx=XM_023253612.1;refGene_gn=RIPOR2;refGene_tx=XM_023253613.1
B2 5193826 5193982
refGene_gn=RIPOR2;refGene_tx=XM_023253604.1;refGene_gn=RIPOR2;refGene_tx=XM_023253605.1;refGene_gn=RIPOR2;refGene_tx=XM_023253607.1;refGene_gn=RIPOR2;refGene_tx=XM_023253608.1;refGene_gn=RIPOR2;refGene_tx=XM_023253609.1;refGene_gn=RIPOR2;refGene_tx=XM_023253610.1;refGene_gn=RIPOR2;refGene_tx=XM_023253611.1;refGene_gn=RIPOR2;refGene_tx=XM_023253612.1;refGene_gn=RIPOR2;refGene_tx=XM_023253613.1
B2 5194799 5194878
refGene_gn=RIPOR2;refGene_tx=XM_023253604.1;refGene_gn=RIPOR2;refGene_tx=XM_023253605.1;refGene_gn=RIPOR2;refGene_tx=XM_023253607.1;refGene_gn=RIPOR2;refGene_tx=XM_023253608.1;refGene_gn=RIPOR2;refGene_tx=XM_023253609.1;refGene_gn=RIPOR2;refGene_tx=XM_023253610.1;refGene_gn=RIPOR2;refGene_tx=XM_023253611.1;refGene_gn=RIPOR2;refGene_tx=XM_023253612.1;refGene_gn=RIPOR2;refGene_tx=XM_023253613.1
B2 5197824 5197848
refGene_gn=RIPOR2;refGene_tx=XM_023253604.1;refGene_gn=RIPOR2;refGene_tx=XM_023253605.1;refGene_gn=RIPOR2;refGene_tx=XM_023253607.1;refGene_gn=RIPOR2;refGene_tx=XM_023253608.1;refGene_gn=RIPOR2;refGene_tx=XM_023253609.1;refGene_gn=RIPOR2;refGene_tx=XM_023253610.1;refGene_gn=RIPOR2;refGene_tx=XM_023253611.1;refGene_gn=RIPOR2;refGene_tx=XM_023253612.1;refGene_gn=RIPOR2;refGene_tx=XM_023253613.1
B2 5198919 5198973
refGene_gn=RIPOR2;refGene_tx=XM_023253604.1;refGene_gn=RIPOR2;refGene_tx=XM_023253605.1;refGene_gn=RIPOR2;refGene_tx=XM_023253607.1;refGene_gn=RIPOR2;refGene_tx=XM_023253608.1;refGene_gn=RIPOR2;refGene_tx=XM_023253609.1;refGene_gn=RIPOR2;refGene_tx=XM_023253610.1;refGene_gn=RIPOR

2;refGene_tx=XM_023253611.1;refGene_gn=RIPOR2;refGene_tx=XM_023253612.1;refGene_gn=RIPOR2;refGene_tx=XM_023253613.1
B2 5202384 5202534
refGene_gn=RIPOR2;refGene_tx=XM_023253604.1;refGene_gn=RIPOR2;refGene_tx=XM_023253605.1;refGene_gn=RIPOR2;refGene_tx=XM_023253607.1;refGene_gn=RIPOR2;refGene_tx=XM_023253608.1;refGene_gn=RIPOR2;refGene_tx=XM_023253609.1;refGene_gn=RIPOR2;refGene_tx=XM_023253610.1;refGene_gn=RIPOR2;refGene_tx=XM_023253611.1;refGene_gn=RIPOR2;refGene_tx=XM_023253612.1;refGene_gn=RIPOR2;refGene_tx=XM_023253613.1
B2 5206242 5206306
refGene_gn=RIPOR2;refGene_tx=XM_023253604.1;refGene_gn=RIPOR2;refGene_tx=XM_023253605.1;refGene_gn=RIPOR2;refGene_tx=XM_023253607.1;refGene_gn=RIPOR2;refGene_tx=XM_023253608.1;refGene_gn=RIPOR2;refGene_tx=XM_023253609.1;refGene_gn=RIPOR2;refGene_tx=XM_023253610.1;refGene_gn=RIPOR2;refGene_tx=XM_023253611.1;refGene_gn=RIPOR2;refGene_tx=XM_023253612.1;refGene_gn=RIPOR2;refGene_tx=XM_023253613.1
B2 5214563 5214607
refGene_gn=RIPOR2;refGene_tx=XM_023253604.1;refGene_gn=RIPOR2;refGene_tx=XM_023253605.1;refGene_gn=RIPOR2;refGene_tx=XM_023253607.1;refGene_gn=RIPOR2;refGene_tx=XM_023253608.1;refGene_gn=RIPOR2;refGene_tx=XM_023253609.1;refGene_gn=RIPOR2;refGene_tx=XM_023253610.1;refGene_gn=RIPOR2;refGene_tx=XM_023253611.1;refGene_gn=RIPOR2;refGene_tx=XM_023253612.1;refGene_gn=RIPOR2;refGene_tx=XM_023253613.1
B2 5216606 5216732
refGene_gn=RIPOR2;refGene_tx=XM_023253604.1;refGene_gn=RIPOR2;refGene_tx=XM_023253605.1;refGene_gn=RIPOR2;refGene_tx=XM_023253607.1;refGene_gn=RIPOR2;refGene_tx=XM_023253608.1;refGene_gn=RIPOR2;refGene_tx=XM_023253609.1;refGene_gn=RIPOR2;refGene_tx=XM_023253610.1;refGene_gn=RIPOR2;refGene_tx=XM_023253611.1;refGene_gn=RIPOR2;refGene_tx=XM_023253612.1;refGene_gn=RIPOR2;refGene_tx=XM_023253613.1
B2 5216982 5217131
refGene_gn=RIPOR2;refGene_tx=XM_023253604.1;refGene_gn=RIPOR2;refGene_tx=XM_023253605.1;refGene_gn=RIPOR2;refGene_tx=XM_023253607.1;refGene_gn=RIPOR2;refGene_tx=XM_023253608.1;refGene_gn=RIPOR2;refGene_tx=XM_023253609.1;refGene_gn=RIPOR2;refGene_tx=XM_023253610.1;refGene_gn=RIPOR2;refGene_tx=XM_023253611.1;refGene_gn=RIPOR2;refGene_tx=XM_023253612.1;refGene_gn=RIPOR2;refGene_tx=XM_023253613.1
B2 5218641 5218771
refGene_gn=RIPOR2;refGene_tx=XM_023253604.1;refGene_gn=RIPOR2;refGene_tx=XM_023253605.1;refGene_gn=RIPOR2;refGene_tx=XM_023253607.1;refGene_gn=RIPOR2;refGene_tx=XM_023253608.1;refGene_gn=RIPOR2;refGene_tx=XM_023253609.1;refGene_gn=RIPOR2;refGene_tx=XM_023253610.1;refGene_gn=RIPOR2;refGene_tx=XM_023253611.1;refGene_gn=RIPOR2;refGene_tx=XM_023253612.1;refGene_gn=RIPOR2;refGene_tx=XM_023253613.1
B2 5219138 5219288 refGene_gn=RIPOR2;refGene_tx=XM_023253612.1
B2 5224284 5224971
refGene_gn=RIPOR2;refGene_tx=XM_023253604.1;refGene_gn=RIPOR2;refGene_tx=XM_023253605.1;refGene_gn=RIPOR2;refGene_tx=XM_023253607.1;refGene_gn=RIPOR2;refGene_tx=XM_023253608.1;refGene_gn=RIPOR2;refGene_tx=XM_023253609.1;refGene_gn=RIPOR2;refGene_tx=XM_023253610.1;refGene_gn=RIPOR

2;refGene_tx=XM_023253611.1;refGene_gn=RIPOR2;refGene_tx=XM_023253612.1;refGene_gn=RIPOR2;refGene_tx=XM_023253613.1
B2 5226784 5226983
refGene_gn=RIPOR2;refGene_tx=XM_023253611.1;refGene_gn=RIPOR2;refGene_tx=XM_023253613.1
B2 5228217 5228399
refGene_gn=RIPOR2;refGene_tx=XM_023253604.1;refGene_gn=RIPOR2;refGene_tx=XM_023253605.1;refGene_gn=RIPOR2;refGene_tx=XM_023253607.1;refGene_gn=RIPOR2;refGene_tx=XM_023253608.1;refGene_gn=RIPOR2;refGene_tx=XM_023253609.1;refGene_gn=RIPOR2;refGene_tx=XM_023253610.1;refGene_gn=RIPOR2;refGene_tx=XM_023253612.1
B2 5231107 5231279
refGene_gn=RIPOR2;refGene_tx=XM_023253604.1;refGene_gn=RIPOR2;refGene_tx=XM_023253605.1;refGene_gn=RIPOR2;refGene_tx=XM_023253607.1;refGene_gn=RIPOR2;refGene_tx=XM_023253608.1;refGene_gn=RIPOR2;refGene_tx=XM_023253609.1;refGene_gn=RIPOR2;refGene_tx=XM_023253610.1;refGene_gn=RIPOR2;refGene_tx=XM_023253612.1
B2 5233973 5234109
refGene_gn=RIPOR2;refGene_tx=XM_023253604.1;refGene_gn=RIPOR2;refGene_tx=XM_023253605.1;refGene_gn=RIPOR2;refGene_tx=XM_023253607.1;refGene_gn=RIPOR2;refGene_tx=XM_023253608.1;refGene_gn=RIPOR2;refGene_tx=XM_023253609.1;refGene_gn=RIPOR2;refGene_tx=XM_023253610.1;refGene_gn=RIPOR2;refGene_tx=XM_023253612.1
B2 5236157 5236319
refGene_gn=RIPOR2;refGene_tx=XM_023253604.1;refGene_gn=RIPOR2;refGene_tx=XM_023253605.1;refGene_gn=RIPOR2;refGene_tx=XM_023253607.1;refGene_gn=RIPOR2;refGene_tx=XM_023253608.1;refGene_gn=RIPOR2;refGene_tx=XM_023253609.1;refGene_gn=RIPOR2;refGene_tx=XM_023253610.1;refGene_gn=RIPOR2;refGene_tx=XM_023253612.1
B2 5237334 5237493
refGene_gn=RIPOR2;refGene_tx=XM_023253604.1;refGene_gn=RIPOR2;refGene_tx=XM_023253605.1;refGene_gn=RIPOR2;refGene_tx=XM_023253607.1;refGene_gn=RIPOR2;refGene_tx=XM_023253608.1;refGene_gn=RIPOR2;refGene_tx=XM_023253609.1;refGene_gn=RIPOR2;refGene_tx=XM_023253610.1;refGene_gn=RIPOR2;refGene_tx=XM_023253612.1
B2 5239752 5239955
refGene_gn=RIPOR2;refGene_tx=XM_023253604.1;refGene_gn=RIPOR2;refGene_tx=XM_023253605.1;refGene_gn=RIPOR2;refGene_tx=XM_023253607.1;refGene_gn=RIPOR2;refGene_tx=XM_023253608.1;refGene_gn=RIPOR2;refGene_tx=XM_023253609.1;refGene_gn=RIPOR2;refGene_tx=XM_023253610.1;refGene_gn=RIPOR2;refGene_tx=XM_023253612.1
B2 5245193 5245277
refGene_gn=RIPOR2;refGene_tx=XM_023253604.1;refGene_gn=RIPOR2;refGene_tx=XM_023253605.1;refGene_gn=RIPOR2;refGene_tx=XM_023253608.1;refGene_gn=RIPOR2;refGene_tx=XM_023253610.1;refGene_gn=RIPOR2;refGene_tx=XM_023253612.1
B2 5249275 5249366
refGene_gn=RIPOR2;refGene_tx=XM_023253604.1;refGene_gn=RIPOR2;refGene_tx=XM_023253605.1;refGene_gn=RIPOR2;refGene_tx=XM_023253607.1;refGene_gn=RIPOR2;refGene_tx=XM_023253608.1;refGene_gn=RIPOR2;refGene_tx=XM_02

3253609.1;refGene_gn=RIPOR2;refGene_tx=XM_023253610.1;refGene_gn=RIPOR2;refGene_tx=XM_023253612.1
B2 5251573 5251767
refGene_gn=RIPOR2;refGene_tx=XM_023253604.1;refGene_gn=RIPOR2;refGene_tx=XM_023253605.1;refGene_gn=RIPOR2;refGene_tx=XM_023253607.1;refGene_gn=RIPOR2;refGene_tx=XM_023253610.1
B2 5254044 5254145
refGene_gn=RIPOR2;refGene_tx=XM_023253608.1;refGene_gn=RIPOR2;refGene_tx=XM_023253609.1;refGene_gn=RIPOR2;refGene_tx=XM_023253612.1
B2 5259813 5260078
refGene_gn=CB2H6orf229;refGene_tx=XM_006931403.3
B2 5262393 5262821
refGene_gn=CB2H6orf229;refGene_tx=XM_006931403.3
B2 5273394 5273556
refGene_gn=GMNN;refGene_tx=XM_003985741.5;refGene_gn=GMNN;refGene_tx=XM_006931404.4
B2 5275233 5275344
refGene_gn=GMNN;refGene_tx=XM_003985741.5;refGene_gn=GMNN;refGene_tx=XM_006931404.4
B2 5275650 5275733
refGene_gn=GMNN;refGene_tx=XM_003985741.5;refGene_gn=GMNN;refGene_tx=XM_006931404.4
B2 5276253 5276401
refGene_gn=GMNN;refGene_tx=XM_003985741.5;refGene_gn=GMNN;refGene_tx=XM_006931404.4
B2 5276959 5277037
refGene_gn=GMNN;refGene_tx=XM_003985741.5;refGene_gn=GMNN;refGene_tx=XM_006931404.4
B2 5280976 5281027
refGene_gn=GMNN;refGene_tx=XM_003985741.5;refGene_gn=GMNN;refGene_tx=XM_006931404.4
B2 5338198 5338240
refGene_gn=CB2H6orf62;refGene_tx=XM_006931406.4
B2 5339473 5339602
refGene_gn=CB2H6orf62;refGene_tx=XM_003985742.4
B2 5341859 5342036
refGene_gn=CB2H6orf62;refGene_tx=XM_003985742.4;refGene_gn=CB2H6orf62;refGene_tx=XM_006931406.4
B2 5342862 5342985
refGene_gn=CB2H6orf62;refGene_tx=XM_003985742.4;refGene_gn=CB2H6orf62;refGene_tx=XM_006931406.4
B2 5347299 5347434
refGene_gn=CB2H6orf62;refGene_tx=XM_003985742.4;refGene_gn=CB2H6orf62;refGene_tx=XM_006931406.4
B2 5351742 5351868
refGene_gn=CB2H6orf62;refGene_tx=XM_003985742.4;refGene_gn=CB2H6orf62;refGene_tx=XM_006931406.4
B2 5356063 5356235 refGene_gn=ACOT13;refGene_tx=XM_006931407.4
B2 5358722 5358907 refGene_gn=ACOT13;refGene_tx=XM_006931407.4
B2 5371440 5371521 refGene_gn=ACOT13;refGene_tx=XM_006931407.4

B2 5371904 5372069 refGene_gn=TDP2;refGene_tx=XM_011282089.3
B2 5372148 5372234 refGene_gn=TDP2;refGene_tx=XM_011282089.3
B2 5375662 5375874
refGene_gn=TDP2;refGene_tx=XM_011282089.3;refGene_gn=TDP2;refGene_tx=XM_019830141.1
B2 5376539 5376631
refGene_gn=TDP2;refGene_tx=XM_011282089.3;refGene_gn=TDP2;refGene_tx=XM_019830141.1
B2 5378343 5378462
refGene_gn=TDP2;refGene_tx=XM_011282089.3;refGene_gn=TDP2;refGene_tx=XM_019830141.1
B2 5380118 5380301
refGene_gn=TDP2;refGene_tx=XM_011282089.3;refGene_gn=TDP2;refGene_tx=XM_019830141.1
B2 5385574 5385856
refGene_gn=TDP2;refGene_tx=XM_011282089.3;refGene_gn=TDP2;refGene_tx=XM_019830141.1
B2 5429056 5429111 refGene_gn=KIAA0319;refGene_tx=XM_023254653.1
B2 5432508 5432556 refGene_gn=KIAA0319;refGene_tx=XM_023254653.1
B2 5432558 5432591 refGene_gn=KIAA0319;refGene_tx=XM_023254653.1
B2 5432593 5432655 refGene_gn=KIAA0319;refGene_tx=XM_023254653.1
B2 5432656 5432691 refGene_gn=KIAA0319;refGene_tx=XM_023254653.1
B2 5432929 5433150 refGene_gn=KIAA0319;refGene_tx=XM_023254653.1
B2 5434367 5434515 refGene_gn=KIAA0319;refGene_tx=XM_023254653.1
B2 5434649 5434848 refGene_gn=KIAA0319;refGene_tx=XM_023254653.1
B2 5439484 5439620 refGene_gn=KIAA0319;refGene_tx=XM_023254653.1
B2 5440746 5440932 refGene_gn=KIAA0319;refGene_tx=XM_023254653.1
B2 5444415 5444564 refGene_gn=KIAA0319;refGene_tx=XM_023254653.1
B2 5445154 5445439 refGene_gn=KIAA0319;refGene_tx=XM_023254653.1
B2 5449725 5449912 refGene_gn=KIAA0319;refGene_tx=XM_023254653.1
B2 5452972 5453065 refGene_gn=KIAA0319;refGene_tx=XM_023254653.1
B2 5454542 5454587 refGene_gn=KIAA0319;refGene_tx=XM_023254653.1
B2 5454589 5454643 refGene_gn=KIAA0319;refGene_tx=XM_023254653.1
B2 5455748 5455829 refGene_gn=KIAA0319;refGene_tx=XM_023254653.1
B2 5456906 5457003 refGene_gn=KIAA0319;refGene_tx=XM_023254653.1
B2 5458445 5458531 refGene_gn=KIAA0319;refGene_tx=XM_023254653.1
B2 5458533 5458570 refGene_gn=KIAA0319;refGene_tx=XM_023254653.1
B2 5478972 5479201 refGene_gn=KIAA0319;refGene_tx=XM_023254653.1
B2 5482184 5482308 refGene_gn=KIAA0319;refGene_tx=XM_023254653.1
B2 5484388 5484500 refGene_gn=KIAA0319;refGene_tx=XM_023254653.1
B2 5485157 5485306 refGene_gn=KIAA0319;refGene_tx=XM_023254653.1
B2 5489322 5489407 refGene_gn=KIAA0319;refGene_tx=XM_023254653.1
B2 5489408 5489454 refGene_gn=KIAA0319;refGene_tx=XM_023254653.1
B2 5491080 5491219 refGene_gn=KIAA0319;refGene_tx=XM_023254653.1
B2 5491786 5491925 refGene_gn=KIAA0319;refGene_tx=XM_023254653.1
B2 5495081 5495224 refGene_gn=KIAA0319;refGene_tx=XM_023254653.1
B2 5496898 5497021 refGene_gn=KIAA0319;refGene_tx=XM_023254653.1
B2 5499399 5499490 refGene_gn=KIAA0319;refGene_tx=XM_023254653.1
B2 5502081 5502173 refGene_gn=KIAA0319;refGene_tx=XM_023254653.1
B2 5504565 5504709 refGene_gn=KIAA0319;refGene_tx=XM_023254653.1

B2	5504711	5504743	refGene_gn=KIAA0319;refGene_tx=XM_023254653.1
B2	5514982	5515188
refGene_gn=ALDH5A1;refGene_tx=XM_003985776.5;refGene_gn=ALDH5A1;refGen
e_tx=XM_023253619.1
B2	5516159	5516218
refGene_gn=ALDH5A1;refGene_tx=XM_003985776.5;refGene_gn=ALDH5A1;refGen
e_tx=XM_023253619.1
B2	5517798	5517968
refGene_gn=ALDH5A1;refGene_tx=XM_003985776.5;refGene_gn=ALDH5A1;refGen
e_tx=XM_023253619.1
B2	5522355	5522514
refGene_gn=ALDH5A1;refGene_tx=XM_003985776.5;refGene_gn=ALDH5A1;refGen
e_tx=XM_023253619.1
B2	5524550	5524694
refGene_gn=ALDH5A1;refGene_tx=XM_003985776.5;refGene_gn=ALDH5A1;refGen
e_tx=XM_023253619.1
B2	5526582	5526726
refGene_gn=ALDH5A1;refGene_tx=XM_003985776.5;refGene_gn=ALDH5A1;refGen
e_tx=XM_023253619.1
B2	5532120	5532237	refGene_gn=ALDH5A1;refGene_tx=XM_003985776.5
B2	5533858	5534029
refGene_gn=ALDH5A1;refGene_tx=XM_003985776.5;refGene_gn=ALDH5A1;refGen
e_tx=XM_023253619.1
B2	5535411	5535495
refGene_gn=ALDH5A1;refGene_tx=XM_003985776.5;refGene_gn=ALDH5A1;refGen
e_tx=XM_023253619.1
B2	5538847	5539165
refGene_gn=ALDH5A1;refGene_tx=XM_003985776.5;refGene_gn=ALDH5A1;refGen
e_tx=XM_023253619.1
B2	5542735	5542832
refGene_gn=GPLD1;refGene_tx=XM_019830142.2;refGene_gn=GPLD1;refGene_tx
=XM_019830143.2;refGene_gn=GPLD1;refGene_tx=XM_019830144.2;refGene_gn=
GPLD1;refGene_tx=XM_019830145.2;refGene_gn=GPLD1;refGene_tx=XM_0198301
46.2;refGene_gn=GPLD1;refGene_tx=XM_019830147.2;refGene_gn=GPLD1;refGe
ne_tx=XM_023253615.1
B2	5545385	5545441
refGene_gn=GPLD1;refGene_tx=XM_019830142.2;refGene_gn=GPLD1;refGene_tx
=XM_019830143.2;refGene_gn=GPLD1;refGene_tx=XM_019830144.2;refGene_gn=
GPLD1;refGene_tx=XM_019830145.2;refGene_gn=GPLD1;refGene_tx=XM_0198301
47.2;refGene_gn=GPLD1;refGene_tx=XM_023253615.1
B2	5556967	5557046
refGene_gn=GPLD1;refGene_tx=XM_019830142.2;refGene_gn=GPLD1;refGene_tx
=XM_019830143.2;refGene_gn=GPLD1;refGene_tx=XM_019830144.2;refGene_gn=
GPLD1;refGene_tx=XM_019830145.2;refGene_gn=GPLD1;refGene_tx=XM_0198301
47.2;refGene_gn=GPLD1;refGene_tx=XM_023253615.1
B2	5560150	5560248
refGene_gn=GPLD1;refGene_tx=XM_019830142.2;refGene_gn=GPLD1;refGene_tx
=XM_019830143.2;refGene_gn=GPLD1;refGene_tx=XM_019830144.2;refGene_gn=
GPLD1;refGene_tx=XM_019830145.2;refGene_gn=GPLD1;refGene_tx=XM_0198301
46.2;refGene_gn=GPLD1;refGene_tx=XM_019830147.2;refGene_gn=GPLD1;refGe

ne_tx=XM_023253615.1
B2	5560671	5560782
refGene_gn=GPLD1;refGene_tx=XM_019830142.2;refGene_gn=GPLD1;refGene_tx=XM_019830143.2;refGene_gn=GPLD1;refGene_tx=XM_019830144.2;refGene_gn=GPLD1;refGene_tx=XM_019830145.2;refGene_gn=GPLD1;refGene_tx=XM_019830146.2;refGene_gn=GPLD1;refGene_tx=XM_019830147.2;refGene_gn=GPLD1;refGene_tx=XM_019830148.1;refGene_gn=GPLD1;refGene_tx=XM_023253615.1
B2	5562371	5562420
refGene_gn=GPLD1;refGene_tx=XM_019830142.2;refGene_gn=GPLD1;refGene_tx=XM_019830143.2;refGene_gn=GPLD1;refGene_tx=XM_019830144.2;refGene_gn=GPLD1;refGene_tx=XM_019830145.2;refGene_gn=GPLD1;refGene_tx=XM_019830146.2;refGene_gn=GPLD1;refGene_tx=XM_019830147.2;refGene_gn=GPLD1;refGene_tx=XM_019830148.1;refGene_gn=GPLD1;refGene_tx=XM_023253615.1
B2	5563370	5563425
refGene_gn=GPLD1;refGene_tx=XM_019830142.2;refGene_gn=GPLD1;refGene_tx=XM_019830143.2;refGene_gn=GPLD1;refGene_tx=XM_019830144.2;refGene_gn=GPLD1;refGene_tx=XM_019830145.2;refGene_gn=GPLD1;refGene_tx=XM_019830146.2;refGene_gn=GPLD1;refGene_tx=XM_019830147.2;refGene_gn=GPLD1;refGene_tx=XM_019830148.1;refGene_gn=GPLD1;refGene_tx=XM_023253615.1
B2	5572598	5572706
refGene_gn=GPLD1;refGene_tx=XM_019830142.2;refGene_gn=GPLD1;refGene_tx=XM_019830143.2;refGene_gn=GPLD1;refGene_tx=XM_019830144.2;refGene_gn=GPLD1;refGene_tx=XM_019830145.2;refGene_gn=GPLD1;refGene_tx=XM_019830146.2;refGene_gn=GPLD1;refGene_tx=XM_019830147.2;refGene_gn=GPLD1;refGene_tx=XM_019830148.1;refGene_gn=GPLD1;refGene_tx=XM_023253615.1
B2	5572933	5572961
refGene_gn=GPLD1;refGene_tx=XM_019830142.2;refGene_gn=GPLD1;refGene_tx=XM_019830143.2;refGene_gn=GPLD1;refGene_tx=XM_019830144.2;refGene_gn=GPLD1;refGene_tx=XM_019830145.2;refGene_gn=GPLD1;refGene_tx=XM_019830146.2;refGene_gn=GPLD1;refGene_tx=XM_019830147.2;refGene_gn=GPLD1;refGene_tx=XM_019830148.1;refGene_gn=GPLD1;refGene_tx=XM_023253615.1
B2	5573054	5573194
refGene_gn=GPLD1;refGene_tx=XM_019830142.2;refGene_gn=GPLD1;refGene_tx=XM_019830143.2;refGene_gn=GPLD1;refGene_tx=XM_019830144.2;refGene_gn=GPLD1;refGene_tx=XM_019830145.2;refGene_gn=GPLD1;refGene_tx=XM_019830146.2;refGene_gn=GPLD1;refGene_tx=XM_019830147.2;refGene_gn=GPLD1;refGene_tx=XM_019830148.1;refGene_gn=GPLD1;refGene_tx=XM_023253615.1
B2	5582167	5582233
refGene_gn=GPLD1;refGene_tx=XM_019830142.2;refGene_gn=GPLD1;refGene_tx=XM_019830143.2;refGene_gn=GPLD1;refGene_tx=XM_019830144.2;refGene_gn=GPLD1;refGene_tx=XM_019830145.2;refGene_gn=GPLD1;refGene_tx=XM_019830146.2;refGene_gn=GPLD1;refGene_tx=XM_019830147.2;refGene_gn=GPLD1;refGene_tx=XM_019830148.1;refGene_gn=GPLD1;refGene_tx=XM_023253615.1
B2	5583877	5584001
refGene_gn=GPLD1;refGene_tx=XM_019830142.2;refGene_gn=GPLD1;refGene_tx=XM_019830143.2;refGene_gn=GPLD1;refGene_tx=XM_019830144.2;refGene_gn=GPLD1;refGene_tx=XM_019830145.2;refGene_gn=GPLD1;refGene_tx=XM_019830146.2;refGene_gn=GPLD1;refGene_tx=XM_019830147.2;refGene_gn=GPLD1;refGene_tx=XM_019830148.1;refGene_gn=GPLD1;refGene_tx=XM_023253615.1
B2	5587481	5587621

refGene_gn=GPLD1;refGene_tx=XM_019830142.2;refGene_gn=GPLD1;refGene_tx=XM_019830143.2;refGene_gn=GPLD1;refGene_tx=XM_019830144.2;refGene_gn=GPLD1;refGene_tx=XM_019830145.2;refGene_gn=GPLD1;refGene_tx=XM_019830146.2;refGene_gn=GPLD1;refGene_tx=XM_019830147.2;refGene_gn=GPLD1;refGene_tx=XM_019830148.1;refGene_gn=GPLD1;refGene_tx=XM_023253615.1
B2 5589562 5589749
refGene_gn=GPLD1;refGene_tx=XM_019830142.2;refGene_gn=GPLD1;refGene_tx=XM_019830143.2;refGene_gn=GPLD1;refGene_tx=XM_019830144.2;refGene_gn=GPLD1;refGene_tx=XM_019830145.2;refGene_gn=GPLD1;refGene_tx=XM_019830146.2;refGene_gn=GPLD1;refGene_tx=XM_019830147.2;refGene_gn=GPLD1;refGene_tx=XM_019830148.1;refGene_gn=GPLD1;refGene_tx=XM_023253615.1
B2 5592486 5592597
refGene_gn=GPLD1;refGene_tx=XM_019830142.2;refGene_gn=GPLD1;refGene_tx=XM_019830143.2;refGene_gn=GPLD1;refGene_tx=XM_019830144.2;refGene_gn=GPLD1;refGene_tx=XM_019830145.2;refGene_gn=GPLD1;refGene_tx=XM_019830146.2;refGene_gn=GPLD1;refGene_tx=XM_019830147.2;refGene_gn=GPLD1;refGene_tx=XM_019830148.1;refGene_gn=GPLD1;refGene_tx=XM_023253615.1
B2 5593890 5593965
refGene_gn=GPLD1;refGene_tx=XM_019830142.2;refGene_gn=GPLD1;refGene_tx=XM_019830143.2;refGene_gn=GPLD1;refGene_tx=XM_019830144.2;refGene_gn=GPLD1;refGene_tx=XM_019830145.2;refGene_gn=GPLD1;refGene_tx=XM_019830146.2;refGene_gn=GPLD1;refGene_tx=XM_019830147.2;refGene_gn=GPLD1;refGene_tx=XM_019830148.1;refGene_gn=GPLD1;refGene_tx=XM_023253615.1
B2 5594061 5594218
refGene_gn=GPLD1;refGene_tx=XM_019830142.2;refGene_gn=GPLD1;refGene_tx=XM_019830143.2;refGene_gn=GPLD1;refGene_tx=XM_019830144.2;refGene_gn=GPLD1;refGene_tx=XM_019830145.2;refGene_gn=GPLD1;refGene_tx=XM_019830146.2;refGene_gn=GPLD1;refGene_tx=XM_019830147.2;refGene_gn=GPLD1;refGene_tx=XM_019830148.1;refGene_gn=GPLD1;refGene_tx=XM_023253615.1
B2 5594711 5594853
refGene_gn=GPLD1;refGene_tx=XM_019830142.2;refGene_gn=GPLD1;refGene_tx=XM_019830143.2;refGene_gn=GPLD1;refGene_tx=XM_019830144.2;refGene_gn=GPLD1;refGene_tx=XM_019830145.2;refGene_gn=GPLD1;refGene_tx=XM_019830146.2;refGene_gn=GPLD1;refGene_tx=XM_019830147.2;refGene_gn=GPLD1;refGene_tx=XM_019830148.1;refGene_gn=GPLD1;refGene_tx=XM_023253615.1
B2 5595616 5595722
refGene_gn=GPLD1;refGene_tx=XM_019830142.2;refGene_gn=GPLD1;refGene_tx=XM_019830143.2;refGene_gn=GPLD1;refGene_tx=XM_019830144.2;refGene_gn=GPLD1;refGene_tx=XM_019830145.2;refGene_gn=GPLD1;refGene_tx=XM_019830146.2;refGene_gn=GPLD1;refGene_tx=XM_019830147.2;refGene_gn=GPLD1;refGene_tx=XM_019830148.1;refGene_gn=GPLD1;refGene_tx=XM_023253615.1
B2 5595807 5595901
refGene_gn=GPLD1;refGene_tx=XM_019830142.2;refGene_gn=GPLD1;refGene_tx=XM_019830143.2;refGene_gn=GPLD1;refGene_tx=XM_019830144.2;refGene_gn=GPLD1;refGene_tx=XM_019830145.2;refGene_gn=GPLD1;refGene_tx=XM_019830146.2;refGene_gn=GPLD1;refGene_tx=XM_019830147.2;refGene_gn=GPLD1;refGene_tx=XM_019830148.1;refGene_gn=GPLD1;refGene_tx=XM_023253615.1
B2 5601999 5602176
refGene_gn=GPLD1;refGene_tx=XM_019830142.2;refGene_gn=GPLD1;refGene_tx=XM_019830143.2;refGene_gn=GPLD1;refGene_tx=XM_019830144.2;refGene_gn=

GPLD1;refGene_tx=XM_019830145.2;refGene_gn=GPLD1;refGene_tx=XM_0198301
46.2;refGene_gn=GPLD1;refGene_tx=XM_019830147.2;refGene_gn=GPLD1;refGe
ne_tx=XM_019830148.1;refGene_gn=GPLD1;refGene_tx=XM_023253615.1
B2 5602518 5602706
refGene_gn=GPLD1;refGene_tx=XM_019830142.2;refGene_gn=GPLD1;refGene_tx
=XM_019830143.2;refGene_gn=GPLD1;refGene_tx=XM_019830144.2;refGene_gn=
GPLD1;refGene_tx=XM_019830145.2;refGene_gn=GPLD1;refGene_tx=XM_0198301
46.2;refGene_gn=GPLD1;refGene_tx=XM_019830147.2;refGene_gn=GPLD1;refGe
ne_tx=XM_019830148.1;refGene_gn=GPLD1;refGene_tx=XM_023253615.1
B2 5603667 5603921
refGene_gn=GPLD1;refGene_tx=XM_019830142.2;refGene_gn=GPLD1;refGene_tx
=XM_019830143.2;refGene_gn=GPLD1;refGene_tx=XM_019830144.2;refGene_gn=
GPLD1;refGene_tx=XM_019830145.2;refGene_gn=GPLD1;refGene_tx=XM_0198301
46.2;refGene_gn=GPLD1;refGene_tx=XM_019830148.1;refGene_gn=GPLD1;refGe
ne_tx=XM_023253615.1;refGene_gn=MRS2;refGene_tx=XM_023253616.1
B2 5604028 5604079
refGene_gn=GPLD1;refGene_tx=XM_019830142.2;refGene_gn=GPLD1;refGene_tx
=XM_019830143.2;refGene_gn=GPLD1;refGene_tx=XM_019830144.2;refGene_gn=
GPLD1;refGene_tx=XM_019830146.2;refGene_gn=GPLD1;refGene_tx=XM_0198301
48.1;refGene_gn=GPLD1;refGene_tx=XM_023253615.1
B2 5605256 5605293 refGene_gn=MRS2;refGene_tx=XM_023253617.1
B2 5605831 5605959
refGene_gn=GPLD1;refGene_tx=XM_023253615.1;refGene_gn=MRS2;refGene_tx=
XM_023253616.1;refGene_gn=MRS2;refGene_tx=XM_023253617.1
B2 5606570 5606657
refGene_gn=GPLD1;refGene_tx=XM_019830142.2;refGene_gn=GPLD1;refGene_tx
=XM_019830143.2;refGene_gn=GPLD1;refGene_tx=XM_019830144.2;refGene_gn=
GPLD1;refGene_tx=XM_019830146.2;refGene_gn=GPLD1;refGene_tx=XM_0198301
48.1
B2 5608271 5608296 refGene_gn=MRS2;refGene_tx=XM_023253618.1
B2 5610553 5610660
refGene_gn=MRS2;refGene_tx=XM_023253616.1;refGene_gn=MRS2;refGene_tx=X
M_023253617.1;refGene_gn=MRS2;refGene_tx=XM_023253618.1
B2 5611195 5611309
refGene_gn=MRS2;refGene_tx=XM_023253616.1;refGene_gn=MRS2;refGene_tx=X
M_023253617.1;refGene_gn=MRS2;refGene_tx=XM_023253618.1
B2 5622904 5623022
refGene_gn=MRS2;refGene_tx=XM_023253616.1;refGene_gn=MRS2;refGene_tx=X
M_023253617.1;refGene_gn=MRS2;refGene_tx=XM_023253618.1
B2 5623259 5623412
refGene_gn=MRS2;refGene_tx=XM_023253616.1;refGene_gn=MRS2;refGene_tx=X
M_023253617.1;refGene_gn=MRS2;refGene_tx=XM_023253618.1
B2 5624675 5624792
refGene_gn=MRS2;refGene_tx=XM_023253616.1;refGene_gn=MRS2;refGene_tx=X
M_023253617.1;refGene_gn=MRS2;refGene_tx=XM_023253618.1
B2 5625228 5625359
refGene_gn=MRS2;refGene_tx=XM_023253616.1;refGene_gn=MRS2;refGene_tx=X
M_023253617.1;refGene_gn=MRS2;refGene_tx=XM_023253618.1
B2 5627611 5627785
refGene_gn=MRS2;refGene_tx=XM_023253616.1;refGene_gn=MRS2;refGene_tx=X

M_023253617.1;refGene_gn=MRS2;refGene_tx=XM_023253618.1
B2 5636545 5636658
refGene_gn=MRS2;refGene_tx=XM_023253616.1;refGene_gn=MRS2;refGene_tx=XM_023253617.1;refGene_gn=MRS2;refGene_tx=XM_023253618.1
B2 5637880 5637917
refGene_gn=MRS2;refGene_tx=XM_023253616.1;refGene_gn=MRS2;refGene_tx=XM_023253617.1;refGene_gn=MRS2;refGene_tx=XM_023253618.1
B2 5641432 5641506
refGene_gn=MRS2;refGene_tx=XM_023253616.1;refGene_gn=MRS2;refGene_tx=XM_023253617.1;refGene_gn=MRS2;refGene_tx=XM_023253618.1
B2 5643280 5643452
refGene_gn=MRS2;refGene_tx=XM_023253616.1;refGene_gn=MRS2;refGene_tx=XM_023253617.1;refGene_gn=MRS2;refGene_tx=XM_023253618.1
B2 5679770 5680063 refGene_gn=DCDC2;refGene_tx=XM_023253620.1
B2 5684579 5684634 refGene_gn=DCDC2;refGene_tx=XM_023253620.1
B2 5728441 5728518 refGene_gn=DCDC2;refGene_tx=XM_023253620.1
B2 5728667 5728799 refGene_gn=DCDC2;refGene_tx=XM_023253620.1
B2 5734387 5734534 refGene_gn=DCDC2;refGene_tx=XM_023253620.1
B2 5736559 5736614 refGene_gn=DCDC2;refGene_tx=XM_023253620.1
B2 5742874 5743037 refGene_gn=DCDC2;refGene_tx=XM_023253620.1
B2 5818024 5818125 refGene_gn=DCDC2;refGene_tx=XM_023253620.1
B2 5838974 5839277 refGene_gn=DCDC2;refGene_tx=XM_023253620.1
B2 5843511 5843616 refGene_gn=DCDC2;refGene_tx=XM_023253620.1
B2 5878092 5878491
refGene_gn=NRSN1;refGene_tx=XM_003985744.4;refGene_gn=NRSN1;refGene_tx=XM_019830152.2
B2 5887887 5888100
refGene_gn=NRSN1;refGene_tx=XM_003985744.4;refGene_gn=NRSN1;refGene_tx=XM_019830152.2
B2 7298945 7299818 refGene_gn=HDGFL1;refGene_tx=XM_023253621.1
B2 7554554 7554582 refGene_gn=PRL;refGene_tx=NM_001043341.1
B2 7556546 7556728 refGene_gn=PRL;refGene_tx=NM_001043341.1
B2 7559328 7559436 refGene_gn=PRL;refGene_tx=NM_001043341.1
B2 7561838 7562018 refGene_gn=PRL;refGene_tx=NM_001043341.1
B2 7564231 7564423 refGene_gn=PRL;refGene_tx=NM_001043341.1
B2 8159841 8161281 refGene_gn=SOX4;refGene_tx=XM_023253622.1
B2 8438309 8438327 refGene_gn=CDKAL1;refGene_tx=XM_023253543.1
B2 8489413 8489599
refGene_gn=CDKAL1;refGene_tx=NM_001278832.1;refGene_gn=CDKAL1;refGene_tx=XM_006931361.3;refGene_gn=CDKAL1;refGene_tx=XM_023253541.1;refGene_gn=CDKAL1;refGene_tx=XM_023253542.1
B2 8522228 8522393
refGene_gn=CDKAL1;refGene_tx=NM_001278832.1;refGene_gn=CDKAL1;refGene_tx=XM_006931361.3;refGene_gn=CDKAL1;refGene_tx=XM_023253541.1;refGene_gn=CDKAL1;refGene_tx=XM_023253542.1;refGene_gn=CDKAL1;refGene_tx=XM_023253543.1
B2 8525311 8525395
refGene_gn=CDKAL1;refGene_tx=NM_001278832.1;refGene_gn=CDKAL1;refGene_tx=XM_006931361.3;refGene_gn=CDKAL1;refGene_tx=XM_023253541.1;refGene_gn=CDKAL1;refGene_tx=XM_023253542.1;refGene_gn=CDKAL1;refGene_tx=XM_02

3253543.1
B2	8611548	8611611
refGene_gn=CDKAL1;refGene_tx=NM_001278832.1;refGene_gn=CDKAL1;refGene_tx=XM_006931361.3;refGene_gn=CDKAL1;refGene_tx=XM_023253541.1;refGene_gn=CDKAL1;refGene_tx=XM_023253542.1;refGene_gn=CDKAL1;refGene_tx=XM_023253543.1
B2	8654350	8654531
refGene_gn=CDKAL1;refGene_tx=NM_001278832.1;refGene_gn=CDKAL1;refGene_tx=XM_006931361.3;refGene_gn=CDKAL1;refGene_tx=XM_023253541.1;refGene_gn=CDKAL1;refGene_tx=XM_023253542.1;refGene_gn=CDKAL1;refGene_tx=XM_023253543.1
B2	8714923	8715069
refGene_gn=CDKAL1;refGene_tx=NM_001278832.1;refGene_gn=CDKAL1;refGene_tx=XM_006931361.3;refGene_gn=CDKAL1;refGene_tx=XM_023253541.1;refGene_gn=CDKAL1;refGene_tx=XM_023253542.1;refGene_gn=CDKAL1;refGene_tx=XM_023253543.1
B2	8748606	8748773
refGene_gn=CDKAL1;refGene_tx=NM_001278832.1;refGene_gn=CDKAL1;refGene_tx=XM_006931361.3;refGene_gn=CDKAL1;refGene_tx=XM_023253541.1;refGene_gn=CDKAL1;refGene_tx=XM_023253542.1;refGene_gn=CDKAL1;refGene_tx=XM_023253543.1
B2	8842289	8842393
refGene_gn=CDKAL1;refGene_tx=NM_001278832.1;refGene_gn=CDKAL1;refGene_tx=XM_006931361.3;refGene_gn=CDKAL1;refGene_tx=XM_023253541.1;refGene_gn=CDKAL1;refGene_tx=XM_023253542.1;refGene_gn=CDKAL1;refGene_tx=XM_023253543.1
B2	8920738	8920859
refGene_gn=CDKAL1;refGene_tx=NM_001278832.1;refGene_gn=CDKAL1;refGene_tx=XM_006931361.3;refGene_gn=CDKAL1;refGene_tx=XM_023253541.1;refGene_gn=CDKAL1;refGene_tx=XM_023253542.1;refGene_gn=CDKAL1;refGene_tx=XM_023253543.1
B2	8954643	8954692
refGene_gn=CDKAL1;refGene_tx=NM_001278832.1;refGene_gn=CDKAL1;refGene_tx=XM_006931361.3;refGene_gn=CDKAL1;refGene_tx=XM_023253541.1;refGene_gn=CDKAL1;refGene_tx=XM_023253542.1;refGene_gn=CDKAL1;refGene_tx=XM_023253543.1
B2	8971155	8971252
refGene_gn=CDKAL1;refGene_tx=NM_001278832.1;refGene_gn=CDKAL1;refGene_tx=XM_006931361.3;refGene_gn=CDKAL1;refGene_tx=XM_023253541.1;refGene_gn=CDKAL1;refGene_tx=XM_023253542.1;refGene_gn=CDKAL1;refGene_tx=XM_023253543.1
B2	9038740	9038825
refGene_gn=CDKAL1;refGene_tx=NM_001278832.1;refGene_gn=CDKAL1;refGene_tx=XM_006931361.3;refGene_gn=CDKAL1;refGene_tx=XM_023253541.1;refGene_gn=CDKAL1;refGene_tx=XM_023253542.1;refGene_gn=CDKAL1;refGene_tx=XM_023253543.1
B2	9130228	9130341
refGene_gn=CDKAL1;refGene_tx=NM_001278832.1;refGene_gn=CDKAL1;refGene_tx=XM_006931361.3;refGene_gn=CDKAL1;refGene_tx=XM_023253541.1;refGene_gn=CDKAL1;refGene_tx=XM_023253542.1;refGene_gn=CDKAL1;refGene_tx=XM_02

3253543.1
B2	9131661	9131831
refGene_gn=CDKAL1;refGene_tx=NM_001278832.1;refGene_gn=CDKAL1;refGene_tx=XM_006931361.3;refGene_gn=CDKAL1;refGene_tx=XM_023253541.1;refGene_gn=CDKAL1;refGene_tx=XM_023253542.1;refGene_gn=CDKAL1;refGene_tx=XM_023253543.1
B2	9195196	9195459
refGene_gn=E2F3;refGene_tx=XM_019830153.2;refGene_gn=E2F3;refGene_tx=XM_023253623.1
B2	9196745	9196881
refGene_gn=E2F3;refGene_tx=XM_019830153.2;refGene_gn=E2F3;refGene_tx=XM_023253623.1
B2	9198580	9198695
refGene_gn=E2F3;refGene_tx=XM_019830153.2;refGene_gn=E2F3;refGene_tx=XM_023253623.1
B2	9201682	9201841
refGene_gn=E2F3;refGene_tx=XM_019830153.2;refGene_gn=E2F3;refGene_tx=XM_023253623.1
B2	9203393	9203613
refGene_gn=E2F3;refGene_tx=XM_019830153.2;refGene_gn=E2F3;refGene_tx=XM_023253623.1
B2	9205154	9205266
refGene_gn=E2F3;refGene_tx=XM_019830153.2;refGene_gn=E2F3;refGene_tx=XM_023253623.1
B2	9267413	9267431
refGene_gn=E2F3;refGene_tx=XM_019830153.2;refGene_gn=E2F3;refGene_tx=XM_023253623.1
B2	9447662	9447758
refGene_gn=MBOAT1;refGene_tx=XM_023253624.1;refGene_gn=MBOAT1;refGene_tx=XM_023253625.1
B2	9506424	9506570
refGene_gn=MBOAT1;refGene_tx=XM_023253624.1;refGene_gn=MBOAT1;refGene_tx=XM_023253625.1
B2	9507752	9507830
refGene_gn=MBOAT1;refGene_tx=XM_019830159.2;refGene_gn=MBOAT1;refGene_tx=XM_023253624.1;refGene_gn=MBOAT1;refGene_tx=XM_023253625.1
B2	9513442	9513538
refGene_gn=MBOAT1;refGene_tx=XM_019830159.2;refGene_gn=MBOAT1;refGene_tx=XM_023253624.1;refGene_gn=MBOAT1;refGene_tx=XM_023253625.1
B2	9526973	9527029
refGene_gn=MBOAT1;refGene_tx=XM_019830159.2;refGene_gn=MBOAT1;refGene_tx=XM_023253624.1;refGene_gn=MBOAT1;refGene_tx=XM_023253625.1
B2	9529718	9529773
refGene_gn=MBOAT1;refGene_tx=XM_019830159.2;refGene_gn=MBOAT1;refGene_tx=XM_023253624.1;refGene_gn=MBOAT1;refGene_tx=XM_023253625.1
B2	9531248	9531432
refGene_gn=MBOAT1;refGene_tx=XM_019830159.2;refGene_gn=MBOAT1;refGene_tx=XM_023253624.1;refGene_gn=MBOAT1;refGene_tx=XM_023253625.1
B2	9533571	9533764
refGene_gn=MBOAT1;refGene_tx=XM_019830159.2;refGene_gn=MBOAT1;refGene_

tx=XM_023253624.1;refGene_gn=MBOAT1;refGene_tx=XM_023253625.1
B2 9539140 9539244
refGene_gn=MBOAT1;refGene_tx=XM_019830159.2;refGene_gn=MBOAT1;refGene_tx=XM_023253624.1
B2 9541639 9541704
refGene_gn=MBOAT1;refGene_tx=XM_019830159.2;refGene_gn=MBOAT1;refGene_tx=XM_023253624.1;refGene_gn=MBOAT1;refGene_tx=XM_023253625.1
B2 9543928 9544061
refGene_gn=MBOAT1;refGene_tx=XM_019830159.2;refGene_gn=MBOAT1;refGene_tx=XM_023253624.1
B2 9546212 9546364
refGene_gn=MBOAT1;refGene_tx=XM_019830159.2;refGene_gn=MBOAT1;refGene_tx=XM_023253624.1;refGene_gn=MBOAT1;refGene_tx=XM_023253625.1
B2 9551281 9551408
refGene_gn=MBOAT1;refGene_tx=XM_019830159.2;refGene_gn=MBOAT1;refGene_tx=XM_023253624.1;refGene_gn=MBOAT1;refGene_tx=XM_023253625.1
B2 10002196 10002241 refGene_gn=ID4;refGene_tx=XM_023253626.1
B2 10002634 10003075 refGene_gn=ID4;refGene_tx=XM_023253626.1
B2 11180442 11180583
refGene_gn=RNF144B;refGene_tx=XM_003985746.5;refGene_gn=RNF144B;refGene_tx=XM_006931411.4;refGene_gn=RNF144B;refGene_tx=XM_019830163.2;refGene_gn=RNF144B;refGene_tx=XM_023253627.1;refGene_gn=RNF144B;refGene_tx=XM_023253628.1
B2 11181734 11181824
refGene_gn=RNF144B;refGene_tx=XM_003985746.5;refGene_gn=RNF144B;refGene_tx=XM_006931411.4;refGene_gn=RNF144B;refGene_tx=XM_019830163.2;refGene_gn=RNF144B;refGene_tx=XM_023253627.1;refGene_gn=RNF144B;refGene_tx=XM_023253628.1
B2 11185202 11185347
refGene_gn=RNF144B;refGene_tx=XM_003985746.5;refGene_gn=RNF144B;refGene_tx=XM_006931411.4;refGene_gn=RNF144B;refGene_tx=XM_019830163.2;refGene_gn=RNF144B;refGene_tx=XM_023253627.1;refGene_gn=RNF144B;refGene_tx=XM_023253628.1
B2 11187376 11187581
refGene_gn=RNF144B;refGene_tx=XM_003985746.5;refGene_gn=RNF144B;refGene_tx=XM_006931411.4;refGene_gn=RNF144B;refGene_tx=XM_019830163.2;refGene_gn=RNF144B;refGene_tx=XM_023253627.1;refGene_gn=RNF144B;refGene_tx=XM_023253628.1
B2 11205815 11205876
refGene_gn=RNF144B;refGene_tx=XM_003985746.5;refGene_gn=RNF144B;refGene_tx=XM_006931411.4;refGene_gn=RNF144B;refGene_tx=XM_019830163.2;refGene_gn=RNF144B;refGene_tx=XM_023253627.1;refGene_gn=RNF144B;refGene_tx=XM_023253628.1
B2 11218118 11218223
refGene_gn=RNF144B;refGene_tx=XM_003985746.5;refGene_gn=RNF144B;refGene_tx=XM_006931411.4;refGene_gn=RNF144B;refGene_tx=XM_019830163.2;refGene_gn=RNF144B;refGene_tx=XM_023253627.1;refGene_gn=RNF144B;refGene_tx=XM_023253628.1
B2 11245358 11245523
refGene_gn=RNF144B;refGene_tx=XM_003985746.5;refGene_gn=RNF144B;refGen

e_tx=XM_006931411.4;refGene_gn=RNF144B;refGene_tx=XM_019830163.2;refGene_gn=RNF144B;refGene_tx=XM_023253627.1;refGene_gn=RNF144B;refGene_tx=XM_023253628.1
B2 11377644 11377795
refGene_gn=DEK;refGene_tx=XM_023253632.1;refGene_gn=DEK;refGene_tx=XM_023253633.1;refGene_gn=DEK;refGene_tx=XM_023253634.1;refGene_gn=DEK;refGene_tx=XM_023253635.1
B2 11381237 11381339
refGene_gn=DEK;refGene_tx=XM_023253632.1;refGene_gn=DEK;refGene_tx=XM_023253633.1;refGene_gn=DEK;refGene_tx=XM_023253634.1;refGene_gn=DEK;refGene_tx=XM_023253635.1
B2 11381587 11381697
refGene_gn=DEK;refGene_tx=XM_023253632.1;refGene_gn=DEK;refGene_tx=XM_023253633.1;refGene_gn=DEK;refGene_tx=XM_023253634.1;refGene_gn=DEK;refGene_tx=XM_023253635.1
B2 11382657 11382752
refGene_gn=DEK;refGene_tx=XM_023253632.1;refGene_gn=DEK;refGene_tx=XM_023253633.1;refGene_gn=DEK;refGene_tx=XM_023253634.1;refGene_gn=DEK;refGene_tx=XM_023253635.1
B2 11384602 11384723
refGene_gn=DEK;refGene_tx=XM_023253632.1;refGene_gn=DEK;refGene_tx=XM_023253633.1;refGene_gn=DEK;refGene_tx=XM_023253634.1;refGene_gn=DEK;refGene_tx=XM_023253635.1
B2 11389147 11389336
refGene_gn=DEK;refGene_tx=XM_023253632.1;refGene_gn=DEK;refGene_tx=XM_023253633.1;refGene_gn=DEK;refGene_tx=XM_023253634.1;refGene_gn=DEK;refGene_tx=XM_023253635.1
B2 11401668 11401804
refGene_gn=DEK;refGene_tx=XM_023253632.1;refGene_gn=DEK;refGene_tx=XM_023253633.1;refGene_gn=DEK;refGene_tx=XM_023253634.1;refGene_gn=DEK;refGene_tx=XM_023253635.1
B2 11402060 11402209
refGene_gn=DEK;refGene_tx=XM_023253632.1;refGene_gn=DEK;refGene_tx=XM_023253633.1;refGene_gn=DEK;refGene_tx=XM_023253634.1;refGene_gn=DEK;refGene_tx=XM_023253635.1
B2 11404131 11404152 refGene_gn=DEK;refGene_tx=XM_023253635.1
B2 11406004 11406073
refGene_gn=DEK;refGene_tx=XM_023253632.1;refGene_gn=DEK;refGene_tx=XM_023253633.1;refGene_gn=DEK;refGene_tx=XM_023253634.1
B2 11406521 11406533
refGene_gn=DEK;refGene_tx=XM_023253632.1;refGene_gn=DEK;refGene_tx=XM_023253633.1;refGene_gn=DEK;refGene_tx=XM_023253634.1
B2 11410296 11410380
refGene_gn=KDM1B;refGene_tx=XM_011282094.2;refGene_gn=KDM1B;refGene_tx=XM_019830164.1;refGene_gn=KDM1B;refGene_tx=XM_023253629.1;refGene_gn=KDM1B;refGene_tx=XM_023253630.1;refGene_gn=KDM1B;refGene_tx=XM_023253631.1
B2 11414341 11414494
refGene_gn=KDM1B;refGene_tx=XM_011282094.2;refGene_gn=KDM1B;refGene_tx=XM_019830164.1;refGene_gn=KDM1B;refGene_tx=XM_023253629.1;refGene_gn=

KDM1B;refGene_tx=XM_023253630.1;refGene_gn=KDM1B;refGene_tx=XM_0232536
31.1
B2 11416328 11416451
refGene_gn=KDM1B;refGene_tx=XM_011282094.2;refGene_gn=KDM1B;refGene_tx
=XM_019830164.1;refGene_gn=KDM1B;refGene_tx=XM_023253629.1;refGene_gn=
KDM1B;refGene_tx=XM_023253630.1;refGene_gn=KDM1B;refGene_tx=XM_0232536
31.1
B2 11417758 11417884
refGene_gn=KDM1B;refGene_tx=XM_011282094.2;refGene_gn=KDM1B;refGene_tx
=XM_019830164.1;refGene_gn=KDM1B;refGene_tx=XM_023253629.1;refGene_gn=
KDM1B;refGene_tx=XM_023253630.1;refGene_gn=KDM1B;refGene_tx=XM_0232536
31.1
B2 11418609 11418726
refGene_gn=KDM1B;refGene_tx=XM_011282094.2;refGene_gn=KDM1B;refGene_tx
=XM_019830164.1;refGene_gn=KDM1B;refGene_tx=XM_023253629.1;refGene_gn=
KDM1B;refGene_tx=XM_023253630.1;refGene_gn=KDM1B;refGene_tx=XM_0232536
31.1
B2 11423043 11423118
refGene_gn=KDM1B;refGene_tx=XM_011282094.2;refGene_gn=KDM1B;refGene_tx
=XM_019830164.1;refGene_gn=KDM1B;refGene_tx=XM_023253629.1;refGene_gn=
KDM1B;refGene_tx=XM_023253630.1;refGene_gn=KDM1B;refGene_tx=XM_0232536
31.1
B2 11423661 11423793
refGene_gn=KDM1B;refGene_tx=XM_011282094.2;refGene_gn=KDM1B;refGene_tx
=XM_019830164.1;refGene_gn=KDM1B;refGene_tx=XM_023253629.1;refGene_gn=
KDM1B;refGene_tx=XM_023253630.1;refGene_gn=KDM1B;refGene_tx=XM_0232536
31.1
B2 11424786 11424914
refGene_gn=KDM1B;refGene_tx=XM_011282094.2;refGene_gn=KDM1B;refGene_tx
=XM_019830164.1;refGene_gn=KDM1B;refGene_tx=XM_023253629.1;refGene_gn=
KDM1B;refGene_tx=XM_023253630.1;refGene_gn=KDM1B;refGene_tx=XM_0232536
31.1
B2 11427769 11427941
refGene_gn=KDM1B;refGene_tx=XM_011282094.2;refGene_gn=KDM1B;refGene_tx
=XM_019830164.1;refGene_gn=KDM1B;refGene_tx=XM_023253629.1;refGene_gn=
KDM1B;refGene_tx=XM_023253630.1;refGene_gn=KDM1B;refGene_tx=XM_0232536
31.1
B2 11428901 11429039
refGene_gn=KDM1B;refGene_tx=XM_011282094.2;refGene_gn=KDM1B;refGene_tx
=XM_019830164.1;refGene_gn=KDM1B;refGene_tx=XM_023253629.1;refGene_gn=
KDM1B;refGene_tx=XM_023253630.1;refGene_gn=KDM1B;refGene_tx=XM_0232536
31.1
B2 11432180 11432255
refGene_gn=KDM1B;refGene_tx=XM_011282094.2;refGene_gn=KDM1B;refGene_tx
=XM_019830164.1;refGene_gn=KDM1B;refGene_tx=XM_023253629.1;refGene_gn=
KDM1B;refGene_tx=XM_023253630.1;refGene_gn=KDM1B;refGene_tx=XM_0232536
31.1
B2 11432649 11432826
refGene_gn=KDM1B;refGene_tx=XM_011282094.2;refGene_gn=KDM1B;refGene_tx
=XM_019830164.1;refGene_gn=KDM1B;refGene_tx=XM_023253629.1;refGene_gn=

KDM1B;refGene_tx=XM_023253630.1;refGene_gn=KDM1B;refGene_tx=XM_0232536
31.1
B2 11436160 11436345
refGene_gn=KDM1B;refGene_tx=XM_011282094.2;refGene_gn=KDM1B;refGene_tx
=XM_019830164.1;refGene_gn=KDM1B;refGene_tx=XM_023253629.1;refGene_gn=
KDM1B;refGene_tx=XM_023253630.1;refGene_gn=KDM1B;refGene_tx=XM_0232536
31.1
B2 11438604 11438809
refGene_gn=KDM1B;refGene_tx=XM_011282094.2;refGene_gn=KDM1B;refGene_tx
=XM_019830164.1;refGene_gn=KDM1B;refGene_tx=XM_023253629.1;refGene_gn=
KDM1B;refGene_tx=XM_023253630.1;refGene_gn=KDM1B;refGene_tx=XM_0232536
31.1
B2 11442967 11443006
refGene_gn=KDM1B;refGene_tx=XM_011282094.2;refGene_gn=KDM1B;refGene_tx
=XM_019830164.1;refGene_gn=KDM1B;refGene_tx=XM_023253629.1;refGene_gn=
KDM1B;refGene_tx=XM_023253630.1;refGene_gn=KDM1B;refGene_tx=XM_0232536
31.1
B2 11454862 11454982
refGene_gn=KDM1B;refGene_tx=XM_011282094.2;refGene_gn=KDM1B;refGene_tx
=XM_019830164.1;refGene_gn=KDM1B;refGene_tx=XM_023253629.1;refGene_gn=
KDM1B;refGene_tx=XM_023253630.1;refGene_gn=KDM1B;refGene_tx=XM_0232536
31.1
B2 11460390 11460502
refGene_gn=KDM1B;refGene_tx=XM_011282094.2;refGene_gn=KDM1B;refGene_tx
=XM_019830164.1;refGene_gn=KDM1B;refGene_tx=XM_023253629.1;refGene_gn=
KDM1B;refGene_tx=XM_023253630.1;refGene_gn=KDM1B;refGene_tx=XM_0232536
31.1
B2 11461055 11461145
refGene_gn=KDM1B;refGene_tx=XM_011282094.2;refGene_gn=KDM1B;refGene_tx
=XM_019830164.1;refGene_gn=KDM1B;refGene_tx=XM_023253629.1;refGene_gn=
KDM1B;refGene_tx=XM_023253630.1;refGene_gn=KDM1B;refGene_tx=XM_0232536
31.1
B2 11462171 11462299
refGene_gn=KDM1B;refGene_tx=XM_011282094.2;refGene_gn=KDM1B;refGene_tx
=XM_019830164.1;refGene_gn=KDM1B;refGene_tx=XM_023253629.1;refGene_gn=
KDM1B;refGene_tx=XM_023253630.1;refGene_gn=KDM1B;refGene_tx=XM_0232536
31.1
B2 11463535 11463622
refGene_gn=KDM1B;refGene_tx=XM_011282094.2;refGene_gn=KDM1B;refGene_tx
=XM_019830164.1;refGene_gn=KDM1B;refGene_tx=XM_023253629.1;refGene_gn=
KDM1B;refGene_tx=XM_023253630.1;refGene_gn=KDM1B;refGene_tx=XM_0232536
31.1
B2 11469362 11469364 refGene_gn=TPMT;refGene_tx=XM_011282109.3
B2 11472305 11472488
refGene_gn=TPMT;refGene_tx=NM_001009836.1;refGene_gn=TPMT;refGene_tx=X
M_011282109.3
B2 11473704 11473797
refGene_gn=TPMT;refGene_tx=NM_001009836.1;refGene_gn=TPMT;refGene_tx=X
M_011282109.3
B2 11478605 11478738

refGene_gn=TPMT;refGene_tx=NM_001009836.1;refGene_gn=TPMT;refGene_tx=XM_011282109.3
B2 11482942 11482995
refGene_gn=TPMT;refGene_tx=NM_001009836.1;refGene_gn=TPMT;refGene_tx=XM_011282109.3
B2 11483816 11483891
refGene_gn=TPMT;refGene_tx=NM_001009836.1;refGene_gn=TPMT;refGene_tx=XM_011282109.3
B2 11489824 11489910
refGene_gn=TPMT;refGene_tx=NM_001009836.1;refGene_gn=TPMT;refGene_tx=XM_011282109.3
B2 11492224 11492269
refGene_gn=TPMT;refGene_tx=NM_001009836.1;refGene_gn=TPMT;refGene_tx=XM_011282109.3
B2 11492892 11493005
refGene_gn=TPMT;refGene_tx=NM_001009836.1;refGene_gn=TPMT;refGene_tx=XM_011282109.3
B2 11502647 11503835 refGene_gn=NHLRC1;refGene_tx=XM_003985749.4
B2 11602665 11602720
refGene_gn=KIF13A;refGene_tx=XM_023253636.1;refGene_gn=KIF13A;refGene_tx=XM_023253637.1;refGene_gn=KIF13A;refGene_tx=XM_023253638.1;refGene_gn=KIF13A;refGene_tx=XM_023253639.1;refGene_gn=KIF13A;refGene_tx=XM_023253640.1;refGene_gn=KIF13A;refGene_tx=XM_023253641.1;refGene_gn=KIF13A;refGene_tx=XM_023253642.1;refGene_gn=KIF13A;refGene_tx=XM_023253643.1
B2 11603009 11603100
refGene_gn=KIF13A;refGene_tx=XM_023253636.1;refGene_gn=KIF13A;refGene_tx=XM_023253637.1;refGene_gn=KIF13A;refGene_tx=XM_023253638.1;refGene_gn=KIF13A;refGene_tx=XM_023253639.1;refGene_gn=KIF13A;refGene_tx=XM_023253640.1;refGene_gn=KIF13A;refGene_tx=XM_023253641.1;refGene_gn=KIF13A;refGene_tx=XM_023253642.1;refGene_gn=KIF13A;refGene_tx=XM_023253643.1
B2 11685744 11685757
refGene_gn=KIF13A;refGene_tx=XM_023253636.1;refGene_gn=KIF13A;refGene_tx=XM_023253637.1;refGene_gn=KIF13A;refGene_tx=XM_023253638.1;refGene_gn=KIF13A;refGene_tx=XM_023253639.1;refGene_gn=KIF13A;refGene_tx=XM_023253640.1;refGene_gn=KIF13A;refGene_tx=XM_023253641.1;refGene_gn=KIF13A;refGene_tx=XM_023253642.1;refGene_gn=KIF13A;refGene_tx=XM_023253643.1
B2 11712960 11713021
refGene_gn=KIF13A;refGene_tx=XM_023253636.1;refGene_gn=KIF13A;refGene_tx=XM_023253637.1;refGene_gn=KIF13A;refGene_tx=XM_023253638.1;refGene_gn=KIF13A;refGene_tx=XM_023253639.1;refGene_gn=KIF13A;refGene_tx=XM_023253640.1;refGene_gn=KIF13A;refGene_tx=XM_023253641.1;refGene_gn=KIF13A;refGene_tx=XM_023253642.1;refGene_gn=KIF13A;refGene_tx=XM_023253643.1
B2 11732284 11732377
refGene_gn=KIF13A;refGene_tx=XM_023253636.1;refGene_gn=KIF13A;refGene_tx=XM_023253637.1;refGene_gn=KIF13A;refGene_tx=XM_023253638.1;refGene_gn=KIF13A;refGene_tx=XM_023253639.1;refGene_gn=KIF13A;refGene_tx=XM_02

3253640.1;refGene_gn=KIF13A;refGene_tx=XM_023253641.1;refGene_gn=KIF13
A;refGene_tx=XM_023253642.1;refGene_gn=KIF13A;refGene_tx=XM_023253643.
1
B2 11733470 11733651
refGene_gn=KIF13A;refGene_tx=XM_023253636.1;refGene_gn=KIF13A;refGene_
tx=XM_023253637.1;refGene_gn=KIF13A;refGene_tx=XM_023253638.1;refGene_
gn=KIF13A;refGene_tx=XM_023253639.1;refGene_gn=KIF13A;refGene_tx=XM_02
3253640.1;refGene_gn=KIF13A;refGene_tx=XM_023253641.1;refGene_gn=KIF13
A;refGene_tx=XM_023253642.1;refGene_gn=KIF13A;refGene_tx=XM_023253643.
1
B2 11735499 11735587
refGene_gn=KIF13A;refGene_tx=XM_023253636.1;refGene_gn=KIF13A;refGene_
tx=XM_023253637.1;refGene_gn=KIF13A;refGene_tx=XM_023253638.1;refGene_
gn=KIF13A;refGene_tx=XM_023253639.1;refGene_gn=KIF13A;refGene_tx=XM_02
3253640.1;refGene_gn=KIF13A;refGene_tx=XM_023253641.1;refGene_gn=KIF13
A;refGene_tx=XM_023253642.1;refGene_gn=KIF13A;refGene_tx=XM_023253643.
1
B2 11737074 11737209
refGene_gn=KIF13A;refGene_tx=XM_023253636.1;refGene_gn=KIF13A;refGene_
tx=XM_023253637.1;refGene_gn=KIF13A;refGene_tx=XM_023253638.1;refGene_
gn=KIF13A;refGene_tx=XM_023253639.1;refGene_gn=KIF13A;refGene_tx=XM_02
3253640.1;refGene_gn=KIF13A;refGene_tx=XM_023253641.1;refGene_gn=KIF13
A;refGene_tx=XM_023253642.1;refGene_gn=KIF13A;refGene_tx=XM_023253643.
1
B2 11738147 11738260
refGene_gn=KIF13A;refGene_tx=XM_023253636.1;refGene_gn=KIF13A;refGene_
tx=XM_023253637.1;refGene_gn=KIF13A;refGene_tx=XM_023253638.1;refGene_
gn=KIF13A;refGene_tx=XM_023253639.1;refGene_gn=KIF13A;refGene_tx=XM_02
3253640.1;refGene_gn=KIF13A;refGene_tx=XM_023253641.1;refGene_gn=KIF13
A;refGene_tx=XM_023253642.1;refGene_gn=KIF13A;refGene_tx=XM_023253643.
1
B2 11751606 11751718
refGene_gn=KIF13A;refGene_tx=XM_023253636.1;refGene_gn=KIF13A;refGene_
tx=XM_023253637.1;refGene_gn=KIF13A;refGene_tx=XM_023253638.1;refGene_
gn=KIF13A;refGene_tx=XM_023253639.1;refGene_gn=KIF13A;refGene_tx=XM_02
3253640.1;refGene_gn=KIF13A;refGene_tx=XM_023253641.1;refGene_gn=KIF13
A;refGene_tx=XM_023253642.1;refGene_gn=KIF13A;refGene_tx=XM_023253643.
1
B2 11752112 11752325
refGene_gn=KIF13A;refGene_tx=XM_023253636.1;refGene_gn=KIF13A;refGene_
tx=XM_023253637.1;refGene_gn=KIF13A;refGene_tx=XM_023253638.1;refGene_
gn=KIF13A;refGene_tx=XM_023253639.1;refGene_gn=KIF13A;refGene_tx=XM_02
3253640.1;refGene_gn=KIF13A;refGene_tx=XM_023253641.1;refGene_gn=KIF13
A;refGene_tx=XM_023253642.1;refGene_gn=KIF13A;refGene_tx=XM_023253643.
1
B2 11755410 11755521
refGene_gn=KIF13A;refGene_tx=XM_023253636.1;refGene_gn=KIF13A;refGene_
tx=XM_023253637.1;refGene_gn=KIF13A;refGene_tx=XM_023253638.1;refGene_
gn=KIF13A;refGene_tx=XM_023253639.1;refGene_gn=KIF13A;refGene_tx=XM_02
3253640.1;refGene_gn=KIF13A;refGene_tx=XM_023253641.1;refGene_gn=KIF13

A;refGene_tx=XM_023253642.1;refGene_gn=KIF13A;refGene_tx=XM_023253643.
1
B2 11756866 11757001
refGene_gn=KIF13A;refGene_tx=XM_023253636.1;refGene_gn=KIF13A;refGene_
tx=XM_023253637.1;refGene_gn=KIF13A;refGene_tx=XM_023253638.1;refGene_
gn=KIF13A;refGene_tx=XM_023253639.1;refGene_gn=KIF13A;refGene_tx=XM_02
3253640.1;refGene_gn=KIF13A;refGene_tx=XM_023253641.1;refGene_gn=KIF13
A;refGene_tx=XM_023253642.1;refGene_gn=KIF13A;refGene_tx=XM_023253643.
1
B2 11758904 11759035
refGene_gn=KIF13A;refGene_tx=XM_023253636.1;refGene_gn=KIF13A;refGene_
tx=XM_023253637.1;refGene_gn=KIF13A;refGene_tx=XM_023253638.1;refGene_
gn=KIF13A;refGene_tx=XM_023253639.1;refGene_gn=KIF13A;refGene_tx=XM_02
3253640.1;refGene_gn=KIF13A;refGene_tx=XM_023253641.1;refGene_gn=KIF13
A;refGene_tx=XM_023253642.1;refGene_gn=KIF13A;refGene_tx=XM_023253643.
1
B2 11763464 11763551
refGene_gn=KIF13A;refGene_tx=XM_023253636.1;refGene_gn=KIF13A;refGene_
tx=XM_023253637.1;refGene_gn=KIF13A;refGene_tx=XM_023253638.1;refGene_
gn=KIF13A;refGene_tx=XM_023253639.1;refGene_gn=KIF13A;refGene_tx=XM_02
3253640.1;refGene_gn=KIF13A;refGene_tx=XM_023253641.1;refGene_gn=KIF13
A;refGene_tx=XM_023253642.1;refGene_gn=KIF13A;refGene_tx=XM_023253643.
1
B2 11763662 11763829
refGene_gn=KIF13A;refGene_tx=XM_023253636.1;refGene_gn=KIF13A;refGene_
tx=XM_023253637.1;refGene_gn=KIF13A;refGene_tx=XM_023253638.1;refGene_
gn=KIF13A;refGene_tx=XM_023253639.1;refGene_gn=KIF13A;refGene_tx=XM_02
3253640.1;refGene_gn=KIF13A;refGene_tx=XM_023253641.1;refGene_gn=KIF13
A;refGene_tx=XM_023253642.1;refGene_gn=KIF13A;refGene_tx=XM_023253643.
1
B2 11765858 11765978
refGene_gn=KIF13A;refGene_tx=XM_023253636.1;refGene_gn=KIF13A;refGene_
tx=XM_023253637.1;refGene_gn=KIF13A;refGene_tx=XM_023253638.1;refGene_
gn=KIF13A;refGene_tx=XM_023253639.1
B2 11769909 11770123
refGene_gn=KIF13A;refGene_tx=XM_023253636.1;refGene_gn=KIF13A;refGene_
tx=XM_023253637.1;refGene_gn=KIF13A;refGene_tx=XM_023253638.1;refGene_
gn=KIF13A;refGene_tx=XM_023253639.1;refGene_gn=KIF13A;refGene_tx=XM_02
3253640.1;refGene_gn=KIF13A;refGene_tx=XM_023253641.1;refGene_gn=KIF13
A;refGene_tx=XM_023253642.1;refGene_gn=KIF13A;refGene_tx=XM_023253643.
1
B2 11773252 11773415
refGene_gn=KIF13A;refGene_tx=XM_023253636.1;refGene_gn=KIF13A;refGene_
tx=XM_023253637.1;refGene_gn=KIF13A;refGene_tx=XM_023253638.1;refGene_
gn=KIF13A;refGene_tx=XM_023253639.1;refGene_gn=KIF13A;refGene_tx=XM_02
3253640.1;refGene_gn=KIF13A;refGene_tx=XM_023253641.1;refGene_gn=KIF13
A;refGene_tx=XM_023253642.1;refGene_gn=KIF13A;refGene_tx=XM_023253643.
1
B2 11775147 11775288
refGene_gn=KIF13A;refGene_tx=XM_023253636.1;refGene_gn=KIF13A;refGene_

tx=XM_023253637.1;refGene_gn=KIF13A;refGene_tx=XM_023253638.1;refGene_gn=KIF13A;refGene_tx=XM_023253639.1;refGene_gn=KIF13A;refGene_tx=XM_023253640.1;refGene_gn=KIF13A;refGene_tx=XM_023253641.1;refGene_gn=KIF13A;refGene_tx=XM_023253642.1;refGene_gn=KIF13A;refGene_tx=XM_023253643.1
B2 11776173 11776323
refGene_gn=KIF13A;refGene_tx=XM_023253636.1;refGene_gn=KIF13A;refGene_tx=XM_023253637.1;refGene_gn=KIF13A;refGene_tx=XM_023253638.1;refGene_gn=KIF13A;refGene_tx=XM_023253639.1;refGene_gn=KIF13A;refGene_tx=XM_023253640.1;refGene_gn=KIF13A;refGene_tx=XM_023253641.1;refGene_gn=KIF13A;refGene_tx=XM_023253642.1;refGene_gn=KIF13A;refGene_tx=XM_023253643.1
B2 11777756 11777918
refGene_gn=KIF13A;refGene_tx=XM_023253636.1;refGene_gn=KIF13A;refGene_tx=XM_023253637.1;refGene_gn=KIF13A;refGene_tx=XM_023253638.1;refGene_gn=KIF13A;refGene_tx=XM_023253639.1;refGene_gn=KIF13A;refGene_tx=XM_023253640.1;refGene_gn=KIF13A;refGene_tx=XM_023253641.1;refGene_gn=KIF13A;refGene_tx=XM_023253642.1;refGene_gn=KIF13A;refGene_tx=XM_023253643.1
B2 11778638 11778812
refGene_gn=KIF13A;refGene_tx=XM_023253636.1;refGene_gn=KIF13A;refGene_tx=XM_023253637.1;refGene_gn=KIF13A;refGene_tx=XM_023253638.1;refGene_gn=KIF13A;refGene_tx=XM_023253639.1;refGene_gn=KIF13A;refGene_tx=XM_023253640.1;refGene_gn=KIF13A;refGene_tx=XM_023253641.1;refGene_gn=KIF13A;refGene_tx=XM_023253642.1;refGene_gn=KIF13A;refGene_tx=XM_023253643.1
B2 11781289 11781441
refGene_gn=KIF13A;refGene_tx=XM_023253636.1;refGene_gn=KIF13A;refGene_tx=XM_023253637.1;refGene_gn=KIF13A;refGene_tx=XM_023253638.1;refGene_gn=KIF13A;refGene_tx=XM_023253639.1;refGene_gn=KIF13A;refGene_tx=XM_023253640.1;refGene_gn=KIF13A;refGene_tx=XM_023253641.1;refGene_gn=KIF13A;refGene_tx=XM_023253642.1;refGene_gn=KIF13A;refGene_tx=XM_023253643.1
B2 11781972 11782105
refGene_gn=KIF13A;refGene_tx=XM_023253636.1;refGene_gn=KIF13A;refGene_tx=XM_023253637.1;refGene_gn=KIF13A;refGene_tx=XM_023253638.1;refGene_gn=KIF13A;refGene_tx=XM_023253639.1;refGene_gn=KIF13A;refGene_tx=XM_023253640.1;refGene_gn=KIF13A;refGene_tx=XM_023253641.1;refGene_gn=KIF13A;refGene_tx=XM_023253642.1;refGene_gn=KIF13A;refGene_tx=XM_023253643.1
B2 11782292 11782439
refGene_gn=KIF13A;refGene_tx=XM_023253636.1;refGene_gn=KIF13A;refGene_tx=XM_023253637.1;refGene_gn=KIF13A;refGene_tx=XM_023253638.1;refGene_gn=KIF13A;refGene_tx=XM_023253639.1;refGene_gn=KIF13A;refGene_tx=XM_023253640.1;refGene_gn=KIF13A;refGene_tx=XM_023253641.1;refGene_gn=KIF13A;refGene_tx=XM_023253642.1;refGene_gn=KIF13A;refGene_tx=XM_023253643.1
B2 11786756 11786795
refGene_gn=KIF13A;refGene_tx=XM_023253636.1;refGene_gn=KIF13A;refGene_tx=XM_023253638.1;refGene_gn=KIF13A;refGene_tx=XM_023253639.1;refGene_

gn=KIF13A;refGene_tx=XM_023253640.1;refGene_gn=KIF13A;refGene_tx=XM_023253642.1
B2 11788425 11788525
refGene_gn=KIF13A;refGene_tx=XM_023253636.1;refGene_gn=KIF13A;refGene_tx=XM_023253637.1;refGene_gn=KIF13A;refGene_tx=XM_023253638.1;refGene_gn=KIF13A;refGene_tx=XM_023253639.1;refGene_gn=KIF13A;refGene_tx=XM_023253640.1;refGene_gn=KIF13A;refGene_tx=XM_023253641.1;refGene_gn=KIF13A;refGene_tx=XM_023253642.1;refGene_gn=KIF13A;refGene_tx=XM_023253643.1
B2 11790610 11790737
refGene_gn=KIF13A;refGene_tx=XM_023253636.1;refGene_gn=KIF13A;refGene_tx=XM_023253637.1;refGene_gn=KIF13A;refGene_tx=XM_023253638.1;refGene_gn=KIF13A;refGene_tx=XM_023253639.1;refGene_gn=KIF13A;refGene_tx=XM_023253640.1;refGene_gn=KIF13A;refGene_tx=XM_023253641.1;refGene_gn=KIF13A;refGene_tx=XM_023253642.1;refGene_gn=KIF13A;refGene_tx=XM_023253643.1
B2 11793085 11793141
refGene_gn=KIF13A;refGene_tx=XM_023253636.1;refGene_gn=KIF13A;refGene_tx=XM_023253637.1;refGene_gn=KIF13A;refGene_tx=XM_023253638.1;refGene_gn=KIF13A;refGene_tx=XM_023253639.1;refGene_gn=KIF13A;refGene_tx=XM_023253640.1;refGene_gn=KIF13A;refGene_tx=XM_023253641.1;refGene_gn=KIF13A;refGene_tx=XM_023253642.1;refGene_gn=KIF13A;refGene_tx=XM_023253643.1
B2 11794913 11795038
refGene_gn=KIF13A;refGene_tx=XM_023253636.1;refGene_gn=KIF13A;refGene_tx=XM_023253637.1;refGene_gn=KIF13A;refGene_tx=XM_023253638.1;refGene_gn=KIF13A;refGene_tx=XM_023253639.1;refGene_gn=KIF13A;refGene_tx=XM_023253640.1;refGene_gn=KIF13A;refGene_tx=XM_023253641.1;refGene_gn=KIF13A;refGene_tx=XM_023253642.1;refGene_gn=KIF13A;refGene_tx=XM_023253643.1
B2 11795294 11795471
refGene_gn=KIF13A;refGene_tx=XM_023253636.1;refGene_gn=KIF13A;refGene_tx=XM_023253637.1;refGene_gn=KIF13A;refGene_tx=XM_023253638.1;refGene_gn=KIF13A;refGene_tx=XM_023253639.1;refGene_gn=KIF13A;refGene_tx=XM_023253640.1;refGene_gn=KIF13A;refGene_tx=XM_023253641.1;refGene_gn=KIF13A;refGene_tx=XM_023253642.1;refGene_gn=KIF13A;refGene_tx=XM_023253643.1
B2 11796178 11796271
refGene_gn=KIF13A;refGene_tx=XM_023253636.1;refGene_gn=KIF13A;refGene_tx=XM_023253637.1;refGene_gn=KIF13A;refGene_tx=XM_023253638.1;refGene_gn=KIF13A;refGene_tx=XM_023253639.1;refGene_gn=KIF13A;refGene_tx=XM_023253640.1;refGene_gn=KIF13A;refGene_tx=XM_023253641.1;refGene_gn=KIF13A;refGene_tx=XM_023253642.1;refGene_gn=KIF13A;refGene_tx=XM_023253643.1
B2 11796585 11796738
refGene_gn=KIF13A;refGene_tx=XM_023253636.1;refGene_gn=KIF13A;refGene_tx=XM_023253637.1;refGene_gn=KIF13A;refGene_tx=XM_023253638.1;refGene_gn=KIF13A;refGene_tx=XM_023253639.1;refGene_gn=KIF13A;refGene_tx=XM_023253640.1;refGene_gn=KIF13A;refGene_tx=XM_023253641.1;refGene_gn=KIF13A;refGene_tx=XM_023253642.1;refGene_gn=KIF13A;refGene_tx=XM_023253643.

1
B2 11798244 11798322
refGene_gn=KIF13A;refGene_tx=XM_023253636.1;refGene_gn=KIF13A;refGene_tx=XM_023253637.1;refGene_gn=KIF13A;refGene_tx=XM_023253638.1;refGene_gn=KIF13A;refGene_tx=XM_023253639.1;refGene_gn=KIF13A;refGene_tx=XM_023253640.1;refGene_gn=KIF13A;refGene_tx=XM_023253641.1;refGene_gn=KIF13A;refGene_tx=XM_023253642.1;refGene_gn=KIF13A;refGene_tx=XM_023253643.1
B2 11801621 11801669
refGene_gn=KIF13A;refGene_tx=XM_023253636.1;refGene_gn=KIF13A;refGene_tx=XM_023253637.1;refGene_gn=KIF13A;refGene_tx=XM_023253638.1;refGene_gn=KIF13A;refGene_tx=XM_023253639.1;refGene_gn=KIF13A;refGene_tx=XM_023253640.1;refGene_gn=KIF13A;refGene_tx=XM_023253641.1;refGene_gn=KIF13A;refGene_tx=XM_023253642.1;refGene_gn=KIF13A;refGene_tx=XM_023253643.1
B2 11802742 11802854
refGene_gn=KIF13A;refGene_tx=XM_023253636.1;refGene_gn=KIF13A;refGene_tx=XM_023253637.1;refGene_gn=KIF13A;refGene_tx=XM_023253638.1;refGene_gn=KIF13A;refGene_tx=XM_023253639.1;refGene_gn=KIF13A;refGene_tx=XM_023253640.1;refGene_gn=KIF13A;refGene_tx=XM_023253641.1;refGene_gn=KIF13A;refGene_tx=XM_023253642.1;refGene_gn=KIF13A;refGene_tx=XM_023253643.1
B2 11804032 11804184
refGene_gn=KIF13A;refGene_tx=XM_023253636.1;refGene_gn=KIF13A;refGene_tx=XM_023253637.1;refGene_gn=KIF13A;refGene_tx=XM_023253638.1;refGene_gn=KIF13A;refGene_tx=XM_023253639.1;refGene_gn=KIF13A;refGene_tx=XM_023253640.1;refGene_gn=KIF13A;refGene_tx=XM_023253641.1;refGene_gn=KIF13A;refGene_tx=XM_023253642.1;refGene_gn=KIF13A;refGene_tx=XM_023253643.1
B2 11804672 11804777
refGene_gn=KIF13A;refGene_tx=XM_023253636.1;refGene_gn=KIF13A;refGene_tx=XM_023253637.1;refGene_gn=KIF13A;refGene_tx=XM_023253640.1;refGene_gn=KIF13A;refGene_tx=XM_023253641.1
B2 11805183 11805225 refGene_gn=KIF13A;refGene_tx=XM_023253638.1
B2 11812340 11813129
refGene_gn=KIF13A;refGene_tx=XM_023253636.1;refGene_gn=KIF13A;refGene_tx=XM_023253637.1;refGene_gn=KIF13A;refGene_tx=XM_023253638.1;refGene_gn=KIF13A;refGene_tx=XM_023253639.1;refGene_gn=KIF13A;refGene_tx=XM_023253640.1;refGene_gn=KIF13A;refGene_tx=XM_023253641.1;refGene_gn=KIF13A;refGene_tx=XM_023253642.1;refGene_gn=KIF13A;refGene_tx=XM_023253643.1
B2 11815904 11815925
refGene_gn=KIF13A;refGene_tx=XM_023253636.1;refGene_gn=KIF13A;refGene_tx=XM_023253637.1;refGene_gn=KIF13A;refGene_tx=XM_023253638.1;refGene_gn=KIF13A;refGene_tx=XM_023253639.1;refGene_gn=KIF13A;refGene_tx=XM_023253640.1;refGene_gn=KIF13A;refGene_tx=XM_023253641.1;refGene_gn=KIF13A;refGene_tx=XM_023253642.1;refGene_gn=KIF13A;refGene_tx=XM_023253643.1
B2 11852470 11852527
refGene_gn=NUP153;refGene_tx=XM_006931414.4;refGene_gn=NUP153;refGene_

tx=XM_019830197.2
B2 11854613 11854724
refGene_gn=NUP153;refGene_tx=XM_003985750.5;refGene_gn=NUP153;refGene_tx=XM_006931416.4;refGene_gn=NUP153;refGene_tx=XM_019830198.2
B2 11869884 11870110
refGene_gn=NUP153;refGene_tx=XM_003985750.5;refGene_gn=NUP153;refGene_tx=XM_006931414.4;refGene_gn=NUP153;refGene_tx=XM_006931416.4;refGene_gn=NUP153;refGene_tx=XM_019830197.2;refGene_gn=NUP153;refGene_tx=XM_019830198.2
B2 11881921 11882170
refGene_gn=NUP153;refGene_tx=XM_003985750.5;refGene_gn=NUP153;refGene_tx=XM_006931414.4;refGene_gn=NUP153;refGene_tx=XM_006931415.4;refGene_gn=NUP153;refGene_tx=XM_006931416.4;refGene_gn=NUP153;refGene_tx=XM_019830197.2;refGene_gn=NUP153;refGene_tx=XM_019830198.2
B2 11882308 11882448
refGene_gn=NUP153;refGene_tx=XM_003985750.5;refGene_gn=NUP153;refGene_tx=XM_006931414.4;refGene_gn=NUP153;refGene_tx=XM_006931415.4;refGene_gn=NUP153;refGene_tx=XM_006931416.4;refGene_gn=NUP153;refGene_tx=XM_019830197.2;refGene_gn=NUP153;refGene_tx=XM_019830198.2
B2 11882529 11882658
refGene_gn=NUP153;refGene_tx=XM_003985750.5;refGene_gn=NUP153;refGene_tx=XM_006931414.4;refGene_gn=NUP153;refGene_tx=XM_006931415.4;refGene_gn=NUP153;refGene_tx=XM_006931416.4;refGene_gn=NUP153;refGene_tx=XM_019830197.2;refGene_gn=NUP153;refGene_tx=XM_019830198.2
B2 11887888 11888005
refGene_gn=NUP153;refGene_tx=XM_003985750.5;refGene_gn=NUP153;refGene_tx=XM_006931414.4;refGene_gn=NUP153;refGene_tx=XM_006931415.4;refGene_gn=NUP153;refGene_tx=XM_006931416.4;refGene_gn=NUP153;refGene_tx=XM_019830197.2;refGene_gn=NUP153;refGene_tx=XM_019830198.2
B2 11888098 11888143
refGene_gn=NUP153;refGene_tx=XM_003985750.5;refGene_gn=NUP153;refGene_tx=XM_006931414.4;refGene_gn=NUP153;refGene_tx=XM_006931415.4;refGene_gn=NUP153;refGene_tx=XM_006931416.4;refGene_gn=NUP153;refGene_tx=XM_019830197.2;refGene_gn=NUP153;refGene_tx=XM_019830198.2
B2 11888253 11888310
refGene_gn=NUP153;refGene_tx=XM_003985750.5;refGene_gn=NUP153;refGene_tx=XM_006931414.4;refGene_gn=NUP153;refGene_tx=XM_006931415.4;refGene_gn=NUP153;refGene_tx=XM_006931416.4;refGene_gn=NUP153;refGene_tx=XM_019830197.2
B2 11890457 11890604
refGene_gn=NUP153;refGene_tx=XM_003985750.5;refGene_gn=NUP153;refGene_tx=XM_006931414.4;refGene_gn=NUP153;refGene_tx=XM_006931415.4;refGene_gn=NUP153;refGene_tx=XM_006931416.4;refGene_gn=NUP153;refGene_tx=XM_019830197.2;refGene_gn=NUP153;refGene_tx=XM_019830198.2
B2 11893495 11893548
refGene_gn=NUP153;refGene_tx=XM_003985750.5;refGene_gn=NUP153;refGene_tx=XM_006931414.4;refGene_gn=NUP153;refGene_tx=XM_006931415.4;refGene_gn=NUP153;refGene_tx=XM_006931416.4;refGene_gn=NUP153;refGene_tx=XM_019830197.2;refGene_gn=NUP153;refGene_tx=XM_019830198.2
B2 11894038 11894165

refGene_gn=NUP153;refGene_tx=XM_003985750.5;refGene_gn=NUP153;refGene_tx=XM_006931414.4;refGene_gn=NUP153;refGene_tx=XM_006931415.4;refGene_gn=NUP153;refGene_tx=XM_006931416.4;refGene_gn=NUP153;refGene_tx=XM_019830197.2;refGene_gn=NUP153;refGene_tx=XM_019830198.2
B2 11902249 11902387
refGene_gn=NUP153;refGene_tx=XM_003985750.5;refGene_gn=NUP153;refGene_tx=XM_006931414.4;refGene_gn=NUP153;refGene_tx=XM_006931415.4;refGene_gn=NUP153;refGene_tx=XM_006931416.4;refGene_gn=NUP153;refGene_tx=XM_019830197.2;refGene_gn=NUP153;refGene_tx=XM_019830198.2
B2 11903143 11903242
refGene_gn=NUP153;refGene_tx=XM_003985750.5;refGene_gn=NUP153;refGene_tx=XM_006931414.4;refGene_gn=NUP153;refGene_tx=XM_006931415.4;refGene_gn=NUP153;refGene_tx=XM_006931416.4;refGene_gn=NUP153;refGene_tx=XM_019830197.2;refGene_gn=NUP153;refGene_tx=XM_019830198.2
B2 11904465 11904556
refGene_gn=NUP153;refGene_tx=XM_003985750.5;refGene_gn=NUP153;refGene_tx=XM_006931414.4;refGene_gn=NUP153;refGene_tx=XM_006931415.4;refGene_gn=NUP153;refGene_tx=XM_006931416.4;refGene_gn=NUP153;refGene_tx=XM_019830197.2;refGene_gn=NUP153;refGene_tx=XM_019830198.2
B2 11908816 11908942
refGene_gn=NUP153;refGene_tx=XM_003985750.5;refGene_gn=NUP153;refGene_tx=XM_006931414.4;refGene_gn=NUP153;refGene_tx=XM_006931415.4;refGene_gn=NUP153;refGene_tx=XM_006931416.4;refGene_gn=NUP153;refGene_tx=XM_019830197.2;refGene_gn=NUP153;refGene_tx=XM_019830198.2
B2 11911940 11912570
refGene_gn=NUP153;refGene_tx=XM_003985750.5;refGene_gn=NUP153;refGene_tx=XM_006931414.4;refGene_gn=NUP153;refGene_tx=XM_006931415.4;refGene_gn=NUP153;refGene_tx=XM_006931416.4;refGene_gn=NUP153;refGene_tx=XM_019830197.2;refGene_gn=NUP153;refGene_tx=XM_019830198.2
B2 11916682 11916877
refGene_gn=NUP153;refGene_tx=XM_003985750.5;refGene_gn=NUP153;refGene_tx=XM_006931414.4;refGene_gn=NUP153;refGene_tx=XM_006931415.4;refGene_gn=NUP153;refGene_tx=XM_006931416.4;refGene_gn=NUP153;refGene_tx=XM_019830197.2;refGene_gn=NUP153;refGene_tx=XM_019830198.2
B2 11919985 11920870
refGene_gn=NUP153;refGene_tx=XM_003985750.5;refGene_gn=NUP153;refGene_tx=XM_006931414.4;refGene_gn=NUP153;refGene_tx=XM_006931415.4;refGene_gn=NUP153;refGene_tx=XM_006931416.4;refGene_gn=NUP153;refGene_tx=XM_019830197.2;refGene_gn=NUP153;refGene_tx=XM_019830198.2
B2 11921230 11921352 refGene_gn=NUP153;refGene_tx=XM_006931416.4
B2 11924053 11924410
refGene_gn=NUP153;refGene_tx=XM_003985750.5;refGene_gn=NUP153;refGene_tx=XM_006931414.4;refGene_gn=NUP153;refGene_tx=XM_006931415.4;refGene_gn=NUP153;refGene_tx=XM_019830197.2;refGene_gn=NUP153;refGene_tx=XM_019830198.2
B2 11925054 11925327
refGene_gn=NUP153;refGene_tx=XM_003985750.5;refGene_gn=NUP153;refGene_tx=XM_006931414.4;refGene_gn=NUP153;refGene_tx=XM_006931415.4;refGene_gn=NUP153;refGene_tx=XM_019830197.2;refGene_gn=NUP153;refGene_tx=XM_019830198.2

B2 11930169 11930338
refGene_gn=NUP153;refGene_tx=XM_003985750.5;refGene_gn=NUP153;refGene_tx=XM_006931414.4;refGene_gn=NUP153;refGene_tx=XM_006931415.4;refGene_gn=NUP153;refGene_tx=XM_019830197.2;refGene_gn=NUP153;refGene_tx=XM_019830198.2
B2 11930599 11930684
refGene_gn=NUP153;refGene_tx=XM_003985750.5;refGene_gn=NUP153;refGene_tx=XM_006931414.4;refGene_gn=NUP153;refGene_tx=XM_006931415.4;refGene_gn=NUP153;refGene_tx=XM_019830197.2;refGene_gn=NUP153;refGene_tx=XM_019830198.2
B2 11937780 11937925 refGene_gn=FAM8A1;refGene_tx=XM_019831563.2
B2 11939338 11939478 refGene_gn=FAM8A1;refGene_tx=XM_019831563.2
B2 11940449 11940573 refGene_gn=FAM8A1;refGene_tx=XM_019831563.2
B2 11942211 11942332 refGene_gn=FAM8A1;refGene_tx=XM_019831563.2
B2 11943618 11944336 refGene_gn=FAM8A1;refGene_tx=XM_019831563.2
B2 11976046 11976130
refGene_gn=CAP2;refGene_tx=XM_003985751.5;refGene_gn=CAP2;refGene_tx=XM_011282140.3;refGene_gn=CAP2;refGene_tx=XM_011282143.3;refGene_gn=CAP2;refGene_tx=XM_019830199.2
B2 11980423 11980564
refGene_gn=CAP2;refGene_tx=XM_003985751.5;refGene_gn=CAP2;refGene_tx=XM_011282140.3;refGene_gn=CAP2;refGene_tx=XM_011282143.3;refGene_gn=CAP2;refGene_tx=XM_019830199.2
B2 11987045 11987128
refGene_gn=CAP2;refGene_tx=XM_003985751.5;refGene_gn=CAP2;refGene_tx=XM_011282140.3;refGene_gn=CAP2;refGene_tx=XM_011282143.3;refGene_gn=CAP2;refGene_tx=XM_019830199.2
B2 11987225 11987349
refGene_gn=CAP2;refGene_tx=XM_003985751.5;refGene_gn=CAP2;refGene_tx=XM_011282140.3;refGene_gn=CAP2;refGene_tx=XM_011282143.3;refGene_gn=CAP2;refGene_tx=XM_019830199.2
B2 11988465 11988641
refGene_gn=CAP2;refGene_tx=XM_003985751.5;refGene_gn=CAP2;refGene_tx=XM_011282140.3;refGene_gn=CAP2;refGene_tx=XM_011282143.3;refGene_gn=CAP2;refGene_tx=XM_019830199.2
B2 11989603 11989793
refGene_gn=CAP2;refGene_tx=XM_003985751.5;refGene_gn=CAP2;refGene_tx=XM_011282140.3;refGene_gn=CAP2;refGene_tx=XM_011282143.3;refGene_gn=CAP2;refGene_tx=XM_019830199.2
B2 12009571 12009677
refGene_gn=CAP2;refGene_tx=XM_003985751.5;refGene_gn=CAP2;refGene_tx=XM_011282140.3;refGene_gn=CAP2;refGene_tx=XM_011282143.3;refGene_gn=CAP2;refGene_tx=XM_019830199.2
B2 12012627 12012713
refGene_gn=CAP2;refGene_tx=XM_003985751.5;refGene_gn=CAP2;refGene_tx=XM_011282140.3;refGene_gn=CAP2;refGene_tx=XM_011282143.3;refGene_gn=CAP2;refGene_tx=XM_019830199.2
B2 12013012 12013156
refGene_gn=CAP2;refGene_tx=XM_003985751.5;refGene_gn=CAP2;refGene_tx=XM_011282140.3;refGene_gn=CAP2;refGene_tx=XM_011282143.3;refGene_gn=CAP

2;refGene_tx=XM_019830199.2
B2 12066632 12066710
refGene_gn=CAP2;refGene_tx=XM_003985751.5;refGene_gn=CAP2;refGene_tx=XM_011282140.3;refGene_gn=CAP2;refGene_tx=XM_019830199.2
B2 12097414 12097515
refGene_gn=CAP2;refGene_tx=XM_003985751.5;refGene_gn=CAP2;refGene_tx=XM_011282140.3;refGene_gn=CAP2;refGene_tx=XM_011282143.3;refGene_gn=CAP2;refGene_tx=XM_019830199.2
B2 12101892 12102013
refGene_gn=CAP2;refGene_tx=XM_003985751.5;refGene_gn=CAP2;refGene_tx=XM_011282140.3;refGene_gn=CAP2;refGene_tx=XM_011282143.3
B2 12196750 12197114
refGene_gn=RBM24;refGene_tx=XM_011282139.3;refGene_gn=RBM24;refGene_tx=XM_019830201.2
B2 12203973 12204028
refGene_gn=RBM24;refGene_tx=XM_011282139.3;refGene_gn=RBM24;refGene_tx=XM_019830201.2
B2 12205435 12205559
refGene_gn=RBM24;refGene_tx=XM_011282139.3;refGene_gn=RBM24;refGene_tx=XM_019830201.2
B2 12206726 12206894 refGene_gn=RBM24;refGene_tx=XM_011282139.3
B2 12300744 12301032 refGene_gn=STMND1;refGene_tx=XM_006931422.3
B2 12301933 12302065 refGene_gn=STMND1;refGene_tx=XM_006931422.3
B2 12310494 12310646 refGene_gn=STMND1;refGene_tx=XM_006931422.3
B2 12315091 12315272 refGene_gn=STMND1;refGene_tx=XM_006931422.3
B2 12327060 12327141 refGene_gn=STMND1;refGene_tx=XM_006931422.3
B2 13006673 13008560
refGene_gn=ATXN1;refGene_tx=XM_019830207.2;refGene_gn=ATXN1;refGene_tx=XM_023253646.1;refGene_gn=ATXN1;refGene_tx=XM_023253647.1;refGene_gn=ATXN1;refGene_tx=XM_023253649.1;refGene_gn=ATXN1;refGene_tx=XM_023253650.1;refGene_gn=ATXN1;refGene_tx=XM_023253651.1;refGene_gn=ATXN1;refGene_tx=XM_023253652.1;refGene_gn=ATXN1;refGene_tx=XM_023253653.1;refGene_gn=ATXN1;refGene_tx=XM_023253654.1;refGene_gn=ATXN1;refGene_tx=XM_023253655.1
B2 13024827 13025358
refGene_gn=ATXN1;refGene_tx=XM_019830207.2;refGene_gn=ATXN1;refGene_tx=XM_023253646.1;refGene_gn=ATXN1;refGene_tx=XM_023253647.1;refGene_gn=ATXN1;refGene_tx=XM_023253649.1;refGene_gn=ATXN1;refGene_tx=XM_023253650.1;refGene_gn=ATXN1;refGene_tx=XM_023253651.1;refGene_gn=ATXN1;refGene_tx=XM_023253652.1;refGene_gn=ATXN1;refGene_tx=XM_023253653.1;refGene_gn=ATXN1;refGene_tx=XM_023253654.1;refGene_gn=ATXN1;refGene_tx=XM_023253655.1
B2 13035978 13036159
refGene_gn=GMPR;refGene_tx=XM_003985789.4;refGene_gn=GMPR;refGene_tx=XM_011282135.3
B2 13039558 13039718
refGene_gn=GMPR;refGene_tx=XM_003985789.4;refGene_gn=GMPR;refGene_tx=XM_011282135.3
B2 13042147 13042190
refGene_gn=GMPR;refGene_tx=XM_003985789.4;refGene_gn=GMPR;refGene_tx=X

M_011282135.3
B2	13052136 13052243
refGene_gn=GMPR;refGene_tx=XM_003985789.4;refGene_gn=GMPR;refGene_tx=XM_011282135.3
B2	13055776 13055858
refGene_gn=GMPR;refGene_tx=XM_003985789.4;refGene_gn=GMPR;refGene_tx=XM_011282135.3
B2	13065239 13065413
refGene_gn=GMPR;refGene_tx=XM_003985789.4;refGene_gn=GMPR;refGene_tx=XM_011282135.3
B2	13067806 13067890
refGene_gn=GMPR;refGene_tx=XM_003985789.4;refGene_gn=GMPR;refGene_tx=XM_011282135.3
B2	13070311 13070431
refGene_gn=GMPR;refGene_tx=XM_003985789.4;refGene_gn=GMPR;refGene_tx=XM_011282135.3
B2	13079130 13079217 refGene_gn=GMPR;refGene_tx=XM_003985789.4
B2	13201827 13202187
refGene_gn=LOC111560420;refGene_tx=XM_023253657.1
B2	13202386 13202448
refGene_gn=LOC111560420;refGene_tx=XM_023253657.1
B2	13202719 13202918
refGene_gn=LOC111560420;refGene_tx=XM_023253657.1
B2	13207593 13207683 refGene_gn=MYLIP;refGene_tx=XM_023253658.1
B2	13209590 13210011 refGene_gn=MYLIP;refGene_tx=XM_023253658.1
B2	13210964 13211129 refGene_gn=MYLIP;refGene_tx=XM_023253658.1
B2	13211633 13211831 refGene_gn=MYLIP;refGene_tx=XM_023253658.1
B2	13213568 13213754 refGene_gn=MYLIP;refGene_tx=XM_023253658.1
B2	13225209 13225400 refGene_gn=MYLIP;refGene_tx=XM_023253658.1
B2	13226528 13226615 refGene_gn=MYLIP;refGene_tx=XM_023253658.1
B2	13658157 13658213
refGene_gn=DTNBP1;refGene_tx=XM_023253667.1;refGene_gn=DTNBP1;refGene_tx=XM_023253668.1
B2	13668859 13668913
refGene_gn=DTNBP1;refGene_tx=XM_023253667.1;refGene_gn=DTNBP1;refGene_tx=XM_023253668.1
B2	13670150 13670201
refGene_gn=DTNBP1;refGene_tx=XM_023253667.1;refGene_gn=DTNBP1;refGene_tx=XM_023253668.1
B2	13681763 13681824
refGene_gn=DTNBP1;refGene_tx=XM_023253667.1;refGene_gn=DTNBP1;refGene_tx=XM_023253668.1
B2	13691619 13691752
refGene_gn=DTNBP1;refGene_tx=XM_023253667.1;refGene_gn=DTNBP1;refGene_tx=XM_023253668.1
B2	13702999 13703132
refGene_gn=DTNBP1;refGene_tx=XM_023253667.1;refGene_gn=DTNBP1;refGene_tx=XM_023253668.1
B2	13730420 13730443
refGene_gn=DTNBP1;refGene_tx=XM_023253667.1;refGene_gn=DTNBP1;refGene_

tx=XM_023253668.1
B2 13780028 13780184 refGene_gn=DTNBP1;refGene_tx=XM_023253667.1
B2 13788899 13789043
refGene_gn=DTNBP1;refGene_tx=XM_023253667.1;refGene_gn=DTNBP1;refGene_tx=XM_023253668.1
B2 13790217 13790417
refGene_gn=DTNBP1;refGene_tx=XM_023253667.1;refGene_gn=DTNBP1;refGene_tx=XM_023253668.1
B2 13792771 13792954
refGene_gn=JARID2;refGene_tx=XM_023253659.1;refGene_gn=JARID2;refGene_tx=XM_023253660.1;refGene_gn=JARID2;refGene_tx=XM_023253662.1;refGene_gn=JARID2;refGene_tx=XM_023253663.1;refGene_gn=JARID2;refGene_tx=XM_023253664.1;refGene_gn=JARID2;refGene_tx=XM_023253665.1;refGene_gn=JARID2;refGene_tx=XM_023253666.1
B2 13796525 13796633
refGene_gn=JARID2;refGene_tx=XM_023253659.1;refGene_gn=JARID2;refGene_tx=XM_023253660.1;refGene_gn=JARID2;refGene_tx=XM_023253662.1;refGene_gn=JARID2;refGene_tx=XM_023253663.1;refGene_gn=JARID2;refGene_tx=XM_023253664.1;refGene_gn=JARID2;refGene_tx=XM_023253665.1;refGene_gn=JARID2;refGene_tx=XM_023253666.1
B2 13799738 13799922
refGene_gn=JARID2;refGene_tx=XM_023253659.1;refGene_gn=JARID2;refGene_tx=XM_023253660.1;refGene_gn=JARID2;refGene_tx=XM_023253662.1;refGene_gn=JARID2;refGene_tx=XM_023253663.1;refGene_gn=JARID2;refGene_tx=XM_023253664.1;refGene_gn=JARID2;refGene_tx=XM_023253665.1;refGene_gn=JARID2;refGene_tx=XM_023253666.1
B2 13800516 13800647
refGene_gn=JARID2;refGene_tx=XM_023253659.1;refGene_gn=JARID2;refGene_tx=XM_023253660.1;refGene_gn=JARID2;refGene_tx=XM_023253662.1;refGene_gn=JARID2;refGene_tx=XM_023253663.1;refGene_gn=JARID2;refGene_tx=XM_023253664.1;refGene_gn=JARID2;refGene_tx=XM_023253665.1;refGene_gn=JARID2;refGene_tx=XM_023253666.1
B2 13801320 13801503
refGene_gn=JARID2;refGene_tx=XM_023253659.1;refGene_gn=JARID2;refGene_tx=XM_023253660.1;refGene_gn=JARID2;refGene_tx=XM_023253662.1;refGene_gn=JARID2;refGene_tx=XM_023253663.1;refGene_gn=JARID2;refGene_tx=XM_023253664.1;refGene_gn=JARID2;refGene_tx=XM_023253665.1;refGene_gn=JARID2;refGene_tx=XM_023253666.1
B2 13803191 13803297
refGene_gn=JARID2;refGene_tx=XM_023253659.1;refGene_gn=JARID2;refGene_tx=XM_023253660.1;refGene_gn=JARID2;refGene_tx=XM_023253662.1;refGene_gn=JARID2;refGene_tx=XM_023253663.1;refGene_gn=JARID2;refGene_tx=XM_023253664.1;refGene_gn=JARID2;refGene_tx=XM_023253665.1;refGene_gn=JARID2;refGene_tx=XM_023253666.1
B2 13805729 13805844
refGene_gn=JARID2;refGene_tx=XM_023253659.1;refGene_gn=JARID2;refGene_tx=XM_023253660.1;refGene_gn=JARID2;refGene_tx=XM_023253662.1;refGene_gn=JARID2;refGene_tx=XM_023253663.1;refGene_gn=JARID2;refGene_tx=XM_023253664.1;refGene_gn=JARID2;refGene_tx=XM_023253665.1;refGene_gn=JARID2;refGene_tx=XM_023253666.1

B2 13806676 13806747
refGene_gn=JARID2;refGene_tx=XM_023253659.1;refGene_gn=JARID2;refGene_tx=XM_023253660.1;refGene_gn=JARID2;refGene_tx=XM_023253662.1;refGene_gn=JARID2;refGene_tx=XM_023253663.1;refGene_gn=JARID2;refGene_tx=XM_023253664.1;refGene_gn=JARID2;refGene_tx=XM_023253665.1;refGene_gn=JARID2;refGene_tx=XM_023253666.1
B2 13806838 13806957
refGene_gn=JARID2;refGene_tx=XM_023253659.1;refGene_gn=JARID2;refGene_tx=XM_023253660.1;refGene_gn=JARID2;refGene_tx=XM_023253662.1;refGene_gn=JARID2;refGene_tx=XM_023253663.1;refGene_gn=JARID2;refGene_tx=XM_023253664.1;refGene_gn=JARID2;refGene_tx=XM_023253665.1;refGene_gn=JARID2;refGene_tx=XM_023253666.1
B2 13809863 13809956
refGene_gn=JARID2;refGene_tx=XM_023253659.1;refGene_gn=JARID2;refGene_tx=XM_023253660.1;refGene_gn=JARID2;refGene_tx=XM_023253662.1;refGene_gn=JARID2;refGene_tx=XM_023253663.1;refGene_gn=JARID2;refGene_tx=XM_023253664.1;refGene_gn=JARID2;refGene_tx=XM_023253665.1;refGene_gn=JARID2;refGene_tx=XM_023253666.1
B2 13812642 13813145
refGene_gn=JARID2;refGene_tx=XM_023253659.1;refGene_gn=JARID2;refGene_tx=XM_023253660.1;refGene_gn=JARID2;refGene_tx=XM_023253662.1;refGene_gn=JARID2;refGene_tx=XM_023253663.1;refGene_gn=JARID2;refGene_tx=XM_023253664.1;refGene_gn=JARID2;refGene_tx=XM_023253665.1;refGene_gn=JARID2;refGene_tx=XM_023253666.1
B2 13815937 13816991
refGene_gn=JARID2;refGene_tx=XM_023253659.1;refGene_gn=JARID2;refGene_tx=XM_023253660.1;refGene_gn=JARID2;refGene_tx=XM_023253662.1;refGene_gn=JARID2;refGene_tx=XM_023253663.1;refGene_gn=JARID2;refGene_tx=XM_023253664.1;refGene_gn=JARID2;refGene_tx=XM_023253665.1;refGene_gn=JARID2;refGene_tx=XM_023253666.1
B2 13825480 13825716
refGene_gn=JARID2;refGene_tx=XM_023253659.1;refGene_gn=JARID2;refGene_tx=XM_023253660.1;refGene_gn=JARID2;refGene_tx=XM_023253662.1;refGene_gn=JARID2;refGene_tx=XM_023253663.1;refGene_gn=JARID2;refGene_tx=XM_023253664.1;refGene_gn=JARID2;refGene_tx=XM_023253665.1;refGene_gn=JARID2;refGene_tx=XM_023253666.1
B2 13849172 13849349
refGene_gn=JARID2;refGene_tx=XM_023253659.1;refGene_gn=JARID2;refGene_tx=XM_023253660.1;refGene_gn=JARID2;refGene_tx=XM_023253662.1;refGene_gn=JARID2;refGene_tx=XM_023253663.1;refGene_gn=JARID2;refGene_tx=XM_023253664.1;refGene_gn=JARID2;refGene_tx=XM_023253665.1;refGene_gn=JARID2;refGene_tx=XM_023253666.1
B2 13865507 13865677
refGene_gn=JARID2;refGene_tx=XM_023253660.1;refGene_gn=JARID2;refGene_tx=XM_023253662.1;refGene_gn=JARID2;refGene_tx=XM_023253663.1;refGene_gn=JARID2;refGene_tx=XM_023253664.1;refGene_gn=JARID2;refGene_tx=XM_023253665.1;refGene_gn=JARID2;refGene_tx=XM_023253666.1
B2 13904830 13904972 refGene_gn=JARID2;refGene_tx=XM_023253660.1
B2 13914310 13914377 refGene_gn=JARID2;refGene_tx=XM_023253660.1
B2 13939241 13939357

refGene_gn=JARID2;refGene_tx=XM_023253662.1;refGene_gn=JARID2;refGene_tx=XM_023253663.1;refGene_gn=JARID2;refGene_tx=XM_023253664.1;refGene_gn=JARID2;refGene_tx=XM_023253665.1;refGene_gn=JARID2;refGene_tx=XM_023253666.1
B2 15069692 15069821
refGene_gn=CD83;refGene_tx=XM_023253670.1;refGene_gn=CD83;refGene_tx=XM_023253671.1
B2 15071122 15071229
refGene_gn=CD83;refGene_tx=XM_023253670.1;refGene_gn=CD83;refGene_tx=XM_023253671.1
B2 15074827 15074998
refGene_gn=CD83;refGene_tx=XM_023253670.1;refGene_gn=CD83;refGene_tx=XM_023253671.1
B2 15079772 15079777
refGene_gn=CD83;refGene_tx=XM_023253670.1;refGene_gn=CD83;refGene_tx=XM_023253671.1
B2 15081871 15081917
refGene_gn=CD83;refGene_tx=XM_023253670.1;refGene_gn=CD83;refGene_tx=XM_023253671.1
B2 15088183 15088303
refGene_gn=CD83;refGene_tx=XM_023253670.1;refGene_gn=CD83;refGene_tx=XM_023253671.1
B2 15088411 15088448
refGene_gn=CD83;refGene_tx=XM_023253670.1;refGene_gn=CD83;refGene_tx=XM_023253671.1
B2 15229662 15230406 refGene_gn=RNF182;refGene_tx=XM_003985792.4
B2 15352026 15352459
refGene_gn=MCUR1;refGene_tx=XM_023253672.1;refGene_gn=MCUR1;refGene_tx=XM_023253673.1;refGene_gn=MCUR1;refGene_tx=XM_023253674.1;refGene_gn=MCUR1;refGene_tx=XM_023253676.1;refGene_gn=MCUR1;refGene_tx=XM_023253677.1;refGene_gn=MCUR1;refGene_tx=XM_023253678.1
B2 15360421 15360559
refGene_gn=MCUR1;refGene_tx=XM_023253672.1;refGene_gn=MCUR1;refGene_tx=XM_023253673.1;refGene_gn=MCUR1;refGene_tx=XM_023253674.1;refGene_gn=MCUR1;refGene_tx=XM_023253676.1;refGene_gn=MCUR1;refGene_tx=XM_023253678.1
B2 15364018 15364122
refGene_gn=MCUR1;refGene_tx=XM_023253672.1;refGene_gn=MCUR1;refGene_tx=XM_023253673.1;refGene_gn=MCUR1;refGene_tx=XM_023253674.1;refGene_gn=MCUR1;refGene_tx=XM_023253676.1;refGene_gn=MCUR1;refGene_tx=XM_023253677.1;refGene_gn=MCUR1;refGene_tx=XM_023253678.1
B2 15364865 15364967
refGene_gn=MCUR1;refGene_tx=XM_023253672.1;refGene_gn=MCUR1;refGene_tx=XM_023253673.1;refGene_gn=MCUR1;refGene_tx=XM_023253674.1;refGene_gn=MCUR1;refGene_tx=XM_023253676.1;refGene_gn=MCUR1;refGene_tx=XM_023253677.1;refGene_gn=MCUR1;refGene_tx=XM_023253678.1
B2 15366483 15366525
refGene_gn=MCUR1;refGene_tx=XM_023253672.1;refGene_gn=MCUR1;refGene_tx=XM_023253673.1;refGene_gn=MCUR1;refGene_tx=XM_023253674.1;refGene_gn=MCUR1;refGene_tx=XM_023253676.1;refGene_gn=MCUR1;refGene_tx=XM_0232536

77.1
B2 15367323 15367395
refGene_gn=MCUR1;refGene_tx=XM_023253672.1;refGene_gn=MCUR1;refGene_tx=XM_023253673.1;refGene_gn=MCUR1;refGene_tx=XM_023253674.1;refGene_gn=MCUR1;refGene_tx=XM_023253676.1;refGene_gn=MCUR1;refGene_tx=XM_023253677.1
B2 15371350 15371404
refGene_gn=MCUR1;refGene_tx=XM_023253672.1;refGene_gn=MCUR1;refGene_tx=XM_023253673.1;refGene_gn=MCUR1;refGene_tx=XM_023253674.1;refGene_gn=MCUR1;refGene_tx=XM_023253677.1
B2 15372710 15372825
refGene_gn=MCUR1;refGene_tx=XM_023253672.1;refGene_gn=MCUR1;refGene_tx=XM_023253673.1;refGene_gn=MCUR1;refGene_tx=XM_023253674.1;refGene_gn=MCUR1;refGene_tx=XM_023253676.1;refGene_gn=MCUR1;refGene_tx=XM_023253677.1;refGene_gn=MCUR1;refGene_tx=XM_023253678.1
B2 15373896 15373952 refGene_gn=MCUR1;refGene_tx=XM_023253673.1
B2 15374154 15374219
refGene_gn=MCUR1;refGene_tx=XM_023253672.1;refGene_gn=MCUR1;refGene_tx=XM_023253676.1;refGene_gn=MCUR1;refGene_tx=XM_023253677.1;refGene_gn=MCUR1;refGene_tx=XM_023253678.1
B2 15377610 15377645 refGene_gn=MCUR1;refGene_tx=XM_023253674.1
B2 15424417 15425000 refGene_gn=RANBP9;refGene_tx=XM_023253679.1
B2 15439315 15439427 refGene_gn=RANBP9;refGene_tx=XM_023253679.1
B2 15477240 15477293 refGene_gn=RANBP9;refGene_tx=XM_023253679.1
B2 15478186 15478354 refGene_gn=RANBP9;refGene_tx=XM_023253679.1
B2 15482572 15482595 refGene_gn=RANBP9;refGene_tx=XM_023253679.1
B2 15490299 15490484 refGene_gn=RANBP9;refGene_tx=XM_023253679.1
B2 15493132 15493245 refGene_gn=RANBP9;refGene_tx=XM_023253679.1
B2 15494595 15494704 refGene_gn=RANBP9;refGene_tx=XM_023253679.1
B2 15497216 15497407 refGene_gn=RANBP9;refGene_tx=XM_023253679.1
B2 15498559 15498704 refGene_gn=RANBP9;refGene_tx=XM_023253679.1
B2 15502032 15502154 refGene_gn=RANBP9;refGene_tx=XM_023253679.1
B2 15503807 15503959 refGene_gn=RANBP9;refGene_tx=XM_023253679.1
B2 15510502 15510614 refGene_gn=RANBP9;refGene_tx=XM_023253679.1
B2 15513960 15514091 refGene_gn=RANBP9;refGene_tx=XM_023253679.1
B2 15515736 15515813 refGene_gn=NOL7;refGene_tx=XM_003985794.5
B2 15516073 15516151 refGene_gn=NOL7;refGene_tx=XM_003985794.5
B2 15516229 15516351 refGene_gn=NOL7;refGene_tx=XM_003985794.5
B2 15516919 15517001 refGene_gn=NOL7;refGene_tx=XM_003985794.5
B2 15517241 15517273 refGene_gn=NOL7;refGene_tx=XM_003985794.5
B2 15518939 15518998 refGene_gn=NOL7;refGene_tx=XM_003985794.5
B2 15519642 15519703 refGene_gn=NOL7;refGene_tx=XM_003985794.5
B2 15519793 15520059 refGene_gn=NOL7;refGene_tx=XM_003985794.5
B2 15524092 15524253
refGene_gn=SIRT5;refGene_tx=XM_003985795.5;refGene_gn=SIRT5;refGene_tx=XM_006931445.4;refGene_gn=SIRT5;refGene_tx=XM_006931448.3;refGene_gn=SIRT5;refGene_tx=XM_019830216.2;refGene_gn=SIRT5;refGene_tx=XM_019830219.1;refGene_gn=SIRT5;refGene_tx=XM_019830220.2;refGene_gn=SIRT5;refGene_tx=XM_023253680.1;refGene_gn=SIRT5;refGene_tx=XM_023253681.1
B2 15536275 15536391

refGene_gn=SIRT5;refGene_tx=XM_003985795.5;refGene_gn=SIRT5;refGene_tx=XM_006931445.4;refGene_gn=SIRT5;refGene_tx=XM_006931448.3;refGene_gn=SIRT5;refGene_tx=XM_019830216.2;refGene_gn=SIRT5;refGene_tx=XM_019830219.1;refGene_gn=SIRT5;refGene_tx=XM_019830220.2;refGene_gn=SIRT5;refGene_tx=XM_023253680.1;refGene_gn=SIRT5;refGene_tx=XM_023253681.1
B2 15538438 15538562
refGene_gn=SIRT5;refGene_tx=XM_003985795.5;refGene_gn=SIRT5;refGene_tx=XM_006931445.4;refGene_gn=SIRT5;refGene_tx=XM_006931448.3;refGene_gn=SIRT5;refGene_tx=XM_019830216.2;refGene_gn=SIRT5;refGene_tx=XM_019830219.1;refGene_gn=SIRT5;refGene_tx=XM_023253680.1;refGene_gn=SIRT5;refGene_tx=XM_023253681.1
B2 15540566 15540620
refGene_gn=SIRT5;refGene_tx=XM_003985795.5;refGene_gn=SIRT5;refGene_tx=XM_006931445.4;refGene_gn=SIRT5;refGene_tx=XM_006931448.3;refGene_gn=SIRT5;refGene_tx=XM_019830216.2;refGene_gn=SIRT5;refGene_tx=XM_019830219.1;refGene_gn=SIRT5;refGene_tx=XM_019830220.2;refGene_gn=SIRT5;refGene_tx=XM_023253680.1;refGene_gn=SIRT5;refGene_tx=XM_023253681.1
B2 15541713 15541801
refGene_gn=SIRT5;refGene_tx=XM_003985795.5;refGene_gn=SIRT5;refGene_tx=XM_006931445.4;refGene_gn=SIRT5;refGene_tx=XM_006931448.3;refGene_gn=SIRT5;refGene_tx=XM_019830216.2;refGene_gn=SIRT5;refGene_tx=XM_019830219.1;refGene_gn=SIRT5;refGene_tx=XM_019830220.2;refGene_gn=SIRT5;refGene_tx=XM_023253680.1;refGene_gn=SIRT5;refGene_tx=XM_023253681.1
B2 15544681 15544907
refGene_gn=SIRT5;refGene_tx=XM_003985795.5;refGene_gn=SIRT5;refGene_tx=XM_006931445.4;refGene_gn=SIRT5;refGene_tx=XM_006931448.3;refGene_gn=SIRT5;refGene_tx=XM_019830216.2;refGene_gn=SIRT5;refGene_tx=XM_019830219.1;refGene_gn=SIRT5;refGene_tx=XM_019830220.2;refGene_gn=SIRT5;refGene_tx=XM_023253680.1;refGene_gn=SIRT5;refGene_tx=XM_023253681.1
B2 15551834 15551968
refGene_gn=SIRT5;refGene_tx=XM_003985795.5;refGene_gn=SIRT5;refGene_tx=XM_006931445.4;refGene_gn=SIRT5;refGene_tx=XM_006931448.3;refGene_gn=SIRT5;refGene_tx=XM_019830216.2;refGene_gn=SIRT5;refGene_tx=XM_019830219.1;refGene_gn=SIRT5;refGene_tx=XM_019830220.2;refGene_gn=SIRT5;refGene_tx=XM_023253680.1;refGene_gn=SIRT5;refGene_tx=XM_023253681.1
B2 15555765 15555880
refGene_gn=SIRT5;refGene_tx=XM_003985795.5;refGene_gn=SIRT5;refGene_tx=XM_006931445.4;refGene_gn=SIRT5;refGene_tx=XM_006931448.3;refGene_gn=SIRT5;refGene_tx=XM_019830216.2;refGene_gn=SIRT5;refGene_tx=XM_019830219.1;refGene_gn=SIRT5;refGene_tx=XM_019830220.2;refGene_gn=SIRT5;refGene_tx=XM_023253680.1;refGene_gn=SIRT5;refGene_tx=XM_023253681.1
B2 15661693 15661946 refGene_gn=GFOD1;refGene_tx=XM_003985798.5
B2 15724942 15725087 refGene_gn=GFOD1;refGene_tx=XM_019830221.2
B2 15774333 15775253
refGene_gn=GFOD1;refGene_tx=XM_003985798.5;refGene_gn=GFOD1;refGene_tx=XM_006931450.4;refGene_gn=GFOD1;refGene_tx=XM_006931451.4;refGene_gn=GFOD1;refGene_tx=XM_019830221.2;refGene_gn=GFOD1;refGene_tx=XM_019830222.2;refGene_gn=GFOD1;refGene_tx=XM_019830223.2
B2 15808222 15808334
refGene_gn=LOC101090689;refGene_tx=XM_003985799.5;refGene_gn=LOC101090

689;refGene_tx=XM_019830224.2;refGene_gn=LOC101090689;refGene_tx=XM_019830225.2;refGene_gn=LOC101090689;refGene_tx=XM_019830226.2;refGene_gn=LOC101090689;refGene_tx=XM_019830227.2
B2 15809333 15809414
refGene_gn=LOC101090689;refGene_tx=XM_003985799.5;refGene_gn=LOC101090689;refGene_tx=XM_019830224.2;refGene_gn=LOC101090689;refGene_tx=XM_019830225.2;refGene_gn=LOC101090689;refGene_tx=XM_019830226.2
B2 15812779 15812967
refGene_gn=LOC101090689;refGene_tx=XM_003985799.5;refGene_gn=LOC101090689;refGene_tx=XM_019830224.2;refGene_gn=LOC101090689;refGene_tx=XM_019830225.2;refGene_gn=LOC101090689;refGene_tx=XM_019830226.2;refGene_gn=LOC101090689;refGene_tx=XM_019830227.2
B2 15818748 15818886
refGene_gn=LOC101090689;refGene_tx=XM_003985799.5;refGene_gn=LOC101090689;refGene_tx=XM_019830224.2;refGene_gn=LOC101090689;refGene_tx=XM_019830225.2;refGene_gn=LOC101090689;refGene_tx=XM_019830226.2;refGene_gn=LOC101090689;refGene_tx=XM_019830227.2
B2 15827174 15827320
refGene_gn=LOC101090689;refGene_tx=XM_003985799.5;refGene_gn=LOC101090689;refGene_tx=XM_019830224.2;refGene_gn=LOC101090689;refGene_tx=XM_019830225.2;refGene_gn=LOC101090689;refGene_tx=XM_019830226.2;refGene_gn=LOC101090689;refGene_tx=XM_019830227.2
B2 15828895 15829025
refGene_gn=LOC101090689;refGene_tx=XM_003985799.5;refGene_gn=LOC101090689;refGene_tx=XM_019830224.2;refGene_gn=LOC101090689;refGene_tx=XM_019830225.2;refGene_gn=LOC101090689;refGene_tx=XM_019830226.2;refGene_gn=LOC101090689;refGene_tx=XM_019830227.2
B2 15830076 15830163
refGene_gn=LOC101090689;refGene_tx=XM_003985799.5;refGene_gn=LOC101090689;refGene_tx=XM_019830224.2;refGene_gn=LOC101090689;refGene_tx=XM_019830225.2;refGene_gn=LOC101090689;refGene_tx=XM_019830226.2;refGene_gn=LOC101090689;refGene_tx=XM_019830227.2
B2 15853016 15853032
refGene_gn=PHACTR1;refGene_tx=XM_011282184.3;refGene_gn=PHACTR1;refGene_tx=XM_011282187.3;refGene_gn=PHACTR1;refGene_tx=XM_011282188.3;refGene_gn=PHACTR1;refGene_tx=XM_019830228.2;refGene_gn=PHACTR1;refGene_tx=XM_019830229.2;refGene_gn=PHACTR1;refGene_tx=XM_019830230.2;refGene_gn=PHACTR1;refGene_tx=XM_019830231.2;refGene_gn=PHACTR1;refGene_tx=XM_019830232.2;refGene_gn=PHACTR1;refGene_tx=XM_019830233.2;refGene_gn=PHACTR1;refGene_tx=XM_019830234.2;refGene_gn=PHACTR1;refGene_tx=XM_023253684.1
B2 15853841 15853918
refGene_gn=PHACTR1;refGene_tx=XM_011282184.3;refGene_gn=PHACTR1;refGene_tx=XM_011282187.3;refGene_gn=PHACTR1;refGene_tx=XM_011282188.3;refGene_gn=PHACTR1;refGene_tx=XM_019830228.2;refGene_gn=PHACTR1;refGene_tx=XM_019830229.2;refGene_gn=PHACTR1;refGene_tx=XM_019830230.2;refGene_gn=PHACTR1;refGene_tx=XM_019830231.2;refGene_gn=PHACTR1;refGene_tx=XM_019830232.2;refGene_gn=PHACTR1;refGene_tx=XM_019830233.2;refGene_gn=PHACTR1;refGene_tx=XM_019830234.2;refGene_gn=PHACTR1;refGene_tx=XM_023253684.1

B2 15856705 15856846
refGene_gn=PHACTR1;refGene_tx=XM_011282184.3;refGene_gn=PHACTR1;refGene_tx=XM_011282187.3;refGene_gn=PHACTR1;refGene_tx=XM_011282188.3;refGene_gn=PHACTR1;refGene_tx=XM_019830228.2;refGene_gn=PHACTR1;refGene_tx=XM_019830229.2;refGene_gn=PHACTR1;refGene_tx=XM_019830230.2;refGene_gn=PHACTR1;refGene_tx=XM_019830231.2;refGene_gn=PHACTR1;refGene_tx=XM_019830232.2;refGene_gn=PHACTR1;refGene_tx=XM_019830233.2;refGene_gn=PHACTR1;refGene_tx=XM_019830234.2;refGene_gn=PHACTR1;refGene_tx=XM_023253684.1
B2 15861209 15861271
refGene_gn=PHACTR1;refGene_tx=XM_011282184.3;refGene_gn=PHACTR1;refGene_tx=XM_011282187.3;refGene_gn=PHACTR1;refGene_tx=XM_011282188.3;refGene_gn=PHACTR1;refGene_tx=XM_019830228.2;refGene_gn=PHACTR1;refGene_tx=XM_019830229.2;refGene_gn=PHACTR1;refGene_tx=XM_019830230.2;refGene_gn=PHACTR1;refGene_tx=XM_019830231.2;refGene_gn=PHACTR1;refGene_tx=XM_019830232.2;refGene_gn=PHACTR1;refGene_tx=XM_019830233.2;refGene_gn=PHACTR1;refGene_tx=XM_019830234.2;refGene_gn=PHACTR1;refGene_tx=XM_023253684.1
B2 15866609 15866665
refGene_gn=PHACTR1;refGene_tx=XM_011282184.3;refGene_gn=PHACTR1;refGene_tx=XM_011282187.3;refGene_gn=PHACTR1;refGene_tx=XM_011282188.3;refGene_gn=PHACTR1;refGene_tx=XM_019830228.2;refGene_gn=PHACTR1;refGene_tx=XM_019830229.2;refGene_gn=PHACTR1;refGene_tx=XM_019830230.2;refGene_gn=PHACTR1;refGene_tx=XM_019830231.2;refGene_gn=PHACTR1;refGene_tx=XM_019830232.2;refGene_gn=PHACTR1;refGene_tx=XM_019830233.2;refGene_gn=PHACTR1;refGene_tx=XM_019830234.2;refGene_gn=PHACTR1;refGene_tx=XM_023253684.1
B2 15897742 15897899
refGene_gn=PHACTR1;refGene_tx=XM_011282184.3;refGene_gn=PHACTR1;refGene_tx=XM_011282187.3;refGene_gn=PHACTR1;refGene_tx=XM_011282188.3;refGene_gn=PHACTR1;refGene_tx=XM_019830228.2;refGene_gn=PHACTR1;refGene_tx=XM_019830229.2;refGene_gn=PHACTR1;refGene_tx=XM_019830230.2;refGene_gn=PHACTR1;refGene_tx=XM_019830231.2;refGene_gn=PHACTR1;refGene_tx=XM_019830232.2;refGene_gn=PHACTR1;refGene_tx=XM_019830233.2;refGene_gn=PHACTR1;refGene_tx=XM_019830234.2;refGene_gn=PHACTR1;refGene_tx=XM_023253684.1
B2 15901022 15901270
refGene_gn=PHACTR1;refGene_tx=XM_011282184.3;refGene_gn=PHACTR1;refGene_tx=XM_011282187.3;refGene_gn=PHACTR1;refGene_tx=XM_011282188.3;refGene_gn=PHACTR1;refGene_tx=XM_019830228.2;refGene_gn=PHACTR1;refGene_tx=XM_019830229.2;refGene_gn=PHACTR1;refGene_tx=XM_019830230.2;refGene_gn=PHACTR1;refGene_tx=XM_019830231.2;refGene_gn=PHACTR1;refGene_tx=XM_019830232.2;refGene_gn=PHACTR1;refGene_tx=XM_019830233.2;refGene_gn=PHACTR1;refGene_tx=XM_019830234.2;refGene_gn=PHACTR1;refGene_tx=XM_023253684.1
B2 15921347 15921669
refGene_gn=PHACTR1;refGene_tx=XM_011282184.3;refGene_gn=PHACTR1;refGene_tx=XM_011282187.3;refGene_gn=PHACTR1;refGene_tx=XM_011282188.3;refGene_gn=PHACTR1;refGene_tx=XM_019830228.2;refGene_gn=PHACTR1;refGene_tx=XM_019830229.2;refGene_gn=PHACTR1;refGene_tx=XM_019830230.2;refGene_gn

=PHACTR1;refGene_tx=XM_019830231.2;refGene_gn=PHACTR1;refGene_tx=XM_019830232.2;refGene_gn=PHACTR1;refGene_tx=XM_019830233.2;refGene_gn=PHACTR1;refGene_tx=XM_019830234.2;refGene_gn=PHACTR1;refGene_tx=XM_023253684.1
B2 15938527 15938734
refGene_gn=PHACTR1;refGene_tx=XM_011282184.3;refGene_gn=PHACTR1;refGene_tx=XM_011282187.3;refGene_gn=PHACTR1;refGene_tx=XM_011282188.3;refGene_gn=PHACTR1;refGene_tx=XM_019830228.2;refGene_gn=PHACTR1;refGene_tx=XM_019830229.2;refGene_gn=PHACTR1;refGene_tx=XM_019830230.2;refGene_gn=PHACTR1;refGene_tx=XM_019830231.2;refGene_gn=PHACTR1;refGene_tx=XM_019830234.2;refGene_gn=PHACTR1;refGene_tx=XM_023253684.1
B2 15940625 15940793
refGene_gn=PHACTR1;refGene_tx=XM_011282184.3;refGene_gn=PHACTR1;refGene_tx=XM_011282187.3;refGene_gn=PHACTR1;refGene_tx=XM_011282188.3;refGene_gn=PHACTR1;refGene_tx=XM_019830228.2;refGene_gn=PHACTR1;refGene_tx=XM_019830229.2;refGene_gn=PHACTR1;refGene_tx=XM_019830230.2;refGene_gn=PHACTR1;refGene_tx=XM_019830231.2;refGene_gn=PHACTR1;refGene_tx=XM_019830232.2;refGene_gn=PHACTR1;refGene_tx=XM_019830233.2;refGene_gn=PHACTR1;refGene_tx=XM_019830234.2;refGene_gn=PHACTR1;refGene_tx=XM_023253684.1
B2 15962685 15962766
refGene_gn=PHACTR1;refGene_tx=XM_011282184.3;refGene_gn=PHACTR1;refGene_tx=XM_011282187.3;refGene_gn=PHACTR1;refGene_tx=XM_011282188.3;refGene_gn=PHACTR1;refGene_tx=XM_019830228.2;refGene_gn=PHACTR1;refGene_tx=XM_019830229.2;refGene_gn=PHACTR1;refGene_tx=XM_019830230.2;refGene_gn=PHACTR1;refGene_tx=XM_019830231.2;refGene_gn=PHACTR1;refGene_tx=XM_019830232.2;refGene_gn=PHACTR1;refGene_tx=XM_019830233.2;refGene_gn=PHACTR1;refGene_tx=XM_019830234.2;refGene_gn=PHACTR1;refGene_tx=XM_023253684.1
B2 16067665 16067830
refGene_gn=PHACTR1;refGene_tx=XM_011282184.3;refGene_gn=PHACTR1;refGene_tx=XM_011282187.3;refGene_gn=PHACTR1;refGene_tx=XM_011282188.3;refGene_gn=PHACTR1;refGene_tx=XM_019830228.2;refGene_gn=PHACTR1;refGene_tx=XM_019830229.2;refGene_gn=PHACTR1;refGene_tx=XM_019830230.2;refGene_gn=PHACTR1;refGene_tx=XM_019830231.2;refGene_gn=PHACTR1;refGene_tx=XM_019830232.2;refGene_gn=PHACTR1;refGene_tx=XM_019830233.2;refGene_gn=PHACTR1;refGene_tx=XM_019830234.2;refGene_gn=PHACTR1;refGene_tx=XM_023253684.1
B2 16113085 16113335 refGene_gn=PHACTR1;refGene_tx=XM_023253684.1
B2 16387789 16387936
refGene_gn=PHACTR1;refGene_tx=XM_011282184.3;refGene_gn=PHACTR1;refGene_tx=XM_019830228.2;refGene_gn=PHACTR1;refGene_tx=XM_019830229.2;refGene_gn=PHACTR1;refGene_tx=XM_019830230.2;refGene_gn=PHACTR1;refGene_tx=XM_019830231.2;refGene_gn=PHACTR1;refGene_tx=XM_019830232.2;refGene_gn=PHACTR1;refGene_tx=XM_019830233.2
B2 16423310 16423413
refGene_gn=PHACTR1;refGene_tx=XM_011282184.3;refGene_gn=PHACTR1;refGene_tx=XM_019830228.2;refGene_gn=PHACTR1;refGene_tx=XM_019830229.2;refGene_gn=PHACTR1;refGene_tx=XM_019830230.2;refGene_gn=PHACTR1;refGene_tx=XM_019830231.2;refGene_gn=PHACTR1;refGene_tx=XM_019830232.2;refGene_gn

=PHACTR1;refGene_tx=XM_019830233.2
B2 16828017 16828120
refGene_gn=EDN1;refGene_tx=NM_001009386.1;refGene_gn=EDN1;refGene_tx=XM_011282148.3
B2 16829749 16829863
refGene_gn=EDN1;refGene_tx=NM_001009386.1;refGene_gn=EDN1;refGene_tx=XM_011282148.3
B2 16830020 16830179
refGene_gn=EDN1;refGene_tx=NM_001009386.1;refGene_gn=EDN1;refGene_tx=XM_011282148.3
B2 16831604 16831773
refGene_gn=EDN1;refGene_tx=NM_001009386.1;refGene_gn=EDN1;refGene_tx=XM_011282148.3
B2 16833303 16833367
refGene_gn=EDN1;refGene_tx=NM_001009386.1;refGene_gn=EDN1;refGene_tx=XM_011282148.3
B2 16952299 16952584 refGene_gn=HIVEP1;refGene_tx=XM_023253685.1
B2 16952585 16952855 refGene_gn=HIVEP1;refGene_tx=XM_023253685.1
B2 16952856 16953480 refGene_gn=HIVEP1;refGene_tx=XM_023253685.1
B2 16954745 16955257 refGene_gn=HIVEP1;refGene_tx=XM_023253685.1
B2 16976278 16976380 refGene_gn=HIVEP1;refGene_tx=XM_023253685.1
B2 16980987 16981163 refGene_gn=HIVEP1;refGene_tx=XM_023253685.1
B2 16981879 16982013 refGene_gn=HIVEP1;refGene_tx=XM_023253685.1
B2 16985314 16991268 refGene_gn=HIVEP1;refGene_tx=XM_023253685.1
B2 17022863 17022917 refGene_gn=HIVEP1;refGene_tx=XM_023253685.1
B2 17095019 17095059 refGene_gn=HIVEP1;refGene_tx=XM_023253685.1
B2 17314267 17314420 refGene_gn=ADTRP;refGene_tx=XM_006931454.3
B2 17324723 17324858 refGene_gn=ADTRP;refGene_tx=XM_006931454.3
B2 17326253 17326355 refGene_gn=ADTRP;refGene_tx=XM_006931454.3
B2 17357500 17357616 refGene_gn=ADTRP;refGene_tx=XM_006931454.3
B2 17371137 17371289 refGene_gn=ADTRP;refGene_tx=XM_006931454.3
B2 17379974 17380009 refGene_gn=ADTRP;refGene_tx=XM_006931454.3
B2 17480005 17480096
refGene_gn=LOC109500091;refGene_tx=XM_019831565.1
B2 17480435 17480580
refGene_gn=LOC109500091;refGene_tx=XM_019831565.1
B2 17481364 17481509
refGene_gn=LOC109500091;refGene_tx=XM_019831565.1
B2 17481698 17481808
refGene_gn=LOC109500091;refGene_tx=XM_019831565.1
B2 17482492 17482649
refGene_gn=LOC109500091;refGene_tx=XM_019831565.1
B2 17509768 17509899
refGene_gn=TMEM170B;refGene_tx=XM_003985801.5;refGene_gn=TMEM170B;refGene_tx=XM_023253686.1
B2 17520542 17520713 refGene_gn=TMEM170B;refGene_tx=XM_003985801.5
B2 17538643 17538962
refGene_gn=TMEM170B;refGene_tx=XM_003985801.5;refGene_gn=TMEM170B;refGene_tx=XM_023253686.1
B2 17732058 17732070 refGene_gn=NEDD9;refGene_tx=XM_019830242.2

B2 17801544 17801556 refGene_gn=NEDD9;refGene_tx=XM_003985802.3
B2 17820834 17821281
refGene_gn=NEDD9;refGene_tx=XM_003985802.3;refGene_gn=NEDD9;refGene_tx
=XM_019830242.2
B2 17841747 17841849
refGene_gn=NEDD9;refGene_tx=XM_003985802.3;refGene_gn=NEDD9;refGene_tx
=XM_019830242.2
B2 17842598 17842700
refGene_gn=NEDD9;refGene_tx=XM_003985802.3;refGene_gn=NEDD9;refGene_tx
=XM_019830242.2
B2 17843653 17844889
refGene_gn=NEDD9;refGene_tx=XM_003985802.3;refGene_gn=NEDD9;refGene_tx
=XM_019830242.2
B2 17846182 17846272
refGene_gn=NEDD9;refGene_tx=XM_003985802.3;refGene_gn=NEDD9;refGene_tx
=XM_019830242.2
B2 17848345 17848855
refGene_gn=NEDD9;refGene_tx=XM_003985802.3;refGene_gn=NEDD9;refGene_tx
=XM_019830242.2
B2 17897759 17897947 refGene_gn=SMIM13;refGene_tx=XM_006931455.4
B2 17920854 17920930 refGene_gn=SMIM13;refGene_tx=XM_006931455.4
B2 17965616 17965619 refGene_gn=ELOVL2;refGene_tx=XM_023253687.1
B2 18010838 18010868 refGene_gn=ELOVL2;refGene_tx=XM_011282182.3
B2 18010982 18011046
refGene_gn=ELOVL2;refGene_tx=XM_011282182.3;refGene_gn=ELOVL2;refGene_
tx=XM_023253687.1
B2 18015657 18015845
refGene_gn=ELOVL2;refGene_tx=XM_011282182.3;refGene_gn=ELOVL2;refGene_
tx=XM_023253687.1
B2 18020156 18020234
refGene_gn=ELOVL2;refGene_tx=XM_011282182.3;refGene_gn=ELOVL2;refGene_
tx=XM_023253687.1
B2 18023844 18024016
refGene_gn=ELOVL2;refGene_tx=XM_011282182.3;refGene_gn=ELOVL2;refGene_
tx=XM_023253687.1
B2 18026888 18027013
refGene_gn=ELOVL2;refGene_tx=XM_011282182.3;refGene_gn=ELOVL2;refGene_
tx=XM_023253687.1
B2 18027489 18027624
refGene_gn=ELOVL2;refGene_tx=XM_011282182.3;refGene_gn=ELOVL2;refGene_
tx=XM_023253687.1
B2 18029410 18029536
refGene_gn=ELOVL2;refGene_tx=XM_011282182.3;refGene_gn=ELOVL2;refGene_
tx=XM_023253687.1
B2 18049581 18049606 refGene_gn=SYCP2L;refGene_tx=XM_019830249.2
B2 18056578 18056637 refGene_gn=SYCP2L;refGene_tx=XM_019830249.2
B2 18056731 18056831 refGene_gn=SYCP2L;refGene_tx=XM_019830249.2
B2 18058846 18058938 refGene_gn=SYCP2L;refGene_tx=XM_019830249.2
B2 18059832 18059939 refGene_gn=SYCP2L;refGene_tx=XM_019830249.2
B2 18061023 18061125 refGene_gn=SYCP2L;refGene_tx=XM_019830249.2

B2 18075012 18075082 refGene_gn=SYCP2L;refGene_tx=XM_019830249.2
B2 18075234 18075308 refGene_gn=SYCP2L;refGene_tx=XM_019830249.2
B2 18076322 18076452 refGene_gn=SYCP2L;refGene_tx=XM_019830249.2
B2 18079036 18079086 refGene_gn=SYCP2L;refGene_tx=XM_019830249.2
B2 18079705 18079853 refGene_gn=SYCP2L;refGene_tx=XM_019830249.2
B2 18081361 18081410 refGene_gn=SYCP2L;refGene_tx=XM_019830249.2
B2 18108195 18108274 refGene_gn=SYCP2L;refGene_tx=XM_019830249.2
B2 18108371 18108429 refGene_gn=SYCP2L;refGene_tx=XM_019830249.2
B2 18109170 18109271 refGene_gn=SYCP2L;refGene_tx=XM_019830249.2
B2 18109779 18109873 refGene_gn=SYCP2L;refGene_tx=XM_019830249.2
B2 18112009 18112155 refGene_gn=SYCP2L;refGene_tx=XM_019830249.2
B2 18117404 18117465 refGene_gn=SYCP2L;refGene_tx=XM_019830249.2
B2 18117563 18117656 refGene_gn=SYCP2L;refGene_tx=XM_019830249.2
B2 18119555 18119601 refGene_gn=SYCP2L;refGene_tx=XM_019830249.2
B2 18120205 18120258 refGene_gn=SYCP2L;refGene_tx=XM_019830249.2
B2 18122523 18122666 refGene_gn=SYCP2L;refGene_tx=XM_019830249.2
B2 18123516 18123551 refGene_gn=SYCP2L;refGene_tx=XM_019830249.2
B2 18125980 18126069 refGene_gn=SYCP2L;refGene_tx=XM_019830249.2
B2 18126186 18126272 refGene_gn=SYCP2L;refGene_tx=XM_019830249.2
B2 18137584 18137609 refGene_gn=SYCP2L;refGene_tx=XM_019830249.2
B2 18138878 18138983 refGene_gn=SYCP2L;refGene_tx=XM_019830249.2
B2 18150814 18150904 refGene_gn=GCM2;refGene_tx=XM_003985808.2
B2 18154967 18155220 refGene_gn=GCM2;refGene_tx=XM_003985808.2
B2 18155333 18155446 refGene_gn=GCM2;refGene_tx=XM_003985808.2
B2 18155842 18155968 refGene_gn=GCM2;refGene_tx=XM_003985808.2
B2 18157625 18158561 refGene_gn=GCM2;refGene_tx=XM_003985808.2
B2 18195344 18195445
refGene_gn=MAK;refGene_tx=XM_006931459.3;refGene_gn=MAK;refGene_tx=XM_019830251.2;refGene_gn=MAK;refGene_tx=XM_019830252.1;refGene_gn=MAK;refGene_tx=XM_019830253.2;refGene_gn=MAK;refGene_tx=XM_019830254.1;refGene_gn=MAK;refGene_tx=XM_019830255.2;refGene_gn=MAK;refGene_tx=XM_019830256.2;refGene_gn=MAK;refGene_tx=XM_019830257.2;refGene_gn=MAK;refGene_tx=XM_019830258.2;refGene_gn=MAK;refGene_tx=XM_019830259.2;refGene_gn=MAK;refGene_tx=XM_019830261.2;refGene_gn=MAK;refGene_tx=XM_023253688.1
B2 18203528 18203583
refGene_gn=MAK;refGene_tx=XM_006931459.3;refGene_gn=MAK;refGene_tx=XM_019830251.2;refGene_gn=MAK;refGene_tx=XM_019830252.1;refGene_gn=MAK;refGene_tx=XM_019830253.2;refGene_gn=MAK;refGene_tx=XM_019830254.1;refGene_gn=MAK;refGene_tx=XM_019830255.2;refGene_gn=MAK;refGene_tx=XM_019830256.2;refGene_gn=MAK;refGene_tx=XM_019830257.2;refGene_gn=MAK;refGene_tx=XM_019830258.2;refGene_gn=MAK;refGene_tx=XM_019830259.2;refGene_gn=MAK;refGene_tx=XM_019830261.2;refGene_gn=MAK;refGene_tx=XM_023253688.1
B2 18204245 18204367
refGene_gn=MAK;refGene_tx=XM_006931459.3;refGene_gn=MAK;refGene_tx=XM_019830251.2;refGene_gn=MAK;refGene_tx=XM_019830252.1;refGene_gn=MAK;refGene_tx=XM_019830253.2;refGene_gn=MAK;refGene_tx=XM_019830254.1;refGene_gn=MAK;refGene_tx=XM_019830255.2;refGene_gn=MAK;refGene_tx=XM_019830256.2;refGene_gn=MAK;refGene_tx=XM_019830257.2;refGene_gn=MAK;refGene

_tx=XM_019830258.2;refGene_gn=MAK;refGene_tx=XM_019830259.2;refGene_gn=MAK;refGene_tx=XM_019830261.2;refGene_gn=MAK;refGene_tx=XM_023253688.1
B2 18208728 18208808
refGene_gn=MAK;refGene_tx=XM_006931459.3;refGene_gn=MAK;refGene_tx=XM_019830251.2;refGene_gn=MAK;refGene_tx=XM_019830252.1;refGene_gn=MAK;refGene_tx=XM_019830253.2;refGene_gn=MAK;refGene_tx=XM_019830254.1;refGene_gn=MAK;refGene_tx=XM_019830255.2;refGene_gn=MAK;refGene_tx=XM_019830256.2;refGene_gn=MAK;refGene_tx=XM_019830257.2;refGene_gn=MAK;refGene_tx=XM_019830258.2;refGene_gn=MAK;refGene_tx=XM_019830259.2;refGene_gn=MAK;refGene_tx=XM_019830261.2;refGene_gn=MAK;refGene_tx=XM_023253688.1
B2 18211636 18211769
refGene_gn=MAK;refGene_tx=XM_006931459.3;refGene_gn=MAK;refGene_tx=XM_019830251.2;refGene_gn=MAK;refGene_tx=XM_019830252.1;refGene_gn=MAK;refGene_tx=XM_019830253.2;refGene_gn=MAK;refGene_tx=XM_019830254.1;refGene_gn=MAK;refGene_tx=XM_019830255.2;refGene_gn=MAK;refGene_tx=XM_019830256.2;refGene_gn=MAK;refGene_tx=XM_019830257.2;refGene_gn=MAK;refGene_tx=XM_019830258.2;refGene_gn=MAK;refGene_tx=XM_019830259.2;refGene_gn=MAK;refGene_tx=XM_019830261.2;refGene_gn=MAK;refGene_tx=XM_023253688.1
B2 18215274 18215446
refGene_gn=MAK;refGene_tx=XM_006931459.3;refGene_gn=MAK;refGene_tx=XM_019830251.2;refGene_gn=MAK;refGene_tx=XM_019830252.1;refGene_gn=MAK;refGene_tx=XM_019830253.2;refGene_gn=MAK;refGene_tx=XM_019830254.1;refGene_gn=MAK;refGene_tx=XM_019830255.2;refGene_gn=MAK;refGene_tx=XM_019830256.2;refGene_gn=MAK;refGene_tx=XM_019830257.2;refGene_gn=MAK;refGene_tx=XM_019830258.2;refGene_gn=MAK;refGene_tx=XM_019830259.2;refGene_gn=MAK;refGene_tx=XM_019830261.2;refGene_gn=MAK;refGene_tx=XM_023253688.1
B2 18216435 18216603
refGene_gn=MAK;refGene_tx=XM_006931459.3;refGene_gn=MAK;refGene_tx=XM_019830251.2;refGene_gn=MAK;refGene_tx=XM_019830252.1;refGene_gn=MAK;refGene_tx=XM_019830253.2;refGene_gn=MAK;refGene_tx=XM_019830254.1;refGene_gn=MAK;refGene_tx=XM_019830255.2;refGene_gn=MAK;refGene_tx=XM_019830256.2;refGene_gn=MAK;refGene_tx=XM_019830257.2;refGene_gn=MAK;refGene_tx=XM_019830258.2;refGene_gn=MAK;refGene_tx=XM_019830259.2;refGene_gn=MAK;refGene_tx=XM_019830261.2;refGene_gn=MAK;refGene_tx=XM_023253688.1
B2 18222621 18222927
refGene_gn=MAK;refGene_tx=XM_006931459.3;refGene_gn=MAK;refGene_tx=XM_019830251.2;refGene_gn=MAK;refGene_tx=XM_019830252.1;refGene_gn=MAK;refGene_tx=XM_019830253.2;refGene_gn=MAK;refGene_tx=XM_019830254.1;refGene_gn=MAK;refGene_tx=XM_019830255.2;refGene_gn=MAK;refGene_tx=XM_019830256.2;refGene_gn=MAK;refGene_tx=XM_019830257.2;refGene_gn=MAK;refGene_tx=XM_019830258.2;refGene_gn=MAK;refGene_tx=XM_019830259.2;refGene_gn=MAK;refGene_tx=XM_019830261.2;refGene_gn=MAK;refGene_tx=XM_023253688.1
B2 18227305 18227478
refGene_gn=MAK;refGene_tx=XM_006931459.3;refGene_gn=MAK;refGene_tx=XM_

019830251.2;refGene_gn=MAK;refGene_tx=XM_019830252.1;refGene_gn=MAK;refGene_tx=XM_019830253.2;refGene_gn=MAK;refGene_tx=XM_019830254.1;refGene_gn=MAK;refGene_tx=XM_019830255.2;refGene_gn=MAK;refGene_tx=XM_019830256.2;refGene_gn=MAK;refGene_tx=XM_019830257.2;refGene_gn=MAK;refGene_tx=XM_019830258.2;refGene_gn=MAK;refGene_tx=XM_019830259.2;refGene_gn=MAK;refGene_tx=XM_019830261.2;refGene_gn=MAK;refGene_tx=XM_023253688.1
B2 18230360 18230509
refGene_gn=MAK;refGene_tx=XM_006931459.3;refGene_gn=MAK;refGene_tx=XM_019830251.2;refGene_gn=MAK;refGene_tx=XM_019830252.1;refGene_gn=MAK;refGene_tx=XM_019830253.2;refGene_gn=MAK;refGene_tx=XM_019830254.1;refGene_gn=MAK;refGene_tx=XM_019830255.2;refGene_gn=MAK;refGene_tx=XM_019830256.2;refGene_gn=MAK;refGene_tx=XM_019830257.2;refGene_gn=MAK;refGene_tx=XM_019830258.2;refGene_gn=MAK;refGene_tx=XM_019830259.2;refGene_gn=MAK;refGene_tx=XM_019830261.2;refGene_gn=MAK;refGene_tx=XM_023253688.1
B2 18235544 18235676
refGene_gn=MAK;refGene_tx=XM_006931459.3;refGene_gn=MAK;refGene_tx=XM_019830251.2;refGene_gn=MAK;refGene_tx=XM_019830252.1;refGene_gn=MAK;refGene_tx=XM_019830253.2;refGene_gn=MAK;refGene_tx=XM_019830254.1;refGene_gn=MAK;refGene_tx=XM_019830255.2;refGene_gn=MAK;refGene_tx=XM_019830256.2;refGene_gn=MAK;refGene_tx=XM_019830257.2;refGene_gn=MAK;refGene_tx=XM_019830258.2;refGene_gn=MAK;refGene_tx=XM_019830259.2;refGene_gn=MAK;refGene_tx=XM_019830261.2;refGene_gn=MAK;refGene_tx=XM_023253688.1
B2 18238342 18238417
refGene_gn=MAK;refGene_tx=XM_019830251.2;refGene_gn=MAK;refGene_tx=XM_019830252.1;refGene_gn=MAK;refGene_tx=XM_019830253.2;refGene_gn=MAK;refGene_tx=XM_019830254.1;refGene_gn=MAK;refGene_tx=XM_019830255.2;refGene_gn=MAK;refGene_tx=XM_023253688.1
B2 18238511 18238543 refGene_gn=MAK;refGene_tx=XM_019830259.2
B2 18240896 18241016
refGene_gn=MAK;refGene_tx=XM_006931459.3;refGene_gn=MAK;refGene_tx=XM_019830251.2;refGene_gn=MAK;refGene_tx=XM_019830252.1;refGene_gn=MAK;refGene_tx=XM_019830253.2;refGene_gn=MAK;refGene_tx=XM_019830254.1;refGene_gn=MAK;refGene_tx=XM_019830255.2;refGene_gn=MAK;refGene_tx=XM_019830256.2;refGene_gn=MAK;refGene_tx=XM_023253688.1
B2 18242331 18242361
refGene_gn=MAK;refGene_tx=XM_019830251.2;refGene_gn=MAK;refGene_tx=XM_019830252.1;refGene_gn=MAK;refGene_tx=XM_019830253.2;refGene_gn=MAK;refGene_tx=XM_019830254.1;refGene_gn=MAK;refGene_tx=XM_019830256.2;refGene_gn=MAK;refGene_tx=XM_023253688.1
B2 18244828 18244983
refGene_gn=MAK;refGene_tx=XM_006931459.3;refGene_gn=MAK;refGene_tx=XM_019830251.2;refGene_gn=MAK;refGene_tx=XM_019830252.1;refGene_gn=MAK;refGene_tx=XM_019830253.2;refGene_gn=MAK;refGene_tx=XM_019830254.1;refGene_gn=MAK;refGene_tx=XM_019830255.2;refGene_gn=MAK;refGene_tx=XM_019830256.2;refGene_gn=MAK;refGene_tx=XM_019830257.2;refGene_gn=MAK;refGene_tx=XM_023253688.1
B2 18260763 18260855 refGene_gn=MAK;refGene_tx=XM_019830258.2

B2 18263176 18263231 refGene_gn=TMEM14C;refGene_tx=XM_003985812.3
B2 18264175 18264263 refGene_gn=TMEM14C;refGene_tx=XM_003985812.3
B2 18267350 18267452 refGene_gn=TMEM14C;refGene_tx=XM_003985812.3
B2 18268298 18268375 refGene_gn=TMEM14C;refGene_tx=XM_003985812.3
B2 18268693 18268716 refGene_gn=TMEM14C;refGene_tx=XM_003985812.3
B2 18277735 18277935
refGene_gn=PAK1IP1;refGene_tx=XM_003985813.5;refGene_gn=PAK1IP1;refGene_tx=XM_023253689.1
B2 18278316 18278437
refGene_gn=PAK1IP1;refGene_tx=XM_003985813.5;refGene_gn=PAK1IP1;refGene_tx=XM_023253689.1
B2 18279420 18279520
refGene_gn=PAK1IP1;refGene_tx=XM_003985813.5;refGene_gn=PAK1IP1;refGene_tx=XM_023253689.1
B2 18281862 18281960
refGene_gn=PAK1IP1;refGene_tx=XM_003985813.5;refGene_gn=PAK1IP1;refGene_tx=XM_023253689.1
B2 18282054 18282200
refGene_gn=PAK1IP1;refGene_tx=XM_003985813.5;refGene_gn=PAK1IP1;refGene_tx=XM_023253689.1
B2 18282604 18282657
refGene_gn=PAK1IP1;refGene_tx=XM_003985813.5;refGene_gn=PAK1IP1;refGene_tx=XM_023253689.1
B2 18283190 18283265
refGene_gn=PAK1IP1;refGene_tx=XM_003985813.5;refGene_gn=PAK1IP1;refGene_tx=XM_023253689.1
B2 18283340 18283461
refGene_gn=PAK1IP1;refGene_tx=XM_003985813.5;refGene_gn=PAK1IP1;refGene_tx=XM_023253689.1
B2 18286952 18287118
refGene_gn=PAK1IP1;refGene_tx=XM_003985813.5;refGene_gn=PAK1IP1;refGene_tx=XM_023253689.1
B2 18289380 18289464 refGene_gn=PAK1IP1;refGene_tx=XM_003985813.5
B2 18294667 18294738
refGene_gn=CB2H6orf52;refGene_tx=XM_003985816.3
B2 18294958 18295136
refGene_gn=CB2H6orf52;refGene_tx=XM_003985816.3
B2 18299546 18299592
refGene_gn=CB2H6orf52;refGene_tx=XM_003985816.3
B2 18302970 18303119
refGene_gn=CB2H6orf52;refGene_tx=XM_003985816.3
B2 18343365 18343556
refGene_gn=GCNT2;refGene_tx=XM_003985814.5;refGene_gn=GCNT2;refGene_tx=XM_023253690.1
B2 18348653 18348746
refGene_gn=GCNT2;refGene_tx=XM_003985814.5;refGene_gn=GCNT2;refGene_tx=XM_023253690.1
B2 18372057 18372982 refGene_gn=GCNT2;refGene_tx=XM_003985814.5
B2 18396894 18397816 refGene_gn=GCNT2;refGene_tx=XM_023253690.1
B2 18424051 18424122

refGene_gn=LOC109491352;refGene_tx=XM_006931463.3
B2	18424283	18425208
refGene_gn=LOC109491352;refGene_tx=XM_006931463.3
B2	18557335	18557386	refGene_gn=TFAP2A;refGene_tx=XM_023253691.1
B2	18562041	18562355	refGene_gn=TFAP2A;refGene_tx=XM_023253691.1
B2	18562356	18562477	refGene_gn=TFAP2A;refGene_tx=XM_023253691.1
B2	18565605	18565657	refGene_gn=TFAP2A;refGene_tx=XM_023253691.1
B2	18567745	18567977	refGene_gn=TFAP2A;refGene_tx=XM_023253691.1
B2	18569892	18570011	refGene_gn=TFAP2A;refGene_tx=XM_023253691.1
B2	18571946	18572088	refGene_gn=TFAP2A;refGene_tx=XM_023253691.1
B2	18573937	18574226	refGene_gn=TFAP2A;refGene_tx=XM_023253691.1
B2	19003534	19003546
refGene_gn=LOC101096891;refGene_tx=XM_023254704.1
B2	19009518	19009585
refGene_gn=LOC101096891;refGene_tx=XM_023254704.1
B2	19009884	19010061
refGene_gn=LOC101096891;refGene_tx=XM_023254704.1
B2	19030687	19030774
refGene_gn=LOC101096891;refGene_tx=XM_023254704.1
B2	19037658	19037739
refGene_gn=LOC101096891;refGene_tx=XM_023254704.1
B2	19041214	19041360
refGene_gn=LOC101096891;refGene_tx=XM_023254704.1
B2	19079240	19079317
refGene_gn=LOC101096891;refGene_tx=XM_023254704.1
B2	19082382	19082506
refGene_gn=LOC101096891;refGene_tx=XM_023254704.1
B2	19085088	19085322
refGene_gn=LOC101096891;refGene_tx=XM_023254704.1
B2	19110869	19111004
refGene_gn=LOC101096891;refGene_tx=XM_023254704.1
B2	19112011	19112275
refGene_gn=LOC101096891;refGene_tx=XM_023254704.1
B2	19115938	19116116
refGene_gn=LOC101096891;refGene_tx=XM_023254704.1
B2	19161563	19161646
refGene_gn=LOC101096891;refGene_tx=XM_023254704.1
B2	19171750	19171875
refGene_gn=LOC101096891;refGene_tx=XM_023254704.1
B2	19220922	19221141
refGene_gn=LOC101096891;refGene_tx=XM_023254704.1
B2	19223429	19223682
refGene_gn=LOC101096891;refGene_tx=XM_023254704.1
B2	19229390	19229554
refGene_gn=LOC101096891;refGene_tx=XM_023254704.1
B2	19229586	19229665
refGene_gn=LOC101096891;refGene_tx=XM_023254704.1
B2	20520763	20520909	refGene_gn=SLC35B3;refGene_tx=XM_023254705.1
B2	20521049	20521182	refGene_gn=SLC35B3;refGene_tx=XM_023254705.1
B2	20527216	20527510	refGene_gn=SLC35B3;refGene_tx=XM_023254705.1

B2 20529734 20529856 refGene_gn=SLC35B3;refGene_tx=XM_023254705.1
B2 20533395 20533550 refGene_gn=SLC35B3;refGene_tx=XM_023254705.1
B2 20534827 20534935 refGene_gn=SLC35B3;refGene_tx=XM_023254705.1
B2 20539932 20540030 refGene_gn=SLC35B3;refGene_tx=XM_023254705.1
B2 20541995 20542088 refGene_gn=SLC35B3;refGene_tx=XM_023254705.1
B2 20542496 20542608 refGene_gn=SLC35B3;refGene_tx=XM_023254705.1
B2 20545843 20545913 refGene_gn=SLC35B3;refGene_tx=XM_023254705.1
B2 20547231 20547382 refGene_gn=SLC35B3;refGene_tx=XM_023254705.1
B2 20831145 20831232
refGene_gn=EEF1E1;refGene_tx=XM_003985822.5;refGene_gn=EEF1E1;refGene_tx=XM_019830265.2;refGene_gn=EEF1E1;refGene_tx=XM_019830266.2
B2 20832542 20832564 refGene_gn=EEF1E1;refGene_tx=XM_023253692.1
B2 20833562 20833594 refGene_gn=EEF1E1;refGene_tx=XM_023253692.1
B2 20837253 20837454
refGene_gn=EEF1E1;refGene_tx=XM_003985822.5;refGene_gn=EEF1E1;refGene_tx=XM_019830265.2;refGene_gn=EEF1E1;refGene_tx=XM_019830266.2;refGene_gn=EEF1E1;refGene_tx=XM_023253692.1
B2 20844774 20844870
refGene_gn=EEF1E1;refGene_tx=XM_003985822.5;refGene_gn=EEF1E1;refGene_tx=XM_019830265.2;refGene_gn=EEF1E1;refGene_tx=XM_019830266.2;refGene_gn=EEF1E1;refGene_tx=XM_023253692.1
B2 20852206 20852335 refGene_gn=EEF1E1;refGene_tx=XM_019830266.2
B2 20853732 20853873
refGene_gn=EEF1E1;refGene_tx=XM_003985822.5;refGene_gn=EEF1E1;refGene_tx=XM_023253692.1
B2 20866926 20867058 refGene_gn=EEF1E1;refGene_tx=XM_019830265.2
B2 20867143 20867258 refGene_gn=BLOC1S5;refGene_tx=XM_003985823.5
B2 20874021 20874104 refGene_gn=BLOC1S5;refGene_tx=XM_003985823.5
B2 20887864 20887994 refGene_gn=BLOC1S5;refGene_tx=XM_003985823.5
B2 20896007 20896066 refGene_gn=BLOC1S5;refGene_tx=XM_003985823.5
B2 20910645 20910825 refGene_gn=BLOC1S5;refGene_tx=XM_003985823.5
B2 21014881 21015159 refGene_gn=TXNDC5;refGene_tx=XM_023253694.1
B2 21019284 21019434 refGene_gn=TXNDC5;refGene_tx=XM_023253694.1
B2 21024025 21024131 refGene_gn=TXNDC5;refGene_tx=XM_023253694.1
B2 21028089 21028186 refGene_gn=TXNDC5;refGene_tx=XM_023253694.1
B2 21030444 21030560 refGene_gn=TXNDC5;refGene_tx=XM_023253694.1
B2 21032741 21032828 refGene_gn=TXNDC5;refGene_tx=XM_023253694.1
B2 21033758 21033899 refGene_gn=TXNDC5;refGene_tx=XM_023253694.1
B2 21036911 21036994 refGene_gn=TXNDC5;refGene_tx=XM_023253694.1
B2 21038465 21038595 refGene_gn=TXNDC5;refGene_tx=XM_023253694.1
B2 21039830 21039953 refGene_gn=TXNDC5;refGene_tx=XM_023253694.1
B2 21042792 21042942 refGene_gn=BMP6;refGene_tx=XM_023253693.1
B2 21043270 21043381 refGene_gn=BMP6;refGene_tx=XM_023253693.1
B2 21044338 21044415 refGene_gn=BMP6;refGene_tx=XM_023253693.1
B2 21056800 21056998 refGene_gn=BMP6;refGene_tx=XM_023253693.1
B2 21057861 21058010 refGene_gn=BMP6;refGene_tx=XM_023253693.1
B2 21073202 21073395 refGene_gn=BMP6;refGene_tx=XM_023253693.1
B2 21197843 21198474 refGene_gn=BMP6;refGene_tx=XM_023253693.1
B2 21347371 21347420
refGene_gn=SNRNP48;refGene_tx=XM_011282175.3;refGene_gn=SNRNP48;refGen

e_tx=XM_011282176.3;refGene_gn=SNRNP48;refGene_tx=XM_019830269.2;refGene_gn=SNRNP48;refGene_tx=XM_019830270.2;refGene_gn=SNRNP48;refGene_tx=XM_019830271.2
B2 21349663 21349828
refGene_gn=SNRNP48;refGene_tx=XM_011282175.3;refGene_gn=SNRNP48;refGene_tx=XM_011282176.3;refGene_gn=SNRNP48;refGene_tx=XM_019830269.2;refGene_gn=SNRNP48;refGene_tx=XM_019830270.2;refGene_gn=SNRNP48;refGene_tx=XM_019830271.2
B2 21350677 21350766
refGene_gn=SNRNP48;refGene_tx=XM_011282175.3;refGene_gn=SNRNP48;refGene_tx=XM_011282176.3;refGene_gn=SNRNP48;refGene_tx=XM_019830269.2;refGene_gn=SNRNP48;refGene_tx=XM_019830270.2;refGene_gn=SNRNP48;refGene_tx=XM_019830271.2
B2 21354216 21354338
refGene_gn=SNRNP48;refGene_tx=XM_011282175.3;refGene_gn=SNRNP48;refGene_tx=XM_011282176.3;refGene_gn=SNRNP48;refGene_tx=XM_019830269.2;refGene_gn=SNRNP48;refGene_tx=XM_019830270.2;refGene_gn=SNRNP48;refGene_tx=XM_019830271.2
B2 21354782 21354971
refGene_gn=SNRNP48;refGene_tx=XM_011282175.3;refGene_gn=SNRNP48;refGene_tx=XM_011282176.3;refGene_gn=SNRNP48;refGene_tx=XM_019830269.2;refGene_gn=SNRNP48;refGene_tx=XM_019830270.2;refGene_gn=SNRNP48;refGene_tx=XM_019830271.2
B2 21360050 21360125
refGene_gn=SNRNP48;refGene_tx=XM_011282175.3;refGene_gn=SNRNP48;refGene_tx=XM_011282176.3;refGene_gn=SNRNP48;refGene_tx=XM_019830269.2;refGene_gn=SNRNP48;refGene_tx=XM_019830270.2;refGene_gn=SNRNP48;refGene_tx=XM_019830271.2
B2 21361172 21361233
refGene_gn=SNRNP48;refGene_tx=XM_011282175.3;refGene_gn=SNRNP48;refGene_tx=XM_011282176.3;refGene_gn=SNRNP48;refGene_tx=XM_019830269.2;refGene_gn=SNRNP48;refGene_tx=XM_019830270.2;refGene_gn=SNRNP48;refGene_tx=XM_019830271.2
B2 21361484 21361598
refGene_gn=SNRNP48;refGene_tx=XM_011282175.3;refGene_gn=SNRNP48;refGene_tx=XM_011282176.3;refGene_gn=SNRNP48;refGene_tx=XM_019830269.2;refGene_gn=SNRNP48;refGene_tx=XM_019830270.2;refGene_gn=SNRNP48;refGene_tx=XM_019830271.2
B2 21364927 21365164
refGene_gn=SNRNP48;refGene_tx=XM_011282175.3;refGene_gn=SNRNP48;refGene_tx=XM_011282176.3;refGene_gn=SNRNP48;refGene_tx=XM_019830269.2;refGene_gn=SNRNP48;refGene_tx=XM_019830270.2;refGene_gn=SNRNP48;refGene_tx=XM_019830271.2
B2 21370238 21373487
refGene_gn=DSP;refGene_tx=XM_003985824.5;refGene_gn=DSP;refGene_tx=XM_006931468.4
B2 21374478 21376773
refGene_gn=DSP;refGene_tx=XM_003985824.5;refGene_gn=DSP;refGene_tx=XM_006931468.4
B2 21378074 21378173

refGene_gn=DSP;refGene_tx=XM_003985824.5;refGene_gn=DSP;refGene_tx=XM_006931468.4
B2 21378979 21379087
refGene_gn=DSP;refGene_tx=XM_003985824.5;refGene_gn=DSP;refGene_tx=XM_006931468.4
B2 21379643 21379727
refGene_gn=DSP;refGene_tx=XM_003985824.5;refGene_gn=DSP;refGene_tx=XM_006931468.4
B2 21380411 21380574
refGene_gn=DSP;refGene_tx=XM_003985824.5;refGene_gn=DSP;refGene_tx=XM_006931468.4
B2 21381633 21381827
refGene_gn=DSP;refGene_tx=XM_003985824.5;refGene_gn=DSP;refGene_tx=XM_006931468.4
B2 21382360 21382499
refGene_gn=DSP;refGene_tx=XM_003985824.5;refGene_gn=DSP;refGene_tx=XM_006931468.4
B2 21382960 21383127
refGene_gn=DSP;refGene_tx=XM_003985824.5;refGene_gn=DSP;refGene_tx=XM_006931468.4
B2 21384007 21384234
refGene_gn=DSP;refGene_tx=XM_003985824.5;refGene_gn=DSP;refGene_tx=XM_006931468.4
B2 21384501 21384703
refGene_gn=DSP;refGene_tx=XM_003985824.5;refGene_gn=DSP;refGene_tx=XM_006931468.4
B2 21385486 21385613
refGene_gn=DSP;refGene_tx=XM_003985824.5;refGene_gn=DSP;refGene_tx=XM_006931468.4
B2 21388013 21388168
refGene_gn=DSP;refGene_tx=XM_003985824.5;refGene_gn=DSP;refGene_tx=XM_006931468.4
B2 21390761 21390914
refGene_gn=DSP;refGene_tx=XM_003985824.5;refGene_gn=DSP;refGene_tx=XM_006931468.4
B2 21391244 21391370
refGene_gn=DSP;refGene_tx=XM_003985824.5;refGene_gn=DSP;refGene_tx=XM_006931468.4
B2 21391945 21392041
refGene_gn=DSP;refGene_tx=XM_003985824.5;refGene_gn=DSP;refGene_tx=XM_006931468.4
B2 21393096 21393201
refGene_gn=DSP;refGene_tx=XM_003985824.5;refGene_gn=DSP;refGene_tx=XM_006931468.4
B2 21394010 21394172
refGene_gn=DSP;refGene_tx=XM_003985824.5;refGene_gn=DSP;refGene_tx=XM_006931468.4
B2 21395339 21395390
refGene_gn=DSP;refGene_tx=XM_003985824.5;refGene_gn=DSP;refGene_tx=XM_006931468.4

B2 21396580 21396709
refGene_gn=DSP;refGene_tx=XM_003985824.5;refGene_gn=DSP;refGene_tx=XM_006931468.4
B2 21400249 21400424
refGene_gn=DSP;refGene_tx=XM_003985824.5;refGene_gn=DSP;refGene_tx=XM_006931468.4
B2 21401425 21401574
refGene_gn=DSP;refGene_tx=XM_003985824.5;refGene_gn=DSP;refGene_tx=XM_006931468.4
B2 21403925 21404028
refGene_gn=DSP;refGene_tx=XM_003985824.5;refGene_gn=DSP;refGene_tx=XM_006931468.4
B2 21418367 21418564
refGene_gn=DSP;refGene_tx=XM_003985824.5;refGene_gn=DSP;refGene_tx=XM_006931468.4
B2 21528306 21528417
refGene_gn=RIOK1;refGene_tx=XM_011282174.3;refGene_gn=RIOK1;refGene_tx=XM_019830273.2
B2 21531705 21531858
refGene_gn=RIOK1;refGene_tx=XM_011282174.3;refGene_gn=RIOK1;refGene_tx=XM_019830273.2
B2 21533087 21533141
refGene_gn=RIOK1;refGene_tx=XM_011282174.3;refGene_gn=RIOK1;refGene_tx=XM_019830273.2
B2 21533968 21534088
refGene_gn=RIOK1;refGene_tx=XM_011282174.3;refGene_gn=RIOK1;refGene_tx=XM_019830273.2
B2 21536733 21536799
refGene_gn=RIOK1;refGene_tx=XM_011282174.3;refGene_gn=RIOK1;refGene_tx=XM_019830273.2
B2 21539111 21539218
refGene_gn=RIOK1;refGene_tx=XM_011282174.3;refGene_gn=RIOK1;refGene_tx=XM_019830273.2
B2 21539424 21539528
refGene_gn=RIOK1;refGene_tx=XM_011282174.3;refGene_gn=RIOK1;refGene_tx=XM_019830273.2
B2 21539888 21540026
refGene_gn=RIOK1;refGene_tx=XM_011282174.3;refGene_gn=RIOK1;refGene_tx=XM_019830273.2
B2 21540428 21540515
refGene_gn=RIOK1;refGene_tx=XM_011282174.3;refGene_gn=RIOK1;refGene_tx=XM_019830273.2
B2 21543066 21543147
refGene_gn=RIOK1;refGene_tx=XM_011282174.3;refGene_gn=RIOK1;refGene_tx=XM_019830273.2
B2 21543255 21543368
refGene_gn=RIOK1;refGene_tx=XM_011282174.3;refGene_gn=RIOK1;refGene_tx=XM_019830273.2
B2 21544627 21544720
refGene_gn=RIOK1;refGene_tx=XM_011282174.3;refGene_gn=RIOK1;refGene_tx

=XM_019830273.2
B2 21547625 21547668
refGene_gn=RIOK1;refGene_tx=XM_011282174.3;refGene_gn=RIOK1;refGene_tx=XM_019830273.2
B2 21548946 21549016
refGene_gn=RIOK1;refGene_tx=XM_011282174.3;refGene_gn=RIOK1;refGene_tx=XM_019830273.2
B2 21549819 21549907
refGene_gn=RIOK1;refGene_tx=XM_011282174.3;refGene_gn=RIOK1;refGene_tx=XM_019830273.2
B2 21551649 21551863
refGene_gn=RIOK1;refGene_tx=XM_011282174.3;refGene_gn=RIOK1;refGene_tx=XM_019830273.2
B2 21554922 21554993 refGene_gn=RIOK1;refGene_tx=XM_011282174.3
B2 21555497 21555510
refGene_gn=CAGE1;refGene_tx=XM_023253696.1;refGene_gn=CAGE1;refGene_tx=XM_023253697.1;refGene_gn=CAGE1;refGene_tx=XM_023253698.1;refGene_gn=CAGE1;refGene_tx=XM_023253699.1;refGene_gn=CAGE1;refGene_tx=XM_023253700.1;refGene_gn=CAGE1;refGene_tx=XM_023253701.1;refGene_gn=CAGE1;refGene_tx=XM_023253702.1;refGene_gn=CAGE1;refGene_tx=XM_023253703.1;refGene_gn=CAGE1;refGene_tx=XM_023253705.1
B2 21557195 21557413
refGene_gn=CAGE1;refGene_tx=XM_023253696.1;refGene_gn=CAGE1;refGene_tx=XM_023253697.1;refGene_gn=CAGE1;refGene_tx=XM_023253698.1;refGene_gn=CAGE1;refGene_tx=XM_023253699.1;refGene_gn=CAGE1;refGene_tx=XM_023253700.1;refGene_gn=CAGE1;refGene_tx=XM_023253701.1;refGene_gn=CAGE1;refGene_tx=XM_023253702.1;refGene_gn=CAGE1;refGene_tx=XM_023253703.1;refGene_gn=CAGE1;refGene_tx=XM_023253704.1;refGene_gn=CAGE1;refGene_tx=XM_023253705.1
B2 21561050 21561138
refGene_gn=CAGE1;refGene_tx=XM_023253696.1;refGene_gn=CAGE1;refGene_tx=XM_023253697.1;refGene_gn=CAGE1;refGene_tx=XM_023253698.1;refGene_gn=CAGE1;refGene_tx=XM_023253699.1;refGene_gn=CAGE1;refGene_tx=XM_023253700.1;refGene_gn=CAGE1;refGene_tx=XM_023253701.1;refGene_gn=CAGE1;refGene_tx=XM_023253702.1;refGene_gn=CAGE1;refGene_tx=XM_023253703.1;refGene_gn=CAGE1;refGene_tx=XM_023253704.1;refGene_gn=CAGE1;refGene_tx=XM_023253705.1
B2 21564125 21564649
refGene_gn=CAGE1;refGene_tx=XM_023253696.1;refGene_gn=CAGE1;refGene_tx=XM_023253697.1;refGene_gn=CAGE1;refGene_tx=XM_023253698.1;refGene_gn=CAGE1;refGene_tx=XM_023253699.1;refGene_gn=CAGE1;refGene_tx=XM_023253700.1;refGene_gn=CAGE1;refGene_tx=XM_023253701.1;refGene_gn=CAGE1;refGene_tx=XM_023253702.1;refGene_gn=CAGE1;refGene_tx=XM_023253703.1;refGene_gn=CAGE1;refGene_tx=XM_023253704.1;refGene_gn=CAGE1;refGene_tx=XM_023253705.1;refGene_gn=CAGE1;refGene_tx=XM_023253706.1
B2 21567389 21568442
refGene_gn=CAGE1;refGene_tx=XM_023253696.1;refGene_gn=CAGE1;refGene_tx=XM_023253697.1;refGene_gn=CAGE1;refGene_tx=XM_023253698.1;refGene_gn=CAGE1;refGene_tx=XM_023253699.1;refGene_gn=CAGE1;refGene_tx=XM_023253700.1;refGene_gn=CAGE1;refGene_tx=XM_023253701.1;refGene_gn=CAGE1;refGe

ne_tx=XM_023253702.1;refGene_gn=CAGE1;refGene_tx=XM_023253703.1;refGen
e_gn=CAGE1;refGene_tx=XM_023253704.1;refGene_gn=CAGE1;refGene_tx=XM_02
3253705.1;refGene_gn=CAGE1;refGene_tx=XM_023253706.1
B2 21571913 21572063
refGene_gn=CAGE1;refGene_tx=XM_023253696.1;refGene_gn=CAGE1;refGene_tx
=XM_023253697.1;refGene_gn=CAGE1;refGene_tx=XM_023253698.1;refGene_gn=
CAGE1;refGene_tx=XM_023253699.1;refGene_gn=CAGE1;refGene_tx=XM_0232537
00.1;refGene_gn=CAGE1;refGene_tx=XM_023253701.1;refGene_gn=CAGE1;refGe
ne_tx=XM_023253702.1;refGene_gn=CAGE1;refGene_tx=XM_023253703.1;refGen
e_gn=CAGE1;refGene_tx=XM_023253704.1;refGene_gn=CAGE1;refGene_tx=XM_02
3253705.1;refGene_gn=CAGE1;refGene_tx=XM_023253706.1
B2 21573913 21574024
refGene_gn=CAGE1;refGene_tx=XM_023253696.1;refGene_gn=CAGE1;refGene_tx
=XM_023253697.1;refGene_gn=CAGE1;refGene_tx=XM_023253698.1;refGene_gn=
CAGE1;refGene_tx=XM_023253699.1;refGene_gn=CAGE1;refGene_tx=XM_0232537
00.1;refGene_gn=CAGE1;refGene_tx=XM_023253701.1;refGene_gn=CAGE1;refGe
ne_tx=XM_023253702.1;refGene_gn=CAGE1;refGene_tx=XM_023253703.1;refGen
e_gn=CAGE1;refGene_tx=XM_023253704.1;refGene_gn=CAGE1;refGene_tx=XM_02
3253705.1;refGene_gn=CAGE1;refGene_tx=XM_023253706.1
B2 21576431 21576512
refGene_gn=CAGE1;refGene_tx=XM_023253696.1;refGene_gn=CAGE1;refGene_tx
=XM_023253697.1;refGene_gn=CAGE1;refGene_tx=XM_023253698.1;refGene_gn=
CAGE1;refGene_tx=XM_023253699.1;refGene_gn=CAGE1;refGene_tx=XM_0232537
00.1;refGene_gn=CAGE1;refGene_tx=XM_023253701.1;refGene_gn=CAGE1;refGe
ne_tx=XM_023253702.1;refGene_gn=CAGE1;refGene_tx=XM_023253703.1;refGen
e_gn=CAGE1;refGene_tx=XM_023253704.1;refGene_gn=CAGE1;refGene_tx=XM_02
3253705.1;refGene_gn=CAGE1;refGene_tx=XM_023253706.1
B2 21576740 21576848
refGene_gn=CAGE1;refGene_tx=XM_023253696.1;refGene_gn=CAGE1;refGene_tx
=XM_023253697.1;refGene_gn=CAGE1;refGene_tx=XM_023253698.1;refGene_gn=
CAGE1;refGene_tx=XM_023253699.1;refGene_gn=CAGE1;refGene_tx=XM_0232537
00.1;refGene_gn=CAGE1;refGene_tx=XM_023253701.1;refGene_gn=CAGE1;refGe
ne_tx=XM_023253702.1;refGene_gn=CAGE1;refGene_tx=XM_023253703.1;refGen
e_gn=CAGE1;refGene_tx=XM_023253704.1;refGene_gn=CAGE1;refGene_tx=XM_02
3253705.1;refGene_gn=CAGE1;refGene_tx=XM_023253706.1
B2 21591564 21591669
refGene_gn=CAGE1;refGene_tx=XM_023253696.1;refGene_gn=CAGE1;refGene_tx
=XM_023253697.1;refGene_gn=CAGE1;refGene_tx=XM_023253698.1;refGene_gn=
CAGE1;refGene_tx=XM_023253699.1;refGene_gn=CAGE1;refGene_tx=XM_0232537
00.1;refGene_gn=CAGE1;refGene_tx=XM_023253701.1;refGene_gn=CAGE1;refGe
ne_tx=XM_023253704.1;refGene_gn=CAGE1;refGene_tx=XM_023253705.1;refGen
e_gn=CAGE1;refGene_tx=XM_023253706.1
B2 21592291 21592362
refGene_gn=CAGE1;refGene_tx=XM_023253696.1;refGene_gn=CAGE1;refGene_tx
=XM_023253697.1;refGene_gn=CAGE1;refGene_tx=XM_023253698.1;refGene_gn=
CAGE1;refGene_tx=XM_023253699.1;refGene_gn=CAGE1;refGene_tx=XM_0232537
00.1;refGene_gn=CAGE1;refGene_tx=XM_023253701.1;refGene_gn=CAGE1;refGe
ne_tx=XM_023253702.1;refGene_gn=CAGE1;refGene_tx=XM_023253703.1;refGen
e_gn=CAGE1;refGene_tx=XM_023253704.1;refGene_gn=CAGE1;refGene_tx=XM_02
3253705.1;refGene_gn=CAGE1;refGene_tx=XM_023253706.1

B2	21592450	21592475
refGene_gn=CAGE1;refGene_tx=XM_023253700.1;refGene_gn=CAGE1;refGene_tx
=XM_023253703.1
B2	21606897	21606966
refGene_gn=CAGE1;refGene_tx=XM_023253696.1;refGene_gn=CAGE1;refGene_tx
=XM_023253697.1;refGene_gn=CAGE1;refGene_tx=XM_023253698.1;refGene_gn=
CAGE1;refGene_tx=XM_023253699.1;refGene_gn=CAGE1;refGene_tx=XM_0232537
01.1;refGene_gn=CAGE1;refGene_tx=XM_023253702.1;refGene_gn=CAGE1;refGe
ne_tx=XM_023253704.1;refGene_gn=CAGE1;refGene_tx=XM_023253705.1;refGen
e_gn=CAGE1;refGene_tx=XM_023253706.1
B2	21609394	21609410
refGene_gn=CAGE1;refGene_tx=XM_023253696.1;refGene_gn=CAGE1;refGene_tx
=XM_023253697.1;refGene_gn=CAGE1;refGene_tx=XM_023253698.1;refGene_gn=
CAGE1;refGene_tx=XM_023253699.1;refGene_gn=CAGE1;refGene_tx=XM_0232537
01.1;refGene_gn=CAGE1;refGene_tx=XM_023253702.1;refGene_gn=CAGE1;refGe
ne_tx=XM_023253704.1;refGene_gn=CAGE1;refGene_tx=XM_023253705.1;refGen
e_gn=CAGE1;refGene_tx=XM_023253706.1
B2	21620715	21620794
refGene_gn=SSR1;refGene_tx=XM_023253707.1;refGene_gn=SSR1;refGene_tx=X
M_023253708.1;refGene_gn=SSR1;refGene_tx=XM_023253709.1
B2	21623756	21623869
refGene_gn=SSR1;refGene_tx=XM_023253707.1;refGene_gn=SSR1;refGene_tx=X
M_023253708.1;refGene_gn=SSR1;refGene_tx=XM_023253709.1
B2	21627745	21627833
refGene_gn=SSR1;refGene_tx=XM_023253707.1;refGene_gn=SSR1;refGene_tx=X
M_023253708.1;refGene_gn=SSR1;refGene_tx=XM_023253709.1
B2	21629115	21629378
refGene_gn=SSR1;refGene_tx=XM_023253707.1;refGene_gn=SSR1;refGene_tx=X
M_023253708.1;refGene_gn=SSR1;refGene_tx=XM_023253709.1
B2	21631072	21631149
refGene_gn=SSR1;refGene_tx=XM_023253707.1;refGene_gn=SSR1;refGene_tx=X
M_023253708.1;refGene_gn=SSR1;refGene_tx=XM_023253709.1
B2	21631934	21632013
refGene_gn=SSR1;refGene_tx=XM_023253707.1;refGene_gn=SSR1;refGene_tx=X
M_023253708.1;refGene_gn=SSR1;refGene_tx=XM_023253709.1
B2	21634995	21635089
refGene_gn=SSR1;refGene_tx=XM_023253707.1;refGene_gn=SSR1;refGene_tx=X
M_023253708.1;refGene_gn=SSR1;refGene_tx=XM_023253709.1
B2	21635375	21635390	refGene_gn=SSR1;refGene_tx=XM_023253707.1
B2	21645051	21645119
refGene_gn=SSR1;refGene_tx=XM_023253707.1;refGene_gn=SSR1;refGene_tx=X
M_023253708.1;refGene_gn=SSR1;refGene_tx=XM_023253709.1
B2	21660482	21660507
refGene_gn=SSR1;refGene_tx=XM_023253707.1;refGene_gn=SSR1;refGene_tx=X
M_023253708.1
B2	21681425	21681943
refGene_gn=RREB1;refGene_tx=XM_011282167.3;refGene_gn=RREB1;refGene_tx
=XM_011282168.3;refGene_gn=RREB1;refGene_tx=XM_011282169.3;refGene_gn=
RREB1;refGene_tx=XM_011282170.3;refGene_gn=RREB1;refGene_tx=XM_0112821
71.3;refGene_gn=RREB1;refGene_tx=XM_011282172.3

B2	21683311	21684118
refGene_gn=RREB1;refGene_tx=XM_011282167.3;refGene_gn=RREB1;refGene_tx=XM_011282168.3;refGene_gn=RREB1;refGene_tx=XM_011282169.3;refGene_gn=RREB1;refGene_tx=XM_011282170.3;refGene_gn=RREB1;refGene_tx=XM_011282171.3;refGene_gn=RREB1;refGene_tx=XM_011282172.3
B2	21689038	21689203
refGene_gn=RREB1;refGene_tx=XM_011282167.3;refGene_gn=RREB1;refGene_tx=XM_011282168.3;refGene_gn=RREB1;refGene_tx=XM_011282169.3;refGene_gn=RREB1;refGene_tx=XM_011282170.3;refGene_gn=RREB1;refGene_tx=XM_011282171.3
B2	21695789	21698703
refGene_gn=RREB1;refGene_tx=XM_011282167.3;refGene_gn=RREB1;refGene_tx=XM_011282168.3;refGene_gn=RREB1;refGene_tx=XM_011282169.3;refGene_gn=RREB1;refGene_tx=XM_011282170.3;refGene_gn=RREB1;refGene_tx=XM_011282171.3;refGene_gn=RREB1;refGene_tx=XM_011282172.3
B2	21699874	21700064
refGene_gn=RREB1;refGene_tx=XM_011282167.3;refGene_gn=RREB1;refGene_tx=XM_011282168.3;refGene_gn=RREB1;refGene_tx=XM_011282169.3;refGene_gn=RREB1;refGene_tx=XM_011282170.3;refGene_gn=RREB1;refGene_tx=XM_011282171.3;refGene_gn=RREB1;refGene_tx=XM_011282172.3
B2	21715877	21716011
refGene_gn=RREB1;refGene_tx=XM_011282167.3;refGene_gn=RREB1;refGene_tx=XM_011282168.3;refGene_gn=RREB1;refGene_tx=XM_011282169.3;refGene_gn=RREB1;refGene_tx=XM_011282170.3;refGene_gn=RREB1;refGene_tx=XM_011282171.3;refGene_gn=RREB1;refGene_tx=XM_011282172.3
B2	21716556	21716701
refGene_gn=RREB1;refGene_tx=XM_011282167.3;refGene_gn=RREB1;refGene_tx=XM_011282168.3;refGene_gn=RREB1;refGene_tx=XM_011282169.3;refGene_gn=RREB1;refGene_tx=XM_011282170.3;refGene_gn=RREB1;refGene_tx=XM_011282171.3;refGene_gn=RREB1;refGene_tx=XM_011282172.3
B2	21739401	21739565
refGene_gn=RREB1;refGene_tx=XM_011282167.3;refGene_gn=RREB1;refGene_tx=XM_011282168.3;refGene_gn=RREB1;refGene_tx=XM_011282169.3;refGene_gn=RREB1;refGene_tx=XM_011282170.3;refGene_gn=RREB1;refGene_tx=XM_011282171.3;refGene_gn=RREB1;refGene_tx=XM_011282172.3
B2	21740965	21741055
refGene_gn=RREB1;refGene_tx=XM_011282167.3;refGene_gn=RREB1;refGene_tx=XM_011282168.3;refGene_gn=RREB1;refGene_tx=XM_011282169.3;refGene_gn=RREB1;refGene_tx=XM_011282170.3;refGene_gn=RREB1;refGene_tx=XM_011282171.3;refGene_gn=RREB1;refGene_tx=XM_011282172.3
B2	21745937	21746108
refGene_gn=RREB1;refGene_tx=XM_011282167.3;refGene_gn=RREB1;refGene_tx=XM_011282168.3;refGene_gn=RREB1;refGene_tx=XM_011282169.3;refGene_gn=RREB1;refGene_tx=XM_011282170.3;refGene_gn=RREB1;refGene_tx=XM_011282172.3
B2	21746783	21746840	refGene_gn=RREB1;refGene_tx=XM_011282171.3
B2	22263794	22263878	refGene_gn=LY86;refGene_tx=XM_003985832.4
B2	22268331	22268384	refGene_gn=LY86;refGene_tx=XM_003985832.4
B2	22289485	22289614	refGene_gn=LY86;refGene_tx=XM_003985832.4
B2	22291012	22291099	refGene_gn=LY86;refGene_tx=XM_003985832.4

B2 22324589 22324716 refGene_gn=LY86;refGene_tx=XM_003985832.4
B2 22613882 22614012
refGene_gn=F13A1;refGene_tx=XM_006931478.2;refGene_gn=F13A1;refGene_tx
=XM_019830279.1
B2 22624760 22624949
refGene_gn=F13A1;refGene_tx=XM_006931478.2;refGene_gn=F13A1;refGene_tx
=XM_019830279.1
B2 22657753 22658005
refGene_gn=F13A1;refGene_tx=XM_006931478.2;refGene_gn=F13A1;refGene_tx
=XM_019830279.1
B2 22678752 22678871
refGene_gn=F13A1;refGene_tx=XM_006931478.2;refGene_gn=F13A1;refGene_tx
=XM_019830279.1
B2 22681423 22681531
refGene_gn=F13A1;refGene_tx=XM_006931478.2;refGene_gn=F13A1;refGene_tx
=XM_019830279.1
B2 22702929 22703104
refGene_gn=F13A1;refGene_tx=XM_006931478.2;refGene_gn=F13A1;refGene_tx
=XM_019830279.1
B2 22706367 22706506
refGene_gn=F13A1;refGene_tx=XM_006931478.2;refGene_gn=F13A1;refGene_tx
=XM_019830279.1
B2 22727172 22727276
refGene_gn=F13A1;refGene_tx=XM_006931478.2;refGene_gn=F13A1;refGene_tx
=XM_019830279.1
B2 22728656 22728745
refGene_gn=F13A1;refGene_tx=XM_006931478.2;refGene_gn=F13A1;refGene_tx
=XM_019830279.1
B2 22737770 22737924
refGene_gn=F13A1;refGene_tx=XM_006931478.2;refGene_gn=F13A1;refGene_tx
=XM_019830279.1
B2 22745201 22745489
refGene_gn=F13A1;refGene_tx=XM_006931478.2;refGene_gn=F13A1;refGene_tx
=XM_019830279.1
B2 22751172 22751333
refGene_gn=F13A1;refGene_tx=XM_006931478.2;refGene_gn=F13A1;refGene_tx
=XM_019830279.1
B2 22766797 22766934
refGene_gn=F13A1;refGene_tx=XM_006931478.2;refGene_gn=F13A1;refGene_tx
=XM_019830279.1
B2 22773269 22773423 refGene_gn=F13A1;refGene_tx=XM_006931478.2
B2 22859892 22860124 refGene_gn=F13A1;refGene_tx=XM_019830279.1
B2 22897702 22897757 refGene_gn=NRN1;refGene_tx=XM_003985833.5
B2 22901870 22902015 refGene_gn=NRN1;refGene_tx=XM_003985833.5
B2 22905287 22905516 refGene_gn=NRN1;refGene_tx=XM_003985833.5
B2 23023495 23023547 refGene_gn=FARS2;refGene_tx=XM_019830295.2
B2 23064208 23064212 refGene_gn=FARS2;refGene_tx=XM_019830299.2
B2 23081880 23081917 refGene_gn=FARS2;refGene_tx=XM_019830296.2
B2 23104098 23104287
refGene_gn=FARS2;refGene_tx=XM_011282158.3;refGene_gn=FARS2;refGene_tx

=XM_019830292.2;refGene_gn=FARS2;refGene_tx=XM_019830297.2
B2 23171335 23171690
refGene_gn=FARS2;refGene_tx=XM_019830281.2;refGene_gn=FARS2;refGene_tx=XM_019830282.2;refGene_gn=FARS2;refGene_tx=XM_019830283.2;refGene_gn=FARS2;refGene_tx=XM_019830284.2;refGene_gn=FARS2;refGene_tx=XM_019830285.1;refGene_gn=FARS2;refGene_tx=XM_019830286.2;refGene_gn=FARS2;refGene_tx=XM_019830287.2;refGene_gn=FARS2;refGene_tx=XM_019830288.2;refGene_gn=FARS2;refGene_tx=XM_019830290.2;refGene_gn=FARS2;refGene_tx=XM_019830291.2
B2 23171773 23171863
refGene_gn=FARS2;refGene_tx=XM_019830281.2;refGene_gn=FARS2;refGene_tx=XM_019830282.2;refGene_gn=FARS2;refGene_tx=XM_019830283.2;refGene_gn=FARS2;refGene_tx=XM_019830284.2;refGene_gn=FARS2;refGene_tx=XM_019830285.1;refGene_gn=FARS2;refGene_tx=XM_019830286.2;refGene_gn=FARS2;refGene_tx=XM_019830287.2;refGene_gn=FARS2;refGene_tx=XM_019830288.2;refGene_gn=FARS2;refGene_tx=XM_019830290.2;refGene_gn=FARS2;refGene_tx=XM_019830291.2;refGene_gn=FARS2;refGene_tx=XM_019830292.2
B2 23192368 23192435 refGene_gn=FARS2;refGene_tx=XM_019830294.2
B2 23238936 23239030 refGene_gn=FARS2;refGene_tx=XM_023253710.1
B2 23243488 23243507 refGene_gn=FARS2;refGene_tx=XM_023253711.1
B2 23264366 23264518
refGene_gn=FARS2;refGene_tx=XM_011282158.3;refGene_gn=FARS2;refGene_tx=XM_019830281.2;refGene_gn=FARS2;refGene_tx=XM_019830282.2;refGene_gn=FARS2;refGene_tx=XM_019830283.2;refGene_gn=FARS2;refGene_tx=XM_019830284.2;refGene_gn=FARS2;refGene_tx=XM_019830285.1;refGene_gn=FARS2;refGene_tx=XM_019830286.2;refGene_gn=FARS2;refGene_tx=XM_019830287.2;refGene_gn=FARS2;refGene_tx=XM_019830288.2;refGene_gn=FARS2;refGene_tx=XM_019830290.2;refGene_gn=FARS2;refGene_tx=XM_019830291.2;refGene_gn=FARS2;refGene_tx=XM_019830292.2;refGene_gn=FARS2;refGene_tx=XM_019830294.2;refGene_gn=FARS2;refGene_tx=XM_019830295.2;refGene_gn=FARS2;refGene_tx=XM_019830296.2;refGene_gn=FARS2;refGene_tx=XM_019830299.2;refGene_gn=FARS2;refGene_tx=XM_023253710.1;refGene_gn=FARS2;refGene_tx=XM_023253711.1
B2 23315346 23315507
refGene_gn=FARS2;refGene_tx=XM_011282158.3;refGene_gn=FARS2;refGene_tx=XM_019830281.2;refGene_gn=FARS2;refGene_tx=XM_019830282.2;refGene_gn=FARS2;refGene_tx=XM_019830283.2;refGene_gn=FARS2;refGene_tx=XM_019830284.2;refGene_gn=FARS2;refGene_tx=XM_019830285.1;refGene_gn=FARS2;refGene_tx=XM_019830286.2;refGene_gn=FARS2;refGene_tx=XM_019830287.2;refGene_gn=FARS2;refGene_tx=XM_019830288.2;refGene_gn=FARS2;refGene_tx=XM_019830290.2;refGene_gn=FARS2;refGene_tx=XM_019830291.2;refGene_gn=FARS2;refGene_tx=XM_019830292.2;refGene_gn=FARS2;refGene_tx=XM_019830294.2;refGene_gn=FARS2;refGene_tx=XM_019830295.2;refGene_gn=FARS2;refGene_tx=XM_019830296.2;refGene_gn=FARS2;refGene_tx=XM_019830297.2;refGene_gn=FARS2;refGene_tx=XM_019830299.2;refGene_gn=FARS2;refGene_tx=XM_023253710.1;refGene_gn=FARS2;refGene_tx=XM_023253711.1
B2 23441647 23441779
refGene_gn=FARS2;refGene_tx=XM_011282158.3;refGene_gn=FARS2;refGene_tx=XM_019830281.2;refGene_gn=FARS2;refGene_tx=XM_019830282.2;refGene_gn=FARS2;refGene_tx=XM_019830283.2;refGene_gn=FARS2;refGene_tx=XM_0198302

84.2;refGene_gn=FARS2;refGene_tx=XM_019830285.1;refGene_gn=FARS2;refGene_tx=XM_019830286.2;refGene_gn=FARS2;refGene_tx=XM_019830287.2;refGene_gn=FARS2;refGene_tx=XM_019830288.2;refGene_gn=FARS2;refGene_tx=XM_019830290.2;refGene_gn=FARS2;refGene_tx=XM_019830292.2;refGene_gn=FARS2;refGene_tx=XM_019830294.2;refGene_gn=FARS2;refGene_tx=XM_019830295.2;refGene_gn=FARS2;refGene_tx=XM_019830296.2;refGene_gn=FARS2;refGene_tx=XM_019830297.2;refGene_gn=FARS2;refGene_tx=XM_019830299.2;refGene_gn=FARS2;refGene_tx=XM_023253710.1;refGene_gn=FARS2;refGene_tx=XM_023253711.1
B2 23465198 23465358
refGene_gn=FARS2;refGene_tx=XM_011282158.3;refGene_gn=FARS2;refGene_tx=XM_019830281.2;refGene_gn=FARS2;refGene_tx=XM_019830282.2;refGene_gn=FARS2;refGene_tx=XM_019830283.2;refGene_gn=FARS2;refGene_tx=XM_019830284.2;refGene_gn=FARS2;refGene_tx=XM_019830285.1;refGene_gn=FARS2;refGene_tx=XM_019830286.2;refGene_gn=FARS2;refGene_tx=XM_019830287.2;refGene_gn=FARS2;refGene_tx=XM_019830288.2;refGene_gn=FARS2;refGene_tx=XM_019830290.2;refGene_gn=FARS2;refGene_tx=XM_019830291.2;refGene_gn=FARS2;refGene_tx=XM_019830292.2;refGene_gn=FARS2;refGene_tx=XM_019830294.2;refGene_gn=FARS2;refGene_tx=XM_019830295.2;refGene_gn=FARS2;refGene_tx=XM_019830296.2;refGene_gn=FARS2;refGene_tx=XM_019830297.2;refGene_gn=FARS2;refGene_tx=XM_019830299.2;refGene_gn=FARS2;refGene_tx=XM_023253710.1;refGene_gn=FARS2;refGene_tx=XM_023253711.1
B2 23508227 23508839
refGene_gn=FARS2;refGene_tx=XM_011282158.3;refGene_gn=FARS2;refGene_tx=XM_019830281.2;refGene_gn=FARS2;refGene_tx=XM_019830282.2;refGene_gn=FARS2;refGene_tx=XM_019830283.2;refGene_gn=FARS2;refGene_tx=XM_019830284.2;refGene_gn=FARS2;refGene_tx=XM_019830285.1;refGene_gn=FARS2;refGene_tx=XM_019830286.2;refGene_gn=FARS2;refGene_tx=XM_019830287.2;refGene_gn=FARS2;refGene_tx=XM_019830288.2;refGene_gn=FARS2;refGene_tx=XM_019830290.2;refGene_gn=FARS2;refGene_tx=XM_019830291.2;refGene_gn=FARS2;refGene_tx=XM_019830292.2;refGene_gn=FARS2;refGene_tx=XM_019830294.2;refGene_gn=FARS2;refGene_tx=XM_019830295.2;refGene_gn=FARS2;refGene_tx=XM_019830296.2;refGene_gn=FARS2;refGene_tx=XM_019830297.2;refGene_gn=FARS2;refGene_tx=XM_019830299.2;refGene_gn=FARS2;refGene_tx=XM_023253710.1;refGene_gn=FARS2;refGene_tx=XM_023253711.1
B2 23646215 23646301
refGene_gn=LYRM4;refGene_tx=XM_003985837.4;refGene_gn=LYRM4;refGene_tx=XM_006931479.3
B2 23690748 23690783 refGene_gn=LYRM4;refGene_tx=XM_019830300.1
B2 23692620 23692741
refGene_gn=LYRM4;refGene_tx=XM_003985837.4;refGene_gn=LYRM4;refGene_tx=XM_006931479.3;refGene_gn=LYRM4;refGene_tx=XM_019830300.1
B2 23762678 23762786
refGene_gn=LYRM4;refGene_tx=XM_006931479.3;refGene_gn=LYRM4;refGene_tx=XM_019830300.1
B2 23829479 23829548
refGene_gn=LYRM4;refGene_tx=XM_003985837.4;refGene_gn=LYRM4;refGene_tx=XM_006931479.3;refGene_gn=LYRM4;refGene_tx=XM_019830300.1
B2 23850617 23851655 refGene_gn=PPP1R3G;refGene_tx=XM_023253712.1
B2 23935989 23936112 refGene_gn=RPP40;refGene_tx=XM_003985838.4

B2 23937701 23937846 refGene_gn=RPP40;refGene_tx=XM_003985838.4
B2 23939172 23939241 refGene_gn=RPP40;refGene_tx=XM_003985838.4
B2 23940077 23940173 refGene_gn=RPP40;refGene_tx=XM_003985838.4
B2 23940852 23940978 refGene_gn=RPP40;refGene_tx=XM_003985838.4
B2 23942703 23942902 refGene_gn=RPP40;refGene_tx=XM_003985838.4
B2 23943035 23943170 refGene_gn=RPP40;refGene_tx=XM_003985838.4
B2 23944504 23944703 refGene_gn=RPP40;refGene_tx=XM_003985838.4
B2 23982081 23982240
refGene_gn=CDYL;refGene_tx=XM_019830302.2;refGene_gn=CDYL;refGene_tx=XM_019830305.2;refGene_gn=CDYL;refGene_tx=XM_019830306.2;refGene_gn=CDYL;refGene_tx=XM_023253713.1;refGene_gn=CDYL;refGene_tx=XM_023253714.1
B2 23983437 23983581
refGene_gn=CDYL;refGene_tx=XM_019830302.2;refGene_gn=CDYL;refGene_tx=XM_019830305.2;refGene_gn=CDYL;refGene_tx=XM_019830306.2;refGene_gn=CDYL;refGene_tx=XM_023253713.1;refGene_gn=CDYL;refGene_tx=XM_023253714.1
B2 23991316 23991527
refGene_gn=CDYL;refGene_tx=XM_019830302.2;refGene_gn=CDYL;refGene_tx=XM_019830305.2;refGene_gn=CDYL;refGene_tx=XM_019830306.2;refGene_gn=CDYL;refGene_tx=XM_023253713.1;refGene_gn=CDYL;refGene_tx=XM_023253714.1
B2 23997611 23997784
refGene_gn=CDYL;refGene_tx=XM_019830302.2;refGene_gn=CDYL;refGene_tx=XM_019830305.2;refGene_gn=CDYL;refGene_tx=XM_019830306.2;refGene_gn=CDYL;refGene_tx=XM_023253713.1;refGene_gn=CDYL;refGene_tx=XM_023253714.1
B2 23999003 23999260
refGene_gn=CDYL;refGene_tx=XM_019830302.2;refGene_gn=CDYL;refGene_tx=XM_019830305.2;refGene_gn=CDYL;refGene_tx=XM_019830306.2;refGene_gn=CDYL;refGene_tx=XM_023253713.1;refGene_gn=CDYL;refGene_tx=XM_023253714.1
B2 24053264 24053931
refGene_gn=CDYL;refGene_tx=XM_019830302.2;refGene_gn=CDYL;refGene_tx=XM_019830305.2;refGene_gn=CDYL;refGene_tx=XM_019830306.2;refGene_gn=CDYL;refGene_tx=XM_023253713.1;refGene_gn=CDYL;refGene_tx=XM_023253714.1
B2 24057886 24057898 refGene_gn=CDYL;refGene_tx=XM_019830305.2
B2 24164383 24164407 refGene_gn=CDYL;refGene_tx=XM_023253714.1
B2 24197382 24197472
refGene_gn=CDYL;refGene_tx=XM_019830302.2;refGene_gn=CDYL;refGene_tx=XM_019830306.2;refGene_gn=CDYL;refGene_tx=XM_023253713.1
B2 24209941 24209944
refGene_gn=CDYL;refGene_tx=XM_019830302.2;refGene_gn=CDYL;refGene_tx=XM_019830306.2;refGene_gn=CDYL;refGene_tx=XM_023253713.1
B2 24728299 24728355 refGene_gn=ECI2;refGene_tx=XM_003985843.4
B2 24730728 24730879
refGene_gn=ECI2;refGene_tx=XM_003985843.4;refGene_gn=ECI2;refGene_tx=XM_019830318.2
B2 24732962 24733061
refGene_gn=ECI2;refGene_tx=XM_003985843.4;refGene_gn=ECI2;refGene_tx=XM_019830318.2
B2 24733293 24733482
refGene_gn=ECI2;refGene_tx=XM_003985843.4;refGene_gn=ECI2;refGene_tx=XM_019830318.2
B2 24735543 24735613

refGene_gn=ECI2;refGene_tx=XM_003985843.4;refGene_gn=ECI2;refGene_tx=XM_019830318.2
B2 24740540 24740643
refGene_gn=ECI2;refGene_tx=XM_003985843.4;refGene_gn=ECI2;refGene_tx=XM_019830318.2
B2 24741214 24741335
refGene_gn=ECI2;refGene_tx=XM_003985843.4;refGene_gn=ECI2;refGene_tx=XM_019830318.2
B2 24745134 24745224
refGene_gn=ECI2;refGene_tx=XM_003985843.4;refGene_gn=ECI2;refGene_tx=XM_019830318.2
B2 24746625 24746769
refGene_gn=ECI2;refGene_tx=XM_003985843.4;refGene_gn=ECI2;refGene_tx=XM_019830318.2
B2 24747642 24747798
refGene_gn=ECI2;refGene_tx=XM_003985843.4;refGene_gn=ECI2;refGene_tx=XM_019830318.2
B2 24783842 24784046
refGene_gn=PRPF4B;refGene_tx=XM_019830307.1;refGene_gn=PRPF4B;refGene_tx=XM_019830308.1;refGene_gn=PRPF4B;refGene_tx=XM_019830309.1;refGene_gn=PRPF4B;refGene_tx=XM_019830310.1;refGene_gn=PRPF4B;refGene_tx=XM_019830311.1;refGene_gn=PRPF4B;refGene_tx=XM_019830312.1;refGene_gn=PRPF4B;refGene_tx=XM_019830313.1;refGene_gn=PRPF4B;refGene_tx=XM_019830314.1;refGene_gn=PRPF4B;refGene_tx=XM_019830315.1;refGene_gn=PRPF4B;refGene_tx=XM_019830316.1
B2 24785098 24785185
refGene_gn=PRPF4B;refGene_tx=XM_019830307.1;refGene_gn=PRPF4B;refGene_tx=XM_019830308.1;refGene_gn=PRPF4B;refGene_tx=XM_019830309.1;refGene_gn=PRPF4B;refGene_tx=XM_019830310.1;refGene_gn=PRPF4B;refGene_tx=XM_019830311.1;refGene_gn=PRPF4B;refGene_tx=XM_019830312.1;refGene_gn=PRPF4B;refGene_tx=XM_019830313.1;refGene_gn=PRPF4B;refGene_tx=XM_019830314.1;refGene_gn=PRPF4B;refGene_tx=XM_019830315.1;refGene_gn=PRPF4B;refGene_tx=XM_019830316.1
B2 24786523 24786675
refGene_gn=PRPF4B;refGene_tx=XM_019830307.1;refGene_gn=PRPF4B;refGene_tx=XM_019830308.1;refGene_gn=PRPF4B;refGene_tx=XM_019830309.1;refGene_gn=PRPF4B;refGene_tx=XM_019830310.1;refGene_gn=PRPF4B;refGene_tx=XM_019830311.1;refGene_gn=PRPF4B;refGene_tx=XM_019830312.1;refGene_gn=PRPF4B;refGene_tx=XM_019830313.1;refGene_gn=PRPF4B;refGene_tx=XM_019830314.1;refGene_gn=PRPF4B;refGene_tx=XM_019830315.1;refGene_gn=PRPF4B;refGene_tx=XM_019830316.1
B2 24787274 24787389
refGene_gn=PRPF4B;refGene_tx=XM_019830307.1;refGene_gn=PRPF4B;refGene_tx=XM_019830308.1;refGene_gn=PRPF4B;refGene_tx=XM_019830309.1;refGene_gn=PRPF4B;refGene_tx=XM_019830310.1;refGene_gn=PRPF4B;refGene_tx=XM_019830311.1;refGene_gn=PRPF4B;refGene_tx=XM_019830312.1;refGene_gn=PRPF4B;refGene_tx=XM_019830313.1;refGene_gn=PRPF4B;refGene_tx=XM_019830314.1;refGene_gn=PRPF4B;refGene_tx=XM_019830315.1;refGene_gn=PRPF4B;refGene_tx=XM_019830316.1
B2 24788193 24788217 refGene_gn=PRPF4B;refGene_tx=XM_006931480.4

B2	24790031 24790185
refGene_gn=PRPF4B;refGene_tx=XM_006931480.4;refGene_gn=PRPF4B;refGene_tx=XM_019830307.1;refGene_gn=PRPF4B;refGene_tx=XM_019830308.1;refGene_gn=PRPF4B;refGene_tx=XM_019830309.1;refGene_gn=PRPF4B;refGene_tx=XM_019830310.1;refGene_gn=PRPF4B;refGene_tx=XM_019830311.1;refGene_gn=PRPF4B;refGene_tx=XM_019830312.1;refGene_gn=PRPF4B;refGene_tx=XM_019830313.1;refGene_gn=PRPF4B;refGene_tx=XM_019830314.1;refGene_gn=PRPF4B;refGene_tx=XM_019830315.1;refGene_gn=PRPF4B;refGene_tx=XM_019830316.1
B2	24790793 24790931
refGene_gn=PRPF4B;refGene_tx=XM_006931480.4;refGene_gn=PRPF4B;refGene_tx=XM_019830307.1;refGene_gn=PRPF4B;refGene_tx=XM_019830308.1;refGene_gn=PRPF4B;refGene_tx=XM_019830309.1;refGene_gn=PRPF4B;refGene_tx=XM_019830310.1;refGene_gn=PRPF4B;refGene_tx=XM_019830311.1;refGene_gn=PRPF4B;refGene_tx=XM_019830312.1;refGene_gn=PRPF4B;refGene_tx=XM_019830313.1;refGene_gn=PRPF4B;refGene_tx=XM_019830314.1;refGene_gn=PRPF4B;refGene_tx=XM_019830315.1;refGene_gn=PRPF4B;refGene_tx=XM_019830316.1
B2	24792450 24792598
refGene_gn=PRPF4B;refGene_tx=XM_006931480.4;refGene_gn=PRPF4B;refGene_tx=XM_019830307.1;refGene_gn=PRPF4B;refGene_tx=XM_019830308.1;refGene_gn=PRPF4B;refGene_tx=XM_019830309.1;refGene_gn=PRPF4B;refGene_tx=XM_019830310.1;refGene_gn=PRPF4B;refGene_tx=XM_019830311.1;refGene_gn=PRPF4B;refGene_tx=XM_019830312.1;refGene_gn=PRPF4B;refGene_tx=XM_019830313.1;refGene_gn=PRPF4B;refGene_tx=XM_019830314.1;refGene_gn=PRPF4B;refGene_tx=XM_019830315.1;refGene_gn=PRPF4B;refGene_tx=XM_019830316.1
B2	24793249 24793340
refGene_gn=PRPF4B;refGene_tx=XM_006931480.4;refGene_gn=PRPF4B;refGene_tx=XM_019830307.1;refGene_gn=PRPF4B;refGene_tx=XM_019830308.1;refGene_gn=PRPF4B;refGene_tx=XM_019830309.1;refGene_gn=PRPF4B;refGene_tx=XM_019830310.1;refGene_gn=PRPF4B;refGene_tx=XM_019830311.1;refGene_gn=PRPF4B;refGene_tx=XM_019830312.1;refGene_gn=PRPF4B;refGene_tx=XM_019830313.1;refGene_gn=PRPF4B;refGene_tx=XM_019830314.1;refGene_gn=PRPF4B;refGene_tx=XM_019830315.1;refGene_gn=PRPF4B;refGene_tx=XM_019830316.1
B2	24794656 24794730
refGene_gn=PRPF4B;refGene_tx=XM_006931480.4;refGene_gn=PRPF4B;refGene_tx=XM_019830307.1;refGene_gn=PRPF4B;refGene_tx=XM_019830308.1;refGene_gn=PRPF4B;refGene_tx=XM_019830309.1;refGene_gn=PRPF4B;refGene_tx=XM_019830310.1;refGene_gn=PRPF4B;refGene_tx=XM_019830311.1;refGene_gn=PRPF4B;refGene_tx=XM_019830312.1;refGene_gn=PRPF4B;refGene_tx=XM_019830313.1;refGene_gn=PRPF4B;refGene_tx=XM_019830314.1;refGene_gn=PRPF4B;refGene_tx=XM_019830315.1;refGene_gn=PRPF4B;refGene_tx=XM_019830316.1
B2	24798273 24798484
refGene_gn=PRPF4B;refGene_tx=XM_006931480.4;refGene_gn=PRPF4B;refGene_tx=XM_019830307.1;refGene_gn=PRPF4B;refGene_tx=XM_019830308.1;refGene_gn=PRPF4B;refGene_tx=XM_019830309.1;refGene_gn=PRPF4B;refGene_tx=XM_019830310.1;refGene_gn=PRPF4B;refGene_tx=XM_019830311.1;refGene_gn=PRPF4B;refGene_tx=XM_019830312.1;refGene_gn=PRPF4B;refGene_tx=XM_019830313.1;refGene_gn=PRPF4B;refGene_tx=XM_019830314.1;refGene_gn=PRPF4B;refGene_tx=XM_019830315.1;refGene_gn=PRPF4B;refGene_tx=XM_019830316.1
B2	24800012 24800094
refGene_gn=PRPF4B;refGene_tx=XM_006931480.4;refGene_gn=PRPF4B;refGene_

tx=XM_019830307.1;refGene_gn=PRPF4B;refGene_tx=XM_019830308.1;refGene_gn=PRPF4B;refGene_tx=XM_019830309.1;refGene_gn=PRPF4B;refGene_tx=XM_019830310.1;refGene_gn=PRPF4B;refGene_tx=XM_019830311.1;refGene_gn=PRPF4B;refGene_tx=XM_019830312.1;refGene_gn=PRPF4B;refGene_tx=XM_019830313.1;refGene_gn=PRPF4B;refGene_tx=XM_019830314.1;refGene_gn=PRPF4B;refGene_tx=XM_019830315.1;refGene_gn=PRPF4B;refGene_tx=XM_019830316.1
B2 24801488 24801644
refGene_gn=PRPF4B;refGene_tx=XM_006931480.4;refGene_gn=PRPF4B;refGene_tx=XM_019830307.1;refGene_gn=PRPF4B;refGene_tx=XM_019830308.1;refGene_gn=PRPF4B;refGene_tx=XM_019830309.1;refGene_gn=PRPF4B;refGene_tx=XM_019830310.1;refGene_gn=PRPF4B;refGene_tx=XM_019830311.1;refGene_gn=PRPF4B;refGene_tx=XM_019830312.1;refGene_gn=PRPF4B;refGene_tx=XM_019830313.1;refGene_gn=PRPF4B;refGene_tx=XM_019830314.1;refGene_gn=PRPF4B;refGene_tx=XM_019830315.1;refGene_gn=PRPF4B;refGene_tx=XM_019830316.1
B2 24804953 24805130
refGene_gn=PRPF4B;refGene_tx=XM_006931480.4;refGene_gn=PRPF4B;refGene_tx=XM_019830307.1;refGene_gn=PRPF4B;refGene_tx=XM_019830308.1;refGene_gn=PRPF4B;refGene_tx=XM_019830309.1;refGene_gn=PRPF4B;refGene_tx=XM_019830310.1;refGene_gn=PRPF4B;refGene_tx=XM_019830311.1;refGene_gn=PRPF4B;refGene_tx=XM_019830312.1;refGene_gn=PRPF4B;refGene_tx=XM_019830313.1;refGene_gn=PRPF4B;refGene_tx=XM_019830314.1;refGene_gn=PRPF4B;refGene_tx=XM_019830315.1;refGene_gn=PRPF4B;refGene_tx=XM_019830316.1
B2 24807720 24808920
refGene_gn=PRPF4B;refGene_tx=XM_006931480.4;refGene_gn=PRPF4B;refGene_tx=XM_019830307.1;refGene_gn=PRPF4B;refGene_tx=XM_019830308.1;refGene_gn=PRPF4B;refGene_tx=XM_019830309.1;refGene_gn=PRPF4B;refGene_tx=XM_019830310.1;refGene_gn=PRPF4B;refGene_tx=XM_019830311.1;refGene_gn=PRPF4B;refGene_tx=XM_019830312.1;refGene_gn=PRPF4B;refGene_tx=XM_019830313.1;refGene_gn=PRPF4B;refGene_tx=XM_019830314.1;refGene_gn=PRPF4B;refGene_tx=XM_019830315.1;refGene_gn=PRPF4B;refGene_tx=XM_019830316.1
B2 24815916 24816053
refGene_gn=PRPF4B;refGene_tx=XM_006931480.4;refGene_gn=PRPF4B;refGene_tx=XM_019830307.1;refGene_gn=PRPF4B;refGene_tx=XM_019830308.1;refGene_gn=PRPF4B;refGene_tx=XM_019830309.1;refGene_gn=PRPF4B;refGene_tx=XM_019830310.1;refGene_gn=PRPF4B;refGene_tx=XM_019830311.1;refGene_gn=PRPF4B;refGene_tx=XM_019830312.1;refGene_gn=PRPF4B;refGene_tx=XM_019830313.1;refGene_gn=PRPF4B;refGene_tx=XM_019830314.1;refGene_gn=PRPF4B;refGene_tx=XM_019830315.1;refGene_gn=PRPF4B;refGene_tx=XM_019830316.1
B2 25058859 25059870
refGene_gn=FAM50B;refGene_tx=XM_003985841.4;refGene_gn=FAM50B;refGene_tx=XM_006931483.3
B2 25163119 25163375 refGene_gn=PXDC1;refGene_tx=XM_019831569.2
B2 25177535 25177627 refGene_gn=PXDC1;refGene_tx=XM_019831569.2
B2 25178326 25178444 refGene_gn=PXDC1;refGene_tx=XM_019831569.2
B2 25186334 25186446 refGene_gn=PXDC1;refGene_tx=XM_019831569.2
B2 25190533 25190651 refGene_gn=PXDC1;refGene_tx=XM_019831569.2
B2 25482427 25483096
refGene_gn=SLC22A23;refGene_tx=XM_023253717.1;refGene_gn=SLC22A23;refGene_tx=XM_023253718.1
B2 25522422 25522526

refGene_gn=SLC22A23;refGene_tx=XM_023253717.1;refGene_gn=SLC22A23;refGene_tx=XM_023253718.1
B2 25528329 25528484
refGene_gn=SLC22A23;refGene_tx=XM_023253717.1;refGene_gn=SLC22A23;refGene_tx=XM_023253718.1
B2 25612810 25612979
refGene_gn=SLC22A23;refGene_tx=XM_023253717.1;refGene_gn=SLC22A23;refGene_tx=XM_023253718.1
B2 25633557 25633685
refGene_gn=SLC22A23;refGene_tx=XM_023253717.1;refGene_gn=SLC22A23;refGene_tx=XM_023253718.1
B2 25643160 25643263
refGene_gn=SLC22A23;refGene_tx=XM_023253717.1;refGene_gn=SLC22A23;refGene_tx=XM_023253718.1
B2 25645232 25645465
refGene_gn=SLC22A23;refGene_tx=XM_023253717.1;refGene_gn=SLC22A23;refGene_tx=XM_023253718.1
B2 25647122 25647155
refGene_gn=SLC22A23;refGene_tx=XM_023253717.1;refGene_gn=SLC22A23;refGene_tx=XM_023253718.1
B2 25647287 25647311 refGene_gn=SLC22A23;refGene_tx=XM_023253717.1
B2 25648386 25648510
refGene_gn=SLC22A23;refGene_tx=XM_023253717.1;refGene_gn=SLC22A23;refGene_tx=XM_023253718.1
B2 25660203 25660561
refGene_gn=SLC22A23;refGene_tx=XM_023253717.1;refGene_gn=SLC22A23;refGene_tx=XM_023253718.1
B2 25666604 25666726 refGene_gn=PSMG4;refGene_tx=XM_019830319.2
B2 25670342 25670418
refGene_gn=PSMG4;refGene_tx=XM_019830319.2;refGene_gn=PSMG4;refGene_tx=XM_023253719.1
B2 25670527 25670601 refGene_gn=PSMG4;refGene_tx=XM_023253719.1
B2 25676596 25676770
refGene_gn=PSMG4;refGene_tx=XM_019830319.2;refGene_gn=PSMG4;refGene_tx=XM_023253719.1
B2 25709994 25710051 refGene_gn=TUBB2B;refGene_tx=XM_003985847.4
B2 25710858 25710967 refGene_gn=TUBB2B;refGene_tx=XM_003985847.4
B2 25711249 25711360 refGene_gn=TUBB2B;refGene_tx=XM_003985847.4
B2 25711715 25712776 refGene_gn=TUBB2B;refGene_tx=XM_003985847.4
B2 25749434 25749567
refGene_gn=LOC111560610;refGene_tx=XM_023254744.1
B2 25749922 25750041
refGene_gn=LOC111560610;refGene_tx=XM_023254744.1
B2 25761331 25761388 refGene_gn=TUBB2A;refGene_tx=XM_023253720.1
B2 25762674 25762783 refGene_gn=TUBB2A;refGene_tx=XM_023253720.1
B2 25763138 25763249 refGene_gn=TUBB2A;refGene_tx=XM_023253720.1
B2 25763935 25764996 refGene_gn=TUBB2A;refGene_tx=XM_023253720.1
B2 25766314 25766402
refGene_gn=BPHL;refGene_tx=XM_003985848.4;refGene_gn=BPHL;refGene_tx=XM_006931486.3;refGene_gn=BPHL;refGene_tx=XM_006931487.3

B2	25777111 25777235
refGene_gn=BPHL;refGene_tx=XM_003985848.4;refGene_gn=BPHL;refGene_tx=XM_006931486.3;refGene_gn=BPHL;refGene_tx=XM_006931487.3
B2	25780735 25780867
refGene_gn=BPHL;refGene_tx=XM_003985848.4;refGene_gn=BPHL;refGene_tx=XM_006931486.3;refGene_gn=BPHL;refGene_tx=XM_006931487.3
B2	25791548 25791702
refGene_gn=BPHL;refGene_tx=XM_003985848.4;refGene_gn=BPHL;refGene_tx=XM_006931486.3;refGene_gn=BPHL;refGene_tx=XM_006931487.3
B2	25793010 25793177
refGene_gn=BPHL;refGene_tx=XM_003985848.4;refGene_gn=BPHL;refGene_tx=XM_006931486.3;refGene_gn=BPHL;refGene_tx=XM_006931487.3
B2	25796199 25796303
refGene_gn=BPHL;refGene_tx=XM_003985848.4;refGene_gn=BPHL;refGene_tx=XM_006931486.3;refGene_gn=BPHL;refGene_tx=XM_006931487.3
B2	25803628 25803672 refGene_gn=BPHL;refGene_tx=XM_006931486.3
B2	25803958 25804065
refGene_gn=BPHL;refGene_tx=XM_003985848.4;refGene_gn=BPHL;refGene_tx=XM_006931487.3
B2	25809222 25809509
refGene_gn=RIPK1;refGene_tx=XM_006931488.4;refGene_gn=RIPK1;refGene_tx=XM_006931489.4;refGene_gn=RIPK1;refGene_tx=XM_006931490.3;refGene_gn=RIPK1;refGene_tx=XM_023253721.1;refGene_gn=RIPK1;refGene_tx=XM_023253722.1;refGene_gn=RIPK1;refGene_tx=XM_023253723.1
B2	25811106 25811259
refGene_gn=RIPK1;refGene_tx=XM_006931488.4;refGene_gn=RIPK1;refGene_tx=XM_006931489.4;refGene_gn=RIPK1;refGene_tx=XM_006931490.3;refGene_gn=RIPK1;refGene_tx=XM_023253721.1;refGene_gn=RIPK1;refGene_tx=XM_023253722.1;refGene_gn=RIPK1;refGene_tx=XM_023253723.1
B2	25812671 25813238
refGene_gn=RIPK1;refGene_tx=XM_006931488.4;refGene_gn=RIPK1;refGene_tx=XM_006931489.4;refGene_gn=RIPK1;refGene_tx=XM_006931490.3;refGene_gn=RIPK1;refGene_tx=XM_023253721.1;refGene_gn=RIPK1;refGene_tx=XM_023253722.1;refGene_gn=RIPK1;refGene_tx=XM_023253723.1
B2	25814102 25814193
refGene_gn=RIPK1;refGene_tx=XM_006931488.4;refGene_gn=RIPK1;refGene_tx=XM_006931489.4;refGene_gn=RIPK1;refGene_tx=XM_006931490.3;refGene_gn=RIPK1;refGene_tx=XM_023253721.1;refGene_gn=RIPK1;refGene_tx=XM_023253722.1;refGene_gn=RIPK1;refGene_tx=XM_023253723.1
B2	25815562 25815639
refGene_gn=RIPK1;refGene_tx=XM_006931488.4;refGene_gn=RIPK1;refGene_tx=XM_006931489.4;refGene_gn=RIPK1;refGene_tx=XM_006931490.3;refGene_gn=RIPK1;refGene_tx=XM_023253721.1;refGene_gn=RIPK1;refGene_tx=XM_023253722.1;refGene_gn=RIPK1;refGene_tx=XM_023253723.1
B2	25826521 25826671
refGene_gn=RIPK1;refGene_tx=XM_006931488.4;refGene_gn=RIPK1;refGene_tx=XM_006931489.4;refGene_gn=RIPK1;refGene_tx=XM_006931490.3;refGene_gn=RIPK1;refGene_tx=XM_023253721.1;refGene_gn=RIPK1;refGene_tx=XM_023253722.1;refGene_gn=RIPK1;refGene_tx=XM_023253723.1
B2	25828702 25828934

refGene_gn=RIPK1;refGene_tx=XM_006931488.4;refGene_gn=RIPK1;refGene_tx=XM_006931489.4;refGene_gn=RIPK1;refGene_tx=XM_006931490.3;refGene_gn=RIPK1;refGene_tx=XM_023253721.1;refGene_gn=RIPK1;refGene_tx=XM_023253722.1;refGene_gn=RIPK1;refGene_tx=XM_023253723.1
B2 25830272 25830410
refGene_gn=RIPK1;refGene_tx=XM_006931488.4;refGene_gn=RIPK1;refGene_tx=XM_006931489.4;refGene_gn=RIPK1;refGene_tx=XM_006931490.3;refGene_gn=RIPK1;refGene_tx=XM_023253721.1;refGene_gn=RIPK1;refGene_tx=XM_023253722.1
B2 25833773 25833930
refGene_gn=RIPK1;refGene_tx=XM_006931488.4;refGene_gn=RIPK1;refGene_tx=XM_006931489.4;refGene_gn=RIPK1;refGene_tx=XM_006931490.3;refGene_gn=RIPK1;refGene_tx=XM_023253721.1;refGene_gn=RIPK1;refGene_tx=XM_023253722.1
B2 25834973 25835125
refGene_gn=RIPK1;refGene_tx=XM_006931488.4;refGene_gn=RIPK1;refGene_tx=XM_006931489.4;refGene_gn=RIPK1;refGene_tx=XM_006931490.3;refGene_gn=RIPK1;refGene_tx=XM_023253721.1;refGene_gn=RIPK1;refGene_tx=XM_023253722.1
B2 25897355 25897532 refGene_gn=NQO2;refGene_tx=XM_003985849.4
B2 25899238 25899340 refGene_gn=NQO2;refGene_tx=XM_003985849.4
B2 25900313 25900427 refGene_gn=NQO2;refGene_tx=XM_003985849.4
B2 25902836 25902967 refGene_gn=NQO2;refGene_tx=XM_003985849.4
B2 25905086 25905251 refGene_gn=NQO2;refGene_tx=XM_003985849.4
B2 25907240 25907247 refGene_gn=NQO2;refGene_tx=XM_003985849.4
B2 25962872 25963037
refGene_gn=SERPINB6;refGene_tx=XM_006931495.4;refGene_gn=SERPINB6;refGene_tx=XM_011282153.2;refGene_gn=SERPINB6;refGene_tx=XM_011282154.3
B2 25966116 25966263
refGene_gn=SERPINB6;refGene_tx=XM_006931495.4;refGene_gn=SERPINB6;refGene_tx=XM_011282153.2;refGene_gn=SERPINB6;refGene_tx=XM_011282154.3
B2 25969588 25969706
refGene_gn=SERPINB6;refGene_tx=XM_006931495.4;refGene_gn=SERPINB6;refGene_tx=XM_011282153.2;refGene_gn=SERPINB6;refGene_tx=XM_011282154.3
B2 25970685 25970828
refGene_gn=SERPINB6;refGene_tx=XM_006931495.4;refGene_gn=SERPINB6;refGene_tx=XM_011282153.2;refGene_gn=SERPINB6;refGene_tx=XM_011282154.3
B2 25974727 25974883
refGene_gn=SERPINB6;refGene_tx=XM_006931495.4;refGene_gn=SERPINB6;refGene_tx=XM_011282153.2;refGene_gn=SERPINB6;refGene_tx=XM_011282154.3
B2 25975066 25975471
refGene_gn=SERPINB6;refGene_tx=XM_006931495.4;refGene_gn=SERPINB6;refGene_tx=XM_011282153.2;refGene_gn=SERPINB6;refGene_tx=XM_011282154.3
B2 26028382 26028550
refGene_gn=SERPINB9;refGene_tx=XM_003985851.5;refGene_gn=SERPINB9;refGene_tx=XM_006931497.4;refGene_gn=SERPINB9;refGene_tx=XM_006931498.4;refGene_gn=SERPINB9;refGene_tx=XM_011282150.3;refGene_gn=SERPINB9;refGene_tx=XM_011282151.3;refGene_gn=SERPINB9;refGene_tx=XM_019830322.2;refGene_gn=SERPINB9;refGene_tx=XM_019830323.2
B2 26030800 26030938

refGene_gn=SERPINB9;refGene_tx=XM_003985851.5;refGene_gn=SERPINB9;refGene_tx=XM_006931497.4;refGene_gn=SERPINB9;refGene_tx=XM_006931498.4;refGene_gn=SERPINB9;refGene_tx=XM_011282150.3;refGene_gn=SERPINB9;refGene_tx=XM_011282151.3;refGene_gn=SERPINB9;refGene_tx=XM_019830322.2;refGene_gn=SERPINB9;refGene_tx=XM_019830323.2
B2 26031283 26031401
refGene_gn=SERPINB9;refGene_tx=XM_003985851.5;refGene_gn=SERPINB9;refGene_tx=XM_006931497.4;refGene_gn=SERPINB9;refGene_tx=XM_006931498.4;refGene_gn=SERPINB9;refGene_tx=XM_011282150.3;refGene_gn=SERPINB9;refGene_tx=XM_011282151.3;refGene_gn=SERPINB9;refGene_tx=XM_019830322.2;refGene_gn=SERPINB9;refGene_tx=XM_019830323.2
B2 26032927 26033070
refGene_gn=SERPINB9;refGene_tx=XM_003985851.5;refGene_gn=SERPINB9;refGene_tx=XM_006931497.4;refGene_gn=SERPINB9;refGene_tx=XM_006931498.4;refGene_gn=SERPINB9;refGene_tx=XM_011282150.3;refGene_gn=SERPINB9;refGene_tx=XM_011282151.3;refGene_gn=SERPINB9;refGene_tx=XM_019830322.2;refGene_gn=SERPINB9;refGene_tx=XM_019830323.2
B2 26035234 26035390
refGene_gn=SERPINB9;refGene_tx=XM_003985851.5;refGene_gn=SERPINB9;refGene_tx=XM_006931497.4;refGene_gn=SERPINB9;refGene_tx=XM_006931498.4;refGene_gn=SERPINB9;refGene_tx=XM_011282150.3;refGene_gn=SERPINB9;refGene_tx=XM_011282151.3;refGene_gn=SERPINB9;refGene_tx=XM_019830323.2
B2 26036723 26037128
refGene_gn=SERPINB9;refGene_tx=XM_003985851.5;refGene_gn=SERPINB9;refGene_tx=XM_006931497.4;refGene_gn=SERPINB9;refGene_tx=XM_006931498.4;refGene_gn=SERPINB9;refGene_tx=XM_011282150.3;refGene_gn=SERPINB9;refGene_tx=XM_011282151.3;refGene_gn=SERPINB9;refGene_tx=XM_019830322.2;refGene_gn=SERPINB9;refGene_tx=XM_019830323.2
B2 26078290 26078458 refGene_gn=SERPINB1;refGene_tx=XM_003985852.4
B2 26080234 26080372 refGene_gn=SERPINB1;refGene_tx=XM_003985852.4
B2 26081401 26081519 refGene_gn=SERPINB1;refGene_tx=XM_003985852.4
B2 26082538 26082681 refGene_gn=SERPINB1;refGene_tx=XM_003985852.4
B2 26082767 26082935 refGene_gn=SERPINB1;refGene_tx=XM_003985852.4
B2 26084677 26085082 refGene_gn=SERPINB1;refGene_tx=XM_003985852.4
B2 26124848 26125124
refGene_gn=WRNIP1;refGene_tx=XM_023253726.1;refGene_gn=WRNIP1;refGene_tx=XM_023253728.1
B2 26125707 26125787 refGene_gn=WRNIP1;refGene_tx=XM_023253726.1
B2 26126484 26126640
refGene_gn=WRNIP1;refGene_tx=XM_023253726.1;refGene_gn=WRNIP1;refGene_tx=XM_023253728.1
B2 26130874 26131104
refGene_gn=WRNIP1;refGene_tx=XM_023253726.1;refGene_gn=WRNIP1;refGene_tx=XM_023253728.1
B2 26144328 26144570
refGene_gn=WRNIP1;refGene_tx=XM_023253726.1;refGene_gn=WRNIP1;refGene_tx=XM_023253728.1
B2 26145807 26145999
refGene_gn=WRNIP1;refGene_tx=XM_023253726.1;refGene_gn=WRNIP1;refGene_tx=XM_023253728.1

B2 26148182 26148998
refGene_gn=WRNIP1;refGene_tx=XM_023253726.1;refGene_gn=WRNIP1;refGene_tx=XM_023253728.1
B2 26169831 26169848
refGene_gn=MYLK4;refGene_tx=XM_006931500.2;refGene_gn=MYLK4;refGene_tx=XM_019830324.1;refGene_gn=MYLK4;refGene_tx=XM_023253729.1
B2 26171460 26171731
refGene_gn=MYLK4;refGene_tx=XM_006931500.2;refGene_gn=MYLK4;refGene_tx=XM_019830324.1;refGene_gn=MYLK4;refGene_tx=XM_023253729.1
B2 26176033 26176084 refGene_gn=MYLK4;refGene_tx=XM_019830325.1
B2 26230384 26230463
refGene_gn=MYLK4;refGene_tx=XM_006931500.2;refGene_gn=MYLK4;refGene_tx=XM_019830324.1;refGene_gn=MYLK4;refGene_tx=XM_019830325.1;refGene_gn=MYLK4;refGene_tx=XM_023253729.1
B2 26230807 26231258
refGene_gn=MYLK4;refGene_tx=XM_006931500.2;refGene_gn=MYLK4;refGene_tx=XM_019830324.1;refGene_gn=MYLK4;refGene_tx=XM_019830325.1
B2 26232464 26232763
refGene_gn=MYLK4;refGene_tx=XM_006931500.2;refGene_gn=MYLK4;refGene_tx=XM_019830324.1;refGene_gn=MYLK4;refGene_tx=XM_019830325.1
B2 26234150 26234256
refGene_gn=MYLK4;refGene_tx=XM_006931500.2;refGene_gn=MYLK4;refGene_tx=XM_019830324.1;refGene_gn=MYLK4;refGene_tx=XM_019830325.1;refGene_gn=MYLK4;refGene_tx=XM_023253729.1
B2 26236517 26236611
refGene_gn=MYLK4;refGene_tx=XM_006931500.2;refGene_gn=MYLK4;refGene_tx=XM_019830324.1;refGene_gn=MYLK4;refGene_tx=XM_019830325.1;refGene_gn=MYLK4;refGene_tx=XM_023253729.1
B2 26236684 26236794
refGene_gn=MYLK4;refGene_tx=XM_006931500.2;refGene_gn=MYLK4;refGene_tx=XM_019830324.1;refGene_gn=MYLK4;refGene_tx=XM_019830325.1;refGene_gn=MYLK4;refGene_tx=XM_023253729.1
B2 26237747 26237889
refGene_gn=MYLK4;refGene_tx=XM_006931500.2;refGene_gn=MYLK4;refGene_tx=XM_019830324.1;refGene_gn=MYLK4;refGene_tx=XM_019830325.1;refGene_gn=MYLK4;refGene_tx=XM_023253729.1
B2 26241116 26241187
refGene_gn=MYLK4;refGene_tx=XM_006931500.2;refGene_gn=MYLK4;refGene_tx=XM_019830324.1;refGene_gn=MYLK4;refGene_tx=XM_019830325.1;refGene_gn=MYLK4;refGene_tx=XM_023253729.1
B2 26241989 26242118
refGene_gn=MYLK4;refGene_tx=XM_006931500.2;refGene_gn=MYLK4;refGene_tx=XM_019830324.1;refGene_gn=MYLK4;refGene_tx=XM_019830325.1;refGene_gn=MYLK4;refGene_tx=XM_023253729.1
B2 26242733 26242886
refGene_gn=MYLK4;refGene_tx=XM_006931500.2;refGene_gn=MYLK4;refGene_tx=XM_019830324.1;refGene_gn=MYLK4;refGene_tx=XM_019830325.1;refGene_gn=MYLK4;refGene_tx=XM_023253729.1
B2 26246085 26246173
refGene_gn=MYLK4;refGene_tx=XM_006931500.2;refGene_gn=MYLK4;refGene_tx

=XM_019830324.1;refGene_gn=MYLK4;refGene_tx=XM_019830325.1;refGene_gn=MYLK4;refGene_tx=XM_023253729.1
B2 26249930 26249978
refGene_gn=MYLK4;refGene_tx=XM_006931500.2;refGene_gn=MYLK4;refGene_tx=XM_019830324.1;refGene_gn=MYLK4;refGene_tx=XM_019830325.1;refGene_gn=MYLK4;refGene_tx=XM_023253729.1
B2 26679508 26679610 refGene_gn=GMDS;refGene_tx=XM_003985853.4
B2 26681695 26681713 refGene_gn=GMDS;refGene_tx=XM_019830326.1
B2 26829672 26829717
refGene_gn=GMDS;refGene_tx=XM_003985853.4;refGene_gn=GMDS;refGene_tx=XM_019830326.1
B2 26838353 26838441
refGene_gn=GMDS;refGene_tx=XM_003985853.4;refGene_gn=GMDS;refGene_tx=XM_019830326.1
B2 26840300 26840410
refGene_gn=GMDS;refGene_tx=XM_003985853.4;refGene_gn=GMDS;refGene_tx=XM_019830326.1
B2 26968927 26969120
refGene_gn=GMDS;refGene_tx=XM_003985853.4;refGene_gn=GMDS;refGene_tx=XM_019830326.1
B2 26969992 26970097
refGene_gn=GMDS;refGene_tx=XM_003985853.4;refGene_gn=GMDS;refGene_tx=XM_019830326.1
B2 27002787 27002915
refGene_gn=GMDS;refGene_tx=XM_003985853.4;refGene_gn=GMDS;refGene_tx=XM_019830326.1
B2 27184335 27184454
refGene_gn=GMDS;refGene_tx=XM_003985853.4;refGene_gn=GMDS;refGene_tx=XM_019830326.1
B2 27211767 27211864
refGene_gn=GMDS;refGene_tx=XM_003985853.4;refGene_gn=GMDS;refGene_tx=XM_019830326.1
B2 27307492 27307561
refGene_gn=GMDS;refGene_tx=XM_003985853.4;refGene_gn=GMDS;refGene_tx=XM_019830326.1
B2 27307870 27307933
refGene_gn=GMDS;refGene_tx=XM_003985853.4;refGene_gn=GMDS;refGene_tx=XM_019830326.1
B2 27320663 27322322 refGene_gn=FOXC1;refGene_tx=XM_023253730.1
B2 27579067 27579234 refGene_gn=FOXF2;refGene_tx=XM_023253731.1
B2 27582788 27583971 refGene_gn=FOXF2;refGene_tx=XM_023253731.1
B2 27674959 27676180 refGene_gn=FOXQ1;refGene_tx=XM_023253732.1
B2 28139552 28139612
refGene_gn=LOC109500146;refGene_tx=XM_023254706.1
B2 28141071 28141228
refGene_gn=LOC109500146;refGene_tx=XM_023254706.1
B2 28574167 28574263 refGene_gn=HUS1B;refGene_tx=XM_003985865.4
B2 28574265 28574346 refGene_gn=HUS1B;refGene_tx=XM_003985865.4
B2 28574348 28574564 refGene_gn=HUS1B;refGene_tx=XM_003985865.4
B2 28574565 28574799 refGene_gn=HUS1B;refGene_tx=XM_003985865.4

B2 28574800 28574902 refGene_gn=HUS1B;refGene_tx=XM_003985865.4
B2 28574903 28574966 refGene_gn=HUS1B;refGene_tx=XM_003985865.4
B2 28574967 28575029 refGene_gn=HUS1B;refGene_tx=XM_003985865.4
B2 28577541 28577545 refGene_gn=HUS1B;refGene_tx=XM_003985865.4
B2 28601817 28601935
refGene_gn=EXOC2;refGene_tx=XM_003985860.4;refGene_gn=EXOC2;refGene_tx=XM_006931507.4
B2 28606447 28606624
refGene_gn=EXOC2;refGene_tx=XM_003985860.4;refGene_gn=EXOC2;refGene_tx=XM_006931507.4
B2 28608782 28608909
refGene_gn=EXOC2;refGene_tx=XM_003985860.4;refGene_gn=EXOC2;refGene_tx=XM_006931507.4
B2 28619966 28620080
refGene_gn=EXOC2;refGene_tx=XM_003985860.4;refGene_gn=EXOC2;refGene_tx=XM_006931507.4
B2 28621767 28621892
refGene_gn=EXOC2;refGene_tx=XM_003985860.4;refGene_gn=EXOC2;refGene_tx=XM_006931507.4
B2 28637396 28637477
refGene_gn=EXOC2;refGene_tx=XM_003985860.4;refGene_gn=EXOC2;refGene_tx=XM_006931507.4
B2 28652261 28652407
refGene_gn=EXOC2;refGene_tx=XM_003985860.4;refGene_gn=EXOC2;refGene_tx=XM_006931507.4
B2 28652541 28652623
refGene_gn=EXOC2;refGene_tx=XM_003985860.4;refGene_gn=EXOC2;refGene_tx=XM_006931507.4
B2 28653374 28653477
refGene_gn=EXOC2;refGene_tx=XM_003985860.4;refGene_gn=EXOC2;refGene_tx=XM_006931507.4
B2 28659108 28659227
refGene_gn=EXOC2;refGene_tx=XM_003985860.4;refGene_gn=EXOC2;refGene_tx=XM_006931507.4
B2 28675463 28675589
refGene_gn=EXOC2;refGene_tx=XM_003985860.4;refGene_gn=EXOC2;refGene_tx=XM_006931507.4
B2 28679925 28680050
refGene_gn=EXOC2;refGene_tx=XM_003985860.4;refGene_gn=EXOC2;refGene_tx=XM_006931507.4
B2 28699328 28699394
refGene_gn=EXOC2;refGene_tx=XM_003985860.4;refGene_gn=EXOC2;refGene_tx=XM_006931507.4
B2 28699562 28699720
refGene_gn=EXOC2;refGene_tx=XM_003985860.4;refGene_gn=EXOC2;refGene_tx=XM_006931507.4
B2 28700129 28700251
refGene_gn=EXOC2;refGene_tx=XM_003985860.4;refGene_gn=EXOC2;refGene_tx=XM_006931507.4
B2 28701674 28701736

refGene_gn=EXOC2;refGene_tx=XM_003985860.4;refGene_gn=EXOC2;refGene_tx=XM_006931507.4
B2 28712933 28713014
refGene_gn=EXOC2;refGene_tx=XM_003985860.4;refGene_gn=EXOC2;refGene_tx=XM_006931507.4
B2 28713472 28713532
refGene_gn=EXOC2;refGene_tx=XM_003985860.4;refGene_gn=EXOC2;refGene_tx=XM_006931507.4
B2 28714212 28714274
refGene_gn=EXOC2;refGene_tx=XM_003985860.4;refGene_gn=EXOC2;refGene_tx=XM_006931507.4
B2 28718045 28718112
refGene_gn=EXOC2;refGene_tx=XM_003985860.4;refGene_gn=EXOC2;refGene_tx=XM_006931507.4
B2 28722322 28722439
refGene_gn=EXOC2;refGene_tx=XM_003985860.4;refGene_gn=EXOC2;refGene_tx=XM_006931507.4
B2 28758420 28758562
refGene_gn=EXOC2;refGene_tx=XM_003985860.4;refGene_gn=EXOC2;refGene_tx=XM_006931507.4
B2 28811581 28811637
refGene_gn=EXOC2;refGene_tx=XM_003985860.4;refGene_gn=EXOC2;refGene_tx=XM_006931507.4
B2 28813681 28813804
refGene_gn=EXOC2;refGene_tx=XM_003985860.4;refGene_gn=EXOC2;refGene_tx=XM_006931507.4
B2 28827426 28827488
refGene_gn=EXOC2;refGene_tx=XM_003985860.4;refGene_gn=EXOC2;refGene_tx=XM_006931507.4
B2 28829633 28829693
refGene_gn=EXOC2;refGene_tx=XM_003985860.4;refGene_gn=EXOC2;refGene_tx=XM_006931507.4
B2 28831211 28831305
refGene_gn=EXOC2;refGene_tx=XM_003985860.4;refGene_gn=EXOC2;refGene_tx=XM_006931507.4
B2 28940170 28940314
refGene_gn=IRF4;refGene_tx=XM_006931509.2;refGene_gn=IRF4;refGene_tx=XM_006931510.2;refGene_gn=IRF4;refGene_tx=XM_006931511.2
B2 28942222 28942335
refGene_gn=IRF4;refGene_tx=XM_006931509.2;refGene_gn=IRF4;refGene_tx=XM_006931510.2;refGene_gn=IRF4;refGene_tx=XM_006931511.2
B2 28945123 28945477
refGene_gn=IRF4;refGene_tx=XM_006931509.2;refGene_gn=IRF4;refGene_tx=XM_006931510.2;refGene_gn=IRF4;refGene_tx=XM_006931511.2
B2 28947767 28947875
refGene_gn=IRF4;refGene_tx=XM_006931509.2;refGene_gn=IRF4;refGene_tx=XM_006931510.2
B2 28950118 28950263
refGene_gn=IRF4;refGene_tx=XM_006931509.2;refGene_gn=IRF4;refGene_tx=XM_006931510.2;refGene_gn=IRF4;refGene_tx=XM_006931511.2

B2 28950974 28951063
refGene_gn=IRF4;refGene_tx=XM_006931509.2;refGene_gn=IRF4;refGene_tx=XM_006931510.2;refGene_gn=IRF4;refGene_tx=XM_006931511.2
B2 28951916 28952103
refGene_gn=IRF4;refGene_tx=XM_006931509.2;refGene_gn=IRF4;refGene_tx=XM_006931510.2;refGene_gn=IRF4;refGene_tx=XM_006931511.2
B2 28953376 28953592
refGene_gn=IRF4;refGene_tx=XM_006931509.2;refGene_gn=IRF4;refGene_tx=XM_006931510.2;refGene_gn=IRF4;refGene_tx=XM_006931511.2
B2 28992257 28992310 refGene_gn=DUSP22;refGene_tx=XM_011282149.3
B2 28999633 28999680 refGene_gn=DUSP22;refGene_tx=XM_003985862.4
B2 29001448 29001521
refGene_gn=DUSP22;refGene_tx=XM_003985862.4;refGene_gn=DUSP22;refGene_tx=XM_011282149.3
B2 29001962 29002134
refGene_gn=DUSP22;refGene_tx=XM_003985862.4;refGene_gn=DUSP22;refGene_tx=XM_011282149.3
B2 29005004 29005079
refGene_gn=DUSP22;refGene_tx=XM_003985862.4;refGene_gn=DUSP22;refGene_tx=XM_011282149.3
B2 29014921 29014971
refGene_gn=DUSP22;refGene_tx=XM_003985862.4;refGene_gn=DUSP22;refGene_tx=XM_011282149.3
B2 29038932 29039015
refGene_gn=DUSP22;refGene_tx=XM_003985862.4;refGene_gn=DUSP22;refGene_tx=XM_011282149.3
B2 29048025 29048059
refGene_gn=DUSP22;refGene_tx=XM_003985862.4;refGene_gn=DUSP22;refGene_tx=XM_011282149.3
B2 29061824 29061845
refGene_gn=DUSP22;refGene_tx=XM_003985862.4;refGene_gn=DUSP22;refGene_tx=XM_011282149.3
B2 31239611 31240118
refGene_gn=LOC109500088;refGene_tx=XM_019831560.1
B2 31271911 31272364
refGene_gn=LOC102901769;refGene_tx=XM_006931514.4;refGene_gn=LOC102901769;refGene_tx=XM_006931515.4;refGene_gn=LOC102901769;refGene_tx=XM_019830167.2;refGene_gn=LOC102901769;refGene_tx=XM_023253734.1
B2 31273537 31273633
refGene_gn=LOC102901769;refGene_tx=XM_006931514.4;refGene_gn=LOC102901769;refGene_tx=XM_006931515.4;refGene_gn=LOC102901769;refGene_tx=XM_019830167.2;refGene_gn=LOC102901769;refGene_tx=XM_023253734.1
B2 31278278 31278509
refGene_gn=LOC102901769;refGene_tx=XM_006931514.4;refGene_gn=LOC102901769;refGene_tx=XM_006931515.4;refGene_gn=LOC102901769;refGene_tx=XM_019830167.2;refGene_gn=LOC102901769;refGene_tx=XM_023253734.1
B2 31279693 31279716
refGene_gn=LOC102901769;refGene_tx=XM_006931514.4;refGene_gn=LOC102901769;refGene_tx=XM_006931515.4;refGene_gn=LOC102901769;refGene_tx=XM_019830167.2;refGene_gn=LOC102901769;refGene_tx=XM_023253734.1

B2 31280298 31280414
refGene_gn=LOC102901769;refGene_tx=XM_006931514.4;refGene_gn=LOC102901769;refGene_tx=XM_006931515.4;refGene_gn=LOC102901769;refGene_tx=XM_019830167.2;refGene_gn=LOC102901769;refGene_tx=XM_023253734.1
B2 31281123 31281678
refGene_gn=LOC102901769;refGene_tx=XM_006931514.4;refGene_gn=LOC102901769;refGene_tx=XM_006931515.4;refGene_gn=LOC102901769;refGene_tx=XM_019830167.2;refGene_gn=LOC102901769;refGene_tx=XM_023253734.1
B2 31284305 31284362 refGene_gn=RPP21;refGene_tx=XM_003985866.5
B2 31284434 31284535 refGene_gn=RPP21;refGene_tx=XM_003985866.5
B2 31284630 31284713 refGene_gn=RPP21;refGene_tx=XM_003985866.5
B2 31285540 31285666 refGene_gn=RPP21;refGene_tx=XM_003985866.5
B2 31285812 31285910 refGene_gn=RPP21;refGene_tx=XM_003985866.5
B2 31395923 31396165 refGene_gn=GNL1;refGene_tx=XM_003985869.5
B2 31396530 31396671 refGene_gn=GNL1;refGene_tx=XM_003985869.5
B2 31396954 31397117 refGene_gn=GNL1;refGene_tx=XM_003985869.5
B2 31397194 31397373 refGene_gn=GNL1;refGene_tx=XM_003985869.5
B2 31399146 31399341 refGene_gn=GNL1;refGene_tx=XM_003985869.5
B2 31399739 31399835 refGene_gn=GNL1;refGene_tx=XM_003985869.5
B2 31400003 31400211 refGene_gn=GNL1;refGene_tx=XM_003985869.5
B2 31401183 31401255 refGene_gn=GNL1;refGene_tx=XM_003985869.5
B2 31401362 31401514 refGene_gn=GNL1;refGene_tx=XM_003985869.5
B2 31401672 31401809 refGene_gn=GNL1;refGene_tx=XM_003985869.5
B2 31402181 31402347 refGene_gn=GNL1;refGene_tx=XM_003985869.5
B2 31402756 31402829 refGene_gn=GNL1;refGene_tx=XM_003985869.5
B2 31403947 31404053
refGene_gn=PRR3;refGene_tx=XM_006931516.4;refGene_gn=PRR3;refGene_tx=XM_011282130.3;refGene_gn=PRR3;refGene_tx=XM_019830170.1
B2 31404782 31404845
refGene_gn=PRR3;refGene_tx=XM_006931516.4;refGene_gn=PRR3;refGene_tx=XM_011282130.3;refGene_gn=PRR3;refGene_tx=XM_019830169.2
B2 31406753 31406936
refGene_gn=PRR3;refGene_tx=XM_006931516.4;refGene_gn=PRR3;refGene_tx=XM_019830169.2
B2 31407287 31407578
refGene_gn=PRR3;refGene_tx=XM_006931516.4;refGene_gn=PRR3;refGene_tx=XM_011282130.3;refGene_gn=PRR3;refGene_tx=XM_019830169.2;refGene_gn=PRR3;refGene_tx=XM_019830170.1
B2 31407848 31407955
refGene_gn=PRR3;refGene_tx=XM_006931516.4;refGene_gn=PRR3;refGene_tx=XM_011282130.3;refGene_gn=PRR3;refGene_tx=XM_019830169.2;refGene_gn=PRR3;refGene_tx=XM_019830170.1
B2 31415704 31415777
refGene_gn=ABCF1;refGene_tx=XM_003985870.4;refGene_gn=ABCF1;refGene_tx=XM_011282128.2;refGene_gn=ABCF1;refGene_tx=XM_019830171.1
B2 31420108 31420174
refGene_gn=ABCF1;refGene_tx=XM_003985870.4;refGene_gn=ABCF1;refGene_tx=XM_011282128.2;refGene_gn=ABCF1;refGene_tx=XM_019830171.1
B2 31420393 31420483
refGene_gn=ABCF1;refGene_tx=XM_003985870.4;refGene_gn=ABCF1;refGene_tx

=XM_011282128.2;refGene_gn=ABCF1;refGene_tx=XM_019830171.1
B2 31420595 31420725
refGene_gn=ABCF1;refGene_tx=XM_003985870.4;refGene_gn=ABCF1;refGene_tx=XM_011282128.2;refGene_gn=ABCF1;refGene_tx=XM_019830171.1
B2 31420875 31420913
refGene_gn=ABCF1;refGene_tx=XM_003985870.4;refGene_gn=ABCF1;refGene_tx=XM_011282128.2;refGene_gn=ABCF1;refGene_tx=XM_019830171.1
B2 31420998 31421106
refGene_gn=ABCF1;refGene_tx=XM_003985870.4;refGene_gn=ABCF1;refGene_tx=XM_011282128.2;refGene_gn=ABCF1;refGene_tx=XM_019830171.1
B2 31422543 31422618
refGene_gn=ABCF1;refGene_tx=XM_003985870.4;refGene_gn=ABCF1;refGene_tx=XM_011282128.2;refGene_gn=ABCF1;refGene_tx=XM_019830171.1
B2 31422699 31422816
refGene_gn=ABCF1;refGene_tx=XM_003985870.4;refGene_gn=ABCF1;refGene_tx=XM_011282128.2;refGene_gn=ABCF1;refGene_tx=XM_019830171.1
B2 31423657 31423762
refGene_gn=ABCF1;refGene_tx=XM_003985870.4;refGene_gn=ABCF1;refGene_tx=XM_011282128.2;refGene_gn=ABCF1;refGene_tx=XM_019830171.1
B2 31424352 31424475
refGene_gn=ABCF1;refGene_tx=XM_003985870.4;refGene_gn=ABCF1;refGene_tx=XM_011282128.2;refGene_gn=ABCF1;refGene_tx=XM_019830171.1
B2 31424884 31424985
refGene_gn=ABCF1;refGene_tx=XM_003985870.4;refGene_gn=ABCF1;refGene_tx=XM_011282128.2;refGene_gn=ABCF1;refGene_tx=XM_019830171.1
B2 31425078 31425164
refGene_gn=ABCF1;refGene_tx=XM_003985870.4;refGene_gn=ABCF1;refGene_tx=XM_011282128.2;refGene_gn=ABCF1;refGene_tx=XM_019830171.1
B2 31425437 31425577
refGene_gn=ABCF1;refGene_tx=XM_003985870.4;refGene_gn=ABCF1;refGene_tx=XM_011282128.2;refGene_gn=ABCF1;refGene_tx=XM_019830171.1
B2 31425662 31425811
refGene_gn=ABCF1;refGene_tx=XM_003985870.4;refGene_gn=ABCF1;refGene_tx=XM_011282128.2;refGene_gn=ABCF1;refGene_tx=XM_019830171.1
B2 31426135 31426219
refGene_gn=ABCF1;refGene_tx=XM_003985870.4;refGene_gn=ABCF1;refGene_tx=XM_011282128.2;refGene_gn=ABCF1;refGene_tx=XM_019830171.1
B2 31426419 31426544
refGene_gn=ABCF1;refGene_tx=XM_003985870.4;refGene_gn=ABCF1;refGene_tx=XM_011282128.2;refGene_gn=ABCF1;refGene_tx=XM_019830171.1
B2 31426732 31426845
refGene_gn=ABCF1;refGene_tx=XM_003985870.4;refGene_gn=ABCF1;refGene_tx=XM_011282128.2;refGene_gn=ABCF1;refGene_tx=XM_019830171.1
B2 31426964 31427136
refGene_gn=ABCF1;refGene_tx=XM_003985870.4;refGene_gn=ABCF1;refGene_tx=XM_011282128.2;refGene_gn=ABCF1;refGene_tx=XM_019830171.1
B2 31427301 31427376
refGene_gn=ABCF1;refGene_tx=XM_003985870.4;refGene_gn=ABCF1;refGene_tx=XM_011282128.2;refGene_gn=ABCF1;refGene_tx=XM_019830171.1
B2 31427468 31427539

refGene_gn=ABCF1;refGene_tx=XM_003985870.4;refGene_gn=ABCF1;refGene_tx=XM_011282128.2;refGene_gn=ABCF1;refGene_tx=XM_019830171.1
B2 31429050 31429083
refGene_gn=ABCF1;refGene_tx=XM_003985870.4;refGene_gn=ABCF1;refGene_tx=XM_011282128.2;refGene_gn=ABCF1;refGene_tx=XM_019830171.1
B2 31429165 31429334
refGene_gn=ABCF1;refGene_tx=XM_003985870.4;refGene_gn=ABCF1;refGene_tx=XM_011282128.2;refGene_gn=ABCF1;refGene_tx=XM_019830171.1
B2 31429513 31429578
refGene_gn=ABCF1;refGene_tx=XM_003985870.4;refGene_gn=ABCF1;refGene_tx=XM_011282128.2;refGene_gn=ABCF1;refGene_tx=XM_019830171.1
B2 31429664 31429737
refGene_gn=ABCF1;refGene_tx=XM_003985870.4;refGene_gn=ABCF1;refGene_tx=XM_011282128.2;refGene_gn=ABCF1;refGene_tx=XM_019830171.1
B2 31429861 31430028
refGene_gn=ABCF1;refGene_tx=XM_003985870.4;refGene_gn=ABCF1;refGene_tx=XM_011282128.2;refGene_gn=ABCF1;refGene_tx=XM_019830171.1
B2 31436980 31437090
refGene_gn=PPP1R10;refGene_tx=XM_023253736.1;refGene_gn=PPP1R10;refGene_tx=XM_023253737.1
B2 31437385 31438186
refGene_gn=PPP1R10;refGene_tx=XM_023253736.1;refGene_gn=PPP1R10;refGene_tx=XM_023253737.1
B2 31438340 31438454
refGene_gn=PPP1R10;refGene_tx=XM_023253736.1;refGene_gn=PPP1R10;refGene_tx=XM_023253737.1
B2 31438718 31438794
refGene_gn=PPP1R10;refGene_tx=XM_023253736.1;refGene_gn=PPP1R10;refGene_tx=XM_023253737.1
B2 31438962 31439157
refGene_gn=PPP1R10;refGene_tx=XM_023253736.1;refGene_gn=PPP1R10;refGene_tx=XM_023253737.1
B2 31439257 31439321
refGene_gn=PPP1R10;refGene_tx=XM_023253736.1;refGene_gn=PPP1R10;refGene_tx=XM_023253737.1
B2 31439505 31439752
refGene_gn=PPP1R10;refGene_tx=XM_023253736.1;refGene_gn=PPP1R10;refGene_tx=XM_023253737.1
B2 31439853 31440012
refGene_gn=PPP1R10;refGene_tx=XM_023253736.1;refGene_gn=PPP1R10;refGene_tx=XM_023253737.1
B2 31440089 31440237
refGene_gn=PPP1R10;refGene_tx=XM_023253736.1;refGene_gn=PPP1R10;refGene_tx=XM_023253737.1
B2 31440496 31440597
refGene_gn=PPP1R10;refGene_tx=XM_023253736.1;refGene_gn=PPP1R10;refGene_tx=XM_023253737.1
B2 31441951 31442064
refGene_gn=PPP1R10;refGene_tx=XM_023253736.1;refGene_gn=PPP1R10;refGene_tx=XM_023253737.1

B2 31442176 31442282
refGene_gn=PPP1R10;refGene_tx=XM_023253736.1;refGene_gn=PPP1R10;refGen
e_tx=XM_023253737.1
B2 31442500 31442674
refGene_gn=PPP1R10;refGene_tx=XM_023253736.1;refGene_gn=PPP1R10;refGen
e_tx=XM_023253737.1
B2 31442812 31442890
refGene_gn=PPP1R10;refGene_tx=XM_023253736.1;refGene_gn=PPP1R10;refGen
e_tx=XM_023253737.1
B2 31445392 31445444 refGene_gn=PPP1R10;refGene_tx=XM_023253736.1
B2 31446680 31446816 refGene_gn=PPP1R10;refGene_tx=XM_023253736.1
B2 31446970 31447057 refGene_gn=PPP1R10;refGene_tx=XM_023253736.1
B2 31447743 31447850 refGene_gn=PPP1R10;refGene_tx=XM_023253736.1
B2 31455441 31455519 refGene_gn=MRPS18B;refGene_tx=XM_003985874.4
B2 31456698 31456807 refGene_gn=MRPS18B;refGene_tx=XM_003985874.4
B2 31456919 31457017 refGene_gn=MRPS18B;refGene_tx=XM_003985874.4
B2 31457144 31457213 refGene_gn=MRPS18B;refGene_tx=XM_003985874.4
B2 31458551 31458618 refGene_gn=MRPS18B;refGene_tx=XM_003985874.4
B2 31460144 31460204 refGene_gn=MRPS18B;refGene_tx=XM_003985874.4
B2 31460777 31461073 refGene_gn=MRPS18B;refGene_tx=XM_003985874.4
B2 31461869 31461976
refGene_gn=ATAT1;refGene_tx=XM_003985875.5;refGene_gn=ATAT1;refGene_tx
=XM_003985876.4;refGene_gn=ATAT1;refGene_tx=XM_003985877.4;refGene_gn=
ATAT1;refGene_tx=XM_006931518.4;refGene_gn=ATAT1;refGene_tx=XM_0198301
72.2
B2 31462409 31462470
refGene_gn=ATAT1;refGene_tx=XM_003985875.5;refGene_gn=ATAT1;refGene_tx
=XM_003985876.4;refGene_gn=ATAT1;refGene_tx=XM_003985877.4;refGene_gn=
ATAT1;refGene_tx=XM_006931518.4;refGene_gn=ATAT1;refGene_tx=XM_0198301
72.2
B2 31462596 31462688
refGene_gn=ATAT1;refGene_tx=XM_003985875.5;refGene_gn=ATAT1;refGene_tx
=XM_003985876.4;refGene_gn=ATAT1;refGene_tx=XM_003985877.4;refGene_gn=
ATAT1;refGene_tx=XM_006931518.4;refGene_gn=ATAT1;refGene_tx=XM_0198301
72.2
B2 31462807 31462868
refGene_gn=ATAT1;refGene_tx=XM_003985875.5;refGene_gn=ATAT1;refGene_tx
=XM_003985876.4;refGene_gn=ATAT1;refGene_tx=XM_003985877.4;refGene_gn=
ATAT1;refGene_tx=XM_006931518.4;refGene_gn=ATAT1;refGene_tx=XM_0198301
72.2
B2 31462985 31463099
refGene_gn=ATAT1;refGene_tx=XM_003985875.5;refGene_gn=ATAT1;refGene_tx
=XM_003985876.4;refGene_gn=ATAT1;refGene_tx=XM_003985877.4;refGene_gn=
ATAT1;refGene_tx=XM_006931518.4;refGene_gn=ATAT1;refGene_tx=XM_0198301
72.2
B2 31463315 31463417
refGene_gn=ATAT1;refGene_tx=XM_003985875.5;refGene_gn=ATAT1;refGene_tx
=XM_003985876.4;refGene_gn=ATAT1;refGene_tx=XM_003985877.4;refGene_gn=
ATAT1;refGene_tx=XM_006931518.4;refGene_gn=ATAT1;refGene_tx=XM_0198301
72.2

B2 31470376 31470422
refGene_gn=ATAT1;refGene_tx=XM_003985875.5;refGene_gn=ATAT1;refGene_tx=XM_003985876.4;refGene_gn=ATAT1;refGene_tx=XM_003985877.4;refGene_gn=ATAT1;refGene_tx=XM_006931518.4;refGene_gn=ATAT1;refGene_tx=XM_019830172.2
B2 31470541 31470610
refGene_gn=ATAT1;refGene_tx=XM_003985875.5;refGene_gn=ATAT1;refGene_tx=XM_003985877.4;refGene_gn=ATAT1;refGene_tx=XM_019830172.2
B2 31472176 31472248
refGene_gn=ATAT1;refGene_tx=XM_003985875.5;refGene_gn=ATAT1;refGene_tx=XM_003985876.4;refGene_gn=ATAT1;refGene_tx=XM_003985877.4;refGene_gn=ATAT1;refGene_tx=XM_006931518.4;refGene_gn=ATAT1;refGene_tx=XM_019830172.2
B2 31472418 31472662
refGene_gn=ATAT1;refGene_tx=XM_003985875.5;refGene_gn=ATAT1;refGene_tx=XM_003985876.4;refGene_gn=ATAT1;refGene_tx=XM_003985877.4;refGene_gn=ATAT1;refGene_tx=XM_006931518.4;refGene_gn=ATAT1;refGene_tx=XM_019830172.2
B2 31473186 31473220
refGene_gn=ATAT1;refGene_tx=XM_003985876.4;refGene_gn=ATAT1;refGene_tx=XM_003985877.4
B2 31475192 31475272
refGene_gn=ATAT1;refGene_tx=XM_003985875.5;refGene_gn=ATAT1;refGene_tx=XM_006931518.4;refGene_gn=ATAT1;refGene_tx=XM_019830172.2
B2 31475363 31475406
refGene_gn=ATAT1;refGene_tx=XM_003985875.5;refGene_gn=ATAT1;refGene_tx=XM_006931518.4;refGene_gn=ATAT1;refGene_tx=XM_019830172.2
B2 31475738 31475913
refGene_gn=ATAT1;refGene_tx=XM_003985875.5;refGene_gn=ATAT1;refGene_tx=XM_006931518.4;refGene_gn=ATAT1;refGene_tx=XM_019830172.2
B2 31476488 31476560
refGene_gn=CB2H6orf136;refGene_tx=XM_003985879.4
B2 31476762 31476793
refGene_gn=CB2H6orf136;refGene_tx=XM_019830173.2;refGene_gn=CB2H6orf136;refGene_tx=XM_019830174.2
B2 31476840 31476865
refGene_gn=CB2H6orf136;refGene_tx=XM_019830173.2;refGene_gn=CB2H6orf136;refGene_tx=XM_019830174.2
B2 31476969 31477138
refGene_gn=CB2H6orf136;refGene_tx=XM_019830173.2;refGene_gn=CB2H6orf136;refGene_tx=XM_019830174.2
B2 31477935 31478334
refGene_gn=CB2H6orf136;refGene_tx=XM_003985879.4;refGene_gn=CB2H6orf136;refGene_tx=XM_019830173.2;refGene_gn=CB2H6orf136;refGene_tx=XM_019830174.2
B2 31478604 31478693
refGene_gn=CB2H6orf136;refGene_tx=XM_003985879.4;refGene_gn=CB2H6orf136;refGene_tx=XM_019830173.2;refGene_gn=CB2H6orf136;refGene_tx=XM_019830174.2
B2 31478875 31479076

refGene_gn=CB2H6orf136;refGene_tx=XM_003985879.4;refGene_gn=CB2H6orf13
6;refGene_tx=XM_019830173.2
B2 31479667 31479737
refGene_gn=CB2H6orf136;refGene_tx=XM_003985879.4;refGene_gn=CB2H6orf13
6;refGene_tx=XM_019830173.2;refGene_gn=CB2H6orf136;refGene_tx=XM_01983
0174.2
B2 31479816 31479930
refGene_gn=CB2H6orf136;refGene_tx=XM_003985879.4;refGene_gn=CB2H6orf13
6;refGene_tx=XM_019830173.2;refGene_gn=CB2H6orf136;refGene_tx=XM_01983
0174.2
B2 31480251 31480380 refGene_gn=DHX16;refGene_tx=XM_003985878.4
B2 31480892 31481066 refGene_gn=DHX16;refGene_tx=XM_003985878.4
B2 31481357 31481519 refGene_gn=DHX16;refGene_tx=XM_003985878.4
B2 31481621 31481784 refGene_gn=DHX16;refGene_tx=XM_003985878.4
B2 31482494 31482562 refGene_gn=DHX16;refGene_tx=XM_003985878.4
B2 31482675 31482794 refGene_gn=DHX16;refGene_tx=XM_003985878.4
B2 31482892 31483055 refGene_gn=DHX16;refGene_tx=XM_003985878.4
B2 31483689 31483830 refGene_gn=DHX16;refGene_tx=XM_003985878.4
B2 31485238 31485391 refGene_gn=DHX16;refGene_tx=XM_003985878.4
B2 31485495 31485594 refGene_gn=DHX16;refGene_tx=XM_003985878.4
B2 31485778 31485989 refGene_gn=DHX16;refGene_tx=XM_003985878.4
B2 31487638 31487754 refGene_gn=DHX16;refGene_tx=XM_003985878.4
B2 31487925 31488036 refGene_gn=DHX16;refGene_tx=XM_003985878.4
B2 31488682 31488874 refGene_gn=DHX16;refGene_tx=XM_003985878.4
B2 31488954 31489158 refGene_gn=DHX16;refGene_tx=XM_003985878.4
B2 31489344 31489599 refGene_gn=DHX16;refGene_tx=XM_003985878.4
B2 31492879 31492936 refGene_gn=DHX16;refGene_tx=XM_003985878.4
B2 31493244 31493407 refGene_gn=DHX16;refGene_tx=XM_003985878.4
B2 31493493 31493741 refGene_gn=DHX16;refGene_tx=XM_003985878.4
B2 31495177 31495384 refGene_gn=DHX16;refGene_tx=XM_003985878.4
B2 31498532 31498552 refGene_gn=PPP1R18;refGene_tx=XM_019830179.2
B2 31499198 31499409 refGene_gn=PPP1R18;refGene_tx=XM_019830179.2
B2 31504100 31505210 refGene_gn=PPP1R18;refGene_tx=XM_019830179.2
B2 31505211 31505727 refGene_gn=PPP1R18;refGene_tx=XM_019830179.2
B2 31508277 31508559
refGene_gn=NRM;refGene_tx=XM_003985882.5;refGene_gn=NRM;refGene_tx=XM_
003985883.5
B2 31508908 31509085 refGene_gn=NRM;refGene_tx=XM_003985882.5
B2 31509700 31509897
refGene_gn=NRM;refGene_tx=XM_003985882.5;refGene_gn=NRM;refGene_tx=XM_
003985883.5
B2 31510470 31510603
refGene_gn=NRM;refGene_tx=XM_003985882.5;refGene_gn=NRM;refGene_tx=XM_
003985883.5
B2 31514779 31514947
refGene_gn=MDC1;refGene_tx=XM_006931521.4;refGene_gn=MDC1;refGene_tx=X
M_006931523.4;refGene_gn=MDC1;refGene_tx=XM_006931524.3;refGene_gn=MDC
1;refGene_tx=XM_006931525.4;refGene_gn=MDC1;refGene_tx=XM_006931526.4;
refGene_gn=MDC1;refGene_tx=XM_006931527.4
B2 31516745 31516856

refGene_gn=MDC1;refGene_tx=XM_006931521.4;refGene_gn=MDC1;refGene_tx=XM_006931523.4;refGene_gn=MDC1;refGene_tx=XM_006931524.3;refGene_gn=MDC1;refGene_tx=XM_006931525.4;refGene_gn=MDC1;refGene_tx=XM_006931526.4;refGene_gn=MDC1;refGene_tx=XM_006931527.4
B2 31516967 31517093
refGene_gn=MDC1;refGene_tx=XM_006931521.4;refGene_gn=MDC1;refGene_tx=XM_006931523.4;refGene_gn=MDC1;refGene_tx=XM_006931524.3;refGene_gn=MDC1;refGene_tx=XM_006931525.4;refGene_gn=MDC1;refGene_tx=XM_006931526.4;refGene_gn=MDC1;refGene_tx=XM_006931527.4
B2 31517323 31517506
refGene_gn=MDC1;refGene_tx=XM_006931521.4;refGene_gn=MDC1;refGene_tx=XM_006931523.4;refGene_gn=MDC1;refGene_tx=XM_006931524.3;refGene_gn=MDC1;refGene_tx=XM_006931525.4;refGene_gn=MDC1;refGene_tx=XM_006931526.4;refGene_gn=MDC1;refGene_tx=XM_006931527.4
B2 31517634 31517754
refGene_gn=MDC1;refGene_tx=XM_006931521.4;refGene_gn=MDC1;refGene_tx=XM_006931523.4;refGene_gn=MDC1;refGene_tx=XM_006931524.3;refGene_gn=MDC1;refGene_tx=XM_006931525.4;refGene_gn=MDC1;refGene_tx=XM_006931526.4;refGene_gn=MDC1;refGene_tx=XM_006931527.4
B2 31517835 31519857
refGene_gn=MDC1;refGene_tx=XM_006931521.4;refGene_gn=MDC1;refGene_tx=XM_006931523.4;refGene_gn=MDC1;refGene_tx=XM_006931524.3;refGene_gn=MDC1;refGene_tx=XM_006931525.4;refGene_gn=MDC1;refGene_tx=XM_006931526.4;refGene_gn=MDC1;refGene_tx=XM_006931527.4
B2 31520561 31520605
refGene_gn=MDC1;refGene_tx=XM_006931521.4;refGene_gn=MDC1;refGene_tx=XM_006931523.4;refGene_gn=MDC1;refGene_tx=XM_006931524.3;refGene_gn=MDC1;refGene_tx=XM_006931525.4;refGene_gn=MDC1;refGene_tx=XM_006931526.4;refGene_gn=MDC1;refGene_tx=XM_006931527.4
B2 31520710 31521487
refGene_gn=MDC1;refGene_tx=XM_006931521.4;refGene_gn=MDC1;refGene_tx=XM_006931523.4;refGene_gn=MDC1;refGene_tx=XM_006931524.3;refGene_gn=MDC1;refGene_tx=XM_006931525.4;refGene_gn=MDC1;refGene_tx=XM_006931526.4
B2 31522436 31522526
refGene_gn=MDC1;refGene_tx=XM_006931521.4;refGene_gn=MDC1;refGene_tx=XM_006931523.4;refGene_gn=MDC1;refGene_tx=XM_006931524.3;refGene_gn=MDC1;refGene_tx=XM_006931525.4;refGene_gn=MDC1;refGene_tx=XM_006931527.4
B2 31522694 31522760
refGene_gn=MDC1;refGene_tx=XM_006931521.4;refGene_gn=MDC1;refGene_tx=XM_006931523.4;refGene_gn=MDC1;refGene_tx=XM_006931524.3;refGene_gn=MDC1;refGene_tx=XM_006931525.4;refGene_gn=MDC1;refGene_tx=XM_006931526.4;refGene_gn=MDC1;refGene_tx=XM_006931527.4
B2 31523485 31524951
refGene_gn=MDC1;refGene_tx=XM_006931521.4;refGene_gn=MDC1;refGene_tx=XM_006931523.4;refGene_gn=MDC1;refGene_tx=XM_006931524.3;refGene_gn=MDC1;refGene_tx=XM_006931525.4;refGene_gn=MDC1;refGene_tx=XM_006931526.4;refGene_gn=MDC1;refGene_tx=XM_006931527.4
B2 31525237 31525307
refGene_gn=MDC1;refGene_tx=XM_006931521.4;refGene_gn=MDC1;refGene_tx=XM_006931523.4;refGene_gn=MDC1;refGene_tx=XM_006931524.3;refGene_gn=MDC

1;refGene_tx=XM_006931525.4;refGene_gn=MDC1;refGene_tx=XM_006931526.4;
refGene_gn=MDC1;refGene_tx=XM_006931527.4
B2	31525392 31525773
refGene_gn=MDC1;refGene_tx=XM_006931521.4;refGene_gn=MDC1;refGene_tx=X
M_006931523.4;refGene_gn=MDC1;refGene_tx=XM_006931524.3;refGene_gn=MDC
1;refGene_tx=XM_006931525.4;refGene_gn=MDC1;refGene_tx=XM_006931526.4;
refGene_gn=MDC1;refGene_tx=XM_006931527.4
B2	31526325 31526464
refGene_gn=MDC1;refGene_tx=XM_006931521.4;refGene_gn=MDC1;refGene_tx=X
M_006931523.4;refGene_gn=MDC1;refGene_tx=XM_006931524.3;refGene_gn=MDC
1;refGene_tx=XM_006931525.4;refGene_gn=MDC1;refGene_tx=XM_006931526.4;
refGene_gn=MDC1;refGene_tx=XM_006931527.4
B2	31528683 31528686
refGene_gn=MDC1;refGene_tx=XM_006931521.4;refGene_gn=MDC1;refGene_tx=X
M_006931526.4;refGene_gn=MDC1;refGene_tx=XM_006931527.4
B2	31530767 31530824 refGene_gn=TUBB;refGene_tx=XM_003985886.4
B2	31532215 31532324 refGene_gn=TUBB;refGene_tx=XM_003985886.4
B2	31532594 31532705 refGene_gn=TUBB;refGene_tx=XM_003985886.4
B2	31533002 31534060 refGene_gn=TUBB;refGene_tx=XM_003985886.4
B2	31536938 31536968 refGene_gn=FLOT1;refGene_tx=XM_011282126.3
B2	31538334 31538499 refGene_gn=FLOT1;refGene_tx=XM_011282126.3
B2	31538758 31538896 refGene_gn=FLOT1;refGene_tx=XM_011282126.3
B2	31539014 31539060 refGene_gn=FLOT1;refGene_tx=XM_011282126.3
B2	31539221 31539403 refGene_gn=FLOT1;refGene_tx=XM_011282126.3
B2	31547628 31547781 refGene_gn=FLOT1;refGene_tx=XM_011282126.3
B2	31548004 31548100 refGene_gn=FLOT1;refGene_tx=XM_011282126.3
B2	31548188 31548308 refGene_gn=FLOT1;refGene_tx=XM_011282126.3
B2	31548444 31548588 refGene_gn=FLOT1;refGene_tx=XM_011282126.3
B2	31548825 31548916 refGene_gn=FLOT1;refGene_tx=XM_011282126.3
B2	31549009 31549085 refGene_gn=FLOT1;refGene_tx=XM_011282126.3
B2	31549379 31549422 refGene_gn=FLOT1;refGene_tx=XM_011282126.3
B2	31550998 31551259 refGene_gn=IER3;refGene_tx=XM_003985890.4
B2	31551358 31551568 refGene_gn=IER3;refGene_tx=XM_003985890.4
B2	31676841 31676932
refGene_gn=DDR1;refGene_tx=XM_006931528.3;refGene_gn=DDR1;refGene_tx=X
M_019830180.2;refGene_gn=DDR1;refGene_tx=XM_019830181.2;refGene_gn=DDR
1;refGene_tx=XM_019830182.2;refGene_gn=DDR1;refGene_tx=XM_023253738.1
B2	31677023 31677126
refGene_gn=DDR1;refGene_tx=XM_006931528.3;refGene_gn=DDR1;refGene_tx=X
M_019830180.2;refGene_gn=DDR1;refGene_tx=XM_019830181.2;refGene_gn=DDR
1;refGene_tx=XM_019830182.2;refGene_gn=DDR1;refGene_tx=XM_023253738.1
B2	31677323 31677552
refGene_gn=DDR1;refGene_tx=XM_006931528.3;refGene_gn=DDR1;refGene_tx=X
M_019830180.2;refGene_gn=DDR1;refGene_tx=XM_019830181.2;refGene_gn=DDR
1;refGene_tx=XM_019830182.2;refGene_gn=DDR1;refGene_tx=XM_023253738.1
B2	31678932 31679080
refGene_gn=DDR1;refGene_tx=XM_006931528.3;refGene_gn=DDR1;refGene_tx=X
M_019830180.2;refGene_gn=DDR1;refGene_tx=XM_019830181.2;refGene_gn=DDR
1;refGene_tx=XM_019830182.2;refGene_gn=DDR1;refGene_tx=XM_023253738.1
B2	31679327 31679436

refGene_gn=DDR1;refGene_tx=XM_006931528.3;refGene_gn=DDR1;refGene_tx=XM_019830180.2;refGene_gn=DDR1;refGene_tx=XM_019830181.2;refGene_gn=DDR1;refGene_tx=XM_019830182.2;refGene_gn=DDR1;refGene_tx=XM_023253738.1
B2 31679907 31680094
refGene_gn=DDR1;refGene_tx=XM_006931528.3;refGene_gn=DDR1;refGene_tx=XM_019830180.2;refGene_gn=DDR1;refGene_tx=XM_019830181.2;refGene_gn=DDR1;refGene_tx=XM_019830182.2;refGene_gn=DDR1;refGene_tx=XM_023253738.1
B2 31680205 31680452
refGene_gn=DDR1;refGene_tx=XM_006931528.3;refGene_gn=DDR1;refGene_tx=XM_019830180.2;refGene_gn=DDR1;refGene_tx=XM_019830181.2;refGene_gn=DDR1;refGene_tx=XM_019830182.2;refGene_gn=DDR1;refGene_tx=XM_023253738.1
B2 31680983 31681079
refGene_gn=DDR1;refGene_tx=XM_006931528.3;refGene_gn=DDR1;refGene_tx=XM_019830180.2;refGene_gn=DDR1;refGene_tx=XM_019830181.2;refGene_gn=DDR1;refGene_tx=XM_019830182.2;refGene_gn=DDR1;refGene_tx=XM_023253738.1
B2 31681206 31681358
refGene_gn=DDR1;refGene_tx=XM_006931528.3;refGene_gn=DDR1;refGene_tx=XM_019830180.2;refGene_gn=DDR1;refGene_tx=XM_019830181.2;refGene_gn=DDR1;refGene_tx=XM_019830182.2;refGene_gn=DDR1;refGene_tx=XM_023253738.1
B2 31682255 31682421
refGene_gn=DDR1;refGene_tx=XM_006931528.3;refGene_gn=DDR1;refGene_tx=XM_019830180.2;refGene_gn=DDR1;refGene_tx=XM_019830181.2;refGene_gn=DDR1;refGene_tx=XM_019830182.2;refGene_gn=DDR1;refGene_tx=XM_023253738.1
B2 31683403 31683514
refGene_gn=DDR1;refGene_tx=XM_006931528.3;refGene_gn=DDR1;refGene_tx=XM_019830180.2;refGene_gn=DDR1;refGene_tx=XM_023253738.1
B2 31684627 31684872
refGene_gn=DDR1;refGene_tx=XM_006931528.3;refGene_gn=DDR1;refGene_tx=XM_019830180.2;refGene_gn=DDR1;refGene_tx=XM_019830181.2;refGene_gn=DDR1;refGene_tx=XM_023253738.1
B2 31685045 31685173
refGene_gn=DDR1;refGene_tx=XM_006931528.3;refGene_gn=DDR1;refGene_tx=XM_019830180.2;refGene_gn=DDR1;refGene_tx=XM_019830181.2;refGene_gn=DDR1;refGene_tx=XM_019830182.2;refGene_gn=DDR1;refGene_tx=XM_023253738.1
B2 31685418 31685637
refGene_gn=DDR1;refGene_tx=XM_006931528.3;refGene_gn=DDR1;refGene_tx=XM_019830180.2;refGene_gn=DDR1;refGene_tx=XM_019830181.2;refGene_gn=DDR1;refGene_tx=XM_019830182.2;refGene_gn=DDR1;refGene_tx=XM_023253738.1
B2 31686116 31686351
refGene_gn=DDR1;refGene_tx=XM_006931528.3;refGene_gn=DDR1;refGene_tx=XM_019830180.2;refGene_gn=DDR1;refGene_tx=XM_019830181.2;refGene_gn=DDR1;refGene_tx=XM_019830182.2;refGene_gn=DDR1;refGene_tx=XM_023253738.1
B2 31686956 31687106
refGene_gn=DDR1;refGene_tx=XM_006931528.3;refGene_gn=DDR1;refGene_tx=XM_019830180.2;refGene_gn=DDR1;refGene_tx=XM_019830181.2;refGene_gn=DDR1;refGene_tx=XM_019830182.2;refGene_gn=DDR1;refGene_tx=XM_023253738.1
B2 31687212 31687353
refGene_gn=DDR1;refGene_tx=XM_006931528.3;refGene_gn=DDR1;refGene_tx=XM_019830180.2;refGene_gn=DDR1;refGene_tx=XM_019830181.2;refGene_gn=DDR1;refGene_tx=XM_019830182.2;refGene_gn=DDR1;refGene_tx=XM_023253738.1

B2 31694556 31694696
refGene_gn=GTF2H4;refGene_tx=XM_003985895.3;refGene_gn=GTF2H4;refGene_tx=XM_006931530.2;refGene_gn=GTF2H4;refGene_tx=XM_006931531.2;refGene_gn=GTF2H4;refGene_tx=XM_023253739.1
B2 31694954 31695059
refGene_gn=GTF2H4;refGene_tx=XM_003985895.3;refGene_gn=GTF2H4;refGene_tx=XM_006931530.2;refGene_gn=GTF2H4;refGene_tx=XM_006931531.2;refGene_gn=GTF2H4;refGene_tx=XM_023253739.1
B2 31695407 31695539
refGene_gn=GTF2H4;refGene_tx=XM_003985895.3;refGene_gn=GTF2H4;refGene_tx=XM_006931530.2;refGene_gn=GTF2H4;refGene_tx=XM_006931531.2;refGene_gn=GTF2H4;refGene_tx=XM_023253739.1
B2 31695817 31695914
refGene_gn=GTF2H4;refGene_tx=XM_003985895.3;refGene_gn=GTF2H4;refGene_tx=XM_006931530.2;refGene_gn=GTF2H4;refGene_tx=XM_006931531.2;refGene_gn=GTF2H4;refGene_tx=XM_023253739.1
B2 31696002 31696091
refGene_gn=GTF2H4;refGene_tx=XM_003985895.3;refGene_gn=GTF2H4;refGene_tx=XM_006931530.2;refGene_gn=GTF2H4;refGene_tx=XM_006931531.2;refGene_gn=GTF2H4;refGene_tx=XM_023253739.1
B2 31696287 31696399
refGene_gn=GTF2H4;refGene_tx=XM_003985895.3;refGene_gn=GTF2H4;refGene_tx=XM_006931530.2;refGene_gn=GTF2H4;refGene_tx=XM_006931531.2;refGene_gn=GTF2H4;refGene_tx=XM_023253739.1
B2 31696534 31696603
refGene_gn=GTF2H4;refGene_tx=XM_003985895.3;refGene_gn=GTF2H4;refGene_tx=XM_006931530.2;refGene_gn=GTF2H4;refGene_tx=XM_006931531.2;refGene_gn=GTF2H4;refGene_tx=XM_023253739.1
B2 31696804 31696888
refGene_gn=GTF2H4;refGene_tx=XM_003985895.3;refGene_gn=GTF2H4;refGene_tx=XM_006931530.2;refGene_gn=GTF2H4;refGene_tx=XM_006931531.2;refGene_gn=GTF2H4;refGene_tx=XM_023253739.1
B2 31697287 31697420
refGene_gn=GTF2H4;refGene_tx=XM_003985895.3;refGene_gn=GTF2H4;refGene_tx=XM_006931530.2;refGene_gn=GTF2H4;refGene_tx=XM_006931531.2;refGene_gn=GTF2H4;refGene_tx=XM_023253739.1
B2 31697606 31697737
refGene_gn=GTF2H4;refGene_tx=XM_003985895.3;refGene_gn=GTF2H4;refGene_tx=XM_006931530.2;refGene_gn=GTF2H4;refGene_tx=XM_006931531.2;refGene_gn=GTF2H4;refGene_tx=XM_023253739.1
B2 31698411 31698459
refGene_gn=GTF2H4;refGene_tx=XM_003985895.3;refGene_gn=GTF2H4;refGene_tx=XM_006931530.2;refGene_gn=GTF2H4;refGene_tx=XM_006931531.2;refGene_gn=GTF2H4;refGene_tx=XM_023253739.1
B2 31698606 31698685
refGene_gn=GTF2H4;refGene_tx=XM_003985895.3;refGene_gn=GTF2H4;refGene_tx=XM_006931530.2;refGene_gn=GTF2H4;refGene_tx=XM_006931531.2;refGene_gn=GTF2H4;refGene_tx=XM_023253739.1
B2 31699155 31699328
refGene_gn=GTF2H4;refGene_tx=XM_003985895.3;refGene_gn=GTF2H4;refGene_

tx=XM_006931530.2;refGene_gn=GTF2H4;refGene_tx=XM_006931531.2;refGene_gn=GTF2H4;refGene_tx=XM_023253739.1
B2 31699551 31699710 refGene_gn=VARS2;refGene_tx=XM_006931532.4
B2 31700167 31700368 refGene_gn=VARS2;refGene_tx=XM_006931533.4;refGene_gn=VARS2;refGene_tx=XM_019830184.2
B2 31700507 31700589 refGene_gn=VARS2;refGene_tx=XM_006931533.4
B2 31700705 31700806 refGene_gn=VARS2;refGene_tx=XM_006931533.4
B2 31701110 31701232 refGene_gn=VARS2;refGene_tx=XM_006931532.4;refGene_gn=VARS2;refGene_tx=XM_006931533.4;refGene_gn=VARS2;refGene_tx=XM_006931534.3;refGene_gn=VARS2;refGene_tx=XM_019830184.2
B2 31701351 31701418 refGene_gn=VARS2;refGene_tx=XM_006931532.4;refGene_gn=VARS2;refGene_tx=XM_006931533.4;refGene_gn=VARS2;refGene_tx=XM_006931534.3;refGene_gn=VARS2;refGene_tx=XM_019830184.2
B2 31701520 31701618 refGene_gn=VARS2;refGene_tx=XM_006931532.4;refGene_gn=VARS2;refGene_tx=XM_006931533.4;refGene_gn=VARS2;refGene_tx=XM_006931534.3;refGene_gn=VARS2;refGene_tx=XM_019830184.2
B2 31702328 31702410 refGene_gn=VARS2;refGene_tx=XM_006931532.4;refGene_gn=VARS2;refGene_tx=XM_006931533.4;refGene_gn=VARS2;refGene_tx=XM_006931534.3;refGene_gn=VARS2;refGene_tx=XM_019830184.2
B2 31702572 31702692 refGene_gn=VARS2;refGene_tx=XM_006931532.4;refGene_gn=VARS2;refGene_tx=XM_006931533.4;refGene_gn=VARS2;refGene_tx=XM_006931534.3;refGene_gn=VARS2;refGene_tx=XM_019830184.2
B2 31702972 31703084 refGene_gn=VARS2;refGene_tx=XM_006931532.4;refGene_gn=VARS2;refGene_tx=XM_006931533.4;refGene_gn=VARS2;refGene_tx=XM_006931534.3;refGene_gn=VARS2;refGene_tx=XM_019830184.2
B2 31703703 31703792 refGene_gn=VARS2;refGene_tx=XM_006931532.4;refGene_gn=VARS2;refGene_tx=XM_006931533.4;refGene_gn=VARS2;refGene_tx=XM_006931534.3;refGene_gn=VARS2;refGene_tx=XM_019830184.2
B2 31704307 31704398 refGene_gn=VARS2;refGene_tx=XM_006931532.4;refGene_gn=VARS2;refGene_tx=XM_006931533.4;refGene_gn=VARS2;refGene_tx=XM_006931534.3;refGene_gn=VARS2;refGene_tx=XM_019830184.2
B2 31704631 31704759 refGene_gn=VARS2;refGene_tx=XM_006931532.4;refGene_gn=VARS2;refGene_tx=XM_006931533.4;refGene_gn=VARS2;refGene_tx=XM_006931534.3;refGene_gn=VARS2;refGene_tx=XM_019830184.2
B2 31704899 31705003 refGene_gn=VARS2;refGene_tx=XM_006931532.4;refGene_gn=VARS2;refGene_tx=XM_006931533.4;refGene_gn=VARS2;refGene_tx=XM_006931534.3;refGene_gn=VARS2;refGene_tx=XM_019830184.2
B2 31705738 31705820 refGene_gn=VARS2;refGene_tx=XM_006931532.4;refGene_gn=VARS2;refGene_tx

=XM_006931533.4;refGene_gn=VARS2;refGene_tx=XM_006931534.3;refGene_gn=VARS2;refGene_tx=XM_019830184.2
B2 31706136 31706213
refGene_gn=VARS2;refGene_tx=XM_006931532.4;refGene_gn=VARS2;refGene_tx=XM_006931533.4;refGene_gn=VARS2;refGene_tx=XM_006931534.3;refGene_gn=VARS2;refGene_tx=XM_019830184.2
B2 31706293 31706369
refGene_gn=VARS2;refGene_tx=XM_006931532.4;refGene_gn=VARS2;refGene_tx=XM_006931533.4;refGene_gn=VARS2;refGene_tx=XM_006931534.3;refGene_gn=VARS2;refGene_tx=XM_019830184.2
B2 31706658 31706764
refGene_gn=VARS2;refGene_tx=XM_006931532.4;refGene_gn=VARS2;refGene_tx=XM_006931533.4;refGene_gn=VARS2;refGene_tx=XM_006931534.3;refGene_gn=VARS2;refGene_tx=XM_019830184.2
B2 31706963 31707034
refGene_gn=VARS2;refGene_tx=XM_006931532.4;refGene_gn=VARS2;refGene_tx=XM_006931533.4;refGene_gn=VARS2;refGene_tx=XM_006931534.3;refGene_gn=VARS2;refGene_tx=XM_019830184.2
B2 31707164 31707290
refGene_gn=VARS2;refGene_tx=XM_006931532.4;refGene_gn=VARS2;refGene_tx=XM_006931533.4;refGene_gn=VARS2;refGene_tx=XM_006931534.3;refGene_gn=VARS2;refGene_tx=XM_019830184.2
B2 31707511 31707616
refGene_gn=VARS2;refGene_tx=XM_006931532.4;refGene_gn=VARS2;refGene_tx=XM_006931533.4;refGene_gn=VARS2;refGene_tx=XM_006931534.3;refGene_gn=VARS2;refGene_tx=XM_019830184.2
B2 31707757 31707826
refGene_gn=VARS2;refGene_tx=XM_006931532.4;refGene_gn=VARS2;refGene_tx=XM_006931533.4;refGene_gn=VARS2;refGene_tx=XM_006931534.3;refGene_gn=VARS2;refGene_tx=XM_019830184.2
B2 31707951 31708030
refGene_gn=VARS2;refGene_tx=XM_006931532.4;refGene_gn=VARS2;refGene_tx=XM_006931533.4;refGene_gn=VARS2;refGene_tx=XM_006931534.3;refGene_gn=VARS2;refGene_tx=XM_019830184.2
B2 31708151 31708279
refGene_gn=VARS2;refGene_tx=XM_006931532.4;refGene_gn=VARS2;refGene_tx=XM_006931533.4;refGene_gn=VARS2;refGene_tx=XM_006931534.3;refGene_gn=VARS2;refGene_tx=XM_019830184.2
B2 31708398 31708551
refGene_gn=VARS2;refGene_tx=XM_006931532.4;refGene_gn=VARS2;refGene_tx=XM_006931533.4;refGene_gn=VARS2;refGene_tx=XM_006931534.3;refGene_gn=VARS2;refGene_tx=XM_019830184.2
B2 31708925 31709132
refGene_gn=VARS2;refGene_tx=XM_006931532.4;refGene_gn=VARS2;refGene_tx=XM_006931533.4;refGene_gn=VARS2;refGene_tx=XM_006931534.3;refGene_gn=VARS2;refGene_tx=XM_019830184.2
B2 31709836 31709976
refGene_gn=SFTA2;refGene_tx=XM_019830185.2;refGene_gn=VARS2;refGene_tx=XM_006931532.4;refGene_gn=VARS2;refGene_tx=XM_006931533.4;refGene_gn=VARS2;refGene_tx=XM_006931534.3;refGene_gn=VARS2;refGene_tx=XM_0198301

84.2
B2 31710106 31710282
refGene_gn=VARS2;refGene_tx=XM_006931532.4;refGene_gn=VARS2;refGene_tx=XM_006931533.4;refGene_gn=VARS2;refGene_tx=XM_006931534.3;refGene_gn=VARS2;refGene_tx=XM_019830184.2
B2 31710453 31710582
refGene_gn=VARS2;refGene_tx=XM_006931532.4;refGene_gn=VARS2;refGene_tx=XM_006931533.4;refGene_gn=VARS2;refGene_tx=XM_006931534.3;refGene_gn=VARS2;refGene_tx=XM_019830184.2
B2 31710676 31710772
refGene_gn=VARS2;refGene_tx=XM_006931532.4;refGene_gn=VARS2;refGene_tx=XM_006931533.4;refGene_gn=VARS2;refGene_tx=XM_006931534.3;refGene_gn=VARS2;refGene_tx=XM_019830184.2
B2 31713602 31713700 refGene_gn=SFTA2;refGene_tx=XM_019830185.2
B2 31714440 31714527 refGene_gn=SFTA2;refGene_tx=XM_003985902.3
B2 31714676 31714765
refGene_gn=SFTA2;refGene_tx=XM_003985902.3;refGene_gn=SFTA2;refGene_tx=XM_019830185.2
B2 31715004 31715065
refGene_gn=SFTA2;refGene_tx=XM_003985902.3;refGene_gn=SFTA2;refGene_tx=XM_019830185.2
B2 31723846 31723928 refGene_gn=DPCR1;refGene_tx=XM_019830183.2
B2 31729900 31731701 refGene_gn=DPCR1;refGene_tx=XM_019830183.2
B2 31731703 31733045 refGene_gn=DPCR1;refGene_tx=XM_019830183.2
B2 31733548 31733695 refGene_gn=DPCR1;refGene_tx=XM_019830183.2
B2 31765661 31765725
refGene_gn=MUC21;refGene_tx=XM_019830186.2;refGene_gn=MUC21;refGene_tx=XM_019830187.2;refGene_gn=MUC21;refGene_tx=XM_023253740.1
B2 31767104 31768246
refGene_gn=MUC21;refGene_tx=XM_019830186.2;refGene_gn=MUC21;refGene_tx=XM_019830187.2;refGene_gn=MUC21;refGene_tx=XM_023253740.1
B2 31768848 31769037
refGene_gn=MUC21;refGene_tx=XM_019830186.2;refGene_gn=MUC21;refGene_tx=XM_019830187.2;refGene_gn=MUC21;refGene_tx=XM_023253740.1
B2 31851969 31852856
refGene_gn=CDSN;refGene_tx=XM_003985896.3;refGene_gn=CDSN;refGene_tx=XM_019830188.1;refGene_gn=CDSN;refGene_tx=XM_019830189.1
B2 31853061 31853149 refGene_gn=CDSN;refGene_tx=XM_003985896.3
B2 31857154 31858650
refGene_gn=CDSN;refGene_tx=XM_003985897.4;refGene_gn=CDSN;refGene_tx=XM_019830188.1;refGene_gn=CDSN;refGene_tx=XM_019830189.1
B2 31860798 31860883
refGene_gn=CDSN;refGene_tx=XM_003985897.4;refGene_gn=CDSN;refGene_tx=XM_019830188.1;refGene_gn=CDSN;refGene_tx=XM_019830189.1
B2 31875033 31875389 refGene_gn=PSORS1C2;refGene_tx=XM_003985898.5
B2 31876581 31876636 refGene_gn=PSORS1C2;refGene_tx=XM_003985898.5
B2 31879589 31879702
refGene_gn=CCHCR1;refGene_tx=XM_011282120.3;refGene_gn=CCHCR1;refGene_tx=XM_011282122.3;refGene_gn=CCHCR1;refGene_tx=XM_011282124.3;refGene_gn=CCHCR1;refGene_tx=XM_011282125.3;refGene_gn=CCHCR1;refGene_tx=XM_02

3253741.1
B2	31879947	31880119
refGene_gn=CCHCR1;refGene_tx=XM_011282120.3;refGene_gn=CCHCR1;refGene_tx=XM_011282122.3;refGene_gn=CCHCR1;refGene_tx=XM_011282124.3;refGene_gn=CCHCR1;refGene_tx=XM_011282125.3;refGene_gn=CCHCR1;refGene_tx=XM_023253741.1
B2	31880232	31880384
refGene_gn=CCHCR1;refGene_tx=XM_011282120.3;refGene_gn=CCHCR1;refGene_tx=XM_011282122.3;refGene_gn=CCHCR1;refGene_tx=XM_011282124.3;refGene_gn=CCHCR1;refGene_tx=XM_011282125.3;refGene_gn=CCHCR1;refGene_tx=XM_023253741.1
B2	31882097	31882199
refGene_gn=CCHCR1;refGene_tx=XM_011282120.3;refGene_gn=CCHCR1;refGene_tx=XM_011282122.3;refGene_gn=CCHCR1;refGene_tx=XM_011282124.3;refGene_gn=CCHCR1;refGene_tx=XM_011282125.3;refGene_gn=CCHCR1;refGene_tx=XM_023253741.1
B2	31882299	31882488
refGene_gn=CCHCR1;refGene_tx=XM_011282120.3;refGene_gn=CCHCR1;refGene_tx=XM_011282122.3;refGene_gn=CCHCR1;refGene_tx=XM_011282124.3;refGene_gn=CCHCR1;refGene_tx=XM_011282125.3;refGene_gn=CCHCR1;refGene_tx=XM_023253741.1
B2	31882610	31882747
refGene_gn=CCHCR1;refGene_tx=XM_011282120.3;refGene_gn=CCHCR1;refGene_tx=XM_011282122.3;refGene_gn=CCHCR1;refGene_tx=XM_011282124.3;refGene_gn=CCHCR1;refGene_tx=XM_011282125.3;refGene_gn=CCHCR1;refGene_tx=XM_023253741.1
B2	31882887	31882933
refGene_gn=CCHCR1;refGene_tx=XM_011282120.3;refGene_gn=CCHCR1;refGene_tx=XM_011282122.3;refGene_gn=CCHCR1;refGene_tx=XM_011282124.3;refGene_gn=CCHCR1;refGene_tx=XM_011282125.3;refGene_gn=CCHCR1;refGene_tx=XM_023253741.1
B2	31883105	31883218
refGene_gn=CCHCR1;refGene_tx=XM_011282120.3;refGene_gn=CCHCR1;refGene_tx=XM_011282122.3;refGene_gn=CCHCR1;refGene_tx=XM_011282124.3;refGene_gn=CCHCR1;refGene_tx=XM_011282125.3;refGene_gn=CCHCR1;refGene_tx=XM_023253741.1
B2	31885990	31886097
refGene_gn=CCHCR1;refGene_tx=XM_011282120.3;refGene_gn=CCHCR1;refGene_tx=XM_011282122.3;refGene_gn=CCHCR1;refGene_tx=XM_011282124.3;refGene_gn=CCHCR1;refGene_tx=XM_011282125.3;refGene_gn=CCHCR1;refGene_tx=XM_023253741.1
B2	31886186	31886297
refGene_gn=CCHCR1;refGene_tx=XM_011282120.3;refGene_gn=CCHCR1;refGene_tx=XM_011282122.3;refGene_gn=CCHCR1;refGene_tx=XM_011282124.3;refGene_gn=CCHCR1;refGene_tx=XM_011282125.3;refGene_gn=CCHCR1;refGene_tx=XM_023253741.1
B2	31886975	31887125
refGene_gn=CCHCR1;refGene_tx=XM_011282120.3;refGene_gn=CCHCR1;refGene_tx=XM_011282122.3;refGene_gn=CCHCR1;refGene_tx=XM_011282124.3;refGene_gn=CCHCR1;refGene_tx=XM_011282125.3;refGene_gn=CCHCR1;refGene_tx=XM_02

3253741.1
B2	31887319 31887430
refGene_gn=CCHCR1;refGene_tx=XM_011282120.3;refGene_gn=CCHCR1;refGene_tx=XM_011282122.3;refGene_gn=CCHCR1;refGene_tx=XM_011282124.3;refGene_gn=CCHCR1;refGene_tx=XM_011282125.3;refGene_gn=CCHCR1;refGene_tx=XM_023253741.1
B2	31887645 31887781
refGene_gn=CCHCR1;refGene_tx=XM_011282120.3;refGene_gn=CCHCR1;refGene_tx=XM_011282122.3;refGene_gn=CCHCR1;refGene_tx=XM_011282124.3;refGene_gn=CCHCR1;refGene_tx=XM_011282125.3;refGene_gn=CCHCR1;refGene_tx=XM_023253741.1
B2	31887882 31888046
refGene_gn=CCHCR1;refGene_tx=XM_011282120.3;refGene_gn=CCHCR1;refGene_tx=XM_011282122.3;refGene_gn=CCHCR1;refGene_tx=XM_011282124.3;refGene_gn=CCHCR1;refGene_tx=XM_011282125.3;refGene_gn=CCHCR1;refGene_tx=XM_023253741.1
B2	31890454 31890758
refGene_gn=CCHCR1;refGene_tx=XM_011282120.3;refGene_gn=CCHCR1;refGene_tx=XM_011282122.3;refGene_gn=CCHCR1;refGene_tx=XM_011282124.3;refGene_gn=CCHCR1;refGene_tx=XM_011282125.3;refGene_gn=CCHCR1;refGene_tx=XM_023253741.1
B2	31892505 31892716
refGene_gn=CCHCR1;refGene_tx=XM_011282120.3;refGene_gn=CCHCR1;refGene_tx=XM_011282122.3;refGene_gn=CCHCR1;refGene_tx=XM_011282124.3;refGene_gn=CCHCR1;refGene_tx=XM_011282125.3;refGene_gn=CCHCR1;refGene_tx=XM_023253741.1
B2	31892798 31892865
refGene_gn=CCHCR1;refGene_tx=XM_011282120.3;refGene_gn=CCHCR1;refGene_tx=XM_011282122.3;refGene_gn=CCHCR1;refGene_tx=XM_011282124.3;refGene_gn=CCHCR1;refGene_tx=XM_011282125.3;refGene_gn=CCHCR1;refGene_tx=XM_023253741.1
B2	31893122 31893272 refGene_gn=CCHCR1;refGene_tx=XM_011282120.3
B2	31895292 31895530
refGene_gn=TCF19;refGene_tx=XM_006931536.3;refGene_gn=TCF19;refGene_tx=XM_006931537.3
B2	31896635 31897209
refGene_gn=TCF19;refGene_tx=XM_006931536.3;refGene_gn=TCF19;refGene_tx=XM_006931537.3
B2	31897476 31897717
refGene_gn=TCF19;refGene_tx=XM_006931536.3;refGene_gn=TCF19;refGene_tx=XM_006931537.3
B2	31899613 31899880 refGene_gn=POU5F1;refGene_tx=NM_001173441.1
B2	31900064 31900223 refGene_gn=POU5F1;refGene_tx=NM_001173441.1
B2	31900545 31900676 refGene_gn=POU5F1;refGene_tx=NM_001173441.1
B2	31900989 31901110 refGene_gn=POU5F1;refGene_tx=NM_001173441.1
B2	31904429 31904834 refGene_gn=POU5F1;refGene_tx=NM_001173441.1
B2	31915802 31915828
refGene_gn=LOC101080285;refGene_tx=XM_023254707.1
B2	31931295 31931571
refGene_gn=LOC101080285;refGene_tx=XM_023254707.1

B2 31932213 31932450
refGene_gn=LOC101080285;refGene_tx=XM_023254707.1
B2 31932724 31932993
refGene_gn=LOC101080285;refGene_tx=XM_023254707.1
B2 31933264 31933473
refGene_gn=LOC101080285;refGene_tx=XM_023254707.1
B2 31998258 31998263
refGene_gn=LOC105260551;refGene_tx=XM_011282114.3
B2 31998392 31998440
refGene_gn=LOC105260551;refGene_tx=XM_011282114.3
B2 31998591 31998624
refGene_gn=LOC105260551;refGene_tx=XM_011282114.3
B2 31999062 31999170
refGene_gn=LOC105260551;refGene_tx=XM_011282114.3
B2 31999286 31999562
refGene_gn=LOC105260551;refGene_tx=XM_011282114.3
B2 32000175 32000407
refGene_gn=LOC105260551;refGene_tx=XM_011282114.3
B2 32000657 32000927
refGene_gn=LOC105260551;refGene_tx=XM_011282114.3
B2 32001208 32001281
refGene_gn=LOC105260551;refGene_tx=XM_011282114.3
B2 32066649 32066691
refGene_gn=LOC101081297;refGene_tx=XM_023253743.1
B2 32066692 32066723
refGene_gn=LOC101081297;refGene_tx=XM_023253743.1
B2 32067004 32067274
refGene_gn=LOC101081297;refGene_tx=XM_023253743.1
B2 32067612 32067650
refGene_gn=LOC101081297;refGene_tx=XM_023253743.1
B2 32067651 32067895
refGene_gn=LOC101081297;refGene_tx=XM_023253743.1
B2 32068503 32068779
refGene_gn=LOC101081297;refGene_tx=XM_023253743.1
B2 32068896 32069004
refGene_gn=LOC101081297;refGene_tx=XM_023253743.1
B2 32069431 32069464
refGene_gn=LOC101081297;refGene_tx=XM_023253743.1
B2 32069608 32069656
refGene_gn=LOC101081297;refGene_tx=XM_023253743.1
B2 32069786 32069791
refGene_gn=LOC101081297;refGene_tx=XM_023253743.1
B2 32105224 32105297
refGene_gn=LOC101080790;refGene_tx=XM_023253744.1
B2 32105581 32105851
refGene_gn=LOC101080790;refGene_tx=XM_023253744.1
B2 32106051 32106374
refGene_gn=LOC101080790;refGene_tx=XM_023253744.1
B2 32497200 32497204
refGene_gn=LOC101099296;refGene_tx=XM_019830338.2

B2 32497341 32497374
refGene_gn=LOC101099296;refGene_tx=XM_019830338.2
B2 32520311 32520447
refGene_gn=LOC101099296;refGene_tx=XM_019830327.2;refGene_gn=LOC101099296;refGene_tx=XM_019830334.2
B2 32522961 32523025
refGene_gn=LOC101099296;refGene_tx=XM_019830335.2;refGene_gn=LOC101099296;refGene_tx=XM_019830336.2
B2 32523223 32523227
refGene_gn=LOC101099296;refGene_tx=XM_019830341.2;refGene_gn=LOC101099296;refGene_tx=XM_019830342.2;refGene_gn=LOC101099296;refGene_tx=XM_019830343.2
B2 32523363 32523396
refGene_gn=LOC101099296;refGene_tx=XM_019830327.2;refGene_gn=LOC101099296;refGene_tx=XM_019830334.2;refGene_gn=LOC101099296;refGene_tx=XM_019830335.2;refGene_gn=LOC101099296;refGene_tx=XM_019830336.2;refGene_gn=LOC101099296;refGene_tx=XM_019830338.2;refGene_gn=LOC101099296;refGene_tx=XM_019830341.2;refGene_gn=LOC101099296;refGene_tx=XM_019830342.2;refGene_gn=LOC101099296;refGene_tx=XM_019830343.2
B2 32523829 32523943
refGene_gn=LOC101099296;refGene_tx=XM_019830327.2;refGene_gn=LOC101099296;refGene_tx=XM_019830328.2;refGene_gn=LOC101099296;refGene_tx=XM_019830329.2;refGene_gn=LOC101099296;refGene_tx=XM_019830330.2;refGene_gn=LOC101099296;refGene_tx=XM_019830331.2;refGene_gn=LOC101099296;refGene_tx=XM_019830332.2;refGene_gn=LOC101099296;refGene_tx=XM_019830333.2;refGene_gn=LOC101099296;refGene_tx=XM_019830334.2;refGene_gn=LOC101099296;refGene_tx=XM_019830335.2;refGene_gn=LOC101099296;refGene_tx=XM_019830336.2;refGene_gn=LOC101099296;refGene_tx=XM_019830338.2;refGene_gn=LOC101099296;refGene_tx=XM_019830339.1;refGene_gn=LOC101099296;refGene_tx=XM_019830341.2;refGene_gn=LOC101099296;refGene_tx=XM_019830342.2;refGene_gn=LOC101099296;refGene_tx=XM_019830343.2;refGene_gn=LOC101099296;refGene_tx=XM_019830344.2;refGene_gn=LOC101099296;refGene_tx=XM_019830345.1;refGene_gn=LOC101099296;refGene_tx=XM_023253745.1
B2 32524070 32524346
refGene_gn=LOC101099296;refGene_tx=XM_019830327.2;refGene_gn=LOC101099296;refGene_tx=XM_019830328.2;refGene_gn=LOC101099296;refGene_tx=XM_019830329.2;refGene_gn=LOC101099296;refGene_tx=XM_019830330.2;refGene_gn=LOC101099296;refGene_tx=XM_019830331.2;refGene_gn=LOC101099296;refGene_tx=XM_019830332.2;refGene_gn=LOC101099296;refGene_tx=XM_019830333.2;refGene_gn=LOC101099296;refGene_tx=XM_019830334.2;refGene_gn=LOC101099296;refGene_tx=XM_019830335.2;refGene_gn=LOC101099296;refGene_tx=XM_019830336.2;refGene_gn=LOC101099296;refGene_tx=XM_019830338.2;refGene_gn=LOC101099296;refGene_tx=XM_019830339.1;refGene_gn=LOC101099296;refGene_tx=XM_019830341.2;refGene_gn=LOC101099296;refGene_tx=XM_019830342.2;refGene_gn=LOC101099296;refGene_tx=XM_019830343.2;refGene_gn=LOC101099296;refGene_tx=XM_023253745.1
B2 32524959 32525235
refGene_gn=LOC101099296;refGene_tx=XM_019830327.2;refGene_gn=LOC101099296;refGene_tx=XM_019830328.2;refGene_gn=LOC101099296;refGene_tx=XM_019830329.2;refGene_gn=LOC101099296;refGene_tx=XM_019830330.2;refGene_gn

=LOC101099296;refGene_tx=XM_019830331.2;refGene_gn=LOC101099296;refGene_tx=XM_019830332.2;refGene_gn=LOC101099296;refGene_tx=XM_019830333.2;refGene_gn=LOC101099296;refGene_tx=XM_019830334.2;refGene_gn=LOC101099296;refGene_tx=XM_019830335.2;refGene_gn=LOC101099296;refGene_tx=XM_019830336.2;refGene_gn=LOC101099296;refGene_tx=XM_019830338.2;refGene_gn=LOC101099296;refGene_tx=XM_019830339.1;refGene_gn=LOC101099296;refGene_tx=XM_019830341.2;refGene_gn=LOC101099296;refGene_tx=XM_019830342.2;refGene_gn=LOC101099296;refGene_tx=XM_019830343.2;refGene_gn=LOC101099296;refGene_tx=XM_019830344.2;refGene_gn=LOC101099296;refGene_tx=XM_019830345.1;refGene_gn=LOC101099296;refGene_tx=XM_023253745.1
B2 32525460 32525730
refGene_gn=LOC101099296;refGene_tx=XM_019830327.2;refGene_gn=LOC101099296;refGene_tx=XM_019830328.2;refGene_gn=LOC101099296;refGene_tx=XM_019830329.2;refGene_gn=LOC101099296;refGene_tx=XM_019830330.2;refGene_gn=LOC101099296;refGene_tx=XM_019830331.2;refGene_gn=LOC101099296;refGene_tx=XM_019830332.2;refGene_gn=LOC101099296;refGene_tx=XM_019830333.2;refGene_gn=LOC101099296;refGene_tx=XM_019830334.2;refGene_gn=LOC101099296;refGene_tx=XM_019830335.2;refGene_gn=LOC101099296;refGene_tx=XM_019830336.2;refGene_gn=LOC101099296;refGene_tx=XM_019830338.2;refGene_gn=LOC101099296;refGene_tx=XM_019830339.1;refGene_gn=LOC101099296;refGene_tx=XM_019830341.2;refGene_gn=LOC101099296;refGene_tx=XM_019830342.2;refGene_gn=LOC101099296;refGene_tx=XM_019830343.2;refGene_gn=LOC101099296;refGene_tx=XM_019830344.2;refGene_gn=LOC101099296;refGene_tx=XM_019830345.1;refGene_gn=LOC101099296;refGene_tx=XM_023253745.1
B2 32525998 32526198
refGene_gn=LOC101099296;refGene_tx=XM_019830327.2;refGene_gn=LOC101099296;refGene_tx=XM_019830328.2;refGene_gn=LOC101099296;refGene_tx=XM_019830329.2;refGene_gn=LOC101099296;refGene_tx=XM_019830330.2;refGene_gn=LOC101099296;refGene_tx=XM_019830331.2;refGene_gn=LOC101099296;refGene_tx=XM_019830332.2;refGene_gn=LOC101099296;refGene_tx=XM_019830333.2;refGene_gn=LOC101099296;refGene_tx=XM_019830334.2;refGene_gn=LOC101099296;refGene_tx=XM_019830335.2;refGene_gn=LOC101099296;refGene_tx=XM_019830336.2;refGene_gn=LOC101099296;refGene_tx=XM_019830338.2;refGene_gn=LOC101099296;refGene_tx=XM_019830339.1;refGene_gn=LOC101099296;refGene_tx=XM_019830341.2;refGene_gn=LOC101099296;refGene_tx=XM_019830342.2;refGene_gn=LOC101099296;refGene_tx=XM_019830343.2;refGene_gn=LOC101099296;refGene_tx=XM_019830344.2;refGene_gn=LOC101099296;refGene_tx=XM_019830345.1;refGene_gn=LOC101099296;refGene_tx=XM_023253745.1
B2 32541545 32541710 refGene_gn=MCCD1;refGene_tx=XM_003985909.3
B2 32542226 32542412 refGene_gn=MCCD1;refGene_tx=XM_003985909.3
B2 32542946 32542963
refGene_gn=DDX39B;refGene_tx=XM_003985910.4;refGene_gn=DDX39B;refGene_tx=XM_019830346.1
B2 32543284 32543432
refGene_gn=DDX39B;refGene_tx=XM_003985910.4;refGene_gn=DDX39B;refGene_tx=XM_019830346.1
B2 32543576 32543721
refGene_gn=DDX39B;refGene_tx=XM_003985910.4;refGene_gn=DDX39B;refGene_tx=XM_019830346.1
B2 32543819 32543929

refGene_gn=DDX39B;refGene_tx=XM_003985910.4;refGene_gn=DDX39B;refGene_tx=XM_019830346.1
B2 32544689 32544821
refGene_gn=DDX39B;refGene_tx=XM_003985910.4;refGene_gn=DDX39B;refGene_tx=XM_019830346.1
B2 32546891 32547010
refGene_gn=DDX39B;refGene_tx=XM_003985910.4;refGene_gn=DDX39B;refGene_tx=XM_019830346.1
B2 32548301 32548485
refGene_gn=DDX39B;refGene_tx=XM_003985910.4;refGene_gn=DDX39B;refGene_tx=XM_019830346.1
B2 32550882 32550975
refGene_gn=DDX39B;refGene_tx=XM_003985910.4;refGene_gn=DDX39B;refGene_tx=XM_019830346.1
B2 32551269 32551397
refGene_gn=DDX39B;refGene_tx=XM_003985910.4;refGene_gn=DDX39B;refGene_tx=XM_019830346.1
B2 32552381 32552592
refGene_gn=DDX39B;refGene_tx=XM_003985910.4;refGene_gn=DDX39B;refGene_tx=XM_019830346.1
B2 32556542 32556716
refGene_gn=ATP6V1G2;refGene_tx=XM_003985913.5;refGene_gn=ATP6V1G2;refGene_tx=XM_019830347.2
B2 32557246 32557347
refGene_gn=ATP6V1G2;refGene_tx=XM_003985913.5;refGene_gn=ATP6V1G2;refGene_tx=XM_019830347.2
B2 32557636 32557718 refGene_gn=ATP6V1G2;refGene_tx=XM_003985913.5
B2 32559024 32559081 refGene_gn=NFKBIL1;refGene_tx=XM_003985912.5
B2 32559475 32559752 refGene_gn=NFKBIL1;refGene_tx=XM_003985912.5
B2 32566775 32566997 refGene_gn=NFKBIL1;refGene_tx=XM_003985912.5
B2 32567177 32567761 refGene_gn=NFKBIL1;refGene_tx=XM_003985912.5
B2 32580230 32580329 refGene_gn=LTA;refGene_tx=XM_023253748.1
B2 32580420 32580526 refGene_gn=LTA;refGene_tx=XM_023253748.1
B2 32580771 32581184 refGene_gn=LTA;refGene_tx=XM_023253748.1
B2 32583571 32583757 refGene_gn=TNF;refGene_tx=NM_001009835.1
B2 32584310 32584356 refGene_gn=TNF;refGene_tx=NM_001009835.1
B2 32584540 32584588 refGene_gn=TNF;refGene_tx=NM_001009835.1
B2 32584917 32585339 refGene_gn=TNF;refGene_tx=NM_001009835.1
B2 32588297 32588752 refGene_gn=LTB;refGene_tx=XM_003985916.5
B2 32589177 32589249
refGene_gn=LTB;refGene_tx=XM_003985916.5;refGene_gn=LTB;refGene_tx=XM_019830349.2
B2 32589447 32589499 refGene_gn=LTB;refGene_tx=XM_003985916.5
B2 32589923 32590085
refGene_gn=LTB;refGene_tx=XM_003985916.5;refGene_gn=LTB;refGene_tx=XM_019830349.2
B2 32593292 32593320 refGene_gn=LST1;refGene_tx=XM_006931549.4
B2 32593686 32593788 refGene_gn=LST1;refGene_tx=XM_006931549.4
B2 32594075 32594098 refGene_gn=LST1;refGene_tx=XM_006931549.4
B2 32594582 32594717 refGene_gn=LST1;refGene_tx=XM_006931549.4

B2 32595322 32595369
refGene_gn=NCR3;refGene_tx=XM_003985976.4;refGene_gn=NCR3;refGene_tx=XM_019830350.2
B2 32595478 32595586 refGene_gn=NCR3;refGene_tx=XM_003985976.4
B2 32595726 32596071
refGene_gn=NCR3;refGene_tx=XM_003985976.4;refGene_gn=NCR3;refGene_tx=XM_019830350.2
B2 32597743 32597786
refGene_gn=NCR3;refGene_tx=XM_003985976.4;refGene_gn=NCR3;refGene_tx=XM_019830350.2
B2 32615181 32615206
refGene_gn=AIF1;refGene_tx=XM_003985918.4;refGene_gn=AIF1;refGene_tx=XM_006931550.3;refGene_gn=AIF1;refGene_tx=XM_019830351.1
B2 32615383 32615445
refGene_gn=AIF1;refGene_tx=XM_003985918.4;refGene_gn=AIF1;refGene_tx=XM_006931550.3;refGene_gn=AIF1;refGene_tx=XM_019830351.1;refGene_gn=AIF1;refGene_tx=XM_019830352.2
B2 32615536 32615603
refGene_gn=AIF1;refGene_tx=XM_003985918.4;refGene_gn=AIF1;refGene_tx=XM_006931550.3;refGene_gn=AIF1;refGene_tx=XM_019830352.2
B2 32615872 32616011
refGene_gn=AIF1;refGene_tx=XM_003985918.4;refGene_gn=AIF1;refGene_tx=XM_019830351.1;refGene_gn=AIF1;refGene_tx=XM_019830352.2
B2 32616193 32616356
refGene_gn=AIF1;refGene_tx=XM_003985918.4;refGene_gn=AIF1;refGene_tx=XM_006931550.3;refGene_gn=AIF1;refGene_tx=XM_019830351.1;refGene_gn=AIF1;refGene_tx=XM_019830352.2
B2 32616905 32616990
refGene_gn=AIF1;refGene_tx=XM_003985918.4;refGene_gn=AIF1;refGene_tx=XM_006931550.3;refGene_gn=AIF1;refGene_tx=XM_019830351.1;refGene_gn=AIF1;refGene_tx=XM_019830352.2
B2 32624030 32624142
refGene_gn=PRRC2A;refGene_tx=XM_023253749.1;refGene_gn=PRRC2A;refGene_tx=XM_023253750.1;refGene_gn=PRRC2A;refGene_tx=XM_023253751.1
B2 32624633 32624811
refGene_gn=PRRC2A;refGene_tx=XM_023253749.1;refGene_gn=PRRC2A;refGene_tx=XM_023253750.1;refGene_gn=PRRC2A;refGene_tx=XM_023253751.1
B2 32625163 32625263
refGene_gn=PRRC2A;refGene_tx=XM_023253749.1;refGene_gn=PRRC2A;refGene_tx=XM_023253750.1;refGene_gn=PRRC2A;refGene_tx=XM_023253751.1
B2 32625358 32625431
refGene_gn=PRRC2A;refGene_tx=XM_023253749.1;refGene_gn=PRRC2A;refGene_tx=XM_023253750.1;refGene_gn=PRRC2A;refGene_tx=XM_023253751.1
B2 32625756 32625900
refGene_gn=PRRC2A;refGene_tx=XM_023253749.1;refGene_gn=PRRC2A;refGene_tx=XM_023253750.1;refGene_gn=PRRC2A;refGene_tx=XM_023253751.1
B2 32626060 32626212
refGene_gn=PRRC2A;refGene_tx=XM_023253749.1;refGene_gn=PRRC2A;refGene_tx=XM_023253750.1;refGene_gn=PRRC2A;refGene_tx=XM_023253751.1
B2 32626394 32626474

refGene_gn=PRRC2A;refGene_tx=XM_023253749.1;refGene_gn=PRRC2A;refGene_tx=XM_023253750.1;refGene_gn=PRRC2A;refGene_tx=XM_023253751.1
B2 32626852 32626995
refGene_gn=PRRC2A;refGene_tx=XM_023253749.1;refGene_gn=PRRC2A;refGene_tx=XM_023253750.1;refGene_gn=PRRC2A;refGene_tx=XM_023253751.1
B2 32627295 32627386
refGene_gn=PRRC2A;refGene_tx=XM_023253749.1;refGene_gn=PRRC2A;refGene_tx=XM_023253750.1;refGene_gn=PRRC2A;refGene_tx=XM_023253751.1
B2 32627479 32627696
refGene_gn=PRRC2A;refGene_tx=XM_023253749.1;refGene_gn=PRRC2A;refGene_tx=XM_023253750.1;refGene_gn=PRRC2A;refGene_tx=XM_023253751.1
B2 32628101 32628576
refGene_gn=PRRC2A;refGene_tx=XM_023253749.1;refGene_gn=PRRC2A;refGene_tx=XM_023253750.1;refGene_gn=PRRC2A;refGene_tx=XM_023253751.1
B2 32628948 32629139
refGene_gn=PRRC2A;refGene_tx=XM_023253749.1;refGene_gn=PRRC2A;refGene_tx=XM_023253750.1;refGene_gn=PRRC2A;refGene_tx=XM_023253751.1
B2 32629389 32629687
refGene_gn=PRRC2A;refGene_tx=XM_023253749.1;refGene_gn=PRRC2A;refGene_tx=XM_023253750.1;refGene_gn=PRRC2A;refGene_tx=XM_023253751.1
B2 32630087 32630298
refGene_gn=PRRC2A;refGene_tx=XM_023253749.1;refGene_gn=PRRC2A;refGene_tx=XM_023253750.1;refGene_gn=PRRC2A;refGene_tx=XM_023253751.1
B2 32630489 32632355
refGene_gn=PRRC2A;refGene_tx=XM_023253749.1;refGene_gn=PRRC2A;refGene_tx=XM_023253750.1;refGene_gn=PRRC2A;refGene_tx=XM_023253751.1
B2 32632731 32632997
refGene_gn=PRRC2A;refGene_tx=XM_023253749.1;refGene_gn=PRRC2A;refGene_tx=XM_023253750.1;refGene_gn=PRRC2A;refGene_tx=XM_023253751.1
B2 32633197 32633328
refGene_gn=PRRC2A;refGene_tx=XM_023253749.1;refGene_gn=PRRC2A;refGene_tx=XM_023253750.1;refGene_gn=PRRC2A;refGene_tx=XM_023253751.1
B2 32633556 32633686
refGene_gn=PRRC2A;refGene_tx=XM_023253749.1;refGene_gn=PRRC2A;refGene_tx=XM_023253750.1;refGene_gn=PRRC2A;refGene_tx=XM_023253751.1
B2 32633791 32633877
refGene_gn=PRRC2A;refGene_tx=XM_023253749.1;refGene_gn=PRRC2A;refGene_tx=XM_023253750.1;refGene_gn=PRRC2A;refGene_tx=XM_023253751.1
B2 32634073 32634307
refGene_gn=PRRC2A;refGene_tx=XM_023253749.1;refGene_gn=PRRC2A;refGene_tx=XM_023253750.1;refGene_gn=PRRC2A;refGene_tx=XM_023253751.1
B2 32634446 32634587
refGene_gn=PRRC2A;refGene_tx=XM_023253749.1;refGene_gn=PRRC2A;refGene_tx=XM_023253750.1;refGene_gn=PRRC2A;refGene_tx=XM_023253751.1
B2 32634706 32634775
refGene_gn=PRRC2A;refGene_tx=XM_023253749.1;refGene_gn=PRRC2A;refGene_tx=XM_023253750.1
B2 32634891 32635059
refGene_gn=PRRC2A;refGene_tx=XM_023253749.1;refGene_gn=PRRC2A;refGene_tx=XM_023253750.1;refGene_gn=PRRC2A;refGene_tx=XM_023253751.1

B2	32635249 32635332
refGene_gn=PRRC2A;refGene_tx=XM_023253749.1;refGene_gn=PRRC2A;refGene_tx=XM_023253750.1;refGene_gn=PRRC2A;refGene_tx=XM_023253751.1
B2	32635499 32635710
refGene_gn=PRRC2A;refGene_tx=XM_023253749.1;refGene_gn=PRRC2A;refGene_tx=XM_023253750.1;refGene_gn=PRRC2A;refGene_tx=XM_023253751.1
B2	32635801 32635900
refGene_gn=PRRC2A;refGene_tx=XM_023253749.1;refGene_gn=PRRC2A;refGene_tx=XM_023253750.1;refGene_gn=PRRC2A;refGene_tx=XM_023253751.1
B2	32636032 32636245
refGene_gn=PRRC2A;refGene_tx=XM_023253749.1;refGene_gn=PRRC2A;refGene_tx=XM_023253750.1;refGene_gn=PRRC2A;refGene_tx=XM_023253751.1
B2	32636360 32636455
refGene_gn=PRRC2A;refGene_tx=XM_023253749.1;refGene_gn=PRRC2A;refGene_tx=XM_023253750.1;refGene_gn=PRRC2A;refGene_tx=XM_023253751.1
B2	32636558 32636649
refGene_gn=PRRC2A;refGene_tx=XM_023253749.1;refGene_gn=PRRC2A;refGene_tx=XM_023253750.1;refGene_gn=PRRC2A;refGene_tx=XM_023253751.1
B2	32636760 32636901
refGene_gn=PRRC2A;refGene_tx=XM_023253749.1;refGene_gn=PRRC2A;refGene_tx=XM_023253750.1;refGene_gn=PRRC2A;refGene_tx=XM_023253751.1
B2	32638510 32638606
refGene_gn=BAG6;refGene_tx=XM_006931552.4;refGene_gn=BAG6;refGene_tx=XM_011282315.3;refGene_gn=BAG6;refGene_tx=XM_011282316.3;refGene_gn=BAG6;refGene_tx=XM_011282317.3;refGene_gn=BAG6;refGene_tx=XM_011282318.3;refGene_gn=BAG6;refGene_tx=XM_011282319.3;refGene_gn=BAG6;refGene_tx=XM_019830354.1;refGene_gn=BAG6;refGene_tx=XM_019830355.2;refGene_gn=BAG6;refGene_tx=XM_019830356.2;refGene_gn=BAG6;refGene_tx=XM_019830357.2;refGene_gn=BAG6;refGene_tx=XM_019830358.2;refGene_gn=BAG6;refGene_tx=XM_019830359.2;refGene_gn=BAG6;refGene_tx=XM_019830360.2;refGene_gn=BAG6;refGene_tx=XM_023253752.1
B2	32638893 32639040
refGene_gn=BAG6;refGene_tx=XM_006931552.4;refGene_gn=BAG6;refGene_tx=XM_011282315.3;refGene_gn=BAG6;refGene_tx=XM_011282316.3;refGene_gn=BAG6;refGene_tx=XM_011282318.3;refGene_gn=BAG6;refGene_tx=XM_019830354.1;refGene_gn=BAG6;refGene_tx=XM_019830355.2;refGene_gn=BAG6;refGene_tx=XM_019830356.2;refGene_gn=BAG6;refGene_tx=XM_019830357.2;refGene_gn=BAG6;refGene_tx=XM_019830358.2;refGene_gn=BAG6;refGene_tx=XM_023253752.1
B2	32639571 32639679
refGene_gn=BAG6;refGene_tx=XM_006931552.4;refGene_gn=BAG6;refGene_tx=XM_011282315.3;refGene_gn=BAG6;refGene_tx=XM_011282316.3;refGene_gn=BAG6;refGene_tx=XM_011282317.3;refGene_gn=BAG6;refGene_tx=XM_011282318.3;refGene_gn=BAG6;refGene_tx=XM_011282319.3;refGene_gn=BAG6;refGene_tx=XM_019830354.1;refGene_gn=BAG6;refGene_tx=XM_019830355.2;refGene_gn=BAG6;refGene_tx=XM_019830356.2;refGene_gn=BAG6;refGene_tx=XM_019830357.2;refGene_gn=BAG6;refGene_tx=XM_019830358.2;refGene_gn=BAG6;refGene_tx=XM_019830359.2;refGene_gn=BAG6;refGene_tx=XM_019830360.2;refGene_gn=BAG6;refGene_tx=XM_023253752.1
B2	32639756 32639903
refGene_gn=BAG6;refGene_tx=XM_006931552.4;refGene_gn=BAG6;refGene_tx=X

M_011282315.3;refGene_gn=BAG6;refGene_tx=XM_011282316.3;refGene_gn=BAG6;refGene_tx=XM_011282317.3;refGene_gn=BAG6;refGene_tx=XM_011282318.3;refGene_gn=BAG6;refGene_tx=XM_011282319.3;refGene_gn=BAG6;refGene_tx=XM_019830354.1;refGene_gn=BAG6;refGene_tx=XM_019830355.2;refGene_gn=BAG6;refGene_tx=XM_019830356.2;refGene_gn=BAG6;refGene_tx=XM_019830357.2;refGene_gn=BAG6;refGene_tx=XM_019830358.2;refGene_gn=BAG6;refGene_tx=XM_019830359.2;refGene_gn=BAG6;refGene_tx=XM_019830360.2;refGene_gn=BAG6;refGene_tx=XM_023253752.1
B2 32640012 32640072
refGene_gn=BAG6;refGene_tx=XM_006931552.4;refGene_gn=BAG6;refGene_tx=XM_011282315.3;refGene_gn=BAG6;refGene_tx=XM_011282316.3;refGene_gn=BAG6;refGene_tx=XM_011282317.3;refGene_gn=BAG6;refGene_tx=XM_011282318.3;refGene_gn=BAG6;refGene_tx=XM_011282319.3;refGene_gn=BAG6;refGene_tx=XM_019830354.1;refGene_gn=BAG6;refGene_tx=XM_019830355.2;refGene_gn=BAG6;refGene_tx=XM_019830356.2;refGene_gn=BAG6;refGene_tx=XM_019830357.2;refGene_gn=BAG6;refGene_tx=XM_019830358.2;refGene_gn=BAG6;refGene_tx=XM_019830359.2;refGene_gn=BAG6;refGene_tx=XM_019830360.2;refGene_gn=BAG6;refGene_tx=XM_023253752.1
B2 32640170 32640317
refGene_gn=BAG6;refGene_tx=XM_006931552.4;refGene_gn=BAG6;refGene_tx=XM_011282315.3;refGene_gn=BAG6;refGene_tx=XM_011282316.3;refGene_gn=BAG6;refGene_tx=XM_011282317.3;refGene_gn=BAG6;refGene_tx=XM_011282318.3;refGene_gn=BAG6;refGene_tx=XM_011282319.3;refGene_gn=BAG6;refGene_tx=XM_019830354.1;refGene_gn=BAG6;refGene_tx=XM_019830355.2;refGene_gn=BAG6;refGene_tx=XM_019830356.2;refGene_gn=BAG6;refGene_tx=XM_019830357.2;refGene_gn=BAG6;refGene_tx=XM_019830358.2;refGene_gn=BAG6;refGene_tx=XM_019830359.2;refGene_gn=BAG6;refGene_tx=XM_019830360.2;refGene_gn=BAG6;refGene_tx=XM_023253752.1
B2 32640488 32640613
refGene_gn=BAG6;refGene_tx=XM_006931552.4;refGene_gn=BAG6;refGene_tx=XM_011282315.3;refGene_gn=BAG6;refGene_tx=XM_011282316.3;refGene_gn=BAG6;refGene_tx=XM_011282317.3;refGene_gn=BAG6;refGene_tx=XM_011282318.3;refGene_gn=BAG6;refGene_tx=XM_011282319.3;refGene_gn=BAG6;refGene_tx=XM_019830354.1;refGene_gn=BAG6;refGene_tx=XM_019830355.2;refGene_gn=BAG6;refGene_tx=XM_019830356.2;refGene_gn=BAG6;refGene_tx=XM_019830357.2;refGene_gn=BAG6;refGene_tx=XM_019830358.2;refGene_gn=BAG6;refGene_tx=XM_019830359.2;refGene_gn=BAG6;refGene_tx=XM_019830360.2;refGene_gn=BAG6;refGene_tx=XM_023253752.1
B2 32640701 32640804
refGene_gn=BAG6;refGene_tx=XM_006931552.4;refGene_gn=BAG6;refGene_tx=XM_011282315.3;refGene_gn=BAG6;refGene_tx=XM_011282316.3;refGene_gn=BAG6;refGene_tx=XM_011282317.3;refGene_gn=BAG6;refGene_tx=XM_011282318.3;refGene_gn=BAG6;refGene_tx=XM_011282319.3;refGene_gn=BAG6;refGene_tx=XM_019830354.1;refGene_gn=BAG6;refGene_tx=XM_019830355.2;refGene_gn=BAG6;refGene_tx=XM_019830356.2;refGene_gn=BAG6;refGene_tx=XM_019830357.2;refGene_gn=BAG6;refGene_tx=XM_019830358.2;refGene_gn=BAG6;refGene_tx=XM_019830359.2;refGene_gn=BAG6;refGene_tx=XM_019830360.2;refGene_gn=BAG6;refGene_tx=XM_023253752.1
B2 32640935 32640989
refGene_gn=BAG6;refGene_tx=XM_006931552.4;refGene_gn=BAG6;refGene_tx=X

M_011282315.3;refGene_gn=BAG6;refGene_tx=XM_011282316.3;refGene_gn=BAG6;refGene_tx=XM_011282317.3;refGene_gn=BAG6;refGene_tx=XM_011282318.3;refGene_gn=BAG6;refGene_tx=XM_011282319.3;refGene_gn=BAG6;refGene_tx=XM_019830354.1;refGene_gn=BAG6;refGene_tx=XM_019830355.2;refGene_gn=BAG6;refGene_tx=XM_019830356.2;refGene_gn=BAG6;refGene_tx=XM_019830357.2;refGene_gn=BAG6;refGene_tx=XM_019830358.2;refGene_gn=BAG6;refGene_tx=XM_019830359.2;refGene_gn=BAG6;refGene_tx=XM_019830360.2;refGene_gn=BAG6;refGene_tx=XM_023253752.1
B2 32641178 32641348
refGene_gn=BAG6;refGene_tx=XM_006931552.4;refGene_gn=BAG6;refGene_tx=XM_011282315.3;refGene_gn=BAG6;refGene_tx=XM_011282316.3;refGene_gn=BAG6;refGene_tx=XM_011282317.3;refGene_gn=BAG6;refGene_tx=XM_011282318.3;refGene_gn=BAG6;refGene_tx=XM_011282319.3;refGene_gn=BAG6;refGene_tx=XM_019830354.1;refGene_gn=BAG6;refGene_tx=XM_019830355.2;refGene_gn=BAG6;refGene_tx=XM_019830356.2;refGene_gn=BAG6;refGene_tx=XM_019830357.2;refGene_gn=BAG6;refGene_tx=XM_019830358.2;refGene_gn=BAG6;refGene_tx=XM_019830359.2;refGene_gn=BAG6;refGene_tx=XM_019830360.2;refGene_gn=BAG6;refGene_tx=XM_023253752.1
B2 32641505 32641797
refGene_gn=BAG6;refGene_tx=XM_006931552.4;refGene_gn=BAG6;refGene_tx=XM_011282315.3;refGene_gn=BAG6;refGene_tx=XM_011282316.3;refGene_gn=BAG6;refGene_tx=XM_011282317.3;refGene_gn=BAG6;refGene_tx=XM_011282318.3;refGene_gn=BAG6;refGene_tx=XM_011282319.3;refGene_gn=BAG6;refGene_tx=XM_019830354.1;refGene_gn=BAG6;refGene_tx=XM_019830355.2;refGene_gn=BAG6;refGene_tx=XM_019830356.2;refGene_gn=BAG6;refGene_tx=XM_019830357.2;refGene_gn=BAG6;refGene_tx=XM_019830358.2;refGene_gn=BAG6;refGene_tx=XM_019830359.2;refGene_gn=BAG6;refGene_tx=XM_019830360.2;refGene_gn=BAG6;refGene_tx=XM_023253752.1
B2 32642235 32642519
refGene_gn=BAG6;refGene_tx=XM_006931552.4;refGene_gn=BAG6;refGene_tx=XM_011282315.3;refGene_gn=BAG6;refGene_tx=XM_011282316.3;refGene_gn=BAG6;refGene_tx=XM_011282317.3;refGene_gn=BAG6;refGene_tx=XM_011282318.3;refGene_gn=BAG6;refGene_tx=XM_011282319.3;refGene_gn=BAG6;refGene_tx=XM_019830354.1;refGene_gn=BAG6;refGene_tx=XM_019830355.2;refGene_gn=BAG6;refGene_tx=XM_019830356.2;refGene_gn=BAG6;refGene_tx=XM_019830357.2;refGene_gn=BAG6;refGene_tx=XM_019830358.2;refGene_gn=BAG6;refGene_tx=XM_019830359.2;refGene_gn=BAG6;refGene_tx=XM_019830360.2;refGene_gn=BAG6;refGene_tx=XM_023253752.1
B2 32642716 32642804
refGene_gn=BAG6;refGene_tx=XM_006931552.4;refGene_gn=BAG6;refGene_tx=XM_011282315.3;refGene_gn=BAG6;refGene_tx=XM_011282316.3;refGene_gn=BAG6;refGene_tx=XM_011282317.3;refGene_gn=BAG6;refGene_tx=XM_011282318.3;refGene_gn=BAG6;refGene_tx=XM_011282319.3;refGene_gn=BAG6;refGene_tx=XM_019830354.1;refGene_gn=BAG6;refGene_tx=XM_019830355.2;refGene_gn=BAG6;refGene_tx=XM_019830356.2;refGene_gn=BAG6;refGene_tx=XM_019830357.2;refGene_gn=BAG6;refGene_tx=XM_019830358.2;refGene_gn=BAG6;refGene_tx=XM_019830359.2;refGene_gn=BAG6;refGene_tx=XM_019830360.2;refGene_gn=BAG6;refGene_tx=XM_023253752.1
B2 32642907 32643020
refGene_gn=BAG6;refGene_tx=XM_006931552.4;refGene_gn=BAG6;refGene_tx=X

M_011282315.3;refGene_gn=BAG6;refGene_tx=XM_011282316.3;refGene_gn=BAG6;refGene_tx=XM_011282317.3;refGene_gn=BAG6;refGene_tx=XM_011282318.3;refGene_gn=BAG6;refGene_tx=XM_011282319.3;refGene_gn=BAG6;refGene_tx=XM_019830354.1;refGene_gn=BAG6;refGene_tx=XM_019830355.2;refGene_gn=BAG6;refGene_tx=XM_019830356.2;refGene_gn=BAG6;refGene_tx=XM_019830357.2;refGene_gn=BAG6;refGene_tx=XM_019830358.2;refGene_gn=BAG6;refGene_tx=XM_019830359.2;refGene_gn=BAG6;refGene_tx=XM_019830360.2;refGene_gn=BAG6;refGene_tx=XM_023253752.1
B2 32643142 32643250
refGene_gn=BAG6;refGene_tx=XM_006931552.4;refGene_gn=BAG6;refGene_tx=XM_011282315.3;refGene_gn=BAG6;refGene_tx=XM_011282317.3;refGene_gn=BAG6;refGene_tx=XM_019830354.1;refGene_gn=BAG6;refGene_tx=XM_019830355.2;refGene_gn=BAG6;refGene_tx=XM_019830356.2;refGene_gn=BAG6;refGene_tx=XM_019830359.2;refGene_gn=BAG6;refGene_tx=XM_023253752.1
B2 32643361 32643439
refGene_gn=BAG6;refGene_tx=XM_006931552.4;refGene_gn=BAG6;refGene_tx=XM_011282315.3;refGene_gn=BAG6;refGene_tx=XM_011282316.3;refGene_gn=BAG6;refGene_tx=XM_011282317.3;refGene_gn=BAG6;refGene_tx=XM_011282318.3;refGene_gn=BAG6;refGene_tx=XM_011282319.3;refGene_gn=BAG6;refGene_tx=XM_019830354.1;refGene_gn=BAG6;refGene_tx=XM_019830355.2;refGene_gn=BAG6;refGene_tx=XM_019830357.2;refGene_gn=BAG6;refGene_tx=XM_019830359.2;refGene_gn=BAG6;refGene_tx=XM_019830360.2;refGene_gn=BAG6;refGene_tx=XM_023253752.1
B2 32643792 32644045
refGene_gn=BAG6;refGene_tx=XM_006931552.4;refGene_gn=BAG6;refGene_tx=XM_011282315.3;refGene_gn=BAG6;refGene_tx=XM_011282316.3;refGene_gn=BAG6;refGene_tx=XM_011282317.3;refGene_gn=BAG6;refGene_tx=XM_011282318.3;refGene_gn=BAG6;refGene_tx=XM_011282319.3;refGene_gn=BAG6;refGene_tx=XM_019830354.1;refGene_gn=BAG6;refGene_tx=XM_019830355.2;refGene_gn=BAG6;refGene_tx=XM_019830356.2;refGene_gn=BAG6;refGene_tx=XM_019830357.2;refGene_gn=BAG6;refGene_tx=XM_019830358.2;refGene_gn=BAG6;refGene_tx=XM_019830359.2;refGene_gn=BAG6;refGene_tx=XM_019830360.2;refGene_gn=BAG6;refGene_tx=XM_023253752.1
B2 32644262 32644460
refGene_gn=BAG6;refGene_tx=XM_006931552.4;refGene_gn=BAG6;refGene_tx=XM_011282315.3;refGene_gn=BAG6;refGene_tx=XM_011282316.3;refGene_gn=BAG6;refGene_tx=XM_011282317.3;refGene_gn=BAG6;refGene_tx=XM_011282318.3;refGene_gn=BAG6;refGene_tx=XM_011282319.3;refGene_gn=BAG6;refGene_tx=XM_019830354.1;refGene_gn=BAG6;refGene_tx=XM_019830355.2;refGene_gn=BAG6;refGene_tx=XM_019830356.2;refGene_gn=BAG6;refGene_tx=XM_019830357.2;refGene_gn=BAG6;refGene_tx=XM_019830358.2;refGene_gn=BAG6;refGene_tx=XM_019830359.2;refGene_gn=BAG6;refGene_tx=XM_019830360.2;refGene_gn=BAG6;refGene_tx=XM_023253752.1
B2 32644920 32645050
refGene_gn=BAG6;refGene_tx=XM_006931552.4;refGene_gn=BAG6;refGene_tx=XM_011282315.3;refGene_gn=BAG6;refGene_tx=XM_011282316.3;refGene_gn=BAG6;refGene_tx=XM_011282317.3;refGene_gn=BAG6;refGene_tx=XM_011282318.3;refGene_gn=BAG6;refGene_tx=XM_011282319.3;refGene_gn=BAG6;refGene_tx=XM_019830354.1;refGene_gn=BAG6;refGene_tx=XM_019830355.2;refGene_gn=BAG6;refGene_tx=XM_019830356.2;refGene_gn=BAG6;refGene_tx=XM_019830357.2;

refGene_gn=BAG6;refGene_tx=XM_019830358.2;refGene_gn=BAG6;refGene_tx=XM_019830359.2;refGene_gn=BAG6;refGene_tx=XM_019830360.2;refGene_gn=BAG6;refGene_tx=XM_023253752.1
B2 32645577 32645816
refGene_gn=BAG6;refGene_tx=XM_006931552.4;refGene_gn=BAG6;refGene_tx=XM_011282315.3;refGene_gn=BAG6;refGene_tx=XM_011282316.3;refGene_gn=BAG6;refGene_tx=XM_011282317.3;refGene_gn=BAG6;refGene_tx=XM_011282318.3;refGene_gn=BAG6;refGene_tx=XM_011282319.3;refGene_gn=BAG6;refGene_tx=XM_019830354.1;refGene_gn=BAG6;refGene_tx=XM_019830355.2;refGene_gn=BAG6;refGene_tx=XM_019830356.2;refGene_gn=BAG6;refGene_tx=XM_019830357.2;refGene_gn=BAG6;refGene_tx=XM_019830358.2;refGene_gn=BAG6;refGene_tx=XM_019830359.2;refGene_gn=BAG6;refGene_tx=XM_019830360.2;refGene_gn=BAG6;refGene_tx=XM_023253752.1
B2 32646467 32646542
refGene_gn=BAG6;refGene_tx=XM_006931552.4;refGene_gn=BAG6;refGene_tx=XM_011282315.3;refGene_gn=BAG6;refGene_tx=XM_011282316.3;refGene_gn=BAG6;refGene_tx=XM_011282317.3;refGene_gn=BAG6;refGene_tx=XM_011282318.3;refGene_gn=BAG6;refGene_tx=XM_011282319.3;refGene_gn=BAG6;refGene_tx=XM_019830354.1;refGene_gn=BAG6;refGene_tx=XM_019830355.2;refGene_gn=BAG6;refGene_tx=XM_019830356.2;refGene_gn=BAG6;refGene_tx=XM_019830357.2;refGene_gn=BAG6;refGene_tx=XM_019830358.2;refGene_gn=BAG6;refGene_tx=XM_019830359.2;refGene_gn=BAG6;refGene_tx=XM_019830360.2;refGene_gn=BAG6;refGene_tx=XM_023253752.1
B2 32646703 32646757
refGene_gn=BAG6;refGene_tx=XM_006931552.4;refGene_gn=BAG6;refGene_tx=XM_011282316.3;refGene_gn=BAG6;refGene_tx=XM_011282317.3;refGene_gn=BAG6;refGene_tx=XM_011282319.3;refGene_gn=BAG6;refGene_tx=XM_019830354.1;refGene_gn=BAG6;refGene_tx=XM_019830355.2;refGene_gn=BAG6;refGene_tx=XM_019830356.2;refGene_gn=BAG6;refGene_tx=XM_019830357.2;refGene_gn=BAG6;refGene_tx=XM_019830358.2;refGene_gn=BAG6;refGene_tx=XM_019830360.2;refGene_gn=BAG6;refGene_tx=XM_023253752.1
B2 32646991 32647188
refGene_gn=BAG6;refGene_tx=XM_006931552.4;refGene_gn=BAG6;refGene_tx=XM_011282315.3;refGene_gn=BAG6;refGene_tx=XM_011282316.3;refGene_gn=BAG6;refGene_tx=XM_011282317.3;refGene_gn=BAG6;refGene_tx=XM_011282318.3;refGene_gn=BAG6;refGene_tx=XM_011282319.3;refGene_gn=BAG6;refGene_tx=XM_019830354.1;refGene_gn=BAG6;refGene_tx=XM_019830355.2;refGene_gn=BAG6;refGene_tx=XM_019830356.2;refGene_gn=BAG6;refGene_tx=XM_019830357.2;refGene_gn=BAG6;refGene_tx=XM_019830358.2;refGene_gn=BAG6;refGene_tx=XM_019830359.2;refGene_gn=BAG6;refGene_tx=XM_019830360.2;refGene_gn=BAG6;refGene_tx=XM_023253752.1
B2 32647308 32647426
refGene_gn=BAG6;refGene_tx=XM_006931552.4;refGene_gn=BAG6;refGene_tx=XM_011282315.3;refGene_gn=BAG6;refGene_tx=XM_011282316.3;refGene_gn=BAG6;refGene_tx=XM_011282317.3;refGene_gn=BAG6;refGene_tx=XM_011282318.3;refGene_gn=BAG6;refGene_tx=XM_011282319.3;refGene_gn=BAG6;refGene_tx=XM_019830354.1;refGene_gn=BAG6;refGene_tx=XM_019830355.2;refGene_gn=BAG6;refGene_tx=XM_019830356.2;refGene_gn=BAG6;refGene_tx=XM_019830357.2;refGene_gn=BAG6;refGene_tx=XM_019830358.2;refGene_gn=BAG6;refGene_tx=XM_019830359.2;refGene_gn=BAG6;refGene_tx=XM_019830360.2;refGene_gn=BAG

6;refGene_tx=XM_023253752.1
B2 32648523 32648637
refGene_gn=BAG6;refGene_tx=XM_006931552.4;refGene_gn=BAG6;refGene_tx=XM_011282315.3;refGene_gn=BAG6;refGene_tx=XM_011282316.3;refGene_gn=BAG6;refGene_tx=XM_011282317.3;refGene_gn=BAG6;refGene_tx=XM_011282318.3;refGene_gn=BAG6;refGene_tx=XM_011282319.3;refGene_gn=BAG6;refGene_tx=XM_019830354.1;refGene_gn=BAG6;refGene_tx=XM_019830355.2;refGene_gn=BAG6;refGene_tx=XM_019830356.2;refGene_gn=BAG6;refGene_tx=XM_019830357.2;refGene_gn=BAG6;refGene_tx=XM_019830358.2;refGene_gn=BAG6;refGene_tx=XM_019830359.2;refGene_gn=BAG6;refGene_tx=XM_019830360.2;refGene_gn=BAG6;refGene_tx=XM_023253752.1
B2 32651779 32651893
refGene_gn=APOM;refGene_tx=XM_003985923.5;refGene_gn=APOM;refGene_tx=XM_019830364.2
B2 32652289 32652444
refGene_gn=APOM;refGene_tx=XM_003985923.5;refGene_gn=APOM;refGene_tx=XM_006931559.4;refGene_gn=APOM;refGene_tx=XM_011282320.3;refGene_gn=APOM;refGene_tx=XM_019830364.2
B2 32652729 32652803
refGene_gn=APOM;refGene_tx=XM_003985923.5;refGene_gn=APOM;refGene_tx=XM_006931559.4;refGene_gn=APOM;refGene_tx=XM_011282320.3;refGene_gn=APOM;refGene_tx=XM_019830364.2
B2 32652901 32653000
refGene_gn=APOM;refGene_tx=XM_003985923.5;refGene_gn=APOM;refGene_tx=XM_006931559.4;refGene_gn=APOM;refGene_tx=XM_011282320.3
B2 32653180 32653279
refGene_gn=APOM;refGene_tx=XM_003985923.5;refGene_gn=APOM;refGene_tx=XM_006931559.4;refGene_gn=APOM;refGene_tx=XM_011282320.3;refGene_gn=APOM;refGene_tx=XM_019830364.2
B2 32653698 32653724
refGene_gn=APOM;refGene_tx=XM_003985923.5;refGene_gn=APOM;refGene_tx=XM_006931559.4;refGene_gn=APOM;refGene_tx=XM_011282320.3;refGene_gn=APOM;refGene_tx=XM_019830364.2
B2 32654735 32655629
refGene_gn=CB2H6orf47;refGene_tx=XM_006931560.4
B2 32657696 32658141
refGene_gn=GPANK1;refGene_tx=XM_003985926.5;refGene_gn=GPANK1;refGene_tx=XM_006931561.4;refGene_gn=GPANK1;refGene_tx=XM_006931562.3;refGene_gn=GPANK1;refGene_tx=XM_006931564.4;refGene_gn=GPANK1;refGene_tx=XM_006931565.4;refGene_gn=GPANK1;refGene_tx=XM_019830362.2
B2 32658680 32659291
refGene_gn=GPANK1;refGene_tx=XM_003985926.5;refGene_gn=GPANK1;refGene_tx=XM_006931561.4;refGene_gn=GPANK1;refGene_tx=XM_006931562.3;refGene_gn=GPANK1;refGene_tx=XM_006931564.4;refGene_gn=GPANK1;refGene_tx=XM_006931565.4;refGene_gn=GPANK1;refGene_tx=XM_019830362.2
B2 32661696 32661768 refGene_gn=CSNK2B;refGene_tx=XM_003985927.4
B2 32662484 32662587 refGene_gn=CSNK2B;refGene_tx=XM_003985927.4
B2 32663170 32663286 refGene_gn=CSNK2B;refGene_tx=XM_003985927.4
B2 32663733 32663809 refGene_gn=CSNK2B;refGene_tx=XM_003985927.4
B2 32663956 32664146 refGene_gn=CSNK2B;refGene_tx=XM_003985927.4

B2 32664787 32664878 refGene_gn=CSNK2B;refGene_tx=XM_003985927.4
B2 32665804 32665925 refGene_gn=LY6G5B;refGene_tx=XM_003985977.5
B2 32666073 32666202 refGene_gn=LY6G5B;refGene_tx=XM_003985977.5
B2 32666495 32666896 refGene_gn=LY6G5B;refGene_tx=XM_003985977.5
B2 32670298 32670462 refGene_gn=LY6G5C;refGene_tx=XM_023254708.1
B2 32673248 32673416 refGene_gn=LY6G5C;refGene_tx=XM_023254708.1
B2 32674493 32674605 refGene_gn=LY6G5C;refGene_tx=XM_023254708.1
B2 32674733 32674754 refGene_gn=LY6G5C;refGene_tx=XM_023254708.1
B2 32679109 32679193
refGene_gn=ABHD16A;refGene_tx=XM_003985929.4;refGene_gn=ABHD16A;refGene_tx=XM_019830365.2
B2 32679385 32679432
refGene_gn=ABHD16A;refGene_tx=XM_003985929.4;refGene_gn=ABHD16A;refGene_tx=XM_019830365.2
B2 32679532 32679631
refGene_gn=ABHD16A;refGene_tx=XM_003985929.4;refGene_gn=ABHD16A;refGene_tx=XM_019830365.2
B2 32679715 32679792
refGene_gn=ABHD16A;refGene_tx=XM_003985929.4;refGene_gn=ABHD16A;refGene_tx=XM_019830365.2
B2 32679929 32679992
refGene_gn=ABHD16A;refGene_tx=XM_003985929.4;refGene_gn=ABHD16A;refGene_tx=XM_019830365.2
B2 32680138 32680195
refGene_gn=ABHD16A;refGene_tx=XM_003985929.4;refGene_gn=ABHD16A;refGene_tx=XM_019830365.2
B2 32680448 32680512
refGene_gn=ABHD16A;refGene_tx=XM_003985929.4;refGene_gn=ABHD16A;refGene_tx=XM_019830365.2
B2 32680743 32680848 refGene_gn=ABHD16A;refGene_tx=XM_003985929.4
B2 32681306 32681430
refGene_gn=ABHD16A;refGene_tx=XM_003985929.4;refGene_gn=ABHD16A;refGene_tx=XM_019830365.2
B2 32681754 32681804
refGene_gn=ABHD16A;refGene_tx=XM_003985929.4;refGene_gn=ABHD16A;refGene_tx=XM_019830365.2
B2 32682224 32682288
refGene_gn=ABHD16A;refGene_tx=XM_003985929.4;refGene_gn=ABHD16A;refGene_tx=XM_019830365.2
B2 32683219 32683321
refGene_gn=ABHD16A;refGene_tx=XM_003985929.4;refGene_gn=ABHD16A;refGene_tx=XM_019830365.2
B2 32683414 32683529
refGene_gn=ABHD16A;refGene_tx=XM_003985929.4;refGene_gn=ABHD16A;refGene_tx=XM_019830365.2
B2 32685910 32686033
refGene_gn=ABHD16A;refGene_tx=XM_003985929.4;refGene_gn=ABHD16A;refGene_tx=XM_019830365.2
B2 32686192 32686266
refGene_gn=ABHD16A;refGene_tx=XM_003985929.4;refGene_gn=ABHD16A;refGen

e_tx=XM_019830365.2
B2	32687926 32688012
refGene_gn=ABHD16A;refGene_tx=XM_003985929.4;refGene_gn=ABHD16A;refGen
e_tx=XM_019830365.2
B2	32690807 32690894
refGene_gn=ABHD16A;refGene_tx=XM_003985929.4;refGene_gn=ABHD16A;refGen
e_tx=XM_019830365.2
B2	32691103 32691170
refGene_gn=ABHD16A;refGene_tx=XM_003985929.4;refGene_gn=ABHD16A;refGen
e_tx=XM_019830365.2
B2	32691879 32691936
refGene_gn=ABHD16A;refGene_tx=XM_003985929.4;refGene_gn=ABHD16A;refGen
e_tx=XM_019830365.2
B2	32692924 32693056
refGene_gn=ABHD16A;refGene_tx=XM_003985929.4;refGene_gn=ABHD16A;refGen
e_tx=XM_019830365.2
B2	32695239 32695294
refGene_gn=LY6G6F;refGene_tx=XM_006931567.4;refGene_gn=LY6G6F;refGene_
tx=XM_023253753.1
B2	32695797 32696139
refGene_gn=LY6G6F;refGene_tx=XM_006931567.4;refGene_gn=LY6G6F;refGene_
tx=XM_023253753.1
B2	32696222 32696486
refGene_gn=LY6G6F;refGene_tx=XM_006931567.4;refGene_gn=LY6G6F;refGene_
tx=XM_023253753.1
B2	32697696 32697852
refGene_gn=LY6G6F;refGene_tx=XM_006931567.4;refGene_gn=LY6G6F;refGene_
tx=XM_023253753.1
B2	32698012 32698079 refGene_gn=LY6G6F;refGene_tx=XM_006931567.4
B2	32698212 32698237 refGene_gn=LY6G6F;refGene_tx=XM_006931567.4
B2	32699954 32700178
refGene_gn=LOC102901967;refGene_tx=XM_006931568.3;refGene_gn=LOC102901
967;refGene_tx=XM_019830371.1;refGene_gn=LOC102901967;refGene_tx=XM_01
9830372.2
B2	32700307 32700493
refGene_gn=LOC102901967;refGene_tx=XM_006931568.3;refGene_gn=LOC102901
967;refGene_tx=XM_019830371.1;refGene_gn=LOC102901967;refGene_tx=XM_01
9830372.2
B2	32700814 32700968
refGene_gn=LOC102901967;refGene_tx=XM_019830371.1
B2	32701046 32701098
refGene_gn=LOC102901967;refGene_tx=XM_006931568.3;refGene_gn=LOC102901
967;refGene_tx=XM_019830372.2
B2	32702773 32702920 refGene_gn=LY6G6F;refGene_tx=XM_023253753.1
B2	32704481 32704696 refGene_gn=LY6G6F;refGene_tx=XM_023253753.1
B2	32706056 32706271 refGene_gn=LY6G6C;refGene_tx=XM_003985932.4
B2	32706745 32706856 refGene_gn=LY6G6C;refGene_tx=XM_003985932.4
B2	32708331 32708383 refGene_gn=LY6G6C;refGene_tx=XM_003985932.4
B2	32709927 32709988
refGene_gn=MPIG6B;refGene_tx=XM_003985933.4;refGene_gn=MPIG6B;refGene_

tx=XM_003985934.4;refGene_gn=MPIG6B;refGene_tx=XM_019830368.2;refGene_gn=MPIG6B;refGene_tx=XM_019830369.2;refGene_gn=MPIG6B;refGene_tx=XM_019830370.2
B2 32710191 32710539
refGene_gn=MPIG6B;refGene_tx=XM_003985933.4;refGene_gn=MPIG6B;refGene_tx=XM_003985934.4;refGene_gn=MPIG6B;refGene_tx=XM_019830368.2;refGene_gn=MPIG6B;refGene_tx=XM_019830369.2;refGene_gn=MPIG6B;refGene_tx=XM_019830370.2
B2 32710696 32710787
refGene_gn=MPIG6B;refGene_tx=XM_003985933.4;refGene_gn=MPIG6B;refGene_tx=XM_019830368.2
B2 32711116 32711157
refGene_gn=MPIG6B;refGene_tx=XM_003985933.4;refGene_gn=MPIG6B;refGene_tx=XM_003985934.4;refGene_gn=MPIG6B;refGene_tx=XM_019830368.2
B2 32711275 32711374
refGene_gn=MPIG6B;refGene_tx=XM_003985933.4;refGene_gn=MPIG6B;refGene_tx=XM_003985934.4;refGene_gn=MPIG6B;refGene_tx=XM_019830368.2;refGene_gn=MPIG6B;refGene_tx=XM_019830369.2;refGene_gn=MPIG6B;refGene_tx=XM_019830370.2
B2 32711484 32711589
refGene_gn=MPIG6B;refGene_tx=XM_003985933.4;refGene_gn=MPIG6B;refGene_tx=XM_003985934.4;refGene_gn=MPIG6B;refGene_tx=XM_019830369.2
B2 32713652 32713769 refGene_gn=DDAH2;refGene_tx=XM_003985937.5
B2 32713958 32714108 refGene_gn=DDAH2;refGene_tx=XM_003985937.5
B2 32714595 32714715 refGene_gn=DDAH2;refGene_tx=XM_003985937.5
B2 32714877 32714951 refGene_gn=DDAH2;refGene_tx=XM_003985937.5
B2 32715074 32715174 refGene_gn=DDAH2;refGene_tx=XM_003985937.5
B2 32715285 32715582 refGene_gn=DDAH2;refGene_tx=XM_003985937.5
B2 32717256 32717418 refGene_gn=CLIC1;refGene_tx=XM_003985936.4
B2 32718423 32718605 refGene_gn=CLIC1;refGene_tx=XM_003985936.4
B2 32719223 32719330 refGene_gn=CLIC1;refGene_tx=XM_003985936.4
B2 32719489 32719615 refGene_gn=CLIC1;refGene_tx=XM_003985936.4
B2 32719831 32719941 refGene_gn=CLIC1;refGene_tx=XM_003985936.4
B2 32721996 32722035 refGene_gn=CLIC1;refGene_tx=XM_003985936.4
B2 32725729 32725873
refGene_gn=MSH5;refGene_tx=XM_019830375.2;refGene_gn=MSH5;refGene_tx=XM_019830376.2;refGene_gn=MSH5;refGene_tx=XM_019830378.2;refGene_gn=MSH5;refGene_tx=XM_019830380.2
B2 32726150 32726274
refGene_gn=MSH5;refGene_tx=XM_019830375.2;refGene_gn=MSH5;refGene_tx=XM_019830376.2;refGene_gn=MSH5;refGene_tx=XM_019830378.2;refGene_gn=MSH5;refGene_tx=XM_019830380.2
B2 32726545 32726612 refGene_gn=MSH5;refGene_tx=XM_019830379.2
B2 32726736 32726817
refGene_gn=MSH5;refGene_tx=XM_019830375.2;refGene_gn=MSH5;refGene_tx=XM_019830376.2;refGene_gn=MSH5;refGene_tx=XM_019830378.2;refGene_gn=MSH5;refGene_tx=XM_019830379.2;refGene_gn=MSH5;refGene_tx=XM_019830380.2
B2 32727000 32727063
refGene_gn=MSH5;refGene_tx=XM_019830375.2;refGene_gn=MSH5;refGene_tx=XM_019830376.2;refGene_gn=MSH5;refGene_tx=XM_019830378.2;refGene_gn=MSH

5;refGene_tx=XM_019830379.2;refGene_gn=MSH5;refGene_tx=XM_019830380.2
B2 32727994 32728116
refGene_gn=MSH5;refGene_tx=XM_019830375.2;refGene_gn=MSH5;refGene_tx=XM_019830376.2;refGene_gn=MSH5;refGene_tx=XM_019830378.2;refGene_gn=MSH5;refGene_tx=XM_019830379.2;refGene_gn=MSH5;refGene_tx=XM_019830380.2
B2 32728290 32728400
refGene_gn=MSH5;refGene_tx=XM_019830375.2;refGene_gn=MSH5;refGene_tx=XM_019830376.2;refGene_gn=MSH5;refGene_tx=XM_019830378.2;refGene_gn=MSH5;refGene_tx=XM_019830379.2;refGene_gn=MSH5;refGene_tx=XM_019830380.2
B2 32728673 32728709
refGene_gn=MSH5;refGene_tx=XM_019830375.2;refGene_gn=MSH5;refGene_tx=XM_019830376.2;refGene_gn=MSH5;refGene_tx=XM_019830378.2;refGene_gn=MSH5;refGene_tx=XM_019830379.2;refGene_gn=MSH5;refGene_tx=XM_019830380.2
B2 32729250 32729333
refGene_gn=MSH5;refGene_tx=XM_019830375.2;refGene_gn=MSH5;refGene_tx=XM_019830376.2;refGene_gn=MSH5;refGene_tx=XM_019830378.2;refGene_gn=MSH5;refGene_tx=XM_019830379.2;refGene_gn=MSH5;refGene_tx=XM_019830380.2
B2 32730118 32730164
refGene_gn=MSH5;refGene_tx=XM_019830375.2;refGene_gn=MSH5;refGene_tx=XM_019830376.2;refGene_gn=MSH5;refGene_tx=XM_019830378.2;refGene_gn=MSH5;refGene_tx=XM_019830379.2;refGene_gn=MSH5;refGene_tx=XM_019830380.2
B2 32734158 32734297
refGene_gn=MSH5;refGene_tx=XM_019830375.2;refGene_gn=MSH5;refGene_tx=XM_019830376.2;refGene_gn=MSH5;refGene_tx=XM_019830378.2;refGene_gn=MSH5;refGene_tx=XM_019830379.2;refGene_gn=MSH5;refGene_tx=XM_019830380.2
B2 32734418 32734481
refGene_gn=MSH5;refGene_tx=XM_019830375.2;refGene_gn=MSH5;refGene_tx=XM_019830376.2;refGene_gn=MSH5;refGene_tx=XM_019830378.2;refGene_gn=MSH5;refGene_tx=XM_019830379.2;refGene_gn=MSH5;refGene_tx=XM_019830380.2
B2 32742013 32742142
refGene_gn=MSH5;refGene_tx=XM_019830375.2;refGene_gn=MSH5;refGene_tx=XM_019830376.2;refGene_gn=MSH5;refGene_tx=XM_019830378.2;refGene_gn=MSH5;refGene_tx=XM_019830379.2
B2 32742434 32742507
refGene_gn=MSH5;refGene_tx=XM_019830375.2;refGene_gn=MSH5;refGene_tx=XM_019830376.2;refGene_gn=MSH5;refGene_tx=XM_019830378.2;refGene_gn=MSH5;refGene_tx=XM_019830379.2
B2 32742670 32742780
refGene_gn=MSH5;refGene_tx=XM_019830375.2;refGene_gn=MSH5;refGene_tx=XM_019830376.2;refGene_gn=MSH5;refGene_tx=XM_019830378.2;refGene_gn=MSH5;refGene_tx=XM_019830379.2
B2 32743016 32743097
refGene_gn=MSH5;refGene_tx=XM_019830375.2;refGene_gn=MSH5;refGene_tx=XM_019830376.2;refGene_gn=MSH5;refGene_tx=XM_019830378.2;refGene_gn=MSH5;refGene_tx=XM_019830379.2
B2 32746329 32746417
refGene_gn=MSH5;refGene_tx=XM_019830375.2;refGene_gn=MSH5;refGene_tx=XM_019830376.2;refGene_gn=MSH5;refGene_tx=XM_019830378.2;refGene_gn=MSH5;refGene_tx=XM_019830379.2;refGene_gn=MSH5;refGene_tx=XM_019830380.2
B2 32746646 32746836

refGene_gn=MSH5;refGene_tx=XM_019830375.2;refGene_gn=MSH5;refGene_tx=XM_019830376.2;refGene_gn=MSH5;refGene_tx=XM_019830378.2;refGene_gn=MSH5;refGene_tx=XM_019830379.2;refGene_gn=MSH5;refGene_tx=XM_019830380.2
B2 32746952 32747079
refGene_gn=MSH5;refGene_tx=XM_019830375.2;refGene_gn=MSH5;refGene_tx=XM_019830376.2;refGene_gn=MSH5;refGene_tx=XM_019830378.2;refGene_gn=MSH5;refGene_tx=XM_019830379.2;refGene_gn=MSH5;refGene_tx=XM_019830380.2
B2 32747732 32747882
refGene_gn=MSH5;refGene_tx=XM_019830375.2;refGene_gn=MSH5;refGene_tx=XM_019830376.2;refGene_gn=MSH5;refGene_tx=XM_019830378.2;refGene_gn=MSH5;refGene_tx=XM_019830379.2;refGene_gn=MSH5;refGene_tx=XM_019830380.2
B2 32748516 32748591
refGene_gn=MSH5;refGene_tx=XM_019830375.2;refGene_gn=MSH5;refGene_tx=XM_019830376.2;refGene_gn=MSH5;refGene_tx=XM_019830378.2;refGene_gn=MSH5;refGene_tx=XM_019830379.2;refGene_gn=MSH5;refGene_tx=XM_019830380.2
B2 32748799 32748943
refGene_gn=MSH5;refGene_tx=XM_019830375.2;refGene_gn=MSH5;refGene_tx=XM_019830376.2;refGene_gn=MSH5;refGene_tx=XM_019830378.2;refGene_gn=MSH5;refGene_tx=XM_019830379.2;refGene_gn=MSH5;refGene_tx=XM_019830380.2
B2 32749077 32749287
refGene_gn=MSH5;refGene_tx=XM_019830375.2;refGene_gn=MSH5;refGene_tx=XM_019830376.2;refGene_gn=MSH5;refGene_tx=XM_019830378.2;refGene_gn=MSH5;refGene_tx=XM_019830379.2;refGene_gn=MSH5;refGene_tx=XM_019830380.2
B2 32749448 32749522
refGene_gn=MSH5;refGene_tx=XM_019830375.2;refGene_gn=MSH5;refGene_tx=XM_019830376.2;refGene_gn=MSH5;refGene_tx=XM_019830378.2;refGene_gn=MSH5;refGene_tx=XM_019830379.2;refGene_gn=MSH5;refGene_tx=XM_019830380.2
B2 32749734 32749846
refGene_gn=MSH5;refGene_tx=XM_019830375.2;refGene_gn=MSH5;refGene_tx=XM_019830376.2;refGene_gn=MSH5;refGene_tx=XM_019830378.2;refGene_gn=MSH5;refGene_tx=XM_019830379.2;refGene_gn=MSH5;refGene_tx=XM_019830380.2
B2 32750490 32750604
refGene_gn=SAPCD1;refGene_tx=XM_019830383.1;refGene_gn=SAPCD1;refGene_tx=XM_019830385.1
B2 32750842 32750983
refGene_gn=SAPCD1;refGene_tx=XM_019830383.1;refGene_gn=SAPCD1;refGene_tx=XM_019830385.1
B2 32751475 32751571
refGene_gn=SAPCD1;refGene_tx=XM_019830383.1;refGene_gn=SAPCD1;refGene_tx=XM_019830385.1
B2 32751683 32751773
refGene_gn=SAPCD1;refGene_tx=XM_019830383.1;refGene_gn=SAPCD1;refGene_tx=XM_019830385.1
B2 32751855 32751948
refGene_gn=SAPCD1;refGene_tx=XM_019830383.1;refGene_gn=SAPCD1;refGene_tx=XM_019830385.1
B2 32752640 32752805
refGene_gn=VWA7;refGene_tx=XM_003985940.5;refGene_gn=VWA7;refGene_tx=XM_011282308.3
B2 32752911 32753158

refGene_gn=VWA7;refGene_tx=XM_003985940.5;refGene_gn=VWA7;refGene_tx=XM_011282308.3
B2 32753261 32753401
refGene_gn=VWA7;refGene_tx=XM_003985940.5;refGene_gn=VWA7;refGene_tx=XM_006931571.4;refGene_gn=VWA7;refGene_tx=XM_011282308.3;refGene_gn=VWA7;refGene_tx=XM_023253755.1
B2 32753482 32753784
refGene_gn=VWA7;refGene_tx=XM_003985940.5;refGene_gn=VWA7;refGene_tx=XM_006931571.4;refGene_gn=VWA7;refGene_tx=XM_011282308.3;refGene_gn=VWA7;refGene_tx=XM_023253755.1
B2 32754225 32754318
refGene_gn=VWA7;refGene_tx=XM_003985940.5;refGene_gn=VWA7;refGene_tx=XM_006931571.4;refGene_gn=VWA7;refGene_tx=XM_011282308.3;refGene_gn=VWA7;refGene_tx=XM_023253755.1
B2 32754432 32754585
refGene_gn=VWA7;refGene_tx=XM_003985940.5;refGene_gn=VWA7;refGene_tx=XM_006931571.4;refGene_gn=VWA7;refGene_tx=XM_011282308.3;refGene_gn=VWA7;refGene_tx=XM_023253755.1
B2 32754698 32754831
refGene_gn=VWA7;refGene_tx=XM_003985940.5;refGene_gn=VWA7;refGene_tx=XM_006931571.4;refGene_gn=VWA7;refGene_tx=XM_011282308.3;refGene_gn=VWA7;refGene_tx=XM_023253755.1
B2 32755316 32755502
refGene_gn=VWA7;refGene_tx=XM_003985940.5;refGene_gn=VWA7;refGene_tx=XM_006931571.4;refGene_gn=VWA7;refGene_tx=XM_011282308.3;refGene_gn=VWA7;refGene_tx=XM_023253755.1
B2 32755965 32756082
refGene_gn=VWA7;refGene_tx=XM_003985940.5;refGene_gn=VWA7;refGene_tx=XM_006931571.4;refGene_gn=VWA7;refGene_tx=XM_011282308.3;refGene_gn=VWA7;refGene_tx=XM_023253755.1
B2 32756314 32756427
refGene_gn=VWA7;refGene_tx=XM_003985940.5;refGene_gn=VWA7;refGene_tx=XM_006931571.4;refGene_gn=VWA7;refGene_tx=XM_011282308.3;refGene_gn=VWA7;refGene_tx=XM_023253755.1
B2 32757715 32757885
refGene_gn=VWA7;refGene_tx=XM_003985940.5;refGene_gn=VWA7;refGene_tx=XM_006931571.4;refGene_gn=VWA7;refGene_tx=XM_011282308.3;refGene_gn=VWA7;refGene_tx=XM_023253755.1
B2 32758003 32758199
refGene_gn=VWA7;refGene_tx=XM_003985940.5;refGene_gn=VWA7;refGene_tx=XM_006931571.4;refGene_gn=VWA7;refGene_tx=XM_011282308.3;refGene_gn=VWA7;refGene_tx=XM_023253755.1
B2 32758642 32758753
refGene_gn=VWA7;refGene_tx=XM_003985940.5;refGene_gn=VWA7;refGene_tx=XM_006931571.4;refGene_gn=VWA7;refGene_tx=XM_011282308.3;refGene_gn=VWA7;refGene_tx=XM_023253755.1
B2 32759317 32759414
refGene_gn=VWA7;refGene_tx=XM_003985940.5;refGene_gn=VWA7;refGene_tx=XM_006931571.4;refGene_gn=VWA7;refGene_tx=XM_011282308.3;refGene_gn=VWA7;refGene_tx=XM_023253755.1

B2 32760141 32760423
refGene_gn=VWA7;refGene_tx=XM_003985940.5;refGene_gn=VWA7;refGene_tx=XM_006931571.4;refGene_gn=VWA7;refGene_tx=XM_011282308.3;refGene_gn=VWA7;refGene_tx=XM_023253755.1
B2 32760755 32760992
refGene_gn=VWA7;refGene_tx=XM_003985940.5;refGene_gn=VWA7;refGene_tx=XM_006931571.4;refGene_gn=VWA7;refGene_tx=XM_011282308.3;refGene_gn=VWA7;refGene_tx=XM_023253755.1
B2 32761814 32761883
refGene_gn=VARS;refGene_tx=XM_003985941.4;refGene_gn=VARS;refGene_tx=XM_006931574.3
B2 32762731 32763057
refGene_gn=VARS;refGene_tx=XM_003985941.4;refGene_gn=VARS;refGene_tx=XM_006931574.3
B2 32763170 32763282
refGene_gn=VARS;refGene_tx=XM_003985941.4;refGene_gn=VARS;refGene_tx=XM_006931574.3
B2 32763354 32763561
refGene_gn=VARS;refGene_tx=XM_003985941.4;refGene_gn=VARS;refGene_tx=XM_006931574.3
B2 32763741 32763897
refGene_gn=VARS;refGene_tx=XM_003985941.4;refGene_gn=VARS;refGene_tx=XM_006931574.3
B2 32764265 32764393
refGene_gn=VARS;refGene_tx=XM_003985941.4;refGene_gn=VARS;refGene_tx=XM_006931574.3
B2 32764514 32764593
refGene_gn=VARS;refGene_tx=XM_003985941.4;refGene_gn=VARS;refGene_tx=XM_006931574.3
B2 32764679 32764748
refGene_gn=VARS;refGene_tx=XM_003985941.4;refGene_gn=VARS;refGene_tx=XM_006931574.3
B2 32764830 32764935
refGene_gn=VARS;refGene_tx=XM_003985941.4;refGene_gn=VARS;refGene_tx=XM_006931574.3
B2 32765294 32765420
refGene_gn=VARS;refGene_tx=XM_003985941.4;refGene_gn=VARS;refGene_tx=XM_006931574.3
B2 32765507 32765578
refGene_gn=VARS;refGene_tx=XM_003985941.4;refGene_gn=VARS;refGene_tx=XM_006931574.3
B2 32765668 32765774
refGene_gn=VARS;refGene_tx=XM_003985941.4;refGene_gn=VARS;refGene_tx=XM_006931574.3
B2 32765901 32765992
refGene_gn=VARS;refGene_tx=XM_003985941.4;refGene_gn=VARS;refGene_tx=XM_006931574.3
B2 32766122 32766281
refGene_gn=VARS;refGene_tx=XM_003985941.4;refGene_gn=VARS;refGene_tx=XM_006931574.3

B2 32766366 32766470
refGene_gn=VARS;refGene_tx=XM_003985941.4;refGene_gn=VARS;refGene_tx=XM_006931574.3
B2 32766555 32766680
refGene_gn=VARS;refGene_tx=XM_003985941.4;refGene_gn=VARS;refGene_tx=XM_006931574.3
B2 32766919 32767010
refGene_gn=VARS;refGene_tx=XM_003985941.4;refGene_gn=VARS;refGene_tx=XM_006931574.3
B2 32767287 32767382
refGene_gn=VARS;refGene_tx=XM_003985941.4;refGene_gn=VARS;refGene_tx=XM_006931574.3
B2 32767474 32767583
refGene_gn=VARS;refGene_tx=XM_003985941.4;refGene_gn=VARS;refGene_tx=XM_006931574.3
B2 32767666 32767786
refGene_gn=VARS;refGene_tx=XM_003985941.4;refGene_gn=VARS;refGene_tx=XM_006931574.3
B2 32768283 32768365
refGene_gn=VARS;refGene_tx=XM_003985941.4;refGene_gn=VARS;refGene_tx=XM_006931574.3
B2 32768589 32768754
refGene_gn=VARS;refGene_tx=XM_003985941.4;refGene_gn=VARS;refGene_tx=XM_006931574.3
B2 32773790 32773918
refGene_gn=VARS;refGene_tx=XM_003985941.4;refGene_gn=VARS;refGene_tx=XM_006931574.3
B2 32774054 32774155
refGene_gn=VARS;refGene_tx=XM_003985941.4;refGene_gn=VARS;refGene_tx=XM_006931574.3
B2 32774379 32774467
refGene_gn=VARS;refGene_tx=XM_003985941.4;refGene_gn=VARS;refGene_tx=XM_006931574.3
B2 32774552 32774677
refGene_gn=VARS;refGene_tx=XM_003985941.4;refGene_gn=VARS;refGene_tx=XM_006931574.3
B2 32774918 32775057
refGene_gn=VARS;refGene_tx=XM_003985941.4;refGene_gn=VARS;refGene_tx=XM_006931574.3
B2 32775152 32775287
refGene_gn=VARS;refGene_tx=XM_003985941.4;refGene_gn=VARS;refGene_tx=XM_006931574.3
B2 32776373 32776760
refGene_gn=VARS;refGene_tx=XM_003985941.4;refGene_gn=VARS;refGene_tx=XM_006931574.3
B2 32778480 32778606 refGene_gn=LSM2;refGene_tx=XM_023254709.1
B2 32778705 32778765 refGene_gn=LSM2;refGene_tx=XM_023254709.1
B2 32778882 32778913 refGene_gn=LSM2;refGene_tx=XM_023254709.1
B2 32783494 32783562 refGene_gn=LSM2;refGene_tx=XM_023254709.1
B2 32784206 32784529 refGene_gn=LSM2;refGene_tx=XM_023254709.1

B2 32785013 32785017 refGene_gn=LSM2;refGene_tx=XM_023254709.1
B2 32786137 32788063
refGene_gn=LOC101091022;refGene_tx=XM_003985942.4;refGene_gn=LOC101091022;refGene_tx=XM_011282307.3
B2 32790185 32792102
refGene_gn=LOC105260573;refGene_tx=XM_019830386.2
B2 32802642 32802651
refGene_gn=LOC105260573;refGene_tx=XM_019830386.2
B2 32833776 32834003 refGene_gn=NEU1;refGene_tx=XM_003985943.5
B2 32834111 32834334 refGene_gn=NEU1;refGene_tx=XM_003985943.5
B2 32834465 32834648 refGene_gn=NEU1;refGene_tx=XM_003985943.5
B2 32835159 32835422 refGene_gn=NEU1;refGene_tx=XM_003985943.5
B2 32836143 32836336 refGene_gn=NEU1;refGene_tx=XM_003985943.5
B2 32836782 32836947 refGene_gn=NEU1;refGene_tx=XM_003985943.5
B2 32837506 32837628 refGene_gn=SLC44A4;refGene_tx=XM_003985944.3
B2 32838486 32838571 refGene_gn=SLC44A4;refGene_tx=XM_003985944.3
B2 32838646 32838741 refGene_gn=SLC44A4;refGene_tx=XM_003985944.3
B2 32838848 32838919 refGene_gn=SLC44A4;refGene_tx=XM_003985944.3
B2 32839140 32839214 refGene_gn=SLC44A4;refGene_tx=XM_003985944.3
B2 32839325 32839429 refGene_gn=SLC44A4;refGene_tx=XM_003985944.3
B2 32839516 32839611 refGene_gn=SLC44A4;refGene_tx=XM_003985944.3
B2 32839697 32839951 refGene_gn=SLC44A4;refGene_tx=XM_003985944.3
B2 32842145 32842248 refGene_gn=SLC44A4;refGene_tx=XM_003985944.3
B2 32842515 32842608 refGene_gn=SLC44A4;refGene_tx=XM_003985944.3
B2 32843252 32843352 refGene_gn=SLC44A4;refGene_tx=XM_003985944.3
B2 32843458 32843694 refGene_gn=SLC44A4;refGene_tx=XM_003985944.3
B2 32843944 32844028 refGene_gn=SLC44A4;refGene_tx=XM_003985944.3
B2 32844108 32844196 refGene_gn=SLC44A4;refGene_tx=XM_003985944.3
B2 32845442 32845503 refGene_gn=SLC44A4;refGene_tx=XM_003985944.3
B2 32845661 32845787 refGene_gn=SLC44A4;refGene_tx=XM_003985944.3
B2 32845867 32845967 refGene_gn=SLC44A4;refGene_tx=XM_003985944.3
B2 32846765 32846844 refGene_gn=SLC44A4;refGene_tx=XM_003985944.3
B2 32846974 32847048 refGene_gn=SLC44A4;refGene_tx=XM_003985944.3
B2 32847611 32847660 refGene_gn=SLC44A4;refGene_tx=XM_003985944.3
B2 32849585 32849619 refGene_gn=SLC44A4;refGene_tx=XM_003985944.3
B2 32850400 32850581
refGene_gn=EHMT2;refGene_tx=XM_003985946.5;refGene_gn=EHMT2;refGene_tx=XM_011282304.3;refGene_gn=EHMT2;refGene_tx=XM_011282305.3;refGene_gn=EHMT2;refGene_tx=XM_023253756.1;refGene_gn=EHMT2;refGene_tx=XM_023253757.1
B2 32850925 32851101
refGene_gn=EHMT2;refGene_tx=XM_003985946.5;refGene_gn=EHMT2;refGene_tx=XM_011282304.3;refGene_gn=EHMT2;refGene_tx=XM_011282305.3;refGene_gn=EHMT2;refGene_tx=XM_023253756.1;refGene_gn=EHMT2;refGene_tx=XM_023253757.1
B2 32851242 32851321
refGene_gn=EHMT2;refGene_tx=XM_003985946.5;refGene_gn=EHMT2;refGene_tx=XM_011282304.3;refGene_gn=EHMT2;refGene_tx=XM_011282305.3;refGene_gn=EHMT2;refGene_tx=XM_023253756.1;refGene_gn=EHMT2;refGene_tx=XM_023253757.1

B2 32852729 32852816
refGene_gn=EHMT2;refGene_tx=XM_003985946.5;refGene_gn=EHMT2;refGene_tx=XM_011282304.3;refGene_gn=EHMT2;refGene_tx=XM_011282305.3;refGene_gn=EHMT2;refGene_tx=XM_023253756.1;refGene_gn=EHMT2;refGene_tx=XM_023253757.1
B2 32852925 32853041
refGene_gn=EHMT2;refGene_tx=XM_003985946.5;refGene_gn=EHMT2;refGene_tx=XM_011282304.3;refGene_gn=EHMT2;refGene_tx=XM_011282305.3;refGene_gn=EHMT2;refGene_tx=XM_023253756.1;refGene_gn=EHMT2;refGene_tx=XM_023253757.1
B2 32853293 32853371
refGene_gn=EHMT2;refGene_tx=XM_003985946.5;refGene_gn=EHMT2;refGene_tx=XM_011282304.3;refGene_gn=EHMT2;refGene_tx=XM_011282305.3;refGene_gn=EHMT2;refGene_tx=XM_023253756.1;refGene_gn=EHMT2;refGene_tx=XM_023253757.1
B2 32853718 32853863
refGene_gn=EHMT2;refGene_tx=XM_003985946.5;refGene_gn=EHMT2;refGene_tx=XM_011282304.3;refGene_gn=EHMT2;refGene_tx=XM_011282305.3;refGene_gn=EHMT2;refGene_tx=XM_023253756.1;refGene_gn=EHMT2;refGene_tx=XM_023253757.1
B2 32854307 32854475
refGene_gn=EHMT2;refGene_tx=XM_003985946.5;refGene_gn=EHMT2;refGene_tx=XM_011282304.3;refGene_gn=EHMT2;refGene_tx=XM_011282305.3;refGene_gn=EHMT2;refGene_tx=XM_023253756.1;refGene_gn=EHMT2;refGene_tx=XM_023253757.1
B2 32854558 32854713
refGene_gn=EHMT2;refGene_tx=XM_003985946.5;refGene_gn=EHMT2;refGene_tx=XM_011282304.3;refGene_gn=EHMT2;refGene_tx=XM_011282305.3;refGene_gn=EHMT2;refGene_tx=XM_023253756.1;refGene_gn=EHMT2;refGene_tx=XM_023253757.1
B2 32854828 32854933
refGene_gn=EHMT2;refGene_tx=XM_003985946.5;refGene_gn=EHMT2;refGene_tx=XM_011282304.3;refGene_gn=EHMT2;refGene_tx=XM_011282305.3;refGene_gn=EHMT2;refGene_tx=XM_023253756.1;refGene_gn=EHMT2;refGene_tx=XM_023253757.1
B2 32855692 32855794
refGene_gn=EHMT2;refGene_tx=XM_003985946.5;refGene_gn=EHMT2;refGene_tx=XM_011282304.3;refGene_gn=EHMT2;refGene_tx=XM_011282305.3;refGene_gn=EHMT2;refGene_tx=XM_023253756.1;refGene_gn=EHMT2;refGene_tx=XM_023253757.1
B2 32855879 32856002
refGene_gn=EHMT2;refGene_tx=XM_003985946.5;refGene_gn=EHMT2;refGene_tx=XM_011282304.3;refGene_gn=EHMT2;refGene_tx=XM_011282305.3;refGene_gn=EHMT2;refGene_tx=XM_023253756.1;refGene_gn=EHMT2;refGene_tx=XM_023253757.1
B2 32856103 32856210
refGene_gn=EHMT2;refGene_tx=XM_003985946.5;refGene_gn=EHMT2;refGene_tx=XM_011282304.3;refGene_gn=EHMT2;refGene_tx=XM_011282305.3;refGene_gn=EHMT2;refGene_tx=XM_023253756.1;refGene_gn=EHMT2;refGene_tx=XM_023253757.1

B2 32856537 32856620
refGene_gn=EHMT2;refGene_tx=XM_003985946.5;refGene_gn=EHMT2;refGene_tx=XM_011282304.3;refGene_gn=EHMT2;refGene_tx=XM_011282305.3;refGene_gn=EHMT2;refGene_tx=XM_023253756.1;refGene_gn=EHMT2;refGene_tx=XM_023253757.1
B2 32856737 32856920
refGene_gn=EHMT2;refGene_tx=XM_003985946.5;refGene_gn=EHMT2;refGene_tx=XM_011282304.3;refGene_gn=EHMT2;refGene_tx=XM_011282305.3;refGene_gn=EHMT2;refGene_tx=XM_023253756.1;refGene_gn=EHMT2;refGene_tx=XM_023253757.1
B2 32856999 32857235
refGene_gn=EHMT2;refGene_tx=XM_003985946.5;refGene_gn=EHMT2;refGene_tx=XM_011282304.3;refGene_gn=EHMT2;refGene_tx=XM_011282305.3;refGene_gn=EHMT2;refGene_tx=XM_023253756.1;refGene_gn=EHMT2;refGene_tx=XM_023253757.1
B2 32857314 32857458
refGene_gn=EHMT2;refGene_tx=XM_003985946.5;refGene_gn=EHMT2;refGene_tx=XM_011282304.3;refGene_gn=EHMT2;refGene_tx=XM_011282305.3;refGene_gn=EHMT2;refGene_tx=XM_023253756.1;refGene_gn=EHMT2;refGene_tx=XM_023253757.1
B2 32857558 32857707
refGene_gn=EHMT2;refGene_tx=XM_003985946.5;refGene_gn=EHMT2;refGene_tx=XM_011282304.3;refGene_gn=EHMT2;refGene_tx=XM_011282305.3;refGene_gn=EHMT2;refGene_tx=XM_023253756.1;refGene_gn=EHMT2;refGene_tx=XM_023253757.1
B2 32857931 32858033
refGene_gn=EHMT2;refGene_tx=XM_003985946.5;refGene_gn=EHMT2;refGene_tx=XM_011282305.3;refGene_gn=EHMT2;refGene_tx=XM_023253756.1
B2 32858203 32858318
refGene_gn=EHMT2;refGene_tx=XM_003985946.5;refGene_gn=EHMT2;refGene_tx=XM_011282304.3;refGene_gn=EHMT2;refGene_tx=XM_011282305.3;refGene_gn=EHMT2;refGene_tx=XM_023253756.1;refGene_gn=EHMT2;refGene_tx=XM_023253757.1
B2 32858434 32858572
refGene_gn=EHMT2;refGene_tx=XM_003985946.5;refGene_gn=EHMT2;refGene_tx=XM_011282304.3;refGene_gn=EHMT2;refGene_tx=XM_011282305.3;refGene_gn=EHMT2;refGene_tx=XM_023253756.1;refGene_gn=EHMT2;refGene_tx=XM_023253757.1
B2 32860641 32860797
refGene_gn=EHMT2;refGene_tx=XM_003985946.5;refGene_gn=EHMT2;refGene_tx=XM_011282304.3;refGene_gn=EHMT2;refGene_tx=XM_023253756.1;refGene_gn=EHMT2;refGene_tx=XM_023253757.1
B2 32860915 32860956
refGene_gn=EHMT2;refGene_tx=XM_003985946.5;refGene_gn=EHMT2;refGene_tx=XM_011282304.3;refGene_gn=EHMT2;refGene_tx=XM_011282305.3;refGene_gn=EHMT2;refGene_tx=XM_023253756.1;refGene_gn=EHMT2;refGene_tx=XM_023253757.1
B2 32861051 32861136
refGene_gn=EHMT2;refGene_tx=XM_003985946.5;refGene_gn=EHMT2;refGene_tx=XM_011282304.3;refGene_gn=EHMT2;refGene_tx=XM_011282305.3;refGene_gn=

EHMT2;refGene_tx=XM_023253756.1;refGene_gn=EHMT2;refGene_tx=XM_023253757.1
B2 32861998 32862249
refGene_gn=EHMT2;refGene_tx=XM_003985946.5;refGene_gn=EHMT2;refGene_tx=XM_011282304.3;refGene_gn=EHMT2;refGene_tx=XM_011282305.3;refGene_gn=EHMT2;refGene_tx=XM_023253756.1;refGene_gn=EHMT2;refGene_tx=XM_023253757.1
B2 32862346 32862565
refGene_gn=EHMT2;refGene_tx=XM_003985946.5;refGene_gn=EHMT2;refGene_tx=XM_011282304.3;refGene_gn=EHMT2;refGene_tx=XM_011282305.3;refGene_gn=EHMT2;refGene_tx=XM_023253756.1;refGene_gn=EHMT2;refGene_tx=XM_023253757.1
B2 32862663 32862943
refGene_gn=EHMT2;refGene_tx=XM_003985946.5;refGene_gn=EHMT2;refGene_tx=XM_011282304.3;refGene_gn=EHMT2;refGene_tx=XM_011282305.3;refGene_gn=EHMT2;refGene_tx=XM_023253756.1;refGene_gn=EHMT2;refGene_tx=XM_023253757.1
B2 32863376 32863418
refGene_gn=EHMT2;refGene_tx=XM_023253756.1;refGene_gn=EHMT2;refGene_tx=XM_023253757.1
B2 32865647 32867027 refGene_gn=ZBTB12;refGene_tx=XM_003985947.5
B2 32867881 32868287 refGene_gn=C2;refGene_tx=XM_019830388.2
B2 32899568 32899668 refGene_gn=C2;refGene_tx=XM_011282299.3
B2 32899799 32900006
refGene_gn=C2;refGene_tx=XM_011282299.3;refGene_gn=C2;refGene_tx=XM_019830388.2
B2 32900615 32900801
refGene_gn=C2;refGene_tx=XM_011282299.3;refGene_gn=C2;refGene_tx=XM_019830388.2
B2 32902531 32902705
refGene_gn=C2;refGene_tx=XM_011282299.3;refGene_gn=C2;refGene_tx=XM_019830388.2
B2 32902791 32902890
refGene_gn=C2;refGene_tx=XM_011282299.3;refGene_gn=C2;refGene_tx=XM_019830388.2
B2 32903086 32903220
refGene_gn=C2;refGene_tx=XM_011282299.3;refGene_gn=C2;refGene_tx=XM_019830388.2
B2 32904332 32904471
refGene_gn=C2;refGene_tx=XM_011282299.3;refGene_gn=C2;refGene_tx=XM_019830388.2
B2 32905078 32905219
refGene_gn=C2;refGene_tx=XM_011282299.3;refGene_gn=C2;refGene_tx=XM_019830388.2
B2 32905876 32905966
refGene_gn=C2;refGene_tx=XM_011282299.3;refGene_gn=C2;refGene_tx=XM_019830388.2
B2 32908802 32908943
refGene_gn=C2;refGene_tx=XM_011282299.3;refGene_gn=C2;refGene_tx=XM_019830388.2

B2 32909068 32909163
refGene_gn=C2;refGene_tx=XM_011282299.3;refGene_gn=C2;refGene_tx=XM_01
9830388.2
B2 32909259 32909371
refGene_gn=C2;refGene_tx=XM_011282299.3;refGene_gn=C2;refGene_tx=XM_01
9830388.2
B2 32909488 32909654
refGene_gn=C2;refGene_tx=XM_011282299.3;refGene_gn=C2;refGene_tx=XM_01
9830388.2
B2 32909763 32909840
refGene_gn=C2;refGene_tx=XM_011282299.3;refGene_gn=C2;refGene_tx=XM_01
9830388.2
B2 32909981 32910073
refGene_gn=C2;refGene_tx=XM_011282299.3;refGene_gn=C2;refGene_tx=XM_01
9830388.2
B2 32910277 32910404
refGene_gn=C2;refGene_tx=XM_011282299.3;refGene_gn=C2;refGene_tx=XM_01
9830388.2
B2 32910564 32910614
refGene_gn=C2;refGene_tx=XM_011282299.3;refGene_gn=C2;refGene_tx=XM_01
9830388.2
B2 32910767 32910944
refGene_gn=C2;refGene_tx=XM_011282299.3;refGene_gn=C2;refGene_tx=XM_01
9830388.2
B2 32911671 32911735 refGene_gn=CFB;refGene_tx=XM_003985951.3
B2 32911834 32912068 refGene_gn=CFB;refGene_tx=XM_003985951.3
B2 32912470 32912656 refGene_gn=CFB;refGene_tx=XM_003985951.3
B2 32912801 32912975 refGene_gn=CFB;refGene_tx=XM_003985951.3
B2 32913218 32913320 refGene_gn=CFB;refGene_tx=XM_003985951.3
B2 32913420 32913557 refGene_gn=CFB;refGene_tx=XM_003985951.3
B2 32913855 32913994 refGene_gn=CFB;refGene_tx=XM_003985951.3
B2 32914320 32914452 refGene_gn=CFB;refGene_tx=XM_003985951.3
B2 32915006 32915108 refGene_gn=CFB;refGene_tx=XM_003985951.3
B2 32915196 32915334 refGene_gn=CFB;refGene_tx=XM_003985951.3
B2 32915811 32915909 refGene_gn=CFB;refGene_tx=XM_003985951.3
B2 32916033 32916151 refGene_gn=CFB;refGene_tx=XM_003985951.3
B2 32916396 32916550 refGene_gn=CFB;refGene_tx=XM_003985951.3
B2 32916655 32916732 refGene_gn=CFB;refGene_tx=XM_003985951.3
B2 32916928 32917038 refGene_gn=CFB;refGene_tx=XM_003985951.3
B2 32917142 32917275 refGene_gn=CFB;refGene_tx=XM_003985951.3
B2 32917355 32917405 refGene_gn=CFB;refGene_tx=XM_003985951.3
B2 32917702 32917858 refGene_gn=CFB;refGene_tx=XM_003985951.3
B2 32918128 32918226
refGene_gn=NELFE;refGene_tx=XM_003985953.4;refGene_gn=NELFE;refGene_tx
=XM_023253758.1
B2 32919460 32919563
refGene_gn=NELFE;refGene_tx=XM_003985953.4;refGene_gn=NELFE;refGene_tx
=XM_023253758.1
B2 32919813 32919868
refGene_gn=NELFE;refGene_tx=XM_003985953.4;refGene_gn=NELFE;refGene_tx

=XM_023253758.1
B2 32920022 32920167
refGene_gn=NELFE;refGene_tx=XM_003985953.4;refGene_gn=NELFE;refGene_tx
=XM_023253758.1
B2 32920286 32920624
refGene_gn=NELFE;refGene_tx=XM_003985953.4;refGene_gn=NELFE;refGene_tx
=XM_023253758.1
B2 32920810 32920848
refGene_gn=NELFE;refGene_tx=XM_003985953.4;refGene_gn=NELFE;refGene_tx
=XM_023253758.1
B2 32920970 32921045
refGene_gn=NELFE;refGene_tx=XM_003985953.4;refGene_gn=NELFE;refGene_tx
=XM_023253758.1
B2 32921534 32921680
refGene_gn=NELFE;refGene_tx=XM_003985953.4;refGene_gn=NELFE;refGene_tx
=XM_023253758.1
B2 32921783 32921853
refGene_gn=NELFE;refGene_tx=XM_003985953.4;refGene_gn=NELFE;refGene_tx
=XM_023253758.1
B2 32923254 32923329
refGene_gn=NELFE;refGene_tx=XM_003985953.4;refGene_gn=NELFE;refGene_tx
=XM_023253758.1
B2 32924077 32924099
refGene_gn=SKIV2L;refGene_tx=XM_003985952.4;refGene_gn=SKIV2L;refGene_
tx=XM_019830387.1
B2 32924178 32924282
refGene_gn=SKIV2L;refGene_tx=XM_003985952.4;refGene_gn=SKIV2L;refGene_
tx=XM_019830387.1
B2 32924896 32925006
refGene_gn=SKIV2L;refGene_tx=XM_003985952.4;refGene_gn=SKIV2L;refGene_
tx=XM_019830387.1
B2 32925086 32925222
refGene_gn=SKIV2L;refGene_tx=XM_003985952.4;refGene_gn=SKIV2L;refGene_
tx=XM_019830387.1
B2 32925309 32925424
refGene_gn=SKIV2L;refGene_tx=XM_003985952.4;refGene_gn=SKIV2L;refGene_
tx=XM_019830387.1
B2 32925542 32925617
refGene_gn=SKIV2L;refGene_tx=XM_003985952.4;refGene_gn=SKIV2L;refGene_
tx=XM_019830387.1
B2 32925900 32925954
refGene_gn=SKIV2L;refGene_tx=XM_003985952.4;refGene_gn=SKIV2L;refGene_
tx=XM_019830387.1
B2 32926053 32926244
refGene_gn=SKIV2L;refGene_tx=XM_003985952.4;refGene_gn=SKIV2L;refGene_
tx=XM_019830387.1
B2 32926422 32926551
refGene_gn=SKIV2L;refGene_tx=XM_003985952.4;refGene_gn=SKIV2L;refGene_
tx=XM_019830387.1
B2 32926775 32926921

refGene_gn=SKIV2L;refGene_tx=XM_003985952.4;refGene_gn=SKIV2L;refGene_tx=XM_019830387.1
B2 32927488 32927635
refGene_gn=SKIV2L;refGene_tx=XM_003985952.4;refGene_gn=SKIV2L;refGene_tx=XM_019830387.1
B2 32927766 32927851
refGene_gn=SKIV2L;refGene_tx=XM_003985952.4;refGene_gn=SKIV2L;refGene_tx=XM_019830387.1
B2 32928040 32928147
refGene_gn=SKIV2L;refGene_tx=XM_003985952.4;refGene_gn=SKIV2L;refGene_tx=XM_019830387.1
B2 32928722 32928869
refGene_gn=SKIV2L;refGene_tx=XM_003985952.4;refGene_gn=SKIV2L;refGene_tx=XM_019830387.1
B2 32928946 32929043
refGene_gn=SKIV2L;refGene_tx=XM_003985952.4;refGene_gn=SKIV2L;refGene_tx=XM_019830387.1
B2 32929226 32929439
refGene_gn=SKIV2L;refGene_tx=XM_003985952.4;refGene_gn=SKIV2L;refGene_tx=XM_019830387.1
B2 32929521 32929632
refGene_gn=SKIV2L;refGene_tx=XM_003985952.4;refGene_gn=SKIV2L;refGene_tx=XM_019830387.1
B2 32931375 32931606
refGene_gn=SKIV2L;refGene_tx=XM_003985952.4;refGene_gn=SKIV2L;refGene_tx=XM_019830387.1
B2 32932319 32932457
refGene_gn=SKIV2L;refGene_tx=XM_003985952.4;refGene_gn=SKIV2L;refGene_tx=XM_019830387.1
B2 32932623 32932761
refGene_gn=SKIV2L;refGene_tx=XM_003985952.4;refGene_gn=SKIV2L;refGene_tx=XM_019830387.1
B2 32932905 32933010
refGene_gn=SKIV2L;refGene_tx=XM_003985952.4;refGene_gn=SKIV2L;refGene_tx=XM_019830387.1
B2 32933353 32933501
refGene_gn=SKIV2L;refGene_tx=XM_003985952.4;refGene_gn=SKIV2L;refGene_tx=XM_019830387.1
B2 32933596 32933723
refGene_gn=SKIV2L;refGene_tx=XM_003985952.4;refGene_gn=SKIV2L;refGene_tx=XM_019830387.1
B2 32933873 32934084
refGene_gn=SKIV2L;refGene_tx=XM_003985952.4;refGene_gn=SKIV2L;refGene_tx=XM_019830387.1
B2 32934239 32934350
refGene_gn=SKIV2L;refGene_tx=XM_003985952.4;refGene_gn=SKIV2L;refGene_tx=XM_019830387.1
B2 32934429 32934648
refGene_gn=SKIV2L;refGene_tx=XM_003985952.4;refGene_gn=SKIV2L;refGene_tx=XM_019830387.1

B2 32934840 32934981
refGene_gn=SKIV2L;refGene_tx=XM_003985952.4;refGene_gn=SKIV2L;refGene_tx=XM_019830387.1
B2 32935069 32935270
refGene_gn=SKIV2L;refGene_tx=XM_003985952.4;refGene_gn=SKIV2L;refGene_tx=XM_019830387.1
B2 32935429 32935580
refGene_gn=DXO;refGene_tx=XM_003985956.5;refGene_gn=DXO;refGene_tx=XM_006931581.4;refGene_gn=DXO;refGene_tx=XM_006931582.4
B2 32935662 32935757
refGene_gn=DXO;refGene_tx=XM_003985956.5;refGene_gn=DXO;refGene_tx=XM_006931581.4;refGene_gn=DXO;refGene_tx=XM_006931582.4
B2 32935898 32936034
refGene_gn=DXO;refGene_tx=XM_003985956.5;refGene_gn=DXO;refGene_tx=XM_006931581.4;refGene_gn=DXO;refGene_tx=XM_006931582.4
B2 32936151 32936371
refGene_gn=DXO;refGene_tx=XM_003985956.5;refGene_gn=DXO;refGene_tx=XM_006931581.4;refGene_gn=DXO;refGene_tx=XM_006931582.4
B2 32936459 32936695
refGene_gn=DXO;refGene_tx=XM_003985956.5;refGene_gn=DXO;refGene_tx=XM_006931581.4;refGene_gn=DXO;refGene_tx=XM_006931582.4
B2 32936865 32937224
refGene_gn=DXO;refGene_tx=XM_003985956.5;refGene_gn=DXO;refGene_tx=XM_006931581.4;refGene_gn=DXO;refGene_tx=XM_006931582.4;refGene_gn=STK19;refGene_tx=XM_011282353.1
B2 32937420 32938007 refGene_gn=STK19;refGene_tx=XM_011282353.1
B2 32938138 32938275 refGene_gn=STK19;refGene_tx=XM_011282353.1
B2 32941414 32941510 refGene_gn=STK19;refGene_tx=XM_011282353.1
B2 32941831 32941971 refGene_gn=STK19;refGene_tx=XM_011282353.1
B2 32942784 32942882 refGene_gn=STK19;refGene_tx=XM_011282353.1
B2 32942989 32943137 refGene_gn=STK19;refGene_tx=XM_011282353.1
B2 32943282 32943328 refGene_gn=STK19;refGene_tx=XM_011282353.1
B2 32944352 32944417
refGene_gn=LOC101094346;refGene_tx=XM_006931583.3
B2 32944556 32944755
refGene_gn=LOC101094346;refGene_tx=XM_006931583.3
B2 32945223 32945425
refGene_gn=LOC101094346;refGene_tx=XM_006931583.3
B2 32945634 32945705
refGene_gn=LOC101094346;refGene_tx=XM_006931583.3
B2 32945785 32945874
refGene_gn=LOC101094346;refGene_tx=XM_006931583.3
B2 32946085 32946168
refGene_gn=LOC101094346;refGene_tx=XM_006931583.3
B2 32946349 32946446
refGene_gn=LOC101094346;refGene_tx=XM_006931583.3
B2 32946604 32946710
refGene_gn=LOC101094346;refGene_tx=XM_006931583.3
B2 32946810 32946943
refGene_gn=LOC101094346;refGene_tx=XM_006931583.3

B2	32947412 32947528
refGene_gn=LOC101094346;refGene_tx=XM_006931583.3
B2	32947633 32947810
refGene_gn=LOC101094346;refGene_tx=XM_006931583.3
B2	32947962 32948145
refGene_gn=LOC101094346;refGene_tx=XM_006931583.3
B2	32948278 32948464
refGene_gn=LOC101094346;refGene_tx=XM_006931583.3
B2	32948579 32948738
refGene_gn=LOC101094346;refGene_tx=XM_006931583.3
B2	32949014 32949141
refGene_gn=LOC101094346;refGene_tx=XM_006931583.3
B2	32949317 32949392
refGene_gn=LOC101094346;refGene_tx=XM_006931583.3
B2	32949670 32949868
refGene_gn=LOC101094346;refGene_tx=XM_006931583.3
B2	32949959 32950071
refGene_gn=LOC101094346;refGene_tx=XM_006931583.3
B2	32950162 32950233
refGene_gn=LOC101094346;refGene_tx=XM_006931583.3
B2	32950507 32950647
refGene_gn=LOC101094346;refGene_tx=XM_006931583.3
B2	32950762 32950972
refGene_gn=LOC101094346;refGene_tx=XM_006931583.3
B2	32951219 32951271
refGene_gn=LOC101094346;refGene_tx=XM_006931583.3
B2	32951387 32951477
refGene_gn=LOC101094346;refGene_tx=XM_006931583.3
B2	32951613 32951823
refGene_gn=LOC101094346;refGene_tx=XM_006931583.3
B2	32952002 32952078
refGene_gn=LOC101094346;refGene_tx=XM_006931583.3
B2	32952240 32952397
refGene_gn=LOC101094346;refGene_tx=XM_006931583.3
B2	32952498 32952615
refGene_gn=LOC101094346;refGene_tx=XM_006931583.3
B2	32952721 32952893
refGene_gn=LOC101094346;refGene_tx=XM_006931583.3
B2	32953091 32953324
refGene_gn=LOC101094346;refGene_tx=XM_006931583.3
B2	32953397 32953565
refGene_gn=LOC101094346;refGene_tx=XM_006931583.3
B2	32953937 32953997
refGene_gn=LOC101094346;refGene_tx=XM_006931583.3
B2	32954993 32955087
refGene_gn=LOC101094346;refGene_tx=XM_006931583.3
B2	32955166 32955353
refGene_gn=LOC101094346;refGene_tx=XM_006931583.3
B2	32955475 32955566
refGene_gn=LOC101094346;refGene_tx=XM_006931583.3

B2 32955702 32955777
refGene_gn=LOC101094346;refGene_tx=XM_006931583.3
B2 32957706 32957809
refGene_gn=LOC101094346;refGene_tx=XM_006931583.3
B2 32957969 32958059
refGene_gn=LOC101094346;refGene_tx=XM_006931583.3
B2 32958142 32958241
refGene_gn=LOC101094346;refGene_tx=XM_006931583.3
B2 32958341 32958425
refGene_gn=LOC101094346;refGene_tx=XM_006931583.3
B2 32958706 32958839
refGene_gn=LOC101094346;refGene_tx=XM_006931583.3
B2 32959002 32959144
refGene_gn=LOC101094346;refGene_tx=XM_006931583.3
B2 32962160 32962359
refGene_gn=LOC101094596;refGene_tx=XM_003985958.5
B2 32962443 32962533
refGene_gn=LOC101094596;refGene_tx=XM_003985958.5
B2 32962636 32962791
refGene_gn=LOC101094596;refGene_tx=XM_003985958.5
B2 32962880 32962976
refGene_gn=LOC101094596;refGene_tx=XM_003985958.5
B2 32963090 32963192
refGene_gn=LOC101094596;refGene_tx=XM_003985958.5
B2 32963289 32963376
refGene_gn=LOC101094596;refGene_tx=XM_003985958.5
B2 32963531 32963732
refGene_gn=LOC101094596;refGene_tx=XM_003985958.5
B2 32963947 32964126
refGene_gn=LOC101094596;refGene_tx=XM_003985958.5
B2 32964204 32964308
refGene_gn=LOC101094596;refGene_tx=XM_003985958.5
B2 32964398 32964664
refGene_gn=LOC101094596;refGene_tx=XM_003985958.5
B2 32964882 32964987 refGene_gn=TNXB;refGene_tx=XM_023254711.1
B2 32965345 32965509 refGene_gn=TNXB;refGene_tx=XM_023254711.1
B2 32965588 32965750 refGene_gn=TNXB;refGene_tx=XM_023254711.1
B2 32965841 32965938 refGene_gn=TNXB;refGene_tx=XM_023254711.1
B2 32966038 32966190 refGene_gn=TNXB;refGene_tx=XM_023254711.1
B2 32966286 32966419 refGene_gn=TNXB;refGene_tx=XM_023254711.1
B2 32966551 32966682 refGene_gn=TNXB;refGene_tx=XM_023254711.1
B2 32966779 32966923 refGene_gn=TNXB;refGene_tx=XM_023254711.1
B2 32967026 32967146 refGene_gn=TNXB;refGene_tx=XM_023254711.1
B2 32967286 32967430 refGene_gn=TNXB;refGene_tx=XM_023254711.1
B2 32967548 32967671 refGene_gn=TNXB;refGene_tx=XM_023254711.1
B2 32967958 32968294 refGene_gn=TNXB;refGene_tx=XM_023254711.1
B2 32968593 32968917 refGene_gn=TNXB;refGene_tx=XM_023254711.1
B2 32969709 32969997 refGene_gn=TNXB;refGene_tx=XM_023254711.1
B2 32971236 32971512 refGene_gn=TNXB;refGene_tx=XM_023254711.1
B2 32971900 32972188 refGene_gn=TNXB;refGene_tx=XM_023254711.1

B2 32972780 32972873 refGene_gn=TNXB;refGene_tx=XM_023254711.1
B2 32972875 32973086 refGene_gn=TNXB;refGene_tx=XM_023254711.1
B2 32973502 32973599 refGene_gn=TNXB;refGene_tx=XM_023254711.1
B2 32973600 32973735 refGene_gn=TNXB;refGene_tx=XM_023254711.1
B2 32973736 32973833 refGene_gn=TNXB;refGene_tx=XM_023254711.1
B2 32975470 32975608 refGene_gn=TNXB;refGene_tx=XM_023254711.1
B2 32975610 32975655 refGene_gn=TNXB;refGene_tx=XM_023254711.1
B2 32977722 32978073 refGene_gn=TNXB;refGene_tx=XM_023254711.1
B2 32978466 32978817 refGene_gn=TNXB;refGene_tx=XM_023254711.1
B2 32980770 32981067 refGene_gn=TNXB;refGene_tx=XM_023254711.1
B2 32982681 32983032 refGene_gn=TNXB;refGene_tx=XM_023254711.1
B2 32983425 32983749 refGene_gn=TNXB;refGene_tx=XM_023254711.1
B2 32985277 32985628 refGene_gn=TNXB;refGene_tx=XM_023254711.1
B2 32986021 32986324 refGene_gn=TNXB;refGene_tx=XM_023254711.1
B2 32986483 32986792 refGene_gn=TNXB;refGene_tx=XM_023254711.1
B2 32987324 32987633 refGene_gn=TNXB;refGene_tx=XM_023254711.1
B2 32987895 32988183 refGene_gn=TNXB;refGene_tx=XM_023254711.1
B2 32989639 32989939 refGene_gn=TNXB;refGene_tx=XM_023254711.1
B2 32991180 32991483 refGene_gn=TNXB;refGene_tx=XM_023254711.1
B2 32994492 32994852 refGene_gn=TNXB;refGene_tx=XM_023254711.1
B2 32996731 32997037 refGene_gn=TNXB;refGene_tx=XM_023254711.1
B2 32997391 32997682 refGene_gn=TNXB;refGene_tx=XM_023254711.1
B2 32999462 32999756 refGene_gn=TNXB;refGene_tx=XM_023254711.1
B2 33000829 33001198 refGene_gn=TNXB;refGene_tx=XM_023254711.1
B2 33003962 33004226 refGene_gn=TNXB;refGene_tx=XM_023254711.1
B2 33004408 33004565 refGene_gn=TNXB;refGene_tx=XM_023254711.1
B2 33009129 33009245 refGene_gn=TNXB;refGene_tx=XM_023254711.1
B2 33009728 33011567 refGene_gn=TNXB;refGene_tx=XM_023254711.1
B2 33011930 33012330 refGene_gn=TNXB;refGene_tx=XM_023254711.1
B2 33028131 33028361
refGene_gn=ATF6B;refGene_tx=XM_003985959.4;refGene_gn=ATF6B;refGene_tx
=XM_006931584.4;refGene_gn=ATF6B;refGene_tx=XM_011282294.3;refGene_gn=
ATF6B;refGene_tx=XM_011282295.2;refGene_gn=ATF6B;refGene_tx=XM_0198303
89.2
B2 33028886 33028971
refGene_gn=ATF6B;refGene_tx=XM_003985959.4;refGene_gn=ATF6B;refGene_tx
=XM_006931584.4;refGene_gn=ATF6B;refGene_tx=XM_011282294.3;refGene_gn=
ATF6B;refGene_tx=XM_011282295.2;refGene_gn=ATF6B;refGene_tx=XM_0198303
89.2
B2 33029111 33029223
refGene_gn=ATF6B;refGene_tx=XM_003985959.4;refGene_gn=ATF6B;refGene_tx
=XM_006931584.4;refGene_gn=ATF6B;refGene_tx=XM_011282294.3;refGene_gn=
ATF6B;refGene_tx=XM_011282295.2;refGene_gn=ATF6B;refGene_tx=XM_0198303
89.2
B2 33029446 33029533
refGene_gn=ATF6B;refGene_tx=XM_003985959.4;refGene_gn=ATF6B;refGene_tx
=XM_006931584.4;refGene_gn=ATF6B;refGene_tx=XM_011282294.3;refGene_gn=
ATF6B;refGene_tx=XM_011282295.2;refGene_gn=ATF6B;refGene_tx=XM_0112822
96.3;refGene_gn=ATF6B;refGene_tx=XM_019830389.2
B2 33029725 33029807

refGene_gn=ATF6B;refGene_tx=XM_003985959.4;refGene_gn=ATF6B;refGene_tx=XM_006931584.4;refGene_gn=ATF6B;refGene_tx=XM_011282294.3;refGene_gn=ATF6B;refGene_tx=XM_011282295.2;refGene_gn=ATF6B;refGene_tx=XM_011282296.3;refGene_gn=ATF6B;refGene_tx=XM_019830389.2
B2 33029989 33030100
refGene_gn=ATF6B;refGene_tx=XM_003985959.4;refGene_gn=ATF6B;refGene_tx=XM_006931584.4;refGene_gn=ATF6B;refGene_tx=XM_011282294.3;refGene_gn=ATF6B;refGene_tx=XM_011282295.2;refGene_gn=ATF6B;refGene_tx=XM_011282296.3;refGene_gn=ATF6B;refGene_tx=XM_019830389.2
B2 33030267 33030447
refGene_gn=ATF6B;refGene_tx=XM_003985959.4;refGene_gn=ATF6B;refGene_tx=XM_006931584.4;refGene_gn=ATF6B;refGene_tx=XM_011282294.3;refGene_gn=ATF6B;refGene_tx=XM_011282295.2;refGene_gn=ATF6B;refGene_tx=XM_011282296.3;refGene_gn=ATF6B;refGene_tx=XM_019830389.2
B2 33030865 33030957
refGene_gn=ATF6B;refGene_tx=XM_003985959.4;refGene_gn=ATF6B;refGene_tx=XM_006931584.4;refGene_gn=ATF6B;refGene_tx=XM_011282294.3;refGene_gn=ATF6B;refGene_tx=XM_011282295.2;refGene_gn=ATF6B;refGene_tx=XM_011282296.3;refGene_gn=ATF6B;refGene_tx=XM_019830389.2
B2 33031050 33031236
refGene_gn=ATF6B;refGene_tx=XM_003985959.4;refGene_gn=ATF6B;refGene_tx=XM_006931584.4;refGene_gn=ATF6B;refGene_tx=XM_011282294.3;refGene_gn=ATF6B;refGene_tx=XM_011282295.2;refGene_gn=ATF6B;refGene_tx=XM_011282296.3;refGene_gn=ATF6B;refGene_tx=XM_019830389.2
B2 33031906 33032040
refGene_gn=ATF6B;refGene_tx=XM_003985959.4;refGene_gn=ATF6B;refGene_tx=XM_006931584.4;refGene_gn=ATF6B;refGene_tx=XM_011282294.3;refGene_gn=ATF6B;refGene_tx=XM_011282295.2;refGene_gn=ATF6B;refGene_tx=XM_011282296.3;refGene_gn=ATF6B;refGene_tx=XM_019830389.2
B2 33032185 33032317
refGene_gn=ATF6B;refGene_tx=XM_003985959.4;refGene_gn=ATF6B;refGene_tx=XM_006931584.4;refGene_gn=ATF6B;refGene_tx=XM_011282294.3;refGene_gn=ATF6B;refGene_tx=XM_011282295.2;refGene_gn=ATF6B;refGene_tx=XM_011282296.3;refGene_gn=ATF6B;refGene_tx=XM_019830389.2
B2 33032405 33032541
refGene_gn=ATF6B;refGene_tx=XM_003985959.4;refGene_gn=ATF6B;refGene_tx=XM_006931584.4;refGene_gn=ATF6B;refGene_tx=XM_011282294.3;refGene_gn=ATF6B;refGene_tx=XM_011282295.2;refGene_gn=ATF6B;refGene_tx=XM_011282296.3;refGene_gn=ATF6B;refGene_tx=XM_019830389.2
B2 33032683 33032769
refGene_gn=ATF6B;refGene_tx=XM_003985959.4;refGene_gn=ATF6B;refGene_tx=XM_006931584.4;refGene_gn=ATF6B;refGene_tx=XM_011282294.3;refGene_gn=ATF6B;refGene_tx=XM_011282295.2;refGene_gn=ATF6B;refGene_tx=XM_011282296.3;refGene_gn=ATF6B;refGene_tx=XM_019830389.2
B2 33033445 33033538
refGene_gn=ATF6B;refGene_tx=XM_011282294.3;refGene_gn=ATF6B;refGene_tx=XM_011282295.2;refGene_gn=ATF6B;refGene_tx=XM_011282296.3;refGene_gn=ATF6B;refGene_tx=XM_019830389.2
B2 33035119 33035255
refGene_gn=ATF6B;refGene_tx=XM_003985959.4;refGene_gn=ATF6B;refGene_tx

=XM_006931584.4;refGene_gn=ATF6B;refGene_tx=XM_011282294.3;refGene_gn=
ATF6B;refGene_tx=XM_011282295.2;refGene_gn=ATF6B;refGene_tx=XM_0112822
96.3;refGene_gn=ATF6B;refGene_tx=XM_019830389.2
B2 33035781 33035873
refGene_gn=ATF6B;refGene_tx=XM_003985959.4;refGene_gn=ATF6B;refGene_tx
=XM_006931584.4;refGene_gn=ATF6B;refGene_tx=XM_011282294.3;refGene_gn=
ATF6B;refGene_tx=XM_011282295.2;refGene_gn=ATF6B;refGene_tx=XM_0112822
96.3;refGene_gn=ATF6B;refGene_tx=XM_019830389.2
B2 33036123 33036202
refGene_gn=ATF6B;refGene_tx=XM_003985959.4;refGene_gn=ATF6B;refGene_tx
=XM_006931584.4;refGene_gn=ATF6B;refGene_tx=XM_011282294.3;refGene_gn=
ATF6B;refGene_tx=XM_011282295.2;refGene_gn=ATF6B;refGene_tx=XM_0112822
96.3;refGene_gn=ATF6B;refGene_tx=XM_019830389.2
B2 33036335 33036415
refGene_gn=ATF6B;refGene_tx=XM_003985959.4;refGene_gn=ATF6B;refGene_tx
=XM_006931584.4;refGene_gn=ATF6B;refGene_tx=XM_011282294.3;refGene_gn=
ATF6B;refGene_tx=XM_011282295.2;refGene_gn=ATF6B;refGene_tx=XM_0112822
96.3;refGene_gn=ATF6B;refGene_tx=XM_019830389.2
B2 33036783 33036898
refGene_gn=ATF6B;refGene_tx=XM_003985959.4;refGene_gn=ATF6B;refGene_tx
=XM_006931584.4;refGene_gn=ATF6B;refGene_tx=XM_011282294.3;refGene_gn=
ATF6B;refGene_tx=XM_011282295.2;refGene_gn=ATF6B;refGene_tx=XM_0112822
96.3;refGene_gn=ATF6B;refGene_tx=XM_019830389.2
B2 33037433 33038483 refGene_gn=FKBPL;refGene_tx=XM_003985960.4
B2 33053625 33053802 refGene_gn=PRRT1;refGene_tx=XM_003985961.4
B2 33053944 33054130 refGene_gn=PRRT1;refGene_tx=XM_003985961.4
B2 33054768 33055307 refGene_gn=PRRT1;refGene_tx=XM_003985961.4
B2 33056209 33056228 refGene_gn=PRRT1;refGene_tx=XM_003985961.4
B2 33057941 33057951 refGene_gn=PPT2;refGene_tx=XM_019830396.2
B2 33058882 33059076
refGene_gn=PPT2;refGene_tx=XM_019830396.2;refGene_gn=PPT2;refGene_tx=X
M_023253762.1;refGene_gn=PPT2;refGene_tx=XM_023253763.1
B2 33059328 33059482
refGene_gn=PPT2;refGene_tx=XM_019830396.2;refGene_gn=PPT2;refGene_tx=X
M_023253762.1;refGene_gn=PPT2;refGene_tx=XM_023253763.1
B2 33059943 33060039
refGene_gn=PPT2;refGene_tx=XM_019830396.2;refGene_gn=PPT2;refGene_tx=X
M_023253762.1;refGene_gn=PPT2;refGene_tx=XM_023253763.1
B2 33060127 33060235
refGene_gn=PPT2;refGene_tx=XM_019830396.2;refGene_gn=PPT2;refGene_tx=X
M_023253762.1;refGene_gn=PPT2;refGene_tx=XM_023253763.1
B2 33060990 33061074
refGene_gn=PPT2;refGene_tx=XM_019830396.2;refGene_gn=PPT2;refGene_tx=X
M_023253762.1;refGene_gn=PPT2;refGene_tx=XM_023253763.1
B2 33061190 33061275
refGene_gn=PPT2;refGene_tx=XM_019830396.2;refGene_gn=PPT2;refGene_tx=X
M_023253762.1;refGene_gn=PPT2;refGene_tx=XM_023253763.1
B2 33064100 33064155
refGene_gn=PPT2;refGene_tx=XM_019830396.2;refGene_gn=PPT2;refGene_tx=X
M_023253762.1;refGene_gn=PPT2;refGene_tx=XM_023253763.1

B2 33064328 33064472
refGene_gn=PPT2;refGene_tx=XM_019830396.2;refGene_gn=PPT2;refGene_tx=XM_023253762.1;refGene_gn=PPT2;refGene_tx=XM_023253763.1
B2 33066625 33066783 refGene_gn=EGFL8;refGene_tx=XM_019830395.1
B2 33066994 33067117 refGene_gn=EGFL8;refGene_tx=XM_019830395.1
B2 33067219 33067329 refGene_gn=EGFL8;refGene_tx=XM_019830395.1
B2 33067427 33067523 refGene_gn=EGFL8;refGene_tx=XM_019830395.1
B2 33067630 33067801 refGene_gn=EGFL8;refGene_tx=XM_019830395.1
B2 33067874 33067954 refGene_gn=EGFL8;refGene_tx=XM_019830395.1
B2 33068036 33068190 refGene_gn=EGFL8;refGene_tx=XM_019830395.1
B2 33068459 33068545 refGene_gn=EGFL8;refGene_tx=XM_019830395.1
B2 33069829 33070014
refGene_gn=AGPAT1;refGene_tx=XM_003985964.4;refGene_gn=AGPAT1;refGene_tx=XM_019830394.1;refGene_gn=AGPAT1;refGene_tx=XM_023253761.1
B2 33070528 33070601
refGene_gn=AGPAT1;refGene_tx=XM_003985964.4;refGene_gn=AGPAT1;refGene_tx=XM_019830394.1;refGene_gn=AGPAT1;refGene_tx=XM_023253761.1
B2 33070726 33070822
refGene_gn=AGPAT1;refGene_tx=XM_003985964.4;refGene_gn=AGPAT1;refGene_tx=XM_019830394.1;refGene_gn=AGPAT1;refGene_tx=XM_023253761.1
B2 33070958 33071134
refGene_gn=AGPAT1;refGene_tx=XM_003985964.4;refGene_gn=AGPAT1;refGene_tx=XM_019830394.1;refGene_gn=AGPAT1;refGene_tx=XM_023253761.1
B2 33071427 33071561
refGene_gn=AGPAT1;refGene_tx=XM_003985964.4;refGene_gn=AGPAT1;refGene_tx=XM_019830394.1;refGene_gn=AGPAT1;refGene_tx=XM_023253761.1
B2 33071778 33071978
refGene_gn=AGPAT1;refGene_tx=XM_003985964.4;refGene_gn=AGPAT1;refGene_tx=XM_019830394.1;refGene_gn=AGPAT1;refGene_tx=XM_023253761.1
B2 33078586 33078726
refGene_gn=RNF5;refGene_tx=XM_003985966.5;refGene_gn=RNF5;refGene_tx=XM_019830398.2
B2 33079691 33079710 refGene_gn=RNF5;refGene_tx=XM_003985966.5
B2 33079805 33079918
refGene_gn=RNF5;refGene_tx=XM_003985966.5;refGene_gn=RNF5;refGene_tx=XM_019830398.2
B2 33080028 33080083
refGene_gn=RNF5;refGene_tx=XM_003985966.5;refGene_gn=RNF5;refGene_tx=XM_019830398.2
B2 33080158 33080276
refGene_gn=RNF5;refGene_tx=XM_003985966.5;refGene_gn=RNF5;refGene_tx=XM_019830398.2
B2 33080382 33080480
refGene_gn=RNF5;refGene_tx=XM_003985966.5;refGene_gn=RNF5;refGene_tx=XM_019830398.2
B2 33081301 33081429
refGene_gn=AGER;refGene_tx=XM_003985968.4;refGene_gn=AGER;refGene_tx=XM_006931589.3;refGene_gn=AGER;refGene_tx=XM_019830391.2;refGene_gn=AGER;refGene_tx=XM_019830392.1;refGene_gn=AGER;refGene_tx=XM_023253760.1
B2 33081527 33081654

refGene_gn=AGER;refGene_tx=XM_003985968.4;refGene_gn=AGER;refGene_tx=XM_006931589.3;refGene_gn=AGER;refGene_tx=XM_019830391.2;refGene_gn=AGER;refGene_tx=XM_019830392.1;refGene_gn=AGER;refGene_tx=XM_023253760.1
B2 33081767 33081791
refGene_gn=AGER;refGene_tx=XM_003985968.4;refGene_gn=AGER;refGene_tx=XM_006931589.3;refGene_gn=AGER;refGene_tx=XM_019830391.2;refGene_gn=AGER;refGene_tx=XM_019830392.1;refGene_gn=AGER;refGene_tx=XM_023253760.1
B2 33082055 33082197
refGene_gn=AGER;refGene_tx=XM_003985968.4;refGene_gn=AGER;refGene_tx=XM_006931589.3;refGene_gn=AGER;refGene_tx=XM_019830391.2;refGene_gn=AGER;refGene_tx=XM_019830392.1;refGene_gn=AGER;refGene_tx=XM_023253760.1
B2 33082397 33082525
refGene_gn=AGER;refGene_tx=XM_003985968.4;refGene_gn=AGER;refGene_tx=XM_006931589.3;refGene_gn=AGER;refGene_tx=XM_019830391.2;refGene_gn=AGER;refGene_tx=XM_019830392.1;refGene_gn=AGER;refGene_tx=XM_023253760.1
B2 33082640 33082823
refGene_gn=AGER;refGene_tx=XM_003985968.4;refGene_gn=AGER;refGene_tx=XM_006931589.3;refGene_gn=AGER;refGene_tx=XM_019830391.2;refGene_gn=AGER;refGene_tx=XM_019830392.1;refGene_gn=AGER;refGene_tx=XM_023253760.1
B2 33082922 33083010
refGene_gn=AGER;refGene_tx=XM_003985968.4;refGene_gn=AGER;refGene_tx=XM_006931589.3;refGene_gn=AGER;refGene_tx=XM_019830391.2;refGene_gn=AGER;refGene_tx=XM_019830392.1;refGene_gn=AGER;refGene_tx=XM_023253760.1
B2 33083153 33083218
refGene_gn=AGER;refGene_tx=XM_003985968.4;refGene_gn=AGER;refGene_tx=XM_006931589.3;refGene_gn=AGER;refGene_tx=XM_019830391.2;refGene_gn=AGER;refGene_tx=XM_019830392.1;refGene_gn=AGER;refGene_tx=XM_023253760.1
B2 33083399 33084149
refGene_gn=AGER;refGene_tx=XM_003985968.4;refGene_gn=AGER;refGene_tx=XM_006931589.3;refGene_gn=AGER;refGene_tx=XM_019830391.2;refGene_gn=AGER;refGene_tx=XM_019830392.1;refGene_gn=AGER;refGene_tx=XM_023253760.1
B2 33086251 33086344 refGene_gn=PBX2;refGene_tx=XM_003985970.5
B2 33086470 33086557 refGene_gn=PBX2;refGene_tx=XM_003985970.5
B2 33086663 33086752 refGene_gn=PBX2;refGene_tx=XM_003985970.5
B2 33087146 33087300 refGene_gn=PBX2;refGene_tx=XM_003985970.5
B2 33087415 33087551 refGene_gn=PBX2;refGene_tx=XM_003985970.5
B2 33087732 33087923 refGene_gn=PBX2;refGene_tx=XM_003985970.5
B2 33088021 33088269 refGene_gn=PBX2;refGene_tx=XM_003985970.5
B2 33088506 33088580 refGene_gn=PBX2;refGene_tx=XM_003985970.5
B2 33089519 33089740 refGene_gn=PBX2;refGene_tx=XM_003985970.5
B2 33091228 33091366
refGene_gn=GPSM3;refGene_tx=XM_006931590.4;refGene_gn=GPSM3;refGene_tx=XM_019830397.2
B2 33091589 33091789
refGene_gn=GPSM3;refGene_tx=XM_006931590.4;refGene_gn=GPSM3;refGene_tx=XM_019830397.2
B2 33091985 33092094
refGene_gn=GPSM3;refGene_tx=XM_006931590.4;refGene_gn=GPSM3;refGene_tx=XM_019830397.2
B2 33092303 33092348

refGene_gn=GPSM3;refGene_tx=XM_006931590.4;refGene_gn=GPSM3;refGene_tx=XM_019830397.2
B2	33092574 33092646 refGene_gn=GPSM3;refGene_tx=XM_019830397.2
B2	33094282 33094417 refGene_gn=GPSM3;refGene_tx=XM_006931590.4
B2	33094545 33095250 refGene_gn=NOTCH4;refGene_tx=XM_023253759.1
B2	33095646 33095744 refGene_gn=NOTCH4;refGene_tx=XM_023253759.1
B2	33096266 33096414 refGene_gn=NOTCH4;refGene_tx=XM_023253759.1
B2	33096644 33096830 refGene_gn=NOTCH4;refGene_tx=XM_023253759.1
B2	33096831 33096941 refGene_gn=NOTCH4;refGene_tx=XM_023253759.1
B2	33097468 33097607 refGene_gn=NOTCH4;refGene_tx=XM_023253759.1
B2	33097710 33097783 refGene_gn=NOTCH4;refGene_tx=XM_023253759.1
B2	33097936 33098156 refGene_gn=NOTCH4;refGene_tx=XM_023253759.1
B2	33098692 33098868 refGene_gn=NOTCH4;refGene_tx=XM_023253759.1
B2	33098973 33099357 refGene_gn=NOTCH4;refGene_tx=XM_023253759.1
B2	33099915 33100439 refGene_gn=NOTCH4;refGene_tx=XM_023253759.1
B2	33101266 33101379 refGene_gn=NOTCH4;refGene_tx=XM_023253759.1
B2	33101606 33101859 refGene_gn=NOTCH4;refGene_tx=XM_023253759.1
B2	33103819 33104004 refGene_gn=NOTCH4;refGene_tx=XM_023253759.1
B2	33104807 33104961 refGene_gn=NOTCH4;refGene_tx=XM_023253759.1
B2	33105314 33105402 refGene_gn=NOTCH4;refGene_tx=XM_023253759.1
B2	33105618 33105707 refGene_gn=NOTCH4;refGene_tx=XM_023253759.1
B2	33198715 33198800
refGene_gn=LOC111560582;refGene_tx=XM_023254712.1
B2	33201184 33201532
refGene_gn=LOC111560582;refGene_tx=XM_023254712.1
B2	33205950 33206232
refGene_gn=LOC111560582;refGene_tx=XM_023254712.1
B2	33206347 33206548
refGene_gn=LOC111560582;refGene_tx=XM_023254712.1
B2	33206789 33206810
refGene_gn=LOC111560582;refGene_tx=XM_023254712.1
B2	33206902 33206929
refGene_gn=LOC111560582;refGene_tx=XM_023254712.1
B2	33207936 33207963
refGene_gn=LOC111560582;refGene_tx=XM_023254712.1
B2	33210425 33210452
refGene_gn=LOC111560582;refGene_tx=XM_023254712.1
B2	33210872 33211618
refGene_gn=LOC111560582;refGene_tx=XM_023254712.1
B2	33218278 33218299
refGene_gn=LOC111560582;refGene_tx=XM_023254712.1
B2	33219962 33220016
refGene_gn=LOC111560582;refGene_tx=XM_023254712.1
B2	33226780 33226808
refGene_gn=LOC111560583;refGene_tx=XM_023254713.1
B2	33227065 33227463
refGene_gn=LOC111560583;refGene_tx=XM_023254713.1
B2	33251374 33252367
refGene_gn=LOC111560438;refGene_tx=XM_023253776.1
B2	33392330 33393106

refGene_gn=CB2H6orf10;refGene_tx=XM_023253764.1;refGene_gn=CB2H6orf10;
refGene_tx=XM_023253765.1;refGene_gn=CB2H6orf10;refGene_tx=XM_02325376
7.1;refGene_gn=CB2H6orf10;refGene_tx=XM_023253768.1;refGene_gn=CB2H6or
f10;refGene_tx=XM_023253769.1;refGene_gn=CB2H6orf10;refGene_tx=XM_0232
53770.1;refGene_gn=CB2H6orf10;refGene_tx=XM_023253771.1;refGene_gn=CB2
H6orf10;refGene_tx=XM_023253772.1;refGene_gn=CB2H6orf10;refGene_tx=XM_
023253773.1;refGene_gn=CB2H6orf10;refGene_tx=XM_023253774.1;refGene_gn
=CB2H6orf10;refGene_tx=XM_023253775.1
B2 33409981 33410002
refGene_gn=CB2H6orf10;refGene_tx=XM_023253764.1;refGene_gn=CB2H6orf10;
refGene_tx=XM_023253765.1;refGene_gn=CB2H6orf10;refGene_tx=XM_02325376
7.1;refGene_gn=CB2H6orf10;refGene_tx=XM_023253769.1;refGene_gn=CB2H6or
f10;refGene_tx=XM_023253770.1;refGene_gn=CB2H6orf10;refGene_tx=XM_0232
53771.1;refGene_gn=CB2H6orf10;refGene_tx=XM_023253772.1;refGene_gn=CB2
H6orf10;refGene_tx=XM_023253773.1;refGene_gn=CB2H6orf10;refGene_tx=XM_
023253774.1;refGene_gn=CB2H6orf10;refGene_tx=XM_023253775.1
B2 33418097 33418115
refGene_gn=CB2H6orf10;refGene_tx=XM_023253764.1;refGene_gn=CB2H6orf10;
refGene_tx=XM_023253765.1;refGene_gn=CB2H6orf10;refGene_tx=XM_02325376
7.1;refGene_gn=CB2H6orf10;refGene_tx=XM_023253768.1;refGene_gn=CB2H6or
f10;refGene_tx=XM_023253769.1;refGene_gn=CB2H6orf10;refGene_tx=XM_0232
53770.1;refGene_gn=CB2H6orf10;refGene_tx=XM_023253771.1;refGene_gn=CB2
H6orf10;refGene_tx=XM_023253772.1;refGene_gn=CB2H6orf10;refGene_tx=XM_
023253773.1;refGene_gn=CB2H6orf10;refGene_tx=XM_023253774.1;refGene_gn
=CB2H6orf10;refGene_tx=XM_023253775.1
B2 33419516 33419537
refGene_gn=CB2H6orf10;refGene_tx=XM_023253764.1;refGene_gn=CB2H6orf10;
refGene_tx=XM_023253765.1;refGene_gn=CB2H6orf10;refGene_tx=XM_02325376
7.1;refGene_gn=CB2H6orf10;refGene_tx=XM_023253768.1;refGene_gn=CB2H6or
f10;refGene_tx=XM_023253769.1;refGene_gn=CB2H6orf10;refGene_tx=XM_0232
53770.1;refGene_gn=CB2H6orf10;refGene_tx=XM_023253771.1;refGene_gn=CB2
H6orf10;refGene_tx=XM_023253772.1;refGene_gn=CB2H6orf10;refGene_tx=XM_
023253773.1;refGene_gn=CB2H6orf10;refGene_tx=XM_023253774.1;refGene_gn
=CB2H6orf10;refGene_tx=XM_023253775.1
B2 33432518 33432539
refGene_gn=CB2H6orf10;refGene_tx=XM_023253764.1;refGene_gn=CB2H6orf10;
refGene_tx=XM_023253765.1;refGene_gn=CB2H6orf10;refGene_tx=XM_02325376
7.1;refGene_gn=CB2H6orf10;refGene_tx=XM_023253768.1;refGene_gn=CB2H6or
f10;refGene_tx=XM_023253769.1;refGene_gn=CB2H6orf10;refGene_tx=XM_0232
53770.1;refGene_gn=CB2H6orf10;refGene_tx=XM_023253771.1;refGene_gn=CB2
H6orf10;refGene_tx=XM_023253772.1;refGene_gn=CB2H6orf10;refGene_tx=XM_
023253773.1;refGene_gn=CB2H6orf10;refGene_tx=XM_023253774.1;refGene_gn
=CB2H6orf10;refGene_tx=XM_023253775.1
B2 33432987 33433008
refGene_gn=CB2H6orf10;refGene_tx=XM_023253764.1;refGene_gn=CB2H6orf10;
refGene_tx=XM_023253765.1;refGene_gn=CB2H6orf10;refGene_tx=XM_02325376
7.1;refGene_gn=CB2H6orf10;refGene_tx=XM_023253768.1;refGene_gn=CB2H6or
f10;refGene_tx=XM_023253770.1;refGene_gn=CB2H6orf10;refGene_tx=XM_0232
53771.1;refGene_gn=CB2H6orf10;refGene_tx=XM_023253772.1;refGene_gn=CB2
H6orf10;refGene_tx=XM_023253773.1;refGene_gn=CB2H6orf10;refGene_tx=XM_

023253774.1;refGene_gn=CB2H6orf10;refGene_tx=XM_023253775.1
B2 33433466 33433493
refGene_gn=CB2H6orf10;refGene_tx=XM_023253764.1;refGene_gn=CB2H6orf10;refGene_tx=XM_023253765.1;refGene_gn=CB2H6orf10;refGene_tx=XM_023253767.1;refGene_gn=CB2H6orf10;refGene_tx=XM_023253768.1;refGene_gn=CB2H6orf10;refGene_tx=XM_023253769.1;refGene_gn=CB2H6orf10;refGene_tx=XM_023253770.1;refGene_gn=CB2H6orf10;refGene_tx=XM_023253771.1;refGene_gn=CB2H6orf10;refGene_tx=XM_023253772.1;refGene_gn=CB2H6orf10;refGene_tx=XM_023253773.1;refGene_gn=CB2H6orf10;refGene_tx=XM_023253774.1;refGene_gn=CB2H6orf10;refGene_tx=XM_023253775.1
B2 33433793 33433817
refGene_gn=CB2H6orf10;refGene_tx=XM_023253764.1;refGene_gn=CB2H6orf10;refGene_tx=XM_023253765.1;refGene_gn=CB2H6orf10;refGene_tx=XM_023253767.1;refGene_gn=CB2H6orf10;refGene_tx=XM_023253768.1;refGene_gn=CB2H6orf10;refGene_tx=XM_023253769.1;refGene_gn=CB2H6orf10;refGene_tx=XM_023253770.1;refGene_gn=CB2H6orf10;refGene_tx=XM_023253771.1;refGene_gn=CB2H6orf10;refGene_tx=XM_023253772.1;refGene_gn=CB2H6orf10;refGene_tx=XM_023253773.1;refGene_gn=CB2H6orf10;refGene_tx=XM_023253774.1;refGene_gn=CB2H6orf10;refGene_tx=XM_023253775.1
B2 33434275 33434299
refGene_gn=CB2H6orf10;refGene_tx=XM_023253764.1;refGene_gn=CB2H6orf10;refGene_tx=XM_023253765.1;refGene_gn=CB2H6orf10;refGene_tx=XM_023253767.1;refGene_gn=CB2H6orf10;refGene_tx=XM_023253768.1;refGene_gn=CB2H6orf10;refGene_tx=XM_023253769.1;refGene_gn=CB2H6orf10;refGene_tx=XM_023253770.1;refGene_gn=CB2H6orf10;refGene_tx=XM_023253771.1;refGene_gn=CB2H6orf10;refGene_tx=XM_023253772.1;refGene_gn=CB2H6orf10;refGene_tx=XM_023253773.1;refGene_gn=CB2H6orf10;refGene_tx=XM_023253774.1;refGene_gn=CB2H6orf10;refGene_tx=XM_023253775.1
B2 33434935 33434950
refGene_gn=CB2H6orf10;refGene_tx=XM_023253764.1;refGene_gn=CB2H6orf10;refGene_tx=XM_023253765.1;refGene_gn=CB2H6orf10;refGene_tx=XM_023253768.1;refGene_gn=CB2H6orf10;refGene_tx=XM_023253769.1;refGene_gn=CB2H6orf10;refGene_tx=XM_023253770.1;refGene_gn=CB2H6orf10;refGene_tx=XM_023253771.1;refGene_gn=CB2H6orf10;refGene_tx=XM_023253772.1;refGene_gn=CB2H6orf10;refGene_tx=XM_023253773.1;refGene_gn=CB2H6orf10;refGene_tx=XM_023253774.1;refGene_gn=CB2H6orf10;refGene_tx=XM_023253775.1
B2 33435814 33435835
refGene_gn=CB2H6orf10;refGene_tx=XM_023253764.1;refGene_gn=CB2H6orf10;refGene_tx=XM_023253765.1;refGene_gn=CB2H6orf10;refGene_tx=XM_023253767.1;refGene_gn=CB2H6orf10;refGene_tx=XM_023253768.1;refGene_gn=CB2H6orf10;refGene_tx=XM_023253769.1;refGene_gn=CB2H6orf10;refGene_tx=XM_023253770.1;refGene_gn=CB2H6orf10;refGene_tx=XM_023253772.1;refGene_gn=CB2H6orf10;refGene_tx=XM_023253773.1;refGene_gn=CB2H6orf10;refGene_tx=XM_023253774.1;refGene_gn=CB2H6orf10;refGene_tx=XM_023253775.1
B2 33438003 33438024
refGene_gn=CB2H6orf10;refGene_tx=XM_023253764.1;refGene_gn=CB2H6orf10;refGene_tx=XM_023253765.1;refGene_gn=CB2H6orf10;refGene_tx=XM_023253767.1;refGene_gn=CB2H6orf10;refGene_tx=XM_023253768.1;refGene_gn=CB2H6orf10;refGene_tx=XM_023253769.1;refGene_gn=CB2H6orf10;refGene_tx=XM_023253770.1;refGene_gn=CB2H6orf10;refGene_tx=XM_023253771.1;refGene_gn=CB2

H6orf10;refGene_tx=XM_023253772.1;refGene_gn=CB2H6orf10;refGene_tx=XM_023253773.1;refGene_gn=CB2H6orf10;refGene_tx=XM_023253774.1;refGene_gn=CB2H6orf10;refGene_tx=XM_023253775.1
B2 33438543 33438564
refGene_gn=CB2H6orf10;refGene_tx=XM_023253764.1;refGene_gn=CB2H6orf10;refGene_tx=XM_023253765.1;refGene_gn=CB2H6orf10;refGene_tx=XM_023253767.1;refGene_gn=CB2H6orf10;refGene_tx=XM_023253768.1;refGene_gn=CB2H6orf10;refGene_tx=XM_023253769.1;refGene_gn=CB2H6orf10;refGene_tx=XM_023253770.1;refGene_gn=CB2H6orf10;refGene_tx=XM_023253771.1;refGene_gn=CB2H6orf10;refGene_tx=XM_023253772.1;refGene_gn=CB2H6orf10;refGene_tx=XM_023253773.1;refGene_gn=CB2H6orf10;refGene_tx=XM_023253774.1;refGene_gn=CB2H6orf10;refGene_tx=XM_023253775.1
B2 33441195 33441216
refGene_gn=CB2H6orf10;refGene_tx=XM_023253764.1;refGene_gn=CB2H6orf10;refGene_tx=XM_023253765.1;refGene_gn=CB2H6orf10;refGene_tx=XM_023253767.1;refGene_gn=CB2H6orf10;refGene_tx=XM_023253768.1;refGene_gn=CB2H6orf10;refGene_tx=XM_023253769.1;refGene_gn=CB2H6orf10;refGene_tx=XM_023253770.1;refGene_gn=CB2H6orf10;refGene_tx=XM_023253771.1;refGene_gn=CB2H6orf10;refGene_tx=XM_023253772.1;refGene_gn=CB2H6orf10;refGene_tx=XM_023253773.1;refGene_gn=CB2H6orf10;refGene_tx=XM_023253774.1;refGene_gn=CB2H6orf10;refGene_tx=XM_023253775.1
B2 33441866 33441887
refGene_gn=CB2H6orf10;refGene_tx=XM_023253764.1;refGene_gn=CB2H6orf10;refGene_tx=XM_023253765.1;refGene_gn=CB2H6orf10;refGene_tx=XM_023253767.1;refGene_gn=CB2H6orf10;refGene_tx=XM_023253768.1;refGene_gn=CB2H6orf10;refGene_tx=XM_023253769.1;refGene_gn=CB2H6orf10;refGene_tx=XM_023253770.1;refGene_gn=CB2H6orf10;refGene_tx=XM_023253771.1;refGene_gn=CB2H6orf10;refGene_tx=XM_023253772.1;refGene_gn=CB2H6orf10;refGene_tx=XM_023253773.1;refGene_gn=CB2H6orf10;refGene_tx=XM_023253774.1;refGene_gn=CB2H6orf10;refGene_tx=XM_023253775.1
B2 33442556 33442577
refGene_gn=CB2H6orf10;refGene_tx=XM_023253764.1;refGene_gn=CB2H6orf10;refGene_tx=XM_023253765.1;refGene_gn=CB2H6orf10;refGene_tx=XM_023253767.1;refGene_gn=CB2H6orf10;refGene_tx=XM_023253768.1;refGene_gn=CB2H6orf10;refGene_tx=XM_023253769.1;refGene_gn=CB2H6orf10;refGene_tx=XM_023253770.1;refGene_gn=CB2H6orf10;refGene_tx=XM_023253771.1;refGene_gn=CB2H6orf10;refGene_tx=XM_023253772.1;refGene_gn=CB2H6orf10;refGene_tx=XM_023253773.1;refGene_gn=CB2H6orf10;refGene_tx=XM_023253774.1;refGene_gn=CB2H6orf10;refGene_tx=XM_023253775.1
B2 33443304 33443325
refGene_gn=CB2H6orf10;refGene_tx=XM_023253764.1;refGene_gn=CB2H6orf10;refGene_tx=XM_023253765.1;refGene_gn=CB2H6orf10;refGene_tx=XM_023253767.1;refGene_gn=CB2H6orf10;refGene_tx=XM_023253768.1;refGene_gn=CB2H6orf10;refGene_tx=XM_023253769.1;refGene_gn=CB2H6orf10;refGene_tx=XM_023253770.1;refGene_gn=CB2H6orf10;refGene_tx=XM_023253771.1;refGene_gn=CB2H6orf10;refGene_tx=XM_023253772.1;refGene_gn=CB2H6orf10;refGene_tx=XM_023253773.1;refGene_gn=CB2H6orf10;refGene_tx=XM_023253774.1;refGene_gn=CB2H6orf10;refGene_tx=XM_023253775.1
B2 33445492 33445513
refGene_gn=CB2H6orf10;refGene_tx=XM_023253764.1;refGene_gn=CB2H6orf10;

refGene_tx=XM_023253765.1;refGene_gn=CB2H6orf10;refGene_tx=XM_02325376
7.1;refGene_gn=CB2H6orf10;refGene_tx=XM_023253768.1;refGene_gn=CB2H6or
f10;refGene_tx=XM_023253769.1;refGene_gn=CB2H6orf10;refGene_tx=XM_0232
53770.1;refGene_gn=CB2H6orf10;refGene_tx=XM_023253771.1;refGene_gn=CB2
H6orf10;refGene_tx=XM_023253772.1;refGene_gn=CB2H6orf10;refGene_tx=XM_
023253773.1;refGene_gn=CB2H6orf10;refGene_tx=XM_023253774.1;refGene_gn
=CB2H6orf10;refGene_tx=XM_023253775.1
B2 33446075 33446114
refGene_gn=CB2H6orf10;refGene_tx=XM_023253764.1;refGene_gn=CB2H6orf10;
refGene_tx=XM_023253765.1;refGene_gn=CB2H6orf10;refGene_tx=XM_02325376
7.1;refGene_gn=CB2H6orf10;refGene_tx=XM_023253768.1;refGene_gn=CB2H6or
f10;refGene_tx=XM_023253769.1;refGene_gn=CB2H6orf10;refGene_tx=XM_0232
53770.1;refGene_gn=CB2H6orf10;refGene_tx=XM_023253771.1;refGene_gn=CB2
H6orf10;refGene_tx=XM_023253772.1;refGene_gn=CB2H6orf10;refGene_tx=XM_
023253773.1;refGene_gn=CB2H6orf10;refGene_tx=XM_023253774.1;refGene_gn
=CB2H6orf10;refGene_tx=XM_023253775.1
B2 33453442 33453463
refGene_gn=CB2H6orf10;refGene_tx=XM_023253764.1;refGene_gn=CB2H6orf10;
refGene_tx=XM_023253765.1;refGene_gn=CB2H6orf10;refGene_tx=XM_02325376
7.1;refGene_gn=CB2H6orf10;refGene_tx=XM_023253768.1;refGene_gn=CB2H6or
f10;refGene_tx=XM_023253769.1;refGene_gn=CB2H6orf10;refGene_tx=XM_0232
53771.1;refGene_gn=CB2H6orf10;refGene_tx=XM_023253773.1
B2 33465842 33465863
refGene_gn=CB2H6orf10;refGene_tx=XM_023253764.1;refGene_gn=CB2H6orf10;
refGene_tx=XM_023253765.1;refGene_gn=CB2H6orf10;refGene_tx=XM_02325376
7.1;refGene_gn=CB2H6orf10;refGene_tx=XM_023253768.1;refGene_gn=CB2H6or
f10;refGene_tx=XM_023253769.1;refGene_gn=CB2H6orf10;refGene_tx=XM_0232
53770.1;refGene_gn=CB2H6orf10;refGene_tx=XM_023253771.1;refGene_gn=CB2
H6orf10;refGene_tx=XM_023253772.1;refGene_gn=CB2H6orf10;refGene_tx=XM_
023253773.1;refGene_gn=CB2H6orf10;refGene_tx=XM_023253774.1;refGene_gn
=CB2H6orf10;refGene_tx=XM_023253775.1
B2 33465956 33465986
refGene_gn=CB2H6orf10;refGene_tx=XM_023253764.1;refGene_gn=CB2H6orf10;
refGene_tx=XM_023253765.1;refGene_gn=CB2H6orf10;refGene_tx=XM_02325376
7.1;refGene_gn=CB2H6orf10;refGene_tx=XM_023253768.1;refGene_gn=CB2H6or
f10;refGene_tx=XM_023253769.1;refGene_gn=CB2H6orf10;refGene_tx=XM_0232
53770.1;refGene_gn=CB2H6orf10;refGene_tx=XM_023253771.1;refGene_gn=CB2
H6orf10;refGene_tx=XM_023253772.1;refGene_gn=CB2H6orf10;refGene_tx=XM_
023253773.1;refGene_gn=CB2H6orf10;refGene_tx=XM_023253774.1;refGene_gn
=CB2H6orf10;refGene_tx=XM_023253775.1
B2 33466071 33466104
refGene_gn=CB2H6orf10;refGene_tx=XM_023253764.1;refGene_gn=CB2H6orf10;
refGene_tx=XM_023253765.1;refGene_gn=CB2H6orf10;refGene_tx=XM_02325376
7.1;refGene_gn=CB2H6orf10;refGene_tx=XM_023253768.1;refGene_gn=CB2H6or
f10;refGene_tx=XM_023253769.1;refGene_gn=CB2H6orf10;refGene_tx=XM_0232
53770.1;refGene_gn=CB2H6orf10;refGene_tx=XM_023253771.1;refGene_gn=CB2
H6orf10;refGene_tx=XM_023253772.1
B2 33467245 33467278
refGene_gn=CB2H6orf10;refGene_tx=XM_023253764.1;refGene_gn=CB2H6orf10;
refGene_tx=XM_023253765.1;refGene_gn=CB2H6orf10;refGene_tx=XM_02325376

7.1;refGene_gn=CB2H6orf10;refGene_tx=XM_023253768.1;refGene_gn=CB2H6or
f10;refGene_tx=XM_023253769.1;refGene_gn=CB2H6orf10;refGene_tx=XM_0232
53770.1;refGene_gn=CB2H6orf10;refGene_tx=XM_023253771.1;refGene_gn=CB2
H6orf10;refGene_tx=XM_023253772.1;refGene_gn=CB2H6orf10;refGene_tx=XM_
023253773.1;refGene_gn=CB2H6orf10;refGene_tx=XM_023253774.1;refGene_gn
=CB2H6orf10;refGene_tx=XM_023253775.1
B2 33468089 33468122
refGene_gn=CB2H6orf10;refGene_tx=XM_023253764.1;refGene_gn=CB2H6orf10;
refGene_tx=XM_023253765.1;refGene_gn=CB2H6orf10;refGene_tx=XM_02325376
7.1;refGene_gn=CB2H6orf10;refGene_tx=XM_023253768.1;refGene_gn=CB2H6or
f10;refGene_tx=XM_023253769.1;refGene_gn=CB2H6orf10;refGene_tx=XM_0232
53770.1;refGene_gn=CB2H6orf10;refGene_tx=XM_023253771.1;refGene_gn=CB2
H6orf10;refGene_tx=XM_023253772.1;refGene_gn=CB2H6orf10;refGene_tx=XM_
023253773.1;refGene_gn=CB2H6orf10;refGene_tx=XM_023253774.1;refGene_gn
=CB2H6orf10;refGene_tx=XM_023253775.1
B2 33468872 33468959
refGene_gn=CB2H6orf10;refGene_tx=XM_023253764.1;refGene_gn=CB2H6orf10;
refGene_tx=XM_023253765.1;refGene_gn=CB2H6orf10;refGene_tx=XM_02325376
7.1;refGene_gn=CB2H6orf10;refGene_tx=XM_023253768.1;refGene_gn=CB2H6or
f10;refGene_tx=XM_023253769.1;refGene_gn=CB2H6orf10;refGene_tx=XM_0232
53770.1;refGene_gn=CB2H6orf10;refGene_tx=XM_023253771.1;refGene_gn=CB2
H6orf10;refGene_tx=XM_023253772.1;refGene_gn=CB2H6orf10;refGene_tx=XM_
023253773.1;refGene_gn=CB2H6orf10;refGene_tx=XM_023253774.1;refGene_gn
=CB2H6orf10;refGene_tx=XM_023253775.1
B2 33470673 33470686
refGene_gn=CB2H6orf10;refGene_tx=XM_023253764.1;refGene_gn=CB2H6orf10;
refGene_tx=XM_023253765.1;refGene_gn=CB2H6orf10;refGene_tx=XM_02325376
7.1;refGene_gn=CB2H6orf10;refGene_tx=XM_023253768.1;refGene_gn=CB2H6or
f10;refGene_tx=XM_023253769.1;refGene_gn=CB2H6orf10;refGene_tx=XM_0232
53770.1;refGene_gn=CB2H6orf10;refGene_tx=XM_023253771.1;refGene_gn=CB2
H6orf10;refGene_tx=XM_023253772.1;refGene_gn=CB2H6orf10;refGene_tx=XM_
023253773.1;refGene_gn=CB2H6orf10;refGene_tx=XM_023253774.1;refGene_gn
=CB2H6orf10;refGene_tx=XM_023253775.1
B2 33601518 33602033
refGene_gn=LOC101081137;refGene_tx=XM_023254714.1
B2 33602730 33602757
refGene_gn=LOC101081137;refGene_tx=XM_023254714.1
B2 33604561 33604588
refGene_gn=LOC101081137;refGene_tx=XM_023254714.1
B2 33604771 33604793
refGene_gn=LOC101081137;refGene_tx=XM_023254714.1
B2 33605477 33605707
refGene_gn=LOC101081137;refGene_tx=XM_023254714.1
B2 33605846 33606128
refGene_gn=LOC101081137;refGene_tx=XM_023254714.1
B2 33606730 33607081
refGene_gn=LOC101081137;refGene_tx=XM_023254714.1
B2 33608045 33608124
refGene_gn=LOC101081137;refGene_tx=XM_023254714.1
B2 33615323 33615402

refGene_gn=LOC101081382;refGene_tx=XM_023254745.1
B2 33615956 33616304
refGene_gn=LOC101081382;refGene_tx=XM_023254745.1
B2 33620067 33620349
refGene_gn=LOC101081382;refGene_tx=XM_023254745.1
B2 33620467 33620635
refGene_gn=LOC101081382;refGene_tx=XM_023254745.1
B2 33623505 33623526
refGene_gn=LOC101081382;refGene_tx=XM_023254745.1
B2 33623620 33623647
refGene_gn=LOC101081382;refGene_tx=XM_023254745.1
B2 33624468 33624495
refGene_gn=LOC101081382;refGene_tx=XM_023254745.1
B2 33624941 33624968
refGene_gn=LOC101081382;refGene_tx=XM_023254745.1
B2 33626187 33626828
refGene_gn=LOC101081382;refGene_tx=XM_023254745.1
B2 33634844 33634852 refGene_gn=BTNL2;refGene_tx=XM_019830431.1
B2 33636030 33636126 refGene_gn=BTNL2;refGene_tx=XM_019830431.1
B2 33636820 33637102 refGene_gn=BTNL2;refGene_tx=XM_019830431.1
B2 33638114 33638462 refGene_gn=BTNL2;refGene_tx=XM_019830431.1
B2 33641838 33641859 refGene_gn=BTNL2;refGene_tx=XM_019830431.1
B2 33643515 33643797 refGene_gn=BTNL2;refGene_tx=XM_019830431.1
B2 33645430 33645778 refGene_gn=BTNL2;refGene_tx=XM_019830431.1
B2 33647701 33647780 refGene_gn=BTNL2;refGene_tx=XM_019830431.1
B2 33667483 33667565 refGene_gn=SLC26A3;refGene_tx=NM_001128071.1
B2 33669206 33669452 refGene_gn=SLC26A3;refGene_tx=NM_001128071.1
B2 33669913 33670195 refGene_gn=SLC26A3;refGene_tx=NM_001128071.1
B2 33670500 33670655 refGene_gn=SLC26A3;refGene_tx=NM_001128071.1
B2 33701364 33701446 refGene_gn=DRA;refGene_tx=NM_001309051.1
B2 33702604 33702850 refGene_gn=DRA;refGene_tx=NM_001309051.1
B2 33703312 33703594 refGene_gn=DRA;refGene_tx=NM_001309051.1
B2 33703894 33704049 refGene_gn=DRA;refGene_tx=NM_001309051.1
B2 33718344 33718368
refGene_gn=LOC101098301;refGene_tx=NM_001305030.1
B2 33718822 33718933
refGene_gn=LOC101098301;refGene_tx=NM_001305030.1
B2 33719349 33719631
refGene_gn=LOC101098301;refGene_tx=NM_001305030.1
B2 33722225 33722495
refGene_gn=LOC101098301;refGene_tx=NM_001305030.1
B2 33727594 33727694
refGene_gn=LOC101098301;refGene_tx=NM_001305030.1
B2 33768535 33768617
refGene_gn=LOC101088734;refGene_tx=XM_004001103.4
B2 33769775 33770021
refGene_gn=LOC101088734;refGene_tx=XM_004001103.4
B2 33770483 33770765
refGene_gn=LOC101088734;refGene_tx=XM_004001103.4
B2 33771065 33771220

refGene_gn=LOC101088734;refGene_tx=XM_004001103.4
B2 33786199 33786223
refGene_gn=LOC102899262;refGene_tx=XM_019830433.2
B2 33786677 33786788
refGene_gn=LOC102899262;refGene_tx=XM_019830433.2
B2 33787536 33787818
refGene_gn=LOC102899262;refGene_tx=XM_019830433.2
B2 33790420 33790690
refGene_gn=LOC102899262;refGene_tx=XM_019830433.2
B2 33795792 33795892
refGene_gn=LOC102899262;refGene_tx=XM_019830433.2
B2 33871205 33871305
refGene_gn=LOC101082653;refGene_tx=XM_023253777.1
B2 33877560 33877830
refGene_gn=LOC101082653;refGene_tx=XM_023253777.1
B2 33881848 33882130
refGene_gn=LOC101082653;refGene_tx=XM_023253777.1
B2 33882828 33882939
refGene_gn=LOC101082653;refGene_tx=XM_023253777.1
B2 33883345 33883369
refGene_gn=LOC101082653;refGene_tx=XM_023253777.1
B2 33926487 33926514
refGene_gn=LOC101082906;refGene_tx=XM_003985988.4;refGene_gn=LOC101082906;refGene_tx=XM_019830434.1;refGene_gn=LOC101082906;refGene_tx=XM_019830435.1;refGene_gn=LOC101082906;refGene_tx=XM_019830436.1;refGene_gn=LOC101082906;refGene_tx=XM_019830437.1
B2 33926688 33926720
refGene_gn=LOC101082906;refGene_tx=XM_003985988.4;refGene_gn=LOC101082906;refGene_tx=XM_019830434.1;refGene_gn=LOC101082906;refGene_tx=XM_019830435.1;refGene_gn=LOC101082906;refGene_tx=XM_019830436.1;refGene_gn=LOC101082906;refGene_tx=XM_019830437.1
B2 33927216 33927327
refGene_gn=LOC101082906;refGene_tx=XM_003985988.4;refGene_gn=LOC101082906;refGene_tx=XM_019830434.1;refGene_gn=LOC101082906;refGene_tx=XM_019830435.1;refGene_gn=LOC101082906;refGene_tx=XM_019830436.1;refGene_gn=LOC101082906;refGene_tx=XM_019830437.1
B2 33927800 33928082
refGene_gn=LOC101082906;refGene_tx=XM_003985988.4;refGene_gn=LOC101082906;refGene_tx=XM_019830434.1;refGene_gn=LOC101082906;refGene_tx=XM_019830435.1;refGene_gn=LOC101082906;refGene_tx=XM_019830436.1;refGene_gn=LOC101082906;refGene_tx=XM_019830437.1
B2 33928532 33928814
refGene_gn=LOC101082906;refGene_tx=XM_003985988.4;refGene_gn=LOC101082906;refGene_tx=XM_019830434.1;refGene_gn=LOC101082906;refGene_tx=XM_019830435.1;refGene_gn=LOC101082906;refGene_tx=XM_019830436.1;refGene_gn=LOC101082906;refGene_tx=XM_019830437.1
B2 33932026 33932153
refGene_gn=LOC101082906;refGene_tx=XM_003985988.4;refGene_gn=LOC101082906;refGene_tx=XM_019830434.1;refGene_gn=LOC101082906;refGene_tx=XM_019830435.1;refGene_gn=LOC101082906;refGene_tx=XM_019830436.1;refGene_gn

=LOC101082906;refGene_tx=XM_019830437.1
B2 33946163 33946344
refGene_gn=LOC101089770;refGene_tx=XM_006931604.4
B2 33946345 33946524
refGene_gn=LOC101089770;refGene_tx=XM_006931604.4
B2 33946870 33947007
refGene_gn=LOC101089770;refGene_tx=XM_006931604.4
B2 33947366 33947526
refGene_gn=LOC101089770;refGene_tx=XM_006931604.4
B2 33947939 33948113
refGene_gn=LOC101089770;refGene_tx=XM_006931604.4
B2 33948263 33948452
refGene_gn=LOC101089770;refGene_tx=XM_006931604.4
B2 33949808 33949937
refGene_gn=LOC101089770;refGene_tx=XM_006931604.4
B2 33950435 33950633
refGene_gn=LOC101089770;refGene_tx=XM_006931604.4
B2 33953391 33953597
refGene_gn=LOC101089770;refGene_tx=XM_006931604.4
B2 33953888 33954019
refGene_gn=LOC101089770;refGene_tx=XM_006931604.4
B2 33955503 33955618
refGene_gn=LOC101089770;refGene_tx=XM_006931604.4
B2 33955707 33956197
refGene_gn=LOC101089770;refGene_tx=XM_006931604.4
B2 33958787 33958876
refGene_gn=LOC101083171;refGene_tx=XM_023253778.1
B2 33959123 33959328
refGene_gn=LOC101083171;refGene_tx=XM_023253778.1
B2 33959685 33959815
refGene_gn=LOC101083171;refGene_tx=XM_023253778.1
B2 33960475 33960587
refGene_gn=LOC101083171;refGene_tx=XM_023253778.1
B2 33960735 33960883
refGene_gn=LOC101083171;refGene_tx=XM_023253778.1
B2 33961657 33961804
refGene_gn=LOC101083171;refGene_tx=XM_023253778.1
B2 33963358 33963571
refGene_gn=LOC101083591;refGene_tx=XM_003985991.4
B2 33964341 33964478
refGene_gn=LOC101083591;refGene_tx=XM_003985991.4
B2 33964743 33964906
refGene_gn=LOC101083591;refGene_tx=XM_003985991.4
B2 33965161 33965335
refGene_gn=LOC101083591;refGene_tx=XM_003985991.4
B2 33965915 33966104
refGene_gn=LOC101083591;refGene_tx=XM_003985991.4
B2 33966260 33966389
refGene_gn=LOC101083591;refGene_tx=XM_003985991.4
B2 33967168 33967366

refGene_gn=LOC101083591;refGene_tx=XM_003985991.4
B2 33967803 33968009
refGene_gn=LOC101083591;refGene_tx=XM_003985991.4
B2 33969258 33969389
refGene_gn=LOC101083591;refGene_tx=XM_003985991.4
B2 33969544 33969659
refGene_gn=LOC101083591;refGene_tx=XM_003985991.4
B2 33970137 33970735
refGene_gn=LOC101083591;refGene_tx=XM_003985991.4
B2 33971379 33971409
refGene_gn=LOC101083848;refGene_tx=XM_023253779.1;refGene_gn=LOC101083848;refGene_tx=XM_023253780.1;refGene_gn=LOC101083848;refGene_tx=XM_023253781.1
B2 33973242 33973310
refGene_gn=LOC101083848;refGene_tx=XM_023253779.1;refGene_gn=LOC101083848;refGene_tx=XM_023253780.1;refGene_gn=LOC101083848;refGene_tx=XM_023253781.1
B2 33974316 33974451
refGene_gn=LOC101083848;refGene_tx=XM_023253779.1;refGene_gn=LOC101083848;refGene_tx=XM_023253780.1
B2 33975111 33975241
refGene_gn=LOC101083848;refGene_tx=XM_023253779.1;refGene_gn=LOC101083848;refGene_tx=XM_023253780.1;refGene_gn=LOC101083848;refGene_tx=XM_023253781.1
B2 33975484 33975626
refGene_gn=LOC101083848;refGene_tx=XM_023253779.1;refGene_gn=LOC101083848;refGene_tx=XM_023253780.1;refGene_gn=LOC101083848;refGene_tx=XM_023253781.1
B2 33976181 33976309
refGene_gn=LOC101083848;refGene_tx=XM_023253779.1;refGene_gn=LOC101083848;refGene_tx=XM_023253780.1;refGene_gn=LOC101083848;refGene_tx=XM_023253781.1
B2 34045237 34045254
refGene_gn=LOC101090265;refGene_tx=XM_023253782.1;refGene_gn=LOC101090265;refGene_tx=XM_023253783.1
B2 34045624 34045660
refGene_gn=LOC101090265;refGene_tx=XM_023253782.1
B2 34045808 34045925
refGene_gn=LOC101090265;refGene_tx=XM_023253782.1;refGene_gn=LOC101090265;refGene_tx=XM_023253783.1
B2 34047387 34047672
refGene_gn=LOC101090265;refGene_tx=XM_023253782.1;refGene_gn=LOC101090265;refGene_tx=XM_023253783.1
B2 34048935 34049217
refGene_gn=LOC101090265;refGene_tx=XM_023253782.1;refGene_gn=LOC101090265;refGene_tx=XM_023253783.1
B2 34049899 34049954
refGene_gn=LOC101090265;refGene_tx=XM_023253782.1;refGene_gn=LOC101090265;refGene_tx=XM_023253783.1
B2 34060253 34060258

refGene_gn=LOC101084108;refGene_tx=XM_003985993.5
B2 34060623 34060752
refGene_gn=LOC101084108;refGene_tx=XM_003985993.5
B2 34060985 34061264
refGene_gn=LOC101084108;refGene_tx=XM_003985993.5
B2 34061870 34062155
refGene_gn=LOC101084108;refGene_tx=XM_003985993.5
B2 34063668 34063756
refGene_gn=LOC101084108;refGene_tx=XM_003985993.5
B2 34081156 34081185
refGene_gn=LOC101084362;refGene_tx=XM_023253784.1
B2 34082832 34083136
refGene_gn=LOC101084362;refGene_tx=XM_023253784.1
B2 34083750 34083888
refGene_gn=LOC101084362;refGene_tx=XM_023253784.1
B2 34084406 34084545
refGene_gn=LOC101084362;refGene_tx=XM_023253784.1
B2 34084647 34084685
refGene_gn=LOC101084362;refGene_tx=XM_023253784.1
B2 34084686 34084726
refGene_gn=LOC101084362;refGene_tx=XM_023253784.1
B2 34084728 34084862
refGene_gn=LOC101084362;refGene_tx=XM_023253784.1
B2 34084964 34085339
refGene_gn=LOC101084362;refGene_tx=XM_023253784.1
B2 34085848 34085977
refGene_gn=LOC101084362;refGene_tx=XM_023253784.1
B2 34086275 34086527
refGene_gn=LOC101084362;refGene_tx=XM_023253784.1
B2 34086654 34086920
refGene_gn=LOC101084362;refGene_tx=XM_023253784.1
B2 34087443 34087664
refGene_gn=LOC101084362;refGene_tx=XM_023253784.1
B2 34087666 34087747
refGene_gn=LOC101084362;refGene_tx=XM_023253784.1
B2 34087969 34088092
refGene_gn=LOC101084362;refGene_tx=XM_023253784.1
B2 34088196 34088336
refGene_gn=LOC101084362;refGene_tx=XM_023253784.1
B2 34110938 34110942
refGene_gn=LOC101090518;refGene_tx=XM_006931614.3
B2 34111180 34111316
refGene_gn=LOC101090518;refGene_tx=XM_006931614.3
B2 34111461 34111743
refGene_gn=LOC101090518;refGene_tx=XM_006931614.3
B2 34112176 34112425
refGene_gn=LOC101090518;refGene_tx=XM_006931614.3
B2 34113611 34113693
refGene_gn=LOC101090518;refGene_tx=XM_006931614.3
B2 34142650 34142928

refGene_gn=LOC101090771;refGene_tx=XM_023253786.1
B2 34143149 34143293
refGene_gn=LOC101090771;refGene_tx=XM_023253786.1
B2 34143295 34143430
refGene_gn=LOC101090771;refGene_tx=XM_023253786.1
B2 34143808 34144054
refGene_gn=LOC101090771;refGene_tx=XM_023253786.1
B2 34147101 34147201
refGene_gn=LOC101090771;refGene_tx=XM_023253786.1
B2 34149668 34149768
refGene_gn=LOC101084962;refGene_tx=XM_023253785.1
B2 34158835 34159105
refGene_gn=LOC101084962;refGene_tx=XM_023253785.1
B2 34159779 34160061
refGene_gn=LOC101084962;refGene_tx=XM_023253785.1
B2 34160582 34160692
refGene_gn=LOC101084962;refGene_tx=XM_023253785.1
B2 34180136 34180277
refGene_gn=LOC101085209;refGene_tx=XM_003985999.5;refGene_gn=LOC101085209;refGene_tx=XM_019830441.2;refGene_gn=LOC101085209;refGene_tx=XM_019830443.2;refGene_gn=LOC101085209;refGene_tx=XM_019830444.2;refGene_gn=LOC101085209;refGene_tx=XM_023253787.1;refGene_gn=LOC101085209;refGene_tx=XM_023253788.1
B2 34180984 34181191
refGene_gn=LOC101085209;refGene_tx=XM_003985999.5;refGene_gn=LOC101085209;refGene_tx=XM_019830441.2;refGene_gn=LOC101085209;refGene_tx=XM_019830443.2;refGene_gn=LOC101085209;refGene_tx=XM_019830444.2;refGene_gn=LOC101085209;refGene_tx=XM_023253787.1;refGene_gn=LOC101085209;refGene_tx=XM_023253788.1
B2 34181561 34181674
refGene_gn=LOC101085209;refGene_tx=XM_003985999.5;refGene_gn=LOC101085209;refGene_tx=XM_019830441.2;refGene_gn=LOC101085209;refGene_tx=XM_019830443.2;refGene_gn=LOC101085209;refGene_tx=XM_019830444.2;refGene_gn=LOC101085209;refGene_tx=XM_023253787.1;refGene_gn=LOC101085209;refGene_tx=XM_023253788.1
B2 34182366 34182634
refGene_gn=LOC101085209;refGene_tx=XM_003985999.5;refGene_gn=LOC101085209;refGene_tx=XM_019830441.2;refGene_gn=LOC101085209;refGene_tx=XM_019830443.2;refGene_gn=LOC101085209;refGene_tx=XM_019830444.2;refGene_gn=LOC101085209;refGene_tx=XM_023253787.1;refGene_gn=LOC101085209;refGene_tx=XM_023253788.1
B2 34182745 34182799
refGene_gn=LOC101085209;refGene_tx=XM_003985999.5;refGene_gn=LOC101085209;refGene_tx=XM_019830441.2;refGene_gn=LOC101085209;refGene_tx=XM_019830443.2;refGene_gn=LOC101085209;refGene_tx=XM_019830444.2;refGene_gn=LOC101085209;refGene_tx=XM_023253787.1;refGene_gn=LOC101085209;refGene_tx=XM_023253788.1
B2 34182984 34183020
refGene_gn=LOC101085209;refGene_tx=XM_003985999.5;refGene_gn=LOC101085209;refGene_tx=XM_019830441.2;refGene_gn=LOC101085209;refGene_tx=XM_01

9830443.2;refGene_gn=LOC101085209;refGene_tx=XM_019830444.2;refGene_gn=LOC101085209;refGene_tx=XM_023253787.1;refGene_gn=LOC101085209;refGene_tx=XM_023253788.1
B2 34183330 34183384
refGene_gn=LOC101085209;refGene_tx=XM_003985999.5;refGene_gn=LOC101085209;refGene_tx=XM_019830441.2;refGene_gn=LOC101085209;refGene_tx=XM_019830443.2;refGene_gn=LOC101085209;refGene_tx=XM_019830444.2;refGene_gn=LOC101085209;refGene_tx=XM_023253787.1;refGene_gn=LOC101085209;refGene_tx=XM_023253788.1
B2 34183520 34183628
refGene_gn=LOC101085209;refGene_tx=XM_003985999.5;refGene_gn=LOC101085209;refGene_tx=XM_019830441.2;refGene_gn=LOC101085209;refGene_tx=XM_019830443.2;refGene_gn=LOC101085209;refGene_tx=XM_019830444.2;refGene_gn=LOC101085209;refGene_tx=XM_023253787.1;refGene_gn=LOC101085209;refGene_tx=XM_023253788.1
B2 34183894 34183948
refGene_gn=LOC101085209;refGene_tx=XM_003985999.5;refGene_gn=LOC101085209;refGene_tx=XM_019830441.2;refGene_gn=LOC101085209;refGene_tx=XM_019830443.2;refGene_gn=LOC101085209;refGene_tx=XM_019830444.2;refGene_gn=LOC101085209;refGene_tx=XM_023253787.1;refGene_gn=LOC101085209;refGene_tx=XM_023253788.1
B2 34184105 34184159
refGene_gn=LOC101085209;refGene_tx=XM_003985999.5;refGene_gn=LOC101085209;refGene_tx=XM_019830441.2;refGene_gn=LOC101085209;refGene_tx=XM_019830443.2;refGene_gn=LOC101085209;refGene_tx=XM_019830444.2;refGene_gn=LOC101085209;refGene_tx=XM_023253787.1;refGene_gn=LOC101085209;refGene_tx=XM_023253788.1
B2 34184269 34184377
refGene_gn=LOC101085209;refGene_tx=XM_003985999.5;refGene_gn=LOC101085209;refGene_tx=XM_019830441.2;refGene_gn=LOC101085209;refGene_tx=XM_019830443.2;refGene_gn=LOC101085209;refGene_tx=XM_019830444.2;refGene_gn=LOC101085209;refGene_tx=XM_023253787.1;refGene_gn=LOC101085209;refGene_tx=XM_023253788.1
B2 34184644 34184698
refGene_gn=LOC101085209;refGene_tx=XM_003985999.5;refGene_gn=LOC101085209;refGene_tx=XM_019830441.2;refGene_gn=LOC101085209;refGene_tx=XM_019830443.2;refGene_gn=LOC101085209;refGene_tx=XM_019830444.2;refGene_gn=LOC101085209;refGene_tx=XM_023253787.1;refGene_gn=LOC101085209;refGene_tx=XM_023253788.1
B2 34185268 34185322
refGene_gn=LOC101085209;refGene_tx=XM_003985999.5;refGene_gn=LOC101085209;refGene_tx=XM_019830441.2;refGene_gn=LOC101085209;refGene_tx=XM_019830443.2;refGene_gn=LOC101085209;refGene_tx=XM_019830444.2;refGene_gn=LOC101085209;refGene_tx=XM_023253787.1;refGene_gn=LOC101085209;refGene_tx=XM_023253788.1
B2 34185458 34185512
refGene_gn=LOC101085209;refGene_tx=XM_003985999.5;refGene_gn=LOC101085209;refGene_tx=XM_019830441.2;refGene_gn=LOC101085209;refGene_tx=XM_019830443.2;refGene_gn=LOC101085209;refGene_tx=XM_019830444.2;refGene_gn=LOC101085209;refGene_tx=XM_023253787.1;refGene_gn=LOC101085209;refGen

e_tx=XM_023253788.1
B2 34185699 34185753
refGene_gn=LOC101085209;refGene_tx=XM_003985999.5;refGene_gn=LOC101085209;refGene_tx=XM_019830441.2;refGene_gn=LOC101085209;refGene_tx=XM_019830443.2;refGene_gn=LOC101085209;refGene_tx=XM_019830444.2;refGene_gn=LOC101085209;refGene_tx=XM_023253787.1;refGene_gn=LOC101085209;refGene_tx=XM_023253788.1
B2 34186152 34186260
refGene_gn=LOC101085209;refGene_tx=XM_003985999.5;refGene_gn=LOC101085209;refGene_tx=XM_019830441.2;refGene_gn=LOC101085209;refGene_tx=XM_019830443.2;refGene_gn=LOC101085209;refGene_tx=XM_019830444.2;refGene_gn=LOC101085209;refGene_tx=XM_023253787.1;refGene_gn=LOC101085209;refGene_tx=XM_023253788.1
B2 34186604 34186658
refGene_gn=LOC101085209;refGene_tx=XM_003985999.5;refGene_gn=LOC101085209;refGene_tx=XM_019830441.2;refGene_gn=LOC101085209;refGene_tx=XM_019830443.2;refGene_gn=LOC101085209;refGene_tx=XM_019830444.2;refGene_gn=LOC101085209;refGene_tx=XM_023253787.1;refGene_gn=LOC101085209;refGene_tx=XM_023253788.1
B2 34186817 34186871
refGene_gn=LOC101085209;refGene_tx=XM_003985999.5;refGene_gn=LOC101085209;refGene_tx=XM_019830441.2;refGene_gn=LOC101085209;refGene_tx=XM_019830443.2;refGene_gn=LOC101085209;refGene_tx=XM_019830444.2;refGene_gn=LOC101085209;refGene_tx=XM_023253787.1;refGene_gn=LOC101085209;refGene_tx=XM_023253788.1
B2 34187095 34187149
refGene_gn=LOC101085209;refGene_tx=XM_003985999.5;refGene_gn=LOC101085209;refGene_tx=XM_019830441.2;refGene_gn=LOC101085209;refGene_tx=XM_019830443.2;refGene_gn=LOC101085209;refGene_tx=XM_019830444.2;refGene_gn=LOC101085209;refGene_tx=XM_023253787.1;refGene_gn=LOC101085209;refGene_tx=XM_023253788.1
B2 34187365 34187419
refGene_gn=LOC101085209;refGene_tx=XM_003985999.5;refGene_gn=LOC101085209;refGene_tx=XM_019830441.2;refGene_gn=LOC101085209;refGene_tx=XM_019830443.2;refGene_gn=LOC101085209;refGene_tx=XM_019830444.2;refGene_gn=LOC101085209;refGene_tx=XM_023253787.1;refGene_gn=LOC101085209;refGene_tx=XM_023253788.1
B2 34187609 34187717
refGene_gn=LOC101085209;refGene_tx=XM_003985999.5;refGene_gn=LOC101085209;refGene_tx=XM_019830441.2;refGene_gn=LOC101085209;refGene_tx=XM_019830443.2;refGene_gn=LOC101085209;refGene_tx=XM_019830444.2;refGene_gn=LOC101085209;refGene_tx=XM_023253787.1;refGene_gn=LOC101085209;refGene_tx=XM_023253788.1
B2 34187901 34187955
refGene_gn=LOC101085209;refGene_tx=XM_003985999.5;refGene_gn=LOC101085209;refGene_tx=XM_019830441.2;refGene_gn=LOC101085209;refGene_tx=XM_019830443.2;refGene_gn=LOC101085209;refGene_tx=XM_019830444.2;refGene_gn=LOC101085209;refGene_tx=XM_023253787.1;refGene_gn=LOC101085209;refGene_tx=XM_023253788.1
B2 34188060 34188114

refGene_gn=LOC101085209;refGene_tx=XM_003985999.5;refGene_gn=LOC101085209;refGene_tx=XM_019830441.2;refGene_gn=LOC101085209;refGene_tx=XM_019830443.2;refGene_gn=LOC101085209;refGene_tx=XM_019830444.2;refGene_gn=LOC101085209;refGene_tx=XM_023253787.1;refGene_gn=LOC101085209;refGene_tx=XM_023253788.1
B2 34188256 34188364
refGene_gn=LOC101085209;refGene_tx=XM_003985999.5;refGene_gn=LOC101085209;refGene_tx=XM_019830441.2;refGene_gn=LOC101085209;refGene_tx=XM_019830443.2;refGene_gn=LOC101085209;refGene_tx=XM_019830444.2;refGene_gn=LOC101085209;refGene_tx=XM_023253787.1;refGene_gn=LOC101085209;refGene_tx=XM_023253788.1
B2 34188502 34188610
refGene_gn=LOC101085209;refGene_tx=XM_003985999.5;refGene_gn=LOC101085209;refGene_tx=XM_019830441.2;refGene_gn=LOC101085209;refGene_tx=XM_019830443.2;refGene_gn=LOC101085209;refGene_tx=XM_019830444.2;refGene_gn=LOC101085209;refGene_tx=XM_023253787.1;refGene_gn=LOC101085209;refGene_tx=XM_023253788.1
B2 34188845 34188899
refGene_gn=LOC101085209;refGene_tx=XM_003985999.5;refGene_gn=LOC101085209;refGene_tx=XM_019830441.2;refGene_gn=LOC101085209;refGene_tx=XM_019830443.2;refGene_gn=LOC101085209;refGene_tx=XM_019830444.2;refGene_gn=LOC101085209;refGene_tx=XM_023253787.1;refGene_gn=LOC101085209;refGene_tx=XM_023253788.1
B2 34189102 34189192
refGene_gn=LOC101085209;refGene_tx=XM_003985999.5;refGene_gn=LOC101085209;refGene_tx=XM_019830441.2;refGene_gn=LOC101085209;refGene_tx=XM_019830443.2;refGene_gn=LOC101085209;refGene_tx=XM_019830444.2;refGene_gn=LOC101085209;refGene_tx=XM_023253787.1;refGene_gn=LOC101085209;refGene_tx=XM_023253788.1
B2 34189340 34189448
refGene_gn=LOC101085209;refGene_tx=XM_003985999.5;refGene_gn=LOC101085209;refGene_tx=XM_019830441.2;refGene_gn=LOC101085209;refGene_tx=XM_019830443.2;refGene_gn=LOC101085209;refGene_tx=XM_019830444.2;refGene_gn=LOC101085209;refGene_tx=XM_023253787.1;refGene_gn=LOC101085209;refGene_tx=XM_023253788.1
B2 34189864 34189918
refGene_gn=LOC101085209;refGene_tx=XM_003985999.5;refGene_gn=LOC101085209;refGene_tx=XM_019830441.2;refGene_gn=LOC101085209;refGene_tx=XM_019830443.2;refGene_gn=LOC101085209;refGene_tx=XM_019830444.2;refGene_gn=LOC101085209;refGene_tx=XM_023253787.1;refGene_gn=LOC101085209;refGene_tx=XM_023253788.1
B2 34190138 34190192
refGene_gn=LOC101085209;refGene_tx=XM_003985999.5;refGene_gn=LOC101085209;refGene_tx=XM_019830441.2;refGene_gn=LOC101085209;refGene_tx=XM_019830443.2;refGene_gn=LOC101085209;refGene_tx=XM_019830444.2;refGene_gn=LOC101085209;refGene_tx=XM_023253787.1;refGene_gn=LOC101085209;refGene_tx=XM_023253788.1
B2 34190287 34190341
refGene_gn=LOC101085209;refGene_tx=XM_003985999.5;refGene_gn=LOC101085209;refGene_tx=XM_019830441.2;refGene_gn=LOC101085209;refGene_tx=XM_01

9830443.2;refGene_gn=LOC101085209;refGene_tx=XM_019830444.2;refGene_gn=LOC101085209;refGene_tx=XM_023253787.1;refGene_gn=LOC101085209;refGene_tx=XM_023253788.1
B2 34190472 34190517
refGene_gn=LOC101085209;refGene_tx=XM_003985999.5;refGene_gn=LOC101085209;refGene_tx=XM_019830441.2;refGene_gn=LOC101085209;refGene_tx=XM_019830443.2;refGene_gn=LOC101085209;refGene_tx=XM_019830444.2;refGene_gn=LOC101085209;refGene_tx=XM_023253787.1;refGene_gn=LOC101085209;refGene_tx=XM_023253788.1
B2 34190643 34190697
refGene_gn=LOC101085209;refGene_tx=XM_003985999.5;refGene_gn=LOC101085209;refGene_tx=XM_019830441.2;refGene_gn=LOC101085209;refGene_tx=XM_019830443.2;refGene_gn=LOC101085209;refGene_tx=XM_019830444.2;refGene_gn=LOC101085209;refGene_tx=XM_023253787.1;refGene_gn=LOC101085209;refGene_tx=XM_023253788.1
B2 34190782 34190827
refGene_gn=LOC101085209;refGene_tx=XM_003985999.5;refGene_gn=LOC101085209;refGene_tx=XM_019830441.2;refGene_gn=LOC101085209;refGene_tx=XM_019830443.2;refGene_gn=LOC101085209;refGene_tx=XM_019830444.2;refGene_gn=LOC101085209;refGene_tx=XM_023253787.1;refGene_gn=LOC101085209;refGene_tx=XM_023253788.1
B2 34190924 34190978
refGene_gn=LOC101085209;refGene_tx=XM_003985999.5;refGene_gn=LOC101085209;refGene_tx=XM_019830441.2;refGene_gn=LOC101085209;refGene_tx=XM_019830443.2;refGene_gn=LOC101085209;refGene_tx=XM_019830444.2;refGene_gn=LOC101085209;refGene_tx=XM_023253787.1;refGene_gn=LOC101085209;refGene_tx=XM_023253788.1
B2 34191273 34191327
refGene_gn=LOC101085209;refGene_tx=XM_003985999.5;refGene_gn=LOC101085209;refGene_tx=XM_019830441.2;refGene_gn=LOC101085209;refGene_tx=XM_019830443.2;refGene_gn=LOC101085209;refGene_tx=XM_019830444.2;refGene_gn=LOC101085209;refGene_tx=XM_023253787.1;refGene_gn=LOC101085209;refGene_tx=XM_023253788.1
B2 34192405 34192513
refGene_gn=LOC101085209;refGene_tx=XM_003985999.5;refGene_gn=LOC101085209;refGene_tx=XM_019830441.2;refGene_gn=LOC101085209;refGene_tx=XM_019830443.2;refGene_gn=LOC101085209;refGene_tx=XM_019830444.2;refGene_gn=LOC101085209;refGene_tx=XM_023253787.1;refGene_gn=LOC101085209;refGene_tx=XM_023253788.1
B2 34192829 34192883
refGene_gn=LOC101085209;refGene_tx=XM_003985999.5;refGene_gn=LOC101085209;refGene_tx=XM_019830441.2;refGene_gn=LOC101085209;refGene_tx=XM_019830443.2;refGene_gn=LOC101085209;refGene_tx=XM_019830444.2;refGene_gn=LOC101085209;refGene_tx=XM_023253787.1;refGene_gn=LOC101085209;refGene_tx=XM_023253788.1
B2 34193112 34193157
refGene_gn=LOC101085209;refGene_tx=XM_003985999.5;refGene_gn=LOC101085209;refGene_tx=XM_019830441.2;refGene_gn=LOC101085209;refGene_tx=XM_019830443.2;refGene_gn=LOC101085209;refGene_tx=XM_019830444.2;refGene_gn=LOC101085209;refGene_tx=XM_023253787.1;refGene_gn=LOC101085209;refGen

e_tx=XM_023253788.1
B2 34193282 34193336
refGene_gn=LOC101085209;refGene_tx=XM_003985999.5;refGene_gn=LOC101085209;refGene_tx=XM_019830441.2;refGene_gn=LOC101085209;refGene_tx=XM_019830443.2;refGene_gn=LOC101085209;refGene_tx=XM_019830444.2;refGene_gn=LOC101085209;refGene_tx=XM_023253787.1;refGene_gn=LOC101085209;refGene_tx=XM_023253788.1
B2 34193573 34193618
refGene_gn=LOC101085209;refGene_tx=XM_003985999.5;refGene_gn=LOC101085209;refGene_tx=XM_019830441.2;refGene_gn=LOC101085209;refGene_tx=XM_019830443.2;refGene_gn=LOC101085209;refGene_tx=XM_019830444.2;refGene_gn=LOC101085209;refGene_tx=XM_023253787.1;refGene_gn=LOC101085209;refGene_tx=XM_023253788.1
B2 34193864 34193918
refGene_gn=LOC101085209;refGene_tx=XM_003985999.5;refGene_gn=LOC101085209;refGene_tx=XM_019830441.2;refGene_gn=LOC101085209;refGene_tx=XM_019830443.2;refGene_gn=LOC101085209;refGene_tx=XM_019830444.2;refGene_gn=LOC101085209;refGene_tx=XM_023253787.1;refGene_gn=LOC101085209;refGene_tx=XM_023253788.1
B2 34194055 34194100
refGene_gn=LOC101085209;refGene_tx=XM_003985999.5;refGene_gn=LOC101085209;refGene_tx=XM_019830441.2;refGene_gn=LOC101085209;refGene_tx=XM_019830443.2;refGene_gn=LOC101085209;refGene_tx=XM_019830444.2;refGene_gn=LOC101085209;refGene_tx=XM_023253787.1;refGene_gn=LOC101085209;refGene_tx=XM_023253788.1
B2 34194289 34194343
refGene_gn=LOC101085209;refGene_tx=XM_003985999.5;refGene_gn=LOC101085209;refGene_tx=XM_019830441.2;refGene_gn=LOC101085209;refGene_tx=XM_019830443.2;refGene_gn=LOC101085209;refGene_tx=XM_019830444.2;refGene_gn=LOC101085209;refGene_tx=XM_023253787.1;refGene_gn=LOC101085209;refGene_tx=XM_023253788.1
B2 34194518 34194563
refGene_gn=LOC101085209;refGene_tx=XM_003985999.5;refGene_gn=LOC101085209;refGene_tx=XM_019830441.2;refGene_gn=LOC101085209;refGene_tx=XM_019830443.2;refGene_gn=LOC101085209;refGene_tx=XM_019830444.2;refGene_gn=LOC101085209;refGene_tx=XM_023253787.1;refGene_gn=LOC101085209;refGene_tx=XM_023253788.1
B2 34195000 34195054
refGene_gn=LOC101085209;refGene_tx=XM_003985999.5;refGene_gn=LOC101085209;refGene_tx=XM_019830441.2;refGene_gn=LOC101085209;refGene_tx=XM_019830443.2;refGene_gn=LOC101085209;refGene_tx=XM_019830444.2;refGene_gn=LOC101085209;refGene_tx=XM_023253787.1;refGene_gn=LOC101085209;refGene_tx=XM_023253788.1
B2 34195176 34195221
refGene_gn=LOC101085209;refGene_tx=XM_003985999.5;refGene_gn=LOC101085209;refGene_tx=XM_019830441.2;refGene_gn=LOC101085209;refGene_tx=XM_019830443.2;refGene_gn=LOC101085209;refGene_tx=XM_019830444.2;refGene_gn=LOC101085209;refGene_tx=XM_023253787.1;refGene_gn=LOC101085209;refGene_tx=XM_023253788.1
B2 34195303 34195357

refGene_gn=LOC101085209;refGene_tx=XM_003985999.5;refGene_gn=LOC101085209;refGene_tx=XM_019830441.2;refGene_gn=LOC101085209;refGene_tx=XM_019830443.2;refGene_gn=LOC101085209;refGene_tx=XM_019830444.2;refGene_gn=LOC101085209;refGene_tx=XM_023253787.1;refGene_gn=LOC101085209;refGene_tx=XM_023253788.1
B2 34195562 34195616
refGene_gn=LOC101085209;refGene_tx=XM_003985999.5;refGene_gn=LOC101085209;refGene_tx=XM_019830441.2;refGene_gn=LOC101085209;refGene_tx=XM_019830443.2;refGene_gn=LOC101085209;refGene_tx=XM_019830444.2;refGene_gn=LOC101085209;refGene_tx=XM_023253787.1;refGene_gn=LOC101085209;refGene_tx=XM_023253788.1
B2 34195784 34195838
refGene_gn=LOC101085209;refGene_tx=XM_003985999.5;refGene_gn=LOC101085209;refGene_tx=XM_019830441.2;refGene_gn=LOC101085209;refGene_tx=XM_019830443.2;refGene_gn=LOC101085209;refGene_tx=XM_019830444.2;refGene_gn=LOC101085209;refGene_tx=XM_023253787.1;refGene_gn=LOC101085209;refGene_tx=XM_023253788.1
B2 34195937 34195991
refGene_gn=LOC101085209;refGene_tx=XM_003985999.5;refGene_gn=LOC101085209;refGene_tx=XM_019830441.2;refGene_gn=LOC101085209;refGene_tx=XM_019830443.2;refGene_gn=LOC101085209;refGene_tx=XM_019830444.2;refGene_gn=LOC101085209;refGene_tx=XM_023253787.1;refGene_gn=LOC101085209;refGene_tx=XM_023253788.1
B2 34196079 34196133
refGene_gn=LOC101085209;refGene_tx=XM_003985999.5;refGene_gn=LOC101085209;refGene_tx=XM_019830441.2;refGene_gn=LOC101085209;refGene_tx=XM_019830443.2;refGene_gn=LOC101085209;refGene_tx=XM_019830444.2;refGene_gn=LOC101085209;refGene_tx=XM_023253787.1;refGene_gn=LOC101085209;refGene_tx=XM_023253788.1
B2 34196302 34196359
refGene_gn=LOC101085209;refGene_tx=XM_003985999.5;refGene_gn=LOC101085209;refGene_tx=XM_019830441.2;refGene_gn=LOC101085209;refGene_tx=XM_019830443.2;refGene_gn=LOC101085209;refGene_tx=XM_019830444.2;refGene_gn=LOC101085209;refGene_tx=XM_023253787.1;refGene_gn=LOC101085209;refGene_tx=XM_023253788.1
B2 34196590 34196677
refGene_gn=LOC101085209;refGene_tx=XM_003985999.5;refGene_gn=LOC101085209;refGene_tx=XM_019830441.2;refGene_gn=LOC101085209;refGene_tx=XM_019830443.2;refGene_gn=LOC101085209;refGene_tx=XM_019830444.2;refGene_gn=LOC101085209;refGene_tx=XM_023253787.1;refGene_gn=LOC101085209;refGene_tx=XM_023253788.1
B2 34197150 34197225
refGene_gn=LOC101085209;refGene_tx=XM_003985999.5;refGene_gn=LOC101085209;refGene_tx=XM_019830441.2;refGene_gn=LOC101085209;refGene_tx=XM_019830443.2;refGene_gn=LOC101085209;refGene_tx=XM_019830444.2;refGene_gn=LOC101085209;refGene_tx=XM_023253787.1;refGene_gn=LOC101085209;refGene_tx=XM_023253788.1
B2 34197598 34197661
refGene_gn=LOC101085209;refGene_tx=XM_003985999.5;refGene_gn=LOC101085209;refGene_tx=XM_019830441.2;refGene_gn=LOC101085209;refGene_tx=XM_01

9830443.2;refGene_gn=LOC101085209;refGene_tx=XM_019830444.2;refGene_gn=LOC101085209;refGene_tx=XM_023253787.1;refGene_gn=LOC101085209;refGene_tx=XM_023253788.1
B2 34197887 34197929
refGene_gn=LOC101085209;refGene_tx=XM_003985999.5;refGene_gn=LOC101085209;refGene_tx=XM_019830441.2;refGene_gn=LOC101085209;refGene_tx=XM_019830443.2;refGene_gn=LOC101085209;refGene_tx=XM_019830444.2;refGene_gn=LOC101085209;refGene_tx=XM_023253787.1;refGene_gn=LOC101085209;refGene_tx=XM_023253788.1
B2 34198031 34198091
refGene_gn=LOC101085209;refGene_tx=XM_003985999.5;refGene_gn=LOC101085209;refGene_tx=XM_019830441.2;refGene_gn=LOC101085209;refGene_tx=XM_019830443.2;refGene_gn=LOC101085209;refGene_tx=XM_019830444.2
B2 34200212 34200392
refGene_gn=LOC101085209;refGene_tx=XM_003985999.5;refGene_gn=LOC101085209;refGene_tx=XM_019830441.2;refGene_gn=LOC101085209;refGene_tx=XM_019830450.2
B2 34201065 34201128
refGene_gn=LOC101085209;refGene_tx=XM_003985999.5;refGene_gn=LOC101085209;refGene_tx=XM_019830441.2;refGene_gn=LOC101085209;refGene_tx=XM_019830450.2
B2 34201785 34201863
refGene_gn=LOC101085209;refGene_tx=XM_003985999.5;refGene_gn=LOC101085209;refGene_tx=XM_019830441.2;refGene_gn=LOC101085209;refGene_tx=XM_019830443.2;refGene_gn=LOC101085209;refGene_tx=XM_019830450.2
B2 34202754 34202946
refGene_gn=LOC101085209;refGene_tx=XM_003985999.5;refGene_gn=LOC101085209;refGene_tx=XM_019830441.2;refGene_gn=LOC101085209;refGene_tx=XM_019830443.2;refGene_gn=LOC101085209;refGene_tx=XM_019830444.2;refGene_gn=LOC101085209;refGene_tx=XM_019830450.2
B2 34204378 34204541
refGene_gn=LOC101085209;refGene_tx=XM_003985999.5;refGene_gn=LOC101085209;refGene_tx=XM_019830441.2;refGene_gn=LOC101085209;refGene_tx=XM_019830443.2;refGene_gn=LOC101085209;refGene_tx=XM_019830444.2;refGene_gn=LOC101085209;refGene_tx=XM_019830450.2
B2 34204999 34205210
refGene_gn=LOC101085209;refGene_tx=XM_003985999.5;refGene_gn=LOC101085209;refGene_tx=XM_019830441.2;refGene_gn=LOC101085209;refGene_tx=XM_019830443.2;refGene_gn=LOC101085209;refGene_tx=XM_019830444.2;refGene_gn=LOC101085209;refGene_tx=XM_019830450.2
B2 34205346 34205496
refGene_gn=LOC101085209;refGene_tx=XM_003985999.5;refGene_gn=LOC101085209;refGene_tx=XM_019830441.2;refGene_gn=LOC101085209;refGene_tx=XM_019830443.2;refGene_gn=LOC101085209;refGene_tx=XM_019830444.2;refGene_gn=LOC101085209;refGene_tx=XM_019830450.2
B2 34208082 34208164
refGene_gn=LOC101085209;refGene_tx=XM_003985999.5;refGene_gn=LOC101085209;refGene_tx=XM_019830441.2;refGene_gn=LOC101085209;refGene_tx=XM_019830443.2;refGene_gn=LOC101085209;refGene_tx=XM_019830444.2;refGene_gn=LOC101085209;refGene_tx=XM_019830450.2

B2 34210678 34210826
refGene_gn=LOC101085801;refGene_tx=XM_023253789.1;refGene_gn=LOC101085801;refGene_tx=XM_023253790.1;refGene_gn=LOC101085801;refGene_tx=XM_023253791.1;refGene_gn=LOC101085801;refGene_tx=XM_023253792.1
B2 34210923 34211029
refGene_gn=LOC101085801;refGene_tx=XM_023253789.1;refGene_gn=LOC101085801;refGene_tx=XM_023253790.1;refGene_gn=LOC101085801;refGene_tx=XM_023253791.1;refGene_gn=LOC101085801;refGene_tx=XM_023253792.1
B2 34211319 34211423
refGene_gn=LOC101085801;refGene_tx=XM_023253789.1;refGene_gn=LOC101085801;refGene_tx=XM_023253790.1;refGene_gn=LOC101085801;refGene_tx=XM_023253791.1;refGene_gn=LOC101085801;refGene_tx=XM_023253792.1
B2 34211526 34211659
refGene_gn=LOC101085801;refGene_tx=XM_023253789.1;refGene_gn=LOC101085801;refGene_tx=XM_023253790.1;refGene_gn=LOC101085801;refGene_tx=XM_023253791.1;refGene_gn=LOC101085801;refGene_tx=XM_023253792.1
B2 34211853 34211983
refGene_gn=LOC101085801;refGene_tx=XM_023253789.1;refGene_gn=LOC101085801;refGene_tx=XM_023253790.1;refGene_gn=LOC101085801;refGene_tx=XM_023253791.1;refGene_gn=LOC101085801;refGene_tx=XM_023253792.1
B2 34212384 34212557
refGene_gn=LOC101085801;refGene_tx=XM_023253789.1;refGene_gn=LOC101085801;refGene_tx=XM_023253790.1;refGene_gn=LOC101085801;refGene_tx=XM_023253791.1;refGene_gn=LOC101085801;refGene_tx=XM_023253792.1
B2 34213954 34214134
refGene_gn=LOC101085801;refGene_tx=XM_023253789.1;refGene_gn=LOC101085801;refGene_tx=XM_023253790.1;refGene_gn=LOC101085801;refGene_tx=XM_023253791.1;refGene_gn=LOC101085801;refGene_tx=XM_023253792.1
B2 34214490 34214647
refGene_gn=LOC101085801;refGene_tx=XM_023253789.1;refGene_gn=LOC101085801;refGene_tx=XM_023253790.1;refGene_gn=LOC101085801;refGene_tx=XM_023253791.1;refGene_gn=LOC101085801;refGene_tx=XM_023253792.1
B2 34214999 34215406
refGene_gn=LOC101085801;refGene_tx=XM_023253789.1;refGene_gn=LOC101085801;refGene_tx=XM_023253790.1;refGene_gn=LOC101085801;refGene_tx=XM_023253791.1;refGene_gn=LOC101085801;refGene_tx=XM_023253792.1
B2 34216087 34216319
refGene_gn=LOC101085801;refGene_tx=XM_023253789.1;refGene_gn=LOC101085801;refGene_tx=XM_023253790.1;refGene_gn=LOC101085801;refGene_tx=XM_023253791.1;refGene_gn=LOC101085801;refGene_tx=XM_023253792.1
B2 34217095 34217506
refGene_gn=LOC101086060;refGene_tx=XM_023253793.1;refGene_gn=LOC101086060;refGene_tx=XM_023253794.1
B2 34217596 34217765
refGene_gn=LOC101086060;refGene_tx=XM_023253793.1;refGene_gn=LOC101086060;refGene_tx=XM_023253794.1
B2 34217926 34217980
refGene_gn=LOC101086060;refGene_tx=XM_023253793.1;refGene_gn=LOC101086060;refGene_tx=XM_023253794.1
B2 34218112 34218277

refGene_gn=LOC101086060;refGene_tx=XM_023253793.1;refGene_gn=LOC101086060;refGene_tx=XM_023253794.1
B2 34218422 34218563
refGene_gn=LOC101086060;refGene_tx=XM_023253793.1;refGene_gn=LOC101086060;refGene_tx=XM_023253794.1
B2 34218768 34218965
refGene_gn=LOC101086060;refGene_tx=XM_023253793.1;refGene_gn=LOC101086060;refGene_tx=XM_023253794.1
B2 34219421 34219694
refGene_gn=LOC101086060;refGene_tx=XM_023253793.1;refGene_gn=LOC101086060;refGene_tx=XM_023253794.1
B2 34220546 34220592
refGene_gn=LOC101086309;refGene_tx=XM_023253795.1
B2 34220784 34221002
refGene_gn=LOC101086309;refGene_tx=XM_023253795.1
B2 34221104 34221221
refGene_gn=LOC101086309;refGene_tx=XM_023253795.1
B2 34221331 34221424
refGene_gn=LOC101086309;refGene_tx=XM_023253795.1
B2 34221526 34221612
refGene_gn=LOC101086309;refGene_tx=XM_023253795.1
B2 34221724 34221809
refGene_gn=LOC101086309;refGene_tx=XM_023253795.1
B2 34222016 34222059
refGene_gn=LOC101086309;refGene_tx=XM_023253795.1
B2 34222290 34222365
refGene_gn=LOC101086309;refGene_tx=XM_023253795.1
B2 34222558 34222575
refGene_gn=LOC101086309;refGene_tx=XM_023253795.1
B2 34224771 34224849
refGene_gn=LOC101086561;refGene_tx=XM_023253796.1
B2 34225644 34225805
refGene_gn=LOC101086561;refGene_tx=XM_023253796.1
B2 34225912 34226128
refGene_gn=LOC101086561;refGene_tx=XM_023253796.1
B2 34226839 34227229
refGene_gn=LOC101086561;refGene_tx=XM_023253796.1
B2 34227414 34227688
refGene_gn=LOC101086561;refGene_tx=XM_023253796.1
B2 34227990 34228092
refGene_gn=LOC101086561;refGene_tx=XM_023253796.1
B2 34256169 34256316
refGene_gn=LOC101086807;refGene_tx=XM_003986005.4
B2 34256514 34256633
refGene_gn=LOC101086807;refGene_tx=XM_003986005.4
B2 34256837 34256949
refGene_gn=LOC101086807;refGene_tx=XM_003986005.4
B2 34264584 34264650
refGene_gn=LOC101086807;refGene_tx=XM_003986005.4
B2 34264874 34264982

refGene_gn=LOC101086807;refGene_tx=XM_003986005.4
B2 34265098 34265194
refGene_gn=LOC101086807;refGene_tx=XM_003986005.4
B2 34265462 34265586
refGene_gn=LOC101086807;refGene_tx=XM_003986005.4
B2 34265994 34266113
refGene_gn=LOC101086807;refGene_tx=XM_003986005.4
B2 34266581 34266737
refGene_gn=LOC101086807;refGene_tx=XM_003986005.4
B2 34267212 34267346
refGene_gn=LOC101086807;refGene_tx=XM_003986005.4
B2 34267704 34267762
refGene_gn=LOC101086807;refGene_tx=XM_003986005.4
B2 34267901 34268034
refGene_gn=LOC101086807;refGene_tx=XM_003986005.4
B2 34268145 34268246
refGene_gn=LOC101086807;refGene_tx=XM_003986005.4
B2 34268534 34268685
refGene_gn=LOC101086807;refGene_tx=XM_003986005.4
B2 34269071 34269247
refGene_gn=LOC101086807;refGene_tx=XM_003986005.4
B2 34269547 34269615
refGene_gn=LOC101086807;refGene_tx=XM_003986005.4
B2 34269813 34269889
refGene_gn=LOC101086807;refGene_tx=XM_003986005.4
B2 34270074 34270127
refGene_gn=LOC101086807;refGene_tx=XM_003986005.4
B2 34270272 34270357
refGene_gn=LOC101086807;refGene_tx=XM_003986005.4
B2 34271225 34271315
refGene_gn=LOC101086807;refGene_tx=XM_003986005.4;refGene_gn=LOC101087230;refGene_tx=XM_019830455.1
B2 34271752 34271755
refGene_gn=LOC101087230;refGene_tx=XM_003986007.3
B2 34272226 34272325
refGene_gn=LOC101087230;refGene_tx=XM_003986007.3;refGene_gn=LOC101087230;refGene_tx=XM_019830455.1
B2 34276103 34276190
refGene_gn=LOC101087230;refGene_tx=XM_003986007.3;refGene_gn=LOC101087230;refGene_tx=XM_019830455.1
B2 34276274 34276376
refGene_gn=LOC101087230;refGene_tx=XM_003986007.3;refGene_gn=LOC101087230;refGene_tx=XM_019830455.1
B2 34276474 34276566
refGene_gn=LOC101087230;refGene_tx=XM_003986007.3;refGene_gn=LOC101087230;refGene_tx=XM_019830455.1
B2 34276688 34276764
refGene_gn=LOC101087230;refGene_tx=XM_003986007.3;refGene_gn=LOC101087230;refGene_tx=XM_019830455.1
B2 34277763 34277973

refGene_gn=LOC101087480;refGene_tx=XM_023253798.1
B2	34277974	34278916
refGene_gn=LOC101087480;refGene_tx=XM_023253798.1
B2	34279624	34279726
refGene_gn=LOC101087740;refGene_tx=XM_003986009.4;refGene_gn=LOC101087740;refGene_tx=XM_019830452.1
B2	34279820	34279934
refGene_gn=LOC101087740;refGene_tx=XM_003986009.4;refGene_gn=LOC101087740;refGene_tx=XM_019830452.1
B2	34280082	34280178
refGene_gn=LOC101087740;refGene_tx=XM_003986009.4;refGene_gn=LOC101087740;refGene_tx=XM_019830452.1
B2	34280352	34280447
refGene_gn=LOC101087740;refGene_tx=XM_003986009.4;refGene_gn=LOC101087740;refGene_tx=XM_019830452.1
B2	34280591	34280905
refGene_gn=LOC101087740;refGene_tx=XM_003986009.4;refGene_gn=LOC101087740;refGene_tx=XM_019830452.1
B2	34284825	34284925
refGene_gn=LOC101087740;refGene_tx=XM_003986009.4;refGene_gn=LOC101087740;refGene_tx=XM_019830452.1
B2	34285118	34285254
refGene_gn=LOC101087740;refGene_tx=XM_003986009.4;refGene_gn=LOC101087740;refGene_tx=XM_019830452.1
B2	34285376	34285527
refGene_gn=LOC101087740;refGene_tx=XM_003986009.4;refGene_gn=LOC101087740;refGene_tx=XM_019830452.1
B2	34285616	34285721
refGene_gn=LOC101087740;refGene_tx=XM_003986009.4;refGene_gn=LOC101087740;refGene_tx=XM_019830452.1
B2	34285990	34286052
refGene_gn=LOC101087740;refGene_tx=XM_003986009.4;refGene_gn=LOC101087740;refGene_tx=XM_019830452.1
B2	34286225	34286313
refGene_gn=LOC101087740;refGene_tx=XM_003986009.4;refGene_gn=LOC101087740;refGene_tx=XM_019830452.1
B2	34286433	34286546
refGene_gn=LOC101087740;refGene_tx=XM_003986009.4;refGene_gn=LOC101087740;refGene_tx=XM_019830452.1
B2	34286686	34286767
refGene_gn=LOC101087740;refGene_tx=XM_003986009.4;refGene_gn=LOC101087740;refGene_tx=XM_019830452.1
B2	34286876	34287089
refGene_gn=LOC101087740;refGene_tx=XM_003986009.4;refGene_gn=LOC101087740;refGene_tx=XM_019830452.1
B2	34287194	34287263
refGene_gn=LOC101087740;refGene_tx=XM_003986009.4;refGene_gn=LOC101087740;refGene_tx=XM_019830452.1
B2	34288013	34288077
refGene_gn=LOC101088333;refGene_tx=XM_003986013.4;refGene_gn=LOC101088

333;refGene_tx=XM_006931619.4;refGene_gn=LOC101088333;refGene_tx=XM_019830456.1;refGene_gn=LOC101088333;refGene_tx=XM_019830457.2
B2 34288323 34288394
refGene_gn=LOC101088333;refGene_tx=XM_003986013.4;refGene_gn=LOC101088333;refGene_tx=XM_006931619.4;refGene_gn=LOC101088333;refGene_tx=XM_019830456.1;refGene_gn=LOC101088333;refGene_tx=XM_019830457.2
B2 34288489 34288614
refGene_gn=LOC101088333;refGene_tx=XM_003986013.4;refGene_gn=LOC101088333;refGene_tx=XM_006931619.4;refGene_gn=LOC101088333;refGene_tx=XM_019830456.1;refGene_gn=LOC101088333;refGene_tx=XM_019830457.2
B2 34288849 34288979
refGene_gn=LOC101088333;refGene_tx=XM_003986013.4;refGene_gn=LOC101088333;refGene_tx=XM_006931619.4;refGene_gn=LOC101088333;refGene_tx=XM_019830456.1;refGene_gn=LOC101088333;refGene_tx=XM_019830457.2
B2 34290383 34290595
refGene_gn=LOC101088748;refGene_tx=XM_023253797.1
B2 34290724 34290839
refGene_gn=LOC101088748;refGene_tx=XM_023253797.1
B2 34291511 34291802
refGene_gn=LOC101088748;refGene_tx=XM_023253797.1
B2 34291912 34292024
refGene_gn=LOC101088748;refGene_tx=XM_023253797.1
B2 34292118 34292214
refGene_gn=LOC101088748;refGene_tx=XM_023253797.1
B2 34292311 34292433
refGene_gn=LOC101088748;refGene_tx=XM_023253797.1
B2 34292555 34292588
refGene_gn=LOC101088748;refGene_tx=XM_023253797.1
B2 34293176 34293251
refGene_gn=LOC101088748;refGene_tx=XM_023253797.1
B2 34293447 34293516
refGene_gn=LOC101088748;refGene_tx=XM_023253797.1
B2 34293618 34293703
refGene_gn=LOC101088748;refGene_tx=XM_023253797.1
B2 34293800 34293904
refGene_gn=LOC101088748;refGene_tx=XM_023253797.1
B2 34293989 34294241
refGene_gn=LOC101088748;refGene_tx=XM_023253797.1
B2 34294516 34294814
refGene_gn=LOC101088748;refGene_tx=XM_023253797.1
B2 34294902 34294953
refGene_gn=LOC101088748;refGene_tx=XM_023253797.1
B2 34295071 34295250
refGene_gn=LOC101088748;refGene_tx=XM_023253797.1
B2 34295508 34295592
refGene_gn=LOC101088748;refGene_tx=XM_023253797.1
B2 34296883 34297042
refGene_gn=LOC101088748;refGene_tx=XM_023253797.1
B2 34299201 34299260
refGene_gn=LOC101091023;refGene_tx=XM_003986020.5;refGene_gn=LOC101091

023;refGene_tx=XM_019830454.2
B2 34300276 34300314
refGene_gn=LOC101091023;refGene_tx=XM_003986020.5
B2 34300472 34300562
refGene_gn=LOC101091023;refGene_tx=XM_003986020.5;refGene_gn=LOC101091023;refGene_tx=XM_019830454.2
B2 34300690 34301032
refGene_gn=LOC101091023;refGene_tx=XM_003986020.5;refGene_gn=LOC101091023;refGene_tx=XM_019830454.2
B2 34301426 34301825
refGene_gn=LOC101091023;refGene_tx=XM_003986020.5;refGene_gn=LOC101091023;refGene_tx=XM_019830454.2
B2 34309477 34309738
refGene_gn=LOC101091023;refGene_tx=XM_003986020.5;refGene_gn=LOC101091023;refGene_tx=XM_019830454.2
B2 34309965 34310136
refGene_gn=LOC101091023;refGene_tx=XM_003986020.5;refGene_gn=LOC101091023;refGene_tx=XM_019830454.2
B2 34310274 34310311
refGene_gn=LOC101091023;refGene_tx=XM_003986020.5;refGene_gn=LOC101091023;refGene_tx=XM_019830454.2
B2 34311269 34313177
refGene_gn=LOC101089010;refGene_tx=XM_004001534.3
B2 34314929 34314989
refGene_gn=LOC101089265;refGene_tx=XM_006931620.3;refGene_gn=LOC101089265;refGene_tx=XM_019830458.1;refGene_gn=LOC101089265;refGene_tx=XM_023253800.1
B2 34315194 34315417
refGene_gn=LOC101089265;refGene_tx=XM_006931620.3;refGene_gn=LOC101089265;refGene_tx=XM_019830458.1;refGene_gn=LOC101089265;refGene_tx=XM_023253800.1
B2 34315575 34316053
refGene_gn=LOC101089265;refGene_tx=XM_006931620.3;refGene_gn=LOC101089265;refGene_tx=XM_019830458.1;refGene_gn=LOC101089265;refGene_tx=XM_023253800.1
B2 34316208 34316440
refGene_gn=LOC101089265;refGene_tx=XM_006931620.3;refGene_gn=LOC101089265;refGene_tx=XM_019830458.1;refGene_gn=LOC101089265;refGene_tx=XM_023253800.1
B2 34316596 34316811
refGene_gn=LOC101089265;refGene_tx=XM_006931620.3;refGene_gn=LOC101089265;refGene_tx=XM_019830458.1;refGene_gn=LOC101089265;refGene_tx=XM_023253800.1
B2 34316936 34317768
refGene_gn=LOC101089265;refGene_tx=XM_006931620.3;refGene_gn=LOC101089265;refGene_tx=XM_019830458.1;refGene_gn=LOC101089265;refGene_tx=XM_023253800.1
B2 34317921 34318128
refGene_gn=LOC101089265;refGene_tx=XM_006931620.3;refGene_gn=LOC101089265;refGene_tx=XM_019830458.1;refGene_gn=LOC101089265;refGene_tx=XM_02

3253800.1
B2	34379120	34379132
refGene_gn=LOC101087231;refGene_tx=XM_023253801.1
B2	34385469	34385607
refGene_gn=LOC101087231;refGene_tx=XM_023253801.1;refGene_gn=LOC101087
231;refGene_tx=XM_023253802.1
B2	34385729	34385829
refGene_gn=LOC101087231;refGene_tx=XM_023253801.1;refGene_gn=LOC101087
231;refGene_tx=XM_023253802.1
B2	34386340	34386394
refGene_gn=LOC101087231;refGene_tx=XM_023253801.1;refGene_gn=LOC101087
231;refGene_tx=XM_023253802.1
B2	34386504	34386555
refGene_gn=LOC101087231;refGene_tx=XM_023253801.1;refGene_gn=LOC101087
231;refGene_tx=XM_023253802.1
B2	34386758	34387153
refGene_gn=LOC101087231;refGene_tx=XM_023253801.1;refGene_gn=LOC101087
231;refGene_tx=XM_023253802.1
B2	34388867	34389647
refGene_gn=LOC101087231;refGene_tx=XM_023253801.1;refGene_gn=LOC101087
231;refGene_tx=XM_023253802.1
B2	34390295	34390586
refGene_gn=LOC101087231;refGene_tx=XM_023253801.1;refGene_gn=LOC101087
231;refGene_tx=XM_023253802.1
B2	34390686	34390760
refGene_gn=LOC101087231;refGene_tx=XM_023253801.1;refGene_gn=LOC101087
231;refGene_tx=XM_023253802.1
B2	34391028	34391155
refGene_gn=LOC101087231;refGene_tx=XM_023253801.1;refGene_gn=LOC101087
231;refGene_tx=XM_023253802.1
B2	34402279	34402293
refGene_gn=LOC111560440;refGene_tx=XM_023253803.1
B2	34402593	34402617
refGene_gn=LOC111560440;refGene_tx=XM_023253803.1
B2	34403070	34403181
refGene_gn=LOC111560440;refGene_tx=XM_023253803.1
B2	34403929	34404211
refGene_gn=LOC111560440;refGene_tx=XM_023253803.1
B2	34406988	34407258
refGene_gn=LOC111560440;refGene_tx=XM_023253803.1
B2	34412131	34412231
refGene_gn=LOC111560440;refGene_tx=XM_023253803.1
B2	34487450	34487550
refGene_gn=LOC111556169;refGene_tx=XM_003985987.5
B2	34493704	34493974
refGene_gn=LOC111556169;refGene_tx=XM_003985987.5
B2	34498002	34498284
refGene_gn=LOC111556169;refGene_tx=XM_003985987.5
B2	34498982	34499093
refGene_gn=LOC111556169;refGene_tx=XM_003985987.5

B2 34499499 34499523
refGene_gn=LOC111556169;refGene_tx=XM_003985987.5
B2 34499816 34499830
refGene_gn=LOC111556169;refGene_tx=XM_003985987.5
B2 34542356 34542383
refGene_gn=LOC111560441;refGene_tx=XM_023253804.1;refGene_gn=LOC111560
441;refGene_tx=XM_023253805.1
B2 34542557 34542589
refGene_gn=LOC111560441;refGene_tx=XM_023253804.1;refGene_gn=LOC111560
441;refGene_tx=XM_023253805.1
B2 34542842 34542953
refGene_gn=LOC111560441;refGene_tx=XM_023253804.1;refGene_gn=LOC111560
441;refGene_tx=XM_023253805.1
B2 34543426 34543708
refGene_gn=LOC111560441;refGene_tx=XM_023253804.1;refGene_gn=LOC111560
441;refGene_tx=XM_023253805.1
B2 34544165 34544447
refGene_gn=LOC111560441;refGene_tx=XM_023253804.1;refGene_gn=LOC111560
441;refGene_tx=XM_023253805.1
B2 34547630 34547757
refGene_gn=LOC111560441;refGene_tx=XM_023253804.1;refGene_gn=LOC111560
441;refGene_tx=XM_023253805.1
B2 34562284 34562468
refGene_gn=LOC111556311;refGene_tx=XM_023253806.1
B2 34562469 34562648
refGene_gn=LOC111556311;refGene_tx=XM_023253806.1
B2 34562994 34563131
refGene_gn=LOC111556311;refGene_tx=XM_023253806.1
B2 34563490 34563650
refGene_gn=LOC111556311;refGene_tx=XM_023253806.1
B2 34564063 34564237
refGene_gn=LOC111556311;refGene_tx=XM_023253806.1
B2 34564387 34564576
refGene_gn=LOC111556311;refGene_tx=XM_023253806.1
B2 34565941 34566070
refGene_gn=LOC111556311;refGene_tx=XM_023253806.1
B2 34566566 34566764
refGene_gn=LOC111556311;refGene_tx=XM_023253806.1
B2 34569513 34569719
refGene_gn=LOC111556311;refGene_tx=XM_023253806.1
B2 34570009 34570140
refGene_gn=LOC111556311;refGene_tx=XM_023253806.1
B2 34571627 34571742
refGene_gn=LOC111556311;refGene_tx=XM_023253806.1
B2 34571831 34572321
refGene_gn=LOC111556311;refGene_tx=XM_023253806.1
B2 34574912 34575001
refGene_gn=LOC111556170;refGene_tx=XM_003985989.5
B2 34575248 34575453
refGene_gn=LOC111556170;refGene_tx=XM_003985989.5

B2 34575795 34575925
refGene_gn=LOC111556170;refGene_tx=XM_003985989.5
B2 34576585 34576697
refGene_gn=LOC111556170;refGene_tx=XM_003985989.5
B2 34576845 34576993
refGene_gn=LOC111556170;refGene_tx=XM_003985989.5
B2 34577764 34577911
refGene_gn=LOC111556170;refGene_tx=XM_003985989.5
B2 34579465 34579678
refGene_gn=LOC111560442;refGene_tx=XM_023253807.1
B2 34580448 34580585
refGene_gn=LOC111560442;refGene_tx=XM_023253807.1
B2 34580850 34581013
refGene_gn=LOC111560442;refGene_tx=XM_023253807.1
B2 34581268 34581442
refGene_gn=LOC111560442;refGene_tx=XM_023253807.1
B2 34582022 34582211
refGene_gn=LOC111560442;refGene_tx=XM_023253807.1
B2 34582367 34582496
refGene_gn=LOC111560442;refGene_tx=XM_023253807.1
B2 34583275 34583473
refGene_gn=LOC111560442;refGene_tx=XM_023253807.1
B2 34583910 34584116
refGene_gn=LOC111560442;refGene_tx=XM_023253807.1
B2 34585373 34585504
refGene_gn=LOC111560442;refGene_tx=XM_023253807.1
B2 34585661 34585776
refGene_gn=LOC111560442;refGene_tx=XM_023253807.1
B2 34586258 34586856
refGene_gn=LOC111560442;refGene_tx=XM_023253807.1
B2 34587499 34587529
refGene_gn=LOC111556171;refGene_tx=XM_011282280.3;refGene_gn=LOC111556171;refGene_tx=XM_011282281.3;refGene_gn=LOC111556171;refGene_tx=XM_023253808.1
B2 34589362 34589430
refGene_gn=LOC111556171;refGene_tx=XM_011282280.3;refGene_gn=LOC111556171;refGene_tx=XM_011282281.3;refGene_gn=LOC111556171;refGene_tx=XM_011282282.3;refGene_gn=LOC111556171;refGene_tx=XM_011282283.3;refGene_gn=LOC111556171;refGene_tx=XM_023253808.1
B2 34590436 34590571
refGene_gn=LOC111556171;refGene_tx=XM_011282280.3;refGene_gn=LOC111556171;refGene_tx=XM_011282281.3;refGene_gn=LOC111556171;refGene_tx=XM_011282282.3;refGene_gn=LOC111556171;refGene_tx=XM_011282283.3
B2 34591231 34591361
refGene_gn=LOC111556171;refGene_tx=XM_011282280.3;refGene_gn=LOC111556171;refGene_tx=XM_011282281.3;refGene_gn=LOC111556171;refGene_tx=XM_011282282.3;refGene_gn=LOC111556171;refGene_tx=XM_011282283.3;refGene_gn=LOC111556171;refGene_tx=XM_023253808.1
B2 34591603 34591745
refGene_gn=LOC111556171;refGene_tx=XM_011282280.3;refGene_gn=LOC111556

171;refGene_tx=XM_011282281.3;refGene_gn=LOC111556171;refGene_tx=XM_01
1282282.3;refGene_gn=LOC111556171;refGene_tx=XM_011282283.3;refGene_gn
=LOC111556171;refGene_tx=XM_023253808.1
B2 34592298 34592426
refGene_gn=LOC111556171;refGene_tx=XM_011282280.3;refGene_gn=LOC111556
171;refGene_tx=XM_011282281.3;refGene_gn=LOC111556171;refGene_tx=XM_01
1282282.3;refGene_gn=LOC111556171;refGene_tx=XM_011282283.3;refGene_gn
=LOC111556171;refGene_tx=XM_023253808.1
B2 34661358 34661375
refGene_gn=LOC111556173;refGene_tx=XM_006931606.4;refGene_gn=LOC111556
173;refGene_tx=XM_019830438.2
B2 34661745 34661781
refGene_gn=LOC111556173;refGene_tx=XM_006931606.4
B2 34661929 34662046
refGene_gn=LOC111556173;refGene_tx=XM_006931606.4;refGene_gn=LOC111556
173;refGene_tx=XM_019830438.2
B2 34663508 34663793
refGene_gn=LOC111556173;refGene_tx=XM_006931606.4;refGene_gn=LOC111556
173;refGene_tx=XM_019830438.2
B2 34665056 34665338
refGene_gn=LOC111556173;refGene_tx=XM_006931606.4;refGene_gn=LOC111556
173;refGene_tx=XM_019830438.2
B2 34666019 34666074
refGene_gn=LOC111556173;refGene_tx=XM_006931606.4;refGene_gn=LOC111556
173;refGene_tx=XM_019830438.2
B2 34676286 34676291
refGene_gn=LOC111560443;refGene_tx=XM_023253809.1
B2 34676638 34676767
refGene_gn=LOC111560443;refGene_tx=XM_023253809.1
B2 34677000 34677279
refGene_gn=LOC111560443;refGene_tx=XM_023253809.1
B2 34677885 34678170
refGene_gn=LOC111560443;refGene_tx=XM_023253809.1
B2 34679677 34679765
refGene_gn=LOC111560443;refGene_tx=XM_023253809.1
B2 34697349 34697378
refGene_gn=LOC111556174;refGene_tx=XM_023253810.1;refGene_gn=LOC111556
174;refGene_tx=XM_023253811.1;refGene_gn=LOC111556174;refGene_tx=XM_02
3253812.1
B2 34699028 34699332
refGene_gn=LOC111556174;refGene_tx=XM_023253810.1;refGene_gn=LOC111556
174;refGene_tx=XM_023253811.1;refGene_gn=LOC111556174;refGene_tx=XM_02
3253812.1;refGene_gn=LOC111556174;refGene_tx=XM_023253813.1
B2 34699944 34700082
refGene_gn=LOC111556174;refGene_tx=XM_023253810.1;refGene_gn=LOC111556
174;refGene_tx=XM_023253811.1;refGene_gn=LOC111556174;refGene_tx=XM_02
3253812.1;refGene_gn=LOC111556174;refGene_tx=XM_023253813.1;refGene_gn
=LOC111556174;refGene_tx=XM_023253814.1
B2 34700600 34700739
refGene_gn=LOC111556174;refGene_tx=XM_023253810.1;refGene_gn=LOC111556

174;refGene_tx=XM_023253811.1;refGene_gn=LOC111556174;refGene_tx=XM_023253812.1;refGene_gn=LOC111556174;refGene_tx=XM_023253813.1;refGene_gn=LOC111556174;refGene_tx=XM_023253814.1
B2 34700841 34701056
refGene_gn=LOC111556174;refGene_tx=XM_023253810.1;refGene_gn=LOC111556174;refGene_tx=XM_023253811.1;refGene_gn=LOC111556174;refGene_tx=XM_023253812.1;refGene_gn=LOC111556174;refGene_tx=XM_023253813.1;refGene_gn=LOC111556174;refGene_tx=XM_023253814.1
B2 34701158 34701533
refGene_gn=LOC111556174;refGene_tx=XM_023253810.1;refGene_gn=LOC111556174;refGene_tx=XM_023253811.1;refGene_gn=LOC111556174;refGene_tx=XM_023253812.1;refGene_gn=LOC111556174;refGene_tx=XM_023253813.1;refGene_gn=LOC111556174;refGene_tx=XM_023253814.1
B2 34702043 34702172
refGene_gn=LOC111556174;refGene_tx=XM_023253810.1;refGene_gn=LOC111556174;refGene_tx=XM_023253811.1;refGene_gn=LOC111556174;refGene_tx=XM_023253812.1;refGene_gn=LOC111556174;refGene_tx=XM_023253813.1;refGene_gn=LOC111556174;refGene_tx=XM_023253814.1
B2 34702469 34702721
refGene_gn=LOC111556174;refGene_tx=XM_023253810.1;refGene_gn=LOC111556174;refGene_tx=XM_023253811.1;refGene_gn=LOC111556174;refGene_tx=XM_023253812.1;refGene_gn=LOC111556174;refGene_tx=XM_023253813.1;refGene_gn=LOC111556174;refGene_tx=XM_023253814.1
B2 34702848 34703114
refGene_gn=LOC111556174;refGene_tx=XM_023253810.1;refGene_gn=LOC111556174;refGene_tx=XM_023253811.1;refGene_gn=LOC111556174;refGene_tx=XM_023253812.1;refGene_gn=LOC111556174;refGene_tx=XM_023253813.1;refGene_gn=LOC111556174;refGene_tx=XM_023253814.1
B2 34703637 34703942
refGene_gn=LOC111556174;refGene_tx=XM_023253810.1;refGene_gn=LOC111556174;refGene_tx=XM_023253811.1;refGene_gn=LOC111556174;refGene_tx=XM_023253812.1;refGene_gn=LOC111556174;refGene_tx=XM_023253813.1;refGene_gn=LOC111556174;refGene_tx=XM_023253814.1
B2 34704164 34704287
refGene_gn=LOC111556174;refGene_tx=XM_023253810.1;refGene_gn=LOC111556174;refGene_tx=XM_023253811.1;refGene_gn=LOC111556174;refGene_tx=XM_023253812.1;refGene_gn=LOC111556174;refGene_tx=XM_023253813.1;refGene_gn=LOC111556174;refGene_tx=XM_023253814.1
B2 34704391 34704531
refGene_gn=LOC111556174;refGene_tx=XM_023253810.1;refGene_gn=LOC111556174;refGene_tx=XM_023253811.1;refGene_gn=LOC111556174;refGene_tx=XM_023253812.1;refGene_gn=LOC111556174;refGene_tx=XM_023253813.1;refGene_gn=LOC111556174;refGene_tx=XM_023253814.1
B2 34727094 34727098
refGene_gn=LOC111560444;refGene_tx=XM_023253815.1
B2 34727336 34727472
refGene_gn=LOC111560444;refGene_tx=XM_023253815.1
B2 34727617 34727899
refGene_gn=LOC111560444;refGene_tx=XM_023253815.1
B2 34728313 34728562

refGene_gn=LOC111560444;refGene_tx=XM_023253815.1
B2	34729739	34729821
refGene_gn=LOC111560444;refGene_tx=XM_023253815.1
B2	34758682	34758858
refGene_gn=LOC111560445;refGene_tx=XM_023253816.1
B2	34760483	34760627
refGene_gn=LOC111560445;refGene_tx=XM_023253816.1
B2	34760629	34760764
refGene_gn=LOC111560445;refGene_tx=XM_023253816.1
B2	34761142	34761388
refGene_gn=LOC111560445;refGene_tx=XM_023253816.1
B2	34764435	34764535
refGene_gn=LOC111560445;refGene_tx=XM_023253816.1
B2	34766744	34766844
refGene_gn=LOC111556176;refGene_tx=XM_023253817.1
B2	34776100	34776370
refGene_gn=LOC111556176;refGene_tx=XM_023253817.1
B2	34777045	34777327
refGene_gn=LOC111556176;refGene_tx=XM_023253817.1
B2	34777848	34777958
refGene_gn=LOC111556176;refGene_tx=XM_023253817.1
B2	34797386	34797527
refGene_gn=LOC111556258;refGene_tx=XM_023253819.1;refGene_gn=LOC111556
258;refGene_tx=XM_023253820.1
B2	34798234	34798441
refGene_gn=LOC111556258;refGene_tx=XM_023253819.1;refGene_gn=LOC111556
258;refGene_tx=XM_023253820.1
B2	34798811	34798924
refGene_gn=LOC111556258;refGene_tx=XM_023253819.1;refGene_gn=LOC111556
258;refGene_tx=XM_023253820.1
B2	34799616	34799884
refGene_gn=LOC111556258;refGene_tx=XM_023253819.1;refGene_gn=LOC111556
258;refGene_tx=XM_023253820.1
B2	34799995	34800049
refGene_gn=LOC111556258;refGene_tx=XM_023253819.1;refGene_gn=LOC111556
258;refGene_tx=XM_023253820.1
B2	34800234	34800270
refGene_gn=LOC111556258;refGene_tx=XM_023253819.1;refGene_gn=LOC111556
258;refGene_tx=XM_023253820.1
B2	34800580	34800634
refGene_gn=LOC111556258;refGene_tx=XM_023253819.1;refGene_gn=LOC111556
258;refGene_tx=XM_023253820.1
B2	34800770	34800878
refGene_gn=LOC111556258;refGene_tx=XM_023253819.1;refGene_gn=LOC111556
258;refGene_tx=XM_023253820.1
B2	34801144	34801198
refGene_gn=LOC111556258;refGene_tx=XM_023253819.1;refGene_gn=LOC111556
258;refGene_tx=XM_023253820.1
B2	34801355	34801409
refGene_gn=LOC111556258;refGene_tx=XM_023253819.1;refGene_gn=LOC111556

258;refGene_tx=XM_023253820.1
B2 34801519 34801627
refGene_gn=LOC111556258;refGene_tx=XM_023253819.1;refGene_gn=LOC111556258;refGene_tx=XM_023253820.1
B2 34801894 34801948
refGene_gn=LOC111556258;refGene_tx=XM_023253819.1;refGene_gn=LOC111556258;refGene_tx=XM_023253820.1
B2 34802518 34802572
refGene_gn=LOC111556258;refGene_tx=XM_023253819.1;refGene_gn=LOC111556258;refGene_tx=XM_023253820.1
B2 34802708 34802762
refGene_gn=LOC111556258;refGene_tx=XM_023253819.1;refGene_gn=LOC111556258;refGene_tx=XM_023253820.1
B2 34802949 34803003
refGene_gn=LOC111556258;refGene_tx=XM_023253819.1;refGene_gn=LOC111556258;refGene_tx=XM_023253820.1
B2 34803402 34803510
refGene_gn=LOC111556258;refGene_tx=XM_023253819.1;refGene_gn=LOC111556258;refGene_tx=XM_023253820.1
B2 34803854 34803908
refGene_gn=LOC111556258;refGene_tx=XM_023253819.1;refGene_gn=LOC111556258;refGene_tx=XM_023253820.1
B2 34804067 34804121
refGene_gn=LOC111556258;refGene_tx=XM_023253819.1;refGene_gn=LOC111556258;refGene_tx=XM_023253820.1
B2 34804345 34804399
refGene_gn=LOC111556258;refGene_tx=XM_023253819.1;refGene_gn=LOC111556258;refGene_tx=XM_023253820.1
B2 34804615 34804669
refGene_gn=LOC111556258;refGene_tx=XM_023253819.1;refGene_gn=LOC111556258;refGene_tx=XM_023253820.1
B2 34804859 34804967
refGene_gn=LOC111556258;refGene_tx=XM_023253819.1;refGene_gn=LOC111556258;refGene_tx=XM_023253820.1
B2 34805151 34805205
refGene_gn=LOC111556258;refGene_tx=XM_023253819.1;refGene_gn=LOC111556258;refGene_tx=XM_023253820.1
B2 34805310 34805364
refGene_gn=LOC111556258;refGene_tx=XM_023253819.1;refGene_gn=LOC111556258;refGene_tx=XM_023253820.1
B2 34805506 34805614
refGene_gn=LOC111556258;refGene_tx=XM_023253819.1;refGene_gn=LOC111556258;refGene_tx=XM_023253820.1
B2 34805752 34805860
refGene_gn=LOC111556258;refGene_tx=XM_023253819.1;refGene_gn=LOC111556258;refGene_tx=XM_023253820.1
B2 34806095 34806149
refGene_gn=LOC111556258;refGene_tx=XM_023253819.1;refGene_gn=LOC111556258;refGene_tx=XM_023253820.1
B2 34806352 34806442

refGene_gn=LOC111556258;refGene_tx=XM_023253819.1;refGene_gn=LOC111556258;refGene_tx=XM_023253820.1
B2 34806590 34806698
refGene_gn=LOC111556258;refGene_tx=XM_023253819.1;refGene_gn=LOC111556258;refGene_tx=XM_023253820.1
B2 34807114 34807168
refGene_gn=LOC111556258;refGene_tx=XM_023253819.1;refGene_gn=LOC111556258;refGene_tx=XM_023253820.1
B2 34807388 34807442
refGene_gn=LOC111556258;refGene_tx=XM_023253819.1;refGene_gn=LOC111556258;refGene_tx=XM_023253820.1
B2 34807537 34807591
refGene_gn=LOC111556258;refGene_tx=XM_023253819.1;refGene_gn=LOC111556258;refGene_tx=XM_023253820.1
B2 34807722 34807767
refGene_gn=LOC111556258;refGene_tx=XM_023253819.1;refGene_gn=LOC111556258;refGene_tx=XM_023253820.1
B2 34807893 34807947
refGene_gn=LOC111556258;refGene_tx=XM_023253819.1;refGene_gn=LOC111556258;refGene_tx=XM_023253820.1
B2 34808032 34808077
refGene_gn=LOC111556258;refGene_tx=XM_023253819.1;refGene_gn=LOC111556258;refGene_tx=XM_023253820.1
B2 34808174 34808228
refGene_gn=LOC111556258;refGene_tx=XM_023253819.1;refGene_gn=LOC111556258;refGene_tx=XM_023253820.1
B2 34808523 34808577
refGene_gn=LOC111556258;refGene_tx=XM_023253819.1;refGene_gn=LOC111556258;refGene_tx=XM_023253820.1
B2 34809655 34809763
refGene_gn=LOC111556258;refGene_tx=XM_023253819.1;refGene_gn=LOC111556258;refGene_tx=XM_023253820.1
B2 34810079 34810133
refGene_gn=LOC111556258;refGene_tx=XM_023253819.1;refGene_gn=LOC111556258;refGene_tx=XM_023253820.1
B2 34810362 34810407
refGene_gn=LOC111556258;refGene_tx=XM_023253819.1;refGene_gn=LOC111556258;refGene_tx=XM_023253820.1
B2 34810532 34810586
refGene_gn=LOC111556258;refGene_tx=XM_023253819.1;refGene_gn=LOC111556258;refGene_tx=XM_023253820.1
B2 34810822 34810867
refGene_gn=LOC111556258;refGene_tx=XM_023253819.1;refGene_gn=LOC111556258;refGene_tx=XM_023253820.1
B2 34811105 34811159
refGene_gn=LOC111556258;refGene_tx=XM_023253819.1;refGene_gn=LOC111556258;refGene_tx=XM_023253820.1
B2 34811296 34811341
refGene_gn=LOC111556258;refGene_tx=XM_023253819.1;refGene_gn=LOC111556258;refGene_tx=XM_023253820.1

B2 34811530 34811584
refGene_gn=LOC111556258;refGene_tx=XM_023253819.1;refGene_gn=LOC111556258;refGene_tx=XM_023253820.1
B2 34811759 34811804
refGene_gn=LOC111556258;refGene_tx=XM_023253819.1;refGene_gn=LOC111556258;refGene_tx=XM_023253820.1
B2 34812248 34812302
refGene_gn=LOC111556258;refGene_tx=XM_023253819.1;refGene_gn=LOC111556258;refGene_tx=XM_023253820.1
B2 34812424 34812469
refGene_gn=LOC111556258;refGene_tx=XM_023253819.1;refGene_gn=LOC111556258;refGene_tx=XM_023253820.1
B2 34812551 34812605
refGene_gn=LOC111556258;refGene_tx=XM_023253819.1;refGene_gn=LOC111556258;refGene_tx=XM_023253820.1
B2 34812810 34812864
refGene_gn=LOC111556258;refGene_tx=XM_023253819.1;refGene_gn=LOC111556258;refGene_tx=XM_023253820.1
B2 34813032 34813086
refGene_gn=LOC111556258;refGene_tx=XM_023253819.1;refGene_gn=LOC111556258;refGene_tx=XM_023253820.1
B2 34813184 34813238
refGene_gn=LOC111556258;refGene_tx=XM_023253819.1;refGene_gn=LOC111556258;refGene_tx=XM_023253820.1
B2 34813326 34813380
refGene_gn=LOC111556258;refGene_tx=XM_023253819.1;refGene_gn=LOC111556258;refGene_tx=XM_023253820.1
B2 34813550 34813607
refGene_gn=LOC111556258;refGene_tx=XM_023253819.1;refGene_gn=LOC111556258;refGene_tx=XM_023253820.1
B2 34813838 34813925
refGene_gn=LOC111556258;refGene_tx=XM_023253819.1;refGene_gn=LOC111556258;refGene_tx=XM_023253820.1
B2 34814398 34814473
refGene_gn=LOC111556258;refGene_tx=XM_023253819.1;refGene_gn=LOC111556258;refGene_tx=XM_023253820.1
B2 34814856 34814919
refGene_gn=LOC111556258;refGene_tx=XM_023253819.1;refGene_gn=LOC111556258;refGene_tx=XM_023253820.1
B2 34815145 34815187
refGene_gn=LOC111556258;refGene_tx=XM_023253819.1;refGene_gn=LOC111556258;refGene_tx=XM_023253820.1
B2 34815289 34815349
refGene_gn=LOC111556258;refGene_tx=XM_023253819.1;refGene_gn=LOC111556258;refGene_tx=XM_023253820.1
B2 34817479 34817659
refGene_gn=LOC111556258;refGene_tx=XM_019830449.2;refGene_gn=LOC111556258;refGene_tx=XM_023253818.1;refGene_gn=LOC111556258;refGene_tx=XM_023253819.1
B2 34818332 34818395

refGene_gn=LOC111556258;refGene_tx=XM_019830449.2;refGene_gn=LOC111556258;refGene_tx=XM_023253818.1;refGene_gn=LOC111556258;refGene_tx=XM_023253819.1
B2 34818640 34818892
refGene_gn=LOC111556258;refGene_tx=XM_019830449.2
B2 34819051 34819129
refGene_gn=LOC111556258;refGene_tx=XM_019830449.2;refGene_gn=LOC111556258;refGene_tx=XM_023253818.1;refGene_gn=LOC111556258;refGene_tx=XM_023253819.1
B2 34820020 34820212
refGene_gn=LOC111556258;refGene_tx=XM_019830449.2;refGene_gn=LOC111556258;refGene_tx=XM_023253818.1;refGene_gn=LOC111556258;refGene_tx=XM_023253819.1;refGene_gn=LOC111556258;refGene_tx=XM_023253820.1
B2 34821649 34821812
refGene_gn=LOC111556258;refGene_tx=XM_019830449.2;refGene_gn=LOC111556258;refGene_tx=XM_023253818.1;refGene_gn=LOC111556258;refGene_tx=XM_023253819.1;refGene_gn=LOC111556258;refGene_tx=XM_023253820.1
B2 34822270 34822481
refGene_gn=LOC111556258;refGene_tx=XM_019830449.2;refGene_gn=LOC111556258;refGene_tx=XM_023253818.1;refGene_gn=LOC111556258;refGene_tx=XM_023253819.1;refGene_gn=LOC111556258;refGene_tx=XM_023253820.1
B2 34822617 34822767
refGene_gn=LOC111556258;refGene_tx=XM_019830449.2;refGene_gn=LOC111556258;refGene_tx=XM_023253818.1;refGene_gn=LOC111556258;refGene_tx=XM_023253819.1;refGene_gn=LOC111556258;refGene_tx=XM_023253820.1
B2 34825354 34825436
refGene_gn=LOC111556258;refGene_tx=XM_019830449.2;refGene_gn=LOC111556258;refGene_tx=XM_023253818.1;refGene_gn=LOC111556258;refGene_tx=XM_023253819.1;refGene_gn=LOC111556258;refGene_tx=XM_023253820.1
B2 34827940 34828088
refGene_gn=LOC111556172;refGene_tx=XM_011282271.3;refGene_gn=LOC111556172;refGene_tx=XM_011282272.3;refGene_gn=LOC111556172;refGene_tx=XM_011282273.3;refGene_gn=LOC111556172;refGene_tx=XM_011282275.3
B2 34828185 34828291
refGene_gn=LOC111556172;refGene_tx=XM_011282271.3;refGene_gn=LOC111556172;refGene_tx=XM_011282272.3;refGene_gn=LOC111556172;refGene_tx=XM_011282273.3;refGene_gn=LOC111556172;refGene_tx=XM_011282275.3
B2 34828581 34828685
refGene_gn=LOC111556172;refGene_tx=XM_011282271.3;refGene_gn=LOC111556172;refGene_tx=XM_011282272.3;refGene_gn=LOC111556172;refGene_tx=XM_011282273.3;refGene_gn=LOC111556172;refGene_tx=XM_011282275.3
B2 34828788 34828921
refGene_gn=LOC111556172;refGene_tx=XM_011282271.3;refGene_gn=LOC111556172;refGene_tx=XM_011282272.3;refGene_gn=LOC111556172;refGene_tx=XM_011282273.3;refGene_gn=LOC111556172;refGene_tx=XM_011282275.3
B2 34829115 34829245
refGene_gn=LOC111556172;refGene_tx=XM_011282271.3;refGene_gn=LOC111556172;refGene_tx=XM_011282272.3;refGene_gn=LOC111556172;refGene_tx=XM_011282273.3;refGene_gn=LOC111556172;refGene_tx=XM_011282275.3
B2 34829646 34829819

refGene_gn=LOC111556172;refGene_tx=XM_011282271.3;refGene_gn=LOC111556172;refGene_tx=XM_011282272.3;refGene_gn=LOC111556172;refGene_tx=XM_011282273.3;refGene_gn=LOC111556172;refGene_tx=XM_011282275.3
B2 34831216 34831396
refGene_gn=LOC111556172;refGene_tx=XM_011282271.3;refGene_gn=LOC111556172;refGene_tx=XM_011282272.3;refGene_gn=LOC111556172;refGene_tx=XM_011282273.3;refGene_gn=LOC111556172;refGene_tx=XM_011282275.3
B2 34831752 34831909
refGene_gn=LOC111556172;refGene_tx=XM_011282271.3;refGene_gn=LOC111556172;refGene_tx=XM_011282272.3;refGene_gn=LOC111556172;refGene_tx=XM_011282273.3;refGene_gn=LOC111556172;refGene_tx=XM_011282275.3
B2 34832261 34832668
refGene_gn=LOC111556172;refGene_tx=XM_011282271.3;refGene_gn=LOC111556172;refGene_tx=XM_011282272.3;refGene_gn=LOC111556172;refGene_tx=XM_011282273.3;refGene_gn=LOC111556172;refGene_tx=XM_011282275.3
B2 34833359 34833591
refGene_gn=LOC111556172;refGene_tx=XM_011282271.3;refGene_gn=LOC111556172;refGene_tx=XM_011282272.3;refGene_gn=LOC111556172;refGene_tx=XM_011282273.3;refGene_gn=LOC111556172;refGene_tx=XM_011282275.3
B2 34834367 34834778
refGene_gn=LOC111556168;refGene_tx=XM_006931616.4;refGene_gn=LOC111556168;refGene_tx=XM_011282276.2
B2 34834868 34835037
refGene_gn=LOC111556168;refGene_tx=XM_006931616.4;refGene_gn=LOC111556168;refGene_tx=XM_011282276.2
B2 34835198 34835252
refGene_gn=LOC111556168;refGene_tx=XM_006931616.4;refGene_gn=LOC111556168;refGene_tx=XM_011282276.2
B2 34835384 34835549
refGene_gn=LOC111556168;refGene_tx=XM_006931616.4;refGene_gn=LOC111556168;refGene_tx=XM_011282276.2
B2 34835694 34835835
refGene_gn=LOC111556168;refGene_tx=XM_006931616.4;refGene_gn=LOC111556168;refGene_tx=XM_011282276.2
B2 34836040 34836237
refGene_gn=LOC111556168;refGene_tx=XM_006931616.4;refGene_gn=LOC111556168;refGene_tx=XM_011282276.2
B2 34836693 34836966
refGene_gn=LOC111556168;refGene_tx=XM_006931616.4;refGene_gn=LOC111556168;refGene_tx=XM_011282276.2
B2 34837819 34837865
refGene_gn=LOC111556166;refGene_tx=XM_003986003.4
B2 34838057 34838275
refGene_gn=LOC111556166;refGene_tx=XM_003986003.4
B2 34838377 34838494
refGene_gn=LOC111556166;refGene_tx=XM_003986003.4
B2 34838604 34838697
refGene_gn=LOC111556166;refGene_tx=XM_003986003.4
B2 34838799 34838885
refGene_gn=LOC111556166;refGene_tx=XM_003986003.4

B2 34838997 34839082
refGene_gn=LOC111556166;refGene_tx=XM_003986003.4
B2 34839289 34839332
refGene_gn=LOC111556166;refGene_tx=XM_003986003.4
B2 34839576 34839651
refGene_gn=LOC111556166;refGene_tx=XM_003986003.4
B2 34839844 34839861
refGene_gn=LOC111556166;refGene_tx=XM_003986003.4
B2 34842046 34842124
refGene_gn=LOC111556167;refGene_tx=XM_011282270.3
B2 34842921 34843082
refGene_gn=LOC111556167;refGene_tx=XM_011282270.3
B2 34843189 34843405
refGene_gn=LOC111556167;refGene_tx=XM_011282270.3
B2 34844116 34844506
refGene_gn=LOC111556167;refGene_tx=XM_011282270.3
B2 34844691 34844965
refGene_gn=LOC111556167;refGene_tx=XM_011282270.3
B2 34845267 34845369
refGene_gn=LOC111556167;refGene_tx=XM_011282270.3
B2 34873456 34873603
refGene_gn=VPS52;refGene_tx=XM_023253821.1;refGene_gn=VPS52;refGene_tx=XM_023253822.1
B2 34873802 34873921
refGene_gn=VPS52;refGene_tx=XM_023253821.1;refGene_gn=VPS52;refGene_tx=XM_023253822.1
B2 34874125 34874237
refGene_gn=VPS52;refGene_tx=XM_023253821.1;refGene_gn=VPS52;refGene_tx=XM_023253822.1
B2 34881864 34881930
refGene_gn=VPS52;refGene_tx=XM_023253821.1;refGene_gn=VPS52;refGene_tx=XM_023253822.1
B2 34882154 34882262
refGene_gn=VPS52;refGene_tx=XM_023253821.1;refGene_gn=VPS52;refGene_tx=XM_023253822.1
B2 34882378 34882474
refGene_gn=VPS52;refGene_tx=XM_023253821.1;refGene_gn=VPS52;refGene_tx=XM_023253822.1
B2 34882742 34882866
refGene_gn=VPS52;refGene_tx=XM_023253821.1;refGene_gn=VPS52;refGene_tx=XM_023253822.1
B2 34883272 34883391
refGene_gn=VPS52;refGene_tx=XM_023253821.1;refGene_gn=VPS52;refGene_tx=XM_023253822.1
B2 34883859 34884015
refGene_gn=VPS52;refGene_tx=XM_023253821.1;refGene_gn=VPS52;refGene_tx=XM_023253822.1
B2 34884490 34884624
refGene_gn=VPS52;refGene_tx=XM_023253821.1;refGene_gn=VPS52;refGene_tx=XM_023253822.1

B2 34884982 34885040
refGene_gn=VPS52;refGene_tx=XM_023253821.1;refGene_gn=VPS52;refGene_tx=XM_023253822.1
B2 34885179 34885312
refGene_gn=VPS52;refGene_tx=XM_023253821.1;refGene_gn=VPS52;refGene_tx=XM_023253822.1
B2 34885423 34885524
refGene_gn=VPS52;refGene_tx=XM_023253821.1;refGene_gn=VPS52;refGene_tx=XM_023253822.1
B2 34885812 34885963
refGene_gn=VPS52;refGene_tx=XM_023253821.1;refGene_gn=VPS52;refGene_tx=XM_023253822.1
B2 34886349 34886525
refGene_gn=VPS52;refGene_tx=XM_023253821.1;refGene_gn=VPS52;refGene_tx=XM_023253822.1
B2 34886825 34886893
refGene_gn=VPS52;refGene_tx=XM_023253821.1;refGene_gn=VPS52;refGene_tx=XM_023253822.1
B2 34887091 34887167
refGene_gn=VPS52;refGene_tx=XM_023253821.1;refGene_gn=VPS52;refGene_tx=XM_023253822.1
B2 34887352 34887405 refGene_gn=VPS52;refGene_tx=XM_023253821.1
B2 34887553 34887638
refGene_gn=VPS52;refGene_tx=XM_023253821.1;refGene_gn=VPS52;refGene_tx=XM_023253822.1
B2 34888507 34888597
refGene_gn=RPS18;refGene_tx=XM_023253824.1;refGene_gn=VPS52;refGene_tx=XM_023253821.1
B2 34889034 34889037 refGene_gn=RPS18;refGene_tx=XM_023253823.1
B2 34889507 34889606
refGene_gn=RPS18;refGene_tx=XM_023253823.1;refGene_gn=RPS18;refGene_tx=XM_023253824.1
B2 34893396 34893483
refGene_gn=RPS18;refGene_tx=XM_023253823.1;refGene_gn=RPS18;refGene_tx=XM_023253824.1
B2 34893567 34893669
refGene_gn=RPS18;refGene_tx=XM_023253823.1;refGene_gn=RPS18;refGene_tx=XM_023253824.1
B2 34893767 34893859
refGene_gn=RPS18;refGene_tx=XM_023253823.1;refGene_gn=RPS18;refGene_tx=XM_023253824.1
B2 34893981 34894057
refGene_gn=RPS18;refGene_tx=XM_023253823.1;refGene_gn=RPS18;refGene_tx=XM_023253824.1
B2 34895056 34896208
refGene_gn=B3GALT4;refGene_tx=XM_003986008.5;refGene_gn=B3GALT4;refGene_tx=XM_023253830.1
B2 34896917 34897019
refGene_gn=WDR46;refGene_tx=XM_023253826.1;refGene_gn=WDR46;refGene_tx=XM_023253827.1

B2 34897113 34897227
refGene_gn=WDR46;refGene_tx=XM_023253826.1;refGene_gn=WDR46;refGene_tx=XM_023253827.1
B2 34897375 34897471
refGene_gn=WDR46;refGene_tx=XM_023253826.1;refGene_gn=WDR46;refGene_tx=XM_023253827.1
B2 34897644 34897739
refGene_gn=WDR46;refGene_tx=XM_023253826.1;refGene_gn=WDR46;refGene_tx=XM_023253827.1
B2 34897883 34898197
refGene_gn=WDR46;refGene_tx=XM_023253826.1;refGene_gn=WDR46;refGene_tx=XM_023253827.1
B2 34902121 34902221
refGene_gn=WDR46;refGene_tx=XM_023253826.1;refGene_gn=WDR46;refGene_tx=XM_023253827.1
B2 34902414 34902550
refGene_gn=WDR46;refGene_tx=XM_023253826.1;refGene_gn=WDR46;refGene_tx=XM_023253827.1
B2 34902672 34902823
refGene_gn=WDR46;refGene_tx=XM_023253826.1;refGene_gn=WDR46;refGene_tx=XM_023253827.1
B2 34902912 34903017
refGene_gn=WDR46;refGene_tx=XM_023253826.1;refGene_gn=WDR46;refGene_tx=XM_023253827.1
B2 34903286 34903348
refGene_gn=WDR46;refGene_tx=XM_023253826.1;refGene_gn=WDR46;refGene_tx=XM_023253827.1
B2 34903525 34903613
refGene_gn=WDR46;refGene_tx=XM_023253826.1;refGene_gn=WDR46;refGene_tx=XM_023253827.1
B2 34903732 34903845
refGene_gn=WDR46;refGene_tx=XM_023253826.1;refGene_gn=WDR46;refGene_tx=XM_023253827.1
B2 34903985 34904066
refGene_gn=WDR46;refGene_tx=XM_023253826.1;refGene_gn=WDR46;refGene_tx=XM_023253827.1
B2 34904175 34904388
refGene_gn=WDR46;refGene_tx=XM_023253826.1;refGene_gn=WDR46;refGene_tx=XM_023253827.1
B2 34904493 34904562
refGene_gn=WDR46;refGene_tx=XM_023253826.1;refGene_gn=WDR46;refGene_tx=XM_023253827.1
B2 34905320 34905384
refGene_gn=PFDN6;refGene_tx=XM_023253831.1;refGene_gn=PFDN6;refGene_tx=XM_023253832.1;refGene_gn=PFDN6;refGene_tx=XM_023253833.1;refGene_gn=PFDN6;refGene_tx=XM_023253834.1
B2 34905630 34905701
refGene_gn=PFDN6;refGene_tx=XM_023253831.1;refGene_gn=PFDN6;refGene_tx=XM_023253832.1;refGene_gn=PFDN6;refGene_tx=XM_023253833.1;refGene_gn=PFDN6;refGene_tx=XM_023253834.1

B2 34905798 34905923
refGene_gn=PFDN6;refGene_tx=XM_023253831.1;refGene_gn=PFDN6;refGene_tx
=XM_023253832.1;refGene_gn=PFDN6;refGene_tx=XM_023253833.1;refGene_gn=
PFDN6;refGene_tx=XM_023253834.1
B2 34906158 34906288
refGene_gn=PFDN6;refGene_tx=XM_023253831.1;refGene_gn=PFDN6;refGene_tx
=XM_023253832.1;refGene_gn=PFDN6;refGene_tx=XM_023253833.1;refGene_gn=
PFDN6;refGene_tx=XM_023253834.1
B2 34907692 34907904 refGene_gn=RGL2;refGene_tx=XM_003986014.5
B2 34908033 34908148 refGene_gn=RGL2;refGene_tx=XM_003986014.5
B2 34908820 34909111 refGene_gn=RGL2;refGene_tx=XM_003986014.5
B2 34909221 34909333 refGene_gn=RGL2;refGene_tx=XM_003986014.5
B2 34909430 34909526 refGene_gn=RGL2;refGene_tx=XM_003986014.5
B2 34909623 34909745 refGene_gn=RGL2;refGene_tx=XM_003986014.5
B2 34909867 34909900 refGene_gn=RGL2;refGene_tx=XM_003986014.5
B2 34910488 34910563 refGene_gn=RGL2;refGene_tx=XM_003986014.5
B2 34910759 34910828 refGene_gn=RGL2;refGene_tx=XM_003986014.5
B2 34910931 34911016 refGene_gn=RGL2;refGene_tx=XM_003986014.5
B2 34911113 34911217 refGene_gn=RGL2;refGene_tx=XM_003986014.5
B2 34911302 34911554 refGene_gn=RGL2;refGene_tx=XM_003986014.5
B2 34911828 34912126 refGene_gn=RGL2;refGene_tx=XM_003986014.5
B2 34912214 34912265 refGene_gn=RGL2;refGene_tx=XM_003986014.5
B2 34912383 34912562 refGene_gn=RGL2;refGene_tx=XM_003986014.5
B2 34912820 34912904 refGene_gn=RGL2;refGene_tx=XM_003986014.5
B2 34914196 34914235 refGene_gn=RGL2;refGene_tx=XM_003986014.5
B2 34914236 34914356 refGene_gn=RGL2;refGene_tx=XM_003986014.5
B2 34916513 34916572
refGene_gn=TAPBP;refGene_tx=XM_023253828.1;refGene_gn=TAPBP;refGene_tx
=XM_023253829.1
B2 34917577 34917615 refGene_gn=TAPBP;refGene_tx=XM_023253829.1
B2 34917773 34917863
refGene_gn=TAPBP;refGene_tx=XM_023253828.1;refGene_gn=TAPBP;refGene_tx
=XM_023253829.1
B2 34917991 34918333
refGene_gn=TAPBP;refGene_tx=XM_023253828.1;refGene_gn=TAPBP;refGene_tx
=XM_023253829.1
B2 34918705 34919104
refGene_gn=TAPBP;refGene_tx=XM_023253828.1;refGene_gn=TAPBP;refGene_tx
=XM_023253829.1
B2 34926577 34926838
refGene_gn=TAPBP;refGene_tx=XM_023253828.1;refGene_gn=TAPBP;refGene_tx
=XM_023253829.1
B2 34927065 34927236
refGene_gn=TAPBP;refGene_tx=XM_023253828.1;refGene_gn=TAPBP;refGene_tx
=XM_023253829.1
B2 34927374 34927411
refGene_gn=TAPBP;refGene_tx=XM_023253828.1;refGene_gn=TAPBP;refGene_tx
=XM_023253829.1
B2 34928369 34930277 refGene_gn=ZBTB22;refGene_tx=XM_023253825.1
B2 34932029 34932089

refGene_gn=DAXX;refGene_tx=XM_023253835.1;refGene_gn=DAXX;refGene_tx=XM_023253836.1;refGene_gn=DAXX;refGene_tx=XM_023253837.1
B2 34932294 34932517
refGene_gn=DAXX;refGene_tx=XM_023253835.1;refGene_gn=DAXX;refGene_tx=XM_023253836.1;refGene_gn=DAXX;refGene_tx=XM_023253837.1
B2 34932675 34933153
refGene_gn=DAXX;refGene_tx=XM_023253835.1;refGene_gn=DAXX;refGene_tx=XM_023253836.1;refGene_gn=DAXX;refGene_tx=XM_023253837.1
B2 34933309 34933541
refGene_gn=DAXX;refGene_tx=XM_023253835.1;refGene_gn=DAXX;refGene_tx=XM_023253836.1;refGene_gn=DAXX;refGene_tx=XM_023253837.1
B2 34933698 34933913
refGene_gn=DAXX;refGene_tx=XM_023253835.1;refGene_gn=DAXX;refGene_tx=XM_023253836.1;refGene_gn=DAXX;refGene_tx=XM_023253837.1
B2 34934038 34934870
refGene_gn=DAXX;refGene_tx=XM_023253835.1;refGene_gn=DAXX;refGene_tx=XM_023253836.1;refGene_gn=DAXX;refGene_tx=XM_023253837.1
B2 34935023 34935230
refGene_gn=DAXX;refGene_tx=XM_023253835.1;refGene_gn=DAXX;refGene_tx=XM_023253836.1;refGene_gn=DAXX;refGene_tx=XM_023253837.1
B2 34996119 34996131 refGene_gn=KIFC1;refGene_tx=XM_019830459.2
B2 35002459 35002597
refGene_gn=KIFC1;refGene_tx=XM_019830459.2;refGene_gn=KIFC1;refGene_tx=XM_019830460.2
B2 35002719 35002819
refGene_gn=KIFC1;refGene_tx=XM_019830459.2;refGene_gn=KIFC1;refGene_tx=XM_019830460.2
B2 35003330 35003384
refGene_gn=KIFC1;refGene_tx=XM_019830459.2;refGene_gn=KIFC1;refGene_tx=XM_019830460.2
B2 35003494 35003545
refGene_gn=KIFC1;refGene_tx=XM_019830459.2;refGene_gn=KIFC1;refGene_tx=XM_019830460.2
B2 35003748 35004143
refGene_gn=KIFC1;refGene_tx=XM_019830459.2;refGene_gn=KIFC1;refGene_tx=XM_019830460.2
B2 35005854 35006634
refGene_gn=KIFC1;refGene_tx=XM_019830459.2;refGene_gn=KIFC1;refGene_tx=XM_019830460.2
B2 35007282 35007573
refGene_gn=KIFC1;refGene_tx=XM_019830459.2;refGene_gn=KIFC1;refGene_tx=XM_019830460.2
B2 35007673 35007747
refGene_gn=KIFC1;refGene_tx=XM_019830459.2;refGene_gn=KIFC1;refGene_tx=XM_019830460.2
B2 35008015 35008091
refGene_gn=KIFC1;refGene_tx=XM_019830459.2;refGene_gn=KIFC1;refGene_tx=XM_019830460.2
B2 35010969 35011014
refGene_gn=KIFC1;refGene_tx=XM_019830459.2;refGene_gn=KIFC1;refGene_tx

=XM_019830460.2
B2 35013177 35013336
refGene_gn=PHF1;refGene_tx=XM_003986021.5;refGene_gn=PHF1;refGene_tx=XM_006931622.4;refGene_gn=PHF1;refGene_tx=XM_011282267.3;refGene_gn=PHF1;refGene_tx=XM_011282268.3;refGene_gn=PHF1;refGene_tx=XM_019830463.2
B2 35013430 35013512
refGene_gn=PHF1;refGene_tx=XM_003986021.5;refGene_gn=PHF1;refGene_tx=XM_006931622.4;refGene_gn=PHF1;refGene_tx=XM_011282267.3;refGene_gn=PHF1;refGene_tx=XM_011282268.3;refGene_gn=PHF1;refGene_tx=XM_019830463.2
B2 35013621 35013717
refGene_gn=PHF1;refGene_tx=XM_003986021.5;refGene_gn=PHF1;refGene_tx=XM_006931622.4;refGene_gn=PHF1;refGene_tx=XM_011282267.3;refGene_gn=PHF1;refGene_tx=XM_011282268.3;refGene_gn=PHF1;refGene_tx=XM_019830463.2
B2 35014136 35014237
refGene_gn=PHF1;refGene_tx=XM_003986021.5;refGene_gn=PHF1;refGene_tx=XM_006931622.4;refGene_gn=PHF1;refGene_tx=XM_011282267.3;refGene_gn=PHF1;refGene_tx=XM_011282268.3;refGene_gn=PHF1;refGene_tx=XM_019830463.2
B2 35014323 35014472
refGene_gn=PHF1;refGene_tx=XM_003986021.5;refGene_gn=PHF1;refGene_tx=XM_006931622.4;refGene_gn=PHF1;refGene_tx=XM_011282267.3;refGene_gn=PHF1;refGene_tx=XM_011282268.3;refGene_gn=PHF1;refGene_tx=XM_019830463.2
B2 35014661 35014757
refGene_gn=PHF1;refGene_tx=XM_003986021.5;refGene_gn=PHF1;refGene_tx=XM_006931622.4;refGene_gn=PHF1;refGene_tx=XM_011282267.3;refGene_gn=PHF1;refGene_tx=XM_011282268.3;refGene_gn=PHF1;refGene_tx=XM_019830463.2
B2 35014967 35015036
refGene_gn=PHF1;refGene_tx=XM_003986021.5;refGene_gn=PHF1;refGene_tx=XM_006931622.4;refGene_gn=PHF1;refGene_tx=XM_011282267.3;refGene_gn=PHF1;refGene_tx=XM_011282268.3;refGene_gn=PHF1;refGene_tx=XM_019830463.2
B2 35015172 35015296
refGene_gn=PHF1;refGene_tx=XM_003986021.5;refGene_gn=PHF1;refGene_tx=XM_006931622.4;refGene_gn=PHF1;refGene_tx=XM_011282267.3;refGene_gn=PHF1;refGene_tx=XM_011282268.3;refGene_gn=PHF1;refGene_tx=XM_019830463.2
B2 35015405 35015473
refGene_gn=PHF1;refGene_tx=XM_003986021.5;refGene_gn=PHF1;refGene_tx=XM_006931622.4;refGene_gn=PHF1;refGene_tx=XM_011282267.3;refGene_gn=PHF1;refGene_tx=XM_011282268.3;refGene_gn=PHF1;refGene_tx=XM_019830463.2
B2 35015638 35015743
refGene_gn=PHF1;refGene_tx=XM_003986021.5;refGene_gn=PHF1;refGene_tx=XM_006931622.4;refGene_gn=PHF1;refGene_tx=XM_011282267.3;refGene_gn=PHF1;refGene_tx=XM_011282268.3;refGene_gn=PHF1;refGene_tx=XM_019830463.2
B2 35015867 35016057
refGene_gn=PHF1;refGene_tx=XM_003986021.5;refGene_gn=PHF1;refGene_tx=XM_006931622.4;refGene_gn=PHF1;refGene_tx=XM_011282267.3
B2 35016145 35016240
refGene_gn=PHF1;refGene_tx=XM_003986021.5;refGene_gn=PHF1;refGene_tx=XM_006931622.4;refGene_gn=PHF1;refGene_tx=XM_011282267.3;refGene_gn=PHF1;refGene_tx=XM_011282268.3;refGene_gn=PHF1;refGene_tx=XM_019830463.2
B2 35016509 35016590
refGene_gn=PHF1;refGene_tx=XM_003986021.5;refGene_gn=PHF1;refGene_tx=X

M_011282267.3;refGene_gn=PHF1;refGene_tx=XM_011282268.3;refGene_gn=PHF1;refGene_tx=XM_019830463.2
B2 35016751 35017040
refGene_gn=PHF1;refGene_tx=XM_003986021.5;refGene_gn=PHF1;refGene_tx=XM_006931622.4;refGene_gn=PHF1;refGene_tx=XM_011282267.3;refGene_gn=PHF1;refGene_tx=XM_011282268.3;refGene_gn=PHF1;refGene_tx=XM_019830463.2
B2 35017594 35017721
refGene_gn=CUTA;refGene_tx=XM_003986023.4;refGene_gn=CUTA;refGene_tx=XM_006931624.2;refGene_gn=CUTA;refGene_tx=XM_006931625.3
B2 35017851 35017901
refGene_gn=CUTA;refGene_tx=XM_003986023.4;refGene_gn=CUTA;refGene_tx=XM_006931624.2;refGene_gn=CUTA;refGene_tx=XM_006931625.3
B2 35018008 35018054
refGene_gn=CUTA;refGene_tx=XM_003986023.4;refGene_gn=CUTA;refGene_tx=XM_006931624.2;refGene_gn=CUTA;refGene_tx=XM_006931625.3
B2 35018165 35018225
refGene_gn=CUTA;refGene_tx=XM_003986023.4;refGene_gn=CUTA;refGene_tx=XM_006931624.2;refGene_gn=CUTA;refGene_tx=XM_006931625.3
B2 35018406 35018615
refGene_gn=CUTA;refGene_tx=XM_003986023.4;refGene_gn=CUTA;refGene_tx=XM_006931624.2;refGene_gn=CUTA;refGene_tx=XM_006931625.3
B2 35018877 35018919 refGene_gn=CUTA;refGene_tx=XM_003986023.4
B2 35019013 35019079 refGene_gn=CUTA;refGene_tx=XM_006931625.3
B2 35021282 35021349
refGene_gn=SYNGAP1;refGene_tx=XM_019830461.2;refGene_gn=SYNGAP1;refGene_tx=XM_023253838.1;refGene_gn=SYNGAP1;refGene_tx=XM_023253839.1;refGene_gn=SYNGAP1;refGene_tx=XM_023253840.1;refGene_gn=SYNGAP1;refGene_tx=XM_023253841.1;refGene_gn=SYNGAP1;refGene_tx=XM_023253842.1;refGene_gn=SYNGAP1;refGene_tx=XM_023253845.1;refGene_gn=SYNGAP1;refGene_tx=XM_023253846.1;refGene_gn=SYNGAP1;refGene_tx=XM_023253847.1;refGene_gn=SYNGAP1;refGene_tx=XM_023253848.1;refGene_gn=SYNGAP1;refGene_tx=XM_023253849.1;refGene_gn=SYNGAP1;refGene_tx=XM_023253850.1
B2 35024776 35024898
refGene_gn=SYNGAP1;refGene_tx=XM_019830461.2;refGene_gn=SYNGAP1;refGene_tx=XM_023253838.1;refGene_gn=SYNGAP1;refGene_tx=XM_023253839.1;refGene_gn=SYNGAP1;refGene_tx=XM_023253840.1;refGene_gn=SYNGAP1;refGene_tx=XM_023253841.1;refGene_gn=SYNGAP1;refGene_tx=XM_023253842.1;refGene_gn=SYNGAP1;refGene_tx=XM_023253843.1;refGene_gn=SYNGAP1;refGene_tx=XM_023253844.1;refGene_gn=SYNGAP1;refGene_tx=XM_023253845.1;refGene_gn=SYNGAP1;refGene_tx=XM_023253846.1;refGene_gn=SYNGAP1;refGene_tx=XM_023253847.1;refGene_gn=SYNGAP1;refGene_tx=XM_023253848.1;refGene_gn=SYNGAP1;refGene_tx=XM_023253849.1;refGene_gn=SYNGAP1;refGene_tx=XM_023253850.1
B2 35027052 35027158
refGene_gn=SYNGAP1;refGene_tx=XM_019830461.2;refGene_gn=SYNGAP1;refGene_tx=XM_023253838.1;refGene_gn=SYNGAP1;refGene_tx=XM_023253839.1;refGene_gn=SYNGAP1;refGene_tx=XM_023253840.1;refGene_gn=SYNGAP1;refGene_tx=XM_023253841.1;refGene_gn=SYNGAP1;refGene_tx=XM_023253842.1;refGene_gn=SYNGAP1;refGene_tx=XM_023253843.1;refGene_gn=SYNGAP1;refGene_tx=XM_023253844.1;refGene_gn=SYNGAP1;refGene_tx=XM_023253845.1;refGene_gn=SYNGAP1;refGene_tx=XM_023253846.1;refGene_gn=SYNGAP1;refGene_tx=XM_0232538

47.1;refGene_gn=SYNGAP1;refGene_tx=XM_023253848.1;refGene_gn=SYNGAP1;refGene_tx=XM_023253849.1;refGene_gn=SYNGAP1;refGene_tx=XM_023253850.1
B2 35029594 35029631 refGene_gn=SYNGAP1;refGene_tx=XM_023253852.1
B2 35033187 35033305 refGene_gn=SYNGAP1;refGene_tx=XM_023253851.1
B2 35033427 35033519
refGene_gn=SYNGAP1;refGene_tx=XM_019830461.2;refGene_gn=SYNGAP1;refGene_tx=XM_023253838.1;refGene_gn=SYNGAP1;refGene_tx=XM_023253839.1;refGene_gn=SYNGAP1;refGene_tx=XM_023253840.1;refGene_gn=SYNGAP1;refGene_tx=XM_023253841.1;refGene_gn=SYNGAP1;refGene_tx=XM_023253842.1;refGene_gn=SYNGAP1;refGene_tx=XM_023253843.1;refGene_gn=SYNGAP1;refGene_tx=XM_023253844.1;refGene_gn=SYNGAP1;refGene_tx=XM_023253845.1;refGene_gn=SYNGAP1;refGene_tx=XM_023253846.1;refGene_gn=SYNGAP1;refGene_tx=XM_023253847.1;refGene_gn=SYNGAP1;refGene_tx=XM_023253848.1;refGene_gn=SYNGAP1;refGene_tx=XM_023253849.1;refGene_gn=SYNGAP1;refGene_tx=XM_023253850.1;refGene_gn=SYNGAP1;refGene_tx=XM_023253851.1;refGene_gn=SYNGAP1;refGene_tx=XM_023253852.1;refGene_gn=SYNGAP1;refGene_tx=XM_023253853.1;refGene_gn=SYNGAP1;refGene_tx=XM_023253854.1;refGene_gn=SYNGAP1;refGene_tx=XM_023253855.1
B2 35033978 35034100
refGene_gn=SYNGAP1;refGene_tx=XM_019830461.2;refGene_gn=SYNGAP1;refGene_tx=XM_023253838.1;refGene_gn=SYNGAP1;refGene_tx=XM_023253839.1;refGene_gn=SYNGAP1;refGene_tx=XM_023253840.1;refGene_gn=SYNGAP1;refGene_tx=XM_023253841.1;refGene_gn=SYNGAP1;refGene_tx=XM_023253842.1;refGene_gn=SYNGAP1;refGene_tx=XM_023253843.1;refGene_gn=SYNGAP1;refGene_tx=XM_023253844.1;refGene_gn=SYNGAP1;refGene_tx=XM_023253845.1;refGene_gn=SYNGAP1;refGene_tx=XM_023253846.1;refGene_gn=SYNGAP1;refGene_tx=XM_023253847.1;refGene_gn=SYNGAP1;refGene_tx=XM_023253848.1;refGene_gn=SYNGAP1;refGene_tx=XM_023253849.1;refGene_gn=SYNGAP1;refGene_tx=XM_023253850.1;refGene_gn=SYNGAP1;refGene_tx=XM_023253851.1;refGene_gn=SYNGAP1;refGene_tx=XM_023253852.1;refGene_gn=SYNGAP1;refGene_tx=XM_023253853.1;refGene_gn=SYNGAP1;refGene_tx=XM_023253854.1;refGene_gn=SYNGAP1;refGene_tx=XM_023253855.1
B2 35035882 35036036
refGene_gn=SYNGAP1;refGene_tx=XM_019830461.2;refGene_gn=SYNGAP1;refGene_tx=XM_023253838.1;refGene_gn=SYNGAP1;refGene_tx=XM_023253839.1;refGene_gn=SYNGAP1;refGene_tx=XM_023253840.1;refGene_gn=SYNGAP1;refGene_tx=XM_023253841.1;refGene_gn=SYNGAP1;refGene_tx=XM_023253842.1;refGene_gn=SYNGAP1;refGene_tx=XM_023253843.1;refGene_gn=SYNGAP1;refGene_tx=XM_023253844.1;refGene_gn=SYNGAP1;refGene_tx=XM_023253845.1;refGene_gn=SYNGAP1;refGene_tx=XM_023253846.1;refGene_gn=SYNGAP1;refGene_tx=XM_023253847.1;refGene_gn=SYNGAP1;refGene_tx=XM_023253848.1;refGene_gn=SYNGAP1;refGene_tx=XM_023253849.1;refGene_gn=SYNGAP1;refGene_tx=XM_023253850.1;refGene_gn=SYNGAP1;refGene_tx=XM_023253851.1;refGene_gn=SYNGAP1;refGene_tx=XM_023253852.1;refGene_gn=SYNGAP1;refGene_tx=XM_023253853.1;refGene_gn=SYNGAP1;refGene_tx=XM_023253854.1;refGene_gn=SYNGAP1;refGene_tx=XM_023253855.1
B2 35036252 35036351
refGene_gn=SYNGAP1;refGene_tx=XM_019830461.2;refGene_gn=SYNGAP1;refGene_tx=XM_023253838.1;refGene_gn=SYNGAP1;refGene_tx=XM_023253839.1;refGene_gn=SYNGAP1;refGene_tx=XM_023253840.1;refGene_gn=SYNGAP1;refGene_tx=

XM_023253841.1;refGene_gn=SYNGAP1;refGene_tx=XM_023253842.1;refGene_gn=SYNGAP1;refGene_tx=XM_023253843.1;refGene_gn=SYNGAP1;refGene_tx=XM_023253844.1;refGene_gn=SYNGAP1;refGene_tx=XM_023253845.1;refGene_gn=SYNGAP1;refGene_tx=XM_023253846.1;refGene_gn=SYNGAP1;refGene_tx=XM_023253847.1;refGene_gn=SYNGAP1;refGene_tx=XM_023253848.1;refGene_gn=SYNGAP1;refGene_tx=XM_023253849.1;refGene_gn=SYNGAP1;refGene_tx=XM_023253850.1;refGene_gn=SYNGAP1;refGene_tx=XM_023253851.1;refGene_gn=SYNGAP1;refGene_tx=XM_023253852.1;refGene_gn=SYNGAP1;refGene_tx=XM_023253853.1;refGene_gn=SYNGAP1;refGene_tx=XM_023253854.1;refGene_gn=SYNGAP1;refGene_tx=XM_023253855.1
B2 35037895 35038522
refGene_gn=SYNGAP1;refGene_tx=XM_019830461.2;refGene_gn=SYNGAP1;refGene_tx=XM_023253838.1;refGene_gn=SYNGAP1;refGene_tx=XM_023253839.1;refGene_gn=SYNGAP1;refGene_tx=XM_023253840.1;refGene_gn=SYNGAP1;refGene_tx=XM_023253841.1;refGene_gn=SYNGAP1;refGene_tx=XM_023253842.1;refGene_gn=SYNGAP1;refGene_tx=XM_023253843.1;refGene_gn=SYNGAP1;refGene_tx=XM_023253844.1;refGene_gn=SYNGAP1;refGene_tx=XM_023253845.1;refGene_gn=SYNGAP1;refGene_tx=XM_023253846.1;refGene_gn=SYNGAP1;refGene_tx=XM_023253847.1;refGene_gn=SYNGAP1;refGene_tx=XM_023253848.1;refGene_gn=SYNGAP1;refGene_tx=XM_023253849.1;refGene_gn=SYNGAP1;refGene_tx=XM_023253850.1;refGene_gn=SYNGAP1;refGene_tx=XM_023253851.1;refGene_gn=SYNGAP1;refGene_tx=XM_023253852.1;refGene_gn=SYNGAP1;refGene_tx=XM_023253853.1;refGene_gn=SYNGAP1;refGene_tx=XM_023253854.1;refGene_gn=SYNGAP1;refGene_tx=XM_023253855.1
B2 35038664 35038809
refGene_gn=SYNGAP1;refGene_tx=XM_019830461.2;refGene_gn=SYNGAP1;refGene_tx=XM_023253838.1;refGene_gn=SYNGAP1;refGene_tx=XM_023253839.1;refGene_gn=SYNGAP1;refGene_tx=XM_023253840.1;refGene_gn=SYNGAP1;refGene_tx=XM_023253841.1;refGene_gn=SYNGAP1;refGene_tx=XM_023253842.1;refGene_gn=SYNGAP1;refGene_tx=XM_023253843.1;refGene_gn=SYNGAP1;refGene_tx=XM_023253844.1;refGene_gn=SYNGAP1;refGene_tx=XM_023253845.1;refGene_gn=SYNGAP1;refGene_tx=XM_023253846.1;refGene_gn=SYNGAP1;refGene_tx=XM_023253847.1;refGene_gn=SYNGAP1;refGene_tx=XM_023253848.1;refGene_gn=SYNGAP1;refGene_tx=XM_023253849.1;refGene_gn=SYNGAP1;refGene_tx=XM_023253850.1;refGene_gn=SYNGAP1;refGene_tx=XM_023253851.1;refGene_gn=SYNGAP1;refGene_tx=XM_023253852.1;refGene_gn=SYNGAP1;refGene_tx=XM_023253853.1;refGene_gn=SYNGAP1;refGene_tx=XM_023253854.1;refGene_gn=SYNGAP1;refGene_tx=XM_023253855.1
B2 35039036 35039181
refGene_gn=SYNGAP1;refGene_tx=XM_019830461.2;refGene_gn=SYNGAP1;refGene_tx=XM_023253838.1;refGene_gn=SYNGAP1;refGene_tx=XM_023253839.1;refGene_gn=SYNGAP1;refGene_tx=XM_023253840.1;refGene_gn=SYNGAP1;refGene_tx=XM_023253841.1;refGene_gn=SYNGAP1;refGene_tx=XM_023253842.1;refGene_gn=SYNGAP1;refGene_tx=XM_023253843.1;refGene_gn=SYNGAP1;refGene_tx=XM_023253844.1;refGene_gn=SYNGAP1;refGene_tx=XM_023253845.1;refGene_gn=SYNGAP1;refGene_tx=XM_023253846.1;refGene_gn=SYNGAP1;refGene_tx=XM_023253847.1;refGene_gn=SYNGAP1;refGene_tx=XM_023253848.1;refGene_gn=SYNGAP1;refGene_tx=XM_023253849.1;refGene_gn=SYNGAP1;refGene_tx=XM_023253850.1;refGene_gn=SYNGAP1;refGene_tx=XM_023253851.1;refGene_gn=SYNGAP1;refGene_tx=XM_023253852.1;refGene_gn=SYNGAP1;refGene_tx=XM_023253853.1;refGe

ne_gn=SYNGAP1;refGene_tx=XM_023253854.1;refGene_gn=SYNGAP1;refGene_tx=
XM_023253855.1
B2 35040717 35040954
refGene_gn=SYNGAP1;refGene_tx=XM_019830461.2;refGene_gn=SYNGAP1;refGen
e_tx=XM_023253838.1;refGene_gn=SYNGAP1;refGene_tx=XM_023253839.1;refGe
ne_gn=SYNGAP1;refGene_tx=XM_023253840.1;refGene_gn=SYNGAP1;refGene_tx=
XM_023253841.1;refGene_gn=SYNGAP1;refGene_tx=XM_023253842.1;refGene_gn
=SYNGAP1;refGene_tx=XM_023253843.1;refGene_gn=SYNGAP1;refGene_tx=XM_02
3253844.1;refGene_gn=SYNGAP1;refGene_tx=XM_023253845.1;refGene_gn=SYNG
AP1;refGene_tx=XM_023253846.1;refGene_gn=SYNGAP1;refGene_tx=XM_0232538
47.1;refGene_gn=SYNGAP1;refGene_tx=XM_023253848.1;refGene_gn=SYNGAP1;r
efGene_tx=XM_023253849.1;refGene_gn=SYNGAP1;refGene_tx=XM_023253850.1;
refGene_gn=SYNGAP1;refGene_tx=XM_023253851.1;refGene_gn=SYNGAP1;refGen
e_tx=XM_023253852.1;refGene_gn=SYNGAP1;refGene_tx=XM_023253853.1;refGe
ne_gn=SYNGAP1;refGene_tx=XM_023253854.1;refGene_gn=SYNGAP1;refGene_tx=
XM_023253855.1
B2 35041094 35041296
refGene_gn=SYNGAP1;refGene_tx=XM_019830461.2;refGene_gn=SYNGAP1;refGen
e_tx=XM_023253838.1;refGene_gn=SYNGAP1;refGene_tx=XM_023253839.1;refGe
ne_gn=SYNGAP1;refGene_tx=XM_023253840.1;refGene_gn=SYNGAP1;refGene_tx=
XM_023253841.1;refGene_gn=SYNGAP1;refGene_tx=XM_023253842.1;refGene_gn
=SYNGAP1;refGene_tx=XM_023253843.1;refGene_gn=SYNGAP1;refGene_tx=XM_02
3253844.1;refGene_gn=SYNGAP1;refGene_tx=XM_023253845.1;refGene_gn=SYNG
AP1;refGene_tx=XM_023253846.1;refGene_gn=SYNGAP1;refGene_tx=XM_0232538
47.1;refGene_gn=SYNGAP1;refGene_tx=XM_023253848.1;refGene_gn=SYNGAP1;r
efGene_tx=XM_023253849.1;refGene_gn=SYNGAP1;refGene_tx=XM_023253850.1;
refGene_gn=SYNGAP1;refGene_tx=XM_023253851.1;refGene_gn=SYNGAP1;refGen
e_tx=XM_023253852.1;refGene_gn=SYNGAP1;refGene_tx=XM_023253853.1;refGe
ne_gn=SYNGAP1;refGene_tx=XM_023253854.1;refGene_gn=SYNGAP1;refGene_tx=
XM_023253855.1
B2 35041525 35041704
refGene_gn=SYNGAP1;refGene_tx=XM_019830461.2;refGene_gn=SYNGAP1;refGen
e_tx=XM_023253838.1;refGene_gn=SYNGAP1;refGene_tx=XM_023253839.1;refGe
ne_gn=SYNGAP1;refGene_tx=XM_023253840.1;refGene_gn=SYNGAP1;refGene_tx=
XM_023253841.1;refGene_gn=SYNGAP1;refGene_tx=XM_023253842.1;refGene_gn
=SYNGAP1;refGene_tx=XM_023253843.1;refGene_gn=SYNGAP1;refGene_tx=XM_02
3253844.1;refGene_gn=SYNGAP1;refGene_tx=XM_023253845.1;refGene_gn=SYNG
AP1;refGene_tx=XM_023253846.1;refGene_gn=SYNGAP1;refGene_tx=XM_0232538
47.1;refGene_gn=SYNGAP1;refGene_tx=XM_023253848.1;refGene_gn=SYNGAP1;r
efGene_tx=XM_023253849.1;refGene_gn=SYNGAP1;refGene_tx=XM_023253850.1;
refGene_gn=SYNGAP1;refGene_tx=XM_023253851.1;refGene_gn=SYNGAP1;refGen
e_tx=XM_023253852.1;refGene_gn=SYNGAP1;refGene_tx=XM_023253853.1;refGe
ne_gn=SYNGAP1;refGene_tx=XM_023253854.1;refGene_gn=SYNGAP1;refGene_tx=
XM_023253855.1
B2 35042407 35042449
refGene_gn=SYNGAP1;refGene_tx=XM_019830461.2;refGene_gn=SYNGAP1;refGen
e_tx=XM_023253838.1;refGene_gn=SYNGAP1;refGene_tx=XM_023253839.1;refGe
ne_gn=SYNGAP1;refGene_tx=XM_023253841.1;refGene_gn=SYNGAP1;refGene_tx=
XM_023253843.1;refGene_gn=SYNGAP1;refGene_tx=XM_023253844.1;refGene_gn
=SYNGAP1;refGene_tx=XM_023253845.1;refGene_gn=SYNGAP1;refGene_tx=XM_02

3253846.1;refGene_gn=SYNGAP1;refGene_tx=XM_023253847.1;refGene_gn=SYNGAP1;refGene_tx=XM_023253848.1;refGene_gn=SYNGAP1;refGene_tx=XM_023253849.1;refGene_gn=SYNGAP1;refGene_tx=XM_023253850.1;refGene_gn=SYNGAP1;refGene_tx=XM_023253851.1;refGene_gn=SYNGAP1;refGene_tx=XM_023253852.1;refGene_gn=SYNGAP1;refGene_tx=XM_023253853.1;refGene_gn=SYNGAP1;refGene_tx=XM_023253854.1;refGene_gn=SYNGAP1;refGene_tx=XM_023253855.1
B2 35042842 35043917
refGene_gn=SYNGAP1;refGene_tx=XM_019830461.2;refGene_gn=SYNGAP1;refGene_tx=XM_023253838.1;refGene_gn=SYNGAP1;refGene_tx=XM_023253839.1;refGene_gn=SYNGAP1;refGene_tx=XM_023253840.1;refGene_gn=SYNGAP1;refGene_tx=XM_023253841.1;refGene_gn=SYNGAP1;refGene_tx=XM_023253842.1;refGene_gn=SYNGAP1;refGene_tx=XM_023253843.1;refGene_gn=SYNGAP1;refGene_tx=XM_023253844.1;refGene_gn=SYNGAP1;refGene_tx=XM_023253845.1;refGene_gn=SYNGAP1;refGene_tx=XM_023253846.1;refGene_gn=SYNGAP1;refGene_tx=XM_023253847.1;refGene_gn=SYNGAP1;refGene_tx=XM_023253848.1;refGene_gn=SYNGAP1;refGene_tx=XM_023253849.1;refGene_gn=SYNGAP1;refGene_tx=XM_023253850.1;refGene_gn=SYNGAP1;refGene_tx=XM_023253851.1;refGene_gn=SYNGAP1;refGene_tx=XM_023253852.1;refGene_gn=SYNGAP1;refGene_tx=XM_023253853.1;refGene_gn=SYNGAP1;refGene_tx=XM_023253854.1;refGene_gn=SYNGAP1;refGene_tx=XM_023253855.1
B2 35044239 35044413
refGene_gn=SYNGAP1;refGene_tx=XM_019830461.2;refGene_gn=SYNGAP1;refGene_tx=XM_023253838.1;refGene_gn=SYNGAP1;refGene_tx=XM_023253839.1;refGene_gn=SYNGAP1;refGene_tx=XM_023253840.1;refGene_gn=SYNGAP1;refGene_tx=XM_023253841.1;refGene_gn=SYNGAP1;refGene_tx=XM_023253842.1;refGene_gn=SYNGAP1;refGene_tx=XM_023253843.1;refGene_gn=SYNGAP1;refGene_tx=XM_023253844.1;refGene_gn=SYNGAP1;refGene_tx=XM_023253845.1;refGene_gn=SYNGAP1;refGene_tx=XM_023253846.1;refGene_gn=SYNGAP1;refGene_tx=XM_023253847.1;refGene_gn=SYNGAP1;refGene_tx=XM_023253848.1;refGene_gn=SYNGAP1;refGene_tx=XM_023253849.1;refGene_gn=SYNGAP1;refGene_tx=XM_023253850.1;refGene_gn=SYNGAP1;refGene_tx=XM_023253851.1;refGene_gn=SYNGAP1;refGene_tx=XM_023253852.1;refGene_gn=SYNGAP1;refGene_tx=XM_023253853.1;refGene_gn=SYNGAP1;refGene_tx=XM_023253854.1;refGene_gn=SYNGAP1;refGene_tx=XM_023253855.1
B2 35046056 35046268
refGene_gn=SYNGAP1;refGene_tx=XM_019830461.2;refGene_gn=SYNGAP1;refGene_tx=XM_023253838.1;refGene_gn=SYNGAP1;refGene_tx=XM_023253839.1;refGene_gn=SYNGAP1;refGene_tx=XM_023253840.1;refGene_gn=SYNGAP1;refGene_tx=XM_023253841.1;refGene_gn=SYNGAP1;refGene_tx=XM_023253842.1;refGene_gn=SYNGAP1;refGene_tx=XM_023253843.1;refGene_gn=SYNGAP1;refGene_tx=XM_023253844.1;refGene_gn=SYNGAP1;refGene_tx=XM_023253845.1;refGene_gn=SYNGAP1;refGene_tx=XM_023253846.1;refGene_gn=SYNGAP1;refGene_tx=XM_023253847.1;refGene_gn=SYNGAP1;refGene_tx=XM_023253848.1;refGene_gn=SYNGAP1;refGene_tx=XM_023253849.1;refGene_gn=SYNGAP1;refGene_tx=XM_023253850.1;refGene_gn=SYNGAP1;refGene_tx=XM_023253851.1;refGene_gn=SYNGAP1;refGene_tx=XM_023253852.1;refGene_gn=SYNGAP1;refGene_tx=XM_023253853.1;refGene_gn=SYNGAP1;refGene_tx=XM_023253854.1;refGene_gn=SYNGAP1;refGene_tx=XM_023253855.1
B2 35047375 35047482
refGene_gn=SYNGAP1;refGene_tx=XM_019830461.2;refGene_gn=SYNGAP1;refGen

e_tx=XM_023253838.1;refGene_gn=SYNGAP1;refGene_tx=XM_023253839.1;refGene_gn=SYNGAP1;refGene_tx=XM_023253840.1;refGene_gn=SYNGAP1;refGene_tx=XM_023253841.1;refGene_gn=SYNGAP1;refGene_tx=XM_023253842.1;refGene_gn=SYNGAP1;refGene_tx=XM_023253843.1;refGene_gn=SYNGAP1;refGene_tx=XM_023253844.1;refGene_gn=SYNGAP1;refGene_tx=XM_023253845.1;refGene_gn=SYNGAP1;refGene_tx=XM_023253846.1;refGene_gn=SYNGAP1;refGene_tx=XM_023253847.1;refGene_gn=SYNGAP1;refGene_tx=XM_023253848.1;refGene_gn=SYNGAP1;refGene_tx=XM_023253849.1;refGene_gn=SYNGAP1;refGene_tx=XM_023253850.1;refGene_gn=SYNGAP1;refGene_tx=XM_023253851.1;refGene_gn=SYNGAP1;refGene_tx=XM_023253852.1;refGene_gn=SYNGAP1;refGene_tx=XM_023253853.1;refGene_gn=SYNGAP1;refGene_tx=XM_023253854.1;refGene_gn=SYNGAP1;refGene_tx=XM_023253855.1
B2 35047754 35047859 refGene_gn=SYNGAP1;refGene_tx=XM_023253841.1
B2 35048224 35048263 refGene_gn=SYNGAP1;refGene_tx=XM_023253845.1
B2 35051384 35051531
refGene_gn=SYNGAP1;refGene_tx=XM_019830461.2;refGene_gn=SYNGAP1;refGene_tx=XM_023253838.1;refGene_gn=SYNGAP1;refGene_tx=XM_023253839.1;refGene_gn=SYNGAP1;refGene_tx=XM_023253840.1;refGene_gn=SYNGAP1;refGene_tx=XM_023253842.1;refGene_gn=SYNGAP1;refGene_tx=XM_023253843.1;refGene_gn=SYNGAP1;refGene_tx=XM_023253844.1;refGene_gn=SYNGAP1;refGene_tx=XM_023253851.1;refGene_gn=SYNGAP1;refGene_tx=XM_023253852.1;refGene_gn=SYNGAP1;refGene_tx=XM_023253853.1;refGene_gn=SYNGAP1;refGene_tx=XM_023253854.1;refGene_gn=SYNGAP1;refGene_tx=XM_023253855.1
B2 35054715 35056143 refGene_gn=ZBTB9;refGene_tx=XM_003986025.5
B2 35146567 35146672
refGene_gn=BAK1;refGene_tx=XM_003986026.4;refGene_gn=BAK1;refGene_tx=XM_019830465.2
B2 35146792 35146973
refGene_gn=BAK1;refGene_tx=XM_003986026.4;refGene_gn=BAK1;refGene_tx=XM_019830465.2
B2 35148062 35148206
refGene_gn=BAK1;refGene_tx=XM_003986026.4;refGene_gn=BAK1;refGene_tx=XM_019830465.2
B2 35148551 35148687
refGene_gn=BAK1;refGene_tx=XM_003986026.4;refGene_gn=BAK1;refGene_tx=XM_019830465.2
B2 35149684 35149754
refGene_gn=BAK1;refGene_tx=XM_003986026.4;refGene_gn=BAK1;refGene_tx=XM_019830465.2
B2 35153321 35153788
refGene_gn=LOC101087741;refGene_tx=XM_006931627.4;refGene_gn=LOC101087741;refGene_tx=XM_023253856.1
B2 35154450 35154569
refGene_gn=LOC101087741;refGene_tx=XM_006931627.4;refGene_gn=LOC101087741;refGene_tx=XM_023253856.1
B2 35155029 35155127
refGene_gn=LOC101087741;refGene_tx=XM_006931627.4;refGene_gn=LOC101087741;refGene_tx=XM_023253856.1
B2 35155335 35155483
refGene_gn=LOC101087741;refGene_tx=XM_006931627.4;refGene_gn=LOC101087

741;refGene_tx=XM_023253856.1
B2 35156398 35156538
refGene_gn=LOC101087741;refGene_tx=XM_006931627.4
B2 35158745 35158876
refGene_gn=LOC101087741;refGene_tx=XM_006931627.4;refGene_gn=LOC101087
741;refGene_tx=XM_023253856.1
B2 35178576 35178665 refGene_gn=ITPR3;refGene_tx=XM_023253857.1
B2 35194892 35194963 refGene_gn=ITPR3;refGene_tx=XM_023253857.1
B2 35206657 35206779 refGene_gn=ITPR3;refGene_tx=XM_023253857.1
B2 35208686 35208773 refGene_gn=ITPR3;refGene_tx=XM_023253857.1
B2 35209364 35209523 refGene_gn=ITPR3;refGene_tx=XM_023253857.1
B2 35209714 35209813 refGene_gn=ITPR3;refGene_tx=XM_023253857.1
B2 35210152 35210236 refGene_gn=ITPR3;refGene_tx=XM_023253857.1
B2 35212554 35212701 refGene_gn=ITPR3;refGene_tx=XM_023253857.1
B2 35212910 35213006 refGene_gn=ITPR3;refGene_tx=XM_023253857.1
B2 35213604 35213655 refGene_gn=ITPR3;refGene_tx=XM_023253857.1
B2 35213828 35213971 refGene_gn=ITPR3;refGene_tx=XM_023253857.1
B2 35214979 35215079 refGene_gn=ITPR3;refGene_tx=XM_023253857.1
B2 35215160 35215321 refGene_gn=ITPR3;refGene_tx=XM_023253857.1
B2 35215943 35216085 refGene_gn=ITPR3;refGene_tx=XM_023253857.1
B2 35217549 35217711 refGene_gn=ITPR3;refGene_tx=XM_023253857.1
B2 35218272 35218445 refGene_gn=ITPR3;refGene_tx=XM_023253857.1
B2 35219017 35219137 refGene_gn=ITPR3;refGene_tx=XM_023253857.1
B2 35219491 35219674 refGene_gn=ITPR3;refGene_tx=XM_023253857.1
B2 35220584 35220836 refGene_gn=ITPR3;refGene_tx=XM_023253857.1
B2 35220927 35221072 refGene_gn=ITPR3;refGene_tx=XM_023253857.1
B2 35221630 35221772 refGene_gn=ITPR3;refGene_tx=XM_023253857.1
B2 35222225 35222425 refGene_gn=ITPR3;refGene_tx=XM_023253857.1
B2 35223614 35223744 refGene_gn=ITPR3;refGene_tx=XM_023253857.1
B2 35224262 35224320 refGene_gn=ITPR3;refGene_tx=XM_023253857.1
B2 35225411 35225577 refGene_gn=ITPR3;refGene_tx=XM_023253857.1
B2 35226547 35226712 refGene_gn=ITPR3;refGene_tx=XM_023253857.1
B2 35226795 35226870 refGene_gn=ITPR3;refGene_tx=XM_023253857.1
B2 35227295 35227409 refGene_gn=ITPR3;refGene_tx=XM_023253857.1
B2 35227410 35227422 refGene_gn=ITPR3;refGene_tx=XM_023253857.1
B2 35228069 35228192 refGene_gn=ITPR3;refGene_tx=XM_023253857.1
B2 35228268 35228469 refGene_gn=ITPR3;refGene_tx=XM_023253857.1
B2 35229422 35229674 refGene_gn=ITPR3;refGene_tx=XM_023253857.1
B2 35229945 35230071 refGene_gn=ITPR3;refGene_tx=XM_023253857.1
B2 35230207 35230333 refGene_gn=ITPR3;refGene_tx=XM_023253857.1
B2 35231552 35231673 refGene_gn=ITPR3;refGene_tx=XM_023253857.1
B2 35232215 35232406 refGene_gn=ITPR3;refGene_tx=XM_023253857.1
B2 35233009 35233158 refGene_gn=ITPR3;refGene_tx=XM_023253857.1
B2 35233356 35233465 refGene_gn=ITPR3;refGene_tx=XM_023253857.1
B2 35233595 35233686 refGene_gn=ITPR3;refGene_tx=XM_023253857.1
B2 35233801 35233971 refGene_gn=ITPR3;refGene_tx=XM_023253857.1
B2 35235146 35235321 refGene_gn=ITPR3;refGene_tx=XM_023253857.1
B2 35235489 35235674 refGene_gn=ITPR3;refGene_tx=XM_023253857.1
B2 35235819 35236020 refGene_gn=ITPR3;refGene_tx=XM_023253857.1
B2 35236167 35236278 refGene_gn=ITPR3;refGene_tx=XM_023253857.1

B2	35236762 35236858 refGene_gn=ITPR3;refGene_tx=XM_023253857.1
B2	35236975 35237077 refGene_gn=ITPR3;refGene_tx=XM_023253857.1
B2	35237171 35237258 refGene_gn=ITPR3;refGene_tx=XM_023253857.1
B2	35237809 35237920 refGene_gn=ITPR3;refGene_tx=XM_023253857.1
B2	35238020 35238213 refGene_gn=ITPR3;refGene_tx=XM_023253857.1
B2	35238431 35238557 refGene_gn=ITPR3;refGene_tx=XM_023253857.1
B2	35239018 35239191 refGene_gn=ITPR3;refGene_tx=XM_023253857.1
B2	35239888 35240053 refGene_gn=ITPR3;refGene_tx=XM_023253857.1
B2	35240731 35240924 refGene_gn=ITPR3;refGene_tx=XM_023253857.1
B2	35241380 35241485 refGene_gn=ITPR3;refGene_tx=XM_023253857.1
B2	35241571 35241699 refGene_gn=ITPR3;refGene_tx=XM_023253857.1
B2	35242589 35242755 refGene_gn=ITPR3;refGene_tx=XM_023253857.1
B2	35243193 35243354 refGene_gn=ITPR3;refGene_tx=XM_023253857.1
B2	35244576 35244738 refGene_gn=ITPR3;refGene_tx=XM_023253857.1
B2	35245436 35245505 refGene_gn=ITPR3;refGene_tx=XM_023253857.1
B2	35247344 35247472
refGene_gn=LOC101093686;refGene_tx=XM_003986027.5
B2	35250164 35250234
refGene_gn=LOC101093686;refGene_tx=XM_003986027.5
B2	35251138 35251213
refGene_gn=LOC101093686;refGene_tx=XM_003986027.5
B2	35261159 35261297
refGene_gn=LOC101093686;refGene_tx=XM_003986027.5
B2	35270056 35270524 refGene_gn=IP6K3;refGene_tx=XM_003986028.5
B2	35272446 35272622 refGene_gn=IP6K3;refGene_tx=XM_003986028.5
B2	35273623 35273799 refGene_gn=IP6K3;refGene_tx=XM_003986028.5
B2	35274944 35275158 refGene_gn=IP6K3;refGene_tx=XM_003986028.5
B2	35281500 35281699 refGene_gn=IP6K3;refGene_tx=XM_003986028.5
B2	35312467 35312618 refGene_gn=LEMD2;refGene_tx=XM_019830466.2
B2	35315822 35315925 refGene_gn=LEMD2;refGene_tx=XM_019830466.2
B2	35316007 35316109 refGene_gn=LEMD2;refGene_tx=XM_019830466.2
B2	35317136 35317282 refGene_gn=LEMD2;refGene_tx=XM_019830466.2
B2	35318906 35318986 refGene_gn=LEMD2;refGene_tx=XM_019830466.2
B2	35319897 35319974 refGene_gn=LEMD2;refGene_tx=XM_019830466.2
B2	35323080 35323156 refGene_gn=LEMD2;refGene_tx=XM_019830466.2
B2	35326220 35326261 refGene_gn=LEMD2;refGene_tx=XM_019830466.2
B2	35327879 35328609 refGene_gn=LEMD2;refGene_tx=XM_019830466.2
B2	35334299 35334307 refGene_gn=MLN;refGene_tx=NM_001009278.1
B2	35335107 35335157 refGene_gn=MLN;refGene_tx=NM_001009278.1
B2	35335158 35335214 refGene_gn=MLN;refGene_tx=NM_001009278.1
B2	35338428 35338545 refGene_gn=MLN;refGene_tx=NM_001009278.1
B2	35340616 35340733 refGene_gn=MLN;refGene_tx=NM_001009278.1
B2	35513974 35514075 refGene_gn=GRM4;refGene_tx=XM_023253862.1
B2	35550569 35550619 refGene_gn=GRM4;refGene_tx=XM_023253863.1
B2	35551508 35551690
refGene_gn=GRM4;refGene_tx=XM_019830473.1;refGene_gn=GRM4;refGene_tx=XM_019830474.1;refGene_gn=GRM4;refGene_tx=XM_023253858.1;refGene_gn=GRM4;refGene_tx=XM_023253859.1;refGene_gn=GRM4;refGene_tx=XM_023253860.1;refGene_gn=GRM4;refGene_tx=XM_023253861.1;refGene_gn=GRM4;refGene_tx=XM_023253865.1

B2 35553212 35553262 refGene_gn=GRM4;refGene_tx=XM_023253864.1
B2 35556567 35556814
refGene_gn=GRM4;refGene_tx=XM_019830473.1;refGene_gn=GRM4;refGene_tx=XM_019830474.1;refGene_gn=GRM4;refGene_tx=XM_023253858.1;refGene_gn=GRM4;refGene_tx=XM_023253859.1;refGene_gn=GRM4;refGene_tx=XM_023253860.1;refGene_gn=GRM4;refGene_tx=XM_023253861.1;refGene_gn=GRM4;refGene_tx=XM_023253862.1;refGene_gn=GRM4;refGene_tx=XM_023253863.1;refGene_gn=GRM4;refGene_tx=XM_023253864.1;refGene_gn=GRM4;refGene_tx=XM_023253865.1
B2 35562981 35563917
refGene_gn=GRM4;refGene_tx=XM_019830473.1;refGene_gn=GRM4;refGene_tx=XM_019830474.1;refGene_gn=GRM4;refGene_tx=XM_023253858.1;refGene_gn=GRM4;refGene_tx=XM_023253859.1;refGene_gn=GRM4;refGene_tx=XM_023253860.1;refGene_gn=GRM4;refGene_tx=XM_023253861.1;refGene_gn=GRM4;refGene_tx=XM_023253862.1;refGene_gn=GRM4;refGene_tx=XM_023253863.1;refGene_gn=GRM4;refGene_tx=XM_023253864.1;refGene_gn=GRM4;refGene_tx=XM_023253865.1
B2 35567224 35567361
refGene_gn=GRM4;refGene_tx=XM_019830473.1;refGene_gn=GRM4;refGene_tx=XM_019830474.1;refGene_gn=GRM4;refGene_tx=XM_023253858.1;refGene_gn=GRM4;refGene_tx=XM_023253859.1;refGene_gn=GRM4;refGene_tx=XM_023253860.1;refGene_gn=GRM4;refGene_tx=XM_023253861.1;refGene_gn=GRM4;refGene_tx=XM_023253862.1;refGene_gn=GRM4;refGene_tx=XM_023253863.1;refGene_gn=GRM4;refGene_tx=XM_023253864.1;refGene_gn=GRM4;refGene_tx=XM_023253865.1
B2 35567589 35567790
refGene_gn=GRM4;refGene_tx=XM_019830473.1;refGene_gn=GRM4;refGene_tx=XM_019830474.1;refGene_gn=GRM4;refGene_tx=XM_023253858.1;refGene_gn=GRM4;refGene_tx=XM_023253859.1;refGene_gn=GRM4;refGene_tx=XM_023253860.1;refGene_gn=GRM4;refGene_tx=XM_023253861.1;refGene_gn=GRM4;refGene_tx=XM_023253862.1;refGene_gn=GRM4;refGene_tx=XM_023253863.1;refGene_gn=GRM4;refGene_tx=XM_023253864.1;refGene_gn=GRM4;refGene_tx=XM_023253865.1
B2 35580909 35581050
refGene_gn=GRM4;refGene_tx=XM_019830473.1;refGene_gn=GRM4;refGene_tx=XM_019830474.1;refGene_gn=GRM4;refGene_tx=XM_023253858.1;refGene_gn=GRM4;refGene_tx=XM_023253859.1;refGene_gn=GRM4;refGene_tx=XM_023253860.1;refGene_gn=GRM4;refGene_tx=XM_023253861.1;refGene_gn=GRM4;refGene_tx=XM_023253862.1;refGene_gn=GRM4;refGene_tx=XM_023253863.1;refGene_gn=GRM4;refGene_tx=XM_023253864.1;refGene_gn=GRM4;refGene_tx=XM_023253865.1
B2 35583276 35583431
refGene_gn=GRM4;refGene_tx=XM_019830473.1;refGene_gn=GRM4;refGene_tx=XM_019830474.1;refGene_gn=GRM4;refGene_tx=XM_023253858.1;refGene_gn=GRM4;refGene_tx=XM_023253859.1;refGene_gn=GRM4;refGene_tx=XM_023253860.1;refGene_gn=GRM4;refGene_tx=XM_023253861.1;refGene_gn=GRM4;refGene_tx=XM_023253862.1;refGene_gn=GRM4;refGene_tx=XM_023253863.1;refGene_gn=GRM4;refGene_tx=XM_023253864.1;refGene_gn=GRM4;refGene_tx=XM_023253865.1
B2 35586367 35586503
refGene_gn=GRM4;refGene_tx=XM_019830473.1;refGene_gn=GRM4;refGene_tx=XM_019830474.1;refGene_gn=GRM4;refGene_tx=XM_023253858.1;refGene_gn=GRM4;refGene_tx=XM_023253859.1;refGene_gn=GRM4;refGene_tx=XM_023253860.1;refGene_gn=GRM4;refGene_tx=XM_023253861.1;refGene_gn=GRM4;refGene_tx=XM_023253862.1;refGene_gn=GRM4;refGene_tx=XM_023253863.1;refGene_gn=GRM4;refGene_tx=XM_023253864.1;refGene_gn=GRM4;refGene_tx=XM_023253865.1

B2 35591549 35591667 refGene_gn=GRM4;refGene_tx=XM_019830473.1
B2 35612636 35612853
refGene_gn=GRM4;refGene_tx=XM_019830474.1;refGene_gn=GRM4;refGene_tx=XM_023253858.1;refGene_gn=GRM4;refGene_tx=XM_023253859.1;refGene_gn=GRM4;refGene_tx=XM_023253860.1;refGene_gn=GRM4;refGene_tx=XM_023253861.1;refGene_gn=GRM4;refGene_tx=XM_023253862.1;refGene_gn=GRM4;refGene_tx=XM_023253863.1;refGene_gn=GRM4;refGene_tx=XM_023253864.1;refGene_gn=GRM4;refGene_tx=XM_023253865.1
B2 35654026 35654545
refGene_gn=GRM4;refGene_tx=XM_023253858.1;refGene_gn=GRM4;refGene_tx=XM_023253859.1;refGene_gn=GRM4;refGene_tx=XM_023253860.1;refGene_gn=GRM4;refGene_tx=XM_023253861.1;refGene_gn=GRM4;refGene_tx=XM_023253862.1;refGene_gn=GRM4;refGene_tx=XM_023253863.1;refGene_gn=GRM4;refGene_tx=XM_023253864.1
B2 35742326 35742461
refGene_gn=HMGA1;refGene_tx=XM_023253866.1;refGene_gn=HMGA1;refGene_tx=XM_023253867.1
B2 35744180 35744264
refGene_gn=HMGA1;refGene_tx=XM_023253866.1;refGene_gn=HMGA1;refGene_tx=XM_023253867.1
B2 35744930 35744981
refGene_gn=HMGA1;refGene_tx=XM_023253866.1;refGene_gn=HMGA1;refGene_tx=XM_023253867.1
B2 35746257 35746311
refGene_gn=HMGA1;refGene_tx=XM_023253866.1;refGene_gn=HMGA1;refGene_tx=XM_023253867.1
B2 35748166 35748251
refGene_gn=SMIM29;refGene_tx=XM_003986034.4;refGene_gn=SMIM29;refGene_tx=XM_006931629.3;refGene_gn=SMIM29;refGene_tx=XM_011282262.2
B2 35748441 35748606
refGene_gn=SMIM29;refGene_tx=XM_003986034.4;refGene_gn=SMIM29;refGene_tx=XM_006931629.3;refGene_gn=SMIM29;refGene_tx=XM_011282262.2
B2 35748870 35748896
refGene_gn=SMIM29;refGene_tx=XM_003986034.4;refGene_gn=SMIM29;refGene_tx=XM_006931629.3;refGene_gn=SMIM29;refGene_tx=XM_011282262.2
B2 35749093 35749204
refGene_gn=SMIM29;refGene_tx=XM_003986034.4;refGene_gn=SMIM29;refGene_tx=XM_006931629.3;refGene_gn=SMIM29;refGene_tx=XM_011282262.2
B2 35780671 35780850
refGene_gn=LOC101088487;refGene_tx=XM_023253868.1
B2 35783650 35783735
refGene_gn=LOC101088487;refGene_tx=XM_023253868.1
B2 35788546 35788591
refGene_gn=LOC101088487;refGene_tx=XM_023253868.1
B2 35840858 35840969
refGene_gn=LOC101088487;refGene_tx=XM_023253868.1
B2 35906315 35906414
refGene_gn=LOC101088487;refGene_tx=XM_023253868.1
B2 35934261 35934303 refGene_gn=RPS10;refGene_tx=XM_006931630.2
B2 35935059 35935115 refGene_gn=RPS10;refGene_tx=XM_006931630.2

B2	35937413 35937491 refGene_gn=RPS10;refGene_tx=XM_006931630.2
B2	35939610 35939782 refGene_gn=RPS10;refGene_tx=XM_006931630.2
B2	35940007 35940157 refGene_gn=RPS10;refGene_tx=XM_006931630.2
B2	36014977 36015040
refGene_gn=PACSIN1;refGene_tx=XM_011282261.3;refGene_gn=PACSIN1;refGene_tx=XM_019830476.2;refGene_gn=PACSIN1;refGene_tx=XM_023253869.1;refGene_gn=PACSIN1;refGene_tx=XM_023253870.1
B2	36015958 36016115
refGene_gn=PACSIN1;refGene_tx=XM_011282261.3;refGene_gn=PACSIN1;refGene_tx=XM_019830476.2;refGene_gn=PACSIN1;refGene_tx=XM_023253869.1;refGene_gn=PACSIN1;refGene_tx=XM_023253870.1
B2	36017289 36017525
refGene_gn=PACSIN1;refGene_tx=XM_011282261.3;refGene_gn=PACSIN1;refGene_tx=XM_019830476.2;refGene_gn=PACSIN1;refGene_tx=XM_023253869.1;refGene_gn=PACSIN1;refGene_tx=XM_023253870.1
B2	36017918 36018074
refGene_gn=PACSIN1;refGene_tx=XM_011282261.3;refGene_gn=PACSIN1;refGene_tx=XM_019830476.2;refGene_gn=PACSIN1;refGene_tx=XM_023253869.1;refGene_gn=PACSIN1;refGene_tx=XM_023253870.1
B2	36018166 36018342
refGene_gn=PACSIN1;refGene_tx=XM_011282261.3;refGene_gn=PACSIN1;refGene_tx=XM_019830476.2;refGene_gn=PACSIN1;refGene_tx=XM_023253869.1;refGene_gn=PACSIN1;refGene_tx=XM_023253870.1
B2	36018779 36018900
refGene_gn=PACSIN1;refGene_tx=XM_011282261.3;refGene_gn=PACSIN1;refGene_tx=XM_019830476.2;refGene_gn=PACSIN1;refGene_tx=XM_023253869.1;refGene_gn=PACSIN1;refGene_tx=XM_023253870.1
B2	36018998 36019126
refGene_gn=PACSIN1;refGene_tx=XM_011282261.3;refGene_gn=PACSIN1;refGene_tx=XM_019830476.2;refGene_gn=PACSIN1;refGene_tx=XM_023253869.1;refGene_gn=PACSIN1;refGene_tx=XM_023253870.1
B2	36019728 36019916
refGene_gn=PACSIN1;refGene_tx=XM_011282261.3;refGene_gn=PACSIN1;refGene_tx=XM_019830476.2;refGene_gn=PACSIN1;refGene_tx=XM_023253869.1;refGene_gn=PACSIN1;refGene_tx=XM_023253870.1
B2	36020452 36020562
refGene_gn=PACSIN1;refGene_tx=XM_011282261.3;refGene_gn=PACSIN1;refGene_tx=XM_019830476.2;refGene_gn=PACSIN1;refGene_tx=XM_023253869.1;refGene_gn=PACSIN1;refGene_tx=XM_023253870.1
B2	36026812 36026991 refGene_gn=SPDEF;refGene_tx=XM_003986039.2
B2	36029442 36029589 refGene_gn=SPDEF;refGene_tx=XM_003986039.2
B2	36029681 36029729 refGene_gn=SPDEF;refGene_tx=XM_003986039.2
B2	36030738 36030936 refGene_gn=SPDEF;refGene_tx=XM_003986039.2
B2	36033029 36033465 refGene_gn=SPDEF;refGene_tx=XM_003986039.2
B2	36075752 36075788
refGene_gn=CB2H6orf106;refGene_tx=XM_023253871.1;refGene_gn=CB2H6orf106;refGene_tx=XM_023253872.1
B2	36092227 36092556
refGene_gn=CB2H6orf106;refGene_tx=XM_023253871.1;refGene_gn=CB2H6orf106;refGene_tx=XM_023253872.1

B2 36125463 36125661
refGene_gn=CB2H6orf106;refGene_tx=XM_023253871.1
B2 36131281 36131436
refGene_gn=CB2H6orf106;refGene_tx=XM_023253871.1;refGene_gn=CB2H6orf10
6;refGene_tx=XM_023253872.1
B2 36168681 36168839
refGene_gn=CB2H6orf106;refGene_tx=XM_023253871.1;refGene_gn=CB2H6orf10
6;refGene_tx=XM_023253872.1
B2 36213020 36213028 refGene_gn=SNRPC;refGene_tx=XM_003986041.5
B2 36213365 36213408 refGene_gn=SNRPC;refGene_tx=XM_003986041.5
B2 36217212 36217321 refGene_gn=SNRPC;refGene_tx=XM_003986041.5
B2 36218875 36218965 refGene_gn=SNRPC;refGene_tx=XM_003986041.5
B2 36220534 36220639 refGene_gn=SNRPC;refGene_tx=XM_003986041.5
B2 36223406 36223531 refGene_gn=SNRPC;refGene_tx=XM_003986041.5
B2 36240522 36240566 refGene_gn=UHRF1BP1;refGene_tx=XM_011282259.2
B2 36268577 36268740 refGene_gn=UHRF1BP1;refGene_tx=XM_011282259.2
B2 36268841 36268880 refGene_gn=UHRF1BP1;refGene_tx=XM_011282259.2
B2 36269584 36269676 refGene_gn=UHRF1BP1;refGene_tx=XM_011282259.2
B2 36278526 36278716 refGene_gn=UHRF1BP1;refGene_tx=XM_011282259.2
B2 36279024 36279165 refGene_gn=UHRF1BP1;refGene_tx=XM_011282259.2
B2 36279595 36279766 refGene_gn=UHRF1BP1;refGene_tx=XM_011282259.2
B2 36280444 36280637 refGene_gn=UHRF1BP1;refGene_tx=XM_011282259.2
B2 36287661 36287748 refGene_gn=UHRF1BP1;refGene_tx=XM_011282259.2
B2 36288282 36288453 refGene_gn=UHRF1BP1;refGene_tx=XM_011282259.2
B2 36289086 36289226 refGene_gn=UHRF1BP1;refGene_tx=XM_011282259.2
B2 36289628 36289770 refGene_gn=UHRF1BP1;refGene_tx=XM_011282259.2
B2 36290026 36290220 refGene_gn=UHRF1BP1;refGene_tx=XM_011282259.2
B2 36290421 36291878 refGene_gn=UHRF1BP1;refGene_tx=XM_011282259.2
B2 36296612 36296809 refGene_gn=UHRF1BP1;refGene_tx=XM_011282259.2
B2 36298253 36298408 refGene_gn=UHRF1BP1;refGene_tx=XM_011282259.2
B2 36298496 36298647 refGene_gn=UHRF1BP1;refGene_tx=XM_011282259.2
B2 36301070 36301333 refGene_gn=UHRF1BP1;refGene_tx=XM_011282259.2
B2 36302017 36302126 refGene_gn=UHRF1BP1;refGene_tx=XM_011282259.2
B2 36302251 36302358 refGene_gn=UHRF1BP1;refGene_tx=XM_011282259.2
B2 36302754 36302880 refGene_gn=UHRF1BP1;refGene_tx=XM_011282259.2
B2 36308313 36308444
refGene_gn=TAF11;refGene_tx=XM_003986042.5;refGene_gn=TAF11;refGene_tx
=XM_003986043.5
B2 36309088 36309185 refGene_gn=TAF11;refGene_tx=XM_003986042.5
B2 36309800 36309888
refGene_gn=TAF11;refGene_tx=XM_003986042.5;refGene_gn=TAF11;refGene_tx
=XM_003986043.5
B2 36311336 36311485
refGene_gn=TAF11;refGene_tx=XM_003986042.5;refGene_gn=TAF11;refGene_tx
=XM_003986043.5
B2 36314931 36315102
refGene_gn=TAF11;refGene_tx=XM_003986042.5;refGene_gn=TAF11;refGene_tx
=XM_003986043.5
B2 36316549 36316743
refGene_gn=ANKS1A;refGene_tx=XM_023253873.1;refGene_gn=ANKS1A;refGene_

tx=XM_023253874.1;refGene_gn=ANKS1A;refGene_tx=XM_023253875.1;refGene_gn=ANKS1A;refGene_tx=XM_023253876.1;refGene_gn=ANKS1A;refGene_tx=XM_023253877.1;refGene_gn=ANKS1A;refGene_tx=XM_023253878.1
B2 36387115 36387196
refGene_gn=ANKS1A;refGene_tx=XM_023253873.1;refGene_gn=ANKS1A;refGene_tx=XM_023253874.1;refGene_gn=ANKS1A;refGene_tx=XM_023253875.1;refGene_gn=ANKS1A;refGene_tx=XM_023253876.1;refGene_gn=ANKS1A;refGene_tx=XM_023253877.1;refGene_gn=ANKS1A;refGene_tx=XM_023253878.1
B2 36387889 36387894 refGene_gn=ANKS1A;refGene_tx=XM_023253879.1
B2 36388943 36389100
refGene_gn=ANKS1A;refGene_tx=XM_023253873.1;refGene_gn=ANKS1A;refGene_tx=XM_023253874.1;refGene_gn=ANKS1A;refGene_tx=XM_023253875.1;refGene_gn=ANKS1A;refGene_tx=XM_023253876.1;refGene_gn=ANKS1A;refGene_tx=XM_023253877.1;refGene_gn=ANKS1A;refGene_tx=XM_023253878.1;refGene_gn=ANKS1A;refGene_tx=XM_023253879.1
B2 36390918 36390981
refGene_gn=ANKS1A;refGene_tx=XM_023253873.1;refGene_gn=ANKS1A;refGene_tx=XM_023253874.1;refGene_gn=ANKS1A;refGene_tx=XM_023253876.1;refGene_gn=ANKS1A;refGene_tx=XM_023253877.1;refGene_gn=ANKS1A;refGene_tx=XM_023253879.1
B2 36397509 36397806
refGene_gn=ANKS1A;refGene_tx=XM_023253873.1;refGene_gn=ANKS1A;refGene_tx=XM_023253874.1;refGene_gn=ANKS1A;refGene_tx=XM_023253875.1;refGene_gn=ANKS1A;refGene_tx=XM_023253876.1;refGene_gn=ANKS1A;refGene_tx=XM_023253877.1;refGene_gn=ANKS1A;refGene_tx=XM_023253878.1;refGene_gn=ANKS1A;refGene_tx=XM_023253879.1;refGene_gn=ANKS1A;refGene_tx=XM_023253880.1
B2 36398608 36398684
refGene_gn=ANKS1A;refGene_tx=XM_023253873.1;refGene_gn=ANKS1A;refGene_tx=XM_023253874.1;refGene_gn=ANKS1A;refGene_tx=XM_023253875.1;refGene_gn=ANKS1A;refGene_tx=XM_023253876.1;refGene_gn=ANKS1A;refGene_tx=XM_023253877.1;refGene_gn=ANKS1A;refGene_tx=XM_023253878.1;refGene_gn=ANKS1A;refGene_tx=XM_023253879.1;refGene_gn=ANKS1A;refGene_tx=XM_023253880.1
B2 36398972 36399074
refGene_gn=ANKS1A;refGene_tx=XM_023253873.1;refGene_gn=ANKS1A;refGene_tx=XM_023253874.1;refGene_gn=ANKS1A;refGene_tx=XM_023253875.1;refGene_gn=ANKS1A;refGene_tx=XM_023253876.1;refGene_gn=ANKS1A;refGene_tx=XM_023253877.1;refGene_gn=ANKS1A;refGene_tx=XM_023253878.1;refGene_gn=ANKS1A;refGene_tx=XM_023253879.1;refGene_gn=ANKS1A;refGene_tx=XM_023253880.1
B2 36399178 36399280
refGene_gn=ANKS1A;refGene_tx=XM_023253873.1;refGene_gn=ANKS1A;refGene_tx=XM_023253874.1;refGene_gn=ANKS1A;refGene_tx=XM_023253875.1;refGene_gn=ANKS1A;refGene_tx=XM_023253876.1;refGene_gn=ANKS1A;refGene_tx=XM_023253877.1;refGene_gn=ANKS1A;refGene_tx=XM_023253878.1;refGene_gn=ANKS1A;refGene_tx=XM_023253879.1;refGene_gn=ANKS1A;refGene_tx=XM_023253880.1
B2 36400675 36400872
refGene_gn=ANKS1A;refGene_tx=XM_023253873.1;refGene_gn=ANKS1A;refGene_

tx=XM_023253874.1;refGene_gn=ANKS1A;refGene_tx=XM_023253875.1;refGene_gn=ANKS1A;refGene_tx=XM_023253876.1;refGene_gn=ANKS1A;refGene_tx=XM_023253877.1;refGene_gn=ANKS1A;refGene_tx=XM_023253878.1;refGene_gn=ANKS1A;refGene_tx=XM_023253879.1;refGene_gn=ANKS1A;refGene_tx=XM_023253880.1

B2 36404663 36404756
refGene_gn=ANKS1A;refGene_tx=XM_023253873.1;refGene_gn=ANKS1A;refGene_tx=XM_023253874.1;refGene_gn=ANKS1A;refGene_tx=XM_023253875.1;refGene_gn=ANKS1A;refGene_tx=XM_023253876.1;refGene_gn=ANKS1A;refGene_tx=XM_023253877.1;refGene_gn=ANKS1A;refGene_tx=XM_023253878.1;refGene_gn=ANKS1A;refGene_tx=XM_023253879.1;refGene_gn=ANKS1A;refGene_tx=XM_023253880.1

B2 36409283 36409404
refGene_gn=ANKS1A;refGene_tx=XM_023253873.1;refGene_gn=ANKS1A;refGene_tx=XM_023253874.1;refGene_gn=ANKS1A;refGene_tx=XM_023253875.1;refGene_gn=ANKS1A;refGene_tx=XM_023253876.1;refGene_gn=ANKS1A;refGene_tx=XM_023253877.1;refGene_gn=ANKS1A;refGene_tx=XM_023253878.1;refGene_gn=ANKS1A;refGene_tx=XM_023253879.1;refGene_gn=ANKS1A;refGene_tx=XM_023253880.1

B2 36423189 36423779
refGene_gn=ANKS1A;refGene_tx=XM_023253873.1;refGene_gn=ANKS1A;refGene_tx=XM_023253874.1;refGene_gn=ANKS1A;refGene_tx=XM_023253875.1;refGene_gn=ANKS1A;refGene_tx=XM_023253876.1;refGene_gn=ANKS1A;refGene_tx=XM_023253877.1;refGene_gn=ANKS1A;refGene_tx=XM_023253878.1;refGene_gn=ANKS1A;refGene_tx=XM_023253879.1;refGene_gn=ANKS1A;refGene_tx=XM_023253880.1

B2 36462406 36462473
refGene_gn=ANKS1A;refGene_tx=XM_023253873.1;refGene_gn=ANKS1A;refGene_tx=XM_023253874.1;refGene_gn=ANKS1A;refGene_tx=XM_023253875.1;refGene_gn=ANKS1A;refGene_tx=XM_023253876.1;refGene_gn=ANKS1A;refGene_tx=XM_023253877.1;refGene_gn=ANKS1A;refGene_tx=XM_023253878.1;refGene_gn=ANKS1A;refGene_tx=XM_023253879.1;refGene_gn=ANKS1A;refGene_tx=XM_023253880.1

B2 36467681 36467788
refGene_gn=ANKS1A;refGene_tx=XM_023253873.1;refGene_gn=ANKS1A;refGene_tx=XM_023253874.1;refGene_gn=ANKS1A;refGene_tx=XM_023253875.1;refGene_gn=ANKS1A;refGene_tx=XM_023253876.1;refGene_gn=ANKS1A;refGene_tx=XM_023253877.1;refGene_gn=ANKS1A;refGene_tx=XM_023253878.1;refGene_gn=ANKS1A;refGene_tx=XM_023253879.1;refGene_gn=ANKS1A;refGene_tx=XM_023253880.1

B2 36482607 36482706
refGene_gn=ANKS1A;refGene_tx=XM_023253873.1;refGene_gn=ANKS1A;refGene_tx=XM_023253874.1;refGene_gn=ANKS1A;refGene_tx=XM_023253875.1;refGene_gn=ANKS1A;refGene_tx=XM_023253876.1;refGene_gn=ANKS1A;refGene_tx=XM_023253877.1;refGene_gn=ANKS1A;refGene_tx=XM_023253878.1;refGene_gn=ANKS1A;refGene_tx=XM_023253879.1;refGene_gn=ANKS1A;refGene_tx=XM_023253880.1

B2 36483520 36483673
refGene_gn=ANKS1A;refGene_tx=XM_023253873.1;refGene_gn=ANKS1A;refGene_tx=XM_023253874.1;refGene_gn=ANKS1A;refGene_tx=XM_023253875.1;refGene_

gn=ANKS1A;refGene_tx=XM_023253876.1;refGene_gn=ANKS1A;refGene_tx=XM_023253877.1;refGene_gn=ANKS1A;refGene_tx=XM_023253878.1;refGene_gn=ANKS1A;refGene_tx=XM_023253879.1;refGene_gn=ANKS1A;refGene_tx=XM_023253880.1
B2 36483788 36483896
refGene_gn=ANKS1A;refGene_tx=XM_023253873.1;refGene_gn=ANKS1A;refGene_tx=XM_023253874.1;refGene_gn=ANKS1A;refGene_tx=XM_023253875.1;refGene_gn=ANKS1A;refGene_tx=XM_023253876.1;refGene_gn=ANKS1A;refGene_tx=XM_023253877.1;refGene_gn=ANKS1A;refGene_tx=XM_023253878.1;refGene_gn=ANKS1A;refGene_tx=XM_023253879.1;refGene_gn=ANKS1A;refGene_tx=XM_023253880.1
B2 36485596 36485761
refGene_gn=ANKS1A;refGene_tx=XM_023253873.1;refGene_gn=ANKS1A;refGene_tx=XM_023253874.1;refGene_gn=ANKS1A;refGene_tx=XM_023253875.1;refGene_gn=ANKS1A;refGene_tx=XM_023253876.1;refGene_gn=ANKS1A;refGene_tx=XM_023253877.1;refGene_gn=ANKS1A;refGene_tx=XM_023253878.1;refGene_gn=ANKS1A;refGene_tx=XM_023253879.1;refGene_gn=ANKS1A;refGene_tx=XM_023253880.1
B2 36487454 36487580
refGene_gn=ANKS1A;refGene_tx=XM_023253873.1;refGene_gn=ANKS1A;refGene_tx=XM_023253874.1;refGene_gn=ANKS1A;refGene_tx=XM_023253875.1;refGene_gn=ANKS1A;refGene_tx=XM_023253876.1;refGene_gn=ANKS1A;refGene_tx=XM_023253877.1;refGene_gn=ANKS1A;refGene_tx=XM_023253878.1;refGene_gn=ANKS1A;refGene_tx=XM_023253879.1;refGene_gn=ANKS1A;refGene_tx=XM_023253880.1
B2 36487803 36487875
refGene_gn=ANKS1A;refGene_tx=XM_023253873.1;refGene_gn=ANKS1A;refGene_tx=XM_023253874.1;refGene_gn=ANKS1A;refGene_tx=XM_023253875.1;refGene_gn=ANKS1A;refGene_tx=XM_023253876.1;refGene_gn=ANKS1A;refGene_tx=XM_023253877.1;refGene_gn=ANKS1A;refGene_tx=XM_023253878.1;refGene_gn=ANKS1A;refGene_tx=XM_023253879.1;refGene_gn=ANKS1A;refGene_tx=XM_023253880.1
B2 36488047 36488134
refGene_gn=ANKS1A;refGene_tx=XM_023253873.1;refGene_gn=ANKS1A;refGene_tx=XM_023253874.1;refGene_gn=ANKS1A;refGene_tx=XM_023253875.1;refGene_gn=ANKS1A;refGene_tx=XM_023253876.1;refGene_gn=ANKS1A;refGene_tx=XM_023253877.1;refGene_gn=ANKS1A;refGene_tx=XM_023253878.1;refGene_gn=ANKS1A;refGene_tx=XM_023253879.1;refGene_gn=ANKS1A;refGene_tx=XM_023253880.1
B2 36488725 36488863
refGene_gn=ANKS1A;refGene_tx=XM_023253873.1;refGene_gn=ANKS1A;refGene_tx=XM_023253874.1;refGene_gn=ANKS1A;refGene_tx=XM_023253875.1;refGene_gn=ANKS1A;refGene_tx=XM_023253876.1;refGene_gn=ANKS1A;refGene_tx=XM_023253877.1;refGene_gn=ANKS1A;refGene_tx=XM_023253878.1;refGene_gn=ANKS1A;refGene_tx=XM_023253879.1;refGene_gn=ANKS1A;refGene_tx=XM_023253880.1
B2 36490345 36490516
refGene_gn=ANKS1A;refGene_tx=XM_023253873.1;refGene_gn=ANKS1A;refGene_tx=XM_023253874.1;refGene_gn=ANKS1A;refGene_tx=XM_023253875.1;refGene_gn=ANKS1A;refGene_tx=XM_023253876.1;refGene_gn=ANKS1A;refGene_tx=XM_02

3253877.1;refGene_gn=ANKS1A;refGene_tx=XM_023253878.1;refGene_gn=ANKS1A;refGene_tx=XM_023253879.1;refGene_gn=ANKS1A;refGene_tx=XM_023253880.1
B2 36490823 36490922
refGene_gn=ANKS1A;refGene_tx=XM_023253873.1;refGene_gn=ANKS1A;refGene_tx=XM_023253874.1;refGene_gn=ANKS1A;refGene_tx=XM_023253875.1;refGene_gn=ANKS1A;refGene_tx=XM_023253879.1;refGene_gn=ANKS1A;refGene_tx=XM_023253880.1
B2 36491536 36491656
refGene_gn=ANKS1A;refGene_tx=XM_023253873.1;refGene_gn=ANKS1A;refGene_tx=XM_023253874.1;refGene_gn=ANKS1A;refGene_tx=XM_023253875.1;refGene_gn=ANKS1A;refGene_tx=XM_023253876.1;refGene_gn=ANKS1A;refGene_tx=XM_023253877.1;refGene_gn=ANKS1A;refGene_tx=XM_023253878.1;refGene_gn=ANKS1A;refGene_tx=XM_023253879.1;refGene_gn=ANKS1A;refGene_tx=XM_023253880.1
B2 36493138 36493142
refGene_gn=ANKS1A;refGene_tx=XM_023253874.1;refGene_gn=ANKS1A;refGene_tx=XM_023253877.1
B2 36508677 36508910
refGene_gn=TCP11;refGene_tx=XM_006931635.4;refGene_gn=TCP11;refGene_tx=XM_011282258.3;refGene_gn=TCP11;refGene_tx=XM_019830479.2;refGene_gn=TCP11;refGene_tx=XM_019830480.1;refGene_gn=TCP11;refGene_tx=XM_023253881.1
B2 36509663 36509827
refGene_gn=TCP11;refGene_tx=XM_006931635.4;refGene_gn=TCP11;refGene_tx=XM_011282258.3;refGene_gn=TCP11;refGene_tx=XM_019830479.2;refGene_gn=TCP11;refGene_tx=XM_019830480.1;refGene_gn=TCP11;refGene_tx=XM_023253881.1
B2 36510582 36510764
refGene_gn=TCP11;refGene_tx=XM_006931635.4;refGene_gn=TCP11;refGene_tx=XM_011282258.3;refGene_gn=TCP11;refGene_tx=XM_019830479.2;refGene_gn=TCP11;refGene_tx=XM_019830480.1;refGene_gn=TCP11;refGene_tx=XM_023253881.1
B2 36510843 36511061
refGene_gn=TCP11;refGene_tx=XM_006931635.4;refGene_gn=TCP11;refGene_tx=XM_011282258.3;refGene_gn=TCP11;refGene_tx=XM_019830479.2;refGene_gn=TCP11;refGene_tx=XM_019830480.1;refGene_gn=TCP11;refGene_tx=XM_023253881.1
B2 36511343 36511480
refGene_gn=TCP11;refGene_tx=XM_006931635.4;refGene_gn=TCP11;refGene_tx=XM_011282258.3;refGene_gn=TCP11;refGene_tx=XM_019830479.2;refGene_gn=TCP11;refGene_tx=XM_019830480.1;refGene_gn=TCP11;refGene_tx=XM_023253881.1
B2 36512440 36512661
refGene_gn=TCP11;refGene_tx=XM_006931635.4;refGene_gn=TCP11;refGene_tx=XM_011282258.3;refGene_gn=TCP11;refGene_tx=XM_019830479.2;refGene_gn=TCP11;refGene_tx=XM_019830480.1;refGene_gn=TCP11;refGene_tx=XM_023253881.1
B2 36519659 36519780
refGene_gn=TCP11;refGene_tx=XM_006931635.4;refGene_gn=TCP11;refGene_tx

=XM_011282258.3;refGene_gn=TCP11;refGene_tx=XM_019830479.2;refGene_gn=TCP11;refGene_tx=XM_019830480.1;refGene_gn=TCP11;refGene_tx=XM_023253881.1
B2 36525814 36525911
refGene_gn=TCP11;refGene_tx=XM_006931635.4;refGene_gn=TCP11;refGene_tx=XM_011282258.3;refGene_gn=TCP11;refGene_tx=XM_019830479.2;refGene_gn=TCP11;refGene_tx=XM_019830480.1;refGene_gn=TCP11;refGene_tx=XM_023253881.1
B2 36528806 36528930
refGene_gn=TCP11;refGene_tx=XM_006931635.4;refGene_gn=TCP11;refGene_tx=XM_011282258.3;refGene_gn=TCP11;refGene_tx=XM_019830479.2;refGene_gn=TCP11;refGene_tx=XM_019830480.1
B2 36606802 36606887
refGene_gn=SCUBE3;refGene_tx=XM_019831576.2;refGene_gn=SCUBE3;refGene_tx=XM_023253882.1;refGene_gn=SCUBE3;refGene_tx=XM_023253883.1;refGene_gn=SCUBE3;refGene_tx=XM_023253884.1;refGene_gn=SCUBE3;refGene_tx=XM_023253885.1;refGene_gn=SCUBE3;refGene_tx=XM_023253886.1;refGene_gn=SCUBE3;refGene_tx=XM_023253887.1;refGene_gn=SCUBE3;refGene_tx=XM_023253888.1;refGene_gn=SCUBE3;refGene_tx=XM_023253889.1;refGene_gn=SCUBE3;refGene_tx=XM_023253890.1;refGene_gn=SCUBE3;refGene_tx=XM_023253891.1
B2 36619835 36619958
refGene_gn=SCUBE3;refGene_tx=XM_019831576.2;refGene_gn=SCUBE3;refGene_tx=XM_023253882.1;refGene_gn=SCUBE3;refGene_tx=XM_023253883.1;refGene_gn=SCUBE3;refGene_tx=XM_023253884.1;refGene_gn=SCUBE3;refGene_tx=XM_023253885.1;refGene_gn=SCUBE3;refGene_tx=XM_023253886.1;refGene_gn=SCUBE3;refGene_tx=XM_023253887.1;refGene_gn=SCUBE3;refGene_tx=XM_023253888.1;refGene_gn=SCUBE3;refGene_tx=XM_023253889.1;refGene_gn=SCUBE3;refGene_tx=XM_023253890.1;refGene_gn=SCUBE3;refGene_tx=XM_023253891.1
B2 36620952 36621078
refGene_gn=SCUBE3;refGene_tx=XM_019831576.2;refGene_gn=SCUBE3;refGene_tx=XM_023253882.1;refGene_gn=SCUBE3;refGene_tx=XM_023253883.1;refGene_gn=SCUBE3;refGene_tx=XM_023253884.1;refGene_gn=SCUBE3;refGene_tx=XM_023253885.1;refGene_gn=SCUBE3;refGene_tx=XM_023253886.1;refGene_gn=SCUBE3;refGene_tx=XM_023253887.1;refGene_gn=SCUBE3;refGene_tx=XM_023253888.1;refGene_gn=SCUBE3;refGene_tx=XM_023253889.1;refGene_gn=SCUBE3;refGene_tx=XM_023253890.1;refGene_gn=SCUBE3;refGene_tx=XM_023253891.1
B2 36623740 36623875
refGene_gn=SCUBE3;refGene_tx=XM_019831576.2;refGene_gn=SCUBE3;refGene_tx=XM_023253882.1;refGene_gn=SCUBE3;refGene_tx=XM_023253883.1;refGene_gn=SCUBE3;refGene_tx=XM_023253884.1;refGene_gn=SCUBE3;refGene_tx=XM_023253885.1;refGene_gn=SCUBE3;refGene_tx=XM_023253886.1;refGene_gn=SCUBE3;refGene_tx=XM_023253887.1;refGene_gn=SCUBE3;refGene_tx=XM_023253888.1;refGene_gn=SCUBE3;refGene_tx=XM_023253889.1;refGene_gn=SCUBE3;refGene_tx=XM_023253890.1;refGene_gn=SCUBE3;refGene_tx=XM_023253891.1
B2 36625512 36625638
refGene_gn=SCUBE3;refGene_tx=XM_019831576.2;refGene_gn=SCUBE3;refGene_tx=XM_023253882.1;refGene_gn=SCUBE3;refGene_tx=XM_023253883.1;refGene_gn=SCUBE3;refGene_tx=XM_023253884.1;refGene_gn=SCUBE3;refGene_tx=XM_023253885.1;refGene_gn=SCUBE3;refGene_tx=XM_023253886.1;refGene_gn=SCUBE3;refGene_tx=XM_023253887.1;refGene_gn=SCUBE3;refGene_tx=XM_023253888.

1;refGene_gn=SCUBE3;refGene_tx=XM_023253889.1;refGene_gn=SCUBE3;refGene_tx=XM_023253890.1;refGene_gn=SCUBE3;refGene_tx=XM_023253891.1
B2 36625825 36625942
refGene_gn=SCUBE3;refGene_tx=XM_019831576.2;refGene_gn=SCUBE3;refGene_tx=XM_023253882.1;refGene_gn=SCUBE3;refGene_tx=XM_023253883.1;refGene_gn=SCUBE3;refGene_tx=XM_023253884.1;refGene_gn=SCUBE3;refGene_tx=XM_023253885.1;refGene_gn=SCUBE3;refGene_tx=XM_023253886.1;refGene_gn=SCUBE3;refGene_tx=XM_023253887.1;refGene_gn=SCUBE3;refGene_tx=XM_023253888.1;refGene_gn=SCUBE3;refGene_tx=XM_023253889.1;refGene_gn=SCUBE3;refGene_tx=XM_023253890.1;refGene_gn=SCUBE3;refGene_tx=XM_023253891.1
B2 36628322 36628370
refGene_gn=SCUBE3;refGene_tx=XM_019831576.2;refGene_gn=SCUBE3;refGene_tx=XM_023253882.1;refGene_gn=SCUBE3;refGene_tx=XM_023253885.1;refGene_gn=SCUBE3;refGene_tx=XM_023253886.1;refGene_gn=SCUBE3;refGene_tx=XM_023253887.1;refGene_gn=SCUBE3;refGene_tx=XM_023253888.1;refGene_gn=SCUBE3;refGene_tx=XM_023253889.1;refGene_gn=SCUBE3;refGene_tx=XM_023253890.1;refGene_gn=SCUBE3;refGene_tx=XM_023253891.1
B2 36630752 36630869
refGene_gn=SCUBE3;refGene_tx=XM_019831576.2;refGene_gn=SCUBE3;refGene_tx=XM_023253882.1;refGene_gn=SCUBE3;refGene_tx=XM_023253883.1;refGene_gn=SCUBE3;refGene_tx=XM_023253884.1;refGene_gn=SCUBE3;refGene_tx=XM_023253885.1;refGene_gn=SCUBE3;refGene_tx=XM_023253886.1;refGene_gn=SCUBE3;refGene_tx=XM_023253887.1;refGene_gn=SCUBE3;refGene_tx=XM_023253888.1;refGene_gn=SCUBE3;refGene_tx=XM_023253889.1;refGene_gn=SCUBE3;refGene_tx=XM_023253890.1;refGene_gn=SCUBE3;refGene_tx=XM_023253891.1
B2 36632557 36632680
refGene_gn=SCUBE3;refGene_tx=XM_019831576.2;refGene_gn=SCUBE3;refGene_tx=XM_023253882.1;refGene_gn=SCUBE3;refGene_tx=XM_023253883.1;refGene_gn=SCUBE3;refGene_tx=XM_023253884.1;refGene_gn=SCUBE3;refGene_tx=XM_023253885.1;refGene_gn=SCUBE3;refGene_tx=XM_023253886.1;refGene_gn=SCUBE3;refGene_tx=XM_023253887.1;refGene_gn=SCUBE3;refGene_tx=XM_023253888.1;refGene_gn=SCUBE3;refGene_tx=XM_023253889.1;refGene_gn=SCUBE3;refGene_tx=XM_023253890.1;refGene_gn=SCUBE3;refGene_tx=XM_023253891.1
B2 36633154 36633271
refGene_gn=SCUBE3;refGene_tx=XM_019831576.2;refGene_gn=SCUBE3;refGene_tx=XM_023253882.1;refGene_gn=SCUBE3;refGene_tx=XM_023253883.1;refGene_gn=SCUBE3;refGene_tx=XM_023253884.1;refGene_gn=SCUBE3;refGene_tx=XM_023253885.1;refGene_gn=SCUBE3;refGene_tx=XM_023253886.1;refGene_gn=SCUBE3;refGene_tx=XM_023253887.1;refGene_gn=SCUBE3;refGene_tx=XM_023253888.1;refGene_gn=SCUBE3;refGene_tx=XM_023253889.1;refGene_gn=SCUBE3;refGene_tx=XM_023253890.1;refGene_gn=SCUBE3;refGene_tx=XM_023253891.1
B2 36633764 36633890
refGene_gn=SCUBE3;refGene_tx=XM_019831576.2;refGene_gn=SCUBE3;refGene_tx=XM_023253882.1;refGene_gn=SCUBE3;refGene_tx=XM_023253883.1;refGene_gn=SCUBE3;refGene_tx=XM_023253884.1;refGene_gn=SCUBE3;refGene_tx=XM_023253885.1;refGene_gn=SCUBE3;refGene_tx=XM_023253886.1;refGene_gn=SCUBE3;refGene_tx=XM_023253887.1;refGene_gn=SCUBE3;refGene_tx=XM_023253888.1;refGene_gn=SCUBE3;refGene_tx=XM_023253889.1;refGene_gn=SCUBE3;refGene_tx=XM_023253890.1;refGene_gn=SCUBE3;refGene_tx=XM_023253891.1
B2 36634175 36634292

refGene_gn=SCUBE3;refGene_tx=XM_019831576.2;refGene_gn=SCUBE3;refGene_tx=XM_023253882.1;refGene_gn=SCUBE3;refGene_tx=XM_023253883.1;refGene_gn=SCUBE3;refGene_tx=XM_023253884.1;refGene_gn=SCUBE3;refGene_tx=XM_023253885.1;refGene_gn=SCUBE3;refGene_tx=XM_023253886.1;refGene_gn=SCUBE3;refGene_tx=XM_023253887.1;refGene_gn=SCUBE3;refGene_tx=XM_023253888.1;refGene_gn=SCUBE3;refGene_tx=XM_023253889.1;refGene_gn=SCUBE3;refGene_tx=XM_023253890.1;refGene_gn=SCUBE3;refGene_tx=XM_023253891.1
B2 36634432 36634537
refGene_gn=SCUBE3;refGene_tx=XM_019831576.2;refGene_gn=SCUBE3;refGene_tx=XM_023253882.1;refGene_gn=SCUBE3;refGene_tx=XM_023253883.1;refGene_gn=SCUBE3;refGene_tx=XM_023253884.1;refGene_gn=SCUBE3;refGene_tx=XM_023253885.1;refGene_gn=SCUBE3;refGene_tx=XM_023253887.1;refGene_gn=SCUBE3;refGene_tx=XM_023253888.1;refGene_gn=SCUBE3;refGene_tx=XM_023253889.1;refGene_gn=SCUBE3;refGene_tx=XM_023253890.1;refGene_gn=SCUBE3;refGene_tx=XM_023253891.1
B2 36634827 36634944
refGene_gn=SCUBE3;refGene_tx=XM_019831576.2;refGene_gn=SCUBE3;refGene_tx=XM_023253882.1;refGene_gn=SCUBE3;refGene_tx=XM_023253883.1;refGene_gn=SCUBE3;refGene_tx=XM_023253884.1;refGene_gn=SCUBE3;refGene_tx=XM_023253885.1;refGene_gn=SCUBE3;refGene_tx=XM_023253886.1;refGene_gn=SCUBE3;refGene_tx=XM_023253887.1;refGene_gn=SCUBE3;refGene_tx=XM_023253888.1;refGene_gn=SCUBE3;refGene_tx=XM_023253889.1;refGene_gn=SCUBE3;refGene_tx=XM_023253890.1;refGene_gn=SCUBE3;refGene_tx=XM_023253891.1
B2 36635281 36635440
refGene_gn=SCUBE3;refGene_tx=XM_019831576.2;refGene_gn=SCUBE3;refGene_tx=XM_023253882.1;refGene_gn=SCUBE3;refGene_tx=XM_023253883.1;refGene_gn=SCUBE3;refGene_tx=XM_023253884.1;refGene_gn=SCUBE3;refGene_tx=XM_023253885.1;refGene_gn=SCUBE3;refGene_tx=XM_023253886.1;refGene_gn=SCUBE3;refGene_tx=XM_023253887.1;refGene_gn=SCUBE3;refGene_tx=XM_023253888.1;refGene_gn=SCUBE3;refGene_tx=XM_023253889.1;refGene_gn=SCUBE3;refGene_tx=XM_023253890.1;refGene_gn=SCUBE3;refGene_tx=XM_023253891.1
B2 36635682 36635898
refGene_gn=SCUBE3;refGene_tx=XM_019831576.2;refGene_gn=SCUBE3;refGene_tx=XM_023253882.1;refGene_gn=SCUBE3;refGene_tx=XM_023253883.1;refGene_gn=SCUBE3;refGene_tx=XM_023253884.1;refGene_gn=SCUBE3;refGene_tx=XM_023253885.1;refGene_gn=SCUBE3;refGene_tx=XM_023253886.1;refGene_gn=SCUBE3;refGene_tx=XM_023253887.1;refGene_gn=SCUBE3;refGene_tx=XM_023253888.1;refGene_gn=SCUBE3;refGene_tx=XM_023253889.1;refGene_gn=SCUBE3;refGene_tx=XM_023253890.1;refGene_gn=SCUBE3;refGene_tx=XM_023253891.1
B2 36636260 36636422
refGene_gn=SCUBE3;refGene_tx=XM_019831576.2;refGene_gn=SCUBE3;refGene_tx=XM_023253882.1;refGene_gn=SCUBE3;refGene_tx=XM_023253883.1;refGene_gn=SCUBE3;refGene_tx=XM_023253884.1;refGene_gn=SCUBE3;refGene_tx=XM_023253885.1;refGene_gn=SCUBE3;refGene_tx=XM_023253886.1;refGene_gn=SCUBE3;refGene_tx=XM_023253889.1;refGene_gn=SCUBE3;refGene_tx=XM_023253890.1;refGene_gn=SCUBE3;refGene_tx=XM_023253891.1
B2 36636625 36636793
refGene_gn=SCUBE3;refGene_tx=XM_019831576.2;refGene_gn=SCUBE3;refGene_tx=XM_023253882.1;refGene_gn=SCUBE3;refGene_tx=XM_023253883.1;refGene_gn=SCUBE3;refGene_tx=XM_023253884.1;refGene_gn=SCUBE3;refGene_tx=XM_02

3253885.1;refGene_gn=SCUBE3;refGene_tx=XM_023253886.1;refGene_gn=SCUBE3;refGene_tx=XM_023253887.1;refGene_gn=SCUBE3;refGene_tx=XM_023253888.1;refGene_gn=SCUBE3;refGene_tx=XM_023253889.1;refGene_gn=SCUBE3;refGene_tx=XM_023253890.1;refGene_gn=SCUBE3;refGene_tx=XM_023253891.1
B2 36637294 36637495
refGene_gn=SCUBE3;refGene_tx=XM_019831576.2;refGene_gn=SCUBE3;refGene_tx=XM_023253882.1;refGene_gn=SCUBE3;refGene_tx=XM_023253883.1;refGene_gn=SCUBE3;refGene_tx=XM_023253884.1;refGene_gn=SCUBE3;refGene_tx=XM_023253885.1;refGene_gn=SCUBE3;refGene_tx=XM_023253886.1;refGene_gn=SCUBE3;refGene_tx=XM_023253887.1;refGene_gn=SCUBE3;refGene_tx=XM_023253888.1;refGene_gn=SCUBE3;refGene_tx=XM_023253889.1;refGene_gn=SCUBE3;refGene_tx=XM_023253890.1;refGene_gn=SCUBE3;refGene_tx=XM_023253891.1
B2 36637875 36638073
refGene_gn=SCUBE3;refGene_tx=XM_019831576.2;refGene_gn=SCUBE3;refGene_tx=XM_023253882.1;refGene_gn=SCUBE3;refGene_tx=XM_023253883.1;refGene_gn=SCUBE3;refGene_tx=XM_023253884.1;refGene_gn=SCUBE3;refGene_tx=XM_023253885.1;refGene_gn=SCUBE3;refGene_tx=XM_023253886.1;refGene_gn=SCUBE3;refGene_tx=XM_023253887.1;refGene_gn=SCUBE3;refGene_tx=XM_023253888.1;refGene_gn=SCUBE3;refGene_tx=XM_023253889.1;refGene_gn=SCUBE3;refGene_tx=XM_023253890.1;refGene_gn=SCUBE3;refGene_tx=XM_023253891.1
B2 36638564 36638717
refGene_gn=SCUBE3;refGene_tx=XM_019831576.2;refGene_gn=SCUBE3;refGene_tx=XM_023253882.1;refGene_gn=SCUBE3;refGene_tx=XM_023253883.1;refGene_gn=SCUBE3;refGene_tx=XM_023253884.1;refGene_gn=SCUBE3;refGene_tx=XM_023253885.1;refGene_gn=SCUBE3;refGene_tx=XM_023253886.1;refGene_gn=SCUBE3;refGene_tx=XM_023253887.1;refGene_gn=SCUBE3;refGene_tx=XM_023253888.1
B2 36638839 36638919
refGene_gn=SCUBE3;refGene_tx=XM_019831576.2;refGene_gn=SCUBE3;refGene_tx=XM_023253882.1;refGene_gn=SCUBE3;refGene_tx=XM_023253883.1;refGene_gn=SCUBE3;refGene_tx=XM_023253884.1;refGene_gn=SCUBE3;refGene_tx=XM_023253885.1;refGene_gn=SCUBE3;refGene_tx=XM_023253886.1;refGene_gn=SCUBE3;refGene_tx=XM_023253887.1;refGene_gn=SCUBE3;refGene_tx=XM_023253888.1
B2 36640546 36640696
refGene_gn=SCUBE3;refGene_tx=XM_019831576.2;refGene_gn=SCUBE3;refGene_tx=XM_023253882.1;refGene_gn=SCUBE3;refGene_tx=XM_023253883.1;refGene_gn=SCUBE3;refGene_tx=XM_023253884.1;refGene_gn=SCUBE3;refGene_tx=XM_023253885.1;refGene_gn=SCUBE3;refGene_tx=XM_023253886.1;refGene_gn=SCUBE3;refGene_tx=XM_023253887.1;refGene_gn=SCUBE3;refGene_tx=XM_023253888.1
B2 36669645 36669718
refGene_gn=ZNF76;refGene_tx=XM_011282254.2;refGene_gn=ZNF76;refGene_tx=XM_011282255.2;refGene_gn=ZNF76;refGene_tx=XM_011282256.3;refGene_gn=ZNF76;refGene_tx=XM_011282257.2
B2 36671076 36671149 refGene_gn=ZNF76;refGene_tx=XM_023253892.1
B2 36675916 36675997
refGene_gn=ZNF76;refGene_tx=XM_011282254.2;refGene_gn=ZNF76;refGene_tx=XM_011282255.2;refGene_gn=ZNF76;refGene_tx=XM_011282256.3;refGene_gn=ZNF76;refGene_tx=XM_011282257.2;refGene_gn=ZNF76;refGene_tx=XM_0232538

92.1
B2 36676111 36676189
refGene_gn=ZNF76;refGene_tx=XM_011282254.2;refGene_gn=ZNF76;refGene_tx=XM_011282255.2;refGene_gn=ZNF76;refGene_tx=XM_011282256.3;refGene_gn=ZNF76;refGene_tx=XM_011282257.2;refGene_gn=ZNF76;refGene_tx=XM_023253892.1
B2 36677603 36677803
refGene_gn=ZNF76;refGene_tx=XM_011282254.2;refGene_gn=ZNF76;refGene_tx=XM_011282255.2;refGene_gn=ZNF76;refGene_tx=XM_011282256.3;refGene_gn=ZNF76;refGene_tx=XM_011282257.2;refGene_gn=ZNF76;refGene_tx=XM_023253892.1
B2 36679862 36679979
refGene_gn=ZNF76;refGene_tx=XM_011282254.2;refGene_gn=ZNF76;refGene_tx=XM_011282255.2;refGene_gn=ZNF76;refGene_tx=XM_011282256.3;refGene_gn=ZNF76;refGene_tx=XM_011282257.2;refGene_gn=ZNF76;refGene_tx=XM_023253892.1
B2 36680299 36680375
refGene_gn=ZNF76;refGene_tx=XM_011282254.2;refGene_gn=ZNF76;refGene_tx=XM_011282255.2;refGene_gn=ZNF76;refGene_tx=XM_011282256.3;refGene_gn=ZNF76;refGene_tx=XM_011282257.2;refGene_gn=ZNF76;refGene_tx=XM_023253892.1
B2 36680901 36681027
refGene_gn=ZNF76;refGene_tx=XM_011282254.2;refGene_gn=ZNF76;refGene_tx=XM_011282255.2;refGene_gn=ZNF76;refGene_tx=XM_011282256.3;refGene_gn=ZNF76;refGene_tx=XM_011282257.2;refGene_gn=ZNF76;refGene_tx=XM_023253892.1
B2 36681128 36681308
refGene_gn=ZNF76;refGene_tx=XM_011282254.2;refGene_gn=ZNF76;refGene_tx=XM_011282255.2;refGene_gn=ZNF76;refGene_tx=XM_011282256.3;refGene_gn=ZNF76;refGene_tx=XM_011282257.2;refGene_gn=ZNF76;refGene_tx=XM_023253892.1
B2 36682102 36682336
refGene_gn=ZNF76;refGene_tx=XM_011282254.2;refGene_gn=ZNF76;refGene_tx=XM_011282255.2;refGene_gn=ZNF76;refGene_tx=XM_011282256.3;refGene_gn=ZNF76;refGene_tx=XM_011282257.2;refGene_gn=ZNF76;refGene_tx=XM_023253892.1
B2 36682429 36682593
refGene_gn=ZNF76;refGene_tx=XM_011282254.2;refGene_gn=ZNF76;refGene_tx=XM_011282255.2;refGene_gn=ZNF76;refGene_tx=XM_011282256.3;refGene_gn=ZNF76;refGene_tx=XM_011282257.2;refGene_gn=ZNF76;refGene_tx=XM_023253892.1
B2 36683289 36683460
refGene_gn=ZNF76;refGene_tx=XM_011282254.2;refGene_gn=ZNF76;refGene_tx=XM_011282255.2;refGene_gn=ZNF76;refGene_tx=XM_011282256.3;refGene_gn=ZNF76;refGene_tx=XM_011282257.2;refGene_gn=ZNF76;refGene_tx=XM_023253892.1
B2 36683977 36684091
refGene_gn=ZNF76;refGene_tx=XM_011282254.2;refGene_gn=ZNF76;refGene_tx=XM_011282255.2;refGene_gn=ZNF76;refGene_tx=XM_011282256.3;refGene_gn=ZNF76;refGene_tx=XM_023253892.1

B2	36684655	36684760
refGene_gn=ZNF76;refGene_tx=XM_011282254.2;refGene_gn=ZNF76;refGene_tx=XM_011282255.2;refGene_gn=ZNF76;refGene_tx=XM_011282256.3;refGene_gn=ZNF76;refGene_tx=XM_023253892.1
B2	36686805	36686880	refGene_gn=DEF6;refGene_tx=XM_006931639.3
B2	36687363	36687459
refGene_gn=DEF6;refGene_tx=XM_006931638.3;refGene_gn=DEF6;refGene_tx=XM_023253893.1
B2	36696985	36697126
refGene_gn=DEF6;refGene_tx=XM_006931638.3;refGene_gn=DEF6;refGene_tx=XM_006931639.3;refGene_gn=DEF6;refGene_tx=XM_023253893.1
B2	36697754	36697940
refGene_gn=DEF6;refGene_tx=XM_006931638.3;refGene_gn=DEF6;refGene_tx=XM_006931639.3;refGene_gn=DEF6;refGene_tx=XM_023253893.1
B2	36699514	36699751
refGene_gn=DEF6;refGene_tx=XM_006931638.3;refGene_gn=DEF6;refGene_tx=XM_006931639.3;refGene_gn=DEF6;refGene_tx=XM_023253893.1
B2	36699840	36699987
refGene_gn=DEF6;refGene_tx=XM_006931638.3;refGene_gn=DEF6;refGene_tx=XM_006931639.3;refGene_gn=DEF6;refGene_tx=XM_023253893.1
B2	36701953	36702062
refGene_gn=DEF6;refGene_tx=XM_006931638.3;refGene_gn=DEF6;refGene_tx=XM_006931639.3;refGene_gn=DEF6;refGene_tx=XM_023253893.1
B2	36702251	36702550
refGene_gn=DEF6;refGene_tx=XM_006931638.3;refGene_gn=DEF6;refGene_tx=XM_006931639.3;refGene_gn=DEF6;refGene_tx=XM_023253893.1
B2	36703599	36703766
refGene_gn=DEF6;refGene_tx=XM_006931638.3;refGene_gn=DEF6;refGene_tx=XM_006931639.3;refGene_gn=DEF6;refGene_tx=XM_023253893.1
B2	36703879	36704078
refGene_gn=DEF6;refGene_tx=XM_006931638.3;refGene_gn=DEF6;refGene_tx=XM_006931639.3;refGene_gn=DEF6;refGene_tx=XM_023253893.1
B2	36705039	36705130
refGene_gn=DEF6;refGene_tx=XM_006931638.3;refGene_gn=DEF6;refGene_tx=XM_006931639.3;refGene_gn=DEF6;refGene_tx=XM_023253893.1
B2	36705333	36705557
refGene_gn=DEF6;refGene_tx=XM_006931638.3;refGene_gn=DEF6;refGene_tx=XM_006931639.3;refGene_gn=DEF6;refGene_tx=XM_023253893.1
B2	36781271	36781442
refGene_gn=PPARD;refGene_tx=XM_011282251.3;refGene_gn=PPARD;refGene_tx=XM_011282253.3;refGene_gn=PPARD;refGene_tx=XM_019830482.2;refGene_gn=PPARD;refGene_tx=XM_019830483.2;refGene_gn=PPARD;refGene_tx=XM_019830484.2
B2	36791230	36791454
refGene_gn=PPARD;refGene_tx=XM_011282251.3;refGene_gn=PPARD;refGene_tx=XM_011282253.3
B2	36792596	36792735
refGene_gn=PPARD;refGene_tx=XM_011282251.3;refGene_gn=PPARD;refGene_tx=XM_011282253.3;refGene_gn=PPARD;refGene_tx=XM_019830482.2;refGene_gn=PPARD;refGene_tx=XM_019830483.2;refGene_gn=PPARD;refGene_tx=XM_0198304

84.2
B2 36794131 36794334
refGene_gn=PPARD;refGene_tx=XM_011282251.3;refGene_gn=PPARD;refGene_tx
=XM_011282253.3;refGene_gn=PPARD;refGene_tx=XM_019830482.2;refGene_gn=
PPARD;refGene_tx=XM_019830483.2;refGene_gn=PPARD;refGene_tx=XM_0198304
84.2
B2 36794511 36794962
refGene_gn=PPARD;refGene_tx=XM_011282251.3;refGene_gn=PPARD;refGene_tx
=XM_011282253.3;refGene_gn=PPARD;refGene_tx=XM_019830482.2;refGene_gn=
PPARD;refGene_tx=XM_019830483.2;refGene_gn=PPARD;refGene_tx=XM_0198304
84.2
B2 36795993 36796241
refGene_gn=PPARD;refGene_tx=XM_011282251.3;refGene_gn=PPARD;refGene_tx
=XM_011282253.3;refGene_gn=PPARD;refGene_tx=XM_019830482.2;refGene_gn=
PPARD;refGene_tx=XM_019830483.2;refGene_gn=PPARD;refGene_tx=XM_0198304
84.2
B2 36806297 36806368
refGene_gn=LOC109499639;refGene_tx=XM_019830485.2;refGene_gn=LOC109499
639;refGene_tx=XM_019830486.2
B2 36811573 36811693
refGene_gn=LOC109499639;refGene_tx=XM_019830485.2;refGene_gn=LOC109499
639;refGene_tx=XM_019830486.2
B2 36811875 36812120
refGene_gn=LOC109499639;refGene_tx=XM_019830485.2;refGene_gn=LOC109499
639;refGene_tx=XM_019830486.2
B2 36813959 36814063
refGene_gn=LOC109499639;refGene_tx=XM_019830485.2;refGene_gn=LOC109499
639;refGene_tx=XM_019830486.2
B2 36814230 36814403
refGene_gn=LOC109499639;refGene_tx=XM_019830485.2;refGene_gn=LOC109499
639;refGene_tx=XM_019830486.2
B2 36814493 36814604
refGene_gn=LOC109499639;refGene_tx=XM_019830485.2;refGene_gn=LOC109499
639;refGene_tx=XM_019830486.2
B2 36816290 36816435
refGene_gn=LOC109499639;refGene_tx=XM_019830485.2;refGene_gn=LOC109499
639;refGene_tx=XM_019830486.2
B2 36816671 36816809
refGene_gn=LOC109499639;refGene_tx=XM_019830485.2;refGene_gn=LOC109499
639;refGene_tx=XM_019830486.2
B2 36819475 36819723
refGene_gn=FANCE;refGene_tx=XM_019831577.2;refGene_gn=FANCE;refGene_tx
=XM_023253894.1
B2 36822916 36823538
refGene_gn=FANCE;refGene_tx=XM_019831577.2;refGene_gn=FANCE;refGene_tx
=XM_023253894.1
B2 36824114 36824159 refGene_gn=FANCE;refGene_tx=XM_019831577.2
B2 36824468 36824528
refGene_gn=FANCE;refGene_tx=XM_019831577.2;refGene_gn=FANCE;refGene_tx
=XM_023253894.1

B2 36824825 36824969
refGene_gn=FANCE;refGene_tx=XM_019831577.2;refGene_gn=FANCE;refGene_tx=XM_023253894.1
B2 36825333 36825457
refGene_gn=FANCE;refGene_tx=XM_019831577.2;refGene_gn=FANCE;refGene_tx=XM_023253894.1
B2 36825668 36825747
refGene_gn=FANCE;refGene_tx=XM_019831577.2;refGene_gn=FANCE;refGene_tx=XM_023253894.1
B2 36827322 36827389
refGene_gn=FANCE;refGene_tx=XM_019831577.2;refGene_gn=FANCE;refGene_tx=XM_023253894.1
B2 36828397 36828523
refGene_gn=FANCE;refGene_tx=XM_019831577.2;refGene_gn=FANCE;refGene_tx=XM_023253894.1
B2 36830582 36830684
refGene_gn=FANCE;refGene_tx=XM_019831577.2;refGene_gn=FANCE;refGene_tx=XM_023253894.1
B2 36832124 36832129 refGene_gn=RPL10A;refGene_tx=XM_003986099.5
B2 36832492 36832567 refGene_gn=RPL10A;refGene_tx=XM_003986099.5
B2 36832658 36832739 refGene_gn=RPL10A;refGene_tx=XM_003986099.5
B2 36833096 36833245 refGene_gn=RPL10A;refGene_tx=XM_003986099.5
B2 36834364 36834537 refGene_gn=RPL10A;refGene_tx=XM_003986099.5
B2 36834842 36835013 refGene_gn=RPL10A;refGene_tx=XM_003986099.5
B2 36840027 36840141 refGene_gn=TEAD3;refGene_tx=XM_023254715.1
B2 36840319 36840472 refGene_gn=TEAD3;refGene_tx=XM_023254715.1
B2 36840550 36840691 refGene_gn=TEAD3;refGene_tx=XM_023254715.1
B2 36840905 36841079 refGene_gn=TEAD3;refGene_tx=XM_023254715.1
B2 36841266 36841400 refGene_gn=TEAD3;refGene_tx=XM_023254715.1
B2 36842154 36842216 refGene_gn=TEAD3;refGene_tx=XM_023254715.1
B2 36842796 36842846 refGene_gn=TEAD3;refGene_tx=XM_023254715.1
B2 36842956 36843094 refGene_gn=TEAD3;refGene_tx=XM_023254715.1
B2 36843672 36843684 refGene_gn=TEAD3;refGene_tx=XM_023254715.1
B2 36844502 36844565 refGene_gn=TEAD3;refGene_tx=XM_023254715.1
B2 36849795 36849860 refGene_gn=TEAD3;refGene_tx=XM_023254715.1
B2 36851652 36851904 refGene_gn=TEAD3;refGene_tx=XM_023254715.1
B2 36861314 36861522 refGene_gn=TEAD3;refGene_tx=XM_023254715.1
B2 36862366 36862384 refGene_gn=TEAD3;refGene_tx=XM_023254715.1
B2 36862788 36862922
refGene_gn=TULP1;refGene_tx=XM_019830487.2;refGene_gn=TULP1;refGene_tx=XM_023253895.1;refGene_gn=TULP1;refGene_tx=XM_023253896.1;refGene_gn=TULP1;refGene_tx=XM_023253897.1;refGene_gn=TULP1;refGene_tx=XM_023253898.1;refGene_gn=TULP1;refGene_tx=XM_023253899.1
B2 36864573 36864745
refGene_gn=TULP1;refGene_tx=XM_019830487.2;refGene_gn=TULP1;refGene_tx=XM_023253895.1;refGene_gn=TULP1;refGene_tx=XM_023253896.1;refGene_gn=TULP1;refGene_tx=XM_023253897.1;refGene_gn=TULP1;refGene_tx=XM_023253898.1;refGene_gn=TULP1;refGene_tx=XM_023253899.1
B2 36866824 36866923
refGene_gn=TULP1;refGene_tx=XM_019830487.2;refGene_gn=TULP1;refGene_tx

=XM_023253895.1;refGene_gn=TULP1;refGene_tx=XM_023253896.1;refGene_gn=TULP1;refGene_tx=XM_023253897.1;refGene_gn=TULP1;refGene_tx=XM_023253898.1;refGene_gn=TULP1;refGene_tx=XM_023253899.1
B2 36866999 36867114
refGene_gn=TULP1;refGene_tx=XM_019830487.2;refGene_gn=TULP1;refGene_tx=XM_023253895.1;refGene_gn=TULP1;refGene_tx=XM_023253896.1;refGene_gn=TULP1;refGene_tx=XM_023253897.1;refGene_gn=TULP1;refGene_tx=XM_023253898.1;refGene_gn=TULP1;refGene_tx=XM_023253899.1
B2 36868416 36868529
refGene_gn=TULP1;refGene_tx=XM_019830487.2;refGene_gn=TULP1;refGene_tx=XM_023253895.1;refGene_gn=TULP1;refGene_tx=XM_023253896.1;refGene_gn=TULP1;refGene_tx=XM_023253897.1;refGene_gn=TULP1;refGene_tx=XM_023253898.1;refGene_gn=TULP1;refGene_tx=XM_023253899.1
B2 36868732 36868903
refGene_gn=TULP1;refGene_tx=XM_019830487.2;refGene_gn=TULP1;refGene_tx=XM_023253895.1;refGene_gn=TULP1;refGene_tx=XM_023253896.1;refGene_gn=TULP1;refGene_tx=XM_023253897.1;refGene_gn=TULP1;refGene_tx=XM_023253898.1;refGene_gn=TULP1;refGene_tx=XM_023253899.1
B2 36869004 36869010
refGene_gn=TULP1;refGene_tx=XM_019830487.2;refGene_gn=TULP1;refGene_tx=XM_023253895.1;refGene_gn=TULP1;refGene_tx=XM_023253896.1;refGene_gn=TULP1;refGene_tx=XM_023253898.1;refGene_gn=TULP1;refGene_tx=XM_023253899.1
B2 36872252 36872359
refGene_gn=TULP1;refGene_tx=XM_019830487.2;refGene_gn=TULP1;refGene_tx=XM_023253895.1;refGene_gn=TULP1;refGene_tx=XM_023253896.1;refGene_gn=TULP1;refGene_tx=XM_023253897.1;refGene_gn=TULP1;refGene_tx=XM_023253898.1;refGene_gn=TULP1;refGene_tx=XM_023253899.1
B2 36872634 36872754
refGene_gn=TULP1;refGene_tx=XM_019830487.2;refGene_gn=TULP1;refGene_tx=XM_023253895.1;refGene_gn=TULP1;refGene_tx=XM_023253896.1;refGene_gn=TULP1;refGene_tx=XM_023253897.1;refGene_gn=TULP1;refGene_tx=XM_023253898.1;refGene_gn=TULP1;refGene_tx=XM_023253899.1
B2 36872828 36872930
refGene_gn=TULP1;refGene_tx=XM_019830487.2;refGene_gn=TULP1;refGene_tx=XM_023253895.1;refGene_gn=TULP1;refGene_tx=XM_023253896.1;refGene_gn=TULP1;refGene_tx=XM_023253897.1;refGene_gn=TULP1;refGene_tx=XM_023253898.1;refGene_gn=TULP1;refGene_tx=XM_023253899.1
B2 36873623 36873886
refGene_gn=TULP1;refGene_tx=XM_019830487.2;refGene_gn=TULP1;refGene_tx=XM_023253895.1;refGene_gn=TULP1;refGene_tx=XM_023253896.1;refGene_gn=TULP1;refGene_tx=XM_023253897.1;refGene_gn=TULP1;refGene_tx=XM_023253898.1;refGene_gn=TULP1;refGene_tx=XM_023253899.1
B2 36874368 36874637
refGene_gn=TULP1;refGene_tx=XM_023253895.1;refGene_gn=TULP1;refGene_tx=XM_023253896.1;refGene_gn=TULP1;refGene_tx=XM_023253897.1;refGene_gn=TULP1;refGene_tx=XM_023253898.1;refGene_gn=TULP1;refGene_tx=XM_023253899.1
B2 36874755 36874888
refGene_gn=TULP1;refGene_tx=XM_019830487.2;refGene_gn=TULP1;refGene_tx

=XM_023253895.1;refGene_gn=TULP1;refGene_tx=XM_023253896.1;refGene_gn=TULP1;refGene_tx=XM_023253897.1
B2 36875300 36875346
refGene_gn=TULP1;refGene_tx=XM_019830487.2;refGene_gn=TULP1;refGene_tx=XM_023253895.1;refGene_gn=TULP1;refGene_tx=XM_023253896.1;refGene_gn=TULP1;refGene_tx=XM_023253897.1
B2 36875462 36875509
refGene_gn=TULP1;refGene_tx=XM_019830487.2;refGene_gn=TULP1;refGene_tx=XM_023253895.1;refGene_gn=TULP1;refGene_tx=XM_023253896.1;refGene_gn=TULP1;refGene_tx=XM_023253897.1
B2 36958666 36958774
refGene_gn=FKBP5;refGene_tx=XM_011282248.3;refGene_gn=FKBP5;refGene_tx=XM_023253900.1;refGene_gn=FKBP5;refGene_tx=XM_023253901.1;refGene_gn=FKBP5;refGene_tx=XM_023253902.1;refGene_gn=FKBP5;refGene_tx=XM_023253903.1
B2 36959689 36959929
refGene_gn=FKBP5;refGene_tx=XM_011282248.3;refGene_gn=FKBP5;refGene_tx=XM_023253900.1;refGene_gn=FKBP5;refGene_tx=XM_023253901.1;refGene_gn=FKBP5;refGene_tx=XM_023253902.1;refGene_gn=FKBP5;refGene_tx=XM_023253903.1
B2 36961639 36961825
refGene_gn=FKBP5;refGene_tx=XM_011282248.3;refGene_gn=FKBP5;refGene_tx=XM_023253900.1;refGene_gn=FKBP5;refGene_tx=XM_023253901.1;refGene_gn=FKBP5;refGene_tx=XM_023253902.1;refGene_gn=FKBP5;refGene_tx=XM_023253903.1
B2 36969155 36969239
refGene_gn=FKBP5;refGene_tx=XM_011282248.3;refGene_gn=FKBP5;refGene_tx=XM_023253900.1;refGene_gn=FKBP5;refGene_tx=XM_023253901.1;refGene_gn=FKBP5;refGene_tx=XM_023253902.1;refGene_gn=FKBP5;refGene_tx=XM_023253903.1
B2 36971688 36971779
refGene_gn=FKBP5;refGene_tx=XM_011282248.3;refGene_gn=FKBP5;refGene_tx=XM_023253900.1;refGene_gn=FKBP5;refGene_tx=XM_023253901.1;refGene_gn=FKBP5;refGene_tx=XM_023253902.1;refGene_gn=FKBP5;refGene_tx=XM_023253903.1
B2 36978284 36978441
refGene_gn=FKBP5;refGene_tx=XM_011282248.3;refGene_gn=FKBP5;refGene_tx=XM_023253900.1;refGene_gn=FKBP5;refGene_tx=XM_023253901.1;refGene_gn=FKBP5;refGene_tx=XM_023253902.1;refGene_gn=FKBP5;refGene_tx=XM_023253903.1
B2 36994346 36994461
refGene_gn=FKBP5;refGene_tx=XM_011282248.3;refGene_gn=FKBP5;refGene_tx=XM_023253900.1;refGene_gn=FKBP5;refGene_tx=XM_023253901.1;refGene_gn=FKBP5;refGene_tx=XM_023253902.1;refGene_gn=FKBP5;refGene_tx=XM_023253903.1
B2 36995429 36995572
refGene_gn=FKBP5;refGene_tx=XM_011282248.3;refGene_gn=FKBP5;refGene_tx=XM_023253900.1;refGene_gn=FKBP5;refGene_tx=XM_023253901.1;refGene_gn=FKBP5;refGene_tx=XM_023253902.1;refGene_gn=FKBP5;refGene_tx=XM_023253903.1

B2	37011025	37011170
refGene_gn=FKBP5;refGene_tx=XM_011282248.3;refGene_gn=FKBP5;refGene_tx=XM_023253900.1;refGene_gn=FKBP5;refGene_tx=XM_023253901.1;refGene_gn=FKBP5;refGene_tx=XM_023253902.1;refGene_gn=FKBP5;refGene_tx=XM_023253903.1
B2	37015363	37015468
refGene_gn=FKBP5;refGene_tx=XM_011282248.3;refGene_gn=FKBP5;refGene_tx=XM_023253900.1;refGene_gn=FKBP5;refGene_tx=XM_023253901.1;refGene_gn=FKBP5;refGene_tx=XM_023253902.1;refGene_gn=FKBP5;refGene_tx=XM_023253903.1
B2	37109481	37109725
refGene_gn=ARMC12;refGene_tx=XM_003986056.4;refGene_gn=ARMC12;refGene_tx=XM_006931643.3
B2	37110310	37110456
refGene_gn=ARMC12;refGene_tx=XM_003986056.4;refGene_gn=ARMC12;refGene_tx=XM_006931643.3
B2	37110672	37110807
refGene_gn=ARMC12;refGene_tx=XM_003986056.4;refGene_gn=ARMC12;refGene_tx=XM_006931643.3
B2	37117570	37117744
refGene_gn=ARMC12;refGene_tx=XM_003986056.4;refGene_gn=ARMC12;refGene_tx=XM_006931643.3
B2	37117886	37117958
refGene_gn=ARMC12;refGene_tx=XM_003986056.4;refGene_gn=ARMC12;refGene_tx=XM_006931643.3
B2	37120508	37120841
refGene_gn=ARMC12;refGene_tx=XM_003986056.4;refGene_gn=ARMC12;refGene_tx=XM_006931643.3
B2	37139299	37139398	refGene_gn=CLPSL2;refGene_tx=XM_003986101.2
B2	37140051	37140174	refGene_gn=CLPSL2;refGene_tx=XM_003986101.2
B2	37141258	37141354	refGene_gn=CLPSL2;refGene_tx=XM_003986101.2
B2	37145824	37145941
refGene_gn=LOC102900686;refGene_tx=XM_019831578.2
B2	37146455	37146575
refGene_gn=LOC102900686;refGene_tx=XM_019831578.2
B2	37148565	37148872
refGene_gn=LOC102900686;refGene_tx=XM_019831578.2
B2	37149183	37149274
refGene_gn=LOC102900686;refGene_tx=XM_019831578.2
B2	37149514	37149800
refGene_gn=LOC102900686;refGene_tx=XM_019831578.2
B2	37154972	37155104	refGene_gn=CLPS;refGene_tx=XM_003986058.3
B2	37155567	37155690	refGene_gn=CLPS;refGene_tx=XM_003986058.3
B2	37157499	37157583	refGene_gn=CLPS;refGene_tx=XM_003986058.3
B2	37166551	37166963
refGene_gn=LHFPL5;refGene_tx=XM_003986102.3;refGene_gn=LHFPL5;refGene_tx=XM_019830498.2;refGene_gn=LHFPL5;refGene_tx=XM_019830500.2;refGene_gn=LHFPL5;refGene_tx=XM_023253904.1
B2	37173608	37173845
refGene_gn=LHFPL5;refGene_tx=XM_003986102.3;refGene_gn=LHFPL5;refGene_

tx=XM_019830498.2;refGene_gn=LHFPL5;refGene_tx=XM_019830500.2;refGene_gn=LHFPL5;refGene_tx=XM_023253904.1
B2 37179836 37179847 refGene_gn=LHFPL5;refGene_tx=XM_019830500.2
B2 37183369 37183464
refGene_gn=LHFPL5;refGene_tx=XM_003986102.3;refGene_gn=LHFPL5;refGene_tx=XM_019830498.2;refGene_gn=LHFPL5;refGene_tx=XM_023253904.1
B2 37191470 37191655 refGene_gn=SRPK1;refGene_tx=XM_023253905.1
B2 37193164 37193257 refGene_gn=SRPK1;refGene_tx=XM_023253905.1
B2 37197933 37198003 refGene_gn=SRPK1;refGene_tx=XM_023253905.1
B2 37202448 37202556 refGene_gn=SRPK1;refGene_tx=XM_023253905.1
B2 37211672 37211773 refGene_gn=SRPK1;refGene_tx=XM_023253905.1
B2 37212143 37212563 refGene_gn=SRPK1;refGene_tx=XM_023253905.1
B2 37212940 37213154 refGene_gn=SRPK1;refGene_tx=XM_023253905.1
B2 37213580 37213606 refGene_gn=SRPK1;refGene_tx=XM_023253905.1
B2 37215343 37215509 refGene_gn=SRPK1;refGene_tx=XM_023253905.1
B2 37216361 37216468 refGene_gn=SRPK1;refGene_tx=XM_023253905.1
B2 37220457 37220545 refGene_gn=SRPK1;refGene_tx=XM_023253905.1
B2 37221578 37221666 refGene_gn=SRPK1;refGene_tx=XM_023253905.1
B2 37222605 37222714 refGene_gn=SRPK1;refGene_tx=XM_023253905.1
B2 37224141 37224260 refGene_gn=SRPK1;refGene_tx=XM_023253905.1
B2 37270601 37271124 refGene_gn=SRPK1;refGene_tx=XM_023253905.1
B2 37271126 37271202 refGene_gn=SRPK1;refGene_tx=XM_023253905.1
B2 37297636 37298119
refGene_gn=SLC26A8;refGene_tx=XM_023253906.1;refGene_gn=SLC26A8;refGene_tx=XM_023253907.1;refGene_gn=SLC26A8;refGene_tx=XM_023253908.1
B2 37303960 37304145
refGene_gn=SLC26A8;refGene_tx=XM_023253906.1;refGene_gn=SLC26A8;refGene_tx=XM_023253907.1;refGene_gn=SLC26A8;refGene_tx=XM_023253908.1
B2 37304241 37304296
refGene_gn=SLC26A8;refGene_tx=XM_023253906.1;refGene_gn=SLC26A8;refGene_tx=XM_023253907.1;refGene_gn=SLC26A8;refGene_tx=XM_023253908.1
B2 37309334 37309706
refGene_gn=SLC26A8;refGene_tx=XM_023253906.1;refGene_gn=SLC26A8;refGene_tx=XM_023253907.1;refGene_gn=SLC26A8;refGene_tx=XM_023253908.1
B2 37313713 37313845
refGene_gn=SLC26A8;refGene_tx=XM_023253906.1;refGene_gn=SLC26A8;refGene_tx=XM_023253907.1;refGene_gn=SLC26A8;refGene_tx=XM_023253908.1
B2 37313960 37314053
refGene_gn=SLC26A8;refGene_tx=XM_023253906.1;refGene_gn=SLC26A8;refGene_tx=XM_023253907.1;refGene_gn=SLC26A8;refGene_tx=XM_023253908.1
B2 37314486 37314556
refGene_gn=SLC26A8;refGene_tx=XM_023253906.1;refGene_gn=SLC26A8;refGene_tx=XM_023253907.1;refGene_gn=SLC26A8;refGene_tx=XM_023253908.1
B2 37314637 37314744
refGene_gn=SLC26A8;refGene_tx=XM_023253906.1;refGene_gn=SLC26A8;refGene_tx=XM_023253907.1;refGene_gn=SLC26A8;refGene_tx=XM_023253908.1
B2 37315906 37316002
refGene_gn=SLC26A8;refGene_tx=XM_023253906.1;refGene_gn=SLC26A8;refGene_tx=XM_023253907.1;refGene_gn=SLC26A8;refGene_tx=XM_023253908.1
B2 37318533 37318611

refGene_gn=SLC26A8;refGene_tx=XM_023253906.1;refGene_gn=SLC26A8;refGene_tx=XM_023253907.1;refGene_gn=SLC26A8;refGene_tx=XM_023253908.1
B2 37324130 37324244
refGene_gn=SLC26A8;refGene_tx=XM_023253906.1;refGene_gn=SLC26A8;refGene_tx=XM_023253907.1;refGene_gn=SLC26A8;refGene_tx=XM_023253908.1
B2 37325539 37325687
refGene_gn=SLC26A8;refGene_tx=XM_023253906.1;refGene_gn=SLC26A8;refGene_tx=XM_023253907.1;refGene_gn=SLC26A8;refGene_tx=XM_023253908.1
B2 37336019 37336102
refGene_gn=SLC26A8;refGene_tx=XM_023253906.1;refGene_gn=SLC26A8;refGene_tx=XM_023253907.1;refGene_gn=SLC26A8;refGene_tx=XM_023253908.1
B2 37341582 37341732
refGene_gn=SLC26A8;refGene_tx=XM_023253906.1;refGene_gn=SLC26A8;refGene_tx=XM_023253907.1
B2 37342434 37342599
refGene_gn=SLC26A8;refGene_tx=XM_023253906.1;refGene_gn=SLC26A8;refGene_tx=XM_023253907.1
B2 37350128 37350310
refGene_gn=SLC26A8;refGene_tx=XM_023253906.1;refGene_gn=SLC26A8;refGene_tx=XM_023253907.1;refGene_gn=SLC26A8;refGene_tx=XM_023253908.1
B2 37351997 37352114
refGene_gn=SLC26A8;refGene_tx=XM_023253906.1;refGene_gn=SLC26A8;refGene_tx=XM_023253907.1;refGene_gn=SLC26A8;refGene_tx=XM_023253908.1
B2 37363818 37363958
refGene_gn=SLC26A8;refGene_tx=XM_023253906.1;refGene_gn=SLC26A8;refGene_tx=XM_023253907.1;refGene_gn=SLC26A8;refGene_tx=XM_023253908.1
B2 37370331 37370516
refGene_gn=SLC26A8;refGene_tx=XM_023253906.1;refGene_gn=SLC26A8;refGene_tx=XM_023253907.1;refGene_gn=SLC26A8;refGene_tx=XM_023253908.1
B2 37383607 37383723
refGene_gn=MAPK14;refGene_tx=XM_003986061.3;refGene_gn=MAPK14;refGene_tx=XM_003986062.3;refGene_gn=MAPK14;refGene_tx=XM_011282246.2
B2 37408174 37408304
refGene_gn=MAPK14;refGene_tx=XM_003986061.3;refGene_gn=MAPK14;refGene_tx=XM_003986062.3;refGene_gn=MAPK14;refGene_tx=XM_011282246.2
B2 37415235 37415294
refGene_gn=MAPK14;refGene_tx=XM_003986061.3;refGene_gn=MAPK14;refGene_tx=XM_003986062.3;refGene_gn=MAPK14;refGene_tx=XM_011282246.2
B2 37429230 37429342
refGene_gn=MAPK14;refGene_tx=XM_003986061.3;refGene_gn=MAPK14;refGene_tx=XM_003986062.3;refGene_gn=MAPK14;refGene_tx=XM_011282246.2
B2 37430065 37430095
refGene_gn=MAPK14;refGene_tx=XM_003986061.3;refGene_gn=MAPK14;refGene_tx=XM_003986062.3;refGene_gn=MAPK14;refGene_tx=XM_011282246.2
B2 37430424 37430472
refGene_gn=MAPK14;refGene_tx=XM_003986061.3;refGene_gn=MAPK14;refGene_tx=XM_003986062.3;refGene_gn=MAPK14;refGene_tx=XM_011282246.2
B2 37431600 37431715
refGene_gn=MAPK14;refGene_tx=XM_003986061.3;refGene_gn=MAPK14;refGene_tx=XM_003986062.3;refGene_gn=MAPK14;refGene_tx=XM_011282246.2

B2 37432287 37432359
refGene_gn=MAPK14;refGene_tx=XM_003986061.3;refGene_gn=MAPK14;refGene_tx=XM_003986062.3;refGene_gn=MAPK14;refGene_tx=XM_011282246.2
B2 37445197 37445277
refGene_gn=MAPK14;refGene_tx=XM_003986061.3;refGene_gn=MAPK14;refGene_tx=XM_011282246.2
B2 37449274 37449354 refGene_gn=MAPK14;refGene_tx=XM_003986062.3
B2 37451693 37451772
refGene_gn=MAPK14;refGene_tx=XM_003986061.3;refGene_gn=MAPK14;refGene_tx=XM_003986062.3
B2 37454610 37454784
refGene_gn=MAPK14;refGene_tx=XM_003986061.3;refGene_gn=MAPK14;refGene_tx=XM_003986062.3;refGene_gn=MAPK14;refGene_tx=XM_011282246.2
B2 37455599 37455667
refGene_gn=MAPK14;refGene_tx=XM_003986061.3;refGene_gn=MAPK14;refGene_tx=XM_003986062.3
B2 37476775 37476894
refGene_gn=MAPK13;refGene_tx=XM_023253909.1;refGene_gn=MAPK13;refGene_tx=XM_023253910.1
B2 37477759 37477889
refGene_gn=MAPK13;refGene_tx=XM_023253909.1;refGene_gn=MAPK13;refGene_tx=XM_023253910.1
B2 37478877 37478936
refGene_gn=MAPK13;refGene_tx=XM_023253909.1;refGene_gn=MAPK13;refGene_tx=XM_023253910.1
B2 37481890 37481999
refGene_gn=MAPK13;refGene_tx=XM_023253909.1;refGene_gn=MAPK13;refGene_tx=XM_023253910.1
B2 37482194 37482224 refGene_gn=MAPK13;refGene_tx=XM_023253909.1
B2 37482612 37482660
refGene_gn=MAPK13;refGene_tx=XM_023253909.1;refGene_gn=MAPK13;refGene_tx=XM_023253910.1
B2 37482776 37482891
refGene_gn=MAPK13;refGene_tx=XM_023253909.1;refGene_gn=MAPK13;refGene_tx=XM_023253910.1
B2 37483034 37483106
refGene_gn=MAPK13;refGene_tx=XM_023253909.1;refGene_gn=MAPK13;refGene_tx=XM_023253910.1
B2 37483918 37483998
refGene_gn=MAPK13;refGene_tx=XM_023253909.1;refGene_gn=MAPK13;refGene_tx=XM_023253910.1
B2 37484265 37484344
refGene_gn=MAPK13;refGene_tx=XM_023253909.1;refGene_gn=MAPK13;refGene_tx=XM_023253910.1
B2 37484454 37484631
refGene_gn=MAPK13;refGene_tx=XM_023253909.1;refGene_gn=MAPK13;refGene_tx=XM_023253910.1
B2 37484910 37484993
refGene_gn=MAPK13;refGene_tx=XM_023253909.1;refGene_gn=MAPK13;refGene_tx=XM_023253910.1

B2	37544515	37545966
refGene_gn=BRPF3;refGene_tx=XM_011282244.3;refGene_gn=BRPF3;refGene_tx=XM_023253911.1;refGene_gn=BRPF3;refGene_tx=XM_023253912.1;refGene_gn=BRPF3;refGene_tx=XM_023253913.1;refGene_gn=BRPF3;refGene_tx=XM_023253914.1
B2	37549285	37549442
refGene_gn=BRPF3;refGene_tx=XM_011282244.3;refGene_gn=BRPF3;refGene_tx=XM_023253911.1;refGene_gn=BRPF3;refGene_tx=XM_023253912.1;refGene_gn=BRPF3;refGene_tx=XM_023253913.1;refGene_gn=BRPF3;refGene_tx=XM_023253914.1
B2	37551823	37551955
refGene_gn=BRPF3;refGene_tx=XM_011282244.3;refGene_gn=BRPF3;refGene_tx=XM_023253911.1;refGene_gn=BRPF3;refGene_tx=XM_023253912.1;refGene_gn=BRPF3;refGene_tx=XM_023253913.1;refGene_gn=BRPF3;refGene_tx=XM_023253914.1
B2	37554200	37554329
refGene_gn=BRPF3;refGene_tx=XM_011282244.3;refGene_gn=BRPF3;refGene_tx=XM_023253911.1;refGene_gn=BRPF3;refGene_tx=XM_023253912.1;refGene_gn=BRPF3;refGene_tx=XM_023253913.1;refGene_gn=BRPF3;refGene_tx=XM_023253914.1
B2	37554619	37554932
refGene_gn=BRPF3;refGene_tx=XM_011282244.3;refGene_gn=BRPF3;refGene_tx=XM_023253911.1;refGene_gn=BRPF3;refGene_tx=XM_023253912.1;refGene_gn=BRPF3;refGene_tx=XM_023253913.1;refGene_gn=BRPF3;refGene_tx=XM_023253914.1
B2	37555662	37555965
refGene_gn=BRPF3;refGene_tx=XM_011282244.3;refGene_gn=BRPF3;refGene_tx=XM_023253911.1;refGene_gn=BRPF3;refGene_tx=XM_023253912.1;refGene_gn=BRPF3;refGene_tx=XM_023253913.1
B2	37557594	37558101
refGene_gn=BRPF3;refGene_tx=XM_011282244.3;refGene_gn=BRPF3;refGene_tx=XM_023253911.1;refGene_gn=BRPF3;refGene_tx=XM_023253912.1
B2	37560936	37561030
refGene_gn=BRPF3;refGene_tx=XM_011282244.3;refGene_gn=BRPF3;refGene_tx=XM_023253911.1;refGene_gn=BRPF3;refGene_tx=XM_023253912.1;refGene_gn=BRPF3;refGene_tx=XM_023253913.1;refGene_gn=BRPF3;refGene_tx=XM_023253914.1
B2	37564770	37564868
refGene_gn=BRPF3;refGene_tx=XM_011282244.3;refGene_gn=BRPF3;refGene_tx=XM_023253911.1;refGene_gn=BRPF3;refGene_tx=XM_023253912.1;refGene_gn=BRPF3;refGene_tx=XM_023253913.1;refGene_gn=BRPF3;refGene_tx=XM_023253914.1
B2	37567783	37567881
refGene_gn=BRPF3;refGene_tx=XM_011282244.3;refGene_gn=BRPF3;refGene_tx=XM_023253911.1;refGene_gn=BRPF3;refGene_tx=XM_023253913.1;refGene_gn=BRPF3;refGene_tx=XM_023253914.1
B2	37571515	37571670
refGene_gn=BRPF3;refGene_tx=XM_011282244.3;refGene_gn=BRPF3;refGene_tx=XM_023253911.1;refGene_gn=BRPF3;refGene_tx=XM_023253912.1;refGene_gn=BRPF3;refGene_tx=XM_023253913.1;refGene_gn=BRPF3;refGene_tx=XM_0232539

14.1
B2 37572928 37573112
refGene_gn=BRPF3;refGene_tx=XM_011282244.3;refGene_gn=BRPF3;refGene_tx=XM_023253911.1;refGene_gn=BRPF3;refGene_tx=XM_023253913.1;refGene_gn=BRPF3;refGene_tx=XM_023253914.1
B2 37612352 37612557
refGene_gn=PNPLA1;refGene_tx=XM_003986105.4;refGene_gn=PNPLA1;refGene_tx=XM_019830508.1;refGene_gn=PNPLA1;refGene_tx=XM_023253915.1
B2 37637773 37638006
refGene_gn=PNPLA1;refGene_tx=XM_003986105.4;refGene_gn=PNPLA1;refGene_tx=XM_019830508.1;refGene_gn=PNPLA1;refGene_tx=XM_023253915.1
B2 37639110 37639176
refGene_gn=PNPLA1;refGene_tx=XM_003986105.4;refGene_gn=PNPLA1;refGene_tx=XM_019830508.1;refGene_gn=PNPLA1;refGene_tx=XM_023253915.1
B2 37640375 37640585
refGene_gn=PNPLA1;refGene_tx=XM_003986105.4;refGene_gn=PNPLA1;refGene_tx=XM_019830508.1;refGene_gn=PNPLA1;refGene_tx=XM_023253915.1
B2 37643920 37643981
refGene_gn=PNPLA1;refGene_tx=XM_003986105.4;refGene_gn=PNPLA1;refGene_tx=XM_019830508.1;refGene_gn=PNPLA1;refGene_tx=XM_023253915.1
B2 37652061 37652619
refGene_gn=PNPLA1;refGene_tx=XM_003986105.4;refGene_gn=PNPLA1;refGene_tx=XM_019830508.1;refGene_gn=PNPLA1;refGene_tx=XM_023253915.1
B2 37655511 37655590
refGene_gn=PNPLA1;refGene_tx=XM_003986105.4;refGene_gn=PNPLA1;refGene_tx=XM_019830508.1;refGene_gn=PNPLA1;refGene_tx=XM_023253915.1
B2 37656947 37657077
refGene_gn=PNPLA1;refGene_tx=XM_003986105.4;refGene_gn=PNPLA1;refGene_tx=XM_019830508.1;refGene_gn=PNPLA1;refGene_tx=XM_023253915.1
B2 37660177 37660304 refGene_gn=PNPLA1;refGene_tx=XM_003986105.4
B2 37661052 37661103
refGene_gn=PNPLA1;refGene_tx=XM_003986105.4;refGene_gn=PNPLA1;refGene_tx=XM_019830508.1
B2 37665634 37665699
refGene_gn=CB2H6orf222;refGene_tx=XM_023254716.1
B2 37666115 37666303
refGene_gn=CB2H6orf222;refGene_tx=XM_023254716.1
B2 37666956 37667306
refGene_gn=CB2H6orf222;refGene_tx=XM_023254716.1
B2 37668707 37668772
refGene_gn=CB2H6orf222;refGene_tx=XM_023254716.1
B2 37669651 37669783
refGene_gn=CB2H6orf222;refGene_tx=XM_023254716.1
B2 37670788 37671050
refGene_gn=CB2H6orf222;refGene_tx=XM_023254716.1
B2 37671343 37671402
refGene_gn=CB2H6orf222;refGene_tx=XM_023254716.1
B2 37672914 37673046
refGene_gn=CB2H6orf222;refGene_tx=XM_023254716.1
B2 37674065 37674363

refGene_gn=CB2H6orf222;refGene_tx=XM_023254716.1
B2	37674556	37674621
refGene_gn=CB2H6orf222;refGene_tx=XM_023254716.1
B2	37676133	37676253
refGene_gn=CB2H6orf222;refGene_tx=XM_023254716.1
B2	37677011	37677015
refGene_gn=CB2H6orf222;refGene_tx=XM_023254716.1
B2	37677166	37677823
refGene_gn=CB2H6orf222;refGene_tx=XM_023254716.1
B2	37727021	37727031	refGene_gn=ETV7;refGene_tx=XM_023253917.1
B2	37729375	37729481
refGene_gn=ETV7;refGene_tx=XM_023253916.1;refGene_gn=ETV7;refGene_tx=XM_023253918.1;refGene_gn=ETV7;refGene_tx=XM_023253919.1;refGene_gn=ETV7;refGene_tx=XM_023253920.1
B2	37729594	37729695
refGene_gn=ETV7;refGene_tx=XM_023253916.1;refGene_gn=ETV7;refGene_tx=XM_023253917.1;refGene_gn=ETV7;refGene_tx=XM_023253918.1;refGene_gn=ETV7;refGene_tx=XM_023253919.1;refGene_gn=ETV7;refGene_tx=XM_023253920.1
B2	37732494	37732637
refGene_gn=ETV7;refGene_tx=XM_023253916.1;refGene_gn=ETV7;refGene_tx=XM_023253917.1;refGene_gn=ETV7;refGene_tx=XM_023253918.1;refGene_gn=ETV7;refGene_tx=XM_023253919.1;refGene_gn=ETV7;refGene_tx=XM_023253920.1
B2	37736541	37736757
refGene_gn=ETV7;refGene_tx=XM_023253916.1;refGene_gn=ETV7;refGene_tx=XM_023253917.1;refGene_gn=ETV7;refGene_tx=XM_023253918.1;refGene_gn=ETV7;refGene_tx=XM_023253919.1;refGene_gn=ETV7;refGene_tx=XM_023253920.1
B2	37738840	37738966
refGene_gn=ETV7;refGene_tx=XM_023253916.1;refGene_gn=ETV7;refGene_tx=XM_023253917.1;refGene_gn=ETV7;refGene_tx=XM_023253918.1;refGene_gn=ETV7;refGene_tx=XM_023253919.1;refGene_gn=ETV7;refGene_tx=XM_023253920.1
B2	37739328	37739493
refGene_gn=ETV7;refGene_tx=XM_023253916.1;refGene_gn=ETV7;refGene_tx=XM_023253917.1;refGene_gn=ETV7;refGene_tx=XM_023253918.1;refGene_gn=ETV7;refGene_tx=XM_023253919.1;refGene_gn=ETV7;refGene_tx=XM_023253920.1
B2	37753220	37753350
refGene_gn=ETV7;refGene_tx=XM_023253916.1;refGene_gn=ETV7;refGene_tx=XM_023253917.1;refGene_gn=ETV7;refGene_tx=XM_023253918.1;refGene_gn=ETV7;refGene_tx=XM_023253919.1;refGene_gn=ETV7;refGene_tx=XM_023253920.1
B2	37754001	37754157
refGene_gn=ETV7;refGene_tx=XM_023253916.1;refGene_gn=ETV7;refGene_tx=XM_023253917.1;refGene_gn=ETV7;refGene_tx=XM_023253919.1
B2	37754900	37754906	refGene_gn=ETV7;refGene_tx=XM_023253918.1
B2	37757471	37757579	refGene_gn=PXT1;refGene_tx=XM_023253921.1
B2	37762960	37763017	refGene_gn=PXT1;refGene_tx=XM_023253922.1
B2	37763904	37764035
refGene_gn=PXT1;refGene_tx=XM_023253921.1;refGene_gn=PXT1;refGene_tx=XM_023253922.1
B2	37785254	37785306
refGene_gn=PXT1;refGene_tx=XM_023253921.1;refGene_gn=PXT1;refGene_tx=XM_023253922.1

B2	37828997 37829157
refGene_gn=KCTD20;refGene_tx=XM_006931651.2;refGene_gn=KCTD20;refGene_tx=XM_023253923.1
B2	37831785 37832059
refGene_gn=KCTD20;refGene_tx=XM_006931651.2;refGene_gn=KCTD20;refGene_tx=XM_023253923.1
B2	37834859 37834962
refGene_gn=KCTD20;refGene_tx=XM_006931651.2;refGene_gn=KCTD20;refGene_tx=XM_023253923.1
B2	37835320 37835441
refGene_gn=KCTD20;refGene_tx=XM_006931651.2;refGene_gn=KCTD20;refGene_tx=XM_023253923.1
B2	37836944 37837142
refGene_gn=KCTD20;refGene_tx=XM_006931651.2;refGene_gn=KCTD20;refGene_tx=XM_023253923.1
B2	37838941 37839052
refGene_gn=KCTD20;refGene_tx=XM_006931651.2;refGene_gn=KCTD20;refGene_tx=XM_023253923.1
B2	37840350 37840643
refGene_gn=KCTD20;refGene_tx=XM_006931651.2;refGene_gn=KCTD20;refGene_tx=XM_023253923.1
B2	37849631 37849762 refGene_gn=STK38;refGene_tx=XM_023253924.1
B2	37850243 37850338 refGene_gn=STK38;refGene_tx=XM_023253924.1
B2	37851403 37851499 refGene_gn=STK38;refGene_tx=XM_023253924.1
B2	37851687 37851811 refGene_gn=STK38;refGene_tx=XM_023253924.1
B2	37853338 37853456 refGene_gn=STK38;refGene_tx=XM_023253924.1
B2	37859206 37859268 refGene_gn=STK38;refGene_tx=XM_023253924.1
B2	37860078 37860181 refGene_gn=STK38;refGene_tx=XM_023253924.1
B2	37864658 37864813 refGene_gn=STK38;refGene_tx=XM_023253924.1
B2	37867028 37867152 refGene_gn=STK38;refGene_tx=XM_023253924.1
B2	37871161 37871245 refGene_gn=STK38;refGene_tx=XM_023253924.1
B2	37873055 37873178 refGene_gn=STK38;refGene_tx=XM_023253924.1
B2	37874090 37874142 refGene_gn=STK38;refGene_tx=XM_023253924.1
B2	37884139 37884218 refGene_gn=STK38;refGene_tx=XM_023253924.1
B2	37884220 37884269 refGene_gn=STK38;refGene_tx=XM_023253924.1
B2	37931075 37931281 refGene_gn=SRSF3;refGene_tx=XM_003986068.5
B2	37932540 37932675 refGene_gn=SRSF3;refGene_tx=XM_003986068.5
B2	37934885 37934924 refGene_gn=SRSF3;refGene_tx=XM_003986068.5
B2	37935161 37935248 refGene_gn=SRSF3;refGene_tx=XM_003986068.5
B2	37935403 37935431 refGene_gn=SRSF3;refGene_tx=XM_003986068.5
B2	38006745 38007190
refGene_gn=CDKN1A;refGene_tx=NM_001009865.1;refGene_gn=CDKN1A;refGene_tx=XM_006931621.4
B2	38008303 38008353
refGene_gn=CDKN1A;refGene_tx=NM_001009865.1;refGene_gn=CDKN1A;refGene_tx=XM_006931621.4
B2	38022571 38022778
refGene_gn=RAB44;refGene_tx=XM_003986108.3;refGene_gn=RAB44;refGene_tx=XM_019830515.2;refGene_gn=RAB44;refGene_tx=XM_019830516.1
B2	38032705 38032817

refGene_gn=RAB44;refGene_tx=XM_003986108.3;refGene_gn=RAB44;refGene_tx=XM_019830515.2;refGene_gn=RAB44;refGene_tx=XM_019830516.1
B2 38033805 38033974 refGene_gn=RAB44;refGene_tx=XM_003986108.3;refGene_gn=RAB44;refGene_tx=XM_019830515.2;refGene_gn=RAB44;refGene_tx=XM_019830516.1
B2 38035230 38035377 refGene_gn=RAB44;refGene_tx=XM_003986108.3;refGene_gn=RAB44;refGene_tx=XM_019830515.2;refGene_gn=RAB44;refGene_tx=XM_019830516.1
B2 38036025 38036116 refGene_gn=RAB44;refGene_tx=XM_003986108.3;refGene_gn=RAB44;refGene_tx=XM_019830515.2;refGene_gn=RAB44;refGene_tx=XM_019830516.1
B2 38036509 38036605 refGene_gn=RAB44;refGene_tx=XM_003986108.3;refGene_gn=RAB44;refGene_tx=XM_019830515.2;refGene_gn=RAB44;refGene_tx=XM_019830516.1
B2 38038986 38039174 refGene_gn=RAB44;refGene_tx=XM_003986108.3;refGene_gn=RAB44;refGene_tx=XM_019830515.2;refGene_gn=RAB44;refGene_tx=XM_019830516.1
B2 38039745 38041223 refGene_gn=RAB44;refGene_tx=XM_003986108.3;refGene_gn=RAB44;refGene_tx=XM_019830515.2;refGene_gn=RAB44;refGene_tx=XM_019830516.1
B2 38045096 38045178 refGene_gn=RAB44;refGene_tx=XM_003986108.3;refGene_gn=RAB44;refGene_tx=XM_019830515.2;refGene_gn=RAB44;refGene_tx=XM_019830516.1
B2 38046365 38046480 refGene_gn=RAB44;refGene_tx=XM_003986108.3;refGene_gn=RAB44;refGene_tx=XM_019830515.2;refGene_gn=RAB44;refGene_tx=XM_019830516.1
B2 38047552 38047654 refGene_gn=RAB44;refGene_tx=XM_003986108.3;refGene_gn=RAB44;refGene_tx=XM_019830515.2;refGene_gn=RAB44;refGene_tx=XM_019830516.1
B2 38048064 38048406 refGene_gn=RAB44;refGene_tx=XM_019830515.2
B2 38049881 38049958 refGene_gn=RAB44;refGene_tx=XM_003986108.3
B2 38051904 38052177 refGene_gn=RAB44;refGene_tx=XM_003986108.3;refGene_gn=RAB44;refGene_tx=XM_019830516.1
B2 38056763 38057822 refGene_gn=LOC101093019;refGene_tx=XM_011282339.1
B2 38061825 38062044 refGene_gn=CPNE5;refGene_tx=XM_023253925.1;refGene_gn=CPNE5;refGene_tx=XM_023253926.1;refGene_gn=CPNE5;refGene_tx=XM_023253927.1
B2 38063282 38063356 refGene_gn=CPNE5;refGene_tx=XM_023253925.1;refGene_gn=CPNE5;refGene_tx=XM_023253926.1;refGene_gn=CPNE5;refGene_tx=XM_023253927.1
B2 38063801 38063859 refGene_gn=CPNE5;refGene_tx=XM_023253925.1;refGene_gn=CPNE5;refGene_tx=XM_023253926.1;refGene_gn=CPNE5;refGene_tx=XM_023253927.1
B2 38064485 38064588 refGene_gn=CPNE5;refGene_tx=XM_023253925.1;refGene_gn=CPNE5;refGene_tx=XM_023253926.1;refGene_gn=CPNE5;refGene_tx=XM_023253927.1
B2 38064821 38064949 refGene_gn=CPNE5;refGene_tx=XM_023253925.1;refGene_gn=CPNE5;refGene_tx

=XM_023253926.1;refGene_gn=CPNE5;refGene_tx=XM_023253927.1
B2 38065781 38065963
refGene_gn=CPNE5;refGene_tx=XM_023253925.1;refGene_gn=CPNE5;refGene_tx
=XM_023253926.1;refGene_gn=CPNE5;refGene_tx=XM_023253927.1
B2 38067053 38067100
refGene_gn=CPNE5;refGene_tx=XM_023253925.1;refGene_gn=CPNE5;refGene_tx
=XM_023253926.1;refGene_gn=CPNE5;refGene_tx=XM_023253927.1
B2 38071970 38072032
refGene_gn=CPNE5;refGene_tx=XM_023253925.1;refGene_gn=CPNE5;refGene_tx
=XM_023253926.1;refGene_gn=CPNE5;refGene_tx=XM_023253927.1
B2 38075284 38075338
refGene_gn=CPNE5;refGene_tx=XM_023253925.1;refGene_gn=CPNE5;refGene_tx
=XM_023253926.1;refGene_gn=CPNE5;refGene_tx=XM_023253927.1
B2 38080946 38081022
refGene_gn=CPNE5;refGene_tx=XM_023253925.1;refGene_gn=CPNE5;refGene_tx
=XM_023253926.1
B2 38083140 38083182
refGene_gn=CPNE5;refGene_tx=XM_023253925.1;refGene_gn=CPNE5;refGene_tx
=XM_023253926.1
B2 38091301 38091406
refGene_gn=CPNE5;refGene_tx=XM_023253925.1;refGene_gn=CPNE5;refGene_tx
=XM_023253926.1
B2 38096274 38096378
refGene_gn=CPNE5;refGene_tx=XM_023253925.1;refGene_gn=CPNE5;refGene_tx
=XM_023253926.1
B2 38108862 38108926
refGene_gn=CPNE5;refGene_tx=XM_023253925.1;refGene_gn=CPNE5;refGene_tx
=XM_023253926.1
B2 38109227 38109278 refGene_gn=CPNE5;refGene_tx=XM_023253925.1
B2 38111342 38111402
refGene_gn=CPNE5;refGene_tx=XM_023253925.1;refGene_gn=CPNE5;refGene_tx
=XM_023253926.1
B2 38112368 38112445
refGene_gn=CPNE5;refGene_tx=XM_023253925.1;refGene_gn=CPNE5;refGene_tx
=XM_023253926.1
B2 38112662 38112702
refGene_gn=CPNE5;refGene_tx=XM_023253925.1;refGene_gn=CPNE5;refGene_tx
=XM_023253926.1
B2 38114075 38114179
refGene_gn=CPNE5;refGene_tx=XM_023253925.1;refGene_gn=CPNE5;refGene_tx
=XM_023253926.1
B2 38139529 38139576
refGene_gn=CPNE5;refGene_tx=XM_023253925.1;refGene_gn=CPNE5;refGene_tx
=XM_023253926.1
B2 38140344 38140385
refGene_gn=CPNE5;refGene_tx=XM_023253925.1;refGene_gn=CPNE5;refGene_tx
=XM_023253926.1
B2 38155139 38155234
refGene_gn=CPNE5;refGene_tx=XM_023253925.1;refGene_gn=CPNE5;refGene_tx
=XM_023253926.1

B2 38169980 38170201 refGene_gn=PPIL1;refGene_tx=XM_003986069.4
B2 38170802 38170871 refGene_gn=PPIL1;refGene_tx=XM_003986069.4
B2 38185455 38185610 refGene_gn=PPIL1;refGene_tx=XM_003986069.4
B2 38188885 38188941 refGene_gn=PPIL1;refGene_tx=XM_003986069.4
B2 38195559 38195561
refGene_gn=CB2H6orf89;refGene_tx=XM_019830523.2;refGene_gn=CB2H6orf89;
refGene_tx=XM_019830524.2
B2 38206810 38206894
refGene_gn=CB2H6orf89;refGene_tx=XM_019830523.2
B2 38212128 38212336
refGene_gn=CB2H6orf89;refGene_tx=XM_019830521.2;refGene_gn=CB2H6orf89;
refGene_tx=XM_019830522.2;refGene_gn=CB2H6orf89;refGene_tx=XM_01983052
3.2;refGene_gn=CB2H6orf89;refGene_tx=XM_019830524.2;refGene_gn=CB2H6or
f89;refGene_tx=XM_019830525.2;refGene_gn=CB2H6orf89;refGene_tx=XM_0198
30526.2;refGene_gn=CB2H6orf89;refGene_tx=XM_019830527.2;refGene_gn=CB2
H6orf89;refGene_tx=XM_019830528.2
B2 38213023 38213237
refGene_gn=CB2H6orf89;refGene_tx=XM_019830521.2;refGene_gn=CB2H6orf89;
refGene_tx=XM_019830522.2;refGene_gn=CB2H6orf89;refGene_tx=XM_01983052
3.2;refGene_gn=CB2H6orf89;refGene_tx=XM_019830524.2;refGene_gn=CB2H6or
f89;refGene_tx=XM_019830525.2;refGene_gn=CB2H6orf89;refGene_tx=XM_0198
30526.2;refGene_gn=CB2H6orf89;refGene_tx=XM_019830527.2;refGene_gn=CB2
H6orf89;refGene_tx=XM_019830528.2
B2 38228051 38228206
refGene_gn=CB2H6orf89;refGene_tx=XM_019830521.2;refGene_gn=CB2H6orf89;
refGene_tx=XM_019830522.2;refGene_gn=CB2H6orf89;refGene_tx=XM_01983052
3.2;refGene_gn=CB2H6orf89;refGene_tx=XM_019830524.2;refGene_gn=CB2H6or
f89;refGene_tx=XM_019830525.2;refGene_gn=CB2H6orf89;refGene_tx=XM_0198
30526.2;refGene_gn=CB2H6orf89;refGene_tx=XM_019830527.2;refGene_gn=CB2
H6orf89;refGene_tx=XM_019830528.2
B2 38228333 38228473
refGene_gn=CB2H6orf89;refGene_tx=XM_019830521.2;refGene_gn=CB2H6orf89;
refGene_tx=XM_019830522.2;refGene_gn=CB2H6orf89;refGene_tx=XM_01983052
3.2;refGene_gn=CB2H6orf89;refGene_tx=XM_019830524.2;refGene_gn=CB2H6or
f89;refGene_tx=XM_019830525.2;refGene_gn=CB2H6orf89;refGene_tx=XM_0198
30526.2;refGene_gn=CB2H6orf89;refGene_tx=XM_019830527.2;refGene_gn=CB2
H6orf89;refGene_tx=XM_019830528.2
B2 38229641 38229771
refGene_gn=CB2H6orf89;refGene_tx=XM_019830521.2;refGene_gn=CB2H6orf89;
refGene_tx=XM_019830522.2;refGene_gn=CB2H6orf89;refGene_tx=XM_01983052
3.2;refGene_gn=CB2H6orf89;refGene_tx=XM_019830524.2;refGene_gn=CB2H6or
f89;refGene_tx=XM_019830525.2;refGene_gn=CB2H6orf89;refGene_tx=XM_0198
30526.2;refGene_gn=CB2H6orf89;refGene_tx=XM_019830527.2;refGene_gn=CB2
H6orf89;refGene_tx=XM_019830528.2
B2 38232842 38232990
refGene_gn=CB2H6orf89;refGene_tx=XM_019830521.2;refGene_gn=CB2H6orf89;
refGene_tx=XM_019830523.2;refGene_gn=CB2H6orf89;refGene_tx=XM_01983052
4.2;refGene_gn=CB2H6orf89;refGene_tx=XM_019830525.2;refGene_gn=CB2H6or
f89;refGene_tx=XM_019830526.2;refGene_gn=CB2H6orf89;refGene_tx=XM_0198
30527.2;refGene_gn=CB2H6orf89;refGene_tx=XM_019830528.2

B2	38236019 38236217
refGene_gn=CB2H6orf89;refGene_tx=XM_019830521.2;refGene_gn=CB2H6orf89;
refGene_tx=XM_019830522.2;refGene_gn=CB2H6orf89;refGene_tx=XM_01983052
3.2;refGene_gn=CB2H6orf89;refGene_tx=XM_019830524.2;refGene_gn=CB2H6or
f89;refGene_tx=XM_019830525.2;refGene_gn=CB2H6orf89;refGene_tx=XM_0198
30526.2;refGene_gn=CB2H6orf89;refGene_tx=XM_019830527.2;refGene_gn=CB2
H6orf89;refGene_tx=XM_019830528.2
B2	38270004 38270160
refGene_gn=PI16;refGene_tx=XM_006931660.4;refGene_gn=PI16;refGene_tx=X
M_006931661.4;refGene_gn=PI16;refGene_tx=XM_011282241.2
B2	38274450 38274672
refGene_gn=PI16;refGene_tx=XM_006931660.4;refGene_gn=PI16;refGene_tx=X
M_006931661.4;refGene_gn=PI16;refGene_tx=XM_011282241.2
B2	38276471 38276581
refGene_gn=PI16;refGene_tx=XM_006931660.4;refGene_gn=PI16;refGene_tx=X
M_006931661.4;refGene_gn=PI16;refGene_tx=XM_011282241.2
B2	38276852 38276941
refGene_gn=PI16;refGene_tx=XM_006931660.4;refGene_gn=PI16;refGene_tx=X
M_006931661.4;refGene_gn=PI16;refGene_tx=XM_011282241.2
B2	38277822 38278500
refGene_gn=PI16;refGene_tx=XM_006931660.4;refGene_gn=PI16;refGene_tx=X
M_006931661.4;refGene_gn=PI16;refGene_tx=XM_011282241.2
B2	38278711 38278833
refGene_gn=PI16;refGene_tx=XM_006931660.4;refGene_gn=PI16;refGene_tx=X
M_006931661.4;refGene_gn=PI16;refGene_tx=XM_011282241.2
B2	38285414 38286021
refGene_gn=MTCH1;refGene_tx=XM_019830529.2;refGene_gn=MTCH1;refGene_tx
=XM_019830530.2;refGene_gn=MTCH1;refGene_tx=XM_019830531.2;refGene_gn=
MTCH1;refGene_tx=XM_023253929.1
B2	38287083 38287159
refGene_gn=MTCH1;refGene_tx=XM_019830531.2;refGene_gn=MTCH1;refGene_tx
=XM_023253929.1
B2	38287435 38287503
refGene_gn=MTCH1;refGene_tx=XM_019830529.2;refGene_gn=MTCH1;refGene_tx
=XM_019830530.2;refGene_gn=MTCH1;refGene_tx=XM_019830531.2;refGene_gn=
MTCH1;refGene_tx=XM_023253929.1
B2	38287704 38287752
refGene_gn=MTCH1;refGene_tx=XM_019830529.2;refGene_gn=MTCH1;refGene_tx
=XM_019830530.2;refGene_gn=MTCH1;refGene_tx=XM_019830531.2;refGene_gn=
MTCH1;refGene_tx=XM_023253929.1
B2	38289792 38289937
refGene_gn=MTCH1;refGene_tx=XM_019830529.2;refGene_gn=MTCH1;refGene_tx
=XM_019830530.2;refGene_gn=MTCH1;refGene_tx=XM_019830531.2;refGene_gn=
MTCH1;refGene_tx=XM_023253929.1
B2	38293160 38293220
refGene_gn=MTCH1;refGene_tx=XM_019830529.2;refGene_gn=MTCH1;refGene_tx
=XM_019830530.2;refGene_gn=MTCH1;refGene_tx=XM_019830531.2;refGene_gn=
MTCH1;refGene_tx=XM_023253929.1
B2	38294772 38294824
refGene_gn=MTCH1;refGene_tx=XM_019830529.2;refGene_gn=MTCH1;refGene_tx

=XM_019830530.2;refGene_gn=MTCH1;refGene_tx=XM_019830531.2;refGene_gn=MTCH1;refGene_tx=XM_023253929.1
B2 38295171 38295229
refGene_gn=MTCH1;refGene_tx=XM_019830529.2;refGene_gn=MTCH1;refGene_tx=XM_019830530.2;refGene_gn=MTCH1;refGene_tx=XM_019830531.2;refGene_gn=MTCH1;refGene_tx=XM_023253929.1
B2 38295609 38295687
refGene_gn=MTCH1;refGene_tx=XM_019830529.2;refGene_gn=MTCH1;refGene_tx=XM_019830530.2;refGene_gn=MTCH1;refGene_tx=XM_019830531.2;refGene_gn=MTCH1;refGene_tx=XM_023253929.1
B2 38296234 38296341
refGene_gn=MTCH1;refGene_tx=XM_019830529.2;refGene_gn=MTCH1;refGene_tx=XM_019830530.2;refGene_gn=MTCH1;refGene_tx=XM_019830531.2;refGene_gn=MTCH1;refGene_tx=XM_023253929.1
B2 38300439 38300524
refGene_gn=MTCH1;refGene_tx=XM_019830529.2;refGene_gn=MTCH1;refGene_tx=XM_019830530.2;refGene_gn=MTCH1;refGene_tx=XM_019830531.2;refGene_gn=MTCH1;refGene_tx=XM_023253929.1
B2 38304575 38304896
refGene_gn=MTCH1;refGene_tx=XM_019830529.2;refGene_gn=MTCH1;refGene_tx=XM_019830530.2;refGene_gn=MTCH1;refGene_tx=XM_019830531.2;refGene_gn=MTCH1;refGene_tx=XM_023253929.1
B2 38321693 38321761
refGene_gn=FGD2;refGene_tx=XM_003986073.4;refGene_gn=FGD2;refGene_tx=XM_006931662.3;refGene_gn=FGD2;refGene_tx=XM_019830532.1
B2 38325002 38325234
refGene_gn=FGD2;refGene_tx=XM_003986073.4;refGene_gn=FGD2;refGene_tx=XM_006931662.3;refGene_gn=FGD2;refGene_tx=XM_019830532.1
B2 38326991 38327072
refGene_gn=FGD2;refGene_tx=XM_003986073.4;refGene_gn=FGD2;refGene_tx=XM_006931662.3;refGene_gn=FGD2;refGene_tx=XM_019830532.1
B2 38328217 38328366
refGene_gn=FGD2;refGene_tx=XM_003986073.4;refGene_gn=FGD2;refGene_tx=XM_006931662.3;refGene_gn=FGD2;refGene_tx=XM_019830532.1
B2 38330056 38330213
refGene_gn=FGD2;refGene_tx=XM_003986073.4;refGene_gn=FGD2;refGene_tx=XM_006931662.3;refGene_gn=FGD2;refGene_tx=XM_019830532.1;refGene_gn=FGD2;refGene_tx=XM_019830534.1
B2 38330428 38330567
refGene_gn=FGD2;refGene_tx=XM_003986073.4;refGene_gn=FGD2;refGene_tx=XM_006931662.3;refGene_gn=FGD2;refGene_tx=XM_019830532.1;refGene_gn=FGD2;refGene_tx=XM_019830534.1;refGene_gn=FGD2;refGene_tx=XM_019830535.1
B2 38331341 38331400
refGene_gn=FGD2;refGene_tx=XM_003986073.4;refGene_gn=FGD2;refGene_tx=XM_006931662.3;refGene_gn=FGD2;refGene_tx=XM_019830532.1;refGene_gn=FGD2;refGene_tx=XM_019830534.1;refGene_gn=FGD2;refGene_tx=XM_019830535.1
B2 38331638 38331785
refGene_gn=FGD2;refGene_tx=XM_003986073.4;refGene_gn=FGD2;refGene_tx=XM_006931662.3;refGene_gn=FGD2;refGene_tx=XM_019830532.1;refGene_gn=FGD2;refGene_tx=XM_019830534.1;refGene_gn=FGD2;refGene_tx=XM_019830535.1

B2 38332463 38332556
refGene_gn=FGD2;refGene_tx=XM_003986073.4;refGene_gn=FGD2;refGene_tx=XM_006931662.3;refGene_gn=FGD2;refGene_tx=XM_019830532.1;refGene_gn=FGD2;refGene_tx=XM_019830534.1;refGene_gn=FGD2;refGene_tx=XM_019830535.1
B2 38339139 38339219
refGene_gn=FGD2;refGene_tx=XM_003986073.4;refGene_gn=FGD2;refGene_tx=XM_006931662.3;refGene_gn=FGD2;refGene_tx=XM_019830532.1;refGene_gn=FGD2;refGene_tx=XM_019830534.1;refGene_gn=FGD2;refGene_tx=XM_019830535.1
B2 38339303 38339334
refGene_gn=FGD2;refGene_tx=XM_003986073.4;refGene_gn=FGD2;refGene_tx=XM_006931662.3;refGene_gn=FGD2;refGene_tx=XM_019830532.1;refGene_gn=FGD2;refGene_tx=XM_019830534.1;refGene_gn=FGD2;refGene_tx=XM_019830535.1
B2 38339799 38339892
refGene_gn=FGD2;refGene_tx=XM_003986073.4;refGene_gn=FGD2;refGene_tx=XM_006931662.3;refGene_gn=FGD2;refGene_tx=XM_019830532.1;refGene_gn=FGD2;refGene_tx=XM_019830534.1;refGene_gn=FGD2;refGene_tx=XM_019830535.1
B2 38340987 38341119
refGene_gn=FGD2;refGene_tx=XM_003986073.4;refGene_gn=FGD2;refGene_tx=XM_006931662.3;refGene_gn=FGD2;refGene_tx=XM_019830532.1;refGene_gn=FGD2;refGene_tx=XM_019830534.1;refGene_gn=FGD2;refGene_tx=XM_019830535.1
B2 38344077 38344224
refGene_gn=FGD2;refGene_tx=XM_003986073.4;refGene_gn=FGD2;refGene_tx=XM_006931662.3;refGene_gn=FGD2;refGene_tx=XM_019830532.1;refGene_gn=FGD2;refGene_tx=XM_019830534.1;refGene_gn=FGD2;refGene_tx=XM_019830535.1
B2 38345364 38345511
refGene_gn=FGD2;refGene_tx=XM_003986073.4;refGene_gn=FGD2;refGene_tx=XM_006931662.3;refGene_gn=FGD2;refGene_tx=XM_019830532.1;refGene_gn=FGD2;refGene_tx=XM_019830534.1;refGene_gn=FGD2;refGene_tx=XM_019830535.1
B2 38345848 38346064
refGene_gn=FGD2;refGene_tx=XM_003986073.4;refGene_gn=FGD2;refGene_tx=XM_006931662.3;refGene_gn=FGD2;refGene_tx=XM_019830534.1;refGene_gn=FGD2;refGene_tx=XM_019830535.1
B2 38350042 38350125 refGene_gn=FGD2;refGene_tx=XM_019830532.1
B2 38352022 38352113 refGene_gn=FGD2;refGene_tx=XM_019830532.1
B2 38465793 38465875
refGene_gn=PIM1;refGene_tx=NM_001009322.1;refGene_gn=PIM1;refGene_tx=XM_023253551.1
B2 38466001 38466108
refGene_gn=PIM1;refGene_tx=NM_001009322.1;refGene_gn=PIM1;refGene_tx=XM_023253551.1
B2 38466215 38466281
refGene_gn=PIM1;refGene_tx=NM_001009322.1;refGene_gn=PIM1;refGene_tx=XM_023253551.1
B2 38466360 38466727
refGene_gn=PIM1;refGene_tx=NM_001009322.1;refGene_gn=PIM1;refGene_tx=XM_023253551.1
B2 38468294 38468471
refGene_gn=PIM1;refGene_tx=NM_001009322.1;refGene_gn=PIM1;refGene_tx=XM_023253551.1
B2 38469251 38469409

refGene_gn=PIM1;refGene_tx=NM_001009322.1;refGene_gn=PIM1;refGene_tx=XM_023253551.1
B2 38499377 38499941
refGene_gn=TMEM217;refGene_tx=XM_011282240.3;refGene_gn=TMEM217;refGene_tx=XM_019830536.2
B2 38517094 38517673 refGene_gn=TMEM217;refGene_tx=XM_003986112.5
B2 38546372 38546428 refGene_gn=TBC1D22B;refGene_tx=XM_006931682.4
B2 38563906 38563963 refGene_gn=TBC1D22B;refGene_tx=XM_006931682.4
B2 38574581 38574889 refGene_gn=TBC1D22B;refGene_tx=XM_006931682.4
B2 38578342 38578522 refGene_gn=TBC1D22B;refGene_tx=XM_006931682.4
B2 38579055 38579126 refGene_gn=TBC1D22B;refGene_tx=XM_006931682.4
B2 38580527 38580656 refGene_gn=TBC1D22B;refGene_tx=XM_006931682.4
B2 38581926 38581992 refGene_gn=TBC1D22B;refGene_tx=XM_006931682.4
B2 38586120 38586235 refGene_gn=TBC1D22B;refGene_tx=XM_006931682.4
B2 38603938 38604045 refGene_gn=TBC1D22B;refGene_tx=XM_006931682.4
B2 38604366 38604442 refGene_gn=TBC1D22B;refGene_tx=XM_006931682.4
B2 38606743 38606871 refGene_gn=TBC1D22B;refGene_tx=XM_006931682.4
B2 38607155 38607251 refGene_gn=TBC1D22B;refGene_tx=XM_006931682.4
B2 38624843 38624972 refGene_gn=TBC1D22B;refGene_tx=XM_006931682.4
B2 38642968 38643079
refGene_gn=RNF8;refGene_tx=XM_023253930.1;refGene_gn=RNF8;refGene_tx=XM_023253931.1
B2 38649219 38649348
refGene_gn=RNF8;refGene_tx=XM_023253930.1;refGene_gn=RNF8;refGene_tx=XM_023253931.1
B2 38658348 38659089
refGene_gn=RNF8;refGene_tx=XM_023253930.1;refGene_gn=RNF8;refGene_tx=XM_023253931.1
B2 38661521 38661584
refGene_gn=RNF8;refGene_tx=XM_023253930.1;refGene_gn=RNF8;refGene_tx=XM_023253931.1
B2 38663751 38663841
refGene_gn=RNF8;refGene_tx=XM_023253930.1;refGene_gn=RNF8;refGene_tx=XM_023253931.1
B2 38666200 38666308
refGene_gn=RNF8;refGene_tx=XM_023253930.1;refGene_gn=RNF8;refGene_tx=XM_023253931.1
B2 38668244 38668449 refGene_gn=RNF8;refGene_tx=XM_023253930.1
B2 38675625 38675709
refGene_gn=RNF8;refGene_tx=XM_023253930.1;refGene_gn=RNF8;refGene_tx=XM_023253931.1
B2 38723163 38723311
refGene_gn=CMTR1;refGene_tx=XM_003986075.5;refGene_gn=CMTR1;refGene_tx=XM_019830537.2;refGene_gn=CMTR1;refGene_tx=XM_019830538.2;refGene_gn=CMTR1;refGene_tx=XM_019830539.2
B2 38732456 38732608
refGene_gn=CMTR1;refGene_tx=XM_003986075.5;refGene_gn=CMTR1;refGene_tx=XM_019830537.2;refGene_gn=CMTR1;refGene_tx=XM_019830538.2;refGene_gn=CMTR1;refGene_tx=XM_019830539.2
B2 38734173 38734332

refGene_gn=CMTR1;refGene_tx=XM_003986075.5;refGene_gn=CMTR1;refGene_tx=XM_019830537.2;refGene_gn=CMTR1;refGene_tx=XM_019830538.2;refGene_gn=CMTR1;refGene_tx=XM_019830539.2
B2 38737234 38737327
refGene_gn=CMTR1;refGene_tx=XM_003986075.5;refGene_gn=CMTR1;refGene_tx=XM_019830537.2;refGene_gn=CMTR1;refGene_tx=XM_019830538.2
B2 38738728 38738800
refGene_gn=CMTR1;refGene_tx=XM_003986075.5;refGene_gn=CMTR1;refGene_tx=XM_019830537.2;refGene_gn=CMTR1;refGene_tx=XM_019830538.2
B2 38739897 38739992
refGene_gn=CMTR1;refGene_tx=XM_003986075.5;refGene_gn=CMTR1;refGene_tx=XM_019830537.2;refGene_gn=CMTR1;refGene_tx=XM_019830538.2
B2 38740105 38740178
refGene_gn=CMTR1;refGene_tx=XM_003986075.5;refGene_gn=CMTR1;refGene_tx=XM_019830537.2;refGene_gn=CMTR1;refGene_tx=XM_019830538.2
B2 38745286 38745485
refGene_gn=CMTR1;refGene_tx=XM_003986075.5;refGene_gn=CMTR1;refGene_tx=XM_019830537.2;refGene_gn=CMTR1;refGene_tx=XM_019830538.2;refGene_gn=CMTR1;refGene_tx=XM_019830539.2
B2 38746273 38746392
refGene_gn=CMTR1;refGene_tx=XM_003986075.5;refGene_gn=CMTR1;refGene_tx=XM_019830537.2;refGene_gn=CMTR1;refGene_tx=XM_019830538.2;refGene_gn=CMTR1;refGene_tx=XM_019830539.2
B2 38748669 38748766
refGene_gn=CMTR1;refGene_tx=XM_003986075.5;refGene_gn=CMTR1;refGene_tx=XM_019830537.2;refGene_gn=CMTR1;refGene_tx=XM_019830538.2;refGene_gn=CMTR1;refGene_tx=XM_019830539.2
B2 38749083 38749216
refGene_gn=CMTR1;refGene_tx=XM_003986075.5;refGene_gn=CMTR1;refGene_tx=XM_019830537.2;refGene_gn=CMTR1;refGene_tx=XM_019830538.2;refGene_gn=CMTR1;refGene_tx=XM_019830539.2
B2 38749927 38750107
refGene_gn=CMTR1;refGene_tx=XM_003986075.5;refGene_gn=CMTR1;refGene_tx=XM_019830537.2;refGene_gn=CMTR1;refGene_tx=XM_019830539.2
B2 38760739 38760796
refGene_gn=CMTR1;refGene_tx=XM_003986075.5;refGene_gn=CMTR1;refGene_tx=XM_019830537.2;refGene_gn=CMTR1;refGene_tx=XM_019830538.2;refGene_gn=CMTR1;refGene_tx=XM_019830539.2
B2 38761560 38761618
refGene_gn=CMTR1;refGene_tx=XM_003986075.5;refGene_gn=CMTR1;refGene_tx=XM_019830537.2;refGene_gn=CMTR1;refGene_tx=XM_019830538.2;refGene_gn=CMTR1;refGene_tx=XM_019830539.2
B2 38762119 38762188
refGene_gn=CMTR1;refGene_tx=XM_003986075.5;refGene_gn=CMTR1;refGene_tx=XM_019830537.2;refGene_gn=CMTR1;refGene_tx=XM_019830538.2;refGene_gn=CMTR1;refGene_tx=XM_019830539.2
B2 38762820 38762952
refGene_gn=CMTR1;refGene_tx=XM_003986075.5;refGene_gn=CMTR1;refGene_tx=XM_019830537.2;refGene_gn=CMTR1;refGene_tx=XM_019830538.2;refGene_gn=CMTR1;refGene_tx=XM_019830539.2

B2 38763930 38764053
refGene_gn=CMTR1;refGene_tx=XM_003986075.5;refGene_gn=CMTR1;refGene_tx=XM_019830537.2;refGene_gn=CMTR1;refGene_tx=XM_019830538.2;refGene_gn=CMTR1;refGene_tx=XM_019830539.2
B2 38764709 38764801
refGene_gn=CMTR1;refGene_tx=XM_003986075.5;refGene_gn=CMTR1;refGene_tx=XM_019830537.2;refGene_gn=CMTR1;refGene_tx=XM_019830538.2;refGene_gn=CMTR1;refGene_tx=XM_019830539.2
B2 38765632 38765701
refGene_gn=CMTR1;refGene_tx=XM_003986075.5;refGene_gn=CMTR1;refGene_tx=XM_019830537.2;refGene_gn=CMTR1;refGene_tx=XM_019830538.2;refGene_gn=CMTR1;refGene_tx=XM_019830539.2
B2 38766883 38766931
refGene_gn=CMTR1;refGene_tx=XM_003986075.5;refGene_gn=CMTR1;refGene_tx=XM_019830537.2;refGene_gn=CMTR1;refGene_tx=XM_019830538.2;refGene_gn=CMTR1;refGene_tx=XM_019830539.2
B2 38767775 38767888
refGene_gn=CMTR1;refGene_tx=XM_003986075.5;refGene_gn=CMTR1;refGene_tx=XM_019830537.2;refGene_gn=CMTR1;refGene_tx=XM_019830538.2;refGene_gn=CMTR1;refGene_tx=XM_019830539.2
B2 38768504 38768613
refGene_gn=CMTR1;refGene_tx=XM_003986075.5;refGene_gn=CMTR1;refGene_tx=XM_019830537.2;refGene_gn=CMTR1;refGene_tx=XM_019830538.2;refGene_gn=CMTR1;refGene_tx=XM_019830539.2
B2 38769404 38769537
refGene_gn=CMTR1;refGene_tx=XM_003986075.5;refGene_gn=CMTR1;refGene_tx=XM_019830537.2;refGene_gn=CMTR1;refGene_tx=XM_019830538.2;refGene_gn=CMTR1;refGene_tx=XM_019830539.2
B2 38772503 38772607 refGene_gn=CCDC167;refGene_tx=XM_023253932.1
B2 38774248 38774301 refGene_gn=CCDC167;refGene_tx=XM_023253932.1
B2 38774481 38774576 refGene_gn=CCDC167;refGene_tx=XM_023253932.1
B2 38788478 38788520 refGene_gn=CCDC167;refGene_tx=XM_023253932.1
B2 38920188 38920280
refGene_gn=MDGA1;refGene_tx=XM_003986077.4;refGene_gn=MDGA1;refGene_tx=XM_019830542.2;refGene_gn=MDGA1;refGene_tx=XM_019830543.2;refGene_gn=MDGA1;refGene_tx=XM_019830544.2
B2 38921021 38921130
refGene_gn=MDGA1;refGene_tx=XM_003986077.4;refGene_gn=MDGA1;refGene_tx=XM_019830542.2;refGene_gn=MDGA1;refGene_tx=XM_019830543.2;refGene_gn=MDGA1;refGene_tx=XM_019830544.2
B2 38921343 38921474
refGene_gn=MDGA1;refGene_tx=XM_003986077.4;refGene_gn=MDGA1;refGene_tx=XM_019830542.2;refGene_gn=MDGA1;refGene_tx=XM_019830543.2;refGene_gn=MDGA1;refGene_tx=XM_019830544.2
B2 38925665 38925800
refGene_gn=MDGA1;refGene_tx=XM_003986077.4;refGene_gn=MDGA1;refGene_tx=XM_019830542.2;refGene_gn=MDGA1;refGene_tx=XM_019830543.2;refGene_gn=MDGA1;refGene_tx=XM_019830544.2
B2 38926167 38926323
refGene_gn=MDGA1;refGene_tx=XM_003986077.4;refGene_gn=MDGA1;refGene_tx

=XM_019830542.2;refGene_gn=MDGA1;refGene_tx=XM_019830543.2;refGene_gn=MDGA1;refGene_tx=XM_019830544.2
B2 38927372 38927396
refGene_gn=MDGA1;refGene_tx=XM_003986077.4;refGene_gn=MDGA1;refGene_tx=XM_019830542.2;refGene_gn=MDGA1;refGene_tx=XM_019830543.2;refGene_gn=MDGA1;refGene_tx=XM_019830544.2
B2 38927633 38927887
refGene_gn=MDGA1;refGene_tx=XM_003986077.4;refGene_gn=MDGA1;refGene_tx=XM_019830542.2;refGene_gn=MDGA1;refGene_tx=XM_019830543.2;refGene_gn=MDGA1;refGene_tx=XM_019830544.2
B2 38928692 38928844
refGene_gn=MDGA1;refGene_tx=XM_003986077.4;refGene_gn=MDGA1;refGene_tx=XM_019830542.2;refGene_gn=MDGA1;refGene_tx=XM_019830543.2;refGene_gn=MDGA1;refGene_tx=XM_019830544.2
B2 38930562 38930847
refGene_gn=MDGA1;refGene_tx=XM_003986077.4;refGene_gn=MDGA1;refGene_tx=XM_019830542.2;refGene_gn=MDGA1;refGene_tx=XM_019830543.2;refGene_gn=MDGA1;refGene_tx=XM_019830544.2
B2 38931689 38931986
refGene_gn=MDGA1;refGene_tx=XM_003986077.4;refGene_gn=MDGA1;refGene_tx=XM_019830542.2;refGene_gn=MDGA1;refGene_tx=XM_019830543.2;refGene_gn=MDGA1;refGene_tx=XM_019830544.2
B2 38934365 38934695
refGene_gn=MDGA1;refGene_tx=XM_003986077.4;refGene_gn=MDGA1;refGene_tx=XM_019830542.2;refGene_gn=MDGA1;refGene_tx=XM_019830543.2;refGene_gn=MDGA1;refGene_tx=XM_019830544.2
B2 38936832 38937102
refGene_gn=MDGA1;refGene_tx=XM_003986077.4;refGene_gn=MDGA1;refGene_tx=XM_019830542.2;refGene_gn=MDGA1;refGene_tx=XM_019830543.2;refGene_gn=MDGA1;refGene_tx=XM_019830544.2
B2 38937352 38937485
refGene_gn=MDGA1;refGene_tx=XM_003986077.4;refGene_gn=MDGA1;refGene_tx=XM_019830542.2;refGene_gn=MDGA1;refGene_tx=XM_019830543.2;refGene_gn=MDGA1;refGene_tx=XM_019830544.2
B2 38938300 38938497
refGene_gn=MDGA1;refGene_tx=XM_003986077.4;refGene_gn=MDGA1;refGene_tx=XM_019830542.2;refGene_gn=MDGA1;refGene_tx=XM_019830543.2;refGene_gn=MDGA1;refGene_tx=XM_019830544.2
B2 38941114 38941289
refGene_gn=MDGA1;refGene_tx=XM_003986077.4;refGene_gn=MDGA1;refGene_tx=XM_019830542.2;refGene_gn=MDGA1;refGene_tx=XM_019830543.2;refGene_gn=MDGA1;refGene_tx=XM_019830544.2
B2 38945790 38945930
refGene_gn=MDGA1;refGene_tx=XM_003986077.4;refGene_gn=MDGA1;refGene_tx=XM_019830542.2;refGene_gn=MDGA1;refGene_tx=XM_019830543.2;refGene_gn=MDGA1;refGene_tx=XM_019830544.2
B2 38976472 38976539
refGene_gn=MDGA1;refGene_tx=XM_003986077.4;refGene_gn=MDGA1;refGene_tx=XM_019830542.2;refGene_gn=MDGA1;refGene_tx=XM_019830543.2;refGene_gn=MDGA1;refGene_tx=XM_019830544.2

B2 39085517 39085588
refGene_gn=ZFAND3;refGene_tx=XM_023253933.1;refGene_gn=ZFAND3;refGene_tx=XM_023253936.1
B2 39192338 39192379
refGene_gn=ZFAND3;refGene_tx=XM_023253933.1;refGene_gn=ZFAND3;refGene_tx=XM_023253936.1
B2 39194039 39194063 refGene_gn=ZFAND3;refGene_tx=XM_023253933.1
B2 39227560 39227678 refGene_gn=ZFAND3;refGene_tx=XM_023253935.1
B2 39267558 39267593 refGene_gn=ZFAND3;refGene_tx=XM_023253934.1
B2 39268002 39268085 refGene_gn=ZFAND3;refGene_tx=XM_023253934.1
B2 39331697 39331880
refGene_gn=ZFAND3;refGene_tx=XM_023253933.1;refGene_gn=ZFAND3;refGene_tx=XM_023253934.1;refGene_gn=ZFAND3;refGene_tx=XM_023253935.1;refGene_gn=ZFAND3;refGene_tx=XM_023253936.1
B2 39353019 39353085
refGene_gn=ZFAND3;refGene_tx=XM_023253933.1;refGene_gn=ZFAND3;refGene_tx=XM_023253934.1;refGene_gn=ZFAND3;refGene_tx=XM_023253935.1;refGene_gn=ZFAND3;refGene_tx=XM_023253936.1
B2 39379580 39379748
refGene_gn=ZFAND3;refGene_tx=XM_023253933.1;refGene_gn=ZFAND3;refGene_tx=XM_023253934.1;refGene_gn=ZFAND3;refGene_tx=XM_023253935.1;refGene_gn=ZFAND3;refGene_tx=XM_023253936.1
B2 39407913 39408068
refGene_gn=ZFAND3;refGene_tx=XM_023253933.1;refGene_gn=ZFAND3;refGene_tx=XM_023253934.1;refGene_gn=ZFAND3;refGene_tx=XM_023253935.1;refGene_gn=ZFAND3;refGene_tx=XM_023253936.1
B2 39432201 39432399 refGene_gn=BTBD9;refGene_tx=XM_019830546.2
B2 39452923 39453002 refGene_gn=BTBD9;refGene_tx=XM_019830546.2
B2 39510680 39510788 refGene_gn=BTBD9;refGene_tx=XM_019830546.2
B2 39540903 39541093 refGene_gn=BTBD9;refGene_tx=XM_019830546.2
B2 39575731 39575841 refGene_gn=BTBD9;refGene_tx=XM_019830546.2
B2 39789734 39789854 refGene_gn=BTBD9;refGene_tx=XM_019830546.2
B2 39792875 39793095 refGene_gn=BTBD9;refGene_tx=XM_019830546.2
B2 39801597 39801862 refGene_gn=BTBD9;refGene_tx=XM_019830546.2
B2 39803174 39803538 refGene_gn=BTBD9;refGene_tx=XM_019830546.2
B2 39807093 39807278 refGene_gn=BTBD9;refGene_tx=XM_019830546.2
B2 39867432 39867521 refGene_gn=GLO1;refGene_tx=XM_003986083.5
B2 39870985 39871075 refGene_gn=GLO1;refGene_tx=XM_003986083.5
B2 39871751 39871819 refGene_gn=GLO1;refGene_tx=XM_003986083.5
B2 39872643 39872784 refGene_gn=GLO1;refGene_tx=XM_003986083.5
B2 39882696 39882779 refGene_gn=GLO1;refGene_tx=XM_003986083.5
B2 39903060 39903144 refGene_gn=GLO1;refGene_tx=XM_003986083.5
B2 39922382 39922805 refGene_gn=DNAH8;refGene_tx=XM_019830547.2
B2 39922913 39923048 refGene_gn=DNAH8;refGene_tx=XM_019830547.2
B2 39931293 39931378 refGene_gn=DNAH8;refGene_tx=XM_019830547.2
B2 39934626 39934778 refGene_gn=DNAH8;refGene_tx=XM_019830547.2
B2 39944528 39944718 refGene_gn=DNAH8;refGene_tx=XM_019830547.2
B2 39945193 39945357 refGene_gn=DNAH8;refGene_tx=XM_019830547.2
B2 39950722 39950899 refGene_gn=DNAH8;refGene_tx=XM_019830547.2
B2 39953237 39953351 refGene_gn=DNAH8;refGene_tx=XM_019830547.2

| | | | |
|---|---|---|---|
| B2 | 39959159 | 39959267 | refGene_gn=DNAH8;refGene_tx=XM_019830547.2 |
| B2 | 39961838 | 39961940 | refGene_gn=DNAH8;refGene_tx=XM_019830547.2 |
| B2 | 39969665 | 39969812 | refGene_gn=DNAH8;refGene_tx=XM_019830547.2 |
| B2 | 39974782 | 39974980 | refGene_gn=DNAH8;refGene_tx=XM_019830547.2 |
| B2 | 39980778 | 39980855 | refGene_gn=DNAH8;refGene_tx=XM_019830547.2 |
| B2 | 39982166 | 39982266 | refGene_gn=DNAH8;refGene_tx=XM_019830547.2 |
| B2 | 39982868 | 39982988 | refGene_gn=DNAH8;refGene_tx=XM_019830547.2 |
| B2 | 39984561 | 39984697 | refGene_gn=DNAH8;refGene_tx=XM_019830547.2 |
| B2 | 39987874 | 39988062 | refGene_gn=DNAH8;refGene_tx=XM_019830547.2 |
| B2 | 39990541 | 39990622 | refGene_gn=DNAH8;refGene_tx=XM_019830547.2 |
| B2 | 39991030 | 39991147 | refGene_gn=DNAH8;refGene_tx=XM_019830547.2 |
| B2 | 39991938 | 39992058 | refGene_gn=DNAH8;refGene_tx=XM_019830547.2 |
| B2 | 39999147 | 39999283 | refGene_gn=DNAH8;refGene_tx=XM_019830547.2 |
| B2 | 40001092 | 40001208 | refGene_gn=DNAH8;refGene_tx=XM_019830547.2 |
| B2 | 40007144 | 40007251 | refGene_gn=DNAH8;refGene_tx=XM_019830547.2 |
| B2 | 40012725 | 40012801 | refGene_gn=DNAH8;refGene_tx=XM_019830547.2 |
| B2 | 40013961 | 40014154 | refGene_gn=DNAH8;refGene_tx=XM_019830547.2 |
| B2 | 40021494 | 40021691 | refGene_gn=DNAH8;refGene_tx=XM_019830547.2 |
| B2 | 40022271 | 40022398 | refGene_gn=DNAH8;refGene_tx=XM_019830547.2 |
| B2 | 40024764 | 40025000 | refGene_gn=DNAH8;refGene_tx=XM_019830547.2 |
| B2 | 40027083 | 40027188 | refGene_gn=DNAH8;refGene_tx=XM_019830547.2 |
| B2 | 40032756 | 40032870 | refGene_gn=DNAH8;refGene_tx=XM_019830547.2 |
| B2 | 40034428 | 40034491 | refGene_gn=DNAH8;refGene_tx=XM_019830547.2 |
| B2 | 40038751 | 40038852 | refGene_gn=DNAH8;refGene_tx=XM_019830547.2 |
| B2 | 40042286 | 40042424 | refGene_gn=DNAH8;refGene_tx=XM_019830547.2 |
| B2 | 40042652 | 40042893 | refGene_gn=DNAH8;refGene_tx=XM_019830547.2 |
| B2 | 40048581 | 40048781 | refGene_gn=DNAH8;refGene_tx=XM_019830547.2 |
| B2 | 40051662 | 40051816 | refGene_gn=DNAH8;refGene_tx=XM_019830547.2 |
| B2 | 40054335 | 40054499 | refGene_gn=DNAH8;refGene_tx=XM_019830547.2 |
| B2 | 40055603 | 40055706 | refGene_gn=DNAH8;refGene_tx=XM_019830547.2 |
| B2 | 40056708 | 40056813 | refGene_gn=DNAH8;refGene_tx=XM_019830547.2 |
| B2 | 40057621 | 40057783 | refGene_gn=DNAH8;refGene_tx=XM_019830547.2 |
| B2 | 40060606 | 40060831 | refGene_gn=DNAH8;refGene_tx=XM_019830547.2 |
| B2 | 40063564 | 40063737 | refGene_gn=DNAH8;refGene_tx=XM_019830547.2 |
| B2 | 40066472 | 40066651 | refGene_gn=DNAH8;refGene_tx=XM_019830547.2 |
| B2 | 40069698 | 40069886 | refGene_gn=DNAH8;refGene_tx=XM_019830547.2 |
| B2 | 40072305 | 40072392 | refGene_gn=DNAH8;refGene_tx=XM_019830547.2 |
| B2 | 40072479 | 40072587 | refGene_gn=DNAH8;refGene_tx=XM_019830547.2 |
| B2 | 40073592 | 40073727 | refGene_gn=DNAH8;refGene_tx=XM_019830547.2 |
| B2 | 40075457 | 40075619 | refGene_gn=DNAH8;refGene_tx=XM_019830547.2 |
| B2 | 40078155 | 40078402 | refGene_gn=DNAH8;refGene_tx=XM_019830547.2 |
| B2 | 40078517 | 40078546 | refGene_gn=DNAH8;refGene_tx=XM_019830547.2 |
| B2 | 40078547 | 40078656 | refGene_gn=DNAH8;refGene_tx=XM_019830547.2 |
| B2 | 40078658 | 40078759 | refGene_gn=DNAH8;refGene_tx=XM_019830547.2 |
| B2 | 40078849 | 40078990 | refGene_gn=DNAH8;refGene_tx=XM_019830547.2 |
| B2 | 40081973 | 40082211 | refGene_gn=DNAH8;refGene_tx=XM_019830547.2 |
| B2 | 40088593 | 40088736 | refGene_gn=DNAH8;refGene_tx=XM_019830547.2 |
| B2 | 40089195 | 40089330 | refGene_gn=DNAH8;refGene_tx=XM_019830547.2 |
| B2 | 40089989 | 40090112 | refGene_gn=DNAH8;refGene_tx=XM_019830547.2 |
| B2 | 40095200 | 40095414 | refGene_gn=DNAH8;refGene_tx=XM_019830547.2 |

| | | | | |
|---|---|---|---|---|
| B2 | 40097776 | 40097886 | refGene_gn=DNAH8; | refGene_tx=XM_019830547.2 |
| B2 | 40102102 | 40102266 | refGene_gn=DNAH8; | refGene_tx=XM_019830547.2 |
| B2 | 40102972 | 40103165 | refGene_gn=DNAH8; | refGene_tx=XM_019830547.2 |
| B2 | 40104549 | 40104672 | refGene_gn=DNAH8; | refGene_tx=XM_019830547.2 |
| B2 | 40106195 | 40106326 | refGene_gn=DNAH8; | refGene_tx=XM_019830547.2 |
| B2 | 40110853 | 40111007 | refGene_gn=DNAH8; | refGene_tx=XM_019830547.2 |
| B2 | 40120452 | 40120617 | refGene_gn=DNAH8; | refGene_tx=XM_019830547.2 |
| B2 | 40121954 | 40122181 | refGene_gn=DNAH8; | refGene_tx=XM_019830547.2 |
| B2 | 40124231 | 40124350 | refGene_gn=DNAH8; | refGene_tx=XM_019830547.2 |
| B2 | 40126153 | 40126257 | refGene_gn=DNAH8; | refGene_tx=XM_019830547.2 |
| B2 | 40127782 | 40127959 | refGene_gn=DNAH8; | refGene_tx=XM_019830547.2 |
| B2 | 40129927 | 40130095 | refGene_gn=DNAH8; | refGene_tx=XM_019830547.2 |
| B2 | 40130614 | 40130830 | refGene_gn=DNAH8; | refGene_tx=XM_019830547.2 |
| B2 | 40134747 | 40134768 | refGene_gn=DNAH8; | refGene_tx=XM_019830547.2 |
| B2 | 40134769 | 40134886 | refGene_gn=DNAH8; | refGene_tx=XM_019830547.2 |
| B2 | 40136351 | 40136479 | refGene_gn=DNAH8; | refGene_tx=XM_019830547.2 |
| B2 | 40137280 | 40137452 | refGene_gn=DNAH8; | refGene_tx=XM_019830547.2 |
| B2 | 40139420 | 40139576 | refGene_gn=DNAH8; | refGene_tx=XM_019830547.2 |
| B2 | 40143940 | 40144096 | refGene_gn=DNAH8; | refGene_tx=XM_019830547.2 |
| B2 | 40145943 | 40146126 | refGene_gn=DNAH8; | refGene_tx=XM_019830547.2 |
| B2 | 40149954 | 40150060 | refGene_gn=DNAH8; | refGene_tx=XM_019830547.2 |
| B2 | 40151799 | 40152052 | refGene_gn=DNAH8; | refGene_tx=XM_019830547.2 |
| B2 | 40152172 | 40152363 | refGene_gn=DNAH8; | refGene_tx=XM_019830547.2 |
| B2 | 40158906 | 40159028 | refGene_gn=DNAH8; | refGene_tx=XM_019830547.2 |
| B2 | 40162366 | 40162485 | refGene_gn=DNAH8; | refGene_tx=XM_019830547.2 |
| B2 | 40164624 | 40164827 | refGene_gn=DNAH8; | refGene_tx=XM_019830547.2 |
| B2 | 40177157 | 40177231 | refGene_gn=DNAH8; | refGene_tx=XM_019830547.2 |
| B2 | 40179557 | 40179710 | refGene_gn=DNAH8; | refGene_tx=XM_019830547.2 |
| B2 | 40180083 | 40180239 | refGene_gn=DNAH8; | refGene_tx=XM_019830547.2 |
| B2 | 40188977 | 40189094 | refGene_gn=DNAH8; | refGene_tx=XM_019830547.2 |
| B2 | 40190889 | 40190991 | refGene_gn=DNAH8; | refGene_tx=XM_019830547.2 |
| B2 | 40199046 | 40199207 | refGene_gn=DNAH8; | refGene_tx=XM_019830547.2 |
| B2 | 40216805 | 40216962 | refGene_gn=DNAH8; | refGene_tx=XM_019830547.2 |
| B2 | 40220877 | 40221030 | refGene_gn=DNAH8; | refGene_tx=XM_019830547.2 |
| B2 | 40221108 | 40221298 | refGene_gn=DNAH8; | refGene_tx=XM_019830547.2 |
| B2 | 40238625 | 40238747 | refGene_gn=DNAH8; | refGene_tx=XM_019830547.2 |
| B2 | 40241917 | 40242205 | refGene_gn=DNAH8; | refGene_tx=XM_019830547.2 |
| B2 | 40262862 | 40262940 | refGene_gn=GLP1R; | refGene_tx=XM_023253938.1 |
| B2 | 40271060 | 40271157 | refGene_gn=GLP1R; | refGene_tx=XM_023253938.1 |
| B2 | 40272080 | 40272188 | refGene_gn=GLP1R; | refGene_tx=XM_023253938.1 |
| B2 | 40279919 | 40280038 | refGene_gn=GLP1R; | refGene_tx=XM_023253938.1 |
| B2 | 40280397 | 40280504 | refGene_gn=GLP1R; | refGene_tx=XM_023253938.1 |
| B2 | 40284191 | 40284345 | refGene_gn=GLP1R; | refGene_tx=XM_023253938.1 |
| B2 | 40284873 | 40285033 | refGene_gn=GLP1R; | refGene_tx=XM_023253938.1 |
| B2 | 40289534 | 40289595 | refGene_gn=GLP1R; | refGene_tx=XM_023253938.1 |
| B2 | 40290150 | 40290220 | refGene_gn=GLP1R; | refGene_tx=XM_023253938.1 |
| B2 | 40290343 | 40290432 | refGene_gn=GLP1R; | refGene_tx=XM_023253938.1 |
| B2 | 40290755 | 40290894 | refGene_gn=GLP1R; | refGene_tx=XM_023253938.1 |
| B2 | 40291799 | 40291841 | refGene_gn=GLP1R; | refGene_tx=XM_023253938.1 |
| B2 | 40296355 | 40296523 | refGene_gn=GLP1R; | refGene_tx=XM_023253938.1 |

B2 40313892 40314237 refGene_gn=SAYSD1;refGene_tx=XM_023253939.1
B2 40319768 40319975 refGene_gn=SAYSD1;refGene_tx=XM_023253939.1
B2 40386139 40387005 refGene_gn=KCNK5;refGene_tx=XM_003986084.5
B2 40389167 40389336 refGene_gn=KCNK5;refGene_tx=XM_003986084.5
B2 40389605 40389772 refGene_gn=KCNK5;refGene_tx=XM_003986084.5
B2 40390937 40391049 refGene_gn=KCNK5;refGene_tx=XM_003986084.5
B2 40421680 40421866 refGene_gn=KCNK5;refGene_tx=XM_003986084.5
B2 40475560 40475871 refGene_gn=KCNK17;refGene_tx=XM_019830556.1
B2 40479993 40480168 refGene_gn=KCNK17;refGene_tx=XM_019830556.1
B2 40480584 40480745 refGene_gn=KCNK17;refGene_tx=XM_019830556.1
B2 40485576 40485691 refGene_gn=KCNK17;refGene_tx=XM_019830556.1
B2 40488800 40489073 refGene_gn=KCNK17;refGene_tx=XM_019830556.1
B2 40490904 40491128 refGene_gn=KCNK16;refGene_tx=XM_003986085.4
B2 40491402 40491568 refGene_gn=KCNK16;refGene_tx=XM_003986085.4
B2 40492418 40492585 refGene_gn=KCNK16;refGene_tx=XM_003986085.4
B2 40493403 40493518 refGene_gn=KCNK16;refGene_tx=XM_003986085.4
B2 40496529 40496742 refGene_gn=KCNK16;refGene_tx=XM_003986085.4
B2 40509147 40509158
refGene_gn=KIF6;refGene_tx=XM_006931669.4;refGene_gn=KIF6;refGene_tx=XM_019830554.2;refGene_gn=KIF6;refGene_tx=XM_019830555.2
B2 40514710 40514817
refGene_gn=KIF6;refGene_tx=XM_006931669.4;refGene_gn=KIF6;refGene_tx=XM_019830554.2;refGene_gn=KIF6;refGene_tx=XM_019830555.2
B2 40517226 40517316
refGene_gn=KIF6;refGene_tx=XM_006931669.4;refGene_gn=KIF6;refGene_tx=XM_019830554.2;refGene_gn=KIF6;refGene_tx=XM_019830555.2
B2 40524743 40524841
refGene_gn=KIF6;refGene_tx=XM_006931669.4;refGene_gn=KIF6;refGene_tx=XM_019830554.2;refGene_gn=KIF6;refGene_tx=XM_019830555.2
B2 40527682 40527818
refGene_gn=KIF6;refGene_tx=XM_006931669.4;refGene_gn=KIF6;refGene_tx=XM_019830554.2;refGene_gn=KIF6;refGene_tx=XM_019830555.2
B2 40529504 40529589
refGene_gn=KIF6;refGene_tx=XM_006931669.4;refGene_gn=KIF6;refGene_tx=XM_019830554.2;refGene_gn=KIF6;refGene_tx=XM_019830555.2
B2 40551341 40551392
refGene_gn=KIF6;refGene_tx=XM_006931669.4;refGene_gn=KIF6;refGene_tx=XM_019830554.2;refGene_gn=KIF6;refGene_tx=XM_019830555.2
B2 40595135 40595191
refGene_gn=KIF6;refGene_tx=XM_006931669.4;refGene_gn=KIF6;refGene_tx=XM_019830554.2;refGene_gn=KIF6;refGene_tx=XM_019830555.2
B2 40604444 40604553
refGene_gn=KIF6;refGene_tx=XM_006931669.4;refGene_gn=KIF6;refGene_tx=XM_019830554.2;refGene_gn=KIF6;refGene_tx=XM_019830555.2
B2 40707991 40708207
refGene_gn=KIF6;refGene_tx=XM_006931669.4;refGene_gn=KIF6;refGene_tx=XM_019830554.2;refGene_gn=KIF6;refGene_tx=XM_019830555.2
B2 40712754 40712893
refGene_gn=KIF6;refGene_tx=XM_006931669.4;refGene_gn=KIF6;refGene_tx=XM_019830554.2;refGene_gn=KIF6;refGene_tx=XM_019830555.2

B2 40713990 40714096
refGene_gn=KIF6;refGene_tx=XM_006931669.4;refGene_gn=KIF6;refGene_tx=XM_019830554.2;refGene_gn=KIF6;refGene_tx=XM_019830555.2
B2 40751182 40751286
refGene_gn=KIF6;refGene_tx=XM_006931669.4;refGene_gn=KIF6;refGene_tx=XM_019830554.2;refGene_gn=KIF6;refGene_tx=XM_019830555.2
B2 40757318 40757405
refGene_gn=KIF6;refGene_tx=XM_006931669.4;refGene_gn=KIF6;refGene_tx=XM_019830554.2;refGene_gn=KIF6;refGene_tx=XM_019830555.2
B2 40758349 40758493
refGene_gn=KIF6;refGene_tx=XM_006931669.4;refGene_gn=KIF6;refGene_tx=XM_019830554.2;refGene_gn=KIF6;refGene_tx=XM_019830555.2
B2 40766368 40766575
refGene_gn=KIF6;refGene_tx=XM_006931669.4;refGene_gn=KIF6;refGene_tx=XM_019830554.2;refGene_gn=KIF6;refGene_tx=XM_019830555.2
B2 40786815 40786945
refGene_gn=KIF6;refGene_tx=XM_006931669.4;refGene_gn=KIF6;refGene_tx=XM_019830554.2;refGene_gn=KIF6;refGene_tx=XM_019830555.2
B2 40807515 40807625
refGene_gn=KIF6;refGene_tx=XM_006931669.4;refGene_gn=KIF6;refGene_tx=XM_019830554.2;refGene_gn=KIF6;refGene_tx=XM_019830555.2
B2 40812683 40812831
refGene_gn=KIF6;refGene_tx=XM_006931669.4;refGene_gn=KIF6;refGene_tx=XM_019830554.2;refGene_gn=KIF6;refGene_tx=XM_019830555.2
B2 40843798 40843824 refGene_gn=KIF6;refGene_tx=XM_019830555.2
B2 40882199 40882274 refGene_gn=KIF6;refGene_tx=XM_006931669.4
B2 40888352 40888462
refGene_gn=KIF6;refGene_tx=XM_006931669.4;refGene_gn=KIF6;refGene_tx=XM_019830554.2
B2 40892629 40892695
refGene_gn=KIF6;refGene_tx=XM_006931669.4;refGene_gn=KIF6;refGene_tx=XM_019830554.2
B2 41025594 41025762
refGene_gn=DAAM2;refGene_tx=XM_006931672.4;refGene_gn=DAAM2;refGene_tx=XM_019830558.2;refGene_gn=DAAM2;refGene_tx=XM_019830559.2;refGene_gn=DAAM2;refGene_tx=XM_019830560.2;refGene_gn=DAAM2;refGene_tx=XM_019830561.2;refGene_gn=DAAM2;refGene_tx=XM_019830562.2
B2 41031197 41031287
refGene_gn=DAAM2;refGene_tx=XM_006931672.4;refGene_gn=DAAM2;refGene_tx=XM_019830558.2;refGene_gn=DAAM2;refGene_tx=XM_019830559.2;refGene_gn=DAAM2;refGene_tx=XM_019830560.2;refGene_gn=DAAM2;refGene_tx=XM_019830561.2;refGene_gn=DAAM2;refGene_tx=XM_019830562.2
B2 41034575 41034650
refGene_gn=DAAM2;refGene_tx=XM_006931672.4;refGene_gn=DAAM2;refGene_tx=XM_019830558.2;refGene_gn=DAAM2;refGene_tx=XM_019830559.2;refGene_gn=DAAM2;refGene_tx=XM_019830560.2;refGene_gn=DAAM2;refGene_tx=XM_019830561.2;refGene_gn=DAAM2;refGene_tx=XM_019830562.2
B2 41035394 41035489
refGene_gn=DAAM2;refGene_tx=XM_006931672.4;refGene_gn=DAAM2;refGene_tx=XM_019830558.2;refGene_gn=DAAM2;refGene_tx=XM_019830559.2;refGene_gn=

DAAM2;refGene_tx=XM_019830560.2;refGene_gn=DAAM2;refGene_tx=XM_019830561.2;refGene_gn=DAAM2;refGene_tx=XM_019830562.2
B2 41038328 41038662
refGene_gn=DAAM2;refGene_tx=XM_006931672.4;refGene_gn=DAAM2;refGene_tx=XM_019830558.2;refGene_gn=DAAM2;refGene_tx=XM_019830559.2;refGene_gn=DAAM2;refGene_tx=XM_019830560.2;refGene_gn=DAAM2;refGene_tx=XM_019830561.2;refGene_gn=DAAM2;refGene_tx=XM_019830562.2
B2 41039523 41039634
refGene_gn=DAAM2;refGene_tx=XM_006931672.4;refGene_gn=DAAM2;refGene_tx=XM_019830558.2;refGene_gn=DAAM2;refGene_tx=XM_019830559.2;refGene_gn=DAAM2;refGene_tx=XM_019830560.2;refGene_gn=DAAM2;refGene_tx=XM_019830561.2;refGene_gn=DAAM2;refGene_tx=XM_019830562.2
B2 41040827 41040931
refGene_gn=DAAM2;refGene_tx=XM_006931672.4;refGene_gn=DAAM2;refGene_tx=XM_019830558.2;refGene_gn=DAAM2;refGene_tx=XM_019830559.2;refGene_gn=DAAM2;refGene_tx=XM_019830560.2;refGene_gn=DAAM2;refGene_tx=XM_019830561.2;refGene_gn=DAAM2;refGene_tx=XM_019830562.2
B2 41042046 41042113
refGene_gn=DAAM2;refGene_tx=XM_006931672.4;refGene_gn=DAAM2;refGene_tx=XM_019830558.2;refGene_gn=DAAM2;refGene_tx=XM_019830559.2;refGene_gn=DAAM2;refGene_tx=XM_019830560.2;refGene_gn=DAAM2;refGene_tx=XM_019830561.2;refGene_gn=DAAM2;refGene_tx=XM_019830562.2
B2 41043571 41043689
refGene_gn=DAAM2;refGene_tx=XM_006931672.4;refGene_gn=DAAM2;refGene_tx=XM_019830558.2;refGene_gn=DAAM2;refGene_tx=XM_019830559.2;refGene_gn=DAAM2;refGene_tx=XM_019830560.2;refGene_gn=DAAM2;refGene_tx=XM_019830561.2;refGene_gn=DAAM2;refGene_tx=XM_019830562.2
B2 41045690 41045829
refGene_gn=DAAM2;refGene_tx=XM_006931672.4;refGene_gn=DAAM2;refGene_tx=XM_019830558.2;refGene_gn=DAAM2;refGene_tx=XM_019830559.2;refGene_gn=DAAM2;refGene_tx=XM_019830560.2;refGene_gn=DAAM2;refGene_tx=XM_019830561.2;refGene_gn=DAAM2;refGene_tx=XM_019830562.2
B2 41049262 41049321
refGene_gn=DAAM2;refGene_tx=XM_006931672.4;refGene_gn=DAAM2;refGene_tx=XM_019830558.2;refGene_gn=DAAM2;refGene_tx=XM_019830559.2;refGene_gn=DAAM2;refGene_tx=XM_019830560.2;refGene_gn=DAAM2;refGene_tx=XM_019830561.2;refGene_gn=DAAM2;refGene_tx=XM_019830562.2
B2 41049456 41049641
refGene_gn=DAAM2;refGene_tx=XM_006931672.4;refGene_gn=DAAM2;refGene_tx=XM_019830558.2;refGene_gn=DAAM2;refGene_tx=XM_019830559.2;refGene_gn=DAAM2;refGene_tx=XM_019830560.2;refGene_gn=DAAM2;refGene_tx=XM_019830561.2;refGene_gn=DAAM2;refGene_tx=XM_019830562.2
B2 41050246 41050546
refGene_gn=DAAM2;refGene_tx=XM_006931672.4;refGene_gn=DAAM2;refGene_tx=XM_019830558.2;refGene_gn=DAAM2;refGene_tx=XM_019830559.2;refGene_gn=DAAM2;refGene_tx=XM_019830560.2;refGene_gn=DAAM2;refGene_tx=XM_019830561.2;refGene_gn=DAAM2;refGene_tx=XM_019830562.2
B2 41055375 41055483
refGene_gn=DAAM2;refGene_tx=XM_006931672.4;refGene_gn=DAAM2;refGene_tx=XM_019830558.2;refGene_gn=DAAM2;refGene_tx=XM_019830559.2;refGene_gn=

DAAM2;refGene_tx=XM_019830560.2;refGene_gn=DAAM2;refGene_tx=XM_0198305
61.2;refGene_gn=DAAM2;refGene_tx=XM_019830562.2
B2 41057787 41057811
refGene_gn=DAAM2;refGene_tx=XM_019830558.2;refGene_gn=DAAM2;refGene_tx
=XM_019830559.2;refGene_gn=DAAM2;refGene_tx=XM_019830560.2;refGene_gn=
DAAM2;refGene_tx=XM_019830561.2;refGene_gn=DAAM2;refGene_tx=XM_0198305
62.2
B2 41058816 41058923
refGene_gn=DAAM2;refGene_tx=XM_006931672.4;refGene_gn=DAAM2;refGene_tx
=XM_019830558.2;refGene_gn=DAAM2;refGene_tx=XM_019830559.2;refGene_gn=
DAAM2;refGene_tx=XM_019830560.2;refGene_gn=DAAM2;refGene_tx=XM_0198305
61.2;refGene_gn=DAAM2;refGene_tx=XM_019830562.2
B2 41060126 41060211
refGene_gn=DAAM2;refGene_tx=XM_006931672.4;refGene_gn=DAAM2;refGene_tx
=XM_019830558.2;refGene_gn=DAAM2;refGene_tx=XM_019830559.2;refGene_gn=
DAAM2;refGene_tx=XM_019830560.2;refGene_gn=DAAM2;refGene_tx=XM_0198305
61.2;refGene_gn=DAAM2;refGene_tx=XM_019830562.2
B2 41061774 41061881
refGene_gn=DAAM2;refGene_tx=XM_006931672.4;refGene_gn=DAAM2;refGene_tx
=XM_019830558.2;refGene_gn=DAAM2;refGene_tx=XM_019830559.2;refGene_gn=
DAAM2;refGene_tx=XM_019830560.2;refGene_gn=DAAM2;refGene_tx=XM_0198305
61.2;refGene_gn=DAAM2;refGene_tx=XM_019830562.2
B2 41061996 41062085
refGene_gn=DAAM2;refGene_tx=XM_006931672.4;refGene_gn=DAAM2;refGene_tx
=XM_019830558.2;refGene_gn=DAAM2;refGene_tx=XM_019830559.2;refGene_gn=
DAAM2;refGene_tx=XM_019830560.2;refGene_gn=DAAM2;refGene_tx=XM_0198305
61.2;refGene_gn=DAAM2;refGene_tx=XM_019830562.2
B2 41071954 41072123
refGene_gn=DAAM2;refGene_tx=XM_006931672.4;refGene_gn=DAAM2;refGene_tx
=XM_019830558.2;refGene_gn=DAAM2;refGene_tx=XM_019830559.2;refGene_gn=
DAAM2;refGene_tx=XM_019830560.2;refGene_gn=DAAM2;refGene_tx=XM_0198305
61.2;refGene_gn=DAAM2;refGene_tx=XM_019830562.2
B2 41072296 41072404
refGene_gn=DAAM2;refGene_tx=XM_006931672.4;refGene_gn=DAAM2;refGene_tx
=XM_019830558.2;refGene_gn=DAAM2;refGene_tx=XM_019830559.2;refGene_gn=
DAAM2;refGene_tx=XM_019830560.2;refGene_gn=DAAM2;refGene_tx=XM_0198305
61.2;refGene_gn=DAAM2;refGene_tx=XM_019830562.2
B2 41073360 41073421
refGene_gn=DAAM2;refGene_tx=XM_006931672.4;refGene_gn=DAAM2;refGene_tx
=XM_019830558.2;refGene_gn=DAAM2;refGene_tx=XM_019830559.2;refGene_gn=
DAAM2;refGene_tx=XM_019830560.2;refGene_gn=DAAM2;refGene_tx=XM_0198305
61.2;refGene_gn=DAAM2;refGene_tx=XM_019830562.2
B2 41074637 41074769
refGene_gn=DAAM2;refGene_tx=XM_006931672.4;refGene_gn=DAAM2;refGene_tx
=XM_019830558.2;refGene_gn=DAAM2;refGene_tx=XM_019830559.2;refGene_gn=
DAAM2;refGene_tx=XM_019830560.2;refGene_gn=DAAM2;refGene_tx=XM_0198305
61.2;refGene_gn=DAAM2;refGene_tx=XM_019830562.2
B2 41075828 41075999
refGene_gn=DAAM2;refGene_tx=XM_006931672.4;refGene_gn=DAAM2;refGene_tx
=XM_019830558.2;refGene_gn=DAAM2;refGene_tx=XM_019830559.2;refGene_gn=

DAAM2;refGene_tx=XM_019830560.2;refGene_gn=DAAM2;refGene_tx=XM_0198305
61.2;refGene_gn=DAAM2;refGene_tx=XM_019830562.2
B2 41076125 41076347
refGene_gn=DAAM2;refGene_tx=XM_006931672.4;refGene_gn=DAAM2;refGene_tx
=XM_019830558.2;refGene_gn=DAAM2;refGene_tx=XM_019830559.2;refGene_gn=
DAAM2;refGene_tx=XM_019830560.2;refGene_gn=DAAM2;refGene_tx=XM_0198305
61.2;refGene_gn=DAAM2;refGene_tx=XM_019830562.2
B2 41080640 41081401
refGene_gn=MOCS1;refGene_tx=XM_011282235.3;refGene_gn=MOCS1;refGene_tx
=XM_011282236.3;refGene_gn=MOCS1;refGene_tx=XM_023253940.1
B2 41083430 41083486
refGene_gn=MOCS1;refGene_tx=XM_006931670.3;refGene_gn=MOCS1;refGene_tx
=XM_011282235.3;refGene_gn=MOCS1;refGene_tx=XM_023253940.1
B2 41083979 41084100
refGene_gn=MOCS1;refGene_tx=XM_006931670.3;refGene_gn=MOCS1;refGene_tx
=XM_011282235.3;refGene_gn=MOCS1;refGene_tx=XM_011282236.3;refGene_gn=
MOCS1;refGene_tx=XM_023253940.1
B2 41086348 41086459
refGene_gn=MOCS1;refGene_tx=XM_006931670.3;refGene_gn=MOCS1;refGene_tx
=XM_011282235.3;refGene_gn=MOCS1;refGene_tx=XM_011282236.3;refGene_gn=
MOCS1;refGene_tx=XM_023253940.1
B2 41086895 41087008
refGene_gn=MOCS1;refGene_tx=XM_006931670.3;refGene_gn=MOCS1;refGene_tx
=XM_011282235.3;refGene_gn=MOCS1;refGene_tx=XM_011282236.3;refGene_gn=
MOCS1;refGene_tx=XM_023253940.1
B2 41087428 41087540
refGene_gn=MOCS1;refGene_tx=XM_006931670.3;refGene_gn=MOCS1;refGene_tx
=XM_011282235.3;refGene_gn=MOCS1;refGene_tx=XM_011282236.3;refGene_gn=
MOCS1;refGene_tx=XM_023253940.1
B2 41087885 41087947
refGene_gn=MOCS1;refGene_tx=XM_006931670.3;refGene_gn=MOCS1;refGene_tx
=XM_011282235.3;refGene_gn=MOCS1;refGene_tx=XM_011282236.3;refGene_gn=
MOCS1;refGene_tx=XM_023253940.1
B2 41090107 41090272
refGene_gn=MOCS1;refGene_tx=XM_006931670.3;refGene_gn=MOCS1;refGene_tx
=XM_011282235.3;refGene_gn=MOCS1;refGene_tx=XM_011282236.3;refGene_gn=
MOCS1;refGene_tx=XM_023253940.1
B2 41107484 41107652
refGene_gn=MOCS1;refGene_tx=XM_006931670.3;refGene_gn=MOCS1;refGene_tx
=XM_011282235.3;refGene_gn=MOCS1;refGene_tx=XM_011282236.3;refGene_gn=
MOCS1;refGene_tx=XM_023253940.1
B2 41109053 41109180
refGene_gn=MOCS1;refGene_tx=XM_006931670.3;refGene_gn=MOCS1;refGene_tx
=XM_011282235.3;refGene_gn=MOCS1;refGene_tx=XM_011282236.3
B2 41116570 41116693
refGene_gn=MOCS1;refGene_tx=XM_006931670.3;refGene_gn=MOCS1;refGene_tx
=XM_011282235.3;refGene_gn=MOCS1;refGene_tx=XM_011282236.3
B2 41494750 41495729 refGene_gn=LRFN2;refGene_tx=XM_003986090.3
B2 41533175 41534575 refGene_gn=LRFN2;refGene_tx=XM_003986090.3
B2 42083458 42083663 refGene_gn=UNC5CL;refGene_tx=XM_003986122.5

B2 42084788 42084902 refGene_gn=UNC5CL;refGene_tx=XM_003986122.5
B2 42085051 42085152 refGene_gn=UNC5CL;refGene_tx=XM_003986122.5
B2 42085984 42086052 refGene_gn=UNC5CL;refGene_tx=XM_003986122.5
B2 42086305 42086407 refGene_gn=UNC5CL;refGene_tx=XM_003986122.5
B2 42087033 42087296 refGene_gn=UNC5CL;refGene_tx=XM_003986122.5
B2 42088019 42088320 refGene_gn=UNC5CL;refGene_tx=XM_003986122.5
B2 42088842 42089288 refGene_gn=UNC5CL;refGene_tx=XM_003986122.5
B2 42093197 42093235 refGene_gn=UNC5CL;refGene_tx=XM_003986122.5
B2 42096716 42096886 refGene_gn=TSPO2;refGene_tx=XM_003986123.3
B2 42097258 42097400 refGene_gn=TSPO2;refGene_tx=XM_003986123.3
B2 42097658 42097859 refGene_gn=TSPO2;refGene_tx=XM_003986123.3
B2 42107717 42107875 refGene_gn=APOBEC2;refGene_tx=XM_003986124.5
B2 42116121 42116665 refGene_gn=APOBEC2;refGene_tx=XM_003986124.5
B2 42122469 42122572
refGene_gn=OARD1;refGene_tx=XM_003986125.4;refGene_gn=OARD1;refGene_tx=XM_006931686.4;refGene_gn=OARD1;refGene_tx=XM_019830580.2;refGene_gn=OARD1;refGene_tx=XM_019830581.2
B2 42123932 42124045
refGene_gn=OARD1;refGene_tx=XM_003986125.4;refGene_gn=OARD1;refGene_tx=XM_006931686.4;refGene_gn=OARD1;refGene_tx=XM_019830580.2
B2 42125070 42125129
refGene_gn=OARD1;refGene_tx=XM_003986125.4;refGene_gn=OARD1;refGene_tx=XM_006931686.4;refGene_gn=OARD1;refGene_tx=XM_019830580.2;refGene_gn=OARD1;refGene_tx=XM_019830581.2
B2 42126055 42126200
refGene_gn=OARD1;refGene_tx=XM_003986125.4;refGene_gn=OARD1;refGene_tx=XM_006931686.4;refGene_gn=OARD1;refGene_tx=XM_019830580.2;refGene_gn=OARD1;refGene_tx=XM_019830581.2
B2 42126724 42126763
refGene_gn=OARD1;refGene_tx=XM_003986125.4;refGene_gn=OARD1;refGene_tx=XM_006931686.4;refGene_gn=OARD1;refGene_tx=XM_019830580.2;refGene_gn=OARD1;refGene_tx=XM_019830581.2
B2 42134924 42134999
refGene_gn=NFYA;refGene_tx=XM_003986126.4;refGene_gn=NFYA;refGene_tx=XM_006931687.3;refGene_gn=NFYA;refGene_tx=XM_006931688.3;refGene_gn=NFYA;refGene_tx=XM_006931689.3;refGene_gn=NFYA;refGene_tx=XM_006931690.3;refGene_gn=NFYA;refGene_tx=XM_006931691.3;refGene_gn=NFYA;refGene_tx=XM_006931692.3;refGene_gn=NFYA;refGene_tx=XM_011282232.3;refGene_gn=NFYA;refGene_tx=XM_019830579.2;refGene_gn=NFYA;refGene_tx=XM_023253949.1
B2 42136552 42136639
refGene_gn=NFYA;refGene_tx=XM_006931687.3;refGene_gn=NFYA;refGene_tx=XM_006931688.3;refGene_gn=NFYA;refGene_tx=XM_006931689.3;refGene_gn=NFYA;refGene_tx=XM_006931690.3;refGene_gn=NFYA;refGene_tx=XM_006931691.3;refGene_gn=NFYA;refGene_tx=XM_011282232.3;refGene_gn=NFYA;refGene_tx=XM_019830579.2;refGene_gn=NFYA;refGene_tx=XM_023253949.1
B2 42139457 42139604
refGene_gn=NFYA;refGene_tx=XM_003986126.4;refGene_gn=NFYA;refGene_tx=XM_006931687.3;refGene_gn=NFYA;refGene_tx=XM_006931688.3;refGene_gn=NFYA;refGene_tx=XM_006931689.3;refGene_gn=NFYA;refGene_tx=XM_006931690.3;refGene_gn=NFYA;refGene_tx=XM_006931691.3;refGene_gn=NFYA;refGene_tx=X

M_006931692.3;refGene_gn=NFYA;refGene_tx=XM_011282232.3;refGene_gn=NFYA;refGene_tx=XM_019830579.2;refGene_gn=NFYA;refGene_tx=XM_023253949.1;refGene_gn=NFYA;refGene_tx=XM_023253950.1
B2 42151970 42152102
refGene_gn=NFYA;refGene_tx=XM_003986126.4;refGene_gn=NFYA;refGene_tx=XM_006931687.3;refGene_gn=NFYA;refGene_tx=XM_006931688.3;refGene_gn=NFYA;refGene_tx=XM_006931689.3;refGene_gn=NFYA;refGene_tx=XM_006931690.3;refGene_gn=NFYA;refGene_tx=XM_006931691.3;refGene_gn=NFYA;refGene_tx=XM_006931692.3;refGene_gn=NFYA;refGene_tx=XM_011282232.3;refGene_gn=NFYA;refGene_tx=XM_019830579.2;refGene_gn=NFYA;refGene_tx=XM_023253949.1;refGene_gn=NFYA;refGene_tx=XM_023253950.1
B2 42152307 42152413
refGene_gn=NFYA;refGene_tx=XM_003986126.4;refGene_gn=NFYA;refGene_tx=XM_006931687.3;refGene_gn=NFYA;refGene_tx=XM_006931688.3;refGene_gn=NFYA;refGene_tx=XM_006931689.3;refGene_gn=NFYA;refGene_tx=XM_006931690.3;refGene_gn=NFYA;refGene_tx=XM_006931691.3;refGene_gn=NFYA;refGene_tx=XM_006931692.3;refGene_gn=NFYA;refGene_tx=XM_011282232.3;refGene_gn=NFYA;refGene_tx=XM_019830579.2;refGene_gn=NFYA;refGene_tx=XM_023253949.1;refGene_gn=NFYA;refGene_tx=XM_023253950.1
B2 42153769 42153936
refGene_gn=NFYA;refGene_tx=XM_003986126.4;refGene_gn=NFYA;refGene_tx=XM_006931687.3;refGene_gn=NFYA;refGene_tx=XM_006931688.3;refGene_gn=NFYA;refGene_tx=XM_006931689.3;refGene_gn=NFYA;refGene_tx=XM_006931690.3;refGene_gn=NFYA;refGene_tx=XM_006931691.3;refGene_gn=NFYA;refGene_tx=XM_006931692.3;refGene_gn=NFYA;refGene_tx=XM_011282232.3;refGene_gn=NFYA;refGene_tx=XM_019830579.2;refGene_gn=NFYA;refGene_tx=XM_023253949.1;refGene_gn=NFYA;refGene_tx=XM_023253950.1
B2 42154999 42155173
refGene_gn=NFYA;refGene_tx=XM_003986126.4;refGene_gn=NFYA;refGene_tx=XM_006931687.3;refGene_gn=NFYA;refGene_tx=XM_006931688.3;refGene_gn=NFYA;refGene_tx=XM_006931689.3;refGene_gn=NFYA;refGene_tx=XM_006931690.3;refGene_gn=NFYA;refGene_tx=XM_006931691.3;refGene_gn=NFYA;refGene_tx=XM_006931692.3;refGene_gn=NFYA;refGene_tx=XM_011282232.3;refGene_gn=NFYA;refGene_tx=XM_019830579.2;refGene_gn=NFYA;refGene_tx=XM_023253949.1;refGene_gn=NFYA;refGene_tx=XM_023253950.1
B2 42156157 42156259
refGene_gn=NFYA;refGene_tx=XM_003986126.4;refGene_gn=NFYA;refGene_tx=XM_006931687.3;refGene_gn=NFYA;refGene_tx=XM_006931688.3;refGene_gn=NFYA;refGene_tx=XM_006931689.3;refGene_gn=NFYA;refGene_tx=XM_006931690.3;refGene_gn=NFYA;refGene_tx=XM_006931691.3;refGene_gn=NFYA;refGene_tx=XM_006931692.3;refGene_gn=NFYA;refGene_tx=XM_011282232.3;refGene_gn=NFYA;refGene_tx=XM_019830579.2;refGene_gn=NFYA;refGene_tx=XM_023253949.1;refGene_gn=NFYA;refGene_tx=XM_023253950.1
B2 42159248 42159302
refGene_gn=NFYA;refGene_tx=XM_003986126.4;refGene_gn=NFYA;refGene_tx=XM_006931687.3;refGene_gn=NFYA;refGene_tx=XM_006931688.3;refGene_gn=NFYA;refGene_tx=XM_006931689.3;refGene_gn=NFYA;refGene_tx=XM_006931690.3;refGene_gn=NFYA;refGene_tx=XM_006931691.3;refGene_gn=NFYA;refGene_tx=XM_006931692.3;refGene_gn=NFYA;refGene_tx=XM_011282232.3;refGene_gn=NFYA;refGene_tx=XM_019830579.2;refGene_gn=NFYA;refGene_tx=XM_023253949.1;

refGene_gn=NFYA;refGene_tx=XM_023253950.1
B2 42163117 42163274
refGene_gn=LOC101090105;refGene_tx=XM_006931757.4;refGene_gn=LOC101090105;refGene_tx=XM_023253942.1;refGene_gn=LOC101090105;refGene_tx=XM_023253943.1;refGene_gn=LOC101090105;refGene_tx=XM_023253944.1;refGene_gn=LOC101090105;refGene_tx=XM_023253946.1;refGene_gn=LOC101090105;refGene_tx=XM_023253947.1;refGene_gn=LOC101090105;refGene_tx=XM_023253948.1
B2 42163490 42163521
refGene_gn=LOC101090105;refGene_tx=XM_023253941.1
B2 42164064 42164169
refGene_gn=LOC101090105;refGene_tx=XM_006931757.4;refGene_gn=LOC101090105;refGene_tx=XM_023253941.1;refGene_gn=LOC101090105;refGene_tx=XM_023253942.1;refGene_gn=LOC101090105;refGene_tx=XM_023253943.1;refGene_gn=LOC101090105;refGene_tx=XM_023253944.1;refGene_gn=LOC101090105;refGene_tx=XM_023253946.1;refGene_gn=LOC101090105;refGene_tx=XM_023253947.1;refGene_gn=LOC101090105;refGene_tx=XM_023253948.1
B2 42167455 42167494
refGene_gn=LOC101090105;refGene_tx=XM_006931757.4;refGene_gn=LOC101090105;refGene_tx=XM_023253941.1;refGene_gn=LOC101090105;refGene_tx=XM_023253942.1;refGene_gn=LOC101090105;refGene_tx=XM_023253943.1;refGene_gn=LOC101090105;refGene_tx=XM_023253944.1;refGene_gn=LOC101090105;refGene_tx=XM_023253946.1;refGene_gn=LOC101090105;refGene_tx=XM_023253947.1;refGene_gn=LOC101090105;refGene_tx=XM_023253948.1
B2 42168265 42168409
refGene_gn=LOC101090105;refGene_tx=XM_006931757.4;refGene_gn=LOC101090105;refGene_tx=XM_023253941.1;refGene_gn=LOC101090105;refGene_tx=XM_023253943.1;refGene_gn=LOC101090105;refGene_tx=XM_023253944.1;refGene_gn=LOC101090105;refGene_tx=XM_023253946.1;refGene_gn=LOC101090105;refGene_tx=XM_023253947.1;refGene_gn=LOC101090105;refGene_tx=XM_023253948.1
B2 42179539 42179745
refGene_gn=LOC101090105;refGene_tx=XM_006931757.4;refGene_gn=LOC101090105;refGene_tx=XM_023253941.1;refGene_gn=LOC101090105;refGene_tx=XM_023253942.1;refGene_gn=LOC101090105;refGene_tx=XM_023253943.1;refGene_gn=LOC101090105;refGene_tx=XM_023253944.1;refGene_gn=LOC101090105;refGene_tx=XM_023253946.1;refGene_gn=LOC101090105;refGene_tx=XM_023253947.1;refGene_gn=LOC101090105;refGene_tx=XM_023253948.1
B2 42180001 42180092
refGene_gn=LOC101090105;refGene_tx=XM_006931757.4;refGene_gn=LOC101090105;refGene_tx=XM_023253941.1;refGene_gn=LOC101090105;refGene_tx=XM_023253942.1;refGene_gn=LOC101090105;refGene_tx=XM_023253943.1;refGene_gn=LOC101090105;refGene_tx=XM_023253944.1;refGene_gn=LOC101090105;refGene_tx=XM_023253946.1;refGene_gn=LOC101090105;refGene_tx=XM_023253947.1;refGene_gn=LOC101090105;refGene_tx=XM_023253948.1
B2 42180182 42180271
refGene_gn=LOC101090105;refGene_tx=XM_006931757.4;refGene_gn=LOC101090105;refGene_tx=XM_023253941.1;refGene_gn=LOC101090105;refGene_tx=XM_023253942.1;refGene_gn=LOC101090105;refGene_tx=XM_023253943.1;refGene_gn=LOC101090105;refGene_tx=XM_023253944.1;refGene_gn=LOC101090105;refGene_tx=XM_023253945.1;refGene_gn=LOC101090105;refGene_tx=XM_023253946.1;refGene_gn=LOC101090105;refGene_tx=XM_023253947.1;refGene_gn=LOC101090

105;refGene_tx=XM_023253948.1
B2 42180874 42181066
refGene_gn=LOC101090105;refGene_tx=XM_006931757.4;refGene_gn=LOC101090105;refGene_tx=XM_023253941.1;refGene_gn=LOC101090105;refGene_tx=XM_023253942.1;refGene_gn=LOC101090105;refGene_tx=XM_023253943.1;refGene_gn=LOC101090105;refGene_tx=XM_023253944.1;refGene_gn=LOC101090105;refGene_tx=XM_023253945.1;refGene_gn=LOC101090105;refGene_tx=XM_023253946.1;refGene_gn=LOC101090105;refGene_tx=XM_023253947.1;refGene_gn=LOC101090105;refGene_tx=XM_023253948.1
B2 42181746 42181868
refGene_gn=LOC101090105;refGene_tx=XM_006931757.4;refGene_gn=LOC101090105;refGene_tx=XM_023253941.1;refGene_gn=LOC101090105;refGene_tx=XM_023253942.1;refGene_gn=LOC101090105;refGene_tx=XM_023253943.1;refGene_gn=LOC101090105;refGene_tx=XM_023253944.1;refGene_gn=LOC101090105;refGene_tx=XM_023253945.1;refGene_gn=LOC101090105;refGene_tx=XM_023253946.1;refGene_gn=LOC101090105;refGene_tx=XM_023253947.1;refGene_gn=LOC101090105;refGene_tx=XM_023253948.1
B2 42186135 42186212
refGene_gn=LOC101090105;refGene_tx=XM_006931757.4;refGene_gn=LOC101090105;refGene_tx=XM_023253941.1;refGene_gn=LOC101090105;refGene_tx=XM_023253942.1;refGene_gn=LOC101090105;refGene_tx=XM_023253943.1;refGene_gn=LOC101090105;refGene_tx=XM_023253944.1;refGene_gn=LOC101090105;refGene_tx=XM_023253945.1;refGene_gn=LOC101090105;refGene_tx=XM_023253946.1;refGene_gn=LOC101090105;refGene_tx=XM_023253947.1;refGene_gn=LOC101090105;refGene_tx=XM_023253948.1
B2 42186311 42186501
refGene_gn=LOC101090105;refGene_tx=XM_006931757.4;refGene_gn=LOC101090105;refGene_tx=XM_023253941.1;refGene_gn=LOC101090105;refGene_tx=XM_023253942.1;refGene_gn=LOC101090105;refGene_tx=XM_023253943.1;refGene_gn=LOC101090105;refGene_tx=XM_023253944.1;refGene_gn=LOC101090105;refGene_tx=XM_023253945.1;refGene_gn=LOC101090105;refGene_tx=XM_023253946.1;refGene_gn=LOC101090105;refGene_tx=XM_023253947.1;refGene_gn=LOC101090105;refGene_tx=XM_023253948.1
B2 42187000 42187056
refGene_gn=LOC101090105;refGene_tx=XM_006931757.4;refGene_gn=LOC101090105;refGene_tx=XM_023253941.1;refGene_gn=LOC101090105;refGene_tx=XM_023253942.1;refGene_gn=LOC101090105;refGene_tx=XM_023253943.1;refGene_gn=LOC101090105;refGene_tx=XM_023253944.1;refGene_gn=LOC101090105;refGene_tx=XM_023253945.1;refGene_gn=LOC101090105;refGene_tx=XM_023253946.1;refGene_gn=LOC101090105;refGene_tx=XM_023253947.1;refGene_gn=LOC101090105;refGene_tx=XM_023253948.1
B2 42187799 42187953
refGene_gn=LOC101090105;refGene_tx=XM_006931757.4;refGene_gn=LOC101090105;refGene_tx=XM_023253941.1;refGene_gn=LOC101090105;refGene_tx=XM_023253942.1;refGene_gn=LOC101090105;refGene_tx=XM_023253943.1;refGene_gn=LOC101090105;refGene_tx=XM_023253944.1;refGene_gn=LOC101090105;refGene_tx=XM_023253945.1;refGene_gn=LOC101090105;refGene_tx=XM_023253946.1;refGene_gn=LOC101090105;refGene_tx=XM_023253947.1;refGene_gn=LOC101090105;refGene_tx=XM_023253948.1
B2 42188115 42188305

refGene_gn=LOC101090105;refGene_tx=XM_006931757.4;refGene_gn=LOC101090105;refGene_tx=XM_023253941.1;refGene_gn=LOC101090105;refGene_tx=XM_023253942.1;refGene_gn=LOC101090105;refGene_tx=XM_023253943.1;refGene_gn=LOC101090105;refGene_tx=XM_023253944.1;refGene_gn=LOC101090105;refGene_tx=XM_023253945.1;refGene_gn=LOC101090105;refGene_tx=XM_023253946.1;refGene_gn=LOC101090105;refGene_tx=XM_023253947.1;refGene_gn=LOC101090105;refGene_tx=XM_023253948.1
B2 42189479 42189566
refGene_gn=LOC101090105;refGene_tx=XM_006931757.4;refGene_gn=LOC101090105;refGene_tx=XM_023253941.1;refGene_gn=LOC101090105;refGene_tx=XM_023253942.1;refGene_gn=LOC101090105;refGene_tx=XM_023253943.1;refGene_gn=LOC101090105;refGene_tx=XM_023253944.1;refGene_gn=LOC101090105;refGene_tx=XM_023253945.1;refGene_gn=LOC101090105;refGene_tx=XM_023253946.1;refGene_gn=LOC101090105;refGene_tx=XM_023253947.1;refGene_gn=LOC101090105;refGene_tx=XM_023253948.1
B2 42190759 42191031
refGene_gn=LOC101090105;refGene_tx=XM_006931757.4;refGene_gn=LOC101090105;refGene_tx=XM_023253941.1;refGene_gn=LOC101090105;refGene_tx=XM_023253942.1;refGene_gn=LOC101090105;refGene_tx=XM_023253943.1;refGene_gn=LOC101090105;refGene_tx=XM_023253944.1;refGene_gn=LOC101090105;refGene_tx=XM_023253945.1;refGene_gn=LOC101090105;refGene_tx=XM_023253946.1;refGene_gn=LOC101090105;refGene_tx=XM_023253947.1;refGene_gn=LOC101090105;refGene_tx=XM_023253948.1
B2 42191551 42191685
refGene_gn=LOC101090105;refGene_tx=XM_006931757.4;refGene_gn=LOC101090105;refGene_tx=XM_023253941.1;refGene_gn=LOC101090105;refGene_tx=XM_023253942.1;refGene_gn=LOC101090105;refGene_tx=XM_023253943.1;refGene_gn=LOC101090105;refGene_tx=XM_023253944.1;refGene_gn=LOC101090105;refGene_tx=XM_023253945.1;refGene_gn=LOC101090105;refGene_tx=XM_023253946.1;refGene_gn=LOC101090105;refGene_tx=XM_023253947.1;refGene_gn=LOC101090105;refGene_tx=XM_023253948.1
B2 42192300 42192426
refGene_gn=LOC101090105;refGene_tx=XM_006931757.4;refGene_gn=LOC101090105;refGene_tx=XM_023253941.1;refGene_gn=LOC101090105;refGene_tx=XM_023253942.1;refGene_gn=LOC101090105;refGene_tx=XM_023253943.1;refGene_gn=LOC101090105;refGene_tx=XM_023253944.1;refGene_gn=LOC101090105;refGene_tx=XM_023253945.1;refGene_gn=LOC101090105;refGene_tx=XM_023253946.1;refGene_gn=LOC101090105;refGene_tx=XM_023253947.1;refGene_gn=LOC101090105;refGene_tx=XM_023253948.1
B2 42194663 42194939
refGene_gn=LOC101090105;refGene_tx=XM_006931757.4;refGene_gn=LOC101090105;refGene_tx=XM_023253941.1;refGene_gn=LOC101090105;refGene_tx=XM_023253942.1;refGene_gn=LOC101090105;refGene_tx=XM_023253943.1;refGene_gn=LOC101090105;refGene_tx=XM_023253944.1;refGene_gn=LOC101090105;refGene_tx=XM_023253945.1;refGene_gn=LOC101090105;refGene_tx=XM_023253946.1;refGene_gn=LOC101090105;refGene_tx=XM_023253947.1;refGene_gn=LOC101090105;refGene_tx=XM_023253948.1
B2 42195772 42196102
refGene_gn=LOC101090105;refGene_tx=XM_006931757.4;refGene_gn=LOC101090105;refGene_tx=XM_023253941.1;refGene_gn=LOC101090105;refGene_tx=XM_02

3253942.1;refGene_gn=LOC101090105;refGene_tx=XM_023253943.1;refGene_gn=LOC101090105;refGene_tx=XM_023253944.1;refGene_gn=LOC101090105;refGene_tx=XM_023253945.1;refGene_gn=LOC101090105;refGene_tx=XM_023253946.1;refGene_gn=LOC101090105;refGene_tx=XM_023253947.1;refGene_gn=LOC101090105;refGene_tx=XM_023253948.1
B2 42201180 42201244
refGene_gn=LOC101090105;refGene_tx=XM_006931757.4;refGene_gn=LOC101090105;refGene_tx=XM_023253941.1;refGene_gn=LOC101090105;refGene_tx=XM_023253942.1;refGene_gn=LOC101090105;refGene_tx=XM_023253943.1;refGene_gn=LOC101090105;refGene_tx=XM_023253944.1;refGene_gn=LOC101090105;refGene_tx=XM_023253945.1;refGene_gn=LOC101090105;refGene_tx=XM_023253946.1;refGene_gn=LOC101090105;refGene_tx=XM_023253947.1;refGene_gn=LOC101090105;refGene_tx=XM_023253948.1
B2 42202448 42202668
refGene_gn=LOC101090105;refGene_tx=XM_006931757.4;refGene_gn=LOC101090105;refGene_tx=XM_023253941.1;refGene_gn=LOC101090105;refGene_tx=XM_023253942.1;refGene_gn=LOC101090105;refGene_tx=XM_023253943.1;refGene_gn=LOC101090105;refGene_tx=XM_023253944.1;refGene_gn=LOC101090105;refGene_tx=XM_023253945.1;refGene_gn=LOC101090105;refGene_tx=XM_023253946.1;refGene_gn=LOC101090105;refGene_tx=XM_023253947.1;refGene_gn=LOC101090105;refGene_tx=XM_023253948.1
B2 42203026 42203152
refGene_gn=LOC101090105;refGene_tx=XM_006931757.4;refGene_gn=LOC101090105;refGene_tx=XM_023253941.1;refGene_gn=LOC101090105;refGene_tx=XM_023253942.1;refGene_gn=LOC101090105;refGene_tx=XM_023253943.1;refGene_gn=LOC101090105;refGene_tx=XM_023253944.1;refGene_gn=LOC101090105;refGene_tx=XM_023253945.1;refGene_gn=LOC101090105;refGene_tx=XM_023253946.1;refGene_gn=LOC101090105;refGene_tx=XM_023253947.1;refGene_gn=LOC101090105;refGene_tx=XM_023253948.1
B2 42206426 42206599
refGene_gn=LOC101090105;refGene_tx=XM_006931757.4;refGene_gn=LOC101090105;refGene_tx=XM_023253941.1;refGene_gn=LOC101090105;refGene_tx=XM_023253942.1;refGene_gn=LOC101090105;refGene_tx=XM_023253943.1;refGene_gn=LOC101090105;refGene_tx=XM_023253944.1;refGene_gn=LOC101090105;refGene_tx=XM_023253945.1;refGene_gn=LOC101090105;refGene_tx=XM_023253946.1;refGene_gn=LOC101090105;refGene_tx=XM_023253947.1;refGene_gn=LOC101090105;refGene_tx=XM_023253948.1
B2 42207520 42207677
refGene_gn=LOC101090105;refGene_tx=XM_006931757.4;refGene_gn=LOC101090105;refGene_tx=XM_023253941.1;refGene_gn=LOC101090105;refGene_tx=XM_023253942.1;refGene_gn=LOC101090105;refGene_tx=XM_023253943.1;refGene_gn=LOC101090105;refGene_tx=XM_023253944.1;refGene_gn=LOC101090105;refGene_tx=XM_023253945.1;refGene_gn=LOC101090105;refGene_tx=XM_023253946.1;refGene_gn=LOC101090105;refGene_tx=XM_023253947.1;refGene_gn=LOC101090105;refGene_tx=XM_023253948.1
B2 42209624 42209829
refGene_gn=LOC101090105;refGene_tx=XM_006931757.4;refGene_gn=LOC101090105;refGene_tx=XM_023253941.1;refGene_gn=LOC101090105;refGene_tx=XM_023253942.1;refGene_gn=LOC101090105;refGene_tx=XM_023253943.1;refGene_gn=LOC101090105;refGene_tx=XM_023253944.1;refGene_gn=LOC101090105;refGen

e_tx=XM_023253945.1
B2 42209918 42210015
refGene_gn=LOC101090105;refGene_tx=XM_006931757.4;refGene_gn=LOC101090105;refGene_tx=XM_023253941.1;refGene_gn=LOC101090105;refGene_tx=XM_023253942.1;refGene_gn=LOC101090105;refGene_tx=XM_023253943.1;refGene_gn=LOC101090105;refGene_tx=XM_023253944.1;refGene_gn=LOC101090105;refGene_tx=XM_023253945.1;refGene_gn=LOC101090105;refGene_tx=XM_023253946.1;refGene_gn=LOC101090105;refGene_tx=XM_023253947.1
B2 42211253 42211372
refGene_gn=LOC101090105;refGene_tx=XM_006931757.4;refGene_gn=LOC101090105;refGene_tx=XM_023253941.1;refGene_gn=LOC101090105;refGene_tx=XM_023253942.1;refGene_gn=LOC101090105;refGene_tx=XM_023253943.1;refGene_gn=LOC101090105;refGene_tx=XM_023253944.1;refGene_gn=LOC101090105;refGene_tx=XM_023253945.1
B2 42211634 42211756
refGene_gn=LOC101090105;refGene_tx=XM_006931757.4;refGene_gn=LOC101090105;refGene_tx=XM_023253941.1;refGene_gn=LOC101090105;refGene_tx=XM_023253942.1;refGene_gn=LOC101090105;refGene_tx=XM_023253943.1;refGene_gn=LOC101090105;refGene_tx=XM_023253944.1;refGene_gn=LOC101090105;refGene_tx=XM_023253945.1;refGene_gn=LOC101090105;refGene_tx=XM_023253948.1
B2 42212169 42212368
refGene_gn=LOC101090105;refGene_tx=XM_006931757.4;refGene_gn=LOC101090105;refGene_tx=XM_023253941.1;refGene_gn=LOC101090105;refGene_tx=XM_023253942.1;refGene_gn=LOC101090105;refGene_tx=XM_023253943.1;refGene_gn=LOC101090105;refGene_tx=XM_023253945.1
B2 42212891 42213105
refGene_gn=LOC101090105;refGene_tx=XM_006931757.4;refGene_gn=LOC101090105;refGene_tx=XM_023253941.1;refGene_gn=LOC101090105;refGene_tx=XM_023253942.1;refGene_gn=LOC101090105;refGene_tx=XM_023253945.1
B2 42213615 42213723
refGene_gn=LOC101090105;refGene_tx=XM_006931757.4;refGene_gn=LOC101090105;refGene_tx=XM_023253941.1;refGene_gn=LOC101090105;refGene_tx=XM_023253942.1;refGene_gn=LOC101090105;refGene_tx=XM_023253943.1;refGene_gn=LOC101090105;refGene_tx=XM_023253945.1
B2 42222764 42223079
refGene_gn=TREML1;refGene_tx=XM_006931693.4;refGene_gn=TREML1;refGene_tx=XM_019830582.2
B2 42223427 42223480
refGene_gn=TREML1;refGene_tx=XM_006931693.4;refGene_gn=TREML1;refGene_tx=XM_019830582.2
B2 42223960 42224046
refGene_gn=TREML1;refGene_tx=XM_006931693.4;refGene_gn=TREML1;refGene_tx=XM_019830582.2
B2 42224463 42224578
refGene_gn=TREML1;refGene_tx=XM_006931693.4;refGene_gn=TREML1;refGene_tx=XM_019830582.2
B2 42227093 42227423
refGene_gn=TREML1;refGene_tx=XM_006931693.4;refGene_gn=TREML1;refGene_tx=XM_019830582.2
B2 42227622 42227665

refGene_gn=TREML1;refGene_tx=XM_006931693.4;refGene_gn=TREML1;refGene_tx=XM_019830582.2
B2 42231751 42231768 refGene_gn=TREM2;refGene_tx=XM_003986128.5
B2 42231864 42232058 refGene_gn=TREM2;refGene_tx=XM_003986128.5
B2 42232390 42232565 refGene_gn=TREM2;refGene_tx=XM_019830583.2
B2 42232767 42232858
refGene_gn=TREM2;refGene_tx=XM_003986128.5;refGene_gn=TREM2;refGene_tx=XM_019830583.2
B2 42234240 42234591
refGene_gn=TREM2;refGene_tx=XM_003986128.5;refGene_gn=TREM2;refGene_tx=XM_019830583.2
B2 42236013 42236053
refGene_gn=TREM2;refGene_tx=XM_003986128.5;refGene_gn=TREM2;refGene_tx=XM_019830583.2
B2 42248172 42248421
refGene_gn=LOC101090615;refGene_tx=XM_006931696.3;refGene_gn=LOC101090615;refGene_tx=XM_006931697.3
B2 42249293 42249400
refGene_gn=LOC101090615;refGene_tx=XM_006931696.3;refGene_gn=LOC101090615;refGene_tx=XM_006931697.3
B2 42251353 42251472
refGene_gn=LOC101090615;refGene_tx=XM_006931696.3;refGene_gn=LOC101090615;refGene_tx=XM_006931697.3
B2 42252774 42252850
refGene_gn=LOC101090615;refGene_tx=XM_006931696.3;refGene_gn=LOC101090615;refGene_tx=XM_006931697.3
B2 42254513 42254846
refGene_gn=LOC101090615;refGene_tx=XM_006931696.3;refGene_gn=LOC101090615;refGene_tx=XM_006931697.3
B2 42255039 42255403
refGene_gn=LOC101090615;refGene_tx=XM_006931696.3;refGene_gn=LOC101090615;refGene_tx=XM_006931697.3
B2 42256366 42256468
refGene_gn=LOC101090615;refGene_tx=XM_006931696.3;refGene_gn=LOC101090615;refGene_tx=XM_006931697.3
B2 42258873 42258968
refGene_gn=TREML2;refGene_tx=XM_023253951.1;refGene_gn=TREML2;refGene_tx=XM_023253952.1
B2 42259272 42259379 refGene_gn=TREML2;refGene_tx=XM_023253951.1
B2 42260398 42260816
refGene_gn=TREML2;refGene_tx=XM_023253951.1;refGene_gn=TREML2;refGene_tx=XM_023253952.1
B2 42264379 42264700
refGene_gn=TREML2;refGene_tx=XM_023253951.1;refGene_gn=TREML2;refGene_tx=XM_023253952.1
B2 42267027 42267088
refGene_gn=TREML2;refGene_tx=XM_023253951.1;refGene_gn=TREML2;refGene_tx=XM_023253952.1
B2 42293246 42293352 refGene_gn=TREM1;refGene_tx=XM_003986129.4
B2 42297553 42297764 refGene_gn=TREM1;refGene_tx=XM_003986129.4

B2 42299132 42299486 refGene_gn=TREM1;refGene_tx=XM_003986129.4
B2 42302767 42302816 refGene_gn=TREM1;refGene_tx=XM_003986129.4
B2 42356432 42356487 refGene_gn=NCR2;refGene_tx=XM_003986182.2
B2 42356633 42356975 refGene_gn=NCR2;refGene_tx=XM_003986182.2
B2 42363024 42363154 refGene_gn=NCR2;refGene_tx=XM_003986182.2
B2 42363261 42363374 refGene_gn=NCR2;refGene_tx=XM_003986182.2
B2 42363961 42364092 refGene_gn=NCR2;refGene_tx=XM_003986182.2
B2 42566128 42566341
refGene_gn=FOXP4;refGene_tx=XM_006931700.3;refGene_gn=FOXP4;refGene_tx=XM_006931702.3;refGene_gn=FOXP4;refGene_tx=XM_006931703.3;refGene_gn=FOXP4;refGene_tx=XM_006931705.4;refGene_gn=FOXP4;refGene_tx=XM_006931706.4;refGene_gn=FOXP4;refGene_tx=XM_023253953.1;refGene_gn=FOXP4;refGene_tx=XM_023253954.1;refGene_gn=FOXP4;refGene_tx=XM_023253955.1
B2 42578781 42578871
refGene_gn=FOXP4;refGene_tx=XM_006931700.3;refGene_gn=FOXP4;refGene_tx=XM_006931702.3;refGene_gn=FOXP4;refGene_tx=XM_006931703.3;refGene_gn=FOXP4;refGene_tx=XM_006931705.4;refGene_gn=FOXP4;refGene_tx=XM_006931706.4;refGene_gn=FOXP4;refGene_tx=XM_023253953.1;refGene_gn=FOXP4;refGene_tx=XM_023253954.1;refGene_gn=FOXP4;refGene_tx=XM_023253955.1
B2 42584737 42584860
refGene_gn=FOXP4;refGene_tx=XM_006931700.3;refGene_gn=FOXP4;refGene_tx=XM_006931702.3;refGene_gn=FOXP4;refGene_tx=XM_006931703.3;refGene_gn=FOXP4;refGene_tx=XM_006931705.4;refGene_gn=FOXP4;refGene_tx=XM_006931706.4;refGene_gn=FOXP4;refGene_tx=XM_023253953.1;refGene_gn=FOXP4;refGene_tx=XM_023253954.1;refGene_gn=FOXP4;refGene_tx=XM_023253955.1
B2 42585474 42585561
refGene_gn=FOXP4;refGene_tx=XM_006931700.3;refGene_gn=FOXP4;refGene_tx=XM_006931702.3;refGene_gn=FOXP4;refGene_tx=XM_006931703.3;refGene_gn=FOXP4;refGene_tx=XM_006931705.4;refGene_gn=FOXP4;refGene_tx=XM_006931706.4;refGene_gn=FOXP4;refGene_tx=XM_023253953.1;refGene_gn=FOXP4;refGene_tx=XM_023253954.1;refGene_gn=FOXP4;refGene_tx=XM_023253955.1
B2 42586861 42587009
refGene_gn=FOXP4;refGene_tx=XM_006931700.3;refGene_gn=FOXP4;refGene_tx=XM_006931702.3;refGene_gn=FOXP4;refGene_tx=XM_006931703.3;refGene_gn=FOXP4;refGene_tx=XM_006931705.4;refGene_gn=FOXP4;refGene_tx=XM_006931706.4;refGene_gn=FOXP4;refGene_tx=XM_023253953.1;refGene_gn=FOXP4;refGene_tx=XM_023253954.1;refGene_gn=FOXP4;refGene_tx=XM_023253955.1
B2 42587130 42587344
refGene_gn=FOXP4;refGene_tx=XM_006931700.3;refGene_gn=FOXP4;refGene_tx=XM_006931702.3;refGene_gn=FOXP4;refGene_tx=XM_006931703.3;refGene_gn=FOXP4;refGene_tx=XM_006931705.4;refGene_gn=FOXP4;refGene_tx=XM_006931706.4;refGene_gn=FOXP4;refGene_tx=XM_023253953.1;refGene_gn=FOXP4;refGene_tx=XM_023253954.1;refGene_gn=FOXP4;refGene_tx=XM_023253955.1
B2 42587591 42587696
refGene_gn=FOXP4;refGene_tx=XM_006931700.3;refGene_gn=FOXP4;refGene_tx=XM_006931702.3;refGene_gn=FOXP4;refGene_tx=XM_006931703.3;refGene_gn=FOXP4;refGene_tx=XM_006931705.4;refGene_gn=FOXP4;refGene_tx=XM_006931706.4;refGene_gn=FOXP4;refGene_tx=XM_023253953.1;refGene_gn=FOXP4;refGene_tx=XM_023253954.1;refGene_gn=FOXP4;refGene_tx=XM_023253955.1
B2 42588497 42588585

refGene_gn=FOXP4;refGene_tx=XM_006931700.3;refGene_gn=FOXP4;refGene_tx
=XM_006931702.3;refGene_gn=FOXP4;refGene_tx=XM_006931703.3;refGene_gn=
FOXP4;refGene_tx=XM_006931705.4;refGene_gn=FOXP4;refGene_tx=XM_0069317
06.4;refGene_gn=FOXP4;refGene_tx=XM_023253953.1;refGene_gn=FOXP4;refGe
ne_tx=XM_023253954.1;refGene_gn=FOXP4;refGene_tx=XM_023253955.1
B2 42589582 42589666
refGene_gn=FOXP4;refGene_tx=XM_006931700.3;refGene_gn=FOXP4;refGene_tx
=XM_006931702.3;refGene_gn=FOXP4;refGene_tx=XM_006931703.3;refGene_gn=
FOXP4;refGene_tx=XM_006931705.4;refGene_gn=FOXP4;refGene_tx=XM_0069317
06.4;refGene_gn=FOXP4;refGene_tx=XM_023253953.1;refGene_gn=FOXP4;refGe
ne_tx=XM_023253954.1;refGene_gn=FOXP4;refGene_tx=XM_023253955.1
B2 42589748 42589956
refGene_gn=FOXP4;refGene_tx=XM_006931700.3;refGene_gn=FOXP4;refGene_tx
=XM_006931702.3;refGene_gn=FOXP4;refGene_tx=XM_006931703.3;refGene_gn=
FOXP4;refGene_tx=XM_006931705.4;refGene_gn=FOXP4;refGene_tx=XM_0069317
06.4;refGene_gn=FOXP4;refGene_tx=XM_023253953.1;refGene_gn=FOXP4;refGe
ne_tx=XM_023253954.1;refGene_gn=FOXP4;refGene_tx=XM_023253955.1
B2 42590038 42590115
refGene_gn=FOXP4;refGene_tx=XM_006931700.3;refGene_gn=FOXP4;refGene_tx
=XM_006931702.3;refGene_gn=FOXP4;refGene_tx=XM_006931703.3;refGene_gn=
FOXP4;refGene_tx=XM_006931705.4;refGene_gn=FOXP4;refGene_tx=XM_0069317
06.4;refGene_gn=FOXP4;refGene_tx=XM_023253953.1;refGene_gn=FOXP4;refGe
ne_tx=XM_023253954.1;refGene_gn=FOXP4;refGene_tx=XM_023253955.1
B2 42590995 42591097
refGene_gn=FOXP4;refGene_tx=XM_006931700.3;refGene_gn=FOXP4;refGene_tx
=XM_006931702.3;refGene_gn=FOXP4;refGene_tx=XM_006931703.3;refGene_gn=
FOXP4;refGene_tx=XM_006931705.4;refGene_gn=FOXP4;refGene_tx=XM_0069317
06.4;refGene_gn=FOXP4;refGene_tx=XM_023253953.1;refGene_gn=FOXP4;refGe
ne_tx=XM_023253954.1;refGene_gn=FOXP4;refGene_tx=XM_023253955.1
B2 42593912 42593963
refGene_gn=FOXP4;refGene_tx=XM_006931700.3;refGene_gn=FOXP4;refGene_tx
=XM_006931702.3;refGene_gn=FOXP4;refGene_tx=XM_023253953.1;refGene_gn=
FOXP4;refGene_tx=XM_023253954.1
B2 42594600 42594722
refGene_gn=FOXP4;refGene_tx=XM_006931700.3;refGene_gn=FOXP4;refGene_tx
=XM_006931702.3;refGene_gn=FOXP4;refGene_tx=XM_006931703.3;refGene_gn=
FOXP4;refGene_tx=XM_006931705.4;refGene_gn=FOXP4;refGene_tx=XM_0069317
06.4;refGene_gn=FOXP4;refGene_tx=XM_023253953.1;refGene_gn=FOXP4;refGe
ne_tx=XM_023253954.1;refGene_gn=FOXP4;refGene_tx=XM_023253955.1
B2 42596228 42596295
refGene_gn=FOXP4;refGene_tx=XM_006931700.3;refGene_gn=FOXP4;refGene_tx
=XM_006931702.3;refGene_gn=FOXP4;refGene_tx=XM_006931703.3;refGene_gn=
FOXP4;refGene_tx=XM_006931705.4;refGene_gn=FOXP4;refGene_tx=XM_0069317
06.4;refGene_gn=FOXP4;refGene_tx=XM_023253953.1;refGene_gn=FOXP4;refGe
ne_tx=XM_023253954.1;refGene_gn=FOXP4;refGene_tx=XM_023253955.1
B2 42596784 42596954
refGene_gn=FOXP4;refGene_tx=XM_006931700.3;refGene_gn=FOXP4;refGene_tx
=XM_006931702.3;refGene_gn=FOXP4;refGene_tx=XM_006931703.3;refGene_gn=
FOXP4;refGene_tx=XM_006931705.4;refGene_gn=FOXP4;refGene_tx=XM_0069317
06.4;refGene_gn=FOXP4;refGene_tx=XM_023253953.1;refGene_gn=FOXP4;refGe

ne_tx=XM_023253954.1;refGene_gn=FOXP4;refGene_tx=XM_023253955.1
B2 42597428 42597576
refGene_gn=FOXP4;refGene_tx=XM_006931700.3;refGene_gn=FOXP4;refGene_tx=XM_006931702.3;refGene_gn=FOXP4;refGene_tx=XM_006931703.3;refGene_gn=FOXP4;refGene_tx=XM_006931705.4;refGene_gn=FOXP4;refGene_tx=XM_006931706.4;refGene_gn=FOXP4;refGene_tx=XM_023253953.1;refGene_gn=FOXP4;refGene_tx=XM_023253954.1;refGene_gn=FOXP4;refGene_tx=XM_023253955.1
B2 42635478 42635554
refGene_gn=MDFI;refGene_tx=XM_023253956.1;refGene_gn=MDFI;refGene_tx=XM_023253957.1;refGene_gn=MDFI;refGene_tx=XM_023253958.1
B2 42643097 42643280
refGene_gn=MDFI;refGene_tx=XM_023253956.1;refGene_gn=MDFI;refGene_tx=XM_023253957.1;refGene_gn=MDFI;refGene_tx=XM_023253958.1
B2 42646202 42646427
refGene_gn=MDFI;refGene_tx=XM_023253956.1;refGene_gn=MDFI;refGene_tx=XM_023253957.1;refGene_gn=MDFI;refGene_tx=XM_023253958.1
B2 42648812 42649069
refGene_gn=MDFI;refGene_tx=XM_023253956.1;refGene_gn=MDFI;refGene_tx=XM_023253957.1;refGene_gn=MDFI;refGene_tx=XM_023253958.1
B2 42683459 42683948 refGene_gn=TFEB;refGene_tx=XM_006931710.4
B2 42685108 42685256 refGene_gn=TFEB;refGene_tx=XM_006931710.4
B2 42685837 42685913 refGene_gn=TFEB;refGene_tx=XM_006931710.4
B2 42686530 42686587 refGene_gn=TFEB;refGene_tx=XM_006931710.4
B2 42686681 42686802 refGene_gn=TFEB;refGene_tx=XM_006931710.4
B2 42688668 42688746 refGene_gn=TFEB;refGene_tx=XM_006931710.4
B2 42689369 42689483 refGene_gn=TFEB;refGene_tx=XM_006931710.4
B2 42689484 42689625 refGene_gn=TFEB;refGene_tx=XM_006931710.4
B2 42689710 42689986 refGene_gn=TFEB;refGene_tx=XM_006931710.4
B2 42739302 42739455 refGene_gn=PGC;refGene_tx=XM_003986131.5
B2 42740147 42740246 refGene_gn=PGC;refGene_tx=XM_003986131.5
B2 42740656 42740804 refGene_gn=PGC;refGene_tx=XM_003986131.5
B2 42741439 42741562 refGene_gn=PGC;refGene_tx=XM_003986131.5
B2 42743175 42743375 refGene_gn=PGC;refGene_tx=XM_003986131.5
B2 42743852 42743971 refGene_gn=PGC;refGene_tx=XM_003986131.5
B2 42745086 42745204 refGene_gn=PGC;refGene_tx=XM_003986131.5
B2 42745490 42745641 refGene_gn=PGC;refGene_tx=XM_003986131.5
B2 42747563 42747622 refGene_gn=PGC;refGene_tx=XM_003986131.5
B2 42768841 42769756 refGene_gn=FRS3;refGene_tx=XM_003986132.5
B2 42770032 42770190 refGene_gn=FRS3;refGene_tx=XM_003986132.5
B2 42771041 42771203 refGene_gn=FRS3;refGene_tx=XM_003986132.5
B2 42773642 42773829 refGene_gn=FRS3;refGene_tx=XM_003986132.5
B2 42774908 42774974 refGene_gn=FRS3;refGene_tx=XM_003986132.5
B2 42781220 42781352
refGene_gn=PRICKLE4;refGene_tx=XM_023253964.1;refGene_gn=PRICKLE4;refGene_tx=XM_023253965.1;refGene_gn=PRICKLE4;refGene_tx=XM_023253966.1;refGene_gn=PRICKLE4;refGene_tx=XM_023253967.1
B2 42781879 42781987
refGene_gn=PRICKLE4;refGene_tx=XM_023253964.1;refGene_gn=PRICKLE4;refGene_tx=XM_023253965.1;refGene_gn=PRICKLE4;refGene_tx=XM_023253966.1;refGene_gn=PRICKLE4;refGene_tx=XM_023253967.1

B2 42782716 42782854
refGene_gn=PRICKLE4;refGene_tx=XM_023253964.1;refGene_gn=PRICKLE4;refGene_tx=XM_023253965.1;refGene_gn=PRICKLE4;refGene_tx=XM_023253966.1;refGene_gn=PRICKLE4;refGene_tx=XM_023253967.1
B2 42783127 42783331
refGene_gn=PRICKLE4;refGene_tx=XM_023253964.1;refGene_gn=PRICKLE4;refGene_tx=XM_023253965.1;refGene_gn=PRICKLE4;refGene_tx=XM_023253966.1;refGene_gn=PRICKLE4;refGene_tx=XM_023253967.1
B2 42783938 42784128
refGene_gn=PRICKLE4;refGene_tx=XM_023253964.1;refGene_gn=PRICKLE4;refGene_tx=XM_023253965.1;refGene_gn=PRICKLE4;refGene_tx=XM_023253966.1;refGene_gn=PRICKLE4;refGene_tx=XM_023253967.1
B2 42784634 42784993
refGene_gn=PRICKLE4;refGene_tx=XM_023253964.1;refGene_gn=PRICKLE4;refGene_tx=XM_023253965.1;refGene_gn=PRICKLE4;refGene_tx=XM_023253966.1;refGene_gn=PRICKLE4;refGene_tx=XM_023253967.1
B2 42785581 42785709 refGene_gn=TOMM6;refGene_tx=XM_003986133.5
B2 42786222 42786319 refGene_gn=TOMM6;refGene_tx=XM_003986133.5
B2 42793262 42793453
refGene_gn=USP49;refGene_tx=XM_023253959.1;refGene_gn=USP49;refGene_tx=XM_023253960.1;refGene_gn=USP49;refGene_tx=XM_023253961.1;refGene_gn=USP49;refGene_tx=XM_023253962.1;refGene_gn=USP49;refGene_tx=XM_023253963.1
B2 42794646 42794852
refGene_gn=USP49;refGene_tx=XM_023253959.1;refGene_gn=USP49;refGene_tx=XM_023253960.1;refGene_gn=USP49;refGene_tx=XM_023253961.1;refGene_gn=USP49;refGene_tx=XM_023253962.1;refGene_gn=USP49;refGene_tx=XM_023253963.1
B2 42795347 42795456
refGene_gn=USP49;refGene_tx=XM_023253959.1;refGene_gn=USP49;refGene_tx=XM_023253960.1;refGene_gn=USP49;refGene_tx=XM_023253961.1;refGene_gn=USP49;refGene_tx=XM_023253962.1;refGene_gn=USP49;refGene_tx=XM_023253963.1
B2 42799854 42800059
refGene_gn=USP49;refGene_tx=XM_023253959.1;refGene_gn=USP49;refGene_tx=XM_023253960.1;refGene_gn=USP49;refGene_tx=XM_023253961.1;refGene_gn=USP49;refGene_tx=XM_023253962.1;refGene_gn=USP49;refGene_tx=XM_023253963.1
B2 42801380 42802718
refGene_gn=USP49;refGene_tx=XM_023253959.1;refGene_gn=USP49;refGene_tx=XM_023253960.1;refGene_gn=USP49;refGene_tx=XM_023253961.1;refGene_gn=USP49;refGene_tx=XM_023253962.1;refGene_gn=USP49;refGene_tx=XM_023253963.1
B2 42879376 42879592
refGene_gn=MED20;refGene_tx=XM_003986134.4;refGene_gn=MED20;refGene_tx=XM_019830588.2
B2 42881430 42881684
refGene_gn=MED20;refGene_tx=XM_003986134.4;refGene_gn=MED20;refGene_tx=XM_019830588.2
B2 42885785 42885940

refGene_gn=MED20;refGene_tx=XM_003986134.4;refGene_gn=MED20;refGene_tx=XM_019830588.2
B2 42888544 42888558 refGene_gn=MED20;refGene_tx=XM_003986134.4
B2 42889100 42889365 refGene_gn=BYSL;refGene_tx=XM_003986135.5
B2 42892812 42892975 refGene_gn=BYSL;refGene_tx=XM_003986135.5
B2 42895047 42895186 refGene_gn=BYSL;refGene_tx=XM_003986135.5
B2 42895534 42895668 refGene_gn=BYSL;refGene_tx=XM_003986135.5
B2 42896238 42896399 refGene_gn=BYSL;refGene_tx=XM_003986135.5
B2 42896566 42896669 refGene_gn=BYSL;refGene_tx=XM_003986135.5
B2 42897190 42897536 refGene_gn=BYSL;refGene_tx=XM_003986135.5
B2 42900576 42900744
refGene_gn=CCND3;refGene_tx=XM_011282227.3;refGene_gn=CCND3;refGene_tx=XM_023253968.1;refGene_gn=CCND3;refGene_tx=XM_023253969.1;refGene_gn=CCND3;refGene_tx=XM_023253970.1;refGene_gn=CCND3;refGene_tx=XM_023253971.1
B2 42901243 42901380
refGene_gn=CCND3;refGene_tx=XM_011282227.3;refGene_gn=CCND3;refGene_tx=XM_023253968.1;refGene_gn=CCND3;refGene_tx=XM_023253969.1;refGene_gn=CCND3;refGene_tx=XM_023253970.1;refGene_gn=CCND3;refGene_tx=XM_023253971.1
B2 42902130 42902290
refGene_gn=CCND3;refGene_tx=XM_011282227.3;refGene_gn=CCND3;refGene_tx=XM_023253968.1;refGene_gn=CCND3;refGene_tx=XM_023253969.1;refGene_gn=CCND3;refGene_tx=XM_023253970.1
B2 42905318 42905534
refGene_gn=CCND3;refGene_tx=XM_011282227.3;refGene_gn=CCND3;refGene_tx=XM_023253968.1;refGene_gn=CCND3;refGene_tx=XM_023253969.1
B2 42906378 42906576 refGene_gn=CCND3;refGene_tx=XM_011282227.3
B2 42997564 42997570
refGene_gn=CCND3;refGene_tx=XM_023253968.1;refGene_gn=CCND3;refGene_tx=XM_023253970.1
B2 42998983 42999028
refGene_gn=TAF8;refGene_tx=XM_003986137.4;refGene_gn=TAF8;refGene_tx=XM_006931715.3;refGene_gn=TAF8;refGene_tx=XM_019830589.2;refGene_gn=TAF8;refGene_tx=XM_019830590.2;refGene_gn=TAF8;refGene_tx=XM_019830592.2
B2 42999818 42999975
refGene_gn=TAF8;refGene_tx=XM_003986137.4;refGene_gn=TAF8;refGene_tx=XM_006931715.3;refGene_gn=TAF8;refGene_tx=XM_019830589.2;refGene_gn=TAF8;refGene_tx=XM_019830590.2;refGene_gn=TAF8;refGene_tx=XM_019830592.2
B2 43003968 43004067
refGene_gn=TAF8;refGene_tx=XM_003986137.4;refGene_gn=TAF8;refGene_tx=XM_006931715.3;refGene_gn=TAF8;refGene_tx=XM_019830589.2;refGene_gn=TAF8;refGene_tx=XM_019830590.2;refGene_gn=TAF8;refGene_tx=XM_019830592.2
B2 43004411 43004474
refGene_gn=TAF8;refGene_tx=XM_003986137.4;refGene_gn=TAF8;refGene_tx=XM_006931715.3;refGene_gn=TAF8;refGene_tx=XM_019830589.2;refGene_gn=TAF8;refGene_tx=XM_019830590.2;refGene_gn=TAF8;refGene_tx=XM_019830592.2
B2 43005513 43005638
refGene_gn=TAF8;refGene_tx=XM_003986137.4;refGene_gn=TAF8;refGene_tx=XM_006931715.3;refGene_gn=TAF8;refGene_tx=XM_019830589.2;refGene_gn=TAF

8;refGene_tx=XM_019830590.2;refGene_gn=TAF8;refGene_tx=XM_019830592.2
B2 43010364 43010512
refGene_gn=TAF8;refGene_tx=XM_003986137.4;refGene_gn=TAF8;refGene_tx=XM_006931715.3;refGene_gn=TAF8;refGene_tx=XM_019830589.2;refGene_gn=TAF8;refGene_tx=XM_019830590.2;refGene_gn=TAF8;refGene_tx=XM_019830592.2
B2 43011915 43012058
refGene_gn=TAF8;refGene_tx=XM_003986137.4;refGene_gn=TAF8;refGene_tx=XM_006931715.3;refGene_gn=TAF8;refGene_tx=XM_019830589.2;refGene_gn=TAF8;refGene_tx=XM_019830590.2;refGene_gn=TAF8;refGene_tx=XM_019830592.2
B2 43013651 43013744
refGene_gn=TAF8;refGene_tx=XM_019830589.2;refGene_gn=TAF8;refGene_tx=XM_019830590.2
B2 43014283 43014289 refGene_gn=TAF8;refGene_tx=XM_019830592.2
B2 43018446 43018590
refGene_gn=TAF8;refGene_tx=XM_003986137.4;refGene_gn=TAF8;refGene_tx=XM_006931715.3
B2 43018891 43018904 refGene_gn=TAF8;refGene_tx=XM_006931715.3
B2 43043975 43044093
refGene_gn=CB2H6orf132;refGene_tx=XM_023253972.1;refGene_gn=CB2H6orf132;refGene_tx=XM_023253973.1
B2 43044748 43048202
refGene_gn=CB2H6orf132;refGene_tx=XM_023253972.1;refGene_gn=CB2H6orf132;refGene_tx=XM_023253973.1
B2 43050642 43050718
refGene_gn=CB2H6orf132;refGene_tx=XM_023253972.1;refGene_gn=CB2H6orf132;refGene_tx=XM_023253973.1
B2 43066045 43066152
refGene_gn=CB2H6orf132;refGene_tx=XM_023253972.1;refGene_gn=CB2H6orf132;refGene_tx=XM_023253973.1
B2 43076435 43076580
refGene_gn=CB2H6orf132;refGene_tx=XM_023253972.1;refGene_gn=CB2H6orf132;refGene_tx=XM_023253973.1
B2 43088442 43088451
refGene_gn=LOC105260566;refGene_tx=XM_011282225.3
B2 43093268 43093277
refGene_gn=LOC105260566;refGene_tx=XM_011282224.3
B2 43093990 43094320
refGene_gn=LOC105260566;refGene_tx=XM_011282224.3;refGene_gn=LOC105260566;refGene_tx=XM_011282225.3
B2 43102508 43102709 refGene_gn=GUCA1A;refGene_tx=XM_003986138.4
B2 43107907 43108057 refGene_gn=GUCA1A;refGene_tx=XM_003986138.4
B2 43108428 43108522 refGene_gn=GUCA1A;refGene_tx=XM_003986138.4
B2 43108799 43108969 refGene_gn=GUCA1A;refGene_tx=XM_003986138.4
B2 43111512 43111640 refGene_gn=GUCA1B;refGene_tx=XM_003986139.3
B2 43112386 43112507 refGene_gn=GUCA1B;refGene_tx=XM_003986139.3
B2 43114290 43114440 refGene_gn=GUCA1B;refGene_tx=XM_003986139.3
B2 43121131 43121338 refGene_gn=GUCA1B;refGene_tx=XM_003986139.3
B2 43130681 43130765
refGene_gn=MRPS10;refGene_tx=XM_003986140.4;refGene_gn=MRPS10;refGene_tx=XM_006931716.3;refGene_gn=MRPS10;refGene_tx=XM_023253974.1;refGene_

gn=MRPS10;refGene_tx=XM_023253975.1
B2	43131237 43131327
refGene_gn=MRPS10;refGene_tx=XM_003986140.4;refGene_gn=MRPS10;refGene_tx=XM_006931716.3;refGene_gn=MRPS10;refGene_tx=XM_023253974.1;refGene_gn=MRPS10;refGene_tx=XM_023253975.1
B2	43133040 43133149
refGene_gn=MRPS10;refGene_tx=XM_003986140.4;refGene_gn=MRPS10;refGene_tx=XM_006931716.3;refGene_gn=MRPS10;refGene_tx=XM_023253974.1;refGene_gn=MRPS10;refGene_tx=XM_023253975.1
B2	43134991 43135128
refGene_gn=MRPS10;refGene_tx=XM_003986140.4;refGene_gn=MRPS10;refGene_tx=XM_006931716.3;refGene_gn=MRPS10;refGene_tx=XM_023253974.1;refGene_gn=MRPS10;refGene_tx=XM_023253975.1
B2	43137118 43137191
refGene_gn=MRPS10;refGene_tx=XM_003986140.4;refGene_gn=MRPS10;refGene_tx=XM_006931716.3;refGene_gn=MRPS10;refGene_tx=XM_023253974.1;refGene_gn=MRPS10;refGene_tx=XM_023253975.1
B2	43137282 43137347
refGene_gn=MRPS10;refGene_tx=XM_003986140.4;refGene_gn=MRPS10;refGene_tx=XM_006931716.3;refGene_gn=MRPS10;refGene_tx=XM_023253974.1;refGene_gn=MRPS10;refGene_tx=XM_023253975.1
B2	43148835 43148995
refGene_gn=MRPS10;refGene_tx=XM_003986140.4;refGene_gn=MRPS10;refGene_tx=XM_006931716.3;refGene_gn=MRPS10;refGene_tx=XM_023253974.1;refGene_gn=MRPS10;refGene_tx=XM_023253975.1
B2	43156856 43157181 refGene_gn=TRERF1;refGene_tx=XM_011282223.3
B2	43160244 43160456 refGene_gn=TRERF1;refGene_tx=XM_011282223.3
B2	43163028 43163232 refGene_gn=TRERF1;refGene_tx=XM_011282223.3
B2	43169543 43169654 refGene_gn=TRERF1;refGene_tx=XM_011282223.3
B2	43171550 43171639 refGene_gn=TRERF1;refGene_tx=XM_011282223.3
B2	43179718 43179794 refGene_gn=TRERF1;refGene_tx=XM_011282223.3
B2	43181315 43181419 refGene_gn=TRERF1;refGene_tx=XM_011282223.3
B2	43181551 43181691 refGene_gn=TRERF1;refGene_tx=XM_011282223.3
B2	43183649 43183716 refGene_gn=TRERF1;refGene_tx=XM_011282223.3
B2	43184407 43184657 refGene_gn=TRERF1;refGene_tx=XM_011282223.3
B2	43184658 43184796 refGene_gn=TRERF1;refGene_tx=XM_011282223.3
B2	43188330 43188570 refGene_gn=TRERF1;refGene_tx=XM_011282223.3
B2	43189626 43189777 refGene_gn=TRERF1;refGene_tx=XM_011282223.3
B2	43190634 43190681 refGene_gn=TRERF1;refGene_tx=XM_011282223.3
B2	43192133 43192768 refGene_gn=TRERF1;refGene_tx=XM_011282223.3
B2	43192770 43193575 refGene_gn=TRERF1;refGene_tx=XM_011282223.3
B2	43479718 43479796
refGene_gn=UBR2;refGene_tx=XM_011282222.3;refGene_gn=UBR2;refGene_tx=XM_023253976.1;refGene_gn=UBR2;refGene_tx=XM_023253977.1;refGene_gn=UBR2;refGene_tx=XM_023253978.1;refGene_gn=UBR2;refGene_tx=XM_023253979.1;refGene_gn=UBR2;refGene_tx=XM_023253980.1;refGene_gn=UBR2;refGene_tx=XM_023253981.1
B2	43488618 43488878
refGene_gn=UBR2;refGene_tx=XM_011282222.3;refGene_gn=UBR2;refGene_tx=XM_023253976.1;refGene_gn=UBR2;refGene_tx=XM_023253977.1;refGene_gn=UBR

2;refGene_tx=XM_023253978.1;refGene_gn=UBR2;refGene_tx=XM_023253979.1;
refGene_gn=UBR2;refGene_tx=XM_023253980.1;refGene_gn=UBR2;refGene_tx=X
M_023253981.1
B2 43517900 43517979
refGene_gn=UBR2;refGene_tx=XM_011282222.3;refGene_gn=UBR2;refGene_tx=X
M_023253976.1;refGene_gn=UBR2;refGene_tx=XM_023253977.1;refGene_gn=UBR
2;refGene_tx=XM_023253978.1;refGene_gn=UBR2;refGene_tx=XM_023253979.1;
refGene_gn=UBR2;refGene_tx=XM_023253980.1;refGene_gn=UBR2;refGene_tx=X
M_023253981.1
B2 43519853 43519967
refGene_gn=UBR2;refGene_tx=XM_011282222.3;refGene_gn=UBR2;refGene_tx=X
M_023253976.1;refGene_gn=UBR2;refGene_tx=XM_023253977.1;refGene_gn=UBR
2;refGene_tx=XM_023253978.1;refGene_gn=UBR2;refGene_tx=XM_023253979.1;
refGene_gn=UBR2;refGene_tx=XM_023253980.1;refGene_gn=UBR2;refGene_tx=X
M_023253981.1
B2 43529055 43529186
refGene_gn=UBR2;refGene_tx=XM_011282222.3;refGene_gn=UBR2;refGene_tx=X
M_023253976.1;refGene_gn=UBR2;refGene_tx=XM_023253977.1;refGene_gn=UBR
2;refGene_tx=XM_023253978.1;refGene_gn=UBR2;refGene_tx=XM_023253979.1;
refGene_gn=UBR2;refGene_tx=XM_023253980.1;refGene_gn=UBR2;refGene_tx=X
M_023253981.1
B2 43532015 43532154
refGene_gn=UBR2;refGene_tx=XM_011282222.3;refGene_gn=UBR2;refGene_tx=X
M_023253976.1;refGene_gn=UBR2;refGene_tx=XM_023253977.1;refGene_gn=UBR
2;refGene_tx=XM_023253978.1;refGene_gn=UBR2;refGene_tx=XM_023253979.1;
refGene_gn=UBR2;refGene_tx=XM_023253980.1;refGene_gn=UBR2;refGene_tx=X
M_023253981.1
B2 43532657 43532720
refGene_gn=UBR2;refGene_tx=XM_011282222.3;refGene_gn=UBR2;refGene_tx=X
M_023253976.1;refGene_gn=UBR2;refGene_tx=XM_023253977.1;refGene_gn=UBR
2;refGene_tx=XM_023253978.1;refGene_gn=UBR2;refGene_tx=XM_023253979.1;
refGene_gn=UBR2;refGene_tx=XM_023253980.1;refGene_gn=UBR2;refGene_tx=X
M_023253981.1
B2 43536598 43536719
refGene_gn=UBR2;refGene_tx=XM_011282222.3;refGene_gn=UBR2;refGene_tx=X
M_023253976.1;refGene_gn=UBR2;refGene_tx=XM_023253977.1;refGene_gn=UBR
2;refGene_tx=XM_023253978.1;refGene_gn=UBR2;refGene_tx=XM_023253979.1;
refGene_gn=UBR2;refGene_tx=XM_023253980.1;refGene_gn=UBR2;refGene_tx=X
M_023253981.1
B2 43538452 43538560
refGene_gn=UBR2;refGene_tx=XM_011282222.3;refGene_gn=UBR2;refGene_tx=X
M_023253976.1;refGene_gn=UBR2;refGene_tx=XM_023253977.1;refGene_gn=UBR
2;refGene_tx=XM_023253978.1;refGene_gn=UBR2;refGene_tx=XM_023253979.1;
refGene_gn=UBR2;refGene_tx=XM_023253980.1;refGene_gn=UBR2;refGene_tx=X
M_023253981.1
B2 43539147 43539236
refGene_gn=UBR2;refGene_tx=XM_011282222.3;refGene_gn=UBR2;refGene_tx=X
M_023253976.1;refGene_gn=UBR2;refGene_tx=XM_023253977.1;refGene_gn=UBR
2;refGene_tx=XM_023253978.1;refGene_gn=UBR2;refGene_tx=XM_023253979.1;
refGene_gn=UBR2;refGene_tx=XM_023253980.1;refGene_gn=UBR2;refGene_tx=X

M_023253981.1
B2 43540309 43540408 refGene_gn=UBR2;refGene_tx=XM_011282222.3
B2 43540478 43540577
refGene_gn=UBR2;refGene_tx=XM_023253976.1;refGene_gn=UBR2;refGene_tx=XM_023253977.1;refGene_gn=UBR2;refGene_tx=XM_023253978.1;refGene_gn=UBR2;refGene_tx=XM_023253979.1;refGene_gn=UBR2;refGene_tx=XM_023253980.1;refGene_gn=UBR2;refGene_tx=XM_023253981.1
B2 43551548 43551712
refGene_gn=UBR2;refGene_tx=XM_011282222.3;refGene_gn=UBR2;refGene_tx=XM_023253976.1;refGene_gn=UBR2;refGene_tx=XM_023253977.1;refGene_gn=UBR2;refGene_tx=XM_023253978.1;refGene_gn=UBR2;refGene_tx=XM_023253979.1;refGene_gn=UBR2;refGene_tx=XM_023253980.1;refGene_gn=UBR2;refGene_tx=XM_023253981.1
B2 43551801 43551901
refGene_gn=UBR2;refGene_tx=XM_011282222.3;refGene_gn=UBR2;refGene_tx=XM_023253976.1;refGene_gn=UBR2;refGene_tx=XM_023253977.1;refGene_gn=UBR2;refGene_tx=XM_023253978.1;refGene_gn=UBR2;refGene_tx=XM_023253979.1;refGene_gn=UBR2;refGene_tx=XM_023253980.1;refGene_gn=UBR2;refGene_tx=XM_023253981.1
B2 43554423 43554552
refGene_gn=UBR2;refGene_tx=XM_011282222.3;refGene_gn=UBR2;refGene_tx=XM_023253976.1;refGene_gn=UBR2;refGene_tx=XM_023253977.1;refGene_gn=UBR2;refGene_tx=XM_023253978.1;refGene_gn=UBR2;refGene_tx=XM_023253979.1;refGene_gn=UBR2;refGene_tx=XM_023253980.1;refGene_gn=UBR2;refGene_tx=XM_023253981.1
B2 43555629 43555813
refGene_gn=UBR2;refGene_tx=XM_011282222.3;refGene_gn=UBR2;refGene_tx=XM_023253976.1;refGene_gn=UBR2;refGene_tx=XM_023253977.1;refGene_gn=UBR2;refGene_tx=XM_023253978.1;refGene_gn=UBR2;refGene_tx=XM_023253979.1;refGene_gn=UBR2;refGene_tx=XM_023253980.1;refGene_gn=UBR2;refGene_tx=XM_023253981.1
B2 43558730 43558792
refGene_gn=UBR2;refGene_tx=XM_011282222.3;refGene_gn=UBR2;refGene_tx=XM_023253976.1;refGene_gn=UBR2;refGene_tx=XM_023253977.1;refGene_gn=UBR2;refGene_tx=XM_023253978.1;refGene_gn=UBR2;refGene_tx=XM_023253979.1;refGene_gn=UBR2;refGene_tx=XM_023253980.1;refGene_gn=UBR2;refGene_tx=XM_023253981.1
B2 43559451 43559562
refGene_gn=UBR2;refGene_tx=XM_011282222.3;refGene_gn=UBR2;refGene_tx=XM_023253976.1;refGene_gn=UBR2;refGene_tx=XM_023253977.1;refGene_gn=UBR2;refGene_tx=XM_023253978.1;refGene_gn=UBR2;refGene_tx=XM_023253979.1;refGene_gn=UBR2;refGene_tx=XM_023253980.1;refGene_gn=UBR2;refGene_tx=XM_023253981.1
B2 43560501 43560567
refGene_gn=UBR2;refGene_tx=XM_011282222.3;refGene_gn=UBR2;refGene_tx=XM_023253976.1;refGene_gn=UBR2;refGene_tx=XM_023253977.1;refGene_gn=UBR2;refGene_tx=XM_023253978.1;refGene_gn=UBR2;refGene_tx=XM_023253979.1;refGene_gn=UBR2;refGene_tx=XM_023253980.1;refGene_gn=UBR2;refGene_tx=XM_023253981.1
B2 43561952 43562075

refGene_gn=UBR2;refGene_tx=XM_011282222.3;refGene_gn=UBR2;refGene_tx=XM_023253976.1;refGene_gn=UBR2;refGene_tx=XM_023253977.1;refGene_gn=UBR2;refGene_tx=XM_023253978.1;refGene_gn=UBR2;refGene_tx=XM_023253979.1;refGene_gn=UBR2;refGene_tx=XM_023253980.1;refGene_gn=UBR2;refGene_tx=XM_023253981.1
B2 43562215 43562279
refGene_gn=UBR2;refGene_tx=XM_011282222.3;refGene_gn=UBR2;refGene_tx=XM_023253976.1;refGene_gn=UBR2;refGene_tx=XM_023253977.1;refGene_gn=UBR2;refGene_tx=XM_023253978.1;refGene_gn=UBR2;refGene_tx=XM_023253979.1;refGene_gn=UBR2;refGene_tx=XM_023253980.1
B2 43563252 43563377
refGene_gn=UBR2;refGene_tx=XM_011282222.3;refGene_gn=UBR2;refGene_tx=XM_023253976.1;refGene_gn=UBR2;refGene_tx=XM_023253977.1;refGene_gn=UBR2;refGene_tx=XM_023253978.1;refGene_gn=UBR2;refGene_tx=XM_023253979.1;refGene_gn=UBR2;refGene_tx=XM_023253980.1
B2 43566391 43566444
refGene_gn=UBR2;refGene_tx=XM_011282222.3;refGene_gn=UBR2;refGene_tx=XM_023253976.1;refGene_gn=UBR2;refGene_tx=XM_023253977.1;refGene_gn=UBR2;refGene_tx=XM_023253978.1;refGene_gn=UBR2;refGene_tx=XM_023253979.1;refGene_gn=UBR2;refGene_tx=XM_023253980.1
B2 43567748 43567851
refGene_gn=UBR2;refGene_tx=XM_011282222.3;refGene_gn=UBR2;refGene_tx=XM_023253976.1;refGene_gn=UBR2;refGene_tx=XM_023253977.1;refGene_gn=UBR2;refGene_tx=XM_023253978.1;refGene_gn=UBR2;refGene_tx=XM_023253979.1;refGene_gn=UBR2;refGene_tx=XM_023253980.1
B2 43568859 43568908
refGene_gn=UBR2;refGene_tx=XM_011282222.3;refGene_gn=UBR2;refGene_tx=XM_023253976.1;refGene_gn=UBR2;refGene_tx=XM_023253977.1;refGene_gn=UBR2;refGene_tx=XM_023253978.1;refGene_gn=UBR2;refGene_tx=XM_023253979.1;refGene_gn=UBR2;refGene_tx=XM_023253980.1
B2 43569345 43569500
refGene_gn=UBR2;refGene_tx=XM_011282222.3;refGene_gn=UBR2;refGene_tx=XM_023253976.1;refGene_gn=UBR2;refGene_tx=XM_023253977.1;refGene_gn=UBR2;refGene_tx=XM_023253978.1;refGene_gn=UBR2;refGene_tx=XM_023253979.1;refGene_gn=UBR2;refGene_tx=XM_023253980.1
B2 43571393 43571496
refGene_gn=UBR2;refGene_tx=XM_011282222.3;refGene_gn=UBR2;refGene_tx=XM_023253976.1;refGene_gn=UBR2;refGene_tx=XM_023253977.1;refGene_gn=UBR2;refGene_tx=XM_023253978.1;refGene_gn=UBR2;refGene_tx=XM_023253979.1
B2 43574452 43574562
refGene_gn=UBR2;refGene_tx=XM_011282222.3;refGene_gn=UBR2;refGene_tx=XM_023253976.1;refGene_gn=UBR2;refGene_tx=XM_023253977.1;refGene_gn=UBR2;refGene_tx=XM_023253978.1;refGene_gn=UBR2;refGene_tx=XM_023253979.1;refGene_gn=UBR2;refGene_tx=XM_023253980.1
B2 43574673 43574754
refGene_gn=UBR2;refGene_tx=XM_011282222.3;refGene_gn=UBR2;refGene_tx=XM_023253976.1;refGene_gn=UBR2;refGene_tx=XM_023253977.1;refGene_gn=UBR2;refGene_tx=XM_023253978.1;refGene_gn=UBR2;refGene_tx=XM_023253979.1
B2 43575080 43575259
refGene_gn=UBR2;refGene_tx=XM_011282222.3;refGene_gn=UBR2;refGene_tx=X

M_023253976.1;refGene_gn=UBR2;refGene_tx=XM_023253977.1;refGene_gn=UBR2;refGene_tx=XM_023253978.1;refGene_gn=UBR2;refGene_tx=XM_023253979.1
B2 43576221 43576421
refGene_gn=UBR2;refGene_tx=XM_011282222.3;refGene_gn=UBR2;refGene_tx=XM_023253976.1;refGene_gn=UBR2;refGene_tx=XM_023253977.1;refGene_gn=UBR2;refGene_tx=XM_023253978.1;refGene_gn=UBR2;refGene_tx=XM_023253979.1
B2 43578431 43578525
refGene_gn=UBR2;refGene_tx=XM_011282222.3;refGene_gn=UBR2;refGene_tx=XM_023253976.1;refGene_gn=UBR2;refGene_tx=XM_023253977.1;refGene_gn=UBR2;refGene_tx=XM_023253978.1;refGene_gn=UBR2;refGene_tx=XM_023253979.1
B2 43579190 43579352
refGene_gn=UBR2;refGene_tx=XM_011282222.3;refGene_gn=UBR2;refGene_tx=XM_023253976.1;refGene_gn=UBR2;refGene_tx=XM_023253977.1;refGene_gn=UBR2;refGene_tx=XM_023253978.1;refGene_gn=UBR2;refGene_tx=XM_023253979.1
B2 43580808 43580912
refGene_gn=UBR2;refGene_tx=XM_011282222.3;refGene_gn=UBR2;refGene_tx=XM_023253976.1;refGene_gn=UBR2;refGene_tx=XM_023253977.1;refGene_gn=UBR2;refGene_tx=XM_023253978.1
B2 43582069 43582148
refGene_gn=UBR2;refGene_tx=XM_011282222.3;refGene_gn=UBR2;refGene_tx=XM_023253976.1;refGene_gn=UBR2;refGene_tx=XM_023253977.1;refGene_gn=UBR2;refGene_tx=XM_023253978.1
B2 43585134 43585283
refGene_gn=UBR2;refGene_tx=XM_011282222.3;refGene_gn=UBR2;refGene_tx=XM_023253976.1;refGene_gn=UBR2;refGene_tx=XM_023253977.1;refGene_gn=UBR2;refGene_tx=XM_023253978.1
B2 43585655 43585711
refGene_gn=UBR2;refGene_tx=XM_011282222.3;refGene_gn=UBR2;refGene_tx=XM_023253976.1;refGene_gn=UBR2;refGene_tx=XM_023253977.1;refGene_gn=UBR2;refGene_tx=XM_023253978.1
B2 43587996 43588093
refGene_gn=UBR2;refGene_tx=XM_011282222.3;refGene_gn=UBR2;refGene_tx=XM_023253976.1;refGene_gn=UBR2;refGene_tx=XM_023253977.1;refGene_gn=UBR2;refGene_tx=XM_023253978.1
B2 43588346 43588414
refGene_gn=UBR2;refGene_tx=XM_011282222.3;refGene_gn=UBR2;refGene_tx=XM_023253976.1;refGene_gn=UBR2;refGene_tx=XM_023253977.1;refGene_gn=UBR2;refGene_tx=XM_023253978.1
B2 43589654 43589790
refGene_gn=UBR2;refGene_tx=XM_011282222.3;refGene_gn=UBR2;refGene_tx=XM_023253976.1;refGene_gn=UBR2;refGene_tx=XM_023253977.1;refGene_gn=UBR2;refGene_tx=XM_023253978.1
B2 43590294 43590385
refGene_gn=UBR2;refGene_tx=XM_011282222.3;refGene_gn=UBR2;refGene_tx=XM_023253976.1;refGene_gn=UBR2;refGene_tx=XM_023253977.1;refGene_gn=UBR2;refGene_tx=XM_023253978.1
B2 43592454 43592585
refGene_gn=UBR2;refGene_tx=XM_011282222.3;refGene_gn=UBR2;refGene_tx=XM_023253976.1;refGene_gn=UBR2;refGene_tx=XM_023253977.1;refGene_gn=UBR2;refGene_tx=XM_023253978.1

B2	43593627	43593736
refGene_gn=UBR2;refGene_tx=XM_011282222.3;refGene_gn=UBR2;refGene_tx=XM_023253976.1;refGene_gn=UBR2;refGene_tx=XM_023253977.1;refGene_gn=UBR2;refGene_tx=XM_023253978.1
B2	43596098	43596155
refGene_gn=UBR2;refGene_tx=XM_011282222.3;refGene_gn=UBR2;refGene_tx=XM_023253976.1;refGene_gn=UBR2;refGene_tx=XM_023253977.1;refGene_gn=UBR2;refGene_tx=XM_023253978.1
B2	43598680	43598758
refGene_gn=UBR2;refGene_tx=XM_011282222.3;refGene_gn=UBR2;refGene_tx=XM_023253976.1;refGene_gn=UBR2;refGene_tx=XM_023253977.1;refGene_gn=UBR2;refGene_tx=XM_023253978.1
B2	43599652	43599823
refGene_gn=UBR2;refGene_tx=XM_011282222.3;refGene_gn=UBR2;refGene_tx=XM_023253976.1;refGene_gn=UBR2;refGene_tx=XM_023253977.1;refGene_gn=UBR2;refGene_tx=XM_023253978.1
B2	43601679	43601781
refGene_gn=UBR2;refGene_tx=XM_011282222.3;refGene_gn=UBR2;refGene_tx=XM_023253976.1;refGene_gn=UBR2;refGene_tx=XM_023253977.1;refGene_gn=UBR2;refGene_tx=XM_023253978.1
B2	43603255	43603397
refGene_gn=UBR2;refGene_tx=XM_011282222.3;refGene_gn=UBR2;refGene_tx=XM_023253976.1;refGene_gn=UBR2;refGene_tx=XM_023253977.1;refGene_gn=UBR2;refGene_tx=XM_023253978.1
B2	43609575	43609788	refGene_gn=PRPH2;refGene_tx=NM_001042568.1
B2	43612697	43612944	refGene_gn=PRPH2;refGene_tx=NM_001042568.1
B2	43625902	43626483	refGene_gn=PRPH2;refGene_tx=NM_001042568.1
B2	43643036	43644164	refGene_gn=TBCC;refGene_tx=XM_003986144.5
B2	43712959	43713107	refGene_gn=BICRAL;refGene_tx=XM_019830597.2
B2	43713605	43713651
refGene_gn=BICRAL;refGene_tx=XM_003986145.5;refGene_gn=BICRAL;refGene_tx=XM_019830596.2;refGene_gn=BICRAL;refGene_tx=XM_019830597.2;refGene_gn=BICRAL;refGene_tx=XM_023253982.1
B2	43715587	43715636
refGene_gn=BICRAL;refGene_tx=XM_003986145.5;refGene_gn=BICRAL;refGene_tx=XM_019830596.2;refGene_gn=BICRAL;refGene_tx=XM_019830597.2;refGene_gn=BICRAL;refGene_tx=XM_023253982.1
B2	43715724	43715793
refGene_gn=BICRAL;refGene_tx=XM_003986145.5;refGene_gn=BICRAL;refGene_tx=XM_019830596.2;refGene_gn=BICRAL;refGene_tx=XM_019830597.2;refGene_gn=BICRAL;refGene_tx=XM_023253982.1
B2	43718609	43720289
refGene_gn=BICRAL;refGene_tx=XM_003986145.5;refGene_gn=BICRAL;refGene_tx=XM_019830596.2;refGene_gn=BICRAL;refGene_tx=XM_019830597.2;refGene_gn=BICRAL;refGene_tx=XM_023253982.1
B2	43737895	43738001
refGene_gn=BICRAL;refGene_tx=XM_003986145.5;refGene_gn=BICRAL;refGene_tx=XM_019830596.2;refGene_gn=BICRAL;refGene_tx=XM_023253982.1
B2	43738871	43738972
refGene_gn=BICRAL;refGene_tx=XM_003986145.5;refGene_gn=BICRAL;refGene_

tx=XM_019830596.2;refGene_gn=BICRAL;refGene_tx=XM_023253982.1
B2	43740730	43740792
refGene_gn=BICRAL;refGene_tx=XM_003986145.5;refGene_gn=BICRAL;refGene_tx=XM_019830596.2;refGene_gn=BICRAL;refGene_tx=XM_019830597.2;refGene_gn=BICRAL;refGene_tx=XM_023253982.1
B2	43741585	43741731
refGene_gn=BICRAL;refGene_tx=XM_003986145.5;refGene_gn=BICRAL;refGene_tx=XM_019830596.2;refGene_gn=BICRAL;refGene_tx=XM_019830597.2;refGene_gn=BICRAL;refGene_tx=XM_023253982.1
B2	43743043	43743138
refGene_gn=BICRAL;refGene_tx=XM_003986145.5;refGene_gn=BICRAL;refGene_tx=XM_019830596.2;refGene_gn=BICRAL;refGene_tx=XM_019830597.2;refGene_gn=BICRAL;refGene_tx=XM_023253982.1
B2	43745406	43745509
refGene_gn=BICRAL;refGene_tx=XM_003986145.5;refGene_gn=BICRAL;refGene_tx=XM_019830596.2;refGene_gn=BICRAL;refGene_tx=XM_019830597.2;refGene_gn=BICRAL;refGene_tx=XM_023253982.1
B2	43750216	43750980
refGene_gn=BICRAL;refGene_tx=XM_003986145.5;refGene_gn=BICRAL;refGene_tx=XM_019830596.2;refGene_gn=BICRAL;refGene_tx=XM_019830597.2;refGene_gn=BICRAL;refGene_tx=XM_023253982.1
B2	43759933	43759947
refGene_gn=RPL7L1;refGene_tx=XM_003986146.5;refGene_gn=RPL7L1;refGene_tx=XM_006931718.4;refGene_gn=RPL7L1;refGene_tx=XM_019830598.2
B2	43760557	43760666
refGene_gn=RPL7L1;refGene_tx=XM_003986146.5;refGene_gn=RPL7L1;refGene_tx=XM_006931718.4;refGene_gn=RPL7L1;refGene_tx=XM_019830598.2
B2	43761340	43761504
refGene_gn=RPL7L1;refGene_tx=XM_003986146.5;refGene_gn=RPL7L1;refGene_tx=XM_006931718.4;refGene_gn=RPL7L1;refGene_tx=XM_019830598.2
B2	43762487	43762625
refGene_gn=RPL7L1;refGene_tx=XM_003986146.5;refGene_gn=RPL7L1;refGene_tx=XM_006931718.4;refGene_gn=RPL7L1;refGene_tx=XM_019830598.2
B2	43763401	43763511
refGene_gn=RPL7L1;refGene_tx=XM_003986146.5;refGene_gn=RPL7L1;refGene_tx=XM_006931718.4;refGene_gn=RPL7L1;refGene_tx=XM_019830598.2
B2	43763703	43763912
refGene_gn=RPL7L1;refGene_tx=XM_003986146.5;refGene_gn=RPL7L1;refGene_tx=XM_006931718.4;refGene_gn=RPL7L1;refGene_tx=XM_019830598.2
B2	43767392	43767728
refGene_gn=CB2H6orf226;refGene_tx=XM_006931761.2
B2	43777154	43777212
refGene_gn=PTCRA;refGene_tx=XM_019830599.2;refGene_gn=PTCRA;refGene_tx=XM_019830600.2
B2	43780021	43780081	refGene_gn=PTCRA;refGene_tx=XM_019830599.2
B2	43780495	43780816
refGene_gn=PTCRA;refGene_tx=XM_019830599.2;refGene_gn=PTCRA;refGene_tx=XM_019830600.2
B2	43781906	43781951
refGene_gn=PTCRA;refGene_tx=XM_019830599.2;refGene_gn=PTCRA;refGene_tx

=XM_019830600.2
B2 43782598 43782951
refGene_gn=PTCRA;refGene_tx=XM_019830599.2;refGene_gn=PTCRA;refGene_tx=XM_019830600.2
B2 43786274 43786437
refGene_gn=CNPY3;refGene_tx=XM_003986147.5;refGene_gn=CNPY3;refGene_tx=XM_006931720.4
B2 43789618 43789619 refGene_gn=CNPY3;refGene_tx=XM_019830604.2
B2 43791063 43791187
refGene_gn=CNPY3;refGene_tx=XM_003986147.5;refGene_gn=CNPY3;refGene_tx=XM_006931720.4;refGene_gn=CNPY3;refGene_tx=XM_019830604.2
B2 43791860 43791918
refGene_gn=CNPY3;refGene_tx=XM_006931720.4;refGene_gn=CNPY3;refGene_tx=XM_019830604.2
B2 43791995 43792118
refGene_gn=CNPY3;refGene_tx=XM_003986147.5;refGene_gn=CNPY3;refGene_tx=XM_006931720.4;refGene_gn=CNPY3;refGene_tx=XM_019830604.2
B2 43794282 43794405
refGene_gn=CNPY3;refGene_tx=XM_003986147.5;refGene_gn=CNPY3;refGene_tx=XM_006931720.4;refGene_gn=CNPY3;refGene_tx=XM_019830604.2
B2 43794653 43794771
refGene_gn=CNPY3;refGene_tx=XM_003986147.5;refGene_gn=CNPY3;refGene_tx=XM_006931720.4;refGene_gn=CNPY3;refGene_tx=XM_019830604.2
B2 43795109 43795324
refGene_gn=CNPY3;refGene_tx=XM_003986147.5;refGene_gn=CNPY3;refGene_tx=XM_006931720.4;refGene_gn=CNPY3;refGene_tx=XM_019830604.2
B2 43806820 43807065 refGene_gn=GNMT;refGene_tx=XM_003986148.4
B2 43807914 43808042 refGene_gn=GNMT;refGene_tx=XM_003986148.4
B2 43808464 43808581 refGene_gn=GNMT;refGene_tx=XM_003986148.4
B2 43808792 43808935 refGene_gn=GNMT;refGene_tx=XM_003986148.4
B2 43809048 43809170 refGene_gn=GNMT;refGene_tx=XM_003986148.4
B2 43809252 43809424 refGene_gn=GNMT;refGene_tx=XM_003986148.4
B2 43810083 43810220
refGene_gn=PEX6;refGene_tx=XM_019830601.2;refGene_gn=PEX6;refGene_tx=XM_023253984.1;refGene_gn=PEX6;refGene_tx=XM_023253985.1
B2 43810402 43810542
refGene_gn=PEX6;refGene_tx=XM_019830601.2;refGene_gn=PEX6;refGene_tx=XM_023253984.1;refGene_gn=PEX6;refGene_tx=XM_023253985.1
B2 43810699 43810777
refGene_gn=PEX6;refGene_tx=XM_019830601.2;refGene_gn=PEX6;refGene_tx=XM_023253985.1
B2 43810901 43811018
refGene_gn=PEX6;refGene_tx=XM_019830601.2;refGene_gn=PEX6;refGene_tx=XM_023253984.1;refGene_gn=PEX6;refGene_tx=XM_023253985.1
B2 43811293 43811402
refGene_gn=PEX6;refGene_tx=XM_019830601.2;refGene_gn=PEX6;refGene_tx=XM_023253984.1;refGene_gn=PEX6;refGene_tx=XM_023253985.1
B2 43811648 43811710
refGene_gn=PEX6;refGene_tx=XM_019830601.2;refGene_gn=PEX6;refGene_tx=XM_023253984.1;refGene_gn=PEX6;refGene_tx=XM_023253985.1

B2 43811836 43812042
refGene_gn=PEX6;refGene_tx=XM_019830601.2;refGene_gn=PEX6;refGene_tx=XM_023253984.1
B2 43812100 43812256
refGene_gn=PEX6;refGene_tx=XM_019830601.2;refGene_gn=PEX6;refGene_tx=XM_023253984.1;refGene_gn=PEX6;refGene_tx=XM_023253985.1
B2 43812417 43812494
refGene_gn=PEX6;refGene_tx=XM_019830601.2;refGene_gn=PEX6;refGene_tx=XM_023253984.1;refGene_gn=PEX6;refGene_tx=XM_023253985.1
B2 43812683 43812879
refGene_gn=PEX6;refGene_tx=XM_019830601.2;refGene_gn=PEX6;refGene_tx=XM_023253984.1;refGene_gn=PEX6;refGene_tx=XM_023253985.1
B2 43813309 43813518
refGene_gn=PEX6;refGene_tx=XM_019830601.2;refGene_gn=PEX6;refGene_tx=XM_023253984.1;refGene_gn=PEX6;refGene_tx=XM_023253985.1
B2 43813872 43813984
refGene_gn=PEX6;refGene_tx=XM_019830601.2;refGene_gn=PEX6;refGene_tx=XM_023253984.1;refGene_gn=PEX6;refGene_tx=XM_023253985.1
B2 43814633 43814767
refGene_gn=PEX6;refGene_tx=XM_019830601.2;refGene_gn=PEX6;refGene_tx=XM_023253984.1;refGene_gn=PEX6;refGene_tx=XM_023253985.1
B2 43814851 43814954
refGene_gn=PEX6;refGene_tx=XM_019830601.2;refGene_gn=PEX6;refGene_tx=XM_023253984.1;refGene_gn=PEX6;refGene_tx=XM_023253985.1
B2 43818079 43818163
refGene_gn=PEX6;refGene_tx=XM_019830601.2;refGene_gn=PEX6;refGene_tx=XM_023253984.1;refGene_gn=PEX6;refGene_tx=XM_023253985.1
B2 43818646 43818810
refGene_gn=PEX6;refGene_tx=XM_019830601.2;refGene_gn=PEX6;refGene_tx=XM_023253984.1;refGene_gn=PEX6;refGene_tx=XM_023253985.1
B2 43820287 43821169
refGene_gn=PEX6;refGene_tx=XM_019830601.2;refGene_gn=PEX6;refGene_tx=XM_023253984.1;refGene_gn=PEX6;refGene_tx=XM_023253985.1
B2 43825774 43825801 refGene_gn=PPP2R5D;refGene_tx=XM_003986149.5
B2 43830431 43830509 refGene_gn=PPP2R5D;refGene_tx=XM_003986149.5
B2 43841758 43841975 refGene_gn=PPP2R5D;refGene_tx=XM_003986149.5
B2 43842224 43842424 refGene_gn=PPP2R5D;refGene_tx=XM_003986149.5
B2 43842511 43842622 refGene_gn=PPP2R5D;refGene_tx=XM_003986149.5
B2 43842743 43842836 refGene_gn=PPP2R5D;refGene_tx=XM_003986149.5
B2 43843276 43843407 refGene_gn=PPP2R5D;refGene_tx=XM_003986149.5
B2 43843540 43843600 refGene_gn=PPP2R5D;refGene_tx=XM_003986149.5
B2 43843704 43843813 refGene_gn=PPP2R5D;refGene_tx=XM_003986149.5
B2 43844027 43844081 refGene_gn=PPP2R5D;refGene_tx=XM_003986149.5
B2 43844372 43844543 refGene_gn=PPP2R5D;refGene_tx=XM_003986149.5
B2 43844648 43844776 refGene_gn=PPP2R5D;refGene_tx=XM_003986149.5
B2 43845430 43845532 refGene_gn=PPP2R5D;refGene_tx=XM_003986149.5
B2 43845627 43845700 refGene_gn=PPP2R5D;refGene_tx=XM_003986149.5
B2 43845826 43845943 refGene_gn=PPP2R5D;refGene_tx=XM_003986149.5
B2 43846094 43846232 refGene_gn=PPP2R5D;refGene_tx=XM_003986149.5
B2 43847425 43847580

refGene_gn=MEA1;refGene_tx=XM_006931722.4;refGene_gn=MEA1;refGene_tx=XM_011282219.3;refGene_gn=MEA1;refGene_tx=XM_011282220.3;refGene_gn=MEA1;refGene_tx=XM_011282221.3
B2 43847883 43847986
refGene_gn=MEA1;refGene_tx=XM_006931722.4;refGene_gn=MEA1;refGene_tx=XM_011282219.3;refGene_gn=MEA1;refGene_tx=XM_011282220.3;refGene_gn=MEA1;refGene_tx=XM_011282221.3
B2 43848073 43848354
refGene_gn=MEA1;refGene_tx=XM_006931722.4;refGene_gn=MEA1;refGene_tx=XM_011282219.3;refGene_gn=MEA1;refGene_tx=XM_011282220.3;refGene_gn=MEA1;refGene_tx=XM_011282221.3
B2 43848864 43848928
refGene_gn=MEA1;refGene_tx=XM_006931722.4;refGene_gn=MEA1;refGene_tx=XM_011282219.3
B2 43851505 43851659
refGene_gn=KLHDC3;refGene_tx=XM_003986152.5;refGene_gn=KLHDC3;refGene_tx=XM_006931725.4;refGene_gn=KLHDC3;refGene_tx=XM_006931727.4;refGene_gn=KLHDC3;refGene_tx=XM_006931728.4;refGene_gn=KLHDC3;refGene_tx=XM_019830603.2
B2 43851836 43852013
refGene_gn=KLHDC3;refGene_tx=XM_003986152.5;refGene_gn=KLHDC3;refGene_tx=XM_006931725.4;refGene_gn=KLHDC3;refGene_tx=XM_006931727.4;refGene_gn=KLHDC3;refGene_tx=XM_006931728.4;refGene_gn=KLHDC3;refGene_tx=XM_019830603.2
B2 43852160 43852276
refGene_gn=KLHDC3;refGene_tx=XM_003986152.5;refGene_gn=KLHDC3;refGene_tx=XM_006931725.4;refGene_gn=KLHDC3;refGene_tx=XM_006931727.4;refGene_gn=KLHDC3;refGene_tx=XM_006931728.4;refGene_gn=KLHDC3;refGene_tx=XM_019830603.2
B2 43852454 43852526
refGene_gn=KLHDC3;refGene_tx=XM_003986152.5;refGene_gn=KLHDC3;refGene_tx=XM_006931725.4;refGene_gn=KLHDC3;refGene_tx=XM_006931727.4;refGene_gn=KLHDC3;refGene_tx=XM_006931728.4;refGene_gn=KLHDC3;refGene_tx=XM_019830603.2
B2 43852655 43852869
refGene_gn=KLHDC3;refGene_tx=XM_003986152.5;refGene_gn=KLHDC3;refGene_tx=XM_006931725.4;refGene_gn=KLHDC3;refGene_tx=XM_006931727.4;refGene_gn=KLHDC3;refGene_tx=XM_006931728.4;refGene_gn=KLHDC3;refGene_tx=XM_019830603.2
B2 43852949 43853036
refGene_gn=KLHDC3;refGene_tx=XM_003986152.5;refGene_gn=KLHDC3;refGene_tx=XM_006931725.4;refGene_gn=KLHDC3;refGene_tx=XM_006931727.4;refGene_gn=KLHDC3;refGene_tx=XM_006931728.4;refGene_gn=KLHDC3;refGene_tx=XM_019830603.2
B2 43853421 43853530
refGene_gn=KLHDC3;refGene_tx=XM_003986152.5;refGene_gn=KLHDC3;refGene_tx=XM_006931725.4;refGene_gn=KLHDC3;refGene_tx=XM_006931727.4;refGene_gn=KLHDC3;refGene_tx=XM_006931728.4;refGene_gn=KLHDC3;refGene_tx=XM_019830603.2
B2 43853649 43853723

refGene_gn=KLHDC3;refGene_tx=XM_003986152.5;refGene_gn=KLHDC3;refGene_tx=XM_006931725.4;refGene_gn=KLHDC3;refGene_tx=XM_006931727.4;refGene_gn=KLHDC3;refGene_tx=XM_006931728.4;refGene_gn=KLHDC3;refGene_tx=XM_019830603.2
B2 43853834 43853913 refGene_gn=KLHDC3;refGene_tx=XM_003986152.5;refGene_gn=KLHDC3;refGene_tx=XM_006931725.4;refGene_gn=KLHDC3;refGene_tx=XM_006931727.4;refGene_gn=KLHDC3;refGene_tx=XM_006931728.4
B2 43855046 43855113 refGene_gn=KLHDC3;refGene_tx=XM_003986152.5;refGene_gn=KLHDC3;refGene_tx=XM_006931725.4;refGene_gn=KLHDC3;refGene_tx=XM_006931727.4;refGene_gn=KLHDC3;refGene_tx=XM_006931728.4
B2 43857608 43857756 refGene_gn=KLHDC3;refGene_tx=XM_019830603.2
B2 43858698 43858765 refGene_gn=KLHDC3;refGene_tx=XM_019830603.2
B2 43858845 43858950 refGene_gn=KLHDC3;refGene_tx=XM_019830603.2
B2 43859310 43859385 refGene_gn=KLHDC3;refGene_tx=XM_019830603.2
B2 43859495 43859613 refGene_gn=KLHDC3;refGene_tx=XM_019830603.2
B2 43860058 43860195 refGene_gn=KLHDC3;refGene_tx=XM_019830603.2
B2 43865093 43865417 refGene_gn=CUL7;refGene_tx=XM_023253988.1
B2 43865661 43865867 refGene_gn=CUL7;refGene_tx=XM_023253988.1
B2 43865944 43866071 refGene_gn=CUL7;refGene_tx=XM_023253988.1
B2 43866220 43866366 refGene_gn=CUL7;refGene_tx=XM_023253988.1
B2 43867118 43867401 refGene_gn=CUL7;refGene_tx=XM_023253988.1
B2 43867507 43867724 refGene_gn=CUL7;refGene_tx=XM_023253988.1
B2 43867878 43868039 refGene_gn=CUL7;refGene_tx=XM_023253988.1
B2 43869533 43869716 refGene_gn=CUL7;refGene_tx=XM_023253988.1
B2 43869805 43869912 refGene_gn=CUL7;refGene_tx=XM_023253988.1
B2 43870129 43870312 refGene_gn=CUL7;refGene_tx=XM_023253988.1
B2 43870608 43870742 refGene_gn=CUL7;refGene_tx=XM_023253988.1
B2 43871070 43871246 refGene_gn=CUL7;refGene_tx=XM_023253988.1
B2 43871435 43871531 refGene_gn=CUL7;refGene_tx=XM_023253988.1
B2 43871824 43871930 refGene_gn=CUL7;refGene_tx=XM_023253988.1
B2 43872112 43872284 refGene_gn=CUL7;refGene_tx=XM_023253988.1
B2 43872382 43872473 refGene_gn=CUL7;refGene_tx=XM_023253988.1
B2 43872767 43872995 refGene_gn=CUL7;refGene_tx=XM_023253988.1
B2 43873973 43874079 refGene_gn=CUL7;refGene_tx=XM_023253988.1
B2 43874162 43874400 refGene_gn=CUL7;refGene_tx=XM_023253988.1
B2 43875104 43875333 refGene_gn=CUL7;refGene_tx=XM_023253988.1
B2 43875625 43875828 refGene_gn=CUL7;refGene_tx=XM_023253988.1
B2 43875921 43876060 refGene_gn=CUL7;refGene_tx=XM_023253988.1
B2 43876587 43877088 refGene_gn=CUL7;refGene_tx=XM_023253988.1
B2 43877226 43877378 refGene_gn=CUL7;refGene_tx=XM_023253988.1
B2 43877911 43878491 refGene_gn=CUL7;refGene_tx=XM_023253988.1
B2 43880081 43880215 refGene_gn=MRPL2;refGene_tx=XM_011282215.3
B2 43880217 43880293 refGene_gn=MRPL2;refGene_tx=XM_011282215.3
B2 43880646 43880720 refGene_gn=MRPL2;refGene_tx=XM_011282215.3
B2 43881022 43881133 refGene_gn=MRPL2;refGene_tx=XM_011282215.3
B2 43881209 43881325 refGene_gn=MRPL2;refGene_tx=XM_011282215.3
B2 43881437 43881579 refGene_gn=MRPL2;refGene_tx=XM_011282215.3
B2 43882634 43882803 refGene_gn=MRPL2;refGene_tx=XM_011282215.3

B2 43884071 43884167 refGene_gn=MRPL2;refGene_tx=XM_011282215.3
B2 43886137 43886395 refGene_gn=KLC4;refGene_tx=XM_023253989.1
B2 43888053 43888284 refGene_gn=KLC4;refGene_tx=XM_023253989.1
B2 43890165 43890247 refGene_gn=KLC4;refGene_tx=XM_023253989.1
B2 43890849 43891060 refGene_gn=KLC4;refGene_tx=XM_023253989.1
B2 43891536 43891624 refGene_gn=KLC4;refGene_tx=XM_023253989.1
B2 43893222 43893324 refGene_gn=KLC4;refGene_tx=XM_023253989.1
B2 43893546 43893720 refGene_gn=KLC4;refGene_tx=XM_023253989.1
B2 43894026 43894126 refGene_gn=KLC4;refGene_tx=XM_023253989.1
B2 43894339 43894392 refGene_gn=KLC4;refGene_tx=XM_023253989.1
B2 43894598 43894669 refGene_gn=KLC4;refGene_tx=XM_023253989.1
B2 43894862 43894971 refGene_gn=KLC4;refGene_tx=XM_023253989.1
B2 43896176 43896317 refGene_gn=KLC4;refGene_tx=XM_023253989.1
B2 43896604 43896719 refGene_gn=KLC4;refGene_tx=XM_023253989.1
B2 43897053 43897145 refGene_gn=KLC4;refGene_tx=XM_023253989.1
B2 43898050 43898101 refGene_gn=KLC4;refGene_tx=XM_023253989.1
B2 43899918 43899997 refGene_gn=PTK7;refGene_tx=XM_003986191.4
B2 43937005 43937293 refGene_gn=PTK7;refGene_tx=XM_003986191.4
B2 43937765 43937868 refGene_gn=PTK7;refGene_tx=XM_003986191.4
B2 43938239 43938430 refGene_gn=PTK7;refGene_tx=XM_003986191.4
B2 43938523 43938674 refGene_gn=PTK7;refGene_tx=XM_003986191.4
B2 43939684 43939833 refGene_gn=PTK7;refGene_tx=XM_003986191.4
B2 43940081 43940348 refGene_gn=PTK7;refGene_tx=XM_003986191.4
B2 43942553 43942687 refGene_gn=PTK7;refGene_tx=XM_003986191.4
B2 43942821 43942957 refGene_gn=PTK7;refGene_tx=XM_003986191.4
B2 43943117 43943237 refGene_gn=PTK7;refGene_tx=XM_003986191.4
B2 43944728 43944878 refGene_gn=PTK7;refGene_tx=XM_003986191.4
B2 43944993 43945144 refGene_gn=PTK7;refGene_tx=XM_003986191.4
B2 43945242 43945370 refGene_gn=PTK7;refGene_tx=XM_003986191.4
B2 43946590 43946794 refGene_gn=PTK7;refGene_tx=XM_003986191.4
B2 43947545 43947701 refGene_gn=PTK7;refGene_tx=XM_003986191.4
B2 43948326 43948559 refGene_gn=PTK7;refGene_tx=XM_003986191.4
B2 43949511 43949592 refGene_gn=PTK7;refGene_tx=XM_003986191.4
B2 43959636 43959788 refGene_gn=PTK7;refGene_tx=XM_003986191.4
B2 43960852 43961031 refGene_gn=PTK7;refGene_tx=XM_003986191.4
B2 43961529 43961690 refGene_gn=PTK7;refGene_tx=XM_003986191.4
B2 43968834 43969359 refGene_gn=SRF;refGene_tx=XM_006931762.3
B2 43971003 43971270 refGene_gn=SRF;refGene_tx=XM_006931762.3
B2 43973070 43973332 refGene_gn=SRF;refGene_tx=XM_006931762.3
B2 43973894 43974014 refGene_gn=SRF;refGene_tx=XM_006931762.3
B2 43975031 43975223 refGene_gn=SRF;refGene_tx=XM_006931762.3
B2 43975627 43975704 refGene_gn=SRF;refGene_tx=XM_006931762.3
B2 43975934 43976030 refGene_gn=SRF;refGene_tx=XM_006931762.3
B2 43979870 43979948
refGene_gn=CUL9;refGene_tx=XM_006931735.4;refGene_gn=CUL9;refGene_tx=XM_019830608.2;refGene_gn=CUL9;refGene_tx=XM_019830609.2;refGene_gn=CUL9;refGene_tx=XM_019830610.2;refGene_gn=CUL9;refGene_tx=XM_019830611.2;refGene_gn=CUL9;refGene_tx=XM_019830612.2;refGene_gn=CUL9;refGene_tx=XM_019830614.2;refGene_gn=CUL9;refGene_tx=XM_019830615.2;refGene_gn=CUL9;refGene_tx=XM_019830616.1;refGene_gn=CUL9;refGene_tx=XM_019830617.2;

refGene_gn=CUL9;refGene_tx=XM_019830618.2
B2	43980746	43981350
refGene_gn=CUL9;refGene_tx=XM_006931735.4;refGene_gn=CUL9;refGene_tx=XM_019830608.2;refGene_gn=CUL9;refGene_tx=XM_019830609.2;refGene_gn=CUL9;refGene_tx=XM_019830610.2;refGene_gn=CUL9;refGene_tx=XM_019830611.2;refGene_gn=CUL9;refGene_tx=XM_019830612.2;refGene_gn=CUL9;refGene_tx=XM_019830613.1;refGene_gn=CUL9;refGene_tx=XM_019830614.2;refGene_gn=CUL9;refGene_tx=XM_019830615.2;refGene_gn=CUL9;refGene_tx=XM_019830616.1;refGene_gn=CUL9;refGene_tx=XM_019830617.2;refGene_gn=CUL9;refGene_tx=XM_019830618.2
B2	43981914	43982069
refGene_gn=CUL9;refGene_tx=XM_006931735.4;refGene_gn=CUL9;refGene_tx=XM_019830608.2;refGene_gn=CUL9;refGene_tx=XM_019830609.2;refGene_gn=CUL9;refGene_tx=XM_019830610.2;refGene_gn=CUL9;refGene_tx=XM_019830611.2;refGene_gn=CUL9;refGene_tx=XM_019830612.2;refGene_gn=CUL9;refGene_tx=XM_019830613.1;refGene_gn=CUL9;refGene_tx=XM_019830614.2;refGene_gn=CUL9;refGene_tx=XM_019830615.2;refGene_gn=CUL9;refGene_tx=XM_019830616.1;refGene_gn=CUL9;refGene_tx=XM_019830617.2;refGene_gn=CUL9;refGene_tx=XM_019830618.2
B2	43982385	43982895
refGene_gn=CUL9;refGene_tx=XM_006931735.4;refGene_gn=CUL9;refGene_tx=XM_019830608.2;refGene_gn=CUL9;refGene_tx=XM_019830609.2;refGene_gn=CUL9;refGene_tx=XM_019830610.2;refGene_gn=CUL9;refGene_tx=XM_019830611.2;refGene_gn=CUL9;refGene_tx=XM_019830612.2;refGene_gn=CUL9;refGene_tx=XM_019830613.1;refGene_gn=CUL9;refGene_tx=XM_019830614.2;refGene_gn=CUL9;refGene_tx=XM_019830615.2;refGene_gn=CUL9;refGene_tx=XM_019830616.1;refGene_gn=CUL9;refGene_tx=XM_019830617.2;refGene_gn=CUL9;refGene_tx=XM_019830618.2
B2	43983378	43983514
refGene_gn=CUL9;refGene_tx=XM_006931735.4;refGene_gn=CUL9;refGene_tx=XM_019830608.2;refGene_gn=CUL9;refGene_tx=XM_019830609.2;refGene_gn=CUL9;refGene_tx=XM_019830610.2;refGene_gn=CUL9;refGene_tx=XM_019830611.2;refGene_gn=CUL9;refGene_tx=XM_019830612.2;refGene_gn=CUL9;refGene_tx=XM_019830613.1;refGene_gn=CUL9;refGene_tx=XM_019830614.2;refGene_gn=CUL9;refGene_tx=XM_019830615.2;refGene_gn=CUL9;refGene_tx=XM_019830616.1;refGene_gn=CUL9;refGene_tx=XM_019830617.2;refGene_gn=CUL9;refGene_tx=XM_019830618.2
B2	43983662	43983856
refGene_gn=CUL9;refGene_tx=XM_006931735.4;refGene_gn=CUL9;refGene_tx=XM_019830608.2;refGene_gn=CUL9;refGene_tx=XM_019830609.2;refGene_gn=CUL9;refGene_tx=XM_019830610.2;refGene_gn=CUL9;refGene_tx=XM_019830611.2;refGene_gn=CUL9;refGene_tx=XM_019830612.2;refGene_gn=CUL9;refGene_tx=XM_019830613.1;refGene_gn=CUL9;refGene_tx=XM_019830614.2;refGene_gn=CUL9;refGene_tx=XM_019830615.2;refGene_gn=CUL9;refGene_tx=XM_019830616.1;refGene_gn=CUL9;refGene_tx=XM_019830617.2;refGene_gn=CUL9;refGene_tx=XM_019830618.2
B2	43984123	43984529
refGene_gn=CUL9;refGene_tx=XM_006931735.4;refGene_gn=CUL9;refGene_tx=XM_019830608.2;refGene_gn=CUL9;refGene_tx=XM_019830609.2;refGene_gn=CUL9;refGene_tx=XM_019830610.2;refGene_gn=CUL9;refGene_tx=XM_019830611.2;

refGene_gn=CUL9;refGene_tx=XM_019830612.2;refGene_gn=CUL9;refGene_tx=XM_019830613.1;refGene_gn=CUL9;refGene_tx=XM_019830614.2;refGene_gn=CUL9;refGene_tx=XM_019830615.2;refGene_gn=CUL9;refGene_tx=XM_019830616.1;refGene_gn=CUL9;refGene_tx=XM_019830617.2;refGene_gn=CUL9;refGene_tx=XM_019830618.2
B2 43984941 43985134
refGene_gn=CUL9;refGene_tx=XM_006931735.4;refGene_gn=CUL9;refGene_tx=XM_019830608.2;refGene_gn=CUL9;refGene_tx=XM_019830609.2;refGene_gn=CUL9;refGene_tx=XM_019830610.2;refGene_gn=CUL9;refGene_tx=XM_019830611.2;refGene_gn=CUL9;refGene_tx=XM_019830612.2;refGene_gn=CUL9;refGene_tx=XM_019830613.1;refGene_gn=CUL9;refGene_tx=XM_019830614.2;refGene_gn=CUL9;refGene_tx=XM_019830615.2;refGene_gn=CUL9;refGene_tx=XM_019830616.1;refGene_gn=CUL9;refGene_tx=XM_019830617.2;refGene_gn=CUL9;refGene_tx=XM_019830618.2
B2 43988013 43988221
refGene_gn=CUL9;refGene_tx=XM_006931735.4;refGene_gn=CUL9;refGene_tx=XM_019830608.2;refGene_gn=CUL9;refGene_tx=XM_019830609.2;refGene_gn=CUL9;refGene_tx=XM_019830610.2;refGene_gn=CUL9;refGene_tx=XM_019830611.2;refGene_gn=CUL9;refGene_tx=XM_019830612.2;refGene_gn=CUL9;refGene_tx=XM_019830613.1;refGene_gn=CUL9;refGene_tx=XM_019830614.2;refGene_gn=CUL9;refGene_tx=XM_019830615.2;refGene_gn=CUL9;refGene_tx=XM_019830616.1;refGene_gn=CUL9;refGene_tx=XM_019830617.2;refGene_gn=CUL9;refGene_tx=XM_019830618.2
B2 43990055 43990252
refGene_gn=CUL9;refGene_tx=XM_006931735.4;refGene_gn=CUL9;refGene_tx=XM_019830608.2;refGene_gn=CUL9;refGene_tx=XM_019830609.2;refGene_gn=CUL9;refGene_tx=XM_019830610.2;refGene_gn=CUL9;refGene_tx=XM_019830611.2;refGene_gn=CUL9;refGene_tx=XM_019830612.2;refGene_gn=CUL9;refGene_tx=XM_019830613.1;refGene_gn=CUL9;refGene_tx=XM_019830614.2;refGene_gn=CUL9;refGene_tx=XM_019830615.2;refGene_gn=CUL9;refGene_tx=XM_019830616.1;refGene_gn=CUL9;refGene_tx=XM_019830617.2;refGene_gn=CUL9;refGene_tx=XM_019830618.2
B2 43990632 43990850
refGene_gn=CUL9;refGene_tx=XM_006931735.4;refGene_gn=CUL9;refGene_tx=XM_019830608.2;refGene_gn=CUL9;refGene_tx=XM_019830609.2;refGene_gn=CUL9;refGene_tx=XM_019830610.2;refGene_gn=CUL9;refGene_tx=XM_019830611.2;refGene_gn=CUL9;refGene_tx=XM_019830612.2;refGene_gn=CUL9;refGene_tx=XM_019830613.1;refGene_gn=CUL9;refGene_tx=XM_019830614.2;refGene_gn=CUL9;refGene_tx=XM_019830615.2;refGene_gn=CUL9;refGene_tx=XM_019830616.1;refGene_gn=CUL9;refGene_tx=XM_019830617.2;refGene_gn=CUL9;refGene_tx=XM_019830618.2
B2 43992001 43992251
refGene_gn=CUL9;refGene_tx=XM_006931735.4;refGene_gn=CUL9;refGene_tx=XM_019830608.2;refGene_gn=CUL9;refGene_tx=XM_019830609.2;refGene_gn=CUL9;refGene_tx=XM_019830610.2;refGene_gn=CUL9;refGene_tx=XM_019830611.2;refGene_gn=CUL9;refGene_tx=XM_019830612.2;refGene_gn=CUL9;refGene_tx=XM_019830613.1;refGene_gn=CUL9;refGene_tx=XM_019830614.2;refGene_gn=CUL9;refGene_tx=XM_019830615.2;refGene_gn=CUL9;refGene_tx=XM_019830616.1;refGene_gn=CUL9;refGene_tx=XM_019830617.2;refGene_gn=CUL9;refGene_tx=XM_019830618.2

B2 43992337 43992443
refGene_gn=CUL9;refGene_tx=XM_006931735.4;refGene_gn=CUL9;refGene_tx=XM_019830608.2;refGene_gn=CUL9;refGene_tx=XM_019830609.2;refGene_gn=CUL9;refGene_tx=XM_019830610.2;refGene_gn=CUL9;refGene_tx=XM_019830611.2;refGene_gn=CUL9;refGene_tx=XM_019830612.2;refGene_gn=CUL9;refGene_tx=XM_019830613.1;refGene_gn=CUL9;refGene_tx=XM_019830614.2;refGene_gn=CUL9;refGene_tx=XM_019830615.2;refGene_gn=CUL9;refGene_tx=XM_019830616.1;refGene_gn=CUL9;refGene_tx=XM_019830617.2;refGene_gn=CUL9;refGene_tx=XM_019830618.2
B2 43992995 43993223
refGene_gn=CUL9;refGene_tx=XM_006931735.4;refGene_gn=CUL9;refGene_tx=XM_019830608.2;refGene_gn=CUL9;refGene_tx=XM_019830609.2;refGene_gn=CUL9;refGene_tx=XM_019830610.2;refGene_gn=CUL9;refGene_tx=XM_019830611.2;refGene_gn=CUL9;refGene_tx=XM_019830612.2;refGene_gn=CUL9;refGene_tx=XM_019830613.1;refGene_gn=CUL9;refGene_tx=XM_019830614.2;refGene_gn=CUL9;refGene_tx=XM_019830615.2;refGene_gn=CUL9;refGene_tx=XM_019830616.1;refGene_gn=CUL9;refGene_tx=XM_019830617.2;refGene_gn=CUL9;refGene_tx=XM_019830618.2
B2 43993514 43993605
refGene_gn=CUL9;refGene_tx=XM_006931735.4;refGene_gn=CUL9;refGene_tx=XM_019830608.2;refGene_gn=CUL9;refGene_tx=XM_019830609.2;refGene_gn=CUL9;refGene_tx=XM_019830610.2;refGene_gn=CUL9;refGene_tx=XM_019830611.2;refGene_gn=CUL9;refGene_tx=XM_019830612.2;refGene_gn=CUL9;refGene_tx=XM_019830613.1;refGene_gn=CUL9;refGene_tx=XM_019830614.2;refGene_gn=CUL9;refGene_tx=XM_019830615.2;refGene_gn=CUL9;refGene_tx=XM_019830616.1;refGene_gn=CUL9;refGene_tx=XM_019830617.2;refGene_gn=CUL9;refGene_tx=XM_019830618.2
B2 43993706 43993878
refGene_gn=CUL9;refGene_tx=XM_006931735.4;refGene_gn=CUL9;refGene_tx=XM_019830608.2;refGene_gn=CUL9;refGene_tx=XM_019830609.2;refGene_gn=CUL9;refGene_tx=XM_019830610.2;refGene_gn=CUL9;refGene_tx=XM_019830611.2;refGene_gn=CUL9;refGene_tx=XM_019830612.2;refGene_gn=CUL9;refGene_tx=XM_019830613.1;refGene_gn=CUL9;refGene_tx=XM_019830614.2;refGene_gn=CUL9;refGene_tx=XM_019830615.2;refGene_gn=CUL9;refGene_tx=XM_019830616.1;refGene_gn=CUL9;refGene_tx=XM_019830617.2;refGene_gn=CUL9;refGene_tx=XM_019830618.2
B2 43995340 43995446
refGene_gn=CUL9;refGene_tx=XM_006931735.4;refGene_gn=CUL9;refGene_tx=XM_019830608.2;refGene_gn=CUL9;refGene_tx=XM_019830609.2;refGene_gn=CUL9;refGene_tx=XM_019830610.2;refGene_gn=CUL9;refGene_tx=XM_019830611.2;refGene_gn=CUL9;refGene_tx=XM_019830612.2;refGene_gn=CUL9;refGene_tx=XM_019830613.1;refGene_gn=CUL9;refGene_tx=XM_019830614.2;refGene_gn=CUL9;refGene_tx=XM_019830615.2;refGene_gn=CUL9;refGene_tx=XM_019830616.1;refGene_gn=CUL9;refGene_tx=XM_019830617.2;refGene_gn=CUL9;refGene_tx=XM_019830618.2
B2 43995684 43995780
refGene_gn=CUL9;refGene_tx=XM_006931735.4;refGene_gn=CUL9;refGene_tx=XM_019830608.2;refGene_gn=CUL9;refGene_tx=XM_019830609.2;refGene_gn=CUL9;refGene_tx=XM_019830610.2;refGene_gn=CUL9;refGene_tx=XM_019830611.2;refGene_gn=CUL9;refGene_tx=XM_019830612.2;refGene_gn=CUL9;refGene_tx=X

M_019830613.1;refGene_gn=CUL9;refGene_tx=XM_019830614.2;refGene_gn=CUL9;refGene_tx=XM_019830615.2;refGene_gn=CUL9;refGene_tx=XM_019830616.1;refGene_gn=CUL9;refGene_tx=XM_019830617.2;refGene_gn=CUL9;refGene_tx=XM_019830618.2
B2 43996038 43996214
refGene_gn=CUL9;refGene_tx=XM_006931735.4;refGene_gn=CUL9;refGene_tx=XM_019830608.2;refGene_gn=CUL9;refGene_tx=XM_019830609.2;refGene_gn=CUL9;refGene_tx=XM_019830610.2;refGene_gn=CUL9;refGene_tx=XM_019830611.2;refGene_gn=CUL9;refGene_tx=XM_019830612.2;refGene_gn=CUL9;refGene_tx=XM_019830613.1;refGene_gn=CUL9;refGene_tx=XM_019830614.2;refGene_gn=CUL9;refGene_tx=XM_019830615.2;refGene_gn=CUL9;refGene_tx=XM_019830616.1;refGene_gn=CUL9;refGene_tx=XM_019830617.2;refGene_gn=CUL9;refGene_tx=XM_019830618.2
B2 43996499 43996633
refGene_gn=CUL9;refGene_tx=XM_006931735.4;refGene_gn=CUL9;refGene_tx=XM_019830608.2;refGene_gn=CUL9;refGene_tx=XM_019830609.2;refGene_gn=CUL9;refGene_tx=XM_019830610.2;refGene_gn=CUL9;refGene_tx=XM_019830611.2;refGene_gn=CUL9;refGene_tx=XM_019830612.2;refGene_gn=CUL9;refGene_tx=XM_019830613.1;refGene_gn=CUL9;refGene_tx=XM_019830614.2;refGene_gn=CUL9;refGene_tx=XM_019830615.2;refGene_gn=CUL9;refGene_tx=XM_019830616.1;refGene_gn=CUL9;refGene_tx=XM_019830617.2;refGene_gn=CUL9;refGene_tx=XM_019830618.2
B2 43996990 43997170
refGene_gn=CUL9;refGene_tx=XM_006931735.4;refGene_gn=CUL9;refGene_tx=XM_019830608.2;refGene_gn=CUL9;refGene_tx=XM_019830609.2;refGene_gn=CUL9;refGene_tx=XM_019830610.2;refGene_gn=CUL9;refGene_tx=XM_019830611.2;refGene_gn=CUL9;refGene_tx=XM_019830612.2;refGene_gn=CUL9;refGene_tx=XM_019830613.1;refGene_gn=CUL9;refGene_tx=XM_019830614.2;refGene_gn=CUL9;refGene_tx=XM_019830615.2;refGene_gn=CUL9;refGene_tx=XM_019830616.1;refGene_gn=CUL9;refGene_tx=XM_019830617.2;refGene_gn=CUL9;refGene_tx=XM_019830618.2
B2 43997356 43997466
refGene_gn=CUL9;refGene_tx=XM_006931735.4;refGene_gn=CUL9;refGene_tx=XM_019830608.2;refGene_gn=CUL9;refGene_tx=XM_019830609.2;refGene_gn=CUL9;refGene_tx=XM_019830610.2;refGene_gn=CUL9;refGene_tx=XM_019830611.2;refGene_gn=CUL9;refGene_tx=XM_019830612.2;refGene_gn=CUL9;refGene_tx=XM_019830613.1;refGene_gn=CUL9;refGene_tx=XM_019830614.2;refGene_gn=CUL9;refGene_tx=XM_019830615.2;refGene_gn=CUL9;refGene_tx=XM_019830616.1;refGene_gn=CUL9;refGene_tx=XM_019830617.2;refGene_gn=CUL9;refGene_tx=XM_019830618.2
B2 43997561 43997744
refGene_gn=CUL9;refGene_tx=XM_006931735.4;refGene_gn=CUL9;refGene_tx=XM_019830608.2;refGene_gn=CUL9;refGene_tx=XM_019830609.2;refGene_gn=CUL9;refGene_tx=XM_019830610.2;refGene_gn=CUL9;refGene_tx=XM_019830611.2;refGene_gn=CUL9;refGene_tx=XM_019830612.2;refGene_gn=CUL9;refGene_tx=XM_019830613.1;refGene_gn=CUL9;refGene_tx=XM_019830614.2;refGene_gn=CUL9;refGene_tx=XM_019830615.2;refGene_gn=CUL9;refGene_tx=XM_019830616.1;refGene_gn=CUL9;refGene_tx=XM_019830617.2;refGene_gn=CUL9;refGene_tx=XM_019830618.2
B2 43997875 43998036

refGene_gn=CUL9;refGene_tx=XM_006931735.4;refGene_gn=CUL9;refGene_tx=XM_019830608.2;refGene_gn=CUL9;refGene_tx=XM_019830609.2;refGene_gn=CUL9;refGene_tx=XM_019830610.2;refGene_gn=CUL9;refGene_tx=XM_019830611.2;refGene_gn=CUL9;refGene_tx=XM_019830612.2;refGene_gn=CUL9;refGene_tx=XM_019830613.1;refGene_gn=CUL9;refGene_tx=XM_019830614.2;refGene_gn=CUL9;refGene_tx=XM_019830615.2;refGene_gn=CUL9;refGene_tx=XM_019830616.1;refGene_gn=CUL9;refGene_tx=XM_019830617.2;refGene_gn=CUL9;refGene_tx=XM_019830618.2
B2 43998240 43998469
refGene_gn=CUL9;refGene_tx=XM_006931735.4;refGene_gn=CUL9;refGene_tx=XM_019830608.2;refGene_gn=CUL9;refGene_tx=XM_019830609.2;refGene_gn=CUL9;refGene_tx=XM_019830610.2;refGene_gn=CUL9;refGene_tx=XM_019830611.2;refGene_gn=CUL9;refGene_tx=XM_019830612.2;refGene_gn=CUL9;refGene_tx=XM_019830613.1;refGene_gn=CUL9;refGene_tx=XM_019830614.2;refGene_gn=CUL9;refGene_tx=XM_019830615.2;refGene_gn=CUL9;refGene_tx=XM_019830616.1;refGene_gn=CUL9;refGene_tx=XM_019830617.2;refGene_gn=CUL9;refGene_tx=XM_019830618.2
B2 43998563 43998753
refGene_gn=CUL9;refGene_tx=XM_006931735.4;refGene_gn=CUL9;refGene_tx=XM_019830608.2;refGene_gn=CUL9;refGene_tx=XM_019830609.2;refGene_gn=CUL9;refGene_tx=XM_019830610.2;refGene_gn=CUL9;refGene_tx=XM_019830611.2;refGene_gn=CUL9;refGene_tx=XM_019830612.2;refGene_gn=CUL9;refGene_tx=XM_019830613.1;refGene_gn=CUL9;refGene_tx=XM_019830614.2;refGene_gn=CUL9;refGene_tx=XM_019830615.2;refGene_gn=CUL9;refGene_tx=XM_019830616.1;refGene_gn=CUL9;refGene_tx=XM_019830617.2;refGene_gn=CUL9;refGene_tx=XM_019830618.2
B2 44005147 44005293
refGene_gn=CUL9;refGene_tx=XM_006931735.4;refGene_gn=CUL9;refGene_tx=XM_019830608.2;refGene_gn=CUL9;refGene_tx=XM_019830609.2;refGene_gn=CUL9;refGene_tx=XM_019830610.2;refGene_gn=CUL9;refGene_tx=XM_019830611.2;refGene_gn=CUL9;refGene_tx=XM_019830612.2;refGene_gn=CUL9;refGene_tx=XM_019830613.1;refGene_gn=CUL9;refGene_tx=XM_019830614.2;refGene_gn=CUL9;refGene_tx=XM_019830615.2;refGene_gn=CUL9;refGene_tx=XM_019830616.1;refGene_gn=CUL9;refGene_tx=XM_019830617.2;refGene_gn=CUL9;refGene_tx=XM_019830618.2
B2 44005424 44005554
refGene_gn=CUL9;refGene_tx=XM_006931735.4;refGene_gn=CUL9;refGene_tx=XM_019830608.2;refGene_gn=CUL9;refGene_tx=XM_019830609.2;refGene_gn=CUL9;refGene_tx=XM_019830610.2;refGene_gn=CUL9;refGene_tx=XM_019830611.2;refGene_gn=CUL9;refGene_tx=XM_019830612.2;refGene_gn=CUL9;refGene_tx=XM_019830613.1;refGene_gn=CUL9;refGene_tx=XM_019830614.2;refGene_gn=CUL9;refGene_tx=XM_019830615.2;refGene_gn=CUL9;refGene_tx=XM_019830616.1;refGene_gn=CUL9;refGene_tx=XM_019830617.2;refGene_gn=CUL9;refGene_tx=XM_019830618.2
B2 44005695 44005895
refGene_gn=CUL9;refGene_tx=XM_006931735.4;refGene_gn=CUL9;refGene_tx=XM_019830608.2;refGene_gn=CUL9;refGene_tx=XM_019830609.2;refGene_gn=CUL9;refGene_tx=XM_019830610.2;refGene_gn=CUL9;refGene_tx=XM_019830611.2;refGene_gn=CUL9;refGene_tx=XM_019830612.2;refGene_gn=CUL9;refGene_tx=XM_019830613.1;refGene_gn=CUL9;refGene_tx=XM_019830614.2;refGene_gn=CUL

9;refGene_tx=XM_019830615.2;refGene_gn=CUL9;refGene_tx=XM_019830616.1;
refGene_gn=CUL9;refGene_tx=XM_019830617.2;refGene_gn=CUL9;refGene_tx=X
M_019830618.2
B2 44006370 44006621
refGene_gn=CUL9;refGene_tx=XM_006931735.4;refGene_gn=CUL9;refGene_tx=X
M_019830608.2;refGene_gn=CUL9;refGene_tx=XM_019830609.2;refGene_gn=CUL
9;refGene_tx=XM_019830610.2;refGene_gn=CUL9;refGene_tx=XM_019830611.2;
refGene_gn=CUL9;refGene_tx=XM_019830612.2;refGene_gn=CUL9;refGene_tx=X
M_019830613.1;refGene_gn=CUL9;refGene_tx=XM_019830614.2;refGene_gn=CUL
9;refGene_tx=XM_019830615.2;refGene_gn=CUL9;refGene_tx=XM_019830616.1;
refGene_gn=CUL9;refGene_tx=XM_019830617.2;refGene_gn=CUL9;refGene_tx=X
M_019830618.2
B2 44007652 44007998
refGene_gn=CUL9;refGene_tx=XM_006931735.4;refGene_gn=CUL9;refGene_tx=X
M_019830608.2;refGene_gn=CUL9;refGene_tx=XM_019830609.2;refGene_gn=CUL
9;refGene_tx=XM_019830610.2;refGene_gn=CUL9;refGene_tx=XM_019830611.2;
refGene_gn=CUL9;refGene_tx=XM_019830612.2;refGene_gn=CUL9;refGene_tx=X
M_019830613.1;refGene_gn=CUL9;refGene_tx=XM_019830614.2;refGene_gn=CUL
9;refGene_tx=XM_019830615.2;refGene_gn=CUL9;refGene_tx=XM_019830616.1;
refGene_gn=CUL9;refGene_tx=XM_019830617.2;refGene_gn=CUL9;refGene_tx=X
M_019830618.2
B2 44011370 44011511
refGene_gn=CUL9;refGene_tx=XM_006931735.4;refGene_gn=CUL9;refGene_tx=X
M_019830608.2;refGene_gn=CUL9;refGene_tx=XM_019830609.2;refGene_gn=CUL
9;refGene_tx=XM_019830610.2;refGene_gn=CUL9;refGene_tx=XM_019830611.2;
refGene_gn=CUL9;refGene_tx=XM_019830612.2;refGene_gn=CUL9;refGene_tx=X
M_019830613.1;refGene_gn=CUL9;refGene_tx=XM_019830614.2;refGene_gn=CUL
9;refGene_tx=XM_019830615.2;refGene_gn=CUL9;refGene_tx=XM_019830616.1;
refGene_gn=CUL9;refGene_tx=XM_019830617.2;refGene_gn=CUL9;refGene_tx=X
M_019830618.2
B2 44011672 44011837
refGene_gn=CUL9;refGene_tx=XM_006931735.4;refGene_gn=CUL9;refGene_tx=X
M_019830608.2;refGene_gn=CUL9;refGene_tx=XM_019830609.2;refGene_gn=CUL
9;refGene_tx=XM_019830610.2;refGene_gn=CUL9;refGene_tx=XM_019830611.2;
refGene_gn=CUL9;refGene_tx=XM_019830612.2;refGene_gn=CUL9;refGene_tx=X
M_019830613.1;refGene_gn=CUL9;refGene_tx=XM_019830614.2;refGene_gn=CUL
9;refGene_tx=XM_019830615.2;refGene_gn=CUL9;refGene_tx=XM_019830616.1;
refGene_gn=CUL9;refGene_tx=XM_019830618.2
B2 44012062 44012226
refGene_gn=CUL9;refGene_tx=XM_006931735.4;refGene_gn=CUL9;refGene_tx=X
M_019830608.2;refGene_gn=CUL9;refGene_tx=XM_019830609.2;refGene_gn=CUL
9;refGene_tx=XM_019830610.2;refGene_gn=CUL9;refGene_tx=XM_019830611.2;
refGene_gn=CUL9;refGene_tx=XM_019830612.2;refGene_gn=CUL9;refGene_tx=X
M_019830613.1;refGene_gn=CUL9;refGene_tx=XM_019830614.2;refGene_gn=CUL
9;refGene_tx=XM_019830615.2;refGene_gn=CUL9;refGene_tx=XM_019830616.1;
refGene_gn=CUL9;refGene_tx=XM_019830617.2;refGene_gn=CUL9;refGene_tx=X
M_019830618.2
B2 44012992 44013086
refGene_gn=CUL9;refGene_tx=XM_006931735.4;refGene_gn=CUL9;refGene_tx=X
M_019830608.2;refGene_gn=CUL9;refGene_tx=XM_019830609.2;refGene_gn=CUL

9;refGene_tx=XM_019830610.2;refGene_gn=CUL9;refGene_tx=XM_019830611.2;refGene_gn=CUL9;refGene_tx=XM_019830612.2;refGene_gn=CUL9;refGene_tx=XM_019830613.1;refGene_gn=CUL9;refGene_tx=XM_019830614.2;refGene_gn=CUL9;refGene_tx=XM_019830615.2;refGene_gn=CUL9;refGene_tx=XM_019830616.1;refGene_gn=CUL9;refGene_tx=XM_019830617.2
B2 44013503 44013578
refGene_gn=CUL9;refGene_tx=XM_006931735.4;refGene_gn=CUL9;refGene_tx=XM_019830608.2;refGene_gn=CUL9;refGene_tx=XM_019830609.2;refGene_gn=CUL9;refGene_tx=XM_019830610.2;refGene_gn=CUL9;refGene_tx=XM_019830612.2;refGene_gn=CUL9;refGene_tx=XM_019830613.1;refGene_gn=CUL9;refGene_tx=XM_019830614.2;refGene_gn=CUL9;refGene_tx=XM_019830615.2;refGene_gn=CUL9;refGene_tx=XM_019830616.1;refGene_gn=CUL9;refGene_tx=XM_019830617.2;refGene_gn=CUL9;refGene_tx=XM_019830618.2
B2 44013732 44013843
refGene_gn=CUL9;refGene_tx=XM_006931735.4;refGene_gn=CUL9;refGene_tx=XM_019830608.2;refGene_gn=CUL9;refGene_tx=XM_019830609.2;refGene_gn=CUL9;refGene_tx=XM_019830610.2;refGene_gn=CUL9;refGene_tx=XM_019830611.2;refGene_gn=CUL9;refGene_tx=XM_019830612.2;refGene_gn=CUL9;refGene_tx=XM_019830613.1;refGene_gn=CUL9;refGene_tx=XM_019830614.2;refGene_gn=CUL9;refGene_tx=XM_019830615.2;refGene_gn=CUL9;refGene_tx=XM_019830616.1;refGene_gn=CUL9;refGene_tx=XM_019830617.2
B2 44014000 44014118
refGene_gn=CUL9;refGene_tx=XM_006931735.4;refGene_gn=CUL9;refGene_tx=XM_019830608.2;refGene_gn=CUL9;refGene_tx=XM_019830609.2;refGene_gn=CUL9;refGene_tx=XM_019830610.2;refGene_gn=CUL9;refGene_tx=XM_019830611.2;refGene_gn=CUL9;refGene_tx=XM_019830612.2;refGene_gn=CUL9;refGene_tx=XM_019830613.1;refGene_gn=CUL9;refGene_tx=XM_019830614.2;refGene_gn=CUL9;refGene_tx=XM_019830615.2;refGene_gn=CUL9;refGene_tx=XM_019830616.1;refGene_gn=CUL9;refGene_tx=XM_019830617.2
B2 44014483 44014824
refGene_gn=CUL9;refGene_tx=XM_006931735.4;refGene_gn=CUL9;refGene_tx=XM_019830608.2;refGene_gn=CUL9;refGene_tx=XM_019830609.2;refGene_gn=CUL9;refGene_tx=XM_019830610.2;refGene_gn=CUL9;refGene_tx=XM_019830611.2;refGene_gn=CUL9;refGene_tx=XM_019830612.2;refGene_gn=CUL9;refGene_tx=XM_019830613.1;refGene_gn=CUL9;refGene_tx=XM_019830614.2;refGene_gn=CUL9;refGene_tx=XM_019830615.2;refGene_gn=CUL9;refGene_tx=XM_019830616.1;refGene_gn=CUL9;refGene_tx=XM_019830617.2
B2 44015044 44015184
refGene_gn=CUL9;refGene_tx=XM_019830608.2;refGene_gn=CUL9;refGene_tx=XM_019830609.2;refGene_gn=CUL9;refGene_tx=XM_019830611.2;refGene_gn=CUL9;refGene_tx=XM_019830613.1;refGene_gn=CUL9;refGene_tx=XM_019830614.2;refGene_gn=CUL9;refGene_tx=XM_019830615.2;refGene_gn=CUL9;refGene_tx=XM_019830617.2
B2 44015359 44015433
refGene_gn=CUL9;refGene_tx=XM_006931735.4;refGene_gn=CUL9;refGene_tx=XM_019830608.2;refGene_gn=CUL9;refGene_tx=XM_019830609.2;refGene_gn=CUL9;refGene_tx=XM_019830610.2;refGene_gn=CUL9;refGene_tx=XM_019830611.2;refGene_gn=CUL9;refGene_tx=XM_019830613.1;refGene_gn=CUL9;refGene_tx=XM_019830614.2;refGene_gn=CUL9;refGene_tx=XM_019830615.2;refGene_gn=CUL9;refGene_tx=XM_019830616.1;refGene_gn=CUL9;refGene_tx=XM_019830617.2

B2 44015518 44015720
refGene_gn=CUL9;refGene_tx=XM_006931735.4;refGene_gn=CUL9;refGene_tx=XM_019830608.2;refGene_gn=CUL9;refGene_tx=XM_019830609.2;refGene_gn=CUL9;refGene_tx=XM_019830610.2;refGene_gn=CUL9;refGene_tx=XM_019830611.2;refGene_gn=CUL9;refGene_tx=XM_019830613.1;refGene_gn=CUL9;refGene_tx=XM_019830615.2;refGene_gn=CUL9;refGene_tx=XM_019830616.1;refGene_gn=CUL9;refGene_tx=XM_019830617.2
B2 44016421 44016558 refGene_gn=DNPH1;refGene_tx=XM_003986161.4
B2 44016738 44016849 refGene_gn=DNPH1;refGene_tx=XM_003986161.4
B2 44017021 44017090 refGene_gn=DNPH1;refGene_tx=XM_003986161.4
B2 44018599 44018792 refGene_gn=DNPH1;refGene_tx=XM_003986161.4
B2 44037851 44037959
refGene_gn=TTBK1;refGene_tx=XM_023253991.1;refGene_gn=TTBK1;refGene_tx=XM_023253992.1
B2 44042562 44042710
refGene_gn=TTBK1;refGene_tx=XM_023253991.1;refGene_gn=TTBK1;refGene_tx=XM_023253992.1
B2 44042948 44043022
refGene_gn=TTBK1;refGene_tx=XM_023253991.1;refGene_gn=TTBK1;refGene_tx=XM_023253992.1
B2 44043191 44043332
refGene_gn=TTBK1;refGene_tx=XM_023253991.1;refGene_gn=TTBK1;refGene_tx=XM_023253992.1
B2 44044539 44044644
refGene_gn=TTBK1;refGene_tx=XM_023253991.1;refGene_gn=TTBK1;refGene_tx=XM_023253992.1
B2 44045065 44045131
refGene_gn=TTBK1;refGene_tx=XM_023253991.1;refGene_gn=TTBK1;refGene_tx=XM_023253992.1
B2 44045543 44045636
refGene_gn=TTBK1;refGene_tx=XM_023253991.1;refGene_gn=TTBK1;refGene_tx=XM_023253992.1
B2 44045721 44045847
refGene_gn=TTBK1;refGene_tx=XM_023253991.1;refGene_gn=TTBK1;refGene_tx=XM_023253992.1
B2 44047457 44047612
refGene_gn=TTBK1;refGene_tx=XM_023253991.1;refGene_gn=TTBK1;refGene_tx=XM_023253992.1
B2 44048540 44048772
refGene_gn=TTBK1;refGene_tx=XM_023253991.1;refGene_gn=TTBK1;refGene_tx=XM_023253992.1
B2 44049024 44049203
refGene_gn=TTBK1;refGene_tx=XM_023253991.1;refGene_gn=TTBK1;refGene_tx=XM_023253992.1
B2 44051912 44052474
refGene_gn=TTBK1;refGene_tx=XM_023253991.1;refGene_gn=TTBK1;refGene_tx=XM_023253992.1
B2 44069697 44071289
refGene_gn=TTBK1;refGene_tx=XM_023253991.1;refGene_gn=TTBK1;refGene_tx=XM_023253992.1

B2 44071957 44072348
refGene_gn=TTBK1;refGene_tx=XM_023253991.1;refGene_gn=TTBK1;refGene_tx=XM_023253992.1
B2 44084342 44084735 refGene_gn=SLC22A7;refGene_tx=XM_003986162.4
B2 44085378 44085482 refGene_gn=SLC22A7;refGene_tx=XM_003986162.4
B2 44085615 44085770 refGene_gn=SLC22A7;refGene_tx=XM_003986162.4
B2 44085894 44086063 refGene_gn=SLC22A7;refGene_tx=XM_003986162.4
B2 44086865 44086989 refGene_gn=SLC22A7;refGene_tx=XM_003986162.4
B2 44087353 44087463 refGene_gn=SLC22A7;refGene_tx=XM_003986162.4
B2 44087927 44088142 refGene_gn=SLC22A7;refGene_tx=XM_003986162.4
B2 44088315 44088424 refGene_gn=SLC22A7;refGene_tx=XM_003986162.4
B2 44089126 44089330 refGene_gn=SLC22A7;refGene_tx=XM_003986162.4
B2 44089758 44089813 refGene_gn=SLC22A7;refGene_tx=XM_003986162.4
B2 44090880 44090942
refGene_gn=CRIP3;refGene_tx=XM_006931739.4;refGene_gn=CRIP3;refGene_tx=XM_019830621.2
B2 44091116 44091174
refGene_gn=CRIP3;refGene_tx=XM_006931739.4;refGene_gn=CRIP3;refGene_tx=XM_019830621.2
B2 44091263 44091367
refGene_gn=CRIP3;refGene_tx=XM_006931739.4;refGene_gn=CRIP3;refGene_tx=XM_019830621.2
B2 44091494 44091566
refGene_gn=CRIP3;refGene_tx=XM_006931739.4;refGene_gn=CRIP3;refGene_tx=XM_019830621.2
B2 44092349 44092481
refGene_gn=CRIP3;refGene_tx=XM_006931739.4;refGene_gn=CRIP3;refGene_tx=XM_019830621.2
B2 44092577 44092635
refGene_gn=CRIP3;refGene_tx=XM_006931739.4;refGene_gn=CRIP3;refGene_tx=XM_019830621.2
B2 44093064 44093159
refGene_gn=CRIP3;refGene_tx=XM_006931739.4;refGene_gn=CRIP3;refGene_tx=XM_019830621.2
B2 44093410 44093453
refGene_gn=CRIP3;refGene_tx=XM_006931739.4;refGene_gn=CRIP3;refGene_tx=XM_019830621.2
B2 44111700 44111823
refGene_gn=ZNF318;refGene_tx=XM_023253994.1;refGene_gn=ZNF318;refGene_tx=XM_023253995.1;refGene_gn=ZNF318;refGene_tx=XM_023253996.1
B2 44117750 44121059 refGene_gn=ZNF318;refGene_tx=XM_023253993.1
B2 44121349 44121468 refGene_gn=ZNF318;refGene_tx=XM_023253993.1
B2 44122423 44122523 refGene_gn=ZNF318;refGene_tx=XM_023253993.1
B2 44122789 44122873 refGene_gn=ZNF318;refGene_tx=XM_023253997.1
B2 44122982 44123186
refGene_gn=ZNF318;refGene_tx=XM_023253993.1;refGene_gn=ZNF318;refGene_tx=XM_023253994.1;refGene_gn=ZNF318;refGene_tx=XM_023253995.1;refGene_gn=ZNF318;refGene_tx=XM_023253996.1;refGene_gn=ZNF318;refGene_tx=XM_023253997.1
B2 44127811 44128116

refGene_gn=ZNF318;refGene_tx=XM_023253993.1;refGene_gn=ZNF318;refGene_tx=XM_023253994.1;refGene_gn=ZNF318;refGene_tx=XM_023253995.1;refGene_gn=ZNF318;refGene_tx=XM_023253996.1;refGene_gn=ZNF318;refGene_tx=XM_023253997.1
B2 44130011 44130111
refGene_gn=ZNF318;refGene_tx=XM_023253993.1;refGene_gn=ZNF318;refGene_tx=XM_023253994.1;refGene_gn=ZNF318;refGene_tx=XM_023253995.1;refGene_gn=ZNF318;refGene_tx=XM_023253996.1;refGene_gn=ZNF318;refGene_tx=XM_023253997.1
B2 44131784 44133227
refGene_gn=ZNF318;refGene_tx=XM_023253993.1;refGene_gn=ZNF318;refGene_tx=XM_023253994.1;refGene_gn=ZNF318;refGene_tx=XM_023253995.1;refGene_gn=ZNF318;refGene_tx=XM_023253996.1;refGene_gn=ZNF318;refGene_tx=XM_023253997.1
B2 44134194 44134834
refGene_gn=ZNF318;refGene_tx=XM_023253993.1;refGene_gn=ZNF318;refGene_tx=XM_023253994.1;refGene_gn=ZNF318;refGene_tx=XM_023253995.1;refGene_gn=ZNF318;refGene_tx=XM_023253996.1;refGene_gn=ZNF318;refGene_tx=XM_023253997.1
B2 44140840 44140992
refGene_gn=ZNF318;refGene_tx=XM_023253993.1;refGene_gn=ZNF318;refGene_tx=XM_023253994.1;refGene_gn=ZNF318;refGene_tx=XM_023253995.1;refGene_gn=ZNF318;refGene_tx=XM_023253996.1;refGene_gn=ZNF318;refGene_tx=XM_023253997.1
B2 44143519 44143975
refGene_gn=ZNF318;refGene_tx=XM_023253993.1;refGene_gn=ZNF318;refGene_tx=XM_023253994.1;refGene_gn=ZNF318;refGene_tx=XM_023253995.1;refGene_gn=ZNF318;refGene_tx=XM_023253996.1;refGene_gn=ZNF318;refGene_tx=XM_023253997.1
B2 44202145 44202306 refGene_gn=ABCC10;refGene_tx=XM_006931742.4
B2 44204963 44205304 refGene_gn=ABCC10;refGene_tx=XM_006931742.4
B2 44205306 44206181 refGene_gn=ABCC10;refGene_tx=XM_006931742.4
B2 44207508 44207736 refGene_gn=ABCC10;refGene_tx=XM_006931742.4
B2 44208361 44208518 refGene_gn=ABCC10;refGene_tx=XM_006931742.4
B2 44208801 44208911 refGene_gn=ABCC10;refGene_tx=XM_006931742.4
B2 44210534 44210614 refGene_gn=ABCC10;refGene_tx=XM_006931742.4
B2 44211265 44211437 refGene_gn=ABCC10;refGene_tx=XM_006931742.4
B2 44212818 44212917 refGene_gn=ABCC10;refGene_tx=XM_006931742.4
B2 44213313 44213503 refGene_gn=ABCC10;refGene_tx=XM_006931742.4
B2 44214275 44214353 refGene_gn=ABCC10;refGene_tx=XM_006931742.4
B2 44214512 44214707 refGene_gn=ABCC10;refGene_tx=XM_006931742.4
B2 44215108 44215262 refGene_gn=ABCC10;refGene_tx=XM_006931742.4
B2 44215447 44215637 refGene_gn=ABCC10;refGene_tx=XM_006931742.4
B2 44217598 44217942 refGene_gn=ABCC10;refGene_tx=XM_006931742.4
B2 44218229 44218399 refGene_gn=ABCC10;refGene_tx=XM_006931742.4
B2 44218888 44219049 refGene_gn=ABCC10;refGene_tx=XM_006931742.4
B2 44219379 44219633 refGene_gn=ABCC10;refGene_tx=XM_006931742.4
B2 44220801 44220947 refGene_gn=ABCC10;refGene_tx=XM_006931742.4
B2 44221042 44221140 refGene_gn=ABCC10;refGene_tx=XM_006931742.4
B2 44221378 44221491 refGene_gn=ABCC10;refGene_tx=XM_006931742.4

B2	44221794 44221948 refGene_gn=ABCC10;refGene_tx=XM_006931742.4
B2	44222392 44223170
refGene_gn=DLK2;refGene_tx=XM_006931740.4;refGene_gn=DLK2;refGene_tx=XM_023253998.1;refGene_gn=DLK2;refGene_tx=XM_023253999.1
B2	44223927 44224072
refGene_gn=DLK2;refGene_tx=XM_006931740.4;refGene_gn=DLK2;refGene_tx=XM_023253998.1;refGene_gn=DLK2;refGene_tx=XM_023253999.1
B2	44224523 44224654
refGene_gn=DLK2;refGene_tx=XM_006931740.4;refGene_gn=DLK2;refGene_tx=XM_023253998.1;refGene_gn=DLK2;refGene_tx=XM_023253999.1
B2	44225643 44225707
refGene_gn=DLK2;refGene_tx=XM_006931740.4;refGene_gn=DLK2;refGene_tx=XM_023253998.1;refGene_gn=DLK2;refGene_tx=XM_023253999.1
B2	44225976 44226052
refGene_gn=DLK2;refGene_tx=XM_006931740.4;refGene_gn=DLK2;refGene_tx=XM_023253998.1;refGene_gn=DLK2;refGene_tx=XM_023253999.1
B2	44260673 44260772
refGene_gn=TJAP1;refGene_tx=XM_019830627.2;refGene_gn=TJAP1;refGene_tx=XM_023254001.1;refGene_gn=TJAP1;refGene_tx=XM_023254002.1;refGene_gn=TJAP1;refGene_tx=XM_023254003.1;refGene_gn=TJAP1;refGene_tx=XM_023254004.1;refGene_gn=TJAP1;refGene_tx=XM_023254005.1;refGene_gn=TJAP1;refGene_tx=XM_023254006.1;refGene_gn=TJAP1;refGene_tx=XM_023254007.1;refGene_gn=TJAP1;refGene_tx=XM_023254008.1;refGene_gn=TJAP1;refGene_tx=XM_023254009.1;refGene_gn=TJAP1;refGene_tx=XM_023254010.1;refGene_gn=TJAP1;refGene_tx=XM_023254011.1
B2	44262366 44262395
refGene_gn=TJAP1;refGene_tx=XM_019830627.2;refGene_gn=TJAP1;refGene_tx=XM_023254001.1;refGene_gn=TJAP1;refGene_tx=XM_023254002.1;refGene_gn=TJAP1;refGene_tx=XM_023254003.1;refGene_gn=TJAP1;refGene_tx=XM_023254004.1;refGene_gn=TJAP1;refGene_tx=XM_023254005.1;refGene_gn=TJAP1;refGene_tx=XM_023254006.1;refGene_gn=TJAP1;refGene_tx=XM_023254007.1;refGene_gn=TJAP1;refGene_tx=XM_023254008.1;refGene_gn=TJAP1;refGene_tx=XM_023254009.1;refGene_gn=TJAP1;refGene_tx=XM_023254010.1;refGene_gn=TJAP1;refGene_tx=XM_023254011.1
B2	44263135 44263297
refGene_gn=TJAP1;refGene_tx=XM_019830627.2;refGene_gn=TJAP1;refGene_tx=XM_023254001.1;refGene_gn=TJAP1;refGene_tx=XM_023254002.1;refGene_gn=TJAP1;refGene_tx=XM_023254003.1;refGene_gn=TJAP1;refGene_tx=XM_023254004.1;refGene_gn=TJAP1;refGene_tx=XM_023254005.1;refGene_gn=TJAP1;refGene_tx=XM_023254006.1;refGene_gn=TJAP1;refGene_tx=XM_023254007.1;refGene_gn=TJAP1;refGene_tx=XM_023254008.1;refGene_gn=TJAP1;refGene_tx=XM_023254009.1;refGene_gn=TJAP1;refGene_tx=XM_023254010.1;refGene_gn=TJAP1;refGene_tx=XM_023254011.1
B2	44263511 44263578
refGene_gn=TJAP1;refGene_tx=XM_019830627.2;refGene_gn=TJAP1;refGene_tx=XM_023254001.1;refGene_gn=TJAP1;refGene_tx=XM_023254002.1;refGene_gn=TJAP1;refGene_tx=XM_023254003.1;refGene_gn=TJAP1;refGene_tx=XM_023254004.1;refGene_gn=TJAP1;refGene_tx=XM_023254005.1;refGene_gn=TJAP1;refGene_tx=XM_023254006.1;refGene_gn=TJAP1;refGene_tx=XM_023254007.1;refGene_gn=TJAP1;refGene_tx=XM_023254008.1;refGene_gn=TJAP1;refGene_tx=XM_02

3254009.1;refGene_gn=TJAP1;refGene_tx=XM_023254010.1;refGene_gn=TJAP1;
refGene_tx=XM_023254011.1
B2 44263812 44263842
refGene_gn=TJAP1;refGene_tx=XM_019830627.2;refGene_gn=TJAP1;refGene_tx
=XM_023254001.1;refGene_gn=TJAP1;refGene_tx=XM_023254002.1;refGene_gn=
TJAP1;refGene_tx=XM_023254003.1;refGene_gn=TJAP1;refGene_tx=XM_0232540
04.1;refGene_gn=TJAP1;refGene_tx=XM_023254005.1;refGene_gn=TJAP1;refGe
ne_tx=XM_023254006.1;refGene_gn=TJAP1;refGene_tx=XM_023254007.1;refGen
e_gn=TJAP1;refGene_tx=XM_023254008.1;refGene_gn=TJAP1;refGene_tx=XM_02
3254009.1;refGene_gn=TJAP1;refGene_tx=XM_023254010.1
B2 44264561 44264669
refGene_gn=TJAP1;refGene_tx=XM_019830627.2;refGene_gn=TJAP1;refGene_tx
=XM_023254001.1;refGene_gn=TJAP1;refGene_tx=XM_023254002.1;refGene_gn=
TJAP1;refGene_tx=XM_023254003.1;refGene_gn=TJAP1;refGene_tx=XM_0232540
04.1;refGene_gn=TJAP1;refGene_tx=XM_023254005.1;refGene_gn=TJAP1;refGe
ne_tx=XM_023254006.1;refGene_gn=TJAP1;refGene_tx=XM_023254007.1;refGen
e_gn=TJAP1;refGene_tx=XM_023254008.1;refGene_gn=TJAP1;refGene_tx=XM_02
3254009.1;refGene_gn=TJAP1;refGene_tx=XM_023254010.1;refGene_gn=TJAP1;
refGene_tx=XM_023254011.1
B2 44264770 44264854
refGene_gn=TJAP1;refGene_tx=XM_019830627.2;refGene_gn=TJAP1;refGene_tx
=XM_023254001.1;refGene_gn=TJAP1;refGene_tx=XM_023254002.1;refGene_gn=
TJAP1;refGene_tx=XM_023254003.1;refGene_gn=TJAP1;refGene_tx=XM_0232540
04.1;refGene_gn=TJAP1;refGene_tx=XM_023254005.1;refGene_gn=TJAP1;refGe
ne_tx=XM_023254006.1;refGene_gn=TJAP1;refGene_tx=XM_023254007.1;refGen
e_gn=TJAP1;refGene_tx=XM_023254008.1;refGene_gn=TJAP1;refGene_tx=XM_02
3254009.1;refGene_gn=TJAP1;refGene_tx=XM_023254010.1;refGene_gn=TJAP1;
refGene_tx=XM_023254011.1
B2 44265612 44266710
refGene_gn=TJAP1;refGene_tx=XM_019830627.2;refGene_gn=TJAP1;refGene_tx
=XM_023254001.1;refGene_gn=TJAP1;refGene_tx=XM_023254002.1;refGene_gn=
TJAP1;refGene_tx=XM_023254003.1;refGene_gn=TJAP1;refGene_tx=XM_0232540
04.1;refGene_gn=TJAP1;refGene_tx=XM_023254005.1;refGene_gn=TJAP1;refGe
ne_tx=XM_023254006.1;refGene_gn=TJAP1;refGene_tx=XM_023254007.1;refGen
e_gn=TJAP1;refGene_tx=XM_023254008.1;refGene_gn=TJAP1;refGene_tx=XM_02
3254009.1;refGene_gn=TJAP1;refGene_tx=XM_023254010.1;refGene_gn=TJAP1;
refGene_tx=XM_023254011.1
B2 44268065 44268136 refGene_gn=LRRC73;refGene_tx=XM_019831588.2
B2 44268284 44268507 refGene_gn=LRRC73;refGene_tx=XM_019831588.2
B2 44268654 44268755 refGene_gn=LRRC73;refGene_tx=XM_019831588.2
B2 44269145 44269268 refGene_gn=LRRC73;refGene_tx=XM_019831588.2
B2 44269609 44269770 refGene_gn=LRRC73;refGene_tx=XM_019831588.2
B2 44270398 44270670 refGene_gn=LRRC73;refGene_tx=XM_019831588.2
B2 44272655 44272722
refGene_gn=YIPF3;refGene_tx=XM_006931747.4;refGene_gn=YIPF3;refGene_tx
=XM_006931748.4;refGene_gn=YIPF3;refGene_tx=XM_019830629.2;refGene_gn=
YIPF3;refGene_tx=XM_019830630.2
B2 44273169 44273318
refGene_gn=YIPF3;refGene_tx=XM_003986169.5;refGene_gn=YIPF3;refGene_tx
=XM_006931747.4;refGene_gn=YIPF3;refGene_tx=XM_006931748.4;refGene_gn=

YIPF3;refGene_tx=XM_019830629.2;refGene_gn=YIPF3;refGene_tx=XM_0198306
30.2
B2 44273410 44273534
refGene_gn=YIPF3;refGene_tx=XM_003986169.5;refGene_gn=YIPF3;refGene_tx
=XM_006931747.4;refGene_gn=YIPF3;refGene_tx=XM_006931748.4
B2 44273732 44273846
refGene_gn=YIPF3;refGene_tx=XM_003986169.5;refGene_gn=YIPF3;refGene_tx
=XM_006931747.4;refGene_gn=YIPF3;refGene_tx=XM_006931748.4;refGene_gn=
YIPF3;refGene_tx=XM_019830629.2;refGene_gn=YIPF3;refGene_tx=XM_0198306
30.2
B2 44274041 44274173
refGene_gn=YIPF3;refGene_tx=XM_003986169.5;refGene_gn=YIPF3;refGene_tx
=XM_006931747.4;refGene_gn=YIPF3;refGene_tx=XM_006931748.4;refGene_gn=
YIPF3;refGene_tx=XM_019830629.2;refGene_gn=YIPF3;refGene_tx=XM_0198306
30.2
B2 44274321 44274414
refGene_gn=YIPF3;refGene_tx=XM_003986169.5;refGene_gn=YIPF3;refGene_tx
=XM_006931747.4;refGene_gn=YIPF3;refGene_tx=XM_006931748.4;refGene_gn=
YIPF3;refGene_tx=XM_019830629.2;refGene_gn=YIPF3;refGene_tx=XM_0198306
30.2
B2 44274553 44274599
refGene_gn=YIPF3;refGene_tx=XM_003986169.5;refGene_gn=YIPF3;refGene_tx
=XM_006931747.4;refGene_gn=YIPF3;refGene_tx=XM_006931748.4;refGene_gn=
YIPF3;refGene_tx=XM_019830629.2;refGene_gn=YIPF3;refGene_tx=XM_0198306
30.2
B2 44275929 44276036
refGene_gn=YIPF3;refGene_tx=XM_003986169.5;refGene_gn=YIPF3;refGene_tx
=XM_006931747.4;refGene_gn=YIPF3;refGene_tx=XM_006931748.4;refGene_gn=
YIPF3;refGene_tx=XM_019830629.2;refGene_gn=YIPF3;refGene_tx=XM_0198306
30.2
B2 44276232 44276430
refGene_gn=YIPF3;refGene_tx=XM_003986169.5;refGene_gn=YIPF3;refGene_tx
=XM_006931747.4;refGene_gn=YIPF3;refGene_tx=XM_006931748.4;refGene_gn=
YIPF3;refGene_tx=XM_019830629.2;refGene_gn=YIPF3;refGene_tx=XM_0198306
30.2
B2 44277043 44277124
refGene_gn=YIPF3;refGene_tx=XM_003986169.5;refGene_gn=YIPF3;refGene_tx
=XM_006931747.4;refGene_gn=YIPF3;refGene_tx=XM_006931748.4;refGene_gn=
YIPF3;refGene_tx=XM_019830629.2;refGene_gn=YIPF3;refGene_tx=XM_0198306
30.2
B2 44277415 44277484 refGene_gn=POLR1C;refGene_tx=XM_003986171.5
B2 44277598 44277670 refGene_gn=POLR1C;refGene_tx=XM_003986171.5
B2 44279131 44279239
refGene_gn=POLR1C;refGene_tx=XM_003986171.5;refGene_gn=POLR1C;refGene_
tx=XM_006931749.4
B2 44279475 44279608
refGene_gn=POLR1C;refGene_tx=XM_003986171.5;refGene_gn=POLR1C;refGene_
tx=XM_006931749.4
B2 44279896 44280016
refGene_gn=POLR1C;refGene_tx=XM_003986171.5;refGene_gn=POLR1C;refGene_

tx=XM_006931749.4
B2	44280106	44280259
refGene_gn=POLR1C;refGene_tx=XM_003986171.5;refGene_gn=POLR1C;refGene_tx=XM_006931749.4
B2	44280433	44280583
refGene_gn=POLR1C;refGene_tx=XM_003986171.5;refGene_gn=POLR1C;refGene_tx=XM_006931749.4
B2	44280696	44280813
refGene_gn=POLR1C;refGene_tx=XM_003986171.5;refGene_gn=POLR1C;refGene_tx=XM_006931749.4
B2	44280961	44281080
refGene_gn=POLR1C;refGene_tx=XM_003986171.5;refGene_gn=POLR1C;refGene_tx=XM_006931749.4
B2	44283923	44284061
refGene_gn=XPO5;refGene_tx=XM_006931756.4;refGene_gn=XPO5;refGene_tx=XM_019830631.2
B2	44284224	44284389
refGene_gn=XPO5;refGene_tx=XM_006931756.4;refGene_gn=XPO5;refGene_tx=XM_019830631.2
B2	44284570	44284708
refGene_gn=XPO5;refGene_tx=XM_006931756.4;refGene_gn=XPO5;refGene_tx=XM_019830631.2
B2	44284842	44284950
refGene_gn=XPO5;refGene_tx=XM_006931756.4;refGene_gn=XPO5;refGene_tx=XM_019830631.2
B2	44285411	44285494
refGene_gn=XPO5;refGene_tx=XM_006931756.4;refGene_gn=XPO5;refGene_tx=XM_019830631.2
B2	44286634	44286697
refGene_gn=XPO5;refGene_tx=XM_006931756.4;refGene_gn=XPO5;refGene_tx=XM_019830631.2
B2	44287242	44287340
refGene_gn=XPO5;refGene_tx=XM_006931756.4;refGene_gn=XPO5;refGene_tx=XM_019830631.2
B2	44287790	44287837
refGene_gn=XPO5;refGene_tx=XM_006931756.4;refGene_gn=XPO5;refGene_tx=XM_019830631.2
B2	44288392	44288490
refGene_gn=XPO5;refGene_tx=XM_006931756.4;refGene_gn=XPO5;refGene_tx=XM_019830631.2
B2	44289138	44289275
refGene_gn=XPO5;refGene_tx=XM_006931756.4;refGene_gn=XPO5;refGene_tx=XM_019830631.2
B2	44290974	44291071
refGene_gn=XPO5;refGene_tx=XM_006931756.4;refGene_gn=XPO5;refGene_tx=XM_019830631.2
B2	44292844	44292945
refGene_gn=XPO5;refGene_tx=XM_006931756.4;refGene_gn=XPO5;refGene_tx=XM_019830631.2
B2	44302469	44302651

refGene_gn=XPO5;refGene_tx=XM_006931756.4;refGene_gn=XPO5;refGene_tx=XM_019830631.2
B2 44303520 44303620
refGene_gn=XPO5;refGene_tx=XM_006931756.4;refGene_gn=XPO5;refGene_tx=XM_019830631.2
B2 44304417 44304617
refGene_gn=XPO5;refGene_tx=XM_006931756.4;refGene_gn=XPO5;refGene_tx=XM_019830631.2
B2 44305357 44305447
refGene_gn=XPO5;refGene_tx=XM_006931756.4;refGene_gn=XPO5;refGene_tx=XM_019830631.2
B2 44305770 44305812
refGene_gn=XPO5;refGene_tx=XM_006931756.4;refGene_gn=XPO5;refGene_tx=XM_019830631.2
B2 44306977 44307133
refGene_gn=XPO5;refGene_tx=XM_006931756.4;refGene_gn=XPO5;refGene_tx=XM_019830631.2
B2 44310618 44310737
refGene_gn=XPO5;refGene_tx=XM_006931756.4;refGene_gn=XPO5;refGene_tx=XM_019830631.2
B2 44311988 44312117
refGene_gn=XPO5;refGene_tx=XM_006931756.4;refGene_gn=XPO5;refGene_tx=XM_019830631.2
B2 44313514 44313605
refGene_gn=XPO5;refGene_tx=XM_006931756.4;refGene_gn=XPO5;refGene_tx=XM_019830631.2
B2 44314581 44314707
refGene_gn=XPO5;refGene_tx=XM_006931756.4;refGene_gn=XPO5;refGene_tx=XM_019830631.2
B2 44315442 44315526
refGene_gn=XPO5;refGene_tx=XM_006931756.4;refGene_gn=XPO5;refGene_tx=XM_019830631.2
B2 44316713 44316813
refGene_gn=XPO5;refGene_tx=XM_006931756.4;refGene_gn=XPO5;refGene_tx=XM_019830631.2
B2 44320375 44320452
refGene_gn=XPO5;refGene_tx=XM_006931756.4;refGene_gn=XPO5;refGene_tx=XM_019830631.2
B2 44321448 44321634
refGene_gn=XPO5;refGene_tx=XM_006931756.4;refGene_gn=XPO5;refGene_tx=XM_019830631.2
B2 44322026 44322053 refGene_gn=XPO5;refGene_tx=XM_006931756.4
B2 44325539 44325722
refGene_gn=XPO5;refGene_tx=XM_006931756.4;refGene_gn=XPO5;refGene_tx=XM_019830631.2
B2 44325866 44326004
refGene_gn=XPO5;refGene_tx=XM_006931756.4;refGene_gn=XPO5;refGene_tx=XM_019830631.2
B2 44327782 44327855
refGene_gn=XPO5;refGene_tx=XM_006931756.4;refGene_gn=XPO5;refGene_tx=X

M_019830631.2
B2	44328786	44328908
refGene_gn=XPO5;refGene_tx=XM_006931756.4;refGene_gn=XPO5;refGene_tx=XM_019830631.2
B2	44330906	44331011
refGene_gn=XPO5;refGene_tx=XM_006931756.4;refGene_gn=XPO5;refGene_tx=XM_019830631.2
B2	44338359	44338496	refGene_gn=POLH;refGene_tx=XM_019830633.2
B2	44338728	44338863
refGene_gn=POLH;refGene_tx=XM_011282210.2;refGene_gn=POLH;refGene_tx=XM_019830633.2
B2	44344438	44344656	refGene_gn=POLH;refGene_tx=XM_019830633.2
B2	44356517	44356646	refGene_gn=GTPBP2;refGene_tx=XM_023254013.1
B2	44358131	44358301
refGene_gn=POLH;refGene_tx=XM_011282210.2;refGene_gn=POLH;refGene_tx=XM_019830633.2
B2	44360307	44360411
refGene_gn=POLH;refGene_tx=XM_011282210.2;refGene_gn=POLH;refGene_tx=XM_019830633.2
B2	44361581	44361701
refGene_gn=POLH;refGene_tx=XM_011282210.2;refGene_gn=POLH;refGene_tx=XM_019830633.2
B2	44362459	44362583
refGene_gn=POLH;refGene_tx=XM_011282210.2;refGene_gn=POLH;refGene_tx=XM_019830633.2
B2	44363119	44363185
refGene_gn=POLH;refGene_tx=XM_011282210.2;refGene_gn=POLH;refGene_tx=XM_019830633.2
B2	44366540	44366710
refGene_gn=POLH;refGene_tx=XM_011282210.2;refGene_gn=POLH;refGene_tx=XM_019830633.2
B2	44368423	44369318
refGene_gn=POLH;refGene_tx=XM_011282210.2;refGene_gn=POLH;refGene_tx=XM_019830633.2
B2	44373614	44373791
refGene_gn=GTPBP2;refGene_tx=XM_003986174.5;refGene_gn=GTPBP2;refGene_tx=XM_023254014.1
B2	44374026	44374191
refGene_gn=GTPBP2;refGene_tx=XM_003986174.5;refGene_gn=GTPBP2;refGene_tx=XM_023254013.1;refGene_gn=GTPBP2;refGene_tx=XM_023254014.1
B2	44374618	44374790
refGene_gn=GTPBP2;refGene_tx=XM_003986174.5;refGene_gn=GTPBP2;refGene_tx=XM_023254013.1;refGene_gn=GTPBP2;refGene_tx=XM_023254014.1
B2	44375401	44375460
refGene_gn=GTPBP2;refGene_tx=XM_003986174.5;refGene_gn=GTPBP2;refGene_tx=XM_023254013.1;refGene_gn=GTPBP2;refGene_tx=XM_023254014.1
B2	44375793	44375929
refGene_gn=GTPBP2;refGene_tx=XM_003986174.5;refGene_gn=GTPBP2;refGene_tx=XM_023254013.1;refGene_gn=GTPBP2;refGene_tx=XM_023254014.1
B2	44376348	44376568

refGene_gn=GTPBP2;refGene_tx=XM_003986174.5;refGene_gn=GTPBP2;refGene_tx=XM_023254013.1;refGene_gn=GTPBP2;refGene_tx=XM_023254014.1
B2 44376727 44376902
refGene_gn=GTPBP2;refGene_tx=XM_003986174.5;refGene_gn=GTPBP2;refGene_tx=XM_023254013.1;refGene_gn=GTPBP2;refGene_tx=XM_023254014.1
B2 44377200 44377398
refGene_gn=GTPBP2;refGene_tx=XM_003986174.5;refGene_gn=GTPBP2;refGene_tx=XM_023254013.1
B2 44377594 44377703
refGene_gn=GTPBP2;refGene_tx=XM_003986174.5;refGene_gn=GTPBP2;refGene_tx=XM_023254013.1;refGene_gn=GTPBP2;refGene_tx=XM_023254014.1
B2 44378056 44378241
refGene_gn=GTPBP2;refGene_tx=XM_003986174.5;refGene_gn=GTPBP2;refGene_tx=XM_023254013.1;refGene_gn=GTPBP2;refGene_tx=XM_023254014.1
B2 44378525 44378552
refGene_gn=GTPBP2;refGene_tx=XM_003986174.5;refGene_gn=GTPBP2;refGene_tx=XM_023254013.1;refGene_gn=GTPBP2;refGene_tx=XM_023254014.1
B2 44380318 44380504
refGene_gn=GTPBP2;refGene_tx=XM_003986174.5;refGene_gn=GTPBP2;refGene_tx=XM_023254013.1;refGene_gn=GTPBP2;refGene_tx=XM_023254014.1
B2 44383702 44383733
refGene_gn=MAD2L1BP;refGene_tx=XM_019830641.2;refGene_gn=MAD2L1BP;refGene_tx=XM_019830642.2
B2 44384287 44384333 refGene_gn=MAD2L1BP;refGene_tx=XM_011282211.3
B2 44384769 44385102
refGene_gn=MAD2L1BP;refGene_tx=XM_006931753.3;refGene_gn=MAD2L1BP;refGene_tx=XM_011282211.3;refGene_gn=MAD2L1BP;refGene_tx=XM_019830638.2;refGene_gn=MAD2L1BP;refGene_tx=XM_019830640.2;refGene_gn=MAD2L1BP;refGene_tx=XM_019830641.2;refGene_gn=MAD2L1BP;refGene_tx=XM_019830642.2;refGene_gn=MAD2L1BP;refGene_tx=XM_019830643.2
B2 44388644 44389157
refGene_gn=MAD2L1BP;refGene_tx=XM_006931753.3;refGene_gn=MAD2L1BP;refGene_tx=XM_011282211.3;refGene_gn=MAD2L1BP;refGene_tx=XM_019830638.2;refGene_gn=MAD2L1BP;refGene_tx=XM_019830640.2;refGene_gn=MAD2L1BP;refGene_tx=XM_019830641.2;refGene_gn=MAD2L1BP;refGene_tx=XM_019830642.2;refGene_gn=MAD2L1BP;refGene_tx=XM_019830643.2
B2 44392205 44392432
refGene_gn=RSPH9;refGene_tx=XM_003986176.4;refGene_gn=RSPH9;refGene_tx=XM_006931751.3
B2 44393244 44393345 refGene_gn=RSPH9;refGene_tx=XM_019830636.2
B2 44395749 44395915
refGene_gn=RSPH9;refGene_tx=XM_003986176.4;refGene_gn=RSPH9;refGene_tx=XM_006931751.3;refGene_gn=RSPH9;refGene_tx=XM_019830636.2
B2 44398106 44398236
refGene_gn=RSPH9;refGene_tx=XM_003986176.4;refGene_gn=RSPH9;refGene_tx=XM_006931751.3;refGene_gn=RSPH9;refGene_tx=XM_019830636.2
B2 44398694 44398841
refGene_gn=RSPH9;refGene_tx=XM_003986176.4;refGene_gn=RSPH9;refGene_tx=XM_006931751.3;refGene_gn=RSPH9;refGene_tx=XM_019830636.2
B2 44399067 44399114 refGene_gn=RSPH9;refGene_tx=XM_006931751.3

B2 44404448 44404609
refGene_gn=RSPH9;refGene_tx=XM_003986176.4;refGene_gn=RSPH9;refGene_tx=XM_019830636.2
B2 44405323 44405468 refGene_gn=MRPS18A;refGene_tx=XM_003986177.3
B2 44408388 44408519
refGene_gn=MRPS18A;refGene_tx=XM_003986177.3;refGene_gn=MRPS18A;refGene_tx=XM_006931750.2
B2 44408697 44408821
refGene_gn=MRPS18A;refGene_tx=XM_003986177.3;refGene_gn=MRPS18A;refGene_tx=XM_006931750.2
B2 44409392 44409639 refGene_gn=MRPS18A;refGene_tx=XM_019830644.1
B2 44410105 44410167 refGene_gn=MRPS18A;refGene_tx=XM_019830644.1
B2 44411495 44411603
refGene_gn=MRPS18A;refGene_tx=XM_003986177.3;refGene_gn=MRPS18A;refGene_tx=XM_006931750.2;refGene_gn=MRPS18A;refGene_tx=XM_019830644.1
B2 44413992 44414024
refGene_gn=MRPS18A;refGene_tx=XM_003986177.3;refGene_gn=MRPS18A;refGene_tx=XM_006931750.2;refGene_gn=MRPS18A;refGene_tx=XM_019830644.1
B2 44421626 44421738
refGene_gn=MRPS18A;refGene_tx=XM_003986177.3;refGene_gn=MRPS18A;refGene_tx=XM_006931750.2;refGene_gn=MRPS18A;refGene_tx=XM_019830644.1
B2 44506725 44506779 refGene_gn=VEGFA;refGene_tx=XM_023253550.1
B2 44507291 44508068
refGene_gn=VEGFA;refGene_tx=NM_001009854.1;refGene_gn=VEGFA;refGene_tx=XM_023253550.1
B2 44511159 44511208
refGene_gn=VEGFA;refGene_tx=NM_001009854.1;refGene_gn=VEGFA;refGene_tx=XM_023253550.1
B2 44514413 44514610
refGene_gn=VEGFA;refGene_tx=NM_001009854.1;refGene_gn=VEGFA;refGene_tx=XM_023253550.1
B2 44515354 44515431
refGene_gn=VEGFA;refGene_tx=NM_001009854.1;refGene_gn=VEGFA;refGene_tx=XM_023253550.1
B2 44515757 44515784
refGene_gn=VEGFA;refGene_tx=NM_001009854.1;refGene_gn=VEGFA;refGene_tx=XM_023253550.1
B2 44517682 44517754 refGene_gn=VEGFA;refGene_tx=XM_023253550.1
B2 44518878 44519010
refGene_gn=VEGFA;refGene_tx=NM_001009854.1;refGene_gn=VEGFA;refGene_tx=XM_023253550.1
B2 44521453 44521475
refGene_gn=VEGFA;refGene_tx=NM_001009854.1;refGene_gn=VEGFA;refGene_tx=XM_023253550.1
B2 44738723 44738755
refGene_gn=CB2H6orf223;refGene_tx=XM_023254717.1
B2 44738904 44738992
refGene_gn=CB2H6orf223;refGene_tx=XM_023254717.1
B2 44738994 44739144
refGene_gn=CB2H6orf223;refGene_tx=XM_023254717.1

B2 44860468 44860835
refGene_gn=MRPL14;refGene_tx=XM_006931766.2;refGene_gn=MRPL14;refGene_tx=XM_011282209.2
B2 44863086 44863157
refGene_gn=MRPL14;refGene_tx=XM_006931766.2;refGene_gn=MRPL14;refGene_tx=XM_011282209.2
B2 44880828 44880987
refGene_gn=TMEM63B;refGene_tx=XM_019830647.2;refGene_gn=TMEM63B;refGene_tx=XM_019830648.2
B2 44881273 44881353
refGene_gn=TMEM63B;refGene_tx=XM_019830647.2;refGene_gn=TMEM63B;refGene_tx=XM_019830648.2
B2 44881614 44881656 refGene_gn=TMEM63B;refGene_tx=XM_019830647.2
B2 44882634 44882725
refGene_gn=TMEM63B;refGene_tx=XM_019830647.2;refGene_gn=TMEM63B;refGene_tx=XM_019830648.2
B2 44884153 44884191
refGene_gn=TMEM63B;refGene_tx=XM_019830647.2;refGene_gn=TMEM63B;refGene_tx=XM_019830648.2
B2 44885113 44885256
refGene_gn=TMEM63B;refGene_tx=XM_019830647.2;refGene_gn=TMEM63B;refGene_tx=XM_019830648.2
B2 44885368 44885420
refGene_gn=TMEM63B;refGene_tx=XM_019830647.2;refGene_gn=TMEM63B;refGene_tx=XM_019830648.2
B2 44885878 44885987
refGene_gn=TMEM63B;refGene_tx=XM_019830647.2;refGene_gn=TMEM63B;refGene_tx=XM_019830648.2
B2 44886634 44886705
refGene_gn=TMEM63B;refGene_tx=XM_019830647.2;refGene_gn=TMEM63B;refGene_tx=XM_019830648.2
B2 44889586 44889667
refGene_gn=TMEM63B;refGene_tx=XM_019830647.2;refGene_gn=TMEM63B;refGene_tx=XM_019830648.2
B2 44890139 44890263
refGene_gn=TMEM63B;refGene_tx=XM_019830647.2;refGene_gn=TMEM63B;refGene_tx=XM_019830648.2
B2 44891106 44891240
refGene_gn=TMEM63B;refGene_tx=XM_019830647.2;refGene_gn=TMEM63B;refGene_tx=XM_019830648.2
B2 44891382 44891520
refGene_gn=TMEM63B;refGene_tx=XM_019830647.2;refGene_gn=TMEM63B;refGene_tx=XM_019830648.2
B2 44891644 44891798
refGene_gn=TMEM63B;refGene_tx=XM_019830647.2;refGene_gn=TMEM63B;refGene_tx=XM_019830648.2
B2 44892704 44892811
refGene_gn=TMEM63B;refGene_tx=XM_019830647.2;refGene_gn=TMEM63B;refGene_tx=XM_019830648.2
B2 44893080 44893167

refGene_gn=TMEM63B;refGene_tx=XM_019830647.2;refGene_gn=TMEM63B;refGene_tx=XM_019830648.2
B2 44893370 44893436
refGene_gn=TMEM63B;refGene_tx=XM_019830647.2;refGene_gn=TMEM63B;refGene_tx=XM_019830648.2
B2 44894737 44894900
refGene_gn=TMEM63B;refGene_tx=XM_019830647.2;refGene_gn=TMEM63B;refGene_tx=XM_019830648.2
B2 44895464 44895570
refGene_gn=TMEM63B;refGene_tx=XM_019830647.2;refGene_gn=TMEM63B;refGene_tx=XM_019830648.2
B2 44896546 44896714
refGene_gn=TMEM63B;refGene_tx=XM_019830647.2;refGene_gn=TMEM63B;refGene_tx=XM_019830648.2
B2 44896946 44897062
refGene_gn=TMEM63B;refGene_tx=XM_019830647.2;refGene_gn=TMEM63B;refGene_tx=XM_019830648.2
B2 44897217 44897298
refGene_gn=TMEM63B;refGene_tx=XM_019830647.2;refGene_gn=TMEM63B;refGene_tx=XM_019830648.2
B2 44897569 44897761
refGene_gn=TMEM63B;refGene_tx=XM_019830647.2;refGene_gn=TMEM63B;refGene_tx=XM_019830648.2
B2 44904100 44904131
refGene_gn=CAPN11;refGene_tx=XM_006931767.4;refGene_gn=CAPN11;refGene_tx=XM_019830649.2;refGene_gn=CAPN11;refGene_tx=XM_019830650.2
B2 44905744 44905760 refGene_gn=CAPN11;refGene_tx=XM_019830651.2
B2 44910338 44910592
refGene_gn=CAPN11;refGene_tx=XM_006931767.4;refGene_gn=CAPN11;refGene_tx=XM_019830649.2;refGene_gn=CAPN11;refGene_tx=XM_019830650.2;refGene_gn=CAPN11;refGene_tx=XM_019830651.2
B2 44910970 44911040
refGene_gn=CAPN11;refGene_tx=XM_006931767.4;refGene_gn=CAPN11;refGene_tx=XM_019830649.2;refGene_gn=CAPN11;refGene_tx=XM_019830650.2;refGene_gn=CAPN11;refGene_tx=XM_019830651.2
B2 44912844 44912963
refGene_gn=CAPN11;refGene_tx=XM_006931767.4;refGene_gn=CAPN11;refGene_tx=XM_019830649.2;refGene_gn=CAPN11;refGene_tx=XM_019830650.2;refGene_gn=CAPN11;refGene_tx=XM_019830651.2
B2 44913774 44913908
refGene_gn=CAPN11;refGene_tx=XM_006931767.4;refGene_gn=CAPN11;refGene_tx=XM_019830649.2;refGene_gn=CAPN11;refGene_tx=XM_019830650.2;refGene_gn=CAPN11;refGene_tx=XM_019830651.2
B2 44914092 44914261
refGene_gn=CAPN11;refGene_tx=XM_006931767.4;refGene_gn=CAPN11;refGene_tx=XM_019830649.2;refGene_gn=CAPN11;refGene_tx=XM_019830650.2;refGene_gn=CAPN11;refGene_tx=XM_019830651.2
B2 44915487 44915571
refGene_gn=CAPN11;refGene_tx=XM_006931767.4;refGene_gn=CAPN11;refGene_tx=XM_019830649.2;refGene_gn=CAPN11;refGene_tx=XM_019830650.2;refGene_

gn=CAPN11;refGene_tx=XM_019830651.2
B2 44915677 44915763
refGene_gn=CAPN11;refGene_tx=XM_006931767.4;refGene_gn=CAPN11;refGene_tx=XM_019830649.2;refGene_gn=CAPN11;refGene_tx=XM_019830650.2;refGene_gn=CAPN11;refGene_tx=XM_019830651.2
B2 44916002 44916077
refGene_gn=CAPN11;refGene_tx=XM_006931767.4;refGene_gn=CAPN11;refGene_tx=XM_019830649.2;refGene_gn=CAPN11;refGene_tx=XM_019830650.2;refGene_gn=CAPN11;refGene_tx=XM_019830651.2
B2 44916233 44916394
refGene_gn=CAPN11;refGene_tx=XM_006931767.4;refGene_gn=CAPN11;refGene_tx=XM_019830649.2;refGene_gn=CAPN11;refGene_tx=XM_019830650.2;refGene_gn=CAPN11;refGene_tx=XM_019830651.2
B2 44916535 44916705
refGene_gn=CAPN11;refGene_tx=XM_006931767.4;refGene_gn=CAPN11;refGene_tx=XM_019830649.2;refGene_gn=CAPN11;refGene_tx=XM_019830650.2;refGene_gn=CAPN11;refGene_tx=XM_019830651.2
B2 44918548 44918560
refGene_gn=CAPN11;refGene_tx=XM_006931767.4;refGene_gn=CAPN11;refGene_tx=XM_019830649.2;refGene_gn=CAPN11;refGene_tx=XM_019830651.2
B2 44918896 44919108
refGene_gn=CAPN11;refGene_tx=XM_006931767.4;refGene_gn=CAPN11;refGene_tx=XM_019830649.2;refGene_gn=CAPN11;refGene_tx=XM_019830650.2;refGene_gn=CAPN11;refGene_tx=XM_019830651.2
B2 44919399 44919439
refGene_gn=CAPN11;refGene_tx=XM_006931767.4;refGene_gn=CAPN11;refGene_tx=XM_019830649.2;refGene_gn=CAPN11;refGene_tx=XM_019830650.2;refGene_gn=CAPN11;refGene_tx=XM_019830651.2
B2 44919537 44919606
refGene_gn=CAPN11;refGene_tx=XM_006931767.4;refGene_gn=CAPN11;refGene_tx=XM_019830649.2;refGene_gn=CAPN11;refGene_tx=XM_019830650.2;refGene_gn=CAPN11;refGene_tx=XM_019830651.2
B2 44919688 44919746
refGene_gn=CAPN11;refGene_tx=XM_006931767.4;refGene_gn=CAPN11;refGene_tx=XM_019830649.2;refGene_gn=CAPN11;refGene_tx=XM_019830650.2;refGene_gn=CAPN11;refGene_tx=XM_019830651.2
B2 44919882 44919947
refGene_gn=CAPN11;refGene_tx=XM_006931767.4;refGene_gn=CAPN11;refGene_tx=XM_019830649.2;refGene_gn=CAPN11;refGene_tx=XM_019830650.2;refGene_gn=CAPN11;refGene_tx=XM_019830651.2
B2 44920220 44920289
refGene_gn=CAPN11;refGene_tx=XM_006931767.4;refGene_gn=CAPN11;refGene_tx=XM_019830649.2;refGene_gn=CAPN11;refGene_tx=XM_019830650.2;refGene_gn=CAPN11;refGene_tx=XM_019830651.2
B2 44920747 44920826
refGene_gn=CAPN11;refGene_tx=XM_006931767.4;refGene_gn=CAPN11;refGene_tx=XM_019830649.2;refGene_gn=CAPN11;refGene_tx=XM_019830650.2;refGene_gn=CAPN11;refGene_tx=XM_019830651.2
B2 44920922 44921039
refGene_gn=CAPN11;refGene_tx=XM_006931767.4;refGene_gn=CAPN11;refGene_

tx=XM_019830649.2;refGene_gn=CAPN11;refGene_tx=XM_019830650.2;refGene_gn=CAPN11;refGene_tx=XM_019830651.2
B2 44921509 44921568
refGene_gn=CAPN11;refGene_tx=XM_006931767.4;refGene_gn=CAPN11;refGene_tx=XM_019830649.2;refGene_gn=CAPN11;refGene_tx=XM_019830650.2;refGene_gn=CAPN11;refGene_tx=XM_019830651.2
B2 44921658 44921951
refGene_gn=CAPN11;refGene_tx=XM_006931767.4;refGene_gn=CAPN11;refGene_tx=XM_019830649.2;refGene_gn=CAPN11;refGene_tx=XM_019830650.2;refGene_gn=CAPN11;refGene_tx=XM_019830651.2
B2 44955411 44955666
refGene_gn=MYMX;refGene_tx=XM_019830654.1;refGene_gn=MYMX;refGene_tx=XM_019830655.1
B2 44963977 44964006
refGene_gn=SLC29A1;refGene_tx=XM_006931770.4;refGene_gn=SLC29A1;refGene_tx=XM_006931771.4;refGene_gn=SLC29A1;refGene_tx=XM_011282207.3;refGene_gn=SLC29A1;refGene_tx=XM_019830652.2
B2 44965990 44966072
refGene_gn=SLC29A1;refGene_tx=XM_003986198.5;refGene_gn=SLC29A1;refGene_tx=XM_006931770.4;refGene_gn=SLC29A1;refGene_tx=XM_006931771.4;refGene_gn=SLC29A1;refGene_tx=XM_011282207.3;refGene_gn=SLC29A1;refGene_tx=XM_019830652.2
B2 44966189 44966392
refGene_gn=SLC29A1;refGene_tx=XM_003986198.5;refGene_gn=SLC29A1;refGene_tx=XM_006931770.4;refGene_gn=SLC29A1;refGene_tx=XM_006931771.4;refGene_gn=SLC29A1;refGene_tx=XM_011282207.3;refGene_gn=SLC29A1;refGene_tx=XM_019830652.2
B2 44966504 44966644
refGene_gn=SLC29A1;refGene_tx=XM_003986198.5;refGene_gn=SLC29A1;refGene_tx=XM_006931770.4;refGene_gn=SLC29A1;refGene_tx=XM_006931771.4;refGene_gn=SLC29A1;refGene_tx=XM_011282207.3;refGene_gn=SLC29A1;refGene_tx=XM_019830652.2
B2 44966948 44967083
refGene_gn=SLC29A1;refGene_tx=XM_003986198.5;refGene_gn=SLC29A1;refGene_tx=XM_006931770.4;refGene_gn=SLC29A1;refGene_tx=XM_006931771.4;refGene_gn=SLC29A1;refGene_tx=XM_011282207.3;refGene_gn=SLC29A1;refGene_tx=XM_019830652.2
B2 44967177 44967275
refGene_gn=SLC29A1;refGene_tx=XM_003986198.5;refGene_gn=SLC29A1;refGene_tx=XM_006931770.4;refGene_gn=SLC29A1;refGene_tx=XM_006931771.4;refGene_gn=SLC29A1;refGene_tx=XM_011282207.3;refGene_gn=SLC29A1;refGene_tx=XM_019830652.2
B2 44967425 44967504
refGene_gn=SLC29A1;refGene_tx=XM_003986198.5;refGene_gn=SLC29A1;refGene_tx=XM_006931770.4;refGene_gn=SLC29A1;refGene_tx=XM_006931771.4;refGene_gn=SLC29A1;refGene_tx=XM_011282207.3;refGene_gn=SLC29A1;refGene_tx=XM_019830652.2
B2 44967935 44968033
refGene_gn=SLC29A1;refGene_tx=XM_003986198.5;refGene_gn=SLC29A1;refGene_tx=XM_006931770.4;refGene_gn=SLC29A1;refGene_tx=XM_006931771.4;refGe

ne_gn=SLC29A1;refGene_tx=XM_011282207.3;refGene_gn=SLC29A1;refGene_tx=XM_019830652.2
B2 44968278 44968387
refGene_gn=SLC29A1;refGene_tx=XM_003986198.5;refGene_gn=SLC29A1;refGene_tx=XM_006931770.4;refGene_gn=SLC29A1;refGene_tx=XM_006931771.4;refGene_gn=SLC29A1;refGene_tx=XM_011282207.3;refGene_gn=SLC29A1;refGene_tx=XM_019830652.2
B2 44968634 44968720
refGene_gn=SLC29A1;refGene_tx=XM_003986198.5;refGene_gn=SLC29A1;refGene_tx=XM_006931770.4;refGene_gn=SLC29A1;refGene_tx=XM_006931771.4;refGene_gn=SLC29A1;refGene_tx=XM_011282207.3;refGene_gn=SLC29A1;refGene_tx=XM_019830652.2
B2 44969094 44969294
refGene_gn=SLC29A1;refGene_tx=XM_003986198.5;refGene_gn=SLC29A1;refGene_tx=XM_006931770.4;refGene_gn=SLC29A1;refGene_tx=XM_006931771.4;refGene_gn=SLC29A1;refGene_tx=XM_011282207.3;refGene_gn=SLC29A1;refGene_tx=XM_019830652.2
B2 44969654 44969766
refGene_gn=SLC29A1;refGene_tx=XM_003986198.5;refGene_gn=SLC29A1;refGene_tx=XM_006931770.4;refGene_gn=SLC29A1;refGene_tx=XM_006931771.4;refGene_gn=SLC29A1;refGene_tx=XM_011282207.3;refGene_gn=SLC29A1;refGene_tx=XM_019830652.2
B2 44983469 44983562 refGene_gn=HSP90AB1;refGene_tx=XM_023254019.1
B2 44983563 44983617 refGene_gn=HSP90AB1;refGene_tx=XM_023254019.1
B2 44984062 44984269 refGene_gn=HSP90AB1;refGene_tx=XM_023254019.1
B2 44984363 44984523 refGene_gn=HSP90AB1;refGene_tx=XM_023254019.1
B2 44984665 44984799 refGene_gn=HSP90AB1;refGene_tx=XM_023254019.1
B2 44984909 44985218 refGene_gn=HSP90AB1;refGene_tx=XM_023254019.1
B2 44985641 44985807 refGene_gn=HSP90AB1;refGene_tx=XM_023254019.1
B2 44985993 44986184 refGene_gn=HSP90AB1;refGene_tx=XM_023254019.1
B2 44986292 44986440 refGene_gn=HSP90AB1;refGene_tx=XM_023254019.1
B2 44986559 44986828 refGene_gn=HSP90AB1;refGene_tx=XM_023254019.1
B2 44987110 44987444 refGene_gn=HSP90AB1;refGene_tx=XM_023254019.1
B2 44987564 44987674 refGene_gn=HSP90AB1;refGene_tx=XM_023254019.1
B2 44988806 44989745
refGene_gn=SLC35B2;refGene_tx=XM_006931774.4;refGene_gn=SLC35B2;refGene_tx=XM_023254020.1
B2 44990466 44990621 refGene_gn=SLC35B2;refGene_tx=XM_023254020.1
B2 44990815 44991009
refGene_gn=SLC35B2;refGene_tx=XM_006931774.4;refGene_gn=SLC35B2;refGene_tx=XM_023254020.1
B2 44991541 44991552 refGene_gn=SLC35B2;refGene_tx=XM_023254020.1
B2 44993314 44993380 refGene_gn=NFKBIE;refGene_tx=XM_003986200.4
B2 44994271 44994511 refGene_gn=NFKBIE;refGene_tx=XM_003986200.4
B2 44994678 44994767 refGene_gn=NFKBIE;refGene_tx=XM_003986200.4
B2 44995832 44996055 refGene_gn=NFKBIE;refGene_tx=XM_003986200.4
B2 44996720 44996823 refGene_gn=NFKBIE;refGene_tx=XM_003986200.4
B2 44999258 44999901 refGene_gn=NFKBIE;refGene_tx=XM_003986200.4
B2 45001563 45001630 refGene_gn=NFKBIE;refGene_tx=XM_003986200.4
B2 45005558 45005681 refGene_gn=TMEM151B;refGene_tx=XM_023254021.1

B2 45007851 45008292 refGene_gn=TMEM151B;refGene_tx=XM_023254021.1
B2 45009477 45010605 refGene_gn=TMEM151B;refGene_tx=XM_023254021.1
B2 45014263 45014473
refGene_gn=TCTE1;refGene_tx=XM_003986201.5;refGene_gn=TCTE1;refGene_tx=XM_019830670.2;refGene_gn=TCTE1;refGene_tx=XM_023254022.1
B2 45016351 45016801
refGene_gn=TCTE1;refGene_tx=XM_003986201.5;refGene_gn=TCTE1;refGene_tx=XM_019830670.2;refGene_gn=TCTE1;refGene_tx=XM_023254022.1
B2 45019621 45020190
refGene_gn=TCTE1;refGene_tx=XM_003986201.5;refGene_gn=TCTE1;refGene_tx=XM_019830670.2;refGene_gn=TCTE1;refGene_tx=XM_023254022.1
B2 45021159 45021454
refGene_gn=TCTE1;refGene_tx=XM_003986201.5;refGene_gn=TCTE1;refGene_tx=XM_019830670.2;refGene_gn=TCTE1;refGene_tx=XM_023254022.1
B2 45034150 45034315
refGene_gn=AARS2;refGene_tx=XM_003986202.5;refGene_gn=AARS2;refGene_tx=XM_019830660.2;refGene_gn=AARS2;refGene_tx=XM_019830662.2
B2 45034685 45034796
refGene_gn=AARS2;refGene_tx=XM_003986202.5;refGene_gn=AARS2;refGene_tx=XM_019830662.2
B2 45034909 45034993
refGene_gn=AARS2;refGene_tx=XM_003986202.5;refGene_gn=AARS2;refGene_tx=XM_019830660.2;refGene_gn=AARS2;refGene_tx=XM_019830662.2
B2 45035564 45035675
refGene_gn=AARS2;refGene_tx=XM_003986202.5;refGene_gn=AARS2;refGene_tx=XM_019830660.2;refGene_gn=AARS2;refGene_tx=XM_019830662.2
B2 45035886 45036009
refGene_gn=AARS2;refGene_tx=XM_003986202.5;refGene_gn=AARS2;refGene_tx=XM_019830660.2;refGene_gn=AARS2;refGene_tx=XM_019830662.2
B2 45036303 45036412
refGene_gn=AARS2;refGene_tx=XM_003986202.5;refGene_gn=AARS2;refGene_tx=XM_019830660.2;refGene_gn=AARS2;refGene_tx=XM_019830662.2
B2 45036565 45036675
refGene_gn=AARS2;refGene_tx=XM_003986202.5;refGene_gn=AARS2;refGene_tx=XM_019830660.2;refGene_gn=AARS2;refGene_tx=XM_019830662.2
B2 45036753 45036891
refGene_gn=AARS2;refGene_tx=XM_003986202.5;refGene_gn=AARS2;refGene_tx=XM_019830660.2;refGene_gn=AARS2;refGene_tx=XM_019830662.2
B2 45037501 45037642
refGene_gn=AARS2;refGene_tx=XM_003986202.5;refGene_gn=AARS2;refGene_tx=XM_019830660.2;refGene_gn=AARS2;refGene_tx=XM_019830662.2
B2 45037731 45037845
refGene_gn=AARS2;refGene_tx=XM_003986202.5;refGene_gn=AARS2;refGene_tx=XM_019830660.2;refGene_gn=AARS2;refGene_tx=XM_019830662.2
B2 45037982 45038155
refGene_gn=AARS2;refGene_tx=XM_003986202.5;refGene_gn=AARS2;refGene_tx=XM_019830660.2;refGene_gn=AARS2;refGene_tx=XM_019830662.2
B2 45038434 45038579
refGene_gn=AARS2;refGene_tx=XM_003986202.5;refGene_gn=AARS2;refGene_tx=XM_019830660.2;refGene_gn=AARS2;refGene_tx=XM_019830662.2

B2 45039012 45039146
refGene_gn=AARS2;refGene_tx=XM_003986202.5;refGene_gn=AARS2;refGene_tx=XM_019830660.2;refGene_gn=AARS2;refGene_tx=XM_019830662.2
B2 45039643 45039755
refGene_gn=AARS2;refGene_tx=XM_003986202.5;refGene_gn=AARS2;refGene_tx=XM_019830660.2;refGene_gn=AARS2;refGene_tx=XM_019830662.2
B2 45039859 45039898
refGene_gn=AARS2;refGene_tx=XM_003986202.5;refGene_gn=AARS2;refGene_tx=XM_019830660.2;refGene_gn=AARS2;refGene_tx=XM_019830662.2
B2 45040240 45040349
refGene_gn=AARS2;refGene_tx=XM_003986202.5;refGene_gn=AARS2;refGene_tx=XM_019830660.2;refGene_gn=AARS2;refGene_tx=XM_019830662.2
B2 45040537 45040683
refGene_gn=AARS2;refGene_tx=XM_003986202.5;refGene_gn=AARS2;refGene_tx=XM_019830660.2;refGene_gn=AARS2;refGene_tx=XM_019830662.2
B2 45042696 45042841
refGene_gn=AARS2;refGene_tx=XM_003986202.5;refGene_gn=AARS2;refGene_tx=XM_019830660.2;refGene_gn=AARS2;refGene_tx=XM_019830662.2
B2 45043403 45043571
refGene_gn=AARS2;refGene_tx=XM_003986202.5;refGene_gn=AARS2;refGene_tx=XM_019830660.2;refGene_gn=AARS2;refGene_tx=XM_019830662.2
B2 45043796 45043942
refGene_gn=AARS2;refGene_tx=XM_003986202.5;refGene_gn=AARS2;refGene_tx=XM_019830660.2;refGene_gn=AARS2;refGene_tx=XM_019830662.2
B2 45044477 45044669
refGene_gn=AARS2;refGene_tx=XM_003986202.5;refGene_gn=AARS2;refGene_tx=XM_019830660.2;refGene_gn=AARS2;refGene_tx=XM_019830662.2
B2 45045922 45046165
refGene_gn=AARS2;refGene_tx=XM_003986202.5;refGene_gn=AARS2;refGene_tx=XM_019830660.2
B2 45074167 45074294
refGene_gn=SPATS1;refGene_tx=XM_019830672.2;refGene_gn=SPATS1;refGene_tx=XM_023254023.1;refGene_gn=SPATS1;refGene_tx=XM_023254025.1;refGene_gn=SPATS1;refGene_tx=XM_023254026.1;refGene_gn=SPATS1;refGene_tx=XM_023254027.1;refGene_gn=SPATS1;refGene_tx=XM_023254028.1
B2 45082668 45082804
refGene_gn=SPATS1;refGene_tx=XM_019830672.2;refGene_gn=SPATS1;refGene_tx=XM_023254023.1;refGene_gn=SPATS1;refGene_tx=XM_023254025.1;refGene_gn=SPATS1;refGene_tx=XM_023254026.1;refGene_gn=SPATS1;refGene_tx=XM_023254027.1;refGene_gn=SPATS1;refGene_tx=XM_023254028.1
B2 45085625 45085750
refGene_gn=SPATS1;refGene_tx=XM_019830672.2;refGene_gn=SPATS1;refGene_tx=XM_023254023.1;refGene_gn=SPATS1;refGene_tx=XM_023254025.1;refGene_gn=SPATS1;refGene_tx=XM_023254026.1;refGene_gn=SPATS1;refGene_tx=XM_023254027.1;refGene_gn=SPATS1;refGene_tx=XM_023254028.1
B2 45087878 45088040
refGene_gn=SPATS1;refGene_tx=XM_019830672.2;refGene_gn=SPATS1;refGene_tx=XM_023254023.1;refGene_gn=SPATS1;refGene_tx=XM_023254025.1;refGene_gn=SPATS1;refGene_tx=XM_023254026.1;refGene_gn=SPATS1;refGene_tx=XM_023254027.1

B2 45093525 45093646
refGene_gn=SPATS1;refGene_tx=XM_019830672.2;refGene_gn=SPATS1;refGene_tx=XM_023254023.1;refGene_gn=SPATS1;refGene_tx=XM_023254025.1;refGene_gn=SPATS1;refGene_tx=XM_023254026.1;refGene_gn=SPATS1;refGene_tx=XM_023254027.1;refGene_gn=SPATS1;refGene_tx=XM_023254028.1
B2 45094467 45094530
refGene_gn=SPATS1;refGene_tx=XM_019830672.2;refGene_gn=SPATS1;refGene_tx=XM_023254023.1;refGene_gn=SPATS1;refGene_tx=XM_023254025.1;refGene_gn=SPATS1;refGene_tx=XM_023254026.1;refGene_gn=SPATS1;refGene_tx=XM_023254027.1;refGene_gn=SPATS1;refGene_tx=XM_023254028.1
B2 45100954 45101102
refGene_gn=SPATS1;refGene_tx=XM_019830672.2;refGene_gn=SPATS1;refGene_tx=XM_023254023.1;refGene_gn=SPATS1;refGene_tx=XM_023254025.1;refGene_gn=SPATS1;refGene_tx=XM_023254026.1;refGene_gn=SPATS1;refGene_tx=XM_023254027.1;refGene_gn=SPATS1;refGene_tx=XM_023254028.1
B2 45101320 45101349 refGene_gn=SPATS1;refGene_tx=XM_019830672.2
B2 45103663 45103728
refGene_gn=SPATS1;refGene_tx=XM_023254023.1;refGene_gn=SPATS1;refGene_tx=XM_023254025.1;refGene_gn=SPATS1;refGene_tx=XM_023254027.1;refGene_gn=SPATS1;refGene_tx=XM_023254028.1
B2 45108882 45108927 refGene_gn=CDC5L;refGene_tx=XM_019830671.2
B2 45111242 45111346 refGene_gn=CDC5L;refGene_tx=XM_019830671.2
B2 45112832 45112994 refGene_gn=CDC5L;refGene_tx=XM_019830671.2
B2 45113588 45113716 refGene_gn=CDC5L;refGene_tx=XM_019830671.2
B2 45115091 45115191 refGene_gn=CDC5L;refGene_tx=XM_019830671.2
B2 45122980 45123199 refGene_gn=CDC5L;refGene_tx=XM_019830671.2
B2 45124578 45124723 refGene_gn=CDC5L;refGene_tx=XM_019830671.2
B2 45126321 45126510 refGene_gn=CDC5L;refGene_tx=XM_019830671.2
B2 45131007 45131156 refGene_gn=CDC5L;refGene_tx=XM_019830671.2
B2 45133212 45133375 refGene_gn=CDC5L;refGene_tx=XM_019830671.2
B2 45135656 45135821 refGene_gn=CDC5L;refGene_tx=XM_019830671.2
B2 45138631 45138712 refGene_gn=CDC5L;refGene_tx=XM_019830671.2
B2 45139001 45139244 refGene_gn=CDC5L;refGene_tx=XM_019830671.2
B2 45142726 45142924 refGene_gn=CDC5L;refGene_tx=XM_019830671.2
B2 45155010 45155223 refGene_gn=CDC5L;refGene_tx=XM_019830671.2
B2 45155977 45156082 refGene_gn=CDC5L;refGene_tx=XM_019830671.2
B2 45502271 45502313
refGene_gn=SUPT3H;refGene_tx=XM_023254038.1;refGene_gn=SUPT3H;refGene_tx=XM_023254039.1
B2 45591575 45591686
refGene_gn=SUPT3H;refGene_tx=XM_023254038.1;refGene_gn=SUPT3H;refGene_tx=XM_023254039.1
B2 45602018 45602186
refGene_gn=SUPT3H;refGene_tx=XM_023254038.1;refGene_gn=SUPT3H;refGene_tx=XM_023254039.1
B2 45605837 45605945
refGene_gn=SUPT3H;refGene_tx=XM_023254038.1;refGene_gn=SUPT3H;refGene_tx=XM_023254039.1
B2 45607225 45607338
refGene_gn=SUPT3H;refGene_tx=XM_023254038.1;refGene_gn=SUPT3H;refGene_

tx=XM_023254039.1
B2	45612491	45612567
refGene_gn=SUPT3H;refGene_tx=XM_023254038.1;refGene_gn=SUPT3H;refGene_tx=XM_023254039.1
B2	45652129	45652269
refGene_gn=SUPT3H;refGene_tx=XM_023254038.1;refGene_gn=SUPT3H;refGene_tx=XM_023254039.1
B2	45660074	45660165
refGene_gn=SUPT3H;refGene_tx=XM_023254038.1;refGene_gn=SUPT3H;refGene_tx=XM_023254039.1
B2	45663141	45663228
refGene_gn=SUPT3H;refGene_tx=XM_023254038.1;refGene_gn=SUPT3H;refGene_tx=XM_023254039.1
B2	45760387	45760472
refGene_gn=SUPT3H;refGene_tx=XM_023254038.1;refGene_gn=SUPT3H;refGene_tx=XM_023254039.1
B2	45989372	45989510
refGene_gn=RUNX2;refGene_tx=XM_023254029.1;refGene_gn=RUNX2;refGene_tx=XM_023254030.1;refGene_gn=RUNX2;refGene_tx=XM_023254031.1;refGene_gn=RUNX2;refGene_tx=XM_023254033.1;refGene_gn=RUNX2;refGene_tx=XM_023254034.1;refGene_gn=RUNX2;refGene_tx=XM_023254035.1;refGene_gn=RUNX2;refGene_tx=XM_023254036.1;refGene_gn=RUNX2;refGene_tx=XM_023254037.1
B2	45989708	45989832
refGene_gn=RUNX2;refGene_tx=XM_023254029.1;refGene_gn=RUNX2;refGene_tx=XM_023254030.1;refGene_gn=RUNX2;refGene_tx=XM_023254031.1;refGene_gn=RUNX2;refGene_tx=XM_023254033.1;refGene_gn=RUNX2;refGene_tx=XM_023254034.1;refGene_gn=RUNX2;refGene_tx=XM_023254035.1;refGene_gn=RUNX2;refGene_tx=XM_023254036.1;refGene_gn=RUNX2;refGene_tx=XM_023254037.1
B2	46032260	46032361
refGene_gn=SUPT3H;refGene_tx=XM_023254038.1;refGene_gn=SUPT3H;refGene_tx=XM_023254039.1
B2	46083769	46084119
refGene_gn=RUNX2;refGene_tx=XM_023254029.1;refGene_gn=RUNX2;refGene_tx=XM_023254030.1;refGene_gn=RUNX2;refGene_tx=XM_023254031.1;refGene_gn=RUNX2;refGene_tx=XM_023254033.1;refGene_gn=RUNX2;refGene_tx=XM_023254034.1;refGene_gn=RUNX2;refGene_tx=XM_023254035.1;refGene_gn=RUNX2;refGene_tx=XM_023254036.1;refGene_gn=RUNX2;refGene_tx=XM_023254037.1
B2	46092969	46093126
refGene_gn=RUNX2;refGene_tx=XM_023254029.1;refGene_gn=RUNX2;refGene_tx=XM_023254030.1;refGene_gn=RUNX2;refGene_tx=XM_023254031.1;refGene_gn=RUNX2;refGene_tx=XM_023254033.1;refGene_gn=RUNX2;refGene_tx=XM_023254034.1;refGene_gn=RUNX2;refGene_tx=XM_023254035.1;refGene_gn=RUNX2;refGene_tx=XM_023254036.1;refGene_gn=RUNX2;refGene_tx=XM_023254037.1
B2	46098999	46099104
refGene_gn=RUNX2;refGene_tx=XM_023254029.1;refGene_gn=RUNX2;refGene_tx=XM_023254030.1;refGene_gn=RUNX2;refGene_tx=XM_023254031.1;refGene_gn=RUNX2;refGene_tx=XM_023254033.1;refGene_gn=RUNX2;refGene_tx=XM_023254034.1;refGene_gn=RUNX2;refGene_tx=XM_023254035.1;refGene_gn=RUNX2;refGene_tx=XM_023254036.1;refGene_gn=RUNX2;refGene_tx=XM_023254037.1
B2	46155770	46155944

refGene_gn=RUNX2;refGene_tx=XM_023254029.1;refGene_gn=RUNX2;refGene_tx=XM_023254030.1;refGene_gn=RUNX2;refGene_tx=XM_023254031.1;refGene_gn=RUNX2;refGene_tx=XM_023254034.1;refGene_gn=RUNX2;refGene_tx=XM_023254035.1;refGene_gn=RUNX2;refGene_tx=XM_023254036.1;refGene_gn=RUNX2;refGene_tx=XM_023254037.1
B2 46177062 46177224
refGene_gn=RUNX2;refGene_tx=XM_023254029.1;refGene_gn=RUNX2;refGene_tx=XM_023254030.1;refGene_gn=RUNX2;refGene_tx=XM_023254031.1;refGene_gn=RUNX2;refGene_tx=XM_023254033.1;refGene_gn=RUNX2;refGene_tx=XM_023254034.1;refGene_gn=RUNX2;refGene_tx=XM_023254035.1;refGene_gn=RUNX2;refGene_tx=XM_023254036.1;refGene_gn=RUNX2;refGene_tx=XM_023254037.1
B2 46178269 46178295
refGene_gn=RUNX2;refGene_tx=XM_023254036.1;refGene_gn=RUNX2;refGene_tx=XM_023254037.1
B2 46208130 46208196
refGene_gn=RUNX2;refGene_tx=XM_023254029.1;refGene_gn=RUNX2;refGene_tx=XM_023254031.1;refGene_gn=RUNX2;refGene_tx=XM_023254033.1
B2 46209811 46210290
refGene_gn=RUNX2;refGene_tx=XM_023254029.1;refGene_gn=RUNX2;refGene_tx=XM_023254030.1;refGene_gn=RUNX2;refGene_tx=XM_023254031.1;refGene_gn=RUNX2;refGene_tx=XM_023254033.1
B2 46213287 46213325 refGene_gn=RUNX2;refGene_tx=XM_023254035.1
B2 46250751 46250795 refGene_gn=RUNX2;refGene_tx=XM_023254034.1
B2 46546729 46546897 refGene_gn=CLIC5;refGene_tx=XM_011282370.3
B2 46553440 46553625 refGene_gn=CLIC5;refGene_tx=XM_011282370.3
B2 46577917 46578024 refGene_gn=CLIC5;refGene_tx=XM_011282370.3
B2 46587070 46587196 refGene_gn=CLIC5;refGene_tx=XM_011282370.3
B2 46590950 46591060 refGene_gn=CLIC5;refGene_tx=XM_011282370.3
B2 46646770 46646833 refGene_gn=CLIC5;refGene_tx=XM_011282370.3
B2 46745075 46745901
refGene_gn=ENPP4;refGene_tx=XM_006931779.4;refGene_gn=ENPP4;refGene_tx=XM_006931781.4
B2 46746539 46746710
refGene_gn=ENPP4;refGene_tx=XM_006931779.4;refGene_gn=ENPP4;refGene_tx=XM_006931781.4
B2 46748688 46749053
refGene_gn=ENPP4;refGene_tx=XM_006931779.4;refGene_gn=ENPP4;refGene_tx=XM_006931781.4
B2 46761506 46761934 refGene_gn=ENPP5;refGene_tx=XM_003986211.5
B2 46766547 46766724 refGene_gn=ENPP5;refGene_tx=XM_003986211.5
B2 46768919 46769748 refGene_gn=ENPP5;refGene_tx=XM_003986211.5
B2 46818001 46818162
refGene_gn=RCAN2;refGene_tx=XM_003986213.5;refGene_gn=RCAN2;refGene_tx=XM_011282368.3;refGene_gn=RCAN2;refGene_tx=XM_023254041.1
B2 46837915 46838087
refGene_gn=RCAN2;refGene_tx=XM_003986213.5;refGene_gn=RCAN2;refGene_tx=XM_011282368.3;refGene_gn=RCAN2;refGene_tx=XM_023254041.1
B2 46842608 46842782
refGene_gn=RCAN2;refGene_tx=XM_003986213.5;refGene_gn=RCAN2;refGene_tx=XM_011282368.3;refGene_gn=RCAN2;refGene_tx=XM_023254041.1

B2 46915681 46915768 refGene_gn=RCAN2;refGene_tx=XM_011282368.3
B2 47047200 47047425
refGene_gn=RCAN2;refGene_tx=XM_003986213.5;refGene_gn=RCAN2;refGene_tx=XM_023254041.1
B2 47136676 47136748
refGene_gn=LOC101100564;refGene_tx=XM_003986216.5
B2 47139343 47139431
refGene_gn=LOC101100564;refGene_tx=XM_003986216.5
B2 47158090 47158179
refGene_gn=LOC101100564;refGene_tx=XM_003986216.5
B2 47159045 47159141
refGene_gn=LOC101100564;refGene_tx=XM_003986216.5
B2 47169401 47169535
refGene_gn=LOC101100564;refGene_tx=XM_003986216.5
B2 47196591 47196682
refGene_gn=LOC101100564;refGene_tx=XM_003986216.5
B2 47201919 47202027
refGene_gn=LOC101100564;refGene_tx=XM_003986216.5
B2 47210372 47210466
refGene_gn=LOC101100564;refGene_tx=XM_003986216.5
B2 47211382 47211532
refGene_gn=LOC101100564;refGene_tx=XM_003986216.5
B2 47212842 47213017
refGene_gn=LOC101100564;refGene_tx=XM_003986216.5
B2 47215473 47215609
refGene_gn=LOC101100564;refGene_tx=XM_003986216.5
B2 47229946 47230123
refGene_gn=LOC101100564;refGene_tx=XM_003986216.5
B2 47230766 47230916
refGene_gn=SLC25A27;refGene_tx=XM_003986215.5;refGene_gn=SLC25A27;refGene_tx=XM_006931783.4;refGene_gn=SLC25A27;refGene_tx=XM_006931784.4;refGene_gn=SLC25A27;refGene_tx=XM_019830679.2;refGene_gn=SLC25A27;refGene_tx=XM_019830680.2;refGene_gn=SLC25A27;refGene_tx=XM_019830683.2;refGene_gn=SLC25A27;refGene_tx=XM_019830684.2;refGene_gn=SLC25A27;refGene_tx=XM_019830685.2
B2 47232031 47232056 refGene_gn=SLC25A27;refGene_tx=XM_019830682.2
B2 47232341 47232432 refGene_gn=SLC25A27;refGene_tx=XM_019830681.2
B2 47233389 47233581
refGene_gn=SLC25A27;refGene_tx=XM_003986215.5;refGene_gn=SLC25A27;refGene_tx=XM_006931783.4;refGene_gn=SLC25A27;refGene_tx=XM_006931784.4;refGene_gn=SLC25A27;refGene_tx=XM_019830679.2;refGene_gn=SLC25A27;refGene_tx=XM_019830680.2;refGene_gn=SLC25A27;refGene_tx=XM_019830681.2;refGene_gn=SLC25A27;refGene_tx=XM_019830682.2;refGene_gn=SLC25A27;refGene_tx=XM_019830683.2;refGene_gn=SLC25A27;refGene_tx=XM_019830684.2;refGene_gn=SLC25A27;refGene_tx=XM_019830685.2
B2 47235765 47235850
refGene_gn=SLC25A27;refGene_tx=XM_003986215.5;refGene_gn=SLC25A27;refGene_tx=XM_006931783.4;refGene_gn=SLC25A27;refGene_tx=XM_006931784.4;refGene_gn=SLC25A27;refGene_tx=XM_019830679.2;refGene_gn=SLC25A27;refGene_tx=XM_019830680.2;refGene_gn=SLC25A27;refGene_tx=XM_019830681.2;refG

ene_gn=SLC25A27;refGene_tx=XM_019830682.2;refGene_gn=SLC25A27;refGene_tx=XM_019830683.2;refGene_gn=SLC25A27;refGene_tx=XM_019830684.2;refGene_gn=SLC25A27;refGene_tx=XM_019830685.2
B2 47239781 47239904
refGene_gn=SLC25A27;refGene_tx=XM_003986215.5;refGene_gn=SLC25A27;refGene_tx=XM_006931783.4;refGene_gn=SLC25A27;refGene_tx=XM_006931784.4;refGene_gn=SLC25A27;refGene_tx=XM_019830679.2;refGene_gn=SLC25A27;refGene_tx=XM_019830680.2;refGene_gn=SLC25A27;refGene_tx=XM_019830681.2;refGene_gn=SLC25A27;refGene_tx=XM_019830682.2;refGene_gn=SLC25A27;refGene_tx=XM_019830683.2;refGene_gn=SLC25A27;refGene_tx=XM_019830684.2;refGene_gn=SLC25A27;refGene_tx=XM_019830685.2
B2 47242134 47242247
refGene_gn=SLC25A27;refGene_tx=XM_003986215.5;refGene_gn=SLC25A27;refGene_tx=XM_006931783.4;refGene_gn=SLC25A27;refGene_tx=XM_006931784.4;refGene_gn=SLC25A27;refGene_tx=XM_019830679.2;refGene_gn=SLC25A27;refGene_tx=XM_019830680.2;refGene_gn=SLC25A27;refGene_tx=XM_019830681.2;refGene_gn=SLC25A27;refGene_tx=XM_019830682.2;refGene_gn=SLC25A27;refGene_tx=XM_019830683.2;refGene_gn=SLC25A27;refGene_tx=XM_019830684.2;refGene_gn=SLC25A27;refGene_tx=XM_019830685.2
B2 47245332 47245417
refGene_gn=SLC25A27;refGene_tx=XM_003986215.5;refGene_gn=SLC25A27;refGene_tx=XM_006931783.4;refGene_gn=SLC25A27;refGene_tx=XM_006931784.4;refGene_gn=SLC25A27;refGene_tx=XM_019830679.2;refGene_gn=SLC25A27;refGene_tx=XM_019830680.2;refGene_gn=SLC25A27;refGene_tx=XM_019830681.2;refGene_gn=SLC25A27;refGene_tx=XM_019830682.2;refGene_gn=SLC25A27;refGene_tx=XM_019830683.2;refGene_gn=SLC25A27;refGene_tx=XM_019830684.2;refGene_gn=SLC25A27;refGene_tx=XM_019830685.2
B2 47251225 47251318
refGene_gn=SLC25A27;refGene_tx=XM_003986215.5;refGene_gn=SLC25A27;refGene_tx=XM_006931783.4;refGene_gn=SLC25A27;refGene_tx=XM_006931784.4;refGene_gn=SLC25A27;refGene_tx=XM_019830679.2;refGene_gn=SLC25A27;refGene_tx=XM_019830680.2;refGene_gn=SLC25A27;refGene_tx=XM_019830681.2;refGene_gn=SLC25A27;refGene_tx=XM_019830682.2;refGene_gn=SLC25A27;refGene_tx=XM_019830683.2;refGene_gn=SLC25A27;refGene_tx=XM_019830684.2;refGene_gn=SLC25A27;refGene_tx=XM_019830685.2
B2 47251569 47251594
refGene_gn=SLC25A27;refGene_tx=XM_006931784.4;refGene_gn=SLC25A27;refGene_tx=XM_019830685.2
B2 47251949 47252241
refGene_gn=SLC25A27;refGene_tx=XM_003986215.5;refGene_gn=SLC25A27;refGene_tx=XM_006931783.4;refGene_gn=SLC25A27;refGene_tx=XM_019830679.2;refGene_gn=SLC25A27;refGene_tx=XM_019830680.2;refGene_gn=SLC25A27;refGene_tx=XM_019830681.2;refGene_gn=SLC25A27;refGene_tx=XM_019830682.2;refGene_gn=SLC25A27;refGene_tx=XM_019830683.2;refGene_gn=SLC25A27;refGene_tx=XM_019830684.2
B2 47257092 47257164 refGene_gn=SLC25A27;refGene_tx=XM_006931783.4
B2 47257258 47257294 refGene_gn=SLC25A27;refGene_tx=XM_019830684.2
B2 47257358 47257463 refGene_gn=SLC25A27;refGene_tx=XM_003986215.5
B2 47261782 47261851 refGene_gn=SLC25A27;refGene_tx=XM_019830683.2
B2 47266739 47269047 refGene_gn=TDRD6;refGene_tx=XM_023254042.1

B2 47269049 47272844 refGene_gn=TDRD6;refGene_tx=XM_023254042.1
B2 47275123 47275248 refGene_gn=TDRD6;refGene_tx=XM_023254042.1
B2 47277226 47277316 refGene_gn=TDRD6;refGene_tx=XM_023254042.1
B2 47281657 47281681 refGene_gn=TDRD6;refGene_tx=XM_023254042.1
B2 47290265 47290396
refGene_gn=PLA2G7;refGene_tx=XM_023254043.1;refGene_gn=PLA2G7;refGene_tx=XM_023254044.1
B2 47290939 47291088
refGene_gn=PLA2G7;refGene_tx=XM_023254043.1;refGene_gn=PLA2G7;refGene_tx=XM_023254044.1
B2 47293566 47293737
refGene_gn=PLA2G7;refGene_tx=XM_023254043.1;refGene_gn=PLA2G7;refGene_tx=XM_023254044.1
B2 47295141 47295233
refGene_gn=PLA2G7;refGene_tx=XM_023254043.1;refGene_gn=PLA2G7;refGene_tx=XM_023254044.1
B2 47296389 47296503
refGene_gn=PLA2G7;refGene_tx=XM_023254043.1;refGene_gn=PLA2G7;refGene_tx=XM_023254044.1
B2 47297655 47297782
refGene_gn=PLA2G7;refGene_tx=XM_023254043.1;refGene_gn=PLA2G7;refGene_tx=XM_023254044.1
B2 47298639 47298708
refGene_gn=PLA2G7;refGene_tx=XM_023254043.1;refGene_gn=PLA2G7;refGene_tx=XM_023254044.1
B2 47300772 47300866
refGene_gn=PLA2G7;refGene_tx=XM_023254043.1;refGene_gn=PLA2G7;refGene_tx=XM_023254044.1
B2 47302328 47302473
refGene_gn=PLA2G7;refGene_tx=XM_023254043.1;refGene_gn=PLA2G7;refGene_tx=XM_023254044.1
B2 47302911 47303033
refGene_gn=PLA2G7;refGene_tx=XM_023254043.1;refGene_gn=PLA2G7;refGene_tx=XM_023254044.1
B2 47307986 47308095
refGene_gn=PLA2G7;refGene_tx=XM_023254043.1;refGene_gn=PLA2G7;refGene_tx=XM_023254044.1
B2 47345501 47345664 refGene_gn=ANKRD66;refGene_tx=XM_023254045.1
B2 47347659 47347888 refGene_gn=ANKRD66;refGene_tx=XM_023254045.1
B2 47352369 47352571 refGene_gn=ANKRD66;refGene_tx=XM_023254045.1
B2 47383904 47383973 refGene_gn=MEP1A;refGene_tx=XM_003986218.5
B2 47384058 47384083 refGene_gn=MEP1A;refGene_tx=XM_003986218.5
B2 47384171 47384222 refGene_gn=MEP1A;refGene_tx=XM_003986218.5
B2 47388729 47388770 refGene_gn=MEP1A;refGene_tx=XM_003986218.5
B2 47389497 47389573 refGene_gn=MEP1A;refGene_tx=XM_003986218.5
B2 47400427 47400545 refGene_gn=MEP1A;refGene_tx=XM_003986218.5
B2 47404712 47404888 refGene_gn=MEP1A;refGene_tx=XM_003986218.5
B2 47409154 47409376 refGene_gn=MEP1A;refGene_tx=XM_003986218.5
B2 47410145 47410295 refGene_gn=MEP1A;refGene_tx=XM_003986218.5
B2 47413072 47413288 refGene_gn=MEP1A;refGene_tx=XM_003986218.5

B2 47414588 47415062 refGene_gn=MEP1A;refGene_tx=XM_003986218.5
B2 47415786 47415960 refGene_gn=MEP1A;refGene_tx=XM_003986218.5
B2 47416475 47416776 refGene_gn=MEP1A;refGene_tx=XM_003986218.5
B2 47420043 47420200 refGene_gn=MEP1A;refGene_tx=XM_003986218.5
B2 47436002 47436082
refGene_gn=ADGRF5;refGene_tx=XM_003986219.5;refGene_gn=ADGRF5;refGene_tx=XM_006931786.4;refGene_gn=ADGRF5;refGene_tx=XM_006931787.4;refGene_gn=ADGRF5;refGene_tx=XM_006931788.4;refGene_gn=ADGRF5;refGene_tx=XM_006931790.4;refGene_gn=ADGRF5;refGene_tx=XM_023254046.1
B2 47436737 47436799 refGene_gn=ADGRF5;refGene_tx=XM_019830688.2
B2 47438057 47438142
refGene_gn=ADGRF5;refGene_tx=XM_003986219.5;refGene_gn=ADGRF5;refGene_tx=XM_006931786.4;refGene_gn=ADGRF5;refGene_tx=XM_006931787.4;refGene_gn=ADGRF5;refGene_tx=XM_006931788.4;refGene_gn=ADGRF5;refGene_tx=XM_006931790.4;refGene_gn=ADGRF5;refGene_tx=XM_019830688.2;refGene_gn=ADGRF5;refGene_tx=XM_023254046.1
B2 47438983 47439043
refGene_gn=ADGRF5;refGene_tx=XM_003986219.5;refGene_gn=ADGRF5;refGene_tx=XM_006931786.4;refGene_gn=ADGRF5;refGene_tx=XM_006931787.4;refGene_gn=ADGRF5;refGene_tx=XM_006931788.4;refGene_gn=ADGRF5;refGene_tx=XM_006931790.4;refGene_gn=ADGRF5;refGene_tx=XM_019830688.2;refGene_gn=ADGRF5;refGene_tx=XM_023254046.1
B2 47439131 47439173
refGene_gn=ADGRF5;refGene_tx=XM_003986219.5;refGene_gn=ADGRF5;refGene_tx=XM_006931786.4;refGene_gn=ADGRF5;refGene_tx=XM_006931787.4;refGene_gn=ADGRF5;refGene_tx=XM_006931788.4;refGene_gn=ADGRF5;refGene_tx=XM_006931790.4;refGene_gn=ADGRF5;refGene_tx=XM_019830688.2;refGene_gn=ADGRF5;refGene_tx=XM_023254046.1
B2 47440716 47442108
refGene_gn=ADGRF5;refGene_tx=XM_003986219.5;refGene_gn=ADGRF5;refGene_tx=XM_006931786.4;refGene_gn=ADGRF5;refGene_tx=XM_006931787.4;refGene_gn=ADGRF5;refGene_tx=XM_006931788.4;refGene_gn=ADGRF5;refGene_tx=XM_006931790.4;refGene_gn=ADGRF5;refGene_tx=XM_019830688.2;refGene_gn=ADGRF5;refGene_tx=XM_023254046.1
B2 47443907 47444087
refGene_gn=ADGRF5;refGene_tx=XM_003986219.5;refGene_gn=ADGRF5;refGene_tx=XM_006931786.4;refGene_gn=ADGRF5;refGene_tx=XM_006931787.4;refGene_gn=ADGRF5;refGene_tx=XM_006931788.4;refGene_gn=ADGRF5;refGene_tx=XM_006931790.4;refGene_gn=ADGRF5;refGene_tx=XM_019830688.2;refGene_gn=ADGRF5;refGene_tx=XM_023254046.1
B2 47446113 47446322
refGene_gn=ADGRF5;refGene_tx=XM_003986219.5;refGene_gn=ADGRF5;refGene_tx=XM_006931786.4;refGene_gn=ADGRF5;refGene_tx=XM_006931787.4;refGene_gn=ADGRF5;refGene_tx=XM_006931788.4;refGene_gn=ADGRF5;refGene_tx=XM_006931790.4;refGene_gn=ADGRF5;refGene_tx=XM_019830688.2;refGene_gn=ADGRF5;refGene_tx=XM_023254046.1
B2 47448147 47448303
refGene_gn=ADGRF5;refGene_tx=XM_003986219.5;refGene_gn=ADGRF5;refGene_tx=XM_006931786.4;refGene_gn=ADGRF5;refGene_tx=XM_006931787.4;refGene_gn=ADGRF5;refGene_tx=XM_006931788.4;refGene_gn=ADGRF5;refGene_tx=XM_00

6931790.4;refGene_gn=ADGRF5;refGene_tx=XM_019830688.2;refGene_gn=ADGRF5;refGene_tx=XM_023254046.1
B2 47450052 47450265
refGene_gn=ADGRF5;refGene_tx=XM_003986219.5;refGene_gn=ADGRF5;refGene_tx=XM_006931786.4;refGene_gn=ADGRF5;refGene_tx=XM_006931787.4;refGene_gn=ADGRF5;refGene_tx=XM_006931788.4;refGene_gn=ADGRF5;refGene_tx=XM_006931790.4;refGene_gn=ADGRF5;refGene_tx=XM_019830688.2;refGene_gn=ADGRF5;refGene_tx=XM_023254046.1
B2 47452515 47452725
refGene_gn=ADGRF5;refGene_tx=XM_003986219.5;refGene_gn=ADGRF5;refGene_tx=XM_006931786.4;refGene_gn=ADGRF5;refGene_tx=XM_006931787.4;refGene_gn=ADGRF5;refGene_tx=XM_006931788.4;refGene_gn=ADGRF5;refGene_tx=XM_006931790.4;refGene_gn=ADGRF5;refGene_tx=XM_019830688.2;refGene_gn=ADGRF5;refGene_tx=XM_023254046.1
B2 47452878 47452998
refGene_gn=ADGRF5;refGene_tx=XM_006931786.4;refGene_gn=ADGRF5;refGene_tx=XM_006931787.4;refGene_gn=ADGRF5;refGene_tx=XM_006931788.4;refGene_gn=ADGRF5;refGene_tx=XM_019830688.2;refGene_gn=ADGRF5;refGene_tx=XM_023254046.1
B2 47454893 47455064
refGene_gn=ADGRF5;refGene_tx=XM_003986219.5;refGene_gn=ADGRF5;refGene_tx=XM_006931786.4;refGene_gn=ADGRF5;refGene_tx=XM_006931787.4;refGene_gn=ADGRF5;refGene_tx=XM_006931788.4;refGene_gn=ADGRF5;refGene_tx=XM_006931790.4;refGene_gn=ADGRF5;refGene_tx=XM_019830688.2;refGene_gn=ADGRF5;refGene_tx=XM_023254046.1
B2 47460529 47460733
refGene_gn=ADGRF5;refGene_tx=XM_003986219.5;refGene_gn=ADGRF5;refGene_tx=XM_006931786.4;refGene_gn=ADGRF5;refGene_tx=XM_006931787.4;refGene_gn=ADGRF5;refGene_tx=XM_006931788.4;refGene_gn=ADGRF5;refGene_tx=XM_006931790.4;refGene_gn=ADGRF5;refGene_tx=XM_019830688.2;refGene_gn=ADGRF5;refGene_tx=XM_023254046.1
B2 47461682 47461904
refGene_gn=ADGRF5;refGene_tx=XM_003986219.5;refGene_gn=ADGRF5;refGene_tx=XM_006931786.4;refGene_gn=ADGRF5;refGene_tx=XM_006931787.4;refGene_gn=ADGRF5;refGene_tx=XM_006931788.4;refGene_gn=ADGRF5;refGene_tx=XM_006931790.4;refGene_gn=ADGRF5;refGene_tx=XM_019830688.2;refGene_gn=ADGRF5;refGene_tx=XM_023254046.1
B2 47466125 47466268
refGene_gn=ADGRF5;refGene_tx=XM_003986219.5;refGene_gn=ADGRF5;refGene_tx=XM_006931786.4;refGene_gn=ADGRF5;refGene_tx=XM_006931787.4;refGene_gn=ADGRF5;refGene_tx=XM_006931788.4;refGene_gn=ADGRF5;refGene_tx=XM_006931790.4;refGene_gn=ADGRF5;refGene_tx=XM_019830688.2;refGene_gn=ADGRF5;refGene_tx=XM_023254046.1
B2 47467095 47467154
refGene_gn=ADGRF5;refGene_tx=XM_003986219.5;refGene_gn=ADGRF5;refGene_tx=XM_006931786.4;refGene_gn=ADGRF5;refGene_tx=XM_006931787.4;refGene_gn=ADGRF5;refGene_tx=XM_006931788.4;refGene_gn=ADGRF5;refGene_tx=XM_006931790.4;refGene_gn=ADGRF5;refGene_tx=XM_019830688.2;refGene_gn=ADGRF5;refGene_tx=XM_023254046.1
B2 47468389 47468499

refGene_gn=ADGRF5;refGene_tx=XM_003986219.5;refGene_gn=ADGRF5;refGene_tx=XM_006931786.4;refGene_gn=ADGRF5;refGene_tx=XM_006931787.4;refGene_gn=ADGRF5;refGene_tx=XM_006931788.4;refGene_gn=ADGRF5;refGene_tx=XM_006931790.4;refGene_gn=ADGRF5;refGene_tx=XM_019830688.2;refGene_gn=ADGRF5;refGene_tx=XM_023254046.1
B2 47468640 47468817
refGene_gn=ADGRF5;refGene_tx=XM_003986219.5;refGene_gn=ADGRF5;refGene_tx=XM_006931786.4;refGene_gn=ADGRF5;refGene_tx=XM_006931787.4;refGene_gn=ADGRF5;refGene_tx=XM_006931788.4;refGene_gn=ADGRF5;refGene_tx=XM_006931790.4;refGene_gn=ADGRF5;refGene_tx=XM_019830688.2;refGene_gn=ADGRF5;refGene_tx=XM_023254046.1
B2 47470780 47470954
refGene_gn=ADGRF5;refGene_tx=XM_003986219.5;refGene_gn=ADGRF5;refGene_tx=XM_006931786.4;refGene_gn=ADGRF5;refGene_tx=XM_006931787.4;refGene_gn=ADGRF5;refGene_tx=XM_006931788.4;refGene_gn=ADGRF5;refGene_tx=XM_006931790.4;refGene_gn=ADGRF5;refGene_tx=XM_019830688.2;refGene_gn=ADGRF5;refGene_tx=XM_023254046.1
B2 47481664 47481719
refGene_gn=ADGRF5;refGene_tx=XM_003986219.5;refGene_gn=ADGRF5;refGene_tx=XM_006931786.4;refGene_gn=ADGRF5;refGene_tx=XM_006931787.4;refGene_gn=ADGRF5;refGene_tx=XM_006931788.4;refGene_gn=ADGRF5;refGene_tx=XM_006931790.4;refGene_gn=ADGRF5;refGene_tx=XM_019830688.2;refGene_gn=ADGRF5;refGene_tx=XM_023254046.1
B2 47488712 47488814
refGene_gn=ADGRF5;refGene_tx=XM_003986219.5;refGene_gn=ADGRF5;refGene_tx=XM_006931786.4;refGene_gn=ADGRF5;refGene_tx=XM_006931787.4;refGene_gn=ADGRF5;refGene_tx=XM_006931788.4;refGene_gn=ADGRF5;refGene_tx=XM_006931790.4;refGene_gn=ADGRF5;refGene_tx=XM_019830688.2;refGene_gn=ADGRF5;refGene_tx=XM_023254046.1
B2 47581713 47581787 refGene_gn=ADGRF1;refGene_tx=XM_006931791.4
B2 47583017 47583084 refGene_gn=ADGRF1;refGene_tx=XM_006931791.4
B2 47587682 47587742 refGene_gn=ADGRF1;refGene_tx=XM_006931791.4
B2 47588992 47589034 refGene_gn=ADGRF1;refGene_tx=XM_006931791.4
B2 47590888 47592049 refGene_gn=ADGRF1;refGene_tx=XM_006931791.4
B2 47592050 47592263 refGene_gn=ADGRF1;refGene_tx=XM_006931791.4
B2 47594962 47595151 refGene_gn=ADGRF1;refGene_tx=XM_006931791.4
B2 47596557 47596721 refGene_gn=ADGRF1;refGene_tx=XM_006931791.4
B2 47597436 47597597 refGene_gn=ADGRF1;refGene_tx=XM_006931791.4
B2 47601711 47601770 refGene_gn=ADGRF1;refGene_tx=XM_006931791.4
B2 47602627 47602728 refGene_gn=ADGRF1;refGene_tx=XM_006931791.4
B2 47604643 47604817 refGene_gn=ADGRF1;refGene_tx=XM_006931791.4
B2 47606064 47606214 refGene_gn=ADGRF1;refGene_tx=XM_006931791.4
B2 47608168 47608226 refGene_gn=ADGRF1;refGene_tx=XM_006931791.4
B2 47609543 47609612 refGene_gn=ADGRF1;refGene_tx=XM_006931791.4
B2 47781370 47781600 refGene_gn=TNFRSF21;refGene_tx=XM_023254047.1
B2 47782928 47783157 refGene_gn=TNFRSF21;refGene_tx=XM_023254047.1
B2 47803357 47803623 refGene_gn=TNFRSF21;refGene_tx=XM_023254047.1
B2 47827773 47828268 refGene_gn=TNFRSF21;refGene_tx=XM_023254047.1
B2 47830227 47830876 refGene_gn=TNFRSF21;refGene_tx=XM_023254047.1
B2 47849742 47849829 refGene_gn=TNFRSF21;refGene_tx=XM_023254047.1

B2 47979979 47979983 refGene_gn=CD2AP;refGene_tx=XM_011282366.2
B2 48018354 48018515 refGene_gn=CD2AP;refGene_tx=XM_011282366.2
B2 48036911 48037065 refGene_gn=CD2AP;refGene_tx=XM_011282366.2
B2 48046404 48046505 refGene_gn=CD2AP;refGene_tx=XM_011282366.2
B2 48049736 48049857 refGene_gn=CD2AP;refGene_tx=XM_011282366.2
B2 48068654 48068842 refGene_gn=CD2AP;refGene_tx=XM_011282366.2
B2 48071171 48071250 refGene_gn=CD2AP;refGene_tx=XM_011282366.2
B2 48071648 48071743 refGene_gn=CD2AP;refGene_tx=XM_011282366.2
B2 48075035 48075140 refGene_gn=CD2AP;refGene_tx=XM_011282366.2
B2 48080363 48080400 refGene_gn=CD2AP;refGene_tx=XM_011282366.2
B2 48082311 48082374 refGene_gn=CD2AP;refGene_tx=XM_011282366.2
B2 48098506 48098672 refGene_gn=CD2AP;refGene_tx=XM_011282366.2
B2 48099059 48099202 refGene_gn=CD2AP;refGene_tx=XM_011282366.2
B2 48108700 48108813 refGene_gn=CD2AP;refGene_tx=XM_011282366.2
B2 48109523 48109625 refGene_gn=CD2AP;refGene_tx=XM_011282366.2
B2 48111598 48111780 refGene_gn=CD2AP;refGene_tx=XM_011282366.2
B2 48116152 48116216 refGene_gn=CD2AP;refGene_tx=XM_011282366.2
B2 48120320 48120362 refGene_gn=CD2AP;refGene_tx=XM_011282366.2
B2 48173170 48173242 refGene_gn=ADGRF2;refGene_tx=XM_023254048.1
B2 48178027 48178067 refGene_gn=ADGRF2;refGene_tx=XM_023254048.1
B2 48178771 48178923 refGene_gn=ADGRF2;refGene_tx=XM_023254048.1
B2 48181367 48181601 refGene_gn=ADGRF2;refGene_tx=XM_023254048.1
B2 48183259 48183751 refGene_gn=ADGRF2;refGene_tx=XM_023254048.1
B2 48183752 48184275 refGene_gn=ADGRF2;refGene_tx=XM_023254048.1
B2 48184277 48184621 refGene_gn=ADGRF2;refGene_tx=XM_023254048.1
B2 48190573 48190615 refGene_gn=ADGRF2;refGene_tx=XM_023254048.1
B2 48193472 48193532 refGene_gn=ADGRF2;refGene_tx=XM_023254048.1
B2 48195959 48196007 refGene_gn=ADGRF2;refGene_tx=XM_023254048.1
B2 48209795 48209882
refGene_gn=ADGRF4;refGene_tx=XM_006931815.4;refGene_gn=ADGRF4;refGene_tx=XM_019830695.2;refGene_gn=ADGRF4;refGene_tx=XM_019830696.2;refGene_gn=ADGRF4;refGene_tx=XM_019830697.2;refGene_gn=ADGRF4;refGene_tx=XM_019830698.2
B2 48211051 48211106
refGene_gn=ADGRF4;refGene_tx=XM_006931815.4;refGene_gn=ADGRF4;refGene_tx=XM_019830695.2;refGene_gn=ADGRF4;refGene_tx=XM_019830696.2;refGene_gn=ADGRF4;refGene_tx=XM_019830697.2;refGene_gn=ADGRF4;refGene_tx=XM_019830698.2
B2 48213802 48213954
refGene_gn=ADGRF4;refGene_tx=XM_006931815.4;refGene_gn=ADGRF4;refGene_tx=XM_019830695.2;refGene_gn=ADGRF4;refGene_tx=XM_019830696.2;refGene_gn=ADGRF4;refGene_tx=XM_019830697.2;refGene_gn=ADGRF4;refGene_tx=XM_019830698.2
B2 48214715 48214967
refGene_gn=ADGRF4;refGene_tx=XM_006931815.4;refGene_gn=ADGRF4;refGene_tx=XM_019830695.2;refGene_gn=ADGRF4;refGene_tx=XM_019830696.2;refGene_gn=ADGRF4;refGene_tx=XM_019830697.2;refGene_gn=ADGRF4;refGene_tx=XM_019830698.2
B2 48215816 48217196
refGene_gn=ADGRF4;refGene_tx=XM_006931815.4;refGene_gn=ADGRF4;refGene_

tx=XM_019830695.2;refGene_gn=ADGRF4;refGene_tx=XM_019830696.2;refGene_gn=ADGRF4;refGene_tx=XM_019830697.2;refGene_gn=ADGRF4;refGene_tx=XM_019830698.2
B2 48219353 48219426
refGene_gn=ADGRF4;refGene_tx=XM_006931815.4;refGene_gn=ADGRF4;refGene_tx=XM_019830695.2;refGene_gn=ADGRF4;refGene_tx=XM_019830696.2;refGene_gn=ADGRF4;refGene_tx=XM_019830697.2;refGene_gn=ADGRF4;refGene_tx=XM_019830698.2
B2 48219813 48219873
refGene_gn=ADGRF4;refGene_tx=XM_006931815.4;refGene_gn=ADGRF4;refGene_tx=XM_019830695.2;refGene_gn=ADGRF4;refGene_tx=XM_019830696.2;refGene_gn=ADGRF4;refGene_tx=XM_019830697.2;refGene_gn=ADGRF4;refGene_tx=XM_019830698.2
B2 48220897 48220945
refGene_gn=ADGRF4;refGene_tx=XM_006931815.4;refGene_gn=ADGRF4;refGene_tx=XM_019830695.2;refGene_gn=ADGRF4;refGene_tx=XM_019830696.2;refGene_gn=ADGRF4;refGene_tx=XM_019830697.2;refGene_gn=ADGRF4;refGene_tx=XM_019830698.2
B2 48295219 48295349 refGene_gn=OPN5;refGene_tx=XM_023254746.1
B2 48299275 48299395 refGene_gn=OPN5;refGene_tx=XM_023254746.1
B2 48303362 48303533 refGene_gn=OPN5;refGene_tx=XM_023254746.1
B2 48307397 48307732 refGene_gn=OPN5;refGene_tx=XM_023254746.1
B2 48318703 48318945 refGene_gn=OPN5;refGene_tx=XM_023254746.1
B2 48322817 48322881 refGene_gn=OPN5;refGene_tx=XM_023254746.1
B2 48360939 48361046 refGene_gn=PTCHD4;refGene_tx=XM_023254051.1
B2 48382678 48384312
refGene_gn=PTCHD4;refGene_tx=XM_003986223.4;refGene_gn=PTCHD4;refGene_tx=XM_023254049.1
B2 48457081 48457212 refGene_gn=PTCHD4;refGene_tx=XM_023254050.1
B2 48506836 48507317
refGene_gn=PTCHD4;refGene_tx=XM_003986223.4;refGene_gn=PTCHD4;refGene_tx=XM_023254049.1;refGene_gn=PTCHD4;refGene_tx=XM_023254050.1;refGene_gn=PTCHD4;refGene_tx=XM_023254051.1
B2 48563231 48563657
refGene_gn=PTCHD4;refGene_tx=XM_003986223.4;refGene_gn=PTCHD4;refGene_tx=XM_023254049.1;refGene_gn=PTCHD4;refGene_tx=XM_023254050.1;refGene_gn=PTCHD4;refGene_tx=XM_023254051.1
B2 48563949 48563954 refGene_gn=PTCHD4;refGene_tx=XM_023254049.1
B2 49896129 49896258 refGene_gn=MUT;refGene_tx=XM_006931792.2
B2 49901818 49901986 refGene_gn=MUT;refGene_tx=XM_006931792.2
B2 49906019 49906167 refGene_gn=MUT;refGene_tx=XM_006931792.2
B2 49907341 49907473 refGene_gn=MUT;refGene_tx=XM_006931792.2
B2 49910768 49910884 refGene_gn=MUT;refGene_tx=XM_006931792.2
B2 49914345 49914461 refGene_gn=MUT;refGene_tx=XM_006931792.2
B2 49915532 49915644 refGene_gn=MUT;refGene_tx=XM_006931792.2
B2 49918420 49918669 refGene_gn=MUT;refGene_tx=XM_006931792.2
B2 49919892 49920064 refGene_gn=MUT;refGene_tx=XM_006931792.2
B2 49922051 49922209 refGene_gn=MUT;refGene_tx=XM_006931792.2
B2 49925737 49926105 refGene_gn=MUT;refGene_tx=XM_006931792.2
B2 49927637 49928022 refGene_gn=MUT;refGene_tx=XM_006931792.2

B2	49936412	49936514
refGene_gn=CENPQ;refGene_tx=XM_003986263.4;refGene_gn=CENPQ;refGene_tx=XM_011282362.2;refGene_gn=CENPQ;refGene_tx=XM_011282364.3;refGene_gn=CENPQ;refGene_tx=XM_011282365.3;refGene_gn=CENPQ;refGene_tx=XM_019830701.1
B2	49937252	49937301
refGene_gn=CENPQ;refGene_tx=XM_003986263.4;refGene_gn=CENPQ;refGene_tx=XM_011282362.2;refGene_gn=CENPQ;refGene_tx=XM_011282364.3;refGene_gn=CENPQ;refGene_tx=XM_011282365.3;refGene_gn=CENPQ;refGene_tx=XM_019830701.1
B2	49938352	49938479
refGene_gn=CENPQ;refGene_tx=XM_003986263.4;refGene_gn=CENPQ;refGene_tx=XM_011282362.2;refGene_gn=CENPQ;refGene_tx=XM_011282364.3;refGene_gn=CENPQ;refGene_tx=XM_011282365.3;refGene_gn=CENPQ;refGene_tx=XM_019830701.1
B2	49939046	49939115
refGene_gn=CENPQ;refGene_tx=XM_003986263.4;refGene_gn=CENPQ;refGene_tx=XM_011282362.2;refGene_gn=CENPQ;refGene_tx=XM_011282364.3;refGene_gn=CENPQ;refGene_tx=XM_011282365.3;refGene_gn=CENPQ;refGene_tx=XM_019830701.1
B2	49940707	49940837
refGene_gn=CENPQ;refGene_tx=XM_003986263.4;refGene_gn=CENPQ;refGene_tx=XM_011282362.2;refGene_gn=CENPQ;refGene_tx=XM_011282364.3;refGene_gn=CENPQ;refGene_tx=XM_011282365.3;refGene_gn=CENPQ;refGene_tx=XM_019830701.1
B2	49945693	49945813
refGene_gn=CENPQ;refGene_tx=XM_003986263.4;refGene_gn=CENPQ;refGene_tx=XM_011282362.2;refGene_gn=CENPQ;refGene_tx=XM_011282364.3;refGene_gn=CENPQ;refGene_tx=XM_019830701.1
B2	49945948	49946023
refGene_gn=CENPQ;refGene_tx=XM_003986263.4;refGene_gn=CENPQ;refGene_tx=XM_011282362.2;refGene_gn=CENPQ;refGene_tx=XM_011282364.3;refGene_gn=CENPQ;refGene_tx=XM_011282365.3;refGene_gn=CENPQ;refGene_tx=XM_019830701.1
B2	49947021	49947153
refGene_gn=CENPQ;refGene_tx=XM_003986263.4;refGene_gn=CENPQ;refGene_tx=XM_011282362.2;refGene_gn=CENPQ;refGene_tx=XM_011282364.3;refGene_gn=CENPQ;refGene_tx=XM_011282365.3;refGene_gn=CENPQ;refGene_tx=XM_019830701.1
B2	49957466	49957544	refGene_gn=GLYATL3;refGene_tx=XM_023254052.1
B2	49961701	49961809	refGene_gn=GLYATL3;refGene_tx=XM_023254052.1
B2	49962546	49962673	refGene_gn=GLYATL3;refGene_tx=XM_023254052.1
B2	49967085	49967212	refGene_gn=GLYATL3;refGene_tx=XM_023254052.1
B2	49971223	49971650	refGene_gn=GLYATL3;refGene_tx=XM_023254052.1
B2	49989441	49990041
refGene_gn=CB2H6orf141;refGene_tx=XM_023254053.1
B2	50038655	50038688	refGene_gn=RHAG;refGene_tx=XM_006931794.3
B2	50040448	50040522	refGene_gn=RHAG;refGene_tx=XM_006931794.3
B2	50040896	50040970	refGene_gn=RHAG;refGene_tx=XM_006931794.3
B2	50045301	50045423	refGene_gn=RHAG;refGene_tx=XM_006931794.3

B2 50046016 50046154 refGene_gn=RHAG;refGene_tx=XM_006931794.3
B2 50047869 50048036 refGene_gn=RHAG;refGene_tx=XM_006931794.3
B2 50048140 50048288 refGene_gn=RHAG;refGene_tx=XM_006931794.3
B2 50050291 50050442 refGene_gn=RHAG;refGene_tx=XM_006931794.3
B2 50051041 50051225 refGene_gn=RHAG;refGene_tx=XM_006931794.3
B2 50059590 50059747 refGene_gn=RHAG;refGene_tx=XM_006931794.3
B2 50129072 50129200
refGene_gn=CRISP2;refGene_tx=XM_006931795.4;refGene_gn=CRISP2;refGene_tx=XM_019830704.2;refGene_gn=CRISP2;refGene_tx=XM_023254054.1;refGene_gn=CRISP2;refGene_tx=XM_023254055.1
B2 50131185 50131274
refGene_gn=CRISP2;refGene_tx=XM_006931795.4;refGene_gn=CRISP2;refGene_tx=XM_019830704.2;refGene_gn=CRISP2;refGene_tx=XM_023254054.1;refGene_gn=CRISP2;refGene_tx=XM_023254055.1
B2 50133625 50133723
refGene_gn=CRISP2;refGene_tx=XM_006931795.4;refGene_gn=CRISP2;refGene_tx=XM_019830704.2;refGene_gn=CRISP2;refGene_tx=XM_023254054.1;refGene_gn=CRISP2;refGene_tx=XM_023254055.1
B2 50134373 50134519
refGene_gn=CRISP2;refGene_tx=XM_006931795.4;refGene_gn=CRISP2;refGene_tx=XM_019830704.2;refGene_gn=CRISP2;refGene_tx=XM_023254054.1;refGene_gn=CRISP2;refGene_tx=XM_023254055.1
B2 50135342 50135523 refGene_gn=CRISP2;refGene_tx=XM_023254055.1
B2 50138232 50138320
refGene_gn=CRISP2;refGene_tx=XM_006931795.4;refGene_gn=CRISP2;refGene_tx=XM_019830704.2;refGene_gn=CRISP2;refGene_tx=XM_023254054.1;refGene_gn=CRISP2;refGene_tx=XM_023254055.1
B2 50139261 50139378
refGene_gn=CRISP2;refGene_tx=XM_006931795.4;refGene_gn=CRISP2;refGene_tx=XM_019830704.2;refGene_gn=CRISP2;refGene_tx=XM_023254054.1
B2 50145216 50145285
refGene_gn=CRISP2;refGene_tx=XM_006931795.4;refGene_gn=CRISP2;refGene_tx=XM_019830704.2;refGene_gn=CRISP2;refGene_tx=XM_023254054.1
B2 50148988 50149133
refGene_gn=LOC109500100;refGene_tx=XM_023254719.1
B2 50149176 50149544
refGene_gn=LOC109500100;refGene_tx=XM_023254719.1
B2 50158096 50158224
refGene_gn=LOC101081299;refGene_tx=XM_003986225.5
B2 50160623 50160712
refGene_gn=LOC101081299;refGene_tx=XM_003986225.5
B2 50162627 50162725
refGene_gn=LOC101081299;refGene_tx=XM_003986225.5
B2 50163135 50163281
refGene_gn=LOC101081299;refGene_tx=XM_003986225.5
B2 50168324 50168412
refGene_gn=LOC101081299;refGene_tx=XM_003986225.5
B2 50169195 50169312
refGene_gn=LOC101081299;refGene_tx=XM_003986225.5
B2 50174531 50174602

refGene_gn=LOC101081299;refGene_tx=XM_003986225.5
B2	50184750	50184778
refGene_gn=LOC101081299;refGene_tx=XM_003986225.5
B2	50205725	50205959
refGene_gn=LOC109500100;refGene_tx=XM_023254719.1
B2	50222306	50223560	refGene_gn=PGK2;refGene_tx=XM_003986226.4
B2	50306780	50306908	refGene_gn=CRISP1;refGene_tx=XM_019830703.2
B2	50312505	50312594	refGene_gn=CRISP1;refGene_tx=XM_019830703.2
B2	50314808	50314906	refGene_gn=CRISP1;refGene_tx=XM_019830703.2
B2	50318097	50318246	refGene_gn=CRISP1;refGene_tx=XM_019830703.2
B2	50319329	50319420	refGene_gn=CRISP1;refGene_tx=XM_019830703.2
B2	50324050	50324179	refGene_gn=CRISP1;refGene_tx=XM_019830703.2
B2	50330687	50330747	refGene_gn=CRISP1;refGene_tx=XM_019830703.2
B2	50394303	50394468	refGene_gn=DEFB133;refGene_tx=XM_023254720.1
B2	50407219	50407401	refGene_gn=DEFB114;refGene_tx=XM_019831593.1
B2	50437997	50438007	refGene_gn=DEFB114;refGene_tx=XM_019831593.1
B2	50438097	50438324	refGene_gn=DEFB113;refGene_tx=XM_023254721.1
B2	50439082	50439154	refGene_gn=DEFB113;refGene_tx=XM_023254721.1
B2	50457362	50457515
refGene_gn=LOC109500102;refGene_tx=XM_023254747.1
B2	50469317	50469466
refGene_gn=LOC101082313;refGene_tx=XM_003986228.3
B2	50474121	50474176
refGene_gn=LOC101082313;refGene_tx=XM_003986228.3
B2	50500292	50500453	refGene_gn=DEFB112;refGene_tx=XM_023254748.1
B2	50504015	50504159	refGene_gn=DEFB112;refGene_tx=XM_023254748.1
B2	50505334	50505389	refGene_gn=DEFB112;refGene_tx=XM_023254748.1
B2	51137745	51137784	refGene_gn=TFAP2D;refGene_tx=XM_003986229.3
B2	51138822	51139143	refGene_gn=TFAP2D;refGene_tx=XM_003986229.3
B2	51139144	51139321	refGene_gn=TFAP2D;refGene_tx=XM_003986229.3
B2	51142972	51143033	refGene_gn=TFAP2D;refGene_tx=XM_003986229.3
B2	51153105	51153271	refGene_gn=TFAP2D;refGene_tx=XM_003986229.3
B2	51153429	51153548	refGene_gn=TFAP2D;refGene_tx=XM_003986229.3
B2	51169807	51169949	refGene_gn=TFAP2D;refGene_tx=XM_003986229.3
B2	51176158	51176272	refGene_gn=TFAP2D;refGene_tx=XM_003986229.3
B2	51194978	51195198	refGene_gn=TFAP2D;refGene_tx=XM_003986229.3
B2	51239316	51239397
refGene_gn=TFAP2B;refGene_tx=XM_003986230.4;refGene_gn=TFAP2B;refGene_tx=XM_019830707.2;refGene_gn=TFAP2B;refGene_tx=XM_019830708.2;refGene_gn=TFAP2B;refGene_tx=XM_019830709.2;refGene_gn=TFAP2B;refGene_tx=XM_019830710.2
B2	51242536	51242563
refGene_gn=TFAP2B;refGene_tx=XM_019830707.2;refGene_gn=TFAP2B;refGene_tx=XM_019830708.2;refGene_gn=TFAP2B;refGene_tx=XM_019830709.2;refGene_gn=TFAP2B;refGene_tx=XM_019830711.2;refGene_gn=TFAP2B;refGene_tx=XM_019830712.1
B2	51243846	51244305
refGene_gn=TFAP2B;refGene_tx=XM_003986230.4;refGene_gn=TFAP2B;refGene_tx=XM_019830707.2;refGene_gn=TFAP2B;refGene_tx=XM_019830708.2;refGene_gn=TFAP2B;refGene_tx=XM_019830709.2;refGene_gn=TFAP2B;refGene_tx=XM_01

9830710.2;refGene_gn=TFAP2B;refGene_tx=XM_019830711.2;refGene_gn=TFAP2B;refGene_tx=XM_019830712.1
B2 51249155 51249216
refGene_gn=TFAP2B;refGene_tx=XM_003986230.4;refGene_gn=TFAP2B;refGene_tx=XM_019830707.2;refGene_gn=TFAP2B;refGene_tx=XM_019830708.2;refGene_gn=TFAP2B;refGene_tx=XM_019830709.2;refGene_gn=TFAP2B;refGene_tx=XM_019830710.2;refGene_gn=TFAP2B;refGene_tx=XM_019830711.2;refGene_gn=TFAP2B;refGene_tx=XM_019830712.1
B2 51257237 51257457
refGene_gn=TFAP2B;refGene_tx=XM_003986230.4;refGene_gn=TFAP2B;refGene_tx=XM_019830707.2;refGene_gn=TFAP2B;refGene_tx=XM_019830708.2;refGene_gn=TFAP2B;refGene_tx=XM_019830709.2;refGene_gn=TFAP2B;refGene_tx=XM_019830710.2;refGene_gn=TFAP2B;refGene_tx=XM_019830711.2;refGene_gn=TFAP2B;refGene_tx=XM_019830712.1
B2 51259106 51259225
refGene_gn=TFAP2B;refGene_tx=XM_003986230.4;refGene_gn=TFAP2B;refGene_tx=XM_019830707.2;refGene_gn=TFAP2B;refGene_tx=XM_019830708.2;refGene_gn=TFAP2B;refGene_tx=XM_019830709.2;refGene_gn=TFAP2B;refGene_tx=XM_019830710.2;refGene_gn=TFAP2B;refGene_tx=XM_019830711.2;refGene_gn=TFAP2B;refGene_tx=XM_019830712.1
B2 51261394 51261536
refGene_gn=TFAP2B;refGene_tx=XM_003986230.4;refGene_gn=TFAP2B;refGene_tx=XM_019830707.2;refGene_gn=TFAP2B;refGene_tx=XM_019830708.2;refGene_gn=TFAP2B;refGene_tx=XM_019830709.2;refGene_gn=TFAP2B;refGene_tx=XM_019830710.2;refGene_gn=TFAP2B;refGene_tx=XM_019830711.2;refGene_gn=TFAP2B;refGene_tx=XM_019830712.1
B2 51264389 51264690
refGene_gn=TFAP2B;refGene_tx=XM_003986230.4;refGene_gn=TFAP2B;refGene_tx=XM_019830707.2;refGene_gn=TFAP2B;refGene_tx=XM_019830708.2;refGene_gn=TFAP2B;refGene_tx=XM_019830709.2;refGene_gn=TFAP2B;refGene_tx=XM_019830710.2;refGene_gn=TFAP2B;refGene_tx=XM_019830711.2;refGene_gn=TFAP2B;refGene_tx=XM_019830712.1
B2 51265711 51265773
refGene_gn=TFAP2B;refGene_tx=XM_003986230.4;refGene_gn=TFAP2B;refGene_tx=XM_019830707.2;refGene_gn=TFAP2B;refGene_tx=XM_019830711.2;refGene_gn=TFAP2B;refGene_tx=XM_019830712.1
B2 51266996 51267019
refGene_gn=TFAP2B;refGene_tx=XM_019830709.2;refGene_gn=TFAP2B;refGene_tx=XM_019830710.2
B2 52014305 52014760 refGene_gn=PKHD1;refGene_tx=XM_011282361.3
B2 52041856 52041979 refGene_gn=PKHD1;refGene_tx=XM_011282361.3
B2 52046648 52046807 refGene_gn=PKHD1;refGene_tx=XM_011282361.3
B2 52053534 52053642 refGene_gn=PKHD1;refGene_tx=XM_011282361.3
B2 52061507 52061595 refGene_gn=PKHD1;refGene_tx=XM_011282361.3
B2 52062579 52062715 refGene_gn=PKHD1;refGene_tx=XM_011282361.3
B2 52072875 52073890 refGene_gn=PKHD1;refGene_tx=XM_011282361.3
B2 52148953 52149111 refGene_gn=PKHD1;refGene_tx=XM_011282361.3
B2 52150922 52151091 refGene_gn=PKHD1;refGene_tx=XM_011282361.3
B2 52152222 52153101 refGene_gn=PKHD1;refGene_tx=XM_011282361.3
B2 52157492 52157645 refGene_gn=PKHD1;refGene_tx=XM_011282361.3

| | | | |
|---|---|---|---|
| B2 | 52159023 | 52159178 | refGene_gn=PKHD1;refGene_tx=XM_011282361.3 |
| B2 | 52178284 | 52178372 | refGene_gn=PKHD1;refGene_tx=XM_011282361.3 |
| B2 | 52180805 | 52180919 | refGene_gn=PKHD1;refGene_tx=XM_011282361.3 |
| B2 | 52199834 | 52199972 | refGene_gn=PKHD1;refGene_tx=XM_011282361.3 |
| B2 | 52236278 | 52236407 | refGene_gn=PKHD1;refGene_tx=XM_011282361.3 |
| B2 | 52242445 | 52242511 | refGene_gn=PKHD1;refGene_tx=XM_011282361.3 |
| B2 | 52253709 | 52253905 | refGene_gn=PKHD1;refGene_tx=XM_011282361.3 |
| B2 | 52255358 | 52255536 | refGene_gn=PKHD1;refGene_tx=XM_011282361.3 |
| B2 | 52269216 | 52269463 | refGene_gn=PKHD1;refGene_tx=XM_011282361.3 |
| B2 | 52271987 | 52272123 | refGene_gn=PKHD1;refGene_tx=XM_011282361.3 |
| B2 | 52278168 | 52278303 | refGene_gn=PKHD1;refGene_tx=XM_011282361.3 |
| B2 | 52282279 | 52282385 | refGene_gn=PKHD1;refGene_tx=XM_011282361.3 |
| B2 | 52283634 | 52283747 | refGene_gn=PKHD1;refGene_tx=XM_011282361.3 |
| B2 | 52296380 | 52296511 | refGene_gn=PKHD1;refGene_tx=XM_011282361.3 |
| B2 | 52296877 | 52296934 | refGene_gn=PKHD1;refGene_tx=XM_011282361.3 |
| B2 | 52299592 | 52299718 | refGene_gn=PKHD1;refGene_tx=XM_011282361.3 |
| B2 | 52302945 | 52303137 | refGene_gn=PKHD1;refGene_tx=XM_011282361.3 |
| B2 | 52305377 | 52305535 | refGene_gn=PKHD1;refGene_tx=XM_011282361.3 |
| B2 | 52305934 | 52306145 | refGene_gn=PKHD1;refGene_tx=XM_011282361.3 |
| B2 | 52326816 | 52327029 | refGene_gn=PKHD1;refGene_tx=XM_011282361.3 |
| B2 | 52347325 | 52347482 | refGene_gn=PKHD1;refGene_tx=XM_011282361.3 |
| B2 | 52405276 | 52405427 | refGene_gn=PKHD1;refGene_tx=XM_011282361.3 |
| B2 | 52412056 | 52412276 | refGene_gn=PKHD1;refGene_tx=XM_011282361.3 |
| B2 | 52417890 | 52418034 | refGene_gn=PKHD1;refGene_tx=XM_011282361.3 |
| B2 | 52420099 | 52421707 | refGene_gn=PKHD1;refGene_tx=XM_011282361.3 |
| B2 | 52423443 | 52423511 | refGene_gn=PKHD1;refGene_tx=XM_011282361.3 |
| B2 | 52424184 | 52424380 | refGene_gn=PKHD1;refGene_tx=XM_011282361.3 |
| B2 | 52428821 | 52428957 | refGene_gn=PKHD1;refGene_tx=XM_011282361.3 |
| B2 | 52431195 | 52431326 | refGene_gn=PKHD1;refGene_tx=XM_011282361.3 |
| B2 | 52446645 | 52446921 | refGene_gn=PKHD1;refGene_tx=XM_011282361.3 |
| B2 | 52447405 | 52447511 | refGene_gn=PKHD1;refGene_tx=XM_011282361.3 |
| B2 | 52449289 | 52449412 | refGene_gn=PKHD1;refGene_tx=XM_011282361.3 |
| B2 | 52450315 | 52450500 | refGene_gn=PKHD1;refGene_tx=XM_011282361.3 |
| B2 | 52452995 | 52453123 | refGene_gn=PKHD1;refGene_tx=XM_011282361.3 |
| B2 | 52455849 | 52455988 | refGene_gn=PKHD1;refGene_tx=XM_011282361.3 |
| B2 | 52458648 | 52458824 | refGene_gn=PKHD1;refGene_tx=XM_011282361.3 |
| B2 | 52460081 | 52460212 | refGene_gn=PKHD1;refGene_tx=XM_011282361.3 |
| B2 | 52462172 | 52462315 | refGene_gn=PKHD1;refGene_tx=XM_011282361.3 |
| B2 | 52463196 | 52463287 | refGene_gn=PKHD1;refGene_tx=XM_011282361.3 |
| B2 | 52463611 | 52463701 | refGene_gn=PKHD1;refGene_tx=XM_011282361.3 |
| B2 | 52464711 | 52464990 | refGene_gn=PKHD1;refGene_tx=XM_011282361.3 |
| B2 | 52466208 | 52466323 | refGene_gn=PKHD1;refGene_tx=XM_011282361.3 |
| B2 | 52469422 | 52469564 | refGene_gn=PKHD1;refGene_tx=XM_011282361.3 |
| B2 | 52472178 | 52472274 | refGene_gn=PKHD1;refGene_tx=XM_011282361.3 |
| B2 | 52473748 | 52473850 | refGene_gn=PKHD1;refGene_tx=XM_011282361.3 |
| B2 | 52477053 | 52477124 | refGene_gn=PKHD1;refGene_tx=XM_011282361.3 |
| B2 | 52477965 | 52478005 | refGene_gn=PKHD1;refGene_tx=XM_011282361.3 |
| B2 | 52478490 | 52478555 | refGene_gn=PKHD1;refGene_tx=XM_011282361.3 |
| B2 | 52479456 | 52479531 | refGene_gn=PKHD1;refGene_tx=XM_011282361.3 |
| B2 | 52480809 | 52480888 | refGene_gn=PKHD1;refGene_tx=XM_011282361.3 |

| | | | |
|---|---|---|---|
| B2 | 52484249 | 52484307 | refGene_gn=PKHD1;refGene_tx=XM_011282361.3 |
| B2 | 52488608 | 52488717 | refGene_gn=PKHD1;refGene_tx=XM_011282361.3 |
| B2 | 52492517 | 52492662 | refGene_gn=PKHD1;refGene_tx=XM_011282361.3 |
| B2 | 52493493 | 52493571 | refGene_gn=PKHD1;refGene_tx=XM_011282361.3 |
| B2 | 52495185 | 52495237 | refGene_gn=PKHD1;refGene_tx=XM_011282361.3 |
| B2 | 52598739 | 52598766 | refGene_gn=IL17A;refGene_tx=XM_006931816.2 |
| B2 | 52599892 | 52600095 | refGene_gn=IL17A;refGene_tx=XM_006931816.2 |
| B2 | 52601568 | 52601806 | refGene_gn=IL17A;refGene_tx=XM_006931816.2 |
| B2 | 52661389 | 52661630 | refGene_gn=IL17F;refGene_tx=XM_023254749.1 |
| B2 | 52662755 | 52662943 | refGene_gn=IL17F;refGene_tx=XM_023254749.1 |
| B2 | 52684943 | 52685142 | refGene_gn=MCM3;refGene_tx=XM_003986232.4 |
| B2 | 52686503 | 52686573 | refGene_gn=MCM3;refGene_tx=XM_003986232.4 |
| B2 | 52686984 | 52687070 | refGene_gn=MCM3;refGene_tx=XM_003986232.4 |
| B2 | 52687597 | 52687701 | refGene_gn=MCM3;refGene_tx=XM_003986232.4 |
| B2 | 52689849 | 52689990 | refGene_gn=MCM3;refGene_tx=XM_003986232.4 |
| B2 | 52692801 | 52692952 | refGene_gn=MCM3;refGene_tx=XM_003986232.4 |
| B2 | 52693742 | 52693869 | refGene_gn=MCM3;refGene_tx=XM_003986232.4 |
| B2 | 52694201 | 52694376 | refGene_gn=MCM3;refGene_tx=XM_003986232.4 |
| B2 | 52695232 | 52695441 | refGene_gn=MCM3;refGene_tx=XM_003986232.4 |
| B2 | 52696018 | 52696150 | refGene_gn=MCM3;refGene_tx=XM_003986232.4 |
| B2 | 52696486 | 52696640 | refGene_gn=MCM3;refGene_tx=XM_003986232.4 |
| B2 | 52697556 | 52697665 | refGene_gn=MCM3;refGene_tx=XM_003986232.4 |
| B2 | 52698140 | 52698379 | refGene_gn=MCM3;refGene_tx=XM_003986232.4 |
| B2 | 52700337 | 52700468 | refGene_gn=MCM3;refGene_tx=XM_003986232.4 |
| B2 | 52700884 | 52701093 | refGene_gn=MCM3;refGene_tx=XM_003986232.4 |
| B2 | 52701515 | 52701628 | refGene_gn=MCM3;refGene_tx=XM_003986232.4 |
| B2 | 52702988 | 52703066 | refGene_gn=MCM3;refGene_tx=XM_003986232.4 |
| B2 | 52820555 | 52821620 | refGene_gn=PAQR8;refGene_tx=XM_023254057.1 |
| B2 | 52833967 | 52834030 | refGene_gn=EFHC1;refGene_tx=XM_003986234.5 |
| B2 | 52843156 | 52843378 | refGene_gn=EFHC1;refGene_tx=XM_003986234.5 |
| B2 | 52846593 | 52846881 | refGene_gn=EFHC1;refGene_tx=XM_003986234.5 |
| B2 | 52851693 | 52851843 | refGene_gn=EFHC1;refGene_tx=XM_003986234.5 |
| B2 | 52855427 | 52855620 | refGene_gn=EFHC1;refGene_tx=XM_003986234.5 |
| B2 | 52862521 | 52862742 | refGene_gn=EFHC1;refGene_tx=XM_003986234.5 |
| B2 | 52866843 | 52866984 | refGene_gn=EFHC1;refGene_tx=XM_003986234.5 |
| B2 | 52875600 | 52875814 | refGene_gn=EFHC1;refGene_tx=XM_003986234.5 |
| B2 | 52876207 | 52876355 | refGene_gn=EFHC1;refGene_tx=XM_003986234.5 |
| B2 | 52900155 | 52900366 | refGene_gn=EFHC1;refGene_tx=XM_003986234.5 |
| B2 | 52901783 | 52901864 | refGene_gn=EFHC1;refGene_tx=XM_003986234.5 |
| B2 | 52910976 | 52911050 | refGene_gn=TRAM2;refGene_tx=XM_003986235.5 |
| B2 | 52912271 | 52912435 | refGene_gn=TRAM2;refGene_tx=XM_003986235.5 |
| B2 | 52913275 | 52913419 | refGene_gn=TRAM2;refGene_tx=XM_003986235.5 |
| B2 | 52913682 | 52913787 | refGene_gn=TRAM2;refGene_tx=XM_003986235.5 |
| B2 | 52916552 | 52916623 | refGene_gn=TRAM2;refGene_tx=XM_003986235.5 |
| B2 | 52917193 | 52917278 | refGene_gn=TRAM2;refGene_tx=XM_003986235.5 |
| B2 | 52918447 | 52918506 | refGene_gn=TRAM2;refGene_tx=XM_003986235.5 |
| B2 | 52924847 | 52924964 | refGene_gn=TRAM2;refGene_tx=XM_003986235.5 |
| B2 | 52925471 | 52925581 | refGene_gn=TRAM2;refGene_tx=XM_003986235.5 |
| B2 | 52945905 | 52945969 | refGene_gn=TRAM2;refGene_tx=XM_003986235.5 |
| B2 | 52988820 | 52988940 | refGene_gn=TRAM2;refGene_tx=XM_003986235.5 |

B2 53088834 53088904 refGene_gn=TMEM14A;refGene_tx=XM_003986236.5
B2 53093285 53093387 refGene_gn=TMEM14A;refGene_tx=XM_003986236.5
B2 53095570 53095658 refGene_gn=TMEM14A;refGene_tx=XM_003986236.5
B2 53097618 53097658 refGene_gn=TMEM14A;refGene_tx=XM_003986236.5
B2 53134860 53134986
refGene_gn=LOC101096317;refGene_tx=XM_011282429.3;refGene_gn=LOC101096317;refGene_tx=XM_023254059.1
B2 53135851 53135983
refGene_gn=LOC101096317;refGene_tx=XM_011282429.3;refGene_gn=LOC101096317;refGene_tx=XM_023254059.1
B2 53137487 53137629
refGene_gn=LOC101096317;refGene_tx=XM_011282429.3;refGene_gn=LOC101096317;refGene_tx=XM_023254059.1
B2 53139528 53139661
refGene_gn=LOC101096317;refGene_tx=XM_011282429.3
B2 53140903 53140955
refGene_gn=LOC101096317;refGene_tx=XM_011282429.3
B2 53141336 53141423
refGene_gn=LOC101096317;refGene_tx=XM_011282429.3
B2 53151106 53151232
refGene_gn=LOC111560471;refGene_tx=XM_023254060.1
B2 53152096 53152228
refGene_gn=LOC111560471;refGene_tx=XM_023254060.1
B2 53153877 53154019
refGene_gn=LOC111560471;refGene_tx=XM_023254060.1
B2 53156500 53156520
refGene_gn=LOC111560471;refGene_tx=XM_023254060.1
B2 53158001 53158004
refGene_gn=LOC111560471;refGene_tx=XM_023254060.1
B2 53171383 53171506
refGene_gn=LOC101084793;refGene_tx=XM_023254061.1
B2 53172716 53172848
refGene_gn=LOC101084793;refGene_tx=XM_023254061.1
B2 53177471 53177613
refGene_gn=LOC101084793;refGene_tx=XM_023254061.1
B2 53179697 53179830
refGene_gn=LOC101084793;refGene_tx=XM_023254061.1
B2 53182399 53182451
refGene_gn=LOC101084793;refGene_tx=XM_023254061.1
B2 53184019 53184106
refGene_gn=LOC101084793;refGene_tx=XM_023254061.1
B2 53197801 53197924 refGene_gn=GSTA4;refGene_tx=XM_011282427.2
B2 53201492 53201624 refGene_gn=GSTA4;refGene_tx=XM_011282427.2
B2 53203994 53204136 refGene_gn=GSTA4;refGene_tx=XM_011282427.2
B2 53205012 53205145 refGene_gn=GSTA4;refGene_tx=XM_011282427.2
B2 53206498 53206550 refGene_gn=GSTA4;refGene_tx=XM_011282427.2
B2 53210761 53210848 refGene_gn=GSTA4;refGene_tx=XM_011282427.2
B2 53224136 53224291
refGene_gn=ICK;refGene_tx=XM_003986239.4;refGene_gn=ICK;refGene_tx=XM_019830723.1;refGene_gn=ICK;refGene_tx=XM_019830724.1

B2 53225522 53225645
refGene_gn=ICK;refGene_tx=XM_003986239.4;refGene_gn=ICK;refGene_tx=XM_019830723.1;refGene_gn=ICK;refGene_tx=XM_019830724.1
B2 53227867 53227996
refGene_gn=ICK;refGene_tx=XM_003986239.4;refGene_gn=ICK;refGene_tx=XM_019830723.1;refGene_gn=ICK;refGene_tx=XM_019830724.1
B2 53229929 53230078
refGene_gn=ICK;refGene_tx=XM_003986239.4;refGene_gn=ICK;refGene_tx=XM_019830723.1;refGene_gn=ICK;refGene_tx=XM_019830724.1
B2 53230195 53230386
refGene_gn=ICK;refGene_tx=XM_003986239.4;refGene_gn=ICK;refGene_tx=XM_019830723.1;refGene_gn=ICK;refGene_tx=XM_019830724.1
B2 53232127 53232448
refGene_gn=ICK;refGene_tx=XM_003986239.4;refGene_gn=ICK;refGene_tx=XM_019830723.1;refGene_gn=ICK;refGene_tx=XM_019830724.1
B2 53234949 53235117
refGene_gn=ICK;refGene_tx=XM_003986239.4;refGene_gn=ICK;refGene_tx=XM_019830723.1;refGene_gn=ICK;refGene_tx=XM_019830724.1
B2 53237116 53237288
refGene_gn=ICK;refGene_tx=XM_003986239.4;refGene_gn=ICK;refGene_tx=XM_019830723.1;refGene_gn=ICK;refGene_tx=XM_019830724.1
B2 53238644 53238777
refGene_gn=ICK;refGene_tx=XM_003986239.4;refGene_gn=ICK;refGene_tx=XM_019830723.1;refGene_gn=ICK;refGene_tx=XM_019830724.1
B2 53248281 53248361
refGene_gn=ICK;refGene_tx=XM_003986239.4;refGene_gn=ICK;refGene_tx=XM_019830723.1;refGene_gn=ICK;refGene_tx=XM_019830724.1
B2 53249363 53249485
refGene_gn=ICK;refGene_tx=XM_003986239.4;refGene_gn=ICK;refGene_tx=XM_019830723.1;refGene_gn=ICK;refGene_tx=XM_019830724.1
B2 53254315 53254370
refGene_gn=ICK;refGene_tx=XM_003986239.4;refGene_gn=ICK;refGene_tx=XM_019830723.1
B2 53258028 53258129
refGene_gn=ICK;refGene_tx=XM_003986239.4;refGene_gn=ICK;refGene_tx=XM_019830723.1
B2 53283033 53283036 refGene_gn=FBXO9;refGene_tx=XM_019830726.1
B2 53286849 53286933 refGene_gn=FBXO9;refGene_tx=XM_019830726.1
B2 53301678 53301834 refGene_gn=FBXO9;refGene_tx=XM_019830726.1
B2 53305035 53305093 refGene_gn=FBXO9;refGene_tx=XM_019830726.1
B2 53306456 53306556 refGene_gn=FBXO9;refGene_tx=XM_019830726.1
B2 53309701 53309832 refGene_gn=FBXO9;refGene_tx=XM_019830726.1
B2 53310585 53310700 refGene_gn=FBXO9;refGene_tx=XM_019830726.1
B2 53313368 53313487 refGene_gn=FBXO9;refGene_tx=XM_019830726.1
B2 53313697 53313788 refGene_gn=FBXO9;refGene_tx=XM_019830726.1
B2 53314894 53314990 refGene_gn=FBXO9;refGene_tx=XM_019830726.1
B2 53315318 53315412 refGene_gn=FBXO9;refGene_tx=XM_019830726.1
B2 53317624 53317776 refGene_gn=FBXO9;refGene_tx=XM_019830726.1
B2 53328044 53328153 refGene_gn=FBXO9;refGene_tx=XM_019830726.1
B2 53382343 53383081

refGene_gn=GCM1;refGene_tx=XM_003986242.2;refGene_gn=GCM1;refGene_tx=XM_023254062.1
B2 53384839 53384968
refGene_gn=GCM1;refGene_tx=XM_003986242.2;refGene_gn=GCM1;refGene_tx=XM_023254062.1
B2 53385943 53386056
refGene_gn=GCM1;refGene_tx=XM_003986242.2;refGene_gn=GCM1;refGene_tx=XM_023254062.1
B2 53387097 53387350
refGene_gn=GCM1;refGene_tx=XM_003986242.2;refGene_gn=GCM1;refGene_tx=XM_023254062.1
B2 53398745 53398820
refGene_gn=GCM1;refGene_tx=XM_003986242.2;refGene_gn=GCM1;refGene_tx=XM_023254062.1
B2 53490183 53490327
refGene_gn=ELOVL5;refGene_tx=XM_003986243.5;refGene_gn=ELOVL5;refGene_tx=XM_011282426.3;refGene_gn=ELOVL5;refGene_tx=XM_019830732.2
B2 53491447 53491582
refGene_gn=ELOVL5;refGene_tx=XM_003986243.5;refGene_gn=ELOVL5;refGene_tx=XM_011282426.3;refGene_gn=ELOVL5;refGene_tx=XM_019830732.2
B2 53494958 53495083
refGene_gn=ELOVL5;refGene_tx=XM_003986243.5;refGene_gn=ELOVL5;refGene_tx=XM_011282426.3;refGene_gn=ELOVL5;refGene_tx=XM_019830732.2
B2 53496847 53497019
refGene_gn=ELOVL5;refGene_tx=XM_003986243.5;refGene_gn=ELOVL5;refGene_tx=XM_011282426.3;refGene_gn=ELOVL5;refGene_tx=XM_019830732.2
B2 53497758 53497836
refGene_gn=ELOVL5;refGene_tx=XM_003986243.5;refGene_gn=ELOVL5;refGene_tx=XM_011282426.3;refGene_gn=ELOVL5;refGene_tx=XM_019830732.2
B2 53512435 53512623
refGene_gn=ELOVL5;refGene_tx=XM_003986243.5;refGene_gn=ELOVL5;refGene_tx=XM_011282426.3;refGene_gn=ELOVL5;refGene_tx=XM_019830732.2
B2 53515665 53515723
refGene_gn=ELOVL5;refGene_tx=XM_003986243.5;refGene_gn=ELOVL5;refGene_tx=XM_019830732.2
B2 53559246 53559291
refGene_gn=LOC105260579;refGene_tx=XM_019831595.1
B2 53561487 53561566
refGene_gn=LOC105260579;refGene_tx=XM_019831595.1
B2 53562098 53562855
refGene_gn=LOC105260579;refGene_tx=XM_019831595.1
B2 53563530 53563762
refGene_gn=LOC105260579;refGene_tx=XM_019831595.1
B2 53764279 53764491 refGene_gn=GCLC;refGene_tx=XM_023254063.1
B2 53765729 53765850 refGene_gn=GCLC;refGene_tx=XM_023254063.1
B2 53765926 53766030 refGene_gn=GCLC;refGene_tx=XM_023254063.1
B2 53766330 53766412 refGene_gn=GCLC;refGene_tx=XM_023254063.1
B2 53773417 53773522 refGene_gn=GCLC;refGene_tx=XM_023254063.1
B2 53773820 53773913 refGene_gn=GCLC;refGene_tx=XM_023254063.1
B2 53774645 53774758 refGene_gn=GCLC;refGene_tx=XM_023254063.1

B2 53775200 53775339 refGene_gn=GCLC;refGene_tx=XM_023254063.1
B2 53776369 53776486 refGene_gn=GCLC;refGene_tx=XM_023254063.1
B2 53777180 53777255 refGene_gn=GCLC;refGene_tx=XM_023254063.1
B2 53781504 53781638 refGene_gn=GCLC;refGene_tx=XM_023254063.1
B2 53781747 53781806 refGene_gn=GCLC;refGene_tx=XM_023254063.1
B2 53783457 53783571 refGene_gn=GCLC;refGene_tx=XM_023254063.1
B2 53787075 53787258 refGene_gn=GCLC;refGene_tx=XM_023254063.1
B2 53788119 53788232 refGene_gn=GCLC;refGene_tx=XM_023254063.1
B2 53812001 53812151 refGene_gn=GCLC;refGene_tx=XM_023254063.1
B2 53930902 53931635 refGene_gn=KLHL31;refGene_tx=XM_006931802.3
B2 53933320 53934492 refGene_gn=KLHL31;refGene_tx=XM_006931802.3
B2 54082153 54082312
refGene_gn=LRRC1;refGene_tx=XM_023254064.1;refGene_gn=LRRC1;refGene_tx=XM_023254065.1
B2 54135122 54135240
refGene_gn=LRRC1;refGene_tx=XM_023254064.1;refGene_gn=LRRC1;refGene_tx=XM_023254065.1
B2 54166853 54166932
refGene_gn=LRRC1;refGene_tx=XM_023254064.1;refGene_gn=LRRC1;refGene_tx=XM_023254065.1
B2 54171212 54171302
refGene_gn=LRRC1;refGene_tx=XM_023254064.1;refGene_gn=LRRC1;refGene_tx=XM_023254065.1
B2 54178493 54178517 refGene_gn=LRRC1;refGene_tx=XM_023254064.1
B2 54188100 54188157
refGene_gn=LRRC1;refGene_tx=XM_023254064.1;refGene_gn=LRRC1;refGene_tx=XM_023254065.1
B2 54188443 54188507
refGene_gn=LRRC1;refGene_tx=XM_023254064.1;refGene_gn=LRRC1;refGene_tx=XM_023254065.1
B2 54189139 54189214
refGene_gn=LRRC1;refGene_tx=XM_023254064.1;refGene_gn=LRRC1;refGene_tx=XM_023254065.1
B2 54192404 54192549
refGene_gn=LRRC1;refGene_tx=XM_023254064.1;refGene_gn=LRRC1;refGene_tx=XM_023254065.1
B2 54194692 54194811
refGene_gn=LRRC1;refGene_tx=XM_023254064.1;refGene_gn=LRRC1;refGene_tx=XM_023254065.1
B2 54196310 54196394
refGene_gn=LRRC1;refGene_tx=XM_023254064.1;refGene_gn=LRRC1;refGene_tx=XM_023254065.1
B2 54206260 54206376
refGene_gn=LRRC1;refGene_tx=XM_023254064.1;refGene_gn=LRRC1;refGene_tx=XM_023254065.1
B2 54214117 54214290
refGene_gn=LRRC1;refGene_tx=XM_023254064.1;refGene_gn=LRRC1;refGene_tx=XM_023254065.1
B2 54215164 54215301
refGene_gn=LRRC1;refGene_tx=XM_023254064.1;refGene_gn=LRRC1;refGene_tx

=XM_023254065.1
B2 54216943 54217102
refGene_gn=LRRC1;refGene_tx=XM_023254064.1;refGene_gn=LRRC1;refGene_tx
=XM_023254065.1
B2 54304913 54304976 refGene_gn=MLIP;refGene_tx=XM_023254067.1
B2 54410148 54410244
refGene_gn=MLIP;refGene_tx=XM_023254068.1;refGene_gn=MLIP;refGene_tx=X
M_023254069.1
B2 54417723 54417879
refGene_gn=MLIP;refGene_tx=XM_023254067.1;refGene_gn=MLIP;refGene_tx=X
M_023254068.1;refGene_gn=MLIP;refGene_tx=XM_023254069.1
B2 54421607 54422000
refGene_gn=MLIP;refGene_tx=XM_023254067.1;refGene_gn=MLIP;refGene_tx=X
M_023254068.1;refGene_gn=MLIP;refGene_tx=XM_023254069.1
B2 54438081 54439647
refGene_gn=MLIP;refGene_tx=XM_023254067.1;refGene_gn=MLIP;refGene_tx=X
M_023254068.1;refGene_gn=MLIP;refGene_tx=XM_023254069.1
B2 54451792 54451864
refGene_gn=MLIP;refGene_tx=XM_023254067.1;refGene_gn=MLIP;refGene_tx=X
M_023254068.1;refGene_gn=MLIP;refGene_tx=XM_023254069.1
B2 54466500 54466566
refGene_gn=MLIP;refGene_tx=XM_023254067.1;refGene_gn=MLIP;refGene_tx=X
M_023254068.1;refGene_gn=MLIP;refGene_tx=XM_023254069.1
B2 54466649 54466733
refGene_gn=MLIP;refGene_tx=XM_023254067.1;refGene_gn=MLIP;refGene_tx=X
M_023254068.1;refGene_gn=MLIP;refGene_tx=XM_023254069.1
B2 54466845 54466905
refGene_gn=MLIP;refGene_tx=XM_023254067.1;refGene_gn=MLIP;refGene_tx=X
M_023254068.1;refGene_gn=MLIP;refGene_tx=XM_023254069.1
B2 54481109 54481154
refGene_gn=MLIP;refGene_tx=XM_023254067.1;refGene_gn=MLIP;refGene_tx=X
M_023254068.1;refGene_gn=MLIP;refGene_tx=XM_023254069.1
B2 54490977 54491022
refGene_gn=MLIP;refGene_tx=XM_023254067.1;refGene_gn=MLIP;refGene_tx=X
M_023254068.1;refGene_gn=MLIP;refGene_tx=XM_023254069.1
B2 54502167 54502296
refGene_gn=MLIP;refGene_tx=XM_023254067.1;refGene_gn=MLIP;refGene_tx=X
M_023254068.1;refGene_gn=MLIP;refGene_tx=XM_023254069.1
B2 54517974 54518181
refGene_gn=MLIP;refGene_tx=XM_023254067.1;refGene_gn=MLIP;refGene_tx=X
M_023254068.1
B2 54541038 54541092 refGene_gn=MLIP;refGene_tx=XM_023254067.1
B2 54552502 54552508
refGene_gn=MLIP;refGene_tx=XM_023254067.1;refGene_gn=MLIP;refGene_tx=X
M_023254068.1;refGene_gn=MLIP;refGene_tx=XM_023254069.1
B2 54585605 54585951
refGene_gn=TINAG;refGene_tx=XM_006931811.2;refGene_gn=TINAG;refGene_tx
=XM_019830738.1
B2 54590862 54590926
refGene_gn=TINAG;refGene_tx=XM_006931811.2;refGene_gn=TINAG;refGene_tx

=XM_019830738.1
B2	54591594	54591684
refGene_gn=TINAG;refGene_tx=XM_006931811.2;refGene_gn=TINAG;refGene_tx
=XM_019830738.1
B2	54598110	54598225
refGene_gn=TINAG;refGene_tx=XM_006931811.2;refGene_gn=TINAG;refGene_tx
=XM_019830738.1
B2	54615904	54616028
refGene_gn=TINAG;refGene_tx=XM_006931811.2;refGene_gn=TINAG;refGene_tx
=XM_019830738.1
B2	54619582	54619733
refGene_gn=TINAG;refGene_tx=XM_006931811.2;refGene_gn=TINAG;refGene_tx
=XM_019830738.1
B2	54621824	54622005
refGene_gn=TINAG;refGene_tx=XM_006931811.2;refGene_gn=TINAG;refGene_tx
=XM_019830738.1
B2	54623033	54623120	refGene_gn=TINAG;refGene_tx=XM_019830738.1
B2	54623691	54623737	refGene_gn=TINAG;refGene_tx=XM_006931811.2
B2	54626850	54626977	refGene_gn=TINAG;refGene_tx=XM_006931811.2
B2	54658281	54658327	refGene_gn=TINAG;refGene_tx=XM_006931811.2
B2	54667844	54667979	refGene_gn=TINAG;refGene_tx=XM_006931811.2
B2	55037929	55038373	refGene_gn=FAM83B;refGene_tx=XM_003986249.4
B2	55097362	55097527	refGene_gn=FAM83B;refGene_tx=XM_003986249.4
B2	55098304	55098429	refGene_gn=FAM83B;refGene_tx=XM_003986249.4
B2	55105315	55107617	refGene_gn=FAM83B;refGene_tx=XM_003986249.4
B2	55292507	55292730
refGene_gn=HCRTR2;refGene_tx=XM_003986250.4;refGene_gn=HCRTR2;refGene_
tx=XM_019830741.1
B2	55363338	55363517
refGene_gn=HCRTR2;refGene_tx=XM_003986250.4;refGene_gn=HCRTR2;refGene_
tx=XM_019830741.1
B2	55369040	55369284
refGene_gn=HCRTR2;refGene_tx=XM_003986250.4;refGene_gn=HCRTR2;refGene_
tx=XM_019830741.1
B2	55376703	55376819
refGene_gn=HCRTR2;refGene_tx=XM_003986250.4;refGene_gn=HCRTR2;refGene_
tx=XM_019830741.1
B2	55395654	55395875
refGene_gn=HCRTR2;refGene_tx=XM_003986250.4;refGene_gn=HCRTR2;refGene_
tx=XM_019830741.1
B2	55398886	55399008
refGene_gn=HCRTR2;refGene_tx=XM_003986250.4;refGene_gn=HCRTR2;refGene_
tx=XM_019830741.1
B2	55401087	55401317
refGene_gn=HCRTR2;refGene_tx=XM_003986250.4;refGene_gn=HCRTR2;refGene_
tx=XM_019830741.1
B2	55403485	55403486	refGene_gn=HCRTR2;refGene_tx=XM_019830741.1
B2	55427621	55427643	refGene_gn=GFRAL;refGene_tx=XM_023254070.1
B2	55432349	55432484	refGene_gn=GFRAL;refGene_tx=XM_023254070.1
B2	55434986	55435139	refGene_gn=GFRAL;refGene_tx=XM_023254070.1

B2 55447764 55447821 refGene_gn=GFRAL;refGene_tx=XM_023254070.1
B2 55448672 55449003 refGene_gn=GFRAL;refGene_tx=XM_023254070.1
B2 55454491 55454742 refGene_gn=GFRAL;refGene_tx=XM_023254070.1
B2 55484490 55484586 refGene_gn=GFRAL;refGene_tx=XM_023254070.1
B2 55484673 55484746 refGene_gn=GFRAL;refGene_tx=XM_023254070.1
B2 55487159 55487223 refGene_gn=GFRAL;refGene_tx=XM_023254070.1
B2 55523262 55523364
refGene_gn=HMGCLL1;refGene_tx=XM_003986251.4;refGene_gn=HMGCLL1;refGene_tx=XM_011282424.3
B2 55527224 55527350
refGene_gn=HMGCLL1;refGene_tx=XM_003986251.4;refGene_gn=HMGCLL1;refGene_tx=XM_011282424.3
B2 55547096 55547141 refGene_gn=HMGCLL1;refGene_tx=XM_019830747.2
B2 55560488 55560539 refGene_gn=HMGCLL1;refGene_tx=XM_019830745.2
B2 55593519 55593564 refGene_gn=HMGCLL1;refGene_tx=XM_019830746.2
B2 55622413 55622602
refGene_gn=HMGCLL1;refGene_tx=XM_003986251.4;refGene_gn=HMGCLL1;refGene_tx=XM_011282424.3;refGene_gn=HMGCLL1;refGene_tx=XM_019830745.2;refGene_gn=HMGCLL1;refGene_tx=XM_019830746.2;refGene_gn=HMGCLL1;refGene_tx=XM_019830747.2
B2 55624185 55624249
refGene_gn=HMGCLL1;refGene_tx=XM_003986251.4;refGene_gn=HMGCLL1;refGene_tx=XM_011282424.3;refGene_gn=HMGCLL1;refGene_tx=XM_019830745.2;refGene_gn=HMGCLL1;refGene_tx=XM_019830746.2;refGene_gn=HMGCLL1;refGene_tx=XM_019830747.2
B2 55634891 55635040
refGene_gn=HMGCLL1;refGene_tx=XM_003986251.4;refGene_gn=HMGCLL1;refGene_tx=XM_011282424.3;refGene_gn=HMGCLL1;refGene_tx=XM_019830745.2;refGene_gn=HMGCLL1;refGene_tx=XM_019830746.2;refGene_gn=HMGCLL1;refGene_tx=XM_019830747.2
B2 55637012 55637108
refGene_gn=HMGCLL1;refGene_tx=XM_003986251.4;refGene_gn=HMGCLL1;refGene_tx=XM_011282424.3;refGene_gn=HMGCLL1;refGene_tx=XM_019830745.2;refGene_gn=HMGCLL1;refGene_tx=XM_019830746.2;refGene_gn=HMGCLL1;refGene_tx=XM_019830747.2
B2 55668403 55668511
refGene_gn=HMGCLL1;refGene_tx=XM_003986251.4;refGene_gn=HMGCLL1;refGene_tx=XM_019830745.2;refGene_gn=HMGCLL1;refGene_tx=XM_019830746.2;refGene_gn=HMGCLL1;refGene_tx=XM_019830747.2
B2 55668743 55668824
refGene_gn=HMGCLL1;refGene_tx=XM_003986251.4;refGene_gn=HMGCLL1;refGene_tx=XM_019830745.2;refGene_gn=HMGCLL1;refGene_tx=XM_019830746.2;refGene_gn=HMGCLL1;refGene_tx=XM_019830747.2
B2 55715183 55715291
refGene_gn=HMGCLL1;refGene_tx=XM_003986251.4;refGene_gn=HMGCLL1;refGene_tx=XM_011282424.3;refGene_gn=HMGCLL1;refGene_tx=XM_019830745.2;refGene_gn=HMGCLL1;refGene_tx=XM_019830746.2;refGene_gn=HMGCLL1;refGene_tx=XM_019830747.2
B2 55859393 55859543 refGene_gn=BMP5;refGene_tx=XM_003986274.4
B2 55862962 55863073 refGene_gn=BMP5;refGene_tx=XM_003986274.4

B2 55864676 55864753 refGene_gn=BMP5;refGene_tx=XM_003986274.4
B2 55875941 55876136 refGene_gn=BMP5;refGene_tx=XM_003986274.4
B2 55895741 55895890 refGene_gn=BMP5;refGene_tx=XM_003986274.4
B2 55915578 55915771 refGene_gn=BMP5;refGene_tx=XM_003986274.4
B2 55972814 55973304 refGene_gn=BMP5;refGene_tx=XM_003986274.4
B2 56131430 56131618 refGene_gn=COL21A1;refGene_tx=XM_023254071.1
B2 56132942 56133020 refGene_gn=COL21A1;refGene_tx=XM_023254071.1
B2 56134217 56134418 refGene_gn=COL21A1;refGene_tx=XM_023254071.1
B2 56134915 56134970 refGene_gn=COL21A1;refGene_tx=XM_023254071.1
B2 56135060 56135207 refGene_gn=COL21A1;refGene_tx=XM_023254071.1
B2 56135791 56135824 refGene_gn=COL21A1;refGene_tx=XM_023254071.1
B2 56138687 56138737 refGene_gn=COL21A1;refGene_tx=XM_023254071.1
B2 56141600 56141631 refGene_gn=COL21A1;refGene_tx=XM_023254071.1
B2 56143547 56143619 refGene_gn=COL21A1;refGene_tx=XM_023254071.1
B2 56145215 56145269 refGene_gn=COL21A1;refGene_tx=XM_023254071.1
B2 56148206 56148260 refGene_gn=COL21A1;refGene_tx=XM_023254071.1
B2 56149430 56149484 refGene_gn=COL21A1;refGene_tx=XM_023254071.1
B2 56151514 56151559 refGene_gn=COL21A1;refGene_tx=XM_023254071.1
B2 56170752 56170806 refGene_gn=COL21A1;refGene_tx=XM_023254071.1
B2 56187614 56187668 refGene_gn=COL21A1;refGene_tx=XM_023254071.1
B2 56187785 56187839 refGene_gn=COL21A1;refGene_tx=XM_023254071.1
B2 56188872 56188926 refGene_gn=COL21A1;refGene_tx=XM_023254071.1
B2 56190037 56190091 refGene_gn=COL21A1;refGene_tx=XM_023254071.1
B2 56206504 56206558 refGene_gn=COL21A1;refGene_tx=XM_023254071.1
B2 56206648 56206702 refGene_gn=COL21A1;refGene_tx=XM_023254071.1
B2 56221203 56221266 refGene_gn=COL21A1;refGene_tx=XM_023254071.1
B2 56228550 56228634 refGene_gn=COL21A1;refGene_tx=XM_023254071.1
B2 56228944 56228953 refGene_gn=COL21A1;refGene_tx=XM_023254071.1
B2 56230767 56230845 refGene_gn=COL21A1;refGene_tx=XM_023254071.1
B2 56232004 56232178 refGene_gn=COL21A1;refGene_tx=XM_023254071.1
B2 56234733 56234950 refGene_gn=COL21A1;refGene_tx=XM_023254071.1
B2 56235062 56235231 refGene_gn=COL21A1;refGene_tx=XM_023254071.1
B2 56250635 56251187 refGene_gn=COL21A1;refGene_tx=XM_023254071.1
B2 56254524 56254612 refGene_gn=COL21A1;refGene_tx=XM_023254071.1
B2 56519557 56519820
refGene_gn=DST;refGene_tx=XM_011282416.2;refGene_gn=DST;refGene_tx=XM_011282421.2;refGene_gn=DST;refGene_tx=XM_011282422.3;refGene_gn=DST;refGene_tx=XM_019830750.2;refGene_gn=DST;refGene_tx=XM_019830751.2;refGene_gn=DST;refGene_tx=XM_019830753.2;refGene_gn=DST;refGene_tx=XM_019830754.2;refGene_gn=DST;refGene_tx=XM_019830755.2;refGene_gn=DST;refGene_tx=XM_019830756.2;refGene_gn=DST;refGene_tx=XM_019830757.2;refGene_gn=DST;refGene_tx=XM_019830758.2;refGene_gn=DST;refGene_tx=XM_019830759.2;refGene_gn=DST;refGene_tx=XM_019830760.2;refGene_gn=DST;refGene_tx=XM_019830761.2;refGene_gn=DST;refGene_tx=XM_019830762.2;refGene_gn=DST;refGene_tx=XM_019830763.2;refGene_gn=DST;refGene_tx=XM_019830764.1;refGene_gn=DST;refGene_tx=XM_019830765.2;refGene_gn=DST;refGene_tx=XM_019830766.2;refGene_gn=DST;refGene_tx=XM_019830768.1;refGene_gn=DST;refGene_tx=XM_019830770.2;refGene_gn=DST;refGene_tx=XM_019830771.2;refGene_gn=DST;refGene_tx=XM_023254073.1
B2 56520638 56520762

refGene_gn=DST;refGene_tx=XM_011282416.2;refGene_gn=DST;refGene_tx=XM_011282421.2;refGene_gn=DST;refGene_tx=XM_011282422.3;refGene_gn=DST;refGene_tx=XM_019830750.2;refGene_gn=DST;refGene_tx=XM_019830751.2;refGene_gn=DST;refGene_tx=XM_019830753.2;refGene_gn=DST;refGene_tx=XM_019830754.2;refGene_gn=DST;refGene_tx=XM_019830755.2;refGene_gn=DST;refGene_tx=XM_019830756.2;refGene_gn=DST;refGene_tx=XM_019830757.2;refGene_gn=DST;refGene_tx=XM_019830758.2;refGene_gn=DST;refGene_tx=XM_019830759.2;refGene_gn=DST;refGene_tx=XM_019830760.2;refGene_gn=DST;refGene_tx=XM_019830761.2;refGene_gn=DST;refGene_tx=XM_019830762.2;refGene_gn=DST;refGene_tx=XM_019830763.2;refGene_gn=DST;refGene_tx=XM_019830764.1;refGene_gn=DST;refGene_tx=XM_019830765.2;refGene_gn=DST;refGene_tx=XM_019830766.2;refGene_gn=DST;refGene_tx=XM_019830768.1;refGene_gn=DST;refGene_tx=XM_019830770.2;refGene_gn=DST;refGene_tx=XM_019830771.2;refGene_gn=DST;refGene_tx=XM_023254073.1
B2 56524098 56524209
refGene_gn=DST;refGene_tx=XM_011282416.2;refGene_gn=DST;refGene_tx=XM_011282421.2;refGene_gn=DST;refGene_tx=XM_011282422.3;refGene_gn=DST;refGene_tx=XM_019830750.2;refGene_gn=DST;refGene_tx=XM_019830751.2;refGene_gn=DST;refGene_tx=XM_019830753.2;refGene_gn=DST;refGene_tx=XM_019830754.2;refGene_gn=DST;refGene_tx=XM_019830755.2;refGene_gn=DST;refGene_tx=XM_019830756.2;refGene_gn=DST;refGene_tx=XM_019830758.2;refGene_gn=DST;refGene_tx=XM_019830759.2;refGene_gn=DST;refGene_tx=XM_019830760.2;refGene_gn=DST;refGene_tx=XM_019830762.2;refGene_gn=DST;refGene_tx=XM_019830763.2;refGene_gn=DST;refGene_tx=XM_019830765.2;refGene_gn=DST;refGene_tx=XM_019830766.2;refGene_gn=DST;refGene_tx=XM_019830768.1;refGene_gn=DST;refGene_tx=XM_019830771.2
B2 56524628 56524828
refGene_gn=DST;refGene_tx=XM_011282416.2;refGene_gn=DST;refGene_tx=XM_011282421.2;refGene_gn=DST;refGene_tx=XM_011282422.3;refGene_gn=DST;refGene_tx=XM_019830750.2;refGene_gn=DST;refGene_tx=XM_019830751.2;refGene_gn=DST;refGene_tx=XM_019830753.2;refGene_gn=DST;refGene_tx=XM_019830754.2;refGene_gn=DST;refGene_tx=XM_019830755.2;refGene_gn=DST;refGene_tx=XM_019830756.2;refGene_gn=DST;refGene_tx=XM_019830757.2;refGene_gn=DST;refGene_tx=XM_019830758.2;refGene_gn=DST;refGene_tx=XM_019830759.2;refGene_gn=DST;refGene_tx=XM_019830760.2;refGene_gn=DST;refGene_tx=XM_019830761.2;refGene_gn=DST;refGene_tx=XM_019830762.2;refGene_gn=DST;refGene_tx=XM_019830763.2;refGene_gn=DST;refGene_tx=XM_019830764.1;refGene_gn=DST;refGene_tx=XM_019830765.2;refGene_gn=DST;refGene_tx=XM_019830766.2;refGene_gn=DST;refGene_tx=XM_019830768.1;refGene_gn=DST;refGene_tx=XM_019830770.2;refGene_gn=DST;refGene_tx=XM_019830771.2;refGene_gn=DST;refGene_tx=XM_023254073.1
B2 56525760 56525832
refGene_gn=DST;refGene_tx=XM_011282422.3;refGene_gn=DST;refGene_tx=XM_019830750.2;refGene_gn=DST;refGene_tx=XM_019830751.2;refGene_gn=DST;refGene_tx=XM_019830753.2;refGene_gn=DST;refGene_tx=XM_019830756.2;refGene_gn=DST;refGene_tx=XM_019830757.2;refGene_gn=DST;refGene_tx=XM_019830758.2;refGene_gn=DST;refGene_tx=XM_019830759.2;refGene_gn=DST;refGene_tx=XM_019830760.2;refGene_gn=DST;refGene_tx=XM_019830761.2;refGene_gn=DST;refGene_tx=XM_019830762.2;refGene_gn=DST;refGene_tx=XM_019830763.2;refGene_gn=DST;refGene_tx=XM_019830764.1;refGene_gn=DST;refGene_tx=X

M_019830765.2;refGene_gn=DST;refGene_tx=XM_019830770.2;refGene_gn=DST;
refGene_tx=XM_023254073.1
B2 56527137 56527255
refGene_gn=DST;refGene_tx=XM_011282416.2;refGene_gn=DST;refGene_tx=XM_
011282421.2;refGene_gn=DST;refGene_tx=XM_011282422.3;refGene_gn=DST;re
fGene_tx=XM_019830750.2;refGene_gn=DST;refGene_tx=XM_019830751.2;refGe
ne_gn=DST;refGene_tx=XM_019830753.2;refGene_gn=DST;refGene_tx=XM_01983
0754.2;refGene_gn=DST;refGene_tx=XM_019830755.2;refGene_gn=DST;refGene
_tx=XM_019830756.2;refGene_gn=DST;refGene_tx=XM_019830757.2;refGene_gn
=DST;refGene_tx=XM_019830758.2;refGene_gn=DST;refGene_tx=XM_019830759.
2;refGene_gn=DST;refGene_tx=XM_019830760.2;refGene_gn=DST;refGene_tx=X
M_019830761.2;refGene_gn=DST;refGene_tx=XM_019830762.2;refGene_gn=DST;
refGene_tx=XM_019830763.2;refGene_gn=DST;refGene_tx=XM_019830764.1;ref
Gene_gn=DST;refGene_tx=XM_019830765.2;refGene_gn=DST;refGene_tx=XM_019
830766.2;refGene_gn=DST;refGene_tx=XM_019830768.1;refGene_gn=DST;refGe
ne_tx=XM_019830770.2;refGene_gn=DST;refGene_tx=XM_019830771.2;refGene_
gn=DST;refGene_tx=XM_023254073.1
B2 56529976 56529994
refGene_gn=DST;refGene_tx=XM_011282416.2;refGene_gn=DST;refGene_tx=XM_
011282421.2;refGene_gn=DST;refGene_tx=XM_011282422.3;refGene_gn=DST;re
fGene_tx=XM_019830750.2;refGene_gn=DST;refGene_tx=XM_019830753.2;refGe
ne_gn=DST;refGene_tx=XM_019830754.2;refGene_gn=DST;refGene_tx=XM_01983
0756.2;refGene_gn=DST;refGene_tx=XM_019830757.2;refGene_gn=DST;refGene
_tx=XM_019830758.2;refGene_gn=DST;refGene_tx=XM_019830759.2;refGene_gn
=DST;refGene_tx=XM_019830760.2;refGene_gn=DST;refGene_tx=XM_019830761.
2;refGene_gn=DST;refGene_tx=XM_019830762.2;refGene_gn=DST;refGene_tx=X
M_019830763.2;refGene_gn=DST;refGene_tx=XM_019830764.1;refGene_gn=DST;
refGene_tx=XM_019830765.2;refGene_gn=DST;refGene_tx=XM_019830766.2;ref
Gene_gn=DST;refGene_tx=XM_019830768.1;refGene_gn=DST;refGene_tx=XM_019
830770.2;refGene_gn=DST;refGene_tx=XM_023254073.1
B2 56530893 56530968
refGene_gn=DST;refGene_tx=XM_011282416.2;refGene_gn=DST;refGene_tx=XM_
011282421.2;refGene_gn=DST;refGene_tx=XM_011282422.3;refGene_gn=DST;re
fGene_tx=XM_019830750.2;refGene_gn=DST;refGene_tx=XM_019830751.2;refGe
ne_gn=DST;refGene_tx=XM_019830753.2;refGene_gn=DST;refGene_tx=XM_01983
0754.2;refGene_gn=DST;refGene_tx=XM_019830755.2;refGene_gn=DST;refGene
_tx=XM_019830756.2;refGene_gn=DST;refGene_tx=XM_019830757.2;refGene_gn
=DST;refGene_tx=XM_019830758.2;refGene_gn=DST;refGene_tx=XM_019830759.
2;refGene_gn=DST;refGene_tx=XM_019830760.2;refGene_gn=DST;refGene_tx=X
M_019830761.2;refGene_gn=DST;refGene_tx=XM_019830762.2;refGene_gn=DST;
refGene_tx=XM_019830763.2;refGene_gn=DST;refGene_tx=XM_019830764.1;ref
Gene_gn=DST;refGene_tx=XM_019830765.2;refGene_gn=DST;refGene_tx=XM_019
830766.2;refGene_gn=DST;refGene_tx=XM_019830768.1;refGene_gn=DST;refGe
ne_tx=XM_019830770.2;refGene_gn=DST;refGene_tx=XM_019830771.2;refGene_
gn=DST;refGene_tx=XM_023254073.1
B2 56531137 56531292
refGene_gn=DST;refGene_tx=XM_011282416.2;refGene_gn=DST;refGene_tx=XM_
011282421.2;refGene_gn=DST;refGene_tx=XM_011282422.3;refGene_gn=DST;re
fGene_tx=XM_019830750.2;refGene_gn=DST;refGene_tx=XM_019830751.2;refGe
ne_gn=DST;refGene_tx=XM_019830753.2;refGene_gn=DST;refGene_tx=XM_01983

0754.2;refGene_gn=DST;refGene_tx=XM_019830755.2;refGene_gn=DST;refGene_tx=XM_019830756.2;refGene_gn=DST;refGene_tx=XM_019830757.2;refGene_gn=DST;refGene_tx=XM_019830758.2;refGene_gn=DST;refGene_tx=XM_019830759.2;refGene_gn=DST;refGene_tx=XM_019830760.2;refGene_gn=DST;refGene_tx=XM_019830761.2;refGene_gn=DST;refGene_tx=XM_019830762.2;refGene_gn=DST;refGene_tx=XM_019830763.2;refGene_gn=DST;refGene_tx=XM_019830764.1;refGene_gn=DST;refGene_tx=XM_019830765.2;refGene_gn=DST;refGene_tx=XM_019830766.2;refGene_gn=DST;refGene_tx=XM_019830768.1;refGene_gn=DST;refGene_tx=XM_019830770.2;refGene_gn=DST;refGene_tx=XM_019830771.2;refGene_gn=DST;refGene_tx=XM_023254073.1
B2 56532605 56532768
refGene_gn=DST;refGene_tx=XM_011282416.2;refGene_gn=DST;refGene_tx=XM_011282421.2;refGene_gn=DST;refGene_tx=XM_011282422.3;refGene_gn=DST;refGene_tx=XM_019830750.2;refGene_gn=DST;refGene_tx=XM_019830751.2;refGene_gn=DST;refGene_tx=XM_019830753.2;refGene_gn=DST;refGene_tx=XM_019830754.2;refGene_gn=DST;refGene_tx=XM_019830755.2;refGene_gn=DST;refGene_tx=XM_019830756.2;refGene_gn=DST;refGene_tx=XM_019830757.2;refGene_gn=DST;refGene_tx=XM_019830758.2;refGene_gn=DST;refGene_tx=XM_019830759.2;refGene_gn=DST;refGene_tx=XM_019830760.2;refGene_gn=DST;refGene_tx=XM_019830761.2;refGene_gn=DST;refGene_tx=XM_019830762.2;refGene_gn=DST;refGene_tx=XM_019830763.2;refGene_gn=DST;refGene_tx=XM_019830764.1;refGene_gn=DST;refGene_tx=XM_019830765.2;refGene_gn=DST;refGene_tx=XM_019830766.2;refGene_gn=DST;refGene_tx=XM_019830768.1;refGene_gn=DST;refGene_tx=XM_019830770.2;refGene_gn=DST;refGene_tx=XM_019830771.2;refGene_gn=DST;refGene_tx=XM_023254073.1
B2 56534244 56534408
refGene_gn=DST;refGene_tx=XM_011282416.2;refGene_gn=DST;refGene_tx=XM_011282421.2;refGene_gn=DST;refGene_tx=XM_011282422.3;refGene_gn=DST;refGene_tx=XM_019830750.2;refGene_gn=DST;refGene_tx=XM_019830751.2;refGene_gn=DST;refGene_tx=XM_019830753.2;refGene_gn=DST;refGene_tx=XM_019830754.2;refGene_gn=DST;refGene_tx=XM_019830755.2;refGene_gn=DST;refGene_tx=XM_019830756.2;refGene_gn=DST;refGene_tx=XM_019830757.2;refGene_gn=DST;refGene_tx=XM_019830758.2;refGene_gn=DST;refGene_tx=XM_019830759.2;refGene_gn=DST;refGene_tx=XM_019830760.2;refGene_gn=DST;refGene_tx=XM_019830761.2;refGene_gn=DST;refGene_tx=XM_019830762.2;refGene_gn=DST;refGene_tx=XM_019830763.2;refGene_gn=DST;refGene_tx=XM_019830764.1;refGene_gn=DST;refGene_tx=XM_019830765.2;refGene_gn=DST;refGene_tx=XM_019830766.2;refGene_gn=DST;refGene_tx=XM_019830768.1;refGene_gn=DST;refGene_tx=XM_019830770.2;refGene_gn=DST;refGene_tx=XM_019830771.2;refGene_gn=DST;refGene_tx=XM_023254073.1
B2 56536266 56536396
refGene_gn=DST;refGene_tx=XM_011282416.2;refGene_gn=DST;refGene_tx=XM_011282421.2;refGene_gn=DST;refGene_tx=XM_011282422.3;refGene_gn=DST;refGene_tx=XM_019830750.2;refGene_gn=DST;refGene_tx=XM_019830751.2;refGene_gn=DST;refGene_tx=XM_019830753.2;refGene_gn=DST;refGene_tx=XM_019830754.2;refGene_gn=DST;refGene_tx=XM_019830755.2;refGene_gn=DST;refGene_tx=XM_019830756.2;refGene_gn=DST;refGene_tx=XM_019830757.2;refGene_gn=DST;refGene_tx=XM_019830758.2;refGene_gn=DST;refGene_tx=XM_019830759.2;refGene_gn=DST;refGene_tx=XM_019830760.2;refGene_gn=DST;refGene_tx=XM_019830761.2;refGene_gn=DST;refGene_tx=XM_019830762.2;refGene_gn=DST;

refGene_tx=XM_019830763.2;refGene_gn=DST;refGene_tx=XM_019830764.1;refGene_gn=DST;refGene_tx=XM_019830765.2;refGene_gn=DST;refGene_tx=XM_019830766.2;refGene_gn=DST;refGene_tx=XM_019830768.1;refGene_gn=DST;refGene_tx=XM_019830770.2;refGene_gn=DST;refGene_tx=XM_019830771.2;refGene_gn=DST;refGene_tx=XM_023254073.1
B2 56537626 56537815
refGene_gn=DST;refGene_tx=XM_011282416.2;refGene_gn=DST;refGene_tx=XM_011282421.2;refGene_gn=DST;refGene_tx=XM_011282422.3;refGene_gn=DST;refGene_tx=XM_019830750.2;refGene_gn=DST;refGene_tx=XM_019830751.2;refGene_gn=DST;refGene_tx=XM_019830753.2;refGene_gn=DST;refGene_tx=XM_019830754.2;refGene_gn=DST;refGene_tx=XM_019830755.2;refGene_gn=DST;refGene_tx=XM_019830756.2;refGene_gn=DST;refGene_tx=XM_019830757.2;refGene_gn=DST;refGene_tx=XM_019830758.2;refGene_gn=DST;refGene_tx=XM_019830759.2;refGene_gn=DST;refGene_tx=XM_019830760.2;refGene_gn=DST;refGene_tx=XM_019830761.2;refGene_gn=DST;refGene_tx=XM_019830762.2;refGene_gn=DST;refGene_tx=XM_019830763.2;refGene_gn=DST;refGene_tx=XM_019830764.1;refGene_gn=DST;refGene_tx=XM_019830765.2;refGene_gn=DST;refGene_tx=XM_019830766.2;refGene_gn=DST;refGene_tx=XM_019830768.1;refGene_gn=DST;refGene_tx=XM_019830770.2;refGene_gn=DST;refGene_tx=XM_019830771.2;refGene_gn=DST;refGene_tx=XM_023254073.1
B2 56538918 56539062
refGene_gn=DST;refGene_tx=XM_011282416.2;refGene_gn=DST;refGene_tx=XM_011282421.2;refGene_gn=DST;refGene_tx=XM_011282422.3;refGene_gn=DST;refGene_tx=XM_019830750.2;refGene_gn=DST;refGene_tx=XM_019830751.2;refGene_gn=DST;refGene_tx=XM_019830753.2;refGene_gn=DST;refGene_tx=XM_019830754.2;refGene_gn=DST;refGene_tx=XM_019830755.2;refGene_gn=DST;refGene_tx=XM_019830756.2;refGene_gn=DST;refGene_tx=XM_019830757.2;refGene_gn=DST;refGene_tx=XM_019830758.2;refGene_gn=DST;refGene_tx=XM_019830759.2;refGene_gn=DST;refGene_tx=XM_019830760.2;refGene_gn=DST;refGene_tx=XM_019830761.2;refGene_gn=DST;refGene_tx=XM_019830762.2;refGene_gn=DST;refGene_tx=XM_019830763.2;refGene_gn=DST;refGene_tx=XM_019830764.1;refGene_gn=DST;refGene_tx=XM_019830765.2;refGene_gn=DST;refGene_tx=XM_019830766.2;refGene_gn=DST;refGene_tx=XM_019830768.1;refGene_gn=DST;refGene_tx=XM_019830770.2;refGene_gn=DST;refGene_tx=XM_019830771.2;refGene_gn=DST;refGene_tx=XM_023254073.1
B2 56542232 56542361
refGene_gn=DST;refGene_tx=XM_011282416.2;refGene_gn=DST;refGene_tx=XM_011282421.2;refGene_gn=DST;refGene_tx=XM_011282422.3;refGene_gn=DST;refGene_tx=XM_019830750.2;refGene_gn=DST;refGene_tx=XM_019830751.2;refGene_gn=DST;refGene_tx=XM_019830753.2;refGene_gn=DST;refGene_tx=XM_019830754.2;refGene_gn=DST;refGene_tx=XM_019830755.2;refGene_gn=DST;refGene_tx=XM_019830756.2;refGene_gn=DST;refGene_tx=XM_019830757.2;refGene_gn=DST;refGene_tx=XM_019830758.2;refGene_gn=DST;refGene_tx=XM_019830759.2;refGene_gn=DST;refGene_tx=XM_019830760.2;refGene_gn=DST;refGene_tx=XM_019830761.2;refGene_gn=DST;refGene_tx=XM_019830762.2;refGene_gn=DST;refGene_tx=XM_019830763.2;refGene_gn=DST;refGene_tx=XM_019830764.1;refGene_gn=DST;refGene_tx=XM_019830765.2;refGene_gn=DST;refGene_tx=XM_019830766.2;refGene_gn=DST;refGene_tx=XM_019830768.1;refGene_gn=DST;refGene_tx=XM_019830770.2;refGene_gn=DST;refGene_tx=XM_019830771.2;refGene_gn=DST;refGene_tx=XM_023254073.1

B2 56542858 56543053
refGene_gn=DST;refGene_tx=XM_011282416.2;refGene_gn=DST;refGene_tx=XM_011282421.2;refGene_gn=DST;refGene_tx=XM_011282422.3;refGene_gn=DST;refGene_tx=XM_019830750.2;refGene_gn=DST;refGene_tx=XM_019830751.2;refGene_gn=DST;refGene_tx=XM_019830753.2;refGene_gn=DST;refGene_tx=XM_019830754.2;refGene_gn=DST;refGene_tx=XM_019830755.2;refGene_gn=DST;refGene_tx=XM_019830756.2;refGene_gn=DST;refGene_tx=XM_019830757.2;refGene_gn=DST;refGene_tx=XM_019830758.2;refGene_gn=DST;refGene_tx=XM_019830759.2;refGene_gn=DST;refGene_tx=XM_019830760.2;refGene_gn=DST;refGene_tx=XM_019830761.2;refGene_gn=DST;refGene_tx=XM_019830762.2;refGene_gn=DST;refGene_tx=XM_019830763.2;refGene_gn=DST;refGene_tx=XM_019830764.1;refGene_gn=DST;refGene_tx=XM_019830765.2;refGene_gn=DST;refGene_tx=XM_019830766.2;refGene_gn=DST;refGene_tx=XM_019830768.1;refGene_gn=DST;refGene_tx=XM_019830770.2;refGene_gn=DST;refGene_tx=XM_019830771.2;refGene_gn=DST;refGene_tx=XM_023254073.1
B2 56545455 56545615
refGene_gn=DST;refGene_tx=XM_011282416.2;refGene_gn=DST;refGene_tx=XM_011282421.2;refGene_gn=DST;refGene_tx=XM_011282422.3;refGene_gn=DST;refGene_tx=XM_019830750.2;refGene_gn=DST;refGene_tx=XM_019830751.2;refGene_gn=DST;refGene_tx=XM_019830753.2;refGene_gn=DST;refGene_tx=XM_019830754.2;refGene_gn=DST;refGene_tx=XM_019830755.2;refGene_gn=DST;refGene_tx=XM_019830756.2;refGene_gn=DST;refGene_tx=XM_019830757.2;refGene_gn=DST;refGene_tx=XM_019830758.2;refGene_gn=DST;refGene_tx=XM_019830759.2;refGene_gn=DST;refGene_tx=XM_019830760.2;refGene_gn=DST;refGene_tx=XM_019830761.2;refGene_gn=DST;refGene_tx=XM_019830762.2;refGene_gn=DST;refGene_tx=XM_019830763.2;refGene_gn=DST;refGene_tx=XM_019830764.1;refGene_gn=DST;refGene_tx=XM_019830765.2;refGene_gn=DST;refGene_tx=XM_019830766.2;refGene_gn=DST;refGene_tx=XM_019830768.1;refGene_gn=DST;refGene_tx=XM_019830770.2;refGene_gn=DST;refGene_tx=XM_019830771.2;refGene_gn=DST;refGene_tx=XM_023254073.1
B2 56547267 56547437
refGene_gn=DST;refGene_tx=XM_011282416.2;refGene_gn=DST;refGene_tx=XM_011282421.2;refGene_gn=DST;refGene_tx=XM_011282422.3;refGene_gn=DST;refGene_tx=XM_019830750.2;refGene_gn=DST;refGene_tx=XM_019830751.2;refGene_gn=DST;refGene_tx=XM_019830753.2;refGene_gn=DST;refGene_tx=XM_019830754.2;refGene_gn=DST;refGene_tx=XM_019830755.2;refGene_gn=DST;refGene_tx=XM_019830756.2;refGene_gn=DST;refGene_tx=XM_019830757.2;refGene_gn=DST;refGene_tx=XM_019830758.2;refGene_gn=DST;refGene_tx=XM_019830759.2;refGene_gn=DST;refGene_tx=XM_019830760.2;refGene_gn=DST;refGene_tx=XM_019830761.2;refGene_gn=DST;refGene_tx=XM_019830762.2;refGene_gn=DST;refGene_tx=XM_019830763.2;refGene_gn=DST;refGene_tx=XM_019830764.1;refGene_gn=DST;refGene_tx=XM_019830765.2;refGene_gn=DST;refGene_tx=XM_019830766.2;refGene_gn=DST;refGene_tx=XM_019830768.1;refGene_gn=DST;refGene_tx=XM_019830770.2;refGene_gn=DST;refGene_tx=XM_019830771.2;refGene_gn=DST;refGene_tx=XM_023254073.1
B2 56550228 56550348
refGene_gn=DST;refGene_tx=XM_011282416.2;refGene_gn=DST;refGene_tx=XM_011282421.2;refGene_gn=DST;refGene_tx=XM_011282422.3;refGene_gn=DST;refGene_tx=XM_019830750.2;refGene_gn=DST;refGene_tx=XM_019830751.2;refGene_gn=DST;refGene_tx=XM_019830753.2;refGene_gn=DST;refGene_tx=XM_01983

0754.2;refGene_gn=DST;refGene_tx=XM_019830755.2;refGene_gn=DST;refGene_tx=XM_019830756.2;refGene_gn=DST;refGene_tx=XM_019830757.2;refGene_gn=DST;refGene_tx=XM_019830758.2;refGene_gn=DST;refGene_tx=XM_019830759.2;refGene_gn=DST;refGene_tx=XM_019830760.2;refGene_gn=DST;refGene_tx=XM_019830761.2;refGene_gn=DST;refGene_tx=XM_019830762.2;refGene_gn=DST;refGene_tx=XM_019830763.2;refGene_gn=DST;refGene_tx=XM_019830764.1;refGene_gn=DST;refGene_tx=XM_019830765.2;refGene_gn=DST;refGene_tx=XM_019830766.2;refGene_gn=DST;refGene_tx=XM_019830768.1;refGene_gn=DST;refGene_tx=XM_019830770.2;refGene_gn=DST;refGene_tx=XM_019830771.2;refGene_gn=DST;refGene_tx=XM_023254073.1
B2 56553659 56553866
refGene_gn=DST;refGene_tx=XM_011282416.2;refGene_gn=DST;refGene_tx=XM_011282421.2;refGene_gn=DST;refGene_tx=XM_011282422.3;refGene_gn=DST;refGene_tx=XM_019830750.2;refGene_gn=DST;refGene_tx=XM_019830751.2;refGene_gn=DST;refGene_tx=XM_019830753.2;refGene_gn=DST;refGene_tx=XM_019830754.2;refGene_gn=DST;refGene_tx=XM_019830755.2;refGene_gn=DST;refGene_tx=XM_019830756.2;refGene_gn=DST;refGene_tx=XM_019830757.2;refGene_gn=DST;refGene_tx=XM_019830758.2;refGene_gn=DST;refGene_tx=XM_019830759.2;refGene_gn=DST;refGene_tx=XM_019830760.2;refGene_gn=DST;refGene_tx=XM_019830761.2;refGene_gn=DST;refGene_tx=XM_019830762.2;refGene_gn=DST;refGene_tx=XM_019830763.2;refGene_gn=DST;refGene_tx=XM_019830764.1;refGene_gn=DST;refGene_tx=XM_019830765.2;refGene_gn=DST;refGene_tx=XM_019830766.2;refGene_gn=DST;refGene_tx=XM_019830768.1;refGene_gn=DST;refGene_tx=XM_019830770.2;refGene_gn=DST;refGene_tx=XM_019830771.2;refGene_gn=DST;refGene_tx=XM_023254073.1
B2 56554067 56554223
refGene_gn=DST;refGene_tx=XM_011282416.2;refGene_gn=DST;refGene_tx=XM_011282421.2;refGene_gn=DST;refGene_tx=XM_011282422.3;refGene_gn=DST;refGene_tx=XM_019830750.2;refGene_gn=DST;refGene_tx=XM_019830751.2;refGene_gn=DST;refGene_tx=XM_019830753.2;refGene_gn=DST;refGene_tx=XM_019830754.2;refGene_gn=DST;refGene_tx=XM_019830755.2;refGene_gn=DST;refGene_tx=XM_019830756.2;refGene_gn=DST;refGene_tx=XM_019830757.2;refGene_gn=DST;refGene_tx=XM_019830758.2;refGene_gn=DST;refGene_tx=XM_019830759.2;refGene_gn=DST;refGene_tx=XM_019830760.2;refGene_gn=DST;refGene_tx=XM_019830761.2;refGene_gn=DST;refGene_tx=XM_019830762.2;refGene_gn=DST;refGene_tx=XM_019830763.2;refGene_gn=DST;refGene_tx=XM_019830764.1;refGene_gn=DST;refGene_tx=XM_019830765.2;refGene_gn=DST;refGene_tx=XM_019830766.2;refGene_gn=DST;refGene_tx=XM_019830768.1;refGene_gn=DST;refGene_tx=XM_019830770.2;refGene_gn=DST;refGene_tx=XM_019830771.2;refGene_gn=DST;refGene_tx=XM_023254073.1
B2 56555058 56555226
refGene_gn=DST;refGene_tx=XM_011282416.2;refGene_gn=DST;refGene_tx=XM_011282421.2;refGene_gn=DST;refGene_tx=XM_011282422.3;refGene_gn=DST;refGene_tx=XM_019830750.2;refGene_gn=DST;refGene_tx=XM_019830751.2;refGene_gn=DST;refGene_tx=XM_019830753.2;refGene_gn=DST;refGene_tx=XM_019830754.2;refGene_gn=DST;refGene_tx=XM_019830755.2;refGene_gn=DST;refGene_tx=XM_019830756.2;refGene_gn=DST;refGene_tx=XM_019830757.2;refGene_gn=DST;refGene_tx=XM_019830758.2;refGene_gn=DST;refGene_tx=XM_019830759.2;refGene_gn=DST;refGene_tx=XM_019830760.2;refGene_gn=DST;refGene_tx=XM_019830761.2;refGene_gn=DST;refGene_tx=XM_019830762.2;refGene_gn=DST;

refGene_tx=XM_019830763.2;refGene_gn=DST;refGene_tx=XM_019830764.1;refGene_gn=DST;refGene_tx=XM_019830765.2;refGene_gn=DST;refGene_tx=XM_019830766.2;refGene_gn=DST;refGene_tx=XM_019830768.1;refGene_gn=DST;refGene_tx=XM_019830770.2;refGene_gn=DST;refGene_tx=XM_019830771.2;refGene_gn=DST;refGene_tx=XM_023254073.1
B2 56557232 56557361
refGene_gn=DST;refGene_tx=XM_011282416.2;refGene_gn=DST;refGene_tx=XM_011282421.2;refGene_gn=DST;refGene_tx=XM_011282422.3;refGene_gn=DST;refGene_tx=XM_019830750.2;refGene_gn=DST;refGene_tx=XM_019830751.2;refGene_gn=DST;refGene_tx=XM_019830753.2;refGene_gn=DST;refGene_tx=XM_019830754.2;refGene_gn=DST;refGene_tx=XM_019830755.2;refGene_gn=DST;refGene_tx=XM_019830756.2;refGene_gn=DST;refGene_tx=XM_019830757.2;refGene_gn=DST;refGene_tx=XM_019830758.2;refGene_gn=DST;refGene_tx=XM_019830759.2;refGene_gn=DST;refGene_tx=XM_019830760.2;refGene_gn=DST;refGene_tx=XM_019830761.2;refGene_gn=DST;refGene_tx=XM_019830762.2;refGene_gn=DST;refGene_tx=XM_019830763.2;refGene_gn=DST;refGene_tx=XM_019830764.1;refGene_gn=DST;refGene_tx=XM_019830765.2;refGene_gn=DST;refGene_tx=XM_019830766.2;refGene_gn=DST;refGene_tx=XM_019830768.1;refGene_gn=DST;refGene_tx=XM_019830770.2;refGene_gn=DST;refGene_tx=XM_019830771.2;refGene_gn=DST;refGene_tx=XM_023254073.1
B2 56557710 56557908
refGene_gn=DST;refGene_tx=XM_011282416.2;refGene_gn=DST;refGene_tx=XM_011282421.2;refGene_gn=DST;refGene_tx=XM_011282422.3;refGene_gn=DST;refGene_tx=XM_019830750.2;refGene_gn=DST;refGene_tx=XM_019830751.2;refGene_gn=DST;refGene_tx=XM_019830753.2;refGene_gn=DST;refGene_tx=XM_019830754.2;refGene_gn=DST;refGene_tx=XM_019830755.2;refGene_gn=DST;refGene_tx=XM_019830756.2;refGene_gn=DST;refGene_tx=XM_019830757.2;refGene_gn=DST;refGene_tx=XM_019830758.2;refGene_gn=DST;refGene_tx=XM_019830759.2;refGene_gn=DST;refGene_tx=XM_019830760.2;refGene_gn=DST;refGene_tx=XM_019830761.2;refGene_gn=DST;refGene_tx=XM_019830762.2;refGene_gn=DST;refGene_tx=XM_019830763.2;refGene_gn=DST;refGene_tx=XM_019830764.1;refGene_gn=DST;refGene_tx=XM_019830765.2;refGene_gn=DST;refGene_tx=XM_019830766.2;refGene_gn=DST;refGene_tx=XM_019830768.1;refGene_gn=DST;refGene_tx=XM_019830770.2;refGene_gn=DST;refGene_tx=XM_019830771.2;refGene_gn=DST;refGene_tx=XM_023254073.1
B2 56561366 56561522
refGene_gn=DST;refGene_tx=XM_011282416.2;refGene_gn=DST;refGene_tx=XM_011282421.2;refGene_gn=DST;refGene_tx=XM_011282422.3;refGene_gn=DST;refGene_tx=XM_019830750.2;refGene_gn=DST;refGene_tx=XM_019830751.2;refGene_gn=DST;refGene_tx=XM_019830753.2;refGene_gn=DST;refGene_tx=XM_019830754.2;refGene_gn=DST;refGene_tx=XM_019830755.2;refGene_gn=DST;refGene_tx=XM_019830756.2;refGene_gn=DST;refGene_tx=XM_019830757.2;refGene_gn=DST;refGene_tx=XM_019830758.2;refGene_gn=DST;refGene_tx=XM_019830759.2;refGene_gn=DST;refGene_tx=XM_019830760.2;refGene_gn=DST;refGene_tx=XM_019830761.2;refGene_gn=DST;refGene_tx=XM_019830762.2;refGene_gn=DST;refGene_tx=XM_019830763.2;refGene_gn=DST;refGene_tx=XM_019830764.1;refGene_gn=DST;refGene_tx=XM_019830765.2;refGene_gn=DST;refGene_tx=XM_019830766.2;refGene_gn=DST;refGene_tx=XM_019830768.1;refGene_gn=DST;refGene_tx=XM_019830770.2;refGene_gn=DST;refGene_tx=XM_019830771.2;refGene_gn=DST;refGene_tx=XM_023254073.1

B2 56561855 56562029
refGene_gn=DST;refGene_tx=XM_011282416.2;refGene_gn=DST;refGene_tx=XM_011282421.2;refGene_gn=DST;refGene_tx=XM_011282422.3;refGene_gn=DST;refGene_tx=XM_019830750.2;refGene_gn=DST;refGene_tx=XM_019830751.2;refGene_gn=DST;refGene_tx=XM_019830753.2;refGene_gn=DST;refGene_tx=XM_019830754.2;refGene_gn=DST;refGene_tx=XM_019830755.2;refGene_gn=DST;refGene_tx=XM_019830756.2;refGene_gn=DST;refGene_tx=XM_019830757.2;refGene_gn=DST;refGene_tx=XM_019830758.2;refGene_gn=DST;refGene_tx=XM_019830759.2;refGene_gn=DST;refGene_tx=XM_019830760.2;refGene_gn=DST;refGene_tx=XM_019830761.2;refGene_gn=DST;refGene_tx=XM_019830762.2;refGene_gn=DST;refGene_tx=XM_019830763.2;refGene_gn=DST;refGene_tx=XM_019830764.1;refGene_gn=DST;refGene_tx=XM_019830765.2;refGene_gn=DST;refGene_tx=XM_019830766.2;refGene_gn=DST;refGene_tx=XM_019830768.1;refGene_gn=DST;refGene_tx=XM_019830770.2;refGene_gn=DST;refGene_tx=XM_019830771.2;refGene_gn=DST;refGene_tx=XM_023254073.1
B2 56564174 56564276
refGene_gn=DST;refGene_tx=XM_011282416.2;refGene_gn=DST;refGene_tx=XM_011282421.2;refGene_gn=DST;refGene_tx=XM_011282422.3;refGene_gn=DST;refGene_tx=XM_019830750.2;refGene_gn=DST;refGene_tx=XM_019830751.2;refGene_gn=DST;refGene_tx=XM_019830753.2;refGene_gn=DST;refGene_tx=XM_019830754.2;refGene_gn=DST;refGene_tx=XM_019830755.2;refGene_gn=DST;refGene_tx=XM_019830756.2;refGene_gn=DST;refGene_tx=XM_019830757.2;refGene_gn=DST;refGene_tx=XM_019830758.2;refGene_gn=DST;refGene_tx=XM_019830759.2;refGene_gn=DST;refGene_tx=XM_019830760.2;refGene_gn=DST;refGene_tx=XM_019830761.2;refGene_gn=DST;refGene_tx=XM_019830762.2;refGene_gn=DST;refGene_tx=XM_019830763.2;refGene_gn=DST;refGene_tx=XM_019830764.1;refGene_gn=DST;refGene_tx=XM_019830765.2;refGene_gn=DST;refGene_tx=XM_019830766.2;refGene_gn=DST;refGene_tx=XM_019830768.1;refGene_gn=DST;refGene_tx=XM_019830770.2;refGene_gn=DST;refGene_tx=XM_019830771.2;refGene_gn=DST;refGene_tx=XM_023254073.1
B2 56566905 56567007
refGene_gn=DST;refGene_tx=XM_011282416.2;refGene_gn=DST;refGene_tx=XM_011282421.2;refGene_gn=DST;refGene_tx=XM_011282422.3;refGene_gn=DST;refGene_tx=XM_019830750.2;refGene_gn=DST;refGene_tx=XM_019830751.2;refGene_gn=DST;refGene_tx=XM_019830753.2;refGene_gn=DST;refGene_tx=XM_019830754.2;refGene_gn=DST;refGene_tx=XM_019830755.2;refGene_gn=DST;refGene_tx=XM_019830756.2;refGene_gn=DST;refGene_tx=XM_019830757.2;refGene_gn=DST;refGene_tx=XM_019830758.2;refGene_gn=DST;refGene_tx=XM_019830759.2;refGene_gn=DST;refGene_tx=XM_019830760.2;refGene_gn=DST;refGene_tx=XM_019830761.2;refGene_gn=DST;refGene_tx=XM_019830762.2;refGene_gn=DST;refGene_tx=XM_019830763.2;refGene_gn=DST;refGene_tx=XM_019830764.1;refGene_gn=DST;refGene_tx=XM_019830765.2;refGene_gn=DST;refGene_tx=XM_019830766.2;refGene_gn=DST;refGene_tx=XM_019830768.1;refGene_gn=DST;refGene_tx=XM_019830770.2;refGene_gn=DST;refGene_tx=XM_019830771.2;refGene_gn=DST;refGene_tx=XM_023254073.1
B2 56567137 56567260
refGene_gn=DST;refGene_tx=XM_011282416.2;refGene_gn=DST;refGene_tx=XM_011282421.2;refGene_gn=DST;refGene_tx=XM_011282422.3;refGene_gn=DST;refGene_tx=XM_019830750.2;refGene_gn=DST;refGene_tx=XM_019830751.2;refGene_gn=DST;refGene_tx=XM_019830753.2;refGene_gn=DST;refGene_tx=XM_01983

0754.2;refGene_gn=DST;refGene_tx=XM_019830755.2;refGene_gn=DST;refGene_tx=XM_019830756.2;refGene_gn=DST;refGene_tx=XM_019830757.2;refGene_gn=DST;refGene_tx=XM_019830758.2;refGene_gn=DST;refGene_tx=XM_019830759.2;refGene_gn=DST;refGene_tx=XM_019830760.2;refGene_gn=DST;refGene_tx=XM_019830761.2;refGene_gn=DST;refGene_tx=XM_019830762.2;refGene_gn=DST;refGene_tx=XM_019830763.2;refGene_gn=DST;refGene_tx=XM_019830764.1;refGene_gn=DST;refGene_tx=XM_019830765.2;refGene_gn=DST;refGene_tx=XM_019830766.2;refGene_gn=DST;refGene_tx=XM_019830768.1;refGene_gn=DST;refGene_tx=XM_019830770.2;refGene_gn=DST;refGene_tx=XM_019830771.2;refGene_gn=DST;refGene_tx=XM_023254073.1
B2 56569149 56569376
refGene_gn=DST;refGene_tx=XM_011282416.2;refGene_gn=DST;refGene_tx=XM_011282421.2;refGene_gn=DST;refGene_tx=XM_011282422.3;refGene_gn=DST;refGene_tx=XM_019830750.2;refGene_gn=DST;refGene_tx=XM_019830751.2;refGene_gn=DST;refGene_tx=XM_019830753.2;refGene_gn=DST;refGene_tx=XM_019830754.2;refGene_gn=DST;refGene_tx=XM_019830755.2;refGene_gn=DST;refGene_tx=XM_019830756.2;refGene_gn=DST;refGene_tx=XM_019830757.2;refGene_gn=DST;refGene_tx=XM_019830758.2;refGene_gn=DST;refGene_tx=XM_019830759.2;refGene_gn=DST;refGene_tx=XM_019830760.2;refGene_gn=DST;refGene_tx=XM_019830761.2;refGene_gn=DST;refGene_tx=XM_019830762.2;refGene_gn=DST;refGene_tx=XM_019830763.2;refGene_gn=DST;refGene_tx=XM_019830764.1;refGene_gn=DST;refGene_tx=XM_019830765.2;refGene_gn=DST;refGene_tx=XM_019830766.2;refGene_gn=DST;refGene_tx=XM_019830768.1;refGene_gn=DST;refGene_tx=XM_019830770.2;refGene_gn=DST;refGene_tx=XM_019830771.2;refGene_gn=DST;refGene_tx=XM_023254073.1
B2 56570261 56570493
refGene_gn=DST;refGene_tx=XM_011282416.2;refGene_gn=DST;refGene_tx=XM_011282421.2;refGene_gn=DST;refGene_tx=XM_011282422.3;refGene_gn=DST;refGene_tx=XM_019830750.2;refGene_gn=DST;refGene_tx=XM_019830751.2;refGene_gn=DST;refGene_tx=XM_019830753.2;refGene_gn=DST;refGene_tx=XM_019830754.2;refGene_gn=DST;refGene_tx=XM_019830755.2;refGene_gn=DST;refGene_tx=XM_019830756.2;refGene_gn=DST;refGene_tx=XM_019830757.2;refGene_gn=DST;refGene_tx=XM_019830758.2;refGene_gn=DST;refGene_tx=XM_019830759.2;refGene_gn=DST;refGene_tx=XM_019830760.2;refGene_gn=DST;refGene_tx=XM_019830761.2;refGene_gn=DST;refGene_tx=XM_019830762.2;refGene_gn=DST;refGene_tx=XM_019830763.2;refGene_gn=DST;refGene_tx=XM_019830764.1;refGene_gn=DST;refGene_tx=XM_019830765.2;refGene_gn=DST;refGene_tx=XM_019830766.2;refGene_gn=DST;refGene_tx=XM_019830768.1;refGene_gn=DST;refGene_tx=XM_019830770.2;refGene_gn=DST;refGene_tx=XM_019830771.2;refGene_gn=DST;refGene_tx=XM_023254073.1
B2 56571683 56571887
refGene_gn=DST;refGene_tx=XM_011282416.2;refGene_gn=DST;refGene_tx=XM_011282421.2;refGene_gn=DST;refGene_tx=XM_011282422.3;refGene_gn=DST;refGene_tx=XM_019830750.2;refGene_gn=DST;refGene_tx=XM_019830751.2;refGene_gn=DST;refGene_tx=XM_019830753.2;refGene_gn=DST;refGene_tx=XM_019830754.2;refGene_gn=DST;refGene_tx=XM_019830755.2;refGene_gn=DST;refGene_tx=XM_019830756.2;refGene_gn=DST;refGene_tx=XM_019830757.2;refGene_gn=DST;refGene_tx=XM_019830758.2;refGene_gn=DST;refGene_tx=XM_019830759.2;refGene_gn=DST;refGene_tx=XM_019830760.2;refGene_gn=DST;refGene_tx=XM_019830761.2;refGene_gn=DST;refGene_tx=XM_019830762.2;refGene_gn=DST;

refGene_tx=XM_019830763.2;refGene_gn=DST;refGene_tx=XM_019830764.1;refGene_gn=DST;refGene_tx=XM_019830765.2;refGene_gn=DST;refGene_tx=XM_019830766.2;refGene_gn=DST;refGene_tx=XM_019830768.1;refGene_gn=DST;refGene_tx=XM_019830770.2;refGene_gn=DST;refGene_tx=XM_019830771.2;refGene_gn=DST;refGene_tx=XM_023254073.1
B2 56575531 56575750
refGene_gn=DST;refGene_tx=XM_011282416.2;refGene_gn=DST;refGene_tx=XM_011282421.2;refGene_gn=DST;refGene_tx=XM_011282422.3;refGene_gn=DST;refGene_tx=XM_019830750.2;refGene_gn=DST;refGene_tx=XM_019830751.2;refGene_gn=DST;refGene_tx=XM_019830753.2;refGene_gn=DST;refGene_tx=XM_019830754.2;refGene_gn=DST;refGene_tx=XM_019830755.2;refGene_gn=DST;refGene_tx=XM_019830756.2;refGene_gn=DST;refGene_tx=XM_019830757.2;refGene_gn=DST;refGene_tx=XM_019830758.2;refGene_gn=DST;refGene_tx=XM_019830759.2;refGene_gn=DST;refGene_tx=XM_019830760.2;refGene_gn=DST;refGene_tx=XM_019830761.2;refGene_gn=DST;refGene_tx=XM_019830762.2;refGene_gn=DST;refGene_tx=XM_019830763.2;refGene_gn=DST;refGene_tx=XM_019830764.1;refGene_gn=DST;refGene_tx=XM_019830765.2;refGene_gn=DST;refGene_tx=XM_019830766.2;refGene_gn=DST;refGene_tx=XM_019830768.1;refGene_gn=DST;refGene_tx=XM_019830770.2;refGene_gn=DST;refGene_tx=XM_019830771.2;refGene_gn=DST;refGene_tx=XM_023254073.1
B2 56578011 56578119
refGene_gn=DST;refGene_tx=XM_011282416.2;refGene_gn=DST;refGene_tx=XM_011282421.2;refGene_gn=DST;refGene_tx=XM_011282422.3;refGene_gn=DST;refGene_tx=XM_019830750.2;refGene_gn=DST;refGene_tx=XM_019830751.2;refGene_gn=DST;refGene_tx=XM_019830753.2;refGene_gn=DST;refGene_tx=XM_019830754.2;refGene_gn=DST;refGene_tx=XM_019830755.2;refGene_gn=DST;refGene_tx=XM_019830756.2;refGene_gn=DST;refGene_tx=XM_019830757.2;refGene_gn=DST;refGene_tx=XM_019830758.2;refGene_gn=DST;refGene_tx=XM_019830759.2;refGene_gn=DST;refGene_tx=XM_019830760.2;refGene_gn=DST;refGene_tx=XM_019830761.2;refGene_gn=DST;refGene_tx=XM_019830762.2;refGene_gn=DST;refGene_tx=XM_019830763.2;refGene_gn=DST;refGene_tx=XM_019830764.1;refGene_gn=DST;refGene_tx=XM_019830765.2;refGene_gn=DST;refGene_tx=XM_019830766.2;refGene_gn=DST;refGene_tx=XM_019830768.1;refGene_gn=DST;refGene_tx=XM_019830770.2;refGene_gn=DST;refGene_tx=XM_019830771.2;refGene_gn=DST;refGene_tx=XM_023254073.1
B2 56578315 56578435
refGene_gn=DST;refGene_tx=XM_011282416.2;refGene_gn=DST;refGene_tx=XM_011282421.2;refGene_gn=DST;refGene_tx=XM_011282422.3;refGene_gn=DST;refGene_tx=XM_019830750.2;refGene_gn=DST;refGene_tx=XM_019830751.2;refGene_gn=DST;refGene_tx=XM_019830753.2;refGene_gn=DST;refGene_tx=XM_019830754.2;refGene_gn=DST;refGene_tx=XM_019830755.2;refGene_gn=DST;refGene_tx=XM_019830756.2;refGene_gn=DST;refGene_tx=XM_019830757.2;refGene_gn=DST;refGene_tx=XM_019830758.2;refGene_gn=DST;refGene_tx=XM_019830759.2;refGene_gn=DST;refGene_tx=XM_019830760.2;refGene_gn=DST;refGene_tx=XM_019830761.2;refGene_gn=DST;refGene_tx=XM_019830762.2;refGene_gn=DST;refGene_tx=XM_019830763.2;refGene_gn=DST;refGene_tx=XM_019830764.1;refGene_gn=DST;refGene_tx=XM_019830765.2;refGene_gn=DST;refGene_tx=XM_019830766.2;refGene_gn=DST;refGene_tx=XM_019830768.1;refGene_gn=DST;refGene_tx=XM_019830770.2;refGene_gn=DST;refGene_tx=XM_019830771.2;refGene_gn=DST;refGene_tx=XM_023254073.1

B2 56588327 56588534
refGene_gn=DST;refGene_tx=XM_011282416.2;refGene_gn=DST;refGene_tx=XM_011282421.2;refGene_gn=DST;refGene_tx=XM_011282422.3;refGene_gn=DST;refGene_tx=XM_019830750.2;refGene_gn=DST;refGene_tx=XM_019830751.2;refGene_gn=DST;refGene_tx=XM_019830753.2;refGene_gn=DST;refGene_tx=XM_019830754.2;refGene_gn=DST;refGene_tx=XM_019830755.2;refGene_gn=DST;refGene_tx=XM_019830756.2;refGene_gn=DST;refGene_tx=XM_019830757.2;refGene_gn=DST;refGene_tx=XM_019830758.2;refGene_gn=DST;refGene_tx=XM_019830759.2;refGene_gn=DST;refGene_tx=XM_019830760.2;refGene_gn=DST;refGene_tx=XM_019830761.2;refGene_gn=DST;refGene_tx=XM_019830762.2;refGene_gn=DST;refGene_tx=XM_019830763.2;refGene_gn=DST;refGene_tx=XM_019830764.1;refGene_gn=DST;refGene_tx=XM_019830765.2;refGene_gn=DST;refGene_tx=XM_019830766.2;refGene_gn=DST;refGene_tx=XM_019830768.1;refGene_gn=DST;refGene_tx=XM_019830770.2;refGene_gn=DST;refGene_tx=XM_019830771.2;refGene_gn=DST;refGene_tx=XM_023254073.1
B2 56589432 56589674
refGene_gn=DST;refGene_tx=XM_011282416.2;refGene_gn=DST;refGene_tx=XM_011282421.2;refGene_gn=DST;refGene_tx=XM_011282422.3;refGene_gn=DST;refGene_tx=XM_019830750.2;refGene_gn=DST;refGene_tx=XM_019830751.2;refGene_gn=DST;refGene_tx=XM_019830753.2;refGene_gn=DST;refGene_tx=XM_019830754.2;refGene_gn=DST;refGene_tx=XM_019830755.2;refGene_gn=DST;refGene_tx=XM_019830756.2;refGene_gn=DST;refGene_tx=XM_019830757.2;refGene_gn=DST;refGene_tx=XM_019830758.2;refGene_gn=DST;refGene_tx=XM_019830759.2;refGene_gn=DST;refGene_tx=XM_019830760.2;refGene_gn=DST;refGene_tx=XM_019830761.2;refGene_gn=DST;refGene_tx=XM_019830762.2;refGene_gn=DST;refGene_tx=XM_019830763.2;refGene_gn=DST;refGene_tx=XM_019830764.1;refGene_gn=DST;refGene_tx=XM_019830765.2;refGene_gn=DST;refGene_tx=XM_019830766.2;refGene_gn=DST;refGene_tx=XM_019830768.1;refGene_gn=DST;refGene_tx=XM_019830770.2;refGene_gn=DST;refGene_tx=XM_019830771.2;refGene_gn=DST;refGene_tx=XM_023254073.1
B2 56590886 56590971
refGene_gn=DST;refGene_tx=XM_011282416.2;refGene_gn=DST;refGene_tx=XM_011282421.2;refGene_gn=DST;refGene_tx=XM_011282422.3;refGene_gn=DST;refGene_tx=XM_019830750.2;refGene_gn=DST;refGene_tx=XM_019830751.2;refGene_gn=DST;refGene_tx=XM_019830753.2;refGene_gn=DST;refGene_tx=XM_019830754.2;refGene_gn=DST;refGene_tx=XM_019830755.2;refGene_gn=DST;refGene_tx=XM_019830756.2;refGene_gn=DST;refGene_tx=XM_019830757.2;refGene_gn=DST;refGene_tx=XM_019830758.2;refGene_gn=DST;refGene_tx=XM_019830759.2;refGene_gn=DST;refGene_tx=XM_019830760.2;refGene_gn=DST;refGene_tx=XM_019830761.2;refGene_gn=DST;refGene_tx=XM_019830762.2;refGene_gn=DST;refGene_tx=XM_019830763.2;refGene_gn=DST;refGene_tx=XM_019830764.1;refGene_gn=DST;refGene_tx=XM_019830765.2;refGene_gn=DST;refGene_tx=XM_019830766.2;refGene_gn=DST;refGene_tx=XM_019830768.1;refGene_gn=DST;refGene_tx=XM_019830770.2;refGene_gn=DST;refGene_tx=XM_019830771.2;refGene_gn=DST;refGene_tx=XM_023254073.1
B2 56591522 56591849
refGene_gn=DST;refGene_tx=XM_011282416.2;refGene_gn=DST;refGene_tx=XM_011282421.2;refGene_gn=DST;refGene_tx=XM_011282422.3;refGene_gn=DST;refGene_tx=XM_019830750.2;refGene_gn=DST;refGene_tx=XM_019830751.2;refGene_gn=DST;refGene_tx=XM_019830753.2;refGene_gn=DST;refGene_tx=XM_01983

0754.2;refGene_gn=DST;refGene_tx=XM_019830755.2;refGene_gn=DST;refGene_tx=XM_019830756.2;refGene_gn=DST;refGene_tx=XM_019830757.2;refGene_gn=DST;refGene_tx=XM_019830758.2;refGene_gn=DST;refGene_tx=XM_019830759.2;refGene_gn=DST;refGene_tx=XM_019830762.2;refGene_gn=DST;refGene_tx=XM_019830763.2;refGene_gn=DST;refGene_tx=XM_019830764.1;refGene_gn=DST;refGene_tx=XM_019830765.2;refGene_gn=DST;refGene_tx=XM_019830766.2;refGene_gn=DST;refGene_tx=XM_019830768.1;refGene_gn=DST;refGene_tx=XM_019830771.2
B2 56592049 56592209
refGene_gn=DST;refGene_tx=XM_011282416.2;refGene_gn=DST;refGene_tx=XM_011282421.2;refGene_gn=DST;refGene_tx=XM_011282422.3;refGene_gn=DST;refGene_tx=XM_019830750.2;refGene_gn=DST;refGene_tx=XM_019830751.2;refGene_gn=DST;refGene_tx=XM_019830753.2;refGene_gn=DST;refGene_tx=XM_019830754.2;refGene_gn=DST;refGene_tx=XM_019830755.2;refGene_gn=DST;refGene_tx=XM_019830756.2;refGene_gn=DST;refGene_tx=XM_019830757.2;refGene_gn=DST;refGene_tx=XM_019830758.2;refGene_gn=DST;refGene_tx=XM_019830759.2;refGene_gn=DST;refGene_tx=XM_019830760.2;refGene_gn=DST;refGene_tx=XM_019830761.2;refGene_gn=DST;refGene_tx=XM_019830762.2;refGene_gn=DST;refGene_tx=XM_019830763.2;refGene_gn=DST;refGene_tx=XM_019830764.1;refGene_gn=DST;refGene_tx=XM_019830765.2;refGene_gn=DST;refGene_tx=XM_019830766.2;refGene_gn=DST;refGene_tx=XM_019830768.1;refGene_gn=DST;refGene_tx=XM_019830770.2;refGene_gn=DST;refGene_tx=XM_019830771.2;refGene_gn=DST;refGene_tx=XM_023254073.1
B2 56594725 56594892
refGene_gn=DST;refGene_tx=XM_011282416.2;refGene_gn=DST;refGene_tx=XM_011282421.2;refGene_gn=DST;refGene_tx=XM_011282422.3;refGene_gn=DST;refGene_tx=XM_019830750.2;refGene_gn=DST;refGene_tx=XM_019830751.2;refGene_gn=DST;refGene_tx=XM_019830753.2;refGene_gn=DST;refGene_tx=XM_019830754.2;refGene_gn=DST;refGene_tx=XM_019830755.2;refGene_gn=DST;refGene_tx=XM_019830756.2;refGene_gn=DST;refGene_tx=XM_019830757.2;refGene_gn=DST;refGene_tx=XM_019830758.2;refGene_gn=DST;refGene_tx=XM_019830759.2;refGene_gn=DST;refGene_tx=XM_019830760.2;refGene_gn=DST;refGene_tx=XM_019830761.2;refGene_gn=DST;refGene_tx=XM_019830762.2;refGene_gn=DST;refGene_tx=XM_019830763.2;refGene_gn=DST;refGene_tx=XM_019830764.1;refGene_gn=DST;refGene_tx=XM_019830765.2;refGene_gn=DST;refGene_tx=XM_019830766.2;refGene_gn=DST;refGene_tx=XM_019830768.1;refGene_gn=DST;refGene_tx=XM_019830770.2;refGene_gn=DST;refGene_tx=XM_019830771.2;refGene_gn=DST;refGene_tx=XM_023254073.1
B2 56597447 56597618
refGene_gn=DST;refGene_tx=XM_011282416.2;refGene_gn=DST;refGene_tx=XM_011282421.2;refGene_gn=DST;refGene_tx=XM_011282422.3;refGene_gn=DST;refGene_tx=XM_019830750.2;refGene_gn=DST;refGene_tx=XM_019830751.2;refGene_gn=DST;refGene_tx=XM_019830753.2;refGene_gn=DST;refGene_tx=XM_019830754.2;refGene_gn=DST;refGene_tx=XM_019830755.2;refGene_gn=DST;refGene_tx=XM_019830756.2;refGene_gn=DST;refGene_tx=XM_019830757.2;refGene_gn=DST;refGene_tx=XM_019830758.2;refGene_gn=DST;refGene_tx=XM_019830759.2;refGene_gn=DST;refGene_tx=XM_019830760.2;refGene_gn=DST;refGene_tx=XM_019830761.2;refGene_gn=DST;refGene_tx=XM_019830762.2;refGene_gn=DST;refGene_tx=XM_019830763.2;refGene_gn=DST;refGene_tx=XM_019830764.1;refGene_gn=DST;refGene_tx=XM_019830765.2;refGene_gn=DST;refGene_tx=XM_019

830766.2;refGene_gn=DST;refGene_tx=XM_019830768.1;refGene_gn=DST;refGene_tx=XM_019830770.2;refGene_gn=DST;refGene_tx=XM_019830771.2;refGene_gn=DST;refGene_tx=XM_023254073.1
B2 56599349 56599511
refGene_gn=DST;refGene_tx=XM_011282416.2;refGene_gn=DST;refGene_tx=XM_011282421.2;refGene_gn=DST;refGene_tx=XM_011282422.3;refGene_gn=DST;refGene_tx=XM_019830750.2;refGene_gn=DST;refGene_tx=XM_019830751.2;refGene_gn=DST;refGene_tx=XM_019830753.2;refGene_gn=DST;refGene_tx=XM_019830754.2;refGene_gn=DST;refGene_tx=XM_019830755.2;refGene_gn=DST;refGene_tx=XM_019830756.2;refGene_gn=DST;refGene_tx=XM_019830757.2;refGene_gn=DST;refGene_tx=XM_019830758.2;refGene_gn=DST;refGene_tx=XM_019830759.2;refGene_gn=DST;refGene_tx=XM_019830760.2;refGene_gn=DST;refGene_tx=XM_019830761.2;refGene_gn=DST;refGene_tx=XM_019830762.2;refGene_gn=DST;refGene_tx=XM_019830763.2;refGene_gn=DST;refGene_tx=XM_019830764.1;refGene_gn=DST;refGene_tx=XM_019830765.2;refGene_gn=DST;refGene_tx=XM_019830766.2;refGene_gn=DST;refGene_tx=XM_019830768.1;refGene_gn=DST;refGene_tx=XM_019830770.2;refGene_gn=DST;refGene_tx=XM_019830771.2;refGene_gn=DST;refGene_tx=XM_023254073.1
B2 56615726 56617198
refGene_gn=DST;refGene_tx=XM_011282416.2;refGene_gn=DST;refGene_tx=XM_011282421.2;refGene_gn=DST;refGene_tx=XM_011282422.3;refGene_gn=DST;refGene_tx=XM_019830750.2;refGene_gn=DST;refGene_tx=XM_019830751.2;refGene_gn=DST;refGene_tx=XM_019830753.2;refGene_gn=DST;refGene_tx=XM_019830754.2;refGene_gn=DST;refGene_tx=XM_019830755.2;refGene_gn=DST;refGene_tx=XM_019830756.2;refGene_gn=DST;refGene_tx=XM_019830757.2;refGene_gn=DST;refGene_tx=XM_019830758.2;refGene_gn=DST;refGene_tx=XM_019830759.2;refGene_gn=DST;refGene_tx=XM_019830760.2;refGene_gn=DST;refGene_tx=XM_019830761.2;refGene_gn=DST;refGene_tx=XM_019830762.2;refGene_gn=DST;refGene_tx=XM_019830763.2;refGene_gn=DST;refGene_tx=XM_019830764.1;refGene_gn=DST;refGene_tx=XM_019830765.2;refGene_gn=DST;refGene_tx=XM_019830766.2;refGene_gn=DST;refGene_tx=XM_019830768.1;refGene_gn=DST;refGene_tx=XM_019830770.2;refGene_gn=DST;refGene_tx=XM_019830771.2;refGene_gn=DST;refGene_tx=XM_023254073.1
B2 56618722 56619218
refGene_gn=DST;refGene_tx=XM_011282416.2;refGene_gn=DST;refGene_tx=XM_011282421.2;refGene_gn=DST;refGene_tx=XM_011282422.3;refGene_gn=DST;refGene_tx=XM_019830750.2;refGene_gn=DST;refGene_tx=XM_019830751.2;refGene_gn=DST;refGene_tx=XM_019830753.2;refGene_gn=DST;refGene_tx=XM_019830754.2;refGene_gn=DST;refGene_tx=XM_019830755.2;refGene_gn=DST;refGene_tx=XM_019830756.2;refGene_gn=DST;refGene_tx=XM_019830757.2;refGene_gn=DST;refGene_tx=XM_019830758.2;refGene_gn=DST;refGene_tx=XM_019830759.2;refGene_gn=DST;refGene_tx=XM_019830760.2;refGene_gn=DST;refGene_tx=XM_019830761.2;refGene_gn=DST;refGene_tx=XM_019830762.2;refGene_gn=DST;refGene_tx=XM_019830763.2;refGene_gn=DST;refGene_tx=XM_019830764.1;refGene_gn=DST;refGene_tx=XM_019830765.2;refGene_gn=DST;refGene_tx=XM_019830766.2;refGene_gn=DST;refGene_tx=XM_019830768.1;refGene_gn=DST;refGene_tx=XM_019830770.2;refGene_gn=DST;refGene_tx=XM_019830771.2;refGene_gn=DST;refGene_tx=XM_023254073.1
B2 56620920 56621120
refGene_gn=DST;refGene_tx=XM_011282416.2;refGene_gn=DST;refGene_tx=XM_

011282421.2;refGene_gn=DST;refGene_tx=XM_011282422.3;refGene_gn=DST;refGene_tx=XM_019830750.2;refGene_gn=DST;refGene_tx=XM_019830751.2;refGene_gn=DST;refGene_tx=XM_019830753.2;refGene_gn=DST;refGene_tx=XM_019830754.2;refGene_gn=DST;refGene_tx=XM_019830755.2;refGene_gn=DST;refGene_tx=XM_019830756.2;refGene_gn=DST;refGene_tx=XM_019830757.2;refGene_gn=DST;refGene_tx=XM_019830758.2;refGene_gn=DST;refGene_tx=XM_019830759.2;refGene_gn=DST;refGene_tx=XM_019830760.2;refGene_gn=DST;refGene_tx=XM_019830761.2;refGene_gn=DST;refGene_tx=XM_019830762.2;refGene_gn=DST;refGene_tx=XM_019830763.2;refGene_gn=DST;refGene_tx=XM_019830764.1;refGene_gn=DST;refGene_tx=XM_019830765.2;refGene_gn=DST;refGene_tx=XM_019830766.2;refGene_gn=DST;refGene_tx=XM_019830768.1;refGene_gn=DST;refGene_tx=XM_019830770.2;refGene_gn=DST;refGene_tx=XM_019830771.2;refGene_gn=DST;refGene_tx=XM_023254073.1
B2 56623936 56624066
refGene_gn=DST;refGene_tx=XM_011282416.2;refGene_gn=DST;refGene_tx=XM_011282421.2;refGene_gn=DST;refGene_tx=XM_011282422.3;refGene_gn=DST;refGene_tx=XM_019830750.2;refGene_gn=DST;refGene_tx=XM_019830751.2;refGene_gn=DST;refGene_tx=XM_019830753.2;refGene_gn=DST;refGene_tx=XM_019830754.2;refGene_gn=DST;refGene_tx=XM_019830755.2;refGene_gn=DST;refGene_tx=XM_019830756.2;refGene_gn=DST;refGene_tx=XM_019830757.2;refGene_gn=DST;refGene_tx=XM_019830758.2;refGene_gn=DST;refGene_tx=XM_019830759.2;refGene_gn=DST;refGene_tx=XM_019830760.2;refGene_gn=DST;refGene_tx=XM_019830761.2;refGene_gn=DST;refGene_tx=XM_019830762.2;refGene_gn=DST;refGene_tx=XM_019830763.2;refGene_gn=DST;refGene_tx=XM_019830764.1;refGene_gn=DST;refGene_tx=XM_019830765.2;refGene_gn=DST;refGene_tx=XM_019830766.2;refGene_gn=DST;refGene_tx=XM_019830768.1;refGene_gn=DST;refGene_tx=XM_019830770.2;refGene_gn=DST;refGene_tx=XM_019830771.2;refGene_gn=DST;refGene_tx=XM_023254073.1
B2 56625273 56625515
refGene_gn=DST;refGene_tx=XM_011282416.2;refGene_gn=DST;refGene_tx=XM_011282421.2;refGene_gn=DST;refGene_tx=XM_011282422.3;refGene_gn=DST;refGene_tx=XM_019830750.2;refGene_gn=DST;refGene_tx=XM_019830751.2;refGene_gn=DST;refGene_tx=XM_019830753.2;refGene_gn=DST;refGene_tx=XM_019830754.2;refGene_gn=DST;refGene_tx=XM_019830755.2;refGene_gn=DST;refGene_tx=XM_019830756.2;refGene_gn=DST;refGene_tx=XM_019830757.2;refGene_gn=DST;refGene_tx=XM_019830758.2;refGene_gn=DST;refGene_tx=XM_019830759.2;refGene_gn=DST;refGene_tx=XM_019830760.2;refGene_gn=DST;refGene_tx=XM_019830761.2;refGene_gn=DST;refGene_tx=XM_019830762.2;refGene_gn=DST;refGene_tx=XM_019830763.2;refGene_gn=DST;refGene_tx=XM_019830764.1;refGene_gn=DST;refGene_tx=XM_019830765.2;refGene_gn=DST;refGene_tx=XM_019830766.2;refGene_gn=DST;refGene_tx=XM_019830768.1;refGene_gn=DST;refGene_tx=XM_019830770.2;refGene_gn=DST;refGene_tx=XM_019830771.2;refGene_gn=DST;refGene_tx=XM_023254073.1
B2 56626333 56626393
refGene_gn=DST;refGene_tx=XM_011282416.2;refGene_gn=DST;refGene_tx=XM_011282421.2;refGene_gn=DST;refGene_tx=XM_011282422.3;refGene_gn=DST;refGene_tx=XM_019830750.2;refGene_gn=DST;refGene_tx=XM_019830751.2;refGene_gn=DST;refGene_tx=XM_019830753.2;refGene_gn=DST;refGene_tx=XM_019830754.2;refGene_gn=DST;refGene_tx=XM_019830755.2;refGene_gn=DST;refGene_tx=XM_019830756.2;refGene_gn=DST;refGene_tx=XM_019830757.2;refGene_gn

=DST;refGene_tx=XM_019830758.2;refGene_gn=DST;refGene_tx=XM_019830759.2;refGene_gn=DST;refGene_tx=XM_019830760.2;refGene_gn=DST;refGene_tx=XM_019830761.2;refGene_gn=DST;refGene_tx=XM_019830762.2;refGene_gn=DST;refGene_tx=XM_019830763.2;refGene_gn=DST;refGene_tx=XM_019830764.1;refGene_gn=DST;refGene_tx=XM_019830765.2;refGene_gn=DST;refGene_tx=XM_019830766.2;refGene_gn=DST;refGene_tx=XM_019830768.1;refGene_gn=DST;refGene_tx=XM_019830770.2;refGene_gn=DST;refGene_tx=XM_019830771.2;refGene_gn=DST;refGene_tx=XM_023254073.1
B2 56628880 56629007
refGene_gn=DST;refGene_tx=XM_011282416.2;refGene_gn=DST;refGene_tx=XM_011282421.2;refGene_gn=DST;refGene_tx=XM_011282422.3;refGene_gn=DST;refGene_tx=XM_019830750.2;refGene_gn=DST;refGene_tx=XM_019830751.2;refGene_gn=DST;refGene_tx=XM_019830753.2;refGene_gn=DST;refGene_tx=XM_019830754.2;refGene_gn=DST;refGene_tx=XM_019830755.2;refGene_gn=DST;refGene_tx=XM_019830756.2;refGene_gn=DST;refGene_tx=XM_019830757.2;refGene_gn=DST;refGene_tx=XM_019830758.2;refGene_gn=DST;refGene_tx=XM_019830759.2;refGene_gn=DST;refGene_tx=XM_019830760.2;refGene_gn=DST;refGene_tx=XM_019830761.2;refGene_gn=DST;refGene_tx=XM_019830762.2;refGene_gn=DST;refGene_tx=XM_019830763.2;refGene_gn=DST;refGene_tx=XM_019830764.1;refGene_gn=DST;refGene_tx=XM_019830765.2;refGene_gn=DST;refGene_tx=XM_019830766.2;refGene_gn=DST;refGene_tx=XM_019830768.1;refGene_gn=DST;refGene_tx=XM_019830770.2;refGene_gn=DST;refGene_tx=XM_019830771.2;refGene_gn=DST;refGene_tx=XM_023254073.1
B2 56629921 56630078
refGene_gn=DST;refGene_tx=XM_011282416.2;refGene_gn=DST;refGene_tx=XM_011282421.2;refGene_gn=DST;refGene_tx=XM_011282422.3;refGene_gn=DST;refGene_tx=XM_019830750.2;refGene_gn=DST;refGene_tx=XM_019830751.2;refGene_gn=DST;refGene_tx=XM_019830753.2;refGene_gn=DST;refGene_tx=XM_019830754.2;refGene_gn=DST;refGene_tx=XM_019830755.2;refGene_gn=DST;refGene_tx=XM_019830756.2;refGene_gn=DST;refGene_tx=XM_019830757.2;refGene_gn=DST;refGene_tx=XM_019830758.2;refGene_gn=DST;refGene_tx=XM_019830759.2;refGene_gn=DST;refGene_tx=XM_019830760.2;refGene_gn=DST;refGene_tx=XM_019830761.2;refGene_gn=DST;refGene_tx=XM_019830762.2;refGene_gn=DST;refGene_tx=XM_019830763.2;refGene_gn=DST;refGene_tx=XM_019830764.1;refGene_gn=DST;refGene_tx=XM_019830765.2;refGene_gn=DST;refGene_tx=XM_019830766.2;refGene_gn=DST;refGene_tx=XM_019830768.1;refGene_gn=DST;refGene_tx=XM_019830770.2;refGene_gn=DST;refGene_tx=XM_019830771.2;refGene_gn=DST;refGene_tx=XM_023254073.1
B2 56631669 56631836
refGene_gn=DST;refGene_tx=XM_011282416.2;refGene_gn=DST;refGene_tx=XM_011282421.2;refGene_gn=DST;refGene_tx=XM_011282422.3;refGene_gn=DST;refGene_tx=XM_019830750.2;refGene_gn=DST;refGene_tx=XM_019830751.2;refGene_gn=DST;refGene_tx=XM_019830753.2;refGene_gn=DST;refGene_tx=XM_019830754.2;refGene_gn=DST;refGene_tx=XM_019830755.2;refGene_gn=DST;refGene_tx=XM_019830756.2;refGene_gn=DST;refGene_tx=XM_019830757.2;refGene_gn=DST;refGene_tx=XM_019830758.2;refGene_gn=DST;refGene_tx=XM_019830759.2;refGene_gn=DST;refGene_tx=XM_019830760.2;refGene_gn=DST;refGene_tx=XM_019830761.2;refGene_gn=DST;refGene_tx=XM_019830762.2;refGene_gn=DST;refGene_tx=XM_019830763.2;refGene_gn=DST;refGene_tx=XM_019830764.1;refGene_gn=DST;refGene_tx=XM_019830765.2;refGene_gn=DST;refGene_tx=XM_019

830766.2;refGene_gn=DST;refGene_tx=XM_019830768.1;refGene_gn=DST;refGene_tx=XM_019830770.2;refGene_gn=DST;refGene_tx=XM_019830771.2;refGene_gn=DST;refGene_tx=XM_023254073.1
B2 56632360 56632678
refGene_gn=DST;refGene_tx=XM_011282416.2;refGene_gn=DST;refGene_tx=XM_011282421.2;refGene_gn=DST;refGene_tx=XM_011282422.3;refGene_gn=DST;refGene_tx=XM_019830750.2;refGene_gn=DST;refGene_tx=XM_019830751.2;refGene_gn=DST;refGene_tx=XM_019830753.2;refGene_gn=DST;refGene_tx=XM_019830754.2;refGene_gn=DST;refGene_tx=XM_019830755.2;refGene_gn=DST;refGene_tx=XM_019830756.2;refGene_gn=DST;refGene_tx=XM_019830757.2;refGene_gn=DST;refGene_tx=XM_019830758.2;refGene_gn=DST;refGene_tx=XM_019830759.2;refGene_gn=DST;refGene_tx=XM_019830760.2;refGene_gn=DST;refGene_tx=XM_019830761.2;refGene_gn=DST;refGene_tx=XM_019830762.2;refGene_gn=DST;refGene_tx=XM_019830763.2;refGene_gn=DST;refGene_tx=XM_019830764.1;refGene_gn=DST;refGene_tx=XM_019830765.2;refGene_gn=DST;refGene_tx=XM_019830766.2;refGene_gn=DST;refGene_tx=XM_019830768.1;refGene_gn=DST;refGene_tx=XM_019830770.2;refGene_gn=DST;refGene_tx=XM_019830771.2;refGene_gn=DST;refGene_tx=XM_023254073.1
B2 56633299 56633508
refGene_gn=DST;refGene_tx=XM_011282416.2;refGene_gn=DST;refGene_tx=XM_011282421.2;refGene_gn=DST;refGene_tx=XM_011282422.3;refGene_gn=DST;refGene_tx=XM_019830750.2;refGene_gn=DST;refGene_tx=XM_019830751.2;refGene_gn=DST;refGene_tx=XM_019830753.2;refGene_gn=DST;refGene_tx=XM_019830754.2;refGene_gn=DST;refGene_tx=XM_019830755.2;refGene_gn=DST;refGene_tx=XM_019830756.2;refGene_gn=DST;refGene_tx=XM_019830757.2;refGene_gn=DST;refGene_tx=XM_019830758.2;refGene_gn=DST;refGene_tx=XM_019830759.2;refGene_gn=DST;refGene_tx=XM_019830760.2;refGene_gn=DST;refGene_tx=XM_019830761.2;refGene_gn=DST;refGene_tx=XM_019830762.2;refGene_gn=DST;refGene_tx=XM_019830763.2;refGene_gn=DST;refGene_tx=XM_019830764.1;refGene_gn=DST;refGene_tx=XM_019830765.2;refGene_gn=DST;refGene_tx=XM_019830766.2;refGene_gn=DST;refGene_tx=XM_019830768.1;refGene_gn=DST;refGene_tx=XM_019830770.2;refGene_gn=DST;refGene_tx=XM_019830771.2;refGene_gn=DST;refGene_tx=XM_023254073.1
B2 56636604 56636728
refGene_gn=DST;refGene_tx=XM_011282416.2;refGene_gn=DST;refGene_tx=XM_011282421.2;refGene_gn=DST;refGene_tx=XM_011282422.3;refGene_gn=DST;refGene_tx=XM_019830750.2;refGene_gn=DST;refGene_tx=XM_019830751.2;refGene_gn=DST;refGene_tx=XM_019830753.2;refGene_gn=DST;refGene_tx=XM_019830754.2;refGene_gn=DST;refGene_tx=XM_019830755.2;refGene_gn=DST;refGene_tx=XM_019830756.2;refGene_gn=DST;refGene_tx=XM_019830757.2;refGene_gn=DST;refGene_tx=XM_019830758.2;refGene_gn=DST;refGene_tx=XM_019830759.2;refGene_gn=DST;refGene_tx=XM_019830760.2;refGene_gn=DST;refGene_tx=XM_019830761.2;refGene_gn=DST;refGene_tx=XM_019830762.2;refGene_gn=DST;refGene_tx=XM_019830763.2;refGene_gn=DST;refGene_tx=XM_019830764.1;refGene_gn=DST;refGene_tx=XM_019830765.2;refGene_gn=DST;refGene_tx=XM_019830766.2;refGene_gn=DST;refGene_tx=XM_019830768.1;refGene_gn=DST;refGene_tx=XM_019830770.2;refGene_gn=DST;refGene_tx=XM_019830771.2;refGene_gn=DST;refGene_tx=XM_023254073.1
B2 56643419 56643596
refGene_gn=DST;refGene_tx=XM_011282416.2;refGene_gn=DST;refGene_tx=XM_

011282421.2;refGene_gn=DST;refGene_tx=XM_011282422.3;refGene_gn=DST;refGene_tx=XM_019830750.2;refGene_gn=DST;refGene_tx=XM_019830751.2;refGene_gn=DST;refGene_tx=XM_019830753.2;refGene_gn=DST;refGene_tx=XM_019830754.2;refGene_gn=DST;refGene_tx=XM_019830755.2;refGene_gn=DST;refGene_tx=XM_019830756.2;refGene_gn=DST;refGene_tx=XM_019830757.2;refGene_gn=DST;refGene_tx=XM_019830758.2;refGene_gn=DST;refGene_tx=XM_019830759.2;refGene_gn=DST;refGene_tx=XM_019830760.2;refGene_gn=DST;refGene_tx=XM_019830761.2;refGene_gn=DST;refGene_tx=XM_019830762.2;refGene_gn=DST;refGene_tx=XM_019830763.2;refGene_gn=DST;refGene_tx=XM_019830764.1;refGene_gn=DST;refGene_tx=XM_019830765.2;refGene_gn=DST;refGene_tx=XM_019830766.2;refGene_gn=DST;refGene_tx=XM_019830768.1;refGene_gn=DST;refGene_tx=XM_019830770.2;refGene_gn=DST;refGene_tx=XM_019830771.2;refGene_gn=DST;refGene_tx=XM_023254073.1
B2 56644656 56645205
refGene_gn=DST;refGene_tx=XM_011282416.2;refGene_gn=DST;refGene_tx=XM_011282421.2;refGene_gn=DST;refGene_tx=XM_011282422.3;refGene_gn=DST;refGene_tx=XM_019830750.2;refGene_gn=DST;refGene_tx=XM_019830751.2;refGene_gn=DST;refGene_tx=XM_019830753.2;refGene_gn=DST;refGene_tx=XM_019830754.2;refGene_gn=DST;refGene_tx=XM_019830755.2;refGene_gn=DST;refGene_tx=XM_019830756.2;refGene_gn=DST;refGene_tx=XM_019830757.2;refGene_gn=DST;refGene_tx=XM_019830758.2;refGene_gn=DST;refGene_tx=XM_019830759.2;refGene_gn=DST;refGene_tx=XM_019830760.2;refGene_gn=DST;refGene_tx=XM_019830761.2;refGene_gn=DST;refGene_tx=XM_019830762.2;refGene_gn=DST;refGene_tx=XM_019830763.2;refGene_gn=DST;refGene_tx=XM_019830764.1;refGene_gn=DST;refGene_tx=XM_019830765.2;refGene_gn=DST;refGene_tx=XM_019830766.2;refGene_gn=DST;refGene_tx=XM_019830768.1;refGene_gn=DST;refGene_tx=XM_019830770.2;refGene_gn=DST;refGene_tx=XM_019830771.2;refGene_gn=DST;refGene_tx=XM_023254073.1
B2 56648504 56648771
refGene_gn=DST;refGene_tx=XM_011282416.2;refGene_gn=DST;refGene_tx=XM_011282421.2;refGene_gn=DST;refGene_tx=XM_011282422.3;refGene_gn=DST;refGene_tx=XM_019830750.2;refGene_gn=DST;refGene_tx=XM_019830751.2;refGene_gn=DST;refGene_tx=XM_019830753.2;refGene_gn=DST;refGene_tx=XM_019830754.2;refGene_gn=DST;refGene_tx=XM_019830755.2;refGene_gn=DST;refGene_tx=XM_019830756.2;refGene_gn=DST;refGene_tx=XM_019830757.2;refGene_gn=DST;refGene_tx=XM_019830758.2;refGene_gn=DST;refGene_tx=XM_019830759.2;refGene_gn=DST;refGene_tx=XM_019830760.2;refGene_gn=DST;refGene_tx=XM_019830761.2;refGene_gn=DST;refGene_tx=XM_019830762.2;refGene_gn=DST;refGene_tx=XM_019830763.2;refGene_gn=DST;refGene_tx=XM_019830764.1;refGene_gn=DST;refGene_tx=XM_019830765.2;refGene_gn=DST;refGene_tx=XM_019830766.2;refGene_gn=DST;refGene_tx=XM_019830768.1;refGene_gn=DST;refGene_tx=XM_019830770.2;refGene_gn=DST;refGene_tx=XM_019830771.2;refGene_gn=DST;refGene_tx=XM_023254073.1
B2 56649210 56649444
refGene_gn=DST;refGene_tx=XM_011282416.2;refGene_gn=DST;refGene_tx=XM_011282421.2;refGene_gn=DST;refGene_tx=XM_011282422.3;refGene_gn=DST;refGene_tx=XM_019830750.2;refGene_gn=DST;refGene_tx=XM_019830751.2;refGene_gn=DST;refGene_tx=XM_019830753.2;refGene_gn=DST;refGene_tx=XM_019830754.2;refGene_gn=DST;refGene_tx=XM_019830755.2;refGene_gn=DST;refGene_tx=XM_019830756.2;refGene_gn=DST;refGene_tx=XM_019830757.2;refGene_gn

=DST;refGene_tx=XM_019830758.2;refGene_gn=DST;refGene_tx=XM_019830759.2;refGene_gn=DST;refGene_tx=XM_019830760.2;refGene_gn=DST;refGene_tx=XM_019830761.2;refGene_gn=DST;refGene_tx=XM_019830762.2;refGene_gn=DST;refGene_tx=XM_019830763.2;refGene_gn=DST;refGene_tx=XM_019830764.1;refGene_gn=DST;refGene_tx=XM_019830765.2;refGene_gn=DST;refGene_tx=XM_019830766.2;refGene_gn=DST;refGene_tx=XM_019830768.1;refGene_gn=DST;refGene_tx=XM_019830770.2;refGene_gn=DST;refGene_tx=XM_019830771.2;refGene_gn=DST;refGene_tx=XM_023254073.1

B2 56651120 56651273

refGene_gn=DST;refGene_tx=XM_011282416.2;refGene_gn=DST;refGene_tx=XM_011282421.2;refGene_gn=DST;refGene_tx=XM_011282422.3;refGene_gn=DST;refGene_tx=XM_019830750.2;refGene_gn=DST;refGene_tx=XM_019830751.2;refGene_gn=DST;refGene_tx=XM_019830753.2;refGene_gn=DST;refGene_tx=XM_019830754.2;refGene_gn=DST;refGene_tx=XM_019830755.2;refGene_gn=DST;refGene_tx=XM_019830756.2;refGene_gn=DST;refGene_tx=XM_019830757.2;refGene_gn=DST;refGene_tx=XM_019830758.2;refGene_gn=DST;refGene_tx=XM_019830759.2;refGene_gn=DST;refGene_tx=XM_019830760.2;refGene_gn=DST;refGene_tx=XM_019830761.2;refGene_gn=DST;refGene_tx=XM_019830762.2;refGene_gn=DST;refGene_tx=XM_019830763.2;refGene_gn=DST;refGene_tx=XM_019830764.1;refGene_gn=DST;refGene_tx=XM_019830765.2;refGene_gn=DST;refGene_tx=XM_019830766.2;refGene_gn=DST;refGene_tx=XM_019830768.1;refGene_gn=DST;refGene_tx=XM_019830770.2;refGene_gn=DST;refGene_tx=XM_019830771.2;refGene_gn=DST;refGene_tx=XM_023254073.1

B2 56652498 56652732

refGene_gn=DST;refGene_tx=XM_019830750.2;refGene_gn=DST;refGene_tx=XM_019830751.2;refGene_gn=DST;refGene_tx=XM_019830753.2;refGene_gn=DST;refGene_tx=XM_019830754.2;refGene_gn=DST;refGene_tx=XM_019830755.2;refGene_gn=DST;refGene_tx=XM_019830756.2;refGene_gn=DST;refGene_tx=XM_019830757.2;refGene_gn=DST;refGene_tx=XM_019830758.2;refGene_gn=DST;refGene_tx=XM_019830759.2;refGene_gn=DST;refGene_tx=XM_019830760.2;refGene_gn=DST;refGene_tx=XM_019830761.2;refGene_gn=DST;refGene_tx=XM_019830762.2;refGene_gn=DST;refGene_tx=XM_019830763.2;refGene_gn=DST;refGene_tx=XM_019830764.1;refGene_gn=DST;refGene_tx=XM_019830765.2;refGene_gn=DST;refGene_tx=XM_019830770.2;refGene_gn=DST;refGene_tx=XM_019830771.2

B2 56654052 56654202

refGene_gn=DST;refGene_tx=XM_019830750.2;refGene_gn=DST;refGene_tx=XM_019830751.2;refGene_gn=DST;refGene_tx=XM_019830753.2;refGene_gn=DST;refGene_tx=XM_019830754.2;refGene_gn=DST;refGene_tx=XM_019830755.2;refGene_gn=DST;refGene_tx=XM_019830756.2;refGene_gn=DST;refGene_tx=XM_019830757.2;refGene_gn=DST;refGene_tx=XM_019830758.2;refGene_gn=DST;refGene_tx=XM_019830759.2;refGene_gn=DST;refGene_tx=XM_019830760.2;refGene_gn=DST;refGene_tx=XM_019830761.2;refGene_gn=DST;refGene_tx=XM_019830762.2;refGene_gn=DST;refGene_tx=XM_019830763.2;refGene_gn=DST;refGene_tx=XM_019830764.1;refGene_gn=DST;refGene_tx=XM_019830765.2;refGene_gn=DST;refGene_tx=XM_019830770.2;refGene_gn=DST;refGene_tx=XM_019830771.2

B2 56654389 56654605

refGene_gn=DST;refGene_tx=XM_019830750.2;refGene_gn=DST;refGene_tx=XM_019830751.2;refGene_gn=DST;refGene_tx=XM_019830753.2;refGene_gn=DST;refGene_tx=XM_019830754.2;refGene_gn=DST;refGene_tx=XM_019830755.2;refGene_gn=DST;refGene_tx=XM_019830756.2;refGene_gn=DST;refGene_tx=XM_01983

0757.2;refGene_gn=DST;refGene_tx=XM_019830758.2;refGene_gn=DST;refGene_tx=XM_019830759.2;refGene_gn=DST;refGene_tx=XM_019830760.2;refGene_gn=DST;refGene_tx=XM_019830761.2;refGene_gn=DST;refGene_tx=XM_019830762.2;refGene_gn=DST;refGene_tx=XM_019830763.2;refGene_gn=DST;refGene_tx=XM_019830764.1;refGene_gn=DST;refGene_tx=XM_019830765.2;refGene_gn=DST;refGene_tx=XM_019830770.2;refGene_gn=DST;refGene_tx=XM_019830771.2
B2 56654791 56654941
refGene_gn=DST;refGene_tx=XM_019830750.2;refGene_gn=DST;refGene_tx=XM_019830751.2;refGene_gn=DST;refGene_tx=XM_019830753.2;refGene_gn=DST;refGene_tx=XM_019830754.2;refGene_gn=DST;refGene_tx=XM_019830755.2;refGene_gn=DST;refGene_tx=XM_019830756.2;refGene_gn=DST;refGene_tx=XM_019830757.2;refGene_gn=DST;refGene_tx=XM_019830758.2;refGene_gn=DST;refGene_tx=XM_019830759.2;refGene_gn=DST;refGene_tx=XM_019830760.2;refGene_gn=DST;refGene_tx=XM_019830761.2;refGene_gn=DST;refGene_tx=XM_019830762.2;refGene_gn=DST;refGene_tx=XM_019830763.2;refGene_gn=DST;refGene_tx=XM_019830764.1;refGene_gn=DST;refGene_tx=XM_019830765.2;refGene_gn=DST;refGene_tx=XM_019830770.2;refGene_gn=DST;refGene_tx=XM_019830771.2
B2 56655068 56660552
refGene_gn=DST;refGene_tx=XM_019830750.2;refGene_gn=DST;refGene_tx=XM_019830751.2;refGene_gn=DST;refGene_tx=XM_019830753.2;refGene_gn=DST;refGene_tx=XM_019830754.2;refGene_gn=DST;refGene_tx=XM_019830755.2;refGene_gn=DST;refGene_tx=XM_019830756.2;refGene_gn=DST;refGene_tx=XM_019830757.2;refGene_gn=DST;refGene_tx=XM_019830758.2;refGene_gn=DST;refGene_tx=XM_019830759.2;refGene_gn=DST;refGene_tx=XM_019830760.2;refGene_gn=DST;refGene_tx=XM_019830761.2;refGene_gn=DST;refGene_tx=XM_019830762.2;refGene_gn=DST;refGene_tx=XM_019830763.2;refGene_gn=DST;refGene_tx=XM_019830764.1;refGene_gn=DST;refGene_tx=XM_019830765.2;refGene_gn=DST;refGene_tx=XM_019830770.2;refGene_gn=DST;refGene_tx=XM_019830771.2
B2 56661632 56661768
refGene_gn=DST;refGene_tx=XM_011282416.2;refGene_gn=DST;refGene_tx=XM_011282421.2;refGene_gn=DST;refGene_tx=XM_011282422.3;refGene_gn=DST;refGene_tx=XM_019830750.2;refGene_gn=DST;refGene_tx=XM_019830751.2;refGene_gn=DST;refGene_tx=XM_019830753.2;refGene_gn=DST;refGene_tx=XM_019830754.2;refGene_gn=DST;refGene_tx=XM_019830755.2;refGene_gn=DST;refGene_tx=XM_019830756.2;refGene_gn=DST;refGene_tx=XM_019830757.2;refGene_gn=DST;refGene_tx=XM_019830758.2;refGene_gn=DST;refGene_tx=XM_019830759.2;refGene_gn=DST;refGene_tx=XM_019830760.2;refGene_gn=DST;refGene_tx=XM_019830761.2;refGene_gn=DST;refGene_tx=XM_019830762.2;refGene_gn=DST;refGene_tx=XM_019830763.2;refGene_gn=DST;refGene_tx=XM_019830764.1;refGene_gn=DST;refGene_tx=XM_019830765.2;refGene_gn=DST;refGene_tx=XM_019830766.2;refGene_gn=DST;refGene_tx=XM_019830768.1;refGene_gn=DST;refGene_tx=XM_019830770.2;refGene_gn=DST;refGene_tx=XM_019830771.2;refGene_gn=DST;refGene_tx=XM_023254073.1
B2 56662742 56662831
refGene_gn=DST;refGene_tx=XM_011282416.2;refGene_gn=DST;refGene_tx=XM_011282421.2;refGene_gn=DST;refGene_tx=XM_011282422.3;refGene_gn=DST;refGene_tx=XM_019830750.2;refGene_gn=DST;refGene_tx=XM_019830751.2;refGene_gn=DST;refGene_tx=XM_019830753.2;refGene_gn=DST;refGene_tx=XM_019830754.2;refGene_gn=DST;refGene_tx=XM_019830755.2;refGene_gn=DST;refGene_tx=XM_019830756.2;refGene_gn=DST;refGene_tx=XM_019830757.2;refGene_gn

=DST;refGene_tx=XM_019830758.2;refGene_gn=DST;refGene_tx=XM_019830759.2;refGene_gn=DST;refGene_tx=XM_019830760.2;refGene_gn=DST;refGene_tx=XM_019830761.2;refGene_gn=DST;refGene_tx=XM_019830762.2;refGene_gn=DST;refGene_tx=XM_019830763.2;refGene_gn=DST;refGene_tx=XM_019830764.1;refGene_gn=DST;refGene_tx=XM_019830765.2;refGene_gn=DST;refGene_tx=XM_019830766.2;refGene_gn=DST;refGene_tx=XM_019830768.1;refGene_gn=DST;refGene_tx=XM_019830770.2;refGene_gn=DST;refGene_tx=XM_019830771.2;refGene_gn=DST;refGene_tx=XM_023254073.1
B2 56665015 56665144
refGene_gn=DST;refGene_tx=XM_011282416.2;refGene_gn=DST;refGene_tx=XM_011282421.2;refGene_gn=DST;refGene_tx=XM_011282422.3;refGene_gn=DST;refGene_tx=XM_019830750.2;refGene_gn=DST;refGene_tx=XM_019830751.2;refGene_gn=DST;refGene_tx=XM_019830753.2;refGene_gn=DST;refGene_tx=XM_019830754.2;refGene_gn=DST;refGene_tx=XM_019830755.2;refGene_gn=DST;refGene_tx=XM_019830756.2;refGene_gn=DST;refGene_tx=XM_019830757.2;refGene_gn=DST;refGene_tx=XM_019830758.2;refGene_gn=DST;refGene_tx=XM_019830759.2;refGene_gn=DST;refGene_tx=XM_019830760.2;refGene_gn=DST;refGene_tx=XM_019830761.2;refGene_gn=DST;refGene_tx=XM_019830762.2;refGene_gn=DST;refGene_tx=XM_019830763.2;refGene_gn=DST;refGene_tx=XM_019830764.1;refGene_gn=DST;refGene_tx=XM_019830765.2;refGene_gn=DST;refGene_tx=XM_019830766.2;refGene_gn=DST;refGene_tx=XM_019830768.1;refGene_gn=DST;refGene_tx=XM_019830770.2;refGene_gn=DST;refGene_tx=XM_019830771.2;refGene_gn=DST;refGene_tx=XM_023254073.1
B2 56666199 56668098 refGene_gn=DST;refGene_tx=XM_019830769.1
B2 56668998 56671716 refGene_gn=DST;refGene_tx=XM_019830769.1
B2 56675204 56675303
refGene_gn=DST;refGene_tx=XM_011282416.2;refGene_gn=DST;refGene_tx=XM_011282421.2;refGene_gn=DST;refGene_tx=XM_011282422.3;refGene_gn=DST;refGene_tx=XM_019830750.2;refGene_gn=DST;refGene_tx=XM_019830751.2;refGene_gn=DST;refGene_tx=XM_019830753.2;refGene_gn=DST;refGene_tx=XM_019830754.2;refGene_gn=DST;refGene_tx=XM_019830755.2;refGene_gn=DST;refGene_tx=XM_019830756.2;refGene_gn=DST;refGene_tx=XM_019830757.2;refGene_gn=DST;refGene_tx=XM_019830758.2;refGene_gn=DST;refGene_tx=XM_019830759.2;refGene_gn=DST;refGene_tx=XM_019830760.2;refGene_gn=DST;refGene_tx=XM_019830761.2;refGene_gn=DST;refGene_tx=XM_019830762.2;refGene_gn=DST;refGene_tx=XM_019830763.2;refGene_gn=DST;refGene_tx=XM_019830764.1;refGene_gn=DST;refGene_tx=XM_019830765.2;refGene_gn=DST;refGene_tx=XM_019830766.2;refGene_gn=DST;refGene_tx=XM_019830768.1;refGene_gn=DST;refGene_tx=XM_019830769.1;refGene_gn=DST;refGene_tx=XM_019830770.2;refGene_gn=DST;refGene_tx=XM_019830771.2;refGene_gn=DST;refGene_tx=XM_023254073.1
B2 56675830 56675938
refGene_gn=DST;refGene_tx=XM_011282416.2;refGene_gn=DST;refGene_tx=XM_011282421.2;refGene_gn=DST;refGene_tx=XM_011282422.3;refGene_gn=DST;refGene_tx=XM_019830750.2;refGene_gn=DST;refGene_tx=XM_019830751.2;refGene_gn=DST;refGene_tx=XM_019830753.2;refGene_gn=DST;refGene_tx=XM_019830754.2;refGene_gn=DST;refGene_tx=XM_019830755.2;refGene_gn=DST;refGene_tx=XM_019830756.2;refGene_gn=DST;refGene_tx=XM_019830757.2;refGene_gn=DST;refGene_tx=XM_019830758.2;refGene_gn=DST;refGene_tx=XM_019830759.2;refGene_gn=DST;refGene_tx=XM_019830760.2;refGene_gn=DST;refGene_tx=X

M_019830761.2;refGene_gn=DST;refGene_tx=XM_019830762.2;refGene_gn=DST;refGene_tx=XM_019830763.2;refGene_gn=DST;refGene_tx=XM_019830764.1;refGene_gn=DST;refGene_tx=XM_019830765.2;refGene_gn=DST;refGene_tx=XM_019830766.2;refGene_gn=DST;refGene_tx=XM_019830768.1;refGene_gn=DST;refGene_tx=XM_019830769.1;refGene_gn=DST;refGene_tx=XM_019830770.2;refGene_gn=DST;refGene_tx=XM_019830771.2;refGene_gn=DST;refGene_tx=XM_023254073.1
B2 56676676 56676760
refGene_gn=DST;refGene_tx=XM_011282416.2;refGene_gn=DST;refGene_tx=XM_011282421.2;refGene_gn=DST;refGene_tx=XM_011282422.3;refGene_gn=DST;refGene_tx=XM_019830750.2;refGene_gn=DST;refGene_tx=XM_019830751.2;refGene_gn=DST;refGene_tx=XM_019830753.2;refGene_gn=DST;refGene_tx=XM_019830754.2;refGene_gn=DST;refGene_tx=XM_019830755.2;refGene_gn=DST;refGene_tx=XM_019830756.2;refGene_gn=DST;refGene_tx=XM_019830757.2;refGene_gn=DST;refGene_tx=XM_019830758.2;refGene_gn=DST;refGene_tx=XM_019830759.2;refGene_gn=DST;refGene_tx=XM_019830760.2;refGene_gn=DST;refGene_tx=XM_019830761.2;refGene_gn=DST;refGene_tx=XM_019830762.2;refGene_gn=DST;refGene_tx=XM_019830763.2;refGene_gn=DST;refGene_tx=XM_019830764.1;refGene_gn=DST;refGene_tx=XM_019830765.2;refGene_gn=DST;refGene_tx=XM_019830766.2;refGene_gn=DST;refGene_tx=XM_019830768.1;refGene_gn=DST;refGene_tx=XM_019830769.1;refGene_gn=DST;refGene_tx=XM_019830770.2;refGene_gn=DST;refGene_tx=XM_019830771.2;refGene_gn=DST;refGene_tx=XM_023254073.1
B2 56677476 56677639
refGene_gn=DST;refGene_tx=XM_011282416.2;refGene_gn=DST;refGene_tx=XM_011282421.2;refGene_gn=DST;refGene_tx=XM_011282422.3;refGene_gn=DST;refGene_tx=XM_019830750.2;refGene_gn=DST;refGene_tx=XM_019830751.2;refGene_gn=DST;refGene_tx=XM_019830753.2;refGene_gn=DST;refGene_tx=XM_019830754.2;refGene_gn=DST;refGene_tx=XM_019830755.2;refGene_gn=DST;refGene_tx=XM_019830756.2;refGene_gn=DST;refGene_tx=XM_019830757.2;refGene_gn=DST;refGene_tx=XM_019830758.2;refGene_gn=DST;refGene_tx=XM_019830759.2;refGene_gn=DST;refGene_tx=XM_019830760.2;refGene_gn=DST;refGene_tx=XM_019830761.2;refGene_gn=DST;refGene_tx=XM_019830762.2;refGene_gn=DST;refGene_tx=XM_019830763.2;refGene_gn=DST;refGene_tx=XM_019830764.1;refGene_gn=DST;refGene_tx=XM_019830765.2;refGene_gn=DST;refGene_tx=XM_019830766.2;refGene_gn=DST;refGene_tx=XM_019830768.1;refGene_gn=DST;refGene_tx=XM_019830769.1;refGene_gn=DST;refGene_tx=XM_019830770.2;refGene_gn=DST;refGene_tx=XM_019830771.2;refGene_gn=DST;refGene_tx=XM_023254073.1
B2 56679262 56679456
refGene_gn=DST;refGene_tx=XM_011282416.2;refGene_gn=DST;refGene_tx=XM_011282421.2;refGene_gn=DST;refGene_tx=XM_011282422.3;refGene_gn=DST;refGene_tx=XM_019830750.2;refGene_gn=DST;refGene_tx=XM_019830751.2;refGene_gn=DST;refGene_tx=XM_019830753.2;refGene_gn=DST;refGene_tx=XM_019830754.2;refGene_gn=DST;refGene_tx=XM_019830755.2;refGene_gn=DST;refGene_tx=XM_019830756.2;refGene_gn=DST;refGene_tx=XM_019830757.2;refGene_gn=DST;refGene_tx=XM_019830758.2;refGene_gn=DST;refGene_tx=XM_019830759.2;refGene_gn=DST;refGene_tx=XM_019830760.2;refGene_gn=DST;refGene_tx=XM_019830761.2;refGene_gn=DST;refGene_tx=XM_019830762.2;refGene_gn=DST;refGene_tx=XM_019830763.2;refGene_gn=DST;refGene_tx=XM_019830764.1;ref

Gene_gn=DST;refGene_tx=XM_019830765.2;refGene_gn=DST;refGene_tx=XM_019830766.2;refGene_gn=DST;refGene_tx=XM_019830768.1;refGene_gn=DST;refGene_tx=XM_019830769.1;refGene_gn=DST;refGene_tx=XM_019830770.2;refGene_gn=DST;refGene_tx=XM_019830771.2;refGene_gn=DST;refGene_tx=XM_023254073.1
B2 56680210 56680349
refGene_gn=DST;refGene_tx=XM_011282416.2;refGene_gn=DST;refGene_tx=XM_011282421.2;refGene_gn=DST;refGene_tx=XM_011282422.3;refGene_gn=DST;refGene_tx=XM_019830750.2;refGene_gn=DST;refGene_tx=XM_019830751.2;refGene_gn=DST;refGene_tx=XM_019830753.2;refGene_gn=DST;refGene_tx=XM_019830754.2;refGene_gn=DST;refGene_tx=XM_019830755.2;refGene_gn=DST;refGene_tx=XM_019830756.2;refGene_gn=DST;refGene_tx=XM_019830757.2;refGene_gn=DST;refGene_tx=XM_019830758.2;refGene_gn=DST;refGene_tx=XM_019830759.2;refGene_gn=DST;refGene_tx=XM_019830760.2;refGene_gn=DST;refGene_tx=XM_019830761.2;refGene_gn=DST;refGene_tx=XM_019830762.2;refGene_gn=DST;refGene_tx=XM_019830763.2;refGene_gn=DST;refGene_tx=XM_019830764.1;refGene_gn=DST;refGene_tx=XM_019830765.2;refGene_gn=DST;refGene_tx=XM_019830766.2;refGene_gn=DST;refGene_tx=XM_019830768.1;refGene_gn=DST;refGene_tx=XM_019830769.1;refGene_gn=DST;refGene_tx=XM_019830770.2;refGene_gn=DST;refGene_tx=XM_019830771.2;refGene_gn=DST;refGene_tx=XM_023254073.1
B2 56681338 56681517
refGene_gn=DST;refGene_tx=XM_011282416.2;refGene_gn=DST;refGene_tx=XM_011282421.2;refGene_gn=DST;refGene_tx=XM_011282422.3;refGene_gn=DST;refGene_tx=XM_019830750.2;refGene_gn=DST;refGene_tx=XM_019830751.2;refGene_gn=DST;refGene_tx=XM_019830753.2;refGene_gn=DST;refGene_tx=XM_019830754.2;refGene_gn=DST;refGene_tx=XM_019830755.2;refGene_gn=DST;refGene_tx=XM_019830756.2;refGene_gn=DST;refGene_tx=XM_019830757.2;refGene_gn=DST;refGene_tx=XM_019830758.2;refGene_gn=DST;refGene_tx=XM_019830759.2;refGene_gn=DST;refGene_tx=XM_019830760.2;refGene_gn=DST;refGene_tx=XM_019830761.2;refGene_gn=DST;refGene_tx=XM_019830762.2;refGene_gn=DST;refGene_tx=XM_019830763.2;refGene_gn=DST;refGene_tx=XM_019830764.1;refGene_gn=DST;refGene_tx=XM_019830765.2;refGene_gn=DST;refGene_tx=XM_019830766.2;refGene_gn=DST;refGene_tx=XM_019830768.1;refGene_gn=DST;refGene_tx=XM_019830769.1;refGene_gn=DST;refGene_tx=XM_019830770.2;refGene_gn=DST;refGene_tx=XM_019830771.2;refGene_gn=DST;refGene_tx=XM_023254073.1
B2 56682008 56682166
refGene_gn=DST;refGene_tx=XM_011282416.2;refGene_gn=DST;refGene_tx=XM_011282421.2;refGene_gn=DST;refGene_tx=XM_011282422.3;refGene_gn=DST;refGene_tx=XM_019830750.2;refGene_gn=DST;refGene_tx=XM_019830751.2;refGene_gn=DST;refGene_tx=XM_019830753.2;refGene_gn=DST;refGene_tx=XM_019830754.2;refGene_gn=DST;refGene_tx=XM_019830755.2;refGene_gn=DST;refGene_tx=XM_019830756.2;refGene_gn=DST;refGene_tx=XM_019830757.2;refGene_gn=DST;refGene_tx=XM_019830758.2;refGene_gn=DST;refGene_tx=XM_019830759.2;refGene_gn=DST;refGene_tx=XM_019830760.2;refGene_gn=DST;refGene_tx=XM_019830761.2;refGene_gn=DST;refGene_tx=XM_019830762.2;refGene_gn=DST;refGene_tx=XM_019830763.2;refGene_gn=DST;refGene_tx=XM_019830764.1;refGene_gn=DST;refGene_tx=XM_019830765.2;refGene_gn=DST;refGene_tx=XM_019830766.2;refGene_gn=DST;refGene_tx=XM_019830768.1;refGene_gn=DST;refGe

ne_tx=XM_019830769.1;refGene_gn=DST;refGene_tx=XM_019830770.2;refGene_gn=DST;refGene_tx=XM_019830771.2;refGene_gn=DST;refGene_tx=XM_023254073.1
B2 56683560 56683744
refGene_gn=DST;refGene_tx=XM_011282416.2;refGene_gn=DST;refGene_tx=XM_011282421.2;refGene_gn=DST;refGene_tx=XM_011282422.3;refGene_gn=DST;refGene_tx=XM_019830750.2;refGene_gn=DST;refGene_tx=XM_019830751.2;refGene_gn=DST;refGene_tx=XM_019830753.2;refGene_gn=DST;refGene_tx=XM_019830754.2;refGene_gn=DST;refGene_tx=XM_019830755.2;refGene_gn=DST;refGene_tx=XM_019830756.2;refGene_gn=DST;refGene_tx=XM_019830757.2;refGene_gn=DST;refGene_tx=XM_019830758.2;refGene_gn=DST;refGene_tx=XM_019830759.2;refGene_gn=DST;refGene_tx=XM_019830760.2;refGene_gn=DST;refGene_tx=XM_019830761.2;refGene_gn=DST;refGene_tx=XM_019830762.2;refGene_gn=DST;refGene_tx=XM_019830763.2;refGene_gn=DST;refGene_tx=XM_019830764.1;refGene_gn=DST;refGene_tx=XM_019830765.2;refGene_gn=DST;refGene_tx=XM_019830766.2;refGene_gn=DST;refGene_tx=XM_019830768.1;refGene_gn=DST;refGene_tx=XM_019830769.1;refGene_gn=DST;refGene_tx=XM_019830770.2;refGene_gn=DST;refGene_tx=XM_019830771.2;refGene_gn=DST;refGene_tx=XM_023254073.1
B2 56684230 56684357
refGene_gn=DST;refGene_tx=XM_011282416.2;refGene_gn=DST;refGene_tx=XM_011282421.2;refGene_gn=DST;refGene_tx=XM_011282422.3;refGene_gn=DST;refGene_tx=XM_019830750.2;refGene_gn=DST;refGene_tx=XM_019830751.2;refGene_gn=DST;refGene_tx=XM_019830753.2;refGene_gn=DST;refGene_tx=XM_019830754.2;refGene_gn=DST;refGene_tx=XM_019830755.2;refGene_gn=DST;refGene_tx=XM_019830756.2;refGene_gn=DST;refGene_tx=XM_019830757.2;refGene_gn=DST;refGene_tx=XM_019830758.2;refGene_gn=DST;refGene_tx=XM_019830759.2;refGene_gn=DST;refGene_tx=XM_019830760.2;refGene_gn=DST;refGene_tx=XM_019830761.2;refGene_gn=DST;refGene_tx=XM_019830762.2;refGene_gn=DST;refGene_tx=XM_019830763.2;refGene_gn=DST;refGene_tx=XM_019830764.1;refGene_gn=DST;refGene_tx=XM_019830765.2;refGene_gn=DST;refGene_tx=XM_019830766.2;refGene_gn=DST;refGene_tx=XM_019830768.1;refGene_gn=DST;refGene_tx=XM_019830769.1;refGene_gn=DST;refGene_tx=XM_019830770.2;refGene_gn=DST;refGene_tx=XM_019830771.2;refGene_gn=DST;refGene_tx=XM_023254073.1
B2 56684562 56684717
refGene_gn=DST;refGene_tx=XM_011282416.2;refGene_gn=DST;refGene_tx=XM_011282421.2;refGene_gn=DST;refGene_tx=XM_011282422.3;refGene_gn=DST;refGene_tx=XM_019830750.2;refGene_gn=DST;refGene_tx=XM_019830751.2;refGene_gn=DST;refGene_tx=XM_019830753.2;refGene_gn=DST;refGene_tx=XM_019830754.2;refGene_gn=DST;refGene_tx=XM_019830755.2;refGene_gn=DST;refGene_tx=XM_019830756.2;refGene_gn=DST;refGene_tx=XM_019830757.2;refGene_gn=DST;refGene_tx=XM_019830758.2;refGene_gn=DST;refGene_tx=XM_019830759.2;refGene_gn=DST;refGene_tx=XM_019830760.2;refGene_gn=DST;refGene_tx=XM_019830761.2;refGene_gn=DST;refGene_tx=XM_019830762.2;refGene_gn=DST;refGene_tx=XM_019830763.2;refGene_gn=DST;refGene_tx=XM_019830764.1;refGene_gn=DST;refGene_tx=XM_019830765.2;refGene_gn=DST;refGene_tx=XM_019830766.2;refGene_gn=DST;refGene_tx=XM_019830768.1;refGene_gn=DST;refGene_tx=XM_019830769.1;refGene_gn=DST;refGene_tx=XM_019830770.2;refGene_gn=DST;refGene_tx=XM_019830771.2;refGene_gn=DST;refGene_tx=XM_02325407

3.1
B2	56685110 56685263
refGene_gn=DST;refGene_tx=XM_011282416.2;refGene_gn=DST;refGene_tx=XM_011282421.2;refGene_gn=DST;refGene_tx=XM_011282422.3;refGene_gn=DST;refGene_tx=XM_019830750.2;refGene_gn=DST;refGene_tx=XM_019830751.2;refGene_gn=DST;refGene_tx=XM_019830753.2;refGene_gn=DST;refGene_tx=XM_019830754.2;refGene_gn=DST;refGene_tx=XM_019830755.2;refGene_gn=DST;refGene_tx=XM_019830756.2;refGene_gn=DST;refGene_tx=XM_019830757.2;refGene_gn=DST;refGene_tx=XM_019830758.2;refGene_gn=DST;refGene_tx=XM_019830759.2;refGene_gn=DST;refGene_tx=XM_019830760.2;refGene_gn=DST;refGene_tx=XM_019830761.2;refGene_gn=DST;refGene_tx=XM_019830762.2;refGene_gn=DST;refGene_tx=XM_019830763.2;refGene_gn=DST;refGene_tx=XM_019830764.1;refGene_gn=DST;refGene_tx=XM_019830765.2;refGene_gn=DST;refGene_tx=XM_019830766.2;refGene_gn=DST;refGene_tx=XM_019830768.1;refGene_gn=DST;refGene_tx=XM_019830769.1;refGene_gn=DST;refGene_tx=XM_019830770.2;refGene_gn=DST;refGene_tx=XM_019830771.2;refGene_gn=DST;refGene_tx=XM_023254073.1
B2	56685894 56686020
refGene_gn=DST;refGene_tx=XM_011282416.2;refGene_gn=DST;refGene_tx=XM_011282421.2;refGene_gn=DST;refGene_tx=XM_011282422.3;refGene_gn=DST;refGene_tx=XM_019830750.2;refGene_gn=DST;refGene_tx=XM_019830751.2;refGene_gn=DST;refGene_tx=XM_019830753.2;refGene_gn=DST;refGene_tx=XM_019830754.2;refGene_gn=DST;refGene_tx=XM_019830755.2;refGene_gn=DST;refGene_tx=XM_019830756.2;refGene_gn=DST;refGene_tx=XM_019830757.2;refGene_gn=DST;refGene_tx=XM_019830758.2;refGene_gn=DST;refGene_tx=XM_019830759.2;refGene_gn=DST;refGene_tx=XM_019830760.2;refGene_gn=DST;refGene_tx=XM_019830761.2;refGene_gn=DST;refGene_tx=XM_019830762.2;refGene_gn=DST;refGene_tx=XM_019830763.2;refGene_gn=DST;refGene_tx=XM_019830764.1;refGene_gn=DST;refGene_tx=XM_019830765.2;refGene_gn=DST;refGene_tx=XM_019830766.2;refGene_gn=DST;refGene_tx=XM_019830768.1;refGene_gn=DST;refGene_tx=XM_019830769.1;refGene_gn=DST;refGene_tx=XM_019830770.2;refGene_gn=DST;refGene_tx=XM_019830771.2;refGene_gn=DST;refGene_tx=XM_023254073.1
B2	56686623 56686719
refGene_gn=DST;refGene_tx=XM_011282416.2;refGene_gn=DST;refGene_tx=XM_011282421.2;refGene_gn=DST;refGene_tx=XM_011282422.3;refGene_gn=DST;refGene_tx=XM_019830750.2;refGene_gn=DST;refGene_tx=XM_019830751.2;refGene_gn=DST;refGene_tx=XM_019830753.2;refGene_gn=DST;refGene_tx=XM_019830754.2;refGene_gn=DST;refGene_tx=XM_019830755.2;refGene_gn=DST;refGene_tx=XM_019830756.2;refGene_gn=DST;refGene_tx=XM_019830757.2;refGene_gn=DST;refGene_tx=XM_019830758.2;refGene_gn=DST;refGene_tx=XM_019830759.2;refGene_gn=DST;refGene_tx=XM_019830760.2;refGene_gn=DST;refGene_tx=XM_019830761.2;refGene_gn=DST;refGene_tx=XM_019830762.2;refGene_gn=DST;refGene_tx=XM_019830763.2;refGene_gn=DST;refGene_tx=XM_019830764.1;refGene_gn=DST;refGene_tx=XM_019830765.2;refGene_gn=DST;refGene_tx=XM_019830766.2;refGene_gn=DST;refGene_tx=XM_019830768.1;refGene_gn=DST;refGene_tx=XM_019830769.1;refGene_gn=DST;refGene_tx=XM_019830770.2;refGene_gn=DST;refGene_tx=XM_019830771.2;refGene_gn=DST;refGene_tx=XM_023254073.1
B2	56689027 56689132

refGene_gn=DST;refGene_tx=XM_011282416.2;refGene_gn=DST;refGene_tx=XM_011282421.2;refGene_gn=DST;refGene_tx=XM_011282422.3;refGene_gn=DST;refGene_tx=XM_019830750.2;refGene_gn=DST;refGene_tx=XM_019830751.2;refGene_gn=DST;refGene_tx=XM_019830753.2;refGene_gn=DST;refGene_tx=XM_019830754.2;refGene_gn=DST;refGene_tx=XM_019830755.2;refGene_gn=DST;refGene_tx=XM_019830756.2;refGene_gn=DST;refGene_tx=XM_019830757.2;refGene_gn=DST;refGene_tx=XM_019830758.2;refGene_gn=DST;refGene_tx=XM_019830759.2;refGene_gn=DST;refGene_tx=XM_019830760.2;refGene_gn=DST;refGene_tx=XM_019830761.2;refGene_gn=DST;refGene_tx=XM_019830762.2;refGene_gn=DST;refGene_tx=XM_019830763.2;refGene_gn=DST;refGene_tx=XM_019830764.1;refGene_gn=DST;refGene_tx=XM_019830765.2;refGene_gn=DST;refGene_tx=XM_019830766.2;refGene_gn=DST;refGene_tx=XM_019830768.1;refGene_gn=DST;refGene_tx=XM_019830769.1;refGene_gn=DST;refGene_tx=XM_019830770.2;refGene_gn=DST;refGene_tx=XM_019830771.2;refGene_gn=DST;refGene_tx=XM_023254073.1

B2 56689222 56689384

refGene_gn=DST;refGene_tx=XM_011282416.2;refGene_gn=DST;refGene_tx=XM_011282421.2;refGene_gn=DST;refGene_tx=XM_011282422.3;refGene_gn=DST;refGene_tx=XM_019830750.2;refGene_gn=DST;refGene_tx=XM_019830751.2;refGene_gn=DST;refGene_tx=XM_019830753.2;refGene_gn=DST;refGene_tx=XM_019830754.2;refGene_gn=DST;refGene_tx=XM_019830755.2;refGene_gn=DST;refGene_tx=XM_019830756.2;refGene_gn=DST;refGene_tx=XM_019830757.2;refGene_gn=DST;refGene_tx=XM_019830758.2;refGene_gn=DST;refGene_tx=XM_019830759.2;refGene_gn=DST;refGene_tx=XM_019830760.2;refGene_gn=DST;refGene_tx=XM_019830761.2;refGene_gn=DST;refGene_tx=XM_019830762.2;refGene_gn=DST;refGene_tx=XM_019830763.2;refGene_gn=DST;refGene_tx=XM_019830764.1;refGene_gn=DST;refGene_tx=XM_019830765.2;refGene_gn=DST;refGene_tx=XM_019830766.2;refGene_gn=DST;refGene_tx=XM_019830768.1;refGene_gn=DST;refGene_tx=XM_019830769.1;refGene_gn=DST;refGene_tx=XM_019830770.2;refGene_gn=DST;refGene_tx=XM_019830771.2;refGene_gn=DST;refGene_tx=XM_023254073.1

B2 56689503 56689581

refGene_gn=DST;refGene_tx=XM_011282416.2;refGene_gn=DST;refGene_tx=XM_011282421.2;refGene_gn=DST;refGene_tx=XM_011282422.3;refGene_gn=DST;refGene_tx=XM_019830750.2;refGene_gn=DST;refGene_tx=XM_019830751.2;refGene_gn=DST;refGene_tx=XM_019830753.2;refGene_gn=DST;refGene_tx=XM_019830754.2;refGene_gn=DST;refGene_tx=XM_019830755.2;refGene_gn=DST;refGene_tx=XM_019830756.2;refGene_gn=DST;refGene_tx=XM_019830757.2;refGene_gn=DST;refGene_tx=XM_019830758.2;refGene_gn=DST;refGene_tx=XM_019830759.2;refGene_gn=DST;refGene_tx=XM_019830760.2;refGene_gn=DST;refGene_tx=XM_019830761.2;refGene_gn=DST;refGene_tx=XM_019830762.2;refGene_gn=DST;refGene_tx=XM_019830763.2;refGene_gn=DST;refGene_tx=XM_019830764.1;refGene_gn=DST;refGene_tx=XM_019830765.2;refGene_gn=DST;refGene_tx=XM_019830766.2;refGene_gn=DST;refGene_tx=XM_019830768.1;refGene_gn=DST;refGene_tx=XM_019830769.1;refGene_gn=DST;refGene_tx=XM_019830770.2;refGene_gn=DST;refGene_tx=XM_019830771.2;refGene_gn=DST;refGene_tx=XM_023254073.1

B2 56689742 56689871

refGene_gn=DST;refGene_tx=XM_011282416.2;refGene_gn=DST;refGene_tx=XM_011282421.2;refGene_gn=DST;refGene_tx=XM_011282422.3;refGene_gn=DST;re fGene_tx=XM_019830750.2;refGene_gn=DST;refGene_tx=XM_019830751.2;refGene_gn=DST;refGene_tx=XM_019830753.2;refGene_gn=DST;refGene_tx=XM_019830754.2;refGene_gn=DST;refGene_tx=XM_019830755.2;refGene_gn=DST;refGene_tx=XM_019830756.2;refGene_gn=DST;refGene_tx=XM_019830757.2;refGene_gn=DST;refGene_tx=XM_019830758.2;refGene_gn=DST;refGene_tx=XM_019830759.2;refGene_gn=DST;refGene_tx=XM_019830760.2;refGene_gn=DST;refGene_tx=XM_019830761.2;refGene_gn=DST;refGene_tx=XM_019830762.2;refGene_gn=DST;refGene_tx=XM_019830763.2;refGene_gn=DST;refGene_tx=XM_019830764.1;refGene_gn=DST;refGene_tx=XM_019830765.2;refGene_gn=DST;refGene_tx=XM_019830766.2;refGene_gn=DST;refGene_tx=XM_019830768.1;refGene_gn=DST;refGene_tx=XM_019830769.1;refGene_gn=DST;refGene_tx=XM_019830770.2;refGene_gn=DST;refGene_tx=XM_019830771.2;refGene_gn=DST;refGene_tx=XM_023254073.1

B2 56689959 56690425

refGene_gn=DST;refGene_tx=XM_011282416.2;refGene_gn=DST;refGene_tx=XM_011282421.2;refGene_gn=DST;refGene_tx=XM_011282422.3;refGene_gn=DST;refGene_tx=XM_019830750.2;refGene_gn=DST;refGene_tx=XM_019830751.2;refGene_gn=DST;refGene_tx=XM_019830753.2;refGene_gn=DST;refGene_tx=XM_019830754.2;refGene_gn=DST;refGene_tx=XM_019830755.2;refGene_gn=DST;refGene_tx=XM_019830756.2;refGene_gn=DST;refGene_tx=XM_019830757.2;refGene_gn=DST;refGene_tx=XM_019830758.2;refGene_gn=DST;refGene_tx=XM_019830759.2;refGene_gn=DST;refGene_tx=XM_019830760.2;refGene_gn=DST;refGene_tx=XM_019830761.2;refGene_gn=DST;refGene_tx=XM_019830762.2;refGene_gn=DST;refGene_tx=XM_019830763.2;refGene_gn=DST;refGene_tx=XM_019830764.1;refGene_gn=DST;refGene_tx=XM_019830765.2;refGene_gn=DST;refGene_tx=XM_019830766.2;refGene_gn=DST;refGene_tx=XM_019830768.1;refGene_gn=DST;refGene_tx=XM_019830769.1;refGene_gn=DST;refGene_tx=XM_019830770.2;refGene_gn=DST;refGene_tx=XM_019830771.2;refGene_gn=DST;refGene_tx=XM_023254073.1

B2 56691691 56691846

refGene_gn=DST;refGene_tx=XM_011282416.2;refGene_gn=DST;refGene_tx=XM_011282421.2;refGene_gn=DST;refGene_tx=XM_011282422.3;refGene_gn=DST;refGene_tx=XM_019830750.2;refGene_gn=DST;refGene_tx=XM_019830751.2;refGene_gn=DST;refGene_tx=XM_019830753.2;refGene_gn=DST;refGene_tx=XM_019830754.2;refGene_gn=DST;refGene_tx=XM_019830755.2;refGene_gn=DST;refGene_tx=XM_019830756.2;refGene_gn=DST;refGene_tx=XM_019830757.2;refGene_gn=DST;refGene_tx=XM_019830758.2;refGene_gn=DST;refGene_tx=XM_019830759.2;refGene_gn=DST;refGene_tx=XM_019830760.2;refGene_gn=DST;refGene_tx=XM_019830761.2;refGene_gn=DST;refGene_tx=XM_019830762.2;refGene_gn=DST;refGene_tx=XM_019830763.2;refGene_gn=DST;refGene_tx=XM_019830764.1;refGene_gn=DST;refGene_tx=XM_019830765.2;refGene_gn=DST;refGene_tx=XM_019830766.2;refGene_gn=DST;refGene_tx=XM_019830768.1;refGene_gn=DST;refGene_tx=XM_019830769.1;refGene_gn=DST;refGene_tx=XM_019830770.2;refGene_gn=DST;refGene_tx=XM_019830771.2;refGene_gn=DST;refGene_tx=XM_023254073.1

B2 56692154 56692248

refGene_gn=DST;refGene_tx=XM_011282416.2;refGene_gn=DST;refGene_tx=XM_011282421.2;refGene_gn=DST;refGene_tx=XM_011282422.3;refGene_gn=DST;refGene_tx=XM_019830750.2;refGene_gn=DST;refGene_tx=XM_019830751.2;refGene_gn=DST;refGene_tx=XM_019830753.2;refGene_gn=DST;refGene_tx=XM_01983

0754.2;refGene_gn=DST;refGene_tx=XM_019830755.2;refGene_gn=DST;refGene_tx=XM_019830756.2;refGene_gn=DST;refGene_tx=XM_019830757.2;refGene_gn=DST;refGene_tx=XM_019830758.2;refGene_gn=DST;refGene_tx=XM_019830759.2;refGene_gn=DST;refGene_tx=XM_019830760.2;refGene_gn=DST;refGene_tx=XM_019830761.2;refGene_gn=DST;refGene_tx=XM_019830762.2;refGene_gn=DST;refGene_tx=XM_019830763.2;refGene_gn=DST;refGene_tx=XM_019830764.1;refGene_gn=DST;refGene_tx=XM_019830765.2;refGene_gn=DST;refGene_tx=XM_019830766.2;refGene_gn=DST;refGene_tx=XM_019830768.1;refGene_gn=DST;refGene_tx=XM_019830769.1;refGene_gn=DST;refGene_tx=XM_019830770.2;refGene_gn=DST;refGene_tx=XM_019830771.2;refGene_gn=DST;refGene_tx=XM_023254073.1

B2 56692365 56692532 refGene_gn=DST;refGene_tx=XM_019830765.2;refGene_gn=DST;refGene_tx=XM_019830768.1;refGene_gn=DST;refGene_tx=XM_019830769.1

B2 56695603 56695731 refGene_gn=DST;refGene_tx=XM_011282416.2;refGene_gn=DST;refGene_tx=XM_011282421.2;refGene_gn=DST;refGene_tx=XM_011282422.3;refGene_gn=DST;refGene_tx=XM_019830750.2;refGene_gn=DST;refGene_tx=XM_019830751.2;refGene_gn=DST;refGene_tx=XM_019830753.2;refGene_gn=DST;refGene_tx=XM_019830754.2;refGene_gn=DST;refGene_tx=XM_019830755.2;refGene_gn=DST;refGene_tx=XM_019830756.2;refGene_gn=DST;refGene_tx=XM_019830757.2;refGene_gn=DST;refGene_tx=XM_019830758.2;refGene_gn=DST;refGene_tx=XM_019830759.2;refGene_gn=DST;refGene_tx=XM_019830760.2;refGene_gn=DST;refGene_tx=XM_019830761.2;refGene_gn=DST;refGene_tx=XM_019830762.2;refGene_gn=DST;refGene_tx=XM_019830763.2;refGene_gn=DST;refGene_tx=XM_019830764.1;refGene_gn=DST;refGene_tx=XM_019830766.2;refGene_gn=DST;refGene_tx=XM_019830770.2;refGene_gn=DST;refGene_tx=XM_019830771.2;refGene_gn=DST;refGene_tx=XM_023254073.1

B2 56695828 56695924 refGene_gn=DST;refGene_tx=XM_011282416.2;refGene_gn=DST;refGene_tx=XM_011282421.2;refGene_gn=DST;refGene_tx=XM_011282422.3;refGene_gn=DST;refGene_tx=XM_019830750.2;refGene_gn=DST;refGene_tx=XM_019830751.2;refGene_gn=DST;refGene_tx=XM_019830753.2;refGene_gn=DST;refGene_tx=XM_019830754.2;refGene_gn=DST;refGene_tx=XM_019830755.2;refGene_gn=DST;refGene_tx=XM_019830756.2;refGene_gn=DST;refGene_tx=XM_019830757.2;refGene_gn=DST;refGene_tx=XM_019830758.2;refGene_gn=DST;refGene_tx=XM_019830759.2;refGene_gn=DST;refGene_tx=XM_019830760.2;refGene_gn=DST;refGene_tx=XM_019830761.2;refGene_gn=DST;refGene_tx=XM_019830762.2;refGene_gn=DST;refGene_tx=XM_019830763.2;refGene_gn=DST;refGene_tx=XM_019830764.1;refGene_gn=DST;refGene_tx=XM_019830766.2;refGene_gn=DST;refGene_tx=XM_019830770.2;refGene_gn=DST;refGene_tx=XM_019830771.2;refGene_gn=DST;refGene_tx=XM_023254073.1

B2 56698030 56698150 refGene_gn=DST;refGene_tx=XM_011282416.2;refGene_gn=DST;refGene_tx=XM_011282421.2;refGene_gn=DST;refGene_tx=XM_011282422.3;refGene_gn=DST;refGene_tx=XM_019830750.2;refGene_gn=DST;refGene_tx=XM_019830751.2;refGene_gn=DST;refGene_tx=XM_019830753.2;refGene_gn=DST;refGene_tx=XM_019830754.2;refGene_gn=DST;refGene_tx=XM_019830755.2;refGene_gn=DST;refGene_tx=XM_019830756.2;refGene_gn=DST;refGene_tx=XM_019830757.2;refGene_gn=DST;refGene_tx=XM_019830758.2;refGene_gn=DST;refGene_tx=XM_019830759.

2;refGene_gn=DST;refGene_tx=XM_019830760.2;refGene_gn=DST;refGene_tx=XM_019830761.2;refGene_gn=DST;refGene_tx=XM_019830762.2;refGene_gn=DST;refGene_tx=XM_019830763.2;refGene_gn=DST;refGene_tx=XM_019830764.1;refGene_gn=DST;refGene_tx=XM_019830766.2;refGene_gn=DST;refGene_tx=XM_019830770.2;refGene_gn=DST;refGene_tx=XM_019830771.2;refGene_gn=DST;refGene_tx=XM_023254073.1
B2 56700308 56700415
refGene_gn=DST;refGene_tx=XM_011282416.2;refGene_gn=DST;refGene_tx=XM_011282421.2;refGene_gn=DST;refGene_tx=XM_011282422.3;refGene_gn=DST;refGene_tx=XM_019830750.2;refGene_gn=DST;refGene_tx=XM_019830751.2;refGene_gn=DST;refGene_tx=XM_019830753.2;refGene_gn=DST;refGene_tx=XM_019830754.2;refGene_gn=DST;refGene_tx=XM_019830755.2;refGene_gn=DST;refGene_tx=XM_019830756.2;refGene_gn=DST;refGene_tx=XM_019830757.2;refGene_gn=DST;refGene_tx=XM_019830758.2;refGene_gn=DST;refGene_tx=XM_019830759.2;refGene_gn=DST;refGene_tx=XM_019830760.2;refGene_gn=DST;refGene_tx=XM_019830761.2;refGene_gn=DST;refGene_tx=XM_019830762.2;refGene_gn=DST;refGene_tx=XM_019830763.2;refGene_gn=DST;refGene_tx=XM_019830764.1;refGene_gn=DST;refGene_tx=XM_019830766.2;refGene_gn=DST;refGene_tx=XM_019830770.2;refGene_gn=DST;refGene_tx=XM_019830771.2;refGene_gn=DST;refGene_tx=XM_023254073.1
B2 56700507 56700620
refGene_gn=DST;refGene_tx=XM_011282416.2;refGene_gn=DST;refGene_tx=XM_011282421.2;refGene_gn=DST;refGene_tx=XM_011282422.3;refGene_gn=DST;refGene_tx=XM_019830750.2;refGene_gn=DST;refGene_tx=XM_019830751.2;refGene_gn=DST;refGene_tx=XM_019830753.2;refGene_gn=DST;refGene_tx=XM_019830754.2;refGene_gn=DST;refGene_tx=XM_019830755.2;refGene_gn=DST;refGene_tx=XM_019830756.2;refGene_gn=DST;refGene_tx=XM_019830757.2;refGene_gn=DST;refGene_tx=XM_019830758.2;refGene_gn=DST;refGene_tx=XM_019830759.2;refGene_gn=DST;refGene_tx=XM_019830760.2;refGene_gn=DST;refGene_tx=XM_019830761.2;refGene_gn=DST;refGene_tx=XM_019830762.2;refGene_gn=DST;refGene_tx=XM_019830763.2;refGene_gn=DST;refGene_tx=XM_019830764.1;refGene_gn=DST;refGene_tx=XM_019830766.2;refGene_gn=DST;refGene_tx=XM_019830770.2;refGene_gn=DST;refGene_tx=XM_019830771.2;refGene_gn=DST;refGene_tx=XM_023254073.1
B2 56718236 56718403
refGene_gn=DST;refGene_tx=XM_011282416.2;refGene_gn=DST;refGene_tx=XM_011282421.2;refGene_gn=DST;refGene_tx=XM_011282422.3;refGene_gn=DST;refGene_tx=XM_019830750.2;refGene_gn=DST;refGene_tx=XM_019830751.2;refGene_gn=DST;refGene_tx=XM_019830753.2;refGene_gn=DST;refGene_tx=XM_019830754.2;refGene_gn=DST;refGene_tx=XM_019830755.2;refGene_gn=DST;refGene_tx=XM_019830756.2;refGene_gn=DST;refGene_tx=XM_019830757.2;refGene_gn=DST;refGene_tx=XM_019830758.2;refGene_gn=DST;refGene_tx=XM_019830759.2;refGene_gn=DST;refGene_tx=XM_019830760.2;refGene_gn=DST;refGene_tx=XM_019830761.2;refGene_gn=DST;refGene_tx=XM_019830762.2;refGene_gn=DST;refGene_tx=XM_019830763.2;refGene_gn=DST;refGene_tx=XM_019830764.1;refGene_gn=DST;refGene_tx=XM_019830766.2;refGene_gn=DST;refGene_tx=XM_019830770.2;refGene_gn=DST;refGene_tx=XM_019830771.2;refGene_gn=DST;refGene_tx=XM_023254073.1
B2 56736366 56737125 refGene_gn=DST;refGene_tx=XM_019830759.2
B2 56742759 56742852

refGene_gn=DST;refGene_tx=XM_011282416.2;refGene_gn=DST;refGene_tx=XM_
011282421.2;refGene_gn=DST;refGene_tx=XM_011282422.3;refGene_gn=DST;re
fGene_tx=XM_019830750.2;refGene_gn=DST;refGene_tx=XM_019830751.2;refGe
ne_gn=DST;refGene_tx=XM_019830753.2;refGene_gn=DST;refGene_tx=XM_01983
0754.2;refGene_gn=DST;refGene_tx=XM_019830755.2;refGene_gn=DST;refGene
_tx=XM_019830756.2;refGene_gn=DST;refGene_tx=XM_019830757.2;refGene_gn
=DST;refGene_tx=XM_019830758.2;refGene_gn=DST;refGene_tx=XM_019830760.
2;refGene_gn=DST;refGene_tx=XM_019830761.2;refGene_gn=DST;refGene_tx=X
M_019830762.2;refGene_gn=DST;refGene_tx=XM_019830763.2;refGene_gn=DST;
refGene_tx=XM_019830764.1;refGene_gn=DST;refGene_tx=XM_019830766.2;ref
Gene_gn=DST;refGene_tx=XM_019830770.2;refGene_gn=DST;refGene_tx=XM_019
830771.2;refGene_gn=DST;refGene_tx=XM_023254073.1
B2 56744766 56744844
refGene_gn=DST;refGene_tx=XM_011282416.2;refGene_gn=DST;refGene_tx=XM_
011282421.2;refGene_gn=DST;refGene_tx=XM_011282422.3;refGene_gn=DST;re
fGene_tx=XM_019830750.2;refGene_gn=DST;refGene_tx=XM_019830751.2;refGe
ne_gn=DST;refGene_tx=XM_019830753.2;refGene_gn=DST;refGene_tx=XM_01983
0754.2;refGene_gn=DST;refGene_tx=XM_019830755.2;refGene_gn=DST;refGene
_tx=XM_019830756.2;refGene_gn=DST;refGene_tx=XM_019830757.2;refGene_gn
=DST;refGene_tx=XM_019830758.2;refGene_gn=DST;refGene_tx=XM_019830760.
2;refGene_gn=DST;refGene_tx=XM_019830761.2;refGene_gn=DST;refGene_tx=X
M_019830762.2;refGene_gn=DST;refGene_tx=XM_019830763.2;refGene_gn=DST;
refGene_tx=XM_019830764.1;refGene_gn=DST;refGene_tx=XM_019830766.2;ref
Gene_gn=DST;refGene_tx=XM_019830770.2;refGene_gn=DST;refGene_tx=XM_019
830771.2;refGene_gn=DST;refGene_tx=XM_023254073.1
B2 56746460 56746559
refGene_gn=DST;refGene_tx=XM_011282422.3;refGene_gn=DST;refGene_tx=XM_
019830750.2;refGene_gn=DST;refGene_tx=XM_019830751.2;refGene_gn=DST;re
fGene_tx=XM_019830753.2;refGene_gn=DST;refGene_tx=XM_019830754.2;refGe
ne_gn=DST;refGene_tx=XM_019830755.2;refGene_gn=DST;refGene_tx=XM_01983
0757.2;refGene_gn=DST;refGene_tx=XM_019830760.2;refGene_gn=DST;refGene
_tx=XM_019830761.2;refGene_gn=DST;refGene_tx=XM_019830762.2
B2 56747090 56747180
refGene_gn=DST;refGene_tx=XM_011282416.2;refGene_gn=DST;refGene_tx=XM_
011282421.2;refGene_gn=DST;refGene_tx=XM_011282422.3;refGene_gn=DST;re
fGene_tx=XM_019830750.2;refGene_gn=DST;refGene_tx=XM_019830751.2;refGe
ne_gn=DST;refGene_tx=XM_019830753.2;refGene_gn=DST;refGene_tx=XM_01983
0754.2;refGene_gn=DST;refGene_tx=XM_019830755.2;refGene_gn=DST;refGene
_tx=XM_019830756.2;refGene_gn=DST;refGene_tx=XM_019830757.2;refGene_gn
=DST;refGene_tx=XM_019830758.2;refGene_gn=DST;refGene_tx=XM_019830760.
2;refGene_gn=DST;refGene_tx=XM_019830761.2;refGene_gn=DST;refGene_tx=X
M_019830762.2;refGene_gn=DST;refGene_tx=XM_019830763.2;refGene_gn=DST;
refGene_tx=XM_019830764.1;refGene_gn=DST;refGene_tx=XM_019830766.2;ref
Gene_gn=DST;refGene_tx=XM_019830770.2;refGene_gn=DST;refGene_tx=XM_019
830771.2;refGene_gn=DST;refGene_tx=XM_023254073.1
B2 56783054 56783116
refGene_gn=DST;refGene_tx=XM_011282416.2;refGene_gn=DST;refGene_tx=XM_
011282421.2;refGene_gn=DST;refGene_tx=XM_011282422.3;refGene_gn=DST;re
fGene_tx=XM_019830750.2;refGene_gn=DST;refGene_tx=XM_019830751.2;refGe
ne_gn=DST;refGene_tx=XM_019830753.2;refGene_gn=DST;refGene_tx=XM_01983

0754.2;refGene_gn=DST;refGene_tx=XM_019830755.2;refGene_gn=DST;refGene_tx=XM_019830756.2;refGene_gn=DST;refGene_tx=XM_019830757.2;refGene_gn=DST;refGene_tx=XM_019830758.2;refGene_gn=DST;refGene_tx=XM_019830760.2;refGene_gn=DST;refGene_tx=XM_019830761.2;refGene_gn=DST;refGene_tx=XM_019830762.2;refGene_gn=DST;refGene_tx=XM_019830763.2;refGene_gn=DST;refGene_tx=XM_019830764.1;refGene_gn=DST;refGene_tx=XM_019830766.2;refGene_gn=DST;refGene_tx=XM_019830770.2;refGene_gn=DST;refGene_tx=XM_019830771.2;refGene_gn=DST;refGene_tx=XM_023254073.1
B2 56884055 56884146
refGene_gn=DST;refGene_tx=XM_011282421.2;refGene_gn=DST;refGene_tx=XM_019830762.2;refGene_gn=DST;refGene_tx=XM_019830763.2;refGene_gn=DST;refGene_tx=XM_019830764.1;refGene_gn=DST;refGene_tx=XM_019830770.2;refGene_gn=DST;refGene_tx=XM_023254073.1
B2 56890648 56890856
refGene_gn=DST;refGene_tx=XM_011282416.2;refGene_gn=DST;refGene_tx=XM_011282422.3;refGene_gn=DST;refGene_tx=XM_019830750.2;refGene_gn=DST;refGene_tx=XM_019830751.2;refGene_gn=DST;refGene_tx=XM_019830753.2;refGene_gn=DST;refGene_tx=XM_019830754.2;refGene_gn=DST;refGene_tx=XM_019830755.2;refGene_gn=DST;refGene_tx=XM_019830756.2;refGene_gn=DST;refGene_tx=XM_019830757.2;refGene_gn=DST;refGene_tx=XM_019830758.2;refGene_gn=DST;refGene_tx=XM_019830760.2;refGene_gn=DST;refGene_tx=XM_019830761.2;refGene_gn=DST;refGene_tx=XM_019830766.2;refGene_gn=DST;refGene_tx=XM_019830771.2
B2 56947498 56947699
refGene_gn=DST;refGene_tx=XM_011282416.2;refGene_gn=DST;refGene_tx=XM_011282422.3;refGene_gn=DST;refGene_tx=XM_019830750.2;refGene_gn=DST;refGene_tx=XM_019830751.2;refGene_gn=DST;refGene_tx=XM_019830753.2;refGene_gn=DST;refGene_tx=XM_019830754.2;refGene_gn=DST;refGene_tx=XM_019830755.2;refGene_gn=DST;refGene_tx=XM_019830756.2;refGene_gn=DST;refGene_tx=XM_019830757.2;refGene_gn=DST;refGene_tx=XM_019830758.2;refGene_gn=DST;refGene_tx=XM_019830760.2;refGene_gn=DST;refGene_tx=XM_019830761.2;refGene_gn=DST;refGene_tx=XM_019830766.2;refGene_gn=DST;refGene_tx=XM_019830771.2
B2 57021313 57021375
refGene_gn=DST;refGene_tx=XM_011282416.2;refGene_gn=DST;refGene_tx=XM_011282422.3;refGene_gn=DST;refGene_tx=XM_019830750.2;refGene_gn=DST;refGene_tx=XM_019830751.2;refGene_gn=DST;refGene_tx=XM_019830753.2;refGene_gn=DST;refGene_tx=XM_019830754.2;refGene_gn=DST;refGene_tx=XM_019830755.2;refGene_gn=DST;refGene_tx=XM_019830756.2;refGene_gn=DST;refGene_tx=XM_019830757.2;refGene_gn=DST;refGene_tx=XM_019830758.2;refGene_gn=DST;refGene_tx=XM_019830760.2;refGene_gn=DST;refGene_tx=XM_019830761.2;refGene_gn=DST;refGene_tx=XM_019830766.2;refGene_gn=DST;refGene_tx=XM_019830771.2
B2 57021972 57022153
refGene_gn=DST;refGene_tx=XM_011282416.2;refGene_gn=DST;refGene_tx=XM_011282422.3;refGene_gn=DST;refGene_tx=XM_019830750.2;refGene_gn=DST;refGene_tx=XM_019830751.2;refGene_gn=DST;refGene_tx=XM_019830753.2;refGene_gn=DST;refGene_tx=XM_019830754.2;refGene_gn=DST;refGene_tx=XM_019830755.2;refGene_gn=DST;refGene_tx=XM_019830756.2;refGene_gn=DST;refGene_tx=XM_019830757.2;refGene_gn=DST;refGene_tx=XM_019830758.2;refGene_gn

=DST;refGene_tx=XM_019830760.2;refGene_gn=DST;refGene_tx=XM_019830761.2;refGene_gn=DST;refGene_tx=XM_019830766.2;refGene_gn=DST;refGene_tx=XM_019830771.2
B2 57040137 57040257 refGene_gn=BEND6;refGene_tx=XM_011282412.2;refGene_gn=BEND6;refGene_tx=XM_011282414.2
B2 57052344 57052380 refGene_gn=BEND6;refGene_tx=XM_011282412.2
B2 57054201 57054379 refGene_gn=BEND6;refGene_tx=XM_011282412.2;refGene_gn=BEND6;refGene_tx=XM_011282414.2
B2 57068110 57068331 refGene_gn=BEND6;refGene_tx=XM_011282412.2;refGene_gn=BEND6;refGene_tx=XM_011282414.2
B2 57070362 57070555 refGene_gn=BEND6;refGene_tx=XM_011282412.2;refGene_gn=BEND6;refGene_tx=XM_011282414.2
B2 57071410 57071538 refGene_gn=BEND6;refGene_tx=XM_011282412.2;refGene_gn=BEND6;refGene_tx=XM_011282414.2
B2 57091418 57091439 refGene_gn=KIAA1586;refGene_tx=XM_023254075.1
B2 57091989 57092073 refGene_gn=KIAA1586;refGene_tx=XM_023254075.1
B2 57096769 57096850 refGene_gn=KIAA1586;refGene_tx=XM_023254075.1
B2 57098609 57099128 refGene_gn=KIAA1586;refGene_tx=XM_023254075.1
B2 57099129 57099473 refGene_gn=KIAA1586;refGene_tx=XM_023254075.1
B2 57099474 57099668 refGene_gn=KIAA1586;refGene_tx=XM_023254075.1
B2 57099670 57100785 refGene_gn=KIAA1586;refGene_tx=XM_023254075.1
B2 57135288 57135309 refGene_gn=LOC101099964;refGene_tx=XM_023254076.1
B2 57135859 57135943 refGene_gn=LOC101099964;refGene_tx=XM_023254076.1
B2 57145082 57145163 refGene_gn=LOC101099964;refGene_tx=XM_023254076.1
B2 57168390 57168516 refGene_gn=LOC101099964;refGene_tx=XM_023254076.1
B2 57173568 57173680 refGene_gn=LOC101099964;refGene_tx=XM_023254076.1
B2 57178034 57178185 refGene_gn=LOC101099964;refGene_tx=XM_023254076.1
B2 57179426 57179553 refGene_gn=LOC101099964;refGene_tx=XM_023254076.1
B2 57189298 57189452 refGene_gn=LOC101099964;refGene_tx=XM_023254076.1
B2 57189959 57190107 refGene_gn=LOC101099964;refGene_tx=XM_023254076.1
B2 57193392 57194092 refGene_gn=LOC101099964;refGene_tx=XM_023254076.1
B2 57194093 57195003 refGene_gn=LOC101099964;refGene_tx=XM_023254076.1
B2 57199514 57199658 refGene_gn=LOC101099964;refGene_tx=XM_023254076.1

B2 57200979 57201110
refGene_gn=LOC101099964;refGene_tx=XM_023254076.1
B2 57202142 57202329
refGene_gn=LOC101099964;refGene_tx=XM_023254076.1
B2 57211507 57211576
refGene_gn=LOC101099964;refGene_tx=XM_023254076.1
B2 57219820 57219850
refGene_gn=LOC101099964;refGene_tx=XM_023254076.1
B2 57222292 57222339
refGene_gn=LOC101099964;refGene_tx=XM_023254076.1
B2 57225932 57226045 refGene_gn=BAG2;refGene_tx=XM_023254077.1
B2 57237527 57237637 refGene_gn=BAG2;refGene_tx=XM_023254077.1
B2 57239495 57239908 refGene_gn=BAG2;refGene_tx=XM_023254077.1
B2 57269558 57269713 refGene_gn=RAB23;refGene_tx=XM_023254078.1
B2 57275910 57275996 refGene_gn=RAB23;refGene_tx=XM_023254078.1
B2 57287489 57287646 refGene_gn=RAB23;refGene_tx=XM_023254078.1
B2 57289172 57289255 refGene_gn=RAB23;refGene_tx=XM_023254078.1
B2 57291450 57291543 refGene_gn=RAB23;refGene_tx=XM_023254078.1
B2 57294703 57294843 refGene_gn=RAB23;refGene_tx=XM_023254078.1
B2 57382769 57382923 refGene_gn=PRIM2;refGene_tx=XM_006931825.3
B2 57385863 57385967 refGene_gn=PRIM2;refGene_tx=XM_006931825.3
B2 57388833 57388913 refGene_gn=PRIM2;refGene_tx=XM_006931825.3
B2 57390424 57390545 refGene_gn=PRIM2;refGene_tx=XM_006931825.3
B2 57446573 57446675 refGene_gn=PRIM2;refGene_tx=XM_006931825.3
B2 57449028 57449166 refGene_gn=PRIM2;refGene_tx=XM_006931825.3
B2 57557531 57557599 refGene_gn=PRIM2;refGene_tx=XM_006931825.3
B2 57581821 57581894 refGene_gn=PRIM2;refGene_tx=XM_006931825.3
B2 57587459 57587645 refGene_gn=PRIM2;refGene_tx=XM_006931825.3
B2 57651185 57651312 refGene_gn=PRIM2;refGene_tx=XM_006931825.3
B2 57657217 57657300 refGene_gn=PRIM2;refGene_tx=XM_006931825.3
B2 57713456 57713525 refGene_gn=PRIM2;refGene_tx=XM_006931825.3
B2 57724136 57724361 refGene_gn=PRIM2;refGene_tx=XM_006931825.3
B2 57810990 57811146
refGene_gn=LOC101098952;refGene_tx=XM_011282410.2
B2 57811148 57811343
refGene_gn=LOC101098952;refGene_tx=XM_011282410.2
B2 58588827 58588925 refGene_gn=KHDRBS2;refGene_tx=XM_019830780.2
B2 58606617 58606676 refGene_gn=KHDRBS2;refGene_tx=XM_019830780.2
B2 58632278 58632361 refGene_gn=KHDRBS2;refGene_tx=XM_019830780.2
B2 58780837 58781036 refGene_gn=KHDRBS2;refGene_tx=XM_019830780.2
B2 58786712 58786840 refGene_gn=KHDRBS2;refGene_tx=XM_019830780.2
B2 58870896 58871043 refGene_gn=KHDRBS2;refGene_tx=XM_019830780.2
B2 58944515 58944632 refGene_gn=KHDRBS2;refGene_tx=XM_019830780.2
B2 59069485 59069613 refGene_gn=KHDRBS2;refGene_tx=XM_019830780.2
B2 59172321 59172412 refGene_gn=KHDRBS2;refGene_tx=XM_019830780.2
B2 59966815 59968015
refGene_gn=LGSN;refGene_tx=XM_023254081.1;refGene_gn=LGSN;refGene_tx=X
M_023254082.1
B2 59971583 59971750
refGene_gn=LGSN;refGene_tx=XM_023254081.1;refGene_gn=LGSN;refGene_tx=X

M_023254082.1
B2 59979493 59979691 refGene_gn=LGSN;refGene_tx=XM_023254081.1
B2 59980424 59980554
refGene_gn=LGSN;refGene_tx=XM_023254081.1;refGene_gn=LGSN;refGene_tx=XM_023254082.1
B2 59994885 59994915
refGene_gn=LGSN;refGene_tx=XM_023254081.1;refGene_gn=LGSN;refGene_tx=XM_023254082.1
B2 60160671 60160776
refGene_gn=PTP4A1;refGene_tx=XM_003986286.5;refGene_gn=PTP4A1;refGene_tx=XM_023254083.1;refGene_gn=PTP4A1;refGene_tx=XM_023254084.1
B2 60162059 60162152
refGene_gn=PTP4A1;refGene_tx=XM_003986286.5;refGene_gn=PTP4A1;refGene_tx=XM_023254083.1;refGene_gn=PTP4A1;refGene_tx=XM_023254084.1
B2 60162480 60162611
refGene_gn=PTP4A1;refGene_tx=XM_003986286.5;refGene_gn=PTP4A1;refGene_tx=XM_023254083.1;refGene_gn=PTP4A1;refGene_tx=XM_023254084.1
B2 60163798 60163873
refGene_gn=PTP4A1;refGene_tx=XM_003986286.5;refGene_gn=PTP4A1;refGene_tx=XM_023254083.1;refGene_gn=PTP4A1;refGene_tx=XM_023254084.1
B2 60164316 60164434
refGene_gn=PTP4A1;refGene_tx=XM_003986286.5;refGene_gn=PTP4A1;refGene_tx=XM_023254083.1;refGene_gn=PTP4A1;refGene_tx=XM_023254084.1
B2 60268874 60269016 refGene_gn=PHF3;refGene_tx=XM_011282408.3
B2 60273783 60273804 refGene_gn=PHF3;refGene_tx=XM_011282408.3
B2 60273806 60275565 refGene_gn=PHF3;refGene_tx=XM_011282408.3
B2 60281181 60281491 refGene_gn=PHF3;refGene_tx=XM_011282408.3
B2 60284447 60284631 refGene_gn=PHF3;refGene_tx=XM_011282408.3
B2 60286580 60286728 refGene_gn=PHF3;refGene_tx=XM_011282408.3
B2 60286824 60286981 refGene_gn=PHF3;refGene_tx=XM_011282408.3
B2 60289256 60289373 refGene_gn=PHF3;refGene_tx=XM_011282408.3
B2 60291333 60291465 refGene_gn=PHF3;refGene_tx=XM_011282408.3
B2 60292039 60292175 refGene_gn=PHF3;refGene_tx=XM_011282408.3
B2 60295253 60295449 refGene_gn=PHF3;refGene_tx=XM_011282408.3
B2 60295937 60296085 refGene_gn=PHF3;refGene_tx=XM_011282408.3
B2 60298054 60298144 refGene_gn=PHF3;refGene_tx=XM_011282408.3
B2 60300174 60300370 refGene_gn=PHF3;refGene_tx=XM_011282408.3
B2 60300596 60302713 refGene_gn=PHF3;refGene_tx=XM_011282408.3
B2 60310644 60311516
refGene_gn=LOC111560589;refGene_tx=XM_023254723.1
B2 60317468 60317630
refGene_gn=LOC111560589;refGene_tx=XM_023254723.1
B2 60350257 60350430
refGene_gn=LOC111560589;refGene_tx=XM_023254723.1
B2 60373080 60373255
refGene_gn=LOC111560589;refGene_tx=XM_023254723.1
B2 60375310 60375395
refGene_gn=LOC111560589;refGene_tx=XM_023254723.1
B2 60377708 60377783
refGene_gn=LOC111560589;refGene_tx=XM_023254723.1

B2 60380263 60380413
refGene_gn=LOC109499722;refGene_tx=XM_023254086.1
B2 60381243 60381410
refGene_gn=LOC109499722;refGene_tx=XM_023254086.1
B2 60395956 60396139
refGene_gn=LOC109499722;refGene_tx=XM_023254086.1
B2 60441989 60442162
refGene_gn=LOC109499722;refGene_tx=XM_023254086.1
B2 60533043 60533264
refGene_gn=LOC109499722;refGene_tx=XM_023254086.1
B2 60542670 60542779
refGene_gn=LOC109499722;refGene_tx=XM_023254086.1
B2 60606499 60606653
refGene_gn=LOC109499722;refGene_tx=XM_023254086.1
B2 60621741 60621906
refGene_gn=LOC109499722;refGene_tx=XM_023254086.1
B2 60671353 60671402
refGene_gn=LOC109499722;refGene_tx=XM_023254086.1
B2 60763305 60763600
refGene_gn=LOC111560475;refGene_tx=XM_023254087.1
B2 60831070 60831141
refGene_gn=LOC111560475;refGene_tx=XM_023254087.1
B2 60889707 60889879
refGene_gn=LOC111560590;refGene_tx=XM_023254724.1
B2 60930089 60930181
refGene_gn=LOC111560590;refGene_tx=XM_023254724.1
B2 60934373 60934573
refGene_gn=LOC111560590;refGene_tx=XM_023254724.1
B2 60943084 60943112
refGene_gn=LOC111560590;refGene_tx=XM_023254724.1
B2 61066012 61066218
refGene_gn=LOC111560591;refGene_tx=XM_023254725.1
B2 61066274 61066481
refGene_gn=LOC111560591;refGene_tx=XM_023254725.1
B2 61071020 61071213
refGene_gn=LOC111560591;refGene_tx=XM_023254725.1
B2 61090747 61090872
refGene_gn=LOC111560591;refGene_tx=XM_023254725.1
B2 61101517 61101642
refGene_gn=LOC111560591;refGene_tx=XM_023254725.1
B2 61103482 61103502
refGene_gn=LOC111560591;refGene_tx=XM_023254725.1
B2 61373335 61373528
refGene_gn=LOC111560592;refGene_tx=XM_023254727.1
B2 61388590 61388698
refGene_gn=LOC111560592;refGene_tx=XM_023254727.1
B2 61388890 61388987
refGene_gn=LOC111560592;refGene_tx=XM_023254727.1
B2 61398729 61398989
refGene_gn=LOC111560592;refGene_tx=XM_023254727.1

B2 61437749 61437871
refGene_gn=LOC111560592;refGene_tx=XM_023254727.1
B2 61499861 61499983
refGene_gn=LOC111560592;refGene_tx=XM_023254727.1
B2 61504438 61504565
refGene_gn=LOC111560592;refGene_tx=XM_023254727.1
B2 61748183 61748451
refGene_gn=LOC101082908;refGene_tx=XM_023254085.1
B2 61757391 61757558
refGene_gn=LOC101082908;refGene_tx=XM_023254085.1
B2 61769403 61769543
refGene_gn=LOC101082908;refGene_tx=XM_023254085.1
B2 61779171 61779337
refGene_gn=LOC101082908;refGene_tx=XM_023254085.1
B2 61814301 61814416
refGene_gn=LOC101082908;refGene_tx=XM_023254085.1
B2 61838160 61838291
refGene_gn=LOC101082908;refGene_tx=XM_023254085.1
B2 61839511 61839705
refGene_gn=LOC101082908;refGene_tx=XM_023254085.1
B2 61915770 61915884
refGene_gn=LOC101082908;refGene_tx=XM_023254085.1
B2 61920383 61921176
refGene_gn=LOC101082908;refGene_tx=XM_023254085.1
B2 64708054 64708811
refGene_gn=ADGRB3;refGene_tx=XM_006931831.4;refGene_gn=ADGRB3;refGene_tx=XM_011282404.3;refGene_gn=ADGRB3;refGene_tx=XM_011282405.3
B2 65001874 65001985
refGene_gn=ADGRB3;refGene_tx=XM_006931831.4;refGene_gn=ADGRB3;refGene_tx=XM_011282404.3;refGene_gn=ADGRB3;refGene_tx=XM_011282405.3
B2 65007962 65008124
refGene_gn=ADGRB3;refGene_tx=XM_006931831.4;refGene_gn=ADGRB3;refGene_tx=XM_011282404.3;refGene_gn=ADGRB3;refGene_tx=XM_011282405.3
B2 65013810 65013975
refGene_gn=ADGRB3;refGene_tx=XM_006931831.4;refGene_gn=ADGRB3;refGene_tx=XM_011282404.3;refGene_gn=ADGRB3;refGene_tx=XM_011282405.3
B2 65025288 65025453
refGene_gn=ADGRB3;refGene_tx=XM_006931831.4;refGene_gn=ADGRB3;refGene_tx=XM_011282404.3;refGene_gn=ADGRB3;refGene_tx=XM_011282405.3
B2 65025901 65026066
refGene_gn=ADGRB3;refGene_tx=XM_006931831.4;refGene_gn=ADGRB3;refGene_tx=XM_011282404.3;refGene_gn=ADGRB3;refGene_tx=XM_011282405.3
B2 65042804 65042906
refGene_gn=ADGRB3;refGene_tx=XM_006931831.4;refGene_gn=ADGRB3;refGene_tx=XM_011282404.3;refGene_gn=ADGRB3;refGene_tx=XM_011282405.3
B2 65043272 65043379
refGene_gn=ADGRB3;refGene_tx=XM_006931831.4;refGene_gn=ADGRB3;refGene_tx=XM_011282404.3;refGene_gn=ADGRB3;refGene_tx=XM_011282405.3
B2 65067750 65067945
refGene_gn=ADGRB3;refGene_tx=XM_006931831.4;refGene_gn=ADGRB3;refGene_

tx=XM_011282404.3;refGene_gn=ADGRB3;refGene_tx=XM_011282405.3
B2 65084845 65084914
refGene_gn=ADGRB3;refGene_tx=XM_006931831.4;refGene_gn=ADGRB3;refGene_tx=XM_011282404.3;refGene_gn=ADGRB3;refGene_tx=XM_011282405.3
B2 65088592 65088701
refGene_gn=ADGRB3;refGene_tx=XM_006931831.4;refGene_gn=ADGRB3;refGene_tx=XM_011282404.3;refGene_gn=ADGRB3;refGene_tx=XM_011282405.3
B2 65116613 65116763
refGene_gn=ADGRB3;refGene_tx=XM_006931831.4;refGene_gn=ADGRB3;refGene_tx=XM_011282404.3;refGene_gn=ADGRB3;refGene_tx=XM_011282405.3
B2 65117550 65117626
refGene_gn=ADGRB3;refGene_tx=XM_006931831.4;refGene_gn=ADGRB3;refGene_tx=XM_011282404.3;refGene_gn=ADGRB3;refGene_tx=XM_011282405.3
B2 65131528 65131631
refGene_gn=ADGRB3;refGene_tx=XM_006931831.4;refGene_gn=ADGRB3;refGene_tx=XM_011282404.3;refGene_gn=ADGRB3;refGene_tx=XM_011282405.3
B2 65144402 65144446
refGene_gn=ADGRB3;refGene_tx=XM_006931831.4;refGene_gn=ADGRB3;refGene_tx=XM_011282404.3;refGene_gn=ADGRB3;refGene_tx=XM_011282405.3
B2 65269121 65269248
refGene_gn=ADGRB3;refGene_tx=XM_006931831.4;refGene_gn=ADGRB3;refGene_tx=XM_011282404.3;refGene_gn=ADGRB3;refGene_tx=XM_011282405.3
B2 65270865 65270969
refGene_gn=ADGRB3;refGene_tx=XM_006931831.4;refGene_gn=ADGRB3;refGene_tx=XM_011282404.3;refGene_gn=ADGRB3;refGene_tx=XM_011282405.3
B2 65274866 65274969
refGene_gn=ADGRB3;refGene_tx=XM_006931831.4;refGene_gn=ADGRB3;refGene_tx=XM_011282404.3;refGene_gn=ADGRB3;refGene_tx=XM_011282405.3
B2 65364386 65364537
refGene_gn=ADGRB3;refGene_tx=XM_006931831.4;refGene_gn=ADGRB3;refGene_tx=XM_011282404.3;refGene_gn=ADGRB3;refGene_tx=XM_011282405.3
B2 65367116 65367186
refGene_gn=ADGRB3;refGene_tx=XM_006931831.4;refGene_gn=ADGRB3;refGene_tx=XM_011282404.3;refGene_gn=ADGRB3;refGene_tx=XM_011282405.3
B2 65369722 65369789
refGene_gn=ADGRB3;refGene_tx=XM_006931831.4;refGene_gn=ADGRB3;refGene_tx=XM_011282404.3;refGene_gn=ADGRB3;refGene_tx=XM_011282405.3
B2 65372286 65372372
refGene_gn=ADGRB3;refGene_tx=XM_006931831.4;refGene_gn=ADGRB3;refGene_tx=XM_011282404.3;refGene_gn=ADGRB3;refGene_tx=XM_011282405.3
B2 65379426 65379525
refGene_gn=ADGRB3;refGene_tx=XM_006931831.4;refGene_gn=ADGRB3;refGene_tx=XM_011282404.3;refGene_gn=ADGRB3;refGene_tx=XM_011282405.3
B2 65379847 65380019
refGene_gn=ADGRB3;refGene_tx=XM_006931831.4;refGene_gn=ADGRB3;refGene_tx=XM_011282404.3;refGene_gn=ADGRB3;refGene_tx=XM_011282405.3
B2 65390312 65390408
refGene_gn=ADGRB3;refGene_tx=XM_006931831.4;refGene_gn=ADGRB3;refGene_tx=XM_011282404.3;refGene_gn=ADGRB3;refGene_tx=XM_011282405.3
B2 65391828 65391868

refGene_gn=ADGRB3;refGene_tx=XM_006931831.4;refGene_gn=ADGRB3;refGene_tx=XM_011282404.3;refGene_gn=ADGRB3;refGene_tx=XM_011282405.3
B2 65396742 65397386 refGene_gn=ADGRB3;refGene_tx=XM_006931831.4;refGene_gn=ADGRB3;refGene_tx=XM_011282404.3;refGene_gn=ADGRB3;refGene_tx=XM_011282405.3
B2 65408413 65408449 refGene_gn=ADGRB3;refGene_tx=XM_006931831.4;refGene_gn=ADGRB3;refGene_tx=XM_011282404.3
B2 65422728 65422833 refGene_gn=ADGRB3;refGene_tx=XM_006931831.4;refGene_gn=ADGRB3;refGene_tx=XM_011282404.3;refGene_gn=ADGRB3;refGene_tx=XM_011282405.3
B2 65432891 65433080 refGene_gn=ADGRB3;refGene_tx=XM_006931831.4;refGene_gn=ADGRB3;refGene_tx=XM_011282404.3;refGene_gn=ADGRB3;refGene_tx=XM_011282405.3
B2 65684900 65685017 refGene_gn=LMBRD1;refGene_tx=XM_023254088.1
B2 65685159 65685251 refGene_gn=LMBRD1;refGene_tx=XM_023254088.1
B2 65712894 65712973 refGene_gn=LMBRD1;refGene_tx=XM_023254088.1
B2 65714748 65714898 refGene_gn=LMBRD1;refGene_tx=XM_023254088.1
B2 65716827 65716932 refGene_gn=LMBRD1;refGene_tx=XM_023254088.1
B2 65718165 65718268 refGene_gn=LMBRD1;refGene_tx=XM_023254088.1
B2 65718374 65718439 refGene_gn=LMBRD1;refGene_tx=XM_023254088.1
B2 65738048 65738201 refGene_gn=LMBRD1;refGene_tx=XM_023254088.1
B2 65743434 65743560 refGene_gn=LMBRD1;refGene_tx=XM_023254088.1
B2 65754163 65754237 refGene_gn=LMBRD1;refGene_tx=XM_023254088.1
B2 65757438 65757527 refGene_gn=LMBRD1;refGene_tx=XM_023254088.1
B2 65769708 65769776 refGene_gn=LMBRD1;refGene_tx=XM_023254088.1
B2 65771891 65771989 refGene_gn=LMBRD1;refGene_tx=XM_023254088.1
B2 65787682 65787743 refGene_gn=LMBRD1;refGene_tx=XM_023254088.1
B2 65792936 65793113 refGene_gn=LMBRD1;refGene_tx=XM_023254088.1
B2 65799301 65799379 refGene_gn=LMBRD1;refGene_tx=XM_023254088.1
B2 65871615 65871810 refGene_gn=COL19A1;refGene_tx=XM_023254089.1
B2 65885198 65885325 refGene_gn=COL19A1;refGene_tx=XM_023254089.1
B2 65898134 65898209 refGene_gn=COL19A1;refGene_tx=XM_023254089.1
B2 65899294 65899394 refGene_gn=COL19A1;refGene_tx=XM_023254089.1
B2 65922056 65922180 refGene_gn=COL19A1;refGene_tx=XM_023254089.1
B2 65923495 65923771 refGene_gn=COL19A1;refGene_tx=XM_023254089.1
B2 65934196 65934277 refGene_gn=COL19A1;refGene_tx=XM_023254089.1
B2 65939756 65939882 refGene_gn=COL19A1;refGene_tx=XM_023254089.1
B2 65940976 65941039 refGene_gn=COL19A1;refGene_tx=XM_023254089.1
B2 65949425 65949479 refGene_gn=COL19A1;refGene_tx=XM_023254089.1
B2 65966724 65966769 refGene_gn=COL19A1;refGene_tx=XM_023254089.1
B2 65969135 65969180 refGene_gn=COL19A1;refGene_tx=XM_023254089.1
B2 66032523 66032577 refGene_gn=COL19A1;refGene_tx=XM_023254089.1
B2 66041694 66041748 refGene_gn=COL19A1;refGene_tx=XM_023254089.1
B2 66043387 66043423 refGene_gn=COL19A1;refGene_tx=XM_023254089.1
B2 66070910 66070964 refGene_gn=COL19A1;refGene_tx=XM_023254089.1
B2 66092032 66092049 refGene_gn=COL19A1;refGene_tx=XM_023254089.1
B2 66111299 66111342 refGene_gn=COL19A1;refGene_tx=XM_023254089.1
B2 66132779 66132830 refGene_gn=COL19A1;refGene_tx=XM_023254089.1
B2 66139835 66139880 refGene_gn=COL19A1;refGene_tx=XM_023254089.1

| | | | |
|---|---|---|---|
| B2 | 66145099 | 66145162 | refGene_gn=COL19A1;refGene_tx=XM_023254089.1 |
| B2 | 66150652 | 66150688 | refGene_gn=COL19A1;refGene_tx=XM_023254089.1 |
| B2 | 66151608 | 66151644 | refGene_gn=COL19A1;refGene_tx=XM_023254089.1 |
| B2 | 66151738 | 66151792 | refGene_gn=COL19A1;refGene_tx=XM_023254089.1 |
| B2 | 66152468 | 66152522 | refGene_gn=COL19A1;refGene_tx=XM_023254089.1 |
| B2 | 66153973 | 66154027 | refGene_gn=COL19A1;refGene_tx=XM_023254089.1 |
| B2 | 66154517 | 66154607 | refGene_gn=COL19A1;refGene_tx=XM_023254089.1 |
| B2 | 66155573 | 66155618 | refGene_gn=COL19A1;refGene_tx=XM_023254089.1 |
| B2 | 66155722 | 66155800 | refGene_gn=COL19A1;refGene_tx=XM_023254089.1 |
| B2 | 66157890 | 66157926 | refGene_gn=COL19A1;refGene_tx=XM_023254089.1 |
| B2 | 66158032 | 66158086 | refGene_gn=COL19A1;refGene_tx=XM_023254089.1 |
| B2 | 66158177 | 66158231 | refGene_gn=COL19A1;refGene_tx=XM_023254089.1 |
| B2 | 66159598 | 66159640 | refGene_gn=COL19A1;refGene_tx=XM_023254089.1 |
| B2 | 66163049 | 66163154 | refGene_gn=COL19A1;refGene_tx=XM_023254089.1 |
| B2 | 66163237 | 66163291 | refGene_gn=COL19A1;refGene_tx=XM_023254089.1 |
| B2 | 66163754 | 66163808 | refGene_gn=COL19A1;refGene_tx=XM_023254089.1 |
| B2 | 66169945 | 66169999 | refGene_gn=COL19A1;refGene_tx=XM_023254089.1 |
| B2 | 66171457 | 66171511 | refGene_gn=COL19A1;refGene_tx=XM_023254089.1 |
| B2 | 66173702 | 66173747 | refGene_gn=COL19A1;refGene_tx=XM_023254089.1 |
| B2 | 66176310 | 66176361 | refGene_gn=COL19A1;refGene_tx=XM_023254089.1 |
| B2 | 66176446 | 66176491 | refGene_gn=COL19A1;refGene_tx=XM_023254089.1 |
| B2 | 66176924 | 66176951 | refGene_gn=COL19A1;refGene_tx=XM_023254089.1 |
| B2 | 66180149 | 66180203 | refGene_gn=COL19A1;refGene_tx=XM_023254089.1 |
| B2 | 66184525 | 66184570 | refGene_gn=COL19A1;refGene_tx=XM_023254089.1 |
| B2 | 66187774 | 66187819 | refGene_gn=COL19A1;refGene_tx=XM_023254089.1 |
| B2 | 66187906 | 66187969 | refGene_gn=COL19A1;refGene_tx=XM_023254089.1 |
| B2 | 66192228 | 66192264 | refGene_gn=COL19A1;refGene_tx=XM_023254089.1 |
| B2 | 66192400 | 66192445 | refGene_gn=COL19A1;refGene_tx=XM_023254089.1 |
| B2 | 66195242 | 66195413 | refGene_gn=COL19A1;refGene_tx=XM_023254089.1 |
| B2 | 66197412 | 66197479 | refGene_gn=COL19A1;refGene_tx=XM_023254089.1 |
| B2 | 66206365 | 66206494 | refGene_gn=COL19A1;refGene_tx=XM_023254089.1 |
| B2 | 66213349 | 66213427 | refGene_gn=COL19A1;refGene_tx=XM_023254089.1 |
| B2 | 66213604 | 66213732 | refGene_gn=COL19A1;refGene_tx=XM_023254089.1 |
| B2 | 66223204 | 66223389 | refGene_gn=COL9A1;refGene_tx=XM_003986294.4 |
| B2 | 66231795 | 66231873 | refGene_gn=COL9A1;refGene_tx=XM_003986294.4 |
| B2 | 66238582 | 66238771 | refGene_gn=COL9A1;refGene_tx=XM_003986294.4 |
| B2 | 66240370 | 66240425 | refGene_gn=COL9A1;refGene_tx=XM_003986294.4 |
| B2 | 66240648 | 66240795 | refGene_gn=COL9A1;refGene_tx=XM_003986294.4 |
| B2 | 66244756 | 66244789 | refGene_gn=COL9A1;refGene_tx=XM_003986294.4 |
| B2 | 66246059 | 66246104 | refGene_gn=COL9A1;refGene_tx=XM_003986294.4 |
| B2 | 66247030 | 66247066 | refGene_gn=COL9A1;refGene_tx=XM_003986294.4 |
| B2 | 66247571 | 66247643 | refGene_gn=COL9A1;refGene_tx=XM_003986294.4 |
| B2 | 66248267 | 66248321 | refGene_gn=COL9A1;refGene_tx=XM_003986294.4 |
| B2 | 66257038 | 66257092 | refGene_gn=COL9A1;refGene_tx=XM_003986294.4 |
| B2 | 66257197 | 66257251 | refGene_gn=COL9A1;refGene_tx=XM_003986294.4 |
| B2 | 66258248 | 66258293 | refGene_gn=COL9A1;refGene_tx=XM_003986294.4 |
| B2 | 66259288 | 66259342 | refGene_gn=COL9A1;refGene_tx=XM_003986294.4 |
| B2 | 66259744 | 66259798 | refGene_gn=COL9A1;refGene_tx=XM_003986294.4 |
| B2 | 66259933 | 66259987 | refGene_gn=COL9A1;refGene_tx=XM_003986294.4 |
| B2 | 66260123 | 66260177 | refGene_gn=COL9A1;refGene_tx=XM_003986294.4 |

B2 66261785 66261839 refGene_gn=COL9A1;refGene_tx=XM_003986294.4
B2 66265615 66265669 refGene_gn=COL9A1;refGene_tx=XM_003986294.4
B2 66268199 66268253 refGene_gn=COL9A1;refGene_tx=XM_003986294.4
B2 66271825 66271879 refGene_gn=COL9A1;refGene_tx=XM_003986294.4
B2 66273305 66273362 refGene_gn=COL9A1;refGene_tx=XM_003986294.4
B2 66274085 66274118 refGene_gn=COL9A1;refGene_tx=XM_003986294.4
B2 66274771 66274825 refGene_gn=COL9A1;refGene_tx=XM_003986294.4
B2 66276111 66276165 refGene_gn=COL9A1;refGene_tx=XM_003986294.4
B2 66277049 66277073 refGene_gn=COL9A1;refGene_tx=XM_003986294.4
B2 66279391 66279427 refGene_gn=COL9A1;refGene_tx=XM_003986294.4
B2 66279687 66279741 refGene_gn=COL9A1;refGene_tx=XM_003986294.4
B2 66285912 66285975 refGene_gn=COL9A1;refGene_tx=XM_003986294.4
B2 66286097 66286133 refGene_gn=COL9A1;refGene_tx=XM_003986294.4
B2 66286469 66286544 refGene_gn=COL9A1;refGene_tx=XM_003986294.4
B2 66288029 66288050 refGene_gn=COL9A1;refGene_tx=XM_003986294.4
B2 66288821 66288905 refGene_gn=COL9A1;refGene_tx=XM_003986294.4
B2 66301897 66302294 refGene_gn=COL9A1;refGene_tx=XM_003986294.4
B2 66308329 66308462 refGene_gn=COL9A1;refGene_tx=XM_003986294.4
B2 66308601 66308679 refGene_gn=COL9A1;refGene_tx=XM_003986294.4
B2 66309883 66309957 refGene_gn=COL9A1;refGene_tx=XM_003986294.4
B2 66310496 66310510 refGene_gn=COL9A1;refGene_tx=XM_003986294.4
B2 66416857 66416934
refGene_gn=FAM135A;refGene_tx=XM_011282396.3;refGene_gn=FAM135A;refGene_tx=XM_011282397.3;refGene_gn=FAM135A;refGene_tx=XM_011282398.3;refGene_gn=FAM135A;refGene_tx=XM_011282399.3;refGene_gn=FAM135A;refGene_tx=XM_019830808.2;refGene_gn=FAM135A;refGene_tx=XM_019830809.2;refGene_gn=FAM135A;refGene_tx=XM_019830810.2;refGene_gn=FAM135A;refGene_tx=XM_019830811.2;refGene_gn=FAM135A;refGene_tx=XM_019830815.2;refGene_gn=FAM135A;refGene_tx=XM_019830816.2
B2 66419546 66419626
refGene_gn=FAM135A;refGene_tx=XM_011282396.3;refGene_gn=FAM135A;refGene_tx=XM_011282397.3;refGene_gn=FAM135A;refGene_tx=XM_011282398.3;refGene_gn=FAM135A;refGene_tx=XM_011282399.3;refGene_gn=FAM135A;refGene_tx=XM_019830808.2;refGene_gn=FAM135A;refGene_tx=XM_019830809.2;refGene_gn=FAM135A;refGene_tx=XM_019830810.2;refGene_gn=FAM135A;refGene_tx=XM_019830811.2;refGene_gn=FAM135A;refGene_tx=XM_019830815.2;refGene_gn=FAM135A;refGene_tx=XM_019830816.2
B2 66436114 66436254
refGene_gn=FAM135A;refGene_tx=XM_011282396.3;refGene_gn=FAM135A;refGene_tx=XM_011282397.3;refGene_gn=FAM135A;refGene_tx=XM_011282398.3;refGene_gn=FAM135A;refGene_tx=XM_011282399.3;refGene_gn=FAM135A;refGene_tx=XM_019830808.2;refGene_gn=FAM135A;refGene_tx=XM_019830809.2;refGene_gn=FAM135A;refGene_tx=XM_019830810.2;refGene_gn=FAM135A;refGene_tx=XM_019830811.2;refGene_gn=FAM135A;refGene_tx=XM_019830815.2;refGene_gn=FAM135A;refGene_tx=XM_019830816.2
B2 66436381 66436452
refGene_gn=FAM135A;refGene_tx=XM_011282396.3;refGene_gn=FAM135A;refGene_tx=XM_011282397.3;refGene_gn=FAM135A;refGene_tx=XM_011282398.3;refGene_gn=FAM135A;refGene_tx=XM_011282399.3;refGene_gn=FAM135A;refGene_tx=XM_019830808.2;refGene_gn=FAM135A;refGene_tx=XM_019830809.2;refGene_gn

=FAM135A;refGene_tx=XM_019830810.2;refGene_gn=FAM135A;refGene_tx=XM_01
9830811.2;refGene_gn=FAM135A;refGene_tx=XM_019830815.2;refGene_gn=FAM1
35A;refGene_tx=XM_019830816.2
B2 66438135 66438309
refGene_gn=FAM135A;refGene_tx=XM_011282396.3;refGene_gn=FAM135A;refGen
e_tx=XM_011282397.3;refGene_gn=FAM135A;refGene_tx=XM_011282398.3;refGe
ne_gn=FAM135A;refGene_tx=XM_011282399.3;refGene_gn=FAM135A;refGene_tx=
XM_019830808.2;refGene_gn=FAM135A;refGene_tx=XM_019830809.2;refGene_gn
=FAM135A;refGene_tx=XM_019830810.2;refGene_gn=FAM135A;refGene_tx=XM_01
9830811.2;refGene_gn=FAM135A;refGene_tx=XM_019830815.2;refGene_gn=FAM1
35A;refGene_tx=XM_019830816.2
B2 66442244 66442371
refGene_gn=FAM135A;refGene_tx=XM_011282396.3;refGene_gn=FAM135A;refGen
e_tx=XM_011282397.3;refGene_gn=FAM135A;refGene_tx=XM_011282398.3;refGe
ne_gn=FAM135A;refGene_tx=XM_011282399.3;refGene_gn=FAM135A;refGene_tx=
XM_019830808.2;refGene_gn=FAM135A;refGene_tx=XM_019830809.2;refGene_gn
=FAM135A;refGene_tx=XM_019830810.2;refGene_gn=FAM135A;refGene_tx=XM_01
9830811.2;refGene_gn=FAM135A;refGene_tx=XM_019830815.2;refGene_gn=FAM1
35A;refGene_tx=XM_019830816.2
B2 66443051 66443205
refGene_gn=FAM135A;refGene_tx=XM_011282396.3;refGene_gn=FAM135A;refGen
e_tx=XM_011282397.3;refGene_gn=FAM135A;refGene_tx=XM_011282398.3;refGe
ne_gn=FAM135A;refGene_tx=XM_011282399.3;refGene_gn=FAM135A;refGene_tx=
XM_019830808.2;refGene_gn=FAM135A;refGene_tx=XM_019830809.2;refGene_gn
=FAM135A;refGene_tx=XM_019830810.2;refGene_gn=FAM135A;refGene_tx=XM_01
9830811.2;refGene_gn=FAM135A;refGene_tx=XM_019830815.2;refGene_gn=FAM1
35A;refGene_tx=XM_019830816.2
B2 66446895 66446973
refGene_gn=FAM135A;refGene_tx=XM_011282396.3;refGene_gn=FAM135A;refGen
e_tx=XM_011282398.3;refGene_gn=FAM135A;refGene_tx=XM_019830808.2;refGe
ne_gn=FAM135A;refGene_tx=XM_019830809.2;refGene_gn=FAM135A;refGene_tx=
XM_019830811.2;refGene_gn=FAM135A;refGene_tx=XM_019830812.1;refGene_gn
=FAM135A;refGene_tx=XM_019830815.2
B2 66463531 66463581
refGene_gn=FAM135A;refGene_tx=XM_011282396.3;refGene_gn=FAM135A;refGen
e_tx=XM_011282397.3;refGene_gn=FAM135A;refGene_tx=XM_011282398.3;refGe
ne_gn=FAM135A;refGene_tx=XM_011282399.3;refGene_gn=FAM135A;refGene_tx=
XM_019830808.2;refGene_gn=FAM135A;refGene_tx=XM_019830809.2;refGene_gn
=FAM135A;refGene_tx=XM_019830810.2;refGene_gn=FAM135A;refGene_tx=XM_01
9830811.2;refGene_gn=FAM135A;refGene_tx=XM_019830812.1;refGene_gn=FAM1
35A;refGene_tx=XM_019830813.2;refGene_gn=FAM135A;refGene_tx=XM_0198308
15.2;refGene_gn=FAM135A;refGene_tx=XM_019830816.2
B2 66468740 66468896
refGene_gn=FAM135A;refGene_tx=XM_011282396.3;refGene_gn=FAM135A;refGen
e_tx=XM_011282397.3;refGene_gn=FAM135A;refGene_tx=XM_011282398.3;refGe
ne_gn=FAM135A;refGene_tx=XM_011282399.3;refGene_gn=FAM135A;refGene_tx=
XM_019830808.2;refGene_gn=FAM135A;refGene_tx=XM_019830811.2;refGene_gn
=FAM135A;refGene_tx=XM_019830812.1;refGene_gn=FAM135A;refGene_tx=XM_01
9830813.2;refGene_gn=FAM135A;refGene_tx=XM_019830814.2;refGene_gn=FAM1
35A;refGene_tx=XM_019830815.2;refGene_gn=FAM135A;refGene_tx=XM_0198308

16.2
B2 66495228 66495302
refGene_gn=FAM135A;refGene_tx=XM_011282396.3;refGene_gn=FAM135A;refGen
e_tx=XM_011282397.3;refGene_gn=FAM135A;refGene_tx=XM_011282398.3;refGe
ne_gn=FAM135A;refGene_tx=XM_011282399.3;refGene_gn=FAM135A;refGene_tx=
XM_019830808.2;refGene_gn=FAM135A;refGene_tx=XM_019830809.2;refGene_gn
=FAM135A;refGene_tx=XM_019830810.2;refGene_gn=FAM135A;refGene_tx=XM_01
9830811.2;refGene_gn=FAM135A;refGene_tx=XM_019830812.1;refGene_gn=FAM1
35A;refGene_tx=XM_019830813.2;refGene_gn=FAM135A;refGene_tx=XM_0198308
14.2;refGene_gn=FAM135A;refGene_tx=XM_019830815.2;refGene_gn=FAM135A;r
efGene_tx=XM_019830816.2
B2 66496642 66496797
refGene_gn=FAM135A;refGene_tx=XM_011282396.3;refGene_gn=FAM135A;refGen
e_tx=XM_011282397.3;refGene_gn=FAM135A;refGene_tx=XM_011282398.3;refGe
ne_gn=FAM135A;refGene_tx=XM_011282399.3;refGene_gn=FAM135A;refGene_tx=
XM_019830808.2;refGene_gn=FAM135A;refGene_tx=XM_019830809.2;refGene_gn
=FAM135A;refGene_tx=XM_019830810.2;refGene_gn=FAM135A;refGene_tx=XM_01
9830811.2;refGene_gn=FAM135A;refGene_tx=XM_019830812.1;refGene_gn=FAM1
35A;refGene_tx=XM_019830813.2;refGene_gn=FAM135A;refGene_tx=XM_0198308
14.2;refGene_gn=FAM135A;refGene_tx=XM_019830815.2;refGene_gn=FAM135A;r
efGene_tx=XM_019830816.2
B2 66497014 66499373
refGene_gn=FAM135A;refGene_tx=XM_011282396.3;refGene_gn=FAM135A;refGen
e_tx=XM_011282397.3;refGene_gn=FAM135A;refGene_tx=XM_011282398.3;refGe
ne_gn=FAM135A;refGene_tx=XM_011282399.3;refGene_gn=FAM135A;refGene_tx=
XM_019830808.2;refGene_gn=FAM135A;refGene_tx=XM_019830809.2;refGene_gn
=FAM135A;refGene_tx=XM_019830810.2;refGene_gn=FAM135A;refGene_tx=XM_01
9830811.2;refGene_gn=FAM135A;refGene_tx=XM_019830812.1;refGene_gn=FAM1
35A;refGene_tx=XM_019830813.2;refGene_gn=FAM135A;refGene_tx=XM_0198308
14.2;refGene_gn=FAM135A;refGene_tx=XM_019830815.2;refGene_gn=FAM135A;r
efGene_tx=XM_019830816.2
B2 66500828 66500989
refGene_gn=FAM135A;refGene_tx=XM_011282396.3;refGene_gn=FAM135A;refGen
e_tx=XM_011282397.3;refGene_gn=FAM135A;refGene_tx=XM_011282398.3;refGe
ne_gn=FAM135A;refGene_tx=XM_011282399.3;refGene_gn=FAM135A;refGene_tx=
XM_019830808.2;refGene_gn=FAM135A;refGene_tx=XM_019830809.2;refGene_gn
=FAM135A;refGene_tx=XM_019830810.2;refGene_gn=FAM135A;refGene_tx=XM_01
9830811.2;refGene_gn=FAM135A;refGene_tx=XM_019830812.1;refGene_gn=FAM1
35A;refGene_tx=XM_019830813.2;refGene_gn=FAM135A;refGene_tx=XM_0198308
14.2;refGene_gn=FAM135A;refGene_tx=XM_019830815.2;refGene_gn=FAM135A;r
efGene_tx=XM_019830816.2
B2 66503057 66503149
refGene_gn=FAM135A;refGene_tx=XM_011282396.3;refGene_gn=FAM135A;refGen
e_tx=XM_011282397.3;refGene_gn=FAM135A;refGene_tx=XM_011282398.3;refGe
ne_gn=FAM135A;refGene_tx=XM_011282399.3;refGene_gn=FAM135A;refGene_tx=
XM_019830808.2;refGene_gn=FAM135A;refGene_tx=XM_019830809.2;refGene_gn
=FAM135A;refGene_tx=XM_019830810.2;refGene_gn=FAM135A;refGene_tx=XM_01
9830811.2;refGene_gn=FAM135A;refGene_tx=XM_019830812.1;refGene_gn=FAM1
35A;refGene_tx=XM_019830813.2;refGene_gn=FAM135A;refGene_tx=XM_0198308
14.2;refGene_gn=FAM135A;refGene_tx=XM_019830815.2;refGene_gn=FAM135A;r

efGene_tx=XM_019830816.2
B2 66503656 66503754
refGene_gn=FAM135A;refGene_tx=XM_011282396.3;refGene_gn=FAM135A;refGene_tx=XM_011282397.3;refGene_gn=FAM135A;refGene_tx=XM_011282398.3;refGene_gn=FAM135A;refGene_tx=XM_011282399.3;refGene_gn=FAM135A;refGene_tx=XM_019830808.2;refGene_gn=FAM135A;refGene_tx=XM_019830809.2;refGene_gn=FAM135A;refGene_tx=XM_019830810.2;refGene_gn=FAM135A;refGene_tx=XM_019830811.2;refGene_gn=FAM135A;refGene_tx=XM_019830812.1;refGene_gn=FAM135A;refGene_tx=XM_019830813.2;refGene_gn=FAM135A;refGene_tx=XM_019830814.2;refGene_gn=FAM135A;refGene_tx=XM_019830815.2;refGene_gn=FAM135A;refGene_tx=XM_019830816.2
B2 66505408 66505560
refGene_gn=FAM135A;refGene_tx=XM_011282396.3;refGene_gn=FAM135A;refGene_tx=XM_011282397.3;refGene_gn=FAM135A;refGene_tx=XM_011282398.3;refGene_gn=FAM135A;refGene_tx=XM_011282399.3;refGene_gn=FAM135A;refGene_tx=XM_019830808.2;refGene_gn=FAM135A;refGene_tx=XM_019830809.2;refGene_gn=FAM135A;refGene_tx=XM_019830810.2;refGene_gn=FAM135A;refGene_tx=XM_019830811.2;refGene_gn=FAM135A;refGene_tx=XM_019830812.1;refGene_gn=FAM135A;refGene_tx=XM_019830813.2;refGene_gn=FAM135A;refGene_tx=XM_019830814.2;refGene_gn=FAM135A;refGene_tx=XM_019830815.2;refGene_gn=FAM135A;refGene_tx=XM_019830816.2
B2 66507362 66507473
refGene_gn=FAM135A;refGene_tx=XM_011282396.3;refGene_gn=FAM135A;refGene_tx=XM_011282397.3;refGene_gn=FAM135A;refGene_tx=XM_011282398.3;refGene_gn=FAM135A;refGene_tx=XM_011282399.3;refGene_gn=FAM135A;refGene_tx=XM_019830808.2;refGene_gn=FAM135A;refGene_tx=XM_019830809.2;refGene_gn=FAM135A;refGene_tx=XM_019830810.2;refGene_gn=FAM135A;refGene_tx=XM_019830811.2;refGene_gn=FAM135A;refGene_tx=XM_019830812.1;refGene_gn=FAM135A;refGene_tx=XM_019830813.2;refGene_gn=FAM135A;refGene_tx=XM_019830814.2;refGene_gn=FAM135A;refGene_tx=XM_019830815.2;refGene_gn=FAM135A;refGene_tx=XM_019830816.2
B2 66516313 66516427
refGene_gn=FAM135A;refGene_tx=XM_011282396.3;refGene_gn=FAM135A;refGene_tx=XM_011282397.3;refGene_gn=FAM135A;refGene_tx=XM_011282398.3;refGene_gn=FAM135A;refGene_tx=XM_011282399.3;refGene_gn=FAM135A;refGene_tx=XM_019830808.2;refGene_gn=FAM135A;refGene_tx=XM_019830809.2;refGene_gn=FAM135A;refGene_tx=XM_019830810.2;refGene_gn=FAM135A;refGene_tx=XM_019830811.2;refGene_gn=FAM135A;refGene_tx=XM_019830812.1;refGene_gn=FAM135A;refGene_tx=XM_019830813.2;refGene_gn=FAM135A;refGene_tx=XM_019830814.2;refGene_gn=FAM135A;refGene_tx=XM_019830815.2;refGene_gn=FAM135A;refGene_tx=XM_019830816.2
B2 66519058 66519264
refGene_gn=FAM135A;refGene_tx=XM_011282396.3;refGene_gn=FAM135A;refGene_tx=XM_011282397.3;refGene_gn=FAM135A;refGene_tx=XM_011282398.3;refGene_gn=FAM135A;refGene_tx=XM_011282399.3;refGene_gn=FAM135A;refGene_tx=XM_019830808.2;refGene_gn=FAM135A;refGene_tx=XM_019830809.2;refGene_gn=FAM135A;refGene_tx=XM_019830810.2;refGene_gn=FAM135A;refGene_tx=XM_019830811.2;refGene_gn=FAM135A;refGene_tx=XM_019830812.1;refGene_gn=FAM135A;refGene_tx=XM_019830813.2;refGene_gn=FAM135A;refGene_tx=XM_019830814.2;refGene_gn=FAM135A;refGene_tx=XM_019830815.2;refGene_gn=FAM135A;r

efGene_tx=XM_019830816.2
B2 66526301 66526365 refGene_gn=SDHAF4;refGene_tx=XM_006931837.3
B2 66540901 66541051 refGene_gn=SDHAF4;refGene_tx=XM_006931837.3
B2 66559132 66559242 refGene_gn=SDHAF4;refGene_tx=XM_006931837.3
B2 66592779 66592888
refGene_gn=LOC109499732;refGene_tx=XM_019830817.1
B2 66595929 66596033
refGene_gn=LOC109499732;refGene_tx=XM_019830817.1
B2 66598395 66598508
refGene_gn=LOC109499732;refGene_tx=XM_019830817.1
B2 66598696 66599156
refGene_gn=LOC109499732;refGene_tx=XM_019830817.1
B2 66618827 66618945
refGene_gn=SMAP1;refGene_tx=XM_011282395.3;refGene_gn=SMAP1;refGene_tx=XM_019830818.2;refGene_gn=SMAP1;refGene_tx=XM_019830819.2;refGene_gn=SMAP1;refGene_tx=XM_019830821.2
B2 66666037 66666171
refGene_gn=SMAP1;refGene_tx=XM_011282395.3;refGene_gn=SMAP1;refGene_tx=XM_019830818.2;refGene_gn=SMAP1;refGene_tx=XM_019830819.2;refGene_gn=SMAP1;refGene_tx=XM_019830821.2
B2 66682316 66682388 refGene_gn=SMAP1;refGene_tx=XM_019830820.2
B2 66690323 66690409
refGene_gn=SMAP1;refGene_tx=XM_011282395.3;refGene_gn=SMAP1;refGene_tx=XM_019830818.2;refGene_gn=SMAP1;refGene_tx=XM_019830819.2;refGene_gn=SMAP1;refGene_tx=XM_019830820.2;refGene_gn=SMAP1;refGene_tx=XM_019830821.2
B2 66698440 66698516
refGene_gn=SMAP1;refGene_tx=XM_011282395.3;refGene_gn=SMAP1;refGene_tx=XM_019830818.2;refGene_gn=SMAP1;refGene_tx=XM_019830819.2;refGene_gn=SMAP1;refGene_tx=XM_019830820.2;refGene_gn=SMAP1;refGene_tx=XM_019830821.2
B2 66710458 66710539
refGene_gn=SMAP1;refGene_tx=XM_011282395.3;refGene_gn=SMAP1;refGene_tx=XM_019830820.2;refGene_gn=SMAP1;refGene_tx=XM_019830821.2
B2 66737046 66737127
refGene_gn=SMAP1;refGene_tx=XM_011282395.3;refGene_gn=SMAP1;refGene_tx=XM_019830818.2;refGene_gn=SMAP1;refGene_tx=XM_019830820.2;refGene_gn=SMAP1;refGene_tx=XM_019830821.2
B2 66790637 66790728
refGene_gn=SMAP1;refGene_tx=XM_011282395.3;refGene_gn=SMAP1;refGene_tx=XM_019830818.2;refGene_gn=SMAP1;refGene_tx=XM_019830819.2;refGene_gn=SMAP1;refGene_tx=XM_019830820.2;refGene_gn=SMAP1;refGene_tx=XM_019830821.2
B2 66813886 66814014
refGene_gn=SMAP1;refGene_tx=XM_011282395.3;refGene_gn=SMAP1;refGene_tx=XM_019830818.2;refGene_gn=SMAP1;refGene_tx=XM_019830819.2;refGene_gn=SMAP1;refGene_tx=XM_019830820.2;refGene_gn=SMAP1;refGene_tx=XM_019830821.2
B2 66818748 66818920
refGene_gn=SMAP1;refGene_tx=XM_011282395.3;refGene_gn=SMAP1;refGene_tx

=XM_019830818.2;refGene_gn=SMAP1;refGene_tx=XM_019830819.2;refGene_gn=SMAP1;refGene_tx=XM_019830820.2;refGene_gn=SMAP1;refGene_tx=XM_019830821.2
B2 66819754 66820068
refGene_gn=SMAP1;refGene_tx=XM_011282395.3;refGene_gn=SMAP1;refGene_tx=XM_019830818.2;refGene_gn=SMAP1;refGene_tx=XM_019830819.2;refGene_gn=SMAP1;refGene_tx=XM_019830820.2
B2 66822146 66822319
refGene_gn=SMAP1;refGene_tx=XM_011282395.3;refGene_gn=SMAP1;refGene_tx=XM_019830818.2;refGene_gn=SMAP1;refGene_tx=XM_019830819.2;refGene_gn=SMAP1;refGene_tx=XM_019830820.2;refGene_gn=SMAP1;refGene_tx=XM_019830821.2
B2 66823587 66823674 refGene_gn=B3GAT2;refGene_tx=XM_003986299.5
B2 66823759 66823908 refGene_gn=B3GAT2;refGene_tx=XM_003986299.5
B2 66849408 66849553 refGene_gn=B3GAT2;refGene_tx=XM_003986299.5
B2 66914506 66915115 refGene_gn=B3GAT2;refGene_tx=XM_003986299.5
B2 67200231 67200471
refGene_gn=OGFRL1;refGene_tx=XM_023254091.1;refGene_gn=OGFRL1;refGene_tx=XM_023254092.1
B2 67204561 67204648
refGene_gn=OGFRL1;refGene_tx=XM_023254091.1;refGene_gn=OGFRL1;refGene_tx=XM_023254092.1
B2 67204798 67204880
refGene_gn=OGFRL1;refGene_tx=XM_023254091.1;refGene_gn=OGFRL1;refGene_tx=XM_023254092.1
B2 67208170 67208249
refGene_gn=OGFRL1;refGene_tx=XM_023254091.1;refGene_gn=OGFRL1;refGene_tx=XM_023254092.1
B2 67208345 67208412
refGene_gn=OGFRL1;refGene_tx=XM_023254091.1;refGene_gn=OGFRL1;refGene_tx=XM_023254092.1
B2 67208518 67208664
refGene_gn=OGFRL1;refGene_tx=XM_023254091.1;refGene_gn=OGFRL1;refGene_tx=XM_023254092.1
B2 67212756 67213417
refGene_gn=OGFRL1;refGene_tx=XM_023254091.1;refGene_gn=OGFRL1;refGene_tx=XM_023254092.1
B2 67731994 67732158
refGene_gn=RIMS1;refGene_tx=XM_019830824.2;refGene_gn=RIMS1;refGene_tx=XM_019830825.2;refGene_gn=RIMS1;refGene_tx=XM_019830833.2;refGene_gn=RIMS1;refGene_tx=XM_019830841.2;refGene_gn=RIMS1;refGene_tx=XM_019830845.2;refGene_gn=RIMS1;refGene_tx=XM_019830850.2;refGene_gn=RIMS1;refGene_tx=XM_019830852.2;refGene_gn=RIMS1;refGene_tx=XM_019830859.2;refGene_gn=RIMS1;refGene_tx=XM_023254093.1;refGene_gn=RIMS1;refGene_tx=XM_023254094.1;refGene_gn=RIMS1;refGene_tx=XM_023254095.1;refGene_gn=RIMS1;refGene_tx=XM_023254096.1;refGene_gn=RIMS1;refGene_tx=XM_023254097.1;refGene_gn=RIMS1;refGene_tx=XM_023254098.1;refGene_gn=RIMS1;refGene_tx=XM_023254099.1;refGene_gn=RIMS1;refGene_tx=XM_023254100.1;refGene_gn=RIMS1;refGene_tx=XM_023254101.1;refGene_gn=RIMS1;refGene_tx=XM_023254102.1;refGene_gn=RIMS1;refGene_tx=XM_023254103.1;refGene_gn=RIMS1;refGen

e_tx=XM_023254104.1;refGene_gn=RIMS1;refGene_tx=XM_023254105.1;refGene_gn=RIMS1;refGene_tx=XM_023254106.1;refGene_gn=RIMS1;refGene_tx=XM_023254107.1
B2 67811381 67811462
refGene_gn=RIMS1;refGene_tx=XM_019830824.2;refGene_gn=RIMS1;refGene_tx=XM_019830825.2;refGene_gn=RIMS1;refGene_tx=XM_019830833.2;refGene_gn=RIMS1;refGene_tx=XM_019830841.2;refGene_gn=RIMS1;refGene_tx=XM_019830845.2;refGene_gn=RIMS1;refGene_tx=XM_019830850.2;refGene_gn=RIMS1;refGene_tx=XM_019830852.2;refGene_gn=RIMS1;refGene_tx=XM_019830859.2;refGene_gn=RIMS1;refGene_tx=XM_023254093.1;refGene_gn=RIMS1;refGene_tx=XM_023254094.1;refGene_gn=RIMS1;refGene_tx=XM_023254095.1;refGene_gn=RIMS1;refGene_tx=XM_023254096.1;refGene_gn=RIMS1;refGene_tx=XM_023254098.1;refGene_gn=RIMS1;refGene_tx=XM_023254099.1;refGene_gn=RIMS1;refGene_tx=XM_023254100.1;refGene_gn=RIMS1;refGene_tx=XM_023254101.1;refGene_gn=RIMS1;refGene_tx=XM_023254102.1;refGene_gn=RIMS1;refGene_tx=XM_023254103.1;refGene_gn=RIMS1;refGene_tx=XM_023254104.1;refGene_gn=RIMS1;refGene_tx=XM_023254105.1;refGene_gn=RIMS1;refGene_tx=XM_023254106.1;refGene_gn=RIMS1;refGene_tx=XM_023254107.1
B2 67932426 67932640
refGene_gn=RIMS1;refGene_tx=XM_019830824.2;refGene_gn=RIMS1;refGene_tx=XM_019830825.2;refGene_gn=RIMS1;refGene_tx=XM_019830833.2;refGene_gn=RIMS1;refGene_tx=XM_019830841.2;refGene_gn=RIMS1;refGene_tx=XM_019830845.2;refGene_gn=RIMS1;refGene_tx=XM_019830850.2;refGene_gn=RIMS1;refGene_tx=XM_019830852.2;refGene_gn=RIMS1;refGene_tx=XM_019830859.2;refGene_gn=RIMS1;refGene_tx=XM_023254093.1;refGene_gn=RIMS1;refGene_tx=XM_023254094.1;refGene_gn=RIMS1;refGene_tx=XM_023254095.1;refGene_gn=RIMS1;refGene_tx=XM_023254096.1;refGene_gn=RIMS1;refGene_tx=XM_023254097.1;refGene_gn=RIMS1;refGene_tx=XM_023254098.1;refGene_gn=RIMS1;refGene_tx=XM_023254099.1;refGene_gn=RIMS1;refGene_tx=XM_023254100.1;refGene_gn=RIMS1;refGene_tx=XM_023254101.1;refGene_gn=RIMS1;refGene_tx=XM_023254102.1;refGene_gn=RIMS1;refGene_tx=XM_023254103.1;refGene_gn=RIMS1;refGene_tx=XM_023254104.1;refGene_gn=RIMS1;refGene_tx=XM_023254105.1;refGene_gn=RIMS1;refGene_tx=XM_023254106.1;refGene_gn=RIMS1;refGene_tx=XM_023254107.1
B2 67935145 67935157
refGene_gn=RIMS1;refGene_tx=XM_019830824.2;refGene_gn=RIMS1;refGene_tx=XM_019830825.2;refGene_gn=RIMS1;refGene_tx=XM_019830833.2;refGene_gn=RIMS1;refGene_tx=XM_019830841.2;refGene_gn=RIMS1;refGene_tx=XM_019830845.2;refGene_gn=RIMS1;refGene_tx=XM_019830850.2;refGene_gn=RIMS1;refGene_tx=XM_019830852.2;refGene_gn=RIMS1;refGene_tx=XM_019830859.2;refGene_gn=RIMS1;refGene_tx=XM_023254093.1;refGene_gn=RIMS1;refGene_tx=XM_023254094.1;refGene_gn=RIMS1;refGene_tx=XM_023254095.1;refGene_gn=RIMS1;refGene_tx=XM_023254096.1;refGene_gn=RIMS1;refGene_tx=XM_023254097.1;refGene_gn=RIMS1;refGene_tx=XM_023254098.1;refGene_gn=RIMS1;refGene_tx=XM_023254099.1;refGene_gn=RIMS1;refGene_tx=XM_023254100.1;refGene_gn=RIMS1;refGene_tx=XM_023254101.1;refGene_gn=RIMS1;refGene_tx=XM_023254102.1;refGene_gn=RIMS1;refGene_tx=XM_023254103.1;refGene_gn=RIMS1;refGene_tx=XM_023254104.1;refGene_gn=RIMS1;refGene_tx=XM_023254105.1;refGene_gn=RIMS1;refGene_tx=XM_023254106.1;refGene_gn=RIMS1;refGene_tx=XM_023254107.1

B2 67995468 67995809
refGene_gn=RIMS1;refGene_tx=XM_019830824.2;refGene_gn=RIMS1;refGene_tx=XM_019830825.2;refGene_gn=RIMS1;refGene_tx=XM_019830833.2;refGene_gn=RIMS1;refGene_tx=XM_019830841.2;refGene_gn=RIMS1;refGene_tx=XM_019830845.2;refGene_gn=RIMS1;refGene_tx=XM_019830850.2;refGene_gn=RIMS1;refGene_tx=XM_019830852.2;refGene_gn=RIMS1;refGene_tx=XM_019830859.2;refGene_gn=RIMS1;refGene_tx=XM_023254093.1;refGene_gn=RIMS1;refGene_tx=XM_023254094.1;refGene_gn=RIMS1;refGene_tx=XM_023254095.1;refGene_gn=RIMS1;refGene_tx=XM_023254096.1;refGene_gn=RIMS1;refGene_tx=XM_023254097.1;refGene_gn=RIMS1;refGene_tx=XM_023254098.1;refGene_gn=RIMS1;refGene_tx=XM_023254099.1;refGene_gn=RIMS1;refGene_tx=XM_023254100.1;refGene_gn=RIMS1;refGene_tx=XM_023254101.1;refGene_gn=RIMS1;refGene_tx=XM_023254102.1;refGene_gn=RIMS1;refGene_tx=XM_023254103.1;refGene_gn=RIMS1;refGene_tx=XM_023254104.1;refGene_gn=RIMS1;refGene_tx=XM_023254105.1;refGene_gn=RIMS1;refGene_tx=XM_023254106.1;refGene_gn=RIMS1;refGene_tx=XM_023254107.1
B2 67999239 68000105
refGene_gn=RIMS1;refGene_tx=XM_019830824.2;refGene_gn=RIMS1;refGene_tx=XM_019830825.2;refGene_gn=RIMS1;refGene_tx=XM_019830833.2;refGene_gn=RIMS1;refGene_tx=XM_019830841.2;refGene_gn=RIMS1;refGene_tx=XM_019830845.2;refGene_gn=RIMS1;refGene_tx=XM_019830850.2;refGene_gn=RIMS1;refGene_tx=XM_019830852.2;refGene_gn=RIMS1;refGene_tx=XM_019830859.2;refGene_gn=RIMS1;refGene_tx=XM_023254093.1;refGene_gn=RIMS1;refGene_tx=XM_023254094.1;refGene_gn=RIMS1;refGene_tx=XM_023254095.1;refGene_gn=RIMS1;refGene_tx=XM_023254096.1;refGene_gn=RIMS1;refGene_tx=XM_023254097.1;refGene_gn=RIMS1;refGene_tx=XM_023254098.1;refGene_gn=RIMS1;refGene_tx=XM_023254099.1;refGene_gn=RIMS1;refGene_tx=XM_023254100.1;refGene_gn=RIMS1;refGene_tx=XM_023254101.1;refGene_gn=RIMS1;refGene_tx=XM_023254102.1;refGene_gn=RIMS1;refGene_tx=XM_023254103.1;refGene_gn=RIMS1;refGene_tx=XM_023254104.1;refGene_gn=RIMS1;refGene_tx=XM_023254105.1;refGene_gn=RIMS1;refGene_tx=XM_023254106.1;refGene_gn=RIMS1;refGene_tx=XM_023254107.1
B2 68025962 68026062
refGene_gn=RIMS1;refGene_tx=XM_019830848.1;refGene_gn=RIMS1;refGene_tx=XM_019830858.1;refGene_gn=RIMS1;refGene_tx=XM_023254108.1;refGene_gn=RIMS1;refGene_tx=XM_023254109.1;refGene_gn=RIMS1;refGene_tx=XM_023254110.1;refGene_gn=RIMS1;refGene_tx=XM_023254112.1;refGene_gn=RIMS1;refGene_tx=XM_023254113.1;refGene_gn=RIMS1;refGene_tx=XM_023254117.1;refGene_gn=RIMS1;refGene_tx=XM_023254118.1;refGene_gn=RIMS1;refGene_tx=XM_023254120.1;refGene_gn=RIMS1;refGene_tx=XM_023254121.1;refGene_gn=RIMS1;refGene_tx=XM_023254122.1;refGene_gn=RIMS1;refGene_tx=XM_023254123.1;refGene_gn=RIMS1;refGene_tx=XM_023254126.1;refGene_gn=RIMS1;refGene_tx=XM_023254132.1;refGene_gn=RIMS1;refGene_tx=XM_023254133.1
B2 68029324 68029379
refGene_gn=RIMS1;refGene_tx=XM_019830857.1;refGene_gn=RIMS1;refGene_tx=XM_023254119.1;refGene_gn=RIMS1;refGene_tx=XM_023254127.1;refGene_gn=RIMS1;refGene_tx=XM_023254128.1
B2 68038637 68038668
refGene_gn=RIMS1;refGene_tx=XM_023254111.1;refGene_gn=RIMS1;refGene_tx=XM_023254114.1;refGene_gn=RIMS1;refGene_tx=XM_023254115.1;refGene_gn=

RIMS1;refGene_tx=XM_023254124.1;refGene_gn=RIMS1;refGene_tx=XM_0232541
25.1;refGene_gn=RIMS1;refGene_tx=XM_023254129.1;refGene_gn=RIMS1;refGe
ne_tx=XM_023254131.1
B2 68046548 68046616
refGene_gn=RIMS1;refGene_tx=XM_019830824.2;refGene_gn=RIMS1;refGene_tx
=XM_019830825.2;refGene_gn=RIMS1;refGene_tx=XM_019830833.2;refGene_gn=
RIMS1;refGene_tx=XM_019830841.2;refGene_gn=RIMS1;refGene_tx=XM_0198308
45.2;refGene_gn=RIMS1;refGene_tx=XM_019830848.1;refGene_gn=RIMS1;refGe
ne_tx=XM_019830850.2;refGene_gn=RIMS1;refGene_tx=XM_019830852.2;refGen
e_gn=RIMS1;refGene_tx=XM_019830857.1;refGene_gn=RIMS1;refGene_tx=XM_01
9830858.1;refGene_gn=RIMS1;refGene_tx=XM_019830859.2;refGene_gn=RIMS1;
refGene_tx=XM_023254093.1;refGene_gn=RIMS1;refGene_tx=XM_023254094.1;r
efGene_gn=RIMS1;refGene_tx=XM_023254095.1;refGene_gn=RIMS1;refGene_tx=
XM_023254096.1;refGene_gn=RIMS1;refGene_tx=XM_023254097.1;refGene_gn=R
IMS1;refGene_tx=XM_023254098.1;refGene_gn=RIMS1;refGene_tx=XM_02325409
9.1;refGene_gn=RIMS1;refGene_tx=XM_023254100.1;refGene_gn=RIMS1;refGen
e_tx=XM_023254101.1;refGene_gn=RIMS1;refGene_tx=XM_023254102.1;refGene
_gn=RIMS1;refGene_tx=XM_023254103.1;refGene_gn=RIMS1;refGene_tx=XM_023
254104.1;refGene_gn=RIMS1;refGene_tx=XM_023254105.1;refGene_gn=RIMS1;r
efGene_tx=XM_023254106.1;refGene_gn=RIMS1;refGene_tx=XM_023254107.1;re
fGene_gn=RIMS1;refGene_tx=XM_023254108.1;refGene_gn=RIMS1;refGene_tx=X
M_023254109.1;refGene_gn=RIMS1;refGene_tx=XM_023254110.1;refGene_gn=RI
MS1;refGene_tx=XM_023254111.1;refGene_gn=RIMS1;refGene_tx=XM_023254112
.
1;refGene_gn=RIMS1;refGene_tx=XM_023254113.1;refGene_gn=RIMS1;refGene_
tx=XM_023254114.1;refGene_gn=RIMS1;refGene_tx=XM_023254115.1;refGene_g
n=RIMS1;refGene_tx=XM_023254117.1;refGene_gn=RIMS1;refGene_tx=XM_02325
4118.1;refGene_gn=RIMS1;refGene_tx=XM_023254119.1;refGene_gn=RIMS1;ref
Gene_tx=XM_023254120.1;refGene_gn=RIMS1;refGene_tx=XM_023254121.1;refG
ene_gn=RIMS1;refGene_tx=XM_023254122.1;refGene_gn=RIMS1;refGene_tx=XM_
023254123.1;refGene_gn=RIMS1;refGene_tx=XM_023254124.1;refGene_gn=RIMS
1;refGene_tx=XM_023254125.1;refGene_gn=RIMS1;refGene_tx=XM_023254126.1
;refGene_gn=RIMS1;refGene_tx=XM_023254127.1;refGene_gn=RIMS1;refGene_t
x=XM_023254128.1;refGene_gn=RIMS1;refGene_tx=XM_023254129.1;refGene_gn
=RIMS1;refGene_tx=XM_023254131.1;refGene_gn=RIMS1;refGene_tx=XM_023254
132.1;refGene_gn=RIMS1;refGene_tx=XM_023254133.1
B2 68047668 68047779
refGene_gn=RIMS1;refGene_tx=XM_019830824.2;refGene_gn=RIMS1;refGene_tx
=XM_019830825.2;refGene_gn=RIMS1;refGene_tx=XM_019830833.2;refGene_gn=
RIMS1;refGene_tx=XM_019830841.2;refGene_gn=RIMS1;refGene_tx=XM_0198308
45.2;refGene_gn=RIMS1;refGene_tx=XM_019830848.1;refGene_gn=RIMS1;refGe
ne_tx=XM_019830850.2;refGene_gn=RIMS1;refGene_tx=XM_019830852.2;refGen
e_gn=RIMS1;refGene_tx=XM_019830857.1;refGene_gn=RIMS1;refGene_tx=XM_01
9830858.1;refGene_gn=RIMS1;refGene_tx=XM_019830859.2;refGene_gn=RIMS1;
refGene_tx=XM_023254093.1;refGene_gn=RIMS1;refGene_tx=XM_023254094.1;r
efGene_gn=RIMS1;refGene_tx=XM_023254095.1;refGene_gn=RIMS1;refGene_tx=
XM_023254096.1;refGene_gn=RIMS1;refGene_tx=XM_023254097.1;refGene_gn=R
IMS1;refGene_tx=XM_023254098.1;refGene_gn=RIMS1;refGene_tx=XM_02325409
9.1;refGene_gn=RIMS1;refGene_tx=XM_023254100.1;refGene_gn=RIMS1;refGen
e_tx=XM_023254101.1;refGene_gn=RIMS1;refGene_tx=XM_023254102.1;refGene

_gn=RIMS1;refGene_tx=XM_023254103.1;refGene_gn=RIMS1;refGene_tx=XM_023254104.1;refGene_gn=RIMS1;refGene_tx=XM_023254105.1;refGene_gn=RIMS1;refGene_tx=XM_023254106.1;refGene_gn=RIMS1;refGene_tx=XM_023254107.1;refGene_gn=RIMS1;refGene_tx=XM_023254108.1;refGene_gn=RIMS1;refGene_tx=XM_023254109.1;refGene_gn=RIMS1;refGene_tx=XM_023254110.1;refGene_gn=RIMS1;refGene_tx=XM_023254111.1;refGene_gn=RIMS1;refGene_tx=XM_023254112.
1;refGene_gn=RIMS1;refGene_tx=XM_023254113.1;refGene_gn=RIMS1;refGene_tx=XM_023254114.1;refGene_gn=RIMS1;refGene_tx=XM_023254115.1;refGene_gn=RIMS1;refGene_tx=XM_023254117.1;refGene_gn=RIMS1;refGene_tx=XM_023254118.1;refGene_gn=RIMS1;refGene_tx=XM_023254119.1;refGene_gn=RIMS1;refGene_tx=XM_023254120.1;refGene_gn=RIMS1;refGene_tx=XM_023254121.1;refGene_gn=RIMS1;refGene_tx=XM_023254122.1;refGene_gn=RIMS1;refGene_tx=XM_023254123.1;refGene_gn=RIMS1;refGene_tx=XM_023254124.1;refGene_gn=RIMS1;refGene_tx=XM_023254125.1;refGene_gn=RIMS1;refGene_tx=XM_023254126.1;refGene_gn=RIMS1;refGene_tx=XM_023254127.1;refGene_gn=RIMS1;refGene_tx=XM_023254128.1;refGene_gn=RIMS1;refGene_tx=XM_023254129.1;refGene_gn=RIMS1;refGene_tx=XM_023254131.1;refGene_gn=RIMS1;refGene_tx=XM_023254132.1;refGene_gn=RIMS1;refGene_tx=XM_023254133.1

B2 68049385 68049485

refGene_gn=RIMS1;refGene_tx=XM_019830824.2;refGene_gn=RIMS1;refGene_tx=XM_019830825.2;refGene_gn=RIMS1;refGene_tx=XM_019830833.2;refGene_gn=RIMS1;refGene_tx=XM_019830841.2;refGene_gn=RIMS1;refGene_tx=XM_019830845.2;refGene_gn=RIMS1;refGene_tx=XM_019830848.1;refGene_gn=RIMS1;refGene_tx=XM_019830850.2;refGene_gn=RIMS1;refGene_tx=XM_019830852.2;refGene_gn=RIMS1;refGene_tx=XM_019830857.1;refGene_gn=RIMS1;refGene_tx=XM_019830858.1;refGene_gn=RIMS1;refGene_tx=XM_019830859.2;refGene_gn=RIMS1;refGene_tx=XM_023254093.1;refGene_gn=RIMS1;refGene_tx=XM_023254094.1;refGene_gn=RIMS1;refGene_tx=XM_023254095.1;refGene_gn=RIMS1;refGene_tx=XM_023254096.1;refGene_gn=RIMS1;refGene_tx=XM_023254097.1;refGene_gn=RIMS1;refGene_tx=XM_023254098.1;refGene_gn=RIMS1;refGene_tx=XM_023254099.1;refGene_gn=RIMS1;refGene_tx=XM_023254100.1;refGene_gn=RIMS1;refGene_tx=XM_023254101.1;refGene_gn=RIMS1;refGene_tx=XM_023254102.1;refGene_gn=RIMS1;refGene_tx=XM_023254103.1;refGene_gn=RIMS1;refGene_tx=XM_023254104.1;refGene_gn=RIMS1;refGene_tx=XM_023254105.1;refGene_gn=RIMS1;refGene_tx=XM_023254106.1;refGene_gn=RIMS1;refGene_tx=XM_023254107.1;refGene_gn=RIMS1;refGene_tx=XM_023254108.1;refGene_gn=RIMS1;refGene_tx=XM_023254109.1;refGene_gn=RIMS1;refGene_tx=XM_023254110.1;refGene_gn=RIMS1;refGene_tx=XM_023254111.1;refGene_gn=RIMS1;refGene_tx=XM_023254112.
1;refGene_gn=RIMS1;refGene_tx=XM_023254113.1;refGene_gn=RIMS1;refGene_tx=XM_023254114.1;refGene_gn=RIMS1;refGene_tx=XM_023254115.1;refGene_gn=RIMS1;refGene_tx=XM_023254117.1;refGene_gn=RIMS1;refGene_tx=XM_023254118.1;refGene_gn=RIMS1;refGene_tx=XM_023254119.1;refGene_gn=RIMS1;refGene_tx=XM_023254120.1;refGene_gn=RIMS1;refGene_tx=XM_023254121.1;refGene_gn=RIMS1;refGene_tx=XM_023254122.1;refGene_gn=RIMS1;refGene_tx=XM_023254123.1;refGene_gn=RIMS1;refGene_tx=XM_023254124.1;refGene_gn=RIMS1;refGene_tx=XM_023254125.1;refGene_gn=RIMS1;refGene_tx=XM_023254126.1;refGene_gn=RIMS1;refGene_tx=XM_023254127.1;refGene_gn=RIMS1;refGene_tx=XM_023254128.1;refGene_gn=RIMS1;refGene_tx=XM_023254129.1;refGene_gn

=RIMS1;refGene_tx=XM_023254131.1;refGene_gn=RIMS1;refGene_tx=XM_023254
132.1;refGene_gn=RIMS1;refGene_tx=XM_023254133.1
B2 68053601 68053725
refGene_gn=RIMS1;refGene_tx=XM_019830824.2;refGene_gn=RIMS1;refGene_tx
=XM_019830825.2;refGene_gn=RIMS1;refGene_tx=XM_019830833.2;refGene_gn=
RIMS1;refGene_tx=XM_019830841.2;refGene_gn=RIMS1;refGene_tx=XM_0198308
45.2;refGene_gn=RIMS1;refGene_tx=XM_019830848.1;refGene_gn=RIMS1;refGe
ne_tx=XM_019830850.2;refGene_gn=RIMS1;refGene_tx=XM_019830852.2;refGen
e_gn=RIMS1;refGene_tx=XM_019830857.1;refGene_gn=RIMS1;refGene_tx=XM_01
9830858.1;refGene_gn=RIMS1;refGene_tx=XM_019830859.2;refGene_gn=RIMS1;
refGene_tx=XM_023254093.1;refGene_gn=RIMS1;refGene_tx=XM_023254094.1;r
efGene_gn=RIMS1;refGene_tx=XM_023254095.1;refGene_gn=RIMS1;refGene_tx=
XM_023254096.1;refGene_gn=RIMS1;refGene_tx=XM_023254097.1;refGene_gn=R
IMS1;refGene_tx=XM_023254098.1;refGene_gn=RIMS1;refGene_tx=XM_02325409
9.1;refGene_gn=RIMS1;refGene_tx=XM_023254100.1;refGene_gn=RIMS1;refGen
e_tx=XM_023254101.1;refGene_gn=RIMS1;refGene_tx=XM_023254102.1;refGene
_gn=RIMS1;refGene_tx=XM_023254103.1;refGene_gn=RIMS1;refGene_tx=XM_023
254104.1;refGene_gn=RIMS1;refGene_tx=XM_023254105.1;refGene_gn=RIMS1;r
efGene_tx=XM_023254106.1;refGene_gn=RIMS1;refGene_tx=XM_023254107.1;re
fGene_gn=RIMS1;refGene_tx=XM_023254108.1;refGene_gn=RIMS1;refGene_tx=X
M_023254109.1;refGene_gn=RIMS1;refGene_tx=XM_023254110.1;refGene_gn=RI
MS1;refGene_tx=XM_023254111.1;refGene_gn=RIMS1;refGene_tx=XM_023254112
.
1;refGene_gn=RIMS1;refGene_tx=XM_023254113.1;refGene_gn=RIMS1;refGene_
tx=XM_023254114.1;refGene_gn=RIMS1;refGene_tx=XM_023254115.1;refGene_g
n=RIMS1;refGene_tx=XM_023254117.1;refGene_gn=RIMS1;refGene_tx=XM_02325
4118.1;refGene_gn=RIMS1;refGene_tx=XM_023254119.1;refGene_gn=RIMS1;ref
Gene_tx=XM_023254120.1;refGene_gn=RIMS1;refGene_tx=XM_023254121.1;refG
ene_gn=RIMS1;refGene_tx=XM_023254122.1;refGene_gn=RIMS1;refGene_tx=XM_
023254123.1;refGene_gn=RIMS1;refGene_tx=XM_023254124.1;refGene_gn=RIMS
1;refGene_tx=XM_023254125.1;refGene_gn=RIMS1;refGene_tx=XM_023254126.1
;refGene_gn=RIMS1;refGene_tx=XM_023254127.1;refGene_gn=RIMS1;refGene_t
x=XM_023254128.1;refGene_gn=RIMS1;refGene_tx=XM_023254129.1;refGene_gn
=RIMS1;refGene_tx=XM_023254131.1;refGene_gn=RIMS1;refGene_tx=XM_023254
132.1;refGene_gn=RIMS1;refGene_tx=XM_023254133.1
B2 68057032 68057079
refGene_gn=RIMS1;refGene_tx=XM_019830824.2;refGene_gn=RIMS1;refGene_tx
=XM_019830825.2;refGene_gn=RIMS1;refGene_tx=XM_019830833.2;refGene_gn=
RIMS1;refGene_tx=XM_019830841.2;refGene_gn=RIMS1;refGene_tx=XM_0198308
45.2;refGene_gn=RIMS1;refGene_tx=XM_019830848.1;refGene_gn=RIMS1;refGe
ne_tx=XM_019830850.2;refGene_gn=RIMS1;refGene_tx=XM_019830852.2;refGen
e_gn=RIMS1;refGene_tx=XM_019830857.1;refGene_gn=RIMS1;refGene_tx=XM_01
9830858.1;refGene_gn=RIMS1;refGene_tx=XM_019830859.2;refGene_gn=RIMS1;
refGene_tx=XM_023254093.1;refGene_gn=RIMS1;refGene_tx=XM_023254094.1;r
efGene_gn=RIMS1;refGene_tx=XM_023254095.1;refGene_gn=RIMS1;refGene_tx=
XM_023254096.1;refGene_gn=RIMS1;refGene_tx=XM_023254097.1;refGene_gn=R
IMS1;refGene_tx=XM_023254098.1;refGene_gn=RIMS1;refGene_tx=XM_02325409
9.1;refGene_gn=RIMS1;refGene_tx=XM_023254100.1;refGene_gn=RIMS1;refGen
e_tx=XM_023254101.1;refGene_gn=RIMS1;refGene_tx=XM_023254102.1;refGene
_gn=RIMS1;refGene_tx=XM_023254103.1;refGene_gn=RIMS1;refGene_tx=XM_023

254104.1;refGene_gn=RIMS1;refGene_tx=XM_023254105.1;refGene_gn=RIMS1;refGene_tx=XM_023254106.1;refGene_gn=RIMS1;refGene_tx=XM_023254107.1;refGene_gn=RIMS1;refGene_tx=XM_023254108.1;refGene_gn=RIMS1;refGene_tx=XM_023254109.1;refGene_gn=RIMS1;refGene_tx=XM_023254110.1;refGene_gn=RIMS1;refGene_tx=XM_023254111.1;refGene_gn=RIMS1;refGene_tx=XM_023254112
.
1;refGene_gn=RIMS1;refGene_tx=XM_023254113.1;refGene_gn=RIMS1;refGene_tx=XM_023254114.1;refGene_gn=RIMS1;refGene_tx=XM_023254115.1;refGene_gn=RIMS1;refGene_tx=XM_023254117.1;refGene_gn=RIMS1;refGene_tx=XM_023254118.1;refGene_gn=RIMS1;refGene_tx=XM_023254119.1;refGene_gn=RIMS1;refGene_tx=XM_023254120.1;refGene_gn=RIMS1;refGene_tx=XM_023254121.1;refGene_gn=RIMS1;refGene_tx=XM_023254122.1;refGene_gn=RIMS1;refGene_tx=XM_023254123.1;refGene_gn=RIMS1;refGene_tx=XM_023254124.1;refGene_gn=RIMS1;refGene_tx=XM_023254125.1;refGene_gn=RIMS1;refGene_tx=XM_023254126.1;refGene_gn=RIMS1;refGene_tx=XM_023254127.1;refGene_gn=RIMS1;refGene_tx=XM_023254128.1;refGene_gn=RIMS1;refGene_tx=XM_023254129.1;refGene_gn=RIMS1;refGene_tx=XM_023254131.1;refGene_gn=RIMS1;refGene_tx=XM_023254132.1;refGene_gn=RIMS1;refGene_tx=XM_023254133.1

B2 68059086 68059199
refGene_gn=RIMS1;refGene_tx=XM_019830824.2;refGene_gn=RIMS1;refGene_tx=XM_019830825.2;refGene_gn=RIMS1;refGene_tx=XM_019830833.2;refGene_gn=RIMS1;refGene_tx=XM_019830841.2;refGene_gn=RIMS1;refGene_tx=XM_019830845.2;refGene_gn=RIMS1;refGene_tx=XM_019830848.1;refGene_gn=RIMS1;refGene_tx=XM_019830850.2;refGene_gn=RIMS1;refGene_tx=XM_019830852.2;refGene_gn=RIMS1;refGene_tx=XM_019830857.1;refGene_gn=RIMS1;refGene_tx=XM_019830858.1;refGene_gn=RIMS1;refGene_tx=XM_019830859.2;refGene_gn=RIMS1;refGene_tx=XM_023254093.1;refGene_gn=RIMS1;refGene_tx=XM_023254094.1;refGene_gn=RIMS1;refGene_tx=XM_023254095.1;refGene_gn=RIMS1;refGene_tx=XM_023254096.1;refGene_gn=RIMS1;refGene_tx=XM_023254097.1;refGene_gn=RIMS1;refGene_tx=XM_023254098.1;refGene_gn=RIMS1;refGene_tx=XM_023254099.1;refGene_gn=RIMS1;refGene_tx=XM_023254100.1;refGene_gn=RIMS1;refGene_tx=XM_023254101.1;refGene_gn=RIMS1;refGene_tx=XM_023254102.1;refGene_gn=RIMS1;refGene_tx=XM_023254103.1;refGene_gn=RIMS1;refGene_tx=XM_023254104.1;refGene_gn=RIMS1;refGene_tx=XM_023254105.1;refGene_gn=RIMS1;refGene_tx=XM_023254106.1;refGene_gn=RIMS1;refGene_tx=XM_023254107.1;refGene_gn=RIMS1;refGene_tx=XM_023254108.1;refGene_gn=RIMS1;refGene_tx=XM_023254109.1;refGene_gn=RIMS1;refGene_tx=XM_023254110.1;refGene_gn=RIMS1;refGene_tx=XM_023254111.1;refGene_gn=RIMS1;refGene_tx=XM_023254112
.
1;refGene_gn=RIMS1;refGene_tx=XM_023254113.1;refGene_gn=RIMS1;refGene_tx=XM_023254114.1;refGene_gn=RIMS1;refGene_tx=XM_023254115.1;refGene_gn=RIMS1;refGene_tx=XM_023254117.1;refGene_gn=RIMS1;refGene_tx=XM_023254118.1;refGene_gn=RIMS1;refGene_tx=XM_023254119.1;refGene_gn=RIMS1;refGene_tx=XM_023254120.1;refGene_gn=RIMS1;refGene_tx=XM_023254121.1;refGene_gn=RIMS1;refGene_tx=XM_023254122.1;refGene_gn=RIMS1;refGene_tx=XM_023254123.1;refGene_gn=RIMS1;refGene_tx=XM_023254124.1;refGene_gn=RIMS1;refGene_tx=XM_023254125.1;refGene_gn=RIMS1;refGene_tx=XM_023254126.1;refGene_gn=RIMS1;refGene_tx=XM_023254127.1;refGene_gn=RIMS1;refGene_tx=XM_023254128.1;refGene_gn=RIMS1;refGene_tx=XM_023254129.1;refGene_gn=RIMS1;refGene_tx=XM_023254131.1;refGene_gn=RIMS1;refGene_tx=XM_023254

132.1;refGene_gn=RIMS1;refGene_tx=XM_023254133.1
B2 68061077 68061208
refGene_gn=RIMS1;refGene_tx=XM_019830824.2;refGene_gn=RIMS1;refGene_tx=XM_019830825.2;refGene_gn=RIMS1;refGene_tx=XM_019830833.2;refGene_gn=RIMS1;refGene_tx=XM_019830841.2;refGene_gn=RIMS1;refGene_tx=XM_019830845.2;refGene_gn=RIMS1;refGene_tx=XM_019830848.1;refGene_gn=RIMS1;refGene_tx=XM_019830850.2;refGene_gn=RIMS1;refGene_tx=XM_019830852.2;refGene_gn=RIMS1;refGene_tx=XM_019830857.1;refGene_gn=RIMS1;refGene_tx=XM_019830858.1;refGene_gn=RIMS1;refGene_tx=XM_019830859.2;refGene_gn=RIMS1;refGene_tx=XM_023254093.1;refGene_gn=RIMS1;refGene_tx=XM_023254094.1;refGene_gn=RIMS1;refGene_tx=XM_023254095.1;refGene_gn=RIMS1;refGene_tx=XM_023254096.1;refGene_gn=RIMS1;refGene_tx=XM_023254097.1;refGene_gn=RIMS1;refGene_tx=XM_023254098.1;refGene_gn=RIMS1;refGene_tx=XM_023254099.1;refGene_gn=RIMS1;refGene_tx=XM_023254100.1;refGene_gn=RIMS1;refGene_tx=XM_023254101.1;refGene_gn=RIMS1;refGene_tx=XM_023254102.1;refGene_gn=RIMS1;refGene_tx=XM_023254103.1;refGene_gn=RIMS1;refGene_tx=XM_023254104.1;refGene_gn=RIMS1;refGene_tx=XM_023254105.1;refGene_gn=RIMS1;refGene_tx=XM_023254106.1;refGene_gn=RIMS1;refGene_tx=XM_023254107.1;refGene_gn=RIMS1;refGene_tx=XM_023254108.1;refGene_gn=RIMS1;refGene_tx=XM_023254109.1;refGene_gn=RIMS1;refGene_tx=XM_023254110.1;refGene_gn=RIMS1;refGene_tx=XM_023254111.1;refGene_gn=RIMS1;refGene_tx=XM_023254112.
1;refGene_gn=RIMS1;refGene_tx=XM_023254113.1;refGene_gn=RIMS1;refGene_tx=XM_023254114.1;refGene_gn=RIMS1;refGene_tx=XM_023254115.1;refGene_gn=RIMS1;refGene_tx=XM_023254117.1;refGene_gn=RIMS1;refGene_tx=XM_023254118.1;refGene_gn=RIMS1;refGene_tx=XM_023254119.1;refGene_gn=RIMS1;refGene_tx=XM_023254120.1;refGene_gn=RIMS1;refGene_tx=XM_023254121.1;refGene_gn=RIMS1;refGene_tx=XM_023254122.1;refGene_gn=RIMS1;refGene_tx=XM_023254123.1;refGene_gn=RIMS1;refGene_tx=XM_023254124.1;refGene_gn=RIMS1;refGene_tx=XM_023254125.1;refGene_gn=RIMS1;refGene_tx=XM_023254126.1;refGene_gn=RIMS1;refGene_tx=XM_023254127.1;refGene_gn=RIMS1;refGene_tx=XM_023254128.1;refGene_gn=RIMS1;refGene_tx=XM_023254129.1;refGene_gn=RIMS1;refGene_tx=XM_023254131.1;refGene_gn=RIMS1;refGene_tx=XM_023254132.1;refGene_gn=RIMS1;refGene_tx=XM_023254133.1
B2 68061763 68061935
refGene_gn=RIMS1;refGene_tx=XM_019830824.2;refGene_gn=RIMS1;refGene_tx=XM_019830825.2;refGene_gn=RIMS1;refGene_tx=XM_019830833.2;refGene_gn=RIMS1;refGene_tx=XM_019830841.2;refGene_gn=RIMS1;refGene_tx=XM_019830845.2;refGene_gn=RIMS1;refGene_tx=XM_019830848.1;refGene_gn=RIMS1;refGene_tx=XM_019830850.2;refGene_gn=RIMS1;refGene_tx=XM_019830852.2;refGene_gn=RIMS1;refGene_tx=XM_019830857.1;refGene_gn=RIMS1;refGene_tx=XM_019830858.1;refGene_gn=RIMS1;refGene_tx=XM_019830859.2;refGene_gn=RIMS1;refGene_tx=XM_023254093.1;refGene_gn=RIMS1;refGene_tx=XM_023254094.1;refGene_gn=RIMS1;refGene_tx=XM_023254095.1;refGene_gn=RIMS1;refGene_tx=XM_023254096.1;refGene_gn=RIMS1;refGene_tx=XM_023254097.1;refGene_gn=RIMS1;refGene_tx=XM_023254098.1;refGene_gn=RIMS1;refGene_tx=XM_023254099.1;refGene_gn=RIMS1;refGene_tx=XM_023254100.1;refGene_gn=RIMS1;refGene_tx=XM_023254101.1;refGene_gn=RIMS1;refGene_tx=XM_023254102.1;refGene_gn=RIMS1;refGene_tx=XM_023254103.1;refGene_gn=RIMS1;refGene_tx=XM_023254104.1;refGene_gn=RIMS1;refGene_tx=XM_023254105.1;refGene_gn=RIMS1;r

efGene_tx=XM_023254106.1;refGene_gn=RIMS1;refGene_tx=XM_023254107.1;re
fGene_gn=RIMS1;refGene_tx=XM_023254108.1;refGene_gn=RIMS1;refGene_tx=X
M_023254109.1;refGene_gn=RIMS1;refGene_tx=XM_023254110.1;refGene_gn=RI
MS1;refGene_tx=XM_023254111.1;refGene_gn=RIMS1;refGene_tx=XM_023254112
.
1;refGene_gn=RIMS1;refGene_tx=XM_023254113.1;refGene_gn=RIMS1;refGene_
tx=XM_023254114.1;refGene_gn=RIMS1;refGene_tx=XM_023254115.1;refGene_g
n=RIMS1;refGene_tx=XM_023254117.1;refGene_gn=RIMS1;refGene_tx=XM_02325
4118.1;refGene_gn=RIMS1;refGene_tx=XM_023254119.1;refGene_gn=RIMS1;ref
Gene_tx=XM_023254120.1;refGene_gn=RIMS1;refGene_tx=XM_023254121.1;refG
ene_gn=RIMS1;refGene_tx=XM_023254122.1;refGene_gn=RIMS1;refGene_tx=XM_
023254123.1;refGene_gn=RIMS1;refGene_tx=XM_023254124.1;refGene_gn=RIMS
1;refGene_tx=XM_023254125.1;refGene_gn=RIMS1;refGene_tx=XM_023254126.1
;refGene_gn=RIMS1;refGene_tx=XM_023254127.1;refGene_gn=RIMS1;refGene_t
x=XM_023254128.1;refGene_gn=RIMS1;refGene_tx=XM_023254129.1;refGene_gn
=RIMS1;refGene_tx=XM_023254131.1;refGene_gn=RIMS1;refGene_tx=XM_023254
132.1;refGene_gn=RIMS1;refGene_tx=XM_023254133.1
B2 68062061 68062215
refGene_gn=RIMS1;refGene_tx=XM_019830824.2;refGene_gn=RIMS1;refGene_tx
=XM_019830825.2;refGene_gn=RIMS1;refGene_tx=XM_019830833.2;refGene_gn=
RIMS1;refGene_tx=XM_019830841.2;refGene_gn=RIMS1;refGene_tx=XM_0198308
45.2;refGene_gn=RIMS1;refGene_tx=XM_019830848.1;refGene_gn=RIMS1;refGe
ne_tx=XM_019830850.2;refGene_gn=RIMS1;refGene_tx=XM_019830852.2;refGen
e_gn=RIMS1;refGene_tx=XM_019830857.1;refGene_gn=RIMS1;refGene_tx=XM_01
9830858.1;refGene_gn=RIMS1;refGene_tx=XM_019830859.2;refGene_gn=RIMS1;
refGene_tx=XM_023254093.1;refGene_gn=RIMS1;refGene_tx=XM_023254094.1;r
efGene_gn=RIMS1;refGene_tx=XM_023254095.1;refGene_gn=RIMS1;refGene_tx=
XM_023254096.1;refGene_gn=RIMS1;refGene_tx=XM_023254097.1;refGene_gn=R
IMS1;refGene_tx=XM_023254098.1;refGene_gn=RIMS1;refGene_tx=XM_02325409
9.1;refGene_gn=RIMS1;refGene_tx=XM_023254100.1;refGene_gn=RIMS1;refGen
e_tx=XM_023254101.1;refGene_gn=RIMS1;refGene_tx=XM_023254102.1;refGene
_gn=RIMS1;refGene_tx=XM_023254103.1;refGene_gn=RIMS1;refGene_tx=XM_023
254104.1;refGene_gn=RIMS1;refGene_tx=XM_023254105.1;refGene_gn=RIMS1;r
efGene_tx=XM_023254106.1;refGene_gn=RIMS1;refGene_tx=XM_023254107.1;re
fGene_gn=RIMS1;refGene_tx=XM_023254108.1;refGene_gn=RIMS1;refGene_tx=X
M_023254109.1;refGene_gn=RIMS1;refGene_tx=XM_023254110.1;refGene_gn=RI
MS1;refGene_tx=XM_023254111.1;refGene_gn=RIMS1;refGene_tx=XM_023254112
.
1;refGene_gn=RIMS1;refGene_tx=XM_023254113.1;refGene_gn=RIMS1;refGene_
tx=XM_023254114.1;refGene_gn=RIMS1;refGene_tx=XM_023254115.1;refGene_g
n=RIMS1;refGene_tx=XM_023254117.1;refGene_gn=RIMS1;refGene_tx=XM_02325
4118.1;refGene_gn=RIMS1;refGene_tx=XM_023254119.1;refGene_gn=RIMS1;ref
Gene_tx=XM_023254120.1;refGene_gn=RIMS1;refGene_tx=XM_023254121.1;refG
ene_gn=RIMS1;refGene_tx=XM_023254122.1;refGene_gn=RIMS1;refGene_tx=XM_
023254123.1;refGene_gn=RIMS1;refGene_tx=XM_023254124.1;refGene_gn=RIMS
1;refGene_tx=XM_023254125.1;refGene_gn=RIMS1;refGene_tx=XM_023254126.1
;refGene_gn=RIMS1;refGene_tx=XM_023254127.1;refGene_gn=RIMS1;refGene_t
x=XM_023254128.1;refGene_gn=RIMS1;refGene_tx=XM_023254129.1;refGene_gn
=RIMS1;refGene_tx=XM_023254131.1;refGene_gn=RIMS1;refGene_tx=XM_023254
132.1;refGene_gn=RIMS1;refGene_tx=XM_023254133.1

B2 68063268 68063340
refGene_gn=RIMS1;refGene_tx=XM_019830824.2;refGene_gn=RIMS1;refGene_tx=XM_019830825.2;refGene_gn=RIMS1;refGene_tx=XM_019830833.2;refGene_gn=RIMS1;refGene_tx=XM_019830841.2;refGene_gn=RIMS1;refGene_tx=XM_019830845.2;refGene_gn=RIMS1;refGene_tx=XM_019830848.1;refGene_gn=RIMS1;refGene_tx=XM_019830850.2;refGene_gn=RIMS1;refGene_tx=XM_019830852.2;refGene_gn=RIMS1;refGene_tx=XM_019830857.1;refGene_gn=RIMS1;refGene_tx=XM_019830858.1;refGene_gn=RIMS1;refGene_tx=XM_019830859.2;refGene_gn=RIMS1;refGene_tx=XM_023254093.1;refGene_gn=RIMS1;refGene_tx=XM_023254094.1;refGene_gn=RIMS1;refGene_tx=XM_023254096.1;refGene_gn=RIMS1;refGene_tx=XM_023254097.1;refGene_gn=RIMS1;refGene_tx=XM_023254098.1;refGene_gn=RIMS1;refGene_tx=XM_023254099.1;refGene_gn=RIMS1;refGene_tx=XM_023254100.1;refGene_gn=RIMS1;refGene_tx=XM_023254101.1;refGene_gn=RIMS1;refGene_tx=XM_023254102.1;refGene_gn=RIMS1;refGene_tx=XM_023254103.1;refGene_gn=RIMS1;refGene_tx=XM_023254104.1;refGene_gn=RIMS1;refGene_tx=XM_023254105.1;refGene_gn=RIMS1;refGene_tx=XM_023254106.1;refGene_gn=RIMS1;refGene_tx=XM_023254107.1;refGene_gn=RIMS1;refGene_tx=XM_023254108.1;refGene_gn=RIMS1;refGene_tx=XM_023254109.1;refGene_gn=RIMS1;refGene_tx=XM_023254110.1;refGene_gn=RIMS1;refGene_tx=XM_023254111.1;refGene_gn=RIMS1;refGene_tx=XM_023254112.1;refGene_gn=RIMS1;refGene_tx=XM_023254113.
1;refGene_gn=RIMS1;refGene_tx=XM_023254114.1;refGene_gn=RIMS1;refGene_tx=XM_023254115.1;refGene_gn=RIMS1;refGene_tx=XM_023254117.1;refGene_gn=RIMS1;refGene_tx=XM_023254118.1;refGene_gn=RIMS1;refGene_tx=XM_023254119.1;refGene_gn=RIMS1;refGene_tx=XM_023254120.1;refGene_gn=RIMS1;refGene_tx=XM_023254121.1;refGene_gn=RIMS1;refGene_tx=XM_023254122.1;refGene_gn=RIMS1;refGene_tx=XM_023254123.1;refGene_gn=RIMS1;refGene_tx=XM_023254124.1;refGene_gn=RIMS1;refGene_tx=XM_023254125.1;refGene_gn=RIMS1;refGene_tx=XM_023254126.1;refGene_gn=RIMS1;refGene_tx=XM_023254127.1;refGene_gn=RIMS1;refGene_tx=XM_023254128.1;refGene_gn=RIMS1;refGene_tx=XM_023254129.1;refGene_gn=RIMS1;refGene_tx=XM_023254131.1;refGene_gn=RIMS1;refGene_tx=XM_023254132.1;refGene_gn=RIMS1;refGene_tx=XM_023254133.1
B2 68068600 68068757
refGene_gn=RIMS1;refGene_tx=XM_019830824.2;refGene_gn=RIMS1;refGene_tx=XM_019830825.2;refGene_gn=RIMS1;refGene_tx=XM_019830833.2;refGene_gn=RIMS1;refGene_tx=XM_019830841.2;refGene_gn=RIMS1;refGene_tx=XM_019830845.2;refGene_gn=RIMS1;refGene_tx=XM_019830848.1;refGene_gn=RIMS1;refGene_tx=XM_019830850.2;refGene_gn=RIMS1;refGene_tx=XM_019830852.2;refGene_gn=RIMS1;refGene_tx=XM_019830857.1;refGene_gn=RIMS1;refGene_tx=XM_019830858.1;refGene_gn=RIMS1;refGene_tx=XM_019830859.2;refGene_gn=RIMS1;refGene_tx=XM_023254093.1;refGene_gn=RIMS1;refGene_tx=XM_023254094.1;refGene_gn=RIMS1;refGene_tx=XM_023254095.1;refGene_gn=RIMS1;refGene_tx=XM_023254096.1;refGene_gn=RIMS1;refGene_tx=XM_023254097.1;refGene_gn=RIMS1;refGene_tx=XM_023254098.1;refGene_gn=RIMS1;refGene_tx=XM_023254099.1;refGene_gn=RIMS1;refGene_tx=XM_023254100.1;refGene_gn=RIMS1;refGene_tx=XM_023254101.1;refGene_gn=RIMS1;refGene_tx=XM_023254102.1;refGene_gn=RIMS1;refGene_tx=XM_023254103.1;refGene_gn=RIMS1;refGene_tx=XM_023254104.1;refGene_gn=RIMS1;refGene_tx=XM_023254105.1;refGene_gn=RIMS1;refGene_tx=XM_023254106.1;refGene_gn=RIMS1;refGene_tx=XM_023254107.1;re

fGene_gn=RIMS1;refGene_tx=XM_023254108.1;refGene_gn=RIMS1;refGene_tx=XM_023254109.1;refGene_gn=RIMS1;refGene_tx=XM_023254110.1;refGene_gn=RIMS1;refGene_tx=XM_023254111.1;refGene_gn=RIMS1;refGene_tx=XM_023254112.
1;refGene_gn=RIMS1;refGene_tx=XM_023254113.1;refGene_gn=RIMS1;refGene_tx=XM_023254114.1;refGene_gn=RIMS1;refGene_tx=XM_023254115.1;refGene_gn=RIMS1;refGene_tx=XM_023254117.1;refGene_gn=RIMS1;refGene_tx=XM_023254118.1;refGene_gn=RIMS1;refGene_tx=XM_023254119.1;refGene_gn=RIMS1;refGene_tx=XM_023254120.1;refGene_gn=RIMS1;refGene_tx=XM_023254121.1;refGene_gn=RIMS1;refGene_tx=XM_023254122.1;refGene_gn=RIMS1;refGene_tx=XM_023254123.1;refGene_gn=RIMS1;refGene_tx=XM_023254124.1;refGene_gn=RIMS1;refGene_tx=XM_023254125.1;refGene_gn=RIMS1;refGene_tx=XM_023254126.1;refGene_gn=RIMS1;refGene_tx=XM_023254127.1;refGene_gn=RIMS1;refGene_tx=XM_023254128.1;refGene_gn=RIMS1;refGene_tx=XM_023254129.1;refGene_gn=RIMS1;refGene_tx=XM_023254131.1;refGene_gn=RIMS1;refGene_tx=XM_023254132.1;refGene_gn=RIMS1;refGene_tx=XM_023254133.1
B2 68069741 68069867
refGene_gn=RIMS1;refGene_tx=XM_019830824.2;refGene_gn=RIMS1;refGene_tx=XM_019830825.2;refGene_gn=RIMS1;refGene_tx=XM_019830833.2;refGene_gn=RIMS1;refGene_tx=XM_019830841.2;refGene_gn=RIMS1;refGene_tx=XM_019830845.2;refGene_gn=RIMS1;refGene_tx=XM_019830848.1;refGene_gn=RIMS1;refGene_tx=XM_019830850.2;refGene_gn=RIMS1;refGene_tx=XM_019830852.2;refGene_gn=RIMS1;refGene_tx=XM_019830857.1;refGene_gn=RIMS1;refGene_tx=XM_019830858.1;refGene_gn=RIMS1;refGene_tx=XM_019830859.2;refGene_gn=RIMS1;refGene_tx=XM_023254093.1;refGene_gn=RIMS1;refGene_tx=XM_023254094.1;refGene_gn=RIMS1;refGene_tx=XM_023254095.1;refGene_gn=RIMS1;refGene_tx=XM_023254096.1;refGene_gn=RIMS1;refGene_tx=XM_023254097.1;refGene_gn=RIMS1;refGene_tx=XM_023254098.1;refGene_gn=RIMS1;refGene_tx=XM_023254099.1;refGene_gn=RIMS1;refGene_tx=XM_023254100.1;refGene_gn=RIMS1;refGene_tx=XM_023254101.1;refGene_gn=RIMS1;refGene_tx=XM_023254102.1;refGene_gn=RIMS1;refGene_tx=XM_023254103.1;refGene_gn=RIMS1;refGene_tx=XM_023254104.1;refGene_gn=RIMS1;refGene_tx=XM_023254105.1;refGene_gn=RIMS1;refGene_tx=XM_023254106.1;refGene_gn=RIMS1;refGene_tx=XM_023254107.1;refGene_gn=RIMS1;refGene_tx=XM_023254108.1;refGene_gn=RIMS1;refGene_tx=XM_023254109.1;refGene_gn=RIMS1;refGene_tx=XM_023254110.1;refGene_gn=RIMS1;refGene_tx=XM_023254111.1;refGene_gn=RIMS1;refGene_tx=XM_023254112.
1;refGene_gn=RIMS1;refGene_tx=XM_023254113.1;refGene_gn=RIMS1;refGene_tx=XM_023254114.1;refGene_gn=RIMS1;refGene_tx=XM_023254115.1;refGene_gn=RIMS1;refGene_tx=XM_023254117.1;refGene_gn=RIMS1;refGene_tx=XM_023254118.1;refGene_gn=RIMS1;refGene_tx=XM_023254119.1;refGene_gn=RIMS1;refGene_tx=XM_023254120.1;refGene_gn=RIMS1;refGene_tx=XM_023254121.1;refGene_gn=RIMS1;refGene_tx=XM_023254122.1;refGene_gn=RIMS1;refGene_tx=XM_023254123.1;refGene_gn=RIMS1;refGene_tx=XM_023254124.1;refGene_gn=RIMS1;refGene_tx=XM_023254125.1;refGene_gn=RIMS1;refGene_tx=XM_023254126.1;refGene_gn=RIMS1;refGene_tx=XM_023254127.1;refGene_gn=RIMS1;refGene_tx=XM_023254128.1;refGene_gn=RIMS1;refGene_tx=XM_023254129.1;refGene_gn=RIMS1;refGene_tx=XM_023254131.1;refGene_gn=RIMS1;refGene_tx=XM_023254132.1;refGene_gn=RIMS1;refGene_tx=XM_023254133.1
B2 68071412 68071475

refGene_gn=RIMS1;refGene_tx=XM_019830824.2;refGene_gn=RIMS1;refGene_tx
=XM_019830825.2;refGene_gn=RIMS1;refGene_tx=XM_019830833.2;refGene_gn=
RIMS1;refGene_tx=XM_019830845.2;refGene_gn=RIMS1;refGene_tx=XM_0198308
48.1;refGene_gn=RIMS1;refGene_tx=XM_019830857.1;refGene_gn=RIMS1;refGe
ne_tx=XM_019830858.1;refGene_gn=RIMS1;refGene_tx=XM_023254093.1;refGen
e_gn=RIMS1;refGene_tx=XM_023254094.1;refGene_gn=RIMS1;refGene_tx=XM_02
3254095.1;refGene_gn=RIMS1;refGene_tx=XM_023254096.1;refGene_gn=RIMS1;
refGene_tx=XM_023254097.1;refGene_gn=RIMS1;refGene_tx=XM_023254098.1;r
efGene_gn=RIMS1;refGene_tx=XM_023254099.1;refGene_gn=RIMS1;refGene_tx=
XM_023254100.1;refGene_gn=RIMS1;refGene_tx=XM_023254101.1;refGene_gn=R
IMS1;refGene_tx=XM_023254102.1;refGene_gn=RIMS1;refGene_tx=XM_02325410
3.1;refGene_gn=RIMS1;refGene_tx=XM_023254104.1;refGene_gn=RIMS1;refGen
e_tx=XM_023254106.1;refGene_gn=RIMS1;refGene_tx=XM_023254107.1;refGene
_gn=RIMS1;refGene_tx=XM_023254108.1;refGene_gn=RIMS1;refGene_tx=XM_023
254109.1;refGene_gn=RIMS1;refGene_tx=XM_023254110.1;refGene_gn=RIMS1;r
efGene_tx=XM_023254111.1;refGene_gn=RIMS1;refGene_tx=XM_023254112.1;re
fGene_gn=RIMS1;refGene_tx=XM_023254113.1;refGene_gn=RIMS1;refGene_tx=X
M_023254114.1;refGene_gn=RIMS1;refGene_tx=XM_023254115.1;refGene_gn=RI
MS1;refGene_tx=XM_023254120.1;refGene_gn=RIMS1;refGene_tx=XM_023254121
.
1;refGene_gn=RIMS1;refGene_tx=XM_023254122.1;refGene_gn=RIMS1;refGene_
tx=XM_023254123.1;refGene_gn=RIMS1;refGene_tx=XM_023254124.1;refGene_g
n=RIMS1;refGene_tx=XM_023254125.1;refGene_gn=RIMS1;refGene_tx=XM_02325
4126.1;refGene_gn=RIMS1;refGene_tx=XM_023254127.1;refGene_gn=RIMS1;ref
Gene_tx=XM_023254128.1;refGene_gn=RIMS1;refGene_tx=XM_023254129.1;refG
ene_gn=RIMS1;refGene_tx=XM_023254131.1
B2 68075085 68075163
refGene_gn=RIMS1;refGene_tx=XM_019830824.2;refGene_gn=RIMS1;refGene_tx
=XM_019830825.2;refGene_gn=RIMS1;refGene_tx=XM_023254093.1;refGene_gn=
RIMS1;refGene_tx=XM_023254094.1;refGene_gn=RIMS1;refGene_tx=XM_0232540
95.1;refGene_gn=RIMS1;refGene_tx=XM_023254097.1;refGene_gn=RIMS1;refGe
ne_tx=XM_023254098.1;refGene_gn=RIMS1;refGene_tx=XM_023254099.1
B2 68076058 68076148
refGene_gn=RIMS1;refGene_tx=XM_019830824.2;refGene_gn=RIMS1;refGene_tx
=XM_019830825.2;refGene_gn=RIMS1;refGene_tx=XM_019830833.2;refGene_gn=
RIMS1;refGene_tx=XM_019830845.2;refGene_gn=RIMS1;refGene_tx=XM_0198308
48.1;refGene_gn=RIMS1;refGene_tx=XM_019830857.1;refGene_gn=RIMS1;refGe
ne_tx=XM_019830858.1;refGene_gn=RIMS1;refGene_tx=XM_023254093.1;refGen
e_gn=RIMS1;refGene_tx=XM_023254094.1;refGene_gn=RIMS1;refGene_tx=XM_02
3254095.1;refGene_gn=RIMS1;refGene_tx=XM_023254096.1;refGene_gn=RIMS1;
refGene_tx=XM_023254097.1;refGene_gn=RIMS1;refGene_tx=XM_023254098.1;r
efGene_gn=RIMS1;refGene_tx=XM_023254099.1;refGene_gn=RIMS1;refGene_tx=
XM_023254100.1;refGene_gn=RIMS1;refGene_tx=XM_023254101.1;refGene_gn=R
IMS1;refGene_tx=XM_023254102.1;refGene_gn=RIMS1;refGene_tx=XM_02325410
3.1;refGene_gn=RIMS1;refGene_tx=XM_023254104.1;refGene_gn=RIMS1;refGen
e_tx=XM_023254106.1;refGene_gn=RIMS1;refGene_tx=XM_023254107.1;refGene
_gn=RIMS1;refGene_tx=XM_023254108.1;refGene_gn=RIMS1;refGene_tx=XM_023
254109.1;refGene_gn=RIMS1;refGene_tx=XM_023254110.1;refGene_gn=RIMS1;r
efGene_tx=XM_023254111.1;refGene_gn=RIMS1;refGene_tx=XM_023254112.1;re
fGene_gn=RIMS1;refGene_tx=XM_023254113.1;refGene_gn=RIMS1;refGene_tx=X

M_023254114.1;refGene_gn=RIMS1;refGene_tx=XM_023254115.1;refGene_gn=RIMS1;refGene_tx=XM_023254120.1;refGene_gn=RIMS1;refGene_tx=XM_023254121.
1;refGene_gn=RIMS1;refGene_tx=XM_023254122.1;refGene_gn=RIMS1;refGene_tx=XM_023254123.1;refGene_gn=RIMS1;refGene_tx=XM_023254124.1;refGene_gn=RIMS1;refGene_tx=XM_023254125.1;refGene_gn=RIMS1;refGene_tx=XM_023254126.1;refGene_gn=RIMS1;refGene_tx=XM_023254127.1;refGene_gn=RIMS1;refGene_tx=XM_023254128.1;refGene_gn=RIMS1;refGene_tx=XM_023254129.1;refGene_gn=RIMS1;refGene_tx=XM_023254131.1
B2 68084683 68084767
refGene_gn=RIMS1;refGene_tx=XM_019830824.2;refGene_gn=RIMS1;refGene_tx=XM_019830825.2;refGene_gn=RIMS1;refGene_tx=XM_023254093.1;refGene_gn=RIMS1;refGene_tx=XM_023254094.1;refGene_gn=RIMS1;refGene_tx=XM_023254095.1;refGene_gn=RIMS1;refGene_tx=XM_023254096.1;refGene_gn=RIMS1;refGene_tx=XM_023254097.1;refGene_gn=RIMS1;refGene_tx=XM_023254099.1;refGene_gn=RIMS1;refGene_tx=XM_023254111.1
B2 68093795 68093867
refGene_gn=RIMS1;refGene_tx=XM_019830824.2;refGene_gn=RIMS1;refGene_tx=XM_019830825.2;refGene_gn=RIMS1;refGene_tx=XM_019830833.2;refGene_gn=RIMS1;refGene_tx=XM_019830841.2;refGene_gn=RIMS1;refGene_tx=XM_019830848.1;refGene_gn=RIMS1;refGene_tx=XM_019830850.2;refGene_gn=RIMS1;refGene_tx=XM_019830857.1;refGene_gn=RIMS1;refGene_tx=XM_023254093.1;refGene_gn=RIMS1;refGene_tx=XM_023254094.1;refGene_gn=RIMS1;refGene_tx=XM_023254095.1;refGene_gn=RIMS1;refGene_tx=XM_023254096.1;refGene_gn=RIMS1;refGene_tx=XM_023254097.1;refGene_gn=RIMS1;refGene_tx=XM_023254098.1;refGene_gn=RIMS1;refGene_tx=XM_023254099.1;refGene_gn=RIMS1;refGene_tx=XM_023254100.1;refGene_gn=RIMS1;refGene_tx=XM_023254101.1;refGene_gn=RIMS1;refGene_tx=XM_023254102.1;refGene_gn=RIMS1;refGene_tx=XM_023254106.1;refGene_gn=RIMS1;refGene_tx=XM_023254108.1;refGene_gn=RIMS1;refGene_tx=XM_023254109.1;refGene_gn=RIMS1;refGene_tx=XM_023254110.1;refGene_gn=RIMS1;refGene_tx=XM_023254120.1;refGene_gn=RIMS1;refGene_tx=XM_023254121.1;refGene_gn=RIMS1;refGene_tx=XM_023254122.1;refGene_gn=RIMS1;refGene_tx=XM_023254123.1;refGene_gn=RIMS1;refGene_tx=XM_023254124.1;refGene_gn=RIMS1;refGene_tx=XM_023254125.1
B2 68099755 68099938
refGene_gn=RIMS1;refGene_tx=XM_019830824.2;refGene_gn=RIMS1;refGene_tx=XM_019830825.2;refGene_gn=RIMS1;refGene_tx=XM_019830833.2;refGene_gn=RIMS1;refGene_tx=XM_019830841.2;refGene_gn=RIMS1;refGene_tx=XM_023254093.1;refGene_gn=RIMS1;refGene_tx=XM_023254094.1;refGene_gn=RIMS1;refGene_tx=XM_023254095.1;refGene_gn=RIMS1;refGene_tx=XM_023254096.1;refGene_gn=RIMS1;refGene_tx=XM_023254097.1;refGene_gn=RIMS1;refGene_tx=XM_023254098.1;refGene_gn=RIMS1;refGene_tx=XM_023254100.1;refGene_gn=RIMS1;refGene_tx=XM_023254108.1;refGene_gn=RIMS1;refGene_tx=XM_023254109.1;refGene_gn=RIMS1;refGene_tx=XM_023254120.1;refGene_gn=RIMS1;refGene_tx=XM_023254121.1
B2 68101042 68101155
refGene_gn=RIMS1;refGene_tx=XM_019830824.2;refGene_gn=RIMS1;refGene_tx=XM_019830825.2;refGene_gn=RIMS1;refGene_tx=XM_019830833.2;refGene_gn=RIMS1;refGene_tx=XM_019830841.2;refGene_gn=RIMS1;refGene_tx=XM_019830845.2;refGene_gn=RIMS1;refGene_tx=XM_019830848.1;refGene_gn=RIMS1;refGe

ne_tx=XM_019830850.2;refGene_gn=RIMS1;refGene_tx=XM_019830852.2;refGene_gn=RIMS1;refGene_tx=XM_019830857.1;refGene_gn=RIMS1;refGene_tx=XM_019830858.1;refGene_gn=RIMS1;refGene_tx=XM_019830859.2;refGene_gn=RIMS1;refGene_tx=XM_023254093.1;refGene_gn=RIMS1;refGene_tx=XM_023254094.1;refGene_gn=RIMS1;refGene_tx=XM_023254095.1;refGene_gn=RIMS1;refGene_tx=XM_023254096.1;refGene_gn=RIMS1;refGene_tx=XM_023254097.1;refGene_gn=RIMS1;refGene_tx=XM_023254098.1;refGene_gn=RIMS1;refGene_tx=XM_023254099.1;refGene_gn=RIMS1;refGene_tx=XM_023254100.1;refGene_gn=RIMS1;refGene_tx=XM_023254101.1;refGene_gn=RIMS1;refGene_tx=XM_023254102.1;refGene_gn=RIMS1;refGene_tx=XM_023254103.1;refGene_gn=RIMS1;refGene_tx=XM_023254104.1;refGene_gn=RIMS1;refGene_tx=XM_023254105.1;refGene_gn=RIMS1;refGene_tx=XM_023254106.1;refGene_gn=RIMS1;refGene_tx=XM_023254107.1;refGene_gn=RIMS1;refGene_tx=XM_023254108.1;refGene_gn=RIMS1;refGene_tx=XM_023254109.1;refGene_gn=RIMS1;refGene_tx=XM_023254110.1;refGene_gn=RIMS1;refGene_tx=XM_023254111.1;refGene_gn=RIMS1;refGene_tx=XM_023254112.
1;refGene_gn=RIMS1;refGene_tx=XM_023254113.1;refGene_gn=RIMS1;refGene_tx=XM_023254114.1;refGene_gn=RIMS1;refGene_tx=XM_023254115.1;refGene_gn=RIMS1;refGene_tx=XM_023254117.1;refGene_gn=RIMS1;refGene_tx=XM_023254118.1;refGene_gn=RIMS1;refGene_tx=XM_023254119.1;refGene_gn=RIMS1;refGene_tx=XM_023254120.1;refGene_gn=RIMS1;refGene_tx=XM_023254121.1;refGene_gn=RIMS1;refGene_tx=XM_023254122.1;refGene_gn=RIMS1;refGene_tx=XM_023254123.1;refGene_gn=RIMS1;refGene_tx=XM_023254124.1;refGene_gn=RIMS1;refGene_tx=XM_023254125.1;refGene_gn=RIMS1;refGene_tx=XM_023254126.1;refGene_gn=RIMS1;refGene_tx=XM_023254127.1;refGene_gn=RIMS1;refGene_tx=XM_023254128.1;refGene_gn=RIMS1;refGene_tx=XM_023254129.1;refGene_gn=RIMS1;refGene_tx=XM_023254131.1;refGene_gn=RIMS1;refGene_tx=XM_023254132.1;refGene_gn=RIMS1;refGene_tx=XM_023254133.1

B2 68105115 68105145

refGene_gn=RIMS1;refGene_tx=XM_019830824.2;refGene_gn=RIMS1;refGene_tx=XM_019830825.2;refGene_gn=RIMS1;refGene_tx=XM_019830833.2;refGene_gn=RIMS1;refGene_tx=XM_019830841.2;refGene_gn=RIMS1;refGene_tx=XM_019830845.2;refGene_gn=RIMS1;refGene_tx=XM_019830850.2;refGene_gn=RIMS1;refGene_tx=XM_019830852.2;refGene_gn=RIMS1;refGene_tx=XM_023254093.1;refGene_gn=RIMS1;refGene_tx=XM_023254095.1;refGene_gn=RIMS1;refGene_tx=XM_023254096.1;refGene_gn=RIMS1;refGene_tx=XM_023254097.1;refGene_gn=RIMS1;refGene_tx=XM_023254098.1;refGene_gn=RIMS1;refGene_tx=XM_023254099.1

B2 68116226 68116339

refGene_gn=RIMS1;refGene_tx=XM_019830824.2;refGene_gn=RIMS1;refGene_tx=XM_019830825.2;refGene_gn=RIMS1;refGene_tx=XM_019830833.2;refGene_gn=RIMS1;refGene_tx=XM_019830841.2;refGene_gn=RIMS1;refGene_tx=XM_019830845.2;refGene_gn=RIMS1;refGene_tx=XM_019830848.1;refGene_gn=RIMS1;refGene_tx=XM_019830850.2;refGene_gn=RIMS1;refGene_tx=XM_019830852.2;refGene_gn=RIMS1;refGene_tx=XM_023254093.1;refGene_gn=RIMS1;refGene_tx=XM_023254094.1;refGene_gn=RIMS1;refGene_tx=XM_023254095.1;refGene_gn=RIMS1;refGene_tx=XM_023254096.1;refGene_gn=RIMS1;refGene_tx=XM_023254097.1;refGene_gn=RIMS1;refGene_tx=XM_023254098.1;refGene_gn=RIMS1;refGene_tx=XM_023254099.1;refGene_gn=RIMS1;refGene_tx=XM_023254100.1;refGene_gn=RIMS1;refGene_tx=XM_023254101.1;refGene_gn=RIMS1;refGene_tx=XM_023254102.1;refGene_gn=RIMS1;refGene_tx=XM_023254103.1;refGene_gn=RIMS1;refGen

e_tx=XM_023254104.1;refGene_gn=RIMS1;refGene_tx=XM_023254105.1;refGene_gn=RIMS1;refGene_tx=XM_023254108.1;refGene_gn=RIMS1;refGene_tx=XM_023254109.1;refGene_gn=RIMS1;refGene_tx=XM_023254110.1;refGene_gn=RIMS1;refGene_tx=XM_023254111.1;refGene_gn=RIMS1;refGene_tx=XM_023254112.1;refGene_gn=RIMS1;refGene_tx=XM_023254113.1;refGene_gn=RIMS1;refGene_tx=XM_023254114.1;refGene_gn=RIMS1;refGene_tx=XM_023254115.1;refGene_gn=RIMS1;refGene_tx=XM_023254117.1;refGene_gn=RIMS1;refGene_tx=XM_023254118.1;refGene_gn=RIMS1;refGene_tx=XM_023254119.1
B2 68122089 68122256
refGene_gn=RIMS1;refGene_tx=XM_019830824.2;refGene_gn=RIMS1;refGene_tx=XM_019830825.2;refGene_gn=RIMS1;refGene_tx=XM_019830833.2;refGene_gn=RIMS1;refGene_tx=XM_019830841.2;refGene_gn=RIMS1;refGene_tx=XM_019830845.2;refGene_gn=RIMS1;refGene_tx=XM_019830848.1;refGene_gn=RIMS1;refGene_tx=XM_019830850.2;refGene_gn=RIMS1;refGene_tx=XM_019830852.2;refGene_gn=RIMS1;refGene_tx=XM_023254093.1;refGene_gn=RIMS1;refGene_tx=XM_023254094.1;refGene_gn=RIMS1;refGene_tx=XM_023254095.1;refGene_gn=RIMS1;refGene_tx=XM_023254096.1;refGene_gn=RIMS1;refGene_tx=XM_023254097.1;refGene_gn=RIMS1;refGene_tx=XM_023254098.1;refGene_gn=RIMS1;refGene_tx=XM_023254099.1;refGene_gn=RIMS1;refGene_tx=XM_023254100.1;refGene_gn=RIMS1;refGene_tx=XM_023254101.1;refGene_gn=RIMS1;refGene_tx=XM_023254102.1;refGene_gn=RIMS1;refGene_tx=XM_023254103.1;refGene_gn=RIMS1;refGene_tx=XM_023254104.1;refGene_gn=RIMS1;refGene_tx=XM_023254105.1;refGene_gn=RIMS1;refGene_tx=XM_023254108.1;refGene_gn=RIMS1;refGene_tx=XM_023254109.1;refGene_gn=RIMS1;refGene_tx=XM_023254110.1;refGene_gn=RIMS1;refGene_tx=XM_023254111.1;refGene_gn=RIMS1;refGene_tx=XM_023254112.1;refGene_gn=RIMS1;refGene_tx=XM_023254113.1;refGene_gn=RIMS1;refGene_tx=XM_023254114.1;refGene_gn=RIMS1;refGene_tx=XM_023254115.1;refGene_gn=RIMS1;refGene_tx=XM_023254117.1;refGene_gn=RIMS1;refGene_tx=XM_023254118.1;refGene_gn=RIMS1;refGene_tx=XM_023254119.1
B2 68147338 68147574
refGene_gn=RIMS1;refGene_tx=XM_019830824.2;refGene_gn=RIMS1;refGene_tx=XM_019830825.2;refGene_gn=RIMS1;refGene_tx=XM_019830833.2;refGene_gn=RIMS1;refGene_tx=XM_019830841.2;refGene_gn=RIMS1;refGene_tx=XM_019830845.2;refGene_gn=RIMS1;refGene_tx=XM_019830848.1;refGene_gn=RIMS1;refGene_tx=XM_019830850.2;refGene_gn=RIMS1;refGene_tx=XM_019830852.2;refGene_gn=RIMS1;refGene_tx=XM_023254093.1;refGene_gn=RIMS1;refGene_tx=XM_023254094.1;refGene_gn=RIMS1;refGene_tx=XM_023254095.1;refGene_gn=RIMS1;refGene_tx=XM_023254096.1;refGene_gn=RIMS1;refGene_tx=XM_023254097.1;refGene_gn=RIMS1;refGene_tx=XM_023254098.1;refGene_gn=RIMS1;refGene_tx=XM_023254099.1;refGene_gn=RIMS1;refGene_tx=XM_023254100.1;refGene_gn=RIMS1;refGene_tx=XM_023254101.1;refGene_gn=RIMS1;refGene_tx=XM_023254102.1;refGene_gn=RIMS1;refGene_tx=XM_023254103.1;refGene_gn=RIMS1;refGene_tx=XM_023254104.1;refGene_gn=RIMS1;refGene_tx=XM_023254105.1;refGene_gn=RIMS1;refGene_tx=XM_023254108.1;refGene_gn=RIMS1;refGene_tx=XM_023254109.1;refGene_gn=RIMS1;refGene_tx=XM_023254110.1;refGene_gn=RIMS1;refGene_tx=XM_023254111.1;refGene_gn=RIMS1;refGene_tx=XM_023254112.1;refGene_gn=RIMS1;refGene_tx=XM_023254113.1;refGene_gn=RIMS1;refGene_tx=XM_023254114.1;refGene_gn=RIMS1;refGene_tx=XM_023254115.1;refGene_gn=RIMS1;refGene_tx=XM_023254117.1;refGene_gn=RIMS1;refGene_tx=XM_023254118.1;refGene_gn=RIMS1;refGene_tx=XM_023254119.1

B2 68214873 68215012
refGene_gn=RIMS1;refGene_tx=XM_019830824.2;refGene_gn=RIMS1;refGene_tx
=XM_019830825.2;refGene_gn=RIMS1;refGene_tx=XM_019830833.2;refGene_gn=
RIMS1;refGene_tx=XM_019830841.2;refGene_gn=RIMS1;refGene_tx=XM_0198308
45.2;refGene_gn=RIMS1;refGene_tx=XM_019830848.1;refGene_gn=RIMS1;refGe
ne_tx=XM_019830850.2;refGene_gn=RIMS1;refGene_tx=XM_019830852.2;refGen
e_gn=RIMS1;refGene_tx=XM_019830857.1;refGene_gn=RIMS1;refGene_tx=XM_01
9830858.1;refGene_gn=RIMS1;refGene_tx=XM_019830859.2;refGene_gn=RIMS1;
refGene_tx=XM_023254093.1;refGene_gn=RIMS1;refGene_tx=XM_023254094.1;r
efGene_gn=RIMS1;refGene_tx=XM_023254095.1;refGene_gn=RIMS1;refGene_tx=
XM_023254096.1;refGene_gn=RIMS1;refGene_tx=XM_023254097.1;refGene_gn=R
IMS1;refGene_tx=XM_023254098.1;refGene_gn=RIMS1;refGene_tx=XM_02325409
9.1;refGene_gn=RIMS1;refGene_tx=XM_023254100.1;refGene_gn=RIMS1;refGen
e_tx=XM_023254101.1;refGene_gn=RIMS1;refGene_tx=XM_023254102.1;refGene
_gn=RIMS1;refGene_tx=XM_023254103.1;refGene_gn=RIMS1;refGene_tx=XM_023
254104.1;refGene_gn=RIMS1;refGene_tx=XM_023254105.1;refGene_gn=RIMS1;r
efGene_tx=XM_023254106.1;refGene_gn=RIMS1;refGene_tx=XM_023254107.1;re
fGene_gn=RIMS1;refGene_tx=XM_023254108.1;refGene_gn=RIMS1;refGene_tx=X
M_023254109.1;refGene_gn=RIMS1;refGene_tx=XM_023254110.1;refGene_gn=RI
MS1;refGene_tx=XM_023254111.1;refGene_gn=RIMS1;refGene_tx=XM_023254112
.
1;refGene_gn=RIMS1;refGene_tx=XM_023254113.1;refGene_gn=RIMS1;refGene_
tx=XM_023254114.1;refGene_gn=RIMS1;refGene_tx=XM_023254115.1;refGene_g
n=RIMS1;refGene_tx=XM_023254117.1;refGene_gn=RIMS1;refGene_tx=XM_02325
4118.1;refGene_gn=RIMS1;refGene_tx=XM_023254119.1;refGene_gn=RIMS1;ref
Gene_tx=XM_023254120.1;refGene_gn=RIMS1;refGene_tx=XM_023254121.1;refG
ene_gn=RIMS1;refGene_tx=XM_023254122.1;refGene_gn=RIMS1;refGene_tx=XM_
023254123.1;refGene_gn=RIMS1;refGene_tx=XM_023254124.1;refGene_gn=RIMS
1;refGene_tx=XM_023254125.1;refGene_gn=RIMS1;refGene_tx=XM_023254126.1
;refGene_gn=RIMS1;refGene_tx=XM_023254127.1;refGene_gn=RIMS1;refGene_t
x=XM_023254128.1;refGene_gn=RIMS1;refGene_tx=XM_023254129.1;refGene_gn
=RIMS1;refGene_tx=XM_023254131.1;refGene_gn=RIMS1;refGene_tx=XM_023254
132.1;refGene_gn=RIMS1;refGene_tx=XM_023254133.1
B2 68217474 68217587
refGene_gn=RIMS1;refGene_tx=XM_019830824.2;refGene_gn=RIMS1;refGene_tx
=XM_019830825.2;refGene_gn=RIMS1;refGene_tx=XM_019830833.2;refGene_gn=
RIMS1;refGene_tx=XM_019830841.2;refGene_gn=RIMS1;refGene_tx=XM_0198308
45.2;refGene_gn=RIMS1;refGene_tx=XM_019830848.1;refGene_gn=RIMS1;refGe
ne_tx=XM_019830850.2;refGene_gn=RIMS1;refGene_tx=XM_019830852.2;refGen
e_gn=RIMS1;refGene_tx=XM_019830857.1;refGene_gn=RIMS1;refGene_tx=XM_01
9830858.1;refGene_gn=RIMS1;refGene_tx=XM_019830859.2;refGene_gn=RIMS1;
refGene_tx=XM_023254093.1;refGene_gn=RIMS1;refGene_tx=XM_023254094.1;r
efGene_gn=RIMS1;refGene_tx=XM_023254095.1;refGene_gn=RIMS1;refGene_tx=
XM_023254096.1;refGene_gn=RIMS1;refGene_tx=XM_023254097.1;refGene_gn=R
IMS1;refGene_tx=XM_023254098.1;refGene_gn=RIMS1;refGene_tx=XM_02325409
9.1;refGene_gn=RIMS1;refGene_tx=XM_023254100.1;refGene_gn=RIMS1;refGen
e_tx=XM_023254101.1;refGene_gn=RIMS1;refGene_tx=XM_023254102.1;refGene
_gn=RIMS1;refGene_tx=XM_023254103.1;refGene_gn=RIMS1;refGene_tx=XM_023
254104.1;refGene_gn=RIMS1;refGene_tx=XM_023254105.1;refGene_gn=RIMS1;r
efGene_tx=XM_023254106.1;refGene_gn=RIMS1;refGene_tx=XM_023254107.1;re

fGene_gn=RIMS1;refGene_tx=XM_023254108.1;refGene_gn=RIMS1;refGene_tx=X
M_023254109.1;refGene_gn=RIMS1;refGene_tx=XM_023254110.1;refGene_gn=RI
MS1;refGene_tx=XM_023254111.1;refGene_gn=RIMS1;refGene_tx=XM_023254112
.
1;refGene_gn=RIMS1;refGene_tx=XM_023254113.1;refGene_gn=RIMS1;refGene_
tx=XM_023254114.1;refGene_gn=RIMS1;refGene_tx=XM_023254115.1;refGene_g
n=RIMS1;refGene_tx=XM_023254117.1;refGene_gn=RIMS1;refGene_tx=XM_02325
4118.1;refGene_gn=RIMS1;refGene_tx=XM_023254119.1;refGene_gn=RIMS1;ref
Gene_tx=XM_023254120.1;refGene_gn=RIMS1;refGene_tx=XM_023254121.1;refG
ene_gn=RIMS1;refGene_tx=XM_023254122.1;refGene_gn=RIMS1;refGene_tx=XM_
023254123.1;refGene_gn=RIMS1;refGene_tx=XM_023254124.1;refGene_gn=RIMS
1;refGene_tx=XM_023254125.1;refGene_gn=RIMS1;refGene_tx=XM_023254126.1
;refGene_gn=RIMS1;refGene_tx=XM_023254127.1;refGene_gn=RIMS1;refGene_t
x=XM_023254128.1;refGene_gn=RIMS1;refGene_tx=XM_023254129.1;refGene_gn
=RIMS1;refGene_tx=XM_023254131.1;refGene_gn=RIMS1;refGene_tx=XM_023254
132.1;refGene_gn=RIMS1;refGene_tx=XM_023254133.1
B2 68220860 68220962
refGene_gn=RIMS1;refGene_tx=XM_019830824.2;refGene_gn=RIMS1;refGene_tx
=XM_019830825.2;refGene_gn=RIMS1;refGene_tx=XM_019830833.2;refGene_gn=
RIMS1;refGene_tx=XM_019830841.2;refGene_gn=RIMS1;refGene_tx=XM_0198308
45.2;refGene_gn=RIMS1;refGene_tx=XM_019830848.1;refGene_gn=RIMS1;refGe
ne_tx=XM_019830850.2;refGene_gn=RIMS1;refGene_tx=XM_019830852.2;refGen
e_gn=RIMS1;refGene_tx=XM_019830857.1;refGene_gn=RIMS1;refGene_tx=XM_01
9830858.1;refGene_gn=RIMS1;refGene_tx=XM_019830859.2;refGene_gn=RIMS1;
refGene_tx=XM_023254093.1;refGene_gn=RIMS1;refGene_tx=XM_023254094.1;r
efGene_gn=RIMS1;refGene_tx=XM_023254095.1;refGene_gn=RIMS1;refGene_tx=
XM_023254096.1;refGene_gn=RIMS1;refGene_tx=XM_023254097.1;refGene_gn=R
IMS1;refGene_tx=XM_023254098.1;refGene_gn=RIMS1;refGene_tx=XM_02325409
9.1;refGene_gn=RIMS1;refGene_tx=XM_023254100.1;refGene_gn=RIMS1;refGen
e_tx=XM_023254101.1;refGene_gn=RIMS1;refGene_tx=XM_023254102.1;refGene
_gn=RIMS1;refGene_tx=XM_023254103.1;refGene_gn=RIMS1;refGene_tx=XM_023
254104.1;refGene_gn=RIMS1;refGene_tx=XM_023254105.1;refGene_gn=RIMS1;r
efGene_tx=XM_023254106.1;refGene_gn=RIMS1;refGene_tx=XM_023254107.1;re
fGene_gn=RIMS1;refGene_tx=XM_023254108.1;refGene_gn=RIMS1;refGene_tx=X
M_023254109.1;refGene_gn=RIMS1;refGene_tx=XM_023254110.1;refGene_gn=RI
MS1;refGene_tx=XM_023254111.1;refGene_gn=RIMS1;refGene_tx=XM_023254112
.
1;refGene_gn=RIMS1;refGene_tx=XM_023254113.1;refGene_gn=RIMS1;refGene_
tx=XM_023254114.1;refGene_gn=RIMS1;refGene_tx=XM_023254115.1;refGene_g
n=RIMS1;refGene_tx=XM_023254117.1;refGene_gn=RIMS1;refGene_tx=XM_02325
4118.1;refGene_gn=RIMS1;refGene_tx=XM_023254119.1;refGene_gn=RIMS1;ref
Gene_tx=XM_023254120.1;refGene_gn=RIMS1;refGene_tx=XM_023254121.1;refG
ene_gn=RIMS1;refGene_tx=XM_023254122.1;refGene_gn=RIMS1;refGene_tx=XM_
023254123.1;refGene_gn=RIMS1;refGene_tx=XM_023254124.1;refGene_gn=RIMS
1;refGene_tx=XM_023254125.1;refGene_gn=RIMS1;refGene_tx=XM_023254126.1
;refGene_gn=RIMS1;refGene_tx=XM_023254127.1;refGene_gn=RIMS1;refGene_t
x=XM_023254128.1;refGene_gn=RIMS1;refGene_tx=XM_023254129.1;refGene_gn
=RIMS1;refGene_tx=XM_023254131.1;refGene_gn=RIMS1;refGene_tx=XM_023254
132.1;refGene_gn=RIMS1;refGene_tx=XM_023254133.1
B2 68221773 68221913

refGene_gn=RIMS1;refGene_tx=XM_019830824.2;refGene_gn=RIMS1;refGene_tx
=XM_019830825.2;refGene_gn=RIMS1;refGene_tx=XM_019830833.2;refGene_gn=
RIMS1;refGene_tx=XM_019830841.2;refGene_gn=RIMS1;refGene_tx=XM_0198308
45.2;refGene_gn=RIMS1;refGene_tx=XM_019830848.1;refGene_gn=RIMS1;refGe
ne_tx=XM_019830850.2;refGene_gn=RIMS1;refGene_tx=XM_019830852.2;refGen
e_gn=RIMS1;refGene_tx=XM_019830857.1;refGene_gn=RIMS1;refGene_tx=XM_01
9830858.1;refGene_gn=RIMS1;refGene_tx=XM_019830859.2;refGene_gn=RIMS1;
refGene_tx=XM_023254093.1;refGene_gn=RIMS1;refGene_tx=XM_023254094.1;r
efGene_gn=RIMS1;refGene_tx=XM_023254095.1;refGene_gn=RIMS1;refGene_tx=
XM_023254096.1;refGene_gn=RIMS1;refGene_tx=XM_023254097.1;refGene_gn=R
IMS1;refGene_tx=XM_023254098.1;refGene_gn=RIMS1;refGene_tx=XM_02325409
9.1;refGene_gn=RIMS1;refGene_tx=XM_023254100.1;refGene_gn=RIMS1;refGen
e_tx=XM_023254101.1;refGene_gn=RIMS1;refGene_tx=XM_023254102.1;refGene
_gn=RIMS1;refGene_tx=XM_023254103.1;refGene_gn=RIMS1;refGene_tx=XM_023
254104.1;refGene_gn=RIMS1;refGene_tx=XM_023254105.1;refGene_gn=RIMS1;r
efGene_tx=XM_023254106.1;refGene_gn=RIMS1;refGene_tx=XM_023254107.1;re
fGene_gn=RIMS1;refGene_tx=XM_023254108.1;refGene_gn=RIMS1;refGene_tx=X
M_023254109.1;refGene_gn=RIMS1;refGene_tx=XM_023254110.1;refGene_gn=RI
MS1;refGene_tx=XM_023254111.1;refGene_gn=RIMS1;refGene_tx=XM_023254112
.
1;refGene_gn=RIMS1;refGene_tx=XM_023254113.1;refGene_gn=RIMS1;refGene_
tx=XM_023254114.1;refGene_gn=RIMS1;refGene_tx=XM_023254115.1;refGene_g
n=RIMS1;refGene_tx=XM_023254117.1;refGene_gn=RIMS1;refGene_tx=XM_02325
4118.1;refGene_gn=RIMS1;refGene_tx=XM_023254119.1;refGene_gn=RIMS1;ref
Gene_tx=XM_023254120.1;refGene_gn=RIMS1;refGene_tx=XM_023254121.1;refG
ene_gn=RIMS1;refGene_tx=XM_023254122.1;refGene_gn=RIMS1;refGene_tx=XM_
023254123.1;refGene_gn=RIMS1;refGene_tx=XM_023254124.1;refGene_gn=RIMS
1;refGene_tx=XM_023254125.1;refGene_gn=RIMS1;refGene_tx=XM_023254126.1
;refGene_gn=RIMS1;refGene_tx=XM_023254127.1;refGene_gn=RIMS1;refGene_t
x=XM_023254128.1;refGene_gn=RIMS1;refGene_tx=XM_023254129.1;refGene_gn
=RIMS1;refGene_tx=XM_023254131.1;refGene_gn=RIMS1;refGene_tx=XM_023254
132.1;refGene_gn=RIMS1;refGene_tx=XM_023254133.1
B2 68223218 68223437
refGene_gn=RIMS1;refGene_tx=XM_019830824.2;refGene_gn=RIMS1;refGene_tx
=XM_019830825.2;refGene_gn=RIMS1;refGene_tx=XM_019830833.2;refGene_gn=
RIMS1;refGene_tx=XM_019830841.2;refGene_gn=RIMS1;refGene_tx=XM_0198308
45.2;refGene_gn=RIMS1;refGene_tx=XM_019830848.1;refGene_gn=RIMS1;refGe
ne_tx=XM_019830850.2;refGene_gn=RIMS1;refGene_tx=XM_019830852.2;refGen
e_gn=RIMS1;refGene_tx=XM_019830857.1;refGene_gn=RIMS1;refGene_tx=XM_01
9830858.1;refGene_gn=RIMS1;refGene_tx=XM_019830859.2;refGene_gn=RIMS1;
refGene_tx=XM_023254093.1;refGene_gn=RIMS1;refGene_tx=XM_023254094.1;r
efGene_gn=RIMS1;refGene_tx=XM_023254095.1;refGene_gn=RIMS1;refGene_tx=
XM_023254096.1;refGene_gn=RIMS1;refGene_tx=XM_023254097.1;refGene_gn=R
IMS1;refGene_tx=XM_023254098.1;refGene_gn=RIMS1;refGene_tx=XM_02325409
9.1;refGene_gn=RIMS1;refGene_tx=XM_023254100.1;refGene_gn=RIMS1;refGen
e_tx=XM_023254101.1;refGene_gn=RIMS1;refGene_tx=XM_023254102.1;refGene
_gn=RIMS1;refGene_tx=XM_023254103.1;refGene_gn=RIMS1;refGene_tx=XM_023
254104.1;refGene_gn=RIMS1;refGene_tx=XM_023254105.1;refGene_gn=RIMS1;r
efGene_tx=XM_023254106.1;refGene_gn=RIMS1;refGene_tx=XM_023254107.1;re
fGene_gn=RIMS1;refGene_tx=XM_023254108.1;refGene_gn=RIMS1;refGene_tx=X

M_023254109.1;refGene_gn=RIMS1;refGene_tx=XM_023254110.1;refGene_gn=RIMS1;refGene_tx=XM_023254111.1;refGene_gn=RIMS1;refGene_tx=XM_023254112.
1;refGene_gn=RIMS1;refGene_tx=XM_023254113.1;refGene_gn=RIMS1;refGene_tx=XM_023254114.1;refGene_gn=RIMS1;refGene_tx=XM_023254115.1;refGene_gn=RIMS1;refGene_tx=XM_023254117.1;refGene_gn=RIMS1;refGene_tx=XM_023254118.1;refGene_gn=RIMS1;refGene_tx=XM_023254119.1;refGene_gn=RIMS1;refGene_tx=XM_023254120.1;refGene_gn=RIMS1;refGene_tx=XM_023254121.1;refGene_gn=RIMS1;refGene_tx=XM_023254122.1;refGene_gn=RIMS1;refGene_tx=XM_023254123.1;refGene_gn=RIMS1;refGene_tx=XM_023254124.1;refGene_gn=RIMS1;refGene_tx=XM_023254125.1;refGene_gn=RIMS1;refGene_tx=XM_023254126.1;refGene_gn=RIMS1;refGene_tx=XM_023254127.1;refGene_gn=RIMS1;refGene_tx=XM_023254128.1;refGene_gn=RIMS1;refGene_tx=XM_023254129.1;refGene_gn=RIMS1;refGene_tx=XM_023254131.1;refGene_gn=RIMS1;refGene_tx=XM_023254132.1;refGene_gn=RIMS1;refGene_tx=XM_023254133.1
B2 68439019 68439438
refGene_gn=KCNQ5;refGene_tx=XM_011282435.2;refGene_gn=KCNQ5;refGene_tx=XM_023254134.1;refGene_gn=KCNQ5;refGene_tx=XM_023254135.1
B2 68762555 68762581 refGene_gn=KCNQ5;refGene_tx=XM_023254136.1
B2 68779428 68779519
refGene_gn=KCNQ5;refGene_tx=XM_011282435.2;refGene_gn=KCNQ5;refGene_tx=XM_023254134.1;refGene_gn=KCNQ5;refGene_tx=XM_023254135.1;refGene_gn=KCNQ5;refGene_tx=XM_023254136.1
B2 68811865 68811992
refGene_gn=KCNQ5;refGene_tx=XM_011282435.2;refGene_gn=KCNQ5;refGene_tx=XM_023254134.1;refGene_gn=KCNQ5;refGene_tx=XM_023254135.1;refGene_gn=KCNQ5;refGene_tx=XM_023254136.1
B2 68846731 68846907
refGene_gn=KCNQ5;refGene_tx=XM_011282435.2;refGene_gn=KCNQ5;refGene_tx=XM_023254134.1;refGene_gn=KCNQ5;refGene_tx=XM_023254135.1;refGene_gn=KCNQ5;refGene_tx=XM_023254136.1
B2 68847787 68847913
refGene_gn=KCNQ5;refGene_tx=XM_011282435.2;refGene_gn=KCNQ5;refGene_tx=XM_023254134.1;refGene_gn=KCNQ5;refGene_tx=XM_023254135.1;refGene_gn=KCNQ5;refGene_tx=XM_023254136.1
B2 68861266 68861377
refGene_gn=KCNQ5;refGene_tx=XM_011282435.2;refGene_gn=KCNQ5;refGene_tx=XM_023254134.1;refGene_gn=KCNQ5;refGene_tx=XM_023254135.1;refGene_gn=KCNQ5;refGene_tx=XM_023254136.1
B2 68867618 68867714
refGene_gn=KCNQ5;refGene_tx=XM_011282435.2;refGene_gn=KCNQ5;refGene_tx=XM_023254134.1;refGene_gn=KCNQ5;refGene_tx=XM_023254135.1;refGene_gn=KCNQ5;refGene_tx=XM_023254136.1
B2 68879831 68879926
refGene_gn=KCNQ5;refGene_tx=XM_011282435.2;refGene_gn=KCNQ5;refGene_tx=XM_023254134.1;refGene_gn=KCNQ5;refGene_tx=XM_023254135.1;refGene_gn=KCNQ5;refGene_tx=XM_023254136.1
B2 68883909 68883936
refGene_gn=KCNQ5;refGene_tx=XM_011282435.2;refGene_gn=KCNQ5;refGene_tx=XM_023254134.1;refGene_gn=KCNQ5;refGene_tx=XM_023254136.1

B2 68888748 68888805
refGene_gn=KCNQ5;refGene_tx=XM_011282435.2;refGene_gn=KCNQ5;refGene_tx=XM_023254136.1
B2 68892380 68892601
refGene_gn=KCNQ5;refGene_tx=XM_011282435.2;refGene_gn=KCNQ5;refGene_tx=XM_023254134.1;refGene_gn=KCNQ5;refGene_tx=XM_023254135.1;refGene_gn=KCNQ5;refGene_tx=XM_023254136.1
B2 68927582 68927691
refGene_gn=KCNQ5;refGene_tx=XM_011282435.2;refGene_gn=KCNQ5;refGene_tx=XM_023254134.1;refGene_gn=KCNQ5;refGene_tx=XM_023254135.1;refGene_gn=KCNQ5;refGene_tx=XM_023254136.1
B2 68946517 68946649
refGene_gn=KCNQ5;refGene_tx=XM_011282435.2;refGene_gn=KCNQ5;refGene_tx=XM_023254134.1;refGene_gn=KCNQ5;refGene_tx=XM_023254135.1;refGene_gn=KCNQ5;refGene_tx=XM_023254136.1
B2 68948476 68948603
refGene_gn=KCNQ5;refGene_tx=XM_011282435.2;refGene_gn=KCNQ5;refGene_tx=XM_023254134.1;refGene_gn=KCNQ5;refGene_tx=XM_023254135.1;refGene_gn=KCNQ5;refGene_tx=XM_023254136.1
B2 68950129 68951092
refGene_gn=KCNQ5;refGene_tx=XM_011282435.2;refGene_gn=KCNQ5;refGene_tx=XM_023254134.1;refGene_gn=KCNQ5;refGene_tx=XM_023254135.1;refGene_gn=KCNQ5;refGene_tx=XM_023254136.1
B2 69029309 69029368 refGene_gn=DPPA5;refGene_tx=XM_003986310.4
B2 69029792 69029972 refGene_gn=DPPA5;refGene_tx=XM_003986310.4
B2 69030045 69030157 refGene_gn=DPPA5;refGene_tx=XM_003986310.4
B2 69045865 69046034 refGene_gn=KHDC3L;refGene_tx=XM_023254750.1
B2 69046225 69046461 refGene_gn=KHDC3L;refGene_tx=XM_023254750.1
B2 69053280 69053360 refGene_gn=OOEP;refGene_tx=NM_001077245.1
B2 69053728 69053908 refGene_gn=OOEP;refGene_tx=NM_001077245.1
B2 69054112 69054302 refGene_gn=OOEP;refGene_tx=NM_001077245.1
B2 69083732 69083982
refGene_gn=DDX43;refGene_tx=XM_003986327.4;refGene_gn=DDX43;refGene_tx=XM_019830862.2
B2 69085689 69085745
refGene_gn=DDX43;refGene_tx=XM_003986327.4;refGene_gn=DDX43;refGene_tx=XM_019830862.2
B2 69087684 69087811
refGene_gn=DDX43;refGene_tx=XM_003986327.4;refGene_gn=DDX43;refGene_tx=XM_019830862.2
B2 69088877 69089009 refGene_gn=DDX43;refGene_tx=XM_003986327.4
B2 69090806 69090888
refGene_gn=DDX43;refGene_tx=XM_003986327.4;refGene_gn=DDX43;refGene_tx=XM_019830862.2
B2 69091750 69091907
refGene_gn=DDX43;refGene_tx=XM_003986327.4;refGene_gn=DDX43;refGene_tx=XM_019830862.2
B2 69092119 69092238
refGene_gn=DDX43;refGene_tx=XM_003986327.4;refGene_gn=DDX43;refGene_tx=XM_019830862.2

B2	69092691 69092802
refGene_gn=DDX43;refGene_tx=XM_003986327.4;refGene_gn=DDX43;refGene_tx=XM_019830862.2
B2	69093170 69093312
refGene_gn=DDX43;refGene_tx=XM_003986327.4;refGene_gn=DDX43;refGene_tx=XM_019830862.2
B2	69094231 69094332
refGene_gn=DDX43;refGene_tx=XM_003986327.4;refGene_gn=DDX43;refGene_tx=XM_019830862.2
B2	69095504 69095592
refGene_gn=DDX43;refGene_tx=XM_003986327.4;refGene_gn=DDX43;refGene_tx=XM_019830862.2
B2	69096584 69096712
refGene_gn=DDX43;refGene_tx=XM_003986327.4;refGene_gn=DDX43;refGene_tx=XM_019830862.2
B2	69096942 69097052
refGene_gn=DDX43;refGene_tx=XM_003986327.4;refGene_gn=DDX43;refGene_tx=XM_019830862.2
B2	69097818 69097957
refGene_gn=DDX43;refGene_tx=XM_003986327.4;refGene_gn=DDX43;refGene_tx=XM_019830862.2
B2	69098600 69098688
refGene_gn=DDX43;refGene_tx=XM_003986327.4;refGene_gn=DDX43;refGene_tx=XM_019830862.2
B2	69099371 69099485
refGene_gn=DDX43;refGene_tx=XM_003986327.4;refGene_gn=DDX43;refGene_tx=XM_019830862.2
B2	69105920 69106272 refGene_gn=CGAS;refGene_tx=XM_006931843.3
B2	69108576 69108679 refGene_gn=CGAS;refGene_tx=XM_006931843.3
B2	69116659 69116896 refGene_gn=CGAS;refGene_tx=XM_006931843.3
B2	69122232 69122452 refGene_gn=CGAS;refGene_tx=XM_006931843.3
B2	69132401 69133034 refGene_gn=CGAS;refGene_tx=XM_006931843.3
B2	69142203 69142353
refGene_gn=LOC101092856;refGene_tx=XM_003986311.5;refGene_gn=LOC101092856;refGene_tx=XM_006931845.4;refGene_gn=LOC101092856;refGene_tx=XM_006931846.4;refGene_gn=LOC101092856;refGene_tx=XM_019830864.2
B2	69142433 69142551
refGene_gn=LOC101092856;refGene_tx=XM_003986311.5;refGene_gn=LOC101092856;refGene_tx=XM_006931845.4;refGene_gn=LOC101092856;refGene_tx=XM_006931846.4;refGene_gn=LOC101092856;refGene_tx=XM_019830864.2
B2	69144425 69144715
refGene_gn=LOC101092856;refGene_tx=XM_003986311.5;refGene_gn=LOC101092856;refGene_tx=XM_006931845.4;refGene_gn=LOC101092856;refGene_tx=XM_006931846.4;refGene_gn=LOC101092856;refGene_tx=XM_019830864.2
B2	69146658 69146771
refGene_gn=LOC101092856;refGene_tx=XM_003986311.5;refGene_gn=LOC101092856;refGene_tx=XM_006931845.4;refGene_gn=LOC101092856;refGene_tx=XM_006931846.4;refGene_gn=LOC101092856;refGene_tx=XM_019830864.2
B2	69146858 69147049
refGene_gn=LOC101092856;refGene_tx=XM_003986311.5;refGene_gn=LOC101092

856;refGene_tx=XM_006931845.4;refGene_gn=LOC101092856;refGene_tx=XM_006931846.4;refGene_gn=LOC101092856;refGene_tx=XM_019830864.2
B2 69147285 69147416
refGene_gn=LOC101092856;refGene_tx=XM_003986311.5;refGene_gn=LOC101092856;refGene_tx=XM_006931845.4;refGene_gn=LOC101092856;refGene_tx=XM_006931846.4;refGene_gn=LOC101092856;refGene_tx=XM_019830864.2
B2 69148358 69148563
refGene_gn=LOC101092856;refGene_tx=XM_003986311.5;refGene_gn=LOC101092856;refGene_tx=XM_006931845.4;refGene_gn=LOC101092856;refGene_tx=XM_006931846.4;refGene_gn=LOC101092856;refGene_tx=XM_019830864.2
B2 69148642 69148814
refGene_gn=LOC101092856;refGene_tx=XM_003986311.5;refGene_gn=LOC101092856;refGene_tx=XM_006931845.4;refGene_gn=LOC101092856;refGene_tx=XM_006931846.4;refGene_gn=LOC101092856;refGene_tx=XM_019830864.2
B2 69153760 69153879
refGene_gn=LOC101092856;refGene_tx=XM_003986311.5;refGene_gn=LOC101092856;refGene_tx=XM_006931845.4;refGene_gn=LOC101092856;refGene_tx=XM_006931846.4;refGene_gn=LOC101092856;refGene_tx=XM_019830864.2
B2 69154842 69154967
refGene_gn=LOC101092856;refGene_tx=XM_019830864.2
B2 69157440 69157601
refGene_gn=LOC101092856;refGene_tx=XM_003986311.5;refGene_gn=LOC101092856;refGene_tx=XM_006931845.4
B2 69159378 69159540
refGene_gn=LOC101092856;refGene_tx=XM_003986311.5;refGene_gn=LOC101092856;refGene_tx=XM_006931846.4
B2 69174169 69174294
refGene_gn=EEF1A1;refGene_tx=NM_001009326.1;refGene_gn=EEF1A1;refGene_tx=XM_006931840.4;refGene_gn=EEF1A1;refGene_tx=XM_019830069.2
B2 69174388 69174623
refGene_gn=EEF1A1;refGene_tx=NM_001009326.1;refGene_gn=EEF1A1;refGene_tx=XM_006931840.4;refGene_gn=EEF1A1;refGene_tx=XM_019830069.2
B2 69174709 69174966
refGene_gn=EEF1A1;refGene_tx=NM_001009326.1;refGene_gn=EEF1A1;refGene_tx=XM_006931840.4;refGene_gn=EEF1A1;refGene_tx=XM_019830069.2
B2 69175058 69175209
refGene_gn=EEF1A1;refGene_tx=NM_001009326.1;refGene_gn=EEF1A1;refGene_tx=XM_006931840.4;refGene_gn=EEF1A1;refGene_tx=XM_019830069.2
B2 69175309 69175606
refGene_gn=EEF1A1;refGene_tx=NM_001009326.1;refGene_gn=EEF1A1;refGene_tx=XM_006931840.4;refGene_gn=EEF1A1;refGene_tx=XM_019830069.2
B2 69175724 69175904
refGene_gn=EEF1A1;refGene_tx=NM_001009326.1;refGene_gn=EEF1A1;refGene_tx=XM_006931840.4;refGene_gn=EEF1A1;refGene_tx=XM_019830069.2
B2 69176177 69176321
refGene_gn=EEF1A1;refGene_tx=NM_001009326.1;refGene_gn=EEF1A1;refGene_tx=XM_006931840.4;refGene_gn=EEF1A1;refGene_tx=XM_019830069.2
B2 69239636 69239762 refGene_gn=SLC17A5;refGene_tx=XM_023254143.1
B2 69243496 69243634
refGene_gn=SLC17A5;refGene_tx=XM_003986312.4;refGene_gn=SLC17A5;refGen

e_tx=XM_019830866.2
B2 69247453 69247544
refGene_gn=SLC17A5;refGene_tx=XM_003986312.4;refGene_gn=SLC17A5;refGene_tx=XM_019830866.2;refGene_gn=SLC17A5;refGene_tx=XM_023254143.1
B2 69264976 69265124
refGene_gn=SLC17A5;refGene_tx=XM_003986312.4;refGene_gn=SLC17A5;refGene_tx=XM_019830866.2;refGene_gn=SLC17A5;refGene_tx=XM_023254143.1
B2 69266758 69266891
refGene_gn=SLC17A5;refGene_tx=XM_003986312.4;refGene_gn=SLC17A5;refGene_tx=XM_019830866.2;refGene_gn=SLC17A5;refGene_tx=XM_023254143.1
B2 69271621 69271780
refGene_gn=SLC17A5;refGene_tx=XM_003986312.4;refGene_gn=SLC17A5;refGene_tx=XM_019830866.2;refGene_gn=SLC17A5;refGene_tx=XM_023254143.1
B2 69276214 69276333
refGene_gn=SLC17A5;refGene_tx=XM_003986312.4;refGene_gn=SLC17A5;refGene_tx=XM_019830866.2;refGene_gn=SLC17A5;refGene_tx=XM_023254143.1
B2 69277121 69277208
refGene_gn=SLC17A5;refGene_tx=XM_003986312.4;refGene_gn=SLC17A5;refGene_tx=XM_019830866.2;refGene_gn=SLC17A5;refGene_tx=XM_023254143.1
B2 69279591 69279679
refGene_gn=SLC17A5;refGene_tx=XM_003986312.4;refGene_gn=SLC17A5;refGene_tx=XM_019830866.2;refGene_gn=SLC17A5;refGene_tx=XM_023254143.1
B2 69283103 69283337
refGene_gn=SLC17A5;refGene_tx=XM_003986312.4;refGene_gn=SLC17A5;refGene_tx=XM_019830866.2;refGene_gn=SLC17A5;refGene_tx=XM_023254143.1
B2 69285063 69285260
refGene_gn=SLC17A5;refGene_tx=XM_003986312.4;refGene_gn=SLC17A5;refGene_tx=XM_019830866.2;refGene_gn=SLC17A5;refGene_tx=XM_023254143.1
B2 69285783 69285826 refGene_gn=SLC17A5;refGene_tx=XM_019830866.2
B2 69292262 69292356
refGene_gn=SLC17A5;refGene_tx=XM_003986312.4;refGene_gn=SLC17A5;refGene_tx=XM_023254143.1
B2 69315952 69315977 refGene_gn=CD109;refGene_tx=XM_023254142.1
B2 69321997 69322071
refGene_gn=CD109;refGene_tx=XM_023254138.1;refGene_gn=CD109;refGene_tx=XM_023254139.1
B2 69323161 69323334
refGene_gn=CD109;refGene_tx=XM_023254138.1;refGene_gn=CD109;refGene_tx=XM_023254139.1;refGene_gn=CD109;refGene_tx=XM_023254140.1;refGene_gn=CD109;refGene_tx=XM_023254141.1
B2 69343507 69343536
refGene_gn=CD109;refGene_tx=XM_023254138.1;refGene_gn=CD109;refGene_tx=XM_023254139.1;refGene_gn=CD109;refGene_tx=XM_023254140.1;refGene_gn=CD109;refGene_tx=XM_023254141.1;refGene_gn=CD109;refGene_tx=XM_023254142.1
B2 69350995 69351226
refGene_gn=CD109;refGene_tx=XM_023254138.1;refGene_gn=CD109;refGene_tx=XM_023254139.1;refGene_gn=CD109;refGene_tx=XM_023254140.1;refGene_gn=CD109;refGene_tx=XM_023254141.1;refGene_gn=CD109;refGene_tx=XM_023254142.1

B2 69356578 69356704
refGene_gn=CD109;refGene_tx=XM_023254138.1;refGene_gn=CD109;refGene_tx=XM_023254139.1;refGene_gn=CD109;refGene_tx=XM_023254140.1;refGene_gn=CD109;refGene_tx=XM_023254141.1;refGene_gn=CD109;refGene_tx=XM_023254142.1
B2 69382348 69382388
refGene_gn=CD109;refGene_tx=XM_023254138.1;refGene_gn=CD109;refGene_tx=XM_023254139.1;refGene_gn=CD109;refGene_tx=XM_023254140.1;refGene_gn=CD109;refGene_tx=XM_023254141.1;refGene_gn=CD109;refGene_tx=XM_023254142.1
B2 69385343 69385428
refGene_gn=CD109;refGene_tx=XM_023254138.1;refGene_gn=CD109;refGene_tx=XM_023254139.1;refGene_gn=CD109;refGene_tx=XM_023254140.1;refGene_gn=CD109;refGene_tx=XM_023254141.1;refGene_gn=CD109;refGene_tx=XM_023254142.1
B2 69389893 69389990
refGene_gn=CD109;refGene_tx=XM_023254138.1;refGene_gn=CD109;refGene_tx=XM_023254139.1;refGene_gn=CD109;refGene_tx=XM_023254140.1;refGene_gn=CD109;refGene_tx=XM_023254141.1;refGene_gn=CD109;refGene_tx=XM_023254142.1
B2 69390258 69390400
refGene_gn=CD109;refGene_tx=XM_023254138.1;refGene_gn=CD109;refGene_tx=XM_023254139.1;refGene_gn=CD109;refGene_tx=XM_023254140.1;refGene_gn=CD109;refGene_tx=XM_023254141.1;refGene_gn=CD109;refGene_tx=XM_023254142.1
B2 69391099 69391209
refGene_gn=CD109;refGene_tx=XM_023254138.1;refGene_gn=CD109;refGene_tx=XM_023254139.1;refGene_gn=CD109;refGene_tx=XM_023254140.1;refGene_gn=CD109;refGene_tx=XM_023254141.1;refGene_gn=CD109;refGene_tx=XM_023254142.1
B2 69393762 69393987
refGene_gn=CD109;refGene_tx=XM_023254138.1;refGene_gn=CD109;refGene_tx=XM_023254139.1;refGene_gn=CD109;refGene_tx=XM_023254140.1;refGene_gn=CD109;refGene_tx=XM_023254141.1;refGene_gn=CD109;refGene_tx=XM_023254142.1
B2 69394843 69394945
refGene_gn=CD109;refGene_tx=XM_023254138.1;refGene_gn=CD109;refGene_tx=XM_023254139.1;refGene_gn=CD109;refGene_tx=XM_023254140.1;refGene_gn=CD109;refGene_tx=XM_023254141.1;refGene_gn=CD109;refGene_tx=XM_023254142.1
B2 69395033 69395099
refGene_gn=CD109;refGene_tx=XM_023254138.1;refGene_gn=CD109;refGene_tx=XM_023254139.1;refGene_gn=CD109;refGene_tx=XM_023254140.1;refGene_gn=CD109;refGene_tx=XM_023254141.1;refGene_gn=CD109;refGene_tx=XM_023254142.1
B2 69395969 69396146
refGene_gn=CD109;refGene_tx=XM_023254138.1;refGene_gn=CD109;refGene_tx=XM_023254139.1;refGene_gn=CD109;refGene_tx=XM_023254140.1;refGene_gn=CD109;refGene_tx=XM_023254141.1;refGene_gn=CD109;refGene_tx=XM_023254142.1

B2 69399364 69399517
refGene_gn=CD109;refGene_tx=XM_023254138.1;refGene_gn=CD109;refGene_tx=XM_023254139.1;refGene_gn=CD109;refGene_tx=XM_023254140.1;refGene_gn=CD109;refGene_tx=XM_023254141.1;refGene_gn=CD109;refGene_tx=XM_023254142.1
B2 69405734 69405809
refGene_gn=CD109;refGene_tx=XM_023254138.1;refGene_gn=CD109;refGene_tx=XM_023254139.1;refGene_gn=CD109;refGene_tx=XM_023254140.1;refGene_gn=CD109;refGene_tx=XM_023254141.1;refGene_gn=CD109;refGene_tx=XM_023254142.1
B2 69406239 69406300
refGene_gn=CD109;refGene_tx=XM_023254138.1;refGene_gn=CD109;refGene_tx=XM_023254139.1;refGene_gn=CD109;refGene_tx=XM_023254140.1;refGene_gn=CD109;refGene_tx=XM_023254141.1;refGene_gn=CD109;refGene_tx=XM_023254142.1
B2 69409749 69409891
refGene_gn=CD109;refGene_tx=XM_023254138.1;refGene_gn=CD109;refGene_tx=XM_023254139.1;refGene_gn=CD109;refGene_tx=XM_023254140.1;refGene_gn=CD109;refGene_tx=XM_023254141.1;refGene_gn=CD109;refGene_tx=XM_023254142.1
B2 69410795 69410913
refGene_gn=CD109;refGene_tx=XM_023254138.1;refGene_gn=CD109;refGene_tx=XM_023254139.1;refGene_gn=CD109;refGene_tx=XM_023254140.1;refGene_gn=CD109;refGene_tx=XM_023254141.1;refGene_gn=CD109;refGene_tx=XM_023254142.1
B2 69413683 69413797
refGene_gn=CD109;refGene_tx=XM_023254138.1;refGene_gn=CD109;refGene_tx=XM_023254139.1;refGene_gn=CD109;refGene_tx=XM_023254140.1;refGene_gn=CD109;refGene_tx=XM_023254141.1;refGene_gn=CD109;refGene_tx=XM_023254142.1
B2 69415207 69415426
refGene_gn=CD109;refGene_tx=XM_023254138.1;refGene_gn=CD109;refGene_tx=XM_023254139.1;refGene_gn=CD109;refGene_tx=XM_023254140.1;refGene_gn=CD109;refGene_tx=XM_023254141.1;refGene_gn=CD109;refGene_tx=XM_023254142.1
B2 69416680 69416825
refGene_gn=CD109;refGene_tx=XM_023254138.1;refGene_gn=CD109;refGene_tx=XM_023254139.1;refGene_gn=CD109;refGene_tx=XM_023254140.1;refGene_gn=CD109;refGene_tx=XM_023254141.1;refGene_gn=CD109;refGene_tx=XM_023254142.1
B2 69420188 69420365
refGene_gn=CD109;refGene_tx=XM_023254138.1;refGene_gn=CD109;refGene_tx=XM_023254139.1;refGene_gn=CD109;refGene_tx=XM_023254140.1;refGene_gn=CD109;refGene_tx=XM_023254141.1;refGene_gn=CD109;refGene_tx=XM_023254142.1
B2 69429884 69429966
refGene_gn=CD109;refGene_tx=XM_023254138.1;refGene_gn=CD109;refGene_tx=XM_023254139.1;refGene_gn=CD109;refGene_tx=XM_023254140.1;refGene_gn=CD109;refGene_tx=XM_023254141.1;refGene_gn=CD109;refGene_tx=XM_023254142.1

B2 69432136 69432365
refGene_gn=CD109;refGene_tx=XM_023254138.1;refGene_gn=CD109;refGene_tx=XM_023254139.1;refGene_gn=CD109;refGene_tx=XM_023254140.1;refGene_gn=CD109;refGene_tx=XM_023254141.1;refGene_gn=CD109;refGene_tx=XM_023254142.1
B2 69435429 69435595
refGene_gn=CD109;refGene_tx=XM_023254138.1;refGene_gn=CD109;refGene_tx=XM_023254139.1;refGene_gn=CD109;refGene_tx=XM_023254140.1;refGene_gn=CD109;refGene_tx=XM_023254141.1;refGene_gn=CD109;refGene_tx=XM_023254142.1
B2 69437360 69437551
refGene_gn=CD109;refGene_tx=XM_023254138.1;refGene_gn=CD109;refGene_tx=XM_023254139.1;refGene_gn=CD109;refGene_tx=XM_023254140.1;refGene_gn=CD109;refGene_tx=XM_023254141.1;refGene_gn=CD109;refGene_tx=XM_023254142.1
B2 69438222 69438378
refGene_gn=CD109;refGene_tx=XM_023254138.1;refGene_gn=CD109;refGene_tx=XM_023254139.1;refGene_gn=CD109;refGene_tx=XM_023254140.1;refGene_gn=CD109;refGene_tx=XM_023254141.1;refGene_gn=CD109;refGene_tx=XM_023254142.1
B2 69439427 69439493
refGene_gn=CD109;refGene_tx=XM_023254138.1;refGene_gn=CD109;refGene_tx=XM_023254139.1;refGene_gn=CD109;refGene_tx=XM_023254140.1;refGene_gn=CD109;refGene_tx=XM_023254141.1;refGene_gn=CD109;refGene_tx=XM_023254142.1
B2 69443050 69443193
refGene_gn=CD109;refGene_tx=XM_023254138.1;refGene_gn=CD109;refGene_tx=XM_023254139.1;refGene_gn=CD109;refGene_tx=XM_023254140.1;refGene_gn=CD109;refGene_tx=XM_023254141.1;refGene_gn=CD109;refGene_tx=XM_023254142.1
B2 69446580 69446728
refGene_gn=CD109;refGene_tx=XM_023254138.1;refGene_gn=CD109;refGene_tx=XM_023254139.1;refGene_gn=CD109;refGene_tx=XM_023254140.1;refGene_gn=CD109;refGene_tx=XM_023254141.1;refGene_gn=CD109;refGene_tx=XM_023254142.1
B2 69448384 69448487
refGene_gn=CD109;refGene_tx=XM_023254138.1;refGene_gn=CD109;refGene_tx=XM_023254139.1;refGene_gn=CD109;refGene_tx=XM_023254140.1;refGene_gn=CD109;refGene_tx=XM_023254141.1;refGene_gn=CD109;refGene_tx=XM_023254142.1
B2 69450633 69450809
refGene_gn=CD109;refGene_tx=XM_023254138.1;refGene_gn=CD109;refGene_tx=XM_023254140.1;refGene_gn=CD109;refGene_tx=XM_023254141.1;refGene_gn=CD109;refGene_tx=XM_023254142.1
B2 69452350 69452370 refGene_gn=CD109;refGene_tx=XM_023254139.1
B2 70408498 70408674
refGene_gn=COL12A1;refGene_tx=XM_006931849.4;refGene_gn=COL12A1;refGene_tx=XM_006931850.4;refGene_gn=COL12A1;refGene_tx=XM_019830867.2;refGene_gn=COL12A1;refGene_tx=XM_019830868.2
B2 70409847 70409858 refGene_gn=COL12A1;refGene_tx=XM_019830870.1

B2 70410877 70411048
refGene_gn=COL12A1;refGene_tx=XM_006931849.4;refGene_gn=COL12A1;refGene_tx=XM_006931850.4;refGene_gn=COL12A1;refGene_tx=XM_019830867.2;refGene_gn=COL12A1;refGene_tx=XM_019830868.2;refGene_gn=COL12A1;refGene_tx=XM_019830870.1
B2 70412067 70412136
refGene_gn=COL12A1;refGene_tx=XM_006931849.4;refGene_gn=COL12A1;refGene_tx=XM_006931850.4;refGene_gn=COL12A1;refGene_tx=XM_019830867.2;refGene_gn=COL12A1;refGene_tx=XM_019830868.2;refGene_gn=COL12A1;refGene_tx=XM_019830870.1
B2 70413107 70413296
refGene_gn=COL12A1;refGene_tx=XM_006931849.4;refGene_gn=COL12A1;refGene_tx=XM_006931850.4;refGene_gn=COL12A1;refGene_tx=XM_019830867.2;refGene_gn=COL12A1;refGene_tx=XM_019830868.2;refGene_gn=COL12A1;refGene_tx=XM_019830870.1
B2 70414396 70414463
refGene_gn=COL12A1;refGene_tx=XM_006931849.4;refGene_gn=COL12A1;refGene_tx=XM_006931850.4;refGene_gn=COL12A1;refGene_tx=XM_019830867.2;refGene_gn=COL12A1;refGene_tx=XM_019830868.2;refGene_gn=COL12A1;refGene_tx=XM_019830870.1
B2 70414572 70414608
refGene_gn=COL12A1;refGene_tx=XM_006931849.4;refGene_gn=COL12A1;refGene_tx=XM_006931850.4;refGene_gn=COL12A1;refGene_tx=XM_019830867.2;refGene_gn=COL12A1;refGene_tx=XM_019830868.2;refGene_gn=COL12A1;refGene_tx=XM_019830870.1
B2 70417808 70417880
refGene_gn=COL12A1;refGene_tx=XM_006931849.4;refGene_gn=COL12A1;refGene_tx=XM_006931850.4;refGene_gn=COL12A1;refGene_tx=XM_019830867.2;refGene_gn=COL12A1;refGene_tx=XM_019830868.2;refGene_gn=COL12A1;refGene_tx=XM_019830870.1
B2 70419289 70419343
refGene_gn=COL12A1;refGene_tx=XM_006931849.4;refGene_gn=COL12A1;refGene_tx=XM_006931850.4;refGene_gn=COL12A1;refGene_tx=XM_019830867.2;refGene_gn=COL12A1;refGene_tx=XM_019830868.2;refGene_gn=COL12A1;refGene_tx=XM_019830870.1
B2 70422330 70422384
refGene_gn=COL12A1;refGene_tx=XM_006931849.4;refGene_gn=COL12A1;refGene_tx=XM_006931850.4;refGene_gn=COL12A1;refGene_tx=XM_019830867.2;refGene_gn=COL12A1;refGene_tx=XM_019830868.2;refGene_gn=COL12A1;refGene_tx=XM_019830870.1
B2 70422741 70422795
refGene_gn=COL12A1;refGene_tx=XM_006931849.4;refGene_gn=COL12A1;refGene_tx=XM_006931850.4;refGene_gn=COL12A1;refGene_tx=XM_019830867.2;refGene_gn=COL12A1;refGene_tx=XM_019830868.2;refGene_gn=COL12A1;refGene_tx=XM_019830870.1
B2 70424236 70424332
refGene_gn=COL12A1;refGene_tx=XM_006931849.4;refGene_gn=COL12A1;refGene_tx=XM_006931850.4;refGene_gn=COL12A1;refGene_tx=XM_019830867.2;refGene_gn=COL12A1;refGene_tx=XM_019830868.2;refGene_gn=COL12A1;refGene_tx=XM_019830870.1

B2 70425425 70425479
refGene_gn=COL12A1;refGene_tx=XM_006931849.4;refGene_gn=COL12A1;refGene_tx=XM_006931850.4;refGene_gn=COL12A1;refGene_tx=XM_019830867.2;refGene_gn=COL12A1;refGene_tx=XM_019830868.2;refGene_gn=COL12A1;refGene_tx=XM_019830870.1
B2 70426881 70426968
refGene_gn=COL12A1;refGene_tx=XM_006931849.4;refGene_gn=COL12A1;refGene_tx=XM_006931850.4;refGene_gn=COL12A1;refGene_tx=XM_019830867.2;refGene_gn=COL12A1;refGene_tx=XM_019830868.2;refGene_gn=COL12A1;refGene_tx=XM_019830870.1
B2 70428286 70428364
refGene_gn=COL12A1;refGene_tx=XM_006931849.4;refGene_gn=COL12A1;refGene_tx=XM_006931850.4;refGene_gn=COL12A1;refGene_tx=XM_019830867.2;refGene_gn=COL12A1;refGene_tx=XM_019830868.2;refGene_gn=COL12A1;refGene_tx=XM_019830870.1
B2 70430428 70430578
refGene_gn=COL12A1;refGene_tx=XM_006931849.4;refGene_gn=COL12A1;refGene_tx=XM_006931850.4;refGene_gn=COL12A1;refGene_tx=XM_019830867.2;refGene_gn=COL12A1;refGene_tx=XM_019830868.2;refGene_gn=COL12A1;refGene_tx=XM_019830870.1
B2 70434193 70434303
refGene_gn=COL12A1;refGene_tx=XM_006931849.4;refGene_gn=COL12A1;refGene_tx=XM_006931850.4;refGene_gn=COL12A1;refGene_tx=XM_019830867.2;refGene_gn=COL12A1;refGene_tx=XM_019830868.2;refGene_gn=COL12A1;refGene_tx=XM_019830870.1
B2 70434579 70434722
refGene_gn=COL12A1;refGene_tx=XM_006931849.4;refGene_gn=COL12A1;refGene_tx=XM_006931850.4;refGene_gn=COL12A1;refGene_tx=XM_019830867.2;refGene_gn=COL12A1;refGene_tx=XM_019830868.2;refGene_gn=COL12A1;refGene_tx=XM_019830870.1
B2 70436811 70436950
refGene_gn=COL12A1;refGene_tx=XM_006931849.4;refGene_gn=COL12A1;refGene_tx=XM_006931850.4;refGene_gn=COL12A1;refGene_tx=XM_019830867.2;refGene_gn=COL12A1;refGene_tx=XM_019830868.2;refGene_gn=COL12A1;refGene_tx=XM_019830870.1
B2 70437046 70437085
refGene_gn=COL12A1;refGene_tx=XM_006931849.4;refGene_gn=COL12A1;refGene_tx=XM_006931850.4;refGene_gn=COL12A1;refGene_tx=XM_019830867.2;refGene_gn=COL12A1;refGene_tx=XM_019830868.2;refGene_gn=COL12A1;refGene_tx=XM_019830870.1
B2 70438414 70438579
refGene_gn=COL12A1;refGene_tx=XM_006931849.4;refGene_gn=COL12A1;refGene_tx=XM_006931850.4;refGene_gn=COL12A1;refGene_tx=XM_019830867.2;refGene_gn=COL12A1;refGene_tx=XM_019830868.2;refGene_gn=COL12A1;refGene_tx=XM_019830870.1
B2 70440517 70440661
refGene_gn=COL12A1;refGene_tx=XM_006931849.4;refGene_gn=COL12A1;refGene_tx=XM_006931850.4;refGene_gn=COL12A1;refGene_tx=XM_019830867.2;refGene_gn=COL12A1;refGene_tx=XM_019830868.2;refGene_gn=COL12A1;refGene_tx=XM_019830870.1

B2 70440835 70440959
refGene_gn=COL12A1;refGene_tx=XM_006931849.4;refGene_gn=COL12A1;refGene_tx=XM_006931850.4;refGene_gn=COL12A1;refGene_tx=XM_019830867.2;refGene_gn=COL12A1;refGene_tx=XM_019830868.2;refGene_gn=COL12A1;refGene_tx=XM_019830870.1
B2 70442287 70442427
refGene_gn=COL12A1;refGene_tx=XM_006931849.4;refGene_gn=COL12A1;refGene_tx=XM_006931850.4;refGene_gn=COL12A1;refGene_tx=XM_019830867.2;refGene_gn=COL12A1;refGene_tx=XM_019830868.2;refGene_gn=COL12A1;refGene_tx=XM_019830870.1
B2 70444261 70444336
refGene_gn=COL12A1;refGene_tx=XM_006931849.4;refGene_gn=COL12A1;refGene_tx=XM_006931850.4;refGene_gn=COL12A1;refGene_tx=XM_019830867.2;refGene_gn=COL12A1;refGene_tx=XM_019830868.2;refGene_gn=COL12A1;refGene_tx=XM_019830870.1
B2 70444937 70445084
refGene_gn=COL12A1;refGene_tx=XM_006931849.4;refGene_gn=COL12A1;refGene_tx=XM_006931850.4;refGene_gn=COL12A1;refGene_tx=XM_019830867.2;refGene_gn=COL12A1;refGene_tx=XM_019830868.2;refGene_gn=COL12A1;refGene_tx=XM_019830870.1
B2 70445277 70445394
refGene_gn=COL12A1;refGene_tx=XM_006931849.4;refGene_gn=COL12A1;refGene_tx=XM_006931850.4;refGene_gn=COL12A1;refGene_tx=XM_019830867.2;refGene_gn=COL12A1;refGene_tx=XM_019830868.2;refGene_gn=COL12A1;refGene_tx=XM_019830870.1
B2 70446195 70446342
refGene_gn=COL12A1;refGene_tx=XM_006931849.4;refGene_gn=COL12A1;refGene_tx=XM_006931850.4;refGene_gn=COL12A1;refGene_tx=XM_019830867.2;refGene_gn=COL12A1;refGene_tx=XM_019830868.2;refGene_gn=COL12A1;refGene_tx=XM_019830870.1
B2 70447625 70447745
refGene_gn=COL12A1;refGene_tx=XM_006931849.4;refGene_gn=COL12A1;refGene_tx=XM_006931850.4;refGene_gn=COL12A1;refGene_tx=XM_019830867.2;refGene_gn=COL12A1;refGene_tx=XM_019830868.2;refGene_gn=COL12A1;refGene_tx=XM_019830870.1
B2 70450118 70450248
refGene_gn=COL12A1;refGene_tx=XM_006931849.4;refGene_gn=COL12A1;refGene_tx=XM_006931850.4;refGene_gn=COL12A1;refGene_tx=XM_019830867.2;refGene_gn=COL12A1;refGene_tx=XM_019830868.2;refGene_gn=COL12A1;refGene_tx=XM_019830870.1
B2 70451947 70452090
refGene_gn=COL12A1;refGene_tx=XM_006931849.4;refGene_gn=COL12A1;refGene_tx=XM_006931850.4;refGene_gn=COL12A1;refGene_tx=XM_019830867.2;refGene_gn=COL12A1;refGene_tx=XM_019830868.2;refGene_gn=COL12A1;refGene_tx=XM_019830870.1
B2 70452664 70452794
refGene_gn=COL12A1;refGene_tx=XM_006931849.4;refGene_gn=COL12A1;refGene_tx=XM_006931850.4;refGene_gn=COL12A1;refGene_tx=XM_019830867.2;refGene_gn=COL12A1;refGene_tx=XM_019830868.2;refGene_gn=COL12A1;refGene_tx=XM_019830870.1

B2 70454112 70454255
refGene_gn=COL12A1;refGene_tx=XM_006931849.4;refGene_gn=COL12A1;refGene_tx=XM_006931850.4;refGene_gn=COL12A1;refGene_tx=XM_019830867.2;refGene_gn=COL12A1;refGene_tx=XM_019830868.2;refGene_gn=COL12A1;refGene_tx=XM_019830870.1
B2 70455504 70455634
refGene_gn=COL12A1;refGene_tx=XM_006931849.4;refGene_gn=COL12A1;refGene_tx=XM_006931850.4;refGene_gn=COL12A1;refGene_tx=XM_019830867.2;refGene_gn=COL12A1;refGene_tx=XM_019830868.2;refGene_gn=COL12A1;refGene_tx=XM_019830870.1
B2 70456066 70456206
refGene_gn=COL12A1;refGene_tx=XM_006931849.4;refGene_gn=COL12A1;refGene_tx=XM_006931850.4;refGene_gn=COL12A1;refGene_tx=XM_019830867.2;refGene_gn=COL12A1;refGene_tx=XM_019830868.2;refGene_gn=COL12A1;refGene_tx=XM_019830870.1
B2 70456890 70457020
refGene_gn=COL12A1;refGene_tx=XM_006931849.4;refGene_gn=COL12A1;refGene_tx=XM_006931850.4;refGene_gn=COL12A1;refGene_tx=XM_019830867.2;refGene_gn=COL12A1;refGene_tx=XM_019830868.2;refGene_gn=COL12A1;refGene_tx=XM_019830870.1
B2 70459700 70459843
refGene_gn=COL12A1;refGene_tx=XM_006931849.4;refGene_gn=COL12A1;refGene_tx=XM_006931850.4;refGene_gn=COL12A1;refGene_tx=XM_019830867.2;refGene_gn=COL12A1;refGene_tx=XM_019830868.2;refGene_gn=COL12A1;refGene_tx=XM_019830870.1
B2 70460183 70460210
refGene_gn=COL12A1;refGene_tx=XM_006931849.4;refGene_gn=COL12A1;refGene_tx=XM_006931850.4;refGene_gn=COL12A1;refGene_tx=XM_019830867.2;refGene_gn=COL12A1;refGene_tx=XM_019830870.1
B2 70460339 70460472
refGene_gn=COL12A1;refGene_tx=XM_006931849.4;refGene_gn=COL12A1;refGene_tx=XM_006931850.4;refGene_gn=COL12A1;refGene_tx=XM_019830867.2;refGene_gn=COL12A1;refGene_tx=XM_019830868.2;refGene_gn=COL12A1;refGene_tx=XM_019830870.1
B2 70460740 70460880
refGene_gn=COL12A1;refGene_tx=XM_006931849.4;refGene_gn=COL12A1;refGene_tx=XM_006931850.4;refGene_gn=COL12A1;refGene_tx=XM_019830867.2;refGene_gn=COL12A1;refGene_tx=XM_019830868.2;refGene_gn=COL12A1;refGene_tx=XM_019830870.1
B2 70464193 70464323
refGene_gn=COL12A1;refGene_tx=XM_006931849.4;refGene_gn=COL12A1;refGene_tx=XM_006931850.4;refGene_gn=COL12A1;refGene_tx=XM_019830867.2;refGene_gn=COL12A1;refGene_tx=XM_019830868.2;refGene_gn=COL12A1;refGene_tx=XM_019830870.1
B2 70465411 70465548
refGene_gn=COL12A1;refGene_tx=XM_006931849.4;refGene_gn=COL12A1;refGene_tx=XM_006931850.4;refGene_gn=COL12A1;refGene_tx=XM_019830867.2;refGene_gn=COL12A1;refGene_tx=XM_019830868.2;refGene_gn=COL12A1;refGene_tx=XM_019830870.1
B2 70467694 70467824

refGene_gn=COL12A1;refGene_tx=XM_006931849.4;refGene_gn=COL12A1;refGene_tx=XM_006931850.4;refGene_gn=COL12A1;refGene_tx=XM_019830867.2;refGene_gn=COL12A1;refGene_tx=XM_019830868.2;refGene_gn=COL12A1;refGene_tx=XM_019830870.1
B2 70468135 70468278
refGene_gn=COL12A1;refGene_tx=XM_006931849.4;refGene_gn=COL12A1;refGene_tx=XM_006931850.4;refGene_gn=COL12A1;refGene_tx=XM_019830867.2;refGene_gn=COL12A1;refGene_tx=XM_019830868.2;refGene_gn=COL12A1;refGene_tx=XM_019830870.1
B2 70475414 70475544
refGene_gn=COL12A1;refGene_tx=XM_006931849.4;refGene_gn=COL12A1;refGene_tx=XM_006931850.4;refGene_gn=COL12A1;refGene_tx=XM_019830867.2;refGene_gn=COL12A1;refGene_tx=XM_019830868.2;refGene_gn=COL12A1;refGene_tx=XM_019830870.1
B2 70476092 70476232
refGene_gn=COL12A1;refGene_tx=XM_006931849.4;refGene_gn=COL12A1;refGene_tx=XM_006931850.4;refGene_gn=COL12A1;refGene_tx=XM_019830867.2;refGene_gn=COL12A1;refGene_tx=XM_019830868.2;refGene_gn=COL12A1;refGene_tx=XM_019830870.1
B2 70477776 70477923
refGene_gn=COL12A1;refGene_tx=XM_006931849.4;refGene_gn=COL12A1;refGene_tx=XM_006931850.4;refGene_gn=COL12A1;refGene_tx=XM_019830867.2;refGene_gn=COL12A1;refGene_tx=XM_019830868.2;refGene_gn=COL12A1;refGene_tx=XM_019830870.1
B2 70478478 70478643
refGene_gn=COL12A1;refGene_tx=XM_006931849.4;refGene_gn=COL12A1;refGene_tx=XM_006931850.4;refGene_gn=COL12A1;refGene_tx=XM_019830867.2;refGene_gn=COL12A1;refGene_tx=XM_019830868.2;refGene_gn=COL12A1;refGene_tx=XM_019830870.1
B2 70478743 70478863
refGene_gn=COL12A1;refGene_tx=XM_006931849.4;refGene_gn=COL12A1;refGene_tx=XM_006931850.4;refGene_gn=COL12A1;refGene_tx=XM_019830867.2;refGene_gn=COL12A1;refGene_tx=XM_019830868.2;refGene_gn=COL12A1;refGene_tx=XM_019830870.1
B2 70478941 70479091
refGene_gn=COL12A1;refGene_tx=XM_006931849.4;refGene_gn=COL12A1;refGene_tx=XM_006931850.4;refGene_gn=COL12A1;refGene_tx=XM_019830867.2;refGene_gn=COL12A1;refGene_tx=XM_019830868.2;refGene_gn=COL12A1;refGene_tx=XM_019830870.1
B2 70480828 70480950
refGene_gn=COL12A1;refGene_tx=XM_006931849.4;refGene_gn=COL12A1;refGene_tx=XM_019830867.2;refGene_gn=COL12A1;refGene_tx=XM_019830868.2;refGene_gn=COL12A1;refGene_tx=XM_019830870.1
B2 70481823 70482016
refGene_gn=COL12A1;refGene_tx=XM_006931849.4;refGene_gn=COL12A1;refGene_tx=XM_019830867.2;refGene_gn=COL12A1;refGene_tx=XM_019830868.2;refGene_gn=COL12A1;refGene_tx=XM_019830870.1
B2 70482453 70482720
refGene_gn=COL12A1;refGene_tx=XM_006931849.4;refGene_gn=COL12A1;refGene_tx=XM_019830867.2;refGene_gn=COL12A1;refGene_tx=XM_019830868.2;refGe

ne_gn=COL12A1;refGene_tx=XM_019830870.1
B2 70488879 70489152
refGene_gn=COL12A1;refGene_tx=XM_006931849.4;refGene_gn=COL12A1;refGene_tx=XM_019830867.2;refGene_gn=COL12A1;refGene_tx=XM_019830868.2;refGene_gn=COL12A1;refGene_tx=XM_019830870.1
B2 70497864 70498137
refGene_gn=COL12A1;refGene_tx=XM_006931849.4;refGene_gn=COL12A1;refGene_tx=XM_019830867.2;refGene_gn=COL12A1;refGene_tx=XM_019830868.2;refGene_gn=COL12A1;refGene_tx=XM_019830870.1
B2 70499817 70500090
refGene_gn=COL12A1;refGene_tx=XM_006931849.4;refGene_gn=COL12A1;refGene_tx=XM_019830867.2;refGene_gn=COL12A1;refGene_tx=XM_019830868.2;refGene_gn=COL12A1;refGene_tx=XM_019830870.1
B2 70503334 70503607
refGene_gn=COL12A1;refGene_tx=XM_006931849.4;refGene_gn=COL12A1;refGene_tx=XM_019830867.2;refGene_gn=COL12A1;refGene_tx=XM_019830868.2;refGene_gn=COL12A1;refGene_tx=XM_019830870.1
B2 70505023 70505626
refGene_gn=COL12A1;refGene_tx=XM_006931849.4;refGene_gn=COL12A1;refGene_tx=XM_019830867.2;refGene_gn=COL12A1;refGene_tx=XM_019830868.2;refGene_gn=COL12A1;refGene_tx=XM_019830870.1
B2 70506069 70506360
refGene_gn=COL12A1;refGene_tx=XM_006931849.4;refGene_gn=COL12A1;refGene_tx=XM_019830867.2;refGene_gn=COL12A1;refGene_tx=XM_019830868.2;refGene_gn=COL12A1;refGene_tx=XM_019830870.1
B2 70508617 70508791
refGene_gn=COL12A1;refGene_tx=XM_006931849.4;refGene_gn=COL12A1;refGene_tx=XM_019830867.2;refGene_gn=COL12A1;refGene_tx=XM_019830868.2;refGene_gn=COL12A1;refGene_tx=XM_019830870.1
B2 70509480 70509645
refGene_gn=COL12A1;refGene_tx=XM_006931849.4;refGene_gn=COL12A1;refGene_tx=XM_019830867.2;refGene_gn=COL12A1;refGene_tx=XM_019830868.2;refGene_gn=COL12A1;refGene_tx=XM_019830870.1
B2 70509821 70510085
refGene_gn=COL12A1;refGene_tx=XM_006931849.4;refGene_gn=COL12A1;refGene_tx=XM_019830867.2;refGene_gn=COL12A1;refGene_tx=XM_019830868.2;refGene_gn=COL12A1;refGene_tx=XM_019830870.1
B2 70512316 70512376
refGene_gn=COL12A1;refGene_tx=XM_006931849.4;refGene_gn=COL12A1;refGene_tx=XM_019830867.2;refGene_gn=COL12A1;refGene_tx=XM_019830868.2;refGene_gn=COL12A1;refGene_tx=XM_019830870.1
B2 70513034 70513178
refGene_gn=COL12A1;refGene_tx=XM_006931849.4;refGene_gn=COL12A1;refGene_tx=XM_019830867.2;refGene_gn=COL12A1;refGene_tx=XM_019830868.2;refGene_gn=COL12A1;refGene_tx=XM_019830870.1
B2 70515194 70515311
refGene_gn=COL12A1;refGene_tx=XM_006931849.4;refGene_gn=COL12A1;refGene_tx=XM_019830867.2;refGene_gn=COL12A1;refGene_tx=XM_019830868.2;refGene_gn=COL12A1;refGene_tx=XM_019830870.1
B2 70522130 70522203

refGene_gn=COL12A1;refGene_tx=XM_006931849.4;refGene_gn=COL12A1;refGene_tx=XM_006931850.4;refGene_gn=COL12A1;refGene_tx=XM_019830867.2;refGene_gn=COL12A1;refGene_tx=XM_019830868.2;refGene_gn=COL12A1;refGene_tx=XM_019830870.1
B2 70561409 70561468
refGene_gn=LOC101093608;refGene_tx=XM_023254729.1
B2 70563011 70563096
refGene_gn=LOC101093608;refGene_tx=XM_023254729.1
B2 70566353 70566443
refGene_gn=LOC101093608;refGene_tx=XM_023254729.1
B2 70567905 70567929
refGene_gn=LOC101093608;refGene_tx=XM_023254729.1
B2 70567931 70568153
refGene_gn=LOC101093608;refGene_tx=XM_023254729.1
B2 70568453 70568653
refGene_gn=LOC101093608;refGene_tx=XM_023254729.1
B2 70570083 70570219
refGene_gn=LOC101093608;refGene_tx=XM_023254729.1
B2 70579861 70580055 refGene_gn=TMEM30A;refGene_tx=XM_003986315.4
B2 70582505 70582712 refGene_gn=TMEM30A;refGene_tx=XM_003986315.4
B2 70583377 70583521 refGene_gn=TMEM30A;refGene_tx=XM_003986315.4
B2 70584650 70584738 refGene_gn=TMEM30A;refGene_tx=XM_003986315.4
B2 70589687 70589795 refGene_gn=TMEM30A;refGene_tx=XM_003986315.4
B2 70592388 70592496 refGene_gn=TMEM30A;refGene_tx=XM_003986315.4
B2 70616189 70616543 refGene_gn=TMEM30A;refGene_tx=XM_003986315.4
B2 70616544 70616715 refGene_gn=TMEM30A;refGene_tx=XM_003986315.4
B2 70629259 70629300 refGene_gn=FILIP1;refGene_tx=XM_003986317.5
B2 70643115 70643156 refGene_gn=FILIP1;refGene_tx=XM_023254144.1
B2 70643635 70643842
refGene_gn=FILIP1;refGene_tx=XM_003986317.5;refGene_gn=FILIP1;refGene_tx=XM_006931852.4;refGene_gn=FILIP1;refGene_tx=XM_023254144.1
B2 70647734 70650540
refGene_gn=FILIP1;refGene_tx=XM_003986317.5;refGene_gn=FILIP1;refGene_tx=XM_006931852.4;refGene_gn=FILIP1;refGene_tx=XM_023254144.1
B2 70687800 70687979
refGene_gn=FILIP1;refGene_tx=XM_003986317.5;refGene_gn=FILIP1;refGene_tx=XM_006931852.4;refGene_gn=FILIP1;refGene_tx=XM_023254144.1
B2 70695312 70695486
refGene_gn=FILIP1;refGene_tx=XM_003986317.5;refGene_gn=FILIP1;refGene_tx=XM_006931852.4;refGene_gn=FILIP1;refGene_tx=XM_023254144.1
B2 70735512 70735788
refGene_gn=FILIP1;refGene_tx=XM_003986317.5;refGene_gn=FILIP1;refGene_tx=XM_006931852.4;refGene_gn=FILIP1;refGene_tx=XM_023254144.1
B2 70933782 70933834 refGene_gn=SENP6;refGene_tx=XM_006931853.4
B2 70933978 70934096
refGene_gn=SENP6;refGene_tx=XM_023254145.1;refGene_gn=SENP6;refGene_tx=XM_023254146.1
B2 70956470 70956564
refGene_gn=SENP6;refGene_tx=XM_006931853.4;refGene_gn=SENP6;refGene_tx=XM_023254145.1;refGene_gn=SENP6;refGene_tx=XM_023254146.1

B2 70958506 70958567
refGene_gn=SENP6;refGene_tx=XM_006931853.4;refGene_gn=SENP6;refGene_tx=XM_023254145.1;refGene_gn=SENP6;refGene_tx=XM_023254146.1
B2 70967413 70967556
refGene_gn=SENP6;refGene_tx=XM_006931853.4;refGene_gn=SENP6;refGene_tx=XM_019830873.1;refGene_gn=SENP6;refGene_tx=XM_023254145.1;refGene_gn=SENP6;refGene_tx=XM_023254146.1
B2 70968593 70968698
refGene_gn=SENP6;refGene_tx=XM_006931853.4;refGene_gn=SENP6;refGene_tx=XM_019830873.1;refGene_gn=SENP6;refGene_tx=XM_023254145.1;refGene_gn=SENP6;refGene_tx=XM_023254146.1
B2 70973365 70973386
refGene_gn=SENP6;refGene_tx=XM_006931853.4;refGene_gn=SENP6;refGene_tx=XM_019830873.1;refGene_gn=SENP6;refGene_tx=XM_023254145.1
B2 70985998 70986069
refGene_gn=SENP6;refGene_tx=XM_006931853.4;refGene_gn=SENP6;refGene_tx=XM_019830873.1;refGene_gn=SENP6;refGene_tx=XM_023254145.1;refGene_gn=SENP6;refGene_tx=XM_023254146.1
B2 70999668 70999814
refGene_gn=SENP6;refGene_tx=XM_006931853.4;refGene_gn=SENP6;refGene_tx=XM_019830873.1;refGene_gn=SENP6;refGene_tx=XM_023254145.1;refGene_gn=SENP6;refGene_tx=XM_023254146.1
B2 71006055 71006353
refGene_gn=SENP6;refGene_tx=XM_006931853.4;refGene_gn=SENP6;refGene_tx=XM_019830873.1;refGene_gn=SENP6;refGene_tx=XM_023254145.1;refGene_gn=SENP6;refGene_tx=XM_023254146.1
B2 71012348 71012581
refGene_gn=SENP6;refGene_tx=XM_006931853.4;refGene_gn=SENP6;refGene_tx=XM_019830873.1;refGene_gn=SENP6;refGene_tx=XM_023254145.1;refGene_gn=SENP6;refGene_tx=XM_023254146.1
B2 71015327 71015498
refGene_gn=SENP6;refGene_tx=XM_006931853.4;refGene_gn=SENP6;refGene_tx=XM_019830873.1;refGene_gn=SENP6;refGene_tx=XM_023254145.1;refGene_gn=SENP6;refGene_tx=XM_023254146.1
B2 71020540 71020574
refGene_gn=SENP6;refGene_tx=XM_006931853.4;refGene_gn=SENP6;refGene_tx=XM_019830873.1;refGene_gn=SENP6;refGene_tx=XM_023254145.1;refGene_gn=SENP6;refGene_tx=XM_023254146.1
B2 71020982 71021177
refGene_gn=SENP6;refGene_tx=XM_006931853.4;refGene_gn=SENP6;refGene_tx=XM_019830873.1;refGene_gn=SENP6;refGene_tx=XM_023254145.1;refGene_gn=SENP6;refGene_tx=XM_023254146.1
B2 71022115 71022342
refGene_gn=SENP6;refGene_tx=XM_006931853.4;refGene_gn=SENP6;refGene_tx=XM_019830873.1;refGene_gn=SENP6;refGene_tx=XM_023254145.1;refGene_gn=SENP6;refGene_tx=XM_023254146.1
B2 71023900 71024010
refGene_gn=SENP6;refGene_tx=XM_006931853.4;refGene_gn=SENP6;refGene_tx=XM_019830873.1;refGene_gn=SENP6;refGene_tx=XM_023254145.1;refGene_gn=SENP6;refGene_tx=XM_023254146.1

B2	71024146	71024263
refGene_gn=SENP6;refGene_tx=XM_006931853.4;refGene_gn=SENP6;refGene_tx
=XM_019830873.1;refGene_gn=SENP6;refGene_tx=XM_023254145.1;refGene_gn=
SENP6;refGene_tx=XM_023254146.1
B2	71035555	71035678
refGene_gn=SENP6;refGene_tx=XM_006931853.4;refGene_gn=SENP6;refGene_tx
=XM_019830873.1;refGene_gn=SENP6;refGene_tx=XM_023254145.1;refGene_gn=
SENP6;refGene_tx=XM_023254146.1
B2	71036824	71036917
refGene_gn=SENP6;refGene_tx=XM_006931853.4;refGene_gn=SENP6;refGene_tx
=XM_019830873.1;refGene_gn=SENP6;refGene_tx=XM_023254145.1;refGene_gn=
SENP6;refGene_tx=XM_023254146.1
B2	71039593	71040021
refGene_gn=SENP6;refGene_tx=XM_006931853.4;refGene_gn=SENP6;refGene_tx
=XM_019830873.1;refGene_gn=SENP6;refGene_tx=XM_023254145.1;refGene_gn=
SENP6;refGene_tx=XM_023254146.1
B2	71046122	71046226
refGene_gn=SENP6;refGene_tx=XM_006931853.4;refGene_gn=SENP6;refGene_tx
=XM_019830873.1;refGene_gn=SENP6;refGene_tx=XM_023254145.1;refGene_gn=
SENP6;refGene_tx=XM_023254146.1
B2	71047786	71047875
refGene_gn=SENP6;refGene_tx=XM_006931853.4;refGene_gn=SENP6;refGene_tx
=XM_019830873.1;refGene_gn=SENP6;refGene_tx=XM_023254145.1;refGene_gn=
SENP6;refGene_tx=XM_023254146.1
B2	71049505	71049574
refGene_gn=SENP6;refGene_tx=XM_006931853.4;refGene_gn=SENP6;refGene_tx
=XM_019830873.1;refGene_gn=SENP6;refGene_tx=XM_023254145.1;refGene_gn=
SENP6;refGene_tx=XM_023254146.1
B2	71049673	71049824
refGene_gn=SENP6;refGene_tx=XM_006931853.4;refGene_gn=SENP6;refGene_tx
=XM_019830873.1;refGene_gn=SENP6;refGene_tx=XM_023254145.1;refGene_gn=
SENP6;refGene_tx=XM_023254146.1
B2	71051672	71051882
refGene_gn=SENP6;refGene_tx=XM_006931853.4;refGene_gn=SENP6;refGene_tx
=XM_019830873.1;refGene_gn=SENP6;refGene_tx=XM_023254145.1;refGene_gn=
SENP6;refGene_tx=XM_023254146.1
B2	71135756	71135873
refGene_gn=MYO6;refGene_tx=XM_023254147.1;refGene_gn=MYO6;refGene_tx=X
M_023254148.1;refGene_gn=MYO6;refGene_tx=XM_023254149.1;refGene_gn=MYO
6;refGene_tx=XM_023254150.1;refGene_gn=MYO6;refGene_tx=XM_023254151.1;
refGene_gn=MYO6;refGene_tx=XM_023254152.1
B2	71140896	71140966
refGene_gn=MYO6;refGene_tx=XM_023254147.1;refGene_gn=MYO6;refGene_tx=X
M_023254148.1;refGene_gn=MYO6;refGene_tx=XM_023254149.1;refGene_gn=MYO
6;refGene_tx=XM_023254150.1;refGene_gn=MYO6;refGene_tx=XM_023254151.1;
refGene_gn=MYO6;refGene_tx=XM_023254152.1
B2	71145199	71145273
refGene_gn=MYO6;refGene_tx=XM_023254147.1;refGene_gn=MYO6;refGene_tx=X
M_023254148.1;refGene_gn=MYO6;refGene_tx=XM_023254149.1;refGene_gn=MYO
6;refGene_tx=XM_023254150.1;refGene_gn=MYO6;refGene_tx=XM_023254151.1;

refGene_gn=MYO6;refGene_tx=XM_023254152.1
B2 71150018 71150148
refGene_gn=MYO6;refGene_tx=XM_023254147.1;refGene_gn=MYO6;refGene_tx=XM_023254148.1;refGene_gn=MYO6;refGene_tx=XM_023254149.1;refGene_gn=MYO6;refGene_tx=XM_023254150.1;refGene_gn=MYO6;refGene_tx=XM_023254151.1;refGene_gn=MYO6;refGene_tx=XM_023254152.1
B2 71152535 71152641
refGene_gn=MYO6;refGene_tx=XM_023254147.1;refGene_gn=MYO6;refGene_tx=XM_023254148.1;refGene_gn=MYO6;refGene_tx=XM_023254149.1;refGene_gn=MYO6;refGene_tx=XM_023254150.1;refGene_gn=MYO6;refGene_tx=XM_023254151.1;refGene_gn=MYO6;refGene_tx=XM_023254152.1
B2 71153824 71153880
refGene_gn=MYO6;refGene_tx=XM_023254147.1;refGene_gn=MYO6;refGene_tx=XM_023254148.1;refGene_gn=MYO6;refGene_tx=XM_023254149.1;refGene_gn=MYO6;refGene_tx=XM_023254150.1;refGene_gn=MYO6;refGene_tx=XM_023254151.1;refGene_gn=MYO6;refGene_tx=XM_023254152.1
B2 71158389 71158487
refGene_gn=MYO6;refGene_tx=XM_023254147.1;refGene_gn=MYO6;refGene_tx=XM_023254148.1;refGene_gn=MYO6;refGene_tx=XM_023254149.1;refGene_gn=MYO6;refGene_tx=XM_023254150.1;refGene_gn=MYO6;refGene_tx=XM_023254151.1;refGene_gn=MYO6;refGene_tx=XM_023254152.1
B2 71158797 71158962
refGene_gn=MYO6;refGene_tx=XM_023254147.1;refGene_gn=MYO6;refGene_tx=XM_023254148.1;refGene_gn=MYO6;refGene_tx=XM_023254149.1;refGene_gn=MYO6;refGene_tx=XM_023254150.1;refGene_gn=MYO6;refGene_tx=XM_023254151.1;refGene_gn=MYO6;refGene_tx=XM_023254152.1
B2 71161348 71161429
refGene_gn=MYO6;refGene_tx=XM_023254147.1;refGene_gn=MYO6;refGene_tx=XM_023254148.1;refGene_gn=MYO6;refGene_tx=XM_023254149.1;refGene_gn=MYO6;refGene_tx=XM_023254150.1;refGene_gn=MYO6;refGene_tx=XM_023254151.1;refGene_gn=MYO6;refGene_tx=XM_023254152.1
B2 71162826 71162835
refGene_gn=MYO6;refGene_tx=XM_023254147.1;refGene_gn=MYO6;refGene_tx=XM_023254149.1
B2 71164098 71164279
refGene_gn=MYO6;refGene_tx=XM_023254147.1;refGene_gn=MYO6;refGene_tx=XM_023254148.1;refGene_gn=MYO6;refGene_tx=XM_023254149.1;refGene_gn=MYO6;refGene_tx=XM_023254150.1;refGene_gn=MYO6;refGene_tx=XM_023254151.1;refGene_gn=MYO6;refGene_tx=XM_023254152.1
B2 71170152 71170297
refGene_gn=MYO6;refGene_tx=XM_023254147.1;refGene_gn=MYO6;refGene_tx=XM_023254148.1;refGene_gn=MYO6;refGene_tx=XM_023254149.1;refGene_gn=MYO6;refGene_tx=XM_023254150.1;refGene_gn=MYO6;refGene_tx=XM_023254151.1;refGene_gn=MYO6;refGene_tx=XM_023254152.1
B2 71172376 71172534
refGene_gn=MYO6;refGene_tx=XM_023254147.1;refGene_gn=MYO6;refGene_tx=XM_023254148.1;refGene_gn=MYO6;refGene_tx=XM_023254149.1;refGene_gn=MYO6;refGene_tx=XM_023254150.1;refGene_gn=MYO6;refGene_tx=XM_023254151.1;refGene_gn=MYO6;refGene_tx=XM_023254152.1
B2 71173974 71174066

refGene_gn=MYO6;refGene_tx=XM_023254147.1;refGene_gn=MYO6;refGene_tx=XM_023254148.1;refGene_gn=MYO6;refGene_tx=XM_023254149.1;refGene_gn=MYO6;refGene_tx=XM_023254150.1;refGene_gn=MYO6;refGene_tx=XM_023254151.1;refGene_gn=MYO6;refGene_tx=XM_023254152.1
B2 71177101 71177174
refGene_gn=MYO6;refGene_tx=XM_023254147.1;refGene_gn=MYO6;refGene_tx=XM_023254148.1;refGene_gn=MYO6;refGene_tx=XM_023254149.1;refGene_gn=MYO6;refGene_tx=XM_023254150.1;refGene_gn=MYO6;refGene_tx=XM_023254151.1;refGene_gn=MYO6;refGene_tx=XM_023254152.1
B2 71179114 71179242
refGene_gn=MYO6;refGene_tx=XM_023254147.1;refGene_gn=MYO6;refGene_tx=XM_023254148.1;refGene_gn=MYO6;refGene_tx=XM_023254149.1;refGene_gn=MYO6;refGene_tx=XM_023254150.1;refGene_gn=MYO6;refGene_tx=XM_023254151.1;refGene_gn=MYO6;refGene_tx=XM_023254152.1
B2 71181718 71181814
refGene_gn=MYO6;refGene_tx=XM_023254147.1;refGene_gn=MYO6;refGene_tx=XM_023254148.1;refGene_gn=MYO6;refGene_tx=XM_023254149.1;refGene_gn=MYO6;refGene_tx=XM_023254150.1;refGene_gn=MYO6;refGene_tx=XM_023254151.1;refGene_gn=MYO6;refGene_tx=XM_023254152.1
B2 71182090 71182264
refGene_gn=MYO6;refGene_tx=XM_023254147.1;refGene_gn=MYO6;refGene_tx=XM_023254148.1;refGene_gn=MYO6;refGene_tx=XM_023254149.1;refGene_gn=MYO6;refGene_tx=XM_023254150.1;refGene_gn=MYO6;refGene_tx=XM_023254151.1;refGene_gn=MYO6;refGene_tx=XM_023254152.1
B2 71185801 71185840
refGene_gn=MYO6;refGene_tx=XM_023254147.1;refGene_gn=MYO6;refGene_tx=XM_023254148.1;refGene_gn=MYO6;refGene_tx=XM_023254149.1;refGene_gn=MYO6;refGene_tx=XM_023254150.1;refGene_gn=MYO6;refGene_tx=XM_023254151.1;refGene_gn=MYO6;refGene_tx=XM_023254152.1
B2 71187697 71187791
refGene_gn=MYO6;refGene_tx=XM_023254147.1;refGene_gn=MYO6;refGene_tx=XM_023254148.1;refGene_gn=MYO6;refGene_tx=XM_023254149.1;refGene_gn=MYO6;refGene_tx=XM_023254150.1;refGene_gn=MYO6;refGene_tx=XM_023254151.1;refGene_gn=MYO6;refGene_tx=XM_023254152.1
B2 71194186 71194317
refGene_gn=MYO6;refGene_tx=XM_023254147.1;refGene_gn=MYO6;refGene_tx=XM_023254148.1;refGene_gn=MYO6;refGene_tx=XM_023254149.1;refGene_gn=MYO6;refGene_tx=XM_023254150.1;refGene_gn=MYO6;refGene_tx=XM_023254151.1;refGene_gn=MYO6;refGene_tx=XM_023254152.1
B2 71194412 71194490
refGene_gn=MYO6;refGene_tx=XM_023254147.1;refGene_gn=MYO6;refGene_tx=XM_023254148.1;refGene_gn=MYO6;refGene_tx=XM_023254149.1;refGene_gn=MYO6;refGene_tx=XM_023254150.1;refGene_gn=MYO6;refGene_tx=XM_023254151.1;refGene_gn=MYO6;refGene_tx=XM_023254152.1
B2 71196291 71196421
refGene_gn=MYO6;refGene_tx=XM_023254147.1;refGene_gn=MYO6;refGene_tx=XM_023254148.1;refGene_gn=MYO6;refGene_tx=XM_023254149.1;refGene_gn=MYO6;refGene_tx=XM_023254150.1;refGene_gn=MYO6;refGene_tx=XM_023254151.1;refGene_gn=MYO6;refGene_tx=XM_023254152.1
B2 71198259 71198350

refGene_gn=MYO6;refGene_tx=XM_023254147.1;refGene_gn=MYO6;refGene_tx=XM_023254148.1;refGene_gn=MYO6;refGene_tx=XM_023254149.1;refGene_gn=MYO6;refGene_tx=XM_023254150.1;refGene_gn=MYO6;refGene_tx=XM_023254151.1;refGene_gn=MYO6;refGene_tx=XM_023254152.1
B2 71199507 71199658
refGene_gn=MYO6;refGene_tx=XM_023254147.1;refGene_gn=MYO6;refGene_tx=XM_023254148.1;refGene_gn=MYO6;refGene_tx=XM_023254149.1;refGene_gn=MYO6;refGene_tx=XM_023254150.1;refGene_gn=MYO6;refGene_tx=XM_023254151.1;refGene_gn=MYO6;refGene_tx=XM_023254152.1
B2 71201962 71202171
refGene_gn=MYO6;refGene_tx=XM_023254147.1;refGene_gn=MYO6;refGene_tx=XM_023254148.1;refGene_gn=MYO6;refGene_tx=XM_023254149.1;refGene_gn=MYO6;refGene_tx=XM_023254150.1;refGene_gn=MYO6;refGene_tx=XM_023254151.1;refGene_gn=MYO6;refGene_tx=XM_023254152.1
B2 71202828 71202907
refGene_gn=MYO6;refGene_tx=XM_023254147.1;refGene_gn=MYO6;refGene_tx=XM_023254148.1;refGene_gn=MYO6;refGene_tx=XM_023254149.1;refGene_gn=MYO6;refGene_tx=XM_023254150.1;refGene_gn=MYO6;refGene_tx=XM_023254151.1;refGene_gn=MYO6;refGene_tx=XM_023254152.1
B2 71204018 71204179
refGene_gn=MYO6;refGene_tx=XM_023254147.1;refGene_gn=MYO6;refGene_tx=XM_023254148.1;refGene_gn=MYO6;refGene_tx=XM_023254149.1;refGene_gn=MYO6;refGene_tx=XM_023254150.1;refGene_gn=MYO6;refGene_tx=XM_023254151.1;refGene_gn=MYO6;refGene_tx=XM_023254152.1
B2 71206481 71206508
refGene_gn=MYO6;refGene_tx=XM_023254147.1;refGene_gn=MYO6;refGene_tx=XM_023254148.1;refGene_gn=MYO6;refGene_tx=XM_023254152.1
B2 71206912 71206942
refGene_gn=MYO6;refGene_tx=XM_023254147.1;refGene_gn=MYO6;refGene_tx=XM_023254148.1;refGene_gn=MYO6;refGene_tx=XM_023254152.1
B2 71209082 71209121
refGene_gn=MYO6;refGene_tx=XM_023254147.1;refGene_gn=MYO6;refGene_tx=XM_023254148.1;refGene_gn=MYO6;refGene_tx=XM_023254152.1
B2 71213132 71213236
refGene_gn=MYO6;refGene_tx=XM_023254147.1;refGene_gn=MYO6;refGene_tx=XM_023254148.1;refGene_gn=MYO6;refGene_tx=XM_023254149.1;refGene_gn=MYO6;refGene_tx=XM_023254150.1;refGene_gn=MYO6;refGene_tx=XM_023254151.1;refGene_gn=MYO6;refGene_tx=XM_023254152.1
B2 71214160 71214292
refGene_gn=MYO6;refGene_tx=XM_023254147.1;refGene_gn=MYO6;refGene_tx=XM_023254148.1;refGene_gn=MYO6;refGene_tx=XM_023254149.1;refGene_gn=MYO6;refGene_tx=XM_023254150.1;refGene_gn=MYO6;refGene_tx=XM_023254151.1;refGene_gn=MYO6;refGene_tx=XM_023254152.1
B2 71216635 71216662
refGene_gn=MYO6;refGene_tx=XM_023254147.1;refGene_gn=MYO6;refGene_tx=XM_023254148.1;refGene_gn=MYO6;refGene_tx=XM_023254149.1;refGene_gn=MYO6;refGene_tx=XM_023254150.1;refGene_gn=MYO6;refGene_tx=XM_023254152.1
B2 71219208 71219433
refGene_gn=MYO6;refGene_tx=XM_023254147.1;refGene_gn=MYO6;refGene_tx=XM_023254148.1;refGene_gn=MYO6;refGene_tx=XM_023254149.1;refGene_gn=MYO

6;refGene_tx=XM_023254150.1;refGene_gn=MYO6;refGene_tx=XM_023254151.1;
refGene_gn=MYO6;refGene_tx=XM_023254152.1
B2 71219948 71220148
refGene_gn=MYO6;refGene_tx=XM_023254147.1;refGene_gn=MYO6;refGene_tx=X
M_023254148.1;refGene_gn=MYO6;refGene_tx=XM_023254149.1;refGene_gn=MYO
6;refGene_tx=XM_023254150.1;refGene_gn=MYO6;refGene_tx=XM_023254151.1;
refGene_gn=MYO6;refGene_tx=XM_023254152.1
B2 71227460 71227538
refGene_gn=IMPG1;refGene_tx=XM_019830876.2;refGene_gn=IMPG1;refGene_tx
=XM_019830877.1;refGene_gn=IMPG1;refGene_tx=XM_019830878.1
B2 71228906 71228979
refGene_gn=IMPG1;refGene_tx=XM_019830876.2;refGene_gn=IMPG1;refGene_tx
=XM_019830877.1;refGene_gn=IMPG1;refGene_tx=XM_019830878.1
B2 71236581 71236777
refGene_gn=IMPG1;refGene_tx=XM_019830876.2;refGene_gn=IMPG1;refGene_tx
=XM_019830877.1;refGene_gn=IMPG1;refGene_tx=XM_019830878.1
B2 71256566 71256786
refGene_gn=IMPG1;refGene_tx=XM_019830876.2;refGene_gn=IMPG1;refGene_tx
=XM_019830877.1;refGene_gn=IMPG1;refGene_tx=XM_019830878.1
B2 71259740 71260252
refGene_gn=IMPG1;refGene_tx=XM_019830876.2;refGene_gn=IMPG1;refGene_tx
=XM_019830877.1;refGene_gn=IMPG1;refGene_tx=XM_019830878.1
B2 71302005 71302084
refGene_gn=IMPG1;refGene_tx=XM_019830876.2;refGene_gn=IMPG1;refGene_tx
=XM_019830877.1;refGene_gn=IMPG1;refGene_tx=XM_019830878.1
B2 71302955 71303032
refGene_gn=IMPG1;refGene_tx=XM_019830876.2;refGene_gn=IMPG1;refGene_tx
=XM_019830877.1;refGene_gn=IMPG1;refGene_tx=XM_019830878.1
B2 71304418 71304666
refGene_gn=IMPG1;refGene_tx=XM_019830876.2;refGene_gn=IMPG1;refGene_tx
=XM_019830877.1;refGene_gn=IMPG1;refGene_tx=XM_019830878.1
B2 71306328 71306349
refGene_gn=IMPG1;refGene_tx=XM_019830876.2;refGene_gn=IMPG1;refGene_tx
=XM_019830877.1;refGene_gn=IMPG1;refGene_tx=XM_019830878.1
B2 71310902 71310961
refGene_gn=IMPG1;refGene_tx=XM_019830876.2;refGene_gn=IMPG1;refGene_tx
=XM_019830877.1;refGene_gn=IMPG1;refGene_tx=XM_019830878.1
B2 71316686 71316839
refGene_gn=IMPG1;refGene_tx=XM_019830876.2;refGene_gn=IMPG1;refGene_tx
=XM_019830877.1;refGene_gn=IMPG1;refGene_tx=XM_019830878.1
B2 71322498 71322578
refGene_gn=IMPG1;refGene_tx=XM_019830876.2;refGene_gn=IMPG1;refGene_tx
=XM_019830877.1;refGene_gn=IMPG1;refGene_tx=XM_019830878.1
B2 71323174 71323239
refGene_gn=IMPG1;refGene_tx=XM_019830876.2;refGene_gn=IMPG1;refGene_tx
=XM_019830877.1;refGene_gn=IMPG1;refGene_tx=XM_019830878.1
B2 71327936 71327965
refGene_gn=IMPG1;refGene_tx=XM_019830876.2;refGene_gn=IMPG1;refGene_tx
=XM_019830877.1;refGene_gn=IMPG1;refGene_tx=XM_019830878.1
B2 71328232 71328399

refGene_gn=IMPG1;refGene_tx=XM_019830876.2;refGene_gn=IMPG1;refGene_tx=XM_019830877.1;refGene_gn=IMPG1;refGene_tx=XM_019830878.1
B2 71337540 71337774
refGene_gn=IMPG1;refGene_tx=XM_019830876.2;refGene_gn=IMPG1;refGene_tx=XM_019830877.1
B2 71340306 71340382 refGene_gn=IMPG1;refGene_tx=XM_019830876.2
B2 71362761 71362828
refGene_gn=IMPG1;refGene_tx=XM_019830877.1;refGene_gn=IMPG1;refGene_tx=XM_019830878.1
B2 72454237 72455407
refGene_gn=HTR1B;refGene_tx=NM_001122872.1;refGene_gn=HTR1B;refGene_tx=XM_019830057.1
B2 72662399 72662631 refGene_gn=MEI4;refGene_tx=XM_006931862.2
B2 72692990 72693526 refGene_gn=MEI4;refGene_tx=XM_006931862.2
B2 72764650 72764782 refGene_gn=MEI4;refGene_tx=XM_006931862.2
B2 72835218 72835479 refGene_gn=MEI4;refGene_tx=XM_006931862.2
B2 73680103 73680418 refGene_gn=IRAK1BP1;refGene_tx=XM_003986321.5
B2 73686764 73686830 refGene_gn=IRAK1BP1;refGene_tx=XM_003986321.5
B2 73694823 73694954 refGene_gn=IRAK1BP1;refGene_tx=XM_003986321.5
B2 73695057 73695328 refGene_gn=IRAK1BP1;refGene_tx=XM_003986321.5
B2 73728870 73729508
refGene_gn=PHIP;refGene_tx=XM_023254153.1;refGene_gn=PHIP;refGene_tx=XM_023254154.1;refGene_gn=PHIP;refGene_tx=XM_023254155.1;refGene_gn=PHIP;refGene_tx=XM_023254156.1
B2 73733476 73733674
refGene_gn=PHIP;refGene_tx=XM_023254153.1;refGene_gn=PHIP;refGene_tx=XM_023254154.1;refGene_gn=PHIP;refGene_tx=XM_023254155.1;refGene_gn=PHIP;refGene_tx=XM_023254156.1
B2 73734211 73734471
refGene_gn=PHIP;refGene_tx=XM_023254153.1;refGene_gn=PHIP;refGene_tx=XM_023254154.1;refGene_gn=PHIP;refGene_tx=XM_023254155.1;refGene_gn=PHIP;refGene_tx=XM_023254156.1
B2 73734925 73735089
refGene_gn=PHIP;refGene_tx=XM_023254153.1;refGene_gn=PHIP;refGene_tx=XM_023254154.1;refGene_gn=PHIP;refGene_tx=XM_023254155.1;refGene_gn=PHIP;refGene_tx=XM_023254156.1
B2 73736460 73736613
refGene_gn=PHIP;refGene_tx=XM_023254153.1;refGene_gn=PHIP;refGene_tx=XM_023254154.1;refGene_gn=PHIP;refGene_tx=XM_023254155.1;refGene_gn=PHIP;refGene_tx=XM_023254156.1
B2 73743809 73743959
refGene_gn=PHIP;refGene_tx=XM_023254153.1;refGene_gn=PHIP;refGene_tx=XM_023254154.1;refGene_gn=PHIP;refGene_tx=XM_023254155.1;refGene_gn=PHIP;refGene_tx=XM_023254156.1
B2 73744272 73744323
refGene_gn=PHIP;refGene_tx=XM_023254153.1;refGene_gn=PHIP;refGene_tx=XM_023254154.1;refGene_gn=PHIP;refGene_tx=XM_023254155.1;refGene_gn=PHIP;refGene_tx=XM_023254156.1
B2 73744697 73744767
refGene_gn=PHIP;refGene_tx=XM_023254153.1;refGene_gn=PHIP;refGene_tx=X

M_023254154.1;refGene_gn=PHIP;refGene_tx=XM_023254155.1;refGene_gn=PHIP;refGene_tx=XM_023254156.1
B2 73747568 73747694
refGene_gn=PHIP;refGene_tx=XM_023254153.1;refGene_gn=PHIP;refGene_tx=XM_023254154.1;refGene_gn=PHIP;refGene_tx=XM_023254155.1;refGene_gn=PHIP;refGene_tx=XM_023254156.1
B2 73750137 73750258
refGene_gn=PHIP;refGene_tx=XM_023254153.1;refGene_gn=PHIP;refGene_tx=XM_023254154.1;refGene_gn=PHIP;refGene_tx=XM_023254155.1;refGene_gn=PHIP;refGene_tx=XM_023254156.1
B2 73751443 73751599
refGene_gn=PHIP;refGene_tx=XM_023254153.1;refGene_gn=PHIP;refGene_tx=XM_023254154.1;refGene_gn=PHIP;refGene_tx=XM_023254155.1;refGene_gn=PHIP;refGene_tx=XM_023254156.1
B2 73753965 73754027
refGene_gn=PHIP;refGene_tx=XM_023254153.1;refGene_gn=PHIP;refGene_tx=XM_023254154.1;refGene_gn=PHIP;refGene_tx=XM_023254155.1;refGene_gn=PHIP;refGene_tx=XM_023254156.1
B2 73754190 73754302
refGene_gn=PHIP;refGene_tx=XM_023254153.1;refGene_gn=PHIP;refGene_tx=XM_023254154.1;refGene_gn=PHIP;refGene_tx=XM_023254155.1;refGene_gn=PHIP;refGene_tx=XM_023254156.1
B2 73757836 73757919
refGene_gn=PHIP;refGene_tx=XM_023254153.1;refGene_gn=PHIP;refGene_tx=XM_023254154.1;refGene_gn=PHIP;refGene_tx=XM_023254155.1;refGene_gn=PHIP;refGene_tx=XM_023254156.1
B2 73758046 73758171
refGene_gn=PHIP;refGene_tx=XM_023254153.1;refGene_gn=PHIP;refGene_tx=XM_023254154.1;refGene_gn=PHIP;refGene_tx=XM_023254155.1;refGene_gn=PHIP;refGene_tx=XM_023254156.1
B2 73758923 73759031
refGene_gn=PHIP;refGene_tx=XM_023254153.1;refGene_gn=PHIP;refGene_tx=XM_023254154.1;refGene_gn=PHIP;refGene_tx=XM_023254155.1;refGene_gn=PHIP;refGene_tx=XM_023254156.1
B2 73761006 73761126
refGene_gn=PHIP;refGene_tx=XM_023254153.1;refGene_gn=PHIP;refGene_tx=XM_023254154.1;refGene_gn=PHIP;refGene_tx=XM_023254155.1;refGene_gn=PHIP;refGene_tx=XM_023254156.1
B2 73764622 73764854
refGene_gn=PHIP;refGene_tx=XM_023254153.1;refGene_gn=PHIP;refGene_tx=XM_023254154.1;refGene_gn=PHIP;refGene_tx=XM_023254155.1;refGene_gn=PHIP;refGene_tx=XM_023254156.1
B2 73767394 73767471
refGene_gn=PHIP;refGene_tx=XM_023254153.1;refGene_gn=PHIP;refGene_tx=XM_023254154.1;refGene_gn=PHIP;refGene_tx=XM_023254155.1;refGene_gn=PHIP;refGene_tx=XM_023254156.1
B2 73770139 73770280
refGene_gn=PHIP;refGene_tx=XM_023254153.1;refGene_gn=PHIP;refGene_tx=XM_023254154.1;refGene_gn=PHIP;refGene_tx=XM_023254155.1;refGene_gn=PHIP;refGene_tx=XM_023254156.1

B2 73781006 73781124
refGene_gn=PHIP;refGene_tx=XM_023254153.1;refGene_gn=PHIP;refGene_tx=XM_023254154.1;refGene_gn=PHIP;refGene_tx=XM_023254155.1;refGene_gn=PHIP;refGene_tx=XM_023254156.1
B2 73786081 73786265
refGene_gn=PHIP;refGene_tx=XM_023254153.1;refGene_gn=PHIP;refGene_tx=XM_023254154.1;refGene_gn=PHIP;refGene_tx=XM_023254155.1;refGene_gn=PHIP;refGene_tx=XM_023254156.1
B2 73786932 73787070
refGene_gn=PHIP;refGene_tx=XM_023254153.1;refGene_gn=PHIP;refGene_tx=XM_023254154.1;refGene_gn=PHIP;refGene_tx=XM_023254155.1;refGene_gn=PHIP;refGene_tx=XM_023254156.1
B2 73790672 73790898
refGene_gn=PHIP;refGene_tx=XM_023254153.1;refGene_gn=PHIP;refGene_tx=XM_023254154.1;refGene_gn=PHIP;refGene_tx=XM_023254155.1;refGene_gn=PHIP;refGene_tx=XM_023254156.1
B2 73792060 73792189
refGene_gn=PHIP;refGene_tx=XM_023254153.1;refGene_gn=PHIP;refGene_tx=XM_023254154.1;refGene_gn=PHIP;refGene_tx=XM_023254155.1;refGene_gn=PHIP;refGene_tx=XM_023254156.1
B2 73804030 73804165
refGene_gn=PHIP;refGene_tx=XM_023254153.1;refGene_gn=PHIP;refGene_tx=XM_023254154.1;refGene_gn=PHIP;refGene_tx=XM_023254155.1;refGene_gn=PHIP;refGene_tx=XM_023254156.1
B2 73804595 73804749
refGene_gn=PHIP;refGene_tx=XM_023254153.1;refGene_gn=PHIP;refGene_tx=XM_023254154.1;refGene_gn=PHIP;refGene_tx=XM_023254155.1;refGene_gn=PHIP;refGene_tx=XM_023254156.1
B2 73805541 73805640
refGene_gn=PHIP;refGene_tx=XM_023254153.1;refGene_gn=PHIP;refGene_tx=XM_023254154.1;refGene_gn=PHIP;refGene_tx=XM_023254155.1;refGene_gn=PHIP;refGene_tx=XM_023254156.1
B2 73806404 73806445
refGene_gn=PHIP;refGene_tx=XM_023254153.1;refGene_gn=PHIP;refGene_tx=XM_023254154.1;refGene_gn=PHIP;refGene_tx=XM_023254155.1;refGene_gn=PHIP;refGene_tx=XM_023254156.1
B2 73806534 73806635
refGene_gn=PHIP;refGene_tx=XM_023254153.1;refGene_gn=PHIP;refGene_tx=XM_023254154.1;refGene_gn=PHIP;refGene_tx=XM_023254155.1;refGene_gn=PHIP;refGene_tx=XM_023254156.1
B2 73808030 73808101
refGene_gn=PHIP;refGene_tx=XM_023254153.1;refGene_gn=PHIP;refGene_tx=XM_023254154.1;refGene_gn=PHIP;refGene_tx=XM_023254155.1;refGene_gn=PHIP;refGene_tx=XM_023254156.1
B2 73812448 73812549
refGene_gn=PHIP;refGene_tx=XM_023254153.1;refGene_gn=PHIP;refGene_tx=XM_023254154.1;refGene_gn=PHIP;refGene_tx=XM_023254155.1;refGene_gn=PHIP;refGene_tx=XM_023254156.1
B2 73812824 73813046
refGene_gn=PHIP;refGene_tx=XM_023254153.1;refGene_gn=PHIP;refGene_tx=X

M_023254154.1;refGene_gn=PHIP;refGene_tx=XM_023254155.1;refGene_gn=PHIP;refGene_tx=XM_023254156.1
B2 73828932 73829093
refGene_gn=PHIP;refGene_tx=XM_023254153.1;refGene_gn=PHIP;refGene_tx=XM_023254154.1;refGene_gn=PHIP;refGene_tx=XM_023254155.1;refGene_gn=PHIP;refGene_tx=XM_023254156.1
B2 73847483 73847582
refGene_gn=PHIP;refGene_tx=XM_023254153.1;refGene_gn=PHIP;refGene_tx=XM_023254154.1;refGene_gn=PHIP;refGene_tx=XM_023254155.1
B2 73847669 73847820
refGene_gn=PHIP;refGene_tx=XM_023254153.1;refGene_gn=PHIP;refGene_tx=XM_023254154.1;refGene_gn=PHIP;refGene_tx=XM_023254155.1
B2 73862475 73862535
refGene_gn=PHIP;refGene_tx=XM_023254153.1;refGene_gn=PHIP;refGene_tx=XM_023254154.1;refGene_gn=PHIP;refGene_tx=XM_023254155.1
B2 73862724 73862754
refGene_gn=PHIP;refGene_tx=XM_023254153.1;refGene_gn=PHIP;refGene_tx=XM_023254154.1;refGene_gn=PHIP;refGene_tx=XM_023254155.1
B2 73862886 73862945
refGene_gn=PHIP;refGene_tx=XM_023254153.1;refGene_gn=PHIP;refGene_tx=XM_023254154.1;refGene_gn=PHIP;refGene_tx=XM_023254155.1
B2 73863060 73863100
refGene_gn=PHIP;refGene_tx=XM_023254153.1;refGene_gn=PHIP;refGene_tx=XM_023254154.1;refGene_gn=PHIP;refGene_tx=XM_023254155.1
B2 73976446 73976485
refGene_gn=HMGN3;refGene_tx=XM_003986322.4;refGene_gn=HMGN3;refGene_tx=XM_006931858.3;refGene_gn=HMGN3;refGene_tx=XM_006931859.3;refGene_gn=HMGN3;refGene_tx=XM_019830883.1
B2 73976769 73976901
refGene_gn=HMGN3;refGene_tx=XM_006931856.2;refGene_gn=HMGN3;refGene_tx=XM_006931858.3;refGene_gn=HMGN3;refGene_tx=XM_006931859.3;refGene_gn=HMGN3;refGene_tx=XM_019830882.1
B2 73977021 73977135
refGene_gn=HMGN3;refGene_tx=XM_003986322.4;refGene_gn=HMGN3;refGene_tx=XM_006931856.2;refGene_gn=HMGN3;refGene_tx=XM_006931858.3;refGene_gn=HMGN3;refGene_tx=XM_006931859.3;refGene_gn=HMGN3;refGene_tx=XM_019830882.1;refGene_gn=HMGN3;refGene_tx=XM_019830883.1
B2 73978516 73978567
refGene_gn=HMGN3;refGene_tx=XM_003986322.4;refGene_gn=HMGN3;refGene_tx=XM_006931856.2;refGene_gn=HMGN3;refGene_tx=XM_006931858.3;refGene_gn=HMGN3;refGene_tx=XM_006931859.3;refGene_gn=HMGN3;refGene_tx=XM_019830882.1;refGene_gn=HMGN3;refGene_tx=XM_019830883.1
B2 73981307 73981337
refGene_gn=HMGN3;refGene_tx=XM_003986322.4;refGene_gn=HMGN3;refGene_tx=XM_006931856.2;refGene_gn=HMGN3;refGene_tx=XM_006931858.3;refGene_gn=HMGN3;refGene_tx=XM_006931859.3;refGene_gn=HMGN3;refGene_tx=XM_019830882.1;refGene_gn=HMGN3;refGene_tx=XM_019830883.1
B2 73988053 73988104
refGene_gn=HMGN3;refGene_tx=XM_003986322.4;refGene_gn=HMGN3;refGene_tx=XM_006931856.2;refGene_gn=HMGN3;refGene_tx=XM_006931858.3;refGene_gn=

HMGN3;refGene_tx=XM_006931859.3;refGene_gn=HMGN3;refGene_tx=XM_0198308
82.1;refGene_gn=HMGN3;refGene_tx=XM_019830883.1
B2	74007012	74007027
refGene_gn=HMGN3;refGene_tx=XM_003986322.4;refGene_gn=HMGN3;refGene_tx
=XM_006931856.2;refGene_gn=HMGN3;refGene_tx=XM_006931858.3;refGene_gn=
HMGN3;refGene_tx=XM_006931859.3;refGene_gn=HMGN3;refGene_tx=XM_0198308
82.1;refGene_gn=HMGN3;refGene_tx=XM_019830883.1
B2	74267775	74268647	refGene_gn=LCA5;refGene_tx=XM_003986325.4
B2	74270374	74270507	refGene_gn=LCA5;refGene_tx=XM_003986325.4
B2	74272622	74272765	refGene_gn=LCA5;refGene_tx=XM_003986325.4
B2	74273471	74273568	refGene_gn=LCA5;refGene_tx=XM_003986325.4
B2	74282647	74282785	refGene_gn=LCA5;refGene_tx=XM_003986325.4
B2	74319716	74319856	refGene_gn=LCA5;refGene_tx=XM_003986325.4
B2	74319858	74320153	refGene_gn=LCA5;refGene_tx=XM_003986325.4
B2	74320154	74320246	refGene_gn=LCA5;refGene_tx=XM_003986325.4
B2	74325198	74325397	refGene_gn=LCA5;refGene_tx=XM_003986325.4
B2	74446133	74446580	refGene_gn=SH3BGRL2;refGene_tx=XM_019831599.2
B2	74481674	74481860	refGene_gn=SH3BGRL2;refGene_tx=XM_019831599.2
B2	74499495	74499576	refGene_gn=SH3BGRL2;refGene_tx=XM_019831599.2
B2	74502529	74502541	refGene_gn=SH3BGRL2;refGene_tx=XM_019831599.2
B2	74686581	74686857	refGene_gn=ELOVL4;refGene_tx=XM_003986334.5
B2	74690091	74690219	refGene_gn=ELOVL4;refGene_tx=XM_003986334.5
B2	74691233	74691405	refGene_gn=ELOVL4;refGene_tx=XM_003986334.5
B2	74692753	74692834	refGene_gn=ELOVL4;refGene_tx=XM_003986334.5
B2	74694001	74694189	refGene_gn=ELOVL4;refGene_tx=XM_003986334.5
B2	74717903	74718003	refGene_gn=ELOVL4;refGene_tx=XM_003986334.5
B2	74754611	74754750
refGene_gn=TTK;refGene_tx=XM_019830885.2;refGene_gn=TTK;refGene_tx=XM_
019830886.2;refGene_gn=TTK;refGene_tx=XM_019830887.2;refGene_gn=TTK;re
fGene_tx=XM_019830888.2;refGene_gn=TTK;refGene_tx=XM_019830889.2;refGe
ne_gn=TTK;refGene_tx=XM_019830890.2;refGene_gn=TTK;refGene_tx=XM_01983
0891.2;refGene_gn=TTK;refGene_tx=XM_019830892.2
B2	74758132	74758355
refGene_gn=TTK;refGene_tx=XM_019830885.2;refGene_gn=TTK;refGene_tx=XM_
019830886.2;refGene_gn=TTK;refGene_tx=XM_019830887.2;refGene_gn=TTK;re
fGene_tx=XM_019830888.2;refGene_gn=TTK;refGene_tx=XM_019830889.2;refGe
ne_gn=TTK;refGene_tx=XM_019830890.2;refGene_gn=TTK;refGene_tx=XM_01983
0891.2;refGene_gn=TTK;refGene_tx=XM_019830892.2
B2	74758711	74758818
refGene_gn=TTK;refGene_tx=XM_019830885.2;refGene_gn=TTK;refGene_tx=XM_
019830886.2;refGene_gn=TTK;refGene_tx=XM_019830887.2;refGene_gn=TTK;re
fGene_tx=XM_019830888.2;refGene_gn=TTK;refGene_tx=XM_019830889.2;refGe
ne_gn=TTK;refGene_tx=XM_019830890.2;refGene_gn=TTK;refGene_tx=XM_01983
0891.2;refGene_gn=TTK;refGene_tx=XM_019830892.2
B2	74761239	74761383
refGene_gn=TTK;refGene_tx=XM_019830885.2;refGene_gn=TTK;refGene_tx=XM_
019830886.2;refGene_gn=TTK;refGene_tx=XM_019830887.2;refGene_gn=TTK;re
fGene_tx=XM_019830888.2;refGene_gn=TTK;refGene_tx=XM_019830889.2;refGe
ne_gn=TTK;refGene_tx=XM_019830890.2;refGene_gn=TTK;refGene_tx=XM_01983
0891.2;refGene_gn=TTK;refGene_tx=XM_019830892.2

B2 74762765 74762880
refGene_gn=TTK;refGene_tx=XM_019830885.2;refGene_gn=TTK;refGene_tx=XM_019830886.2;refGene_gn=TTK;refGene_tx=XM_019830887.2;refGene_gn=TTK;refGene_tx=XM_019830888.2;refGene_gn=TTK;refGene_tx=XM_019830889.2;refGene_gn=TTK;refGene_tx=XM_019830890.2;refGene_gn=TTK;refGene_tx=XM_019830891.2;refGene_gn=TTK;refGene_tx=XM_019830892.2
B2 74763041 74763114
refGene_gn=TTK;refGene_tx=XM_019830885.2;refGene_gn=TTK;refGene_tx=XM_019830886.2;refGene_gn=TTK;refGene_tx=XM_019830887.2;refGene_gn=TTK;refGene_tx=XM_019830888.2;refGene_gn=TTK;refGene_tx=XM_019830889.2;refGene_gn=TTK;refGene_tx=XM_019830890.2;refGene_gn=TTK;refGene_tx=XM_019830891.2;refGene_gn=TTK;refGene_tx=XM_019830892.2
B2 74763453 74763545
refGene_gn=TTK;refGene_tx=XM_019830885.2;refGene_gn=TTK;refGene_tx=XM_019830886.2;refGene_gn=TTK;refGene_tx=XM_019830887.2;refGene_gn=TTK;refGene_tx=XM_019830888.2;refGene_gn=TTK;refGene_tx=XM_019830889.2;refGene_gn=TTK;refGene_tx=XM_019830890.2;refGene_gn=TTK;refGene_tx=XM_019830891.2;refGene_gn=TTK;refGene_tx=XM_019830892.2
B2 74765252 74765343
refGene_gn=TTK;refGene_tx=XM_019830885.2;refGene_gn=TTK;refGene_tx=XM_019830886.2;refGene_gn=TTK;refGene_tx=XM_019830887.2;refGene_gn=TTK;refGene_tx=XM_019830888.2;refGene_gn=TTK;refGene_tx=XM_019830889.2;refGene_gn=TTK;refGene_tx=XM_019830890.2;refGene_gn=TTK;refGene_tx=XM_019830891.2;refGene_gn=TTK;refGene_tx=XM_019830892.2
B2 74766322 74766443
refGene_gn=TTK;refGene_tx=XM_019830885.2;refGene_gn=TTK;refGene_tx=XM_019830886.2;refGene_gn=TTK;refGene_tx=XM_019830887.2;refGene_gn=TTK;refGene_tx=XM_019830888.2;refGene_gn=TTK;refGene_tx=XM_019830889.2;refGene_gn=TTK;refGene_tx=XM_019830890.2;refGene_gn=TTK;refGene_tx=XM_019830891.2;refGene_gn=TTK;refGene_tx=XM_019830892.2
B2 74770786 74770935
refGene_gn=TTK;refGene_tx=XM_019830885.2;refGene_gn=TTK;refGene_tx=XM_019830886.2;refGene_gn=TTK;refGene_tx=XM_019830887.2;refGene_gn=TTK;refGene_tx=XM_019830888.2;refGene_gn=TTK;refGene_tx=XM_019830889.2;refGene_gn=TTK;refGene_tx=XM_019830890.2;refGene_gn=TTK;refGene_tx=XM_019830891.2;refGene_gn=TTK;refGene_tx=XM_019830892.2
B2 74775085 74775222
refGene_gn=TTK;refGene_tx=XM_019830885.2;refGene_gn=TTK;refGene_tx=XM_019830886.2;refGene_gn=TTK;refGene_tx=XM_019830887.2;refGene_gn=TTK;refGene_tx=XM_019830888.2;refGene_gn=TTK;refGene_tx=XM_019830889.2;refGene_gn=TTK;refGene_tx=XM_019830890.2;refGene_gn=TTK;refGene_tx=XM_019830891.2;refGene_gn=TTK;refGene_tx=XM_019830892.2
B2 74776786 74776910
refGene_gn=TTK;refGene_tx=XM_019830885.2;refGene_gn=TTK;refGene_tx=XM_019830886.2;refGene_gn=TTK;refGene_tx=XM_019830887.2;refGene_gn=TTK;refGene_tx=XM_019830888.2;refGene_gn=TTK;refGene_tx=XM_019830889.2;refGene_gn=TTK;refGene_tx=XM_019830890.2;refGene_gn=TTK;refGene_tx=XM_019830891.2;refGene_gn=TTK;refGene_tx=XM_019830892.2
B2 74779704 74779797
refGene_gn=TTK;refGene_tx=XM_019830885.2;refGene_gn=TTK;refGene_tx=XM_

019830886.2;refGene_gn=TTK;refGene_tx=XM_019830887.2;refGene_gn=TTK;refGene_tx=XM_019830888.2;refGene_gn=TTK;refGene_tx=XM_019830889.2;refGene_gn=TTK;refGene_tx=XM_019830890.2;refGene_gn=TTK;refGene_tx=XM_019830891.2;refGene_gn=TTK;refGene_tx=XM_019830892.2
B2 74784042 74784200
refGene_gn=TTK;refGene_tx=XM_019830885.2;refGene_gn=TTK;refGene_tx=XM_019830886.2;refGene_gn=TTK;refGene_tx=XM_019830887.2;refGene_gn=TTK;refGene_tx=XM_019830888.2;refGene_gn=TTK;refGene_tx=XM_019830889.2;refGene_gn=TTK;refGene_tx=XM_019830890.2;refGene_gn=TTK;refGene_tx=XM_019830891.2;refGene_gn=TTK;refGene_tx=XM_019830892.2
B2 74784566 74784718
refGene_gn=TTK;refGene_tx=XM_019830885.2;refGene_gn=TTK;refGene_tx=XM_019830886.2;refGene_gn=TTK;refGene_tx=XM_019830887.2;refGene_gn=TTK;refGene_tx=XM_019830888.2;refGene_gn=TTK;refGene_tx=XM_019830889.2;refGene_gn=TTK;refGene_tx=XM_019830890.2;refGene_gn=TTK;refGene_tx=XM_019830891.2;refGene_gn=TTK;refGene_tx=XM_019830892.2
B2 74786157 74786282
refGene_gn=TTK;refGene_tx=XM_019830885.2;refGene_gn=TTK;refGene_tx=XM_019830886.2;refGene_gn=TTK;refGene_tx=XM_019830887.2;refGene_gn=TTK;refGene_tx=XM_019830888.2;refGene_gn=TTK;refGene_tx=XM_019830889.2;refGene_gn=TTK;refGene_tx=XM_019830890.2;refGene_gn=TTK;refGene_tx=XM_019830891.2;refGene_gn=TTK;refGene_tx=XM_019830892.2
B2 74787629 74787710
refGene_gn=TTK;refGene_tx=XM_019830885.2;refGene_gn=TTK;refGene_tx=XM_019830886.2;refGene_gn=TTK;refGene_tx=XM_019830887.2;refGene_gn=TTK;refGene_tx=XM_019830888.2;refGene_gn=TTK;refGene_tx=XM_019830889.2;refGene_gn=TTK;refGene_tx=XM_019830890.2;refGene_gn=TTK;refGene_tx=XM_019830891.2;refGene_gn=TTK;refGene_tx=XM_019830892.2
B2 74791181 74791358
refGene_gn=TTK;refGene_tx=XM_019830885.2;refGene_gn=TTK;refGene_tx=XM_019830886.2;refGene_gn=TTK;refGene_tx=XM_019830887.2;refGene_gn=TTK;refGene_tx=XM_019830888.2;refGene_gn=TTK;refGene_tx=XM_019830889.2;refGene_gn=TTK;refGene_tx=XM_019830890.2;refGene_gn=TTK;refGene_tx=XM_019830891.2;refGene_gn=TTK;refGene_tx=XM_019830892.2
B2 74791702 74791787
refGene_gn=TTK;refGene_tx=XM_019830885.2;refGene_gn=TTK;refGene_tx=XM_019830886.2;refGene_gn=TTK;refGene_tx=XM_019830887.2;refGene_gn=TTK;refGene_tx=XM_019830888.2;refGene_gn=TTK;refGene_tx=XM_019830889.2;refGene_gn=TTK;refGene_tx=XM_019830890.2;refGene_gn=TTK;refGene_tx=XM_019830891.2;refGene_gn=TTK;refGene_tx=XM_019830892.2
B2 74792157 74792255
refGene_gn=TTK;refGene_tx=XM_019830885.2;refGene_gn=TTK;refGene_tx=XM_019830886.2;refGene_gn=TTK;refGene_tx=XM_019830887.2;refGene_gn=TTK;refGene_tx=XM_019830888.2;refGene_gn=TTK;refGene_tx=XM_019830889.2;refGene_gn=TTK;refGene_tx=XM_019830890.2;refGene_gn=TTK;refGene_tx=XM_019830891.2;refGene_gn=TTK;refGene_tx=XM_019830892.2
B2 74793454 74793541
refGene_gn=TTK;refGene_tx=XM_019830885.2;refGene_gn=TTK;refGene_tx=XM_019830886.2;refGene_gn=TTK;refGene_tx=XM_019830887.2;refGene_gn=TTK;refGene_tx=XM_019830888.2;refGene_gn=TTK;refGene_tx=XM_019830889.2;refGe

ne_gn=TTK;refGene_tx=XM_019830890.2;refGene_gn=TTK;refGene_tx=XM_01983
0891.2;refGene_gn=TTK;refGene_tx=XM_019830892.2
B2 74855516 74855712
refGene_gn=BCKDHB;refGene_tx=XM_003986336.4;refGene_gn=BCKDHB;refGene_
tx=XM_006931864.3
B2 74868095 74868173
refGene_gn=BCKDHB;refGene_tx=XM_003986336.4;refGene_gn=BCKDHB;refGene_
tx=XM_006931864.3
B2 74874374 74874443
refGene_gn=BCKDHB;refGene_tx=XM_003986336.4;refGene_gn=BCKDHB;refGene_
tx=XM_006931864.3
B2 74908103 74908237
refGene_gn=BCKDHB;refGene_tx=XM_003986336.4;refGene_gn=BCKDHB;refGene_
tx=XM_006931864.3
B2 74908481 74908637
refGene_gn=BCKDHB;refGene_tx=XM_003986336.4;refGene_gn=BCKDHB;refGene_
tx=XM_006931864.3
B2 74911480 74911589
refGene_gn=BCKDHB;refGene_tx=XM_003986336.4;refGene_gn=BCKDHB;refGene_
tx=XM_006931864.3
B2 74924752 74924850
refGene_gn=BCKDHB;refGene_tx=XM_003986336.4;refGene_gn=BCKDHB;refGene_
tx=XM_006931864.3
B2 74926908 74927019
refGene_gn=BCKDHB;refGene_tx=XM_003986336.4;refGene_gn=BCKDHB;refGene_
tx=XM_006931864.3
B2 75000609 75000696
refGene_gn=BCKDHB;refGene_tx=XM_003986336.4;refGene_gn=BCKDHB;refGene_
tx=XM_006931864.3
B2 75061500 75061641
refGene_gn=BCKDHB;refGene_tx=XM_003986336.4;refGene_gn=BCKDHB;refGene_
tx=XM_006931864.3
B2 76231132 76231909
refGene_gn=FAM46A;refGene_tx=XM_003986337.4;refGene_gn=FAM46A;refGene_
tx=XM_006931865.3
B2 76233008 76233558
refGene_gn=FAM46A;refGene_tx=XM_003986337.4;refGene_gn=FAM46A;refGene_
tx=XM_006931865.3
B2 76233740 76233760 refGene_gn=FAM46A;refGene_tx=XM_006931865.3
B2 76636266 76636344
refGene_gn=IBTK;refGene_tx=XM_011282486.3;refGene_gn=IBTK;refGene_tx=X
M_011282487.3;refGene_gn=IBTK;refGene_tx=XM_011282488.3;refGene_gn=IBT
K;refGene_tx=XM_011282490.3
B2 76637114 76637247
refGene_gn=IBTK;refGene_tx=XM_011282486.3;refGene_gn=IBTK;refGene_tx=X
M_011282487.3;refGene_gn=IBTK;refGene_tx=XM_011282488.3;refGene_gn=IBT
K;refGene_tx=XM_011282490.3
B2 76648126 76648276
refGene_gn=IBTK;refGene_tx=XM_011282486.3;refGene_gn=IBTK;refGene_tx=X
M_011282487.3;refGene_gn=IBTK;refGene_tx=XM_011282488.3;refGene_gn=IBT

K;refGene_tx=XM_011282490.3
B2 76660247 76660391
refGene_gn=IBTK;refGene_tx=XM_011282486.3;refGene_gn=IBTK;refGene_tx=XM_011282487.3;refGene_gn=IBTK;refGene_tx=XM_011282488.3;refGene_gn=IBTK;refGene_tx=XM_011282490.3
B2 76660913 76661006
refGene_gn=IBTK;refGene_tx=XM_011282486.3;refGene_gn=IBTK;refGene_tx=XM_011282487.3;refGene_gn=IBTK;refGene_tx=XM_011282488.3;refGene_gn=IBTK;refGene_tx=XM_011282490.3
B2 76662383 76662547
refGene_gn=IBTK;refGene_tx=XM_011282486.3;refGene_gn=IBTK;refGene_tx=XM_011282487.3;refGene_gn=IBTK;refGene_tx=XM_011282488.3;refGene_gn=IBTK;refGene_tx=XM_011282490.3
B2 76663912 76664061
refGene_gn=IBTK;refGene_tx=XM_011282486.3;refGene_gn=IBTK;refGene_tx=XM_011282487.3;refGene_gn=IBTK;refGene_tx=XM_011282488.3;refGene_gn=IBTK;refGene_tx=XM_011282490.3
B2 76668053 76668166
refGene_gn=IBTK;refGene_tx=XM_011282486.3;refGene_gn=IBTK;refGene_tx=XM_011282487.3;refGene_gn=IBTK;refGene_tx=XM_011282488.3;refGene_gn=IBTK;refGene_tx=XM_011282490.3
B2 76669469 76669591
refGene_gn=IBTK;refGene_tx=XM_011282486.3;refGene_gn=IBTK;refGene_tx=XM_011282487.3;refGene_gn=IBTK;refGene_tx=XM_011282488.3;refGene_gn=IBTK;refGene_tx=XM_011282490.3
B2 76670154 76670215
refGene_gn=IBTK;refGene_tx=XM_011282486.3;refGene_gn=IBTK;refGene_tx=XM_011282487.3;refGene_gn=IBTK;refGene_tx=XM_011282488.3;refGene_gn=IBTK;refGene_tx=XM_011282490.3
B2 76672921 76673039
refGene_gn=IBTK;refGene_tx=XM_011282486.3;refGene_gn=IBTK;refGene_tx=XM_011282487.3;refGene_gn=IBTK;refGene_tx=XM_011282488.3;refGene_gn=IBTK;refGene_tx=XM_011282490.3
B2 76676071 76676173
refGene_gn=IBTK;refGene_tx=XM_011282486.3;refGene_gn=IBTK;refGene_tx=XM_011282487.3;refGene_gn=IBTK;refGene_tx=XM_011282488.3;refGene_gn=IBTK;refGene_tx=XM_011282490.3
B2 76684408 76684505
refGene_gn=IBTK;refGene_tx=XM_011282486.3;refGene_gn=IBTK;refGene_tx=XM_011282487.3;refGene_gn=IBTK;refGene_tx=XM_011282488.3;refGene_gn=IBTK;refGene_tx=XM_011282490.3
B2 76684965 76685003
refGene_gn=IBTK;refGene_tx=XM_011282486.3;refGene_gn=IBTK;refGene_tx=XM_011282487.3;refGene_gn=IBTK;refGene_tx=XM_011282488.3;refGene_gn=IBTK;refGene_tx=XM_011282490.3
B2 76685088 76685171
refGene_gn=IBTK;refGene_tx=XM_011282486.3;refGene_gn=IBTK;refGene_tx=XM_011282487.3;refGene_gn=IBTK;refGene_tx=XM_011282488.3;refGene_gn=IBTK;refGene_tx=XM_011282490.3
B2 76686217 76686304

refGene_gn=IBTK;refGene_tx=XM_011282486.3;refGene_gn=IBTK;refGene_tx=XM_011282487.3;refGene_gn=IBTK;refGene_tx=XM_011282488.3;refGene_gn=IBTK;refGene_tx=XM_011282490.3
B2 76687483 76688083
refGene_gn=IBTK;refGene_tx=XM_011282486.3;refGene_gn=IBTK;refGene_tx=XM_011282487.3;refGene_gn=IBTK;refGene_tx=XM_011282488.3;refGene_gn=IBTK;refGene_tx=XM_011282490.3
B2 76689317 76689495
refGene_gn=IBTK;refGene_tx=XM_011282486.3;refGene_gn=IBTK;refGene_tx=XM_011282487.3;refGene_gn=IBTK;refGene_tx=XM_011282488.3;refGene_gn=IBTK;refGene_tx=XM_011282490.3
B2 76692527 76692705
refGene_gn=IBTK;refGene_tx=XM_011282486.3;refGene_gn=IBTK;refGene_tx=XM_011282487.3;refGene_gn=IBTK;refGene_tx=XM_011282488.3;refGene_gn=IBTK;refGene_tx=XM_011282490.3
B2 76694688 76694812
refGene_gn=IBTK;refGene_tx=XM_011282486.3;refGene_gn=IBTK;refGene_tx=XM_011282487.3;refGene_gn=IBTK;refGene_tx=XM_011282488.3;refGene_gn=IBTK;refGene_tx=XM_011282490.3
B2 76697021 76697202
refGene_gn=IBTK;refGene_tx=XM_011282486.3;refGene_gn=IBTK;refGene_tx=XM_011282487.3;refGene_gn=IBTK;refGene_tx=XM_011282488.3;refGene_gn=IBTK;refGene_tx=XM_011282490.3
B2 76697285 76697403
refGene_gn=IBTK;refGene_tx=XM_011282486.3;refGene_gn=IBTK;refGene_tx=XM_011282487.3;refGene_gn=IBTK;refGene_tx=XM_011282488.3;refGene_gn=IBTK;refGene_tx=XM_011282490.3
B2 76700879 76701050
refGene_gn=IBTK;refGene_tx=XM_011282486.3;refGene_gn=IBTK;refGene_tx=XM_011282487.3;refGene_gn=IBTK;refGene_tx=XM_011282488.3;refGene_gn=IBTK;refGene_tx=XM_011282490.3
B2 76702163 76702274
refGene_gn=IBTK;refGene_tx=XM_011282486.3;refGene_gn=IBTK;refGene_tx=XM_011282487.3;refGene_gn=IBTK;refGene_tx=XM_011282488.3;refGene_gn=IBTK;refGene_tx=XM_011282490.3
B2 76709130 76709255
refGene_gn=IBTK;refGene_tx=XM_011282486.3;refGene_gn=IBTK;refGene_tx=XM_011282487.3;refGene_gn=IBTK;refGene_tx=XM_011282488.3;refGene_gn=IBTK;refGene_tx=XM_011282490.3
B2 76710579 76710676
refGene_gn=IBTK;refGene_tx=XM_011282486.3;refGene_gn=IBTK;refGene_tx=XM_011282487.3;refGene_gn=IBTK;refGene_tx=XM_011282488.3;refGene_gn=IBTK;refGene_tx=XM_011282490.3
B2 76715901 76716222
refGene_gn=IBTK;refGene_tx=XM_011282486.3;refGene_gn=IBTK;refGene_tx=XM_011282487.3;refGene_gn=IBTK;refGene_tx=XM_011282488.3;refGene_gn=IBTK;refGene_tx=XM_011282490.3
B2 76811289 76812549 refGene_gn=TPBG;refGene_tx=XM_019830894.2
B2 77203775 77203937
refGene_gn=UBE3D;refGene_tx=XM_023254157.1;refGene_gn=UBE3D;refGene_tx

=XM_023254158.1
B2 77228290 77229367
refGene_gn=LOC101085976;refGene_tx=XM_011282500.2
B2 77287362 77287383 refGene_gn=UBE3D;refGene_tx=XM_006931866.4
B2 77327528 77327555 refGene_gn=UBE3D;refGene_tx=XM_011282484.3
B2 77355203 77355342
refGene_gn=UBE3D;refGene_tx=XM_006931866.4;refGene_gn=UBE3D;refGene_tx
=XM_011282484.3;refGene_gn=UBE3D;refGene_tx=XM_023254157.1;refGene_gn=
UBE3D;refGene_tx=XM_023254158.1
B2 77397557 77397721
refGene_gn=UBE3D;refGene_tx=XM_006931866.4;refGene_gn=UBE3D;refGene_tx
=XM_011282484.3;refGene_gn=UBE3D;refGene_tx=XM_023254157.1;refGene_gn=
UBE3D;refGene_tx=XM_023254158.1
B2 77400561 77400670
refGene_gn=UBE3D;refGene_tx=XM_006931866.4;refGene_gn=UBE3D;refGene_tx
=XM_011282484.3;refGene_gn=UBE3D;refGene_tx=XM_023254157.1;refGene_gn=
UBE3D;refGene_tx=XM_023254158.1
B2 77401647 77401717
refGene_gn=UBE3D;refGene_tx=XM_006931866.4;refGene_gn=UBE3D;refGene_tx
=XM_011282484.3;refGene_gn=UBE3D;refGene_tx=XM_023254157.1;refGene_gn=
UBE3D;refGene_tx=XM_023254158.1
B2 77409895 77409965
refGene_gn=UBE3D;refGene_tx=XM_006931866.4;refGene_gn=UBE3D;refGene_tx
=XM_011282484.3;refGene_gn=UBE3D;refGene_tx=XM_023254157.1;refGene_gn=
UBE3D;refGene_tx=XM_023254158.1
B2 77415461 77415693
refGene_gn=UBE3D;refGene_tx=XM_006931866.4;refGene_gn=UBE3D;refGene_tx
=XM_011282484.3;refGene_gn=UBE3D;refGene_tx=XM_023254157.1;refGene_gn=
UBE3D;refGene_tx=XM_023254158.1
B2 77427435 77427526
refGene_gn=UBE3D;refGene_tx=XM_006931866.4;refGene_gn=UBE3D;refGene_tx
=XM_011282484.3;refGene_gn=UBE3D;refGene_tx=XM_023254157.1;refGene_gn=
UBE3D;refGene_tx=XM_023254158.1
B2 77431242 77431439
refGene_gn=UBE3D;refGene_tx=XM_006931866.4;refGene_gn=UBE3D;refGene_tx
=XM_011282484.3;refGene_gn=UBE3D;refGene_tx=XM_023254157.1;refGene_gn=
UBE3D;refGene_tx=XM_023254158.1
B2 77441744 77441821
refGene_gn=UBE3D;refGene_tx=XM_006931866.4;refGene_gn=UBE3D;refGene_tx
=XM_011282484.3;refGene_gn=UBE3D;refGene_tx=XM_023254157.1
B2 77444450 77444488 refGene_gn=UBE3D;refGene_tx=XM_023254158.1
B2 77477057 77477195
refGene_gn=DOPEY1;refGene_tx=XM_011282483.3;refGene_gn=DOPEY1;refGene_
tx=XM_019830895.2;refGene_gn=DOPEY1;refGene_tx=XM_019830896.2;refGene_
gn=DOPEY1;refGene_tx=XM_019830898.2;refGene_gn=DOPEY1;refGene_tx=XM_01
9830899.2
B2 77481689 77481871
refGene_gn=DOPEY1;refGene_tx=XM_011282483.3;refGene_gn=DOPEY1;refGene_
tx=XM_019830895.2;refGene_gn=DOPEY1;refGene_tx=XM_019830896.2;refGene_
gn=DOPEY1;refGene_tx=XM_019830898.2;refGene_gn=DOPEY1;refGene_tx=XM_01

9830899.2
B2	77489563 77489734
refGene_gn=DOPEY1;refGene_tx=XM_011282483.3;refGene_gn=DOPEY1;refGene_tx=XM_019830895.2;refGene_gn=DOPEY1;refGene_tx=XM_019830896.2;refGene_gn=DOPEY1;refGene_tx=XM_019830898.2;refGene_gn=DOPEY1;refGene_tx=XM_019830899.2;refGene_gn=DOPEY1;refGene_tx=XM_019830900.2
B2	77492597 77492787
refGene_gn=DOPEY1;refGene_tx=XM_011282483.3;refGene_gn=DOPEY1;refGene_tx=XM_019830895.2;refGene_gn=DOPEY1;refGene_tx=XM_019830896.2;refGene_gn=DOPEY1;refGene_tx=XM_019830898.2;refGene_gn=DOPEY1;refGene_tx=XM_019830899.2;refGene_gn=DOPEY1;refGene_tx=XM_019830900.2
B2	77495121 77495220
refGene_gn=DOPEY1;refGene_tx=XM_011282483.3;refGene_gn=DOPEY1;refGene_tx=XM_019830895.2;refGene_gn=DOPEY1;refGene_tx=XM_019830896.2;refGene_gn=DOPEY1;refGene_tx=XM_019830898.2;refGene_gn=DOPEY1;refGene_tx=XM_019830899.2;refGene_gn=DOPEY1;refGene_tx=XM_019830900.2
B2	77500521 77500621
refGene_gn=DOPEY1;refGene_tx=XM_011282483.3;refGene_gn=DOPEY1;refGene_tx=XM_019830895.2;refGene_gn=DOPEY1;refGene_tx=XM_019830896.2;refGene_gn=DOPEY1;refGene_tx=XM_019830898.2;refGene_gn=DOPEY1;refGene_tx=XM_019830899.2;refGene_gn=DOPEY1;refGene_tx=XM_019830900.2
B2	77501372 77501482
refGene_gn=DOPEY1;refGene_tx=XM_011282483.3;refGene_gn=DOPEY1;refGene_tx=XM_019830895.2;refGene_gn=DOPEY1;refGene_tx=XM_019830896.2;refGene_gn=DOPEY1;refGene_tx=XM_019830898.2;refGene_gn=DOPEY1;refGene_tx=XM_019830899.2;refGene_gn=DOPEY1;refGene_tx=XM_019830900.2
B2	77502373 77502482
refGene_gn=DOPEY1;refGene_tx=XM_011282483.3;refGene_gn=DOPEY1;refGene_tx=XM_019830895.2;refGene_gn=DOPEY1;refGene_tx=XM_019830896.2;refGene_gn=DOPEY1;refGene_tx=XM_019830898.2;refGene_gn=DOPEY1;refGene_tx=XM_019830899.2;refGene_gn=DOPEY1;refGene_tx=XM_019830900.2;refGene_gn=DOPEY1;refGene_tx=XM_019830901.2
B2	77503813 77503934
refGene_gn=DOPEY1;refGene_tx=XM_011282483.3;refGene_gn=DOPEY1;refGene_tx=XM_019830895.2;refGene_gn=DOPEY1;refGene_tx=XM_019830896.2;refGene_gn=DOPEY1;refGene_tx=XM_019830898.2;refGene_gn=DOPEY1;refGene_tx=XM_019830899.2;refGene_gn=DOPEY1;refGene_tx=XM_019830900.2;refGene_gn=DOPEY1;refGene_tx=XM_019830901.2
B2	77506922 77507042
refGene_gn=DOPEY1;refGene_tx=XM_011282483.3;refGene_gn=DOPEY1;refGene_tx=XM_019830895.2;refGene_gn=DOPEY1;refGene_tx=XM_019830896.2;refGene_gn=DOPEY1;refGene_tx=XM_019830898.2;refGene_gn=DOPEY1;refGene_tx=XM_019830899.2;refGene_gn=DOPEY1;refGene_tx=XM_019830900.2;refGene_gn=DOPEY1;refGene_tx=XM_019830901.2
B2	77510141 77510256
refGene_gn=DOPEY1;refGene_tx=XM_011282483.3;refGene_gn=DOPEY1;refGene_tx=XM_019830895.2;refGene_gn=DOPEY1;refGene_tx=XM_019830896.2;refGene_gn=DOPEY1;refGene_tx=XM_019830898.2;refGene_gn=DOPEY1;refGene_tx=XM_019830899.2;refGene_gn=DOPEY1;refGene_tx=XM_019830900.2;refGene_gn=DOPEY1;refGene_tx=XM_019830901.2

B2 77510598 77510628
refGene_gn=DOPEY1;refGene_tx=XM_011282483.3;refGene_gn=DOPEY1;refGene_tx=XM_019830895.2;refGene_gn=DOPEY1;refGene_tx=XM_019830898.2;refGene_gn=DOPEY1;refGene_tx=XM_019830899.2;refGene_gn=DOPEY1;refGene_tx=XM_019830900.2;refGene_gn=DOPEY1;refGene_tx=XM_019830901.2
B2 77510920 77511154
refGene_gn=DOPEY1;refGene_tx=XM_011282483.3;refGene_gn=DOPEY1;refGene_tx=XM_019830895.2;refGene_gn=DOPEY1;refGene_tx=XM_019830896.2;refGene_gn=DOPEY1;refGene_tx=XM_019830898.2;refGene_gn=DOPEY1;refGene_tx=XM_019830899.2;refGene_gn=DOPEY1;refGene_tx=XM_019830900.2;refGene_gn=DOPEY1;refGene_tx=XM_019830901.2
B2 77516849 77517471
refGene_gn=DOPEY1;refGene_tx=XM_011282483.3;refGene_gn=DOPEY1;refGene_tx=XM_019830895.2;refGene_gn=DOPEY1;refGene_tx=XM_019830896.2;refGene_gn=DOPEY1;refGene_tx=XM_019830898.2;refGene_gn=DOPEY1;refGene_tx=XM_019830899.2;refGene_gn=DOPEY1;refGene_tx=XM_019830900.2;refGene_gn=DOPEY1;refGene_tx=XM_019830901.2
B2 77518337 77518612
refGene_gn=DOPEY1;refGene_tx=XM_011282483.3;refGene_gn=DOPEY1;refGene_tx=XM_019830895.2;refGene_gn=DOPEY1;refGene_tx=XM_019830896.2;refGene_gn=DOPEY1;refGene_tx=XM_019830898.2;refGene_gn=DOPEY1;refGene_tx=XM_019830899.2;refGene_gn=DOPEY1;refGene_tx=XM_019830900.2;refGene_gn=DOPEY1;refGene_tx=XM_019830901.2
B2 77519817 77519970
refGene_gn=DOPEY1;refGene_tx=XM_011282483.3;refGene_gn=DOPEY1;refGene_tx=XM_019830895.2;refGene_gn=DOPEY1;refGene_tx=XM_019830896.2;refGene_gn=DOPEY1;refGene_tx=XM_019830898.2;refGene_gn=DOPEY1;refGene_tx=XM_019830899.2;refGene_gn=DOPEY1;refGene_tx=XM_019830900.2;refGene_gn=DOPEY1;refGene_tx=XM_019830901.2
B2 77522053 77522154
refGene_gn=DOPEY1;refGene_tx=XM_011282483.3;refGene_gn=DOPEY1;refGene_tx=XM_019830895.2;refGene_gn=DOPEY1;refGene_tx=XM_019830896.2;refGene_gn=DOPEY1;refGene_tx=XM_019830898.2;refGene_gn=DOPEY1;refGene_tx=XM_019830899.2;refGene_gn=DOPEY1;refGene_tx=XM_019830900.2;refGene_gn=DOPEY1;refGene_tx=XM_019830901.2
B2 77523470 77523730
refGene_gn=DOPEY1;refGene_tx=XM_011282483.3;refGene_gn=DOPEY1;refGene_tx=XM_019830895.2;refGene_gn=DOPEY1;refGene_tx=XM_019830896.2;refGene_gn=DOPEY1;refGene_tx=XM_019830898.2;refGene_gn=DOPEY1;refGene_tx=XM_019830899.2;refGene_gn=DOPEY1;refGene_tx=XM_019830900.2;refGene_gn=DOPEY1;refGene_tx=XM_019830901.2
B2 77525478 77527468
refGene_gn=DOPEY1;refGene_tx=XM_011282483.3;refGene_gn=DOPEY1;refGene_tx=XM_019830895.2;refGene_gn=DOPEY1;refGene_tx=XM_019830896.2;refGene_gn=DOPEY1;refGene_tx=XM_019830898.2;refGene_gn=DOPEY1;refGene_tx=XM_019830899.2;refGene_gn=DOPEY1;refGene_tx=XM_019830900.2;refGene_gn=DOPEY1;refGene_tx=XM_019830901.2
B2 77528296 77528408
refGene_gn=DOPEY1;refGene_tx=XM_011282483.3;refGene_gn=DOPEY1;refGene_tx=XM_019830895.2;refGene_gn=DOPEY1;refGene_tx=XM_019830896.2;refGene_

gn=DOPEY1;refGene_tx=XM_019830898.2;refGene_gn=DOPEY1;refGene_tx=XM_01
9830899.2;refGene_gn=DOPEY1;refGene_tx=XM_019830900.2;refGene_gn=DOPEY
1;refGene_tx=XM_019830901.2
B2 77528513 77528696
refGene_gn=DOPEY1;refGene_tx=XM_011282483.3;refGene_gn=DOPEY1;refGene_
tx=XM_019830895.2;refGene_gn=DOPEY1;refGene_tx=XM_019830896.2;refGene_
gn=DOPEY1;refGene_tx=XM_019830898.2;refGene_gn=DOPEY1;refGene_tx=XM_01
9830899.2;refGene_gn=DOPEY1;refGene_tx=XM_019830900.2;refGene_gn=DOPEY
1;refGene_tx=XM_019830901.2
B2 77530457 77530583
refGene_gn=DOPEY1;refGene_tx=XM_011282483.3;refGene_gn=DOPEY1;refGene_
tx=XM_019830895.2;refGene_gn=DOPEY1;refGene_tx=XM_019830896.2;refGene_
gn=DOPEY1;refGene_tx=XM_019830898.2;refGene_gn=DOPEY1;refGene_tx=XM_01
9830899.2;refGene_gn=DOPEY1;refGene_tx=XM_019830900.2;refGene_gn=DOPEY
1;refGene_tx=XM_019830901.2
B2 77536860 77536995
refGene_gn=DOPEY1;refGene_tx=XM_011282483.3;refGene_gn=DOPEY1;refGene_
tx=XM_019830895.2;refGene_gn=DOPEY1;refGene_tx=XM_019830896.2;refGene_
gn=DOPEY1;refGene_tx=XM_019830898.2;refGene_gn=DOPEY1;refGene_tx=XM_01
9830899.2;refGene_gn=DOPEY1;refGene_tx=XM_019830900.2;refGene_gn=DOPEY
1;refGene_tx=XM_019830901.2
B2 77538636 77538692
refGene_gn=DOPEY1;refGene_tx=XM_011282483.3;refGene_gn=DOPEY1;refGene_
tx=XM_019830895.2;refGene_gn=DOPEY1;refGene_tx=XM_019830896.2;refGene_
gn=DOPEY1;refGene_tx=XM_019830898.2;refGene_gn=DOPEY1;refGene_tx=XM_01
9830899.2;refGene_gn=DOPEY1;refGene_tx=XM_019830900.2;refGene_gn=DOPEY
1;refGene_tx=XM_019830901.2
B2 77539832 77539937
refGene_gn=DOPEY1;refGene_tx=XM_011282483.3;refGene_gn=DOPEY1;refGene_
tx=XM_019830895.2;refGene_gn=DOPEY1;refGene_tx=XM_019830896.2;refGene_
gn=DOPEY1;refGene_tx=XM_019830898.2;refGene_gn=DOPEY1;refGene_tx=XM_01
9830899.2;refGene_gn=DOPEY1;refGene_tx=XM_019830900.2;refGene_gn=DOPEY
1;refGene_tx=XM_019830901.2
B2 77544078 77544145
refGene_gn=DOPEY1;refGene_tx=XM_011282483.3;refGene_gn=DOPEY1;refGene_
tx=XM_019830895.2;refGene_gn=DOPEY1;refGene_tx=XM_019830896.2;refGene_
gn=DOPEY1;refGene_tx=XM_019830898.2;refGene_gn=DOPEY1;refGene_tx=XM_01
9830899.2;refGene_gn=DOPEY1;refGene_tx=XM_019830900.2;refGene_gn=DOPEY
1;refGene_tx=XM_019830901.2
B2 77544314 77544459
refGene_gn=DOPEY1;refGene_tx=XM_011282483.3;refGene_gn=DOPEY1;refGene_
tx=XM_019830895.2;refGene_gn=DOPEY1;refGene_tx=XM_019830896.2;refGene_
gn=DOPEY1;refGene_tx=XM_019830898.2;refGene_gn=DOPEY1;refGene_tx=XM_01
9830899.2;refGene_gn=DOPEY1;refGene_tx=XM_019830900.2;refGene_gn=DOPEY
1;refGene_tx=XM_019830901.2
B2 77544635 77544715
refGene_gn=DOPEY1;refGene_tx=XM_011282483.3;refGene_gn=DOPEY1;refGene_
tx=XM_019830895.2;refGene_gn=DOPEY1;refGene_tx=XM_019830896.2;refGene_
gn=DOPEY1;refGene_tx=XM_019830898.2;refGene_gn=DOPEY1;refGene_tx=XM_01
9830899.2;refGene_gn=DOPEY1;refGene_tx=XM_019830900.2;refGene_gn=DOPEY

1;refGene_tx=XM_019830901.2
B2 77545784 77545894
refGene_gn=DOPEY1;refGene_tx=XM_011282483.3;refGene_gn=DOPEY1;refGene_tx=XM_019830895.2;refGene_gn=DOPEY1;refGene_tx=XM_019830896.2;refGene_gn=DOPEY1;refGene_tx=XM_019830898.2;refGene_gn=DOPEY1;refGene_tx=XM_019830899.2;refGene_gn=DOPEY1;refGene_tx=XM_019830900.2;refGene_gn=DOPEY1;refGene_tx=XM_019830901.2
B2 77546149 77546299
refGene_gn=DOPEY1;refGene_tx=XM_011282483.3;refGene_gn=DOPEY1;refGene_tx=XM_019830895.2;refGene_gn=DOPEY1;refGene_tx=XM_019830896.2;refGene_gn=DOPEY1;refGene_tx=XM_019830898.2;refGene_gn=DOPEY1;refGene_tx=XM_019830899.2;refGene_gn=DOPEY1;refGene_tx=XM_019830900.2;refGene_gn=DOPEY1;refGene_tx=XM_019830901.2
B2 77546438 77546500
refGene_gn=DOPEY1;refGene_tx=XM_011282483.3;refGene_gn=DOPEY1;refGene_tx=XM_019830895.2;refGene_gn=DOPEY1;refGene_tx=XM_019830896.2;refGene_gn=DOPEY1;refGene_tx=XM_019830898.2;refGene_gn=DOPEY1;refGene_tx=XM_019830899.2;refGene_gn=DOPEY1;refGene_tx=XM_019830900.2;refGene_gn=DOPEY1;refGene_tx=XM_019830901.2
B2 77548078 77548231
refGene_gn=DOPEY1;refGene_tx=XM_011282483.3;refGene_gn=DOPEY1;refGene_tx=XM_019830895.2;refGene_gn=DOPEY1;refGene_tx=XM_019830896.2;refGene_gn=DOPEY1;refGene_tx=XM_019830898.2;refGene_gn=DOPEY1;refGene_tx=XM_019830899.2;refGene_gn=DOPEY1;refGene_tx=XM_019830900.2;refGene_gn=DOPEY1;refGene_tx=XM_019830901.2
B2 77549333 77549470
refGene_gn=DOPEY1;refGene_tx=XM_011282483.3;refGene_gn=DOPEY1;refGene_tx=XM_019830895.2;refGene_gn=DOPEY1;refGene_tx=XM_019830896.2;refGene_gn=DOPEY1;refGene_tx=XM_019830898.2;refGene_gn=DOPEY1;refGene_tx=XM_019830899.2;refGene_gn=DOPEY1;refGene_tx=XM_019830900.2;refGene_gn=DOPEY1;refGene_tx=XM_019830901.2
B2 77550360 77550416
refGene_gn=DOPEY1;refGene_tx=XM_011282483.3;refGene_gn=DOPEY1;refGene_tx=XM_019830895.2;refGene_gn=DOPEY1;refGene_tx=XM_019830896.2;refGene_gn=DOPEY1;refGene_tx=XM_019830898.2;refGene_gn=DOPEY1;refGene_tx=XM_019830899.2;refGene_gn=DOPEY1;refGene_tx=XM_019830900.2;refGene_gn=DOPEY1;refGene_tx=XM_019830901.2
B2 77551269 77551434
refGene_gn=DOPEY1;refGene_tx=XM_011282483.3;refGene_gn=DOPEY1;refGene_tx=XM_019830895.2;refGene_gn=DOPEY1;refGene_tx=XM_019830896.2;refGene_gn=DOPEY1;refGene_tx=XM_019830898.2;refGene_gn=DOPEY1;refGene_tx=XM_019830899.2;refGene_gn=DOPEY1;refGene_tx=XM_019830900.2;refGene_gn=DOPEY1;refGene_tx=XM_019830901.2
B2 77554337 77554467
refGene_gn=DOPEY1;refGene_tx=XM_011282483.3;refGene_gn=DOPEY1;refGene_tx=XM_019830895.2;refGene_gn=DOPEY1;refGene_tx=XM_019830896.2;refGene_gn=DOPEY1;refGene_tx=XM_019830898.2;refGene_gn=DOPEY1;refGene_tx=XM_019830899.2;refGene_gn=DOPEY1;refGene_tx=XM_019830900.2;refGene_gn=DOPEY1;refGene_tx=XM_019830901.2
B2 77555428 77555488

refGene_gn=DOPEY1;refGene_tx=XM_011282483.3;refGene_gn=DOPEY1;refGene_tx=XM_019830895.2;refGene_gn=DOPEY1;refGene_tx=XM_019830896.2;refGene_gn=DOPEY1;refGene_tx=XM_019830900.2;refGene_gn=DOPEY1;refGene_tx=XM_019830901.2
B2 77557476 77557782
refGene_gn=DOPEY1;refGene_tx=XM_011282483.3;refGene_gn=DOPEY1;refGene_tx=XM_019830895.2;refGene_gn=DOPEY1;refGene_tx=XM_019830896.2;refGene_gn=DOPEY1;refGene_tx=XM_019830898.2;refGene_gn=DOPEY1;refGene_tx=XM_019830899.2;refGene_gn=DOPEY1;refGene_tx=XM_019830900.2;refGene_gn=DOPEY1;refGene_tx=XM_019830901.2
B2 77558838 77558916 refGene_gn=PGM3;refGene_tx=XM_003986339.5
B2 77559972 77560146 refGene_gn=PGM3;refGene_tx=XM_003986339.5
B2 77561131 77561254 refGene_gn=PGM3;refGene_tx=XM_003986339.5
B2 77561932 77562049 refGene_gn=PGM3;refGene_tx=XM_003986339.5
B2 77563966 77564065 refGene_gn=PGM3;refGene_tx=XM_003986339.5
B2 77566501 77566585 refGene_gn=PGM3;refGene_tx=XM_003986339.5
B2 77567302 77567460 refGene_gn=PGM3;refGene_tx=XM_003986339.5
B2 77569891 77570087 refGene_gn=PGM3;refGene_tx=XM_003986339.5
B2 77571009 77571143 refGene_gn=PGM3;refGene_tx=XM_003986339.5
B2 77576293 77576361 refGene_gn=PGM3;refGene_tx=XM_003986339.5
B2 77578158 77578343 refGene_gn=PGM3;refGene_tx=XM_003986339.5
B2 77579755 77579959 refGene_gn=PGM3;refGene_tx=XM_003986339.5
B2 77581952 77581965 refGene_gn=RWDD2A;refGene_tx=XM_019830905.2
B2 77582984 77583103 refGene_gn=RWDD2A;refGene_tx=XM_019830905.2
B2 77583104 77583203 refGene_gn=RWDD2A;refGene_tx=XM_019830905.2
B2 77584222 77584900 refGene_gn=RWDD2A;refGene_tx=XM_019830905.2
B2 77595471 77595642 refGene_gn=ME1;refGene_tx=XM_011282499.3
B2 77599530 77599629 refGene_gn=ME1;refGene_tx=XM_011282499.3
B2 77607298 77607472 refGene_gn=ME1;refGene_tx=XM_011282499.3
B2 77609980 77610123 refGene_gn=ME1;refGene_tx=XM_011282499.3
B2 77611493 77611599 refGene_gn=ME1;refGene_tx=XM_011282499.3
B2 77627706 77627820 refGene_gn=ME1;refGene_tx=XM_011282499.3
B2 77629532 77629630 refGene_gn=ME1;refGene_tx=XM_011282499.3
B2 77644801 77644911 refGene_gn=ME1;refGene_tx=XM_011282499.3
B2 77690791 77690895 refGene_gn=ME1;refGene_tx=XM_011282499.3
B2 77702144 77702306 refGene_gn=ME1;refGene_tx=XM_011282499.3
B2 77708331 77708407 refGene_gn=ME1;refGene_tx=XM_011282499.3
B2 77752246 77752396 refGene_gn=ME1;refGene_tx=XM_011282499.3
B2 77756450 77756584 refGene_gn=ME1;refGene_tx=XM_011282499.3
B2 77784188 77784266 refGene_gn=ME1;refGene_tx=XM_011282499.3
B2 77850794 77850969
refGene_gn=LOC101082405;refGene_tx=XM_023254751.1
B2 77850971 77851307
refGene_gn=LOC101082405;refGene_tx=XM_023254751.1
B2 77851309 77851648
refGene_gn=LOC101082405;refGene_tx=XM_023254751.1
B2 77851880 77852068
refGene_gn=LOC101082405;refGene_tx=XM_023254751.1
B2 77852069 77852333
refGene_gn=LOC101082405;refGene_tx=XM_023254751.1

B2 77885713 77885724 refGene_gn=PRSS35;refGene_tx=XM_003986342.5
B2 77896161 77897422 refGene_gn=PRSS35;refGene_tx=XM_003986342.5
B2 77915007 77915100 refGene_gn=SNAP91;refGene_tx=XM_023254159.1
B2 77919041 77919146 refGene_gn=SNAP91;refGene_tx=XM_023254159.1
B2 77920878 77920962 refGene_gn=SNAP91;refGene_tx=XM_023254159.1
B2 77931573 77931685 refGene_gn=SNAP91;refGene_tx=XM_023254159.1
B2 77932649 77932680 refGene_gn=SNAP91;refGene_tx=XM_023254159.1
B2 77936452 77936602 refGene_gn=SNAP91;refGene_tx=XM_023254159.1
B2 77938255 77938390 refGene_gn=SNAP91;refGene_tx=XM_023254159.1
B2 77946951 77947035 refGene_gn=SNAP91;refGene_tx=XM_023254159.1
B2 77948361 77948445 refGene_gn=SNAP91;refGene_tx=XM_023254159.1
B2 77948883 77948955 refGene_gn=SNAP91;refGene_tx=XM_023254159.1
B2 77949129 77949206 refGene_gn=SNAP91;refGene_tx=XM_023254159.1
B2 77949443 77949697 refGene_gn=SNAP91;refGene_tx=XM_023254159.1
B2 77949698 77949709 refGene_gn=SNAP91;refGene_tx=XM_023254159.1
B2 77950340 77950448 refGene_gn=SNAP91;refGene_tx=XM_023254159.1
B2 77956699 77956867 refGene_gn=SNAP91;refGene_tx=XM_023254159.1
B2 77957012 77957027 refGene_gn=SNAP91;refGene_tx=XM_023254159.1
B2 77971759 77971884 refGene_gn=SNAP91;refGene_tx=XM_023254159.1
B2 77973978 77974088 refGene_gn=SNAP91;refGene_tx=XM_023254159.1
B2 77977956 77977984 refGene_gn=SNAP91;refGene_tx=XM_023254159.1
B2 77980824 77980830 refGene_gn=SNAP91;refGene_tx=XM_023254159.1
B2 77982924 77982995 refGene_gn=SNAP91;refGene_tx=XM_023254159.1
B2 77988782 77988824 refGene_gn=SNAP91;refGene_tx=XM_023254159.1
B2 77993225 77993332 refGene_gn=SNAP91;refGene_tx=XM_023254159.1
B2 78003027 78003139 refGene_gn=SNAP91;refGene_tx=XM_023254159.1
B2 78004851 78004945 refGene_gn=SNAP91;refGene_tx=XM_023254159.1
B2 78006618 78006721 refGene_gn=SNAP91;refGene_tx=XM_023254159.1
B2 78007485 78007561 refGene_gn=SNAP91;refGene_tx=XM_023254159.1
B2 78008201 78008344 refGene_gn=SNAP91;refGene_tx=XM_023254159.1
B2 78056259 78056389 refGene_gn=SNAP91;refGene_tx=XM_023254159.1
B2 78179671 78180022 refGene_gn=RIPPLY2;refGene_tx=XM_023254732.1
B2 78180346 78180411 refGene_gn=RIPPLY2;refGene_tx=XM_023254732.1
B2 78183835 78183983 refGene_gn=RIPPLY2;refGene_tx=XM_023254732.1
B2 78186857 78186932
refGene_gn=LOC101083174;refGene_tx=XM_003986344.3;refGene_gn=LOC101083
174;refGene_tx=XM_019830914.2
B2 78190856 78191010
refGene_gn=LOC101083174;refGene_tx=XM_003986344.3;refGene_gn=LOC101083
174;refGene_tx=XM_019830914.2;refGene_gn=LOC101083174;refGene_tx=XM_01
9830915.2
B2 78216656 78216757
refGene_gn=LOC101083174;refGene_tx=XM_003986344.3;refGene_gn=LOC101083
174;refGene_tx=XM_019830914.2;refGene_gn=LOC101083174;refGene_tx=XM_01
9830915.2
B2 78220337 78220419
refGene_gn=LOC101083174;refGene_tx=XM_003986344.3;refGene_gn=LOC101083
174;refGene_tx=XM_019830914.2;refGene_gn=LOC101083174;refGene_tx=XM_01
9830915.2
B2 78230990 78231020

refGene_gn=LOC101083174;refGene_tx=XM_003986344.3;refGene_gn=LOC101083174;refGene_tx=XM_019830914.2;refGene_gn=LOC101083174;refGene_tx=XM_019830915.2
B2 78235118 78235179
refGene_gn=LOC101083174;refGene_tx=XM_003986344.3;refGene_gn=LOC101083174;refGene_tx=XM_019830914.2;refGene_gn=LOC101083174;refGene_tx=XM_019830915.2
B2 78236611 78236669
refGene_gn=LOC101083174;refGene_tx=XM_003986344.3;refGene_gn=LOC101083174;refGene_tx=XM_019830914.2;refGene_gn=LOC101083174;refGene_tx=XM_019830915.2
B2 78238405 78238499
refGene_gn=LOC101083174;refGene_tx=XM_003986344.3;refGene_gn=LOC101083174;refGene_tx=XM_019830914.2;refGene_gn=LOC101083174;refGene_tx=XM_019830915.2
B2 78241904 78241937
refGene_gn=LOC101083174;refGene_tx=XM_003986344.3;refGene_gn=LOC101083174;refGene_tx=XM_019830914.2;refGene_gn=LOC101083174;refGene_tx=XM_019830915.2
B2 78243957 78244080
refGene_gn=LOC101083174;refGene_tx=XM_003986344.3;refGene_gn=LOC101083174;refGene_tx=XM_019830914.2;refGene_gn=LOC101083174;refGene_tx=XM_019830915.2
B2 78259424 78259565
refGene_gn=LOC101083174;refGene_tx=XM_003986344.3;refGene_gn=LOC101083174;refGene_tx=XM_019830914.2;refGene_gn=LOC101083174;refGene_tx=XM_019830915.2
B2 78261098 78261251
refGene_gn=LOC101083174;refGene_tx=XM_003986344.3;refGene_gn=LOC101083174;refGene_tx=XM_019830914.2;refGene_gn=LOC101083174;refGene_tx=XM_019830915.2
B2 78264059 78264210
refGene_gn=LOC101083174;refGene_tx=XM_003986344.3;refGene_gn=LOC101083174;refGene_tx=XM_019830914.2;refGene_gn=LOC101083174;refGene_tx=XM_019830915.2
B2 78264516 78264603
refGene_gn=LOC101083174;refGene_tx=XM_003986344.3;refGene_gn=LOC101083174;refGene_tx=XM_019830914.2;refGene_gn=LOC101083174;refGene_tx=XM_019830915.2
B2 78275488 78275653
refGene_gn=LOC101083174;refGene_tx=XM_003986344.3;refGene_gn=LOC101083174;refGene_tx=XM_019830914.2;refGene_gn=LOC101083174;refGene_tx=XM_019830915.2
B2 78279375 78279499
refGene_gn=LOC101083174;refGene_tx=XM_019830914.2;refGene_gn=LOC101083174;refGene_tx=XM_019830915.2
B2 78279999 78280054
refGene_gn=LOC101083174;refGene_tx=XM_003986344.3;refGene_gn=LOC101083174;refGene_tx=XM_019830914.2;refGene_gn=LOC101083174;refGene_tx=XM_019830915.2

B2 78392653 78392780 refGene_gn=MRAP2;refGene_tx=XM_003986345.5
B2 78405451 78405551 refGene_gn=MRAP2;refGene_tx=XM_003986345.5
B2 78422273 78422667
refGene_gn=MRAP2;refGene_tx=XM_003986345.5;refGene_gn=MRAP2;refGene_tx=XM_019830919.1
B2 78439621 78439831
refGene_gn=CEP162;refGene_tx=XM_011282478.2;refGene_gn=CEP162;refGene_tx=XM_019830920.1;refGene_gn=CEP162;refGene_tx=XM_019830922.1;refGene_gn=CEP162;refGene_tx=XM_019830923.1
B2 78456027 78456162
refGene_gn=CEP162;refGene_tx=XM_011282478.2;refGene_gn=CEP162;refGene_tx=XM_019830920.1;refGene_gn=CEP162;refGene_tx=XM_019830922.1;refGene_gn=CEP162;refGene_tx=XM_019830923.1
B2 78464827 78464926
refGene_gn=CEP162;refGene_tx=XM_011282478.2;refGene_gn=CEP162;refGene_tx=XM_019830920.1;refGene_gn=CEP162;refGene_tx=XM_019830922.1;refGene_gn=CEP162;refGene_tx=XM_019830923.1
B2 78467198 78467340
refGene_gn=CEP162;refGene_tx=XM_011282478.2;refGene_gn=CEP162;refGene_tx=XM_019830920.1;refGene_gn=CEP162;refGene_tx=XM_019830922.1;refGene_gn=CEP162;refGene_tx=XM_019830923.1
B2 78470617 78471270
refGene_gn=CEP162;refGene_tx=XM_011282478.2;refGene_gn=CEP162;refGene_tx=XM_019830920.1;refGene_gn=CEP162;refGene_tx=XM_019830922.1;refGene_gn=CEP162;refGene_tx=XM_019830923.1
B2 78474835 78475048
refGene_gn=CEP162;refGene_tx=XM_011282478.2;refGene_gn=CEP162;refGene_tx=XM_019830920.1;refGene_gn=CEP162;refGene_tx=XM_019830922.1;refGene_gn=CEP162;refGene_tx=XM_019830923.1
B2 78482796 78482901
refGene_gn=CEP162;refGene_tx=XM_011282478.2;refGene_gn=CEP162;refGene_tx=XM_019830920.1;refGene_gn=CEP162;refGene_tx=XM_019830922.1;refGene_gn=CEP162;refGene_tx=XM_019830923.1
B2 78483881 78484045
refGene_gn=CEP162;refGene_tx=XM_011282478.2;refGene_gn=CEP162;refGene_tx=XM_019830920.1;refGene_gn=CEP162;refGene_tx=XM_019830922.1;refGene_gn=CEP162;refGene_tx=XM_019830923.1
B2 78485244 78485371
refGene_gn=CEP162;refGene_tx=XM_011282478.2;refGene_gn=CEP162;refGene_tx=XM_019830920.1;refGene_gn=CEP162;refGene_tx=XM_019830922.1;refGene_gn=CEP162;refGene_tx=XM_019830923.1
B2 78490949 78491055
refGene_gn=CEP162;refGene_tx=XM_011282478.2;refGene_gn=CEP162;refGene_tx=XM_019830920.1;refGene_gn=CEP162;refGene_tx=XM_019830922.1;refGene_gn=CEP162;refGene_tx=XM_019830923.1
B2 78492669 78492782
refGene_gn=CEP162;refGene_tx=XM_011282478.2;refGene_gn=CEP162;refGene_tx=XM_019830920.1;refGene_gn=CEP162;refGene_tx=XM_019830922.1;refGene_gn=CEP162;refGene_tx=XM_019830923.1
B2 78494544 78494685

refGene_gn=CEP162;refGene_tx=XM_011282478.2;refGene_gn=CEP162;refGene_tx=XM_019830920.1;refGene_gn=CEP162;refGene_tx=XM_019830922.1;refGene_gn=CEP162;refGene_tx=XM_019830923.1
B2 78495224 78495452
refGene_gn=CEP162;refGene_tx=XM_011282478.2;refGene_gn=CEP162;refGene_tx=XM_019830920.1;refGene_gn=CEP162;refGene_tx=XM_019830922.1;refGene_gn=CEP162;refGene_tx=XM_019830923.1
B2 78495709 78495843
refGene_gn=CEP162;refGene_tx=XM_011282478.2;refGene_gn=CEP162;refGene_tx=XM_019830920.1;refGene_gn=CEP162;refGene_tx=XM_019830922.1;refGene_gn=CEP162;refGene_tx=XM_019830923.1
B2 78503342 78503604
refGene_gn=CEP162;refGene_tx=XM_011282478.2;refGene_gn=CEP162;refGene_tx=XM_019830920.1;refGene_gn=CEP162;refGene_tx=XM_019830922.1;refGene_gn=CEP162;refGene_tx=XM_019830923.1
B2 78504485 78504768
refGene_gn=CEP162;refGene_tx=XM_011282478.2;refGene_gn=CEP162;refGene_tx=XM_019830920.1;refGene_gn=CEP162;refGene_tx=XM_019830922.1;refGene_gn=CEP162;refGene_tx=XM_019830923.1
B2 78509438 78509520
refGene_gn=CEP162;refGene_tx=XM_011282478.2;refGene_gn=CEP162;refGene_tx=XM_019830920.1;refGene_gn=CEP162;refGene_tx=XM_019830922.1;refGene_gn=CEP162;refGene_tx=XM_019830923.1
B2 78510112 78510304
refGene_gn=CEP162;refGene_tx=XM_011282478.2;refGene_gn=CEP162;refGene_tx=XM_019830920.1;refGene_gn=CEP162;refGene_tx=XM_019830922.1;refGene_gn=CEP162;refGene_tx=XM_019830923.1
B2 78514818 78514932
refGene_gn=CEP162;refGene_tx=XM_011282478.2;refGene_gn=CEP162;refGene_tx=XM_019830920.1;refGene_gn=CEP162;refGene_tx=XM_019830922.1;refGene_gn=CEP162;refGene_tx=XM_019830923.1
B2 78522528 78522559
refGene_gn=CEP162;refGene_tx=XM_011282478.2;refGene_gn=CEP162;refGene_tx=XM_019830920.1;refGene_gn=CEP162;refGene_tx=XM_019830922.1;refGene_gn=CEP162;refGene_tx=XM_019830923.1
B2 78524746 78524862
refGene_gn=CEP162;refGene_tx=XM_011282478.2;refGene_gn=CEP162;refGene_tx=XM_019830920.1;refGene_gn=CEP162;refGene_tx=XM_019830922.1;refGene_gn=CEP162;refGene_tx=XM_019830923.1
B2 78531595 78531660
refGene_gn=CEP162;refGene_tx=XM_011282478.2;refGene_gn=CEP162;refGene_tx=XM_019830920.1;refGene_gn=CEP162;refGene_tx=XM_019830922.1
B2 78534320 78534504
refGene_gn=CEP162;refGene_tx=XM_011282478.2;refGene_gn=CEP162;refGene_tx=XM_019830920.1;refGene_gn=CEP162;refGene_tx=XM_019830922.1;refGene_gn=CEP162;refGene_tx=XM_019830923.1
B2 78534830 78534977
refGene_gn=CEP162;refGene_tx=XM_011282478.2;refGene_gn=CEP162;refGene_tx=XM_019830920.1;refGene_gn=CEP162;refGene_tx=XM_019830922.1
B2 78537938 78538053

refGene_gn=CEP162;refGene_tx=XM_011282478.2;refGene_gn=CEP162;refGene_tx=XM_019830920.1;refGene_gn=CEP162;refGene_tx=XM_019830922.1
B2 78550285 78550342
refGene_gn=CEP162;refGene_tx=XM_011282478.2;refGene_gn=CEP162;refGene_tx=XM_019830920.1;refGene_gn=CEP162;refGene_tx=XM_019830922.1
B2 78969539 78970264 refGene_gn=TBX18;refGene_tx=XM_003986346.5
B2 78971813 78971908 refGene_gn=TBX18;refGene_tx=XM_003986346.5
B2 78977942 78978007 refGene_gn=TBX18;refGene_tx=XM_003986346.5
B2 78981236 78981404 refGene_gn=TBX18;refGene_tx=XM_003986346.5
B2 78989291 78989463 refGene_gn=TBX18;refGene_tx=XM_003986346.5
B2 78992454 78992556 refGene_gn=TBX18;refGene_tx=XM_003986346.5
B2 78994733 78994938 refGene_gn=TBX18;refGene_tx=XM_003986346.5
B2 78996090 78996886 refGene_gn=TBX18;refGene_tx=XM_003986346.5
B2 79580794 79581133
refGene_gn=NT5E;refGene_tx=XM_011282497.2;refGene_gn=NT5E;refGene_tx=XM_023254164.1
B2 79600573 79600796
refGene_gn=NT5E;refGene_tx=XM_011282497.2;refGene_gn=NT5E;refGene_tx=XM_023254164.1
B2 79605730 79605919
refGene_gn=NT5E;refGene_tx=XM_011282497.2;refGene_gn=NT5E;refGene_tx=XM_023254164.1
B2 79616811 79617009
refGene_gn=NT5E;refGene_tx=XM_011282497.2;refGene_gn=NT5E;refGene_tx=XM_023254164.1
B2 79617927 79618082
refGene_gn=NT5E;refGene_tx=XM_011282497.2;refGene_gn=NT5E;refGene_tx=XM_023254164.1
B2 79621575 79621681
refGene_gn=NT5E;refGene_tx=XM_011282497.2;refGene_gn=NT5E;refGene_tx=XM_023254164.1
B2 79622570 79622720
refGene_gn=NT5E;refGene_tx=XM_011282497.2;refGene_gn=NT5E;refGene_tx=XM_023254164.1
B2 79625118 79625319
refGene_gn=NT5E;refGene_tx=XM_011282497.2;refGene_gn=NT5E;refGene_tx=XM_023254164.1
B2 79628068 79628232 refGene_gn=NT5E;refGene_tx=XM_011282497.2
B2 79628287 79628310 refGene_gn=NT5E;refGene_tx=XM_023254164.1
B2 79633783 79633882
refGene_gn=SNX14;refGene_tx=XM_019830924.2;refGene_gn=SNX14;refGene_tx=XM_023254160.1;refGene_gn=SNX14;refGene_tx=XM_023254161.1
B2 79636489 79636528
refGene_gn=SNX14;refGene_tx=XM_011282475.3;refGene_gn=SNX14;refGene_tx=XM_011282477.3;refGene_gn=SNX14;refGene_tx=XM_023254162.1;refGene_gn=SNX14;refGene_tx=XM_023254163.1
B2 79637086 79637143
refGene_gn=SNX14;refGene_tx=XM_011282475.3;refGene_gn=SNX14;refGene_tx=XM_011282477.3;refGene_gn=SNX14;refGene_tx=XM_019830924.2;refGene_gn=SNX14;refGene_tx=XM_023254160.1;refGene_gn=SNX14;refGene_tx=XM_0232541

61.1;refGene_gn=SNX14;refGene_tx=XM_023254162.1;refGene_gn=SNX14;refGene_tx=XM_023254163.1
B2 79637939 79638031
refGene_gn=SNX14;refGene_tx=XM_011282475.3;refGene_gn=SNX14;refGene_tx=XM_011282477.3;refGene_gn=SNX14;refGene_tx=XM_019830924.2;refGene_gn=SNX14;refGene_tx=XM_023254160.1;refGene_gn=SNX14;refGene_tx=XM_023254161.1;refGene_gn=SNX14;refGene_tx=XM_023254162.1;refGene_gn=SNX14;refGene_tx=XM_023254163.1
B2 79640320 79640416
refGene_gn=SNX14;refGene_tx=XM_011282475.3;refGene_gn=SNX14;refGene_tx=XM_011282477.3;refGene_gn=SNX14;refGene_tx=XM_019830924.2;refGene_gn=SNX14;refGene_tx=XM_023254160.1;refGene_gn=SNX14;refGene_tx=XM_023254161.1;refGene_gn=SNX14;refGene_tx=XM_023254162.1;refGene_gn=SNX14;refGene_tx=XM_023254163.1
B2 79640549 79640714
refGene_gn=SNX14;refGene_tx=XM_011282475.3;refGene_gn=SNX14;refGene_tx=XM_011282477.3;refGene_gn=SNX14;refGene_tx=XM_019830924.2;refGene_gn=SNX14;refGene_tx=XM_023254160.1;refGene_gn=SNX14;refGene_tx=XM_023254161.1;refGene_gn=SNX14;refGene_tx=XM_023254162.1;refGene_gn=SNX14;refGene_tx=XM_023254163.1
B2 79640985 79641109
refGene_gn=SNX14;refGene_tx=XM_011282475.3;refGene_gn=SNX14;refGene_tx=XM_011282477.3;refGene_gn=SNX14;refGene_tx=XM_019830924.2;refGene_gn=SNX14;refGene_tx=XM_023254160.1;refGene_gn=SNX14;refGene_tx=XM_023254161.1;refGene_gn=SNX14;refGene_tx=XM_023254162.1;refGene_gn=SNX14;refGene_tx=XM_023254163.1
B2 79643897 79644017
refGene_gn=SNX14;refGene_tx=XM_011282475.3;refGene_gn=SNX14;refGene_tx=XM_011282477.3;refGene_gn=SNX14;refGene_tx=XM_019830924.2;refGene_gn=SNX14;refGene_tx=XM_023254160.1;refGene_gn=SNX14;refGene_tx=XM_023254161.1;refGene_gn=SNX14;refGene_tx=XM_023254162.1;refGene_gn=SNX14;refGene_tx=XM_023254163.1
B2 79644160 79644201
refGene_gn=SNX14;refGene_tx=XM_011282475.3;refGene_gn=SNX14;refGene_tx=XM_011282477.3;refGene_gn=SNX14;refGene_tx=XM_019830924.2;refGene_gn=SNX14;refGene_tx=XM_023254160.1;refGene_gn=SNX14;refGene_tx=XM_023254161.1;refGene_gn=SNX14;refGene_tx=XM_023254162.1;refGene_gn=SNX14;refGene_tx=XM_023254163.1
B2 79647148 79647260
refGene_gn=SNX14;refGene_tx=XM_011282475.3;refGene_gn=SNX14;refGene_tx=XM_011282477.3;refGene_gn=SNX14;refGene_tx=XM_019830924.2;refGene_gn=SNX14;refGene_tx=XM_023254160.1;refGene_gn=SNX14;refGene_tx=XM_023254161.1;refGene_gn=SNX14;refGene_tx=XM_023254162.1;refGene_gn=SNX14;refGene_tx=XM_023254163.1
B2 79649285 79649386
refGene_gn=SNX14;refGene_tx=XM_011282475.3;refGene_gn=SNX14;refGene_tx=XM_011282477.3;refGene_gn=SNX14;refGene_tx=XM_019830924.2;refGene_gn=SNX14;refGene_tx=XM_023254160.1;refGene_gn=SNX14;refGene_tx=XM_023254161.1;refGene_gn=SNX14;refGene_tx=XM_023254162.1;refGene_gn=SNX14;refGene_tx=XM_023254163.1

B2 79651502 79651586
refGene_gn=SNX14;refGene_tx=XM_011282475.3;refGene_gn=SNX14;refGene_tx
=XM_011282477.3;refGene_gn=SNX14;refGene_tx=XM_019830924.2;refGene_gn=
SNX14;refGene_tx=XM_023254160.1;refGene_gn=SNX14;refGene_tx=XM_0232541
61.1;refGene_gn=SNX14;refGene_tx=XM_023254162.1;refGene_gn=SNX14;refGe
ne_tx=XM_023254163.1
B2 79653671 79653873
refGene_gn=SNX14;refGene_tx=XM_011282475.3;refGene_gn=SNX14;refGene_tx
=XM_011282477.3;refGene_gn=SNX14;refGene_tx=XM_019830924.2;refGene_gn=
SNX14;refGene_tx=XM_023254160.1;refGene_gn=SNX14;refGene_tx=XM_0232541
61.1;refGene_gn=SNX14;refGene_tx=XM_023254162.1;refGene_gn=SNX14;refGe
ne_tx=XM_023254163.1
B2 79662833 79662966
refGene_gn=SNX14;refGene_tx=XM_011282475.3;refGene_gn=SNX14;refGene_tx
=XM_011282477.3;refGene_gn=SNX14;refGene_tx=XM_019830924.2;refGene_gn=
SNX14;refGene_tx=XM_023254160.1;refGene_gn=SNX14;refGene_tx=XM_0232541
61.1;refGene_gn=SNX14;refGene_tx=XM_023254162.1;refGene_gn=SNX14;refGe
ne_tx=XM_023254163.1
B2 79666107 79666134
refGene_gn=SNX14;refGene_tx=XM_011282475.3;refGene_gn=SNX14;refGene_tx
=XM_019830924.2;refGene_gn=SNX14;refGene_tx=XM_023254160.1;refGene_gn=
SNX14;refGene_tx=XM_023254162.1
B2 79668549 79668608
refGene_gn=SNX14;refGene_tx=XM_011282475.3;refGene_gn=SNX14;refGene_tx
=XM_011282477.3;refGene_gn=SNX14;refGene_tx=XM_019830924.2;refGene_gn=
SNX14;refGene_tx=XM_023254160.1;refGene_gn=SNX14;refGene_tx=XM_0232541
61.1;refGene_gn=SNX14;refGene_tx=XM_023254162.1;refGene_gn=SNX14;refGe
ne_tx=XM_023254163.1
B2 79669110 79669235
refGene_gn=SNX14;refGene_tx=XM_011282475.3;refGene_gn=SNX14;refGene_tx
=XM_011282477.3;refGene_gn=SNX14;refGene_tx=XM_019830924.2;refGene_gn=
SNX14;refGene_tx=XM_023254160.1;refGene_gn=SNX14;refGene_tx=XM_0232541
61.1;refGene_gn=SNX14;refGene_tx=XM_023254162.1;refGene_gn=SNX14;refGe
ne_tx=XM_023254163.1
B2 79669541 79669697
refGene_gn=SNX14;refGene_tx=XM_011282475.3;refGene_gn=SNX14;refGene_tx
=XM_011282477.3;refGene_gn=SNX14;refGene_tx=XM_019830924.2;refGene_gn=
SNX14;refGene_tx=XM_023254160.1;refGene_gn=SNX14;refGene_tx=XM_0232541
61.1;refGene_gn=SNX14;refGene_tx=XM_023254162.1;refGene_gn=SNX14;refGe
ne_tx=XM_023254163.1
B2 79673417 79673532
refGene_gn=SNX14;refGene_tx=XM_011282475.3;refGene_gn=SNX14;refGene_tx
=XM_011282477.3;refGene_gn=SNX14;refGene_tx=XM_019830924.2;refGene_gn=
SNX14;refGene_tx=XM_023254160.1;refGene_gn=SNX14;refGene_tx=XM_0232541
61.1;refGene_gn=SNX14;refGene_tx=XM_023254162.1;refGene_gn=SNX14;refGe
ne_tx=XM_023254163.1
B2 79673629 79673710
refGene_gn=SNX14;refGene_tx=XM_011282475.3;refGene_gn=SNX14;refGene_tx
=XM_011282477.3;refGene_gn=SNX14;refGene_tx=XM_019830924.2;refGene_gn=
SNX14;refGene_tx=XM_023254160.1;refGene_gn=SNX14;refGene_tx=XM_0232541

61.1;refGene_gn=SNX14;refGene_tx=XM_023254162.1;refGene_gn=SNX14;refGene_tx=XM_023254163.1
B2 79673814 79673859
refGene_gn=SNX14;refGene_tx=XM_011282475.3;refGene_gn=SNX14;refGene_tx=XM_011282477.3;refGene_gn=SNX14;refGene_tx=XM_019830924.2;refGene_gn=SNX14;refGene_tx=XM_023254160.1;refGene_gn=SNX14;refGene_tx=XM_023254161.1;refGene_gn=SNX14;refGene_tx=XM_023254162.1;refGene_gn=SNX14;refGene_tx=XM_023254163.1
B2 79674870 79674946
refGene_gn=SNX14;refGene_tx=XM_011282475.3;refGene_gn=SNX14;refGene_tx=XM_011282477.3;refGene_gn=SNX14;refGene_tx=XM_019830924.2;refGene_gn=SNX14;refGene_tx=XM_023254160.1;refGene_gn=SNX14;refGene_tx=XM_023254161.1;refGene_gn=SNX14;refGene_tx=XM_023254162.1;refGene_gn=SNX14;refGene_tx=XM_023254163.1
B2 79676491 79676648
refGene_gn=SNX14;refGene_tx=XM_011282475.3;refGene_gn=SNX14;refGene_tx=XM_011282477.3;refGene_gn=SNX14;refGene_tx=XM_019830924.2;refGene_gn=SNX14;refGene_tx=XM_023254160.1;refGene_gn=SNX14;refGene_tx=XM_023254161.1;refGene_gn=SNX14;refGene_tx=XM_023254162.1;refGene_gn=SNX14;refGene_tx=XM_023254163.1
B2 79683413 79683498
refGene_gn=SNX14;refGene_tx=XM_011282475.3;refGene_gn=SNX14;refGene_tx=XM_011282477.3;refGene_gn=SNX14;refGene_tx=XM_019830924.2;refGene_gn=SNX14;refGene_tx=XM_023254160.1;refGene_gn=SNX14;refGene_tx=XM_023254161.1;refGene_gn=SNX14;refGene_tx=XM_023254162.1;refGene_gn=SNX14;refGene_tx=XM_023254163.1
B2 79688373 79688461
refGene_gn=SNX14;refGene_tx=XM_011282475.3;refGene_gn=SNX14;refGene_tx=XM_011282477.3;refGene_gn=SNX14;refGene_tx=XM_019830924.2;refGene_gn=SNX14;refGene_tx=XM_023254160.1;refGene_gn=SNX14;refGene_tx=XM_023254161.1;refGene_gn=SNX14;refGene_tx=XM_023254162.1;refGene_gn=SNX14;refGene_tx=XM_023254163.1
B2 79690254 79690298
refGene_gn=SNX14;refGene_tx=XM_011282475.3;refGene_gn=SNX14;refGene_tx=XM_011282477.3;refGene_gn=SNX14;refGene_tx=XM_019830924.2;refGene_gn=SNX14;refGene_tx=XM_023254160.1;refGene_gn=SNX14;refGene_tx=XM_023254161.1;refGene_gn=SNX14;refGene_tx=XM_023254162.1;refGene_gn=SNX14;refGene_tx=XM_023254163.1
B2 79694581 79694660
refGene_gn=SNX14;refGene_tx=XM_011282475.3;refGene_gn=SNX14;refGene_tx=XM_011282477.3;refGene_gn=SNX14;refGene_tx=XM_019830924.2;refGene_gn=SNX14;refGene_tx=XM_023254160.1;refGene_gn=SNX14;refGene_tx=XM_023254161.1;refGene_gn=SNX14;refGene_tx=XM_023254162.1;refGene_gn=SNX14;refGene_tx=XM_023254163.1
B2 79694741 79694818
refGene_gn=SNX14;refGene_tx=XM_011282475.3;refGene_gn=SNX14;refGene_tx=XM_011282477.3;refGene_gn=SNX14;refGene_tx=XM_019830924.2;refGene_gn=SNX14;refGene_tx=XM_023254160.1;refGene_gn=SNX14;refGene_tx=XM_023254161.1;refGene_gn=SNX14;refGene_tx=XM_023254162.1;refGene_gn=SNX14;refGene_tx=XM_023254163.1

B2 79696403 79696524
refGene_gn=SNX14;refGene_tx=XM_011282475.3;refGene_gn=SNX14;refGene_tx=XM_011282477.3;refGene_gn=SNX14;refGene_tx=XM_019830924.2;refGene_gn=SNX14;refGene_tx=XM_023254160.1;refGene_gn=SNX14;refGene_tx=XM_023254161.1;refGene_gn=SNX14;refGene_tx=XM_023254162.1;refGene_gn=SNX14;refGene_tx=XM_023254163.1
B2 79715194 79715394
refGene_gn=SNX14;refGene_tx=XM_011282475.3;refGene_gn=SNX14;refGene_tx=XM_011282477.3;refGene_gn=SNX14;refGene_tx=XM_019830924.2;refGene_gn=SNX14;refGene_tx=XM_023254160.1;refGene_gn=SNX14;refGene_tx=XM_023254161.1;refGene_gn=SNX14;refGene_tx=XM_023254162.1;refGene_gn=SNX14;refGene_tx=XM_023254163.1
B2 79735421 79735463
refGene_gn=SYNCRIP;refGene_tx=XM_011282473.3;refGene_gn=SYNCRIP;refGene_tx=XM_011282474.3
B2 79737361 79737953
refGene_gn=SYNCRIP;refGene_tx=XM_011282473.3;refGene_gn=SYNCRIP;refGene_tx=XM_011282474.3;refGene_gn=SYNCRIP;refGene_tx=XM_023254165.1;refGene_gn=SYNCRIP;refGene_tx=XM_023254166.1
B2 79741351 79741473
refGene_gn=SYNCRIP;refGene_tx=XM_011282473.3;refGene_gn=SYNCRIP;refGene_tx=XM_011282474.3;refGene_gn=SYNCRIP;refGene_tx=XM_023254165.1;refGene_gn=SYNCRIP;refGene_tx=XM_023254166.1
B2 79741790 79741940
refGene_gn=SYNCRIP;refGene_tx=XM_011282473.3;refGene_gn=SYNCRIP;refGene_tx=XM_011282474.3;refGene_gn=SYNCRIP;refGene_tx=XM_023254165.1;refGene_gn=SYNCRIP;refGene_tx=XM_023254166.1
B2 79744322 79744528
refGene_gn=SYNCRIP;refGene_tx=XM_011282473.3;refGene_gn=SYNCRIP;refGene_tx=XM_011282474.3;refGene_gn=SYNCRIP;refGene_tx=XM_023254165.1;refGene_gn=SYNCRIP;refGene_tx=XM_023254166.1
B2 79745730 79745866
refGene_gn=SYNCRIP;refGene_tx=XM_011282473.3;refGene_gn=SYNCRIP;refGene_tx=XM_011282474.3;refGene_gn=SYNCRIP;refGene_tx=XM_023254165.1;refGene_gn=SYNCRIP;refGene_tx=XM_023254166.1
B2 79756330 79756507
refGene_gn=SYNCRIP;refGene_tx=XM_011282473.3;refGene_gn=SYNCRIP;refGene_tx=XM_011282474.3;refGene_gn=SYNCRIP;refGene_tx=XM_023254165.1;refGene_gn=SYNCRIP;refGene_tx=XM_023254166.1
B2 79756607 79756721
refGene_gn=SYNCRIP;refGene_tx=XM_011282473.3;refGene_gn=SYNCRIP;refGene_tx=XM_011282474.3;refGene_gn=SYNCRIP;refGene_tx=XM_023254165.1;refGene_gn=SYNCRIP;refGene_tx=XM_023254166.1
B2 79759154 79759262
refGene_gn=SYNCRIP;refGene_tx=XM_011282473.3;refGene_gn=SYNCRIP;refGene_tx=XM_011282474.3;refGene_gn=SYNCRIP;refGene_tx=XM_023254165.1;refGene_gn=SYNCRIP;refGene_tx=XM_023254166.1
B2 79759378 79759497
refGene_gn=SYNCRIP;refGene_tx=XM_011282473.3;refGene_gn=SYNCRIP;refGene_tx=XM_011282474.3;refGene_gn=SYNCRIP;refGene_tx=XM_023254165.1

B2 79760173 79760321
refGene_gn=SYNCRIP;refGene_tx=XM_011282473.3;refGene_gn=SYNCRIP;refGene_tx=XM_011282474.3;refGene_gn=SYNCRIP;refGene_tx=XM_023254165.1
B2 80774430 80775528 refGene_gn=HTR1E;refGene_tx=XM_006931876.3
B2 80850564 80850642 refGene_gn=CGA;refGene_tx=NM_001048158.1
B2 80851041 80851226 refGene_gn=CGA;refGene_tx=NM_001048158.1
B2 80852388 80852488 refGene_gn=CGA;refGene_tx=NM_001048158.1
B2 80920167 80920335 refGene_gn=ZNF292;refGene_tx=XM_003986354.5
B2 80970238 80970393 refGene_gn=ZNF292;refGene_tx=XM_003986354.5
B2 80970578 80970657 refGene_gn=ZNF292;refGene_tx=XM_003986354.5
B2 80972884 80973020 refGene_gn=ZNF292;refGene_tx=XM_003986354.5
B2 80988709 80988912
refGene_gn=ZNF292;refGene_tx=XM_003986354.5;refGene_gn=ZNF292;refGene_tx=XM_019830931.2
B2 80995219 80995356
refGene_gn=ZNF292;refGene_tx=XM_003986354.5;refGene_gn=ZNF292;refGene_tx=XM_019830931.2
B2 80997653 80997795
refGene_gn=ZNF292;refGene_tx=XM_003986354.5;refGene_gn=ZNF292;refGene_tx=XM_019830931.2
B2 81006460 81013615
refGene_gn=ZNF292;refGene_tx=XM_003986354.5;refGene_gn=ZNF292;refGene_tx=XM_019830931.2
B2 81031776 81032640 refGene_gn=GJB7;refGene_tx=XM_023254167.1
B2 81076585 81076720
refGene_gn=SMIM8;refGene_tx=XM_006931877.2;refGene_gn=SMIM8;refGene_tx=XM_006931878.3;refGene_gn=SMIM8;refGene_tx=XM_006931879.3;refGene_gn=SMIM8;refGene_tx=XM_019830934.2;refGene_gn=SMIM8;refGene_tx=XM_023254168.1
B2 81078828 81078987
refGene_gn=SMIM8;refGene_tx=XM_006931877.2;refGene_gn=SMIM8;refGene_tx=XM_006931878.3;refGene_gn=SMIM8;refGene_tx=XM_006931879.3;refGene_gn=SMIM8;refGene_tx=XM_019830934.2;refGene_gn=SMIM8;refGene_tx=XM_023254168.1
B2 81084494 81084642
refGene_gn=CB2H6orf163;refGene_tx=XM_011282471.2
B2 81088306 81088343
refGene_gn=CB2H6orf163;refGene_tx=XM_023254169.1
B2 81088506 81088601
refGene_gn=CB2H6orf163;refGene_tx=XM_011282471.2;refGene_gn=CB2H6orf163;refGene_tx=XM_023254169.1
B2 81090015 81090123
refGene_gn=CB2H6orf163;refGene_tx=XM_011282471.2;refGene_gn=CB2H6orf163;refGene_tx=XM_023254169.1
B2 81095193 81095396
refGene_gn=CB2H6orf163;refGene_tx=XM_011282471.2;refGene_gn=CB2H6orf163;refGene_tx=XM_023254169.1
B2 81102963 81103399
refGene_gn=CB2H6orf163;refGene_tx=XM_011282471.2;refGene_gn=CB2H6orf163;refGene_tx=XM_023254169.1

B2 81152394 81152410 refGene_gn=CFAP206;refGene_tx=XM_003986356.5
B2 81154973 81155086 refGene_gn=CFAP206;refGene_tx=XM_003986356.5
B2 81155694 81155778 refGene_gn=CFAP206;refGene_tx=XM_003986356.5
B2 81158734 81158825 refGene_gn=CFAP206;refGene_tx=XM_003986356.5
B2 81160281 81160470 refGene_gn=CFAP206;refGene_tx=XM_003986356.5
B2 81161264 81161423 refGene_gn=CFAP206;refGene_tx=XM_003986356.5
B2 81162687 81162896 refGene_gn=CFAP206;refGene_tx=XM_003986356.5
B2 81173630 81173750 refGene_gn=CFAP206;refGene_tx=XM_003986356.5
B2 81175131 81175330 refGene_gn=CFAP206;refGene_tx=XM_003986356.5
B2 81177533 81177628 refGene_gn=CFAP206;refGene_tx=XM_003986356.5
B2 81177629 81177675 refGene_gn=CFAP206;refGene_tx=XM_003986356.5
B2 81180738 81180932 refGene_gn=CFAP206;refGene_tx=XM_003986356.5
B2 81184256 81184400 refGene_gn=CFAP206;refGene_tx=XM_003986356.5
B2 81194458 81194689 refGene_gn=CFAP206;refGene_tx=XM_003986356.5
B2 81202245 81202261 refGene_gn=SLC35A1;refGene_tx=XM_011282468.3
B2 81203827 81204005 refGene_gn=SLC35A1;refGene_tx=XM_011282468.3
B2 81216636 81216707 refGene_gn=SLC35A1;refGene_tx=XM_006931883.4
B2 81219062 81219128
refGene_gn=RARS2;refGene_tx=XM_019830943.1;refGene_gn=RARS2;refGene_tx=XM_023254170.1;refGene_gn=RARS2;refGene_tx=XM_023254173.1
B2 81220324 81220484
refGene_gn=SLC35A1;refGene_tx=XM_006931883.4;refGene_gn=SLC35A1;refGene_tx=XM_011282468.3
B2 81220676 81220829
refGene_gn=SLC35A1;refGene_tx=XM_006931883.4;refGene_gn=SLC35A1;refGene_tx=XM_011282468.3
B2 81224271 81224338
refGene_gn=SLC35A1;refGene_tx=XM_006931883.4;refGene_gn=SLC35A1;refGene_tx=XM_011282468.3
B2 81226200 81226377
refGene_gn=SLC35A1;refGene_tx=XM_006931883.4;refGene_gn=SLC35A1;refGene_tx=XM_011282468.3
B2 81226824 81226959
refGene_gn=SLC35A1;refGene_tx=XM_006931883.4;refGene_gn=SLC35A1;refGene_tx=XM_011282468.3
B2 81228009 81228033 refGene_gn=RARS2;refGene_tx=XM_023254172.1
B2 81229039 81229167
refGene_gn=SLC35A1;refGene_tx=XM_006931883.4;refGene_gn=SLC35A1;refGene_tx=XM_011282468.3
B2 81230049 81230106 refGene_gn=RARS2;refGene_tx=XM_023254171.1
B2 81231805 81231892
refGene_gn=RARS2;refGene_tx=XM_003986357.4;refGene_gn=RARS2;refGene_tx=XM_019830942.1;refGene_gn=RARS2;refGene_tx=XM_019830943.1;refGene_gn=RARS2;refGene_tx=XM_023254170.1;refGene_gn=RARS2;refGene_tx=XM_023254171.1;refGene_gn=RARS2;refGene_tx=XM_023254172.1;refGene_gn=RARS2;refGene_tx=XM_023254173.1
B2 81232355 81232419
refGene_gn=RARS2;refGene_tx=XM_003986357.4;refGene_gn=RARS2;refGene_tx=XM_019830942.1;refGene_gn=RARS2;refGene_tx=XM_019830943.1;refGene_gn=RARS2;refGene_tx=XM_023254170.1;refGene_gn=RARS2;refGene_tx=XM_0232541

71.1;refGene_gn=RARS2;refGene_tx=XM_023254172.1;refGene_gn=RARS2;refGene_tx=XM_023254173.1
B2 81233989 81234064
refGene_gn=RARS2;refGene_tx=XM_003986357.4;refGene_gn=RARS2;refGene_tx=XM_019830942.1;refGene_gn=RARS2;refGene_tx=XM_019830943.1;refGene_gn=RARS2;refGene_tx=XM_023254170.1;refGene_gn=RARS2;refGene_tx=XM_023254171.1;refGene_gn=RARS2;refGene_tx=XM_023254172.1;refGene_gn=RARS2;refGene_tx=XM_023254173.1
B2 81235068 81235164
refGene_gn=RARS2;refGene_tx=XM_003986357.4;refGene_gn=RARS2;refGene_tx=XM_019830942.1;refGene_gn=RARS2;refGene_tx=XM_019830943.1;refGene_gn=RARS2;refGene_tx=XM_023254170.1;refGene_gn=RARS2;refGene_tx=XM_023254171.1;refGene_gn=RARS2;refGene_tx=XM_023254172.1;refGene_gn=RARS2;refGene_tx=XM_023254173.1
B2 81235541 81235651
refGene_gn=RARS2;refGene_tx=XM_003986357.4;refGene_gn=RARS2;refGene_tx=XM_019830942.1;refGene_gn=RARS2;refGene_tx=XM_019830943.1;refGene_gn=RARS2;refGene_tx=XM_023254170.1;refGene_gn=RARS2;refGene_tx=XM_023254171.1;refGene_gn=RARS2;refGene_tx=XM_023254172.1;refGene_gn=RARS2;refGene_tx=XM_023254173.1
B2 81235736 81235804
refGene_gn=RARS2;refGene_tx=XM_003986357.4;refGene_gn=RARS2;refGene_tx=XM_019830942.1;refGene_gn=RARS2;refGene_tx=XM_019830943.1;refGene_gn=RARS2;refGene_tx=XM_023254170.1;refGene_gn=RARS2;refGene_tx=XM_023254171.1;refGene_gn=RARS2;refGene_tx=XM_023254172.1;refGene_gn=RARS2;refGene_tx=XM_023254173.1
B2 81236386 81236511
refGene_gn=RARS2;refGene_tx=XM_003986357.4;refGene_gn=RARS2;refGene_tx=XM_019830942.1;refGene_gn=RARS2;refGene_tx=XM_019830943.1;refGene_gn=RARS2;refGene_tx=XM_023254170.1;refGene_gn=RARS2;refGene_tx=XM_023254171.1;refGene_gn=RARS2;refGene_tx=XM_023254172.1;refGene_gn=RARS2;refGene_tx=XM_023254173.1
B2 81236984 81237061
refGene_gn=RARS2;refGene_tx=XM_003986357.4;refGene_gn=RARS2;refGene_tx=XM_019830942.1;refGene_gn=RARS2;refGene_tx=XM_019830943.1;refGene_gn=RARS2;refGene_tx=XM_023254170.1;refGene_gn=RARS2;refGene_tx=XM_023254171.1;refGene_gn=RARS2;refGene_tx=XM_023254172.1;refGene_gn=RARS2;refGene_tx=XM_023254173.1
B2 81238579 81238640
refGene_gn=RARS2;refGene_tx=XM_003986357.4;refGene_gn=RARS2;refGene_tx=XM_019830942.1;refGene_gn=RARS2;refGene_tx=XM_019830943.1;refGene_gn=RARS2;refGene_tx=XM_023254170.1;refGene_gn=RARS2;refGene_tx=XM_023254171.1;refGene_gn=RARS2;refGene_tx=XM_023254172.1;refGene_gn=RARS2;refGene_tx=XM_023254173.1
B2 81241418 81241514
refGene_gn=RARS2;refGene_tx=XM_003986357.4;refGene_gn=RARS2;refGene_tx=XM_019830942.1;refGene_gn=RARS2;refGene_tx=XM_019830943.1;refGene_gn=RARS2;refGene_tx=XM_023254170.1;refGene_gn=RARS2;refGene_tx=XM_023254171.1;refGene_gn=RARS2;refGene_tx=XM_023254172.1;refGene_gn=RARS2;refGene_tx=XM_023254173.1

B2	81242642	81242749
refGene_gn=RARS2;refGene_tx=XM_003986357.4;refGene_gn=RARS2;refGene_tx=XM_019830942.1;refGene_gn=RARS2;refGene_tx=XM_019830943.1;refGene_gn=RARS2;refGene_tx=XM_023254170.1;refGene_gn=RARS2;refGene_tx=XM_023254171.1;refGene_gn=RARS2;refGene_tx=XM_023254172.1;refGene_gn=RARS2;refGene_tx=XM_023254173.1
B2	81243759	81243918
refGene_gn=RARS2;refGene_tx=XM_003986357.4;refGene_gn=RARS2;refGene_tx=XM_019830942.1;refGene_gn=RARS2;refGene_tx=XM_019830943.1;refGene_gn=RARS2;refGene_tx=XM_023254170.1;refGene_gn=RARS2;refGene_tx=XM_023254171.1;refGene_gn=RARS2;refGene_tx=XM_023254172.1;refGene_gn=RARS2;refGene_tx=XM_023254173.1
B2	81253370	81253447
refGene_gn=RARS2;refGene_tx=XM_003986357.4;refGene_gn=RARS2;refGene_tx=XM_019830942.1;refGene_gn=RARS2;refGene_tx=XM_019830943.1;refGene_gn=RARS2;refGene_tx=XM_023254170.1;refGene_gn=RARS2;refGene_tx=XM_023254171.1;refGene_gn=RARS2;refGene_tx=XM_023254172.1;refGene_gn=RARS2;refGene_tx=XM_023254173.1
B2	81257455	81257539
refGene_gn=RARS2;refGene_tx=XM_003986357.4;refGene_gn=RARS2;refGene_tx=XM_019830942.1;refGene_gn=RARS2;refGene_tx=XM_019830943.1;refGene_gn=RARS2;refGene_tx=XM_023254170.1;refGene_gn=RARS2;refGene_tx=XM_023254171.1;refGene_gn=RARS2;refGene_tx=XM_023254172.1;refGene_gn=RARS2;refGene_tx=XM_023254173.1
B2	81260032	81260088
refGene_gn=RARS2;refGene_tx=XM_003986357.4;refGene_gn=RARS2;refGene_tx=XM_019830942.1;refGene_gn=RARS2;refGene_tx=XM_019830943.1
B2	81265979	81266077
refGene_gn=RARS2;refGene_tx=XM_003986357.4;refGene_gn=RARS2;refGene_tx=XM_019830942.1;refGene_gn=RARS2;refGene_tx=XM_019830943.1
B2	81273046	81273130
refGene_gn=RARS2;refGene_tx=XM_003986357.4;refGene_gn=RARS2;refGene_tx=XM_019830942.1;refGene_gn=RARS2;refGene_tx=XM_019830943.1
B2	81274674	81274777
refGene_gn=RARS2;refGene_tx=XM_003986357.4;refGene_gn=RARS2;refGene_tx=XM_019830942.1;refGene_gn=RARS2;refGene_tx=XM_019830943.1
B2	81276798	81276887	refGene_gn=RARS2;refGene_tx=XM_019830942.1
B2	81286304	81286378
refGene_gn=RARS2;refGene_tx=XM_003986357.4;refGene_gn=RARS2;refGene_tx=XM_019830942.1;refGene_gn=RARS2;refGene_tx=XM_019830943.1
B2	81293949	81293989	refGene_gn=RARS2;refGene_tx=XM_019830942.1
B2	81300376	81300412
refGene_gn=RARS2;refGene_tx=XM_003986357.4;refGene_gn=RARS2;refGene_tx=XM_019830943.1
B2	81300625	81300649
refGene_gn=ORC3;refGene_tx=XM_003986358.4;refGene_gn=ORC3;refGene_tx=XM_006931884.3;refGene_gn=ORC3;refGene_tx=XM_006931885.3;refGene_gn=ORC3;refGene_tx=XM_006931886.3
B2	81304399	81304454
refGene_gn=ORC3;refGene_tx=XM_003986358.4;refGene_gn=ORC3;refGene_tx=X

M_006931884.3;refGene_gn=ORC3;refGene_tx=XM_006931885.3;refGene_gn=ORC3;refGene_tx=XM_006931886.3
B2 81310977 81311075
refGene_gn=ORC3;refGene_tx=XM_003986358.4;refGene_gn=ORC3;refGene_tx=XM_006931884.3;refGene_gn=ORC3;refGene_tx=XM_006931885.3
B2 81312773 81312918
refGene_gn=ORC3;refGene_tx=XM_003986358.4;refGene_gn=ORC3;refGene_tx=XM_006931884.3;refGene_gn=ORC3;refGene_tx=XM_006931885.3
B2 81315789 81315894
refGene_gn=ORC3;refGene_tx=XM_003986358.4;refGene_gn=ORC3;refGene_tx=XM_006931884.3;refGene_gn=ORC3;refGene_tx=XM_006931885.3;refGene_gn=ORC3;refGene_tx=XM_006931886.3
B2 81316921 81317070
refGene_gn=ORC3;refGene_tx=XM_003986358.4;refGene_gn=ORC3;refGene_tx=XM_006931884.3;refGene_gn=ORC3;refGene_tx=XM_006931885.3;refGene_gn=ORC3;refGene_tx=XM_006931886.3
B2 81318340 81318474
refGene_gn=ORC3;refGene_tx=XM_003986358.4;refGene_gn=ORC3;refGene_tx=XM_006931884.3;refGene_gn=ORC3;refGene_tx=XM_006931885.3;refGene_gn=ORC3;refGene_tx=XM_006931886.3
B2 81320120 81320280
refGene_gn=ORC3;refGene_tx=XM_003986358.4;refGene_gn=ORC3;refGene_tx=XM_006931884.3;refGene_gn=ORC3;refGene_tx=XM_006931885.3;refGene_gn=ORC3;refGene_tx=XM_006931886.3
B2 81321759 81321873
refGene_gn=ORC3;refGene_tx=XM_003986358.4;refGene_gn=ORC3;refGene_tx=XM_006931884.3;refGene_gn=ORC3;refGene_tx=XM_006931885.3;refGene_gn=ORC3;refGene_tx=XM_006931886.3
B2 81327598 81327732
refGene_gn=ORC3;refGene_tx=XM_003986358.4;refGene_gn=ORC3;refGene_tx=XM_006931884.3;refGene_gn=ORC3;refGene_tx=XM_006931885.3;refGene_gn=ORC3;refGene_tx=XM_006931886.3
B2 81328211 81328275
refGene_gn=ORC3;refGene_tx=XM_003986358.4;refGene_gn=ORC3;refGene_tx=XM_006931884.3;refGene_gn=ORC3;refGene_tx=XM_006931885.3;refGene_gn=ORC3;refGene_tx=XM_006931886.3
B2 81337549 81337666
refGene_gn=ORC3;refGene_tx=XM_003986358.4;refGene_gn=ORC3;refGene_tx=XM_006931884.3;refGene_gn=ORC3;refGene_tx=XM_006931885.3;refGene_gn=ORC3;refGene_tx=XM_006931886.3
B2 81338789 81338869
refGene_gn=ORC3;refGene_tx=XM_003986358.4;refGene_gn=ORC3;refGene_tx=XM_006931884.3;refGene_gn=ORC3;refGene_tx=XM_006931885.3;refGene_gn=ORC3;refGene_tx=XM_006931886.3
B2 81351142 81351276
refGene_gn=ORC3;refGene_tx=XM_003986358.4;refGene_gn=ORC3;refGene_tx=XM_006931884.3;refGene_gn=ORC3;refGene_tx=XM_006931885.3;refGene_gn=ORC3;refGene_tx=XM_006931886.3
B2 81353773 81353853
refGene_gn=ORC3;refGene_tx=XM_003986358.4;refGene_gn=ORC3;refGene_tx=X

M_006931884.3;refGene_gn=ORC3;refGene_tx=XM_006931885.3;refGene_gn=ORC3;refGene_tx=XM_006931886.3
B2 81355026 81355127 refGene_gn=ORC3;refGene_tx=XM_003986358.4;refGene_gn=ORC3;refGene_tx=XM_006931884.3;refGene_gn=ORC3;refGene_tx=XM_006931885.3;refGene_gn=ORC3;refGene_tx=XM_006931886.3
B2 81358791 81358933 refGene_gn=ORC3;refGene_tx=XM_003986358.4;refGene_gn=ORC3;refGene_tx=XM_006931884.3;refGene_gn=ORC3;refGene_tx=XM_006931885.3;refGene_gn=ORC3;refGene_tx=XM_006931886.3
B2 81360665 81360782 refGene_gn=ORC3;refGene_tx=XM_003986358.4;refGene_gn=ORC3;refGene_tx=XM_006931884.3;refGene_gn=ORC3;refGene_tx=XM_006931885.3;refGene_gn=ORC3;refGene_tx=XM_006931886.3
B2 81361517 81361597 refGene_gn=ORC3;refGene_tx=XM_003986358.4;refGene_gn=ORC3;refGene_tx=XM_006931884.3;refGene_gn=ORC3;refGene_tx=XM_006931885.3;refGene_gn=ORC3;refGene_tx=XM_006931886.3
B2 81362129 81362242 refGene_gn=ORC3;refGene_tx=XM_003986358.4;refGene_gn=ORC3;refGene_tx=XM_006931884.3;refGene_gn=ORC3;refGene_tx=XM_006931885.3;refGene_gn=ORC3;refGene_tx=XM_006931886.3
B2 81369352 81369363 refGene_gn=AKIRIN2;refGene_tx=XM_011282467.3
B2 81369613 81369685 refGene_gn=AKIRIN2;refGene_tx=XM_011282467.3
B2 81370592 81370742 refGene_gn=AKIRIN2;refGene_tx=XM_011282467.3
B2 81372295 81372439 refGene_gn=AKIRIN2;refGene_tx=XM_011282467.3
B2 81387412 81387647 refGene_gn=AKIRIN2;refGene_tx=XM_011282467.3
B2 81710776 81710990 refGene_gn=SPACA1;refGene_tx=XM_003986364.4
B2 81716670 81716727 refGene_gn=SPACA1;refGene_tx=XM_003986364.4
B2 81720311 81720413 refGene_gn=SPACA1;refGene_tx=XM_003986364.4
B2 81721180 81721287 refGene_gn=SPACA1;refGene_tx=XM_003986364.4
B2 81721928 81722064 refGene_gn=SPACA1;refGene_tx=XM_003986364.4
B2 81727450 81727571 refGene_gn=SPACA1;refGene_tx=XM_003986364.4
B2 81728098 81728255 refGene_gn=SPACA1;refGene_tx=XM_003986364.4
B2 81799711 81801130 refGene_gn=CNR1;refGene_tx=NM_001009331.1;refGene_gn=CNR1;refGene_tx=XM_011282446.3;refGene_gn=CNR1;refGene_tx=XM_019830070.2
B2 82210609 82210773 refGene_gn=RNGTT;refGene_tx=XM_011282466.3
B2 82212257 82212381 refGene_gn=RNGTT;refGene_tx=XM_011282466.3
B2 82260313 82260380 refGene_gn=RNGTT;refGene_tx=XM_011282466.3
B2 82343837 82343938 refGene_gn=RNGTT;refGene_tx=XM_011282466.3
B2 82367191 82367260 refGene_gn=RNGTT;refGene_tx=XM_011282466.3
B2 82415029 82415194 refGene_gn=RNGTT;refGene_tx=XM_011282466.3
B2 82420682 82420754 refGene_gn=RNGTT;refGene_tx=XM_011282466.3
B2 82436138 82436274 refGene_gn=RNGTT;refGene_tx=XM_011282466.3
B2 82479000 82479102 refGene_gn=RNGTT;refGene_tx=XM_011282466.3
B2 82480290 82480400 refGene_gn=RNGTT;refGene_tx=XM_011282466.3
B2 82491922 82492163 refGene_gn=RNGTT;refGene_tx=XM_011282466.3
B2 82493696 82493772 refGene_gn=RNGTT;refGene_tx=XM_011282466.3
B2 82508361 82508450 refGene_gn=RNGTT;refGene_tx=XM_011282466.3

B2 82508546 82508650 refGene_gn=RNGTT;refGene_tx=XM_011282466.3
B2 82514270 82514380 refGene_gn=RNGTT;refGene_tx=XM_011282466.3
B2 82523375 82523439 refGene_gn=RNGTT;refGene_tx=XM_011282466.3
B2 82592630 82593170 refGene_gn=PNRC1;refGene_tx=XM_011282465.3
B2 82595169 82595613 refGene_gn=PNRC1;refGene_tx=XM_011282465.3
B2 82610206 82610576
refGene_gn=SRSF12;refGene_tx=XM_006931890.4;refGene_gn=SRSF12;refGene_tx=XM_023254175.1
B2 82615911 82616053
refGene_gn=SRSF12;refGene_tx=XM_006931890.4;refGene_gn=SRSF12;refGene_tx=XM_023254175.1
B2 82616205 82616309
refGene_gn=SRSF12;refGene_tx=XM_006931890.4;refGene_gn=SRSF12;refGene_tx=XM_023254175.1
B2 82617616 82617721
refGene_gn=SRSF12;refGene_tx=XM_006931890.4;refGene_gn=SRSF12;refGene_tx=XM_023254175.1
B2 82625659 82625679 refGene_gn=SRSF12;refGene_tx=XM_006931890.4
B2 82626112 82626177 refGene_gn=SRSF12;refGene_tx=XM_023254175.1
B2 82656475 82656931 refGene_gn=PM20D2;refGene_tx=XM_003986367.5
B2 82659019 82659168 refGene_gn=PM20D2;refGene_tx=XM_003986367.5
B2 82666432 82666575 refGene_gn=PM20D2;refGene_tx=XM_003986367.5
B2 82667870 82668025 refGene_gn=PM20D2;refGene_tx=XM_003986367.5
B2 82670124 82670260 refGene_gn=PM20D2;refGene_tx=XM_003986367.5
B2 82671286 82671394 refGene_gn=PM20D2;refGene_tx=XM_003986367.5
B2 82671606 82671761 refGene_gn=PM20D2;refGene_tx=XM_003986367.5
B2 82686748 82687042 refGene_gn=GABRR1;refGene_tx=XM_003986368.3
B2 82687957 82688154 refGene_gn=GABRR1;refGene_tx=XM_003986368.3
B2 82689822 82689975 refGene_gn=GABRR1;refGene_tx=XM_003986368.3
B2 82691873 82692014 refGene_gn=GABRR1;refGene_tx=XM_003986368.3
B2 82694647 82694730 refGene_gn=GABRR1;refGene_tx=XM_003986368.3
B2 82695540 82695764 refGene_gn=GABRR1;refGene_tx=XM_003986368.3
B2 82696769 82696837 refGene_gn=GABRR1;refGene_tx=XM_003986368.3
B2 82698751 82698858 refGene_gn=GABRR1;refGene_tx=XM_003986368.3
B2 82713732 82713854 refGene_gn=GABRR1;refGene_tx=XM_003986368.3
B2 82739734 82740046
refGene_gn=GABRR2;refGene_tx=XM_003986370.4;refGene_gn=GABRR2;refGene_tx=XM_019830948.1;refGene_gn=GABRR2;refGene_tx=XM_023254177.1
B2 82742186 82742383
refGene_gn=GABRR2;refGene_tx=XM_003986370.4;refGene_gn=GABRR2;refGene_tx=XM_019830948.1;refGene_gn=GABRR2;refGene_tx=XM_023254177.1
B2 82743613 82743766
refGene_gn=GABRR2;refGene_tx=XM_003986370.4;refGene_gn=GABRR2;refGene_tx=XM_019830948.1;refGene_gn=GABRR2;refGene_tx=XM_023254177.1
B2 82746708 82746849
refGene_gn=GABRR2;refGene_tx=XM_003986370.4;refGene_gn=GABRR2;refGene_tx=XM_019830948.1;refGene_gn=GABRR2;refGene_tx=XM_023254177.1
B2 82747047 82747130
refGene_gn=GABRR2;refGene_tx=XM_003986370.4;refGene_gn=GABRR2;refGene_tx=XM_019830948.1;refGene_gn=GABRR2;refGene_tx=XM_023254177.1

B2 82748112 82748336
refGene_gn=GABRR2;refGene_tx=XM_003986370.4;refGene_gn=GABRR2;refGene_tx=XM_019830948.1;refGene_gn=GABRR2;refGene_tx=XM_023254177.1
B2 82750114 82750213
refGene_gn=GABRR2;refGene_tx=XM_003986370.4;refGene_gn=GABRR2;refGene_tx=XM_019830948.1;refGene_gn=GABRR2;refGene_tx=XM_023254177.1
B2 82759858 82759918 refGene_gn=GABRR2;refGene_tx=XM_019830948.1
B2 82772795 82772902 refGene_gn=GABRR2;refGene_tx=XM_003986370.4
B2 82780266 82780454 refGene_gn=GABRR2;refGene_tx=XM_003986370.4
B2 82787532 82787811
refGene_gn=UBE2J1;refGene_tx=XM_003986371.5;refGene_gn=UBE2J1;refGene_tx=XM_019830949.2
B2 82794142 82794262
refGene_gn=UBE2J1;refGene_tx=XM_003986371.5;refGene_gn=UBE2J1;refGene_tx=XM_019830949.2
B2 82795822 82795952
refGene_gn=UBE2J1;refGene_tx=XM_003986371.5;refGene_gn=UBE2J1;refGene_tx=XM_019830949.2
B2 82799333 82799439
refGene_gn=UBE2J1;refGene_tx=XM_003986371.5;refGene_gn=UBE2J1;refGene_tx=XM_019830949.2
B2 82799983 82800068
refGene_gn=UBE2J1;refGene_tx=XM_003986371.5;refGene_gn=UBE2J1;refGene_tx=XM_019830949.2
B2 82802951 82803083
refGene_gn=UBE2J1;refGene_tx=XM_003986371.5;refGene_gn=UBE2J1;refGene_tx=XM_019830949.2
B2 82804491 82804565
refGene_gn=UBE2J1;refGene_tx=XM_003986371.5;refGene_gn=UBE2J1;refGene_tx=XM_019830949.2
B2 82813886 82813917 refGene_gn=UBE2J1;refGene_tx=XM_003986371.5
B2 82829899 82830051 refGene_gn=RRAGD;refGene_tx=XM_011282464.3
B2 82838444 82838593 refGene_gn=RRAGD;refGene_tx=XM_011282464.3
B2 82843228 82843371 refGene_gn=RRAGD;refGene_tx=XM_011282464.3
B2 82844822 82844937 refGene_gn=RRAGD;refGene_tx=XM_011282464.3
B2 82846381 82846581 refGene_gn=RRAGD;refGene_tx=XM_011282464.3
B2 82851884 82852180 refGene_gn=RRAGD;refGene_tx=XM_011282464.3
B2 82868778 82868914 refGene_gn=RRAGD;refGene_tx=XM_011282464.3
B2 83023088 83023208
refGene_gn=ANKRD6;refGene_tx=XM_019830952.1;refGene_gn=ANKRD6;refGene_tx=XM_019830954.2;refGene_gn=ANKRD6;refGene_tx=XM_023254179.1;refGene_gn=ANKRD6;refGene_tx=XM_023254180.1
B2 83066328 83066427
refGene_gn=ANKRD6;refGene_tx=XM_019830952.1;refGene_gn=ANKRD6;refGene_tx=XM_019830954.2;refGene_gn=ANKRD6;refGene_tx=XM_023254179.1;refGene_gn=ANKRD6;refGene_tx=XM_023254180.1
B2 83068680 83068974
refGene_gn=ANKRD6;refGene_tx=XM_019830951.1;refGene_gn=ANKRD6;refGene_tx=XM_019830953.1;refGene_gn=ANKRD6;refGene_tx=XM_019830955.1
B2 83072935 83073034

refGene_gn=ANKRD6;refGene_tx=XM_019830951.1;refGene_gn=ANKRD6;refGene_tx=XM_019830952.1;refGene_gn=ANKRD6;refGene_tx=XM_019830953.1;refGene_gn=ANKRD6;refGene_tx=XM_019830954.2;refGene_gn=ANKRD6;refGene_tx=XM_019830955.1;refGene_gn=ANKRD6;refGene_tx=XM_023254179.1;refGene_gn=ANKRD6;refGene_tx=XM_023254180.1
B2 83075370 83075469
refGene_gn=ANKRD6;refGene_tx=XM_019830951.1;refGene_gn=ANKRD6;refGene_tx=XM_019830952.1;refGene_gn=ANKRD6;refGene_tx=XM_019830953.1;refGene_gn=ANKRD6;refGene_tx=XM_019830954.2;refGene_gn=ANKRD6;refGene_tx=XM_019830955.1;refGene_gn=ANKRD6;refGene_tx=XM_023254179.1;refGene_gn=ANKRD6;refGene_tx=XM_023254180.1
B2 83079620 83079719
refGene_gn=ANKRD6;refGene_tx=XM_019830951.1;refGene_gn=ANKRD6;refGene_tx=XM_019830952.1;refGene_gn=ANKRD6;refGene_tx=XM_019830953.1;refGene_gn=ANKRD6;refGene_tx=XM_019830954.2;refGene_gn=ANKRD6;refGene_tx=XM_023254179.1;refGene_gn=ANKRD6;refGene_tx=XM_023254180.1
B2 83081341 83081440
refGene_gn=ANKRD6;refGene_tx=XM_019830951.1;refGene_gn=ANKRD6;refGene_tx=XM_019830952.1;refGene_gn=ANKRD6;refGene_tx=XM_019830953.1;refGene_gn=ANKRD6;refGene_tx=XM_019830954.2;refGene_gn=ANKRD6;refGene_tx=XM_023254179.1;refGene_gn=ANKRD6;refGene_tx=XM_023254180.1
B2 83083355 83083454
refGene_gn=ANKRD6;refGene_tx=XM_019830951.1;refGene_gn=ANKRD6;refGene_tx=XM_019830952.1;refGene_gn=ANKRD6;refGene_tx=XM_019830953.1;refGene_gn=ANKRD6;refGene_tx=XM_019830954.2;refGene_gn=ANKRD6;refGene_tx=XM_019830955.1;refGene_gn=ANKRD6;refGene_tx=XM_023254179.1;refGene_gn=ANKRD6;refGene_tx=XM_023254180.1
B2 83084669 83084747
refGene_gn=ANKRD6;refGene_tx=XM_019830951.1;refGene_gn=ANKRD6;refGene_tx=XM_019830952.1;refGene_gn=ANKRD6;refGene_tx=XM_019830953.1;refGene_gn=ANKRD6;refGene_tx=XM_019830954.2;refGene_gn=ANKRD6;refGene_tx=XM_019830955.1;refGene_gn=ANKRD6;refGene_tx=XM_023254179.1;refGene_gn=ANKRD6;refGene_tx=XM_023254180.1
B2 83088596 83088701
refGene_gn=ANKRD6;refGene_tx=XM_019830951.1;refGene_gn=ANKRD6;refGene_tx=XM_019830952.1;refGene_gn=ANKRD6;refGene_tx=XM_019830955.1;refGene_gn=ANKRD6;refGene_tx=XM_023254179.1;refGene_gn=ANKRD6;refGene_tx=XM_023254180.1
B2 83090027 83090162
refGene_gn=ANKRD6;refGene_tx=XM_019830951.1;refGene_gn=ANKRD6;refGene_tx=XM_019830952.1;refGene_gn=ANKRD6;refGene_tx=XM_019830953.1;refGene_gn=ANKRD6;refGene_tx=XM_019830954.2;refGene_gn=ANKRD6;refGene_tx=XM_019830955.1;refGene_gn=ANKRD6;refGene_tx=XM_023254179.1;refGene_gn=ANKRD6;refGene_tx=XM_023254180.1
B2 83090474 83090660
refGene_gn=ANKRD6;refGene_tx=XM_019830951.1;refGene_gn=ANKRD6;refGene_tx=XM_019830952.1;refGene_gn=ANKRD6;refGene_tx=XM_019830953.1;refGene_gn=ANKRD6;refGene_tx=XM_019830954.2;refGene_gn=ANKRD6;refGene_tx=XM_019830955.1;refGene_gn=ANKRD6;refGene_tx=XM_023254179.1;refGene_gn=ANKRD6;refGene_tx=XM_023254180.1

B2 83091123 83091276
refGene_gn=ANKRD6;refGene_tx=XM_019830951.1;refGene_gn=ANKRD6;refGene_tx=XM_019830952.1;refGene_gn=ANKRD6;refGene_tx=XM_019830953.1;refGene_gn=ANKRD6;refGene_tx=XM_019830954.2;refGene_gn=ANKRD6;refGene_tx=XM_019830955.1;refGene_gn=ANKRD6;refGene_tx=XM_023254179.1;refGene_gn=ANKRD6;refGene_tx=XM_023254180.1
B2 83099789 83099903
refGene_gn=ANKRD6;refGene_tx=XM_019830951.1;refGene_gn=ANKRD6;refGene_tx=XM_019830952.1;refGene_gn=ANKRD6;refGene_tx=XM_019830953.1;refGene_gn=ANKRD6;refGene_tx=XM_019830954.2;refGene_gn=ANKRD6;refGene_tx=XM_019830955.1;refGene_gn=ANKRD6;refGene_tx=XM_023254179.1;refGene_gn=ANKRD6;refGene_tx=XM_023254180.1
B2 83100684 83100811
refGene_gn=ANKRD6;refGene_tx=XM_019830951.1;refGene_gn=ANKRD6;refGene_tx=XM_019830952.1;refGene_gn=ANKRD6;refGene_tx=XM_019830953.1;refGene_gn=ANKRD6;refGene_tx=XM_019830954.2;refGene_gn=ANKRD6;refGene_tx=XM_019830955.1;refGene_gn=ANKRD6;refGene_tx=XM_023254179.1;refGene_gn=ANKRD6;refGene_tx=XM_023254180.1
B2 83102164 83102736
refGene_gn=ANKRD6;refGene_tx=XM_019830951.1;refGene_gn=ANKRD6;refGene_tx=XM_019830952.1;refGene_gn=ANKRD6;refGene_tx=XM_019830953.1;refGene_gn=ANKRD6;refGene_tx=XM_019830954.2;refGene_gn=ANKRD6;refGene_tx=XM_019830955.1;refGene_gn=ANKRD6;refGene_tx=XM_023254179.1;refGene_gn=ANKRD6;refGene_tx=XM_023254180.1
B2 83108991 83109072 refGene_gn=LYRM2;refGene_tx=XM_003986376.4
B2 83109513 83109654 refGene_gn=LYRM2;refGene_tx=XM_003986376.4
B2 83110279 83110324 refGene_gn=LYRM2;refGene_tx=XM_003986376.4
B2 83115219 83115408 refGene_gn=MDN1;refGene_tx=XM_023254181.1
B2 83116158 83116301 refGene_gn=MDN1;refGene_tx=XM_023254181.1
B2 83118175 83118239 refGene_gn=MDN1;refGene_tx=XM_023254181.1
B2 83119122 83119237 refGene_gn=MDN1;refGene_tx=XM_023254181.1
B2 83119330 83119404 refGene_gn=MDN1;refGene_tx=XM_023254181.1
B2 83121038 83121213 refGene_gn=MDN1;refGene_tx=XM_023254181.1
B2 83121802 83121918 refGene_gn=MDN1;refGene_tx=XM_023254181.1
B2 83122827 83122917 refGene_gn=MDN1;refGene_tx=XM_023254181.1
B2 83123637 83123801 refGene_gn=MDN1;refGene_tx=XM_023254181.1
B2 83125549 83125714 refGene_gn=MDN1;refGene_tx=XM_023254181.1
B2 83126818 83127020 refGene_gn=MDN1;refGene_tx=XM_023254181.1
B2 83127748 83127850 refGene_gn=MDN1;refGene_tx=XM_023254181.1
B2 83128944 83129019 refGene_gn=MDN1;refGene_tx=XM_023254181.1
B2 83129020 83129116 refGene_gn=MDN1;refGene_tx=XM_023254181.1
B2 83129597 83129920 refGene_gn=MDN1;refGene_tx=XM_023254181.1
B2 83132625 83132770 refGene_gn=MDN1;refGene_tx=XM_023254181.1
B2 83133266 83133413 refGene_gn=MDN1;refGene_tx=XM_023254181.1
B2 83133974 83134147 refGene_gn=MDN1;refGene_tx=XM_023254181.1
B2 83135586 83135725 refGene_gn=MDN1;refGene_tx=XM_023254181.1
B2 83139208 83139346 refGene_gn=MDN1;refGene_tx=XM_023254181.1
B2 83141986 83142148 refGene_gn=MDN1;refGene_tx=XM_023254181.1
B2 83142925 83143095 refGene_gn=MDN1;refGene_tx=XM_023254181.1
B2 83143494 83143650 refGene_gn=MDN1;refGene_tx=XM_023254181.1

| | | | |
|---|---|---|---|
| B2 | 83144437 | 83144664 | refGene_gn=MDN1;refGene_tx=XM_023254181.1 |
| B2 | 83144937 | 83145423 | refGene_gn=MDN1;refGene_tx=XM_023254181.1 |
| B2 | 83146328 | 83146444 | refGene_gn=MDN1;refGene_tx=XM_023254181.1 |
| B2 | 83147494 | 83147600 | refGene_gn=MDN1;refGene_tx=XM_023254181.1 |
| B2 | 83148796 | 83148920 | refGene_gn=MDN1;refGene_tx=XM_023254181.1 |
| B2 | 83149459 | 83149606 | refGene_gn=MDN1;refGene_tx=XM_023254181.1 |
| B2 | 83151701 | 83151861 | refGene_gn=MDN1;refGene_tx=XM_023254181.1 |
| B2 | 83153175 | 83153368 | refGene_gn=MDN1;refGene_tx=XM_023254181.1 |
| B2 | 83153850 | 83153924 | refGene_gn=MDN1;refGene_tx=XM_023254181.1 |
| B2 | 83154537 | 83154647 | refGene_gn=MDN1;refGene_tx=XM_023254181.1 |
| B2 | 83155489 | 83155636 | refGene_gn=MDN1;refGene_tx=XM_023254181.1 |
| B2 | 83156160 | 83156282 | refGene_gn=MDN1;refGene_tx=XM_023254181.1 |
| B2 | 83156932 | 83157027 | refGene_gn=MDN1;refGene_tx=XM_023254181.1 |
| B2 | 83157614 | 83157710 | refGene_gn=MDN1;refGene_tx=XM_023254181.1 |
| B2 | 83158454 | 83158690 | refGene_gn=MDN1;refGene_tx=XM_023254181.1 |
| B2 | 83159437 | 83159711 | refGene_gn=MDN1;refGene_tx=XM_023254181.1 |
| B2 | 83159926 | 83160088 | refGene_gn=MDN1;refGene_tx=XM_023254181.1 |
| B2 | 83162000 | 83162706 | refGene_gn=MDN1;refGene_tx=XM_023254181.1 |
| B2 | 83163535 | 83163645 | refGene_gn=MDN1;refGene_tx=XM_023254181.1 |
| B2 | 83164361 | 83164749 | refGene_gn=MDN1;refGene_tx=XM_023254181.1 |
| B2 | 83165120 | 83165335 | refGene_gn=MDN1;refGene_tx=XM_023254181.1 |
| B2 | 83168769 | 83168940 | refGene_gn=MDN1;refGene_tx=XM_023254181.1 |
| B2 | 83170022 | 83170149 | refGene_gn=MDN1;refGene_tx=XM_023254181.1 |
| B2 | 83170491 | 83170723 | refGene_gn=MDN1;refGene_tx=XM_023254181.1 |
| B2 | 83171071 | 83171282 | refGene_gn=MDN1;refGene_tx=XM_023254181.1 |
| B2 | 83171623 | 83171744 | refGene_gn=MDN1;refGene_tx=XM_023254181.1 |
| B2 | 83171963 | 83172121 | refGene_gn=MDN1;refGene_tx=XM_023254181.1 |
| B2 | 83173113 | 83173247 | refGene_gn=MDN1;refGene_tx=XM_023254181.1 |
| B2 | 83174246 | 83174362 | refGene_gn=MDN1;refGene_tx=XM_023254181.1 |
| B2 | 83175510 | 83175643 | refGene_gn=MDN1;refGene_tx=XM_023254181.1 |
| B2 | 83177275 | 83177389 | refGene_gn=MDN1;refGene_tx=XM_023254181.1 |
| B2 | 83178511 | 83178732 | refGene_gn=MDN1;refGene_tx=XM_023254181.1 |
| B2 | 83179068 | 83179280 | refGene_gn=MDN1;refGene_tx=XM_023254181.1 |
| B2 | 83179644 | 83179793 | refGene_gn=MDN1;refGene_tx=XM_023254181.1 |
| B2 | 83183080 | 83183289 | refGene_gn=MDN1;refGene_tx=XM_023254181.1 |
| B2 | 83184522 | 83184639 | refGene_gn=MDN1;refGene_tx=XM_023254181.1 |
| B2 | 83185699 | 83185859 | refGene_gn=MDN1;refGene_tx=XM_023254181.1 |
| B2 | 83197078 | 83197340 | refGene_gn=MDN1;refGene_tx=XM_023254181.1 |
| B2 | 83197630 | 83197894 | refGene_gn=MDN1;refGene_tx=XM_023254181.1 |
| B2 | 83197999 | 83198089 | refGene_gn=MDN1;refGene_tx=XM_023254181.1 |
| B2 | 83202071 | 83202260 | refGene_gn=MDN1;refGene_tx=XM_023254181.1 |
| B2 | 83202607 | 83202715 | refGene_gn=MDN1;refGene_tx=XM_023254181.1 |
| B2 | 83203989 | 83204187 | refGene_gn=MDN1;refGene_tx=XM_023254181.1 |
| B2 | 83205921 | 83206044 | refGene_gn=MDN1;refGene_tx=XM_023254181.1 |
| B2 | 83206730 | 83206939 | refGene_gn=MDN1;refGene_tx=XM_023254181.1 |
| B2 | 83208314 | 83208512 | refGene_gn=MDN1;refGene_tx=XM_023254181.1 |
| B2 | 83209224 | 83209443 | refGene_gn=MDN1;refGene_tx=XM_023254181.1 |
| B2 | 83215253 | 83215383 | refGene_gn=MDN1;refGene_tx=XM_023254181.1 |
| B2 | 83217660 | 83217805 | refGene_gn=MDN1;refGene_tx=XM_023254181.1 |
| B2 | 83219732 | 83219863 | refGene_gn=MDN1;refGene_tx=XM_023254181.1 |

B2 83220126 83220265 refGene_gn=MDN1;refGene_tx=XM_023254181.1
B2 83220865 83221004 refGene_gn=MDN1;refGene_tx=XM_023254181.1
B2 83221082 83221217 refGene_gn=MDN1;refGene_tx=XM_023254181.1
B2 83222594 83222736 refGene_gn=MDN1;refGene_tx=XM_023254181.1
B2 83223711 83223858 refGene_gn=MDN1;refGene_tx=XM_023254181.1
B2 83224018 83224227 refGene_gn=MDN1;refGene_tx=XM_023254181.1
B2 83224824 83225003 refGene_gn=MDN1;refGene_tx=XM_023254181.1
B2 83225512 83225664 refGene_gn=MDN1;refGene_tx=XM_023254181.1
B2 83228621 83228732 refGene_gn=MDN1;refGene_tx=XM_023254181.1
B2 83229544 83229692 refGene_gn=MDN1;refGene_tx=XM_023254181.1
B2 83231763 83231877 refGene_gn=MDN1;refGene_tx=XM_023254181.1
B2 83233367 83233464 refGene_gn=MDN1;refGene_tx=XM_023254181.1
B2 83233938 83234083 refGene_gn=MDN1;refGene_tx=XM_023254181.1
B2 83236332 83236436 refGene_gn=MDN1;refGene_tx=XM_023254181.1
B2 83244417 83244629 refGene_gn=MDN1;refGene_tx=XM_023254181.1
B2 83250125 83250186 refGene_gn=MDN1;refGene_tx=XM_023254181.1
B2 83251064 83251213 refGene_gn=MDN1;refGene_tx=XM_023254181.1
B2 83252427 83252540 refGene_gn=MDN1;refGene_tx=XM_023254181.1
B2 83254109 83254205 refGene_gn=MDN1;refGene_tx=XM_023254181.1
B2 83258919 83259001 refGene_gn=MDN1;refGene_tx=XM_023254181.1
B2 83260552 83260746 refGene_gn=MDN1;refGene_tx=XM_023254181.1
B2 83265115 83265230 refGene_gn=MDN1;refGene_tx=XM_023254181.1
B2 83268645 83268749 refGene_gn=MDN1;refGene_tx=XM_023254181.1
B2 83271435 83271567 refGene_gn=MDN1;refGene_tx=XM_023254181.1
B2 83271808 83272051 refGene_gn=MDN1;refGene_tx=XM_023254181.1
B2 83274565 83274758 refGene_gn=MDN1;refGene_tx=XM_023254181.1
B2 83274838 83274897 refGene_gn=MDN1;refGene_tx=XM_023254181.1
B2 83274898 83274947 refGene_gn=MDN1;refGene_tx=XM_023254181.1
B2 83275458 83275683 refGene_gn=MDN1;refGene_tx=XM_023254181.1
B2 83281775 83282002 refGene_gn=MDN1;refGene_tx=XM_023254181.1
B2 83290352 83290484 refGene_gn=MDN1;refGene_tx=XM_023254181.1
B2 83324244 83324299 refGene_gn=CASP8AP2;refGene_tx=XM_023254733.1
B2 83326682 83326751 refGene_gn=CASP8AP2;refGene_tx=XM_023254733.1
B2 83332153 83332254 refGene_gn=CASP8AP2;refGene_tx=XM_023254733.1
B2 83332800 83332875 refGene_gn=CASP8AP2;refGene_tx=XM_023254733.1
B2 83334307 83334423 refGene_gn=CASP8AP2;refGene_tx=XM_023254733.1
B2 83338124 83339842 refGene_gn=CASP8AP2;refGene_tx=XM_023254733.1
B2 83341092 83341583 refGene_gn=CASP8AP2;refGene_tx=XM_023254733.1
B2 83341584 83344220 refGene_gn=CASP8AP2;refGene_tx=XM_023254733.1
B2 83346506 83346606 refGene_gn=CASP8AP2;refGene_tx=XM_023254733.1
B2 83347393 83347450 refGene_gn=CASP8AP2;refGene_tx=XM_023254733.1
B2 83365833 83367270 refGene_gn=GJA10;refGene_tx=XM_004001506.5
B2 83375853 83375892 refGene_gn=GJA10;refGene_tx=XM_004001506.5
B2 83385266 83385749
refGene_gn=BACH2;refGene_tx=XM_011282461.3;refGene_gn=BACH2;refGene_tx=XM_019830959.2;refGene_gn=BACH2;refGene_tx=XM_019830960.2;refGene_gn=BACH2;refGene_tx=XM_019830962.2
B2 83390269 83390476
refGene_gn=BACH2;refGene_tx=XM_011282461.3;refGene_gn=BACH2;refGene_tx=XM_019830959.2;refGene_gn=BACH2;refGene_tx=XM_019830960.2;refGene_gn=

BACH2;refGene_tx=XM_019830962.2
B2 83401170 83402760
refGene_gn=BACH2;refGene_tx=XM_011282461.3;refGene_gn=BACH2;refGene_tx=XM_019830959.2;refGene_gn=BACH2;refGene_tx=XM_019830960.2;refGene_gn=BACH2;refGene_tx=XM_019830962.2
B2 83458983 83459238
refGene_gn=BACH2;refGene_tx=XM_011282461.3;refGene_gn=BACH2;refGene_tx=XM_019830959.2;refGene_gn=BACH2;refGene_tx=XM_019830960.2;refGene_gn=BACH2;refGene_tx=XM_019830962.2
B2 83641015 83641128
refGene_gn=BACH2;refGene_tx=XM_019830959.2;refGene_gn=BACH2;refGene_tx=XM_019830960.2
B2 83654587 83654894 refGene_gn=BACH2;refGene_tx=XM_019830959.2
B2 83951678 83951859
refGene_gn=MAP3K7;refGene_tx=XM_003986379.5;refGene_gn=MAP3K7;refGene_tx=XM_003986380.5
B2 83954295 83954411
refGene_gn=MAP3K7;refGene_tx=XM_003986379.5;refGene_gn=MAP3K7;refGene_tx=XM_003986380.5
B2 83955149 83955211
refGene_gn=MAP3K7;refGene_tx=XM_003986379.5;refGene_gn=MAP3K7;refGene_tx=XM_003986380.5
B2 83960006 83960112
refGene_gn=MAP3K7;refGene_tx=XM_003986379.5;refGene_gn=MAP3K7;refGene_tx=XM_003986380.5
B2 83973700 83973765
refGene_gn=MAP3K7;refGene_tx=XM_003986379.5;refGene_gn=MAP3K7;refGene_tx=XM_003986380.5
B2 83982578 83982659 refGene_gn=MAP3K7;refGene_tx=XM_003986379.5
B2 83985552 83985682
refGene_gn=MAP3K7;refGene_tx=XM_003986379.5;refGene_gn=MAP3K7;refGene_tx=XM_003986380.5
B2 83986311 83986442
refGene_gn=MAP3K7;refGene_tx=XM_003986379.5;refGene_gn=MAP3K7;refGene_tx=XM_003986380.5
B2 83988971 83989053
refGene_gn=MAP3K7;refGene_tx=XM_003986379.5;refGene_gn=MAP3K7;refGene_tx=XM_003986380.5
B2 83990862 83990993
refGene_gn=MAP3K7;refGene_tx=XM_003986379.5;refGene_gn=MAP3K7;refGene_tx=XM_003986380.5
B2 83992773 83992902
refGene_gn=MAP3K7;refGene_tx=XM_003986379.5;refGene_gn=MAP3K7;refGene_tx=XM_003986380.5
B2 83999005 83999130
refGene_gn=MAP3K7;refGene_tx=XM_003986379.5;refGene_gn=MAP3K7;refGene_tx=XM_003986380.5
B2 84007983 84008122
refGene_gn=MAP3K7;refGene_tx=XM_003986379.5;refGene_gn=MAP3K7;refGene_tx=XM_003986380.5

B2 84009198 84009244
refGene_gn=MAP3K7;refGene_tx=XM_003986379.5;refGene_gn=MAP3K7;refGene_tx=XM_003986380.5
B2 84009735 84009801
refGene_gn=MAP3K7;refGene_tx=XM_003986379.5;refGene_gn=MAP3K7;refGene_tx=XM_003986380.5
B2 84013961 84014072
refGene_gn=MAP3K7;refGene_tx=XM_003986379.5;refGene_gn=MAP3K7;refGene_tx=XM_003986380.5
B2 84029281 84029401
refGene_gn=MAP3K7;refGene_tx=XM_003986379.5;refGene_gn=MAP3K7;refGene_tx=XM_003986380.5
B2 86505811 86505926 refGene_gn=EPHA7;refGene_tx=XM_011282457.3
B2 86507900 86508056 refGene_gn=EPHA7;refGene_tx=XM_011282457.3
B2 86509315 86509509 refGene_gn=EPHA7;refGene_tx=XM_011282457.3
B2 86516967 86517117 refGene_gn=EPHA7;refGene_tx=XM_011282457.3
B2 86517836 86518046 refGene_gn=EPHA7;refGene_tx=XM_011282457.3
B2 86519481 86519543 refGene_gn=EPHA7;refGene_tx=XM_011282457.3
B2 86520133 86520319 refGene_gn=EPHA7;refGene_tx=XM_011282457.3
B2 86521541 86521667 refGene_gn=EPHA7;refGene_tx=XM_011282457.3
B2 86525581 86525637 refGene_gn=EPHA7;refGene_tx=XM_011282457.3
B2 86526279 86526388 refGene_gn=EPHA7;refGene_tx=XM_011282457.3
B2 86532225 86532409 refGene_gn=EPHA7;refGene_tx=XM_011282457.3
B2 86535077 86535202 refGene_gn=EPHA7;refGene_tx=XM_011282457.3
B2 86612654 86612990 refGene_gn=EPHA7;refGene_tx=XM_011282457.3
B2 86614065 86614221 refGene_gn=EPHA7;refGene_tx=XM_011282457.3
B2 86665711 86666381 refGene_gn=EPHA7;refGene_tx=XM_011282457.3
B2 86669991 86670056 refGene_gn=EPHA7;refGene_tx=XM_011282457.3
B2 86674578 86674599 refGene_gn=EPHA7;refGene_tx=XM_011282457.3
B2 86674601 86674674 refGene_gn=EPHA7;refGene_tx=XM_011282457.3
B2 87465394 87465448
refGene_gn=LOC101100814;refGene_tx=XM_019831600.2
B2 87476144 87476624
refGene_gn=LOC101100814;refGene_tx=XM_019831600.2
B2 87855517 87857179
refGene_gn=LOC101099553;refGene_tx=XM_019830965.2;refGene_gn=LOC101099553;refGene_tx=XM_019830966.2
B2 88643947 88644488
refGene_gn=MANEA;refGene_tx=XM_003986390.5;refGene_gn=MANEA;refGene_tx=XM_019830967.2
B2 88668146 88668256
refGene_gn=MANEA;refGene_tx=XM_003986390.5;refGene_gn=MANEA;refGene_tx=XM_019830967.2
B2 88693138 88693215
refGene_gn=MANEA;refGene_tx=XM_003986390.5;refGene_gn=MANEA;refGene_tx=XM_019830967.2
B2 88693991 88694649
refGene_gn=MANEA;refGene_tx=XM_003986390.5;refGene_gn=MANEA;refGene_tx=XM_019830967.2
B2 89187461 89187480

refGene_gn=FUT9;refGene_tx=XM_006944368.4;refGene_gn=FUT9;refGene_tx=XM_019830970.2
B2 89256979 89257052 refGene_gn=FUT9;refGene_tx=XM_019830968.2
B2 89282852 89283940
refGene_gn=FUT9;refGene_tx=XM_006944368.4;refGene_gn=FUT9;refGene_tx=XM_019830968.2;refGene_gn=FUT9;refGene_tx=XM_019830970.2;refGene_gn=FUT9;refGene_tx=XM_019830971.2
B2 89537620 89537697 refGene_gn=UFL1;refGene_tx=XM_006931895.4
B2 89538981 89539127
refGene_gn=UFL1;refGene_tx=XM_006931895.4;refGene_gn=UFL1;refGene_tx=XM_019830973.2
B2 89540203 89540232
refGene_gn=UFL1;refGene_tx=XM_006931895.4;refGene_gn=UFL1;refGene_tx=XM_019830973.2
B2 89541106 89541204
refGene_gn=UFL1;refGene_tx=XM_006931895.4;refGene_gn=UFL1;refGene_tx=XM_019830973.2
B2 89541899 89542014
refGene_gn=UFL1;refGene_tx=XM_006931895.4;refGene_gn=UFL1;refGene_tx=XM_019830973.2
B2 89543584 89543715
refGene_gn=UFL1;refGene_tx=XM_006931895.4;refGene_gn=UFL1;refGene_tx=XM_019830973.2
B2 89550489 89550548
refGene_gn=UFL1;refGene_tx=XM_006931895.4;refGene_gn=UFL1;refGene_tx=XM_019830973.2
B2 89552589 89552736
refGene_gn=UFL1;refGene_tx=XM_006931895.4;refGene_gn=UFL1;refGene_tx=XM_019830973.2
B2 89554058 89554234
refGene_gn=UFL1;refGene_tx=XM_006931895.4;refGene_gn=UFL1;refGene_tx=XM_019830973.2
B2 89555429 89555609
refGene_gn=UFL1;refGene_tx=XM_006931895.4;refGene_gn=UFL1;refGene_tx=XM_019830973.2
B2 89557809 89557930
refGene_gn=UFL1;refGene_tx=XM_006931895.4;refGene_gn=UFL1;refGene_tx=XM_019830973.2
B2 89560142 89560259
refGene_gn=UFL1;refGene_tx=XM_006931895.4;refGene_gn=UFL1;refGene_tx=XM_019830973.2
B2 89563508 89563626
refGene_gn=UFL1;refGene_tx=XM_006931895.4;refGene_gn=UFL1;refGene_tx=XM_019830973.2
B2 89565021 89565188
refGene_gn=UFL1;refGene_tx=XM_006931895.4;refGene_gn=UFL1;refGene_tx=XM_019830973.2
B2 89565269 89565400
refGene_gn=UFL1;refGene_tx=XM_006931895.4;refGene_gn=UFL1;refGene_tx=XM_019830973.2

B2 89566978 89567059
refGene_gn=UFL1;refGene_tx=XM_006931895.4;refGene_gn=UFL1;refGene_tx=XM_019830973.2
B2 89567347 89567433
refGene_gn=UFL1;refGene_tx=XM_006931895.4;refGene_gn=UFL1;refGene_tx=XM_019830973.2
B2 89568201 89568382
refGene_gn=UFL1;refGene_tx=XM_006931895.4;refGene_gn=UFL1;refGene_tx=XM_019830973.2
B2 89569011 89569230
refGene_gn=UFL1;refGene_tx=XM_006931895.4;refGene_gn=UFL1;refGene_tx=XM_019830973.2
B2 89607139 89607298
refGene_gn=FHL5;refGene_tx=XM_003986392.4;refGene_gn=FHL5;refGene_tx=XM_019830974.2;refGene_gn=FHL5;refGene_tx=XM_019830975.2;refGene_gn=FHL5;refGene_tx=XM_019830976.2;refGene_gn=FHL5;refGene_tx=XM_023254185.1
B2 89608131 89608306
refGene_gn=FHL5;refGene_tx=XM_003986392.4;refGene_gn=FHL5;refGene_tx=XM_019830974.2;refGene_gn=FHL5;refGene_tx=XM_019830975.2;refGene_gn=FHL5;refGene_tx=XM_019830976.2;refGene_gn=FHL5;refGene_tx=XM_023254185.1
B2 89609258 89609428
refGene_gn=FHL5;refGene_tx=XM_003986392.4;refGene_gn=FHL5;refGene_tx=XM_019830974.2;refGene_gn=FHL5;refGene_tx=XM_019830975.2;refGene_gn=FHL5;refGene_tx=XM_019830976.2;refGene_gn=FHL5;refGene_tx=XM_023254185.1
B2 89613655 89613842
refGene_gn=FHL5;refGene_tx=XM_003986392.4;refGene_gn=FHL5;refGene_tx=XM_019830974.2;refGene_gn=FHL5;refGene_tx=XM_019830975.2;refGene_gn=FHL5;refGene_tx=XM_019830976.2;refGene_gn=FHL5;refGene_tx=XM_023254185.1
B2 89624406 89624570
refGene_gn=FHL5;refGene_tx=XM_003986392.4;refGene_gn=FHL5;refGene_tx=XM_019830974.2;refGene_gn=FHL5;refGene_tx=XM_019830975.2;refGene_gn=FHL5;refGene_tx=XM_019830976.2;refGene_gn=FHL5;refGene_tx=XM_023254185.1
B2 89762214 89763474
refGene_gn=GPR63;refGene_tx=XM_006931897.4;refGene_gn=GPR63;refGene_tx=XM_019830977.2;refGene_gn=GPR63;refGene_tx=XM_019830978.2
B2 89869126 89869414 refGene_gn=NDUFAF4;refGene_tx=XM_003986394.5
B2 89874293 89874397 refGene_gn=NDUFAF4;refGene_tx=XM_003986394.5
B2 89874907 89875043 refGene_gn=NDUFAF4;refGene_tx=XM_003986394.5
B2 89948409 89948432
refGene_gn=KLHL32;refGene_tx=XM_003986395.5;refGene_gn=KLHL32;refGene_tx=XM_011282456.3;refGene_gn=KLHL32;refGene_tx=XM_019830988.2;refGene_gn=KLHL32;refGene_tx=XM_019830992.2;refGene_gn=KLHL32;refGene_tx=XM_019830993.2;refGene_gn=KLHL32;refGene_tx=XM_019830994.2;refGene_gn=KLHL32;refGene_tx=XM_019830995.2;refGene_gn=KLHL32;refGene_tx=XM_023254187.1;refGene_gn=KLHL32;refGene_tx=XM_023254188.1
B2 89956810 89956991
refGene_gn=KLHL32;refGene_tx=XM_003986395.5;refGene_gn=KLHL32;refGene_tx=XM_011282456.3;refGene_gn=KLHL32;refGene_tx=XM_019830986.2;refGene_gn=KLHL32;refGene_tx=XM_019830988.2;refGene_gn=KLHL32;refGene_tx=XM_019830992.2;refGene_gn=KLHL32;refGene_tx=XM_019830993.2;refGene_gn=KLHL3

2;refGene_tx=XM_019830994.2;refGene_gn=KLHL32;refGene_tx=XM_019830995.2;refGene_gn=KLHL32;refGene_tx=XM_023254187.1;refGene_gn=KLHL32;refGene_tx=XM_023254188.1
B2 89997864 89997972
refGene_gn=KLHL32;refGene_tx=XM_003986395.5;refGene_gn=KLHL32;refGene_tx=XM_011282456.3;refGene_gn=KLHL32;refGene_tx=XM_019830986.2;refGene_gn=KLHL32;refGene_tx=XM_019830988.2;refGene_gn=KLHL32;refGene_tx=XM_019830992.2;refGene_gn=KLHL32;refGene_tx=XM_019830993.2;refGene_gn=KLHL32;refGene_tx=XM_019830994.2;refGene_gn=KLHL32;refGene_tx=XM_019830995.2;refGene_gn=KLHL32;refGene_tx=XM_023254187.1;refGene_gn=KLHL32;refGene_tx=XM_023254188.1
B2 90025686 90025785
refGene_gn=KLHL32;refGene_tx=XM_003986395.5;refGene_gn=KLHL32;refGene_tx=XM_011282456.3;refGene_gn=KLHL32;refGene_tx=XM_019830986.2;refGene_gn=KLHL32;refGene_tx=XM_019830988.2;refGene_gn=KLHL32;refGene_tx=XM_019830992.2;refGene_gn=KLHL32;refGene_tx=XM_019830993.2;refGene_gn=KLHL32;refGene_tx=XM_019830994.2;refGene_gn=KLHL32;refGene_tx=XM_019830995.2;refGene_gn=KLHL32;refGene_tx=XM_023254187.1;refGene_gn=KLHL32;refGene_tx=XM_023254188.1
B2 90053839 90054055
refGene_gn=KLHL32;refGene_tx=XM_003986395.5;refGene_gn=KLHL32;refGene_tx=XM_011282456.3;refGene_gn=KLHL32;refGene_tx=XM_019830986.2;refGene_gn=KLHL32;refGene_tx=XM_019830992.2;refGene_gn=KLHL32;refGene_tx=XM_019830993.2;refGene_gn=KLHL32;refGene_tx=XM_019830994.2;refGene_gn=KLHL32;refGene_tx=XM_019830995.2;refGene_gn=KLHL32;refGene_tx=XM_023254187.1;refGene_gn=KLHL32;refGene_tx=XM_023254188.1
B2 90071341 90071476 refGene_gn=KLHL32;refGene_tx=XM_019830993.2
B2 90081787 90082514
refGene_gn=KLHL32;refGene_tx=XM_003986395.5;refGene_gn=KLHL32;refGene_tx=XM_011282456.3;refGene_gn=KLHL32;refGene_tx=XM_019830986.2;refGene_gn=KLHL32;refGene_tx=XM_019830988.2;refGene_gn=KLHL32;refGene_tx=XM_019830990.2;refGene_gn=KLHL32;refGene_tx=XM_019830991.2;refGene_gn=KLHL32;refGene_tx=XM_019830992.2;refGene_gn=KLHL32;refGene_tx=XM_019830995.2;refGene_gn=KLHL32;refGene_tx=XM_023254187.1;refGene_gn=KLHL32;refGene_tx=XM_023254188.1
B2 90104080 90104139
refGene_gn=KLHL32;refGene_tx=XM_003986395.5;refGene_gn=KLHL32;refGene_tx=XM_011282456.3;refGene_gn=KLHL32;refGene_tx=XM_019830986.2;refGene_gn=KLHL32;refGene_tx=XM_019830988.2;refGene_gn=KLHL32;refGene_tx=XM_019830990.2;refGene_gn=KLHL32;refGene_tx=XM_019830991.2;refGene_gn=KLHL32;refGene_tx=XM_019830992.2;refGene_gn=KLHL32;refGene_tx=XM_019830994.2;refGene_gn=KLHL32;refGene_tx=XM_019830996.2;refGene_gn=KLHL32;refGene_tx=XM_019830997.2;refGene_gn=KLHL32;refGene_tx=XM_023254187.1;refGene_gn=KLHL32;refGene_tx=XM_023254188.1
B2 90107371 90107564
refGene_gn=KLHL32;refGene_tx=XM_003986395.5;refGene_gn=KLHL32;refGene_tx=XM_011282456.3;refGene_gn=KLHL32;refGene_tx=XM_019830986.2;refGene_gn=KLHL32;refGene_tx=XM_019830988.2;refGene_gn=KLHL32;refGene_tx=XM_019830990.2;refGene_gn=KLHL32;refGene_tx=XM_019830991.2;refGene_gn=KLHL32;refGene_tx=XM_019830996.2;refGene_gn=KLHL32;refGene_tx=XM_019830997.

2;refGene_gn=KLHL32;refGene_tx=XM_023254187.1;refGene_gn=KLHL32;refGene_tx=XM_023254188.1
B2 90109273 90109368
refGene_gn=KLHL32;refGene_tx=XM_003986395.5;refGene_gn=KLHL32;refGene_tx=XM_011282456.3;refGene_gn=KLHL32;refGene_tx=XM_019830986.2;refGene_gn=KLHL32;refGene_tx=XM_019830988.2;refGene_gn=KLHL32;refGene_tx=XM_019830990.2;refGene_gn=KLHL32;refGene_tx=XM_019830991.2;refGene_gn=KLHL32;refGene_tx=XM_019830996.2;refGene_gn=KLHL32;refGene_tx=XM_019830997.2;refGene_gn=KLHL32;refGene_tx=XM_023254187.1;refGene_gn=KLHL32;refGene_tx=XM_023254188.1
B2 90142099 90142261
refGene_gn=KLHL32;refGene_tx=XM_003986395.5;refGene_gn=KLHL32;refGene_tx=XM_011282456.3;refGene_gn=KLHL32;refGene_tx=XM_019830986.2;refGene_gn=KLHL32;refGene_tx=XM_019830988.2;refGene_gn=KLHL32;refGene_tx=XM_019830990.2;refGene_gn=KLHL32;refGene_tx=XM_019830991.2;refGene_gn=KLHL32;refGene_tx=XM_019830996.2;refGene_gn=KLHL32;refGene_tx=XM_019830997.2;refGene_gn=KLHL32;refGene_tx=XM_023254187.1;refGene_gn=KLHL32;refGene_tx=XM_023254188.1
B2 90149165 90149247 refGene_gn=MMS22L;refGene_tx=XM_023254186.1
B2 90149981 90150149 refGene_gn=MMS22L;refGene_tx=XM_023254186.1
B2 90151739 90151836 refGene_gn=MMS22L;refGene_tx=XM_023254186.1
B2 90162376 90162540 refGene_gn=MMS22L;refGene_tx=XM_023254186.1
B2 90165765 90165977 refGene_gn=MMS22L;refGene_tx=XM_023254186.1
B2 90168703 90168873 refGene_gn=MMS22L;refGene_tx=XM_023254186.1
B2 90173073 90173233 refGene_gn=MMS22L;refGene_tx=XM_023254186.1
B2 90178496 90178639 refGene_gn=MMS22L;refGene_tx=XM_023254186.1
B2 90179505 90179657 refGene_gn=MMS22L;refGene_tx=XM_023254186.1
B2 90181508 90181659 refGene_gn=MMS22L;refGene_tx=XM_023254186.1
B2 90186920 90187114 refGene_gn=MMS22L;refGene_tx=XM_023254186.1
B2 90229521 90230025 refGene_gn=MMS22L;refGene_tx=XM_023254186.1
B2 90231929 90232156 refGene_gn=MMS22L;refGene_tx=XM_023254186.1
B2 90237920 90238040 refGene_gn=MMS22L;refGene_tx=XM_023254186.1
B2 90246028 90246091 refGene_gn=MMS22L;refGene_tx=XM_023254186.1
B2 90253987 90254164 refGene_gn=MMS22L;refGene_tx=XM_023254186.1
B2 90264094 90264208 refGene_gn=MMS22L;refGene_tx=XM_023254186.1
B2 90265662 90265793 refGene_gn=MMS22L;refGene_tx=XM_023254186.1
B2 90267880 90267971 refGene_gn=MMS22L;refGene_tx=XM_023254186.1
B2 90270965 90271143 refGene_gn=MMS22L;refGene_tx=XM_023254186.1
B2 90271238 90271326 refGene_gn=MMS22L;refGene_tx=XM_023254186.1
B2 90273021 90273071 refGene_gn=MMS22L;refGene_tx=XM_023254186.1
B2 90274374 90274500 refGene_gn=MMS22L;refGene_tx=XM_023254186.1
B2 90275572 90275736 refGene_gn=MMS22L;refGene_tx=XM_023254186.1
B2 91737919 91738408 refGene_gn=POU3F2;refGene_tx=XM_023254189.1
B2 91738409 91739252 refGene_gn=POU3F2;refGene_tx=XM_023254189.1
B2 91773829 91773993
refGene_gn=FBXL4;refGene_tx=XM_006931899.4;refGene_gn=FBXL4;refGene_tx=XM_011282454.3
B2 91786941 91787254
refGene_gn=FBXL4;refGene_tx=XM_006931899.4;refGene_gn=FBXL4;refGene_tx=XM_011282454.3

B2 91791298 91791370
refGene_gn=FBXL4;refGene_tx=XM_006931899.4;refGene_gn=FBXL4;refGene_tx=XM_011282454.3
B2 91812478 91812692
refGene_gn=FBXL4;refGene_tx=XM_006931899.4;refGene_gn=FBXL4;refGene_tx=XM_011282454.3
B2 91821194 91821439
refGene_gn=FBXL4;refGene_tx=XM_006931899.4;refGene_gn=FBXL4;refGene_tx=XM_011282454.3
B2 91829219 91829565
refGene_gn=FBXL4;refGene_tx=XM_006931899.4;refGene_gn=FBXL4;refGene_tx=XM_011282454.3
B2 91839223 91839735
refGene_gn=FBXL4;refGene_tx=XM_006931899.4;refGene_gn=FBXL4;refGene_tx=XM_011282454.3
B2 92160233 92160523
refGene_gn=FAXC;refGene_tx=XM_011282453.3;refGene_gn=FAXC;refGene_tx=XM_019830998.2;refGene_gn=FAXC;refGene_tx=XM_019830999.2;refGene_gn=FAXC;refGene_tx=XM_023254190.1
B2 92168875 92168992
refGene_gn=FAXC;refGene_tx=XM_011282453.3;refGene_gn=FAXC;refGene_tx=XM_019830998.2;refGene_gn=FAXC;refGene_tx=XM_019830999.2;refGene_gn=FAXC;refGene_tx=XM_023254190.1
B2 92201132 92201356
refGene_gn=FAXC;refGene_tx=XM_011282453.3;refGene_gn=FAXC;refGene_tx=XM_019830998.2;refGene_gn=FAXC;refGene_tx=XM_019830999.2;refGene_gn=FAXC;refGene_tx=XM_023254190.1
B2 92209877 92210074
refGene_gn=FAXC;refGene_tx=XM_011282453.3;refGene_gn=FAXC;refGene_tx=XM_019830998.2;refGene_gn=FAXC;refGene_tx=XM_019830999.2;refGene_gn=FAXC;refGene_tx=XM_023254190.1
B2 92220134 92220215
refGene_gn=FAXC;refGene_tx=XM_019830998.2;refGene_gn=FAXC;refGene_tx=XM_019830999.2;refGene_gn=FAXC;refGene_tx=XM_023254190.1
B2 92224635 92224771
refGene_gn=FAXC;refGene_tx=XM_011282453.3;refGene_gn=FAXC;refGene_tx=XM_019830998.2;refGene_gn=FAXC;refGene_tx=XM_019830999.2;refGene_gn=FAXC;refGene_tx=XM_023254190.1
B2 92230538 92230804
refGene_gn=FAXC;refGene_tx=XM_011282453.3;refGene_gn=FAXC;refGene_tx=XM_019830998.2;refGene_gn=FAXC;refGene_tx=XM_023254190.1
B2 92254756 92254977
refGene_gn=COQ3;refGene_tx=XM_003986401.5;refGene_gn=COQ3;refGene_tx=XM_006931900.4;refGene_gn=COQ3;refGene_tx=XM_019831011.2
B2 92256183 92256343
refGene_gn=COQ3;refGene_tx=XM_003986401.5;refGene_gn=COQ3;refGene_tx=XM_006931900.4;refGene_gn=COQ3;refGene_tx=XM_019831011.2
B2 92266123 92266366
refGene_gn=COQ3;refGene_tx=XM_003986401.5;refGene_gn=COQ3;refGene_tx=XM_006931900.4;refGene_gn=COQ3;refGene_tx=XM_019831011.2

B2 92267348 92267448
refGene_gn=COQ3;refGene_tx=XM_003986401.5;refGene_gn=COQ3;refGene_tx=XM_006931900.4;refGene_gn=COQ3;refGene_tx=XM_019831011.2
B2 92271834 92271987
refGene_gn=COQ3;refGene_tx=XM_003986401.5;refGene_gn=COQ3;refGene_tx=XM_006931900.4;refGene_gn=COQ3;refGene_tx=XM_019831011.2
B2 92274237 92274364
refGene_gn=COQ3;refGene_tx=XM_003986401.5;refGene_gn=COQ3;refGene_tx=XM_006931900.4;refGene_gn=COQ3;refGene_tx=XM_019831011.2
B2 92280158 92280267 refGene_gn=COQ3;refGene_tx=XM_003986401.5
B2 92284940 92284971
refGene_gn=PNISR;refGene_tx=XM_006931901.4;refGene_gn=PNISR;refGene_tx=XM_006931903.4;refGene_gn=PNISR;refGene_tx=XM_019831000.2;refGene_gn=PNISR;refGene_tx=XM_019831001.2
B2 92285209 92286300
refGene_gn=PNISR;refGene_tx=XM_006931901.4;refGene_gn=PNISR;refGene_tx=XM_006931903.4;refGene_gn=PNISR;refGene_tx=XM_011282452.3;refGene_gn=PNISR;refGene_tx=XM_019831000.2;refGene_gn=PNISR;refGene_tx=XM_019831001.2
B2 92287537 92287708
refGene_gn=PNISR;refGene_tx=XM_006931901.4;refGene_gn=PNISR;refGene_tx=XM_006931903.4;refGene_gn=PNISR;refGene_tx=XM_011282452.3;refGene_gn=PNISR;refGene_tx=XM_019831000.2;refGene_gn=PNISR;refGene_tx=XM_019831001.2
B2 92288776 92288830
refGene_gn=PNISR;refGene_tx=XM_006931901.4;refGene_gn=PNISR;refGene_tx=XM_006931903.4;refGene_gn=PNISR;refGene_tx=XM_011282452.3;refGene_gn=PNISR;refGene_tx=XM_019831000.2;refGene_gn=PNISR;refGene_tx=XM_019831001.2
B2 92289573 92289673
refGene_gn=PNISR;refGene_tx=XM_006931901.4;refGene_gn=PNISR;refGene_tx=XM_006931903.4;refGene_gn=PNISR;refGene_tx=XM_011282452.3;refGene_gn=PNISR;refGene_tx=XM_019831000.2;refGene_gn=PNISR;refGene_tx=XM_019831001.2
B2 92291071 92291209
refGene_gn=PNISR;refGene_tx=XM_006931901.4;refGene_gn=PNISR;refGene_tx=XM_006931903.4;refGene_gn=PNISR;refGene_tx=XM_011282452.3;refGene_gn=PNISR;refGene_tx=XM_019831000.2;refGene_gn=PNISR;refGene_tx=XM_019831001.2
B2 92293407 92293598
refGene_gn=PNISR;refGene_tx=XM_006931901.4;refGene_gn=PNISR;refGene_tx=XM_006931903.4;refGene_gn=PNISR;refGene_tx=XM_011282452.3;refGene_gn=PNISR;refGene_tx=XM_019831000.2;refGene_gn=PNISR;refGene_tx=XM_019831001.2
B2 92295379 92295551
refGene_gn=PNISR;refGene_tx=XM_006931901.4;refGene_gn=PNISR;refGene_tx=XM_006931903.4;refGene_gn=PNISR;refGene_tx=XM_011282452.3;refGene_gn=PNISR;refGene_tx=XM_019831000.2;refGene_gn=PNISR;refGene_tx=XM_019831001.2;refGene_gn=PNISR;refGene_tx=XM_019831002.2
B2 92297044 92297268

refGene_gn=PNISR;refGene_tx=XM_006931901.4;refGene_gn=PNISR;refGene_tx=XM_006931903.4;refGene_gn=PNISR;refGene_tx=XM_011282452.3;refGene_gn=PNISR;refGene_tx=XM_019831000.2;refGene_gn=PNISR;refGene_tx=XM_019831001.2;refGene_gn=PNISR;refGene_tx=XM_019831002.2
B2 92298417 92298422 refGene_gn=PNISR;refGene_tx=XM_019831003.2
B2 92298653 92298842
refGene_gn=PNISR;refGene_tx=XM_006931901.4;refGene_gn=PNISR;refGene_tx=XM_006931903.4;refGene_gn=PNISR;refGene_tx=XM_011282452.3;refGene_gn=PNISR;refGene_tx=XM_019831000.2;refGene_gn=PNISR;refGene_tx=XM_019831001.2;refGene_gn=PNISR;refGene_tx=XM_019831002.2;refGene_gn=PNISR;refGene_tx=XM_019831003.2
B2 92300323 92300411
refGene_gn=PNISR;refGene_tx=XM_006931901.4;refGene_gn=PNISR;refGene_tx=XM_006931903.4;refGene_gn=PNISR;refGene_tx=XM_011282452.3;refGene_gn=PNISR;refGene_tx=XM_019831000.2;refGene_gn=PNISR;refGene_tx=XM_019831001.2;refGene_gn=PNISR;refGene_tx=XM_019831002.2;refGene_gn=PNISR;refGene_tx=XM_019831003.2
B2 92317841 92317972
refGene_gn=USP45;refGene_tx=XM_011282451.3;refGene_gn=USP45;refGene_tx=XM_019831004.2;refGene_gn=USP45;refGene_tx=XM_019831005.2;refGene_gn=USP45;refGene_tx=XM_019831006.2;refGene_gn=USP45;refGene_tx=XM_019831007.2
B2 92318563 92318717
refGene_gn=USP45;refGene_tx=XM_011282451.3;refGene_gn=USP45;refGene_tx=XM_019831004.2;refGene_gn=USP45;refGene_tx=XM_019831005.2;refGene_gn=USP45;refGene_tx=XM_019831006.2;refGene_gn=USP45;refGene_tx=XM_019831007.2
B2 92320606 92320693
refGene_gn=USP45;refGene_tx=XM_011282451.3;refGene_gn=USP45;refGene_tx=XM_019831004.2;refGene_gn=USP45;refGene_tx=XM_019831005.2;refGene_gn=USP45;refGene_tx=XM_019831006.2;refGene_gn=USP45;refGene_tx=XM_019831007.2
B2 92322303 92322401
refGene_gn=USP45;refGene_tx=XM_011282451.3;refGene_gn=USP45;refGene_tx=XM_019831004.2;refGene_gn=USP45;refGene_tx=XM_019831005.2;refGene_gn=USP45;refGene_tx=XM_019831006.2;refGene_gn=USP45;refGene_tx=XM_019831007.2
B2 92324039 92324700
refGene_gn=USP45;refGene_tx=XM_011282451.3;refGene_gn=USP45;refGene_tx=XM_019831004.2;refGene_gn=USP45;refGene_tx=XM_019831005.2;refGene_gn=USP45;refGene_tx=XM_019831006.2;refGene_gn=USP45;refGene_tx=XM_019831007.2
B2 92325024 92325048 refGene_gn=USP45;refGene_tx=XM_019831008.2
B2 92326705 92326849
refGene_gn=USP45;refGene_tx=XM_011282451.3;refGene_gn=USP45;refGene_tx=XM_019831004.2;refGene_gn=USP45;refGene_tx=XM_019831005.2;refGene_gn=USP45;refGene_tx=XM_019831006.2;refGene_gn=USP45;refGene_tx=XM_019831007.2;refGene_gn=USP45;refGene_tx=XM_019831008.2
B2 92327311 92327368
refGene_gn=USP45;refGene_tx=XM_011282451.3;refGene_gn=USP45;refGene_tx

=XM_019831004.2;refGene_gn=USP45;refGene_tx=XM_019831005.2;refGene_gn=USP45;refGene_tx=XM_019831006.2;refGene_gn=USP45;refGene_tx=XM_019831007.2;refGene_gn=USP45;refGene_tx=XM_019831008.2;refGene_gn=USP45;refGene_tx=XM_019831009.1
B2 92328822 92328914
refGene_gn=USP45;refGene_tx=XM_011282451.3;refGene_gn=USP45;refGene_tx=XM_019831004.2;refGene_gn=USP45;refGene_tx=XM_019831005.2;refGene_gn=USP45;refGene_tx=XM_019831006.2;refGene_gn=USP45;refGene_tx=XM_019831008.2
B2 92331804 92331886
refGene_gn=USP45;refGene_tx=XM_011282451.3;refGene_gn=USP45;refGene_tx=XM_019831004.2;refGene_gn=USP45;refGene_tx=XM_019831005.2;refGene_gn=USP45;refGene_tx=XM_019831006.2;refGene_gn=USP45;refGene_tx=XM_019831007.2;refGene_gn=USP45;refGene_tx=XM_019831008.2;refGene_gn=USP45;refGene_tx=XM_019831009.1
B2 92332899 92332920 refGene_gn=USP45;refGene_tx=XM_019831010.2
B2 92347149 92347237
refGene_gn=USP45;refGene_tx=XM_011282451.3;refGene_gn=USP45;refGene_tx=XM_019831004.2;refGene_gn=USP45;refGene_tx=XM_019831005.2;refGene_gn=USP45;refGene_tx=XM_019831006.2;refGene_gn=USP45;refGene_tx=XM_019831008.2;refGene_gn=USP45;refGene_tx=XM_019831009.1;refGene_gn=USP45;refGene_tx=XM_019831010.2
B2 92355869 92356000
refGene_gn=USP45;refGene_tx=XM_011282451.3;refGene_gn=USP45;refGene_tx=XM_019831004.2;refGene_gn=USP45;refGene_tx=XM_019831005.2;refGene_gn=USP45;refGene_tx=XM_019831006.2;refGene_gn=USP45;refGene_tx=XM_019831008.2;refGene_gn=USP45;refGene_tx=XM_019831009.1;refGene_gn=USP45;refGene_tx=XM_019831010.2
B2 92360093 92360189
refGene_gn=USP45;refGene_tx=XM_011282451.3;refGene_gn=USP45;refGene_tx=XM_019831004.2;refGene_gn=USP45;refGene_tx=XM_019831005.2;refGene_gn=USP45;refGene_tx=XM_019831006.2;refGene_gn=USP45;refGene_tx=XM_019831008.2;refGene_gn=USP45;refGene_tx=XM_019831009.1;refGene_gn=USP45;refGene_tx=XM_019831010.2
B2 92360569 92360712
refGene_gn=USP45;refGene_tx=XM_011282451.3;refGene_gn=USP45;refGene_tx=XM_019831004.2;refGene_gn=USP45;refGene_tx=XM_019831005.2;refGene_gn=USP45;refGene_tx=XM_019831006.2;refGene_gn=USP45;refGene_tx=XM_019831008.2;refGene_gn=USP45;refGene_tx=XM_019831009.1;refGene_gn=USP45;refGene_tx=XM_019831010.2
B2 92375930 92376031
refGene_gn=USP45;refGene_tx=XM_011282451.3;refGene_gn=USP45;refGene_tx=XM_019831004.2;refGene_gn=USP45;refGene_tx=XM_019831005.2;refGene_gn=USP45;refGene_tx=XM_019831006.2;refGene_gn=USP45;refGene_tx=XM_019831008.2;refGene_gn=USP45;refGene_tx=XM_019831009.1;refGene_gn=USP45;refGene_tx=XM_019831010.2
B2 92380208 92380312
refGene_gn=USP45;refGene_tx=XM_011282451.3;refGene_gn=USP45;refGene_tx=XM_019831004.2;refGene_gn=USP45;refGene_tx=XM_019831005.2;refGene_gn=USP45;refGene_tx=XM_019831006.2;refGene_gn=USP45;refGene_tx=XM_0198310

08.2;refGene_gn=USP45;refGene_tx=XM_019831009.1;refGene_gn=USP45;refGene_tx=XM_019831010.2
B2	92382407	92382580
refGene_gn=USP45;refGene_tx=XM_011282451.3;refGene_gn=USP45;refGene_tx=XM_019831004.2;refGene_gn=USP45;refGene_tx=XM_019831005.2;refGene_gn=USP45;refGene_tx=XM_019831006.2;refGene_gn=USP45;refGene_tx=XM_019831008.2;refGene_gn=USP45;refGene_tx=XM_019831009.1;refGene_gn=USP45;refGene_tx=XM_019831010.2
B2	92387594	92387694
refGene_gn=USP45;refGene_tx=XM_011282451.3;refGene_gn=USP45;refGene_tx=XM_019831004.2;refGene_gn=USP45;refGene_tx=XM_019831005.2;refGene_gn=USP45;refGene_tx=XM_019831006.2;refGene_gn=USP45;refGene_tx=XM_019831008.2;refGene_gn=USP45;refGene_tx=XM_019831009.1;refGene_gn=USP45;refGene_tx=XM_019831010.2
B2	92410442	92410497	refGene_gn=CCNC;refGene_tx=XM_006931905.3
B2	92411082	92411134	refGene_gn=CCNC;refGene_tx=XM_011282450.2
B2	92411925	92412044
refGene_gn=CCNC;refGene_tx=XM_006931905.3;refGene_gn=CCNC;refGene_tx=XM_011282450.2
B2	92412941	92413021
refGene_gn=CCNC;refGene_tx=XM_006931905.3;refGene_gn=CCNC;refGene_tx=XM_011282450.2
B2	92416581	92416649
refGene_gn=CCNC;refGene_tx=XM_006931905.3;refGene_gn=CCNC;refGene_tx=XM_011282450.2
B2	92417248	92417340
refGene_gn=CCNC;refGene_tx=XM_006931905.3;refGene_gn=CCNC;refGene_tx=XM_011282450.2
B2	92418015	92418051
refGene_gn=CCNC;refGene_tx=XM_006931905.3;refGene_gn=CCNC;refGene_tx=XM_011282450.2
B2	92418933	92418989
refGene_gn=CCNC;refGene_tx=XM_006931905.3;refGene_gn=CCNC;refGene_tx=XM_011282450.2
B2	92425637	92425689
refGene_gn=CCNC;refGene_tx=XM_006931905.3;refGene_gn=CCNC;refGene_tx=XM_011282450.2
B2	92429786	92429856
refGene_gn=CCNC;refGene_tx=XM_006931905.3;refGene_gn=CCNC;refGene_tx=XM_011282450.2
B2	92430249	92430334
refGene_gn=CCNC;refGene_tx=XM_006931905.3;refGene_gn=CCNC;refGene_tx=XM_011282450.2
B2	92431464	92431571
refGene_gn=CCNC;refGene_tx=XM_006931905.3;refGene_gn=CCNC;refGene_tx=XM_011282450.2
B2	92438079	92438111
refGene_gn=CCNC;refGene_tx=XM_006931905.3;refGene_gn=CCNC;refGene_tx=XM_011282450.2
B2	92475235	92475379

refGene_gn=PRDM13;refGene_tx=XM_023254192.1;refGene_gn=PRDM13;refGene_tx=XM_023254193.1
B2 92476902 92477034 refGene_gn=PRDM13;refGene_tx=XM_023254192.1
B2 92477335 92477456
refGene_gn=PRDM13;refGene_tx=XM_023254192.1;refGene_gn=PRDM13;refGene_tx=XM_023254193.1
B2 92480848 92482599
refGene_gn=PRDM13;refGene_tx=XM_023254192.1;refGene_gn=PRDM13;refGene_tx=XM_023254193.1
B2 92762609 92762925 refGene_gn=MCHR2;refGene_tx=XM_023254194.1
B2 92768673 92768793 refGene_gn=MCHR2;refGene_tx=XM_023254194.1
B2 92771189 92771384 refGene_gn=MCHR2;refGene_tx=XM_023254194.1
B2 92778708 92778918 refGene_gn=MCHR2;refGene_tx=XM_023254194.1
B2 92788000 92788182 refGene_gn=MCHR2;refGene_tx=XM_023254194.1
B2 93172493 93173224
refGene_gn=SIM1;refGene_tx=XM_023254195.1;refGene_gn=SIM1;refGene_tx=XM_023254196.1;refGene_gn=SIM1;refGene_tx=XM_023254197.1
B2 93175674 93176077
refGene_gn=SIM1;refGene_tx=XM_023254195.1;refGene_gn=SIM1;refGene_tx=XM_023254196.1;refGene_gn=SIM1;refGene_tx=XM_023254197.1
B2 93198666 93198835
refGene_gn=SIM1;refGene_tx=XM_023254195.1;refGene_gn=SIM1;refGene_tx=XM_023254196.1;refGene_gn=SIM1;refGene_tx=XM_023254197.1
B2 93221630 93221778
refGene_gn=SIM1;refGene_tx=XM_023254195.1;refGene_gn=SIM1;refGene_tx=XM_023254196.1;refGene_gn=SIM1;refGene_tx=XM_023254197.1
B2 93222441 93222548
refGene_gn=SIM1;refGene_tx=XM_023254195.1;refGene_gn=SIM1;refGene_tx=XM_023254196.1;refGene_gn=SIM1;refGene_tx=XM_023254197.1
B2 93222744 93222944
refGene_gn=SIM1;refGene_tx=XM_023254195.1;refGene_gn=SIM1;refGene_tx=XM_023254196.1;refGene_gn=SIM1;refGene_tx=XM_023254197.1
B2 93223606 93223692
refGene_gn=SIM1;refGene_tx=XM_023254195.1;refGene_gn=SIM1;refGene_tx=XM_023254196.1;refGene_gn=SIM1;refGene_tx=XM_023254197.1
B2 93223878 93223987
refGene_gn=SIM1;refGene_tx=XM_023254195.1;refGene_gn=SIM1;refGene_tx=XM_023254196.1;refGene_gn=SIM1;refGene_tx=XM_023254197.1
B2 93224494 93224584
refGene_gn=SIM1;refGene_tx=XM_023254195.1;refGene_gn=SIM1;refGene_tx=XM_023254196.1;refGene_gn=SIM1;refGene_tx=XM_023254197.1
B2 93227998 93228081 refGene_gn=SIM1;refGene_tx=XM_023254195.1
B2 93237500 93237675
refGene_gn=SIM1;refGene_tx=XM_023254195.1;refGene_gn=SIM1;refGene_tx=XM_023254196.1
B2 93274795 93274943
refGene_gn=ASCC3;refGene_tx=XM_011282448.3;refGene_gn=ASCC3;refGene_tx=XM_019831018.2;refGene_gn=ASCC3;refGene_tx=XM_019831019.2
B2 93275024 93275200
refGene_gn=ASCC3;refGene_tx=XM_011282448.3;refGene_gn=ASCC3;refGene_tx

=XM_019831018.2;refGene_gn=ASCC3;refGene_tx=XM_019831019.2
B2	93277398 93277608
refGene_gn=ASCC3;refGene_tx=XM_011282448.3;refGene_gn=ASCC3;refGene_tx
=XM_019831018.2;refGene_gn=ASCC3;refGene_tx=XM_019831019.2
B2	93281601 93281749
refGene_gn=ASCC3;refGene_tx=XM_011282448.3;refGene_gn=ASCC3;refGene_tx
=XM_019831018.2;refGene_gn=ASCC3;refGene_tx=XM_019831019.2
B2	93283034 93283070
refGene_gn=ASCC3;refGene_tx=XM_019831018.2;refGene_gn=ASCC3;refGene_tx
=XM_019831019.2
B2	93283437 93283589
refGene_gn=ASCC3;refGene_tx=XM_011282448.3;refGene_gn=ASCC3;refGene_tx
=XM_019831018.2;refGene_gn=ASCC3;refGene_tx=XM_019831019.2
B2	93305034 93305259
refGene_gn=ASCC3;refGene_tx=XM_011282448.3;refGene_gn=ASCC3;refGene_tx
=XM_019831018.2;refGene_gn=ASCC3;refGene_tx=XM_019831019.2
B2	93345254 93345269 refGene_gn=ASCC3;refGene_tx=XM_023254198.1
B2	93358044 93358179
refGene_gn=ASCC3;refGene_tx=XM_011282448.3;refGene_gn=ASCC3;refGene_tx
=XM_019831018.2;refGene_gn=ASCC3;refGene_tx=XM_019831019.2;refGene_gn=
ASCC3;refGene_tx=XM_023254198.1
B2	93358375 93358487
refGene_gn=ASCC3;refGene_tx=XM_011282448.3;refGene_gn=ASCC3;refGene_tx
=XM_019831018.2;refGene_gn=ASCC3;refGene_tx=XM_019831019.2;refGene_gn=
ASCC3;refGene_tx=XM_023254198.1
B2	93369211 93369337
refGene_gn=ASCC3;refGene_tx=XM_011282448.3;refGene_gn=ASCC3;refGene_tx
=XM_019831018.2;refGene_gn=ASCC3;refGene_tx=XM_019831019.2;refGene_gn=
ASCC3;refGene_tx=XM_023254198.1
B2	93372301 93372434
refGene_gn=ASCC3;refGene_tx=XM_011282448.3;refGene_gn=ASCC3;refGene_tx
=XM_019831018.2;refGene_gn=ASCC3;refGene_tx=XM_019831019.2;refGene_gn=
ASCC3;refGene_tx=XM_023254198.1
B2	93373407 93373528
refGene_gn=ASCC3;refGene_tx=XM_011282448.3;refGene_gn=ASCC3;refGene_tx
=XM_019831018.2;refGene_gn=ASCC3;refGene_tx=XM_019831019.2;refGene_gn=
ASCC3;refGene_tx=XM_023254198.1
B2	93373605 93373743
refGene_gn=ASCC3;refGene_tx=XM_011282448.3;refGene_gn=ASCC3;refGene_tx
=XM_019831018.2;refGene_gn=ASCC3;refGene_tx=XM_019831019.2;refGene_gn=
ASCC3;refGene_tx=XM_023254198.1
B2	93387268 93387411
refGene_gn=ASCC3;refGene_tx=XM_011282448.3;refGene_gn=ASCC3;refGene_tx
=XM_019831018.2;refGene_gn=ASCC3;refGene_tx=XM_019831019.2;refGene_gn=
ASCC3;refGene_tx=XM_023254198.1
B2	93389864 93389985
refGene_gn=ASCC3;refGene_tx=XM_011282448.3;refGene_gn=ASCC3;refGene_tx
=XM_019831018.2;refGene_gn=ASCC3;refGene_tx=XM_019831019.2;refGene_gn=
ASCC3;refGene_tx=XM_023254198.1
B2	93390108 93390254

refGene_gn=ASCC3;refGene_tx=XM_011282448.3;refGene_gn=ASCC3;refGene_tx=XM_019831018.2;refGene_gn=ASCC3;refGene_tx=XM_019831019.2;refGene_gn=ASCC3;refGene_tx=XM_023254198.1
B2 93390362 93390529
refGene_gn=ASCC3;refGene_tx=XM_011282448.3;refGene_gn=ASCC3;refGene_tx=XM_019831018.2;refGene_gn=ASCC3;refGene_tx=XM_019831019.2;refGene_gn=ASCC3;refGene_tx=XM_023254198.1
B2 93391958 93392044
refGene_gn=ASCC3;refGene_tx=XM_011282448.3;refGene_gn=ASCC3;refGene_tx=XM_019831018.2;refGene_gn=ASCC3;refGene_tx=XM_019831019.2;refGene_gn=ASCC3;refGene_tx=XM_023254198.1
B2 93393510 93393731
refGene_gn=ASCC3;refGene_tx=XM_011282448.3;refGene_gn=ASCC3;refGene_tx=XM_019831018.2;refGene_gn=ASCC3;refGene_tx=XM_019831019.2;refGene_gn=ASCC3;refGene_tx=XM_023254198.1
B2 93396497 93396666
refGene_gn=ASCC3;refGene_tx=XM_011282448.3;refGene_gn=ASCC3;refGene_tx=XM_019831018.2;refGene_gn=ASCC3;refGene_tx=XM_019831019.2;refGene_gn=ASCC3;refGene_tx=XM_023254198.1
B2 93398522 93398621
refGene_gn=ASCC3;refGene_tx=XM_011282448.3;refGene_gn=ASCC3;refGene_tx=XM_019831018.2;refGene_gn=ASCC3;refGene_tx=XM_019831019.2;refGene_gn=ASCC3;refGene_tx=XM_023254198.1
B2 93402352 93402507
refGene_gn=ASCC3;refGene_tx=XM_011282448.3;refGene_gn=ASCC3;refGene_tx=XM_019831018.2;refGene_gn=ASCC3;refGene_tx=XM_019831019.2;refGene_gn=ASCC3;refGene_tx=XM_023254198.1
B2 93403762 93403988
refGene_gn=ASCC3;refGene_tx=XM_011282448.3;refGene_gn=ASCC3;refGene_tx=XM_019831018.2;refGene_gn=ASCC3;refGene_tx=XM_019831019.2;refGene_gn=ASCC3;refGene_tx=XM_023254198.1
B2 93407787 93407964
refGene_gn=ASCC3;refGene_tx=XM_011282448.3;refGene_gn=ASCC3;refGene_tx=XM_019831018.2;refGene_gn=ASCC3;refGene_tx=XM_019831019.2;refGene_gn=ASCC3;refGene_tx=XM_023254198.1
B2 93408825 93408912
refGene_gn=ASCC3;refGene_tx=XM_011282448.3;refGene_gn=ASCC3;refGene_tx=XM_019831018.2;refGene_gn=ASCC3;refGene_tx=XM_019831019.2;refGene_gn=ASCC3;refGene_tx=XM_023254198.1
B2 93410858 93411023
refGene_gn=ASCC3;refGene_tx=XM_011282448.3;refGene_gn=ASCC3;refGene_tx=XM_019831018.2;refGene_gn=ASCC3;refGene_tx=XM_019831019.2;refGene_gn=ASCC3;refGene_tx=XM_023254198.1
B2 93413941 93414061
refGene_gn=ASCC3;refGene_tx=XM_011282448.3;refGene_gn=ASCC3;refGene_tx=XM_019831018.2;refGene_gn=ASCC3;refGene_tx=XM_019831019.2;refGene_gn=ASCC3;refGene_tx=XM_023254198.1
B2 93420519 93420744
refGene_gn=ASCC3;refGene_tx=XM_011282448.3;refGene_gn=ASCC3;refGene_tx=XM_019831018.2;refGene_gn=ASCC3;refGene_tx=XM_019831019.2;refGene_gn=

ASCC3;refGene_tx=XM_023254198.1
B2 93421044 93421236
refGene_gn=ASCC3;refGene_tx=XM_011282448.3;refGene_gn=ASCC3;refGene_tx=XM_019831018.2;refGene_gn=ASCC3;refGene_tx=XM_019831019.2;refGene_gn=ASCC3;refGene_tx=XM_023254198.1
B2 93443516 93443651
refGene_gn=ASCC3;refGene_tx=XM_011282448.3;refGene_gn=ASCC3;refGene_tx=XM_019831018.2;refGene_gn=ASCC3;refGene_tx=XM_019831019.2;refGene_gn=ASCC3;refGene_tx=XM_023254198.1
B2 93481643 93481715
refGene_gn=ASCC3;refGene_tx=XM_011282448.3;refGene_gn=ASCC3;refGene_tx=XM_019831018.2;refGene_gn=ASCC3;refGene_tx=XM_019831019.2;refGene_gn=ASCC3;refGene_tx=XM_023254198.1
B2 93484498 93484675
refGene_gn=ASCC3;refGene_tx=XM_011282448.3;refGene_gn=ASCC3;refGene_tx=XM_019831018.2;refGene_gn=ASCC3;refGene_tx=XM_019831019.2;refGene_gn=ASCC3;refGene_tx=XM_023254198.1
B2 93495445 93495610
refGene_gn=ASCC3;refGene_tx=XM_011282448.3;refGene_gn=ASCC3;refGene_tx=XM_019831018.2;refGene_gn=ASCC3;refGene_tx=XM_019831019.2;refGene_gn=ASCC3;refGene_tx=XM_023254198.1
B2 93535581 93535722
refGene_gn=ASCC3;refGene_tx=XM_011282448.3;refGene_gn=ASCC3;refGene_tx=XM_019831018.2;refGene_gn=ASCC3;refGene_tx=XM_019831019.2;refGene_gn=ASCC3;refGene_tx=XM_023254198.1
B2 93536175 93536376
refGene_gn=ASCC3;refGene_tx=XM_011282448.3;refGene_gn=ASCC3;refGene_tx=XM_019831018.2;refGene_gn=ASCC3;refGene_tx=XM_019831019.2;refGene_gn=ASCC3;refGene_tx=XM_023254198.1
B2 93563035 93563161
refGene_gn=ASCC3;refGene_tx=XM_011282448.3;refGene_gn=ASCC3;refGene_tx=XM_019831018.2;refGene_gn=ASCC3;refGene_tx=XM_019831019.2;refGene_gn=ASCC3;refGene_tx=XM_023254198.1
B2 93563975 93564117
refGene_gn=ASCC3;refGene_tx=XM_011282448.3;refGene_gn=ASCC3;refGene_tx=XM_019831018.2;refGene_gn=ASCC3;refGene_tx=XM_019831019.2;refGene_gn=ASCC3;refGene_tx=XM_023254198.1
B2 93564838 93565043
refGene_gn=ASCC3;refGene_tx=XM_011282448.3;refGene_gn=ASCC3;refGene_tx=XM_019831018.2;refGene_gn=ASCC3;refGene_tx=XM_019831019.2;refGene_gn=ASCC3;refGene_tx=XM_023254198.1
B2 93569732 93569853
refGene_gn=ASCC3;refGene_tx=XM_011282448.3;refGene_gn=ASCC3;refGene_tx=XM_019831018.2;refGene_gn=ASCC3;refGene_tx=XM_019831019.2;refGene_gn=ASCC3;refGene_tx=XM_023254198.1
B2 93596121 93596681
refGene_gn=ASCC3;refGene_tx=XM_011282448.3;refGene_gn=ASCC3;refGene_tx=XM_019831018.2;refGene_gn=ASCC3;refGene_tx=XM_019831019.2;refGene_gn=ASCC3;refGene_tx=XM_023254198.1
B2 93611181 93611332

refGene_gn=ASCC3;refGene_tx=XM_011282448.3;refGene_gn=ASCC3;refGene_tx=XM_019831018.2;refGene_gn=ASCC3;refGene_tx=XM_019831019.2;refGene_gn=ASCC3;refGene_tx=XM_023254198.1
B2 93614018 93614108
refGene_gn=ASCC3;refGene_tx=XM_011282448.3;refGene_gn=ASCC3;refGene_tx=XM_019831018.2;refGene_gn=ASCC3;refGene_tx=XM_019831019.2;refGene_gn=ASCC3;refGene_tx=XM_023254198.1
B2 94086587 94086702
refGene_gn=GRIK2;refGene_tx=XM_011282447.3;refGene_gn=GRIK2;refGene_tx=XM_019831020.2;refGene_gn=GRIK2;refGene_tx=XM_019831021.2;refGene_gn=GRIK2;refGene_tx=XM_019831022.1;refGene_gn=GRIK2;refGene_tx=XM_019831023.2;refGene_gn=GRIK2;refGene_tx=XM_019831024.2;refGene_gn=GRIK2;refGene_tx=XM_019831025.1
B2 94328923 94329091
refGene_gn=GRIK2;refGene_tx=XM_011282447.3;refGene_gn=GRIK2;refGene_tx=XM_019831020.2;refGene_gn=GRIK2;refGene_tx=XM_019831021.2;refGene_gn=GRIK2;refGene_tx=XM_019831022.1;refGene_gn=GRIK2;refGene_tx=XM_019831023.2;refGene_gn=GRIK2;refGene_tx=XM_019831024.2;refGene_gn=GRIK2;refGene_tx=XM_019831025.1
B2 94333135 94333393
refGene_gn=GRIK2;refGene_tx=XM_011282447.3;refGene_gn=GRIK2;refGene_tx=XM_019831020.2;refGene_gn=GRIK2;refGene_tx=XM_019831021.2;refGene_gn=GRIK2;refGene_tx=XM_019831022.1;refGene_gn=GRIK2;refGene_tx=XM_019831023.2;refGene_gn=GRIK2;refGene_tx=XM_019831024.2;refGene_gn=GRIK2;refGene_tx=XM_019831025.1
B2 94367510 94367692
refGene_gn=GRIK2;refGene_tx=XM_011282447.3;refGene_gn=GRIK2;refGene_tx=XM_019831020.2;refGene_gn=GRIK2;refGene_tx=XM_019831021.2;refGene_gn=GRIK2;refGene_tx=XM_019831022.1;refGene_gn=GRIK2;refGene_tx=XM_019831023.2;refGene_gn=GRIK2;refGene_tx=XM_019831024.2;refGene_gn=GRIK2;refGene_tx=XM_019831025.1
B2 94374025 94374079
refGene_gn=GRIK2;refGene_tx=XM_011282447.3;refGene_gn=GRIK2;refGene_tx=XM_019831020.2;refGene_gn=GRIK2;refGene_tx=XM_019831021.2;refGene_gn=GRIK2;refGene_tx=XM_019831022.1;refGene_gn=GRIK2;refGene_tx=XM_019831023.2;refGene_gn=GRIK2;refGene_tx=XM_019831024.2;refGene_gn=GRIK2;refGene_tx=XM_019831025.1
B2 94377353 94377527
refGene_gn=GRIK2;refGene_tx=XM_011282447.3;refGene_gn=GRIK2;refGene_tx=XM_019831020.2;refGene_gn=GRIK2;refGene_tx=XM_019831021.2;refGene_gn=GRIK2;refGene_tx=XM_019831022.1;refGene_gn=GRIK2;refGene_tx=XM_019831023.2;refGene_gn=GRIK2;refGene_tx=XM_019831024.2;refGene_gn=GRIK2;refGene_tx=XM_019831025.1
B2 94475513 94475657
refGene_gn=GRIK2;refGene_tx=XM_011282447.3;refGene_gn=GRIK2;refGene_tx=XM_019831020.2;refGene_gn=GRIK2;refGene_tx=XM_019831021.2;refGene_gn=GRIK2;refGene_tx=XM_019831022.1;refGene_gn=GRIK2;refGene_tx=XM_019831023.2;refGene_gn=GRIK2;refGene_tx=XM_019831024.2;refGene_gn=GRIK2;refGene_tx=XM_019831025.1
B2 94478074 94478182

refGene_gn=GRIK2;refGene_tx=XM_011282447.3;refGene_gn=GRIK2;refGene_tx=XM_019831020.2;refGene_gn=GRIK2;refGene_tx=XM_019831021.2;refGene_gn=GRIK2;refGene_tx=XM_019831022.1;refGene_gn=GRIK2;refGene_tx=XM_019831023.2;refGene_gn=GRIK2;refGene_tx=XM_019831024.2;refGene_gn=GRIK2;refGene_tx=XM_019831025.1
B2 94492910 94493024
refGene_gn=GRIK2;refGene_tx=XM_011282447.3;refGene_gn=GRIK2;refGene_tx=XM_019831020.2;refGene_gn=GRIK2;refGene_tx=XM_019831021.2;refGene_gn=GRIK2;refGene_tx=XM_019831022.1;refGene_gn=GRIK2;refGene_tx=XM_019831023.2;refGene_gn=GRIK2;refGene_tx=XM_019831024.2;refGene_gn=GRIK2;refGene_tx=XM_019831025.1
B2 94525904 94526111
refGene_gn=GRIK2;refGene_tx=XM_011282447.3;refGene_gn=GRIK2;refGene_tx=XM_019831020.2;refGene_gn=GRIK2;refGene_tx=XM_019831021.2;refGene_gn=GRIK2;refGene_tx=XM_019831022.1;refGene_gn=GRIK2;refGene_tx=XM_019831023.2;refGene_gn=GRIK2;refGene_tx=XM_019831024.2;refGene_gn=GRIK2;refGene_tx=XM_019831025.1
B2 94571440 94571668
refGene_gn=GRIK2;refGene_tx=XM_011282447.3;refGene_gn=GRIK2;refGene_tx=XM_019831020.2;refGene_gn=GRIK2;refGene_tx=XM_019831021.2;refGene_gn=GRIK2;refGene_tx=XM_019831022.1;refGene_gn=GRIK2;refGene_tx=XM_019831023.2;refGene_gn=GRIK2;refGene_tx=XM_019831024.2;refGene_gn=GRIK2;refGene_tx=XM_019831025.1
B2 94606867 94606986
refGene_gn=GRIK2;refGene_tx=XM_011282447.3;refGene_gn=GRIK2;refGene_tx=XM_019831020.2;refGene_gn=GRIK2;refGene_tx=XM_019831021.2;refGene_gn=GRIK2;refGene_tx=XM_019831022.1;refGene_gn=GRIK2;refGene_tx=XM_019831023.2;refGene_gn=GRIK2;refGene_tx=XM_019831024.2
B2 94610642 94610860
refGene_gn=GRIK2;refGene_tx=XM_011282447.3;refGene_gn=GRIK2;refGene_tx=XM_019831020.2;refGene_gn=GRIK2;refGene_tx=XM_019831021.2;refGene_gn=GRIK2;refGene_tx=XM_019831022.1;refGene_gn=GRIK2;refGene_tx=XM_019831023.2;refGene_gn=GRIK2;refGene_tx=XM_019831024.2
B2 94703281 94703507
refGene_gn=GRIK2;refGene_tx=XM_011282447.3;refGene_gn=GRIK2;refGene_tx=XM_019831020.2;refGene_gn=GRIK2;refGene_tx=XM_019831021.2;refGene_gn=GRIK2;refGene_tx=XM_019831022.1;refGene_gn=GRIK2;refGene_tx=XM_019831023.2;refGene_gn=GRIK2;refGene_tx=XM_019831024.2
B2 94724869 94725120
refGene_gn=GRIK2;refGene_tx=XM_011282447.3;refGene_gn=GRIK2;refGene_tx=XM_019831020.2;refGene_gn=GRIK2;refGene_tx=XM_019831021.2;refGene_gn=GRIK2;refGene_tx=XM_019831022.1;refGene_gn=GRIK2;refGene_tx=XM_019831023.2;refGene_gn=GRIK2;refGene_tx=XM_019831024.2
B2 94734585 94734633
refGene_gn=GRIK2;refGene_tx=XM_019831023.2;refGene_gn=GRIK2;refGene_tx=XM_019831024.2
B2 94736234 94736354
refGene_gn=GRIK2;refGene_tx=XM_019831021.2;refGene_gn=GRIK2;refGene_tx=XM_019831022.1
B2 94738724 94738889

refGene_gn=GRIK2;refGene_tx=XM_011282447.3;refGene_gn=GRIK2;refGene_tx=XM_019831020.2
B2 97160505 97160608
refGene_gn=HACE1;refGene_tx=XM_019831029.1;refGene_gn=HACE1;refGene_tx=XM_023254199.1;refGene_gn=HACE1;refGene_tx=XM_023254200.1;refGene_gn=HACE1;refGene_tx=XM_023254202.1
B2 97161158 97161272
refGene_gn=HACE1;refGene_tx=XM_019831029.1;refGene_gn=HACE1;refGene_tx=XM_023254199.1;refGene_gn=HACE1;refGene_tx=XM_023254200.1;refGene_gn=HACE1;refGene_tx=XM_023254202.1
B2 97173355 97173426
refGene_gn=HACE1;refGene_tx=XM_019831029.1;refGene_gn=HACE1;refGene_tx=XM_023254199.1;refGene_gn=HACE1;refGene_tx=XM_023254200.1;refGene_gn=HACE1;refGene_tx=XM_023254201.1;refGene_gn=HACE1;refGene_tx=XM_023254202.1
B2 97173721 97173820
refGene_gn=HACE1;refGene_tx=XM_019831029.1;refGene_gn=HACE1;refGene_tx=XM_023254199.1;refGene_gn=HACE1;refGene_tx=XM_023254200.1;refGene_gn=HACE1;refGene_tx=XM_023254201.1;refGene_gn=HACE1;refGene_tx=XM_023254202.1
B2 97182016 97182148
refGene_gn=HACE1;refGene_tx=XM_019831029.1;refGene_gn=HACE1;refGene_tx=XM_023254199.1;refGene_gn=HACE1;refGene_tx=XM_023254200.1;refGene_gn=HACE1;refGene_tx=XM_023254201.1;refGene_gn=HACE1;refGene_tx=XM_023254202.1
B2 97190797 97190994
refGene_gn=HACE1;refGene_tx=XM_019831029.1;refGene_gn=HACE1;refGene_tx=XM_023254199.1;refGene_gn=HACE1;refGene_tx=XM_023254200.1;refGene_gn=HACE1;refGene_tx=XM_023254201.1;refGene_gn=HACE1;refGene_tx=XM_023254202.1
B2 97191522 97191672
refGene_gn=HACE1;refGene_tx=XM_019831029.1;refGene_gn=HACE1;refGene_tx=XM_023254199.1;refGene_gn=HACE1;refGene_tx=XM_023254200.1;refGene_gn=HACE1;refGene_tx=XM_023254201.1;refGene_gn=HACE1;refGene_tx=XM_023254202.1
B2 97195602 97195690
refGene_gn=HACE1;refGene_tx=XM_019831029.1;refGene_gn=HACE1;refGene_tx=XM_023254199.1;refGene_gn=HACE1;refGene_tx=XM_023254200.1;refGene_gn=HACE1;refGene_tx=XM_023254201.1;refGene_gn=HACE1;refGene_tx=XM_023254202.1
B2 97195870 97195968
refGene_gn=HACE1;refGene_tx=XM_019831029.1;refGene_gn=HACE1;refGene_tx=XM_023254199.1;refGene_gn=HACE1;refGene_tx=XM_023254200.1;refGene_gn=HACE1;refGene_tx=XM_023254201.1;refGene_gn=HACE1;refGene_tx=XM_023254202.1
B2 97196061 97196173
refGene_gn=HACE1;refGene_tx=XM_019831029.1;refGene_gn=HACE1;refGene_tx=XM_023254199.1;refGene_gn=HACE1;refGene_tx=XM_023254200.1;refGene_gn=HACE1;refGene_tx=XM_023254201.1;refGene_gn=HACE1;refGene_tx=XM_023254202.1

B2 97216638 97216726
refGene_gn=HACE1;refGene_tx=XM_019831029.1;refGene_gn=HACE1;refGene_tx=XM_023254199.1;refGene_gn=HACE1;refGene_tx=XM_023254200.1;refGene_gn=HACE1;refGene_tx=XM_023254201.1;refGene_gn=HACE1;refGene_tx=XM_023254202.1
B2 97216961 97217030
refGene_gn=HACE1;refGene_tx=XM_019831029.1;refGene_gn=HACE1;refGene_tx=XM_023254199.1;refGene_gn=HACE1;refGene_tx=XM_023254201.1;refGene_gn=HACE1;refGene_tx=XM_023254202.1
B2 97217517 97217852
refGene_gn=HACE1;refGene_tx=XM_019831029.1;refGene_gn=HACE1;refGene_tx=XM_023254199.1;refGene_gn=HACE1;refGene_tx=XM_023254200.1;refGene_gn=HACE1;refGene_tx=XM_023254201.1;refGene_gn=HACE1;refGene_tx=XM_023254202.1
B2 97225999 97226150
refGene_gn=HACE1;refGene_tx=XM_019831029.1;refGene_gn=HACE1;refGene_tx=XM_023254199.1;refGene_gn=HACE1;refGene_tx=XM_023254200.1;refGene_gn=HACE1;refGene_tx=XM_023254201.1;refGene_gn=HACE1;refGene_tx=XM_023254202.1
B2 97232607 97232714
refGene_gn=HACE1;refGene_tx=XM_019831029.1;refGene_gn=HACE1;refGene_tx=XM_023254199.1;refGene_gn=HACE1;refGene_tx=XM_023254200.1;refGene_gn=HACE1;refGene_tx=XM_023254201.1;refGene_gn=HACE1;refGene_tx=XM_023254202.1
B2 97233331 97233433
refGene_gn=HACE1;refGene_tx=XM_019831029.1;refGene_gn=HACE1;refGene_tx=XM_023254199.1;refGene_gn=HACE1;refGene_tx=XM_023254200.1;refGene_gn=HACE1;refGene_tx=XM_023254201.1;refGene_gn=HACE1;refGene_tx=XM_023254202.1
B2 97234462 97234559
refGene_gn=HACE1;refGene_tx=XM_019831029.1;refGene_gn=HACE1;refGene_tx=XM_023254199.1;refGene_gn=HACE1;refGene_tx=XM_023254200.1;refGene_gn=HACE1;refGene_tx=XM_023254201.1;refGene_gn=HACE1;refGene_tx=XM_023254202.1
B2 97242863 97242946
refGene_gn=HACE1;refGene_tx=XM_023254199.1;refGene_gn=HACE1;refGene_tx=XM_023254200.1;refGene_gn=HACE1;refGene_tx=XM_023254201.1;refGene_gn=HACE1;refGene_tx=XM_023254202.1
B2 97255464 97255596
refGene_gn=HACE1;refGene_tx=XM_019831029.1;refGene_gn=HACE1;refGene_tx=XM_023254199.1;refGene_gn=HACE1;refGene_tx=XM_023254200.1;refGene_gn=HACE1;refGene_tx=XM_023254201.1
B2 97263699 97263775
refGene_gn=HACE1;refGene_tx=XM_023254199.1;refGene_gn=HACE1;refGene_tx=XM_023254200.1;refGene_gn=HACE1;refGene_tx=XM_023254201.1;refGene_gn=HACE1;refGene_tx=XM_023254202.1
B2 97270839 97270944
refGene_gn=HACE1;refGene_tx=XM_023254199.1;refGene_gn=HACE1;refGene_tx=XM_023254200.1;refGene_gn=HACE1;refGene_tx=XM_023254201.1;refGene_gn=HACE1;refGene_tx=XM_023254202.1

B2 97272201 97272291
refGene_gn=HACE1;refGene_tx=XM_023254199.1;refGene_gn=HACE1;refGene_tx=XM_023254200.1;refGene_gn=HACE1;refGene_tx=XM_023254201.1;refGene_gn=HACE1;refGene_tx=XM_023254202.1
B2 97273509 97273564
refGene_gn=HACE1;refGene_tx=XM_023254199.1;refGene_gn=HACE1;refGene_tx=XM_023254200.1;refGene_gn=HACE1;refGene_tx=XM_023254201.1;refGene_gn=HACE1;refGene_tx=XM_023254202.1
B2 97281284 97281360
refGene_gn=HACE1;refGene_tx=XM_023254199.1;refGene_gn=HACE1;refGene_tx=XM_023254200.1;refGene_gn=HACE1;refGene_tx=XM_023254201.1;refGene_gn=HACE1;refGene_tx=XM_023254202.1
B2 97289524 97289632
refGene_gn=LOC101095660;refGene_tx=XM_011282536.2
B2 97289633 97289792
refGene_gn=LOC101095660;refGene_tx=XM_011282536.2
B2 97342546 97342580 refGene_gn=LIN28B;refGene_tx=XM_023254203.1
B2 97349292 97349302 refGene_gn=LIN28B;refGene_tx=XM_023254204.1
B2 97350129 97350317
refGene_gn=LIN28B;refGene_tx=XM_023254203.1;refGene_gn=LIN28B;refGene_tx=XM_023254204.1
B2 97403779 97403964
refGene_gn=LIN28B;refGene_tx=XM_023254203.1;refGene_gn=LIN28B;refGene_tx=XM_023254204.1
B2 97460957 97461327
refGene_gn=LIN28B;refGene_tx=XM_023254203.1;refGene_gn=LIN28B;refGene_tx=XM_023254204.1
B2 97483666 97483791
refGene_gn=BVES;refGene_tx=XM_003986413.5;refGene_gn=BVES;refGene_tx=XM_006931912.4;refGene_gn=BVES;refGene_tx=XM_019831030.2
B2 97498787 97498929
refGene_gn=BVES;refGene_tx=XM_003986413.5;refGene_gn=BVES;refGene_tx=XM_006931912.4;refGene_gn=BVES;refGene_tx=XM_019831030.2
B2 97499832 97500000
refGene_gn=BVES;refGene_tx=XM_003986413.5;refGene_gn=BVES;refGene_tx=XM_006931912.4;refGene_gn=BVES;refGene_tx=XM_019831030.2
B2 97505776 97505885
refGene_gn=BVES;refGene_tx=XM_003986413.5;refGene_gn=BVES;refGene_tx=XM_006931912.4;refGene_gn=BVES;refGene_tx=XM_019831030.2
B2 97506631 97506819
refGene_gn=BVES;refGene_tx=XM_003986413.5;refGene_gn=BVES;refGene_tx=XM_006931912.4;refGene_gn=BVES;refGene_tx=XM_019831030.2
B2 97509890 97510015
refGene_gn=BVES;refGene_tx=XM_003986413.5;refGene_gn=BVES;refGene_tx=XM_006931912.4;refGene_gn=BVES;refGene_tx=XM_019831030.2
B2 97512964 97513190
refGene_gn=BVES;refGene_tx=XM_003986413.5;refGene_gn=BVES;refGene_tx=XM_006931912.4;refGene_gn=BVES;refGene_tx=XM_019831030.2
B2 97534330 97534612 refGene_gn=POPDC3;refGene_tx=XM_023254205.1
B2 97535622 97535731 refGene_gn=POPDC3;refGene_tx=XM_023254205.1

B2 97536658 97537143 refGene_gn=POPDC3;refGene_tx=XM_023254205.1
B2 97646934 97647229 refGene_gn=PREP;refGene_tx=XM_011282523.2
B2 97650278 97650435 refGene_gn=PREP;refGene_tx=XM_011282523.2
B2 97654003 97654135 refGene_gn=PREP;refGene_tx=XM_011282523.2
B2 97656918 97657013 refGene_gn=PREP;refGene_tx=XM_011282523.2
B2 97659745 97659882 refGene_gn=PREP;refGene_tx=XM_011282523.2
B2 97689580 97689684 refGene_gn=PREP;refGene_tx=XM_011282523.2
B2 97695494 97695692 refGene_gn=PREP;refGene_tx=XM_011282523.2
B2 97700169 97700361 refGene_gn=PREP;refGene_tx=XM_011282523.2
B2 97714958 97715064 refGene_gn=PREP;refGene_tx=XM_011282523.2
B2 97732900 97733022 refGene_gn=PREP;refGene_tx=XM_011282523.2
B2 97737785 97737995 refGene_gn=PREP;refGene_tx=XM_011282523.2
B2 97740502 97740633 refGene_gn=PREP;refGene_tx=XM_011282523.2
B2 97741823 97741957 refGene_gn=PREP;refGene_tx=XM_011282523.2
B2 97760569 97760644 refGene_gn=PREP;refGene_tx=XM_011282523.2
B2 97766509 97766554 refGene_gn=PREP;refGene_tx=XM_011282523.2
B2 98422678 98422720 refGene_gn=PRDM1;refGene_tx=XM_003986416.5
B2 98424350 98424590
refGene_gn=PRDM1;refGene_tx=XM_003986416.5;refGene_gn=PRDM1;refGene_tx
=XM_019831032.2
B2 98431065 98431185
refGene_gn=PRDM1;refGene_tx=XM_003986416.5;refGene_gn=PRDM1;refGene_tx
=XM_019831032.2
B2 98435215 98435468
refGene_gn=PRDM1;refGene_tx=XM_003986416.5;refGene_gn=PRDM1;refGene_tx
=XM_019831032.2
B2 98440868 98441968
refGene_gn=PRDM1;refGene_tx=XM_003986416.5;refGene_gn=PRDM1;refGene_tx
=XM_019831032.2
B2 98442420 98442549
refGene_gn=PRDM1;refGene_tx=XM_003986416.5;refGene_gn=PRDM1;refGene_tx
=XM_019831032.2
B2 98442924 98443497
refGene_gn=PRDM1;refGene_tx=XM_003986416.5;refGene_gn=PRDM1;refGene_tx
=XM_019831032.2
B2 98516672 98516809
refGene_gn=ATG5;refGene_tx=XM_003986419.5;refGene_gn=ATG5;refGene_tx=X
M_006931914.3;refGene_gn=ATG5;refGene_tx=XM_019831033.2
B2 98536431 98536549
refGene_gn=ATG5;refGene_tx=XM_003986419.5;refGene_gn=ATG5;refGene_tx=X
M_006931914.3;refGene_gn=ATG5;refGene_tx=XM_019831033.2
B2 98571065 98571160
refGene_gn=ATG5;refGene_tx=XM_003986419.5;refGene_gn=ATG5;refGene_tx=X
M_006931914.3;refGene_gn=ATG5;refGene_tx=XM_019831033.2
B2 98594149 98594312
refGene_gn=ATG5;refGene_tx=XM_003986419.5;refGene_gn=ATG5;refGene_tx=X
M_006931914.3;refGene_gn=ATG5;refGene_tx=XM_019831033.2
B2 98616180 98616259
refGene_gn=ATG5;refGene_tx=XM_003986419.5;refGene_gn=ATG5;refGene_tx=X
M_006931914.3;refGene_gn=ATG5;refGene_tx=XM_019831033.2

B2 98625065 98625193
refGene_gn=ATG5;refGene_tx=XM_003986419.5;refGene_gn=ATG5;refGene_tx=XM_019831033.2
B2 98637192 98637300
refGene_gn=ATG5;refGene_tx=XM_003986419.5;refGene_gn=ATG5;refGene_tx=XM_006931914.3
B2 98674005 98674178 refGene_gn=CRYBG1;refGene_tx=XM_023254207.1
B2 98743624 98743763 refGene_gn=CRYBG1;refGene_tx=XM_023254207.1
B2 98754675 98754828 refGene_gn=CRYBG1;refGene_tx=XM_023254208.1
B2 98790151 98790154 refGene_gn=CRYBG1;refGene_tx=XM_023254209.1
B2 98806824 98808440
refGene_gn=CRYBG1;refGene_tx=XM_023254207.1;refGene_gn=CRYBG1;refGene_tx=XM_023254208.1;refGene_gn=CRYBG1;refGene_tx=XM_023254209.1
B2 98813528 98815869
refGene_gn=CRYBG1;refGene_tx=XM_023254207.1;refGene_gn=CRYBG1;refGene_tx=XM_023254208.1;refGene_gn=CRYBG1;refGene_tx=XM_023254209.1
B2 98818922 98818970
refGene_gn=CRYBG1;refGene_tx=XM_023254207.1;refGene_gn=CRYBG1;refGene_tx=XM_023254208.1;refGene_gn=CRYBG1;refGene_tx=XM_023254209.1
B2 98819055 98819174
refGene_gn=CRYBG1;refGene_tx=XM_023254207.1;refGene_gn=CRYBG1;refGene_tx=XM_023254208.1;refGene_gn=CRYBG1;refGene_tx=XM_023254209.1
B2 98820857 98821023
refGene_gn=CRYBG1;refGene_tx=XM_023254207.1;refGene_gn=CRYBG1;refGene_tx=XM_023254208.1;refGene_gn=CRYBG1;refGene_tx=XM_023254209.1
B2 98824038 98824178
refGene_gn=CRYBG1;refGene_tx=XM_023254207.1;refGene_gn=CRYBG1;refGene_tx=XM_023254208.1;refGene_gn=CRYBG1;refGene_tx=XM_023254209.1
B2 98831498 98831625
refGene_gn=CRYBG1;refGene_tx=XM_023254207.1;refGene_gn=CRYBG1;refGene_tx=XM_023254208.1;refGene_gn=CRYBG1;refGene_tx=XM_023254209.1
B2 98835583 98835619
refGene_gn=CRYBG1;refGene_tx=XM_023254207.1;refGene_gn=CRYBG1;refGene_tx=XM_023254208.1;refGene_gn=CRYBG1;refGene_tx=XM_023254209.1
B2 98836616 98836771
refGene_gn=CRYBG1;refGene_tx=XM_023254207.1;refGene_gn=CRYBG1;refGene_tx=XM_023254208.1;refGene_gn=CRYBG1;refGene_tx=XM_023254209.1
B2 98837449 98837576
refGene_gn=CRYBG1;refGene_tx=XM_023254207.1;refGene_gn=CRYBG1;refGene_tx=XM_023254208.1;refGene_gn=CRYBG1;refGene_tx=XM_023254209.1
B2 98837666 98837812
refGene_gn=CRYBG1;refGene_tx=XM_023254207.1;refGene_gn=CRYBG1;refGene_tx=XM_023254208.1;refGene_gn=CRYBG1;refGene_tx=XM_023254209.1
B2 98844537 98844664
refGene_gn=CRYBG1;refGene_tx=XM_023254207.1;refGene_gn=CRYBG1;refGene_tx=XM_023254208.1;refGene_gn=CRYBG1;refGene_tx=XM_023254209.1
B2 98844869 98844902
refGene_gn=CRYBG1;refGene_tx=XM_023254207.1;refGene_gn=CRYBG1;refGene_tx=XM_023254208.1;refGene_gn=CRYBG1;refGene_tx=XM_023254209.1
B2 98845930 98846043

refGene_gn=CRYBG1;refGene_tx=XM_023254207.1;refGene_gn=CRYBG1;refGene_tx=XM_023254208.1;refGene_gn=CRYBG1;refGene_tx=XM_023254209.1
B2 98847666 98847796
refGene_gn=CRYBG1;refGene_tx=XM_023254207.1;refGene_gn=CRYBG1;refGene_tx=XM_023254208.1;refGene_gn=CRYBG1;refGene_tx=XM_023254209.1
B2 98850060 98850200
refGene_gn=CRYBG1;refGene_tx=XM_023254207.1;refGene_gn=CRYBG1;refGene_tx=XM_023254208.1;refGene_gn=CRYBG1;refGene_tx=XM_023254209.1
B2 98852423 98852547
refGene_gn=CRYBG1;refGene_tx=XM_023254207.1;refGene_gn=CRYBG1;refGene_tx=XM_023254208.1;refGene_gn=CRYBG1;refGene_tx=XM_023254209.1
B2 98852980 98853139
refGene_gn=CRYBG1;refGene_tx=XM_023254207.1;refGene_gn=CRYBG1;refGene_tx=XM_023254208.1;refGene_gn=CRYBG1;refGene_tx=XM_023254209.1
B2 98854640 98854803
refGene_gn=CRYBG1;refGene_tx=XM_023254207.1;refGene_gn=CRYBG1;refGene_tx=XM_023254208.1;refGene_gn=CRYBG1;refGene_tx=XM_023254209.1
B2 98857240 98857329
refGene_gn=CRYBG1;refGene_tx=XM_023254207.1;refGene_gn=CRYBG1;refGene_tx=XM_023254208.1;refGene_gn=CRYBG1;refGene_tx=XM_023254209.1
B2 98860319 98860427 refGene_gn=RTN4IP1;refGene_tx=XM_011282520.3
B2 98869033 98869126 refGene_gn=RTN4IP1;refGene_tx=XM_011282520.3
B2 98873234 98873418 refGene_gn=RTN4IP1;refGene_tx=XM_011282520.3
B2 98877044 98877181 refGene_gn=RTN4IP1;refGene_tx=XM_011282520.3
B2 98885583 98885632 refGene_gn=RTN4IP1;refGene_tx=XM_011282520.3
B2 98896558 98896683 refGene_gn=RTN4IP1;refGene_tx=XM_011282520.3
B2 98897584 98897653 refGene_gn=RTN4IP1;refGene_tx=XM_011282520.3
B2 98898910 98899062 refGene_gn=RTN4IP1;refGene_tx=XM_011282520.3
B2 98909079 98909356 refGene_gn=RTN4IP1;refGene_tx=XM_011282520.3
B2 98909983 98910007
refGene_gn=QRSL1;refGene_tx=XM_003986421.5;refGene_gn=QRSL1;refGene_tx=XM_006931915.4
B2 98918740 98918900
refGene_gn=QRSL1;refGene_tx=XM_003986421.5;refGene_gn=QRSL1;refGene_tx=XM_006931915.4
B2 98919221 98919320
refGene_gn=QRSL1;refGene_tx=XM_003986421.5;refGene_gn=QRSL1;refGene_tx=XM_006931915.4
B2 98920339 98920436
refGene_gn=QRSL1;refGene_tx=XM_003986421.5;refGene_gn=QRSL1;refGene_tx=XM_006931915.4
B2 98923166 98923343 refGene_gn=QRSL1;refGene_tx=XM_003986421.5
B2 98927017 98927193
refGene_gn=QRSL1;refGene_tx=XM_003986421.5;refGene_gn=QRSL1;refGene_tx=XM_006931915.4
B2 98927283 98927396
refGene_gn=QRSL1;refGene_tx=XM_003986421.5;refGene_gn=QRSL1;refGene_tx=XM_006931915.4
B2 98929652 98929845
refGene_gn=QRSL1;refGene_tx=XM_003986421.5;refGene_gn=QRSL1;refGene_tx

=XM_006931915.4
B2 98930374 98930492
refGene_gn=QRSL1;refGene_tx=XM_003986421.5;refGene_gn=QRSL1;refGene_tx
=XM_006931915.4
B2 98937684 98937890
refGene_gn=QRSL1;refGene_tx=XM_003986421.5;refGene_gn=QRSL1;refGene_tx
=XM_006931915.4
B2 98940688 98940909
refGene_gn=QRSL1;refGene_tx=XM_003986421.5;refGene_gn=QRSL1;refGene_tx
=XM_006931915.4
B2 99234266 99234733
refGene_gn=CB2H6orf203;refGene_tx=XM_019831037.2;refGene_gn=CB2H6orf20
3;refGene_tx=XM_023254212.1;refGene_gn=CB2H6orf203;refGene_tx=XM_02325
4213.1;refGene_gn=CB2H6orf203;refGene_tx=XM_023254214.1;refGene_gn=CB2
H6orf203;refGene_tx=XM_023254215.1;refGene_gn=CB2H6orf203;refGene_tx=X
M_023254216.1;refGene_gn=CB2H6orf203;refGene_tx=XM_023254217.1
B2 99236339 99236412
refGene_gn=CB2H6orf203;refGene_tx=XM_019831037.2;refGene_gn=CB2H6orf20
3;refGene_tx=XM_023254212.1;refGene_gn=CB2H6orf203;refGene_tx=XM_02325
4213.1;refGene_gn=CB2H6orf203;refGene_tx=XM_023254214.1;refGene_gn=CB2
H6orf203;refGene_tx=XM_023254215.1;refGene_gn=CB2H6orf203;refGene_tx=X
M_023254216.1;refGene_gn=CB2H6orf203;refGene_tx=XM_023254217.1
B2 99238080 99238164
refGene_gn=CB2H6orf203;refGene_tx=XM_023254216.1
B2 99240640 99240820
refGene_gn=CB2H6orf203;refGene_tx=XM_019831037.2;refGene_gn=CB2H6orf20
3;refGene_tx=XM_023254212.1;refGene_gn=CB2H6orf203;refGene_tx=XM_02325
4213.1;refGene_gn=CB2H6orf203;refGene_tx=XM_023254214.1;refGene_gn=CB2
H6orf203;refGene_tx=XM_023254217.1
B2 99248836 99248983
refGene_gn=CB2H6orf203;refGene_tx=XM_019831037.2;refGene_gn=CB2H6orf20
3;refGene_tx=XM_023254212.1;refGene_gn=CB2H6orf203;refGene_tx=XM_02325
4213.1;refGene_gn=CB2H6orf203;refGene_tx=XM_023254215.1;refGene_gn=CB2
H6orf203;refGene_tx=XM_023254217.1
B2 99252226 99254473 refGene_gn=BEND3;refGene_tx=XM_023254210.1
B2 99267355 99267558 refGene_gn=BEND3;refGene_tx=XM_023254210.1
B2 99268055 99268092 refGene_gn=BEND3;refGene_tx=XM_023254210.1
B2 99284135 99284693
refGene_gn=LOC111560599;refGene_tx=XM_023254734.1
B2 99284782 99284944
refGene_gn=LOC111560599;refGene_tx=XM_023254734.1
B2 99316817 99316976 refGene_gn=PDSS2;refGene_tx=XM_011282517.3
B2 99341586 99341619 refGene_gn=PDSS2;refGene_tx=XM_011282517.3
B2 99356767 99356899 refGene_gn=PDSS2;refGene_tx=XM_011282517.3
B2 99358101 99358275 refGene_gn=PDSS2;refGene_tx=XM_011282517.3
B2 99394959 99395034 refGene_gn=PDSS2;refGene_tx=XM_011282517.3
B2 99419491 99419690 refGene_gn=PDSS2;refGene_tx=XM_011282517.3
B2 99469262 99469397 refGene_gn=PDSS2;refGene_tx=XM_011282517.3
B2 99576754 99577050 refGene_gn=PDSS2;refGene_tx=XM_011282517.3
B2 99610662 99610758

refGene_gn=SOBP;refGene_tx=XM_023254219.1;refGene_gn=SOBP;refGene_tx=XM_023254220.1
B2 99624136 99624275
refGene_gn=SOBP;refGene_tx=XM_023254219.1;refGene_gn=SOBP;refGene_tx=XM_023254220.1
B2 99626316 99626502
refGene_gn=SOBP;refGene_tx=XM_023254219.1;refGene_gn=SOBP;refGene_tx=XM_023254220.1
B2 99652865 99653017
refGene_gn=SOBP;refGene_tx=XM_023254219.1;refGene_gn=SOBP;refGene_tx=XM_023254220.1
B2 99707001 99707097 refGene_gn=SOBP;refGene_tx=XM_023254219.1
B2 99752822 99754769
refGene_gn=SOBP;refGene_tx=XM_023254219.1;refGene_gn=SOBP;refGene_tx=XM_023254220.1
B2 99815282 99815408
refGene_gn=SCML4;refGene_tx=XM_006931920.3;refGene_gn=SCML4;refGene_tx=XM_006931921.4;refGene_gn=SCML4;refGene_tx=XM_006931922.2;refGene_gn=SCML4;refGene_tx=XM_019831047.2;refGene_gn=SCML4;refGene_tx=XM_019831049.2
B2 99818165 99818311
refGene_gn=SCML4;refGene_tx=XM_006931920.3;refGene_gn=SCML4;refGene_tx=XM_006931921.4;refGene_gn=SCML4;refGene_tx=XM_006931922.2;refGene_gn=SCML4;refGene_tx=XM_019831047.2;refGene_gn=SCML4;refGene_tx=XM_019831049.2
B2 99827836 99828127
refGene_gn=SCML4;refGene_tx=XM_006931920.3;refGene_gn=SCML4;refGene_tx=XM_006931921.4;refGene_gn=SCML4;refGene_tx=XM_006931922.2;refGene_gn=SCML4;refGene_tx=XM_019831047.2;refGene_gn=SCML4;refGene_tx=XM_019831049.2
B2 99846157 99846352
refGene_gn=SCML4;refGene_tx=XM_019831047.2;refGene_gn=SCML4;refGene_tx=XM_019831049.2
B2 99850516 99850717
refGene_gn=SCML4;refGene_tx=XM_019831047.2;refGene_gn=SCML4;refGene_tx=XM_019831049.2
B2 99852880 99853010
refGene_gn=SCML4;refGene_tx=XM_019831047.2;refGene_gn=SCML4;refGene_tx=XM_019831049.2
B2 99870298 99870454 refGene_gn=SCML4;refGene_tx=XM_019831047.2
B2 99961317 99961461 refGene_gn=SEC63;refGene_tx=XM_003986424.5
B2 99962454 99962559 refGene_gn=SEC63;refGene_tx=XM_003986424.5
B2 99973478 99973577 refGene_gn=SEC63;refGene_tx=XM_003986424.5
B2 99977150 99977252 refGene_gn=SEC63;refGene_tx=XM_003986424.5
B2 99981566 99981725 refGene_gn=SEC63;refGene_tx=XM_003986424.5
B2 99985911 99986085 refGene_gn=SEC63;refGene_tx=XM_003986424.5
B2 99986285 99986345 refGene_gn=SEC63;refGene_tx=XM_003986424.5
B2 99989326 99989409 refGene_gn=SEC63;refGene_tx=XM_003986424.5
B2 99992549 99992697 refGene_gn=SEC63;refGene_tx=XM_003986424.5
B2 99994949 99995104 refGene_gn=SEC63;refGene_tx=XM_003986424.5

B2	99995659	99995752	refGene_gn=SEC63;refGene_tx=XM_003986424.5
B2	99997330	99997463	refGene_gn=SEC63;refGene_tx=XM_003986424.5
B2	99997567	99997662	refGene_gn=SEC63;refGene_tx=XM_003986424.5
B2	99999918	100000027
refGene_gn=SEC63;refGene_tx=XM_003986424.5
B2	100001023	100001074
refGene_gn=SEC63;refGene_tx=XM_003986424.5
B2	100002601	100002660
refGene_gn=SEC63;refGene_tx=XM_003986424.5
B2	100003241	100003303
refGene_gn=SEC63;refGene_tx=XM_003986424.5
B2	100011871	100011984
refGene_gn=SEC63;refGene_tx=XM_003986424.5
B2	100013490	100013605
refGene_gn=SEC63;refGene_tx=XM_003986424.5
B2	100018534	100018634
refGene_gn=SEC63;refGene_tx=XM_003986424.5
B2	100035207	100035331
refGene_gn=SEC63;refGene_tx=XM_003986424.5
B2	100099225	100099281
refGene_gn=OSTM1;refGene_tx=XM_003986433.5
B2	100104309	100104475
refGene_gn=OSTM1;refGene_tx=XM_003986433.5
B2	100106690	100106858
refGene_gn=OSTM1;refGene_tx=XM_003986433.5
B2	100121097	100121195
refGene_gn=OSTM1;refGene_tx=XM_003986433.5
B2	100132275	100132390
refGene_gn=OSTM1;refGene_tx=XM_003986433.5
B2	100137642	100138044
refGene_gn=OSTM1;refGene_tx=XM_003986433.5
B2	100233810	100233835
refGene_gn=NR2E1;refGene_tx=XM_023254222.1;refGene_gn=NR2E1;refGene_tx
=XM_023254223.1
B2	100238509	100238655
refGene_gn=NR2E1;refGene_tx=XM_023254222.1;refGene_gn=NR2E1;refGene_tx
=XM_023254223.1
B2	100242112	100242200
refGene_gn=NR2E1;refGene_tx=XM_023254222.1;refGene_gn=NR2E1;refGene_tx
=XM_023254223.1
B2	100243801	100244037
refGene_gn=NR2E1;refGene_tx=XM_023254222.1;refGene_gn=NR2E1;refGene_tx
=XM_023254223.1
B2	100245373	100245520
refGene_gn=NR2E1;refGene_tx=XM_023254222.1;refGene_gn=NR2E1;refGene_tx
=XM_023254223.1
B2	100247640	100247737
refGene_gn=NR2E1;refGene_tx=XM_023254222.1;refGene_gn=NR2E1;refGene_tx
=XM_023254223.1
B2	100248124	100248274

refGene_gn=NR2E1;refGene_tx=XM_023254222.1;refGene_gn=NR2E1;refGene_tx=XM_023254223.1
B2 100248527 100248633
refGene_gn=NR2E1;refGene_tx=XM_023254222.1;refGene_gn=NR2E1;refGene_tx=XM_023254223.1
B2 100252738 100252901
refGene_gn=NR2E1;refGene_tx=XM_023254222.1
B2 100254297 100254364
refGene_gn=NR2E1;refGene_tx=XM_023254223.1
B2 100276713 100276819
refGene_gn=SNX3;refGene_tx=XM_023254224.1
B2 100278216 100278341
refGene_gn=SNX3;refGene_tx=XM_023254224.1
B2 100287837 100287933
refGene_gn=SNX3;refGene_tx=XM_023254224.1
B2 100324348 100324510
refGene_gn=SNX3;refGene_tx=XM_023254224.1
B2 100341381 100341463
refGene_gn=AFG1L;refGene_tx=XM_006931927.4;refGene_gn=AFG1L;refGene_tx=XM_019831051.2
B2 100341591 100341735
refGene_gn=AFG1L;refGene_tx=XM_006931927.4;refGene_gn=AFG1L;refGene_tx=XM_019831051.2
B2 100376544 100376768
refGene_gn=AFG1L;refGene_tx=XM_006931927.4;refGene_gn=AFG1L;refGene_tx=XM_019831051.2
B2 100398210 100398262
refGene_gn=AFG1L;refGene_tx=XM_006931927.4;refGene_gn=AFG1L;refGene_tx=XM_019831051.2
B2 100409240 100409342
refGene_gn=AFG1L;refGene_tx=XM_006931927.4;refGene_gn=AFG1L;refGene_tx=XM_019831051.2
B2 100410521 100410652
refGene_gn=AFG1L;refGene_tx=XM_006931927.4;refGene_gn=AFG1L;refGene_tx=XM_019831051.2
B2 100420908 100421008
refGene_gn=AFG1L;refGene_tx=XM_006931927.4;refGene_gn=AFG1L;refGene_tx=XM_019831051.2
B2 100446538 100446597
refGene_gn=AFG1L;refGene_tx=XM_006931927.4;refGene_gn=AFG1L;refGene_tx=XM_019831051.2
B2 100483145 100483228
refGene_gn=AFG1L;refGene_tx=XM_006931927.4;refGene_gn=AFG1L;refGene_tx=XM_019831051.2
B2 100506956 100507027
refGene_gn=AFG1L;refGene_tx=XM_006931927.4;refGene_gn=AFG1L;refGene_tx=XM_019831051.2
B2 100507283 100507384
refGene_gn=AFG1L;refGene_tx=XM_006931927.4;refGene_gn=AFG1L;refGene_tx=XM_019831051.2

B2 100525327 100525463
refGene_gn=AFG1L;refGene_tx=XM_019831051.2
B2 100535996 100536137
refGene_gn=AFG1L;refGene_tx=XM_006931927.4;refGene_gn=AFG1L;refGene_tx=XM_019831051.2
B2 100538380 100538494
refGene_gn=AFG1L;refGene_tx=XM_006931927.4
B2 100541294 100541423
refGene_gn=AFG1L;refGene_tx=XM_006931927.4
B2 100580317 100580935
refGene_gn=FOXO3;refGene_tx=XM_003986453.5
B2 100684342 100685743
refGene_gn=FOXO3;refGene_tx=XM_003986453.5
B2 100724923 100725383
refGene_gn=LOC101098872;refGene_tx=XM_003986436.3
B2 100725624 100725707
refGene_gn=LOC101098872;refGene_tx=XM_003986436.3
B2 100872346 100872486
refGene_gn=ARMC2;refGene_tx=XM_003986437.5;refGene_gn=ARMC2;refGene_tx=XM_019831054.2
B2 100880902 100881163
refGene_gn=ARMC2;refGene_tx=XM_003986437.5;refGene_gn=ARMC2;refGene_tx=XM_019831054.2;refGene_gn=ARMC2;refGene_tx=XM_019831055.2
B2 100881642 100881715
refGene_gn=ARMC2;refGene_tx=XM_003986437.5;refGene_gn=ARMC2;refGene_tx=XM_019831054.2;refGene_gn=ARMC2;refGene_tx=XM_019831055.2
B2 100890695 100890870
refGene_gn=ARMC2;refGene_tx=XM_003986437.5;refGene_gn=ARMC2;refGene_tx=XM_019831054.2;refGene_gn=ARMC2;refGene_tx=XM_019831055.2
B2 100897695 100897903
refGene_gn=ARMC2;refGene_tx=XM_003986437.5;refGene_gn=ARMC2;refGene_tx=XM_019831054.2;refGene_gn=ARMC2;refGene_tx=XM_019831055.2
B2 100911916 100911993
refGene_gn=ARMC2;refGene_tx=XM_003986437.5;refGene_gn=ARMC2;refGene_tx=XM_019831054.2;refGene_gn=ARMC2;refGene_tx=XM_019831055.2
B2 100916936 100917035
refGene_gn=ARMC2;refGene_tx=XM_003986437.5;refGene_gn=ARMC2;refGene_tx=XM_019831054.2;refGene_gn=ARMC2;refGene_tx=XM_019831055.2
B2 100921165 100921341
refGene_gn=ARMC2;refGene_tx=XM_003986437.5;refGene_gn=ARMC2;refGene_tx=XM_019831054.2;refGene_gn=ARMC2;refGene_tx=XM_019831055.2
B2 100926838 100926941
refGene_gn=ARMC2;refGene_tx=XM_003986437.5;refGene_gn=ARMC2;refGene_tx=XM_019831054.2;refGene_gn=ARMC2;refGene_tx=XM_019831055.2
B2 100928011 100928235
refGene_gn=ARMC2;refGene_tx=XM_003986437.5;refGene_gn=ARMC2;refGene_tx=XM_019831054.2;refGene_gn=ARMC2;refGene_tx=XM_019831055.2
B2 100946126 100946272
refGene_gn=ARMC2;refGene_tx=XM_003986437.5;refGene_gn=ARMC2;refGene_tx=XM_019831054.2;refGene_gn=ARMC2;refGene_tx=XM_019831055.2

B2 100949438 100949538
refGene_gn=ARMC2;refGene_tx=XM_003986437.5;refGene_gn=ARMC2;refGene_tx=XM_019831054.2;refGene_gn=ARMC2;refGene_tx=XM_019831055.2
B2 100962249 100962565
refGene_gn=ARMC2;refGene_tx=XM_003986437.5;refGene_gn=ARMC2;refGene_tx=XM_019831054.2;refGene_gn=ARMC2;refGene_tx=XM_019831055.2
B2 100965806 100965929
refGene_gn=ARMC2;refGene_tx=XM_003986437.5;refGene_gn=ARMC2;refGene_tx=XM_019831054.2;refGene_gn=ARMC2;refGene_tx=XM_019831055.2
B2 100966237 100966351
refGene_gn=ARMC2;refGene_tx=XM_003986437.5;refGene_gn=ARMC2;refGene_tx=XM_019831054.2;refGene_gn=ARMC2;refGene_tx=XM_019831055.2
B2 100968918 100969051
refGene_gn=ARMC2;refGene_tx=XM_003986437.5;refGene_gn=ARMC2;refGene_tx=XM_019831054.2;refGene_gn=ARMC2;refGene_tx=XM_019831055.2
B2 100969584 100969745
refGene_gn=ARMC2;refGene_tx=XM_003986437.5;refGene_gn=ARMC2;refGene_tx=XM_019831054.2;refGene_gn=ARMC2;refGene_tx=XM_019831055.2
B2 100975457 100975540
refGene_gn=ARMC2;refGene_tx=XM_019831054.2
B2 100976305 100976463
refGene_gn=ARMC2;refGene_tx=XM_003986437.5;refGene_gn=ARMC2;refGene_tx=XM_019831055.2
B2 101002077 101002164
refGene_gn=SESN1;refGene_tx=XM_003986438.4;refGene_gn=SESN1;refGene_tx=XM_003986439.5;refGene_gn=SESN1;refGene_tx=XM_019831056.2;refGene_gn=SESN1;refGene_tx=XM_019831057.2;refGene_gn=SESN1;refGene_tx=XM_023254226.1
B2 101002718 101002863
refGene_gn=SESN1;refGene_tx=XM_003986438.4;refGene_gn=SESN1;refGene_tx=XM_003986439.5;refGene_gn=SESN1;refGene_tx=XM_019831056.2;refGene_gn=SESN1;refGene_tx=XM_019831057.2;refGene_gn=SESN1;refGene_tx=XM_023254226.1
B2 101004113 101004304
refGene_gn=SESN1;refGene_tx=XM_003986438.4;refGene_gn=SESN1;refGene_tx=XM_003986439.5;refGene_gn=SESN1;refGene_tx=XM_019831056.2;refGene_gn=SESN1;refGene_tx=XM_019831057.2;refGene_gn=SESN1;refGene_tx=XM_023254226.1
B2 101006342 101006455
refGene_gn=SESN1;refGene_tx=XM_003986438.4;refGene_gn=SESN1;refGene_tx=XM_003986439.5;refGene_gn=SESN1;refGene_tx=XM_019831056.2;refGene_gn=SESN1;refGene_tx=XM_019831057.2;refGene_gn=SESN1;refGene_tx=XM_023254226.1
B2 101008165 101008313
refGene_gn=SESN1;refGene_tx=XM_003986438.4;refGene_gn=SESN1;refGene_tx=XM_003986439.5;refGene_gn=SESN1;refGene_tx=XM_019831056.2;refGene_gn=SESN1;refGene_tx=XM_019831057.2;refGene_gn=SESN1;refGene_tx=XM_023254226.1
B2 101011895 101012138
refGene_gn=SESN1;refGene_tx=XM_003986438.4;refGene_gn=SESN1;refGene_tx

=XM_003986439.5;refGene_gn=SESN1;refGene_tx=XM_019831056.2;refGene_gn=SESN1;refGene_tx=XM_019831057.2;refGene_gn=SESN1;refGene_tx=XM_023254226.1
B2 101014098 101014281
refGene_gn=SESN1;refGene_tx=XM_003986438.4;refGene_gn=SESN1;refGene_tx=XM_003986439.5;refGene_gn=SESN1;refGene_tx=XM_019831056.2;refGene_gn=SESN1;refGene_tx=XM_019831057.2;refGene_gn=SESN1;refGene_tx=XM_023254226.1
B2 101014908 101015109
refGene_gn=SESN1;refGene_tx=XM_003986438.4;refGene_gn=SESN1;refGene_tx=XM_003986439.5;refGene_gn=SESN1;refGene_tx=XM_019831056.2;refGene_gn=SESN1;refGene_tx=XM_019831057.2;refGene_gn=SESN1;refGene_tx=XM_023254226.1
B2 101015888 101015954
refGene_gn=SESN1;refGene_tx=XM_003986438.4;refGene_gn=SESN1;refGene_tx=XM_003986439.5
B2 101022803 101022905
refGene_gn=SESN1;refGene_tx=XM_003986438.4
B2 101116048 101116327
refGene_gn=SESN1;refGene_tx=XM_003986439.5
B2 101162824 101162827
refGene_gn=CEP57L1;refGene_tx=XM_011282515.3
B2 101164024 101164190
refGene_gn=CEP57L1;refGene_tx=XM_011282515.3;refGene_gn=CEP57L1;refGene_tx=XM_011282516.3;refGene_gn=CEP57L1;refGene_tx=XM_023254227.1;refGene_gn=CEP57L1;refGene_tx=XM_023254228.1
B2 101166636 101166816
refGene_gn=CEP57L1;refGene_tx=XM_011282515.3;refGene_gn=CEP57L1;refGene_tx=XM_011282516.3;refGene_gn=CEP57L1;refGene_tx=XM_023254227.1;refGene_gn=CEP57L1;refGene_tx=XM_023254228.1
B2 101168159 101168281
refGene_gn=CEP57L1;refGene_tx=XM_011282515.3;refGene_gn=CEP57L1;refGene_tx=XM_011282516.3;refGene_gn=CEP57L1;refGene_tx=XM_023254227.1;refGene_gn=CEP57L1;refGene_tx=XM_023254228.1;refGene_gn=CEP57L1;refGene_tx=XM_023254229.1
B2 101182595 101182712
refGene_gn=CEP57L1;refGene_tx=XM_011282515.3;refGene_gn=CEP57L1;refGene_tx=XM_011282516.3;refGene_gn=CEP57L1;refGene_tx=XM_023254227.1;refGene_gn=CEP57L1;refGene_tx=XM_023254228.1;refGene_gn=CEP57L1;refGene_tx=XM_023254229.1
B2 101185916 101185994
refGene_gn=CEP57L1;refGene_tx=XM_011282515.3;refGene_gn=CEP57L1;refGene_tx=XM_011282516.3;refGene_gn=CEP57L1;refGene_tx=XM_023254227.1;refGene_gn=CEP57L1;refGene_tx=XM_023254228.1;refGene_gn=CEP57L1;refGene_tx=XM_023254229.1
B2 101186463 101186550
refGene_gn=CEP57L1;refGene_tx=XM_011282515.3;refGene_gn=CEP57L1;refGene_tx=XM_011282516.3;refGene_gn=CEP57L1;refGene_tx=XM_023254227.1;refGene_gn=CEP57L1;refGene_tx=XM_023254228.1;refGene_gn=CEP57L1;refGene_tx=XM_023254229.1

B2 101189586 101189664
refGene_gn=CEP57L1;refGene_tx=XM_011282515.3;refGene_gn=CEP57L1;refGene_tx=XM_011282516.3;refGene_gn=CEP57L1;refGene_tx=XM_023254227.1;refGene_gn=CEP57L1;refGene_tx=XM_023254228.1;refGene_gn=CEP57L1;refGene_tx=XM_023254229.1
B2 101189780 101190025
refGene_gn=CEP57L1;refGene_tx=XM_011282515.3;refGene_gn=CEP57L1;refGene_tx=XM_011282516.3;refGene_gn=CEP57L1;refGene_tx=XM_023254227.1;refGene_gn=CEP57L1;refGene_tx=XM_023254228.1;refGene_gn=CEP57L1;refGene_tx=XM_023254229.1
B2 101191280 101191425
refGene_gn=CEP57L1;refGene_tx=XM_011282515.3;refGene_gn=CEP57L1;refGene_tx=XM_011282516.3;refGene_gn=CEP57L1;refGene_tx=XM_023254227.1;refGene_gn=CEP57L1;refGene_tx=XM_023254228.1;refGene_gn=CEP57L1;refGene_tx=XM_023254229.1
B2 101193247 101193466
refGene_gn=CEP57L1;refGene_tx=XM_011282515.3;refGene_gn=CEP57L1;refGene_tx=XM_011282516.3;refGene_gn=CEP57L1;refGene_tx=XM_023254227.1;refGene_gn=CEP57L1;refGene_tx=XM_023254228.1;refGene_gn=CEP57L1;refGene_tx=XM_023254229.1
B2 101219532 101219663
refGene_gn=CCDC162P;refGene_tx=XM_023254735.1
B2 101221030 101221103
refGene_gn=CCDC162P;refGene_tx=XM_023254735.1
B2 101221498 101221609
refGene_gn=CCDC162P;refGene_tx=XM_023254735.1
B2 101225622 101225736
refGene_gn=CCDC162P;refGene_tx=XM_023254735.1
B2 101226979 101227174
refGene_gn=CCDC162P;refGene_tx=XM_023254735.1
B2 101228784 101228938
refGene_gn=CCDC162P;refGene_tx=XM_023254735.1
B2 101229977 101230135
refGene_gn=CCDC162P;refGene_tx=XM_023254735.1
B2 101231854 101232007
refGene_gn=CCDC162P;refGene_tx=XM_023254735.1
B2 101233713 101233815
refGene_gn=CCDC162P;refGene_tx=XM_023254735.1
B2 101238058 101238220
refGene_gn=CCDC162P;refGene_tx=XM_023254735.1
B2 101247567 101247680
refGene_gn=CCDC162P;refGene_tx=XM_023254735.1
B2 101248062 101248198
refGene_gn=CCDC162P;refGene_tx=XM_023254735.1
B2 101248453 101248657
refGene_gn=CCDC162P;refGene_tx=XM_023254735.1
B2 101261944 101262137
refGene_gn=CCDC162P;refGene_tx=XM_023254735.1
B2 101264102 101264230
refGene_gn=CCDC162P;refGene_tx=XM_023254735.1

B2 101269990 101270191
refGene_gn=CCDC162P;refGene_tx=XM_023254735.1
B2 101270905 101271027
refGene_gn=CCDC162P;refGene_tx=XM_023254735.1
B2 101273040 101273269
refGene_gn=CCDC162P;refGene_tx=XM_023254735.1
B2 101276911 101277114
refGene_gn=CCDC162P;refGene_tx=XM_023254735.1
B2 101279398 101279554
refGene_gn=CCDC162P;refGene_tx=XM_023254735.1
B2 101281688 101281782
refGene_gn=CCDC162P;refGene_tx=XM_023254735.1
B2 101282477 101282615
refGene_gn=CCDC162P;refGene_tx=XM_023254735.1
B2 101287938 101288096
refGene_gn=CCDC162P;refGene_tx=XM_023254735.1
B2 101289165 101289277
refGene_gn=CCDC162P;refGene_tx=XM_023254735.1
B2 101290658 101290708
refGene_gn=CCDC162P;refGene_tx=XM_023254735.1
B2 101291078 101291255
refGene_gn=CCDC162P;refGene_tx=XM_023254735.1
B2 101299564 101299682
refGene_gn=CCDC162P;refGene_tx=XM_023254735.1
B2 101302145 101302311
refGene_gn=CCDC162P;refGene_tx=XM_023254735.1
B2 101307114 101307253
refGene_gn=CCDC162P;refGene_tx=XM_023254735.1
B2 101315054 101315208
refGene_gn=CCDC162P;refGene_tx=XM_023254735.1
B2 101317618 101317714
refGene_gn=CCDC162P;refGene_tx=XM_023254735.1
B2 101318778 101318982
refGene_gn=CCDC162P;refGene_tx=XM_023254735.1
B2 101319166 101319172
refGene_gn=LOC111560600;refGene_tx=XM_023254736.1
B2 101319701 101319863
refGene_gn=LOC111560600;refGene_tx=XM_023254736.1
B2 101331530 101331760
refGene_gn=LOC111560600;refGene_tx=XM_023254736.1
B2 101332904 101333043
refGene_gn=LOC111560600;refGene_tx=XM_023254736.1
B2 101334489 101334610
refGene_gn=LOC111560600;refGene_tx=XM_023254736.1
B2 101346235 101346345
refGene_gn=LOC111560600;refGene_tx=XM_023254736.1
B2 101353981 101354131
refGene_gn=LOC111560600;refGene_tx=XM_023254736.1
B2 101355022 101355076
refGene_gn=LOC111560600;refGene_tx=XM_023254736.1

B2 101355948 101356116
refGene_gn=LOC111560600;refGene_tx=XM_023254736.1
B2 101357975 101358128
refGene_gn=LOC111560600;refGene_tx=XM_023254736.1
B2 101358888 101359020
refGene_gn=LOC111560600;refGene_tx=XM_023254736.1
B2 101359861 101360014
refGene_gn=LOC111560600;refGene_tx=XM_023254736.1
B2 101362178 101362388
refGene_gn=LOC111560600;refGene_tx=XM_023254736.1
B2 101366095 101366216
refGene_gn=LOC111560600;refGene_tx=XM_023254736.1
B2 101369270 101369398
refGene_gn=LOC111560600;refGene_tx=XM_023254736.1
B2 101371963 101372089
refGene_gn=LOC111560600;refGene_tx=XM_023254736.1
B2 101383563 101383730
refGene_gn=CD164;refGene_tx=XM_011282514.3
B2 101386572 101386629
refGene_gn=CD164;refGene_tx=XM_011282514.3
B2 101390291 101390327
refGene_gn=CD164;refGene_tx=XM_011282514.3
B2 101392223 101392295
refGene_gn=CD164;refGene_tx=XM_011282514.3
B2 101393872 101393956
refGene_gn=CD164;refGene_tx=XM_011282514.3
B2 101396732 101396913
refGene_gn=CD164;refGene_tx=XM_011282514.3
B2 101408516 101408628
refGene_gn=PPIL6;refGene_tx=XM_003986442.4
B2 101413175 101413311
refGene_gn=PPIL6;refGene_tx=XM_003986442.4
B2 101423895 101423952
refGene_gn=PPIL6;refGene_tx=XM_003986442.4
B2 101429686 101429834
refGene_gn=PPIL6;refGene_tx=XM_003986442.4
B2 101429971 101430034
refGene_gn=PPIL6;refGene_tx=XM_003986442.4
B2 101433125 101433269
refGene_gn=PPIL6;refGene_tx=XM_003986442.4
B2 101433270 101433315
refGene_gn=PPIL6;refGene_tx=XM_003986442.4
B2 101436510 101436606
refGene_gn=PPIL6;refGene_tx=XM_003986442.4
B2 101440723 101440858
refGene_gn=PPIL6;refGene_tx=XM_003986442.4
B2 101441440 101441490
refGene_gn=SMPD2;refGene_tx=XM_011282513.3
B2 101441715 101441812
refGene_gn=SMPD2;refGene_tx=XM_011282513.3

B2	101441940	101442017
refGene_gn=SMPD2;refGene_tx=XM_011282513.3
B2	101442371	101442465
refGene_gn=SMPD2;refGene_tx=XM_011282513.3
B2	101442607	101442697
refGene_gn=SMPD2;refGene_tx=XM_011282513.3
B2	101442932	101443015
refGene_gn=SMPD2;refGene_tx=XM_011282513.3
B2	101443144	101443277
refGene_gn=SMPD2;refGene_tx=XM_011282513.3
B2	101443369	101443474
refGene_gn=SMPD2;refGene_tx=XM_011282513.3
B2	101443634	101443791
refGene_gn=SMPD2;refGene_tx=XM_011282513.3
B2	101443891	101444280
refGene_gn=SMPD2;refGene_tx=XM_011282513.3
B2	101444568	101444717
refGene_gn=MICAL1;refGene_tx=XM_003986443.3;refGene_gn=MICAL1;refGene_tx=XM_023254230.1;refGene_gn=MICAL1;refGene_tx=XM_023254231.1;refGene_gn=MICAL1;refGene_tx=XM_023254232.1;refGene_gn=MICAL1;refGene_tx=XM_023254233.1;refGene_gn=MICAL1;refGene_tx=XM_023254234.1
B2	101445337	101445411
refGene_gn=MICAL1;refGene_tx=XM_003986443.3;refGene_gn=MICAL1;refGene_tx=XM_023254230.1;refGene_gn=MICAL1;refGene_tx=XM_023254231.1;refGene_gn=MICAL1;refGene_tx=XM_023254232.1;refGene_gn=MICAL1;refGene_tx=XM_023254233.1;refGene_gn=MICAL1;refGene_tx=XM_023254234.1
B2	101445508	101445608
refGene_gn=MICAL1;refGene_tx=XM_003986443.3;refGene_gn=MICAL1;refGene_tx=XM_023254230.1;refGene_gn=MICAL1;refGene_tx=XM_023254231.1;refGene_gn=MICAL1;refGene_tx=XM_023254232.1;refGene_gn=MICAL1;refGene_tx=XM_023254233.1;refGene_gn=MICAL1;refGene_tx=XM_023254234.1
B2	101445791	101445885
refGene_gn=MICAL1;refGene_tx=XM_003986443.3;refGene_gn=MICAL1;refGene_tx=XM_023254230.1;refGene_gn=MICAL1;refGene_tx=XM_023254231.1;refGene_gn=MICAL1;refGene_tx=XM_023254232.1;refGene_gn=MICAL1;refGene_tx=XM_023254233.1;refGene_gn=MICAL1;refGene_tx=XM_023254234.1
B2	101446011	101446125
refGene_gn=MICAL1;refGene_tx=XM_003986443.3;refGene_gn=MICAL1;refGene_tx=XM_023254230.1;refGene_gn=MICAL1;refGene_tx=XM_023254231.1;refGene_gn=MICAL1;refGene_tx=XM_023254232.1;refGene_gn=MICAL1;refGene_tx=XM_023254233.1;refGene_gn=MICAL1;refGene_tx=XM_023254234.1
B2	101446420	101446521
refGene_gn=MICAL1;refGene_tx=XM_003986443.3;refGene_gn=MICAL1;refGene_tx=XM_023254230.1;refGene_gn=MICAL1;refGene_tx=XM_023254231.1;refGene_gn=MICAL1;refGene_tx=XM_023254232.1;refGene_gn=MICAL1;refGene_tx=XM_023254233.1;refGene_gn=MICAL1;refGene_tx=XM_023254234.1
B2	101446648	101446925
refGene_gn=MICAL1;refGene_tx=XM_003986443.3;refGene_gn=MICAL1;refGene_tx=XM_023254230.1;refGene_gn=MICAL1;refGene_tx=XM_023254231.1;refGene_gn=MICAL1;refGene_tx=XM_023254232.1;refGene_gn=MICAL1;refGene_tx=XM_02

3254233.1;refGene_gn=MICAL1;refGene_tx=XM_023254234.1
B2 101447206 101447283
refGene_gn=MICAL1;refGene_tx=XM_003986443.3;refGene_gn=MICAL1;refGene_tx=XM_023254230.1;refGene_gn=MICAL1;refGene_tx=XM_023254231.1;refGene_gn=MICAL1;refGene_tx=XM_023254232.1;refGene_gn=MICAL1;refGene_tx=XM_023254233.1;refGene_gn=MICAL1;refGene_tx=XM_023254234.1
B2 101447600 101447757
refGene_gn=MICAL1;refGene_tx=XM_003986443.3;refGene_gn=MICAL1;refGene_tx=XM_023254230.1;refGene_gn=MICAL1;refGene_tx=XM_023254231.1;refGene_gn=MICAL1;refGene_tx=XM_023254232.1;refGene_gn=MICAL1;refGene_tx=XM_023254233.1;refGene_gn=MICAL1;refGene_tx=XM_023254234.1
B2 101447884 101447968
refGene_gn=MICAL1;refGene_tx=XM_003986443.3;refGene_gn=MICAL1;refGene_tx=XM_023254230.1;refGene_gn=MICAL1;refGene_tx=XM_023254231.1;refGene_gn=MICAL1;refGene_tx=XM_023254232.1;refGene_gn=MICAL1;refGene_tx=XM_023254233.1;refGene_gn=MICAL1;refGene_tx=XM_023254234.1
B2 101448196 101448250
refGene_gn=MICAL1;refGene_tx=XM_003986443.3;refGene_gn=MICAL1;refGene_tx=XM_023254230.1;refGene_gn=MICAL1;refGene_tx=XM_023254231.1;refGene_gn=MICAL1;refGene_tx=XM_023254232.1;refGene_gn=MICAL1;refGene_tx=XM_023254233.1;refGene_gn=MICAL1;refGene_tx=XM_023254234.1
B2 101448374 101448463
refGene_gn=MICAL1;refGene_tx=XM_003986443.3;refGene_gn=MICAL1;refGene_tx=XM_023254230.1;refGene_gn=MICAL1;refGene_tx=XM_023254231.1;refGene_gn=MICAL1;refGene_tx=XM_023254232.1;refGene_gn=MICAL1;refGene_tx=XM_023254233.1;refGene_gn=MICAL1;refGene_tx=XM_023254234.1
B2 101448598 101448789
refGene_gn=MICAL1;refGene_tx=XM_003986443.3;refGene_gn=MICAL1;refGene_tx=XM_023254230.1;refGene_gn=MICAL1;refGene_tx=XM_023254231.1;refGene_gn=MICAL1;refGene_tx=XM_023254232.1;refGene_gn=MICAL1;refGene_tx=XM_023254233.1;refGene_gn=MICAL1;refGene_tx=XM_023254234.1
B2 101449212 101449360
refGene_gn=MICAL1;refGene_tx=XM_003986443.3;refGene_gn=MICAL1;refGene_tx=XM_023254230.1;refGene_gn=MICAL1;refGene_tx=XM_023254231.1;refGene_gn=MICAL1;refGene_tx=XM_023254232.1;refGene_gn=MICAL1;refGene_tx=XM_023254233.1;refGene_gn=MICAL1;refGene_tx=XM_023254234.1
B2 101449916 101449998
refGene_gn=MICAL1;refGene_tx=XM_003986443.3;refGene_gn=MICAL1;refGene_tx=XM_023254230.1;refGene_gn=MICAL1;refGene_tx=XM_023254231.1;refGene_gn=MICAL1;refGene_tx=XM_023254232.1;refGene_gn=MICAL1;refGene_tx=XM_023254233.1;refGene_gn=MICAL1;refGene_tx=XM_023254234.1
B2 101450139 101450266
refGene_gn=MICAL1;refGene_tx=XM_003986443.3;refGene_gn=MICAL1;refGene_tx=XM_023254230.1;refGene_gn=MICAL1;refGene_tx=XM_023254231.1;refGene_gn=MICAL1;refGene_tx=XM_023254232.1;refGene_gn=MICAL1;refGene_tx=XM_023254233.1;refGene_gn=MICAL1;refGene_tx=XM_023254234.1
B2 101450458 101450574
refGene_gn=MICAL1;refGene_tx=XM_003986443.3;refGene_gn=MICAL1;refGene_tx=XM_023254230.1;refGene_gn=MICAL1;refGene_tx=XM_023254231.1;refGene_gn=MICAL1;refGene_tx=XM_023254232.1;refGene_gn=MICAL1;refGene_tx=XM_02

3254233.1;refGene_gn=MICAL1;refGene_tx=XM_023254234.1
B2	101450822	101451080
refGene_gn=MICAL1;refGene_tx=XM_003986443.3;refGene_gn=MICAL1;refGene_tx=XM_023254230.1;refGene_gn=MICAL1;refGene_tx=XM_023254231.1;refGene_gn=MICAL1;refGene_tx=XM_023254232.1;refGene_gn=MICAL1;refGene_tx=XM_023254233.1;refGene_gn=MICAL1;refGene_tx=XM_023254234.1
B2	101451817	101451918
refGene_gn=MICAL1;refGene_tx=XM_003986443.3;refGene_gn=MICAL1;refGene_tx=XM_023254230.1;refGene_gn=MICAL1;refGene_tx=XM_023254231.1;refGene_gn=MICAL1;refGene_tx=XM_023254232.1;refGene_gn=MICAL1;refGene_tx=XM_023254233.1;refGene_gn=MICAL1;refGene_tx=XM_023254234.1
B2	101452785	101452941
refGene_gn=MICAL1;refGene_tx=XM_003986443.3;refGene_gn=MICAL1;refGene_tx=XM_023254230.1;refGene_gn=MICAL1;refGene_tx=XM_023254231.1;refGene_gn=MICAL1;refGene_tx=XM_023254232.1;refGene_gn=MICAL1;refGene_tx=XM_023254233.1;refGene_gn=MICAL1;refGene_tx=XM_023254234.1
B2	101453061	101453166
refGene_gn=MICAL1;refGene_tx=XM_003986443.3;refGene_gn=MICAL1;refGene_tx=XM_023254230.1;refGene_gn=MICAL1;refGene_tx=XM_023254231.1;refGene_gn=MICAL1;refGene_tx=XM_023254232.1;refGene_gn=MICAL1;refGene_tx=XM_023254233.1;refGene_gn=MICAL1;refGene_tx=XM_023254234.1
B2	101453496	101453601
refGene_gn=MICAL1;refGene_tx=XM_003986443.3;refGene_gn=MICAL1;refGene_tx=XM_023254230.1;refGene_gn=MICAL1;refGene_tx=XM_023254231.1;refGene_gn=MICAL1;refGene_tx=XM_023254232.1;refGene_gn=MICAL1;refGene_tx=XM_023254233.1;refGene_gn=MICAL1;refGene_tx=XM_023254234.1
B2	101453846	101454085
refGene_gn=MICAL1;refGene_tx=XM_003986443.3;refGene_gn=MICAL1;refGene_tx=XM_023254230.1;refGene_gn=MICAL1;refGene_tx=XM_023254231.1;refGene_gn=MICAL1;refGene_tx=XM_023254232.1;refGene_gn=MICAL1;refGene_tx=XM_023254233.1;refGene_gn=MICAL1;refGene_tx=XM_023254234.1
B2	101454188	101454499
refGene_gn=MICAL1;refGene_tx=XM_003986443.3;refGene_gn=MICAL1;refGene_tx=XM_023254230.1;refGene_gn=MICAL1;refGene_tx=XM_023254231.1;refGene_gn=MICAL1;refGene_tx=XM_023254232.1;refGene_gn=MICAL1;refGene_tx=XM_023254233.1
B2	101454999	101455030
refGene_gn=MICAL1;refGene_tx=XM_023254231.1
B2	101465010	101465728
refGene_gn=ZBTB24;refGene_tx=XM_006931931.4;refGene_gn=ZBTB24;refGene_tx=XM_006931932.4
B2	101465888	101465901
refGene_gn=ZBTB24;refGene_tx=XM_006931933.3
B2	101466829	101466911
refGene_gn=ZBTB24;refGene_tx=XM_006931931.4;refGene_gn=ZBTB24;refGene_tx=XM_006931932.4;refGene_gn=ZBTB24;refGene_tx=XM_006931933.3
B2	101470768	101470852
refGene_gn=ZBTB24;refGene_tx=XM_006931931.4;refGene_gn=ZBTB24;refGene_tx=XM_006931932.4;refGene_gn=ZBTB24;refGene_tx=XM_006931933.3
B2	101470910	101470936

refGene_gn=ZBTB24;refGene_tx=XM_006931934.3
B2	101471544	101471628
refGene_gn=ZBTB24;refGene_tx=XM_006931931.4;refGene_gn=ZBTB24;refGene_tx=XM_006931932.4;refGene_gn=ZBTB24;refGene_tx=XM_006931933.3;refGene_gn=ZBTB24;refGene_tx=XM_006931934.3
B2	101472108	101472276
refGene_gn=ZBTB24;refGene_tx=XM_006931931.4;refGene_gn=ZBTB24;refGene_tx=XM_006931932.4;refGene_gn=ZBTB24;refGene_tx=XM_006931933.3;refGene_gn=ZBTB24;refGene_tx=XM_006931934.3
B2	101475294	101476237
refGene_gn=ZBTB24;refGene_tx=XM_006931931.4;refGene_gn=ZBTB24;refGene_tx=XM_006931932.4;refGene_gn=ZBTB24;refGene_tx=XM_006931933.3;refGene_gn=ZBTB24;refGene_tx=XM_006931934.3
B2	101487257	101487460
refGene_gn=AK9;refGene_tx=XM_023254235.1
B2	101487872	101487987
refGene_gn=AK9;refGene_tx=XM_023254235.1
B2	101489137	101489240
refGene_gn=AK9;refGene_tx=XM_023254235.1
B2	101493051	101493150
refGene_gn=AK9;refGene_tx=XM_023254235.1
B2	101493385	101493555
refGene_gn=AK9;refGene_tx=XM_023254235.1
B2	101495118	101495315
refGene_gn=AK9;refGene_tx=XM_023254235.1
B2	101499810	101500031
refGene_gn=AK9;refGene_tx=XM_023254235.1
B2	101500134	101500281
refGene_gn=AK9;refGene_tx=XM_023254235.1
B2	101502247	101502449
refGene_gn=AK9;refGene_tx=XM_023254235.1
B2	101509049	101509263
refGene_gn=AK9;refGene_tx=XM_023254235.1
B2	101512135	101512354
refGene_gn=AK9;refGene_tx=XM_023254235.1
B2	101512931	101513138
refGene_gn=AK9;refGene_tx=XM_023254235.1
B2	101525031	101525097
refGene_gn=AK9;refGene_tx=XM_023254235.1
B2	101529428	101529648
refGene_gn=AK9;refGene_tx=XM_023254235.1
B2	101533307	101533432
refGene_gn=AK9;refGene_tx=XM_023254235.1
B2	101534815	101535076
refGene_gn=AK9;refGene_tx=XM_023254235.1
B2	101536531	101536744
refGene_gn=AK9;refGene_tx=XM_023254235.1
B2	101543226	101543348
refGene_gn=AK9;refGene_tx=XM_023254235.1
B2	101543886	101544087

refGene_gn=AK9;refGene_tx=XM_023254235.1
B2	101544554	101544644
refGene_gn=AK9;refGene_tx=XM_023254235.1
B2	101551042	101551201
refGene_gn=AK9;refGene_tx=XM_023254235.1
B2	101554659	101554733
refGene_gn=AK9;refGene_tx=XM_023254235.1
B2	101558680	101558795
refGene_gn=AK9;refGene_tx=XM_023254235.1
B2	101559603	101559760
refGene_gn=AK9;refGene_tx=XM_023254235.1
B2	101570463	101570612
refGene_gn=AK9;refGene_tx=XM_023254235.1
B2	101575605	101575701
refGene_gn=AK9;refGene_tx=XM_023254235.1
B2	101575787	101575886
refGene_gn=AK9;refGene_tx=XM_023254235.1
B2	101579497	101579642
refGene_gn=AK9;refGene_tx=XM_023254235.1
B2	101583497	101583678
refGene_gn=AK9;refGene_tx=XM_023254235.1
B2	101583758	101583898
refGene_gn=AK9;refGene_tx=XM_023254235.1
B2	101585601	101585700
refGene_gn=AK9;refGene_tx=XM_023254235.1
B2	101588405	101588480
refGene_gn=AK9;refGene_tx=XM_023254235.1
B2	101591332	101591461
refGene_gn=AK9;refGene_tx=XM_023254235.1
B2	101596116	101596305
refGene_gn=AK9;refGene_tx=XM_023254235.1
B2	101599097	101599210
refGene_gn=AK9;refGene_tx=XM_023254235.1
B2	101606741	101606838
refGene_gn=AK9;refGene_tx=XM_023254235.1
B2	101606937	101606990
refGene_gn=AK9;refGene_tx=XM_023254235.1
B2	101609259	101609323
refGene_gn=AK9;refGene_tx=XM_023254235.1
B2	101610613	101610730
refGene_gn=AK9;refGene_tx=XM_023254235.1
B2	101621751	101621817
refGene_gn=FIG4;refGene_tx=XM_023254237.1;refGene_gn=FIG4;refGene_tx=XM_023254238.1
B2	101639328	101639427
refGene_gn=FIG4;refGene_tx=XM_023254237.1;refGene_gn=FIG4;refGene_tx=XM_023254238.1
B2	101640643	101640767
refGene_gn=FIG4;refGene_tx=XM_023254237.1;refGene_gn=FIG4;refGene_tx=XM_023254238.1

B2	101648873	101649030
refGene_gn=FIG4;refGene_tx=XM_023254237.1;refGene_gn=FIG4;refGene_tx=XM_023254238.1
B2	101652979	101653030
refGene_gn=FIG4;refGene_tx=XM_023254237.1;refGene_gn=FIG4;refGene_tx=XM_023254238.1
B2	101658937	101659086
refGene_gn=FIG4;refGene_tx=XM_023254237.1;refGene_gn=FIG4;refGene_tx=XM_023254238.1
B2	101662496	101662625
refGene_gn=FIG4;refGene_tx=XM_023254237.1;refGene_gn=FIG4;refGene_tx=XM_023254238.1
B2	101664927	101665028
refGene_gn=FIG4;refGene_tx=XM_023254237.1;refGene_gn=FIG4;refGene_tx=XM_023254238.1
B2	101666495	101666658
refGene_gn=FIG4;refGene_tx=XM_023254237.1;refGene_gn=FIG4;refGene_tx=XM_023254238.1
B2	101667095	101667193
refGene_gn=FIG4;refGene_tx=XM_023254237.1;refGene_gn=FIG4;refGene_tx=XM_023254238.1
B2	101671304	101671438
refGene_gn=FIG4;refGene_tx=XM_023254237.1;refGene_gn=FIG4;refGene_tx=XM_023254238.1
B2	101673361	101673478
refGene_gn=FIG4;refGene_tx=XM_023254237.1;refGene_gn=FIG4;refGene_tx=XM_023254238.1
B2	101675749	101675795
refGene_gn=FIG4;refGene_tx=XM_023254237.1;refGene_gn=FIG4;refGene_tx=XM_023254238.1
B2	101676460	101676609
refGene_gn=FIG4;refGene_tx=XM_023254237.1;refGene_gn=FIG4;refGene_tx=XM_023254238.1
B2	101677544	101677711
refGene_gn=FIG4;refGene_tx=XM_023254237.1;refGene_gn=FIG4;refGene_tx=XM_023254238.1
B2	101688351	101688490
refGene_gn=FIG4;refGene_tx=XM_023254237.1;refGene_gn=FIG4;refGene_tx=XM_023254238.1
B2	101697697	101697756
refGene_gn=FIG4;refGene_tx=XM_023254237.1;refGene_gn=FIG4;refGene_tx=XM_023254238.1
B2	101698956	101699104
refGene_gn=FIG4;refGene_tx=XM_023254237.1;refGene_gn=FIG4;refGene_tx=XM_023254238.1
B2	101702421	101702505
refGene_gn=FIG4;refGene_tx=XM_023254237.1;refGene_gn=FIG4;refGene_tx=XM_023254238.1
B2	101704242	101704438
refGene_gn=FIG4;refGene_tx=XM_023254237.1;refGene_gn=FIG4;refGene_tx=X

M_023254238.1
B2 101706734 101706817
refGene_gn=FIG4;refGene_tx=XM_023254237.1;refGene_gn=FIG4;refGene_tx=X
M_023254238.1
B2 101711163 101711250
refGene_gn=FIG4;refGene_tx=XM_023254237.1;refGene_gn=FIG4;refGene_tx=X
M_023254238.1
B2 101715775 101715815
refGene_gn=FIG4;refGene_tx=XM_023254238.1
B2 101752246 101752424
refGene_gn=FIG4;refGene_tx=XM_023254237.1
B2 101850254 101851343
refGene_gn=GPR6;refGene_tx=XM_023253555.1
B2 101961135 101961293
refGene_gn=WASF1;refGene_tx=XM_011282505.2;refGene_gn=WASF1;refGene_tx
=XM_019831067.2;refGene_gn=WASF1;refGene_tx=XM_019831068.2;refGene_gn=
WASF1;refGene_tx=XM_019831069.2;refGene_gn=WASF1;refGene_tx=XM_0198310
70.2
B2 101962381 101963010
refGene_gn=WASF1;refGene_tx=XM_011282505.2;refGene_gn=WASF1;refGene_tx
=XM_019831067.2;refGene_gn=WASF1;refGene_tx=XM_019831068.2;refGene_gn=
WASF1;refGene_tx=XM_019831069.2;refGene_gn=WASF1;refGene_tx=XM_0198310
70.2
B2 101964095 101964275
refGene_gn=WASF1;refGene_tx=XM_011282505.2;refGene_gn=WASF1;refGene_tx
=XM_019831067.2;refGene_gn=WASF1;refGene_tx=XM_019831068.2;refGene_gn=
WASF1;refGene_tx=XM_019831069.2;refGene_gn=WASF1;refGene_tx=XM_0198310
70.2
B2 101966112 101966285
refGene_gn=WASF1;refGene_tx=XM_011282505.2;refGene_gn=WASF1;refGene_tx
=XM_019831067.2;refGene_gn=WASF1;refGene_tx=XM_019831068.2;refGene_gn=
WASF1;refGene_tx=XM_019831069.2;refGene_gn=WASF1;refGene_tx=XM_0198310
70.2
B2 101967746 101967891
refGene_gn=WASF1;refGene_tx=XM_011282505.2;refGene_gn=WASF1;refGene_tx
=XM_019831067.2;refGene_gn=WASF1;refGene_tx=XM_019831068.2;refGene_gn=
WASF1;refGene_tx=XM_019831069.2;refGene_gn=WASF1;refGene_tx=XM_0198310
70.2
B2 101968929 101969083
refGene_gn=WASF1;refGene_tx=XM_011282505.2;refGene_gn=WASF1;refGene_tx
=XM_019831067.2;refGene_gn=WASF1;refGene_tx=XM_019831068.2;refGene_gn=
WASF1;refGene_tx=XM_019831069.2;refGene_gn=WASF1;refGene_tx=XM_0198310
70.2
B2 101973981 101974116
refGene_gn=WASF1;refGene_tx=XM_011282505.2;refGene_gn=WASF1;refGene_tx
=XM_019831067.2;refGene_gn=WASF1;refGene_tx=XM_019831068.2;refGene_gn=
WASF1;refGene_tx=XM_019831069.2
B2 101979574 101979707
refGene_gn=WASF1;refGene_tx=XM_011282505.2;refGene_gn=WASF1;refGene_tx
=XM_019831067.2;refGene_gn=WASF1;refGene_tx=XM_019831068.2;refGene_gn=

WASF1;refGene_tx=XM_019831069.2
B2	102033888	102034077
refGene_gn=CDC40;refGene_tx=XM_023254239.1;refGene_gn=CDC40;refGene_tx=XM_023254240.1
B2	102036499	102036526
refGene_gn=METTL24;refGene_tx=XM_023254245.1
B2	102042805	102042859
refGene_gn=METTL24;refGene_tx=XM_023254243.1
B2	102042917	102042956
refGene_gn=METTL24;refGene_tx=XM_023254244.1
B2	102047556	102047643
refGene_gn=CDC40;refGene_tx=XM_023254239.1;refGene_gn=CDC40;refGene_tx=XM_023254240.1
B2	102055342	102055472
refGene_gn=CDC40;refGene_tx=XM_023254239.1;refGene_gn=CDC40;refGene_tx=XM_023254240.1
B2	102066191	102066332
refGene_gn=METTL24;refGene_tx=XM_023254242.1
B2	102068743	102068827
refGene_gn=CDC40;refGene_tx=XM_023254239.1;refGene_gn=CDC40;refGene_tx=XM_023254240.1
B2	102070323	102070463
refGene_gn=CDC40;refGene_tx=XM_023254239.1;refGene_gn=CDC40;refGene_tx=XM_023254240.1
B2	102072074	102072171
refGene_gn=CDC40;refGene_tx=XM_023254239.1;refGene_gn=CDC40;refGene_tx=XM_023254240.1
B2	102074005	102074145
refGene_gn=CDC40;refGene_tx=XM_023254239.1;refGene_gn=CDC40;refGene_tx=XM_023254240.1
B2	102074777	102074852
refGene_gn=CDC40;refGene_tx=XM_023254239.1;refGene_gn=CDC40;refGene_tx=XM_023254240.1
B2	102077295	102077341
refGene_gn=CDC40;refGene_tx=XM_023254239.1
B2	102079173	102079219
refGene_gn=CDC40;refGene_tx=XM_023254240.1
B2	102081811	102081913
refGene_gn=CDC40;refGene_tx=XM_023254239.1;refGene_gn=CDC40;refGene_tx=XM_023254240.1
B2	102082867	102082983
refGene_gn=CDC40;refGene_tx=XM_023254239.1;refGene_gn=CDC40;refGene_tx=XM_023254240.1
B2	102083241	102083375
refGene_gn=CDC40;refGene_tx=XM_023254239.1;refGene_gn=CDC40;refGene_tx=XM_023254240.1
B2	102095172	102095249
refGene_gn=CDC40;refGene_tx=XM_023254239.1;refGene_gn=CDC40;refGene_tx=XM_023254240.1
B2	102096754	102096899

refGene_gn=CDC40;refGene_tx=XM_023254239.1;refGene_gn=CDC40;refGene_tx=XM_023254240.1
B2 102097928 102098106
refGene_gn=CDC40;refGene_tx=XM_023254239.1;refGene_gn=CDC40;refGene_tx=XM_023254240.1
B2 102108742 102109057
refGene_gn=METTL24;refGene_tx=XM_023254241.1;refGene_gn=METTL24;refGene_tx=XM_023254242.1;refGene_gn=METTL24;refGene_tx=XM_023254243.1;refGene_gn=METTL24;refGene_tx=XM_023254244.1;refGene_gn=METTL24;refGene_tx=XM_023254245.1;refGene_gn=METTL24;refGene_tx=XM_023254248.1
B2 102121657 102121753
refGene_gn=METTL24;refGene_tx=XM_023254247.1
B2 102154774 102154873
refGene_gn=METTL24;refGene_tx=XM_023254246.1
B2 102161492 102161721
refGene_gn=METTL24;refGene_tx=XM_023254241.1;refGene_gn=METTL24;refGene_tx=XM_023254242.1;refGene_gn=METTL24;refGene_tx=XM_023254243.1;refGene_gn=METTL24;refGene_tx=XM_023254244.1;refGene_gn=METTL24;refGene_tx=XM_023254245.1;refGene_gn=METTL24;refGene_tx=XM_023254246.1;refGene_gn=METTL24;refGene_tx=XM_023254247.1;refGene_gn=METTL24;refGene_tx=XM_023254248.1
B2 102170421 102170561
refGene_gn=METTL24;refGene_tx=XM_023254241.1;refGene_gn=METTL24;refGene_tx=XM_023254242.1;refGene_gn=METTL24;refGene_tx=XM_023254243.1;refGene_gn=METTL24;refGene_tx=XM_023254244.1;refGene_gn=METTL24;refGene_tx=XM_023254245.1;refGene_gn=METTL24;refGene_tx=XM_023254246.1;refGene_gn=METTL24;refGene_tx=XM_023254247.1
B2 102177837 102177936
refGene_gn=METTL24;refGene_tx=XM_023254241.1;refGene_gn=METTL24;refGene_tx=XM_023254242.1;refGene_gn=METTL24;refGene_tx=XM_023254243.1;refGene_gn=METTL24;refGene_tx=XM_023254244.1;refGene_gn=METTL24;refGene_tx=XM_023254245.1;refGene_gn=METTL24;refGene_tx=XM_023254246.1;refGene_gn=METTL24;refGene_tx=XM_023254247.1
B2 102210748 102211066
refGene_gn=METTL24;refGene_tx=XM_023254241.1;refGene_gn=METTL24;refGene_tx=XM_023254242.1;refGene_gn=METTL24;refGene_tx=XM_023254243.1;refGene_gn=METTL24;refGene_tx=XM_023254244.1;refGene_gn=METTL24;refGene_tx=XM_023254245.1;refGene_gn=METTL24;refGene_tx=XM_023254246.1;refGene_gn=METTL24;refGene_tx=XM_023254247.1
B2 102236102 102236673
refGene_gn=DDO;refGene_tx=XM_003986451.5;refGene_gn=DDO;refGene_tx=XM_019831071.2;refGene_gn=DDO;refGene_tx=XM_019831072.2;refGene_gn=DDO;refGene_tx=XM_019831073.2;refGene_gn=DDO;refGene_tx=XM_019831074.2
B2 102248747 102248924
refGene_gn=DDO;refGene_tx=XM_003986451.5;refGene_gn=DDO;refGene_tx=XM_019831071.2;refGene_gn=DDO;refGene_tx=XM_019831072.2;refGene_gn=DDO;refGene_tx=XM_019831073.2;refGene_gn=DDO;refGene_tx=XM_019831074.2
B2 102253185 102253294
refGene_gn=DDO;refGene_tx=XM_003986451.5;refGene_gn=DDO;refGene_tx=XM_019831071.2;refGene_gn=DDO;refGene_tx=XM_019831072.2;refGene_gn=DDO;re

fGene_tx=XM_019831073.2;refGene_gn=DDO;refGene_tx=XM_019831074.2
B2 102256633 102256809
refGene_gn=DDO;refGene_tx=XM_003986451.5;refGene_gn=DDO;refGene_tx=XM_019831071.2;refGene_gn=DDO;refGene_tx=XM_019831072.2;refGene_gn=DDO;refGene_tx=XM_019831073.2;refGene_gn=DDO;refGene_tx=XM_019831074.2
B2 102258628 102258712
refGene_gn=DDO;refGene_tx=XM_003986451.5;refGene_gn=DDO;refGene_tx=XM_019831071.2
B2 102262756 102262873
refGene_gn=DDO;refGene_tx=XM_003986451.5;refGene_gn=DDO;refGene_tx=XM_019831071.2;refGene_gn=DDO;refGene_tx=XM_019831072.2;refGene_gn=DDO;refGene_tx=XM_019831074.2
B2 102265575 102265785
refGene_gn=SLC22A16;refGene_tx=XM_011282527.3;refGene_gn=SLC22A16;refGene_tx=XM_019831076.2;refGene_gn=SLC22A16;refGene_tx=XM_019831077.2;refGene_gn=SLC22A16;refGene_tx=XM_019831081.2;refGene_gn=SLC22A16;refGene_tx=XM_019831083.2;refGene_gn=SLC22A16;refGene_tx=XM_023254249.1;refGene_gn=SLC22A16;refGene_tx=XM_023254250.1
B2 102269412 102269477
refGene_gn=SLC22A16;refGene_tx=XM_019831078.2;refGene_gn=SLC22A16;refGene_tx=XM_019831080.2
B2 102272088 102272188
refGene_gn=SLC22A16;refGene_tx=XM_011282527.3;refGene_gn=SLC22A16;refGene_tx=XM_019831076.2;refGene_gn=SLC22A16;refGene_tx=XM_019831077.2;refGene_gn=SLC22A16;refGene_tx=XM_019831078.2;refGene_gn=SLC22A16;refGene_tx=XM_019831080.2;refGene_gn=SLC22A16;refGene_tx=XM_019831081.2;refGene_gn=SLC22A16;refGene_tx=XM_023254249.1;refGene_gn=SLC22A16;refGene_tx=XM_023254250.1
B2 102273865 102273975
refGene_gn=SLC22A16;refGene_tx=XM_011282527.3;refGene_gn=SLC22A16;refGene_tx=XM_019831076.2;refGene_gn=SLC22A16;refGene_tx=XM_019831077.2;refGene_gn=SLC22A16;refGene_tx=XM_019831078.2;refGene_gn=SLC22A16;refGene_tx=XM_019831080.2;refGene_gn=SLC22A16;refGene_tx=XM_019831081.2;refGene_gn=SLC22A16;refGene_tx=XM_019831083.2;refGene_gn=SLC22A16;refGene_tx=XM_023254249.1;refGene_gn=SLC22A16;refGene_tx=XM_023254250.1
B2 102276434 102276562
refGene_gn=SLC22A16;refGene_tx=XM_011282527.3;refGene_gn=SLC22A16;refGene_tx=XM_019831076.2;refGene_gn=SLC22A16;refGene_tx=XM_019831077.2;refGene_gn=SLC22A16;refGene_tx=XM_019831078.2;refGene_gn=SLC22A16;refGene_tx=XM_019831080.2;refGene_gn=SLC22A16;refGene_tx=XM_019831081.2;refGene_gn=SLC22A16;refGene_tx=XM_019831083.2;refGene_gn=SLC22A16;refGene_tx=XM_023254249.1;refGene_gn=SLC22A16;refGene_tx=XM_023254250.1
B2 102280035 102280567
refGene_gn=SLC22A16;refGene_tx=XM_011282527.3;refGene_gn=SLC22A16;refGene_tx=XM_019831076.2;refGene_gn=SLC22A16;refGene_tx=XM_019831077.2;refGene_gn=SLC22A16;refGene_tx=XM_019831078.2;refGene_gn=SLC22A16;refGene_tx=XM_019831080.2;refGene_gn=SLC22A16;refGene_tx=XM_019831081.2;refGene_gn=SLC22A16;refGene_tx=XM_019831083.2;refGene_gn=SLC22A16;refGene_tx=XM_023254249.1;refGene_gn=SLC22A16;refGene_tx=XM_023254250.1
B2 102284976 102285094

refGene_gn=SLC22A16;refGene_tx=XM_011282527.3;refGene_gn=SLC22A16;refGene_tx=XM_019831076.2;refGene_gn=SLC22A16;refGene_tx=XM_019831077.2;refGene_gn=SLC22A16;refGene_tx=XM_019831078.2;refGene_gn=SLC22A16;refGene_tx=XM_019831080.2;refGene_gn=SLC22A16;refGene_tx=XM_019831081.2;refGene_gn=SLC22A16;refGene_tx=XM_019831083.2;refGene_gn=SLC22A16;refGene_tx=XM_023254249.1;refGene_gn=SLC22A16;refGene_tx=XM_023254250.1
B2 102293737 102294217
refGene_gn=SLC22A16;refGene_tx=XM_011282527.3;refGene_gn=SLC22A16;refGene_tx=XM_019831076.2;refGene_gn=SLC22A16;refGene_tx=XM_019831077.2;refGene_gn=SLC22A16;refGene_tx=XM_019831078.2;refGene_gn=SLC22A16;refGene_tx=XM_019831080.2;refGene_gn=SLC22A16;refGene_tx=XM_019831081.2;refGene_gn=SLC22A16;refGene_tx=XM_019831083.2;refGene_gn=SLC22A16;refGene_tx=XM_023254249.1;refGene_gn=SLC22A16;refGene_tx=XM_023254250.1
B2 102299117 102299125
refGene_gn=SLC22A16;refGene_tx=XM_019831076.2
B2 102308447 102308500
refGene_gn=SLC22A16;refGene_tx=XM_023254249.1
B2 102308830 102308946
refGene_gn=SLC22A16;refGene_tx=XM_011282527.3;refGene_gn=SLC22A16;refGene_tx=XM_019831077.2;refGene_gn=SLC22A16;refGene_tx=XM_019831078.2;refGene_gn=SLC22A16;refGene_tx=XM_019831080.2;refGene_gn=SLC22A16;refGene_tx=XM_019831083.2
B2 102388651 102388783
refGene_gn=CDK19;refGene_tx=XM_023254251.1;refGene_gn=CDK19;refGene_tx=XM_023254252.1;refGene_gn=CDK19;refGene_tx=XM_023254253.1;refGene_gn=CDK19;refGene_tx=XM_023254254.1;refGene_gn=CDK19;refGene_tx=XM_023254255.1;refGene_gn=CDK19;refGene_tx=XM_023254256.1;refGene_gn=CDK19;refGene_tx=XM_023254257.1;refGene_gn=CDK19;refGene_tx=XM_023254258.1;refGene_gn=CDK19;refGene_tx=XM_023254259.1
B2 102399539 102399806
refGene_gn=CDK19;refGene_tx=XM_023254251.1;refGene_gn=CDK19;refGene_tx=XM_023254252.1;refGene_gn=CDK19;refGene_tx=XM_023254253.1;refGene_gn=CDK19;refGene_tx=XM_023254254.1;refGene_gn=CDK19;refGene_tx=XM_023254255.1;refGene_gn=CDK19;refGene_tx=XM_023254256.1;refGene_gn=CDK19;refGene_tx=XM_023254257.1;refGene_gn=CDK19;refGene_tx=XM_023254258.1;refGene_gn=CDK19;refGene_tx=XM_023254259.1
B2 102400732 102400811
refGene_gn=CDK19;refGene_tx=XM_023254251.1;refGene_gn=CDK19;refGene_tx=XM_023254252.1;refGene_gn=CDK19;refGene_tx=XM_023254253.1;refGene_gn=CDK19;refGene_tx=XM_023254254.1;refGene_gn=CDK19;refGene_tx=XM_023254255.1;refGene_gn=CDK19;refGene_tx=XM_023254256.1;refGene_gn=CDK19;refGene_tx=XM_023254257.1;refGene_gn=CDK19;refGene_tx=XM_023254258.1;refGene_gn=CDK19;refGene_tx=XM_023254259.1
B2 102402043 102402141
refGene_gn=CDK19;refGene_tx=XM_023254251.1;refGene_gn=CDK19;refGene_tx=XM_023254252.1;refGene_gn=CDK19;refGene_tx=XM_023254253.1;refGene_gn=CDK19;refGene_tx=XM_023254254.1;refGene_gn=CDK19;refGene_tx=XM_023254255.1;refGene_gn=CDK19;refGene_tx=XM_023254256.1;refGene_gn=CDK19;refGene_tx=XM_023254257.1;refGene_gn=CDK19;refGene_tx=XM_023254258.1;refGene_gn=CDK19;refGene_tx=XM_023254259.1

B2 102403217 102403290
refGene_gn=CDK19;refGene_tx=XM_023254251.1;refGene_gn=CDK19;refGene_tx=XM_023254252.1;refGene_gn=CDK19;refGene_tx=XM_023254253.1;refGene_gn=CDK19;refGene_tx=XM_023254254.1;refGene_gn=CDK19;refGene_tx=XM_023254255.1;refGene_gn=CDK19;refGene_tx=XM_023254256.1;refGene_gn=CDK19;refGene_tx=XM_023254257.1;refGene_gn=CDK19;refGene_tx=XM_023254258.1;refGene_gn=CDK19;refGene_tx=XM_023254259.1
B2 102408456 102408526
refGene_gn=CDK19;refGene_tx=XM_023254251.1;refGene_gn=CDK19;refGene_tx=XM_023254252.1;refGene_gn=CDK19;refGene_tx=XM_023254253.1;refGene_gn=CDK19;refGene_tx=XM_023254254.1;refGene_gn=CDK19;refGene_tx=XM_023254255.1;refGene_gn=CDK19;refGene_tx=XM_023254256.1;refGene_gn=CDK19;refGene_tx=XM_023254257.1;refGene_gn=CDK19;refGene_tx=XM_023254258.1;refGene_gn=CDK19;refGene_tx=XM_023254259.1
B2 102408688 102408832
refGene_gn=CDK19;refGene_tx=XM_023254251.1;refGene_gn=CDK19;refGene_tx=XM_023254252.1;refGene_gn=CDK19;refGene_tx=XM_023254253.1;refGene_gn=CDK19;refGene_tx=XM_023254254.1;refGene_gn=CDK19;refGene_tx=XM_023254255.1;refGene_gn=CDK19;refGene_tx=XM_023254256.1;refGene_gn=CDK19;refGene_tx=XM_023254257.1;refGene_gn=CDK19;refGene_tx=XM_023254258.1;refGene_gn=CDK19;refGene_tx=XM_023254259.1
B2 102413219 102413351
refGene_gn=CDK19;refGene_tx=XM_023254251.1;refGene_gn=CDK19;refGene_tx=XM_023254252.1;refGene_gn=CDK19;refGene_tx=XM_023254253.1;refGene_gn=CDK19;refGene_tx=XM_023254254.1;refGene_gn=CDK19;refGene_tx=XM_023254255.1;refGene_gn=CDK19;refGene_tx=XM_023254256.1;refGene_gn=CDK19;refGene_tx=XM_023254257.1;refGene_gn=CDK19;refGene_tx=XM_023254258.1;refGene_gn=CDK19;refGene_tx=XM_023254259.1
B2 102418386 102418444
refGene_gn=CDK19;refGene_tx=XM_023254251.1;refGene_gn=CDK19;refGene_tx=XM_023254252.1;refGene_gn=CDK19;refGene_tx=XM_023254253.1;refGene_gn=CDK19;refGene_tx=XM_023254254.1;refGene_gn=CDK19;refGene_tx=XM_023254255.1;refGene_gn=CDK19;refGene_tx=XM_023254256.1;refGene_gn=CDK19;refGene_tx=XM_023254257.1;refGene_gn=CDK19;refGene_tx=XM_023254258.1;refGene_gn=CDK19;refGene_tx=XM_023254259.1
B2 102442619 102442760
refGene_gn=CDK19;refGene_tx=XM_023254251.1;refGene_gn=CDK19;refGene_tx=XM_023254252.1;refGene_gn=CDK19;refGene_tx=XM_023254253.1;refGene_gn=CDK19;refGene_tx=XM_023254254.1;refGene_gn=CDK19;refGene_tx=XM_023254255.1;refGene_gn=CDK19;refGene_tx=XM_023254256.1;refGene_gn=CDK19;refGene_tx=XM_023254257.1;refGene_gn=CDK19;refGene_tx=XM_023254258.1;refGene_gn=CDK19;refGene_tx=XM_023254259.1
B2 102445312 102445423
refGene_gn=CDK19;refGene_tx=XM_023254251.1;refGene_gn=CDK19;refGene_tx=XM_023254252.1;refGene_gn=CDK19;refGene_tx=XM_023254253.1;refGene_gn=CDK19;refGene_tx=XM_023254254.1;refGene_gn=CDK19;refGene_tx=XM_023254255.1;refGene_gn=CDK19;refGene_tx=XM_023254256.1;refGene_gn=CDK19;refGene_tx=XM_023254257.1;refGene_gn=CDK19;refGene_tx=XM_023254258.1;refGene_gn=CDK19;refGene_tx=XM_023254259.1
B2 102505411 102505487

refGene_gn=CDK19;refGene_tx=XM_023254251.1;refGene_gn=CDK19;refGene_tx
=XM_023254252.1;refGene_gn=CDK19;refGene_tx=XM_023254253.1;refGene_gn=
CDK19;refGene_tx=XM_023254254.1;refGene_gn=CDK19;refGene_tx=XM_0232542
55.1;refGene_gn=CDK19;refGene_tx=XM_023254256.1;refGene_gn=CDK19;refGe
ne_tx=XM_023254257.1;refGene_gn=CDK19;refGene_tx=XM_023254258.1;refGen
e_gn=CDK19;refGene_tx=XM_023254259.1
B2 102522308 102522377
refGene_gn=CDK19;refGene_tx=XM_023254251.1;refGene_gn=CDK19;refGene_tx
=XM_023254253.1
B2 102531727 102531789
refGene_gn=CDK19;refGene_tx=XM_023254251.1;refGene_gn=CDK19;refGene_tx
=XM_023254252.1
B2 102577129 102577224
refGene_gn=CDK19;refGene_tx=XM_023254251.1;refGene_gn=CDK19;refGene_tx
=XM_023254252.1
B2 102578152 102578280
refGene_gn=CDK19;refGene_tx=XM_023254251.1;refGene_gn=CDK19;refGene_tx
=XM_023254252.1;refGene_gn=CDK19;refGene_tx=XM_023254253.1;refGene_gn=
CDK19;refGene_tx=XM_023254254.1
B2 102631301 102631411
refGene_gn=AMD1;refGene_tx=XM_023254260.1
B2 102647664 102647751
refGene_gn=AMD1;refGene_tx=XM_023254260.1
B2 102648897 102649024
refGene_gn=AMD1;refGene_tx=XM_023254260.1
B2 102649594 102649697
refGene_gn=AMD1;refGene_tx=XM_023254260.1
B2 102650796 102650839
refGene_gn=AMD1;refGene_tx=XM_023254260.1
B2 102650975 102651120
refGene_gn=AMD1;refGene_tx=XM_023254260.1
B2 102651460 102651553
refGene_gn=AMD1;refGene_tx=XM_023254260.1
B2 102651629 102651785
refGene_gn=AMD1;refGene_tx=XM_023254260.1
B2 102652191 102652332
refGene_gn=AMD1;refGene_tx=XM_023254260.1
B2 102703460 102703574
refGene_gn=GTF3C6;refGene_tx=XM_023254261.1;refGene_gn=GTF3C6;refGene_
tx=XM_023254262.1
B2 102703890 102703971
refGene_gn=GTF3C6;refGene_tx=XM_023254261.1;refGene_gn=GTF3C6;refGene_
tx=XM_023254262.1
B2 102705106 102705170
refGene_gn=GTF3C6;refGene_tx=XM_023254261.1;refGene_gn=GTF3C6;refGene_
tx=XM_023254262.1
B2 102705246 102705312
refGene_gn=GTF3C6;refGene_tx=XM_023254261.1;refGene_gn=GTF3C6;refGene_
tx=XM_023254262.1
B2 102707378 102707492

refGene_gn=GTF3C6;refGene_tx=XM_023254261.1;refGene_gn=GTF3C6;refGene_
tx=XM_023254262.1
B2 102712987 102713286
refGene_gn=GTF3C6;refGene_tx=XM_023254261.1;refGene_gn=GTF3C6;refGene_
tx=XM_023254262.1
B2 102727589 102727612
refGene_gn=RPF2;refGene_tx=XM_003986464.5
B2 102729582 102729715
refGene_gn=RPF2;refGene_tx=XM_003986464.5;refGene_gn=RPF2;refGene_tx=X
M_019831086.2
B2 102732931 102732969
refGene_gn=RPF2;refGene_tx=XM_003986464.5;refGene_gn=RPF2;refGene_tx=X
M_019831086.2
B2 102739480 102739520
refGene_gn=RPF2;refGene_tx=XM_003986464.5;refGene_gn=RPF2;refGene_tx=X
M_019831086.2
B2 102740182 102740264
refGene_gn=RPF2;refGene_tx=XM_003986464.5;refGene_gn=RPF2;refGene_tx=X
M_019831086.2
B2 102740878 102740955
refGene_gn=RPF2;refGene_tx=XM_003986464.5;refGene_gn=RPF2;refGene_tx=X
M_019831086.2
B2 102745539 102745639
refGene_gn=RPF2;refGene_tx=XM_003986464.5;refGene_gn=RPF2;refGene_tx=X
M_019831086.2
B2 102754483 102754586
refGene_gn=RPF2;refGene_tx=XM_003986464.5;refGene_gn=RPF2;refGene_tx=X
M_019831086.2
B2 102760282 102760427
refGene_gn=RPF2;refGene_tx=XM_003986464.5;refGene_gn=RPF2;refGene_tx=X
M_019831086.2
B2 102761210 102761390
refGene_gn=RPF2;refGene_tx=XM_003986464.5;refGene_gn=RPF2;refGene_tx=X
M_019831086.2
B2 102829063 102829363
refGene_gn=SLC16A10;refGene_tx=XM_011282525.3
B2 102829364 102829401
refGene_gn=SLC16A10;refGene_tx=XM_011282525.3
B2 102901545 102901690
refGene_gn=SLC16A10;refGene_tx=XM_011282525.3
B2 102903876 102904342
refGene_gn=SLC16A10;refGene_tx=XM_011282525.3
B2 102931018 102931162
refGene_gn=SLC16A10;refGene_tx=XM_011282525.3
B2 102939136 102939365
refGene_gn=SLC16A10;refGene_tx=XM_011282525.3
B2 102942152 102942385
refGene_gn=SLC16A10;refGene_tx=XM_011282525.3
B2 102971986 102972037
refGene_gn=MFSD4B;refGene_tx=XM_003986474.5

B2 102973650 102973819
refGene_gn=MFSD4B;refGene_tx=XM_003986474.5
B2 102975359 102975476
refGene_gn=MFSD4B;refGene_tx=XM_003986474.5
B2 102976679 102977887
refGene_gn=MFSD4B;refGene_tx=XM_003986474.5
B2 102977888 102977924
refGene_gn=MFSD4B;refGene_tx=XM_003986474.5
B2 103015930 103016071
refGene_gn=REV3L;refGene_tx=XM_019831087.2;refGene_gn=REV3L;refGene_tx
=XM_023254263.1
B2 103019596 103019806
refGene_gn=REV3L;refGene_tx=XM_019831087.2;refGene_gn=REV3L;refGene_tx
=XM_023254263.1
B2 103021416 103021663
refGene_gn=REV3L;refGene_tx=XM_019831087.2;refGene_gn=REV3L;refGene_tx
=XM_023254263.1
B2 103022546 103022737
refGene_gn=REV3L;refGene_tx=XM_019831087.2;refGene_gn=REV3L;refGene_tx
=XM_023254263.1
B2 103023665 103023803
refGene_gn=REV3L;refGene_tx=XM_019831087.2;refGene_gn=REV3L;refGene_tx
=XM_023254263.1
B2 103028074 103028189
refGene_gn=REV3L;refGene_tx=XM_019831087.2;refGene_gn=REV3L;refGene_tx
=XM_023254263.1
B2 103032742 103032852
refGene_gn=REV3L;refGene_tx=XM_019831087.2;refGene_gn=REV3L;refGene_tx
=XM_023254263.1
B2 103041694 103041901
refGene_gn=REV3L;refGene_tx=XM_019831087.2;refGene_gn=REV3L;refGene_tx
=XM_023254263.1
B2 103043721 103043830
refGene_gn=REV3L;refGene_tx=XM_019831087.2;refGene_gn=REV3L;refGene_tx
=XM_023254263.1
B2 103045062 103045307
refGene_gn=REV3L;refGene_tx=XM_019831087.2;refGene_gn=REV3L;refGene_tx
=XM_023254263.1;refGene_gn=REV3L;refGene_tx=XM_023254264.1
B2 103047213 103047355
refGene_gn=REV3L;refGene_tx=XM_019831087.2;refGene_gn=REV3L;refGene_tx
=XM_023254263.1
B2 103056192 103056311
refGene_gn=REV3L;refGene_tx=XM_019831087.2;refGene_gn=REV3L;refGene_tx
=XM_023254263.1;refGene_gn=REV3L;refGene_tx=XM_023254264.1
B2 103063227 103063346
refGene_gn=REV3L;refGene_tx=XM_019831087.2;refGene_gn=REV3L;refGene_tx
=XM_023254263.1;refGene_gn=REV3L;refGene_tx=XM_023254264.1
B2 103066026 103066142
refGene_gn=REV3L;refGene_tx=XM_019831087.2;refGene_gn=REV3L;refGene_tx
=XM_023254263.1;refGene_gn=REV3L;refGene_tx=XM_023254264.1

B2	103072194	103072306
refGene_gn=REV3L;refGene_tx=XM_019831087.2;refGene_gn=REV3L;refGene_tx=XM_023254263.1;refGene_gn=REV3L;refGene_tx=XM_023254264.1
B2	103073768	103073961
refGene_gn=REV3L;refGene_tx=XM_019831087.2;refGene_gn=REV3L;refGene_tx=XM_023254263.1;refGene_gn=REV3L;refGene_tx=XM_023254264.1
B2	103078780	103078906
refGene_gn=REV3L;refGene_tx=XM_019831087.2;refGene_gn=REV3L;refGene_tx=XM_023254263.1;refGene_gn=REV3L;refGene_tx=XM_023254264.1
B2	103080217	103080297
refGene_gn=REV3L;refGene_tx=XM_019831087.2;refGene_gn=REV3L;refGene_tx=XM_023254263.1;refGene_gn=REV3L;refGene_tx=XM_023254264.1
B2	103082062	103082976
refGene_gn=REV3L;refGene_tx=XM_019831087.2;refGene_gn=REV3L;refGene_tx=XM_023254263.1;refGene_gn=REV3L;refGene_tx=XM_023254264.1
B2	103086821	103090986
refGene_gn=REV3L;refGene_tx=XM_019831087.2;refGene_gn=REV3L;refGene_tx=XM_023254263.1;refGene_gn=REV3L;refGene_tx=XM_023254264.1
B2	103091655	103091798
refGene_gn=REV3L;refGene_tx=XM_019831087.2;refGene_gn=REV3L;refGene_tx=XM_023254263.1;refGene_gn=REV3L;refGene_tx=XM_023254264.1
B2	103097558	103097796
refGene_gn=REV3L;refGene_tx=XM_019831087.2;refGene_gn=REV3L;refGene_tx=XM_023254263.1;refGene_gn=REV3L;refGene_tx=XM_023254264.1
B2	103100884	103101004
refGene_gn=REV3L;refGene_tx=XM_019831087.2;refGene_gn=REV3L;refGene_tx=XM_023254263.1;refGene_gn=REV3L;refGene_tx=XM_023254264.1
B2	103110726	103110875
refGene_gn=REV3L;refGene_tx=XM_019831087.2;refGene_gn=REV3L;refGene_tx=XM_023254263.1;refGene_gn=REV3L;refGene_tx=XM_023254264.1
B2	103110962	103111047
refGene_gn=REV3L;refGene_tx=XM_019831087.2;refGene_gn=REV3L;refGene_tx=XM_023254263.1;refGene_gn=REV3L;refGene_tx=XM_023254264.1
B2	103112016	103112121
refGene_gn=REV3L;refGene_tx=XM_019831087.2;refGene_gn=REV3L;refGene_tx=XM_023254263.1;refGene_gn=REV3L;refGene_tx=XM_023254264.1
B2	103112880	103112975
refGene_gn=REV3L;refGene_tx=XM_019831087.2;refGene_gn=REV3L;refGene_tx=XM_023254263.1;refGene_gn=REV3L;refGene_tx=XM_023254264.1
B2	103117723	103117820
refGene_gn=REV3L;refGene_tx=XM_019831087.2;refGene_gn=REV3L;refGene_tx=XM_023254263.1;refGene_gn=REV3L;refGene_tx=XM_023254264.1
B2	103123202	103123363
refGene_gn=REV3L;refGene_tx=XM_019831087.2;refGene_gn=REV3L;refGene_tx=XM_023254263.1;refGene_gn=REV3L;refGene_tx=XM_023254264.1
B2	103128386	103128461
refGene_gn=REV3L;refGene_tx=XM_019831087.2;refGene_gn=REV3L;refGene_tx=XM_023254263.1;refGene_gn=REV3L;refGene_tx=XM_023254264.1
B2	103136084	103136274
refGene_gn=REV3L;refGene_tx=XM_019831087.2;refGene_gn=REV3L;refGene_tx

=XM_023254263.1;refGene_gn=REV3L;refGene_tx=XM_023254264.1
B2	103192837	103192976
refGene_gn=REV3L;refGene_tx=XM_023254263.1;refGene_gn=REV3L;refGene_tx=XM_023254264.1
B2	103273881	103274028
refGene_gn=TRAF3IP2;refGene_tx=XM_003986465.5;refGene_gn=TRAF3IP2;refGene_tx=XM_006931950.2;refGene_gn=TRAF3IP2;refGene_tx=XM_019831088.1;refGene_gn=TRAF3IP2;refGene_tx=XM_019831089.2;refGene_gn=TRAF3IP2;refGene_tx=XM_019831090.1
B2	103276697	103276772
refGene_gn=TRAF3IP2;refGene_tx=XM_003986465.5;refGene_gn=TRAF3IP2;refGene_tx=XM_006931950.2;refGene_gn=TRAF3IP2;refGene_tx=XM_019831088.1;refGene_gn=TRAF3IP2;refGene_tx=XM_019831089.2;refGene_gn=TRAF3IP2;refGene_tx=XM_019831090.1
B2	103280187	103280304
refGene_gn=TRAF3IP2;refGene_tx=XM_003986465.5;refGene_gn=TRAF3IP2;refGene_tx=XM_006931950.2;refGene_gn=TRAF3IP2;refGene_tx=XM_019831088.1;refGene_gn=TRAF3IP2;refGene_tx=XM_019831089.2;refGene_gn=TRAF3IP2;refGene_tx=XM_019831090.1
B2	103281421	103281490
refGene_gn=TRAF3IP2;refGene_tx=XM_003986465.5;refGene_gn=TRAF3IP2;refGene_tx=XM_006931950.2;refGene_gn=TRAF3IP2;refGene_tx=XM_019831088.1;refGene_gn=TRAF3IP2;refGene_tx=XM_019831089.2
B2	103285932	103286021
refGene_gn=TRAF3IP2;refGene_tx=XM_003986465.5;refGene_gn=TRAF3IP2;refGene_tx=XM_019831088.1
B2	103288063	103288242
refGene_gn=TRAF3IP2;refGene_tx=XM_003986465.5;refGene_gn=TRAF3IP2;refGene_tx=XM_019831088.1
B2	103294627	103294805
refGene_gn=TRAF3IP2;refGene_tx=XM_003986465.5;refGene_gn=TRAF3IP2;refGene_tx=XM_019831088.1
B2	103304700	103305523
refGene_gn=TRAF3IP2;refGene_tx=XM_003986465.5;refGene_gn=TRAF3IP2;refGene_tx=XM_019831088.1
B2	103364501	103364710
refGene_gn=FYN;refGene_tx=XM_003986468.5;refGene_gn=FYN;refGene_tx=XM_019831092.2;refGene_gn=FYN;refGene_tx=XM_019831093.2;refGene_gn=FYN;refGene_tx=XM_019831094.2;refGene_gn=FYN;refGene_tx=XM_019831095.2;refGene_gn=FYN;refGene_tx=XM_019831096.2;refGene_gn=FYN;refGene_tx=XM_023254265.1;refGene_gn=FYN;refGene_tx=XM_023254266.1;refGene_gn=FYN;refGene_tx=XM_023254267.1
B2	103376158	103376290
refGene_gn=FYN;refGene_tx=XM_003986468.5;refGene_gn=FYN;refGene_tx=XM_019831092.2;refGene_gn=FYN;refGene_tx=XM_019831093.2;refGene_gn=FYN;refGene_tx=XM_019831094.2;refGene_gn=FYN;refGene_tx=XM_019831095.2;refGene_gn=FYN;refGene_tx=XM_019831096.2;refGene_gn=FYN;refGene_tx=XM_023254265.1;refGene_gn=FYN;refGene_tx=XM_023254266.1;refGene_gn=FYN;refGene_tx=XM_023254267.1
B2	103396227	103396381

refGene_gn=FYN;refGene_tx=XM_003986468.5;refGene_gn=FYN;refGene_tx=XM_019831092.2;refGene_gn=FYN;refGene_tx=XM_019831093.2;refGene_gn=FYN;refGene_tx=XM_019831094.2;refGene_gn=FYN;refGene_tx=XM_019831095.2;refGene_gn=FYN;refGene_tx=XM_019831096.2;refGene_gn=FYN;refGene_tx=XM_023254265.1;refGene_gn=FYN;refGene_tx=XM_023254266.1;refGene_gn=FYN;refGene_tx=XM_023254267.1
B2	103396474	103396551
refGene_gn=FYN;refGene_tx=XM_003986468.5;refGene_gn=FYN;refGene_tx=XM_019831092.2;refGene_gn=FYN;refGene_tx=XM_019831093.2;refGene_gn=FYN;refGene_tx=XM_019831094.2;refGene_gn=FYN;refGene_tx=XM_019831095.2;refGene_gn=FYN;refGene_tx=XM_019831096.2;refGene_gn=FYN;refGene_tx=XM_023254265.1;refGene_gn=FYN;refGene_tx=XM_023254266.1;refGene_gn=FYN;refGene_tx=XM_023254267.1
B2	103398750	103398930
refGene_gn=FYN;refGene_tx=XM_003986468.5;refGene_gn=FYN;refGene_tx=XM_019831092.2;refGene_gn=FYN;refGene_tx=XM_019831093.2;refGene_gn=FYN;refGene_tx=XM_019831094.2;refGene_gn=FYN;refGene_tx=XM_019831095.2;refGene_gn=FYN;refGene_tx=XM_019831096.2;refGene_gn=FYN;refGene_tx=XM_023254265.1;refGene_gn=FYN;refGene_tx=XM_023254266.1;refGene_gn=FYN;refGene_tx=XM_023254267.1
B2	103401885	103402041
refGene_gn=FYN;refGene_tx=XM_023254267.1
B2	103402461	103402626
refGene_gn=FYN;refGene_tx=XM_003986468.5;refGene_gn=FYN;refGene_tx=XM_019831092.2;refGene_gn=FYN;refGene_tx=XM_019831093.2;refGene_gn=FYN;refGene_tx=XM_019831094.2;refGene_gn=FYN;refGene_tx=XM_019831095.2;refGene_gn=FYN;refGene_tx=XM_019831096.2;refGene_gn=FYN;refGene_tx=XM_023254265.1;refGene_gn=FYN;refGene_tx=XM_023254266.1
B2	103405235	103405385
refGene_gn=FYN;refGene_tx=XM_003986468.5;refGene_gn=FYN;refGene_tx=XM_019831092.2;refGene_gn=FYN;refGene_tx=XM_019831093.2;refGene_gn=FYN;refGene_tx=XM_019831094.2;refGene_gn=FYN;refGene_tx=XM_019831095.2;refGene_gn=FYN;refGene_tx=XM_019831096.2;refGene_gn=FYN;refGene_tx=XM_023254265.1;refGene_gn=FYN;refGene_tx=XM_023254266.1;refGene_gn=FYN;refGene_tx=XM_023254267.1
B2	103406359	103406463
refGene_gn=FYN;refGene_tx=XM_003986468.5;refGene_gn=FYN;refGene_tx=XM_019831092.2;refGene_gn=FYN;refGene_tx=XM_019831093.2;refGene_gn=FYN;refGene_tx=XM_019831094.2;refGene_gn=FYN;refGene_tx=XM_019831095.2;refGene_gn=FYN;refGene_tx=XM_019831096.2;refGene_gn=FYN;refGene_tx=XM_023254265.1;refGene_gn=FYN;refGene_tx=XM_023254266.1;refGene_gn=FYN;refGene_tx=XM_023254267.1
B2	103409197	103409296
refGene_gn=FYN;refGene_tx=XM_003986468.5;refGene_gn=FYN;refGene_tx=XM_019831092.2;refGene_gn=FYN;refGene_tx=XM_019831093.2;refGene_gn=FYN;refGene_tx=XM_019831094.2;refGene_gn=FYN;refGene_tx=XM_019831095.2;refGene_gn=FYN;refGene_tx=XM_019831096.2;refGene_gn=FYN;refGene_tx=XM_023254265.1;refGene_gn=FYN;refGene_tx=XM_023254266.1;refGene_gn=FYN;refGene_tx=XM_023254267.1
B2	103418796	103418893

refGene_gn=FYN;refGene_tx=XM_003986468.5;refGene_gn=FYN;refGene_tx=XM_019831092.2;refGene_gn=FYN;refGene_tx=XM_019831093.2;refGene_gn=FYN;refGene_tx=XM_019831094.2;refGene_gn=FYN;refGene_tx=XM_019831095.2;refGene_gn=FYN;refGene_tx=XM_019831096.2;refGene_gn=FYN;refGene_tx=XM_023254265.1;refGene_gn=FYN;refGene_tx=XM_023254266.1;refGene_gn=FYN;refGene_tx=XM_023254267.1
B2 103422805 103423052
refGene_gn=FYN;refGene_tx=XM_003986468.5;refGene_gn=FYN;refGene_tx=XM_019831092.2;refGene_gn=FYN;refGene_tx=XM_019831093.2;refGene_gn=FYN;refGene_tx=XM_019831094.2;refGene_gn=FYN;refGene_tx=XM_019831095.2;refGene_gn=FYN;refGene_tx=XM_019831096.2;refGene_gn=FYN;refGene_tx=XM_023254265.1;refGene_gn=FYN;refGene_tx=XM_023254266.1;refGene_gn=FYN;refGene_tx=XM_023254267.1
B2 103743063 103743111
refGene_gn=WISP3;refGene_tx=XM_003986469.5
B2 103750263 103750567
refGene_gn=WISP3;refGene_tx=XM_003986469.5;refGene_gn=WISP3;refGene_tx=XM_023254271.1
B2 103754440 103754683
refGene_gn=WISP3;refGene_tx=XM_003986469.5;refGene_gn=WISP3;refGene_tx=XM_023254271.1
B2 103758038 103758232
refGene_gn=WISP3;refGene_tx=XM_003986469.5;refGene_gn=WISP3;refGene_tx=XM_023254271.1
B2 103759298 103759580
refGene_gn=WISP3;refGene_tx=XM_003986469.5;refGene_gn=WISP3;refGene_tx=XM_023254271.1
B2 103761280 103761439
refGene_gn=TUBE1;refGene_tx=XM_011282648.3;refGene_gn=TUBE1;refGene_tx=XM_023254268.1;refGene_gn=TUBE1;refGene_tx=XM_023254269.1;refGene_gn=TUBE1;refGene_tx=XM_023254270.1
B2 103761968 103762143
refGene_gn=TUBE1;refGene_tx=XM_011282648.3;refGene_gn=TUBE1;refGene_tx=XM_023254268.1;refGene_gn=TUBE1;refGene_tx=XM_023254269.1;refGene_gn=TUBE1;refGene_tx=XM_023254270.1
B2 103762974 103763115
refGene_gn=TUBE1;refGene_tx=XM_011282648.3;refGene_gn=TUBE1;refGene_tx=XM_023254268.1;refGene_gn=TUBE1;refGene_tx=XM_023254269.1;refGene_gn=TUBE1;refGene_tx=XM_023254270.1
B2 103765685 103765826
refGene_gn=TUBE1;refGene_tx=XM_011282648.3;refGene_gn=TUBE1;refGene_tx=XM_023254268.1;refGene_gn=TUBE1;refGene_tx=XM_023254269.1;refGene_gn=TUBE1;refGene_tx=XM_023254270.1
B2 103766655 103766831
refGene_gn=TUBE1;refGene_tx=XM_011282648.3;refGene_gn=TUBE1;refGene_tx=XM_023254268.1;refGene_gn=TUBE1;refGene_tx=XM_023254269.1;refGene_gn=TUBE1;refGene_tx=XM_023254270.1
B2 103767049 103767237
refGene_gn=TUBE1;refGene_tx=XM_011282648.3;refGene_gn=TUBE1;refGene_tx=XM_023254268.1;refGene_gn=TUBE1;refGene_tx=XM_023254269.1;refGene_gn=

TUBE1;refGene_tx=XM_023254270.1
B2	103769421	103769543
refGene_gn=TUBE1;refGene_tx=XM_011282648.3;refGene_gn=TUBE1;refGene_tx=XM_023254268.1;refGene_gn=TUBE1;refGene_tx=XM_023254269.1;refGene_gn=TUBE1;refGene_tx=XM_023254270.1
B2	103771516	103771632
refGene_gn=TUBE1;refGene_tx=XM_011282648.3;refGene_gn=TUBE1;refGene_tx=XM_023254268.1;refGene_gn=TUBE1;refGene_tx=XM_023254269.1;refGene_gn=TUBE1;refGene_tx=XM_023254270.1
B2	103774143	103774201
refGene_gn=TUBE1;refGene_tx=XM_011282648.3;refGene_gn=TUBE1;refGene_tx=XM_023254268.1
B2	103777147	103777200
refGene_gn=TUBE1;refGene_tx=XM_011282648.3;refGene_gn=TUBE1;refGene_tx=XM_023254268.1
B2	103777862	103777936
refGene_gn=TUBE1;refGene_tx=XM_011282648.3;refGene_gn=TUBE1;refGene_tx=XM_023254268.1
B2	103778037	103778062
refGene_gn=TUBE1;refGene_tx=XM_011282648.3;refGene_gn=TUBE1;refGene_tx=XM_023254268.1
B2	103790688	103790813
refGene_gn=FAM229B;refGene_tx=XM_006931957.3;refGene_gn=FAM229B;refGene_tx=XM_011282647.2;refGene_gn=FAM229B;refGene_tx=XM_019831099.1
B2	103792268	103792386
refGene_gn=FAM229B;refGene_tx=XM_006931957.3;refGene_gn=FAM229B;refGene_tx=XM_011282647.2;refGene_gn=FAM229B;refGene_tx=XM_019831099.1
B2	103802394	103802540
refGene_gn=LAMA4;refGene_tx=XM_003986472.4;refGene_gn=LAMA4;refGene_tx=XM_006931958.3
B2	103806918	103807038
refGene_gn=LAMA4;refGene_tx=XM_003986472.4;refGene_gn=LAMA4;refGene_tx=XM_006931958.3
B2	103807509	103807603
refGene_gn=LAMA4;refGene_tx=XM_003986472.4;refGene_gn=LAMA4;refGene_tx=XM_006931958.3
B2	103808810	103808941
refGene_gn=LAMA4;refGene_tx=XM_003986472.4;refGene_gn=LAMA4;refGene_tx=XM_006931958.3
B2	103810279	103810439
refGene_gn=LAMA4;refGene_tx=XM_003986472.4;refGene_gn=LAMA4;refGene_tx=XM_006931958.3
B2	103812011	103812167
refGene_gn=LAMA4;refGene_tx=XM_003986472.4;refGene_gn=LAMA4;refGene_tx=XM_006931958.3
B2	103813045	103813235
refGene_gn=LAMA4;refGene_tx=XM_003986472.4;refGene_gn=LAMA4;refGene_tx=XM_006931958.3
B2	103814920	103815108
refGene_gn=LAMA4;refGene_tx=XM_003986472.4;refGene_gn=LAMA4;refGene_tx

=XM_006931958.3
B2	103819991	103820145
refGene_gn=LAMA4;refGene_tx=XM_003986472.4;refGene_gn=LAMA4;refGene_tx
=XM_006931958.3
B2	103821566	103821731
refGene_gn=LAMA4;refGene_tx=XM_003986472.4;refGene_gn=LAMA4;refGene_tx
=XM_006931958.3
B2	103822886	103823020
refGene_gn=LAMA4;refGene_tx=XM_003986472.4;refGene_gn=LAMA4;refGene_tx
=XM_006931958.3
B2	103824886	103825024
refGene_gn=LAMA4;refGene_tx=XM_003986472.4;refGene_gn=LAMA4;refGene_tx
=XM_006931958.3
B2	103825665	103825804
refGene_gn=LAMA4;refGene_tx=XM_003986472.4;refGene_gn=LAMA4;refGene_tx
=XM_006931958.3
B2	103827555	103827698
refGene_gn=LAMA4;refGene_tx=XM_003986472.4;refGene_gn=LAMA4;refGene_tx
=XM_006931958.3
B2	103829278	103829410
refGene_gn=LAMA4;refGene_tx=XM_003986472.4;refGene_gn=LAMA4;refGene_tx
=XM_006931958.3
B2	103833896	103834068
refGene_gn=LAMA4;refGene_tx=XM_003986472.4;refGene_gn=LAMA4;refGene_tx
=XM_006931958.3
B2	103835664	103835798
refGene_gn=LAMA4;refGene_tx=XM_003986472.4;refGene_gn=LAMA4;refGene_tx
=XM_006931958.3
B2	103837083	103837246
refGene_gn=LAMA4;refGene_tx=XM_003986472.4;refGene_gn=LAMA4;refGene_tx
=XM_006931958.3
B2	103837557	103837703
refGene_gn=LAMA4;refGene_tx=XM_003986472.4;refGene_gn=LAMA4;refGene_tx
=XM_006931958.3
B2	103838349	103838526
refGene_gn=LAMA4;refGene_tx=XM_003986472.4;refGene_gn=LAMA4;refGene_tx
=XM_006931958.3
B2	103841180	103841320
refGene_gn=LAMA4;refGene_tx=XM_003986472.4;refGene_gn=LAMA4;refGene_tx
=XM_006931958.3
B2	103844774	103844954
refGene_gn=LAMA4;refGene_tx=XM_003986472.4;refGene_gn=LAMA4;refGene_tx
=XM_006931958.3
B2	103846551	103846668
refGene_gn=LAMA4;refGene_tx=XM_003986472.4;refGene_gn=LAMA4;refGene_tx
=XM_006931958.3
B2	103851197	103851294
refGene_gn=LAMA4;refGene_tx=XM_003986472.4;refGene_gn=LAMA4;refGene_tx
=XM_006931958.3
B2	103852046	103852188

refGene_gn=LAMA4;refGene_tx=XM_003986472.4;refGene_gn=LAMA4;refGene_tx=XM_006931958.3
B2 103856575 103856724
refGene_gn=LAMA4;refGene_tx=XM_003986472.4;refGene_gn=LAMA4;refGene_tx=XM_006931958.3
B2 103862141 103862258
refGene_gn=LAMA4;refGene_tx=XM_003986472.4;refGene_gn=LAMA4;refGene_tx=XM_006931958.3
B2 103866700 103866894
refGene_gn=LAMA4;refGene_tx=XM_003986472.4;refGene_gn=LAMA4;refGene_tx=XM_006931958.3
B2 103869343 103869511
refGene_gn=LAMA4;refGene_tx=XM_003986472.4;refGene_gn=LAMA4;refGene_tx=XM_006931958.3
B2 103872597 103872709
refGene_gn=LAMA4;refGene_tx=XM_003986472.4;refGene_gn=LAMA4;refGene_tx=XM_006931958.3
B2 103880040 103880151
refGene_gn=LAMA4;refGene_tx=XM_003986472.4;refGene_gn=LAMA4;refGene_tx=XM_006931958.3
B2 103881914 103882066
refGene_gn=LAMA4;refGene_tx=XM_003986472.4;refGene_gn=LAMA4;refGene_tx=XM_006931958.3
B2 103883793 103883889
refGene_gn=LAMA4;refGene_tx=XM_003986472.4;refGene_gn=LAMA4;refGene_tx=XM_006931958.3
B2 103885786 103886001
refGene_gn=LAMA4;refGene_tx=XM_003986472.4;refGene_gn=LAMA4;refGene_tx=XM_006931958.3
B2 103896045 103896126
refGene_gn=LAMA4;refGene_tx=XM_003986472.4;refGene_gn=LAMA4;refGene_tx=XM_006931958.3
B2 103901633 103901758
refGene_gn=LAMA4;refGene_tx=XM_003986472.4;refGene_gn=LAMA4;refGene_tx=XM_006931958.3
B2 103910934 103911036
refGene_gn=LAMA4;refGene_tx=XM_003986472.4;refGene_gn=LAMA4;refGene_tx=XM_006931958.3
B2 103947712 103948058
refGene_gn=LAMA4;refGene_tx=XM_003986472.4;refGene_gn=LAMA4;refGene_tx=XM_006931958.3
B2 103948350 103948466
refGene_gn=LAMA4;refGene_tx=XM_003986472.4;refGene_gn=LAMA4;refGene_tx=XM_006931958.3
B2 104033917 104034730
refGene_gn=RFPL4B;refGene_tx=XM_023254272.1
B2 105407677 105407779
refGene_gn=MARCKS;refGene_tx=XM_023254273.1
B2 105409616 105410522
refGene_gn=MARCKS;refGene_tx=XM_023254273.1

B2 105487834 105487865
refGene_gn=HDAC2;refGene_tx=XM_003986476.4;refGene_gn=HDAC2;refGene_tx=XM_023254274.1
B2 105488779 105488837
refGene_gn=HDAC2;refGene_tx=XM_003986476.4;refGene_gn=HDAC2;refGene_tx=XM_023254274.1
B2 105492394 105492550
refGene_gn=HDAC2;refGene_tx=XM_003986476.4;refGene_gn=HDAC2;refGene_tx=XM_023254274.1
B2 105493012 105493143
refGene_gn=HDAC2;refGene_tx=XM_003986476.4;refGene_gn=HDAC2;refGene_tx=XM_023254274.1
B2 105493758 105493867
refGene_gn=HDAC2;refGene_tx=XM_003986476.4;refGene_gn=HDAC2;refGene_tx=XM_023254274.1
B2 105494138 105494279
refGene_gn=HDAC2;refGene_tx=XM_003986476.4;refGene_gn=HDAC2;refGene_tx=XM_023254274.1
B2 105497429 105497538
refGene_gn=HDAC2;refGene_tx=XM_003986476.4;refGene_gn=HDAC2;refGene_tx=XM_023254274.1
B2 105497741 105497834
refGene_gn=HDAC2;refGene_tx=XM_003986476.4;refGene_gn=HDAC2;refGene_tx=XM_023254274.1
B2 105501069 105501211
refGene_gn=HDAC2;refGene_tx=XM_003986476.4;refGene_gn=HDAC2;refGene_tx=XM_023254274.1
B2 105503456 105503595
refGene_gn=HDAC2;refGene_tx=XM_003986476.4;refGene_gn=HDAC2;refGene_tx=XM_023254274.1
B2 105504628 105504703
refGene_gn=HDAC2;refGene_tx=XM_003986476.4;refGene_gn=HDAC2;refGene_tx=XM_023254274.1
B2 105506310 105506428
refGene_gn=HDAC2;refGene_tx=XM_003986476.4;refGene_gn=HDAC2;refGene_tx=XM_023254274.1
B2 105507580 105507693
refGene_gn=HDAC2;refGene_tx=XM_003986476.4;refGene_gn=HDAC2;refGene_tx=XM_023254274.1
B2 105517089 105517141
refGene_gn=HDAC2;refGene_tx=XM_023254274.1
B2 105579663 105580594
refGene_gn=HS3ST5;refGene_tx=XM_003986477.5;refGene_gn=HS3ST5;refGene_tx=XM_019831102.2
B2 105584078 105584185
refGene_gn=HS3ST5;refGene_tx=XM_003986477.5;refGene_gn=HS3ST5;refGene_tx=XM_019831102.2
B2 107241725 107241937
refGene_gn=FRK;refGene_tx=XM_003986478.4
B2 107242325 107242491

refGene_gn=FRK;refGene_tx=XM_003986478.4
B2 107243595 107243777
refGene_gn=FRK;refGene_tx=XM_003986478.4
B2 107253499 107253658
refGene_gn=FRK;refGene_tx=XM_003986478.4
B2 107258215 107258384
refGene_gn=FRK;refGene_tx=XM_003986478.4
B2 107260292 107260456
refGene_gn=FRK;refGene_tx=XM_003986478.4
B2 107297104 107297226
refGene_gn=FRK;refGene_tx=XM_003986478.4
B2 107346805 107347176
refGene_gn=FRK;refGene_tx=XM_003986478.4
B2 107385979 107386072
refGene_gn=NT5DC1;refGene_tx=XM_003986479.5;refGene_gn=NT5DC1;refGene_tx=XM_006931961.4
B2 107391491 107391583
refGene_gn=NT5DC1;refGene_tx=XM_003986479.5;refGene_gn=NT5DC1;refGene_tx=XM_006931961.4
B2 107392981 107393053
refGene_gn=NT5DC1;refGene_tx=XM_003986479.5;refGene_gn=NT5DC1;refGene_tx=XM_006931961.4
B2 107394181 107394288
refGene_gn=NT5DC1;refGene_tx=XM_003986479.5;refGene_gn=NT5DC1;refGene_tx=XM_006931961.4
B2 107399451 107399531
refGene_gn=NT5DC1;refGene_tx=XM_003986479.5;refGene_gn=NT5DC1;refGene_tx=XM_006931961.4
B2 107401191 107401276
refGene_gn=NT5DC1;refGene_tx=XM_003986479.5;refGene_gn=NT5DC1;refGene_tx=XM_006931961.4
B2 107403411 107405282
refGene_gn=COL10A1;refGene_tx=XM_003986480.5
B2 107409784 107409938
refGene_gn=COL10A1;refGene_tx=XM_003986480.5
B2 107498809 107498984
refGene_gn=NT5DC1;refGene_tx=XM_003986479.5;refGene_gn=NT5DC1;refGene_tx=XM_006931961.4
B2 107500743 107500841
refGene_gn=NT5DC1;refGene_tx=XM_003986479.5;refGene_gn=NT5DC1;refGene_tx=XM_006931961.4
B2 107511078 107511197
refGene_gn=NT5DC1;refGene_tx=XM_003986479.5;refGene_gn=NT5DC1;refGene_tx=XM_006931961.4
B2 107512291 107512444
refGene_gn=NT5DC1;refGene_tx=XM_003986479.5;refGene_gn=NT5DC1;refGene_tx=XM_006931961.4
B2 107521763 107521932
refGene_gn=NT5DC1;refGene_tx=XM_003986479.5;refGene_gn=NT5DC1;refGene_tx=XM_006931961.4

B2	107525323	107525439
refGene_gn=NT5DC1;refGene_tx=XM_003986479.5;refGene_gn=NT5DC1;refGene_tx=XM_006931961.4
B2	107535704	107536958
refGene_gn=TSPYL4;refGene_tx=XM_006931962.4
B2	107555481	107556822
refGene_gn=TSPYL1;refGene_tx=XM_003986500.5
B2	107632844	107633022
refGene_gn=DSE;refGene_tx=XM_019831105.2
B2	107661647	107662116
refGene_gn=DSE;refGene_tx=XM_003986481.5;refGene_gn=DSE;refGene_tx=XM_019831105.2;refGene_gn=DSE;refGene_tx=XM_023254275.1
B2	107704315	107704569
refGene_gn=DSE;refGene_tx=XM_003986481.5;refGene_gn=DSE;refGene_tx=XM_019831105.2;refGene_gn=DSE;refGene_tx=XM_019831106.2;refGene_gn=DSE;refGene_tx=XM_023254275.1
B2	107709122	107709362
refGene_gn=DSE;refGene_tx=XM_003986481.5;refGene_gn=DSE;refGene_tx=XM_019831105.2;refGene_gn=DSE;refGene_tx=XM_019831106.2;refGene_gn=DSE;refGene_tx=XM_023254275.1
B2	107711735	107711943
refGene_gn=DSE;refGene_tx=XM_003986481.5;refGene_gn=DSE;refGene_tx=XM_019831105.2;refGene_gn=DSE;refGene_tx=XM_019831106.2;refGene_gn=DSE;refGene_tx=XM_023254275.1
B2	107714256	107716006
refGene_gn=DSE;refGene_tx=XM_003986481.5;refGene_gn=DSE;refGene_tx=XM_019831105.2;refGene_gn=DSE;refGene_tx=XM_019831106.2;refGene_gn=DSE;refGene_tx=XM_023254275.1
B2	107745381	107745912
refGene_gn=CALHM6;refGene_tx=XM_023254276.1;refGene_gn=CALHM6;refGene_tx=XM_023254277.1
B2	107746542	107746965
refGene_gn=CALHM6;refGene_tx=XM_023254276.1;refGene_gn=CALHM6;refGene_tx=XM_023254277.1
B2	107752006	107752107
refGene_gn=CALHM6;refGene_tx=XM_023254276.1
B2	107774998	107775118
refGene_gn=TRAPPC3L;refGene_tx=XM_023254737.1
B2	107779577	107779763
refGene_gn=TRAPPC3L;refGene_tx=XM_023254737.1
B2	107782117	107782657
refGene_gn=CALHM5;refGene_tx=XM_003986482.5
B2	107786099	107786489
refGene_gn=CALHM5;refGene_tx=XM_003986482.5
B2	107807820	107807920
refGene_gn=TRAPPC3L;refGene_tx=XM_023254737.1
B2	107811618	107811716
refGene_gn=TRAPPC3L;refGene_tx=XM_023254737.1
B2	107813635	107813665
refGene_gn=TRAPPC3L;refGene_tx=XM_023254737.1

B2 107839025 107839098
refGene_gn=RWDD1;refGene_tx=XM_011282644.2
B2 107847319 107847385
refGene_gn=RWDD1;refGene_tx=XM_011282644.2;refGene_gn=RWDD1;refGene_tx=XM_019831108.2;refGene_gn=RWDD1;refGene_tx=XM_019831109.2;refGene_gn=RWDD1;refGene_tx=XM_019831110.2
B2 107850436 107850567
refGene_gn=RWDD1;refGene_tx=XM_011282644.2
B2 107856060 107856204
refGene_gn=RWDD1;refGene_tx=XM_011282644.2;refGene_gn=RWDD1;refGene_tx=XM_019831108.2;refGene_gn=RWDD1;refGene_tx=XM_019831109.2;refGene_gn=RWDD1;refGene_tx=XM_019831110.2;refGene_gn=RWDD1;refGene_tx=XM_019831111.2;refGene_gn=RWDD1;refGene_tx=XM_019831112.2
B2 107856563 107856696
refGene_gn=RWDD1;refGene_tx=XM_011282644.2;refGene_gn=RWDD1;refGene_tx=XM_019831108.2;refGene_gn=RWDD1;refGene_tx=XM_019831109.2;refGene_gn=RWDD1;refGene_tx=XM_019831110.2;refGene_gn=RWDD1;refGene_tx=XM_019831111.2;refGene_gn=RWDD1;refGene_tx=XM_019831112.2
B2 107857230 107857293
refGene_gn=RWDD1;refGene_tx=XM_011282644.2;refGene_gn=RWDD1;refGene_tx=XM_019831108.2;refGene_gn=RWDD1;refGene_tx=XM_019831109.2;refGene_gn=RWDD1;refGene_tx=XM_019831110.2;refGene_gn=RWDD1;refGene_tx=XM_019831111.2;refGene_gn=RWDD1;refGene_tx=XM_019831112.2
B2 107865153 107865275
refGene_gn=RWDD1;refGene_tx=XM_011282644.2;refGene_gn=RWDD1;refGene_tx=XM_019831108.2;refGene_gn=RWDD1;refGene_tx=XM_019831109.2;refGene_gn=RWDD1;refGene_tx=XM_019831110.2;refGene_gn=RWDD1;refGene_tx=XM_019831111.2;refGene_gn=RWDD1;refGene_tx=XM_019831112.2
B2 107882223 107882365
refGene_gn=LOC101090867;refGene_tx=XM_003986487.3;refGene_gn=LOC101090867;refGene_tx=XM_019831114.1;refGene_gn=LOC101090867;refGene_tx=XM_019831115.1
B2 107903275 107903484
refGene_gn=LOC101090867;refGene_tx=XM_003986487.3;refGene_gn=LOC101090867;refGene_tx=XM_019831114.1;refGene_gn=LOC101090867;refGene_tx=XM_019831115.1
B2 107908390 107908514
refGene_gn=LOC101090867;refGene_tx=XM_003986487.3;refGene_gn=LOC101090867;refGene_tx=XM_019831114.1;refGene_gn=LOC101090867;refGene_tx=XM_019831115.1
B2 107909501 107909596
refGene_gn=LOC101090867;refGene_tx=XM_003986487.3;refGene_gn=LOC101090867;refGene_tx=XM_019831114.1;refGene_gn=LOC101090867;refGene_tx=XM_019831115.1
B2 107912034 107912218
refGene_gn=LOC101090867;refGene_tx=XM_003986487.3;refGene_gn=LOC101090867;refGene_tx=XM_019831114.1;refGene_gn=LOC101090867;refGene_tx=XM_019831115.1
B2 107913994 107914116
refGene_gn=LOC101090867;refGene_tx=XM_003986487.3;refGene_gn=LOC101090

867;refGene_tx=XM_019831114.1;refGene_gn=LOC101090867;refGene_tx=XM_019831115.1
B2 107926829 107926869
refGene_gn=LOC101091113;refGene_tx=XM_023254278.1
B2 107932584 107932737
refGene_gn=LOC101091113;refGene_tx=XM_023254278.1
B2 107937068 107937277
refGene_gn=LOC101091113;refGene_tx=XM_023254278.1
B2 107941699 107941823
refGene_gn=LOC101091113;refGene_tx=XM_023254278.1
B2 107942798 107942893
refGene_gn=LOC101091113;refGene_tx=XM_023254278.1
B2 107945833 107946017
refGene_gn=LOC101091113;refGene_tx=XM_023254278.1
B2 107968382 107968414
refGene_gn=LOC101091113;refGene_tx=XM_023254278.1
B2 108066701 108066843
refGene_gn=LOC101091370;refGene_tx=XM_011282642.3;refGene_gn=LOC101091370;refGene_tx=XM_019831118.2;refGene_gn=LOC101091370;refGene_tx=XM_019831119.1
B2 108077420 108077629
refGene_gn=LOC101091370;refGene_tx=XM_011282642.3;refGene_gn=LOC101091370;refGene_tx=XM_019831118.2;refGene_gn=LOC101091370;refGene_tx=XM_019831119.1
B2 108078515 108078639
refGene_gn=LOC101091370;refGene_tx=XM_011282642.3;refGene_gn=LOC101091370;refGene_tx=XM_019831118.2;refGene_gn=LOC101091370;refGene_tx=XM_019831119.1
B2 108079887 108079982
refGene_gn=LOC101091370;refGene_tx=XM_011282642.3;refGene_gn=LOC101091370;refGene_tx=XM_019831118.2;refGene_gn=LOC101091370;refGene_tx=XM_019831119.1
B2 108081980 108082164
refGene_gn=LOC101091370;refGene_tx=XM_011282642.3;refGene_gn=LOC101091370;refGene_tx=XM_019831118.2;refGene_gn=LOC101091370;refGene_tx=XM_019831119.1
B2 108090561 108090683
refGene_gn=LOC101091370;refGene_tx=XM_011282642.3;refGene_gn=LOC101091370;refGene_tx=XM_019831118.2;refGene_gn=LOC101091370;refGene_tx=XM_019831119.1
B2 108153716 108154393
refGene_gn=RSPH4A;refGene_tx=XM_003986489.5;refGene_gn=RSPH4A;refGene_tx=XM_019831120.2;refGene_gn=RSPH4A;refGene_tx=XM_019831121.2
B2 108157614 108157849
refGene_gn=RSPH4A;refGene_tx=XM_003986489.5;refGene_gn=RSPH4A;refGene_tx=XM_019831120.2;refGene_gn=RSPH4A;refGene_tx=XM_019831121.2
B2 108162764 108163505
refGene_gn=RSPH4A;refGene_tx=XM_003986489.5;refGene_gn=RSPH4A;refGene_tx=XM_019831120.2;refGene_gn=RSPH4A;refGene_tx=XM_019831121.2
B2 108164877 108165028

refGene_gn=RSPH4A;refGene_tx=XM_019831120.2;refGene_gn=RSPH4A;refGene_tx=XM_019831121.2
B2 108165884 108166002
refGene_gn=RSPH4A;refGene_tx=XM_003986489.5;refGene_gn=RSPH4A;refGene_tx=XM_019831120.2;refGene_gn=RSPH4A;refGene_tx=XM_019831121.2
B2 108168237 108168478
refGene_gn=RSPH4A;refGene_tx=XM_003986489.5;refGene_gn=RSPH4A;refGene_tx=XM_019831120.2;refGene_gn=RSPH4A;refGene_tx=XM_019831121.2
B2 108171402 108171450
refGene_gn=ZUFSP;refGene_tx=XM_003986490.5;refGene_gn=ZUFSP;refGene_tx=XM_019831122.2;refGene_gn=ZUFSP;refGene_tx=XM_019831123.2
B2 108174925 108175146
refGene_gn=ZUFSP;refGene_tx=XM_003986490.5;refGene_gn=ZUFSP;refGene_tx=XM_019831122.2
B2 108176219 108176371
refGene_gn=ZUFSP;refGene_tx=XM_003986490.5;refGene_gn=ZUFSP;refGene_tx=XM_019831122.2;refGene_gn=ZUFSP;refGene_tx=XM_019831123.2;refGene_gn=ZUFSP;refGene_tx=XM_019831124.2
B2 108180228 108180394
refGene_gn=ZUFSP;refGene_tx=XM_003986490.5;refGene_gn=ZUFSP;refGene_tx=XM_019831122.2;refGene_gn=ZUFSP;refGene_tx=XM_019831123.2;refGene_gn=ZUFSP;refGene_tx=XM_019831124.2
B2 108181030 108181219
refGene_gn=ZUFSP;refGene_tx=XM_003986490.5;refGene_gn=ZUFSP;refGene_tx=XM_019831122.2;refGene_gn=ZUFSP;refGene_tx=XM_019831123.2;refGene_gn=ZUFSP;refGene_tx=XM_019831124.2
B2 108182723 108182892
refGene_gn=ZUFSP;refGene_tx=XM_003986490.5;refGene_gn=ZUFSP;refGene_tx=XM_019831122.2;refGene_gn=ZUFSP;refGene_tx=XM_019831123.2;refGene_gn=ZUFSP;refGene_tx=XM_019831124.2
B2 108185055 108185177
refGene_gn=ZUFSP;refGene_tx=XM_003986490.5;refGene_gn=ZUFSP;refGene_tx=XM_019831122.2;refGene_gn=ZUFSP;refGene_tx=XM_019831123.2;refGene_gn=ZUFSP;refGene_tx=XM_019831124.2
B2 108186786 108186897
refGene_gn=ZUFSP;refGene_tx=XM_003986490.5;refGene_gn=ZUFSP;refGene_tx=XM_019831122.2;refGene_gn=ZUFSP;refGene_tx=XM_019831123.2;refGene_gn=ZUFSP;refGene_tx=XM_019831124.2
B2 108190101 108190179
refGene_gn=ZUFSP;refGene_tx=XM_019831122.2;refGene_gn=ZUFSP;refGene_tx=XM_019831123.2;refGene_gn=ZUFSP;refGene_tx=XM_019831124.2
B2 108195769 108196331
refGene_gn=ZUFSP;refGene_tx=XM_003986490.5;refGene_gn=ZUFSP;refGene_tx=XM_019831122.2;refGene_gn=ZUFSP;refGene_tx=XM_019831123.2;refGene_gn=ZUFSP;refGene_tx=XM_019831124.2
B2 108200907 108200911
refGene_gn=KPNA5;refGene_tx=XM_023254279.1
B2 108206652 108206786
refGene_gn=KPNA5;refGene_tx=XM_023254279.1
B2 108208207 108208309

refGene_gn=KPNA5;refGene_tx=XM_023254279.1
B2 108208428 108208528
refGene_gn=KPNA5;refGene_tx=XM_023254279.1
B2 108210044 108210139
refGene_gn=KPNA5;refGene_tx=XM_023254279.1
B2 108211115 108211247
refGene_gn=KPNA5;refGene_tx=XM_023254279.1
B2 108213395 108213484
refGene_gn=KPNA5;refGene_tx=XM_023254279.1
B2 108223268 108223368
refGene_gn=KPNA5;refGene_tx=XM_023254279.1
B2 108229037 108229201
refGene_gn=KPNA5;refGene_tx=XM_023254279.1
B2 108231624 108231703
refGene_gn=KPNA5;refGene_tx=XM_023254279.1
B2 108233312 108233438
refGene_gn=KPNA5;refGene_tx=XM_023254279.1
B2 108234032 108234089
refGene_gn=KPNA5;refGene_tx=XM_023254279.1
B2 108234090 108234161
refGene_gn=KPNA5;refGene_tx=XM_023254279.1
B2 108237147 108237326
refGene_gn=KPNA5;refGene_tx=XM_023254279.1
B2 108240419 108240607
refGene_gn=KPNA5;refGene_tx=XM_023254279.1
B2 108259476 108259575
refGene_gn=FAM162B;refGene_tx=XM_023254281.1;refGene_gn=FAM162B;refGene_tx=XM_023254282.1;refGene_gn=FAM162B;refGene_tx=XM_023254283.1
B2 108262539 108262648
refGene_gn=FAM162B;refGene_tx=XM_023254281.1;refGene_gn=FAM162B;refGene_tx=XM_023254282.1;refGene_gn=FAM162B;refGene_tx=XM_023254283.1
B2 108264273 108264316
refGene_gn=FAM162B;refGene_tx=XM_023254282.1
B2 108266211 108266323
refGene_gn=FAM162B;refGene_tx=XM_023254281.1;refGene_gn=FAM162B;refGene_tx=XM_023254282.1;refGene_gn=FAM162B;refGene_tx=XM_023254283.1
B2 108266489 108266655
refGene_gn=FAM162B;refGene_tx=XM_023254281.1;refGene_gn=FAM162B;refGene_tx=XM_023254282.1;refGene_gn=FAM162B;refGene_tx=XM_023254283.1
B2 108288611 108289726
refGene_gn=GPRC6A;refGene_tx=XM_003986492.5
B2 108292249 108292373
refGene_gn=GPRC6A;refGene_tx=XM_003986492.5
B2 108295507 108295720
refGene_gn=GPRC6A;refGene_tx=XM_003986492.5
B2 108302861 108303698
refGene_gn=GPRC6A;refGene_tx=XM_003986492.5
B2 108306011 108306315
refGene_gn=GPRC6A;refGene_tx=XM_003986492.5
B2 108310011 108310205

refGene_gn=GPRC6A;refGene_tx=XM_003986492.5
B2 108340427 108340650
refGene_gn=RFX6;refGene_tx=XM_023254284.1
B2 108340960 108341117
refGene_gn=RFX6;refGene_tx=XM_023254284.1
B2 108343894 108344018
refGene_gn=RFX6;refGene_tx=XM_023254284.1
B2 108346300 108346362
refGene_gn=RFX6;refGene_tx=XM_023254284.1
B2 108356662 108356740
refGene_gn=RFX6;refGene_tx=XM_023254284.1
B2 108357981 108358009
refGene_gn=RFX6;refGene_tx=XM_023254284.1
B2 108372758 108372866
refGene_gn=RFX6;refGene_tx=XM_023254284.1
B2 108377241 108377319
refGene_gn=RFX6;refGene_tx=XM_023254284.1
B2 108377435 108377549
refGene_gn=RFX6;refGene_tx=XM_023254284.1
B2 108380702 108380752
refGene_gn=RFX6;refGene_tx=XM_023254284.1
B2 108383642 108383802
refGene_gn=RFX6;refGene_tx=XM_023254284.1
B2 108384702 108384847
refGene_gn=RFX6;refGene_tx=XM_023254284.1
B2 108386061 108386171
refGene_gn=RFX6;refGene_tx=XM_023254284.1
B2 108387369 108387487
refGene_gn=RFX6;refGene_tx=XM_023254284.1
B2 108388951 108389074
refGene_gn=RFX6;refGene_tx=XM_023254284.1
B2 108389734 108389941
refGene_gn=RFX6;refGene_tx=XM_023254284.1
B2 108391123 108391636
refGene_gn=RFX6;refGene_tx=XM_023254284.1
B2 108392486 108392699
refGene_gn=RFX6;refGene_tx=XM_023254284.1
B2 108395315 108395491
refGene_gn=RFX6;refGene_tx=XM_023254284.1
B2 108630408 108630489
refGene_gn=VGLL2;refGene_tx=XM_011282685.3;refGene_gn=VGLL2;refGene_tx
=XM_023254287.1
B2 108630716 108630869
refGene_gn=VGLL2;refGene_tx=XM_023254285.1;refGene_gn=VGLL2;refGene_tx
=XM_023254286.1
B2 108632745 108633052
refGene_gn=VGLL2;refGene_tx=XM_011282685.3;refGene_gn=VGLL2;refGene_tx
=XM_023254285.1;refGene_gn=VGLL2;refGene_tx=XM_023254286.1;refGene_gn=
VGLL2;refGene_tx=XM_023254287.1;refGene_gn=VGLL2;refGene_tx=XM_0232542
88.1

B2	108635084	108635710
refGene_gn=VGLL2;refGene_tx=XM_011282685.3;refGene_gn=VGLL2;refGene_tx
=XM_023254285.1;refGene_gn=VGLL2;refGene_tx=XM_023254286.1;refGene_gn=
VGLL2;refGene_tx=XM_023254287.1;refGene_gn=VGLL2;refGene_tx=XM_0232542
88.1
B2	108636737	108636781
refGene_gn=VGLL2;refGene_tx=XM_023254285.1
B2	108636880	108636921
refGene_gn=VGLL2;refGene_tx=XM_023254286.1;refGene_gn=VGLL2;refGene_tx
=XM_023254287.1
B2	108655926	108656237
refGene_gn=ROS1;refGene_tx=XM_023254738.1
B2	108665139	108665303
refGene_gn=ROS1;refGene_tx=XM_023254738.1
B2	108672515	108672650
refGene_gn=ROS1;refGene_tx=XM_023254738.1
B2	108673594	108673798
refGene_gn=ROS1;refGene_tx=XM_023254738.1
B2	108674013	108674111
refGene_gn=ROS1;refGene_tx=XM_023254738.1
B2	108680835	108680965
refGene_gn=ROS1;refGene_tx=XM_023254738.1
B2	108681900	108681965
refGene_gn=ROS1;refGene_tx=XM_023254738.1
B2	108683626	108683789
refGene_gn=ROS1;refGene_tx=XM_023254738.1
B2	108685052	108685188
refGene_gn=ROS1;refGene_tx=XM_023254738.1
B2	108688157	108688241
refGene_gn=ROS1;refGene_tx=XM_023254738.1
B2	108690038	108690229
refGene_gn=ROS1;refGene_tx=XM_023254738.1
B2	108693330	108693448
refGene_gn=ROS1;refGene_tx=XM_023254738.1
B2	108694182	108694351
refGene_gn=ROS1;refGene_tx=XM_023254738.1
B2	108698394	108698571
refGene_gn=ROS1;refGene_tx=XM_023254738.1
B2	108698658	108698891
refGene_gn=ROS1;refGene_tx=XM_023254738.1
B2	108699660	108699805
refGene_gn=ROS1;refGene_tx=XM_023254738.1
B2	108701484	108701687
refGene_gn=ROS1;refGene_tx=XM_023254738.1
B2	108703328	108703505
refGene_gn=ROS1;refGene_tx=XM_023254738.1
B2	108706305	108706592
refGene_gn=ROS1;refGene_tx=XM_023254738.1
B2	108707466	108707672
refGene_gn=ROS1;refGene_tx=XM_023254738.1

B2 108709227 108709430
refGene_gn=ROS1;refGene_tx=XM_023254738.1
B2 108709743 108709807
refGene_gn=ROS1;refGene_tx=XM_023254738.1
B2 108713193 108713456
refGene_gn=ROS1;refGene_tx=XM_023254738.1
B2 108715643 108715788
refGene_gn=ROS1;refGene_tx=XM_023254738.1
B2 108716655 108716816
refGene_gn=ROS1;refGene_tx=XM_023254738.1
B2 108717713 108717928
refGene_gn=ROS1;refGene_tx=XM_023254738.1
B2 108724843 108724944
refGene_gn=ROS1;refGene_tx=XM_023254738.1
B2 108728386 108728578
refGene_gn=ROS1;refGene_tx=XM_023254738.1
B2 108731977 108732156
refGene_gn=ROS1;refGene_tx=XM_023254738.1
B2 108733985 108734198
refGene_gn=ROS1;refGene_tx=XM_023254738.1
B2 108735653 108736150
refGene_gn=ROS1;refGene_tx=XM_023254738.1
B2 108740242 108740340
refGene_gn=ROS1;refGene_tx=XM_023254738.1
B2 108741496 108741681
refGene_gn=ROS1;refGene_tx=XM_023254738.1
B2 108741850 108741973
refGene_gn=ROS1;refGene_tx=XM_023254738.1
B2 108743310 108743387
refGene_gn=ROS1;refGene_tx=XM_023254738.1
B2 108744047 108744249
refGene_gn=ROS1;refGene_tx=XM_023254738.1
B2 108747241 108747358
refGene_gn=ROS1;refGene_tx=XM_023254738.1
B2 108747360 108747379
refGene_gn=ROS1;refGene_tx=XM_023254738.1
B2 108748289 108748438
refGene_gn=ROS1;refGene_tx=XM_023254738.1
B2 108756127 108756188
refGene_gn=ROS1;refGene_tx=XM_023254738.1
B2 108762837 108762864
refGene_gn=ROS1;refGene_tx=XM_023254738.1
B2 108764559 108764619
refGene_gn=ROS1;refGene_tx=XM_023254738.1
B2 108765758 108765803
refGene_gn=ROS1;refGene_tx=XM_023254738.1
B2 108771943 108772066
refGene_gn=ROS1;refGene_tx=XM_023254738.1
B2 108804052 108804132
refGene_gn=DCBLD1;refGene_tx=XM_023254739.1

B2 108805050 108805499
refGene_gn=DCBLD1;refGene_tx=XM_023254739.1
B2 108827106 108827319
refGene_gn=DCBLD1;refGene_tx=XM_023254739.1
B2 108843318 108843453
refGene_gn=DCBLD1;refGene_tx=XM_023254739.1
B2 108844899 108844951
refGene_gn=DCBLD1;refGene_tx=XM_023254739.1
B2 108847548 108847621
refGene_gn=DCBLD1;refGene_tx=XM_023254739.1
B2 108856729 108856863
refGene_gn=DCBLD1;refGene_tx=XM_023254739.1
B2 108862692 108862733
refGene_gn=DCBLD1;refGene_tx=XM_023254739.1
B2 108864056 108864266
refGene_gn=DCBLD1;refGene_tx=XM_023254739.1
B2 108864875 108865000
refGene_gn=DCBLD1;refGene_tx=XM_023254739.1
B2 108865272 108865420
refGene_gn=DCBLD1;refGene_tx=XM_023254739.1
B2 108865520 108865628
refGene_gn=DCBLD1;refGene_tx=XM_023254739.1
B2 108868235 108868320
refGene_gn=DCBLD1;refGene_tx=XM_023254739.1
B2 108868555 108868605
refGene_gn=DCBLD1;refGene_tx=XM_023254739.1
B2 108869480 108869600
refGene_gn=DCBLD1;refGene_tx=XM_023254739.1
B2 108872258 108872749
refGene_gn=DCBLD1;refGene_tx=XM_023254739.1
B2 108889821 108889952
refGene_gn=GOPC;refGene_tx=XM_011282638.3;refGene_gn=GOPC;refGene_tx=XM_011282639.3
B2 108891729 108891737
refGene_gn=GOPC;refGene_tx=XM_023254293.1
B2 108893254 108893435
refGene_gn=GOPC;refGene_tx=XM_011282638.3;refGene_gn=GOPC;refGene_tx=XM_011282639.3;refGene_gn=GOPC;refGene_tx=XM_023254293.1
B2 108895542 108895707
refGene_gn=GOPC;refGene_tx=XM_011282638.3;refGene_gn=GOPC;refGene_tx=XM_011282639.3;refGene_gn=GOPC;refGene_tx=XM_023254293.1
B2 108897045 108897141
refGene_gn=GOPC;refGene_tx=XM_011282638.3;refGene_gn=GOPC;refGene_tx=XM_011282639.3;refGene_gn=GOPC;refGene_tx=XM_023254293.1
B2 108899750 108899916
refGene_gn=GOPC;refGene_tx=XM_011282638.3;refGene_gn=GOPC;refGene_tx=XM_011282639.3;refGene_gn=GOPC;refGene_tx=XM_023254293.1
B2 108900952 108901128
refGene_gn=GOPC;refGene_tx=XM_011282638.3;refGene_gn=GOPC;refGene_tx=XM_011282639.3;refGene_gn=GOPC;refGene_tx=XM_023254293.1

B2 108904305 108904329
refGene_gn=GOPC;refGene_tx=XM_011282638.3;refGene_gn=GOPC;refGene_tx=XM_023254293.1
B2 108905865 108906030
refGene_gn=GOPC;refGene_tx=XM_011282638.3;refGene_gn=GOPC;refGene_tx=XM_011282639.3;refGene_gn=GOPC;refGene_tx=XM_023254293.1
B2 108926788 108927076
refGene_gn=GOPC;refGene_tx=XM_011282638.3;refGene_gn=GOPC;refGene_tx=XM_011282639.3;refGene_gn=GOPC;refGene_tx=XM_023254293.1
B2 108936286 108936292
refGene_gn=LOC101093938;refGene_tx=XM_023254292.1
B2 108936947 108937136
refGene_gn=LOC101093938;refGene_tx=XM_023254289.1;refGene_gn=LOC101093938;refGene_tx=XM_023254290.1;refGene_gn=LOC101093938;refGene_tx=XM_023254291.1
B2 108948208 108949110
refGene_gn=LOC101093938;refGene_tx=XM_023254289.1;refGene_gn=LOC101093938;refGene_tx=XM_023254290.1;refGene_gn=LOC101093938;refGene_tx=XM_023254291.1;refGene_gn=LOC101093938;refGene_tx=XM_023254292.1
B2 108958623 108959071
refGene_gn=LOC101093938;refGene_tx=XM_023254289.1;refGene_gn=LOC101093938;refGene_tx=XM_023254290.1;refGene_gn=LOC101093938;refGene_tx=XM_023254291.1;refGene_gn=LOC101093938;refGene_tx=XM_023254292.1
B2 108970145 108970560
refGene_gn=NUS1;refGene_tx=XM_003986497.5
B2 108987023 108987149
refGene_gn=NUS1;refGene_tx=XM_003986497.5
B2 108988066 108988216
refGene_gn=NUS1;refGene_tx=XM_003986497.5
B2 108994843 108994943
refGene_gn=NUS1;refGene_tx=XM_003986497.5
B2 108998963 108999054
refGene_gn=NUS1;refGene_tx=XM_003986497.5
B2 109127918 109128100
refGene_gn=SLC35F1;refGene_tx=XM_011282636.3
B2 109382223 109382399
refGene_gn=SLC35F1;refGene_tx=XM_011282636.3
B2 109459025 109459153
refGene_gn=SLC35F1;refGene_tx=XM_011282636.3
B2 109489320 109489480
refGene_gn=SLC35F1;refGene_tx=XM_011282636.3
B2 109496273 109496430
refGene_gn=SLC35F1;refGene_tx=XM_011282636.3
B2 109498997 109499050
refGene_gn=SLC35F1;refGene_tx=XM_011282636.3
B2 109506571 109506726
refGene_gn=SLC35F1;refGene_tx=XM_011282636.3
B2 109536957 109537182
refGene_gn=SLC35F1;refGene_tx=XM_011282636.3
B2 109640541 109640705

refGene_gn=CEP85L;refGene_tx=XM_023254294.1;refGene_gn=CEP85L;refGene_tx=XM_023254295.1
B2 109645495 109645727
refGene_gn=CEP85L;refGene_tx=XM_023254294.1;refGene_gn=CEP85L;refGene_tx=XM_023254295.1
B2 109647684 109647792
refGene_gn=CEP85L;refGene_tx=XM_023254294.1;refGene_gn=CEP85L;refGene_tx=XM_023254295.1
B2 109664807 109664858
refGene_gn=CEP85L;refGene_tx=XM_023254294.1;refGene_gn=CEP85L;refGene_tx=XM_023254295.1
B2 109665912 109666030
refGene_gn=CEP85L;refGene_tx=XM_023254294.1;refGene_gn=CEP85L;refGene_tx=XM_023254295.1
B2 109670084 109670239
refGene_gn=CEP85L;refGene_tx=XM_023254294.1;refGene_gn=CEP85L;refGene_tx=XM_023254295.1
B2 109672166 109672319
refGene_gn=CEP85L;refGene_tx=XM_023254294.1;refGene_gn=CEP85L;refGene_tx=XM_023254295.1
B2 109679896 109680076
refGene_gn=CEP85L;refGene_tx=XM_023254294.1
B2 109700711 109700829
refGene_gn=CEP85L;refGene_tx=XM_023254294.1;refGene_gn=CEP85L;refGene_tx=XM_023254295.1
B2 109713291 109713413
refGene_gn=CEP85L;refGene_tx=XM_023254294.1;refGene_gn=CEP85L;refGene_tx=XM_023254295.1
B2 109748494 109748653
refGene_gn=PLN;refGene_tx=XM_003986508.5
B2 109756081 109756869
refGene_gn=CEP85L;refGene_tx=XM_023254294.1;refGene_gn=CEP85L;refGene_tx=XM_023254295.1
B2 109788449 109788611
refGene_gn=CEP85L;refGene_tx=XM_023254294.1;refGene_gn=CEP85L;refGene_tx=XM_023254295.1
B2 109806107 109806180
refGene_gn=CEP85L;refGene_tx=XM_023254294.1;refGene_gn=CEP85L;refGene_tx=XM_023254295.1
B2 109949297 109950765
refGene_gn=MCM9;refGene_tx=XM_003986512.5;refGene_gn=MCM9;refGene_tx=XM_023254296.1;refGene_gn=MCM9;refGene_tx=XM_023254297.1
B2 109954846 109954992
refGene_gn=MCM9;refGene_tx=XM_003986512.5;refGene_gn=MCM9;refGene_tx=XM_023254296.1;refGene_gn=MCM9;refGene_tx=XM_023254297.1
B2 109955473 109955556
refGene_gn=MCM9;refGene_tx=XM_003986512.5;refGene_gn=MCM9;refGene_tx=XM_023254296.1;refGene_gn=MCM9;refGene_tx=XM_023254297.1
B2 109957190 109957394
refGene_gn=MCM9;refGene_tx=XM_003986512.5;refGene_gn=MCM9;refGene_tx=X

M_023254296.1;refGene_gn=MCM9;refGene_tx=XM_023254297.1
B2 109958008 109958211
refGene_gn=MCM9;refGene_tx=XM_003986512.5;refGene_gn=MCM9;refGene_tx=XM_023254296.1;refGene_gn=MCM9;refGene_tx=XM_023254297.1
B2 109992190 109992365
refGene_gn=MCM9;refGene_tx=XM_003986512.5;refGene_gn=MCM9;refGene_tx=XM_023254296.1;refGene_gn=MCM9;refGene_tx=XM_023254297.1
B2 110020860 110020969
refGene_gn=ASF1A;refGene_tx=XM_023254298.1
B2 110025434 110025483
refGene_gn=ASF1A;refGene_tx=XM_011282651.3
B2 110026539 110026655
refGene_gn=ASF1A;refGene_tx=XM_011282651.3;refGene_gn=ASF1A;refGene_tx=XM_023254298.1
B2 110031074 110031251
refGene_gn=ASF1A;refGene_tx=XM_011282651.3;refGene_gn=ASF1A;refGene_tx=XM_023254298.1
B2 110032490 110032703
refGene_gn=ASF1A;refGene_tx=XM_011282651.3;refGene_gn=ASF1A;refGene_tx=XM_023254298.1
B2 110036649 110036769
refGene_gn=MCM9;refGene_tx=XM_003986512.5;refGene_gn=MCM9;refGene_tx=XM_023254296.1;refGene_gn=MCM9;refGene_tx=XM_023254297.1
B2 110038194 110038320
refGene_gn=MCM9;refGene_tx=XM_003986512.5;refGene_gn=MCM9;refGene_tx=XM_023254296.1;refGene_gn=MCM9;refGene_tx=XM_023254297.1
B2 110041979 110042180
refGene_gn=MCM9;refGene_tx=XM_003986512.5;refGene_gn=MCM9;refGene_tx=XM_023254297.1
B2 110047212 110047294
refGene_gn=MCM9;refGene_tx=XM_003986512.5;refGene_gn=MCM9;refGene_tx=XM_023254296.1;refGene_gn=MCM9;refGene_tx=XM_023254297.1
B2 110048742 110049059
refGene_gn=MCM9;refGene_tx=XM_003986512.5;refGene_gn=MCM9;refGene_tx=XM_023254296.1
B2 110057936 110058240
refGene_gn=MCM9;refGene_tx=XM_003986512.5;refGene_gn=MCM9;refGene_tx=XM_023254296.1;refGene_gn=MCM9;refGene_tx=XM_023254297.1
B2 110114844 110114926
refGene_gn=FAM184A;refGene_tx=XM_003986510.5;refGene_gn=FAM184A;refGene_tx=XM_019831135.2;refGene_gn=FAM184A;refGene_tx=XM_019831136.2;refGene_gn=FAM184A;refGene_tx=XM_019831137.2;refGene_gn=FAM184A;refGene_tx=XM_019831139.2;refGene_gn=FAM184A;refGene_tx=XM_019831140.2;refGene_gn=FAM184A;refGene_tx=XM_019831142.2;refGene_gn=FAM184A;refGene_tx=XM_019831144.2
B2 110115647 110115657
refGene_gn=FAM184A;refGene_tx=XM_019831138.2
B2 110116493 110116729
refGene_gn=FAM184A;refGene_tx=XM_003986510.5;refGene_gn=FAM184A;refGene_tx=XM_019831135.2;refGene_gn=FAM184A;refGene_tx=XM_019831136.2;refGe

ne_gn=FAM184A;refGene_tx=XM_019831137.2;refGene_gn=FAM184A;refGene_tx=XM_019831138.2;refGene_gn=FAM184A;refGene_tx=XM_019831139.2;refGene_gn=FAM184A;refGene_tx=XM_019831140.2;refGene_gn=FAM184A;refGene_tx=XM_019831142.2;refGene_gn=FAM184A;refGene_tx=XM_019831144.2
B2 110117113 110117134
refGene_gn=FAM184A;refGene_tx=XM_019831143.2
B2 110119152 110119257
refGene_gn=FAM184A;refGene_tx=XM_003986510.5;refGene_gn=FAM184A;refGene_tx=XM_019831135.2;refGene_gn=FAM184A;refGene_tx=XM_019831136.2;refGene_gn=FAM184A;refGene_tx=XM_019831137.2;refGene_gn=FAM184A;refGene_tx=XM_019831138.2;refGene_gn=FAM184A;refGene_tx=XM_019831140.2;refGene_gn=FAM184A;refGene_tx=XM_019831142.2;refGene_gn=FAM184A;refGene_tx=XM_019831143.2
B2 110121862 110121980
refGene_gn=FAM184A;refGene_tx=XM_003986510.5;refGene_gn=FAM184A;refGene_tx=XM_019831135.2;refGene_gn=FAM184A;refGene_tx=XM_019831136.2;refGene_gn=FAM184A;refGene_tx=XM_019831137.2;refGene_gn=FAM184A;refGene_tx=XM_019831138.2;refGene_gn=FAM184A;refGene_tx=XM_019831139.2;refGene_gn=FAM184A;refGene_tx=XM_019831140.2;refGene_gn=FAM184A;refGene_tx=XM_019831142.2;refGene_gn=FAM184A;refGene_tx=XM_019831143.2;refGene_gn=FAM184A;refGene_tx=XM_019831144.2
B2 110128897 110129044
refGene_gn=FAM184A;refGene_tx=XM_003986510.5;refGene_gn=FAM184A;refGene_tx=XM_019831135.2;refGene_gn=FAM184A;refGene_tx=XM_019831136.2;refGene_gn=FAM184A;refGene_tx=XM_019831137.2;refGene_gn=FAM184A;refGene_tx=XM_019831138.2;refGene_gn=FAM184A;refGene_tx=XM_019831139.2;refGene_gn=FAM184A;refGene_tx=XM_019831143.2
B2 110129464 110129649
refGene_gn=FAM184A;refGene_tx=XM_003986510.5;refGene_gn=FAM184A;refGene_tx=XM_019831135.2;refGene_gn=FAM184A;refGene_tx=XM_019831136.2;refGene_gn=FAM184A;refGene_tx=XM_019831137.2;refGene_gn=FAM184A;refGene_tx=XM_019831138.2;refGene_gn=FAM184A;refGene_tx=XM_019831139.2;refGene_gn=FAM184A;refGene_tx=XM_019831140.2;refGene_gn=FAM184A;refGene_tx=XM_019831142.2;refGene_gn=FAM184A;refGene_tx=XM_019831143.2;refGene_gn=FAM184A;refGene_tx=XM_019831144.2
B2 110131107 110131235
refGene_gn=FAM184A;refGene_tx=XM_003986510.5;refGene_gn=FAM184A;refGene_tx=XM_019831135.2;refGene_gn=FAM184A;refGene_tx=XM_019831136.2;refGene_gn=FAM184A;refGene_tx=XM_019831137.2;refGene_gn=FAM184A;refGene_tx=XM_019831138.2;refGene_gn=FAM184A;refGene_tx=XM_019831139.2;refGene_gn=FAM184A;refGene_tx=XM_019831140.2;refGene_gn=FAM184A;refGene_tx=XM_019831142.2;refGene_gn=FAM184A;refGene_tx=XM_019831143.2;refGene_gn=FAM184A;refGene_tx=XM_019831144.2
B2 110135260 110135414
refGene_gn=FAM184A;refGene_tx=XM_003986510.5;refGene_gn=FAM184A;refGene_tx=XM_019831135.2;refGene_gn=FAM184A;refGene_tx=XM_019831136.2;refGene_gn=FAM184A;refGene_tx=XM_019831137.2;refGene_gn=FAM184A;refGene_tx=XM_019831138.2;refGene_gn=FAM184A;refGene_tx=XM_019831139.2;refGene_gn=FAM184A;refGene_tx=XM_019831140.2;refGene_gn=FAM184A;refGene_tx=XM_019831142.2;refGene_gn=FAM184A;refGene_tx=XM_019831143.2;refGene_gn=FAM1

84A;refGene_tx=XM_019831144.2
B2	110144930	110145143
refGene_gn=FAM184A;refGene_tx=XM_003986510.5;refGene_gn=FAM184A;refGene_tx=XM_019831135.2;refGene_gn=FAM184A;refGene_tx=XM_019831136.2;refGene_gn=FAM184A;refGene_tx=XM_019831137.2;refGene_gn=FAM184A;refGene_tx=XM_019831138.2;refGene_gn=FAM184A;refGene_tx=XM_019831139.2;refGene_gn=FAM184A;refGene_tx=XM_019831140.2;refGene_gn=FAM184A;refGene_tx=XM_019831142.2;refGene_gn=FAM184A;refGene_tx=XM_019831143.2;refGene_gn=FAM184A;refGene_tx=XM_019831144.2
B2	110151348	110151390
refGene_gn=FAM184A;refGene_tx=XM_019831135.2;refGene_gn=FAM184A;refGene_tx=XM_019831136.2;refGene_gn=FAM184A;refGene_tx=XM_019831138.2;refGene_gn=FAM184A;refGene_tx=XM_019831139.2;refGene_gn=FAM184A;refGene_tx=XM_019831140.2;refGene_gn=FAM184A;refGene_tx=XM_019831143.2
B2	110164225	110164376
refGene_gn=FAM184A;refGene_tx=XM_003986510.5;refGene_gn=FAM184A;refGene_tx=XM_019831135.2;refGene_gn=FAM184A;refGene_tx=XM_019831136.2;refGene_gn=FAM184A;refGene_tx=XM_019831137.2;refGene_gn=FAM184A;refGene_tx=XM_019831138.2;refGene_gn=FAM184A;refGene_tx=XM_019831139.2;refGene_gn=FAM184A;refGene_tx=XM_019831140.2;refGene_gn=FAM184A;refGene_tx=XM_019831142.2;refGene_gn=FAM184A;refGene_tx=XM_019831143.2;refGene_gn=FAM184A;refGene_tx=XM_019831144.2
B2	110165049	110165171
refGene_gn=FAM184A;refGene_tx=XM_003986510.5;refGene_gn=FAM184A;refGene_tx=XM_019831135.2;refGene_gn=FAM184A;refGene_tx=XM_019831136.2;refGene_gn=FAM184A;refGene_tx=XM_019831137.2;refGene_gn=FAM184A;refGene_tx=XM_019831138.2;refGene_gn=FAM184A;refGene_tx=XM_019831139.2;refGene_gn=FAM184A;refGene_tx=XM_019831140.2;refGene_gn=FAM184A;refGene_tx=XM_019831142.2;refGene_gn=FAM184A;refGene_tx=XM_019831143.2;refGene_gn=FAM184A;refGene_tx=XM_019831144.2
B2	110168213	110168375
refGene_gn=FAM184A;refGene_tx=XM_003986510.5;refGene_gn=FAM184A;refGene_tx=XM_019831135.2;refGene_gn=FAM184A;refGene_tx=XM_019831136.2;refGene_gn=FAM184A;refGene_tx=XM_019831137.2;refGene_gn=FAM184A;refGene_tx=XM_019831138.2;refGene_gn=FAM184A;refGene_tx=XM_019831139.2;refGene_gn=FAM184A;refGene_tx=XM_019831140.2;refGene_gn=FAM184A;refGene_tx=XM_019831142.2;refGene_gn=FAM184A;refGene_tx=XM_019831143.2;refGene_gn=FAM184A;refGene_tx=XM_019831144.2
B2	110180802	110180925
refGene_gn=FAM184A;refGene_tx=XM_003986510.5;refGene_gn=FAM184A;refGene_tx=XM_019831135.2;refGene_gn=FAM184A;refGene_tx=XM_019831136.2;refGene_gn=FAM184A;refGene_tx=XM_019831137.2;refGene_gn=FAM184A;refGene_tx=XM_019831138.2;refGene_gn=FAM184A;refGene_tx=XM_019831139.2;refGene_gn=FAM184A;refGene_tx=XM_019831140.2;refGene_gn=FAM184A;refGene_tx=XM_019831142.2;refGene_gn=FAM184A;refGene_tx=XM_019831143.2;refGene_gn=FAM184A;refGene_tx=XM_019831144.2
B2	110184716	110184914
refGene_gn=FAM184A;refGene_tx=XM_003986510.5;refGene_gn=FAM184A;refGene_tx=XM_019831135.2;refGene_gn=FAM184A;refGene_tx=XM_019831136.2;refGene_gn=FAM184A;refGene_tx=XM_019831137.2;refGene_gn=FAM184A;refGene_tx=

XM_019831138.2;refGene_gn=FAM184A;refGene_tx=XM_019831139.2;refGene_gn=FAM184A;refGene_tx=XM_019831140.2;refGene_gn=FAM184A;refGene_tx=XM_019831142.2;refGene_gn=FAM184A;refGene_tx=XM_019831143.2;refGene_gn=FAM184A;refGene_tx=XM_019831144.2
B2 110188046 110188228
refGene_gn=FAM184A;refGene_tx=XM_003986510.5;refGene_gn=FAM184A;refGene_tx=XM_019831135.2;refGene_gn=FAM184A;refGene_tx=XM_019831136.2;refGene_gn=FAM184A;refGene_tx=XM_019831137.2;refGene_gn=FAM184A;refGene_tx=XM_019831138.2;refGene_gn=FAM184A;refGene_tx=XM_019831139.2;refGene_gn=FAM184A;refGene_tx=XM_019831140.2;refGene_gn=FAM184A;refGene_tx=XM_019831142.2;refGene_gn=FAM184A;refGene_tx=XM_019831143.2;refGene_gn=FAM184A;refGene_tx=XM_019831144.2
B2 110189836 110189972
refGene_gn=FAM184A;refGene_tx=XM_003986510.5;refGene_gn=FAM184A;refGene_tx=XM_019831135.2;refGene_gn=FAM184A;refGene_tx=XM_019831136.2;refGene_gn=FAM184A;refGene_tx=XM_019831137.2;refGene_gn=FAM184A;refGene_tx=XM_019831138.2;refGene_gn=FAM184A;refGene_tx=XM_019831139.2;refGene_gn=FAM184A;refGene_tx=XM_019831140.2;refGene_gn=FAM184A;refGene_tx=XM_019831142.2;refGene_gn=FAM184A;refGene_tx=XM_019831143.2;refGene_gn=FAM184A;refGene_tx=XM_019831144.2
B2 110190626 110191481
refGene_gn=FAM184A;refGene_tx=XM_003986510.5;refGene_gn=FAM184A;refGene_tx=XM_019831135.2;refGene_gn=FAM184A;refGene_tx=XM_019831136.2;refGene_gn=FAM184A;refGene_tx=XM_019831137.2;refGene_gn=FAM184A;refGene_tx=XM_019831138.2;refGene_gn=FAM184A;refGene_tx=XM_019831139.2;refGene_gn=FAM184A;refGene_tx=XM_019831140.2;refGene_gn=FAM184A;refGene_tx=XM_019831142.2;refGene_gn=FAM184A;refGene_tx=XM_019831143.2;refGene_gn=FAM184A;refGene_tx=XM_019831144.2
B2 110236661 110236829
refGene_gn=FAM184A;refGene_tx=XM_003986510.5;refGene_gn=FAM184A;refGene_tx=XM_019831135.2;refGene_gn=FAM184A;refGene_tx=XM_019831136.2;refGene_gn=FAM184A;refGene_tx=XM_019831137.2;refGene_gn=FAM184A;refGene_tx=XM_019831138.2;refGene_gn=FAM184A;refGene_tx=XM_019831139.2;refGene_gn=FAM184A;refGene_tx=XM_019831140.2;refGene_gn=FAM184A;refGene_tx=XM_019831142.2;refGene_gn=FAM184A;refGene_tx=XM_019831143.2;refGene_gn=FAM184A;refGene_tx=XM_019831144.2
B2 110335728 110335855
refGene_gn=MAN1A1;refGene_tx=XM_023254299.1
B2 110336256 110336372
refGene_gn=MAN1A1;refGene_tx=XM_023254299.1
B2 110345062 110345235
refGene_gn=MAN1A1;refGene_tx=XM_023254299.1
B2 110346163 110346383
refGene_gn=MAN1A1;refGene_tx=XM_023254299.1
B2 110352220 110352336
refGene_gn=MAN1A1;refGene_tx=XM_023254299.1
B2 110359556 110359650
refGene_gn=MAN1A1;refGene_tx=XM_023254299.1
B2 110361556 110361680
refGene_gn=MAN1A1;refGene_tx=XM_023254299.1

B2	110402880	110402975
refGene_gn=MAN1A1;refGene_tx=XM_023254299.1
B2	110439168	110439249
refGene_gn=MAN1A1;refGene_tx=XM_023254299.1
B2	110453884	110454000
refGene_gn=MAN1A1;refGene_tx=XM_023254299.1
B2	110456279	110456376
refGene_gn=MAN1A1;refGene_tx=XM_023254299.1
B2	110495610	110496231
refGene_gn=MAN1A1;refGene_tx=XM_023254299.1
B2	111918954	111919074
refGene_gn=TBC1D32;refGene_tx=XM_011282635.3;refGene_gn=TBC1D32;refGene_tx=XM_023254300.1;refGene_gn=TBC1D32;refGene_tx=XM_023254301.1;refGene_gn=TBC1D32;refGene_tx=XM_023254302.1;refGene_gn=TBC1D32;refGene_tx=XM_023254303.1
B2	111930294	111930483
refGene_gn=TBC1D32;refGene_tx=XM_011282635.3;refGene_gn=TBC1D32;refGene_tx=XM_023254300.1;refGene_gn=TBC1D32;refGene_tx=XM_023254301.1;refGene_gn=TBC1D32;refGene_tx=XM_023254302.1;refGene_gn=TBC1D32;refGene_tx=XM_023254303.1
B2	111940670	111940811
refGene_gn=TBC1D32;refGene_tx=XM_011282635.3;refGene_gn=TBC1D32;refGene_tx=XM_023254300.1;refGene_gn=TBC1D32;refGene_tx=XM_023254301.1;refGene_gn=TBC1D32;refGene_tx=XM_023254302.1
B2	111944751	111944906
refGene_gn=TBC1D32;refGene_tx=XM_011282635.3;refGene_gn=TBC1D32;refGene_tx=XM_023254300.1;refGene_gn=TBC1D32;refGene_tx=XM_023254301.1;refGene_gn=TBC1D32;refGene_tx=XM_023254302.1;refGene_gn=TBC1D32;refGene_tx=XM_023254303.1
B2	111945658	111945774
refGene_gn=TBC1D32;refGene_tx=XM_011282635.3;refGene_gn=TBC1D32;refGene_tx=XM_023254300.1;refGene_gn=TBC1D32;refGene_tx=XM_023254301.1;refGene_gn=TBC1D32;refGene_tx=XM_023254302.1;refGene_gn=TBC1D32;refGene_tx=XM_023254303.1
B2	111947706	111947776
refGene_gn=TBC1D32;refGene_tx=XM_011282635.3;refGene_gn=TBC1D32;refGene_tx=XM_023254300.1;refGene_gn=TBC1D32;refGene_tx=XM_023254301.1;refGene_gn=TBC1D32;refGene_tx=XM_023254302.1;refGene_gn=TBC1D32;refGene_tx=XM_023254303.1
B2	111962460	111962544
refGene_gn=TBC1D32;refGene_tx=XM_011282635.3;refGene_gn=TBC1D32;refGene_tx=XM_023254300.1;refGene_gn=TBC1D32;refGene_tx=XM_023254301.1;refGene_gn=TBC1D32;refGene_tx=XM_023254302.1;refGene_gn=TBC1D32;refGene_tx=XM_023254303.1
B2	111968341	111968467
refGene_gn=TBC1D32;refGene_tx=XM_011282635.3;refGene_gn=TBC1D32;refGene_tx=XM_023254300.1;refGene_gn=TBC1D32;refGene_tx=XM_023254301.1;refGene_gn=TBC1D32;refGene_tx=XM_023254302.1;refGene_gn=TBC1D32;refGene_tx=XM_023254303.1
B2	111990464	111990558

refGene_gn=TBC1D32;refGene_tx=XM_011282635.3;refGene_gn=TBC1D32;refGene_tx=XM_023254300.1;refGene_gn=TBC1D32;refGene_tx=XM_023254301.1;refGene_gn=TBC1D32;refGene_tx=XM_023254302.1;refGene_gn=TBC1D32;refGene_tx=XM_023254303.1
B2 111991251 111991360
refGene_gn=TBC1D32;refGene_tx=XM_011282635.3;refGene_gn=TBC1D32;refGene_tx=XM_023254300.1;refGene_gn=TBC1D32;refGene_tx=XM_023254301.1;refGene_gn=TBC1D32;refGene_tx=XM_023254302.1;refGene_gn=TBC1D32;refGene_tx=XM_023254303.1
B2 111992667 111992790
refGene_gn=TBC1D32;refGene_tx=XM_011282635.3;refGene_gn=TBC1D32;refGene_tx=XM_023254300.1;refGene_gn=TBC1D32;refGene_tx=XM_023254301.1;refGene_gn=TBC1D32;refGene_tx=XM_023254302.1;refGene_gn=TBC1D32;refGene_tx=XM_023254303.1
B2 112015786 112015875
refGene_gn=TBC1D32;refGene_tx=XM_011282635.3;refGene_gn=TBC1D32;refGene_tx=XM_023254300.1;refGene_gn=TBC1D32;refGene_tx=XM_023254301.1;refGene_gn=TBC1D32;refGene_tx=XM_023254302.1;refGene_gn=TBC1D32;refGene_tx=XM_023254303.1
B2 112028889 112029006
refGene_gn=TBC1D32;refGene_tx=XM_011282635.3;refGene_gn=TBC1D32;refGene_tx=XM_023254300.1;refGene_gn=TBC1D32;refGene_tx=XM_023254301.1;refGene_gn=TBC1D32;refGene_tx=XM_023254302.1;refGene_gn=TBC1D32;refGene_tx=XM_023254303.1
B2 112040232 112040351
refGene_gn=TBC1D32;refGene_tx=XM_011282635.3;refGene_gn=TBC1D32;refGene_tx=XM_023254300.1;refGene_gn=TBC1D32;refGene_tx=XM_023254301.1;refGene_gn=TBC1D32;refGene_tx=XM_023254302.1;refGene_gn=TBC1D32;refGene_tx=XM_023254303.1
B2 112042268 112042356
refGene_gn=TBC1D32;refGene_tx=XM_011282635.3;refGene_gn=TBC1D32;refGene_tx=XM_023254300.1;refGene_gn=TBC1D32;refGene_tx=XM_023254301.1;refGene_gn=TBC1D32;refGene_tx=XM_023254302.1;refGene_gn=TBC1D32;refGene_tx=XM_023254303.1
B2 112045457 112045596
refGene_gn=TBC1D32;refGene_tx=XM_011282635.3;refGene_gn=TBC1D32;refGene_tx=XM_023254300.1;refGene_gn=TBC1D32;refGene_tx=XM_023254301.1;refGene_gn=TBC1D32;refGene_tx=XM_023254302.1;refGene_gn=TBC1D32;refGene_tx=XM_023254303.1
B2 112053600 112053683
refGene_gn=TBC1D32;refGene_tx=XM_011282635.3;refGene_gn=TBC1D32;refGene_tx=XM_023254300.1;refGene_gn=TBC1D32;refGene_tx=XM_023254301.1;refGene_gn=TBC1D32;refGene_tx=XM_023254302.1;refGene_gn=TBC1D32;refGene_tx=XM_023254303.1
B2 112054250 112054452
refGene_gn=TBC1D32;refGene_tx=XM_011282635.3;refGene_gn=TBC1D32;refGene_tx=XM_023254300.1;refGene_gn=TBC1D32;refGene_tx=XM_023254301.1;refGene_gn=TBC1D32;refGene_tx=XM_023254302.1;refGene_gn=TBC1D32;refGene_tx=XM_023254303.1
B2 112058863 112058991

refGene_gn=TBC1D32;refGene_tx=XM_011282635.3;refGene_gn=TBC1D32;refGene_tx=XM_023254300.1;refGene_gn=TBC1D32;refGene_tx=XM_023254301.1;refGene_gn=TBC1D32;refGene_tx=XM_023254302.1;refGene_gn=TBC1D32;refGene_tx=XM_023254303.1
B2 112062865 112063008
refGene_gn=TBC1D32;refGene_tx=XM_011282635.3;refGene_gn=TBC1D32;refGene_tx=XM_023254300.1;refGene_gn=TBC1D32;refGene_tx=XM_023254301.1;refGene_gn=TBC1D32;refGene_tx=XM_023254302.1;refGene_gn=TBC1D32;refGene_tx=XM_023254303.1
B2 112064796 112064889
refGene_gn=TBC1D32;refGene_tx=XM_011282635.3;refGene_gn=TBC1D32;refGene_tx=XM_023254300.1;refGene_gn=TBC1D32;refGene_tx=XM_023254301.1;refGene_gn=TBC1D32;refGene_tx=XM_023254302.1;refGene_gn=TBC1D32;refGene_tx=XM_023254303.1
B2 112067540 112067681
refGene_gn=TBC1D32;refGene_tx=XM_011282635.3;refGene_gn=TBC1D32;refGene_tx=XM_023254300.1;refGene_gn=TBC1D32;refGene_tx=XM_023254301.1;refGene_gn=TBC1D32;refGene_tx=XM_023254303.1
B2 112071165 112071256
refGene_gn=TBC1D32;refGene_tx=XM_011282635.3;refGene_gn=TBC1D32;refGene_tx=XM_023254300.1;refGene_gn=TBC1D32;refGene_tx=XM_023254301.1;refGene_gn=TBC1D32;refGene_tx=XM_023254302.1;refGene_gn=TBC1D32;refGene_tx=XM_023254303.1
B2 112079366 112079426
refGene_gn=TBC1D32;refGene_tx=XM_011282635.3;refGene_gn=TBC1D32;refGene_tx=XM_023254300.1;refGene_gn=TBC1D32;refGene_tx=XM_023254301.1;refGene_gn=TBC1D32;refGene_tx=XM_023254302.1;refGene_gn=TBC1D32;refGene_tx=XM_023254303.1
B2 112083535 112083680
refGene_gn=TBC1D32;refGene_tx=XM_011282635.3;refGene_gn=TBC1D32;refGene_tx=XM_023254300.1;refGene_gn=TBC1D32;refGene_tx=XM_023254301.1;refGene_gn=TBC1D32;refGene_tx=XM_023254302.1;refGene_gn=TBC1D32;refGene_tx=XM_023254303.1
B2 112084276 112084338
refGene_gn=TBC1D32;refGene_tx=XM_011282635.3;refGene_gn=TBC1D32;refGene_tx=XM_023254300.1;refGene_gn=TBC1D32;refGene_tx=XM_023254301.1;refGene_gn=TBC1D32;refGene_tx=XM_023254302.1;refGene_gn=TBC1D32;refGene_tx=XM_023254303.1
B2 112084430 112084534
refGene_gn=TBC1D32;refGene_tx=XM_011282635.3;refGene_gn=TBC1D32;refGene_tx=XM_023254300.1;refGene_gn=TBC1D32;refGene_tx=XM_023254301.1;refGene_gn=TBC1D32;refGene_tx=XM_023254302.1;refGene_gn=TBC1D32;refGene_tx=XM_023254303.1
B2 112084659 112084738
refGene_gn=TBC1D32;refGene_tx=XM_011282635.3;refGene_gn=TBC1D32;refGene_tx=XM_023254300.1;refGene_gn=TBC1D32;refGene_tx=XM_023254301.1;refGene_gn=TBC1D32;refGene_tx=XM_023254302.1;refGene_gn=TBC1D32;refGene_tx=XM_023254303.1
B2 112087506 112087632
refGene_gn=TBC1D32;refGene_tx=XM_011282635.3;refGene_gn=TBC1D32;refGen

e_tx=XM_023254300.1;refGene_gn=TBC1D32;refGene_tx=XM_023254302.1;refGene_gn=TBC1D32;refGene_tx=XM_023254303.1
B2 112090075 112090144
refGene_gn=TBC1D32;refGene_tx=XM_011282635.3;refGene_gn=TBC1D32;refGene_tx=XM_023254300.1;refGene_gn=TBC1D32;refGene_tx=XM_023254301.1;refGene_gn=TBC1D32;refGene_tx=XM_023254302.1;refGene_gn=TBC1D32;refGene_tx=XM_023254303.1
B2 112102421 112102599
refGene_gn=TBC1D32;refGene_tx=XM_011282635.3;refGene_gn=TBC1D32;refGene_tx=XM_023254300.1;refGene_gn=TBC1D32;refGene_tx=XM_023254301.1;refGene_gn=TBC1D32;refGene_tx=XM_023254302.1;refGene_gn=TBC1D32;refGene_tx=XM_023254303.1
B2 112111920 112112082
refGene_gn=TBC1D32;refGene_tx=XM_011282635.3;refGene_gn=TBC1D32;refGene_tx=XM_023254300.1;refGene_gn=TBC1D32;refGene_tx=XM_023254301.1;refGene_gn=TBC1D32;refGene_tx=XM_023254302.1;refGene_gn=TBC1D32;refGene_tx=XM_023254303.1
B2 112121682 112121837
refGene_gn=TBC1D32;refGene_tx=XM_011282635.3;refGene_gn=TBC1D32;refGene_tx=XM_023254300.1;refGene_gn=TBC1D32;refGene_tx=XM_023254301.1;refGene_gn=TBC1D32;refGene_tx=XM_023254302.1;refGene_gn=TBC1D32;refGene_tx=XM_023254303.1
B2 112198496 112199645
refGene_gn=GJA1;refGene_tx=XM_003986514.5
B2 113063386 113063479
refGene_gn=HSF2;refGene_tx=XM_023254304.1;refGene_gn=HSF2;refGene_tx=XM_023254305.1;refGene_gn=HSF2;refGene_tx=XM_023254306.1;refGene_gn=HSF2;refGene_tx=XM_023254307.1;refGene_gn=HSF2;refGene_tx=XM_023254308.1
B2 113064055 113064100
refGene_gn=HSF2;refGene_tx=XM_011282630.3
B2 113077319 113077428
refGene_gn=HSF2;refGene_tx=XM_011282630.3;refGene_gn=HSF2;refGene_tx=XM_023254304.1;refGene_gn=HSF2;refGene_tx=XM_023254305.1;refGene_gn=HSF2;refGene_tx=XM_023254306.1;refGene_gn=HSF2;refGene_tx=XM_023254307.1;refGene_gn=HSF2;refGene_tx=XM_023254308.1
B2 113077564 113077692
refGene_gn=HSF2;refGene_tx=XM_011282630.3;refGene_gn=HSF2;refGene_tx=XM_023254304.1;refGene_gn=HSF2;refGene_tx=XM_023254305.1;refGene_gn=HSF2;refGene_tx=XM_023254306.1;refGene_gn=HSF2;refGene_tx=XM_023254307.1;refGene_gn=HSF2;refGene_tx=XM_023254308.1
B2 113078561 113078686
refGene_gn=HSF2;refGene_tx=XM_011282630.3;refGene_gn=HSF2;refGene_tx=XM_023254304.1;refGene_gn=HSF2;refGene_tx=XM_023254305.1;refGene_gn=HSF2;refGene_tx=XM_023254306.1;refGene_gn=HSF2;refGene_tx=XM_023254307.1;refGene_gn=HSF2;refGene_tx=XM_023254308.1
B2 113081551 113081627
refGene_gn=HSF2;refGene_tx=XM_011282630.3;refGene_gn=HSF2;refGene_tx=XM_023254304.1;refGene_gn=HSF2;refGene_tx=XM_023254305.1;refGene_gn=HSF2;refGene_tx=XM_023254306.1;refGene_gn=HSF2;refGene_tx=XM_023254307.1;refGene_gn=HSF2;refGene_tx=XM_023254308.1

B2 113085191 113085253
refGene_gn=HSF2;refGene_tx=XM_011282630.3;refGene_gn=HSF2;refGene_tx=XM_023254304.1;refGene_gn=HSF2;refGene_tx=XM_023254305.1;refGene_gn=HSF2;refGene_tx=XM_023254306.1;refGene_gn=HSF2;refGene_tx=XM_023254307.1;refGene_gn=HSF2;refGene_tx=XM_023254308.1
B2 113086399 113086487
refGene_gn=HSF2;refGene_tx=XM_011282630.3;refGene_gn=HSF2;refGene_tx=XM_023254304.1;refGene_gn=HSF2;refGene_tx=XM_023254305.1;refGene_gn=HSF2;refGene_tx=XM_023254306.1;refGene_gn=HSF2;refGene_tx=XM_023254307.1;refGene_gn=HSF2;refGene_tx=XM_023254308.1
B2 113088262 113088411
refGene_gn=HSF2;refGene_tx=XM_011282630.3;refGene_gn=HSF2;refGene_tx=XM_023254304.1;refGene_gn=HSF2;refGene_tx=XM_023254305.1;refGene_gn=HSF2;refGene_tx=XM_023254306.1;refGene_gn=HSF2;refGene_tx=XM_023254307.1;refGene_gn=HSF2;refGene_tx=XM_023254308.1
B2 113088822 113089062
refGene_gn=HSF2;refGene_tx=XM_011282630.3;refGene_gn=HSF2;refGene_tx=XM_023254304.1;refGene_gn=HSF2;refGene_tx=XM_023254305.1;refGene_gn=HSF2;refGene_tx=XM_023254306.1;refGene_gn=HSF2;refGene_tx=XM_023254307.1;refGene_gn=HSF2;refGene_tx=XM_023254308.1
B2 113089348 113089454
refGene_gn=HSF2;refGene_tx=XM_011282630.3;refGene_gn=HSF2;refGene_tx=XM_023254304.1;refGene_gn=HSF2;refGene_tx=XM_023254305.1;refGene_gn=HSF2;refGene_tx=XM_023254306.1;refGene_gn=HSF2;refGene_tx=XM_023254307.1;refGene_gn=HSF2;refGene_tx=XM_023254308.1
B2 113094143 113094197
refGene_gn=HSF2;refGene_tx=XM_011282630.3;refGene_gn=HSF2;refGene_tx=XM_023254304.1;refGene_gn=HSF2;refGene_tx=XM_023254306.1;refGene_gn=HSF2;refGene_tx=XM_023254307.1
B2 113097182 113097267
refGene_gn=HSF2;refGene_tx=XM_011282630.3;refGene_gn=HSF2;refGene_tx=XM_023254304.1;refGene_gn=HSF2;refGene_tx=XM_023254305.1;refGene_gn=HSF2;refGene_tx=XM_023254306.1;refGene_gn=HSF2;refGene_tx=XM_023254307.1;refGene_gn=HSF2;refGene_tx=XM_023254308.1
B2 113097727 113098020
refGene_gn=HSF2;refGene_tx=XM_011282630.3;refGene_gn=HSF2;refGene_tx=XM_023254304.1;refGene_gn=HSF2;refGene_tx=XM_023254305.1;refGene_gn=HSF2;refGene_tx=XM_023254306.1
B2 113100153 113100158
refGene_gn=HSF2;refGene_tx=XM_023254306.1;refGene_gn=HSF2;refGene_tx=XM_023254307.1;refGene_gn=HSF2;refGene_tx=XM_023254308.1
B2 113104603 113104739
refGene_gn=SERINC1;refGene_tx=XM_003986516.5
B2 113106387 113106618
refGene_gn=SERINC1;refGene_tx=XM_003986516.5
B2 113106733 113106878
refGene_gn=SERINC1;refGene_tx=XM_003986516.5
B2 113108474 113108565
refGene_gn=SERINC1;refGene_tx=XM_003986516.5
B2 113108674 113108844

refGene_gn=SERINC1;refGene_tx=XM_003986516.5
B2 113110671 113110809
refGene_gn=SERINC1;refGene_tx=XM_003986516.5
B2 113111041 113111121
refGene_gn=SERINC1;refGene_tx=XM_003986516.5
B2 113119310 113119480
refGene_gn=SERINC1;refGene_tx=XM_003986516.5
B2 113122544 113122706
refGene_gn=SERINC1;refGene_tx=XM_003986516.5
B2 113138987 113139026
refGene_gn=SERINC1;refGene_tx=XM_003986516.5
B2 113173907 113174163
refGene_gn=PKIB;refGene_tx=XM_019831156.2;refGene_gn=PKIB;refGene_tx=XM_019831158.2;refGene_gn=PKIB;refGene_tx=XM_023254309.1
B2 113194807 113194853
refGene_gn=PKIB;refGene_tx=XM_019831156.2;refGene_gn=PKIB;refGene_tx=XM_019831158.2;refGene_gn=PKIB;refGene_tx=XM_023254309.1
B2 113195797 113195843
refGene_gn=PKIB;refGene_tx=XM_019831158.2
B2 113260288 113260386
refGene_gn=PKIB;refGene_tx=XM_019831156.2;refGene_gn=PKIB;refGene_tx=XM_023254309.1
B2 113294936 113295048
refGene_gn=PKIB;refGene_tx=XM_023254309.1
B2 113302734 113302790
refGene_gn=PKIB;refGene_tx=XM_019831156.2;refGene_gn=PKIB;refGene_tx=XM_023254309.1
B2 113317993 113318081
refGene_gn=PKIB;refGene_tx=XM_006931971.4;refGene_gn=PKIB;refGene_tx=XM_019831159.2
B2 113339807 113339984
refGene_gn=PKIB;refGene_tx=XM_006931971.4;refGene_gn=PKIB;refGene_tx=XM_019831155.2;refGene_gn=PKIB;refGene_tx=XM_019831159.2;refGene_gn=PKIB;refGene_tx=XM_019831160.2;refGene_gn=PKIB;refGene_tx=XM_019831161.2;refGene_gn=PKIB;refGene_tx=XM_019831162.2
B2 113344589 113344657
refGene_gn=PKIB;refGene_tx=XM_006931971.4;refGene_gn=PKIB;refGene_tx=XM_019831155.2;refGene_gn=PKIB;refGene_tx=XM_019831159.2;refGene_gn=PKIB;refGene_tx=XM_019831160.2;refGene_gn=PKIB;refGene_tx=XM_019831161.2;refGene_gn=PKIB;refGene_tx=XM_019831162.2
B2 113387708 113387781
refGene_gn=FABP7;refGene_tx=XM_003986517.5
B2 113388225 113388398
refGene_gn=FABP7;refGene_tx=XM_003986517.5
B2 113389017 113389071
refGene_gn=FABP7;refGene_tx=XM_003986517.5
B2 113389072 113389120
refGene_gn=FABP7;refGene_tx=XM_003986517.5
B2 113390939 113390990
refGene_gn=FABP7;refGene_tx=XM_003986517.5

B2 113396413 113396516
refGene_gn=SMPDL3A;refGene_tx=XM_003986518.5
B2 113401505 113401719
refGene_gn=SMPDL3A;refGene_tx=XM_003986518.5
B2 113402450 113402595
refGene_gn=SMPDL3A;refGene_tx=XM_003986518.5;refGene_gn=SMPDL3A;refGene_tx=XM_019831163.2
B2 113406908 113407005
refGene_gn=SMPDL3A;refGene_tx=XM_003986518.5;refGene_gn=SMPDL3A;refGene_tx=XM_019831163.2
B2 113407749 113407919
refGene_gn=SMPDL3A;refGene_tx=XM_003986518.5;refGene_gn=SMPDL3A;refGene_tx=XM_019831163.2
B2 113408388 113408569
refGene_gn=SMPDL3A;refGene_tx=XM_003986518.5;refGene_gn=SMPDL3A;refGene_tx=XM_019831163.2
B2 113409999 113410124
refGene_gn=SMPDL3A;refGene_tx=XM_003986518.5;refGene_gn=SMPDL3A;refGene_tx=XM_019831163.2
B2 113412941 113413259
refGene_gn=SMPDL3A;refGene_tx=XM_003986518.5;refGene_gn=SMPDL3A;refGene_tx=XM_019831163.2
B2 113524443 113524832
refGene_gn=CLVS2;refGene_tx=XM_006931973.4;refGene_gn=CLVS2;refGene_tx=XM_019831164.2
B2 113535911 113536086
refGene_gn=CLVS2;refGene_tx=XM_006931973.4;refGene_gn=CLVS2;refGene_tx=XM_019831164.2
B2 113575768 113575879
refGene_gn=CLVS2;refGene_tx=XM_006931973.4;refGene_gn=CLVS2;refGene_tx=XM_019831164.2
B2 113582164 113582385
refGene_gn=CLVS2;refGene_tx=XM_006931973.4;refGene_gn=CLVS2;refGene_tx=XM_019831164.2
B2 113587444 113587532
refGene_gn=CLVS2;refGene_tx=XM_006931973.4;refGene_gn=CLVS2;refGene_tx=XM_019831164.2
B2 113733402 113733542
refGene_gn=TRDN;refGene_tx=XM_023254312.1
B2 113736949 113736985
refGene_gn=TRDN;refGene_tx=XM_023254312.1
B2 113739060 113739102
refGene_gn=TRDN;refGene_tx=XM_023254312.1
B2 113760012 113760033
refGene_gn=TRDN;refGene_tx=XM_023254312.1
B2 113762890 113762935
refGene_gn=TRDN;refGene_tx=XM_023254312.1
B2 113763701 113763737
refGene_gn=TRDN;refGene_tx=XM_023254312.1
B2 113764108 113764147

refGene_gn=TRDN;refGene_tx=XM_023254312.1
B2 113764916 113764943
refGene_gn=TRDN;refGene_tx=XM_023254312.1
B2 113766674 113766695
refGene_gn=TRDN;refGene_tx=XM_023254312.1
B2 113768726 113768771
refGene_gn=TRDN;refGene_tx=XM_023254312.1
B2 113770632 113770650
refGene_gn=TRDN;refGene_tx=XM_023254312.1
B2 113771970 113772018
refGene_gn=TRDN;refGene_tx=XM_023254312.1
B2 113773841 113773889
refGene_gn=TRDN;refGene_tx=XM_023254312.1
B2 113775493 113775523
refGene_gn=TRDN;refGene_tx=XM_023254312.1
B2 113786093 113786123
refGene_gn=TRDN;refGene_tx=XM_023254312.1
B2 113786615 113786642
refGene_gn=TRDN;refGene_tx=XM_023254312.1
B2 113814039 113814078
refGene_gn=TRDN;refGene_tx=XM_023254312.1
B2 113831435 113831486
refGene_gn=TRDN;refGene_tx=XM_023254312.1
B2 113837561 113837612
refGene_gn=TRDN;refGene_tx=XM_023254312.1
B2 113855578 113855626
refGene_gn=TRDN;refGene_tx=XM_023254312.1
B2 113865814 113865862
refGene_gn=TRDN;refGene_tx=XM_023254312.1
B2 113874189 113874216
refGene_gn=TRDN;refGene_tx=XM_023254312.1
B2 113876573 113876600
refGene_gn=TRDN;refGene_tx=XM_023254312.1
B2 113876721 113876754
refGene_gn=TRDN;refGene_tx=XM_023254312.1
B2 113879910 113879928
refGene_gn=TRDN;refGene_tx=XM_023254312.1
B2 113880639 113880669
refGene_gn=TRDN;refGene_tx=XM_023254312.1
B2 113890670 113890700
refGene_gn=TRDN;refGene_tx=XM_023254312.1
B2 113897961 113898018
refGene_gn=TRDN;refGene_tx=XM_019831165.2;refGene_gn=TRDN;refGene_tx=XM_023254312.1
B2 113957459 113957519
refGene_gn=TRDN;refGene_tx=XM_019831165.2;refGene_gn=TRDN;refGene_tx=XM_023254312.1
B2 113959009 113959069
refGene_gn=TRDN;refGene_tx=XM_019831165.2;refGene_gn=TRDN;refGene_tx=XM_023254312.1

B2 113978504 113978582
refGene_gn=TRDN;refGene_tx=XM_019831165.2;refGene_gn=TRDN;refGene_tx=XM_023254312.1
B2 114004367 114004427
refGene_gn=TRDN;refGene_tx=XM_019831165.2;refGene_gn=TRDN;refGene_tx=XM_023254312.1
B2 114010254 114010425
refGene_gn=TRDN;refGene_tx=XM_019831165.2;refGene_gn=TRDN;refGene_tx=XM_023254312.1
B2 114018661 114018721
refGene_gn=TRDN;refGene_tx=XM_019831165.2;refGene_gn=TRDN;refGene_tx=XM_023254312.1
B2 114022327 114022375
refGene_gn=TRDN;refGene_tx=XM_023254312.1
B2 114036712 114036772
refGene_gn=TRDN;refGene_tx=XM_019831165.2;refGene_gn=TRDN;refGene_tx=XM_023254312.1
B2 114056142 114056175
refGene_gn=TRDN;refGene_tx=XM_019831165.2;refGene_gn=TRDN;refGene_tx=XM_023254312.1
B2 114057250 114057415
refGene_gn=TRDN;refGene_tx=XM_019831165.2;refGene_gn=TRDN;refGene_tx=XM_023254312.1
B2 114078215 114078425
refGene_gn=TRDN;refGene_tx=XM_019831165.2;refGene_gn=TRDN;refGene_tx=XM_023254312.1
B2 114126146 114126168
refGene_gn=TRDN;refGene_tx=XM_019831165.2;refGene_gn=TRDN;refGene_tx=XM_023254312.1
B2 114274678 114274732
refGene_gn=NKAIN2;refGene_tx=XM_003986520.5;refGene_gn=NKAIN2;refGene_tx=XM_019831167.2
B2 114736931 114737069
refGene_gn=NKAIN2;refGene_tx=XM_003986520.5;refGene_gn=NKAIN2;refGene_tx=XM_019831167.2
B2 114808102 114808183
refGene_gn=NKAIN2;refGene_tx=XM_003986520.5;refGene_gn=NKAIN2;refGene_tx=XM_019831167.2
B2 115089207 115089408
refGene_gn=NKAIN2;refGene_tx=XM_003986520.5;refGene_gn=NKAIN2;refGene_tx=XM_019831167.2
B2 115236450 115236511
refGene_gn=NKAIN2;refGene_tx=XM_003986520.5;refGene_gn=NKAIN2;refGene_tx=XM_019831167.2
B2 115257710 115257792
refGene_gn=NKAIN2;refGene_tx=XM_003986520.5;refGene_gn=NKAIN2;refGene_tx=XM_019831167.2
B2 115258478 115258503
refGene_gn=NKAIN2;refGene_tx=XM_019831167.2
B2 115261963 115261973

refGene_gn=NKAIN2;refGene_tx=XM_003986520.5
B2	115370222	115371092
refGene_gn=RNF217;refGene_tx=XM_019831614.2
B2	115450443	115450677
refGene_gn=RNF217;refGene_tx=XM_019831614.2
B2	115464350	115464515
refGene_gn=RNF217;refGene_tx=XM_019831614.2
B2	115486480	115486682
refGene_gn=RNF217;refGene_tx=XM_019831614.2
B2	115491547	115491619
refGene_gn=RNF217;refGene_tx=XM_019831614.2
B2	115493477	115493551
refGene_gn=RNF217;refGene_tx=XM_019831614.2
B2	115532559	115532578
refGene_gn=TPD52L1;refGene_tx=XM_019831169.2;refGene_gn=TPD52L1;refGene_tx=XM_023254314.1;refGene_gn=TPD52L1;refGene_tx=XM_023254315.1;refGene_gn=TPD52L1;refGene_tx=XM_023254318.1;refGene_gn=TPD52L1;refGene_tx=XM_023254319.1;refGene_gn=TPD52L1;refGene_tx=XM_023254320.1
B2	115596467	115596583
refGene_gn=TPD52L1;refGene_tx=XM_019831169.2;refGene_gn=TPD52L1;refGene_tx=XM_023254314.1;refGene_gn=TPD52L1;refGene_tx=XM_023254315.1;refGene_gn=TPD52L1;refGene_tx=XM_023254316.1;refGene_gn=TPD52L1;refGene_tx=XM_023254317.1;refGene_gn=TPD52L1;refGene_tx=XM_023254318.1;refGene_gn=TPD52L1;refGene_tx=XM_023254319.1;refGene_gn=TPD52L1;refGene_tx=XM_023254320.1
B2	115608264	115608413
refGene_gn=TPD52L1;refGene_tx=XM_019831169.2;refGene_gn=TPD52L1;refGene_tx=XM_023254314.1;refGene_gn=TPD52L1;refGene_tx=XM_023254315.1;refGene_gn=TPD52L1;refGene_tx=XM_023254316.1;refGene_gn=TPD52L1;refGene_tx=XM_023254317.1;refGene_gn=TPD52L1;refGene_tx=XM_023254318.1;refGene_gn=TPD52L1;refGene_tx=XM_023254319.1;refGene_gn=TPD52L1;refGene_tx=XM_023254320.1
B2	115619324	115619426
refGene_gn=TPD52L1;refGene_tx=XM_019831169.2;refGene_gn=TPD52L1;refGene_tx=XM_023254314.1;refGene_gn=TPD52L1;refGene_tx=XM_023254315.1;refGene_gn=TPD52L1;refGene_tx=XM_023254316.1;refGene_gn=TPD52L1;refGene_tx=XM_023254317.1;refGene_gn=TPD52L1;refGene_tx=XM_023254318.1;refGene_gn=TPD52L1;refGene_tx=XM_023254319.1;refGene_gn=TPD52L1;refGene_tx=XM_023254320.1
B2	115622824	115622852
refGene_gn=TPD52L1;refGene_tx=XM_019831169.2
B2	115623123	115623138
refGene_gn=TPD52L1;refGene_tx=XM_023254314.1;refGene_gn=TPD52L1;refGene_tx=XM_023254316.1;refGene_gn=TPD52L1;refGene_tx=XM_023254317.1;refGene_gn=TPD52L1;refGene_tx=XM_023254318.1
B2	115625002	115625041
refGene_gn=TPD52L1;refGene_tx=XM_023254314.1;refGene_gn=TPD52L1;refGene_tx=XM_023254315.1;refGene_gn=TPD52L1;refGene_tx=XM_023254316.1;refGene_gn=TPD52L1;refGene_tx=XM_023254317.1;refGene_gn=TPD52L1;refGene_tx=XM_023254318.1;refGene_gn=TPD52L1;refGene_tx=XM_023254319.1

B2 115627960 115628021
refGene_gn=TPD52L1;refGene_tx=XM_023254314.1;refGene_gn=TPD52L1;refGene_tx=XM_023254315.1;refGene_gn=TPD52L1;refGene_tx=XM_023254316.1;refGene_gn=TPD52L1;refGene_tx=XM_023254317.1
B2 115632647 115632776
refGene_gn=TPD52L1;refGene_tx=XM_023254314.1;refGene_gn=TPD52L1;refGene_tx=XM_023254315.1;refGene_gn=TPD52L1;refGene_tx=XM_023254316.1;refGene_gn=TPD52L1;refGene_tx=XM_023254317.1;refGene_gn=TPD52L1;refGene_tx=XM_023254318.1;refGene_gn=TPD52L1;refGene_tx=XM_023254319.1;refGene_gn=TPD52L1;refGene_tx=XM_023254320.1
B2 115637953 115638051
refGene_gn=HDDC2;refGene_tx=XM_023254321.1;refGene_gn=HDDC2;refGene_tx=XM_023254322.1
B2 115641636 115641775
refGene_gn=HDDC2;refGene_tx=XM_023254321.1;refGene_gn=HDDC2;refGene_tx=XM_023254322.1
B2 115654368 115654437
refGene_gn=HDDC2;refGene_tx=XM_023254321.1;refGene_gn=HDDC2;refGene_tx=XM_023254322.1
B2 115659867 115659970
refGene_gn=HDDC2;refGene_tx=XM_023254321.1;refGene_gn=HDDC2;refGene_tx=XM_023254322.1
B2 115661732 115661854
refGene_gn=HDDC2;refGene_tx=XM_023254321.1;refGene_gn=HDDC2;refGene_tx=XM_023254322.1
B2 115663087 115663171
refGene_gn=HDDC2;refGene_tx=XM_023254321.1;refGene_gn=HDDC2;refGene_tx=XM_023254322.1
B2 116063437 116063520
refGene_gn=HEY2;refGene_tx=XM_003986528.5;refGene_gn=HEY2;refGene_tx=XM_023254323.1
B2 116065457 116065536
refGene_gn=HEY2;refGene_tx=XM_003986528.5;refGene_gn=HEY2;refGene_tx=XM_019831170.2;refGene_gn=HEY2;refGene_tx=XM_023254323.1
B2 116065670 116065754
refGene_gn=HEY2;refGene_tx=XM_003986528.5;refGene_gn=HEY2;refGene_tx=XM_019831170.2;refGene_gn=HEY2;refGene_tx=XM_023254323.1
B2 116068028 116068110
refGene_gn=HEY2;refGene_tx=XM_003986528.5;refGene_gn=HEY2;refGene_tx=XM_019831170.2;refGene_gn=HEY2;refGene_tx=XM_023254323.1
B2 116072532 116073218
refGene_gn=HEY2;refGene_tx=XM_003986528.5;refGene_gn=HEY2;refGene_tx=XM_019831170.2;refGene_gn=HEY2;refGene_tx=XM_023254323.1
B2 116073266 116073277
refGene_gn=HEY2;refGene_tx=XM_023254323.1
B2 116187261 116187300
refGene_gn=NCOA7;refGene_tx=XM_023254324.1
B2 116190499 116190607
refGene_gn=NCOA7;refGene_tx=XM_023254324.1
B2 116195419 116195533

refGene_gn=NCOA7;refGene_tx=XM_023254324.1
B2 116196117 116196243
refGene_gn=NCOA7;refGene_tx=XM_023254324.1
B2 116198056 116198241
refGene_gn=NCOA7;refGene_tx=XM_023254324.1
B2 116202395 116203111
refGene_gn=NCOA7;refGene_tx=XM_023254324.1
B2 116203113 116203440
refGene_gn=NCOA7;refGene_tx=XM_023254324.1
B2 116204284 116204453
refGene_gn=NCOA7;refGene_tx=XM_023254324.1
B2 116227163 116227311
refGene_gn=NCOA7;refGene_tx=XM_023254324.1
B2 116236196 116236322
refGene_gn=NCOA7;refGene_tx=XM_023254324.1
B2 116237663 116237816
refGene_gn=NCOA7;refGene_tx=XM_023254324.1
B2 116249493 116249589
refGene_gn=NCOA7;refGene_tx=XM_023254324.1
B2 116250027 116250101
refGene_gn=NCOA7;refGene_tx=XM_023254324.1
B2 116250472 116250608
refGene_gn=NCOA7;refGene_tx=XM_023254324.1
B2 116269531 116269732
refGene_gn=HINT3;refGene_tx=XM_003986531.5
B2 116278154 116278272
refGene_gn=HINT3;refGene_tx=XM_003986531.5
B2 116282299 116282369
refGene_gn=HINT3;refGene_tx=XM_003986531.5
B2 116284334 116284461
refGene_gn=HINT3;refGene_tx=XM_003986531.5
B2 116286614 116286647
refGene_gn=HINT3;refGene_tx=XM_003986531.5
B2 116298006 116298078
refGene_gn=TRMT11;refGene_tx=XM_003986532.5
B2 116298258 116298336
refGene_gn=TRMT11;refGene_tx=XM_011282620.3;refGene_gn=TRMT11;refGene_tx=XM_011282622.2;refGene_gn=TRMT11;refGene_tx=XM_023254326.1
B2 116304826 116304892
refGene_gn=TRMT11;refGene_tx=XM_003986532.5;refGene_gn=TRMT11;refGene_tx=XM_011282620.3;refGene_gn=TRMT11;refGene_tx=XM_011282622.2;refGene_gn=TRMT11;refGene_tx=XM_023254326.1
B2 116306433 116306507
refGene_gn=TRMT11;refGene_tx=XM_003986532.5;refGene_gn=TRMT11;refGene_tx=XM_011282620.3;refGene_gn=TRMT11;refGene_tx=XM_011282622.2;refGene_gn=TRMT11;refGene_tx=XM_023254326.1
B2 116307863 116307945
refGene_gn=TRMT11;refGene_tx=XM_003986532.5;refGene_gn=TRMT11;refGene_tx=XM_011282620.3;refGene_gn=TRMT11;refGene_tx=XM_011282622.2;refGene_gn=TRMT11;refGene_tx=XM_023254326.1

B2 116308034 116308127
refGene_gn=TRMT11;refGene_tx=XM_003986532.5;refGene_gn=TRMT11;refGene_tx=XM_011282620.3
B2 116308352 116308487
refGene_gn=TRMT11;refGene_tx=XM_003986532.5;refGene_gn=TRMT11;refGene_tx=XM_011282620.3;refGene_gn=TRMT11;refGene_tx=XM_011282622.2;refGene_gn=TRMT11;refGene_tx=XM_023254326.1
B2 116309190 116309347
refGene_gn=TRMT11;refGene_tx=XM_003986532.5;refGene_gn=TRMT11;refGene_tx=XM_011282620.3;refGene_gn=TRMT11;refGene_tx=XM_011282622.2;refGene_gn=TRMT11;refGene_tx=XM_023254326.1
B2 116317832 116317913
refGene_gn=TRMT11;refGene_tx=XM_003986532.5;refGene_gn=TRMT11;refGene_tx=XM_011282620.3;refGene_gn=TRMT11;refGene_tx=XM_011282622.2
B2 116320946 116321111
refGene_gn=TRMT11;refGene_tx=XM_003986532.5;refGene_gn=TRMT11;refGene_tx=XM_011282620.3;refGene_gn=TRMT11;refGene_tx=XM_011282622.2;refGene_gn=TRMT11;refGene_tx=XM_023254326.1
B2 116321563 116321636
refGene_gn=TRMT11;refGene_tx=XM_003986532.5;refGene_gn=TRMT11;refGene_tx=XM_011282620.3;refGene_gn=TRMT11;refGene_tx=XM_011282622.2;refGene_gn=TRMT11;refGene_tx=XM_023254326.1
B2 116321757 116321889
refGene_gn=TRMT11;refGene_tx=XM_003986532.5;refGene_gn=TRMT11;refGene_tx=XM_011282620.3;refGene_gn=TRMT11;refGene_tx=XM_011282622.2;refGene_gn=TRMT11;refGene_tx=XM_023254326.1
B2 116336293 116336414
refGene_gn=TRMT11;refGene_tx=XM_003986532.5;refGene_gn=TRMT11;refGene_tx=XM_011282620.3;refGene_gn=TRMT11;refGene_tx=XM_011282622.2;refGene_gn=TRMT11;refGene_tx=XM_023254326.1
B2 116355840 116355972
refGene_gn=TRMT11;refGene_tx=XM_003986532.5;refGene_gn=TRMT11;refGene_tx=XM_011282620.3;refGene_gn=TRMT11;refGene_tx=XM_011282622.2;refGene_gn=TRMT11;refGene_tx=XM_023254326.1
B2 116619943 116620069
refGene_gn=CENPW;refGene_tx=XM_003986533.4
B2 116625787 116625901
refGene_gn=CENPW;refGene_tx=XM_003986533.4
B2 116627913 116627940
refGene_gn=CENPW;refGene_tx=XM_003986533.4
B2 117341540 117341637
refGene_gn=RSPO3;refGene_tx=XM_003986534.4;refGene_gn=RSPO3;refGene_tx=XM_006931980.4
B2 117373598 117373790
refGene_gn=RSPO3;refGene_tx=XM_003986534.4;refGene_gn=RSPO3;refGene_tx=XM_006931980.4
B2 117375378 117375525
refGene_gn=RSPO3;refGene_tx=XM_003986534.4;refGene_gn=RSPO3;refGene_tx=XM_006931980.4
B2 117380795 117380993

refGene_gn=RSPO3;refGene_tx=XM_003986534.4;refGene_gn=RSPO3;refGene_tx=XM_006931980.4
B2 117425415 117425609
refGene_gn=RSPO3;refGene_tx=XM_003986534.4;refGene_gn=RSPO3;refGene_tx=XM_006931980.4
B2 117426973 117427009
refGene_gn=RSPO3;refGene_tx=XM_006931980.4
B2 117486180 117486182
refGene_gn=RNF146;refGene_tx=XM_003986535.3;refGene_gn=RNF146;refGene_tx=XM_003986536.3;refGene_gn=RNF146;refGene_tx=XM_006931983.4;refGene_gn=RNF146;refGene_tx=XM_019831176.2;refGene_gn=RNF146;refGene_tx=XM_023254327.1
B2 117495040 117496109
refGene_gn=RNF146;refGene_tx=XM_003986535.3;refGene_gn=RNF146;refGene_tx=XM_003986536.3;refGene_gn=RNF146;refGene_tx=XM_006931983.4;refGene_gn=RNF146;refGene_tx=XM_019831176.2;refGene_gn=RNF146;refGene_tx=XM_023254327.1
B2 117499021 117499430
refGene_gn=ECHDC1;refGene_tx=XM_011282615.2;refGene_gn=ECHDC1;refGene_tx=XM_011282616.2;refGene_gn=ECHDC1;refGene_tx=XM_011282617.3;refGene_gn=ECHDC1;refGene_tx=XM_019831177.1
B2 117536178 117536259
refGene_gn=ECHDC1;refGene_tx=XM_011282615.2;refGene_gn=ECHDC1;refGene_tx=XM_011282616.2;refGene_gn=ECHDC1;refGene_tx=XM_011282617.3;refGene_gn=ECHDC1;refGene_tx=XM_019831177.1
B2 117537891 117537944
refGene_gn=ECHDC1;refGene_tx=XM_011282615.2;refGene_gn=ECHDC1;refGene_tx=XM_011282616.2;refGene_gn=ECHDC1;refGene_tx=XM_011282617.3;refGene_gn=ECHDC1;refGene_tx=XM_019831177.1
B2 117540618 117540761
refGene_gn=ECHDC1;refGene_tx=XM_011282615.2;refGene_gn=ECHDC1;refGene_tx=XM_011282616.2;refGene_gn=ECHDC1;refGene_tx=XM_011282617.3;refGene_gn=ECHDC1;refGene_tx=XM_019831177.1
B2 117544393 117544615
refGene_gn=ECHDC1;refGene_tx=XM_011282615.2;refGene_gn=ECHDC1;refGene_tx=XM_011282616.2;refGene_gn=ECHDC1;refGene_tx=XM_011282617.3;refGene_gn=ECHDC1;refGene_tx=XM_019831177.1
B2 117551955 117551971
refGene_gn=ECHDC1;refGene_tx=XM_011282615.2
B2 117630321 117630495
refGene_gn=KIAA0408;refGene_tx=XM_019831179.2
B2 117633549 117634870
refGene_gn=KIAA0408;refGene_tx=XM_019831179.2
B2 117636843 117636923
refGene_gn=KIAA0408;refGene_tx=XM_019831179.2
B2 117637012 117637375
refGene_gn=KIAA0408;refGene_tx=XM_019831179.2
B2 117642349 117642484
refGene_gn=KIAA0408;refGene_tx=XM_019831179.2
B2 117661912 117662023

refGene_gn=SOGA3;refGene_tx=XM_019831178.2
B2	117662675	117662687
refGene_gn=SOGA3;refGene_tx=XM_023254328.1
B2	117663449	117664625
refGene_gn=SOGA3;refGene_tx=XM_019831178.2;refGene_gn=SOGA3;refGene_tx
=XM_023254328.1
B2	117670196	117670268
refGene_gn=SOGA3;refGene_tx=XM_019831178.2;refGene_gn=SOGA3;refGene_tx
=XM_023254328.1
B2	117699593	117699752
refGene_gn=SOGA3;refGene_tx=XM_019831178.2;refGene_gn=SOGA3;refGene_tx
=XM_023254328.1
B2	117701791	117702016
refGene_gn=SOGA3;refGene_tx=XM_019831178.2;refGene_gn=SOGA3;refGene_tx
=XM_023254328.1
B2	117702475	117703570
refGene_gn=SOGA3;refGene_tx=XM_019831178.2;refGene_gn=SOGA3;refGene_tx
=XM_023254328.1
B2	117775943	117776241
refGene_gn=CB2H6orf58;refGene_tx=XM_003986539.2
B2	117778147	117778234
refGene_gn=CB2H6orf58;refGene_tx=XM_003986539.2
B2	117779986	117780171
refGene_gn=CB2H6orf58;refGene_tx=XM_003986539.2
B2	117781095	117781196
refGene_gn=CB2H6orf58;refGene_tx=XM_003986539.2
B2	117787539	117787778
refGene_gn=CB2H6orf58;refGene_tx=XM_003986539.2
B2	117790540	117790668
refGene_gn=CB2H6orf58;refGene_tx=XM_003986539.2
B2	117821666	117821767
refGene_gn=LOC101091873;refGene_tx=XM_019831180.1
B2	117825005	117825244
refGene_gn=LOC101091873;refGene_tx=XM_019831180.1
B2	117826888	117826989
refGene_gn=LOC101091873;refGene_tx=XM_019831180.1
B2	117827906	117828091
refGene_gn=LOC101091873;refGene_tx=XM_019831180.1
B2	117829291	117829378
refGene_gn=LOC101091873;refGene_tx=XM_019831180.1
B2	117833862	117834163
refGene_gn=LOC101091873;refGene_tx=XM_019831180.1
B2	117870322	117870354
refGene_gn=THEMIS;refGene_tx=XM_006931988.3
B2	117882194	117882342
refGene_gn=THEMIS;refGene_tx=XM_006931988.3
B2	117956829	117957878
refGene_gn=THEMIS;refGene_tx=XM_006931988.3
B2	117976133	117976592
refGene_gn=THEMIS;refGene_tx=XM_006931988.3

B2	118010256	118010415
refGene_gn=THEMIS;refGene_tx=XM_006931988.3
B2	118057837	118057928
refGene_gn=THEMIS;refGene_tx=XM_006931988.3
B2	118123034	118123088
refGene_gn=PTPRK;refGene_tx=XM_023254330.1;refGene_gn=PTPRK;refGene_tx=XM_023254331.1;refGene_gn=PTPRK;refGene_tx=XM_023254332.1;refGene_gn=PTPRK;refGene_tx=XM_023254333.1;refGene_gn=PTPRK;refGene_tx=XM_023254334.1;refGene_gn=PTPRK;refGene_tx=XM_023254335.1;refGene_gn=PTPRK;refGene_tx=XM_023254336.1;refGene_gn=PTPRK;refGene_tx=XM_023254337.1;refGene_gn=PTPRK;refGene_tx=XM_023254338.1;refGene_gn=PTPRK;refGene_tx=XM_023254339.1;refGene_gn=PTPRK;refGene_tx=XM_023254340.1
B2	118126776	118126912
refGene_gn=PTPRK;refGene_tx=XM_023254330.1;refGene_gn=PTPRK;refGene_tx=XM_023254331.1;refGene_gn=PTPRK;refGene_tx=XM_023254332.1;refGene_gn=PTPRK;refGene_tx=XM_023254333.1;refGene_gn=PTPRK;refGene_tx=XM_023254334.1;refGene_gn=PTPRK;refGene_tx=XM_023254335.1;refGene_gn=PTPRK;refGene_tx=XM_023254336.1;refGene_gn=PTPRK;refGene_tx=XM_023254337.1;refGene_gn=PTPRK;refGene_tx=XM_023254338.1;refGene_gn=PTPRK;refGene_tx=XM_023254339.1;refGene_gn=PTPRK;refGene_tx=XM_023254340.1
B2	118127663	118127827
refGene_gn=PTPRK;refGene_tx=XM_023254330.1;refGene_gn=PTPRK;refGene_tx=XM_023254331.1;refGene_gn=PTPRK;refGene_tx=XM_023254332.1;refGene_gn=PTPRK;refGene_tx=XM_023254333.1;refGene_gn=PTPRK;refGene_tx=XM_023254334.1;refGene_gn=PTPRK;refGene_tx=XM_023254335.1;refGene_gn=PTPRK;refGene_tx=XM_023254336.1;refGene_gn=PTPRK;refGene_tx=XM_023254337.1;refGene_gn=PTPRK;refGene_tx=XM_023254338.1;refGene_gn=PTPRK;refGene_tx=XM_023254339.1;refGene_gn=PTPRK;refGene_tx=XM_023254340.1
B2	118130440	118130566
refGene_gn=PTPRK;refGene_tx=XM_023254330.1;refGene_gn=PTPRK;refGene_tx=XM_023254331.1;refGene_gn=PTPRK;refGene_tx=XM_023254332.1;refGene_gn=PTPRK;refGene_tx=XM_023254333.1;refGene_gn=PTPRK;refGene_tx=XM_023254334.1;refGene_gn=PTPRK;refGene_tx=XM_023254335.1;refGene_gn=PTPRK;refGene_tx=XM_023254336.1;refGene_gn=PTPRK;refGene_tx=XM_023254337.1;refGene_gn=PTPRK;refGene_tx=XM_023254338.1;refGene_gn=PTPRK;refGene_tx=XM_023254339.1;refGene_gn=PTPRK;refGene_tx=XM_023254340.1
B2	118130715	118130847
refGene_gn=PTPRK;refGene_tx=XM_023254330.1;refGene_gn=PTPRK;refGene_tx=XM_023254331.1;refGene_gn=PTPRK;refGene_tx=XM_023254332.1;refGene_gn=PTPRK;refGene_tx=XM_023254333.1;refGene_gn=PTPRK;refGene_tx=XM_023254334.1;refGene_gn=PTPRK;refGene_tx=XM_023254335.1;refGene_gn=PTPRK;refGene_tx=XM_023254336.1;refGene_gn=PTPRK;refGene_tx=XM_023254337.1;refGene_gn=PTPRK;refGene_tx=XM_023254338.1;refGene_gn=PTPRK;refGene_tx=XM_023254339.1;refGene_gn=PTPRK;refGene_tx=XM_023254340.1
B2	118133817	118133991
refGene_gn=PTPRK;refGene_tx=XM_023254330.1;refGene_gn=PTPRK;refGene_tx=XM_023254331.1;refGene_gn=PTPRK;refGene_tx=XM_023254332.1;refGene_gn=PTPRK;refGene_tx=XM_023254333.1;refGene_gn=PTPRK;refGene_tx=XM_023254334.1;refGene_gn=PTPRK;refGene_tx=XM_023254335.1;refGene_gn=PTPRK;refGene_tx=XM_023254336.1;refGene_gn=PTPRK;refGene_tx=XM_023254337.1;refGen

e_gn=PTPRK;refGene_tx=XM_023254338.1;refGene_gn=PTPRK;refGene_tx=XM_023254339.1;refGene_gn=PTPRK;refGene_tx=XM_023254340.1
B2 118134877 118135027
refGene_gn=PTPRK;refGene_tx=XM_023254330.1;refGene_gn=PTPRK;refGene_tx=XM_023254331.1;refGene_gn=PTPRK;refGene_tx=XM_023254332.1;refGene_gn=PTPRK;refGene_tx=XM_023254333.1;refGene_gn=PTPRK;refGene_tx=XM_023254334.1;refGene_gn=PTPRK;refGene_tx=XM_023254335.1;refGene_gn=PTPRK;refGene_tx=XM_023254336.1;refGene_gn=PTPRK;refGene_tx=XM_023254337.1;refGene_gn=PTPRK;refGene_tx=XM_023254338.1;refGene_gn=PTPRK;refGene_tx=XM_023254339.1;refGene_gn=PTPRK;refGene_tx=XM_023254340.1
B2 118135473 118135609
refGene_gn=PTPRK;refGene_tx=XM_023254330.1;refGene_gn=PTPRK;refGene_tx=XM_023254331.1;refGene_gn=PTPRK;refGene_tx=XM_023254332.1;refGene_gn=PTPRK;refGene_tx=XM_023254333.1;refGene_gn=PTPRK;refGene_tx=XM_023254334.1;refGene_gn=PTPRK;refGene_tx=XM_023254335.1;refGene_gn=PTPRK;refGene_tx=XM_023254336.1;refGene_gn=PTPRK;refGene_tx=XM_023254337.1;refGene_gn=PTPRK;refGene_tx=XM_023254338.1;refGene_gn=PTPRK;refGene_tx=XM_023254339.1;refGene_gn=PTPRK;refGene_tx=XM_023254340.1
B2 118137703 118137858
refGene_gn=PTPRK;refGene_tx=XM_023254330.1;refGene_gn=PTPRK;refGene_tx=XM_023254331.1;refGene_gn=PTPRK;refGene_tx=XM_023254332.1;refGene_gn=PTPRK;refGene_tx=XM_023254333.1;refGene_gn=PTPRK;refGene_tx=XM_023254334.1;refGene_gn=PTPRK;refGene_tx=XM_023254335.1;refGene_gn=PTPRK;refGene_tx=XM_023254336.1;refGene_gn=PTPRK;refGene_tx=XM_023254337.1;refGene_gn=PTPRK;refGene_tx=XM_023254338.1;refGene_gn=PTPRK;refGene_tx=XM_023254339.1;refGene_gn=PTPRK;refGene_tx=XM_023254340.1
B2 118142462 118142579
refGene_gn=PTPRK;refGene_tx=XM_023254330.1;refGene_gn=PTPRK;refGene_tx=XM_023254331.1;refGene_gn=PTPRK;refGene_tx=XM_023254332.1;refGene_gn=PTPRK;refGene_tx=XM_023254333.1;refGene_gn=PTPRK;refGene_tx=XM_023254334.1;refGene_gn=PTPRK;refGene_tx=XM_023254335.1;refGene_gn=PTPRK;refGene_tx=XM_023254336.1;refGene_gn=PTPRK;refGene_tx=XM_023254337.1;refGene_gn=PTPRK;refGene_tx=XM_023254338.1;refGene_gn=PTPRK;refGene_tx=XM_023254339.1;refGene_gn=PTPRK;refGene_tx=XM_023254340.1
B2 118142990 118143088
refGene_gn=PTPRK;refGene_tx=XM_023254330.1;refGene_gn=PTPRK;refGene_tx=XM_023254331.1;refGene_gn=PTPRK;refGene_tx=XM_023254332.1;refGene_gn=PTPRK;refGene_tx=XM_023254333.1;refGene_gn=PTPRK;refGene_tx=XM_023254334.1;refGene_gn=PTPRK;refGene_tx=XM_023254335.1;refGene_gn=PTPRK;refGene_tx=XM_023254336.1;refGene_gn=PTPRK;refGene_tx=XM_023254337.1;refGene_gn=PTPRK;refGene_tx=XM_023254338.1;refGene_gn=PTPRK;refGene_tx=XM_023254339.1;refGene_gn=PTPRK;refGene_tx=XM_023254340.1
B2 118144694 118144731
refGene_gn=PTPRK;refGene_tx=XM_023254330.1;refGene_gn=PTPRK;refGene_tx=XM_023254331.1;refGene_gn=PTPRK;refGene_tx=XM_023254332.1;refGene_gn=PTPRK;refGene_tx=XM_023254333.1;refGene_gn=PTPRK;refGene_tx=XM_023254334.1;refGene_gn=PTPRK;refGene_tx=XM_023254335.1;refGene_gn=PTPRK;refGene_tx=XM_023254336.1;refGene_gn=PTPRK;refGene_tx=XM_023254337.1;refGene_gn=PTPRK;refGene_tx=XM_023254338.1;refGene_gn=PTPRK;refGene_tx=XM_023254339.1;refGene_gn=PTPRK;refGene_tx=XM_023254340.1

B2 118147330 118147348
refGene_gn=PTPRK;refGene_tx=XM_023254330.1;refGene_gn=PTPRK;refGene_tx
=XM_023254331.1;refGene_gn=PTPRK;refGene_tx=XM_023254332.1;refGene_gn=
PTPRK;refGene_tx=XM_023254334.1;refGene_gn=PTPRK;refGene_tx=XM_0232543
35.1;refGene_gn=PTPRK;refGene_tx=XM_023254336.1;refGene_gn=PTPRK;refGe
ne_tx=XM_023254337.1;refGene_gn=PTPRK;refGene_tx=XM_023254338.1
B2 118147543 118147620
refGene_gn=PTPRK;refGene_tx=XM_023254330.1;refGene_gn=PTPRK;refGene_tx
=XM_023254331.1;refGene_gn=PTPRK;refGene_tx=XM_023254332.1;refGene_gn=
PTPRK;refGene_tx=XM_023254333.1;refGene_gn=PTPRK;refGene_tx=XM_0232543
34.1;refGene_gn=PTPRK;refGene_tx=XM_023254335.1;refGene_gn=PTPRK;refGe
ne_tx=XM_023254336.1;refGene_gn=PTPRK;refGene_tx=XM_023254337.1;refGen
e_gn=PTPRK;refGene_tx=XM_023254338.1;refGene_gn=PTPRK;refGene_tx=XM_02
3254339.1;refGene_gn=PTPRK;refGene_tx=XM_023254340.1
B2 118148974 118149062
refGene_gn=PTPRK;refGene_tx=XM_023254330.1;refGene_gn=PTPRK;refGene_tx
=XM_023254331.1;refGene_gn=PTPRK;refGene_tx=XM_023254332.1;refGene_gn=
PTPRK;refGene_tx=XM_023254333.1;refGene_gn=PTPRK;refGene_tx=XM_0232543
34.1;refGene_gn=PTPRK;refGene_tx=XM_023254335.1;refGene_gn=PTPRK;refGe
ne_tx=XM_023254336.1;refGene_gn=PTPRK;refGene_tx=XM_023254337.1;refGen
e_gn=PTPRK;refGene_tx=XM_023254338.1;refGene_gn=PTPRK;refGene_tx=XM_02
3254339.1;refGene_gn=PTPRK;refGene_tx=XM_023254340.1
B2 118150735 118150920
refGene_gn=PTPRK;refGene_tx=XM_023254330.1;refGene_gn=PTPRK;refGene_tx
=XM_023254331.1;refGene_gn=PTPRK;refGene_tx=XM_023254332.1;refGene_gn=
PTPRK;refGene_tx=XM_023254333.1;refGene_gn=PTPRK;refGene_tx=XM_0232543
34.1;refGene_gn=PTPRK;refGene_tx=XM_023254335.1;refGene_gn=PTPRK;refGe
ne_tx=XM_023254336.1;refGene_gn=PTPRK;refGene_tx=XM_023254337.1;refGen
e_gn=PTPRK;refGene_tx=XM_023254338.1;refGene_gn=PTPRK;refGene_tx=XM_02
3254339.1;refGene_gn=PTPRK;refGene_tx=XM_023254340.1
B2 118152308 118152367
refGene_gn=PTPRK;refGene_tx=XM_023254341.1
B2 118153228 118153240
refGene_gn=PTPRK;refGene_tx=XM_023254330.1;refGene_gn=PTPRK;refGene_tx
=XM_023254331.1;refGene_gn=PTPRK;refGene_tx=XM_023254333.1;refGene_gn=
PTPRK;refGene_tx=XM_023254334.1;refGene_gn=PTPRK;refGene_tx=XM_0232543
36.1;refGene_gn=PTPRK;refGene_tx=XM_023254341.1
B2 118155191 118155227
refGene_gn=PTPRK;refGene_tx=XM_023254330.1;refGene_gn=PTPRK;refGene_tx
=XM_023254331.1;refGene_gn=PTPRK;refGene_tx=XM_023254332.1;refGene_gn=
PTPRK;refGene_tx=XM_023254333.1;refGene_gn=PTPRK;refGene_tx=XM_0232543
34.1;refGene_gn=PTPRK;refGene_tx=XM_023254336.1;refGene_gn=PTPRK;refGe
ne_tx=XM_023254337.1;refGene_gn=PTPRK;refGene_tx=XM_023254341.1
B2 118156331 118156492
refGene_gn=PTPRK;refGene_tx=XM_023254330.1;refGene_gn=PTPRK;refGene_tx
=XM_023254331.1;refGene_gn=PTPRK;refGene_tx=XM_023254332.1;refGene_gn=
PTPRK;refGene_tx=XM_023254333.1;refGene_gn=PTPRK;refGene_tx=XM_0232543
34.1;refGene_gn=PTPRK;refGene_tx=XM_023254335.1;refGene_gn=PTPRK;refGe
ne_tx=XM_023254336.1;refGene_gn=PTPRK;refGene_tx=XM_023254337.1;refGen
e_gn=PTPRK;refGene_tx=XM_023254338.1;refGene_gn=PTPRK;refGene_tx=XM_02

3254339.1;refGene_gn=PTPRK;refGene_tx=XM_023254340.1;refGene_gn=PTPRK;
refGene_tx=XM_023254341.1
B2 118161415 118161445
refGene_gn=PTPRK;refGene_tx=XM_023254330.1;refGene_gn=PTPRK;refGene_tx
=XM_023254331.1;refGene_gn=PTPRK;refGene_tx=XM_023254332.1;refGene_gn=
PTPRK;refGene_tx=XM_023254333.1;refGene_gn=PTPRK;refGene_tx=XM_0232543
34.1;refGene_gn=PTPRK;refGene_tx=XM_023254335.1;refGene_gn=PTPRK;refGe
ne_tx=XM_023254341.1
B2 118163434 118163476
refGene_gn=PTPRK;refGene_tx=XM_023254330.1;refGene_gn=PTPRK;refGene_tx
=XM_023254331.1;refGene_gn=PTPRK;refGene_tx=XM_023254332.1;refGene_gn=
PTPRK;refGene_tx=XM_023254333.1;refGene_gn=PTPRK;refGene_tx=XM_0232543
35.1
B2 118164499 118164638
refGene_gn=PTPRK;refGene_tx=XM_023254330.1;refGene_gn=PTPRK;refGene_tx
=XM_023254331.1;refGene_gn=PTPRK;refGene_tx=XM_023254332.1;refGene_gn=
PTPRK;refGene_tx=XM_023254333.1;refGene_gn=PTPRK;refGene_tx=XM_0232543
34.1;refGene_gn=PTPRK;refGene_tx=XM_023254335.1;refGene_gn=PTPRK;refGe
ne_tx=XM_023254336.1;refGene_gn=PTPRK;refGene_tx=XM_023254337.1;refGen
e_gn=PTPRK;refGene_tx=XM_023254338.1;refGene_gn=PTPRK;refGene_tx=XM_02
3254339.1;refGene_gn=PTPRK;refGene_tx=XM_023254340.1;refGene_gn=PTPRK;
refGene_tx=XM_023254341.1
B2 118215801 118215838
refGene_gn=PTPRK;refGene_tx=XM_023254330.1;refGene_gn=PTPRK;refGene_tx
=XM_023254331.1;refGene_gn=PTPRK;refGene_tx=XM_023254332.1;refGene_gn=
PTPRK;refGene_tx=XM_023254333.1;refGene_gn=PTPRK;refGene_tx=XM_0232543
34.1;refGene_gn=PTPRK;refGene_tx=XM_023254335.1;refGene_gn=PTPRK;refGe
ne_tx=XM_023254336.1;refGene_gn=PTPRK;refGene_tx=XM_023254337.1;refGen
e_gn=PTPRK;refGene_tx=XM_023254338.1;refGene_gn=PTPRK;refGene_tx=XM_02
3254339.1;refGene_gn=PTPRK;refGene_tx=XM_023254340.1;refGene_gn=PTPRK;
refGene_tx=XM_023254341.1
B2 118218575 118218849
refGene_gn=PTPRK;refGene_tx=XM_023254330.1;refGene_gn=PTPRK;refGene_tx
=XM_023254331.1;refGene_gn=PTPRK;refGene_tx=XM_023254332.1;refGene_gn=
PTPRK;refGene_tx=XM_023254333.1;refGene_gn=PTPRK;refGene_tx=XM_0232543
34.1;refGene_gn=PTPRK;refGene_tx=XM_023254335.1;refGene_gn=PTPRK;refGe
ne_tx=XM_023254336.1;refGene_gn=PTPRK;refGene_tx=XM_023254337.1;refGen
e_gn=PTPRK;refGene_tx=XM_023254338.1;refGene_gn=PTPRK;refGene_tx=XM_02
3254339.1;refGene_gn=PTPRK;refGene_tx=XM_023254340.1;refGene_gn=PTPRK;
refGene_tx=XM_023254341.1
B2 118238343 118238449
refGene_gn=PTPRK;refGene_tx=XM_023254330.1;refGene_gn=PTPRK;refGene_tx
=XM_023254331.1;refGene_gn=PTPRK;refGene_tx=XM_023254332.1;refGene_gn=
PTPRK;refGene_tx=XM_023254333.1;refGene_gn=PTPRK;refGene_tx=XM_0232543
34.1;refGene_gn=PTPRK;refGene_tx=XM_023254335.1;refGene_gn=PTPRK;refGe
ne_tx=XM_023254336.1;refGene_gn=PTPRK;refGene_tx=XM_023254337.1;refGen
e_gn=PTPRK;refGene_tx=XM_023254338.1;refGene_gn=PTPRK;refGene_tx=XM_02
3254339.1;refGene_gn=PTPRK;refGene_tx=XM_023254340.1;refGene_gn=PTPRK;
refGene_tx=XM_023254341.1
B2 118241998 118242200

refGene_gn=PTPRK;refGene_tx=XM_023254330.1;refGene_gn=PTPRK;refGene_tx
=XM_023254331.1;refGene_gn=PTPRK;refGene_tx=XM_023254332.1;refGene_gn=
PTPRK;refGene_tx=XM_023254333.1;refGene_gn=PTPRK;refGene_tx=XM_0232543
34.1;refGene_gn=PTPRK;refGene_tx=XM_023254335.1;refGene_gn=PTPRK;refGe
ne_tx=XM_023254336.1;refGene_gn=PTPRK;refGene_tx=XM_023254337.1;refGen
e_gn=PTPRK;refGene_tx=XM_023254338.1;refGene_gn=PTPRK;refGene_tx=XM_02
3254339.1;refGene_gn=PTPRK;refGene_tx=XM_023254340.1;refGene_gn=PTPRK;
refGene_tx=XM_023254341.1
B2 118243291 118243401
refGene_gn=PTPRK;refGene_tx=XM_023254330.1;refGene_gn=PTPRK;refGene_tx
=XM_023254331.1;refGene_gn=PTPRK;refGene_tx=XM_023254332.1;refGene_gn=
PTPRK;refGene_tx=XM_023254333.1;refGene_gn=PTPRK;refGene_tx=XM_0232543
34.1;refGene_gn=PTPRK;refGene_tx=XM_023254335.1;refGene_gn=PTPRK;refGe
ne_tx=XM_023254336.1;refGene_gn=PTPRK;refGene_tx=XM_023254337.1;refGen
e_gn=PTPRK;refGene_tx=XM_023254338.1;refGene_gn=PTPRK;refGene_tx=XM_02
3254339.1;refGene_gn=PTPRK;refGene_tx=XM_023254340.1;refGene_gn=PTPRK;
refGene_tx=XM_023254341.1
B2 118249129 118249432
refGene_gn=PTPRK;refGene_tx=XM_023254330.1;refGene_gn=PTPRK;refGene_tx
=XM_023254331.1;refGene_gn=PTPRK;refGene_tx=XM_023254332.1;refGene_gn=
PTPRK;refGene_tx=XM_023254333.1;refGene_gn=PTPRK;refGene_tx=XM_0232543
34.1;refGene_gn=PTPRK;refGene_tx=XM_023254335.1;refGene_gn=PTPRK;refGe
ne_tx=XM_023254336.1;refGene_gn=PTPRK;refGene_tx=XM_023254337.1;refGen
e_gn=PTPRK;refGene_tx=XM_023254338.1;refGene_gn=PTPRK;refGene_tx=XM_02
3254339.1;refGene_gn=PTPRK;refGene_tx=XM_023254340.1;refGene_gn=PTPRK;
refGene_tx=XM_023254341.1
B2 118349497 118349791
refGene_gn=PTPRK;refGene_tx=XM_023254330.1;refGene_gn=PTPRK;refGene_tx
=XM_023254331.1;refGene_gn=PTPRK;refGene_tx=XM_023254332.1;refGene_gn=
PTPRK;refGene_tx=XM_023254333.1;refGene_gn=PTPRK;refGene_tx=XM_0232543
34.1;refGene_gn=PTPRK;refGene_tx=XM_023254335.1;refGene_gn=PTPRK;refGe
ne_tx=XM_023254336.1;refGene_gn=PTPRK;refGene_tx=XM_023254337.1;refGen
e_gn=PTPRK;refGene_tx=XM_023254338.1;refGene_gn=PTPRK;refGene_tx=XM_02
3254339.1;refGene_gn=PTPRK;refGene_tx=XM_023254340.1;refGene_gn=PTPRK;
refGene_tx=XM_023254341.1
B2 118377932 118378107
refGene_gn=PTPRK;refGene_tx=XM_023254330.1;refGene_gn=PTPRK;refGene_tx
=XM_023254331.1;refGene_gn=PTPRK;refGene_tx=XM_023254332.1;refGene_gn=
PTPRK;refGene_tx=XM_023254333.1;refGene_gn=PTPRK;refGene_tx=XM_0232543
34.1;refGene_gn=PTPRK;refGene_tx=XM_023254335.1;refGene_gn=PTPRK;refGe
ne_tx=XM_023254336.1;refGene_gn=PTPRK;refGene_tx=XM_023254337.1;refGen
e_gn=PTPRK;refGene_tx=XM_023254338.1;refGene_gn=PTPRK;refGene_tx=XM_02
3254339.1;refGene_gn=PTPRK;refGene_tx=XM_023254340.1;refGene_gn=PTPRK;
refGene_tx=XM_023254341.1
B2 118398774 118398890
refGene_gn=PTPRK;refGene_tx=XM_023254330.1;refGene_gn=PTPRK;refGene_tx
=XM_023254331.1;refGene_gn=PTPRK;refGene_tx=XM_023254332.1;refGene_gn=
PTPRK;refGene_tx=XM_023254333.1;refGene_gn=PTPRK;refGene_tx=XM_0232543
34.1;refGene_gn=PTPRK;refGene_tx=XM_023254335.1;refGene_gn=PTPRK;refGe
ne_tx=XM_023254336.1;refGene_gn=PTPRK;refGene_tx=XM_023254337.1;refGen

e_gn=PTPRK;refGene_tx=XM_023254338.1;refGene_gn=PTPRK;refGene_tx=XM_02
3254339.1;refGene_gn=PTPRK;refGene_tx=XM_023254340.1;refGene_gn=PTPRK;
refGene_tx=XM_023254341.1
B2 118401188 118401270
refGene_gn=PTPRK;refGene_tx=XM_023254330.1;refGene_gn=PTPRK;refGene_tx
=XM_023254331.1;refGene_gn=PTPRK;refGene_tx=XM_023254332.1;refGene_gn=
PTPRK;refGene_tx=XM_023254333.1;refGene_gn=PTPRK;refGene_tx=XM_0232543
34.1;refGene_gn=PTPRK;refGene_tx=XM_023254335.1;refGene_gn=PTPRK;refGe
ne_tx=XM_023254336.1;refGene_gn=PTPRK;refGene_tx=XM_023254337.1;refGen
e_gn=PTPRK;refGene_tx=XM_023254338.1;refGene_gn=PTPRK;refGene_tx=XM_02
3254339.1;refGene_gn=PTPRK;refGene_tx=XM_023254340.1;refGene_gn=PTPRK;
refGene_tx=XM_023254341.1
B2 118475109 118475381
refGene_gn=PTPRK;refGene_tx=XM_023254330.1;refGene_gn=PTPRK;refGene_tx
=XM_023254331.1;refGene_gn=PTPRK;refGene_tx=XM_023254332.1;refGene_gn=
PTPRK;refGene_tx=XM_023254333.1;refGene_gn=PTPRK;refGene_tx=XM_0232543
34.1;refGene_gn=PTPRK;refGene_tx=XM_023254335.1;refGene_gn=PTPRK;refGe
ne_tx=XM_023254336.1;refGene_gn=PTPRK;refGene_tx=XM_023254337.1;refGen
e_gn=PTPRK;refGene_tx=XM_023254338.1;refGene_gn=PTPRK;refGene_tx=XM_02
3254339.1;refGene_gn=PTPRK;refGene_tx=XM_023254340.1;refGene_gn=PTPRK;
refGene_tx=XM_023254341.1
B2 118549163 118549286
refGene_gn=PTPRK;refGene_tx=XM_023254330.1;refGene_gn=PTPRK;refGene_tx
=XM_023254331.1;refGene_gn=PTPRK;refGene_tx=XM_023254332.1;refGene_gn=
PTPRK;refGene_tx=XM_023254333.1;refGene_gn=PTPRK;refGene_tx=XM_0232543
34.1;refGene_gn=PTPRK;refGene_tx=XM_023254335.1;refGene_gn=PTPRK;refGe
ne_tx=XM_023254336.1;refGene_gn=PTPRK;refGene_tx=XM_023254337.1;refGen
e_gn=PTPRK;refGene_tx=XM_023254338.1;refGene_gn=PTPRK;refGene_tx=XM_02
3254339.1;refGene_gn=PTPRK;refGene_tx=XM_023254340.1;refGene_gn=PTPRK;
refGene_tx=XM_023254341.1
B2 118683411 118683508
refGene_gn=PTPRK;refGene_tx=XM_023254330.1;refGene_gn=PTPRK;refGene_tx
=XM_023254331.1;refGene_gn=PTPRK;refGene_tx=XM_023254332.1;refGene_gn=
PTPRK;refGene_tx=XM_023254333.1;refGene_gn=PTPRK;refGene_tx=XM_0232543
34.1;refGene_gn=PTPRK;refGene_tx=XM_023254335.1;refGene_gn=PTPRK;refGe
ne_tx=XM_023254336.1;refGene_gn=PTPRK;refGene_tx=XM_023254337.1;refGen
e_gn=PTPRK;refGene_tx=XM_023254338.1;refGene_gn=PTPRK;refGene_tx=XM_02
3254339.1;refGene_gn=PTPRK;refGene_tx=XM_023254340.1;refGene_gn=PTPRK;
refGene_tx=XM_023254341.1
B2 119066157 119066278
refGene_gn=LAMA2;refGene_tx=XM_011282613.3
B2 119214389 119214560
refGene_gn=LAMA2;refGene_tx=XM_011282613.3
B2 119224354 119224467
refGene_gn=LAMA2;refGene_tx=XM_011282613.3
B2 119264177 119264420
refGene_gn=LAMA2;refGene_tx=XM_011282613.3
B2 119310234 119310414
refGene_gn=LAMA2;refGene_tx=XM_011282613.3
B2 119313323 119313413

refGene_gn=LAMA2;refGene_tx=XM_011282613.3
B2 119314778 119314896
refGene_gn=LAMA2;refGene_tx=XM_011282613.3
B2 119320799 119320978
refGene_gn=LAMA2;refGene_tx=XM_011282613.3
B2 119326669 119326769
refGene_gn=LAMA2;refGene_tx=XM_011282613.3
B2 119331584 119331745
refGene_gn=LAMA2;refGene_tx=XM_011282613.3
B2 119346451 119346592
refGene_gn=LAMA2;refGene_tx=XM_011282613.3
B2 119348921 119349095
refGene_gn=LAMA2;refGene_tx=XM_011282613.3
B2 119404003 119404105
refGene_gn=LAMA2;refGene_tx=XM_011282613.3
B2 119405545 119405757
refGene_gn=LAMA2;refGene_tx=XM_011282613.3
B2 119414203 119414315
refGene_gn=LAMA2;refGene_tx=XM_011282613.3
B2 119419932 119420046
refGene_gn=LAMA2;refGene_tx=XM_011282613.3
B2 119425210 119425338
refGene_gn=LAMA2;refGene_tx=XM_011282613.3
B2 119435840 119435927
refGene_gn=LAMA2;refGene_tx=XM_011282613.3
B2 119443960 119444172
refGene_gn=LAMA2;refGene_tx=XM_011282613.3
B2 119450792 119450899
refGene_gn=LAMA2;refGene_tx=XM_011282613.3
B2 119456885 119457066
refGene_gn=LAMA2;refGene_tx=XM_011282613.3
B2 119459918 119460055
refGene_gn=LAMA2;refGene_tx=XM_011282613.3
B2 119472602 119472839
refGene_gn=LAMA2;refGene_tx=XM_011282613.3
B2 119474068 119474212
refGene_gn=LAMA2;refGene_tx=XM_011282613.3
B2 119474904 119475084
refGene_gn=LAMA2;refGene_tx=XM_011282613.3
B2 119475193 119475211
refGene_gn=LAMA2;refGene_tx=XM_011282613.3
B2 119475212 119475383
refGene_gn=LAMA2;refGene_tx=XM_011282613.3
B2 119475463 119475597
refGene_gn=LAMA2;refGene_tx=XM_011282613.3
B2 119479651 119479769
refGene_gn=LAMA2;refGene_tx=XM_011282613.3
B2 119486891 119487026
refGene_gn=LAMA2;refGene_tx=XM_011282613.3
B2 119499015 119499140

refGene_gn=LAMA2;refGene_tx=XM_011282613.3
B2 119505665 119505752
refGene_gn=LAMA2;refGene_tx=XM_011282613.3
B2 119510075 119510269
refGene_gn=LAMA2;refGene_tx=XM_011282613.3
B2 119526706 119526849
refGene_gn=LAMA2;refGene_tx=XM_011282613.3
B2 119530492 119530591
refGene_gn=LAMA2;refGene_tx=XM_011282613.3
B2 119546996 119547108
refGene_gn=LAMA2;refGene_tx=XM_011282613.3
B2 119552397 119552560
refGene_gn=LAMA2;refGene_tx=XM_011282613.3
B2 119553857 119554068
refGene_gn=LAMA2;refGene_tx=XM_011282613.3
B2 119561084 119561201
refGene_gn=LAMA2;refGene_tx=XM_011282613.3
B2 119561894 119562058
refGene_gn=LAMA2;refGene_tx=XM_011282613.3
B2 119563354 119563493
refGene_gn=LAMA2;refGene_tx=XM_011282613.3
B2 119588458 119588561
refGene_gn=LAMA2;refGene_tx=XM_011282613.3
B2 119600069 119600186
refGene_gn=LAMA2;refGene_tx=XM_011282613.3
B2 119602488 119602671
refGene_gn=LAMA2;refGene_tx=XM_011282613.3
B2 119604283 119604289
refGene_gn=LAMA2;refGene_tx=XM_011282613.3
B2 119606974 119607129
refGene_gn=LAMA2;refGene_tx=XM_011282613.3
B2 119613445 119613589
refGene_gn=LAMA2;refGene_tx=XM_011282613.3
B2 119615231 119615365
refGene_gn=LAMA2;refGene_tx=XM_011282613.3
B2 119617339 119617499
refGene_gn=LAMA2;refGene_tx=XM_011282613.3
B2 119620274 119620399
refGene_gn=LAMA2;refGene_tx=XM_011282613.3
B2 119625867 119626030
refGene_gn=LAMA2;refGene_tx=XM_011282613.3
B2 119626708 119626853
refGene_gn=LAMA2;refGene_tx=XM_011282613.3
B2 119634182 119634321
refGene_gn=LAMA2;refGene_tx=XM_011282613.3
B2 119637119 119637131
refGene_gn=LAMA2;refGene_tx=XM_011282613.3
B2 119639896 119640017
refGene_gn=LAMA2;refGene_tx=XM_011282613.3
B2 119642430 119642607

refGene_gn=LAMA2;refGene_tx=XM_011282613.3
B2 119646431 119646580
refGene_gn=LAMA2;refGene_tx=XM_011282613.3
B2 119651510 119651687
refGene_gn=LAMA2;refGene_tx=XM_011282613.3
B2 119652149 119652318
refGene_gn=LAMA2;refGene_tx=XM_011282613.3
B2 119661933 119662046
refGene_gn=LAMA2;refGene_tx=XM_011282613.3
B2 119662365 119662555
refGene_gn=LAMA2;refGene_tx=XM_011282613.3
B2 119664543 119664699
refGene_gn=LAMA2;refGene_tx=XM_011282613.3
B2 119672890 119673044
refGene_gn=LAMA2;refGene_tx=XM_011282613.3
B2 119675348 119675479
refGene_gn=LAMA2;refGene_tx=XM_011282613.3
B2 119677394 119677617
refGene_gn=LAMA2;refGene_tx=XM_011282613.3
B2 119685296 119685454
refGene_gn=LAMA2;refGene_tx=XM_011282613.3
B2 119747398 119747490
refGene_gn=ARHGAP18;refGene_tx=XM_003986540.3;refGene_gn=ARHGAP18;refGene_tx=XM_019831190.1;refGene_gn=ARHGAP18;refGene_tx=XM_019831191.1;refGene_gn=ARHGAP18;refGene_tx=XM_019831194.2
B2 119748714 119748776
refGene_gn=ARHGAP18;refGene_tx=XM_003986540.3;refGene_gn=ARHGAP18;refGene_tx=XM_019831190.1;refGene_gn=ARHGAP18;refGene_tx=XM_019831191.1;refGene_gn=ARHGAP18;refGene_tx=XM_019831192.1;refGene_gn=ARHGAP18;refGene_tx=XM_019831194.2
B2 119752684 119752809
refGene_gn=ARHGAP18;refGene_tx=XM_003986540.3;refGene_gn=ARHGAP18;refGene_tx=XM_019831190.1;refGene_gn=ARHGAP18;refGene_tx=XM_019831191.1;refGene_gn=ARHGAP18;refGene_tx=XM_019831194.2
B2 119763620 119763761
refGene_gn=ARHGAP18;refGene_tx=XM_003986540.3;refGene_gn=ARHGAP18;refGene_tx=XM_019831190.1;refGene_gn=ARHGAP18;refGene_tx=XM_019831191.1;refGene_gn=ARHGAP18;refGene_tx=XM_019831192.1;refGene_gn=ARHGAP18;refGene_tx=XM_019831194.2
B2 119764545 119764752
refGene_gn=ARHGAP18;refGene_tx=XM_003986540.3;refGene_gn=ARHGAP18;refGene_tx=XM_019831190.1;refGene_gn=ARHGAP18;refGene_tx=XM_019831191.1;refGene_gn=ARHGAP18;refGene_tx=XM_019831192.1;refGene_gn=ARHGAP18;refGene_tx=XM_019831194.2
B2 119769977 119770060
refGene_gn=ARHGAP18;refGene_tx=XM_003986540.3;refGene_gn=ARHGAP18;refGene_tx=XM_019831190.1;refGene_gn=ARHGAP18;refGene_tx=XM_019831191.1;refGene_gn=ARHGAP18;refGene_tx=XM_019831192.1;refGene_gn=ARHGAP18;refGene_tx=XM_019831194.2
B2 119772168 119772328

refGene_gn=ARHGAP18;refGene_tx=XM_003986540.3;refGene_gn=ARHGAP18;refGene_tx=XM_019831190.1;refGene_gn=ARHGAP18;refGene_tx=XM_019831191.1;refGene_gn=ARHGAP18;refGene_tx=XM_019831192.1;refGene_gn=ARHGAP18;refGene_tx=XM_019831194.2
B2	119775996	119776074
refGene_gn=ARHGAP18;refGene_tx=XM_003986540.3;refGene_gn=ARHGAP18;refGene_tx=XM_019831190.1;refGene_gn=ARHGAP18;refGene_tx=XM_019831191.1;refGene_gn=ARHGAP18;refGene_tx=XM_019831192.1;refGene_gn=ARHGAP18;refGene_tx=XM_019831194.2
B2	119780817	119780909
refGene_gn=ARHGAP18;refGene_tx=XM_003986540.3;refGene_gn=ARHGAP18;refGene_tx=XM_019831190.1;refGene_gn=ARHGAP18;refGene_tx=XM_019831192.1;refGene_gn=ARHGAP18;refGene_tx=XM_019831194.2
B2	119783236	119783402
refGene_gn=ARHGAP18;refGene_tx=XM_003986540.3;refGene_gn=ARHGAP18;refGene_tx=XM_019831190.1;refGene_gn=ARHGAP18;refGene_tx=XM_019831192.1;refGene_gn=ARHGAP18;refGene_tx=XM_019831194.2
B2	119793134	119793313
refGene_gn=ARHGAP18;refGene_tx=XM_003986540.3;refGene_gn=ARHGAP18;refGene_tx=XM_019831190.1;refGene_gn=ARHGAP18;refGene_tx=XM_019831191.1;refGene_gn=ARHGAP18;refGene_tx=XM_019831192.1;refGene_gn=ARHGAP18;refGene_tx=XM_019831194.2
B2	119797562	119797626
refGene_gn=ARHGAP18;refGene_tx=XM_003986540.3;refGene_gn=ARHGAP18;refGene_tx=XM_019831190.1;refGene_gn=ARHGAP18;refGene_tx=XM_019831191.1;refGene_gn=ARHGAP18;refGene_tx=XM_019831192.1;refGene_gn=ARHGAP18;refGene_tx=XM_019831194.2
B2	119801180	119801416
refGene_gn=ARHGAP18;refGene_tx=XM_003986540.3;refGene_gn=ARHGAP18;refGene_tx=XM_019831190.1;refGene_gn=ARHGAP18;refGene_tx=XM_019831191.1;refGene_gn=ARHGAP18;refGene_tx=XM_019831192.1;refGene_gn=ARHGAP18;refGene_tx=XM_019831194.2
B2	119804517	119804711
refGene_gn=ARHGAP18;refGene_tx=XM_003986540.3;refGene_gn=ARHGAP18;refGene_tx=XM_019831190.1;refGene_gn=ARHGAP18;refGene_tx=XM_019831191.1;refGene_gn=ARHGAP18;refGene_tx=XM_019831192.1;refGene_gn=ARHGAP18;refGene_tx=XM_019831194.2
B2	119869112	119869225
refGene_gn=ARHGAP18;refGene_tx=XM_003986540.3
B2	119939227	119939721
refGene_gn=ARHGAP18;refGene_tx=XM_019831190.1;refGene_gn=ARHGAP18;refGene_tx=XM_019831191.1;refGene_gn=ARHGAP18;refGene_tx=XM_019831192.1
B2	119994859	119994954
refGene_gn=TMEM244;refGene_tx=XM_006931991.3
B2	119996744	119996789
refGene_gn=TMEM244;refGene_tx=XM_006931991.3
B2	119998325	119998451
refGene_gn=TMEM244;refGene_tx=XM_006931991.3
B2	120001921	120001995
refGene_gn=TMEM244;refGene_tx=XM_006931991.3

B2 120003543 120003629
refGene_gn=TMEM244;refGene_tx=XM_006931991.3
B2 120018276 120018309
refGene_gn=TMEM244;refGene_tx=XM_006931991.3
B2 120197785 120197887
refGene_gn=L3MBTL3;refGene_tx=XM_003986541.5;refGene_gn=L3MBTL3;refGene_tx=XM_023254343.1;refGene_gn=L3MBTL3;refGene_tx=XM_023254344.1;refGene_gn=L3MBTL3;refGene_tx=XM_023254345.1;refGene_gn=L3MBTL3;refGene_tx=XM_023254346.1;refGene_gn=L3MBTL3;refGene_tx=XM_023254347.1;refGene_gn=L3MBTL3;refGene_tx=XM_023254348.1
B2 120204608 120204720
refGene_gn=L3MBTL3;refGene_tx=XM_003986541.5;refGene_gn=L3MBTL3;refGene_tx=XM_023254343.1;refGene_gn=L3MBTL3;refGene_tx=XM_023254344.1;refGene_gn=L3MBTL3;refGene_tx=XM_023254345.1;refGene_gn=L3MBTL3;refGene_tx=XM_023254346.1;refGene_gn=L3MBTL3;refGene_tx=XM_023254347.1;refGene_gn=L3MBTL3;refGene_tx=XM_023254348.1
B2 120205092 120205167
refGene_gn=L3MBTL3;refGene_tx=XM_003986541.5;refGene_gn=L3MBTL3;refGene_tx=XM_023254343.1;refGene_gn=L3MBTL3;refGene_tx=XM_023254344.1;refGene_gn=L3MBTL3;refGene_tx=XM_023254345.1;refGene_gn=L3MBTL3;refGene_tx=XM_023254346.1;refGene_gn=L3MBTL3;refGene_tx=XM_023254347.1;refGene_gn=L3MBTL3;refGene_tx=XM_023254348.1
B2 120206653 120206813
refGene_gn=L3MBTL3;refGene_tx=XM_003986541.5;refGene_gn=L3MBTL3;refGene_tx=XM_023254343.1;refGene_gn=L3MBTL3;refGene_tx=XM_023254344.1;refGene_gn=L3MBTL3;refGene_tx=XM_023254345.1;refGene_gn=L3MBTL3;refGene_tx=XM_023254346.1;refGene_gn=L3MBTL3;refGene_tx=XM_023254347.1;refGene_gn=L3MBTL3;refGene_tx=XM_023254348.1
B2 120208176 120208309
refGene_gn=L3MBTL3;refGene_tx=XM_003986541.5;refGene_gn=L3MBTL3;refGene_tx=XM_023254343.1;refGene_gn=L3MBTL3;refGene_tx=XM_023254344.1;refGene_gn=L3MBTL3;refGene_tx=XM_023254345.1;refGene_gn=L3MBTL3;refGene_tx=XM_023254346.1;refGene_gn=L3MBTL3;refGene_tx=XM_023254347.1;refGene_gn=L3MBTL3;refGene_tx=XM_023254348.1
B2 120210388 120210473
refGene_gn=L3MBTL3;refGene_tx=XM_003986541.5;refGene_gn=L3MBTL3;refGene_tx=XM_023254343.1;refGene_gn=L3MBTL3;refGene_tx=XM_023254344.1;refGene_gn=L3MBTL3;refGene_tx=XM_023254345.1;refGene_gn=L3MBTL3;refGene_tx=XM_023254346.1;refGene_gn=L3MBTL3;refGene_tx=XM_023254347.1;refGene_gn=L3MBTL3;refGene_tx=XM_023254348.1
B2 120212609 120212701
refGene_gn=L3MBTL3;refGene_tx=XM_003986541.5;refGene_gn=L3MBTL3;refGene_tx=XM_023254343.1;refGene_gn=L3MBTL3;refGene_tx=XM_023254344.1;refGene_gn=L3MBTL3;refGene_tx=XM_023254345.1;refGene_gn=L3MBTL3;refGene_tx=XM_023254346.1;refGene_gn=L3MBTL3;refGene_tx=XM_023254347.1;refGene_gn=L3MBTL3;refGene_tx=XM_023254348.1
B2 120214852 120214957
refGene_gn=L3MBTL3;refGene_tx=XM_003986541.5;refGene_gn=L3MBTL3;refGene_tx=XM_023254343.1;refGene_gn=L3MBTL3;refGene_tx=XM_023254344.1;refGene_gn=L3MBTL3;refGene_tx=XM_023254345.1;refGene_gn=L3MBTL3;refGene_tx=

XM_023254346.1;refGene_gn=L3MBTL3;refGene_tx=XM_023254347.1;refGene_gn
=L3MBTL3;refGene_tx=XM_023254348.1
B2 120220488 120220624
refGene_gn=L3MBTL3;refGene_tx=XM_003986541.5;refGene_gn=L3MBTL3;refGen
e_tx=XM_023254343.1;refGene_gn=L3MBTL3;refGene_tx=XM_023254344.1;refGe
ne_gn=L3MBTL3;refGene_tx=XM_023254345.1;refGene_gn=L3MBTL3;refGene_tx=
XM_023254346.1;refGene_gn=L3MBTL3;refGene_tx=XM_023254347.1;refGene_gn
=L3MBTL3;refGene_tx=XM_023254348.1
B2 120222559 120222651
refGene_gn=L3MBTL3;refGene_tx=XM_003986541.5;refGene_gn=L3MBTL3;refGen
e_tx=XM_023254343.1;refGene_gn=L3MBTL3;refGene_tx=XM_023254344.1;refGe
ne_gn=L3MBTL3;refGene_tx=XM_023254345.1;refGene_gn=L3MBTL3;refGene_tx=
XM_023254346.1;refGene_gn=L3MBTL3;refGene_tx=XM_023254347.1;refGene_gn
=L3MBTL3;refGene_tx=XM_023254348.1
B2 120225158 120225310
refGene_gn=L3MBTL3;refGene_tx=XM_003986541.5;refGene_gn=L3MBTL3;refGen
e_tx=XM_023254343.1;refGene_gn=L3MBTL3;refGene_tx=XM_023254344.1;refGe
ne_gn=L3MBTL3;refGene_tx=XM_023254345.1;refGene_gn=L3MBTL3;refGene_tx=
XM_023254346.1;refGene_gn=L3MBTL3;refGene_tx=XM_023254347.1;refGene_gn
=L3MBTL3;refGene_tx=XM_023254348.1
B2 120236623 120236700
refGene_gn=L3MBTL3;refGene_tx=XM_003986541.5;refGene_gn=L3MBTL3;refGen
e_tx=XM_023254343.1;refGene_gn=L3MBTL3;refGene_tx=XM_023254344.1;refGe
ne_gn=L3MBTL3;refGene_tx=XM_023254345.1;refGene_gn=L3MBTL3;refGene_tx=
XM_023254346.1;refGene_gn=L3MBTL3;refGene_tx=XM_023254347.1;refGene_gn
=L3MBTL3;refGene_tx=XM_023254348.1
B2 120241422 120241508
refGene_gn=L3MBTL3;refGene_tx=XM_003986541.5;refGene_gn=L3MBTL3;refGen
e_tx=XM_023254343.1;refGene_gn=L3MBTL3;refGene_tx=XM_023254344.1;refGe
ne_gn=L3MBTL3;refGene_tx=XM_023254345.1;refGene_gn=L3MBTL3;refGene_tx=
XM_023254346.1;refGene_gn=L3MBTL3;refGene_tx=XM_023254347.1;refGene_gn
=L3MBTL3;refGene_tx=XM_023254348.1
B2 120243780 120243891
refGene_gn=L3MBTL3;refGene_tx=XM_003986541.5;refGene_gn=L3MBTL3;refGen
e_tx=XM_023254343.1;refGene_gn=L3MBTL3;refGene_tx=XM_023254344.1;refGe
ne_gn=L3MBTL3;refGene_tx=XM_023254345.1;refGene_gn=L3MBTL3;refGene_tx=
XM_023254346.1;refGene_gn=L3MBTL3;refGene_tx=XM_023254347.1;refGene_gn
=L3MBTL3;refGene_tx=XM_023254348.1
B2 120254570 120254685
refGene_gn=L3MBTL3;refGene_tx=XM_003986541.5;refGene_gn=L3MBTL3;refGen
e_tx=XM_023254343.1;refGene_gn=L3MBTL3;refGene_tx=XM_023254344.1;refGe
ne_gn=L3MBTL3;refGene_tx=XM_023254345.1;refGene_gn=L3MBTL3;refGene_tx=
XM_023254346.1;refGene_gn=L3MBTL3;refGene_tx=XM_023254347.1;refGene_gn
=L3MBTL3;refGene_tx=XM_023254348.1
B2 120256014 120256349
refGene_gn=L3MBTL3;refGene_tx=XM_003986541.5;refGene_gn=L3MBTL3;refGen
e_tx=XM_023254343.1;refGene_gn=L3MBTL3;refGene_tx=XM_023254344.1;refGe
ne_gn=L3MBTL3;refGene_tx=XM_023254345.1;refGene_gn=L3MBTL3;refGene_tx=
XM_023254346.1;refGene_gn=L3MBTL3;refGene_tx=XM_023254347.1;refGene_gn
=L3MBTL3;refGene_tx=XM_023254348.1

B2 120270773 120270923
refGene_gn=L3MBTL3;refGene_tx=XM_003986541.5
B2 120283142 120283222
refGene_gn=L3MBTL3;refGene_tx=XM_003986541.5
B2 120297325 120297495
refGene_gn=L3MBTL3;refGene_tx=XM_003986541.5
B2 120297713 120297776
refGene_gn=L3MBTL3;refGene_tx=XM_003986541.5
B2 120302827 120302971
refGene_gn=L3MBTL3;refGene_tx=XM_003986541.5
B2 120307311 120307596
refGene_gn=SAMD3;refGene_tx=XM_023254349.1
B2 120307823 120307906
refGene_gn=SAMD3;refGene_tx=XM_023254349.1
B2 120308632 120308804
refGene_gn=SAMD3;refGene_tx=XM_023254349.1
B2 120312365 120312566
refGene_gn=SAMD3;refGene_tx=XM_023254349.1
B2 120331790 120331958
refGene_gn=SAMD3;refGene_tx=XM_023254349.1
B2 120340267 120340352
refGene_gn=SAMD3;refGene_tx=XM_023254349.1
B2 120340601 120340787
refGene_gn=SAMD3;refGene_tx=XM_023254349.1
B2 120357909 120358017
refGene_gn=SAMD3;refGene_tx=XM_023254349.1
B2 120361366 120361556
refGene_gn=SAMD3;refGene_tx=XM_023254349.1
B2 120362229 120362308
refGene_gn=SAMD3;refGene_tx=XM_023254349.1
B2 120546418 120547894
refGene_gn=TMEM200A;refGene_tx=XM_003986544.5;refGene_gn=TMEM200A;refGene_tx=XM_019831199.2;refGene_gn=TMEM200A;refGene_tx=XM_019831200.2;refGene_gn=TMEM200A;refGene_tx=XM_019831201.2
B2 120930121 120930189
refGene_gn=EPB41L2;refGene_tx=XM_023254361.1
B2 120938739 120938743
refGene_gn=EPB41L2;refGene_tx=XM_023254361.1
B2 120939926 120940031
refGene_gn=EPB41L2;refGene_tx=XM_006931999.4;refGene_gn=EPB41L2;refGene_tx=XM_006932000.3;refGene_gn=EPB41L2;refGene_tx=XM_019831204.2;refGene_gn=EPB41L2;refGene_tx=XM_019831205.2;refGene_gn=EPB41L2;refGene_tx=XM_023254350.1;refGene_gn=EPB41L2;refGene_tx=XM_023254351.1;refGene_gn=EPB41L2;refGene_tx=XM_023254352.1;refGene_gn=EPB41L2;refGene_tx=XM_023254353.1;refGene_gn=EPB41L2;refGene_tx=XM_023254354.1;refGene_gn=EPB41L2;refGene_tx=XM_023254355.1;refGene_gn=EPB41L2;refGene_tx=XM_023254356.1;refGene_gn=EPB41L2;refGene_tx=XM_023254357.1;refGene_gn=EPB41L2;refGene_tx=XM_023254358.1;refGene_gn=EPB41L2;refGene_tx=XM_023254359.1;refGene_gn=EPB41L2;refGene_tx=XM_023254360.1;refGene_gn=EPB41L2;refGene_tx=XM_023254362.1;refGene_gn=EPB41L2;refGene_tx=XM_023254363.1;refGe

ne_gn=EPB41L2;refGene_tx=XM_023254364.1;refGene_gn=EPB41L2;refGene_tx=XM_023254365.1;refGene_gn=EPB41L2;refGene_tx=XM_023254366.1;refGene_gn=EPB41L2;refGene_tx=XM_023254367.1;refGene_gn=EPB41L2;refGene_tx=XM_023254368.1;refGene_gn=EPB41L2;refGene_tx=XM_023254369.1
B2 120943910 120943991
refGene_gn=EPB41L2;refGene_tx=XM_006931999.4;refGene_gn=EPB41L2;refGene_tx=XM_006932000.3;refGene_gn=EPB41L2;refGene_tx=XM_019831204.2;refGene_gn=EPB41L2;refGene_tx=XM_019831205.2;refGene_gn=EPB41L2;refGene_tx=XM_023254350.1;refGene_gn=EPB41L2;refGene_tx=XM_023254351.1;refGene_gn=EPB41L2;refGene_tx=XM_023254352.1;refGene_gn=EPB41L2;refGene_tx=XM_023254353.1;refGene_gn=EPB41L2;refGene_tx=XM_023254354.1;refGene_gn=EPB41L2;refGene_tx=XM_023254355.1;refGene_gn=EPB41L2;refGene_tx=XM_023254356.1;refGene_gn=EPB41L2;refGene_tx=XM_023254357.1;refGene_gn=EPB41L2;refGene_tx=XM_023254358.1;refGene_gn=EPB41L2;refGene_tx=XM_023254359.1;refGene_gn=EPB41L2;refGene_tx=XM_023254360.1;refGene_gn=EPB41L2;refGene_tx=XM_023254362.1;refGene_gn=EPB41L2;refGene_tx=XM_023254363.1;refGene_gn=EPB41L2;refGene_tx=XM_023254364.1;refGene_gn=EPB41L2;refGene_tx=XM_023254365.1;refGene_gn=EPB41L2;refGene_tx=XM_023254366.1;refGene_gn=EPB41L2;refGene_tx=XM_023254367.1;refGene_gn=EPB41L2;refGene_tx=XM_023254368.1;refGene_gn=EPB41L2;refGene_tx=XM_023254369.1
B2 120945817 120945916
refGene_gn=EPB41L2;refGene_tx=XM_006931999.4;refGene_gn=EPB41L2;refGene_tx=XM_006932000.3;refGene_gn=EPB41L2;refGene_tx=XM_019831204.2;refGene_gn=EPB41L2;refGene_tx=XM_019831205.2;refGene_gn=EPB41L2;refGene_tx=XM_023254350.1;refGene_gn=EPB41L2;refGene_tx=XM_023254351.1;refGene_gn=EPB41L2;refGene_tx=XM_023254352.1;refGene_gn=EPB41L2;refGene_tx=XM_023254353.1;refGene_gn=EPB41L2;refGene_tx=XM_023254354.1;refGene_gn=EPB41L2;refGene_tx=XM_023254355.1;refGene_gn=EPB41L2;refGene_tx=XM_023254356.1;refGene_gn=EPB41L2;refGene_tx=XM_023254357.1;refGene_gn=EPB41L2;refGene_tx=XM_023254358.1;refGene_gn=EPB41L2;refGene_tx=XM_023254359.1;refGene_gn=EPB41L2;refGene_tx=XM_023254360.1;refGene_gn=EPB41L2;refGene_tx=XM_023254362.1;refGene_gn=EPB41L2;refGene_tx=XM_023254363.1;refGene_gn=EPB41L2;refGene_tx=XM_023254364.1;refGene_gn=EPB41L2;refGene_tx=XM_023254365.1;refGene_gn=EPB41L2;refGene_tx=XM_023254366.1;refGene_gn=EPB41L2;refGene_tx=XM_023254367.1;refGene_gn=EPB41L2;refGene_tx=XM_023254368.1;refGene_gn=EPB41L2;refGene_tx=XM_023254369.1
B2 120947637 120947760
refGene_gn=EPB41L2;refGene_tx=XM_006931999.4;refGene_gn=EPB41L2;refGene_tx=XM_006932000.3;refGene_gn=EPB41L2;refGene_tx=XM_019831204.2;refGene_gn=EPB41L2;refGene_tx=XM_019831205.2;refGene_gn=EPB41L2;refGene_tx=XM_023254350.1;refGene_gn=EPB41L2;refGene_tx=XM_023254351.1;refGene_gn=EPB41L2;refGene_tx=XM_023254352.1;refGene_gn=EPB41L2;refGene_tx=XM_023254353.1;refGene_gn=EPB41L2;refGene_tx=XM_023254354.1;refGene_gn=EPB41L2;refGene_tx=XM_023254355.1;refGene_gn=EPB41L2;refGene_tx=XM_023254356.1;refGene_gn=EPB41L2;refGene_tx=XM_023254357.1;refGene_gn=EPB41L2;refGene_tx=XM_023254358.1;refGene_gn=EPB41L2;refGene_tx=XM_023254359.1;refGene_gn=EPB41L2;refGene_tx=XM_023254360.1;refGene_gn=EPB41L2;refGene_tx=XM_023254362.1;refGene_gn=EPB41L2;refGene_tx=XM_023254364.1;refGene_gn=EPB41L2;refGene_tx=XM_023254365.1;refGene_gn=EPB41L2;refGene_tx=XM_023254367.1;refGene_gn=EPB41L2;refGene_tx=XM_023254369.1

B2 120949692 120950262
refGene_gn=EPB41L2;refGene_tx=XM_006931999.4;refGene_gn=EPB41L2;refGene_tx=XM_006932000.3;refGene_gn=EPB41L2;refGene_tx=XM_019831204.2;refGene_gn=EPB41L2;refGene_tx=XM_019831205.2;refGene_gn=EPB41L2;refGene_tx=XM_023254350.1;refGene_gn=EPB41L2;refGene_tx=XM_023254351.1;refGene_gn=EPB41L2;refGene_tx=XM_023254352.1;refGene_gn=EPB41L2;refGene_tx=XM_023254353.1;refGene_gn=EPB41L2;refGene_tx=XM_023254354.1;refGene_gn=EPB41L2;refGene_tx=XM_023254355.1;refGene_gn=EPB41L2;refGene_tx=XM_023254356.1;refGene_gn=EPB41L2;refGene_tx=XM_023254357.1;refGene_gn=EPB41L2;refGene_tx=XM_023254358.1;refGene_gn=EPB41L2;refGene_tx=XM_023254359.1;refGene_gn=EPB41L2;refGene_tx=XM_023254360.1;refGene_gn=EPB41L2;refGene_tx=XM_023254361.1;refGene_gn=EPB41L2;refGene_tx=XM_023254362.1;refGene_gn=EPB41L2;refGene_tx=XM_023254363.1;refGene_gn=EPB41L2;refGene_tx=XM_023254364.1;refGene_gn=EPB41L2;refGene_tx=XM_023254365.1;refGene_gn=EPB41L2;refGene_tx=XM_023254366.1;refGene_gn=EPB41L2;refGene_tx=XM_023254369.1
B2 120950468 120950522
refGene_gn=EPB41L2;refGene_tx=XM_006931999.4;refGene_gn=EPB41L2;refGene_tx=XM_006932000.3;refGene_gn=EPB41L2;refGene_tx=XM_019831204.2;refGene_gn=EPB41L2;refGene_tx=XM_019831205.2;refGene_gn=EPB41L2;refGene_tx=XM_023254350.1;refGene_gn=EPB41L2;refGene_tx=XM_023254351.1;refGene_gn=EPB41L2;refGene_tx=XM_023254352.1;refGene_gn=EPB41L2;refGene_tx=XM_023254354.1;refGene_gn=EPB41L2;refGene_tx=XM_023254355.1;refGene_gn=EPB41L2;refGene_tx=XM_023254357.1;refGene_gn=EPB41L2;refGene_tx=XM_023254359.1;refGene_gn=EPB41L2;refGene_tx=XM_023254360.1;refGene_gn=EPB41L2;refGene_tx=XM_023254361.1;refGene_gn=EPB41L2;refGene_tx=XM_023254363.1;refGene_gn=EPB41L2;refGene_tx=XM_023254364.1;refGene_gn=EPB41L2;refGene_tx=XM_023254369.1
B2 120952568 120952736
refGene_gn=EPB41L2;refGene_tx=XM_023254369.1
B2 120956259 120956361
refGene_gn=EPB41L2;refGene_tx=XM_006931999.4;refGene_gn=EPB41L2;refGene_tx=XM_006932000.3;refGene_gn=EPB41L2;refGene_tx=XM_019831204.2;refGene_gn=EPB41L2;refGene_tx=XM_019831205.2;refGene_gn=EPB41L2;refGene_tx=XM_023254350.1;refGene_gn=EPB41L2;refGene_tx=XM_023254351.1;refGene_gn=EPB41L2;refGene_tx=XM_023254352.1;refGene_gn=EPB41L2;refGene_tx=XM_023254353.1;refGene_gn=EPB41L2;refGene_tx=XM_023254354.1;refGene_gn=EPB41L2;refGene_tx=XM_023254357.1;refGene_gn=EPB41L2;refGene_tx=XM_023254359.1;refGene_gn=EPB41L2;refGene_tx=XM_023254361.1;refGene_gn=EPB41L2;refGene_tx=XM_023254363.1
B2 120957749 120957896
refGene_gn=EPB41L2;refGene_tx=XM_006931999.4;refGene_gn=EPB41L2;refGene_tx=XM_006932000.3;refGene_gn=EPB41L2;refGene_tx=XM_019831204.2;refGene_gn=EPB41L2;refGene_tx=XM_019831205.2;refGene_gn=EPB41L2;refGene_tx=XM_023254350.1;refGene_gn=EPB41L2;refGene_tx=XM_023254351.1;refGene_gn=EPB41L2;refGene_tx=XM_023254352.1;refGene_gn=EPB41L2;refGene_tx=XM_023254353.1;refGene_gn=EPB41L2;refGene_tx=XM_023254354.1;refGene_gn=EPB41L2;refGene_tx=XM_023254355.1;refGene_gn=EPB41L2;refGene_tx=XM_023254356.1;refGene_gn=EPB41L2;refGene_tx=XM_023254358.1;refGene_gn=EPB41L2;refGene_tx=XM_023254359.1;refGene_gn=EPB41L2;refGene_tx=XM_023254361.1;

refGene_gn=EPB41L2;refGene_tx=XM_023254363.1;refGene_gn=EPB41L2;refGene_tx=XM_023254369.1
B2 120959839 120959902
refGene_gn=EPB41L2;refGene_tx=XM_006931999.4;refGene_gn=EPB41L2;refGene_tx=XM_006932000.3;refGene_gn=EPB41L2;refGene_tx=XM_019831204.2;refGene_gn=EPB41L2;refGene_tx=XM_019831205.2;refGene_gn=EPB41L2;refGene_tx=XM_023254350.1;refGene_gn=EPB41L2;refGene_tx=XM_023254351.1;refGene_gn=EPB41L2;refGene_tx=XM_023254352.1;refGene_gn=EPB41L2;refGene_tx=XM_023254353.1;refGene_gn=EPB41L2;refGene_tx=XM_023254355.1;refGene_gn=EPB41L2;refGene_tx=XM_023254356.1;refGene_gn=EPB41L2;refGene_tx=XM_023254359.1;refGene_gn=EPB41L2;refGene_tx=XM_023254361.1;refGene_gn=EPB41L2;refGene_tx=XM_023254363.1;refGene_gn=EPB41L2;refGene_tx=XM_023254369.1
B2 120964717 120964890
refGene_gn=EPB41L2;refGene_tx=XM_006931999.4;refGene_gn=EPB41L2;refGene_tx=XM_006932000.3;refGene_gn=EPB41L2;refGene_tx=XM_019831204.2;refGene_gn=EPB41L2;refGene_tx=XM_019831205.2;refGene_gn=EPB41L2;refGene_tx=XM_023254350.1;refGene_gn=EPB41L2;refGene_tx=XM_023254351.1;refGene_gn=EPB41L2;refGene_tx=XM_023254352.1;refGene_gn=EPB41L2;refGene_tx=XM_023254353.1;refGene_gn=EPB41L2;refGene_tx=XM_023254354.1;refGene_gn=EPB41L2;refGene_tx=XM_023254355.1;refGene_gn=EPB41L2;refGene_tx=XM_023254356.1;refGene_gn=EPB41L2;refGene_tx=XM_023254357.1;refGene_gn=EPB41L2;refGene_tx=XM_023254358.1;refGene_gn=EPB41L2;refGene_tx=XM_023254359.1;refGene_gn=EPB41L2;refGene_tx=XM_023254360.1;refGene_gn=EPB41L2;refGene_tx=XM_023254361.1;refGene_gn=EPB41L2;refGene_tx=XM_023254362.1;refGene_gn=EPB41L2;refGene_tx=XM_023254363.1;refGene_gn=EPB41L2;refGene_tx=XM_023254364.1;refGene_gn=EPB41L2;refGene_tx=XM_023254365.1;refGene_gn=EPB41L2;refGene_tx=XM_023254366.1;refGene_gn=EPB41L2;refGene_tx=XM_023254367.1;refGene_gn=EPB41L2;refGene_tx=XM_023254368.1;refGene_gn=EPB41L2;refGene_tx=XM_023254369.1
B2 120970237 120970410
refGene_gn=EPB41L2;refGene_tx=XM_006931999.4;refGene_gn=EPB41L2;refGene_tx=XM_006932000.3;refGene_gn=EPB41L2;refGene_tx=XM_019831204.2;refGene_gn=EPB41L2;refGene_tx=XM_019831205.2;refGene_gn=EPB41L2;refGene_tx=XM_023254350.1;refGene_gn=EPB41L2;refGene_tx=XM_023254351.1;refGene_gn=EPB41L2;refGene_tx=XM_023254352.1;refGene_gn=EPB41L2;refGene_tx=XM_023254353.1;refGene_gn=EPB41L2;refGene_tx=XM_023254354.1;refGene_gn=EPB41L2;refGene_tx=XM_023254355.1;refGene_gn=EPB41L2;refGene_tx=XM_023254356.1;refGene_gn=EPB41L2;refGene_tx=XM_023254357.1;refGene_gn=EPB41L2;refGene_tx=XM_023254358.1;refGene_gn=EPB41L2;refGene_tx=XM_023254359.1;refGene_gn=EPB41L2;refGene_tx=XM_023254360.1;refGene_gn=EPB41L2;refGene_tx=XM_023254361.1;refGene_gn=EPB41L2;refGene_tx=XM_023254362.1;refGene_gn=EPB41L2;refGene_tx=XM_023254363.1;refGene_gn=EPB41L2;refGene_tx=XM_023254364.1;refGene_gn=EPB41L2;refGene_tx=XM_023254365.1;refGene_gn=EPB41L2;refGene_tx=XM_023254366.1;refGene_gn=EPB41L2;refGene_tx=XM_023254367.1;refGene_gn=EPB41L2;refGene_tx=XM_023254368.1;refGene_gn=EPB41L2;refGene_tx=XM_023254369.1
B2 120979706 120979804
refGene_gn=EPB41L2;refGene_tx=XM_006931999.4;refGene_gn=EPB41L2;refGene_tx=XM_006932000.3;refGene_gn=EPB41L2;refGene_tx=XM_019831204.2;refGene_gn=EPB41L2;refGene_tx=XM_019831205.2;refGene_gn=EPB41L2;refGene_tx=

XM_023254350.1;refGene_gn=EPB41L2;refGene_tx=XM_023254351.1;refGene_gn
=EPB41L2;refGene_tx=XM_023254352.1;refGene_gn=EPB41L2;refGene_tx=XM_02
3254353.1;refGene_gn=EPB41L2;refGene_tx=XM_023254354.1;refGene_gn=EPB4
1L2;refGene_tx=XM_023254355.1;refGene_gn=EPB41L2;refGene_tx=XM_0232543
56.1;refGene_gn=EPB41L2;refGene_tx=XM_023254357.1;refGene_gn=EPB41L2;r
efGene_tx=XM_023254358.1;refGene_gn=EPB41L2;refGene_tx=XM_023254359.1;
refGene_gn=EPB41L2;refGene_tx=XM_023254360.1;refGene_gn=EPB41L2;refGen
e_tx=XM_023254361.1;refGene_gn=EPB41L2;refGene_tx=XM_023254362.1;refGe
ne_gn=EPB41L2;refGene_tx=XM_023254363.1;refGene_gn=EPB41L2;refGene_tx=
XM_023254364.1;refGene_gn=EPB41L2;refGene_tx=XM_023254365.1;refGene_gn
=EPB41L2;refGene_tx=XM_023254366.1;refGene_gn=EPB41L2;refGene_tx=XM_02
3254367.1;refGene_gn=EPB41L2;refGene_tx=XM_023254368.1;refGene_gn=EPB4
1L2;refGene_tx=XM_023254369.1
B2 120980322 120980475
refGene_gn=EPB41L2;refGene_tx=XM_006931999.4;refGene_gn=EPB41L2;refGen
e_tx=XM_006932000.3;refGene_gn=EPB41L2;refGene_tx=XM_019831204.2;refGe
ne_gn=EPB41L2;refGene_tx=XM_019831205.2;refGene_gn=EPB41L2;refGene_tx=
XM_023254350.1;refGene_gn=EPB41L2;refGene_tx=XM_023254351.1;refGene_gn
=EPB41L2;refGene_tx=XM_023254352.1;refGene_gn=EPB41L2;refGene_tx=XM_02
3254353.1;refGene_gn=EPB41L2;refGene_tx=XM_023254354.1;refGene_gn=EPB4
1L2;refGene_tx=XM_023254355.1;refGene_gn=EPB41L2;refGene_tx=XM_0232543
56.1;refGene_gn=EPB41L2;refGene_tx=XM_023254357.1;refGene_gn=EPB41L2;r
efGene_tx=XM_023254358.1;refGene_gn=EPB41L2;refGene_tx=XM_023254359.1;
refGene_gn=EPB41L2;refGene_tx=XM_023254360.1;refGene_gn=EPB41L2;refGen
e_tx=XM_023254361.1;refGene_gn=EPB41L2;refGene_tx=XM_023254362.1;refGe
ne_gn=EPB41L2;refGene_tx=XM_023254363.1;refGene_gn=EPB41L2;refGene_tx=
XM_023254364.1;refGene_gn=EPB41L2;refGene_tx=XM_023254365.1;refGene_gn
=EPB41L2;refGene_tx=XM_023254366.1;refGene_gn=EPB41L2;refGene_tx=XM_02
3254367.1;refGene_gn=EPB41L2;refGene_tx=XM_023254368.1;refGene_gn=EPB4
1L2;refGene_tx=XM_023254369.1
B2 120984698 120984786
refGene_gn=EPB41L2;refGene_tx=XM_006931999.4;refGene_gn=EPB41L2;refGen
e_tx=XM_006932000.3;refGene_gn=EPB41L2;refGene_tx=XM_019831204.2;refGe
ne_gn=EPB41L2;refGene_tx=XM_019831205.2;refGene_gn=EPB41L2;refGene_tx=
XM_023254350.1;refGene_gn=EPB41L2;refGene_tx=XM_023254351.1;refGene_gn
=EPB41L2;refGene_tx=XM_023254352.1;refGene_gn=EPB41L2;refGene_tx=XM_02
3254353.1;refGene_gn=EPB41L2;refGene_tx=XM_023254354.1;refGene_gn=EPB4
1L2;refGene_tx=XM_023254355.1;refGene_gn=EPB41L2;refGene_tx=XM_0232543
56.1;refGene_gn=EPB41L2;refGene_tx=XM_023254357.1;refGene_gn=EPB41L2;r
efGene_tx=XM_023254358.1;refGene_gn=EPB41L2;refGene_tx=XM_023254359.1;
refGene_gn=EPB41L2;refGene_tx=XM_023254360.1;refGene_gn=EPB41L2;refGen
e_tx=XM_023254361.1;refGene_gn=EPB41L2;refGene_tx=XM_023254362.1;refGe
ne_gn=EPB41L2;refGene_tx=XM_023254363.1;refGene_gn=EPB41L2;refGene_tx=
XM_023254364.1;refGene_gn=EPB41L2;refGene_tx=XM_023254365.1;refGene_gn
=EPB41L2;refGene_tx=XM_023254366.1;refGene_gn=EPB41L2;refGene_tx=XM_02
3254367.1;refGene_gn=EPB41L2;refGene_tx=XM_023254368.1;refGene_gn=EPB4
1L2;refGene_tx=XM_023254369.1
B2 120986332 120986551
refGene_gn=EPB41L2;refGene_tx=XM_006931999.4;refGene_gn=EPB41L2;refGen
e_tx=XM_006932000.3;refGene_gn=EPB41L2;refGene_tx=XM_019831204.2;refGe

ne_gn=EPB41L2;refGene_tx=XM_019831205.2;refGene_gn=EPB41L2;refGene_tx=XM_023254350.1;refGene_gn=EPB41L2;refGene_tx=XM_023254351.1;refGene_gn=EPB41L2;refGene_tx=XM_023254352.1;refGene_gn=EPB41L2;refGene_tx=XM_023254353.1;refGene_gn=EPB41L2;refGene_tx=XM_023254354.1;refGene_gn=EPB41L2;refGene_tx=XM_023254355.1;refGene_gn=EPB41L2;refGene_tx=XM_023254356.1;refGene_gn=EPB41L2;refGene_tx=XM_023254357.1;refGene_gn=EPB41L2;refGene_tx=XM_023254358.1;refGene_gn=EPB41L2;refGene_tx=XM_023254359.1;refGene_gn=EPB41L2;refGene_tx=XM_023254360.1;refGene_gn=EPB41L2;refGene_tx=XM_023254361.1;refGene_gn=EPB41L2;refGene_tx=XM_023254362.1;refGene_gn=EPB41L2;refGene_tx=XM_023254363.1;refGene_gn=EPB41L2;refGene_tx=XM_023254364.1;refGene_gn=EPB41L2;refGene_tx=XM_023254365.1;refGene_gn=EPB41L2;refGene_tx=XM_023254366.1;refGene_gn=EPB41L2;refGene_tx=XM_023254367.1;refGene_gn=EPB41L2;refGene_tx=XM_023254368.1;refGene_gn=EPB41L2;refGene_tx=XM_023254369.1
B2 120989585 120989661
refGene_gn=EPB41L2;refGene_tx=XM_006931999.4;refGene_gn=EPB41L2;refGene_tx=XM_006932000.3;refGene_gn=EPB41L2;refGene_tx=XM_019831204.2;refGene_gn=EPB41L2;refGene_tx=XM_019831205.2;refGene_gn=EPB41L2;refGene_tx=XM_023254350.1;refGene_gn=EPB41L2;refGene_tx=XM_023254351.1;refGene_gn=EPB41L2;refGene_tx=XM_023254352.1;refGene_gn=EPB41L2;refGene_tx=XM_023254353.1;refGene_gn=EPB41L2;refGene_tx=XM_023254354.1;refGene_gn=EPB41L2;refGene_tx=XM_023254355.1;refGene_gn=EPB41L2;refGene_tx=XM_023254356.1;refGene_gn=EPB41L2;refGene_tx=XM_023254357.1;refGene_gn=EPB41L2;refGene_tx=XM_023254358.1;refGene_gn=EPB41L2;refGene_tx=XM_023254359.1;refGene_gn=EPB41L2;refGene_tx=XM_023254360.1;refGene_gn=EPB41L2;refGene_tx=XM_023254361.1;refGene_gn=EPB41L2;refGene_tx=XM_023254362.1;refGene_gn=EPB41L2;refGene_tx=XM_023254363.1;refGene_gn=EPB41L2;refGene_tx=XM_023254364.1;refGene_gn=EPB41L2;refGene_tx=XM_023254365.1;refGene_gn=EPB41L2;refGene_tx=XM_023254366.1;refGene_gn=EPB41L2;refGene_tx=XM_023254367.1;refGene_gn=EPB41L2;refGene_tx=XM_023254368.1;refGene_gn=EPB41L2;refGene_tx=XM_023254369.1
B2 120994406 120994449
refGene_gn=EPB41L2;refGene_tx=XM_006931999.4;refGene_gn=EPB41L2;refGene_tx=XM_006932000.3;refGene_gn=EPB41L2;refGene_tx=XM_019831204.2;refGene_gn=EPB41L2;refGene_tx=XM_019831205.2;refGene_gn=EPB41L2;refGene_tx=XM_023254350.1;refGene_gn=EPB41L2;refGene_tx=XM_023254351.1;refGene_gn=EPB41L2;refGene_tx=XM_023254352.1;refGene_gn=EPB41L2;refGene_tx=XM_023254353.1;refGene_gn=EPB41L2;refGene_tx=XM_023254354.1;refGene_gn=EPB41L2;refGene_tx=XM_023254355.1;refGene_gn=EPB41L2;refGene_tx=XM_023254356.1;refGene_gn=EPB41L2;refGene_tx=XM_023254357.1;refGene_gn=EPB41L2;refGene_tx=XM_023254358.1;refGene_gn=EPB41L2;refGene_tx=XM_023254359.1;refGene_gn=EPB41L2;refGene_tx=XM_023254360.1;refGene_gn=EPB41L2;refGene_tx=XM_023254361.1;refGene_gn=EPB41L2;refGene_tx=XM_023254362.1;refGene_gn=EPB41L2;refGene_tx=XM_023254363.1;refGene_gn=EPB41L2;refGene_tx=XM_023254364.1;refGene_gn=EPB41L2;refGene_tx=XM_023254365.1;refGene_gn=EPB41L2;refGene_tx=XM_023254366.1;refGene_gn=EPB41L2;refGene_tx=XM_023254367.1;refGene_gn=EPB41L2;refGene_tx=XM_023254368.1;refGene_gn=EPB41L2;refGene_tx=XM_023254369.1
B2 121009707 121009812
refGene_gn=EPB41L2;refGene_tx=XM_006931999.4;refGene_gn=EPB41L2;refGen

e_tx=XM_006932000.3;refGene_gn=EPB41L2;refGene_tx=XM_019831204.2;refGene_gn=EPB41L2;refGene_tx=XM_019831205.2;refGene_gn=EPB41L2;refGene_tx=XM_023254350.1;refGene_gn=EPB41L2;refGene_tx=XM_023254351.1;refGene_gn=EPB41L2;refGene_tx=XM_023254352.1;refGene_gn=EPB41L2;refGene_tx=XM_023254353.1;refGene_gn=EPB41L2;refGene_tx=XM_023254354.1;refGene_gn=EPB41L2;refGene_tx=XM_023254355.1;refGene_gn=EPB41L2;refGene_tx=XM_023254356.1;refGene_gn=EPB41L2;refGene_tx=XM_023254357.1;refGene_gn=EPB41L2;refGene_tx=XM_023254358.1;refGene_gn=EPB41L2;refGene_tx=XM_023254359.1;refGene_gn=EPB41L2;refGene_tx=XM_023254360.1;refGene_gn=EPB41L2;refGene_tx=XM_023254361.1;refGene_gn=EPB41L2;refGene_tx=XM_023254362.1;refGene_gn=EPB41L2;refGene_tx=XM_023254363.1;refGene_gn=EPB41L2;refGene_tx=XM_023254364.1;refGene_gn=EPB41L2;refGene_tx=XM_023254365.1;refGene_gn=EPB41L2;refGene_tx=XM_023254366.1;refGene_gn=EPB41L2;refGene_tx=XM_023254367.1;refGene_gn=EPB41L2;refGene_tx=XM_023254368.1;refGene_gn=EPB41L2;refGene_tx=XM_023254369.1
B2 121040271 121040478
refGene_gn=EPB41L2;refGene_tx=XM_006931999.4;refGene_gn=EPB41L2;refGene_tx=XM_006932000.3;refGene_gn=EPB41L2;refGene_tx=XM_019831204.2;refGene_gn=EPB41L2;refGene_tx=XM_019831205.2;refGene_gn=EPB41L2;refGene_tx=XM_023254350.1;refGene_gn=EPB41L2;refGene_tx=XM_023254351.1;refGene_gn=EPB41L2;refGene_tx=XM_023254352.1;refGene_gn=EPB41L2;refGene_tx=XM_023254353.1;refGene_gn=EPB41L2;refGene_tx=XM_023254354.1;refGene_gn=EPB41L2;refGene_tx=XM_023254355.1;refGene_gn=EPB41L2;refGene_tx=XM_023254356.1;refGene_gn=EPB41L2;refGene_tx=XM_023254357.1;refGene_gn=EPB41L2;refGene_tx=XM_023254358.1;refGene_gn=EPB41L2;refGene_tx=XM_023254359.1;refGene_gn=EPB41L2;refGene_tx=XM_023254360.1;refGene_gn=EPB41L2;refGene_tx=XM_023254361.1;refGene_gn=EPB41L2;refGene_tx=XM_023254362.1;refGene_gn=EPB41L2;refGene_tx=XM_023254363.1;refGene_gn=EPB41L2;refGene_tx=XM_023254364.1;refGene_gn=EPB41L2;refGene_tx=XM_023254365.1;refGene_gn=EPB41L2;refGene_tx=XM_023254366.1;refGene_gn=EPB41L2;refGene_tx=XM_023254367.1;refGene_gn=EPB41L2;refGene_tx=XM_023254368.1;refGene_gn=EPB41L2;refGene_tx=XM_023254369.1
B2 121041604 121042075
refGene_gn=EPB41L2;refGene_tx=XM_006931999.4;refGene_gn=EPB41L2;refGene_tx=XM_006932000.3;refGene_gn=EPB41L2;refGene_tx=XM_019831204.2;refGene_gn=EPB41L2;refGene_tx=XM_019831205.2;refGene_gn=EPB41L2;refGene_tx=XM_023254350.1;refGene_gn=EPB41L2;refGene_tx=XM_023254351.1;refGene_gn=EPB41L2;refGene_tx=XM_023254352.1;refGene_gn=EPB41L2;refGene_tx=XM_023254353.1;refGene_gn=EPB41L2;refGene_tx=XM_023254354.1;refGene_gn=EPB41L2;refGene_tx=XM_023254355.1;refGene_gn=EPB41L2;refGene_tx=XM_023254356.1;refGene_gn=EPB41L2;refGene_tx=XM_023254357.1;refGene_gn=EPB41L2;refGene_tx=XM_023254358.1;refGene_gn=EPB41L2;refGene_tx=XM_023254359.1;refGene_gn=EPB41L2;refGene_tx=XM_023254360.1;refGene_gn=EPB41L2;refGene_tx=XM_023254361.1;refGene_gn=EPB41L2;refGene_tx=XM_023254362.1;refGene_gn=EPB41L2;refGene_tx=XM_023254363.1;refGene_gn=EPB41L2;refGene_tx=XM_023254364.1;refGene_gn=EPB41L2;refGene_tx=XM_023254365.1;refGene_gn=EPB41L2;refGene_tx=XM_023254366.1;refGene_gn=EPB41L2;refGene_tx=XM_023254367.1;refGene_gn=EPB41L2;refGene_tx=XM_023254368.1;refGene_gn=EPB41L2;refGene_tx=XM_023254369.1
B2 121185752 121185771

refGene_gn=AKAP7;refGene_tx=XM_023254370.1;refGene_gn=AKAP7;refGene_tx=XM_023254371.1;refGene_gn=AKAP7;refGene_tx=XM_023254372.1;refGene_gn=AKAP7;refGene_tx=XM_023254373.1
B2 121191924 121192062
refGene_gn=AKAP7;refGene_tx=XM_019831226.2;refGene_gn=AKAP7;refGene_tx=XM_023254370.1;refGene_gn=AKAP7;refGene_tx=XM_023254371.1;refGene_gn=AKAP7;refGene_tx=XM_023254372.1;refGene_gn=AKAP7;refGene_tx=XM_023254373.1
B2 121207528 121207668
refGene_gn=AKAP7;refGene_tx=XM_023254370.1;refGene_gn=AKAP7;refGene_tx=XM_023254371.1;refGene_gn=AKAP7;refGene_tx=XM_023254372.1;refGene_gn=AKAP7;refGene_tx=XM_023254373.1
B2 121219018 121219155
refGene_gn=AKAP7;refGene_tx=XM_019831226.2;refGene_gn=AKAP7;refGene_tx=XM_023254370.1;refGene_gn=AKAP7;refGene_tx=XM_023254371.1;refGene_gn=AKAP7;refGene_tx=XM_023254372.1;refGene_gn=AKAP7;refGene_tx=XM_023254373.1
B2 121225442 121225603
refGene_gn=AKAP7;refGene_tx=XM_019831226.2;refGene_gn=AKAP7;refGene_tx=XM_023254370.1;refGene_gn=AKAP7;refGene_tx=XM_023254371.1;refGene_gn=AKAP7;refGene_tx=XM_023254372.1;refGene_gn=AKAP7;refGene_tx=XM_023254373.1
B2 121244531 121244644
refGene_gn=AKAP7;refGene_tx=XM_019831226.2;refGene_gn=AKAP7;refGene_tx=XM_023254370.1;refGene_gn=AKAP7;refGene_tx=XM_023254371.1;refGene_gn=AKAP7;refGene_tx=XM_023254372.1;refGene_gn=AKAP7;refGene_tx=XM_023254373.1
B2 121280040 121280188
refGene_gn=AKAP7;refGene_tx=XM_019831226.2;refGene_gn=AKAP7;refGene_tx=XM_023254370.1;refGene_gn=AKAP7;refGene_tx=XM_023254371.1;refGene_gn=AKAP7;refGene_tx=XM_023254372.1;refGene_gn=AKAP7;refGene_tx=XM_023254373.1
B2 121302694 121302743
refGene_gn=AKAP7;refGene_tx=XM_023254374.1;refGene_gn=AKAP7;refGene_tx=XM_023254375.1
B2 121305117 121305186
refGene_gn=AKAP7;refGene_tx=XM_019831226.2;refGene_gn=AKAP7;refGene_tx=XM_023254370.1;refGene_gn=AKAP7;refGene_tx=XM_023254371.1;refGene_gn=AKAP7;refGene_tx=XM_023254374.1
B2 121341699 121341896
refGene_gn=AKAP7;refGene_tx=XM_023254371.1;refGene_gn=AKAP7;refGene_tx=XM_023254373.1;refGene_gn=AKAP7;refGene_tx=XM_023254374.1;refGene_gn=AKAP7;refGene_tx=XM_023254375.1
B2 121342582 121342788
refGene_gn=AKAP7;refGene_tx=XM_019831226.2;refGene_gn=AKAP7;refGene_tx=XM_023254370.1;refGene_gn=AKAP7;refGene_tx=XM_023254372.1
B2 121558484 121558541
refGene_gn=ARG1;refGene_tx=XM_003986549.5
B2 121562050 121562123
refGene_gn=ARG1;refGene_tx=XM_003986549.5

B2 121564614 121564789
refGene_gn=ARG1;refGene_tx=XM_003986549.5
B2 121566349 121566509
refGene_gn=ARG1;refGene_tx=XM_003986549.5
B2 121567705 121567800
refGene_gn=ARG1;refGene_tx=XM_003986549.5
B2 121568121 121568226
refGene_gn=ARG1;refGene_tx=XM_003986549.5
B2 121568668 121568805
refGene_gn=ARG1;refGene_tx=XM_003986549.5
B2 121569026 121569193
refGene_gn=ARG1;refGene_tx=XM_003986549.5
B2 121572629 121572797
refGene_gn=MED23;refGene_tx=XM_011282608.3;refGene_gn=MED23;refGene_tx
=XM_011282609.3
B2 121574639 121574771
refGene_gn=MED23;refGene_tx=XM_011282608.3;refGene_gn=MED23;refGene_tx
=XM_011282609.3
B2 121575515 121575636
refGene_gn=MED23;refGene_tx=XM_011282608.3;refGene_gn=MED23;refGene_tx
=XM_011282609.3
B2 121576585 121576800
refGene_gn=MED23;refGene_tx=XM_011282608.3;refGene_gn=MED23;refGene_tx
=XM_011282609.3
B2 121577801 121577874
refGene_gn=MED23;refGene_tx=XM_011282608.3;refGene_gn=MED23;refGene_tx
=XM_011282609.3
B2 121578688 121578854
refGene_gn=MED23;refGene_tx=XM_011282608.3;refGene_gn=MED23;refGene_tx
=XM_011282609.3
B2 121580164 121580401
refGene_gn=MED23;refGene_tx=XM_011282608.3;refGene_gn=MED23;refGene_tx
=XM_011282609.3
B2 121581952 121582169
refGene_gn=MED23;refGene_tx=XM_011282608.3;refGene_gn=MED23;refGene_tx
=XM_011282609.3
B2 121582605 121582776
refGene_gn=MED23;refGene_tx=XM_011282608.3;refGene_gn=MED23;refGene_tx
=XM_011282609.3
B2 121584271 121584452
refGene_gn=MED23;refGene_tx=XM_011282608.3;refGene_gn=MED23;refGene_tx
=XM_011282609.3
B2 121584541 121584747
refGene_gn=MED23;refGene_tx=XM_011282608.3;refGene_gn=MED23;refGene_tx
=XM_011282609.3
B2 121585780 121585905
refGene_gn=MED23;refGene_tx=XM_011282608.3;refGene_gn=MED23;refGene_tx
=XM_011282609.3
B2 121588558 121588722
refGene_gn=MED23;refGene_tx=XM_011282608.3;refGene_gn=MED23;refGene_tx

=XM_011282609.3
B2	121590063	121590238
refGene_gn=MED23;refGene_tx=XM_011282608.3;refGene_gn=MED23;refGene_tx=XM_011282609.3
B2	121591287	121591430
refGene_gn=MED23;refGene_tx=XM_011282608.3;refGene_gn=MED23;refGene_tx=XM_011282609.3
B2	121593223	121593469
refGene_gn=MED23;refGene_tx=XM_011282608.3;refGene_gn=MED23;refGene_tx=XM_011282609.3
B2	121594467	121594613
refGene_gn=MED23;refGene_tx=XM_011282608.3;refGene_gn=MED23;refGene_tx=XM_011282609.3
B2	121596676	121596820
refGene_gn=MED23;refGene_tx=XM_011282608.3;refGene_gn=MED23;refGene_tx=XM_011282609.3
B2	121598408	121598609
refGene_gn=MED23;refGene_tx=XM_011282608.3;refGene_gn=MED23;refGene_tx=XM_011282609.3
B2	121603693	121603711
refGene_gn=MED23;refGene_tx=XM_011282608.3
B2	121604255	121604351
refGene_gn=MED23;refGene_tx=XM_011282608.3;refGene_gn=MED23;refGene_tx=XM_011282609.3
B2	121605818	121605931
refGene_gn=MED23;refGene_tx=XM_011282608.3;refGene_gn=MED23;refGene_tx=XM_011282609.3
B2	121607408	121607478
refGene_gn=MED23;refGene_tx=XM_011282608.3;refGene_gn=MED23;refGene_tx=XM_011282609.3
B2	121608063	121608165
refGene_gn=MED23;refGene_tx=XM_011282608.3;refGene_gn=MED23;refGene_tx=XM_011282609.3
B2	121608765	121608864
refGene_gn=MED23;refGene_tx=XM_011282608.3;refGene_gn=MED23;refGene_tx=XM_011282609.3
B2	121610595	121610707
refGene_gn=MED23;refGene_tx=XM_011282608.3;refGene_gn=MED23;refGene_tx=XM_011282609.3
B2	121611713	121611838
refGene_gn=MED23;refGene_tx=XM_011282608.3;refGene_gn=MED23;refGene_tx=XM_011282609.3
B2	121614792	121614880
refGene_gn=MED23;refGene_tx=XM_011282608.3;refGene_gn=MED23;refGene_tx=XM_011282609.3
B2	121615042	121615074
refGene_gn=MED23;refGene_tx=XM_011282608.3;refGene_gn=MED23;refGene_tx=XM_011282609.3
B2	121615418	121615466
refGene_gn=MED23;refGene_tx=XM_011282608.3;refGene_gn=MED23;refGene_tx

=XM_011282609.3
B2 121631473 121631551
refGene_gn=ENPP3;refGene_tx=XM_006932012.3
B2 121634219 121634295
refGene_gn=ENPP3;refGene_tx=XM_006932012.3
B2 121640130 121640253
refGene_gn=ENPP3;refGene_tx=XM_006932012.3;refGene_gn=ENPP3;refGene_tx
=XM_019831231.2
B2 121642001 121642127
refGene_gn=ENPP3;refGene_tx=XM_006932012.3;refGene_gn=ENPP3;refGene_tx
=XM_019831231.2
B2 121645107 121645168
refGene_gn=ENPP3;refGene_tx=XM_006932012.3;refGene_gn=ENPP3;refGene_tx
=XM_019831231.2
B2 121646636 121646734
refGene_gn=ENPP3;refGene_tx=XM_006932012.3;refGene_gn=ENPP3;refGene_tx
=XM_019831231.2
B2 121655235 121655315
refGene_gn=ENPP3;refGene_tx=XM_006932012.3;refGene_gn=ENPP3;refGene_tx
=XM_019831231.2
B2 121658185 121658305
refGene_gn=ENPP3;refGene_tx=XM_006932012.3;refGene_gn=ENPP3;refGene_tx
=XM_019831231.2
B2 121658824 121658934
refGene_gn=ENPP3;refGene_tx=XM_006932012.3;refGene_gn=ENPP3;refGene_tx
=XM_019831231.2
B2 121660408 121660474
refGene_gn=ENPP3;refGene_tx=XM_006932012.3;refGene_gn=ENPP3;refGene_tx
=XM_019831231.2
B2 121661177 121661250
refGene_gn=ENPP3;refGene_tx=XM_006932012.3;refGene_gn=ENPP3;refGene_tx
=XM_019831231.2
B2 121668749 121668858
refGene_gn=ENPP3;refGene_tx=XM_006932012.3;refGene_gn=ENPP3;refGene_tx
=XM_019831231.2
B2 121669913 121670042
refGene_gn=ENPP3;refGene_tx=XM_006932012.3;refGene_gn=ENPP3;refGene_tx
=XM_019831231.2
B2 121670791 121670823
refGene_gn=ENPP3;refGene_tx=XM_006932012.3;refGene_gn=ENPP3;refGene_tx
=XM_019831231.2
B2 121673413 121673541
refGene_gn=ENPP3;refGene_tx=XM_006932012.3;refGene_gn=ENPP3;refGene_tx
=XM_019831231.2
B2 121679341 121679408
refGene_gn=ENPP3;refGene_tx=XM_006932012.3;refGene_gn=ENPP3;refGene_tx
=XM_019831231.2
B2 121682048 121682136
refGene_gn=ENPP3;refGene_tx=XM_006932012.3;refGene_gn=ENPP3;refGene_tx
=XM_019831231.2

B2 121683676 121683855
refGene_gn=ENPP3;refGene_tx=XM_006932012.3;refGene_gn=ENPP3;refGene_tx=XM_019831231.2
B2 121685345 121685397
refGene_gn=ENPP3;refGene_tx=XM_006932012.3;refGene_gn=ENPP3;refGene_tx=XM_019831231.2
B2 121686721 121686876
refGene_gn=ENPP3;refGene_tx=XM_006932012.3;refGene_gn=ENPP3;refGene_tx=XM_019831231.2
B2 121693736 121693872
refGene_gn=ENPP3;refGene_tx=XM_006932012.3;refGene_gn=ENPP3;refGene_tx=XM_019831231.2
B2 121696380 121696458
refGene_gn=ENPP3;refGene_tx=XM_006932012.3;refGene_gn=ENPP3;refGene_tx=XM_019831231.2
B2 121697077 121697210
refGene_gn=ENPP3;refGene_tx=XM_006932012.3;refGene_gn=ENPP3;refGene_tx=XM_019831231.2
B2 121698823 121698980
refGene_gn=ENPP3;refGene_tx=XM_006932012.3;refGene_gn=ENPP3;refGene_tx=XM_019831231.2
B2 121705644 121705815
refGene_gn=ENPP3;refGene_tx=XM_006932012.3;refGene_gn=ENPP3;refGene_tx=XM_019831231.2
B2 121769934 121770174
refGene_gn=ENPP1;refGene_tx=XM_023254376.1
B2 121802211 121802284
refGene_gn=ENPP1;refGene_tx=XM_023254376.1
B2 121805237 121805354
refGene_gn=ENPP1;refGene_tx=XM_023254376.1
B2 121806171 121806297
refGene_gn=ENPP1;refGene_tx=XM_023254376.1
B2 121807424 121807485
refGene_gn=ENPP1;refGene_tx=XM_023254376.1
B2 121809958 121810056
refGene_gn=ENPP1;refGene_tx=XM_023254376.1
B2 121813362 121813442
refGene_gn=ENPP1;refGene_tx=XM_023254376.1
B2 121814347 121814467
refGene_gn=ENPP1;refGene_tx=XM_023254376.1
B2 121815523 121815633
refGene_gn=ENPP1;refGene_tx=XM_023254376.1
B2 121816438 121816504
refGene_gn=ENPP1;refGene_tx=XM_023254376.1
B2 121816811 121816884
refGene_gn=ENPP1;refGene_tx=XM_023254376.1
B2 121819917 121820026
refGene_gn=ENPP1;refGene_tx=XM_023254376.1
B2 121820975 121821107
refGene_gn=ENPP1;refGene_tx=XM_023254376.1

B2 121822635 121822667
refGene_gn=ENPP1;refGene_tx=XM_023254376.1
B2 121823618 121823746
refGene_gn=ENPP1;refGene_tx=XM_023254376.1
B2 121824914 121824984
refGene_gn=ENPP1;refGene_tx=XM_023254376.1
B2 121825899 121825987
refGene_gn=ENPP1;refGene_tx=XM_023254376.1
B2 121827417 121827587
refGene_gn=ENPP1;refGene_tx=XM_023254376.1
B2 121829040 121829092
refGene_gn=ENPP1;refGene_tx=XM_023254376.1
B2 121830637 121830792
refGene_gn=ENPP1;refGene_tx=XM_023254376.1
B2 121832127 121832257
refGene_gn=ENPP1;refGene_tx=XM_023254376.1
B2 121833345 121833426
refGene_gn=ENPP1;refGene_tx=XM_023254376.1
B2 121834280 121834413
refGene_gn=ENPP1;refGene_tx=XM_023254376.1
B2 121836523 121836686
refGene_gn=ENPP1;refGene_tx=XM_023254376.1
B2 121839959 121840130
refGene_gn=ENPP1;refGene_tx=XM_023254376.1
B2 121883482 121883779
refGene_gn=CTGF;refGene_tx=XM_023254377.1
B2 121884159 121884371
refGene_gn=CTGF;refGene_tx=XM_023254377.1
B2 121884502 121884754
refGene_gn=CTGF;refGene_tx=XM_023254377.1
B2 121884916 121885139
refGene_gn=CTGF;refGene_tx=XM_023254377.1
B2 121885255 121885321
refGene_gn=CTGF;refGene_tx=XM_023254377.1
B2 122227783 122227954
refGene_gn=MOXD1;refGene_tx=XM_023254379.1
B2 122228452 122228621
refGene_gn=MOXD1;refGene_tx=XM_023254379.1
B2 122239722 122239865
refGene_gn=MOXD1;refGene_tx=XM_023254379.1
B2 122243023 122243083
refGene_gn=MOXD1;refGene_tx=XM_023254379.1
B2 122244253 122244445
refGene_gn=MOXD1;refGene_tx=XM_023254379.1
B2 122245538 122245705
refGene_gn=MOXD1;refGene_tx=XM_023254379.1
B2 122247183 122247286
refGene_gn=MOXD1;refGene_tx=XM_023254379.1
B2 122247816 122247996
refGene_gn=MOXD1;refGene_tx=XM_023254379.1

B2 122256069 122256359
refGene_gn=LOC109499899;refGene_tx=XM_019831234.2
B2 122257692 122257772
refGene_gn=LOC109499899;refGene_tx=XM_019831234.2
B2 122257774 122257851
refGene_gn=LOC109499899;refGene_tx=XM_019831234.2
B2 122292393 122292477
refGene_gn=MOXD1;refGene_tx=XM_023254379.1
B2 122292613 122292781
refGene_gn=MOXD1;refGene_tx=XM_023254379.1
B2 122294597 122294744
refGene_gn=MOXD1;refGene_tx=XM_023254379.1
B2 122317656 122317932
refGene_gn=MOXD1;refGene_tx=XM_023254379.1
B2 122368770 122368863
refGene_gn=STX7;refGene_tx=XM_006932016.4
B2 122369488 122369515
refGene_gn=STX7;refGene_tx=XM_019831236.2
B2 122371255 122371338
refGene_gn=STX7;refGene_tx=XM_006932016.4;refGene_gn=STX7;refGene_tx=X
M_019831236.2
B2 122375777 122375850
refGene_gn=STX7;refGene_tx=XM_006932016.4;refGene_gn=STX7;refGene_tx=X
M_019831236.2
B2 122377010 122377107
refGene_gn=STX7;refGene_tx=XM_006932016.4;refGene_gn=STX7;refGene_tx=X
M_019831236.2
B2 122377922 122377975
refGene_gn=STX7;refGene_tx=XM_006932016.4;refGene_gn=STX7;refGene_tx=X
M_019831236.2
B2 122378753 122378891
refGene_gn=STX7;refGene_tx=XM_006932016.4;refGene_gn=STX7;refGene_tx=X
M_019831236.2
B2 122379585 122379679
refGene_gn=STX7;refGene_tx=XM_006932016.4;refGene_gn=STX7;refGene_tx=X
M_019831236.2
B2 122382093 122382163
refGene_gn=STX7;refGene_tx=XM_006932016.4;refGene_gn=STX7;refGene_tx=X
M_019831236.2
B2 122403193 122403278
refGene_gn=STX7;refGene_tx=XM_006932016.4;refGene_gn=STX7;refGene_tx=X
M_019831236.2
B2 122436579 122437626
refGene_gn=TAAR9;refGene_tx=XM_003986587.1
B2 122444990 122446022
refGene_gn=TAAR8;refGene_tx=XM_003986562.2
B2 122455287 122456325
refGene_gn=LOC111560604;refGene_tx=XM_023254740.1
B2 122466293 122467331
refGene_gn=LOC101095497;refGene_tx=XM_003986563.2

B2 122477880 122478042
refGene_gn=LOC102901115;refGene_tx=XM_019831619.2
B2 122482445 122482685
refGene_gn=LOC102901115;refGene_tx=XM_019831619.2
B2 122482686 122482920
refGene_gn=LOC102901115;refGene_tx=XM_019831619.2
B2 122489601 122490627
refGene_gn=LOC101095743;refGene_tx=XM_003986564.2
B2 122503181 122504192
refGene_gn=TAAR5;refGene_tx=XM_003986565.5
B2 122512200 122513244
refGene_gn=TAAR4;refGene_tx=XM_019831237.2;refGene_gn=TAAR4;refGene_tx=XM_019831238.2;refGene_gn=TAAR4;refGene_tx=XM_023254380.1
B2 122525611 122526643
refGene_gn=LOC101096479;refGene_tx=XM_023253559.1
B2 122538798 122539719
refGene_gn=TAAR2;refGene_tx=XM_011282607.2
B2 122565976 122566981
refGene_gn=TAAR1;refGene_tx=XM_011282606.3
B2 122592989 122593073
refGene_gn=LOC101096984;refGene_tx=XM_023254381.1;refGene_gn=LOC101096984;refGene_tx=XM_023254382.1;refGene_gn=LOC101096984;refGene_tx=XM_023254383.1;refGene_gn=LOC101096984;refGene_tx=XM_023254384.1;refGene_gn=LOC101096984;refGene_tx=XM_023254385.1;refGene_gn=LOC101096984;refGene_tx=XM_023254386.1;refGene_gn=LOC101096984;refGene_tx=XM_023254387.1;refGene_gn=LOC101096984;refGene_tx=XM_023254388.1
B2 122595617 122595796
refGene_gn=LOC101096984;refGene_tx=XM_023254381.1;refGene_gn=LOC101096984;refGene_tx=XM_023254382.1;refGene_gn=LOC101096984;refGene_tx=XM_023254383.1;refGene_gn=LOC101096984;refGene_tx=XM_023254384.1;refGene_gn=LOC101096984;refGene_tx=XM_023254385.1;refGene_gn=LOC101096984;refGene_tx=XM_023254386.1;refGene_gn=LOC101096984;refGene_tx=XM_023254388.1
B2 122597126 122597297
refGene_gn=LOC101096984;refGene_tx=XM_023254381.1;refGene_gn=LOC101096984;refGene_tx=XM_023254382.1;refGene_gn=LOC101096984;refGene_tx=XM_023254383.1;refGene_gn=LOC101096984;refGene_tx=XM_023254384.1;refGene_gn=LOC101096984;refGene_tx=XM_023254385.1;refGene_gn=LOC101096984;refGene_tx=XM_023254386.1;refGene_gn=LOC101096984;refGene_tx=XM_023254387.1;refGene_gn=LOC101096984;refGene_tx=XM_023254388.1
B2 122604989 122605351
refGene_gn=LOC101096984;refGene_tx=XM_023254381.1;refGene_gn=LOC101096984;refGene_tx=XM_023254382.1;refGene_gn=LOC101096984;refGene_tx=XM_023254383.1;refGene_gn=LOC101096984;refGene_tx=XM_023254384.1;refGene_gn=LOC101096984;refGene_tx=XM_023254385.1;refGene_gn=LOC101096984;refGene_tx=XM_023254386.1;refGene_gn=LOC101096984;refGene_tx=XM_023254387.1;refGene_gn=LOC101096984;refGene_tx=XM_023254388.1
B2 122605795 122606087
refGene_gn=LOC101096984;refGene_tx=XM_023254381.1;refGene_gn=LOC101096984;refGene_tx=XM_023254382.1;refGene_gn=LOC101096984;refGene_tx=XM_023254383.1;refGene_gn=LOC101096984;refGene_tx=XM_023254384.1;refGene_gn

=LOC101096984;refGene_tx=XM_023254385.1;refGene_gn=LOC101096984;refGene_tx=XM_023254386.1;refGene_gn=LOC101096984;refGene_tx=XM_023254387.1;refGene_gn=LOC101096984;refGene_tx=XM_023254388.1
B2 122606764 122606957
refGene_gn=LOC101096984;refGene_tx=XM_023254381.1;refGene_gn=LOC101096984;refGene_tx=XM_023254382.1;refGene_gn=LOC101096984;refGene_tx=XM_023254383.1;refGene_gn=LOC101096984;refGene_tx=XM_023254384.1;refGene_gn=LOC101096984;refGene_tx=XM_023254385.1;refGene_gn=LOC101096984;refGene_tx=XM_023254386.1;refGene_gn=LOC101096984;refGene_tx=XM_023254387.1
B2 122621678 122621809
refGene_gn=LOC101096984;refGene_tx=XM_023254381.1;refGene_gn=LOC101096984;refGene_tx=XM_023254382.1;refGene_gn=LOC101096984;refGene_tx=XM_023254383.1;refGene_gn=LOC101096984;refGene_tx=XM_023254384.1;refGene_gn=LOC101096984;refGene_tx=XM_023254385.1;refGene_gn=LOC101096984;refGene_tx=XM_023254386.1;refGene_gn=LOC101096984;refGene_tx=XM_023254387.1
B2 122623138 122623351
refGene_gn=LOC101096984;refGene_tx=XM_023254381.1;refGene_gn=LOC101096984;refGene_tx=XM_023254382.1;refGene_gn=LOC101096984;refGene_tx=XM_023254383.1;refGene_gn=LOC101096984;refGene_tx=XM_023254384.1;refGene_gn=LOC101096984;refGene_tx=XM_023254385.1;refGene_gn=LOC101096984;refGene_tx=XM_023254386.1;refGene_gn=LOC101096984;refGene_tx=XM_023254387.1
B2 122662319 122662448
refGene_gn=LOC101097240;refGene_tx=XM_023254389.1;refGene_gn=LOC101097240;refGene_tx=XM_023254390.1;refGene_gn=LOC101097240;refGene_tx=XM_023254391.1;refGene_gn=LOC101097240;refGene_tx=XM_023254392.1;refGene_gn=LOC101097240;refGene_tx=XM_023254393.1
B2 122663116 122663290
refGene_gn=LOC101097240;refGene_tx=XM_023254389.1;refGene_gn=LOC101097240;refGene_tx=XM_023254390.1;refGene_gn=LOC101097240;refGene_tx=XM_023254391.1;refGene_gn=LOC101097240;refGene_tx=XM_023254392.1;refGene_gn=LOC101097240;refGene_tx=XM_023254393.1
B2 122664542 122664916
refGene_gn=LOC101097240;refGene_tx=XM_023254389.1;refGene_gn=LOC101097240;refGene_tx=XM_023254390.1;refGene_gn=LOC101097240;refGene_tx=XM_023254391.1;refGene_gn=LOC101097240;refGene_tx=XM_023254392.1;refGene_gn=LOC101097240;refGene_tx=XM_023254393.1
B2 122665723 122666015
refGene_gn=LOC101097240;refGene_tx=XM_023254389.1;refGene_gn=LOC101097240;refGene_tx=XM_023254390.1;refGene_gn=LOC101097240;refGene_tx=XM_023254391.1;refGene_gn=LOC101097240;refGene_tx=XM_023254392.1;refGene_gn=LOC101097240;refGene_tx=XM_023254393.1
B2 122667466 122667659
refGene_gn=LOC101097240;refGene_tx=XM_023254389.1;refGene_gn=LOC101097240;refGene_tx=XM_023254390.1;refGene_gn=LOC101097240;refGene_tx=XM_023254391.1;refGene_gn=LOC101097240;refGene_tx=XM_023254392.1;refGene_gn=LOC101097240;refGene_tx=XM_023254393.1
B2 122671768 122671899
refGene_gn=LOC101097240;refGene_tx=XM_023254389.1;refGene_gn=LOC101097240;refGene_tx=XM_023254390.1;refGene_gn=LOC101097240;refGene_tx=XM_023254391.1;refGene_gn=LOC101097240;refGene_tx=XM_023254392.1;refGene_gn

=LOC101097240;refGene_tx=XM_023254393.1
B2 122672013 122672226
refGene_gn=LOC101097240;refGene_tx=XM_023254389.1;refGene_gn=LOC101097240;refGene_tx=XM_023254390.1;refGene_gn=LOC101097240;refGene_tx=XM_023254391.1;refGene_gn=LOC101097240;refGene_tx=XM_023254392.1;refGene_gn=LOC101097240;refGene_tx=XM_023254393.1
B2 122673969 122674004
refGene_gn=LOC101097240;refGene_tx=XM_023254390.1;refGene_gn=LOC101097240;refGene_tx=XM_023254391.1;refGene_gn=LOC101097240;refGene_tx=XM_023254392.1;refGene_gn=LOC101097240;refGene_tx=XM_023254393.1
B2 122700404 122700471
refGene_gn=SLC18B1;refGene_tx=XM_003986571.5;refGene_gn=SLC18B1;refGene_tx=XM_019831249.2;refGene_gn=SLC18B1;refGene_tx=XM_023254394.1;refGene_gn=SLC18B1;refGene_tx=XM_023254395.1
B2 122701153 122701203
refGene_gn=SLC18B1;refGene_tx=XM_003986571.5;refGene_gn=SLC18B1;refGene_tx=XM_019831249.2;refGene_gn=SLC18B1;refGene_tx=XM_023254394.1;refGene_gn=SLC18B1;refGene_tx=XM_023254395.1
B2 122701294 122701388
refGene_gn=SLC18B1;refGene_tx=XM_003986571.5;refGene_gn=SLC18B1;refGene_tx=XM_019831249.2;refGene_gn=SLC18B1;refGene_tx=XM_023254394.1;refGene_gn=SLC18B1;refGene_tx=XM_023254395.1
B2 122702075 122702150
refGene_gn=SLC18B1;refGene_tx=XM_003986571.5;refGene_gn=SLC18B1;refGene_tx=XM_019831249.2;refGene_gn=SLC18B1;refGene_tx=XM_023254394.1;refGene_gn=SLC18B1;refGene_tx=XM_023254395.1
B2 122704821 122704917
refGene_gn=SLC18B1;refGene_tx=XM_003986571.5;refGene_gn=SLC18B1;refGene_tx=XM_019831249.2;refGene_gn=SLC18B1;refGene_tx=XM_023254394.1;refGene_gn=SLC18B1;refGene_tx=XM_023254395.1
B2 122706044 122706136
refGene_gn=SLC18B1;refGene_tx=XM_003986571.5;refGene_gn=SLC18B1;refGene_tx=XM_019831249.2;refGene_gn=SLC18B1;refGene_tx=XM_023254394.1;refGene_gn=SLC18B1;refGene_tx=XM_023254395.1
B2 122708395 122708497
refGene_gn=SLC18B1;refGene_tx=XM_003986571.5;refGene_gn=SLC18B1;refGene_tx=XM_019831249.2;refGene_gn=SLC18B1;refGene_tx=XM_023254394.1;refGene_gn=SLC18B1;refGene_tx=XM_023254395.1
B2 122711506 122711643
refGene_gn=SLC18B1;refGene_tx=XM_003986571.5;refGene_gn=SLC18B1;refGene_tx=XM_019831249.2;refGene_gn=SLC18B1;refGene_tx=XM_023254394.1;refGene_gn=SLC18B1;refGene_tx=XM_023254395.1
B2 122716053 122716210
refGene_gn=SLC18B1;refGene_tx=XM_003986571.5;refGene_gn=SLC18B1;refGene_tx=XM_019831249.2;refGene_gn=SLC18B1;refGene_tx=XM_023254394.1
B2 122719035 122719183
refGene_gn=SLC18B1;refGene_tx=XM_003986571.5;refGene_gn=SLC18B1;refGene_tx=XM_019831249.2;refGene_gn=SLC18B1;refGene_tx=XM_023254395.1
B2 122720472 122720546
refGene_gn=SLC18B1;refGene_tx=XM_003986571.5;refGene_gn=SLC18B1;refGen

e_tx=XM_019831249.2
B2	122721260	122721356
refGene_gn=SLC18B1;refGene_tx=XM_003986571.5;refGene_gn=SLC18B1;refGene_tx=XM_019831249.2;refGene_gn=SLC18B1;refGene_tx=XM_023254394.1;refGene_gn=SLC18B1;refGene_tx=XM_023254395.1
B2	122725133	122725273
refGene_gn=SLC18B1;refGene_tx=XM_003986571.5;refGene_gn=SLC18B1;refGene_tx=XM_019831249.2;refGene_gn=SLC18B1;refGene_tx=XM_023254394.1;refGene_gn=SLC18B1;refGene_tx=XM_023254395.1
B2	122726165	122726181
refGene_gn=SLC18B1;refGene_tx=XM_019831249.2
B2	122726442	122726485
refGene_gn=SLC18B1;refGene_tx=XM_003986571.5;refGene_gn=SLC18B1;refGene_tx=XM_023254394.1;refGene_gn=SLC18B1;refGene_tx=XM_023254395.1
B2	122737557	122737571
refGene_gn=RPS12;refGene_tx=XM_003986572.4
B2	122737755	122737872
refGene_gn=RPS12;refGene_tx=XM_003986572.4
B2	122738924	122739027
refGene_gn=RPS12;refGene_tx=XM_003986572.4
B2	122739452	122739554
refGene_gn=RPS12;refGene_tx=XM_003986572.4
B2	122739967	122740030
refGene_gn=RPS12;refGene_tx=XM_003986572.4
B2	123172105	123172138
refGene_gn=EYA4;refGene_tx=XM_006932029.4;refGene_gn=EYA4;refGene_tx=XM_019831256.2;refGene_gn=EYA4;refGene_tx=XM_023254396.1;refGene_gn=EYA4;refGene_tx=XM_023254397.1;refGene_gn=EYA4;refGene_tx=XM_023254398.1;refGene_gn=EYA4;refGene_tx=XM_023254399.1;refGene_gn=EYA4;refGene_tx=XM_023254400.1;refGene_gn=EYA4;refGene_tx=XM_023254401.1;refGene_gn=EYA4;refGene_tx=XM_023254402.1;refGene_gn=EYA4;refGene_tx=XM_023254403.1;refGene_gn=EYA4;refGene_tx=XM_023254404.1;refGene_gn=EYA4;refGene_tx=XM_023254405.1;refGene_gn=EYA4;refGene_tx=XM_023254406.1;refGene_gn=EYA4;refGene_tx=XM_023254407.1
B2	123290032	123290082
refGene_gn=EYA4;refGene_tx=XM_006932029.4;refGene_gn=EYA4;refGene_tx=XM_019831256.2;refGene_gn=EYA4;refGene_tx=XM_023254396.1;refGene_gn=EYA4;refGene_tx=XM_023254397.1;refGene_gn=EYA4;refGene_tx=XM_023254398.1;refGene_gn=EYA4;refGene_tx=XM_023254399.1;refGene_gn=EYA4;refGene_tx=XM_023254400.1;refGene_gn=EYA4;refGene_tx=XM_023254401.1;refGene_gn=EYA4;refGene_tx=XM_023254402.1;refGene_gn=EYA4;refGene_tx=XM_023254403.1;refGene_gn=EYA4;refGene_tx=XM_023254404.1;refGene_gn=EYA4;refGene_tx=XM_023254405.1;refGene_gn=EYA4;refGene_tx=XM_023254406.1;refGene_gn=EYA4;refGene_tx=XM_023254407.1
B2	123352751	123352876
refGene_gn=EYA4;refGene_tx=XM_006932029.4;refGene_gn=EYA4;refGene_tx=XM_019831256.2;refGene_gn=EYA4;refGene_tx=XM_023254396.1;refGene_gn=EYA4;refGene_tx=XM_023254397.1;refGene_gn=EYA4;refGene_tx=XM_023254398.1;refGene_gn=EYA4;refGene_tx=XM_023254399.1;refGene_gn=EYA4;refGene_tx=XM_023254400.1;refGene_gn=EYA4;refGene_tx=XM_023254401.1;refGene_gn=EYA

4;refGene_tx=XM_023254402.1;refGene_gn=EYA4;refGene_tx=XM_023254403.1;
refGene_gn=EYA4;refGene_tx=XM_023254404.1;refGene_gn=EYA4;refGene_tx=X
M_023254405.1;refGene_gn=EYA4;refGene_tx=XM_023254406.1;refGene_gn=EYA
4;refGene_tx=XM_023254407.1
B2 123354260 123354329
refGene_gn=EYA4;refGene_tx=XM_006932029.4;refGene_gn=EYA4;refGene_tx=X
M_019831256.2;refGene_gn=EYA4;refGene_tx=XM_023254396.1;refGene_gn=EYA
4;refGene_tx=XM_023254397.1;refGene_gn=EYA4;refGene_tx=XM_023254398.1;
refGene_gn=EYA4;refGene_tx=XM_023254399.1;refGene_gn=EYA4;refGene_tx=X
M_023254400.1;refGene_gn=EYA4;refGene_tx=XM_023254401.1
B2 123363516 123363609
refGene_gn=EYA4;refGene_tx=XM_006932029.4;refGene_gn=EYA4;refGene_tx=X
M_019831256.2;refGene_gn=EYA4;refGene_tx=XM_023254396.1;refGene_gn=EYA
4;refGene_tx=XM_023254397.1;refGene_gn=EYA4;refGene_tx=XM_023254398.1;
refGene_gn=EYA4;refGene_tx=XM_023254399.1;refGene_gn=EYA4;refGene_tx=X
M_023254400.1;refGene_gn=EYA4;refGene_tx=XM_023254401.1;refGene_gn=EYA
4;refGene_tx=XM_023254402.1;refGene_gn=EYA4;refGene_tx=XM_023254403.1;
refGene_gn=EYA4;refGene_tx=XM_023254404.1;refGene_gn=EYA4;refGene_tx=X
M_023254405.1
B2 123368093 123368160
refGene_gn=EYA4;refGene_tx=XM_006932029.4;refGene_gn=EYA4;refGene_tx=X
M_019831256.2;refGene_gn=EYA4;refGene_tx=XM_023254396.1;refGene_gn=EYA
4;refGene_tx=XM_023254397.1;refGene_gn=EYA4;refGene_tx=XM_023254398.1;
refGene_gn=EYA4;refGene_tx=XM_023254399.1;refGene_gn=EYA4;refGene_tx=X
M_023254400.1;refGene_gn=EYA4;refGene_tx=XM_023254401.1;refGene_gn=EYA
4;refGene_tx=XM_023254402.1;refGene_gn=EYA4;refGene_tx=XM_023254403.1;
refGene_gn=EYA4;refGene_tx=XM_023254404.1;refGene_gn=EYA4;refGene_tx=X
M_023254405.1;refGene_gn=EYA4;refGene_tx=XM_023254406.1;refGene_gn=EYA
4;refGene_tx=XM_023254407.1
B2 123369421 123369564
refGene_gn=EYA4;refGene_tx=XM_006932029.4;refGene_gn=EYA4;refGene_tx=X
M_019831256.2;refGene_gn=EYA4;refGene_tx=XM_019831258.2;refGene_gn=EYA
4;refGene_tx=XM_019831259.2;refGene_gn=EYA4;refGene_tx=XM_023254396.1;
refGene_gn=EYA4;refGene_tx=XM_023254397.1;refGene_gn=EYA4;refGene_tx=X
M_023254398.1;refGene_gn=EYA4;refGene_tx=XM_023254399.1;refGene_gn=EYA
4;refGene_tx=XM_023254400.1;refGene_gn=EYA4;refGene_tx=XM_023254401.1;
refGene_gn=EYA4;refGene_tx=XM_023254402.1;refGene_gn=EYA4;refGene_tx=X
M_023254403.1;refGene_gn=EYA4;refGene_tx=XM_023254404.1;refGene_gn=EYA
4;refGene_tx=XM_023254405.1;refGene_gn=EYA4;refGene_tx=XM_023254406.1;
refGene_gn=EYA4;refGene_tx=XM_023254407.1
B2 123369706 123369850
refGene_gn=EYA4;refGene_tx=XM_006932029.4;refGene_gn=EYA4;refGene_tx=X
M_019831256.2;refGene_gn=EYA4;refGene_tx=XM_019831258.2;refGene_gn=EYA
4;refGene_tx=XM_019831259.2;refGene_gn=EYA4;refGene_tx=XM_023254396.1;
refGene_gn=EYA4;refGene_tx=XM_023254397.1;refGene_gn=EYA4;refGene_tx=X
M_023254398.1;refGene_gn=EYA4;refGene_tx=XM_023254399.1;refGene_gn=EYA
4;refGene_tx=XM_023254400.1;refGene_gn=EYA4;refGene_tx=XM_023254401.1;
refGene_gn=EYA4;refGene_tx=XM_023254402.1;refGene_gn=EYA4;refGene_tx=X
M_023254403.1;refGene_gn=EYA4;refGene_tx=XM_023254404.1;refGene_gn=EYA
4;refGene_tx=XM_023254405.1;refGene_gn=EYA4;refGene_tx=XM_023254406.1;

refGene_gn=EYA4;refGene_tx=XM_023254407.1
B2 123372648 123372728
refGene_gn=EYA4;refGene_tx=XM_006932029.4;refGene_gn=EYA4;refGene_tx=XM_019831256.2;refGene_gn=EYA4;refGene_tx=XM_019831258.2;refGene_gn=EYA4;refGene_tx=XM_019831259.2;refGene_gn=EYA4;refGene_tx=XM_023254396.1;refGene_gn=EYA4;refGene_tx=XM_023254397.1;refGene_gn=EYA4;refGene_tx=XM_023254398.1;refGene_gn=EYA4;refGene_tx=XM_023254399.1;refGene_gn=EYA4;refGene_tx=XM_023254400.1;refGene_gn=EYA4;refGene_tx=XM_023254401.1;refGene_gn=EYA4;refGene_tx=XM_023254402.1;refGene_gn=EYA4;refGene_tx=XM_023254403.1;refGene_gn=EYA4;refGene_tx=XM_023254404.1;refGene_gn=EYA4;refGene_tx=XM_023254405.1;refGene_gn=EYA4;refGene_tx=XM_023254406.1;refGene_gn=EYA4;refGene_tx=XM_023254407.1
B2 123376627 123376811
refGene_gn=EYA4;refGene_tx=XM_006932029.4;refGene_gn=EYA4;refGene_tx=XM_019831256.2;refGene_gn=EYA4;refGene_tx=XM_019831258.2;refGene_gn=EYA4;refGene_tx=XM_019831259.2;refGene_gn=EYA4;refGene_tx=XM_023254396.1;refGene_gn=EYA4;refGene_tx=XM_023254397.1;refGene_gn=EYA4;refGene_tx=XM_023254398.1;refGene_gn=EYA4;refGene_tx=XM_023254399.1;refGene_gn=EYA4;refGene_tx=XM_023254400.1;refGene_gn=EYA4;refGene_tx=XM_023254401.1;refGene_gn=EYA4;refGene_tx=XM_023254402.1;refGene_gn=EYA4;refGene_tx=XM_023254403.1;refGene_gn=EYA4;refGene_tx=XM_023254404.1;refGene_gn=EYA4;refGene_tx=XM_023254405.1;refGene_gn=EYA4;refGene_tx=XM_023254406.1;refGene_gn=EYA4;refGene_tx=XM_023254407.1
B2 123384946 123385083
refGene_gn=EYA4;refGene_tx=XM_006932029.4;refGene_gn=EYA4;refGene_tx=XM_019831256.2;refGene_gn=EYA4;refGene_tx=XM_019831258.2;refGene_gn=EYA4;refGene_tx=XM_019831259.2;refGene_gn=EYA4;refGene_tx=XM_023254396.1;refGene_gn=EYA4;refGene_tx=XM_023254397.1;refGene_gn=EYA4;refGene_tx=XM_023254398.1;refGene_gn=EYA4;refGene_tx=XM_023254399.1;refGene_gn=EYA4;refGene_tx=XM_023254400.1;refGene_gn=EYA4;refGene_tx=XM_023254401.1;refGene_gn=EYA4;refGene_tx=XM_023254402.1;refGene_gn=EYA4;refGene_tx=XM_023254403.1;refGene_gn=EYA4;refGene_tx=XM_023254404.1;refGene_gn=EYA4;refGene_tx=XM_023254405.1;refGene_gn=EYA4;refGene_tx=XM_023254406.1;refGene_gn=EYA4;refGene_tx=XM_023254407.1
B2 123386455 123386539
refGene_gn=EYA4;refGene_tx=XM_006932029.4;refGene_gn=EYA4;refGene_tx=XM_019831256.2;refGene_gn=EYA4;refGene_tx=XM_019831258.2;refGene_gn=EYA4;refGene_tx=XM_019831259.2;refGene_gn=EYA4;refGene_tx=XM_023254396.1;refGene_gn=EYA4;refGene_tx=XM_023254397.1;refGene_gn=EYA4;refGene_tx=XM_023254398.1;refGene_gn=EYA4;refGene_tx=XM_023254399.1;refGene_gn=EYA4;refGene_tx=XM_023254400.1;refGene_gn=EYA4;refGene_tx=XM_023254401.1;refGene_gn=EYA4;refGene_tx=XM_023254402.1;refGene_gn=EYA4;refGene_tx=XM_023254403.1;refGene_gn=EYA4;refGene_tx=XM_023254404.1;refGene_gn=EYA4;refGene_tx=XM_023254405.1;refGene_gn=EYA4;refGene_tx=XM_023254406.1;refGene_gn=EYA4;refGene_tx=XM_023254407.1
B2 123402626 123402716
refGene_gn=EYA4;refGene_tx=XM_006932029.4;refGene_gn=EYA4;refGene_tx=XM_019831256.2;refGene_gn=EYA4;refGene_tx=XM_019831258.2;refGene_gn=EYA4;refGene_tx=XM_019831259.2;refGene_gn=EYA4;refGene_tx=XM_023254396.1;refGene_gn=EYA4;refGene_tx=XM_023254397.1;refGene_gn=EYA4;refGene_tx=X

M_023254398.1;refGene_gn=EYA4;refGene_tx=XM_023254399.1;refGene_gn=EYA4;refGene_tx=XM_023254400.1;refGene_gn=EYA4;refGene_tx=XM_023254401.1;refGene_gn=EYA4;refGene_tx=XM_023254402.1;refGene_gn=EYA4;refGene_tx=XM_023254403.1;refGene_gn=EYA4;refGene_tx=XM_023254404.1;refGene_gn=EYA4;refGene_tx=XM_023254405.1;refGene_gn=EYA4;refGene_tx=XM_023254406.1;refGene_gn=EYA4;refGene_tx=XM_023254407.1
B2 123408835 123408894
refGene_gn=EYA4;refGene_tx=XM_006932029.4;refGene_gn=EYA4;refGene_tx=XM_019831256.2;refGene_gn=EYA4;refGene_tx=XM_019831258.2;refGene_gn=EYA4;refGene_tx=XM_019831259.2;refGene_gn=EYA4;refGene_tx=XM_023254396.1;refGene_gn=EYA4;refGene_tx=XM_023254397.1;refGene_gn=EYA4;refGene_tx=XM_023254398.1;refGene_gn=EYA4;refGene_tx=XM_023254399.1;refGene_gn=EYA4;refGene_tx=XM_023254400.1;refGene_gn=EYA4;refGene_tx=XM_023254401.1;refGene_gn=EYA4;refGene_tx=XM_023254402.1;refGene_gn=EYA4;refGene_tx=XM_023254403.1;refGene_gn=EYA4;refGene_tx=XM_023254404.1;refGene_gn=EYA4;refGene_tx=XM_023254405.1;refGene_gn=EYA4;refGene_tx=XM_023254406.1;refGene_gn=EYA4;refGene_tx=XM_023254407.1
B2 123409001 123409162
refGene_gn=EYA4;refGene_tx=XM_006932029.4;refGene_gn=EYA4;refGene_tx=XM_019831256.2;refGene_gn=EYA4;refGene_tx=XM_019831258.2;refGene_gn=EYA4;refGene_tx=XM_019831259.2;refGene_gn=EYA4;refGene_tx=XM_023254396.1;refGene_gn=EYA4;refGene_tx=XM_023254397.1;refGene_gn=EYA4;refGene_tx=XM_023254398.1;refGene_gn=EYA4;refGene_tx=XM_023254399.1;refGene_gn=EYA4;refGene_tx=XM_023254400.1;refGene_gn=EYA4;refGene_tx=XM_023254401.1;refGene_gn=EYA4;refGene_tx=XM_023254402.1;refGene_gn=EYA4;refGene_tx=XM_023254403.1;refGene_gn=EYA4;refGene_tx=XM_023254404.1;refGene_gn=EYA4;refGene_tx=XM_023254405.1;refGene_gn=EYA4;refGene_tx=XM_023254406.1;refGene_gn=EYA4;refGene_tx=XM_023254407.1
B2 123412176 123412291
refGene_gn=EYA4;refGene_tx=XM_006932029.4;refGene_gn=EYA4;refGene_tx=XM_019831256.2;refGene_gn=EYA4;refGene_tx=XM_019831258.2;refGene_gn=EYA4;refGene_tx=XM_019831259.2;refGene_gn=EYA4;refGene_tx=XM_023254396.1;refGene_gn=EYA4;refGene_tx=XM_023254397.1;refGene_gn=EYA4;refGene_tx=XM_023254398.1;refGene_gn=EYA4;refGene_tx=XM_023254399.1;refGene_gn=EYA4;refGene_tx=XM_023254400.1;refGene_gn=EYA4;refGene_tx=XM_023254401.1;refGene_gn=EYA4;refGene_tx=XM_023254402.1;refGene_gn=EYA4;refGene_tx=XM_023254403.1;refGene_gn=EYA4;refGene_tx=XM_023254404.1;refGene_gn=EYA4;refGene_tx=XM_023254405.1;refGene_gn=EYA4;refGene_tx=XM_023254406.1;refGene_gn=EYA4;refGene_tx=XM_023254407.1
B2 123416362 123416484
refGene_gn=EYA4;refGene_tx=XM_006932029.4;refGene_gn=EYA4;refGene_tx=XM_019831256.2;refGene_gn=EYA4;refGene_tx=XM_019831258.2;refGene_gn=EYA4;refGene_tx=XM_019831259.2;refGene_gn=EYA4;refGene_tx=XM_023254396.1;refGene_gn=EYA4;refGene_tx=XM_023254397.1;refGene_gn=EYA4;refGene_tx=XM_023254398.1;refGene_gn=EYA4;refGene_tx=XM_023254399.1;refGene_gn=EYA4;refGene_tx=XM_023254400.1;refGene_gn=EYA4;refGene_tx=XM_023254401.1;refGene_gn=EYA4;refGene_tx=XM_023254402.1;refGene_gn=EYA4;refGene_tx=XM_023254403.1;refGene_gn=EYA4;refGene_tx=XM_023254404.1;refGene_gn=EYA4;refGene_tx=XM_023254405.1;refGene_gn=EYA4;refGene_tx=XM_023254406.1;refGene_gn=EYA4;refGene_tx=XM_023254407.1

B2	123418389	123418490
refGene_gn=EYA4;refGene_tx=XM_023254397.1;refGene_gn=EYA4;refGene_tx=XM_023254400.1;refGene_gn=EYA4;refGene_tx=XM_023254401.1;refGene_gn=EYA4;refGene_tx=XM_023254405.1;refGene_gn=EYA4;refGene_tx=XM_023254407.1
B2	123418528	123418629
refGene_gn=EYA4;refGene_tx=XM_006932029.4;refGene_gn=EYA4;refGene_tx=XM_019831256.2;refGene_gn=EYA4;refGene_tx=XM_019831258.2;refGene_gn=EYA4;refGene_tx=XM_019831259.2;refGene_gn=EYA4;refGene_tx=XM_023254396.1;refGene_gn=EYA4;refGene_tx=XM_023254398.1;refGene_gn=EYA4;refGene_tx=XM_023254399.1;refGene_gn=EYA4;refGene_tx=XM_023254402.1;refGene_gn=EYA4;refGene_tx=XM_023254403.1;refGene_gn=EYA4;refGene_tx=XM_023254404.1;refGene_gn=EYA4;refGene_tx=XM_023254406.1
B2	123422035	123422116
refGene_gn=EYA4;refGene_tx=XM_006932029.4;refGene_gn=EYA4;refGene_tx=XM_019831256.2;refGene_gn=EYA4;refGene_tx=XM_019831258.2;refGene_gn=EYA4;refGene_tx=XM_019831259.2;refGene_gn=EYA4;refGene_tx=XM_023254396.1;refGene_gn=EYA4;refGene_tx=XM_023254397.1;refGene_gn=EYA4;refGene_tx=XM_023254398.1;refGene_gn=EYA4;refGene_tx=XM_023254399.1;refGene_gn=EYA4;refGene_tx=XM_023254400.1;refGene_gn=EYA4;refGene_tx=XM_023254402.1;refGene_gn=EYA4;refGene_tx=XM_023254403.1;refGene_gn=EYA4;refGene_tx=XM_023254404.1;refGene_gn=EYA4;refGene_tx=XM_023254405.1;refGene_gn=EYA4;refGene_tx=XM_023254406.1;refGene_gn=EYA4;refGene_tx=XM_023254407.1
B2	123422228	123422264
refGene_gn=EYA4;refGene_tx=XM_023254401.1
B2	123722203	123722653
refGene_gn=TCF21;refGene_tx=XM_006932032.4
B2	123724427	123724517
refGene_gn=TCF21;refGene_tx=XM_006932032.4
B2	123801471	123801606
refGene_gn=TBPL1;refGene_tx=XM_003986575.5;refGene_gn=TBPL1;refGene_tx=XM_023254408.1;refGene_gn=TBPL1;refGene_tx=XM_023254409.1;refGene_gn=TBPL1;refGene_tx=XM_023254410.1;refGene_gn=TBPL1;refGene_tx=XM_023254411.1;refGene_gn=TBPL1;refGene_tx=XM_023254412.1;refGene_gn=TBPL1;refGene_tx=XM_023254413.1;refGene_gn=TBPL1;refGene_tx=XM_023254414.1;refGene_gn=TBPL1;refGene_tx=XM_023254415.1;refGene_gn=TBPL1;refGene_tx=XM_023254416.1;refGene_gn=TBPL1;refGene_tx=XM_023254417.1
B2	123803644	123803727
refGene_gn=TBPL1;refGene_tx=XM_003986575.5;refGene_gn=TBPL1;refGene_tx=XM_023254408.1;refGene_gn=TBPL1;refGene_tx=XM_023254409.1;refGene_gn=TBPL1;refGene_tx=XM_023254410.1;refGene_gn=TBPL1;refGene_tx=XM_023254411.1;refGene_gn=TBPL1;refGene_tx=XM_023254412.1;refGene_gn=TBPL1;refGene_tx=XM_023254413.1;refGene_gn=TBPL1;refGene_tx=XM_023254414.1;refGene_gn=TBPL1;refGene_tx=XM_023254415.1;refGene_gn=TBPL1;refGene_tx=XM_023254416.1;refGene_gn=TBPL1;refGene_tx=XM_023254417.1
B2	123803877	123803941
refGene_gn=TBPL1;refGene_tx=XM_003986575.5;refGene_gn=TBPL1;refGene_tx=XM_023254408.1;refGene_gn=TBPL1;refGene_tx=XM_023254409.1;refGene_gn=TBPL1;refGene_tx=XM_023254410.1;refGene_gn=TBPL1;refGene_tx=XM_023254411.1;refGene_gn=TBPL1;refGene_tx=XM_023254412.1;refGene_gn=TBPL1;refGene_tx=XM_023254413.1;refGene_gn=TBPL1;refGene_tx=XM_023254414.1;refGen

e_gn=TBPL1;refGene_tx=XM_023254415.1;refGene_gn=TBPL1;refGene_tx=XM_023254416.1;refGene_gn=TBPL1;refGene_tx=XM_023254417.1
B2 123806381 123806485
refGene_gn=TBPL1;refGene_tx=XM_003986575.5;refGene_gn=TBPL1;refGene_tx=XM_023254408.1;refGene_gn=TBPL1;refGene_tx=XM_023254409.1;refGene_gn=TBPL1;refGene_tx=XM_023254410.1;refGene_gn=TBPL1;refGene_tx=XM_023254411.1;refGene_gn=TBPL1;refGene_tx=XM_023254412.1;refGene_gn=TBPL1;refGene_tx=XM_023254413.1;refGene_gn=TBPL1;refGene_tx=XM_023254414.1;refGene_gn=TBPL1;refGene_tx=XM_023254415.1;refGene_gn=TBPL1;refGene_tx=XM_023254416.1;refGene_gn=TBPL1;refGene_tx=XM_023254417.1
B2 123806584 123806679
refGene_gn=TBPL1;refGene_tx=XM_003986575.5;refGene_gn=TBPL1;refGene_tx=XM_023254408.1;refGene_gn=TBPL1;refGene_tx=XM_023254409.1;refGene_gn=TBPL1;refGene_tx=XM_023254410.1;refGene_gn=TBPL1;refGene_tx=XM_023254411.1;refGene_gn=TBPL1;refGene_tx=XM_023254412.1;refGene_gn=TBPL1;refGene_tx=XM_023254413.1;refGene_gn=TBPL1;refGene_tx=XM_023254414.1;refGene_gn=TBPL1;refGene_tx=XM_023254415.1;refGene_gn=TBPL1;refGene_tx=XM_023254416.1;refGene_gn=TBPL1;refGene_tx=XM_023254417.1
B2 123807318 123807386
refGene_gn=TBPL1;refGene_tx=XM_023254416.1
B2 123808217 123808270
refGene_gn=TBPL1;refGene_tx=XM_023254417.1
B2 123808648 123808728
refGene_gn=TBPL1;refGene_tx=XM_003986575.5;refGene_gn=TBPL1;refGene_tx=XM_023254408.1;refGene_gn=TBPL1;refGene_tx=XM_023254409.1;refGene_gn=TBPL1;refGene_tx=XM_023254410.1;refGene_gn=TBPL1;refGene_tx=XM_023254411.1;refGene_gn=TBPL1;refGene_tx=XM_023254412.1;refGene_gn=TBPL1;refGene_tx=XM_023254413.1;refGene_gn=TBPL1;refGene_tx=XM_023254414.1;refGene_gn=TBPL1;refGene_tx=XM_023254415.1
B2 123813056 123813162
refGene_gn=SLC2A12;refGene_tx=XM_003986576.5;refGene_gn=SLC2A12;refGene_tx=XM_019831260.2
B2 123825153 123825286
refGene_gn=SLC2A12;refGene_tx=XM_003986576.5;refGene_gn=SLC2A12;refGene_tx=XM_019831260.2
B2 123830023 123830146
refGene_gn=SLC2A12;refGene_tx=XM_003986576.5
B2 123846775 123846819
refGene_gn=SLC2A12;refGene_tx=XM_019831261.2
B2 123849716 123851048
refGene_gn=SLC2A12;refGene_tx=XM_003986576.5;refGene_gn=SLC2A12;refGene_tx=XM_019831260.2;refGene_gn=SLC2A12;refGene_tx=XM_019831261.2
B2 123870358 123870473
refGene_gn=SLC2A12;refGene_tx=XM_003986576.5;refGene_gn=SLC2A12;refGene_tx=XM_019831260.2;refGene_gn=SLC2A12;refGene_tx=XM_019831261.2
B2 123949569 123949737
refGene_gn=SGK1;refGene_tx=XM_003986577.5;refGene_gn=SGK1;refGene_tx=XM_003986579.5;refGene_gn=SGK1;refGene_tx=XM_019831272.2;refGene_gn=SGK1;refGene_tx=XM_019831273.2;refGene_gn=SGK1;refGene_tx=XM_019831274.2
B2 123950156 123950246

refGene_gn=SGK1;refGene_tx=XM_003986577.5;refGene_gn=SGK1;refGene_tx=XM_003986579.5;refGene_gn=SGK1;refGene_tx=XM_019831272.2;refGene_gn=SGK1;refGene_tx=XM_019831273.2;refGene_gn=SGK1;refGene_tx=XM_019831274.2
B2 123950351 123950507
refGene_gn=SGK1;refGene_tx=XM_003986577.5;refGene_gn=SGK1;refGene_tx=XM_003986579.5;refGene_gn=SGK1;refGene_tx=XM_019831272.2;refGene_gn=SGK1;refGene_tx=XM_019831273.2;refGene_gn=SGK1;refGene_tx=XM_019831274.2
B2 123950964 123951060
refGene_gn=SGK1;refGene_tx=XM_003986577.5;refGene_gn=SGK1;refGene_tx=XM_003986579.5;refGene_gn=SGK1;refGene_tx=XM_019831272.2;refGene_gn=SGK1;refGene_tx=XM_019831273.2;refGene_gn=SGK1;refGene_tx=XM_019831274.2
B2 123951516 123951640
refGene_gn=SGK1;refGene_tx=XM_003986577.5;refGene_gn=SGK1;refGene_tx=XM_003986579.5;refGene_gn=SGK1;refGene_tx=XM_019831272.2;refGene_gn=SGK1;refGene_tx=XM_019831273.2;refGene_gn=SGK1;refGene_tx=XM_019831274.2
B2 123951988 123952101
refGene_gn=SGK1;refGene_tx=XM_003986577.5;refGene_gn=SGK1;refGene_tx=XM_003986579.5;refGene_gn=SGK1;refGene_tx=XM_019831272.2;refGene_gn=SGK1;refGene_tx=XM_019831273.2;refGene_gn=SGK1;refGene_tx=XM_019831274.2
B2 123952366 123952498
refGene_gn=SGK1;refGene_tx=XM_003986577.5;refGene_gn=SGK1;refGene_tx=XM_003986579.5;refGene_gn=SGK1;refGene_tx=XM_019831272.2;refGene_gn=SGK1;refGene_tx=XM_019831273.2;refGene_gn=SGK1;refGene_tx=XM_019831274.2
B2 123952612 123952696
refGene_gn=SGK1;refGene_tx=XM_003986577.5;refGene_gn=SGK1;refGene_tx=XM_003986579.5;refGene_gn=SGK1;refGene_tx=XM_019831272.2;refGene_gn=SGK1;refGene_tx=XM_019831273.2;refGene_gn=SGK1;refGene_tx=XM_019831274.2
B2 123952801 123952906
refGene_gn=SGK1;refGene_tx=XM_003986577.5;refGene_gn=SGK1;refGene_tx=XM_003986579.5;refGene_gn=SGK1;refGene_tx=XM_019831272.2;refGene_gn=SGK1;refGene_tx=XM_019831273.2;refGene_gn=SGK1;refGene_tx=XM_019831274.2
B2 123953265 123953341
refGene_gn=SGK1;refGene_tx=XM_003986577.5;refGene_gn=SGK1;refGene_tx=XM_003986579.5;refGene_gn=SGK1;refGene_tx=XM_019831272.2;refGene_gn=SGK1;refGene_tx=XM_019831273.2;refGene_gn=SGK1;refGene_tx=XM_019831274.2
B2 123953795 123953871
refGene_gn=SGK1;refGene_tx=XM_003986577.5;refGene_gn=SGK1;refGene_tx=XM_003986579.5;refGene_gn=SGK1;refGene_tx=XM_019831272.2;refGene_gn=SGK1;refGene_tx=XM_019831273.2;refGene_gn=SGK1;refGene_tx=XM_019831274.2
B2 123954009 123954085
refGene_gn=SGK1;refGene_tx=XM_003986577.5
B2 123954915 123955033
refGene_gn=SGK1;refGene_tx=XM_003986579.5
B2 123956729 123956763
refGene_gn=SGK1;refGene_tx=XM_019831274.2
B2 123982387 123982463
refGene_gn=SGK1;refGene_tx=XM_019831272.2;refGene_gn=SGK1;refGene_tx=XM_019831273.2
B2 124020984 124021200
refGene_gn=SGK1;refGene_tx=XM_019831272.2;refGene_gn=SGK1;refGene_tx=X

M_019831273.2
B2	124059013	124059061
refGene_gn=SGK1;refGene_tx=XM_019831273.2
B2	124063237	124063306
refGene_gn=SGK1;refGene_tx=XM_019831272.2
B2	124574806	124575259
refGene_gn=ALDH8A1;refGene_tx=XM_003986580.4
B2	124580943	124581105
refGene_gn=ALDH8A1;refGene_tx=XM_003986580.4
B2	124584993	124585250
refGene_gn=ALDH8A1;refGene_tx=XM_003986580.4
B2	124590385	124590535
refGene_gn=ALDH8A1;refGene_tx=XM_003986580.4
B2	124592422	124592578
refGene_gn=ALDH8A1;refGene_tx=XM_003986580.4
B2	124593845	124593993
refGene_gn=ALDH8A1;refGene_tx=XM_003986580.4
B2	124597812	124597950
refGene_gn=ALDH8A1;refGene_tx=XM_003986580.4
B2	124607647	124607659
refGene_gn=HBS1L;refGene_tx=XM_006932040.3;refGene_gn=HBS1L;refGene_tx=XM_006932041.3
B2	124609242	124609387
refGene_gn=HBS1L;refGene_tx=XM_006932040.3;refGene_gn=HBS1L;refGene_tx=XM_006932041.3
B2	124612521	124612622
refGene_gn=HBS1L;refGene_tx=XM_006932040.3;refGene_gn=HBS1L;refGene_tx=XM_006932041.3
B2	124620544	124620653
refGene_gn=HBS1L;refGene_tx=XM_006932040.3;refGene_gn=HBS1L;refGene_tx=XM_006932041.3
B2	124621086	124621177
refGene_gn=HBS1L;refGene_tx=XM_006932040.3;refGene_gn=HBS1L;refGene_tx=XM_006932041.3
B2	124624951	124625056
refGene_gn=HBS1L;refGene_tx=XM_006932040.3;refGene_gn=HBS1L;refGene_tx=XM_006932041.3
B2	124627807	124627876
refGene_gn=HBS1L;refGene_tx=XM_006932040.3;refGene_gn=HBS1L;refGene_tx=XM_006932041.3
B2	124628269	124628387
refGene_gn=HBS1L;refGene_tx=XM_006932040.3;refGene_gn=HBS1L;refGene_tx=XM_006932041.3
B2	124629139	124629214
refGene_gn=HBS1L;refGene_tx=XM_006932040.3;refGene_gn=HBS1L;refGene_tx=XM_006932041.3
B2	124631739	124631886
refGene_gn=HBS1L;refGene_tx=XM_006932040.3;refGene_gn=HBS1L;refGene_tx=XM_006932041.3
B2	124632893	124633011

refGene_gn=HBS1L;refGene_tx=XM_006932040.3;refGene_gn=HBS1L;refGene_tx=XM_006932041.3
B2 124635815 124635981
refGene_gn=HBS1L;refGene_tx=XM_006932040.3;refGene_gn=HBS1L;refGene_tx=XM_006932041.3
B2 124636447 124636707
refGene_gn=HBS1L;refGene_tx=XM_006932040.3;refGene_gn=HBS1L;refGene_tx=XM_006932041.3
B2 124641043 124641152
refGene_gn=HBS1L;refGene_tx=XM_006932040.3;refGene_gn=HBS1L;refGene_tx=XM_006932041.3
B2 124655378 124655406
refGene_gn=HBS1L;refGene_tx=XM_006932041.3
B2 124675601 124677073
refGene_gn=HBS1L;refGene_tx=XM_011282599.3
B2 124678299 124678494
refGene_gn=HBS1L;refGene_tx=XM_006932040.3;refGene_gn=HBS1L;refGene_tx=XM_011282599.3
B2 124678992 124679121
refGene_gn=HBS1L;refGene_tx=XM_006932040.3;refGene_gn=HBS1L;refGene_tx=XM_011282599.3
B2 124687921 124687987
refGene_gn=HBS1L;refGene_tx=XM_006932040.3;refGene_gn=HBS1L;refGene_tx=XM_011282599.3
B2 124692403 124692446
refGene_gn=HBS1L;refGene_tx=XM_006932040.3;refGene_gn=HBS1L;refGene_tx=XM_011282599.3
B2 124810342 124810365
refGene_gn=MYB;refGene_tx=XM_023254420.1;refGene_gn=MYB;refGene_tx=XM_023254422.1
B2 124815117 124815235
refGene_gn=MYB;refGene_tx=XM_023254420.1;refGene_gn=MYB;refGene_tx=XM_023254421.1;refGene_gn=MYB;refGene_tx=XM_023254422.1
B2 124817004 124817076
refGene_gn=MYB;refGene_tx=XM_023254420.1;refGene_gn=MYB;refGene_tx=XM_023254421.1;refGene_gn=MYB;refGene_tx=XM_023254422.1
B2 124818512 124818605
refGene_gn=MYB;refGene_tx=XM_023254420.1;refGene_gn=MYB;refGene_tx=XM_023254421.1;refGene_gn=MYB;refGene_tx=XM_023254422.1
B2 124818856 124819077
refGene_gn=MYB;refGene_tx=XM_023254420.1;refGene_gn=MYB;refGene_tx=XM_023254421.1;refGene_gn=MYB;refGene_tx=XM_023254422.1
B2 124820792 124821027
refGene_gn=MYB;refGene_tx=XM_023254420.1;refGene_gn=MYB;refGene_tx=XM_023254421.1;refGene_gn=MYB;refGene_tx=XM_023254422.1
B2 124822289 124822370
refGene_gn=MYB;refGene_tx=XM_023254420.1;refGene_gn=MYB;refGene_tx=XM_023254421.1;refGene_gn=MYB;refGene_tx=XM_023254422.1
B2 124822834 124822939
refGene_gn=MYB;refGene_tx=XM_023254420.1;refGene_gn=MYB;refGene_tx=XM_

023254421.1;refGene_gn=MYB;refGene_tx=XM_023254422.1
B2 124824447 124824702
refGene_gn=MYB;refGene_tx=XM_023254420.1;refGene_gn=MYB;refGene_tx=XM_
023254421.1;refGene_gn=MYB;refGene_tx=XM_023254422.1
B2 124825848 124826211
refGene_gn=MYB;refGene_tx=XM_023254420.1;refGene_gn=MYB;refGene_tx=XM_
023254421.1
B2 124827741 124827884
refGene_gn=MYB;refGene_tx=XM_023254420.1;refGene_gn=MYB;refGene_tx=XM_
023254421.1;refGene_gn=MYB;refGene_tx=XM_023254422.1
B2 124828869 124828984
refGene_gn=MYB;refGene_tx=XM_023254420.1;refGene_gn=MYB;refGene_tx=XM_
023254421.1;refGene_gn=MYB;refGene_tx=XM_023254422.1
B2 124829076 124829202
refGene_gn=MYB;refGene_tx=XM_023254420.1;refGene_gn=MYB;refGene_tx=XM_
023254421.1;refGene_gn=MYB;refGene_tx=XM_023254422.1
B2 124830062 124830173
refGene_gn=MYB;refGene_tx=XM_023254420.1;refGene_gn=MYB;refGene_tx=XM_
023254421.1;refGene_gn=MYB;refGene_tx=XM_023254422.1
B2 124832133 124832241
refGene_gn=MYB;refGene_tx=XM_023254420.1;refGene_gn=MYB;refGene_tx=XM_
023254421.1;refGene_gn=MYB;refGene_tx=XM_023254422.1
B2 124843203 124843320
refGene_gn=MYB;refGene_tx=XM_023254420.1;refGene_gn=MYB;refGene_tx=XM_
023254421.1;refGene_gn=MYB;refGene_tx=XM_023254422.1
B2 124902048 124902051
refGene_gn=AHI1;refGene_tx=XM_019831276.2
B2 124906652 124906755
refGene_gn=AHI1;refGene_tx=XM_019831276.2
B2 124922453 124922512
refGene_gn=AHI1;refGene_tx=XM_019831276.2
B2 124943774 124943875
refGene_gn=AHI1;refGene_tx=XM_019831276.2
B2 124947534 124947697
refGene_gn=AHI1;refGene_tx=XM_019831276.2
B2 124983552 124983608
refGene_gn=AHI1;refGene_tx=XM_019831276.2
B2 125009115 125009233
refGene_gn=AHI1;refGene_tx=XM_019831276.2
B2 125023058 125023085
refGene_gn=AHI1;refGene_tx=XM_019831276.2
B2 125026780 125026977
refGene_gn=AHI1;refGene_tx=XM_019831276.2
B2 125040469 125040610
refGene_gn=AHI1;refGene_tx=XM_019831276.2
B2 125042552 125042564
refGene_gn=AHI1;refGene_tx=XM_019831276.2
B2 125042566 125042682
refGene_gn=AHI1;refGene_tx=XM_019831276.2
B2 125045440 125045475

refGene_gn=AHI1;refGene_tx=XM_019831276.2
B2	125045476	125045560
refGene_gn=AHI1;refGene_tx=XM_019831276.2
B2	125048845	125048952
refGene_gn=AHI1;refGene_tx=XM_019831276.2
B2	125051002	125051232
refGene_gn=AHI1;refGene_tx=XM_019831276.2
B2	125058123	125058247
refGene_gn=AHI1;refGene_tx=XM_019831276.2
B2	125061939	125062072
refGene_gn=AHI1;refGene_tx=XM_019831276.2
B2	125066984	125067137
refGene_gn=AHI1;refGene_tx=XM_019831276.2
B2	125068500	125068686
refGene_gn=AHI1;refGene_tx=XM_019831276.2
B2	125073284	125073380
refGene_gn=AHI1;refGene_tx=XM_019831276.2
B2	125075393	125075586
refGene_gn=AHI1;refGene_tx=XM_019831276.2
B2	125076742	125076962
refGene_gn=AHI1;refGene_tx=XM_019831276.2
B2	125086404	125086586
refGene_gn=AHI1;refGene_tx=XM_019831276.2
B2	125089306	125089905
refGene_gn=AHI1;refGene_tx=XM_019831276.2
B2	125097307	125097432
refGene_gn=AHI1;refGene_tx=XM_019831276.2
B2	125098137	125098147
refGene_gn=AHI1;refGene_tx=XM_019831276.2
B2	125380358	125380379
refGene_gn=PDE7B;refGene_tx=XM_019831281.2
B2	125486409	125486491
refGene_gn=PDE7B;refGene_tx=XM_019831281.2;refGene_gn=PDE7B;refGene_tx
=XM_019831282.1
B2	125556672	125556910
refGene_gn=PDE7B;refGene_tx=XM_011282597.3
B2	125619632	125619716
refGene_gn=PDE7B;refGene_tx=XM_011282597.3;refGene_gn=PDE7B;refGene_tx
=XM_019831281.2;refGene_gn=PDE7B;refGene_tx=XM_019831282.1
B2	125655865	125656017
refGene_gn=PDE7B;refGene_tx=XM_011282597.3;refGene_gn=PDE7B;refGene_tx
=XM_019831281.2;refGene_gn=PDE7B;refGene_tx=XM_019831282.1
B2	125658244	125658308
refGene_gn=PDE7B;refGene_tx=XM_011282597.3;refGene_gn=PDE7B;refGene_tx
=XM_019831281.2;refGene_gn=PDE7B;refGene_tx=XM_019831282.1
B2	125660237	125660333
refGene_gn=PDE7B;refGene_tx=XM_011282597.3;refGene_gn=PDE7B;refGene_tx
=XM_019831281.2;refGene_gn=PDE7B;refGene_tx=XM_019831282.1
B2	125662416	125662517
refGene_gn=PDE7B;refGene_tx=XM_011282597.3;refGene_gn=PDE7B;refGene_tx

=XM_019831281.2;refGene_gn=PDE7B;refGene_tx=XM_019831282.1
B2 125664009 125664141
refGene_gn=PDE7B;refGene_tx=XM_011282597.3;refGene_gn=PDE7B;refGene_tx=XM_019831281.2;refGene_gn=PDE7B;refGene_tx=XM_019831282.1
B2 125679812 125679904
refGene_gn=PDE7B;refGene_tx=XM_011282597.3;refGene_gn=PDE7B;refGene_tx=XM_019831281.2;refGene_gn=PDE7B;refGene_tx=XM_019831282.1
B2 125682482 125682627
refGene_gn=PDE7B;refGene_tx=XM_011282597.3;refGene_gn=PDE7B;refGene_tx=XM_019831281.2;refGene_gn=PDE7B;refGene_tx=XM_019831282.1
B2 125684568 125684665
refGene_gn=PDE7B;refGene_tx=XM_011282597.3;refGene_gn=PDE7B;refGene_tx=XM_019831281.2;refGene_gn=PDE7B;refGene_tx=XM_019831282.1
B2 125689787 125689868
refGene_gn=PDE7B;refGene_tx=XM_011282597.3;refGene_gn=PDE7B;refGene_tx=XM_019831281.2;refGene_gn=PDE7B;refGene_tx=XM_019831282.1
B2 125693961 125694191
refGene_gn=PDE7B;refGene_tx=XM_011282597.3;refGene_gn=PDE7B;refGene_tx=XM_019831281.2;refGene_gn=PDE7B;refGene_tx=XM_019831282.1
B2 125727537 125727745
refGene_gn=MTFR2;refGene_tx=XM_006932043.4;refGene_gn=MTFR2;refGene_tx=XM_011282596.3;refGene_gn=MTFR2;refGene_tx=XM_019831284.1
B2 125730420 125730778
refGene_gn=MTFR2;refGene_tx=XM_006932043.4;refGene_gn=MTFR2;refGene_tx=XM_011282596.3;refGene_gn=MTFR2;refGene_tx=XM_019831284.1
B2 125731913 125732149
refGene_gn=MTFR2;refGene_tx=XM_006932043.4;refGene_gn=MTFR2;refGene_tx=XM_011282596.3;refGene_gn=MTFR2;refGene_tx=XM_019831284.1
B2 125733076 125733189
refGene_gn=MTFR2;refGene_tx=XM_006932043.4;refGene_gn=MTFR2;refGene_tx=XM_011282596.3;refGene_gn=MTFR2;refGene_tx=XM_019831284.1
B2 125733655 125733760
refGene_gn=MTFR2;refGene_tx=XM_006932043.4;refGene_gn=MTFR2;refGene_tx=XM_011282596.3;refGene_gn=MTFR2;refGene_tx=XM_019831284.1
B2 125739214 125739277
refGene_gn=MTFR2;refGene_tx=XM_006932043.4;refGene_gn=MTFR2;refGene_tx=XM_011282596.3;refGene_gn=MTFR2;refGene_tx=XM_019831284.1
B2 125749905 125749911
refGene_gn=BCLAF1;refGene_tx=XM_004001520.4;refGene_gn=BCLAF1;refGene_tx=XM_011282593.2;refGene_gn=BCLAF1;refGene_tx=XM_011282594.3;refGene_gn=BCLAF1;refGene_tx=XM_011282595.3
B2 125750069 125750282
refGene_gn=BCLAF1;refGene_tx=XM_004001520.4;refGene_gn=BCLAF1;refGene_tx=XM_011282593.2;refGene_gn=BCLAF1;refGene_tx=XM_011282594.3;refGene_gn=BCLAF1;refGene_tx=XM_011282595.3
B2 125756240 125756387
refGene_gn=BCLAF1;refGene_tx=XM_004001520.4;refGene_gn=BCLAF1;refGene_tx=XM_011282593.2
B2 125757509 125757690
refGene_gn=BCLAF1;refGene_tx=XM_004001520.4;refGene_gn=BCLAF1;refGene_

tx=XM_011282593.2;refGene_gn=BCLAF1;refGene_tx=XM_011282594.3;refGene_gn=BCLAF1;refGene_tx=XM_011282595.3
B2	125758808	125758984
refGene_gn=BCLAF1;refGene_tx=XM_004001520.4;refGene_gn=BCLAF1;refGene_tx=XM_011282593.2;refGene_gn=BCLAF1;refGene_tx=XM_011282594.3;refGene_gn=BCLAF1;refGene_tx=XM_011282595.3
B2	125761355	125761440
refGene_gn=BCLAF1;refGene_tx=XM_004001520.4;refGene_gn=BCLAF1;refGene_tx=XM_011282593.2;refGene_gn=BCLAF1;refGene_tx=XM_011282594.3;refGene_gn=BCLAF1;refGene_tx=XM_011282595.3
B2	125762378	125762484
refGene_gn=BCLAF1;refGene_tx=XM_004001520.4;refGene_gn=BCLAF1;refGene_tx=XM_011282593.2;refGene_gn=BCLAF1;refGene_tx=XM_011282594.3;refGene_gn=BCLAF1;refGene_tx=XM_011282595.3
B2	125764807	125764977
refGene_gn=BCLAF1;refGene_tx=XM_004001520.4;refGene_gn=BCLAF1;refGene_tx=XM_011282593.2;refGene_gn=BCLAF1;refGene_tx=XM_011282594.3;refGene_gn=BCLAF1;refGene_tx=XM_011282595.3
B2	125765130	125765796
refGene_gn=BCLAF1;refGene_tx=XM_004001520.4;refGene_gn=BCLAF1;refGene_tx=XM_011282593.2;refGene_gn=BCLAF1;refGene_tx=XM_011282594.3;refGene_gn=BCLAF1;refGene_tx=XM_011282595.3
B2	125766885	125767797
refGene_gn=BCLAF1;refGene_tx=XM_004001520.4;refGene_gn=BCLAF1;refGene_tx=XM_011282593.2;refGene_gn=BCLAF1;refGene_tx=XM_011282594.3;refGene_gn=BCLAF1;refGene_tx=XM_011282595.3
B2	125768559	125768663
refGene_gn=BCLAF1;refGene_tx=XM_004001520.4;refGene_gn=BCLAF1;refGene_tx=XM_011282593.2;refGene_gn=BCLAF1;refGene_tx=XM_011282594.3;refGene_gn=BCLAF1;refGene_tx=XM_011282595.3
B2	125826284	125826295
refGene_gn=MAP7;refGene_tx=XM_023254426.1;refGene_gn=MAP7;refGene_tx=XM_023254427.1;refGene_gn=MAP7;refGene_tx=XM_023254428.1;refGene_gn=MAP7;refGene_tx=XM_023254429.1;refGene_gn=MAP7;refGene_tx=XM_023254430.1;refGene_gn=MAP7;refGene_tx=XM_023254431.1;refGene_gn=MAP7;refGene_tx=XM_023254432.1;refGene_gn=MAP7;refGene_tx=XM_023254433.1;refGene_gn=MAP7;refGene_tx=XM_023254434.1;refGene_gn=MAP7;refGene_tx=XM_023254435.1
B2	125828034	125828258
refGene_gn=MAP7;refGene_tx=XM_023254426.1;refGene_gn=MAP7;refGene_tx=XM_023254427.1;refGene_gn=MAP7;refGene_tx=XM_023254428.1;refGene_gn=MAP7;refGene_tx=XM_023254429.1;refGene_gn=MAP7;refGene_tx=XM_023254430.1;refGene_gn=MAP7;refGene_tx=XM_023254431.1;refGene_gn=MAP7;refGene_tx=XM_023254432.1;refGene_gn=MAP7;refGene_tx=XM_023254433.1;refGene_gn=MAP7;refGene_tx=XM_023254434.1;refGene_gn=MAP7;refGene_tx=XM_023254435.1
B2	125837412	125837515
refGene_gn=MAP7;refGene_tx=XM_023254426.1;refGene_gn=MAP7;refGene_tx=XM_023254427.1;refGene_gn=MAP7;refGene_tx=XM_023254428.1;refGene_gn=MAP7;refGene_tx=XM_023254429.1;refGene_gn=MAP7;refGene_tx=XM_023254430.1;refGene_gn=MAP7;refGene_tx=XM_023254431.1;refGene_gn=MAP7;refGene_tx=XM_023254432.1;refGene_gn=MAP7;refGene_tx=XM_023254433.1;refGene_gn=MAP

7;refGene_tx=XM_023254434.1;refGene_gn=MAP7;refGene_tx=XM_023254435.1
B2 125839913 125839971
refGene_gn=MAP7;refGene_tx=XM_023254426.1;refGene_gn=MAP7;refGene_tx=XM_023254427.1;refGene_gn=MAP7;refGene_tx=XM_023254428.1;refGene_gn=MAP7;refGene_tx=XM_023254429.1;refGene_gn=MAP7;refGene_tx=XM_023254430.1;refGene_gn=MAP7;refGene_tx=XM_023254431.1;refGene_gn=MAP7;refGene_tx=XM_023254432.1;refGene_gn=MAP7;refGene_tx=XM_023254433.1;refGene_gn=MAP7;refGene_tx=XM_023254434.1;refGene_gn=MAP7;refGene_tx=XM_023254435.1
B2 125840073 125840124
refGene_gn=MAP7;refGene_tx=XM_023254426.1;refGene_gn=MAP7;refGene_tx=XM_023254427.1;refGene_gn=MAP7;refGene_tx=XM_023254428.1;refGene_gn=MAP7;refGene_tx=XM_023254429.1;refGene_gn=MAP7;refGene_tx=XM_023254430.1;refGene_gn=MAP7;refGene_tx=XM_023254431.1;refGene_gn=MAP7;refGene_tx=XM_023254432.1;refGene_gn=MAP7;refGene_tx=XM_023254433.1;refGene_gn=MAP7;refGene_tx=XM_023254434.1;refGene_gn=MAP7;refGene_tx=XM_023254435.1
B2 125840662 125840764
refGene_gn=MAP7;refGene_tx=XM_023254426.1;refGene_gn=MAP7;refGene_tx=XM_023254427.1;refGene_gn=MAP7;refGene_tx=XM_023254428.1;refGene_gn=MAP7;refGene_tx=XM_023254429.1;refGene_gn=MAP7;refGene_tx=XM_023254430.1;refGene_gn=MAP7;refGene_tx=XM_023254431.1;refGene_gn=MAP7;refGene_tx=XM_023254432.1;refGene_gn=MAP7;refGene_tx=XM_023254433.1;refGene_gn=MAP7;refGene_tx=XM_023254434.1;refGene_gn=MAP7;refGene_tx=XM_023254435.1
B2 125840995 125841170
refGene_gn=MAP7;refGene_tx=XM_023254426.1;refGene_gn=MAP7;refGene_tx=XM_023254427.1;refGene_gn=MAP7;refGene_tx=XM_023254428.1;refGene_gn=MAP7;refGene_tx=XM_023254429.1;refGene_gn=MAP7;refGene_tx=XM_023254430.1;refGene_gn=MAP7;refGene_tx=XM_023254431.1;refGene_gn=MAP7;refGene_tx=XM_023254432.1;refGene_gn=MAP7;refGene_tx=XM_023254433.1;refGene_gn=MAP7;refGene_tx=XM_023254434.1;refGene_gn=MAP7;refGene_tx=XM_023254435.1
B2 125842303 125842535
refGene_gn=MAP7;refGene_tx=XM_023254426.1;refGene_gn=MAP7;refGene_tx=XM_023254427.1;refGene_gn=MAP7;refGene_tx=XM_023254428.1;refGene_gn=MAP7;refGene_tx=XM_023254429.1;refGene_gn=MAP7;refGene_tx=XM_023254430.1;refGene_gn=MAP7;refGene_tx=XM_023254431.1;refGene_gn=MAP7;refGene_tx=XM_023254432.1;refGene_gn=MAP7;refGene_tx=XM_023254433.1;refGene_gn=MAP7;refGene_tx=XM_023254434.1;refGene_gn=MAP7;refGene_tx=XM_023254435.1
B2 125844474 125844758
refGene_gn=MAP7;refGene_tx=XM_023254426.1;refGene_gn=MAP7;refGene_tx=XM_023254427.1;refGene_gn=MAP7;refGene_tx=XM_023254428.1;refGene_gn=MAP7;refGene_tx=XM_023254429.1;refGene_gn=MAP7;refGene_tx=XM_023254430.1;refGene_gn=MAP7;refGene_tx=XM_023254431.1;refGene_gn=MAP7;refGene_tx=XM_023254432.1;refGene_gn=MAP7;refGene_tx=XM_023254433.1;refGene_gn=MAP7;refGene_tx=XM_023254434.1;refGene_gn=MAP7;refGene_tx=XM_023254435.1
B2 125845092 125845208
refGene_gn=MAP7;refGene_tx=XM_023254426.1;refGene_gn=MAP7;refGene_tx=XM_023254427.1;refGene_gn=MAP7;refGene_tx=XM_023254428.1;refGene_gn=MAP7;refGene_tx=XM_023254429.1;refGene_gn=MAP7;refGene_tx=XM_023254430.1;refGene_gn=MAP7;refGene_tx=XM_023254431.1;refGene_gn=MAP7;refGene_tx=XM_023254432.1;refGene_gn=MAP7;refGene_tx=XM_023254433.1;refGene_gn=MAP7;refGene_tx=XM_023254434.1;refGene_gn=MAP7;refGene_tx=XM_023254435.1

B2 125850822 125850974
refGene_gn=MAP7;refGene_tx=XM_023254426.1;refGene_gn=MAP7;refGene_tx=XM_023254427.1;refGene_gn=MAP7;refGene_tx=XM_023254428.1;refGene_gn=MAP7;refGene_tx=XM_023254429.1;refGene_gn=MAP7;refGene_tx=XM_023254430.1;refGene_gn=MAP7;refGene_tx=XM_023254431.1;refGene_gn=MAP7;refGene_tx=XM_023254432.1;refGene_gn=MAP7;refGene_tx=XM_023254433.1;refGene_gn=MAP7;refGene_tx=XM_023254434.1;refGene_gn=MAP7;refGene_tx=XM_023254435.1
B2 125855713 125855827
refGene_gn=MAP7;refGene_tx=XM_023254426.1;refGene_gn=MAP7;refGene_tx=XM_023254427.1;refGene_gn=MAP7;refGene_tx=XM_023254428.1;refGene_gn=MAP7;refGene_tx=XM_023254429.1;refGene_gn=MAP7;refGene_tx=XM_023254430.1;refGene_gn=MAP7;refGene_tx=XM_023254431.1;refGene_gn=MAP7;refGene_tx=XM_023254432.1;refGene_gn=MAP7;refGene_tx=XM_023254433.1;refGene_gn=MAP7;refGene_tx=XM_023254434.1;refGene_gn=MAP7;refGene_tx=XM_023254435.1
B2 125861631 125861742
refGene_gn=MAP7;refGene_tx=XM_023254426.1;refGene_gn=MAP7;refGene_tx=XM_023254427.1;refGene_gn=MAP7;refGene_tx=XM_023254428.1;refGene_gn=MAP7;refGene_tx=XM_023254429.1;refGene_gn=MAP7;refGene_tx=XM_023254430.1;refGene_gn=MAP7;refGene_tx=XM_023254431.1;refGene_gn=MAP7;refGene_tx=XM_023254432.1;refGene_gn=MAP7;refGene_tx=XM_023254433.1
B2 125865134 125865221
refGene_gn=MAP7;refGene_tx=XM_023254426.1;refGene_gn=MAP7;refGene_tx=XM_023254427.1;refGene_gn=MAP7;refGene_tx=XM_023254428.1;refGene_gn=MAP7;refGene_tx=XM_023254429.1;refGene_gn=MAP7;refGene_tx=XM_023254432.1;refGene_gn=MAP7;refGene_tx=XM_023254433.1
B2 125865922 125866040
refGene_gn=MAP7;refGene_tx=XM_023254426.1;refGene_gn=MAP7;refGene_tx=XM_023254427.1;refGene_gn=MAP7;refGene_tx=XM_023254428.1;refGene_gn=MAP7;refGene_tx=XM_023254429.1;refGene_gn=MAP7;refGene_tx=XM_023254430.1;refGene_gn=MAP7;refGene_tx=XM_023254431.1;refGene_gn=MAP7;refGene_tx=XM_023254432.1;refGene_gn=MAP7;refGene_tx=XM_023254433.1;refGene_gn=MAP7;refGene_tx=XM_023254434.1;refGene_gn=MAP7;refGene_tx=XM_023254435.1
B2 125866519 125866683
refGene_gn=MAP7;refGene_tx=XM_023254426.1;refGene_gn=MAP7;refGene_tx=XM_023254427.1;refGene_gn=MAP7;refGene_tx=XM_023254428.1;refGene_gn=MAP7;refGene_tx=XM_023254429.1;refGene_gn=MAP7;refGene_tx=XM_023254430.1;refGene_gn=MAP7;refGene_tx=XM_023254431.1;refGene_gn=MAP7;refGene_tx=XM_023254432.1;refGene_gn=MAP7;refGene_tx=XM_023254433.1;refGene_gn=MAP7;refGene_tx=XM_023254434.1;refGene_gn=MAP7;refGene_tx=XM_023254435.1
B2 125881836 125881914
refGene_gn=MAP7;refGene_tx=XM_023254426.1;refGene_gn=MAP7;refGene_tx=XM_023254427.1;refGene_gn=MAP7;refGene_tx=XM_023254428.1;refGene_gn=MAP7;refGene_tx=XM_023254429.1;refGene_gn=MAP7;refGene_tx=XM_023254430.1;refGene_gn=MAP7;refGene_tx=XM_023254431.1;refGene_gn=MAP7;refGene_tx=XM_023254432.1;refGene_gn=MAP7;refGene_tx=XM_023254433.1;refGene_gn=MAP7;refGene_tx=XM_023254434.1;refGene_gn=MAP7;refGene_tx=XM_023254435.1
B2 125889694 125889790
refGene_gn=MAP7;refGene_tx=XM_023254426.1;refGene_gn=MAP7;refGene_tx=XM_023254427.1;refGene_gn=MAP7;refGene_tx=XM_023254428.1;refGene_gn=MAP7;refGene_tx=XM_023254429.1;refGene_gn=MAP7;refGene_tx=XM_023254430.1;

refGene_gn=MAP7;refGene_tx=XM_023254431.1;refGene_gn=MAP7;refGene_tx=XM_023254432.1;refGene_gn=MAP7;refGene_tx=XM_023254433.1;refGene_gn=MAP7;refGene_tx=XM_023254434.1;refGene_gn=MAP7;refGene_tx=XM_023254435.1
B2 125892965 125892984
refGene_gn=MAP7;refGene_tx=XM_023254433.1
B2 125930377 125930399
refGene_gn=MAP7;refGene_tx=XM_023254432.1
B2 125979434 125979567
refGene_gn=MAP7;refGene_tx=XM_023254426.1;refGene_gn=MAP7;refGene_tx=XM_023254427.1;refGene_gn=MAP7;refGene_tx=XM_023254428.1;refGene_gn=MAP7;refGene_tx=XM_023254430.1;refGene_gn=MAP7;refGene_tx=XM_023254431.1;refGene_gn=MAP7;refGene_tx=XM_023254434.1;refGene_gn=MAP7;refGene_tx=XM_023254435.1
B2 126005023 126005093
refGene_gn=MAP7;refGene_tx=XM_023254429.1
B2 126011634 126011695
refGene_gn=MAP3K5;refGene_tx=XM_023254424.1;refGene_gn=MAP3K5;refGene_tx=XM_023254425.1
B2 126012594 126012671
refGene_gn=MAP3K5;refGene_tx=XM_023254424.1;refGene_gn=MAP3K5;refGene_tx=XM_023254425.1
B2 126014503 126014616
refGene_gn=MAP3K5;refGene_tx=XM_023254424.1;refGene_gn=MAP3K5;refGene_tx=XM_023254425.1
B2 126015352 126015465
refGene_gn=MAP3K5;refGene_tx=XM_023254424.1;refGene_gn=MAP3K5;refGene_tx=XM_023254425.1
B2 126021150 126021394
refGene_gn=MAP3K5;refGene_tx=XM_023254424.1;refGene_gn=MAP3K5;refGene_tx=XM_023254425.1
B2 126028887 126028993
refGene_gn=MAP3K5;refGene_tx=XM_023254424.1;refGene_gn=MAP3K5;refGene_tx=XM_023254425.1
B2 126032275 126032461
refGene_gn=MAP3K5;refGene_tx=XM_023254424.1;refGene_gn=MAP3K5;refGene_tx=XM_023254425.1
B2 126038745 126038914
refGene_gn=MAP3K5;refGene_tx=XM_023254424.1;refGene_gn=MAP3K5;refGene_tx=XM_023254425.1
B2 126039006 126039187
refGene_gn=MAP3K5;refGene_tx=XM_023254424.1;refGene_gn=MAP3K5;refGene_tx=XM_023254425.1
B2 126043203 126043224
refGene_gn=MAP3K5;refGene_tx=XM_023254424.1;refGene_gn=MAP3K5;refGene_tx=XM_023254425.1
B2 126043994 126044172
refGene_gn=MAP3K5;refGene_tx=XM_023254424.1;refGene_gn=MAP3K5;refGene_tx=XM_023254425.1
B2 126046304 126046462
refGene_gn=MAP3K5;refGene_tx=XM_023254424.1;refGene_gn=MAP3K5;refGene_

tx=XM_023254425.1
B2	126051942	126052048
refGene_gn=MAP3K5;refGene_tx=XM_023254424.1;refGene_gn=MAP3K5;refGene_
tx=XM_023254425.1
B2	126053011	126053148
refGene_gn=MAP3K5;refGene_tx=XM_023254424.1;refGene_gn=MAP3K5;refGene_
tx=XM_023254425.1
B2	126054629	126054757
refGene_gn=MAP3K5;refGene_tx=XM_023254424.1;refGene_gn=MAP3K5;refGene_
tx=XM_023254425.1
B2	126059678	126059812
refGene_gn=MAP3K5;refGene_tx=XM_023254424.1;refGene_gn=MAP3K5;refGene_
tx=XM_023254425.1
B2	126076768	126076850
refGene_gn=MAP3K5;refGene_tx=XM_023254424.1;refGene_gn=MAP3K5;refGene_
tx=XM_023254425.1
B2	126079748	126079844
refGene_gn=MAP3K5;refGene_tx=XM_023254424.1;refGene_gn=MAP3K5;refGene_
tx=XM_023254425.1
B2	126082296	126082346
refGene_gn=MAP3K5;refGene_tx=XM_023254424.1;refGene_gn=MAP3K5;refGene_
tx=XM_023254425.1
B2	126089497	126089605
refGene_gn=MAP3K5;refGene_tx=XM_023254424.1;refGene_gn=MAP3K5;refGene_
tx=XM_023254425.1
B2	126096788	126096942
refGene_gn=MAP3K5;refGene_tx=XM_023254424.1;refGene_gn=MAP3K5;refGene_
tx=XM_023254425.1
B2	126099506	126099666
refGene_gn=MAP3K5;refGene_tx=XM_023254424.1;refGene_gn=MAP3K5;refGene_
tx=XM_023254425.1
B2	126106112	126106225
refGene_gn=MAP3K5;refGene_tx=XM_023254424.1;refGene_gn=MAP3K5;refGene_
tx=XM_023254425.1
B2	126125984	126126155
refGene_gn=MAP3K5;refGene_tx=XM_023254424.1;refGene_gn=MAP3K5;refGene_
tx=XM_023254425.1
B2	126129261	126129368
refGene_gn=MAP3K5;refGene_tx=XM_023254424.1;refGene_gn=MAP3K5;refGene_
tx=XM_023254425.1
B2	126130796	126130965
refGene_gn=MAP3K5;refGene_tx=XM_023254424.1;refGene_gn=MAP3K5;refGene_
tx=XM_023254425.1
B2	126131689	126131883
refGene_gn=MAP3K5;refGene_tx=XM_023254424.1;refGene_gn=MAP3K5;refGene_
tx=XM_023254425.1
B2	126137761	126137785
refGene_gn=MAP3K5;refGene_tx=XM_023254424.1;refGene_gn=MAP3K5;refGene_
tx=XM_023254425.1
B2	126150938	126151078

refGene_gn=MAP3K5;refGene_tx=XM_023254424.1;refGene_gn=MAP3K5;refGene_tx=XM_023254425.1
B2 126216913 126217337
refGene_gn=MAP3K5;refGene_tx=XM_023254424.1;refGene_gn=MAP3K5;refGene_tx=XM_023254425.1
B2 126248206 126248336
refGene_gn=PEX7;refGene_tx=XM_023254436.1
B2 126250130 126250188
refGene_gn=PEX7;refGene_tx=XM_023254436.1
B2 126251090 126251241
refGene_gn=PEX7;refGene_tx=XM_023254436.1
B2 126262774 126262852
refGene_gn=PEX7;refGene_tx=XM_023254436.1
B2 126263641 126263750
refGene_gn=PEX7;refGene_tx=XM_023254436.1
B2 126278706 126278813
refGene_gn=PEX7;refGene_tx=XM_023254436.1
B2 126280260 126280374
refGene_gn=PEX7;refGene_tx=XM_023254436.1
B2 126281886 126281942
refGene_gn=PEX7;refGene_tx=XM_023254436.1
B2 126300660 126300760
refGene_gn=PEX7;refGene_tx=XM_023254436.1
B2 126326830 126326899
refGene_gn=PEX7;refGene_tx=XM_023254436.1
B2 126339545 126339984
refGene_gn=SLC35D3;refGene_tx=XM_003986593.4
B2 126341019 126341849
refGene_gn=SLC35D3;refGene_tx=XM_003986593.4
B2 126417764 126418559
refGene_gn=IL20RA;refGene_tx=XM_006932047.4;refGene_gn=IL20RA;refGene_tx=XM_019831286.2;refGene_gn=IL20RA;refGene_tx=XM_019831287.2;refGene_gn=IL20RA;refGene_tx=XM_023254437.1;refGene_gn=IL20RA;refGene_tx=XM_023254438.1
B2 126421037 126421177
refGene_gn=IL20RA;refGene_tx=XM_006932047.4;refGene_gn=IL20RA;refGene_tx=XM_019831286.2;refGene_gn=IL20RA;refGene_tx=XM_019831287.2;refGene_gn=IL20RA;refGene_tx=XM_023254437.1;refGene_gn=IL20RA;refGene_tx=XM_023254438.1
B2 126424185 126424330
refGene_gn=IL20RA;refGene_tx=XM_006932047.4;refGene_gn=IL20RA;refGene_tx=XM_019831286.2;refGene_gn=IL20RA;refGene_tx=XM_019831287.2;refGene_gn=IL20RA;refGene_tx=XM_023254437.1;refGene_gn=IL20RA;refGene_tx=XM_023254438.1
B2 126424892 126425068
refGene_gn=IL20RA;refGene_tx=XM_006932047.4;refGene_gn=IL20RA;refGene_tx=XM_019831286.2;refGene_gn=IL20RA;refGene_tx=XM_019831287.2;refGene_gn=IL20RA;refGene_tx=XM_023254437.1;refGene_gn=IL20RA;refGene_tx=XM_023254438.1
B2 126426504 126426683

refGene_gn=IL20RA;refGene_tx=XM_006932047.4;refGene_gn=IL20RA;refGene_tx=XM_019831286.2;refGene_gn=IL20RA;refGene_tx=XM_019831287.2;refGene_gn=IL20RA;refGene_tx=XM_023254437.1;refGene_gn=IL20RA;refGene_tx=XM_023254438.1
B2 126430309 126430332
refGene_gn=IL20RA;refGene_tx=XM_019831287.2
B2 126432659 126432736
refGene_gn=IL20RA;refGene_tx=XM_006932047.4;refGene_gn=IL20RA;refGene_tx=XM_019831286.2;refGene_gn=IL20RA;refGene_tx=XM_023254437.1;refGene_gn=IL20RA;refGene_tx=XM_023254438.1
B2 126513380 126513530
refGene_gn=IL22RA2;refGene_tx=XM_003986594.2
B2 126514557 126514727
refGene_gn=IL22RA2;refGene_tx=XM_003986594.2
B2 126527407 126527586
refGene_gn=IL22RA2;refGene_tx=XM_003986594.2
B2 126530166 126530302
refGene_gn=IL22RA2;refGene_tx=XM_003986594.2
B2 126536425 126536639
refGene_gn=IL22RA2;refGene_tx=XM_003986594.2
B2 126545980 126546541
refGene_gn=IFNGR1;refGene_tx=XM_003986606.4
B2 126548492 126548620
refGene_gn=IFNGR1;refGene_tx=XM_003986606.4
B2 126551338 126551525
refGene_gn=IFNGR1;refGene_tx=XM_003986606.4
B2 126553665 126553835
refGene_gn=IFNGR1;refGene_tx=XM_003986606.4
B2 126555663 126555827
refGene_gn=IFNGR1;refGene_tx=XM_003986606.4
B2 126556520 126556635
refGene_gn=IFNGR1;refGene_tx=XM_003986606.4
B2 126566983 126567068
refGene_gn=IFNGR1;refGene_tx=XM_003986606.4
B2 126818018 126818837
refGene_gn=OLIG3;refGene_tx=XM_019831653.2
B2 127171511 127171806
refGene_gn=TNFAIP3;refGene_tx=XM_006932048.4;refGene_gn=TNFAIP3;refGene_tx=XM_006932049.3
B2 127175040 127175231
refGene_gn=TNFAIP3;refGene_tx=XM_006932048.4;refGene_gn=TNFAIP3;refGene_tx=XM_006932049.3
B2 127175922 127176070
refGene_gn=TNFAIP3;refGene_tx=XM_006932048.4;refGene_gn=TNFAIP3;refGene_tx=XM_006932049.3
B2 127176237 127176408
refGene_gn=TNFAIP3;refGene_tx=XM_006932048.4;refGene_gn=TNFAIP3;refGene_tx=XM_006932049.3
B2 127177234 127177415
refGene_gn=TNFAIP3;refGene_tx=XM_006932048.4;refGene_gn=TNFAIP3;refGen

e_tx=XM_006932049.3
B2	127178568	127179488
refGene_gn=TNFAIP3;refGene_tx=XM_006932048.4;refGene_gn=TNFAIP3;refGene_tx=XM_006932049.3
B2	127180154	127180336
refGene_gn=TNFAIP3;refGene_tx=XM_006932048.4;refGene_gn=TNFAIP3;refGene_tx=XM_006932049.3
B2	127181550	127181835
refGene_gn=TNFAIP3;refGene_tx=XM_006932048.4;refGene_gn=TNFAIP3;refGene_tx=XM_006932049.3
B2	127346811	127347038
refGene_gn=PERP;refGene_tx=XM_003986596.5
B2	127350715	127350856
refGene_gn=PERP;refGene_tx=XM_003986596.5
B2	127361390	127361604
refGene_gn=PERP;refGene_tx=XM_003986596.5
B2	127416026	127416111
refGene_gn=ARFGEF3;refGene_tx=XM_023254439.1;refGene_gn=ARFGEF3;refGene_tx=XM_023254440.1;refGene_gn=ARFGEF3;refGene_tx=XM_023254441.1
B2	127424995	127425047
refGene_gn=ARFGEF3;refGene_tx=XM_023254439.1;refGene_gn=ARFGEF3;refGene_tx=XM_023254440.1;refGene_gn=ARFGEF3;refGene_tx=XM_023254441.1
B2	127462842	127462924
refGene_gn=ARFGEF3;refGene_tx=XM_023254439.1;refGene_gn=ARFGEF3;refGene_tx=XM_023254440.1;refGene_gn=ARFGEF3;refGene_tx=XM_023254441.1
B2	127466656	127466788
refGene_gn=ARFGEF3;refGene_tx=XM_023254439.1;refGene_gn=ARFGEF3;refGene_tx=XM_023254440.1
B2	127473567	127473695
refGene_gn=PBOV1;refGene_tx=XM_023254635.1
B2	127473697	127473740
refGene_gn=PBOV1;refGene_tx=XM_023254635.1
B2	127473742	127474036
refGene_gn=PBOV1;refGene_tx=XM_023254635.1
B2	127475736	127475756
refGene_gn=PBOV1;refGene_tx=XM_023254635.1
B2	127478155	127478317
refGene_gn=PBOV1;refGene_tx=XM_023254635.1
B2	127480505	127480682
refGene_gn=PBOV1;refGene_tx=XM_023254635.1
B2	127481632	127481669
refGene_gn=PBOV1;refGene_tx=XM_023254635.1
B2	127485711	127485780
refGene_gn=ARFGEF3;refGene_tx=XM_023254439.1;refGene_gn=ARFGEF3;refGene_tx=XM_023254440.1;refGene_gn=ARFGEF3;refGene_tx=XM_023254441.1
B2	127499093	127499216
refGene_gn=ARFGEF3;refGene_tx=XM_023254439.1;refGene_gn=ARFGEF3;refGene_tx=XM_023254441.1
B2	127502906	127502949
refGene_gn=ARFGEF3;refGene_tx=XM_023254439.1;refGene_gn=ARFGEF3;refGen

e_tx=XM_023254440.1;refGene_gn=ARFGEF3;refGene_tx=XM_023254441.1
B2 127505544 127505623
refGene_gn=ARFGEF3;refGene_tx=XM_023254439.1;refGene_gn=ARFGEF3;refGene_tx=XM_023254440.1;refGene_gn=ARFGEF3;refGene_tx=XM_023254441.1
B2 127512433 127512538
refGene_gn=ARFGEF3;refGene_tx=XM_023254439.1;refGene_gn=ARFGEF3;refGene_tx=XM_023254440.1;refGene_gn=ARFGEF3;refGene_tx=XM_023254441.1
B2 127513427 127513761
refGene_gn=ARFGEF3;refGene_tx=XM_023254439.1;refGene_gn=ARFGEF3;refGene_tx=XM_023254440.1;refGene_gn=ARFGEF3;refGene_tx=XM_023254441.1
B2 127525149 127525262
refGene_gn=ARFGEF3;refGene_tx=XM_023254439.1;refGene_gn=ARFGEF3;refGene_tx=XM_023254440.1;refGene_gn=ARFGEF3;refGene_tx=XM_023254441.1
B2 127526046 127526957
refGene_gn=ARFGEF3;refGene_tx=XM_023254439.1;refGene_gn=ARFGEF3;refGene_tx=XM_023254440.1;refGene_gn=ARFGEF3;refGene_tx=XM_023254441.1
B2 127542333 127542500
refGene_gn=ARFGEF3;refGene_tx=XM_023254439.1;refGene_gn=ARFGEF3;refGene_tx=XM_023254440.1;refGene_gn=ARFGEF3;refGene_tx=XM_023254441.1
B2 127543781 127543947
refGene_gn=ARFGEF3;refGene_tx=XM_023254439.1;refGene_gn=ARFGEF3;refGene_tx=XM_023254440.1;refGene_gn=ARFGEF3;refGene_tx=XM_023254441.1
B2 127548158 127548266
refGene_gn=ARFGEF3;refGene_tx=XM_023254439.1;refGene_gn=ARFGEF3;refGene_tx=XM_023254440.1;refGene_gn=ARFGEF3;refGene_tx=XM_023254441.1
B2 127548612 127548828
refGene_gn=ARFGEF3;refGene_tx=XM_023254439.1;refGene_gn=ARFGEF3;refGene_tx=XM_023254440.1;refGene_gn=ARFGEF3;refGene_tx=XM_023254441.1
B2 127548988 127549099
refGene_gn=ARFGEF3;refGene_tx=XM_023254439.1;refGene_gn=ARFGEF3;refGene_tx=XM_023254440.1;refGene_gn=ARFGEF3;refGene_tx=XM_023254441.1
B2 127550836 127550987
refGene_gn=ARFGEF3;refGene_tx=XM_023254439.1;refGene_gn=ARFGEF3;refGene_tx=XM_023254440.1;refGene_gn=ARFGEF3;refGene_tx=XM_023254441.1
B2 127552297 127552618
refGene_gn=ARFGEF3;refGene_tx=XM_023254439.1;refGene_gn=ARFGEF3;refGene_tx=XM_023254440.1;refGene_gn=ARFGEF3;refGene_tx=XM_023254441.1
B2 127555140 127555274
refGene_gn=ARFGEF3;refGene_tx=XM_023254439.1;refGene_gn=ARFGEF3;refGene_tx=XM_023254440.1;refGene_gn=ARFGEF3;refGene_tx=XM_023254441.1
B2 127557279 127557425
refGene_gn=ARFGEF3;refGene_tx=XM_023254439.1;refGene_gn=ARFGEF3;refGene_tx=XM_023254440.1;refGene_gn=ARFGEF3;refGene_tx=XM_023254441.1
B2 127559196 127559376
refGene_gn=ARFGEF3;refGene_tx=XM_023254439.1;refGene_gn=ARFGEF3;refGene_tx=XM_023254440.1;refGene_gn=ARFGEF3;refGene_tx=XM_023254441.1
B2 127568293 127568438
refGene_gn=ARFGEF3;refGene_tx=XM_023254439.1;refGene_gn=ARFGEF3;refGene_tx=XM_023254440.1;refGene_gn=ARFGEF3;refGene_tx=XM_023254441.1
B2 127570024 127570147

refGene_gn=ARFGEF3;refGene_tx=XM_023254439.1;refGene_gn=ARFGEF3;refGene_tx=XM_023254440.1;refGene_gn=ARFGEF3;refGene_tx=XM_023254441.1
B2 127574948 127575052
refGene_gn=ARFGEF3;refGene_tx=XM_023254439.1;refGene_gn=ARFGEF3;refGene_tx=XM_023254440.1;refGene_gn=ARFGEF3;refGene_tx=XM_023254441.1
B2 127576677 127576822
refGene_gn=ARFGEF3;refGene_tx=XM_023254439.1;refGene_gn=ARFGEF3;refGene_tx=XM_023254440.1;refGene_gn=ARFGEF3;refGene_tx=XM_023254441.1
B2 127581432 127581561
refGene_gn=ARFGEF3;refGene_tx=XM_023254439.1;refGene_gn=ARFGEF3;refGene_tx=XM_023254440.1;refGene_gn=ARFGEF3;refGene_tx=XM_023254441.1
B2 127583458 127583635
refGene_gn=ARFGEF3;refGene_tx=XM_023254439.1;refGene_gn=ARFGEF3;refGene_tx=XM_023254440.1;refGene_gn=ARFGEF3;refGene_tx=XM_023254441.1
B2 127585738 127585853
refGene_gn=ARFGEF3;refGene_tx=XM_023254439.1;refGene_gn=ARFGEF3;refGene_tx=XM_023254440.1;refGene_gn=ARFGEF3;refGene_tx=XM_023254441.1
B2 127586741 127586844
refGene_gn=ARFGEF3;refGene_tx=XM_023254439.1;refGene_gn=ARFGEF3;refGene_tx=XM_023254440.1;refGene_gn=ARFGEF3;refGene_tx=XM_023254441.1
B2 127587082 127587214
refGene_gn=ARFGEF3;refGene_tx=XM_023254439.1;refGene_gn=ARFGEF3;refGene_tx=XM_023254440.1;refGene_gn=ARFGEF3;refGene_tx=XM_023254441.1
B2 127589475 127589597
refGene_gn=ARFGEF3;refGene_tx=XM_023254439.1;refGene_gn=ARFGEF3;refGene_tx=XM_023254440.1;refGene_gn=ARFGEF3;refGene_tx=XM_023254441.1
B2 127593309 127594528
refGene_gn=ARFGEF3;refGene_tx=XM_023254439.1;refGene_gn=ARFGEF3;refGene_tx=XM_023254440.1;refGene_gn=ARFGEF3;refGene_tx=XM_023254441.1
B2 127598428 127598620
refGene_gn=ARFGEF3;refGene_tx=XM_023254439.1;refGene_gn=ARFGEF3;refGene_tx=XM_023254440.1;refGene_gn=ARFGEF3;refGene_tx=XM_023254441.1
B2 127664693 127664726
refGene_gn=HEBP2;refGene_tx=XM_023254442.1
B2 127665463 127665565
refGene_gn=HEBP2;refGene_tx=XM_023254443.1
B2 127666096 127666232
refGene_gn=HEBP2;refGene_tx=XM_023254442.1;refGene_gn=HEBP2;refGene_tx=XM_023254443.1
B2 127667225 127667406
refGene_gn=HEBP2;refGene_tx=XM_023254442.1;refGene_gn=HEBP2;refGene_tx=XM_023254443.1
B2 127674281 127674480
refGene_gn=HEBP2;refGene_tx=XM_023254442.1;refGene_gn=HEBP2;refGene_tx=XM_023254443.1
B2 127689705 127690441
refGene_gn=NHSL1;refGene_tx=XM_019831290.2;refGene_gn=NHSL1;refGene_tx=XM_019831295.2;refGene_gn=NHSL1;refGene_tx=XM_019831296.2;refGene_gn=NHSL1;refGene_tx=XM_019831297.2;refGene_gn=NHSL1;refGene_tx=XM_019831298.2;refGene_gn=NHSL1;refGene_tx=XM_019831299.2;refGene_gn=NHSL1;refGe

ne_tx=XM_023254444.1;refGene_gn=NHSL1;refGene_tx=XM_023254445.1;refGen
e_gn=NHSL1;refGene_tx=XM_023254446.1;refGene_gn=NHSL1;refGene_tx=XM_02
3254447.1
B2 127696048 127696181
refGene_gn=NHSL1;refGene_tx=XM_019831290.2;refGene_gn=NHSL1;refGene_tx
=XM_019831295.2;refGene_gn=NHSL1;refGene_tx=XM_019831296.2;refGene_gn=
NHSL1;refGene_tx=XM_019831297.2;refGene_gn=NHSL1;refGene_tx=XM_0198312
98.2;refGene_gn=NHSL1;refGene_tx=XM_019831299.2;refGene_gn=NHSL1;refGe
ne_tx=XM_023254444.1;refGene_gn=NHSL1;refGene_tx=XM_023254445.1;refGen
e_gn=NHSL1;refGene_tx=XM_023254446.1;refGene_gn=NHSL1;refGene_tx=XM_02
3254447.1
B2 127696700 127699925
refGene_gn=NHSL1;refGene_tx=XM_019831290.2;refGene_gn=NHSL1;refGene_tx
=XM_019831295.2;refGene_gn=NHSL1;refGene_tx=XM_019831296.2;refGene_gn=
NHSL1;refGene_tx=XM_019831297.2;refGene_gn=NHSL1;refGene_tx=XM_0198312
98.2;refGene_gn=NHSL1;refGene_tx=XM_019831299.2;refGene_gn=NHSL1;refGe
ne_tx=XM_023254444.1;refGene_gn=NHSL1;refGene_tx=XM_023254445.1;refGen
e_gn=NHSL1;refGene_tx=XM_023254446.1;refGene_gn=NHSL1;refGene_tx=XM_02
3254447.1
B2 127705891 127706023
refGene_gn=NHSL1;refGene_tx=XM_019831290.2;refGene_gn=NHSL1;refGene_tx
=XM_019831295.2;refGene_gn=NHSL1;refGene_tx=XM_019831297.2;refGene_gn=
NHSL1;refGene_tx=XM_019831298.2;refGene_gn=NHSL1;refGene_tx=XM_0198312
99.2;refGene_gn=NHSL1;refGene_tx=XM_023254444.1;refGene_gn=NHSL1;refGe
ne_tx=XM_023254445.1
B2 127709983 127710176
refGene_gn=NHSL1;refGene_tx=XM_019831290.2;refGene_gn=NHSL1;refGene_tx
=XM_019831295.2;refGene_gn=NHSL1;refGene_tx=XM_019831296.2;refGene_gn=
NHSL1;refGene_tx=XM_019831297.2;refGene_gn=NHSL1;refGene_tx=XM_0198312
98.2;refGene_gn=NHSL1;refGene_tx=XM_019831299.2;refGene_gn=NHSL1;refGe
ne_tx=XM_023254444.1;refGene_gn=NHSL1;refGene_tx=XM_023254445.1;refGen
e_gn=NHSL1;refGene_tx=XM_023254446.1;refGene_gn=NHSL1;refGene_tx=XM_02
3254447.1
B2 127727057 127727185
refGene_gn=NHSL1;refGene_tx=XM_019831290.2;refGene_gn=NHSL1;refGene_tx
=XM_019831295.2;refGene_gn=NHSL1;refGene_tx=XM_019831296.2;refGene_gn=
NHSL1;refGene_tx=XM_019831297.2;refGene_gn=NHSL1;refGene_tx=XM_0198312
98.2;refGene_gn=NHSL1;refGene_tx=XM_019831299.2;refGene_gn=NHSL1;refGe
ne_tx=XM_023254444.1;refGene_gn=NHSL1;refGene_tx=XM_023254445.1;refGen
e_gn=NHSL1;refGene_tx=XM_023254446.1;refGene_gn=NHSL1;refGene_tx=XM_02
3254447.1
B2 127751458 127751611
refGene_gn=NHSL1;refGene_tx=XM_019831290.2;refGene_gn=NHSL1;refGene_tx
=XM_019831295.2;refGene_gn=NHSL1;refGene_tx=XM_019831296.2;refGene_gn=
NHSL1;refGene_tx=XM_019831297.2;refGene_gn=NHSL1;refGene_tx=XM_0198312
98.2;refGene_gn=NHSL1;refGene_tx=XM_019831299.2;refGene_gn=NHSL1;refGe
ne_tx=XM_023254444.1;refGene_gn=NHSL1;refGene_tx=XM_023254445.1;refGen
e_gn=NHSL1;refGene_tx=XM_023254446.1;refGene_gn=NHSL1;refGene_tx=XM_02
3254447.1
B2 127754325 127754383

refGene_gn=NHSL1;refGene_tx=XM_019831295.2;refGene_gn=NHSL1;refGene_tx=XM_019831297.2
B2	127788473	127788501
refGene_gn=NHSL1;refGene_tx=XM_019831298.2
B2	127794118	127794134
refGene_gn=NHSL1;refGene_tx=XM_019831299.2
B2	127818845	127819047
refGene_gn=NHSL1;refGene_tx=XM_019831290.2;refGene_gn=NHSL1;refGene_tx=XM_019831296.2
B2	127930554	127930837
refGene_gn=NHSL1;refGene_tx=XM_023254444.1;refGene_gn=NHSL1;refGene_tx=XM_023254445.1;refGene_gn=NHSL1;refGene_tx=XM_023254446.1;refGene_gn=NHSL1;refGene_tx=XM_023254447.1
B2	127997869	127998076
refGene_gn=CCDC28A;refGene_tx=XM_006944343.3
B2	128000118	128000318
refGene_gn=CCDC28A;refGene_tx=XM_006944343.3
B2	128001255	128001419
refGene_gn=CCDC28A;refGene_tx=XM_006944343.3
B2	128007441	128007596
refGene_gn=CCDC28A;refGene_tx=XM_006944343.3
B2	128009484	128009507
refGene_gn=CCDC28A;refGene_tx=XM_006944343.3
B2	128012589	128012644
refGene_gn=CCDC28A;refGene_tx=XM_006944343.3
B2	128029097	128029199
refGene_gn=ECT2L;refGene_tx=XM_019831302.2;refGene_gn=ECT2L;refGene_tx=XM_023254448.1;refGene_gn=ECT2L;refGene_tx=XM_023254449.1;refGene_gn=ECT2L;refGene_tx=XM_023254450.1;refGene_gn=ECT2L;refGene_tx=XM_023254451.1;refGene_gn=ECT2L;refGene_tx=XM_023254452.1;refGene_gn=ECT2L;refGene_tx=XM_023254453.1;refGene_gn=ECT2L;refGene_tx=XM_023254454.1
B2	128030349	128030462
refGene_gn=ECT2L;refGene_tx=XM_019831302.2;refGene_gn=ECT2L;refGene_tx=XM_023254448.1;refGene_gn=ECT2L;refGene_tx=XM_023254449.1;refGene_gn=ECT2L;refGene_tx=XM_023254450.1;refGene_gn=ECT2L;refGene_tx=XM_023254451.1;refGene_gn=ECT2L;refGene_tx=XM_023254452.1;refGene_gn=ECT2L;refGene_tx=XM_023254453.1;refGene_gn=ECT2L;refGene_tx=XM_023254454.1
B2	128049037	128049200
refGene_gn=ECT2L;refGene_tx=XM_019831302.2;refGene_gn=ECT2L;refGene_tx=XM_023254448.1;refGene_gn=ECT2L;refGene_tx=XM_023254449.1;refGene_gn=ECT2L;refGene_tx=XM_023254450.1;refGene_gn=ECT2L;refGene_tx=XM_023254451.1;refGene_gn=ECT2L;refGene_tx=XM_023254452.1;refGene_gn=ECT2L;refGene_tx=XM_023254453.1;refGene_gn=ECT2L;refGene_tx=XM_023254454.1;refGene_gn=ECT2L;refGene_tx=XM_023254455.1
B2	128052033	128052286
refGene_gn=ECT2L;refGene_tx=XM_019831302.2;refGene_gn=ECT2L;refGene_tx=XM_023254448.1;refGene_gn=ECT2L;refGene_tx=XM_023254449.1;refGene_gn=ECT2L;refGene_tx=XM_023254450.1;refGene_gn=ECT2L;refGene_tx=XM_023254451.1;refGene_gn=ECT2L;refGene_tx=XM_023254452.1;refGene_gn=ECT2L;refGene_tx=XM_023254453.1;refGene_gn=ECT2L;refGene_tx=XM_023254454.1;refGen

e_gn=ECT2L;refGene_tx=XM_023254455.1
B2 128053154 128053323
refGene_gn=ECT2L;refGene_tx=XM_019831302.2;refGene_gn=ECT2L;refGene_tx=XM_023254448.1;refGene_gn=ECT2L;refGene_tx=XM_023254449.1;refGene_gn=ECT2L;refGene_tx=XM_023254450.1;refGene_gn=ECT2L;refGene_tx=XM_023254451.1;refGene_gn=ECT2L;refGene_tx=XM_023254452.1;refGene_gn=ECT2L;refGene_tx=XM_023254453.1;refGene_gn=ECT2L;refGene_tx=XM_023254454.1;refGene_gn=ECT2L;refGene_tx=XM_023254455.1
B2 128054064 128054203
refGene_gn=ECT2L;refGene_tx=XM_019831302.2;refGene_gn=ECT2L;refGene_tx=XM_023254448.1;refGene_gn=ECT2L;refGene_tx=XM_023254449.1;refGene_gn=ECT2L;refGene_tx=XM_023254450.1;refGene_gn=ECT2L;refGene_tx=XM_023254451.1;refGene_gn=ECT2L;refGene_tx=XM_023254452.1;refGene_gn=ECT2L;refGene_tx=XM_023254453.1;refGene_gn=ECT2L;refGene_tx=XM_023254454.1;refGene_gn=ECT2L;refGene_tx=XM_023254455.1
B2 128056467 128056636
refGene_gn=ECT2L;refGene_tx=XM_019831302.2;refGene_gn=ECT2L;refGene_tx=XM_023254448.1;refGene_gn=ECT2L;refGene_tx=XM_023254449.1;refGene_gn=ECT2L;refGene_tx=XM_023254450.1;refGene_gn=ECT2L;refGene_tx=XM_023254451.1;refGene_gn=ECT2L;refGene_tx=XM_023254452.1;refGene_gn=ECT2L;refGene_tx=XM_023254453.1;refGene_gn=ECT2L;refGene_tx=XM_023254454.1;refGene_gn=ECT2L;refGene_tx=XM_023254455.1
B2 128060828 128060957
refGene_gn=ECT2L;refGene_tx=XM_019831302.2;refGene_gn=ECT2L;refGene_tx=XM_023254448.1;refGene_gn=ECT2L;refGene_tx=XM_023254449.1;refGene_gn=ECT2L;refGene_tx=XM_023254450.1;refGene_gn=ECT2L;refGene_tx=XM_023254451.1;refGene_gn=ECT2L;refGene_tx=XM_023254452.1;refGene_gn=ECT2L;refGene_tx=XM_023254454.1;refGene_gn=ECT2L;refGene_tx=XM_023254455.1
B2 128067592 128067685
refGene_gn=ECT2L;refGene_tx=XM_019831302.2;refGene_gn=ECT2L;refGene_tx=XM_023254448.1;refGene_gn=ECT2L;refGene_tx=XM_023254449.1;refGene_gn=ECT2L;refGene_tx=XM_023254450.1;refGene_gn=ECT2L;refGene_tx=XM_023254451.1;refGene_gn=ECT2L;refGene_tx=XM_023254452.1;refGene_gn=ECT2L;refGene_tx=XM_023254453.1;refGene_gn=ECT2L;refGene_tx=XM_023254454.1;refGene_gn=ECT2L;refGene_tx=XM_023254455.1
B2 128069718 128069901
refGene_gn=ECT2L;refGene_tx=XM_019831302.2;refGene_gn=ECT2L;refGene_tx=XM_023254448.1;refGene_gn=ECT2L;refGene_tx=XM_023254449.1;refGene_gn=ECT2L;refGene_tx=XM_023254450.1;refGene_gn=ECT2L;refGene_tx=XM_023254451.1;refGene_gn=ECT2L;refGene_tx=XM_023254452.1;refGene_gn=ECT2L;refGene_tx=XM_023254453.1;refGene_gn=ECT2L;refGene_tx=XM_023254454.1;refGene_gn=ECT2L;refGene_tx=XM_023254455.1
B2 128071935 128072039
refGene_gn=ECT2L;refGene_tx=XM_019831302.2
B2 128073293 128073397
refGene_gn=ECT2L;refGene_tx=XM_023254448.1;refGene_gn=ECT2L;refGene_tx=XM_023254449.1;refGene_gn=ECT2L;refGene_tx=XM_023254450.1;refGene_gn=ECT2L;refGene_tx=XM_023254451.1;refGene_gn=ECT2L;refGene_tx=XM_023254452.1;refGene_gn=ECT2L;refGene_tx=XM_023254453.1;refGene_gn=ECT2L;refGene_tx=XM_023254454.1;refGene_gn=ECT2L;refGene_tx=XM_023254455.1

B2 128074261 128074387
refGene_gn=ECT2L;refGene_tx=XM_019831302.2;refGene_gn=ECT2L;refGene_tx=XM_023254448.1;refGene_gn=ECT2L;refGene_tx=XM_023254449.1;refGene_gn=ECT2L;refGene_tx=XM_023254450.1;refGene_gn=ECT2L;refGene_tx=XM_023254451.1;refGene_gn=ECT2L;refGene_tx=XM_023254452.1;refGene_gn=ECT2L;refGene_tx=XM_023254453.1;refGene_gn=ECT2L;refGene_tx=XM_023254454.1;refGene_gn=ECT2L;refGene_tx=XM_023254455.1
B2 128079048 128079135
refGene_gn=ECT2L;refGene_tx=XM_019831302.2;refGene_gn=ECT2L;refGene_tx=XM_023254448.1;refGene_gn=ECT2L;refGene_tx=XM_023254449.1;refGene_gn=ECT2L;refGene_tx=XM_023254450.1;refGene_gn=ECT2L;refGene_tx=XM_023254451.1;refGene_gn=ECT2L;refGene_tx=XM_023254452.1;refGene_gn=ECT2L;refGene_tx=XM_023254453.1;refGene_gn=ECT2L;refGene_tx=XM_023254454.1;refGene_gn=ECT2L;refGene_tx=XM_023254455.1
B2 128081781 128081996
refGene_gn=ECT2L;refGene_tx=XM_019831302.2;refGene_gn=ECT2L;refGene_tx=XM_023254448.1;refGene_gn=ECT2L;refGene_tx=XM_023254449.1;refGene_gn=ECT2L;refGene_tx=XM_023254450.1;refGene_gn=ECT2L;refGene_tx=XM_023254451.1;refGene_gn=ECT2L;refGene_tx=XM_023254452.1;refGene_gn=ECT2L;refGene_tx=XM_023254453.1;refGene_gn=ECT2L;refGene_tx=XM_023254454.1;refGene_gn=ECT2L;refGene_tx=XM_023254455.1
B2 128083299 128083447
refGene_gn=ECT2L;refGene_tx=XM_019831302.2;refGene_gn=ECT2L;refGene_tx=XM_023254448.1;refGene_gn=ECT2L;refGene_tx=XM_023254449.1;refGene_gn=ECT2L;refGene_tx=XM_023254450.1;refGene_gn=ECT2L;refGene_tx=XM_023254451.1;refGene_gn=ECT2L;refGene_tx=XM_023254452.1;refGene_gn=ECT2L;refGene_tx=XM_023254453.1;refGene_gn=ECT2L;refGene_tx=XM_023254454.1;refGene_gn=ECT2L;refGene_tx=XM_023254455.1
B2 128084867 128084941
refGene_gn=ECT2L;refGene_tx=XM_019831302.2;refGene_gn=ECT2L;refGene_tx=XM_023254448.1;refGene_gn=ECT2L;refGene_tx=XM_023254449.1;refGene_gn=ECT2L;refGene_tx=XM_023254450.1;refGene_gn=ECT2L;refGene_tx=XM_023254451.1;refGene_gn=ECT2L;refGene_tx=XM_023254452.1;refGene_gn=ECT2L;refGene_tx=XM_023254453.1;refGene_gn=ECT2L;refGene_tx=XM_023254454.1;refGene_gn=ECT2L;refGene_tx=XM_023254455.1
B2 128085040 128085197
refGene_gn=ECT2L;refGene_tx=XM_019831302.2;refGene_gn=ECT2L;refGene_tx=XM_023254448.1;refGene_gn=ECT2L;refGene_tx=XM_023254449.1;refGene_gn=ECT2L;refGene_tx=XM_023254450.1;refGene_gn=ECT2L;refGene_tx=XM_023254451.1;refGene_gn=ECT2L;refGene_tx=XM_023254452.1;refGene_gn=ECT2L;refGene_tx=XM_023254453.1;refGene_gn=ECT2L;refGene_tx=XM_023254454.1;refGene_gn=ECT2L;refGene_tx=XM_023254455.1
B2 128086353 128086419
refGene_gn=ECT2L;refGene_tx=XM_019831302.2;refGene_gn=ECT2L;refGene_tx=XM_023254448.1;refGene_gn=ECT2L;refGene_tx=XM_023254449.1;refGene_gn=ECT2L;refGene_tx=XM_023254450.1;refGene_gn=ECT2L;refGene_tx=XM_023254451.1;refGene_gn=ECT2L;refGene_tx=XM_023254453.1;refGene_gn=ECT2L;refGene_tx=XM_023254454.1;refGene_gn=ECT2L;refGene_tx=XM_023254455.1
B2 128088371 128088460
refGene_gn=ECT2L;refGene_tx=XM_019831302.2;refGene_gn=ECT2L;refGene_tx

=XM_023254448.1;refGene_gn=ECT2L;refGene_tx=XM_023254449.1;refGene_gn=ECT2L;refGene_tx=XM_023254450.1;refGene_gn=ECT2L;refGene_tx=XM_023254451.1;refGene_gn=ECT2L;refGene_tx=XM_023254452.1;refGene_gn=ECT2L;refGene_tx=XM_023254453.1;refGene_gn=ECT2L;refGene_tx=XM_023254454.1;refGene_gn=ECT2L;refGene_tx=XM_023254455.1
B2 128088631 128088692
refGene_gn=ECT2L;refGene_tx=XM_023254454.1
B2 128097133 128097306
refGene_gn=ECT2L;refGene_tx=XM_019831302.2;refGene_gn=ECT2L;refGene_tx=XM_023254448.1;refGene_gn=ECT2L;refGene_tx=XM_023254449.1;refGene_gn=ECT2L;refGene_tx=XM_023254450.1;refGene_gn=ECT2L;refGene_tx=XM_023254451.1;refGene_gn=ECT2L;refGene_tx=XM_023254452.1;refGene_gn=ECT2L;refGene_tx=XM_023254453.1;refGene_gn=ECT2L;refGene_tx=XM_023254455.1
B2 128098979 128099107
refGene_gn=ECT2L;refGene_tx=XM_019831302.2;refGene_gn=ECT2L;refGene_tx=XM_023254448.1;refGene_gn=ECT2L;refGene_tx=XM_023254449.1;refGene_gn=ECT2L;refGene_tx=XM_023254450.1;refGene_gn=ECT2L;refGene_tx=XM_023254451.1;refGene_gn=ECT2L;refGene_tx=XM_023254452.1;refGene_gn=ECT2L;refGene_tx=XM_023254453.1;refGene_gn=ECT2L;refGene_tx=XM_023254455.1
B2 128101353 128101422
refGene_gn=REPS1;refGene_tx=XM_003986599.4;refGene_gn=REPS1;refGene_tx=XM_003986600.4;refGene_gn=REPS1;refGene_tx=XM_006932050.3;refGene_gn=REPS1;refGene_tx=XM_006932051.3;refGene_gn=REPS1;refGene_tx=XM_006932052.3;refGene_gn=REPS1;refGene_tx=XM_019831316.2;refGene_gn=REPS1;refGene_tx=XM_019831317.2
B2 128103428 128103534
refGene_gn=REPS1;refGene_tx=XM_003986599.4;refGene_gn=REPS1;refGene_tx=XM_003986600.4;refGene_gn=REPS1;refGene_tx=XM_006932050.3;refGene_gn=REPS1;refGene_tx=XM_006932051.3;refGene_gn=REPS1;refGene_tx=XM_006932052.3;refGene_gn=REPS1;refGene_tx=XM_019831316.2;refGene_gn=REPS1;refGene_tx=XM_019831317.2
B2 128104230 128104379
refGene_gn=REPS1;refGene_tx=XM_003986599.4;refGene_gn=REPS1;refGene_tx=XM_003986600.4;refGene_gn=REPS1;refGene_tx=XM_006932050.3;refGene_gn=REPS1;refGene_tx=XM_006932051.3;refGene_gn=REPS1;refGene_tx=XM_006932052.3;refGene_gn=REPS1;refGene_tx=XM_019831316.2;refGene_gn=REPS1;refGene_tx=XM_019831317.2
B2 128106241 128106337
refGene_gn=REPS1;refGene_tx=XM_003986599.4;refGene_gn=REPS1;refGene_tx=XM_003986600.4;refGene_gn=REPS1;refGene_tx=XM_006932050.3;refGene_gn=REPS1;refGene_tx=XM_006932051.3;refGene_gn=REPS1;refGene_tx=XM_006932052.3;refGene_gn=REPS1;refGene_tx=XM_019831316.2
B2 128107318 128107504
refGene_gn=REPS1;refGene_tx=XM_003986599.4;refGene_gn=REPS1;refGene_tx=XM_003986600.4;refGene_gn=REPS1;refGene_tx=XM_006932050.3;refGene_gn=REPS1;refGene_tx=XM_006932051.3;refGene_gn=REPS1;refGene_tx=XM_006932052.3;refGene_gn=REPS1;refGene_tx=XM_019831316.2;refGene_gn=REPS1;refGene_tx=XM_019831317.2
B2 128109416 128109481
refGene_gn=REPS1;refGene_tx=XM_003986599.4;refGene_gn=REPS1;refGene_tx

=XM_003986600.4;refGene_gn=REPS1;refGene_tx=XM_006932050.3;refGene_gn=REPS1;refGene_tx=XM_006932051.3;refGene_gn=REPS1;refGene_tx=XM_006932052.3;refGene_gn=REPS1;refGene_tx=XM_019831316.2;refGene_gn=REPS1;refGene_tx=XM_019831317.2
B2 128110291 128110410
refGene_gn=REPS1;refGene_tx=XM_003986599.4;refGene_gn=REPS1;refGene_tx=XM_003986600.4;refGene_gn=REPS1;refGene_tx=XM_006932050.3;refGene_gn=REPS1;refGene_tx=XM_006932051.3;refGene_gn=REPS1;refGene_tx=XM_006932052.3;refGene_gn=REPS1;refGene_tx=XM_019831316.2;refGene_gn=REPS1;refGene_tx=XM_019831317.2
B2 128111726 128111799
refGene_gn=REPS1;refGene_tx=XM_003986599.4;refGene_gn=REPS1;refGene_tx=XM_003986600.4;refGene_gn=REPS1;refGene_tx=XM_006932050.3;refGene_gn=REPS1;refGene_tx=XM_006932051.3;refGene_gn=REPS1;refGene_tx=XM_006932052.3;refGene_gn=REPS1;refGene_tx=XM_019831316.2;refGene_gn=REPS1;refGene_tx=XM_019831317.2
B2 128113534 128113636
refGene_gn=REPS1;refGene_tx=XM_003986599.4;refGene_gn=REPS1;refGene_tx=XM_003986600.4;refGene_gn=REPS1;refGene_tx=XM_006932050.3;refGene_gn=REPS1;refGene_tx=XM_006932051.3;refGene_gn=REPS1;refGene_tx=XM_006932052.3;refGene_gn=REPS1;refGene_tx=XM_019831316.2;refGene_gn=REPS1;refGene_tx=XM_019831317.2
B2 128114844 128114932
refGene_gn=REPS1;refGene_tx=XM_003986599.4;refGene_gn=REPS1;refGene_tx=XM_003986600.4;refGene_gn=REPS1;refGene_tx=XM_006932050.3;refGene_gn=REPS1;refGene_tx=XM_006932051.3;refGene_gn=REPS1;refGene_tx=XM_006932052.3;refGene_gn=REPS1;refGene_tx=XM_019831316.2;refGene_gn=REPS1;refGene_tx=XM_019831317.2
B2 128120404 128120485
refGene_gn=REPS1;refGene_tx=XM_003986600.4;refGene_gn=REPS1;refGene_tx=XM_006932050.3;refGene_gn=REPS1;refGene_tx=XM_006932051.3;refGene_gn=REPS1;refGene_tx=XM_019831316.2;refGene_gn=REPS1;refGene_tx=XM_019831317.2
B2 128123972 128124094
refGene_gn=REPS1;refGene_tx=XM_003986599.4;refGene_gn=REPS1;refGene_tx=XM_003986600.4;refGene_gn=REPS1;refGene_tx=XM_006932050.3;refGene_gn=REPS1;refGene_tx=XM_006932051.3;refGene_gn=REPS1;refGene_tx=XM_006932052.3;refGene_gn=REPS1;refGene_tx=XM_019831316.2;refGene_gn=REPS1;refGene_tx=XM_019831317.2
B2 128135219 128135374
refGene_gn=REPS1;refGene_tx=XM_003986599.4;refGene_gn=REPS1;refGene_tx=XM_003986600.4;refGene_gn=REPS1;refGene_tx=XM_006932050.3;refGene_gn=REPS1;refGene_tx=XM_006932051.3;refGene_gn=REPS1;refGene_tx=XM_006932052.3;refGene_gn=REPS1;refGene_tx=XM_019831316.2;refGene_gn=REPS1;refGene_tx=XM_019831317.2
B2 128136693 128136757
refGene_gn=REPS1;refGene_tx=XM_003986599.4;refGene_gn=REPS1;refGene_tx=XM_003986600.4;refGene_gn=REPS1;refGene_tx=XM_006932050.3;refGene_gn=REPS1;refGene_tx=XM_006932051.3;refGene_gn=REPS1;refGene_tx=XM_006932052.3;refGene_gn=REPS1;refGene_tx=XM_019831316.2;refGene_gn=REPS1;refGe

ne_tx=XM_019831317.2
B2 128137038 128137201
refGene_gn=REPS1;refGene_tx=XM_003986599.4;refGene_gn=REPS1;refGene_tx=XM_003986600.4;refGene_gn=REPS1;refGene_tx=XM_006932050.3;refGene_gn=REPS1;refGene_tx=XM_006932051.3;refGene_gn=REPS1;refGene_tx=XM_006932052.3;refGene_gn=REPS1;refGene_tx=XM_019831316.2;refGene_gn=REPS1;refGene_tx=XM_019831317.2
B2 128137722 128137847
refGene_gn=REPS1;refGene_tx=XM_003986599.4;refGene_gn=REPS1;refGene_tx=XM_003986600.4;refGene_gn=REPS1;refGene_tx=XM_006932050.3;refGene_gn=REPS1;refGene_tx=XM_006932051.3;refGene_gn=REPS1;refGene_tx=XM_006932052.3;refGene_gn=REPS1;refGene_tx=XM_019831316.2;refGene_gn=REPS1;refGene_tx=XM_019831317.2
B2 128138636 128138790
refGene_gn=REPS1;refGene_tx=XM_003986599.4;refGene_gn=REPS1;refGene_tx=XM_003986600.4;refGene_gn=REPS1;refGene_tx=XM_006932050.3;refGene_gn=REPS1;refGene_tx=XM_006932051.3;refGene_gn=REPS1;refGene_tx=XM_006932052.3;refGene_gn=REPS1;refGene_tx=XM_019831316.2;refGene_gn=REPS1;refGene_tx=XM_019831317.2
B2 128138910 128139107
refGene_gn=REPS1;refGene_tx=XM_003986599.4;refGene_gn=REPS1;refGene_tx=XM_003986600.4;refGene_gn=REPS1;refGene_tx=XM_006932050.3;refGene_gn=REPS1;refGene_tx=XM_006932051.3;refGene_gn=REPS1;refGene_tx=XM_006932052.3;refGene_gn=REPS1;refGene_tx=XM_019831316.2;refGene_gn=REPS1;refGene_tx=XM_019831317.2
B2 128140634 128140758
refGene_gn=REPS1;refGene_tx=XM_003986599.4;refGene_gn=REPS1;refGene_tx=XM_003986600.4;refGene_gn=REPS1;refGene_tx=XM_006932050.3;refGene_gn=REPS1;refGene_tx=XM_006932051.3;refGene_gn=REPS1;refGene_tx=XM_006932052.3;refGene_gn=REPS1;refGene_tx=XM_019831316.2;refGene_gn=REPS1;refGene_tx=XM_019831317.2
B2 128187705 128187858
refGene_gn=REPS1;refGene_tx=XM_003986599.4;refGene_gn=REPS1;refGene_tx=XM_003986600.4;refGene_gn=REPS1;refGene_tx=XM_006932050.3;refGene_gn=REPS1;refGene_tx=XM_006932051.3;refGene_gn=REPS1;refGene_tx=XM_006932052.3;refGene_gn=REPS1;refGene_tx=XM_019831316.2;refGene_gn=REPS1;refGene_tx=XM_019831317.2
B2 128223273 128223334
refGene_gn=ABRACL;refGene_tx=XM_003986602.5;refGene_gn=ABRACL;refGene_tx=XM_006932053.4
B2 128231472 128231657
refGene_gn=ABRACL;refGene_tx=XM_003986602.5;refGene_gn=ABRACL;refGene_tx=XM_006932053.4
B2 128315690 128315961
refGene_gn=HECA;refGene_tx=XM_023254456.1
B2 128353029 128354070
refGene_gn=HECA;refGene_tx=XM_023254456.1
B2 128360130 128360285
refGene_gn=HECA;refGene_tx=XM_023254456.1
B2 128363434 128363599

refGene_gn=HECA;refGene_tx=XM_023254456.1
B2	128405541	128406342
refGene_gn=TXLNB;refGene_tx=XM_003986611.5;refGene_gn=TXLNB;refGene_tx=XM_006932054.4
B2	128407570	128407666
refGene_gn=TXLNB;refGene_tx=XM_003986611.5;refGene_gn=TXLNB;refGene_tx=XM_006932054.4
B2	128417058	128417151
refGene_gn=TXLNB;refGene_tx=XM_003986611.5;refGene_gn=TXLNB;refGene_tx=XM_006932054.4
B2	128423749	128423824
refGene_gn=TXLNB;refGene_tx=XM_003986611.5;refGene_gn=TXLNB;refGene_tx=XM_006932054.4
B2	128430052	128430172
refGene_gn=TXLNB;refGene_tx=XM_003986611.5;refGene_gn=TXLNB;refGene_tx=XM_006932054.4
B2	128432003	128432198
refGene_gn=TXLNB;refGene_tx=XM_003986611.5;refGene_gn=TXLNB;refGene_tx=XM_006932054.4
B2	128439592	128439763
refGene_gn=TXLNB;refGene_tx=XM_003986611.5;refGene_gn=TXLNB;refGene_tx=XM_006932054.4
B2	128445449	128445541
refGene_gn=TXLNB;refGene_tx=XM_003986611.5;refGene_gn=TXLNB;refGene_tx=XM_006932054.4
B2	128454743	128455170
refGene_gn=TXLNB;refGene_tx=XM_003986611.5;refGene_gn=TXLNB;refGene_tx=XM_006932054.4
B2	128543020	128543842
refGene_gn=CITED2;refGene_tx=XM_023254457.1
B2	130900060	130900462
refGene_gn=NMBR;refGene_tx=XM_003986612.4
B2	130903484	130903833
refGene_gn=NMBR;refGene_tx=XM_003986612.4
B2	130916709	130917131
refGene_gn=NMBR;refGene_tx=XM_003986612.4
B2	130943361	130943400
refGene_gn=LOC109500226;refGene_tx=XM_019831654.1
B2	130944086	130944269
refGene_gn=LOC109500226;refGene_tx=XM_019831654.1
B2	130944800	130945196
refGene_gn=LOC109500226;refGene_tx=XM_019831654.1
B2	130954677	130954789
refGene_gn=VTA1;refGene_tx=XM_003986613.5;refGene_gn=VTA1;refGene_tx=XM_006932060.4
B2	130976410	130976505
refGene_gn=VTA1;refGene_tx=XM_003986613.5;refGene_gn=VTA1;refGene_tx=XM_006932060.4
B2	130979318	130979446
refGene_gn=VTA1;refGene_tx=XM_003986613.5;refGene_gn=VTA1;refGene_tx=X

M_006932060.4
B2	130980113	130980189
refGene_gn=VTA1;refGene_tx=XM_003986613.5;refGene_gn=VTA1;refGene_tx=XM_006932060.4
B2	130996872	130996981
refGene_gn=VTA1;refGene_tx=XM_003986613.5;refGene_gn=VTA1;refGene_tx=XM_006932060.4
B2	131009774	131009948
refGene_gn=VTA1;refGene_tx=XM_003986613.5;refGene_gn=VTA1;refGene_tx=XM_006932060.4
B2	131015994	131016075
refGene_gn=VTA1;refGene_tx=XM_003986613.5;refGene_gn=VTA1;refGene_tx=XM_006932060.4
B2	131023689	131023835
refGene_gn=VTA1;refGene_tx=XM_003986613.5;refGene_gn=VTA1;refGene_tx=XM_006932060.4
B2	131100512	131100514
refGene_gn=ADGRG6;refGene_tx=XM_006932061.4;refGene_gn=ADGRG6;refGene_tx=XM_006932062.4;refGene_gn=ADGRG6;refGene_tx=XM_019831320.2;refGene_gn=ADGRG6;refGene_tx=XM_019831321.2;refGene_gn=ADGRG6;refGene_tx=XM_019831322.2
B2	131108454	131108555
refGene_gn=ADGRG6;refGene_tx=XM_006932061.4;refGene_gn=ADGRG6;refGene_tx=XM_006932062.4;refGene_gn=ADGRG6;refGene_tx=XM_019831319.2;refGene_gn=ADGRG6;refGene_tx=XM_019831320.2;refGene_gn=ADGRG6;refGene_tx=XM_019831321.2;refGene_gn=ADGRG6;refGene_tx=XM_019831322.2
B2	131166104	131166446
refGene_gn=ADGRG6;refGene_tx=XM_006932061.4;refGene_gn=ADGRG6;refGene_tx=XM_006932062.4;refGene_gn=ADGRG6;refGene_tx=XM_019831319.2;refGene_gn=ADGRG6;refGene_tx=XM_019831320.2;refGene_gn=ADGRG6;refGene_tx=XM_019831321.2;refGene_gn=ADGRG6;refGene_tx=XM_019831322.2
B2	131170300	131170924
refGene_gn=ADGRG6;refGene_tx=XM_006932061.4;refGene_gn=ADGRG6;refGene_tx=XM_006932062.4;refGene_gn=ADGRG6;refGene_tx=XM_019831319.2;refGene_gn=ADGRG6;refGene_tx=XM_019831320.2;refGene_gn=ADGRG6;refGene_tx=XM_019831321.2;refGene_gn=ADGRG6;refGene_tx=XM_019831322.2
B2	131179587	131179656
refGene_gn=ADGRG6;refGene_tx=XM_006932061.4;refGene_gn=ADGRG6;refGene_tx=XM_006932062.4;refGene_gn=ADGRG6;refGene_tx=XM_019831319.2;refGene_gn=ADGRG6;refGene_tx=XM_019831320.2;refGene_gn=ADGRG6;refGene_tx=XM_019831321.2;refGene_gn=ADGRG6;refGene_tx=XM_019831322.2
B2	131182124	131182208
refGene_gn=ADGRG6;refGene_tx=XM_006932061.4;refGene_gn=ADGRG6;refGene_tx=XM_019831319.2;refGene_gn=ADGRG6;refGene_tx=XM_019831320.2;refGene_gn=ADGRG6;refGene_tx=XM_019831321.2
B2	131187819	131187905
refGene_gn=ADGRG6;refGene_tx=XM_006932061.4;refGene_gn=ADGRG6;refGene_tx=XM_006932062.4;refGene_gn=ADGRG6;refGene_tx=XM_019831319.2;refGene_gn=ADGRG6;refGene_tx=XM_019831320.2;refGene_gn=ADGRG6;refGene_tx=XM_019831321.2;refGene_gn=ADGRG6;refGene_tx=XM_019831322.2

B2	131191970	131192023
refGene_gn=ADGRG6;refGene_tx=XM_006932061.4;refGene_gn=ADGRG6;refGene_tx=XM_006932062.4;refGene_gn=ADGRG6;refGene_tx=XM_019831319.2;refGene_gn=ADGRG6;refGene_tx=XM_019831320.2;refGene_gn=ADGRG6;refGene_tx=XM_019831321.2;refGene_gn=ADGRG6;refGene_tx=XM_019831322.2
B2	131193005	131193068
refGene_gn=ADGRG6;refGene_tx=XM_006932061.4;refGene_gn=ADGRG6;refGene_tx=XM_006932062.4;refGene_gn=ADGRG6;refGene_tx=XM_019831319.2;refGene_gn=ADGRG6;refGene_tx=XM_019831320.2;refGene_gn=ADGRG6;refGene_tx=XM_019831321.2;refGene_gn=ADGRG6;refGene_tx=XM_019831322.2
B2	131196161	131196304
refGene_gn=ADGRG6;refGene_tx=XM_006932061.4;refGene_gn=ADGRG6;refGene_tx=XM_006932062.4;refGene_gn=ADGRG6;refGene_tx=XM_019831319.2;refGene_gn=ADGRG6;refGene_tx=XM_019831320.2;refGene_gn=ADGRG6;refGene_tx=XM_019831321.2;refGene_gn=ADGRG6;refGene_tx=XM_019831322.2
B2	131199528	131199640
refGene_gn=ADGRG6;refGene_tx=XM_006932061.4;refGene_gn=ADGRG6;refGene_tx=XM_006932062.4;refGene_gn=ADGRG6;refGene_tx=XM_019831319.2;refGene_gn=ADGRG6;refGene_tx=XM_019831320.2;refGene_gn=ADGRG6;refGene_tx=XM_019831321.2;refGene_gn=ADGRG6;refGene_tx=XM_019831322.2
B2	131200972	131201037
refGene_gn=ADGRG6;refGene_tx=XM_006932061.4;refGene_gn=ADGRG6;refGene_tx=XM_006932062.4;refGene_gn=ADGRG6;refGene_tx=XM_019831319.2;refGene_gn=ADGRG6;refGene_tx=XM_019831320.2;refGene_gn=ADGRG6;refGene_tx=XM_019831321.2;refGene_gn=ADGRG6;refGene_tx=XM_019831322.2
B2	131201483	131201694
refGene_gn=ADGRG6;refGene_tx=XM_006932061.4;refGene_gn=ADGRG6;refGene_tx=XM_006932062.4;refGene_gn=ADGRG6;refGene_tx=XM_019831319.2;refGene_gn=ADGRG6;refGene_tx=XM_019831320.2;refGene_gn=ADGRG6;refGene_tx=XM_019831321.2;refGene_gn=ADGRG6;refGene_tx=XM_019831322.2
B2	131202928	131203100
refGene_gn=ADGRG6;refGene_tx=XM_006932061.4;refGene_gn=ADGRG6;refGene_tx=XM_006932062.4;refGene_gn=ADGRG6;refGene_tx=XM_019831319.2;refGene_gn=ADGRG6;refGene_tx=XM_019831320.2;refGene_gn=ADGRG6;refGene_tx=XM_019831321.2;refGene_gn=ADGRG6;refGene_tx=XM_019831322.2
B2	131204503	131204644
refGene_gn=ADGRG6;refGene_tx=XM_006932061.4;refGene_gn=ADGRG6;refGene_tx=XM_006932062.4;refGene_gn=ADGRG6;refGene_tx=XM_019831319.2;refGene_gn=ADGRG6;refGene_tx=XM_019831320.2;refGene_gn=ADGRG6;refGene_tx=XM_019831321.2;refGene_gn=ADGRG6;refGene_tx=XM_019831322.2
B2	131207021	131207141
refGene_gn=ADGRG6;refGene_tx=XM_006932061.4;refGene_gn=ADGRG6;refGene_tx=XM_006932062.4;refGene_gn=ADGRG6;refGene_tx=XM_019831319.2;refGene_gn=ADGRG6;refGene_tx=XM_019831320.2;refGene_gn=ADGRG6;refGene_tx=XM_019831321.2;refGene_gn=ADGRG6;refGene_tx=XM_019831322.2
B2	131208801	131208847
refGene_gn=ADGRG6;refGene_tx=XM_006932061.4;refGene_gn=ADGRG6;refGene_tx=XM_006932062.4;refGene_gn=ADGRG6;refGene_tx=XM_019831319.2;refGene_gn=ADGRG6;refGene_tx=XM_019831320.2;refGene_gn=ADGRG6;refGene_tx=XM_019831321.2;refGene_gn=ADGRG6;refGene_tx=XM_019831322.2

B2 131210480 131210587
refGene_gn=ADGRG6;refGene_tx=XM_006932061.4;refGene_gn=ADGRG6;refGene_tx=XM_006932062.4;refGene_gn=ADGRG6;refGene_tx=XM_019831319.2;refGene_gn=ADGRG6;refGene_tx=XM_019831320.2;refGene_gn=ADGRG6;refGene_tx=XM_019831321.2;refGene_gn=ADGRG6;refGene_tx=XM_019831322.2
B2 131214303 131214431
refGene_gn=ADGRG6;refGene_tx=XM_006932061.4;refGene_gn=ADGRG6;refGene_tx=XM_006932062.4;refGene_gn=ADGRG6;refGene_tx=XM_019831319.2;refGene_gn=ADGRG6;refGene_tx=XM_019831320.2;refGene_gn=ADGRG6;refGene_tx=XM_019831321.2;refGene_gn=ADGRG6;refGene_tx=XM_019831322.2
B2 131215590 131215859
refGene_gn=ADGRG6;refGene_tx=XM_006932061.4;refGene_gn=ADGRG6;refGene_tx=XM_006932062.4;refGene_gn=ADGRG6;refGene_tx=XM_019831319.2;refGene_gn=ADGRG6;refGene_tx=XM_019831320.2;refGene_gn=ADGRG6;refGene_tx=XM_019831321.2;refGene_gn=ADGRG6;refGene_tx=XM_019831322.2
B2 131217531 131217628
refGene_gn=ADGRG6;refGene_tx=XM_006932061.4;refGene_gn=ADGRG6;refGene_tx=XM_006932062.4;refGene_gn=ADGRG6;refGene_tx=XM_019831319.2;refGene_gn=ADGRG6;refGene_tx=XM_019831320.2;refGene_gn=ADGRG6;refGene_tx=XM_019831321.2;refGene_gn=ADGRG6;refGene_tx=XM_019831322.2
B2 131220207 131220491
refGene_gn=ADGRG6;refGene_tx=XM_006932061.4;refGene_gn=ADGRG6;refGene_tx=XM_006932062.4;refGene_gn=ADGRG6;refGene_tx=XM_019831319.2;refGene_gn=ADGRG6;refGene_tx=XM_019831320.2;refGene_gn=ADGRG6;refGene_tx=XM_019831321.2;refGene_gn=ADGRG6;refGene_tx=XM_019831322.2
B2 131233981 131234083
refGene_gn=ADGRG6;refGene_tx=XM_006932061.4;refGene_gn=ADGRG6;refGene_tx=XM_006932062.4;refGene_gn=ADGRG6;refGene_tx=XM_019831319.2;refGene_gn=ADGRG6;refGene_tx=XM_019831320.2;refGene_gn=ADGRG6;refGene_tx=XM_019831321.2;refGene_gn=ADGRG6;refGene_tx=XM_019831322.2
B2 131234715 131234868
refGene_gn=ADGRG6;refGene_tx=XM_006932061.4;refGene_gn=ADGRG6;refGene_tx=XM_006932062.4;refGene_gn=ADGRG6;refGene_tx=XM_019831319.2;refGene_gn=ADGRG6;refGene_tx=XM_019831320.2;refGene_gn=ADGRG6;refGene_tx=XM_019831321.2;refGene_gn=ADGRG6;refGene_tx=XM_019831322.2
B2 131237420 131237434
refGene_gn=ADGRG6;refGene_tx=XM_019831321.2;refGene_gn=ADGRG6;refGene_tx=XM_019831322.2
B2 131239831 131240010
refGene_gn=ADGRG6;refGene_tx=XM_006932061.4;refGene_gn=ADGRG6;refGene_tx=XM_006932062.4;refGene_gn=ADGRG6;refGene_tx=XM_019831319.2;refGene_gn=ADGRG6;refGene_tx=XM_019831320.2
B2 131515835 131516660
refGene_gn=HIVEP2;refGene_tx=XM_023254458.1
B2 131522063 131522959
refGene_gn=HIVEP2;refGene_tx=XM_023254458.1
B2 131523744 131523846
refGene_gn=HIVEP2;refGene_tx=XM_023254458.1
B2 131527805 131527981
refGene_gn=HIVEP2;refGene_tx=XM_023254458.1

B2	131531781	131531936
refGene_gn=HIVEP2;refGene_tx=XM_023254458.1
B2	131532912	131538081
refGene_gn=HIVEP2;refGene_tx=XM_023254458.1
B2	131798256	131798397
refGene_gn=AIG1;refGene_tx=XM_003986615.5;refGene_gn=AIG1;refGene_tx=XM_006932063.4;refGene_gn=AIG1;refGene_tx=XM_006932065.4;refGene_gn=AIG1;refGene_tx=XM_019831325.2;refGene_gn=AIG1;refGene_tx=XM_019831327.2;refGene_gn=AIG1;refGene_tx=XM_019831328.2;refGene_gn=AIG1;refGene_tx=XM_019831329.2
B2	131856185	131856341
refGene_gn=AIG1;refGene_tx=XM_003986615.5;refGene_gn=AIG1;refGene_tx=XM_006932063.4;refGene_gn=AIG1;refGene_tx=XM_006932065.4;refGene_gn=AIG1;refGene_tx=XM_019831325.2;refGene_gn=AIG1;refGene_tx=XM_019831327.2;refGene_gn=AIG1;refGene_tx=XM_019831328.2;refGene_gn=AIG1;refGene_tx=XM_019831329.2
B2	131883986	131884088
refGene_gn=AIG1;refGene_tx=XM_003986615.5;refGene_gn=AIG1;refGene_tx=XM_006932063.4;refGene_gn=AIG1;refGene_tx=XM_006932065.4;refGene_gn=AIG1;refGene_tx=XM_019831325.2;refGene_gn=AIG1;refGene_tx=XM_019831328.2;refGene_gn=AIG1;refGene_tx=XM_019831329.2
B2	131914886	131914937
refGene_gn=AIG1;refGene_tx=XM_019831329.2
B2	131917930	131918203
refGene_gn=AIG1;refGene_tx=XM_019831325.2
B2	131996094	131996210
refGene_gn=AIG1;refGene_tx=XM_003986615.5;refGene_gn=AIG1;refGene_tx=XM_006932063.4;refGene_gn=AIG1;refGene_tx=XM_006932065.4;refGene_gn=AIG1;refGene_tx=XM_019831327.2;refGene_gn=AIG1;refGene_tx=XM_019831328.2
B2	132034610	132034774
refGene_gn=AIG1;refGene_tx=XM_003986615.5;refGene_gn=AIG1;refGene_tx=XM_006932063.4;refGene_gn=AIG1;refGene_tx=XM_006932065.4;refGene_gn=AIG1;refGene_tx=XM_019831327.2
B2	132038981	132039052
refGene_gn=AIG1;refGene_tx=XM_006932063.4;refGene_gn=AIG1;refGene_tx=XM_019831327.2
B2	132040477	132040491
refGene_gn=AIG1;refGene_tx=XM_006932065.4
B2	132040885	132040923
refGene_gn=AIG1;refGene_tx=XM_003986615.5
B2	132111483	132111535
refGene_gn=AIG1;refGene_tx=XM_019831328.2
B2	132138060	132138104
refGene_gn=ADAT2;refGene_tx=XM_003986629.5
B2	132138238	132138311
refGene_gn=ADAT2;refGene_tx=XM_003986629.5
B2	132143016	132143123
refGene_gn=ADAT2;refGene_tx=XM_003986629.5
B2	132144614	132144765
refGene_gn=ADAT2;refGene_tx=XM_003986629.5

B2 132147748 132147853
refGene_gn=ADAT2;refGene_tx=XM_003986629.5
B2 132166796 132166895
refGene_gn=ADAT2;refGene_tx=XM_003986629.5
B2 132167300 132167373
refGene_gn=PEX3;refGene_tx=XM_003986617.5
B2 132173061 132173193
refGene_gn=PEX3;refGene_tx=XM_003986617.5
B2 132177403 132177485
refGene_gn=PEX3;refGene_tx=XM_003986617.5
B2 132183347 132183391
refGene_gn=PEX3;refGene_tx=XM_003986617.5
B2 132184935 132185060
refGene_gn=PEX3;refGene_tx=XM_003986617.5
B2 132185454 132185521
refGene_gn=PEX3;refGene_tx=XM_003986617.5
B2 132185628 132185683
refGene_gn=PEX3;refGene_tx=XM_003986617.5
B2 132186265 132186434
refGene_gn=PEX3;refGene_tx=XM_003986617.5
B2 132189523 132189594
refGene_gn=PEX3;refGene_tx=XM_003986617.5
B2 132191660 132191783
refGene_gn=PEX3;refGene_tx=XM_003986617.5
B2 132196696 132196793
refGene_gn=PEX3;refGene_tx=XM_003986617.5
B2 132199122 132199206
refGene_gn=PEX3;refGene_tx=XM_003986617.5
B2 132205408 132205549
refGene_gn=FUCA2;refGene_tx=XM_011282671.3;refGene_gn=FUCA2;refGene_tx=XM_023254460.1
B2 132205690 132205744
refGene_gn=FUCA2;refGene_tx=XM_023254459.1
B2 132207312 132207421
refGene_gn=FUCA2;refGene_tx=XM_011282671.3;refGene_gn=FUCA2;refGene_tx=XM_023254459.1;refGene_gn=FUCA2;refGene_tx=XM_023254460.1
B2 132213772 132213960
refGene_gn=FUCA2;refGene_tx=XM_011282671.3;refGene_gn=FUCA2;refGene_tx=XM_023254459.1;refGene_gn=FUCA2;refGene_tx=XM_023254460.1
B2 132214205 132214416
refGene_gn=FUCA2;refGene_tx=XM_011282671.3;refGene_gn=FUCA2;refGene_tx=XM_023254459.1;refGene_gn=FUCA2;refGene_tx=XM_023254460.1
B2 132216287 132216627
refGene_gn=FUCA2;refGene_tx=XM_011282671.3;refGene_gn=FUCA2;refGene_tx=XM_023254459.1;refGene_gn=FUCA2;refGene_tx=XM_023254460.1
B2 132219870 132220058
refGene_gn=FUCA2;refGene_tx=XM_011282671.3;refGene_gn=FUCA2;refGene_tx=XM_023254459.1;refGene_gn=FUCA2;refGene_tx=XM_023254460.1
B2 132223344 132223574
refGene_gn=FUCA2;refGene_tx=XM_011282671.3;refGene_gn=FUCA2;refGene_tx

=XM_023254459.1
B2 132336696 132336709
refGene_gn=PHACTR2;refGene_tx=XM_019831332.2
B2 132400725 132400771
refGene_gn=PHACTR2;refGene_tx=XM_006932071.4;refGene_gn=PHACTR2;refGene_tx=XM_006932072.4;refGene_gn=PHACTR2;refGene_tx=XM_006932073.4;refGene_gn=PHACTR2;refGene_tx=XM_006932074.3;refGene_gn=PHACTR2;refGene_tx=XM_019831331.2;refGene_gn=PHACTR2;refGene_tx=XM_019831333.2;refGene_gn=PHACTR2;refGene_tx=XM_023254461.1
B2 132431054 132431222
refGene_gn=PHACTR2;refGene_tx=XM_006932071.4;refGene_gn=PHACTR2;refGene_tx=XM_006932072.4;refGene_gn=PHACTR2;refGene_tx=XM_006932073.4;refGene_gn=PHACTR2;refGene_tx=XM_006932074.3;refGene_gn=PHACTR2;refGene_tx=XM_019831331.2;refGene_gn=PHACTR2;refGene_tx=XM_019831332.2;refGene_gn=PHACTR2;refGene_tx=XM_019831333.2;refGene_gn=PHACTR2;refGene_tx=XM_023254461.1
B2 132464198 132464279
refGene_gn=PHACTR2;refGene_tx=XM_006932071.4;refGene_gn=PHACTR2;refGene_tx=XM_006932072.4;refGene_gn=PHACTR2;refGene_tx=XM_006932073.4;refGene_gn=PHACTR2;refGene_tx=XM_006932074.3;refGene_gn=PHACTR2;refGene_tx=XM_019831331.2;refGene_gn=PHACTR2;refGene_tx=XM_019831332.2;refGene_gn=PHACTR2;refGene_tx=XM_023254461.1
B2 132469262 132469421
refGene_gn=PHACTR2;refGene_tx=XM_006932071.4;refGene_gn=PHACTR2;refGene_tx=XM_006932072.4;refGene_gn=PHACTR2;refGene_tx=XM_006932073.4;refGene_gn=PHACTR2;refGene_tx=XM_006932074.3;refGene_gn=PHACTR2;refGene_tx=XM_019831331.2;refGene_gn=PHACTR2;refGene_tx=XM_019831332.2;refGene_gn=PHACTR2;refGene_tx=XM_019831333.2;refGene_gn=PHACTR2;refGene_tx=XM_023254461.1
B2 132476063 132476300
refGene_gn=PHACTR2;refGene_tx=XM_006932071.4;refGene_gn=PHACTR2;refGene_tx=XM_006932072.4;refGene_gn=PHACTR2;refGene_tx=XM_019831331.2;refGene_gn=PHACTR2;refGene_tx=XM_019831332.2;refGene_gn=PHACTR2;refGene_tx=XM_019831333.2;refGene_gn=PHACTR2;refGene_tx=XM_023254461.1
B2 132480203 132480735
refGene_gn=PHACTR2;refGene_tx=XM_006932071.4;refGene_gn=PHACTR2;refGene_tx=XM_006932072.4;refGene_gn=PHACTR2;refGene_tx=XM_006932073.4;refGene_gn=PHACTR2;refGene_tx=XM_006932074.3;refGene_gn=PHACTR2;refGene_tx=XM_019831331.2;refGene_gn=PHACTR2;refGene_tx=XM_019831332.2;refGene_gn=PHACTR2;refGene_tx=XM_019831333.2;refGene_gn=PHACTR2;refGene_tx=XM_023254461.1
B2 132486526 132486735
refGene_gn=PHACTR2;refGene_tx=XM_006932071.4;refGene_gn=PHACTR2;refGene_tx=XM_006932072.4;refGene_gn=PHACTR2;refGene_tx=XM_006932073.4;refGene_gn=PHACTR2;refGene_tx=XM_006932074.3;refGene_gn=PHACTR2;refGene_tx=XM_019831331.2;refGene_gn=PHACTR2;refGene_tx=XM_019831332.2;refGene_gn=PHACTR2;refGene_tx=XM_019831333.2;refGene_gn=PHACTR2;refGene_tx=XM_023254461.1
B2 132488004 132488161
refGene_gn=PHACTR2;refGene_tx=XM_006932071.4;refGene_gn=PHACTR2;refGen

e_tx=XM_006932072.4;refGene_gn=PHACTR2;refGene_tx=XM_006932073.4;refGene_gn=PHACTR2;refGene_tx=XM_006932074.3;refGene_gn=PHACTR2;refGene_tx=XM_019831331.2;refGene_gn=PHACTR2;refGene_tx=XM_019831332.2;refGene_gn=PHACTR2;refGene_tx=XM_019831333.2;refGene_gn=PHACTR2;refGene_tx=XM_023254461.1
B2	132492412	132492468
refGene_gn=PHACTR2;refGene_tx=XM_006932071.4;refGene_gn=PHACTR2;refGene_tx=XM_006932072.4;refGene_gn=PHACTR2;refGene_tx=XM_006932073.4;refGene_gn=PHACTR2;refGene_tx=XM_006932074.3;refGene_gn=PHACTR2;refGene_tx=XM_019831331.2;refGene_gn=PHACTR2;refGene_tx=XM_019831332.2;refGene_gn=PHACTR2;refGene_tx=XM_019831333.2;refGene_gn=PHACTR2;refGene_tx=XM_023254461.1
B2	132498898	132498960
refGene_gn=PHACTR2;refGene_tx=XM_006932071.4;refGene_gn=PHACTR2;refGene_tx=XM_006932072.4;refGene_gn=PHACTR2;refGene_tx=XM_006932073.4;refGene_gn=PHACTR2;refGene_tx=XM_006932074.3;refGene_gn=PHACTR2;refGene_tx=XM_019831331.2;refGene_gn=PHACTR2;refGene_tx=XM_019831332.2;refGene_gn=PHACTR2;refGene_tx=XM_019831333.2;refGene_gn=PHACTR2;refGene_tx=XM_023254461.1
B2	132501695	132501833
refGene_gn=PHACTR2;refGene_tx=XM_006932071.4;refGene_gn=PHACTR2;refGene_tx=XM_006932072.4;refGene_gn=PHACTR2;refGene_tx=XM_006932073.4;refGene_gn=PHACTR2;refGene_tx=XM_006932074.3;refGene_gn=PHACTR2;refGene_tx=XM_019831331.2;refGene_gn=PHACTR2;refGene_tx=XM_019831332.2;refGene_gn=PHACTR2;refGene_tx=XM_019831333.2;refGene_gn=PHACTR2;refGene_tx=XM_023254461.1
B2	132515026	132515107
refGene_gn=PHACTR2;refGene_tx=XM_006932071.4;refGene_gn=PHACTR2;refGene_tx=XM_006932072.4;refGene_gn=PHACTR2;refGene_tx=XM_006932073.4;refGene_gn=PHACTR2;refGene_tx=XM_006932074.3;refGene_gn=PHACTR2;refGene_tx=XM_019831331.2;refGene_gn=PHACTR2;refGene_tx=XM_019831332.2;refGene_gn=PHACTR2;refGene_tx=XM_019831333.2;refGene_gn=PHACTR2;refGene_tx=XM_023254461.1
B2	132528477	132528493
refGene_gn=PHACTR2;refGene_tx=XM_006932071.4;refGene_gn=PHACTR2;refGene_tx=XM_006932073.4;refGene_gn=PHACTR2;refGene_tx=XM_019831331.2;refGene_gn=PHACTR2;refGene_tx=XM_019831332.2;refGene_gn=PHACTR2;refGene_tx=XM_019831333.2
B2	132544985	132544988
refGene_gn=LTV1;refGene_tx=XM_006932075.4
B2	132545989	132546121
refGene_gn=LTV1;refGene_tx=XM_006932075.4
B2	132548284	132548458
refGene_gn=LTV1;refGene_tx=XM_006932075.4
B2	132552897	132552985
refGene_gn=LTV1;refGene_tx=XM_006932075.4
B2	132555023	132555162
refGene_gn=LTV1;refGene_tx=XM_006932075.4;refGene_gn=LTV1;refGene_tx=XM_019831335.1
B2	132555527	132555783

refGene_gn=LTV1;refGene_tx=XM_006932075.4;refGene_gn=LTV1;refGene_tx=XM_019831335.1
B2 132556798 132556926
refGene_gn=LTV1;refGene_tx=XM_006932075.4;refGene_gn=LTV1;refGene_tx=XM_019831335.1
B2 132558621 132558761
refGene_gn=LTV1;refGene_tx=XM_006932075.4;refGene_gn=LTV1;refGene_tx=XM_019831335.1
B2 132559074 132559127
refGene_gn=LTV1;refGene_tx=XM_006932075.4;refGene_gn=LTV1;refGene_tx=XM_019831335.1
B2 132559338 132559539
refGene_gn=LTV1;refGene_tx=XM_006932075.4;refGene_gn=LTV1;refGene_tx=XM_019831335.1
B2 132559668 132559779
refGene_gn=LTV1;refGene_tx=XM_006932075.4;refGene_gn=LTV1;refGene_tx=XM_019831335.1
B2 132560776 132560804
refGene_gn=ZC2HC1B;refGene_tx=XM_003986619.4;refGene_gn=ZC2HC1B;refGene_tx=XM_019831345.1
B2 132565853 132565904
refGene_gn=ZC2HC1B;refGene_tx=XM_023254462.1
B2 132570114 132570176
refGene_gn=ZC2HC1B;refGene_tx=XM_003986619.4;refGene_gn=ZC2HC1B;refGene_tx=XM_019831345.1
B2 132572775 132572895
refGene_gn=ZC2HC1B;refGene_tx=XM_003986619.4;refGene_gn=ZC2HC1B;refGene_tx=XM_019831345.1;refGene_gn=ZC2HC1B;refGene_tx=XM_023254462.1
B2 132573423 132573562
refGene_gn=ZC2HC1B;refGene_tx=XM_003986619.4;refGene_gn=ZC2HC1B;refGene_tx=XM_019831345.1;refGene_gn=ZC2HC1B;refGene_tx=XM_023254462.1
B2 132590478 132590618
refGene_gn=ZC2HC1B;refGene_tx=XM_003986619.4;refGene_gn=ZC2HC1B;refGene_tx=XM_019831345.1;refGene_gn=ZC2HC1B;refGene_tx=XM_023254462.1
B2 132594112 132594221
refGene_gn=ZC2HC1B;refGene_tx=XM_003986619.4;refGene_gn=ZC2HC1B;refGene_tx=XM_019831345.1;refGene_gn=ZC2HC1B;refGene_tx=XM_023254462.1
B2 132600886 132600957
refGene_gn=ZC2HC1B;refGene_tx=XM_003986619.4
B2 132601446 132601526
refGene_gn=ZC2HC1B;refGene_tx=XM_019831345.1;refGene_gn=ZC2HC1B;refGene_tx=XM_023254462.1
B2 132604965 132606211
refGene_gn=PLAGL1;refGene_tx=XM_006932077.3;refGene_gn=PLAGL1;refGene_tx=XM_006932078.4;refGene_gn=PLAGL1;refGene_tx=XM_011282582.2;refGene_gn=PLAGL1;refGene_tx=XM_011282583.2;refGene_gn=PLAGL1;refGene_tx=XM_023254463.1;refGene_gn=PLAGL1;refGene_tx=XM_023254464.1;refGene_gn=PLAGL1;refGene_tx=XM_023254465.1;refGene_gn=PLAGL1;refGene_tx=XM_023254466.1;refGene_gn=PLAGL1;refGene_tx=XM_023254467.1;refGene_gn=PLAGL1;refGene_tx=XM_023254468.1;refGene_gn=PLAGL1;refGene_tx=XM_023254469.1

B2	132610959	132611111
refGene_gn=PLAGL1;refGene_tx=XM_023254463.1;refGene_gn=PLAGL1;refGene_tx=XM_023254464.1;refGene_gn=PLAGL1;refGene_tx=XM_023254465.1;refGene_gn=PLAGL1;refGene_tx=XM_023254466.1;refGene_gn=PLAGL1;refGene_tx=XM_023254467.1;refGene_gn=PLAGL1;refGene_tx=XM_023254468.1;refGene_gn=PLAGL1;refGene_tx=XM_023254469.1
B2	132611294	132611323
refGene_gn=PLAGL1;refGene_tx=XM_006932077.3
B2	132744339	132744600
refGene_gn=SF3B5;refGene_tx=XM_003986622.3
B2	132797983	132798026
refGene_gn=STX11;refGene_tx=XM_019831346.1
B2	132842645	132843514
refGene_gn=STX11;refGene_tx=XM_003986623.5;refGene_gn=STX11;refGene_tx=XM_006932080.4;refGene_gn=STX11;refGene_tx=XM_019831346.1;refGene_gn=STX11;refGene_tx=XM_023254470.1;refGene_gn=STX11;refGene_tx=XM_023254471.1
B2	132975435	132975523
refGene_gn=UTRN;refGene_tx=XM_019831348.2
B2	133027760	133027822
refGene_gn=UTRN;refGene_tx=XM_019831348.2
B2	133044700	133044793
refGene_gn=UTRN;refGene_tx=XM_019831348.2
B2	133046433	133046469
refGene_gn=UTRN;refGene_tx=XM_019831348.2
B2	133046470	133046512
refGene_gn=UTRN;refGene_tx=XM_019831348.2
B2	133046865	133046958
refGene_gn=UTRN;refGene_tx=XM_019831348.2
B2	133049114	133049287
refGene_gn=UTRN;refGene_tx=XM_019831348.2
B2	133052124	133052240
refGene_gn=UTRN;refGene_tx=XM_019831348.2
B2	133053141	133053302
refGene_gn=UTRN;refGene_tx=XM_019831348.2
B2	133057733	133057937
refGene_gn=UTRN;refGene_tx=XM_019831348.2
B2	133058863	133059045
refGene_gn=UTRN;refGene_tx=XM_019831348.2
B2	133059753	133059904
refGene_gn=UTRN;refGene_tx=XM_019831348.2
B2	133061074	133061194
refGene_gn=UTRN;refGene_tx=XM_019831348.2
B2	133064033	133064135
refGene_gn=UTRN;refGene_tx=XM_019831348.2
B2	133067093	133067201
refGene_gn=UTRN;refGene_tx=XM_019831348.2
B2	133067483	133067663
refGene_gn=UTRN;refGene_tx=XM_019831348.2
B2	133068496	133068666

refGene_gn=UTRN;refGene_tx=XM_019831348.2
B2 133072436 133072560
refGene_gn=UTRN;refGene_tx=XM_019831348.2
B2 133075806 133075894
refGene_gn=UTRN;refGene_tx=XM_019831348.2
B2 133081463 133081705
refGene_gn=UTRN;refGene_tx=XM_019831348.2
B2 133081869 133082050
refGene_gn=UTRN;refGene_tx=XM_019831348.2
B2 133084676 133084822
refGene_gn=UTRN;refGene_tx=XM_019831348.2
B2 133085481 133085694
refGene_gn=UTRN;refGene_tx=XM_019831348.2
B2 133097561 133097675
refGene_gn=UTRN;refGene_tx=XM_019831348.2
B2 133098335 133098491
refGene_gn=UTRN;refGene_tx=XM_019831348.2
B2 133102570 133102741
refGene_gn=UTRN;refGene_tx=XM_019831348.2
B2 133104612 133104792
refGene_gn=UTRN;refGene_tx=XM_019831348.2
B2 133110248 133110383
refGene_gn=UTRN;refGene_tx=XM_019831348.2
B2 133112225 133112375
refGene_gn=UTRN;refGene_tx=XM_019831348.2
B2 133113656 133113818
refGene_gn=UTRN;refGene_tx=XM_019831348.2
B2 133114253 133114382
refGene_gn=UTRN;refGene_tx=XM_019831348.2
B2 133115071 133115245
refGene_gn=UTRN;refGene_tx=XM_019831348.2
B2 133116974 133117130
refGene_gn=UTRN;refGene_tx=XM_019831348.2
B2 133123743 133123914
refGene_gn=UTRN;refGene_tx=XM_019831348.2
B2 133129472 133129652
refGene_gn=UTRN;refGene_tx=XM_019831348.2
B2 133131963 133132092
refGene_gn=UTRN;refGene_tx=XM_019831348.2
B2 133132404 133132575
refGene_gn=UTRN;refGene_tx=XM_019831348.2
B2 133135415 133135577
refGene_gn=UTRN;refGene_tx=XM_019831348.2
B2 133136014 133136152
refGene_gn=UTRN;refGene_tx=XM_019831348.2
B2 133140228 133140420
refGene_gn=UTRN;refGene_tx=XM_019831348.2
B2 133141484 133141657
refGene_gn=UTRN;refGene_tx=XM_019831348.2
B2 133153031 133153182

refGene_gn=UTRN;refGene_tx=XM_019831348.2
B2 133154431 133154607
refGene_gn=UTRN;refGene_tx=XM_019831348.2
B2 133164745 133164881
refGene_gn=UTRN;refGene_tx=XM_019831348.2
B2 133166949 133167099
refGene_gn=UTRN;refGene_tx=XM_019831348.2
B2 133170187 133170263
refGene_gn=UTRN;refGene_tx=XM_019831348.2
B2 133177684 133177899
refGene_gn=UTRN;refGene_tx=XM_019831348.2
B2 133180499 133180617
refGene_gn=UTRN;refGene_tx=XM_019831348.2
B2 133184660 133184866
refGene_gn=UTRN;refGene_tx=XM_019831348.2
B2 133186677 133186832
refGene_gn=UTRN;refGene_tx=XM_019831348.2
B2 133200905 133201095
refGene_gn=UTRN;refGene_tx=XM_019831348.2
B2 133284512 133284685
refGene_gn=UTRN;refGene_tx=XM_019831348.2
B2 133297745 133297902
refGene_gn=UTRN;refGene_tx=XM_019831348.2
B2 133323310 133323440
refGene_gn=UTRN;refGene_tx=XM_019831348.2
B2 133340690 133340959
refGene_gn=UTRN;refGene_tx=XM_019831348.2
B2 133347062 133347209
refGene_gn=UTRN;refGene_tx=XM_019831348.2
B2 133350295 133350374
refGene_gn=UTRN;refGene_tx=XM_019831348.2
B2 133353513 133353574
refGene_gn=UTRN;refGene_tx=XM_019831348.2
B2 133367701 133367763
refGene_gn=UTRN;refGene_tx=XM_019831348.2
B2 133369839 133369914
refGene_gn=UTRN;refGene_tx=XM_019831348.2
B2 133378381 133378583
refGene_gn=UTRN;refGene_tx=XM_019831348.2
B2 133385116 133385202
refGene_gn=UTRN;refGene_tx=XM_019831348.2
B2 133389147 133389305
refGene_gn=UTRN;refGene_tx=XM_019831348.2
B2 133390758 133390925
refGene_gn=UTRN;refGene_tx=XM_019831348.2
B2 133392011 133392123
refGene_gn=UTRN;refGene_tx=XM_019831348.2
B2 133408901 133409038
refGene_gn=UTRN;refGene_tx=XM_019831348.2
B2 133414886 133414925

refGene_gn=UTRN;refGene_tx=XM_019831348.2
B2 133415150 133415216
refGene_gn=UTRN;refGene_tx=XM_019831348.2
B2 133416292 133416358
refGene_gn=UTRN;refGene_tx=XM_019831348.2
B2 133425259 133425418
refGene_gn=UTRN;refGene_tx=XM_019831348.2
B2 133425777 133426018
refGene_gn=UTRN;refGene_tx=XM_019831348.2
B2 133428988 133429097
refGene_gn=UTRN;refGene_tx=XM_019831348.2
B2 133430322 133430415
refGene_gn=UTRN;refGene_tx=XM_019831348.2
B2 133435196 133435219
refGene_gn=UTRN;refGene_tx=XM_019831348.2
B2 133437596 133437605
refGene_gn=UTRN;refGene_tx=XM_019831348.2
B2 134133263 134133541
refGene_gn=EPM2A;refGene_tx=XM_023254472.1
B2 134141886 134142128
refGene_gn=EPM2A;refGene_tx=XM_023254472.1
B2 134199210 134199385
refGene_gn=EPM2A;refGene_tx=XM_023254472.1
B2 134239863 134240164
refGene_gn=EPM2A;refGene_tx=XM_023254472.1
B2 134282497 134282701
refGene_gn=FBXO30;refGene_tx=XM_023254473.1
B2 134287351 134289373
refGene_gn=FBXO30;refGene_tx=XM_023254473.1
B2 134350367 134350464
refGene_gn=SHPRH;refGene_tx=XM_006932081.4;refGene_gn=SHPRH;refGene_tx=XM_019831355.2;refGene_gn=SHPRH;refGene_tx=XM_019831356.2
B2 134351656 134351737
refGene_gn=SHPRH;refGene_tx=XM_006932081.4;refGene_gn=SHPRH;refGene_tx=XM_019831355.2;refGene_gn=SHPRH;refGene_tx=XM_019831356.2
B2 134356025 134356204
refGene_gn=SHPRH;refGene_tx=XM_006932081.4;refGene_gn=SHPRH;refGene_tx=XM_019831355.2;refGene_gn=SHPRH;refGene_tx=XM_019831356.2
B2 134356958 134357045
refGene_gn=SHPRH;refGene_tx=XM_006932081.4;refGene_gn=SHPRH;refGene_tx=XM_019831355.2;refGene_gn=SHPRH;refGene_tx=XM_019831356.2
B2 134357842 134357935
refGene_gn=SHPRH;refGene_tx=XM_006932081.4;refGene_gn=SHPRH;refGene_tx=XM_019831355.2;refGene_gn=SHPRH;refGene_tx=XM_019831356.2;refGene_gn=SHPRH;refGene_tx=XM_019831357.2
B2 134373672 134373861
refGene_gn=SHPRH;refGene_tx=XM_006932081.4;refGene_gn=SHPRH;refGene_tx=XM_019831355.2;refGene_gn=SHPRH;refGene_tx=XM_019831356.2;refGene_gn=SHPRH;refGene_tx=XM_019831357.2
B2 134376905 134376977

refGene_gn=SHPRH;refGene_tx=XM_006932081.4;refGene_gn=SHPRH;refGene_tx=XM_019831355.2;refGene_gn=SHPRH;refGene_tx=XM_019831356.2;refGene_gn=SHPRH;refGene_tx=XM_019831357.2
B2 134384709 134384811
refGene_gn=SHPRH;refGene_tx=XM_006932081.4;refGene_gn=SHPRH;refGene_tx=XM_019831355.2;refGene_gn=SHPRH;refGene_tx=XM_019831356.2;refGene_gn=SHPRH;refGene_tx=XM_019831357.2
B2 134386162 134386306
refGene_gn=SHPRH;refGene_tx=XM_006932081.4;refGene_gn=SHPRH;refGene_tx=XM_019831355.2;refGene_gn=SHPRH;refGene_tx=XM_019831356.2;refGene_gn=SHPRH;refGene_tx=XM_019831357.2
B2 134388075 134388301
refGene_gn=SHPRH;refGene_tx=XM_006932081.4;refGene_gn=SHPRH;refGene_tx=XM_019831355.2;refGene_gn=SHPRH;refGene_tx=XM_019831356.2;refGene_gn=SHPRH;refGene_tx=XM_019831357.2
B2 134389274 134389337
refGene_gn=SHPRH;refGene_tx=XM_006932081.4;refGene_gn=SHPRH;refGene_tx=XM_019831355.2;refGene_gn=SHPRH;refGene_tx=XM_019831356.2;refGene_gn=SHPRH;refGene_tx=XM_019831357.2
B2 134389812 134389986
refGene_gn=SHPRH;refGene_tx=XM_006932081.4;refGene_gn=SHPRH;refGene_tx=XM_019831355.2;refGene_gn=SHPRH;refGene_tx=XM_019831356.2;refGene_gn=SHPRH;refGene_tx=XM_019831357.2
B2 134393468 134393611
refGene_gn=SHPRH;refGene_tx=XM_006932081.4;refGene_gn=SHPRH;refGene_tx=XM_019831355.2;refGene_gn=SHPRH;refGene_tx=XM_019831356.2;refGene_gn=SHPRH;refGene_tx=XM_019831357.2
B2 134394801 134394909
refGene_gn=SHPRH;refGene_tx=XM_006932081.4;refGene_gn=SHPRH;refGene_tx=XM_019831355.2;refGene_gn=SHPRH;refGene_tx=XM_019831356.2;refGene_gn=SHPRH;refGene_tx=XM_019831357.2
B2 134397237 134397330
refGene_gn=SHPRH;refGene_tx=XM_006932081.4;refGene_gn=SHPRH;refGene_tx=XM_019831355.2;refGene_gn=SHPRH;refGene_tx=XM_019831356.2;refGene_gn=SHPRH;refGene_tx=XM_019831357.2
B2 134399401 134399490
refGene_gn=SHPRH;refGene_tx=XM_006932081.4;refGene_gn=SHPRH;refGene_tx=XM_019831355.2;refGene_gn=SHPRH;refGene_tx=XM_019831356.2;refGene_gn=SHPRH;refGene_tx=XM_019831357.2
B2 134412488 134412610
refGene_gn=SHPRH;refGene_tx=XM_006932081.4;refGene_gn=SHPRH;refGene_tx=XM_019831355.2;refGene_gn=SHPRH;refGene_tx=XM_019831356.2;refGene_gn=SHPRH;refGene_tx=XM_019831357.2
B2 134413767 134414024
refGene_gn=SHPRH;refGene_tx=XM_006932081.4;refGene_gn=SHPRH;refGene_tx=XM_019831355.2;refGene_gn=SHPRH;refGene_tx=XM_019831356.2;refGene_gn=SHPRH;refGene_tx=XM_019831357.2
B2 134414134 134414298
refGene_gn=SHPRH;refGene_tx=XM_006932081.4;refGene_gn=SHPRH;refGene_tx=XM_019831355.2;refGene_gn=SHPRH;refGene_tx=XM_019831356.2;refGene_gn=

SHPRH;refGene_tx=XM_019831357.2
B2 134417843 134417922
refGene_gn=SHPRH;refGene_tx=XM_006932081.4;refGene_gn=SHPRH;refGene_tx=XM_019831355.2;refGene_gn=SHPRH;refGene_tx=XM_019831356.2;refGene_gn=SHPRH;refGene_tx=XM_019831357.2
B2 134418747 134418999
refGene_gn=SHPRH;refGene_tx=XM_006932081.4;refGene_gn=SHPRH;refGene_tx=XM_019831355.2;refGene_gn=SHPRH;refGene_tx=XM_019831356.2;refGene_gn=SHPRH;refGene_tx=XM_019831357.2
B2 134422556 134423216
refGene_gn=SHPRH;refGene_tx=XM_006932081.4;refGene_gn=SHPRH;refGene_tx=XM_019831355.2;refGene_gn=SHPRH;refGene_tx=XM_019831356.2;refGene_gn=SHPRH;refGene_tx=XM_019831357.2
B2 134425291 134425548
refGene_gn=SHPRH;refGene_tx=XM_006932081.4;refGene_gn=SHPRH;refGene_tx=XM_019831355.2;refGene_gn=SHPRH;refGene_tx=XM_019831356.2;refGene_gn=SHPRH;refGene_tx=XM_019831357.2
B2 134426182 134426291
refGene_gn=SHPRH;refGene_tx=XM_006932081.4;refGene_gn=SHPRH;refGene_tx=XM_019831355.2;refGene_gn=SHPRH;refGene_tx=XM_019831356.2;refGene_gn=SHPRH;refGene_tx=XM_019831357.2
B2 134427817 134427968
refGene_gn=SHPRH;refGene_tx=XM_006932081.4;refGene_gn=SHPRH;refGene_tx=XM_019831355.2;refGene_gn=SHPRH;refGene_tx=XM_019831356.2;refGene_gn=SHPRH;refGene_tx=XM_019831357.2
B2 134428647 134428726
refGene_gn=SHPRH;refGene_tx=XM_006932081.4;refGene_gn=SHPRH;refGene_tx=XM_019831355.2;refGene_gn=SHPRH;refGene_tx=XM_019831356.2;refGene_gn=SHPRH;refGene_tx=XM_019831357.2
B2 134431224 134431443
refGene_gn=SHPRH;refGene_tx=XM_006932081.4;refGene_gn=SHPRH;refGene_tx=XM_019831355.2;refGene_gn=SHPRH;refGene_tx=XM_019831356.2;refGene_gn=SHPRH;refGene_tx=XM_019831357.2
B2 134433580 134433710
refGene_gn=SHPRH;refGene_tx=XM_006932081.4;refGene_gn=SHPRH;refGene_tx=XM_019831355.2;refGene_gn=SHPRH;refGene_tx=XM_019831356.2;refGene_gn=SHPRH;refGene_tx=XM_019831357.2
B2 134436224 134436863
refGene_gn=SHPRH;refGene_tx=XM_006932081.4;refGene_gn=SHPRH;refGene_tx=XM_019831355.2;refGene_gn=SHPRH;refGene_tx=XM_019831356.2;refGene_gn=SHPRH;refGene_tx=XM_019831357.2
B2 134488128 134488828
refGene_gn=GRM1;refGene_tx=XM_003986626.4;refGene_gn=GRM1;refGene_tx=XM_011282579.3;refGene_gn=GRM1;refGene_tx=XM_019831358.2;refGene_gn=GRM1;refGene_tx=XM_019831359.2;refGene_gn=GRM1;refGene_tx=XM_019831361.2;refGene_gn=GRM1;refGene_tx=XM_019831362.2
B2 134603508 134603758
refGene_gn=GRM1;refGene_tx=XM_003986626.4;refGene_gn=GRM1;refGene_tx=XM_011282579.3;refGene_gn=GRM1;refGene_tx=XM_019831358.2;refGene_gn=GRM1;refGene_tx=XM_019831359.2;refGene_gn=GRM1;refGene_tx=XM_019831361.2;

refGene_gn=GRM1;refGene_tx=XM_019831362.2
B2 134794167 134794403
refGene_gn=GRM1;refGene_tx=XM_003986626.4;refGene_gn=GRM1;refGene_tx=XM_011282579.3;refGene_gn=GRM1;refGene_tx=XM_019831358.2;refGene_gn=GRM1;refGene_tx=XM_019831359.2;refGene_gn=GRM1;refGene_tx=XM_019831361.2;refGene_gn=GRM1;refGene_tx=XM_019831362.2
B2 134834178 134834425
refGene_gn=GRM1;refGene_tx=XM_003986626.4;refGene_gn=GRM1;refGene_tx=XM_011282579.3;refGene_gn=GRM1;refGene_tx=XM_019831358.2;refGene_gn=GRM1;refGene_tx=XM_019831359.2;refGene_gn=GRM1;refGene_tx=XM_019831361.2;refGene_gn=GRM1;refGene_tx=XM_019831362.2
B2 134839770 134839939
refGene_gn=GRM1;refGene_tx=XM_003986626.4;refGene_gn=GRM1;refGene_tx=XM_011282579.3;refGene_gn=GRM1;refGene_tx=XM_019831358.2;refGene_gn=GRM1;refGene_tx=XM_019831359.2;refGene_gn=GRM1;refGene_tx=XM_019831361.2;refGene_gn=GRM1;refGene_tx=XM_019831362.2
B2 134854806 134854933
refGene_gn=GRM1;refGene_tx=XM_003986626.4;refGene_gn=GRM1;refGene_tx=XM_011282579.3;refGene_gn=GRM1;refGene_tx=XM_019831358.2;refGene_gn=GRM1;refGene_tx=XM_019831359.2;refGene_gn=GRM1;refGene_tx=XM_019831361.2;refGene_gn=GRM1;refGene_tx=XM_019831362.2
B2 134867854 134868785
refGene_gn=GRM1;refGene_tx=XM_003986626.4;refGene_gn=GRM1;refGene_tx=XM_011282579.3;refGene_gn=GRM1;refGene_tx=XM_019831358.2;refGene_gn=GRM1;refGene_tx=XM_019831359.2;refGene_gn=GRM1;refGene_tx=XM_019831361.2;refGene_gn=GRM1;refGene_tx=XM_019831362.2
B2 134896229 134896290
refGene_gn=GRM1;refGene_tx=XM_019831361.2;refGene_gn=GRM1;refGene_tx=XM_019831362.2
B2 134903623 134904536
refGene_gn=GRM1;refGene_tx=XM_003986626.4;refGene_gn=GRM1;refGene_tx=XM_011282579.3;refGene_gn=GRM1;refGene_tx=XM_019831358.2;refGene_gn=GRM1;refGene_tx=XM_019831359.2
B2 135011086 135011336
refGene_gn=RAB32;refGene_tx=XM_003986627.5
B2 135025549 135025827
refGene_gn=RAB32;refGene_tx=XM_003986627.5
B2 135029895 135030045
refGene_gn=RAB32;refGene_tx=XM_003986627.5
B2 135068896 135068970
refGene_gn=ADGB;refGene_tx=XM_006932082.3
B2 135100212 135100375
refGene_gn=ADGB;refGene_tx=XM_006932082.3
B2 135104913 135105006
refGene_gn=ADGB;refGene_tx=XM_006932082.3
B2 135108582 135108654
refGene_gn=ADGB;refGene_tx=XM_006932082.3
B2 135110502 135110712
refGene_gn=ADGB;refGene_tx=XM_006932082.3
B2 135123029 135123169

refGene_gn=ADGB;refGene_tx=XM_006932082.3
B2 135125718 135125805
refGene_gn=ADGB;refGene_tx=XM_006932082.3
B2 135133076 135133306
refGene_gn=ADGB;refGene_tx=XM_006932082.3
B2 135135010 135135151
refGene_gn=ADGB;refGene_tx=XM_006932082.3
B2 135136862 135136957
refGene_gn=ADGB;refGene_tx=XM_006932082.3
B2 135145466 135145592
refGene_gn=ADGB;refGene_tx=XM_006932082.3
B2 135145593 135145642
refGene_gn=ADGB;refGene_tx=XM_006932082.3
B2 135149336 135149427
refGene_gn=ADGB;refGene_tx=XM_006932082.3
B2 135154181 135154311
refGene_gn=ADGB;refGene_tx=XM_006932082.3
B2 135161175 135161209
refGene_gn=ADGB;refGene_tx=XM_006932082.3
B2 135162104 135162288
refGene_gn=ADGB;refGene_tx=XM_006932082.3
B2 135163234 135163298
refGene_gn=ADGB;refGene_tx=XM_006932082.3
B2 135166700 135166803
refGene_gn=ADGB;refGene_tx=XM_006932082.3
B2 135169626 135169768
refGene_gn=ADGB;refGene_tx=XM_006932082.3
B2 135171114 135171229
refGene_gn=ADGB;refGene_tx=XM_006932082.3
B2 135173772 135173940
refGene_gn=ADGB;refGene_tx=XM_006932082.3
B2 135178572 135178708
refGene_gn=ADGB;refGene_tx=XM_006932082.3
B2 135179199 135179337
refGene_gn=ADGB;refGene_tx=XM_006932082.3
B2 135180749 135180843
refGene_gn=ADGB;refGene_tx=XM_006932082.3
B2 135185804 135185939
refGene_gn=ADGB;refGene_tx=XM_006932082.3
B2 135186453 135186607
refGene_gn=ADGB;refGene_tx=XM_006932082.3
B2 135190237 135190425
refGene_gn=ADGB;refGene_tx=XM_006932082.3
B2 135193433 135193618
refGene_gn=ADGB;refGene_tx=XM_006932082.3
B2 135196287 135196487
refGene_gn=ADGB;refGene_tx=XM_006932082.3
B2 135199334 135199446
refGene_gn=ADGB;refGene_tx=XM_006932082.3
B2 135207920 135208090

refGene_gn=ADGB;refGene_tx=XM_006932082.3
B2 135214867 135215044
refGene_gn=ADGB;refGene_tx=XM_006932082.3
B2 135215778 135215881
refGene_gn=ADGB;refGene_tx=XM_006932082.3
B2 135218658 135218868
refGene_gn=ADGB;refGene_tx=XM_006932082.3
B2 135225079 135225170
refGene_gn=ADGB;refGene_tx=XM_006932082.3
B2 135225897 135226084
refGene_gn=ADGB;refGene_tx=XM_006932082.3
B2 135232067 135232253
refGene_gn=ADGB;refGene_tx=XM_006932082.3
B2 135622725 135622878
refGene_gn=STXBP5;refGene_tx=XM_011282577.3;refGene_gn=STXBP5;refGene_tx=XM_011282578.3;refGene_gn=STXBP5;refGene_tx=XM_019831369.2;refGene_gn=STXBP5;refGene_tx=XM_019831370.2
B2 135624202 135624300
refGene_gn=STXBP5;refGene_tx=XM_011282577.3;refGene_gn=STXBP5;refGene_tx=XM_011282578.3;refGene_gn=STXBP5;refGene_tx=XM_019831369.2;refGene_gn=STXBP5;refGene_tx=XM_019831370.2
B2 135649351 135649433
refGene_gn=STXBP5;refGene_tx=XM_011282577.3;refGene_gn=STXBP5;refGene_tx=XM_011282578.3;refGene_gn=STXBP5;refGene_tx=XM_019831369.2;refGene_gn=STXBP5;refGene_tx=XM_019831370.2
B2 135653230 135653331
refGene_gn=STXBP5;refGene_tx=XM_011282577.3;refGene_gn=STXBP5;refGene_tx=XM_011282578.3;refGene_gn=STXBP5;refGene_tx=XM_019831369.2;refGene_gn=STXBP5;refGene_tx=XM_019831370.2
B2 135676235 135676370
refGene_gn=STXBP5;refGene_tx=XM_011282577.3;refGene_gn=STXBP5;refGene_tx=XM_011282578.3;refGene_gn=STXBP5;refGene_tx=XM_019831369.2;refGene_gn=STXBP5;refGene_tx=XM_019831370.2
B2 135678782 135678846
refGene_gn=STXBP5;refGene_tx=XM_011282577.3;refGene_gn=STXBP5;refGene_tx=XM_011282578.3;refGene_gn=STXBP5;refGene_tx=XM_019831369.2;refGene_gn=STXBP5;refGene_tx=XM_019831370.2
B2 135683534 135683618
refGene_gn=STXBP5;refGene_tx=XM_011282577.3;refGene_gn=STXBP5;refGene_tx=XM_011282578.3;refGene_gn=STXBP5;refGene_tx=XM_019831369.2;refGene_gn=STXBP5;refGene_tx=XM_019831370.2
B2 135687234 135687358
refGene_gn=STXBP5;refGene_tx=XM_011282577.3;refGene_gn=STXBP5;refGene_tx=XM_011282578.3;refGene_gn=STXBP5;refGene_tx=XM_019831369.2;refGene_gn=STXBP5;refGene_tx=XM_019831370.2
B2 135698149 135698228
refGene_gn=STXBP5;refGene_tx=XM_011282577.3;refGene_gn=STXBP5;refGene_tx=XM_011282578.3;refGene_gn=STXBP5;refGene_tx=XM_019831369.2;refGene_gn=STXBP5;refGene_tx=XM_019831370.2
B2 135715045 135715200

refGene_gn=STXBP5;refGene_tx=XM_011282577.3;refGene_gn=STXBP5;refGene_
tx=XM_011282578.3;refGene_gn=STXBP5;refGene_tx=XM_019831369.2;refGene_
gn=STXBP5;refGene_tx=XM_019831370.2
B2	135716377	135716450
refGene_gn=STXBP5;refGene_tx=XM_011282577.3;refGene_gn=STXBP5;refGene_
tx=XM_011282578.3;refGene_gn=STXBP5;refGene_tx=XM_019831369.2;refGene_
gn=STXBP5;refGene_tx=XM_019831370.2
B2	135719471	135719619
refGene_gn=STXBP5;refGene_tx=XM_011282577.3;refGene_gn=STXBP5;refGene_
tx=XM_011282578.3;refGene_gn=STXBP5;refGene_tx=XM_019831369.2;refGene_
gn=STXBP5;refGene_tx=XM_019831370.2
B2	135719823	135719891
refGene_gn=STXBP5;refGene_tx=XM_011282577.3;refGene_gn=STXBP5;refGene_
tx=XM_011282578.3;refGene_gn=STXBP5;refGene_tx=XM_019831369.2;refGene_
gn=STXBP5;refGene_tx=XM_019831370.2
B2	135720210	135720251
refGene_gn=STXBP5;refGene_tx=XM_011282577.3;refGene_gn=STXBP5;refGene_
tx=XM_011282578.3;refGene_gn=STXBP5;refGene_tx=XM_019831369.2;refGene_
gn=STXBP5;refGene_tx=XM_019831370.2
B2	135721131	135721352
refGene_gn=STXBP5;refGene_tx=XM_011282577.3;refGene_gn=STXBP5;refGene_
tx=XM_011282578.3;refGene_gn=STXBP5;refGene_tx=XM_019831369.2;refGene_
gn=STXBP5;refGene_tx=XM_019831370.2
B2	135721850	135722029
refGene_gn=STXBP5;refGene_tx=XM_011282577.3;refGene_gn=STXBP5;refGene_
tx=XM_011282578.3;refGene_gn=STXBP5;refGene_tx=XM_019831369.2;refGene_
gn=STXBP5;refGene_tx=XM_019831370.2
B2	135727546	135727672
refGene_gn=STXBP5;refGene_tx=XM_011282577.3;refGene_gn=STXBP5;refGene_
tx=XM_011282578.3;refGene_gn=STXBP5;refGene_tx=XM_019831369.2;refGene_
gn=STXBP5;refGene_tx=XM_019831370.2
B2	135729324	135729476
refGene_gn=STXBP5;refGene_tx=XM_011282577.3;refGene_gn=STXBP5;refGene_
tx=XM_011282578.3;refGene_gn=STXBP5;refGene_tx=XM_019831369.2;refGene_
gn=STXBP5;refGene_tx=XM_019831370.2
B2	135735987	135736053
refGene_gn=STXBP5;refGene_tx=XM_011282577.3;refGene_gn=STXBP5;refGene_
tx=XM_011282578.3;refGene_gn=STXBP5;refGene_tx=XM_019831369.2;refGene_
gn=STXBP5;refGene_tx=XM_019831370.2
B2	135740692	135740752
refGene_gn=STXBP5;refGene_tx=XM_011282577.3;refGene_gn=STXBP5;refGene_
tx=XM_019831369.2
B2	135740851	135740899
refGene_gn=STXBP5;refGene_tx=XM_011282577.3
B2	135753196	135753247
refGene_gn=STXBP5;refGene_tx=XM_011282577.3;refGene_gn=STXBP5;refGene_
tx=XM_011282578.3;refGene_gn=STXBP5;refGene_tx=XM_019831369.2
B2	135758650	135758890
refGene_gn=STXBP5;refGene_tx=XM_011282577.3;refGene_gn=STXBP5;refGene_
tx=XM_011282578.3;refGene_gn=STXBP5;refGene_tx=XM_019831369.2;refGene_

gn=STXBP5;refGene_tx=XM_019831370.2
B2 135760316 135760686
refGene_gn=STXBP5;refGene_tx=XM_011282577.3;refGene_gn=STXBP5;refGene_tx=XM_011282578.3;refGene_gn=STXBP5;refGene_tx=XM_019831369.2;refGene_gn=STXBP5;refGene_tx=XM_019831370.2
B2 135761665 135761831
refGene_gn=STXBP5;refGene_tx=XM_011282577.3;refGene_gn=STXBP5;refGene_tx=XM_011282578.3;refGene_gn=STXBP5;refGene_tx=XM_019831369.2;refGene_gn=STXBP5;refGene_tx=XM_019831370.2
B2 135771074 135771186
refGene_gn=STXBP5;refGene_tx=XM_011282577.3;refGene_gn=STXBP5;refGene_tx=XM_011282578.3;refGene_gn=STXBP5;refGene_tx=XM_019831369.2;refGene_gn=STXBP5;refGene_tx=XM_019831370.2
B2 135792021 135792242
refGene_gn=STXBP5;refGene_tx=XM_011282577.3;refGene_gn=STXBP5;refGene_tx=XM_011282578.3;refGene_gn=STXBP5;refGene_tx=XM_019831369.2;refGene_gn=STXBP5;refGene_tx=XM_019831370.2
B2 135793994 135794036
refGene_gn=STXBP5;refGene_tx=XM_011282577.3;refGene_gn=STXBP5;refGene_tx=XM_011282578.3;refGene_gn=STXBP5;refGene_tx=XM_019831369.2;refGene_gn=STXBP5;refGene_tx=XM_019831370.2
B2 135872085 135872541
refGene_gn=SAMD5;refGene_tx=XM_023254486.1;refGene_gn=SAMD5;refGene_tx=XM_023254487.1;refGene_gn=SAMD5;refGene_tx=XM_023254488.1;refGene_gn=SAMD5;refGene_tx=XM_023254489.1
B2 135921719 135921818
refGene_gn=SAMD5;refGene_tx=XM_023254486.1;refGene_gn=SAMD5;refGene_tx=XM_023254487.1;refGene_gn=SAMD5;refGene_tx=XM_023254488.1
B2 135924358 135924421
refGene_gn=SAMD5;refGene_tx=XM_023254489.1
B2 136501249 136501294
refGene_gn=SASH1;refGene_tx=XM_023254477.1
B2 136578804 136578825
refGene_gn=SASH1;refGene_tx=XM_023254478.1
B2 136642227 136642347
refGene_gn=SASH1;refGene_tx=XM_023254476.1;refGene_gn=SASH1;refGene_tx=XM_023254484.1
B2 136687927 136688056
refGene_gn=SASH1;refGene_tx=XM_023254476.1;refGene_gn=SASH1;refGene_tx=XM_023254477.1;refGene_gn=SASH1;refGene_tx=XM_023254478.1;refGene_gn=SASH1;refGene_tx=XM_023254479.1;refGene_gn=SASH1;refGene_tx=XM_023254480.1;refGene_gn=SASH1;refGene_tx=XM_023254481.1;refGene_gn=SASH1;refGene_tx=XM_023254482.1;refGene_gn=SASH1;refGene_tx=XM_023254483.1;refGene_gn=SASH1;refGene_tx=XM_023254484.1
B2 136729043 136729094
refGene_gn=SASH1;refGene_tx=XM_023254476.1;refGene_gn=SASH1;refGene_tx=XM_023254477.1;refGene_gn=SASH1;refGene_tx=XM_023254478.1;refGene_gn=SASH1;refGene_tx=XM_023254479.1;refGene_gn=SASH1;refGene_tx=XM_023254484.1
B2 136729220 136729270

refGene_gn=SASH1;refGene_tx=XM_023254476.1;refGene_gn=SASH1;refGene_tx
=XM_023254477.1;refGene_gn=SASH1;refGene_tx=XM_023254478.1;refGene_gn=
SASH1;refGene_tx=XM_023254479.1;refGene_gn=SASH1;refGene_tx=XM_0232544
84.1
B2 136753940 136753981
refGene_gn=SASH1;refGene_tx=XM_023254476.1;refGene_gn=SASH1;refGene_tx
=XM_023254477.1;refGene_gn=SASH1;refGene_tx=XM_023254478.1;refGene_gn=
SASH1;refGene_tx=XM_023254479.1;refGene_gn=SASH1;refGene_tx=XM_0232544
80.1;refGene_gn=SASH1;refGene_tx=XM_023254481.1;refGene_gn=SASH1;refGe
ne_tx=XM_023254482.1;refGene_gn=SASH1;refGene_tx=XM_023254483.1;refGen
e_gn=SASH1;refGene_tx=XM_023254484.1
B2 136756269 136756356
refGene_gn=SASH1;refGene_tx=XM_023254476.1;refGene_gn=SASH1;refGene_tx
=XM_023254477.1;refGene_gn=SASH1;refGene_tx=XM_023254478.1;refGene_gn=
SASH1;refGene_tx=XM_023254479.1;refGene_gn=SASH1;refGene_tx=XM_0232544
80.1;refGene_gn=SASH1;refGene_tx=XM_023254481.1;refGene_gn=SASH1;refGe
ne_tx=XM_023254482.1;refGene_gn=SASH1;refGene_tx=XM_023254483.1;refGen
e_gn=SASH1;refGene_tx=XM_023254484.1
B2 136758691 136758804
refGene_gn=SASH1;refGene_tx=XM_023254476.1;refGene_gn=SASH1;refGene_tx
=XM_023254477.1;refGene_gn=SASH1;refGene_tx=XM_023254478.1;refGene_gn=
SASH1;refGene_tx=XM_023254479.1;refGene_gn=SASH1;refGene_tx=XM_0232544
80.1;refGene_gn=SASH1;refGene_tx=XM_023254481.1;refGene_gn=SASH1;refGe
ne_tx=XM_023254482.1;refGene_gn=SASH1;refGene_tx=XM_023254483.1;refGen
e_gn=SASH1;refGene_tx=XM_023254484.1
B2 136766387 136766489
refGene_gn=SASH1;refGene_tx=XM_023254476.1;refGene_gn=SASH1;refGene_tx
=XM_023254477.1;refGene_gn=SASH1;refGene_tx=XM_023254478.1;refGene_gn=
SASH1;refGene_tx=XM_023254479.1;refGene_gn=SASH1;refGene_tx=XM_0232544
80.1;refGene_gn=SASH1;refGene_tx=XM_023254481.1;refGene_gn=SASH1;refGe
ne_tx=XM_023254482.1;refGene_gn=SASH1;refGene_tx=XM_023254483.1;refGen
e_gn=SASH1;refGene_tx=XM_023254484.1
B2 136780682 136780706
refGene_gn=SASH1;refGene_tx=XM_023254485.1
B2 136786126 136786138
refGene_gn=SASH1;refGene_tx=XM_019831373.2;refGene_gn=SASH1;refGene_tx
=XM_019831375.2
B2 136791388 136791391
refGene_gn=SASH1;refGene_tx=XM_019831374.2
B2 136791556 136791689
refGene_gn=SASH1;refGene_tx=XM_019831373.2;refGene_gn=SASH1;refGene_tx
=XM_019831374.2;refGene_gn=SASH1;refGene_tx=XM_019831375.2;refGene_gn=
SASH1;refGene_tx=XM_023254476.1;refGene_gn=SASH1;refGene_tx=XM_0232544
77.1;refGene_gn=SASH1;refGene_tx=XM_023254478.1;refGene_gn=SASH1;refGe
ne_tx=XM_023254479.1;refGene_gn=SASH1;refGene_tx=XM_023254480.1;refGen
e_gn=SASH1;refGene_tx=XM_023254481.1;refGene_gn=SASH1;refGene_tx=XM_02
3254482.1;refGene_gn=SASH1;refGene_tx=XM_023254483.1;refGene_gn=SASH1;
refGene_tx=XM_023254484.1;refGene_gn=SASH1;refGene_tx=XM_023254485.1
B2 136797742 136798572
refGene_gn=SASH1;refGene_tx=XM_019831373.2;refGene_gn=SASH1;refGene_tx

=XM_019831374.2;refGene_gn=SASH1;refGene_tx=XM_019831375.2;refGene_gn=SASH1;refGene_tx=XM_023254476.1;refGene_gn=SASH1;refGene_tx=XM_023254477.1;refGene_gn=SASH1;refGene_tx=XM_023254478.1;refGene_gn=SASH1;refGene_tx=XM_023254479.1;refGene_gn=SASH1;refGene_tx=XM_023254480.1;refGene_gn=SASH1;refGene_tx=XM_023254481.1;refGene_gn=SASH1;refGene_tx=XM_023254482.1;refGene_gn=SASH1;refGene_tx=XM_023254483.1;refGene_gn=SASH1;refGene_tx=XM_023254484.1;refGene_gn=SASH1;refGene_tx=XM_023254485.1
B2 136803550 136803625
refGene_gn=SASH1;refGene_tx=XM_019831373.2;refGene_gn=SASH1;refGene_tx=XM_019831374.2;refGene_gn=SASH1;refGene_tx=XM_019831375.2;refGene_gn=SASH1;refGene_tx=XM_023254476.1;refGene_gn=SASH1;refGene_tx=XM_023254477.1;refGene_gn=SASH1;refGene_tx=XM_023254478.1;refGene_gn=SASH1;refGene_tx=XM_023254479.1;refGene_gn=SASH1;refGene_tx=XM_023254480.1;refGene_gn=SASH1;refGene_tx=XM_023254481.1;refGene_gn=SASH1;refGene_tx=XM_023254482.1;refGene_gn=SASH1;refGene_tx=XM_023254483.1;refGene_gn=SASH1;refGene_tx=XM_023254484.1;refGene_gn=SASH1;refGene_tx=XM_023254485.1
B2 136805054 136805198
refGene_gn=SASH1;refGene_tx=XM_019831373.2;refGene_gn=SASH1;refGene_tx=XM_019831374.2;refGene_gn=SASH1;refGene_tx=XM_019831375.2;refGene_gn=SASH1;refGene_tx=XM_023254476.1;refGene_gn=SASH1;refGene_tx=XM_023254477.1;refGene_gn=SASH1;refGene_tx=XM_023254478.1;refGene_gn=SASH1;refGene_tx=XM_023254479.1;refGene_gn=SASH1;refGene_tx=XM_023254480.1;refGene_gn=SASH1;refGene_tx=XM_023254481.1;refGene_gn=SASH1;refGene_tx=XM_023254482.1;refGene_gn=SASH1;refGene_tx=XM_023254483.1;refGene_gn=SASH1;refGene_tx=XM_023254484.1;refGene_gn=SASH1;refGene_tx=XM_023254485.1
B2 136808463 136808599
refGene_gn=SASH1;refGene_tx=XM_019831373.2;refGene_gn=SASH1;refGene_tx=XM_019831374.2;refGene_gn=SASH1;refGene_tx=XM_019831375.2;refGene_gn=SASH1;refGene_tx=XM_023254476.1;refGene_gn=SASH1;refGene_tx=XM_023254477.1;refGene_gn=SASH1;refGene_tx=XM_023254478.1;refGene_gn=SASH1;refGene_tx=XM_023254479.1;refGene_gn=SASH1;refGene_tx=XM_023254480.1;refGene_gn=SASH1;refGene_tx=XM_023254481.1;refGene_gn=SASH1;refGene_tx=XM_023254482.1;refGene_gn=SASH1;refGene_tx=XM_023254483.1;refGene_gn=SASH1;refGene_tx=XM_023254484.1;refGene_gn=SASH1;refGene_tx=XM_023254485.1
B2 136809656 136809826
refGene_gn=SASH1;refGene_tx=XM_019831373.2;refGene_gn=SASH1;refGene_tx=XM_019831374.2;refGene_gn=SASH1;refGene_tx=XM_019831375.2;refGene_gn=SASH1;refGene_tx=XM_023254476.1;refGene_gn=SASH1;refGene_tx=XM_023254477.1;refGene_gn=SASH1;refGene_tx=XM_023254478.1;refGene_gn=SASH1;refGene_tx=XM_023254479.1;refGene_gn=SASH1;refGene_tx=XM_023254480.1;refGene_gn=SASH1;refGene_tx=XM_023254481.1;refGene_gn=SASH1;refGene_tx=XM_023254482.1;refGene_gn=SASH1;refGene_tx=XM_023254483.1;refGene_gn=SASH1;refGene_tx=XM_023254484.1;refGene_gn=SASH1;refGene_tx=XM_023254485.1
B2 136810589 136810799
refGene_gn=SASH1;refGene_tx=XM_019831373.2;refGene_gn=SASH1;refGene_tx=XM_019831374.2;refGene_gn=SASH1;refGene_tx=XM_019831375.2;refGene_gn=SASH1;refGene_tx=XM_023254476.1;refGene_gn=SASH1;refGene_tx=XM_023254477.1;refGene_gn=SASH1;refGene_tx=XM_023254478.1;refGene_gn=SASH1;refGene_tx=XM_023254479.1;refGene_gn=SASH1;refGene_tx=XM_023254480.1;refGene_gn=SASH1;refGene_tx=XM_023254481.1;refGene_gn=SASH1;refGene_tx=XM_02

3254482.1;refGene_gn=SASH1;refGene_tx=XM_023254483.1;refGene_gn=SASH1;
refGene_tx=XM_023254484.1;refGene_gn=SASH1;refGene_tx=XM_023254485.1
B2 136811592 136811743
refGene_gn=SASH1;refGene_tx=XM_019831373.2;refGene_gn=SASH1;refGene_tx
=XM_019831374.2;refGene_gn=SASH1;refGene_tx=XM_019831375.2;refGene_gn=
SASH1;refGene_tx=XM_023254476.1;refGene_gn=SASH1;refGene_tx=XM_0232544
77.1;refGene_gn=SASH1;refGene_tx=XM_023254478.1;refGene_gn=SASH1;refGe
ne_tx=XM_023254479.1;refGene_gn=SASH1;refGene_tx=XM_023254480.1;refGen
e_gn=SASH1;refGene_tx=XM_023254481.1;refGene_gn=SASH1;refGene_tx=XM_02
3254482.1;refGene_gn=SASH1;refGene_tx=XM_023254483.1;refGene_gn=SASH1;
refGene_tx=XM_023254484.1;refGene_gn=SASH1;refGene_tx=XM_023254485.1
B2 136819106 136819220
refGene_gn=SASH1;refGene_tx=XM_019831373.2;refGene_gn=SASH1;refGene_tx
=XM_019831374.2;refGene_gn=SASH1;refGene_tx=XM_019831375.2;refGene_gn=
SASH1;refGene_tx=XM_023254476.1;refGene_gn=SASH1;refGene_tx=XM_0232544
77.1;refGene_gn=SASH1;refGene_tx=XM_023254478.1;refGene_gn=SASH1;refGe
ne_tx=XM_023254479.1;refGene_gn=SASH1;refGene_tx=XM_023254480.1;refGen
e_gn=SASH1;refGene_tx=XM_023254481.1;refGene_gn=SASH1;refGene_tx=XM_02
3254482.1;refGene_gn=SASH1;refGene_tx=XM_023254483.1;refGene_gn=SASH1;
refGene_tx=XM_023254484.1;refGene_gn=SASH1;refGene_tx=XM_023254485.1
B2 136832009 136833118
refGene_gn=SASH1;refGene_tx=XM_019831373.2;refGene_gn=SASH1;refGene_tx
=XM_019831374.2;refGene_gn=SASH1;refGene_tx=XM_019831375.2;refGene_gn=
SASH1;refGene_tx=XM_023254476.1;refGene_gn=SASH1;refGene_tx=XM_0232544
77.1;refGene_gn=SASH1;refGene_tx=XM_023254478.1;refGene_gn=SASH1;refGe
ne_tx=XM_023254479.1;refGene_gn=SASH1;refGene_tx=XM_023254480.1;refGen
e_gn=SASH1;refGene_tx=XM_023254481.1;refGene_gn=SASH1;refGene_tx=XM_02
3254482.1;refGene_gn=SASH1;refGene_tx=XM_023254483.1;refGene_gn=SASH1;
refGene_tx=XM_023254484.1;refGene_gn=SASH1;refGene_tx=XM_023254485.1
B2 136834405 136834537
refGene_gn=SASH1;refGene_tx=XM_019831373.2;refGene_gn=SASH1;refGene_tx
=XM_019831374.2;refGene_gn=SASH1;refGene_tx=XM_019831375.2;refGene_gn=
SASH1;refGene_tx=XM_023254476.1;refGene_gn=SASH1;refGene_tx=XM_0232544
77.1;refGene_gn=SASH1;refGene_tx=XM_023254478.1;refGene_gn=SASH1;refGe
ne_tx=XM_023254479.1;refGene_gn=SASH1;refGene_tx=XM_023254480.1;refGen
e_gn=SASH1;refGene_tx=XM_023254481.1;refGene_gn=SASH1;refGene_tx=XM_02
3254482.1;refGene_gn=SASH1;refGene_tx=XM_023254483.1;refGene_gn=SASH1;
refGene_tx=XM_023254484.1;refGene_gn=SASH1;refGene_tx=XM_023254485.1
B2 136836177 136836441
refGene_gn=SASH1;refGene_tx=XM_019831373.2;refGene_gn=SASH1;refGene_tx
=XM_019831374.2;refGene_gn=SASH1;refGene_tx=XM_019831375.2;refGene_gn=
SASH1;refGene_tx=XM_023254476.1;refGene_gn=SASH1;refGene_tx=XM_0232544
77.1;refGene_gn=SASH1;refGene_tx=XM_023254478.1;refGene_gn=SASH1;refGe
ne_tx=XM_023254479.1;refGene_gn=SASH1;refGene_tx=XM_023254480.1;refGen
e_gn=SASH1;refGene_tx=XM_023254481.1;refGene_gn=SASH1;refGene_tx=XM_02
3254482.1;refGene_gn=SASH1;refGene_tx=XM_023254483.1;refGene_gn=SASH1;
refGene_tx=XM_023254484.1;refGene_gn=SASH1;refGene_tx=XM_023254485.1
B2 136996115 136996362
refGene_gn=UST;refGene_tx=XM_023254490.1;refGene_gn=UST;refGene_tx=XM_
023254491.1;refGene_gn=UST;refGene_tx=XM_023254492.1;refGene_gn=UST;re

fGene_tx=XM_023254493.1
B2	137115808	137115852
refGene_gn=UST;refGene_tx=XM_023254490.1;refGene_gn=UST;refGene_tx=XM_
023254491.1;refGene_gn=UST;refGene_tx=XM_023254492.1;refGene_gn=UST;re
fGene_tx=XM_023254493.1
B2	137171184	137171340
refGene_gn=UST;refGene_tx=XM_023254490.1;refGene_gn=UST;refGene_tx=XM_
023254491.1;refGene_gn=UST;refGene_tx=XM_023254492.1;refGene_gn=UST;re
fGene_tx=XM_023254493.1
B2	137183839	137183919
refGene_gn=UST;refGene_tx=XM_023254490.1;refGene_gn=UST;refGene_tx=XM_
023254491.1;refGene_gn=UST;refGene_tx=XM_023254492.1;refGene_gn=UST;re
fGene_tx=XM_023254493.1
B2	137195279	137195433
refGene_gn=UST;refGene_tx=XM_023254490.1;refGene_gn=UST;refGene_tx=XM_
023254491.1;refGene_gn=UST;refGene_tx=XM_023254492.1;refGene_gn=UST;re
fGene_tx=XM_023254493.1
B2	137248857	137248955
refGene_gn=UST;refGene_tx=XM_023254490.1;refGene_gn=UST;refGene_tx=XM_
023254491.1;refGene_gn=UST;refGene_tx=XM_023254492.1;refGene_gn=UST;re
fGene_tx=XM_023254493.1
B2	137250914	137251072
refGene_gn=UST;refGene_tx=XM_023254490.1;refGene_gn=UST;refGene_tx=XM_
023254491.1;refGene_gn=UST;refGene_tx=XM_023254492.1;refGene_gn=UST;re
fGene_tx=XM_023254493.1
B2	137300142	137300426
refGene_gn=UST;refGene_tx=XM_023254490.1;refGene_gn=UST;refGene_tx=XM_
023254491.1;refGene_gn=UST;refGene_tx=XM_023254492.1
B2	137301957	137302057
refGene_gn=UST;refGene_tx=XM_023254490.1
B2	137316235	137316348
refGene_gn=UST;refGene_tx=XM_023254491.1;refGene_gn=UST;refGene_tx=XM_
023254493.1
B2	137354204	137354381
refGene_gn=LOC101098462;refGene_tx=XM_011282576.2
B2	137354604	137354772
refGene_gn=LOC101098462;refGene_tx=XM_011282576.2
B2	137566861	137566963
refGene_gn=TAB2;refGene_tx=XM_006932085.4
B2	137576735	137578236
refGene_gn=TAB2;refGene_tx=XM_006932085.4
B2	137599976	137600137
refGene_gn=TAB2;refGene_tx=XM_006932085.4
B2	137600353	137600447
refGene_gn=TAB2;refGene_tx=XM_006932085.4
B2	137601770	137601851
refGene_gn=TAB2;refGene_tx=XM_006932085.4
B2	137613082	137613225
refGene_gn=TAB2;refGene_tx=XM_006932085.4
B2	137649529	137650314

refGene_gn=ZC3H12D;refGene_tx=XM_023254494.1
B2 137651461 137651568
refGene_gn=ZC3H12D;refGene_tx=XM_023254494.1
B2 137654848 137655083
refGene_gn=ZC3H12D;refGene_tx=XM_023254494.1
B2 137661199 137661339
refGene_gn=ZC3H12D;refGene_tx=XM_023254494.1
B2 137671113 137671418
refGene_gn=ZC3H12D;refGene_tx=XM_023254494.1
B2 137699681 137699933
refGene_gn=PPIL4;refGene_tx=XM_003986636.5
B2 137705907 137706055
refGene_gn=PPIL4;refGene_tx=XM_003986636.5
B2 137710464 137710561
refGene_gn=PPIL4;refGene_tx=XM_003986636.5
B2 137714827 137714939
refGene_gn=PPIL4;refGene_tx=XM_003986636.5
B2 137716595 137716662
refGene_gn=PPIL4;refGene_tx=XM_003986636.5
B2 137719060 137719185
refGene_gn=PPIL4;refGene_tx=XM_003986636.5
B2 137724459 137724576
refGene_gn=PPIL4;refGene_tx=XM_003986636.5
B2 137725313 137725410
refGene_gn=PPIL4;refGene_tx=XM_003986636.5
B2 137725905 137726048
refGene_gn=PPIL4;refGene_tx=XM_003986636.5
B2 137726745 137726863
refGene_gn=PPIL4;refGene_tx=XM_003986636.5
B2 137727122 137727187
refGene_gn=PPIL4;refGene_tx=XM_003986636.5
B2 137727271 137727339
refGene_gn=PPIL4;refGene_tx=XM_003986636.5
B2 137733252 137733322
refGene_gn=PPIL4;refGene_tx=XM_003986636.5
B2 137740181 137740301
refGene_gn=GINM1;refGene_tx=XM_003986637.5;refGene_gn=GINM1;refGene_tx
=XM_023254495.1
B2 137745409 137745469
refGene_gn=GINM1;refGene_tx=XM_003986637.5;refGene_gn=GINM1;refGene_tx
=XM_023254495.1
B2 137745639 137745736
refGene_gn=GINM1;refGene_tx=XM_003986637.5;refGene_gn=GINM1;refGene_tx
=XM_023254495.1
B2 137753829 137753981
refGene_gn=GINM1;refGene_tx=XM_003986637.5;refGene_gn=GINM1;refGene_tx
=XM_023254495.1
B2 137754527 137754684
refGene_gn=GINM1;refGene_tx=XM_003986637.5;refGene_gn=GINM1;refGene_tx
=XM_023254495.1

B2 137755493 137755627
refGene_gn=GINM1;refGene_tx=XM_003986637.5;refGene_gn=GINM1;refGene_tx=XM_023254495.1
B2 137757215 137757379
refGene_gn=GINM1;refGene_tx=XM_003986637.5;refGene_gn=GINM1;refGene_tx=XM_023254495.1
B2 137761496 137761608
refGene_gn=GINM1;refGene_tx=XM_003986637.5;refGene_gn=GINM1;refGene_tx=XM_023254495.1
B2 137764827 137765026
refGene_gn=KATNA1;refGene_tx=XM_003986638.5;refGene_gn=KATNA1;refGene_tx=XM_019831380.2
B2 137766271 137766398
refGene_gn=KATNA1;refGene_tx=XM_003986638.5;refGene_gn=KATNA1;refGene_tx=XM_019831380.2
B2 137766711 137766846
refGene_gn=KATNA1;refGene_tx=XM_003986638.5;refGene_gn=KATNA1;refGene_tx=XM_019831380.2
B2 137767366 137767493
refGene_gn=KATNA1;refGene_tx=XM_003986638.5;refGene_gn=KATNA1;refGene_tx=XM_019831380.2
B2 137771972 137772131
refGene_gn=KATNA1;refGene_tx=XM_003986638.5;refGene_gn=KATNA1;refGene_tx=XM_019831380.2
B2 137772950 137773056
refGene_gn=KATNA1;refGene_tx=XM_003986638.5;refGene_gn=KATNA1;refGene_tx=XM_019831380.2
B2 137774226 137774348
refGene_gn=KATNA1;refGene_tx=XM_003986638.5;refGene_gn=KATNA1;refGene_tx=XM_019831380.2
B2 137785700 137785881
refGene_gn=KATNA1;refGene_tx=XM_003986638.5;refGene_gn=KATNA1;refGene_tx=XM_019831380.2
B2 137794745 137794903
refGene_gn=KATNA1;refGene_tx=XM_003986638.5;refGene_gn=KATNA1;refGene_tx=XM_019831380.2
B2 137796387 137796549
refGene_gn=KATNA1;refGene_tx=XM_003986638.5;refGene_gn=KATNA1;refGene_tx=XM_019831380.2
B2 137819442 137819952
refGene_gn=LATS1;refGene_tx=XM_011282574.3
B2 137824848 137824955
refGene_gn=LATS1;refGene_tx=XM_011282574.3
B2 137825154 137825337
refGene_gn=LATS1;refGene_tx=XM_011282574.3;refGene_gn=LATS1;refGene_tx=XM_019831381.2
B2 137827044 137827108
refGene_gn=LATS1;refGene_tx=XM_019831381.2
B2 137827926 137828509
refGene_gn=LATS1;refGene_tx=XM_011282574.3;refGene_gn=LATS1;refGene_tx

=XM_019831381.2
B2 137829570 137829579
refGene_gn=LATS1;refGene_tx=XM_023254496.1
B2 137830776 137832287
refGene_gn=LATS1;refGene_tx=XM_011282574.3;refGene_gn=LATS1;refGene_tx
=XM_019831381.2;refGene_gn=LATS1;refGene_tx=XM_023254496.1
B2 137836081 137836229
refGene_gn=LATS1;refGene_tx=XM_011282574.3;refGene_gn=LATS1;refGene_tx
=XM_019831381.2;refGene_gn=LATS1;refGene_tx=XM_023254496.1
B2 137849122 137849464
refGene_gn=LATS1;refGene_tx=XM_011282574.3;refGene_gn=LATS1;refGene_tx
=XM_019831381.2;refGene_gn=LATS1;refGene_tx=XM_023254496.1
B2 137862490 137862536
refGene_gn=LOC102899086;refGene_tx=XM_006932100.3
B2 137863920 137863984
refGene_gn=LOC102899086;refGene_tx=XM_006932100.3
B2 137869418 137870107
refGene_gn=LOC102899086;refGene_tx=XM_006932100.3
B2 137870183 137870239
refGene_gn=LOC102899086;refGene_tx=XM_006932100.3
B2 137878308 137878538
refGene_gn=NUP43;refGene_tx=XM_003986640.5
B2 137879060 137879183
refGene_gn=NUP43;refGene_tx=XM_003986640.5
B2 137881452 137881604
refGene_gn=NUP43;refGene_tx=XM_003986640.5
B2 137883062 137883198
refGene_gn=NUP43;refGene_tx=XM_003986640.5
B2 137885969 137886150
refGene_gn=NUP43;refGene_tx=XM_003986640.5
B2 137888314 137888392
refGene_gn=NUP43;refGene_tx=XM_003986640.5
B2 137889472 137889595
refGene_gn=NUP43;refGene_tx=XM_003986640.5
B2 137889912 137890032
refGene_gn=NUP43;refGene_tx=XM_003986640.5
B2 137893466 137893695
refGene_gn=PCMT1;refGene_tx=XM_023254497.1;refGene_gn=PCMT1;refGene_tx
=XM_023254498.1
B2 137914992 137915097
refGene_gn=PCMT1;refGene_tx=XM_023254497.1;refGene_gn=PCMT1;refGene_tx
=XM_023254498.1
B2 137916307 137916339
refGene_gn=PCMT1;refGene_tx=XM_023254497.1;refGene_gn=PCMT1;refGene_tx
=XM_023254498.1
B2 137927466 137927571
refGene_gn=PCMT1;refGene_tx=XM_023254497.1;refGene_gn=PCMT1;refGene_tx
=XM_023254498.1
B2 137930156 137930277
refGene_gn=PCMT1;refGene_tx=XM_023254497.1;refGene_gn=PCMT1;refGene_tx

=XM_023254498.1
B2 137932425 137932511
refGene_gn=PCMT1;refGene_tx=XM_023254497.1;refGene_gn=PCMT1;refGene_tx=XM_023254498.1
B2 137937127 137937307
refGene_gn=PCMT1;refGene_tx=XM_023254497.1;refGene_gn=PCMT1;refGene_tx=XM_023254498.1
B2 137944990 137945003
refGene_gn=PCMT1;refGene_tx=XM_023254497.1
B2 137953837 137953992
refGene_gn=LRP11;refGene_tx=XM_006932087.4;refGene_gn=LRP11;refGene_tx=XM_023254499.1;refGene_gn=LRP11;refGene_tx=XM_023254500.1;refGene_gn=LRP11;refGene_tx=XM_023254501.1
B2 137959134 137959230
refGene_gn=LRP11;refGene_tx=XM_006932087.4;refGene_gn=LRP11;refGene_tx=XM_023254499.1;refGene_gn=LRP11;refGene_tx=XM_023254501.1
B2 137971768 137971975
refGene_gn=LRP11;refGene_tx=XM_006932087.4;refGene_gn=LRP11;refGene_tx=XM_023254499.1;refGene_gn=LRP11;refGene_tx=XM_023254500.1;refGene_gn=LRP11;refGene_tx=XM_023254501.1
B2 137973366 137973492
refGene_gn=LRP11;refGene_tx=XM_006932087.4;refGene_gn=LRP11;refGene_tx=XM_023254499.1;refGene_gn=LRP11;refGene_tx=XM_023254500.1
B2 137979265 137979473
refGene_gn=LRP11;refGene_tx=XM_006932087.4;refGene_gn=LRP11;refGene_tx=XM_023254499.1;refGene_gn=LRP11;refGene_tx=XM_023254500.1;refGene_gn=LRP11;refGene_tx=XM_023254501.1
B2 137997406 137997564
refGene_gn=LRP11;refGene_tx=XM_006932087.4;refGene_gn=LRP11;refGene_tx=XM_023254499.1;refGene_gn=LRP11;refGene_tx=XM_023254500.1;refGene_gn=LRP11;refGene_tx=XM_023254501.1
B2 138004416 138005026
refGene_gn=LRP11;refGene_tx=XM_006932087.4;refGene_gn=LRP11;refGene_tx=XM_023254499.1;refGene_gn=LRP11;refGene_tx=XM_023254500.1;refGene_gn=LRP11;refGene_tx=XM_023254501.1
B2 138016592 138016672
refGene_gn=LOC101082237;refGene_tx=XM_019831629.1
B2 138020028 138020157
refGene_gn=LOC101082237;refGene_tx=XM_019831629.1
B2 138020966 138021239
refGene_gn=LOC101082237;refGene_tx=XM_019831629.1
B2 138021423 138021687
refGene_gn=LOC101082237;refGene_tx=XM_019831629.1
B2 138022271 138022380
refGene_gn=LOC101082237;refGene_tx=XM_019831629.1
B2 138058591 138058682
refGene_gn=LOC102899212;refGene_tx=XM_011282572.3;refGene_gn=LOC102899212;refGene_tx=XM_011282573.3;refGene_gn=LOC102899212;refGene_tx=XM_019831384.2
B2 138064586 138064850

refGene_gn=LOC102899212;refGene_tx=XM_011282572.3;refGene_gn=LOC102899212;refGene_tx=XM_011282573.3;refGene_gn=LOC102899212;refGene_tx=XM_019831384.2
B2 138065086 138065359
refGene_gn=LOC102899212;refGene_tx=XM_011282572.3;refGene_gn=LOC102899212;refGene_tx=XM_011282573.3;refGene_gn=LOC102899212;refGene_tx=XM_019831384.2
B2 138066105 138066230
refGene_gn=LOC102899212;refGene_tx=XM_011282572.3;refGene_gn=LOC102899212;refGene_tx=XM_011282573.3;refGene_gn=LOC102899212;refGene_tx=XM_019831384.2
B2 138067795 138067803
refGene_gn=LOC102899212;refGene_tx=XM_011282573.3
B2 138067891 138067947
refGene_gn=LOC102899212;refGene_tx=XM_019831384.2
B2 138081923 138081931
refGene_gn=LOC105260590;refGene_tx=XM_023254502.1
B2 138083329 138083440
refGene_gn=LOC105260590;refGene_tx=XM_023254502.1
B2 138084179 138084452
refGene_gn=LOC105260590;refGene_tx=XM_023254502.1
B2 138084688 138084952
refGene_gn=LOC105260590;refGene_tx=XM_023254502.1
B2 138090480 138090601
refGene_gn=LOC105260590;refGene_tx=XM_023254502.1
B2 138159137 138159476
refGene_gn=PPP1R14C;refGene_tx=XM_019831630.2
B2 138218236 138218320
refGene_gn=PPP1R14C;refGene_tx=XM_019831630.2
B2 138220390 138220423
refGene_gn=PPP1R14C;refGene_tx=XM_019831630.2
B2 138246668 138246743
refGene_gn=PPP1R14C;refGene_tx=XM_019831630.2
B2 138343863 138344041
refGene_gn=IYD;refGene_tx=XM_003986641.4
B2 138353578 138353770
refGene_gn=IYD;refGene_tx=XM_003986641.4
B2 138356046 138356206
refGene_gn=IYD;refGene_tx=XM_003986641.4
B2 138359388 138359545
refGene_gn=IYD;refGene_tx=XM_003986641.4
B2 138361459 138361642
refGene_gn=IYD;refGene_tx=XM_003986641.4
B2 138621293 138621498
refGene_gn=PLEKHG1;refGene_tx=XM_019831385.2
B2 138623688 138623806
refGene_gn=PLEKHG1;refGene_tx=XM_019831386.2
B2 138654602 138654630
refGene_gn=PLEKHG1;refGene_tx=XM_019831387.2
B2 138664589 138665095

refGene_gn=PLEKHG1;refGene_tx=XM_019831385.2;refGene_gn=PLEKHG1;refGene_tx=XM_019831386.2;refGene_gn=PLEKHG1;refGene_tx=XM_019831387.2;refGene_gn=PLEKHG1;refGene_tx=XM_019831388.2;refGene_gn=PLEKHG1;refGene_tx=XM_023254503.1;refGene_gn=PLEKHG1;refGene_tx=XM_023254504.1
B2 138700487 138700588
refGene_gn=PLEKHG1;refGene_tx=XM_019831385.2;refGene_gn=PLEKHG1;refGene_tx=XM_019831386.2;refGene_gn=PLEKHG1;refGene_tx=XM_019831387.2;refGene_gn=PLEKHG1;refGene_tx=XM_019831388.2;refGene_gn=PLEKHG1;refGene_tx=XM_023254503.1;refGene_gn=PLEKHG1;refGene_tx=XM_023254504.1
B2 138718676 138718746
refGene_gn=PLEKHG1;refGene_tx=XM_019831385.2;refGene_gn=PLEKHG1;refGene_tx=XM_019831386.2;refGene_gn=PLEKHG1;refGene_tx=XM_019831387.2;refGene_gn=PLEKHG1;refGene_tx=XM_019831388.2;refGene_gn=PLEKHG1;refGene_tx=XM_023254503.1;refGene_gn=PLEKHG1;refGene_tx=XM_023254504.1
B2 138728389 138728436
refGene_gn=PLEKHG1;refGene_tx=XM_019831385.2;refGene_gn=PLEKHG1;refGene_tx=XM_019831386.2;refGene_gn=PLEKHG1;refGene_tx=XM_019831387.2;refGene_gn=PLEKHG1;refGene_tx=XM_019831388.2;refGene_gn=PLEKHG1;refGene_tx=XM_023254503.1;refGene_gn=PLEKHG1;refGene_tx=XM_023254504.1
B2 138732975 138733126
refGene_gn=PLEKHG1;refGene_tx=XM_019831385.2;refGene_gn=PLEKHG1;refGene_tx=XM_019831386.2;refGene_gn=PLEKHG1;refGene_tx=XM_019831387.2;refGene_gn=PLEKHG1;refGene_tx=XM_019831388.2;refGene_gn=PLEKHG1;refGene_tx=XM_023254503.1;refGene_gn=PLEKHG1;refGene_tx=XM_023254504.1
B2 138737839 138737971
refGene_gn=PLEKHG1;refGene_tx=XM_019831385.2;refGene_gn=PLEKHG1;refGene_tx=XM_019831386.2;refGene_gn=PLEKHG1;refGene_tx=XM_019831387.2;refGene_gn=PLEKHG1;refGene_tx=XM_019831388.2;refGene_gn=PLEKHG1;refGene_tx=XM_023254503.1;refGene_gn=PLEKHG1;refGene_tx=XM_023254504.1
B2 138740766 138740949
refGene_gn=PLEKHG1;refGene_tx=XM_019831385.2;refGene_gn=PLEKHG1;refGene_tx=XM_019831386.2;refGene_gn=PLEKHG1;refGene_tx=XM_019831387.2;refGene_gn=PLEKHG1;refGene_tx=XM_019831388.2;refGene_gn=PLEKHG1;refGene_tx=XM_023254503.1;refGene_gn=PLEKHG1;refGene_tx=XM_023254504.1
B2 138741042 138741138
refGene_gn=PLEKHG1;refGene_tx=XM_019831385.2;refGene_gn=PLEKHG1;refGene_tx=XM_019831386.2;refGene_gn=PLEKHG1;refGene_tx=XM_019831387.2;refGene_gn=PLEKHG1;refGene_tx=XM_019831388.2;refGene_gn=PLEKHG1;refGene_tx=XM_023254503.1;refGene_gn=PLEKHG1;refGene_tx=XM_023254504.1
B2 138741331 138741418
refGene_gn=PLEKHG1;refGene_tx=XM_019831385.2;refGene_gn=PLEKHG1;refGene_tx=XM_019831386.2;refGene_gn=PLEKHG1;refGene_tx=XM_019831387.2;refGene_gn=PLEKHG1;refGene_tx=XM_019831388.2;refGene_gn=PLEKHG1;refGene_tx=XM_023254503.1;refGene_gn=PLEKHG1;refGene_tx=XM_023254504.1
B2 138747386 138747420
refGene_gn=PLEKHG1;refGene_tx=XM_019831385.2;refGene_gn=PLEKHG1;refGene_tx=XM_019831386.2;refGene_gn=PLEKHG1;refGene_tx=XM_019831387.2;refGene_gn=PLEKHG1;refGene_tx=XM_019831388.2;refGene_gn=PLEKHG1;refGene_tx=XM_023254503.1;refGene_gn=PLEKHG1;refGene_tx=XM_023254504.1
B2 138750433 138750529

refGene_gn=PLEKHG1;refGene_tx=XM_019831385.2;refGene_gn=PLEKHG1;refGene_tx=XM_019831386.2;refGene_gn=PLEKHG1;refGene_tx=XM_019831387.2;refGene_gn=PLEKHG1;refGene_tx=XM_019831388.2;refGene_gn=PLEKHG1;refGene_tx=XM_023254503.1;refGene_gn=PLEKHG1;refGene_tx=XM_023254504.1
B2 138752766 138752805
refGene_gn=PLEKHG1;refGene_tx=XM_019831385.2;refGene_gn=PLEKHG1;refGene_tx=XM_019831386.2;refGene_gn=PLEKHG1;refGene_tx=XM_019831387.2;refGene_gn=PLEKHG1;refGene_tx=XM_019831388.2;refGene_gn=PLEKHG1;refGene_tx=XM_023254503.1;refGene_gn=PLEKHG1;refGene_tx=XM_023254504.1
B2 138754757 138754780
refGene_gn=PLEKHG1;refGene_tx=XM_019831385.2;refGene_gn=PLEKHG1;refGene_tx=XM_019831386.2;refGene_gn=PLEKHG1;refGene_tx=XM_019831387.2;refGene_gn=PLEKHG1;refGene_tx=XM_019831388.2;refGene_gn=PLEKHG1;refGene_tx=XM_023254503.1;refGene_gn=PLEKHG1;refGene_tx=XM_023254504.1
B2 138759425 138761067
refGene_gn=PLEKHG1;refGene_tx=XM_019831385.2;refGene_gn=PLEKHG1;refGene_tx=XM_019831386.2;refGene_gn=PLEKHG1;refGene_tx=XM_019831387.2;refGene_gn=PLEKHG1;refGene_tx=XM_019831388.2;refGene_gn=PLEKHG1;refGene_tx=XM_023254503.1;refGene_gn=PLEKHG1;refGene_tx=XM_023254504.1
B2 138767083 138768147
refGene_gn=PLEKHG1;refGene_tx=XM_019831385.2;refGene_gn=PLEKHG1;refGene_tx=XM_019831386.2;refGene_gn=PLEKHG1;refGene_tx=XM_019831387.2;refGene_gn=PLEKHG1;refGene_tx=XM_019831388.2;refGene_gn=PLEKHG1;refGene_tx=XM_023254503.1;refGene_gn=PLEKHG1;refGene_tx=XM_023254504.1
B2 138781419 138781646
refGene_gn=MTHFD1L;refGene_tx=XM_023254505.1;refGene_gn=MTHFD1L;refGene_tx=XM_023254506.1;refGene_gn=MTHFD1L;refGene_tx=XM_023254507.1;refGene_gn=MTHFD1L;refGene_tx=XM_023254509.1
B2 138788149 138788234
refGene_gn=MTHFD1L;refGene_tx=XM_023254505.1;refGene_gn=MTHFD1L;refGene_tx=XM_023254506.1;refGene_gn=MTHFD1L;refGene_tx=XM_023254507.1;refGene_gn=MTHFD1L;refGene_tx=XM_023254509.1
B2 138789620 138789671
refGene_gn=MTHFD1L;refGene_tx=XM_023254505.1;refGene_gn=MTHFD1L;refGene_tx=XM_023254506.1;refGene_gn=MTHFD1L;refGene_tx=XM_023254507.1;refGene_gn=MTHFD1L;refGene_tx=XM_023254509.1
B2 138789761 138789815
refGene_gn=MTHFD1L;refGene_tx=XM_023254505.1;refGene_gn=MTHFD1L;refGene_tx=XM_023254506.1;refGene_gn=MTHFD1L;refGene_tx=XM_023254507.1;refGene_gn=MTHFD1L;refGene_tx=XM_023254509.1
B2 138792402 138792527
refGene_gn=MTHFD1L;refGene_tx=XM_023254505.1;refGene_gn=MTHFD1L;refGene_tx=XM_023254506.1;refGene_gn=MTHFD1L;refGene_tx=XM_023254507.1;refGene_gn=MTHFD1L;refGene_tx=XM_023254509.1
B2 138794377 138794478
refGene_gn=MTHFD1L;refGene_tx=XM_023254505.1;refGene_gn=MTHFD1L;refGene_tx=XM_023254506.1;refGene_gn=MTHFD1L;refGene_tx=XM_023254507.1;refGene_gn=MTHFD1L;refGene_tx=XM_023254509.1
B2 138796438 138796575
refGene_gn=MTHFD1L;refGene_tx=XM_023254505.1;refGene_gn=MTHFD1L;refGen

e_tx=XM_023254506.1;refGene_gn=MTHFD1L;refGene_tx=XM_023254507.1;refGene_gn=MTHFD1L;refGene_tx=XM_023254509.1
B2 138811390 138811502
refGene_gn=MTHFD1L;refGene_tx=XM_023254505.1;refGene_gn=MTHFD1L;refGene_tx=XM_023254506.1;refGene_gn=MTHFD1L;refGene_tx=XM_023254507.1;refGene_gn=MTHFD1L;refGene_tx=XM_023254509.1
B2 138819126 138819218
refGene_gn=MTHFD1L;refGene_tx=XM_023254505.1;refGene_gn=MTHFD1L;refGene_tx=XM_023254506.1;refGene_gn=MTHFD1L;refGene_tx=XM_023254507.1;refGene_gn=MTHFD1L;refGene_tx=XM_023254509.1
B2 138820983 138821081
refGene_gn=MTHFD1L;refGene_tx=XM_023254505.1;refGene_gn=MTHFD1L;refGene_tx=XM_023254506.1;refGene_gn=MTHFD1L;refGene_tx=XM_023254507.1;refGene_gn=MTHFD1L;refGene_tx=XM_023254508.1;refGene_gn=MTHFD1L;refGene_tx=XM_023254509.1
B2 138823675 138823849
refGene_gn=MTHFD1L;refGene_tx=XM_023254505.1;refGene_gn=MTHFD1L;refGene_tx=XM_023254506.1;refGene_gn=MTHFD1L;refGene_tx=XM_023254507.1;refGene_gn=MTHFD1L;refGene_tx=XM_023254508.1;refGene_gn=MTHFD1L;refGene_tx=XM_023254509.1
B2 138832126 138832263
refGene_gn=MTHFD1L;refGene_tx=XM_023254505.1;refGene_gn=MTHFD1L;refGene_tx=XM_023254506.1;refGene_gn=MTHFD1L;refGene_tx=XM_023254507.1;refGene_gn=MTHFD1L;refGene_tx=XM_023254508.1;refGene_gn=MTHFD1L;refGene_tx=XM_023254509.1
B2 138833994 138834041
refGene_gn=MTHFD1L;refGene_tx=XM_023254505.1;refGene_gn=MTHFD1L;refGene_tx=XM_023254506.1;refGene_gn=MTHFD1L;refGene_tx=XM_023254507.1;refGene_gn=MTHFD1L;refGene_tx=XM_023254508.1;refGene_gn=MTHFD1L;refGene_tx=XM_023254509.1
B2 138836788 138836896
refGene_gn=MTHFD1L;refGene_tx=XM_023254505.1;refGene_gn=MTHFD1L;refGene_tx=XM_023254506.1;refGene_gn=MTHFD1L;refGene_tx=XM_023254507.1;refGene_gn=MTHFD1L;refGene_tx=XM_023254508.1;refGene_gn=MTHFD1L;refGene_tx=XM_023254509.1
B2 138837987 138838062
refGene_gn=MTHFD1L;refGene_tx=XM_023254505.1;refGene_gn=MTHFD1L;refGene_tx=XM_023254506.1;refGene_gn=MTHFD1L;refGene_tx=XM_023254507.1;refGene_gn=MTHFD1L;refGene_tx=XM_023254508.1;refGene_gn=MTHFD1L;refGene_tx=XM_023254509.1
B2 138838870 138838973
refGene_gn=MTHFD1L;refGene_tx=XM_023254505.1;refGene_gn=MTHFD1L;refGene_tx=XM_023254506.1;refGene_gn=MTHFD1L;refGene_tx=XM_023254507.1;refGene_gn=MTHFD1L;refGene_tx=XM_023254508.1;refGene_gn=MTHFD1L;refGene_tx=XM_023254509.1
B2 138842268 138842345
refGene_gn=MTHFD1L;refGene_tx=XM_023254505.1;refGene_gn=MTHFD1L;refGene_tx=XM_023254506.1;refGene_gn=MTHFD1L;refGene_tx=XM_023254507.1;refGene_gn=MTHFD1L;refGene_tx=XM_023254508.1;refGene_gn=MTHFD1L;refGene_tx=XM_023254509.1

B2	138845446	138845575
refGene_gn=MTHFD1L;refGene_tx=XM_023254509.1
B2	138847199	138847340
refGene_gn=MTHFD1L;refGene_tx=XM_023254505.1;refGene_gn=MTHFD1L;refGene_tx=XM_023254506.1;refGene_gn=MTHFD1L;refGene_tx=XM_023254507.1;refGene_gn=MTHFD1L;refGene_tx=XM_023254508.1
B2	138850673	138850742
refGene_gn=MTHFD1L;refGene_tx=XM_023254505.1;refGene_gn=MTHFD1L;refGene_tx=XM_023254506.1;refGene_gn=MTHFD1L;refGene_tx=XM_023254507.1;refGene_gn=MTHFD1L;refGene_tx=XM_023254508.1
B2	138857680	138857792
refGene_gn=MTHFD1L;refGene_tx=XM_023254505.1;refGene_gn=MTHFD1L;refGene_tx=XM_023254506.1;refGene_gn=MTHFD1L;refGene_tx=XM_023254507.1;refGene_gn=MTHFD1L;refGene_tx=XM_023254508.1
B2	138890268	138890408
refGene_gn=MTHFD1L;refGene_tx=XM_023254505.1;refGene_gn=MTHFD1L;refGene_tx=XM_023254506.1;refGene_gn=MTHFD1L;refGene_tx=XM_023254507.1;refGene_gn=MTHFD1L;refGene_tx=XM_023254508.1
B2	138897335	138897377
refGene_gn=MTHFD1L;refGene_tx=XM_023254505.1;refGene_gn=MTHFD1L;refGene_tx=XM_023254507.1;refGene_gn=MTHFD1L;refGene_tx=XM_023254508.1
B2	138898071	138898273
refGene_gn=MTHFD1L;refGene_tx=XM_023254505.1;refGene_gn=MTHFD1L;refGene_tx=XM_023254507.1;refGene_gn=MTHFD1L;refGene_tx=XM_023254508.1
B2	138898815	138898993
refGene_gn=MTHFD1L;refGene_tx=XM_023254505.1;refGene_gn=MTHFD1L;refGene_tx=XM_023254508.1
B2	138914862	138914970
refGene_gn=MTHFD1L;refGene_tx=XM_023254505.1;refGene_gn=MTHFD1L;refGene_tx=XM_023254506.1;refGene_gn=MTHFD1L;refGene_tx=XM_023254508.1
B2	138917605	138917758
refGene_gn=MTHFD1L;refGene_tx=XM_023254505.1;refGene_gn=MTHFD1L;refGene_tx=XM_023254506.1;refGene_gn=MTHFD1L;refGene_tx=XM_023254508.1
B2	138964158	138964248
refGene_gn=MTHFD1L;refGene_tx=XM_023254505.1;refGene_gn=MTHFD1L;refGene_tx=XM_023254506.1;refGene_gn=MTHFD1L;refGene_tx=XM_023254508.1
B2	139087994	139088156
refGene_gn=AKAP12;refGene_tx=XM_011282570.3
B2	139144381	139144496
refGene_gn=AKAP12;refGene_tx=XM_011282570.3
B2	139183349	139187498
refGene_gn=AKAP12;refGene_tx=XM_011282570.3
B2	139187500	139187623
refGene_gn=AKAP12;refGene_tx=XM_011282570.3
B2	139187625	139188347
refGene_gn=AKAP12;refGene_tx=XM_011282570.3
B2	139201243	139201383
refGene_gn=ZBTB2;refGene_tx=XM_023254510.1
B2	139201385	139201932
refGene_gn=ZBTB2;refGene_tx=XM_023254510.1

B2 139201933 139202615
refGene_gn=ZBTB2;refGene_tx=XM_023254510.1
B2 139208049 139208222
refGene_gn=ZBTB2;refGene_tx=XM_023254510.1
B2 139235075 139235117
refGene_gn=RMND1;refGene_tx=XM_003986644.4
B2 139235584 139235701
refGene_gn=RMND1;refGene_tx=XM_003986644.4
B2 139239897 139240038
refGene_gn=RMND1;refGene_tx=XM_011282569.3
B2 139249443 139249564
refGene_gn=RMND1;refGene_tx=XM_003986644.4;refGene_gn=RMND1;refGene_tx=XM_011282569.3
B2 139251901 139251978
refGene_gn=RMND1;refGene_tx=XM_003986644.4;refGene_gn=RMND1;refGene_tx=XM_011282569.3
B2 139252692 139252757
refGene_gn=RMND1;refGene_tx=XM_003986644.4;refGene_gn=RMND1;refGene_tx=XM_011282569.3
B2 139253662 139253769
refGene_gn=RMND1;refGene_tx=XM_003986644.4;refGene_gn=RMND1;refGene_tx=XM_011282569.3
B2 139256673 139256774
refGene_gn=RMND1;refGene_tx=XM_003986644.4;refGene_gn=RMND1;refGene_tx=XM_011282569.3
B2 139258692 139258732
refGene_gn=RMND1;refGene_tx=XM_003986644.4;refGene_gn=RMND1;refGene_tx=XM_011282569.3
B2 139262123 139262199
refGene_gn=RMND1;refGene_tx=XM_003986644.4;refGene_gn=RMND1;refGene_tx=XM_011282569.3
B2 139263950 139264059
refGene_gn=RMND1;refGene_tx=XM_003986644.4;refGene_gn=RMND1;refGene_tx=XM_011282569.3
B2 139272929 139273433
refGene_gn=RMND1;refGene_tx=XM_003986644.4;refGene_gn=RMND1;refGene_tx=XM_011282569.3
B2 139279655 139279699
refGene_gn=ARMT1;refGene_tx=XM_003986645.5
B2 139281705 139281811
refGene_gn=ARMT1;refGene_tx=XM_003986645.5
B2 139282219 139282464
refGene_gn=ARMT1;refGene_tx=XM_003986645.5;refGene_gn=ARMT1;refGene_tx=XM_006932093.4
B2 139287961 139288127
refGene_gn=ARMT1;refGene_tx=XM_003986645.5;refGene_gn=ARMT1;refGene_tx=XM_006932093.4
B2 139291433 139292201
refGene_gn=ARMT1;refGene_tx=XM_003986645.5;refGene_gn=ARMT1;refGene_tx=XM_006932093.4

B2	139325590	139325596
refGene_gn=CCDC170;refGene_tx=XM_023254511.1
B2	139357171	139357300
refGene_gn=CCDC170;refGene_tx=XM_019831393.2;refGene_gn=CCDC170;refGene_tx=XM_019831394.2;refGene_gn=CCDC170;refGene_tx=XM_023254511.1;refGene_gn=CCDC170;refGene_tx=XM_023254512.1
B2	139359120	139359377
refGene_gn=CCDC170;refGene_tx=XM_023254511.1
B2	139363540	139363685
refGene_gn=CCDC170;refGene_tx=XM_019831393.2;refGene_gn=CCDC170;refGene_tx=XM_019831394.2;refGene_gn=CCDC170;refGene_tx=XM_023254511.1;refGene_gn=CCDC170;refGene_tx=XM_023254512.1
B2	139368415	139368601
refGene_gn=CCDC170;refGene_tx=XM_019831393.2;refGene_gn=CCDC170;refGene_tx=XM_019831394.2;refGene_gn=CCDC170;refGene_tx=XM_023254511.1;refGene_gn=CCDC170;refGene_tx=XM_023254512.1
B2	139379976	139380294
refGene_gn=CCDC170;refGene_tx=XM_019831393.2;refGene_gn=CCDC170;refGene_tx=XM_019831394.2;refGene_gn=CCDC170;refGene_tx=XM_023254511.1;refGene_gn=CCDC170;refGene_tx=XM_023254512.1
B2	139388841	139389042
refGene_gn=CCDC170;refGene_tx=XM_019831393.2;refGene_gn=CCDC170;refGene_tx=XM_019831394.2;refGene_gn=CCDC170;refGene_tx=XM_023254511.1;refGene_gn=CCDC170;refGene_tx=XM_023254512.1
B2	139394306	139394480
refGene_gn=CCDC170;refGene_tx=XM_019831393.2;refGene_gn=CCDC170;refGene_tx=XM_019831394.2;refGene_gn=CCDC170;refGene_tx=XM_023254511.1;refGene_gn=CCDC170;refGene_tx=XM_023254512.1
B2	139397099	139397342
refGene_gn=CCDC170;refGene_tx=XM_019831393.2;refGene_gn=CCDC170;refGene_tx=XM_019831394.2;refGene_gn=CCDC170;refGene_tx=XM_023254511.1;refGene_gn=CCDC170;refGene_tx=XM_023254512.1
B2	139413914	139414151
refGene_gn=CCDC170;refGene_tx=XM_019831393.2;refGene_gn=CCDC170;refGene_tx=XM_019831394.2;refGene_gn=CCDC170;refGene_tx=XM_023254511.1;refGene_gn=CCDC170;refGene_tx=XM_023254512.1
B2	139415855	139416017
refGene_gn=CCDC170;refGene_tx=XM_019831393.2;refGene_gn=CCDC170;refGene_tx=XM_019831394.2;refGene_gn=CCDC170;refGene_tx=XM_023254511.1;refGene_gn=CCDC170;refGene_tx=XM_023254512.1
B2	139615674	139616126
refGene_gn=ESR1;refGene_tx=NM_001024231.1
B2	139656168	139656359
refGene_gn=ESR1;refGene_tx=NM_001024231.1
B2	139686746	139686863
refGene_gn=ESR1;refGene_tx=NM_001024231.1
B2	139730466	139730802
refGene_gn=ESR1;refGene_tx=NM_001024231.1
B2	139810183	139810322
refGene_gn=ESR1;refGene_tx=NM_001024231.1

B2 139855364 139855498
refGene_gn=ESR1;refGene_tx=NM_001024231.1
B2 139882203 139882387
refGene_gn=ESR1;refGene_tx=NM_001024231.1
B2 139886493 139886728
refGene_gn=ESR1;refGene_tx=NM_001024231.1
B2 139907360 139907601
refGene_gn=SYNE1;refGene_tx=XM_023254513.1
B2 139915153 139915212
refGene_gn=SYNE1;refGene_tx=XM_023254513.1
B2 139916594 139916687
refGene_gn=SYNE1;refGene_tx=XM_023254513.1
B2 139917707 139917920
refGene_gn=SYNE1;refGene_tx=XM_023254513.1
B2 139919465 139919594
refGene_gn=SYNE1;refGene_tx=XM_023254513.1
B2 139920907 139921108
refGene_gn=SYNE1;refGene_tx=XM_023254513.1
B2 139923473 139923685
refGene_gn=SYNE1;refGene_tx=XM_023254513.1
B2 139924783 139924910
refGene_gn=SYNE1;refGene_tx=XM_023254513.1
B2 139926756 139926899
refGene_gn=SYNE1;refGene_tx=XM_023254513.1
B2 139928567 139928636
refGene_gn=SYNE1;refGene_tx=XM_023254513.1
B2 139930804 139931138
refGene_gn=SYNE1;refGene_tx=XM_023254513.1
B2 139932028 139932220
refGene_gn=SYNE1;refGene_tx=XM_023254513.1
B2 139934194 139934332
refGene_gn=SYNE1;refGene_tx=XM_023254513.1
B2 139934591 139934774
refGene_gn=SYNE1;refGene_tx=XM_023254513.1
B2 139938662 139938813
refGene_gn=SYNE1;refGene_tx=XM_023254513.1
B2 139940174 139940362
refGene_gn=SYNE1;refGene_tx=XM_023254513.1
B2 139951639 139951802
refGene_gn=SYNE1;refGene_tx=XM_023254513.1
B2 139966420 139966587
refGene_gn=SYNE1;refGene_tx=XM_023254513.1
B2 139969435 139969549
refGene_gn=SYNE1;refGene_tx=XM_023254513.1
B2 139969550 139969595
refGene_gn=SYNE1;refGene_tx=XM_023254513.1
B2 139977572 139977728
refGene_gn=SYNE1;refGene_tx=XM_023254513.1
B2 139980315 139980441
refGene_gn=SYNE1;refGene_tx=XM_023254513.1

B2 139984020 139984215
refGene_gn=SYNE1;refGene_tx=XM_023254513.1
B2 139986152 139986387
refGene_gn=SYNE1;refGene_tx=XM_023254513.1
B2 139989330 139989425
refGene_gn=SYNE1;refGene_tx=XM_023254513.1
B2 139991840 139991988
refGene_gn=SYNE1;refGene_tx=XM_023254513.1
B2 139992411 139992566
refGene_gn=SYNE1;refGene_tx=XM_023254513.1
B2 139994571 139994718
refGene_gn=SYNE1;refGene_tx=XM_023254513.1
B2 139995265 139995448
refGene_gn=SYNE1;refGene_tx=XM_023254513.1
B2 139997611 139997816
refGene_gn=SYNE1;refGene_tx=XM_023254513.1
B2 139998204 139998338
refGene_gn=SYNE1;refGene_tx=XM_023254513.1
B2 140000058 140000229
refGene_gn=SYNE1;refGene_tx=XM_023254513.1
B2 140003081 140003219
refGene_gn=SYNE1;refGene_tx=XM_023254513.1
B2 140003220 140003238
refGene_gn=SYNE1;refGene_tx=XM_023254513.1
B2 140007376 140007532
refGene_gn=SYNE1;refGene_tx=XM_023254513.1
B2 140007990 140008167
refGene_gn=SYNE1;refGene_tx=XM_023254513.1
B2 140009019 140009169
refGene_gn=SYNE1;refGene_tx=XM_023254513.1
B2 140010383 140010566
refGene_gn=SYNE1;refGene_tx=XM_023254513.1
B2 140010913 140011046
refGene_gn=SYNE1;refGene_tx=XM_023254513.1
B2 140011730 140011927
refGene_gn=SYNE1;refGene_tx=XM_023254513.1
B2 140012636 140012768
refGene_gn=SYNE1;refGene_tx=XM_023254513.1
B2 140014418 140014592
refGene_gn=SYNE1;refGene_tx=XM_023254513.1
B2 140016993 140017194
refGene_gn=SYNE1;refGene_tx=XM_023254513.1
B2 140018833 140018953
refGene_gn=SYNE1;refGene_tx=XM_023254513.1
B2 140023524 140023626
refGene_gn=SYNE1;refGene_tx=XM_023254513.1
B2 140028437 140028647
refGene_gn=SYNE1;refGene_tx=XM_023254513.1
B2 140029232 140029388
refGene_gn=SYNE1;refGene_tx=XM_023254513.1

B2 140029584 140029716
refGene_gn=SYNE1;refGene_tx=XM_023254513.1
B2 140032676 140032833
refGene_gn=SYNE1;refGene_tx=XM_023254513.1
B2 140037565 140037675
refGene_gn=SYNE1;refGene_tx=XM_023254513.1
B2 140038613 140038745
refGene_gn=SYNE1;refGene_tx=XM_023254513.1
B2 140047063 140047255
refGene_gn=SYNE1;refGene_tx=XM_023254513.1
B2 140049997 140050171
refGene_gn=SYNE1;refGene_tx=XM_023254513.1
B2 140052283 140052478
refGene_gn=SYNE1;refGene_tx=XM_023254513.1
B2 140061182 140061344
refGene_gn=SYNE1;refGene_tx=XM_023254513.1
B2 140061567 140061735
refGene_gn=SYNE1;refGene_tx=XM_023254513.1
B2 140069211 140069352
refGene_gn=SYNE1;refGene_tx=XM_023254513.1
B2 140070475 140070670
refGene_gn=SYNE1;refGene_tx=XM_023254513.1
B2 140076084 140076228
refGene_gn=SYNE1;refGene_tx=XM_023254513.1
B2 140077312 140077495
refGene_gn=SYNE1;refGene_tx=XM_023254513.1
B2 140077801 140077924
refGene_gn=SYNE1;refGene_tx=XM_023254513.1
B2 140078107 140078293
refGene_gn=SYNE1;refGene_tx=XM_023254513.1
B2 140082052 140082190
refGene_gn=SYNE1;refGene_tx=XM_023254513.1
B2 140083149 140083332
refGene_gn=SYNE1;refGene_tx=XM_023254513.1
B2 140083997 140084150
refGene_gn=SYNE1;refGene_tx=XM_023254513.1
B2 140086330 140086486
refGene_gn=SYNE1;refGene_tx=XM_023254513.1
B2 140087075 140087241
refGene_gn=SYNE1;refGene_tx=XM_023254513.1
B2 140088566 140088826
refGene_gn=SYNE1;refGene_tx=XM_023254513.1
B2 140089590 140089809
refGene_gn=SYNE1;refGene_tx=XM_023254513.1
B2 140090777 140090922
refGene_gn=SYNE1;refGene_tx=XM_023254513.1
B2 140091112 140091450
refGene_gn=SYNE1;refGene_tx=XM_023254513.1
B2 140096093 140098254
refGene_gn=SYNE1;refGene_tx=XM_023254513.1

B2 140100621 140100887
refGene_gn=SYNE1;refGene_tx=XM_023254513.1
B2 140101835 140102012
refGene_gn=SYNE1;refGene_tx=XM_023254513.1
B2 140103422 140103548
refGene_gn=SYNE1;refGene_tx=XM_023254513.1
B2 140105121 140105268
refGene_gn=SYNE1;refGene_tx=XM_023254513.1
B2 140108111 140108288
refGene_gn=SYNE1;refGene_tx=XM_023254513.1
B2 140110191 140110359
refGene_gn=SYNE1;refGene_tx=XM_023254513.1
B2 140111137 140111290
refGene_gn=SYNE1;refGene_tx=XM_023254513.1
B2 140112546 140112873
refGene_gn=SYNE1;refGene_tx=XM_023254513.1
B2 140113476 140113647
refGene_gn=SYNE1;refGene_tx=XM_023254513.1
B2 140113802 140113958
refGene_gn=SYNE1;refGene_tx=XM_023254513.1
B2 140115113 140115431
refGene_gn=SYNE1;refGene_tx=XM_023254513.1
B2 140118687 140118852
refGene_gn=SYNE1;refGene_tx=XM_023254513.1
B2 140119635 140119779
refGene_gn=SYNE1;refGene_tx=XM_023254513.1
B2 140122920 140123074
refGene_gn=SYNE1;refGene_tx=XM_023254513.1
B2 140125315 140125488
refGene_gn=SYNE1;refGene_tx=XM_023254513.1
B2 140126868 140127033
refGene_gn=SYNE1;refGene_tx=XM_023254513.1
B2 140128549 140128705
refGene_gn=SYNE1;refGene_tx=XM_023254513.1
B2 140129034 140129178
refGene_gn=SYNE1;refGene_tx=XM_023254513.1
B2 140132560 140132743
refGene_gn=SYNE1;refGene_tx=XM_023254513.1
B2 140133235 140133413
refGene_gn=SYNE1;refGene_tx=XM_023254513.1
B2 140133612 140133749
refGene_gn=SYNE1;refGene_tx=XM_023254513.1
B2 140138143 140138500
refGene_gn=SYNE1;refGene_tx=XM_023254513.1
B2 140142592 140142757
refGene_gn=SYNE1;refGene_tx=XM_023254513.1
B2 140143959 140144269
refGene_gn=SYNE1;refGene_tx=XM_023254513.1
B2 140146611 140146784
refGene_gn=SYNE1;refGene_tx=XM_023254513.1

B2 140147790 140148082
refGene_gn=SYNE1;refGene_tx=XM_023254513.1
B2 140151244 140151400
refGene_gn=SYNE1;refGene_tx=XM_023254513.1
B2 140152296 140152502
refGene_gn=SYNE1;refGene_tx=XM_023254513.1
B2 140155119 140155232
refGene_gn=SYNE1;refGene_tx=XM_023254513.1
B2 140156059 140156267
refGene_gn=SYNE1;refGene_tx=XM_023254513.1
B2 140157292 140157496
refGene_gn=SYNE1;refGene_tx=XM_023254513.1
B2 140159831 140159933
refGene_gn=SYNE1;refGene_tx=XM_023254513.1
B2 140162728 140162911
refGene_gn=SYNE1;refGene_tx=XM_023254513.1
B2 140164716 140164875
refGene_gn=SYNE1;refGene_tx=XM_023254513.1
B2 140165709 140165860
refGene_gn=SYNE1;refGene_tx=XM_023254513.1
B2 140167801 140167981
refGene_gn=SYNE1;refGene_tx=XM_023254513.1
B2 140171101 140171730
refGene_gn=SYNE1;refGene_tx=XM_023254513.1
B2 140175066 140175220
refGene_gn=SYNE1;refGene_tx=XM_023254513.1
B2 140177428 140177595
refGene_gn=SYNE1;refGene_tx=XM_023254513.1
B2 140178805 140178929
refGene_gn=SYNE1;refGene_tx=XM_023254513.1
B2 140179335 140179526
refGene_gn=SYNE1;refGene_tx=XM_023254513.1
B2 140181058 140181157
refGene_gn=SYNE1;refGene_tx=XM_023254513.1
B2 140181558 140181786
refGene_gn=SYNE1;refGene_tx=XM_023254513.1
B2 140184898 140185049
refGene_gn=SYNE1;refGene_tx=XM_023254513.1
B2 140186999 140187160
refGene_gn=SYNE1;refGene_tx=XM_023254513.1
B2 140190940 140191081
refGene_gn=SYNE1;refGene_tx=XM_023254513.1
B2 140191852 140192023
refGene_gn=SYNE1;refGene_tx=XM_023254513.1
B2 140193641 140193809
refGene_gn=SYNE1;refGene_tx=XM_023254513.1
B2 140197052 140197217
refGene_gn=SYNE1;refGene_tx=XM_023254513.1
B2 140199107 140199216
refGene_gn=SYNE1;refGene_tx=XM_023254513.1

B2 140199837 140200046
refGene_gn=SYNE1;refGene_tx=XM_023254513.1
B2 140200247 140200406
refGene_gn=SYNE1;refGene_tx=XM_023254513.1
B2 140202589 140202724
refGene_gn=SYNE1;refGene_tx=XM_023254513.1
B2 140204322 140204487
refGene_gn=SYNE1;refGene_tx=XM_023254513.1
B2 140204786 140204945
refGene_gn=SYNE1;refGene_tx=XM_023254513.1
B2 140207142 140207316
refGene_gn=SYNE1;refGene_tx=XM_023254513.1
B2 140210217 140210361
refGene_gn=SYNE1;refGene_tx=XM_023254513.1
B2 140211613 140211766
refGene_gn=SYNE1;refGene_tx=XM_023254513.1
B2 140212200 140212365
refGene_gn=SYNE1;refGene_tx=XM_023254513.1
B2 140214039 140214242
refGene_gn=SYNE1;refGene_tx=XM_023254513.1
B2 140215037 140215134
refGene_gn=SYNE1;refGene_tx=XM_023254513.1
B2 140220388 140220557
refGene_gn=SYNE1;refGene_tx=XM_023254513.1
B2 140221113 140221226
refGene_gn=SYNE1;refGene_tx=XM_023254513.1
B2 140229764 140229929
refGene_gn=SYNE1;refGene_tx=XM_023254513.1
B2 140231591 140231729
refGene_gn=SYNE1;refGene_tx=XM_023254513.1
B2 140235166 140235274
refGene_gn=SYNE1;refGene_tx=XM_023254513.1
B2 140238319 140238370
refGene_gn=SYNE1;refGene_tx=XM_023254513.1
B2 140243719 140243829
refGene_gn=SYNE1;refGene_tx=XM_023254513.1
B2 140246349 140246546
refGene_gn=SYNE1;refGene_tx=XM_023254513.1
B2 140251950 140252129
refGene_gn=SYNE1;refGene_tx=XM_023254513.1
B2 140253022 140253115
refGene_gn=SYNE1;refGene_tx=XM_023254513.1
B2 140253593 140253614
refGene_gn=SYNE1;refGene_tx=XM_023254513.1
B2 140266966 140267050
refGene_gn=SYNE1;refGene_tx=XM_023254513.1
B2 140273703 140273799
refGene_gn=SYNE1;refGene_tx=XM_023254513.1
B2 140296459 140296521
refGene_gn=SYNE1;refGene_tx=XM_023254513.1

B2 140370442 140370509
refGene_gn=SYNE1;refGene_tx=XM_023254513.1
B2 140433512 140433620
refGene_gn=MYCT1;refGene_tx=XM_019831396.1
B2 140433621 140433709
refGene_gn=MYCT1;refGene_tx=XM_019831396.1
B2 140458489 140458998
refGene_gn=MYCT1;refGene_tx=XM_019831396.1
B2 140501597 140501704
refGene_gn=VIP;refGene_tx=XM_011282568.3;refGene_gn=VIP;refGene_tx=XM_019831397.2
B2 140503531 140503654
refGene_gn=VIP;refGene_tx=XM_011282568.3;refGene_gn=VIP;refGene_tx=XM_019831397.2
B2 140504512 140504617
refGene_gn=VIP;refGene_tx=XM_011282568.3;refGene_gn=VIP;refGene_tx=XM_019831397.2
B2 140505589 140505721
refGene_gn=VIP;refGene_tx=XM_011282568.3;refGene_gn=VIP;refGene_tx=XM_019831397.2
B2 140506596 140506642
refGene_gn=VIP;refGene_tx=XM_011282568.3;refGene_gn=VIP;refGene_tx=XM_019831397.2
B2 140692373 140692625
refGene_gn=FBXO5;refGene_tx=XM_011282567.2
B2 140693754 140693937
refGene_gn=FBXO5;refGene_tx=XM_011282567.2
B2 140695129 140695220
refGene_gn=FBXO5;refGene_tx=XM_011282567.2
B2 140696407 140697128
refGene_gn=FBXO5;refGene_tx=XM_011282567.2
B2 140702709 140702812
refGene_gn=FBXO5;refGene_tx=XM_011282567.2
B2 140712115 140712313
refGene_gn=MTRF1L;refGene_tx=XM_003986660.4;refGene_gn=MTRF1L;refGene_tx=XM_011282565.2;refGene_gn=MTRF1L;refGene_tx=XM_023254514.1
B2 140713519 140713656
refGene_gn=MTRF1L;refGene_tx=XM_003986660.4;refGene_gn=MTRF1L;refGene_tx=XM_011282565.2
B2 140714869 140714874
refGene_gn=MTRF1L;refGene_tx=XM_019831398.2
B2 140715012 140715130
refGene_gn=MTRF1L;refGene_tx=XM_003986660.4;refGene_gn=MTRF1L;refGene_tx=XM_011282565.2;refGene_gn=MTRF1L;refGene_tx=XM_019831398.2
B2 140716830 140716994
refGene_gn=MTRF1L;refGene_tx=XM_003986660.4;refGene_gn=MTRF1L;refGene_tx=XM_011282565.2;refGene_gn=MTRF1L;refGene_tx=XM_019831398.2;refGene_gn=MTRF1L;refGene_tx=XM_023254514.1
B2 140717480 140717664
refGene_gn=MTRF1L;refGene_tx=XM_003986660.4;refGene_gn=MTRF1L;refGene_

tx=XM_011282565.2;refGene_gn=MTRF1L;refGene_tx=XM_019831398.2;refGene_gn=MTRF1L;refGene_tx=XM_023254514.1
B2 140719954 140720034
refGene_gn=MTRF1L;refGene_tx=XM_003986660.4;refGene_gn=MTRF1L;refGene_tx=XM_011282565.2;refGene_gn=MTRF1L;refGene_tx=XM_019831398.2;refGene_gn=MTRF1L;refGene_tx=XM_023254514.1
B2 140725512 140725771
refGene_gn=MTRF1L;refGene_tx=XM_003986660.4;refGene_gn=MTRF1L;refGene_tx=XM_019831398.2;refGene_gn=MTRF1L;refGene_tx=XM_023254514.1
B2 140734183 140734372
refGene_gn=RGS17;refGene_tx=XM_003986661.5;refGene_gn=RGS17;refGene_tx=XM_019831399.2
B2 140743309 140743544
refGene_gn=RGS17;refGene_tx=XM_003986661.5;refGene_gn=RGS17;refGene_tx=XM_019831399.2
B2 140745871 140745961
refGene_gn=RGS17;refGene_tx=XM_003986661.5;refGene_gn=RGS17;refGene_tx=XM_019831399.2
B2 140763749 140763868
refGene_gn=RGS17;refGene_tx=XM_003986661.5;refGene_gn=RGS17;refGene_tx=XM_019831399.2
B2 141598083 141598376
refGene_gn=OPRM1;refGene_tx=XM_019831412.1
B2 141637902 141638255
refGene_gn=OPRM1;refGene_tx=XM_019831412.1
B2 141639218 141639739
refGene_gn=OPRM1;refGene_tx=XM_019831412.1
B2 141652530 141652569
refGene_gn=OPRM1;refGene_tx=XM_019831412.1
B2 141696523 141696730
refGene_gn=IPCEF1;refGene_tx=XM_019831406.1;refGene_gn=IPCEF1;refGene_tx=XM_019831408.1;refGene_gn=IPCEF1;refGene_tx=XM_019831409.2;refGene_gn=IPCEF1;refGene_tx=XM_019831410.1
B2 141705220 141705414
refGene_gn=IPCEF1;refGene_tx=XM_019831406.1;refGene_gn=IPCEF1;refGene_tx=XM_019831408.1;refGene_gn=IPCEF1;refGene_tx=XM_019831409.2;refGene_gn=IPCEF1;refGene_tx=XM_019831410.1
B2 141722919 141723292
refGene_gn=IPCEF1;refGene_tx=XM_019831406.1;refGene_gn=IPCEF1;refGene_tx=XM_019831408.1;refGene_gn=IPCEF1;refGene_tx=XM_019831409.2;refGene_gn=IPCEF1;refGene_tx=XM_019831410.1
B2 141734896 141734985
refGene_gn=IPCEF1;refGene_tx=XM_019831406.1;refGene_gn=IPCEF1;refGene_tx=XM_019831408.1;refGene_gn=IPCEF1;refGene_tx=XM_019831409.2;refGene_gn=IPCEF1;refGene_tx=XM_019831410.1
B2 141736333 141736392
refGene_gn=IPCEF1;refGene_tx=XM_019831406.1;refGene_gn=IPCEF1;refGene_tx=XM_019831408.1;refGene_gn=IPCEF1;refGene_tx=XM_019831409.2;refGene_gn=IPCEF1;refGene_tx=XM_019831410.1
B2 141741590 141741662

refGene_gn=IPCEF1;refGene_tx=XM_019831406.1;refGene_gn=IPCEF1;refGene_tx=XM_019831408.1;refGene_gn=IPCEF1;refGene_tx=XM_019831409.2;refGene_gn=IPCEF1;refGene_tx=XM_019831410.1
B2 141743408 141743482
refGene_gn=IPCEF1;refGene_tx=XM_019831406.1;refGene_gn=IPCEF1;refGene_tx=XM_019831408.1;refGene_gn=IPCEF1;refGene_tx=XM_019831409.2;refGene_gn=IPCEF1;refGene_tx=XM_019831410.1
B2 141760901 141760958
refGene_gn=IPCEF1;refGene_tx=XM_019831406.1;refGene_gn=IPCEF1;refGene_tx=XM_019831408.1;refGene_gn=IPCEF1;refGene_tx=XM_019831410.1
B2 141767648 141767818
refGene_gn=IPCEF1;refGene_tx=XM_019831406.1;refGene_gn=IPCEF1;refGene_tx=XM_019831408.1;refGene_gn=IPCEF1;refGene_tx=XM_019831409.2;refGene_gn=IPCEF1;refGene_tx=XM_019831410.1
B2 141768243 141768280
refGene_gn=IPCEF1;refGene_tx=XM_019831408.1
B2 141768453 141768493
refGene_gn=IPCEF1;refGene_tx=XM_019831406.1;refGene_gn=IPCEF1;refGene_tx=XM_019831409.2
B2 141782149 141782173
refGene_gn=IPCEF1;refGene_tx=XM_019831406.1;refGene_gn=IPCEF1;refGene_tx=XM_019831409.2
B2 141920426 141920725
refGene_gn=CNKSR3;refGene_tx=XM_019831402.2;refGene_gn=CNKSR3;refGene_tx=XM_019831403.2;refGene_gn=CNKSR3;refGene_tx=XM_019831404.2;refGene_gn=CNKSR3;refGene_tx=XM_019831405.2;refGene_gn=CNKSR3;refGene_tx=XM_023254515.1;refGene_gn=CNKSR3;refGene_tx=XM_023254516.1;refGene_gn=CNKSR3;refGene_tx=XM_023254517.1
B2 141923501 141923591
refGene_gn=CNKSR3;refGene_tx=XM_019831402.2;refGene_gn=CNKSR3;refGene_tx=XM_019831403.2;refGene_gn=CNKSR3;refGene_tx=XM_019831404.2;refGene_gn=CNKSR3;refGene_tx=XM_019831405.2;refGene_gn=CNKSR3;refGene_tx=XM_023254515.1;refGene_gn=CNKSR3;refGene_tx=XM_023254516.1;refGene_gn=CNKSR3;refGene_tx=XM_023254517.1
B2 141924090 141924299
refGene_gn=CNKSR3;refGene_tx=XM_019831402.2;refGene_gn=CNKSR3;refGene_tx=XM_019831403.2;refGene_gn=CNKSR3;refGene_tx=XM_019831404.2;refGene_gn=CNKSR3;refGene_tx=XM_019831405.2;refGene_gn=CNKSR3;refGene_tx=XM_023254515.1;refGene_gn=CNKSR3;refGene_tx=XM_023254516.1;refGene_gn=CNKSR3;refGene_tx=XM_023254517.1
B2 141926882 141927007
refGene_gn=CNKSR3;refGene_tx=XM_019831402.2;refGene_gn=CNKSR3;refGene_tx=XM_019831403.2;refGene_gn=CNKSR3;refGene_tx=XM_019831404.2;refGene_gn=CNKSR3;refGene_tx=XM_019831405.2;refGene_gn=CNKSR3;refGene_tx=XM_023254515.1;refGene_gn=CNKSR3;refGene_tx=XM_023254516.1;refGene_gn=CNKSR3;refGene_tx=XM_023254517.1
B2 141927712 141927946
refGene_gn=CNKSR3;refGene_tx=XM_019831402.2;refGene_gn=CNKSR3;refGene_tx=XM_019831403.2;refGene_gn=CNKSR3;refGene_tx=XM_019831404.2;refGene_gn=CNKSR3;refGene_tx=XM_019831405.2;refGene_gn=CNKSR3;refGene_tx=XM_02

3254515.1;refGene_gn=CNKSR3;refGene_tx=XM_023254516.1;refGene_gn=CNKSR3;refGene_tx=XM_023254517.1
B2 141928194 141928263
refGene_gn=CNKSR3;refGene_tx=XM_019831402.2;refGene_gn=CNKSR3;refGene_tx=XM_019831403.2;refGene_gn=CNKSR3;refGene_tx=XM_019831404.2;refGene_gn=CNKSR3;refGene_tx=XM_019831405.2;refGene_gn=CNKSR3;refGene_tx=XM_023254515.1;refGene_gn=CNKSR3;refGene_tx=XM_023254516.1;refGene_gn=CNKSR3;refGene_tx=XM_023254517.1
B2 141933102 141933162
refGene_gn=CNKSR3;refGene_tx=XM_019831402.2;refGene_gn=CNKSR3;refGene_tx=XM_019831403.2;refGene_gn=CNKSR3;refGene_tx=XM_019831404.2;refGene_gn=CNKSR3;refGene_tx=XM_019831405.2;refGene_gn=CNKSR3;refGene_tx=XM_023254515.1;refGene_gn=CNKSR3;refGene_tx=XM_023254516.1;refGene_gn=CNKSR3;refGene_tx=XM_023254517.1
B2 141935140 141935260
refGene_gn=CNKSR3;refGene_tx=XM_019831402.2;refGene_gn=CNKSR3;refGene_tx=XM_019831403.2;refGene_gn=CNKSR3;refGene_tx=XM_019831404.2;refGene_gn=CNKSR3;refGene_tx=XM_019831405.2;refGene_gn=CNKSR3;refGene_tx=XM_023254515.1;refGene_gn=CNKSR3;refGene_tx=XM_023254516.1;refGene_gn=CNKSR3;refGene_tx=XM_023254517.1
B2 141937726 141937768
refGene_gn=CNKSR3;refGene_tx=XM_019831402.2;refGene_gn=CNKSR3;refGene_tx=XM_019831403.2;refGene_gn=CNKSR3;refGene_tx=XM_019831404.2;refGene_gn=CNKSR3;refGene_tx=XM_019831405.2;refGene_gn=CNKSR3;refGene_tx=XM_023254515.1;refGene_gn=CNKSR3;refGene_tx=XM_023254516.1;refGene_gn=CNKSR3;refGene_tx=XM_023254517.1
B2 141942759 141942847
refGene_gn=CNKSR3;refGene_tx=XM_019831402.2;refGene_gn=CNKSR3;refGene_tx=XM_019831403.2;refGene_gn=CNKSR3;refGene_tx=XM_019831404.2;refGene_gn=CNKSR3;refGene_tx=XM_019831405.2;refGene_gn=CNKSR3;refGene_tx=XM_023254515.1;refGene_gn=CNKSR3;refGene_tx=XM_023254516.1;refGene_gn=CNKSR3;refGene_tx=XM_023254517.1
B2 141943341 141943544
refGene_gn=CNKSR3;refGene_tx=XM_019831402.2;refGene_gn=CNKSR3;refGene_tx=XM_019831403.2;refGene_gn=CNKSR3;refGene_tx=XM_019831404.2;refGene_gn=CNKSR3;refGene_tx=XM_019831405.2;refGene_gn=CNKSR3;refGene_tx=XM_023254515.1;refGene_gn=CNKSR3;refGene_tx=XM_023254516.1;refGene_gn=CNKSR3;refGene_tx=XM_023254517.1
B2 141949761 141949925
refGene_gn=CNKSR3;refGene_tx=XM_023254515.1;refGene_gn=CNKSR3;refGene_tx=XM_023254516.1
B2 142001398 142001450
refGene_gn=CNKSR3;refGene_tx=XM_023254515.1;refGene_gn=CNKSR3;refGene_tx=XM_023254516.1
B2 142192894 142192975
refGene_gn=SCAF8;refGene_tx=XM_023254520.1
B2 142193318 142193402
refGene_gn=SCAF8;refGene_tx=XM_023254520.1
B2 142199577 142199610
refGene_gn=SCAF8;refGene_tx=XM_023254520.1

B2	142199611	142199641
refGene_gn=SCAF8;refGene_tx=XM_023254520.1
B2	142228282	142228366
refGene_gn=SCAF8;refGene_tx=XM_023254520.1
B2	142232532	142232577
refGene_gn=SCAF8;refGene_tx=XM_023254520.1
B2	142245759	142245921
refGene_gn=SCAF8;refGene_tx=XM_023254520.1
B2	142246950	142247104
refGene_gn=SCAF8;refGene_tx=XM_023254520.1
B2	142248163	142248294
refGene_gn=SCAF8;refGene_tx=XM_023254520.1
B2	142251102	142251279
refGene_gn=SCAF8;refGene_tx=XM_023254520.1
B2	142252359	142252439
refGene_gn=SCAF8;refGene_tx=XM_023254520.1
B2	142254188	142254306
refGene_gn=SCAF8;refGene_tx=XM_023254520.1
B2	142261242	142261374
refGene_gn=SCAF8;refGene_tx=XM_023254520.1
B2	142261642	142261755
refGene_gn=SCAF8;refGene_tx=XM_023254520.1
B2	142263197	142263391
refGene_gn=SCAF8;refGene_tx=XM_023254520.1
B2	142266702	142266803
refGene_gn=SCAF8;refGene_tx=XM_023254520.1
B2	142269328	142269442
refGene_gn=SCAF8;refGene_tx=XM_023254520.1
B2	142271290	142271447
refGene_gn=SCAF8;refGene_tx=XM_023254520.1
B2	142273543	142273677
refGene_gn=SCAF8;refGene_tx=XM_023254520.1
B2	142275262	142275407
refGene_gn=SCAF8;refGene_tx=XM_023254520.1
B2	142278015	142278084
refGene_gn=SCAF8;refGene_tx=XM_023254520.1
B2	142281378	142281597
refGene_gn=SCAF8;refGene_tx=XM_023254520.1
B2	142282580	142284034
refGene_gn=SCAF8;refGene_tx=XM_023254520.1
B2	142548693	142549827
refGene_gn=TIAM2;refGene_tx=XM_006932109.4;refGene_gn=TIAM2;refGene_tx=XM_019831414.2;refGene_gn=TIAM2;refGene_tx=XM_019831415.2;refGene_gn=TIAM2;refGene_tx=XM_019831416.2;refGene_gn=TIAM2;refGene_tx=XM_019831417.2;refGene_gn=TIAM2;refGene_tx=XM_019831418.2;refGene_gn=TIAM2;refGene_tx=XM_023254518.1;refGene_gn=TIAM2;refGene_tx=XM_023254519.1
B2	142555928	142556364
refGene_gn=TIAM2;refGene_tx=XM_006932109.4;refGene_gn=TIAM2;refGene_tx=XM_019831414.2;refGene_gn=TIAM2;refGene_tx=XM_019831415.2;refGene_gn=TIAM2;refGene_tx=XM_019831416.2;refGene_gn=TIAM2;refGene_tx=XM_0198314

17.2;refGene_gn=TIAM2;refGene_tx=XM_019831418.2;refGene_gn=TIAM2;refGene_tx=XM_023254518.1;refGene_gn=TIAM2;refGene_tx=XM_023254519.1
B2 142562289 142562462
refGene_gn=TIAM2;refGene_tx=XM_006932109.4;refGene_gn=TIAM2;refGene_tx=XM_019831414.2;refGene_gn=TIAM2;refGene_tx=XM_019831415.2;refGene_gn=TIAM2;refGene_tx=XM_019831416.2;refGene_gn=TIAM2;refGene_tx=XM_019831417.2;refGene_gn=TIAM2;refGene_tx=XM_019831418.2;refGene_gn=TIAM2;refGene_tx=XM_023254518.1;refGene_gn=TIAM2;refGene_tx=XM_023254519.1
B2 142564526 142564751
refGene_gn=TIAM2;refGene_tx=XM_006932109.4;refGene_gn=TIAM2;refGene_tx=XM_019831414.2;refGene_gn=TIAM2;refGene_tx=XM_019831415.2;refGene_gn=TIAM2;refGene_tx=XM_019831416.2;refGene_gn=TIAM2;refGene_tx=XM_019831417.2;refGene_gn=TIAM2;refGene_tx=XM_019831418.2;refGene_gn=TIAM2;refGene_tx=XM_023254518.1;refGene_gn=TIAM2;refGene_tx=XM_023254519.1
B2 142575165 142575351
refGene_gn=TIAM2;refGene_tx=XM_006932109.4;refGene_gn=TIAM2;refGene_tx=XM_019831414.2;refGene_gn=TIAM2;refGene_tx=XM_019831415.2;refGene_gn=TIAM2;refGene_tx=XM_019831416.2;refGene_gn=TIAM2;refGene_tx=XM_019831417.2;refGene_gn=TIAM2;refGene_tx=XM_019831418.2;refGene_gn=TIAM2;refGene_tx=XM_023254518.1;refGene_gn=TIAM2;refGene_tx=XM_023254519.1
B2 142576036 142576195
refGene_gn=TIAM2;refGene_tx=XM_006932109.4;refGene_gn=TIAM2;refGene_tx=XM_019831414.2;refGene_gn=TIAM2;refGene_tx=XM_019831415.2;refGene_gn=TIAM2;refGene_tx=XM_019831416.2;refGene_gn=TIAM2;refGene_tx=XM_019831417.2;refGene_gn=TIAM2;refGene_tx=XM_019831418.2;refGene_gn=TIAM2;refGene_tx=XM_023254518.1;refGene_gn=TIAM2;refGene_tx=XM_023254519.1
B2 142587705 142587867
refGene_gn=TIAM2;refGene_tx=XM_006932109.4;refGene_gn=TIAM2;refGene_tx=XM_019831414.2;refGene_gn=TIAM2;refGene_tx=XM_019831415.2;refGene_gn=TIAM2;refGene_tx=XM_019831416.2;refGene_gn=TIAM2;refGene_tx=XM_019831417.2;refGene_gn=TIAM2;refGene_tx=XM_019831418.2;refGene_gn=TIAM2;refGene_tx=XM_023254518.1;refGene_gn=TIAM2;refGene_tx=XM_023254519.1
B2 142589474 142589579
refGene_gn=TIAM2;refGene_tx=XM_006932109.4;refGene_gn=TIAM2;refGene_tx=XM_019831414.2;refGene_gn=TIAM2;refGene_tx=XM_019831415.2;refGene_gn=TIAM2;refGene_tx=XM_019831416.2;refGene_gn=TIAM2;refGene_tx=XM_019831417.2;refGene_gn=TIAM2;refGene_tx=XM_019831418.2;refGene_gn=TIAM2;refGene_tx=XM_023254518.1;refGene_gn=TIAM2;refGene_tx=XM_023254519.1
B2 142589784 142589863
refGene_gn=TIAM2;refGene_tx=XM_006932109.4;refGene_gn=TIAM2;refGene_tx=XM_019831414.2;refGene_gn=TIAM2;refGene_tx=XM_019831415.2;refGene_gn=TIAM2;refGene_tx=XM_019831416.2;refGene_gn=TIAM2;refGene_tx=XM_019831417.2;refGene_gn=TIAM2;refGene_tx=XM_019831418.2;refGene_gn=TIAM2;refGene_tx=XM_023254518.1;refGene_gn=TIAM2;refGene_tx=XM_023254519.1
B2 142592376 142592469
refGene_gn=TIAM2;refGene_tx=XM_006932109.4;refGene_gn=TIAM2;refGene_tx=XM_019831414.2;refGene_gn=TIAM2;refGene_tx=XM_019831415.2;refGene_gn=TIAM2;refGene_tx=XM_019831416.2;refGene_gn=TIAM2;refGene_tx=XM_019831417.2;refGene_gn=TIAM2;refGene_tx=XM_019831418.2;refGene_gn=TIAM2;refGene_tx=XM_023254518.1;refGene_gn=TIAM2;refGene_tx=XM_023254519.1

B2 142593590 142593854
refGene_gn=TIAM2;refGene_tx=XM_006932109.4;refGene_gn=TIAM2;refGene_tx=XM_019831414.2;refGene_gn=TIAM2;refGene_tx=XM_019831415.2;refGene_gn=TIAM2;refGene_tx=XM_019831416.2;refGene_gn=TIAM2;refGene_tx=XM_019831417.2;refGene_gn=TIAM2;refGene_tx=XM_019831418.2;refGene_gn=TIAM2;refGene_tx=XM_019831421.2;refGene_gn=TIAM2;refGene_tx=XM_023254518.1;refGene_gn=TIAM2;refGene_tx=XM_023254519.1
B2 142618253 142618357
refGene_gn=TIAM2;refGene_tx=XM_006932109.4;refGene_gn=TIAM2;refGene_tx=XM_019831414.2;refGene_gn=TIAM2;refGene_tx=XM_019831415.2;refGene_gn=TIAM2;refGene_tx=XM_019831416.2;refGene_gn=TIAM2;refGene_tx=XM_019831417.2;refGene_gn=TIAM2;refGene_tx=XM_019831418.2;refGene_gn=TIAM2;refGene_tx=XM_023254518.1;refGene_gn=TIAM2;refGene_tx=XM_023254519.1
B2 142642175 142642355
refGene_gn=TIAM2;refGene_tx=XM_006932109.4;refGene_gn=TIAM2;refGene_tx=XM_006932110.4;refGene_gn=TIAM2;refGene_tx=XM_019831414.2;refGene_gn=TIAM2;refGene_tx=XM_019831415.2;refGene_gn=TIAM2;refGene_tx=XM_019831416.2;refGene_gn=TIAM2;refGene_tx=XM_019831417.2;refGene_gn=TIAM2;refGene_tx=XM_019831418.2;refGene_gn=TIAM2;refGene_tx=XM_019831421.2;refGene_gn=TIAM2;refGene_tx=XM_019831423.2;refGene_gn=TIAM2;refGene_tx=XM_023254518.1;refGene_gn=TIAM2;refGene_tx=XM_023254519.1
B2 142645379 142645448
refGene_gn=TIAM2;refGene_tx=XM_006932109.4;refGene_gn=TIAM2;refGene_tx=XM_006932110.4;refGene_gn=TIAM2;refGene_tx=XM_019831414.2;refGene_gn=TIAM2;refGene_tx=XM_019831415.2;refGene_gn=TIAM2;refGene_tx=XM_019831416.2;refGene_gn=TIAM2;refGene_tx=XM_019831417.2;refGene_gn=TIAM2;refGene_tx=XM_019831418.2;refGene_gn=TIAM2;refGene_tx=XM_019831421.2;refGene_gn=TIAM2;refGene_tx=XM_019831423.2;refGene_gn=TIAM2;refGene_tx=XM_023254518.1;refGene_gn=TIAM2;refGene_tx=XM_023254519.1
B2 142646005 142646131
refGene_gn=TIAM2;refGene_tx=XM_006932109.4;refGene_gn=TIAM2;refGene_tx=XM_006932110.4;refGene_gn=TIAM2;refGene_tx=XM_019831414.2;refGene_gn=TIAM2;refGene_tx=XM_019831415.2;refGene_gn=TIAM2;refGene_tx=XM_019831416.2;refGene_gn=TIAM2;refGene_tx=XM_019831417.2;refGene_gn=TIAM2;refGene_tx=XM_019831418.2;refGene_gn=TIAM2;refGene_tx=XM_019831421.2;refGene_gn=TIAM2;refGene_tx=XM_019831423.2;refGene_gn=TIAM2;refGene_tx=XM_023254518.1;refGene_gn=TIAM2;refGene_tx=XM_023254519.1
B2 142646987 142647096
refGene_gn=TIAM2;refGene_tx=XM_006932109.4;refGene_gn=TIAM2;refGene_tx=XM_006932110.4;refGene_gn=TIAM2;refGene_tx=XM_019831414.2;refGene_gn=TIAM2;refGene_tx=XM_019831415.2;refGene_gn=TIAM2;refGene_tx=XM_019831416.2;refGene_gn=TIAM2;refGene_tx=XM_019831417.2;refGene_gn=TIAM2;refGene_tx=XM_019831418.2;refGene_gn=TIAM2;refGene_tx=XM_019831421.2;refGene_gn=TIAM2;refGene_tx=XM_019831423.2;refGene_gn=TIAM2;refGene_tx=XM_023254518.1;refGene_gn=TIAM2;refGene_tx=XM_023254519.1
B2 142650245 142650425
refGene_gn=TIAM2;refGene_tx=XM_006932109.4;refGene_gn=TIAM2;refGene_tx=XM_006932110.4;refGene_gn=TIAM2;refGene_tx=XM_019831414.2;refGene_gn=TIAM2;refGene_tx=XM_019831415.2;refGene_gn=TIAM2;refGene_tx=XM_019831416.2;refGene_gn=TIAM2;refGene_tx=XM_019831417.2;refGene_gn=TIAM2;refGe

ne_tx=XM_019831418.2;refGene_gn=TIAM2;refGene_tx=XM_019831421.2;refGene_gn=TIAM2;refGene_tx=XM_019831423.2;refGene_gn=TIAM2;refGene_tx=XM_023254518.1;refGene_gn=TIAM2;refGene_tx=XM_023254519.1
B2 142650845 142650964
refGene_gn=TIAM2;refGene_tx=XM_006932109.4;refGene_gn=TIAM2;refGene_tx=XM_006932110.4;refGene_gn=TIAM2;refGene_tx=XM_019831414.2;refGene_gn=TIAM2;refGene_tx=XM_019831415.2;refGene_gn=TIAM2;refGene_tx=XM_019831416.2;refGene_gn=TIAM2;refGene_tx=XM_019831417.2;refGene_gn=TIAM2;refGene_tx=XM_019831418.2;refGene_gn=TIAM2;refGene_tx=XM_019831421.2;refGene_gn=TIAM2;refGene_tx=XM_019831423.2;refGene_gn=TIAM2;refGene_tx=XM_023254518.1;refGene_gn=TIAM2;refGene_tx=XM_023254519.1
B2 142651987 142652096
refGene_gn=TIAM2;refGene_tx=XM_006932109.4;refGene_gn=TIAM2;refGene_tx=XM_006932110.4;refGene_gn=TIAM2;refGene_tx=XM_019831414.2;refGene_gn=TIAM2;refGene_tx=XM_019831415.2;refGene_gn=TIAM2;refGene_tx=XM_019831416.2;refGene_gn=TIAM2;refGene_tx=XM_019831417.2;refGene_gn=TIAM2;refGene_tx=XM_019831418.2;refGene_gn=TIAM2;refGene_tx=XM_019831421.2;refGene_gn=TIAM2;refGene_tx=XM_019831423.2;refGene_gn=TIAM2;refGene_tx=XM_023254518.1;refGene_gn=TIAM2;refGene_tx=XM_023254519.1
B2 142653166 142653225
refGene_gn=TIAM2;refGene_tx=XM_006932109.4;refGene_gn=TIAM2;refGene_tx=XM_006932110.4;refGene_gn=TIAM2;refGene_tx=XM_019831414.2;refGene_gn=TIAM2;refGene_tx=XM_019831415.2;refGene_gn=TIAM2;refGene_tx=XM_019831416.2;refGene_gn=TIAM2;refGene_tx=XM_019831417.2;refGene_gn=TIAM2;refGene_tx=XM_019831418.2;refGene_gn=TIAM2;refGene_tx=XM_019831421.2;refGene_gn=TIAM2;refGene_tx=XM_019831423.2;refGene_gn=TIAM2;refGene_tx=XM_023254518.1;refGene_gn=TIAM2;refGene_tx=XM_023254519.1
B2 142653890 142653996
refGene_gn=TIAM2;refGene_tx=XM_006932109.4;refGene_gn=TIAM2;refGene_tx=XM_006932110.4;refGene_gn=TIAM2;refGene_tx=XM_019831414.2;refGene_gn=TIAM2;refGene_tx=XM_019831415.2;refGene_gn=TIAM2;refGene_tx=XM_019831416.2;refGene_gn=TIAM2;refGene_tx=XM_019831417.2;refGene_gn=TIAM2;refGene_tx=XM_019831418.2;refGene_gn=TIAM2;refGene_tx=XM_019831421.2;refGene_gn=TIAM2;refGene_tx=XM_019831423.2;refGene_gn=TIAM2;refGene_tx=XM_023254518.1;refGene_gn=TIAM2;refGene_tx=XM_023254519.1
B2 142654903 142654991
refGene_gn=TIAM2;refGene_tx=XM_006932109.4;refGene_gn=TIAM2;refGene_tx=XM_006932110.4;refGene_gn=TIAM2;refGene_tx=XM_019831414.2;refGene_gn=TIAM2;refGene_tx=XM_019831415.2;refGene_gn=TIAM2;refGene_tx=XM_019831416.2;refGene_gn=TIAM2;refGene_tx=XM_019831417.2;refGene_gn=TIAM2;refGene_tx=XM_019831418.2;refGene_gn=TIAM2;refGene_tx=XM_019831421.2;refGene_gn=TIAM2;refGene_tx=XM_019831423.2;refGene_gn=TIAM2;refGene_tx=XM_023254518.1;refGene_gn=TIAM2;refGene_tx=XM_023254519.1
B2 142655367 142655522
refGene_gn=TIAM2;refGene_tx=XM_006932109.4;refGene_gn=TIAM2;refGene_tx=XM_006932110.4;refGene_gn=TIAM2;refGene_tx=XM_019831414.2;refGene_gn=TIAM2;refGene_tx=XM_019831415.2;refGene_gn=TIAM2;refGene_tx=XM_019831416.2;refGene_gn=TIAM2;refGene_tx=XM_019831417.2;refGene_gn=TIAM2;refGene_tx=XM_019831418.2;refGene_gn=TIAM2;refGene_tx=XM_019831421.2;refGene_gn=TIAM2;refGene_tx=XM_019831423.2;refGene_gn=TIAM2;refGene_tx=XM_02

3254518.1;refGene_gn=TIAM2;refGene_tx=XM_023254519.1
B2 142657085 142657678
refGene_gn=TIAM2;refGene_tx=XM_006932109.4;refGene_gn=TIAM2;refGene_tx=XM_006932110.4;refGene_gn=TIAM2;refGene_tx=XM_019831414.2;refGene_gn=TIAM2;refGene_tx=XM_019831415.2;refGene_gn=TIAM2;refGene_tx=XM_019831416.2;refGene_gn=TIAM2;refGene_tx=XM_019831417.2;refGene_gn=TIAM2;refGene_tx=XM_019831418.2;refGene_gn=TIAM2;refGene_tx=XM_019831421.2;refGene_gn=TIAM2;refGene_tx=XM_019831423.2;refGene_gn=TIAM2;refGene_tx=XM_023254518.1;refGene_gn=TIAM2;refGene_tx=XM_023254519.1
B2 142658356 142658600
refGene_gn=TFB1M;refGene_tx=XM_003986680.4
B2 142661608 142661736
refGene_gn=TFB1M;refGene_tx=XM_003986680.4
B2 142674979 142675639
refGene_gn=CLDN20;refGene_tx=XM_003986666.4;refGene_gn=CLDN20;refGene_tx=XM_019831428.2
B2 142687562 142687682
refGene_gn=TFB1M;refGene_tx=XM_003986680.4
B2 142703805 142703957
refGene_gn=TFB1M;refGene_tx=XM_003986680.4
B2 142705287 142705396
refGene_gn=TFB1M;refGene_tx=XM_003986680.4
B2 142720512 142720664
refGene_gn=TFB1M;refGene_tx=XM_003986680.4
B2 142723653 142723786
refGene_gn=TFB1M;refGene_tx=XM_003986680.4
B2 142782952 142783079
refGene_gn=NOX3;refGene_tx=XM_023254521.1
B2 142793683 142793808
refGene_gn=NOX3;refGene_tx=XM_023254521.1
B2 142797955 142798102
refGene_gn=NOX3;refGene_tx=XM_023254521.1
B2 142811127 142811287
refGene_gn=NOX3;refGene_tx=XM_023254521.1
B2 142818655 142818909
refGene_gn=NOX3;refGene_tx=XM_023254521.1
B2 142821613 142821706
refGene_gn=NOX3;refGene_tx=XM_023254521.1
B2 142826838 142826968
refGene_gn=NOX3;refGene_tx=XM_023254521.1
B2 142830186 142830368
refGene_gn=NOX3;refGene_tx=XM_023254521.1
B2 142833655 142833801
refGene_gn=NOX3;refGene_tx=XM_023254521.1
B2 142842881 142842966
refGene_gn=NOX3;refGene_tx=XM_023254521.1
B2 142844009 142844120
refGene_gn=NOX3;refGene_tx=XM_023254521.1
B2 142844224 142844320
refGene_gn=NOX3;refGene_tx=XM_023254521.1

B2 142844951 142844996
refGene_gn=NOX3;refGene_tx=XM_023254521.1
B2 144036134 144036823
refGene_gn=ARID1B;refGene_tx=XM_023254522.1
B2 144037350 144037657
refGene_gn=ARID1B;refGene_tx=XM_023254522.1
B2 144037659 144038256
refGene_gn=ARID1B;refGene_tx=XM_023254522.1
B2 144088495 144088690
refGene_gn=ARID1B;refGene_tx=XM_023254522.1
B2 144172633 144172783
refGene_gn=ARID1B;refGene_tx=XM_023254522.1
B2 144203060 144203171
refGene_gn=ARID1B;refGene_tx=XM_023254522.1
B2 144359179 144359423
refGene_gn=ARID1B;refGene_tx=XM_023254522.1
B2 144394277 144394367
refGene_gn=ARID1B;refGene_tx=XM_023254522.1
B2 144414170 144414350
refGene_gn=ARID1B;refGene_tx=XM_023254522.1
B2 144428751 144429079
refGene_gn=ARID1B;refGene_tx=XM_023254522.1
B2 144445976 144446122
refGene_gn=ARID1B;refGene_tx=XM_023254522.1
B2 144453603 144453713
refGene_gn=ARID1B;refGene_tx=XM_023254522.1
B2 144454446 144454605
refGene_gn=ARID1B;refGene_tx=XM_023254522.1
B2 144460223 144460433
refGene_gn=ARID1B;refGene_tx=XM_023254522.1
B2 144463220 144463425
refGene_gn=ARID1B;refGene_tx=XM_023254522.1
B2 144468903 144469042
refGene_gn=ARID1B;refGene_tx=XM_023254522.1
B2 144469292 144469465
refGene_gn=ARID1B;refGene_tx=XM_023254522.1
B2 144475393 144475541
refGene_gn=ARID1B;refGene_tx=XM_023254522.1
B2 144477986 144478083
refGene_gn=ARID1B;refGene_tx=XM_023254522.1
B2 144479548 144480332
refGene_gn=ARID1B;refGene_tx=XM_023254522.1
B2 144482561 144482692
refGene_gn=ARID1B;refGene_tx=XM_023254522.1
B2 144484854 144486591
refGene_gn=ARID1B;refGene_tx=XM_023254522.1
B2 144496424 144496674
refGene_gn=LOC109500119;refGene_tx=XM_023254524.1
B2 144506181 144506500
refGene_gn=LOC109500119;refGene_tx=XM_023254523.1

B2 144506826 144506982
refGene_gn=LOC109500119;refGene_tx=XM_023254523.1
B2 144507229 144507236
refGene_gn=LOC109500119;refGene_tx=XM_023254525.1
B2 144509365 144509471
refGene_gn=LOC109500119;refGene_tx=XM_023254523.1;refGene_gn=LOC109500119;refGene_tx=XM_023254524.1;refGene_gn=LOC109500119;refGene_tx=XM_023254525.1
B2 144515779 144515885
refGene_gn=LOC109500119;refGene_tx=XM_023254523.1;refGene_gn=LOC109500119;refGene_tx=XM_023254524.1;refGene_gn=LOC109500119;refGene_tx=XM_023254525.1
B2 144521677 144521794
refGene_gn=LOC109500119;refGene_tx=XM_023254523.1;refGene_gn=LOC109500119;refGene_tx=XM_023254524.1;refGene_gn=LOC109500119;refGene_tx=XM_023254525.1
B2 144561969 144562068
refGene_gn=TMEM242;refGene_tx=XM_003986667.5
B2 144567968 144569114
refGene_gn=LDHAL6B;refGene_tx=XM_011282559.3
B2 144580317 144580365
refGene_gn=TMEM242;refGene_tx=XM_006932111.4
B2 144585061 144585199
refGene_gn=TMEM242;refGene_tx=XM_003986667.5;refGene_gn=TMEM242;refGene_tx=XM_006932111.4
B2 144590039 144590140
refGene_gn=TMEM242;refGene_tx=XM_003986667.5;refGene_gn=TMEM242;refGene_tx=XM_006932111.4
B2 144591093 144591181
refGene_gn=TMEM242;refGene_tx=XM_003986667.5;refGene_gn=TMEM242;refGene_tx=XM_006932111.4
B2 144638265 144638510
refGene_gn=ZDHHC14;refGene_tx=XM_003986669.3
B2 144809205 144809366
refGene_gn=ZDHHC14;refGene_tx=XM_003986669.3
B2 144854265 144854424
refGene_gn=ZDHHC14;refGene_tx=XM_003986669.3
B2 144888459 144888597
refGene_gn=ZDHHC14;refGene_tx=XM_003986669.3
B2 144892043 144892092
refGene_gn=ZDHHC14;refGene_tx=XM_003986669.3
B2 144906752 144906855
refGene_gn=ZDHHC14;refGene_tx=XM_003986669.3
B2 144907966 144908002
refGene_gn=ZDHHC14;refGene_tx=XM_003986669.3
B2 144908003 144908077
refGene_gn=ZDHHC14;refGene_tx=XM_003986669.3
B2 144913345 144913448
refGene_gn=ZDHHC14;refGene_tx=XM_003986669.3
B2 144928375 144928774

refGene_gn=ZDHHC14;refGene_tx=XM_003986669.3
B2 145071594 145071630
refGene_gn=SNX9;refGene_tx=XM_019831438.2;refGene_gn=SNX9;refGene_tx=XM_023254527.1
B2 145109044 145109131
refGene_gn=SNX9;refGene_tx=XM_019831438.2;refGene_gn=SNX9;refGene_tx=XM_023254527.1
B2 145111870 145111945
refGene_gn=SNX9;refGene_tx=XM_019831438.2;refGene_gn=SNX9;refGene_tx=XM_023254527.1
B2 145113221 145113347
refGene_gn=SNX9;refGene_tx=XM_019831438.2;refGene_gn=SNX9;refGene_tx=XM_023254527.1
B2 145126288 145126375
refGene_gn=SNX9;refGene_tx=XM_023254527.1
B2 145135680 145135846
refGene_gn=SNX9;refGene_tx=XM_019831438.2;refGene_gn=SNX9;refGene_tx=XM_023254527.1
B2 145137392 145137546
refGene_gn=SNX9;refGene_tx=XM_019831438.2;refGene_gn=SNX9;refGene_tx=XM_023254527.1
B2 145141604 145141689
refGene_gn=SNX9;refGene_tx=XM_019831438.2;refGene_gn=SNX9;refGene_tx=XM_023254527.1
B2 145143753 145143879
refGene_gn=SNX9;refGene_tx=XM_019831438.2;refGene_gn=SNX9;refGene_tx=XM_023254527.1
B2 145143999 145144117
refGene_gn=SNX9;refGene_tx=XM_019831438.2;refGene_gn=SNX9;refGene_tx=XM_023254527.1
B2 145153555 145153686
refGene_gn=SNX9;refGene_tx=XM_019831438.2;refGene_gn=SNX9;refGene_tx=XM_023254527.1
B2 145158570 145158674
refGene_gn=SNX9;refGene_tx=XM_019831438.2;refGene_gn=SNX9;refGene_tx=XM_023254527.1
B2 145160553 145160657
refGene_gn=SNX9;refGene_tx=XM_019831438.2;refGene_gn=SNX9;refGene_tx=XM_023254527.1
B2 145164336 145164414
refGene_gn=SNX9;refGene_tx=XM_019831438.2;refGene_gn=SNX9;refGene_tx=XM_023254527.1
B2 145168624 145168701
refGene_gn=SNX9;refGene_tx=XM_019831438.2;refGene_gn=SNX9;refGene_tx=XM_023254527.1
B2 145170021 145170111
refGene_gn=SNX9;refGene_tx=XM_019831438.2;refGene_gn=SNX9;refGene_tx=XM_023254527.1
B2 145171947 145172062
refGene_gn=SNX9;refGene_tx=XM_019831438.2;refGene_gn=SNX9;refGene_tx=X

M_023254527.1
B2	145173933	145174025
refGene_gn=SNX9;refGene_tx=XM_019831438.2;refGene_gn=SNX9;refGene_tx=XM_023254527.1
B2	145175977	145176025
refGene_gn=SNX9;refGene_tx=XM_019831438.2;refGene_gn=SNX9;refGene_tx=XM_023254527.1
B2	145201423	145201550
refGene_gn=SYNJ2;refGene_tx=XM_023254528.1;refGene_gn=SYNJ2;refGene_tx=XM_023254529.1;refGene_gn=SYNJ2;refGene_tx=XM_023254530.1;refGene_gn=SYNJ2;refGene_tx=XM_023254531.1
B2	145228469	145228556
refGene_gn=SYNJ2;refGene_tx=XM_023254528.1;refGene_gn=SYNJ2;refGene_tx=XM_023254529.1;refGene_gn=SYNJ2;refGene_tx=XM_023254530.1;refGene_gn=SYNJ2;refGene_tx=XM_023254531.1
B2	145238448	145238719
refGene_gn=SYNJ2;refGene_tx=XM_023254528.1;refGene_gn=SYNJ2;refGene_tx=XM_023254529.1;refGene_gn=SYNJ2;refGene_tx=XM_023254530.1;refGene_gn=SYNJ2;refGene_tx=XM_023254531.1
B2	145242584	145242810
refGene_gn=SYNJ2;refGene_tx=XM_023254528.1;refGene_gn=SYNJ2;refGene_tx=XM_023254529.1;refGene_gn=SYNJ2;refGene_tx=XM_023254530.1;refGene_gn=SYNJ2;refGene_tx=XM_023254531.1
B2	145249924	145250008
refGene_gn=SYNJ2;refGene_tx=XM_023254528.1;refGene_gn=SYNJ2;refGene_tx=XM_023254529.1;refGene_gn=SYNJ2;refGene_tx=XM_023254530.1;refGene_gn=SYNJ2;refGene_tx=XM_023254531.1
B2	145260444	145260506
refGene_gn=SYNJ2;refGene_tx=XM_023254528.1;refGene_gn=SYNJ2;refGene_tx=XM_023254529.1;refGene_gn=SYNJ2;refGene_tx=XM_023254530.1;refGene_gn=SYNJ2;refGene_tx=XM_023254531.1
B2	145265275	145265372
refGene_gn=SYNJ2;refGene_tx=XM_023254528.1;refGene_gn=SYNJ2;refGene_tx=XM_023254529.1;refGene_gn=SYNJ2;refGene_tx=XM_023254530.1;refGene_gn=SYNJ2;refGene_tx=XM_023254531.1
B2	145267536	145267641
refGene_gn=SYNJ2;refGene_tx=XM_023254533.1
B2	145267828	145268001
refGene_gn=SYNJ2;refGene_tx=XM_023254528.1;refGene_gn=SYNJ2;refGene_tx=XM_023254529.1;refGene_gn=SYNJ2;refGene_tx=XM_023254530.1;refGene_gn=SYNJ2;refGene_tx=XM_023254531.1;refGene_gn=SYNJ2;refGene_tx=XM_023254533.1
B2	145269480	145269562
refGene_gn=SYNJ2;refGene_tx=XM_023254528.1;refGene_gn=SYNJ2;refGene_tx=XM_023254529.1;refGene_gn=SYNJ2;refGene_tx=XM_023254530.1;refGene_gn=SYNJ2;refGene_tx=XM_023254531.1;refGene_gn=SYNJ2;refGene_tx=XM_023254533.1
B2	145270314	145270464
refGene_gn=SYNJ2;refGene_tx=XM_023254528.1;refGene_gn=SYNJ2;refGene_tx=XM_023254529.1;refGene_gn=SYNJ2;refGene_tx=XM_023254530.1;refGene_gn=

SYNJ2;refGene_tx=XM_023254531.1;refGene_gn=SYNJ2;refGene_tx=XM_0232545
33.1
B2 145270539 145270705
refGene_gn=SYNJ2;refGene_tx=XM_023254528.1;refGene_gn=SYNJ2;refGene_tx
=XM_023254529.1;refGene_gn=SYNJ2;refGene_tx=XM_023254530.1;refGene_gn=
SYNJ2;refGene_tx=XM_023254531.1;refGene_gn=SYNJ2;refGene_tx=XM_0232545
33.1
B2 145277750 145277942
refGene_gn=SYNJ2;refGene_tx=XM_023254528.1;refGene_gn=SYNJ2;refGene_tx
=XM_023254529.1;refGene_gn=SYNJ2;refGene_tx=XM_023254530.1;refGene_gn=
SYNJ2;refGene_tx=XM_023254531.1;refGene_gn=SYNJ2;refGene_tx=XM_0232545
33.1
B2 145279900 145279982
refGene_gn=SYNJ2;refGene_tx=XM_023254528.1;refGene_gn=SYNJ2;refGene_tx
=XM_023254529.1;refGene_gn=SYNJ2;refGene_tx=XM_023254530.1;refGene_gn=
SYNJ2;refGene_tx=XM_023254531.1;refGene_gn=SYNJ2;refGene_tx=XM_0232545
33.1
B2 145280745 145280886
refGene_gn=SYNJ2;refGene_tx=XM_023254528.1;refGene_gn=SYNJ2;refGene_tx
=XM_023254529.1;refGene_gn=SYNJ2;refGene_tx=XM_023254530.1;refGene_gn=
SYNJ2;refGene_tx=XM_023254531.1;refGene_gn=SYNJ2;refGene_tx=XM_0232545
33.1
B2 145282495 145282688
refGene_gn=SYNJ2;refGene_tx=XM_023254528.1;refGene_gn=SYNJ2;refGene_tx
=XM_023254529.1;refGene_gn=SYNJ2;refGene_tx=XM_023254530.1;refGene_gn=
SYNJ2;refGene_tx=XM_023254531.1;refGene_gn=SYNJ2;refGene_tx=XM_0232545
33.1
B2 145285552 145285711
refGene_gn=SYNJ2;refGene_tx=XM_023254528.1;refGene_gn=SYNJ2;refGene_tx
=XM_023254529.1;refGene_gn=SYNJ2;refGene_tx=XM_023254530.1;refGene_gn=
SYNJ2;refGene_tx=XM_023254531.1;refGene_gn=SYNJ2;refGene_tx=XM_0232545
33.1
B2 145287472 145287629
refGene_gn=SYNJ2;refGene_tx=XM_023254528.1;refGene_gn=SYNJ2;refGene_tx
=XM_023254529.1;refGene_gn=SYNJ2;refGene_tx=XM_023254530.1;refGene_gn=
SYNJ2;refGene_tx=XM_023254531.1;refGene_gn=SYNJ2;refGene_tx=XM_0232545
33.1
B2 145288578 145288696
refGene_gn=SYNJ2;refGene_tx=XM_023254528.1;refGene_gn=SYNJ2;refGene_tx
=XM_023254529.1;refGene_gn=SYNJ2;refGene_tx=XM_023254530.1;refGene_gn=
SYNJ2;refGene_tx=XM_023254531.1;refGene_gn=SYNJ2;refGene_tx=XM_0232545
33.1
B2 145292921 145293140
refGene_gn=SYNJ2;refGene_tx=XM_023254528.1;refGene_gn=SYNJ2;refGene_tx
=XM_023254529.1;refGene_gn=SYNJ2;refGene_tx=XM_023254530.1;refGene_gn=
SYNJ2;refGene_tx=XM_023254531.1;refGene_gn=SYNJ2;refGene_tx=XM_0232545
33.1
B2 145293251 145293330
refGene_gn=SYNJ2;refGene_tx=XM_023254528.1;refGene_gn=SYNJ2;refGene_tx
=XM_023254529.1;refGene_gn=SYNJ2;refGene_tx=XM_023254530.1;refGene_gn=

SYNJ2;refGene_tx=XM_023254531.1;refGene_gn=SYNJ2;refGene_tx=XM_0232545
33.1
B2	145293896	145294065
refGene_gn=SYNJ2;refGene_tx=XM_023254528.1;refGene_gn=SYNJ2;refGene_tx
=XM_023254529.1;refGene_gn=SYNJ2;refGene_tx=XM_023254530.1;refGene_gn=
SYNJ2;refGene_tx=XM_023254531.1;refGene_gn=SYNJ2;refGene_tx=XM_0232545
33.1
B2	145294503	145294677
refGene_gn=SYNJ2;refGene_tx=XM_023254528.1;refGene_gn=SYNJ2;refGene_tx
=XM_023254529.1;refGene_gn=SYNJ2;refGene_tx=XM_023254530.1;refGene_gn=
SYNJ2;refGene_tx=XM_023254531.1;refGene_gn=SYNJ2;refGene_tx=XM_0232545
33.1
B2	145297459	145297594
refGene_gn=SYNJ2;refGene_tx=XM_023254528.1;refGene_gn=SYNJ2;refGene_tx
=XM_023254529.1;refGene_gn=SYNJ2;refGene_tx=XM_023254531.1;refGene_gn=
SYNJ2;refGene_tx=XM_023254533.1
B2	145299048	145299161
refGene_gn=SYNJ2;refGene_tx=XM_023254528.1;refGene_gn=SYNJ2;refGene_tx
=XM_023254529.1;refGene_gn=SYNJ2;refGene_tx=XM_023254530.1;refGene_gn=
SYNJ2;refGene_tx=XM_023254531.1;refGene_gn=SYNJ2;refGene_tx=XM_0232545
33.1
B2	145300226	145300335
refGene_gn=SYNJ2;refGene_tx=XM_023254528.1;refGene_gn=SYNJ2;refGene_tx
=XM_023254529.1;refGene_gn=SYNJ2;refGene_tx=XM_023254530.1;refGene_gn=
SYNJ2;refGene_tx=XM_023254531.1;refGene_gn=SYNJ2;refGene_tx=XM_0232545
33.1
B2	145301991	145302143
refGene_gn=SYNJ2;refGene_tx=XM_023254528.1;refGene_gn=SYNJ2;refGene_tx
=XM_023254529.1;refGene_gn=SYNJ2;refGene_tx=XM_023254530.1;refGene_gn=
SYNJ2;refGene_tx=XM_023254531.1;refGene_gn=SYNJ2;refGene_tx=XM_0232545
33.1
B2	145302558	145302681
refGene_gn=SYNJ2;refGene_tx=XM_023254531.1
B2	145303896	145304634
refGene_gn=SYNJ2;refGene_tx=XM_023254528.1;refGene_gn=SYNJ2;refGene_tx
=XM_023254529.1;refGene_gn=SYNJ2;refGene_tx=XM_023254530.1;refGene_gn=
SYNJ2;refGene_tx=XM_023254533.1
B2	145325986	145326123
refGene_gn=SERAC1;refGene_tx=XM_023254534.1;refGene_gn=SERAC1;refGene_
tx=XM_023254535.1;refGene_gn=SERAC1;refGene_tx=XM_023254536.1
B2	145327534	145327678
refGene_gn=SERAC1;refGene_tx=XM_023254534.1;refGene_gn=SERAC1;refGene_
tx=XM_023254535.1;refGene_gn=SERAC1;refGene_tx=XM_023254536.1
B2	145328836	145329019
refGene_gn=SERAC1;refGene_tx=XM_023254534.1;refGene_gn=SERAC1;refGene_
tx=XM_023254535.1;refGene_gn=SERAC1;refGene_tx=XM_023254536.1
B2	145329817	145329936
refGene_gn=SERAC1;refGene_tx=XM_023254537.1
B2	145330429	145330527
refGene_gn=SERAC1;refGene_tx=XM_023254534.1;refGene_gn=SERAC1;refGene_

tx=XM_023254535.1;refGene_gn=SERAC1;refGene_tx=XM_023254536.1;refGene_gn=SERAC1;refGene_tx=XM_023254537.1
B2 145332036 145332131
refGene_gn=SERAC1;refGene_tx=XM_023254534.1;refGene_gn=SERAC1;refGene_tx=XM_023254535.1;refGene_gn=SERAC1;refGene_tx=XM_023254536.1;refGene_gn=SERAC1;refGene_tx=XM_023254537.1
B2 145333137 145333279
refGene_gn=SERAC1;refGene_tx=XM_023254534.1;refGene_gn=SERAC1;refGene_tx=XM_023254535.1;refGene_gn=SERAC1;refGene_tx=XM_023254536.1;refGene_gn=SERAC1;refGene_tx=XM_023254537.1
B2 145334945 145335096
refGene_gn=SERAC1;refGene_tx=XM_023254534.1;refGene_gn=SERAC1;refGene_tx=XM_023254535.1;refGene_gn=SERAC1;refGene_tx=XM_023254536.1;refGene_gn=SERAC1;refGene_tx=XM_023254537.1
B2 145349033 145349196
refGene_gn=SERAC1;refGene_tx=XM_023254534.1;refGene_gn=SERAC1;refGene_tx=XM_023254535.1;refGene_gn=SERAC1;refGene_tx=XM_023254536.1;refGene_gn=SERAC1;refGene_tx=XM_023254537.1
B2 145351192 145351306
refGene_gn=SERAC1;refGene_tx=XM_023254534.1;refGene_gn=SERAC1;refGene_tx=XM_023254535.1;refGene_gn=SERAC1;refGene_tx=XM_023254536.1;refGene_gn=SERAC1;refGene_tx=XM_023254537.1
B2 145362733 145362862
refGene_gn=SERAC1;refGene_tx=XM_023254534.1;refGene_gn=SERAC1;refGene_tx=XM_023254535.1;refGene_gn=SERAC1;refGene_tx=XM_023254536.1;refGene_gn=SERAC1;refGene_tx=XM_023254537.1
B2 145363927 145364049
refGene_gn=SERAC1;refGene_tx=XM_023254534.1;refGene_gn=SERAC1;refGene_tx=XM_023254535.1;refGene_gn=SERAC1;refGene_tx=XM_023254536.1;refGene_gn=SERAC1;refGene_tx=XM_023254537.1
B2 145365980 145366112
refGene_gn=SERAC1;refGene_tx=XM_023254534.1;refGene_gn=SERAC1;refGene_tx=XM_023254535.1;refGene_gn=SERAC1;refGene_tx=XM_023254536.1;refGene_gn=SERAC1;refGene_tx=XM_023254537.1
B2 145367419 145367509
refGene_gn=SERAC1;refGene_tx=XM_023254534.1;refGene_gn=SERAC1;refGene_tx=XM_023254535.1;refGene_gn=SERAC1;refGene_tx=XM_023254536.1;refGene_gn=SERAC1;refGene_tx=XM_023254537.1
B2 145368685 145368822
refGene_gn=SERAC1;refGene_tx=XM_023254534.1;refGene_gn=SERAC1;refGene_tx=XM_023254535.1;refGene_gn=SERAC1;refGene_tx=XM_023254537.1
B2 145371662 145371699
refGene_gn=SERAC1;refGene_tx=XM_023254534.1;refGene_gn=SERAC1;refGene_tx=XM_023254535.1;refGene_gn=SERAC1;refGene_tx=XM_023254537.1
B2 145374127 145374218
refGene_gn=SERAC1;refGene_tx=XM_023254534.1;refGene_gn=SERAC1;refGene_tx=XM_023254535.1;refGene_gn=SERAC1;refGene_tx=XM_023254536.1;refGene_gn=SERAC1;refGene_tx=XM_023254537.1
B2 145392309 145392344
refGene_gn=GTF2H5;refGene_tx=XM_003986673.5;refGene_gn=GTF2H5;refGene_

tx=XM_006932114.4;refGene_gn=GTF2H5;refGene_tx=XM_006932115.4
B2	145409196	145409377
refGene_gn=GTF2H5;refGene_tx=XM_003986673.5;refGene_gn=GTF2H5;refGene_tx=XM_006932114.4;refGene_gn=GTF2H5;refGene_tx=XM_006932115.4
B2	145515135	145515387
refGene_gn=TULP4;refGene_tx=XM_023254538.1;refGene_gn=TULP4;refGene_tx=XM_023254539.1;refGene_gn=TULP4;refGene_tx=XM_023254540.1
B2	145579591	145579594
refGene_gn=TULP4;refGene_tx=XM_023254541.1
B2	145607242	145607371
refGene_gn=TULP4;refGene_tx=XM_023254538.1;refGene_gn=TULP4;refGene_tx=XM_023254539.1;refGene_gn=TULP4;refGene_tx=XM_023254540.1;refGene_gn=TULP4;refGene_tx=XM_023254541.1
B2	145621668	145621830
refGene_gn=TULP4;refGene_tx=XM_023254538.1;refGene_gn=TULP4;refGene_tx=XM_023254539.1;refGene_gn=TULP4;refGene_tx=XM_023254540.1;refGene_gn=TULP4;refGene_tx=XM_023254541.1
B2	145647883	145648064
refGene_gn=TULP4;refGene_tx=XM_023254538.1;refGene_gn=TULP4;refGene_tx=XM_023254539.1;refGene_gn=TULP4;refGene_tx=XM_023254540.1;refGene_gn=TULP4;refGene_tx=XM_023254541.1
B2	145650941	145651076
refGene_gn=TULP4;refGene_tx=XM_023254538.1;refGene_gn=TULP4;refGene_tx=XM_023254539.1;refGene_gn=TULP4;refGene_tx=XM_023254540.1;refGene_gn=TULP4;refGene_tx=XM_023254541.1
B2	145657507	145657674
refGene_gn=TULP4;refGene_tx=XM_023254538.1;refGene_gn=TULP4;refGene_tx=XM_023254539.1;refGene_gn=TULP4;refGene_tx=XM_023254540.1;refGene_gn=TULP4;refGene_tx=XM_023254541.1
B2	145665178	145665403
refGene_gn=TULP4;refGene_tx=XM_023254538.1;refGene_gn=TULP4;refGene_tx=XM_023254539.1;refGene_gn=TULP4;refGene_tx=XM_023254540.1;refGene_gn=TULP4;refGene_tx=XM_023254541.1
B2	145666389	145666624
refGene_gn=TULP4;refGene_tx=XM_023254538.1;refGene_gn=TULP4;refGene_tx=XM_023254539.1;refGene_gn=TULP4;refGene_tx=XM_023254540.1;refGene_gn=TULP4;refGene_tx=XM_023254541.1
B2	145671134	145671279
refGene_gn=TULP4;refGene_tx=XM_023254538.1;refGene_gn=TULP4;refGene_tx=XM_023254539.1;refGene_gn=TULP4;refGene_tx=XM_023254540.1;refGene_gn=TULP4;refGene_tx=XM_023254541.1
B2	145673903	145674048
refGene_gn=TULP4;refGene_tx=XM_023254538.1;refGene_gn=TULP4;refGene_tx=XM_023254539.1;refGene_gn=TULP4;refGene_tx=XM_023254540.1;refGene_gn=TULP4;refGene_tx=XM_023254541.1
B2	145675327	145675421
refGene_gn=TULP4;refGene_tx=XM_023254538.1;refGene_gn=TULP4;refGene_tx=XM_023254539.1;refGene_gn=TULP4;refGene_tx=XM_023254540.1;refGene_gn=TULP4;refGene_tx=XM_023254541.1
B2	145678591	145678735

refGene_gn=TULP4;refGene_tx=XM_023254538.1;refGene_gn=TULP4;refGene_tx=XM_023254539.1;refGene_gn=TULP4;refGene_tx=XM_023254540.1;refGene_gn=TULP4;refGene_tx=XM_023254541.1
B2 145681302 145683821
refGene_gn=TULP4;refGene_tx=XM_023254538.1;refGene_gn=TULP4;refGene_tx=XM_023254539.1;refGene_gn=TULP4;refGene_tx=XM_023254540.1;refGene_gn=TULP4;refGene_tx=XM_023254541.1
B2 145685569 145685686
refGene_gn=TULP4;refGene_tx=XM_023254538.1;refGene_gn=TULP4;refGene_tx=XM_023254539.1;refGene_gn=TULP4;refGene_tx=XM_023254540.1;refGene_gn=TULP4;refGene_tx=XM_023254541.1
B2 145719274 145719405
refGene_gn=TMEM181;refGene_tx=XM_003986675.4
B2 145753161 145753265
refGene_gn=TMEM181;refGene_tx=XM_003986675.4;refGene_gn=TMEM181;refGene_tx=XM_023254542.1
B2 145760046 145760102
refGene_gn=TMEM181;refGene_tx=XM_003986675.4;refGene_gn=TMEM181;refGene_tx=XM_023254542.1
B2 145761891 145761982
refGene_gn=TMEM181;refGene_tx=XM_003986675.4;refGene_gn=TMEM181;refGene_tx=XM_023254542.1
B2 145764344 145764466
refGene_gn=TMEM181;refGene_tx=XM_003986675.4;refGene_gn=TMEM181;refGene_tx=XM_023254542.1
B2 145767654 145767765
refGene_gn=TMEM181;refGene_tx=XM_003986675.4;refGene_gn=TMEM181;refGene_tx=XM_023254542.1
B2 145778058 145778139
refGene_gn=TMEM181;refGene_tx=XM_003986675.4;refGene_gn=TMEM181;refGene_tx=XM_023254542.1
B2 145779884 145779984
refGene_gn=TMEM181;refGene_tx=XM_003986675.4;refGene_gn=TMEM181;refGene_tx=XM_023254542.1
B2 145780561 145780692
refGene_gn=TMEM181;refGene_tx=XM_003986675.4;refGene_gn=TMEM181;refGene_tx=XM_023254542.1
B2 145780898 145780990
refGene_gn=TMEM181;refGene_tx=XM_003986675.4;refGene_gn=TMEM181;refGene_tx=XM_023254542.1
B2 145782190 145782248
refGene_gn=TMEM181;refGene_tx=XM_003986675.4;refGene_gn=TMEM181;refGene_tx=XM_023254542.1
B2 145784541 145784644
refGene_gn=TMEM181;refGene_tx=XM_003986675.4;refGene_gn=TMEM181;refGene_tx=XM_023254542.1
B2 145785252 145785304
refGene_gn=TMEM181;refGene_tx=XM_003986675.4;refGene_gn=TMEM181;refGene_tx=XM_023254542.1
B2 145787433 145787516

refGene_gn=TMEM181;refGene_tx=XM_003986675.4;refGene_gn=TMEM181;refGene_tx=XM_023254542.1
B2 145788610 145788700
refGene_gn=TMEM181;refGene_tx=XM_003986675.4;refGene_gn=TMEM181;refGene_tx=XM_023254542.1
B2 145790065 145790132
refGene_gn=TMEM181;refGene_tx=XM_003986675.4;refGene_gn=TMEM181;refGene_tx=XM_023254542.1
B2 145790552 145790631
refGene_gn=TMEM181;refGene_tx=XM_003986675.4;refGene_gn=TMEM181;refGene_tx=XM_023254542.1
B2 145794807 145794878
refGene_gn=DYNLT1;refGene_tx=XM_003986676.5
B2 145795384 145795462
refGene_gn=DYNLT1;refGene_tx=XM_003986676.5
B2 145795856 145795980
refGene_gn=DYNLT1;refGene_tx=XM_003986676.5
B2 145799168 145799210
refGene_gn=DYNLT1;refGene_tx=XM_003986676.5
B2 145802089 145802116
refGene_gn=DYNLT1;refGene_tx=XM_003986676.5
B2 145820445 145820555
refGene_gn=SYTL3;refGene_tx=XM_019831440.2;refGene_gn=SYTL3;refGene_tx=XM_019831441.2;refGene_gn=SYTL3;refGene_tx=XM_019831443.2
B2 145821999 145822218
refGene_gn=SYTL3;refGene_tx=XM_019831440.2;refGene_gn=SYTL3;refGene_tx=XM_019831441.2;refGene_gn=SYTL3;refGene_tx=XM_019831443.2
B2 145839051 145839116
refGene_gn=SYTL3;refGene_tx=XM_019831440.2;refGene_gn=SYTL3;refGene_tx=XM_019831441.2;refGene_gn=SYTL3;refGene_tx=XM_019831443.2
B2 145858268 145858320
refGene_gn=SYTL3;refGene_tx=XM_019831440.2;refGene_gn=SYTL3;refGene_tx=XM_019831441.2;refGene_gn=SYTL3;refGene_tx=XM_019831443.2
B2 145859581 145859651
refGene_gn=SYTL3;refGene_tx=XM_019831440.2;refGene_gn=SYTL3;refGene_tx=XM_019831441.2;refGene_gn=SYTL3;refGene_tx=XM_019831443.2
B2 145862422 145862501
refGene_gn=SYTL3;refGene_tx=XM_019831440.2;refGene_gn=SYTL3;refGene_tx=XM_019831441.2;refGene_gn=SYTL3;refGene_tx=XM_019831443.2
B2 145866477 145866602
refGene_gn=SYTL3;refGene_tx=XM_019831440.2;refGene_gn=SYTL3;refGene_tx=XM_019831441.2;refGene_gn=SYTL3;refGene_tx=XM_019831443.2
B2 145873801 145873936
refGene_gn=SYTL3;refGene_tx=XM_019831440.2;refGene_gn=SYTL3;refGene_tx=XM_019831441.2;refGene_gn=SYTL3;refGene_tx=XM_019831443.2
B2 145885388 145885567
refGene_gn=SYTL3;refGene_tx=XM_019831440.2;refGene_gn=SYTL3;refGene_tx=XM_019831441.2;refGene_gn=SYTL3;refGene_tx=XM_019831443.2
B2 145891308 145891411
refGene_gn=SYTL3;refGene_tx=XM_019831440.2;refGene_gn=SYTL3;refGene_tx

=XM_019831441.2;refGene_gn=SYTL3;refGene_tx=XM_019831443.2
B2 145895362 145895533
refGene_gn=SYTL3;refGene_tx=XM_019831440.2;refGene_gn=SYTL3;refGene_tx
=XM_019831441.2;refGene_gn=SYTL3;refGene_tx=XM_019831443.2
B2 145897292 145897398
refGene_gn=SYTL3;refGene_tx=XM_019831440.2;refGene_gn=SYTL3;refGene_tx
=XM_019831441.2;refGene_gn=SYTL3;refGene_tx=XM_019831443.2
B2 145898389 145898495
refGene_gn=SYTL3;refGene_tx=XM_019831440.2;refGene_gn=SYTL3;refGene_tx
=XM_019831441.2;refGene_gn=SYTL3;refGene_tx=XM_019831443.2
B2 145899415 145899636
refGene_gn=SYTL3;refGene_tx=XM_019831440.2;refGene_gn=SYTL3;refGene_tx
=XM_019831441.2;refGene_gn=SYTL3;refGene_tx=XM_019831443.2
B2 145900961 145901071
refGene_gn=SYTL3;refGene_tx=XM_019831440.2;refGene_gn=SYTL3;refGene_tx
=XM_019831441.2;refGene_gn=SYTL3;refGene_tx=XM_019831443.2
B2 145902743 145902908
refGene_gn=EZR;refGene_tx=XM_023254543.1;refGene_gn=EZR;refGene_tx=XM_
023254544.1
B2 145903027 145903279
refGene_gn=EZR;refGene_tx=XM_023254543.1;refGene_gn=EZR;refGene_tx=XM_
023254544.1
B2 145904837 145904930
refGene_gn=EZR;refGene_tx=XM_023254543.1;refGene_gn=EZR;refGene_tx=XM_
023254544.1
B2 145905280 145905441
refGene_gn=EZR;refGene_tx=XM_023254543.1;refGene_gn=EZR;refGene_tx=XM_
023254544.1
B2 145906091 145906222
refGene_gn=EZR;refGene_tx=XM_023254543.1;refGene_gn=EZR;refGene_tx=XM_
023254544.1
B2 145906516 145906680
refGene_gn=EZR;refGene_tx=XM_023254543.1;refGene_gn=EZR;refGene_tx=XM_
023254544.1
B2 145911552 145911649
refGene_gn=EZR;refGene_tx=XM_023254543.1;refGene_gn=EZR;refGene_tx=XM_
023254544.1
B2 145918987 145919134
refGene_gn=EZR;refGene_tx=XM_023254543.1;refGene_gn=EZR;refGene_tx=XM_
023254544.1
B2 145920323 145920407
refGene_gn=EZR;refGene_tx=XM_023254543.1;refGene_gn=EZR;refGene_tx=XM_
023254544.1
B2 145920998 145921273
refGene_gn=EZR;refGene_tx=XM_023254543.1
B2 145922188 145922284
refGene_gn=EZR;refGene_tx=XM_023254543.1
B2 145926942 145927026
refGene_gn=EZR;refGene_tx=XM_023254543.1;refGene_gn=EZR;refGene_tx=XM_
023254544.1

B2 145951516 145951528
refGene_gn=EZR;refGene_tx=XM_023254543.1
B2 146065735 146066070
refGene_gn=RSPH3;refGene_tx=XM_006932135.4
B2 146066607 146066694
refGene_gn=RSPH3;refGene_tx=XM_006932135.4
B2 146068778 146068941
refGene_gn=RSPH3;refGene_tx=XM_006932135.4
B2 146070498 146070702
refGene_gn=RSPH3;refGene_tx=XM_006932135.4
B2 146071718 146071864
refGene_gn=RSPH3;refGene_tx=XM_006932135.4
B2 146073641 146073783
refGene_gn=RSPH3;refGene_tx=XM_006932135.4
B2 146085867 146085955
refGene_gn=RSPH3;refGene_tx=XM_006932135.4
B2 146091660 146092127
refGene_gn=RSPH3;refGene_tx=XM_006932135.4
B2 146129241 146130509
refGene_gn=TAGAP;refGene_tx=XM_003986684.5
B2 146131113 146131228
refGene_gn=TAGAP;refGene_tx=XM_003986684.5
B2 146132024 146132220
refGene_gn=TAGAP;refGene_tx=XM_003986684.5
B2 146133626 146133736
refGene_gn=TAGAP;refGene_tx=XM_003986684.5
B2 146134220 146134382
refGene_gn=TAGAP;refGene_tx=XM_003986684.5
B2 146135050 146135217
refGene_gn=TAGAP;refGene_tx=XM_003986684.5
B2 146136707 146136774
refGene_gn=TAGAP;refGene_tx=XM_003986684.5
B2 146137080 146137134
refGene_gn=TAGAP;refGene_tx=XM_003986684.5
B2 146137221 146137248
refGene_gn=TAGAP;refGene_tx=XM_003986684.5
B2 146244762 146244871
refGene_gn=FNDC1;refGene_tx=XM_023254545.1
B2 146272756 146272951
refGene_gn=FNDC1;refGene_tx=XM_023254545.1
B2 146275288 146275375
refGene_gn=FNDC1;refGene_tx=XM_023254545.1
B2 146275817 146275886
refGene_gn=FNDC1;refGene_tx=XM_023254545.1
B2 146291843 146292050
refGene_gn=FNDC1;refGene_tx=XM_023254545.1
B2 146297406 146297505
refGene_gn=FNDC1;refGene_tx=XM_023254545.1
B2 146299058 146299176
refGene_gn=FNDC1;refGene_tx=XM_023254545.1

B2	146301594	146301782
refGene_gn=FNDC1;refGene_tx=XM_023254545.1
B2	146302647	146302755
refGene_gn=FNDC1;refGene_tx=XM_023254545.1
B2	146305681	146305870
refGene_gn=FNDC1;refGene_tx=XM_023254545.1
B2	146307260	146309849
refGene_gn=FNDC1;refGene_tx=XM_023254545.1
B2	146311536	146311637
refGene_gn=FNDC1;refGene_tx=XM_023254545.1
B2	146313812	146313924
refGene_gn=FNDC1;refGene_tx=XM_023254545.1
B2	146315025	146315421
refGene_gn=FNDC1;refGene_tx=XM_023254545.1
B2	146330227	146330296
refGene_gn=FNDC1;refGene_tx=XM_023254545.1
B2	146331868	146332042
refGene_gn=FNDC1;refGene_tx=XM_023254545.1
B2	146334116	146334347
refGene_gn=FNDC1;refGene_tx=XM_023254545.1
B2	146337813	146337922
refGene_gn=FNDC1;refGene_tx=XM_023254545.1
B2	146343510	146343590
refGene_gn=FNDC1;refGene_tx=XM_023254545.1
B2	146346914	146346944
refGene_gn=FNDC1;refGene_tx=XM_023254545.1
B2	146347893	146348055
refGene_gn=FNDC1;refGene_tx=XM_023254545.1
B2	146349394	146349517
refGene_gn=FNDC1;refGene_tx=XM_023254545.1
B2	146352107	146352223
refGene_gn=FNDC1;refGene_tx=XM_023254545.1
B2	146705144	146705290
refGene_gn=SOD2;refGene_tx=XM_023254547.1
B2	146707811	146707991
refGene_gn=SOD2;refGene_tx=XM_023254547.1
B2	146709715	146709832
refGene_gn=SOD2;refGene_tx=XM_023254547.1
B2	146714518	146714721
refGene_gn=SOD2;refGene_tx=XM_023254547.1
B2	146715067	146715090
refGene_gn=SOD2;refGene_tx=XM_023254547.1
B2	146744365	146744395
refGene_gn=WTAP;refGene_tx=XM_023254548.1;refGene_gn=WTAP;refGene_tx=XM_023254549.1;refGene_gn=WTAP;refGene_tx=XM_023254551.1
B2	146746641	146746697
refGene_gn=WTAP;refGene_tx=XM_023254548.1;refGene_gn=WTAP;refGene_tx=XM_023254549.1;refGene_gn=WTAP;refGene_tx=XM_023254551.1
B2	146749475	146749534
refGene_gn=WTAP;refGene_tx=XM_023254548.1;refGene_gn=WTAP;refGene_tx=X

M_023254549.1;refGene_gn=WTAP;refGene_tx=XM_023254551.1
B2 146751241 146751369
refGene_gn=WTAP;refGene_tx=XM_023254548.1;refGene_gn=WTAP;refGene_tx=XM_023254549.1;refGene_gn=WTAP;refGene_tx=XM_023254551.1
B2 146755527 146755706
refGene_gn=WTAP;refGene_tx=XM_023254548.1;refGene_gn=WTAP;refGene_tx=XM_023254549.1;refGene_gn=WTAP;refGene_tx=XM_023254551.1
B2 146755925 146755986
refGene_gn=WTAP;refGene_tx=XM_023254551.1
B2 146760790 146760945
refGene_gn=WTAP;refGene_tx=XM_023254548.1;refGene_gn=WTAP;refGene_tx=XM_023254549.1
B2 146762632 146763216
refGene_gn=WTAP;refGene_tx=XM_023254548.1;refGene_gn=WTAP;refGene_tx=XM_023254549.1
B2 146770360 146770415
refGene_gn=ACAT2;refGene_tx=XM_019831446.2
B2 146771015 146771150
refGene_gn=ACAT2;refGene_tx=XM_019831446.2
B2 146774481 146774663
refGene_gn=ACAT2;refGene_tx=XM_019831446.2
B2 146776106 146776224
refGene_gn=ACAT2;refGene_tx=XM_019831446.2
B2 146783164 146783308
refGene_gn=ACAT2;refGene_tx=XM_019831446.2
B2 146784152 146784275
refGene_gn=ACAT2;refGene_tx=XM_019831446.2
B2 146784851 146785006
refGene_gn=ACAT2;refGene_tx=XM_019831446.2
B2 146785575 146785686
refGene_gn=ACAT2;refGene_tx=XM_019831446.2
B2 146786313 146786484
refGene_gn=ACAT2;refGene_tx=XM_019831446.2
B2 146786706 146786923
refGene_gn=TCP1;refGene_tx=XM_003986687.5
B2 146787422 146787586
refGene_gn=TCP1;refGene_tx=XM_003986687.5
B2 146787943 146788136
refGene_gn=TCP1;refGene_tx=XM_003986687.5
B2 146788442 146788566
refGene_gn=TCP1;refGene_tx=XM_003986687.5
B2 146788933 146789109
refGene_gn=TCP1;refGene_tx=XM_003986687.5
B2 146792979 146793106
refGene_gn=TCP1;refGene_tx=XM_003986687.5
B2 146793489 146793671
refGene_gn=TCP1;refGene_tx=XM_003986687.5
B2 146794220 146794331
refGene_gn=TCP1;refGene_tx=XM_003986687.5
B2 146794700 146794798

refGene_gn=TCP1;refGene_tx=XM_003986687.5
B2 146796279 146796408
refGene_gn=TCP1;refGene_tx=XM_003986687.5
B2 146796607 146796693
refGene_gn=TCP1;refGene_tx=XM_003986687.5
B2 146797536 146797600
refGene_gn=TCP1;refGene_tx=XM_003986687.5
B2 146798805 146798857
refGene_gn=MRPL18;refGene_tx=XM_003986690.5;refGene_gn=MRPL18;refGene_tx=XM_019831447.2
B2 146799146 146799333
refGene_gn=MRPL18;refGene_tx=XM_003986690.5;refGene_gn=MRPL18;refGene_tx=XM_019831447.2
B2 146802326 146802558
refGene_gn=MRPL18;refGene_tx=XM_003986690.5;refGene_gn=MRPL18;refGene_tx=XM_019831447.2
B2 146802971 146802973
refGene_gn=MRPL18;refGene_tx=XM_019831447.2
B2 146803046 146803118
refGene_gn=MRPL18;refGene_tx=XM_003986690.5;refGene_gn=MRPL18;refGene_tx=XM_019831447.2
B2 146804823 146804899
refGene_gn=PNLDC1;refGene_tx=XM_011282552.3;refGene_gn=PNLDC1;refGene_tx=XM_019831448.2;refGene_gn=PNLDC1;refGene_tx=XM_023254552.1
B2 146805322 146805380
refGene_gn=PNLDC1;refGene_tx=XM_011282552.3;refGene_gn=PNLDC1;refGene_tx=XM_019831448.2;refGene_gn=PNLDC1;refGene_tx=XM_023254552.1
B2 146805681 146805755
refGene_gn=PNLDC1;refGene_tx=XM_011282552.3;refGene_gn=PNLDC1;refGene_tx=XM_019831448.2;refGene_gn=PNLDC1;refGene_tx=XM_023254552.1
B2 146807375 146807415
refGene_gn=PNLDC1;refGene_tx=XM_011282552.3;refGene_gn=PNLDC1;refGene_tx=XM_019831448.2;refGene_gn=PNLDC1;refGene_tx=XM_023254552.1
B2 146807947 146808071
refGene_gn=PNLDC1;refGene_tx=XM_011282552.3;refGene_gn=PNLDC1;refGene_tx=XM_019831448.2;refGene_gn=PNLDC1;refGene_tx=XM_023254552.1
B2 146808470 146808559
refGene_gn=PNLDC1;refGene_tx=XM_011282552.3;refGene_gn=PNLDC1;refGene_tx=XM_019831448.2;refGene_gn=PNLDC1;refGene_tx=XM_023254552.1
B2 146810005 146810106
refGene_gn=PNLDC1;refGene_tx=XM_011282552.3;refGene_gn=PNLDC1;refGene_tx=XM_019831448.2;refGene_gn=PNLDC1;refGene_tx=XM_023254552.1
B2 146811000 146811077
refGene_gn=PNLDC1;refGene_tx=XM_011282552.3;refGene_gn=PNLDC1;refGene_tx=XM_019831448.2;refGene_gn=PNLDC1;refGene_tx=XM_023254552.1
B2 146811238 146811382
refGene_gn=PNLDC1;refGene_tx=XM_011282552.3;refGene_gn=PNLDC1;refGene_tx=XM_019831448.2;refGene_gn=PNLDC1;refGene_tx=XM_023254552.1
B2 146811871 146811944
refGene_gn=PNLDC1;refGene_tx=XM_011282552.3;refGene_gn=PNLDC1;refGene_

tx=XM_019831448.2;refGene_gn=PNLDC1;refGene_tx=XM_023254552.1
B2 146813562 146813648
refGene_gn=PNLDC1;refGene_tx=XM_011282552.3;refGene_gn=PNLDC1;refGene_tx=XM_019831448.2;refGene_gn=PNLDC1;refGene_tx=XM_023254552.1
B2 146814985 146815041
refGene_gn=PNLDC1;refGene_tx=XM_011282552.3;refGene_gn=PNLDC1;refGene_tx=XM_019831448.2;refGene_gn=PNLDC1;refGene_tx=XM_023254552.1
B2 146818607 146818672
refGene_gn=PNLDC1;refGene_tx=XM_011282552.3;refGene_gn=PNLDC1;refGene_tx=XM_019831448.2;refGene_gn=PNLDC1;refGene_tx=XM_023254552.1
B2 146819120 146819174
refGene_gn=PNLDC1;refGene_tx=XM_011282552.3;refGene_gn=PNLDC1;refGene_tx=XM_019831448.2;refGene_gn=PNLDC1;refGene_tx=XM_023254552.1
B2 146819377 146819420
refGene_gn=PNLDC1;refGene_tx=XM_011282552.3;refGene_gn=PNLDC1;refGene_tx=XM_019831448.2;refGene_gn=PNLDC1;refGene_tx=XM_023254552.1
B2 146820871 146820971
refGene_gn=PNLDC1;refGene_tx=XM_011282552.3;refGene_gn=PNLDC1;refGene_tx=XM_019831448.2;refGene_gn=PNLDC1;refGene_tx=XM_023254552.1
B2 146821238 146821414
refGene_gn=PNLDC1;refGene_tx=XM_011282552.3;refGene_gn=PNLDC1;refGene_tx=XM_019831448.2;refGene_gn=PNLDC1;refGene_tx=XM_023254552.1
B2 146821559 146821658
refGene_gn=PNLDC1;refGene_tx=XM_011282552.3;refGene_gn=PNLDC1;refGene_tx=XM_019831448.2;refGene_gn=PNLDC1;refGene_tx=XM_023254552.1
B2 146822723 146822787
refGene_gn=PNLDC1;refGene_tx=XM_011282552.3;refGene_gn=PNLDC1;refGene_tx=XM_019831448.2;refGene_gn=PNLDC1;refGene_tx=XM_023254552.1
B2 146861646 146862624
refGene_gn=MAS1;refGene_tx=XM_019831449.2
B2 146923721 146923876
refGene_gn=IGF2R;refGene_tx=XM_023254553.1
B2 146944255 146944395
refGene_gn=IGF2R;refGene_tx=XM_023254553.1
B2 146954298 146954420
refGene_gn=IGF2R;refGene_tx=XM_023254553.1
B2 146956089 146956188
refGene_gn=IGF2R;refGene_tx=XM_023254553.1
B2 146960667 146960800
refGene_gn=IGF2R;refGene_tx=XM_023254553.1
B2 146963980 146964110
refGene_gn=IGF2R;refGene_tx=XM_023254553.1
B2 146965791 146965897
refGene_gn=IGF2R;refGene_tx=XM_023254553.1
B2 146968469 146968632
refGene_gn=IGF2R;refGene_tx=XM_023254553.1
B2 146968807 146968982
refGene_gn=IGF2R;refGene_tx=XM_023254553.1
B2 146969813 146969917
refGene_gn=IGF2R;refGene_tx=XM_023254553.1

B2 146975621 146975786
refGene_gn=IGF2R;refGene_tx=XM_023254553.1
B2 146978446 146978587
refGene_gn=IGF2R;refGene_tx=XM_023254553.1
B2 146979809 146979950
refGene_gn=IGF2R;refGene_tx=XM_023254553.1
B2 146981300 146981438
refGene_gn=IGF2R;refGene_tx=XM_023254553.1
B2 146982549 146982697
refGene_gn=IGF2R;refGene_tx=XM_023254553.1
B2 146983209 146983387
refGene_gn=IGF2R;refGene_tx=XM_023254553.1
B2 146984018 146984134
refGene_gn=IGF2R;refGene_tx=XM_023254553.1
B2 146984614 146984783
refGene_gn=IGF2R;refGene_tx=XM_023254553.1
B2 146986009 146986189
refGene_gn=IGF2R;refGene_tx=XM_023254553.1
B2 146988916 146989018
refGene_gn=IGF2R;refGene_tx=XM_023254553.1
B2 146990314 146990416
refGene_gn=IGF2R;refGene_tx=XM_023254553.1
B2 146992489 146992685
refGene_gn=IGF2R;refGene_tx=XM_023254553.1
B2 146994003 146994174
refGene_gn=IGF2R;refGene_tx=XM_023254553.1
B2 146994714 146994858
refGene_gn=IGF2R;refGene_tx=XM_023254553.1
B2 146994941 146995117
refGene_gn=IGF2R;refGene_tx=XM_023254553.1
B2 146995400 146995488
refGene_gn=IGF2R;refGene_tx=XM_023254553.1
B2 146996206 146996422
refGene_gn=IGF2R;refGene_tx=XM_023254553.1
B2 146997271 146997402
refGene_gn=IGF2R;refGene_tx=XM_023254553.1
B2 146997653 146997751
refGene_gn=IGF2R;refGene_tx=XM_023254553.1
B2 147000194 147000331
refGene_gn=IGF2R;refGene_tx=XM_023254553.1
B2 147001146 147001337
refGene_gn=IGF2R;refGene_tx=XM_023254553.1
B2 147002828 147002955
refGene_gn=IGF2R;refGene_tx=XM_023254553.1
B2 147004742 147004862
refGene_gn=IGF2R;refGene_tx=XM_023254553.1
B2 147005184 147005441
refGene_gn=IGF2R;refGene_tx=XM_023254553.1
B2 147005778 147005997
refGene_gn=IGF2R;refGene_tx=XM_023254553.1

B2 147007996 147008146
refGene_gn=IGF2R;refGene_tx=XM_023254553.1
B2 147010496 147010658
refGene_gn=IGF2R;refGene_tx=XM_023254553.1
B2 147011874 147012082
refGene_gn=IGF2R;refGene_tx=XM_023254553.1
B2 147012422 147012569
refGene_gn=IGF2R;refGene_tx=XM_023254553.1
B2 147013577 147013812
refGene_gn=IGF2R;refGene_tx=XM_023254553.1
B2 147014537 147014674
refGene_gn=IGF2R;refGene_tx=XM_023254553.1
B2 147017682 147017797
refGene_gn=IGF2R;refGene_tx=XM_023254553.1
B2 147018747 147018894
refGene_gn=IGF2R;refGene_tx=XM_023254553.1
B2 147019559 147019747
refGene_gn=IGF2R;refGene_tx=XM_023254553.1
B2 147021391 147021578
refGene_gn=IGF2R;refGene_tx=XM_023254553.1
B2 147024019 147024172
refGene_gn=IGF2R;refGene_tx=XM_023254553.1
B2 147026700 147026770
refGene_gn=IGF2R;refGene_tx=XM_023254553.1
B2 147027448 147027865
refGene_gn=IGF2R;refGene_tx=XM_023254553.1
B2 147038121 147038532
refGene_gn=SLC22A1;refGene_tx=XM_003986693.5;refGene_gn=SLC22A1;refGene_tx=XM_006932128.4
B2 147043379 147043483
refGene_gn=SLC22A1;refGene_tx=XM_003986693.5;refGene_gn=SLC22A1;refGene_tx=XM_006932128.4
B2 147045280 147045435
refGene_gn=SLC22A1;refGene_tx=XM_003986693.5;refGene_gn=SLC22A1;refGene_tx=XM_006932128.4
B2 147047785 147047954
refGene_gn=SLC22A1;refGene_tx=XM_003986693.5;refGene_gn=SLC22A1;refGene_tx=XM_006932128.4
B2 147049208 147049323
refGene_gn=SLC22A1;refGene_tx=XM_003986693.5;refGene_gn=SLC22A1;refGene_tx=XM_006932128.4
B2 147049555 147049662
refGene_gn=SLC22A1;refGene_tx=XM_003986693.5;refGene_gn=SLC22A1;refGene_tx=XM_006932128.4
B2 147052864 147053079
refGene_gn=SLC22A1;refGene_tx=XM_003986693.5
B2 147055943 147056052
refGene_gn=SLC22A1;refGene_tx=XM_003986693.5
B2 147063778 147063891
refGene_gn=SLC22A1;refGene_tx=XM_003986693.5;refGene_gn=SLC22A1;refGen

e_tx=XM_006932128.4
B2	147065211	147065311
refGene_gn=SLC22A1;refGene_tx=XM_003986693.5;refGene_gn=SLC22A1;refGene_tx=XM_006932128.4
B2	147067358	147067425
refGene_gn=SLC22A1;refGene_tx=XM_003986693.5;refGene_gn=SLC22A1;refGene_tx=XM_006932128.4
B2	147099774	147099841
refGene_gn=LOC101095017;refGene_tx=XM_003986694.5
B2	147106630	147106730
refGene_gn=LOC101095017;refGene_tx=XM_003986694.5
B2	147110016	147110129
refGene_gn=LOC101095017;refGene_tx=XM_003986694.5
B2	147110665	147110774
refGene_gn=LOC101095017;refGene_tx=XM_003986694.5
B2	147111654	147111869
refGene_gn=LOC101095017;refGene_tx=XM_003986694.5
B2	147113203	147113232
refGene_gn=LOC101095017;refGene_tx=XM_003986694.5
B2	147113233	147113311
refGene_gn=LOC101095017;refGene_tx=XM_003986694.5
B2	147115396	147115511
refGene_gn=LOC101095017;refGene_tx=XM_003986694.5
B2	147119468	147119637
refGene_gn=LOC101095017;refGene_tx=XM_003986694.5
B2	147122026	147122181
refGene_gn=LOC101095017;refGene_tx=XM_003986694.5
B2	147127553	147127657
refGene_gn=LOC101095017;refGene_tx=XM_003986694.5
B2	147129498	147129848
refGene_gn=LOC101095017;refGene_tx=XM_003986694.5
B2	147129850	147129908
refGene_gn=LOC101095017;refGene_tx=XM_003986694.5
B2	147196094	147196505
refGene_gn=SLC22A3;refGene_tx=XM_023254554.1;refGene_gn=SLC22A3;refGene_tx=XM_023254555.1
B2	147242269	147242373
refGene_gn=SLC22A3;refGene_tx=XM_023254554.1;refGene_gn=SLC22A3;refGene_tx=XM_023254555.1
B2	147244753	147244908
refGene_gn=SLC22A3;refGene_tx=XM_023254554.1
B2	147246634	147246803
refGene_gn=SLC22A3;refGene_tx=XM_023254554.1
B2	147247681	147247799
refGene_gn=SLC22A3;refGene_tx=XM_023254554.1
B2	147276702	147276800
refGene_gn=SLC22A3;refGene_tx=XM_023254554.1
B2	147276919	147277134
refGene_gn=SLC22A3;refGene_tx=XM_023254554.1;refGene_gn=SLC22A3;refGene_tx=XM_023254555.1

B2 147286077 147286186
refGene_gn=SLC22A3;refGene_tx=XM_023254554.1;refGene_gn=SLC22A3;refGene_tx=XM_023254555.1
B2 147286948 147287061
refGene_gn=SLC22A3;refGene_tx=XM_023254554.1;refGene_gn=SLC22A3;refGene_tx=XM_023254555.1
B2 147290538 147290638
refGene_gn=SLC22A3;refGene_tx=XM_023254554.1;refGene_gn=SLC22A3;refGene_tx=XM_023254555.1
B2 147293367 147293428
refGene_gn=SLC22A3;refGene_tx=XM_023254554.1;refGene_gn=SLC22A3;refGene_tx=XM_023254555.1
B2 147295767 147295929
refGene_gn=PLG;refGene_tx=XM_006932130.4
B2 147296577 147296723
refGene_gn=PLG;refGene_tx=XM_006932130.4
B2 147300958 147301065
refGene_gn=PLG;refGene_tx=XM_006932130.4
B2 147302966 147303107
refGene_gn=PLG;refGene_tx=XM_006932130.4
B2 147303773 147303848
refGene_gn=PLG;refGene_tx=XM_006932130.4
B2 147305255 147305376
refGene_gn=PLG;refGene_tx=XM_006932130.4
B2 147307542 147307636
refGene_gn=PLG;refGene_tx=XM_006932130.4
B2 147309189 147309338
refGene_gn=PLG;refGene_tx=XM_006932130.4
B2 147310895 147311077
refGene_gn=PLG;refGene_tx=XM_006932130.4
B2 147316484 147316650
refGene_gn=PLG;refGene_tx=XM_006932130.4
B2 147320491 147320637
refGene_gn=PLG;refGene_tx=XM_006932130.4
B2 147321326 147321489
refGene_gn=PLG;refGene_tx=XM_006932130.4
B2 147322664 147322783
refGene_gn=PLG;refGene_tx=XM_006932130.4
B2 147324238 147324359
refGene_gn=PLG;refGene_tx=XM_006932130.4
B2 147326743 147326883
refGene_gn=PLG;refGene_tx=XM_006932130.4
B2 147329105 147329220
refGene_gn=PLG;refGene_tx=XM_006932130.4
B2 147332897 147333004
refGene_gn=PLG;refGene_tx=XM_006932130.4
B2 147334329 147334465
refGene_gn=PLG;refGene_tx=XM_006932130.4
B2 147339226 147339275
refGene_gn=PLG;refGene_tx=XM_006932130.4

B2	147396057	147396197
refGene_gn=MAP3K4;refGene_tx=XM_023254556.1;refGene_gn=MAP3K4;refGene_tx=XM_023254557.1;refGene_gn=MAP3K4;refGene_tx=XM_023254558.1
B2	147422283	147422294
refGene_gn=MAP3K4;refGene_tx=XM_019831456.2
B2	147436088	147436279
refGene_gn=MAP3K4;refGene_tx=XM_019831456.2;refGene_gn=MAP3K4;refGene_tx=XM_023254556.1;refGene_gn=MAP3K4;refGene_tx=XM_023254557.1;refGene_gn=MAP3K4;refGene_tx=XM_023254558.1
B2	147448648	147450012
refGene_gn=MAP3K4;refGene_tx=XM_019831456.2;refGene_gn=MAP3K4;refGene_tx=XM_023254556.1;refGene_gn=MAP3K4;refGene_tx=XM_023254557.1;refGene_gn=MAP3K4;refGene_tx=XM_023254558.1
B2	147468157	147468400
refGene_gn=MAP3K4;refGene_tx=XM_019831456.2;refGene_gn=MAP3K4;refGene_tx=XM_023254556.1;refGene_gn=MAP3K4;refGene_tx=XM_023254557.1;refGene_gn=MAP3K4;refGene_tx=XM_023254558.1
B2	147470662	147470809
refGene_gn=MAP3K4;refGene_tx=XM_019831456.2;refGene_gn=MAP3K4;refGene_tx=XM_023254556.1;refGene_gn=MAP3K4;refGene_tx=XM_023254557.1;refGene_gn=MAP3K4;refGene_tx=XM_023254558.1
B2	147476194	147476352
refGene_gn=MAP3K4;refGene_tx=XM_019831456.2;refGene_gn=MAP3K4;refGene_tx=XM_023254556.1;refGene_gn=MAP3K4;refGene_tx=XM_023254557.1;refGene_gn=MAP3K4;refGene_tx=XM_023254558.1
B2	147479637	147479754
refGene_gn=MAP3K4;refGene_tx=XM_019831456.2;refGene_gn=MAP3K4;refGene_tx=XM_023254556.1;refGene_gn=MAP3K4;refGene_tx=XM_023254557.1;refGene_gn=MAP3K4;refGene_tx=XM_023254558.1
B2	147481239	147481339
refGene_gn=MAP3K4;refGene_tx=XM_019831456.2;refGene_gn=MAP3K4;refGene_tx=XM_023254556.1;refGene_gn=MAP3K4;refGene_tx=XM_023254557.1;refGene_gn=MAP3K4;refGene_tx=XM_023254558.1
B2	147481536	147481620
refGene_gn=MAP3K4;refGene_tx=XM_019831456.2;refGene_gn=MAP3K4;refGene_tx=XM_023254556.1;refGene_gn=MAP3K4;refGene_tx=XM_023254557.1;refGene_gn=MAP3K4;refGene_tx=XM_023254558.1
B2	147483267	147483531
refGene_gn=MAP3K4;refGene_tx=XM_019831456.2;refGene_gn=MAP3K4;refGene_tx=XM_023254556.1;refGene_gn=MAP3K4;refGene_tx=XM_023254557.1;refGene_gn=MAP3K4;refGene_tx=XM_023254558.1
B2	147484835	147484985
refGene_gn=MAP3K4;refGene_tx=XM_019831456.2;refGene_gn=MAP3K4;refGene_tx=XM_023254556.1;refGene_gn=MAP3K4;refGene_tx=XM_023254557.1;refGene_gn=MAP3K4;refGene_tx=XM_023254558.1
B2	147487243	147487405
refGene_gn=MAP3K4;refGene_tx=XM_019831456.2;refGene_gn=MAP3K4;refGene_tx=XM_023254556.1;refGene_gn=MAP3K4;refGene_tx=XM_023254557.1;refGene_gn=MAP3K4;refGene_tx=XM_023254558.1
B2	147487860	147487994

refGene_gn=MAP3K4;refGene_tx=XM_019831456.2;refGene_gn=MAP3K4;refGene_tx=XM_023254556.1;refGene_gn=MAP3K4;refGene_tx=XM_023254557.1;refGene_gn=MAP3K4;refGene_tx=XM_023254558.1
B2 147488687 147488766
refGene_gn=MAP3K4;refGene_tx=XM_019831456.2;refGene_gn=MAP3K4;refGene_tx=XM_023254556.1;refGene_gn=MAP3K4;refGene_tx=XM_023254557.1;refGene_gn=MAP3K4;refGene_tx=XM_023254558.1
B2 147489482 147489561
refGene_gn=MAP3K4;refGene_tx=XM_019831456.2;refGene_gn=MAP3K4;refGene_tx=XM_023254556.1;refGene_gn=MAP3K4;refGene_tx=XM_023254557.1;refGene_gn=MAP3K4;refGene_tx=XM_023254558.1
B2 147492993 147493090
refGene_gn=MAP3K4;refGene_tx=XM_019831456.2;refGene_gn=MAP3K4;refGene_tx=XM_023254556.1;refGene_gn=MAP3K4;refGene_tx=XM_023254557.1;refGene_gn=MAP3K4;refGene_tx=XM_023254558.1
B2 147493883 147494021
refGene_gn=MAP3K4;refGene_tx=XM_019831456.2;refGene_gn=MAP3K4;refGene_tx=XM_023254556.1;refGene_gn=MAP3K4;refGene_tx=XM_023254558.1
B2 147497153 147497254
refGene_gn=MAP3K4;refGene_tx=XM_019831456.2;refGene_gn=MAP3K4;refGene_tx=XM_023254556.1;refGene_gn=MAP3K4;refGene_tx=XM_023254557.1;refGene_gn=MAP3K4;refGene_tx=XM_023254558.1
B2 147497881 147497962
refGene_gn=MAP3K4;refGene_tx=XM_019831456.2;refGene_gn=MAP3K4;refGene_tx=XM_023254556.1;refGene_gn=MAP3K4;refGene_tx=XM_023254557.1;refGene_gn=MAP3K4;refGene_tx=XM_023254558.1
B2 147500847 147501039
refGene_gn=MAP3K4;refGene_tx=XM_019831456.2;refGene_gn=MAP3K4;refGene_tx=XM_023254556.1;refGene_gn=MAP3K4;refGene_tx=XM_023254557.1;refGene_gn=MAP3K4;refGene_tx=XM_023254558.1
B2 147502036 147502107
refGene_gn=MAP3K4;refGene_tx=XM_019831456.2;refGene_gn=MAP3K4;refGene_tx=XM_023254556.1;refGene_gn=MAP3K4;refGene_tx=XM_023254557.1;refGene_gn=MAP3K4;refGene_tx=XM_023254558.1
B2 147503087 147503204
refGene_gn=MAP3K4;refGene_tx=XM_019831456.2;refGene_gn=MAP3K4;refGene_tx=XM_023254556.1;refGene_gn=MAP3K4;refGene_tx=XM_023254557.1;refGene_gn=MAP3K4;refGene_tx=XM_023254558.1
B2 147504087 147504247
refGene_gn=MAP3K4;refGene_tx=XM_019831456.2;refGene_gn=MAP3K4;refGene_tx=XM_023254556.1;refGene_gn=MAP3K4;refGene_tx=XM_023254557.1;refGene_gn=MAP3K4;refGene_tx=XM_023254558.1
B2 147506427 147506550
refGene_gn=MAP3K4;refGene_tx=XM_019831456.2;refGene_gn=MAP3K4;refGene_tx=XM_023254556.1;refGene_gn=MAP3K4;refGene_tx=XM_023254557.1;refGene_gn=MAP3K4;refGene_tx=XM_023254558.1
B2 147507420 147507527
refGene_gn=MAP3K4;refGene_tx=XM_019831456.2;refGene_gn=MAP3K4;refGene_tx=XM_023254556.1;refGene_gn=MAP3K4;refGene_tx=XM_023254557.1;refGene_gn=MAP3K4;refGene_tx=XM_023254558.1

B2 147509383 147509563
refGene_gn=MAP3K4;refGene_tx=XM_019831456.2;refGene_gn=MAP3K4;refGene_tx=XM_023254556.1;refGene_gn=MAP3K4;refGene_tx=XM_023254557.1;refGene_gn=MAP3K4;refGene_tx=XM_023254558.1
B2 147511212 147511233
refGene_gn=MAP3K4;refGene_tx=XM_019831456.2;refGene_gn=MAP3K4;refGene_tx=XM_023254556.1;refGene_gn=MAP3K4;refGene_tx=XM_023254557.1;refGene_gn=MAP3K4;refGene_tx=XM_023254558.1
B2 147519308 147519489
refGene_gn=AGPAT4;refGene_tx=XM_003986696.5;refGene_gn=AGPAT4;refGene_tx=XM_006932131.4;refGene_gn=AGPAT4;refGene_tx=XM_006932132.4;refGene_gn=AGPAT4;refGene_tx=XM_011282550.3;refGene_gn=AGPAT4;refGene_tx=XM_019831458.2;refGene_gn=AGPAT4;refGene_tx=XM_019831459.2
B2 147521936 147522135
refGene_gn=AGPAT4;refGene_tx=XM_003986696.5;refGene_gn=AGPAT4;refGene_tx=XM_006932131.4;refGene_gn=AGPAT4;refGene_tx=XM_006932132.4;refGene_gn=AGPAT4;refGene_tx=XM_011282550.3;refGene_gn=AGPAT4;refGene_tx=XM_019831458.2;refGene_gn=AGPAT4;refGene_tx=XM_019831459.2
B2 147528677 147528753
refGene_gn=AGPAT4;refGene_tx=XM_003986696.5;refGene_gn=AGPAT4;refGene_tx=XM_006932131.4;refGene_gn=AGPAT4;refGene_tx=XM_006932132.4;refGene_gn=AGPAT4;refGene_tx=XM_011282550.3;refGene_gn=AGPAT4;refGene_tx=XM_019831459.2
B2 147531237 147531340
refGene_gn=AGPAT4;refGene_tx=XM_003986696.5;refGene_gn=AGPAT4;refGene_tx=XM_006932131.4;refGene_gn=AGPAT4;refGene_tx=XM_006932132.4;refGene_gn=AGPAT4;refGene_tx=XM_011282550.3;refGene_gn=AGPAT4;refGene_tx=XM_019831459.2
B2 147534746 147534900
refGene_gn=AGPAT4;refGene_tx=XM_003986696.5;refGene_gn=AGPAT4;refGene_tx=XM_006932131.4;refGene_gn=AGPAT4;refGene_tx=XM_006932132.4;refGene_gn=AGPAT4;refGene_tx=XM_011282550.3;refGene_gn=AGPAT4;refGene_tx=XM_019831458.2;refGene_gn=AGPAT4;refGene_tx=XM_019831459.2
B2 147535374 147535536
refGene_gn=AGPAT4;refGene_tx=XM_003986696.5;refGene_gn=AGPAT4;refGene_tx=XM_006932131.4;refGene_gn=AGPAT4;refGene_tx=XM_006932132.4;refGene_gn=AGPAT4;refGene_tx=XM_011282550.3;refGene_gn=AGPAT4;refGene_tx=XM_019831458.2;refGene_gn=AGPAT4;refGene_tx=XM_019831459.2
B2 147549410 147549580
refGene_gn=AGPAT4;refGene_tx=XM_003986696.5;refGene_gn=AGPAT4;refGene_tx=XM_006932131.4;refGene_gn=AGPAT4;refGene_tx=XM_006932132.4;refGene_gn=AGPAT4;refGene_tx=XM_019831458.2
B2 147608165 147608430
refGene_gn=AGPAT4;refGene_tx=XM_003986696.5;refGene_gn=AGPAT4;refGene_tx=XM_006932131.4;refGene_gn=AGPAT4;refGene_tx=XM_006932132.4;refGene_gn=AGPAT4;refGene_tx=XM_011282550.3;refGene_gn=AGPAT4;refGene_tx=XM_019831458.2;refGene_gn=AGPAT4;refGene_tx=XM_019831459.2
B2 147645386 147645473
refGene_gn=AGPAT4;refGene_tx=XM_006932132.4;refGene_gn=AGPAT4;refGene_tx=XM_011282550.3;refGene_gn=AGPAT4;refGene_tx=XM_019831458.2;refGene_

gn=AGPAT4;refGene_tx=XM_019831459.2
B2 147693902 147694015
refGene_gn=PRKN;refGene_tx=XM_003986705.5;refGene_gn=PRKN;refGene_tx=XM_019831462.2
B2 147700895 147701013
refGene_gn=PRKN;refGene_tx=XM_003986705.5;refGene_gn=PRKN;refGene_tx=XM_019831462.2
B2 147730930 147731014
refGene_gn=PRKN;refGene_tx=XM_003986705.5;refGene_gn=PRKN;refGene_tx=XM_019831462.2
B2 147883247 147883397
refGene_gn=PRKN;refGene_tx=XM_003986705.5;refGene_gn=PRKN;refGene_tx=XM_019831462.2
B2 147898286 147898348
refGene_gn=PRKN;refGene_tx=XM_003986705.5;refGene_gn=PRKN;refGene_tx=XM_019831462.2
B2 148121231 148121368
refGene_gn=PRKN;refGene_tx=XM_003986705.5;refGene_gn=PRKN;refGene_tx=XM_019831462.2
B2 148295194 148295310
refGene_gn=PRKN;refGene_tx=XM_003986705.5;refGene_gn=PRKN;refGene_tx=XM_019831462.2
B2 148384291 148384375
refGene_gn=PRKN;refGene_tx=XM_003986705.5;refGene_gn=PRKN;refGene_tx=XM_019831462.2
B2 148542638 148542760
refGene_gn=PRKN;refGene_tx=XM_003986705.5;refGene_gn=PRKN;refGene_tx=XM_019831462.2
B2 148607584 148607816
refGene_gn=PRKN;refGene_tx=XM_003986705.5;refGene_gn=PRKN;refGene_tx=XM_019831462.2
B2 148774862 148775026
refGene_gn=PRKN;refGene_tx=XM_003986705.5;refGene_gn=PRKN;refGene_tx=XM_019831462.2
B2 149043081 149043088
refGene_gn=PRKN;refGene_tx=XM_003986705.5;refGene_gn=PRKN;refGene_tx=XM_019831462.2
B2 149043681 149043837
refGene_gn=PACRG;refGene_tx=XM_003986697.5
B2 149111394 149111529
refGene_gn=PACRG;refGene_tx=XM_003986697.5
B2 149327092 149327264
refGene_gn=PACRG;refGene_tx=XM_003986697.5
B2 149349181 149349331
refGene_gn=PACRG;refGene_tx=XM_003986697.5
B2 149557630 149557791
refGene_gn=PACRG;refGene_tx=XM_003986697.5
B2 149648350 149648492
refGene_gn=QKI;refGene_tx=NM_001009232.1;refGene_gn=QKI;refGene_tx=XM_023253544.1;refGene_gn=QKI;refGene_tx=XM_023253545.1;refGene_gn=QKI;re

fGene_tx=XM_023253546.1;refGene_gn=QKI;refGene_tx=XM_023253547.1;refGene_gn=QKI;refGene_tx=XM_023253548.1
B2 149686524 149686667
refGene_gn=QKI;refGene_tx=NM_001009232.1;refGene_gn=QKI;refGene_tx=XM_023253544.1;refGene_gn=QKI;refGene_tx=XM_023253545.1;refGene_gn=QKI;refGene_tx=XM_023253546.1;refGene_gn=QKI;refGene_tx=XM_023253547.1;refGene_gn=QKI;refGene_tx=XM_023253548.1
B2 149708858 149708975
refGene_gn=QKI;refGene_tx=NM_001009232.1;refGene_gn=QKI;refGene_tx=XM_023253544.1;refGene_gn=QKI;refGene_tx=XM_023253545.1;refGene_gn=QKI;refGene_tx=XM_023253546.1;refGene_gn=QKI;refGene_tx=XM_023253547.1;refGene_gn=QKI;refGene_tx=XM_023253548.1
B2 149760545 149760689
refGene_gn=QKI;refGene_tx=NM_001009232.1;refGene_gn=QKI;refGene_tx=XM_023253544.1;refGene_gn=QKI;refGene_tx=XM_023253545.1;refGene_gn=QKI;refGene_tx=XM_023253546.1;refGene_gn=QKI;refGene_tx=XM_023253547.1;refGene_gn=QKI;refGene_tx=XM_023253548.1
B2 149793115 149793203
refGene_gn=QKI;refGene_tx=NM_001009232.1;refGene_gn=QKI;refGene_tx=XM_023253544.1;refGene_gn=QKI;refGene_tx=XM_023253545.1;refGene_gn=QKI;refGene_tx=XM_023253546.1;refGene_gn=QKI;refGene_tx=XM_023253547.1;refGene_gn=QKI;refGene_tx=XM_023253548.1
B2 149794559 149794859
refGene_gn=QKI;refGene_tx=NM_001009232.1;refGene_gn=QKI;refGene_tx=XM_023253544.1;refGene_gn=QKI;refGene_tx=XM_023253545.1;refGene_gn=QKI;refGene_tx=XM_023253546.1;refGene_gn=QKI;refGene_tx=XM_023253547.1;refGene_gn=QKI;refGene_tx=XM_023253548.1
B2 149795563 149795607
refGene_gn=QKI;refGene_tx=XM_023253546.1;refGene_gn=QKI;refGene_tx=XM_023253548.1
B2 149796833 149796859
refGene_gn=QKI;refGene_tx=XM_023253545.1
B2 149797620 149797695
refGene_gn=QKI;refGene_tx=NM_001009232.1;refGene_gn=QKI;refGene_tx=XM_023253544.1
B2 149802191 149802208
refGene_gn=QKI;refGene_tx=NM_001009232.1;refGene_gn=QKI;refGene_tx=XM_023253544.1;refGene_gn=QKI;refGene_tx=XM_023253547.1
B2 150813037 150813499
refGene_gn=LOC109500035;refGene_tx=XM_023254742.1
B2 150814050 150814152
refGene_gn=LOC109500035;refGene_tx=XM_023254742.1
B2 150814392 150814491
refGene_gn=LOC109500035;refGene_tx=XM_023254742.1
B2 151235345 151235454
refGene_gn=CB2H6orf118;refGene_tx=XM_023254559.1;refGene_gn=CB2H6orf118;refGene_tx=XM_023254560.1
B2 151237738 151237800
refGene_gn=CB2H6orf118;refGene_tx=XM_023254559.1;refGene_gn=CB2H6orf118;refGene_tx=XM_023254560.1

B2 151238380 151238434
refGene_gn=CB2H6orf118;refGene_tx=XM_023254559.1;refGene_gn=CB2H6orf11
8;refGene_tx=XM_023254560.1
B2 151241703 151241885
refGene_gn=CB2H6orf118;refGene_tx=XM_023254559.1;refGene_gn=CB2H6orf11
8;refGene_tx=XM_023254560.1
B2 151246051 151246110
refGene_gn=CB2H6orf118;refGene_tx=XM_023254559.1;refGene_gn=CB2H6orf11
8;refGene_tx=XM_023254560.1
B2 151252465 151252590
refGene_gn=CB2H6orf118;refGene_tx=XM_023254559.1;refGene_gn=CB2H6orf11
8;refGene_tx=XM_023254560.1
B2 151254262 151254322
refGene_gn=CB2H6orf118;refGene_tx=XM_023254559.1;refGene_gn=CB2H6orf11
8;refGene_tx=XM_023254560.1
B2 151255369 151255492
refGene_gn=CB2H6orf118;refGene_tx=XM_023254559.1;refGene_gn=CB2H6orf11
8;refGene_tx=XM_023254560.1
B2 151256327 151257010
refGene_gn=CB2H6orf118;refGene_tx=XM_023254559.1;refGene_gn=CB2H6orf11
8;refGene_tx=XM_023254560.1
B2 151263421 151263446
refGene_gn=CB2H6orf118;refGene_tx=XM_023254559.1;refGene_gn=CB2H6orf11
8;refGene_tx=XM_023254560.1
B2 151284195 151284313
refGene_gn=PDE10A;refGene_tx=XM_006932134.4;refGene_gn=PDE10A;refGene_
tx=XM_019831471.2;refGene_gn=PDE10A;refGene_tx=XM_023254561.1;refGene_
gn=PDE10A;refGene_tx=XM_023254562.1;refGene_gn=PDE10A;refGene_tx=XM_02
3254563.1;refGene_gn=PDE10A;refGene_tx=XM_023254564.1;refGene_gn=PDE10
A;refGene_tx=XM_023254565.1
B2 151286760 151286849
refGene_gn=PDE10A;refGene_tx=XM_006932134.4;refGene_gn=PDE10A;refGene_
tx=XM_019831471.2;refGene_gn=PDE10A;refGene_tx=XM_023254561.1;refGene_
gn=PDE10A;refGene_tx=XM_023254562.1;refGene_gn=PDE10A;refGene_tx=XM_02
3254563.1;refGene_gn=PDE10A;refGene_tx=XM_023254564.1;refGene_gn=PDE10
A;refGene_tx=XM_023254565.1
B2 151290446 151290527
refGene_gn=PDE10A;refGene_tx=XM_006932134.4;refGene_gn=PDE10A;refGene_
tx=XM_019831471.2;refGene_gn=PDE10A;refGene_tx=XM_023254561.1;refGene_
gn=PDE10A;refGene_tx=XM_023254562.1;refGene_gn=PDE10A;refGene_tx=XM_02
3254563.1;refGene_gn=PDE10A;refGene_tx=XM_023254564.1
B2 151293267 151293379
refGene_gn=PDE10A;refGene_tx=XM_006932134.4;refGene_gn=PDE10A;refGene_
tx=XM_019831471.2;refGene_gn=PDE10A;refGene_tx=XM_023254561.1;refGene_
gn=PDE10A;refGene_tx=XM_023254562.1;refGene_gn=PDE10A;refGene_tx=XM_02
3254563.1;refGene_gn=PDE10A;refGene_tx=XM_023254564.1
B2 151313590 151313763
refGene_gn=PDE10A;refGene_tx=XM_006932134.4;refGene_gn=PDE10A;refGene_
tx=XM_019831471.2;refGene_gn=PDE10A;refGene_tx=XM_023254561.1;refGene_
gn=PDE10A;refGene_tx=XM_023254562.1;refGene_gn=PDE10A;refGene_tx=XM_02

3254563.1;refGene_gn=PDE10A;refGene_tx=XM_023254564.1;refGene_gn=PDE10A;refGene_tx=XM_023254565.1
B2 151319565 151319721
refGene_gn=PDE10A;refGene_tx=XM_006932134.4;refGene_gn=PDE10A;refGene_tx=XM_019831471.2;refGene_gn=PDE10A;refGene_tx=XM_023254561.1;refGene_gn=PDE10A;refGene_tx=XM_023254562.1;refGene_gn=PDE10A;refGene_tx=XM_023254563.1;refGene_gn=PDE10A;refGene_tx=XM_023254564.1;refGene_gn=PDE10A;refGene_tx=XM_023254565.1
B2 151322947 151323098
refGene_gn=PDE10A;refGene_tx=XM_006932134.4;refGene_gn=PDE10A;refGene_tx=XM_019831471.2;refGene_gn=PDE10A;refGene_tx=XM_023254561.1;refGene_gn=PDE10A;refGene_tx=XM_023254562.1;refGene_gn=PDE10A;refGene_tx=XM_023254563.1;refGene_gn=PDE10A;refGene_tx=XM_023254564.1;refGene_gn=PDE10A;refGene_tx=XM_023254565.1
B2 151325477 151325561
refGene_gn=PDE10A;refGene_tx=XM_006932134.4;refGene_gn=PDE10A;refGene_tx=XM_019831471.2;refGene_gn=PDE10A;refGene_tx=XM_023254561.1;refGene_gn=PDE10A;refGene_tx=XM_023254562.1;refGene_gn=PDE10A;refGene_tx=XM_023254563.1;refGene_gn=PDE10A;refGene_tx=XM_023254564.1;refGene_gn=PDE10A;refGene_tx=XM_023254565.1
B2 151327856 151327999
refGene_gn=PDE10A;refGene_tx=XM_006932134.4;refGene_gn=PDE10A;refGene_tx=XM_019831471.2;refGene_gn=PDE10A;refGene_tx=XM_023254561.1;refGene_gn=PDE10A;refGene_tx=XM_023254562.1;refGene_gn=PDE10A;refGene_tx=XM_023254563.1;refGene_gn=PDE10A;refGene_tx=XM_023254564.1;refGene_gn=PDE10A;refGene_tx=XM_023254565.1
B2 151347710 151347897
refGene_gn=PDE10A;refGene_tx=XM_006932134.4;refGene_gn=PDE10A;refGene_tx=XM_019831471.2;refGene_gn=PDE10A;refGene_tx=XM_023254561.1;refGene_gn=PDE10A;refGene_tx=XM_023254562.1;refGene_gn=PDE10A;refGene_tx=XM_023254563.1;refGene_gn=PDE10A;refGene_tx=XM_023254564.1;refGene_gn=PDE10A;refGene_tx=XM_023254565.1
B2 151350678 151350771
refGene_gn=PDE10A;refGene_tx=XM_006932134.4;refGene_gn=PDE10A;refGene_tx=XM_019831471.2;refGene_gn=PDE10A;refGene_tx=XM_023254561.1;refGene_gn=PDE10A;refGene_tx=XM_023254562.1;refGene_gn=PDE10A;refGene_tx=XM_023254563.1;refGene_gn=PDE10A;refGene_tx=XM_023254564.1;refGene_gn=PDE10A;refGene_tx=XM_023254565.1
B2 151353259 151353402
refGene_gn=PDE10A;refGene_tx=XM_006932134.4;refGene_gn=PDE10A;refGene_tx=XM_019831471.2;refGene_gn=PDE10A;refGene_tx=XM_023254561.1;refGene_gn=PDE10A;refGene_tx=XM_023254562.1;refGene_gn=PDE10A;refGene_tx=XM_023254563.1;refGene_gn=PDE10A;refGene_tx=XM_023254564.1;refGene_gn=PDE10A;refGene_tx=XM_023254565.1
B2 151361141 151361193
refGene_gn=PDE10A;refGene_tx=XM_006932134.4;refGene_gn=PDE10A;refGene_tx=XM_019831471.2;refGene_gn=PDE10A;refGene_tx=XM_023254561.1;refGene_gn=PDE10A;refGene_tx=XM_023254562.1;refGene_gn=PDE10A;refGene_tx=XM_023254563.1;refGene_gn=PDE10A;refGene_tx=XM_023254564.1;refGene_gn=PDE10A;refGene_tx=XM_023254565.1

B2 151363242 151363301
refGene_gn=PDE10A;refGene_tx=XM_006932134.4;refGene_gn=PDE10A;refGene_tx=XM_019831471.2;refGene_gn=PDE10A;refGene_tx=XM_023254561.1;refGene_gn=PDE10A;refGene_tx=XM_023254562.1;refGene_gn=PDE10A;refGene_tx=XM_023254563.1;refGene_gn=PDE10A;refGene_tx=XM_023254564.1;refGene_gn=PDE10A;refGene_tx=XM_023254565.1
B2 151364331 151364382
refGene_gn=PDE10A;refGene_tx=XM_006932134.4;refGene_gn=PDE10A;refGene_tx=XM_019831471.2;refGene_gn=PDE10A;refGene_tx=XM_023254561.1;refGene_gn=PDE10A;refGene_tx=XM_023254562.1;refGene_gn=PDE10A;refGene_tx=XM_023254563.1;refGene_gn=PDE10A;refGene_tx=XM_023254564.1;refGene_gn=PDE10A;refGene_tx=XM_023254565.1
B2 151365941 151366097
refGene_gn=PDE10A;refGene_tx=XM_006932134.4;refGene_gn=PDE10A;refGene_tx=XM_019831471.2;refGene_gn=PDE10A;refGene_tx=XM_023254561.1;refGene_gn=PDE10A;refGene_tx=XM_023254562.1;refGene_gn=PDE10A;refGene_tx=XM_023254563.1;refGene_gn=PDE10A;refGene_tx=XM_023254564.1;refGene_gn=PDE10A;refGene_tx=XM_023254565.1
B2 151368982 151369123
refGene_gn=PDE10A;refGene_tx=XM_006932134.4;refGene_gn=PDE10A;refGene_tx=XM_019831471.2;refGene_gn=PDE10A;refGene_tx=XM_023254561.1;refGene_gn=PDE10A;refGene_tx=XM_023254562.1;refGene_gn=PDE10A;refGene_tx=XM_023254563.1;refGene_gn=PDE10A;refGene_tx=XM_023254564.1;refGene_gn=PDE10A;refGene_tx=XM_023254565.1
B2 151380333 151380383
refGene_gn=PDE10A;refGene_tx=XM_006932134.4;refGene_gn=PDE10A;refGene_tx=XM_019831471.2;refGene_gn=PDE10A;refGene_tx=XM_023254561.1;refGene_gn=PDE10A;refGene_tx=XM_023254562.1;refGene_gn=PDE10A;refGene_tx=XM_023254563.1;refGene_gn=PDE10A;refGene_tx=XM_023254564.1;refGene_gn=PDE10A;refGene_tx=XM_023254565.1
B2 151381398 151381519
refGene_gn=PDE10A;refGene_tx=XM_006932134.4;refGene_gn=PDE10A;refGene_tx=XM_019831471.2;refGene_gn=PDE10A;refGene_tx=XM_023254561.1;refGene_gn=PDE10A;refGene_tx=XM_023254562.1;refGene_gn=PDE10A;refGene_tx=XM_023254563.1;refGene_gn=PDE10A;refGene_tx=XM_023254564.1;refGene_gn=PDE10A;refGene_tx=XM_023254565.1
B2 151400533 151400562
refGene_gn=PDE10A;refGene_tx=XM_006932134.4;refGene_gn=PDE10A;refGene_tx=XM_019831471.2;refGene_gn=PDE10A;refGene_tx=XM_023254561.1;refGene_gn=PDE10A;refGene_tx=XM_023254562.1;refGene_gn=PDE10A;refGene_tx=XM_023254563.1;refGene_gn=PDE10A;refGene_tx=XM_023254564.1;refGene_gn=PDE10A;refGene_tx=XM_023254565.1
B2 151401494 151401534
refGene_gn=PDE10A;refGene_tx=XM_019831471.2
B2 151467843 151467972
refGene_gn=PDE10A;refGene_tx=XM_006932134.4;refGene_gn=PDE10A;refGene_tx=XM_023254561.1;refGene_gn=PDE10A;refGene_tx=XM_023254562.1;refGene_gn=PDE10A;refGene_tx=XM_023254563.1;refGene_gn=PDE10A;refGene_tx=XM_023254564.1;refGene_gn=PDE10A;refGene_tx=XM_023254565.1
B2 151532962 151532978

refGene_gn=PDE10A;refGene_tx=XM_006932134.4
B2 151533155 151533297
refGene_gn=PDE10A;refGene_tx=XM_023254561.1;refGene_gn=PDE10A;refGene_tx=XM_023254562.1;refGene_gn=PDE10A;refGene_tx=XM_023254565.1
B2 151588211 151588278
refGene_gn=PDE10A;refGene_tx=XM_023254563.1
B2 151637483 151637532
refGene_gn=PDE10A;refGene_tx=XM_023254564.1
B2 152019279 152019553
refGene_gn=T;refGene_tx=XM_003986708.3
B2 152021339 152021469
refGene_gn=T;refGene_tx=XM_003986708.3
B2 152022896 152023070
refGene_gn=T;refGene_tx=XM_003986708.3
B2 152024606 152024668
refGene_gn=T;refGene_tx=XM_003986708.3
B2 152024789 152024851
refGene_gn=T;refGene_tx=XM_003986708.3
B2 152025632 152025767
refGene_gn=T;refGene_tx=XM_003986708.3
B2 152026521 152026786
refGene_gn=T;refGene_tx=XM_003986708.3
B2 152027191 152027397
refGene_gn=T;refGene_tx=XM_003986708.3
B2 152171446 152172101
refGene_gn=PRR18;refGene_tx=XM_023254743.1
B2 152172854 152172898
refGene_gn=PRR18;refGene_tx=XM_023254743.1
B2 152180015 152180055
refGene_gn=SFT2D1;refGene_tx=XM_023254566.1
B2 152181650 152181680
refGene_gn=SFT2D1;refGene_tx=XM_023254566.1
B2 152183390 152183449
refGene_gn=SFT2D1;refGene_tx=XM_023254566.1
B2 152184976 152185012
refGene_gn=SFT2D1;refGene_tx=XM_023254566.1
B2 152187101 152187183
refGene_gn=SFT2D1;refGene_tx=XM_023254566.1
B2 152188356 152188439
refGene_gn=SFT2D1;refGene_tx=XM_023254566.1
B2 152189221 152189308
refGene_gn=SFT2D1;refGene_tx=XM_023254566.1
B2 152195177 152195240
refGene_gn=SFT2D1;refGene_tx=XM_023254566.1
B2 152216346 152216371
refGene_gn=MPC1;refGene_tx=XM_023254567.1;refGene_gn=MPC1;refGene_tx=XM_023254568.1
B2 152216739 152216872
refGene_gn=MPC1;refGene_tx=XM_023254567.1;refGene_gn=MPC1;refGene_tx=XM_023254568.1

B2 152217489 152217586
refGene_gn=MPC1;refGene_tx=XM_023254567.1;refGene_gn=MPC1;refGene_tx=XM_023254568.1
B2 152219611 152219615
refGene_gn=MPC1;refGene_tx=XM_023254567.1
B2 152223795 152223799
refGene_gn=MPC1;refGene_tx=XM_023254568.1
B2 152225130 152225201
refGene_gn=MPC1;refGene_tx=XM_023254567.1;refGene_gn=MPC1;refGene_tx=XM_023254568.1
B2 152249882 152250008
refGene_gn=RPS6KA2;refGene_tx=XM_023254569.1;refGene_gn=RPS6KA2;refGene_tx=XM_023254570.1
B2 152251737 152251875
refGene_gn=RPS6KA2;refGene_tx=XM_023254569.1;refGene_gn=RPS6KA2;refGene_tx=XM_023254570.1
B2 152256917 152257035
refGene_gn=RPS6KA2;refGene_tx=XM_023254569.1;refGene_gn=RPS6KA2;refGene_tx=XM_023254570.1
B2 152259029 152259108
refGene_gn=RPS6KA2;refGene_tx=XM_023254569.1;refGene_gn=RPS6KA2;refGene_tx=XM_023254570.1
B2 152261569 152261729
refGene_gn=RPS6KA2;refGene_tx=XM_023254569.1;refGene_gn=RPS6KA2;refGene_tx=XM_023254570.1
B2 152263202 152263361
refGene_gn=RPS6KA2;refGene_tx=XM_023254569.1;refGene_gn=RPS6KA2;refGene_tx=XM_023254570.1
B2 152265387 152265477
refGene_gn=RPS6KA2;refGene_tx=XM_023254569.1;refGene_gn=RPS6KA2;refGene_tx=XM_023254570.1
B2 152274162 152274288
refGene_gn=RPS6KA2;refGene_tx=XM_023254569.1;refGene_gn=RPS6KA2;refGene_tx=XM_023254570.1
B2 152275317 152275448
refGene_gn=RPS6KA2;refGene_tx=XM_023254569.1;refGene_gn=RPS6KA2;refGene_tx=XM_023254570.1
B2 152277635 152277738
refGene_gn=RPS6KA2;refGene_tx=XM_023254569.1;refGene_gn=RPS6KA2;refGene_tx=XM_023254570.1
B2 152286728 152286793
refGene_gn=RPS6KA2;refGene_tx=XM_023254569.1;refGene_gn=RPS6KA2;refGene_tx=XM_023254570.1
B2 152300038 152300127
refGene_gn=RPS6KA2;refGene_tx=XM_023254569.1;refGene_gn=RPS6KA2;refGene_tx=XM_023254570.1
B2 152302825 152302896
refGene_gn=RPS6KA2;refGene_tx=XM_023254569.1;refGene_gn=RPS6KA2;refGene_tx=XM_023254570.1
B2 152308894 152309037

refGene_gn=RPS6KA2;refGene_tx=XM_023254569.1;refGene_gn=RPS6KA2;refGene_tx=XM_023254570.1
B2 152312108 152312146
refGene_gn=RPS6KA2;refGene_tx=XM_023254569.1;refGene_gn=RPS6KA2;refGene_tx=XM_023254570.1
B2 152314350 152314457
refGene_gn=RPS6KA2;refGene_tx=XM_023254569.1;refGene_gn=RPS6KA2;refGene_tx=XM_023254570.1
B2 152316935 152317015
refGene_gn=RPS6KA2;refGene_tx=XM_023254569.1;refGene_gn=RPS6KA2;refGene_tx=XM_023254570.1
B2 152320652 152320733
refGene_gn=RPS6KA2;refGene_tx=XM_023254569.1;refGene_gn=RPS6KA2;refGene_tx=XM_023254570.1
B2 152335385 152335467
refGene_gn=RPS6KA2;refGene_tx=XM_023254569.1;refGene_gn=RPS6KA2;refGene_tx=XM_023254570.1
B2 152342291 152342408
refGene_gn=RPS6KA2;refGene_tx=XM_023254569.1;refGene_gn=RPS6KA2;refGene_tx=XM_023254570.1
B2 152415929 152416028
refGene_gn=RPS6KA2;refGene_tx=XM_023254570.1
B2 152547051 152547102
refGene_gn=RPS6KA2;refGene_tx=XM_023254569.1
B2 152598007 152598067
refGene_gn=RPS6KA2;refGene_tx=XM_023254569.1
B2 152602390 152602453
refGene_gn=RPS6KA2;refGene_tx=XM_023254569.1
B2 152647828 152648020
refGene_gn=RNASET2;refGene_tx=XM_023254571.1
B2 152649276 152649351
refGene_gn=RNASET2;refGene_tx=XM_023254571.1
B2 152651046 152651092
refGene_gn=RNASET2;refGene_tx=XM_023254571.1
B2 152653197 152653311
refGene_gn=RNASET2;refGene_tx=XM_023254571.1
B2 152656507 152656575
refGene_gn=RNASET2;refGene_tx=XM_023254571.1
B2 152659316 152659374
refGene_gn=RNASET2;refGene_tx=XM_023254571.1
B2 152660919 152660975
refGene_gn=RNASET2;refGene_tx=XM_023254571.1
B2 152662760 152662821
refGene_gn=RNASET2;refGene_tx=XM_023254571.1
B2 152663927 152664004
refGene_gn=RNASET2;refGene_tx=XM_023254571.1
B2 152690073 152690175
refGene_gn=FGFR1OP;refGene_tx=XM_023254572.1;refGene_gn=FGFR1OP;refGene_tx=XM_023254573.1
B2 152690753 152690807

refGene_gn=FGFR1OP;refGene_tx=XM_023254572.1;refGene_gn=FGFR1OP;refGene_tx=XM_023254573.1
B2 152694468 152694523
refGene_gn=FGFR1OP;refGene_tx=XM_023254572.1;refGene_gn=FGFR1OP;refGene_tx=XM_023254573.1
B2 152694993 152695082
refGene_gn=FGFR1OP;refGene_tx=XM_023254572.1;refGene_gn=FGFR1OP;refGene_tx=XM_023254573.1
B2 152695963 152696101
refGene_gn=FGFR1OP;refGene_tx=XM_023254572.1;refGene_gn=FGFR1OP;refGene_tx=XM_023254573.1
B2 152700597 152700678
refGene_gn=FGFR1OP;refGene_tx=XM_023254572.1;refGene_gn=FGFR1OP;refGene_tx=XM_023254573.1
B2 152702771 152702831
refGene_gn=FGFR1OP;refGene_tx=XM_023254572.1
B2 152711312 152711539
refGene_gn=FGFR1OP;refGene_tx=XM_023254572.1;refGene_gn=FGFR1OP;refGene_tx=XM_023254573.1
B2 152713387 152713518
refGene_gn=FGFR1OP;refGene_tx=XM_023254572.1;refGene_gn=FGFR1OP;refGene_tx=XM_023254573.1
B2 152715704 152715773
refGene_gn=FGFR1OP;refGene_tx=XM_023254572.1;refGene_gn=FGFR1OP;refGene_tx=XM_023254573.1
B2 152718386 152718426
refGene_gn=FGFR1OP;refGene_tx=XM_023254572.1;refGene_gn=FGFR1OP;refGene_tx=XM_023254573.1
B2 152719222 152719319
refGene_gn=FGFR1OP;refGene_tx=XM_023254572.1;refGene_gn=FGFR1OP;refGene_tx=XM_023254573.1
B2 152720477 152720552
refGene_gn=FGFR1OP;refGene_tx=XM_023254572.1;refGene_gn=FGFR1OP;refGene_tx=XM_023254573.1
B2 152797044 152797053
refGene_gn=CCR6;refGene_tx=XM_011282547.3;refGene_gn=CCR6;refGene_tx=XM_023254574.1;refGene_gn=CCR6;refGene_tx=XM_023254575.1
B2 152797151 152798273
refGene_gn=CCR6;refGene_tx=XM_011282547.3;refGene_gn=CCR6;refGene_tx=XM_023254574.1;refGene_gn=CCR6;refGene_tx=XM_023254575.1
B2 152812957 152814088
refGene_gn=GPR31;refGene_tx=XM_003986719.3
B2 152864413 152864827
refGene_gn=TTLL2;refGene_tx=XM_023253561.1
B2 152884904 152885170
refGene_gn=UNC93A;refGene_tx=XM_003986720.5;refGene_gn=UNC93A;refGene_tx=XM_023254577.1
B2 152893899 152894031
refGene_gn=UNC93A;refGene_tx=XM_003986720.5;refGene_gn=UNC93A;refGene_tx=XM_023254577.1

B2 152895006 152895142
refGene_gn=UNC93A;refGene_tx=XM_003986720.5;refGene_gn=UNC93A;refGene_tx=XM_023254577.1
B2 152896943 152897155
refGene_gn=UNC93A;refGene_tx=XM_003986720.5;refGene_gn=UNC93A;refGene_tx=XM_023254577.1
B2 152901280 152901406
refGene_gn=UNC93A;refGene_tx=XM_003986720.5;refGene_gn=UNC93A;refGene_tx=XM_023254577.1
B2 152902901 152903131
refGene_gn=UNC93A;refGene_tx=XM_003986720.5;refGene_gn=UNC93A;refGene_tx=XM_023254577.1
B2 152904135 152904317
refGene_gn=UNC93A;refGene_tx=XM_003986720.5;refGene_gn=UNC93A;refGene_tx=XM_023254577.1
B2 152910313 152910451
refGene_gn=UNC93A;refGene_tx=XM_003986720.5;refGene_gn=UNC93A;refGene_tx=XM_023254577.1
B2 152921683 152921815
refGene_gn=UNC93A;refGene_tx=XM_023254577.1
B2 153222394 153222499
refGene_gn=AFDN;refGene_tx=XM_023254578.1;refGene_gn=AFDN;refGene_tx=XM_023254579.1;refGene_gn=AFDN;refGene_tx=XM_023254580.1;refGene_gn=AFDN;refGene_tx=XM_023254581.1;refGene_gn=AFDN;refGene_tx=XM_023254582.1;refGene_gn=AFDN;refGene_tx=XM_023254583.1;refGene_gn=AFDN;refGene_tx=XM_023254584.1;refGene_gn=AFDN;refGene_tx=XM_023254585.1;refGene_gn=AFDN;refGene_tx=XM_023254586.1;refGene_gn=AFDN;refGene_tx=XM_023254587.1;refGene_gn=AFDN;refGene_tx=XM_023254588.1;refGene_gn=AFDN;refGene_tx=XM_023254589.1;refGene_gn=AFDN;refGene_tx=XM_023254590.1;refGene_gn=AFDN;refGene_tx=XM_023254591.1;refGene_gn=AFDN;refGene_tx=XM_023254592.1
B2 153259316 153259512
refGene_gn=AFDN;refGene_tx=XM_023254578.1;refGene_gn=AFDN;refGene_tx=XM_023254579.1;refGene_gn=AFDN;refGene_tx=XM_023254580.1;refGene_gn=AFDN;refGene_tx=XM_023254581.1;refGene_gn=AFDN;refGene_tx=XM_023254582.1;refGene_gn=AFDN;refGene_tx=XM_023254583.1;refGene_gn=AFDN;refGene_tx=XM_023254584.1;refGene_gn=AFDN;refGene_tx=XM_023254585.1;refGene_gn=AFDN;refGene_tx=XM_023254586.1;refGene_gn=AFDN;refGene_tx=XM_023254587.1;refGene_gn=AFDN;refGene_tx=XM_023254588.1;refGene_gn=AFDN;refGene_tx=XM_023254589.1;refGene_gn=AFDN;refGene_tx=XM_023254590.1;refGene_gn=AFDN;refGene_tx=XM_023254591.1;refGene_gn=AFDN;refGene_tx=XM_023254592.1
B2 153262928 153263044
refGene_gn=AFDN;refGene_tx=XM_023254578.1;refGene_gn=AFDN;refGene_tx=XM_023254579.1;refGene_gn=AFDN;refGene_tx=XM_023254580.1;refGene_gn=AFDN;refGene_tx=XM_023254581.1;refGene_gn=AFDN;refGene_tx=XM_023254582.1;refGene_gn=AFDN;refGene_tx=XM_023254583.1;refGene_gn=AFDN;refGene_tx=XM_023254584.1;refGene_gn=AFDN;refGene_tx=XM_023254585.1;refGene_gn=AFDN;refGene_tx=XM_023254586.1;refGene_gn=AFDN;refGene_tx=XM_023254587.1;refGene_gn=AFDN;refGene_tx=XM_023254588.1;refGene_gn=AFDN;refGene_tx=XM_023254589.1;refGene_gn=AFDN;refGene_tx=XM_023254590.1;refGene_gn=AFDN;refGene_tx=XM_023254591.1;refGene_gn=AFDN;refGene_tx=XM_023254592.1

B2 153264689 153264853
refGene_gn=AFDN;refGene_tx=XM_023254578.1;refGene_gn=AFDN;refGene_tx=XM_023254579.1;refGene_gn=AFDN;refGene_tx=XM_023254580.1;refGene_gn=AFDN;refGene_tx=XM_023254581.1;refGene_gn=AFDN;refGene_tx=XM_023254582.1;refGene_gn=AFDN;refGene_tx=XM_023254583.1;refGene_gn=AFDN;refGene_tx=XM_023254584.1;refGene_gn=AFDN;refGene_tx=XM_023254585.1;refGene_gn=AFDN;refGene_tx=XM_023254586.1;refGene_gn=AFDN;refGene_tx=XM_023254587.1;refGene_gn=AFDN;refGene_tx=XM_023254588.1;refGene_gn=AFDN;refGene_tx=XM_023254589.1;refGene_gn=AFDN;refGene_tx=XM_023254590.1;refGene_gn=AFDN;refGene_tx=XM_023254591.1;refGene_gn=AFDN;refGene_tx=XM_023254592.1
B2 153268147 153268308
refGene_gn=AFDN;refGene_tx=XM_023254578.1;refGene_gn=AFDN;refGene_tx=XM_023254579.1;refGene_gn=AFDN;refGene_tx=XM_023254580.1;refGene_gn=AFDN;refGene_tx=XM_023254581.1;refGene_gn=AFDN;refGene_tx=XM_023254582.1;refGene_gn=AFDN;refGene_tx=XM_023254583.1;refGene_gn=AFDN;refGene_tx=XM_023254584.1;refGene_gn=AFDN;refGene_tx=XM_023254585.1;refGene_gn=AFDN;refGene_tx=XM_023254586.1;refGene_gn=AFDN;refGene_tx=XM_023254587.1;refGene_gn=AFDN;refGene_tx=XM_023254588.1;refGene_gn=AFDN;refGene_tx=XM_023254589.1;refGene_gn=AFDN;refGene_tx=XM_023254590.1;refGene_gn=AFDN;refGene_tx=XM_023254591.1;refGene_gn=AFDN;refGene_tx=XM_023254592.1
B2 153274429 153274587
refGene_gn=AFDN;refGene_tx=XM_023254578.1;refGene_gn=AFDN;refGene_tx=XM_023254579.1;refGene_gn=AFDN;refGene_tx=XM_023254580.1;refGene_gn=AFDN;refGene_tx=XM_023254581.1;refGene_gn=AFDN;refGene_tx=XM_023254582.1;refGene_gn=AFDN;refGene_tx=XM_023254583.1;refGene_gn=AFDN;refGene_tx=XM_023254584.1;refGene_gn=AFDN;refGene_tx=XM_023254585.1;refGene_gn=AFDN;refGene_tx=XM_023254586.1;refGene_gn=AFDN;refGene_tx=XM_023254587.1;refGene_gn=AFDN;refGene_tx=XM_023254588.1;refGene_gn=AFDN;refGene_tx=XM_023254589.1;refGene_gn=AFDN;refGene_tx=XM_023254590.1;refGene_gn=AFDN;refGene_tx=XM_023254591.1;refGene_gn=AFDN;refGene_tx=XM_023254592.1
B2 153282460 153282572
refGene_gn=AFDN;refGene_tx=XM_023254578.1;refGene_gn=AFDN;refGene_tx=XM_023254579.1;refGene_gn=AFDN;refGene_tx=XM_023254580.1;refGene_gn=AFDN;refGene_tx=XM_023254581.1;refGene_gn=AFDN;refGene_tx=XM_023254582.1;refGene_gn=AFDN;refGene_tx=XM_023254583.1;refGene_gn=AFDN;refGene_tx=XM_023254584.1;refGene_gn=AFDN;refGene_tx=XM_023254585.1;refGene_gn=AFDN;refGene_tx=XM_023254586.1;refGene_gn=AFDN;refGene_tx=XM_023254587.1;refGene_gn=AFDN;refGene_tx=XM_023254588.1;refGene_gn=AFDN;refGene_tx=XM_023254589.1;refGene_gn=AFDN;refGene_tx=XM_023254590.1;refGene_gn=AFDN;refGene_tx=XM_023254591.1;refGene_gn=AFDN;refGene_tx=XM_023254592.1
B2 153283827 153283995
refGene_gn=AFDN;refGene_tx=XM_023254578.1;refGene_gn=AFDN;refGene_tx=XM_023254579.1;refGene_gn=AFDN;refGene_tx=XM_023254580.1;refGene_gn=AFDN;refGene_tx=XM_023254581.1;refGene_gn=AFDN;refGene_tx=XM_023254582.1;refGene_gn=AFDN;refGene_tx=XM_023254583.1;refGene_gn=AFDN;refGene_tx=XM_023254584.1;refGene_gn=AFDN;refGene_tx=XM_023254585.1;refGene_gn=AFDN;refGene_tx=XM_023254586.1;refGene_gn=AFDN;refGene_tx=XM_023254587.1;refGene_gn=AFDN;refGene_tx=XM_023254588.1;refGene_gn=AFDN;refGene_tx=XM_023254589.1;refGene_gn=AFDN;refGene_tx=XM_023254590.1;refGene_gn=AFDN;refGene_tx=XM_023254591.1;refGene_gn=AFDN;refGene_tx=XM_023254592.1

B2	153286861	153286912
refGene_gn=AFDN;refGene_tx=XM_023254578.1;refGene_gn=AFDN;refGene_tx=XM_023254579.1;refGene_gn=AFDN;refGene_tx=XM_023254580.1;refGene_gn=AFDN;refGene_tx=XM_023254581.1;refGene_gn=AFDN;refGene_tx=XM_023254582.1;refGene_gn=AFDN;refGene_tx=XM_023254583.1;refGene_gn=AFDN;refGene_tx=XM_023254584.1;refGene_gn=AFDN;refGene_tx=XM_023254585.1;refGene_gn=AFDN;refGene_tx=XM_023254588.1;refGene_gn=AFDN;refGene_tx=XM_023254589.1;refGene_gn=AFDN;refGene_tx=XM_023254590.1;refGene_gn=AFDN;refGene_tx=XM_023254592.1
B2	153291456	153291551
refGene_gn=AFDN;refGene_tx=XM_023254578.1;refGene_gn=AFDN;refGene_tx=XM_023254579.1;refGene_gn=AFDN;refGene_tx=XM_023254580.1;refGene_gn=AFDN;refGene_tx=XM_023254581.1;refGene_gn=AFDN;refGene_tx=XM_023254582.1;refGene_gn=AFDN;refGene_tx=XM_023254583.1;refGene_gn=AFDN;refGene_tx=XM_023254584.1;refGene_gn=AFDN;refGene_tx=XM_023254585.1;refGene_gn=AFDN;refGene_tx=XM_023254586.1;refGene_gn=AFDN;refGene_tx=XM_023254587.1;refGene_gn=AFDN;refGene_tx=XM_023254588.1;refGene_gn=AFDN;refGene_tx=XM_023254589.1;refGene_gn=AFDN;refGene_tx=XM_023254590.1;refGene_gn=AFDN;refGene_tx=XM_023254591.1;refGene_gn=AFDN;refGene_tx=XM_023254592.1
B2	153292369	153292632
refGene_gn=AFDN;refGene_tx=XM_023254578.1;refGene_gn=AFDN;refGene_tx=XM_023254579.1;refGene_gn=AFDN;refGene_tx=XM_023254580.1;refGene_gn=AFDN;refGene_tx=XM_023254581.1;refGene_gn=AFDN;refGene_tx=XM_023254582.1;refGene_gn=AFDN;refGene_tx=XM_023254583.1;refGene_gn=AFDN;refGene_tx=XM_023254584.1;refGene_gn=AFDN;refGene_tx=XM_023254585.1;refGene_gn=AFDN;refGene_tx=XM_023254586.1;refGene_gn=AFDN;refGene_tx=XM_023254587.1;refGene_gn=AFDN;refGene_tx=XM_023254588.1;refGene_gn=AFDN;refGene_tx=XM_023254589.1;refGene_gn=AFDN;refGene_tx=XM_023254590.1;refGene_gn=AFDN;refGene_tx=XM_023254591.1;refGene_gn=AFDN;refGene_tx=XM_023254592.1
B2	153298061	153298131
refGene_gn=AFDN;refGene_tx=XM_023254578.1;refGene_gn=AFDN;refGene_tx=XM_023254579.1;refGene_gn=AFDN;refGene_tx=XM_023254580.1;refGene_gn=AFDN;refGene_tx=XM_023254581.1;refGene_gn=AFDN;refGene_tx=XM_023254582.1;refGene_gn=AFDN;refGene_tx=XM_023254583.1;refGene_gn=AFDN;refGene_tx=XM_023254584.1;refGene_gn=AFDN;refGene_tx=XM_023254585.1;refGene_gn=AFDN;refGene_tx=XM_023254586.1;refGene_gn=AFDN;refGene_tx=XM_023254587.1;refGene_gn=AFDN;refGene_tx=XM_023254588.1;refGene_gn=AFDN;refGene_tx=XM_023254589.1;refGene_gn=AFDN;refGene_tx=XM_023254590.1;refGene_gn=AFDN;refGene_tx=XM_023254591.1;refGene_gn=AFDN;refGene_tx=XM_023254592.1
B2	153301958	153302074
refGene_gn=AFDN;refGene_tx=XM_023254578.1;refGene_gn=AFDN;refGene_tx=XM_023254579.1;refGene_gn=AFDN;refGene_tx=XM_023254580.1;refGene_gn=AFDN;refGene_tx=XM_023254581.1;refGene_gn=AFDN;refGene_tx=XM_023254582.1;refGene_gn=AFDN;refGene_tx=XM_023254583.1;refGene_gn=AFDN;refGene_tx=XM_023254584.1;refGene_gn=AFDN;refGene_tx=XM_023254585.1;refGene_gn=AFDN;refGene_tx=XM_023254586.1;refGene_gn=AFDN;refGene_tx=XM_023254587.1;refGene_gn=AFDN;refGene_tx=XM_023254588.1;refGene_gn=AFDN;refGene_tx=XM_023254589.1;refGene_gn=AFDN;refGene_tx=XM_023254590.1;refGene_gn=AFDN;refGene_tx=XM_023254591.1;refGene_gn=AFDN;refGene_tx=XM_023254592.1
B2	153305621	153305683

refGene_gn=AFDN;refGene_tx=XM_023254578.1;refGene_gn=AFDN;refGene_tx=X
M_023254579.1;refGene_gn=AFDN;refGene_tx=XM_023254580.1;refGene_gn=AFD
N;refGene_tx=XM_023254581.1;refGene_gn=AFDN;refGene_tx=XM_023254582.1;
refGene_gn=AFDN;refGene_tx=XM_023254583.1;refGene_gn=AFDN;refGene_tx=X
M_023254584.1;refGene_gn=AFDN;refGene_tx=XM_023254585.1;refGene_gn=AFD
N;refGene_tx=XM_023254586.1;refGene_gn=AFDN;refGene_tx=XM_023254587.1;
refGene_gn=AFDN;refGene_tx=XM_023254588.1;refGene_gn=AFDN;refGene_tx=X
M_023254589.1;refGene_gn=AFDN;refGene_tx=XM_023254590.1;refGene_gn=AFD
N;refGene_tx=XM_023254591.1;refGene_gn=AFDN;refGene_tx=XM_023254592.1
B2 153305807 153306013
refGene_gn=AFDN;refGene_tx=XM_023254578.1;refGene_gn=AFDN;refGene_tx=X
M_023254579.1;refGene_gn=AFDN;refGene_tx=XM_023254580.1;refGene_gn=AFD
N;refGene_tx=XM_023254581.1;refGene_gn=AFDN;refGene_tx=XM_023254582.1;
refGene_gn=AFDN;refGene_tx=XM_023254583.1;refGene_gn=AFDN;refGene_tx=X
M_023254584.1;refGene_gn=AFDN;refGene_tx=XM_023254585.1;refGene_gn=AFD
N;refGene_tx=XM_023254586.1;refGene_gn=AFDN;refGene_tx=XM_023254587.1;
refGene_gn=AFDN;refGene_tx=XM_023254588.1;refGene_gn=AFDN;refGene_tx=X
M_023254589.1;refGene_gn=AFDN;refGene_tx=XM_023254590.1;refGene_gn=AFD
N;refGene_tx=XM_023254591.1;refGene_gn=AFDN;refGene_tx=XM_023254592.1
B2 153307907 153307928
refGene_gn=AFDN;refGene_tx=XM_023254578.1;refGene_gn=AFDN;refGene_tx=X
M_023254579.1;refGene_gn=AFDN;refGene_tx=XM_023254580.1;refGene_gn=AFD
N;refGene_tx=XM_023254581.1;refGene_gn=AFDN;refGene_tx=XM_023254583.1;
refGene_gn=AFDN;refGene_tx=XM_023254584.1;refGene_gn=AFDN;refGene_tx=X
M_023254585.1;refGene_gn=AFDN;refGene_tx=XM_023254586.1
B2 153308652 153308798
refGene_gn=AFDN;refGene_tx=XM_023254578.1;refGene_gn=AFDN;refGene_tx=X
M_023254579.1;refGene_gn=AFDN;refGene_tx=XM_023254580.1;refGene_gn=AFD
N;refGene_tx=XM_023254581.1;refGene_gn=AFDN;refGene_tx=XM_023254582.1;
refGene_gn=AFDN;refGene_tx=XM_023254583.1;refGene_gn=AFDN;refGene_tx=X
M_023254584.1;refGene_gn=AFDN;refGene_tx=XM_023254585.1;refGene_gn=AFD
N;refGene_tx=XM_023254586.1;refGene_gn=AFDN;refGene_tx=XM_023254587.1;
refGene_gn=AFDN;refGene_tx=XM_023254588.1;refGene_gn=AFDN;refGene_tx=X
M_023254589.1;refGene_gn=AFDN;refGene_tx=XM_023254590.1;refGene_gn=AFD
N;refGene_tx=XM_023254591.1;refGene_gn=AFDN;refGene_tx=XM_023254592.1
B2 153309078 153309173
refGene_gn=AFDN;refGene_tx=XM_023254578.1;refGene_gn=AFDN;refGene_tx=X
M_023254579.1;refGene_gn=AFDN;refGene_tx=XM_023254580.1;refGene_gn=AFD
N;refGene_tx=XM_023254581.1;refGene_gn=AFDN;refGene_tx=XM_023254582.1;
refGene_gn=AFDN;refGene_tx=XM_023254583.1;refGene_gn=AFDN;refGene_tx=X
M_023254584.1;refGene_gn=AFDN;refGene_tx=XM_023254585.1;refGene_gn=AFD
N;refGene_tx=XM_023254586.1;refGene_gn=AFDN;refGene_tx=XM_023254587.1;
refGene_gn=AFDN;refGene_tx=XM_023254588.1;refGene_gn=AFDN;refGene_tx=X
M_023254589.1;refGene_gn=AFDN;refGene_tx=XM_023254590.1;refGene_gn=AFD
N;refGene_tx=XM_023254591.1;refGene_gn=AFDN;refGene_tx=XM_023254592.1
B2 153316597 153316863
refGene_gn=AFDN;refGene_tx=XM_023254578.1;refGene_gn=AFDN;refGene_tx=X
M_023254579.1;refGene_gn=AFDN;refGene_tx=XM_023254580.1;refGene_gn=AFD
N;refGene_tx=XM_023254581.1;refGene_gn=AFDN;refGene_tx=XM_023254582.1;
refGene_gn=AFDN;refGene_tx=XM_023254583.1;refGene_gn=AFDN;refGene_tx=X

M_023254584.1;refGene_gn=AFDN;refGene_tx=XM_023254585.1;refGene_gn=AFDN;refGene_tx=XM_023254586.1;refGene_gn=AFDN;refGene_tx=XM_023254587.1;refGene_gn=AFDN;refGene_tx=XM_023254588.1;refGene_gn=AFDN;refGene_tx=XM_023254589.1;refGene_gn=AFDN;refGene_tx=XM_023254590.1;refGene_gn=AFDN;refGene_tx=XM_023254591.1;refGene_gn=AFDN;refGene_tx=XM_023254592.1
B2 153318166 153318310
refGene_gn=AFDN;refGene_tx=XM_023254578.1;refGene_gn=AFDN;refGene_tx=XM_023254579.1;refGene_gn=AFDN;refGene_tx=XM_023254580.1;refGene_gn=AFDN;refGene_tx=XM_023254581.1;refGene_gn=AFDN;refGene_tx=XM_023254582.1;refGene_gn=AFDN;refGene_tx=XM_023254583.1;refGene_gn=AFDN;refGene_tx=XM_023254584.1;refGene_gn=AFDN;refGene_tx=XM_023254585.1;refGene_gn=AFDN;refGene_tx=XM_023254586.1;refGene_gn=AFDN;refGene_tx=XM_023254587.1;refGene_gn=AFDN;refGene_tx=XM_023254588.1;refGene_gn=AFDN;refGene_tx=XM_023254589.1;refGene_gn=AFDN;refGene_tx=XM_023254590.1;refGene_gn=AFDN;refGene_tx=XM_023254591.1;refGene_gn=AFDN;refGene_tx=XM_023254592.1
B2 153319811 153320010
refGene_gn=AFDN;refGene_tx=XM_023254578.1;refGene_gn=AFDN;refGene_tx=XM_023254579.1;refGene_gn=AFDN;refGene_tx=XM_023254580.1;refGene_gn=AFDN;refGene_tx=XM_023254581.1;refGene_gn=AFDN;refGene_tx=XM_023254582.1;refGene_gn=AFDN;refGene_tx=XM_023254583.1;refGene_gn=AFDN;refGene_tx=XM_023254584.1;refGene_gn=AFDN;refGene_tx=XM_023254585.1;refGene_gn=AFDN;refGene_tx=XM_023254586.1;refGene_gn=AFDN;refGene_tx=XM_023254587.1;refGene_gn=AFDN;refGene_tx=XM_023254588.1;refGene_gn=AFDN;refGene_tx=XM_023254589.1;refGene_gn=AFDN;refGene_tx=XM_023254590.1;refGene_gn=AFDN;refGene_tx=XM_023254591.1;refGene_gn=AFDN;refGene_tx=XM_023254592.1
B2 153323733 153323837
refGene_gn=AFDN;refGene_tx=XM_023254578.1;refGene_gn=AFDN;refGene_tx=XM_023254579.1;refGene_gn=AFDN;refGene_tx=XM_023254580.1;refGene_gn=AFDN;refGene_tx=XM_023254581.1;refGene_gn=AFDN;refGene_tx=XM_023254582.1;refGene_gn=AFDN;refGene_tx=XM_023254583.1;refGene_gn=AFDN;refGene_tx=XM_023254584.1;refGene_gn=AFDN;refGene_tx=XM_023254585.1;refGene_gn=AFDN;refGene_tx=XM_023254586.1;refGene_gn=AFDN;refGene_tx=XM_023254587.1;refGene_gn=AFDN;refGene_tx=XM_023254588.1;refGene_gn=AFDN;refGene_tx=XM_023254589.1;refGene_gn=AFDN;refGene_tx=XM_023254590.1;refGene_gn=AFDN;refGene_tx=XM_023254591.1;refGene_gn=AFDN;refGene_tx=XM_023254592.1
B2 153326146 153326233
refGene_gn=AFDN;refGene_tx=XM_023254578.1;refGene_gn=AFDN;refGene_tx=XM_023254579.1;refGene_gn=AFDN;refGene_tx=XM_023254580.1;refGene_gn=AFDN;refGene_tx=XM_023254581.1;refGene_gn=AFDN;refGene_tx=XM_023254582.1;refGene_gn=AFDN;refGene_tx=XM_023254583.1;refGene_gn=AFDN;refGene_tx=XM_023254584.1;refGene_gn=AFDN;refGene_tx=XM_023254585.1;refGene_gn=AFDN;refGene_tx=XM_023254586.1;refGene_gn=AFDN;refGene_tx=XM_023254587.1;refGene_gn=AFDN;refGene_tx=XM_023254588.1;refGene_gn=AFDN;refGene_tx=XM_023254589.1;refGene_gn=AFDN;refGene_tx=XM_023254590.1;refGene_gn=AFDN;refGene_tx=XM_023254591.1;refGene_gn=AFDN;refGene_tx=XM_023254592.1
B2 153339928 153339994
refGene_gn=AFDN;refGene_tx=XM_023254578.1;refGene_gn=AFDN;refGene_tx=XM_023254579.1;refGene_gn=AFDN;refGene_tx=XM_023254580.1;refGene_gn=AFDN;refGene_tx=XM_023254581.1;refGene_gn=AFDN;refGene_tx=XM_023254582.1;refGene_gn=AFDN;refGene_tx=XM_023254583.1;refGene_gn=AFDN;refGene_tx=X

M_023254584.1;refGene_gn=AFDN;refGene_tx=XM_023254585.1;refGene_gn=AFDN;refGene_tx=XM_023254586.1;refGene_gn=AFDN;refGene_tx=XM_023254587.1;refGene_gn=AFDN;refGene_tx=XM_023254588.1;refGene_gn=AFDN;refGene_tx=XM_023254589.1;refGene_gn=AFDN;refGene_tx=XM_023254590.1;refGene_gn=AFDN;refGene_tx=XM_023254591.1;refGene_gn=AFDN;refGene_tx=XM_023254592.1
B2 153340203 153340277
refGene_gn=AFDN;refGene_tx=XM_023254578.1;refGene_gn=AFDN;refGene_tx=XM_023254579.1;refGene_gn=AFDN;refGene_tx=XM_023254580.1;refGene_gn=AFDN;refGene_tx=XM_023254581.1;refGene_gn=AFDN;refGene_tx=XM_023254582.1;refGene_gn=AFDN;refGene_tx=XM_023254583.1;refGene_gn=AFDN;refGene_tx=XM_023254584.1;refGene_gn=AFDN;refGene_tx=XM_023254585.1;refGene_gn=AFDN;refGene_tx=XM_023254586.1;refGene_gn=AFDN;refGene_tx=XM_023254587.1;refGene_gn=AFDN;refGene_tx=XM_023254588.1;refGene_gn=AFDN;refGene_tx=XM_023254589.1;refGene_gn=AFDN;refGene_tx=XM_023254590.1;refGene_gn=AFDN;refGene_tx=XM_023254591.1;refGene_gn=AFDN;refGene_tx=XM_023254592.1
B2 153340949 153341068
refGene_gn=AFDN;refGene_tx=XM_023254578.1;refGene_gn=AFDN;refGene_tx=XM_023254579.1;refGene_gn=AFDN;refGene_tx=XM_023254580.1;refGene_gn=AFDN;refGene_tx=XM_023254581.1;refGene_gn=AFDN;refGene_tx=XM_023254582.1;refGene_gn=AFDN;refGene_tx=XM_023254583.1;refGene_gn=AFDN;refGene_tx=XM_023254584.1;refGene_gn=AFDN;refGene_tx=XM_023254585.1;refGene_gn=AFDN;refGene_tx=XM_023254586.1;refGene_gn=AFDN;refGene_tx=XM_023254587.1;refGene_gn=AFDN;refGene_tx=XM_023254588.1;refGene_gn=AFDN;refGene_tx=XM_023254589.1;refGene_gn=AFDN;refGene_tx=XM_023254590.1;refGene_gn=AFDN;refGene_tx=XM_023254591.1;refGene_gn=AFDN;refGene_tx=XM_023254592.1
B2 153342410 153342605
refGene_gn=AFDN;refGene_tx=XM_023254578.1;refGene_gn=AFDN;refGene_tx=XM_023254579.1;refGene_gn=AFDN;refGene_tx=XM_023254580.1;refGene_gn=AFDN;refGene_tx=XM_023254581.1;refGene_gn=AFDN;refGene_tx=XM_023254582.1;refGene_gn=AFDN;refGene_tx=XM_023254583.1;refGene_gn=AFDN;refGene_tx=XM_023254584.1;refGene_gn=AFDN;refGene_tx=XM_023254585.1;refGene_gn=AFDN;refGene_tx=XM_023254586.1;refGene_gn=AFDN;refGene_tx=XM_023254587.1;refGene_gn=AFDN;refGene_tx=XM_023254588.1;refGene_gn=AFDN;refGene_tx=XM_023254589.1;refGene_gn=AFDN;refGene_tx=XM_023254590.1;refGene_gn=AFDN;refGene_tx=XM_023254591.1;refGene_gn=AFDN;refGene_tx=XM_023254592.1
B2 153343252 153343344
refGene_gn=AFDN;refGene_tx=XM_023254578.1;refGene_gn=AFDN;refGene_tx=XM_023254579.1;refGene_gn=AFDN;refGene_tx=XM_023254580.1;refGene_gn=AFDN;refGene_tx=XM_023254581.1;refGene_gn=AFDN;refGene_tx=XM_023254582.1;refGene_gn=AFDN;refGene_tx=XM_023254583.1;refGene_gn=AFDN;refGene_tx=XM_023254584.1;refGene_gn=AFDN;refGene_tx=XM_023254585.1;refGene_gn=AFDN;refGene_tx=XM_023254586.1;refGene_gn=AFDN;refGene_tx=XM_023254587.1;refGene_gn=AFDN;refGene_tx=XM_023254588.1;refGene_gn=AFDN;refGene_tx=XM_023254589.1;refGene_gn=AFDN;refGene_tx=XM_023254590.1;refGene_gn=AFDN;refGene_tx=XM_023254591.1;refGene_gn=AFDN;refGene_tx=XM_023254592.1
B2 153343661 153343847
refGene_gn=AFDN;refGene_tx=XM_023254578.1;refGene_gn=AFDN;refGene_tx=XM_023254579.1;refGene_gn=AFDN;refGene_tx=XM_023254580.1;refGene_gn=AFDN;refGene_tx=XM_023254581.1;refGene_gn=AFDN;refGene_tx=XM_023254582.1;refGene_gn=AFDN;refGene_tx=XM_023254583.1;refGene_gn=AFDN;refGene_tx=X

M_023254584.1;refGene_gn=AFDN;refGene_tx=XM_023254585.1;refGene_gn=AFD
N;refGene_tx=XM_023254586.1;refGene_gn=AFDN;refGene_tx=XM_023254587.1;
refGene_gn=AFDN;refGene_tx=XM_023254588.1;refGene_gn=AFDN;refGene_tx=X
M_023254589.1;refGene_gn=AFDN;refGene_tx=XM_023254590.1;refGene_gn=AFD
N;refGene_tx=XM_023254591.1;refGene_gn=AFDN;refGene_tx=XM_023254592.1
B2 153346536 153347508
refGene_gn=AFDN;refGene_tx=XM_023254578.1;refGene_gn=AFDN;refGene_tx=X
M_023254579.1;refGene_gn=AFDN;refGene_tx=XM_023254580.1;refGene_gn=AFD
N;refGene_tx=XM_023254581.1;refGene_gn=AFDN;refGene_tx=XM_023254582.1;
refGene_gn=AFDN;refGene_tx=XM_023254583.1;refGene_gn=AFDN;refGene_tx=X
M_023254584.1;refGene_gn=AFDN;refGene_tx=XM_023254585.1;refGene_gn=AFD
N;refGene_tx=XM_023254586.1;refGene_gn=AFDN;refGene_tx=XM_023254587.1;
refGene_gn=AFDN;refGene_tx=XM_023254588.1;refGene_gn=AFDN;refGene_tx=X
M_023254589.1;refGene_gn=AFDN;refGene_tx=XM_023254590.1;refGene_gn=AFD
N;refGene_tx=XM_023254591.1;refGene_gn=AFDN;refGene_tx=XM_023254592.1
B2 153349565 153349601
refGene_gn=AFDN;refGene_tx=XM_023254578.1;refGene_gn=AFDN;refGene_tx=X
M_023254579.1;refGene_gn=AFDN;refGene_tx=XM_023254580.1;refGene_gn=AFD
N;refGene_tx=XM_023254581.1;refGene_gn=AFDN;refGene_tx=XM_023254582.1;
refGene_gn=AFDN;refGene_tx=XM_023254583.1;refGene_gn=AFDN;refGene_tx=X
M_023254586.1;refGene_gn=AFDN;refGene_tx=XM_023254587.1
B2 153356166 153356295
refGene_gn=AFDN;refGene_tx=XM_023254578.1;refGene_gn=AFDN;refGene_tx=X
M_023254579.1;refGene_gn=AFDN;refGene_tx=XM_023254580.1;refGene_gn=AFD
N;refGene_tx=XM_023254581.1;refGene_gn=AFDN;refGene_tx=XM_023254582.1;
refGene_gn=AFDN;refGene_tx=XM_023254583.1;refGene_gn=AFDN;refGene_tx=X
M_023254584.1;refGene_gn=AFDN;refGene_tx=XM_023254585.1;refGene_gn=AFD
N;refGene_tx=XM_023254586.1;refGene_gn=AFDN;refGene_tx=XM_023254587.1;
refGene_gn=AFDN;refGene_tx=XM_023254588.1;refGene_gn=AFDN;refGene_tx=X
M_023254589.1;refGene_gn=AFDN;refGene_tx=XM_023254590.1;refGene_gn=AFD
N;refGene_tx=XM_023254591.1;refGene_gn=AFDN;refGene_tx=XM_023254592.1
B2 153359097 153359410
refGene_gn=AFDN;refGene_tx=XM_023254578.1;refGene_gn=AFDN;refGene_tx=X
M_023254579.1;refGene_gn=AFDN;refGene_tx=XM_023254580.1;refGene_gn=AFD
N;refGene_tx=XM_023254581.1;refGene_gn=AFDN;refGene_tx=XM_023254582.1;
refGene_gn=AFDN;refGene_tx=XM_023254583.1;refGene_gn=AFDN;refGene_tx=X
M_023254584.1;refGene_gn=AFDN;refGene_tx=XM_023254585.1;refGene_gn=AFD
N;refGene_tx=XM_023254586.1;refGene_gn=AFDN;refGene_tx=XM_023254587.1;
refGene_gn=AFDN;refGene_tx=XM_023254588.1;refGene_gn=AFDN;refGene_tx=X
M_023254589.1;refGene_gn=AFDN;refGene_tx=XM_023254590.1;refGene_gn=AFD
N;refGene_tx=XM_023254591.1
B2 153362644 153362729
refGene_gn=AFDN;refGene_tx=XM_023254578.1;refGene_gn=AFDN;refGene_tx=X
M_023254579.1;refGene_gn=AFDN;refGene_tx=XM_023254580.1;refGene_gn=AFD
N;refGene_tx=XM_023254581.1;refGene_gn=AFDN;refGene_tx=XM_023254582.1;
refGene_gn=AFDN;refGene_tx=XM_023254583.1;refGene_gn=AFDN;refGene_tx=X
M_023254584.1;refGene_gn=AFDN;refGene_tx=XM_023254585.1;refGene_gn=AFD
N;refGene_tx=XM_023254586.1;refGene_gn=AFDN;refGene_tx=XM_023254587.1;
refGene_gn=AFDN;refGene_tx=XM_023254588.1;refGene_gn=AFDN;refGene_tx=X
M_023254589.1;refGene_gn=AFDN;refGene_tx=XM_023254590.1;refGene_gn=AFD

N;refGene_tx=XM_023254591.1
B2	153363509	153363663
refGene_gn=AFDN;refGene_tx=XM_023254578.1;refGene_gn=AFDN;refGene_tx=XM_023254579.1;refGene_gn=AFDN;refGene_tx=XM_023254580.1;refGene_gn=AFDN;refGene_tx=XM_023254581.1;refGene_gn=AFDN;refGene_tx=XM_023254582.1;refGene_gn=AFDN;refGene_tx=XM_023254583.1;refGene_gn=AFDN;refGene_tx=XM_023254584.1;refGene_gn=AFDN;refGene_tx=XM_023254585.1;refGene_gn=AFDN;refGene_tx=XM_023254586.1;refGene_gn=AFDN;refGene_tx=XM_023254587.1;refGene_gn=AFDN;refGene_tx=XM_023254588.1;refGene_gn=AFDN;refGene_tx=XM_023254589.1;refGene_gn=AFDN;refGene_tx=XM_023254590.1;refGene_gn=AFDN;refGene_tx=XM_023254591.1
B2	153370211	153370286
refGene_gn=KIF25;refGene_tx=XM_023254593.1;refGene_gn=KIF25;refGene_tx=XM_023254594.1;refGene_gn=KIF25;refGene_tx=XM_023254595.1
B2	153379678	153379744
refGene_gn=KIF25;refGene_tx=XM_023254593.1;refGene_gn=KIF25;refGene_tx=XM_023254594.1;refGene_gn=KIF25;refGene_tx=XM_023254595.1
B2	153385539	153385662
refGene_gn=KIF25;refGene_tx=XM_023254593.1;refGene_gn=KIF25;refGene_tx=XM_023254594.1;refGene_gn=KIF25;refGene_tx=XM_023254595.1
B2	153387782	153387921
refGene_gn=KIF25;refGene_tx=XM_023254593.1;refGene_gn=KIF25;refGene_tx=XM_023254594.1;refGene_gn=KIF25;refGene_tx=XM_023254595.1
B2	153388860	153388964
refGene_gn=KIF25;refGene_tx=XM_023254593.1;refGene_gn=KIF25;refGene_tx=XM_023254594.1;refGene_gn=KIF25;refGene_tx=XM_023254595.1
B2	153390394	153390483
refGene_gn=KIF25;refGene_tx=XM_023254593.1;refGene_gn=KIF25;refGene_tx=XM_023254594.1;refGene_gn=KIF25;refGene_tx=XM_023254595.1
B2	153392354	153392391
refGene_gn=KIF25;refGene_tx=XM_023254593.1;refGene_gn=KIF25;refGene_tx=XM_023254594.1;refGene_gn=KIF25;refGene_tx=XM_023254595.1
B2	153400576	153400644
refGene_gn=KIF25;refGene_tx=XM_023254593.1;refGene_gn=KIF25;refGene_tx=XM_023254594.1;refGene_gn=KIF25;refGene_tx=XM_023254595.1
B2	153407773	153407848
refGene_gn=KIF25;refGene_tx=XM_023254593.1;refGene_gn=KIF25;refGene_tx=XM_023254594.1;refGene_gn=KIF25;refGene_tx=XM_023254595.1
B2	153410016	153410166
refGene_gn=KIF25;refGene_tx=XM_023254593.1;refGene_gn=KIF25;refGene_tx=XM_023254595.1
B2	153413785	153413962
refGene_gn=KIF25;refGene_tx=XM_023254593.1;refGene_gn=KIF25;refGene_tx=XM_023254594.1;refGene_gn=KIF25;refGene_tx=XM_023254595.1
B2	153414928	153415080
refGene_gn=KIF25;refGene_tx=XM_023254593.1;refGene_gn=KIF25;refGene_tx=XM_023254594.1;refGene_gn=KIF25;refGene_tx=XM_023254595.1
B2	153416213	153416387
refGene_gn=KIF25;refGene_tx=XM_023254593.1;refGene_gn=KIF25;refGene_tx=XM_023254594.1;refGene_gn=KIF25;refGene_tx=XM_023254595.1

B2 153416720 153417331
refGene_gn=KIF25;refGene_tx=XM_023254593.1;refGene_gn=KIF25;refGene_tx=XM_023254594.1;refGene_gn=KIF25;refGene_tx=XM_023254595.1
B2 153417333 153417488
refGene_gn=KIF25;refGene_tx=XM_023254595.1
B2 153425025 153425157
refGene_gn=FRMD1;refGene_tx=XM_019831640.2;refGene_gn=FRMD1;refGene_tx=XM_023254596.1;refGene_gn=FRMD1;refGene_tx=XM_023254597.1;refGene_gn=FRMD1;refGene_tx=XM_023254598.1;refGene_gn=FRMD1;refGene_tx=XM_023254599.1;refGene_gn=FRMD1;refGene_tx=XM_023254600.1;refGene_gn=FRMD1;refGene_tx=XM_023254601.1;refGene_gn=FRMD1;refGene_tx=XM_023254602.1
B2 153426336 153426416
refGene_gn=FRMD1;refGene_tx=XM_019831640.2;refGene_gn=FRMD1;refGene_tx=XM_023254596.1;refGene_gn=FRMD1;refGene_tx=XM_023254597.1;refGene_gn=FRMD1;refGene_tx=XM_023254598.1;refGene_gn=FRMD1;refGene_tx=XM_023254599.1;refGene_gn=FRMD1;refGene_tx=XM_023254600.1;refGene_gn=FRMD1;refGene_tx=XM_023254601.1;refGene_gn=FRMD1;refGene_tx=XM_023254602.1
B2 153430113 153430461
refGene_gn=FRMD1;refGene_tx=XM_019831640.2;refGene_gn=FRMD1;refGene_tx=XM_023254596.1;refGene_gn=FRMD1;refGene_tx=XM_023254597.1;refGene_gn=FRMD1;refGene_tx=XM_023254598.1;refGene_gn=FRMD1;refGene_tx=XM_023254599.1;refGene_gn=FRMD1;refGene_tx=XM_023254600.1;refGene_gn=FRMD1;refGene_tx=XM_023254601.1;refGene_gn=FRMD1;refGene_tx=XM_023254602.1
B2 153431008 153431183
refGene_gn=FRMD1;refGene_tx=XM_019831640.2;refGene_gn=FRMD1;refGene_tx=XM_023254596.1;refGene_gn=FRMD1;refGene_tx=XM_023254597.1;refGene_gn=FRMD1;refGene_tx=XM_023254598.1;refGene_gn=FRMD1;refGene_tx=XM_023254599.1;refGene_gn=FRMD1;refGene_tx=XM_023254600.1;refGene_gn=FRMD1;refGene_tx=XM_023254601.1;refGene_gn=FRMD1;refGene_tx=XM_023254602.1
B2 153431792 153431861
refGene_gn=FRMD1;refGene_tx=XM_019831640.2;refGene_gn=FRMD1;refGene_tx=XM_023254596.1;refGene_gn=FRMD1;refGene_tx=XM_023254597.1;refGene_gn=FRMD1;refGene_tx=XM_023254598.1;refGene_gn=FRMD1;refGene_tx=XM_023254599.1;refGene_gn=FRMD1;refGene_tx=XM_023254600.1;refGene_gn=FRMD1;refGene_tx=XM_023254601.1;refGene_gn=FRMD1;refGene_tx=XM_023254602.1
B2 153432198 153432264
refGene_gn=FRMD1;refGene_tx=XM_019831640.2;refGene_gn=FRMD1;refGene_tx=XM_023254596.1;refGene_gn=FRMD1;refGene_tx=XM_023254597.1;refGene_gn=FRMD1;refGene_tx=XM_023254598.1;refGene_gn=FRMD1;refGene_tx=XM_023254599.1;refGene_gn=FRMD1;refGene_tx=XM_023254600.1;refGene_gn=FRMD1;refGene_tx=XM_023254601.1;refGene_gn=FRMD1;refGene_tx=XM_023254602.1
B2 153432600 153432765
refGene_gn=FRMD1;refGene_tx=XM_019831640.2;refGene_gn=FRMD1;refGene_tx=XM_023254596.1;refGene_gn=FRMD1;refGene_tx=XM_023254597.1;refGene_gn=FRMD1;refGene_tx=XM_023254598.1;refGene_gn=FRMD1;refGene_tx=XM_023254599.1;refGene_gn=FRMD1;refGene_tx=XM_023254600.1;refGene_gn=FRMD1;refGene_tx=XM_023254601.1;refGene_gn=FRMD1;refGene_tx=XM_023254602.1
B2 153432908 153433095
refGene_gn=FRMD1;refGene_tx=XM_019831640.2;refGene_gn=FRMD1;refGene_tx=XM_023254596.1;refGene_gn=FRMD1;refGene_tx=XM_023254597.1;refGene_gn=

FRMD1;refGene_tx=XM_023254598.1;refGene_gn=FRMD1;refGene_tx=XM_023254599.1;refGene_gn=FRMD1;refGene_tx=XM_023254600.1;refGene_gn=FRMD1;refGene_tx=XM_023254601.1;refGene_gn=FRMD1;refGene_tx=XM_023254602.1
B2 153433426 153433500
refGene_gn=FRMD1;refGene_tx=XM_019831640.2;refGene_gn=FRMD1;refGene_tx=XM_023254596.1;refGene_gn=FRMD1;refGene_tx=XM_023254597.1;refGene_gn=FRMD1;refGene_tx=XM_023254598.1;refGene_gn=FRMD1;refGene_tx=XM_023254599.1;refGene_gn=FRMD1;refGene_tx=XM_023254600.1;refGene_gn=FRMD1;refGene_tx=XM_023254601.1;refGene_gn=FRMD1;refGene_tx=XM_023254602.1
B2 153433853 153433933
refGene_gn=FRMD1;refGene_tx=XM_019831640.2;refGene_gn=FRMD1;refGene_tx=XM_023254596.1;refGene_gn=FRMD1;refGene_tx=XM_023254597.1;refGene_gn=FRMD1;refGene_tx=XM_023254598.1;refGene_gn=FRMD1;refGene_tx=XM_023254599.1;refGene_gn=FRMD1;refGene_tx=XM_023254600.1;refGene_gn=FRMD1;refGene_tx=XM_023254601.1;refGene_gn=FRMD1;refGene_tx=XM_023254602.1
B2 153434492 153435012
refGene_gn=FRMD1;refGene_tx=XM_023254596.1;refGene_gn=FRMD1;refGene_tx=XM_023254597.1;refGene_gn=FRMD1;refGene_tx=XM_023254598.1;refGene_gn=FRMD1;refGene_tx=XM_023254599.1
B2 153435342 153435433
refGene_gn=FRMD1;refGene_tx=XM_019831640.2;refGene_gn=FRMD1;refGene_tx=XM_023254600.1;refGene_gn=FRMD1;refGene_tx=XM_023254601.1
B2 153437691 153438957
refGene_gn=FRMD1;refGene_tx=XM_023254603.1
B2 153439099 153439168
refGene_gn=FRMD1;refGene_tx=XM_019831640.2;refGene_gn=FRMD1;refGene_tx=XM_023254600.1;refGene_gn=FRMD1;refGene_tx=XM_023254601.1
B2 153567681 153568626
refGene_gn=LOC111560556;refGene_tx=XM_023254604.1
B2 153569263 153570822
refGene_gn=DACT2;refGene_tx=XM_023254605.1;refGene_gn=DACT2;refGene_tx=XM_023254606.1;refGene_gn=DACT2;refGene_tx=XM_023254607.1;refGene_gn=DACT2;refGene_tx=XM_023254608.1
B2 153571648 153571867
refGene_gn=DACT2;refGene_tx=XM_023254605.1;refGene_gn=DACT2;refGene_tx=XM_023254606.1;refGene_gn=DACT2;refGene_tx=XM_023254607.1;refGene_gn=DACT2;refGene_tx=XM_023254608.1
B2 153572757 153572863
refGene_gn=DACT2;refGene_tx=XM_023254605.1;refGene_gn=DACT2;refGene_tx=XM_023254606.1;refGene_gn=DACT2;refGene_tx=XM_023254607.1;refGene_gn=DACT2;refGene_tx=XM_023254608.1
B2 153574488 153574515
refGene_gn=DACT2;refGene_tx=XM_023254608.1
B2 153575082 153575202
refGene_gn=DACT2;refGene_tx=XM_023254607.1
B2 153575457 153575658
refGene_gn=DACT2;refGene_tx=XM_023254606.1
B2 153578380 153578626
refGene_gn=DACT2;refGene_tx=XM_023254605.1
B2 153659374 153659458

refGene_gn=SMOC2;refGene_tx=XM_023254609.1;refGene_gn=SMOC2;refGene_tx
=XM_023254610.1;refGene_gn=SMOC2;refGene_tx=XM_023254611.1;refGene_gn=
SMOC2;refGene_tx=XM_023254612.1;refGene_gn=SMOC2;refGene_tx=XM_0232546
13.1
B2	153704551	153704723
refGene_gn=SMOC2;refGene_tx=XM_023254609.1;refGene_gn=SMOC2;refGene_tx
=XM_023254610.1;refGene_gn=SMOC2;refGene_tx=XM_023254611.1;refGene_gn=
SMOC2;refGene_tx=XM_023254612.1;refGene_gn=SMOC2;refGene_tx=XM_0232546
13.1
B2	153715228	153715335
refGene_gn=SMOC2;refGene_tx=XM_023254609.1;refGene_gn=SMOC2;refGene_tx
=XM_023254610.1;refGene_gn=SMOC2;refGene_tx=XM_023254611.1;refGene_gn=
SMOC2;refGene_tx=XM_023254612.1;refGene_gn=SMOC2;refGene_tx=XM_0232546
13.1
B2	153721513	153721613
refGene_gn=SMOC2;refGene_tx=XM_023254609.1;refGene_gn=SMOC2;refGene_tx
=XM_023254610.1;refGene_gn=SMOC2;refGene_tx=XM_023254611.1;refGene_gn=
SMOC2;refGene_tx=XM_023254612.1;refGene_gn=SMOC2;refGene_tx=XM_0232546
13.1
B2	153735638	153735686
refGene_gn=SMOC2;refGene_tx=XM_023254609.1;refGene_gn=SMOC2;refGene_tx
=XM_023254610.1;refGene_gn=SMOC2;refGene_tx=XM_023254611.1;refGene_gn=
SMOC2;refGene_tx=XM_023254612.1
B2	153738597	153738681
refGene_gn=SMOC2;refGene_tx=XM_023254609.1;refGene_gn=SMOC2;refGene_tx
=XM_023254610.1;refGene_gn=SMOC2;refGene_tx=XM_023254611.1;refGene_gn=
SMOC2;refGene_tx=XM_023254612.1
B2	153739832	153739907
refGene_gn=SMOC2;refGene_tx=XM_023254609.1;refGene_gn=SMOC2;refGene_tx
=XM_023254610.1;refGene_gn=SMOC2;refGene_tx=XM_023254611.1;refGene_gn=
SMOC2;refGene_tx=XM_023254612.1
B2	153780343	153780530
refGene_gn=SMOC2;refGene_tx=XM_023254609.1;refGene_gn=SMOC2;refGene_tx
=XM_023254610.1;refGene_gn=SMOC2;refGene_tx=XM_023254611.1;refGene_gn=
SMOC2;refGene_tx=XM_023254612.1;refGene_gn=SMOC2;refGene_tx=XM_0232546
13.1
B2	153789013	153789096
refGene_gn=SMOC2;refGene_tx=XM_023254609.1;refGene_gn=SMOC2;refGene_tx
=XM_023254610.1;refGene_gn=SMOC2;refGene_tx=XM_023254611.1;refGene_gn=
SMOC2;refGene_tx=XM_023254612.1;refGene_gn=SMOC2;refGene_tx=XM_0232546
13.1
B2	153819028	153819125
refGene_gn=SMOC2;refGene_tx=XM_023254609.1;refGene_gn=SMOC2;refGene_tx
=XM_023254610.1;refGene_gn=SMOC2;refGene_tx=XM_023254611.1;refGene_gn=
SMOC2;refGene_tx=XM_023254612.1;refGene_gn=SMOC2;refGene_tx=XM_0232546
13.1
B2	153821705	153821980
refGene_gn=SMOC2;refGene_tx=XM_023254609.1;refGene_gn=SMOC2;refGene_tx
=XM_023254610.1;refGene_gn=SMOC2;refGene_tx=XM_023254611.1;refGene_gn=
SMOC2;refGene_tx=XM_023254612.1;refGene_gn=SMOC2;refGene_tx=XM_0232546

13.1
B2	153828331	153828369
refGene_gn=SMOC2;refGene_tx=XM_023254609.1;refGene_gn=SMOC2;refGene_tx=XM_023254610.1;refGene_gn=SMOC2;refGene_tx=XM_023254611.1;refGene_gn=SMOC2;refGene_tx=XM_023254612.1;refGene_gn=SMOC2;refGene_tx=XM_023254613.1
B2	153831131	153831149
refGene_gn=SMOC2;refGene_tx=XM_023254609.1;refGene_gn=SMOC2;refGene_tx=XM_023254610.1;refGene_gn=SMOC2;refGene_tx=XM_023254611.1;refGene_gn=SMOC2;refGene_tx=XM_023254612.1;refGene_gn=SMOC2;refGene_tx=XM_023254613.1
B2	154198934	154198942
refGene_gn=THBS2;refGene_tx=XM_006932146.4;refGene_gn=THBS2;refGene_tx=XM_006932147.4
B2	154200006	154200154
refGene_gn=THBS2;refGene_tx=XM_006932146.4;refGene_gn=THBS2;refGene_tx=XM_006932147.4;refGene_gn=THBS2;refGene_tx=XM_019831507.2
B2	154200555	154200653
refGene_gn=THBS2;refGene_tx=XM_006932146.4;refGene_gn=THBS2;refGene_tx=XM_006932147.4;refGene_gn=THBS2;refGene_tx=XM_019831507.2
B2	154201110	154201382
refGene_gn=THBS2;refGene_tx=XM_006932146.4;refGene_gn=THBS2;refGene_tx=XM_006932147.4;refGene_gn=THBS2;refGene_tx=XM_019831507.2
B2	154201830	154202058
refGene_gn=THBS2;refGene_tx=XM_006932146.4;refGene_gn=THBS2;refGene_tx=XM_006932147.4;refGene_gn=THBS2;refGene_tx=XM_019831507.2
B2	154203547	154203782
refGene_gn=THBS2;refGene_tx=XM_006932146.4;refGene_gn=THBS2;refGene_tx=XM_006932147.4;refGene_gn=THBS2;refGene_tx=XM_019831507.2
B2	154204350	154204469
refGene_gn=THBS2;refGene_tx=XM_006932146.4;refGene_gn=THBS2;refGene_tx=XM_006932147.4;refGene_gn=THBS2;refGene_tx=XM_019831507.2
B2	154205711	154205871
refGene_gn=THBS2;refGene_tx=XM_006932146.4;refGene_gn=THBS2;refGene_tx=XM_006932147.4;refGene_gn=THBS2;refGene_tx=XM_019831507.2
B2	154206482	154206590
refGene_gn=THBS2;refGene_tx=XM_006932146.4;refGene_gn=THBS2;refGene_tx=XM_006932147.4;refGene_gn=THBS2;refGene_tx=XM_019831507.2
B2	154207521	154207740
refGene_gn=THBS2;refGene_tx=XM_006932146.4;refGene_gn=THBS2;refGene_tx=XM_006932147.4;refGene_gn=THBS2;refGene_tx=XM_019831507.2
B2	154208195	154208348
refGene_gn=THBS2;refGene_tx=XM_006932146.4;refGene_gn=THBS2;refGene_tx=XM_006932147.4;refGene_gn=THBS2;refGene_tx=XM_019831507.2
B2	154208418	154208546
refGene_gn=THBS2;refGene_tx=XM_006932146.4;refGene_gn=THBS2;refGene_tx=XM_006932147.4;refGene_gn=THBS2;refGene_tx=XM_019831507.2
B2	154209227	154209401
refGene_gn=THBS2;refGene_tx=XM_006932146.4;refGene_gn=THBS2;refGene_tx=XM_006932147.4;refGene_gn=THBS2;refGene_tx=XM_019831507.2

B2 154210068 154210245
refGene_gn=THBS2;refGene_tx=XM_006932146.4;refGene_gn=THBS2;refGene_tx=XM_006932147.4;refGene_gn=THBS2;refGene_tx=XM_019831507.2
B2 154210384 154210555
refGene_gn=THBS2;refGene_tx=XM_006932146.4;refGene_gn=THBS2;refGene_tx=XM_006932147.4;refGene_gn=THBS2;refGene_tx=XM_019831507.2
B2 154211008 154211105
refGene_gn=THBS2;refGene_tx=XM_006932146.4;refGene_gn=THBS2;refGene_tx=XM_006932147.4;refGene_gn=THBS2;refGene_tx=XM_019831507.2
B2 154211637 154211778
refGene_gn=THBS2;refGene_tx=XM_006932146.4;refGene_gn=THBS2;refGene_tx=XM_006932147.4;refGene_gn=THBS2;refGene_tx=XM_019831507.2
B2 154212796 154212993
refGene_gn=THBS2;refGene_tx=XM_006932146.4;refGene_gn=THBS2;refGene_tx=XM_006932147.4;refGene_gn=THBS2;refGene_tx=XM_019831507.2
B2 154215457 154215542
refGene_gn=THBS2;refGene_tx=XM_006932146.4;refGene_gn=THBS2;refGene_tx=XM_006932147.4;refGene_gn=THBS2;refGene_tx=XM_019831507.2
B2 154216704 154217261
refGene_gn=THBS2;refGene_tx=XM_006932146.4;refGene_gn=THBS2;refGene_tx=XM_006932147.4;refGene_gn=THBS2;refGene_tx=XM_019831507.2
B2 154218886 154218938
refGene_gn=THBS2;refGene_tx=XM_006932146.4;refGene_gn=THBS2;refGene_tx=XM_006932147.4;refGene_gn=THBS2;refGene_tx=XM_019831507.2
B2 154346788 154346854
refGene_gn=WDR27;refGene_tx=XM_019831511.2
B2 154369801 154370024
refGene_gn=WDR27;refGene_tx=XM_019831508.2;refGene_gn=WDR27;refGene_tx=XM_019831510.2;refGene_gn=WDR27;refGene_tx=XM_019831517.2;refGene_gn=WDR27;refGene_tx=XM_019831520.2;refGene_gn=WDR27;refGene_tx=XM_023254615.1
B2 154371868 154372783
refGene_gn=LOC111560558;refGene_tx=XM_023254622.1
B2 154440593 154440599
refGene_gn=WDR27;refGene_tx=XM_019831519.2
B2 154445144 154445287
refGene_gn=WDR27;refGene_tx=XM_019831508.2;refGene_gn=WDR27;refGene_tx=XM_019831510.2;refGene_gn=WDR27;refGene_tx=XM_023254615.1
B2 154445415 154445514
refGene_gn=WDR27;refGene_tx=XM_019831508.2;refGene_gn=WDR27;refGene_tx=XM_019831510.2;refGene_gn=WDR27;refGene_tx=XM_019831511.2;refGene_gn=WDR27;refGene_tx=XM_019831517.2;refGene_gn=WDR27;refGene_tx=XM_019831519.2;refGene_gn=WDR27;refGene_tx=XM_023254615.1
B2 154459332 154459435
refGene_gn=WDR27;refGene_tx=XM_019831508.2;refGene_gn=WDR27;refGene_tx=XM_019831510.2;refGene_gn=WDR27;refGene_tx=XM_019831511.2;refGene_gn=WDR27;refGene_tx=XM_019831517.2;refGene_gn=WDR27;refGene_tx=XM_019831519.2;refGene_gn=WDR27;refGene_tx=XM_019831520.2;refGene_gn=WDR27;refGene_tx=XM_023254615.1
B2 154460414 154460512

refGene_gn=WDR27;refGene_tx=XM_019831508.2;refGene_gn=WDR27;refGene_tx
=XM_019831510.2;refGene_gn=WDR27;refGene_tx=XM_019831511.2;refGene_gn=
WDR27;refGene_tx=XM_019831517.2;refGene_gn=WDR27;refGene_tx=XM_0198315
19.2;refGene_gn=WDR27;refGene_tx=XM_019831520.2;refGene_gn=WDR27;refGe
ne_tx=XM_023254615.1
B2 154470649 154470820
refGene_gn=WDR27;refGene_tx=XM_019831522.2
B2 154475632 154475754
refGene_gn=WDR27;refGene_tx=XM_019831508.2;refGene_gn=WDR27;refGene_tx
=XM_019831510.2;refGene_gn=WDR27;refGene_tx=XM_019831511.2;refGene_gn=
WDR27;refGene_tx=XM_019831517.2;refGene_gn=WDR27;refGene_tx=XM_0198315
19.2;refGene_gn=WDR27;refGene_tx=XM_019831520.2;refGene_gn=WDR27;refGe
ne_tx=XM_019831522.2;refGene_gn=WDR27;refGene_tx=XM_023254615.1
B2 154476849 154476947
refGene_gn=WDR27;refGene_tx=XM_019831508.2;refGene_gn=WDR27;refGene_tx
=XM_019831510.2;refGene_gn=WDR27;refGene_tx=XM_019831511.2;refGene_gn=
WDR27;refGene_tx=XM_019831517.2;refGene_gn=WDR27;refGene_tx=XM_0198315
19.2;refGene_gn=WDR27;refGene_tx=XM_019831520.2;refGene_gn=WDR27;refGe
ne_tx=XM_019831522.2;refGene_gn=WDR27;refGene_tx=XM_023254615.1
B2 154481494 154481628
refGene_gn=WDR27;refGene_tx=XM_019831508.2;refGene_gn=WDR27;refGene_tx
=XM_019831510.2;refGene_gn=WDR27;refGene_tx=XM_019831511.2;refGene_gn=
WDR27;refGene_tx=XM_019831517.2;refGene_gn=WDR27;refGene_tx=XM_0198315
19.2;refGene_gn=WDR27;refGene_tx=XM_019831520.2;refGene_gn=WDR27;refGe
ne_tx=XM_019831522.2;refGene_gn=WDR27;refGene_tx=XM_023254615.1
B2 154482466 154482588
refGene_gn=WDR27;refGene_tx=XM_019831508.2;refGene_gn=WDR27;refGene_tx
=XM_019831510.2;refGene_gn=WDR27;refGene_tx=XM_019831511.2;refGene_gn=
WDR27;refGene_tx=XM_019831517.2;refGene_gn=WDR27;refGene_tx=XM_0198315
19.2;refGene_gn=WDR27;refGene_tx=XM_019831520.2;refGene_gn=WDR27;refGe
ne_tx=XM_019831522.2;refGene_gn=WDR27;refGene_tx=XM_023254615.1
B2 154486858 154486948
refGene_gn=WDR27;refGene_tx=XM_019831508.2;refGene_gn=WDR27;refGene_tx
=XM_019831510.2;refGene_gn=WDR27;refGene_tx=XM_019831511.2;refGene_gn=
WDR27;refGene_tx=XM_019831517.2;refGene_gn=WDR27;refGene_tx=XM_0198315
19.2;refGene_gn=WDR27;refGene_tx=XM_019831520.2;refGene_gn=WDR27;refGe
ne_tx=XM_019831522.2;refGene_gn=WDR27;refGene_tx=XM_023254615.1
B2 154488153 154488251
refGene_gn=WDR27;refGene_tx=XM_019831508.2;refGene_gn=WDR27;refGene_tx
=XM_019831510.2;refGene_gn=WDR27;refGene_tx=XM_019831511.2;refGene_gn=
WDR27;refGene_tx=XM_019831517.2;refGene_gn=WDR27;refGene_tx=XM_0198315
19.2;refGene_gn=WDR27;refGene_tx=XM_019831520.2;refGene_gn=WDR27;refGe
ne_tx=XM_019831522.2;refGene_gn=WDR27;refGene_tx=XM_023254615.1
B2 154491300 154491378
refGene_gn=WDR27;refGene_tx=XM_019831508.2;refGene_gn=WDR27;refGene_tx
=XM_019831510.2;refGene_gn=WDR27;refGene_tx=XM_019831511.2;refGene_gn=
WDR27;refGene_tx=XM_019831517.2;refGene_gn=WDR27;refGene_tx=XM_0198315
19.2;refGene_gn=WDR27;refGene_tx=XM_019831520.2;refGene_gn=WDR27;refGe
ne_tx=XM_019831522.2;refGene_gn=WDR27;refGene_tx=XM_023254615.1
B2 154493751 154493830

refGene_gn=WDR27;refGene_tx=XM_019831508.2;refGene_gn=WDR27;refGene_tx=XM_019831510.2;refGene_gn=WDR27;refGene_tx=XM_019831511.2;refGene_gn=WDR27;refGene_tx=XM_019831517.2;refGene_gn=WDR27;refGene_tx=XM_019831519.2;refGene_gn=WDR27;refGene_tx=XM_019831520.2;refGene_gn=WDR27;refGene_tx=XM_019831522.2;refGene_gn=WDR27;refGene_tx=XM_023254615.1
B2 154499083 154499166
refGene_gn=WDR27;refGene_tx=XM_019831508.2;refGene_gn=WDR27;refGene_tx=XM_019831510.2;refGene_gn=WDR27;refGene_tx=XM_019831511.2;refGene_gn=WDR27;refGene_tx=XM_019831517.2;refGene_gn=WDR27;refGene_tx=XM_019831519.2;refGene_gn=WDR27;refGene_tx=XM_019831520.2;refGene_gn=WDR27;refGene_tx=XM_019831522.2;refGene_gn=WDR27;refGene_tx=XM_023254615.1
B2 154499783 154499905
refGene_gn=WDR27;refGene_tx=XM_019831508.2;refGene_gn=WDR27;refGene_tx=XM_019831510.2;refGene_gn=WDR27;refGene_tx=XM_019831511.2;refGene_gn=WDR27;refGene_tx=XM_019831517.2;refGene_gn=WDR27;refGene_tx=XM_019831519.2;refGene_gn=WDR27;refGene_tx=XM_019831520.2;refGene_gn=WDR27;refGene_tx=XM_019831522.2;refGene_gn=WDR27;refGene_tx=XM_023254615.1
B2 154500103 154500171
refGene_gn=WDR27;refGene_tx=XM_019831508.2;refGene_gn=WDR27;refGene_tx=XM_019831510.2;refGene_gn=WDR27;refGene_tx=XM_019831511.2;refGene_gn=WDR27;refGene_tx=XM_019831517.2;refGene_gn=WDR27;refGene_tx=XM_019831519.2;refGene_gn=WDR27;refGene_tx=XM_019831520.2;refGene_gn=WDR27;refGene_tx=XM_019831522.2;refGene_gn=WDR27;refGene_tx=XM_023254615.1
B2 154501184 154501288
refGene_gn=WDR27;refGene_tx=XM_019831508.2;refGene_gn=WDR27;refGene_tx=XM_019831510.2;refGene_gn=WDR27;refGene_tx=XM_019831511.2;refGene_gn=WDR27;refGene_tx=XM_019831517.2;refGene_gn=WDR27;refGene_tx=XM_019831519.2;refGene_gn=WDR27;refGene_tx=XM_019831520.2;refGene_gn=WDR27;refGene_tx=XM_019831522.2;refGene_gn=WDR27;refGene_tx=XM_023254615.1
B2 154502658 154502791
refGene_gn=WDR27;refGene_tx=XM_019831508.2;refGene_gn=WDR27;refGene_tx=XM_019831510.2;refGene_gn=WDR27;refGene_tx=XM_019831511.2;refGene_gn=WDR27;refGene_tx=XM_019831517.2;refGene_gn=WDR27;refGene_tx=XM_019831519.2;refGene_gn=WDR27;refGene_tx=XM_019831520.2;refGene_gn=WDR27;refGene_tx=XM_019831522.2;refGene_gn=WDR27;refGene_tx=XM_023254615.1
B2 154504426 154504544
refGene_gn=WDR27;refGene_tx=XM_019831508.2;refGene_gn=WDR27;refGene_tx=XM_019831510.2;refGene_gn=WDR27;refGene_tx=XM_019831511.2;refGene_gn=WDR27;refGene_tx=XM_019831517.2;refGene_gn=WDR27;refGene_tx=XM_019831519.2;refGene_gn=WDR27;refGene_tx=XM_019831520.2;refGene_gn=WDR27;refGene_tx=XM_019831522.2;refGene_gn=WDR27;refGene_tx=XM_023254615.1
B2 154506363 154506434
refGene_gn=WDR27;refGene_tx=XM_019831508.2;refGene_gn=WDR27;refGene_tx=XM_019831510.2;refGene_gn=WDR27;refGene_tx=XM_019831511.2;refGene_gn=WDR27;refGene_tx=XM_019831517.2;refGene_gn=WDR27;refGene_tx=XM_019831519.2;refGene_gn=WDR27;refGene_tx=XM_019831520.2;refGene_gn=WDR27;refGene_tx=XM_019831522.2;refGene_gn=WDR27;refGene_tx=XM_023254615.1
B2 154508048 154508100
refGene_gn=WDR27;refGene_tx=XM_019831508.2;refGene_gn=WDR27;refGene_tx=XM_019831510.2;refGene_gn=WDR27;refGene_tx=XM_019831511.2;refGene_gn=

WDR27;refGene_tx=XM_019831517.2;refGene_gn=WDR27;refGene_tx=XM_019831519.2;refGene_gn=WDR27;refGene_tx=XM_019831520.2;refGene_gn=WDR27;refGene_tx=XM_019831522.2;refGene_gn=WDR27;refGene_tx=XM_023254615.1
B2 154509310 154509424
refGene_gn=WDR27;refGene_tx=XM_019831508.2;refGene_gn=WDR27;refGene_tx=XM_019831510.2;refGene_gn=WDR27;refGene_tx=XM_019831511.2;refGene_gn=WDR27;refGene_tx=XM_019831517.2;refGene_gn=WDR27;refGene_tx=XM_019831519.2;refGene_gn=WDR27;refGene_tx=XM_019831520.2;refGene_gn=WDR27;refGene_tx=XM_019831522.2;refGene_gn=WDR27;refGene_tx=XM_023254615.1
B2 154511494 154511619
refGene_gn=WDR27;refGene_tx=XM_019831508.2;refGene_gn=WDR27;refGene_tx=XM_019831510.2;refGene_gn=WDR27;refGene_tx=XM_019831511.2;refGene_gn=WDR27;refGene_tx=XM_019831517.2;refGene_gn=WDR27;refGene_tx=XM_019831519.2;refGene_gn=WDR27;refGene_tx=XM_019831520.2;refGene_gn=WDR27;refGene_tx=XM_019831522.2;refGene_gn=WDR27;refGene_tx=XM_023254615.1
B2 154515927 154516069
refGene_gn=WDR27;refGene_tx=XM_019831508.2;refGene_gn=WDR27;refGene_tx=XM_019831510.2;refGene_gn=WDR27;refGene_tx=XM_019831511.2;refGene_gn=WDR27;refGene_tx=XM_019831517.2;refGene_gn=WDR27;refGene_tx=XM_019831519.2;refGene_gn=WDR27;refGene_tx=XM_019831520.2;refGene_gn=WDR27;refGene_tx=XM_019831522.2;refGene_gn=WDR27;refGene_tx=XM_023254615.1
B2 154521870 154522066
refGene_gn=WDR27;refGene_tx=XM_019831508.2;refGene_gn=WDR27;refGene_tx=XM_019831510.2;refGene_gn=WDR27;refGene_tx=XM_019831511.2;refGene_gn=WDR27;refGene_tx=XM_019831517.2;refGene_gn=WDR27;refGene_tx=XM_019831519.2;refGene_gn=WDR27;refGene_tx=XM_019831520.2;refGene_gn=WDR27;refGene_tx=XM_019831522.2;refGene_gn=WDR27;refGene_tx=XM_023254615.1
B2 154522288 154522433
refGene_gn=WDR27;refGene_tx=XM_019831508.2;refGene_gn=WDR27;refGene_tx=XM_019831511.2;refGene_gn=WDR27;refGene_tx=XM_019831517.2;refGene_gn=WDR27;refGene_tx=XM_019831519.2;refGene_gn=WDR27;refGene_tx=XM_019831520.2;refGene_gn=WDR27;refGene_tx=XM_019831522.2
B2 154524359 154524375
refGene_gn=WDR27;refGene_tx=XM_019831508.2;refGene_gn=WDR27;refGene_tx=XM_019831511.2;refGene_gn=WDR27;refGene_tx=XM_019831517.2;refGene_gn=WDR27;refGene_tx=XM_019831519.2;refGene_gn=WDR27;refGene_tx=XM_019831520.2;refGene_gn=WDR27;refGene_tx=XM_019831522.2
B2 154532676 154533231
refGene_gn=CB2H6orf120;refGene_tx=XM_023254624.1
B2 154534149 154534235
refGene_gn=PHF10;refGene_tx=XM_023254618.1;refGene_gn=PHF10;refGene_tx=XM_023254619.1;refGene_gn=PHF10;refGene_tx=XM_023254620.1;refGene_gn=PHF10;refGene_tx=XM_023254621.1
B2 154535302 154535491
refGene_gn=PHF10;refGene_tx=XM_023254618.1;refGene_gn=PHF10;refGene_tx=XM_023254619.1;refGene_gn=PHF10;refGene_tx=XM_023254620.1;refGene_gn=PHF10;refGene_tx=XM_023254621.1
B2 154535759 154535868
refGene_gn=PHF10;refGene_tx=XM_023254618.1;refGene_gn=PHF10;refGene_tx=XM_023254619.1;refGene_gn=PHF10;refGene_tx=XM_023254620.1;refGene_gn=

PHF10;refGene_tx=XM_023254621.1
B2 154540315 154540471
refGene_gn=PHF10;refGene_tx=XM_023254618.1;refGene_gn=PHF10;refGene_tx=XM_023254619.1;refGene_gn=PHF10;refGene_tx=XM_023254620.1;refGene_gn=PHF10;refGene_tx=XM_023254621.1
B2 154542691 154542845
refGene_gn=PHF10;refGene_tx=XM_023254618.1;refGene_gn=PHF10;refGene_tx=XM_023254619.1;refGene_gn=PHF10;refGene_tx=XM_023254620.1;refGene_gn=PHF10;refGene_tx=XM_023254621.1
B2 154544612 154544722
refGene_gn=PHF10;refGene_tx=XM_023254618.1;refGene_gn=PHF10;refGene_tx=XM_023254619.1;refGene_gn=PHF10;refGene_tx=XM_023254620.1;refGene_gn=PHF10;refGene_tx=XM_023254621.1
B2 154545360 154545510
refGene_gn=PHF10;refGene_tx=XM_023254618.1;refGene_gn=PHF10;refGene_tx=XM_023254619.1;refGene_gn=PHF10;refGene_tx=XM_023254620.1;refGene_gn=PHF10;refGene_tx=XM_023254621.1
B2 154545606 154545740
refGene_gn=PHF10;refGene_tx=XM_023254618.1;refGene_gn=PHF10;refGene_tx=XM_023254619.1;refGene_gn=PHF10;refGene_tx=XM_023254620.1;refGene_gn=PHF10;refGene_tx=XM_023254621.1
B2 154547829 154547913
refGene_gn=PHF10;refGene_tx=XM_023254618.1;refGene_gn=PHF10;refGene_tx=XM_023254619.1;refGene_gn=PHF10;refGene_tx=XM_023254620.1;refGene_gn=PHF10;refGene_tx=XM_023254621.1
B2 154548507 154548638
refGene_gn=PHF10;refGene_tx=XM_023254618.1;refGene_gn=PHF10;refGene_tx=XM_023254619.1
B2 154550973 154551080
refGene_gn=PHF10;refGene_tx=XM_023254618.1;refGene_gn=PHF10;refGene_tx=XM_023254619.1;refGene_gn=PHF10;refGene_tx=XM_023254620.1;refGene_gn=PHF10;refGene_tx=XM_023254621.1
B2 154553177 154553264
refGene_gn=PHF10;refGene_tx=XM_023254618.1;refGene_gn=PHF10;refGene_tx=XM_023254619.1
B2 154564161 154564272
refGene_gn=TCTE3;refGene_tx=XM_023254623.1
B2 154566600 154566759
refGene_gn=TCTE3;refGene_tx=XM_023254623.1
B2 154569166 154569373
refGene_gn=TCTE3;refGene_tx=XM_023254623.1
B2 154581848 154581956
refGene_gn=TCTE3;refGene_tx=XM_023254623.1
B2 154583516 154583588
refGene_gn=ERMARD;refGene_tx=XM_019831525.2;refGene_gn=ERMARD;refGene_tx=XM_019831527.2;refGene_gn=ERMARD;refGene_tx=XM_019831529.2;refGene_gn=ERMARD;refGene_tx=XM_019831530.2;refGene_gn=ERMARD;refGene_tx=XM_023254617.1
B2 154585378 154585547
refGene_gn=ERMARD;refGene_tx=XM_019831525.2;refGene_gn=ERMARD;refGene_

tx=XM_019831527.2;refGene_gn=ERMARD;refGene_tx=XM_019831528.2;refGene_gn=ERMARD;refGene_tx=XM_019831529.2;refGene_gn=ERMARD;refGene_tx=XM_019831530.2;refGene_gn=ERMARD;refGene_tx=XM_023254617.1
B2 154586060 154586200
refGene_gn=ERMARD;refGene_tx=XM_019831525.2;refGene_gn=ERMARD;refGene_tx=XM_019831527.2;refGene_gn=ERMARD;refGene_tx=XM_019831528.2;refGene_gn=ERMARD;refGene_tx=XM_019831529.2;refGene_gn=ERMARD;refGene_tx=XM_019831530.2;refGene_gn=ERMARD;refGene_tx=XM_023254617.1
B2 154587629 154587749
refGene_gn=ERMARD;refGene_tx=XM_019831525.2;refGene_gn=ERMARD;refGene_tx=XM_019831527.2;refGene_gn=ERMARD;refGene_tx=XM_019831528.2;refGene_gn=ERMARD;refGene_tx=XM_019831529.2;refGene_gn=ERMARD;refGene_tx=XM_019831530.2;refGene_gn=ERMARD;refGene_tx=XM_019831531.2;refGene_gn=ERMARD;refGene_tx=XM_023254617.1
B2 154588260 154588350
refGene_gn=ERMARD;refGene_tx=XM_019831525.2;refGene_gn=ERMARD;refGene_tx=XM_019831527.2;refGene_gn=ERMARD;refGene_tx=XM_019831528.2;refGene_gn=ERMARD;refGene_tx=XM_019831529.2;refGene_gn=ERMARD;refGene_tx=XM_019831530.2;refGene_gn=ERMARD;refGene_tx=XM_019831531.2;refGene_gn=ERMARD;refGene_tx=XM_023254617.1
B2 154589674 154589785
refGene_gn=ERMARD;refGene_tx=XM_019831532.2
B2 154590322 154590420
refGene_gn=ERMARD;refGene_tx=XM_019831525.2;refGene_gn=ERMARD;refGene_tx=XM_019831527.2;refGene_gn=ERMARD;refGene_tx=XM_019831528.2;refGene_gn=ERMARD;refGene_tx=XM_019831529.2;refGene_gn=ERMARD;refGene_tx=XM_019831530.2;refGene_gn=ERMARD;refGene_tx=XM_019831531.2;refGene_gn=ERMARD;refGene_tx=XM_019831532.2;refGene_gn=ERMARD;refGene_tx=XM_023254617.1
B2 154590837 154590974
refGene_gn=ERMARD;refGene_tx=XM_019831525.2;refGene_gn=ERMARD;refGene_tx=XM_019831527.2;refGene_gn=ERMARD;refGene_tx=XM_019831528.2;refGene_gn=ERMARD;refGene_tx=XM_019831529.2;refGene_gn=ERMARD;refGene_tx=XM_019831530.2;refGene_gn=ERMARD;refGene_tx=XM_019831531.2;refGene_gn=ERMARD;refGene_tx=XM_019831532.2;refGene_gn=ERMARD;refGene_tx=XM_023254617.1
B2 154592920 154593035
refGene_gn=ERMARD;refGene_tx=XM_019831525.2;refGene_gn=ERMARD;refGene_tx=XM_019831527.2;refGene_gn=ERMARD;refGene_tx=XM_019831528.2;refGene_gn=ERMARD;refGene_tx=XM_019831529.2;refGene_gn=ERMARD;refGene_tx=XM_019831530.2;refGene_gn=ERMARD;refGene_tx=XM_019831531.2;refGene_gn=ERMARD;refGene_tx=XM_019831532.2;refGene_gn=ERMARD;refGene_tx=XM_023254617.1
B2 154593776 154593879
refGene_gn=ERMARD;refGene_tx=XM_019831525.2;refGene_gn=ERMARD;refGene_tx=XM_019831527.2;refGene_gn=ERMARD;refGene_tx=XM_019831528.2;refGene_gn=ERMARD;refGene_tx=XM_019831529.2;refGene_gn=ERMARD;refGene_tx=XM_019831530.2;refGene_gn=ERMARD;refGene_tx=XM_019831531.2;refGene_gn=ERMARD;refGene_tx=XM_019831532.2;refGene_gn=ERMARD;refGene_tx=XM_023254617.1

B2 154594746 154595544
refGene_gn=ERMARD;refGene_tx=XM_019831530.2
B2 154596723 154596753
refGene_gn=ERMARD;refGene_tx=XM_019831525.2;refGene_gn=ERMARD;refGene_tx=XM_019831527.2;refGene_gn=ERMARD;refGene_tx=XM_019831528.2;refGene_gn=ERMARD;refGene_tx=XM_019831529.2;refGene_gn=ERMARD;refGene_tx=XM_019831531.2;refGene_gn=ERMARD;refGene_tx=XM_019831532.2
B2 154598486 154598555
refGene_gn=ERMARD;refGene_tx=XM_019831525.2;refGene_gn=ERMARD;refGene_tx=XM_019831527.2;refGene_gn=ERMARD;refGene_tx=XM_019831528.2;refGene_gn=ERMARD;refGene_tx=XM_019831529.2;refGene_gn=ERMARD;refGene_tx=XM_019831531.2;refGene_gn=ERMARD;refGene_tx=XM_019831532.2;refGene_gn=ERMARD;refGene_tx=XM_023254617.1
B2 154599914 154600088
refGene_gn=ERMARD;refGene_tx=XM_019831525.2;refGene_gn=ERMARD;refGene_tx=XM_019831527.2;refGene_gn=ERMARD;refGene_tx=XM_019831528.2;refGene_gn=ERMARD;refGene_tx=XM_019831529.2;refGene_gn=ERMARD;refGene_tx=XM_019831531.2;refGene_gn=ERMARD;refGene_tx=XM_019831532.2;refGene_gn=ERMARD;refGene_tx=XM_023254617.1
B2 154603154 154603238
refGene_gn=ERMARD;refGene_tx=XM_019831525.2;refGene_gn=ERMARD;refGene_tx=XM_019831527.2;refGene_gn=ERMARD;refGene_tx=XM_019831528.2;refGene_gn=ERMARD;refGene_tx=XM_019831529.2;refGene_gn=ERMARD;refGene_tx=XM_019831531.2;refGene_gn=ERMARD;refGene_tx=XM_019831532.2;refGene_gn=ERMARD;refGene_tx=XM_023254617.1
B2 154604903 154604977
refGene_gn=ERMARD;refGene_tx=XM_019831525.2;refGene_gn=ERMARD;refGene_tx=XM_019831527.2;refGene_gn=ERMARD;refGene_tx=XM_019831528.2;refGene_gn=ERMARD;refGene_tx=XM_019831529.2;refGene_gn=ERMARD;refGene_tx=XM_019831531.2;refGene_gn=ERMARD;refGene_tx=XM_019831532.2;refGene_gn=ERMARD;refGene_tx=XM_023254617.1
B2 154605564 154605690
refGene_gn=ERMARD;refGene_tx=XM_019831525.2;refGene_gn=ERMARD;refGene_tx=XM_019831527.2;refGene_gn=ERMARD;refGene_tx=XM_019831528.2;refGene_gn=ERMARD;refGene_tx=XM_019831529.2;refGene_gn=ERMARD;refGene_tx=XM_019831531.2;refGene_gn=ERMARD;refGene_tx=XM_019831532.2;refGene_gn=ERMARD;refGene_tx=XM_023254617.1
B2 154606017 154606236
refGene_gn=ERMARD;refGene_tx=XM_019831525.2;refGene_gn=ERMARD;refGene_tx=XM_019831527.2;refGene_gn=ERMARD;refGene_tx=XM_019831528.2;refGene_gn=ERMARD;refGene_tx=XM_019831529.2;refGene_gn=ERMARD;refGene_tx=XM_019831531.2;refGene_gn=ERMARD;refGene_tx=XM_019831532.2;refGene_gn=ERMARD;refGene_tx=XM_023254617.1
B2 154607137 154607251
refGene_gn=ERMARD;refGene_tx=XM_019831525.2;refGene_gn=ERMARD;refGene_tx=XM_019831527.2;refGene_gn=ERMARD;refGene_tx=XM_019831528.2;refGene_gn=ERMARD;refGene_tx=XM_019831529.2;refGene_gn=ERMARD;refGene_tx=XM_019831531.2;refGene_gn=ERMARD;refGene_tx=XM_019831532.2;refGene_gn=ERMARD;refGene_tx=XM_023254617.1
B2 154608694 154608878

refGene_gn=ERMARD;refGene_tx=XM_019831525.2;refGene_gn=ERMARD;refGene_tx=XM_019831527.2;refGene_gn=ERMARD;refGene_tx=XM_019831528.2;refGene_gn=ERMARD;refGene_tx=XM_019831531.2;refGene_gn=ERMARD;refGene_tx=XM_019831532.2;refGene_gn=ERMARD;refGene_tx=XM_023254617.1
B2 154624958 154624986
refGene_gn=ERMARD;refGene_tx=XM_019831529.2
B2 154982645 154982651
refGene_gn=DLL1;refGene_tx=XM_023254631.1
B2 154982760 154982878
refGene_gn=DLL1;refGene_tx=XM_023254631.1
B2 154983077 154983882
refGene_gn=DLL1;refGene_tx=XM_023254631.1
B2 154985430 154985647
refGene_gn=DLL1;refGene_tx=XM_023254631.1
B2 154985760 154985930
refGene_gn=DLL1;refGene_tx=XM_023254631.1
B2 154986053 154986184
refGene_gn=DLL1;refGene_tx=XM_023254631.1
B2 154986707 154986768
refGene_gn=DLL1;refGene_tx=XM_023254631.1
B2 154988611 154988869
refGene_gn=DLL1;refGene_tx=XM_023254631.1
B2 154989109 154989170
refGene_gn=DLL1;refGene_tx=XM_023254631.1
B2 154989868 154990165
refGene_gn=DLL1;refGene_tx=XM_023254631.1
B2 154990373 154990427
refGene_gn=DLL1;refGene_tx=XM_023254631.1
B2 155008158 155008239
refGene_gn=FAM120B;refGene_tx=XM_019831534.2;refGene_gn=FAM120B;refGene_tx=XM_019831535.2;refGene_gn=FAM120B;refGene_tx=XM_019831536.2;refGene_gn=FAM120B;refGene_tx=XM_019831540.2;refGene_gn=FAM120B;refGene_tx=XM_019831541.2;refGene_gn=FAM120B;refGene_tx=XM_019831542.2;refGene_gn=FAM120B;refGene_tx=XM_023254626.1;refGene_gn=FAM120B;refGene_tx=XM_023254627.1;refGene_gn=FAM120B;refGene_tx=XM_023254628.1;refGene_gn=FAM120B;refGene_tx=XM_023254629.1;refGene_gn=FAM120B;refGene_tx=XM_023254630.1
B2 155019369 155021091
refGene_gn=FAM120B;refGene_tx=XM_019831534.2;refGene_gn=FAM120B;refGene_tx=XM_019831535.2;refGene_gn=FAM120B;refGene_tx=XM_019831536.2;refGene_gn=FAM120B;refGene_tx=XM_019831537.1;refGene_gn=FAM120B;refGene_tx=XM_019831538.1;refGene_gn=FAM120B;refGene_tx=XM_019831540.2;refGene_gn=FAM120B;refGene_tx=XM_019831541.2;refGene_gn=FAM120B;refGene_tx=XM_019831542.2;refGene_gn=FAM120B;refGene_tx=XM_023254626.1;refGene_gn=FAM120B;refGene_tx=XM_023254627.1;refGene_gn=FAM120B;refGene_tx=XM_023254628.1;refGene_gn=FAM120B;refGene_tx=XM_023254629.1;refGene_gn=FAM120B;refGene_tx=XM_023254630.1
B2 155021715 155021736
refGene_gn=FAM120B;refGene_tx=XM_023254630.1
B2 155024082 155024263

refGene_gn=FAM120B;refGene_tx=XM_019831534.2;refGene_gn=FAM120B;refGene_tx=XM_019831535.2;refGene_gn=FAM120B;refGene_tx=XM_019831536.2;refGene_gn=FAM120B;refGene_tx=XM_019831537.1;refGene_gn=FAM120B;refGene_tx=XM_019831538.1;refGene_gn=FAM120B;refGene_tx=XM_019831540.2;refGene_gn=FAM120B;refGene_tx=XM_019831541.2;refGene_gn=FAM120B;refGene_tx=XM_019831542.2;refGene_gn=FAM120B;refGene_tx=XM_019831545.1;refGene_gn=FAM120B;refGene_tx=XM_023254626.1;refGene_gn=FAM120B;refGene_tx=XM_023254627.1;refGene_gn=FAM120B;refGene_tx=XM_023254628.1;refGene_gn=FAM120B;refGene_tx=XM_023254629.1
B2 155030360 155030462
refGene_gn=FAM120B;refGene_tx=XM_019831534.2;refGene_gn=FAM120B;refGene_tx=XM_019831535.2;refGene_gn=FAM120B;refGene_tx=XM_019831537.1;refGene_gn=FAM120B;refGene_tx=XM_019831538.1;refGene_gn=FAM120B;refGene_tx=XM_019831540.2;refGene_gn=FAM120B;refGene_tx=XM_019831541.2;refGene_gn=FAM120B;refGene_tx=XM_019831542.2;refGene_gn=FAM120B;refGene_tx=XM_019831545.1;refGene_gn=FAM120B;refGene_tx=XM_023254626.1;refGene_gn=FAM120B;refGene_tx=XM_023254627.1;refGene_gn=FAM120B;refGene_tx=XM_023254628.1;refGene_gn=FAM120B;refGene_tx=XM_023254629.1
B2 155039259 155039432
refGene_gn=FAM120B;refGene_tx=XM_019831534.2;refGene_gn=FAM120B;refGene_tx=XM_019831535.2;refGene_gn=FAM120B;refGene_tx=XM_019831536.2;refGene_gn=FAM120B;refGene_tx=XM_019831537.1;refGene_gn=FAM120B;refGene_tx=XM_019831538.1;refGene_gn=FAM120B;refGene_tx=XM_019831540.2;refGene_gn=FAM120B;refGene_tx=XM_019831541.2;refGene_gn=FAM120B;refGene_tx=XM_019831542.2;refGene_gn=FAM120B;refGene_tx=XM_019831545.1;refGene_gn=FAM120B;refGene_tx=XM_023254626.1;refGene_gn=FAM120B;refGene_tx=XM_023254627.1;refGene_gn=FAM120B;refGene_tx=XM_023254628.1;refGene_gn=FAM120B;refGene_tx=XM_023254629.1
B2 155044015 155044108
refGene_gn=FAM120B;refGene_tx=XM_019831534.2;refGene_gn=FAM120B;refGene_tx=XM_019831535.2;refGene_gn=FAM120B;refGene_tx=XM_019831536.2;refGene_gn=FAM120B;refGene_tx=XM_019831537.1;refGene_gn=FAM120B;refGene_tx=XM_019831538.1;refGene_gn=FAM120B;refGene_tx=XM_019831540.2;refGene_gn=FAM120B;refGene_tx=XM_019831541.2;refGene_gn=FAM120B;refGene_tx=XM_019831542.2;refGene_gn=FAM120B;refGene_tx=XM_019831545.1;refGene_gn=FAM120B;refGene_tx=XM_023254626.1;refGene_gn=FAM120B;refGene_tx=XM_023254627.1;refGene_gn=FAM120B;refGene_tx=XM_023254628.1;refGene_gn=FAM120B;refGene_tx=XM_023254629.1
B2 155064648 155064717
refGene_gn=FAM120B;refGene_tx=XM_019831542.2
B2 155072324 155072531
refGene_gn=FAM120B;refGene_tx=XM_019831534.2;refGene_gn=FAM120B;refGene_tx=XM_019831535.2;refGene_gn=FAM120B;refGene_tx=XM_019831536.2;refGene_gn=FAM120B;refGene_tx=XM_019831537.1;refGene_gn=FAM120B;refGene_tx=XM_019831538.1;refGene_gn=FAM120B;refGene_tx=XM_019831540.2;refGene_gn=FAM120B;refGene_tx=XM_019831541.2;refGene_gn=FAM120B;refGene_tx=XM_019831545.1;refGene_gn=FAM120B;refGene_tx=XM_023254626.1;refGene_gn=FAM120B;refGene_tx=XM_023254627.1;refGene_gn=FAM120B;refGene_tx=XM_023254628.1;refGene_gn=FAM120B;refGene_tx=XM_023254629.1
B2 155073862 155073971

refGene_gn=FAM120B;refGene_tx=XM_019831534.2;refGene_gn=FAM120B;refGen
e_tx=XM_019831535.2;refGene_gn=FAM120B;refGene_tx=XM_019831536.2;refGe
ne_gn=FAM120B;refGene_tx=XM_019831537.1;refGene_gn=FAM120B;refGene_tx=
XM_019831538.1;refGene_gn=FAM120B;refGene_tx=XM_019831545.1;refGene_gn
=FAM120B;refGene_tx=XM_023254626.1;refGene_gn=FAM120B;refGene_tx=XM_02
3254627.1;refGene_gn=FAM120B;refGene_tx=XM_023254628.1
B2 155075830 155075923
refGene_gn=FAM120B;refGene_tx=XM_019831534.2;refGene_gn=FAM120B;refGen
e_tx=XM_019831536.2;refGene_gn=FAM120B;refGene_tx=XM_019831537.1;refGe
ne_gn=FAM120B;refGene_tx=XM_019831538.1;refGene_gn=FAM120B;refGene_tx=
XM_019831545.1;refGene_gn=FAM120B;refGene_tx=XM_023254626.1;refGene_gn
=FAM120B;refGene_tx=XM_023254627.1
B2 155080748 155080797
refGene_gn=FAM120B;refGene_tx=XM_019831534.2;refGene_gn=FAM120B;refGen
e_tx=XM_019831535.2;refGene_gn=FAM120B;refGene_tx=XM_019831536.2;refGe
ne_gn=FAM120B;refGene_tx=XM_019831537.1;refGene_gn=FAM120B;refGene_tx=
XM_019831538.1;refGene_gn=FAM120B;refGene_tx=XM_019831540.2;refGene_gn
=FAM120B;refGene_tx=XM_019831541.2;refGene_gn=FAM120B;refGene_tx=XM_01
9831545.1;refGene_gn=FAM120B;refGene_tx=XM_023254626.1;refGene_gn=FAM1
20B;refGene_tx=XM_023254627.1;refGene_gn=FAM120B;refGene_tx=XM_0232546
28.1;refGene_gn=FAM120B;refGene_tx=XM_023254629.1
B2 155081331 155081372
refGene_gn=FAM120B;refGene_tx=XM_023254627.1;refGene_gn=FAM120B;refGen
e_tx=XM_023254628.1;refGene_gn=FAM120B;refGene_tx=XM_023254629.1
B2 155093032 155093306
refGene_gn=FAM120B;refGene_tx=XM_019831534.2;refGene_gn=FAM120B;refGen
e_tx=XM_019831535.2;refGene_gn=FAM120B;refGene_tx=XM_019831536.2;refGe
ne_gn=FAM120B;refGene_tx=XM_019831538.1;refGene_gn=FAM120B;refGene_tx=
XM_019831540.2;refGene_gn=FAM120B;refGene_tx=XM_019831545.1;refGene_gn
=FAM120B;refGene_tx=XM_023254626.1
B2 155095871 155095924
refGene_gn=FAM120B;refGene_tx=XM_019831537.1;refGene_gn=FAM120B;refGen
e_tx=XM_019831541.2
B2 155180922 155181108
refGene_gn=PSMB1;refGene_tx=XM_003986715.4
B2 155182112 155182219
refGene_gn=PSMB1;refGene_tx=XM_003986715.4
B2 155187880 155188010
refGene_gn=PSMB1;refGene_tx=XM_003986715.4
B2 155190307 155190389
refGene_gn=PSMB1;refGene_tx=XM_003986715.4
B2 155193901 155194009
refGene_gn=PSMB1;refGene_tx=XM_003986715.4
B2 155197801 155197914
refGene_gn=PSMB1;refGene_tx=XM_003986715.4
B2 155211936 155211990
refGene_gn=TBP;refGene_tx=XM_011282689.3;refGene_gn=TBP;refGene_tx=XM_
011282690.3;refGene_gn=TBP;refGene_tx=XM_019831546.2
B2 155214668 155215075
refGene_gn=TBP;refGene_tx=XM_011282689.3;refGene_gn=TBP;refGene_tx=XM_

011282690.3;refGene_gn=TBP;refGene_tx=XM_019831546.2
B2	155220721	155220809
refGene_gn=TBP;refGene_tx=XM_011282689.3;refGene_gn=TBP;refGene_tx=XM_
011282690.3;refGene_gn=TBP;refGene_tx=XM_019831546.2
B2	155226326	155226418
refGene_gn=TBP;refGene_tx=XM_011282689.3;refGene_gn=TBP;refGene_tx=XM_
011282690.3;refGene_gn=TBP;refGene_tx=XM_019831546.2
B2	155228032	155228200
refGene_gn=TBP;refGene_tx=XM_011282689.3;refGene_gn=TBP;refGene_tx=XM_
011282690.3;refGene_gn=TBP;refGene_tx=XM_019831546.2
B2	155229101	155229196
refGene_gn=TBP;refGene_tx=XM_011282689.3;refGene_gn=TBP;refGene_tx=XM_
011282690.3;refGene_gn=TBP;refGene_tx=XM_019831546.2
B2	155229856	155229936
refGene_gn=TBP;refGene_tx=XM_011282689.3;refGene_gn=TBP;refGene_tx=XM_
011282690.3;refGene_gn=TBP;refGene_tx=XM_019831546.2
B2	155233431	155233590
refGene_gn=PDCD2;refGene_tx=XM_023254632.1
B2	155234356	155234470
refGene_gn=PDCD2;refGene_tx=XM_023254632.1
B2	155235837	155235941
refGene_gn=PDCD2;refGene_tx=XM_023254632.1
B2	155237276	155237408
refGene_gn=PDCD2;refGene_tx=XM_023254632.1
B2	155237766	155238009
refGene_gn=PDCD2;refGene_tx=XM_023254632.1
B2	155238546	155238829
refGene_gn=PDCD2;refGene_tx=XM_023254632.1
B3	169687	169742	refGene_gn=CHRNB4;refGene_tx=XM_011282697.3
B3	173653	173802	refGene_gn=CHRNB4;refGene_tx=XM_011282697.3
B3	180080	180125
refGene_gn=CHRNB4;refGene_tx=XM_011282697.3;refGene_gn=CHRNB4;refGene_
tx=XM_019831740.2
B3	180227	180337
refGene_gn=CHRNB4;refGene_tx=XM_011282697.3;refGene_gn=CHRNB4;refGene_
tx=XM_019831740.2
B3	181828	182777
refGene_gn=CHRNB4;refGene_tx=XM_011282697.3;refGene_gn=CHRNB4;refGene_
tx=XM_019831740.2
B3	185944	186085
refGene_gn=CHRNB4;refGene_tx=XM_011282697.3;refGene_gn=CHRNB4;refGene_
tx=XM_019831740.2
B3	188896	188993	refGene_gn=CHRNA3;refGene_tx=XM_023254770.1
B3	190409	190549	refGene_gn=CHRNA3;refGene_tx=XM_023254770.1
B3	190661	190706	refGene_gn=CHRNA3;refGene_tx=XM_023254770.1
B3	191809	191919	refGene_gn=CHRNA3;refGene_tx=XM_023254770.1
B3	196707	197719	refGene_gn=CHRNA3;refGene_tx=XM_023254770.1
B3	203473	203602	refGene_gn=CHRNA3;refGene_tx=XM_023254770.1
B3	207159	207192	refGene_gn=CHRNA5;refGene_tx=XM_023254772.1
B3	207341	207503

refGene_gn=CHRNA5;refGene_tx=XM_019831741.2;refGene_gn=CHRNA5;refGene_tx=XM_023254771.1;refGene_gn=CHRNA5;refGene_tx=XM_023254772.1
B3 211015 211847
refGene_gn=CHRNA5;refGene_tx=XM_019831741.2;refGene_gn=CHRNA5;refGene_tx=XM_023254771.1;refGene_gn=CHRNA5;refGene_tx=XM_023254772.1;refGene_gn=CHRNA5;refGene_tx=XM_023254773.1;refGene_gn=CHRNA5;refGene_tx=XM_023254774.1
B3 213682 213792
refGene_gn=CHRNA5;refGene_tx=XM_019831741.2;refGene_gn=CHRNA5;refGene_tx=XM_023254771.1;refGene_gn=CHRNA5;refGene_tx=XM_023254772.1;refGene_gn=CHRNA5;refGene_tx=XM_023254773.1;refGene_gn=CHRNA5;refGene_tx=XM_023254774.1
B3 214896 214941
refGene_gn=CHRNA5;refGene_tx=XM_019831741.2;refGene_gn=CHRNA5;refGene_tx=XM_023254771.1;refGene_gn=CHRNA5;refGene_tx=XM_023254772.1;refGene_gn=CHRNA5;refGene_tx=XM_023254773.1;refGene_gn=CHRNA5;refGene_tx=XM_023254774.1
B3 220467 220619
refGene_gn=CHRNA5;refGene_tx=XM_019831741.2;refGene_gn=CHRNA5;refGene_tx=XM_023254771.1;refGene_gn=CHRNA5;refGene_tx=XM_023254772.1;refGene_gn=CHRNA5;refGene_tx=XM_023254773.1;refGene_gn=CHRNA5;refGene_tx=XM_023254774.1
B3 256044 256066 refGene_gn=CHRNA5;refGene_tx=XM_019831741.2
B3 256706 256833
refGene_gn=CHRNA5;refGene_tx=XM_023254771.1;refGene_gn=CHRNA5;refGene_tx=XM_023254772.1;refGene_gn=CHRNA5;refGene_tx=XM_023254773.1;refGene_gn=CHRNA5;refGene_tx=XM_023254774.1
B3 272677 272832 refGene_gn=PSMA4;refGene_tx=XM_011282699.3
B3 273942 274066 refGene_gn=PSMA4;refGene_tx=XM_011282699.3
B3 274788 274919 refGene_gn=PSMA4;refGene_tx=XM_011282699.3
B3 275362 275451 refGene_gn=PSMA4;refGene_tx=XM_011282699.3
B3 276723 276801 refGene_gn=PSMA4;refGene_tx=XM_011282699.3
B3 278323 278486 refGene_gn=PSMA4;refGene_tx=XM_011282699.3
B3 278876 278919 refGene_gn=PSMA4;refGene_tx=XM_011282699.3
B3 279161 279164 refGene_gn=PSMA4;refGene_tx=XM_011282699.3
B3 285177 285638
refGene_gn=HYKK;refGene_tx=XM_003986734.5;refGene_gn=HYKK;refGene_tx=XM_006932166.4
B3 289187 289371
refGene_gn=HYKK;refGene_tx=XM_003986734.5;refGene_gn=HYKK;refGene_tx=XM_006932166.4
B3 294427 294567
refGene_gn=HYKK;refGene_tx=XM_003986734.5;refGene_gn=HYKK;refGene_tx=XM_006932166.4
B3 295608 295954
refGene_gn=HYKK;refGene_tx=XM_003986734.5;refGene_gn=HYKK;refGene_tx=XM_006932166.4
B3 306541 306652 refGene_gn=IREB2;refGene_tx=XM_023254775.1
B3 307749 307935 refGene_gn=IREB2;refGene_tx=XM_023254775.1
B3 309892 310015 refGene_gn=IREB2;refGene_tx=XM_023254775.1

B3 310101 310249 refGene_gn=IREB2;refGene_tx=XM_023254775.1
B3 312725 312868 refGene_gn=IREB2;refGene_tx=XM_023254775.1
B3 312962 313067 refGene_gn=IREB2;refGene_tx=XM_023254775.1
B3 314104 314229 refGene_gn=IREB2;refGene_tx=XM_023254775.1
B3 314890 315047 refGene_gn=IREB2;refGene_tx=XM_023254775.1
B3 315421 315506 refGene_gn=IREB2;refGene_tx=XM_023254775.1
B3 316623 316759 refGene_gn=IREB2;refGene_tx=XM_023254775.1
B3 317756 317916 refGene_gn=IREB2;refGene_tx=XM_023254775.1
B3 319002 319119 refGene_gn=IREB2;refGene_tx=XM_023254775.1
B3 322592 322631 refGene_gn=IREB2;refGene_tx=XM_023254775.1
B3 322632 322694 refGene_gn=IREB2;refGene_tx=XM_023254775.1
B3 324523 324695 refGene_gn=IREB2;refGene_tx=XM_023254775.1
B3 327654 327794 refGene_gn=IREB2;refGene_tx=XM_023254775.1
B3 328513 328697 refGene_gn=IREB2;refGene_tx=XM_023254775.1
B3 329137 329207 refGene_gn=IREB2;refGene_tx=XM_023254775.1
B3 332273 332492 refGene_gn=IREB2;refGene_tx=XM_023254775.1
B3 333422 333560 refGene_gn=IREB2;refGene_tx=XM_023254775.1
B3 335104 335270 refGene_gn=IREB2;refGene_tx=XM_023254775.1
B3 347765 347855 refGene_gn=IREB2;refGene_tx=XM_023254775.1
B3 349082 349101 refGene_gn=IREB2;refGene_tx=XM_023254775.1
B3 399405 399456 refGene_gn=CRABP1;refGene_tx=XM_023254776.1
B3 402590 402704 refGene_gn=CRABP1;refGene_tx=XM_023254776.1
B3 404584 404763 refGene_gn=CRABP1;refGene_tx=XM_023254776.1
B3 405285 405355 refGene_gn=CRABP1;refGene_tx=XM_023254776.1
B3 433873 433885
refGene_gn=WDR61;refGene_tx=XM_003986735.5;refGene_gn=WDR61;refGene_tx
=XM_006932168.4
B3 434602 434630
refGene_gn=WDR61;refGene_tx=XM_003986735.5;refGene_gn=WDR61;refGene_tx
=XM_006932168.4
B3 436126 436232
refGene_gn=WDR61;refGene_tx=XM_003986735.5;refGene_gn=WDR61;refGene_tx
=XM_006932168.4
B3 436609 436782
refGene_gn=WDR61;refGene_tx=XM_003986735.5;refGene_gn=WDR61;refGene_tx
=XM_006932168.4
B3 443227 443376
refGene_gn=WDR61;refGene_tx=XM_003986735.5;refGene_gn=WDR61;refGene_tx
=XM_006932168.4
B3 443564 443658
refGene_gn=WDR61;refGene_tx=XM_003986735.5;refGene_gn=WDR61;refGene_tx
=XM_006932168.4
B3 444746 444840
refGene_gn=WDR61;refGene_tx=XM_003986735.5;refGene_gn=WDR61;refGene_tx
=XM_006932168.4
B3 446132 446225
refGene_gn=WDR61;refGene_tx=XM_003986735.5;refGene_gn=WDR61;refGene_tx
=XM_006932168.4
B3 446929 447008
refGene_gn=WDR61;refGene_tx=XM_003986735.5;refGene_gn=WDR61;refGene_tx

=XM_006932168.4
B3 448693 448783
refGene_gn=WDR61;refGene_tx=XM_003986735.5;refGene_gn=WDR61;refGene_tx
=XM_006932168.4
B3 451264 451480 refGene_gn=DNAJA4;refGene_tx=XM_023254777.1
B3 451584 451685 refGene_gn=DNAJA4;refGene_tx=XM_023254777.1
B3 453832 454063 refGene_gn=DNAJA4;refGene_tx=XM_023254777.1
B3 454933 455161 refGene_gn=DNAJA4;refGene_tx=XM_023254777.1
B3 456303 456408 refGene_gn=DNAJA4;refGene_tx=XM_023254777.1
B3 457969 458150 refGene_gn=DNAJA4;refGene_tx=XM_023254777.1
B3 463303 463442 refGene_gn=DNAJA4;refGene_tx=XM_023254777.1
B3 464047 464127 refGene_gn=DNAJA4;refGene_tx=XM_023254777.1
B3 506651 506701
refGene_gn=ACSBG1;refGene_tx=XM_019831747.2;refGene_gn=ACSBG1;refGene_
tx=XM_019831748.2;refGene_gn=ACSBG1;refGene_tx=XM_019831749.2
B3 509103 509202
refGene_gn=ACSBG1;refGene_tx=XM_019831747.2;refGene_gn=ACSBG1;refGene_
tx=XM_019831748.2;refGene_gn=ACSBG1;refGene_tx=XM_019831749.2
B3 510064 510165
refGene_gn=ACSBG1;refGene_tx=XM_019831747.2;refGene_gn=ACSBG1;refGene_
tx=XM_019831748.2;refGene_gn=ACSBG1;refGene_tx=XM_019831749.2;refGene_
gn=ACSBG1;refGene_tx=XM_019831750.1
B3 517490 517711
refGene_gn=ACSBG1;refGene_tx=XM_019831747.2;refGene_gn=ACSBG1;refGene_
tx=XM_019831748.2;refGene_gn=ACSBG1;refGene_tx=XM_019831749.2
B3 518090 518179
refGene_gn=ACSBG1;refGene_tx=XM_019831747.2;refGene_gn=ACSBG1;refGene_
tx=XM_019831748.2;refGene_gn=ACSBG1;refGene_tx=XM_019831749.2;refGene_
gn=ACSBG1;refGene_tx=XM_019831750.1
B3 518367 518488
refGene_gn=ACSBG1;refGene_tx=XM_019831747.2;refGene_gn=ACSBG1;refGene_
tx=XM_019831748.2;refGene_gn=ACSBG1;refGene_tx=XM_019831749.2;refGene_
gn=ACSBG1;refGene_tx=XM_019831750.1
B3 523362 523443
refGene_gn=ACSBG1;refGene_tx=XM_019831747.2;refGene_gn=ACSBG1;refGene_
tx=XM_019831748.2;refGene_gn=ACSBG1;refGene_tx=XM_019831749.2;refGene_
gn=ACSBG1;refGene_tx=XM_019831750.1
B3 523526 523676
refGene_gn=ACSBG1;refGene_tx=XM_019831747.2;refGene_gn=ACSBG1;refGene_
tx=XM_019831748.2;refGene_gn=ACSBG1;refGene_tx=XM_019831749.2;refGene_
gn=ACSBG1;refGene_tx=XM_019831750.1
B3 523935 524112
refGene_gn=ACSBG1;refGene_tx=XM_019831747.2;refGene_gn=ACSBG1;refGene_
tx=XM_019831748.2;refGene_gn=ACSBG1;refGene_tx=XM_019831749.2;refGene_
gn=ACSBG1;refGene_tx=XM_019831750.1
B3 524713 524895
refGene_gn=ACSBG1;refGene_tx=XM_019831747.2;refGene_gn=ACSBG1;refGene_
tx=XM_019831748.2;refGene_gn=ACSBG1;refGene_tx=XM_019831749.2;refGene_
gn=ACSBG1;refGene_tx=XM_019831750.1
B3 525574 525895

refGene_gn=ACSBG1;refGene_tx=XM_019831747.2;refGene_gn=ACSBG1;refGene_tx=XM_019831748.2;refGene_gn=ACSBG1;refGene_tx=XM_019831749.2;refGene_gn=ACSBG1;refGene_tx=XM_019831750.1
B3 526376 526594
refGene_gn=ACSBG1;refGene_tx=XM_019831747.2;refGene_gn=ACSBG1;refGene_tx=XM_019831748.2;refGene_gn=ACSBG1;refGene_tx=XM_019831749.2;refGene_gn=ACSBG1;refGene_tx=XM_019831750.1
B3 529194 529292 refGene_gn=ACSBG1;refGene_tx=XM_019831749.2
B3 530058 530198
refGene_gn=ACSBG1;refGene_tx=XM_019831747.2;refGene_gn=ACSBG1;refGene_tx=XM_019831748.2;refGene_gn=ACSBG1;refGene_tx=XM_019831749.2;refGene_gn=ACSBG1;refGene_tx=XM_019831750.1
B3 530707 530954
refGene_gn=ACSBG1;refGene_tx=XM_019831747.2;refGene_gn=ACSBG1;refGene_tx=XM_019831748.2;refGene_gn=ACSBG1;refGene_tx=XM_019831750.1
B3 531488 531574
refGene_gn=ACSBG1;refGene_tx=XM_019831747.2;refGene_gn=ACSBG1;refGene_tx=XM_019831748.2;refGene_gn=ACSBG1;refGene_tx=XM_019831750.1
B3 533604 533688
refGene_gn=IDH3A;refGene_tx=XM_003986736.5;refGene_gn=IDH3A;refGene_tx=XM_006932170.4;refGene_gn=IDH3A;refGene_tx=XM_019831751.2
B3 534204 534357
refGene_gn=IDH3A;refGene_tx=XM_003986736.5;refGene_gn=IDH3A;refGene_tx=XM_006932170.4;refGene_gn=IDH3A;refGene_tx=XM_019831751.2
B3 534798 534883
refGene_gn=IDH3A;refGene_tx=XM_003986736.5;refGene_gn=IDH3A;refGene_tx=XM_006932170.4;refGene_gn=IDH3A;refGene_tx=XM_019831751.2
B3 536137 536202
refGene_gn=IDH3A;refGene_tx=XM_003986736.5;refGene_gn=IDH3A;refGene_tx=XM_006932170.4;refGene_gn=IDH3A;refGene_tx=XM_019831751.2
B3 536299 536402
refGene_gn=IDH3A;refGene_tx=XM_003986736.5;refGene_gn=IDH3A;refGene_tx=XM_006932170.4;refGene_gn=IDH3A;refGene_tx=XM_019831751.2
B3 537124 537258
refGene_gn=IDH3A;refGene_tx=XM_003986736.5;refGene_gn=IDH3A;refGene_tx=XM_006932170.4;refGene_gn=IDH3A;refGene_tx=XM_019831751.2
B3 537707 537895
refGene_gn=IDH3A;refGene_tx=XM_003986736.5;refGene_gn=IDH3A;refGene_tx=XM_006932170.4;refGene_gn=IDH3A;refGene_tx=XM_019831751.2
B3 539022 539137
refGene_gn=IDH3A;refGene_tx=XM_003986736.5;refGene_gn=IDH3A;refGene_tx=XM_006932170.4;refGene_gn=IDH3A;refGene_tx=XM_019831751.2
B3 540323 540407
refGene_gn=IDH3A;refGene_tx=XM_003986736.5;refGene_gn=IDH3A;refGene_tx=XM_006932170.4;refGene_gn=IDH3A;refGene_tx=XM_019831751.2
B3 542490 542553
refGene_gn=IDH3A;refGene_tx=XM_003986736.5;refGene_gn=IDH3A;refGene_tx=XM_006932170.4
B3 545952 545979
refGene_gn=IDH3A;refGene_tx=XM_003986736.5;refGene_gn=IDH3A;refGene_tx

=XM_006932170.4
B3 554893 554944
refGene_gn=LOC101085978;refGene_tx=XM_003986737.5;refGene_gn=LOC101085978;refGene_tx=XM_006932171.4;refGene_gn=LOC101085978;refGene_tx=XM_006932172.4;refGene_gn=LOC101085978;refGene_tx=XM_006932174.4
B3 559319 559354
refGene_gn=LOC101085978;refGene_tx=XM_003986737.5;refGene_gn=LOC101085978;refGene_tx=XM_006932171.4;refGene_gn=LOC101085978;refGene_tx=XM_006932172.4
B3 560537 560654
refGene_gn=LOC101085978;refGene_tx=XM_006932171.4;refGene_gn=LOC101085978;refGene_tx=XM_006932172.4
B3 567269 567381
refGene_gn=LOC101085978;refGene_tx=XM_003986737.5;refGene_gn=LOC101085978;refGene_tx=XM_006932171.4;refGene_gn=LOC101085978;refGene_tx=XM_006932172.4;refGene_gn=LOC101085978;refGene_tx=XM_006932173.4
B3 569166 569314
refGene_gn=LOC101085978;refGene_tx=XM_003986737.5;refGene_gn=LOC101085978;refGene_tx=XM_006932171.4;refGene_gn=LOC101085978;refGene_tx=XM_006932172.4;refGene_gn=LOC101085978;refGene_tx=XM_006932173.4;refGene_gn=LOC101085978;refGene_tx=XM_006932174.4
B3 573478 573674
refGene_gn=LOC101085978;refGene_tx=XM_003986737.5;refGene_gn=LOC101085978;refGene_tx=XM_006932171.4;refGene_gn=LOC101085978;refGene_tx=XM_006932172.4;refGene_gn=LOC101085978;refGene_tx=XM_006932173.4;refGene_gn=LOC101085978;refGene_tx=XM_006932174.4
B3 573863 573885
refGene_gn=LOC101085978;refGene_tx=XM_003986737.5;refGene_gn=LOC101085978;refGene_tx=XM_006932171.4;refGene_gn=LOC101085978;refGene_tx=XM_006932172.4;refGene_gn=LOC101085978;refGene_tx=XM_006932173.4;refGene_gn=LOC101085978;refGene_tx=XM_006932174.4
B3 574116 574135
refGene_gn=LOC111560761;refGene_tx=XM_023254764.1
B3 574534 574784
refGene_gn=LOC111560761;refGene_tx=XM_023254764.1
B3 574973 574995
refGene_gn=LOC111560761;refGene_tx=XM_023254764.1
B3 576793 576844 refGene_gn=SH2D7;refGene_tx=XM_019831752.1
B3 577518 578205 refGene_gn=SH2D7;refGene_tx=XM_019831752.1
B3 579551 579740 refGene_gn=SH2D7;refGene_tx=XM_019831752.1
B3 580014 580180 refGene_gn=SH2D7;refGene_tx=XM_019831752.1
B3 582978 583068 refGene_gn=SH2D7;refGene_tx=XM_019831752.1
B3 584456 584632 refGene_gn=SH2D7;refGene_tx=XM_019831752.1
B3 594457 594775 refGene_gn=TBC1D2B;refGene_tx=XM_023254782.1
B3 596913 596949 refGene_gn=TBC1D2B;refGene_tx=XM_006932176.3
B3 608765 608786 refGene_gn=TBC1D2B;refGene_tx=XM_006932177.3
B3 609094 609248
refGene_gn=TBC1D2B;refGene_tx=XM_006932176.3;refGene_gn=TBC1D2B;refGene_tx=XM_006932177.3;refGene_gn=TBC1D2B;refGene_tx=XM_006932178.4;refGene_gn=TBC1D2B;refGene_tx=XM_006932180.4;refGene_gn=TBC1D2B;refGene_tx=

XM_023254782.1
B3 617343 617503
refGene_gn=TBC1D2B;refGene_tx=XM_006932176.3;refGene_gn=TBC1D2B;refGene_tx=XM_006932177.3;refGene_gn=TBC1D2B;refGene_tx=XM_006932178.4;refGene_gn=TBC1D2B;refGene_tx=XM_006932180.4;refGene_gn=TBC1D2B;refGene_tx=XM_023254782.1
B3 632409 632573
refGene_gn=TBC1D2B;refGene_tx=XM_006932176.3;refGene_gn=TBC1D2B;refGene_tx=XM_006932177.3;refGene_gn=TBC1D2B;refGene_tx=XM_006932178.4;refGene_gn=TBC1D2B;refGene_tx=XM_006932180.4;refGene_gn=TBC1D2B;refGene_tx=XM_023254782.1
B3 635411 635638
refGene_gn=TBC1D2B;refGene_tx=XM_006932176.3;refGene_gn=TBC1D2B;refGene_tx=XM_006932177.3;refGene_gn=TBC1D2B;refGene_tx=XM_006932178.4;refGene_gn=TBC1D2B;refGene_tx=XM_006932180.4;refGene_gn=TBC1D2B;refGene_tx=XM_023254782.1
B3 636206 636590
refGene_gn=TBC1D2B;refGene_tx=XM_006932176.3;refGene_gn=TBC1D2B;refGene_tx=XM_006932177.3;refGene_gn=TBC1D2B;refGene_tx=XM_006932178.4;refGene_gn=TBC1D2B;refGene_tx=XM_006932180.4;refGene_gn=TBC1D2B;refGene_tx=XM_023254782.1
B3 641666 641777
refGene_gn=TBC1D2B;refGene_tx=XM_006932176.3;refGene_gn=TBC1D2B;refGene_tx=XM_006932177.3;refGene_gn=TBC1D2B;refGene_tx=XM_006932178.4;refGene_gn=TBC1D2B;refGene_tx=XM_006932180.4;refGene_gn=TBC1D2B;refGene_tx=XM_023254782.1
B3 642598 642792
refGene_gn=TBC1D2B;refGene_tx=XM_006932176.3;refGene_gn=TBC1D2B;refGene_tx=XM_006932177.3;refGene_gn=TBC1D2B;refGene_tx=XM_006932178.4;refGene_gn=TBC1D2B;refGene_tx=XM_006932180.4;refGene_gn=TBC1D2B;refGene_tx=XM_023254782.1
B3 645187 645682
refGene_gn=TBC1D2B;refGene_tx=XM_006932176.3;refGene_gn=TBC1D2B;refGene_tx=XM_006932177.3;refGene_gn=TBC1D2B;refGene_tx=XM_006932178.4;refGene_gn=TBC1D2B;refGene_tx=XM_006932180.4;refGene_gn=TBC1D2B;refGene_tx=XM_023254782.1
B3 648356 648474
refGene_gn=TBC1D2B;refGene_tx=XM_006932176.3;refGene_gn=TBC1D2B;refGene_tx=XM_006932177.3;refGene_gn=TBC1D2B;refGene_tx=XM_006932178.4;refGene_gn=TBC1D2B;refGene_tx=XM_006932180.4;refGene_gn=TBC1D2B;refGene_tx=XM_023254782.1
B3 653080 653266
refGene_gn=TBC1D2B;refGene_tx=XM_006932176.3;refGene_gn=TBC1D2B;refGene_tx=XM_006932177.3;refGene_gn=TBC1D2B;refGene_tx=XM_006932178.4;refGene_gn=TBC1D2B;refGene_tx=XM_006932180.4;refGene_gn=TBC1D2B;refGene_tx=XM_023254782.1
B3 653414 653536
refGene_gn=TBC1D2B;refGene_tx=XM_006932176.3;refGene_gn=TBC1D2B;refGene_tx=XM_006932177.3;refGene_gn=TBC1D2B;refGene_tx=XM_006932178.4;refGene_gn=TBC1D2B;refGene_tx=XM_006932180.4;refGene_gn=TBC1D2B;refGene_tx=

XM_023254782.1
B3	656737	656930
refGene_gn=TBC1D2B;refGene_tx=XM_006932176.3;refGene_gn=TBC1D2B;refGene_tx=XM_006932177.3;refGene_gn=TBC1D2B;refGene_tx=XM_006932178.4;refGene_gn=TBC1D2B;refGene_tx=XM_006932180.4;refGene_gn=TBC1D2B;refGene_tx=XM_023254782.1
B3	661664	661822	refGene_gn=ADAMTS7;refGene_tx=XM_023254780.1
B3	663958	664085	refGene_gn=ADAMTS7;refGene_tx=XM_023254780.1
B3	664086	664122	refGene_gn=ADAMTS7;refGene_tx=XM_023254780.1
B3	666636	666783	refGene_gn=ADAMTS7;refGene_tx=XM_023254780.1
B3	666869	667043	refGene_gn=ADAMTS7;refGene_tx=XM_023254780.1
B3	667303	667456	refGene_gn=ADAMTS7;refGene_tx=XM_023254780.1
B3	668163	669453	refGene_gn=ADAMTS7;refGene_tx=XM_023254780.1
B3	669729	669943	refGene_gn=ADAMTS7;refGene_tx=XM_023254780.1
B3	670505	670632	refGene_gn=ADAMTS7;refGene_tx=XM_023254780.1
B3	672564	672706	refGene_gn=ADAMTS7;refGene_tx=XM_023254780.1
B3	673519	673764	refGene_gn=ADAMTS7;refGene_tx=XM_023254780.1
B3	674796	674917	refGene_gn=ADAMTS7;refGene_tx=XM_023254780.1
B3	675715	675849	refGene_gn=ADAMTS7;refGene_tx=XM_023254780.1
B3	676058	676228	refGene_gn=ADAMTS7;refGene_tx=XM_023254780.1
B3	676314	676460	refGene_gn=ADAMTS7;refGene_tx=XM_023254780.1
B3	676771	676864	refGene_gn=ADAMTS7;refGene_tx=XM_023254780.1
B3	677706	677851	refGene_gn=ADAMTS7;refGene_tx=XM_023254780.1
B3	686650	686794	refGene_gn=ADAMTS7;refGene_tx=XM_023254780.1
B3	687854	688004	refGene_gn=ADAMTS7;refGene_tx=XM_023254780.1
B3	688822	688947	refGene_gn=ADAMTS7;refGene_tx=XM_023254780.1
B3	689264	689348	refGene_gn=ADAMTS7;refGene_tx=XM_023254780.1
B3	692731	692928	refGene_gn=ADAMTS7;refGene_tx=XM_023254780.1
B3	693849	694015	refGene_gn=ADAMTS7;refGene_tx=XM_023254780.1
B3	695529	695885	refGene_gn=ADAMTS7;refGene_tx=XM_023254780.1
B3	703176	703267	refGene_gn=ADAMTS7;refGene_tx=XM_023254780.1
B3	738087	738127
refGene_gn=MORF4L1;refGene_tx=XM_023254783.1;refGene_gn=MORF4L1;refGene_tx=XM_023254784.1;refGene_gn=MORF4L1;refGene_tx=XM_023254785.1
B3	743820	743867
refGene_gn=MORF4L1;refGene_tx=XM_023254783.1;refGene_gn=MORF4L1;refGene_tx=XM_023254784.1;refGene_gn=MORF4L1;refGene_tx=XM_023254785.1
B3	745528	745596
refGene_gn=MORF4L1;refGene_tx=XM_023254783.1;refGene_gn=MORF4L1;refGene_tx=XM_023254784.1;refGene_gn=MORF4L1;refGene_tx=XM_023254785.1
B3	748951	749068
refGene_gn=MORF4L1;refGene_tx=XM_023254783.1;refGene_gn=MORF4L1;refGene_tx=XM_023254784.1
B3	750558	750645
refGene_gn=MORF4L1;refGene_tx=XM_023254783.1;refGene_gn=MORF4L1;refGene_tx=XM_023254784.1;refGene_gn=MORF4L1;refGene_tx=XM_023254785.1
B3	751071	751152
refGene_gn=MORF4L1;refGene_tx=XM_023254783.1;refGene_gn=MORF4L1;refGene_tx=XM_023254784.1;refGene_gn=MORF4L1;refGene_tx=XM_023254785.1
B3	753577	753603

refGene_gn=MORF4L1;refGene_tx=XM_023254783.1;refGene_gn=MORF4L1;refGene_tx=XM_023254784.1;refGene_gn=MORF4L1;refGene_tx=XM_023254785.1
B3 754317 754406
refGene_gn=MORF4L1;refGene_tx=XM_023254783.1;refGene_gn=MORF4L1;refGene_tx=XM_023254784.1;refGene_gn=MORF4L1;refGene_tx=XM_023254785.1
B3 755546 755648
refGene_gn=MORF4L1;refGene_tx=XM_023254783.1;refGene_gn=MORF4L1;refGene_tx=XM_023254784.1;refGene_gn=MORF4L1;refGene_tx=XM_023254785.1
B3 756754 756843
refGene_gn=MORF4L1;refGene_tx=XM_023254783.1;refGene_gn=MORF4L1;refGene_tx=XM_023254784.1;refGene_gn=MORF4L1;refGene_tx=XM_023254785.1
B3 757334 757507
refGene_gn=MORF4L1;refGene_tx=XM_023254783.1;refGene_gn=MORF4L1;refGene_tx=XM_023254784.1;refGene_gn=MORF4L1;refGene_tx=XM_023254785.1
B3 757997 758082
refGene_gn=MORF4L1;refGene_tx=XM_023254783.1;refGene_gn=MORF4L1;refGene_tx=XM_023254784.1;refGene_gn=MORF4L1;refGene_tx=XM_023254785.1
B3 761481 761566
refGene_gn=MORF4L1;refGene_tx=XM_023254783.1;refGene_gn=MORF4L1;refGene_tx=XM_023254784.1;refGene_gn=MORF4L1;refGene_tx=XM_023254785.1
B3 774591 774625 refGene_gn=CTSH;refGene_tx=XM_023254788.1
B3 775905 775981 refGene_gn=CTSH;refGene_tx=XM_023254787.1
B3 776711 776837
refGene_gn=CTSH;refGene_tx=XM_023254787.1;refGene_gn=CTSH;refGene_tx=XM_023254788.1
B3 778323 778430
refGene_gn=CTSH;refGene_tx=XM_023254787.1;refGene_gn=CTSH;refGene_tx=XM_023254788.1
B3 779318 779387
refGene_gn=CTSH;refGene_tx=XM_023254787.1;refGene_gn=CTSH;refGene_tx=XM_023254788.1
B3 780117 780199
refGene_gn=CTSH;refGene_tx=XM_023254787.1;refGene_gn=CTSH;refGene_tx=XM_023254788.1
B3 780849 780905
refGene_gn=CTSH;refGene_tx=XM_023254787.1;refGene_gn=CTSH;refGene_tx=XM_023254788.1
B3 781562 781649
refGene_gn=CTSH;refGene_tx=XM_023254787.1;refGene_gn=CTSH;refGene_tx=XM_023254788.1
B3 783115 783220
refGene_gn=CTSH;refGene_tx=XM_023254787.1;refGene_gn=CTSH;refGene_tx=XM_023254788.1
B3 783726 783797
refGene_gn=CTSH;refGene_tx=XM_023254787.1;refGene_gn=CTSH;refGene_tx=XM_023254788.1
B3 784737 784843
refGene_gn=CTSH;refGene_tx=XM_023254787.1;refGene_gn=CTSH;refGene_tx=XM_023254788.1
B3 785412 785444

refGene_gn=CTSH;refGene_tx=XM_023254787.1;refGene_gn=CTSH;refGene_tx=XM_023254788.1
B3 788757 788848
refGene_gn=CTSH;refGene_tx=XM_023254787.1;refGene_gn=CTSH;refGene_tx=XM_023254788.1
B3 804467 804560
refGene_gn=RASGRF1;refGene_tx=XM_011282702.3;refGene_gn=RASGRF1;refGene_tx=XM_019831755.2;refGene_gn=RASGRF1;refGene_tx=XM_023254789.1
B3 805547 805616
refGene_gn=RASGRF1;refGene_tx=XM_011282702.3;refGene_gn=RASGRF1;refGene_tx=XM_019831755.2;refGene_gn=RASGRF1;refGene_tx=XM_023254789.1
B3 806471 806589
refGene_gn=RASGRF1;refGene_tx=XM_011282702.3;refGene_gn=RASGRF1;refGene_tx=XM_019831755.2;refGene_gn=RASGRF1;refGene_tx=XM_023254789.1
B3 812032 812112
refGene_gn=RASGRF1;refGene_tx=XM_011282702.3;refGene_gn=RASGRF1;refGene_tx=XM_019831755.2;refGene_gn=RASGRF1;refGene_tx=XM_023254789.1
B3 815900 816098
refGene_gn=RASGRF1;refGene_tx=XM_011282702.3;refGene_gn=RASGRF1;refGene_tx=XM_019831755.2;refGene_gn=RASGRF1;refGene_tx=XM_023254789.1
B3 818919 819004
refGene_gn=RASGRF1;refGene_tx=XM_011282702.3;refGene_gn=RASGRF1;refGene_tx=XM_019831755.2;refGene_gn=RASGRF1;refGene_tx=XM_023254789.1
B3 820136 820240
refGene_gn=RASGRF1;refGene_tx=XM_011282702.3;refGene_gn=RASGRF1;refGene_tx=XM_019831755.2;refGene_gn=RASGRF1;refGene_tx=XM_023254789.1
B3 823270 823331
refGene_gn=RASGRF1;refGene_tx=XM_011282702.3;refGene_gn=RASGRF1;refGene_tx=XM_019831755.2;refGene_gn=RASGRF1;refGene_tx=XM_023254789.1
B3 825097 825210
refGene_gn=RASGRF1;refGene_tx=XM_011282702.3;refGene_gn=RASGRF1;refGene_tx=XM_019831755.2;refGene_gn=RASGRF1;refGene_tx=XM_023254789.1
B3 825469 825576
refGene_gn=RASGRF1;refGene_tx=XM_011282702.3;refGene_gn=RASGRF1;refGene_tx=XM_019831755.2;refGene_gn=RASGRF1;refGene_tx=XM_023254789.1
B3 826052 826223
refGene_gn=RASGRF1;refGene_tx=XM_011282702.3;refGene_gn=RASGRF1;refGene_tx=XM_019831755.2;refGene_gn=RASGRF1;refGene_tx=XM_023254789.1
B3 827540 827666
refGene_gn=RASGRF1;refGene_tx=XM_011282702.3;refGene_gn=RASGRF1;refGene_tx=XM_019831755.2;refGene_gn=RASGRF1;refGene_tx=XM_023254789.1
B3 829493 829867
refGene_gn=RASGRF1;refGene_tx=XM_011282702.3;refGene_gn=RASGRF1;refGene_tx=XM_019831755.2;refGene_gn=RASGRF1;refGene_tx=XM_023254789.1
B3 832828 833077
refGene_gn=RASGRF1;refGene_tx=XM_011282702.3;refGene_gn=RASGRF1;refGene_tx=XM_019831755.2;refGene_gn=RASGRF1;refGene_tx=XM_023254789.1
B3 839202 839285
refGene_gn=RASGRF1;refGene_tx=XM_011282702.3;refGene_gn=RASGRF1;refGene_tx=XM_019831755.2;refGene_gn=RASGRF1;refGene_tx=XM_023254789.1

B3 841365 841502
refGene_gn=RASGRF1;refGene_tx=XM_011282702.3;refGene_gn=RASGRF1;refGene_tx=XM_019831755.2;refGene_gn=RASGRF1;refGene_tx=XM_023254789.1
B3 843301 843365
refGene_gn=RASGRF1;refGene_tx=XM_011282702.3;refGene_gn=RASGRF1;refGene_tx=XM_019831755.2;refGene_gn=RASGRF1;refGene_tx=XM_023254789.1
B3 846429 846590
refGene_gn=RASGRF1;refGene_tx=XM_011282702.3;refGene_gn=RASGRF1;refGene_tx=XM_019831755.2;refGene_gn=RASGRF1;refGene_tx=XM_023254789.1
B3 848269 848388
refGene_gn=RASGRF1;refGene_tx=XM_011282702.3;refGene_gn=RASGRF1;refGene_tx=XM_019831755.2;refGene_gn=RASGRF1;refGene_tx=XM_023254789.1
B3 851393 851503
refGene_gn=RASGRF1;refGene_tx=XM_011282702.3;refGene_gn=RASGRF1;refGene_tx=XM_019831755.2;refGene_gn=RASGRF1;refGene_tx=XM_023254789.1
B3 852491 852500 refGene_gn=RASGRF1;refGene_tx=XM_023254789.1
B3 852969 853163
refGene_gn=RASGRF1;refGene_tx=XM_011282702.3;refGene_gn=RASGRF1;refGene_tx=XM_019831755.2
B3 854807 854887
refGene_gn=RASGRF1;refGene_tx=XM_011282702.3;refGene_gn=RASGRF1;refGene_tx=XM_019831755.2
B3 861061 861315
refGene_gn=RASGRF1;refGene_tx=XM_011282702.3;refGene_gn=RASGRF1;refGene_tx=XM_019831755.2
B3 863432 863525
refGene_gn=RASGRF1;refGene_tx=XM_011282702.3;refGene_gn=RASGRF1;refGene_tx=XM_019831755.2
B3 871438 871586
refGene_gn=RASGRF1;refGene_tx=XM_011282702.3;refGene_gn=RASGRF1;refGene_tx=XM_019831755.2
B3 877551 877658
refGene_gn=RASGRF1;refGene_tx=XM_011282702.3;refGene_gn=RASGRF1;refGene_tx=XM_019831755.2
B3 901783 902059 refGene_gn=RASGRF1;refGene_tx=XM_011282702.3
B3 903130 903169 refGene_gn=RASGRF1;refGene_tx=XM_019831755.2
B3 1047347 1048928
refGene_gn=ANKRD34C;refGene_tx=XM_006932185.3;refGene_gn=ANKRD34C;refGene_tx=XM_006932186.3;refGene_gn=ANKRD34C;refGene_tx=XM_011282704.3;refGene_gn=ANKRD34C;refGene_tx=XM_019831757.2;refGene_gn=ANKRD34C;refGene_tx=XM_023254791.1;refGene_gn=ANKRD34C;refGene_tx=XM_023254792.1
B3 1056965 1057151
refGene_gn=TMED3;refGene_tx=XM_003986765.3;refGene_gn=TMED3;refGene_tx=XM_023254795.1
B3 1058673 1058922
refGene_gn=TMED3;refGene_tx=XM_003986765.3;refGene_gn=TMED3;refGene_tx=XM_023254795.1
B3 1064227 1064464 refGene_gn=TMED3;refGene_tx=XM_003986765.3
B3 1076339 1076675 refGene_gn=TMED3;refGene_tx=XM_023254795.1
B3 1155581 1157870

refGene_gn=KIAA1024;refGene_tx=XM_023254793.1;refGene_gn=KIAA1024;refGene_tx=XM_023254794.1
B3 1160020 1160275
refGene_gn=KIAA1024;refGene_tx=XM_023254793.1;refGene_gn=KIAA1024;refGene_tx=XM_023254794.1
B3 1164193 1164391
refGene_gn=KIAA1024;refGene_tx=XM_023254793.1;refGene_gn=KIAA1024;refGene_tx=XM_023254794.1
B3 1311954 1312370
refGene_gn=LOC109500452;refGene_tx=XM_023254796.1
B3 1330581 1330726
refGene_gn=LOC109500452;refGene_tx=XM_023254796.1
B3 1350279 1350381
refGene_gn=LOC109500452;refGene_tx=XM_023254796.1
B3 1360506 1360559
refGene_gn=LOC111560765;refGene_tx=XM_023254797.1
B3 1361140 1361894
refGene_gn=LOC111560765;refGene_tx=XM_023254797.1
B3 1388271 1388498
refGene_gn=MTHFS;refGene_tx=XM_011282707.3;refGene_gn=MTHFS;refGene_tx=XM_023254798.1
B3 1413643 1413905
refGene_gn=MTHFS;refGene_tx=XM_011282707.3;refGene_gn=MTHFS;refGene_tx=XM_023254798.1
B3 1420279 1420399 refGene_gn=MTHFS;refGene_tx=XM_023254798.1
B3 1436578 1436686 refGene_gn=BCL2A1;refGene_tx=XM_003986739.4
B3 1445295 1445712 refGene_gn=BCL2A1;refGene_tx=XM_003986739.4
B3 1450370 1451098
refGene_gn=BCL2A1;refGene_tx=XM_019831763.2;refGene_gn=BCL2A1;refGene_tx=XM_019831764.2
B3 1475501 1475585
refGene_gn=BCL2A1;refGene_tx=XM_019831763.2;refGene_gn=BCL2A1;refGene_tx=XM_019831764.2
B3 1486330 1486442
refGene_gn=BCL2A1;refGene_tx=XM_019831763.2;refGene_gn=BCL2A1;refGene_tx=XM_019831764.2
B3 1489053 1489104
refGene_gn=BCL2A1;refGene_tx=XM_019831763.2;refGene_gn=BCL2A1;refGene_tx=XM_019831764.2
B3 1555230 1555384
refGene_gn=ZFAND6;refGene_tx=XM_003986740.4;refGene_gn=ZFAND6;refGene_tx=XM_006932188.3;refGene_gn=ZFAND6;refGene_tx=XM_011282708.1;refGene_gn=ZFAND6;refGene_tx=XM_019831768.1;refGene_gn=ZFAND6;refGene_tx=XM_023254799.1;refGene_gn=ZFAND6;refGene_tx=XM_023254800.1
B3 1557030 1557139
refGene_gn=ZFAND6;refGene_tx=XM_003986740.4;refGene_gn=ZFAND6;refGene_tx=XM_006932188.3;refGene_gn=ZFAND6;refGene_tx=XM_011282708.1;refGene_gn=ZFAND6;refGene_tx=XM_019831768.1;refGene_gn=ZFAND6;refGene_tx=XM_023254799.1;refGene_gn=ZFAND6;refGene_tx=XM_023254800.1
B3 1557871 1557972

refGene_gn=ZFAND6;refGene_tx=XM_003986740.4;refGene_gn=ZFAND6;refGene_tx=XM_006932188.3;refGene_gn=ZFAND6;refGene_tx=XM_011282708.1;refGene_gn=ZFAND6;refGene_tx=XM_019831768.1;refGene_gn=ZFAND6;refGene_tx=XM_023254799.1;refGene_gn=ZFAND6;refGene_tx=XM_023254800.1
B3 1559426 1559605 refGene_gn=ZFAND6;refGene_tx=XM_019831768.1
B3 1566504 1566618
refGene_gn=ZFAND6;refGene_tx=XM_003986740.4;refGene_gn=ZFAND6;refGene_tx=XM_006932188.3;refGene_gn=ZFAND6;refGene_tx=XM_011282708.1;refGene_gn=ZFAND6;refGene_tx=XM_023254799.1;refGene_gn=ZFAND6;refGene_tx=XM_023254800.1
B3 1568639 1568788
refGene_gn=ZFAND6;refGene_tx=XM_003986740.4;refGene_gn=ZFAND6;refGene_tx=XM_006932188.3;refGene_gn=ZFAND6;refGene_tx=XM_011282708.1;refGene_gn=ZFAND6;refGene_tx=XM_023254799.1;refGene_gn=ZFAND6;refGene_tx=XM_023254800.1
B3 1581995 1582076
refGene_gn=FAH;refGene_tx=XM_003986744.3;refGene_gn=FAH;refGene_tx=XM_019831770.2;refGene_gn=FAH;refGene_tx=XM_019831771.2;refGene_gn=FAH;refGene_tx=XM_019831772.1;refGene_gn=FAH;refGene_tx=XM_019831773.2
B3 1584635 1584746
refGene_gn=FAH;refGene_tx=XM_003986744.3;refGene_gn=FAH;refGene_tx=XM_019831770.2;refGene_gn=FAH;refGene_tx=XM_019831771.2;refGene_gn=FAH;refGene_tx=XM_019831772.1;refGene_gn=FAH;refGene_tx=XM_019831773.2
B3 1585605 1585727
refGene_gn=FAH;refGene_tx=XM_003986744.3;refGene_gn=FAH;refGene_tx=XM_019831770.2;refGene_gn=FAH;refGene_tx=XM_019831771.2;refGene_gn=FAH;refGene_tx=XM_019831772.1;refGene_gn=FAH;refGene_tx=XM_019831773.2
B3 1586184 1586234
refGene_gn=FAH;refGene_tx=XM_003986744.3;refGene_gn=FAH;refGene_tx=XM_019831770.2;refGene_gn=FAH;refGene_tx=XM_019831771.2;refGene_gn=FAH;refGene_tx=XM_019831772.1;refGene_gn=FAH;refGene_tx=XM_019831773.2
B3 1588417 1588508
refGene_gn=FAH;refGene_tx=XM_003986744.3;refGene_gn=FAH;refGene_tx=XM_019831770.2;refGene_gn=FAH;refGene_tx=XM_019831771.2;refGene_gn=FAH;refGene_tx=XM_019831772.1;refGene_gn=FAH;refGene_tx=XM_019831773.2
B3 1590148 1590246
refGene_gn=FAH;refGene_tx=XM_003986744.3;refGene_gn=FAH;refGene_tx=XM_019831770.2;refGene_gn=FAH;refGene_tx=XM_019831771.2;refGene_gn=FAH;refGene_tx=XM_019831772.1;refGene_gn=FAH;refGene_tx=XM_019831773.2
B3 1590341 1590394
refGene_gn=FAH;refGene_tx=XM_003986744.3;refGene_gn=FAH;refGene_tx=XM_019831770.2;refGene_gn=FAH;refGene_tx=XM_019831771.2;refGene_gn=FAH;refGene_tx=XM_019831772.1;refGene_gn=FAH;refGene_tx=XM_019831773.2
B3 1592607 1592707
refGene_gn=FAH;refGene_tx=XM_003986744.3;refGene_gn=FAH;refGene_tx=XM_019831770.2;refGene_gn=FAH;refGene_tx=XM_019831771.2;refGene_gn=FAH;refGene_tx=XM_019831772.1;refGene_gn=FAH;refGene_tx=XM_019831773.2
B3 1593379 1593510
refGene_gn=FAH;refGene_tx=XM_003986744.3;refGene_gn=FAH;refGene_tx=XM_019831770.2;refGene_gn=FAH;refGene_tx=XM_019831771.2;refGene_gn=FAH;re

fGene_tx=XM_019831772.1;refGene_gn=FAH;refGene_tx=XM_019831773.2
B3 1594991 1595067
refGene_gn=FAH;refGene_tx=XM_003986744.3;refGene_gn=FAH;refGene_tx=XM_019831770.2;refGene_gn=FAH;refGene_tx=XM_019831771.2;refGene_gn=FAH;refGene_tx=XM_019831772.1;refGene_gn=FAH;refGene_tx=XM_019831773.2
B3 1596909 1596956
refGene_gn=FAH;refGene_tx=XM_003986744.3;refGene_gn=FAH;refGene_tx=XM_019831770.2;refGene_gn=FAH;refGene_tx=XM_019831771.2;refGene_gn=FAH;refGene_tx=XM_019831772.1;refGene_gn=FAH;refGene_tx=XM_019831773.2
B3 1600137 1600239
refGene_gn=FAH;refGene_tx=XM_003986744.3;refGene_gn=FAH;refGene_tx=XM_019831770.2;refGene_gn=FAH;refGene_tx=XM_019831771.2;refGene_gn=FAH;refGene_tx=XM_019831772.1;refGene_gn=FAH;refGene_tx=XM_019831773.2
B3 1600380 1600498
refGene_gn=FAH;refGene_tx=XM_003986744.3;refGene_gn=FAH;refGene_tx=XM_019831770.2;refGene_gn=FAH;refGene_tx=XM_019831771.2;refGene_gn=FAH;refGene_tx=XM_019831772.1;refGene_gn=FAH;refGene_tx=XM_019831773.2
B3 1602205 1602255 refGene_gn=FAH;refGene_tx=XM_019831772.1
B3 1602837 1602917 refGene_gn=FAH;refGene_tx=XM_003986744.3
B3 1607357 1607458 refGene_gn=FAH;refGene_tx=XM_019831770.2
B3 1607558 1607563 refGene_gn=FAH;refGene_tx=XM_019831773.2
B3 1615333 1615513
refGene_gn=LOC111560766;refGene_tx=XM_023254801.1
B3 1617018 1617077 refGene_gn=FAH;refGene_tx=XM_019831771.2
B3 1642419 1643262
refGene_gn=LOC111560762;refGene_tx=XM_023254765.1
B3 1714984 1715015
refGene_gn=ARNT2;refGene_tx=XM_023254803.1;refGene_gn=ARNT2;refGene_tx=XM_023254804.1;refGene_gn=ARNT2;refGene_tx=XM_023254805.1
B3 1732370 1732401 refGene_gn=ARNT2;refGene_tx=XM_019831774.2
B3 1750718 1750833
refGene_gn=ARNT2;refGene_tx=XM_003986745.5;refGene_gn=ARNT2;refGene_tx=XM_006932194.4;refGene_gn=ARNT2;refGene_tx=XM_019831774.2;refGene_gn=ARNT2;refGene_tx=XM_019831776.2;refGene_gn=ARNT2;refGene_tx=XM_019831777.2;refGene_gn=ARNT2;refGene_tx=XM_023254803.1;refGene_gn=ARNT2;refGene_tx=XM_023254804.1;refGene_gn=ARNT2;refGene_tx=XM_023254805.1
B3 1756935 1756983
refGene_gn=ARNT2;refGene_tx=XM_003986745.5;refGene_gn=ARNT2;refGene_tx=XM_006932194.4;refGene_gn=ARNT2;refGene_tx=XM_019831774.2;refGene_gn=ARNT2;refGene_tx=XM_019831776.2;refGene_gn=ARNT2;refGene_tx=XM_019831777.2;refGene_gn=ARNT2;refGene_tx=XM_023254803.1;refGene_gn=ARNT2;refGene_tx=XM_023254804.1;refGene_gn=ARNT2;refGene_tx=XM_023254805.1
B3 1764073 1764287
refGene_gn=ARNT2;refGene_tx=XM_003986745.5;refGene_gn=ARNT2;refGene_tx=XM_006932194.4;refGene_gn=ARNT2;refGene_tx=XM_019831774.2;refGene_gn=ARNT2;refGene_tx=XM_019831776.2;refGene_gn=ARNT2;refGene_tx=XM_019831777.2;refGene_gn=ARNT2;refGene_tx=XM_023254803.1;refGene_gn=ARNT2;refGene_tx=XM_023254804.1;refGene_gn=ARNT2;refGene_tx=XM_023254805.1
B3 1767849 1768063
refGene_gn=ARNT2;refGene_tx=XM_003986745.5;refGene_gn=ARNT2;refGene_tx

=XM_006932194.4;refGene_gn=ARNT2;refGene_tx=XM_019831774.2;refGene_gn=
ARNT2;refGene_tx=XM_019831776.2;refGene_gn=ARNT2;refGene_tx=XM_0198317
77.2;refGene_gn=ARNT2;refGene_tx=XM_023254803.1;refGene_gn=ARNT2;refGe
ne_tx=XM_023254804.1;refGene_gn=ARNT2;refGene_tx=XM_023254805.1
B3 1792010 1792113
refGene_gn=ARNT2;refGene_tx=XM_003986745.5;refGene_gn=ARNT2;refGene_tx
=XM_006932194.4;refGene_gn=ARNT2;refGene_tx=XM_019831774.2;refGene_gn=
ARNT2;refGene_tx=XM_019831776.2;refGene_gn=ARNT2;refGene_tx=XM_0198317
77.2;refGene_gn=ARNT2;refGene_tx=XM_023254803.1;refGene_gn=ARNT2;refGe
ne_tx=XM_023254804.1
B3 1795613 1795679
refGene_gn=ARNT2;refGene_tx=XM_003986745.5;refGene_gn=ARNT2;refGene_tx
=XM_006932194.4;refGene_gn=ARNT2;refGene_tx=XM_019831774.2;refGene_gn=
ARNT2;refGene_tx=XM_019831776.2;refGene_gn=ARNT2;refGene_tx=XM_0198317
77.2;refGene_gn=ARNT2;refGene_tx=XM_023254803.1;refGene_gn=ARNT2;refGe
ne_tx=XM_023254804.1
B3 1795960 1796046
refGene_gn=ARNT2;refGene_tx=XM_003986745.5;refGene_gn=ARNT2;refGene_tx
=XM_006932194.4;refGene_gn=ARNT2;refGene_tx=XM_019831774.2;refGene_gn=
ARNT2;refGene_tx=XM_019831776.2;refGene_gn=ARNT2;refGene_tx=XM_0198317
77.2;refGene_gn=ARNT2;refGene_tx=XM_023254803.1;refGene_gn=ARNT2;refGe
ne_tx=XM_023254804.1
B3 1829078 1829155
refGene_gn=ARNT2;refGene_tx=XM_003986745.5;refGene_gn=ARNT2;refGene_tx
=XM_006932194.4;refGene_gn=ARNT2;refGene_tx=XM_019831774.2;refGene_gn=
ARNT2;refGene_tx=XM_019831776.2;refGene_gn=ARNT2;refGene_tx=XM_0198317
77.2;refGene_gn=ARNT2;refGene_tx=XM_023254803.1;refGene_gn=ARNT2;refGe
ne_tx=XM_023254804.1;refGene_gn=ARNT2;refGene_tx=XM_023254805.1
B3 1830359 1830494
refGene_gn=ARNT2;refGene_tx=XM_003986745.5;refGene_gn=ARNT2;refGene_tx
=XM_006932194.4;refGene_gn=ARNT2;refGene_tx=XM_019831774.2;refGene_gn=
ARNT2;refGene_tx=XM_019831776.2;refGene_gn=ARNT2;refGene_tx=XM_0198317
77.2;refGene_gn=ARNT2;refGene_tx=XM_023254803.1;refGene_gn=ARNT2;refGe
ne_tx=XM_023254804.1;refGene_gn=ARNT2;refGene_tx=XM_023254805.1
B3 1832645 1832720
refGene_gn=ARNT2;refGene_tx=XM_003986745.5;refGene_gn=ARNT2;refGene_tx
=XM_006932194.4;refGene_gn=ARNT2;refGene_tx=XM_019831774.2;refGene_gn=
ARNT2;refGene_tx=XM_019831776.2;refGene_gn=ARNT2;refGene_tx=XM_0198317
77.2;refGene_gn=ARNT2;refGene_tx=XM_023254803.1;refGene_gn=ARNT2;refGe
ne_tx=XM_023254804.1;refGene_gn=ARNT2;refGene_tx=XM_023254805.1
B3 1837653 1837805
refGene_gn=ARNT2;refGene_tx=XM_003986745.5;refGene_gn=ARNT2;refGene_tx
=XM_006932194.4;refGene_gn=ARNT2;refGene_tx=XM_019831774.2;refGene_gn=
ARNT2;refGene_tx=XM_019831776.2;refGene_gn=ARNT2;refGene_tx=XM_0198317
77.2;refGene_gn=ARNT2;refGene_tx=XM_023254803.1;refGene_gn=ARNT2;refGe
ne_tx=XM_023254804.1;refGene_gn=ARNT2;refGene_tx=XM_023254805.1
B3 1845670 1845743
refGene_gn=ARNT2;refGene_tx=XM_003986745.5;refGene_gn=ARNT2;refGene_tx
=XM_006932194.4;refGene_gn=ARNT2;refGene_tx=XM_019831774.2;refGene_gn=
ARNT2;refGene_tx=XM_019831776.2;refGene_gn=ARNT2;refGene_tx=XM_0198317

77.2;refGene_gn=ARNT2;refGene_tx=XM_023254803.1;refGene_gn=ARNT2;refGene_tx=XM_023254804.1;refGene_gn=ARNT2;refGene_tx=XM_023254805.1
B3 1846517 1846641
refGene_gn=ARNT2;refGene_tx=XM_003986745.5;refGene_gn=ARNT2;refGene_tx=XM_006932194.4;refGene_gn=ARNT2;refGene_tx=XM_019831774.2;refGene_gn=ARNT2;refGene_tx=XM_019831776.2;refGene_gn=ARNT2;refGene_tx=XM_019831777.2;refGene_gn=ARNT2;refGene_tx=XM_023254803.1;refGene_gn=ARNT2;refGene_tx=XM_023254804.1;refGene_gn=ARNT2;refGene_tx=XM_023254805.1
B3 1849700 1849800
refGene_gn=ARNT2;refGene_tx=XM_003986745.5;refGene_gn=ARNT2;refGene_tx=XM_006932194.4;refGene_gn=ARNT2;refGene_tx=XM_019831774.2;refGene_gn=ARNT2;refGene_tx=XM_019831776.2;refGene_gn=ARNT2;refGene_tx=XM_019831777.2;refGene_gn=ARNT2;refGene_tx=XM_023254803.1;refGene_gn=ARNT2;refGene_tx=XM_023254804.1;refGene_gn=ARNT2;refGene_tx=XM_023254805.1
B3 1853245 1853384
refGene_gn=ARNT2;refGene_tx=XM_003986745.5;refGene_gn=ARNT2;refGene_tx=XM_006932194.4;refGene_gn=ARNT2;refGene_tx=XM_019831774.2;refGene_gn=ARNT2;refGene_tx=XM_019831776.2;refGene_gn=ARNT2;refGene_tx=XM_019831777.2;refGene_gn=ARNT2;refGene_tx=XM_023254803.1;refGene_gn=ARNT2;refGene_tx=XM_023254804.1;refGene_gn=ARNT2;refGene_tx=XM_023254805.1
B3 1853744 1853910
refGene_gn=ARNT2;refGene_tx=XM_003986745.5;refGene_gn=ARNT2;refGene_tx=XM_006932194.4;refGene_gn=ARNT2;refGene_tx=XM_019831774.2;refGene_gn=ARNT2;refGene_tx=XM_019831776.2;refGene_gn=ARNT2;refGene_tx=XM_019831777.2;refGene_gn=ARNT2;refGene_tx=XM_023254803.1;refGene_gn=ARNT2;refGene_tx=XM_023254804.1;refGene_gn=ARNT2;refGene_tx=XM_023254805.1
B3 1859211 1859378
refGene_gn=ARNT2;refGene_tx=XM_003986745.5;refGene_gn=ARNT2;refGene_tx=XM_006932194.4;refGene_gn=ARNT2;refGene_tx=XM_019831774.2;refGene_gn=ARNT2;refGene_tx=XM_019831776.2;refGene_gn=ARNT2;refGene_tx=XM_019831777.2;refGene_gn=ARNT2;refGene_tx=XM_023254803.1;refGene_gn=ARNT2;refGene_tx=XM_023254804.1;refGene_gn=ARNT2;refGene_tx=XM_023254805.1
B3 1860703 1860802
refGene_gn=ARNT2;refGene_tx=XM_003986745.5;refGene_gn=ARNT2;refGene_tx=XM_006932194.4;refGene_gn=ARNT2;refGene_tx=XM_019831774.2;refGene_gn=ARNT2;refGene_tx=XM_019831776.2;refGene_gn=ARNT2;refGene_tx=XM_019831777.2;refGene_gn=ARNT2;refGene_tx=XM_023254803.1;refGene_gn=ARNT2;refGene_tx=XM_023254805.1
B3 1929557 1930156 refGene_gn=ABHD17C;refGene_tx=XM_023254806.1
B3 1972668 1972848 refGene_gn=ABHD17C;refGene_tx=XM_023254806.1
B3 1977833 1978053 refGene_gn=ABHD17C;refGene_tx=XM_023254806.1
B3 2066314 2066504
refGene_gn=CEMIP;refGene_tx=XM_019831781.2;refGene_gn=CEMIP;refGene_tx=XM_023254807.1;refGene_gn=CEMIP;refGene_tx=XM_023254808.1;refGene_gn=CEMIP;refGene_tx=XM_023254809.1
B3 2070298 2070448
refGene_gn=CEMIP;refGene_tx=XM_019831781.2;refGene_gn=CEMIP;refGene_tx=XM_023254807.1;refGene_gn=CEMIP;refGene_tx=XM_023254808.1;refGene_gn=CEMIP;refGene_tx=XM_023254809.1
B3 2070939 2071078

refGene_gn=CEMIP;refGene_tx=XM_019831781.2;refGene_gn=CEMIP;refGene_tx=XM_023254807.1;refGene_gn=CEMIP;refGene_tx=XM_023254808.1;refGene_gn=CEMIP;refGene_tx=XM_023254809.1
B3 2071737 2071974
refGene_gn=CEMIP;refGene_tx=XM_019831781.2;refGene_gn=CEMIP;refGene_tx=XM_023254807.1;refGene_gn=CEMIP;refGene_tx=XM_023254808.1;refGene_gn=CEMIP;refGene_tx=XM_023254809.1
B3 2074409 2074589
refGene_gn=CEMIP;refGene_tx=XM_019831781.2;refGene_gn=CEMIP;refGene_tx=XM_023254807.1;refGene_gn=CEMIP;refGene_tx=XM_023254808.1;refGene_gn=CEMIP;refGene_tx=XM_023254809.1
B3 2077002 2077073
refGene_gn=CEMIP;refGene_tx=XM_019831781.2;refGene_gn=CEMIP;refGene_tx=XM_023254807.1;refGene_gn=CEMIP;refGene_tx=XM_023254808.1;refGene_gn=CEMIP;refGene_tx=XM_023254809.1
B3 2077851 2077947
refGene_gn=CEMIP;refGene_tx=XM_019831781.2;refGene_gn=CEMIP;refGene_tx=XM_023254807.1;refGene_gn=CEMIP;refGene_tx=XM_023254808.1;refGene_gn=CEMIP;refGene_tx=XM_023254809.1
B3 2078339 2078461
refGene_gn=CEMIP;refGene_tx=XM_019831781.2;refGene_gn=CEMIP;refGene_tx=XM_023254807.1;refGene_gn=CEMIP;refGene_tx=XM_023254808.1;refGene_gn=CEMIP;refGene_tx=XM_023254809.1
B3 2080243 2080376
refGene_gn=CEMIP;refGene_tx=XM_019831781.2;refGene_gn=CEMIP;refGene_tx=XM_023254807.1;refGene_gn=CEMIP;refGene_tx=XM_023254808.1;refGene_gn=CEMIP;refGene_tx=XM_023254809.1
B3 2081120 2081312
refGene_gn=CEMIP;refGene_tx=XM_019831781.2;refGene_gn=CEMIP;refGene_tx=XM_023254807.1;refGene_gn=CEMIP;refGene_tx=XM_023254808.1;refGene_gn=CEMIP;refGene_tx=XM_023254809.1
B3 2087293 2087469
refGene_gn=CEMIP;refGene_tx=XM_019831781.2;refGene_gn=CEMIP;refGene_tx=XM_023254807.1;refGene_gn=CEMIP;refGene_tx=XM_023254808.1;refGene_gn=CEMIP;refGene_tx=XM_023254809.1
B3 2089105 2089315
refGene_gn=CEMIP;refGene_tx=XM_019831781.2;refGene_gn=CEMIP;refGene_tx=XM_023254807.1;refGene_gn=CEMIP;refGene_tx=XM_023254808.1;refGene_gn=CEMIP;refGene_tx=XM_023254809.1
B3 2097044 2097250
refGene_gn=CEMIP;refGene_tx=XM_019831781.2;refGene_gn=CEMIP;refGene_tx=XM_023254807.1;refGene_gn=CEMIP;refGene_tx=XM_023254808.1;refGene_gn=CEMIP;refGene_tx=XM_023254809.1
B3 2097838 2097908
refGene_gn=CEMIP;refGene_tx=XM_019831781.2;refGene_gn=CEMIP;refGene_tx=XM_023254807.1;refGene_gn=CEMIP;refGene_tx=XM_023254808.1;refGene_gn=CEMIP;refGene_tx=XM_023254809.1
B3 2098681 2098810
refGene_gn=CEMIP;refGene_tx=XM_019831781.2;refGene_gn=CEMIP;refGene_tx=XM_023254807.1;refGene_gn=CEMIP;refGene_tx=XM_023254808.1;refGene_gn=

CEMIP;refGene_tx=XM_023254809.1
B3	2100999	2101085
refGene_gn=CEMIP;refGene_tx=XM_019831781.2;refGene_gn=CEMIP;refGene_tx=XM_023254807.1;refGene_gn=CEMIP;refGene_tx=XM_023254808.1;refGene_gn=CEMIP;refGene_tx=XM_023254809.1
B3	2101978	2102110
refGene_gn=CEMIP;refGene_tx=XM_019831781.2;refGene_gn=CEMIP;refGene_tx=XM_023254807.1;refGene_gn=CEMIP;refGene_tx=XM_023254808.1;refGene_gn=CEMIP;refGene_tx=XM_023254809.1
B3	2103044	2103080
refGene_gn=CEMIP;refGene_tx=XM_019831781.2;refGene_gn=CEMIP;refGene_tx=XM_023254807.1;refGene_gn=CEMIP;refGene_tx=XM_023254808.1;refGene_gn=CEMIP;refGene_tx=XM_023254809.1
B3	2103161	2103317
refGene_gn=CEMIP;refGene_tx=XM_019831781.2;refGene_gn=CEMIP;refGene_tx=XM_023254807.1;refGene_gn=CEMIP;refGene_tx=XM_023254808.1;refGene_gn=CEMIP;refGene_tx=XM_023254809.1
B3	2106008	2106189
refGene_gn=CEMIP;refGene_tx=XM_019831781.2;refGene_gn=CEMIP;refGene_tx=XM_023254807.1;refGene_gn=CEMIP;refGene_tx=XM_023254808.1;refGene_gn=CEMIP;refGene_tx=XM_023254809.1
B3	2107220	2107601
refGene_gn=CEMIP;refGene_tx=XM_019831781.2;refGene_gn=CEMIP;refGene_tx=XM_023254807.1;refGene_gn=CEMIP;refGene_tx=XM_023254808.1;refGene_gn=CEMIP;refGene_tx=XM_023254809.1
B3	2107813	2107902
refGene_gn=LOC102899053;refGene_tx=XM_006932199.3;refGene_gn=LOC102899053;refGene_tx=XM_011282712.3
B3	2109391	2109504
refGene_gn=LOC102899053;refGene_tx=XM_019831786.2
B3	2109511	2109723
refGene_gn=CEMIP;refGene_tx=XM_019831781.2;refGene_gn=CEMIP;refGene_tx=XM_023254807.1;refGene_gn=CEMIP;refGene_tx=XM_023254808.1
B3	2110366	2110552
refGene_gn=CEMIP;refGene_tx=XM_019831781.2;refGene_gn=CEMIP;refGene_tx=XM_023254807.1;refGene_gn=CEMIP;refGene_tx=XM_023254808.1
B3	2110635	2110721
refGene_gn=LOC102899053;refGene_tx=XM_006932199.3;refGene_gn=LOC102899053;refGene_tx=XM_011282712.3;refGene_gn=LOC102899053;refGene_tx=XM_019831786.2
B3	2111018	2111107
refGene_gn=LOC102899053;refGene_tx=XM_006932199.3;refGene_gn=LOC102899053;refGene_tx=XM_011282712.3;refGene_gn=LOC102899053;refGene_tx=XM_019831786.2
B3	2111715	2111927
refGene_gn=LOC102899053;refGene_tx=XM_006932199.3;refGene_gn=LOC102899053;refGene_tx=XM_011282712.3;refGene_gn=LOC102899053;refGene_tx=XM_019831786.2
B3	2112078	2112250
refGene_gn=LOC102899053;refGene_tx=XM_006932199.3;refGene_gn=LOC102899

053;refGene_tx=XM_011282712.3;refGene_gn=LOC102899053;refGene_tx=XM_019831786.2
B3	2112472	2112677	refGene_gn=CEMIP;refGene_tx=XM_019831781.2;refGene_gn=CEMIP;refGene_tx=XM_023254807.1;refGene_gn=CEMIP;refGene_tx=XM_023254808.1
B3	2112845	2112932	refGene_gn=CEMIP;refGene_tx=XM_019831781.2;refGene_gn=CEMIP;refGene_tx=XM_023254807.1;refGene_gn=CEMIP;refGene_tx=XM_023254808.1
B3	2113505	2113663	refGene_gn=CEMIP;refGene_tx=XM_019831781.2;refGene_gn=CEMIP;refGene_tx=XM_023254807.1;refGene_gn=CEMIP;refGene_tx=XM_023254808.1
B3	2115839	2115940	refGene_gn=CEMIP;refGene_tx=XM_019831781.2;refGene_gn=CEMIP;refGene_tx=XM_023254807.1;refGene_gn=CEMIP;refGene_tx=XM_023254808.1
B3	2117544	2117672	refGene_gn=CEMIP;refGene_tx=XM_019831781.2;refGene_gn=CEMIP;refGene_tx=XM_023254807.1;refGene_gn=CEMIP;refGene_tx=XM_023254808.1
B3	2140087	2140352	refGene_gn=MESD;refGene_tx=XM_023254810.1
B3	2141585	2141818	refGene_gn=MESD;refGene_tx=XM_023254810.1
B3	2147666	2147870	refGene_gn=MESD;refGene_tx=XM_023254810.1
B3	2158727	2159819	refGene_gn=TLNRD1;refGene_tx=XM_023254811.1
B3	2234918	2234987	refGene_gn=CFAP161;refGene_tx=XM_019831789.1;refGene_gn=CFAP161;refGene_tx=XM_023254814.1
B3	2235370	2235493	refGene_gn=CFAP161;refGene_tx=XM_023254812.1;refGene_gn=CFAP161;refGene_tx=XM_023254813.1;refGene_gn=CFAP161;refGene_tx=XM_023254815.1
B3	2235740	2235830	refGene_gn=CFAP161;refGene_tx=XM_019831789.1;refGene_gn=CFAP161;refGene_tx=XM_023254812.1;refGene_gn=CFAP161;refGene_tx=XM_023254813.1;refGene_gn=CFAP161;refGene_tx=XM_023254814.1;refGene_gn=CFAP161;refGene_tx=XM_023254815.1
B3	2237097	2237330	refGene_gn=CFAP161;refGene_tx=XM_019831789.1;refGene_gn=CFAP161;refGene_tx=XM_023254812.1;refGene_gn=CFAP161;refGene_tx=XM_023254813.1;refGene_gn=CFAP161;refGene_tx=XM_023254814.1;refGene_gn=CFAP161;refGene_tx=XM_023254815.1
B3	2238287	2238372	refGene_gn=CFAP161;refGene_tx=XM_019831789.1;refGene_gn=CFAP161;refGene_tx=XM_023254812.1;refGene_gn=CFAP161;refGene_tx=XM_023254813.1;refGene_gn=CFAP161;refGene_tx=XM_023254814.1;refGene_gn=CFAP161;refGene_tx=XM_023254815.1
B3	2242593	2242752	refGene_gn=CFAP161;refGene_tx=XM_019831789.1;refGene_gn=CFAP161;refGene_tx=XM_023254812.1;refGene_gn=CFAP161;refGene_tx=XM_023254813.1;refGene_gn=CFAP161;refGene_tx=XM_023254814.1
B3	2244609	2244683	refGene_gn=CFAP161;refGene_tx=XM_019831789.1;refGene_gn=CFAP161;refGene_tx=XM_023254812.1;refGene_gn=CFAP161;refGene_tx=XM_023254813.1;refGene_gn=CFAP161;refGene_tx=XM_023254814.1;refGene_gn=CFAP161;refGene_tx=

XM_023254815.1
B3 2245270 2245466
refGene_gn=CFAP161;refGene_tx=XM_019831789.1;refGene_gn=CFAP161;refGene_tx=XM_023254812.1;refGene_gn=CFAP161;refGene_tx=XM_023254813.1;refGene_gn=CFAP161;refGene_tx=XM_023254814.1;refGene_gn=CFAP161;refGene_tx=XM_023254815.1
B3 2270584 2270624 refGene_gn=IL16;refGene_tx=XM_023254819.1
B3 2276281 2276525
refGene_gn=IL16;refGene_tx=XM_023254817.1;refGene_gn=IL16;refGene_tx=XM_023254818.1;refGene_gn=IL16;refGene_tx=XM_023254821.1
B3 2295185 2295598
refGene_gn=IL16;refGene_tx=XM_023254817.1;refGene_gn=IL16;refGene_tx=XM_023254818.1;refGene_gn=IL16;refGene_tx=XM_023254819.1;refGene_gn=IL16;refGene_tx=XM_023254820.1;refGene_gn=IL16;refGene_tx=XM_023254821.1;refGene_gn=IL16;refGene_tx=XM_023254822.1;refGene_gn=IL16;refGene_tx=XM_023254824.1
B3 2324221 2324330
refGene_gn=IL16;refGene_tx=XM_023254817.1;refGene_gn=IL16;refGene_tx=XM_023254818.1;refGene_gn=IL16;refGene_tx=XM_023254819.1;refGene_gn=IL16;refGene_tx=XM_023254820.1;refGene_gn=IL16;refGene_tx=XM_023254822.1;refGene_gn=IL16;refGene_tx=XM_023254824.1
B3 2324928 2325019 refGene_gn=IL16;refGene_tx=XM_023254823.1
B3 2327913 2328056
refGene_gn=IL16;refGene_tx=XM_023254817.1;refGene_gn=IL16;refGene_tx=XM_023254818.1;refGene_gn=IL16;refGene_tx=XM_023254819.1;refGene_gn=IL16;refGene_tx=XM_023254820.1;refGene_gn=IL16;refGene_tx=XM_023254822.1;refGene_gn=IL16;refGene_tx=XM_023254823.1;refGene_gn=IL16;refGene_tx=XM_023254824.1
B3 2329462 2329573
refGene_gn=IL16;refGene_tx=XM_023254817.1;refGene_gn=IL16;refGene_tx=XM_023254818.1;refGene_gn=IL16;refGene_tx=XM_023254819.1;refGene_gn=IL16;refGene_tx=XM_023254820.1;refGene_gn=IL16;refGene_tx=XM_023254822.1;refGene_gn=IL16;refGene_tx=XM_023254823.1;refGene_gn=IL16;refGene_tx=XM_023254824.1
B3 2332102 2332217
refGene_gn=IL16;refGene_tx=XM_023254817.1;refGene_gn=IL16;refGene_tx=XM_023254818.1;refGene_gn=IL16;refGene_tx=XM_023254819.1;refGene_gn=IL16;refGene_tx=XM_023254820.1;refGene_gn=IL16;refGene_tx=XM_023254821.1;refGene_gn=IL16;refGene_tx=XM_023254822.1;refGene_gn=IL16;refGene_tx=XM_023254823.1;refGene_gn=IL16;refGene_tx=XM_023254824.1
B3 2339003 2339077
refGene_gn=IL16;refGene_tx=XM_023254817.1;refGene_gn=IL16;refGene_tx=XM_023254818.1;refGene_gn=IL16;refGene_tx=XM_023254819.1;refGene_gn=IL16;refGene_tx=XM_023254820.1;refGene_gn=IL16;refGene_tx=XM_023254821.1;refGene_gn=IL16;refGene_tx=XM_023254822.1;refGene_gn=IL16;refGene_tx=XM_023254823.1;refGene_gn=IL16;refGene_tx=XM_023254824.1;refGene_gn=IL16;refGene_tx=XM_023254825.1
B3 2339809 2340026
refGene_gn=IL16;refGene_tx=XM_023254817.1;refGene_gn=IL16;refGene_tx=XM_023254818.1;refGene_gn=IL16;refGene_tx=XM_023254819.1;refGene_gn=IL1

6;refGene_tx=XM_023254820.1;refGene_gn=IL16;refGene_tx=XM_023254821.1;
refGene_gn=IL16;refGene_tx=XM_023254822.1;refGene_gn=IL16;refGene_tx=X
M_023254823.1;refGene_gn=IL16;refGene_tx=XM_023254824.1;refGene_gn=IL1
6;refGene_tx=XM_023254825.1
B3 2340415 2340533
refGene_gn=IL16;refGene_tx=XM_023254817.1;refGene_gn=IL16;refGene_tx=X
M_023254818.1;refGene_gn=IL16;refGene_tx=XM_023254819.1;refGene_gn=IL1
6;refGene_tx=XM_023254820.1;refGene_gn=IL16;refGene_tx=XM_023254821.1;
refGene_gn=IL16;refGene_tx=XM_023254822.1;refGene_gn=IL16;refGene_tx=X
M_023254823.1;refGene_gn=IL16;refGene_tx=XM_023254824.1;refGene_gn=IL1
6;refGene_tx=XM_023254825.1
B3 2343280 2343413
refGene_gn=IL16;refGene_tx=XM_023254817.1;refGene_gn=IL16;refGene_tx=X
M_023254818.1;refGene_gn=IL16;refGene_tx=XM_023254819.1;refGene_gn=IL1
6;refGene_tx=XM_023254820.1;refGene_gn=IL16;refGene_tx=XM_023254821.1;
refGene_gn=IL16;refGene_tx=XM_023254822.1;refGene_gn=IL16;refGene_tx=X
M_023254823.1;refGene_gn=IL16;refGene_tx=XM_023254824.1;refGene_gn=IL1
6;refGene_tx=XM_023254825.1
B3 2348166 2348254
refGene_gn=IL16;refGene_tx=XM_023254817.1;refGene_gn=IL16;refGene_tx=X
M_023254818.1;refGene_gn=IL16;refGene_tx=XM_023254819.1;refGene_gn=IL1
6;refGene_tx=XM_023254820.1;refGene_gn=IL16;refGene_tx=XM_023254821.1;
refGene_gn=IL16;refGene_tx=XM_023254822.1;refGene_gn=IL16;refGene_tx=X
M_023254823.1;refGene_gn=IL16;refGene_tx=XM_023254824.1;refGene_gn=IL1
6;refGene_tx=XM_023254825.1
B3 2350248 2350727
refGene_gn=IL16;refGene_tx=XM_023254817.1;refGene_gn=IL16;refGene_tx=X
M_023254818.1;refGene_gn=IL16;refGene_tx=XM_023254819.1;refGene_gn=IL1
6;refGene_tx=XM_023254820.1;refGene_gn=IL16;refGene_tx=XM_023254821.1;
refGene_gn=IL16;refGene_tx=XM_023254822.1;refGene_gn=IL16;refGene_tx=X
M_023254823.1;refGene_gn=IL16;refGene_tx=XM_023254824.1;refGene_gn=IL1
6;refGene_tx=XM_023254825.1
B3 2354121 2354266
refGene_gn=IL16;refGene_tx=XM_023254817.1;refGene_gn=IL16;refGene_tx=X
M_023254818.1;refGene_gn=IL16;refGene_tx=XM_023254819.1;refGene_gn=IL1
6;refGene_tx=XM_023254820.1;refGene_gn=IL16;refGene_tx=XM_023254821.1;
refGene_gn=IL16;refGene_tx=XM_023254822.1;refGene_gn=IL16;refGene_tx=X
M_023254823.1;refGene_gn=IL16;refGene_tx=XM_023254824.1;refGene_gn=IL1
6;refGene_tx=XM_023254825.1
B3 2356400 2357496
refGene_gn=IL16;refGene_tx=XM_023254817.1;refGene_gn=IL16;refGene_tx=X
M_023254818.1;refGene_gn=IL16;refGene_tx=XM_023254819.1;refGene_gn=IL1
6;refGene_tx=XM_023254820.1;refGene_gn=IL16;refGene_tx=XM_023254821.1;
refGene_gn=IL16;refGene_tx=XM_023254822.1;refGene_gn=IL16;refGene_tx=X
M_023254823.1;refGene_gn=IL16;refGene_tx=XM_023254824.1;refGene_gn=IL1
6;refGene_tx=XM_023254825.1;refGene_gn=IL16;refGene_tx=XM_023254826.1
B3 2358870 2359039
refGene_gn=IL16;refGene_tx=XM_023254817.1;refGene_gn=IL16;refGene_tx=X
M_023254818.1;refGene_gn=IL16;refGene_tx=XM_023254819.1;refGene_gn=IL1
6;refGene_tx=XM_023254820.1;refGene_gn=IL16;refGene_tx=XM_023254821.1;

refGene_gn=IL16;refGene_tx=XM_023254822.1;refGene_gn=IL16;refGene_tx=XM_023254823.1;refGene_gn=IL16;refGene_tx=XM_023254824.1;refGene_gn=IL16;refGene_tx=XM_023254825.1;refGene_gn=IL16;refGene_tx=XM_023254826.1
B3 2361568 2361670
refGene_gn=IL16;refGene_tx=XM_023254817.1;refGene_gn=IL16;refGene_tx=XM_023254818.1;refGene_gn=IL16;refGene_tx=XM_023254819.1;refGene_gn=IL16;refGene_tx=XM_023254820.1;refGene_gn=IL16;refGene_tx=XM_023254821.1;refGene_gn=IL16;refGene_tx=XM_023254822.1;refGene_gn=IL16;refGene_tx=XM_023254823.1;refGene_gn=IL16;refGene_tx=XM_023254824.1;refGene_gn=IL16;refGene_tx=XM_023254825.1;refGene_gn=IL16;refGene_tx=XM_023254826.1
B3 2363857 2364116
refGene_gn=IL16;refGene_tx=XM_023254817.1;refGene_gn=IL16;refGene_tx=XM_023254818.1;refGene_gn=IL16;refGene_tx=XM_023254819.1;refGene_gn=IL16;refGene_tx=XM_023254820.1;refGene_gn=IL16;refGene_tx=XM_023254821.1;refGene_gn=IL16;refGene_tx=XM_023254822.1;refGene_gn=IL16;refGene_tx=XM_023254823.1;refGene_gn=IL16;refGene_tx=XM_023254824.1;refGene_gn=IL16;refGene_tx=XM_023254825.1;refGene_gn=IL16;refGene_tx=XM_023254826.1
B3 2364334 2364457
refGene_gn=IL16;refGene_tx=XM_023254817.1;refGene_gn=IL16;refGene_tx=XM_023254818.1;refGene_gn=IL16;refGene_tx=XM_023254819.1;refGene_gn=IL16;refGene_tx=XM_023254820.1;refGene_gn=IL16;refGene_tx=XM_023254821.1;refGene_gn=IL16;refGene_tx=XM_023254822.1;refGene_gn=IL16;refGene_tx=XM_023254823.1;refGene_gn=IL16;refGene_tx=XM_023254824.1;refGene_gn=IL16;refGene_tx=XM_023254825.1;refGene_gn=IL16;refGene_tx=XM_023254826.1
B3 2366474 2366668
refGene_gn=IL16;refGene_tx=XM_023254817.1;refGene_gn=IL16;refGene_tx=XM_023254818.1;refGene_gn=IL16;refGene_tx=XM_023254819.1;refGene_gn=IL16;refGene_tx=XM_023254820.1;refGene_gn=IL16;refGene_tx=XM_023254821.1;refGene_gn=IL16;refGene_tx=XM_023254822.1;refGene_gn=IL16;refGene_tx=XM_023254823.1;refGene_gn=IL16;refGene_tx=XM_023254824.1;refGene_gn=IL16;refGene_tx=XM_023254825.1;refGene_gn=IL16;refGene_tx=XM_023254826.1
B3 2370319 2370467 refGene_gn=STARD5;refGene_tx=XM_011282714.3
B3 2371923 2372017 refGene_gn=STARD5;refGene_tx=XM_011282714.3
B3 2372930 2373048 refGene_gn=STARD5;refGene_tx=XM_011282714.3
B3 2376534 2376667 refGene_gn=STARD5;refGene_tx=XM_011282714.3
B3 2377017 2377067 refGene_gn=STARD5;refGene_tx=XM_011282714.3
B3 2377717 2377816 refGene_gn=STARD5;refGene_tx=XM_011282714.3
B3 2385910 2386754 refGene_gn=TMC3;refGene_tx=XM_019831793.2
B3 2387775 2388031 refGene_gn=TMC3;refGene_tx=XM_019831793.2
B3 2389609 2389652 refGene_gn=TMC3;refGene_tx=XM_019831793.2
B3 2390794 2390873 refGene_gn=TMC3;refGene_tx=XM_019831793.2
B3 2391242 2391368 refGene_gn=TMC3;refGene_tx=XM_019831793.2
B3 2392025 2392136 refGene_gn=TMC3;refGene_tx=XM_019831793.2
B3 2393263 2393392 refGene_gn=TMC3;refGene_tx=XM_019831793.2
B3 2395217 2395285 refGene_gn=TMC3;refGene_tx=XM_019831793.2
B3 2395789 2395918 refGene_gn=TMC3;refGene_tx=XM_019831793.2
B3 2396582 2396828 refGene_gn=TMC3;refGene_tx=XM_019831793.2
B3 2398155 2398234 refGene_gn=TMC3;refGene_tx=XM_019831793.2
B3 2401677 2401787 refGene_gn=TMC3;refGene_tx=XM_019831793.2
B3 2403390 2403538 refGene_gn=TMC3;refGene_tx=XM_019831793.2

B3 2406793 2406837 refGene_gn=TMC3;refGene_tx=XM_019831793.2
B3 2407704 2407852 refGene_gn=TMC3;refGene_tx=XM_019831793.2
B3 2409086 2409229 refGene_gn=TMC3;refGene_tx=XM_019831793.2
B3 2409329 2409428 refGene_gn=TMC3;refGene_tx=XM_019831793.2
B3 2410637 2410744 refGene_gn=TMC3;refGene_tx=XM_019831793.2
B3 2414014 2414096 refGene_gn=TMC3;refGene_tx=XM_019831793.2
B3 2419724 2419800 refGene_gn=TMC3;refGene_tx=XM_019831793.2
B3 2423209 2423356 refGene_gn=TMC3;refGene_tx=XM_019831793.2
B3 2424612 2424701 refGene_gn=TMC3;refGene_tx=XM_019831793.2
B3 2894372 2895820 refGene_gn=MEX3B;refGene_tx=XM_003986747.4
B3 2896669 2896952 refGene_gn=MEX3B;refGene_tx=XM_003986747.4
B3 2964161 2964350
refGene_gn=EFL1;refGene_tx=XM_023254827.1;refGene_gn=EFL1;refGene_tx=XM_023254828.1
B3 2971595 2971780
refGene_gn=EFL1;refGene_tx=XM_023254827.1;refGene_gn=EFL1;refGene_tx=XM_023254828.1
B3 2979289 2980272
refGene_gn=EFL1;refGene_tx=XM_023254827.1;refGene_gn=EFL1;refGene_tx=XM_023254828.1
B3 2988930 2989078
refGene_gn=EFL1;refGene_tx=XM_023254827.1;refGene_gn=EFL1;refGene_tx=XM_023254828.1
B3 2993544 2993676
refGene_gn=EFL1;refGene_tx=XM_023254827.1;refGene_gn=EFL1;refGene_tx=XM_023254828.1
B3 3039433 3039572
refGene_gn=EFL1;refGene_tx=XM_023254827.1;refGene_gn=EFL1;refGene_tx=XM_023254828.1
B3 3044525 3044692
refGene_gn=EFL1;refGene_tx=XM_023254827.1;refGene_gn=EFL1;refGene_tx=XM_023254828.1
B3 3044928 3045080
refGene_gn=EFL1;refGene_tx=XM_023254827.1;refGene_gn=EFL1;refGene_tx=XM_023254828.1
B3 3051959 3052059
refGene_gn=EFL1;refGene_tx=XM_023254827.1;refGene_gn=EFL1;refGene_tx=XM_023254828.1
B3 3054182 3054305
refGene_gn=EFL1;refGene_tx=XM_023254827.1;refGene_gn=EFL1;refGene_tx=XM_023254828.1
B3 3056044 3056181
refGene_gn=EFL1;refGene_tx=XM_023254827.1;refGene_gn=EFL1;refGene_tx=XM_023254828.1
B3 3057154 3057231
refGene_gn=EFL1;refGene_tx=XM_023254827.1;refGene_gn=EFL1;refGene_tx=XM_023254828.1
B3 3058512 3058636
refGene_gn=EFL1;refGene_tx=XM_023254827.1;refGene_gn=EFL1;refGene_tx=XM_023254828.1

B3 3067334 3067549
refGene_gn=EFL1;refGene_tx=XM_023254827.1;refGene_gn=EFL1;refGene_tx=XM_023254828.1
B3 3069048 3069186
refGene_gn=EFL1;refGene_tx=XM_023254827.1;refGene_gn=EFL1;refGene_tx=XM_023254828.1
B3 3070817 3070951
refGene_gn=EFL1;refGene_tx=XM_023254827.1;refGene_gn=EFL1;refGene_tx=XM_023254828.1
B3 3080869 3080954 refGene_gn=EFL1;refGene_tx=XM_023254827.1
B3 3082495 3082563 refGene_gn=EFL1;refGene_tx=XM_023254827.1
B3 3085461 3085552
refGene_gn=EFL1;refGene_tx=XM_023254827.1;refGene_gn=EFL1;refGene_tx=XM_023254828.1
B3 3086612 3086650 refGene_gn=SAXO2;refGene_tx=XM_019831795.2
B3 3089327 3089507 refGene_gn=SAXO2;refGene_tx=XM_019831795.2
B3 3094606 3094809 refGene_gn=SAXO2;refGene_tx=XM_019831795.2
B3 3102671 3102699 refGene_gn=SAXO2;refGene_tx=XM_023254829.1
B3 3102931 3103875
refGene_gn=SAXO2;refGene_tx=XM_019831795.2;refGene_gn=SAXO2;refGene_tx=XM_023254829.1
B3 3108956 3109063 refGene_gn=ADAMTSL3;refGene_tx=XM_023254830.1
B3 3110626 3110841 refGene_gn=ADAMTSL3;refGene_tx=XM_023254830.1
B3 3115788 3115889 refGene_gn=ADAMTSL3;refGene_tx=XM_023254830.1
B3 3121187 3121392 refGene_gn=ADAMTSL3;refGene_tx=XM_023254830.1
B3 3124880 3125064 refGene_gn=ADAMTSL3;refGene_tx=XM_023254830.1
B3 3131382 3131499 refGene_gn=ADAMTSL3;refGene_tx=XM_023254830.1
B3 3135095 3135278 refGene_gn=ADAMTSL3;refGene_tx=XM_023254830.1
B3 3164503 3164632 refGene_gn=ADAMTSL3;refGene_tx=XM_023254830.1
B3 3166593 3166721 refGene_gn=ADAMTSL3;refGene_tx=XM_023254830.1
B3 3172381 3173453 refGene_gn=ADAMTSL3;refGene_tx=XM_023254830.1
B3 3183655 3183809 refGene_gn=ADAMTSL3;refGene_tx=XM_023254830.1
B3 3195208 3195388 refGene_gn=ADAMTSL3;refGene_tx=XM_023254830.1
B3 3195467 3195660 refGene_gn=ADAMTSL3;refGene_tx=XM_023254830.1
B3 3209727 3209857 refGene_gn=ADAMTSL3;refGene_tx=XM_023254830.1
B3 3222093 3222380 refGene_gn=ADAMTSL3;refGene_tx=XM_023254830.1
B3 3236712 3236797 refGene_gn=ADAMTSL3;refGene_tx=XM_023254830.1
B3 3238457 3238605 refGene_gn=ADAMTSL3;refGene_tx=XM_023254830.1
B3 3243218 3243423 refGene_gn=ADAMTSL3;refGene_tx=XM_023254830.1
B3 3243892 3243943 refGene_gn=ADAMTSL3;refGene_tx=XM_023254830.1
B3 3245031 3245170 refGene_gn=ADAMTSL3;refGene_tx=XM_023254830.1
B3 3250435 3250547 refGene_gn=ADAMTSL3;refGene_tx=XM_023254830.1
B3 3268528 3268686 refGene_gn=ADAMTSL3;refGene_tx=XM_023254830.1
B3 3276069 3276144 refGene_gn=ADAMTSL3;refGene_tx=XM_023254830.1
B3 3304799 3304926 refGene_gn=ADAMTSL3;refGene_tx=XM_023254830.1
B3 3313137 3313374 refGene_gn=ADAMTSL3;refGene_tx=XM_023254830.1
B3 3322455 3322501 refGene_gn=ADAMTSL3;refGene_tx=XM_023254830.1
B3 3349804 3349932 refGene_gn=ADAMTSL3;refGene_tx=XM_023254830.1
B3 3403418 3403484 refGene_gn=ADAMTSL3;refGene_tx=XM_023254830.1
B3 3403485 3403539 refGene_gn=ADAMTSL3;refGene_tx=XM_023254830.1

B3	3450781	3450850	refGene_gn=ADAMTSL3;refGene_tx=XM_023254830.1
B3	3451759	3452656
refGene_gn=LOC111560763;refGene_tx=XM_023254766.1
B3	3468481	3468627	refGene_gn=SH3GL3;refGene_tx=XM_023254842.1
B3	3470192	3470398
refGene_gn=SH3GL3;refGene_tx=XM_006932204.3;refGene_gn=SH3GL3;refGene_tx=XM_011282785.2;refGene_gn=SH3GL3;refGene_tx=XM_019831808.2;refGene_gn=SH3GL3;refGene_tx=XM_023254841.1;refGene_gn=SH3GL3;refGene_tx=XM_023254844.1
B3	3478604	3478633	refGene_gn=SH3GL3;refGene_tx=XM_023254843.1
B3	3497429	3497539
refGene_gn=SH3GL3;refGene_tx=XM_006932204.3;refGene_gn=SH3GL3;refGene_tx=XM_011282785.2;refGene_gn=SH3GL3;refGene_tx=XM_019831808.2;refGene_gn=SH3GL3;refGene_tx=XM_023254841.1;refGene_gn=SH3GL3;refGene_tx=XM_023254842.1;refGene_gn=SH3GL3;refGene_tx=XM_023254843.1;refGene_gn=SH3GL3;refGene_tx=XM_023254844.1
B3	3498624	3498728
refGene_gn=SH3GL3;refGene_tx=XM_006932204.3;refGene_gn=SH3GL3;refGene_tx=XM_011282785.2;refGene_gn=SH3GL3;refGene_tx=XM_019831808.2;refGene_gn=SH3GL3;refGene_tx=XM_023254841.1;refGene_gn=SH3GL3;refGene_tx=XM_023254842.1;refGene_gn=SH3GL3;refGene_tx=XM_023254843.1;refGene_gn=SH3GL3;refGene_tx=XM_023254844.1
B3	3505198	3505357
refGene_gn=SH3GL3;refGene_tx=XM_006932204.3;refGene_gn=SH3GL3;refGene_tx=XM_011282785.2;refGene_gn=SH3GL3;refGene_tx=XM_019831808.2;refGene_gn=SH3GL3;refGene_tx=XM_023254841.1;refGene_gn=SH3GL3;refGene_tx=XM_023254842.1;refGene_gn=SH3GL3;refGene_tx=XM_023254843.1;refGene_gn=SH3GL3;refGene_tx=XM_023254844.1
B3	3509024	3509158
refGene_gn=SH3GL3;refGene_tx=XM_006932204.3;refGene_gn=SH3GL3;refGene_tx=XM_011282785.2;refGene_gn=SH3GL3;refGene_tx=XM_019831808.2;refGene_gn=SH3GL3;refGene_tx=XM_023254841.1;refGene_gn=SH3GL3;refGene_tx=XM_023254842.1;refGene_gn=SH3GL3;refGene_tx=XM_023254843.1;refGene_gn=SH3GL3;refGene_tx=XM_023254844.1
B3	3513029	3513173
refGene_gn=SH3GL3;refGene_tx=XM_006932204.3;refGene_gn=SH3GL3;refGene_tx=XM_011282785.2;refGene_gn=SH3GL3;refGene_tx=XM_019831808.2;refGene_gn=SH3GL3;refGene_tx=XM_023254841.1;refGene_gn=SH3GL3;refGene_tx=XM_023254842.1;refGene_gn=SH3GL3;refGene_tx=XM_023254843.1;refGene_gn=SH3GL3;refGene_tx=XM_023254844.1
B3	3515879	3515952
refGene_gn=SH3GL3;refGene_tx=XM_023254841.1;refGene_gn=SH3GL3;refGene_tx=XM_023254842.1;refGene_gn=SH3GL3;refGene_tx=XM_023254843.1
B3	3521310	3521379
refGene_gn=SH3GL3;refGene_tx=XM_023254841.1;refGene_gn=SH3GL3;refGene_tx=XM_023254842.1;refGene_gn=SH3GL3;refGene_tx=XM_023254843.1
B3	3607060	3607105
refGene_gn=SH3GL3;refGene_tx=XM_023254841.1;refGene_gn=SH3GL3;refGene_tx=XM_023254842.1;refGene_gn=SH3GL3;refGene_tx=XM_023254843.1
B3	3726426	3726522

refGene_gn=BNC1;refGene_tx=XM_023254831.1;refGene_gn=BNC1;refGene_tx=XM_023254832.1;refGene_gn=BNC1;refGene_tx=XM_023254833.1;refGene_gn=BNC1;refGene_tx=XM_023254834.1;refGene_gn=BNC1;refGene_tx=XM_023254835.1;refGene_gn=BNC1;refGene_tx=XM_023254836.1;refGene_gn=BNC1;refGene_tx=XM_023254837.1
B3 3749636 3749681
refGene_gn=BNC1;refGene_tx=XM_019831801.1;refGene_gn=BNC1;refGene_tx=XM_019831802.1
B3 3750822 3750831 refGene_gn=BNC1;refGene_tx=XM_023254839.1
B3 3751812 3751890 refGene_gn=BNC1;refGene_tx=XM_023254840.1
B3 3769795 3769895
refGene_gn=BNC1;refGene_tx=XM_019831801.1;refGene_gn=BNC1;refGene_tx=XM_019831802.1;refGene_gn=BNC1;refGene_tx=XM_023254831.1;refGene_gn=BNC1;refGene_tx=XM_023254832.1;refGene_gn=BNC1;refGene_tx=XM_023254833.1;refGene_gn=BNC1;refGene_tx=XM_023254834.1;refGene_gn=BNC1;refGene_tx=XM_023254835.1;refGene_gn=BNC1;refGene_tx=XM_023254836.1;refGene_gn=BNC1;refGene_tx=XM_023254837.1;refGene_gn=BNC1;refGene_tx=XM_023254839.1;refGene_gn=BNC1;refGene_tx=XM_023254840.1
B3 3771162 3771398
refGene_gn=BNC1;refGene_tx=XM_019831801.1;refGene_gn=BNC1;refGene_tx=XM_019831802.1;refGene_gn=BNC1;refGene_tx=XM_023254831.1;refGene_gn=BNC1;refGene_tx=XM_023254832.1;refGene_gn=BNC1;refGene_tx=XM_023254833.1;refGene_gn=BNC1;refGene_tx=XM_023254834.1;refGene_gn=BNC1;refGene_tx=XM_023254835.1;refGene_gn=BNC1;refGene_tx=XM_023254836.1;refGene_gn=BNC1;refGene_tx=XM_023254837.1;refGene_gn=BNC1;refGene_tx=XM_023254839.1;refGene_gn=BNC1;refGene_tx=XM_023254840.1
B3 3773754 3775583
refGene_gn=BNC1;refGene_tx=XM_019831801.1;refGene_gn=BNC1;refGene_tx=XM_019831802.1;refGene_gn=BNC1;refGene_tx=XM_023254831.1;refGene_gn=BNC1;refGene_tx=XM_023254832.1;refGene_gn=BNC1;refGene_tx=XM_023254833.1;refGene_gn=BNC1;refGene_tx=XM_023254834.1;refGene_gn=BNC1;refGene_tx=XM_023254835.1;refGene_gn=BNC1;refGene_tx=XM_023254836.1;refGene_gn=BNC1;refGene_tx=XM_023254837.1;refGene_gn=BNC1;refGene_tx=XM_023254839.1;refGene_gn=BNC1;refGene_tx=XM_023254840.1
B3 3780824 3781509
refGene_gn=BNC1;refGene_tx=XM_019831801.1;refGene_gn=BNC1;refGene_tx=XM_019831802.1;refGene_gn=BNC1;refGene_tx=XM_023254831.1;refGene_gn=BNC1;refGene_tx=XM_023254832.1;refGene_gn=BNC1;refGene_tx=XM_023254833.1;refGene_gn=BNC1;refGene_tx=XM_023254834.1;refGene_gn=BNC1;refGene_tx=XM_023254835.1;refGene_gn=BNC1;refGene_tx=XM_023254836.1;refGene_gn=BNC1;refGene_tx=XM_023254837.1;refGene_gn=BNC1;refGene_tx=XM_023254839.1;refGene_gn=BNC1;refGene_tx=XM_023254840.1
B3 3843113 3843197 refGene_gn=HDGFL3;refGene_tx=XM_023254853.1
B3 3889704 3889872 refGene_gn=TM6SF1;refGene_tx=XM_023254846.1
B3 3890988 3891065 refGene_gn=HDGFL3;refGene_tx=XM_023254853.1
B3 3896719 3896858 refGene_gn=HDGFL3;refGene_tx=XM_023254853.1
B3 3897182 3897341 refGene_gn=HDGFL3;refGene_tx=XM_023254853.1
B3 3901244 3901391 refGene_gn=HDGFL3;refGene_tx=XM_023254853.1
B3 3914938 3914944 refGene_gn=HDGFL3;refGene_tx=XM_023254853.1
B3 3917577 3917769

refGene_gn=TM6SF1;refGene_tx=XM_023254845.1;refGene_gn=TM6SF1;refGene_
tx=XM_023254847.1;refGene_gn=TM6SF1;refGene_tx=XM_023254848.1;refGene_
gn=TM6SF1;refGene_tx=XM_023254849.1;refGene_gn=TM6SF1;refGene_tx=XM_02
3254851.1;refGene_gn=TM6SF1;refGene_tx=XM_023254852.1
B3 3925069 3925189
refGene_gn=TM6SF1;refGene_tx=XM_023254845.1;refGene_gn=TM6SF1;refGene_
tx=XM_023254846.1;refGene_gn=TM6SF1;refGene_tx=XM_023254847.1;refGene_
gn=TM6SF1;refGene_tx=XM_023254848.1;refGene_gn=TM6SF1;refGene_tx=XM_02
3254849.1;refGene_gn=TM6SF1;refGene_tx=XM_023254851.1;refGene_gn=TM6SF
1;refGene_tx=XM_023254852.1
B3 3925838 3925931
refGene_gn=TM6SF1;refGene_tx=XM_023254845.1;refGene_gn=TM6SF1;refGene_
tx=XM_023254846.1;refGene_gn=TM6SF1;refGene_tx=XM_023254848.1;refGene_
gn=TM6SF1;refGene_tx=XM_023254849.1;refGene_gn=TM6SF1;refGene_tx=XM_02
3254851.1;refGene_gn=TM6SF1;refGene_tx=XM_023254852.1
B3 3927794 3927899
refGene_gn=TM6SF1;refGene_tx=XM_023254845.1;refGene_gn=TM6SF1;refGene_
tx=XM_023254846.1;refGene_gn=TM6SF1;refGene_tx=XM_023254847.1;refGene_
gn=TM6SF1;refGene_tx=XM_023254848.1;refGene_gn=TM6SF1;refGene_tx=XM_02
3254849.1;refGene_gn=TM6SF1;refGene_tx=XM_023254851.1;refGene_gn=TM6SF
1;refGene_tx=XM_023254852.1
B3 3929676 3929798
refGene_gn=TM6SF1;refGene_tx=XM_023254845.1;refGene_gn=TM6SF1;refGene_
tx=XM_023254846.1;refGene_gn=TM6SF1;refGene_tx=XM_023254847.1;refGene_
gn=TM6SF1;refGene_tx=XM_023254848.1;refGene_gn=TM6SF1;refGene_tx=XM_02
3254851.1
B3 3930436 3930519
refGene_gn=TM6SF1;refGene_tx=XM_023254845.1;refGene_gn=TM6SF1;refGene_
tx=XM_023254846.1;refGene_gn=TM6SF1;refGene_tx=XM_023254847.1;refGene_
gn=TM6SF1;refGene_tx=XM_023254848.1;refGene_gn=TM6SF1;refGene_tx=XM_02
3254851.1
B3 3932209 3932313
refGene_gn=TM6SF1;refGene_tx=XM_023254845.1;refGene_gn=TM6SF1;refGene_
tx=XM_023254846.1;refGene_gn=TM6SF1;refGene_tx=XM_023254847.1;refGene_
gn=TM6SF1;refGene_tx=XM_023254848.1
B3 3937157 3937255
refGene_gn=TM6SF1;refGene_tx=XM_023254845.1;refGene_gn=TM6SF1;refGene_
tx=XM_023254846.1;refGene_gn=TM6SF1;refGene_tx=XM_023254847.1;refGene_
gn=TM6SF1;refGene_tx=XM_023254848.1;refGene_gn=TM6SF1;refGene_tx=XM_02
3254849.1;refGene_gn=TM6SF1;refGene_tx=XM_023254852.1
B3 3941313 3941417
refGene_gn=TM6SF1;refGene_tx=XM_023254845.1;refGene_gn=TM6SF1;refGene_
tx=XM_023254846.1;refGene_gn=TM6SF1;refGene_tx=XM_023254847.1;refGene_
gn=TM6SF1;refGene_tx=XM_023254849.1
B3 3946450 3946542
refGene_gn=TM6SF1;refGene_tx=XM_023254845.1;refGene_gn=TM6SF1;refGene_
tx=XM_023254846.1;refGene_gn=TM6SF1;refGene_tx=XM_023254847.1;refGene_
gn=TM6SF1;refGene_tx=XM_023254848.1;refGene_gn=TM6SF1;refGene_tx=XM_02
3254849.1;refGene_gn=TM6SF1;refGene_tx=XM_023254852.1
B3 3983526 3983936 refGene_gn=BTBD1;refGene_tx=XM_023254854.1

B3 3994974 3995131 refGene_gn=BTBD1;refGene_tx=XM_023254854.1
B3 4000016 4000122 refGene_gn=BTBD1;refGene_tx=XM_023254854.1
B3 4006837 4007035 refGene_gn=BTBD1;refGene_tx=XM_023254854.1
B3 4014386 4014579 refGene_gn=BTBD1;refGene_tx=XM_023254854.1
B3 4020298 4020386 refGene_gn=BTBD1;refGene_tx=XM_023254854.1
B3 4022687 4022834 refGene_gn=BTBD1;refGene_tx=XM_023254854.1
B3 4023324 4023483 refGene_gn=BTBD1;refGene_tx=XM_023254854.1
B3 4026309 4026423
refGene_gn=CB3H15orf40;refGene_tx=XM_023254855.1;refGene_gn=CB3H15orf40;refGene_tx=XM_023254856.1;refGene_gn=CB3H15orf40;refGene_tx=XM_023254857.1
B3 4027170 4027297
refGene_gn=CB3H15orf40;refGene_tx=XM_023254855.1;refGene_gn=CB3H15orf40;refGene_tx=XM_023254856.1;refGene_gn=CB3H15orf40;refGene_tx=XM_023254857.1
B3 4029293 4029421
refGene_gn=CB3H15orf40;refGene_tx=XM_023254855.1;refGene_gn=CB3H15orf40;refGene_tx=XM_023254856.1;refGene_gn=CB3H15orf40;refGene_tx=XM_023254857.1
B3 4030692 4030788
refGene_gn=CB3H15orf40;refGene_tx=XM_023254856.1
B3 4032940 4032964
refGene_gn=CB3H15orf40;refGene_tx=XM_023254857.1
B3 4036578 4036722
refGene_gn=CB3H15orf40;refGene_tx=XM_023254855.1
B3 4039896 4040083
refGene_gn=FAM103A1;refGene_tx=XM_023254858.1;refGene_gn=FAM103A1;refGene_tx=XM_023254859.1;refGene_gn=FAM103A1;refGene_tx=XM_023254860.1;refGene_gn=FAM103A1;refGene_tx=XM_023254861.1;refGene_gn=FAM103A1;refGene_tx=XM_023254862.1
B3 4040795 4040965
refGene_gn=FAM103A1;refGene_tx=XM_023254858.1;refGene_gn=FAM103A1;refGene_tx=XM_023254859.1;refGene_gn=FAM103A1;refGene_tx=XM_023254860.1;refGene_gn=FAM103A1;refGene_tx=XM_023254861.1;refGene_gn=FAM103A1;refGene_tx=XM_023254862.1
B3 4077266 4077361
refGene_gn=HOMER2;refGene_tx=XM_023254863.1;refGene_gn=HOMER2;refGene_tx=XM_023254864.1
B3 4081687 4081692 refGene_gn=HOMER2;refGene_tx=XM_023254865.1
B3 4136613 4136770
refGene_gn=HOMER2;refGene_tx=XM_023254863.1;refGene_gn=HOMER2;refGene_tx=XM_023254864.1;refGene_gn=HOMER2;refGene_tx=XM_023254865.1
B3 4151525 4151657
refGene_gn=HOMER2;refGene_tx=XM_023254863.1;refGene_gn=HOMER2;refGene_tx=XM_023254864.1;refGene_gn=HOMER2;refGene_tx=XM_023254865.1;refGene_gn=HOMER2;refGene_tx=XM_023254866.1
B3 4164792 4164885
refGene_gn=HOMER2;refGene_tx=XM_023254863.1;refGene_gn=HOMER2;refGene_tx=XM_023254864.1;refGene_gn=HOMER2;refGene_tx=XM_023254865.1;refGene_gn=HOMER2;refGene_tx=XM_023254866.1

B3	4169132	4169272
refGene_gn=HOMER2;refGene_tx=XM_023254863.1;refGene_gn=HOMER2;refGene_tx=XM_023254864.1;refGene_gn=HOMER2;refGene_tx=XM_023254865.1;refGene_gn=HOMER2;refGene_tx=XM_023254866.1
B3	4172047	4172204
refGene_gn=HOMER2;refGene_tx=XM_023254863.1;refGene_gn=HOMER2;refGene_tx=XM_023254864.1;refGene_gn=HOMER2;refGene_tx=XM_023254865.1;refGene_gn=HOMER2;refGene_tx=XM_023254866.1
B3	4173502	4173613
refGene_gn=HOMER2;refGene_tx=XM_023254863.1;refGene_gn=HOMER2;refGene_tx=XM_023254864.1;refGene_gn=HOMER2;refGene_tx=XM_023254865.1;refGene_gn=HOMER2;refGene_tx=XM_023254866.1
B3	4174196	4174277
refGene_gn=HOMER2;refGene_tx=XM_023254863.1;refGene_gn=HOMER2;refGene_tx=XM_023254864.1;refGene_gn=HOMER2;refGene_tx=XM_023254865.1;refGene_gn=HOMER2;refGene_tx=XM_023254866.1
B3	4175425	4175614
refGene_gn=HOMER2;refGene_tx=XM_023254863.1;refGene_gn=HOMER2;refGene_tx=XM_023254864.1;refGene_gn=HOMER2;refGene_tx=XM_023254865.1;refGene_gn=HOMER2;refGene_tx=XM_023254866.1
B3	4189982	4190172	refGene_gn=WHAMM;refGene_tx=XM_023254878.1
B3	4192435	4192860	refGene_gn=WHAMM;refGene_tx=XM_023254878.1
B3	4197158	4197254	refGene_gn=WHAMM;refGene_tx=XM_023254878.1
B3	4197510	4197597	refGene_gn=WHAMM;refGene_tx=XM_023254878.1
B3	4198987	4199175	refGene_gn=WHAMM;refGene_tx=XM_023254878.1
B3	4201332	4201498	refGene_gn=WHAMM;refGene_tx=XM_023254878.1
B3	4202326	4202496	refGene_gn=WHAMM;refGene_tx=XM_023254878.1
B3	4203558	4203709	refGene_gn=WHAMM;refGene_tx=XM_023254878.1
B3	4204514	4204688	refGene_gn=WHAMM;refGene_tx=XM_023254878.1
B3	4206246	4206837	refGene_gn=WHAMM;refGene_tx=XM_023254878.1
B3	4212779	4212789
refGene_gn=FSD2;refGene_tx=XM_019831817.2;refGene_gn=FSD2;refGene_tx=XM_023254877.1
B3	4227407	4228078
refGene_gn=FSD2;refGene_tx=XM_019831817.2;refGene_gn=FSD2;refGene_tx=XM_023254868.1;refGene_gn=FSD2;refGene_tx=XM_023254869.1;refGene_gn=FSD2;refGene_tx=XM_023254870.1;refGene_gn=FSD2;refGene_tx=XM_023254871.1;refGene_gn=FSD2;refGene_tx=XM_023254872.1;refGene_gn=FSD2;refGene_tx=XM_023254873.1;refGene_gn=FSD2;refGene_tx=XM_023254874.1;refGene_gn=FSD2;refGene_tx=XM_023254875.1;refGene_gn=FSD2;refGene_tx=XM_023254876.1;refGene_gn=FSD2;refGene_tx=XM_023254877.1
B3	4228211	4228307
refGene_gn=FSD2;refGene_tx=XM_019831817.2;refGene_gn=FSD2;refGene_tx=XM_023254868.1;refGene_gn=FSD2;refGene_tx=XM_023254869.1;refGene_gn=FSD2;refGene_tx=XM_023254870.1;refGene_gn=FSD2;refGene_tx=XM_023254871.1;refGene_gn=FSD2;refGene_tx=XM_023254872.1;refGene_gn=FSD2;refGene_tx=XM_023254873.1;refGene_gn=FSD2;refGene_tx=XM_023254874.1;refGene_gn=FSD2;refGene_tx=XM_023254875.1;refGene_gn=FSD2;refGene_tx=XM_023254876.1;refGene_gn=FSD2;refGene_tx=XM_023254877.1
B3	4230314	4230545

refGene_gn=FSD2;refGene_tx=XM_019831817.2;refGene_gn=FSD2;refGene_tx=XM_023254868.1;refGene_gn=FSD2;refGene_tx=XM_023254869.1;refGene_gn=FSD2;refGene_tx=XM_023254870.1;refGene_gn=FSD2;refGene_tx=XM_023254871.1;refGene_gn=FSD2;refGene_tx=XM_023254872.1;refGene_gn=FSD2;refGene_tx=XM_023254873.1;refGene_gn=FSD2;refGene_tx=XM_023254874.1;refGene_gn=FSD2;refGene_tx=XM_023254875.1;refGene_gn=FSD2;refGene_tx=XM_023254876.1;refGene_gn=FSD2;refGene_tx=XM_023254877.1
B3 4231795 4231818
refGene_gn=FSD2;refGene_tx=XM_019831817.2;refGene_gn=FSD2;refGene_tx=XM_023254868.1;refGene_gn=FSD2;refGene_tx=XM_023254869.1;refGene_gn=FSD2;refGene_tx=XM_023254870.1;refGene_gn=FSD2;refGene_tx=XM_023254871.1;refGene_gn=FSD2;refGene_tx=XM_023254872.1;refGene_gn=FSD2;refGene_tx=XM_023254873.1;refGene_gn=FSD2;refGene_tx=XM_023254874.1;refGene_gn=FSD2;refGene_tx=XM_023254875.1;refGene_gn=FSD2;refGene_tx=XM_023254876.1;refGene_gn=FSD2;refGene_tx=XM_023254877.1
B3 4232797 4232919
refGene_gn=FSD2;refGene_tx=XM_019831817.2;refGene_gn=FSD2;refGene_tx=XM_023254868.1;refGene_gn=FSD2;refGene_tx=XM_023254869.1;refGene_gn=FSD2;refGene_tx=XM_023254870.1;refGene_gn=FSD2;refGene_tx=XM_023254871.1;refGene_gn=FSD2;refGene_tx=XM_023254872.1;refGene_gn=FSD2;refGene_tx=XM_023254873.1;refGene_gn=FSD2;refGene_tx=XM_023254874.1;refGene_gn=FSD2;refGene_tx=XM_023254875.1;refGene_gn=FSD2;refGene_tx=XM_023254876.1;refGene_gn=FSD2;refGene_tx=XM_023254877.1
B3 4235811 4235967
refGene_gn=FSD2;refGene_tx=XM_019831817.2;refGene_gn=FSD2;refGene_tx=XM_023254868.1;refGene_gn=FSD2;refGene_tx=XM_023254869.1;refGene_gn=FSD2;refGene_tx=XM_023254870.1;refGene_gn=FSD2;refGene_tx=XM_023254871.1;refGene_gn=FSD2;refGene_tx=XM_023254872.1;refGene_gn=FSD2;refGene_tx=XM_023254873.1;refGene_gn=FSD2;refGene_tx=XM_023254874.1;refGene_gn=FSD2;refGene_tx=XM_023254875.1;refGene_gn=FSD2;refGene_tx=XM_023254876.1;refGene_gn=FSD2;refGene_tx=XM_023254877.1
B3 4238246 4238381
refGene_gn=FSD2;refGene_tx=XM_019831817.2;refGene_gn=FSD2;refGene_tx=XM_023254868.1;refGene_gn=FSD2;refGene_tx=XM_023254869.1;refGene_gn=FSD2;refGene_tx=XM_023254870.1;refGene_gn=FSD2;refGene_tx=XM_023254871.1;refGene_gn=FSD2;refGene_tx=XM_023254872.1;refGene_gn=FSD2;refGene_tx=XM_023254873.1;refGene_gn=FSD2;refGene_tx=XM_023254874.1;refGene_gn=FSD2;refGene_tx=XM_023254875.1;refGene_gn=FSD2;refGene_tx=XM_023254876.1;refGene_gn=FSD2;refGene_tx=XM_023254877.1
B3 4239097 4239248
refGene_gn=FSD2;refGene_tx=XM_019831817.2;refGene_gn=FSD2;refGene_tx=XM_023254868.1;refGene_gn=FSD2;refGene_tx=XM_023254869.1;refGene_gn=FSD2;refGene_tx=XM_023254870.1;refGene_gn=FSD2;refGene_tx=XM_023254871.1;refGene_gn=FSD2;refGene_tx=XM_023254872.1;refGene_gn=FSD2;refGene_tx=XM_023254873.1;refGene_gn=FSD2;refGene_tx=XM_023254874.1;refGene_gn=FSD2;refGene_tx=XM_023254875.1;refGene_gn=FSD2;refGene_tx=XM_023254876.1;refGene_gn=FSD2;refGene_tx=XM_023254877.1
B3 4241201 4241335
refGene_gn=FSD2;refGene_tx=XM_019831817.2;refGene_gn=FSD2;refGene_tx=XM_023254868.1;refGene_gn=FSD2;refGene_tx=XM_023254869.1;refGene_gn=FSD

2;refGene_tx=XM_023254870.1;refGene_gn=FSD2;refGene_tx=XM_023254871.1;
refGene_gn=FSD2;refGene_tx=XM_023254872.1;refGene_gn=FSD2;refGene_tx=X
M_023254873.1;refGene_gn=FSD2;refGene_tx=XM_023254874.1;refGene_gn=FSD
2;refGene_tx=XM_023254875.1;refGene_gn=FSD2;refGene_tx=XM_023254876.1;
refGene_gn=FSD2;refGene_tx=XM_023254877.1
B3 4244047 4244180
refGene_gn=FSD2;refGene_tx=XM_019831817.2;refGene_gn=FSD2;refGene_tx=X
M_023254868.1;refGene_gn=FSD2;refGene_tx=XM_023254869.1;refGene_gn=FSD
2;refGene_tx=XM_023254870.1;refGene_gn=FSD2;refGene_tx=XM_023254871.1;
refGene_gn=FSD2;refGene_tx=XM_023254872.1;refGene_gn=FSD2;refGene_tx=X
M_023254873.1;refGene_gn=FSD2;refGene_tx=XM_023254874.1;refGene_gn=FSD
2;refGene_tx=XM_023254875.1;refGene_gn=FSD2;refGene_tx=XM_023254876.1;
refGene_gn=FSD2;refGene_tx=XM_023254877.1
B3 4248505 4248682
refGene_gn=FSD2;refGene_tx=XM_019831817.2;refGene_gn=FSD2;refGene_tx=X
M_023254868.1;refGene_gn=FSD2;refGene_tx=XM_023254869.1;refGene_gn=FSD
2;refGene_tx=XM_023254870.1;refGene_gn=FSD2;refGene_tx=XM_023254871.1;
refGene_gn=FSD2;refGene_tx=XM_023254872.1;refGene_gn=FSD2;refGene_tx=X
M_023254873.1;refGene_gn=FSD2;refGene_tx=XM_023254874.1;refGene_gn=FSD
2;refGene_tx=XM_023254875.1;refGene_gn=FSD2;refGene_tx=XM_023254876.1
B3 4250668 4250921
refGene_gn=FSD2;refGene_tx=XM_019831817.2;refGene_gn=FSD2;refGene_tx=X
M_023254868.1;refGene_gn=FSD2;refGene_tx=XM_023254869.1;refGene_gn=FSD
2;refGene_tx=XM_023254870.1;refGene_gn=FSD2;refGene_tx=XM_023254871.1;
refGene_gn=FSD2;refGene_tx=XM_023254872.1;refGene_gn=FSD2;refGene_tx=X
M_023254873.1;refGene_gn=FSD2;refGene_tx=XM_023254874.1;refGene_gn=FSD
2;refGene_tx=XM_023254875.1;refGene_gn=FSD2;refGene_tx=XM_023254876.1;
refGene_gn=FSD2;refGene_tx=XM_023254877.1
B3 4291319 4291432 refGene_gn=AP3B2;refGene_tx=XM_023254867.1
B3 4294313 4294450 refGene_gn=AP3B2;refGene_tx=XM_019831815.2
B3 4295026 4295286 refGene_gn=AP3B2;refGene_tx=XM_011282781.3
B3 4300628 4300704
refGene_gn=AP3B2;refGene_tx=XM_011282781.3;refGene_gn=AP3B2;refGene_tx
=XM_019831815.2;refGene_gn=AP3B2;refGene_tx=XM_023254867.1
B3 4300834 4300909
refGene_gn=AP3B2;refGene_tx=XM_011282781.3;refGene_gn=AP3B2;refGene_tx
=XM_019831815.2;refGene_gn=AP3B2;refGene_tx=XM_023254867.1
B3 4301181 4301277
refGene_gn=AP3B2;refGene_tx=XM_011282781.3;refGene_gn=AP3B2;refGene_tx
=XM_019831815.2;refGene_gn=AP3B2;refGene_tx=XM_023254867.1
B3 4306740 4306901
refGene_gn=AP3B2;refGene_tx=XM_011282781.3;refGene_gn=AP3B2;refGene_tx
=XM_019831815.2;refGene_gn=AP3B2;refGene_tx=XM_023254867.1
B3 4307177 4307244
refGene_gn=AP3B2;refGene_tx=XM_011282781.3;refGene_gn=AP3B2;refGene_tx
=XM_019831815.2;refGene_gn=AP3B2;refGene_tx=XM_023254867.1
B3 4307342 4307525
refGene_gn=AP3B2;refGene_tx=XM_011282781.3;refGene_gn=AP3B2;refGene_tx
=XM_019831815.2;refGene_gn=AP3B2;refGene_tx=XM_023254867.1
B3 4307632 4307916

refGene_gn=AP3B2;refGene_tx=XM_011282781.3;refGene_gn=AP3B2;refGene_tx=XM_019831815.2;refGene_gn=AP3B2;refGene_tx=XM_023254867.1
B3 4308376 4308431
refGene_gn=AP3B2;refGene_tx=XM_011282781.3;refGene_gn=AP3B2;refGene_tx=XM_019831815.2;refGene_gn=AP3B2;refGene_tx=XM_023254867.1
B3 4308787 4308859
refGene_gn=AP3B2;refGene_tx=XM_011282781.3;refGene_gn=AP3B2;refGene_tx=XM_019831815.2;refGene_gn=AP3B2;refGene_tx=XM_023254867.1
B3 4310159 4310222
refGene_gn=AP3B2;refGene_tx=XM_011282781.3;refGene_gn=AP3B2;refGene_tx=XM_019831815.2;refGene_gn=AP3B2;refGene_tx=XM_023254867.1
B3 4310601 4310734
refGene_gn=AP3B2;refGene_tx=XM_011282781.3;refGene_gn=AP3B2;refGene_tx=XM_019831815.2;refGene_gn=AP3B2;refGene_tx=XM_023254867.1
B3 4310987 4311097
refGene_gn=AP3B2;refGene_tx=XM_011282781.3;refGene_gn=AP3B2;refGene_tx=XM_019831815.2;refGene_gn=AP3B2;refGene_tx=XM_023254867.1
B3 4311754 4311931
refGene_gn=AP3B2;refGene_tx=XM_011282781.3;refGene_gn=AP3B2;refGene_tx=XM_019831815.2;refGene_gn=AP3B2;refGene_tx=XM_023254867.1
B3 4317078 4317265
refGene_gn=AP3B2;refGene_tx=XM_011282781.3;refGene_gn=AP3B2;refGene_tx=XM_019831815.2;refGene_gn=AP3B2;refGene_tx=XM_023254867.1
B3 4318503 4318622
refGene_gn=AP3B2;refGene_tx=XM_011282781.3;refGene_gn=AP3B2;refGene_tx=XM_019831815.2;refGene_gn=AP3B2;refGene_tx=XM_023254867.1
B3 4319005 4319114
refGene_gn=AP3B2;refGene_tx=XM_011282781.3;refGene_gn=AP3B2;refGene_tx=XM_019831815.2;refGene_gn=AP3B2;refGene_tx=XM_023254867.1
B3 4319401 4319519
refGene_gn=AP3B2;refGene_tx=XM_011282781.3;refGene_gn=AP3B2;refGene_tx=XM_019831815.2;refGene_gn=AP3B2;refGene_tx=XM_023254867.1
B3 4319914 4320098
refGene_gn=AP3B2;refGene_tx=XM_011282781.3;refGene_gn=AP3B2;refGene_tx=XM_019831815.2;refGene_gn=AP3B2;refGene_tx=XM_023254867.1
B3 4320274 4320335
refGene_gn=AP3B2;refGene_tx=XM_011282781.3;refGene_gn=AP3B2;refGene_tx=XM_019831815.2;refGene_gn=AP3B2;refGene_tx=XM_023254867.1
B3 4320684 4320791
refGene_gn=AP3B2;refGene_tx=XM_011282781.3;refGene_gn=AP3B2;refGene_tx=XM_019831815.2;refGene_gn=AP3B2;refGene_tx=XM_023254867.1
B3 4320979 4321208
refGene_gn=AP3B2;refGene_tx=XM_011282781.3;refGene_gn=AP3B2;refGene_tx=XM_019831815.2;refGene_gn=AP3B2;refGene_tx=XM_023254867.1
B3 4321636 4321721
refGene_gn=AP3B2;refGene_tx=XM_011282781.3;refGene_gn=AP3B2;refGene_tx=XM_019831815.2;refGene_gn=AP3B2;refGene_tx=XM_023254867.1
B3 4321923 4322021
refGene_gn=AP3B2;refGene_tx=XM_011282781.3;refGene_gn=AP3B2;refGene_tx=XM_019831815.2;refGene_gn=AP3B2;refGene_tx=XM_023254867.1

B3 4324473 4324612
refGene_gn=AP3B2;refGene_tx=XM_011282781.3;refGene_gn=AP3B2;refGene_tx=XM_019831815.2;refGene_gn=AP3B2;refGene_tx=XM_023254867.1
B3 4324716 4324867
refGene_gn=AP3B2;refGene_tx=XM_011282781.3;refGene_gn=AP3B2;refGene_tx=XM_019831815.2;refGene_gn=AP3B2;refGene_tx=XM_023254867.1
B3 4333956 4333971 refGene_gn=CPEB1;refGene_tx=XM_023254879.1
B3 4334295 4334457
refGene_gn=CPEB1;refGene_tx=XM_011282774.3;refGene_gn=CPEB1;refGene_tx=XM_011282775.3;refGene_gn=CPEB1;refGene_tx=XM_011282776.3;refGene_gn=CPEB1;refGene_tx=XM_011282777.3;refGene_gn=CPEB1;refGene_tx=XM_019831819.2;refGene_gn=CPEB1;refGene_tx=XM_023254883.1
B3 4350173 4350348
refGene_gn=CPEB1;refGene_tx=XM_011282774.3;refGene_gn=CPEB1;refGene_tx=XM_011282775.3;refGene_gn=CPEB1;refGene_tx=XM_011282776.3;refGene_gn=CPEB1;refGene_tx=XM_011282777.3;refGene_gn=CPEB1;refGene_tx=XM_019831819.2;refGene_gn=CPEB1;refGene_tx=XM_023254879.1;refGene_gn=CPEB1;refGene_tx=XM_023254883.1
B3 4399344 4399536
refGene_gn=CPEB1;refGene_tx=XM_006932227.4;refGene_gn=CPEB1;refGene_tx=XM_011282774.3;refGene_gn=CPEB1;refGene_tx=XM_011282775.3;refGene_gn=CPEB1;refGene_tx=XM_011282776.3;refGene_gn=CPEB1;refGene_tx=XM_011282777.3;refGene_gn=CPEB1;refGene_tx=XM_011282780.3;refGene_gn=CPEB1;refGene_tx=XM_019831820.2;refGene_gn=CPEB1;refGene_tx=XM_019831821.2;refGene_gn=CPEB1;refGene_tx=XM_019831822.2;refGene_gn=CPEB1;refGene_tx=XM_019831823.2;refGene_gn=CPEB1;refGene_tx=XM_023254879.1;refGene_gn=CPEB1;refGene_tx=XM_023254880.1;refGene_gn=CPEB1;refGene_tx=XM_023254881.1;refGene_gn=CPEB1;refGene_tx=XM_023254882.1;refGene_gn=CPEB1;refGene_tx=XM_023254883.1;refGene_gn=CPEB1;refGene_tx=XM_023254884.1
B3 4411476 4411703
refGene_gn=CPEB1;refGene_tx=XM_006932227.4;refGene_gn=CPEB1;refGene_tx=XM_011282774.3;refGene_gn=CPEB1;refGene_tx=XM_011282775.3;refGene_gn=CPEB1;refGene_tx=XM_011282776.3;refGene_gn=CPEB1;refGene_tx=XM_011282777.3;refGene_gn=CPEB1;refGene_tx=XM_011282780.3;refGene_gn=CPEB1;refGene_tx=XM_019831819.2;refGene_gn=CPEB1;refGene_tx=XM_019831820.2;refGene_gn=CPEB1;refGene_tx=XM_019831821.2;refGene_gn=CPEB1;refGene_tx=XM_019831822.2;refGene_gn=CPEB1;refGene_tx=XM_019831823.2;refGene_gn=CPEB1;refGene_tx=XM_023254879.1;refGene_gn=CPEB1;refGene_tx=XM_023254880.1;refGene_gn=CPEB1;refGene_tx=XM_023254881.1;refGene_gn=CPEB1;refGene_tx=XM_023254882.1;refGene_gn=CPEB1;refGene_tx=XM_023254883.1;refGene_gn=CPEB1;refGene_tx=XM_023254884.1
B3 4413411 4413664
refGene_gn=CPEB1;refGene_tx=XM_006932227.4;refGene_gn=CPEB1;refGene_tx=XM_011282774.3;refGene_gn=CPEB1;refGene_tx=XM_011282775.3;refGene_gn=CPEB1;refGene_tx=XM_011282776.3;refGene_gn=CPEB1;refGene_tx=XM_011282777.3;refGene_gn=CPEB1;refGene_tx=XM_011282780.3;refGene_gn=CPEB1;refGene_tx=XM_019831819.2;refGene_gn=CPEB1;refGene_tx=XM_019831820.2;refGene_gn=CPEB1;refGene_tx=XM_019831821.2;refGene_gn=CPEB1;refGene_tx=XM_019831822.2;refGene_gn=CPEB1;refGene_tx=XM_019831823.2;refGene_gn=CPEB1;refGene_tx=XM_023254879.1;refGene_gn=CPEB1;refGene_tx=XM_023254880.1;r

efGene_gn=CPEB1;refGene_tx=XM_023254881.1;refGene_gn=CPEB1;refGene_tx=XM_023254882.1;refGene_gn=CPEB1;refGene_tx=XM_023254883.1;refGene_gn=CPEB1;refGene_tx=XM_023254884.1
B3 4415620 4415734
refGene_gn=CPEB1;refGene_tx=XM_006932227.4;refGene_gn=CPEB1;refGene_tx=XM_011282774.3;refGene_gn=CPEB1;refGene_tx=XM_011282775.3;refGene_gn=CPEB1;refGene_tx=XM_011282776.3;refGene_gn=CPEB1;refGene_tx=XM_011282777.3;refGene_gn=CPEB1;refGene_tx=XM_011282780.3;refGene_gn=CPEB1;refGene_tx=XM_019831819.2;refGene_gn=CPEB1;refGene_tx=XM_019831820.2;refGene_gn=CPEB1;refGene_tx=XM_019831821.2;refGene_gn=CPEB1;refGene_tx=XM_019831822.2;refGene_gn=CPEB1;refGene_tx=XM_019831823.2;refGene_gn=CPEB1;refGene_tx=XM_023254879.1;refGene_gn=CPEB1;refGene_tx=XM_023254880.1;refGene_gn=CPEB1;refGene_tx=XM_023254881.1;refGene_gn=CPEB1;refGene_tx=XM_023254882.1;refGene_gn=CPEB1;refGene_tx=XM_023254883.1;refGene_gn=CPEB1;refGene_tx=XM_023254884.1
B3 4415983 4416088
refGene_gn=CPEB1;refGene_tx=XM_006932227.4;refGene_gn=CPEB1;refGene_tx=XM_011282774.3;refGene_gn=CPEB1;refGene_tx=XM_011282775.3;refGene_gn=CPEB1;refGene_tx=XM_011282776.3;refGene_gn=CPEB1;refGene_tx=XM_011282777.3;refGene_gn=CPEB1;refGene_tx=XM_011282780.3;refGene_gn=CPEB1;refGene_tx=XM_019831819.2;refGene_gn=CPEB1;refGene_tx=XM_019831820.2;refGene_gn=CPEB1;refGene_tx=XM_019831821.2;refGene_gn=CPEB1;refGene_tx=XM_019831822.2;refGene_gn=CPEB1;refGene_tx=XM_019831823.2;refGene_gn=CPEB1;refGene_tx=XM_023254879.1;refGene_gn=CPEB1;refGene_tx=XM_023254880.1;refGene_gn=CPEB1;refGene_tx=XM_023254881.1;refGene_gn=CPEB1;refGene_tx=XM_023254882.1;refGene_gn=CPEB1;refGene_tx=XM_023254883.1;refGene_gn=CPEB1;refGene_tx=XM_023254884.1
B3 4416978 4417115
refGene_gn=CPEB1;refGene_tx=XM_006932227.4;refGene_gn=CPEB1;refGene_tx=XM_011282774.3;refGene_gn=CPEB1;refGene_tx=XM_011282775.3;refGene_gn=CPEB1;refGene_tx=XM_011282776.3;refGene_gn=CPEB1;refGene_tx=XM_011282777.3;refGene_gn=CPEB1;refGene_tx=XM_011282780.3;refGene_gn=CPEB1;refGene_tx=XM_019831819.2;refGene_gn=CPEB1;refGene_tx=XM_019831820.2;refGene_gn=CPEB1;refGene_tx=XM_019831821.2;refGene_gn=CPEB1;refGene_tx=XM_019831822.2;refGene_gn=CPEB1;refGene_tx=XM_019831823.2;refGene_gn=CPEB1;refGene_tx=XM_023254879.1;refGene_gn=CPEB1;refGene_tx=XM_023254880.1;refGene_gn=CPEB1;refGene_tx=XM_023254881.1;refGene_gn=CPEB1;refGene_tx=XM_023254882.1;refGene_gn=CPEB1;refGene_tx=XM_023254883.1;refGene_gn=CPEB1;refGene_tx=XM_023254884.1
B3 4418406 4418436 refGene_gn=CPEB1;refGene_tx=XM_023254883.1
B3 4419976 4420175
refGene_gn=CPEB1;refGene_tx=XM_006932227.4;refGene_gn=CPEB1;refGene_tx=XM_011282774.3;refGene_gn=CPEB1;refGene_tx=XM_011282775.3;refGene_gn=CPEB1;refGene_tx=XM_011282776.3;refGene_gn=CPEB1;refGene_tx=XM_011282777.3;refGene_gn=CPEB1;refGene_tx=XM_011282780.3;refGene_gn=CPEB1;refGene_tx=XM_019831819.2;refGene_gn=CPEB1;refGene_tx=XM_019831820.2;refGene_gn=CPEB1;refGene_tx=XM_019831821.2;refGene_gn=CPEB1;refGene_tx=XM_019831822.2;refGene_gn=CPEB1;refGene_tx=XM_019831823.2;refGene_gn=CPEB1;refGene_tx=XM_023254879.1;refGene_gn=CPEB1;refGene_tx=XM_023254880.1;refGene_gn=CPEB1;refGene_tx=XM_023254881.1;refGene_gn=CPEB1;refGene_tx=

XM_023254882.1;refGene_gn=CPEB1;refGene_tx=XM_023254884.1
B3 4421597 4421692
refGene_gn=CPEB1;refGene_tx=XM_006932227.4;refGene_gn=CPEB1;refGene_tx
=XM_011282774.3;refGene_gn=CPEB1;refGene_tx=XM_011282775.3;refGene_gn=
CPEB1;refGene_tx=XM_011282776.3;refGene_gn=CPEB1;refGene_tx=XM_0112827
77.3;refGene_gn=CPEB1;refGene_tx=XM_011282780.3;refGene_gn=CPEB1;refGe
ne_tx=XM_019831819.2;refGene_gn=CPEB1;refGene_tx=XM_019831820.2;refGen
e_gn=CPEB1;refGene_tx=XM_019831821.2;refGene_gn=CPEB1;refGene_tx=XM_01
9831822.2;refGene_gn=CPEB1;refGene_tx=XM_019831823.2;refGene_gn=CPEB1;
refGene_tx=XM_023254879.1;refGene_gn=CPEB1;refGene_tx=XM_023254880.1;r
efGene_gn=CPEB1;refGene_tx=XM_023254881.1;refGene_gn=CPEB1;refGene_tx=
XM_023254882.1;refGene_gn=CPEB1;refGene_tx=XM_023254884.1
B3 4422236 4422317
refGene_gn=CPEB1;refGene_tx=XM_006932227.4;refGene_gn=CPEB1;refGene_tx
=XM_011282774.3;refGene_gn=CPEB1;refGene_tx=XM_011282775.3;refGene_gn=
CPEB1;refGene_tx=XM_011282776.3;refGene_gn=CPEB1;refGene_tx=XM_0112827
77.3;refGene_gn=CPEB1;refGene_tx=XM_011282780.3;refGene_gn=CPEB1;refGe
ne_tx=XM_019831819.2;refGene_gn=CPEB1;refGene_tx=XM_019831820.2;refGen
e_gn=CPEB1;refGene_tx=XM_019831821.2;refGene_gn=CPEB1;refGene_tx=XM_01
9831822.2;refGene_gn=CPEB1;refGene_tx=XM_019831823.2;refGene_gn=CPEB1;
refGene_tx=XM_023254879.1;refGene_gn=CPEB1;refGene_tx=XM_023254880.1;r
efGene_gn=CPEB1;refGene_tx=XM_023254881.1;refGene_gn=CPEB1;refGene_tx=
XM_023254882.1;refGene_gn=CPEB1;refGene_tx=XM_023254884.1
B3 4424058 4424169
refGene_gn=CPEB1;refGene_tx=XM_006932227.4;refGene_gn=CPEB1;refGene_tx
=XM_011282774.3;refGene_gn=CPEB1;refGene_tx=XM_011282775.3;refGene_gn=
CPEB1;refGene_tx=XM_011282776.3;refGene_gn=CPEB1;refGene_tx=XM_0112827
77.3;refGene_gn=CPEB1;refGene_tx=XM_011282780.3;refGene_gn=CPEB1;refGe
ne_tx=XM_019831819.2;refGene_gn=CPEB1;refGene_tx=XM_019831820.2;refGen
e_gn=CPEB1;refGene_tx=XM_019831821.2;refGene_gn=CPEB1;refGene_tx=XM_01
9831822.2;refGene_gn=CPEB1;refGene_tx=XM_019831823.2;refGene_gn=CPEB1;
refGene_tx=XM_023254879.1;refGene_gn=CPEB1;refGene_tx=XM_023254880.1;r
efGene_gn=CPEB1;refGene_tx=XM_023254881.1;refGene_gn=CPEB1;refGene_tx=
XM_023254882.1;refGene_gn=CPEB1;refGene_tx=XM_023254884.1
B3 4427407 4427410 refGene_gn=RPS17;refGene_tx=XM_003986784.3
B3 4427700 4427852 refGene_gn=RPS17;refGene_tx=XM_003986784.3
B3 4428364 4428470 refGene_gn=RPS17;refGene_tx=XM_003986784.3
B3 4428988 4429054 refGene_gn=RPS17;refGene_tx=XM_003986784.3
B3 4430307 4430388 refGene_gn=RPS17;refGene_tx=XM_003986784.3
B3 4474947 4475054
refGene_gn=PDE8A;refGene_tx=XM_023254885.1;refGene_gn=PDE8A;refGene_tx
=XM_023254886.1
B3 4475705 4475835
refGene_gn=PDE8A;refGene_tx=XM_023254885.1;refGene_gn=PDE8A;refGene_tx
=XM_023254886.1
B3 4491979 4492147
refGene_gn=PDE8A;refGene_tx=XM_023254885.1;refGene_gn=PDE8A;refGene_tx
=XM_023254886.1
B3 4494546 4494679
refGene_gn=PDE8A;refGene_tx=XM_023254885.1;refGene_gn=PDE8A;refGene_tx

=XM_023254886.1
B3	4495965	4496183
refGene_gn=PDE8A;refGene_tx=XM_023254885.1;refGene_gn=PDE8A;refGene_tx=XM_023254886.1
B3	4498762	4498961
refGene_gn=PDE8A;refGene_tx=XM_023254885.1;refGene_gn=PDE8A;refGene_tx=XM_023254886.1
B3	4500414	4500550
refGene_gn=PDE8A;refGene_tx=XM_023254885.1;refGene_gn=PDE8A;refGene_tx=XM_023254886.1
B3	4500965	4501014
refGene_gn=PDE8A;refGene_tx=XM_023254885.1;refGene_gn=PDE8A;refGene_tx=XM_023254886.1
B3	4502430	4502595
refGene_gn=PDE8A;refGene_tx=XM_023254885.1;refGene_gn=PDE8A;refGene_tx=XM_023254886.1
B3	4502861	4502932
refGene_gn=PDE8A;refGene_tx=XM_023254885.1;refGene_gn=PDE8A;refGene_tx=XM_023254886.1
B3	4509061	4509139	refGene_gn=PDE8A;refGene_tx=XM_023254885.1
B3	4516550	4516593
refGene_gn=PDE8A;refGene_tx=XM_023254885.1;refGene_gn=PDE8A;refGene_tx=XM_023254886.1
B3	4516693	4516745
refGene_gn=PDE8A;refGene_tx=XM_023254885.1;refGene_gn=PDE8A;refGene_tx=XM_023254886.1
B3	4518379	4518468
refGene_gn=PDE8A;refGene_tx=XM_023254885.1;refGene_gn=PDE8A;refGene_tx=XM_023254886.1
B3	4523488	4523626
refGene_gn=PDE8A;refGene_tx=XM_023254885.1;refGene_gn=PDE8A;refGene_tx=XM_023254886.1
B3	4525470	4525549
refGene_gn=PDE8A;refGene_tx=XM_023254885.1;refGene_gn=PDE8A;refGene_tx=XM_023254886.1
B3	4540985	4541074
refGene_gn=PDE8A;refGene_tx=XM_023254885.1;refGene_gn=PDE8A;refGene_tx=XM_023254886.1
B3	4547924	4547979
refGene_gn=PDE8A;refGene_tx=XM_023254885.1;refGene_gn=PDE8A;refGene_tx=XM_023254886.1
B3	4548807	4548864
refGene_gn=PDE8A;refGene_tx=XM_023254885.1;refGene_gn=PDE8A;refGene_tx=XM_023254886.1
B3	4555544	4555735
refGene_gn=PDE8A;refGene_tx=XM_023254885.1;refGene_gn=PDE8A;refGene_tx=XM_023254886.1
B3	4557404	4557461
refGene_gn=PDE8A;refGene_tx=XM_023254885.1;refGene_gn=PDE8A;refGene_tx=XM_023254886.1

B3 4629212 4629419
refGene_gn=PDE8A;refGene_tx=XM_023254885.1;refGene_gn=PDE8A;refGene_tx
=XM_023254886.1
B3 4664643 4664716
refGene_gn=SLC28A1;refGene_tx=XM_003986785.4;refGene_gn=SLC28A1;refGen
e_tx=XM_019831824.2;refGene_gn=SLC28A1;refGene_tx=XM_019831825.2;refGe
ne_gn=SLC28A1;refGene_tx=XM_019831826.2
B3 4664986 4665098
refGene_gn=SLC28A1;refGene_tx=XM_003986785.4;refGene_gn=SLC28A1;refGen
e_tx=XM_019831824.2;refGene_gn=SLC28A1;refGene_tx=XM_019831825.2;refGe
ne_gn=SLC28A1;refGene_tx=XM_019831826.2
B3 4665241 4665340
refGene_gn=SLC28A1;refGene_tx=XM_003986785.4;refGene_gn=SLC28A1;refGen
e_tx=XM_019831824.2;refGene_gn=SLC28A1;refGene_tx=XM_019831825.2
B3 4666043 4666125
refGene_gn=SLC28A1;refGene_tx=XM_003986785.4;refGene_gn=SLC28A1;refGen
e_tx=XM_019831824.2;refGene_gn=SLC28A1;refGene_tx=XM_019831825.2;refGe
ne_gn=SLC28A1;refGene_tx=XM_019831826.2
B3 4669634 4669832
refGene_gn=SLC28A1;refGene_tx=XM_003986785.4;refGene_gn=SLC28A1;refGen
e_tx=XM_019831824.2;refGene_gn=SLC28A1;refGene_tx=XM_019831825.2;refGe
ne_gn=SLC28A1;refGene_tx=XM_019831826.2
B3 4669968 4670137
refGene_gn=SLC28A1;refGene_tx=XM_003986785.4;refGene_gn=SLC28A1;refGen
e_tx=XM_019831824.2;refGene_gn=SLC28A1;refGene_tx=XM_019831825.2;refGe
ne_gn=SLC28A1;refGene_tx=XM_019831826.2
B3 4671457 4671588
refGene_gn=SLC28A1;refGene_tx=XM_003986785.4;refGene_gn=SLC28A1;refGen
e_tx=XM_019831824.2;refGene_gn=SLC28A1;refGene_tx=XM_019831825.2;refGe
ne_gn=SLC28A1;refGene_tx=XM_019831826.2
B3 4679656 4679782
refGene_gn=SLC28A1;refGene_tx=XM_003986785.4;refGene_gn=SLC28A1;refGen
e_tx=XM_019831824.2;refGene_gn=SLC28A1;refGene_tx=XM_019831825.2;refGe
ne_gn=SLC28A1;refGene_tx=XM_019831826.2
B3 4683829 4683910
refGene_gn=SLC28A1;refGene_tx=XM_003986785.4;refGene_gn=SLC28A1;refGen
e_tx=XM_019831824.2;refGene_gn=SLC28A1;refGene_tx=XM_019831825.2;refGe
ne_gn=SLC28A1;refGene_tx=XM_019831826.2
B3 4684590 4684671
refGene_gn=SLC28A1;refGene_tx=XM_003986785.4;refGene_gn=SLC28A1;refGen
e_tx=XM_019831824.2;refGene_gn=SLC28A1;refGene_tx=XM_019831825.2;refGe
ne_gn=SLC28A1;refGene_tx=XM_019831826.2
B3 4697123 4697201
refGene_gn=SLC28A1;refGene_tx=XM_003986785.4;refGene_gn=SLC28A1;refGen
e_tx=XM_019831824.2;refGene_gn=SLC28A1;refGene_tx=XM_019831826.2
B3 4698612 4698726
refGene_gn=SLC28A1;refGene_tx=XM_003986785.4;refGene_gn=SLC28A1;refGen
e_tx=XM_019831824.2;refGene_gn=SLC28A1;refGene_tx=XM_019831825.2;refGe
ne_gn=SLC28A1;refGene_tx=XM_019831826.2
B3 4699380 4699522

refGene_gn=SLC28A1;refGene_tx=XM_003986785.4;refGene_gn=SLC28A1;refGene_tx=XM_019831824.2;refGene_gn=SLC28A1;refGene_tx=XM_019831825.2;refGene_gn=SLC28A1;refGene_tx=XM_019831826.2
B3	4708028	4708212	refGene_gn=SLC28A1;refGene_tx=XM_003986785.4;refGene_gn=SLC28A1;refGene_tx=XM_019831824.2;refGene_gn=SLC28A1;refGene_tx=XM_019831825.2;refGene_gn=SLC28A1;refGene_tx=XM_019831826.2
B3	4712104	4712196	refGene_gn=SLC28A1;refGene_tx=XM_003986785.4;refGene_gn=SLC28A1;refGene_tx=XM_019831824.2;refGene_gn=SLC28A1;refGene_tx=XM_019831825.2;refGene_gn=SLC28A1;refGene_tx=XM_019831826.2
B3	4713380	4713469	refGene_gn=SLC28A1;refGene_tx=XM_003986785.4;refGene_gn=SLC28A1;refGene_tx=XM_019831824.2;refGene_gn=SLC28A1;refGene_tx=XM_019831825.2;refGene_gn=SLC28A1;refGene_tx=XM_019831826.2
B3	4714331	4714427	refGene_gn=SLC28A1;refGene_tx=XM_003986785.4;refGene_gn=SLC28A1;refGene_tx=XM_019831824.2;refGene_gn=SLC28A1;refGene_tx=XM_019831825.2;refGene_gn=SLC28A1;refGene_tx=XM_019831826.2
B3	4728850	4729196	refGene_gn=ALPK3;refGene_tx=XM_023254767.1
B3	4730041	4730090	refGene_gn=ALPK3;refGene_tx=XM_023254767.1
B3	4732958	4733194	refGene_gn=ALPK3;refGene_tx=XM_023254767.1
B3	4733981	4734070	refGene_gn=ALPK3;refGene_tx=XM_023254767.1
B3	4734657	4734937	refGene_gn=ALPK3;refGene_tx=XM_023254767.1
B3	4737301	4737344	refGene_gn=ALPK3;refGene_tx=XM_023254767.1
B3	4737469	4737597	refGene_gn=ALPK3;refGene_tx=XM_023254767.1
B3	4738006	4738154	refGene_gn=ALPK3;refGene_tx=XM_023254767.1
B3	4739956	4741976	refGene_gn=ALPK3;refGene_tx=XM_023254767.1
B3	4758346	4759559	refGene_gn=ALPK3;refGene_tx=XM_023254767.1
B3	4760149	4760267	refGene_gn=ALPK3;refGene_tx=XM_023254767.1
B3	4770641	4770763	refGene_gn=ALPK3;refGene_tx=XM_023254767.1
B3	4774163	4774202	refGene_gn=ALPK3;refGene_tx=XM_023254767.1
B3	4779211	4779376	refGene_gn=ALPK3;refGene_tx=XM_023254767.1
B3	4779377	4779451	refGene_gn=ALPK3;refGene_tx=XM_023254767.1
B3	4779452	4779776	refGene_gn=ALPK3;refGene_tx=XM_023254767.1
B3	4781551	4781578	refGene_gn=ALPK3;refGene_tx=XM_023254767.1
B3	4782619	4782706	refGene_gn=ZNF592;refGene_tx=XM_011282770.2
B3	4790050	4790131	refGene_gn=ZNF592;refGene_tx=XM_011282771.2
B3	4792916	4793456	refGene_gn=ZNF592;refGene_tx=XM_006932235.4;refGene_gn=ZNF592;refGene_tx=XM_011282770.2;refGene_gn=ZNF592;refGene_tx=XM_011282771.2;refGene_gn=ZNF592;refGene_tx=XM_023254887.1;refGene_gn=ZNF592;refGene_tx=XM_023254888.1
B3	4795042	4795178	refGene_gn=ZNF592;refGene_tx=XM_006932235.4;refGene_gn=ZNF592;refGene_tx=XM_011282770.2;refGene_gn=ZNF592;refGene_tx=XM_011282771.2;refGene_gn=ZNF592;refGene_tx=XM_023254887.1;refGene_gn=ZNF592;refGene_tx=XM_023254888.1
B3	4795738	4795851	refGene_gn=ZNF592;refGene_tx=XM_006932235.4;refGene_gn=ZNF592;refGene_

tx=XM_011282770.2;refGene_gn=ZNF592;refGene_tx=XM_011282771.2;refGene_gn=ZNF592;refGene_tx=XM_023254887.1;refGene_gn=ZNF592;refGene_tx=XM_023254888.1
B3 4796082 4796370
refGene_gn=ZNF592;refGene_tx=XM_006932235.4;refGene_gn=ZNF592;refGene_tx=XM_011282770.2;refGene_gn=ZNF592;refGene_tx=XM_011282771.2;refGene_gn=ZNF592;refGene_tx=XM_023254887.1;refGene_gn=ZNF592;refGene_tx=XM_023254888.1
B3 4796484 4796644
refGene_gn=ZNF592;refGene_tx=XM_006932235.4;refGene_gn=ZNF592;refGene_tx=XM_011282770.2;refGene_gn=ZNF592;refGene_tx=XM_011282771.2;refGene_gn=ZNF592;refGene_tx=XM_023254887.1;refGene_gn=ZNF592;refGene_tx=XM_023254888.1
B3 4796921 4797098
refGene_gn=ZNF592;refGene_tx=XM_006932235.4;refGene_gn=ZNF592;refGene_tx=XM_011282770.2;refGene_gn=ZNF592;refGene_tx=XM_011282771.2;refGene_gn=ZNF592;refGene_tx=XM_023254887.1;refGene_gn=ZNF592;refGene_tx=XM_023254888.1
B3 4802616 4802795
refGene_gn=ZNF592;refGene_tx=XM_006932235.4;refGene_gn=ZNF592;refGene_tx=XM_011282770.2;refGene_gn=ZNF592;refGene_tx=XM_011282771.2;refGene_gn=ZNF592;refGene_tx=XM_023254887.1;refGene_gn=ZNF592;refGene_tx=XM_023254888.1
B3 4809531 4811784
refGene_gn=ZNF592;refGene_tx=XM_006932235.4;refGene_gn=ZNF592;refGene_tx=XM_011282770.2;refGene_gn=ZNF592;refGene_tx=XM_011282771.2;refGene_gn=ZNF592;refGene_tx=XM_023254887.1;refGene_gn=ZNF592;refGene_tx=XM_023254888.1
B3 4869096 4869147
refGene_gn=SEC11A;refGene_tx=XM_003986787.4;refGene_gn=SEC11A;refGene_tx=XM_019831830.2;refGene_gn=SEC11A;refGene_tx=XM_019831831.2;refGene_gn=SEC11A;refGene_tx=XM_019831832.2
B3 4890479 4890589
refGene_gn=SEC11A;refGene_tx=XM_003986787.4;refGene_gn=SEC11A;refGene_tx=XM_019831830.2;refGene_gn=SEC11A;refGene_tx=XM_019831831.2;refGene_gn=SEC11A;refGene_tx=XM_019831832.2
B3 4896165 4896315
refGene_gn=SEC11A;refGene_tx=XM_003986787.4;refGene_gn=SEC11A;refGene_tx=XM_019831830.2;refGene_gn=SEC11A;refGene_tx=XM_019831831.2
B3 4902891 4903011
refGene_gn=SEC11A;refGene_tx=XM_003986787.4;refGene_gn=SEC11A;refGene_tx=XM_019831830.2;refGene_gn=SEC11A;refGene_tx=XM_019831831.2;refGene_gn=SEC11A;refGene_tx=XM_019831832.2
B3 4903525 4903563 refGene_gn=SEC11A;refGene_tx=XM_019831830.2
B3 4903736 4903740 refGene_gn=SEC11A;refGene_tx=XM_019831831.2
B3 4908977 4909035
refGene_gn=SEC11A;refGene_tx=XM_003986787.4;refGene_gn=SEC11A;refGene_tx=XM_019831830.2;refGene_gn=SEC11A;refGene_tx=XM_019831832.2
B3 4909444 4909495
refGene_gn=SEC11A;refGene_tx=XM_003986787.4;refGene_gn=SEC11A;refGene_

tx=XM_019831832.2
B3	4918809	4918966	refGene_gn=NMB;refGene_tx=XM_003986824.4
B3	4919577	4919750	refGene_gn=NMB;refGene_tx=XM_003986824.4
B3	4921684	4921849	refGene_gn=NMB;refGene_tx=XM_003986824.4
B3	4922888	4922929	refGene_gn=WDR73;refGene_tx=XM_011282768.3
B3	4923479	4923547	refGene_gn=WDR73;refGene_tx=XM_011282768.3
B3	4923865	4923954	refGene_gn=WDR73;refGene_tx=XM_011282768.3
B3	4925469	4925558	refGene_gn=WDR73;refGene_tx=XM_011282768.3
B3	4926134	4926199	refGene_gn=WDR73;refGene_tx=XM_011282768.3
B3	4932567	4932732	refGene_gn=WDR73;refGene_tx=XM_011282768.3
B3	4933033	4933399	refGene_gn=WDR73;refGene_tx=XM_011282768.3
B3	4934881	4935117	refGene_gn=WDR73;refGene_tx=XM_011282768.3
B3	4947815	4948837
refGene_gn=LOC101087322;refGene_tx=XM_006932242.4;refGene_gn=LOC101087
322;refGene_tx=XM_019831833.2
B3	4950393	4950495
refGene_gn=LOC101087322;refGene_tx=XM_006932242.4;refGene_gn=LOC101087
322;refGene_tx=XM_019831833.2
B3	4951194	4951647
refGene_gn=LOC101087322;refGene_tx=XM_006932242.4;refGene_gn=LOC101087
322;refGene_tx=XM_019831833.2
B3	4952046	4952814
refGene_gn=LOC101087322;refGene_tx=XM_006932242.4;refGene_gn=LOC101087
322;refGene_tx=XM_019831833.2
B3	4953290	4953414
refGene_gn=LOC101087322;refGene_tx=XM_006932242.4;refGene_gn=LOC101087
322;refGene_tx=XM_019831833.2
B3	4954095	4954172
refGene_gn=LOC101087322;refGene_tx=XM_006932242.4;refGene_gn=LOC101087
322;refGene_tx=XM_019831833.2
B3	4954727	4954812
refGene_gn=LOC101087322;refGene_tx=XM_006932242.4;refGene_gn=LOC101087
322;refGene_tx=XM_019831833.2
B3	4955569	4955968
refGene_gn=LOC101087322;refGene_tx=XM_006932242.4;refGene_gn=LOC101087
322;refGene_tx=XM_019831833.2
B3	4965525	4966053	refGene_gn=ZSCAN2;refGene_tx=XM_006932238.3
B3	4966055	4966969	refGene_gn=ZSCAN2;refGene_tx=XM_006932238.3
B3	4982938	4983329	refGene_gn=ZSCAN2;refGene_tx=XM_006932238.3
B3	5020512	5020697	refGene_gn=ZNF774;refGene_tx=XM_019831836.2
B3	5022285	5022392
refGene_gn=ZNF774;refGene_tx=XM_003986827.4;refGene_gn=ZNF774;refGene_
tx=XM_006932246.3
B3	5025668	5025775
refGene_gn=ZNF774;refGene_tx=XM_003986827.4;refGene_gn=ZNF774;refGene_
tx=XM_006932246.3;refGene_gn=ZNF774;refGene_tx=XM_019831836.2
B3	5026563	5027804
refGene_gn=ZNF774;refGene_tx=XM_003986827.4;refGene_gn=ZNF774;refGene_
tx=XM_006932246.3;refGene_gn=ZNF774;refGene_tx=XM_019831836.2
B3	5055579	5055634	refGene_gn=IQGAP1;refGene_tx=XM_023254889.1

| | | | |
|---|---|---|---|
| B3 | 5057638 | 5057738 | refGene_gn=IQGAP1;refGene_tx=XM_023254889.1 |
| B3 | 5088111 | 5088268 | refGene_gn=IQGAP1;refGene_tx=XM_023254889.1 |
| B3 | 5090001 | 5090079 | refGene_gn=IQGAP1;refGene_tx=XM_023254889.1 |
| B3 | 5094598 | 5094675 | refGene_gn=IQGAP1;refGene_tx=XM_023254889.1 |
| B3 | 5100545 | 5100613 | refGene_gn=IQGAP1;refGene_tx=XM_023254889.1 |
| B3 | 5101973 | 5102087 | refGene_gn=IQGAP1;refGene_tx=XM_023254889.1 |
| B3 | 5102839 | 5103018 | refGene_gn=IQGAP1;refGene_tx=XM_023254889.1 |
| B3 | 5103745 | 5103830 | refGene_gn=IQGAP1;refGene_tx=XM_023254889.1 |
| B3 | 5108163 | 5108327 | refGene_gn=IQGAP1;refGene_tx=XM_023254889.1 |
| B3 | 5110009 | 5110094 | refGene_gn=IQGAP1;refGene_tx=XM_023254889.1 |
| B3 | 5110860 | 5111024 | refGene_gn=IQGAP1;refGene_tx=XM_023254889.1 |
| B3 | 5111237 | 5111398 | refGene_gn=IQGAP1;refGene_tx=XM_023254889.1 |
| B3 | 5113084 | 5113209 | refGene_gn=IQGAP1;refGene_tx=XM_023254889.1 |
| B3 | 5114804 | 5114968 | refGene_gn=IQGAP1;refGene_tx=XM_023254889.1 |
| B3 | 5126700 | 5126791 | refGene_gn=IQGAP1;refGene_tx=XM_023254889.1 |
| B3 | 5126995 | 5127163 | refGene_gn=IQGAP1;refGene_tx=XM_023254889.1 |
| B3 | 5127882 | 5128025 | refGene_gn=IQGAP1;refGene_tx=XM_023254889.1 |
| B3 | 5132711 | 5132882 | refGene_gn=IQGAP1;refGene_tx=XM_023254889.1 |
| B3 | 5133574 | 5133658 | refGene_gn=IQGAP1;refGene_tx=XM_023254889.1 |
| B3 | 5133751 | 5133823 | refGene_gn=IQGAP1;refGene_tx=XM_023254889.1 |
| B3 | 5133917 | 5133987 | refGene_gn=IQGAP1;refGene_tx=XM_023254889.1 |
| B3 | 5134360 | 5134569 | refGene_gn=IQGAP1;refGene_tx=XM_023254889.1 |
| B3 | 5136318 | 5136474 | refGene_gn=IQGAP1;refGene_tx=XM_023254889.1 |
| B3 | 5136698 | 5136862 | refGene_gn=IQGAP1;refGene_tx=XM_023254889.1 |
| B3 | 5137813 | 5138038 | refGene_gn=IQGAP1;refGene_tx=XM_023254889.1 |
| B3 | 5141338 | 5141479 | refGene_gn=IQGAP1;refGene_tx=XM_023254889.1 |
| B3 | 5141569 | 5141654 | refGene_gn=IQGAP1;refGene_tx=XM_023254889.1 |
| B3 | 5142744 | 5142977 | refGene_gn=IQGAP1;refGene_tx=XM_023254889.1 |
| B3 | 5143583 | 5143716 | refGene_gn=IQGAP1;refGene_tx=XM_023254889.1 |
| B3 | 5144742 | 5144845 | refGene_gn=IQGAP1;refGene_tx=XM_023254889.1 |
| B3 | 5145584 | 5145720 | refGene_gn=IQGAP1;refGene_tx=XM_023254889.1 |
| B3 | 5146120 | 5146208 | refGene_gn=IQGAP1;refGene_tx=XM_023254889.1 |
| B3 | 5148652 | 5148865 | refGene_gn=IQGAP1;refGene_tx=XM_023254889.1 |
| B3 | 5149824 | 5149991 | refGene_gn=IQGAP1;refGene_tx=XM_023254889.1 |
| B3 | 5151506 | 5151629 | refGene_gn=IQGAP1;refGene_tx=XM_023254889.1 |
| B3 | 5155384 | 5155493 | refGene_gn=IQGAP1;refGene_tx=XM_023254889.1 |
| B3 | 5157424 | 5157538 | refGene_gn=IQGAP1;refGene_tx=XM_023254889.1 |
| B3 | 5179963 | 5180095 | refGene_gn=CRTC3;refGene_tx=XM_023254890.1 |
| B3 | 5191335 | 5191434 | refGene_gn=CRTC3;refGene_tx=XM_023254890.1 |
| B3 | 5236555 | 5236675 | refGene_gn=CRTC3;refGene_tx=XM_023254890.1 |
| B3 | 5244312 | 5244374 | refGene_gn=CRTC3;refGene_tx=XM_023254890.1 |
| B3 | 5245996 | 5246059 | refGene_gn=CRTC3;refGene_tx=XM_023254890.1 |
| B3 | 5248134 | 5248235 | refGene_gn=CRTC3;refGene_tx=XM_023254890.1 |
| B3 | 5253998 | 5254034 | refGene_gn=CRTC3;refGene_tx=XM_023254890.1 |
| B3 | 5255931 | 5256017 | refGene_gn=CRTC3;refGene_tx=XM_023254890.1 |
| B3 | 5258039 | 5258089 | refGene_gn=CRTC3;refGene_tx=XM_023254890.1 |
| B3 | 5263566 | 5263784 | refGene_gn=CRTC3;refGene_tx=XM_023254890.1 |
| B3 | 5267280 | 5267579 | refGene_gn=CRTC3;refGene_tx=XM_023254890.1 |
| B3 | 5274079 | 5274280 | refGene_gn=CRTC3;refGene_tx=XM_023254890.1 |
| B3 | 5274363 | 5274444 | refGene_gn=CRTC3;refGene_tx=XM_023254890.1 |

B3 5276853 5276953 refGene_gn=CRTC3;refGene_tx=XM_023254890.1
B3 5277386 5277595 refGene_gn=CRTC3;refGene_tx=XM_023254890.1
B3 5367389 5367487
refGene_gn=BLM;refGene_tx=XM_019831837.2;refGene_gn=BLM;refGene_tx=XM_019831839.2;refGene_gn=BLM;refGene_tx=XM_023254891.1
B3 5368652 5369353
refGene_gn=BLM;refGene_tx=XM_019831837.2;refGene_gn=BLM;refGene_tx=XM_019831839.2;refGene_gn=BLM;refGene_tx=XM_023254891.1
B3 5370434 5370591
refGene_gn=BLM;refGene_tx=XM_019831837.2;refGene_gn=BLM;refGene_tx=XM_019831839.2;refGene_gn=BLM;refGene_tx=XM_023254891.1
B3 5373462 5373584
refGene_gn=BLM;refGene_tx=XM_019831837.2;refGene_gn=BLM;refGene_tx=XM_019831839.2;refGene_gn=BLM;refGene_tx=XM_023254891.1
B3 5378728 5378861
refGene_gn=BLM;refGene_tx=XM_019831837.2;refGene_gn=BLM;refGene_tx=XM_019831839.2;refGene_gn=BLM;refGene_tx=XM_019831840.1;refGene_gn=BLM;refGene_tx=XM_023254891.1
B3 5379163 5379867
refGene_gn=BLM;refGene_tx=XM_019831837.2;refGene_gn=BLM;refGene_tx=XM_019831839.2;refGene_gn=BLM;refGene_tx=XM_019831840.1;refGene_gn=BLM;refGene_tx=XM_023254891.1
B3 5381600 5381792
refGene_gn=BLM;refGene_tx=XM_019831837.2;refGene_gn=BLM;refGene_tx=XM_019831839.2;refGene_gn=BLM;refGene_tx=XM_019831840.1;refGene_gn=BLM;refGene_tx=XM_023254891.1
B3 5383877 5383996
refGene_gn=BLM;refGene_tx=XM_019831837.2;refGene_gn=BLM;refGene_tx=XM_019831839.2;refGene_gn=BLM;refGene_tx=XM_019831840.1;refGene_gn=BLM;refGene_tx=XM_023254891.1
B3 5384877 5384991
refGene_gn=BLM;refGene_tx=XM_019831837.2;refGene_gn=BLM;refGene_tx=XM_019831839.2;refGene_gn=BLM;refGene_tx=XM_019831840.1;refGene_gn=BLM;refGene_tx=XM_023254891.1
B3 5386496 5386595
refGene_gn=BLM;refGene_tx=XM_019831837.2;refGene_gn=BLM;refGene_tx=XM_019831839.2;refGene_gn=BLM;refGene_tx=XM_019831840.1;refGene_gn=BLM;refGene_tx=XM_023254891.1
B3 5386795 5386944
refGene_gn=BLM;refGene_tx=XM_019831837.2;refGene_gn=BLM;refGene_tx=XM_019831839.2;refGene_gn=BLM;refGene_tx=XM_019831840.1;refGene_gn=BLM;refGene_tx=XM_023254891.1
B3 5396198 5396305
refGene_gn=BLM;refGene_tx=XM_019831837.2;refGene_gn=BLM;refGene_tx=XM_019831839.2;refGene_gn=BLM;refGene_tx=XM_019831840.1;refGene_gn=BLM;refGene_tx=XM_023254891.1
B3 5397676 5397837
refGene_gn=BLM;refGene_tx=XM_019831837.2;refGene_gn=BLM;refGene_tx=XM_019831839.2;refGene_gn=BLM;refGene_tx=XM_019831840.1;refGene_gn=BLM;refGene_tx=XM_023254891.1

B3 5405085 5405281
refGene_gn=BLM;refGene_tx=XM_019831837.2;refGene_gn=BLM;refGene_tx=XM_019831839.2;refGene_gn=BLM;refGene_tx=XM_019831840.1;refGene_gn=BLM;refGene_tx=XM_023254891.1
B3 5407497 5407688
refGene_gn=BLM;refGene_tx=XM_019831837.2;refGene_gn=BLM;refGene_tx=XM_019831839.2;refGene_gn=BLM;refGene_tx=XM_019831840.1;refGene_gn=BLM;refGene_tx=XM_023254891.1
B3 5411662 5411810
refGene_gn=BLM;refGene_tx=XM_019831837.2;refGene_gn=BLM;refGene_tx=XM_019831839.2;refGene_gn=BLM;refGene_tx=XM_019831840.1;refGene_gn=BLM;refGene_tx=XM_023254891.1
B3 5418346 5418546
refGene_gn=BLM;refGene_tx=XM_019831837.2;refGene_gn=BLM;refGene_tx=XM_019831839.2;refGene_gn=BLM;refGene_tx=XM_019831840.1;refGene_gn=BLM;refGene_tx=XM_023254891.1
B3 5419011 5419204
refGene_gn=BLM;refGene_tx=XM_019831837.2;refGene_gn=BLM;refGene_tx=XM_019831839.2;refGene_gn=BLM;refGene_tx=XM_019831840.1;refGene_gn=BLM;refGene_tx=XM_023254891.1
B3 5421912 5422035
refGene_gn=BLM;refGene_tx=XM_019831837.2;refGene_gn=BLM;refGene_tx=XM_019831840.1;refGene_gn=BLM;refGene_tx=XM_023254891.1
B3 5424334 5424536
refGene_gn=BLM;refGene_tx=XM_019831837.2;refGene_gn=BLM;refGene_tx=XM_019831840.1;refGene_gn=BLM;refGene_tx=XM_023254891.1
B3 5427582 5427760
refGene_gn=BLM;refGene_tx=XM_019831837.2;refGene_gn=BLM;refGene_tx=XM_019831840.1;refGene_gn=BLM;refGene_tx=XM_023254891.1
B3 5474827 5475004 refGene_gn=FURIN;refGene_tx=XM_023254894.1
B3 5475393 5475492 refGene_gn=FURIN;refGene_tx=XM_023254894.1
B3 5475603 5475699 refGene_gn=FURIN;refGene_tx=XM_023254894.1
B3 5476055 5476184 refGene_gn=FURIN;refGene_tx=XM_023254894.1
B3 5476293 5476370 refGene_gn=FURIN;refGene_tx=XM_023254894.1
B3 5476641 5476730 refGene_gn=FURIN;refGene_tx=XM_023254894.1
B3 5477246 5477419 refGene_gn=FURIN;refGene_tx=XM_023254894.1
B3 5478109 5478322 refGene_gn=FURIN;refGene_tx=XM_023254894.1
B3 5478721 5478822 refGene_gn=FURIN;refGene_tx=XM_023254894.1
B3 5478942 5479046 refGene_gn=FURIN;refGene_tx=XM_023254894.1
B3 5479134 5479252 refGene_gn=FURIN;refGene_tx=XM_023254894.1
B3 5479343 5479523 refGene_gn=FURIN;refGene_tx=XM_023254894.1
B3 5479958 5480083 refGene_gn=FURIN;refGene_tx=XM_023254894.1
B3 5480201 5480312 refGene_gn=FURIN;refGene_tx=XM_023254894.1
B3 5480525 5481115 refGene_gn=FURIN;refGene_tx=XM_023254894.1
B3 5484180 5484387
refGene_gn=FES;refGene_tx=XM_023254892.1;refGene_gn=FES;refGene_tx=XM_023254893.1
B3 5484580 5484754
refGene_gn=FES;refGene_tx=XM_023254892.1;refGene_gn=FES;refGene_tx=XM_023254893.1

B3 5486141 5486238
refGene_gn=FES;refGene_tx=XM_023254892.1;refGene_gn=FES;refGene_tx=XM_023254893.1
B3 5486366 5486547
refGene_gn=FES;refGene_tx=XM_023254892.1;refGene_gn=FES;refGene_tx=XM_023254893.1
B3 5487960 5488098
refGene_gn=FES;refGene_tx=XM_023254892.1;refGene_gn=FES;refGene_tx=XM_023254893.1
B3 5488172 5488292
refGene_gn=FES;refGene_tx=XM_023254892.1;refGene_gn=FES;refGene_tx=XM_023254893.1
B3 5488445 5488568
refGene_gn=FES;refGene_tx=XM_023254892.1;refGene_gn=FES;refGene_tx=XM_023254893.1
B3 5488681 5488868
refGene_gn=FES;refGene_tx=XM_023254892.1;refGene_gn=FES;refGene_tx=XM_023254893.1
B3 5488989 5489073
refGene_gn=FES;refGene_tx=XM_023254892.1;refGene_gn=FES;refGene_tx=XM_023254893.1
B3 5489844 5490054
refGene_gn=FES;refGene_tx=XM_023254892.1;refGene_gn=FES;refGene_tx=XM_023254893.1
B3 5490316 5490439 refGene_gn=FES;refGene_tx=XM_023254892.1
B3 5490807 5490861
refGene_gn=FES;refGene_tx=XM_023254892.1;refGene_gn=FES;refGene_tx=XM_023254893.1
B3 5491551 5491670
refGene_gn=FES;refGene_tx=XM_023254892.1;refGene_gn=FES;refGene_tx=XM_023254893.1
B3 5491992 5492087
refGene_gn=FES;refGene_tx=XM_023254892.1;refGene_gn=FES;refGene_tx=XM_023254893.1
B3 5492167 5492291
refGene_gn=FES;refGene_tx=XM_023254892.1;refGene_gn=FES;refGene_tx=XM_023254893.1
B3 5492524 5492682
refGene_gn=FES;refGene_tx=XM_023254892.1;refGene_gn=FES;refGene_tx=XM_023254893.1
B3 5492789 5492912
refGene_gn=FES;refGene_tx=XM_023254892.1;refGene_gn=FES;refGene_tx=XM_023254893.1
B3 5493766 5493909
refGene_gn=FES;refGene_tx=XM_023254892.1;refGene_gn=FES;refGene_tx=XM_023254893.1
B3 5507014 5507146
refGene_gn=MAN2A2;refGene_tx=XM_019831841.1;refGene_gn=MAN2A2;refGene_tx=XM_019831842.1;refGene_gn=MAN2A2;refGene_tx=XM_023254895.1;refGene_gn=MAN2A2;refGene_tx=XM_023254896.1;refGene_gn=MAN2A2;refGene_tx=XM_02

3254898.1
B3	5507694	5507952
refGene_gn=MAN2A2;refGene_tx=XM_019831841.1;refGene_gn=MAN2A2;refGene_tx=XM_019831842.1;refGene_gn=MAN2A2;refGene_tx=XM_023254895.1;refGene_gn=MAN2A2;refGene_tx=XM_023254896.1;refGene_gn=MAN2A2;refGene_tx=XM_023254898.1
B3	5508023	5508168
refGene_gn=MAN2A2;refGene_tx=XM_019831841.1;refGene_gn=MAN2A2;refGene_tx=XM_019831842.1;refGene_gn=MAN2A2;refGene_tx=XM_023254895.1;refGene_gn=MAN2A2;refGene_tx=XM_023254896.1;refGene_gn=MAN2A2;refGene_tx=XM_023254898.1
B3	5508284	5508456
refGene_gn=MAN2A2;refGene_tx=XM_019831841.1;refGene_gn=MAN2A2;refGene_tx=XM_019831842.1;refGene_gn=MAN2A2;refGene_tx=XM_023254895.1;refGene_gn=MAN2A2;refGene_tx=XM_023254896.1;refGene_gn=MAN2A2;refGene_tx=XM_023254898.1
B3	5508899	5509027
refGene_gn=MAN2A2;refGene_tx=XM_019831841.1;refGene_gn=MAN2A2;refGene_tx=XM_019831842.1;refGene_gn=MAN2A2;refGene_tx=XM_023254895.1;refGene_gn=MAN2A2;refGene_tx=XM_023254896.1;refGene_gn=MAN2A2;refGene_tx=XM_023254898.1
B3	5509230	5509404
refGene_gn=MAN2A2;refGene_tx=XM_019831841.1;refGene_gn=MAN2A2;refGene_tx=XM_019831842.1;refGene_gn=MAN2A2;refGene_tx=XM_023254895.1;refGene_gn=MAN2A2;refGene_tx=XM_023254896.1;refGene_gn=MAN2A2;refGene_tx=XM_023254898.1
B3	5509810	5509997
refGene_gn=MAN2A2;refGene_tx=XM_019831841.1;refGene_gn=MAN2A2;refGene_tx=XM_019831842.1;refGene_gn=MAN2A2;refGene_tx=XM_023254895.1;refGene_gn=MAN2A2;refGene_tx=XM_023254896.1;refGene_gn=MAN2A2;refGene_tx=XM_023254898.1
B3	5512245	5512423
refGene_gn=MAN2A2;refGene_tx=XM_019831841.1;refGene_gn=MAN2A2;refGene_tx=XM_019831842.1;refGene_gn=MAN2A2;refGene_tx=XM_023254895.1;refGene_gn=MAN2A2;refGene_tx=XM_023254896.1;refGene_gn=MAN2A2;refGene_tx=XM_023254898.1
B3	5512739	5512942
refGene_gn=MAN2A2;refGene_tx=XM_019831841.1;refGene_gn=MAN2A2;refGene_tx=XM_019831842.1;refGene_gn=MAN2A2;refGene_tx=XM_023254895.1;refGene_gn=MAN2A2;refGene_tx=XM_023254896.1;refGene_gn=MAN2A2;refGene_tx=XM_023254898.1
B3	5513150	5513333
refGene_gn=MAN2A2;refGene_tx=XM_019831841.1;refGene_gn=MAN2A2;refGene_tx=XM_019831842.1;refGene_gn=MAN2A2;refGene_tx=XM_023254895.1;refGene_gn=MAN2A2;refGene_tx=XM_023254896.1;refGene_gn=MAN2A2;refGene_tx=XM_023254898.1
B3	5513487	5513602
refGene_gn=MAN2A2;refGene_tx=XM_019831841.1;refGene_gn=MAN2A2;refGene_tx=XM_019831842.1;refGene_gn=MAN2A2;refGene_tx=XM_023254895.1;refGene_gn=MAN2A2;refGene_tx=XM_023254896.1;refGene_gn=MAN2A2;refGene_tx=XM_02

3254898.1
B3	5513829	5513897
refGene_gn=MAN2A2;refGene_tx=XM_019831841.1;refGene_gn=MAN2A2;refGene_tx=XM_019831842.1;refGene_gn=MAN2A2;refGene_tx=XM_023254895.1;refGene_gn=MAN2A2;refGene_tx=XM_023254896.1;refGene_gn=MAN2A2;refGene_tx=XM_023254898.1
B3	5514046	5514212
refGene_gn=MAN2A2;refGene_tx=XM_019831841.1;refGene_gn=MAN2A2;refGene_tx=XM_019831842.1;refGene_gn=MAN2A2;refGene_tx=XM_023254895.1;refGene_gn=MAN2A2;refGene_tx=XM_023254896.1;refGene_gn=MAN2A2;refGene_tx=XM_023254898.1
B3	5514733	5514970
refGene_gn=MAN2A2;refGene_tx=XM_019831841.1;refGene_gn=MAN2A2;refGene_tx=XM_019831842.1;refGene_gn=MAN2A2;refGene_tx=XM_023254895.1;refGene_gn=MAN2A2;refGene_tx=XM_023254896.1;refGene_gn=MAN2A2;refGene_tx=XM_023254898.1
B3	5515194	5515317
refGene_gn=MAN2A2;refGene_tx=XM_019831841.1;refGene_gn=MAN2A2;refGene_tx=XM_019831842.1;refGene_gn=MAN2A2;refGene_tx=XM_023254895.1;refGene_gn=MAN2A2;refGene_tx=XM_023254896.1;refGene_gn=MAN2A2;refGene_tx=XM_023254898.1
B3	5515636	5515751
refGene_gn=MAN2A2;refGene_tx=XM_019831841.1;refGene_gn=MAN2A2;refGene_tx=XM_019831842.1;refGene_gn=MAN2A2;refGene_tx=XM_023254895.1;refGene_gn=MAN2A2;refGene_tx=XM_023254896.1;refGene_gn=MAN2A2;refGene_tx=XM_023254898.1
B3	5515994	5516128
refGene_gn=MAN2A2;refGene_tx=XM_019831841.1;refGene_gn=MAN2A2;refGene_tx=XM_019831842.1;refGene_gn=MAN2A2;refGene_tx=XM_023254895.1;refGene_gn=MAN2A2;refGene_tx=XM_023254896.1;refGene_gn=MAN2A2;refGene_tx=XM_023254898.1
B3	5516297	5516439
refGene_gn=MAN2A2;refGene_tx=XM_019831841.1;refGene_gn=MAN2A2;refGene_tx=XM_019831842.1;refGene_gn=MAN2A2;refGene_tx=XM_023254895.1;refGene_gn=MAN2A2;refGene_tx=XM_023254896.1;refGene_gn=MAN2A2;refGene_tx=XM_023254898.1
B3	5517997	5518131
refGene_gn=MAN2A2;refGene_tx=XM_019831841.1;refGene_gn=MAN2A2;refGene_tx=XM_019831842.1;refGene_gn=MAN2A2;refGene_tx=XM_023254895.1;refGene_gn=MAN2A2;refGene_tx=XM_023254896.1;refGene_gn=MAN2A2;refGene_tx=XM_023254898.1
B3	5518425	5518500
refGene_gn=MAN2A2;refGene_tx=XM_019831841.1;refGene_gn=MAN2A2;refGene_tx=XM_023254895.1;refGene_gn=MAN2A2;refGene_tx=XM_023254896.1;refGene_gn=MAN2A2;refGene_tx=XM_023254898.1
B3	5520337	5520532
refGene_gn=MAN2A2;refGene_tx=XM_019831841.1;refGene_gn=MAN2A2;refGene_tx=XM_019831842.1;refGene_gn=MAN2A2;refGene_tx=XM_023254895.1;refGene_gn=MAN2A2;refGene_tx=XM_023254896.1;refGene_gn=MAN2A2;refGene_tx=XM_023254898.1

B3 5520697 5520808
refGene_gn=MAN2A2;refGene_tx=XM_019831841.1;refGene_gn=MAN2A2;refGene_tx=XM_019831842.1;refGene_gn=MAN2A2;refGene_tx=XM_023254895.1;refGene_gn=MAN2A2;refGene_tx=XM_023254896.1;refGene_gn=MAN2A2;refGene_tx=XM_023254898.1
B3 5521635 5521788
refGene_gn=MAN2A2;refGene_tx=XM_019831841.1;refGene_gn=MAN2A2;refGene_tx=XM_019831842.1;refGene_gn=MAN2A2;refGene_tx=XM_023254895.1;refGene_gn=MAN2A2;refGene_tx=XM_023254896.1;refGene_gn=MAN2A2;refGene_tx=XM_023254898.1
B3 5531041 5531172 refGene_gn=HDDC3;refGene_tx=XM_003986796.5
B3 5531478 5531719 refGene_gn=HDDC3;refGene_tx=XM_003986796.5
B3 5531829 5531885 refGene_gn=HDDC3;refGene_tx=XM_003986796.5
B3 5532196 5532308 refGene_gn=HDDC3;refGene_tx=XM_003986796.5
B3 5535182 5535233
refGene_gn=UNC45A;refGene_tx=XM_003986797.4;refGene_gn=UNC45A;refGene_tx=XM_003986798.4
B3 5535414 5535576
refGene_gn=UNC45A;refGene_tx=XM_003986797.4;refGene_gn=UNC45A;refGene_tx=XM_003986798.4
B3 5535809 5535846
refGene_gn=UNC45A;refGene_tx=XM_003986797.4;refGene_gn=UNC45A;refGene_tx=XM_003986798.4
B3 5536148 5536324
refGene_gn=UNC45A;refGene_tx=XM_003986797.4;refGene_gn=UNC45A;refGene_tx=XM_003986798.4
B3 5538874 5538967
refGene_gn=UNC45A;refGene_tx=XM_003986797.4;refGene_gn=UNC45A;refGene_tx=XM_003986798.4
B3 5539424 5539592
refGene_gn=UNC45A;refGene_tx=XM_003986797.4;refGene_gn=UNC45A;refGene_tx=XM_003986798.4
B3 5540820 5540989
refGene_gn=UNC45A;refGene_tx=XM_003986797.4;refGene_gn=UNC45A;refGene_tx=XM_003986798.4
B3 5541276 5541447
refGene_gn=UNC45A;refGene_tx=XM_003986797.4;refGene_gn=UNC45A;refGene_tx=XM_003986798.4
B3 5542681 5542853
refGene_gn=UNC45A;refGene_tx=XM_003986797.4;refGene_gn=UNC45A;refGene_tx=XM_003986798.4
B3 5543458 5543759
refGene_gn=UNC45A;refGene_tx=XM_003986797.4;refGene_gn=UNC45A;refGene_tx=XM_003986798.4
B3 5544562 5544657
refGene_gn=UNC45A;refGene_tx=XM_003986797.4;refGene_gn=UNC45A;refGene_tx=XM_003986798.4
B3 5544887 5545029
refGene_gn=UNC45A;refGene_tx=XM_003986797.4;refGene_gn=UNC45A;refGene_tx=XM_003986798.4

B3 5545395 5545536
refGene_gn=UNC45A;refGene_tx=XM_003986797.4;refGene_gn=UNC45A;refGene_tx=XM_003986798.4
B3 5545738 5545866
refGene_gn=UNC45A;refGene_tx=XM_003986797.4;refGene_gn=UNC45A;refGene_tx=XM_003986798.4
B3 5546052 5546119
refGene_gn=UNC45A;refGene_tx=XM_003986797.4;refGene_gn=UNC45A;refGene_tx=XM_003986798.4
B3 5546532 5546646
refGene_gn=UNC45A;refGene_tx=XM_003986797.4;refGene_gn=UNC45A;refGene_tx=XM_003986798.4
B3 5546863 5546979
refGene_gn=UNC45A;refGene_tx=XM_003986797.4;refGene_gn=UNC45A;refGene_tx=XM_003986798.4
B3 5547828 5547946
refGene_gn=UNC45A;refGene_tx=XM_003986797.4;refGene_gn=UNC45A;refGene_tx=XM_003986798.4
B3 5548052 5548208
refGene_gn=UNC45A;refGene_tx=XM_003986797.4;refGene_gn=UNC45A;refGene_tx=XM_003986798.4
B3 5548359 5548617
refGene_gn=UNC45A;refGene_tx=XM_003986797.4;refGene_gn=UNC45A;refGene_tx=XM_003986798.4
B3 5551042 5551223
refGene_gn=RCCD1;refGene_tx=XM_019831852.2;refGene_gn=RCCD1;refGene_tx=XM_023254902.1;refGene_gn=RCCD1;refGene_tx=XM_023254903.1;refGene_gn=RCCD1;refGene_tx=XM_023254904.1;refGene_gn=RCCD1;refGene_tx=XM_023254905.1;refGene_gn=RCCD1;refGene_tx=XM_023254906.1
B3 5551313 5551710
refGene_gn=RCCD1;refGene_tx=XM_019831852.2;refGene_gn=RCCD1;refGene_tx=XM_023254902.1;refGene_gn=RCCD1;refGene_tx=XM_023254903.1;refGene_gn=RCCD1;refGene_tx=XM_023254904.1;refGene_gn=RCCD1;refGene_tx=XM_023254905.1;refGene_gn=RCCD1;refGene_tx=XM_023254906.1
B3 5551869 5551991
refGene_gn=RCCD1;refGene_tx=XM_019831852.2;refGene_gn=RCCD1;refGene_tx=XM_023254902.1;refGene_gn=RCCD1;refGene_tx=XM_023254903.1;refGene_gn=RCCD1;refGene_tx=XM_023254904.1;refGene_gn=RCCD1;refGene_tx=XM_023254905.1;refGene_gn=RCCD1;refGene_tx=XM_023254906.1
B3 5554158 5554257
refGene_gn=RCCD1;refGene_tx=XM_019831852.2;refGene_gn=RCCD1;refGene_tx=XM_023254902.1;refGene_gn=RCCD1;refGene_tx=XM_023254903.1;refGene_gn=RCCD1;refGene_tx=XM_023254904.1;refGene_gn=RCCD1;refGene_tx=XM_023254905.1;refGene_gn=RCCD1;refGene_tx=XM_023254906.1
B3 5554595 5554763
refGene_gn=RCCD1;refGene_tx=XM_019831852.2;refGene_gn=RCCD1;refGene_tx=XM_023254902.1;refGene_gn=RCCD1;refGene_tx=XM_023254903.1;refGene_gn=RCCD1;refGene_tx=XM_023254904.1;refGene_gn=RCCD1;refGene_tx=XM_023254905.1
B3 5555276 5555306

refGene_gn=RCCD1;refGene_tx=XM_019831852.2;refGene_gn=RCCD1;refGene_tx
=XM_023254902.1;refGene_gn=RCCD1;refGene_tx=XM_023254903.1;refGene_gn=
RCCD1;refGene_tx=XM_023254904.1;refGene_gn=RCCD1;refGene_tx=XM_0232549
05.1;refGene_gn=RCCD1;refGene_tx=XM_023254906.1
B3 5555898 5556050
refGene_gn=RCCD1;refGene_tx=XM_019831852.2;refGene_gn=RCCD1;refGene_tx
=XM_023254902.1;refGene_gn=RCCD1;refGene_tx=XM_023254903.1;refGene_gn=
RCCD1;refGene_tx=XM_023254904.1;refGene_gn=RCCD1;refGene_tx=XM_0232549
05.1;refGene_gn=RCCD1;refGene_tx=XM_023254906.1
B3 5575058 5575136
refGene_gn=RCCD1;refGene_tx=XM_019831852.2;refGene_gn=RCCD1;refGene_tx
=XM_023254902.1;refGene_gn=RCCD1;refGene_tx=XM_023254905.1;refGene_gn=
RCCD1;refGene_tx=XM_023254906.1
B3 5575194 5575248 refGene_gn=RCCD1;refGene_tx=XM_023254904.1
B3 5576195 5576267
refGene_gn=PRC1;refGene_tx=XM_023254899.1;refGene_gn=PRC1;refGene_tx=X
M_023254900.1;refGene_gn=PRC1;refGene_tx=XM_023254901.1
B3 5577804 5577846 refGene_gn=PRC1;refGene_tx=XM_023254899.1
B3 5578164 5578341
refGene_gn=PRC1;refGene_tx=XM_023254899.1;refGene_gn=PRC1;refGene_tx=X
M_023254900.1;refGene_gn=PRC1;refGene_tx=XM_023254901.1
B3 5578914 5579025
refGene_gn=PRC1;refGene_tx=XM_023254899.1;refGene_gn=PRC1;refGene_tx=X
M_023254900.1;refGene_gn=PRC1;refGene_tx=XM_023254901.1
B3 5582481 5582592
refGene_gn=PRC1;refGene_tx=XM_023254899.1;refGene_gn=PRC1;refGene_tx=X
M_023254900.1;refGene_gn=PRC1;refGene_tx=XM_023254901.1
B3 5582805 5582952
refGene_gn=PRC1;refGene_tx=XM_023254899.1;refGene_gn=PRC1;refGene_tx=X
M_023254900.1;refGene_gn=PRC1;refGene_tx=XM_023254901.1
B3 5585094 5585190
refGene_gn=PRC1;refGene_tx=XM_023254899.1;refGene_gn=PRC1;refGene_tx=X
M_023254900.1;refGene_gn=PRC1;refGene_tx=XM_023254901.1
B3 5587117 5587254
refGene_gn=PRC1;refGene_tx=XM_023254899.1;refGene_gn=PRC1;refGene_tx=X
M_023254900.1;refGene_gn=PRC1;refGene_tx=XM_023254901.1
B3 5588011 5588159
refGene_gn=PRC1;refGene_tx=XM_023254899.1;refGene_gn=PRC1;refGene_tx=X
M_023254900.1;refGene_gn=PRC1;refGene_tx=XM_023254901.1
B3 5588952 5589102
refGene_gn=PRC1;refGene_tx=XM_023254899.1;refGene_gn=PRC1;refGene_tx=X
M_023254900.1;refGene_gn=PRC1;refGene_tx=XM_023254901.1
B3 5589550 5589721
refGene_gn=PRC1;refGene_tx=XM_023254899.1;refGene_gn=PRC1;refGene_tx=X
M_023254900.1;refGene_gn=PRC1;refGene_tx=XM_023254901.1
B3 5589798 5590032
refGene_gn=PRC1;refGene_tx=XM_023254899.1;refGene_gn=PRC1;refGene_tx=X
M_023254900.1;refGene_gn=PRC1;refGene_tx=XM_023254901.1
B3 5591798 5591921
refGene_gn=PRC1;refGene_tx=XM_023254899.1;refGene_gn=PRC1;refGene_tx=X

M_023254900.1;refGene_gn=PRC1;refGene_tx=XM_023254901.1
B3	5592391	5592524
refGene_gn=PRC1;refGene_tx=XM_023254899.1;refGene_gn=PRC1;refGene_tx=XM_023254900.1;refGene_gn=PRC1;refGene_tx=XM_023254901.1
B3	5601682	5601693
refGene_gn=PRC1;refGene_tx=XM_023254899.1;refGene_gn=PRC1;refGene_tx=XM_023254900.1;refGene_gn=PRC1;refGene_tx=XM_023254901.1
B3	5606727	5606807
refGene_gn=VPS33B;refGene_tx=XM_003986800.4;refGene_gn=VPS33B;refGene_tx=XM_006932253.4;refGene_gn=VPS33B;refGene_tx=XM_019831854.2;refGene_gn=VPS33B;refGene_tx=XM_019831855.2
B3	5607081	5607198
refGene_gn=VPS33B;refGene_tx=XM_003986800.4;refGene_gn=VPS33B;refGene_tx=XM_006932253.4;refGene_gn=VPS33B;refGene_tx=XM_019831854.2;refGene_gn=VPS33B;refGene_tx=XM_019831855.2
B3	5607292	5607368
refGene_gn=VPS33B;refGene_tx=XM_003986800.4;refGene_gn=VPS33B;refGene_tx=XM_006932253.4;refGene_gn=VPS33B;refGene_tx=XM_019831854.2;refGene_gn=VPS33B;refGene_tx=XM_019831855.2
B3	5607561	5607663
refGene_gn=VPS33B;refGene_tx=XM_003986800.4;refGene_gn=VPS33B;refGene_tx=XM_006932253.4;refGene_gn=VPS33B;refGene_tx=XM_019831854.2;refGene_gn=VPS33B;refGene_tx=XM_019831855.2
B3	5608417	5608491
refGene_gn=VPS33B;refGene_tx=XM_003986800.4;refGene_gn=VPS33B;refGene_tx=XM_006932253.4;refGene_gn=VPS33B;refGene_tx=XM_019831854.2;refGene_gn=VPS33B;refGene_tx=XM_019831855.2
B3	5609082	5609215
refGene_gn=VPS33B;refGene_tx=XM_003986800.4;refGene_gn=VPS33B;refGene_tx=XM_006932253.4;refGene_gn=VPS33B;refGene_tx=XM_019831854.2;refGene_gn=VPS33B;refGene_tx=XM_019831855.2
B3	5609757	5609804
refGene_gn=VPS33B;refGene_tx=XM_003986800.4;refGene_gn=VPS33B;refGene_tx=XM_006932253.4;refGene_gn=VPS33B;refGene_tx=XM_019831854.2;refGene_gn=VPS33B;refGene_tx=XM_019831855.2
B3	5611041	5611096
refGene_gn=VPS33B;refGene_tx=XM_003986800.4;refGene_gn=VPS33B;refGene_tx=XM_006932253.4;refGene_gn=VPS33B;refGene_tx=XM_019831854.2;refGene_gn=VPS33B;refGene_tx=XM_019831855.2
B3	5611218	5611283
refGene_gn=VPS33B;refGene_tx=XM_003986800.4;refGene_gn=VPS33B;refGene_tx=XM_006932253.4;refGene_gn=VPS33B;refGene_tx=XM_019831854.2;refGene_gn=VPS33B;refGene_tx=XM_019831855.2
B3	5611559	5611634
refGene_gn=VPS33B;refGene_tx=XM_003986800.4;refGene_gn=VPS33B;refGene_tx=XM_006932253.4;refGene_gn=VPS33B;refGene_tx=XM_019831854.2;refGene_gn=VPS33B;refGene_tx=XM_019831855.2
B3	5612018	5612109
refGene_gn=VPS33B;refGene_tx=XM_003986800.4;refGene_gn=VPS33B;refGene_tx=XM_006932253.4;refGene_gn=VPS33B;refGene_tx=XM_019831854.2;refGene_

gn=VPS33B;refGene_tx=XM_019831855.2
B3 5612298 5612385
refGene_gn=VPS33B;refGene_tx=XM_003986800.4;refGene_gn=VPS33B;refGene_tx=XM_006932253.4;refGene_gn=VPS33B;refGene_tx=XM_019831854.2;refGene_gn=VPS33B;refGene_tx=XM_019831855.2
B3 5612663 5612737
refGene_gn=VPS33B;refGene_tx=XM_003986800.4;refGene_gn=VPS33B;refGene_tx=XM_006932253.4;refGene_gn=VPS33B;refGene_tx=XM_019831854.2;refGene_gn=VPS33B;refGene_tx=XM_019831855.2
B3 5612939 5613017
refGene_gn=VPS33B;refGene_tx=XM_003986800.4;refGene_gn=VPS33B;refGene_tx=XM_006932253.4;refGene_gn=VPS33B;refGene_tx=XM_019831854.2;refGene_gn=VPS33B;refGene_tx=XM_019831855.2
B3 5613238 5613335
refGene_gn=VPS33B;refGene_tx=XM_003986800.4;refGene_gn=VPS33B;refGene_tx=XM_006932253.4;refGene_gn=VPS33B;refGene_tx=XM_019831854.2;refGene_gn=VPS33B;refGene_tx=XM_019831855.2
B3 5613742 5613847
refGene_gn=VPS33B;refGene_tx=XM_003986800.4;refGene_gn=VPS33B;refGene_tx=XM_006932253.4;refGene_gn=VPS33B;refGene_tx=XM_019831854.2;refGene_gn=VPS33B;refGene_tx=XM_019831855.2
B3 5614126 5614221
refGene_gn=VPS33B;refGene_tx=XM_003986800.4;refGene_gn=VPS33B;refGene_tx=XM_006932253.4;refGene_gn=VPS33B;refGene_tx=XM_019831854.2;refGene_gn=VPS33B;refGene_tx=XM_019831855.2
B3 5615235 5615281
refGene_gn=VPS33B;refGene_tx=XM_003986800.4;refGene_gn=VPS33B;refGene_tx=XM_006932253.4;refGene_gn=VPS33B;refGene_tx=XM_019831854.2;refGene_gn=VPS33B;refGene_tx=XM_019831855.2
B3 5619850 5619918
refGene_gn=VPS33B;refGene_tx=XM_003986800.4;refGene_gn=VPS33B;refGene_tx=XM_006932253.4;refGene_gn=VPS33B;refGene_tx=XM_019831854.2;refGene_gn=VPS33B;refGene_tx=XM_019831855.2
B3 5620813 5620863
refGene_gn=VPS33B;refGene_tx=XM_003986800.4;refGene_gn=VPS33B;refGene_tx=XM_006932253.4;refGene_gn=VPS33B;refGene_tx=XM_019831854.2;refGene_gn=VPS33B;refGene_tx=XM_019831855.2
B3 5621968 5622030
refGene_gn=VPS33B;refGene_tx=XM_003986800.4;refGene_gn=VPS33B;refGene_tx=XM_006932253.4;refGene_gn=VPS33B;refGene_tx=XM_019831854.2;refGene_gn=VPS33B;refGene_tx=XM_019831855.2
B3 5622390 5622471
refGene_gn=VPS33B;refGene_tx=XM_006932253.4;refGene_gn=VPS33B;refGene_tx=XM_019831854.2;refGene_gn=VPS33B;refGene_tx=XM_019831855.2
B3 5623780 5623876
refGene_gn=VPS33B;refGene_tx=XM_003986800.4;refGene_gn=VPS33B;refGene_tx=XM_006932253.4
B3 5629503 5630107 refGene_gn=NGRN;refGene_tx=XM_003986802.5
B3 5634157 5634268 refGene_gn=NGRN;refGene_tx=XM_003986802.5
B3 5634632 5634796 refGene_gn=NGRN;refGene_tx=XM_003986802.5

B3	5635544	5635801
refGene_gn=LOC101081807;refGene_tx=XM_003986803.5;refGene_gn=LOC101081807;refGene_tx=XM_006932254.4;refGene_gn=LOC101081807;refGene_tx=XM_019831856.2;refGene_gn=LOC101081807;refGene_tx=XM_019831857.2;refGene_gn=LOC101081807;refGene_tx=XM_019831858.2
B3	5636038	5636467
refGene_gn=LOC101081807;refGene_tx=XM_003986803.5;refGene_gn=LOC101081807;refGene_tx=XM_006932254.4;refGene_gn=LOC101081807;refGene_tx=XM_019831856.2;refGene_gn=LOC101081807;refGene_tx=XM_019831857.2;refGene_gn=LOC101081807;refGene_tx=XM_019831858.2;refGene_gn=LOC101081807;refGene_tx=XM_023254907.1
B3	5637051	5637231
refGene_gn=LOC101081807;refGene_tx=XM_003986803.5;refGene_gn=LOC101081807;refGene_tx=XM_006932254.4;refGene_gn=LOC101081807;refGene_tx=XM_019831856.2;refGene_gn=LOC101081807;refGene_tx=XM_019831857.2;refGene_gn=LOC101081807;refGene_tx=XM_019831858.2;refGene_gn=LOC101081807;refGene_tx=XM_023254907.1
B3	5637508	5637689
refGene_gn=LOC101081807;refGene_tx=XM_003986803.5;refGene_gn=LOC101081807;refGene_tx=XM_006932254.4;refGene_gn=LOC101081807;refGene_tx=XM_019831856.2;refGene_gn=LOC101081807;refGene_tx=XM_019831857.2;refGene_gn=LOC101081807;refGene_tx=XM_019831858.2;refGene_gn=LOC101081807;refGene_tx=XM_023254907.1
B3	5637984	5638170
refGene_gn=LOC101081807;refGene_tx=XM_003986803.5;refGene_gn=LOC101081807;refGene_tx=XM_006932254.4;refGene_gn=LOC101081807;refGene_tx=XM_019831856.2;refGene_gn=LOC101081807;refGene_tx=XM_019831857.2;refGene_gn=LOC101081807;refGene_tx=XM_019831858.2;refGene_gn=LOC101081807;refGene_tx=XM_023254907.1
B3	5639764	5639949
refGene_gn=LOC101081807;refGene_tx=XM_003986803.5;refGene_gn=LOC101081807;refGene_tx=XM_006932254.4;refGene_gn=LOC101081807;refGene_tx=XM_019831856.2;refGene_gn=LOC101081807;refGene_tx=XM_019831857.2;refGene_gn=LOC101081807;refGene_tx=XM_019831858.2;refGene_gn=LOC101081807;refGene_tx=XM_023254907.1
B3	5640433	5640665
refGene_gn=LOC101081807;refGene_tx=XM_003986803.5;refGene_gn=LOC101081807;refGene_tx=XM_006932254.4;refGene_gn=LOC101081807;refGene_tx=XM_019831856.2;refGene_gn=LOC101081807;refGene_tx=XM_019831857.2;refGene_gn=LOC101081807;refGene_tx=XM_019831858.2;refGene_gn=LOC101081807;refGene_tx=XM_023254907.1
B3	5640965	5641051
refGene_gn=LOC101081807;refGene_tx=XM_003986803.5;refGene_gn=LOC101081807;refGene_tx=XM_006932254.4;refGene_gn=LOC101081807;refGene_tx=XM_019831856.2;refGene_gn=LOC101081807;refGene_tx=XM_019831857.2;refGene_gn=LOC101081807;refGene_tx=XM_019831858.2;refGene_gn=LOC101081807;refGene_tx=XM_023254907.1
B3	5641410	5641554
refGene_gn=LOC101081807;refGene_tx=XM_003986803.5;refGene_gn=LOC101081807;refGene_tx=XM_006932254.4;refGene_gn=LOC101081807;refGene_tx=XM_01

9831856.2;refGene_gn=LOC101081807;refGene_tx=XM_019831857.2;refGene_gn
=LOC101081807;refGene_tx=XM_019831858.2;refGene_gn=LOC101081807;refGen
e_tx=XM_023254907.1
B3 5642400 5642557
refGene_gn=LOC101081807;refGene_tx=XM_003986803.5;refGene_gn=LOC101081
807;refGene_tx=XM_006932254.4;refGene_gn=LOC101081807;refGene_tx=XM_01
9831856.2;refGene_gn=LOC101081807;refGene_tx=XM_019831857.2;refGene_gn
=LOC101081807;refGene_tx=XM_019831858.2;refGene_gn=LOC101081807;refGen
e_tx=XM_023254907.1
B3 5642787 5642918
refGene_gn=LOC101081807;refGene_tx=XM_003986803.5;refGene_gn=LOC101081
807;refGene_tx=XM_006932254.4;refGene_gn=LOC101081807;refGene_tx=XM_01
9831856.2;refGene_gn=LOC101081807;refGene_tx=XM_019831857.2;refGene_gn
=LOC101081807;refGene_tx=XM_019831858.2;refGene_gn=LOC101081807;refGen
e_tx=XM_023254907.1
B3 5644406 5644525
refGene_gn=LOC101081807;refGene_tx=XM_003986803.5;refGene_gn=LOC101081
807;refGene_tx=XM_006932254.4;refGene_gn=LOC101081807;refGene_tx=XM_01
9831856.2;refGene_gn=LOC101081807;refGene_tx=XM_019831857.2;refGene_gn
=LOC101081807;refGene_tx=XM_019831858.2;refGene_gn=LOC101081807;refGen
e_tx=XM_023254907.1
B3 5645237 5645278
refGene_gn=LOC101081807;refGene_tx=XM_003986803.5;refGene_gn=LOC101081
807;refGene_tx=XM_006932254.4;refGene_gn=LOC101081807;refGene_tx=XM_01
9831856.2;refGene_gn=LOC101081807;refGene_tx=XM_019831857.2;refGene_gn
=LOC101081807;refGene_tx=XM_019831858.2;refGene_gn=LOC101081807;refGen
e_tx=XM_023254907.1
B3 5645627 5645887
refGene_gn=LOC101081807;refGene_tx=XM_003986803.5;refGene_gn=LOC101081
807;refGene_tx=XM_006932254.4;refGene_gn=LOC101081807;refGene_tx=XM_01
9831856.2;refGene_gn=LOC101081807;refGene_tx=XM_019831857.2;refGene_gn
=LOC101081807;refGene_tx=XM_019831858.2;refGene_gn=LOC101081807;refGen
e_tx=XM_023254907.1
B3 5649704 5650874
refGene_gn=GDPGP1;refGene_tx=XM_006932255.4;refGene_gn=GDPGP1;refGene_
tx=XM_011282759.3;refGene_gn=GDPGP1;refGene_tx=XM_019831860.2;refGene_
gn=GDPGP1;refGene_tx=XM_019831861.2;refGene_gn=GDPGP1;refGene_tx=XM_01
9831862.2;refGene_gn=GDPGP1;refGene_tx=XM_019831864.2;refGene_gn=GDPGP
1;refGene_tx=XM_019831865.2;refGene_gn=GDPGP1;refGene_tx=XM_019831866.
2;refGene_gn=GDPGP1;refGene_tx=XM_019831868.2;refGene_gn=GDPGP1;refGen
e_tx=XM_019831869.2
B3 5651599 5651638
refGene_gn=GDPGP1;refGene_tx=XM_019831860.2;refGene_gn=GDPGP1;refGene_
tx=XM_019831861.2;refGene_gn=GDPGP1;refGene_tx=XM_019831862.2;refGene_
gn=GDPGP1;refGene_tx=XM_019831864.2;refGene_gn=GDPGP1;refGene_tx=XM_01
9831865.2;refGene_gn=GDPGP1;refGene_tx=XM_019831866.2
B3 5659390 5659441 refGene_gn=CIB1;refGene_tx=XM_023254908.1
B3 5659582 5659617 refGene_gn=CIB1;refGene_tx=XM_023254908.1
B3 5660641 5660750 refGene_gn=CIB1;refGene_tx=XM_023254908.1
B3 5661876 5662027 refGene_gn=CIB1;refGene_tx=XM_023254908.1

B3 5662173 5662292 refGene_gn=CIB1;refGene_tx=XM_023254908.1
B3 5662368 5662457 refGene_gn=CIB1;refGene_tx=XM_023254908.1
B3 5662722 5662744 refGene_gn=CIB1;refGene_tx=XM_023254908.1
B3 5666937 5667680 refGene_gn=SEMA4B;refGene_tx=XM_023254909.1
B3 5667905 5667991 refGene_gn=SEMA4B;refGene_tx=XM_023254909.1
B3 5669781 5669948 refGene_gn=SEMA4B;refGene_tx=XM_023254909.1
B3 5670210 5670326 refGene_gn=SEMA4B;refGene_tx=XM_023254909.1
B3 5670412 5670623 refGene_gn=SEMA4B;refGene_tx=XM_023254909.1
B3 5671528 5671679 refGene_gn=SEMA4B;refGene_tx=XM_023254909.1
B3 5673107 5673259 refGene_gn=SEMA4B;refGene_tx=XM_023254909.1
B3 5673394 5673508 refGene_gn=SEMA4B;refGene_tx=XM_023254909.1
B3 5673805 5673917 refGene_gn=SEMA4B;refGene_tx=XM_023254909.1
B3 5674521 5674620 refGene_gn=SEMA4B;refGene_tx=XM_023254909.1
B3 5677105 5677168 refGene_gn=SEMA4B;refGene_tx=XM_023254909.1
B3 5677342 5677431 refGene_gn=SEMA4B;refGene_tx=XM_023254909.1
B3 5770932 5771047 refGene_gn=IDH2;refGene_tx=XM_023254911.1
B3 5780877 5780969 refGene_gn=IDH2;refGene_tx=XM_023254911.1
B3 5781071 5781237 refGene_gn=IDH2;refGene_tx=XM_023254911.1
B3 5782806 5782967
refGene_gn=IDH2;refGene_tx=XM_023254911.1;refGene_gn=IDH2;refGene_tx=X
M_023254912.1
B3 5783058 5783202
refGene_gn=IDH2;refGene_tx=XM_023254911.1;refGene_gn=IDH2;refGene_tx=X
M_023254912.1
B3 5783882 5784019
refGene_gn=IDH2;refGene_tx=XM_023254911.1;refGene_gn=IDH2;refGene_tx=X
M_023254912.1
B3 5784195 5784347
refGene_gn=IDH2;refGene_tx=XM_023254911.1;refGene_gn=IDH2;refGene_tx=X
M_023254912.1
B3 5786174 5786287
refGene_gn=IDH2;refGene_tx=XM_023254911.1;refGene_gn=IDH2;refGene_tx=X
M_023254912.1
B3 5786461 5786559
refGene_gn=IDH2;refGene_tx=XM_023254911.1;refGene_gn=IDH2;refGene_tx=X
M_023254912.1
B3 5786651 5786744
refGene_gn=IDH2;refGene_tx=XM_023254911.1;refGene_gn=IDH2;refGene_tx=X
M_023254912.1
B3 5787143 5787228
refGene_gn=IDH2;refGene_tx=XM_023254911.1;refGene_gn=IDH2;refGene_tx=X
M_023254912.1
B3 5791339 5791506
refGene_gn=ZNF710;refGene_tx=XM_003986806.5;refGene_gn=ZNF710;refGene_
tx=XM_006932256.4;refGene_gn=ZNF710;refGene_tx=XM_019831872.2;refGene_
gn=ZNF710;refGene_tx=XM_019831873.2;refGene_gn=ZNF710;refGene_tx=XM_02
3254910.1
B3 5795092 5795267
refGene_gn=ZNF710;refGene_tx=XM_003986806.5;refGene_gn=ZNF710;refGene_
tx=XM_006932256.4;refGene_gn=ZNF710;refGene_tx=XM_019831872.2;refGene_

gn=ZNF710;refGene_tx=XM_019831873.2;refGene_gn=ZNF710;refGene_tx=XM_023254910.1
B3	5795746	5795938	refGene_gn=ZNF710;refGene_tx=XM_003986806.5;refGene_gn=ZNF710;refGene_tx=XM_006932256.4;refGene_gn=ZNF710;refGene_tx=XM_019831872.2;refGene_gn=ZNF710;refGene_tx=XM_019831873.2;refGene_gn=ZNF710;refGene_tx=XM_023254910.1
B3	5799830	5801483	refGene_gn=ZNF710;refGene_tx=XM_003986806.5;refGene_gn=ZNF710;refGene_tx=XM_006932256.4;refGene_gn=ZNF710;refGene_tx=XM_019831872.2;refGene_gn=ZNF710;refGene_tx=XM_019831873.2;refGene_gn=ZNF710;refGene_tx=XM_023254910.1
B3	5805702	5805776	refGene_gn=ZNF710;refGene_tx=XM_003986806.5
B3	5974966	5975058	refGene_gn=LOC102901344;refGene_tx=XM_023254913.1
B3	5976318	5976394	refGene_gn=LOC102901344;refGene_tx=XM_023254913.1
B3	5979426	5979559	refGene_gn=LOC102901344;refGene_tx=XM_023254913.1
B3	5982753	5982960	refGene_gn=LOC102901344;refGene_tx=XM_023254913.1
B3	5983350	5983514	refGene_gn=LOC102901344;refGene_tx=XM_023254913.1
B3	5984412	5984421	refGene_gn=LOC102901344;refGene_tx=XM_023254913.1
B3	5990800	5990869	refGene_gn=AP3S2;refGene_tx=XM_003986807.4;refGene_gn=AP3S2;refGene_tx=XM_006932257.3;refGene_gn=AP3S2;refGene_tx=XM_019831875.2
B3	5994002	5994094	refGene_gn=AP3S2;refGene_tx=XM_003986807.4;refGene_gn=AP3S2;refGene_tx=XM_006932257.3;refGene_gn=AP3S2;refGene_tx=XM_019831875.2
B3	5994458	5994570	refGene_gn=AP3S2;refGene_tx=XM_003986807.4;refGene_gn=AP3S2;refGene_tx=XM_006932257.3;refGene_gn=AP3S2;refGene_tx=XM_019831875.2
B3	6013664	6013736	refGene_gn=AP3S2;refGene_tx=XM_003986807.4;refGene_gn=AP3S2;refGene_tx=XM_019831875.2
B3	6021524	6021686	refGene_gn=AP3S2;refGene_tx=XM_019831875.2
B3	6043202	6043310	refGene_gn=AP3S2;refGene_tx=XM_003986807.4;refGene_gn=AP3S2;refGene_tx=XM_006932257.3
B3	6045050	6045179	refGene_gn=AP3S2;refGene_tx=XM_003986807.4;refGene_gn=AP3S2;refGene_tx=XM_006932257.3
B3	6074208	6074819	refGene_gn=ANPEP;refGene_tx=NM_001009252.2
B3	6075380	6075523	refGene_gn=ANPEP;refGene_tx=NM_001009252.2
B3	6075638	6075778	refGene_gn=ANPEP;refGene_tx=NM_001009252.2
B3	6076185	6076312	refGene_gn=ANPEP;refGene_tx=NM_001009252.2
B3	6076403	6076558	refGene_gn=ANPEP;refGene_tx=NM_001009252.2
B3	6076774	6076888	refGene_gn=ANPEP;refGene_tx=NM_001009252.2

B3 6076999 6077143 refGene_gn=ANPEP;refGene_tx=NM_001009252.2
B3 6077300 6077366 refGene_gn=ANPEP;refGene_tx=NM_001009252.2
B3 6077507 6077573 refGene_gn=ANPEP;refGene_tx=NM_001009252.2
B3 6078971 6079147 refGene_gn=ANPEP;refGene_tx=NM_001009252.2
B3 6079418 6079498 refGene_gn=ANPEP;refGene_tx=NM_001009252.2
B3 6080756 6080890 refGene_gn=ANPEP;refGene_tx=NM_001009252.2
B3 6081006 6081062 refGene_gn=ANPEP;refGene_tx=NM_001009252.2
B3 6081972 6082120 refGene_gn=ANPEP;refGene_tx=NM_001009252.2
B3 6086346 6086438 refGene_gn=ANPEP;refGene_tx=NM_001009252.2
B3 6086917 6087028 refGene_gn=ANPEP;refGene_tx=NM_001009252.2
B3 6087145 6087313 refGene_gn=ANPEP;refGene_tx=NM_001009252.2
B3 6088081 6088222 refGene_gn=ANPEP;refGene_tx=NM_001009252.2
B3 6088682 6088764 refGene_gn=ANPEP;refGene_tx=NM_001009252.2
B3 6092474 6092624 refGene_gn=ANPEP;refGene_tx=NM_001009252.2
B3 6098351 6098621 refGene_gn=MESP2;refGene_tx=XM_019833746.2
B3 6099438 6100290 refGene_gn=MESP2;refGene_tx=XM_019833746.2
B3 6118087 6118798 refGene_gn=MESP1;refGene_tx=XM_019831877.2
B3 6119088 6119163 refGene_gn=MESP1;refGene_tx=XM_019831877.2
B3 6128485 6128582 refGene_gn=WDR93;refGene_tx=XM_003986836.4
B3 6135598 6135728 refGene_gn=WDR93;refGene_tx=XM_006932258.3
B3 6135902 6136101
refGene_gn=WDR93;refGene_tx=XM_003986836.4;refGene_gn=WDR93;refGene_tx=XM_006932258.3
B3 6136423 6136580
refGene_gn=WDR93;refGene_tx=XM_003986836.4;refGene_gn=WDR93;refGene_tx=XM_006932258.3
B3 6136616 6137466
refGene_gn=WDR93;refGene_tx=XM_003986836.4;refGene_gn=WDR93;refGene_tx=XM_006932258.3;refGene_gn=WDR93;refGene_tx=XM_019831878.1;refGene_gn=WDR93;refGene_tx=XM_023254915.1
B3 6139168 6139382
refGene_gn=WDR93;refGene_tx=XM_003986836.4;refGene_gn=WDR93;refGene_tx=XM_006932258.3;refGene_gn=WDR93;refGene_tx=XM_019831878.1;refGene_gn=WDR93;refGene_tx=XM_023254915.1
B3 6140784 6140904
refGene_gn=WDR93;refGene_tx=XM_003986836.4;refGene_gn=WDR93;refGene_tx=XM_006932258.3;refGene_gn=WDR93;refGene_tx=XM_019831878.1;refGene_gn=WDR93;refGene_tx=XM_023254915.1
B3 6142184 6142271
refGene_gn=WDR93;refGene_tx=XM_003986836.4;refGene_gn=WDR93;refGene_tx=XM_006932258.3;refGene_gn=WDR93;refGene_tx=XM_019831878.1;refGene_gn=WDR93;refGene_tx=XM_023254915.1
B3 6142798 6142869
refGene_gn=WDR93;refGene_tx=XM_003986836.4;refGene_gn=WDR93;refGene_tx=XM_006932258.3;refGene_gn=WDR93;refGene_tx=XM_019831878.1;refGene_gn=WDR93;refGene_tx=XM_023254915.1
B3 6144332 6144504
refGene_gn=WDR93;refGene_tx=XM_003986836.4;refGene_gn=WDR93;refGene_tx=XM_006932258.3;refGene_gn=WDR93;refGene_tx=XM_019831878.1;refGene_gn=WDR93;refGene_tx=XM_023254915.1

B3 6148433 6148518
refGene_gn=WDR93;refGene_tx=XM_003986836.4;refGene_gn=WDR93;refGene_tx=XM_006932258.3;refGene_gn=WDR93;refGene_tx=XM_019831878.1;refGene_gn=WDR93;refGene_tx=XM_023254915.1
B3 6156069 6156108
refGene_gn=WDR93;refGene_tx=XM_003986836.4;refGene_gn=WDR93;refGene_tx=XM_006932258.3;refGene_gn=WDR93;refGene_tx=XM_019831878.1;refGene_gn=WDR93;refGene_tx=XM_023254915.1
B3 6156128 6156209 refGene_gn=WDR93;refGene_tx=XM_011282758.2
B3 6157726 6157842
refGene_gn=WDR93;refGene_tx=XM_003986836.4;refGene_gn=WDR93;refGene_tx=XM_006932258.3;refGene_gn=WDR93;refGene_tx=XM_011282758.2;refGene_gn=WDR93;refGene_tx=XM_019831878.1;refGene_gn=WDR93;refGene_tx=XM_023254915.1
B3 6159553 6159632
refGene_gn=WDR93;refGene_tx=XM_003986836.4;refGene_gn=WDR93;refGene_tx=XM_006932258.3;refGene_gn=WDR93;refGene_tx=XM_011282758.2;refGene_gn=WDR93;refGene_tx=XM_019831878.1;refGene_gn=WDR93;refGene_tx=XM_023254915.1
B3 6166587 6166652
refGene_gn=WDR93;refGene_tx=XM_003986836.4;refGene_gn=WDR93;refGene_tx=XM_006932258.3;refGene_gn=WDR93;refGene_tx=XM_011282758.2;refGene_gn=WDR93;refGene_tx=XM_019831878.1;refGene_gn=WDR93;refGene_tx=XM_023254915.1
B3 6168266 6168459
refGene_gn=WDR93;refGene_tx=XM_003986836.4;refGene_gn=WDR93;refGene_tx=XM_006932258.3;refGene_gn=WDR93;refGene_tx=XM_011282758.2;refGene_gn=WDR93;refGene_tx=XM_019831878.1;refGene_gn=WDR93;refGene_tx=XM_023254915.1
B3 6169066 6169369
refGene_gn=WDR93;refGene_tx=XM_003986836.4;refGene_gn=WDR93;refGene_tx=XM_006932258.3;refGene_gn=WDR93;refGene_tx=XM_011282758.2;refGene_gn=WDR93;refGene_tx=XM_019831878.1;refGene_gn=WDR93;refGene_tx=XM_023254915.1
B3 6177557 6177613
refGene_gn=PEX11A;refGene_tx=XM_003986808.4;refGene_gn=PEX11A;refGene_tx=XM_019831881.2
B3 6187778 6187894 refGene_gn=PEX11A;refGene_tx=XM_003986808.4
B3 6189846 6190418
refGene_gn=PEX11A;refGene_tx=XM_003986808.4;refGene_gn=PEX11A;refGene_tx=XM_019831881.2
B3 6195502 6195550
refGene_gn=PLIN1;refGene_tx=XM_023254916.1;refGene_gn=PLIN1;refGene_tx=XM_023254918.1;refGene_gn=PLIN1;refGene_tx=XM_023254919.1;refGene_gn=PLIN1;refGene_tx=XM_023254920.1
B3 6198318 6198523
refGene_gn=PLIN1;refGene_tx=XM_023254916.1;refGene_gn=PLIN1;refGene_tx=XM_023254918.1;refGene_gn=PLIN1;refGene_tx=XM_023254919.1;refGene_gn=PLIN1;refGene_tx=XM_023254920.1;refGene_gn=PLIN1;refGene_tx=XM_023254921.1

B3 6200233 6200316
refGene_gn=PLIN1;refGene_tx=XM_023254916.1;refGene_gn=PLIN1;refGene_tx=XM_023254918.1;refGene_gn=PLIN1;refGene_tx=XM_023254919.1;refGene_gn=PLIN1;refGene_tx=XM_023254920.1
B3 6201633 6201898
refGene_gn=PLIN1;refGene_tx=XM_023254916.1;refGene_gn=PLIN1;refGene_tx=XM_023254918.1;refGene_gn=PLIN1;refGene_tx=XM_023254919.1;refGene_gn=PLIN1;refGene_tx=XM_023254920.1;refGene_gn=PLIN1;refGene_tx=XM_023254921.1
B3 6202230 6202403
refGene_gn=PLIN1;refGene_tx=XM_023254916.1;refGene_gn=PLIN1;refGene_tx=XM_023254918.1;refGene_gn=PLIN1;refGene_tx=XM_023254919.1;refGene_gn=PLIN1;refGene_tx=XM_023254920.1;refGene_gn=PLIN1;refGene_tx=XM_023254921.1
B3 6204095 6204284
refGene_gn=PLIN1;refGene_tx=XM_023254916.1;refGene_gn=PLIN1;refGene_tx=XM_023254918.1;refGene_gn=PLIN1;refGene_tx=XM_023254919.1;refGene_gn=PLIN1;refGene_tx=XM_023254920.1;refGene_gn=PLIN1;refGene_tx=XM_023254921.1
B3 6204714 6204960
refGene_gn=PLIN1;refGene_tx=XM_023254916.1;refGene_gn=PLIN1;refGene_tx=XM_023254918.1;refGene_gn=PLIN1;refGene_tx=XM_023254919.1;refGene_gn=PLIN1;refGene_tx=XM_023254920.1;refGene_gn=PLIN1;refGene_tx=XM_023254921.1
B3 6205968 6206328
refGene_gn=PLIN1;refGene_tx=XM_023254916.1;refGene_gn=PLIN1;refGene_tx=XM_023254918.1;refGene_gn=PLIN1;refGene_tx=XM_023254919.1;refGene_gn=PLIN1;refGene_tx=XM_023254920.1;refGene_gn=PLIN1;refGene_tx=XM_023254921.1
B3 6221177 6221505
refGene_gn=KIF7;refGene_tx=XM_023254924.1;refGene_gn=KIF7;refGene_tx=XM_023254925.1;refGene_gn=KIF7;refGene_tx=XM_023254926.1;refGene_gn=KIF7;refGene_tx=XM_023254927.1;refGene_gn=KIF7;refGene_tx=XM_023254928.1
B3 6223169 6223370
refGene_gn=KIF7;refGene_tx=XM_023254924.1;refGene_gn=KIF7;refGene_tx=XM_023254925.1;refGene_gn=KIF7;refGene_tx=XM_023254926.1;refGene_gn=KIF7;refGene_tx=XM_023254927.1;refGene_gn=KIF7;refGene_tx=XM_023254928.1
B3 6223755 6224149
refGene_gn=KIF7;refGene_tx=XM_023254924.1;refGene_gn=KIF7;refGene_tx=XM_023254925.1;refGene_gn=KIF7;refGene_tx=XM_023254926.1;refGene_gn=KIF7;refGene_tx=XM_023254927.1;refGene_gn=KIF7;refGene_tx=XM_023254928.1
B3 6224377 6224918
refGene_gn=KIF7;refGene_tx=XM_023254924.1;refGene_gn=KIF7;refGene_tx=XM_023254925.1;refGene_gn=KIF7;refGene_tx=XM_023254926.1;refGene_gn=KIF7;refGene_tx=XM_023254927.1;refGene_gn=KIF7;refGene_tx=XM_023254928.1
B3 6225438 6225552
refGene_gn=KIF7;refGene_tx=XM_023254924.1;refGene_gn=KIF7;refGene_tx=XM_023254925.1;refGene_gn=KIF7;refGene_tx=XM_023254926.1;refGene_gn=KIF7;refGene_tx=XM_023254927.1;refGene_gn=KIF7;refGene_tx=XM_023254928.1
B3 6226092 6226320

refGene_gn=KIF7;refGene_tx=XM_023254924.1;refGene_gn=KIF7;refGene_tx=XM_023254925.1;refGene_gn=KIF7;refGene_tx=XM_023254926.1;refGene_gn=KIF7;refGene_tx=XM_023254927.1;refGene_gn=KIF7;refGene_tx=XM_023254928.1
B3 6227080 6227217
refGene_gn=KIF7;refGene_tx=XM_023254924.1;refGene_gn=KIF7;refGene_tx=XM_023254925.1;refGene_gn=KIF7;refGene_tx=XM_023254926.1;refGene_gn=KIF7;refGene_tx=XM_023254927.1;refGene_gn=KIF7;refGene_tx=XM_023254928.1
B3 6227561 6227677
refGene_gn=KIF7;refGene_tx=XM_023254924.1;refGene_gn=KIF7;refGene_tx=XM_023254925.1;refGene_gn=KIF7;refGene_tx=XM_023254926.1;refGene_gn=KIF7;refGene_tx=XM_023254927.1;refGene_gn=KIF7;refGene_tx=XM_023254928.1
B3 6227853 6228006
refGene_gn=KIF7;refGene_tx=XM_023254924.1;refGene_gn=KIF7;refGene_tx=XM_023254925.1;refGene_gn=KIF7;refGene_tx=XM_023254926.1;refGene_gn=KIF7;refGene_tx=XM_023254927.1;refGene_gn=KIF7;refGene_tx=XM_023254928.1
B3 6229682 6229885
refGene_gn=KIF7;refGene_tx=XM_023254924.1;refGene_gn=KIF7;refGene_tx=XM_023254925.1;refGene_gn=KIF7;refGene_tx=XM_023254926.1;refGene_gn=KIF7;refGene_tx=XM_023254927.1;refGene_gn=KIF7;refGene_tx=XM_023254928.1
B3 6241229 6241427
refGene_gn=KIF7;refGene_tx=XM_023254924.1;refGene_gn=KIF7;refGene_tx=XM_023254925.1;refGene_gn=KIF7;refGene_tx=XM_023254926.1;refGene_gn=KIF7;refGene_tx=XM_023254927.1;refGene_gn=KIF7;refGene_tx=XM_023254928.1
B3 6241723 6241849
refGene_gn=KIF7;refGene_tx=XM_023254924.1;refGene_gn=KIF7;refGene_tx=XM_023254925.1;refGene_gn=KIF7;refGene_tx=XM_023254926.1;refGene_gn=KIF7;refGene_tx=XM_023254927.1;refGene_gn=KIF7;refGene_tx=XM_023254928.1
B3 6242023 6242200
refGene_gn=KIF7;refGene_tx=XM_023254924.1;refGene_gn=KIF7;refGene_tx=XM_023254925.1;refGene_gn=KIF7;refGene_tx=XM_023254926.1;refGene_gn=KIF7;refGene_tx=XM_023254927.1;refGene_gn=KIF7;refGene_tx=XM_023254928.1
B3 6243014 6243230
refGene_gn=KIF7;refGene_tx=XM_023254924.1;refGene_gn=KIF7;refGene_tx=XM_023254925.1;refGene_gn=KIF7;refGene_tx=XM_023254926.1;refGene_gn=KIF7;refGene_tx=XM_023254927.1;refGene_gn=KIF7;refGene_tx=XM_023254928.1
B3 6243887 6244094
refGene_gn=KIF7;refGene_tx=XM_023254924.1;refGene_gn=KIF7;refGene_tx=XM_023254925.1;refGene_gn=KIF7;refGene_tx=XM_023254926.1;refGene_gn=KIF7;refGene_tx=XM_023254927.1;refGene_gn=KIF7;refGene_tx=XM_023254928.1
B3 6244818 6245017
refGene_gn=KIF7;refGene_tx=XM_023254924.1;refGene_gn=KIF7;refGene_tx=XM_023254925.1;refGene_gn=KIF7;refGene_tx=XM_023254926.1;refGene_gn=KIF7;refGene_tx=XM_023254927.1;refGene_gn=KIF7;refGene_tx=XM_023254928.1
B3 6245100 6245247
refGene_gn=KIF7;refGene_tx=XM_023254924.1;refGene_gn=KIF7;refGene_tx=XM_023254925.1;refGene_gn=KIF7;refGene_tx=XM_023254926.1;refGene_gn=KIF7;refGene_tx=XM_023254927.1;refGene_gn=KIF7;refGene_tx=XM_023254928.1
B3 6245385 6245729
refGene_gn=KIF7;refGene_tx=XM_023254924.1;refGene_gn=KIF7;refGene_tx=XM_023254925.1;refGene_gn=KIF7;refGene_tx=XM_023254926.1;refGene_gn=KIF

7;refGene_tx=XM_023254927.1;refGene_gn=KIF7;refGene_tx=XM_023254928.1
B3	6247051	6247161	refGene_gn=TICRR;refGene_tx=XM_023254923.1
B3	6247900	6248026	refGene_gn=TICRR;refGene_tx=XM_023254923.1
B3	6248146	6250289	refGene_gn=TICRR;refGene_tx=XM_023254923.1
B3	6252395	6252553	refGene_gn=TICRR;refGene_tx=XM_023254923.1
B3	6253137	6253267	refGene_gn=TICRR;refGene_tx=XM_023254923.1
B3	6253887	6253892	refGene_gn=TICRR;refGene_tx=XM_023254923.1
B3	6254560	6254622	refGene_gn=TICRR;refGene_tx=XM_023254923.1
B3	6257101	6257192	refGene_gn=TICRR;refGene_tx=XM_023254923.1
B3	6267746	6267896	refGene_gn=TICRR;refGene_tx=XM_023254923.1
B3	6269490	6269548	refGene_gn=TICRR;refGene_tx=XM_023254923.1
B3	6272182	6272279	refGene_gn=TICRR;refGene_tx=XM_023254923.1
B3	6272767	6273007	refGene_gn=TICRR;refGene_tx=XM_023254923.1
B3	6273169	6273249	refGene_gn=TICRR;refGene_tx=XM_023254923.1
B3	6273344	6273438	refGene_gn=TICRR;refGene_tx=XM_023254923.1
B3	6273748	6273816	refGene_gn=TICRR;refGene_tx=XM_023254923.1
B3	6273817	6273847	refGene_gn=TICRR;refGene_tx=XM_023254923.1
B3	6274838	6274990	refGene_gn=TICRR;refGene_tx=XM_023254923.1
B3	6277379	6277598	refGene_gn=TICRR;refGene_tx=XM_023254923.1
B3	6278394	6278534	refGene_gn=TICRR;refGene_tx=XM_023254923.1
B3	6282497	6282627	refGene_gn=TICRR;refGene_tx=XM_023254923.1
B3	6283594	6283829	refGene_gn=TICRR;refGene_tx=XM_023254923.1
B3	6284965	6285204	refGene_gn=TICRR;refGene_tx=XM_023254923.1
B3	6287081	6287361	refGene_gn=TICRR;refGene_tx=XM_023254923.1
B3	6293376	6294039	refGene_gn=TICRR;refGene_tx=XM_023254923.1
B3	6342232	6342419
refGene_gn=RHCG;refGene_tx=XM_003986810.5;refGene_gn=RHCG;refGene_tx=XM_006932260.4;refGene_gn=RHCG;refGene_tx=XM_019831884.2
B3	6350194	6350381
refGene_gn=RHCG;refGene_tx=XM_003986810.5;refGene_gn=RHCG;refGene_tx=XM_006932260.4;refGene_gn=RHCG;refGene_tx=XM_019831884.2
B3	6352582	6352733
refGene_gn=RHCG;refGene_tx=XM_003986810.5;refGene_gn=RHCG;refGene_tx=XM_006932260.4;refGene_gn=RHCG;refGene_tx=XM_019831884.2
B3	6356611	6356759
refGene_gn=RHCG;refGene_tx=XM_003986810.5;refGene_gn=RHCG;refGene_tx=XM_006932260.4;refGene_gn=RHCG;refGene_tx=XM_019831884.2
B3	6357560	6357727
refGene_gn=RHCG;refGene_tx=XM_003986810.5;refGene_gn=RHCG;refGene_tx=XM_006932260.4;refGene_gn=RHCG;refGene_tx=XM_019831884.2
B3	6359584	6359722
refGene_gn=RHCG;refGene_tx=XM_003986810.5;refGene_gn=RHCG;refGene_tx=XM_006932260.4;refGene_gn=RHCG;refGene_tx=XM_019831884.2
B3	6359911	6360048
refGene_gn=RHCG;refGene_tx=XM_003986810.5;refGene_gn=RHCG;refGene_tx=XM_006932260.4;refGene_gn=RHCG;refGene_tx=XM_019831884.2
B3	6360361	6360486
refGene_gn=RHCG;refGene_tx=XM_003986810.5;refGene_gn=RHCG;refGene_tx=XM_006932260.4;refGene_gn=RHCG;refGene_tx=XM_019831884.2
B3	6360708	6360782

refGene_gn=RHCG;refGene_tx=XM_003986810.5;refGene_gn=RHCG;refGene_tx=XM_006932260.4;refGene_gn=RHCG;refGene_tx=XM_019831884.2
B3 6363762 6363831 refGene_gn=RHCG;refGene_tx=XM_003986810.5;refGene_gn=RHCG;refGene_tx=XM_006932260.4;refGene_gn=RHCG;refGene_tx=XM_019831884.2
B3 6488745 6489365 refGene_gn=POLG;refGene_tx=XM_019831892.2
B3 6492118 6492314 refGene_gn=POLG;refGene_tx=XM_019831892.2
B3 6493211 6493376 refGene_gn=POLG;refGene_tx=XM_019831892.2
B3 6493502 6493649 refGene_gn=POLG;refGene_tx=XM_019831892.2
B3 6493800 6493880 refGene_gn=POLG;refGene_tx=XM_019831892.2
B3 6494679 6494862 refGene_gn=POLG;refGene_tx=XM_019831892.2
B3 6494961 6495116 refGene_gn=POLG;refGene_tx=XM_019831892.2
B3 6495293 6495420 refGene_gn=POLG;refGene_tx=XM_019831892.2
B3 6496297 6496534 refGene_gn=POLG;refGene_tx=XM_019831892.2
B3 6497375 6497496 refGene_gn=POLG;refGene_tx=XM_019831892.2
B3 6497716 6497803 refGene_gn=POLG;refGene_tx=XM_019831892.2
B3 6498112 6498220 refGene_gn=POLG;refGene_tx=XM_019831892.2
B3 6499008 6499169 refGene_gn=POLG;refGene_tx=XM_019831892.2
B3 6499854 6499908 refGene_gn=POLG;refGene_tx=XM_019831892.2
B3 6500010 6500128 refGene_gn=POLG;refGene_tx=XM_019831892.2
B3 6500593 6500729 refGene_gn=POLG;refGene_tx=XM_019831892.2
B3 6500823 6501070 refGene_gn=POLG;refGene_tx=XM_019831892.2
B3 6502296 6502419 refGene_gn=POLG;refGene_tx=XM_019831892.2
B3 6502548 6502717 refGene_gn=POLG;refGene_tx=XM_019831892.2
B3 6502806 6503015 refGene_gn=POLG;refGene_tx=XM_019831892.2
B3 6503754 6503915 refGene_gn=POLG;refGene_tx=XM_019831892.2
B3 6504368 6504427 refGene_gn=POLG;refGene_tx=XM_019831892.2
B3 6504731 6504794 refGene_gn=FANCI;refGene_tx=XM_023254929.1
B3 6505604 6505718 refGene_gn=FANCI;refGene_tx=XM_023254929.1
B3 6505992 6506088 refGene_gn=FANCI;refGene_tx=XM_023254929.1
B3 6507464 6507539 refGene_gn=FANCI;refGene_tx=XM_023254929.1
B3 6511175 6511235 refGene_gn=FANCI;refGene_tx=XM_023254929.1
B3 6511316 6511370 refGene_gn=FANCI;refGene_tx=XM_023254929.1
B3 6513176 6513364 refGene_gn=FANCI;refGene_tx=XM_023254929.1
B3 6513659 6513753 refGene_gn=FANCI;refGene_tx=XM_023254929.1
B3 6513926 6513995 refGene_gn=FANCI;refGene_tx=XM_023254929.1
B3 6514093 6514221 refGene_gn=FANCI;refGene_tx=XM_023254929.1
B3 6515009 6515061 refGene_gn=FANCI;refGene_tx=XM_023254929.1
B3 6517576 6517693 refGene_gn=FANCI;refGene_tx=XM_023254929.1
B3 6518784 6518861 refGene_gn=FANCI;refGene_tx=XM_023254929.1
B3 6519224 6519391 refGene_gn=FANCI;refGene_tx=XM_023254929.1
B3 6522436 6522616 refGene_gn=FANCI;refGene_tx=XM_023254929.1
B3 6523257 6523422 refGene_gn=FANCI;refGene_tx=XM_023254929.1
B3 6523934 6524056 refGene_gn=FANCI;refGene_tx=XM_023254929.1
B3 6524932 6525106 refGene_gn=FANCI;refGene_tx=XM_023254929.1
B3 6526091 6526186 refGene_gn=FANCI;refGene_tx=XM_023254929.1
B3 6528869 6528945 refGene_gn=FANCI;refGene_tx=XM_023254929.1
B3 6534190 6534313 refGene_gn=FANCI;refGene_tx=XM_023254929.1
B3 6535765 6535880 refGene_gn=FANCI;refGene_tx=XM_023254929.1
B3 6537601 6537672 refGene_gn=FANCI;refGene_tx=XM_023254929.1

B3	6538145	6538276	refGene_gn=FANCI;refGene_tx=XM_023254929.1
B3	6551979	6552067	refGene_gn=FANCI;refGene_tx=XM_023254929.1
B3	6553936	6554117	refGene_gn=FANCI;refGene_tx=XM_023254929.1
B3	6559870	6560007	refGene_gn=FANCI;refGene_tx=XM_023254929.1
B3	6560621	6560714	refGene_gn=FANCI;refGene_tx=XM_023254929.1
B3	6566020	6566147	refGene_gn=FANCI;refGene_tx=XM_023254929.1
B3	6568353	6568439	refGene_gn=FANCI;refGene_tx=XM_023254929.1
B3	6569140	6569264	refGene_gn=FANCI;refGene_tx=XM_023254929.1
B3	6570048	6570090	refGene_gn=FANCI;refGene_tx=XM_023254929.1
B3	6571705	6571763	refGene_gn=FANCI;refGene_tx=XM_023254929.1
B3	6571842	6571999	refGene_gn=FANCI;refGene_tx=XM_023254929.1
B3	6572406	6572537	refGene_gn=FANCI;refGene_tx=XM_023254929.1
B3	6580861	6580934	refGene_gn=FANCI;refGene_tx=XM_023254929.1
B3	6592062	6592164	refGene_gn=FANCI;refGene_tx=XM_023254929.1
B3	6610272	6610284
refGene_gn=RLBP1;refGene_tx=XM_011282757.2;refGene_gn=RLBP1;refGene_tx
=XM_019831893.1;refGene_gn=RLBP1;refGene_tx=XM_019831894.1
B3	6610562	6610691
refGene_gn=RLBP1;refGene_tx=XM_011282757.2;refGene_gn=RLBP1;refGene_tx
=XM_019831893.1
B3	6611130	6611235	refGene_gn=FANCI;refGene_tx=XM_023254929.1
B3	6611899	6612104
refGene_gn=RLBP1;refGene_tx=XM_011282757.2;refGene_gn=RLBP1;refGene_tx
=XM_019831893.1;refGene_gn=RLBP1;refGene_tx=XM_019831894.1
B3	6613808	6613987
refGene_gn=RLBP1;refGene_tx=XM_011282757.2;refGene_gn=RLBP1;refGene_tx
=XM_019831893.1;refGene_gn=RLBP1;refGene_tx=XM_019831894.1
B3	6617310	6617469
refGene_gn=RLBP1;refGene_tx=XM_011282757.2;refGene_gn=RLBP1;refGene_tx
=XM_019831893.1;refGene_gn=RLBP1;refGene_tx=XM_019831894.1
B3	6618646	6618757
refGene_gn=RLBP1;refGene_tx=XM_011282757.2;refGene_gn=RLBP1;refGene_tx
=XM_019831893.1;refGene_gn=RLBP1;refGene_tx=XM_019831894.1
B3	6619078	6619237
refGene_gn=RLBP1;refGene_tx=XM_011282757.2;refGene_gn=RLBP1;refGene_tx
=XM_019831893.1;refGene_gn=RLBP1;refGene_tx=XM_019831894.1
B3	6625504	6625515	refGene_gn=ABHD2;refGene_tx=XM_019831896.2
B3	6633133	6633330	refGene_gn=ABHD2;refGene_tx=XM_006932261.4
B3	6633479	6633631	refGene_gn=ABHD2;refGene_tx=XM_019831895.2
B3	6635405	6635490
refGene_gn=ABHD2;refGene_tx=XM_006932261.4;refGene_gn=ABHD2;refGene_tx
=XM_019831895.2;refGene_gn=ABHD2;refGene_tx=XM_019831896.2
B3	6636974	6637044
refGene_gn=ABHD2;refGene_tx=XM_006932261.4;refGene_gn=ABHD2;refGene_tx
=XM_019831895.2;refGene_gn=ABHD2;refGene_tx=XM_019831896.2
B3	6639084	6639195
refGene_gn=ABHD2;refGene_tx=XM_006932261.4;refGene_gn=ABHD2;refGene_tx
=XM_019831895.2;refGene_gn=ABHD2;refGene_tx=XM_019831896.2
B3	6642197	6642290
refGene_gn=ABHD2;refGene_tx=XM_006932261.4;refGene_gn=ABHD2;refGene_tx

=XM_019831895.2;refGene_gn=ABHD2;refGene_tx=XM_019831896.2
B3 6652718 6652902
refGene_gn=ABHD2;refGene_tx=XM_006932261.4;refGene_gn=ABHD2;refGene_tx
=XM_019831895.2;refGene_gn=ABHD2;refGene_tx=XM_019831896.2
B3 6667035 6667203
refGene_gn=ABHD2;refGene_tx=XM_006932261.4;refGene_gn=ABHD2;refGene_tx
=XM_019831895.2;refGene_gn=ABHD2;refGene_tx=XM_019831896.2
B3 6670134 6670310
refGene_gn=ABHD2;refGene_tx=XM_006932261.4;refGene_gn=ABHD2;refGene_tx
=XM_019831895.2;refGene_gn=ABHD2;refGene_tx=XM_019831896.2
B3 6706336 6706530
refGene_gn=ABHD2;refGene_tx=XM_006932261.4;refGene_gn=ABHD2;refGene_tx
=XM_019831895.2;refGene_gn=ABHD2;refGene_tx=XM_019831896.2
B3 6839739 6839812
refGene_gn=MFGE8;refGene_tx=XM_011282749.2;refGene_gn=MFGE8;refGene_tx
=XM_011282750.2
B3 6842342 6842468 refGene_gn=MFGE8;refGene_tx=XM_011282749.2
B3 6842829 6842961
refGene_gn=MFGE8;refGene_tx=XM_011282749.2;refGene_gn=MFGE8;refGene_tx
=XM_011282750.2
B3 6845455 6845637
refGene_gn=MFGE8;refGene_tx=XM_011282749.2;refGene_gn=MFGE8;refGene_tx
=XM_011282750.2
B3 6846105 6846261
refGene_gn=MFGE8;refGene_tx=XM_011282749.2;refGene_gn=MFGE8;refGene_tx
=XM_011282750.2
B3 6846963 6847108
refGene_gn=MFGE8;refGene_tx=XM_011282749.2;refGene_gn=MFGE8;refGene_tx
=XM_011282750.2
B3 6851088 6851273
refGene_gn=MFGE8;refGene_tx=XM_011282749.2;refGene_gn=MFGE8;refGene_tx
=XM_011282750.2
B3 6852346 6852502
refGene_gn=MFGE8;refGene_tx=XM_011282749.2;refGene_gn=MFGE8;refGene_tx
=XM_011282750.2
B3 6852629 6852767
refGene_gn=MFGE8;refGene_tx=XM_011282749.2;refGene_gn=MFGE8;refGene_tx
=XM_011282750.2
B3 6856021 6856055 refGene_gn=HAPLN3;refGene_tx=XM_006932263.3
B3 6856188 6856501
refGene_gn=HAPLN3;refGene_tx=XM_003986842.4;refGene_gn=HAPLN3;refGene_
tx=XM_006932263.3
B3 6864684 6864858
refGene_gn=HAPLN3;refGene_tx=XM_003986842.4;refGene_gn=HAPLN3;refGene_
tx=XM_006932263.3
B3 6868063 6868432
refGene_gn=HAPLN3;refGene_tx=XM_003986842.4;refGene_gn=HAPLN3;refGene_
tx=XM_006932263.3
B3 6869746 6870049
refGene_gn=HAPLN3;refGene_tx=XM_003986842.4;refGene_gn=HAPLN3;refGene_

tx=XM_006932263.3
B3	6870743	6871030
refGene_gn=HAPLN3;refGene_tx=XM_003986842.4;refGene_gn=HAPLN3;refGene_tx=XM_006932263.3
B3	6873841	6873918	refGene_gn=ACAN;refGene_tx=XM_023254934.1
B3	6874347	6874530	refGene_gn=ACAN;refGene_tx=XM_023254934.1
B3	6875460	6875605	refGene_gn=ACAN;refGene_tx=XM_023254934.1
B3	6876184	6876267	refGene_gn=ACAN;refGene_tx=XM_023254934.1
B3	6876645	6876804	refGene_gn=ACAN;refGene_tx=XM_023254934.1
B3	6880305	6880419	refGene_gn=ACAN;refGene_tx=XM_023254934.1
B3	6890814	6893344	refGene_gn=ACAN;refGene_tx=XM_023254934.1
B3	6894053	6894847	refGene_gn=ACAN;refGene_tx=XM_023254934.1
B3	6897935	6898145	refGene_gn=ACAN;refGene_tx=XM_023254934.1
B3	6899968	6900262	refGene_gn=ACAN;refGene_tx=XM_023254934.1
B3	6901266	6901394	refGene_gn=ACAN;refGene_tx=XM_023254934.1
B3	6901915	6902090	refGene_gn=ACAN;refGene_tx=XM_023254934.1
B3	6904162	6904564	refGene_gn=ACAN;refGene_tx=XM_023254934.1
B3	6906247	6906541	refGene_gn=ACAN;refGene_tx=XM_023254934.1
B3	6907830	6907958	refGene_gn=ACAN;refGene_tx=XM_023254934.1
B3	6910047	6910222	refGene_gn=ACAN;refGene_tx=XM_023254934.1
B3	6911607	6911991	refGene_gn=ACAN;refGene_tx=XM_023254934.1
B3	6914342	6914412	refGene_gn=ACAN;refGene_tx=XM_023254934.1
B3	7082624	7082711	refGene_gn=ISG20;refGene_tx=XM_019831901.2
B3	7084267	7084273	refGene_gn=ISG20;refGene_tx=XM_023254933.1
B3	7084420	7084621
refGene_gn=ISG20;refGene_tx=XM_019831901.2;refGene_gn=ISG20;refGene_tx=XM_023254933.1
B3	7085835	7086318
refGene_gn=ISG20;refGene_tx=XM_023254931.1;refGene_gn=ISG20;refGene_tx=XM_023254932.1
B3	7092756	7092984
refGene_gn=ISG20;refGene_tx=XM_019831901.2;refGene_gn=ISG20;refGene_tx=XM_023254931.1;refGene_gn=ISG20;refGene_tx=XM_023254932.1;refGene_gn=ISG20;refGene_tx=XM_023254933.1
B3	7101169	7101406
refGene_gn=AEN;refGene_tx=XM_006932265.2;refGene_gn=AEN;refGene_tx=XM_006932266.3;refGene_gn=AEN;refGene_tx=XM_019831903.2
B3	7101966	7102167
refGene_gn=AEN;refGene_tx=XM_006932265.2;refGene_gn=AEN;refGene_tx=XM_006932266.3;refGene_gn=AEN;refGene_tx=XM_019831903.2
B3	7102612	7103152
refGene_gn=AEN;refGene_tx=XM_006932265.2;refGene_gn=AEN;refGene_tx=XM_006932266.3;refGene_gn=AEN;refGene_tx=XM_019831903.2
B3	7164898	7165981
refGene_gn=DET1;refGene_tx=XM_003986815.4;refGene_gn=DET1;refGene_tx=XM_006932269.4;refGene_gn=DET1;refGene_tx=XM_006932270.4;refGene_gn=DET1;refGene_tx=XM_006932271.3;refGene_gn=DET1;refGene_tx=XM_019831905.2;refGene_gn=DET1;refGene_tx=XM_023254936.1
B3	7168783	7168971
refGene_gn=DET1;refGene_tx=XM_003986815.4;refGene_gn=DET1;refGene_tx=X

M_006932269.4;refGene_gn=DET1;refGene_tx=XM_006932270.4;refGene_gn=DET1;refGene_tx=XM_006932271.3;refGene_gn=DET1;refGene_tx=XM_019831905.2;refGene_gn=DET1;refGene_tx=XM_023254936.1
B3 7177653 7177845
refGene_gn=DET1;refGene_tx=XM_003986815.4;refGene_gn=DET1;refGene_tx=XM_006932269.4;refGene_gn=DET1;refGene_tx=XM_006932270.4;refGene_gn=DET1;refGene_tx=XM_006932271.3;refGene_gn=DET1;refGene_tx=XM_019831905.2;refGene_gn=DET1;refGene_tx=XM_023254936.1
B3 7180849 7181039 refGene_gn=DET1;refGene_tx=XM_003986815.4
B3 7187835 7188028
refGene_gn=DET1;refGene_tx=XM_006932269.4;refGene_gn=DET1;refGene_tx=XM_019831905.2
B3 7190279 7190346 refGene_gn=DET1;refGene_tx=XM_023254936.1
B3 7191847 7191863 refGene_gn=DET1;refGene_tx=XM_006932271.3
B3 7216027 7216135
refGene_gn=MRPS11;refGene_tx=XM_003986816.4;refGene_gn=MRPS11;refGene_tx=XM_023254937.1
B3 7216317 7216363 refGene_gn=DET1;refGene_tx=XM_006932270.4
B3 7216831 7216897
refGene_gn=MRPS11;refGene_tx=XM_003986816.4;refGene_gn=MRPS11;refGene_tx=XM_023254937.1
B3 7217913 7218043
refGene_gn=MRPS11;refGene_tx=XM_003986816.4;refGene_gn=MRPS11;refGene_tx=XM_023254937.1
B3 7221059 7221158
refGene_gn=MRPS11;refGene_tx=XM_003986816.4;refGene_gn=MRPS11;refGene_tx=XM_023254937.1
B3 7226654 7226777
refGene_gn=MRPS11;refGene_tx=XM_003986816.4;refGene_gn=MRPS11;refGene_tx=XM_023254937.1
B3 7226903 7227081
refGene_gn=MRPS11;refGene_tx=XM_003986816.4;refGene_gn=MRPS11;refGene_tx=XM_023254937.1
B3 7227361 7227589 refGene_gn=MRPL46;refGene_tx=XM_003986817.5
B3 7228882 7229072 refGene_gn=MRPL46;refGene_tx=XM_003986817.5
B3 7229844 7230018 refGene_gn=MRPL46;refGene_tx=XM_003986817.5
B3 7236507 7236761 refGene_gn=MRPL46;refGene_tx=XM_003986817.5
B3 7429337 7429585
refGene_gn=NTRK3;refGene_tx=XM_011282744.3;refGene_gn=NTRK3;refGene_tx=XM_011282745.2;refGene_gn=NTRK3;refGene_tx=XM_019831907.2;refGene_gn=NTRK3;refGene_tx=XM_019831909.2;refGene_gn=NTRK3;refGene_tx=XM_023254939.1
B3 7503945 7504020
refGene_gn=NTRK3;refGene_tx=XM_011282744.3;refGene_gn=NTRK3;refGene_tx=XM_011282745.2;refGene_gn=NTRK3;refGene_tx=XM_019831907.2;refGene_gn=NTRK3;refGene_tx=XM_019831909.2;refGene_gn=NTRK3;refGene_tx=XM_023254939.1
B3 7505335 7505407
refGene_gn=NTRK3;refGene_tx=XM_011282744.3;refGene_gn=NTRK3;refGene_tx=XM_011282745.2;refGene_gn=NTRK3;refGene_tx=XM_019831907.2;refGene_gn=

NTRK3;refGene_tx=XM_019831909.2;refGene_gn=NTRK3;refGene_tx=XM_0232549
39.1
B3 7542068 7542137
refGene_gn=NTRK3;refGene_tx=XM_011282744.3;refGene_gn=NTRK3;refGene_tx
=XM_011282745.2;refGene_gn=NTRK3;refGene_tx=XM_019831907.2;refGene_gn=
NTRK3;refGene_tx=XM_019831909.2;refGene_gn=NTRK3;refGene_tx=XM_0232549
39.1
B3 7551588 7551746
refGene_gn=NTRK3;refGene_tx=XM_011282744.3;refGene_gn=NTRK3;refGene_tx
=XM_011282745.2;refGene_gn=NTRK3;refGene_tx=XM_019831907.2;refGene_gn=
NTRK3;refGene_tx=XM_019831909.2;refGene_gn=NTRK3;refGene_tx=XM_0232549
39.1
B3 7552423 7552566
refGene_gn=NTRK3;refGene_tx=XM_011282744.3;refGene_gn=NTRK3;refGene_tx
=XM_011282745.2;refGene_gn=NTRK3;refGene_tx=XM_019831907.2;refGene_gn=
NTRK3;refGene_tx=XM_019831909.2;refGene_gn=NTRK3;refGene_tx=XM_0232549
39.1
B3 7552997 7553139
refGene_gn=NTRK3;refGene_tx=XM_011282744.3;refGene_gn=NTRK3;refGene_tx
=XM_011282745.2;refGene_gn=NTRK3;refGene_tx=XM_019831907.2;refGene_gn=
NTRK3;refGene_tx=XM_019831909.2;refGene_gn=NTRK3;refGene_tx=XM_0232549
39.1
B3 7553567 7553864
refGene_gn=NTRK3;refGene_tx=XM_011282744.3;refGene_gn=NTRK3;refGene_tx
=XM_011282745.2;refGene_gn=NTRK3;refGene_tx=XM_019831907.2;refGene_gn=
NTRK3;refGene_tx=XM_019831909.2;refGene_gn=NTRK3;refGene_tx=XM_0232549
39.1
B3 7562497 7562521
refGene_gn=NTRK3;refGene_tx=XM_011282744.3;refGene_gn=NTRK3;refGene_tx
=XM_011282745.2;refGene_gn=NTRK3;refGene_tx=XM_019831907.2;refGene_gn=
NTRK3;refGene_tx=XM_019831909.2;refGene_gn=NTRK3;refGene_tx=XM_0232549
39.1
B3 7563994 7564056
refGene_gn=NTRK3;refGene_tx=XM_011282744.3;refGene_gn=NTRK3;refGene_tx
=XM_011282745.2;refGene_gn=NTRK3;refGene_tx=XM_019831907.2;refGene_gn=
NTRK3;refGene_tx=XM_019831909.2;refGene_gn=NTRK3;refGene_tx=XM_0232549
39.1
B3 7565133 7565236
refGene_gn=NTRK3;refGene_tx=XM_011282744.3;refGene_gn=NTRK3;refGene_tx
=XM_011282745.2;refGene_gn=NTRK3;refGene_tx=XM_019831907.2;refGene_gn=
NTRK3;refGene_tx=XM_019831909.2;refGene_gn=NTRK3;refGene_tx=XM_0232549
39.1
B3 7597355 7597865
refGene_gn=LOC109500501;refGene_tx=XM_019831910.2
B3 7597867 7598476
refGene_gn=LOC109500501;refGene_tx=XM_019831910.2
B3 7659586 7659775
refGene_gn=NTRK3;refGene_tx=XM_011282744.3;refGene_gn=NTRK3;refGene_tx
=XM_011282745.2;refGene_gn=NTRK3;refGene_tx=XM_019831907.2;refGene_gn=
NTRK3;refGene_tx=XM_019831909.2;refGene_gn=NTRK3;refGene_tx=XM_0232549

39.1
B3 7708787 7708922 refGene_gn=NTRK3;refGene_tx=XM_011282745.2
B3 7710480 7710599
refGene_gn=NTRK3;refGene_tx=XM_011282745.2;refGene_gn=NTRK3;refGene_tx
=XM_019831909.2
B3 7747278 7747409
refGene_gn=NTRK3;refGene_tx=XM_011282744.3;refGene_gn=NTRK3;refGene_tx
=XM_019831907.2;refGene_gn=NTRK3;refGene_tx=XM_023254939.1
B3 7756129 7756302
refGene_gn=NTRK3;refGene_tx=XM_011282744.3;refGene_gn=NTRK3;refGene_tx
=XM_019831907.2;refGene_gn=NTRK3;refGene_tx=XM_023254939.1
B3 7759395 7759639
refGene_gn=NTRK3;refGene_tx=XM_011282744.3;refGene_gn=NTRK3;refGene_tx
=XM_019831907.2;refGene_gn=NTRK3;refGene_tx=XM_023254939.1
B3 7779718 7779793
refGene_gn=NTRK3;refGene_tx=XM_019831907.2;refGene_gn=NTRK3;refGene_tx
=XM_023254939.1
B3 7806085 7806244
refGene_gn=NTRK3;refGene_tx=XM_011282744.3;refGene_gn=NTRK3;refGene_tx
=XM_019831907.2;refGene_gn=NTRK3;refGene_tx=XM_023254939.1
B3 7810236 7810422
refGene_gn=NTRK3;refGene_tx=XM_011282744.3;refGene_gn=NTRK3;refGene_tx
=XM_019831907.2;refGene_gn=NTRK3;refGene_tx=XM_023254939.1
B3 8633001 8633212 refGene_gn=AGBL1;refGene_tx=XM_011282742.3
B3 8851184 8851348 refGene_gn=AGBL1;refGene_tx=XM_011282742.3
B3 8957142 8957319 refGene_gn=AGBL1;refGene_tx=XM_011282742.3
B3 8970121 8970253 refGene_gn=AGBL1;refGene_tx=XM_011282742.3
B3 8986797 8986927 refGene_gn=AGBL1;refGene_tx=XM_011282742.3
B3 9087599 9087780 refGene_gn=AGBL1;refGene_tx=XM_011282742.3
B3 9187512 9187666 refGene_gn=AGBL1;refGene_tx=XM_011282742.3
B3 9200614 9200759 refGene_gn=AGBL1;refGene_tx=XM_011282742.3
B3 9216545 9216633 refGene_gn=AGBL1;refGene_tx=XM_011282742.3
B3 9218109 9218258 refGene_gn=AGBL1;refGene_tx=XM_011282742.3
B3 9221330 9221417 refGene_gn=AGBL1;refGene_tx=XM_011282742.3
B3 9221650 9221734 refGene_gn=AGBL1;refGene_tx=XM_011282742.3
B3 9223260 9223841 refGene_gn=AGBL1;refGene_tx=XM_011282742.3
B3 9224953 9225070 refGene_gn=AGBL1;refGene_tx=XM_011282742.3
B3 9230551 9230619 refGene_gn=AGBL1;refGene_tx=XM_011282742.3
B3 9231546 9231712 refGene_gn=AGBL1;refGene_tx=XM_011282742.3
B3 9239361 9239570 refGene_gn=AGBL1;refGene_tx=XM_011282742.3
B3 9260705 9260743 refGene_gn=AGBL1;refGene_tx=XM_011282742.3
B3 9320098 9320192 refGene_gn=AGBL1;refGene_tx=XM_011282742.3
B3 9323818 9323950 refGene_gn=AGBL1;refGene_tx=XM_011282742.3
B3 9327806 9327953 refGene_gn=AGBL1;refGene_tx=XM_011282742.3
B3 9329920 9329984 refGene_gn=AGBL1;refGene_tx=XM_011282742.3
B3 9389182 9389233 refGene_gn=AGBL1;refGene_tx=XM_011282742.3
B3 9704035 9704132 refGene_gn=KLHL25;refGene_tx=XM_006932282.4
B3 9705488 9705561 refGene_gn=KLHL25;refGene_tx=XM_019831912.2
B3 9706391 9708172
refGene_gn=KLHL25;refGene_tx=XM_006932282.4;refGene_gn=KLHL25;refGene_

tx=XM_019831912.2;refGene_gn=KLHL25;refGene_tx=XM_023254940.1
B3	9760531	9760581	refGene_gn=AKAP13;refGene_tx=XM_023254941.1
B3	9761415	9761749	refGene_gn=AKAP13;refGene_tx=XM_023254941.1
B3	9762638	9763088	refGene_gn=AKAP13;refGene_tx=XM_023254941.1
B3	9763721	9763772	refGene_gn=AKAP13;refGene_tx=XM_023254941.1
B3	9767045	9767090	refGene_gn=AKAP13;refGene_tx=XM_023254941.1
B3	9767534	9767605	refGene_gn=AKAP13;refGene_tx=XM_023254941.1
B3	9768152	9768314	refGene_gn=AKAP13;refGene_tx=XM_023254941.1
B3	9770828	9771023	refGene_gn=AKAP13;refGene_tx=XM_023254941.1
B3	9773016	9773099	refGene_gn=AKAP13;refGene_tx=XM_023254941.1
B3	9773213	9773395	refGene_gn=AKAP13;refGene_tx=XM_023254941.1
B3	9774375	9774452	refGene_gn=AKAP13;refGene_tx=XM_023254941.1
B3	9779133	9779382	refGene_gn=AKAP13;refGene_tx=XM_023254941.1
B3	9780088	9780206	refGene_gn=AKAP13;refGene_tx=XM_023254941.1
B3	9780318	9780444	refGene_gn=AKAP13;refGene_tx=XM_023254941.1
B3	9783957	9783996	refGene_gn=AKAP13;refGene_tx=XM_023254941.1
B3	9783997	9784209	refGene_gn=AKAP13;refGene_tx=XM_023254941.1
B3	9784820	9784973	refGene_gn=AKAP13;refGene_tx=XM_023254941.1
B3	9785577	9785690	refGene_gn=AKAP13;refGene_tx=XM_023254941.1
B3	9787071	9787207	refGene_gn=AKAP13;refGene_tx=XM_023254941.1
B3	9791549	9791616	refGene_gn=AKAP13;refGene_tx=XM_023254941.1
B3	9793749	9793817	refGene_gn=AKAP13;refGene_tx=XM_023254941.1
B3	9808761	9808936	refGene_gn=AKAP13;refGene_tx=XM_023254941.1
B3	9815387	9815520	refGene_gn=AKAP13;refGene_tx=XM_023254941.1
B3	9817258	9817301	refGene_gn=AKAP13;refGene_tx=XM_023254941.1
B3	9829966	9830075	refGene_gn=AKAP13;refGene_tx=XM_023254941.1
B3	9833630	9833823	refGene_gn=AKAP13;refGene_tx=XM_023254941.1
B3	9835665	9835731	refGene_gn=AKAP13;refGene_tx=XM_023254941.1
B3	9838693	9838747	refGene_gn=AKAP13;refGene_tx=XM_023254941.1
B3	9841465	9841836	refGene_gn=AKAP13;refGene_tx=XM_023254941.1
B3	9853471	9853608	refGene_gn=AKAP13;refGene_tx=XM_023254941.1
B3	9859042	9859118	refGene_gn=AKAP13;refGene_tx=XM_023254941.1
B3	9906674	9906808	refGene_gn=AKAP13;refGene_tx=XM_023254941.1
B3	9910733	9913809	refGene_gn=AKAP13;refGene_tx=XM_023254941.1
B3	9916418	9916617	refGene_gn=AKAP13;refGene_tx=XM_023254941.1
B3	9941949	9942133	refGene_gn=AKAP13;refGene_tx=XM_023254941.1
B3	9951044	9951341	refGene_gn=AKAP13;refGene_tx=XM_023254941.1
B3	9965371	9965519	refGene_gn=AKAP13;refGene_tx=XM_023254941.1
B3	10000908	10000941	refGene_gn=AKAP13;refGene_tx=XM_023254941.1
B3	10342416	10342867
refGene_gn=SV2B;refGene_tx=XM_003986847.5;refGene_gn=SV2B;refGene_tx=XM_019831915.2;refGene_gn=SV2B;refGene_tx=XM_019831917.2;refGene_gn=SV2B;refGene_tx=XM_019831918.2
B3	10372369	10372550
refGene_gn=SV2B;refGene_tx=XM_003986847.5;refGene_gn=SV2B;refGene_tx=XM_019831915.2;refGene_gn=SV2B;refGene_tx=XM_019831917.2;refGene_gn=SV2B;refGene_tx=XM_019831918.2
B3	10372889	10373041
refGene_gn=SV2B;refGene_tx=XM_003986847.5;refGene_gn=SV2B;refGene_tx=XM_019831915.2;refGene_gn=SV2B;refGene_tx=XM_019831917.2;refGene_gn=SV2

B;refGene_tx=XM_019831918.2
B3	10378968	10379102
refGene_gn=SV2B;refGene_tx=XM_003986847.5;refGene_gn=SV2B;refGene_tx=XM_019831915.2;refGene_gn=SV2B;refGene_tx=XM_019831917.2;refGene_gn=SV2B;refGene_tx=XM_019831918.2
B3	10381191	10381281
refGene_gn=SV2B;refGene_tx=XM_003986847.5;refGene_gn=SV2B;refGene_tx=XM_019831915.2;refGene_gn=SV2B;refGene_tx=XM_019831917.2;refGene_gn=SV2B;refGene_tx=XM_019831918.2
B3	10386910	10387021
refGene_gn=SV2B;refGene_tx=XM_003986847.5;refGene_gn=SV2B;refGene_tx=XM_019831915.2;refGene_gn=SV2B;refGene_tx=XM_019831917.2;refGene_gn=SV2B;refGene_tx=XM_019831918.2
B3	10387833	10387922
refGene_gn=SV2B;refGene_tx=XM_003986847.5;refGene_gn=SV2B;refGene_tx=XM_019831915.2;refGene_gn=SV2B;refGene_tx=XM_019831917.2;refGene_gn=SV2B;refGene_tx=XM_019831918.2
B3	10390198	10390363
refGene_gn=SV2B;refGene_tx=XM_003986847.5;refGene_gn=SV2B;refGene_tx=XM_019831915.2;refGene_gn=SV2B;refGene_tx=XM_019831917.2;refGene_gn=SV2B;refGene_tx=XM_019831918.2
B3	10408370	10408504
refGene_gn=SV2B;refGene_tx=XM_003986847.5;refGene_gn=SV2B;refGene_tx=XM_019831915.2;refGene_gn=SV2B;refGene_tx=XM_019831917.2;refGene_gn=SV2B;refGene_tx=XM_019831918.2
B3	10412042	10412243
refGene_gn=SV2B;refGene_tx=XM_003986847.5;refGene_gn=SV2B;refGene_tx=XM_019831915.2;refGene_gn=SV2B;refGene_tx=XM_019831917.2;refGene_gn=SV2B;refGene_tx=XM_019831918.2
B3	10416670	10416830
refGene_gn=SV2B;refGene_tx=XM_003986847.5;refGene_gn=SV2B;refGene_tx=XM_019831915.2;refGene_gn=SV2B;refGene_tx=XM_019831917.2;refGene_gn=SV2B;refGene_tx=XM_019831918.2
B3	10419454	10419638
refGene_gn=SV2B;refGene_tx=XM_003986847.5;refGene_gn=SV2B;refGene_tx=XM_019831915.2;refGene_gn=SV2B;refGene_tx=XM_019831917.2;refGene_gn=SV2B;refGene_tx=XM_019831918.2
B3	10922277	10922457
refGene_gn=SLCO3A1;refGene_tx=XM_023254947.1;refGene_gn=SLCO3A1;refGene_tx=XM_023254948.1;refGene_gn=SLCO3A1;refGene_tx=XM_023254949.1
B3	10986376	10986842
refGene_gn=SLCO3A1;refGene_tx=XM_023254947.1;refGene_gn=SLCO3A1;refGene_tx=XM_023254948.1;refGene_gn=SLCO3A1;refGene_tx=XM_023254949.1
B3	11156202	11156301
refGene_gn=SLCO3A1;refGene_tx=XM_023254947.1;refGene_gn=SLCO3A1;refGene_tx=XM_023254948.1;refGene_gn=SLCO3A1;refGene_tx=XM_023254949.1
B3	11165892	11166156
refGene_gn=SLCO3A1;refGene_tx=XM_023254947.1;refGene_gn=SLCO3A1;refGene_tx=XM_023254948.1
B3	11181935	11182100

refGene_gn=SLCO3A1;refGene_tx=XM_023254947.1;refGene_gn=SLCO3A1;refGene_tx=XM_023254948.1;refGene_gn=SLCO3A1;refGene_tx=XM_023254949.1
B3 11187085 11187284
refGene_gn=SLCO3A1;refGene_tx=XM_023254947.1;refGene_gn=SLCO3A1;refGene_tx=XM_023254948.1;refGene_gn=SLCO3A1;refGene_tx=XM_023254949.1
B3 11189555 11189694
refGene_gn=SLCO3A1;refGene_tx=XM_023254947.1;refGene_gn=SLCO3A1;refGene_tx=XM_023254948.1;refGene_gn=SLCO3A1;refGene_tx=XM_023254949.1
B3 11202859 11203035
refGene_gn=SLCO3A1;refGene_tx=XM_023254947.1;refGene_gn=SLCO3A1;refGene_tx=XM_023254948.1;refGene_gn=SLCO3A1;refGene_tx=XM_023254949.1
B3 11207578 11207594
refGene_gn=SLCO3A1;refGene_tx=XM_023254947.1;refGene_gn=SLCO3A1;refGene_tx=XM_023254948.1;refGene_gn=SLCO3A1;refGene_tx=XM_023254949.1
B3 11513845 11513943 refGene_gn=ST8SIA2;refGene_tx=XM_011282740.3
B3 11547812 11547875 refGene_gn=ST8SIA2;refGene_tx=XM_011282740.3
B3 11551647 11551776 refGene_gn=ST8SIA2;refGene_tx=XM_011282740.3
B3 11555687 11555945 refGene_gn=ST8SIA2;refGene_tx=XM_011282740.3
B3 11561669 11561963 refGene_gn=ST8SIA2;refGene_tx=XM_011282740.3
B3 11577692 11577978 refGene_gn=ST8SIA2;refGene_tx=XM_011282740.3
B3 11583610 11583616
refGene_gn=CB3H15orf32;refGene_tx=XM_006944362.1
B3 11586111 11586205
refGene_gn=CB3H15orf32;refGene_tx=XM_006944362.1
B3 11586847 11587019
refGene_gn=CB3H15orf32;refGene_tx=XM_006944362.1
B3 11587409 11587479
refGene_gn=CB3H15orf32;refGene_tx=XM_006944362.1
B3 11727572 11727576 refGene_gn=FAM174B;refGene_tx=XM_023254951.1
B3 11738714 11738846 refGene_gn=FAM174B;refGene_tx=XM_023254951.1
B3 11761273 11761641 refGene_gn=FAM174B;refGene_tx=XM_023254951.1
B3 11862870 11863191
refGene_gn=LOC109500522;refGene_tx=XM_023254768.1
B3 11872121 11872411
refGene_gn=LOC109500522;refGene_tx=XM_023254768.1
B3 11872543 11872715
refGene_gn=LOC109500522;refGene_tx=XM_023254768.1
B3 11873372 11873447
refGene_gn=LOC109500522;refGene_tx=XM_023254768.1
B3 11950893 11950955 refGene_gn=CHD2;refGene_tx=XM_011282738.3
B3 11954626 11954733
refGene_gn=CHD2;refGene_tx=XM_011282739.3;refGene_gn=CHD2;refGene_tx=XM_019831921.2;refGene_gn=CHD2;refGene_tx=XM_019831922.2
B3 11972647 11972879
refGene_gn=CHD2;refGene_tx=XM_011282738.3;refGene_gn=CHD2;refGene_tx=XM_011282739.3;refGene_gn=CHD2;refGene_tx=XM_019831921.2;refGene_gn=CHD2;refGene_tx=XM_019831922.2
B3 11976621 11976708
refGene_gn=CHD2;refGene_tx=XM_011282738.3;refGene_gn=CHD2;refGene_tx=XM_011282739.3;refGene_gn=CHD2;refGene_tx=XM_019831921.2;refGene_gn=CHD

2;refGene_tx=XM_019831922.2
B3 11978713 11978775
refGene_gn=CHD2;refGene_tx=XM_011282738.3;refGene_gn=CHD2;refGene_tx=XM_011282739.3;refGene_gn=CHD2;refGene_tx=XM_019831921.2;refGene_gn=CHD2;refGene_tx=XM_019831922.2
B3 11984615 11984723
refGene_gn=CHD2;refGene_tx=XM_011282738.3;refGene_gn=CHD2;refGene_tx=XM_011282739.3;refGene_gn=CHD2;refGene_tx=XM_019831922.2
B3 11987178 11987319
refGene_gn=CHD2;refGene_tx=XM_011282738.3;refGene_gn=CHD2;refGene_tx=XM_011282739.3;refGene_gn=CHD2;refGene_tx=XM_019831921.2;refGene_gn=CHD2;refGene_tx=XM_019831922.2
B3 11988809 11988943
refGene_gn=CHD2;refGene_tx=XM_011282738.3;refGene_gn=CHD2;refGene_tx=XM_011282739.3;refGene_gn=CHD2;refGene_tx=XM_019831921.2;refGene_gn=CHD2;refGene_tx=XM_019831922.2
B3 11990042 11990268
refGene_gn=CHD2;refGene_tx=XM_011282738.3;refGene_gn=CHD2;refGene_tx=XM_011282739.3;refGene_gn=CHD2;refGene_tx=XM_019831921.2;refGene_gn=CHD2;refGene_tx=XM_019831922.2
B3 11991582 11991683
refGene_gn=CHD2;refGene_tx=XM_011282738.3;refGene_gn=CHD2;refGene_tx=XM_011282739.3;refGene_gn=CHD2;refGene_tx=XM_019831921.2;refGene_gn=CHD2;refGene_tx=XM_019831922.2
B3 11992672 11992717
refGene_gn=CHD2;refGene_tx=XM_011282738.3;refGene_gn=CHD2;refGene_tx=XM_011282739.3;refGene_gn=CHD2;refGene_tx=XM_019831921.2;refGene_gn=CHD2;refGene_tx=XM_019831922.2
B3 11992887 11993066
refGene_gn=CHD2;refGene_tx=XM_011282738.3;refGene_gn=CHD2;refGene_tx=XM_011282739.3;refGene_gn=CHD2;refGene_tx=XM_019831921.2;refGene_gn=CHD2;refGene_tx=XM_019831922.2
B3 11995274 11995403
refGene_gn=CHD2;refGene_tx=XM_011282738.3;refGene_gn=CHD2;refGene_tx=XM_011282739.3;refGene_gn=CHD2;refGene_tx=XM_019831921.2;refGene_gn=CHD2;refGene_tx=XM_019831922.2
B3 11998285 11998502
refGene_gn=CHD2;refGene_tx=XM_011282738.3;refGene_gn=CHD2;refGene_tx=XM_019831921.2;refGene_gn=CHD2;refGene_tx=XM_019831922.2
B3 12000714 12000804
refGene_gn=CHD2;refGene_tx=XM_011282738.3;refGene_gn=CHD2;refGene_tx=XM_019831921.2;refGene_gn=CHD2;refGene_tx=XM_019831922.2
B3 12001656 12001847
refGene_gn=CHD2;refGene_tx=XM_011282738.3;refGene_gn=CHD2;refGene_tx=XM_019831921.2;refGene_gn=CHD2;refGene_tx=XM_019831922.2
B3 12013044 12013233
refGene_gn=CHD2;refGene_tx=XM_011282738.3;refGene_gn=CHD2;refGene_tx=XM_019831921.2;refGene_gn=CHD2;refGene_tx=XM_019831922.2
B3 12018982 12019145
refGene_gn=CHD2;refGene_tx=XM_011282738.3;refGene_gn=CHD2;refGene_tx=X

M_019831921.2;refGene_gn=CHD2;refGene_tx=XM_019831922.2
B3 12019916 12020069
refGene_gn=CHD2;refGene_tx=XM_011282738.3;refGene_gn=CHD2;refGene_tx=XM_019831921.2;refGene_gn=CHD2;refGene_tx=XM_019831922.2
B3 12022237 12022309
refGene_gn=CHD2;refGene_tx=XM_011282738.3;refGene_gn=CHD2;refGene_tx=XM_019831921.2;refGene_gn=CHD2;refGene_tx=XM_019831922.2
B3 12025738 12025888
refGene_gn=CHD2;refGene_tx=XM_011282738.3;refGene_gn=CHD2;refGene_tx=XM_019831921.2;refGene_gn=CHD2;refGene_tx=XM_019831922.2
B3 12026753 12026902
refGene_gn=CHD2;refGene_tx=XM_011282738.3;refGene_gn=CHD2;refGene_tx=XM_019831921.2;refGene_gn=CHD2;refGene_tx=XM_019831922.2
B3 12027900 12027997
refGene_gn=CHD2;refGene_tx=XM_011282738.3;refGene_gn=CHD2;refGene_tx=XM_019831921.2;refGene_gn=CHD2;refGene_tx=XM_019831922.2
B3 12028425 12028518
refGene_gn=CHD2;refGene_tx=XM_011282738.3;refGene_gn=CHD2;refGene_tx=XM_019831921.2;refGene_gn=CHD2;refGene_tx=XM_019831922.2
B3 12031651 12031822
refGene_gn=CHD2;refGene_tx=XM_011282738.3;refGene_gn=CHD2;refGene_tx=XM_019831921.2;refGene_gn=CHD2;refGene_tx=XM_019831922.2
B3 12032806 12032982
refGene_gn=CHD2;refGene_tx=XM_011282738.3;refGene_gn=CHD2;refGene_tx=XM_019831921.2;refGene_gn=CHD2;refGene_tx=XM_019831922.2
B3 12036345 12036387
refGene_gn=CHD2;refGene_tx=XM_011282738.3;refGene_gn=CHD2;refGene_tx=XM_019831921.2;refGene_gn=CHD2;refGene_tx=XM_019831922.2
B3 12037694 12037834
refGene_gn=CHD2;refGene_tx=XM_011282738.3;refGene_gn=CHD2;refGene_tx=XM_019831921.2;refGene_gn=CHD2;refGene_tx=XM_019831922.2
B3 12042033 12042172
refGene_gn=CHD2;refGene_tx=XM_011282738.3;refGene_gn=CHD2;refGene_tx=XM_019831921.2;refGene_gn=CHD2;refGene_tx=XM_019831922.2
B3 12042365 12042516
refGene_gn=CHD2;refGene_tx=XM_011282738.3;refGene_gn=CHD2;refGene_tx=XM_019831921.2;refGene_gn=CHD2;refGene_tx=XM_019831922.2
B3 12043654 12043777
refGene_gn=CHD2;refGene_tx=XM_011282738.3;refGene_gn=CHD2;refGene_tx=XM_019831921.2;refGene_gn=CHD2;refGene_tx=XM_019831922.2
B3 12045010 12045139
refGene_gn=CHD2;refGene_tx=XM_011282738.3;refGene_gn=CHD2;refGene_tx=XM_019831921.2;refGene_gn=CHD2;refGene_tx=XM_019831922.2
B3 12046074 12046215
refGene_gn=CHD2;refGene_tx=XM_011282738.3;refGene_gn=CHD2;refGene_tx=XM_019831921.2
B3 12047957 12048092
refGene_gn=CHD2;refGene_tx=XM_011282738.3;refGene_gn=CHD2;refGene_tx=XM_019831921.2;refGene_gn=CHD2;refGene_tx=XM_019831922.2
B3 12053172 12053351

refGene_gn=CHD2;refGene_tx=XM_011282738.3;refGene_gn=CHD2;refGene_tx=XM_019831921.2;refGene_gn=CHD2;refGene_tx=XM_019831922.2
B3 12055284 12055384
refGene_gn=CHD2;refGene_tx=XM_011282738.3;refGene_gn=CHD2;refGene_tx=XM_019831921.2;refGene_gn=CHD2;refGene_tx=XM_019831922.2
B3 12057629 12057843
refGene_gn=CHD2;refGene_tx=XM_011282738.3;refGene_gn=CHD2;refGene_tx=XM_019831921.2;refGene_gn=CHD2;refGene_tx=XM_019831922.2
B3 12065400 12065647
refGene_gn=CHD2;refGene_tx=XM_011282738.3;refGene_gn=CHD2;refGene_tx=XM_019831921.2;refGene_gn=CHD2;refGene_tx=XM_019831922.2
B3 12070144 12070457
refGene_gn=CHD2;refGene_tx=XM_011282738.3;refGene_gn=CHD2;refGene_tx=XM_019831921.2;refGene_gn=CHD2;refGene_tx=XM_019831922.2
B3 12087191 12087914 refGene_gn=RGMA;refGene_tx=XM_023254952.1
B3 12097200 12097715 refGene_gn=RGMA;refGene_tx=XM_023254952.1
B3 12115816 12115932 refGene_gn=RGMA;refGene_tx=XM_023254952.1
B3 12133369 12133383 refGene_gn=RGMA;refGene_tx=XM_023254952.1
B3 12320073 12320088
refGene_gn=LOC111556300;refGene_tx=XM_023254955.1
B3 12340151 12340250
refGene_gn=LOC111556300;refGene_tx=XM_023254953.1;refGene_gn=LOC111556300;refGene_tx=XM_023254954.1;refGene_gn=LOC111556300;refGene_tx=XM_023254956.1
B3 12341175 12341247
refGene_gn=LOC111556300;refGene_tx=XM_023254953.1;refGene_gn=LOC111556300;refGene_tx=XM_023254954.1;refGene_gn=LOC111556300;refGene_tx=XM_023254956.1
B3 12367502 12367647
refGene_gn=LOC111556300;refGene_tx=XM_023254953.1;refGene_gn=LOC111556300;refGene_tx=XM_023254954.1;refGene_gn=LOC111556300;refGene_tx=XM_023254955.1;refGene_gn=LOC111556300;refGene_tx=XM_023254956.1
B3 12385791 12385912
refGene_gn=LOC111556300;refGene_tx=XM_023254953.1;refGene_gn=LOC111556300;refGene_tx=XM_023254954.1;refGene_gn=LOC111556300;refGene_tx=XM_023254955.1
B3 12388468 12388664
refGene_gn=LOC111556300;refGene_tx=XM_023254953.1;refGene_gn=LOC111556300;refGene_tx=XM_023254954.1;refGene_gn=LOC111556300;refGene_tx=XM_023254955.1;refGene_gn=LOC111556300;refGene_tx=XM_023254956.1
B3 12395341 12395343
refGene_gn=LOC111556300;refGene_tx=XM_023254954.1
B3 13140464 13140932 refGene_gn=MCTP2;refGene_tx=XM_011282736.3
B3 13155270 13155333 refGene_gn=MCTP2;refGene_tx=XM_011282736.3
B3 13156797 13156906 refGene_gn=MCTP2;refGene_tx=XM_011282736.3
B3 13182473 13182616 refGene_gn=MCTP2;refGene_tx=XM_011282736.3
B3 13183407 13183484 refGene_gn=MCTP2;refGene_tx=XM_011282736.3
B3 13183998 13184110 refGene_gn=MCTP2;refGene_tx=XM_011282736.3
B3 13188442 13188478 refGene_gn=MCTP2;refGene_tx=XM_011282736.3
B3 13200023 13200188 refGene_gn=MCTP2;refGene_tx=XM_011282736.3

B3	13202384	13202515
refGene_gn=MCTP2;refGene_tx=XM_011282736.3;refGene_gn=MCTP2;refGene_tx=XM_023254957.1
B3	13214528	13214715
refGene_gn=MCTP2;refGene_tx=XM_011282736.3;refGene_gn=MCTP2;refGene_tx=XM_023254957.1
B3	13217527	13217621
refGene_gn=MCTP2;refGene_tx=XM_011282736.3;refGene_gn=MCTP2;refGene_tx=XM_023254957.1
B3	13232376	13232479
refGene_gn=MCTP2;refGene_tx=XM_011282736.3;refGene_gn=MCTP2;refGene_tx=XM_023254957.1
B3	13234828	13234931
refGene_gn=MCTP2;refGene_tx=XM_011282736.3;refGene_gn=MCTP2;refGene_tx=XM_023254957.1
B3	13250996	13251098
refGene_gn=MCTP2;refGene_tx=XM_011282736.3;refGene_gn=MCTP2;refGene_tx=XM_023254957.1
B3	13253979	13254054
refGene_gn=MCTP2;refGene_tx=XM_011282736.3;refGene_gn=MCTP2;refGene_tx=XM_023254957.1
B3	13256154	13256274
refGene_gn=MCTP2;refGene_tx=XM_011282736.3;refGene_gn=MCTP2;refGene_tx=XM_023254957.1
B3	13286253	13286715
refGene_gn=LOC102899291;refGene_tx=XM_019831928.2
B3	13294663	13294786
refGene_gn=MCTP2;refGene_tx=XM_011282736.3;refGene_gn=MCTP2;refGene_tx=XM_023254957.1
B3	13298330	13298372
refGene_gn=MCTP2;refGene_tx=XM_011282736.3;refGene_gn=MCTP2;refGene_tx=XM_023254957.1
B3	13313604	13313714
refGene_gn=MCTP2;refGene_tx=XM_011282736.3;refGene_gn=MCTP2;refGene_tx=XM_023254957.1
B3	13321497	13321607
refGene_gn=MCTP2;refGene_tx=XM_011282736.3;refGene_gn=MCTP2;refGene_tx=XM_023254957.1
B3	13327460	13327558
refGene_gn=MCTP2;refGene_tx=XM_011282736.3;refGene_gn=MCTP2;refGene_tx=XM_023254957.1
B3	13329749	13329818
refGene_gn=MCTP2;refGene_tx=XM_011282736.3;refGene_gn=MCTP2;refGene_tx=XM_023254957.1
B3	15114088	15114131	refGene_gn=NR2F2;refGene_tx=XM_023254959.1
B3	15119926	15120368	refGene_gn=NR2F2;refGene_tx=XM_023254958.1
B3	15121958	15122486
refGene_gn=NR2F2;refGene_tx=XM_023254958.1;refGene_gn=NR2F2;refGene_tx=XM_023254959.1;refGene_gn=NR2F2;refGene_tx=XM_023254960.1
B3	15125305	15125580

refGene_gn=NR2F2;refGene_tx=XM_023254958.1;refGene_gn=NR2F2;refGene_tx=XM_023254959.1;refGene_gn=NR2F2;refGene_tx=XM_023254960.1
B3 16694800 16695089 refGene_gn=ARRDC4;refGene_tx=XM_023254963.1
B3 16699783 16699850 refGene_gn=ARRDC4;refGene_tx=XM_023254963.1
B3 16700107 16700255 refGene_gn=ARRDC4;refGene_tx=XM_023254963.1
B3 16702470 16702573 refGene_gn=ARRDC4;refGene_tx=XM_023254963.1
B3 16703538 16703795 refGene_gn=ARRDC4;refGene_tx=XM_023254963.1
B3 16704650 16704813 refGene_gn=ARRDC4;refGene_tx=XM_023254963.1
B3 16705386 16705541 refGene_gn=ARRDC4;refGene_tx=XM_023254963.1
B3 16705920 16705977 refGene_gn=ARRDC4;refGene_tx=XM_023254963.1
B3 17133923 17134106 refGene_gn=FAM169B;refGene_tx=XM_023254964.1;refGene_gn=FAM169B;refGene_tx=XM_023254965.1
B3 17135408 17135503 refGene_gn=FAM169B;refGene_tx=XM_023254964.1;refGene_gn=FAM169B;refGene_tx=XM_023254965.1
B3 17140699 17140831 refGene_gn=FAM169B;refGene_tx=XM_023254964.1
B3 17144592 17144775 refGene_gn=FAM169B;refGene_tx=XM_023254964.1;refGene_gn=FAM169B;refGene_tx=XM_023254965.1
B3 17166268 17166446 refGene_gn=FAM169B;refGene_tx=XM_023254964.1;refGene_gn=FAM169B;refGene_tx=XM_023254965.1
B3 17172386 17172472 refGene_gn=FAM169B;refGene_tx=XM_023254964.1;refGene_gn=FAM169B;refGene_tx=XM_023254965.1
B3 17196042 17196142 refGene_gn=FAM169B;refGene_tx=XM_023254964.1;refGene_gn=FAM169B;refGene_tx=XM_023254965.1
B3 17200186 17200317 refGene_gn=FAM169B;refGene_tx=XM_023254964.1;refGene_gn=FAM169B;refGene_tx=XM_023254965.1
B3 17221498 17221529 refGene_gn=FAM169B;refGene_tx=XM_023254964.1;refGene_gn=FAM169B;refGene_tx=XM_023254965.1
B3 17266335 17266383 refGene_gn=PGPEP1L;refGene_tx=XM_023254769.1
B3 17267392 17267536 refGene_gn=PGPEP1L;refGene_tx=XM_023254769.1
B3 17269201 17269350 refGene_gn=PGPEP1L;refGene_tx=XM_023254769.1
B3 17270203 17270486 refGene_gn=PGPEP1L;refGene_tx=XM_023254769.1
B3 17270488 17270610 refGene_gn=PGPEP1L;refGene_tx=XM_023254769.1
B3 17271359 17271452 refGene_gn=PGPEP1L;refGene_tx=XM_023254769.1
B3 17271453 17271562 refGene_gn=PGPEP1L;refGene_tx=XM_023254769.1
B3 17281905 17282287 refGene_gn=IGF1R;refGene_tx=XM_023254966.1;refGene_gn=IGF1R;refGene_tx=XM_023254968.1
B3 17291410 17291545 refGene_gn=IGF1R;refGene_tx=XM_023254966.1;refGene_gn=IGF1R;refGene_tx=XM_023254968.1
B3 17297628 17297758 refGene_gn=IGF1R;refGene_tx=XM_023254966.1;refGene_gn=IGF1R;refGene_tx

=XM_023254968.1
B3	17301773	17301933
refGene_gn=IGF1R;refGene_tx=XM_023254966.1;refGene_gn=IGF1R;refGene_tx=XM_023254968.1
B3	17305870	17305981
refGene_gn=IGF1R;refGene_tx=XM_023254966.1;refGene_gn=IGF1R;refGene_tx=XM_023254968.1
B3	17306293	17306523
refGene_gn=IGF1R;refGene_tx=XM_023254966.1;refGene_gn=IGF1R;refGene_tx=XM_023254968.1
B3	17310933	17311004
refGene_gn=IGF1R;refGene_tx=XM_023254966.1;refGene_gn=IGF1R;refGene_tx=XM_023254968.1
B3	17311778	17311881
refGene_gn=IGF1R;refGene_tx=XM_023254966.1;refGene_gn=IGF1R;refGene_tx=XM_023254968.1
B3	17316727	17316887
refGene_gn=IGF1R;refGene_tx=XM_023254966.1;refGene_gn=IGF1R;refGene_tx=XM_023254968.1
B3	17317368	17317505
refGene_gn=IGF1R;refGene_tx=XM_023254966.1;refGene_gn=IGF1R;refGene_tx=XM_023254968.1
B3	17318984	17319268
refGene_gn=IGF1R;refGene_tx=XM_023254966.1;refGene_gn=IGF1R;refGene_tx=XM_023254968.1
B3	17324418	17324623
refGene_gn=IGF1R;refGene_tx=XM_023254966.1;refGene_gn=IGF1R;refGene_tx=XM_023254968.1
B3	17324779	17324947
refGene_gn=IGF1R;refGene_tx=XM_023254966.1;refGene_gn=IGF1R;refGene_tx=XM_023254968.1
B3	17327392	17327631
refGene_gn=IGF1R;refGene_tx=XM_023254966.1;refGene_gn=IGF1R;refGene_tx=XM_023254968.1
B3	17329647	17329774
refGene_gn=IGF1R;refGene_tx=XM_023254966.1;refGene_gn=IGF1R;refGene_tx=XM_023254968.1
B3	17331963	17332178
refGene_gn=IGF1R;refGene_tx=XM_023254966.1;refGene_gn=IGF1R;refGene_tx=XM_023254968.1
B3	17341096	17341241
refGene_gn=IGF1R;refGene_tx=XM_023254966.1;refGene_gn=IGF1R;refGene_tx=XM_023254968.1
B3	17343617	17343766
refGene_gn=IGF1R;refGene_tx=XM_023254966.1;refGene_gn=IGF1R;refGene_tx=XM_023254968.1
B3	17347956	17348269
refGene_gn=IGF1R;refGene_tx=XM_023254966.1;refGene_gn=IGF1R;refGene_tx=XM_023254968.1
B3	17441878	17441945	refGene_gn=IGF1R;refGene_tx=XM_023254968.1

B3 17530258 17530804 refGene_gn=IGF1R;refGene_tx=XM_023254966.1
B3 17585013 17585107 refGene_gn=IGF1R;refGene_tx=XM_023254966.1
B3 17762842 17763652 refGene_gn=SYNM;refGene_tx=XM_023254969.1
B3 17770857 17770982 refGene_gn=SYNM;refGene_tx=XM_023254969.1
B3 17780730 17780801 refGene_gn=SYNM;refGene_tx=XM_023254969.1
B3 17784219 17786678 refGene_gn=SYNM;refGene_tx=XM_023254969.1
B3 17787594 17787933 refGene_gn=SYNM;refGene_tx=XM_023254969.1
B3 17793371 17793471
refGene_gn=TTC23;refGene_tx=XM_006932290.4;refGene_gn=TTC23;refGene_tx=XM_006932291.4;refGene_gn=TTC23;refGene_tx=XM_006932292.4;refGene_gn=TTC23;refGene_tx=XM_006932293.4;refGene_gn=TTC23;refGene_tx=XM_019831931.2;refGene_gn=TTC23;refGene_tx=XM_019831932.2
B3 17794316 17794399
refGene_gn=TTC23;refGene_tx=XM_006932290.4;refGene_gn=TTC23;refGene_tx=XM_006932291.4;refGene_gn=TTC23;refGene_tx=XM_006932292.4;refGene_gn=TTC23;refGene_tx=XM_006932293.4;refGene_gn=TTC23;refGene_tx=XM_019831931.2;refGene_gn=TTC23;refGene_tx=XM_019831932.2
B3 17794534 17794576
refGene_gn=TTC23;refGene_tx=XM_019831933.2;refGene_gn=TTC23;refGene_tx=XM_023254971.1
B3 17799446 17799452 refGene_gn=TTC23;refGene_tx=XM_006932294.4
B3 17801464 17801587
refGene_gn=TTC23;refGene_tx=XM_006932290.4;refGene_gn=TTC23;refGene_tx=XM_006932291.4;refGene_gn=TTC23;refGene_tx=XM_006932292.4;refGene_gn=TTC23;refGene_tx=XM_006932294.4;refGene_gn=TTC23;refGene_tx=XM_019831931.2;refGene_gn=TTC23;refGene_tx=XM_019831932.2;refGene_gn=TTC23;refGene_tx=XM_019831933.2
B3 17810133 17810283
refGene_gn=TTC23;refGene_tx=XM_006932290.4;refGene_gn=TTC23;refGene_tx=XM_006932291.4;refGene_gn=TTC23;refGene_tx=XM_006932292.4;refGene_gn=TTC23;refGene_tx=XM_006932293.4;refGene_gn=TTC23;refGene_tx=XM_006932294.4;refGene_gn=TTC23;refGene_tx=XM_019831931.2;refGene_gn=TTC23;refGene_tx=XM_019831932.2;refGene_gn=TTC23;refGene_tx=XM_019831933.2;refGene_gn=TTC23;refGene_tx=XM_023254971.1
B3 17814497 17814625
refGene_gn=TTC23;refGene_tx=XM_006932290.4;refGene_gn=TTC23;refGene_tx=XM_006932291.4;refGene_gn=TTC23;refGene_tx=XM_006932292.4;refGene_gn=TTC23;refGene_tx=XM_006932293.4;refGene_gn=TTC23;refGene_tx=XM_006932294.4;refGene_gn=TTC23;refGene_tx=XM_019831931.2;refGene_gn=TTC23;refGene_tx=XM_019831932.2;refGene_gn=TTC23;refGene_tx=XM_019831933.2;refGene_gn=TTC23;refGene_tx=XM_023254971.1
B3 17826977 17827083
refGene_gn=TTC23;refGene_tx=XM_006932290.4;refGene_gn=TTC23;refGene_tx=XM_006932291.4;refGene_gn=TTC23;refGene_tx=XM_006932292.4;refGene_gn=TTC23;refGene_tx=XM_006932293.4;refGene_gn=TTC23;refGene_tx=XM_006932294.4;refGene_gn=TTC23;refGene_tx=XM_019831931.2;refGene_gn=TTC23;refGene_tx=XM_019831932.2;refGene_gn=TTC23;refGene_tx=XM_019831933.2;refGene_gn=TTC23;refGene_tx=XM_023254971.1
B3 17844174 17844352
refGene_gn=TTC23;refGene_tx=XM_006932290.4;refGene_gn=TTC23;refGene_tx

=XM_006932291.4;refGene_gn=TTC23;refGene_tx=XM_006932292.4;refGene_gn=TTC23;refGene_tx=XM_006932293.4;refGene_gn=TTC23;refGene_tx=XM_006932294.4;refGene_gn=TTC23;refGene_tx=XM_019831931.2;refGene_gn=TTC23;refGene_tx=XM_019831932.2;refGene_gn=TTC23;refGene_tx=XM_019831933.2;refGene_gn=TTC23;refGene_tx=XM_023254971.1
B3 17864384 17864510
refGene_gn=TTC23;refGene_tx=XM_006932290.4;refGene_gn=TTC23;refGene_tx=XM_006932291.4;refGene_gn=TTC23;refGene_tx=XM_006932292.4;refGene_gn=TTC23;refGene_tx=XM_006932293.4;refGene_gn=TTC23;refGene_tx=XM_006932294.4;refGene_gn=TTC23;refGene_tx=XM_019831931.2;refGene_gn=TTC23;refGene_tx=XM_019831932.2;refGene_gn=TTC23;refGene_tx=XM_019831933.2;refGene_gn=TTC23;refGene_tx=XM_023254971.1
B3 17864701 17864852
refGene_gn=TTC23;refGene_tx=XM_006932290.4;refGene_gn=TTC23;refGene_tx=XM_006932291.4;refGene_gn=TTC23;refGene_tx=XM_006932292.4;refGene_gn=TTC23;refGene_tx=XM_006932293.4;refGene_gn=TTC23;refGene_tx=XM_006932294.4;refGene_gn=TTC23;refGene_tx=XM_019831931.2;refGene_gn=TTC23;refGene_tx=XM_019831932.2;refGene_gn=TTC23;refGene_tx=XM_019831933.2;refGene_gn=TTC23;refGene_tx=XM_023254971.1
B3 17867490 17867614
refGene_gn=TTC23;refGene_tx=XM_006932290.4;refGene_gn=TTC23;refGene_tx=XM_006932291.4;refGene_gn=TTC23;refGene_tx=XM_006932292.4;refGene_gn=TTC23;refGene_tx=XM_006932293.4;refGene_gn=TTC23;refGene_tx=XM_006932294.4;refGene_gn=TTC23;refGene_tx=XM_019831931.2;refGene_gn=TTC23;refGene_tx=XM_019831932.2;refGene_gn=TTC23;refGene_tx=XM_019831933.2;refGene_gn=TTC23;refGene_tx=XM_023254971.1
B3 17873809 17874009
refGene_gn=TTC23;refGene_tx=XM_006932290.4;refGene_gn=TTC23;refGene_tx=XM_006932291.4;refGene_gn=TTC23;refGene_tx=XM_006932292.4;refGene_gn=TTC23;refGene_tx=XM_006932293.4;refGene_gn=TTC23;refGene_tx=XM_006932294.4;refGene_gn=TTC23;refGene_tx=XM_019831931.2;refGene_gn=TTC23;refGene_tx=XM_019831932.2;refGene_gn=TTC23;refGene_tx=XM_019831933.2;refGene_gn=TTC23;refGene_tx=XM_023254971.1
B3 17888925 17888977
refGene_gn=TTC23;refGene_tx=XM_006932290.4;refGene_gn=TTC23;refGene_tx=XM_006932293.4;refGene_gn=TTC23;refGene_tx=XM_006932294.4;refGene_gn=TTC23;refGene_tx=XM_019831933.2;refGene_gn=TTC23;refGene_tx=XM_023254971.1
B3 17905405 17905573
refGene_gn=LRRC28;refGene_tx=XM_003986861.5;refGene_gn=LRRC28;refGene_tx=XM_006932295.4;refGene_gn=LRRC28;refGene_tx=XM_006932296.4
B3 17919509 17919550
refGene_gn=LRRC28;refGene_tx=XM_003986861.5;refGene_gn=LRRC28;refGene_tx=XM_006932295.4;refGene_gn=LRRC28;refGene_tx=XM_006932296.4
B3 17935105 17935143
refGene_gn=LRRC28;refGene_tx=XM_003986861.5;refGene_gn=LRRC28;refGene_tx=XM_006932295.4;refGene_gn=LRRC28;refGene_tx=XM_006932296.4
B3 17935467 17935605
refGene_gn=LRRC28;refGene_tx=XM_003986861.5;refGene_gn=LRRC28;refGene_tx=XM_006932295.4;refGene_gn=LRRC28;refGene_tx=XM_006932296.4

B3 17994377 17994584
refGene_gn=LRRC28;refGene_tx=XM_003986861.5;refGene_gn=LRRC28;refGene_tx=XM_006932295.4
B3 18012422 18012525
refGene_gn=LRRC28;refGene_tx=XM_003986861.5;refGene_gn=LRRC28;refGene_tx=XM_006932295.4;refGene_gn=LRRC28;refGene_tx=XM_006932296.4
B3 18030652 18030828
refGene_gn=LRRC28;refGene_tx=XM_003986861.5;refGene_gn=LRRC28;refGene_tx=XM_006932295.4;refGene_gn=LRRC28;refGene_tx=XM_006932296.4
B3 18032906 18033066
refGene_gn=LRRC28;refGene_tx=XM_003986861.5;refGene_gn=LRRC28;refGene_tx=XM_006932295.4;refGene_gn=LRRC28;refGene_tx=XM_006932296.4
B3 18053940 18054013
refGene_gn=LRRC28;refGene_tx=XM_003986861.5;refGene_gn=LRRC28;refGene_tx=XM_006932295.4;refGene_gn=LRRC28;refGene_tx=XM_006932296.4
B3 18292799 18292853
refGene_gn=MEF2A;refGene_tx=XM_023254972.1;refGene_gn=MEF2A;refGene_tx=XM_023254973.1;refGene_gn=MEF2A;refGene_tx=XM_023254974.1;refGene_gn=MEF2A;refGene_tx=XM_023254975.1;refGene_gn=MEF2A;refGene_tx=XM_023254976.1;refGene_gn=MEF2A;refGene_tx=XM_023254977.1
B3 18320155 18320359
refGene_gn=MEF2A;refGene_tx=XM_023254972.1;refGene_gn=MEF2A;refGene_tx=XM_023254973.1;refGene_gn=MEF2A;refGene_tx=XM_023254974.1;refGene_gn=MEF2A;refGene_tx=XM_023254975.1;refGene_gn=MEF2A;refGene_tx=XM_023254976.1;refGene_gn=MEF2A;refGene_tx=XM_023254977.1
B3 18347460 18347592
refGene_gn=MEF2A;refGene_tx=XM_023254975.1;refGene_gn=MEF2A;refGene_tx=XM_023254977.1
B3 18347657 18347795
refGene_gn=MEF2A;refGene_tx=XM_023254972.1;refGene_gn=MEF2A;refGene_tx=XM_023254973.1;refGene_gn=MEF2A;refGene_tx=XM_023254974.1;refGene_gn=MEF2A;refGene_tx=XM_023254976.1
B3 18350231 18350451
refGene_gn=MEF2A;refGene_tx=XM_023254972.1;refGene_gn=MEF2A;refGene_tx=XM_023254973.1;refGene_gn=MEF2A;refGene_tx=XM_023254974.1;refGene_gn=MEF2A;refGene_tx=XM_023254975.1;refGene_gn=MEF2A;refGene_tx=XM_023254976.1;refGene_gn=MEF2A;refGene_tx=XM_023254977.1
B3 18351175 18351235
refGene_gn=MEF2A;refGene_tx=XM_023254972.1;refGene_gn=MEF2A;refGene_tx=XM_023254973.1;refGene_gn=MEF2A;refGene_tx=XM_023254974.1;refGene_gn=MEF2A;refGene_tx=XM_023254975.1;refGene_gn=MEF2A;refGene_tx=XM_023254976.1;refGene_gn=MEF2A;refGene_tx=XM_023254977.1
B3 18361789 18361977
refGene_gn=MEF2A;refGene_tx=XM_023254972.1;refGene_gn=MEF2A;refGene_tx=XM_023254973.1;refGene_gn=MEF2A;refGene_tx=XM_023254974.1;refGene_gn=MEF2A;refGene_tx=XM_023254975.1;refGene_gn=MEF2A;refGene_tx=XM_023254976.1;refGene_gn=MEF2A;refGene_tx=XM_023254977.1
B3 18371490 18371514
refGene_gn=MEF2A;refGene_tx=XM_023254972.1;refGene_gn=MEF2A;refGene_tx=XM_023254973.1;refGene_gn=MEF2A;refGene_tx=XM_023254974.1;refGene_gn=

MEF2A;refGene_tx=XM_023254975.1
B3 18375795 18375922
refGene_gn=MEF2A;refGene_tx=XM_023254972.1;refGene_gn=MEF2A;refGene_tx=XM_023254973.1;refGene_gn=MEF2A;refGene_tx=XM_023254974.1;refGene_gn=MEF2A;refGene_tx=XM_023254975.1;refGene_gn=MEF2A;refGene_tx=XM_023254976.1;refGene_gn=MEF2A;refGene_tx=XM_023254977.1
B3 18379571 18379698
refGene_gn=MEF2A;refGene_tx=XM_023254972.1;refGene_gn=MEF2A;refGene_tx=XM_023254973.1;refGene_gn=MEF2A;refGene_tx=XM_023254974.1;refGene_gn=MEF2A;refGene_tx=XM_023254975.1;refGene_gn=MEF2A;refGene_tx=XM_023254976.1;refGene_gn=MEF2A;refGene_tx=XM_023254977.1
B3 18381034 18381389
refGene_gn=MEF2A;refGene_tx=XM_023254972.1;refGene_gn=MEF2A;refGene_tx=XM_023254973.1;refGene_gn=MEF2A;refGene_tx=XM_023254974.1;refGene_gn=MEF2A;refGene_tx=XM_023254975.1;refGene_gn=MEF2A;refGene_tx=XM_023254976.1;refGene_gn=MEF2A;refGene_tx=XM_023254977.1
B3 18410242 18410854
refGene_gn=LYSMD4;refGene_tx=XM_003986865.5;refGene_gn=LYSMD4;refGene_tx=XM_006932297.4;refGene_gn=LYSMD4;refGene_tx=XM_019831934.2
B3 18412988 18413270
refGene_gn=LYSMD4;refGene_tx=XM_003986865.5;refGene_gn=LYSMD4;refGene_tx=XM_006932297.4;refGene_gn=LYSMD4;refGene_tx=XM_019831934.2
B3 18475206 18475376
refGene_gn=ADAMTS17;refGene_tx=XM_023254978.1;refGene_gn=ADAMTS17;refGene_tx=XM_023254979.1;refGene_gn=ADAMTS17;refGene_tx=XM_023254980.1;refGene_gn=ADAMTS17;refGene_tx=XM_023254981.1
B3 18476507 18476685
refGene_gn=ADAMTS17;refGene_tx=XM_023254978.1;refGene_gn=ADAMTS17;refGene_tx=XM_023254979.1;refGene_gn=ADAMTS17;refGene_tx=XM_023254980.1;refGene_gn=ADAMTS17;refGene_tx=XM_023254981.1
B3 18490655 18490808
refGene_gn=ADAMTS17;refGene_tx=XM_023254978.1;refGene_gn=ADAMTS17;refGene_tx=XM_023254979.1;refGene_gn=ADAMTS17;refGene_tx=XM_023254980.1;refGene_gn=ADAMTS17;refGene_tx=XM_023254981.1
B3 18495561 18495766
refGene_gn=ADAMTS17;refGene_tx=XM_023254978.1;refGene_gn=ADAMTS17;refGene_tx=XM_023254979.1;refGene_gn=ADAMTS17;refGene_tx=XM_023254980.1;refGene_gn=ADAMTS17;refGene_tx=XM_023254981.1
B3 18543516 18543652
refGene_gn=ADAMTS17;refGene_tx=XM_023254978.1;refGene_gn=ADAMTS17;refGene_tx=XM_023254979.1;refGene_gn=ADAMTS17;refGene_tx=XM_023254980.1;refGene_gn=ADAMTS17;refGene_tx=XM_023254981.1
B3 18546223 18546383
refGene_gn=ADAMTS17;refGene_tx=XM_023254978.1;refGene_gn=ADAMTS17;refGene_tx=XM_023254979.1;refGene_gn=ADAMTS17;refGene_tx=XM_023254980.1;refGene_gn=ADAMTS17;refGene_tx=XM_023254981.1
B3 18549273 18549431
refGene_gn=ADAMTS17;refGene_tx=XM_023254978.1;refGene_gn=ADAMTS17;refGene_tx=XM_023254979.1;refGene_gn=ADAMTS17;refGene_tx=XM_023254980.1;refGene_gn=ADAMTS17;refGene_tx=XM_023254981.1

B3 18559160 18559241
refGene_gn=ADAMTS17;refGene_tx=XM_023254978.1;refGene_gn=ADAMTS17;refGene_tx=XM_023254979.1;refGene_gn=ADAMTS17;refGene_tx=XM_023254980.1;refGene_gn=ADAMTS17;refGene_tx=XM_023254981.1
B3 18572025 18572094
refGene_gn=ADAMTS17;refGene_tx=XM_023254978.1;refGene_gn=ADAMTS17;refGene_tx=XM_023254979.1
B3 18576831 18576952
refGene_gn=ADAMTS17;refGene_tx=XM_023254978.1;refGene_gn=ADAMTS17;refGene_tx=XM_023254979.1;refGene_gn=ADAMTS17;refGene_tx=XM_023254980.1;refGene_gn=ADAMTS17;refGene_tx=XM_023254981.1
B3 18589621 18589749
refGene_gn=ADAMTS17;refGene_tx=XM_023254978.1;refGene_gn=ADAMTS17;refGene_tx=XM_023254979.1;refGene_gn=ADAMTS17;refGene_tx=XM_023254980.1;refGene_gn=ADAMTS17;refGene_tx=XM_023254981.1
B3 18597804 18597971
refGene_gn=ADAMTS17;refGene_tx=XM_023254978.1;refGene_gn=ADAMTS17;refGene_tx=XM_023254979.1;refGene_gn=ADAMTS17;refGene_tx=XM_023254980.1;refGene_gn=ADAMTS17;refGene_tx=XM_023254981.1
B3 18613011 18613157
refGene_gn=ADAMTS17;refGene_tx=XM_023254978.1;refGene_gn=ADAMTS17;refGene_tx=XM_023254979.1;refGene_gn=ADAMTS17;refGene_tx=XM_023254980.1;refGene_gn=ADAMTS17;refGene_tx=XM_023254981.1
B3 18614232 18614337
refGene_gn=ADAMTS17;refGene_tx=XM_023254978.1;refGene_gn=ADAMTS17;refGene_tx=XM_023254979.1;refGene_gn=ADAMTS17;refGene_tx=XM_023254980.1;refGene_gn=ADAMTS17;refGene_tx=XM_023254981.1
B3 18641650 18641801
refGene_gn=ADAMTS17;refGene_tx=XM_023254978.1;refGene_gn=ADAMTS17;refGene_tx=XM_023254979.1;refGene_gn=ADAMTS17;refGene_tx=XM_023254980.1;refGene_gn=ADAMTS17;refGene_tx=XM_023254981.1
B3 18644008 18644149
refGene_gn=ADAMTS17;refGene_tx=XM_023254978.1;refGene_gn=ADAMTS17;refGene_tx=XM_023254979.1;refGene_gn=ADAMTS17;refGene_tx=XM_023254980.1;refGene_gn=ADAMTS17;refGene_tx=XM_023254981.1
B3 18682879 18682985
refGene_gn=ADAMTS17;refGene_tx=XM_023254978.1;refGene_gn=ADAMTS17;refGene_tx=XM_023254979.1;refGene_gn=ADAMTS17;refGene_tx=XM_023254980.1;refGene_gn=ADAMTS17;refGene_tx=XM_023254981.1
B3 18735864 18735908
refGene_gn=ADAMTS17;refGene_tx=XM_023254978.1;refGene_gn=ADAMTS17;refGene_tx=XM_023254979.1;refGene_gn=ADAMTS17;refGene_tx=XM_023254980.1;refGene_gn=ADAMTS17;refGene_tx=XM_023254981.1
B3 18742921 18743079
refGene_gn=ADAMTS17;refGene_tx=XM_023254978.1;refGene_gn=ADAMTS17;refGene_tx=XM_023254979.1;refGene_gn=ADAMTS17;refGene_tx=XM_023254980.1;refGene_gn=ADAMTS17;refGene_tx=XM_023254981.1
B3 18743799 18743883
refGene_gn=ADAMTS17;refGene_tx=XM_023254978.1;refGene_gn=ADAMTS17;refGene_tx=XM_023254979.1;refGene_gn=ADAMTS17;refGene_tx=XM_023254980.1;re

fGene_gn=ADAMTS17;refGene_tx=XM_023254981.1
B3 18754284 18754457
refGene_gn=ADAMTS17;refGene_tx=XM_023254978.1;refGene_gn=ADAMTS17;refGene_tx=XM_023254979.1;refGene_gn=ADAMTS17;refGene_tx=XM_023254980.1;refGene_gn=ADAMTS17;refGene_tx=XM_023254981.1
B3 18805399 18805565
refGene_gn=ADAMTS17;refGene_tx=XM_023254978.1;refGene_gn=ADAMTS17;refGene_tx=XM_023254979.1;refGene_gn=ADAMTS17;refGene_tx=XM_023254980.1;refGene_gn=ADAMTS17;refGene_tx=XM_023254981.1
B3 18817383 18817754
refGene_gn=ADAMTS17;refGene_tx=XM_023254978.1;refGene_gn=ADAMTS17;refGene_tx=XM_023254979.1;refGene_gn=ADAMTS17;refGene_tx=XM_023254980.1;refGene_gn=ADAMTS17;refGene_tx=XM_023254981.1
B3 18818242 18818321
refGene_gn=ADAMTS17;refGene_tx=XM_023254978.1;refGene_gn=ADAMTS17;refGene_tx=XM_023254979.1;refGene_gn=ADAMTS17;refGene_tx=XM_023254980.1;refGene_gn=ADAMTS17;refGene_tx=XM_023254981.1
B3 18863158 18863320 refGene_gn=CERS3;refGene_tx=XM_003986866.5
B3 18893983 18894137 refGene_gn=CERS3;refGene_tx=XM_003986866.5
B3 18905475 18905582 refGene_gn=CERS3;refGene_tx=XM_003986866.5
B3 18907766 18907895 refGene_gn=CERS3;refGene_tx=XM_003986866.5
B3 18910775 18910868 refGene_gn=CERS3;refGene_tx=XM_003986866.5
B3 18915038 18915089 refGene_gn=CERS3;refGene_tx=XM_003986866.5
B3 18915982 18916040 refGene_gn=CERS3;refGene_tx=XM_003986866.5
B3 18923233 18923352 refGene_gn=CERS3;refGene_tx=XM_003986866.5
B3 18928716 18928834 refGene_gn=CERS3;refGene_tx=XM_003986866.5
B3 18940583 18940756 refGene_gn=CERS3;refGene_tx=XM_003986866.5
B3 18988893 18989776
refGene_gn=LINS1;refGene_tx=XM_006932298.4;refGene_gn=LINS1;refGene_tx=XM_006932300.4;refGene_gn=LINS1;refGene_tx=XM_006932301.4;refGene_gn=LINS1;refGene_tx=XM_006932304.4;refGene_gn=LINS1;refGene_tx=XM_006932305.4;refGene_gn=LINS1;refGene_tx=XM_019831935.2;refGene_gn=LINS1;refGene_tx=XM_019831936.2;refGene_gn=LINS1;refGene_tx=XM_019831937.2;refGene_gn=LINS1;refGene_tx=XM_019831938.2;refGene_gn=LINS1;refGene_tx=XM_023254982.1;refGene_gn=LINS1;refGene_tx=XM_023254983.1;refGene_gn=LINS1;refGene_tx=XM_023254984.1;refGene_gn=LINS1;refGene_tx=XM_023254985.1;refGene_gn=LINS1;refGene_tx=XM_023254986.1;refGene_gn=LINS1;refGene_tx=XM_023254987.1;refGene_gn=LINS1;refGene_tx=XM_023254988.1
B3 18991615 18991787
refGene_gn=LINS1;refGene_tx=XM_006932298.4;refGene_gn=LINS1;refGene_tx=XM_006932300.4;refGene_gn=LINS1;refGene_tx=XM_006932301.4;refGene_gn=LINS1;refGene_tx=XM_006932304.4;refGene_gn=LINS1;refGene_tx=XM_006932305.4;refGene_gn=LINS1;refGene_tx=XM_019831935.2;refGene_gn=LINS1;refGene_tx=XM_019831936.2;refGene_gn=LINS1;refGene_tx=XM_019831937.2;refGene_gn=LINS1;refGene_tx=XM_019831938.2;refGene_gn=LINS1;refGene_tx=XM_023254982.1;refGene_gn=LINS1;refGene_tx=XM_023254983.1;refGene_gn=LINS1;refGene_tx=XM_023254984.1;refGene_gn=LINS1;refGene_tx=XM_023254985.1;refGene_gn=LINS1;refGene_tx=XM_023254986.1;refGene_gn=LINS1;refGene_tx=XM_023254987.1;refGene_gn=LINS1;refGene_tx=XM_023254988.1
B3 18993402 18993993

refGene_gn=LINS1;refGene_tx=XM_006932298.4;refGene_gn=LINS1;refGene_tx=XM_006932300.4;refGene_gn=LINS1;refGene_tx=XM_006932301.4;refGene_gn=LINS1;refGene_tx=XM_006932304.4;refGene_gn=LINS1;refGene_tx=XM_006932305.4;refGene_gn=LINS1;refGene_tx=XM_019831935.2;refGene_gn=LINS1;refGene_tx=XM_019831936.2;refGene_gn=LINS1;refGene_tx=XM_019831937.2;refGene_gn=LINS1;refGene_tx=XM_019831938.2;refGene_gn=LINS1;refGene_tx=XM_023254982.1;refGene_gn=LINS1;refGene_tx=XM_023254983.1;refGene_gn=LINS1;refGene_tx=XM_023254984.1;refGene_gn=LINS1;refGene_tx=XM_023254985.1;refGene_gn=LINS1;refGene_tx=XM_023254986.1;refGene_gn=LINS1;refGene_tx=XM_023254987.1;refGene_gn=LINS1;refGene_tx=XM_023254988.1
B3 18994870 18995012
refGene_gn=LINS1;refGene_tx=XM_006932298.4;refGene_gn=LINS1;refGene_tx=XM_006932300.4;refGene_gn=LINS1;refGene_tx=XM_006932301.4;refGene_gn=LINS1;refGene_tx=XM_006932304.4;refGene_gn=LINS1;refGene_tx=XM_006932305.4;refGene_gn=LINS1;refGene_tx=XM_019831935.2;refGene_gn=LINS1;refGene_tx=XM_019831936.2;refGene_gn=LINS1;refGene_tx=XM_019831937.2;refGene_gn=LINS1;refGene_tx=XM_019831938.2;refGene_gn=LINS1;refGene_tx=XM_023254982.1;refGene_gn=LINS1;refGene_tx=XM_023254983.1;refGene_gn=LINS1;refGene_tx=XM_023254984.1;refGene_gn=LINS1;refGene_tx=XM_023254985.1;refGene_gn=LINS1;refGene_tx=XM_023254986.1;refGene_gn=LINS1;refGene_tx=XM_023254987.1;refGene_gn=LINS1;refGene_tx=XM_023254988.1
B3 18995859 18995949
refGene_gn=LINS1;refGene_tx=XM_006932298.4;refGene_gn=LINS1;refGene_tx=XM_006932300.4;refGene_gn=LINS1;refGene_tx=XM_006932301.4;refGene_gn=LINS1;refGene_tx=XM_006932304.4;refGene_gn=LINS1;refGene_tx=XM_006932305.4;refGene_gn=LINS1;refGene_tx=XM_019831935.2;refGene_gn=LINS1;refGene_tx=XM_019831936.2;refGene_gn=LINS1;refGene_tx=XM_019831937.2;refGene_gn=LINS1;refGene_tx=XM_019831938.2;refGene_gn=LINS1;refGene_tx=XM_023254982.1;refGene_gn=LINS1;refGene_tx=XM_023254983.1;refGene_gn=LINS1;refGene_tx=XM_023254984.1;refGene_gn=LINS1;refGene_tx=XM_023254985.1;refGene_gn=LINS1;refGene_tx=XM_023254986.1;refGene_gn=LINS1;refGene_tx=XM_023254987.1;refGene_gn=LINS1;refGene_tx=XM_023254988.1
B3 18996062 18996485
refGene_gn=LINS1;refGene_tx=XM_006932298.4;refGene_gn=LINS1;refGene_tx=XM_006932300.4;refGene_gn=LINS1;refGene_tx=XM_006932301.4;refGene_gn=LINS1;refGene_tx=XM_006932304.4;refGene_gn=LINS1;refGene_tx=XM_006932305.4;refGene_gn=LINS1;refGene_tx=XM_019831935.2;refGene_gn=LINS1;refGene_tx=XM_019831936.2;refGene_gn=LINS1;refGene_tx=XM_019831937.2;refGene_gn=LINS1;refGene_tx=XM_023254982.1;refGene_gn=LINS1;refGene_tx=XM_023254983.1;refGene_gn=LINS1;refGene_tx=XM_023254984.1;refGene_gn=LINS1;refGene_tx=XM_023254985.1;refGene_gn=LINS1;refGene_tx=XM_023254986.1;refGene_gn=LINS1;refGene_tx=XM_023254987.1;refGene_gn=LINS1;refGene_tx=XM_023254988.1
B3 19021720 19021931
refGene_gn=ASB7;refGene_tx=XM_003986867.5;refGene_gn=ASB7;refGene_tx=XM_019831939.2
B3 19043756 19044362
refGene_gn=ASB7;refGene_tx=XM_003986867.5;refGene_gn=ASB7;refGene_tx=XM_019831939.2
B3 19046729 19046749 refGene_gn=ASB7;refGene_tx=XM_019831939.2

B3 19059823 19059963 refGene_gn=ASB7;refGene_tx=XM_003986867.5
B3 19296766 19296865 refGene_gn=ALDH1A3;refGene_tx=XM_023254989.1
B3 19301936 19302041 refGene_gn=ALDH1A3;refGene_tx=XM_023254989.1
B3 19304021 19304162 refGene_gn=ALDH1A3;refGene_tx=XM_023254989.1
B3 19308602 19308732 refGene_gn=ALDH1A3;refGene_tx=XM_023254989.1
B3 19309487 19309549 refGene_gn=ALDH1A3;refGene_tx=XM_023254989.1
B3 19311785 19311914 refGene_gn=ALDH1A3;refGene_tx=XM_023254989.1
B3 19313481 19313595 refGene_gn=ALDH1A3;refGene_tx=XM_023254989.1
B3 19315788 19315891 refGene_gn=ALDH1A3;refGene_tx=XM_023254989.1
B3 19318533 19318718 refGene_gn=ALDH1A3;refGene_tx=XM_023254989.1
B3 19324596 19324761 refGene_gn=ALDH1A3;refGene_tx=XM_023254989.1
B3 19325969 19326127 refGene_gn=ALDH1A3;refGene_tx=XM_023254989.1
B3 19327442 19327517 refGene_gn=ALDH1A3;refGene_tx=XM_023254989.1
B3 19333885 19333958 refGene_gn=ALDH1A3;refGene_tx=XM_023254989.1
B3 19344758 19344855 refGene_gn=LRRK1;refGene_tx=XM_023254990.1
B3 19352267 19352316 refGene_gn=LRRK1;refGene_tx=XM_019831943.2
B3 19384870 19385034
refGene_gn=LRRK1;refGene_tx=XM_019831943.2;refGene_gn=LRRK1;refGene_tx
=XM_023254990.1
B3 19393082 19393254
refGene_gn=LRRK1;refGene_tx=XM_019831943.2;refGene_gn=LRRK1;refGene_tx
=XM_023254990.1
B3 19395882 19396062
refGene_gn=LRRK1;refGene_tx=XM_019831943.2;refGene_gn=LRRK1;refGene_tx
=XM_023254990.1
B3 19396487 19396636
refGene_gn=LRRK1;refGene_tx=XM_019831943.2;refGene_gn=LRRK1;refGene_tx
=XM_023254990.1
B3 19418093 19418320
refGene_gn=LRRK1;refGene_tx=XM_019831943.2;refGene_gn=LRRK1;refGene_tx
=XM_023254990.1
B3 19419522 19419650
refGene_gn=LRRK1;refGene_tx=XM_019831943.2;refGene_gn=LRRK1;refGene_tx
=XM_023254990.1
B3 19419772 19419936
refGene_gn=LRRK1;refGene_tx=XM_019831943.2;refGene_gn=LRRK1;refGene_tx
=XM_023254990.1
B3 19420758 19420896
refGene_gn=LRRK1;refGene_tx=XM_019831943.2;refGene_gn=LRRK1;refGene_tx
=XM_023254990.1
B3 19423511 19423624
refGene_gn=LRRK1;refGene_tx=XM_019831943.2;refGene_gn=LRRK1;refGene_tx
=XM_023254990.1
B3 19424710 19424787
refGene_gn=LRRK1;refGene_tx=XM_019831943.2;refGene_gn=LRRK1;refGene_tx
=XM_023254990.1
B3 19429612 19429742
refGene_gn=LRRK1;refGene_tx=XM_019831943.2;refGene_gn=LRRK1;refGene_tx
=XM_023254990.1
B3 19430424 19430537

refGene_gn=LRRK1;refGene_tx=XM_019831943.2;refGene_gn=LRRK1;refGene_tx=XM_023254990.1
B3 19430970 19431185
refGene_gn=LRRK1;refGene_tx=XM_019831943.2;refGene_gn=LRRK1;refGene_tx=XM_023254990.1
B3 19433225 19433390
refGene_gn=LRRK1;refGene_tx=XM_019831943.2;refGene_gn=LRRK1;refGene_tx=XM_023254990.1
B3 19434100 19434273
refGene_gn=LRRK1;refGene_tx=XM_019831943.2;refGene_gn=LRRK1;refGene_tx=XM_023254990.1
B3 19435310 19435431
refGene_gn=LRRK1;refGene_tx=XM_019831943.2;refGene_gn=LRRK1;refGene_tx=XM_023254990.1
B3 19435942 19436102
refGene_gn=LRRK1;refGene_tx=XM_019831943.2;refGene_gn=LRRK1;refGene_tx=XM_023254990.1
B3 19436494 19436771
refGene_gn=LRRK1;refGene_tx=XM_019831943.2;refGene_gn=LRRK1;refGene_tx=XM_023254990.1
B3 19439980 19440152
refGene_gn=LRRK1;refGene_tx=XM_019831943.2;refGene_gn=LRRK1;refGene_tx=XM_023254990.1
B3 19442924 19443088
refGene_gn=LRRK1;refGene_tx=XM_019831943.2;refGene_gn=LRRK1;refGene_tx=XM_023254990.1
B3 19444064 19444204
refGene_gn=LRRK1;refGene_tx=XM_019831943.2;refGene_gn=LRRK1;refGene_tx=XM_023254990.1
B3 19445992 19446242
refGene_gn=LRRK1;refGene_tx=XM_019831943.2;refGene_gn=LRRK1;refGene_tx=XM_023254990.1
B3 19449274 19449441
refGene_gn=LRRK1;refGene_tx=XM_019831943.2;refGene_gn=LRRK1;refGene_tx=XM_023254990.1
B3 19451770 19451968
refGene_gn=LRRK1;refGene_tx=XM_019831943.2;refGene_gn=LRRK1;refGene_tx=XM_023254990.1
B3 19453820 19454098
refGene_gn=LRRK1;refGene_tx=XM_019831943.2;refGene_gn=LRRK1;refGene_tx=XM_023254990.1
B3 19455713 19455908
refGene_gn=LRRK1;refGene_tx=XM_019831943.2;refGene_gn=LRRK1;refGene_tx=XM_023254990.1
B3 19456871 19457023
refGene_gn=LRRK1;refGene_tx=XM_019831943.2;refGene_gn=LRRK1;refGene_tx=XM_023254990.1
B3 19462442 19462560
refGene_gn=LRRK1;refGene_tx=XM_019831943.2;refGene_gn=LRRK1;refGene_tx=XM_023254990.1

B3 19463612 19463729
refGene_gn=LRRK1;refGene_tx=XM_019831943.2;refGene_gn=LRRK1;refGene_tx=XM_023254990.1
B3 19466326 19467174
refGene_gn=LRRK1;refGene_tx=XM_019831943.2;refGene_gn=LRRK1;refGene_tx=XM_023254990.1
B3 19467554 19467656
refGene_gn=LRRK1;refGene_tx=XM_019831943.2;refGene_gn=LRRK1;refGene_tx=XM_023254990.1
B3 19469456 19469634
refGene_gn=LRRK1;refGene_tx=XM_019831943.2;refGene_gn=LRRK1;refGene_tx=XM_023254990.1
B3 19560807 19562400 refGene_gn=CHSY1;refGene_tx=XM_019831944.2
B3 19614303 19614799 refGene_gn=CHSY1;refGene_tx=XM_019831944.2
B3 19629895 19630215 refGene_gn=CHSY1;refGene_tx=XM_019831944.2
B3 19667095 19667175
refGene_gn=SELENOS;refGene_tx=XM_003986878.5;refGene_gn=SELENOS;refGene_tx=XM_019831946.2
B3 19668641 19668810
refGene_gn=SELENOS;refGene_tx=XM_003986878.5;refGene_gn=SELENOS;refGene_tx=XM_019831946.2
B3 19673554 19673661
refGene_gn=SELENOS;refGene_tx=XM_003986878.5;refGene_gn=SELENOS;refGene_tx=XM_019831946.2
B3 19676504 19676639
refGene_gn=SELENOS;refGene_tx=XM_003986878.5;refGene_gn=SELENOS;refGene_tx=XM_019831946.2
B3 19677269 19677276 refGene_gn=SELENOS;refGene_tx=XM_019831946.2
B3 19677328 19677404 refGene_gn=SELENOS;refGene_tx=XM_003986878.5
B3 19683078 19683137 refGene_gn=SNRPA1;refGene_tx=XM_011282792.3
B3 19685010 19685104 refGene_gn=SNRPA1;refGene_tx=XM_011282792.3
B3 19685681 19685757 refGene_gn=SNRPA1;refGene_tx=XM_011282792.3
B3 19686169 19686249 refGene_gn=SNRPA1;refGene_tx=XM_011282792.3
B3 19686900 19687003 refGene_gn=SNRPA1;refGene_tx=XM_011282792.3
B3 19687289 19687336 refGene_gn=SNRPA1;refGene_tx=XM_011282792.3
B3 19692537 19692616 refGene_gn=SNRPA1;refGene_tx=XM_011282792.3
B3 19693588 19693736 refGene_gn=SNRPA1;refGene_tx=XM_011282792.3
B3 19695637 19695719 refGene_gn=SNRPA1;refGene_tx=XM_011282792.3
B3 19706586 19706684
refGene_gn=PCSK6;refGene_tx=XM_023254991.1;refGene_gn=PCSK6;refGene_tx=XM_023254992.1
B3 19708380 19708493
refGene_gn=PCSK6;refGene_tx=XM_023254991.1;refGene_gn=PCSK6;refGene_tx=XM_023254992.1
B3 19713100 19713230
refGene_gn=PCSK6;refGene_tx=XM_023254991.1;refGene_gn=PCSK6;refGene_tx=XM_023254992.1
B3 19718555 19718659
refGene_gn=PCSK6;refGene_tx=XM_023254991.1;refGene_gn=PCSK6;refGene_tx=XM_023254992.1

B3 19722148 19722236
refGene_gn=PCSK6;refGene_tx=XM_023254991.1;refGene_gn=PCSK6;refGene_tx
=XM_023254992.1
B3 19723905 19724102
refGene_gn=PCSK6;refGene_tx=XM_023254991.1;refGene_gn=PCSK6;refGene_tx
=XM_023254992.1
B3 19725061 19725164
refGene_gn=PCSK6;refGene_tx=XM_023254991.1;refGene_gn=PCSK6;refGene_tx
=XM_023254992.1
B3 19731121 19731160 refGene_gn=PCSK6;refGene_tx=XM_023254992.1
B3 19731326 19731506
refGene_gn=PCSK6;refGene_tx=XM_023254991.1;refGene_gn=PCSK6;refGene_tx
=XM_023254992.1
B3 19763932 19764069
refGene_gn=PCSK6;refGene_tx=XM_023254991.1;refGene_gn=PCSK6;refGene_tx
=XM_023254992.1
B3 19767916 19768105
refGene_gn=PCSK6;refGene_tx=XM_023254991.1;refGene_gn=PCSK6;refGene_tx
=XM_023254992.1
B3 19780525 19780643
refGene_gn=PCSK6;refGene_tx=XM_023254991.1;refGene_gn=PCSK6;refGene_tx
=XM_023254992.1
B3 19782569 19782673
refGene_gn=PCSK6;refGene_tx=XM_023254991.1;refGene_gn=PCSK6;refGene_tx
=XM_023254992.1
B3 19787163 19787264
refGene_gn=PCSK6;refGene_tx=XM_023254991.1;refGene_gn=PCSK6;refGene_tx
=XM_023254992.1
B3 19791033 19791246
refGene_gn=PCSK6;refGene_tx=XM_023254991.1;refGene_gn=PCSK6;refGene_tx
=XM_023254992.1
B3 19797168 19797341
refGene_gn=PCSK6;refGene_tx=XM_023254991.1;refGene_gn=PCSK6;refGene_tx
=XM_023254992.1
B3 19827985 19828074
refGene_gn=PCSK6;refGene_tx=XM_023254991.1;refGene_gn=PCSK6;refGene_tx
=XM_023254992.1
B3 19830158 19830235
refGene_gn=PCSK6;refGene_tx=XM_023254991.1;refGene_gn=PCSK6;refGene_tx
=XM_023254992.1
B3 19831840 19831984
refGene_gn=PCSK6;refGene_tx=XM_023254991.1;refGene_gn=PCSK6;refGene_tx
=XM_023254992.1
B3 19832488 19832599
refGene_gn=PCSK6;refGene_tx=XM_023254991.1;refGene_gn=PCSK6;refGene_tx
=XM_023254992.1
B3 19848110 19848215
refGene_gn=PCSK6;refGene_tx=XM_023254991.1;refGene_gn=PCSK6;refGene_tx
=XM_023254992.1
B3 19898460 19898739

refGene_gn=PCSK6;refGene_tx=XM_023254991.1;refGene_gn=PCSK6;refGene_tx=XM_023254992.1
B3 20050571 20050737
refGene_gn=TM2D3;refGene_tx=XM_003986879.3;refGene_gn=TM2D3;refGene_tx=XM_006932310.2
B3 20050765 20050841 refGene_gn=TM2D3;refGene_tx=XM_019831948.2
B3 20053134 20053210
refGene_gn=TM2D3;refGene_tx=XM_003986879.3;refGene_gn=TM2D3;refGene_tx=XM_006932310.2;refGene_gn=TM2D3;refGene_tx=XM_019831948.2
B3 20055234 20055409
refGene_gn=TM2D3;refGene_tx=XM_003986879.3;refGene_gn=TM2D3;refGene_tx=XM_006932310.2;refGene_gn=TM2D3;refGene_tx=XM_019831948.2
B3 20060790 20060948
refGene_gn=TM2D3;refGene_tx=XM_003986879.3;refGene_gn=TM2D3;refGene_tx=XM_006932310.2;refGene_gn=TM2D3;refGene_tx=XM_019831948.2
B3 20062528 20062621
refGene_gn=TM2D3;refGene_tx=XM_006932310.2;refGene_gn=TM2D3;refGene_tx=XM_019831948.2
B3 20063056 20063147
refGene_gn=TM2D3;refGene_tx=XM_003986879.3;refGene_gn=TM2D3;refGene_tx=XM_006932310.2;refGene_gn=TM2D3;refGene_tx=XM_019831948.2
B3 20065942 20066091 refGene_gn=TARSL2;refGene_tx=XM_011282731.3
B3 20067359 20067474 refGene_gn=TARSL2;refGene_tx=XM_011282731.3
B3 20068453 20068526 refGene_gn=TARSL2;refGene_tx=XM_011282731.3
B3 20072586 20072691 refGene_gn=TARSL2;refGene_tx=XM_011282731.3
B3 20075217 20075318 refGene_gn=TARSL2;refGene_tx=XM_011282731.3
B3 20075402 20075480 refGene_gn=TARSL2;refGene_tx=XM_011282731.3
B3 20079052 20079190 refGene_gn=TARSL2;refGene_tx=XM_011282731.3
B3 20085260 20085423 refGene_gn=TARSL2;refGene_tx=XM_011282731.3
B3 20086277 20086444 refGene_gn=TARSL2;refGene_tx=XM_011282731.3
B3 20101146 20101245 refGene_gn=TARSL2;refGene_tx=XM_011282731.3
B3 20102080 20102227 refGene_gn=TARSL2;refGene_tx=XM_011282731.3
B3 20103101 20103180 refGene_gn=TARSL2;refGene_tx=XM_011282731.3
B3 20106067 20106132 refGene_gn=TARSL2;refGene_tx=XM_011282731.3
B3 20109331 20109706
refGene_gn=LOC101081984;refGene_tx=XM_019831949.2
B3 20111902 20112020 refGene_gn=TARSL2;refGene_tx=XM_011282731.3
B3 20113912 20114034 refGene_gn=TARSL2;refGene_tx=XM_011282731.3
B3 20119006 20119130 refGene_gn=TARSL2;refGene_tx=XM_011282731.3
B3 20126065 20126262 refGene_gn=TARSL2;refGene_tx=XM_011282731.3
B3 20126898 20126967 refGene_gn=TARSL2;refGene_tx=XM_011282731.3
B3 20127969 20128287 refGene_gn=TARSL2;refGene_tx=XM_011282731.3
B3 20144442 20144785
refGene_gn=TJP1;refGene_tx=XM_011282728.3;refGene_gn=TJP1;refGene_tx=XM_019831950.2;refGene_gn=TJP1;refGene_tx=XM_019831951.2;refGene_gn=TJP1;refGene_tx=XM_019831953.2;refGene_gn=TJP1;refGene_tx=XM_019831954.2
B3 20156698 20156825
refGene_gn=TJP1;refGene_tx=XM_011282728.3;refGene_gn=TJP1;refGene_tx=XM_019831950.2;refGene_gn=TJP1;refGene_tx=XM_019831951.2;refGene_gn=TJP1;refGene_tx=XM_019831953.2;refGene_gn=TJP1;refGene_tx=XM_019831954.2

B3 20267943 20268116
refGene_gn=TJP1;refGene_tx=XM_023254994.1;refGene_gn=TJP1;refGene_tx=X
M_023254995.1
B3 20282712 20282845
refGene_gn=TJP1;refGene_tx=XM_011282728.3;refGene_gn=TJP1;refGene_tx=X
M_019831950.2;refGene_gn=TJP1;refGene_tx=XM_019831951.2;refGene_gn=TJP
1;refGene_tx=XM_019831953.2;refGene_gn=TJP1;refGene_tx=XM_019831954.2;
refGene_gn=TJP1;refGene_tx=XM_023254994.1;refGene_gn=TJP1;refGene_tx=X
M_023254995.1
B3 20410337 20410364 refGene_gn=TJP1;refGene_tx=XM_023254996.1
B3 20435659 20435716
refGene_gn=TJP1;refGene_tx=XM_011282728.3;refGene_gn=TJP1;refGene_tx=X
M_019831950.2;refGene_gn=TJP1;refGene_tx=XM_019831951.2;refGene_gn=TJP
1;refGene_tx=XM_019831953.2;refGene_gn=TJP1;refGene_tx=XM_019831954.2;
refGene_gn=TJP1;refGene_tx=XM_023254994.1;refGene_gn=TJP1;refGene_tx=X
M_023254995.1;refGene_gn=TJP1;refGene_tx=XM_023254996.1
B3 20453727 20453852
refGene_gn=TJP1;refGene_tx=XM_011282728.3;refGene_gn=TJP1;refGene_tx=X
M_019831950.2;refGene_gn=TJP1;refGene_tx=XM_019831951.2;refGene_gn=TJP
1;refGene_tx=XM_019831953.2;refGene_gn=TJP1;refGene_tx=XM_019831954.2;
refGene_gn=TJP1;refGene_tx=XM_023254994.1;refGene_gn=TJP1;refGene_tx=X
M_023254995.1;refGene_gn=TJP1;refGene_tx=XM_023254996.1
B3 20454483 20454586
refGene_gn=TJP1;refGene_tx=XM_011282728.3;refGene_gn=TJP1;refGene_tx=X
M_019831950.2;refGene_gn=TJP1;refGene_tx=XM_019831951.2;refGene_gn=TJP
1;refGene_tx=XM_019831953.2;refGene_gn=TJP1;refGene_tx=XM_019831954.2;
refGene_gn=TJP1;refGene_tx=XM_023254994.1;refGene_gn=TJP1;refGene_tx=X
M_023254995.1;refGene_gn=TJP1;refGene_tx=XM_023254996.1
B3 20459758 20460035
refGene_gn=TJP1;refGene_tx=XM_011282728.3;refGene_gn=TJP1;refGene_tx=X
M_019831950.2;refGene_gn=TJP1;refGene_tx=XM_019831951.2;refGene_gn=TJP
1;refGene_tx=XM_019831953.2;refGene_gn=TJP1;refGene_tx=XM_019831954.2;
refGene_gn=TJP1;refGene_tx=XM_023254994.1;refGene_gn=TJP1;refGene_tx=X
M_023254995.1;refGene_gn=TJP1;refGene_tx=XM_023254996.1
B3 20462668 20462772
refGene_gn=TJP1;refGene_tx=XM_011282728.3;refGene_gn=TJP1;refGene_tx=X
M_019831950.2;refGene_gn=TJP1;refGene_tx=XM_019831951.2;refGene_gn=TJP
1;refGene_tx=XM_019831953.2;refGene_gn=TJP1;refGene_tx=XM_019831954.2;
refGene_gn=TJP1;refGene_tx=XM_023254994.1;refGene_gn=TJP1;refGene_tx=X
M_023254995.1;refGene_gn=TJP1;refGene_tx=XM_023254996.1
B3 20463374 20463543
refGene_gn=TJP1;refGene_tx=XM_011282728.3;refGene_gn=TJP1;refGene_tx=X
M_019831950.2;refGene_gn=TJP1;refGene_tx=XM_019831951.2;refGene_gn=TJP
1;refGene_tx=XM_019831953.2;refGene_gn=TJP1;refGene_tx=XM_019831954.2;
refGene_gn=TJP1;refGene_tx=XM_023254994.1;refGene_gn=TJP1;refGene_tx=X
M_023254995.1;refGene_gn=TJP1;refGene_tx=XM_023254996.1
B3 20464356 20464504
refGene_gn=TJP1;refGene_tx=XM_011282728.3;refGene_gn=TJP1;refGene_tx=X
M_019831950.2;refGene_gn=TJP1;refGene_tx=XM_019831951.2;refGene_gn=TJP
1;refGene_tx=XM_019831953.2;refGene_gn=TJP1;refGene_tx=XM_019831954.2;

refGene_gn=TJP1;refGene_tx=XM_023254994.1;refGene_gn=TJP1;refGene_tx=XM_023254995.1;refGene_gn=TJP1;refGene_tx=XM_023254996.1
B3 20477627 20477764
refGene_gn=TJP1;refGene_tx=XM_011282728.3;refGene_gn=TJP1;refGene_tx=XM_019831950.2;refGene_gn=TJP1;refGene_tx=XM_019831951.2;refGene_gn=TJP1;refGene_tx=XM_019831953.2;refGene_gn=TJP1;refGene_tx=XM_019831954.2;refGene_gn=TJP1;refGene_tx=XM_023254994.1;refGene_gn=TJP1;refGene_tx=XM_023254995.1;refGene_gn=TJP1;refGene_tx=XM_023254996.1
B3 20478865 20478971
refGene_gn=TJP1;refGene_tx=XM_011282728.3;refGene_gn=TJP1;refGene_tx=XM_019831950.2;refGene_gn=TJP1;refGene_tx=XM_019831951.2;refGene_gn=TJP1;refGene_tx=XM_019831953.2;refGene_gn=TJP1;refGene_tx=XM_019831954.2;refGene_gn=TJP1;refGene_tx=XM_023254994.1;refGene_gn=TJP1;refGene_tx=XM_023254995.1;refGene_gn=TJP1;refGene_tx=XM_023254996.1
B3 20481704 20481855
refGene_gn=TJP1;refGene_tx=XM_011282728.3;refGene_gn=TJP1;refGene_tx=XM_019831950.2;refGene_gn=TJP1;refGene_tx=XM_019831951.2;refGene_gn=TJP1;refGene_tx=XM_019831953.2;refGene_gn=TJP1;refGene_tx=XM_019831954.2;refGene_gn=TJP1;refGene_tx=XM_023254994.1;refGene_gn=TJP1;refGene_tx=XM_023254995.1;refGene_gn=TJP1;refGene_tx=XM_023254996.1
B3 20484335 20484444
refGene_gn=TJP1;refGene_tx=XM_011282728.3;refGene_gn=TJP1;refGene_tx=XM_019831950.2;refGene_gn=TJP1;refGene_tx=XM_019831951.2;refGene_gn=TJP1;refGene_tx=XM_019831953.2;refGene_gn=TJP1;refGene_tx=XM_019831954.2;refGene_gn=TJP1;refGene_tx=XM_023254994.1;refGene_gn=TJP1;refGene_tx=XM_023254995.1;refGene_gn=TJP1;refGene_tx=XM_023254996.1
B3 20485659 20485879
refGene_gn=TJP1;refGene_tx=XM_011282728.3;refGene_gn=TJP1;refGene_tx=XM_019831950.2;refGene_gn=TJP1;refGene_tx=XM_019831951.2;refGene_gn=TJP1;refGene_tx=XM_019831953.2;refGene_gn=TJP1;refGene_tx=XM_019831954.2;refGene_gn=TJP1;refGene_tx=XM_023254994.1;refGene_gn=TJP1;refGene_tx=XM_023254995.1;refGene_gn=TJP1;refGene_tx=XM_023254996.1
B3 20486324 20486509
refGene_gn=TJP1;refGene_tx=XM_011282728.3;refGene_gn=TJP1;refGene_tx=XM_019831950.2;refGene_gn=TJP1;refGene_tx=XM_019831951.2;refGene_gn=TJP1;refGene_tx=XM_019831953.2;refGene_gn=TJP1;refGene_tx=XM_019831954.2;refGene_gn=TJP1;refGene_tx=XM_023254994.1;refGene_gn=TJP1;refGene_tx=XM_023254995.1;refGene_gn=TJP1;refGene_tx=XM_023254996.1
B3 20486608 20486704
refGene_gn=TJP1;refGene_tx=XM_011282728.3;refGene_gn=TJP1;refGene_tx=XM_019831950.2;refGene_gn=TJP1;refGene_tx=XM_019831951.2;refGene_gn=TJP1;refGene_tx=XM_019831953.2;refGene_gn=TJP1;refGene_tx=XM_019831954.2;refGene_gn=TJP1;refGene_tx=XM_023254994.1;refGene_gn=TJP1;refGene_tx=XM_023254995.1;refGene_gn=TJP1;refGene_tx=XM_023254996.1
B3 20490605 20490688
refGene_gn=TJP1;refGene_tx=XM_011282728.3;refGene_gn=TJP1;refGene_tx=XM_019831950.2;refGene_gn=TJP1;refGene_tx=XM_019831951.2;refGene_gn=TJP1;refGene_tx=XM_019831953.2;refGene_gn=TJP1;refGene_tx=XM_019831954.2;refGene_gn=TJP1;refGene_tx=XM_023254994.1;refGene_gn=TJP1;refGene_tx=XM_023254995.1;refGene_gn=TJP1;refGene_tx=XM_023254996.1

B3 20491239 20491450
refGene_gn=TJP1;refGene_tx=XM_011282728.3;refGene_gn=TJP1;refGene_tx=XM_019831950.2;refGene_gn=TJP1;refGene_tx=XM_019831951.2;refGene_gn=TJP1;refGene_tx=XM_019831953.2;refGene_gn=TJP1;refGene_tx=XM_019831954.2;refGene_gn=TJP1;refGene_tx=XM_023254994.1;refGene_gn=TJP1;refGene_tx=XM_023254995.1;refGene_gn=TJP1;refGene_tx=XM_023254996.1
B3 20491986 20492087
refGene_gn=TJP1;refGene_tx=XM_011282728.3;refGene_gn=TJP1;refGene_tx=XM_019831950.2;refGene_gn=TJP1;refGene_tx=XM_019831951.2;refGene_gn=TJP1;refGene_tx=XM_019831953.2;refGene_gn=TJP1;refGene_tx=XM_019831954.2;refGene_gn=TJP1;refGene_tx=XM_023254994.1;refGene_gn=TJP1;refGene_tx=XM_023254995.1;refGene_gn=TJP1;refGene_tx=XM_023254996.1
B3 20495531 20495882
refGene_gn=TJP1;refGene_tx=XM_011282728.3;refGene_gn=TJP1;refGene_tx=XM_019831950.2;refGene_gn=TJP1;refGene_tx=XM_019831951.2;refGene_gn=TJP1;refGene_tx=XM_019831953.2;refGene_gn=TJP1;refGene_tx=XM_019831954.2;refGene_gn=TJP1;refGene_tx=XM_023254994.1;refGene_gn=TJP1;refGene_tx=XM_023254995.1;refGene_gn=TJP1;refGene_tx=XM_023254996.1
B3 20496216 20496456
refGene_gn=TJP1;refGene_tx=XM_011282728.3;refGene_gn=TJP1;refGene_tx=XM_019831950.2;refGene_gn=TJP1;refGene_tx=XM_019831951.2;refGene_gn=TJP1;refGene_tx=XM_023254994.1;refGene_gn=TJP1;refGene_tx=XM_023254995.1;refGene_gn=TJP1;refGene_tx=XM_023254996.1
B3 20497104 20497977
refGene_gn=TJP1;refGene_tx=XM_011282728.3;refGene_gn=TJP1;refGene_tx=XM_019831950.2;refGene_gn=TJP1;refGene_tx=XM_019831951.2;refGene_gn=TJP1;refGene_tx=XM_019831953.2;refGene_gn=TJP1;refGene_tx=XM_019831954.2;refGene_gn=TJP1;refGene_tx=XM_023254994.1;refGene_gn=TJP1;refGene_tx=XM_023254995.1;refGene_gn=TJP1;refGene_tx=XM_023254996.1
B3 20498131 20498229
refGene_gn=TJP1;refGene_tx=XM_011282728.3;refGene_gn=TJP1;refGene_tx=XM_019831950.2;refGene_gn=TJP1;refGene_tx=XM_019831951.2;refGene_gn=TJP1;refGene_tx=XM_019831953.2;refGene_gn=TJP1;refGene_tx=XM_019831954.2;refGene_gn=TJP1;refGene_tx=XM_023254994.1;refGene_gn=TJP1;refGene_tx=XM_023254995.1;refGene_gn=TJP1;refGene_tx=XM_023254996.1
B3 20500314 20500542
refGene_gn=TJP1;refGene_tx=XM_011282728.3;refGene_gn=TJP1;refGene_tx=XM_019831950.2;refGene_gn=TJP1;refGene_tx=XM_019831951.2;refGene_gn=TJP1;refGene_tx=XM_019831953.2;refGene_gn=TJP1;refGene_tx=XM_019831954.2;refGene_gn=TJP1;refGene_tx=XM_023254994.1;refGene_gn=TJP1;refGene_tx=XM_023254995.1;refGene_gn=TJP1;refGene_tx=XM_023254996.1
B3 20505560 20505721
refGene_gn=TJP1;refGene_tx=XM_011282728.3;refGene_gn=TJP1;refGene_tx=XM_019831950.2;refGene_gn=TJP1;refGene_tx=XM_019831951.2;refGene_gn=TJP1;refGene_tx=XM_019831953.2;refGene_gn=TJP1;refGene_tx=XM_019831954.2;refGene_gn=TJP1;refGene_tx=XM_023254994.1;refGene_gn=TJP1;refGene_tx=XM_023254995.1;refGene_gn=TJP1;refGene_tx=XM_023254996.1
B3 20508153 20508631
refGene_gn=TJP1;refGene_tx=XM_011282728.3;refGene_gn=TJP1;refGene_tx=XM_019831950.2;refGene_gn=TJP1;refGene_tx=XM_019831951.2;refGene_gn=TJP

1;refGene_tx=XM_019831953.2;refGene_gn=TJP1;refGene_tx=XM_019831954.2;
refGene_gn=TJP1;refGene_tx=XM_023254994.1;refGene_gn=TJP1;refGene_tx=X
M_023254995.1;refGene_gn=TJP1;refGene_tx=XM_023254996.1
B3 20510367 20510585
refGene_gn=TJP1;refGene_tx=XM_011282728.3;refGene_gn=TJP1;refGene_tx=X
M_019831950.2;refGene_gn=TJP1;refGene_tx=XM_019831951.2;refGene_gn=TJP
1;refGene_tx=XM_019831953.2;refGene_gn=TJP1;refGene_tx=XM_019831954.2;
refGene_gn=TJP1;refGene_tx=XM_023254994.1;refGene_gn=TJP1;refGene_tx=X
M_023254995.1;refGene_gn=TJP1;refGene_tx=XM_023254996.1
B3 20511790 20511934
refGene_gn=TJP1;refGene_tx=XM_011282728.3;refGene_gn=TJP1;refGene_tx=X
M_019831950.2;refGene_gn=TJP1;refGene_tx=XM_019831951.2;refGene_gn=TJP
1;refGene_tx=XM_019831953.2;refGene_gn=TJP1;refGene_tx=XM_019831954.2;
refGene_gn=TJP1;refGene_tx=XM_023254994.1;refGene_gn=TJP1;refGene_tx=X
M_023254995.1;refGene_gn=TJP1;refGene_tx=XM_023254996.1
B3 20514621 20514716
refGene_gn=TJP1;refGene_tx=XM_011282728.3;refGene_gn=TJP1;refGene_tx=X
M_019831950.2;refGene_gn=TJP1;refGene_tx=XM_019831951.2;refGene_gn=TJP
1;refGene_tx=XM_019831953.2;refGene_gn=TJP1;refGene_tx=XM_019831954.2;
refGene_gn=TJP1;refGene_tx=XM_023254994.1;refGene_gn=TJP1;refGene_tx=X
M_023254995.1;refGene_gn=TJP1;refGene_tx=XM_023254996.1
B3 20636191 20636410 refGene_gn=FAM189A1;refGene_tx=XM_023255006.1
B3 20829031 20829088 refGene_gn=FAM189A1;refGene_tx=XM_023255006.1
B3 20940304 20941165 refGene_gn=NSMCE3;refGene_tx=XM_006932311.4
B3 20958370 20958481 refGene_gn=FAM189A1;refGene_tx=XM_023255006.1
B3 21012527 21012662 refGene_gn=FAM189A1;refGene_tx=XM_023255006.1
B3 21057342 21057450 refGene_gn=FAM189A1;refGene_tx=XM_023255006.1
B3 21066809 21066970 refGene_gn=FAM189A1;refGene_tx=XM_023255006.1
B3 21067568 21067704 refGene_gn=FAM189A1;refGene_tx=XM_023255006.1
B3 21074494 21074527 refGene_gn=APBA2;refGene_tx=XM_023255003.1
B3 21076118 21076194 refGene_gn=FAM189A1;refGene_tx=XM_023255006.1
B3 21081841 21082048 refGene_gn=FAM189A1;refGene_tx=XM_023255006.1
B3 21083638 21083713 refGene_gn=FAM189A1;refGene_tx=XM_023255006.1
B3 21084838 21085161 refGene_gn=FAM189A1;refGene_tx=XM_023255006.1
B3 21090460 21090532
refGene_gn=APBA2;refGene_tx=XM_023254998.1;refGene_gn=APBA2;refGene_tx
=XM_023254999.1;refGene_gn=APBA2;refGene_tx=XM_023255000.1;refGene_gn=
APBA2;refGene_tx=XM_023255001.1;refGene_gn=APBA2;refGene_tx=XM_0232550
02.1;refGene_gn=APBA2;refGene_tx=XM_023255004.1;refGene_gn=APBA2;refGe
ne_tx=XM_023255005.1
B3 21092926 21093067
refGene_gn=APBA2;refGene_tx=XM_023254998.1;refGene_gn=APBA2;refGene_tx
=XM_023254999.1;refGene_gn=APBA2;refGene_tx=XM_023255000.1;refGene_gn=
APBA2;refGene_tx=XM_023255001.1;refGene_gn=APBA2;refGene_tx=XM_0232550
02.1;refGene_gn=APBA2;refGene_tx=XM_023255003.1;refGene_gn=APBA2;refGe
ne_tx=XM_023255004.1
B3 21098350 21098470
refGene_gn=APBA2;refGene_tx=XM_023254998.1;refGene_gn=APBA2;refGene_tx
=XM_023254999.1;refGene_gn=APBA2;refGene_tx=XM_023255000.1;refGene_gn=
APBA2;refGene_tx=XM_023255001.1;refGene_gn=APBA2;refGene_tx=XM_0232550

02.1;refGene_gn=APBA2;refGene_tx=XM_023255003.1;refGene_gn=APBA2;refGene_tx=XM_023255004.1;refGene_gn=APBA2;refGene_tx=XM_023255005.1
B3 21100046 21100259
refGene_gn=APBA2;refGene_tx=XM_023254998.1;refGene_gn=APBA2;refGene_tx=XM_023254999.1;refGene_gn=APBA2;refGene_tx=XM_023255000.1;refGene_gn=APBA2;refGene_tx=XM_023255001.1;refGene_gn=APBA2;refGene_tx=XM_023255002.1;refGene_gn=APBA2;refGene_tx=XM_023255003.1;refGene_gn=APBA2;refGene_tx=XM_023255004.1;refGene_gn=APBA2;refGene_tx=XM_023255005.1
B3 21101305 21101485
refGene_gn=APBA2;refGene_tx=XM_023254998.1;refGene_gn=APBA2;refGene_tx=XM_023254999.1;refGene_gn=APBA2;refGene_tx=XM_023255000.1;refGene_gn=APBA2;refGene_tx=XM_023255001.1;refGene_gn=APBA2;refGene_tx=XM_023255002.1;refGene_gn=APBA2;refGene_tx=XM_023255003.1;refGene_gn=APBA2;refGene_tx=XM_023255004.1;refGene_gn=APBA2;refGene_tx=XM_023255005.1
B3 21105598 21105784
refGene_gn=APBA2;refGene_tx=XM_023254998.1;refGene_gn=APBA2;refGene_tx=XM_023254999.1;refGene_gn=APBA2;refGene_tx=XM_023255000.1;refGene_gn=APBA2;refGene_tx=XM_023255001.1;refGene_gn=APBA2;refGene_tx=XM_023255002.1;refGene_gn=APBA2;refGene_tx=XM_023255003.1;refGene_gn=APBA2;refGene_tx=XM_023255004.1;refGene_gn=APBA2;refGene_tx=XM_023255005.1
B3 21108753 21108840
refGene_gn=APBA2;refGene_tx=XM_023254998.1;refGene_gn=APBA2;refGene_tx=XM_023254999.1;refGene_gn=APBA2;refGene_tx=XM_023255000.1;refGene_gn=APBA2;refGene_tx=XM_023255001.1;refGene_gn=APBA2;refGene_tx=XM_023255002.1;refGene_gn=APBA2;refGene_tx=XM_023255003.1;refGene_gn=APBA2;refGene_tx=XM_023255004.1;refGene_gn=APBA2;refGene_tx=XM_023255005.1
B3 21111896 21111941
refGene_gn=APBA2;refGene_tx=XM_023254998.1;refGene_gn=APBA2;refGene_tx=XM_023254999.1;refGene_gn=APBA2;refGene_tx=XM_023255000.1;refGene_gn=APBA2;refGene_tx=XM_023255001.1;refGene_gn=APBA2;refGene_tx=XM_023255002.1;refGene_gn=APBA2;refGene_tx=XM_023255003.1;refGene_gn=APBA2;refGene_tx=XM_023255005.1
B3 21112912 21113058
refGene_gn=APBA2;refGene_tx=XM_023254998.1;refGene_gn=APBA2;refGene_tx=XM_023254999.1;refGene_gn=APBA2;refGene_tx=XM_023255000.1;refGene_gn=APBA2;refGene_tx=XM_023255001.1;refGene_gn=APBA2;refGene_tx=XM_023255002.1;refGene_gn=APBA2;refGene_tx=XM_023255003.1;refGene_gn=APBA2;refGene_tx=XM_023255004.1;refGene_gn=APBA2;refGene_tx=XM_023255005.1
B3 21132671 21132708
refGene_gn=APBA2;refGene_tx=XM_023254998.1;refGene_gn=APBA2;refGene_tx=XM_023254999.1;refGene_gn=APBA2;refGene_tx=XM_023255000.1;refGene_gn=APBA2;refGene_tx=XM_023255001.1;refGene_gn=APBA2;refGene_tx=XM_023255002.1;refGene_gn=APBA2;refGene_tx=XM_023255003.1;refGene_gn=APBA2;refGene_tx=XM_023255004.1;refGene_gn=APBA2;refGene_tx=XM_023255005.1
B3 21133834 21133915
refGene_gn=APBA2;refGene_tx=XM_023254998.1;refGene_gn=APBA2;refGene_tx=XM_023254999.1;refGene_gn=APBA2;refGene_tx=XM_023255000.1;refGene_gn=APBA2;refGene_tx=XM_023255001.1;refGene_gn=APBA2;refGene_tx=XM_023255002.1;refGene_gn=APBA2;refGene_tx=XM_023255003.1;refGene_gn=APBA2;refGene_tx=XM_023255004.1;refGene_gn=APBA2;refGene_tx=XM_023255005.1

B3 21156247 21157213
refGene_gn=APBA2;refGene_tx=XM_023254998.1;refGene_gn=APBA2;refGene_tx=XM_023254999.1;refGene_gn=APBA2;refGene_tx=XM_023255000.1;refGene_gn=APBA2;refGene_tx=XM_023255001.1;refGene_gn=APBA2;refGene_tx=XM_023255002.1;refGene_gn=APBA2;refGene_tx=XM_023255003.1;refGene_gn=APBA2;refGene_tx=XM_023255004.1;refGene_gn=APBA2;refGene_tx=XM_023255005.1
B3 21410282 21410435 refGene_gn=MCEE;refGene_tx=XM_003986891.5
B3 21429475 21429813 refGene_gn=MCEE;refGene_tx=XM_003986891.5
B3 21438967 21439013 refGene_gn=MCEE;refGene_tx=XM_003986891.5
B3 21439462 21439551
refGene_gn=MPHOSPH10;refGene_tx=XM_003986892.5;refGene_gn=MPHOSPH10;refGene_tx=XM_019831959.2
B3 21441175 21441848
refGene_gn=MPHOSPH10;refGene_tx=XM_003986892.5;refGene_gn=MPHOSPH10;refGene_tx=XM_019831959.2
B3 21442929 21443081
refGene_gn=MPHOSPH10;refGene_tx=XM_003986892.5;refGene_gn=MPHOSPH10;refGene_tx=XM_019831959.2
B3 21443578 21443750
refGene_gn=MPHOSPH10;refGene_tx=XM_003986892.5;refGene_gn=MPHOSPH10;refGene_tx=XM_019831959.2
B3 21446681 21446823
refGene_gn=MPHOSPH10;refGene_tx=XM_003986892.5;refGene_gn=MPHOSPH10;refGene_tx=XM_019831959.2
B3 21447679 21447747
refGene_gn=MPHOSPH10;refGene_tx=XM_003986892.5;refGene_gn=MPHOSPH10;refGene_tx=XM_019831959.2
B3 21448670 21448808
refGene_gn=MPHOSPH10;refGene_tx=XM_003986892.5;refGene_gn=MPHOSPH10;refGene_tx=XM_019831959.2
B3 21451101 21451212
refGene_gn=MPHOSPH10;refGene_tx=XM_003986892.5;refGene_gn=MPHOSPH10;refGene_tx=XM_019831959.2
B3 21457064 21457173
refGene_gn=MPHOSPH10;refGene_tx=XM_003986892.5;refGene_gn=MPHOSPH10;refGene_tx=XM_019831959.2
B3 21458163 21458394
refGene_gn=MPHOSPH10;refGene_tx=XM_003986892.5;refGene_gn=MPHOSPH10;refGene_tx=XM_019831959.2
B3 21458786 21458936
refGene_gn=MPHOSPH10;refGene_tx=XM_003986892.5
B3 21471243 21472507
refGene_gn=FAN1;refGene_tx=XM_006932318.3;refGene_gn=FAN1;refGene_tx=XM_019831972.2;refGene_gn=FAN1;refGene_tx=XM_019831973.1;refGene_gn=FAN1;refGene_tx=XM_019831974.1
B3 21474447 21474588
refGene_gn=FAN1;refGene_tx=XM_006932318.3;refGene_gn=FAN1;refGene_tx=XM_019831972.2;refGene_gn=FAN1;refGene_tx=XM_019831973.1;refGene_gn=FAN1;refGene_tx=XM_019831974.1
B3 21476706 21476908

refGene_gn=FAN1;refGene_tx=XM_006932318.3;refGene_gn=FAN1;refGene_tx=XM_019831972.2;refGene_gn=FAN1;refGene_tx=XM_019831973.1;refGene_gn=FAN1;refGene_tx=XM_019831974.1
B3 21482356 21482590
refGene_gn=FAN1;refGene_tx=XM_006932318.3;refGene_gn=FAN1;refGene_tx=XM_019831972.2;refGene_gn=FAN1;refGene_tx=XM_019831973.1;refGene_gn=FAN1;refGene_tx=XM_019831974.1
B3 21487707 21487839
refGene_gn=FAN1;refGene_tx=XM_006932318.3;refGene_gn=FAN1;refGene_tx=XM_019831972.2;refGene_gn=FAN1;refGene_tx=XM_019831973.1;refGene_gn=FAN1;refGene_tx=XM_019831974.1
B3 21490738 21490847
refGene_gn=FAN1;refGene_tx=XM_006932318.3;refGene_gn=FAN1;refGene_tx=XM_019831972.2;refGene_gn=FAN1;refGene_tx=XM_019831973.1;refGene_gn=FAN1;refGene_tx=XM_019831974.1
B3 21491985 21492105
refGene_gn=FAN1;refGene_tx=XM_006932318.3;refGene_gn=FAN1;refGene_tx=XM_019831972.2;refGene_gn=FAN1;refGene_tx=XM_019831973.1;refGene_gn=FAN1;refGene_tx=XM_019831974.1
B3 21495111 21495276
refGene_gn=FAN1;refGene_tx=XM_006932318.3;refGene_gn=FAN1;refGene_tx=XM_019831972.2;refGene_gn=FAN1;refGene_tx=XM_019831973.1;refGene_gn=FAN1;refGene_tx=XM_019831974.1
B3 21495789 21495940
refGene_gn=FAN1;refGene_tx=XM_006932318.3;refGene_gn=FAN1;refGene_tx=XM_019831972.2;refGene_gn=FAN1;refGene_tx=XM_019831973.1;refGene_gn=FAN1;refGene_tx=XM_019831974.1
B3 21499503 21499607
refGene_gn=FAN1;refGene_tx=XM_006932318.3;refGene_gn=FAN1;refGene_tx=XM_019831972.2;refGene_gn=FAN1;refGene_tx=XM_019831973.1;refGene_gn=FAN1;refGene_tx=XM_019831974.1
B3 21500181 21500376
refGene_gn=FAN1;refGene_tx=XM_006932318.3;refGene_gn=FAN1;refGene_tx=XM_019831972.2;refGene_gn=FAN1;refGene_tx=XM_019831973.1;refGene_gn=FAN1;refGene_tx=XM_019831974.1
B3 21500995 21501124
refGene_gn=FAN1;refGene_tx=XM_006932318.3;refGene_gn=FAN1;refGene_tx=XM_019831972.2;refGene_gn=FAN1;refGene_tx=XM_019831973.1;refGene_gn=FAN1;refGene_tx=XM_019831974.1
B3 21502844 21502910
refGene_gn=FAN1;refGene_tx=XM_019831973.1;refGene_gn=FAN1;refGene_tx=XM_019831974.1
B3 21504138 21504276
refGene_gn=FAN1;refGene_tx=XM_006932318.3;refGene_gn=FAN1;refGene_tx=XM_019831972.2
B3 21508298 21508892
refGene_gn=MTMR10;refGene_tx=XM_006932321.3;refGene_gn=MTMR10;refGene_tx=XM_023255007.1
B3 21509494 21509677
refGene_gn=MTMR10;refGene_tx=XM_006932321.3;refGene_gn=MTMR10;refGene_

tx=XM_023255007.1
B3 21513669 21513840
refGene_gn=MTMR10;refGene_tx=XM_006932321.3;refGene_gn=MTMR10;refGene_
tx=XM_023255007.1
B3 21515598 21515768
refGene_gn=MTMR10;refGene_tx=XM_006932321.3;refGene_gn=MTMR10;refGene_
tx=XM_023255007.1
B3 21518026 21518097
refGene_gn=MTMR10;refGene_tx=XM_006932321.3;refGene_gn=MTMR10;refGene_
tx=XM_023255007.1
B3 21518749 21518819
refGene_gn=MTMR10;refGene_tx=XM_006932321.3;refGene_gn=MTMR10;refGene_
tx=XM_023255007.1
B3 21520063 21520194
refGene_gn=MTMR10;refGene_tx=XM_006932321.3;refGene_gn=MTMR10;refGene_
tx=XM_023255007.1
B3 21522537 21522626
refGene_gn=MTMR10;refGene_tx=XM_006932321.3;refGene_gn=MTMR10;refGene_
tx=XM_023255007.1
B3 21522707 21522795
refGene_gn=MTMR10;refGene_tx=XM_006932321.3;refGene_gn=MTMR10;refGene_
tx=XM_023255007.1
B3 21524076 21524248
refGene_gn=MTMR10;refGene_tx=XM_006932321.3;refGene_gn=MTMR10;refGene_
tx=XM_023255007.1
B3 21530391 21530482
refGene_gn=MTMR10;refGene_tx=XM_006932321.3;refGene_gn=MTMR10;refGene_
tx=XM_023255007.1
B3 21537557 21537700
refGene_gn=MTMR10;refGene_tx=XM_006932321.3;refGene_gn=MTMR10;refGene_
tx=XM_023255007.1
B3 21538362 21538435
refGene_gn=MTMR10;refGene_tx=XM_006932321.3;refGene_gn=MTMR10;refGene_
tx=XM_023255007.1
B3 21540229 21540366
refGene_gn=MTMR10;refGene_tx=XM_006932321.3;refGene_gn=MTMR10;refGene_
tx=XM_023255007.1
B3 21560637 21560698 refGene_gn=MTMR10;refGene_tx=XM_006932321.3
B3 21561277 21561337 refGene_gn=MTMR10;refGene_tx=XM_006932321.3
B3 21572853 21574120
refGene_gn=TRPM1;refGene_tx=XM_019831960.2;refGene_gn=TRPM1;refGene_tx
=XM_019831961.2;refGene_gn=TRPM1;refGene_tx=XM_019831962.2;refGene_gn=
TRPM1;refGene_tx=XM_019831963.2;refGene_gn=TRPM1;refGene_tx=XM_0198319
64.2;refGene_gn=TRPM1;refGene_tx=XM_019831965.2;refGene_gn=TRPM1;refGe
ne_tx=XM_019831966.2;refGene_gn=TRPM1;refGene_tx=XM_019831967.2;refGen
e_gn=TRPM1;refGene_tx=XM_019831968.2;refGene_gn=TRPM1;refGene_tx=XM_01
9831969.2;refGene_gn=TRPM1;refGene_tx=XM_019831971.2
B3 21577296 21577330 refGene_gn=TRPM1;refGene_tx=XM_023255008.1
B3 21594824 21594957
refGene_gn=TRPM1;refGene_tx=XM_019831960.2;refGene_gn=TRPM1;refGene_tx

=XM_019831961.2;refGene_gn=TRPM1;refGene_tx=XM_019831962.2;refGene_gn=TRPM1;refGene_tx=XM_019831963.2;refGene_gn=TRPM1;refGene_tx=XM_019831964.2;refGene_gn=TRPM1;refGene_tx=XM_019831965.2;refGene_gn=TRPM1;refGene_tx=XM_019831966.2;refGene_gn=TRPM1;refGene_tx=XM_019831967.2;refGene_gn=TRPM1;refGene_tx=XM_019831968.2;refGene_gn=TRPM1;refGene_tx=XM_019831969.2;refGene_gn=TRPM1;refGene_tx=XM_019831971.2;refGene_gn=TRPM1;refGene_tx=XM_023255008.1
B3 21595583 21595786
refGene_gn=TRPM1;refGene_tx=XM_019831960.2;refGene_gn=TRPM1;refGene_tx=XM_019831961.2;refGene_gn=TRPM1;refGene_tx=XM_019831962.2;refGene_gn=TRPM1;refGene_tx=XM_019831963.2;refGene_gn=TRPM1;refGene_tx=XM_019831964.2;refGene_gn=TRPM1;refGene_tx=XM_019831965.2;refGene_gn=TRPM1;refGene_tx=XM_019831966.2;refGene_gn=TRPM1;refGene_tx=XM_019831967.2;refGene_gn=TRPM1;refGene_tx=XM_019831968.2;refGene_gn=TRPM1;refGene_tx=XM_019831969.2;refGene_gn=TRPM1;refGene_tx=XM_019831971.2;refGene_gn=TRPM1;refGene_tx=XM_023255008.1
B3 21597303 21597448
refGene_gn=TRPM1;refGene_tx=XM_019831960.2;refGene_gn=TRPM1;refGene_tx=XM_019831961.2;refGene_gn=TRPM1;refGene_tx=XM_019831962.2;refGene_gn=TRPM1;refGene_tx=XM_019831963.2;refGene_gn=TRPM1;refGene_tx=XM_019831964.2;refGene_gn=TRPM1;refGene_tx=XM_019831965.2;refGene_gn=TRPM1;refGene_tx=XM_019831966.2;refGene_gn=TRPM1;refGene_tx=XM_019831967.2;refGene_gn=TRPM1;refGene_tx=XM_019831968.2;refGene_gn=TRPM1;refGene_tx=XM_019831969.2;refGene_gn=TRPM1;refGene_tx=XM_019831971.2;refGene_gn=TRPM1;refGene_tx=XM_023255008.1
B3 21598188 21598209
refGene_gn=TRPM1;refGene_tx=XM_019831960.2;refGene_gn=TRPM1;refGene_tx=XM_019831961.2;refGene_gn=TRPM1;refGene_tx=XM_019831963.2;refGene_gn=TRPM1;refGene_tx=XM_019831965.2;refGene_gn=TRPM1;refGene_tx=XM_019831966.2;refGene_gn=TRPM1;refGene_tx=XM_019831967.2;refGene_gn=TRPM1;refGene_tx=XM_019831968.2;refGene_gn=TRPM1;refGene_tx=XM_019831969.2;refGene_gn=TRPM1;refGene_tx=XM_023255008.1
B3 21599696 21599889
refGene_gn=TRPM1;refGene_tx=XM_019831960.2;refGene_gn=TRPM1;refGene_tx=XM_019831961.2;refGene_gn=TRPM1;refGene_tx=XM_019831962.2;refGene_gn=TRPM1;refGene_tx=XM_019831963.2;refGene_gn=TRPM1;refGene_tx=XM_019831964.2;refGene_gn=TRPM1;refGene_tx=XM_019831965.2;refGene_gn=TRPM1;refGene_tx=XM_019831966.2;refGene_gn=TRPM1;refGene_tx=XM_019831967.2;refGene_gn=TRPM1;refGene_tx=XM_019831968.2;refGene_gn=TRPM1;refGene_tx=XM_019831969.2;refGene_gn=TRPM1;refGene_tx=XM_019831971.2;refGene_gn=TRPM1;refGene_tx=XM_023255008.1
B3 21601008 21601260
refGene_gn=TRPM1;refGene_tx=XM_019831960.2;refGene_gn=TRPM1;refGene_tx=XM_019831961.2;refGene_gn=TRPM1;refGene_tx=XM_019831962.2;refGene_gn=TRPM1;refGene_tx=XM_019831963.2;refGene_gn=TRPM1;refGene_tx=XM_019831964.2;refGene_gn=TRPM1;refGene_tx=XM_019831965.2;refGene_gn=TRPM1;refGene_tx=XM_019831966.2;refGene_gn=TRPM1;refGene_tx=XM_019831967.2;refGene_gn=TRPM1;refGene_tx=XM_019831968.2;refGene_gn=TRPM1;refGene_tx=XM_019831969.2;refGene_gn=TRPM1;refGene_tx=XM_019831971.2;refGene_gn=TRPM1;refGene_tx=XM_023255008.1

B3 21603150 21603279
refGene_gn=TRPM1;refGene_tx=XM_019831960.2;refGene_gn=TRPM1;refGene_tx=XM_019831961.2;refGene_gn=TRPM1;refGene_tx=XM_019831962.2;refGene_gn=TRPM1;refGene_tx=XM_019831963.2;refGene_gn=TRPM1;refGene_tx=XM_019831964.2;refGene_gn=TRPM1;refGene_tx=XM_019831965.2;refGene_gn=TRPM1;refGene_tx=XM_019831966.2;refGene_gn=TRPM1;refGene_tx=XM_019831967.2;refGene_gn=TRPM1;refGene_tx=XM_019831968.2;refGene_gn=TRPM1;refGene_tx=XM_019831969.2;refGene_gn=TRPM1;refGene_tx=XM_019831971.2;refGene_gn=TRPM1;refGene_tx=XM_023255008.1
B3 21605276 21605408
refGene_gn=TRPM1;refGene_tx=XM_019831960.2;refGene_gn=TRPM1;refGene_tx=XM_019831961.2;refGene_gn=TRPM1;refGene_tx=XM_019831962.2;refGene_gn=TRPM1;refGene_tx=XM_019831963.2;refGene_gn=TRPM1;refGene_tx=XM_019831964.2;refGene_gn=TRPM1;refGene_tx=XM_019831965.2;refGene_gn=TRPM1;refGene_tx=XM_019831966.2;refGene_gn=TRPM1;refGene_tx=XM_019831967.2;refGene_gn=TRPM1;refGene_tx=XM_019831968.2;refGene_gn=TRPM1;refGene_tx=XM_019831969.2;refGene_gn=TRPM1;refGene_tx=XM_019831971.2;refGene_gn=TRPM1;refGene_tx=XM_023255008.1
B3 21605607 21605730
refGene_gn=TRPM1;refGene_tx=XM_019831960.2;refGene_gn=TRPM1;refGene_tx=XM_019831961.2;refGene_gn=TRPM1;refGene_tx=XM_019831962.2;refGene_gn=TRPM1;refGene_tx=XM_019831963.2;refGene_gn=TRPM1;refGene_tx=XM_019831964.2;refGene_gn=TRPM1;refGene_tx=XM_019831965.2;refGene_gn=TRPM1;refGene_tx=XM_019831966.2;refGene_gn=TRPM1;refGene_tx=XM_019831967.2;refGene_gn=TRPM1;refGene_tx=XM_019831968.2;refGene_gn=TRPM1;refGene_tx=XM_019831969.2;refGene_gn=TRPM1;refGene_tx=XM_019831971.2;refGene_gn=TRPM1;refGene_tx=XM_023255008.1
B3 21618227 21618456
refGene_gn=TRPM1;refGene_tx=XM_019831960.2;refGene_gn=TRPM1;refGene_tx=XM_019831961.2;refGene_gn=TRPM1;refGene_tx=XM_019831962.2;refGene_gn=TRPM1;refGene_tx=XM_019831963.2;refGene_gn=TRPM1;refGene_tx=XM_019831964.2;refGene_gn=TRPM1;refGene_tx=XM_019831965.2;refGene_gn=TRPM1;refGene_tx=XM_019831966.2;refGene_gn=TRPM1;refGene_tx=XM_019831967.2;refGene_gn=TRPM1;refGene_tx=XM_019831968.2;refGene_gn=TRPM1;refGene_tx=XM_019831969.2;refGene_gn=TRPM1;refGene_tx=XM_019831971.2;refGene_gn=TRPM1;refGene_tx=XM_023255008.1
B3 21620739 21621032
refGene_gn=TRPM1;refGene_tx=XM_019831960.2;refGene_gn=TRPM1;refGene_tx=XM_019831961.2;refGene_gn=TRPM1;refGene_tx=XM_019831962.2;refGene_gn=TRPM1;refGene_tx=XM_019831963.2;refGene_gn=TRPM1;refGene_tx=XM_019831964.2;refGene_gn=TRPM1;refGene_tx=XM_019831965.2;refGene_gn=TRPM1;refGene_tx=XM_019831966.2;refGene_gn=TRPM1;refGene_tx=XM_019831967.2;refGene_gn=TRPM1;refGene_tx=XM_019831968.2;refGene_gn=TRPM1;refGene_tx=XM_019831969.2;refGene_gn=TRPM1;refGene_tx=XM_019831971.2;refGene_gn=TRPM1;refGene_tx=XM_023255008.1
B3 21622994 21623024
refGene_gn=TRPM1;refGene_tx=XM_019831960.2;refGene_gn=TRPM1;refGene_tx=XM_019831961.2;refGene_gn=TRPM1;refGene_tx=XM_019831962.2;refGene_gn=TRPM1;refGene_tx=XM_019831964.2;refGene_gn=TRPM1;refGene_tx=XM_019831965.2;refGene_gn=TRPM1;refGene_tx=XM_019831967.2;refGene_gn=TRPM1;refGe

ne_tx=XM_019831968.2;refGene_gn=TRPM1;refGene_tx=XM_019831969.2;refGen
e_gn=TRPM1;refGene_tx=XM_023255008.1
B3 21623945 21624086
refGene_gn=TRPM1;refGene_tx=XM_019831960.2;refGene_gn=TRPM1;refGene_tx
=XM_019831961.2;refGene_gn=TRPM1;refGene_tx=XM_019831962.2;refGene_gn=
TRPM1;refGene_tx=XM_019831963.2;refGene_gn=TRPM1;refGene_tx=XM_0198319
64.2;refGene_gn=TRPM1;refGene_tx=XM_019831965.2;refGene_gn=TRPM1;refGe
ne_tx=XM_019831967.2;refGene_gn=TRPM1;refGene_tx=XM_019831968.2;refGen
e_gn=TRPM1;refGene_tx=XM_019831969.2;refGene_gn=TRPM1;refGene_tx=XM_01
9831971.2;refGene_gn=TRPM1;refGene_tx=XM_023255008.1
B3 21627151 21627202
refGene_gn=TRPM1;refGene_tx=XM_019831960.2;refGene_gn=TRPM1;refGene_tx
=XM_019831961.2;refGene_gn=TRPM1;refGene_tx=XM_019831962.2;refGene_gn=
TRPM1;refGene_tx=XM_019831963.2;refGene_gn=TRPM1;refGene_tx=XM_0198319
64.2;refGene_gn=TRPM1;refGene_tx=XM_019831965.2;refGene_gn=TRPM1;refGe
ne_tx=XM_019831966.2;refGene_gn=TRPM1;refGene_tx=XM_019831967.2;refGen
e_gn=TRPM1;refGene_tx=XM_019831968.2;refGene_gn=TRPM1;refGene_tx=XM_01
9831969.2;refGene_gn=TRPM1;refGene_tx=XM_019831971.2;refGene_gn=TRPM1;
refGene_tx=XM_023255008.1
B3 21627417 21627552
refGene_gn=TRPM1;refGene_tx=XM_019831960.2;refGene_gn=TRPM1;refGene_tx
=XM_019831961.2;refGene_gn=TRPM1;refGene_tx=XM_019831962.2;refGene_gn=
TRPM1;refGene_tx=XM_019831963.2;refGene_gn=TRPM1;refGene_tx=XM_0198319
64.2;refGene_gn=TRPM1;refGene_tx=XM_019831965.2;refGene_gn=TRPM1;refGe
ne_tx=XM_019831966.2;refGene_gn=TRPM1;refGene_tx=XM_019831967.2;refGen
e_gn=TRPM1;refGene_tx=XM_019831968.2;refGene_gn=TRPM1;refGene_tx=XM_01
9831969.2;refGene_gn=TRPM1;refGene_tx=XM_019831971.2;refGene_gn=TRPM1;
refGene_tx=XM_023255008.1
B3 21628813 21628987
refGene_gn=TRPM1;refGene_tx=XM_019831960.2;refGene_gn=TRPM1;refGene_tx
=XM_019831961.2;refGene_gn=TRPM1;refGene_tx=XM_019831962.2;refGene_gn=
TRPM1;refGene_tx=XM_019831963.2;refGene_gn=TRPM1;refGene_tx=XM_0198319
64.2;refGene_gn=TRPM1;refGene_tx=XM_019831965.2;refGene_gn=TRPM1;refGe
ne_tx=XM_019831966.2;refGene_gn=TRPM1;refGene_tx=XM_019831967.2;refGen
e_gn=TRPM1;refGene_tx=XM_019831968.2;refGene_gn=TRPM1;refGene_tx=XM_01
9831969.2;refGene_gn=TRPM1;refGene_tx=XM_019831971.2;refGene_gn=TRPM1;
refGene_tx=XM_023255008.1
B3 21637360 21637461
refGene_gn=TRPM1;refGene_tx=XM_019831960.2;refGene_gn=TRPM1;refGene_tx
=XM_019831961.2;refGene_gn=TRPM1;refGene_tx=XM_019831962.2;refGene_gn=
TRPM1;refGene_tx=XM_019831963.2;refGene_gn=TRPM1;refGene_tx=XM_0198319
64.2;refGene_gn=TRPM1;refGene_tx=XM_019831965.2;refGene_gn=TRPM1;refGe
ne_tx=XM_019831966.2;refGene_gn=TRPM1;refGene_tx=XM_019831967.2;refGen
e_gn=TRPM1;refGene_tx=XM_019831968.2;refGene_gn=TRPM1;refGene_tx=XM_01
9831969.2;refGene_gn=TRPM1;refGene_tx=XM_019831971.2;refGene_gn=TRPM1;
refGene_tx=XM_023255008.1
B3 21638365 21638438
refGene_gn=TRPM1;refGene_tx=XM_019831960.2;refGene_gn=TRPM1;refGene_tx
=XM_019831961.2;refGene_gn=TRPM1;refGene_tx=XM_019831962.2;refGene_gn=
TRPM1;refGene_tx=XM_019831963.2;refGene_gn=TRPM1;refGene_tx=XM_0198319

64.2;refGene_gn=TRPM1;refGene_tx=XM_019831965.2;refGene_gn=TRPM1;refGene_tx=XM_019831966.2;refGene_gn=TRPM1;refGene_tx=XM_019831967.2;refGene_gn=TRPM1;refGene_tx=XM_019831968.2;refGene_gn=TRPM1;refGene_tx=XM_019831969.2;refGene_gn=TRPM1;refGene_tx=XM_019831971.2;refGene_gn=TRPM1;refGene_tx=XM_023255008.1
B3 21639444 21639568
refGene_gn=TRPM1;refGene_tx=XM_019831960.2;refGene_gn=TRPM1;refGene_tx=XM_019831961.2;refGene_gn=TRPM1;refGene_tx=XM_019831962.2;refGene_gn=TRPM1;refGene_tx=XM_019831963.2;refGene_gn=TRPM1;refGene_tx=XM_019831964.2;refGene_gn=TRPM1;refGene_tx=XM_019831965.2;refGene_gn=TRPM1;refGene_tx=XM_019831966.2;refGene_gn=TRPM1;refGene_tx=XM_019831967.2;refGene_gn=TRPM1;refGene_tx=XM_019831968.2;refGene_gn=TRPM1;refGene_tx=XM_019831969.2;refGene_gn=TRPM1;refGene_tx=XM_019831971.2;refGene_gn=TRPM1;refGene_tx=XM_023255008.1
B3 21639999 21640178
refGene_gn=TRPM1;refGene_tx=XM_019831960.2;refGene_gn=TRPM1;refGene_tx=XM_019831961.2;refGene_gn=TRPM1;refGene_tx=XM_019831962.2;refGene_gn=TRPM1;refGene_tx=XM_019831963.2;refGene_gn=TRPM1;refGene_tx=XM_019831964.2;refGene_gn=TRPM1;refGene_tx=XM_019831965.2;refGene_gn=TRPM1;refGene_tx=XM_019831966.2;refGene_gn=TRPM1;refGene_tx=XM_019831967.2;refGene_gn=TRPM1;refGene_tx=XM_019831968.2;refGene_gn=TRPM1;refGene_tx=XM_019831969.2;refGene_gn=TRPM1;refGene_tx=XM_019831970.1;refGene_gn=TRPM1;refGene_tx=XM_019831971.2;refGene_gn=TRPM1;refGene_tx=XM_023255008.1
B3 21642031 21642203
refGene_gn=TRPM1;refGene_tx=XM_019831960.2;refGene_gn=TRPM1;refGene_tx=XM_019831961.2;refGene_gn=TRPM1;refGene_tx=XM_019831962.2;refGene_gn=TRPM1;refGene_tx=XM_019831963.2;refGene_gn=TRPM1;refGene_tx=XM_019831964.2;refGene_gn=TRPM1;refGene_tx=XM_019831965.2;refGene_gn=TRPM1;refGene_tx=XM_019831966.2;refGene_gn=TRPM1;refGene_tx=XM_019831967.2;refGene_gn=TRPM1;refGene_tx=XM_019831968.2;refGene_gn=TRPM1;refGene_tx=XM_019831969.2;refGene_gn=TRPM1;refGene_tx=XM_019831970.1;refGene_gn=TRPM1;refGene_tx=XM_019831971.2;refGene_gn=TRPM1;refGene_tx=XM_023255008.1
B3 21643087 21643212
refGene_gn=TRPM1;refGene_tx=XM_019831960.2;refGene_gn=TRPM1;refGene_tx=XM_019831961.2;refGene_gn=TRPM1;refGene_tx=XM_019831962.2;refGene_gn=TRPM1;refGene_tx=XM_019831963.2;refGene_gn=TRPM1;refGene_tx=XM_019831964.2;refGene_gn=TRPM1;refGene_tx=XM_019831965.2;refGene_gn=TRPM1;refGene_tx=XM_019831966.2;refGene_gn=TRPM1;refGene_tx=XM_019831967.2;refGene_gn=TRPM1;refGene_tx=XM_019831968.2;refGene_gn=TRPM1;refGene_tx=XM_019831969.2;refGene_gn=TRPM1;refGene_tx=XM_019831970.1;refGene_gn=TRPM1;refGene_tx=XM_019831971.2;refGene_gn=TRPM1;refGene_tx=XM_023255008.1
B3 21643861 21644075
refGene_gn=TRPM1;refGene_tx=XM_019831960.2;refGene_gn=TRPM1;refGene_tx=XM_019831961.2;refGene_gn=TRPM1;refGene_tx=XM_019831962.2;refGene_gn=TRPM1;refGene_tx=XM_019831963.2;refGene_gn=TRPM1;refGene_tx=XM_019831964.2;refGene_gn=TRPM1;refGene_tx=XM_019831965.2;refGene_gn=TRPM1;refGene_tx=XM_019831966.2;refGene_gn=TRPM1;refGene_tx=XM_019831967.2;refGene_gn=TRPM1;refGene_tx=XM_019831968.2;refGene_gn=TRPM1;refGene_tx=XM_019831969.2;refGene_gn=TRPM1;refGene_tx=XM_019831970.1;refGene_gn=TRPM1;refGene_tx=XM_019831971.2;refGene_gn=TRPM1;refGene_tx=XM_023255008.1

B3 21645335 21645531
refGene_gn=TRPM1;refGene_tx=XM_019831960.2;refGene_gn=TRPM1;refGene_tx=XM_019831961.2;refGene_gn=TRPM1;refGene_tx=XM_019831962.2;refGene_gn=TRPM1;refGene_tx=XM_019831963.2;refGene_gn=TRPM1;refGene_tx=XM_019831964.2;refGene_gn=TRPM1;refGene_tx=XM_019831965.2;refGene_gn=TRPM1;refGene_tx=XM_019831966.2;refGene_gn=TRPM1;refGene_tx=XM_019831967.2;refGene_gn=TRPM1;refGene_tx=XM_019831968.2;refGene_gn=TRPM1;refGene_tx=XM_019831969.2;refGene_gn=TRPM1;refGene_tx=XM_019831970.1;refGene_gn=TRPM1;refGene_tx=XM_019831971.2;refGene_gn=TRPM1;refGene_tx=XM_023255008.1
B3 21650178 21650258
refGene_gn=TRPM1;refGene_tx=XM_019831960.2;refGene_gn=TRPM1;refGene_tx=XM_019831961.2;refGene_gn=TRPM1;refGene_tx=XM_019831962.2;refGene_gn=TRPM1;refGene_tx=XM_019831963.2;refGene_gn=TRPM1;refGene_tx=XM_019831964.2;refGene_gn=TRPM1;refGene_tx=XM_019831965.2;refGene_gn=TRPM1;refGene_tx=XM_019831966.2;refGene_gn=TRPM1;refGene_tx=XM_019831967.2;refGene_gn=TRPM1;refGene_tx=XM_019831968.2;refGene_gn=TRPM1;refGene_tx=XM_019831969.2;refGene_gn=TRPM1;refGene_tx=XM_019831970.1;refGene_gn=TRPM1;refGene_tx=XM_019831971.2;refGene_gn=TRPM1;refGene_tx=XM_023255008.1
B3 21673323 21673460
refGene_gn=TRPM1;refGene_tx=XM_019831960.2;refGene_gn=TRPM1;refGene_tx=XM_019831961.2;refGene_gn=TRPM1;refGene_tx=XM_019831962.2;refGene_gn=TRPM1;refGene_tx=XM_019831963.2;refGene_gn=TRPM1;refGene_tx=XM_019831964.2;refGene_gn=TRPM1;refGene_tx=XM_019831965.2;refGene_gn=TRPM1;refGene_tx=XM_019831966.2;refGene_gn=TRPM1;refGene_tx=XM_019831970.1;refGene_gn=TRPM1;refGene_tx=XM_019831971.2;refGene_gn=TRPM1;refGene_tx=XM_023255008.1
B3 21676807 21676837 refGene_gn=TRPM1;refGene_tx=XM_019831969.2
B3 21677030 21677175
refGene_gn=TRPM1;refGene_tx=XM_019831960.2;refGene_gn=TRPM1;refGene_tx=XM_019831961.2;refGene_gn=TRPM1;refGene_tx=XM_019831962.2;refGene_gn=TRPM1;refGene_tx=XM_019831963.2;refGene_gn=TRPM1;refGene_tx=XM_019831964.2;refGene_gn=TRPM1;refGene_tx=XM_019831965.2;refGene_gn=TRPM1;refGene_tx=XM_019831966.2;refGene_gn=TRPM1;refGene_tx=XM_019831970.1;refGene_gn=TRPM1;refGene_tx=XM_019831971.2;refGene_gn=TRPM1;refGene_tx=XM_023255008.1
B3 21717846 21717921
refGene_gn=TRPM1;refGene_tx=XM_019831967.2;refGene_gn=TRPM1;refGene_tx=XM_019831968.2
B3 21867125 21867626 refGene_gn=KLF13;refGene_tx=XM_023255009.1
B3 21867627 21867718 refGene_gn=KLF13;refGene_tx=XM_023255009.1
B3 21911481 21911771 refGene_gn=KLF13;refGene_tx=XM_023255009.1
B3 22004440 22005892 refGene_gn=OTUD7A;refGene_tx=XM_023255010.1
B3 22008231 22008316 refGene_gn=OTUD7A;refGene_tx=XM_023255010.1
B3 22008497 22008612 refGene_gn=OTUD7A;refGene_tx=XM_023255010.1
B3 22025659 22025809 refGene_gn=OTUD7A;refGene_tx=XM_023255010.1
B3 22027747 22027875 refGene_gn=OTUD7A;refGene_tx=XM_023255010.1
B3 22047891 22048004 refGene_gn=OTUD7A;refGene_tx=XM_023255010.1
B3 22048822 22048950 refGene_gn=OTUD7A;refGene_tx=XM_023255010.1
B3 22052501 22052603 refGene_gn=OTUD7A;refGene_tx=XM_023255010.1
B3 22079627 22079846 refGene_gn=OTUD7A;refGene_tx=XM_023255010.1

B3 22089338 22089521 refGene_gn=OTUD7A;refGene_tx=XM_023255010.1
B3 22155991 22156142 refGene_gn=OTUD7A;refGene_tx=XM_023255010.1
B3 22551500 22551546
refGene_gn=CHRNA7;refGene_tx=XM_011282725.3;refGene_gn=CHRNA7;refGene_tx=XM_019831978.2
B3 22551811 22551951
refGene_gn=CHRNA7;refGene_tx=XM_011282725.3;refGene_gn=CHRNA7;refGene_tx=XM_019831978.2
B3 22601467 22601512
refGene_gn=CHRNA7;refGene_tx=XM_011282725.3;refGene_gn=CHRNA7;refGene_tx=XM_019831978.2
B3 22613932 22614042
refGene_gn=CHRNA7;refGene_tx=XM_011282725.3;refGene_gn=CHRNA7;refGene_tx=XM_019831978.2
B3 22649503 22649583
refGene_gn=CHRNA7;refGene_tx=XM_011282725.3;refGene_gn=CHRNA7;refGene_tx=XM_019831978.2
B3 22654969 22655137
refGene_gn=CHRNA7;refGene_tx=XM_011282725.3;refGene_gn=CHRNA7;refGene_tx=XM_019831978.2
B3 22655915 22656110
refGene_gn=CHRNA7;refGene_tx=XM_011282725.3;refGene_gn=CHRNA7;refGene_tx=XM_019831978.2
B3 22657106 22657193
refGene_gn=CHRNA7;refGene_tx=XM_011282725.3;refGene_gn=CHRNA7;refGene_tx=XM_019831978.2
B3 22660063 22660173 refGene_gn=CHRNA7;refGene_tx=XM_011282725.3
B3 22664478 22664997
refGene_gn=CHRNA7;refGene_tx=XM_011282725.3;refGene_gn=CHRNA7;refGene_tx=XM_019831978.2
B3 23158969 23160460 refGene_gn=MKRN3;refGene_tx=XM_023256161.1
B3 23203261 23206822 refGene_gn=MAGEL2;refGene_tx=XM_019831979.2
B3 23250388 23251363 refGene_gn=NDN;refGene_tx=XM_011282821.3
B3 24041165 24041179
refGene_gn=SNURF;refGene_tx=XM_003986901.4;refGene_gn=SNURF;refGene_tx=XM_006932326.3;refGene_gn=SNURF;refGene_tx=XM_006932327.2;refGene_gn=SNURF;refGene_tx=XM_006932328.2
B3 24048124 24048220
refGene_gn=SNURF;refGene_tx=XM_003986901.4;refGene_gn=SNURF;refGene_tx=XM_006932326.3;refGene_gn=SNURF;refGene_tx=XM_006932327.2;refGene_gn=SNURF;refGene_tx=XM_006932328.2
B3 24053671 24053777
refGene_gn=SNURF;refGene_tx=XM_003986901.4;refGene_gn=SNURF;refGene_tx=XM_006932326.3;refGene_gn=SNURF;refGene_tx=XM_006932327.2;refGene_gn=SNURF;refGene_tx=XM_006932328.2
B3 24060273 24060276 refGene_gn=SNRPN;refGene_tx=XM_004001550.4
B3 24062440 24062592 refGene_gn=SNRPN;refGene_tx=XM_004001550.4
B3 24064627 24064739 refGene_gn=SNRPN;refGene_tx=XM_004001550.4
B3 24065164 24065317 refGene_gn=SNRPN;refGene_tx=XM_004001550.4
B3 24065758 24065897 refGene_gn=SNRPN;refGene_tx=XM_004001550.4

B3 24066266 24066392 refGene_gn=SNRPN;refGene_tx=XM_004001550.4
B3 24066498 24066536 refGene_gn=SNRPN;refGene_tx=XM_004001550.4
B3 24153294 24153363
refGene_gn=LOC111556287;refGene_tx=XM_023255011.1;refGene_gn=LOC111556287;refGene_tx=XM_023255012.1;refGene_gn=LOC111556287;refGene_tx=XM_023255013.1;refGene_gn=LOC111556287;refGene_tx=XM_023255014.1;refGene_gn=LOC111556287;refGene_tx=XM_023255015.1;refGene_gn=LOC111556287;refGene_tx=XM_023255016.1;refGene_gn=LOC111556287;refGene_tx=XM_023255017.1;refGene_gn=LOC111556287;refGene_tx=XM_023255018.1;refGene_gn=LOC111556287;refGene_tx=XM_023255019.1;refGene_gn=LOC111556287;refGene_tx=XM_023255020.1;refGene_gn=LOC111556287;refGene_tx=XM_023255021.1;refGene_gn=LOC111556287;refGene_tx=XM_023255022.1;refGene_gn=LOC111556287;refGene_tx=XM_023255023.1
B3 24155888 24155998
refGene_gn=LOC111556287;refGene_tx=XM_023255024.1;refGene_gn=LOC111556287;refGene_tx=XM_023255025.1
B3 24158164 24158299
refGene_gn=LOC111556287;refGene_tx=XM_023255024.1;refGene_gn=LOC111556287;refGene_tx=XM_023255025.1
B3 24159215 24159350
refGene_gn=LOC111556287;refGene_tx=XM_023255011.1;refGene_gn=LOC111556287;refGene_tx=XM_023255012.1;refGene_gn=LOC111556287;refGene_tx=XM_023255013.1;refGene_gn=LOC111556287;refGene_tx=XM_023255014.1;refGene_gn=LOC111556287;refGene_tx=XM_023255015.1;refGene_gn=LOC111556287;refGene_tx=XM_023255016.1;refGene_gn=LOC111556287;refGene_tx=XM_023255017.1;refGene_gn=LOC111556287;refGene_tx=XM_023255018.1;refGene_gn=LOC111556287;refGene_tx=XM_023255019.1;refGene_gn=LOC111556287;refGene_tx=XM_023255020.1;refGene_gn=LOC111556287;refGene_tx=XM_023255021.1;refGene_gn=LOC111556287;refGene_tx=XM_023255022.1;refGene_gn=LOC111556287;refGene_tx=XM_023255023.1;refGene_gn=LOC111556287;refGene_tx=XM_023255024.1;refGene_gn=LOC111556287;refGene_tx=XM_023255025.1
B3 24159470 24159563
refGene_gn=LOC111556287;refGene_tx=XM_023255011.1;refGene_gn=LOC111556287;refGene_tx=XM_023255012.1;refGene_gn=LOC111556287;refGene_tx=XM_023255013.1;refGene_gn=LOC111556287;refGene_tx=XM_023255014.1;refGene_gn=LOC111556287;refGene_tx=XM_023255015.1;refGene_gn=LOC111556287;refGene_tx=XM_023255016.1;refGene_gn=LOC111556287;refGene_tx=XM_023255017.1;refGene_gn=LOC111556287;refGene_tx=XM_023255018.1;refGene_gn=LOC111556287;refGene_tx=XM_023255019.1;refGene_gn=LOC111556287;refGene_tx=XM_023255020.1;refGene_gn=LOC111556287;refGene_tx=XM_023255021.1;refGene_gn=LOC111556287;refGene_tx=XM_023255023.1
B3 24161892 24162027
refGene_gn=LOC111556287;refGene_tx=XM_023255011.1;refGene_gn=LOC111556287;refGene_tx=XM_023255012.1;refGene_gn=LOC111556287;refGene_tx=XM_023255013.1;refGene_gn=LOC111556287;refGene_tx=XM_023255014.1;refGene_gn=LOC111556287;refGene_tx=XM_023255015.1;refGene_gn=LOC111556287;refGene_tx=XM_023255016.1;refGene_gn=LOC111556287;refGene_tx=XM_023255017.1;refGene_gn=LOC111556287;refGene_tx=XM_023255018.1;refGene_gn=LOC111556287;refGene_tx=XM_023255019.1;refGene_gn=LOC111556287;refGene_tx=XM_023255020.1;refGene_gn=LOC111556287;refGene_tx=XM_023255021.1;refGene_gn

=LOC111556287;refGene_tx=XM_023255022.1;refGene_gn=LOC111556287;refGene_tx=XM_023255023.1;refGene_gn=LOC111556287;refGene_tx=XM_023255024.1;refGene_gn=LOC111556287;refGene_tx=XM_023255025.1
B3 24162302 24162395
refGene_gn=LOC111556287;refGene_tx=XM_023255011.1;refGene_gn=LOC111556287;refGene_tx=XM_023255012.1;refGene_gn=LOC111556287;refGene_tx=XM_023255013.1;refGene_gn=LOC111556287;refGene_tx=XM_023255014.1;refGene_gn=LOC111556287;refGene_tx=XM_023255015.1;refGene_gn=LOC111556287;refGene_tx=XM_023255016.1;refGene_gn=LOC111556287;refGene_tx=XM_023255017.1;refGene_gn=LOC111556287;refGene_tx=XM_023255018.1;refGene_gn=LOC111556287;refGene_tx=XM_023255019.1;refGene_gn=LOC111556287;refGene_tx=XM_023255020.1;refGene_gn=LOC111556287;refGene_tx=XM_023255022.1;refGene_gn=LOC111556287;refGene_tx=XM_023255023.1
B3 24164724 24164859
refGene_gn=LOC111556287;refGene_tx=XM_023255011.1;refGene_gn=LOC111556287;refGene_tx=XM_023255012.1;refGene_gn=LOC111556287;refGene_tx=XM_023255013.1;refGene_gn=LOC111556287;refGene_tx=XM_023255014.1;refGene_gn=LOC111556287;refGene_tx=XM_023255015.1;refGene_gn=LOC111556287;refGene_tx=XM_023255016.1;refGene_gn=LOC111556287;refGene_tx=XM_023255017.1;refGene_gn=LOC111556287;refGene_tx=XM_023255018.1;refGene_gn=LOC111556287;refGene_tx=XM_023255019.1;refGene_gn=LOC111556287;refGene_tx=XM_023255020.1;refGene_gn=LOC111556287;refGene_tx=XM_023255021.1;refGene_gn=LOC111556287;refGene_tx=XM_023255022.1;refGene_gn=LOC111556287;refGene_tx=XM_023255023.1;refGene_gn=LOC111556287;refGene_tx=XM_023255024.1;refGene_gn=LOC111556287;refGene_tx=XM_023255025.1
B3 24165134 24165227
refGene_gn=LOC111556287;refGene_tx=XM_023255011.1;refGene_gn=LOC111556287;refGene_tx=XM_023255012.1;refGene_gn=LOC111556287;refGene_tx=XM_023255013.1;refGene_gn=LOC111556287;refGene_tx=XM_023255014.1;refGene_gn=LOC111556287;refGene_tx=XM_023255015.1;refGene_gn=LOC111556287;refGene_tx=XM_023255016.1;refGene_gn=LOC111556287;refGene_tx=XM_023255017.1;refGene_gn=LOC111556287;refGene_tx=XM_023255018.1;refGene_gn=LOC111556287;refGene_tx=XM_023255019.1;refGene_gn=LOC111556287;refGene_tx=XM_023255021.1;refGene_gn=LOC111556287;refGene_tx=XM_023255022.1;refGene_gn=LOC111556287;refGene_tx=XM_023255023.1
B3 24167557 24167692
refGene_gn=LOC111556287;refGene_tx=XM_023255011.1;refGene_gn=LOC111556287;refGene_tx=XM_023255012.1;refGene_gn=LOC111556287;refGene_tx=XM_023255013.1;refGene_gn=LOC111556287;refGene_tx=XM_023255014.1;refGene_gn=LOC111556287;refGene_tx=XM_023255015.1;refGene_gn=LOC111556287;refGene_tx=XM_023255016.1;refGene_gn=LOC111556287;refGene_tx=XM_023255017.1;refGene_gn=LOC111556287;refGene_tx=XM_023255018.1;refGene_gn=LOC111556287;refGene_tx=XM_023255019.1;refGene_gn=LOC111556287;refGene_tx=XM_023255020.1;refGene_gn=LOC111556287;refGene_tx=XM_023255021.1;refGene_gn=LOC111556287;refGene_tx=XM_023255022.1;refGene_gn=LOC111556287;refGene_tx=XM_023255023.1;refGene_gn=LOC111556287;refGene_tx=XM_023255024.1;refGene_gn=LOC111556287;refGene_tx=XM_023255025.1
B3 24167967 24168183
refGene_gn=LOC111556287;refGene_tx=XM_023255011.1;refGene_gn=LOC111556287;refGene_tx=XM_023255012.1;refGene_gn=LOC111556287;refGene_tx=XM_02

3255013.1;refGene_gn=LOC111556287;refGene_tx=XM_023255014.1;refGene_gn=LOC111556287;refGene_tx=XM_023255015.1;refGene_gn=LOC111556287;refGene_tx=XM_023255016.1;refGene_gn=LOC111556287;refGene_tx=XM_023255017.1;refGene_gn=LOC111556287;refGene_tx=XM_023255018.1;refGene_gn=LOC111556287;refGene_tx=XM_023255019.1;refGene_gn=LOC111556287;refGene_tx=XM_023255020.1;refGene_gn=LOC111556287;refGene_tx=XM_023255021.1;refGene_gn=LOC111556287;refGene_tx=XM_023255022.1;refGene_gn=LOC111556287;refGene_tx=XM_023255024.1
B3 24185207 24185240
refGene_gn=LOC111556287;refGene_tx=XM_023255012.1;refGene_gn=LOC111556287;refGene_tx=XM_023255013.1;refGene_gn=LOC111556287;refGene_tx=XM_023255014.1;refGene_gn=LOC111556287;refGene_tx=XM_023255015.1;refGene_gn=LOC111556287;refGene_tx=XM_023255016.1;refGene_gn=LOC111556287;refGene_tx=XM_023255017.1;refGene_gn=LOC111556287;refGene_tx=XM_023255018.1;refGene_gn=LOC111556287;refGene_tx=XM_023255019.1;refGene_gn=LOC111556287;refGene_tx=XM_023255023.1;refGene_gn=LOC111556287;refGene_tx=XM_023255025.1
B3 24284267 24284430
refGene_gn=LOC111560794;refGene_tx=XM_023255026.1
B3 24284797 24284875
refGene_gn=LOC111560794;refGene_tx=XM_023255026.1;refGene_gn=LOC111560794;refGene_tx=XM_023255027.1
B3 24286114 24286231
refGene_gn=LOC111560794;refGene_tx=XM_023255026.1;refGene_gn=LOC111560794;refGene_tx=XM_023255027.1
B3 24287921 24288038
refGene_gn=LOC111560794;refGene_tx=XM_023255026.1;refGene_gn=LOC111560794;refGene_tx=XM_023255027.1
B3 24291565 24291683
refGene_gn=LOC111560794;refGene_tx=XM_023255026.1;refGene_gn=LOC111560794;refGene_tx=XM_023255027.1
B3 24292062 24292129
refGene_gn=LOC111560794;refGene_tx=XM_023255026.1;refGene_gn=LOC111560794;refGene_tx=XM_023255027.1
B3 24296765 24296889
refGene_gn=LOC111560794;refGene_tx=XM_023255027.1
B3 24421026 24421147 refGene_gn=UBE3A;refGene_tx=XM_003986902.5
B3 24421908 24422052 refGene_gn=UBE3A;refGene_tx=XM_003986902.5
B3 24436753 24436827 refGene_gn=UBE3A;refGene_tx=XM_003986902.5
B3 24436918 24437074 refGene_gn=UBE3A;refGene_tx=XM_003986902.5
B3 24438381 24438546 refGene_gn=UBE3A;refGene_tx=XM_003986902.5
B3 24439153 24439359 refGene_gn=UBE3A;refGene_tx=XM_003986902.5
B3 24442407 24442552 refGene_gn=UBE3A;refGene_tx=XM_003986902.5
B3 24451679 24452929 refGene_gn=UBE3A;refGene_tx=XM_003986902.5
B3 24458947 24459086 refGene_gn=UBE3A;refGene_tx=XM_003986902.5
B3 24459087 24459256 refGene_gn=UBE3A;refGene_tx=XM_003986902.5
B3 24488801 24488846 refGene_gn=UBE3A;refGene_tx=XM_003986902.5
B3 24492067 24492087 refGene_gn=UBE3A;refGene_tx=XM_003986902.5
B3 24992322 24992941
refGene_gn=ATP10A;refGene_tx=XM_023255028.1;refGene_gn=ATP10A;refGene_

tx=XM_023255029.1;refGene_gn=ATP10A;refGene_tx=XM_023255030.1;refGene_gn=ATP10A;refGene_tx=XM_023255031.1
B3 24993109 24993297
refGene_gn=ATP10A;refGene_tx=XM_023255028.1;refGene_gn=ATP10A;refGene_tx=XM_023255029.1;refGene_gn=ATP10A;refGene_tx=XM_023255030.1;refGene_gn=ATP10A;refGene_tx=XM_023255031.1
B3 24993778 24993883
refGene_gn=ATP10A;refGene_tx=XM_023255028.1;refGene_gn=ATP10A;refGene_tx=XM_023255029.1;refGene_gn=ATP10A;refGene_tx=XM_023255030.1;refGene_gn=ATP10A;refGene_tx=XM_023255031.1
B3 24993960 24994041
refGene_gn=ATP10A;refGene_tx=XM_023255028.1;refGene_gn=ATP10A;refGene_tx=XM_023255029.1;refGene_gn=ATP10A;refGene_tx=XM_023255030.1;refGene_gn=ATP10A;refGene_tx=XM_023255031.1
B3 24996598 24996799
refGene_gn=ATP10A;refGene_tx=XM_023255028.1;refGene_gn=ATP10A;refGene_tx=XM_023255029.1;refGene_gn=ATP10A;refGene_tx=XM_023255030.1;refGene_gn=ATP10A;refGene_tx=XM_023255031.1
B3 25000417 25000543
refGene_gn=ATP10A;refGene_tx=XM_023255028.1;refGene_gn=ATP10A;refGene_tx=XM_023255029.1;refGene_gn=ATP10A;refGene_tx=XM_023255030.1;refGene_gn=ATP10A;refGene_tx=XM_023255031.1
B3 25003564 25003641
refGene_gn=ATP10A;refGene_tx=XM_023255028.1;refGene_gn=ATP10A;refGene_tx=XM_023255029.1;refGene_gn=ATP10A;refGene_tx=XM_023255030.1;refGene_gn=ATP10A;refGene_tx=XM_023255031.1
B3 25006238 25006566
refGene_gn=ATP10A;refGene_tx=XM_023255028.1;refGene_gn=ATP10A;refGene_tx=XM_023255029.1;refGene_gn=ATP10A;refGene_tx=XM_023255030.1;refGene_gn=ATP10A;refGene_tx=XM_023255031.1
B3 25008062 25008247
refGene_gn=ATP10A;refGene_tx=XM_023255028.1;refGene_gn=ATP10A;refGene_tx=XM_023255029.1;refGene_gn=ATP10A;refGene_tx=XM_023255030.1;refGene_gn=ATP10A;refGene_tx=XM_023255031.1
B3 25011006 25011133
refGene_gn=ATP10A;refGene_tx=XM_023255028.1;refGene_gn=ATP10A;refGene_tx=XM_023255029.1;refGene_gn=ATP10A;refGene_tx=XM_023255030.1;refGene_gn=ATP10A;refGene_tx=XM_023255031.1
B3 25011226 25011330
refGene_gn=ATP10A;refGene_tx=XM_023255028.1;refGene_gn=ATP10A;refGene_tx=XM_023255029.1;refGene_gn=ATP10A;refGene_tx=XM_023255030.1;refGene_gn=ATP10A;refGene_tx=XM_023255031.1
B3 25019063 25019646
refGene_gn=ATP10A;refGene_tx=XM_023255028.1;refGene_gn=ATP10A;refGene_tx=XM_023255029.1;refGene_gn=ATP10A;refGene_tx=XM_023255030.1;refGene_gn=ATP10A;refGene_tx=XM_023255031.1
B3 25022059 25022254
refGene_gn=ATP10A;refGene_tx=XM_023255028.1;refGene_gn=ATP10A;refGene_tx=XM_023255029.1;refGene_gn=ATP10A;refGene_tx=XM_023255030.1;refGene_gn=ATP10A;refGene_tx=XM_023255031.1

B3 25024291 25024509
refGene_gn=ATP10A;refGene_tx=XM_023255028.1;refGene_gn=ATP10A;refGene_tx=XM_023255029.1;refGene_gn=ATP10A;refGene_tx=XM_023255030.1;refGene_gn=ATP10A;refGene_tx=XM_023255031.1
B3 25027933 25028186
refGene_gn=ATP10A;refGene_tx=XM_023255028.1;refGene_gn=ATP10A;refGene_tx=XM_023255029.1;refGene_gn=ATP10A;refGene_tx=XM_023255030.1;refGene_gn=ATP10A;refGene_tx=XM_023255031.1
B3 25029665 25029796
refGene_gn=ATP10A;refGene_tx=XM_023255028.1;refGene_gn=ATP10A;refGene_tx=XM_023255029.1;refGene_gn=ATP10A;refGene_tx=XM_023255030.1;refGene_gn=ATP10A;refGene_tx=XM_023255031.1
B3 25030854 25030986
refGene_gn=ATP10A;refGene_tx=XM_023255028.1;refGene_gn=ATP10A;refGene_tx=XM_023255029.1;refGene_gn=ATP10A;refGene_tx=XM_023255030.1;refGene_gn=ATP10A;refGene_tx=XM_023255031.1
B3 25031378 25031485
refGene_gn=ATP10A;refGene_tx=XM_023255028.1;refGene_gn=ATP10A;refGene_tx=XM_023255029.1;refGene_gn=ATP10A;refGene_tx=XM_023255030.1;refGene_gn=ATP10A;refGene_tx=XM_023255031.1
B3 25046446 25046532
refGene_gn=ATP10A;refGene_tx=XM_023255028.1;refGene_gn=ATP10A;refGene_tx=XM_023255029.1;refGene_gn=ATP10A;refGene_tx=XM_023255030.1;refGene_gn=ATP10A;refGene_tx=XM_023255031.1
B3 25097111 25097316
refGene_gn=ATP10A;refGene_tx=XM_023255028.1;refGene_gn=ATP10A;refGene_tx=XM_023255029.1;refGene_gn=ATP10A;refGene_tx=XM_023255030.1;refGene_gn=ATP10A;refGene_tx=XM_023255031.1
B3 25185026 25185493
refGene_gn=ATP10A;refGene_tx=XM_023255028.1;refGene_gn=ATP10A;refGene_tx=XM_023255029.1;refGene_gn=ATP10A;refGene_tx=XM_023255030.1;refGene_gn=ATP10A;refGene_tx=XM_023255031.1
B3 25913390 25913732
refGene_gn=GABRB3;refGene_tx=XM_011282816.3;refGene_gn=GABRB3;refGene_tx=XM_019831992.2;refGene_gn=GABRB3;refGene_tx=XM_019831993.2
B3 25922569 25922814
refGene_gn=GABRB3;refGene_tx=XM_011282816.3;refGene_gn=GABRB3;refGene_tx=XM_019831992.2;refGene_gn=GABRB3;refGene_tx=XM_019831993.2
B3 25928657 25928810
refGene_gn=GABRB3;refGene_tx=XM_011282816.3;refGene_gn=GABRB3;refGene_tx=XM_019831992.2;refGene_gn=GABRB3;refGene_tx=XM_019831993.2
B3 25949715 25949853
refGene_gn=GABRB3;refGene_tx=XM_011282816.3;refGene_gn=GABRB3;refGene_tx=XM_019831992.2;refGene_gn=GABRB3;refGene_tx=XM_019831993.2
B3 25952512 25952595
refGene_gn=GABRB3;refGene_tx=XM_011282816.3;refGene_gn=GABRB3;refGene_tx=XM_019831992.2;refGene_gn=GABRB3;refGene_tx=XM_019831993.2
B3 25982625 25982846
refGene_gn=GABRB3;refGene_tx=XM_011282816.3;refGene_gn=GABRB3;refGene_tx=XM_019831992.2;refGene_gn=GABRB3;refGene_tx=XM_019831993.2

B3 26157548 26157616
refGene_gn=GABRB3;refGene_tx=XM_011282816.3;refGene_gn=GABRB3;refGene_tx=XM_019831992.2;refGene_gn=GABRB3;refGene_tx=XM_019831993.2
B3 26157841 26157933
refGene_gn=GABRB3;refGene_tx=XM_011282816.3;refGene_gn=GABRB3;refGene_tx=XM_019831992.2;refGene_gn=GABRB3;refGene_tx=XM_019831993.2
B3 26158062 26158142 refGene_gn=GABRB3;refGene_tx=XM_011282816.3
B3 26158787 26158867 refGene_gn=GABRB3;refGene_tx=XM_019831993.2
B3 26219736 26219784 refGene_gn=GABRB3;refGene_tx=XM_019831992.2
B3 26306200 26306286
refGene_gn=GABRA5;refGene_tx=XM_019831994.2;refGene_gn=GABRA5;refGene_tx=XM_023255032.1
B3 26319534 26319656
refGene_gn=GABRA5;refGene_tx=XM_019831994.2;refGene_gn=GABRA5;refGene_tx=XM_019831996.2;refGene_gn=GABRA5;refGene_tx=XM_023255032.1
B3 26322004 26322072
refGene_gn=GABRA5;refGene_tx=XM_019831994.2;refGene_gn=GABRA5;refGene_tx=XM_019831996.2;refGene_gn=GABRA5;refGene_tx=XM_023255032.1
B3 26322182 26322403
refGene_gn=GABRA5;refGene_tx=XM_019831994.2;refGene_gn=GABRA5;refGene_tx=XM_019831996.2;refGene_gn=GABRA5;refGene_tx=XM_023255032.1
B3 26356133 26356216
refGene_gn=GABRA5;refGene_tx=XM_019831994.2;refGene_gn=GABRA5;refGene_tx=XM_019831996.2;refGene_gn=GABRA5;refGene_tx=XM_023255032.1
B3 26375625 26375769
refGene_gn=GABRA5;refGene_tx=XM_019831994.2;refGene_gn=GABRA5;refGene_tx=XM_019831996.2;refGene_gn=GABRA5;refGene_tx=XM_023255032.1
B3 26378250 26378348 refGene_gn=GABRB3;refGene_tx=XM_019831992.2
B3 26378923 26379076
refGene_gn=GABRA5;refGene_tx=XM_019831994.2;refGene_gn=GABRA5;refGene_tx=XM_019831996.2;refGene_gn=GABRA5;refGene_tx=XM_023255032.1
B3 26381995 26382207
refGene_gn=GABRA5;refGene_tx=XM_019831994.2;refGene_gn=GABRA5;refGene_tx=XM_019831996.2;refGene_gn=GABRA5;refGene_tx=XM_023255032.1
B3 26386942 26387245
refGene_gn=GABRA5;refGene_tx=XM_019831994.2;refGene_gn=GABRA5;refGene_tx=XM_019831996.2;refGene_gn=GABRA5;refGene_tx=XM_023255032.1
B3 26410508 26410561
refGene_gn=GABRG3;refGene_tx=XM_011282834.3;refGene_gn=GABRG3;refGene_tx=XM_019831997.2;refGene_gn=GABRG3;refGene_tx=XM_019831998.2;refGene_gn=GABRG3;refGene_tx=XM_019832001.2
B3 26415673 26415822
refGene_gn=GABRG3;refGene_tx=XM_011282834.3;refGene_gn=GABRG3;refGene_tx=XM_019831997.2;refGene_gn=GABRG3;refGene_tx=XM_019831998.2;refGene_gn=GABRG3;refGene_tx=XM_019832001.2
B3 26459959 26460027
refGene_gn=GABRG3;refGene_tx=XM_011282834.3;refGene_gn=GABRG3;refGene_tx=XM_019831997.2;refGene_gn=GABRG3;refGene_tx=XM_019831998.2;refGene_gn=GABRG3;refGene_tx=XM_019832001.2
B3 26788027 26788248

refGene_gn=GABRG3;refGene_tx=XM_011282834.3;refGene_gn=GABRG3;refGene_tx=XM_019831997.2;refGene_gn=GABRG3;refGene_tx=XM_019831998.2;refGene_gn=GABRG3;refGene_tx=XM_019832001.2
B3 26795353 26795436 refGene_gn=GABRG3;refGene_tx=XM_011282834.3;refGene_gn=GABRG3;refGene_tx=XM_019831997.2;refGene_gn=GABRG3;refGene_tx=XM_019831998.2;refGene_gn=GABRG3;refGene_tx=XM_019832001.2
B3 27016162 27016300 refGene_gn=GABRG3;refGene_tx=XM_011282834.3;refGene_gn=GABRG3;refGene_tx=XM_019831997.2;refGene_gn=GABRG3;refGene_tx=XM_019831998.2;refGene_gn=GABRG3;refGene_tx=XM_019832001.2
B3 27064483 27064572 refGene_gn=GABRG3;refGene_tx=XM_019832001.2
B3 27095639 27095792 refGene_gn=GABRG3;refGene_tx=XM_011282834.3;refGene_gn=GABRG3;refGene_tx=XM_019831997.2;refGene_gn=GABRG3;refGene_tx=XM_019831998.2
B3 27099432 27099629 refGene_gn=GABRG3;refGene_tx=XM_011282834.3;refGene_gn=GABRG3;refGene_tx=XM_019831997.2;refGene_gn=GABRG3;refGene_tx=XM_019831998.2
B3 27099925 27100003 refGene_gn=GABRG3;refGene_tx=XM_011282834.3;refGene_gn=GABRG3;refGene_tx=XM_019831997.2;refGene_gn=GABRG3;refGene_tx=XM_019831998.2
B3 27101779 27101848 refGene_gn=GABRG3;refGene_tx=XM_019831997.2
B3 27104950 27105232 refGene_gn=GABRG3;refGene_tx=XM_011282834.3;refGene_gn=GABRG3;refGene_tx=XM_019831997.2;refGene_gn=GABRG3;refGene_tx=XM_019831998.2
B3 27381292 27381377 refGene_gn=OCA2;refGene_tx=XM_003986906.4
B3 27532072 27532166 refGene_gn=OCA2;refGene_tx=XM_003986906.4
B3 27545196 27545290 refGene_gn=OCA2;refGene_tx=XM_003986906.4
B3 27571804 27571909 refGene_gn=OCA2;refGene_tx=XM_003986906.4
B3 27572523 27572583 refGene_gn=OCA2;refGene_tx=XM_003986906.4
B3 27615821 27615949 refGene_gn=OCA2;refGene_tx=XM_003986906.4
B3 27660225 27660334 refGene_gn=OCA2;refGene_tx=XM_003986906.4
B3 27662859 27662917 refGene_gn=OCA2;refGene_tx=XM_003986906.4
B3 27664859 27665007 refGene_gn=OCA2;refGene_tx=XM_003986906.4
B3 27687576 27687709 refGene_gn=OCA2;refGene_tx=XM_003986906.4
B3 27695675 27695814 refGene_gn=OCA2;refGene_tx=XM_003986906.4
B3 27698986 27699111 refGene_gn=OCA2;refGene_tx=XM_003986906.4
B3 27700199 27700256 refGene_gn=OCA2;refGene_tx=XM_003986906.4
B3 27702645 27702711 refGene_gn=OCA2;refGene_tx=XM_003986906.4
B3 27704610 27704682 refGene_gn=OCA2;refGene_tx=XM_003986906.4
B3 27750284 27750438 refGene_gn=OCA2;refGene_tx=XM_003986906.4
B3 27751709 27751792 refGene_gn=OCA2;refGene_tx=XM_003986906.4
B3 27753952 27754113 refGene_gn=OCA2;refGene_tx=XM_003986906.4
B3 27762491 27762564 refGene_gn=OCA2;refGene_tx=XM_003986906.4
B3 27764596 27764654 refGene_gn=OCA2;refGene_tx=XM_003986906.4
B3 27771299 27771488 refGene_gn=OCA2;refGene_tx=XM_003986906.4
B3 27774484 27774583 refGene_gn=OCA2;refGene_tx=XM_003986906.4
B3 27847565 27847813 refGene_gn=OCA2;refGene_tx=XM_003986906.4
B3 27887082 27887355 refGene_gn=HERC2;refGene_tx=XM_023255033.1
B3 27888457 27888670 refGene_gn=HERC2;refGene_tx=XM_023255033.1

| | | | |
|---|---|---|---|
| B3 | 27889062 | 27889168 | refGene_gn=HERC2;refGene_tx=XM_023255033.1 |
| B3 | 27890078 | 27890269 | refGene_gn=HERC2;refGene_tx=XM_023255033.1 |
| B3 | 27890882 | 27890995 | refGene_gn=HERC2;refGene_tx=XM_023255033.1 |
| B3 | 27891882 | 27892077 | refGene_gn=HERC2;refGene_tx=XM_023255033.1 |
| B3 | 27892241 | 27892383 | refGene_gn=HERC2;refGene_tx=XM_023255033.1 |
| B3 | 27896531 | 27896615 | refGene_gn=HERC2;refGene_tx=XM_023255033.1 |
| B3 | 27899137 | 27899335 | refGene_gn=HERC2;refGene_tx=XM_023255033.1 |
| B3 | 27899840 | 27900028 | refGene_gn=HERC2;refGene_tx=XM_023255033.1 |
| B3 | 27903954 | 27904094 | refGene_gn=HERC2;refGene_tx=XM_023255033.1 |
| B3 | 27904298 | 27904390 | refGene_gn=HERC2;refGene_tx=XM_023255033.1 |
| B3 | 27906971 | 27907133 | refGene_gn=HERC2;refGene_tx=XM_023255033.1 |
| B3 | 27907745 | 27907923 | refGene_gn=HERC2;refGene_tx=XM_023255033.1 |
| B3 | 27913201 | 27913416 | refGene_gn=HERC2;refGene_tx=XM_023255033.1 |
| B3 | 27917525 | 27917724 | refGene_gn=HERC2;refGene_tx=XM_023255033.1 |
| B3 | 27917823 | 27917939 | refGene_gn=HERC2;refGene_tx=XM_023255033.1 |
| B3 | 27918336 | 27918492 | refGene_gn=HERC2;refGene_tx=XM_023255033.1 |
| B3 | 27920508 | 27920634 | refGene_gn=HERC2;refGene_tx=XM_023255033.1 |
| B3 | 27921451 | 27921570 | refGene_gn=HERC2;refGene_tx=XM_023255033.1 |
| B3 | 27921655 | 27921814 | refGene_gn=HERC2;refGene_tx=XM_023255033.1 |
| B3 | 27922252 | 27922384 | refGene_gn=HERC2;refGene_tx=XM_023255033.1 |
| B3 | 27923182 | 27923290 | refGene_gn=HERC2;refGene_tx=XM_023255033.1 |
| B3 | 27938076 | 27938230 | refGene_gn=HERC2;refGene_tx=XM_023255033.1 |
| B3 | 27941236 | 27941428 | refGene_gn=HERC2;refGene_tx=XM_023255033.1 |
| B3 | 27945722 | 27945863 | refGene_gn=HERC2;refGene_tx=XM_023255033.1 |
| B3 | 27946403 | 27946593 | refGene_gn=HERC2;refGene_tx=XM_023255033.1 |
| B3 | 27947463 | 27947635 | refGene_gn=HERC2;refGene_tx=XM_023255033.1 |
| B3 | 27954152 | 27954378 | refGene_gn=HERC2;refGene_tx=XM_023255033.1 |
| B3 | 27955094 | 27955239 | refGene_gn=HERC2;refGene_tx=XM_023255033.1 |
| B3 | 27957407 | 27957579 | refGene_gn=HERC2;refGene_tx=XM_023255033.1 |
| B3 | 27957662 | 27957744 | refGene_gn=HERC2;refGene_tx=XM_023255033.1 |
| B3 | 27957930 | 27958108 | refGene_gn=HERC2;refGene_tx=XM_023255033.1 |
| B3 | 27958407 | 27958498 | refGene_gn=HERC2;refGene_tx=XM_023255033.1 |
| B3 | 27960244 | 27960388 | refGene_gn=HERC2;refGene_tx=XM_023255033.1 |
| B3 | 27960494 | 27960576 | refGene_gn=HERC2;refGene_tx=XM_023255033.1 |
| B3 | 27962009 | 27962121 | refGene_gn=HERC2;refGene_tx=XM_023255033.1 |
| B3 | 27969917 | 27970093 | refGene_gn=HERC2;refGene_tx=XM_023255033.1 |
| B3 | 27973351 | 27973443 | refGene_gn=HERC2;refGene_tx=XM_023255033.1 |
| B3 | 27973532 | 27973638 | refGene_gn=HERC2;refGene_tx=XM_023255033.1 |
| B3 | 27974564 | 27974755 | refGene_gn=HERC2;refGene_tx=XM_023255033.1 |
| B3 | 27982869 | 27983009 | refGene_gn=HERC2;refGene_tx=XM_023255033.1 |
| B3 | 27983124 | 27983233 | refGene_gn=HERC2;refGene_tx=XM_023255033.1 |
| B3 | 27987031 | 27987157 | refGene_gn=HERC2;refGene_tx=XM_023255033.1 |
| B3 | 27987257 | 27987426 | refGene_gn=HERC2;refGene_tx=XM_023255033.1 |
| B3 | 27989321 | 27989420 | refGene_gn=HERC2;refGene_tx=XM_023255033.1 |
| B3 | 27989906 | 27990043 | refGene_gn=HERC2;refGene_tx=XM_023255033.1 |
| B3 | 27990189 | 27990457 | refGene_gn=HERC2;refGene_tx=XM_023255033.1 |
| B3 | 28002004 | 28002147 | refGene_gn=HERC2;refGene_tx=XM_023255033.1 |
| B3 | 28005166 | 28005310 | refGene_gn=HERC2;refGene_tx=XM_023255033.1 |
| B3 | 28006850 | 28006989 | refGene_gn=HERC2;refGene_tx=XM_023255033.1 |
| B3 | 28007857 | 28008088 | refGene_gn=HERC2;refGene_tx=XM_023255033.1 |

B3 28008321 28008518 refGene_gn=HERC2;refGene_tx=XM_023255033.1
B3 28008596 28008744 refGene_gn=HERC2;refGene_tx=XM_023255033.1
B3 28009311 28009493 refGene_gn=HERC2;refGene_tx=XM_023255033.1
B3 28011675 28011858 refGene_gn=HERC2;refGene_tx=XM_023255033.1
B3 28013273 28013466 refGene_gn=HERC2;refGene_tx=XM_023255033.1
B3 28014559 28014747 refGene_gn=HERC2;refGene_tx=XM_023255033.1
B3 28026205 28026397 refGene_gn=HERC2;refGene_tx=XM_023255033.1
B3 28027342 28027494 refGene_gn=HERC2;refGene_tx=XM_023255033.1
B3 28027866 28028003 refGene_gn=HERC2;refGene_tx=XM_023255033.1
B3 28028081 28028255 refGene_gn=HERC2;refGene_tx=XM_023255033.1
B3 28028805 28028939 refGene_gn=HERC2;refGene_tx=XM_023255033.1
B3 28031333 28031501 refGene_gn=HERC2;refGene_tx=XM_023255033.1
B3 28031503 28031528 refGene_gn=HERC2;refGene_tx=XM_023255033.1
B3 28031617 28031745 refGene_gn=HERC2;refGene_tx=XM_023255033.1
B3 28031844 28031977 refGene_gn=HERC2;refGene_tx=XM_023255033.1
B3 28032453 28032668 refGene_gn=HERC2;refGene_tx=XM_023255033.1
B3 28034822 28034973 refGene_gn=HERC2;refGene_tx=XM_023255033.1
B3 28036338 28036442 refGene_gn=HERC2;refGene_tx=XM_023255033.1
B3 28036830 28037001 refGene_gn=HERC2;refGene_tx=XM_023255033.1
B3 28043213 28043399 refGene_gn=HERC2;refGene_tx=XM_023255033.1
B3 28044065 28044221 refGene_gn=HERC2;refGene_tx=XM_023255033.1
B3 28045445 28045630 refGene_gn=HERC2;refGene_tx=XM_023255033.1
B3 28051883 28052062 refGene_gn=HERC2;refGene_tx=XM_023255033.1
B3 28056262 28056387 refGene_gn=HERC2;refGene_tx=XM_023255033.1
B3 28056483 28056712 refGene_gn=HERC2;refGene_tx=XM_023255033.1
B3 28062097 28062298 refGene_gn=HERC2;refGene_tx=XM_023255033.1
B3 28068387 28068581 refGene_gn=HERC2;refGene_tx=XM_023255033.1
B3 28074049 28074301 refGene_gn=HERC2;refGene_tx=XM_023255033.1
B3 28081609 28081723 refGene_gn=HERC2;refGene_tx=XM_023255033.1
B3 28081812 28081970 refGene_gn=HERC2;refGene_tx=XM_023255033.1
B3 28083995 28084147 refGene_gn=HERC2;refGene_tx=XM_023255033.1
B3 28084750 28084939 refGene_gn=HERC2;refGene_tx=XM_023255033.1
B3 28086525 28086699 refGene_gn=HERC2;refGene_tx=XM_023255033.1
B3 28088606 28088778 refGene_gn=HERC2;refGene_tx=XM_023255033.1
B3 28089560 28089671 refGene_gn=HERC2;refGene_tx=XM_023255033.1
B3 28091503 28091660 refGene_gn=HERC2;refGene_tx=XM_023255033.1
B3 28092412 28092513 refGene_gn=HERC2;refGene_tx=XM_023255033.1
B3 28096360 28096580 refGene_gn=HERC2;refGene_tx=XM_023255033.1
B3 28132369 28132507 refGene_gn=HERC2;refGene_tx=XM_023255033.1
B3 28155775 28155890 refGene_gn=HERC2;refGene_tx=XM_023255033.1
B3 28167480 28167552 refGene_gn=HERC2;refGene_tx=XM_023255033.1
B3 28201061 28201224 refGene_gn=NIPA1;refGene_tx=XM_023255034.1
B3 28244258 28244306
refGene_gn=NIPA1;refGene_tx=XM_019832006.2;refGene_gn=NIPA1;refGene_tx
=XM_023255034.1;refGene_gn=NIPA1;refGene_tx=XM_023255035.1
B3 28245285 28245376
refGene_gn=NIPA1;refGene_tx=XM_019832006.2;refGene_gn=NIPA1;refGene_tx
=XM_023255034.1;refGene_gn=NIPA1;refGene_tx=XM_023255035.1
B3 28291642 28291803
refGene_gn=NIPA1;refGene_tx=XM_019832006.2;refGene_gn=NIPA1;refGene_tx

=XM_023255034.1;refGene_gn=NIPA1;refGene_tx=XM_023255035.1
B3 28294643 28295155
refGene_gn=NIPA1;refGene_tx=XM_019832006.2;refGene_gn=NIPA1;refGene_tx=XM_023255034.1;refGene_gn=NIPA1;refGene_tx=XM_023255035.1
B3 28315441 28315580
refGene_gn=NIPA2;refGene_tx=XM_006944165.4;refGene_gn=NIPA2;refGene_tx=XM_023255036.1
B3 28316890 28316947
refGene_gn=NIPA2;refGene_tx=XM_006944165.4;refGene_gn=NIPA2;refGene_tx=XM_023255036.1
B3 28325776 28325867
refGene_gn=NIPA2;refGene_tx=XM_006944165.4;refGene_gn=NIPA2;refGene_tx=XM_023255036.1
B3 28327254 28327415
refGene_gn=NIPA2;refGene_tx=XM_006944165.4;refGene_gn=NIPA2;refGene_tx=XM_023255036.1
B3 28330789 28331424
refGene_gn=NIPA2;refGene_tx=XM_006944165.4;refGene_gn=NIPA2;refGene_tx=XM_023255036.1
B3 28345943 28346108
refGene_gn=CYFIP1;refGene_tx=XM_019832007.1;refGene_gn=CYFIP1;refGene_tx=XM_019832008.1;refGene_gn=CYFIP1;refGene_tx=XM_019832009.2
B3 28348433 28348581
refGene_gn=CYFIP1;refGene_tx=XM_019832007.1;refGene_gn=CYFIP1;refGene_tx=XM_019832008.1;refGene_gn=CYFIP1;refGene_tx=XM_019832009.2;refGene_gn=CYFIP1;refGene_tx=XM_019832010.1
B3 28349059 28349298
refGene_gn=CYFIP1;refGene_tx=XM_019832007.1;refGene_gn=CYFIP1;refGene_tx=XM_019832008.1;refGene_gn=CYFIP1;refGene_tx=XM_019832009.2
B3 28353714 28353809
refGene_gn=CYFIP1;refGene_tx=XM_019832007.1;refGene_gn=CYFIP1;refGene_tx=XM_019832008.1;refGene_gn=CYFIP1;refGene_tx=XM_019832010.1
B3 28354367 28354440
refGene_gn=CYFIP1;refGene_tx=XM_019832007.1;refGene_gn=CYFIP1;refGene_tx=XM_019832008.1;refGene_gn=CYFIP1;refGene_tx=XM_019832010.1
B3 28368759 28368890
refGene_gn=CYFIP1;refGene_tx=XM_019832007.1;refGene_gn=CYFIP1;refGene_tx=XM_019832008.1;refGene_gn=CYFIP1;refGene_tx=XM_019832009.2;refGene_gn=CYFIP1;refGene_tx=XM_019832010.1
B3 28370002 28370093
refGene_gn=CYFIP1;refGene_tx=XM_019832007.1;refGene_gn=CYFIP1;refGene_tx=XM_019832008.1;refGene_gn=CYFIP1;refGene_tx=XM_019832009.2;refGene_gn=CYFIP1;refGene_tx=XM_019832010.1
B3 28373406 28373550
refGene_gn=CYFIP1;refGene_tx=XM_019832007.1;refGene_gn=CYFIP1;refGene_tx=XM_019832008.1;refGene_gn=CYFIP1;refGene_tx=XM_019832009.2;refGene_gn=CYFIP1;refGene_tx=XM_019832010.1
B3 28384685 28384773
refGene_gn=CYFIP1;refGene_tx=XM_019832007.1;refGene_gn=CYFIP1;refGene_tx=XM_019832008.1;refGene_gn=CYFIP1;refGene_tx=XM_019832009.2;refGene_

gn=CYFIP1;refGene_tx=XM_019832010.1
B3 28391488 28391688
refGene_gn=CYFIP1;refGene_tx=XM_019832007.1;refGene_gn=CYFIP1;refGene_tx=XM_019832008.1;refGene_gn=CYFIP1;refGene_tx=XM_019832009.2;refGene_gn=CYFIP1;refGene_tx=XM_019832010.1
B3 28393689 28393809
refGene_gn=CYFIP1;refGene_tx=XM_019832007.1;refGene_gn=CYFIP1;refGene_tx=XM_019832008.1;refGene_gn=CYFIP1;refGene_tx=XM_019832009.2;refGene_gn=CYFIP1;refGene_tx=XM_019832010.1
B3 28394828 28394937
refGene_gn=CYFIP1;refGene_tx=XM_019832007.1;refGene_gn=CYFIP1;refGene_tx=XM_019832008.1;refGene_gn=CYFIP1;refGene_tx=XM_019832009.2;refGene_gn=CYFIP1;refGene_tx=XM_019832010.1
B3 28395035 28395112
refGene_gn=CYFIP1;refGene_tx=XM_019832007.1;refGene_gn=CYFIP1;refGene_tx=XM_019832008.1;refGene_gn=CYFIP1;refGene_tx=XM_019832009.2;refGene_gn=CYFIP1;refGene_tx=XM_019832010.1
B3 28397242 28397339
refGene_gn=CYFIP1;refGene_tx=XM_019832007.1;refGene_gn=CYFIP1;refGene_tx=XM_019832008.1;refGene_gn=CYFIP1;refGene_tx=XM_019832009.2;refGene_gn=CYFIP1;refGene_tx=XM_019832010.1
B3 28398499 28398656
refGene_gn=CYFIP1;refGene_tx=XM_019832007.1;refGene_gn=CYFIP1;refGene_tx=XM_019832008.1;refGene_gn=CYFIP1;refGene_tx=XM_019832009.2;refGene_gn=CYFIP1;refGene_tx=XM_019832010.1
B3 28400072 28400226
refGene_gn=CYFIP1;refGene_tx=XM_019832007.1;refGene_gn=CYFIP1;refGene_tx=XM_019832009.2;refGene_gn=CYFIP1;refGene_tx=XM_019832010.1
B3 28400498 28400646 refGene_gn=CYFIP1;refGene_tx=XM_019832008.1
B3 28401460 28401608
refGene_gn=CYFIP1;refGene_tx=XM_019832007.1;refGene_gn=CYFIP1;refGene_tx=XM_019832008.1;refGene_gn=CYFIP1;refGene_tx=XM_019832009.2;refGene_gn=CYFIP1;refGene_tx=XM_019832010.1
B3 28402642 28402809
refGene_gn=CYFIP1;refGene_tx=XM_019832007.1;refGene_gn=CYFIP1;refGene_tx=XM_019832008.1;refGene_gn=CYFIP1;refGene_tx=XM_019832009.2;refGene_gn=CYFIP1;refGene_tx=XM_019832010.1
B3 28409843 28409969
refGene_gn=CYFIP1;refGene_tx=XM_019832007.1;refGene_gn=CYFIP1;refGene_tx=XM_019832008.1;refGene_gn=CYFIP1;refGene_tx=XM_019832009.2;refGene_gn=CYFIP1;refGene_tx=XM_019832010.1
B3 28411037 28411160
refGene_gn=CYFIP1;refGene_tx=XM_019832007.1;refGene_gn=CYFIP1;refGene_tx=XM_019832008.1;refGene_gn=CYFIP1;refGene_tx=XM_019832009.2;refGene_gn=CYFIP1;refGene_tx=XM_019832010.1
B3 28415875 28415993
refGene_gn=CYFIP1;refGene_tx=XM_019832007.1;refGene_gn=CYFIP1;refGene_tx=XM_019832008.1;refGene_gn=CYFIP1;refGene_tx=XM_019832009.2;refGene_gn=CYFIP1;refGene_tx=XM_019832010.1
B3 28420591 28420683

refGene_gn=CYFIP1;refGene_tx=XM_019832007.1;refGene_gn=CYFIP1;refGene_tx=XM_019832008.1;refGene_gn=CYFIP1;refGene_tx=XM_019832009.2;refGene_gn=CYFIP1;refGene_tx=XM_019832010.1
B3 28423007 28423112
refGene_gn=CYFIP1;refGene_tx=XM_019832007.1;refGene_gn=CYFIP1;refGene_tx=XM_019832008.1;refGene_gn=CYFIP1;refGene_tx=XM_019832009.2;refGene_gn=CYFIP1;refGene_tx=XM_019832010.1
B3 28424381 28424510
refGene_gn=CYFIP1;refGene_tx=XM_019832007.1;refGene_gn=CYFIP1;refGene_tx=XM_019832008.1;refGene_gn=CYFIP1;refGene_tx=XM_019832009.2;refGene_gn=CYFIP1;refGene_tx=XM_019832010.1
B3 28424621 28424718
refGene_gn=CYFIP1;refGene_tx=XM_019832007.1;refGene_gn=CYFIP1;refGene_tx=XM_019832008.1;refGene_gn=CYFIP1;refGene_tx=XM_019832009.2;refGene_gn=CYFIP1;refGene_tx=XM_019832010.1
B3 28427839 28428021
refGene_gn=CYFIP1;refGene_tx=XM_019832007.1;refGene_gn=CYFIP1;refGene_tx=XM_019832008.1;refGene_gn=CYFIP1;refGene_tx=XM_019832009.2;refGene_gn=CYFIP1;refGene_tx=XM_019832010.1
B3 28428902 28429004
refGene_gn=CYFIP1;refGene_tx=XM_019832007.1;refGene_gn=CYFIP1;refGene_tx=XM_019832008.1;refGene_gn=CYFIP1;refGene_tx=XM_019832009.2;refGene_gn=CYFIP1;refGene_tx=XM_019832010.1
B3 28429256 28429334
refGene_gn=CYFIP1;refGene_tx=XM_019832007.1;refGene_gn=CYFIP1;refGene_tx=XM_019832008.1;refGene_gn=CYFIP1;refGene_tx=XM_019832009.2;refGene_gn=CYFIP1;refGene_tx=XM_019832010.1
B3 28430711 28430801
refGene_gn=CYFIP1;refGene_tx=XM_019832007.1;refGene_gn=CYFIP1;refGene_tx=XM_019832008.1;refGene_gn=CYFIP1;refGene_tx=XM_019832009.2;refGene_gn=CYFIP1;refGene_tx=XM_019832010.1
B3 28430876 28430993
refGene_gn=CYFIP1;refGene_tx=XM_019832007.1;refGene_gn=CYFIP1;refGene_tx=XM_019832008.1;refGene_gn=CYFIP1;refGene_tx=XM_019832009.2;refGene_gn=CYFIP1;refGene_tx=XM_019832010.1
B3 28527670 28527717
refGene_gn=TUBGCP5;refGene_tx=XM_011282825.3;refGene_gn=TUBGCP5;refGene_tx=XM_011282826.3;refGene_gn=TUBGCP5;refGene_tx=XM_011282827.3;refGene_gn=TUBGCP5;refGene_tx=XM_011282828.3;refGene_gn=TUBGCP5;refGene_tx=XM_011282829.3;refGene_gn=TUBGCP5;refGene_tx=XM_011282830.3;refGene_gn=TUBGCP5;refGene_tx=XM_011282831.3;refGene_gn=TUBGCP5;refGene_tx=XM_019832011.2;refGene_gn=TUBGCP5;refGene_tx=XM_023255037.1
B3 28529644 28529745
refGene_gn=TUBGCP5;refGene_tx=XM_011282825.3;refGene_gn=TUBGCP5;refGene_tx=XM_011282826.3;refGene_gn=TUBGCP5;refGene_tx=XM_011282827.3;refGene_gn=TUBGCP5;refGene_tx=XM_011282828.3;refGene_gn=TUBGCP5;refGene_tx=XM_011282829.3;refGene_gn=TUBGCP5;refGene_tx=XM_011282830.3;refGene_gn=TUBGCP5;refGene_tx=XM_011282831.3;refGene_gn=TUBGCP5;refGene_tx=XM_019832011.2;refGene_gn=TUBGCP5;refGene_tx=XM_023255037.1
B3 28531822 28531911

refGene_gn=TUBGCP5;refGene_tx=XM_011282825.3;refGene_gn=TUBGCP5;refGen
e_tx=XM_011282826.3;refGene_gn=TUBGCP5;refGene_tx=XM_011282827.3;refGe
ne_gn=TUBGCP5;refGene_tx=XM_011282828.3;refGene_gn=TUBGCP5;refGene_tx=
XM_011282829.3;refGene_gn=TUBGCP5;refGene_tx=XM_011282830.3;refGene_gn
=TUBGCP5;refGene_tx=XM_011282831.3;refGene_gn=TUBGCP5;refGene_tx=XM_01
9832011.2;refGene_gn=TUBGCP5;refGene_tx=XM_023255037.1
B3 28532492 28532618
refGene_gn=TUBGCP5;refGene_tx=XM_011282825.3;refGene_gn=TUBGCP5;refGen
e_tx=XM_011282826.3;refGene_gn=TUBGCP5;refGene_tx=XM_011282827.3;refGe
ne_gn=TUBGCP5;refGene_tx=XM_011282828.3;refGene_gn=TUBGCP5;refGene_tx=
XM_011282829.3;refGene_gn=TUBGCP5;refGene_tx=XM_011282830.3;refGene_gn
=TUBGCP5;refGene_tx=XM_011282831.3;refGene_gn=TUBGCP5;refGene_tx=XM_01
9832011.2;refGene_gn=TUBGCP5;refGene_tx=XM_023255037.1
B3 28532842 28532866 refGene_gn=TUBGCP5;refGene_tx=XM_011282832.3
B3 28534251 28534430
refGene_gn=TUBGCP5;refGene_tx=XM_011282825.3;refGene_gn=TUBGCP5;refGen
e_tx=XM_011282826.3;refGene_gn=TUBGCP5;refGene_tx=XM_011282827.3;refGe
ne_gn=TUBGCP5;refGene_tx=XM_011282828.3;refGene_gn=TUBGCP5;refGene_tx=
XM_011282829.3;refGene_gn=TUBGCP5;refGene_tx=XM_011282830.3;refGene_gn
=TUBGCP5;refGene_tx=XM_011282831.3;refGene_gn=TUBGCP5;refGene_tx=XM_01
1282832.3;refGene_gn=TUBGCP5;refGene_tx=XM_019832011.2;refGene_gn=TUBG
CP5;refGene_tx=XM_023255037.1
B3 28534925 28535046
refGene_gn=TUBGCP5;refGene_tx=XM_011282825.3;refGene_gn=TUBGCP5;refGen
e_tx=XM_011282826.3;refGene_gn=TUBGCP5;refGene_tx=XM_011282827.3;refGe
ne_gn=TUBGCP5;refGene_tx=XM_011282828.3;refGene_gn=TUBGCP5;refGene_tx=
XM_011282829.3;refGene_gn=TUBGCP5;refGene_tx=XM_011282830.3;refGene_gn
=TUBGCP5;refGene_tx=XM_011282831.3;refGene_gn=TUBGCP5;refGene_tx=XM_01
1282832.3;refGene_gn=TUBGCP5;refGene_tx=XM_019832011.2;refGene_gn=TUBG
CP5;refGene_tx=XM_023255037.1
B3 28535149 28535234
refGene_gn=TUBGCP5;refGene_tx=XM_011282825.3;refGene_gn=TUBGCP5;refGen
e_tx=XM_011282826.3;refGene_gn=TUBGCP5;refGene_tx=XM_011282827.3;refGe
ne_gn=TUBGCP5;refGene_tx=XM_011282828.3;refGene_gn=TUBGCP5;refGene_tx=
XM_011282829.3;refGene_gn=TUBGCP5;refGene_tx=XM_011282830.3;refGene_gn
=TUBGCP5;refGene_tx=XM_011282831.3;refGene_gn=TUBGCP5;refGene_tx=XM_01
1282832.3;refGene_gn=TUBGCP5;refGene_tx=XM_019832011.2;refGene_gn=TUBG
CP5;refGene_tx=XM_023255037.1
B3 28538122 28538305
refGene_gn=TUBGCP5;refGene_tx=XM_011282825.3;refGene_gn=TUBGCP5;refGen
e_tx=XM_011282826.3;refGene_gn=TUBGCP5;refGene_tx=XM_011282827.3;refGe
ne_gn=TUBGCP5;refGene_tx=XM_011282828.3;refGene_gn=TUBGCP5;refGene_tx=
XM_011282829.3;refGene_gn=TUBGCP5;refGene_tx=XM_011282830.3;refGene_gn
=TUBGCP5;refGene_tx=XM_011282831.3;refGene_gn=TUBGCP5;refGene_tx=XM_01
1282832.3;refGene_gn=TUBGCP5;refGene_tx=XM_019832011.2;refGene_gn=TUBG
CP5;refGene_tx=XM_023255037.1
B3 28539636 28539825
refGene_gn=TUBGCP5;refGene_tx=XM_011282825.3;refGene_gn=TUBGCP5;refGen
e_tx=XM_011282826.3;refGene_gn=TUBGCP5;refGene_tx=XM_011282827.3;refGe
ne_gn=TUBGCP5;refGene_tx=XM_011282828.3;refGene_gn=TUBGCP5;refGene_tx=

XM_011282829.3;refGene_gn=TUBGCP5;refGene_tx=XM_011282830.3;refGene_gn=TUBGCP5;refGene_tx=XM_011282831.3;refGene_gn=TUBGCP5;refGene_tx=XM_011282832.3;refGene_gn=TUBGCP5;refGene_tx=XM_019832011.2;refGene_gn=TUBGCP5;refGene_tx=XM_023255037.1
B3 28540471 28540670
refGene_gn=TUBGCP5;refGene_tx=XM_011282825.3;refGene_gn=TUBGCP5;refGene_tx=XM_011282826.3;refGene_gn=TUBGCP5;refGene_tx=XM_011282827.3;refGene_gn=TUBGCP5;refGene_tx=XM_011282828.3;refGene_gn=TUBGCP5;refGene_tx=XM_011282829.3;refGene_gn=TUBGCP5;refGene_tx=XM_011282830.3;refGene_gn=TUBGCP5;refGene_tx=XM_011282831.3;refGene_gn=TUBGCP5;refGene_tx=XM_011282832.3;refGene_gn=TUBGCP5;refGene_tx=XM_019832011.2;refGene_gn=TUBGCP5;refGene_tx=XM_023255037.1
B3 28541618 28541887
refGene_gn=TUBGCP5;refGene_tx=XM_011282825.3;refGene_gn=TUBGCP5;refGene_tx=XM_011282826.3;refGene_gn=TUBGCP5;refGene_tx=XM_011282827.3;refGene_gn=TUBGCP5;refGene_tx=XM_011282828.3;refGene_gn=TUBGCP5;refGene_tx=XM_011282829.3;refGene_gn=TUBGCP5;refGene_tx=XM_011282830.3;refGene_gn=TUBGCP5;refGene_tx=XM_011282831.3;refGene_gn=TUBGCP5;refGene_tx=XM_011282832.3;refGene_gn=TUBGCP5;refGene_tx=XM_019832011.2;refGene_gn=TUBGCP5;refGene_tx=XM_023255037.1
B3 28543808 28543924
refGene_gn=TUBGCP5;refGene_tx=XM_011282825.3;refGene_gn=TUBGCP5;refGene_tx=XM_011282826.3;refGene_gn=TUBGCP5;refGene_tx=XM_011282827.3;refGene_gn=TUBGCP5;refGene_tx=XM_011282828.3;refGene_gn=TUBGCP5;refGene_tx=XM_011282829.3;refGene_gn=TUBGCP5;refGene_tx=XM_011282830.3;refGene_gn=TUBGCP5;refGene_tx=XM_011282831.3;refGene_gn=TUBGCP5;refGene_tx=XM_011282832.3;refGene_gn=TUBGCP5;refGene_tx=XM_019832011.2;refGene_gn=TUBGCP5;refGene_tx=XM_023255037.1
B3 28545243 28545446
refGene_gn=TUBGCP5;refGene_tx=XM_011282825.3;refGene_gn=TUBGCP5;refGene_tx=XM_011282826.3;refGene_gn=TUBGCP5;refGene_tx=XM_011282827.3;refGene_gn=TUBGCP5;refGene_tx=XM_011282828.3;refGene_gn=TUBGCP5;refGene_tx=XM_011282829.3;refGene_gn=TUBGCP5;refGene_tx=XM_011282830.3;refGene_gn=TUBGCP5;refGene_tx=XM_011282831.3;refGene_gn=TUBGCP5;refGene_tx=XM_011282832.3;refGene_gn=TUBGCP5;refGene_tx=XM_019832011.2;refGene_gn=TUBGCP5;refGene_tx=XM_023255037.1
B3 28547109 28547356
refGene_gn=TUBGCP5;refGene_tx=XM_011282825.3;refGene_gn=TUBGCP5;refGene_tx=XM_011282826.3;refGene_gn=TUBGCP5;refGene_tx=XM_011282827.3;refGene_gn=TUBGCP5;refGene_tx=XM_011282828.3;refGene_gn=TUBGCP5;refGene_tx=XM_011282829.3;refGene_gn=TUBGCP5;refGene_tx=XM_011282830.3;refGene_gn=TUBGCP5;refGene_tx=XM_011282831.3;refGene_gn=TUBGCP5;refGene_tx=XM_011282832.3;refGene_gn=TUBGCP5;refGene_tx=XM_019832011.2;refGene_gn=TUBGCP5;refGene_tx=XM_023255037.1
B3 28547846 28547940
refGene_gn=TUBGCP5;refGene_tx=XM_011282825.3;refGene_gn=TUBGCP5;refGene_tx=XM_011282826.3;refGene_gn=TUBGCP5;refGene_tx=XM_011282827.3;refGene_gn=TUBGCP5;refGene_tx=XM_011282828.3;refGene_gn=TUBGCP5;refGene_tx=XM_011282829.3;refGene_gn=TUBGCP5;refGene_tx=XM_011282830.3;refGene_gn=TUBGCP5;refGene_tx=XM_011282831.3;refGene_gn=TUBGCP5;refGene_tx=XM_01

1282832.3;refGene_gn=TUBGCP5;refGene_tx=XM_019832011.2;refGene_gn=TUBGCP5;refGene_tx=XM_023255037.1
B3 28548710 28548800
refGene_gn=TUBGCP5;refGene_tx=XM_011282825.3;refGene_gn=TUBGCP5;refGene_tx=XM_011282826.3;refGene_gn=TUBGCP5;refGene_tx=XM_011282827.3;refGene_gn=TUBGCP5;refGene_tx=XM_011282828.3;refGene_gn=TUBGCP5;refGene_tx=XM_011282829.3;refGene_gn=TUBGCP5;refGene_tx=XM_011282830.3;refGene_gn=TUBGCP5;refGene_tx=XM_011282831.3;refGene_gn=TUBGCP5;refGene_tx=XM_011282832.3;refGene_gn=TUBGCP5;refGene_tx=XM_019832011.2;refGene_gn=TUBGCP5;refGene_tx=XM_023255037.1
B3 28549555 28549673
refGene_gn=TUBGCP5;refGene_tx=XM_011282825.3;refGene_gn=TUBGCP5;refGene_tx=XM_011282826.3;refGene_gn=TUBGCP5;refGene_tx=XM_011282827.3;refGene_gn=TUBGCP5;refGene_tx=XM_011282828.3;refGene_gn=TUBGCP5;refGene_tx=XM_011282829.3;refGene_gn=TUBGCP5;refGene_tx=XM_011282830.3;refGene_gn=TUBGCP5;refGene_tx=XM_011282831.3;refGene_gn=TUBGCP5;refGene_tx=XM_011282832.3;refGene_gn=TUBGCP5;refGene_tx=XM_019832011.2;refGene_gn=TUBGCP5;refGene_tx=XM_023255037.1
B3 28551604 28551712
refGene_gn=TUBGCP5;refGene_tx=XM_011282825.3;refGene_gn=TUBGCP5;refGene_tx=XM_011282826.3;refGene_gn=TUBGCP5;refGene_tx=XM_011282827.3;refGene_gn=TUBGCP5;refGene_tx=XM_011282828.3;refGene_gn=TUBGCP5;refGene_tx=XM_011282829.3;refGene_gn=TUBGCP5;refGene_tx=XM_011282832.3;refGene_gn=TUBGCP5;refGene_tx=XM_019832011.2;refGene_gn=TUBGCP5;refGene_tx=XM_023255037.1
B3 28576624 28576760
refGene_gn=TUBGCP5;refGene_tx=XM_011282825.3;refGene_gn=TUBGCP5;refGene_tx=XM_011282826.3;refGene_gn=TUBGCP5;refGene_tx=XM_011282827.3;refGene_gn=TUBGCP5;refGene_tx=XM_011282828.3;refGene_gn=TUBGCP5;refGene_tx=XM_011282829.3;refGene_gn=TUBGCP5;refGene_tx=XM_011282830.3;refGene_gn=TUBGCP5;refGene_tx=XM_011282831.3;refGene_gn=TUBGCP5;refGene_tx=XM_011282832.3;refGene_gn=TUBGCP5;refGene_tx=XM_023255037.1
B3 28577392 28577472
refGene_gn=TUBGCP5;refGene_tx=XM_011282825.3;refGene_gn=TUBGCP5;refGene_tx=XM_011282826.3;refGene_gn=TUBGCP5;refGene_tx=XM_011282827.3;refGene_gn=TUBGCP5;refGene_tx=XM_011282828.3;refGene_gn=TUBGCP5;refGene_tx=XM_011282829.3;refGene_gn=TUBGCP5;refGene_tx=XM_011282830.3;refGene_gn=TUBGCP5;refGene_tx=XM_011282831.3;refGene_gn=TUBGCP5;refGene_tx=XM_011282832.3;refGene_gn=TUBGCP5;refGene_tx=XM_023255037.1
B3 28577890 28577987
refGene_gn=TUBGCP5;refGene_tx=XM_011282825.3;refGene_gn=TUBGCP5;refGene_tx=XM_011282826.3;refGene_gn=TUBGCP5;refGene_tx=XM_011282827.3;refGene_gn=TUBGCP5;refGene_tx=XM_011282828.3;refGene_gn=TUBGCP5;refGene_tx=XM_011282829.3;refGene_gn=TUBGCP5;refGene_tx=XM_011282830.3;refGene_gn=TUBGCP5;refGene_tx=XM_011282831.3;refGene_gn=TUBGCP5;refGene_tx=XM_011282832.3
B3 28596307 28596416
refGene_gn=TUBGCP5;refGene_tx=XM_011282825.3;refGene_gn=TUBGCP5;refGene_tx=XM_011282826.3;refGene_gn=TUBGCP5;refGene_tx=XM_011282827.3;refGene_gn=TUBGCP5;refGene_tx=XM_011282828.3;refGene_gn=TUBGCP5;refGene_tx=

XM_011282829.3;refGene_gn=TUBGCP5;refGene_tx=XM_011282830.3;refGene_gn=TUBGCP5;refGene_tx=XM_011282831.3;refGene_gn=TUBGCP5;refGene_tx=XM_011282832.3
B3 28596526 28596580
refGene_gn=TUBGCP5;refGene_tx=XM_011282825.3;refGene_gn=TUBGCP5;refGene_tx=XM_011282826.3;refGene_gn=TUBGCP5;refGene_tx=XM_011282827.3;refGene_gn=TUBGCP5;refGene_tx=XM_011282828.3;refGene_gn=TUBGCP5;refGene_tx=XM_011282829.3;refGene_gn=TUBGCP5;refGene_tx=XM_011282830.3;refGene_gn=TUBGCP5;refGene_tx=XM_011282831.3;refGene_gn=TUBGCP5;refGene_tx=XM_011282832.3
B3 28599107 28599253
refGene_gn=TUBGCP5;refGene_tx=XM_011282825.3;refGene_gn=TUBGCP5;refGene_tx=XM_011282826.3;refGene_gn=TUBGCP5;refGene_tx=XM_011282827.3;refGene_gn=TUBGCP5;refGene_tx=XM_011282828.3;refGene_gn=TUBGCP5;refGene_tx=XM_011282829.3;refGene_gn=TUBGCP5;refGene_tx=XM_011282830.3;refGene_gn=TUBGCP5;refGene_tx=XM_011282831.3;refGene_gn=TUBGCP5;refGene_tx=XM_011282832.3
B3 30728537 30728729
refGene_gn=UBE2Q2;refGene_tx=XM_003986912.5;refGene_gn=UBE2Q2;refGene_tx=XM_011282991.3;refGene_gn=UBE2Q2;refGene_tx=XM_019832013.2;refGene_gn=UBE2Q2;refGene_tx=XM_019832014.2;refGene_gn=UBE2Q2;refGene_tx=XM_019832015.2
B3 30740928 30741030
refGene_gn=UBE2Q2;refGene_tx=XM_003986912.5;refGene_gn=UBE2Q2;refGene_tx=XM_011282991.3;refGene_gn=UBE2Q2;refGene_tx=XM_019832013.2;refGene_gn=UBE2Q2;refGene_tx=XM_019832014.2;refGene_gn=UBE2Q2;refGene_tx=XM_019832015.2
B3 30745496 30745601
refGene_gn=UBE2Q2;refGene_tx=XM_003986912.5;refGene_gn=UBE2Q2;refGene_tx=XM_019832013.2;refGene_gn=UBE2Q2;refGene_tx=XM_019832015.2
B3 30752933 30752990
refGene_gn=UBE2Q2;refGene_tx=XM_003986912.5;refGene_gn=UBE2Q2;refGene_tx=XM_011282991.3;refGene_gn=UBE2Q2;refGene_tx=XM_019832013.2;refGene_gn=UBE2Q2;refGene_tx=XM_019832014.2;refGene_gn=UBE2Q2;refGene_tx=XM_019832015.2
B3 30756529 30756670
refGene_gn=UBE2Q2;refGene_tx=XM_003986912.5;refGene_gn=UBE2Q2;refGene_tx=XM_011282991.3;refGene_gn=UBE2Q2;refGene_tx=XM_019832013.2;refGene_gn=UBE2Q2;refGene_tx=XM_019832014.2;refGene_gn=UBE2Q2;refGene_tx=XM_019832015.2
B3 30759564 30759649
refGene_gn=UBE2Q2;refGene_tx=XM_003986912.5;refGene_gn=UBE2Q2;refGene_tx=XM_011282991.3;refGene_gn=UBE2Q2;refGene_tx=XM_019832013.2;refGene_gn=UBE2Q2;refGene_tx=XM_019832014.2
B3 30761053 30761114
refGene_gn=UBE2Q2;refGene_tx=XM_003986912.5;refGene_gn=UBE2Q2;refGene_tx=XM_011282991.3;refGene_gn=UBE2Q2;refGene_tx=XM_019832013.2;refGene_gn=UBE2Q2;refGene_tx=XM_019832014.2
B3 30761872 30761963
refGene_gn=UBE2Q2;refGene_tx=XM_003986912.5;refGene_gn=UBE2Q2;refGene_

tx=XM_011282991.3;refGene_gn=UBE2Q2;refGene_tx=XM_019832013.2;refGene_gn=UBE2Q2;refGene_tx=XM_019832014.2
B3 30769361 30769420
refGene_gn=UBE2Q2;refGene_tx=XM_003986912.5;refGene_gn=UBE2Q2;refGene_tx=XM_011282991.3;refGene_gn=UBE2Q2;refGene_tx=XM_019832013.2;refGene_gn=UBE2Q2;refGene_tx=XM_019832014.2;refGene_gn=UBE2Q2;refGene_tx=XM_019832015.2
B3 30789472 30789521
refGene_gn=UBE2Q2;refGene_tx=XM_003986912.5;refGene_gn=UBE2Q2;refGene_tx=XM_011282991.3;refGene_gn=UBE2Q2;refGene_tx=XM_019832013.2;refGene_gn=UBE2Q2;refGene_tx=XM_019832014.2;refGene_gn=UBE2Q2;refGene_tx=XM_019832015.2
B3 30789970 30790066
refGene_gn=UBE2Q2;refGene_tx=XM_003986912.5;refGene_gn=UBE2Q2;refGene_tx=XM_011282991.3;refGene_gn=UBE2Q2;refGene_tx=XM_019832013.2;refGene_gn=UBE2Q2;refGene_tx=XM_019832014.2;refGene_gn=UBE2Q2;refGene_tx=XM_019832015.2
B3 30799617 30799684
refGene_gn=UBE2Q2;refGene_tx=XM_003986912.5;refGene_gn=UBE2Q2;refGene_tx=XM_011282991.3
B3 30801497 30801659
refGene_gn=UBE2Q2;refGene_tx=XM_003986912.5;refGene_gn=UBE2Q2;refGene_tx=XM_011282991.3;refGene_gn=UBE2Q2;refGene_tx=XM_019832013.2;refGene_gn=UBE2Q2;refGene_tx=XM_019832014.2;refGene_gn=UBE2Q2;refGene_tx=XM_019832015.2
B3 30805998 30806153
refGene_gn=FBXO22;refGene_tx=XM_003986913.5;refGene_gn=FBXO22;refGene_tx=XM_006932335.4;refGene_gn=FBXO22;refGene_tx=XM_019832012.2
B3 30806516 30806655
refGene_gn=FBXO22;refGene_tx=XM_003986913.5;refGene_gn=FBXO22;refGene_tx=XM_006932335.4;refGene_gn=FBXO22;refGene_tx=XM_019832012.2
B3 30811269 30811360
refGene_gn=FBXO22;refGene_tx=XM_003986913.5;refGene_gn=FBXO22;refGene_tx=XM_006932335.4;refGene_gn=FBXO22;refGene_tx=XM_019832012.2
B3 30812153 30812249
refGene_gn=FBXO22;refGene_tx=XM_003986913.5;refGene_gn=FBXO22;refGene_tx=XM_006932335.4;refGene_gn=FBXO22;refGene_tx=XM_019832012.2
B3 30818216 30818381
refGene_gn=FBXO22;refGene_tx=XM_003986913.5;refGene_gn=FBXO22;refGene_tx=XM_006932335.4
B3 30832533 30832699
refGene_gn=FBXO22;refGene_tx=XM_003986913.5;refGene_gn=FBXO22;refGene_tx=XM_006932335.4;refGene_gn=FBXO22;refGene_tx=XM_019832012.2
B3 30835319 30835737
refGene_gn=FBXO22;refGene_tx=XM_003986913.5;refGene_gn=FBXO22;refGene_tx=XM_006932335.4;refGene_gn=FBXO22;refGene_tx=XM_019832012.2
B3 30853999 30854016
refGene_gn=NRG4;refGene_tx=XM_023255038.1;refGene_gn=NRG4;refGene_tx=XM_023255039.1;refGene_gn=NRG4;refGene_tx=XM_023255040.1;refGene_gn=NRG4;refGene_tx=XM_023255041.1;refGene_gn=NRG4;refGene_tx=XM_023255042.1

B3 30874710 30874790
refGene_gn=NRG4;refGene_tx=XM_023255038.1;refGene_gn=NRG4;refGene_tx=XM_023255039.1;refGene_gn=NRG4;refGene_tx=XM_023255040.1;refGene_gn=NRG4;refGene_tx=XM_023255041.1;refGene_gn=NRG4;refGene_tx=XM_023255042.1
B3 30878114 30878264
refGene_gn=NRG4;refGene_tx=XM_023255038.1;refGene_gn=NRG4;refGene_tx=XM_023255039.1;refGene_gn=NRG4;refGene_tx=XM_023255040.1;refGene_gn=NRG4;refGene_tx=XM_023255041.1;refGene_gn=NRG4;refGene_tx=XM_023255042.1
B3 30953565 30953659
refGene_gn=NRG4;refGene_tx=XM_023255038.1;refGene_gn=NRG4;refGene_tx=XM_023255039.1;refGene_gn=NRG4;refGene_tx=XM_023255040.1;refGene_gn=NRG4;refGene_tx=XM_023255041.1;refGene_gn=NRG4;refGene_tx=XM_023255042.1
B3 30956157 30956167
refGene_gn=NRG4;refGene_tx=XM_023255038.1;refGene_gn=NRG4;refGene_tx=XM_023255039.1;refGene_gn=NRG4;refGene_tx=XM_023255040.1;refGene_gn=NRG4;refGene_tx=XM_023255041.1;refGene_gn=NRG4;refGene_tx=XM_023255042.1
B3 31105214 31105252 refGene_gn=TMEM266;refGene_tx=XM_011282990.3
B3 31109064 31109253 refGene_gn=TMEM266;refGene_tx=XM_011282990.3
B3 31127257 31127412 refGene_gn=TMEM266;refGene_tx=XM_011282990.3
B3 31130391 31130465 refGene_gn=TMEM266;refGene_tx=XM_011282990.3
B3 31144068 31144125 refGene_gn=TMEM266;refGene_tx=XM_011282990.3
B3 31145258 31145397 refGene_gn=TMEM266;refGene_tx=XM_011282990.3
B3 31149143 31149259 refGene_gn=TMEM266;refGene_tx=XM_011282990.3
B3 31162877 31163067 refGene_gn=TMEM266;refGene_tx=XM_011282990.3
B3 31177533 31177596 refGene_gn=TMEM266;refGene_tx=XM_011282990.3
B3 31179096 31179692 refGene_gn=TMEM266;refGene_tx=XM_011282990.3
B3 31190976 31191015
refGene_gn=ETFA;refGene_tx=NM_001287554.1;refGene_gn=ETFA;refGene_tx=XM_006932340.4;refGene_gn=ETFA;refGene_tx=XM_006932341.4
B3 31198094 31198175
refGene_gn=ETFA;refGene_tx=NM_001287554.1;refGene_gn=ETFA;refGene_tx=XM_006932340.4
B3 31205836 31205902
refGene_gn=ETFA;refGene_tx=NM_001287554.1;refGene_gn=ETFA;refGene_tx=XM_006932340.4;refGene_gn=ETFA;refGene_tx=XM_006932341.4
B3 31233722 31233805
refGene_gn=ETFA;refGene_tx=NM_001287554.1;refGene_gn=ETFA;refGene_tx=XM_006932340.4;refGene_gn=ETFA;refGene_tx=XM_006932341.4
B3 31238591 31238660
refGene_gn=ETFA;refGene_tx=NM_001287554.1;refGene_gn=ETFA;refGene_tx=XM_006932340.4;refGene_gn=ETFA;refGene_tx=XM_006932341.4
B3 31239001 31239103
refGene_gn=ETFA;refGene_tx=NM_001287554.1;refGene_gn=ETFA;refGene_tx=XM_006932340.4;refGene_gn=ETFA;refGene_tx=XM_006932341.4
B3 31239983 31240094
refGene_gn=ETFA;refGene_tx=NM_001287554.1;refGene_gn=ETFA;refGene_tx=XM_006932340.4;refGene_gn=ETFA;refGene_tx=XM_006932341.4
B3 31241851 31241951
refGene_gn=ETFA;refGene_tx=NM_001287554.1;refGene_gn=ETFA;refGene_tx=XM_006932340.4;refGene_gn=ETFA;refGene_tx=XM_006932341.4

B3 31247899 31247982
refGene_gn=ETFA;refGene_tx=NM_001287554.1;refGene_gn=ETFA;refGene_tx=XM_006932340.4;refGene_gn=ETFA;refGene_tx=XM_006932341.4
B3 31248085 31248167
refGene_gn=ETFA;refGene_tx=NM_001287554.1;refGene_gn=ETFA;refGene_tx=XM_006932340.4;refGene_gn=ETFA;refGene_tx=XM_006932341.4
B3 31251114 31251261
refGene_gn=ETFA;refGene_tx=NM_001287554.1;refGene_gn=ETFA;refGene_tx=XM_006932340.4;refGene_gn=ETFA;refGene_tx=XM_006932341.4
B3 31266919 31266949 refGene_gn=ETFA;refGene_tx=XM_006932340.4
B3 31266978 31267017
refGene_gn=ETFA;refGene_tx=NM_001287554.1;refGene_gn=ETFA;refGene_tx=XM_006932341.4
B3 31309649 31309707 refGene_gn=ISL2;refGene_tx=XM_023255043.1
B3 31310555 31310745 refGene_gn=ISL2;refGene_tx=XM_023255043.1
B3 31311025 31311288 refGene_gn=ISL2;refGene_tx=XM_023255043.1
B3 31313079 31313363 refGene_gn=ISL2;refGene_tx=XM_023255043.1
B3 31313945 31314113 refGene_gn=ISL2;refGene_tx=XM_023255043.1
B3 31314515 31314632 refGene_gn=ISL2;refGene_tx=XM_023255043.1
B3 31321801 31321905
refGene_gn=SCAPER;refGene_tx=XM_023255044.1;refGene_gn=SCAPER;refGene_tx=XM_023255045.1;refGene_gn=SCAPER;refGene_tx=XM_023255046.1
B3 31324535 31324587
refGene_gn=SCAPER;refGene_tx=XM_023255044.1;refGene_gn=SCAPER;refGene_tx=XM_023255045.1;refGene_gn=SCAPER;refGene_tx=XM_023255046.1
B3 31327157 31327349
refGene_gn=SCAPER;refGene_tx=XM_023255044.1;refGene_gn=SCAPER;refGene_tx=XM_023255045.1;refGene_gn=SCAPER;refGene_tx=XM_023255046.1
B3 31352943 31353093
refGene_gn=SCAPER;refGene_tx=XM_023255044.1;refGene_gn=SCAPER;refGene_tx=XM_023255045.1;refGene_gn=SCAPER;refGene_tx=XM_023255046.1
B3 31358618 31358856
refGene_gn=SCAPER;refGene_tx=XM_023255044.1;refGene_gn=SCAPER;refGene_tx=XM_023255045.1;refGene_gn=SCAPER;refGene_tx=XM_023255046.1
B3 31383939 31384095
refGene_gn=SCAPER;refGene_tx=XM_023255044.1;refGene_gn=SCAPER;refGene_tx=XM_023255045.1;refGene_gn=SCAPER;refGene_tx=XM_023255046.1
B3 31409349 31409582
refGene_gn=SCAPER;refGene_tx=XM_023255044.1;refGene_gn=SCAPER;refGene_tx=XM_023255045.1;refGene_gn=SCAPER;refGene_tx=XM_023255046.1
B3 31461868 31461992
refGene_gn=SCAPER;refGene_tx=XM_023255044.1;refGene_gn=SCAPER;refGene_tx=XM_023255045.1;refGene_gn=SCAPER;refGene_tx=XM_023255046.1
B3 31487988 31488104
refGene_gn=SCAPER;refGene_tx=XM_023255044.1;refGene_gn=SCAPER;refGene_tx=XM_023255045.1;refGene_gn=SCAPER;refGene_tx=XM_023255046.1
B3 31572507 31572634
refGene_gn=SCAPER;refGene_tx=XM_023255044.1;refGene_gn=SCAPER;refGene_tx=XM_023255045.1;refGene_gn=SCAPER;refGene_tx=XM_023255046.1
B3 31583653 31583719

refGene_gn=SCAPER;refGene_tx=XM_023255044.1;refGene_gn=SCAPER;refGene_tx=XM_023255045.1;refGene_gn=SCAPER;refGene_tx=XM_023255046.1
B3 31620109 31620246
refGene_gn=SCAPER;refGene_tx=XM_023255044.1;refGene_gn=SCAPER;refGene_tx=XM_023255045.1;refGene_gn=SCAPER;refGene_tx=XM_023255046.1
B3 31657679 31657787
refGene_gn=SCAPER;refGene_tx=XM_023255044.1;refGene_gn=SCAPER;refGene_tx=XM_023255045.1;refGene_gn=SCAPER;refGene_tx=XM_023255046.1
B3 31658319 31658472
refGene_gn=SCAPER;refGene_tx=XM_023255044.1;refGene_gn=SCAPER;refGene_tx=XM_023255045.1;refGene_gn=SCAPER;refGene_tx=XM_023255046.1
B3 31660522 31660604
refGene_gn=SCAPER;refGene_tx=XM_023255044.1;refGene_gn=SCAPER;refGene_tx=XM_023255045.1;refGene_gn=SCAPER;refGene_tx=XM_023255046.1
B3 31688232 31688375
refGene_gn=SCAPER;refGene_tx=XM_023255044.1;refGene_gn=SCAPER;refGene_tx=XM_023255045.1;refGene_gn=SCAPER;refGene_tx=XM_023255046.1
B3 31697207 31697363
refGene_gn=SCAPER;refGene_tx=XM_023255044.1;refGene_gn=SCAPER;refGene_tx=XM_023255045.1;refGene_gn=SCAPER;refGene_tx=XM_023255046.1
B3 31710447 31710588
refGene_gn=SCAPER;refGene_tx=XM_023255044.1;refGene_gn=SCAPER;refGene_tx=XM_023255045.1;refGene_gn=SCAPER;refGene_tx=XM_023255046.1
B3 31714115 31714227
refGene_gn=SCAPER;refGene_tx=XM_023255044.1;refGene_gn=SCAPER;refGene_tx=XM_023255045.1;refGene_gn=SCAPER;refGene_tx=XM_023255046.1
B3 31714488 31714606
refGene_gn=SCAPER;refGene_tx=XM_023255044.1;refGene_gn=SCAPER;refGene_tx=XM_023255045.1;refGene_gn=SCAPER;refGene_tx=XM_023255046.1
B3 31714811 31714887
refGene_gn=SCAPER;refGene_tx=XM_023255044.1;refGene_gn=SCAPER;refGene_tx=XM_023255045.1;refGene_gn=SCAPER;refGene_tx=XM_023255046.1
B3 31719954 31720125
refGene_gn=SCAPER;refGene_tx=XM_023255044.1;refGene_gn=SCAPER;refGene_tx=XM_023255045.1;refGene_gn=SCAPER;refGene_tx=XM_023255046.1
B3 31722657 31722891
refGene_gn=SCAPER;refGene_tx=XM_023255044.1;refGene_gn=SCAPER;refGene_tx=XM_023255045.1;refGene_gn=SCAPER;refGene_tx=XM_023255046.1
B3 31725933 31726196
refGene_gn=SCAPER;refGene_tx=XM_023255044.1;refGene_gn=SCAPER;refGene_tx=XM_023255045.1;refGene_gn=SCAPER;refGene_tx=XM_023255046.1
B3 31750913 31751074
refGene_gn=SCAPER;refGene_tx=XM_023255044.1;refGene_gn=SCAPER;refGene_tx=XM_023255045.1;refGene_gn=SCAPER;refGene_tx=XM_023255046.1
B3 31753213 31753330
refGene_gn=SCAPER;refGene_tx=XM_023255044.1;refGene_gn=SCAPER;refGene_tx=XM_023255045.1;refGene_gn=SCAPER;refGene_tx=XM_023255046.1
B3 31763293 31763394
refGene_gn=SCAPER;refGene_tx=XM_023255044.1;refGene_gn=SCAPER;refGene_tx=XM_023255045.1;refGene_gn=SCAPER;refGene_tx=XM_023255046.1

B3 31795466 31795664
refGene_gn=SCAPER;refGene_tx=XM_023255044.1;refGene_gn=SCAPER;refGene_tx=XM_023255045.1;refGene_gn=SCAPER;refGene_tx=XM_023255046.1
B3 31808064 31808135
refGene_gn=SCAPER;refGene_tx=XM_023255044.1;refGene_gn=SCAPER;refGene_tx=XM_023255045.1;refGene_gn=SCAPER;refGene_tx=XM_023255046.1
B3 31820196 31820314
refGene_gn=SCAPER;refGene_tx=XM_023255044.1;refGene_gn=SCAPER;refGene_tx=XM_023255045.1;refGene_gn=SCAPER;refGene_tx=XM_023255046.1
B3 31841840 31841846
refGene_gn=SCAPER;refGene_tx=XM_023255044.1;refGene_gn=SCAPER;refGene_tx=XM_023255045.1;refGene_gn=SCAPER;refGene_tx=XM_023255046.1
B3 31873926 31874070
refGene_gn=RCN2;refGene_tx=XM_023255047.1;refGene_gn=RCN2;refGene_tx=XM_023255048.1
B3 31874456 31874562
refGene_gn=RCN2;refGene_tx=XM_023255047.1;refGene_gn=RCN2;refGene_tx=XM_023255048.1
B3 31877628 31877825
refGene_gn=RCN2;refGene_tx=XM_023255047.1;refGene_gn=RCN2;refGene_tx=XM_023255048.1
B3 31884011 31884125
refGene_gn=RCN2;refGene_tx=XM_023255047.1;refGene_gn=RCN2;refGene_tx=XM_023255048.1
B3 31887739 31887836
refGene_gn=RCN2;refGene_tx=XM_023255047.1;refGene_gn=RCN2;refGene_tx=XM_023255048.1
B3 31888978 31889121
refGene_gn=RCN2;refGene_tx=XM_023255047.1;refGene_gn=RCN2;refGene_tx=XM_023255048.1
B3 31889655 31889808
refGene_gn=RCN2;refGene_tx=XM_023255047.1;refGene_gn=RCN2;refGene_tx=XM_023255048.1
B3 31929357 31929393
refGene_gn=PSTPIP1;refGene_tx=XM_011282985.2;refGene_gn=PSTPIP1;refGene_tx=XM_011282986.2
B3 31949233 31949334
refGene_gn=PSTPIP1;refGene_tx=XM_011282985.2;refGene_gn=PSTPIP1;refGene_tx=XM_011282986.2
B3 31949681 31949756 refGene_gn=PSTPIP1;refGene_tx=XM_011282985.2
B3 31958298 31958333
refGene_gn=PSTPIP1;refGene_tx=XM_011282985.2;refGene_gn=PSTPIP1;refGene_tx=XM_011282986.2
B3 31958566 31958673
refGene_gn=PSTPIP1;refGene_tx=XM_011282985.2;refGene_gn=PSTPIP1;refGene_tx=XM_011282986.2
B3 31960887 31960950
refGene_gn=PSTPIP1;refGene_tx=XM_011282985.2;refGene_gn=PSTPIP1;refGene_tx=XM_011282986.2
B3 31961578 31961677

refGene_gn=PSTPIP1;refGene_tx=XM_011282985.2;refGene_gn=PSTPIP1;refGene_tx=XM_011282986.2
B3 31962480 31962526
refGene_gn=PSTPIP1;refGene_tx=XM_011282985.2;refGene_gn=PSTPIP1;refGene_tx=XM_011282986.2
B3 31969459 31969539
refGene_gn=PSTPIP1;refGene_tx=XM_011282985.2;refGene_gn=PSTPIP1;refGene_tx=XM_011282986.2
B3 31970149 31970248
refGene_gn=PSTPIP1;refGene_tx=XM_011282985.2;refGene_gn=PSTPIP1;refGene_tx=XM_011282986.2
B3 31971311 31971408
refGene_gn=PSTPIP1;refGene_tx=XM_011282985.2;refGene_gn=PSTPIP1;refGene_tx=XM_011282986.2
B3 31971922 31972013
refGene_gn=PSTPIP1;refGene_tx=XM_011282985.2;refGene_gn=PSTPIP1;refGene_tx=XM_011282986.2
B3 31974276 31974332
refGene_gn=PSTPIP1;refGene_tx=XM_011282985.2;refGene_gn=PSTPIP1;refGene_tx=XM_011282986.2
B3 31974532 31974657
refGene_gn=PSTPIP1;refGene_tx=XM_011282985.2;refGene_gn=PSTPIP1;refGene_tx=XM_011282986.2
B3 31975614 31975746
refGene_gn=PSTPIP1;refGene_tx=XM_011282985.2;refGene_gn=PSTPIP1;refGene_tx=XM_011282986.2
B3 31985460 31985553 refGene_gn=TSPAN3;refGene_tx=XM_023255049.1
B3 31993696 31993780 refGene_gn=TSPAN3;refGene_tx=XM_023255049.1
B3 31994044 31994197 refGene_gn=TSPAN3;refGene_tx=XM_023255049.1
B3 31995075 31995177 refGene_gn=TSPAN3;refGene_tx=XM_023255049.1
B3 32000036 32000111 refGene_gn=TSPAN3;refGene_tx=XM_023255049.1
B3 32000329 32000521 refGene_gn=TSPAN3;refGene_tx=XM_023255049.1
B3 32013793 32013856 refGene_gn=TSPAN3;refGene_tx=XM_023255049.1
B3 32056014 32057178
refGene_gn=PEAK1;refGene_tx=XM_011282984.3;refGene_gn=PEAK1;refGene_tx=XM_019832021.2;refGene_gn=PEAK1;refGene_tx=XM_019832022.2
B3 32080237 32080983
refGene_gn=PEAK1;refGene_tx=XM_011282984.3;refGene_gn=PEAK1;refGene_tx=XM_019832021.2;refGene_gn=PEAK1;refGene_tx=XM_019832022.2
B3 32105516 32105710
refGene_gn=PEAK1;refGene_tx=XM_011282984.3;refGene_gn=PEAK1;refGene_tx=XM_019832021.2;refGene_gn=PEAK1;refGene_tx=XM_019832022.2
B3 32127153 32130287
refGene_gn=PEAK1;refGene_tx=XM_011282984.3;refGene_gn=PEAK1;refGene_tx=XM_019832021.2;refGene_gn=PEAK1;refGene_tx=XM_019832022.2
B3 32407134 32407223
refGene_gn=HMG20A;refGene_tx=XM_011282981.3;refGene_gn=HMG20A;refGene_tx=XM_011282982.3;refGene_gn=HMG20A;refGene_tx=XM_019832023.2
B3 32412656 32412804
refGene_gn=HMG20A;refGene_tx=XM_011282981.3;refGene_gn=HMG20A;refGene_

tx=XM_011282982.3;refGene_gn=HMG20A;refGene_tx=XM_019832023.2
B3	32414849	32415062
refGene_gn=HMG20A;refGene_tx=XM_011282981.3;refGene_gn=HMG20A;refGene_tx=XM_011282982.3;refGene_gn=HMG20A;refGene_tx=XM_019832023.2
B3	32418276	32418409
refGene_gn=HMG20A;refGene_tx=XM_011282981.3;refGene_gn=HMG20A;refGene_tx=XM_011282982.3;refGene_gn=HMG20A;refGene_tx=XM_019832023.2
B3	32419281	32419313
refGene_gn=HMG20A;refGene_tx=XM_011282981.3;refGene_gn=HMG20A;refGene_tx=XM_011282982.3;refGene_gn=HMG20A;refGene_tx=XM_019832023.2
B3	32426537	32426613
refGene_gn=HMG20A;refGene_tx=XM_011282981.3;refGene_gn=HMG20A;refGene_tx=XM_011282982.3;refGene_gn=HMG20A;refGene_tx=XM_019832023.2
B3	32427323	32427539
refGene_gn=HMG20A;refGene_tx=XM_011282981.3;refGene_gn=HMG20A;refGene_tx=XM_011282982.3;refGene_gn=HMG20A;refGene_tx=XM_019832023.2
B3	32428228	32428365
refGene_gn=HMG20A;refGene_tx=XM_011282981.3;refGene_gn=HMG20A;refGene_tx=XM_011282982.3;refGene_gn=HMG20A;refGene_tx=XM_019832023.2
B3	32547615	32549472
refGene_gn=LINGO1;refGene_tx=XM_003986920.5;refGene_gn=LINGO1;refGene_tx=XM_019832024.2
B3	32563909	32563915	refGene_gn=LINGO1;refGene_tx=XM_003986920.5
B3	32792384	32792866	refGene_gn=ODF3L1;refGene_tx=XM_003986935.5
B3	32793474	32793588	refGene_gn=ODF3L1;refGene_tx=XM_003986935.5
B3	32794488	32794599	refGene_gn=ODF3L1;refGene_tx=XM_003986935.5
B3	32795509	32795630	refGene_gn=ODF3L1;refGene_tx=XM_003986935.5
B3	32806632	32806726	refGene_gn=CSPG4;refGene_tx=XM_003986922.5
B3	32824850	32825014	refGene_gn=CSPG4;refGene_tx=XM_003986922.5
B3	32826427	32829970	refGene_gn=CSPG4;refGene_tx=XM_003986922.5
B3	32831317	32831800	refGene_gn=CSPG4;refGene_tx=XM_003986922.5
B3	32832161	32832338	refGene_gn=CSPG4;refGene_tx=XM_003986922.5
B3	32834004	32834203	refGene_gn=CSPG4;refGene_tx=XM_003986922.5
B3	32834283	32834418	refGene_gn=CSPG4;refGene_tx=XM_003986922.5
B3	32834545	32834712	refGene_gn=CSPG4;refGene_tx=XM_003986922.5
B3	32837843	32838027	refGene_gn=CSPG4;refGene_tx=XM_003986922.5
B3	32838325	32840160	refGene_gn=CSPG4;refGene_tx=XM_003986922.5
B3	32857395	32857649	refGene_gn=SNX33;refGene_tx=XM_003986923.5
B3	32860557	32860571	refGene_gn=SNX33;refGene_tx=XM_019832026.2
B3	32863571	32865039
refGene_gn=SNX33;refGene_tx=XM_003986923.5;refGene_gn=SNX33;refGene_tx=XM_019832026.2
B3	32872601	32873156	refGene_gn=IMP3;refGene_tx=XM_006932352.4
B3	32886593	32886751
refGene_gn=SNUPN;refGene_tx=XM_003986925.5;refGene_gn=SNUPN;refGene_tx=XM_006932345.4;refGene_gn=SNUPN;refGene_tx=XM_006932346.4;refGene_gn=SNUPN;refGene_tx=XM_006932347.4;refGene_gn=SNUPN;refGene_tx=XM_006932348.4;refGene_gn=SNUPN;refGene_tx=XM_006932349.4;refGene_gn=SNUPN;refGene_tx=XM_006932350.4;refGene_gn=SNUPN;refGene_tx=XM_006932351.4;refGene_gn=SNUPN;refGene_tx=XM_019832027.2

B3 32888662 32888810
refGene_gn=SNUPN;refGene_tx=XM_003986925.5;refGene_gn=SNUPN;refGene_tx=XM_006932345.4;refGene_gn=SNUPN;refGene_tx=XM_006932346.4;refGene_gn=SNUPN;refGene_tx=XM_006932347.4;refGene_gn=SNUPN;refGene_tx=XM_006932348.4;refGene_gn=SNUPN;refGene_tx=XM_006932349.4;refGene_gn=SNUPN;refGene_tx=XM_006932350.4;refGene_gn=SNUPN;refGene_tx=XM_006932351.4;refGene_gn=SNUPN;refGene_tx=XM_019832027.2
B3 32894328 32894433
refGene_gn=SNUPN;refGene_tx=XM_003986925.5;refGene_gn=SNUPN;refGene_tx=XM_006932345.4;refGene_gn=SNUPN;refGene_tx=XM_006932346.4;refGene_gn=SNUPN;refGene_tx=XM_006932347.4;refGene_gn=SNUPN;refGene_tx=XM_006932348.4;refGene_gn=SNUPN;refGene_tx=XM_006932349.4;refGene_gn=SNUPN;refGene_tx=XM_006932350.4;refGene_gn=SNUPN;refGene_tx=XM_006932351.4;refGene_gn=SNUPN;refGene_tx=XM_019832027.2
B3 32894651 32894748
refGene_gn=SNUPN;refGene_tx=XM_003986925.5;refGene_gn=SNUPN;refGene_tx=XM_006932345.4;refGene_gn=SNUPN;refGene_tx=XM_006932346.4;refGene_gn=SNUPN;refGene_tx=XM_006932347.4;refGene_gn=SNUPN;refGene_tx=XM_006932348.4;refGene_gn=SNUPN;refGene_tx=XM_006932349.4;refGene_gn=SNUPN;refGene_tx=XM_006932350.4;refGene_gn=SNUPN;refGene_tx=XM_006932351.4;refGene_gn=SNUPN;refGene_tx=XM_019832027.2
B3 32896874 32896972
refGene_gn=SNUPN;refGene_tx=XM_003986925.5;refGene_gn=SNUPN;refGene_tx=XM_006932345.4;refGene_gn=SNUPN;refGene_tx=XM_006932346.4;refGene_gn=SNUPN;refGene_tx=XM_006932347.4;refGene_gn=SNUPN;refGene_tx=XM_006932348.4;refGene_gn=SNUPN;refGene_tx=XM_006932349.4;refGene_gn=SNUPN;refGene_tx=XM_006932350.4;refGene_gn=SNUPN;refGene_tx=XM_006932351.4;refGene_gn=SNUPN;refGene_tx=XM_019832027.2
B3 32898804 32898882
refGene_gn=SNUPN;refGene_tx=XM_003986925.5;refGene_gn=SNUPN;refGene_tx=XM_006932345.4;refGene_gn=SNUPN;refGene_tx=XM_006932346.4;refGene_gn=SNUPN;refGene_tx=XM_006932347.4;refGene_gn=SNUPN;refGene_tx=XM_006932348.4;refGene_gn=SNUPN;refGene_tx=XM_006932349.4;refGene_gn=SNUPN;refGene_tx=XM_006932350.4;refGene_gn=SNUPN;refGene_tx=XM_006932351.4;refGene_gn=SNUPN;refGene_tx=XM_019832027.2
B3 32900476 32900557
refGene_gn=SNUPN;refGene_tx=XM_003986925.5;refGene_gn=SNUPN;refGene_tx=XM_006932345.4;refGene_gn=SNUPN;refGene_tx=XM_006932346.4;refGene_gn=SNUPN;refGene_tx=XM_006932347.4;refGene_gn=SNUPN;refGene_tx=XM_006932348.4;refGene_gn=SNUPN;refGene_tx=XM_006932349.4;refGene_gn=SNUPN;refGene_tx=XM_006932350.4;refGene_gn=SNUPN;refGene_tx=XM_006932351.4;refGene_gn=SNUPN;refGene_tx=XM_019832027.2
B3 32902005 32902326
refGene_gn=SNUPN;refGene_tx=XM_003986925.5;refGene_gn=SNUPN;refGene_tx=XM_006932345.4;refGene_gn=SNUPN;refGene_tx=XM_006932346.4;refGene_gn=SNUPN;refGene_tx=XM_006932347.4;refGene_gn=SNUPN;refGene_tx=XM_006932348.4;refGene_gn=SNUPN;refGene_tx=XM_006932349.4;refGene_gn=SNUPN;refGene_tx=XM_006932350.4;refGene_gn=SNUPN;refGene_tx=XM_006932351.4;refGene_gn=SNUPN;refGene_tx=XM_019832027.2
B3 32914293 32914356 refGene_gn=PTPN9;refGene_tx=XM_023255050.1

B3 32950257 32950401 refGene_gn=PTPN9;refGene_tx=XM_023255050.1
B3 32953117 32953207 refGene_gn=PTPN9;refGene_tx=XM_023255050.1
B3 32953931 32954056 refGene_gn=PTPN9;refGene_tx=XM_023255050.1
B3 32959108 32959214 refGene_gn=PTPN9;refGene_tx=XM_023255050.1
B3 32967497 32967608 refGene_gn=PTPN9;refGene_tx=XM_023255050.1
B3 32969883 32970212 refGene_gn=PTPN9;refGene_tx=XM_023255050.1
B3 32980275 32980369 refGene_gn=PTPN9;refGene_tx=XM_023255050.1
B3 32986037 32986104 refGene_gn=PTPN9;refGene_tx=XM_023255050.1
B3 32990842 32990921 refGene_gn=PTPN9;refGene_tx=XM_023255050.1
B3 32992087 32992238 refGene_gn=PTPN9;refGene_tx=XM_023255050.1
B3 32992662 32992870 refGene_gn=PTPN9;refGene_tx=XM_023255050.1
B3 32993659 32993874 refGene_gn=PTPN9;refGene_tx=XM_023255050.1
B3 33025417 33025606
refGene_gn=SIN3A;refGene_tx=XM_003986928.5;refGene_gn=SIN3A;refGene_tx
=XM_019832029.2
B3 33029459 33029636
refGene_gn=SIN3A;refGene_tx=XM_003986928.5;refGene_gn=SIN3A;refGene_tx
=XM_019832029.2
B3 33034855 33034962
refGene_gn=SIN3A;refGene_tx=XM_003986928.5;refGene_gn=SIN3A;refGene_tx
=XM_019832029.2
B3 33035666 33035949
refGene_gn=SIN3A;refGene_tx=XM_003986928.5;refGene_gn=SIN3A;refGene_tx
=XM_019832029.2
B3 33036999 33037251
refGene_gn=SIN3A;refGene_tx=XM_003986928.5;refGene_gn=SIN3A;refGene_tx
=XM_019832029.2
B3 33038180 33038333
refGene_gn=SIN3A;refGene_tx=XM_003986928.5;refGene_gn=SIN3A;refGene_tx
=XM_019832029.2
B3 33038474 33038633
refGene_gn=SIN3A;refGene_tx=XM_003986928.5;refGene_gn=SIN3A;refGene_tx
=XM_019832029.2
B3 33040380 33040470
refGene_gn=SIN3A;refGene_tx=XM_003986928.5;refGene_gn=SIN3A;refGene_tx
=XM_019832029.2
B3 33043391 33043510
refGene_gn=SIN3A;refGene_tx=XM_003986928.5;refGene_gn=SIN3A;refGene_tx
=XM_019832029.2
B3 33044055 33044266
refGene_gn=SIN3A;refGene_tx=XM_003986928.5;refGene_gn=SIN3A;refGene_tx
=XM_019832029.2
B3 33044559 33044676
refGene_gn=SIN3A;refGene_tx=XM_003986928.5;refGene_gn=SIN3A;refGene_tx
=XM_019832029.2
B3 33047437 33047676
refGene_gn=SIN3A;refGene_tx=XM_003986928.5;refGene_gn=SIN3A;refGene_tx
=XM_019832029.2
B3 33049223 33049407
refGene_gn=SIN3A;refGene_tx=XM_003986928.5;refGene_gn=SIN3A;refGene_tx

=XM_019832029.2
B3 33050791 33051365
refGene_gn=SIN3A;refGene_tx=XM_003986928.5;refGene_gn=SIN3A;refGene_tx
=XM_019832029.2
B3 33053917 33054087
refGene_gn=SIN3A;refGene_tx=XM_003986928.5;refGene_gn=SIN3A;refGene_tx
=XM_019832029.2
B3 33057822 33057996
refGene_gn=SIN3A;refGene_tx=XM_003986928.5;refGene_gn=SIN3A;refGene_tx
=XM_019832029.2
B3 33066833 33066926
refGene_gn=SIN3A;refGene_tx=XM_003986928.5;refGene_gn=SIN3A;refGene_tx
=XM_019832029.2
B3 33067820 33067915
refGene_gn=SIN3A;refGene_tx=XM_003986928.5;refGene_gn=SIN3A;refGene_tx
=XM_019832029.2
B3 33074849 33075057
refGene_gn=SIN3A;refGene_tx=XM_003986928.5;refGene_gn=SIN3A;refGene_tx
=XM_019832029.2
B3 33079208 33079439
refGene_gn=SIN3A;refGene_tx=XM_003986928.5;refGene_gn=SIN3A;refGene_tx
=XM_019832029.2
B3 33082737 33082835 refGene_gn=MAN2C1;refGene_tx=XM_003986929.5
B3 33083109 33083235 refGene_gn=MAN2C1;refGene_tx=XM_003986929.5
B3 33083668 33083792 refGene_gn=MAN2C1;refGene_tx=XM_003986929.5
B3 33084956 33085027 refGene_gn=MAN2C1;refGene_tx=XM_003986929.5
B3 33086530 33086708 refGene_gn=MAN2C1;refGene_tx=XM_003986929.5
B3 33087024 33087214 refGene_gn=MAN2C1;refGene_tx=XM_003986929.5
B3 33088279 33088386 refGene_gn=MAN2C1;refGene_tx=XM_003986929.5
B3 33088562 33088673 refGene_gn=MAN2C1;refGene_tx=XM_003986929.5
B3 33089064 33089157 refGene_gn=MAN2C1;refGene_tx=XM_003986929.5
B3 33089280 33089397 refGene_gn=MAN2C1;refGene_tx=XM_003986929.5
B3 33089578 33089674 refGene_gn=MAN2C1;refGene_tx=XM_003986929.5
B3 33089787 33089933 refGene_gn=MAN2C1;refGene_tx=XM_003986929.5
B3 33090297 33090421 refGene_gn=MAN2C1;refGene_tx=XM_003986929.5
B3 33090761 33090883 refGene_gn=MAN2C1;refGene_tx=XM_003986929.5
B3 33090985 33091071 refGene_gn=MAN2C1;refGene_tx=XM_003986929.5
B3 33091197 33091353 refGene_gn=MAN2C1;refGene_tx=XM_003986929.5
B3 33091558 33091650 refGene_gn=MAN2C1;refGene_tx=XM_003986929.5
B3 33091820 33091915 refGene_gn=MAN2C1;refGene_tx=XM_003986929.5
B3 33092170 33092275 refGene_gn=MAN2C1;refGene_tx=XM_003986929.5
B3 33092357 33092514 refGene_gn=MAN2C1;refGene_tx=XM_003986929.5
B3 33092607 33092751 refGene_gn=MAN2C1;refGene_tx=XM_003986929.5
B3 33093111 33093221 refGene_gn=MAN2C1;refGene_tx=XM_003986929.5
B3 33093323 33093403 refGene_gn=MAN2C1;refGene_tx=XM_003986929.5
B3 33093508 33093657 refGene_gn=MAN2C1;refGene_tx=XM_003986929.5
B3 33093739 33093849 refGene_gn=MAN2C1;refGene_tx=XM_003986929.5
B3 33093921 33094048 refGene_gn=MAN2C1;refGene_tx=XM_003986929.5
B3 33094721 33094913
refGene_gn=NEIL1;refGene_tx=XM_006932356.4;refGene_gn=NEIL1;refGene_tx

=XM_019832030.2;refGene_gn=NEIL1;refGene_tx=XM_019832031.2;refGene_gn=NEIL1;refGene_tx=XM_019832032.2
B3 33095064 33095230 refGene_gn=NEIL1;refGene_tx=XM_006932356.4;refGene_gn=NEIL1;refGene_tx=XM_019832030.2;refGene_gn=NEIL1;refGene_tx=XM_019832031.2
B3 33095362 33095424 refGene_gn=NEIL1;refGene_tx=XM_006932356.4;refGene_gn=NEIL1;refGene_tx=XM_019832030.2;refGene_gn=NEIL1;refGene_tx=XM_019832031.2;refGene_gn=NEIL1;refGene_tx=XM_019832032.2
B3 33095595 33095623 refGene_gn=NEIL1;refGene_tx=XM_006932356.4;refGene_gn=NEIL1;refGene_tx=XM_019832030.2;refGene_gn=NEIL1;refGene_tx=XM_019832031.2;refGene_gn=NEIL1;refGene_tx=XM_019832032.2
B3 33096015 33096143 refGene_gn=NEIL1;refGene_tx=XM_006932356.4;refGene_gn=NEIL1;refGene_tx=XM_019832030.2;refGene_gn=NEIL1;refGene_tx=XM_019832031.2;refGene_gn=NEIL1;refGene_tx=XM_019832032.2
B3 33096522 33096622 refGene_gn=NEIL1;refGene_tx=XM_006932356.4;refGene_gn=NEIL1;refGene_tx=XM_019832030.2;refGene_gn=NEIL1;refGene_tx=XM_019832031.2;refGene_gn=NEIL1;refGene_tx=XM_019832032.2
B3 33096905 33096969 refGene_gn=NEIL1;refGene_tx=XM_006932356.4;refGene_gn=NEIL1;refGene_tx=XM_019832030.2;refGene_gn=NEIL1;refGene_tx=XM_019832031.2;refGene_gn=NEIL1;refGene_tx=XM_019832032.2
B3 33097058 33097178 refGene_gn=NEIL1;refGene_tx=XM_006932356.4;refGene_gn=NEIL1;refGene_tx=XM_019832030.2;refGene_gn=NEIL1;refGene_tx=XM_019832031.2;refGene_gn=NEIL1;refGene_tx=XM_019832032.2
B3 33099095 33099529 refGene_gn=NEIL1;refGene_tx=XM_006932356.4;refGene_gn=NEIL1;refGene_tx=XM_019832030.2;refGene_gn=NEIL1;refGene_tx=XM_019832031.2;refGene_gn=NEIL1;refGene_tx=XM_019832032.2
B3 33107196 33107237 refGene_gn=COMMD4;refGene_tx=XM_006932357.4
B3 33107326 33107503 refGene_gn=COMMD4;refGene_tx=XM_006932357.4
B3 33107830 33107911 refGene_gn=COMMD4;refGene_tx=XM_006932357.4
B3 33108054 33108174 refGene_gn=COMMD4;refGene_tx=XM_006932357.4
B3 33108480 33108520 refGene_gn=COMMD4;refGene_tx=XM_006932357.4
B3 33108682 33108748 refGene_gn=COMMD4;refGene_tx=XM_006932357.4
B3 33108965 33109037 refGene_gn=COMMD4;refGene_tx=XM_006932357.4
B3 33112973 33112976 refGene_gn=COMMD4;refGene_tx=XM_006932357.4
B3 33152995 33153363 refGene_gn=CB3H15orf39;refGene_tx=XM_011282977.3;refGene_gn=CB3H15orf39;refGene_tx=XM_019832033.2;refGene_gn=CB3H15orf39;refGene_tx=XM_023255051.1
B3 33155262 33157977 refGene_gn=CB3H15orf39;refGene_tx=XM_011282977.3;refGene_gn=CB3H15orf39;refGene_tx=XM_011282978.3;refGene_gn=CB3H15orf39;refGene_tx=XM_019832033.2;refGene_gn=CB3H15orf39;refGene_tx=XM_023255051.1
B3 33289679 33289765

refGene_gn=PPCDC;refGene_tx=XM_006932363.4;refGene_gn=PPCDC;refGene_tx=XM_011282973.3;refGene_gn=PPCDC;refGene_tx=XM_011282974.3;refGene_gn=PPCDC;refGene_tx=XM_011282975.3;refGene_gn=PPCDC;refGene_tx=XM_011282976.3;refGene_gn=PPCDC;refGene_tx=XM_019832035.2;refGene_gn=PPCDC;refGene_tx=XM_019832036.2;refGene_gn=PPCDC;refGene_tx=XM_019832037.2
B3 33290187 33290356
refGene_gn=PPCDC;refGene_tx=XM_006932363.4;refGene_gn=PPCDC;refGene_tx=XM_011282973.3;refGene_gn=PPCDC;refGene_tx=XM_011282974.3;refGene_gn=PPCDC;refGene_tx=XM_011282975.3;refGene_gn=PPCDC;refGene_tx=XM_011282976.3;refGene_gn=PPCDC;refGene_tx=XM_019832035.2;refGene_gn=PPCDC;refGene_tx=XM_019832036.2;refGene_gn=PPCDC;refGene_tx=XM_019832037.2
B3 33294038 33294167
refGene_gn=PPCDC;refGene_tx=XM_006932363.4;refGene_gn=PPCDC;refGene_tx=XM_011282973.3;refGene_gn=PPCDC;refGene_tx=XM_011282976.3;refGene_gn=PPCDC;refGene_tx=XM_019832035.2;refGene_gn=PPCDC;refGene_tx=XM_019832036.2;refGene_gn=PPCDC;refGene_tx=XM_019832037.2
B3 33295043 33295139
refGene_gn=PPCDC;refGene_tx=XM_006932363.4;refGene_gn=PPCDC;refGene_tx=XM_011282973.3;refGene_gn=PPCDC;refGene_tx=XM_011282974.3;refGene_gn=PPCDC;refGene_tx=XM_019832035.2;refGene_gn=PPCDC;refGene_tx=XM_019832036.2
B3 33306790 33306859 refGene_gn=PPCDC;refGene_tx=XM_019832036.2
B3 33309006 33309144
refGene_gn=PPCDC;refGene_tx=XM_006932363.4;refGene_gn=PPCDC;refGene_tx=XM_011282973.3;refGene_gn=PPCDC;refGene_tx=XM_011282974.3;refGene_gn=PPCDC;refGene_tx=XM_011282975.3;refGene_gn=PPCDC;refGene_tx=XM_019832035.2
B3 33317006 33317201 refGene_gn=SCAMP5;refGene_tx=XM_019832034.2
B3 33317465 33317583 refGene_gn=SCAMP5;refGene_tx=XM_019832034.2
B3 33318010 33318112 refGene_gn=SCAMP5;refGene_tx=XM_019832034.2
B3 33319262 33319419 refGene_gn=SCAMP5;refGene_tx=XM_019832034.2
B3 33322671 33322800 refGene_gn=SCAMP5;refGene_tx=XM_019832034.2
B3 33323603 33323610 refGene_gn=SCAMP5;refGene_tx=XM_019832034.2
B3 33358052 33358694 refGene_gn=RPP25;refGene_tx=XM_003986952.5
B3 33370322 33370428
refGene_gn=LOC101081808;refGene_tx=XM_003986941.4
B3 33376344 33376461
refGene_gn=LOC101081808;refGene_tx=XM_003986941.4
B3 33378587 33378709
refGene_gn=LOC101081808;refGene_tx=XM_003986941.4
B3 33379893 33380007
refGene_gn=LOC101081808;refGene_tx=XM_003986941.4
B3 33400590 33400804
refGene_gn=FAM219B;refGene_tx=XM_003986942.5;refGene_gn=FAM219B;refGene_tx=XM_011282972.3
B3 33401016 33401104
refGene_gn=FAM219B;refGene_tx=XM_003986942.5;refGene_gn=FAM219B;refGene_tx=XM_011282972.3
B3 33401795 33401873
refGene_gn=FAM219B;refGene_tx=XM_003986942.5;refGene_gn=FAM219B;refGen

e_tx=XM_011282972.3
B3 33402259 33402308
refGene_gn=FAM219B;refGene_tx=XM_003986942.5;refGene_gn=FAM219B;refGen
e_tx=XM_011282972.3
B3 33403836 33404004
refGene_gn=FAM219B;refGene_tx=XM_003986942.5;refGene_gn=FAM219B;refGen
e_tx=XM_011282972.3
B3 33407934 33408153
refGene_gn=MPI;refGene_tx=XM_003986943.5;refGene_gn=MPI;refGene_tx=XM_
006932364.4;refGene_gn=MPI;refGene_tx=XM_011282971.3
B3 33408685 33408888
refGene_gn=MPI;refGene_tx=XM_003986943.5;refGene_gn=MPI;refGene_tx=XM_
006932364.4;refGene_gn=MPI;refGene_tx=XM_011282971.3
B3 33408912 33408962 refGene_gn=MPI;refGene_tx=XM_019832039.1
B3 33409654 33409828
refGene_gn=MPI;refGene_tx=XM_003986943.5;refGene_gn=MPI;refGene_tx=XM_
006932364.4;refGene_gn=MPI;refGene_tx=XM_011282971.3;refGene_gn=MPI;re
fGene_tx=XM_019832039.1
B3 33411814 33411997
refGene_gn=MPI;refGene_tx=XM_003986943.5;refGene_gn=MPI;refGene_tx=XM_
006932364.4;refGene_gn=MPI;refGene_tx=XM_011282971.3;refGene_gn=MPI;re
fGene_tx=XM_019832039.1
B3 33412560 33412702
refGene_gn=MPI;refGene_tx=XM_003986943.5;refGene_gn=MPI;refGene_tx=XM_
006932364.4;refGene_gn=MPI;refGene_tx=XM_011282971.3;refGene_gn=MPI;re
fGene_tx=XM_019832039.1
B3 33413365 33413566
refGene_gn=MPI;refGene_tx=XM_003986943.5;refGene_gn=MPI;refGene_tx=XM_
006932364.4;refGene_gn=MPI;refGene_tx=XM_011282971.3;refGene_gn=MPI;re
fGene_tx=XM_019832039.1
B3 33414272 33414400
refGene_gn=MPI;refGene_tx=XM_003986943.5;refGene_gn=MPI;refGene_tx=XM_
006932364.4;refGene_gn=MPI;refGene_tx=XM_019832039.1
B3 33414803 33414810 refGene_gn=MPI;refGene_tx=XM_003986943.5
B3 33414885 33414901
refGene_gn=MPI;refGene_tx=XM_006932364.4;refGene_gn=MPI;refGene_tx=XM_
019832039.1
B3 33421172 33421229
refGene_gn=SCAMP2;refGene_tx=XM_003986944.5;refGene_gn=SCAMP2;refGene_
tx=XM_019832043.2
B3 33433898 33433967
refGene_gn=SCAMP2;refGene_tx=XM_003986944.5;refGene_gn=SCAMP2;refGene_
tx=XM_019832043.2
B3 33434399 33434498
refGene_gn=SCAMP2;refGene_tx=XM_003986944.5;refGene_gn=SCAMP2;refGene_
tx=XM_019832043.2
B3 33435070 33435199 refGene_gn=SCAMP2;refGene_tx=XM_019832043.2
B3 33436332 33436450
refGene_gn=SCAMP2;refGene_tx=XM_003986944.5;refGene_gn=SCAMP2;refGene_
tx=XM_019832043.2

B3 33437003 33437132
refGene_gn=SCAMP2;refGene_tx=XM_003986944.5;refGene_gn=SCAMP2;refGene_tx=XM_019832043.2
B3 33437808 33437968
refGene_gn=SCAMP2;refGene_tx=XM_003986944.5;refGene_gn=SCAMP2;refGene_tx=XM_019832043.2
B3 33439031 33439133
refGene_gn=SCAMP2;refGene_tx=XM_003986944.5;refGene_gn=SCAMP2;refGene_tx=XM_019832043.2
B3 33443337 33443458
refGene_gn=SCAMP2;refGene_tx=XM_003986944.5;refGene_gn=SCAMP2;refGene_tx=XM_019832043.2
B3 33443648 33443783
refGene_gn=SCAMP2;refGene_tx=XM_003986944.5;refGene_gn=SCAMP2;refGene_tx=XM_019832043.2
B3 33445802 33445904
refGene_gn=ULK3;refGene_tx=XM_003986945.5;refGene_gn=ULK3;refGene_tx=XM_019832041.2
B3 33446567 33446708
refGene_gn=ULK3;refGene_tx=XM_003986945.5;refGene_gn=ULK3;refGene_tx=XM_019832041.2
B3 33446794 33446915
refGene_gn=ULK3;refGene_tx=XM_003986945.5;refGene_gn=ULK3;refGene_tx=XM_006932369.4;refGene_gn=ULK3;refGene_tx=XM_019832041.2
B3 33447389 33447494
refGene_gn=ULK3;refGene_tx=XM_003986945.5;refGene_gn=ULK3;refGene_tx=XM_006932369.4;refGene_gn=ULK3;refGene_tx=XM_019832041.2
B3 33448231 33448375
refGene_gn=ULK3;refGene_tx=XM_003986945.5;refGene_gn=ULK3;refGene_tx=XM_006932369.4;refGene_gn=ULK3;refGene_tx=XM_019832041.2
B3 33448583 33448666
refGene_gn=ULK3;refGene_tx=XM_003986945.5;refGene_gn=ULK3;refGene_tx=XM_006932369.4;refGene_gn=ULK3;refGene_tx=XM_019832041.2
B3 33449016 33449172
refGene_gn=ULK3;refGene_tx=XM_003986945.5;refGene_gn=ULK3;refGene_tx=XM_006932369.4;refGene_gn=ULK3;refGene_tx=XM_019832041.2
B3 33449316 33449422
refGene_gn=ULK3;refGene_tx=XM_003986945.5;refGene_gn=ULK3;refGene_tx=XM_006932369.4;refGene_gn=ULK3;refGene_tx=XM_019832041.2
B3 33449621 33449662
refGene_gn=ULK3;refGene_tx=XM_003986945.5;refGene_gn=ULK3;refGene_tx=XM_006932369.4;refGene_gn=ULK3;refGene_tx=XM_019832041.2
B3 33449890 33449993
refGene_gn=ULK3;refGene_tx=XM_003986945.5;refGene_gn=ULK3;refGene_tx=XM_006932369.4;refGene_gn=ULK3;refGene_tx=XM_019832041.2
B3 33450122 33450187
refGene_gn=ULK3;refGene_tx=XM_003986945.5;refGene_gn=ULK3;refGene_tx=XM_006932369.4;refGene_gn=ULK3;refGene_tx=XM_019832041.2
B3 33450272 33450351
refGene_gn=ULK3;refGene_tx=XM_003986945.5;refGene_gn=ULK3;refGene_tx=X

M_006932369.4;refGene_gn=ULK3;refGene_tx=XM_019832041.2
B3 33450427 33450468
refGene_gn=ULK3;refGene_tx=XM_003986945.5;refGene_gn=ULK3;refGene_tx=XM_006932369.4;refGene_gn=ULK3;refGene_tx=XM_019832041.2
B3 33450856 33450904
refGene_gn=ULK3;refGene_tx=XM_003986945.5;refGene_gn=ULK3;refGene_tx=XM_006932369.4;refGene_gn=ULK3;refGene_tx=XM_019832041.2
B3 33451092 33451159
refGene_gn=ULK3;refGene_tx=XM_003986945.5;refGene_gn=ULK3;refGene_tx=XM_006932369.4;refGene_gn=ULK3;refGene_tx=XM_019832041.2
B3 33451277 33451294
refGene_gn=ULK3;refGene_tx=XM_003986945.5;refGene_gn=ULK3;refGene_tx=XM_006932369.4;refGene_gn=ULK3;refGene_tx=XM_019832041.2
B3 33457697 33457922 refGene_gn=CPLX3;refGene_tx=XM_003986946.4
B3 33459592 33459680 refGene_gn=CPLX3;refGene_tx=XM_003986946.4
B3 33460856 33461020 refGene_gn=CPLX3;refGene_tx=XM_003986946.4
B3 33462083 33462213 refGene_gn=LMAN1L;refGene_tx=XM_023255053.1
B3 33463280 33463408 refGene_gn=LMAN1L;refGene_tx=XM_023255053.1
B3 33464033 33464157 refGene_gn=LMAN1L;refGene_tx=XM_023255053.1
B3 33465008 33465092 refGene_gn=LMAN1L;refGene_tx=XM_023255053.1
B3 33465465 33465524 refGene_gn=LMAN1L;refGene_tx=XM_023255053.1
B3 33466191 33466343 refGene_gn=LMAN1L;refGene_tx=XM_023255053.1
B3 33466647 33466780 refGene_gn=LMAN1L;refGene_tx=XM_023255053.1
B3 33467381 33467437 refGene_gn=LMAN1L;refGene_tx=XM_023255053.1
B3 33468284 33468405 refGene_gn=LMAN1L;refGene_tx=XM_023255053.1
B3 33468799 33468899 refGene_gn=LMAN1L;refGene_tx=XM_023255053.1
B3 33470229 33470288 refGene_gn=LMAN1L;refGene_tx=XM_023255053.1
B3 33470389 33470497 refGene_gn=LMAN1L;refGene_tx=XM_023255053.1
B3 33470615 33470770 refGene_gn=LMAN1L;refGene_tx=XM_023255053.1
B3 33473587 33473765 refGene_gn=LMAN1L;refGene_tx=XM_023255053.1
B3 33479679 33479862 refGene_gn=CSK;refGene_tx=XM_003986947.4
B3 33480125 33480212 refGene_gn=CSK;refGene_tx=XM_003986947.4
B3 33480324 33480520 refGene_gn=CSK;refGene_tx=XM_003986947.4
B3 33480615 33480689 refGene_gn=CSK;refGene_tx=XM_003986947.4
B3 33481149 33481240 refGene_gn=CSK;refGene_tx=XM_003986947.4
B3 33481341 33481441 refGene_gn=CSK;refGene_tx=XM_003986947.4
B3 33481524 33481590 refGene_gn=CSK;refGene_tx=XM_003986947.4
B3 33481777 33481871 refGene_gn=CSK;refGene_tx=XM_003986947.4
B3 33482734 33482954 refGene_gn=CSK;refGene_tx=XM_003986947.4
B3 33483329 33483442 refGene_gn=CSK;refGene_tx=XM_003986947.4
B3 33483564 33483678 refGene_gn=CSK;refGene_tx=XM_003986947.4
B3 33484019 33484034 refGene_gn=CSK;refGene_tx=XM_003986947.4
B3 33525579 33525877
refGene_gn=CYP1A2;refGene_tx=NM_001048013.1;refGene_gn=CYP1A2;refGene_tx=XM_019831732.2
B3 33526602 33526689
refGene_gn=CYP1A2;refGene_tx=NM_001048013.1;refGene_gn=CYP1A2;refGene_tx=XM_019831732.2
B3 33527606 33527730
refGene_gn=CYP1A2;refGene_tx=NM_001048013.1;refGene_gn=CYP1A2;refGene_

tx=XM_019831732.2
B3 33527909 33527999
refGene_gn=CYP1A2;refGene_tx=NM_001048013.1;refGene_gn=CYP1A2;refGene_tx=XM_019831732.2
B3 33528475 33528596
refGene_gn=CYP1A2;refGene_tx=NM_001048013.1;refGene_gn=CYP1A2;refGene_tx=XM_019831732.2
B3 33529142 33529961
refGene_gn=CYP1A2;refGene_tx=NM_001048013.1;refGene_gn=CYP1A2;refGene_tx=XM_019831732.2
B3 33549489 33550329
refGene_gn=CYP1A1;refGene_tx=NM_001024859.1;refGene_gn=CYP1A1;refGene_tx=XM_011282840.2;refGene_gn=CYP1A1;refGene_tx=XM_019831733.1;refGene_gn=CYP1A1;refGene_tx=XM_019831734.1
B3 33550878 33551005
refGene_gn=CYP1A1;refGene_tx=NM_001024859.1;refGene_gn=CYP1A1;refGene_tx=XM_011282840.2;refGene_gn=CYP1A1;refGene_tx=XM_019831733.1;refGene_gn=CYP1A1;refGene_tx=XM_019831734.1
B3 33551096 33551186
refGene_gn=CYP1A1;refGene_tx=NM_001024859.1;refGene_gn=CYP1A1;refGene_tx=XM_011282840.2;refGene_gn=CYP1A1;refGene_tx=XM_019831733.1;refGene_gn=CYP1A1;refGene_tx=XM_019831734.1
B3 33551270 33551394
refGene_gn=CYP1A1;refGene_tx=NM_001024859.1;refGene_gn=CYP1A1;refGene_tx=XM_011282840.2;refGene_gn=CYP1A1;refGene_tx=XM_019831733.1;refGene_gn=CYP1A1;refGene_tx=XM_019831734.1
B3 33551539 33551626
refGene_gn=CYP1A1;refGene_tx=NM_001024859.1;refGene_gn=CYP1A1;refGene_tx=XM_011282840.2;refGene_gn=CYP1A1;refGene_tx=XM_019831733.1;refGene_gn=CYP1A1;refGene_tx=XM_019831734.1
B3 33551777 33552063
refGene_gn=CYP1A1;refGene_tx=NM_001024859.1;refGene_gn=CYP1A1;refGene_tx=XM_011282840.2;refGene_gn=CYP1A1;refGene_tx=XM_019831733.1;refGene_gn=CYP1A1;refGene_tx=XM_019831734.1
B3 33567154 33567241 refGene_gn=EDC3;refGene_tx=XM_011282968.3
B3 33585008 33585190
refGene_gn=EDC3;refGene_tx=XM_011282968.3;refGene_gn=EDC3;refGene_tx=XM_011282969.3;refGene_gn=EDC3;refGene_tx=XM_019832045.2;refGene_gn=EDC3;refGene_tx=XM_023255055.1
B3 33588422 33588742
refGene_gn=EDC3;refGene_tx=XM_011282968.3;refGene_gn=EDC3;refGene_tx=XM_011282969.3;refGene_gn=EDC3;refGene_tx=XM_019832045.2;refGene_gn=EDC3;refGene_tx=XM_023255055.1
B3 33605312 33605648
refGene_gn=EDC3;refGene_tx=XM_011282968.3;refGene_gn=EDC3;refGene_tx=XM_011282969.3;refGene_gn=EDC3;refGene_tx=XM_019832045.2;refGene_gn=EDC3;refGene_tx=XM_023255055.1
B3 33619963 33620117
refGene_gn=EDC3;refGene_tx=XM_011282968.3;refGene_gn=EDC3;refGene_tx=XM_011282969.3;refGene_gn=EDC3;refGene_tx=XM_019832045.2;refGene_gn=EDC

3;refGene_tx=XM_023255055.1
B3	33622564	33622782
refGene_gn=EDC3;refGene_tx=XM_011282968.3;refGene_gn=EDC3;refGene_tx=XM_011282969.3;refGene_gn=EDC3;refGene_tx=XM_019832045.2;refGene_gn=EDC3;refGene_tx=XM_023255055.1
B3	33625436	33625771
refGene_gn=EDC3;refGene_tx=XM_011282968.3;refGene_gn=EDC3;refGene_tx=XM_011282969.3;refGene_gn=EDC3;refGene_tx=XM_019832045.2;refGene_gn=EDC3;refGene_tx=XM_023255055.1
B3	33628589	33628766
refGene_gn=CLK3;refGene_tx=XM_003986949.4;refGene_gn=CLK3;refGene_tx=XM_006932371.4;refGene_gn=CLK3;refGene_tx=XM_019832047.1
B3	33629404	33629495
refGene_gn=CLK3;refGene_tx=XM_003986949.4;refGene_gn=CLK3;refGene_tx=XM_006932371.4;refGene_gn=CLK3;refGene_tx=XM_019832047.1
B3	33629792	33629872
refGene_gn=CLK3;refGene_tx=XM_003986949.4;refGene_gn=CLK3;refGene_tx=XM_006932371.4;refGene_gn=CLK3;refGene_tx=XM_019832047.1
B3	33630286	33630369
refGene_gn=CLK3;refGene_tx=XM_003986949.4;refGene_gn=CLK3;refGene_tx=XM_006932371.4;refGene_gn=CLK3;refGene_tx=XM_019832047.1
B3	33630621	33630751
refGene_gn=CLK3;refGene_tx=XM_003986949.4;refGene_gn=CLK3;refGene_tx=XM_006932371.4;refGene_gn=CLK3;refGene_tx=XM_019832047.1
B3	33630841	33630936
refGene_gn=CLK3;refGene_tx=XM_003986949.4;refGene_gn=CLK3;refGene_tx=XM_006932371.4;refGene_gn=CLK3;refGene_tx=XM_019832047.1
B3	33632133	33632300
refGene_gn=CLK3;refGene_tx=XM_003986949.4;refGene_gn=CLK3;refGene_tx=XM_006932371.4;refGene_gn=CLK3;refGene_tx=XM_019832047.1
B3	33632992	33633109
refGene_gn=CLK3;refGene_tx=XM_003986949.4;refGene_gn=CLK3;refGene_tx=XM_006932371.4;refGene_gn=CLK3;refGene_tx=XM_019832047.1
B3	33635436	33635503
refGene_gn=CLK3;refGene_tx=XM_003986949.4;refGene_gn=CLK3;refGene_tx=XM_006932371.4;refGene_gn=CLK3;refGene_tx=XM_019832047.1
B3	33635780	33635877
refGene_gn=CLK3;refGene_tx=XM_003986949.4;refGene_gn=CLK3;refGene_tx=XM_006932371.4;refGene_gn=CLK3;refGene_tx=XM_019832047.1
B3	33637740	33637957
refGene_gn=CLK3;refGene_tx=XM_003986949.4;refGene_gn=CLK3;refGene_tx=XM_006932371.4;refGene_gn=CLK3;refGene_tx=XM_019832047.1
B3	33638574	33638726
refGene_gn=CLK3;refGene_tx=XM_003986949.4;refGene_gn=CLK3;refGene_tx=XM_006932371.4;refGene_gn=CLK3;refGene_tx=XM_019832047.1
B3	33664405	33664563
refGene_gn=ARID3B;refGene_tx=XM_019832048.2;refGene_gn=ARID3B;refGene_tx=XM_019832049.2;refGene_gn=ARID3B;refGene_tx=XM_019832050.2;refGene_gn=ARID3B;refGene_tx=XM_019832052.2;refGene_gn=ARID3B;refGene_tx=XM_023255056.1;refGene_gn=ARID3B;refGene_tx=XM_023255058.1;refGene_gn=ARID3

B;refGene_tx=XM_023255059.1;refGene_gn=ARID3B;refGene_tx=XM_023255060.
1
B3 33666432 33666534
refGene_gn=ARID3B;refGene_tx=XM_019832048.2;refGene_gn=ARID3B;refGene_
tx=XM_019832049.2;refGene_gn=ARID3B;refGene_tx=XM_019832050.2;refGene_
gn=ARID3B;refGene_tx=XM_019832052.2;refGene_gn=ARID3B;refGene_tx=XM_02
3255056.1;refGene_gn=ARID3B;refGene_tx=XM_023255058.1;refGene_gn=ARID3
B;refGene_tx=XM_023255059.1;refGene_gn=ARID3B;refGene_tx=XM_023255060.
1
B3 33667914 33668187
refGene_gn=ARID3B;refGene_tx=XM_019832048.2;refGene_gn=ARID3B;refGene_
tx=XM_019832049.2;refGene_gn=ARID3B;refGene_tx=XM_019832050.2;refGene_
gn=ARID3B;refGene_tx=XM_019832052.2;refGene_gn=ARID3B;refGene_tx=XM_02
3255056.1;refGene_gn=ARID3B;refGene_tx=XM_023255058.1;refGene_gn=ARID3
B;refGene_tx=XM_023255059.1;refGene_gn=ARID3B;refGene_tx=XM_023255060.
1
B3 33668270 33668551
refGene_gn=ARID3B;refGene_tx=XM_019832048.2;refGene_gn=ARID3B;refGene_
tx=XM_019832049.2;refGene_gn=ARID3B;refGene_tx=XM_019832050.2;refGene_
gn=ARID3B;refGene_tx=XM_019832052.2;refGene_gn=ARID3B;refGene_tx=XM_02
3255056.1;refGene_gn=ARID3B;refGene_tx=XM_023255058.1;refGene_gn=ARID3
B;refGene_tx=XM_023255059.1;refGene_gn=ARID3B;refGene_tx=XM_023255060.
1
B3 33669638 33669822
refGene_gn=ARID3B;refGene_tx=XM_019832048.2;refGene_gn=ARID3B;refGene_
tx=XM_019832049.2;refGene_gn=ARID3B;refGene_tx=XM_019832050.2;refGene_
gn=ARID3B;refGene_tx=XM_019832052.2;refGene_gn=ARID3B;refGene_tx=XM_02
3255056.1;refGene_gn=ARID3B;refGene_tx=XM_023255058.1;refGene_gn=ARID3
B;refGene_tx=XM_023255059.1;refGene_gn=ARID3B;refGene_tx=XM_023255060.
1
B3 33682838 33682911
refGene_gn=ARID3B;refGene_tx=XM_019832048.2;refGene_gn=ARID3B;refGene_
tx=XM_019832049.2;refGene_gn=ARID3B;refGene_tx=XM_019832050.2;refGene_
gn=ARID3B;refGene_tx=XM_019832052.2;refGene_gn=ARID3B;refGene_tx=XM_02
3255056.1;refGene_gn=ARID3B;refGene_tx=XM_023255058.1;refGene_gn=ARID3
B;refGene_tx=XM_023255059.1;refGene_gn=ARID3B;refGene_tx=XM_023255060.
1
B3 33683114 33683186
refGene_gn=ARID3B;refGene_tx=XM_019832048.2;refGene_gn=ARID3B;refGene_
tx=XM_019832049.2;refGene_gn=ARID3B;refGene_tx=XM_019832050.2;refGene_
gn=ARID3B;refGene_tx=XM_019832052.2;refGene_gn=ARID3B;refGene_tx=XM_02
3255056.1;refGene_gn=ARID3B;refGene_tx=XM_023255058.1;refGene_gn=ARID3
B;refGene_tx=XM_023255059.1;refGene_gn=ARID3B;refGene_tx=XM_023255060.
1
B3 33716678 33717233
refGene_gn=ARID3B;refGene_tx=XM_019832048.2;refGene_gn=ARID3B;refGene_
tx=XM_019832049.2;refGene_gn=ARID3B;refGene_tx=XM_019832050.2;refGene_
gn=ARID3B;refGene_tx=XM_019832052.2;refGene_gn=ARID3B;refGene_tx=XM_02
3255056.1;refGene_gn=ARID3B;refGene_tx=XM_023255058.1;refGene_gn=ARID3
B;refGene_tx=XM_023255059.1;refGene_gn=ARID3B;refGene_tx=XM_023255060.

1
B3	33787895	33787983
refGene_gn=UBL7;refGene_tx=XM_003986954.5;refGene_gn=UBL7;refGene_tx=XM_019832056.2
B3	33789561	33789774
refGene_gn=UBL7;refGene_tx=XM_003986954.5;refGene_gn=UBL7;refGene_tx=XM_019832056.2
B3	33790573	33790705
refGene_gn=UBL7;refGene_tx=XM_003986954.5;refGene_gn=UBL7;refGene_tx=XM_019832056.2
B3	33793757	33793840
refGene_gn=UBL7;refGene_tx=XM_003986954.5;refGene_gn=UBL7;refGene_tx=XM_019832056.2
B3	33794671	33794756
refGene_gn=UBL7;refGene_tx=XM_003986954.5;refGene_gn=UBL7;refGene_tx=XM_019832056.2
B3	33795282	33795340
refGene_gn=UBL7;refGene_tx=XM_003986954.5;refGene_gn=UBL7;refGene_tx=XM_019832056.2
B3	33796422	33796556
refGene_gn=UBL7;refGene_tx=XM_003986954.5;refGene_gn=UBL7;refGene_tx=XM_019832056.2
B3	33796815	33796865
refGene_gn=UBL7;refGene_tx=XM_003986954.5;refGene_gn=UBL7;refGene_tx=XM_019832056.2
B3	33797106	33797274
refGene_gn=UBL7;refGene_tx=XM_003986954.5;refGene_gn=UBL7;refGene_tx=XM_019832056.2
B3	33797843	33797966
refGene_gn=UBL7;refGene_tx=XM_003986954.5;refGene_gn=UBL7;refGene_tx=XM_019832056.2
B3	33800651	33800789
refGene_gn=UBL7;refGene_tx=XM_003986954.5;refGene_gn=UBL7;refGene_tx=XM_019832056.2
B3	33811678	33811847	refGene_gn=SEMA7A;refGene_tx=XM_011283023.3
B3	33825045	33825197	refGene_gn=SEMA7A;refGene_tx=XM_011283023.3
B3	33825739	33825781	refGene_gn=SEMA7A;refGene_tx=XM_011283023.3
B3	33826106	33826199	refGene_gn=SEMA7A;refGene_tx=XM_011283023.3
B3	33826421	33826506	refGene_gn=SEMA7A;refGene_tx=XM_011283023.3
B3	33826625	33826736	refGene_gn=SEMA7A;refGene_tx=XM_011283023.3
B3	33827387	33827527	refGene_gn=SEMA7A;refGene_tx=XM_011283023.3
B3	33828101	33828286	refGene_gn=SEMA7A;refGene_tx=XM_011283023.3
B3	33829125	33829234	refGene_gn=SEMA7A;refGene_tx=XM_011283023.3
B3	33829318	33829517	refGene_gn=SEMA7A;refGene_tx=XM_011283023.3
B3	33831938	33832066	refGene_gn=SEMA7A;refGene_tx=XM_011283023.3
B3	33832266	33832421	refGene_gn=SEMA7A;refGene_tx=XM_011283023.3
B3	33832620	33832682	refGene_gn=SEMA7A;refGene_tx=XM_011283023.3
B3	33832990	33833352	refGene_gn=SEMA7A;refGene_tx=XM_011283023.3
B3	33880521	33880790
refGene_gn=LOC101085980;refGene_tx=XM_003986957.3

B3 33884428 33884584
refGene_gn=LOC101085980;refGene_tx=XM_003986957.3;refGene_gn=LOC101085980;refGene_tx=XM_019832057.2
B3 33887270 33887470
refGene_gn=LOC101085980;refGene_tx=XM_003986957.3;refGene_gn=LOC101085980;refGene_tx=XM_019832057.2
B3 33888325 33888529
refGene_gn=LOC101085980;refGene_tx=XM_003986957.3;refGene_gn=LOC101085980;refGene_tx=XM_019832057.2
B3 33889198 33889356
refGene_gn=LOC101085980;refGene_tx=XM_003986957.3;refGene_gn=LOC101085980;refGene_tx=XM_019832057.2
B3 33892529 33892696
refGene_gn=LOC101085980;refGene_tx=XM_003986957.3;refGene_gn=LOC101085980;refGene_tx=XM_019832057.2
B3 33893177 33893256
refGene_gn=LOC101085980;refGene_tx=XM_003986957.3;refGene_gn=LOC101085980;refGene_tx=XM_019832057.2
B3 33893745 33893943
refGene_gn=LOC101085980;refGene_tx=XM_003986957.3;refGene_gn=LOC101085980;refGene_tx=XM_019832057.2
B3 33894389 33894521
refGene_gn=LOC101085980;refGene_tx=XM_003986957.3;refGene_gn=LOC101085980;refGene_tx=XM_019832057.2
B3 33896374 33896482
refGene_gn=CCDC33;refGene_tx=XM_019832060.2;refGene_gn=CCDC33;refGene_tx=XM_023255062.1;refGene_gn=CCDC33;refGene_tx=XM_023255063.1
B3 33897321 33897435
refGene_gn=CCDC33;refGene_tx=XM_019832060.2;refGene_gn=CCDC33;refGene_tx=XM_023255062.1;refGene_gn=CCDC33;refGene_tx=XM_023255063.1
B3 33897790 33897877
refGene_gn=CCDC33;refGene_tx=XM_019832060.2;refGene_gn=CCDC33;refGene_tx=XM_023255062.1;refGene_gn=CCDC33;refGene_tx=XM_023255063.1
B3 33898442 33898544 refGene_gn=CCDC33;refGene_tx=XM_019832060.2
B3 33898902 33899069
refGene_gn=CCDC33;refGene_tx=XM_019832060.2;refGene_gn=CCDC33;refGene_tx=XM_023255062.1;refGene_gn=CCDC33;refGene_tx=XM_023255063.1
B3 33900023 33900114
refGene_gn=CCDC33;refGene_tx=XM_019832060.2;refGene_gn=CCDC33;refGene_tx=XM_023255062.1;refGene_gn=CCDC33;refGene_tx=XM_023255063.1
B3 33900209 33900341
refGene_gn=CCDC33;refGene_tx=XM_019832060.2;refGene_gn=CCDC33;refGene_tx=XM_023255062.1;refGene_gn=CCDC33;refGene_tx=XM_023255063.1
B3 33900507 33900596
refGene_gn=CCDC33;refGene_tx=XM_019832060.2;refGene_gn=CCDC33;refGene_tx=XM_023255062.1;refGene_gn=CCDC33;refGene_tx=XM_023255063.1
B3 33900860 33901026
refGene_gn=CCDC33;refGene_tx=XM_019832060.2;refGene_gn=CCDC33;refGene_tx=XM_023255062.1;refGene_gn=CCDC33;refGene_tx=XM_023255063.1
B3 33936933 33937122

refGene_gn=CCDC33;refGene_tx=XM_019832060.2;refGene_gn=CCDC33;refGene_tx=XM_023255062.1;refGene_gn=CCDC33;refGene_tx=XM_023255063.1
B3 33948917 33948989
refGene_gn=CCDC33;refGene_tx=XM_019832060.2;refGene_gn=CCDC33;refGene_tx=XM_023255062.1;refGene_gn=CCDC33;refGene_tx=XM_023255063.1
B3 33949934 33950068
refGene_gn=CCDC33;refGene_tx=XM_019832060.2;refGene_gn=CCDC33;refGene_tx=XM_023255062.1;refGene_gn=CCDC33;refGene_tx=XM_023255063.1
B3 33950697 33950827
refGene_gn=CCDC33;refGene_tx=XM_019832060.2;refGene_gn=CCDC33;refGene_tx=XM_023255062.1;refGene_gn=CCDC33;refGene_tx=XM_023255063.1
B3 33956824 33956945
refGene_gn=CCDC33;refGene_tx=XM_019832060.2;refGene_gn=CCDC33;refGene_tx=XM_023255062.1;refGene_gn=CCDC33;refGene_tx=XM_023255063.1
B3 33957793 33957885
refGene_gn=CCDC33;refGene_tx=XM_019832060.2;refGene_gn=CCDC33;refGene_tx=XM_023255062.1;refGene_gn=CCDC33;refGene_tx=XM_023255063.1
B3 33961067 33961184
refGene_gn=CCDC33;refGene_tx=XM_019832060.2;refGene_gn=CCDC33;refGene_tx=XM_023255062.1;refGene_gn=CCDC33;refGene_tx=XM_023255063.1
B3 33962747 33962857
refGene_gn=CCDC33;refGene_tx=XM_019832060.2;refGene_gn=CCDC33;refGene_tx=XM_023255062.1;refGene_gn=CCDC33;refGene_tx=XM_023255063.1
B3 33966931 33967065
refGene_gn=CCDC33;refGene_tx=XM_019832060.2;refGene_gn=CCDC33;refGene_tx=XM_023255062.1;refGene_gn=CCDC33;refGene_tx=XM_023255063.1
B3 33980766 33980930
refGene_gn=CCDC33;refGene_tx=XM_019832060.2;refGene_gn=CCDC33;refGene_tx=XM_023255062.1;refGene_gn=CCDC33;refGene_tx=XM_023255063.1
B3 33986073 33986094
refGene_gn=CCDC33;refGene_tx=XM_019832060.2;refGene_gn=CCDC33;refGene_tx=XM_023255062.1
B3 33999808 33999811 refGene_gn=CCDC33;refGene_tx=XM_023255063.1
B3 34002010 34002336 refGene_gn=CCDC33;refGene_tx=XM_023255063.1
B3 34003224 34003534 refGene_gn=CCDC33;refGene_tx=XM_023255063.1
B3 34017188 34017301 refGene_gn=STRA6;refGene_tx=XM_019832074.2
B3 34020463 34020627 refGene_gn=STRA6;refGene_tx=XM_019832075.2
B3 34020778 34020845
refGene_gn=STRA6;refGene_tx=XM_019832074.2;refGene_gn=STRA6;refGene_tx=XM_019832075.2
B3 34021159 34021245
refGene_gn=STRA6;refGene_tx=XM_019832074.2;refGene_gn=STRA6;refGene_tx=XM_019832075.2
B3 34022437 34022577
refGene_gn=STRA6;refGene_tx=XM_019832074.2;refGene_gn=STRA6;refGene_tx=XM_019832075.2
B3 34023214 34023238
refGene_gn=STRA6;refGene_tx=XM_019832074.2;refGene_gn=STRA6;refGene_tx=XM_019832075.2
B3 34023430 34023597

refGene_gn=STRA6;refGene_tx=XM_019832074.2;refGene_gn=STRA6;refGene_tx=XM_019832075.2
B3 34024917 34025040
refGene_gn=STRA6;refGene_tx=XM_019832074.2;refGene_gn=STRA6;refGene_tx=XM_019832075.2
B3 34027208 34027276
refGene_gn=STRA6;refGene_tx=XM_019832074.2;refGene_gn=STRA6;refGene_tx=XM_019832075.2
B3 34027454 34027531
refGene_gn=STRA6;refGene_tx=XM_019832074.2;refGene_gn=STRA6;refGene_tx=XM_019832075.2
B3 34027802 34027864
refGene_gn=STRA6;refGene_tx=XM_019832074.2;refGene_gn=STRA6;refGene_tx=XM_019832075.2
B3 34028807 34028970
refGene_gn=STRA6;refGene_tx=XM_019832074.2;refGene_gn=STRA6;refGene_tx=XM_019832075.2
B3 34032100 34032176
refGene_gn=STRA6;refGene_tx=XM_019832074.2;refGene_gn=STRA6;refGene_tx=XM_019832075.2
B3 34033182 34033316
refGene_gn=STRA6;refGene_tx=XM_019832074.2;refGene_gn=STRA6;refGene_tx=XM_019832075.2
B3 34034444 34034562
refGene_gn=STRA6;refGene_tx=XM_019832074.2;refGene_gn=STRA6;refGene_tx=XM_019832075.2
B3 34034645 34034747
refGene_gn=STRA6;refGene_tx=XM_019832074.2;refGene_gn=STRA6;refGene_tx=XM_019832075.2
B3 34035453 34035617
refGene_gn=STRA6;refGene_tx=XM_019832074.2;refGene_gn=STRA6;refGene_tx=XM_019832075.2
B3 34035873 34036023
refGene_gn=STRA6;refGene_tx=XM_019832074.2;refGene_gn=STRA6;refGene_tx=XM_019832075.2
B3 34036673 34036822
refGene_gn=STRA6;refGene_tx=XM_019832074.2;refGene_gn=STRA6;refGene_tx=XM_019832075.2
B3 34040724 34042011
refGene_gn=ISLR;refGene_tx=XM_003986959.3;refGene_gn=ISLR;refGene_tx=XM_006932377.4
B3 34076470 34078722
refGene_gn=ISLR2;refGene_tx=XM_006932378.4;refGene_gn=ISLR2;refGene_tx=XM_006932380.4;refGene_gn=ISLR2;refGene_tx=XM_006932381.4;refGene_gn=ISLR2;refGene_tx=XM_019832076.2
B3 34080154 34080251
refGene_gn=ISLR2;refGene_tx=XM_006932378.4;refGene_gn=ISLR2;refGene_tx=XM_006932380.4;refGene_gn=ISLR2;refGene_tx=XM_006932381.4;refGene_gn=ISLR2;refGene_tx=XM_019832076.2
B3 34096833 34097609

refGene_gn=PML;refGene_tx=XM_003986961.3;refGene_gn=PML;refGene_tx=XM_006932386.2;refGene_gn=PML;refGene_tx=XM_006932390.3
B3 34098664 34098815
refGene_gn=PML;refGene_tx=XM_003986961.3;refGene_gn=PML;refGene_tx=XM_006932386.2
B3 34105893 34106515
refGene_gn=PML;refGene_tx=XM_006932384.2;refGene_gn=PML;refGene_tx=XM_006932385.3
B3 34107128 34107181
refGene_gn=PML;refGene_tx=XM_003986961.3;refGene_gn=PML;refGene_tx=XM_006932384.2;refGene_gn=PML;refGene_tx=XM_006932385.3;refGene_gn=PML;refGene_tx=XM_006932386.2;refGene_gn=PML;refGene_tx=XM_006932388.3;refGene_gn=PML;refGene_tx=XM_006932389.2;refGene_gn=PML;refGene_tx=XM_006932390.3;refGene_gn=PML;refGene_tx=XM_011282964.2;refGene_gn=PML;refGene_tx=XM_019832077.1
B3 34107578 34107649
refGene_gn=PML;refGene_tx=XM_006932388.3;refGene_gn=PML;refGene_tx=XM_006932389.2;refGene_gn=PML;refGene_tx=XM_011282964.2;refGene_gn=PML;refGene_tx=XM_019832077.1
B3 34108235 34108494
refGene_gn=PML;refGene_tx=XM_003986961.3;refGene_gn=PML;refGene_tx=XM_006932384.2;refGene_gn=PML;refGene_tx=XM_006932385.3;refGene_gn=PML;refGene_tx=XM_006932386.2;refGene_gn=PML;refGene_tx=XM_006932388.3;refGene_gn=PML;refGene_tx=XM_006932389.2;refGene_gn=PML;refGene_tx=XM_006932390.3;refGene_gn=PML;refGene_tx=XM_011282964.2;refGene_gn=PML;refGene_tx=XM_019832077.1
B3 34108888 34109026
refGene_gn=PML;refGene_tx=XM_003986961.3;refGene_gn=PML;refGene_tx=XM_006932384.2;refGene_gn=PML;refGene_tx=XM_006932385.3;refGene_gn=PML;refGene_tx=XM_006932386.2;refGene_gn=PML;refGene_tx=XM_006932388.3;refGene_gn=PML;refGene_tx=XM_006932389.2;refGene_gn=PML;refGene_tx=XM_006932390.3;refGene_gn=PML;refGene_tx=XM_011282964.2;refGene_gn=PML;refGene_tx=XM_019832077.1
B3 34115153 34115224
refGene_gn=PML;refGene_tx=XM_003986961.3;refGene_gn=PML;refGene_tx=XM_006932384.2;refGene_gn=PML;refGene_tx=XM_006932385.3;refGene_gn=PML;refGene_tx=XM_006932386.2;refGene_gn=PML;refGene_tx=XM_006932388.3;refGene_gn=PML;refGene_tx=XM_006932389.2;refGene_gn=PML;refGene_tx=XM_006932390.3;refGene_gn=PML;refGene_tx=XM_011282964.2;refGene_gn=PML;refGene_tx=XM_019832077.1
B3 34116452 34117033
refGene_gn=PML;refGene_tx=XM_003986961.3;refGene_gn=PML;refGene_tx=XM_006932384.2;refGene_gn=PML;refGene_tx=XM_006932385.3;refGene_gn=PML;refGene_tx=XM_006932386.2;refGene_gn=PML;refGene_tx=XM_006932388.3;refGene_gn=PML;refGene_tx=XM_006932389.2;refGene_gn=PML;refGene_tx=XM_006932390.3;refGene_gn=PML;refGene_tx=XM_011282964.2;refGene_gn=PML;refGene_tx=XM_019832077.1
B3 34131464 34131931
refGene_gn=PML;refGene_tx=XM_003986961.3;refGene_gn=PML;refGene_tx=XM_006932384.2;refGene_gn=PML;refGene_tx=XM_006932385.3;refGene_gn=PML;re

fGene_tx=XM_006932386.2;refGene_gn=PML;refGene_tx=XM_006932388.3;refGene_gn=PML;refGene_tx=XM_006932389.2;refGene_gn=PML;refGene_tx=XM_006932390.3;refGene_gn=PML;refGene_tx=XM_011282964.2;refGene_gn=PML;refGene_tx=XM_019832077.1
B3 34134047 34134182
refGene_gn=PML;refGene_tx=XM_003986961.3;refGene_gn=PML;refGene_tx=XM_006932384.2;refGene_gn=PML;refGene_tx=XM_006932385.3;refGene_gn=PML;refGene_tx=XM_006932386.2;refGene_gn=PML;refGene_tx=XM_006932388.3;refGene_gn=PML;refGene_tx=XM_006932389.2;refGene_gn=PML;refGene_tx=XM_006932390.3;refGene_gn=PML;refGene_tx=XM_011282964.2;refGene_gn=PML;refGene_tx=XM_019832077.1
B3 34136605 34136738
refGene_gn=STOML1;refGene_tx=XM_011282963.3;refGene_gn=STOML1;refGene_tx=XM_023255066.1
B3 34138245 34138352
refGene_gn=STOML1;refGene_tx=XM_011282963.3;refGene_gn=STOML1;refGene_tx=XM_023255066.1
B3 34139280 34139430
refGene_gn=STOML1;refGene_tx=XM_006932391.4;refGene_gn=STOML1;refGene_tx=XM_011282963.3;refGene_gn=STOML1;refGene_tx=XM_023255066.1
B3 34139738 34139966
refGene_gn=STOML1;refGene_tx=XM_006932391.4;refGene_gn=STOML1;refGene_tx=XM_011282963.3;refGene_gn=STOML1;refGene_tx=XM_023255066.1
B3 34143966 34144162
refGene_gn=STOML1;refGene_tx=XM_006932391.4;refGene_gn=STOML1;refGene_tx=XM_011282963.3;refGene_gn=STOML1;refGene_tx=XM_023255066.1
B3 34144632 34144845
refGene_gn=STOML1;refGene_tx=XM_006932391.4;refGene_gn=STOML1;refGene_tx=XM_011282963.3;refGene_gn=STOML1;refGene_tx=XM_023255066.1
B3 34145318 34145512
refGene_gn=STOML1;refGene_tx=XM_006932391.4;refGene_gn=STOML1;refGene_tx=XM_011282963.3;refGene_gn=STOML1;refGene_tx=XM_023255066.1
B3 34166897 34166904 refGene_gn=LOXL1;refGene_tx=XM_023255067.1
B3 34169064 34169180 refGene_gn=LOXL1;refGene_tx=XM_023255067.1
B3 34170837 34170933 refGene_gn=LOXL1;refGene_tx=XM_023255067.1
B3 34171745 34171902 refGene_gn=LOXL1;refGene_tx=XM_023255067.1
B3 34172382 34172520 refGene_gn=LOXL1;refGene_tx=XM_023255067.1
B3 34175412 34175521 refGene_gn=LOXL1;refGene_tx=XM_023255067.1
B3 34189322 34190472 refGene_gn=LOXL1;refGene_tx=XM_023255067.1
B3 34222813 34222846
refGene_gn=TBC1D21;refGene_tx=XM_003986962.4;refGene_gn=TBC1D21;refGene_tx=XM_019832081.2
B3 34222953 34223046
refGene_gn=TBC1D21;refGene_tx=XM_006932393.4;refGene_gn=TBC1D21;refGene_tx=XM_019832079.2;refGene_gn=TBC1D21;refGene_tx=XM_019832080.2
B3 34223184 34223268
refGene_gn=TBC1D21;refGene_tx=XM_003986962.4;refGene_gn=TBC1D21;refGene_tx=XM_006932393.4;refGene_gn=TBC1D21;refGene_tx=XM_019832079.2;refGene_gn=TBC1D21;refGene_tx=XM_019832080.2;refGene_gn=TBC1D21;refGene_tx=XM_019832081.2

B3 34223858 34223975
refGene_gn=TBC1D21;refGene_tx=XM_003986962.4;refGene_gn=TBC1D21;refGene_tx=XM_006932393.4;refGene_gn=TBC1D21;refGene_tx=XM_019832079.2;refGene_gn=TBC1D21;refGene_tx=XM_019832080.2;refGene_gn=TBC1D21;refGene_tx=XM_019832081.2
B3 34224943 34225044
refGene_gn=TBC1D21;refGene_tx=XM_003986962.4;refGene_gn=TBC1D21;refGene_tx=XM_006932393.4;refGene_gn=TBC1D21;refGene_tx=XM_019832079.2;refGene_gn=TBC1D21;refGene_tx=XM_019832080.2;refGene_gn=TBC1D21;refGene_tx=XM_019832081.2
B3 34225421 34225518
refGene_gn=TBC1D21;refGene_tx=XM_003986962.4;refGene_gn=TBC1D21;refGene_tx=XM_006932393.4;refGene_gn=TBC1D21;refGene_tx=XM_019832079.2;refGene_gn=TBC1D21;refGene_tx=XM_019832080.2;refGene_gn=TBC1D21;refGene_tx=XM_019832081.2
B3 34226472 34226573
refGene_gn=TBC1D21;refGene_tx=XM_003986962.4;refGene_gn=TBC1D21;refGene_tx=XM_006932393.4;refGene_gn=TBC1D21;refGene_tx=XM_019832079.2;refGene_gn=TBC1D21;refGene_tx=XM_019832080.2;refGene_gn=TBC1D21;refGene_tx=XM_019832081.2
B3 34226683 34226794
refGene_gn=TBC1D21;refGene_tx=XM_003986962.4;refGene_gn=TBC1D21;refGene_tx=XM_006932393.4;refGene_gn=TBC1D21;refGene_tx=XM_019832079.2;refGene_gn=TBC1D21;refGene_tx=XM_019832080.2;refGene_gn=TBC1D21;refGene_tx=XM_019832081.2
B3 34227330 34227425
refGene_gn=TBC1D21;refGene_tx=XM_003986962.4;refGene_gn=TBC1D21;refGene_tx=XM_006932393.4;refGene_gn=TBC1D21;refGene_tx=XM_019832079.2;refGene_gn=TBC1D21;refGene_tx=XM_019832080.2;refGene_gn=TBC1D21;refGene_tx=XM_019832081.2
B3 34228655 34228759
refGene_gn=TBC1D21;refGene_tx=XM_003986962.4;refGene_gn=TBC1D21;refGene_tx=XM_006932393.4;refGene_gn=TBC1D21;refGene_tx=XM_019832079.2;refGene_gn=TBC1D21;refGene_tx=XM_019832080.2;refGene_gn=TBC1D21;refGene_tx=XM_019832081.2
B3 34228891 34228999
refGene_gn=TBC1D21;refGene_tx=XM_003986962.4;refGene_gn=TBC1D21;refGene_tx=XM_006932393.4;refGene_gn=TBC1D21;refGene_tx=XM_019832079.2
B3 34235315 34235375
refGene_gn=TBC1D21;refGene_tx=XM_003986962.4;refGene_gn=TBC1D21;refGene_tx=XM_006932393.4;refGene_gn=TBC1D21;refGene_tx=XM_019832079.2;refGene_gn=TBC1D21;refGene_tx=XM_019832080.2;refGene_gn=TBC1D21;refGene_tx=XM_019832081.2
B3 34350855 34351011
refGene_gn=CB3H15orf59;refGene_tx=XM_003986964.5
B3 34360214 34360937
refGene_gn=CB3H15orf59;refGene_tx=XM_003986964.5
B3 34384612 34384635
refGene_gn=CD276;refGene_tx=XM_023255069.1;refGene_gn=CD276;refGene_tx=XM_023255071.1

B3 34386723 34386759
refGene_gn=CD276;refGene_tx=XM_023255069.1;refGene_gn=CD276;refGene_tx=XM_023255071.1
B3 34388211 34388253
refGene_gn=CD276;refGene_tx=XM_023255069.1;refGene_gn=CD276;refGene_tx=XM_023255071.1
B3 34389160 34389295
refGene_gn=CD276;refGene_tx=XM_023255069.1;refGene_gn=CD276;refGene_tx=XM_023255071.1
B3 34392385 34392682
refGene_gn=CD276;refGene_tx=XM_023255069.1;refGene_gn=CD276;refGene_tx=XM_023255071.1
B3 34392889 34393228
refGene_gn=CD276;refGene_tx=XM_023255069.1;refGene_gn=CD276;refGene_tx=XM_023255071.1
B3 34393846 34394161
refGene_gn=CD276;refGene_tx=XM_023255069.1;refGene_gn=CD276;refGene_tx=XM_023255071.1
B3 34394368 34394707
refGene_gn=CD276;refGene_tx=XM_023255069.1;refGene_gn=CD276;refGene_tx=XM_023255071.1
B3 34397491 34397576
refGene_gn=CD276;refGene_tx=XM_023255069.1;refGene_gn=CD276;refGene_tx=XM_023255071.1
B3 34454390 34454481
refGene_gn=NPTN;refGene_tx=XM_019832082.2;refGene_gn=NPTN;refGene_tx=XM_019832083.2
B3 34493861 34494209
refGene_gn=NPTN;refGene_tx=XM_019832082.2;refGene_gn=NPTN;refGene_tx=XM_019832083.2
B3 34499094 34499266
refGene_gn=NPTN;refGene_tx=XM_019832082.2;refGene_gn=NPTN;refGene_tx=XM_019832083.2
B3 34502885 34502980
refGene_gn=NPTN;refGene_tx=XM_019832082.2;refGene_gn=NPTN;refGene_tx=XM_019832083.2
B3 34515514 34515648
refGene_gn=NPTN;refGene_tx=XM_019832082.2;refGene_gn=NPTN;refGene_tx=XM_019832083.2
B3 34520789 34521063
refGene_gn=NPTN;refGene_tx=XM_019832082.2;refGene_gn=NPTN;refGene_tx=XM_019832083.2
B3 34527962 34527984
refGene_gn=NPTN;refGene_tx=XM_019832082.2;refGene_gn=NPTN;refGene_tx=XM_019832083.2
B3 34528814 34528875
refGene_gn=NPTN;refGene_tx=XM_019832082.2;refGene_gn=NPTN;refGene_tx=XM_019832083.2
B3 34530557 34530722 refGene_gn=REC114;refGene_tx=XM_019832084.2
B3 34533470 34533560 refGene_gn=REC114;refGene_tx=XM_019832084.2

B3 34539244 34539457 refGene_gn=REC114;refGene_tx=XM_019832084.2
B3 34547368 34547452 refGene_gn=REC114;refGene_tx=XM_019832084.2
B3 34614059 34614149 refGene_gn=REC114;refGene_tx=XM_019832084.2
B3 34636752 34636911 refGene_gn=REC114;refGene_tx=XM_019832084.2
B3 34712898 34713686 refGene_gn=HCN4;refGene_tx=XM_023255072.1
B3 34737520 34737944 refGene_gn=HCN4;refGene_tx=XM_023255072.1
B3 34746853 34747015 refGene_gn=HCN4;refGene_tx=XM_023255072.1
B3 34749094 34749313 refGene_gn=HCN4;refGene_tx=XM_023255072.1
B3 34752893 34753040 refGene_gn=HCN4;refGene_tx=XM_023255072.1
B3 34753138 34753379 refGene_gn=HCN4;refGene_tx=XM_023255072.1
B3 34753667 34753832 refGene_gn=HCN4;refGene_tx=XM_023255072.1
B3 34753932 34755395 refGene_gn=HCN4;refGene_tx=XM_023255072.1
B3 34783555 34783639
refGene_gn=NEO1;refGene_tx=XM_023255073.1;refGene_gn=NEO1;refGene_tx=XM_023255074.1;refGene_gn=NEO1;refGene_tx=XM_023255075.1;refGene_gn=NEO1;refGene_tx=XM_023255077.1;refGene_gn=NEO1;refGene_tx=XM_023255078.1;refGene_gn=NEO1;refGene_tx=XM_023255079.1;refGene_gn=NEO1;refGene_tx=XM_023255080.1;refGene_gn=NEO1;refGene_tx=XM_023255081.1
B3 34783838 34783975
refGene_gn=NEO1;refGene_tx=XM_023255073.1;refGene_gn=NEO1;refGene_tx=XM_023255074.1;refGene_gn=NEO1;refGene_tx=XM_023255075.1;refGene_gn=NEO1;refGene_tx=XM_023255077.1;refGene_gn=NEO1;refGene_tx=XM_023255078.1;refGene_gn=NEO1;refGene_tx=XM_023255079.1;refGene_gn=NEO1;refGene_tx=XM_023255080.1;refGene_gn=NEO1;refGene_tx=XM_023255081.1
B3 34786293 34786557
refGene_gn=NEO1;refGene_tx=XM_023255073.1;refGene_gn=NEO1;refGene_tx=XM_023255074.1;refGene_gn=NEO1;refGene_tx=XM_023255075.1;refGene_gn=NEO1;refGene_tx=XM_023255077.1;refGene_gn=NEO1;refGene_tx=XM_023255078.1;refGene_gn=NEO1;refGene_tx=XM_023255079.1;refGene_gn=NEO1;refGene_tx=XM_023255080.1;refGene_gn=NEO1;refGene_tx=XM_023255081.1
B3 34790530 34790689
refGene_gn=NEO1;refGene_tx=XM_023255073.1;refGene_gn=NEO1;refGene_tx=XM_023255074.1;refGene_gn=NEO1;refGene_tx=XM_023255075.1;refGene_gn=NEO1;refGene_tx=XM_023255077.1;refGene_gn=NEO1;refGene_tx=XM_023255078.1;refGene_gn=NEO1;refGene_tx=XM_023255079.1
B3 34794001 34794094
refGene_gn=NEO1;refGene_tx=XM_023255073.1;refGene_gn=NEO1;refGene_tx=XM_023255074.1;refGene_gn=NEO1;refGene_tx=XM_023255075.1;refGene_gn=NEO1;refGene_tx=XM_023255077.1;refGene_gn=NEO1;refGene_tx=XM_023255078.1;refGene_gn=NEO1;refGene_tx=XM_023255079.1;refGene_gn=NEO1;refGene_tx=XM_023255080.1;refGene_gn=NEO1;refGene_tx=XM_023255081.1
B3 34794711 34794950
refGene_gn=NEO1;refGene_tx=XM_023255073.1;refGene_gn=NEO1;refGene_tx=XM_023255074.1;refGene_gn=NEO1;refGene_tx=XM_023255075.1;refGene_gn=NEO1;refGene_tx=XM_023255077.1;refGene_gn=NEO1;refGene_tx=XM_023255078.1;refGene_gn=NEO1;refGene_tx=XM_023255079.1;refGene_gn=NEO1;refGene_tx=XM_023255080.1;refGene_gn=NEO1;refGene_tx=XM_023255081.1
B3 34800535 34800683
refGene_gn=NEO1;refGene_tx=XM_023255073.1;refGene_gn=NEO1;refGene_tx=XM_023255074.1;refGene_gn=NEO1;refGene_tx=XM_023255075.1;refGene_gn=NEO

1;refGene_tx=XM_023255077.1;refGene_gn=NEO1;refGene_tx=XM_023255078.1;
refGene_gn=NEO1;refGene_tx=XM_023255079.1;refGene_gn=NEO1;refGene_tx=X
M_023255080.1;refGene_gn=NEO1;refGene_tx=XM_023255081.1
B3 34804119 34804188
refGene_gn=NEO1;refGene_tx=XM_023255073.1;refGene_gn=NEO1;refGene_tx=X
M_023255074.1;refGene_gn=NEO1;refGene_tx=XM_023255075.1;refGene_gn=NEO
1;refGene_tx=XM_023255077.1;refGene_gn=NEO1;refGene_tx=XM_023255078.1;
refGene_gn=NEO1;refGene_tx=XM_023255079.1;refGene_gn=NEO1;refGene_tx=X
M_023255080.1;refGene_gn=NEO1;refGene_tx=XM_023255081.1
B3 34807141 34807174
refGene_gn=NEO1;refGene_tx=XM_023255073.1;refGene_gn=NEO1;refGene_tx=X
M_023255075.1;refGene_gn=NEO1;refGene_tx=XM_023255077.1;refGene_gn=NEO
1;refGene_tx=XM_023255080.1;refGene_gn=NEO1;refGene_tx=XM_023255081.1
B3 34807807 34808002
refGene_gn=NEO1;refGene_tx=XM_023255073.1;refGene_gn=NEO1;refGene_tx=X
M_023255074.1;refGene_gn=NEO1;refGene_tx=XM_023255075.1;refGene_gn=NEO
1;refGene_tx=XM_023255077.1;refGene_gn=NEO1;refGene_tx=XM_023255078.1;
refGene_gn=NEO1;refGene_tx=XM_023255079.1;refGene_gn=NEO1;refGene_tx=X
M_023255080.1;refGene_gn=NEO1;refGene_tx=XM_023255081.1
B3 34810359 34810467
refGene_gn=NEO1;refGene_tx=XM_023255073.1;refGene_gn=NEO1;refGene_tx=X
M_023255074.1;refGene_gn=NEO1;refGene_tx=XM_023255075.1;refGene_gn=NEO
1;refGene_tx=XM_023255077.1;refGene_gn=NEO1;refGene_tx=XM_023255078.1;
refGene_gn=NEO1;refGene_tx=XM_023255079.1;refGene_gn=NEO1;refGene_tx=X
M_023255080.1;refGene_gn=NEO1;refGene_tx=XM_023255081.1
B3 34812681 34812820
refGene_gn=NEO1;refGene_tx=XM_023255073.1;refGene_gn=NEO1;refGene_tx=X
M_023255074.1;refGene_gn=NEO1;refGene_tx=XM_023255075.1;refGene_gn=NEO
1;refGene_tx=XM_023255077.1;refGene_gn=NEO1;refGene_tx=XM_023255078.1;
refGene_gn=NEO1;refGene_tx=XM_023255079.1;refGene_gn=NEO1;refGene_tx=X
M_023255080.1;refGene_gn=NEO1;refGene_tx=XM_023255081.1
B3 34812906 34813130
refGene_gn=NEO1;refGene_tx=XM_023255073.1;refGene_gn=NEO1;refGene_tx=X
M_023255074.1;refGene_gn=NEO1;refGene_tx=XM_023255075.1;refGene_gn=NEO
1;refGene_tx=XM_023255077.1;refGene_gn=NEO1;refGene_tx=XM_023255078.1;
refGene_gn=NEO1;refGene_tx=XM_023255079.1;refGene_gn=NEO1;refGene_tx=X
M_023255080.1;refGene_gn=NEO1;refGene_tx=XM_023255081.1
B3 34815581 34815677
refGene_gn=NEO1;refGene_tx=XM_023255073.1;refGene_gn=NEO1;refGene_tx=X
M_023255074.1;refGene_gn=NEO1;refGene_tx=XM_023255075.1;refGene_gn=NEO
1;refGene_tx=XM_023255077.1;refGene_gn=NEO1;refGene_tx=XM_023255078.1;
refGene_gn=NEO1;refGene_tx=XM_023255079.1;refGene_gn=NEO1;refGene_tx=X
M_023255080.1;refGene_gn=NEO1;refGene_tx=XM_023255081.1
B3 34821647 34821842
refGene_gn=NEO1;refGene_tx=XM_023255073.1;refGene_gn=NEO1;refGene_tx=X
M_023255074.1;refGene_gn=NEO1;refGene_tx=XM_023255075.1;refGene_gn=NEO
1;refGene_tx=XM_023255077.1;refGene_gn=NEO1;refGene_tx=XM_023255078.1;
refGene_gn=NEO1;refGene_tx=XM_023255079.1;refGene_gn=NEO1;refGene_tx=X
M_023255080.1;refGene_gn=NEO1;refGene_tx=XM_023255081.1
B3 34822958 34823069

refGene_gn=NEO1;refGene_tx=XM_023255073.1;refGene_gn=NEO1;refGene_tx=XM_023255074.1;refGene_gn=NEO1;refGene_tx=XM_023255075.1;refGene_gn=NEO1;refGene_tx=XM_023255077.1;refGene_gn=NEO1;refGene_tx=XM_023255078.1;refGene_gn=NEO1;refGene_tx=XM_023255079.1;refGene_gn=NEO1;refGene_tx=XM_023255080.1;refGene_gn=NEO1;refGene_tx=XM_023255081.1
B3 34827190 34827338
refGene_gn=NEO1;refGene_tx=XM_023255073.1;refGene_gn=NEO1;refGene_tx=XM_023255074.1;refGene_gn=NEO1;refGene_tx=XM_023255075.1;refGene_gn=NEO1;refGene_tx=XM_023255077.1;refGene_gn=NEO1;refGene_tx=XM_023255078.1;refGene_gn=NEO1;refGene_tx=XM_023255079.1;refGene_gn=NEO1;refGene_tx=XM_023255080.1;refGene_gn=NEO1;refGene_tx=XM_023255081.1
B3 34828259 34828309
refGene_gn=NEO1;refGene_tx=XM_023255073.1;refGene_gn=NEO1;refGene_tx=XM_023255074.1;refGene_gn=NEO1;refGene_tx=XM_023255075.1;refGene_gn=NEO1;refGene_tx=XM_023255077.1;refGene_gn=NEO1;refGene_tx=XM_023255078.1;refGene_gn=NEO1;refGene_tx=XM_023255079.1;refGene_gn=NEO1;refGene_tx=XM_023255080.1;refGene_gn=NEO1;refGene_tx=XM_023255081.1
B3 34833364 34833503
refGene_gn=NEO1;refGene_tx=XM_023255073.1;refGene_gn=NEO1;refGene_tx=XM_023255074.1;refGene_gn=NEO1;refGene_tx=XM_023255075.1;refGene_gn=NEO1;refGene_tx=XM_023255077.1;refGene_gn=NEO1;refGene_tx=XM_023255078.1;refGene_gn=NEO1;refGene_tx=XM_023255079.1;refGene_gn=NEO1;refGene_tx=XM_023255080.1;refGene_gn=NEO1;refGene_tx=XM_023255081.1
B3 34833875 34834024
refGene_gn=NEO1;refGene_tx=XM_023255073.1;refGene_gn=NEO1;refGene_tx=XM_023255074.1;refGene_gn=NEO1;refGene_tx=XM_023255075.1;refGene_gn=NEO1;refGene_tx=XM_023255077.1;refGene_gn=NEO1;refGene_tx=XM_023255078.1;refGene_gn=NEO1;refGene_tx=XM_023255079.1;refGene_gn=NEO1;refGene_tx=XM_023255080.1;refGene_gn=NEO1;refGene_tx=XM_023255081.1
B3 34836478 34836633
refGene_gn=NEO1;refGene_tx=XM_023255073.1;refGene_gn=NEO1;refGene_tx=XM_023255074.1;refGene_gn=NEO1;refGene_tx=XM_023255075.1;refGene_gn=NEO1;refGene_tx=XM_023255077.1;refGene_gn=NEO1;refGene_tx=XM_023255078.1;refGene_gn=NEO1;refGene_tx=XM_023255079.1;refGene_gn=NEO1;refGene_tx=XM_023255080.1;refGene_gn=NEO1;refGene_tx=XM_023255081.1
B3 34843215 34843435
refGene_gn=NEO1;refGene_tx=XM_023255073.1;refGene_gn=NEO1;refGene_tx=XM_023255074.1;refGene_gn=NEO1;refGene_tx=XM_023255075.1;refGene_gn=NEO1;refGene_tx=XM_023255077.1;refGene_gn=NEO1;refGene_tx=XM_023255078.1;refGene_gn=NEO1;refGene_tx=XM_023255079.1;refGene_gn=NEO1;refGene_tx=XM_023255080.1;refGene_gn=NEO1;refGene_tx=XM_023255081.1
B3 34889328 34889449
refGene_gn=NEO1;refGene_tx=XM_023255073.1;refGene_gn=NEO1;refGene_tx=XM_023255074.1;refGene_gn=NEO1;refGene_tx=XM_023255075.1;refGene_gn=NEO1;refGene_tx=XM_023255077.1;refGene_gn=NEO1;refGene_tx=XM_023255078.1;refGene_gn=NEO1;refGene_tx=XM_023255079.1;refGene_gn=NEO1;refGene_tx=XM_023255080.1;refGene_gn=NEO1;refGene_tx=XM_023255081.1
B3 34891012 34891167
refGene_gn=NEO1;refGene_tx=XM_023255073.1;refGene_gn=NEO1;refGene_tx=XM_023255074.1;refGene_gn=NEO1;refGene_tx=XM_023255075.1;refGene_gn=NEO

1;refGene_tx=XM_023255077.1;refGene_gn=NEO1;refGene_tx=XM_023255078.1;
refGene_gn=NEO1;refGene_tx=XM_023255079.1;refGene_gn=NEO1;refGene_tx=X
M_023255080.1;refGene_gn=NEO1;refGene_tx=XM_023255081.1
B3 34924315 34924452
refGene_gn=NEO1;refGene_tx=XM_023255073.1;refGene_gn=NEO1;refGene_tx=X
M_023255074.1;refGene_gn=NEO1;refGene_tx=XM_023255075.1;refGene_gn=NEO
1;refGene_tx=XM_023255077.1;refGene_gn=NEO1;refGene_tx=XM_023255078.1;
refGene_gn=NEO1;refGene_tx=XM_023255079.1;refGene_gn=NEO1;refGene_tx=X
M_023255080.1;refGene_gn=NEO1;refGene_tx=XM_023255081.1
B3 34944008 34944162
refGene_gn=NEO1;refGene_tx=XM_023255073.1;refGene_gn=NEO1;refGene_tx=X
M_023255074.1;refGene_gn=NEO1;refGene_tx=XM_023255075.1;refGene_gn=NEO
1;refGene_tx=XM_023255077.1;refGene_gn=NEO1;refGene_tx=XM_023255078.1;
refGene_gn=NEO1;refGene_tx=XM_023255079.1;refGene_gn=NEO1;refGene_tx=X
M_023255080.1;refGene_gn=NEO1;refGene_tx=XM_023255081.1
B3 34947810 34948086
refGene_gn=NEO1;refGene_tx=XM_023255073.1;refGene_gn=NEO1;refGene_tx=X
M_023255074.1;refGene_gn=NEO1;refGene_tx=XM_023255075.1;refGene_gn=NEO
1;refGene_tx=XM_023255077.1;refGene_gn=NEO1;refGene_tx=XM_023255078.1;
refGene_gn=NEO1;refGene_tx=XM_023255079.1;refGene_gn=NEO1;refGene_tx=X
M_023255080.1;refGene_gn=NEO1;refGene_tx=XM_023255081.1
B3 34953779 34954097
refGene_gn=NEO1;refGene_tx=XM_023255073.1;refGene_gn=NEO1;refGene_tx=X
M_023255074.1;refGene_gn=NEO1;refGene_tx=XM_023255075.1;refGene_gn=NEO
1;refGene_tx=XM_023255077.1;refGene_gn=NEO1;refGene_tx=XM_023255078.1;
refGene_gn=NEO1;refGene_tx=XM_023255079.1;refGene_gn=NEO1;refGene_tx=X
M_023255080.1;refGene_gn=NEO1;refGene_tx=XM_023255081.1
B3 35022068 35022195
refGene_gn=NEO1;refGene_tx=XM_023255073.1;refGene_gn=NEO1;refGene_tx=X
M_023255074.1;refGene_gn=NEO1;refGene_tx=XM_023255075.1;refGene_gn=NEO
1;refGene_tx=XM_023255077.1;refGene_gn=NEO1;refGene_tx=XM_023255078.1;
refGene_gn=NEO1;refGene_tx=XM_023255079.1;refGene_gn=NEO1;refGene_tx=X
M_023255080.1;refGene_gn=NEO1;refGene_tx=XM_023255081.1
B3 35285986 35286213 refGene_gn=ADPGK;refGene_tx=XM_023255082.1
B3 35292768 35293012 refGene_gn=ADPGK;refGene_tx=XM_023255082.1
B3 35294288 35294351 refGene_gn=ADPGK;refGene_tx=XM_023255082.1
B3 35304769 35304890 refGene_gn=ADPGK;refGene_tx=XM_023255082.1
B3 35309257 35309454 refGene_gn=ADPGK;refGene_tx=XM_023255082.1
B3 35310062 35310161 refGene_gn=ADPGK;refGene_tx=XM_023255082.1
B3 35312871 35313423 refGene_gn=ADPGK;refGene_tx=XM_023255082.1
B3 35322436 35322546
refGene_gn=BBS4;refGene_tx=XM_011282954.3;refGene_gn=BBS4;refGene_tx=X
M_011282956.3;refGene_gn=BBS4;refGene_tx=XM_011282957.3;refGene_gn=BBS
4;refGene_tx=XM_019832085.2
B3 35323053 35323255
refGene_gn=BBS4;refGene_tx=XM_011282954.3;refGene_gn=BBS4;refGene_tx=X
M_011282956.3;refGene_gn=BBS4;refGene_tx=XM_011282957.3;refGene_gn=BBS
4;refGene_tx=XM_019832085.2
B3 35323776 35323918
refGene_gn=BBS4;refGene_tx=XM_011282954.3;refGene_gn=BBS4;refGene_tx=X

M_011282956.3;refGene_gn=BBS4;refGene_tx=XM_011282957.3;refGene_gn=BBS
4;refGene_tx=XM_019832085.2
B3 35324531 35324601
refGene_gn=BBS4;refGene_tx=XM_011282954.3;refGene_gn=BBS4;refGene_tx=X
M_011282956.3;refGene_gn=BBS4;refGene_tx=XM_011282957.3;refGene_gn=BBS
4;refGene_tx=XM_019832085.2
B3 35326182 35326354
refGene_gn=BBS4;refGene_tx=XM_011282954.3;refGene_gn=BBS4;refGene_tx=X
M_011282956.3;refGene_gn=BBS4;refGene_tx=XM_011282957.3;refGene_gn=BBS
4;refGene_tx=XM_019832085.2
B3 35326456 35326609
refGene_gn=BBS4;refGene_tx=XM_011282954.3;refGene_gn=BBS4;refGene_tx=X
M_011282956.3;refGene_gn=BBS4;refGene_tx=XM_011282957.3;refGene_gn=BBS
4;refGene_tx=XM_019832085.2
B3 35327580 35327649
refGene_gn=BBS4;refGene_tx=XM_011282954.3;refGene_gn=BBS4;refGene_tx=X
M_011282956.3;refGene_gn=BBS4;refGene_tx=XM_011282957.3;refGene_gn=BBS
4;refGene_tx=XM_019832085.2
B3 35328594 35328649
refGene_gn=BBS4;refGene_tx=XM_011282954.3;refGene_gn=BBS4;refGene_tx=X
M_011282956.3;refGene_gn=BBS4;refGene_tx=XM_011282957.3;refGene_gn=BBS
4;refGene_tx=XM_019832085.2
B3 35331468 35331596
refGene_gn=BBS4;refGene_tx=XM_011282954.3;refGene_gn=BBS4;refGene_tx=X
M_011282956.3;refGene_gn=BBS4;refGene_tx=XM_011282957.3;refGene_gn=BBS
4;refGene_tx=XM_019832085.2
B3 35332897 35332951
refGene_gn=BBS4;refGene_tx=XM_011282954.3;refGene_gn=BBS4;refGene_tx=X
M_011282956.3;refGene_gn=BBS4;refGene_tx=XM_019832085.2
B3 35339429 35339502
refGene_gn=BBS4;refGene_tx=XM_011282954.3;refGene_gn=BBS4;refGene_tx=X
M_011282956.3;refGene_gn=BBS4;refGene_tx=XM_019832085.2
B3 35340486 35340598
refGene_gn=BBS4;refGene_tx=XM_011282954.3;refGene_gn=BBS4;refGene_tx=X
M_011282956.3;refGene_gn=BBS4;refGene_tx=XM_019832085.2
B3 35341983 35342047
refGene_gn=BBS4;refGene_tx=XM_011282954.3;refGene_gn=BBS4;refGene_tx=X
M_011282956.3;refGene_gn=BBS4;refGene_tx=XM_019832085.2
B3 35344482 35344562
refGene_gn=BBS4;refGene_tx=XM_011282954.3;refGene_gn=BBS4;refGene_tx=X
M_011282956.3;refGene_gn=BBS4;refGene_tx=XM_019832085.2
B3 35347161 35347201 refGene_gn=BBS4;refGene_tx=XM_019832085.2
B3 35370667 35370719
refGene_gn=BBS4;refGene_tx=XM_011282954.3;refGene_gn=BBS4;refGene_tx=X
M_011282956.3
B3 35377665 35377707 refGene_gn=BBS4;refGene_tx=XM_011282954.3
B3 35378104 35378128 refGene_gn=BBS4;refGene_tx=XM_011282956.3
B3 35395107 35395192 refGene_gn=ARIH1;refGene_tx=XM_023255083.1
B3 35396763 35396876 refGene_gn=ARIH1;refGene_tx=XM_023255083.1
B3 35399133 35399394 refGene_gn=ARIH1;refGene_tx=XM_023255083.1

B3	35403901 35403959 refGene_gn=ARIH1;refGene_tx=XM_023255083.1
B3	35407506 35407637 refGene_gn=ARIH1;refGene_tx=XM_023255083.1
B3	35411710 35411782 refGene_gn=ARIH1;refGene_tx=XM_023255083.1
B3	35412300 35412343 refGene_gn=ARIH1;refGene_tx=XM_023255083.1
B3	35414330 35414437 refGene_gn=ARIH1;refGene_tx=XM_023255083.1
B3	35416226 35416293 refGene_gn=ARIH1;refGene_tx=XM_023255083.1
B3	35428745 35428801 refGene_gn=ARIH1;refGene_tx=XM_023255083.1
B3	35429287 35429380 refGene_gn=ARIH1;refGene_tx=XM_023255083.1
B3	35436714 35436859 refGene_gn=ARIH1;refGene_tx=XM_023255083.1
B3	35461204 35461272 refGene_gn=ARIH1;refGene_tx=XM_023255083.1
B3	35509755 35510124 refGene_gn=ARIH1;refGene_tx=XM_023255083.1
B3	35532705 35533241
refGene_gn=TMEM202;refGene_tx=XM_006932398.4;refGene_gn=TMEM202;refGene_tx=XM_019832089.2;refGene_gn=TMEM202;refGene_tx=XM_019832093.2;refGene_gn=TMEM202;refGene_tx=XM_023255084.1;refGene_gn=TMEM202;refGene_tx=XM_023255085.1;refGene_gn=TMEM202;refGene_tx=XM_023255086.1
B3	35534424 35534556
refGene_gn=TMEM202;refGene_tx=XM_019832089.2;refGene_gn=TMEM202;refGene_tx=XM_023255084.1;refGene_gn=TMEM202;refGene_tx=XM_023255085.1;refGene_gn=TMEM202;refGene_tx=XM_023255086.1
B3	35535140 35535287
refGene_gn=TMEM202;refGene_tx=XM_019832089.2;refGene_gn=TMEM202;refGene_tx=XM_019832093.2;refGene_gn=TMEM202;refGene_tx=XM_023255084.1;refGene_gn=TMEM202;refGene_tx=XM_023255086.1
B3	35535914 35536167
refGene_gn=TMEM202;refGene_tx=XM_006932398.4;refGene_gn=TMEM202;refGene_tx=XM_019832089.2;refGene_gn=TMEM202;refGene_tx=XM_019832093.2;refGene_gn=TMEM202;refGene_tx=XM_023255084.1;refGene_gn=TMEM202;refGene_tx=XM_023255085.1;refGene_gn=TMEM202;refGene_tx=XM_023255086.1
B3	35536474 35536588
refGene_gn=TMEM202;refGene_tx=XM_006932398.4;refGene_gn=TMEM202;refGene_tx=XM_019832089.2;refGene_gn=TMEM202;refGene_tx=XM_019832093.2;refGene_gn=TMEM202;refGene_tx=XM_023255085.1;refGene_gn=TMEM202;refGene_tx=XM_023255086.1
B3	35559993 35560190
refGene_gn=TMEM202;refGene_tx=XM_003986978.4;refGene_gn=TMEM202;refGene_tx=XM_023255087.1
B3	35560660 35560792
refGene_gn=TMEM202;refGene_tx=XM_003986978.4;refGene_gn=TMEM202;refGene_tx=XM_023255087.1
B3	35561132 35561282
refGene_gn=TMEM202;refGene_tx=XM_003986978.4;refGene_gn=TMEM202;refGene_tx=XM_023255087.1
B3	35568322 35568578 refGene_gn=TMEM202;refGene_tx=XM_003986978.4
B3	35568798 35568879 refGene_gn=TMEM202;refGene_tx=XM_003986978.4
B3	35586670 35586773 refGene_gn=TMEM202;refGene_tx=XM_023255087.1
B3	35586947 35587076 refGene_gn=TMEM202;refGene_tx=XM_023255087.1
B3	35588613 35588866 refGene_gn=HEXA;refGene_tx=NM_001278835.1
B3	35604864 35604957 refGene_gn=HEXA;refGene_tx=NM_001278835.1
B3	35605852 35605918

refGene_gn=HEXA;refGene_tx=NM_001278835.1;refGene_gn=HEXA;refGene_tx=XM_019831719.2
B3 35607565 35607612 refGene_gn=HEXA;refGene_tx=NM_001278835.1
B3 35608387 35608498
refGene_gn=HEXA;refGene_tx=NM_001278835.1;refGene_gn=HEXA;refGene_tx=XM_019831719.2
B3 35609383 35609485
refGene_gn=HEXA;refGene_tx=NM_001278835.1;refGene_gn=HEXA;refGene_tx=XM_019831719.2
B3 35610176 35610309
refGene_gn=HEXA;refGene_tx=NM_001278835.1;refGene_gn=HEXA;refGene_tx=XM_019831719.2
B3 35610935 35611116
refGene_gn=HEXA;refGene_tx=NM_001278835.1;refGene_gn=HEXA;refGene_tx=XM_019831719.2
B3 35612119 35612206
refGene_gn=HEXA;refGene_tx=NM_001278835.1;refGene_gn=HEXA;refGene_tx=XM_019831719.2
B3 35612474 35612547
refGene_gn=HEXA;refGene_tx=NM_001278835.1;refGene_gn=HEXA;refGene_tx=XM_019831719.2
B3 35613308 35613492
refGene_gn=HEXA;refGene_tx=NM_001278835.1;refGene_gn=HEXA;refGene_tx=XM_019831719.2
B3 35613683 35613774
refGene_gn=HEXA;refGene_tx=NM_001278835.1;refGene_gn=HEXA;refGene_tx=XM_019831719.2
B3 35614401 35614506
refGene_gn=HEXA;refGene_tx=NM_001278835.1;refGene_gn=HEXA;refGene_tx=XM_019831719.2
B3 35615578 35615642
refGene_gn=HEXA;refGene_tx=NM_001278835.1;refGene_gn=HEXA;refGene_tx=XM_019831719.2
B3 35639129 35639391
refGene_gn=CELF6;refGene_tx=XM_023255088.1;refGene_gn=CELF6;refGene_tx=XM_023255089.1;refGene_gn=CELF6;refGene_tx=XM_023255090.1
B3 35643105 35643257
refGene_gn=CELF6;refGene_tx=XM_019832099.2;refGene_gn=CELF6;refGene_tx=XM_023255088.1;refGene_gn=CELF6;refGene_tx=XM_023255089.1;refGene_gn=CELF6;refGene_tx=XM_023255090.1
B3 35650154 35650230 refGene_gn=CELF6;refGene_tx=XM_019832099.2
B3 35651861 35651910
refGene_gn=CELF6;refGene_tx=XM_019832099.2;refGene_gn=CELF6;refGene_tx=XM_023255088.1;refGene_gn=CELF6;refGene_tx=XM_023255089.1;refGene_gn=CELF6;refGene_tx=XM_023255090.1
B3 35664578 35664707
refGene_gn=CELF6;refGene_tx=XM_019832099.2;refGene_gn=CELF6;refGene_tx=XM_023255088.1;refGene_gn=CELF6;refGene_tx=XM_023255089.1;refGene_gn=CELF6;refGene_tx=XM_023255090.1
B3 35664805 35664885

refGene_gn=CELF6;refGene_tx=XM_019832099.2;refGene_gn=CELF6;refGene_tx=XM_023255088.1;refGene_gn=CELF6;refGene_tx=XM_023255089.1;refGene_gn=CELF6;refGene_tx=XM_023255090.1
B3 35665052 35665196
refGene_gn=CELF6;refGene_tx=XM_019832099.2;refGene_gn=CELF6;refGene_tx=XM_023255088.1;refGene_gn=CELF6;refGene_tx=XM_023255089.1;refGene_gn=CELF6;refGene_tx=XM_023255090.1
B3 35665319 35665452
refGene_gn=CELF6;refGene_tx=XM_019832099.2;refGene_gn=CELF6;refGene_tx=XM_023255088.1;refGene_gn=CELF6;refGene_tx=XM_023255089.1;refGene_gn=CELF6;refGene_tx=XM_023255090.1
B3 35665534 35665684
refGene_gn=CELF6;refGene_tx=XM_019832099.2;refGene_gn=CELF6;refGene_tx=XM_023255088.1;refGene_gn=CELF6;refGene_tx=XM_023255089.1;refGene_gn=CELF6;refGene_tx=XM_023255090.1
B3 35665861 35665924
refGene_gn=CELF6;refGene_tx=XM_019832099.2;refGene_gn=CELF6;refGene_tx=XM_023255088.1;refGene_gn=CELF6;refGene_tx=XM_023255089.1;refGene_gn=CELF6;refGene_tx=XM_023255090.1
B3 35666190 35666271
refGene_gn=CELF6;refGene_tx=XM_019832099.2;refGene_gn=CELF6;refGene_tx=XM_023255088.1;refGene_gn=CELF6;refGene_tx=XM_023255089.1;refGene_gn=CELF6;refGene_tx=XM_023255090.1
B3 35666357 35666501
refGene_gn=CELF6;refGene_tx=XM_019832099.2;refGene_gn=CELF6;refGene_tx=XM_023255088.1;refGene_gn=CELF6;refGene_tx=XM_023255089.1;refGene_gn=CELF6;refGene_tx=XM_023255090.1
B3 35667140 35667268
refGene_gn=CELF6;refGene_tx=XM_019832099.2;refGene_gn=CELF6;refGene_tx=XM_023255088.1;refGene_gn=CELF6;refGene_tx=XM_023255089.1;refGene_gn=CELF6;refGene_tx=XM_023255090.1
B3 35683584 35683587
refGene_gn=PARP6;refGene_tx=XM_006932407.3;refGene_gn=PARP6;refGene_tx=XM_019832104.2;refGene_gn=PARP6;refGene_tx=XM_019832105.2;refGene_gn=PARP6;refGene_tx=XM_019832107.2;refGene_gn=PARP6;refGene_tx=XM_019832108.2;refGene_gn=PARP6;refGene_tx=XM_019832109.2;refGene_gn=PARP6;refGene_tx=XM_019832110.2;refGene_gn=PARP6;refGene_tx=XM_019832111.2;refGene_gn=PARP6;refGene_tx=XM_019832114.1;refGene_gn=PARP6;refGene_tx=XM_019832115.1;refGene_gn=PARP6;refGene_tx=XM_019832116.1
B3 35684206 35684284
refGene_gn=PARP6;refGene_tx=XM_006932407.3;refGene_gn=PARP6;refGene_tx=XM_019832104.2;refGene_gn=PARP6;refGene_tx=XM_019832105.2;refGene_gn=PARP6;refGene_tx=XM_019832107.2;refGene_gn=PARP6;refGene_tx=XM_019832108.2;refGene_gn=PARP6;refGene_tx=XM_019832109.2;refGene_gn=PARP6;refGene_tx=XM_019832110.2;refGene_gn=PARP6;refGene_tx=XM_019832111.2;refGene_gn=PARP6;refGene_tx=XM_019832114.1;refGene_gn=PARP6;refGene_tx=XM_019832115.1;refGene_gn=PARP6;refGene_tx=XM_019832116.1
B3 35685043 35685138
refGene_gn=PARP6;refGene_tx=XM_006932407.3;refGene_gn=PARP6;refGene_tx=XM_019832104.2;refGene_gn=PARP6;refGene_tx=XM_019832105.2;refGene_gn=

PARP6;refGene_tx=XM_019832107.2;refGene_gn=PARP6;refGene_tx=XM_019832108.2;refGene_gn=PARP6;refGene_tx=XM_019832109.2;refGene_gn=PARP6;refGene_tx=XM_019832110.2;refGene_gn=PARP6;refGene_tx=XM_019832111.2;refGene_gn=PARP6;refGene_tx=XM_019832114.1;refGene_gn=PARP6;refGene_tx=XM_019832115.1;refGene_gn=PARP6;refGene_tx=XM_019832116.1
B3 35685565 35685626
refGene_gn=PARP6;refGene_tx=XM_006932407.3;refGene_gn=PARP6;refGene_tx=XM_019832104.2;refGene_gn=PARP6;refGene_tx=XM_019832105.2;refGene_gn=PARP6;refGene_tx=XM_019832107.2;refGene_gn=PARP6;refGene_tx=XM_019832108.2;refGene_gn=PARP6;refGene_tx=XM_019832109.2;refGene_gn=PARP6;refGene_tx=XM_019832110.2;refGene_gn=PARP6;refGene_tx=XM_019832111.2;refGene_gn=PARP6;refGene_tx=XM_019832114.1;refGene_gn=PARP6;refGene_tx=XM_019832115.1;refGene_gn=PARP6;refGene_tx=XM_019832116.1
B3 35685866 35685957
refGene_gn=PARP6;refGene_tx=XM_006932407.3;refGene_gn=PARP6;refGene_tx=XM_019832104.2;refGene_gn=PARP6;refGene_tx=XM_019832105.2;refGene_gn=PARP6;refGene_tx=XM_019832107.2;refGene_gn=PARP6;refGene_tx=XM_019832108.2;refGene_gn=PARP6;refGene_tx=XM_019832109.2;refGene_gn=PARP6;refGene_tx=XM_019832110.2;refGene_gn=PARP6;refGene_tx=XM_019832111.2;refGene_gn=PARP6;refGene_tx=XM_019832114.1;refGene_gn=PARP6;refGene_tx=XM_019832115.1;refGene_gn=PARP6;refGene_tx=XM_019832116.1
B3 35686408 35686475
refGene_gn=PARP6;refGene_tx=XM_006932407.3;refGene_gn=PARP6;refGene_tx=XM_019832104.2;refGene_gn=PARP6;refGene_tx=XM_019832105.2;refGene_gn=PARP6;refGene_tx=XM_019832107.2;refGene_gn=PARP6;refGene_tx=XM_019832108.2;refGene_gn=PARP6;refGene_tx=XM_019832109.2;refGene_gn=PARP6;refGene_tx=XM_019832110.2;refGene_gn=PARP6;refGene_tx=XM_019832111.2;refGene_gn=PARP6;refGene_tx=XM_019832114.1;refGene_gn=PARP6;refGene_tx=XM_019832115.1;refGene_gn=PARP6;refGene_tx=XM_019832116.1
B3 35689029 35689179
refGene_gn=PARP6;refGene_tx=XM_006932407.3;refGene_gn=PARP6;refGene_tx=XM_019832104.2;refGene_gn=PARP6;refGene_tx=XM_019832105.2;refGene_gn=PARP6;refGene_tx=XM_019832107.2;refGene_gn=PARP6;refGene_tx=XM_019832108.2;refGene_gn=PARP6;refGene_tx=XM_019832109.2;refGene_gn=PARP6;refGene_tx=XM_019832110.2;refGene_gn=PARP6;refGene_tx=XM_019832111.2;refGene_gn=PARP6;refGene_tx=XM_019832112.2;refGene_gn=PARP6;refGene_tx=XM_019832113.2;refGene_gn=PARP6;refGene_tx=XM_019832114.1;refGene_gn=PARP6;refGene_tx=XM_019832115.1;refGene_gn=PARP6;refGene_tx=XM_019832116.1
B3 35690315 35690526
refGene_gn=PARP6;refGene_tx=XM_006932407.3;refGene_gn=PARP6;refGene_tx=XM_019832104.2;refGene_gn=PARP6;refGene_tx=XM_019832105.2;refGene_gn=PARP6;refGene_tx=XM_019832107.2;refGene_gn=PARP6;refGene_tx=XM_019832108.2;refGene_gn=PARP6;refGene_tx=XM_019832109.2;refGene_gn=PARP6;refGene_tx=XM_019832110.2;refGene_gn=PARP6;refGene_tx=XM_019832111.2;refGene_gn=PARP6;refGene_tx=XM_019832112.2;refGene_gn=PARP6;refGene_tx=XM_019832113.2;refGene_gn=PARP6;refGene_tx=XM_019832114.1;refGene_gn=PARP6;refGene_tx=XM_019832115.1;refGene_gn=PARP6;refGene_tx=XM_019832116.1
B3 35691045 35691099
refGene_gn=PARP6;refGene_tx=XM_006932407.3;refGene_gn=PARP6;refGene_tx=XM_019832104.2;refGene_gn=PARP6;refGene_tx=XM_019832105.2;refGene_gn=

PARP6;refGene_tx=XM_019832107.2;refGene_gn=PARP6;refGene_tx=XM_019832108.2;refGene_gn=PARP6;refGene_tx=XM_019832109.2;refGene_gn=PARP6;refGene_tx=XM_019832110.2;refGene_gn=PARP6;refGene_tx=XM_019832111.2;refGene_gn=PARP6;refGene_tx=XM_019832112.2;refGene_gn=PARP6;refGene_tx=XM_019832113.2;refGene_gn=PARP6;refGene_tx=XM_019832114.1;refGene_gn=PARP6;refGene_tx=XM_019832115.1;refGene_gn=PARP6;refGene_tx=XM_019832116.1
B3 35692705 35692801
refGene_gn=PARP6;refGene_tx=XM_006932407.3;refGene_gn=PARP6;refGene_tx=XM_019832104.2;refGene_gn=PARP6;refGene_tx=XM_019832105.2;refGene_gn=PARP6;refGene_tx=XM_019832107.2;refGene_gn=PARP6;refGene_tx=XM_019832108.2;refGene_gn=PARP6;refGene_tx=XM_019832109.2;refGene_gn=PARP6;refGene_tx=XM_019832110.2;refGene_gn=PARP6;refGene_tx=XM_019832111.2;refGene_gn=PARP6;refGene_tx=XM_019832112.2;refGene_gn=PARP6;refGene_tx=XM_019832113.2;refGene_gn=PARP6;refGene_tx=XM_019832114.1;refGene_gn=PARP6;refGene_tx=XM_019832115.1;refGene_gn=PARP6;refGene_tx=XM_019832116.1
B3 35693718 35693811
refGene_gn=PARP6;refGene_tx=XM_006932407.3;refGene_gn=PARP6;refGene_tx=XM_019832104.2;refGene_gn=PARP6;refGene_tx=XM_019832105.2;refGene_gn=PARP6;refGene_tx=XM_019832107.2;refGene_gn=PARP6;refGene_tx=XM_019832108.2;refGene_gn=PARP6;refGene_tx=XM_019832109.2;refGene_gn=PARP6;refGene_tx=XM_019832110.2;refGene_gn=PARP6;refGene_tx=XM_019832111.2;refGene_gn=PARP6;refGene_tx=XM_019832112.2;refGene_gn=PARP6;refGene_tx=XM_019832113.2;refGene_gn=PARP6;refGene_tx=XM_019832114.1;refGene_gn=PARP6;refGene_tx=XM_019832115.1;refGene_gn=PARP6;refGene_tx=XM_019832116.1
B3 35694451 35694577
refGene_gn=PARP6;refGene_tx=XM_006932407.3;refGene_gn=PARP6;refGene_tx=XM_019832104.2;refGene_gn=PARP6;refGene_tx=XM_019832105.2;refGene_gn=PARP6;refGene_tx=XM_019832107.2;refGene_gn=PARP6;refGene_tx=XM_019832108.2;refGene_gn=PARP6;refGene_tx=XM_019832109.2;refGene_gn=PARP6;refGene_tx=XM_019832110.2;refGene_gn=PARP6;refGene_tx=XM_019832112.2;refGene_gn=PARP6;refGene_tx=XM_019832113.2;refGene_gn=PARP6;refGene_tx=XM_019832114.1;refGene_gn=PARP6;refGene_tx=XM_019832115.1;refGene_gn=PARP6;refGene_tx=XM_019832116.1
B3 35695853 35695919
refGene_gn=PARP6;refGene_tx=XM_006932407.3;refGene_gn=PARP6;refGene_tx=XM_019832104.2;refGene_gn=PARP6;refGene_tx=XM_019832105.2;refGene_gn=PARP6;refGene_tx=XM_019832107.2;refGene_gn=PARP6;refGene_tx=XM_019832108.2;refGene_gn=PARP6;refGene_tx=XM_019832109.2;refGene_gn=PARP6;refGene_tx=XM_019832110.2;refGene_gn=PARP6;refGene_tx=XM_019832111.2;refGene_gn=PARP6;refGene_tx=XM_019832112.2;refGene_gn=PARP6;refGene_tx=XM_019832113.2;refGene_gn=PARP6;refGene_tx=XM_019832114.1;refGene_gn=PARP6;refGene_tx=XM_019832115.1;refGene_gn=PARP6;refGene_tx=XM_019832116.1
B3 35696829 35696897
refGene_gn=PARP6;refGene_tx=XM_006932407.3;refGene_gn=PARP6;refGene_tx=XM_019832104.2;refGene_gn=PARP6;refGene_tx=XM_019832105.2;refGene_gn=PARP6;refGene_tx=XM_019832107.2;refGene_gn=PARP6;refGene_tx=XM_019832108.2;refGene_gn=PARP6;refGene_tx=XM_019832109.2;refGene_gn=PARP6;refGene_tx=XM_019832110.2;refGene_gn=PARP6;refGene_tx=XM_019832111.2;refGene_gn=PARP6;refGene_tx=XM_019832112.2;refGene_gn=PARP6;refGene_tx=XM_019832113.2;refGene_gn=PARP6;refGene_tx=XM_019832114.1;refGene_gn=PARP6;

refGene_tx=XM_019832115.1;refGene_gn=PARP6;refGene_tx=XM_019832116.1
B3 35698393 35698442
refGene_gn=PARP6;refGene_tx=XM_006932407.3;refGene_gn=PARP6;refGene_tx=XM_019832104.2;refGene_gn=PARP6;refGene_tx=XM_019832105.2;refGene_gn=PARP6;refGene_tx=XM_019832107.2;refGene_gn=PARP6;refGene_tx=XM_019832108.2;refGene_gn=PARP6;refGene_tx=XM_019832109.2;refGene_gn=PARP6;refGene_tx=XM_019832110.2;refGene_gn=PARP6;refGene_tx=XM_019832111.2;refGene_gn=PARP6;refGene_tx=XM_019832112.2;refGene_gn=PARP6;refGene_tx=XM_019832113.2;refGene_gn=PARP6;refGene_tx=XM_019832114.1;refGene_gn=PARP6;refGene_tx=XM_019832115.1;refGene_gn=PARP6;refGene_tx=XM_019832116.1
B3 35698691 35698804
refGene_gn=PARP6;refGene_tx=XM_006932407.3;refGene_gn=PARP6;refGene_tx=XM_019832104.2;refGene_gn=PARP6;refGene_tx=XM_019832105.2;refGene_gn=PARP6;refGene_tx=XM_019832107.2;refGene_gn=PARP6;refGene_tx=XM_019832108.2;refGene_gn=PARP6;refGene_tx=XM_019832109.2;refGene_gn=PARP6;refGene_tx=XM_019832110.2;refGene_gn=PARP6;refGene_tx=XM_019832111.2;refGene_gn=PARP6;refGene_tx=XM_019832112.2;refGene_gn=PARP6;refGene_tx=XM_019832113.2;refGene_gn=PARP6;refGene_tx=XM_019832114.1;refGene_gn=PARP6;refGene_tx=XM_019832115.1;refGene_gn=PARP6;refGene_tx=XM_019832116.1
B3 35699558 35699631
refGene_gn=PARP6;refGene_tx=XM_006932407.3;refGene_gn=PARP6;refGene_tx=XM_019832104.2;refGene_gn=PARP6;refGene_tx=XM_019832105.2;refGene_gn=PARP6;refGene_tx=XM_019832107.2;refGene_gn=PARP6;refGene_tx=XM_019832108.2;refGene_gn=PARP6;refGene_tx=XM_019832109.2;refGene_gn=PARP6;refGene_tx=XM_019832110.2;refGene_gn=PARP6;refGene_tx=XM_019832111.2;refGene_gn=PARP6;refGene_tx=XM_019832112.2;refGene_gn=PARP6;refGene_tx=XM_019832113.2;refGene_gn=PARP6;refGene_tx=XM_019832114.1;refGene_gn=PARP6;refGene_tx=XM_019832115.1;refGene_gn=PARP6;refGene_tx=XM_019832116.1
B3 35700338 35700408
refGene_gn=PARP6;refGene_tx=XM_006932407.3;refGene_gn=PARP6;refGene_tx=XM_019832104.2;refGene_gn=PARP6;refGene_tx=XM_019832105.2;refGene_gn=PARP6;refGene_tx=XM_019832107.2;refGene_gn=PARP6;refGene_tx=XM_019832108.2;refGene_gn=PARP6;refGene_tx=XM_019832109.2;refGene_gn=PARP6;refGene_tx=XM_019832110.2;refGene_gn=PARP6;refGene_tx=XM_019832111.2;refGene_gn=PARP6;refGene_tx=XM_019832112.2;refGene_gn=PARP6;refGene_tx=XM_019832113.2;refGene_gn=PARP6;refGene_tx=XM_019832114.1;refGene_gn=PARP6;refGene_tx=XM_019832115.1;refGene_gn=PARP6;refGene_tx=XM_019832116.1
B3 35705502 35705582
refGene_gn=PARP6;refGene_tx=XM_006932407.3;refGene_gn=PARP6;refGene_tx=XM_019832104.2;refGene_gn=PARP6;refGene_tx=XM_019832105.2;refGene_gn=PARP6;refGene_tx=XM_019832107.2;refGene_gn=PARP6;refGene_tx=XM_019832108.2;refGene_gn=PARP6;refGene_tx=XM_019832109.2;refGene_gn=PARP6;refGene_tx=XM_019832110.2;refGene_gn=PARP6;refGene_tx=XM_019832111.2;refGene_gn=PARP6;refGene_tx=XM_019832112.2;refGene_gn=PARP6;refGene_tx=XM_019832113.2;refGene_gn=PARP6;refGene_tx=XM_019832114.1;refGene_gn=PARP6;refGene_tx=XM_019832115.1;refGene_gn=PARP6;refGene_tx=XM_019832116.1
B3 35705978 35706042
refGene_gn=PARP6;refGene_tx=XM_006932407.3;refGene_gn=PARP6;refGene_tx=XM_019832104.2;refGene_gn=PARP6;refGene_tx=XM_019832105.2;refGene_gn=PARP6;refGene_tx=XM_019832107.2;refGene_gn=PARP6;refGene_tx=XM_0198321

08.2;refGene_gn=PARP6;refGene_tx=XM_019832109.2;refGene_gn=PARP6;refGene_tx=XM_019832110.2;refGene_gn=PARP6;refGene_tx=XM_019832111.2;refGene_gn=PARP6;refGene_tx=XM_019832112.2;refGene_gn=PARP6;refGene_tx=XM_019832113.2;refGene_gn=PARP6;refGene_tx=XM_019832114.1;refGene_gn=PARP6;refGene_tx=XM_019832115.1;refGene_gn=PARP6;refGene_tx=XM_019832116.1
B3 35706205 35706290
refGene_gn=PARP6;refGene_tx=XM_006932407.3;refGene_gn=PARP6;refGene_tx=XM_019832104.2;refGene_gn=PARP6;refGene_tx=XM_019832105.2;refGene_gn=PARP6;refGene_tx=XM_019832107.2;refGene_gn=PARP6;refGene_tx=XM_019832108.2;refGene_gn=PARP6;refGene_tx=XM_019832109.2;refGene_gn=PARP6;refGene_tx=XM_019832110.2;refGene_gn=PARP6;refGene_tx=XM_019832111.2;refGene_gn=PARP6;refGene_tx=XM_019832112.2;refGene_gn=PARP6;refGene_tx=XM_019832113.2;refGene_gn=PARP6;refGene_tx=XM_019832114.1;refGene_gn=PARP6;refGene_tx=XM_019832115.1;refGene_gn=PARP6;refGene_tx=XM_019832116.1
B3 35706622 35706725
refGene_gn=PARP6;refGene_tx=XM_006932407.3;refGene_gn=PARP6;refGene_tx=XM_019832114.1;refGene_gn=PARP6;refGene_tx=XM_019832115.1;refGene_gn=PARP6;refGene_tx=XM_019832116.1
B3 35725813 35725995
refGene_gn=PARP6;refGene_tx=XM_019832104.2;refGene_gn=PARP6;refGene_tx=XM_019832105.2;refGene_gn=PARP6;refGene_tx=XM_019832107.2;refGene_gn=PARP6;refGene_tx=XM_019832108.2;refGene_gn=PARP6;refGene_tx=XM_019832109.2;refGene_gn=PARP6;refGene_tx=XM_019832110.2;refGene_gn=PARP6;refGene_tx=XM_019832111.2;refGene_gn=PARP6;refGene_tx=XM_019832112.2;refGene_gn=PARP6;refGene_tx=XM_019832113.2;refGene_gn=PARP6;refGene_tx=XM_019832117.2;refGene_gn=PARP6;refGene_tx=XM_019832118.2
B3 35726792 35726884
refGene_gn=PARP6;refGene_tx=XM_019832104.2;refGene_gn=PARP6;refGene_tx=XM_019832105.2;refGene_gn=PARP6;refGene_tx=XM_019832107.2;refGene_gn=PARP6;refGene_tx=XM_019832108.2;refGene_gn=PARP6;refGene_tx=XM_019832109.2;refGene_gn=PARP6;refGene_tx=XM_019832110.2;refGene_gn=PARP6;refGene_tx=XM_019832111.2;refGene_gn=PARP6;refGene_tx=XM_019832112.2;refGene_gn=PARP6;refGene_tx=XM_019832113.2;refGene_gn=PARP6;refGene_tx=XM_019832117.2;refGene_gn=PARP6;refGene_tx=XM_019832118.2
B3 35731415 35731547
refGene_gn=PARP6;refGene_tx=XM_019832104.2;refGene_gn=PARP6;refGene_tx=XM_019832105.2;refGene_gn=PARP6;refGene_tx=XM_019832107.2;refGene_gn=PARP6;refGene_tx=XM_019832108.2;refGene_gn=PARP6;refGene_tx=XM_019832109.2;refGene_gn=PARP6;refGene_tx=XM_019832110.2;refGene_gn=PARP6;refGene_tx=XM_019832111.2;refGene_gn=PARP6;refGene_tx=XM_019832112.2;refGene_gn=PARP6;refGene_tx=XM_019832113.2;refGene_gn=PARP6;refGene_tx=XM_019832117.2;refGene_gn=PARP6;refGene_tx=XM_019832118.2
B3 35731986 35732173
refGene_gn=PARP6;refGene_tx=XM_019832104.2;refGene_gn=PARP6;refGene_tx=XM_019832105.2;refGene_gn=PARP6;refGene_tx=XM_019832107.2;refGene_gn=PARP6;refGene_tx=XM_019832108.2;refGene_gn=PARP6;refGene_tx=XM_019832109.2;refGene_gn=PARP6;refGene_tx=XM_019832110.2;refGene_gn=PARP6;refGene_tx=XM_019832111.2;refGene_gn=PARP6;refGene_tx=XM_019832112.2;refGene_gn=PARP6;refGene_tx=XM_019832113.2;refGene_gn=PARP6;refGene_tx=XM_019832117.2;refGene_gn=PARP6;refGene_tx=XM_019832118.2

B3 35732953 35733224
refGene_gn=PARP6;refGene_tx=XM_019832104.2;refGene_gn=PARP6;refGene_tx=XM_019832105.2;refGene_gn=PARP6;refGene_tx=XM_019832107.2;refGene_gn=PARP6;refGene_tx=XM_019832108.2;refGene_gn=PARP6;refGene_tx=XM_019832109.2;refGene_gn=PARP6;refGene_tx=XM_019832110.2;refGene_gn=PARP6;refGene_tx=XM_019832111.2;refGene_gn=PARP6;refGene_tx=XM_019832112.2;refGene_gn=PARP6;refGene_tx=XM_019832113.2;refGene_gn=PARP6;refGene_tx=XM_019832117.2;refGene_gn=PARP6;refGene_tx=XM_019832118.2
B3 35734664 35734815
refGene_gn=PARP6;refGene_tx=XM_019832104.2;refGene_gn=PARP6;refGene_tx=XM_019832105.2;refGene_gn=PARP6;refGene_tx=XM_019832107.2;refGene_gn=PARP6;refGene_tx=XM_019832108.2;refGene_gn=PARP6;refGene_tx=XM_019832109.2;refGene_gn=PARP6;refGene_tx=XM_019832110.2;refGene_gn=PARP6;refGene_tx=XM_019832111.2;refGene_gn=PARP6;refGene_tx=XM_019832112.2;refGene_gn=PARP6;refGene_tx=XM_019832113.2;refGene_gn=PARP6;refGene_tx=XM_019832117.2;refGene_gn=PARP6;refGene_tx=XM_019832118.2
B3 35735014 35735167
refGene_gn=PARP6;refGene_tx=XM_019832104.2;refGene_gn=PARP6;refGene_tx=XM_019832105.2;refGene_gn=PARP6;refGene_tx=XM_019832107.2;refGene_gn=PARP6;refGene_tx=XM_019832108.2;refGene_gn=PARP6;refGene_tx=XM_019832109.2;refGene_gn=PARP6;refGene_tx=XM_019832110.2;refGene_gn=PARP6;refGene_tx=XM_019832111.2;refGene_gn=PARP6;refGene_tx=XM_019832112.2;refGene_gn=PARP6;refGene_tx=XM_019832113.2;refGene_gn=PARP6;refGene_tx=XM_019832117.2;refGene_gn=PARP6;refGene_tx=XM_019832118.2
B3 35737677 35737844
refGene_gn=PARP6;refGene_tx=XM_019832104.2;refGene_gn=PARP6;refGene_tx=XM_019832105.2;refGene_gn=PARP6;refGene_tx=XM_019832107.2;refGene_gn=PARP6;refGene_tx=XM_019832109.2;refGene_gn=PARP6;refGene_tx=XM_019832110.2;refGene_gn=PARP6;refGene_tx=XM_019832111.2;refGene_gn=PARP6;refGene_tx=XM_019832112.2;refGene_gn=PARP6;refGene_tx=XM_019832113.2;refGene_gn=PARP6;refGene_tx=XM_019832117.2;refGene_gn=PARP6;refGene_tx=XM_019832118.2
B3 35738221 35738388 refGene_gn=PARP6;refGene_tx=XM_019832108.2
B3 35740158 35740340
refGene_gn=PARP6;refGene_tx=XM_019832104.2;refGene_gn=PARP6;refGene_tx=XM_019832105.2;refGene_gn=PARP6;refGene_tx=XM_019832107.2;refGene_gn=PARP6;refGene_tx=XM_019832108.2;refGene_gn=PARP6;refGene_tx=XM_019832109.2;refGene_gn=PARP6;refGene_tx=XM_019832110.2;refGene_gn=PARP6;refGene_tx=XM_019832111.2;refGene_gn=PARP6;refGene_tx=XM_019832112.2;refGene_gn=PARP6;refGene_tx=XM_019832113.2;refGene_gn=PARP6;refGene_tx=XM_019832117.2;refGene_gn=PARP6;refGene_tx=XM_019832118.2
B3 35741039 35741146
refGene_gn=PARP6;refGene_tx=XM_019832104.2;refGene_gn=PARP6;refGene_tx=XM_019832105.2;refGene_gn=PARP6;refGene_tx=XM_019832107.2;refGene_gn=PARP6;refGene_tx=XM_019832108.2;refGene_gn=PARP6;refGene_tx=XM_019832109.2;refGene_gn=PARP6;refGene_tx=XM_019832110.2;refGene_gn=PARP6;refGene_tx=XM_019832111.2;refGene_gn=PARP6;refGene_tx=XM_019832112.2;refGene_gn=PARP6;refGene_tx=XM_019832113.2;refGene_gn=PARP6;refGene_tx=XM_019832117.2;refGene_gn=PARP6;refGene_tx=XM_019832118.2
B3 35742962 35743003

refGene_gn=GRAMD2A;refGene_tx=XM_006932411.3;refGene_gn=GRAMD2A;refGene_tx=XM_006932420.3
B3	35768299	35768392
refGene_gn=GRAMD2A;refGene_tx=XM_006932411.3;refGene_gn=GRAMD2A;refGene_tx=XM_006932412.4;refGene_gn=GRAMD2A;refGene_tx=XM_006932420.3;refGene_gn=GRAMD2A;refGene_tx=XM_023255092.1
B3	35769203	35769229
refGene_gn=GRAMD2A;refGene_tx=XM_006932419.4;refGene_gn=GRAMD2A;refGene_tx=XM_019832125.2
B3	35769592	35769703
refGene_gn=GRAMD2A;refGene_tx=XM_006932411.3;refGene_gn=GRAMD2A;refGene_tx=XM_006932412.4;refGene_gn=GRAMD2A;refGene_tx=XM_006932419.4;refGene_gn=GRAMD2A;refGene_tx=XM_006932420.3;refGene_gn=GRAMD2A;refGene_tx=XM_019832123.2;refGene_gn=GRAMD2A;refGene_tx=XM_019832125.2;refGene_gn=GRAMD2A;refGene_tx=XM_023255092.1;refGene_gn=GRAMD2A;refGene_tx=XM_023255093.1;refGene_gn=GRAMD2A;refGene_tx=XM_023255094.1;refGene_gn=GRAMD2A;refGene_tx=XM_023255095.1
B3	35770041	35770138
refGene_gn=GRAMD2A;refGene_tx=XM_006932411.3;refGene_gn=GRAMD2A;refGene_tx=XM_006932412.4;refGene_gn=GRAMD2A;refGene_tx=XM_006932419.4;refGene_gn=GRAMD2A;refGene_tx=XM_006932420.3;refGene_gn=GRAMD2A;refGene_tx=XM_019832123.2;refGene_gn=GRAMD2A;refGene_tx=XM_019832125.2;refGene_gn=GRAMD2A;refGene_tx=XM_023255092.1;refGene_gn=GRAMD2A;refGene_tx=XM_023255093.1;refGene_gn=GRAMD2A;refGene_tx=XM_023255094.1;refGene_gn=GRAMD2A;refGene_tx=XM_023255095.1
B3	35770782	35770886
refGene_gn=GRAMD2A;refGene_tx=XM_006932411.3;refGene_gn=GRAMD2A;refGene_tx=XM_006932412.4;refGene_gn=GRAMD2A;refGene_tx=XM_006932419.4;refGene_gn=GRAMD2A;refGene_tx=XM_006932420.3;refGene_gn=GRAMD2A;refGene_tx=XM_019832123.2;refGene_gn=GRAMD2A;refGene_tx=XM_019832125.2;refGene_gn=GRAMD2A;refGene_tx=XM_023255092.1;refGene_gn=GRAMD2A;refGene_tx=XM_023255093.1;refGene_gn=GRAMD2A;refGene_tx=XM_023255094.1;refGene_gn=GRAMD2A;refGene_tx=XM_023255095.1
B3	35771456	35771555
refGene_gn=GRAMD2A;refGene_tx=XM_006932411.3;refGene_gn=GRAMD2A;refGene_tx=XM_006932412.4;refGene_gn=GRAMD2A;refGene_tx=XM_006932419.4;refGene_gn=GRAMD2A;refGene_tx=XM_006932420.3;refGene_gn=GRAMD2A;refGene_tx=XM_019832123.2;refGene_gn=GRAMD2A;refGene_tx=XM_019832125.2;refGene_gn=GRAMD2A;refGene_tx=XM_023255092.1;refGene_gn=GRAMD2A;refGene_tx=XM_023255093.1;refGene_gn=GRAMD2A;refGene_tx=XM_023255094.1;refGene_gn=GRAMD2A;refGene_tx=XM_023255095.1
B3	35771680	35771752
refGene_gn=GRAMD2A;refGene_tx=XM_006932411.3;refGene_gn=GRAMD2A;refGene_tx=XM_006932412.4;refGene_gn=GRAMD2A;refGene_tx=XM_006932419.4;refGene_gn=GRAMD2A;refGene_tx=XM_006932420.3;refGene_gn=GRAMD2A;refGene_tx=XM_019832123.2;refGene_gn=GRAMD2A;refGene_tx=XM_019832125.2;refGene_gn=GRAMD2A;refGene_tx=XM_023255092.1;refGene_gn=GRAMD2A;refGene_tx=XM_023255093.1;refGene_gn=GRAMD2A;refGene_tx=XM_023255094.1;refGene_gn=GRAMD2A;refGene_tx=XM_023255095.1
B3	35772287	35772350

refGene_gn=GRAMD2A;refGene_tx=XM_006932411.3;refGene_gn=GRAMD2A;refGene_tx=XM_006932412.4;refGene_gn=GRAMD2A;refGene_tx=XM_006932419.4;refGene_gn=GRAMD2A;refGene_tx=XM_006932420.3;refGene_gn=GRAMD2A;refGene_tx=XM_019832123.2;refGene_gn=GRAMD2A;refGene_tx=XM_019832125.2;refGene_gn=GRAMD2A;refGene_tx=XM_023255092.1;refGene_gn=GRAMD2A;refGene_tx=XM_023255093.1;refGene_gn=GRAMD2A;refGene_tx=XM_023255094.1;refGene_gn=GRAMD2A;refGene_tx=XM_023255095.1
B3 35773343 35773488
refGene_gn=GRAMD2A;refGene_tx=XM_006932411.3;refGene_gn=GRAMD2A;refGene_tx=XM_006932412.4;refGene_gn=GRAMD2A;refGene_tx=XM_006932419.4;refGene_gn=GRAMD2A;refGene_tx=XM_006932420.3;refGene_gn=GRAMD2A;refGene_tx=XM_019832123.2;refGene_gn=GRAMD2A;refGene_tx=XM_019832125.2;refGene_gn=GRAMD2A;refGene_tx=XM_023255092.1;refGene_gn=GRAMD2A;refGene_tx=XM_023255093.1;refGene_gn=GRAMD2A;refGene_tx=XM_023255094.1;refGene_gn=GRAMD2A;refGene_tx=XM_023255095.1
B3 35773745 35773845
refGene_gn=GRAMD2A;refGene_tx=XM_006932411.3;refGene_gn=GRAMD2A;refGene_tx=XM_006932412.4;refGene_gn=GRAMD2A;refGene_tx=XM_006932419.4;refGene_gn=GRAMD2A;refGene_tx=XM_006932420.3;refGene_gn=GRAMD2A;refGene_tx=XM_019832123.2;refGene_gn=GRAMD2A;refGene_tx=XM_019832125.2;refGene_gn=GRAMD2A;refGene_tx=XM_023255092.1;refGene_gn=GRAMD2A;refGene_tx=XM_023255093.1;refGene_gn=GRAMD2A;refGene_tx=XM_023255094.1;refGene_gn=GRAMD2A;refGene_tx=XM_023255095.1
B3 35774673 35774782
refGene_gn=GRAMD2A;refGene_tx=XM_006932411.3;refGene_gn=GRAMD2A;refGene_tx=XM_006932412.4;refGene_gn=GRAMD2A;refGene_tx=XM_006932419.4;refGene_gn=GRAMD2A;refGene_tx=XM_006932420.3;refGene_gn=GRAMD2A;refGene_tx=XM_019832123.2;refGene_gn=GRAMD2A;refGene_tx=XM_019832125.2;refGene_gn=GRAMD2A;refGene_tx=XM_023255092.1;refGene_gn=GRAMD2A;refGene_tx=XM_023255093.1;refGene_gn=GRAMD2A;refGene_tx=XM_023255094.1;refGene_gn=GRAMD2A;refGene_tx=XM_023255095.1
B3 35775038 35775042
refGene_gn=GRAMD2A;refGene_tx=XM_006932411.3;refGene_gn=GRAMD2A;refGene_tx=XM_006932412.4;refGene_gn=GRAMD2A;refGene_tx=XM_006932419.4;refGene_gn=GRAMD2A;refGene_tx=XM_019832123.2;refGene_gn=GRAMD2A;refGene_tx=XM_019832125.2;refGene_gn=GRAMD2A;refGene_tx=XM_023255092.1;refGene_gn=GRAMD2A;refGene_tx=XM_023255093.1;refGene_gn=GRAMD2A;refGene_tx=XM_023255094.1;refGene_gn=GRAMD2A;refGene_tx=XM_023255095.1
B3 35792604 35793243
refGene_gn=SENP8;refGene_tx=XM_003986998.5;refGene_gn=SENP8;refGene_tx=XM_006932421.4
B3 35806455 35807256
refGene_gn=LOC111560944;refGene_tx=XM_023256162.1
B3 35807304 35807382
refGene_gn=LOC111560944;refGene_tx=XM_023256162.1
B3 35849595 35850435
refGene_gn=MYO9A;refGene_tx=XM_019832127.2;refGene_gn=MYO9A;refGene_tx=XM_019832128.2;refGene_gn=MYO9A;refGene_tx=XM_019832129.2;refGene_gn=MYO9A;refGene_tx=XM_019832130.2;refGene_gn=MYO9A;refGene_tx=XM_019832131.2;refGene_gn=MYO9A;refGene_tx=XM_019832132.2;refGene_gn=MYO9A;refGe

ne_tx=XM_019832133.2;refGene_gn=MYO9A;refGene_tx=XM_019832134.2;refGene_gn=MYO9A;refGene_tx=XM_019832135.2;refGene_gn=MYO9A;refGene_tx=XM_019832136.2
B3 35862644 35862739
refGene_gn=MYO9A;refGene_tx=XM_019832127.2;refGene_gn=MYO9A;refGene_tx=XM_019832128.2;refGene_gn=MYO9A;refGene_tx=XM_019832129.2;refGene_gn=MYO9A;refGene_tx=XM_019832130.2;refGene_gn=MYO9A;refGene_tx=XM_019832131.2;refGene_gn=MYO9A;refGene_tx=XM_019832132.2;refGene_gn=MYO9A;refGene_tx=XM_019832133.2;refGene_gn=MYO9A;refGene_tx=XM_019832134.2;refGene_gn=MYO9A;refGene_tx=XM_019832135.2;refGene_gn=MYO9A;refGene_tx=XM_019832136.2
B3 35869237 35869300
refGene_gn=MYO9A;refGene_tx=XM_019832127.2;refGene_gn=MYO9A;refGene_tx=XM_019832128.2;refGene_gn=MYO9A;refGene_tx=XM_019832129.2;refGene_gn=MYO9A;refGene_tx=XM_019832130.2;refGene_gn=MYO9A;refGene_tx=XM_019832131.2;refGene_gn=MYO9A;refGene_tx=XM_019832132.2;refGene_gn=MYO9A;refGene_tx=XM_019832133.2;refGene_gn=MYO9A;refGene_tx=XM_019832134.2;refGene_gn=MYO9A;refGene_tx=XM_019832135.2;refGene_gn=MYO9A;refGene_tx=XM_019832136.2
B3 35878241 35878341
refGene_gn=MYO9A;refGene_tx=XM_019832127.2;refGene_gn=MYO9A;refGene_tx=XM_019832128.2;refGene_gn=MYO9A;refGene_tx=XM_019832129.2;refGene_gn=MYO9A;refGene_tx=XM_019832130.2;refGene_gn=MYO9A;refGene_tx=XM_019832131.2;refGene_gn=MYO9A;refGene_tx=XM_019832132.2;refGene_gn=MYO9A;refGene_tx=XM_019832133.2;refGene_gn=MYO9A;refGene_tx=XM_019832134.2;refGene_gn=MYO9A;refGene_tx=XM_019832135.2;refGene_gn=MYO9A;refGene_tx=XM_019832136.2
B3 35879814 35879871
refGene_gn=MYO9A;refGene_tx=XM_019832127.2;refGene_gn=MYO9A;refGene_tx=XM_019832128.2;refGene_gn=MYO9A;refGene_tx=XM_019832129.2;refGene_gn=MYO9A;refGene_tx=XM_019832130.2;refGene_gn=MYO9A;refGene_tx=XM_019832131.2;refGene_gn=MYO9A;refGene_tx=XM_019832132.2;refGene_gn=MYO9A;refGene_tx=XM_019832134.2;refGene_gn=MYO9A;refGene_tx=XM_019832135.2;refGene_gn=MYO9A;refGene_tx=XM_019832136.2
B3 35883542 35883640
refGene_gn=MYO9A;refGene_tx=XM_019832127.2;refGene_gn=MYO9A;refGene_tx=XM_019832128.2;refGene_gn=MYO9A;refGene_tx=XM_019832129.2;refGene_gn=MYO9A;refGene_tx=XM_019832130.2;refGene_gn=MYO9A;refGene_tx=XM_019832131.2;refGene_gn=MYO9A;refGene_tx=XM_019832132.2;refGene_gn=MYO9A;refGene_tx=XM_019832133.2;refGene_gn=MYO9A;refGene_tx=XM_019832134.2;refGene_gn=MYO9A;refGene_tx=XM_019832135.2;refGene_gn=MYO9A;refGene_tx=XM_019832136.2
B3 35887907 35888034
refGene_gn=MYO9A;refGene_tx=XM_019832127.2;refGene_gn=MYO9A;refGene_tx=XM_019832128.2;refGene_gn=MYO9A;refGene_tx=XM_019832129.2;refGene_gn=MYO9A;refGene_tx=XM_019832130.2;refGene_gn=MYO9A;refGene_tx=XM_019832131.2;refGene_gn=MYO9A;refGene_tx=XM_019832132.2;refGene_gn=MYO9A;refGene_tx=XM_019832133.2;refGene_gn=MYO9A;refGene_tx=XM_019832134.2;refGene_gn=MYO9A;refGene_tx=XM_019832135.2;refGene_gn=MYO9A;refGene_tx=XM_019832136.2

B3 35895394 35895484
refGene_gn=MYO9A;refGene_tx=XM_019832127.2;refGene_gn=MYO9A;refGene_tx
=XM_019832128.2;refGene_gn=MYO9A;refGene_tx=XM_019832129.2;refGene_gn=
MYO9A;refGene_tx=XM_019832130.2;refGene_gn=MYO9A;refGene_tx=XM_0198321
31.2;refGene_gn=MYO9A;refGene_tx=XM_019832132.2;refGene_gn=MYO9A;refGe
ne_tx=XM_019832133.2;refGene_gn=MYO9A;refGene_tx=XM_019832134.2;refGen
e_gn=MYO9A;refGene_tx=XM_019832135.2;refGene_gn=MYO9A;refGene_tx=XM_01
9832136.2
B3 35900097 35900214
refGene_gn=MYO9A;refGene_tx=XM_019832127.2;refGene_gn=MYO9A;refGene_tx
=XM_019832128.2;refGene_gn=MYO9A;refGene_tx=XM_019832129.2;refGene_gn=
MYO9A;refGene_tx=XM_019832130.2;refGene_gn=MYO9A;refGene_tx=XM_0198321
31.2;refGene_gn=MYO9A;refGene_tx=XM_019832132.2;refGene_gn=MYO9A;refGe
ne_tx=XM_019832133.2;refGene_gn=MYO9A;refGene_tx=XM_019832134.2;refGen
e_gn=MYO9A;refGene_tx=XM_019832135.2;refGene_gn=MYO9A;refGene_tx=XM_01
9832136.2
B3 35903054 35903189
refGene_gn=MYO9A;refGene_tx=XM_019832127.2;refGene_gn=MYO9A;refGene_tx
=XM_019832128.2;refGene_gn=MYO9A;refGene_tx=XM_019832129.2;refGene_gn=
MYO9A;refGene_tx=XM_019832130.2;refGene_gn=MYO9A;refGene_tx=XM_0198321
31.2;refGene_gn=MYO9A;refGene_tx=XM_019832132.2;refGene_gn=MYO9A;refGe
ne_tx=XM_019832133.2;refGene_gn=MYO9A;refGene_tx=XM_019832134.2;refGen
e_gn=MYO9A;refGene_tx=XM_019832135.2;refGene_gn=MYO9A;refGene_tx=XM_01
9832136.2
B3 35912842 35912964
refGene_gn=MYO9A;refGene_tx=XM_019832127.2;refGene_gn=MYO9A;refGene_tx
=XM_019832128.2;refGene_gn=MYO9A;refGene_tx=XM_019832129.2;refGene_gn=
MYO9A;refGene_tx=XM_019832130.2;refGene_gn=MYO9A;refGene_tx=XM_0198321
31.2;refGene_gn=MYO9A;refGene_tx=XM_019832132.2;refGene_gn=MYO9A;refGe
ne_tx=XM_019832133.2;refGene_gn=MYO9A;refGene_tx=XM_019832134.2;refGen
e_gn=MYO9A;refGene_tx=XM_019832135.2;refGene_gn=MYO9A;refGene_tx=XM_01
9832136.2
B3 35918354 35918496
refGene_gn=MYO9A;refGene_tx=XM_019832127.2;refGene_gn=MYO9A;refGene_tx
=XM_019832128.2;refGene_gn=MYO9A;refGene_tx=XM_019832129.2;refGene_gn=
MYO9A;refGene_tx=XM_019832130.2;refGene_gn=MYO9A;refGene_tx=XM_0198321
31.2;refGene_gn=MYO9A;refGene_tx=XM_019832132.2;refGene_gn=MYO9A;refGe
ne_tx=XM_019832133.2;refGene_gn=MYO9A;refGene_tx=XM_019832134.2;refGen
e_gn=MYO9A;refGene_tx=XM_019832135.2;refGene_gn=MYO9A;refGene_tx=XM_01
9832136.2
B3 35924472 35924668
refGene_gn=MYO9A;refGene_tx=XM_019832127.2;refGene_gn=MYO9A;refGene_tx
=XM_019832128.2;refGene_gn=MYO9A;refGene_tx=XM_019832129.2;refGene_gn=
MYO9A;refGene_tx=XM_019832130.2;refGene_gn=MYO9A;refGene_tx=XM_0198321
31.2;refGene_gn=MYO9A;refGene_tx=XM_019832132.2;refGene_gn=MYO9A;refGe
ne_tx=XM_019832133.2;refGene_gn=MYO9A;refGene_tx=XM_019832134.2;refGen
e_gn=MYO9A;refGene_tx=XM_019832135.2;refGene_gn=MYO9A;refGene_tx=XM_01
9832136.2
B3 35935669 35935789
refGene_gn=MYO9A;refGene_tx=XM_019832127.2;refGene_gn=MYO9A;refGene_tx

=XM_019832128.2;refGene_gn=MYO9A;refGene_tx=XM_019832129.2;refGene_gn=MYO9A;refGene_tx=XM_019832130.2;refGene_gn=MYO9A;refGene_tx=XM_019832131.2;refGene_gn=MYO9A;refGene_tx=XM_019832132.2;refGene_gn=MYO9A;refGene_tx=XM_019832133.2;refGene_gn=MYO9A;refGene_tx=XM_019832135.2;refGene_gn=MYO9A;refGene_tx=XM_019832136.2
B3 35957440 35957516
refGene_gn=MYO9A;refGene_tx=XM_019832127.2;refGene_gn=MYO9A;refGene_tx=XM_019832128.2;refGene_gn=MYO9A;refGene_tx=XM_019832129.2;refGene_gn=MYO9A;refGene_tx=XM_019832130.2;refGene_gn=MYO9A;refGene_tx=XM_019832131.2;refGene_gn=MYO9A;refGene_tx=XM_019832132.2;refGene_gn=MYO9A;refGene_tx=XM_019832133.2;refGene_gn=MYO9A;refGene_tx=XM_019832134.2;refGene_gn=MYO9A;refGene_tx=XM_019832135.2;refGene_gn=MYO9A;refGene_tx=XM_019832136.2
B3 35965920 35966064
refGene_gn=MYO9A;refGene_tx=XM_019832127.2;refGene_gn=MYO9A;refGene_tx=XM_019832128.2;refGene_gn=MYO9A;refGene_tx=XM_019832129.2;refGene_gn=MYO9A;refGene_tx=XM_019832130.2;refGene_gn=MYO9A;refGene_tx=XM_019832131.2;refGene_gn=MYO9A;refGene_tx=XM_019832132.2;refGene_gn=MYO9A;refGene_tx=XM_019832133.2;refGene_gn=MYO9A;refGene_tx=XM_019832134.2;refGene_gn=MYO9A;refGene_tx=XM_019832135.2;refGene_gn=MYO9A;refGene_tx=XM_019832136.2
B3 35967426 35967466
refGene_gn=MYO9A;refGene_tx=XM_019832127.2;refGene_gn=MYO9A;refGene_tx=XM_019832128.2;refGene_gn=MYO9A;refGene_tx=XM_019832129.2;refGene_gn=MYO9A;refGene_tx=XM_019832130.2;refGene_gn=MYO9A;refGene_tx=XM_019832131.2;refGene_gn=MYO9A;refGene_tx=XM_019832132.2;refGene_gn=MYO9A;refGene_tx=XM_019832133.2;refGene_gn=MYO9A;refGene_tx=XM_019832134.2;refGene_gn=MYO9A;refGene_tx=XM_019832135.2;refGene_gn=MYO9A;refGene_tx=XM_019832136.2
B3 35973851 35973974
refGene_gn=MYO9A;refGene_tx=XM_019832127.2;refGene_gn=MYO9A;refGene_tx=XM_019832128.2;refGene_gn=MYO9A;refGene_tx=XM_019832129.2;refGene_gn=MYO9A;refGene_tx=XM_019832130.2;refGene_gn=MYO9A;refGene_tx=XM_019832131.2;refGene_gn=MYO9A;refGene_tx=XM_019832132.2;refGene_gn=MYO9A;refGene_tx=XM_019832133.2;refGene_gn=MYO9A;refGene_tx=XM_019832134.2;refGene_gn=MYO9A;refGene_tx=XM_019832135.2;refGene_gn=MYO9A;refGene_tx=XM_019832136.2
B3 35976824 35976905
refGene_gn=MYO9A;refGene_tx=XM_019832127.2;refGene_gn=MYO9A;refGene_tx=XM_019832128.2;refGene_gn=MYO9A;refGene_tx=XM_019832129.2;refGene_gn=MYO9A;refGene_tx=XM_019832130.2;refGene_gn=MYO9A;refGene_tx=XM_019832131.2;refGene_gn=MYO9A;refGene_tx=XM_019832132.2;refGene_gn=MYO9A;refGene_tx=XM_019832133.2;refGene_gn=MYO9A;refGene_tx=XM_019832134.2;refGene_gn=MYO9A;refGene_tx=XM_019832135.2;refGene_gn=MYO9A;refGene_tx=XM_019832136.2
B3 35978680 35978791
refGene_gn=MYO9A;refGene_tx=XM_019832127.2;refGene_gn=MYO9A;refGene_tx=XM_019832128.2;refGene_gn=MYO9A;refGene_tx=XM_019832129.2;refGene_gn=MYO9A;refGene_tx=XM_019832130.2;refGene_gn=MYO9A;refGene_tx=XM_019832131.2;refGene_gn=MYO9A;refGene_tx=XM_019832132.2;refGene_gn=MYO9A;refGe

ne_tx=XM_019832133.2;refGene_gn=MYO9A;refGene_tx=XM_019832134.2;refGene_gn=MYO9A;refGene_tx=XM_019832135.2;refGene_gn=MYO9A;refGene_tx=XM_019832136.2
B3 35979680 35979803
refGene_gn=MYO9A;refGene_tx=XM_019832127.2;refGene_gn=MYO9A;refGene_tx=XM_019832128.2;refGene_gn=MYO9A;refGene_tx=XM_019832129.2;refGene_gn=MYO9A;refGene_tx=XM_019832130.2;refGene_gn=MYO9A;refGene_tx=XM_019832131.2;refGene_gn=MYO9A;refGene_tx=XM_019832132.2;refGene_gn=MYO9A;refGene_tx=XM_019832133.2;refGene_gn=MYO9A;refGene_tx=XM_019832134.2;refGene_gn=MYO9A;refGene_tx=XM_019832135.2;refGene_gn=MYO9A;refGene_tx=XM_019832136.2
B3 35981590 35981740
refGene_gn=MYO9A;refGene_tx=XM_019832127.2;refGene_gn=MYO9A;refGene_tx=XM_019832128.2;refGene_gn=MYO9A;refGene_tx=XM_019832129.2;refGene_gn=MYO9A;refGene_tx=XM_019832130.2;refGene_gn=MYO9A;refGene_tx=XM_019832131.2;refGene_gn=MYO9A;refGene_tx=XM_019832132.2;refGene_gn=MYO9A;refGene_tx=XM_019832133.2;refGene_gn=MYO9A;refGene_tx=XM_019832134.2;refGene_gn=MYO9A;refGene_tx=XM_019832135.2;refGene_gn=MYO9A;refGene_tx=XM_019832136.2
B3 35982558 35982878
refGene_gn=MYO9A;refGene_tx=XM_019832127.2;refGene_gn=MYO9A;refGene_tx=XM_019832128.2;refGene_gn=MYO9A;refGene_tx=XM_019832129.2;refGene_gn=MYO9A;refGene_tx=XM_019832130.2;refGene_gn=MYO9A;refGene_tx=XM_019832131.2;refGene_gn=MYO9A;refGene_tx=XM_019832132.2;refGene_gn=MYO9A;refGene_tx=XM_019832133.2;refGene_gn=MYO9A;refGene_tx=XM_019832134.2;refGene_gn=MYO9A;refGene_tx=XM_019832135.2;refGene_gn=MYO9A;refGene_tx=XM_019832136.2
B3 35983472 35985074
refGene_gn=MYO9A;refGene_tx=XM_019832127.2;refGene_gn=MYO9A;refGene_tx=XM_019832128.2;refGene_gn=MYO9A;refGene_tx=XM_019832129.2;refGene_gn=MYO9A;refGene_tx=XM_019832130.2;refGene_gn=MYO9A;refGene_tx=XM_019832131.2;refGene_gn=MYO9A;refGene_tx=XM_019832132.2;refGene_gn=MYO9A;refGene_tx=XM_019832133.2;refGene_gn=MYO9A;refGene_tx=XM_019832134.2;refGene_gn=MYO9A;refGene_tx=XM_019832135.2;refGene_gn=MYO9A;refGene_tx=XM_019832136.2
B3 35988876 35988976
refGene_gn=MYO9A;refGene_tx=XM_019832127.2;refGene_gn=MYO9A;refGene_tx=XM_019832128.2;refGene_gn=MYO9A;refGene_tx=XM_019832129.2;refGene_gn=MYO9A;refGene_tx=XM_019832130.2;refGene_gn=MYO9A;refGene_tx=XM_019832131.2;refGene_gn=MYO9A;refGene_tx=XM_019832132.2;refGene_gn=MYO9A;refGene_tx=XM_019832133.2;refGene_gn=MYO9A;refGene_tx=XM_019832134.2;refGene_gn=MYO9A;refGene_tx=XM_019832135.2;refGene_gn=MYO9A;refGene_tx=XM_019832136.2
B3 35989484 35989700
refGene_gn=MYO9A;refGene_tx=XM_019832127.2;refGene_gn=MYO9A;refGene_tx=XM_019832128.2;refGene_gn=MYO9A;refGene_tx=XM_019832129.2;refGene_gn=MYO9A;refGene_tx=XM_019832130.2;refGene_gn=MYO9A;refGene_tx=XM_019832131.2;refGene_gn=MYO9A;refGene_tx=XM_019832132.2;refGene_gn=MYO9A;refGene_tx=XM_019832133.2;refGene_gn=MYO9A;refGene_tx=XM_019832134.2
B3 35998731 35998844

refGene_gn=MYO9A;refGene_tx=XM_019832127.2;refGene_gn=MYO9A;refGene_tx=XM_019832128.2;refGene_gn=MYO9A;refGene_tx=XM_019832129.2;refGene_gn=MYO9A;refGene_tx=XM_019832130.2;refGene_gn=MYO9A;refGene_tx=XM_019832131.2;refGene_gn=MYO9A;refGene_tx=XM_019832132.2;refGene_gn=MYO9A;refGene_tx=XM_019832133.2;refGene_gn=MYO9A;refGene_tx=XM_019832134.2;refGene_gn=MYO9A;refGene_tx=XM_019832135.2;refGene_gn=MYO9A;refGene_tx=XM_019832136.2
B3 36004349 36004492
refGene_gn=MYO9A;refGene_tx=XM_019832127.2;refGene_gn=MYO9A;refGene_tx=XM_019832128.2;refGene_gn=MYO9A;refGene_tx=XM_019832129.2;refGene_gn=MYO9A;refGene_tx=XM_019832130.2;refGene_gn=MYO9A;refGene_tx=XM_019832131.2;refGene_gn=MYO9A;refGene_tx=XM_019832132.2;refGene_gn=MYO9A;refGene_tx=XM_019832133.2;refGene_gn=MYO9A;refGene_tx=XM_019832134.2;refGene_gn=MYO9A;refGene_tx=XM_019832135.2;refGene_gn=MYO9A;refGene_tx=XM_019832136.2
B3 36007104 36007328
refGene_gn=MYO9A;refGene_tx=XM_019832127.2;refGene_gn=MYO9A;refGene_tx=XM_019832128.2;refGene_gn=MYO9A;refGene_tx=XM_019832129.2;refGene_gn=MYO9A;refGene_tx=XM_019832130.2;refGene_gn=MYO9A;refGene_tx=XM_019832131.2;refGene_gn=MYO9A;refGene_tx=XM_019832132.2;refGene_gn=MYO9A;refGene_tx=XM_019832133.2;refGene_gn=MYO9A;refGene_tx=XM_019832134.2;refGene_gn=MYO9A;refGene_tx=XM_019832135.2;refGene_gn=MYO9A;refGene_tx=XM_019832136.2
B3 36007840 36007957
refGene_gn=MYO9A;refGene_tx=XM_019832127.2;refGene_gn=MYO9A;refGene_tx=XM_019832128.2;refGene_gn=MYO9A;refGene_tx=XM_019832129.2;refGene_gn=MYO9A;refGene_tx=XM_019832130.2;refGene_gn=MYO9A;refGene_tx=XM_019832131.2;refGene_gn=MYO9A;refGene_tx=XM_019832132.2;refGene_gn=MYO9A;refGene_tx=XM_019832133.2;refGene_gn=MYO9A;refGene_tx=XM_019832134.2;refGene_gn=MYO9A;refGene_tx=XM_019832135.2;refGene_gn=MYO9A;refGene_tx=XM_019832136.2
B3 36009496 36009685
refGene_gn=MYO9A;refGene_tx=XM_019832127.2;refGene_gn=MYO9A;refGene_tx=XM_019832128.2;refGene_gn=MYO9A;refGene_tx=XM_019832129.2;refGene_gn=MYO9A;refGene_tx=XM_019832130.2;refGene_gn=MYO9A;refGene_tx=XM_019832131.2;refGene_gn=MYO9A;refGene_tx=XM_019832132.2;refGene_gn=MYO9A;refGene_tx=XM_019832133.2;refGene_gn=MYO9A;refGene_tx=XM_019832134.2;refGene_gn=MYO9A;refGene_tx=XM_019832135.2;refGene_gn=MYO9A;refGene_tx=XM_019832136.2
B3 36010945 36010993
refGene_gn=MYO9A;refGene_tx=XM_019832127.2;refGene_gn=MYO9A;refGene_tx=XM_019832128.2;refGene_gn=MYO9A;refGene_tx=XM_019832129.2;refGene_gn=MYO9A;refGene_tx=XM_019832130.2;refGene_gn=MYO9A;refGene_tx=XM_019832131.2;refGene_gn=MYO9A;refGene_tx=XM_019832132.2;refGene_gn=MYO9A;refGene_tx=XM_019832133.2;refGene_gn=MYO9A;refGene_tx=XM_019832134.2;refGene_gn=MYO9A;refGene_tx=XM_019832135.2;refGene_gn=MYO9A;refGene_tx=XM_019832136.2
B3 36020336 36020448
refGene_gn=MYO9A;refGene_tx=XM_019832127.2;refGene_gn=MYO9A;refGene_tx=XM_019832128.2;refGene_gn=MYO9A;refGene_tx=XM_019832129.2;refGene_gn=

MYO9A;refGene_tx=XM_019832130.2;refGene_gn=MYO9A;refGene_tx=XM_019832131.2;refGene_gn=MYO9A;refGene_tx=XM_019832132.2;refGene_gn=MYO9A;refGene_tx=XM_019832133.2;refGene_gn=MYO9A;refGene_tx=XM_019832134.2;refGene_gn=MYO9A;refGene_tx=XM_019832135.2;refGene_gn=MYO9A;refGene_tx=XM_019832136.2
B3 36023554 36023616
refGene_gn=MYO9A;refGene_tx=XM_019832127.2;refGene_gn=MYO9A;refGene_tx=XM_019832128.2;refGene_gn=MYO9A;refGene_tx=XM_019832129.2;refGene_gn=MYO9A;refGene_tx=XM_019832130.2;refGene_gn=MYO9A;refGene_tx=XM_019832131.2;refGene_gn=MYO9A;refGene_tx=XM_019832132.2;refGene_gn=MYO9A;refGene_tx=XM_019832133.2;refGene_gn=MYO9A;refGene_tx=XM_019832134.2;refGene_gn=MYO9A;refGene_tx=XM_019832135.2;refGene_gn=MYO9A;refGene_tx=XM_019832136.2
B3 36035706 36035899
refGene_gn=MYO9A;refGene_tx=XM_019832127.2;refGene_gn=MYO9A;refGene_tx=XM_019832128.2;refGene_gn=MYO9A;refGene_tx=XM_019832129.2;refGene_gn=MYO9A;refGene_tx=XM_019832130.2;refGene_gn=MYO9A;refGene_tx=XM_019832131.2;refGene_gn=MYO9A;refGene_tx=XM_019832132.2;refGene_gn=MYO9A;refGene_tx=XM_019832133.2;refGene_gn=MYO9A;refGene_tx=XM_019832134.2;refGene_gn=MYO9A;refGene_tx=XM_019832135.2;refGene_gn=MYO9A;refGene_tx=XM_019832136.2
B3 36037425 36037554
refGene_gn=MYO9A;refGene_tx=XM_019832127.2;refGene_gn=MYO9A;refGene_tx=XM_019832128.2;refGene_gn=MYO9A;refGene_tx=XM_019832129.2;refGene_gn=MYO9A;refGene_tx=XM_019832130.2;refGene_gn=MYO9A;refGene_tx=XM_019832131.2;refGene_gn=MYO9A;refGene_tx=XM_019832132.2;refGene_gn=MYO9A;refGene_tx=XM_019832133.2;refGene_gn=MYO9A;refGene_tx=XM_019832134.2;refGene_gn=MYO9A;refGene_tx=XM_019832135.2;refGene_gn=MYO9A;refGene_tx=XM_019832136.2
B3 36038373 36038479
refGene_gn=MYO9A;refGene_tx=XM_019832127.2;refGene_gn=MYO9A;refGene_tx=XM_019832128.2;refGene_gn=MYO9A;refGene_tx=XM_019832129.2;refGene_gn=MYO9A;refGene_tx=XM_019832130.2;refGene_gn=MYO9A;refGene_tx=XM_019832131.2;refGene_gn=MYO9A;refGene_tx=XM_019832132.2;refGene_gn=MYO9A;refGene_tx=XM_019832133.2;refGene_gn=MYO9A;refGene_tx=XM_019832134.2;refGene_gn=MYO9A;refGene_tx=XM_019832135.2;refGene_gn=MYO9A;refGene_tx=XM_019832136.2
B3 36039288 36039420
refGene_gn=MYO9A;refGene_tx=XM_019832127.2;refGene_gn=MYO9A;refGene_tx=XM_019832128.2;refGene_gn=MYO9A;refGene_tx=XM_019832129.2;refGene_gn=MYO9A;refGene_tx=XM_019832130.2;refGene_gn=MYO9A;refGene_tx=XM_019832131.2;refGene_gn=MYO9A;refGene_tx=XM_019832132.2;refGene_gn=MYO9A;refGene_tx=XM_019832133.2;refGene_gn=MYO9A;refGene_tx=XM_019832134.2;refGene_gn=MYO9A;refGene_tx=XM_019832135.2;refGene_gn=MYO9A;refGene_tx=XM_019832136.2
B3 36040151 36040275
refGene_gn=MYO9A;refGene_tx=XM_019832127.2;refGene_gn=MYO9A;refGene_tx=XM_019832128.2;refGene_gn=MYO9A;refGene_tx=XM_019832129.2;refGene_gn=MYO9A;refGene_tx=XM_019832130.2;refGene_gn=MYO9A;refGene_tx=XM_019832131.2;refGene_gn=MYO9A;refGene_tx=XM_019832132.2;refGene_gn=MYO9A;refGe

ne_tx=XM_019832133.2;refGene_gn=MYO9A;refGene_tx=XM_019832134.2;refGene_gn=MYO9A;refGene_tx=XM_019832135.2;refGene_gn=MYO9A;refGene_tx=XM_019832136.2
B3 36041600 36041654
refGene_gn=MYO9A;refGene_tx=XM_019832127.2;refGene_gn=MYO9A;refGene_tx=XM_019832128.2;refGene_gn=MYO9A;refGene_tx=XM_019832129.2;refGene_gn=MYO9A;refGene_tx=XM_019832130.2;refGene_gn=MYO9A;refGene_tx=XM_019832131.2;refGene_gn=MYO9A;refGene_tx=XM_019832133.2;refGene_gn=MYO9A;refGene_tx=XM_019832134.2;refGene_gn=MYO9A;refGene_tx=XM_019832135.2
B3 36078622 36078825
refGene_gn=MYO9A;refGene_tx=XM_019832127.2;refGene_gn=MYO9A;refGene_tx=XM_019832128.2;refGene_gn=MYO9A;refGene_tx=XM_019832129.2;refGene_gn=MYO9A;refGene_tx=XM_019832130.2;refGene_gn=MYO9A;refGene_tx=XM_019832131.2;refGene_gn=MYO9A;refGene_tx=XM_019832132.2;refGene_gn=MYO9A;refGene_tx=XM_019832133.2;refGene_gn=MYO9A;refGene_tx=XM_019832134.2;refGene_gn=MYO9A;refGene_tx=XM_019832135.2;refGene_gn=MYO9A;refGene_tx=XM_019832136.2
B3 36080623 36080766
refGene_gn=MYO9A;refGene_tx=XM_019832127.2;refGene_gn=MYO9A;refGene_tx=XM_019832128.2;refGene_gn=MYO9A;refGene_tx=XM_019832129.2;refGene_gn=MYO9A;refGene_tx=XM_019832130.2;refGene_gn=MYO9A;refGene_tx=XM_019832131.2;refGene_gn=MYO9A;refGene_tx=XM_019832132.2;refGene_gn=MYO9A;refGene_tx=XM_019832133.2;refGene_gn=MYO9A;refGene_tx=XM_019832134.2;refGene_gn=MYO9A;refGene_tx=XM_019832135.2;refGene_gn=MYO9A;refGene_tx=XM_019832136.2
B3 36081433 36081897
refGene_gn=MYO9A;refGene_tx=XM_019832127.2;refGene_gn=MYO9A;refGene_tx=XM_019832128.2;refGene_gn=MYO9A;refGene_tx=XM_019832129.2;refGene_gn=MYO9A;refGene_tx=XM_019832130.2;refGene_gn=MYO9A;refGene_tx=XM_019832131.2;refGene_gn=MYO9A;refGene_tx=XM_019832132.2;refGene_gn=MYO9A;refGene_tx=XM_019832133.2;refGene_gn=MYO9A;refGene_tx=XM_019832134.2;refGene_gn=MYO9A;refGene_tx=XM_019832135.2;refGene_gn=MYO9A;refGene_tx=XM_019832136.2
B3 36090313 36090446
refGene_gn=NR2E3;refGene_tx=XM_003986989.3;refGene_gn=NR2E3;refGene_tx=XM_019832138.2;refGene_gn=NR2E3;refGene_tx=XM_019832139.2
B3 36092775 36092881
refGene_gn=NR2E3;refGene_tx=XM_003986989.3;refGene_gn=NR2E3;refGene_tx=XM_019832138.2;refGene_gn=NR2E3;refGene_tx=XM_019832139.2
B3 36093147 36093394
refGene_gn=NR2E3;refGene_tx=XM_003986989.3;refGene_gn=NR2E3;refGene_tx=XM_019832138.2;refGene_gn=NR2E3;refGene_tx=XM_019832139.2
B3 36094281 36094499
refGene_gn=NR2E3;refGene_tx=XM_003986989.3;refGene_gn=NR2E3;refGene_tx=XM_019832138.2;refGene_gn=NR2E3;refGene_tx=XM_019832139.2
B3 36094576 36094825
refGene_gn=NR2E3;refGene_tx=XM_003986989.3;refGene_gn=NR2E3;refGene_tx=XM_019832138.2;refGene_gn=NR2E3;refGene_tx=XM_019832139.2
B3 36094917 36095021
refGene_gn=NR2E3;refGene_tx=XM_003986989.3;refGene_gn=NR2E3;refGene_tx

=XM_019832138.2;refGene_gn=NR2E3;refGene_tx=XM_019832139.2
B3 36095191 36095318
refGene_gn=NR2E3;refGene_tx=XM_003986989.3;refGene_gn=NR2E3;refGene_tx
=XM_019832138.2;refGene_gn=NR2E3;refGene_tx=XM_019832139.2
B3 36095853 36095947
refGene_gn=NR2E3;refGene_tx=XM_003986989.3;refGene_gn=NR2E3;refGene_tx
=XM_019832138.2;refGene_gn=NR2E3;refGene_tx=XM_019832139.2
B3 36122560 36122703
refGene_gn=THSD4;refGene_tx=XM_019832142.2;refGene_gn=THSD4;refGene_tx
=XM_023255096.1;refGene_gn=THSD4;refGene_tx=XM_023255097.1
B3 36128425 36128570
refGene_gn=THSD4;refGene_tx=XM_019832142.2;refGene_gn=THSD4;refGene_tx
=XM_023255096.1;refGene_gn=THSD4;refGene_tx=XM_023255097.1
B3 36129614 36129794
refGene_gn=THSD4;refGene_tx=XM_019832142.2;refGene_gn=THSD4;refGene_tx
=XM_023255096.1;refGene_gn=THSD4;refGene_tx=XM_023255097.1
B3 36136474 36136648
refGene_gn=THSD4;refGene_tx=XM_019832142.2;refGene_gn=THSD4;refGene_tx
=XM_023255096.1;refGene_gn=THSD4;refGene_tx=XM_023255097.1
B3 36144057 36144231
refGene_gn=THSD4;refGene_tx=XM_019832142.2;refGene_gn=THSD4;refGene_tx
=XM_023255096.1;refGene_gn=THSD4;refGene_tx=XM_023255097.1
B3 36144814 36145019
refGene_gn=THSD4;refGene_tx=XM_019832142.2;refGene_gn=THSD4;refGene_tx
=XM_023255096.1;refGene_gn=THSD4;refGene_tx=XM_023255097.1
B3 36146057 36146187
refGene_gn=THSD4;refGene_tx=XM_019832142.2;refGene_gn=THSD4;refGene_tx
=XM_023255096.1;refGene_gn=THSD4;refGene_tx=XM_023255097.1
B3 36152455 36152728
refGene_gn=THSD4;refGene_tx=XM_019832142.2;refGene_gn=THSD4;refGene_tx
=XM_023255096.1;refGene_gn=THSD4;refGene_tx=XM_023255097.1
B3 36156949 36157046
refGene_gn=THSD4;refGene_tx=XM_019832142.2;refGene_gn=THSD4;refGene_tx
=XM_023255096.1;refGene_gn=THSD4;refGene_tx=XM_023255097.1
B3 36159439 36159615
refGene_gn=THSD4;refGene_tx=XM_019832142.2;refGene_gn=THSD4;refGene_tx
=XM_023255096.1;refGene_gn=THSD4;refGene_tx=XM_023255097.1
B3 36223365 36223570
refGene_gn=THSD4;refGene_tx=XM_019832142.2;refGene_gn=THSD4;refGene_tx
=XM_023255096.1;refGene_gn=THSD4;refGene_tx=XM_023255097.1
B3 36320286 36320358 refGene_gn=THSD4;refGene_tx=XM_019832142.2
B3 36436685 36436822
refGene_gn=THSD4;refGene_tx=XM_023255096.1;refGene_gn=THSD4;refGene_tx
=XM_023255097.1
B3 36567464 36567567
refGene_gn=THSD4;refGene_tx=XM_023255096.1;refGene_gn=THSD4;refGene_tx
=XM_023255097.1
B3 36582362 36582810
refGene_gn=THSD4;refGene_tx=XM_023255096.1;refGene_gn=THSD4;refGene_tx
=XM_023255097.1

B3 36612525 36612902 refGene_gn=THSD4;refGene_tx=XM_023255096.1
B3 36661684 36661754 refGene_gn=THSD4;refGene_tx=XM_023255096.1
B3 36675046 36675075 refGene_gn=THSD4;refGene_tx=XM_023255096.1
B3 36753214 36753418
refGene_gn=LRRC49;refGene_tx=XM_011282946.3;refGene_gn=LRRC49;refGene_tx=XM_023255098.1;refGene_gn=LRRC49;refGene_tx=XM_023255099.1;refGene_gn=LRRC49;refGene_tx=XM_023255100.1;refGene_gn=LRRC49;refGene_tx=XM_023255101.1
B3 36761553 36761707
refGene_gn=LRRC49;refGene_tx=XM_011282946.3;refGene_gn=LRRC49;refGene_tx=XM_023255098.1;refGene_gn=LRRC49;refGene_tx=XM_023255099.1;refGene_gn=LRRC49;refGene_tx=XM_023255100.1;refGene_gn=LRRC49;refGene_tx=XM_023255101.1
B3 36768985 36769095
refGene_gn=LRRC49;refGene_tx=XM_011282946.3;refGene_gn=LRRC49;refGene_tx=XM_023255098.1;refGene_gn=LRRC49;refGene_tx=XM_023255099.1;refGene_gn=LRRC49;refGene_tx=XM_023255100.1;refGene_gn=LRRC49;refGene_tx=XM_023255101.1
B3 36772049 36772235
refGene_gn=LRRC49;refGene_tx=XM_011282946.3;refGene_gn=LRRC49;refGene_tx=XM_023255098.1;refGene_gn=LRRC49;refGene_tx=XM_023255099.1;refGene_gn=LRRC49;refGene_tx=XM_023255100.1;refGene_gn=LRRC49;refGene_tx=XM_023255101.1
B3 36783158 36783396
refGene_gn=LRRC49;refGene_tx=XM_011282946.3;refGene_gn=LRRC49;refGene_tx=XM_023255098.1;refGene_gn=LRRC49;refGene_tx=XM_023255099.1;refGene_gn=LRRC49;refGene_tx=XM_023255100.1;refGene_gn=LRRC49;refGene_tx=XM_023255101.1
B3 36800687 36800857
refGene_gn=LRRC49;refGene_tx=XM_011282946.3;refGene_gn=LRRC49;refGene_tx=XM_023255098.1;refGene_gn=LRRC49;refGene_tx=XM_023255099.1;refGene_gn=LRRC49;refGene_tx=XM_023255100.1;refGene_gn=LRRC49;refGene_tx=XM_023255101.1
B3 36805347 36805431
refGene_gn=LRRC49;refGene_tx=XM_011282946.3;refGene_gn=LRRC49;refGene_tx=XM_023255098.1;refGene_gn=LRRC49;refGene_tx=XM_023255099.1;refGene_gn=LRRC49;refGene_tx=XM_023255100.1;refGene_gn=LRRC49;refGene_tx=XM_023255101.1
B3 36814867 36815015
refGene_gn=LRRC49;refGene_tx=XM_011282946.3;refGene_gn=LRRC49;refGene_tx=XM_023255098.1;refGene_gn=LRRC49;refGene_tx=XM_023255099.1;refGene_gn=LRRC49;refGene_tx=XM_023255100.1;refGene_gn=LRRC49;refGene_tx=XM_023255101.1
B3 36855008 36855070
refGene_gn=LRRC49;refGene_tx=XM_011282946.3;refGene_gn=LRRC49;refGene_tx=XM_023255098.1;refGene_gn=LRRC49;refGene_tx=XM_023255099.1;refGene_gn=LRRC49;refGene_tx=XM_023255100.1;refGene_gn=LRRC49;refGene_tx=XM_023255101.1
B3 36873377 36873521
refGene_gn=LRRC49;refGene_tx=XM_011282946.3;refGene_gn=LRRC49;refGene_

tx=XM_023255098.1;refGene_gn=LRRC49;refGene_tx=XM_023255099.1;refGene_gn=LRRC49;refGene_tx=XM_023255100.1;refGene_gn=LRRC49;refGene_tx=XM_023255101.1
B3 36881766 36881833 refGene_gn=LRRC49;refGene_tx=XM_011282946.3;refGene_gn=LRRC49;refGene_tx=XM_023255098.1;refGene_gn=LRRC49;refGene_tx=XM_023255099.1;refGene_gn=LRRC49;refGene_tx=XM_023255100.1;refGene_gn=LRRC49;refGene_tx=XM_023255101.1
B3 36890306 36890510 refGene_gn=LRRC49;refGene_tx=XM_011282946.3;refGene_gn=LRRC49;refGene_tx=XM_023255098.1;refGene_gn=LRRC49;refGene_tx=XM_023255099.1;refGene_gn=LRRC49;refGene_tx=XM_023255100.1;refGene_gn=LRRC49;refGene_tx=XM_023255101.1
B3 36894171 36894274 refGene_gn=LRRC49;refGene_tx=XM_011282946.3;refGene_gn=LRRC49;refGene_tx=XM_023255098.1;refGene_gn=LRRC49;refGene_tx=XM_023255099.1;refGene_gn=LRRC49;refGene_tx=XM_023255100.1;refGene_gn=LRRC49;refGene_tx=XM_023255101.1
B3 36894626 36894824 refGene_gn=LRRC49;refGene_tx=XM_011282946.3;refGene_gn=LRRC49;refGene_tx=XM_023255099.1
B3 36896527 36896588 refGene_gn=LRRC49;refGene_tx=XM_023255101.1
B3 36899060 36899148 refGene_gn=LRRC49;refGene_tx=XM_011282946.3;refGene_gn=LRRC49;refGene_tx=XM_023255098.1
B3 36901320 36901377 refGene_gn=LRRC49;refGene_tx=XM_011282946.3;refGene_gn=LRRC49;refGene_tx=XM_023255098.1
B3 36902041 36902089 refGene_gn=LRRC49;refGene_tx=XM_011282946.3;refGene_gn=LRRC49;refGene_tx=XM_023255098.1
B3 36902651 36903080 refGene_gn=THAP10;refGene_tx=XM_023255102.1
B3 36910299 36910447 refGene_gn=THAP10;refGene_tx=XM_023255102.1
B3 36910555 36910752 refGene_gn=THAP10;refGene_tx=XM_023255102.1
B3 36936348 36936548 refGene_gn=LARP6;refGene_tx=XM_023255103.1
B3 36950210 36950421 refGene_gn=LARP6;refGene_tx=XM_023255103.1
B3 36955477 36956545 refGene_gn=LARP6;refGene_tx=XM_023255103.1
B3 37011477 37011561 refGene_gn=UACA;refGene_tx=XM_003986991.5
B3 37060025 37060064 refGene_gn=UACA;refGene_tx=XM_003986992.5
B3 37062241 37062375 refGene_gn=UACA;refGene_tx=XM_003986991.5;refGene_gn=UACA;refGene_tx=XM_003986992.5;refGene_gn=UACA;refGene_tx=XM_019832145.2
B3 37067181 37067270 refGene_gn=UACA;refGene_tx=XM_003986991.5;refGene_gn=UACA;refGene_tx=XM_003986992.5;refGene_gn=UACA;refGene_tx=XM_019832145.2
B3 37070206 37070271 refGene_gn=UACA;refGene_tx=XM_003986991.5;refGene_gn=UACA;refGene_tx=XM_003986992.5;refGene_gn=UACA;refGene_tx=XM_019832145.2
B3 37071039 37071097 refGene_gn=UACA;refGene_tx=XM_003986991.5;refGene_gn=UACA;refGene_tx=X

M_003986992.5;refGene_gn=UACA;refGene_tx=XM_019832145.2
B3 37075044 37075115
refGene_gn=UACA;refGene_tx=XM_003986991.5;refGene_gn=UACA;refGene_tx=XM_003986992.5;refGene_gn=UACA;refGene_tx=XM_019832145.2
B3 37075214 37075321
refGene_gn=UACA;refGene_tx=XM_003986991.5;refGene_gn=UACA;refGene_tx=XM_003986992.5;refGene_gn=UACA;refGene_tx=XM_019832145.2
B3 37078529 37078711
refGene_gn=UACA;refGene_tx=XM_003986991.5;refGene_gn=UACA;refGene_tx=XM_003986992.5;refGene_gn=UACA;refGene_tx=XM_019832145.2
B3 37080097 37080135
refGene_gn=UACA;refGene_tx=XM_003986991.5;refGene_gn=UACA;refGene_tx=XM_003986992.5;refGene_gn=UACA;refGene_tx=XM_019832145.2
B3 37083141 37083210
refGene_gn=UACA;refGene_tx=XM_003986991.5;refGene_gn=UACA;refGene_tx=XM_003986992.5;refGene_gn=UACA;refGene_tx=XM_019832145.2
B3 37086827 37086935
refGene_gn=UACA;refGene_tx=XM_003986991.5;refGene_gn=UACA;refGene_tx=XM_003986992.5;refGene_gn=UACA;refGene_tx=XM_019832145.2
B3 37087464 37087497
refGene_gn=UACA;refGene_tx=XM_003986991.5;refGene_gn=UACA;refGene_tx=XM_003986992.5;refGene_gn=UACA;refGene_tx=XM_019832145.2
B3 37088003 37088102
refGene_gn=UACA;refGene_tx=XM_003986991.5;refGene_gn=UACA;refGene_tx=XM_003986992.5;refGene_gn=UACA;refGene_tx=XM_019832145.2
B3 37093700 37093737
refGene_gn=UACA;refGene_tx=XM_003986991.5;refGene_gn=UACA;refGene_tx=XM_003986992.5;refGene_gn=UACA;refGene_tx=XM_019832145.2
B3 37094856 37094909
refGene_gn=UACA;refGene_tx=XM_003986991.5;refGene_gn=UACA;refGene_tx=XM_003986992.5;refGene_gn=UACA;refGene_tx=XM_019832145.2
B3 37096710 37099449
refGene_gn=UACA;refGene_tx=XM_003986991.5;refGene_gn=UACA;refGene_tx=XM_003986992.5;refGene_gn=UACA;refGene_tx=XM_019832145.2
B3 37101426 37101579
refGene_gn=UACA;refGene_tx=XM_003986991.5;refGene_gn=UACA;refGene_tx=XM_003986992.5;refGene_gn=UACA;refGene_tx=XM_019832145.2
B3 37103687 37103753
refGene_gn=UACA;refGene_tx=XM_003986991.5;refGene_gn=UACA;refGene_tx=XM_003986992.5;refGene_gn=UACA;refGene_tx=XM_019832145.2
B3 37106638 37106710
refGene_gn=UACA;refGene_tx=XM_003986991.5;refGene_gn=UACA;refGene_tx=XM_003986992.5;refGene_gn=UACA;refGene_tx=XM_019832145.2
B3 37581815 37581839
refGene_gn=TLE3;refGene_tx=XM_023255104.1;refGene_gn=TLE3;refGene_tx=XM_023255105.1;refGene_gn=TLE3;refGene_tx=XM_023255106.1;refGene_gn=TLE3;refGene_tx=XM_023255107.1;refGene_gn=TLE3;refGene_tx=XM_023255108.1;refGene_gn=TLE3;refGene_tx=XM_023255109.1;refGene_gn=TLE3;refGene_tx=XM_023255110.1;refGene_gn=TLE3;refGene_tx=XM_023255111.1;refGene_gn=TLE3;refGene_tx=XM_023255112.1;refGene_gn=TLE3;refGene_tx=XM_023255113.1;

refGene_gn=TLE3;refGene_tx=XM_023255114.1
B3 37582422 37582523
refGene_gn=TLE3;refGene_tx=XM_023255104.1;refGene_gn=TLE3;refGene_tx=XM_023255105.1;refGene_gn=TLE3;refGene_tx=XM_023255106.1;refGene_gn=TLE3;refGene_tx=XM_023255107.1;refGene_gn=TLE3;refGene_tx=XM_023255108.1;refGene_gn=TLE3;refGene_tx=XM_023255109.1;refGene_gn=TLE3;refGene_tx=XM_023255110.1;refGene_gn=TLE3;refGene_tx=XM_023255111.1;refGene_gn=TLE3;refGene_tx=XM_023255112.1;refGene_gn=TLE3;refGene_tx=XM_023255113.1;refGene_gn=TLE3;refGene_tx=XM_023255114.1
B3 37583049 37583113
refGene_gn=TLE3;refGene_tx=XM_023255104.1;refGene_gn=TLE3;refGene_tx=XM_023255105.1;refGene_gn=TLE3;refGene_tx=XM_023255106.1;refGene_gn=TLE3;refGene_tx=XM_023255107.1;refGene_gn=TLE3;refGene_tx=XM_023255108.1;refGene_gn=TLE3;refGene_tx=XM_023255109.1;refGene_gn=TLE3;refGene_tx=XM_023255110.1;refGene_gn=TLE3;refGene_tx=XM_023255111.1;refGene_gn=TLE3;refGene_tx=XM_023255112.1;refGene_gn=TLE3;refGene_tx=XM_023255113.1;refGene_gn=TLE3;refGene_tx=XM_023255114.1
B3 37584019 37584064
refGene_gn=TLE3;refGene_tx=XM_023255104.1;refGene_gn=TLE3;refGene_tx=XM_023255105.1;refGene_gn=TLE3;refGene_tx=XM_023255106.1;refGene_gn=TLE3;refGene_tx=XM_023255107.1;refGene_gn=TLE3;refGene_tx=XM_023255108.1;refGene_gn=TLE3;refGene_tx=XM_023255109.1;refGene_gn=TLE3;refGene_tx=XM_023255110.1;refGene_gn=TLE3;refGene_tx=XM_023255111.1;refGene_gn=TLE3;refGene_tx=XM_023255112.1;refGene_gn=TLE3;refGene_tx=XM_023255113.1;refGene_gn=TLE3;refGene_tx=XM_023255114.1
B3 37602528 37602591
refGene_gn=TLE3;refGene_tx=XM_023255104.1;refGene_gn=TLE3;refGene_tx=XM_023255105.1;refGene_gn=TLE3;refGene_tx=XM_023255106.1;refGene_gn=TLE3;refGene_tx=XM_023255107.1;refGene_gn=TLE3;refGene_tx=XM_023255108.1;refGene_gn=TLE3;refGene_tx=XM_023255109.1;refGene_gn=TLE3;refGene_tx=XM_023255110.1;refGene_gn=TLE3;refGene_tx=XM_023255111.1;refGene_gn=TLE3;refGene_tx=XM_023255112.1;refGene_gn=TLE3;refGene_tx=XM_023255113.1;refGene_gn=TLE3;refGene_tx=XM_023255114.1
B3 37604380 37604485
refGene_gn=TLE3;refGene_tx=XM_023255104.1;refGene_gn=TLE3;refGene_tx=XM_023255105.1;refGene_gn=TLE3;refGene_tx=XM_023255106.1;refGene_gn=TLE3;refGene_tx=XM_023255107.1;refGene_gn=TLE3;refGene_tx=XM_023255108.1;refGene_gn=TLE3;refGene_tx=XM_023255109.1;refGene_gn=TLE3;refGene_tx=XM_023255110.1;refGene_gn=TLE3;refGene_tx=XM_023255111.1;refGene_gn=TLE3;refGene_tx=XM_023255112.1;refGene_gn=TLE3;refGene_tx=XM_023255113.1;refGene_gn=TLE3;refGene_tx=XM_023255114.1
B3 37612726 37612931
refGene_gn=TLE3;refGene_tx=XM_023255104.1;refGene_gn=TLE3;refGene_tx=XM_023255105.1;refGene_gn=TLE3;refGene_tx=XM_023255106.1;refGene_gn=TLE3;refGene_tx=XM_023255107.1;refGene_gn=TLE3;refGene_tx=XM_023255108.1;refGene_gn=TLE3;refGene_tx=XM_023255109.1;refGene_gn=TLE3;refGene_tx=XM_023255110.1;refGene_gn=TLE3;refGene_tx=XM_023255111.1;refGene_gn=TLE3;refGene_tx=XM_023255112.1;refGene_gn=TLE3;refGene_tx=XM_023255113.1;refGene_gn=TLE3;refGene_tx=XM_023255114.1
B3 37614509 37614526

refGene_gn=TLE3;refGene_tx=XM_023255104.1;refGene_gn=TLE3;refGene_tx=XM_023255105.1;refGene_gn=TLE3;refGene_tx=XM_023255106.1;refGene_gn=TLE3;refGene_tx=XM_023255107.1;refGene_gn=TLE3;refGene_tx=XM_023255108.1;refGene_gn=TLE3;refGene_tx=XM_023255109.1;refGene_gn=TLE3;refGene_tx=XM_023255110.1;refGene_gn=TLE3;refGene_tx=XM_023255111.1;refGene_gn=TLE3;refGene_tx=XM_023255112.1;refGene_gn=TLE3;refGene_tx=XM_023255113.1;refGene_gn=TLE3;refGene_tx=XM_023255114.1
B3 37618163 37618283
refGene_gn=TLE3;refGene_tx=XM_023255104.1;refGene_gn=TLE3;refGene_tx=XM_023255105.1;refGene_gn=TLE3;refGene_tx=XM_023255106.1;refGene_gn=TLE3;refGene_tx=XM_023255107.1;refGene_gn=TLE3;refGene_tx=XM_023255108.1;refGene_gn=TLE3;refGene_tx=XM_023255109.1;refGene_gn=TLE3;refGene_tx=XM_023255110.1;refGene_gn=TLE3;refGene_tx=XM_023255111.1;refGene_gn=TLE3;refGene_tx=XM_023255112.1;refGene_gn=TLE3;refGene_tx=XM_023255113.1;refGene_gn=TLE3;refGene_tx=XM_023255114.1
B3 37619151 37619202
refGene_gn=TLE3;refGene_tx=XM_023255104.1;refGene_gn=TLE3;refGene_tx=XM_023255105.1;refGene_gn=TLE3;refGene_tx=XM_023255106.1;refGene_gn=TLE3;refGene_tx=XM_023255107.1;refGene_gn=TLE3;refGene_tx=XM_023255108.1;refGene_gn=TLE3;refGene_tx=XM_023255109.1;refGene_gn=TLE3;refGene_tx=XM_023255110.1;refGene_gn=TLE3;refGene_tx=XM_023255111.1;refGene_gn=TLE3;refGene_tx=XM_023255112.1;refGene_gn=TLE3;refGene_tx=XM_023255113.1;refGene_gn=TLE3;refGene_tx=XM_023255114.1
B3 37619864 37620017
refGene_gn=TLE3;refGene_tx=XM_023255104.1;refGene_gn=TLE3;refGene_tx=XM_023255105.1;refGene_gn=TLE3;refGene_tx=XM_023255106.1;refGene_gn=TLE3;refGene_tx=XM_023255107.1;refGene_gn=TLE3;refGene_tx=XM_023255108.1;refGene_gn=TLE3;refGene_tx=XM_023255109.1;refGene_gn=TLE3;refGene_tx=XM_023255110.1;refGene_gn=TLE3;refGene_tx=XM_023255111.1;refGene_gn=TLE3;refGene_tx=XM_023255112.1;refGene_gn=TLE3;refGene_tx=XM_023255113.1;refGene_gn=TLE3;refGene_tx=XM_023255114.1
B3 37620390 37620532
refGene_gn=TLE3;refGene_tx=XM_023255104.1;refGene_gn=TLE3;refGene_tx=XM_023255105.1;refGene_gn=TLE3;refGene_tx=XM_023255106.1;refGene_gn=TLE3;refGene_tx=XM_023255107.1;refGene_gn=TLE3;refGene_tx=XM_023255108.1;refGene_gn=TLE3;refGene_tx=XM_023255109.1;refGene_gn=TLE3;refGene_tx=XM_023255110.1;refGene_gn=TLE3;refGene_tx=XM_023255111.1;refGene_gn=TLE3;refGene_tx=XM_023255112.1;refGene_gn=TLE3;refGene_tx=XM_023255113.1;refGene_gn=TLE3;refGene_tx=XM_023255114.1
B3 37621078 37621278
refGene_gn=TLE3;refGene_tx=XM_023255104.1;refGene_gn=TLE3;refGene_tx=XM_023255105.1;refGene_gn=TLE3;refGene_tx=XM_023255106.1;refGene_gn=TLE3;refGene_tx=XM_023255107.1;refGene_gn=TLE3;refGene_tx=XM_023255108.1;refGene_gn=TLE3;refGene_tx=XM_023255109.1;refGene_gn=TLE3;refGene_tx=XM_023255110.1;refGene_gn=TLE3;refGene_tx=XM_023255111.1;refGene_gn=TLE3;refGene_tx=XM_023255112.1;refGene_gn=TLE3;refGene_tx=XM_023255113.1;refGene_gn=TLE3;refGene_tx=XM_023255114.1
B3 37621990 37622067
refGene_gn=TLE3;refGene_tx=XM_023255104.1;refGene_gn=TLE3;refGene_tx=XM_023255105.1;refGene_gn=TLE3;refGene_tx=XM_023255106.1;refGene_gn=TLE

3;refGene_tx=XM_023255107.1;refGene_gn=TLE3;refGene_tx=XM_023255108.1;
refGene_gn=TLE3;refGene_tx=XM_023255109.1;refGene_gn=TLE3;refGene_tx=X
M_023255110.1;refGene_gn=TLE3;refGene_tx=XM_023255111.1;refGene_gn=TLE
3;refGene_tx=XM_023255112.1;refGene_gn=TLE3;refGene_tx=XM_023255113.1;
refGene_gn=TLE3;refGene_tx=XM_023255114.1
B3 37622670 37622920
refGene_gn=TLE3;refGene_tx=XM_023255104.1;refGene_gn=TLE3;refGene_tx=X
M_023255105.1;refGene_gn=TLE3;refGene_tx=XM_023255106.1;refGene_gn=TLE
3;refGene_tx=XM_023255107.1;refGene_gn=TLE3;refGene_tx=XM_023255108.1;
refGene_gn=TLE3;refGene_tx=XM_023255109.1;refGene_gn=TLE3;refGene_tx=X
M_023255110.1;refGene_gn=TLE3;refGene_tx=XM_023255111.1;refGene_gn=TLE
3;refGene_tx=XM_023255112.1;refGene_gn=TLE3;refGene_tx=XM_023255113.1;
refGene_gn=TLE3;refGene_tx=XM_023255114.1
B3 37623226 37623474
refGene_gn=TLE3;refGene_tx=XM_023255104.1;refGene_gn=TLE3;refGene_tx=X
M_023255105.1;refGene_gn=TLE3;refGene_tx=XM_023255106.1;refGene_gn=TLE
3;refGene_tx=XM_023255107.1;refGene_gn=TLE3;refGene_tx=XM_023255108.1;
refGene_gn=TLE3;refGene_tx=XM_023255109.1;refGene_gn=TLE3;refGene_tx=X
M_023255110.1;refGene_gn=TLE3;refGene_tx=XM_023255111.1;refGene_gn=TLE
3;refGene_tx=XM_023255112.1;refGene_gn=TLE3;refGene_tx=XM_023255113.1;
refGene_gn=TLE3;refGene_tx=XM_023255114.1
B3 37624281 37624429
refGene_gn=TLE3;refGene_tx=XM_023255104.1;refGene_gn=TLE3;refGene_tx=X
M_023255105.1;refGene_gn=TLE3;refGene_tx=XM_023255106.1;refGene_gn=TLE
3;refGene_tx=XM_023255107.1;refGene_gn=TLE3;refGene_tx=XM_023255108.1;
refGene_gn=TLE3;refGene_tx=XM_023255109.1;refGene_gn=TLE3;refGene_tx=X
M_023255110.1;refGene_gn=TLE3;refGene_tx=XM_023255111.1;refGene_gn=TLE
3;refGene_tx=XM_023255112.1;refGene_gn=TLE3;refGene_tx=XM_023255113.1;
refGene_gn=TLE3;refGene_tx=XM_023255114.1
B3 37625425 37625576
refGene_gn=TLE3;refGene_tx=XM_023255104.1;refGene_gn=TLE3;refGene_tx=X
M_023255105.1;refGene_gn=TLE3;refGene_tx=XM_023255106.1;refGene_gn=TLE
3;refGene_tx=XM_023255107.1;refGene_gn=TLE3;refGene_tx=XM_023255108.1;
refGene_gn=TLE3;refGene_tx=XM_023255109.1;refGene_gn=TLE3;refGene_tx=X
M_023255110.1;refGene_gn=TLE3;refGene_tx=XM_023255111.1;refGene_gn=TLE
3;refGene_tx=XM_023255112.1;refGene_gn=TLE3;refGene_tx=XM_023255113.1;
refGene_gn=TLE3;refGene_tx=XM_023255114.1
B3 37626441 37626518
refGene_gn=TLE3;refGene_tx=XM_023255104.1;refGene_gn=TLE3;refGene_tx=X
M_023255105.1;refGene_gn=TLE3;refGene_tx=XM_023255106.1;refGene_gn=TLE
3;refGene_tx=XM_023255107.1;refGene_gn=TLE3;refGene_tx=XM_023255108.1;
refGene_gn=TLE3;refGene_tx=XM_023255109.1;refGene_gn=TLE3;refGene_tx=X
M_023255110.1;refGene_gn=TLE3;refGene_tx=XM_023255111.1;refGene_gn=TLE
3;refGene_tx=XM_023255112.1;refGene_gn=TLE3;refGene_tx=XM_023255113.1;
refGene_gn=TLE3;refGene_tx=XM_023255114.1
B3 37627145 37627271
refGene_gn=TLE3;refGene_tx=XM_023255104.1;refGene_gn=TLE3;refGene_tx=X
M_023255105.1;refGene_gn=TLE3;refGene_tx=XM_023255106.1;refGene_gn=TLE
3;refGene_tx=XM_023255108.1
B3 37627621 37627729

refGene_gn=TLE3;refGene_tx=XM_023255107.1;refGene_gn=TLE3;refGene_tx=XM_023255109.1;refGene_gn=TLE3;refGene_tx=XM_023255110.1;refGene_gn=TLE3;refGene_tx=XM_023255111.1;refGene_gn=TLE3;refGene_tx=XM_023255112.1;refGene_gn=TLE3;refGene_tx=XM_023255113.1;refGene_gn=TLE3;refGene_tx=XM_023255114.1
B3 38150017 38150097 refGene_gn=RPLP1;refGene_tx=XM_003986994.4
B3 38150557 38150675 refGene_gn=RPLP1;refGene_tx=XM_003986994.4
B3 38151711 38151786 refGene_gn=RPLP1;refGene_tx=XM_003986994.4
B3 38152473 38152545 refGene_gn=RPLP1;refGene_tx=XM_003986994.4
B3 38178415 38178431 refGene_gn=KIF23;refGene_tx=XM_023255116.1;refGene_gn=KIF23;refGene_tx=XM_023255117.1;refGene_gn=KIF23;refGene_tx=XM_023255118.1
B3 38179167 38179238 refGene_gn=KIF23;refGene_tx=XM_023255116.1;refGene_gn=KIF23;refGene_tx=XM_023255117.1;refGene_gn=KIF23;refGene_tx=XM_023255118.1
B3 38179855 38179940 refGene_gn=KIF23;refGene_tx=XM_023255116.1;refGene_gn=KIF23;refGene_tx=XM_023255117.1;refGene_gn=KIF23;refGene_tx=XM_023255118.1
B3 38180120 38180203 refGene_gn=KIF23;refGene_tx=XM_023255116.1;refGene_gn=KIF23;refGene_tx=XM_023255117.1;refGene_gn=KIF23;refGene_tx=XM_023255118.1
B3 38181089 38181320 refGene_gn=KIF23;refGene_tx=XM_023255116.1;refGene_gn=KIF23;refGene_tx=XM_023255117.1
B3 38184378 38184690 refGene_gn=KIF23;refGene_tx=XM_023255116.1;refGene_gn=KIF23;refGene_tx=XM_023255117.1;refGene_gn=KIF23;refGene_tx=XM_023255118.1
B3 38184981 38185161 refGene_gn=KIF23;refGene_tx=XM_023255116.1;refGene_gn=KIF23;refGene_tx=XM_023255117.1;refGene_gn=KIF23;refGene_tx=XM_023255118.1
B3 38185399 38185573 refGene_gn=KIF23;refGene_tx=XM_023255116.1;refGene_gn=KIF23;refGene_tx=XM_023255117.1;refGene_gn=KIF23;refGene_tx=XM_023255118.1
B3 38186228 38186386 refGene_gn=KIF23;refGene_tx=XM_023255116.1;refGene_gn=KIF23;refGene_tx=XM_023255117.1;refGene_gn=KIF23;refGene_tx=XM_023255118.1
B3 38187686 38187845 refGene_gn=KIF23;refGene_tx=XM_023255116.1;refGene_gn=KIF23;refGene_tx=XM_023255117.1;refGene_gn=KIF23;refGene_tx=XM_023255118.1
B3 38188182 38188309 refGene_gn=KIF23;refGene_tx=XM_023255116.1;refGene_gn=KIF23;refGene_tx=XM_023255117.1;refGene_gn=KIF23;refGene_tx=XM_023255118.1
B3 38188535 38188655 refGene_gn=KIF23;refGene_tx=XM_023255116.1;refGene_gn=KIF23;refGene_tx=XM_023255117.1;refGene_gn=KIF23;refGene_tx=XM_023255118.1
B3 38188740 38188820 refGene_gn=KIF23;refGene_tx=XM_023255116.1;refGene_gn=KIF23;refGene_tx=XM_023255117.1;refGene_gn=KIF23;refGene_tx=XM_023255118.1
B3 38192717 38192820 refGene_gn=KIF23;refGene_tx=XM_023255116.1;refGene_gn=KIF23;refGene_tx

=XM_023255117.1;refGene_gn=KIF23;refGene_tx=XM_023255118.1
B3 38194083 38194205
refGene_gn=KIF23;refGene_tx=XM_023255116.1;refGene_gn=KIF23;refGene_tx=XM_023255117.1;refGene_gn=KIF23;refGene_tx=XM_023255118.1
B3 38194347 38194460
refGene_gn=KIF23;refGene_tx=XM_023255116.1;refGene_gn=KIF23;refGene_tx=XM_023255117.1;refGene_gn=KIF23;refGene_tx=XM_023255118.1
B3 38196479 38196650
refGene_gn=KIF23;refGene_tx=XM_023255116.1;refGene_gn=KIF23;refGene_tx=XM_023255117.1;refGene_gn=KIF23;refGene_tx=XM_023255118.1
B3 38197084 38197194
refGene_gn=KIF23;refGene_tx=XM_023255116.1;refGene_gn=KIF23;refGene_tx=XM_023255117.1;refGene_gn=KIF23;refGene_tx=XM_023255118.1
B3 38197382 38197519
refGene_gn=KIF23;refGene_tx=XM_023255116.1;refGene_gn=KIF23;refGene_tx=XM_023255117.1;refGene_gn=KIF23;refGene_tx=XM_023255118.1
B3 38197769 38197875
refGene_gn=KIF23;refGene_tx=XM_023255116.1;refGene_gn=KIF23;refGene_tx=XM_023255117.1;refGene_gn=KIF23;refGene_tx=XM_023255118.1
B3 38199684 38199769
refGene_gn=LOC105260622;refGene_tx=XM_023255125.1;refGene_gn=LOC105260622;refGene_tx=XM_023255126.1
B3 38201856 38201985
refGene_gn=KIF23;refGene_tx=XM_023255116.1;refGene_gn=KIF23;refGene_tx=XM_023255117.1;refGene_gn=KIF23;refGene_tx=XM_023255118.1
B3 38203006 38203076
refGene_gn=KIF23;refGene_tx=XM_023255116.1;refGene_gn=KIF23;refGene_tx=XM_023255117.1;refGene_gn=KIF23;refGene_tx=XM_023255118.1
B3 38204112 38204203
refGene_gn=LOC105260622;refGene_tx=XM_023255125.1;refGene_gn=LOC105260622;refGene_tx=XM_023255126.1
B3 38204805 38204816
refGene_gn=KIF23;refGene_tx=XM_023255116.1;refGene_gn=KIF23;refGene_tx=XM_023255117.1;refGene_gn=KIF23;refGene_tx=XM_023255118.1
B3 38204821 38204976
refGene_gn=LOC105260622;refGene_tx=XM_023255125.1;refGene_gn=LOC105260622;refGene_tx=XM_023255126.1
B3 38205088 38205459
refGene_gn=LOC105260622;refGene_tx=XM_023255125.1;refGene_gn=LOC105260622;refGene_tx=XM_023255126.1
B3 38210734 38210878
refGene_gn=LOC105260622;refGene_tx=XM_023255126.1
B3 38213677 38213919
refGene_gn=LOC105260622;refGene_tx=XM_023255126.1;refGene_gn=PAQR5;refGene_tx=XM_023255119.1;refGene_gn=PAQR5;refGene_tx=XM_023255120.1;refGene_gn=PAQR5;refGene_tx=XM_023255121.1;refGene_gn=PAQR5;refGene_tx=XM_023255122.1;refGene_gn=PAQR5;refGene_tx=XM_023255123.1;refGene_gn=PAQR5;refGene_tx=XM_023255124.1
B3 38217193 38217335
refGene_gn=PAQR5;refGene_tx=XM_023255119.1;refGene_gn=PAQR5;refGene_tx

=XM_023255120.1;refGene_gn=PAQR5;refGene_tx=XM_023255121.1;refGene_gn=PAQR5;refGene_tx=XM_023255122.1;refGene_gn=PAQR5;refGene_tx=XM_023255123.1;refGene_gn=PAQR5;refGene_tx=XM_023255124.1
B3 38219638 38219735 refGene_gn=PAQR5;refGene_tx=XM_023255119.1;refGene_gn=PAQR5;refGene_tx=XM_023255120.1;refGene_gn=PAQR5;refGene_tx=XM_023255121.1;refGene_gn=PAQR5;refGene_tx=XM_023255122.1;refGene_gn=PAQR5;refGene_tx=XM_023255123.1;refGene_gn=PAQR5;refGene_tx=XM_023255124.1
B3 38227048 38227175 refGene_gn=PAQR5;refGene_tx=XM_023255119.1;refGene_gn=PAQR5;refGene_tx=XM_023255120.1;refGene_gn=PAQR5;refGene_tx=XM_023255121.1;refGene_gn=PAQR5;refGene_tx=XM_023255122.1;refGene_gn=PAQR5;refGene_tx=XM_023255123.1;refGene_gn=PAQR5;refGene_tx=XM_023255124.1
B3 38231745 38231951 refGene_gn=PAQR5;refGene_tx=XM_023255119.1;refGene_gn=PAQR5;refGene_tx=XM_023255120.1;refGene_gn=PAQR5;refGene_tx=XM_023255121.1;refGene_gn=PAQR5;refGene_tx=XM_023255122.1;refGene_gn=PAQR5;refGene_tx=XM_023255123.1;refGene_gn=PAQR5;refGene_tx=XM_023255124.1
B3 38234147 38234275 refGene_gn=PAQR5;refGene_tx=XM_023255119.1;refGene_gn=PAQR5;refGene_tx=XM_023255120.1;refGene_gn=PAQR5;refGene_tx=XM_023255121.1;refGene_gn=PAQR5;refGene_tx=XM_023255122.1;refGene_gn=PAQR5;refGene_tx=XM_023255123.1;refGene_gn=PAQR5;refGene_tx=XM_023255124.1
B3 38234582 38234591 refGene_gn=PAQR5;refGene_tx=XM_023255124.1
B3 38252247 38252392 refGene_gn=PAQR5;refGene_tx=XM_023255119.1;refGene_gn=PAQR5;refGene_tx=XM_023255120.1;refGene_gn=PAQR5;refGene_tx=XM_023255121.1;refGene_gn=PAQR5;refGene_tx=XM_023255122.1;refGene_gn=PAQR5;refGene_tx=XM_023255123.1
B3 38255679 38255735 refGene_gn=PAQR5;refGene_tx=XM_023255119.1;refGene_gn=PAQR5;refGene_tx=XM_023255120.1;refGene_gn=PAQR5;refGene_tx=XM_023255121.1
B3 38316538 38317563 refGene_gn=GLCE;refGene_tx=XM_023255127.1;refGene_gn=GLCE;refGene_tx=XM_023255128.1;refGene_gn=GLCE;refGene_tx=XM_023255129.1
B3 38325426 38325672 refGene_gn=GLCE;refGene_tx=XM_023255127.1;refGene_gn=GLCE;refGene_tx=XM_023255128.1;refGene_gn=GLCE;refGene_tx=XM_023255129.1
B3 38329451 38330034 refGene_gn=GLCE;refGene_tx=XM_023255127.1;refGene_gn=GLCE;refGene_tx=XM_023255128.1;refGene_gn=GLCE;refGene_tx=XM_023255129.1
B3 38545585 38545717 refGene_gn=NOX5;refGene_tx=XM_019832157.2
B3 38546701 38546868 refGene_gn=NOX5;refGene_tx=XM_019832157.2
B3 38550484 38550584 refGene_gn=NOX5;refGene_tx=XM_019832157.2
B3 38551257 38551342 refGene_gn=NOX5;refGene_tx=XM_019832157.2
B3 38551617 38551742 refGene_gn=NOX5;refGene_tx=XM_019832157.2
B3 38552142 38552187 refGene_gn=NOX5;refGene_tx=XM_019832157.2
B3 38554288 38554480 refGene_gn=NOX5;refGene_tx=XM_019832157.2
B3 38556519 38556656 refGene_gn=NOX5;refGene_tx=XM_019832157.2
B3 38559867 38560000 refGene_gn=NOX5;refGene_tx=XM_019832157.2

B3 38561486 38561669 refGene_gn=NOX5;refGene_tx=XM_019832157.2
B3 38562593 38562754 refGene_gn=NOX5;refGene_tx=XM_019832157.2
B3 38562756 38562771 refGene_gn=NOX5;refGene_tx=XM_019832157.2
B3 38562955 38563109 refGene_gn=NOX5;refGene_tx=XM_019832157.2
B3 38564842 38565077 refGene_gn=NOX5;refGene_tx=XM_019832157.2
B3 38566716 38566927 refGene_gn=NOX5;refGene_tx=XM_019832157.2
B3 38571486 38571637 refGene_gn=NOX5;refGene_tx=XM_019832157.2
B3 38573290 38573414 refGene_gn=NOX5;refGene_tx=XM_019832157.2
B3 38578782 38578832 refGene_gn=NOX5;refGene_tx=XM_019832157.2
B3 38643439 38644458 refGene_gn=SPESP1;refGene_tx=XM_006932437.4
B3 38675834 38675895 refGene_gn=SPESP1;refGene_tx=XM_006932437.4
B3 38792663 38792717 refGene_gn=ANP32A;refGene_tx=XM_003987005.5
B3 38793697 38793835 refGene_gn=ANP32A;refGene_tx=XM_006932438.4
B3 38796604 38796613 refGene_gn=ANP32A;refGene_tx=XM_019832162.2
B3 38821839 38821848 refGene_gn=ANP32A;refGene_tx=XM_023255131.1
B3 38824180 38824330
refGene_gn=ANP32A;refGene_tx=XM_003987005.5;refGene_gn=ANP32A;refGene_tx=XM_006932438.4;refGene_gn=ANP32A;refGene_tx=XM_019832162.2;refGene_gn=ANP32A;refGene_tx=XM_023255131.1
B3 38824539 38824662
refGene_gn=ANP32A;refGene_tx=XM_003987005.5;refGene_gn=ANP32A;refGene_tx=XM_006932438.4;refGene_gn=ANP32A;refGene_tx=XM_019832162.2;refGene_gn=ANP32A;refGene_tx=XM_023255131.1
B3 38826739 38826938
refGene_gn=ANP32A;refGene_tx=XM_003987005.5;refGene_gn=ANP32A;refGene_tx=XM_006932438.4;refGene_gn=ANP32A;refGene_tx=XM_019832162.2;refGene_gn=ANP32A;refGene_tx=XM_023255131.1
B3 38828146 38828244
refGene_gn=ANP32A;refGene_tx=XM_003987005.5;refGene_gn=ANP32A;refGene_tx=XM_006932438.4;refGene_gn=ANP32A;refGene_tx=XM_019832162.2;refGene_gn=ANP32A;refGene_tx=XM_023255131.1
B3 38830805 38830869
refGene_gn=ANP32A;refGene_tx=XM_003987005.5;refGene_gn=ANP32A;refGene_tx=XM_006932438.4;refGene_gn=ANP32A;refGene_tx=XM_019832162.2;refGene_gn=ANP32A;refGene_tx=XM_023255131.1
B3 38831127 38831189
refGene_gn=ANP32A;refGene_tx=XM_003987005.5;refGene_gn=ANP32A;refGene_tx=XM_006932438.4;refGene_gn=ANP32A;refGene_tx=XM_019832162.2;refGene_gn=ANP32A;refGene_tx=XM_023255131.1
B3 38873658 38873790
refGene_gn=CORO2B;refGene_tx=XM_023255132.1;refGene_gn=CORO2B;refGene_tx=XM_023255133.1;refGene_gn=CORO2B;refGene_tx=XM_023255134.1
B3 38877795 38877935
refGene_gn=CORO2B;refGene_tx=XM_023255132.1;refGene_gn=CORO2B;refGene_tx=XM_023255133.1;refGene_gn=CORO2B;refGene_tx=XM_023255134.1
B3 38878137 38878228
refGene_gn=CORO2B;refGene_tx=XM_023255132.1;refGene_gn=CORO2B;refGene_tx=XM_023255133.1;refGene_gn=CORO2B;refGene_tx=XM_023255134.1
B3 38878567 38878680
refGene_gn=CORO2B;refGene_tx=XM_023255132.1;refGene_gn=CORO2B;refGene_

tx=XM_023255133.1;refGene_gn=CORO2B;refGene_tx=XM_023255134.1
B3 38882229 38882326
refGene_gn=CORO2B;refGene_tx=XM_023255132.1;refGene_gn=CORO2B;refGene_
tx=XM_023255133.1;refGene_gn=CORO2B;refGene_tx=XM_023255134.1
B3 38882866 38882971
refGene_gn=CORO2B;refGene_tx=XM_023255132.1;refGene_gn=CORO2B;refGene_
tx=XM_023255133.1;refGene_gn=CORO2B;refGene_tx=XM_023255134.1
B3 38883096 38883213
refGene_gn=CORO2B;refGene_tx=XM_023255132.1;refGene_gn=CORO2B;refGene_
tx=XM_023255133.1;refGene_gn=CORO2B;refGene_tx=XM_023255134.1
B3 38885440 38885605
refGene_gn=CORO2B;refGene_tx=XM_023255132.1;refGene_gn=CORO2B;refGene_
tx=XM_023255133.1;refGene_gn=CORO2B;refGene_tx=XM_023255134.1
B3 38886244 38886394
refGene_gn=CORO2B;refGene_tx=XM_023255132.1;refGene_gn=CORO2B;refGene_
tx=XM_023255133.1;refGene_gn=CORO2B;refGene_tx=XM_023255134.1
B3 38895911 38896028
refGene_gn=CORO2B;refGene_tx=XM_023255132.1;refGene_gn=CORO2B;refGene_
tx=XM_023255133.1;refGene_gn=CORO2B;refGene_tx=XM_023255134.1
B3 38918027 38918205 refGene_gn=CORO2B;refGene_tx=XM_023255132.1
B3 38936891 38937092
refGene_gn=CORO2B;refGene_tx=XM_023255132.1;refGene_gn=CORO2B;refGene_
tx=XM_023255133.1;refGene_gn=CORO2B;refGene_tx=XM_023255134.1
B3 39006575 39006590 refGene_gn=CORO2B;refGene_tx=XM_023255133.1
B3 39130595 39130647 refGene_gn=ITGA11;refGene_tx=XM_023255135.1
B3 39165295 39165407 refGene_gn=ITGA11;refGene_tx=XM_023255135.1
B3 39194695 39194796 refGene_gn=ITGA11;refGene_tx=XM_023255135.1
B3 39198964 39199056 refGene_gn=ITGA11;refGene_tx=XM_023255135.1
B3 39201988 39202103 refGene_gn=ITGA11;refGene_tx=XM_023255135.1
B3 39205020 39205148 refGene_gn=ITGA11;refGene_tx=XM_023255135.1
B3 39206163 39206312 refGene_gn=ITGA11;refGene_tx=XM_023255135.1
B3 39210745 39210890 refGene_gn=ITGA11;refGene_tx=XM_023255135.1
B3 39211360 39211526 refGene_gn=ITGA11;refGene_tx=XM_023255135.1
B3 39212914 39212985 refGene_gn=ITGA11;refGene_tx=XM_023255135.1
B3 39221563 39221708 refGene_gn=ITGA11;refGene_tx=XM_023255135.1
B3 39224394 39224543 refGene_gn=ITGA11;refGene_tx=XM_023255135.1
B3 39228575 39228716 refGene_gn=ITGA11;refGene_tx=XM_023255135.1
B3 39228953 39229157 refGene_gn=ITGA11;refGene_tx=XM_023255135.1
B3 39229899 39230030 refGene_gn=ITGA11;refGene_tx=XM_023255135.1
B3 39233934 39234101 refGene_gn=ITGA11;refGene_tx=XM_023255135.1
B3 39235310 39235453 refGene_gn=ITGA11;refGene_tx=XM_023255135.1
B3 39236831 39236942 refGene_gn=ITGA11;refGene_tx=XM_023255135.1
B3 39240396 39240482 refGene_gn=ITGA11;refGene_tx=XM_023255135.1
B3 39242223 39242431 refGene_gn=ITGA11;refGene_tx=XM_023255135.1
B3 39244567 39244666 refGene_gn=ITGA11;refGene_tx=XM_023255135.1
B3 39246043 39246120 refGene_gn=ITGA11;refGene_tx=XM_023255135.1
B3 39247008 39247098 refGene_gn=ITGA11;refGene_tx=XM_023255135.1
B3 39248000 39248091 refGene_gn=ITGA11;refGene_tx=XM_023255135.1
B3 39249501 39249615 refGene_gn=ITGA11;refGene_tx=XM_023255135.1
B3 39249824 39249911 refGene_gn=ITGA11;refGene_tx=XM_023255135.1

B3 39251285 39251396 refGene_gn=ITGA11;refGene_tx=XM_023255135.1
B3 39251769 39251865 refGene_gn=ITGA11;refGene_tx=XM_023255135.1
B3 39255671 39255785 refGene_gn=ITGA11;refGene_tx=XM_023255135.1
B3 39256509 39256581 refGene_gn=ITGA11;refGene_tx=XM_023255135.1
B3 39269599 39271235 refGene_gn=FEM1B;refGene_tx=XM_011282943.3
B3 39280965 39281213 refGene_gn=FEM1B;refGene_tx=XM_011282943.3
B3 39294628 39296007
refGene_gn=LOC111560804;refGene_tx=XM_023255136.1
B3 39296008 39296903
refGene_gn=LOC111560804;refGene_tx=XM_023255136.1
B3 39320782 39320868 refGene_gn=CLN6;refGene_tx=XM_003987007.5
B3 39327149 39327264 refGene_gn=CLN6;refGene_tx=XM_003987007.5
B3 39331728 39331827 refGene_gn=CLN6;refGene_tx=XM_003987007.5
B3 39332876 39333065 refGene_gn=CLN6;refGene_tx=XM_003987007.5
B3 39333354 39333410 refGene_gn=CLN6;refGene_tx=XM_003987007.5
B3 39334207 39334330 refGene_gn=CLN6;refGene_tx=XM_003987007.5
B3 39335935 39336206 refGene_gn=CLN6;refGene_tx=XM_003987007.5
B3 39338764 39338767 refGene_gn=CALML4;refGene_tx=XM_023255142.1
B3 39340378 39340381 refGene_gn=CALML4;refGene_tx=XM_023255141.1
B3 39340493 39340524
refGene_gn=CALML4;refGene_tx=XM_023255141.1;refGene_gn=CALML4;refGene_tx=XM_023255142.1
B3 39344883 39345024
refGene_gn=CALML4;refGene_tx=XM_023255141.1;refGene_gn=CALML4;refGene_tx=XM_023255142.1
B3 39346185 39346490
refGene_gn=CALML4;refGene_tx=XM_023255141.1;refGene_gn=CALML4;refGene_tx=XM_023255142.1
B3 39347170 39347318
refGene_gn=LOC111556187;refGene_tx=XM_023255143.1
B3 39350079 39350177
refGene_gn=LOC111556187;refGene_tx=XM_023255143.1
B3 39356652 39356946
refGene_gn=PIAS1;refGene_tx=XM_003987009.4;refGene_gn=PIAS1;refGene_tx=XM_019832168.2;refGene_gn=PIAS1;refGene_tx=XM_019832169.2;refGene_gn=PIAS1;refGene_tx=XM_023255137.1;refGene_gn=PIAS1;refGene_tx=XM_023255138.1;refGene_gn=PIAS1;refGene_tx=XM_023255139.1;refGene_gn=PIAS1;refGene_tx=XM_023255140.1
B3 39357567 39357605
refGene_gn=PIAS1;refGene_tx=XM_003987009.4;refGene_gn=PIAS1;refGene_tx=XM_019832168.2;refGene_gn=PIAS1;refGene_tx=XM_019832169.2;refGene_gn=PIAS1;refGene_tx=XM_023255137.1;refGene_gn=PIAS1;refGene_tx=XM_023255138.1;refGene_gn=PIAS1;refGene_tx=XM_023255139.1
B3 39359452 39359595
refGene_gn=PIAS1;refGene_tx=XM_003987009.4;refGene_gn=PIAS1;refGene_tx=XM_019832168.2;refGene_gn=PIAS1;refGene_tx=XM_019832169.2;refGene_gn=PIAS1;refGene_tx=XM_023255137.1;refGene_gn=PIAS1;refGene_tx=XM_023255138.1;refGene_gn=PIAS1;refGene_tx=XM_023255139.1;refGene_gn=PIAS1;refGene_tx=XM_023255140.1
B3 39363385 39363566

refGene_gn=PIAS1;refGene_tx=XM_003987009.4;refGene_gn=PIAS1;refGene_tx=XM_019832168.2;refGene_gn=PIAS1;refGene_tx=XM_019832169.2;refGene_gn=PIAS1;refGene_tx=XM_023255137.1;refGene_gn=PIAS1;refGene_tx=XM_023255138.1;refGene_gn=PIAS1;refGene_tx=XM_023255139.1;refGene_gn=PIAS1;refGene_tx=XM_023255140.1
B3 39364670 39364801
refGene_gn=PIAS1;refGene_tx=XM_003987009.4;refGene_gn=PIAS1;refGene_tx=XM_019832168.2;refGene_gn=PIAS1;refGene_tx=XM_019832169.2;refGene_gn=PIAS1;refGene_tx=XM_023255137.1;refGene_gn=PIAS1;refGene_tx=XM_023255138.1;refGene_gn=PIAS1;refGene_tx=XM_023255139.1;refGene_gn=PIAS1;refGene_tx=XM_023255140.1
B3 39366578 39366739
refGene_gn=PIAS1;refGene_tx=XM_003987009.4;refGene_gn=PIAS1;refGene_tx=XM_019832168.2;refGene_gn=PIAS1;refGene_tx=XM_019832169.2;refGene_gn=PIAS1;refGene_tx=XM_023255137.1;refGene_gn=PIAS1;refGene_tx=XM_023255138.1;refGene_gn=PIAS1;refGene_tx=XM_023255139.1;refGene_gn=PIAS1;refGene_tx=XM_023255140.1
B3 39376945 39377019
refGene_gn=PIAS1;refGene_tx=XM_003987009.4;refGene_gn=PIAS1;refGene_tx=XM_019832168.2;refGene_gn=PIAS1;refGene_tx=XM_019832169.2;refGene_gn=PIAS1;refGene_tx=XM_023255137.1;refGene_gn=PIAS1;refGene_tx=XM_023255138.1;refGene_gn=PIAS1;refGene_tx=XM_023255139.1;refGene_gn=PIAS1;refGene_tx=XM_023255140.1
B3 39387823 39387929
refGene_gn=PIAS1;refGene_tx=XM_003987009.4;refGene_gn=PIAS1;refGene_tx=XM_019832168.2;refGene_gn=PIAS1;refGene_tx=XM_019832169.2;refGene_gn=PIAS1;refGene_tx=XM_023255137.1;refGene_gn=PIAS1;refGene_tx=XM_023255138.1;refGene_gn=PIAS1;refGene_tx=XM_023255139.1;refGene_gn=PIAS1;refGene_tx=XM_023255140.1
B3 39394069 39394204
refGene_gn=PIAS1;refGene_tx=XM_003987009.4;refGene_gn=PIAS1;refGene_tx=XM_019832168.2;refGene_gn=PIAS1;refGene_tx=XM_019832169.2;refGene_gn=PIAS1;refGene_tx=XM_023255137.1;refGene_gn=PIAS1;refGene_tx=XM_023255138.1;refGene_gn=PIAS1;refGene_tx=XM_023255139.1;refGene_gn=PIAS1;refGene_tx=XM_023255140.1
B3 39394831 39394922
refGene_gn=PIAS1;refGene_tx=XM_003987009.4;refGene_gn=PIAS1;refGene_tx=XM_019832168.2;refGene_gn=PIAS1;refGene_tx=XM_019832169.2;refGene_gn=PIAS1;refGene_tx=XM_023255137.1;refGene_gn=PIAS1;refGene_tx=XM_023255138.1;refGene_gn=PIAS1;refGene_tx=XM_023255139.1;refGene_gn=PIAS1;refGene_tx=XM_023255140.1
B3 39398594 39398642
refGene_gn=PIAS1;refGene_tx=XM_003987009.4;refGene_gn=PIAS1;refGene_tx=XM_019832168.2;refGene_gn=PIAS1;refGene_tx=XM_019832169.2;refGene_gn=PIAS1;refGene_tx=XM_023255137.1;refGene_gn=PIAS1;refGene_tx=XM_023255138.1;refGene_gn=PIAS1;refGene_tx=XM_023255139.1;refGene_gn=PIAS1;refGene_tx=XM_023255140.1
B3 39398903 39398988
refGene_gn=PIAS1;refGene_tx=XM_003987009.4;refGene_gn=PIAS1;refGene_tx=XM_019832168.2;refGene_gn=PIAS1;refGene_tx=XM_019832169.2;refGene_gn=

PIAS1;refGene_tx=XM_023255137.1;refGene_gn=PIAS1;refGene_tx=XM_0232551
38.1;refGene_gn=PIAS1;refGene_tx=XM_023255139.1;refGene_gn=PIAS1;refGe
ne_tx=XM_023255140.1
B3 39445707 39446152
refGene_gn=PIAS1;refGene_tx=XM_003987009.4;refGene_gn=PIAS1;refGene_tx
=XM_019832168.2;refGene_gn=PIAS1;refGene_tx=XM_019832169.2;refGene_gn=
PIAS1;refGene_tx=XM_023255137.1;refGene_gn=PIAS1;refGene_tx=XM_0232551
38.1;refGene_gn=PIAS1;refGene_tx=XM_023255139.1;refGene_gn=PIAS1;refGe
ne_tx=XM_023255140.1
B3 39478464 39478467 refGene_gn=PIAS1;refGene_tx=XM_019832169.2
B3 39478471 39478477 refGene_gn=PIAS1;refGene_tx=XM_019832168.2
B3 39479378 39479402 refGene_gn=PIAS1;refGene_tx=XM_023255137.1
B3 39479596 39479599 refGene_gn=PIAS1;refGene_tx=XM_023255139.1
B3 39483376 39483433
refGene_gn=PIAS1;refGene_tx=XM_003987009.4;refGene_gn=PIAS1;refGene_tx
=XM_023255138.1;refGene_gn=PIAS1;refGene_tx=XM_023255140.1
B3 39642320 39642415 refGene_gn=SKOR1;refGene_tx=XM_023256163.1
B3 39642944 39643010 refGene_gn=SKOR1;refGene_tx=XM_023256163.1
B3 39643523 39643598 refGene_gn=SKOR1;refGene_tx=XM_023256163.1
B3 39643837 39643912 refGene_gn=SKOR1;refGene_tx=XM_023256163.1
B3 39645282 39645354 refGene_gn=SKOR1;refGene_tx=XM_023256163.1
B3 39645875 39645983 refGene_gn=SKOR1;refGene_tx=XM_023256163.1
B3 39646918 39647009 refGene_gn=SKOR1;refGene_tx=XM_023256163.1
B3 39648117 39648674 refGene_gn=SKOR1;refGene_tx=XM_023256163.1
B3 39649169 39650320 refGene_gn=SKOR1;refGene_tx=XM_023256163.1
B3 39652636 39652722 refGene_gn=SKOR1;refGene_tx=XM_023256163.1
B3 39652794 39653346 refGene_gn=SKOR1;refGene_tx=XM_023256163.1
B3 39669434 39669539
refGene_gn=MAP2K5;refGene_tx=XM_003987010.5;refGene_gn=MAP2K5;refGene_
tx=XM_003987011.5;refGene_gn=MAP2K5;refGene_tx=XM_019832174.2;refGene_
gn=MAP2K5;refGene_tx=XM_019832175.2
B3 39706654 39706700
refGene_gn=MAP2K5;refGene_tx=XM_003987010.5;refGene_gn=MAP2K5;refGene_
tx=XM_003987011.5;refGene_gn=MAP2K5;refGene_tx=XM_019832176.2;refGene_
gn=MAP2K5;refGene_tx=XM_019832177.2
B3 39709597 39709659
refGene_gn=MAP2K5;refGene_tx=XM_003987010.5;refGene_gn=MAP2K5;refGene_
tx=XM_003987011.5
B3 39733345 39733378
refGene_gn=MAP2K5;refGene_tx=XM_003987010.5;refGene_gn=MAP2K5;refGene_
tx=XM_003987011.5;refGene_gn=MAP2K5;refGene_tx=XM_019832174.2;refGene_
gn=MAP2K5;refGene_tx=XM_019832175.2;refGene_gn=MAP2K5;refGene_tx=XM_01
9832176.2;refGene_gn=MAP2K5;refGene_tx=XM_019832177.2
B3 39733690 39733717
refGene_gn=MAP2K5;refGene_tx=XM_003987010.5;refGene_gn=MAP2K5;refGene_
tx=XM_003987011.5;refGene_gn=MAP2K5;refGene_tx=XM_019832174.2;refGene_
gn=MAP2K5;refGene_tx=XM_019832175.2;refGene_gn=MAP2K5;refGene_tx=XM_01
9832176.2;refGene_gn=MAP2K5;refGene_tx=XM_019832177.2
B3 39755068 39755098
refGene_gn=MAP2K5;refGene_tx=XM_003987010.5;refGene_gn=MAP2K5;refGene_

tx=XM_019832174.2;refGene_gn=MAP2K5;refGene_tx=XM_019832176.2
B3 39777141 39777213
refGene_gn=MAP2K5;refGene_tx=XM_003987010.5;refGene_gn=MAP2K5;refGene_tx=XM_003987011.5;refGene_gn=MAP2K5;refGene_tx=XM_019832174.2;refGene_gn=MAP2K5;refGene_tx=XM_019832175.2;refGene_gn=MAP2K5;refGene_tx=XM_019832176.2;refGene_gn=MAP2K5;refGene_tx=XM_019832177.2
B3 39787185 39787236
refGene_gn=MAP2K5;refGene_tx=XM_003987010.5;refGene_gn=MAP2K5;refGene_tx=XM_003987011.5;refGene_gn=MAP2K5;refGene_tx=XM_019832174.2;refGene_gn=MAP2K5;refGene_tx=XM_019832175.2;refGene_gn=MAP2K5;refGene_tx=XM_019832176.2;refGene_gn=MAP2K5;refGene_tx=XM_019832177.2
B3 39788200 39788274
refGene_gn=MAP2K5;refGene_tx=XM_003987010.5;refGene_gn=MAP2K5;refGene_tx=XM_003987011.5;refGene_gn=MAP2K5;refGene_tx=XM_019832174.2;refGene_gn=MAP2K5;refGene_tx=XM_019832175.2;refGene_gn=MAP2K5;refGene_tx=XM_019832176.2;refGene_gn=MAP2K5;refGene_tx=XM_019832177.2
B3 39813398 39813447
refGene_gn=MAP2K5;refGene_tx=XM_003987010.5;refGene_gn=MAP2K5;refGene_tx=XM_003987011.5;refGene_gn=MAP2K5;refGene_tx=XM_019832174.2;refGene_gn=MAP2K5;refGene_tx=XM_019832175.2;refGene_gn=MAP2K5;refGene_tx=XM_019832176.2;refGene_gn=MAP2K5;refGene_tx=XM_019832177.2
B3 39820190 39820252
refGene_gn=MAP2K5;refGene_tx=XM_003987010.5;refGene_gn=MAP2K5;refGene_tx=XM_003987011.5;refGene_gn=MAP2K5;refGene_tx=XM_019832174.2;refGene_gn=MAP2K5;refGene_tx=XM_019832175.2;refGene_gn=MAP2K5;refGene_tx=XM_019832176.2;refGene_gn=MAP2K5;refGene_tx=XM_019832177.2
B3 39839461 39839543
refGene_gn=MAP2K5;refGene_tx=XM_003987010.5;refGene_gn=MAP2K5;refGene_tx=XM_003987011.5;refGene_gn=MAP2K5;refGene_tx=XM_019832174.2;refGene_gn=MAP2K5;refGene_tx=XM_019832175.2;refGene_gn=MAP2K5;refGene_tx=XM_019832176.2;refGene_gn=MAP2K5;refGene_tx=XM_019832177.2
B3 39839653 39839722
refGene_gn=MAP2K5;refGene_tx=XM_003987010.5;refGene_gn=MAP2K5;refGene_tx=XM_003987011.5;refGene_gn=MAP2K5;refGene_tx=XM_019832174.2;refGene_gn=MAP2K5;refGene_tx=XM_019832175.2;refGene_gn=MAP2K5;refGene_tx=XM_019832176.2;refGene_gn=MAP2K5;refGene_tx=XM_019832177.2
B3 39850590 39850630
refGene_gn=MAP2K5;refGene_tx=XM_003987010.5;refGene_gn=MAP2K5;refGene_tx=XM_003987011.5;refGene_gn=MAP2K5;refGene_tx=XM_019832174.2;refGene_gn=MAP2K5;refGene_tx=XM_019832175.2;refGene_gn=MAP2K5;refGene_tx=XM_019832176.2;refGene_gn=MAP2K5;refGene_tx=XM_019832177.2
B3 39881896 39881961
refGene_gn=MAP2K5;refGene_tx=XM_003987010.5;refGene_gn=MAP2K5;refGene_tx=XM_003987011.5;refGene_gn=MAP2K5;refGene_tx=XM_019832174.2;refGene_gn=MAP2K5;refGene_tx=XM_019832175.2;refGene_gn=MAP2K5;refGene_tx=XM_019832176.2;refGene_gn=MAP2K5;refGene_tx=XM_019832177.2
B3 39889478 39889527
refGene_gn=MAP2K5;refGene_tx=XM_003987010.5;refGene_gn=MAP2K5;refGene_tx=XM_003987011.5;refGene_gn=MAP2K5;refGene_tx=XM_019832174.2;refGene_gn=MAP2K5;refGene_tx=XM_019832175.2;refGene_gn=MAP2K5;refGene_tx=XM_01

9832176.2;refGene_gn=MAP2K5;refGene_tx=XM_019832177.2
B3 39896191 39896259
refGene_gn=MAP2K5;refGene_tx=XM_003987010.5;refGene_gn=MAP2K5;refGene_tx=XM_003987011.5;refGene_gn=MAP2K5;refGene_tx=XM_019832174.2;refGene_gn=MAP2K5;refGene_tx=XM_019832175.2;refGene_gn=MAP2K5;refGene_tx=XM_019832176.2;refGene_gn=MAP2K5;refGene_tx=XM_019832177.2
B3 39897232 39897273
refGene_gn=MAP2K5;refGene_tx=XM_003987010.5;refGene_gn=MAP2K5;refGene_tx=XM_003987011.5;refGene_gn=MAP2K5;refGene_tx=XM_019832174.2;refGene_gn=MAP2K5;refGene_tx=XM_019832175.2;refGene_gn=MAP2K5;refGene_tx=XM_019832176.2;refGene_gn=MAP2K5;refGene_tx=XM_019832177.2
B3 39902020 39902090
refGene_gn=MAP2K5;refGene_tx=XM_003987010.5;refGene_gn=MAP2K5;refGene_tx=XM_003987011.5;refGene_gn=MAP2K5;refGene_tx=XM_019832174.2;refGene_gn=MAP2K5;refGene_tx=XM_019832175.2;refGene_gn=MAP2K5;refGene_tx=XM_019832176.2;refGene_gn=MAP2K5;refGene_tx=XM_019832177.2
B3 39918441 39918509
refGene_gn=MAP2K5;refGene_tx=XM_003987010.5;refGene_gn=MAP2K5;refGene_tx=XM_003987011.5;refGene_gn=MAP2K5;refGene_tx=XM_019832174.2;refGene_gn=MAP2K5;refGene_tx=XM_019832175.2;refGene_gn=MAP2K5;refGene_tx=XM_019832176.2;refGene_gn=MAP2K5;refGene_tx=XM_019832177.2
B3 39933358 39933407
refGene_gn=MAP2K5;refGene_tx=XM_003987010.5;refGene_gn=MAP2K5;refGene_tx=XM_003987011.5;refGene_gn=MAP2K5;refGene_tx=XM_019832174.2;refGene_gn=MAP2K5;refGene_tx=XM_019832175.2;refGene_gn=MAP2K5;refGene_tx=XM_019832176.2;refGene_gn=MAP2K5;refGene_tx=XM_019832177.2
B3 39940066 39940201
refGene_gn=MAP2K5;refGene_tx=XM_003987010.5;refGene_gn=MAP2K5;refGene_tx=XM_003987011.5;refGene_gn=MAP2K5;refGene_tx=XM_019832174.2;refGene_gn=MAP2K5;refGene_tx=XM_019832175.2;refGene_gn=MAP2K5;refGene_tx=XM_019832176.2;refGene_gn=MAP2K5;refGene_tx=XM_019832177.2
B3 39953670 39953798
refGene_gn=CB3H15orf61;refGene_tx=XM_003987072.5
B3 39958160 39958506
refGene_gn=CB3H15orf61;refGene_tx=XM_003987072.5
B3 39978923 39979040
refGene_gn=IQCH;refGene_tx=XM_023255144.1;refGene_gn=IQCH;refGene_tx=XM_023255145.1;refGene_gn=IQCH;refGene_tx=XM_023255146.1;refGene_gn=IQCH;refGene_tx=XM_023255147.1
B3 39983652 39983761
refGene_gn=IQCH;refGene_tx=XM_023255144.1;refGene_gn=IQCH;refGene_tx=XM_023255145.1;refGene_gn=IQCH;refGene_tx=XM_023255146.1;refGene_gn=IQCH;refGene_tx=XM_023255147.1
B3 39988762 39988824
refGene_gn=IQCH;refGene_tx=XM_023255144.1;refGene_gn=IQCH;refGene_tx=XM_023255145.1;refGene_gn=IQCH;refGene_tx=XM_023255146.1;refGene_gn=IQCH;refGene_tx=XM_023255147.1
B3 39996327 39996450
refGene_gn=IQCH;refGene_tx=XM_023255144.1;refGene_gn=IQCH;refGene_tx=XM_023255145.1;refGene_gn=IQCH;refGene_tx=XM_023255146.1;refGene_gn=IQC

H;refGene_tx=XM_023255147.1
B3 40005929 40006100
refGene_gn=IQCH;refGene_tx=XM_023255144.1;refGene_gn=IQCH;refGene_tx=XM_023255145.1;refGene_gn=IQCH;refGene_tx=XM_023255146.1;refGene_gn=IQCH;refGene_tx=XM_023255147.1
B3 40030270 40030303 refGene_gn=IQCH;refGene_tx=XM_023255149.1
B3 40031465 40031752
refGene_gn=IQCH;refGene_tx=XM_023255144.1;refGene_gn=IQCH;refGene_tx=XM_023255145.1;refGene_gn=IQCH;refGene_tx=XM_023255146.1;refGene_gn=IQCH;refGene_tx=XM_023255147.1;refGene_gn=IQCH;refGene_tx=XM_023255149.1
B3 40035869 40035990
refGene_gn=IQCH;refGene_tx=XM_023255144.1;refGene_gn=IQCH;refGene_tx=XM_023255145.1;refGene_gn=IQCH;refGene_tx=XM_023255146.1;refGene_gn=IQCH;refGene_tx=XM_023255147.1;refGene_gn=IQCH;refGene_tx=XM_023255149.1
B3 40045073 40045238 refGene_gn=IQCH;refGene_tx=XM_023255150.1
B3 40052567 40052759
refGene_gn=IQCH;refGene_tx=XM_023255144.1;refGene_gn=IQCH;refGene_tx=XM_023255145.1;refGene_gn=IQCH;refGene_tx=XM_023255149.1;refGene_gn=IQCH;refGene_tx=XM_023255150.1
B3 40056956 40057229
refGene_gn=IQCH;refGene_tx=XM_023255144.1;refGene_gn=IQCH;refGene_tx=XM_023255145.1;refGene_gn=IQCH;refGene_tx=XM_023255146.1;refGene_gn=IQCH;refGene_tx=XM_023255149.1;refGene_gn=IQCH;refGene_tx=XM_023255150.1
B3 40063742 40063918
refGene_gn=IQCH;refGene_tx=XM_023255144.1;refGene_gn=IQCH;refGene_tx=XM_023255145.1;refGene_gn=IQCH;refGene_tx=XM_023255146.1;refGene_gn=IQCH;refGene_tx=XM_023255147.1;refGene_gn=IQCH;refGene_tx=XM_023255149.1;refGene_gn=IQCH;refGene_tx=XM_023255150.1
B3 40068014 40068098
refGene_gn=IQCH;refGene_tx=XM_023255144.1;refGene_gn=IQCH;refGene_tx=XM_023255145.1;refGene_gn=IQCH;refGene_tx=XM_023255146.1;refGene_gn=IQCH;refGene_tx=XM_023255147.1;refGene_gn=IQCH;refGene_tx=XM_023255149.1;refGene_gn=IQCH;refGene_tx=XM_023255150.1
B3 40079696 40079763
refGene_gn=IQCH;refGene_tx=XM_023255144.1;refGene_gn=IQCH;refGene_tx=XM_023255145.1;refGene_gn=IQCH;refGene_tx=XM_023255146.1;refGene_gn=IQCH;refGene_tx=XM_023255147.1;refGene_gn=IQCH;refGene_tx=XM_023255149.1;refGene_gn=IQCH;refGene_tx=XM_023255150.1
B3 40080468 40081020
refGene_gn=IQCH;refGene_tx=XM_023255144.1;refGene_gn=IQCH;refGene_tx=XM_023255145.1;refGene_gn=IQCH;refGene_tx=XM_023255146.1;refGene_gn=IQCH;refGene_tx=XM_023255147.1;refGene_gn=IQCH;refGene_tx=XM_023255149.1;refGene_gn=IQCH;refGene_tx=XM_023255150.1
B3 40094387 40094426
refGene_gn=IQCH;refGene_tx=XM_023255144.1;refGene_gn=IQCH;refGene_tx=XM_023255146.1;refGene_gn=IQCH;refGene_tx=XM_023255147.1;refGene_gn=IQCH;refGene_tx=XM_023255149.1;refGene_gn=IQCH;refGene_tx=XM_023255150.1
B3 40096213 40096293
refGene_gn=IQCH;refGene_tx=XM_023255144.1;refGene_gn=IQCH;refGene_tx=XM_023255145.1;refGene_gn=IQCH;refGene_tx=XM_023255146.1;refGene_gn=IQC

H;refGene_tx=XM_023255147.1;refGene_gn=IQCH;refGene_tx=XM_023255149.1;refGene_gn=IQCH;refGene_tx=XM_023255150.1
B3 40117335 40117464
refGene_gn=IQCH;refGene_tx=XM_023255144.1;refGene_gn=IQCH;refGene_tx=XM_023255145.1;refGene_gn=IQCH;refGene_tx=XM_023255146.1;refGene_gn=IQCH;refGene_tx=XM_023255147.1;refGene_gn=IQCH;refGene_tx=XM_023255149.1;refGene_gn=IQCH;refGene_tx=XM_023255150.1
B3 40125035 40125156
refGene_gn=IQCH;refGene_tx=XM_023255144.1;refGene_gn=IQCH;refGene_tx=XM_023255145.1;refGene_gn=IQCH;refGene_tx=XM_023255146.1;refGene_gn=IQCH;refGene_tx=XM_023255147.1;refGene_gn=IQCH;refGene_tx=XM_023255149.1;refGene_gn=IQCH;refGene_tx=XM_023255150.1
B3 40140271 40140418 refGene_gn=IQCH;refGene_tx=XM_023255151.1
B3 40144354 40144369 refGene_gn=IQCH;refGene_tx=XM_023255152.1
B3 40153643 40153762
refGene_gn=IQCH;refGene_tx=XM_023255144.1;refGene_gn=IQCH;refGene_tx=XM_023255145.1;refGene_gn=IQCH;refGene_tx=XM_023255146.1;refGene_gn=IQCH;refGene_tx=XM_023255147.1;refGene_gn=IQCH;refGene_tx=XM_023255149.1;refGene_gn=IQCH;refGene_tx=XM_023255150.1;refGene_gn=IQCH;refGene_tx=XM_023255151.1;refGene_gn=IQCH;refGene_tx=XM_023255152.1
B3 40169183 40169299
refGene_gn=IQCH;refGene_tx=XM_023255144.1;refGene_gn=IQCH;refGene_tx=XM_023255145.1;refGene_gn=IQCH;refGene_tx=XM_023255146.1;refGene_gn=IQCH;refGene_tx=XM_023255147.1;refGene_gn=IQCH;refGene_tx=XM_023255149.1;refGene_gn=IQCH;refGene_tx=XM_023255150.1;refGene_gn=IQCH;refGene_tx=XM_023255151.1;refGene_gn=IQCH;refGene_tx=XM_023255152.1
B3 40180155 40180250
refGene_gn=IQCH;refGene_tx=XM_023255144.1;refGene_gn=IQCH;refGene_tx=XM_023255145.1;refGene_gn=IQCH;refGene_tx=XM_023255146.1;refGene_gn=IQCH;refGene_tx=XM_023255147.1;refGene_gn=IQCH;refGene_tx=XM_023255149.1;refGene_gn=IQCH;refGene_tx=XM_023255150.1;refGene_gn=IQCH;refGene_tx=XM_023255151.1;refGene_gn=IQCH;refGene_tx=XM_023255152.1
B3 40181450 40181573
refGene_gn=IQCH;refGene_tx=XM_023255144.1;refGene_gn=IQCH;refGene_tx=XM_023255145.1;refGene_gn=IQCH;refGene_tx=XM_023255146.1;refGene_gn=IQCH;refGene_tx=XM_023255147.1;refGene_gn=IQCH;refGene_tx=XM_023255149.1;refGene_gn=IQCH;refGene_tx=XM_023255150.1;refGene_gn=IQCH;refGene_tx=XM_023255151.1;refGene_gn=IQCH;refGene_tx=XM_023255152.1
B3 40191405 40191456
refGene_gn=IQCH;refGene_tx=XM_023255144.1;refGene_gn=IQCH;refGene_tx=XM_023255145.1;refGene_gn=IQCH;refGene_tx=XM_023255146.1;refGene_gn=IQCH;refGene_tx=XM_023255147.1;refGene_gn=IQCH;refGene_tx=XM_023255149.1;refGene_gn=IQCH;refGene_tx=XM_023255150.1;refGene_gn=IQCH;refGene_tx=XM_023255151.1;refGene_gn=IQCH;refGene_tx=XM_023255152.1
B3 40191724 40191797
refGene_gn=AAGAB;refGene_tx=XM_003987012.4;refGene_gn=AAGAB;refGene_tx=XM_019832189.1
B3 40212252 40212443
refGene_gn=AAGAB;refGene_tx=XM_003987012.4;refGene_gn=AAGAB;refGene_tx=XM_019832189.1

B3 40212579 40212676
refGene_gn=AAGAB;refGene_tx=XM_003987012.4;refGene_gn=AAGAB;refGene_tx=XM_019832189.1;refGene_gn=AAGAB;refGene_tx=XM_019832190.2;refGene_gn=AAGAB;refGene_tx=XM_019832191.2
B3 40213226 40213316
refGene_gn=AAGAB;refGene_tx=XM_003987012.4;refGene_gn=AAGAB;refGene_tx=XM_019832189.1;refGene_gn=AAGAB;refGene_tx=XM_019832190.2;refGene_gn=AAGAB;refGene_tx=XM_019832191.2
B3 40216223 40216307
refGene_gn=AAGAB;refGene_tx=XM_003987012.4;refGene_gn=AAGAB;refGene_tx=XM_019832189.1;refGene_gn=AAGAB;refGene_tx=XM_019832190.2;refGene_gn=AAGAB;refGene_tx=XM_019832191.2
B3 40238418 40238501
refGene_gn=AAGAB;refGene_tx=XM_003987012.4;refGene_gn=AAGAB;refGene_tx=XM_019832189.1;refGene_gn=AAGAB;refGene_tx=XM_019832190.2;refGene_gn=AAGAB;refGene_tx=XM_019832191.2
B3 40239359 40239456
refGene_gn=AAGAB;refGene_tx=XM_003987012.4;refGene_gn=AAGAB;refGene_tx=XM_019832189.1;refGene_gn=AAGAB;refGene_tx=XM_019832190.2;refGene_gn=AAGAB;refGene_tx=XM_019832191.2
B3 40244397 40244502
refGene_gn=AAGAB;refGene_tx=XM_003987012.4;refGene_gn=AAGAB;refGene_tx=XM_019832189.1;refGene_gn=AAGAB;refGene_tx=XM_019832190.2;refGene_gn=AAGAB;refGene_tx=XM_019832191.2
B3 40244737 40244760 refGene_gn=AAGAB;refGene_tx=XM_019832189.1
B3 40244968 40245018
refGene_gn=AAGAB;refGene_tx=XM_003987012.4;refGene_gn=AAGAB;refGene_tx=XM_019832190.2;refGene_gn=AAGAB;refGene_tx=XM_019832191.2
B3 40245684 40245762
refGene_gn=AAGAB;refGene_tx=XM_003987012.4;refGene_gn=AAGAB;refGene_tx=XM_019832190.2;refGene_gn=AAGAB;refGene_tx=XM_019832191.2
B3 40257103 40257227 refGene_gn=SMAD3;refGene_tx=XM_023255153.1
B3 40259678 40259823 refGene_gn=SMAD3;refGene_tx=XM_023255153.1
B3 40262642 40262780 refGene_gn=SMAD3;refGene_tx=XM_023255153.1
B3 40265018 40265231 refGene_gn=SMAD3;refGene_tx=XM_023255153.1
B3 40274676 40274727 refGene_gn=SMAD3;refGene_tx=XM_023255153.1
B3 40277493 40277568 refGene_gn=SMAD3;refGene_tx=XM_023255153.1
B3 40278916 40279048 refGene_gn=SMAD3;refGene_tx=XM_023255153.1
B3 40279225 40279419 refGene_gn=SMAD3;refGene_tx=XM_023255153.1
B3 40371835 40372041 refGene_gn=SMAD3;refGene_tx=XM_023255153.1
B3 40599599 40600141 refGene_gn=SMAD6;refGene_tx=XM_023255154.1
B3 40662206 40662284 refGene_gn=SMAD6;refGene_tx=XM_023255154.1
B3 40665949 40666006 refGene_gn=SMAD6;refGene_tx=XM_023255154.1
B3 40673605 40673660 refGene_gn=SMAD6;refGene_tx=XM_023255154.1
B3 40673661 40674423 refGene_gn=SMAD6;refGene_tx=XM_023255154.1
B3 40798199 40798317
refGene_gn=LCTL;refGene_tx=XM_003987014.4;refGene_gn=LCTL;refGene_tx=XM_019832194.1
B3 40798713 40798877
refGene_gn=LCTL;refGene_tx=XM_003987014.4;refGene_gn=LCTL;refGene_tx=X

M_019832194.1
B3 40799491 40799579
refGene_gn=LCTL;refGene_tx=XM_003987014.4;refGene_gn=LCTL;refGene_tx=XM_019832194.1
B3 40799849 40799959
refGene_gn=LCTL;refGene_tx=XM_003987014.4;refGene_gn=LCTL;refGene_tx=XM_019832194.1
B3 40801346 40801475
refGene_gn=LCTL;refGene_tx=XM_003987014.4;refGene_gn=LCTL;refGene_tx=XM_019832193.2;refGene_gn=LCTL;refGene_tx=XM_019832194.1
B3 40801575 40801671
refGene_gn=LCTL;refGene_tx=XM_003987014.4;refGene_gn=LCTL;refGene_tx=XM_019832193.2;refGene_gn=LCTL;refGene_tx=XM_019832194.1
B3 40803029 40803084
refGene_gn=LCTL;refGene_tx=XM_003987014.4;refGene_gn=LCTL;refGene_tx=XM_019832193.2;refGene_gn=LCTL;refGene_tx=XM_019832194.1
B3 40803456 40803618
refGene_gn=LCTL;refGene_tx=XM_003987014.4;refGene_gn=LCTL;refGene_tx=XM_019832193.2;refGene_gn=LCTL;refGene_tx=XM_019832194.1
B3 40807236 40807511
refGene_gn=LCTL;refGene_tx=XM_003987014.4;refGene_gn=LCTL;refGene_tx=XM_019832193.2;refGene_gn=LCTL;refGene_tx=XM_019832194.1
B3 40808051 40808178
refGene_gn=LCTL;refGene_tx=XM_003987014.4;refGene_gn=LCTL;refGene_tx=XM_019832193.2;refGene_gn=LCTL;refGene_tx=XM_019832194.1
B3 40808363 40808563
refGene_gn=LCTL;refGene_tx=XM_003987014.4;refGene_gn=LCTL;refGene_tx=XM_019832193.2;refGene_gn=LCTL;refGene_tx=XM_019832194.1
B3 40810041 40810105
refGene_gn=LCTL;refGene_tx=XM_003987014.4;refGene_gn=LCTL;refGene_tx=XM_019832193.2;refGene_gn=LCTL;refGene_tx=XM_019832194.1
B3 40811540 40811656
refGene_gn=LCTL;refGene_tx=XM_003987014.4;refGene_gn=LCTL;refGene_tx=XM_019832193.2;refGene_gn=LCTL;refGene_tx=XM_019832194.1
B3 40812930 40812986
refGene_gn=ZWILCH;refGene_tx=XM_006932447.4;refGene_gn=ZWILCH;refGene_tx=XM_023255156.1;refGene_gn=ZWILCH;refGene_tx=XM_023255157.1
B3 40816746 40816868
refGene_gn=ZWILCH;refGene_tx=XM_006932447.4;refGene_gn=ZWILCH;refGene_tx=XM_023255156.1;refGene_gn=ZWILCH;refGene_tx=XM_023255157.1
B3 40817313 40817536
refGene_gn=ZWILCH;refGene_tx=XM_023255155.1;refGene_gn=ZWILCH;refGene_tx=XM_023255159.1
B3 40818065 40818161
refGene_gn=ZWILCH;refGene_tx=XM_006932447.4;refGene_gn=ZWILCH;refGene_tx=XM_023255155.1;refGene_gn=ZWILCH;refGene_tx=XM_023255156.1;refGene_gn=ZWILCH;refGene_tx=XM_023255157.1;refGene_gn=ZWILCH;refGene_tx=XM_023255159.1
B3 40819320 40819457
refGene_gn=ZWILCH;refGene_tx=XM_006932447.4;refGene_gn=ZWILCH;refGene_

tx=XM_023255155.1;refGene_gn=ZWILCH;refGene_tx=XM_023255156.1;refGene_gn=ZWILCH;refGene_tx=XM_023255157.1;refGene_gn=ZWILCH;refGene_tx=XM_023255159.1
B3 40822557 40822586
refGene_gn=ZWILCH;refGene_tx=XM_006932447.4;refGene_gn=ZWILCH;refGene_tx=XM_023255155.1;refGene_gn=ZWILCH;refGene_tx=XM_023255156.1;refGene_gn=ZWILCH;refGene_tx=XM_023255157.1;refGene_gn=ZWILCH;refGene_tx=XM_023255159.1
B3 40823096 40823253
refGene_gn=ZWILCH;refGene_tx=XM_006932447.4;refGene_gn=ZWILCH;refGene_tx=XM_023255155.1;refGene_gn=ZWILCH;refGene_tx=XM_023255156.1;refGene_gn=ZWILCH;refGene_tx=XM_023255157.1;refGene_gn=ZWILCH;refGene_tx=XM_023255159.1
B3 40824795 40824875
refGene_gn=ZWILCH;refGene_tx=XM_006932447.4;refGene_gn=ZWILCH;refGene_tx=XM_023255155.1;refGene_gn=ZWILCH;refGene_tx=XM_023255156.1;refGene_gn=ZWILCH;refGene_tx=XM_023255157.1;refGene_gn=ZWILCH;refGene_tx=XM_023255159.1
B3 40825394 40825500
refGene_gn=ZWILCH;refGene_tx=XM_006932447.4;refGene_gn=ZWILCH;refGene_tx=XM_023255155.1;refGene_gn=ZWILCH;refGene_tx=XM_023255156.1;refGene_gn=ZWILCH;refGene_tx=XM_023255157.1;refGene_gn=ZWILCH;refGene_tx=XM_023255159.1
B3 40826291 40826341
refGene_gn=ZWILCH;refGene_tx=XM_006932447.4;refGene_gn=ZWILCH;refGene_tx=XM_023255155.1;refGene_gn=ZWILCH;refGene_tx=XM_023255156.1;refGene_gn=ZWILCH;refGene_tx=XM_023255157.1;refGene_gn=ZWILCH;refGene_tx=XM_023255159.1
B3 40826867 40826961
refGene_gn=ZWILCH;refGene_tx=XM_006932447.4;refGene_gn=ZWILCH;refGene_tx=XM_023255155.1;refGene_gn=ZWILCH;refGene_tx=XM_023255156.1;refGene_gn=ZWILCH;refGene_tx=XM_023255157.1;refGene_gn=ZWILCH;refGene_tx=XM_023255159.1
B3 40830884 40830956
refGene_gn=ZWILCH;refGene_tx=XM_006932447.4;refGene_gn=ZWILCH;refGene_tx=XM_023255155.1;refGene_gn=ZWILCH;refGene_tx=XM_023255156.1;refGene_gn=ZWILCH;refGene_tx=XM_023255157.1;refGene_gn=ZWILCH;refGene_tx=XM_023255159.1
B3 40832228 40832384
refGene_gn=ZWILCH;refGene_tx=XM_006932447.4;refGene_gn=ZWILCH;refGene_tx=XM_023255155.1;refGene_gn=ZWILCH;refGene_tx=XM_023255156.1;refGene_gn=ZWILCH;refGene_tx=XM_023255157.1;refGene_gn=ZWILCH;refGene_tx=XM_023255159.1
B3 40832729 40832800
refGene_gn=ZWILCH;refGene_tx=XM_006932447.4;refGene_gn=ZWILCH;refGene_tx=XM_023255155.1;refGene_gn=ZWILCH;refGene_tx=XM_023255156.1;refGene_gn=ZWILCH;refGene_tx=XM_023255157.1;refGene_gn=ZWILCH;refGene_tx=XM_023255159.1
B3 40834187 40834387
refGene_gn=ZWILCH;refGene_tx=XM_006932447.4;refGene_gn=ZWILCH;refGene_

tx=XM_023255155.1;refGene_gn=ZWILCH;refGene_tx=XM_023255156.1;refGene_gn=ZWILCH;refGene_tx=XM_023255157.1;refGene_gn=ZWILCH;refGene_tx=XM_023255159.1
B3 40835585 40835741
refGene_gn=ZWILCH;refGene_tx=XM_023255155.1;refGene_gn=ZWILCH;refGene_tx=XM_023255156.1;refGene_gn=ZWILCH;refGene_tx=XM_023255157.1
B3 40838083 40838202
refGene_gn=ZWILCH;refGene_tx=XM_006932447.4;refGene_gn=ZWILCH;refGene_tx=XM_023255155.1;refGene_gn=ZWILCH;refGene_tx=XM_023255156.1;refGene_gn=ZWILCH;refGene_tx=XM_023255157.1;refGene_gn=ZWILCH;refGene_tx=XM_023255159.1
B3 40842395 40842491
refGene_gn=ZWILCH;refGene_tx=XM_006932447.4;refGene_gn=ZWILCH;refGene_tx=XM_023255155.1;refGene_gn=ZWILCH;refGene_tx=XM_023255156.1;refGene_gn=ZWILCH;refGene_tx=XM_023255157.1;refGene_gn=ZWILCH;refGene_tx=XM_023255159.1
B3 40846302 40846354
refGene_gn=ZWILCH;refGene_tx=XM_006932447.4;refGene_gn=ZWILCH;refGene_tx=XM_023255155.1;refGene_gn=ZWILCH;refGene_tx=XM_023255156.1;refGene_gn=ZWILCH;refGene_tx=XM_023255157.1;refGene_gn=ZWILCH;refGene_tx=XM_023255159.1
B3 40849365 40849418
refGene_gn=ZWILCH;refGene_tx=XM_006932447.4;refGene_gn=ZWILCH;refGene_tx=XM_023255155.1;refGene_gn=ZWILCH;refGene_tx=XM_023255156.1;refGene_gn=ZWILCH;refGene_tx=XM_023255157.1;refGene_gn=ZWILCH;refGene_tx=XM_023255159.1
B3 40849969 40849972 refGene_gn=RPL4;refGene_tx=XM_019832195.1
B3 40851311 40851483 refGene_gn=RPL4;refGene_tx=XM_019832195.1
B3 40851669 40851776 refGene_gn=RPL4;refGene_tx=XM_019832195.1
B3 40852097 40852236 refGene_gn=RPL4;refGene_tx=XM_019832195.1
B3 40852673 40852798 refGene_gn=RPL4;refGene_tx=XM_019832195.1
B3 40853094 40853224 refGene_gn=RPL4;refGene_tx=XM_019832195.1
B3 40853498 40853655 refGene_gn=RPL4;refGene_tx=XM_019832195.1
B3 40853907 40853991 refGene_gn=RPL4;refGene_tx=XM_019832195.1
B3 40854100 40854221 refGene_gn=RPL4;refGene_tx=XM_019832195.1
B3 40854303 40854531 refGene_gn=RPL4;refGene_tx=XM_019832195.1
B3 40856426 40856516
refGene_gn=SNAPC5;refGene_tx=XM_006932449.3;refGene_gn=SNAPC5;refGene_tx=XM_006932450.3
B3 40857730 40857820
refGene_gn=SNAPC5;refGene_tx=XM_006932449.3;refGene_gn=SNAPC5;refGene_tx=XM_006932450.3;refGene_gn=SNAPC5;refGene_tx=XM_019832196.2
B3 40858824 40858941
refGene_gn=SNAPC5;refGene_tx=XM_006932449.3;refGene_gn=SNAPC5;refGene_tx=XM_006932450.3;refGene_gn=SNAPC5;refGene_tx=XM_019832196.2
B3 40861968 40862082
refGene_gn=MAP2K1;refGene_tx=XM_003987018.5;refGene_gn=MAP2K1;refGene_tx=XM_011282937.3
B3 40862719 40862765
refGene_gn=MAP2K1;refGene_tx=XM_003987018.5;refGene_gn=MAP2K1;refGene_

tx=XM_011282937.3
B3 40863115 40863177
refGene_gn=MAP2K1;refGene_tx=XM_003987018.5;refGene_gn=MAP2K1;refGene_
tx=XM_011282937.3
B3 40865766 40865831
refGene_gn=MAP2K1;refGene_tx=XM_003987018.5;refGene_gn=MAP2K1;refGene_
tx=XM_011282937.3
B3 40867141 40867343
refGene_gn=MAP2K1;refGene_tx=XM_003987018.5;refGene_gn=MAP2K1;refGene_
tx=XM_011282937.3
B3 40869356 40869481
refGene_gn=MAP2K1;refGene_tx=XM_003987018.5;refGene_gn=MAP2K1;refGene_
tx=XM_011282937.3
B3 40893764 40893816
refGene_gn=MAP2K1;refGene_tx=XM_003987018.5;refGene_gn=MAP2K1;refGene_
tx=XM_011282937.3
B3 40894716 40894794
refGene_gn=MAP2K1;refGene_tx=XM_003987018.5;refGene_gn=MAP2K1;refGene_
tx=XM_011282937.3
B3 40902502 40902649
refGene_gn=MAP2K1;refGene_tx=XM_003987018.5;refGene_gn=MAP2K1;refGene_
tx=XM_011282937.3
B3 40903963 40904174
refGene_gn=MAP2K1;refGene_tx=XM_003987018.5;refGene_gn=MAP2K1;refGene_
tx=XM_011282937.3
B3 40930998 40931012 refGene_gn=MAP2K1;refGene_tx=XM_003987018.5
B3 40942389 40942469 refGene_gn=MAP2K1;refGene_tx=XM_011282937.3
B3 40973580 40973710
refGene_gn=TIPIN;refGene_tx=XM_006932451.3;refGene_gn=TIPIN;refGene_tx
=XM_023255161.1
B3 40973834 40973913
refGene_gn=TIPIN;refGene_tx=XM_006932451.3;refGene_gn=TIPIN;refGene_tx
=XM_023255161.1
B3 40974335 40974411
refGene_gn=TIPIN;refGene_tx=XM_006932451.3;refGene_gn=TIPIN;refGene_tx
=XM_023255161.1
B3 40979276 40979399
refGene_gn=TIPIN;refGene_tx=XM_006932451.3;refGene_gn=TIPIN;refGene_tx
=XM_023255161.1
B3 40979633 40979697
refGene_gn=TIPIN;refGene_tx=XM_006932451.3;refGene_gn=TIPIN;refGene_tx
=XM_023255161.1
B3 40985969 40986176
refGene_gn=TIPIN;refGene_tx=XM_006932451.3;refGene_gn=TIPIN;refGene_tx
=XM_023255161.1
B3 40988247 40988438
refGene_gn=TIPIN;refGene_tx=XM_006932451.3;refGene_gn=TIPIN;refGene_tx
=XM_023255161.1
B3 40990435 40990744
refGene_gn=DIS3L;refGene_tx=XM_023255162.1;refGene_gn=DIS3L;refGene_tx

=XM_023255163.1
B3 40990832 40991007
refGene_gn=DIS3L;refGene_tx=XM_023255162.1;refGene_gn=DIS3L;refGene_tx
=XM_023255163.1
B3 40991625 40991771
refGene_gn=DIS3L;refGene_tx=XM_023255162.1;refGene_gn=DIS3L;refGene_tx
=XM_023255163.1
B3 40992142 40992321
refGene_gn=DIS3L;refGene_tx=XM_023255162.1;refGene_gn=DIS3L;refGene_tx
=XM_023255163.1
B3 40992420 40992575
refGene_gn=DIS3L;refGene_tx=XM_023255162.1;refGene_gn=DIS3L;refGene_tx
=XM_023255163.1
B3 40994063 40994597
refGene_gn=DIS3L;refGene_tx=XM_023255162.1;refGene_gn=DIS3L;refGene_tx
=XM_023255163.1
B3 40998209 40998302
refGene_gn=DIS3L;refGene_tx=XM_023255162.1;refGene_gn=DIS3L;refGene_tx
=XM_023255163.1
B3 40998868 40999116
refGene_gn=DIS3L;refGene_tx=XM_023255162.1;refGene_gn=DIS3L;refGene_tx
=XM_023255163.1
B3 41000807 41000969
refGene_gn=DIS3L;refGene_tx=XM_023255162.1;refGene_gn=DIS3L;refGene_tx
=XM_023255163.1
B3 41002307 41002486
refGene_gn=DIS3L;refGene_tx=XM_023255162.1;refGene_gn=DIS3L;refGene_tx
=XM_023255163.1
B3 41004753 41004939
refGene_gn=DIS3L;refGene_tx=XM_023255162.1;refGene_gn=DIS3L;refGene_tx
=XM_023255163.1
B3 41006783 41007003 refGene_gn=DIS3L;refGene_tx=XM_023255163.1
B3 41010062 41010198 refGene_gn=DIS3L;refGene_tx=XM_023255163.1
B3 41011963 41012092 refGene_gn=DIS3L;refGene_tx=XM_023255163.1
B3 41023011 41023165 refGene_gn=DIS3L;refGene_tx=XM_023255163.1
B3 41024455 41024594 refGene_gn=DIS3L;refGene_tx=XM_023255163.1
B3 41196266 41196364
refGene_gn=MEGF11;refGene_tx=XM_019832204.2;refGene_gn=MEGF11;refGene_
tx=XM_019832205.2;refGene_gn=MEGF11;refGene_tx=XM_023255165.1;refGene_
gn=MEGF11;refGene_tx=XM_023255166.1;refGene_gn=MEGF11;refGene_tx=XM_02
3255167.1;refGene_gn=MEGF11;refGene_tx=XM_023255169.1;refGene_gn=MEGF1
1;refGene_tx=XM_023255170.1;refGene_gn=MEGF11;refGene_tx=XM_023255171.
1;refGene_gn=MEGF11;refGene_tx=XM_023255172.1;refGene_gn=MEGF11;refGen
e_tx=XM_023255173.1;refGene_gn=MEGF11;refGene_tx=XM_023255174.1;refGen
e_gn=MEGF11;refGene_tx=XM_023255175.1;refGene_gn=MEGF11;refGene_tx=XM_
023255176.1
B3 41200617 41200719
refGene_gn=MEGF11;refGene_tx=XM_019832204.2;refGene_gn=MEGF11;refGene_
tx=XM_019832205.2;refGene_gn=MEGF11;refGene_tx=XM_023255165.1;refGene_
gn=MEGF11;refGene_tx=XM_023255166.1;refGene_gn=MEGF11;refGene_tx=XM_02

3255167.1;refGene_gn=MEGF11;refGene_tx=XM_023255169.1;refGene_gn=MEGF11;refGene_tx=XM_023255170.1;refGene_gn=MEGF11;refGene_tx=XM_023255171.1;refGene_gn=MEGF11;refGene_tx=XM_023255172.1;refGene_gn=MEGF11;refGene_tx=XM_023255173.1;refGene_gn=MEGF11;refGene_tx=XM_023255174.1;refGene_gn=MEGF11;refGene_tx=XM_023255175.1;refGene_gn=MEGF11;refGene_tx=XM_023255176.1
B3 41205172 41205273
refGene_gn=MEGF11;refGene_tx=XM_019832204.2;refGene_gn=MEGF11;refGene_tx=XM_019832205.2;refGene_gn=MEGF11;refGene_tx=XM_023255165.1;refGene_gn=MEGF11;refGene_tx=XM_023255166.1;refGene_gn=MEGF11;refGene_tx=XM_023255167.1;refGene_gn=MEGF11;refGene_tx=XM_023255169.1;refGene_gn=MEGF11;refGene_tx=XM_023255170.1;refGene_gn=MEGF11;refGene_tx=XM_023255171.1;refGene_gn=MEGF11;refGene_tx=XM_023255172.1;refGene_gn=MEGF11;refGene_tx=XM_023255173.1;refGene_gn=MEGF11;refGene_tx=XM_023255174.1;refGene_gn=MEGF11;refGene_tx=XM_023255175.1;refGene_gn=MEGF11;refGene_tx=XM_023255176.1
B3 41246705 41246798
refGene_gn=MEGF11;refGene_tx=XM_019832204.2;refGene_gn=MEGF11;refGene_tx=XM_019832205.2;refGene_gn=MEGF11;refGene_tx=XM_023255165.1;refGene_gn=MEGF11;refGene_tx=XM_023255166.1;refGene_gn=MEGF11;refGene_tx=XM_023255167.1;refGene_gn=MEGF11;refGene_tx=XM_023255169.1;refGene_gn=MEGF11;refGene_tx=XM_023255170.1;refGene_gn=MEGF11;refGene_tx=XM_023255171.1;refGene_gn=MEGF11;refGene_tx=XM_023255172.1;refGene_gn=MEGF11;refGene_tx=XM_023255173.1;refGene_gn=MEGF11;refGene_tx=XM_023255174.1;refGene_gn=MEGF11;refGene_tx=XM_023255175.1;refGene_gn=MEGF11;refGene_tx=XM_023255176.1
B3 41350593 41350840
refGene_gn=MEGF11;refGene_tx=XM_019832204.2;refGene_gn=MEGF11;refGene_tx=XM_019832205.2;refGene_gn=MEGF11;refGene_tx=XM_023255165.1;refGene_gn=MEGF11;refGene_tx=XM_023255166.1;refGene_gn=MEGF11;refGene_tx=XM_023255167.1;refGene_gn=MEGF11;refGene_tx=XM_023255169.1;refGene_gn=MEGF11;refGene_tx=XM_023255170.1;refGene_gn=MEGF11;refGene_tx=XM_023255171.1;refGene_gn=MEGF11;refGene_tx=XM_023255172.1;refGene_gn=MEGF11;refGene_tx=XM_023255173.1;refGene_gn=MEGF11;refGene_tx=XM_023255174.1;refGene_gn=MEGF11;refGene_tx=XM_023255175.1;refGene_gn=MEGF11;refGene_tx=XM_023255176.1
B3 41352147 41352268
refGene_gn=MEGF11;refGene_tx=XM_019832204.2;refGene_gn=MEGF11;refGene_tx=XM_019832205.2;refGene_gn=MEGF11;refGene_tx=XM_023255165.1;refGene_gn=MEGF11;refGene_tx=XM_023255166.1;refGene_gn=MEGF11;refGene_tx=XM_023255167.1;refGene_gn=MEGF11;refGene_tx=XM_023255169.1;refGene_gn=MEGF11;refGene_tx=XM_023255170.1;refGene_gn=MEGF11;refGene_tx=XM_023255171.1;refGene_gn=MEGF11;refGene_tx=XM_023255172.1;refGene_gn=MEGF11;refGene_tx=XM_023255173.1;refGene_gn=MEGF11;refGene_tx=XM_023255174.1;refGene_gn=MEGF11;refGene_tx=XM_023255175.1;refGene_gn=MEGF11;refGene_tx=XM_023255176.1
B3 41360100 41360237
refGene_gn=MEGF11;refGene_tx=XM_019832204.2;refGene_gn=MEGF11;refGene_tx=XM_019832205.2;refGene_gn=MEGF11;refGene_tx=XM_023255165.1;refGene_gn=MEGF11;refGene_tx=XM_023255166.1;refGene_gn=MEGF11;refGene_tx=XM_02

3255167.1;refGene_gn=MEGF11;refGene_tx=XM_023255169.1;refGene_gn=MEGF11;refGene_tx=XM_023255170.1;refGene_gn=MEGF11;refGene_tx=XM_023255171.1;refGene_gn=MEGF11;refGene_tx=XM_023255172.1;refGene_gn=MEGF11;refGene_tx=XM_023255173.1;refGene_gn=MEGF11;refGene_tx=XM_023255174.1;refGene_gn=MEGF11;refGene_tx=XM_023255175.1;refGene_gn=MEGF11;refGene_tx=XM_023255176.1
B3 41365185 41365398
refGene_gn=MEGF11;refGene_tx=XM_019832204.2;refGene_gn=MEGF11;refGene_tx=XM_019832205.2;refGene_gn=MEGF11;refGene_tx=XM_023255165.1;refGene_gn=MEGF11;refGene_tx=XM_023255166.1;refGene_gn=MEGF11;refGene_tx=XM_023255167.1;refGene_gn=MEGF11;refGene_tx=XM_023255169.1;refGene_gn=MEGF11;refGene_tx=XM_023255170.1;refGene_gn=MEGF11;refGene_tx=XM_023255171.1;refGene_gn=MEGF11;refGene_tx=XM_023255172.1;refGene_gn=MEGF11;refGene_tx=XM_023255173.1;refGene_gn=MEGF11;refGene_tx=XM_023255174.1;refGene_gn=MEGF11;refGene_tx=XM_023255175.1;refGene_gn=MEGF11;refGene_tx=XM_023255176.1
B3 41372445 41372620
refGene_gn=MEGF11;refGene_tx=XM_019832204.2;refGene_gn=MEGF11;refGene_tx=XM_019832205.2;refGene_gn=MEGF11;refGene_tx=XM_023255165.1;refGene_gn=MEGF11;refGene_tx=XM_023255166.1;refGene_gn=MEGF11;refGene_tx=XM_023255167.1;refGene_gn=MEGF11;refGene_tx=XM_023255169.1;refGene_gn=MEGF11;refGene_tx=XM_023255170.1;refGene_gn=MEGF11;refGene_tx=XM_023255171.1;refGene_gn=MEGF11;refGene_tx=XM_023255172.1;refGene_gn=MEGF11;refGene_tx=XM_023255173.1;refGene_gn=MEGF11;refGene_tx=XM_023255174.1;refGene_gn=MEGF11;refGene_tx=XM_023255175.1;refGene_gn=MEGF11;refGene_tx=XM_023255176.1
B3 41396035 41396156
refGene_gn=MEGF11;refGene_tx=XM_019832204.2;refGene_gn=MEGF11;refGene_tx=XM_019832205.2;refGene_gn=MEGF11;refGene_tx=XM_023255165.1;refGene_gn=MEGF11;refGene_tx=XM_023255166.1;refGene_gn=MEGF11;refGene_tx=XM_023255167.1;refGene_gn=MEGF11;refGene_tx=XM_023255169.1;refGene_gn=MEGF11;refGene_tx=XM_023255170.1;refGene_gn=MEGF11;refGene_tx=XM_023255171.1;refGene_gn=MEGF11;refGene_tx=XM_023255172.1;refGene_gn=MEGF11;refGene_tx=XM_023255173.1;refGene_gn=MEGF11;refGene_tx=XM_023255174.1;refGene_gn=MEGF11;refGene_tx=XM_023255175.1;refGene_gn=MEGF11;refGene_tx=XM_023255176.1
B3 41397137 41397301
refGene_gn=MEGF11;refGene_tx=XM_019832204.2;refGene_gn=MEGF11;refGene_tx=XM_019832205.2;refGene_gn=MEGF11;refGene_tx=XM_023255165.1;refGene_gn=MEGF11;refGene_tx=XM_023255166.1;refGene_gn=MEGF11;refGene_tx=XM_023255167.1;refGene_gn=MEGF11;refGene_tx=XM_023255169.1;refGene_gn=MEGF11;refGene_tx=XM_023255170.1;refGene_gn=MEGF11;refGene_tx=XM_023255171.1;refGene_gn=MEGF11;refGene_tx=XM_023255172.1;refGene_gn=MEGF11;refGene_tx=XM_023255173.1;refGene_gn=MEGF11;refGene_tx=XM_023255174.1;refGene_gn=MEGF11;refGene_tx=XM_023255175.1;refGene_gn=MEGF11;refGene_tx=XM_023255176.1
B3 41398368 41398471
refGene_gn=MEGF11;refGene_tx=XM_019832204.2;refGene_gn=MEGF11;refGene_tx=XM_019832205.2;refGene_gn=MEGF11;refGene_tx=XM_023255165.1;refGene_gn=MEGF11;refGene_tx=XM_023255166.1;refGene_gn=MEGF11;refGene_tx=XM_02

3255167.1;refGene_gn=MEGF11;refGene_tx=XM_023255169.1;refGene_gn=MEGF1
1;refGene_tx=XM_023255170.1;refGene_gn=MEGF11;refGene_tx=XM_023255171.
1;refGene_gn=MEGF11;refGene_tx=XM_023255172.1;refGene_gn=MEGF11;refGen
e_tx=XM_023255173.1;refGene_gn=MEGF11;refGene_tx=XM_023255174.1;refGen
e_gn=MEGF11;refGene_tx=XM_023255175.1;refGene_gn=MEGF11;refGene_tx=XM_
023255176.1
B3 41403690 41403837
refGene_gn=MEGF11;refGene_tx=XM_019832204.2;refGene_gn=MEGF11;refGene_
tx=XM_019832205.2;refGene_gn=MEGF11;refGene_tx=XM_023255165.1;refGene_
gn=MEGF11;refGene_tx=XM_023255166.1;refGene_gn=MEGF11;refGene_tx=XM_02
3255167.1;refGene_gn=MEGF11;refGene_tx=XM_023255169.1;refGene_gn=MEGF1
1;refGene_tx=XM_023255170.1;refGene_gn=MEGF11;refGene_tx=XM_023255171.
1;refGene_gn=MEGF11;refGene_tx=XM_023255172.1;refGene_gn=MEGF11;refGen
e_tx=XM_023255173.1;refGene_gn=MEGF11;refGene_tx=XM_023255174.1;refGen
e_gn=MEGF11;refGene_tx=XM_023255175.1;refGene_gn=MEGF11;refGene_tx=XM_
023255176.1
B3 41404185 41404320
refGene_gn=MEGF11;refGene_tx=XM_019832204.2;refGene_gn=MEGF11;refGene_
tx=XM_019832205.2;refGene_gn=MEGF11;refGene_tx=XM_023255165.1;refGene_
gn=MEGF11;refGene_tx=XM_023255166.1;refGene_gn=MEGF11;refGene_tx=XM_02
3255169.1;refGene_gn=MEGF11;refGene_tx=XM_023255170.1;refGene_gn=MEGF1
1;refGene_tx=XM_023255171.1;refGene_gn=MEGF11;refGene_tx=XM_023255172.
1;refGene_gn=MEGF11;refGene_tx=XM_023255173.1;refGene_gn=MEGF11;refGen
e_tx=XM_023255174.1;refGene_gn=MEGF11;refGene_tx=XM_023255175.1;refGen
e_gn=MEGF11;refGene_tx=XM_023255176.1
B3 41409170 41409299
refGene_gn=MEGF11;refGene_tx=XM_019832204.2;refGene_gn=MEGF11;refGene_
tx=XM_019832205.2;refGene_gn=MEGF11;refGene_tx=XM_023255165.1;refGene_
gn=MEGF11;refGene_tx=XM_023255166.1;refGene_gn=MEGF11;refGene_tx=XM_02
3255167.1;refGene_gn=MEGF11;refGene_tx=XM_023255169.1;refGene_gn=MEGF1
1;refGene_tx=XM_023255170.1;refGene_gn=MEGF11;refGene_tx=XM_023255171.
1;refGene_gn=MEGF11;refGene_tx=XM_023255172.1;refGene_gn=MEGF11;refGen
e_tx=XM_023255173.1;refGene_gn=MEGF11;refGene_tx=XM_023255174.1;refGen
e_gn=MEGF11;refGene_tx=XM_023255175.1;refGene_gn=MEGF11;refGene_tx=XM_
023255176.1
B3 41410316 41410445
refGene_gn=MEGF11;refGene_tx=XM_019832204.2;refGene_gn=MEGF11;refGene_
tx=XM_019832205.2;refGene_gn=MEGF11;refGene_tx=XM_023255165.1;refGene_
gn=MEGF11;refGene_tx=XM_023255166.1;refGene_gn=MEGF11;refGene_tx=XM_02
3255167.1;refGene_gn=MEGF11;refGene_tx=XM_023255169.1;refGene_gn=MEGF1
1;refGene_tx=XM_023255170.1;refGene_gn=MEGF11;refGene_tx=XM_023255171.
1;refGene_gn=MEGF11;refGene_tx=XM_023255172.1;refGene_gn=MEGF11;refGen
e_tx=XM_023255173.1;refGene_gn=MEGF11;refGene_tx=XM_023255174.1;refGen
e_gn=MEGF11;refGene_tx=XM_023255175.1;refGene_gn=MEGF11;refGene_tx=XM_
023255176.1
B3 41410945 41411074
refGene_gn=MEGF11;refGene_tx=XM_019832204.2;refGene_gn=MEGF11;refGene_
tx=XM_019832205.2;refGene_gn=MEGF11;refGene_tx=XM_023255165.1;refGene_
gn=MEGF11;refGene_tx=XM_023255166.1;refGene_gn=MEGF11;refGene_tx=XM_02
3255167.1;refGene_gn=MEGF11;refGene_tx=XM_023255169.1;refGene_gn=MEGF1

1;refGene_tx=XM_023255170.1;refGene_gn=MEGF11;refGene_tx=XM_023255171.
1;refGene_gn=MEGF11;refGene_tx=XM_023255172.1;refGene_gn=MEGF11;refGen
e_tx=XM_023255173.1;refGene_gn=MEGF11;refGene_tx=XM_023255174.1;refGen
e_gn=MEGF11;refGene_tx=XM_023255175.1;refGene_gn=MEGF11;refGene_tx=XM_
023255176.1
B3 41411644 41411773
refGene_gn=MEGF11;refGene_tx=XM_019832204.2;refGene_gn=MEGF11;refGene_
tx=XM_019832205.2;refGene_gn=MEGF11;refGene_tx=XM_023255165.1;refGene_
gn=MEGF11;refGene_tx=XM_023255166.1;refGene_gn=MEGF11;refGene_tx=XM_02
3255167.1;refGene_gn=MEGF11;refGene_tx=XM_023255169.1;refGene_gn=MEGF1
1;refGene_tx=XM_023255170.1;refGene_gn=MEGF11;refGene_tx=XM_023255171.
1;refGene_gn=MEGF11;refGene_tx=XM_023255172.1;refGene_gn=MEGF11;refGen
e_tx=XM_023255173.1;refGene_gn=MEGF11;refGene_tx=XM_023255174.1;refGen
e_gn=MEGF11;refGene_tx=XM_023255175.1;refGene_gn=MEGF11;refGene_tx=XM_
023255176.1
B3 41413287 41413524
refGene_gn=MEGF11;refGene_tx=XM_019832204.2;refGene_gn=MEGF11;refGene_
tx=XM_019832205.2;refGene_gn=MEGF11;refGene_tx=XM_023255165.1;refGene_
gn=MEGF11;refGene_tx=XM_023255166.1;refGene_gn=MEGF11;refGene_tx=XM_02
3255167.1;refGene_gn=MEGF11;refGene_tx=XM_023255169.1;refGene_gn=MEGF1
1;refGene_tx=XM_023255170.1;refGene_gn=MEGF11;refGene_tx=XM_023255171.
1;refGene_gn=MEGF11;refGene_tx=XM_023255172.1;refGene_gn=MEGF11;refGen
e_tx=XM_023255173.1;refGene_gn=MEGF11;refGene_tx=XM_023255174.1;refGen
e_gn=MEGF11;refGene_tx=XM_023255175.1;refGene_gn=MEGF11;refGene_tx=XM_
023255176.1
B3 41415115 41415234
refGene_gn=MEGF11;refGene_tx=XM_019832204.2;refGene_gn=MEGF11;refGene_
tx=XM_019832205.2;refGene_gn=MEGF11;refGene_tx=XM_023255165.1;refGene_
gn=MEGF11;refGene_tx=XM_023255166.1;refGene_gn=MEGF11;refGene_tx=XM_02
3255167.1;refGene_gn=MEGF11;refGene_tx=XM_023255169.1;refGene_gn=MEGF1
1;refGene_tx=XM_023255170.1;refGene_gn=MEGF11;refGene_tx=XM_023255171.
1;refGene_gn=MEGF11;refGene_tx=XM_023255173.1;refGene_gn=MEGF11;refGen
e_tx=XM_023255174.1;refGene_gn=MEGF11;refGene_tx=XM_023255175.1
B3 41417111 41417178
refGene_gn=MEGF11;refGene_tx=XM_019832204.2;refGene_gn=MEGF11;refGene_
tx=XM_019832205.2;refGene_gn=MEGF11;refGene_tx=XM_023255165.1;refGene_
gn=MEGF11;refGene_tx=XM_023255166.1;refGene_gn=MEGF11;refGene_tx=XM_02
3255167.1;refGene_gn=MEGF11;refGene_tx=XM_023255170.1;refGene_gn=MEGF1
1;refGene_tx=XM_023255171.1
B3 41417452 41417559
refGene_gn=MEGF11;refGene_tx=XM_019832204.2;refGene_gn=MEGF11;refGene_
tx=XM_019832205.2;refGene_gn=MEGF11;refGene_tx=XM_023255165.1;refGene_
gn=MEGF11;refGene_tx=XM_023255167.1;refGene_gn=MEGF11;refGene_tx=XM_02
3255170.1;refGene_gn=MEGF11;refGene_tx=XM_023255171.1
B3 41417782 41417884
refGene_gn=MEGF11;refGene_tx=XM_019832204.2;refGene_gn=MEGF11;refGene_
tx=XM_019832205.2;refGene_gn=MEGF11;refGene_tx=XM_023255165.1;refGene_
gn=MEGF11;refGene_tx=XM_023255166.1;refGene_gn=MEGF11;refGene_tx=XM_02
3255167.1;refGene_gn=MEGF11;refGene_tx=XM_023255169.1;refGene_gn=MEGF1
1;refGene_tx=XM_023255170.1;refGene_gn=MEGF11;refGene_tx=XM_023255171.

1;refGene_gn=MEGF11;refGene_tx=XM_023255173.1
B3 41420320 41420377
refGene_gn=MEGF11;refGene_tx=XM_019832204.2;refGene_gn=MEGF11;refGene_tx=XM_019832205.2;refGene_gn=MEGF11;refGene_tx=XM_023255165.1;refGene_gn=MEGF11;refGene_tx=XM_023255166.1;refGene_gn=MEGF11;refGene_tx=XM_023255167.1;refGene_gn=MEGF11;refGene_tx=XM_023255169.1;refGene_gn=MEGF11;refGene_tx=XM_023255170.1;refGene_gn=MEGF11;refGene_tx=XM_023255171.1;refGene_gn=MEGF11;refGene_tx=XM_023255172.1;refGene_gn=MEGF11;refGene_tx=XM_023255173.1;refGene_gn=MEGF11;refGene_tx=XM_023255174.1;refGene_gn=MEGF11;refGene_tx=XM_023255175.1;refGene_gn=MEGF11;refGene_tx=XM_023255176.1
B3 41421237 41421306
refGene_gn=MEGF11;refGene_tx=XM_023255170.1;refGene_gn=MEGF11;refGene_tx=XM_023255173.1;refGene_gn=MEGF11;refGene_tx=XM_023255174.1;refGene_gn=MEGF11;refGene_tx=XM_023255176.1
B3 41427067 41427274
refGene_gn=MEGF11;refGene_tx=XM_019832204.2;refGene_gn=MEGF11;refGene_tx=XM_019832205.2;refGene_gn=MEGF11;refGene_tx=XM_023255165.1;refGene_gn=MEGF11;refGene_tx=XM_023255166.1;refGene_gn=MEGF11;refGene_tx=XM_023255167.1;refGene_gn=MEGF11;refGene_tx=XM_023255169.1;refGene_gn=MEGF11;refGene_tx=XM_023255171.1;refGene_gn=MEGF11;refGene_tx=XM_023255172.1;refGene_gn=MEGF11;refGene_tx=XM_023255175.1
B3 41427939 41428100
refGene_gn=MEGF11;refGene_tx=XM_019832204.2;refGene_gn=MEGF11;refGene_tx=XM_019832205.2;refGene_gn=MEGF11;refGene_tx=XM_023255165.1;refGene_gn=MEGF11;refGene_tx=XM_023255166.1;refGene_gn=MEGF11;refGene_tx=XM_023255167.1;refGene_gn=MEGF11;refGene_tx=XM_023255169.1;refGene_gn=MEGF11;refGene_tx=XM_023255172.1
B3 41437147 41437287 refGene_gn=RAB11A;refGene_tx=XM_003987022.5
B3 41445722 41445803 refGene_gn=RAB11A;refGene_tx=XM_003987022.5
B3 41446811 41447005 refGene_gn=RAB11A;refGene_tx=XM_003987022.5
B3 41447256 41447452 refGene_gn=RAB11A;refGene_tx=XM_003987022.5
B3 41454088 41454128 refGene_gn=RAB11A;refGene_tx=XM_003987022.5
B3 41564182 41564493
refGene_gn=DENND4A;refGene_tx=XM_011282934.3;refGene_gn=DENND4A;refGene_tx=XM_019832212.2;refGene_gn=DENND4A;refGene_tx=XM_019832213.2;refGene_gn=DENND4A;refGene_tx=XM_019832214.2;refGene_gn=DENND4A;refGene_tx=XM_019832216.2;refGene_gn=DENND4A;refGene_tx=XM_019832217.2;refGene_gn=DENND4A;refGene_tx=XM_023255178.1
B3 41570036 41570286
refGene_gn=DENND4A;refGene_tx=XM_011282934.3;refGene_gn=DENND4A;refGene_tx=XM_019832212.2;refGene_gn=DENND4A;refGene_tx=XM_019832213.2;refGene_gn=DENND4A;refGene_tx=XM_019832214.2;refGene_gn=DENND4A;refGene_tx=XM_019832216.2;refGene_gn=DENND4A;refGene_tx=XM_019832217.2;refGene_gn=DENND4A;refGene_tx=XM_023255178.1
B3 41580519 41580589
refGene_gn=DENND4A;refGene_tx=XM_011282934.3;refGene_gn=DENND4A;refGene_tx=XM_019832212.2;refGene_gn=DENND4A;refGene_tx=XM_019832213.2;refGene_gn=DENND4A;refGene_tx=XM_019832214.2;refGene_gn=DENND4A;refGene_tx=XM_019832216.2;refGene_gn=DENND4A;refGene_tx=XM_019832217.2;refGene_gn

=DENND4A;refGene_tx=XM_023255178.1
B3 41581146 41581316
refGene_gn=DENND4A;refGene_tx=XM_011282934.3;refGene_gn=DENND4A;refGene_tx=XM_019832212.2;refGene_gn=DENND4A;refGene_tx=XM_019832213.2;refGene_gn=DENND4A;refGene_tx=XM_019832214.2;refGene_gn=DENND4A;refGene_tx=XM_019832216.2;refGene_gn=DENND4A;refGene_tx=XM_019832217.2;refGene_gn=DENND4A;refGene_tx=XM_023255178.1
B3 41582382 41582624
refGene_gn=DENND4A;refGene_tx=XM_011282934.3;refGene_gn=DENND4A;refGene_tx=XM_019832212.2;refGene_gn=DENND4A;refGene_tx=XM_019832213.2;refGene_gn=DENND4A;refGene_tx=XM_019832214.2;refGene_gn=DENND4A;refGene_tx=XM_019832216.2;refGene_gn=DENND4A;refGene_tx=XM_019832217.2;refGene_gn=DENND4A;refGene_tx=XM_023255178.1
B3 41585137 41585204
refGene_gn=DENND4A;refGene_tx=XM_011282934.3;refGene_gn=DENND4A;refGene_tx=XM_019832212.2;refGene_gn=DENND4A;refGene_tx=XM_019832213.2;refGene_gn=DENND4A;refGene_tx=XM_019832214.2;refGene_gn=DENND4A;refGene_tx=XM_019832216.2;refGene_gn=DENND4A;refGene_tx=XM_019832217.2;refGene_gn=DENND4A;refGene_tx=XM_023255178.1
B3 41586594 41586653
refGene_gn=DENND4A;refGene_tx=XM_011282934.3;refGene_gn=DENND4A;refGene_tx=XM_019832212.2;refGene_gn=DENND4A;refGene_tx=XM_019832213.2;refGene_gn=DENND4A;refGene_tx=XM_019832214.2;refGene_gn=DENND4A;refGene_tx=XM_019832216.2;refGene_gn=DENND4A;refGene_tx=XM_019832217.2;refGene_gn=DENND4A;refGene_tx=XM_023255178.1
B3 41588783 41588928
refGene_gn=DENND4A;refGene_tx=XM_011282934.3;refGene_gn=DENND4A;refGene_tx=XM_019832212.2;refGene_gn=DENND4A;refGene_tx=XM_019832213.2;refGene_gn=DENND4A;refGene_tx=XM_019832214.2;refGene_gn=DENND4A;refGene_tx=XM_019832216.2;refGene_gn=DENND4A;refGene_tx=XM_019832217.2;refGene_gn=DENND4A;refGene_tx=XM_023255178.1
B3 41589219 41589395
refGene_gn=DENND4A;refGene_tx=XM_011282934.3;refGene_gn=DENND4A;refGene_tx=XM_019832212.2;refGene_gn=DENND4A;refGene_tx=XM_019832213.2;refGene_gn=DENND4A;refGene_tx=XM_019832214.2;refGene_gn=DENND4A;refGene_tx=XM_019832216.2;refGene_gn=DENND4A;refGene_tx=XM_019832217.2;refGene_gn=DENND4A;refGene_tx=XM_023255178.1
B3 41593281 41593382
refGene_gn=DENND4A;refGene_tx=XM_011282934.3;refGene_gn=DENND4A;refGene_tx=XM_019832212.2;refGene_gn=DENND4A;refGene_tx=XM_019832213.2;refGene_gn=DENND4A;refGene_tx=XM_019832214.2;refGene_gn=DENND4A;refGene_tx=XM_019832216.2;refGene_gn=DENND4A;refGene_tx=XM_019832217.2;refGene_gn=DENND4A;refGene_tx=XM_023255178.1
B3 41598320 41598539
refGene_gn=DENND4A;refGene_tx=XM_011282934.3;refGene_gn=DENND4A;refGene_tx=XM_019832212.2;refGene_gn=DENND4A;refGene_tx=XM_019832213.2;refGene_gn=DENND4A;refGene_tx=XM_019832214.2;refGene_gn=DENND4A;refGene_tx=XM_019832216.2;refGene_gn=DENND4A;refGene_tx=XM_019832217.2;refGene_gn=DENND4A;refGene_tx=XM_023255178.1
B3 41600200 41600346

refGene_gn=DENND4A;refGene_tx=XM_011282934.3;refGene_gn=DENND4A;refGene_tx=XM_019832212.2;refGene_gn=DENND4A;refGene_tx=XM_019832213.2;refGene_gn=DENND4A;refGene_tx=XM_019832214.2;refGene_gn=DENND4A;refGene_tx=XM_019832216.2;refGene_gn=DENND4A;refGene_tx=XM_019832217.2;refGene_gn=DENND4A;refGene_tx=XM_023255178.1
B3 41615466 41615600
refGene_gn=DENND4A;refGene_tx=XM_011282934.3;refGene_gn=DENND4A;refGene_tx=XM_019832212.2;refGene_gn=DENND4A;refGene_tx=XM_019832213.2;refGene_gn=DENND4A;refGene_tx=XM_019832214.2;refGene_gn=DENND4A;refGene_tx=XM_019832216.2;refGene_gn=DENND4A;refGene_tx=XM_019832217.2;refGene_gn=DENND4A;refGene_tx=XM_023255178.1
B3 41619120 41619256
refGene_gn=DENND4A;refGene_tx=XM_011282934.3;refGene_gn=DENND4A;refGene_tx=XM_019832212.2;refGene_gn=DENND4A;refGene_tx=XM_019832213.2;refGene_gn=DENND4A;refGene_tx=XM_019832214.2;refGene_gn=DENND4A;refGene_tx=XM_019832216.2;refGene_gn=DENND4A;refGene_tx=XM_019832217.2;refGene_gn=DENND4A;refGene_tx=XM_023255178.1
B3 41619614 41619821
refGene_gn=DENND4A;refGene_tx=XM_011282934.3;refGene_gn=DENND4A;refGene_tx=XM_019832212.2;refGene_gn=DENND4A;refGene_tx=XM_019832213.2;refGene_gn=DENND4A;refGene_tx=XM_019832214.2;refGene_gn=DENND4A;refGene_tx=XM_019832216.2;refGene_gn=DENND4A;refGene_tx=XM_019832217.2;refGene_gn=DENND4A;refGene_tx=XM_023255178.1
B3 41620178 41620307
refGene_gn=DENND4A;refGene_tx=XM_011282934.3;refGene_gn=DENND4A;refGene_tx=XM_019832212.2;refGene_gn=DENND4A;refGene_tx=XM_019832213.2;refGene_gn=DENND4A;refGene_tx=XM_019832214.2;refGene_gn=DENND4A;refGene_tx=XM_019832216.2;refGene_gn=DENND4A;refGene_tx=XM_019832217.2;refGene_gn=DENND4A;refGene_tx=XM_023255178.1
B3 41620567 41620709
refGene_gn=DENND4A;refGene_tx=XM_011282934.3;refGene_gn=DENND4A;refGene_tx=XM_019832212.2;refGene_gn=DENND4A;refGene_tx=XM_019832213.2;refGene_gn=DENND4A;refGene_tx=XM_019832214.2;refGene_gn=DENND4A;refGene_tx=XM_019832216.2;refGene_gn=DENND4A;refGene_tx=XM_019832217.2;refGene_gn=DENND4A;refGene_tx=XM_023255178.1
B3 41621109 41621241
refGene_gn=DENND4A;refGene_tx=XM_011282934.3;refGene_gn=DENND4A;refGene_tx=XM_019832212.2;refGene_gn=DENND4A;refGene_tx=XM_019832213.2;refGene_gn=DENND4A;refGene_tx=XM_019832214.2;refGene_gn=DENND4A;refGene_tx=XM_023255178.1
B3 41623408 41623525
refGene_gn=DENND4A;refGene_tx=XM_011282934.3;refGene_gn=DENND4A;refGene_tx=XM_019832212.2;refGene_gn=DENND4A;refGene_tx=XM_019832213.2;refGene_gn=DENND4A;refGene_tx=XM_019832214.2;refGene_gn=DENND4A;refGene_tx=XM_019832216.2;refGene_gn=DENND4A;refGene_tx=XM_019832217.2
B3 41624374 41624506
refGene_gn=DENND4A;refGene_tx=XM_011282934.3;refGene_gn=DENND4A;refGene_tx=XM_019832212.2;refGene_gn=DENND4A;refGene_tx=XM_019832213.2;refGene_gn=DENND4A;refGene_tx=XM_019832214.2;refGene_gn=DENND4A;refGene_tx=XM_019832216.2;refGene_gn=DENND4A;refGene_tx=XM_019832217.2;refGene_gn

=DENND4A;refGene_tx=XM_023255178.1
B3 41628626 41629726
refGene_gn=DENND4A;refGene_tx=XM_011282934.3;refGene_gn=DENND4A;refGene_tx=XM_019832212.2;refGene_gn=DENND4A;refGene_tx=XM_019832213.2;refGene_gn=DENND4A;refGene_tx=XM_019832214.2;refGene_gn=DENND4A;refGene_tx=XM_019832216.2;refGene_gn=DENND4A;refGene_tx=XM_019832217.2;refGene_gn=DENND4A;refGene_tx=XM_023255178.1
B3 41634540 41634730
refGene_gn=DENND4A;refGene_tx=XM_011282934.3;refGene_gn=DENND4A;refGene_tx=XM_019832212.2;refGene_gn=DENND4A;refGene_tx=XM_019832213.2;refGene_gn=DENND4A;refGene_tx=XM_019832214.2;refGene_gn=DENND4A;refGene_tx=XM_019832216.2;refGene_gn=DENND4A;refGene_tx=XM_019832217.2;refGene_gn=DENND4A;refGene_tx=XM_023255178.1
B3 41639210 41639305
refGene_gn=DENND4A;refGene_tx=XM_011282934.3;refGene_gn=DENND4A;refGene_tx=XM_019832212.2;refGene_gn=DENND4A;refGene_tx=XM_019832213.2;refGene_gn=DENND4A;refGene_tx=XM_019832214.2;refGene_gn=DENND4A;refGene_tx=XM_019832216.2;refGene_gn=DENND4A;refGene_tx=XM_019832217.2;refGene_gn=DENND4A;refGene_tx=XM_023255178.1
B3 41641234 41641410
refGene_gn=DENND4A;refGene_tx=XM_011282934.3;refGene_gn=DENND4A;refGene_tx=XM_019832212.2;refGene_gn=DENND4A;refGene_tx=XM_019832213.2;refGene_gn=DENND4A;refGene_tx=XM_019832214.2;refGene_gn=DENND4A;refGene_tx=XM_019832216.2;refGene_gn=DENND4A;refGene_tx=XM_019832217.2;refGene_gn=DENND4A;refGene_tx=XM_023255178.1
B3 41641504 41641651
refGene_gn=DENND4A;refGene_tx=XM_011282934.3;refGene_gn=DENND4A;refGene_tx=XM_019832212.2;refGene_gn=DENND4A;refGene_tx=XM_019832213.2;refGene_gn=DENND4A;refGene_tx=XM_019832214.2;refGene_gn=DENND4A;refGene_tx=XM_019832216.2;refGene_gn=DENND4A;refGene_tx=XM_019832217.2;refGene_gn=DENND4A;refGene_tx=XM_023255178.1
B3 41644035 41644234
refGene_gn=DENND4A;refGene_tx=XM_011282934.3;refGene_gn=DENND4A;refGene_tx=XM_019832212.2;refGene_gn=DENND4A;refGene_tx=XM_019832213.2;refGene_gn=DENND4A;refGene_tx=XM_019832214.2;refGene_gn=DENND4A;refGene_tx=XM_019832216.2;refGene_gn=DENND4A;refGene_tx=XM_019832217.2;refGene_gn=DENND4A;refGene_tx=XM_023255178.1
B3 41644482 41644737
refGene_gn=DENND4A;refGene_tx=XM_011282934.3;refGene_gn=DENND4A;refGene_tx=XM_019832212.2;refGene_gn=DENND4A;refGene_tx=XM_019832213.2;refGene_gn=DENND4A;refGene_tx=XM_019832214.2;refGene_gn=DENND4A;refGene_tx=XM_019832216.2;refGene_gn=DENND4A;refGene_tx=XM_019832217.2;refGene_gn=DENND4A;refGene_tx=XM_023255178.1
B3 41645664 41645782
refGene_gn=DENND4A;refGene_tx=XM_011282934.3;refGene_gn=DENND4A;refGene_tx=XM_019832212.2;refGene_gn=DENND4A;refGene_tx=XM_019832213.2;refGene_gn=DENND4A;refGene_tx=XM_019832214.2;refGene_gn=DENND4A;refGene_tx=XM_019832216.2;refGene_gn=DENND4A;refGene_tx=XM_019832217.2;refGene_gn=DENND4A;refGene_tx=XM_023255178.1
B3 41645899 41645926

refGene_gn=DENND4A;refGene_tx=XM_019832212.2;refGene_gn=DENND4A;refGene_tx=XM_019832213.2;refGene_gn=DENND4A;refGene_tx=XM_019832216.2;refGene_gn=DENND4A;refGene_tx=XM_023255178.1
B3 41646388 41646551
refGene_gn=DENND4A;refGene_tx=XM_011282934.3;refGene_gn=DENND4A;refGene_tx=XM_019832212.2;refGene_gn=DENND4A;refGene_tx=XM_019832213.2;refGene_gn=DENND4A;refGene_tx=XM_019832214.2;refGene_gn=DENND4A;refGene_tx=XM_019832216.2;refGene_gn=DENND4A;refGene_tx=XM_019832217.2;refGene_gn=DENND4A;refGene_tx=XM_023255178.1
B3 41646713 41646778
refGene_gn=DENND4A;refGene_tx=XM_011282934.3;refGene_gn=DENND4A;refGene_tx=XM_019832212.2;refGene_gn=DENND4A;refGene_tx=XM_019832213.2;refGene_gn=DENND4A;refGene_tx=XM_019832214.2;refGene_gn=DENND4A;refGene_tx=XM_019832216.2;refGene_gn=DENND4A;refGene_tx=XM_019832217.2;refGene_gn=DENND4A;refGene_tx=XM_023255178.1
B3 41655636 41655773
refGene_gn=DENND4A;refGene_tx=XM_011282934.3;refGene_gn=DENND4A;refGene_tx=XM_019832212.2;refGene_gn=DENND4A;refGene_tx=XM_019832213.2;refGene_gn=DENND4A;refGene_tx=XM_019832214.2;refGene_gn=DENND4A;refGene_tx=XM_019832216.2;refGene_gn=DENND4A;refGene_tx=XM_019832217.2;refGene_gn=DENND4A;refGene_tx=XM_023255178.1
B3 41662471 41662721
refGene_gn=SLC24A1;refGene_tx=XM_019832218.2;refGene_gn=SLC24A1;refGene_tx=XM_019832219.2;refGene_gn=SLC24A1;refGene_tx=XM_019832220.2;refGene_gn=SLC24A1;refGene_tx=XM_019832221.2
B3 41663669 41663836
refGene_gn=SLC24A1;refGene_tx=XM_019832218.2;refGene_gn=SLC24A1;refGene_tx=XM_019832219.2;refGene_gn=SLC24A1;refGene_tx=XM_019832220.2;refGene_gn=SLC24A1;refGene_tx=XM_019832221.2
B3 41664709 41664799
refGene_gn=SLC24A1;refGene_tx=XM_019832218.2;refGene_gn=SLC24A1;refGene_tx=XM_019832219.2;refGene_gn=SLC24A1;refGene_tx=XM_019832220.2
B3 41665521 41666085
refGene_gn=SLC24A1;refGene_tx=XM_019832218.2;refGene_gn=SLC24A1;refGene_tx=XM_019832219.2;refGene_gn=SLC24A1;refGene_tx=XM_019832220.2;refGene_gn=SLC24A1;refGene_tx=XM_019832221.2
B3 41669422 41669514
refGene_gn=SLC24A1;refGene_tx=XM_019832218.2;refGene_gn=SLC24A1;refGene_tx=XM_019832219.2;refGene_gn=SLC24A1;refGene_tx=XM_019832220.2;refGene_gn=SLC24A1;refGene_tx=XM_019832221.2
B3 41670601 41670685
refGene_gn=SLC24A1;refGene_tx=XM_019832218.2;refGene_gn=SLC24A1;refGene_tx=XM_019832219.2;refGene_gn=SLC24A1;refGene_tx=XM_019832220.2;refGene_gn=SLC24A1;refGene_tx=XM_019832221.2
B3 41674637 41674746
refGene_gn=SLC24A1;refGene_tx=XM_019832218.2;refGene_gn=SLC24A1;refGene_tx=XM_019832219.2;refGene_gn=SLC24A1;refGene_tx=XM_019832220.2;refGene_gn=SLC24A1;refGene_tx=XM_019832221.2
B3 41676079 41676133
refGene_gn=SLC24A1;refGene_tx=XM_019832218.2;refGene_gn=SLC24A1;refGen

e_tx=XM_019832219.2;refGene_gn=SLC24A1;refGene_tx=XM_019832221.2
B3 41687170 41689054
refGene_gn=SLC24A1;refGene_tx=XM_019832218.2;refGene_gn=SLC24A1;refGene_tx=XM_019832219.2;refGene_gn=SLC24A1;refGene_tx=XM_019832220.2;refGene_gn=SLC24A1;refGene_tx=XM_019832221.2
B3 41706981 41707203
refGene_gn=INTS14;refGene_tx=XM_003987027.4;refGene_gn=INTS14;refGene_tx=XM_006932455.4
B3 41709432 41709540
refGene_gn=INTS14;refGene_tx=XM_003987027.4;refGene_gn=INTS14;refGene_tx=XM_006932455.4
B3 41713493 41713649
refGene_gn=INTS14;refGene_tx=XM_003987027.4;refGene_gn=INTS14;refGene_tx=XM_006932455.4
B3 41714429 41714548
refGene_gn=INTS14;refGene_tx=XM_003987027.4;refGene_gn=INTS14;refGene_tx=XM_006932455.4
B3 41714988 41715131
refGene_gn=INTS14;refGene_tx=XM_003987027.4;refGene_gn=INTS14;refGene_tx=XM_006932455.4
B3 41717107 41717200
refGene_gn=INTS14;refGene_tx=XM_003987027.4;refGene_gn=INTS14;refGene_tx=XM_006932455.4
B3 41718412 41718557
refGene_gn=INTS14;refGene_tx=XM_003987027.4;refGene_gn=INTS14;refGene_tx=XM_006932455.4
B3 41719797 41719931
refGene_gn=INTS14;refGene_tx=XM_003987027.4;refGene_gn=INTS14;refGene_tx=XM_006932455.4
B3 41729102 41729221
refGene_gn=INTS14;refGene_tx=XM_003987027.4;refGene_gn=INTS14;refGene_tx=XM_006932455.4
B3 41731410 41731476
refGene_gn=INTS14;refGene_tx=XM_003987027.4;refGene_gn=INTS14;refGene_tx=XM_006932455.4
B3 41733161 41733413
refGene_gn=INTS14;refGene_tx=XM_003987027.4;refGene_gn=INTS14;refGene_tx=XM_006932455.4
B3 41736103 41736180 refGene_gn=HACD3;refGene_tx=XM_003987030.5
B3 41738794 41738926 refGene_gn=HACD3;refGene_tx=XM_003987030.5
B3 41739455 41739562 refGene_gn=HACD3;refGene_tx=XM_003987030.5
B3 41740912 41741025 refGene_gn=HACD3;refGene_tx=XM_003987030.5
B3 41742959 41743087 refGene_gn=HACD3;refGene_tx=XM_003987030.5
B3 41749070 41749181 refGene_gn=HACD3;refGene_tx=XM_003987030.5
B3 41750431 41750483 refGene_gn=HACD3;refGene_tx=XM_003987030.5
B3 41752158 41752323 refGene_gn=HACD3;refGene_tx=XM_003987030.5
B3 41753603 41753677 refGene_gn=HACD3;refGene_tx=XM_003987030.5
B3 41755574 41755617 refGene_gn=HACD3;refGene_tx=XM_003987030.5
B3 41773146 41773233 refGene_gn=HACD3;refGene_tx=XM_003987030.5
B3 41786989 41787026

refGene_gn=DPP8;refGene_tx=XM_023255180.1;refGene_gn=DPP8;refGene_tx=XM_023255181.1;refGene_gn=DPP8;refGene_tx=XM_023255182.1;refGene_gn=DPP8;refGene_tx=XM_023255184.1
B3 41789231 41789501
refGene_gn=DPP8;refGene_tx=XM_023255180.1;refGene_gn=DPP8;refGene_tx=XM_023255181.1;refGene_gn=DPP8;refGene_tx=XM_023255182.1;refGene_gn=DPP8;refGene_tx=XM_023255184.1
B3 41794712 41794825
refGene_gn=DPP8;refGene_tx=XM_023255180.1;refGene_gn=DPP8;refGene_tx=XM_023255181.1;refGene_gn=DPP8;refGene_tx=XM_023255182.1;refGene_gn=DPP8;refGene_tx=XM_023255184.1
B3 41811178 41811352
refGene_gn=DPP8;refGene_tx=XM_023255180.1;refGene_gn=DPP8;refGene_tx=XM_023255181.1;refGene_gn=DPP8;refGene_tx=XM_023255182.1;refGene_gn=DPP8;refGene_tx=XM_023255184.1
B3 41813457 41813626
refGene_gn=DPP8;refGene_tx=XM_023255180.1;refGene_gn=DPP8;refGene_tx=XM_023255181.1;refGene_gn=DPP8;refGene_tx=XM_023255182.1;refGene_gn=DPP8;refGene_tx=XM_023255184.1
B3 41819357 41819468
refGene_gn=DPP8;refGene_tx=XM_023255180.1;refGene_gn=DPP8;refGene_tx=XM_023255181.1;refGene_gn=DPP8;refGene_tx=XM_023255182.1;refGene_gn=DPP8;refGene_tx=XM_023255184.1
B3 41821085 41821214
refGene_gn=DPP8;refGene_tx=XM_023255180.1;refGene_gn=DPP8;refGene_tx=XM_023255181.1;refGene_gn=DPP8;refGene_tx=XM_023255182.1;refGene_gn=DPP8;refGene_tx=XM_023255184.1
B3 41822200 41822262
refGene_gn=DPP8;refGene_tx=XM_023255180.1;refGene_gn=DPP8;refGene_tx=XM_023255181.1;refGene_gn=DPP8;refGene_tx=XM_023255182.1;refGene_gn=DPP8;refGene_tx=XM_023255184.1
B3 41824453 41824554
refGene_gn=DPP8;refGene_tx=XM_023255180.1;refGene_gn=DPP8;refGene_tx=XM_023255181.1;refGene_gn=DPP8;refGene_tx=XM_023255182.1;refGene_gn=DPP8;refGene_tx=XM_023255184.1
B3 41825231 41825409
refGene_gn=DPP8;refGene_tx=XM_023255180.1;refGene_gn=DPP8;refGene_tx=XM_023255181.1;refGene_gn=DPP8;refGene_tx=XM_023255182.1;refGene_gn=DPP8;refGene_tx=XM_023255184.1
B3 41826365 41826525
refGene_gn=DPP8;refGene_tx=XM_023255180.1;refGene_gn=DPP8;refGene_tx=XM_023255181.1;refGene_gn=DPP8;refGene_tx=XM_023255182.1;refGene_gn=DPP8;refGene_tx=XM_023255184.1
B3 41831745 41831825
refGene_gn=DPP8;refGene_tx=XM_023255180.1;refGene_gn=DPP8;refGene_tx=XM_023255181.1;refGene_gn=DPP8;refGene_tx=XM_023255182.1;refGene_gn=DPP8;refGene_tx=XM_023255184.1
B3 41834742 41834895
refGene_gn=DPP8;refGene_tx=XM_023255180.1;refGene_gn=DPP8;refGene_tx=XM_023255181.1;refGene_gn=DPP8;refGene_tx=XM_023255182.1;refGene_gn=DPP

8;refGene_tx=XM_023255184.1
B3 41835139 41835275
refGene_gn=DPP8;refGene_tx=XM_023255180.1;refGene_gn=DPP8;refGene_tx=XM_023255181.1;refGene_gn=DPP8;refGene_tx=XM_023255182.1;refGene_gn=DPP8;refGene_tx=XM_023255184.1
B3 41836760 41836906
refGene_gn=DPP8;refGene_tx=XM_023255180.1;refGene_gn=DPP8;refGene_tx=XM_023255181.1;refGene_gn=DPP8;refGene_tx=XM_023255182.1;refGene_gn=DPP8;refGene_tx=XM_023255184.1
B3 41843008 41843155
refGene_gn=DPP8;refGene_tx=XM_023255180.1;refGene_gn=DPP8;refGene_tx=XM_023255181.1;refGene_gn=DPP8;refGene_tx=XM_023255182.1;refGene_gn=DPP8;refGene_tx=XM_023255184.1
B3 41843663 41843687 refGene_gn=DPP8;refGene_tx=XM_023255184.1
B3 41844464 41844617
refGene_gn=DPP8;refGene_tx=XM_023255180.1;refGene_gn=DPP8;refGene_tx=XM_023255181.1;refGene_gn=DPP8;refGene_tx=XM_023255182.1
B3 41845897 41846040
refGene_gn=DPP8;refGene_tx=XM_023255180.1;refGene_gn=DPP8;refGene_tx=XM_023255181.1;refGene_gn=DPP8;refGene_tx=XM_023255182.1
B3 41846699 41846811
refGene_gn=DPP8;refGene_tx=XM_023255180.1;refGene_gn=DPP8;refGene_tx=XM_023255181.1;refGene_gn=DPP8;refGene_tx=XM_023255182.1
B3 41850389 41850512
refGene_gn=DPP8;refGene_tx=XM_023255180.1;refGene_gn=DPP8;refGene_tx=XM_023255181.1;refGene_gn=DPP8;refGene_tx=XM_023255182.1
B3 41871117 41871187
refGene_gn=IGDCC4;refGene_tx=XM_023255185.1;refGene_gn=IGDCC4;refGene_tx=XM_023255186.1;refGene_gn=IGDCC4;refGene_tx=XM_023255187.1;refGene_gn=IGDCC4;refGene_tx=XM_023255188.1
B3 41880150 41880495
refGene_gn=IGDCC4;refGene_tx=XM_023255185.1;refGene_gn=IGDCC4;refGene_tx=XM_023255186.1;refGene_gn=IGDCC4;refGene_tx=XM_023255187.1;refGene_gn=IGDCC4;refGene_tx=XM_023255188.1
B3 41881158 41881300
refGene_gn=IGDCC4;refGene_tx=XM_023255185.1;refGene_gn=IGDCC4;refGene_tx=XM_023255186.1;refGene_gn=IGDCC4;refGene_tx=XM_023255187.1;refGene_gn=IGDCC4;refGene_tx=XM_023255188.1
B3 41888840 41888977
refGene_gn=IGDCC4;refGene_tx=XM_023255185.1;refGene_gn=IGDCC4;refGene_tx=XM_023255186.1;refGene_gn=IGDCC4;refGene_tx=XM_023255187.1;refGene_gn=IGDCC4;refGene_tx=XM_023255188.1
B3 41890281 41890422
refGene_gn=IGDCC4;refGene_tx=XM_023255185.1;refGene_gn=IGDCC4;refGene_tx=XM_023255186.1;refGene_gn=IGDCC4;refGene_tx=XM_023255187.1;refGene_gn=IGDCC4;refGene_tx=XM_023255188.1
B3 41894238 41894394
refGene_gn=IGDCC4;refGene_tx=XM_023255185.1;refGene_gn=IGDCC4;refGene_tx=XM_023255186.1;refGene_gn=IGDCC4;refGene_tx=XM_023255187.1;refGene_gn=IGDCC4;refGene_tx=XM_023255188.1

B3 41895193 41895607
refGene_gn=IGDCC4;refGene_tx=XM_023255185.1;refGene_gn=IGDCC4;refGene_tx=XM_023255186.1;refGene_gn=IGDCC4;refGene_tx=XM_023255187.1;refGene_gn=IGDCC4;refGene_tx=XM_023255188.1
B3 41896131 41896296
refGene_gn=IGDCC4;refGene_tx=XM_023255185.1;refGene_gn=IGDCC4;refGene_tx=XM_023255186.1;refGene_gn=IGDCC4;refGene_tx=XM_023255187.1;refGene_gn=IGDCC4;refGene_tx=XM_023255188.1
B3 41896834 41896972
refGene_gn=IGDCC4;refGene_tx=XM_023255185.1;refGene_gn=IGDCC4;refGene_tx=XM_023255186.1;refGene_gn=IGDCC4;refGene_tx=XM_023255187.1;refGene_gn=IGDCC4;refGene_tx=XM_023255188.1
B3 41897851 41898025
refGene_gn=IGDCC4;refGene_tx=XM_023255185.1;refGene_gn=IGDCC4;refGene_tx=XM_023255186.1;refGene_gn=IGDCC4;refGene_tx=XM_023255187.1;refGene_gn=IGDCC4;refGene_tx=XM_023255188.1
B3 41899028 41899268
refGene_gn=IGDCC4;refGene_tx=XM_023255185.1;refGene_gn=IGDCC4;refGene_tx=XM_023255186.1;refGene_gn=IGDCC4;refGene_tx=XM_023255187.1;refGene_gn=IGDCC4;refGene_tx=XM_023255188.1
B3 41899420 41899522
refGene_gn=IGDCC4;refGene_tx=XM_023255185.1;refGene_gn=IGDCC4;refGene_tx=XM_023255186.1;refGene_gn=IGDCC4;refGene_tx=XM_023255187.1;refGene_gn=IGDCC4;refGene_tx=XM_023255188.1
B3 41901708 41901892
refGene_gn=IGDCC4;refGene_tx=XM_023255185.1;refGene_gn=IGDCC4;refGene_tx=XM_023255186.1;refGene_gn=IGDCC4;refGene_tx=XM_023255187.1;refGene_gn=IGDCC4;refGene_tx=XM_023255188.1
B3 41902655 41902783
refGene_gn=IGDCC4;refGene_tx=XM_023255185.1;refGene_gn=IGDCC4;refGene_tx=XM_023255186.1;refGene_gn=IGDCC4;refGene_tx=XM_023255187.1;refGene_gn=IGDCC4;refGene_tx=XM_023255188.1
B3 41904313 41904484
refGene_gn=IGDCC4;refGene_tx=XM_023255185.1;refGene_gn=IGDCC4;refGene_tx=XM_023255186.1;refGene_gn=IGDCC4;refGene_tx=XM_023255187.1;refGene_gn=IGDCC4;refGene_tx=XM_023255188.1
B3 41904734 41904872
refGene_gn=IGDCC4;refGene_tx=XM_023255185.1;refGene_gn=IGDCC4;refGene_tx=XM_023255186.1;refGene_gn=IGDCC4;refGene_tx=XM_023255187.1;refGene_gn=IGDCC4;refGene_tx=XM_023255188.1
B3 41906005 41906111
refGene_gn=IGDCC4;refGene_tx=XM_023255185.1;refGene_gn=IGDCC4;refGene_tx=XM_023255186.1;refGene_gn=IGDCC4;refGene_tx=XM_023255187.1;refGene_gn=IGDCC4;refGene_tx=XM_023255188.1
B3 41906601 41906830
refGene_gn=IGDCC4;refGene_tx=XM_023255185.1;refGene_gn=IGDCC4;refGene_tx=XM_023255186.1;refGene_gn=IGDCC4;refGene_tx=XM_023255187.1;refGene_gn=IGDCC4;refGene_tx=XM_023255188.1
B3 41907599 41907761
refGene_gn=IGDCC4;refGene_tx=XM_023255185.1;refGene_gn=IGDCC4;refGene_

tx=XM_023255186.1;refGene_gn=IGDCC4;refGene_tx=XM_023255187.1;refGene_gn=IGDCC4;refGene_tx=XM_023255188.1
B3 41908325 41908736
refGene_gn=IGDCC4;refGene_tx=XM_023255185.1;refGene_gn=IGDCC4;refGene_tx=XM_023255186.1;refGene_gn=IGDCC4;refGene_tx=XM_023255187.1;refGene_gn=IGDCC4;refGene_tx=XM_023255188.1
B3 41913863 41913969
refGene_gn=IGDCC3;refGene_tx=XM_023255189.1;refGene_gn=IGDCC3;refGene_tx=XM_023255190.1;refGene_gn=IGDCC3;refGene_tx=XM_023255191.1;refGene_gn=IGDCC3;refGene_tx=XM_023255194.1
B3 41916658 41916964
refGene_gn=IGDCC3;refGene_tx=XM_023255189.1;refGene_gn=IGDCC3;refGene_tx=XM_023255190.1;refGene_gn=IGDCC3;refGene_tx=XM_023255191.1;refGene_gn=IGDCC3;refGene_tx=XM_023255192.1;refGene_gn=IGDCC3;refGene_tx=XM_023255194.1
B3 41947170 41947315
refGene_gn=IGDCC3;refGene_tx=XM_023255189.1;refGene_gn=IGDCC3;refGene_tx=XM_023255190.1;refGene_gn=IGDCC3;refGene_tx=XM_023255191.1;refGene_gn=IGDCC3;refGene_tx=XM_023255192.1;refGene_gn=IGDCC3;refGene_tx=XM_023255194.1
B3 41947678 41947809
refGene_gn=IGDCC3;refGene_tx=XM_023255189.1;refGene_gn=IGDCC3;refGene_tx=XM_023255190.1;refGene_gn=IGDCC3;refGene_tx=XM_023255191.1;refGene_gn=IGDCC3;refGene_tx=XM_023255192.1;refGene_gn=IGDCC3;refGene_tx=XM_023255194.1
B3 41948246 41948384
refGene_gn=IGDCC3;refGene_tx=XM_023255189.1;refGene_gn=IGDCC3;refGene_tx=XM_023255190.1;refGene_gn=IGDCC3;refGene_tx=XM_023255191.1;refGene_gn=IGDCC3;refGene_tx=XM_023255192.1;refGene_gn=IGDCC3;refGene_tx=XM_023255194.1
B3 41949956 41950115
refGene_gn=IGDCC3;refGene_tx=XM_023255189.1;refGene_gn=IGDCC3;refGene_tx=XM_023255190.1;refGene_gn=IGDCC3;refGene_tx=XM_023255192.1;refGene_gn=IGDCC3;refGene_tx=XM_023255194.1
B3 41951257 41951423
refGene_gn=IGDCC3;refGene_tx=XM_023255189.1;refGene_gn=IGDCC3;refGene_tx=XM_023255190.1;refGene_gn=IGDCC3;refGene_tx=XM_023255191.1;refGene_gn=IGDCC3;refGene_tx=XM_023255192.1;refGene_gn=IGDCC3;refGene_tx=XM_023255194.1
B3 41951678 41951926
refGene_gn=IGDCC3;refGene_tx=XM_023255189.1;refGene_gn=IGDCC3;refGene_tx=XM_023255190.1;refGene_gn=IGDCC3;refGene_tx=XM_023255191.1;refGene_gn=IGDCC3;refGene_tx=XM_023255192.1;refGene_gn=IGDCC3;refGene_tx=XM_023255194.1
B3 41952112 41952277
refGene_gn=IGDCC3;refGene_tx=XM_023255189.1;refGene_gn=IGDCC3;refGene_tx=XM_023255190.1;refGene_gn=IGDCC3;refGene_tx=XM_023255191.1;refGene_gn=IGDCC3;refGene_tx=XM_023255192.1;refGene_gn=IGDCC3;refGene_tx=XM_023255194.1
B3 41952530 41952722

refGene_gn=IGDCC3;refGene_tx=XM_023255189.1;refGene_gn=IGDCC3;refGene_tx=XM_023255190.1;refGene_gn=IGDCC3;refGene_tx=XM_023255191.1;refGene_gn=IGDCC3;refGene_tx=XM_023255192.1;refGene_gn=IGDCC3;refGene_tx=XM_023255194.1
B3 41952875 41952980
refGene_gn=IGDCC3;refGene_tx=XM_023255189.1;refGene_gn=IGDCC3;refGene_tx=XM_023255190.1;refGene_gn=IGDCC3;refGene_tx=XM_023255191.1;refGene_gn=IGDCC3;refGene_tx=XM_023255192.1;refGene_gn=IGDCC3;refGene_tx=XM_023255194.1
B3 41953459 41953598
refGene_gn=IGDCC3;refGene_tx=XM_023255189.1;refGene_gn=IGDCC3;refGene_tx=XM_023255190.1;refGene_gn=IGDCC3;refGene_tx=XM_023255191.1;refGene_gn=IGDCC3;refGene_tx=XM_023255192.1;refGene_gn=IGDCC3;refGene_tx=XM_023255194.1
B3 41953747 41953946
refGene_gn=IGDCC3;refGene_tx=XM_023255189.1;refGene_gn=IGDCC3;refGene_tx=XM_023255190.1;refGene_gn=IGDCC3;refGene_tx=XM_023255191.1;refGene_gn=IGDCC3;refGene_tx=XM_023255192.1;refGene_gn=IGDCC3;refGene_tx=XM_023255194.1
B3 41954212 41954437
refGene_gn=IGDCC3;refGene_tx=XM_023255189.1;refGene_gn=IGDCC3;refGene_tx=XM_023255190.1;refGene_gn=IGDCC3;refGene_tx=XM_023255191.1;refGene_gn=IGDCC3;refGene_tx=XM_023255192.1
B3 42011075 42011249
refGene_gn=PARP16;refGene_tx=XM_006932456.3;refGene_gn=PARP16;refGene_tx=XM_011282929.2;refGene_gn=PARP16;refGene_tx=XM_011282930.2;refGene_gn=PARP16;refGene_tx=XM_019832226.2;refGene_gn=PARP16;refGene_tx=XM_023255195.1
B3 42021764 42021902
refGene_gn=PARP16;refGene_tx=XM_006932456.3;refGene_gn=PARP16;refGene_tx=XM_011282929.2;refGene_gn=PARP16;refGene_tx=XM_011282930.2;refGene_gn=PARP16;refGene_tx=XM_019832226.2;refGene_gn=PARP16;refGene_tx=XM_023255195.1
B3 42024187 42024394
refGene_gn=PARP16;refGene_tx=XM_006932456.3;refGene_gn=PARP16;refGene_tx=XM_011282929.2;refGene_gn=PARP16;refGene_tx=XM_011282930.2;refGene_gn=PARP16;refGene_tx=XM_019832226.2;refGene_gn=PARP16;refGene_tx=XM_023255195.1
B3 42027411 42027583
refGene_gn=PARP16;refGene_tx=XM_006932456.3;refGene_gn=PARP16;refGene_tx=XM_011282929.2;refGene_gn=PARP16;refGene_tx=XM_011282930.2;refGene_gn=PARP16;refGene_tx=XM_019832226.2;refGene_gn=PARP16;refGene_tx=XM_023255195.1
B3 42028902 42029044
refGene_gn=PARP16;refGene_tx=XM_006932456.3;refGene_gn=PARP16;refGene_tx=XM_011282929.2;refGene_gn=PARP16;refGene_tx=XM_011282930.2;refGene_gn=PARP16;refGene_tx=XM_019832226.2;refGene_gn=PARP16;refGene_tx=XM_023255195.1
B3 42030384 42030523
refGene_gn=PARP16;refGene_tx=XM_006932456.3;refGene_gn=PARP16;refGene_

tx=XM_011282929.2;refGene_gn=PARP16;refGene_tx=XM_011282930.2;refGene_gn=PARP16;refGene_tx=XM_023255195.1
B3 42031887 42031891 refGene_gn=PARP16;refGene_tx=XM_019832226.2
B3 42069550 42069611 refGene_gn=CILP;refGene_tx=XM_003987033.5;refGene_gn=CILP;refGene_tx=XM_011282927.3;refGene_gn=CILP;refGene_tx=XM_011282928.3
B3 42071569 42071662 refGene_gn=CILP;refGene_tx=XM_003987033.5;refGene_gn=CILP;refGene_tx=XM_011282927.3;refGene_gn=CILP;refGene_tx=XM_011282928.3
B3 42072324 42072594 refGene_gn=CILP;refGene_tx=XM_003987033.5;refGene_gn=CILP;refGene_tx=XM_011282927.3
B3 42073884 42074064 refGene_gn=CILP;refGene_tx=XM_003987033.5;refGene_gn=CILP;refGene_tx=XM_011282927.3;refGene_gn=CILP;refGene_tx=XM_011282928.3
B3 42074459 42074774 refGene_gn=CILP;refGene_tx=XM_003987033.5;refGene_gn=CILP;refGene_tx=XM_011282927.3;refGene_gn=CILP;refGene_tx=XM_011282928.3
B3 42075662 42075771 refGene_gn=CILP;refGene_tx=XM_003987033.5;refGene_gn=CILP;refGene_tx=XM_011282927.3;refGene_gn=CILP;refGene_tx=XM_011282928.3
B3 42076455 42076613 refGene_gn=CILP;refGene_tx=XM_003987033.5;refGene_gn=CILP;refGene_tx=XM_011282927.3;refGene_gn=CILP;refGene_tx=XM_011282928.3
B3 42078607 42080961 refGene_gn=CILP;refGene_tx=XM_003987033.5;refGene_gn=CILP;refGene_tx=XM_011282927.3;refGene_gn=CILP;refGene_tx=XM_011282928.3
B3 42097172 42097251 refGene_gn=CLPX;refGene_tx=XM_003987034.5
B3 42097806 42097906 refGene_gn=CLPX;refGene_tx=XM_019832224.2
B3 42107679 42107840 refGene_gn=CLPX;refGene_tx=XM_003987034.5;refGene_gn=CLPX;refGene_tx=XM_019832224.2
B3 42108828 42108946 refGene_gn=CLPX;refGene_tx=XM_003987034.5;refGene_gn=CLPX;refGene_tx=XM_019832224.2
B3 42115882 42116037 refGene_gn=CLPX;refGene_tx=XM_003987034.5;refGene_gn=CLPX;refGene_tx=XM_019832224.2
B3 42118757 42118917 refGene_gn=CLPX;refGene_tx=XM_003987034.5;refGene_gn=CLPX;refGene_tx=XM_019832224.2
B3 42120184 42120226 refGene_gn=CLPX;refGene_tx=XM_003987034.5;refGene_gn=CLPX;refGene_tx=XM_019832224.2
B3 42124521 42124698 refGene_gn=CLPX;refGene_tx=XM_003987034.5;refGene_gn=CLPX;refGene_tx=XM_019832224.2
B3 42125077 42125242 refGene_gn=CLPX;refGene_tx=XM_003987034.5;refGene_gn=CLPX;refGene_tx=XM_019832224.2

B3 42126034 42126123
refGene_gn=CLPX;refGene_tx=XM_003987034.5;refGene_gn=CLPX;refGene_tx=XM_019832224.2
B3 42127093 42127258
refGene_gn=CLPX;refGene_tx=XM_003987034.5;refGene_gn=CLPX;refGene_tx=XM_019832224.2
B3 42130654 42130954
refGene_gn=CLPX;refGene_tx=XM_003987034.5;refGene_gn=CLPX;refGene_tx=XM_019832224.2
B3 42132544 42132637
refGene_gn=CLPX;refGene_tx=XM_003987034.5;refGene_gn=CLPX;refGene_tx=XM_019832224.2
B3 42133975 42134082
refGene_gn=CLPX;refGene_tx=XM_003987034.5;refGene_gn=CLPX;refGene_tx=XM_019832224.2
B3 42135846 42135937
refGene_gn=CLPX;refGene_tx=XM_003987034.5;refGene_gn=CLPX;refGene_tx=XM_019832224.2
B3 42144307 42145174 refGene_gn=PDCD7;refGene_tx=XM_023255197.1
B3 42147606 42147745 refGene_gn=PDCD7;refGene_tx=XM_023255197.1
B3 42154390 42154627 refGene_gn=PDCD7;refGene_tx=XM_023255197.1
B3 42154886 42154974 refGene_gn=PDCD7;refGene_tx=XM_023255197.1
B3 42156930 42157054 refGene_gn=PDCD7;refGene_tx=XM_023255197.1
B3 42161057 42161210 refGene_gn=UBAP1L;refGene_tx=XM_023255198.1
B3 42168532 42168700
refGene_gn=UBAP1L;refGene_tx=XM_019832229.2;refGene_gn=UBAP1L;refGene_tx=XM_023255198.1;refGene_gn=UBAP1L;refGene_tx=XM_023255199.1
B3 42173064 42173685
refGene_gn=UBAP1L;refGene_tx=XM_019832229.2;refGene_gn=UBAP1L;refGene_tx=XM_023255198.1;refGene_gn=UBAP1L;refGene_tx=XM_023255199.1
B3 42176042 42176249
refGene_gn=UBAP1L;refGene_tx=XM_019832229.2;refGene_gn=UBAP1L;refGene_tx=XM_023255198.1;refGene_gn=UBAP1L;refGene_tx=XM_023255199.1
B3 42181204 42181306
refGene_gn=UBAP1L;refGene_tx=XM_019832229.2;refGene_gn=UBAP1L;refGene_tx=XM_023255198.1;refGene_gn=UBAP1L;refGene_tx=XM_023255199.1
B3 42182243 42182378
refGene_gn=UBAP1L;refGene_tx=XM_019832229.2;refGene_gn=UBAP1L;refGene_tx=XM_023255198.1;refGene_gn=UBAP1L;refGene_tx=XM_023255199.1
B3 42200458 42201838 refGene_gn=KBTBD13;refGene_tx=XM_003987081.4
B3 42207773 42207876 refGene_gn=RASL12;refGene_tx=XM_003987035.5
B3 42209745 42209802 refGene_gn=RASL12;refGene_tx=XM_003987035.5
B3 42216010 42216084 refGene_gn=RASL12;refGene_tx=XM_003987035.5
B3 42216269 42216362 refGene_gn=RASL12;refGene_tx=XM_019832233.1
B3 42216783 42216974
refGene_gn=RASL12;refGene_tx=XM_003987035.5;refGene_gn=RASL12;refGene_tx=XM_019832233.1
B3 42219620 42219996
refGene_gn=RASL12;refGene_tx=XM_003987035.5;refGene_gn=RASL12;refGene_tx=XM_019832233.1

B3 42221863 42222077
refGene_gn=SLC51B;refGene_tx=XM_003987036.5;refGene_gn=SLC51B;refGene_tx=XM_019832234.2
B3 42224278 42224369
refGene_gn=SLC51B;refGene_tx=XM_003987036.5;refGene_gn=SLC51B;refGene_tx=XM_019832234.2
B3 42225396 42225493
refGene_gn=SLC51B;refGene_tx=XM_003987036.5;refGene_gn=SLC51B;refGene_tx=XM_019832234.2
B3 42248802 42249023 refGene_gn=MTFMT;refGene_tx=XM_011282925.2
B3 42250776 42250986 refGene_gn=MTFMT;refGene_tx=XM_011282925.2
B3 42253522 42253645 refGene_gn=MTFMT;refGene_tx=XM_011282925.2
B3 42255145 42255248 refGene_gn=MTFMT;refGene_tx=XM_011282925.2
B3 42256597 42256673 refGene_gn=MTFMT;refGene_tx=XM_011282925.2
B3 42262554 42262646 refGene_gn=MTFMT;refGene_tx=XM_011282925.2
B3 42269597 42269676 refGene_gn=MTFMT;refGene_tx=XM_011282925.2
B3 42270528 42270611 refGene_gn=MTFMT;refGene_tx=XM_011282925.2
B3 42272956 42273154 refGene_gn=MTFMT;refGene_tx=XM_011282925.2
B3 42287119 42287182
refGene_gn=SPG21;refGene_tx=XM_011282922.2;refGene_gn=SPG21;refGene_tx=XM_011282924.2;refGene_gn=SPG21;refGene_tx=XM_019832236.2;refGene_gn=SPG21;refGene_tx=XM_019832237.2
B3 42289881 42290043
refGene_gn=SPG21;refGene_tx=XM_011282922.2;refGene_gn=SPG21;refGene_tx=XM_011282924.2;refGene_gn=SPG21;refGene_tx=XM_019832236.2;refGene_gn=SPG21;refGene_tx=XM_019832237.2
B3 42294079 42294160
refGene_gn=SPG21;refGene_tx=XM_011282922.2;refGene_gn=SPG21;refGene_tx=XM_011282924.2;refGene_gn=SPG21;refGene_tx=XM_019832236.2;refGene_gn=SPG21;refGene_tx=XM_019832237.2
B3 42295444 42295590
refGene_gn=SPG21;refGene_tx=XM_011282922.2;refGene_gn=SPG21;refGene_tx=XM_011282924.2;refGene_gn=SPG21;refGene_tx=XM_019832236.2;refGene_gn=SPG21;refGene_tx=XM_019832237.2
B3 42298036 42298160
refGene_gn=SPG21;refGene_tx=XM_011282922.2;refGene_gn=SPG21;refGene_tx=XM_011282924.2;refGene_gn=SPG21;refGene_tx=XM_019832236.2;refGene_gn=SPG21;refGene_tx=XM_019832237.2
B3 42298962 42299070
refGene_gn=SPG21;refGene_tx=XM_011282922.2;refGene_gn=SPG21;refGene_tx=XM_011282924.2;refGene_gn=SPG21;refGene_tx=XM_019832236.2;refGene_gn=SPG21;refGene_tx=XM_019832237.2
B3 42301112 42301253
refGene_gn=SPG21;refGene_tx=XM_011282922.2;refGene_gn=SPG21;refGene_tx=XM_011282924.2;refGene_gn=SPG21;refGene_tx=XM_019832236.2;refGene_gn=SPG21;refGene_tx=XM_019832237.2
B3 42302585 42302702
refGene_gn=SPG21;refGene_tx=XM_011282922.2;refGene_gn=SPG21;refGene_tx=XM_011282924.2;refGene_gn=SPG21;refGene_tx=XM_019832236.2;refGene_gn=SPG21;refGene_tx=XM_019832237.2

B3 42307759 42307821
refGene_gn=ANKDD1A;refGene_tx=XM_019832238.2;refGene_gn=ANKDD1A;refGene_tx=XM_019832239.2;refGene_gn=ANKDD1A;refGene_tx=XM_019832240.2;refGene_gn=ANKDD1A;refGene_tx=XM_019832241.2;refGene_gn=ANKDD1A;refGene_tx=XM_019832242.2;refGene_gn=ANKDD1A;refGene_tx=XM_019832243.2;refGene_gn=ANKDD1A;refGene_tx=XM_019832245.2;refGene_gn=ANKDD1A;refGene_tx=XM_019832246.2;refGene_gn=ANKDD1A;refGene_tx=XM_019832247.2;refGene_gn=ANKDD1A;refGene_tx=XM_019832248.2;refGene_gn=ANKDD1A;refGene_tx=XM_019832250.2
B3 42308765 42308897
refGene_gn=ANKDD1A;refGene_tx=XM_019832238.2;refGene_gn=ANKDD1A;refGene_tx=XM_019832239.2;refGene_gn=ANKDD1A;refGene_tx=XM_019832240.2;refGene_gn=ANKDD1A;refGene_tx=XM_019832241.2;refGene_gn=ANKDD1A;refGene_tx=XM_019832242.2;refGene_gn=ANKDD1A;refGene_tx=XM_019832243.2;refGene_gn=ANKDD1A;refGene_tx=XM_019832245.2;refGene_gn=ANKDD1A;refGene_tx=XM_019832246.2;refGene_gn=ANKDD1A;refGene_tx=XM_019832247.2;refGene_gn=ANKDD1A;refGene_tx=XM_019832248.2;refGene_gn=ANKDD1A;refGene_tx=XM_019832250.2
B3 42311032 42311237
refGene_gn=ANKDD1A;refGene_tx=XM_019832238.2;refGene_gn=ANKDD1A;refGene_tx=XM_019832239.2;refGene_gn=ANKDD1A;refGene_tx=XM_019832240.2;refGene_gn=ANKDD1A;refGene_tx=XM_019832241.2;refGene_gn=ANKDD1A;refGene_tx=XM_019832242.2;refGene_gn=ANKDD1A;refGene_tx=XM_019832243.2;refGene_gn=ANKDD1A;refGene_tx=XM_019832245.2;refGene_gn=ANKDD1A;refGene_tx=XM_019832246.2;refGene_gn=ANKDD1A;refGene_tx=XM_019832247.2;refGene_gn=ANKDD1A;refGene_tx=XM_019832248.2;refGene_gn=ANKDD1A;refGene_tx=XM_019832250.2
B3 42313418 42313514
refGene_gn=ANKDD1A;refGene_tx=XM_019832238.2;refGene_gn=ANKDD1A;refGene_tx=XM_019832239.2;refGene_gn=ANKDD1A;refGene_tx=XM_019832240.2;refGene_gn=ANKDD1A;refGene_tx=XM_019832241.2;refGene_gn=ANKDD1A;refGene_tx=XM_019832242.2;refGene_gn=ANKDD1A;refGene_tx=XM_019832243.2;refGene_gn=ANKDD1A;refGene_tx=XM_019832246.2;refGene_gn=ANKDD1A;refGene_tx=XM_019832247.2;refGene_gn=ANKDD1A;refGene_tx=XM_019832248.2;refGene_gn=ANKDD1A;refGene_tx=XM_019832250.2
B3 42314545 42314644
refGene_gn=ANKDD1A;refGene_tx=XM_019832238.2;refGene_gn=ANKDD1A;refGene_tx=XM_019832239.2;refGene_gn=ANKDD1A;refGene_tx=XM_019832240.2;refGene_gn=ANKDD1A;refGene_tx=XM_019832241.2;refGene_gn=ANKDD1A;refGene_tx=XM_019832242.2;refGene_gn=ANKDD1A;refGene_tx=XM_019832243.2;refGene_gn=ANKDD1A;refGene_tx=XM_019832245.2;refGene_gn=ANKDD1A;refGene_tx=XM_019832246.2;refGene_gn=ANKDD1A;refGene_tx=XM_019832247.2;refGene_gn=ANKDD1A;refGene_tx=XM_019832248.2;refGene_gn=ANKDD1A;refGene_tx=XM_019832250.2
B3 42315673 42315772
refGene_gn=ANKDD1A;refGene_tx=XM_019832238.2;refGene_gn=ANKDD1A;refGene_tx=XM_019832239.2;refGene_gn=ANKDD1A;refGene_tx=XM_019832240.2;refGene_gn=ANKDD1A;refGene_tx=XM_019832241.2;refGene_gn=ANKDD1A;refGene_tx=XM_019832242.2;refGene_gn=ANKDD1A;refGene_tx=XM_019832243.2;refGene_gn=ANKDD1A;refGene_tx=XM_019832245.2;refGene_gn=ANKDD1A;refGene_tx=XM_01

9832246.2;refGene_gn=ANKDD1A;refGene_tx=XM_019832247.2;refGene_gn=ANKD
D1A;refGene_tx=XM_019832248.2;refGene_gn=ANKDD1A;refGene_tx=XM_0198322
50.2
B3 42322049 42322148
refGene_gn=ANKDD1A;refGene_tx=XM_019832238.2;refGene_gn=ANKDD1A;refGen
e_tx=XM_019832239.2;refGene_gn=ANKDD1A;refGene_tx=XM_019832240.2;refGe
ne_gn=ANKDD1A;refGene_tx=XM_019832241.2;refGene_gn=ANKDD1A;refGene_tx=
XM_019832242.2;refGene_gn=ANKDD1A;refGene_tx=XM_019832243.2;refGene_gn
=ANKDD1A;refGene_tx=XM_019832245.2;refGene_gn=ANKDD1A;refGene_tx=XM_01
9832246.2;refGene_gn=ANKDD1A;refGene_tx=XM_019832247.2;refGene_gn=ANKD
D1A;refGene_tx=XM_019832248.2;refGene_gn=ANKDD1A;refGene_tx=XM_0198322
50.2
B3 42324150 42324249
refGene_gn=ANKDD1A;refGene_tx=XM_019832238.2;refGene_gn=ANKDD1A;refGen
e_tx=XM_019832239.2;refGene_gn=ANKDD1A;refGene_tx=XM_019832240.2;refGe
ne_gn=ANKDD1A;refGene_tx=XM_019832241.2;refGene_gn=ANKDD1A;refGene_tx=
XM_019832242.2;refGene_gn=ANKDD1A;refGene_tx=XM_019832243.2;refGene_gn
=ANKDD1A;refGene_tx=XM_019832245.2;refGene_gn=ANKDD1A;refGene_tx=XM_01
9832248.2;refGene_gn=ANKDD1A;refGene_tx=XM_019832250.2
B3 42324640 42324757
refGene_gn=ANKDD1A;refGene_tx=XM_019832238.2;refGene_gn=ANKDD1A;refGen
e_tx=XM_019832239.2;refGene_gn=ANKDD1A;refGene_tx=XM_019832240.2;refGe
ne_gn=ANKDD1A;refGene_tx=XM_019832241.2;refGene_gn=ANKDD1A;refGene_tx=
XM_019832242.2;refGene_gn=ANKDD1A;refGene_tx=XM_019832243.2;refGene_gn
=ANKDD1A;refGene_tx=XM_019832245.2;refGene_gn=ANKDD1A;refGene_tx=XM_01
9832246.2;refGene_gn=ANKDD1A;refGene_tx=XM_019832247.2;refGene_gn=ANKD
D1A;refGene_tx=XM_019832248.2;refGene_gn=ANKDD1A;refGene_tx=XM_0198322
50.2
B3 42329626 42329956
refGene_gn=ANKDD1A;refGene_tx=XM_019832238.2;refGene_gn=ANKDD1A;refGen
e_tx=XM_019832239.2;refGene_gn=ANKDD1A;refGene_tx=XM_019832240.2;refGe
ne_gn=ANKDD1A;refGene_tx=XM_019832241.2;refGene_gn=ANKDD1A;refGene_tx=
XM_019832242.2;refGene_gn=ANKDD1A;refGene_tx=XM_019832243.2;refGene_gn
=ANKDD1A;refGene_tx=XM_019832245.2;refGene_gn=ANKDD1A;refGene_tx=XM_01
9832246.2;refGene_gn=ANKDD1A;refGene_tx=XM_019832247.2;refGene_gn=ANKD
D1A;refGene_tx=XM_019832250.2
B3 42330462 42330567
refGene_gn=ANKDD1A;refGene_tx=XM_019832238.2;refGene_gn=ANKDD1A;refGen
e_tx=XM_019832239.2;refGene_gn=ANKDD1A;refGene_tx=XM_019832240.2;refGe
ne_gn=ANKDD1A;refGene_tx=XM_019832241.2;refGene_gn=ANKDD1A;refGene_tx=
XM_019832242.2;refGene_gn=ANKDD1A;refGene_tx=XM_019832243.2;refGene_gn
=ANKDD1A;refGene_tx=XM_019832245.2;refGene_gn=ANKDD1A;refGene_tx=XM_01
9832246.2;refGene_gn=ANKDD1A;refGene_tx=XM_019832247.2;refGene_gn=ANKD
D1A;refGene_tx=XM_019832248.2
B3 42335390 42335489
refGene_gn=ANKDD1A;refGene_tx=XM_019832238.2;refGene_gn=ANKDD1A;refGen
e_tx=XM_019832239.2;refGene_gn=ANKDD1A;refGene_tx=XM_019832240.2;refGe
ne_gn=ANKDD1A;refGene_tx=XM_019832241.2;refGene_gn=ANKDD1A;refGene_tx=
XM_019832242.2;refGene_gn=ANKDD1A;refGene_tx=XM_019832243.2;refGene_gn
=ANKDD1A;refGene_tx=XM_019832245.2;refGene_gn=ANKDD1A;refGene_tx=XM_01

9832246.2;refGene_gn=ANKDD1A;refGene_tx=XM_019832247.2;refGene_gn=ANKDD1A;refGene_tx=XM_019832248.2
B3 42338298 42338598
refGene_gn=ANKDD1A;refGene_tx=XM_019832238.2;refGene_gn=ANKDD1A;refGene_tx=XM_019832239.2;refGene_gn=ANKDD1A;refGene_tx=XM_019832240.2;refGene_gn=ANKDD1A;refGene_tx=XM_019832241.2;refGene_gn=ANKDD1A;refGene_tx=XM_019832242.2;refGene_gn=ANKDD1A;refGene_tx=XM_019832243.2;refGene_gn=ANKDD1A;refGene_tx=XM_019832245.2;refGene_gn=ANKDD1A;refGene_tx=XM_019832246.2;refGene_gn=ANKDD1A;refGene_tx=XM_019832247.2;refGene_gn=ANKDD1A;refGene_tx=XM_019832248.2
B3 42340432 42340536 refGene_gn=ANKDD1A;refGene_tx=XM_019832242.2
B3 42343915 42343949 refGene_gn=ANKDD1A;refGene_tx=XM_019832242.2
B3 42388271 42388310 refGene_gn=PLEKHO2;refGene_tx=XM_003987038.3
B3 42388311 42389277 refGene_gn=PLEKHO2;refGene_tx=XM_003987038.3
B3 42391577 42391676 refGene_gn=PLEKHO2;refGene_tx=XM_003987038.3
B3 42393221 42393326 refGene_gn=PLEKHO2;refGene_tx=XM_003987038.3
B3 42400816 42400933 refGene_gn=PLEKHO2;refGene_tx=XM_003987038.3
B3 42405158 42405308 refGene_gn=PLEKHO2;refGene_tx=XM_003987038.3
B3 42411500 42411512 refGene_gn=PLEKHO2;refGene_tx=XM_003987038.3
B3 42422730 42423288
refGene_gn=PIF1;refGene_tx=XM_006932458.4;refGene_gn=PIF1;refGene_tx=XM_023255203.1;refGene_gn=PIF1;refGene_tx=XM_023255204.1
B3 42424233 42424366
refGene_gn=PIF1;refGene_tx=XM_006932458.4;refGene_gn=PIF1;refGene_tx=XM_023255203.1;refGene_gn=PIF1;refGene_tx=XM_023255204.1
B3 42424451 42424577
refGene_gn=PIF1;refGene_tx=XM_006932458.4;refGene_gn=PIF1;refGene_tx=XM_023255203.1;refGene_gn=PIF1;refGene_tx=XM_023255204.1
B3 42424703 42424857
refGene_gn=PIF1;refGene_tx=XM_006932458.4;refGene_gn=PIF1;refGene_tx=XM_023255203.1;refGene_gn=PIF1;refGene_tx=XM_023255204.1
B3 42424948 42425063
refGene_gn=PIF1;refGene_tx=XM_006932458.4;refGene_gn=PIF1;refGene_tx=XM_023255203.1;refGene_gn=PIF1;refGene_tx=XM_023255204.1
B3 42425139 42425246
refGene_gn=PIF1;refGene_tx=XM_006932458.4;refGene_gn=PIF1;refGene_tx=XM_023255203.1;refGene_gn=PIF1;refGene_tx=XM_023255204.1
B3 42426600 42426740
refGene_gn=PIF1;refGene_tx=XM_006932458.4;refGene_gn=PIF1;refGene_tx=XM_023255203.1;refGene_gn=PIF1;refGene_tx=XM_023255204.1
B3 42427092 42427199
refGene_gn=PIF1;refGene_tx=XM_006932458.4;refGene_gn=PIF1;refGene_tx=XM_023255203.1;refGene_gn=PIF1;refGene_tx=XM_023255204.1
B3 42427888 42427976
refGene_gn=PIF1;refGene_tx=XM_006932458.4;refGene_gn=PIF1;refGene_tx=XM_023255203.1;refGene_gn=PIF1;refGene_tx=XM_023255204.1
B3 42428101 42428247
refGene_gn=PIF1;refGene_tx=XM_006932458.4;refGene_gn=PIF1;refGene_tx=XM_023255203.1;refGene_gn=PIF1;refGene_tx=XM_023255204.1
B3 42428924 42429116

refGene_gn=PIF1;refGene_tx=XM_006932458.4;refGene_gn=PIF1;refGene_tx=XM_023255203.1;refGene_gn=PIF1;refGene_tx=XM_023255204.1
B3 42429327 42429387 refGene_gn=PIF1;refGene_tx=XM_006932458.4
B3 42452128 42452335 refGene_gn=PIF1;refGene_tx=XM_023255203.1
B3 42457355 42457445 refGene_gn=PIF1;refGene_tx=XM_023255203.1
B3 42465752 42465839
refGene_gn=RBPMS2;refGene_tx=XM_023255206.1;refGene_gn=RBPMS2;refGene_tx=XM_023255207.1;refGene_gn=RBPMS2;refGene_tx=XM_023255208.1
B3 42482409 42482487
refGene_gn=RBPMS2;refGene_tx=XM_023255206.1;refGene_gn=RBPMS2;refGene_tx=XM_023255207.1;refGene_gn=RBPMS2;refGene_tx=XM_023255208.1
B3 42483086 42483125
refGene_gn=RBPMS2;refGene_tx=XM_023255206.1;refGene_gn=RBPMS2;refGene_tx=XM_023255207.1;refGene_gn=RBPMS2;refGene_tx=XM_023255208.1
B3 42483706 42483769
refGene_gn=RBPMS2;refGene_tx=XM_023255206.1;refGene_gn=RBPMS2;refGene_tx=XM_023255207.1;refGene_gn=RBPMS2;refGene_tx=XM_023255208.1
B3 42484114 42484265
refGene_gn=RBPMS2;refGene_tx=XM_023255206.1;refGene_gn=RBPMS2;refGene_tx=XM_023255207.1;refGene_gn=RBPMS2;refGene_tx=XM_023255208.1
B3 42485063 42485215
refGene_gn=RBPMS2;refGene_tx=XM_023255206.1;refGene_gn=RBPMS2;refGene_tx=XM_023255207.1;refGene_gn=RBPMS2;refGene_tx=XM_023255208.1
B3 42491280 42491301 refGene_gn=RBPMS2;refGene_tx=XM_023255208.1
B3 42491427 42491490 refGene_gn=RBPMS2;refGene_tx=XM_023255206.1
B3 42491766 42491820 refGene_gn=RBPMS2;refGene_tx=XM_023255207.1
B3 42514199 42514219 refGene_gn=OAZ2;refGene_tx=NM_001290005.1
B3 42526250 42526326 refGene_gn=OAZ2;refGene_tx=NM_001290005.1
B3 42526327 42526408 refGene_gn=OAZ2;refGene_tx=NM_001290005.1
B3 42527664 42527840 refGene_gn=OAZ2;refGene_tx=NM_001290005.1
B3 42528725 42528811 refGene_gn=OAZ2;refGene_tx=NM_001290005.1
B3 42529067 42529198 refGene_gn=OAZ2;refGene_tx=NM_001290005.1
B3 42536365 42536439 refGene_gn=ZNF609;refGene_tx=XM_023255209.1
B3 42537051 42537268 refGene_gn=ZNF609;refGene_tx=XM_023255209.1
B3 42537507 42537683 refGene_gn=ZNF609;refGene_tx=XM_023255209.1
B3 42539303 42539670 refGene_gn=ZNF609;refGene_tx=XM_023255209.1
B3 42540659 42543000 refGene_gn=ZNF609;refGene_tx=XM_023255209.1
B3 42543679 42543767 refGene_gn=ZNF609;refGene_tx=XM_023255209.1
B3 42591624 42591850 refGene_gn=ZNF609;refGene_tx=XM_023255209.1
B3 42716813 42717560 refGene_gn=ZNF609;refGene_tx=XM_023255209.1
B3 42749485 42749553
refGene_gn=TRIP4;refGene_tx=XM_023255211.1;refGene_gn=TRIP4;refGene_tx=XM_023255212.1
B3 42756932 42757035
refGene_gn=TRIP4;refGene_tx=XM_023255211.1;refGene_gn=TRIP4;refGene_tx=XM_023255212.1
B3 42772971 42773063
refGene_gn=TRIP4;refGene_tx=XM_023255211.1;refGene_gn=TRIP4;refGene_tx=XM_023255212.1
B3 42774446 42774571

refGene_gn=TRIP4;refGene_tx=XM_023255211.1;refGene_gn=TRIP4;refGene_tx=XM_023255212.1
B3 42778518 42778706
refGene_gn=TRIP4;refGene_tx=XM_023255211.1;refGene_gn=TRIP4;refGene_tx=XM_023255212.1
B3 42782357 42782484
refGene_gn=TRIP4;refGene_tx=XM_023255211.1;refGene_gn=TRIP4;refGene_tx=XM_023255212.1
B3 42787213 42787429
refGene_gn=TRIP4;refGene_tx=XM_023255211.1;refGene_gn=TRIP4;refGene_tx=XM_023255212.1
B3 42790523 42790653
refGene_gn=TRIP4;refGene_tx=XM_023255211.1;refGene_gn=TRIP4;refGene_tx=XM_023255212.1
B3 42797865 42797944
refGene_gn=TRIP4;refGene_tx=XM_023255211.1;refGene_gn=TRIP4;refGene_tx=XM_023255212.1
B3 42799341 42799554 refGene_gn=TRIP4;refGene_tx=XM_023255211.1
B3 42811536 42811670
refGene_gn=TRIP4;refGene_tx=XM_023255211.1;refGene_gn=TRIP4;refGene_tx=XM_023255212.1
B3 42813605 42813775
refGene_gn=TRIP4;refGene_tx=XM_023255211.1;refGene_gn=TRIP4;refGene_tx=XM_023255212.1
B3 42817107 42817208
refGene_gn=TRIP4;refGene_tx=XM_023255211.1;refGene_gn=TRIP4;refGene_tx=XM_023255212.1
B3 42822219 42822265
refGene_gn=PCLAF;refGene_tx=XM_003987042.5;refGene_gn=PCLAF;refGene_tx=XM_019832257.2
B3 42822570 42822651
refGene_gn=PCLAF;refGene_tx=XM_003987042.5;refGene_gn=PCLAF;refGene_tx=XM_019832257.2
B3 42824461 42824624
refGene_gn=PCLAF;refGene_tx=XM_003987042.5;refGene_gn=PCLAF;refGene_tx=XM_019832257.2
B3 42828376 42828389 refGene_gn=PCLAF;refGene_tx=XM_019832257.2
B3 42829166 42829212 refGene_gn=PCLAF;refGene_tx=XM_003987042.5
B3 42891913 42892094
refGene_gn=CSNK1G1;refGene_tx=XM_023255213.1;refGene_gn=CSNK1G1;refGene_tx=XM_023255214.1;refGene_gn=CSNK1G1;refGene_tx=XM_023255215.1;refGene_gn=CSNK1G1;refGene_tx=XM_023255216.1;refGene_gn=CSNK1G1;refGene_tx=XM_023255217.1;refGene_gn=CSNK1G1;refGene_tx=XM_023255218.1;refGene_gn=CSNK1G1;refGene_tx=XM_023255219.1;refGene_gn=CSNK1G1;refGene_tx=XM_023255220.1
B3 42913415 42913456
refGene_gn=CSNK1G1;refGene_tx=XM_023255213.1;refGene_gn=CSNK1G1;refGene_tx=XM_023255214.1;refGene_gn=CSNK1G1;refGene_tx=XM_023255215.1;refGene_gn=CSNK1G1;refGene_tx=XM_023255216.1;refGene_gn=CSNK1G1;refGene_tx=XM_023255217.1;refGene_gn=CSNK1G1;refGene_tx=XM_023255218.1;refGene_gn

=CSNK1G1;refGene_tx=XM_023255219.1;refGene_gn=CSNK1G1;refGene_tx=XM_023255220.1
B3 42920392 42920462
refGene_gn=CSNK1G1;refGene_tx=XM_023255213.1;refGene_gn=CSNK1G1;refGene_tx=XM_023255214.1;refGene_gn=CSNK1G1;refGene_tx=XM_023255215.1;refGene_gn=CSNK1G1;refGene_tx=XM_023255216.1;refGene_gn=CSNK1G1;refGene_tx=XM_023255217.1;refGene_gn=CSNK1G1;refGene_tx=XM_023255218.1;refGene_gn=CSNK1G1;refGene_tx=XM_023255219.1;refGene_gn=CSNK1G1;refGene_tx=XM_023255220.1
B3 42950389 42950541
refGene_gn=CSNK1G1;refGene_tx=XM_023255213.1;refGene_gn=CSNK1G1;refGene_tx=XM_023255214.1;refGene_gn=CSNK1G1;refGene_tx=XM_023255215.1;refGene_gn=CSNK1G1;refGene_tx=XM_023255216.1;refGene_gn=CSNK1G1;refGene_tx=XM_023255217.1;refGene_gn=CSNK1G1;refGene_tx=XM_023255218.1;refGene_gn=CSNK1G1;refGene_tx=XM_023255219.1;refGene_gn=CSNK1G1;refGene_tx=XM_023255220.1
B3 42951886 42952121
refGene_gn=CSNK1G1;refGene_tx=XM_023255213.1;refGene_gn=CSNK1G1;refGene_tx=XM_023255214.1;refGene_gn=CSNK1G1;refGene_tx=XM_023255215.1;refGene_gn=CSNK1G1;refGene_tx=XM_023255216.1;refGene_gn=CSNK1G1;refGene_tx=XM_023255217.1;refGene_gn=CSNK1G1;refGene_tx=XM_023255218.1;refGene_gn=CSNK1G1;refGene_tx=XM_023255219.1;refGene_gn=CSNK1G1;refGene_tx=XM_023255220.1
B3 42958066 42958152
refGene_gn=CSNK1G1;refGene_tx=XM_023255213.1;refGene_gn=CSNK1G1;refGene_tx=XM_023255214.1;refGene_gn=CSNK1G1;refGene_tx=XM_023255215.1;refGene_gn=CSNK1G1;refGene_tx=XM_023255216.1;refGene_gn=CSNK1G1;refGene_tx=XM_023255217.1;refGene_gn=CSNK1G1;refGene_tx=XM_023255218.1;refGene_gn=CSNK1G1;refGene_tx=XM_023255219.1;refGene_gn=CSNK1G1;refGene_tx=XM_023255220.1
B3 42960181 42960266
refGene_gn=CSNK1G1;refGene_tx=XM_023255213.1;refGene_gn=CSNK1G1;refGene_tx=XM_023255214.1;refGene_gn=CSNK1G1;refGene_tx=XM_023255215.1;refGene_gn=CSNK1G1;refGene_tx=XM_023255216.1;refGene_gn=CSNK1G1;refGene_tx=XM_023255217.1;refGene_gn=CSNK1G1;refGene_tx=XM_023255218.1;refGene_gn=CSNK1G1;refGene_tx=XM_023255219.1;refGene_gn=CSNK1G1;refGene_tx=XM_023255220.1
B3 42960520 42960669
refGene_gn=CSNK1G1;refGene_tx=XM_023255213.1;refGene_gn=CSNK1G1;refGene_tx=XM_023255214.1;refGene_gn=CSNK1G1;refGene_tx=XM_023255215.1;refGene_gn=CSNK1G1;refGene_tx=XM_023255216.1;refGene_gn=CSNK1G1;refGene_tx=XM_023255217.1;refGene_gn=CSNK1G1;refGene_tx=XM_023255218.1;refGene_gn=CSNK1G1;refGene_tx=XM_023255219.1;refGene_gn=CSNK1G1;refGene_tx=XM_023255220.1
B3 42961726 42961834
refGene_gn=CSNK1G1;refGene_tx=XM_023255213.1;refGene_gn=CSNK1G1;refGene_tx=XM_023255214.1;refGene_gn=CSNK1G1;refGene_tx=XM_023255215.1;refGene_gn=CSNK1G1;refGene_tx=XM_023255216.1;refGene_gn=CSNK1G1;refGene_tx=XM_023255217.1;refGene_gn=CSNK1G1;refGene_tx=XM_023255218.1;refGene_gn=CSNK1G1;refGene_tx=XM_023255219.1;refGene_gn=CSNK1G1;refGene_tx=XM_02

3255220.1
B3	42964423	42964450	refGene_gn=CSNK1G1;refGene_tx=XM_023255220.1
B3	42971433	42971493	refGene_gn=CSNK1G1;refGene_tx=XM_023255219.1
B3	42974531	42974642
refGene_gn=CSNK1G1;refGene_tx=XM_023255213.1;refGene_gn=CSNK1G1;refGen
e_tx=XM_023255214.1;refGene_gn=CSNK1G1;refGene_tx=XM_023255215.1;refGe
ne_gn=CSNK1G1;refGene_tx=XM_023255216.1
B3	42986479	42986586
refGene_gn=CSNK1G1;refGene_tx=XM_023255213.1;refGene_gn=CSNK1G1;refGen
e_tx=XM_023255214.1;refGene_gn=CSNK1G1;refGene_tx=XM_023255215.1;refGe
ne_gn=CSNK1G1;refGene_tx=XM_023255216.1;refGene_gn=CSNK1G1;refGene_tx=
XM_023255217.1;refGene_gn=CSNK1G1;refGene_tx=XM_023255218.1
B3	42994236	42994291
refGene_gn=CSNK1G1;refGene_tx=XM_023255213.1;refGene_gn=CSNK1G1;refGen
e_tx=XM_023255214.1;refGene_gn=CSNK1G1;refGene_tx=XM_023255215.1;refGe
ne_gn=CSNK1G1;refGene_tx=XM_023255216.1;refGene_gn=CSNK1G1;refGene_tx=
XM_023255217.1;refGene_gn=CSNK1G1;refGene_tx=XM_023255218.1
B3	43002967	43003102	refGene_gn=PPIB;refGene_tx=NM_001278847.1
B3	43004087	43004201	refGene_gn=PPIB;refGene_tx=NM_001278847.1
B3	43005365	43005459	refGene_gn=PPIB;refGene_tx=NM_001278847.1
B3	43007863	43008048	refGene_gn=PPIB;refGene_tx=NM_001278847.1
B3	43008614	43008737	refGene_gn=PPIB;refGene_tx=NM_001278847.1
B3	43009413	43009535
refGene_gn=SNX22;refGene_tx=XM_003987046.5;refGene_gn=SNX22;refGene_tx
=XM_019832267.2;refGene_gn=SNX22;refGene_tx=XM_019832268.2;refGene_gn=
SNX22;refGene_tx=XM_019832269.2
B3	43010009	43010077
refGene_gn=SNX22;refGene_tx=XM_003987046.5;refGene_gn=SNX22;refGene_tx
=XM_019832267.2;refGene_gn=SNX22;refGene_tx=XM_019832268.2;refGene_gn=
SNX22;refGene_tx=XM_019832269.2
B3	43010341	43010410
refGene_gn=SNX22;refGene_tx=XM_003987046.5;refGene_gn=SNX22;refGene_tx
=XM_019832267.2;refGene_gn=SNX22;refGene_tx=XM_019832269.2
B3	43010722	43010817
refGene_gn=SNX22;refGene_tx=XM_003987046.5;refGene_gn=SNX22;refGene_tx
=XM_019832267.2;refGene_gn=SNX22;refGene_tx=XM_019832268.2;refGene_gn=
SNX22;refGene_tx=XM_019832269.2
B3	43011333	43011438
refGene_gn=SNX22;refGene_tx=XM_003987046.5;refGene_gn=SNX22;refGene_tx
=XM_019832267.2;refGene_gn=SNX22;refGene_tx=XM_019832268.2;refGene_gn=
SNX22;refGene_tx=XM_019832269.2
B3	43011816	43011900
refGene_gn=SNX22;refGene_tx=XM_003987046.5;refGene_gn=SNX22;refGene_tx
=XM_019832267.2;refGene_gn=SNX22;refGene_tx=XM_019832268.2
B3	43012285	43012360
refGene_gn=SNX22;refGene_tx=XM_003987046.5;refGene_gn=SNX22;refGene_tx
=XM_019832267.2;refGene_gn=SNX22;refGene_tx=XM_019832268.2;refGene_gn=
SNX22;refGene_tx=XM_019832269.2
B3	43024201	43024288	refGene_gn=SNX1;refGene_tx=XM_023255221.1
B3	43024706	43024757

refGene_gn=SNX1;refGene_tx=XM_003987049.5;refGene_gn=SNX1;refGene_tx=XM_019832270.2
B3 43025344 43025416 refGene_gn=SNX1;refGene_tx=XM_003987049.5;refGene_gn=SNX1;refGene_tx=XM_019832270.2;refGene_gn=SNX1;refGene_tx=XM_023255221.1
B3 43025892 43025973 refGene_gn=SNX1;refGene_tx=XM_003987049.5;refGene_gn=SNX1;refGene_tx=XM_019832270.2;refGene_gn=SNX1;refGene_tx=XM_023255221.1
B3 43026545 43026689 refGene_gn=SNX1;refGene_tx=XM_003987049.5;refGene_gn=SNX1;refGene_tx=XM_019832270.2;refGene_gn=SNX1;refGene_tx=XM_023255221.1
B3 43029439 43029645 refGene_gn=SNX1;refGene_tx=XM_003987049.5;refGene_gn=SNX1;refGene_tx=XM_019832270.2;refGene_gn=SNX1;refGene_tx=XM_023255221.1
B3 43030876 43030970 refGene_gn=SNX1;refGene_tx=XM_003987049.5;refGene_gn=SNX1;refGene_tx=XM_019832270.2;refGene_gn=SNX1;refGene_tx=XM_023255221.1
B3 43031157 43031271 refGene_gn=SNX1;refGene_tx=XM_003987049.5;refGene_gn=SNX1;refGene_tx=XM_019832270.2;refGene_gn=SNX1;refGene_tx=XM_023255221.1
B3 43033889 43033965 refGene_gn=SNX1;refGene_tx=XM_003987049.5;refGene_gn=SNX1;refGene_tx=XM_019832270.2;refGene_gn=SNX1;refGene_tx=XM_023255221.1
B3 43034411 43034490 refGene_gn=SNX1;refGene_tx=XM_003987049.5;refGene_gn=SNX1;refGene_tx=XM_019832270.2;refGene_gn=SNX1;refGene_tx=XM_023255221.1
B3 43035211 43035353 refGene_gn=SNX1;refGene_tx=XM_003987049.5;refGene_gn=SNX1;refGene_tx=XM_019832270.2;refGene_gn=SNX1;refGene_tx=XM_023255221.1
B3 43037316 43037360 refGene_gn=SNX1;refGene_tx=XM_003987049.5;refGene_gn=SNX1;refGene_tx=XM_019832270.2;refGene_gn=SNX1;refGene_tx=XM_023255221.1
B3 43042186 43042253 refGene_gn=SNX1;refGene_tx=XM_003987049.5;refGene_gn=SNX1;refGene_tx=XM_019832270.2;refGene_gn=SNX1;refGene_tx=XM_023255221.1
B3 43042801 43042929 refGene_gn=SNX1;refGene_tx=XM_003987049.5;refGene_gn=SNX1;refGene_tx=XM_019832270.2;refGene_gn=SNX1;refGene_tx=XM_023255221.1
B3 43048557 43048666 refGene_gn=SNX1;refGene_tx=XM_003987049.5;refGene_gn=SNX1;refGene_tx=XM_019832270.2;refGene_gn=SNX1;refGene_tx=XM_023255221.1
B3 43062070 43062229 refGene_gn=SNX1;refGene_tx=XM_003987049.5;refGene_gn=SNX1;refGene_tx=XM_023255221.1
B3 43063804 43063819 refGene_gn=SNX1;refGene_tx=XM_019832270.2
B3 43064221 43064345 refGene_gn=FAM96A;refGene_tx=XM_003987051.5
B3 43069902 43070067 refGene_gn=FAM96A;refGene_tx=XM_003987051.5
B3 43075239 43075289 refGene_gn=FAM96A;refGene_tx=XM_003987051.5
B3 43077221 43077267 refGene_gn=FAM96A;refGene_tx=XM_003987051.5
B3 43078441 43078539 refGene_gn=FAM96A;refGene_tx=XM_003987051.5

B3	43112295	43112387
refGene_gn=DAPK2;refGene_tx=XM_006932468.3;refGene_gn=DAPK2;refGene_tx=XM_006932471.4;refGene_gn=DAPK2;refGene_tx=XM_019832271.2;refGene_gn=DAPK2;refGene_tx=XM_019832272.2;refGene_gn=DAPK2;refGene_tx=XM_019832273.2;refGene_gn=DAPK2;refGene_tx=XM_023255222.1;refGene_gn=DAPK2;refGene_tx=XM_023255223.1
B3	43172557	43172779
refGene_gn=DAPK2;refGene_tx=XM_006932468.3;refGene_gn=DAPK2;refGene_tx=XM_006932471.4;refGene_gn=DAPK2;refGene_tx=XM_019832271.2;refGene_gn=DAPK2;refGene_tx=XM_019832272.2;refGene_gn=DAPK2;refGene_tx=XM_019832273.2;refGene_gn=DAPK2;refGene_tx=XM_023255222.1;refGene_gn=DAPK2;refGene_tx=XM_023255223.1
B3	43181055	43181194
refGene_gn=DAPK2;refGene_tx=XM_006932468.3;refGene_gn=DAPK2;refGene_tx=XM_006932471.4;refGene_gn=DAPK2;refGene_tx=XM_019832271.2;refGene_gn=DAPK2;refGene_tx=XM_019832272.2;refGene_gn=DAPK2;refGene_tx=XM_019832273.2;refGene_gn=DAPK2;refGene_tx=XM_023255222.1;refGene_gn=DAPK2;refGene_tx=XM_023255223.1
B3	43200589	43200719
refGene_gn=DAPK2;refGene_tx=XM_006932468.3;refGene_gn=DAPK2;refGene_tx=XM_006932471.4;refGene_gn=DAPK2;refGene_tx=XM_019832271.2;refGene_gn=DAPK2;refGene_tx=XM_019832272.2;refGene_gn=DAPK2;refGene_tx=XM_019832273.2;refGene_gn=DAPK2;refGene_tx=XM_023255222.1;refGene_gn=DAPK2;refGene_tx=XM_023255223.1;refGene_gn=DAPK2;refGene_tx=XM_023255224.1
B3	43206145	43206194
refGene_gn=DAPK2;refGene_tx=XM_006932468.3;refGene_gn=DAPK2;refGene_tx=XM_006932471.4;refGene_gn=DAPK2;refGene_tx=XM_019832271.2;refGene_gn=DAPK2;refGene_tx=XM_019832272.2;refGene_gn=DAPK2;refGene_tx=XM_019832273.2;refGene_gn=DAPK2;refGene_tx=XM_023255222.1;refGene_gn=DAPK2;refGene_tx=XM_023255223.1;refGene_gn=DAPK2;refGene_tx=XM_023255224.1
B3	43206993	43207020
refGene_gn=DAPK2;refGene_tx=XM_006932468.3;refGene_gn=DAPK2;refGene_tx=XM_006932471.4;refGene_gn=DAPK2;refGene_tx=XM_019832271.2;refGene_gn=DAPK2;refGene_tx=XM_019832272.2;refGene_gn=DAPK2;refGene_tx=XM_019832273.2;refGene_gn=DAPK2;refGene_tx=XM_023255222.1;refGene_gn=DAPK2;refGene_tx=XM_023255223.1;refGene_gn=DAPK2;refGene_tx=XM_023255224.1
B3	43209852	43210005
refGene_gn=DAPK2;refGene_tx=XM_006932468.3;refGene_gn=DAPK2;refGene_tx=XM_006932471.4;refGene_gn=DAPK2;refGene_tx=XM_019832271.2;refGene_gn=DAPK2;refGene_tx=XM_019832272.2;refGene_gn=DAPK2;refGene_tx=XM_019832273.2;refGene_gn=DAPK2;refGene_tx=XM_023255222.1;refGene_gn=DAPK2;refGene_tx=XM_023255223.1;refGene_gn=DAPK2;refGene_tx=XM_023255224.1
B3	43211030	43211076
refGene_gn=DAPK2;refGene_tx=XM_006932468.3;refGene_gn=DAPK2;refGene_tx=XM_006932471.4;refGene_gn=DAPK2;refGene_tx=XM_019832271.2;refGene_gn=DAPK2;refGene_tx=XM_019832272.2;refGene_gn=DAPK2;refGene_tx=XM_019832273.2;refGene_gn=DAPK2;refGene_tx=XM_023255222.1;refGene_gn=DAPK2;refGene_tx=XM_023255223.1;refGene_gn=DAPK2;refGene_tx=XM_023255224.1
B3	43212571	43213180
refGene_gn=DAPK2;refGene_tx=XM_006932468.3;refGene_gn=DAPK2;refGene_tx

=XM_006932471.4;refGene_gn=DAPK2;refGene_tx=XM_019832271.2;refGene_gn=DAPK2;refGene_tx=XM_019832272.2;refGene_gn=DAPK2;refGene_tx=XM_023255222.1;refGene_gn=DAPK2;refGene_tx=XM_023255224.1
B3 43224397 43224487
refGene_gn=DAPK2;refGene_tx=XM_019832273.2;refGene_gn=DAPK2;refGene_tx=XM_023255223.1
B3 43224596 43224684
refGene_gn=DAPK2;refGene_tx=XM_019832273.2;refGene_gn=DAPK2;refGene_tx=XM_023255223.1
B3 43227114 43227195
refGene_gn=DAPK2;refGene_tx=XM_019832273.2;refGene_gn=DAPK2;refGene_tx=XM_023255223.1
B3 43335975 43336905
refGene_gn=HERC1;refGene_tx=XM_023255225.1;refGene_gn=HERC1;refGene_tx=XM_023255226.1;refGene_gn=HERC1;refGene_tx=XM_023255227.1;refGene_gn=HERC1;refGene_tx=XM_023255228.1;refGene_gn=HERC1;refGene_tx=XM_023255229.1;refGene_gn=HERC1;refGene_tx=XM_023255231.1;refGene_gn=HERC1;refGene_tx=XM_023255232.1;refGene_gn=HERC1;refGene_tx=XM_023255233.1;refGene_gn=HERC1;refGene_tx=XM_023255234.1
B3 43341301 43341397
refGene_gn=HERC1;refGene_tx=XM_023255225.1;refGene_gn=HERC1;refGene_tx=XM_023255226.1;refGene_gn=HERC1;refGene_tx=XM_023255227.1;refGene_gn=HERC1;refGene_tx=XM_023255228.1;refGene_gn=HERC1;refGene_tx=XM_023255229.1;refGene_gn=HERC1;refGene_tx=XM_023255231.1;refGene_gn=HERC1;refGene_tx=XM_023255232.1;refGene_gn=HERC1;refGene_tx=XM_023255233.1;refGene_gn=HERC1;refGene_tx=XM_023255234.1
B3 43349052 43349247
refGene_gn=HERC1;refGene_tx=XM_023255225.1;refGene_gn=HERC1;refGene_tx=XM_023255226.1;refGene_gn=HERC1;refGene_tx=XM_023255227.1;refGene_gn=HERC1;refGene_tx=XM_023255228.1;refGene_gn=HERC1;refGene_tx=XM_023255229.1;refGene_gn=HERC1;refGene_tx=XM_023255231.1;refGene_gn=HERC1;refGene_tx=XM_023255232.1;refGene_gn=HERC1;refGene_tx=XM_023255233.1;refGene_gn=HERC1;refGene_tx=XM_023255234.1
B3 43350046 43350358
refGene_gn=HERC1;refGene_tx=XM_023255225.1;refGene_gn=HERC1;refGene_tx=XM_023255226.1;refGene_gn=HERC1;refGene_tx=XM_023255227.1;refGene_gn=HERC1;refGene_tx=XM_023255228.1;refGene_gn=HERC1;refGene_tx=XM_023255229.1;refGene_gn=HERC1;refGene_tx=XM_023255231.1;refGene_gn=HERC1;refGene_tx=XM_023255232.1;refGene_gn=HERC1;refGene_tx=XM_023255233.1;refGene_gn=HERC1;refGene_tx=XM_023255234.1
B3 43352520 43352617
refGene_gn=HERC1;refGene_tx=XM_023255225.1;refGene_gn=HERC1;refGene_tx=XM_023255226.1;refGene_gn=HERC1;refGene_tx=XM_023255227.1;refGene_gn=HERC1;refGene_tx=XM_023255228.1;refGene_gn=HERC1;refGene_tx=XM_023255229.1;refGene_gn=HERC1;refGene_tx=XM_023255231.1;refGene_gn=HERC1;refGene_tx=XM_023255232.1;refGene_gn=HERC1;refGene_tx=XM_023255233.1;refGene_gn=HERC1;refGene_tx=XM_023255234.1
B3 43353208 43353352
refGene_gn=HERC1;refGene_tx=XM_023255225.1;refGene_gn=HERC1;refGene_tx=XM_023255226.1;refGene_gn=HERC1;refGene_tx=XM_023255227.1;refGene_gn=

HERC1;refGene_tx=XM_023255228.1;refGene_gn=HERC1;refGene_tx=XM_023255229.1;refGene_gn=HERC1;refGene_tx=XM_023255231.1;refGene_gn=HERC1;refGene_tx=XM_023255232.1;refGene_gn=HERC1;refGene_tx=XM_023255233.1;refGene_gn=HERC1;refGene_tx=XM_023255234.1
B3 43354602 43354730
refGene_gn=HERC1;refGene_tx=XM_023255225.1;refGene_gn=HERC1;refGene_tx=XM_023255226.1;refGene_gn=HERC1;refGene_tx=XM_023255227.1;refGene_gn=HERC1;refGene_tx=XM_023255228.1;refGene_gn=HERC1;refGene_tx=XM_023255229.1;refGene_gn=HERC1;refGene_tx=XM_023255231.1;refGene_gn=HERC1;refGene_tx=XM_023255232.1;refGene_gn=HERC1;refGene_tx=XM_023255233.1;refGene_gn=HERC1;refGene_tx=XM_023255234.1
B3 43357169 43357314
refGene_gn=HERC1;refGene_tx=XM_023255225.1;refGene_gn=HERC1;refGene_tx=XM_023255226.1;refGene_gn=HERC1;refGene_tx=XM_023255227.1;refGene_gn=HERC1;refGene_tx=XM_023255228.1;refGene_gn=HERC1;refGene_tx=XM_023255229.1;refGene_gn=HERC1;refGene_tx=XM_023255231.1;refGene_gn=HERC1;refGene_tx=XM_023255232.1;refGene_gn=HERC1;refGene_tx=XM_023255233.1;refGene_gn=HERC1;refGene_tx=XM_023255234.1
B3 43357416 43357588
refGene_gn=HERC1;refGene_tx=XM_023255225.1;refGene_gn=HERC1;refGene_tx=XM_023255226.1;refGene_gn=HERC1;refGene_tx=XM_023255227.1;refGene_gn=HERC1;refGene_tx=XM_023255228.1;refGene_gn=HERC1;refGene_tx=XM_023255229.1;refGene_gn=HERC1;refGene_tx=XM_023255231.1;refGene_gn=HERC1;refGene_tx=XM_023255232.1;refGene_gn=HERC1;refGene_tx=XM_023255233.1;refGene_gn=HERC1;refGene_tx=XM_023255234.1
B3 43358774 43358909
refGene_gn=HERC1;refGene_tx=XM_023255225.1;refGene_gn=HERC1;refGene_tx=XM_023255226.1;refGene_gn=HERC1;refGene_tx=XM_023255227.1;refGene_gn=HERC1;refGene_tx=XM_023255228.1;refGene_gn=HERC1;refGene_tx=XM_023255229.1;refGene_gn=HERC1;refGene_tx=XM_023255231.1;refGene_gn=HERC1;refGene_tx=XM_023255232.1;refGene_gn=HERC1;refGene_tx=XM_023255233.1;refGene_gn=HERC1;refGene_tx=XM_023255234.1
B3 43359270 43359436
refGene_gn=HERC1;refGene_tx=XM_023255225.1;refGene_gn=HERC1;refGene_tx=XM_023255226.1;refGene_gn=HERC1;refGene_tx=XM_023255227.1;refGene_gn=HERC1;refGene_tx=XM_023255228.1;refGene_gn=HERC1;refGene_tx=XM_023255229.1;refGene_gn=HERC1;refGene_tx=XM_023255231.1;refGene_gn=HERC1;refGene_tx=XM_023255232.1;refGene_gn=HERC1;refGene_tx=XM_023255233.1;refGene_gn=HERC1;refGene_tx=XM_023255234.1
B3 43364161 43364287
refGene_gn=HERC1;refGene_tx=XM_023255225.1;refGene_gn=HERC1;refGene_tx=XM_023255226.1;refGene_gn=HERC1;refGene_tx=XM_023255227.1;refGene_gn=HERC1;refGene_tx=XM_023255228.1;refGene_gn=HERC1;refGene_tx=XM_023255229.1;refGene_gn=HERC1;refGene_tx=XM_023255231.1;refGene_gn=HERC1;refGene_tx=XM_023255232.1;refGene_gn=HERC1;refGene_tx=XM_023255233.1;refGene_gn=HERC1;refGene_tx=XM_023255234.1
B3 43365387 43365609
refGene_gn=HERC1;refGene_tx=XM_023255225.1;refGene_gn=HERC1;refGene_tx=XM_023255226.1;refGene_gn=HERC1;refGene_tx=XM_023255227.1;refGene_gn=HERC1;refGene_tx=XM_023255228.1;refGene_gn=HERC1;refGene_tx=XM_0232552

29.1;refGene_gn=HERC1;refGene_tx=XM_023255231.1;refGene_gn=HERC1;refGene_tx=XM_023255232.1;refGene_gn=HERC1;refGene_tx=XM_023255233.1;refGene_gn=HERC1;refGene_tx=XM_023255234.1
B3 43369407 43369560
refGene_gn=HERC1;refGene_tx=XM_023255225.1;refGene_gn=HERC1;refGene_tx=XM_023255226.1;refGene_gn=HERC1;refGene_tx=XM_023255227.1;refGene_gn=HERC1;refGene_tx=XM_023255228.1;refGene_gn=HERC1;refGene_tx=XM_023255229.1;refGene_gn=HERC1;refGene_tx=XM_023255231.1;refGene_gn=HERC1;refGene_tx=XM_023255232.1;refGene_gn=HERC1;refGene_tx=XM_023255233.1;refGene_gn=HERC1;refGene_tx=XM_023255234.1
B3 43369686 43369819
refGene_gn=HERC1;refGene_tx=XM_023255225.1;refGene_gn=HERC1;refGene_tx=XM_023255226.1;refGene_gn=HERC1;refGene_tx=XM_023255227.1;refGene_gn=HERC1;refGene_tx=XM_023255228.1;refGene_gn=HERC1;refGene_tx=XM_023255229.1;refGene_gn=HERC1;refGene_tx=XM_023255231.1;refGene_gn=HERC1;refGene_tx=XM_023255232.1;refGene_gn=HERC1;refGene_tx=XM_023255233.1;refGene_gn=HERC1;refGene_tx=XM_023255234.1
B3 43371616 43371808
refGene_gn=HERC1;refGene_tx=XM_023255225.1;refGene_gn=HERC1;refGene_tx=XM_023255226.1;refGene_gn=HERC1;refGene_tx=XM_023255227.1;refGene_gn=HERC1;refGene_tx=XM_023255228.1;refGene_gn=HERC1;refGene_tx=XM_023255229.1;refGene_gn=HERC1;refGene_tx=XM_023255231.1;refGene_gn=HERC1;refGene_tx=XM_023255232.1;refGene_gn=HERC1;refGene_tx=XM_023255233.1;refGene_gn=HERC1;refGene_tx=XM_023255234.1
B3 43374561 43374783
refGene_gn=HERC1;refGene_tx=XM_023255225.1;refGene_gn=HERC1;refGene_tx=XM_023255226.1;refGene_gn=HERC1;refGene_tx=XM_023255227.1;refGene_gn=HERC1;refGene_tx=XM_023255228.1;refGene_gn=HERC1;refGene_tx=XM_023255229.1;refGene_gn=HERC1;refGene_tx=XM_023255231.1;refGene_gn=HERC1;refGene_tx=XM_023255232.1;refGene_gn=HERC1;refGene_tx=XM_023255233.1;refGene_gn=HERC1;refGene_tx=XM_023255234.1
B3 43376861 43377035
refGene_gn=HERC1;refGene_tx=XM_023255225.1;refGene_gn=HERC1;refGene_tx=XM_023255226.1;refGene_gn=HERC1;refGene_tx=XM_023255227.1;refGene_gn=HERC1;refGene_tx=XM_023255228.1;refGene_gn=HERC1;refGene_tx=XM_023255229.1;refGene_gn=HERC1;refGene_tx=XM_023255231.1;refGene_gn=HERC1;refGene_tx=XM_023255232.1;refGene_gn=HERC1;refGene_tx=XM_023255233.1;refGene_gn=HERC1;refGene_tx=XM_023255234.1
B3 43380141 43380256
refGene_gn=HERC1;refGene_tx=XM_023255225.1;refGene_gn=HERC1;refGene_tx=XM_023255226.1;refGene_gn=HERC1;refGene_tx=XM_023255227.1;refGene_gn=HERC1;refGene_tx=XM_023255228.1;refGene_gn=HERC1;refGene_tx=XM_023255229.1;refGene_gn=HERC1;refGene_tx=XM_023255231.1;refGene_gn=HERC1;refGene_tx=XM_023255232.1;refGene_gn=HERC1;refGene_tx=XM_023255233.1;refGene_gn=HERC1;refGene_tx=XM_023255234.1
B3 43380344 43380465
refGene_gn=HERC1;refGene_tx=XM_023255225.1;refGene_gn=HERC1;refGene_tx=XM_023255226.1;refGene_gn=HERC1;refGene_tx=XM_023255227.1;refGene_gn=HERC1;refGene_tx=XM_023255228.1;refGene_gn=HERC1;refGene_tx=XM_023255229.1;refGene_gn=HERC1;refGene_tx=XM_023255231.1;refGene_gn=HERC1;refGe

ne_tx=XM_023255232.1;refGene_gn=HERC1;refGene_tx=XM_023255233.1;refGene_gn=HERC1;refGene_tx=XM_023255234.1
B3 43382319 43382491
refGene_gn=HERC1;refGene_tx=XM_023255225.1;refGene_gn=HERC1;refGene_tx=XM_023255226.1;refGene_gn=HERC1;refGene_tx=XM_023255227.1;refGene_gn=HERC1;refGene_tx=XM_023255228.1;refGene_gn=HERC1;refGene_tx=XM_023255229.1;refGene_gn=HERC1;refGene_tx=XM_023255231.1;refGene_gn=HERC1;refGene_tx=XM_023255232.1;refGene_gn=HERC1;refGene_tx=XM_023255233.1;refGene_gn=HERC1;refGene_tx=XM_023255234.1
B3 43384764 43385104
refGene_gn=HERC1;refGene_tx=XM_023255225.1;refGene_gn=HERC1;refGene_tx=XM_023255226.1;refGene_gn=HERC1;refGene_tx=XM_023255227.1;refGene_gn=HERC1;refGene_tx=XM_023255228.1;refGene_gn=HERC1;refGene_tx=XM_023255229.1;refGene_gn=HERC1;refGene_tx=XM_023255231.1;refGene_gn=HERC1;refGene_tx=XM_023255232.1;refGene_gn=HERC1;refGene_tx=XM_023255233.1;refGene_gn=HERC1;refGene_tx=XM_023255234.1
B3 43385571 43385692
refGene_gn=HERC1;refGene_tx=XM_023255225.1;refGene_gn=HERC1;refGene_tx=XM_023255226.1;refGene_gn=HERC1;refGene_tx=XM_023255227.1;refGene_gn=HERC1;refGene_tx=XM_023255228.1;refGene_gn=HERC1;refGene_tx=XM_023255229.1;refGene_gn=HERC1;refGene_tx=XM_023255231.1;refGene_gn=HERC1;refGene_tx=XM_023255232.1;refGene_gn=HERC1;refGene_tx=XM_023255233.1;refGene_gn=HERC1;refGene_tx=XM_023255234.1
B3 43391843 43391895
refGene_gn=HERC1;refGene_tx=XM_023255225.1;refGene_gn=HERC1;refGene_tx=XM_023255226.1;refGene_gn=HERC1;refGene_tx=XM_023255227.1;refGene_gn=HERC1;refGene_tx=XM_023255228.1;refGene_gn=HERC1;refGene_tx=XM_023255229.1;refGene_gn=HERC1;refGene_tx=XM_023255231.1;refGene_gn=HERC1;refGene_tx=XM_023255232.1;refGene_gn=HERC1;refGene_tx=XM_023255233.1;refGene_gn=HERC1;refGene_tx=XM_023255234.1
B3 43398604 43398873
refGene_gn=HERC1;refGene_tx=XM_023255225.1;refGene_gn=HERC1;refGene_tx=XM_023255226.1;refGene_gn=HERC1;refGene_tx=XM_023255227.1;refGene_gn=HERC1;refGene_tx=XM_023255228.1;refGene_gn=HERC1;refGene_tx=XM_023255229.1;refGene_gn=HERC1;refGene_tx=XM_023255231.1;refGene_gn=HERC1;refGene_tx=XM_023255232.1;refGene_gn=HERC1;refGene_tx=XM_023255233.1;refGene_gn=HERC1;refGene_tx=XM_023255234.1
B3 43400627 43400843
refGene_gn=HERC1;refGene_tx=XM_023255225.1;refGene_gn=HERC1;refGene_tx=XM_023255226.1;refGene_gn=HERC1;refGene_tx=XM_023255227.1;refGene_gn=HERC1;refGene_tx=XM_023255228.1;refGene_gn=HERC1;refGene_tx=XM_023255229.1;refGene_gn=HERC1;refGene_tx=XM_023255231.1;refGene_gn=HERC1;refGene_tx=XM_023255232.1;refGene_gn=HERC1;refGene_tx=XM_023255233.1;refGene_gn=HERC1;refGene_tx=XM_023255234.1
B3 43401483 43401604
refGene_gn=HERC1;refGene_tx=XM_023255225.1;refGene_gn=HERC1;refGene_tx=XM_023255226.1;refGene_gn=HERC1;refGene_tx=XM_023255227.1;refGene_gn=HERC1;refGene_tx=XM_023255228.1;refGene_gn=HERC1;refGene_tx=XM_023255229.1;refGene_gn=HERC1;refGene_tx=XM_023255231.1;refGene_gn=HERC1;refGene_tx=XM_023255232.1;refGene_gn=HERC1;refGene_tx=XM_023255233.1;refGen

e_gn=HERC1;refGene_tx=XM_023255234.1
B3 43401833 43402071
refGene_gn=HERC1;refGene_tx=XM_023255225.1;refGene_gn=HERC1;refGene_tx=XM_023255226.1;refGene_gn=HERC1;refGene_tx=XM_023255227.1;refGene_gn=HERC1;refGene_tx=XM_023255228.1;refGene_gn=HERC1;refGene_tx=XM_023255229.1;refGene_gn=HERC1;refGene_tx=XM_023255231.1;refGene_gn=HERC1;refGene_tx=XM_023255232.1;refGene_gn=HERC1;refGene_tx=XM_023255233.1;refGene_gn=HERC1;refGene_tx=XM_023255234.1
B3 43402220 43402414
refGene_gn=HERC1;refGene_tx=XM_023255225.1;refGene_gn=HERC1;refGene_tx=XM_023255226.1;refGene_gn=HERC1;refGene_tx=XM_023255227.1;refGene_gn=HERC1;refGene_tx=XM_023255228.1;refGene_gn=HERC1;refGene_tx=XM_023255229.1;refGene_gn=HERC1;refGene_tx=XM_023255231.1;refGene_gn=HERC1;refGene_tx=XM_023255232.1;refGene_gn=HERC1;refGene_tx=XM_023255233.1;refGene_gn=HERC1;refGene_tx=XM_023255234.1
B3 43405450 43405606
refGene_gn=HERC1;refGene_tx=XM_023255225.1;refGene_gn=HERC1;refGene_tx=XM_023255226.1;refGene_gn=HERC1;refGene_tx=XM_023255227.1;refGene_gn=HERC1;refGene_tx=XM_023255228.1;refGene_gn=HERC1;refGene_tx=XM_023255229.1;refGene_gn=HERC1;refGene_tx=XM_023255231.1;refGene_gn=HERC1;refGene_tx=XM_023255232.1;refGene_gn=HERC1;refGene_tx=XM_023255233.1;refGene_gn=HERC1;refGene_tx=XM_023255234.1
B3 43407748 43407855
refGene_gn=HERC1;refGene_tx=XM_023255225.1;refGene_gn=HERC1;refGene_tx=XM_023255226.1;refGene_gn=HERC1;refGene_tx=XM_023255227.1;refGene_gn=HERC1;refGene_tx=XM_023255228.1;refGene_gn=HERC1;refGene_tx=XM_023255229.1;refGene_gn=HERC1;refGene_tx=XM_023255231.1;refGene_gn=HERC1;refGene_tx=XM_023255232.1;refGene_gn=HERC1;refGene_tx=XM_023255233.1;refGene_gn=HERC1;refGene_tx=XM_023255234.1
B3 43408388 43408499
refGene_gn=HERC1;refGene_tx=XM_023255225.1;refGene_gn=HERC1;refGene_tx=XM_023255226.1;refGene_gn=HERC1;refGene_tx=XM_023255227.1;refGene_gn=HERC1;refGene_tx=XM_023255228.1;refGene_gn=HERC1;refGene_tx=XM_023255229.1;refGene_gn=HERC1;refGene_tx=XM_023255231.1;refGene_gn=HERC1;refGene_tx=XM_023255232.1;refGene_gn=HERC1;refGene_tx=XM_023255233.1;refGene_gn=HERC1;refGene_tx=XM_023255234.1
B3 43411314 43411491
refGene_gn=HERC1;refGene_tx=XM_023255225.1;refGene_gn=HERC1;refGene_tx=XM_023255226.1;refGene_gn=HERC1;refGene_tx=XM_023255227.1;refGene_gn=HERC1;refGene_tx=XM_023255228.1;refGene_gn=HERC1;refGene_tx=XM_023255229.1;refGene_gn=HERC1;refGene_tx=XM_023255231.1;refGene_gn=HERC1;refGene_tx=XM_023255232.1;refGene_gn=HERC1;refGene_tx=XM_023255233.1;refGene_gn=HERC1;refGene_tx=XM_023255234.1
B3 43415447 43415687
refGene_gn=HERC1;refGene_tx=XM_023255225.1;refGene_gn=HERC1;refGene_tx=XM_023255226.1;refGene_gn=HERC1;refGene_tx=XM_023255227.1;refGene_gn=HERC1;refGene_tx=XM_023255228.1;refGene_gn=HERC1;refGene_tx=XM_023255229.1;refGene_gn=HERC1;refGene_tx=XM_023255231.1;refGene_gn=HERC1;refGene_tx=XM_023255232.1;refGene_gn=HERC1;refGene_tx=XM_023255233.1;refGene_gn=HERC1;refGene_tx=XM_023255234.1

B3 43416052 43416136
refGene_gn=HERC1;refGene_tx=XM_023255225.1;refGene_gn=HERC1;refGene_tx=XM_023255226.1;refGene_gn=HERC1;refGene_tx=XM_023255227.1;refGene_gn=HERC1;refGene_tx=XM_023255228.1;refGene_gn=HERC1;refGene_tx=XM_023255229.1;refGene_gn=HERC1;refGene_tx=XM_023255231.1;refGene_gn=HERC1;refGene_tx=XM_023255232.1;refGene_gn=HERC1;refGene_tx=XM_023255233.1;refGene_gn=HERC1;refGene_tx=XM_023255234.1
B3 43417884 43418405
refGene_gn=HERC1;refGene_tx=XM_023255225.1;refGene_gn=HERC1;refGene_tx=XM_023255226.1;refGene_gn=HERC1;refGene_tx=XM_023255227.1;refGene_gn=HERC1;refGene_tx=XM_023255228.1;refGene_gn=HERC1;refGene_tx=XM_023255229.1;refGene_gn=HERC1;refGene_tx=XM_023255231.1;refGene_gn=HERC1;refGene_tx=XM_023255232.1;refGene_gn=HERC1;refGene_tx=XM_023255233.1;refGene_gn=HERC1;refGene_tx=XM_023255234.1
B3 43420173 43420949
refGene_gn=HERC1;refGene_tx=XM_023255225.1;refGene_gn=HERC1;refGene_tx=XM_023255226.1;refGene_gn=HERC1;refGene_tx=XM_023255227.1;refGene_gn=HERC1;refGene_tx=XM_023255228.1;refGene_gn=HERC1;refGene_tx=XM_023255229.1;refGene_gn=HERC1;refGene_tx=XM_023255231.1;refGene_gn=HERC1;refGene_tx=XM_023255232.1;refGene_gn=HERC1;refGene_tx=XM_023255233.1;refGene_gn=HERC1;refGene_tx=XM_023255234.1
B3 43422981 43423180
refGene_gn=HERC1;refGene_tx=XM_023255225.1;refGene_gn=HERC1;refGene_tx=XM_023255226.1;refGene_gn=HERC1;refGene_tx=XM_023255227.1;refGene_gn=HERC1;refGene_tx=XM_023255228.1;refGene_gn=HERC1;refGene_tx=XM_023255229.1;refGene_gn=HERC1;refGene_tx=XM_023255231.1;refGene_gn=HERC1;refGene_tx=XM_023255232.1;refGene_gn=HERC1;refGene_tx=XM_023255233.1;refGene_gn=HERC1;refGene_tx=XM_023255234.1
B3 43425655 43425816
refGene_gn=HERC1;refGene_tx=XM_023255225.1;refGene_gn=HERC1;refGene_tx=XM_023255226.1;refGene_gn=HERC1;refGene_tx=XM_023255227.1;refGene_gn=HERC1;refGene_tx=XM_023255228.1;refGene_gn=HERC1;refGene_tx=XM_023255229.1;refGene_gn=HERC1;refGene_tx=XM_023255231.1;refGene_gn=HERC1;refGene_tx=XM_023255232.1;refGene_gn=HERC1;refGene_tx=XM_023255233.1;refGene_gn=HERC1;refGene_tx=XM_023255234.1
B3 43431631 43431748
refGene_gn=HERC1;refGene_tx=XM_023255225.1;refGene_gn=HERC1;refGene_tx=XM_023255226.1;refGene_gn=HERC1;refGene_tx=XM_023255227.1;refGene_gn=HERC1;refGene_tx=XM_023255228.1;refGene_gn=HERC1;refGene_tx=XM_023255229.1;refGene_gn=HERC1;refGene_tx=XM_023255231.1;refGene_gn=HERC1;refGene_tx=XM_023255232.1;refGene_gn=HERC1;refGene_tx=XM_023255233.1;refGene_gn=HERC1;refGene_tx=XM_023255234.1
B3 43431949 43432181
refGene_gn=HERC1;refGene_tx=XM_023255225.1;refGene_gn=HERC1;refGene_tx=XM_023255226.1;refGene_gn=HERC1;refGene_tx=XM_023255227.1;refGene_gn=HERC1;refGene_tx=XM_023255228.1;refGene_gn=HERC1;refGene_tx=XM_023255229.1;refGene_gn=HERC1;refGene_tx=XM_023255231.1;refGene_gn=HERC1;refGene_tx=XM_023255232.1;refGene_gn=HERC1;refGene_tx=XM_023255233.1;refGene_gn=HERC1;refGene_tx=XM_023255234.1
B3 43433083 43433208

refGene_gn=HERC1;refGene_tx=XM_023255225.1;refGene_gn=HERC1;refGene_tx=XM_023255226.1;refGene_gn=HERC1;refGene_tx=XM_023255227.1;refGene_gn=HERC1;refGene_tx=XM_023255228.1;refGene_gn=HERC1;refGene_tx=XM_023255229.1;refGene_gn=HERC1;refGene_tx=XM_023255231.1;refGene_gn=HERC1;refGene_tx=XM_023255232.1;refGene_gn=HERC1;refGene_tx=XM_023255233.1;refGene_gn=HERC1;refGene_tx=XM_023255234.1
B3 43434396 43434617
refGene_gn=HERC1;refGene_tx=XM_023255225.1;refGene_gn=HERC1;refGene_tx=XM_023255226.1;refGene_gn=HERC1;refGene_tx=XM_023255227.1;refGene_gn=HERC1;refGene_tx=XM_023255228.1;refGene_gn=HERC1;refGene_tx=XM_023255229.1;refGene_gn=HERC1;refGene_tx=XM_023255231.1;refGene_gn=HERC1;refGene_tx=XM_023255232.1;refGene_gn=HERC1;refGene_tx=XM_023255233.1;refGene_gn=HERC1;refGene_tx=XM_023255234.1
B3 43435566 43435835
refGene_gn=HERC1;refGene_tx=XM_023255225.1;refGene_gn=HERC1;refGene_tx=XM_023255226.1;refGene_gn=HERC1;refGene_tx=XM_023255227.1;refGene_gn=HERC1;refGene_tx=XM_023255228.1;refGene_gn=HERC1;refGene_tx=XM_023255229.1;refGene_gn=HERC1;refGene_tx=XM_023255231.1;refGene_gn=HERC1;refGene_tx=XM_023255232.1;refGene_gn=HERC1;refGene_tx=XM_023255233.1;refGene_gn=HERC1;refGene_tx=XM_023255234.1
B3 43436579 43436632
refGene_gn=HERC1;refGene_tx=XM_023255225.1;refGene_gn=HERC1;refGene_tx=XM_023255226.1;refGene_gn=HERC1;refGene_tx=XM_023255227.1;refGene_gn=HERC1;refGene_tx=XM_023255228.1;refGene_gn=HERC1;refGene_tx=XM_023255229.1;refGene_gn=HERC1;refGene_tx=XM_023255231.1;refGene_gn=HERC1;refGene_tx=XM_023255232.1;refGene_gn=HERC1;refGene_tx=XM_023255233.1;refGene_gn=HERC1;refGene_tx=XM_023255234.1
B3 43437324 43437525
refGene_gn=HERC1;refGene_tx=XM_023255225.1;refGene_gn=HERC1;refGene_tx=XM_023255226.1;refGene_gn=HERC1;refGene_tx=XM_023255227.1;refGene_gn=HERC1;refGene_tx=XM_023255228.1;refGene_gn=HERC1;refGene_tx=XM_023255229.1;refGene_gn=HERC1;refGene_tx=XM_023255231.1;refGene_gn=HERC1;refGene_tx=XM_023255232.1;refGene_gn=HERC1;refGene_tx=XM_023255233.1;refGene_gn=HERC1;refGene_tx=XM_023255234.1
B3 43438146 43438321
refGene_gn=HERC1;refGene_tx=XM_023255225.1;refGene_gn=HERC1;refGene_tx=XM_023255226.1;refGene_gn=HERC1;refGene_tx=XM_023255227.1;refGene_gn=HERC1;refGene_tx=XM_023255228.1;refGene_gn=HERC1;refGene_tx=XM_023255229.1;refGene_gn=HERC1;refGene_tx=XM_023255231.1;refGene_gn=HERC1;refGene_tx=XM_023255232.1;refGene_gn=HERC1;refGene_tx=XM_023255233.1;refGene_gn=HERC1;refGene_tx=XM_023255234.1
B3 43440539 43440810
refGene_gn=HERC1;refGene_tx=XM_023255225.1;refGene_gn=HERC1;refGene_tx=XM_023255226.1;refGene_gn=HERC1;refGene_tx=XM_023255227.1;refGene_gn=HERC1;refGene_tx=XM_023255228.1;refGene_gn=HERC1;refGene_tx=XM_023255229.1;refGene_gn=HERC1;refGene_tx=XM_023255231.1;refGene_gn=HERC1;refGene_tx=XM_023255232.1;refGene_gn=HERC1;refGene_tx=XM_023255233.1;refGene_gn=HERC1;refGene_tx=XM_023255234.1
B3 43440942 43441156
refGene_gn=HERC1;refGene_tx=XM_023255225.1;refGene_gn=HERC1;refGene_tx

=XM_023255226.1;refGene_gn=HERC1;refGene_tx=XM_023255227.1;refGene_gn=HERC1;refGene_tx=XM_023255228.1;refGene_gn=HERC1;refGene_tx=XM_023255229.1;refGene_gn=HERC1;refGene_tx=XM_023255231.1;refGene_gn=HERC1;refGene_tx=XM_023255232.1;refGene_gn=HERC1;refGene_tx=XM_023255233.1;refGene_gn=HERC1;refGene_tx=XM_023255234.1
B3 43441916 43442173
refGene_gn=HERC1;refGene_tx=XM_023255225.1;refGene_gn=HERC1;refGene_tx=XM_023255226.1;refGene_gn=HERC1;refGene_tx=XM_023255227.1;refGene_gn=HERC1;refGene_tx=XM_023255228.1;refGene_gn=HERC1;refGene_tx=XM_023255229.1;refGene_gn=HERC1;refGene_tx=XM_023255231.1;refGene_gn=HERC1;refGene_tx=XM_023255232.1;refGene_gn=HERC1;refGene_tx=XM_023255233.1;refGene_gn=HERC1;refGene_tx=XM_023255234.1
B3 43443272 43443400
refGene_gn=HERC1;refGene_tx=XM_023255225.1;refGene_gn=HERC1;refGene_tx=XM_023255226.1;refGene_gn=HERC1;refGene_tx=XM_023255227.1;refGene_gn=HERC1;refGene_tx=XM_023255228.1;refGene_gn=HERC1;refGene_tx=XM_023255229.1;refGene_gn=HERC1;refGene_tx=XM_023255231.1;refGene_gn=HERC1;refGene_tx=XM_023255232.1;refGene_gn=HERC1;refGene_tx=XM_023255233.1;refGene_gn=HERC1;refGene_tx=XM_023255234.1
B3 43444735 43444863
refGene_gn=HERC1;refGene_tx=XM_023255225.1;refGene_gn=HERC1;refGene_tx=XM_023255226.1;refGene_gn=HERC1;refGene_tx=XM_023255227.1;refGene_gn=HERC1;refGene_tx=XM_023255228.1;refGene_gn=HERC1;refGene_tx=XM_023255229.1;refGene_gn=HERC1;refGene_tx=XM_023255231.1;refGene_gn=HERC1;refGene_tx=XM_023255232.1;refGene_gn=HERC1;refGene_tx=XM_023255233.1;refGene_gn=HERC1;refGene_tx=XM_023255234.1
B3 43446083 43446284
refGene_gn=HERC1;refGene_tx=XM_023255225.1;refGene_gn=HERC1;refGene_tx=XM_023255226.1;refGene_gn=HERC1;refGene_tx=XM_023255227.1;refGene_gn=HERC1;refGene_tx=XM_023255228.1;refGene_gn=HERC1;refGene_tx=XM_023255229.1;refGene_gn=HERC1;refGene_tx=XM_023255231.1;refGene_gn=HERC1;refGene_tx=XM_023255232.1;refGene_gn=HERC1;refGene_tx=XM_023255233.1;refGene_gn=HERC1;refGene_tx=XM_023255234.1
B3 43447477 43447608
refGene_gn=HERC1;refGene_tx=XM_023255225.1;refGene_gn=HERC1;refGene_tx=XM_023255226.1;refGene_gn=HERC1;refGene_tx=XM_023255227.1;refGene_gn=HERC1;refGene_tx=XM_023255228.1;refGene_gn=HERC1;refGene_tx=XM_023255229.1;refGene_gn=HERC1;refGene_tx=XM_023255231.1;refGene_gn=HERC1;refGene_tx=XM_023255232.1;refGene_gn=HERC1;refGene_tx=XM_023255233.1;refGene_gn=HERC1;refGene_tx=XM_023255234.1
B3 43448410 43448610
refGene_gn=HERC1;refGene_tx=XM_023255225.1;refGene_gn=HERC1;refGene_tx=XM_023255226.1;refGene_gn=HERC1;refGene_tx=XM_023255227.1;refGene_gn=HERC1;refGene_tx=XM_023255228.1;refGene_gn=HERC1;refGene_tx=XM_023255229.1;refGene_gn=HERC1;refGene_tx=XM_023255231.1;refGene_gn=HERC1;refGene_tx=XM_023255232.1;refGene_gn=HERC1;refGene_tx=XM_023255233.1;refGene_gn=HERC1;refGene_tx=XM_023255234.1
B3 43448986 43449092
refGene_gn=HERC1;refGene_tx=XM_023255225.1;refGene_gn=HERC1;refGene_tx=XM_023255226.1;refGene_gn=HERC1;refGene_tx=XM_023255227.1;refGene_gn=

HERC1;refGene_tx=XM_023255228.1;refGene_gn=HERC1;refGene_tx=XM_023255229.1;refGene_gn=HERC1;refGene_tx=XM_023255231.1;refGene_gn=HERC1;refGene_tx=XM_023255232.1;refGene_gn=HERC1;refGene_tx=XM_023255233.1;refGene_gn=HERC1;refGene_tx=XM_023255234.1
B3 43451272 43451419
refGene_gn=HERC1;refGene_tx=XM_023255225.1;refGene_gn=HERC1;refGene_tx=XM_023255226.1;refGene_gn=HERC1;refGene_tx=XM_023255227.1;refGene_gn=HERC1;refGene_tx=XM_023255228.1;refGene_gn=HERC1;refGene_tx=XM_023255229.1;refGene_gn=HERC1;refGene_tx=XM_023255231.1;refGene_gn=HERC1;refGene_tx=XM_023255232.1;refGene_gn=HERC1;refGene_tx=XM_023255233.1;refGene_gn=HERC1;refGene_tx=XM_023255234.1
B3 43453421 43453523
refGene_gn=HERC1;refGene_tx=XM_023255225.1;refGene_gn=HERC1;refGene_tx=XM_023255226.1;refGene_gn=HERC1;refGene_tx=XM_023255227.1;refGene_gn=HERC1;refGene_tx=XM_023255228.1;refGene_gn=HERC1;refGene_tx=XM_023255229.1;refGene_gn=HERC1;refGene_tx=XM_023255231.1;refGene_gn=HERC1;refGene_tx=XM_023255232.1;refGene_gn=HERC1;refGene_tx=XM_023255233.1;refGene_gn=HERC1;refGene_tx=XM_023255234.1
B3 43454605 43454779
refGene_gn=HERC1;refGene_tx=XM_023255225.1;refGene_gn=HERC1;refGene_tx=XM_023255226.1;refGene_gn=HERC1;refGene_tx=XM_023255227.1;refGene_gn=HERC1;refGene_tx=XM_023255228.1;refGene_gn=HERC1;refGene_tx=XM_023255229.1;refGene_gn=HERC1;refGene_tx=XM_023255231.1;refGene_gn=HERC1;refGene_tx=XM_023255232.1;refGene_gn=HERC1;refGene_tx=XM_023255233.1;refGene_gn=HERC1;refGene_tx=XM_023255234.1
B3 43456279 43456573
refGene_gn=HERC1;refGene_tx=XM_023255225.1;refGene_gn=HERC1;refGene_tx=XM_023255226.1;refGene_gn=HERC1;refGene_tx=XM_023255227.1;refGene_gn=HERC1;refGene_tx=XM_023255228.1;refGene_gn=HERC1;refGene_tx=XM_023255229.1;refGene_gn=HERC1;refGene_tx=XM_023255231.1;refGene_gn=HERC1;refGene_tx=XM_023255232.1;refGene_gn=HERC1;refGene_tx=XM_023255233.1;refGene_gn=HERC1;refGene_tx=XM_023255234.1
B3 43457838 43457904
refGene_gn=HERC1;refGene_tx=XM_023255225.1;refGene_gn=HERC1;refGene_tx=XM_023255226.1;refGene_gn=HERC1;refGene_tx=XM_023255227.1;refGene_gn=HERC1;refGene_tx=XM_023255228.1;refGene_gn=HERC1;refGene_tx=XM_023255229.1;refGene_gn=HERC1;refGene_tx=XM_023255231.1;refGene_gn=HERC1;refGene_tx=XM_023255232.1;refGene_gn=HERC1;refGene_tx=XM_023255233.1;refGene_gn=HERC1;refGene_tx=XM_023255234.1
B3 43458074 43458200
refGene_gn=HERC1;refGene_tx=XM_023255225.1;refGene_gn=HERC1;refGene_tx=XM_023255226.1;refGene_gn=HERC1;refGene_tx=XM_023255227.1;refGene_gn=HERC1;refGene_tx=XM_023255228.1;refGene_gn=HERC1;refGene_tx=XM_023255229.1;refGene_gn=HERC1;refGene_tx=XM_023255231.1;refGene_gn=HERC1;refGene_tx=XM_023255232.1;refGene_gn=HERC1;refGene_tx=XM_023255233.1;refGene_gn=HERC1;refGene_tx=XM_023255234.1
B3 43458976 43459115
refGene_gn=HERC1;refGene_tx=XM_023255225.1;refGene_gn=HERC1;refGene_tx=XM_023255226.1;refGene_gn=HERC1;refGene_tx=XM_023255227.1;refGene_gn=HERC1;refGene_tx=XM_023255228.1;refGene_gn=HERC1;refGene_tx=XM_0232552

29.1;refGene_gn=HERC1;refGene_tx=XM_023255231.1;refGene_gn=HERC1;refGene_tx=XM_023255232.1;refGene_gn=HERC1;refGene_tx=XM_023255233.1;refGene_gn=HERC1;refGene_tx=XM_023255234.1
B3 43460391 43460573
refGene_gn=HERC1;refGene_tx=XM_023255225.1;refGene_gn=HERC1;refGene_tx=XM_023255226.1;refGene_gn=HERC1;refGene_tx=XM_023255227.1;refGene_gn=HERC1;refGene_tx=XM_023255228.1;refGene_gn=HERC1;refGene_tx=XM_023255229.1;refGene_gn=HERC1;refGene_tx=XM_023255231.1;refGene_gn=HERC1;refGene_tx=XM_023255232.1;refGene_gn=HERC1;refGene_tx=XM_023255233.1;refGene_gn=HERC1;refGene_tx=XM_023255234.1
B3 43461279 43461435
refGene_gn=HERC1;refGene_tx=XM_023255225.1;refGene_gn=HERC1;refGene_tx=XM_023255226.1;refGene_gn=HERC1;refGene_tx=XM_023255227.1;refGene_gn=HERC1;refGene_tx=XM_023255228.1;refGene_gn=HERC1;refGene_tx=XM_023255229.1;refGene_gn=HERC1;refGene_tx=XM_023255231.1;refGene_gn=HERC1;refGene_tx=XM_023255232.1;refGene_gn=HERC1;refGene_tx=XM_023255233.1;refGene_gn=HERC1;refGene_tx=XM_023255234.1
B3 43462149 43462272
refGene_gn=HERC1;refGene_tx=XM_023255225.1;refGene_gn=HERC1;refGene_tx=XM_023255226.1;refGene_gn=HERC1;refGene_tx=XM_023255227.1;refGene_gn=HERC1;refGene_tx=XM_023255228.1;refGene_gn=HERC1;refGene_tx=XM_023255229.1;refGene_gn=HERC1;refGene_tx=XM_023255231.1;refGene_gn=HERC1;refGene_tx=XM_023255232.1;refGene_gn=HERC1;refGene_tx=XM_023255233.1;refGene_gn=HERC1;refGene_tx=XM_023255234.1
B3 43463248 43463351
refGene_gn=HERC1;refGene_tx=XM_023255225.1;refGene_gn=HERC1;refGene_tx=XM_023255226.1;refGene_gn=HERC1;refGene_tx=XM_023255227.1;refGene_gn=HERC1;refGene_tx=XM_023255228.1;refGene_gn=HERC1;refGene_tx=XM_023255229.1;refGene_gn=HERC1;refGene_tx=XM_023255231.1;refGene_gn=HERC1;refGene_tx=XM_023255232.1;refGene_gn=HERC1;refGene_tx=XM_023255233.1;refGene_gn=HERC1;refGene_tx=XM_023255234.1
B3 43464901 43465071
refGene_gn=HERC1;refGene_tx=XM_023255225.1;refGene_gn=HERC1;refGene_tx=XM_023255226.1;refGene_gn=HERC1;refGene_tx=XM_023255227.1;refGene_gn=HERC1;refGene_tx=XM_023255228.1;refGene_gn=HERC1;refGene_tx=XM_023255229.1;refGene_gn=HERC1;refGene_tx=XM_023255231.1;refGene_gn=HERC1;refGene_tx=XM_023255232.1;refGene_gn=HERC1;refGene_tx=XM_023255233.1;refGene_gn=HERC1;refGene_tx=XM_023255234.1
B3 43466439 43466578
refGene_gn=HERC1;refGene_tx=XM_023255225.1;refGene_gn=HERC1;refGene_tx=XM_023255226.1;refGene_gn=HERC1;refGene_tx=XM_023255227.1;refGene_gn=HERC1;refGene_tx=XM_023255228.1;refGene_gn=HERC1;refGene_tx=XM_023255229.1;refGene_gn=HERC1;refGene_tx=XM_023255231.1;refGene_gn=HERC1;refGene_tx=XM_023255232.1;refGene_gn=HERC1;refGene_tx=XM_023255233.1;refGene_gn=HERC1;refGene_tx=XM_023255234.1
B3 43468523 43468693
refGene_gn=HERC1;refGene_tx=XM_023255225.1;refGene_gn=HERC1;refGene_tx=XM_023255226.1;refGene_gn=HERC1;refGene_tx=XM_023255227.1;refGene_gn=HERC1;refGene_tx=XM_023255228.1;refGene_gn=HERC1;refGene_tx=XM_023255229.1;refGene_gn=HERC1;refGene_tx=XM_023255231.1;refGene_gn=HERC1;refGe

ne_tx=XM_023255232.1;refGene_gn=HERC1;refGene_tx=XM_023255233.1;refGene_gn=HERC1;refGene_tx=XM_023255234.1
B3 43469906 43470076
refGene_gn=HERC1;refGene_tx=XM_023255225.1;refGene_gn=HERC1;refGene_tx=XM_023255226.1;refGene_gn=HERC1;refGene_tx=XM_023255227.1;refGene_gn=HERC1;refGene_tx=XM_023255228.1;refGene_gn=HERC1;refGene_tx=XM_023255229.1;refGene_gn=HERC1;refGene_tx=XM_023255231.1;refGene_gn=HERC1;refGene_tx=XM_023255232.1;refGene_gn=HERC1;refGene_tx=XM_023255233.1;refGene_gn=HERC1;refGene_tx=XM_023255234.1
B3 43470356 43470522
refGene_gn=HERC1;refGene_tx=XM_023255225.1;refGene_gn=HERC1;refGene_tx=XM_023255226.1;refGene_gn=HERC1;refGene_tx=XM_023255227.1;refGene_gn=HERC1;refGene_tx=XM_023255228.1;refGene_gn=HERC1;refGene_tx=XM_023255229.1;refGene_gn=HERC1;refGene_tx=XM_023255231.1;refGene_gn=HERC1;refGene_tx=XM_023255232.1;refGene_gn=HERC1;refGene_tx=XM_023255233.1;refGene_gn=HERC1;refGene_tx=XM_023255234.1
B3 43471317 43471394
refGene_gn=HERC1;refGene_tx=XM_023255225.1;refGene_gn=HERC1;refGene_tx=XM_023255226.1;refGene_gn=HERC1;refGene_tx=XM_023255227.1;refGene_gn=HERC1;refGene_tx=XM_023255228.1;refGene_gn=HERC1;refGene_tx=XM_023255229.1;refGene_gn=HERC1;refGene_tx=XM_023255231.1;refGene_gn=HERC1;refGene_tx=XM_023255232.1;refGene_gn=HERC1;refGene_tx=XM_023255233.1;refGene_gn=HERC1;refGene_tx=XM_023255234.1
B3 43472189 43472442
refGene_gn=HERC1;refGene_tx=XM_023255225.1;refGene_gn=HERC1;refGene_tx=XM_023255226.1;refGene_gn=HERC1;refGene_tx=XM_023255227.1;refGene_gn=HERC1;refGene_tx=XM_023255228.1;refGene_gn=HERC1;refGene_tx=XM_023255229.1;refGene_gn=HERC1;refGene_tx=XM_023255231.1;refGene_gn=HERC1;refGene_tx=XM_023255232.1;refGene_gn=HERC1;refGene_tx=XM_023255233.1;refGene_gn=HERC1;refGene_tx=XM_023255234.1
B3 43472945 43473098
refGene_gn=HERC1;refGene_tx=XM_023255225.1;refGene_gn=HERC1;refGene_tx=XM_023255226.1;refGene_gn=HERC1;refGene_tx=XM_023255227.1;refGene_gn=HERC1;refGene_tx=XM_023255228.1;refGene_gn=HERC1;refGene_tx=XM_023255229.1;refGene_gn=HERC1;refGene_tx=XM_023255231.1;refGene_gn=HERC1;refGene_tx=XM_023255232.1;refGene_gn=HERC1;refGene_tx=XM_023255233.1;refGene_gn=HERC1;refGene_tx=XM_023255234.1
B3 43475934 43476240
refGene_gn=HERC1;refGene_tx=XM_023255225.1;refGene_gn=HERC1;refGene_tx=XM_023255226.1;refGene_gn=HERC1;refGene_tx=XM_023255227.1;refGene_gn=HERC1;refGene_tx=XM_023255228.1;refGene_gn=HERC1;refGene_tx=XM_023255229.1;refGene_gn=HERC1;refGene_tx=XM_023255231.1;refGene_gn=HERC1;refGene_tx=XM_023255232.1;refGene_gn=HERC1;refGene_tx=XM_023255233.1;refGene_gn=HERC1;refGene_tx=XM_023255234.1
B3 43478865 43479051
refGene_gn=HERC1;refGene_tx=XM_023255225.1;refGene_gn=HERC1;refGene_tx=XM_023255226.1;refGene_gn=HERC1;refGene_tx=XM_023255227.1;refGene_gn=HERC1;refGene_tx=XM_023255228.1;refGene_gn=HERC1;refGene_tx=XM_023255229.1;refGene_gn=HERC1;refGene_tx=XM_023255231.1;refGene_gn=HERC1;refGene_tx=XM_023255232.1;refGene_gn=HERC1;refGene_tx=XM_023255233.1;refGen

e_gn=HERC1;refGene_tx=XM_023255234.1
B3 43485916 43486605
refGene_gn=FBXL22;refGene_tx=XM_003987085.5;refGene_gn=FBXL22;refGene_tx=XM_019832286.2
B3 43489578 43489913
refGene_gn=FBXL22;refGene_tx=XM_003987085.5;refGene_gn=FBXL22;refGene_tx=XM_019832286.2
B3 43496522 43496688
refGene_gn=USP3;refGene_tx=XM_019832285.2;refGene_gn=USP3;refGene_tx=XM_023255235.1;refGene_gn=USP3;refGene_tx=XM_023255236.1;refGene_gn=USP3;refGene_tx=XM_023255237.1;refGene_gn=USP3;refGene_tx=XM_023255238.1;refGene_gn=USP3;refGene_tx=XM_023255239.1;refGene_gn=USP3;refGene_tx=XM_023255240.1
B3 43498922 43498990
refGene_gn=USP3;refGene_tx=XM_019832285.2;refGene_gn=USP3;refGene_tx=XM_023255235.1;refGene_gn=USP3;refGene_tx=XM_023255236.1;refGene_gn=USP3;refGene_tx=XM_023255237.1;refGene_gn=USP3;refGene_tx=XM_023255238.1;refGene_gn=USP3;refGene_tx=XM_023255239.1;refGene_gn=USP3;refGene_tx=XM_023255240.1
B3 43499130 43499244
refGene_gn=USP3;refGene_tx=XM_019832285.2;refGene_gn=USP3;refGene_tx=XM_023255235.1;refGene_gn=USP3;refGene_tx=XM_023255236.1;refGene_gn=USP3;refGene_tx=XM_023255237.1;refGene_gn=USP3;refGene_tx=XM_023255238.1;refGene_gn=USP3;refGene_tx=XM_023255239.1;refGene_gn=USP3;refGene_tx=XM_023255240.1
B3 43500467 43500586
refGene_gn=USP3;refGene_tx=XM_019832285.2;refGene_gn=USP3;refGene_tx=XM_023255235.1;refGene_gn=USP3;refGene_tx=XM_023255236.1;refGene_gn=USP3;refGene_tx=XM_023255237.1;refGene_gn=USP3;refGene_tx=XM_023255238.1;refGene_gn=USP3;refGene_tx=XM_023255239.1;refGene_gn=USP3;refGene_tx=XM_023255240.1
B3 43524395 43524476
refGene_gn=USP3;refGene_tx=XM_019832285.2;refGene_gn=USP3;refGene_tx=XM_023255235.1;refGene_gn=USP3;refGene_tx=XM_023255236.1;refGene_gn=USP3;refGene_tx=XM_023255237.1;refGene_gn=USP3;refGene_tx=XM_023255238.1;refGene_gn=USP3;refGene_tx=XM_023255239.1;refGene_gn=USP3;refGene_tx=XM_023255240.1
B3 43524618 43524725
refGene_gn=USP3;refGene_tx=XM_019832285.2;refGene_gn=USP3;refGene_tx=XM_023255235.1;refGene_gn=USP3;refGene_tx=XM_023255236.1;refGene_gn=USP3;refGene_tx=XM_023255237.1;refGene_gn=USP3;refGene_tx=XM_023255238.1;refGene_gn=USP3;refGene_tx=XM_023255239.1;refGene_gn=USP3;refGene_tx=XM_023255240.1
B3 43527329 43527374
refGene_gn=USP3;refGene_tx=XM_019832285.2;refGene_gn=USP3;refGene_tx=XM_023255235.1;refGene_gn=USP3;refGene_tx=XM_023255236.1;refGene_gn=USP3;refGene_tx=XM_023255237.1;refGene_gn=USP3;refGene_tx=XM_023255239.1
B3 43528043 43528190
refGene_gn=USP3;refGene_tx=XM_019832285.2;refGene_gn=USP3;refGene_tx=XM_023255235.1;refGene_gn=USP3;refGene_tx=XM_023255236.1;refGene_gn=USP

3;refGene_tx=XM_023255237.1;refGene_gn=USP3;refGene_tx=XM_023255238.1;
refGene_gn=USP3;refGene_tx=XM_023255239.1;refGene_gn=USP3;refGene_tx=X
M_023255240.1
B3 43538602 43538716
refGene_gn=USP3;refGene_tx=XM_019832285.2;refGene_gn=USP3;refGene_tx=X
M_023255235.1;refGene_gn=USP3;refGene_tx=XM_023255236.1;refGene_gn=USP
3;refGene_tx=XM_023255237.1;refGene_gn=USP3;refGene_tx=XM_023255238.1;
refGene_gn=USP3;refGene_tx=XM_023255239.1;refGene_gn=USP3;refGene_tx=X
M_023255240.1
B3 43541423 43541537
refGene_gn=USP3;refGene_tx=XM_019832285.2;refGene_gn=USP3;refGene_tx=X
M_023255235.1;refGene_gn=USP3;refGene_tx=XM_023255236.1;refGene_gn=USP
3;refGene_tx=XM_023255237.1;refGene_gn=USP3;refGene_tx=XM_023255238.1;
refGene_gn=USP3;refGene_tx=XM_023255239.1;refGene_gn=USP3;refGene_tx=X
M_023255240.1
B3 43543749 43543832
refGene_gn=USP3;refGene_tx=XM_019832285.2;refGene_gn=USP3;refGene_tx=X
M_023255235.1;refGene_gn=USP3;refGene_tx=XM_023255236.1;refGene_gn=USP
3;refGene_tx=XM_023255237.1;refGene_gn=USP3;refGene_tx=XM_023255238.1;
refGene_gn=USP3;refGene_tx=XM_023255239.1;refGene_gn=USP3;refGene_tx=X
M_023255240.1
B3 43545052 43545134
refGene_gn=USP3;refGene_tx=XM_019832285.2;refGene_gn=USP3;refGene_tx=X
M_023255235.1;refGene_gn=USP3;refGene_tx=XM_023255236.1;refGene_gn=USP
3;refGene_tx=XM_023255237.1;refGene_gn=USP3;refGene_tx=XM_023255238.1;
refGene_gn=USP3;refGene_tx=XM_023255239.1;refGene_gn=USP3;refGene_tx=X
M_023255240.1
B3 43547024 43547108
refGene_gn=USP3;refGene_tx=XM_019832285.2;refGene_gn=USP3;refGene_tx=X
M_023255235.1;refGene_gn=USP3;refGene_tx=XM_023255236.1;refGene_gn=USP
3;refGene_tx=XM_023255237.1;refGene_gn=USP3;refGene_tx=XM_023255238.1;
refGene_gn=USP3;refGene_tx=XM_023255239.1;refGene_gn=USP3;refGene_tx=X
M_023255240.1
B3 43562052 43562184
refGene_gn=USP3;refGene_tx=XM_019832285.2;refGene_gn=USP3;refGene_tx=X
M_023255235.1;refGene_gn=USP3;refGene_tx=XM_023255236.1;refGene_gn=USP
3;refGene_tx=XM_023255237.1;refGene_gn=USP3;refGene_tx=XM_023255238.1;
refGene_gn=USP3;refGene_tx=XM_023255239.1;refGene_gn=USP3;refGene_tx=X
M_023255240.1
B3 43565812 43565873
refGene_gn=USP3;refGene_tx=XM_023255235.1;refGene_gn=USP3;refGene_tx=X
M_023255236.1;refGene_gn=USP3;refGene_tx=XM_023255237.1;refGene_gn=USP
3;refGene_tx=XM_023255238.1;refGene_gn=USP3;refGene_tx=XM_023255239.1;
refGene_gn=USP3;refGene_tx=XM_023255240.1
B3 43575637 43575703
refGene_gn=USP3;refGene_tx=XM_023255235.1;refGene_gn=USP3;refGene_tx=X
M_023255236.1;refGene_gn=USP3;refGene_tx=XM_023255237.1;refGene_gn=USP
3;refGene_tx=XM_023255238.1
B3 43595649 43595740
refGene_gn=USP3;refGene_tx=XM_023255235.1;refGene_gn=USP3;refGene_tx=X

M_023255236.1;refGene_gn=USP3;refGene_tx=XM_023255237.1;refGene_gn=USP3;refGene_tx=XM_023255238.1;refGene_gn=USP3;refGene_tx=XM_023255239.1;refGene_gn=USP3;refGene_tx=XM_023255240.1
B3 43716975 43717057
refGene_gn=CA12;refGene_tx=XM_023255241.1;refGene_gn=CA12;refGene_tx=XM_023255242.1;refGene_gn=CA12;refGene_tx=XM_023255243.1;refGene_gn=CA12;refGene_tx=XM_023255244.1;refGene_gn=CA12;refGene_tx=XM_023255245.1;refGene_gn=CA12;refGene_tx=XM_023255246.1
B3 43722174 43722195
refGene_gn=CA12;refGene_tx=XM_023255241.1;refGene_gn=CA12;refGene_tx=XM_023255242.1;refGene_gn=CA12;refGene_tx=XM_023255243.1;refGene_gn=CA12;refGene_tx=XM_023255244.1;refGene_gn=CA12;refGene_tx=XM_023255245.1;refGene_gn=CA12;refGene_tx=XM_023255246.1
B3 43748391 43748571
refGene_gn=CA12;refGene_tx=XM_023255241.1;refGene_gn=CA12;refGene_tx=XM_023255242.1;refGene_gn=CA12;refGene_tx=XM_023255243.1;refGene_gn=CA12;refGene_tx=XM_023255244.1;refGene_gn=CA12;refGene_tx=XM_023255245.1;refGene_gn=CA12;refGene_tx=XM_023255246.1
B3 43749445 43749588
refGene_gn=CA12;refGene_tx=XM_023255241.1;refGene_gn=CA12;refGene_tx=XM_023255242.1;refGene_gn=CA12;refGene_tx=XM_023255243.1;refGene_gn=CA12;refGene_tx=XM_023255244.1;refGene_gn=CA12;refGene_tx=XM_023255245.1;refGene_gn=CA12;refGene_tx=XM_023255246.1
B3 43751923 43752019
refGene_gn=CA12;refGene_tx=XM_023255241.1;refGene_gn=CA12;refGene_tx=XM_023255242.1;refGene_gn=CA12;refGene_tx=XM_023255243.1;refGene_gn=CA12;refGene_tx=XM_023255244.1;refGene_gn=CA12;refGene_tx=XM_023255246.1
B3 43752895 43752959
refGene_gn=CA12;refGene_tx=XM_023255241.1;refGene_gn=CA12;refGene_tx=XM_023255242.1;refGene_gn=CA12;refGene_tx=XM_023255243.1;refGene_gn=CA12;refGene_tx=XM_023255244.1;refGene_gn=CA12;refGene_tx=XM_023255245.1;refGene_gn=CA12;refGene_tx=XM_023255246.1
B3 43753142 43753300
refGene_gn=CA12;refGene_tx=XM_023255241.1;refGene_gn=CA12;refGene_tx=XM_023255242.1;refGene_gn=CA12;refGene_tx=XM_023255243.1;refGene_gn=CA12;refGene_tx=XM_023255244.1;refGene_gn=CA12;refGene_tx=XM_023255245.1;refGene_gn=CA12;refGene_tx=XM_023255246.1
B3 43754564 43754691
refGene_gn=CA12;refGene_tx=XM_023255241.1;refGene_gn=CA12;refGene_tx=XM_023255242.1;refGene_gn=CA12;refGene_tx=XM_023255243.1;refGene_gn=CA12;refGene_tx=XM_023255244.1;refGene_gn=CA12;refGene_tx=XM_023255245.1;refGene_gn=CA12;refGene_tx=XM_023255246.1
B3 43755950 43756039 refGene_gn=CA12;refGene_tx=XM_023255246.1
B3 43763788 43763821
refGene_gn=CA12;refGene_tx=XM_023255241.1;refGene_gn=CA12;refGene_tx=XM_023255242.1;refGene_gn=CA12;refGene_tx=XM_023255245.1
B3 43764682 43764770
refGene_gn=CA12;refGene_tx=XM_023255241.1;refGene_gn=CA12;refGene_tx=XM_023255242.1;refGene_gn=CA12;refGene_tx=XM_023255243.1;refGene_gn=CA12;refGene_tx=XM_023255244.1;refGene_gn=CA12;refGene_tx=XM_023255245.1

B3 43765621 43765694
refGene_gn=CA12;refGene_tx=XM_023255241.1;refGene_gn=CA12;refGene_tx=XM_023255242.1;refGene_gn=CA12;refGene_tx=XM_023255243.1;refGene_gn=CA12;refGene_tx=XM_023255244.1;refGene_gn=CA12;refGene_tx=XM_023255245.1
B3 43786966 43787134
refGene_gn=APH1B;refGene_tx=XM_019832294.2;refGene_gn=APH1B;refGene_tx=XM_023255247.1;refGene_gn=APH1B;refGene_tx=XM_023255248.1;refGene_gn=APH1B;refGene_tx=XM_023255249.1;refGene_gn=APH1B;refGene_tx=XM_023255250.1;refGene_gn=APH1B;refGene_tx=XM_023255251.1
B3 43797161 43797289
refGene_gn=APH1B;refGene_tx=XM_019832294.2;refGene_gn=APH1B;refGene_tx=XM_023255247.1;refGene_gn=APH1B;refGene_tx=XM_023255248.1;refGene_gn=APH1B;refGene_tx=XM_023255249.1;refGene_gn=APH1B;refGene_tx=XM_023255250.1;refGene_gn=APH1B;refGene_tx=XM_023255251.1
B3 43815897 43816020
refGene_gn=APH1B;refGene_tx=XM_019832294.2;refGene_gn=APH1B;refGene_tx=XM_023255247.1;refGene_gn=APH1B;refGene_tx=XM_023255248.1;refGene_gn=APH1B;refGene_tx=XM_023255249.1;refGene_gn=APH1B;refGene_tx=XM_023255250.1
B3 43817405 43817476
refGene_gn=APH1B;refGene_tx=XM_019832294.2;refGene_gn=APH1B;refGene_tx=XM_023255247.1;refGene_gn=APH1B;refGene_tx=XM_023255248.1;refGene_gn=APH1B;refGene_tx=XM_023255249.1;refGene_gn=APH1B;refGene_tx=XM_023255250.1;refGene_gn=APH1B;refGene_tx=XM_023255251.1
B3 43823034 43823205
refGene_gn=APH1B;refGene_tx=XM_019832294.2;refGene_gn=APH1B;refGene_tx=XM_023255247.1;refGene_gn=APH1B;refGene_tx=XM_023255248.1;refGene_gn=APH1B;refGene_tx=XM_023255249.1;refGene_gn=APH1B;refGene_tx=XM_023255250.1;refGene_gn=APH1B;refGene_tx=XM_023255251.1
B3 43824615 43824728
refGene_gn=APH1B;refGene_tx=XM_019832294.2;refGene_gn=APH1B;refGene_tx=XM_023255247.1;refGene_gn=APH1B;refGene_tx=XM_023255248.1;refGene_gn=APH1B;refGene_tx=XM_023255249.1;refGene_gn=APH1B;refGene_tx=XM_023255250.1;refGene_gn=APH1B;refGene_tx=XM_023255251.1
B3 43836487 43836580
refGene_gn=RAB8B;refGene_tx=XM_019832295.2;refGene_gn=RAB8B;refGene_tx=XM_019832298.2
B3 43837424 43837475
refGene_gn=RAB8B;refGene_tx=XM_019832295.2;refGene_gn=RAB8B;refGene_tx=XM_019832298.2
B3 43839948 43840014
refGene_gn=RAB8B;refGene_tx=XM_019832295.2;refGene_gn=RAB8B;refGene_tx=XM_019832298.2
B3 43841501 43841591
refGene_gn=RAB8B;refGene_tx=XM_019832295.2;refGene_gn=RAB8B;refGene_tx=XM_019832298.2
B3 43842563 43842641
refGene_gn=RAB8B;refGene_tx=XM_019832295.2;refGene_gn=RAB8B;refGene_tx=XM_019832298.2
B3 43845857 43845918

refGene_gn=RAB8B;refGene_tx=XM_019832295.2;refGene_gn=RAB8B;refGene_tx=XM_019832298.2
B3 43848463 43848825
refGene_gn=RAB8B;refGene_tx=XM_019832295.2;refGene_gn=RAB8B;refGene_tx=XM_019832298.2
B3 43890838 43890962 refGene_gn=RAB8B;refGene_tx=XM_019832298.2
B3 43920565 43920571
refGene_gn=RPS27L;refGene_tx=XM_003987059.5;refGene_gn=RPS27L;refGene_tx=XM_019832299.2
B3 43921463 43921572
refGene_gn=RPS27L;refGene_tx=XM_003987059.5;refGene_gn=RPS27L;refGene_tx=XM_019832299.2;refGene_gn=RPS27L;refGene_tx=XM_023255252.1;refGene_gn=RPS27L;refGene_tx=XM_023255253.1;refGene_gn=RPS27L;refGene_tx=XM_023255254.1
B3 43922247 43922358
refGene_gn=RPS27L;refGene_tx=XM_003987059.5;refGene_gn=RPS27L;refGene_tx=XM_019832299.2;refGene_gn=RPS27L;refGene_tx=XM_023255252.1;refGene_gn=RPS27L;refGene_tx=XM_023255253.1;refGene_gn=RPS27L;refGene_tx=XM_023255254.1
B3 43922739 43922756 refGene_gn=RPS27L;refGene_tx=XM_019832299.2
B3 43923747 43923776
refGene_gn=RPS27L;refGene_tx=XM_003987059.5;refGene_gn=RPS27L;refGene_tx=XM_023255252.1;refGene_gn=RPS27L;refGene_tx=XM_023255253.1;refGene_gn=RPS27L;refGene_tx=XM_023255254.1
B3 43936122 43936654
refGene_gn=LACTB;refGene_tx=XM_023255255.1;refGene_gn=LACTB;refGene_tx=XM_023255256.1
B3 43946203 43946369
refGene_gn=LACTB;refGene_tx=XM_023255255.1;refGene_gn=LACTB;refGene_tx=XM_023255256.1;refGene_gn=LACTB;refGene_tx=XM_023255257.1
B3 43947569 43947921
refGene_gn=LACTB;refGene_tx=XM_023255255.1;refGene_gn=LACTB;refGene_tx=XM_023255256.1;refGene_gn=LACTB;refGene_tx=XM_023255257.1
B3 43948213 43948404
refGene_gn=LACTB;refGene_tx=XM_023255255.1;refGene_gn=LACTB;refGene_tx=XM_023255256.1;refGene_gn=LACTB;refGene_tx=XM_023255257.1
B3 43950533 43950600
refGene_gn=LACTB;refGene_tx=XM_023255255.1;refGene_gn=LACTB;refGene_tx=XM_023255256.1;refGene_gn=LACTB;refGene_tx=XM_023255257.1
B3 43951045 43951408
refGene_gn=LACTB;refGene_tx=XM_023255255.1;refGene_gn=LACTB;refGene_tx=XM_023255256.1;refGene_gn=LACTB;refGene_tx=XM_023255257.1
B3 44009043 44009126
refGene_gn=TPM1;refGene_tx=XM_011282915.2;refGene_gn=TPM1;refGene_tx=XM_019832305.2;refGene_gn=TPM1;refGene_tx=XM_023255260.1;refGene_gn=TPM1;refGene_tx=XM_023255264.1;refGene_gn=TPM1;refGene_tx=XM_023255266.1;refGene_gn=TPM1;refGene_tx=XM_023255267.1
B3 44010272 44010346
refGene_gn=TPM1;refGene_tx=XM_019832306.2;refGene_gn=TPM1;refGene_tx=XM_019832307.2;refGene_gn=TPM1;refGene_tx=XM_023255269.1

B3 44014301 44014305
refGene_gn=TPM1;refGene_tx=XM_019832301.2;refGene_gn=TPM1;refGene_tx=XM_019832303.2;refGene_gn=TPM1;refGene_tx=XM_023255258.1;refGene_gn=TPM1;refGene_tx=XM_023255261.1;refGene_gn=TPM1;refGene_tx=XM_023255262.1
B3 44015721 44015804
refGene_gn=TPM1;refGene_tx=XM_019832301.2;refGene_gn=TPM1;refGene_tx=XM_019832302.2;refGene_gn=TPM1;refGene_tx=XM_019832303.2;refGene_gn=TPM1;refGene_tx=XM_019832304.2;refGene_gn=TPM1;refGene_tx=XM_023255258.1;refGene_gn=TPM1;refGene_tx=XM_023255259.1;refGene_gn=TPM1;refGene_tx=XM_023255261.1;refGene_gn=TPM1;refGene_tx=XM_023255262.1;refGene_gn=TPM1;refGene_tx=XM_023255263.1;refGene_gn=TPM1;refGene_tx=XM_023255265.1
B3 44017214 44017284
refGene_gn=TPM1;refGene_tx=XM_011282915.2;refGene_gn=TPM1;refGene_tx=XM_019832301.2;refGene_gn=TPM1;refGene_tx=XM_019832302.2;refGene_gn=TPM1;refGene_tx=XM_019832303.2;refGene_gn=TPM1;refGene_tx=XM_019832304.2;refGene_gn=TPM1;refGene_tx=XM_019832305.2;refGene_gn=TPM1;refGene_tx=XM_019832306.2;refGene_gn=TPM1;refGene_tx=XM_019832307.2;refGene_gn=TPM1;refGene_tx=XM_023255258.1;refGene_gn=TPM1;refGene_tx=XM_023255259.1;refGene_gn=TPM1;refGene_tx=XM_023255260.1;refGene_gn=TPM1;refGene_tx=XM_023255261.1;refGene_gn=TPM1;refGene_tx=XM_023255262.1;refGene_gn=TPM1;refGene_tx=XM_023255263.1;refGene_gn=TPM1;refGene_tx=XM_023255264.1;refGene_gn=TPM1;refGene_tx=XM_023255265.1;refGene_gn=TPM1;refGene_tx=XM_023255266.1;refGene_gn=TPM1;refGene_tx=XM_023255267.1;refGene_gn=TPM1;refGene_tx=XM_023255269.1
B3 44017565 44017628
refGene_gn=TPM1;refGene_tx=XM_011282915.2;refGene_gn=TPM1;refGene_tx=XM_019832301.2;refGene_gn=TPM1;refGene_tx=XM_019832302.2;refGene_gn=TPM1;refGene_tx=XM_019832303.2;refGene_gn=TPM1;refGene_tx=XM_019832304.2;refGene_gn=TPM1;refGene_tx=XM_019832305.2;refGene_gn=TPM1;refGene_tx=XM_019832306.2;refGene_gn=TPM1;refGene_tx=XM_019832307.2;refGene_gn=TPM1;refGene_tx=XM_023255258.1;refGene_gn=TPM1;refGene_tx=XM_023255259.1;refGene_gn=TPM1;refGene_tx=XM_023255260.1;refGene_gn=TPM1;refGene_tx=XM_023255261.1;refGene_gn=TPM1;refGene_tx=XM_023255262.1;refGene_gn=TPM1;refGene_tx=XM_023255263.1;refGene_gn=TPM1;refGene_tx=XM_023255264.1;refGene_gn=TPM1;refGene_tx=XM_023255265.1;refGene_gn=TPM1;refGene_tx=XM_023255266.1;refGene_gn=TPM1;refGene_tx=XM_023255267.1;refGene_gn=TPM1;refGene_tx=XM_023255269.1
B3 44018069 44018145
refGene_gn=TPM1;refGene_tx=XM_011282915.2;refGene_gn=TPM1;refGene_tx=XM_019832301.2;refGene_gn=TPM1;refGene_tx=XM_019832302.2;refGene_gn=TPM1;refGene_tx=XM_019832306.2;refGene_gn=TPM1;refGene_tx=XM_023255258.1;refGene_gn=TPM1;refGene_tx=XM_023255259.1;refGene_gn=TPM1;refGene_tx=XM_023255260.1;refGene_gn=TPM1;refGene_tx=XM_023255261.1;refGene_gn=TPM1;refGene_tx=XM_023255263.1;refGene_gn=TPM1;refGene_tx=XM_023255264.1;refGene_gn=TPM1;refGene_tx=XM_023255269.1
B3 44018575 44018651
refGene_gn=TPM1;refGene_tx=XM_019832303.2;refGene_gn=TPM1;refGene_tx=XM_019832304.2;refGene_gn=TPM1;refGene_tx=XM_019832305.2;refGene_gn=TPM1;refGene_tx=XM_019832307.2;refGene_gn=TPM1;refGene_tx=XM_023255262.1;refGene_gn=TPM1;refGene_tx=XM_023255265.1;refGene_gn=TPM1;refGene_tx=X

M_023255266.1;refGene_gn=TPM1;refGene_tx=XM_023255267.1
B3 44018906 44018977
refGene_gn=TPM1;refGene_tx=XM_011282915.2;refGene_gn=TPM1;refGene_tx=XM_019832301.2;refGene_gn=TPM1;refGene_tx=XM_019832302.2;refGene_gn=TPM1;refGene_tx=XM_019832303.2;refGene_gn=TPM1;refGene_tx=XM_019832304.2;refGene_gn=TPM1;refGene_tx=XM_019832305.2;refGene_gn=TPM1;refGene_tx=XM_019832306.2;refGene_gn=TPM1;refGene_tx=XM_019832307.2;refGene_gn=TPM1;refGene_tx=XM_023255258.1;refGene_gn=TPM1;refGene_tx=XM_023255259.1;refGene_gn=TPM1;refGene_tx=XM_023255260.1;refGene_gn=TPM1;refGene_tx=XM_023255261.1;refGene_gn=TPM1;refGene_tx=XM_023255262.1;refGene_gn=TPM1;refGene_tx=XM_023255263.1;refGene_gn=TPM1;refGene_tx=XM_023255264.1;refGene_gn=TPM1;refGene_tx=XM_023255265.1;refGene_gn=TPM1;refGene_tx=XM_023255266.1;refGene_gn=TPM1;refGene_tx=XM_023255267.1;refGene_gn=TPM1;refGene_tx=XM_023255269.1
B3 44020284 44020402
refGene_gn=TPM1;refGene_tx=XM_011282915.2;refGene_gn=TPM1;refGene_tx=XM_019832301.2;refGene_gn=TPM1;refGene_tx=XM_019832302.2;refGene_gn=TPM1;refGene_tx=XM_019832303.2;refGene_gn=TPM1;refGene_tx=XM_019832304.2;refGene_gn=TPM1;refGene_tx=XM_019832305.2;refGene_gn=TPM1;refGene_tx=XM_019832306.2;refGene_gn=TPM1;refGene_tx=XM_019832307.2;refGene_gn=TPM1;refGene_tx=XM_023255258.1;refGene_gn=TPM1;refGene_tx=XM_023255259.1;refGene_gn=TPM1;refGene_tx=XM_023255260.1;refGene_gn=TPM1;refGene_tx=XM_023255261.1;refGene_gn=TPM1;refGene_tx=XM_023255262.1;refGene_gn=TPM1;refGene_tx=XM_023255263.1;refGene_gn=TPM1;refGene_tx=XM_023255264.1;refGene_gn=TPM1;refGene_tx=XM_023255265.1;refGene_gn=TPM1;refGene_tx=XM_023255266.1;refGene_gn=TPM1;refGene_tx=XM_023255267.1;refGene_gn=TPM1;refGene_tx=XM_023255269.1
B3 44022492 44022626
refGene_gn=TPM1;refGene_tx=XM_011282915.2;refGene_gn=TPM1;refGene_tx=XM_019832301.2;refGene_gn=TPM1;refGene_tx=XM_019832302.2;refGene_gn=TPM1;refGene_tx=XM_019832303.2;refGene_gn=TPM1;refGene_tx=XM_019832304.2;refGene_gn=TPM1;refGene_tx=XM_019832305.2;refGene_gn=TPM1;refGene_tx=XM_019832306.2;refGene_gn=TPM1;refGene_tx=XM_019832307.2;refGene_gn=TPM1;refGene_tx=XM_023255258.1;refGene_gn=TPM1;refGene_tx=XM_023255259.1;refGene_gn=TPM1;refGene_tx=XM_023255260.1;refGene_gn=TPM1;refGene_tx=XM_023255261.1;refGene_gn=TPM1;refGene_tx=XM_023255262.1;refGene_gn=TPM1;refGene_tx=XM_023255263.1;refGene_gn=TPM1;refGene_tx=XM_023255264.1;refGene_gn=TPM1;refGene_tx=XM_023255265.1;refGene_gn=TPM1;refGene_tx=XM_023255266.1;refGene_gn=TPM1;refGene_tx=XM_023255267.1;refGene_gn=TPM1;refGene_tx=XM_023255269.1
B3 44030200 44030332
refGene_gn=TPM1;refGene_tx=XM_011282915.2;refGene_gn=TPM1;refGene_tx=XM_019832301.2;refGene_gn=TPM1;refGene_tx=XM_019832302.2;refGene_gn=TPM1;refGene_tx=XM_019832303.2;refGene_gn=TPM1;refGene_tx=XM_019832304.2;refGene_gn=TPM1;refGene_tx=XM_019832305.2;refGene_gn=TPM1;refGene_tx=XM_019832306.2;refGene_gn=TPM1;refGene_tx=XM_019832307.2
B3 44034509 44034635
refGene_gn=TPM1;refGene_tx=XM_023255258.1;refGene_gn=TPM1;refGene_tx=XM_023255259.1;refGene_gn=TPM1;refGene_tx=XM_023255260.1;refGene_gn=TPM1;refGene_tx=XM_023255262.1;refGene_gn=TPM1;refGene_tx=XM_023255265.1;

refGene_gn=TPM1;refGene_tx=XM_023255266.1;refGene_gn=TPM1;refGene_tx=X
M_023255269.1
B3 44034836 44034962
refGene_gn=TPM1;refGene_tx=XM_023255261.1;refGene_gn=TPM1;refGene_tx=X
M_023255263.1;refGene_gn=TPM1;refGene_tx=XM_023255264.1;refGene_gn=TPM
1;refGene_tx=XM_023255267.1
B3 44035769 44035883
refGene_gn=TPM1;refGene_tx=XM_023255258.1;refGene_gn=TPM1;refGene_tx=X
M_023255259.1;refGene_gn=TPM1;refGene_tx=XM_023255260.1;refGene_gn=TPM
1;refGene_tx=XM_023255261.1;refGene_gn=TPM1;refGene_tx=XM_023255262.1;
refGene_gn=TPM1;refGene_tx=XM_023255263.1;refGene_gn=TPM1;refGene_tx=X
M_023255264.1;refGene_gn=TPM1;refGene_tx=XM_023255265.1;refGene_gn=TPM
1;refGene_tx=XM_023255266.1;refGene_gn=TPM1;refGene_tx=XM_023255267.1;
refGene_gn=TPM1;refGene_tx=XM_023255269.1
B3 44239219 44239348
refGene_gn=TLN2;refGene_tx=XM_011282911.2;refGene_gn=TLN2;refGene_tx=X
M_019832308.2;refGene_gn=TLN2;refGene_tx=XM_019832309.2;refGene_gn=TLN
2;refGene_tx=XM_019832310.2;refGene_gn=TLN2;refGene_tx=XM_019832311.2
B3 44240859 44240985
refGene_gn=TLN2;refGene_tx=XM_011282911.2;refGene_gn=TLN2;refGene_tx=X
M_019832308.2;refGene_gn=TLN2;refGene_tx=XM_019832309.2;refGene_gn=TLN
2;refGene_tx=XM_019832310.2;refGene_gn=TLN2;refGene_tx=XM_019832311.2
B3 44244291 44244519
refGene_gn=TLN2;refGene_tx=XM_011282911.2;refGene_gn=TLN2;refGene_tx=X
M_019832308.2;refGene_gn=TLN2;refGene_tx=XM_019832309.2;refGene_gn=TLN
2;refGene_tx=XM_019832310.2;refGene_gn=TLN2;refGene_tx=XM_019832311.2
B3 44244610 44244673
refGene_gn=TLN2;refGene_tx=XM_011282911.2;refGene_gn=TLN2;refGene_tx=X
M_019832308.2;refGene_gn=TLN2;refGene_tx=XM_019832309.2;refGene_gn=TLN
2;refGene_tx=XM_019832310.2;refGene_gn=TLN2;refGene_tx=XM_019832311.2
B3 44246743 44246869
refGene_gn=TLN2;refGene_tx=XM_011282911.2;refGene_gn=TLN2;refGene_tx=X
M_019832308.2;refGene_gn=TLN2;refGene_tx=XM_019832309.2;refGene_gn=TLN
2;refGene_tx=XM_019832310.2;refGene_gn=TLN2;refGene_tx=XM_019832311.2
B3 44258434 44258559
refGene_gn=TLN2;refGene_tx=XM_011282911.2;refGene_gn=TLN2;refGene_tx=X
M_019832308.2;refGene_gn=TLN2;refGene_tx=XM_019832309.2;refGene_gn=TLN
2;refGene_tx=XM_019832310.2;refGene_gn=TLN2;refGene_tx=XM_019832311.2
B3 44259442 44259548
refGene_gn=TLN2;refGene_tx=XM_011282911.2;refGene_gn=TLN2;refGene_tx=X
M_019832308.2;refGene_gn=TLN2;refGene_tx=XM_019832309.2;refGene_gn=TLN
2;refGene_tx=XM_019832310.2;refGene_gn=TLN2;refGene_tx=XM_019832311.2
B3 44270390 44270498
refGene_gn=TLN2;refGene_tx=XM_011282911.2;refGene_gn=TLN2;refGene_tx=X
M_019832308.2;refGene_gn=TLN2;refGene_tx=XM_019832309.2;refGene_gn=TLN
2;refGene_tx=XM_019832310.2;refGene_gn=TLN2;refGene_tx=XM_019832311.2
B3 44274036 44274222
refGene_gn=TLN2;refGene_tx=XM_011282911.2;refGene_gn=TLN2;refGene_tx=X
M_019832308.2;refGene_gn=TLN2;refGene_tx=XM_019832309.2;refGene_gn=TLN
2;refGene_tx=XM_019832310.2;refGene_gn=TLN2;refGene_tx=XM_019832311.2

B3 44276478 44276595
refGene_gn=TLN2;refGene_tx=XM_011282911.2;refGene_gn=TLN2;refGene_tx=XM_019832308.2;refGene_gn=TLN2;refGene_tx=XM_019832309.2;refGene_gn=TLN2;refGene_tx=XM_019832310.2;refGene_gn=TLN2;refGene_tx=XM_019832311.2
B3 44276802 44276928
refGene_gn=TLN2;refGene_tx=XM_011282911.2;refGene_gn=TLN2;refGene_tx=XM_019832308.2;refGene_gn=TLN2;refGene_tx=XM_019832309.2;refGene_gn=TLN2;refGene_tx=XM_019832310.2;refGene_gn=TLN2;refGene_tx=XM_019832311.2
B3 44280202 44280386
refGene_gn=TLN2;refGene_tx=XM_011282911.2;refGene_gn=TLN2;refGene_tx=XM_019832308.2;refGene_gn=TLN2;refGene_tx=XM_019832309.2;refGene_gn=TLN2;refGene_tx=XM_019832310.2;refGene_gn=TLN2;refGene_tx=XM_019832311.2
B3 44281242 44281409
refGene_gn=TLN2;refGene_tx=XM_011282911.2;refGene_gn=TLN2;refGene_tx=XM_019832308.2;refGene_gn=TLN2;refGene_tx=XM_019832309.2;refGene_gn=TLN2;refGene_tx=XM_019832310.2;refGene_gn=TLN2;refGene_tx=XM_019832311.2
B3 44287932 44288079
refGene_gn=TLN2;refGene_tx=XM_011282911.2;refGene_gn=TLN2;refGene_tx=XM_019832308.2;refGene_gn=TLN2;refGene_tx=XM_019832309.2;refGene_gn=TLN2;refGene_tx=XM_019832310.2;refGene_gn=TLN2;refGene_tx=XM_019832311.2
B3 44295052 44295172
refGene_gn=TLN2;refGene_tx=XM_011282911.2;refGene_gn=TLN2;refGene_tx=XM_019832308.2;refGene_gn=TLN2;refGene_tx=XM_019832309.2;refGene_gn=TLN2;refGene_tx=XM_019832310.2
B3 44297731 44297833
refGene_gn=TLN2;refGene_tx=XM_011282911.2;refGene_gn=TLN2;refGene_tx=XM_019832308.2;refGene_gn=TLN2;refGene_tx=XM_019832309.2;refGene_gn=TLN2;refGene_tx=XM_019832310.2;refGene_gn=TLN2;refGene_tx=XM_019832311.2
B3 44301883 44302030
refGene_gn=TLN2;refGene_tx=XM_011282911.2;refGene_gn=TLN2;refGene_tx=XM_019832308.2;refGene_gn=TLN2;refGene_tx=XM_019832309.2;refGene_gn=TLN2;refGene_tx=XM_019832310.2;refGene_gn=TLN2;refGene_tx=XM_019832311.2
B3 44312634 44312805
refGene_gn=TLN2;refGene_tx=XM_011282911.2;refGene_gn=TLN2;refGene_tx=XM_019832308.2;refGene_gn=TLN2;refGene_tx=XM_019832309.2;refGene_gn=TLN2;refGene_tx=XM_019832310.2;refGene_gn=TLN2;refGene_tx=XM_019832311.2
B3 44318893 44318995
refGene_gn=TLN2;refGene_tx=XM_011282911.2;refGene_gn=TLN2;refGene_tx=XM_019832308.2;refGene_gn=TLN2;refGene_tx=XM_019832309.2;refGene_gn=TLN2;refGene_tx=XM_019832310.2;refGene_gn=TLN2;refGene_tx=XM_019832311.2
B3 44326467 44326600
refGene_gn=TLN2;refGene_tx=XM_011282911.2;refGene_gn=TLN2;refGene_tx=XM_019832308.2;refGene_gn=TLN2;refGene_tx=XM_019832309.2;refGene_gn=TLN2;refGene_tx=XM_019832310.2;refGene_gn=TLN2;refGene_tx=XM_019832311.2
B3 44327945 44328127
refGene_gn=TLN2;refGene_tx=XM_011282911.2;refGene_gn=TLN2;refGene_tx=XM_019832308.2;refGene_gn=TLN2;refGene_tx=XM_019832309.2;refGene_gn=TLN2;refGene_tx=XM_019832310.2;refGene_gn=TLN2;refGene_tx=XM_019832311.2
B3 44328585 44328726
refGene_gn=TLN2;refGene_tx=XM_011282911.2;refGene_gn=TLN2;refGene_tx=X

M_019832308.2;refGene_gn=TLN2;refGene_tx=XM_019832309.2;refGene_gn=TLN2;refGene_tx=XM_019832310.2;refGene_gn=TLN2;refGene_tx=XM_019832311.2
B3 44334787 44334949
refGene_gn=TLN2;refGene_tx=XM_011282911.2;refGene_gn=TLN2;refGene_tx=XM_019832308.2;refGene_gn=TLN2;refGene_tx=XM_019832309.2;refGene_gn=TLN2;refGene_tx=XM_019832310.2;refGene_gn=TLN2;refGene_tx=XM_019832311.2
B3 44336783 44336927
refGene_gn=TLN2;refGene_tx=XM_011282911.2;refGene_gn=TLN2;refGene_tx=XM_019832308.2;refGene_gn=TLN2;refGene_tx=XM_019832309.2;refGene_gn=TLN2;refGene_tx=XM_019832310.2;refGene_gn=TLN2;refGene_tx=XM_019832311.2
B3 44338415 44338538
refGene_gn=TLN2;refGene_tx=XM_011282911.2;refGene_gn=TLN2;refGene_tx=XM_019832308.2;refGene_gn=TLN2;refGene_tx=XM_019832309.2;refGene_gn=TLN2;refGene_tx=XM_019832310.2;refGene_gn=TLN2;refGene_tx=XM_019832311.2
B3 44340406 44340496
refGene_gn=TLN2;refGene_tx=XM_011282911.2;refGene_gn=TLN2;refGene_tx=XM_019832308.2;refGene_gn=TLN2;refGene_tx=XM_019832309.2;refGene_gn=TLN2;refGene_tx=XM_019832310.2;refGene_gn=TLN2;refGene_tx=XM_019832311.2
B3 44342395 44342489
refGene_gn=TLN2;refGene_tx=XM_011282911.2;refGene_gn=TLN2;refGene_tx=XM_019832308.2;refGene_gn=TLN2;refGene_tx=XM_019832309.2;refGene_gn=TLN2;refGene_tx=XM_019832310.2;refGene_gn=TLN2;refGene_tx=XM_019832311.2
B3 44349396 44349536
refGene_gn=TLN2;refGene_tx=XM_011282911.2;refGene_gn=TLN2;refGene_tx=XM_019832308.2;refGene_gn=TLN2;refGene_tx=XM_019832309.2;refGene_gn=TLN2;refGene_tx=XM_019832310.2;refGene_gn=TLN2;refGene_tx=XM_019832311.2
B3 44350592 44350790
refGene_gn=TLN2;refGene_tx=XM_011282911.2;refGene_gn=TLN2;refGene_tx=XM_019832308.2;refGene_gn=TLN2;refGene_tx=XM_019832309.2;refGene_gn=TLN2;refGene_tx=XM_019832310.2;refGene_gn=TLN2;refGene_tx=XM_019832311.2
B3 44351806 44351926
refGene_gn=TLN2;refGene_tx=XM_011282911.2;refGene_gn=TLN2;refGene_tx=XM_019832308.2;refGene_gn=TLN2;refGene_tx=XM_019832309.2;refGene_gn=TLN2;refGene_tx=XM_019832310.2;refGene_gn=TLN2;refGene_tx=XM_019832311.2
B3 44353002 44353211
refGene_gn=TLN2;refGene_tx=XM_011282911.2;refGene_gn=TLN2;refGene_tx=XM_019832308.2;refGene_gn=TLN2;refGene_tx=XM_019832309.2;refGene_gn=TLN2;refGene_tx=XM_019832310.2;refGene_gn=TLN2;refGene_tx=XM_019832311.2
B3 44363042 44363145
refGene_gn=TLN2;refGene_tx=XM_011282911.2;refGene_gn=TLN2;refGene_tx=XM_019832308.2;refGene_gn=TLN2;refGene_tx=XM_019832309.2;refGene_gn=TLN2;refGene_tx=XM_019832310.2;refGene_gn=TLN2;refGene_tx=XM_019832311.2
B3 44365562 44365691
refGene_gn=TLN2;refGene_tx=XM_011282911.2;refGene_gn=TLN2;refGene_tx=XM_019832308.2;refGene_gn=TLN2;refGene_tx=XM_019832309.2;refGene_gn=TLN2;refGene_tx=XM_019832310.2;refGene_gn=TLN2;refGene_tx=XM_019832311.2
B3 44367624 44367759
refGene_gn=TLN2;refGene_tx=XM_011282911.2;refGene_gn=TLN2;refGene_tx=XM_019832308.2;refGene_gn=TLN2;refGene_tx=XM_019832309.2;refGene_gn=TLN2;refGene_tx=XM_019832310.2;refGene_gn=TLN2;refGene_tx=XM_019832311.2

B3 44371071 44371185
refGene_gn=TLN2;refGene_tx=XM_011282911.2;refGene_gn=TLN2;refGene_tx=XM_019832308.2;refGene_gn=TLN2;refGene_tx=XM_019832309.2;refGene_gn=TLN2;refGene_tx=XM_019832310.2;refGene_gn=TLN2;refGene_tx=XM_019832311.2
B3 44373301 44373415
refGene_gn=TLN2;refGene_tx=XM_011282911.2;refGene_gn=TLN2;refGene_tx=XM_019832308.2;refGene_gn=TLN2;refGene_tx=XM_019832309.2;refGene_gn=TLN2;refGene_tx=XM_019832310.2;refGene_gn=TLN2;refGene_tx=XM_019832311.2
B3 44375097 44375226
refGene_gn=TLN2;refGene_tx=XM_011282911.2;refGene_gn=TLN2;refGene_tx=XM_019832308.2;refGene_gn=TLN2;refGene_tx=XM_019832309.2;refGene_gn=TLN2;refGene_tx=XM_019832310.2;refGene_gn=TLN2;refGene_tx=XM_019832311.2
B3 44379065 44379232
refGene_gn=TLN2;refGene_tx=XM_011282911.2;refGene_gn=TLN2;refGene_tx=XM_019832308.2;refGene_gn=TLN2;refGene_tx=XM_019832309.2;refGene_gn=TLN2;refGene_tx=XM_019832310.2;refGene_gn=TLN2;refGene_tx=XM_019832311.2
B3 44381804 44382099
refGene_gn=TLN2;refGene_tx=XM_011282911.2;refGene_gn=TLN2;refGene_tx=XM_019832308.2;refGene_gn=TLN2;refGene_tx=XM_019832309.2;refGene_gn=TLN2;refGene_tx=XM_019832310.2;refGene_gn=TLN2;refGene_tx=XM_019832311.2
B3 44383318 44383486
refGene_gn=TLN2;refGene_tx=XM_011282911.2;refGene_gn=TLN2;refGene_tx=XM_019832308.2;refGene_gn=TLN2;refGene_tx=XM_019832309.2;refGene_gn=TLN2;refGene_tx=XM_019832310.2;refGene_gn=TLN2;refGene_tx=XM_019832311.2
B3 44387409 44387508
refGene_gn=TLN2;refGene_tx=XM_011282911.2;refGene_gn=TLN2;refGene_tx=XM_019832308.2;refGene_gn=TLN2;refGene_tx=XM_019832309.2;refGene_gn=TLN2;refGene_tx=XM_019832310.2;refGene_gn=TLN2;refGene_tx=XM_019832311.2
B3 44388062 44388271
refGene_gn=TLN2;refGene_tx=XM_011282911.2;refGene_gn=TLN2;refGene_tx=XM_019832308.2;refGene_gn=TLN2;refGene_tx=XM_019832309.2;refGene_gn=TLN2;refGene_tx=XM_019832310.2;refGene_gn=TLN2;refGene_tx=XM_019832311.2
B3 44388982 44389091
refGene_gn=TLN2;refGene_tx=XM_011282911.2;refGene_gn=TLN2;refGene_tx=XM_019832308.2;refGene_gn=TLN2;refGene_tx=XM_019832309.2;refGene_gn=TLN2;refGene_tx=XM_019832310.2;refGene_gn=TLN2;refGene_tx=XM_019832311.2
B3 44391396 44391510
refGene_gn=TLN2;refGene_tx=XM_011282911.2;refGene_gn=TLN2;refGene_tx=XM_019832308.2;refGene_gn=TLN2;refGene_tx=XM_019832309.2;refGene_gn=TLN2;refGene_tx=XM_019832310.2;refGene_gn=TLN2;refGene_tx=XM_019832311.2
B3 44392631 44392812
refGene_gn=TLN2;refGene_tx=XM_011282911.2;refGene_gn=TLN2;refGene_tx=XM_019832308.2;refGene_gn=TLN2;refGene_tx=XM_019832309.2;refGene_gn=TLN2;refGene_tx=XM_019832310.2;refGene_gn=TLN2;refGene_tx=XM_019832311.2
B3 44395608 44395685
refGene_gn=TLN2;refGene_tx=XM_011282911.2;refGene_gn=TLN2;refGene_tx=XM_019832308.2;refGene_gn=TLN2;refGene_tx=XM_019832309.2;refGene_gn=TLN2;refGene_tx=XM_019832310.2;refGene_gn=TLN2;refGene_tx=XM_019832311.2
B3 44396606 44396708
refGene_gn=TLN2;refGene_tx=XM_011282911.2;refGene_gn=TLN2;refGene_tx=X

M_019832308.2;refGene_gn=TLN2;refGene_tx=XM_019832309.2;refGene_gn=TLN2;refGene_tx=XM_019832310.2;refGene_gn=TLN2;refGene_tx=XM_019832311.2
B3 44400876 44401032 refGene_gn=TLN2;refGene_tx=XM_011282911.2;refGene_gn=TLN2;refGene_tx=XM_019832308.2;refGene_gn=TLN2;refGene_tx=XM_019832309.2;refGene_gn=TLN2;refGene_tx=XM_019832310.2;refGene_gn=TLN2;refGene_tx=XM_019832311.2
B3 44408390 44408495 refGene_gn=TLN2;refGene_tx=XM_011282911.2;refGene_gn=TLN2;refGene_tx=XM_019832308.2;refGene_gn=TLN2;refGene_tx=XM_019832309.2;refGene_gn=TLN2;refGene_tx=XM_019832310.2;refGene_gn=TLN2;refGene_tx=XM_019832311.2
B3 44409543 44409607 refGene_gn=TLN2;refGene_tx=XM_011282911.2;refGene_gn=TLN2;refGene_tx=XM_019832308.2;refGene_gn=TLN2;refGene_tx=XM_019832309.2;refGene_gn=TLN2;refGene_tx=XM_019832310.2;refGene_gn=TLN2;refGene_tx=XM_019832311.2
B3 44428356 44428484 refGene_gn=TLN2;refGene_tx=XM_011282911.2;refGene_gn=TLN2;refGene_tx=XM_019832308.2;refGene_gn=TLN2;refGene_tx=XM_019832309.2;refGene_gn=TLN2;refGene_tx=XM_019832310.2;refGene_gn=TLN2;refGene_tx=XM_019832311.2
B3 44429971 44430114 refGene_gn=TLN2;refGene_tx=XM_011282911.2;refGene_gn=TLN2;refGene_tx=XM_019832308.2;refGene_gn=TLN2;refGene_tx=XM_019832309.2;refGene_gn=TLN2;refGene_tx=XM_019832310.2;refGene_gn=TLN2;refGene_tx=XM_019832311.2
B3 44432075 44432228 refGene_gn=TLN2;refGene_tx=XM_011282911.2;refGene_gn=TLN2;refGene_tx=XM_019832308.2;refGene_gn=TLN2;refGene_tx=XM_019832309.2;refGene_gn=TLN2;refGene_tx=XM_019832310.2;refGene_gn=TLN2;refGene_tx=XM_019832311.2
B3 44433300 44433430 refGene_gn=TLN2;refGene_tx=XM_011282911.2;refGene_gn=TLN2;refGene_tx=XM_019832308.2;refGene_gn=TLN2;refGene_tx=XM_019832309.2;refGene_gn=TLN2;refGene_tx=XM_019832310.2;refGene_gn=TLN2;refGene_tx=XM_019832311.2
B3 44434644 44434742 refGene_gn=TLN2;refGene_tx=XM_011282911.2;refGene_gn=TLN2;refGene_tx=XM_019832308.2;refGene_gn=TLN2;refGene_tx=XM_019832309.2;refGene_gn=TLN2;refGene_tx=XM_019832310.2;refGene_gn=TLN2;refGene_tx=XM_019832311.2
B3 44436841 44436977 refGene_gn=TLN2;refGene_tx=XM_011282911.2;refGene_gn=TLN2;refGene_tx=XM_019832308.2;refGene_gn=TLN2;refGene_tx=XM_019832309.2;refGene_gn=TLN2;refGene_tx=XM_019832310.2;refGene_gn=TLN2;refGene_tx=XM_019832311.2
B3 44820655 44821870 refGene_gn=C2CD4B;refGene_tx=XM_023255271.1;refGene_gn=C2CD4B;refGene_tx=XM_023255272.1
B3 44919350 44920421 refGene_gn=C2CD4A;refGene_tx=XM_023255274.1
B3 44929529 44929629 refGene_gn=VPS13C;refGene_tx=XM_023255275.1;refGene_gn=VPS13C;refGene_tx=XM_023255276.1;refGene_gn=VPS13C;refGene_tx=XM_023255277.1;refGene_gn=VPS13C;refGene_tx=XM_023255278.1;refGene_gn=VPS13C;refGene_tx=XM_023255279.1
B3 44943057 44943101 refGene_gn=VPS13C;refGene_tx=XM_023255275.1;refGene_gn=VPS13C;refGene_tx=XM_023255276.1;refGene_gn=VPS13C;refGene_tx=XM_023255277.1;refGene_

gn=VPS13C;refGene_tx=XM_023255278.1;refGene_gn=VPS13C;refGene_tx=XM_02
3255279.1
B3 44946455 44946498
refGene_gn=VPS13C;refGene_tx=XM_023255275.1;refGene_gn=VPS13C;refGene_
tx=XM_023255276.1;refGene_gn=VPS13C;refGene_tx=XM_023255277.1;refGene_
gn=VPS13C;refGene_tx=XM_023255278.1;refGene_gn=VPS13C;refGene_tx=XM_02
3255279.1
B3 44950878 44950974
refGene_gn=VPS13C;refGene_tx=XM_023255275.1;refGene_gn=VPS13C;refGene_
tx=XM_023255276.1;refGene_gn=VPS13C;refGene_tx=XM_023255277.1;refGene_
gn=VPS13C;refGene_tx=XM_023255278.1;refGene_gn=VPS13C;refGene_tx=XM_02
3255279.1
B3 44952567 44952669
refGene_gn=VPS13C;refGene_tx=XM_023255275.1;refGene_gn=VPS13C;refGene_
tx=XM_023255276.1;refGene_gn=VPS13C;refGene_tx=XM_023255277.1;refGene_
gn=VPS13C;refGene_tx=XM_023255278.1;refGene_gn=VPS13C;refGene_tx=XM_02
3255279.1
B3 44958388 44958451
refGene_gn=VPS13C;refGene_tx=XM_023255275.1;refGene_gn=VPS13C;refGene_
tx=XM_023255277.1;refGene_gn=VPS13C;refGene_tx=XM_023255279.1
B3 44963648 44963714
refGene_gn=VPS13C;refGene_tx=XM_023255275.1;refGene_gn=VPS13C;refGene_
tx=XM_023255277.1;refGene_gn=VPS13C;refGene_tx=XM_023255279.1
B3 44963988 44964098
refGene_gn=VPS13C;refGene_tx=XM_023255275.1;refGene_gn=VPS13C;refGene_
tx=XM_023255276.1;refGene_gn=VPS13C;refGene_tx=XM_023255277.1;refGene_
gn=VPS13C;refGene_tx=XM_023255278.1;refGene_gn=VPS13C;refGene_tx=XM_02
3255279.1
B3 44966329 44966389
refGene_gn=VPS13C;refGene_tx=XM_023255275.1;refGene_gn=VPS13C;refGene_
tx=XM_023255276.1;refGene_gn=VPS13C;refGene_tx=XM_023255277.1;refGene_
gn=VPS13C;refGene_tx=XM_023255278.1;refGene_gn=VPS13C;refGene_tx=XM_02
3255279.1
B3 44969496 44969556
refGene_gn=VPS13C;refGene_tx=XM_023255275.1;refGene_gn=VPS13C;refGene_
tx=XM_023255276.1;refGene_gn=VPS13C;refGene_tx=XM_023255277.1;refGene_
gn=VPS13C;refGene_tx=XM_023255278.1;refGene_gn=VPS13C;refGene_tx=XM_02
3255279.1
B3 44970346 44970427
refGene_gn=VPS13C;refGene_tx=XM_023255275.1;refGene_gn=VPS13C;refGene_
tx=XM_023255276.1;refGene_gn=VPS13C;refGene_tx=XM_023255277.1;refGene_
gn=VPS13C;refGene_tx=XM_023255278.1;refGene_gn=VPS13C;refGene_tx=XM_02
3255279.1
B3 44972111 44972169
refGene_gn=VPS13C;refGene_tx=XM_023255275.1;refGene_gn=VPS13C;refGene_
tx=XM_023255276.1;refGene_gn=VPS13C;refGene_tx=XM_023255277.1;refGene_
gn=VPS13C;refGene_tx=XM_023255278.1;refGene_gn=VPS13C;refGene_tx=XM_02
3255279.1
B3 44973897 44974025
refGene_gn=VPS13C;refGene_tx=XM_023255275.1;refGene_gn=VPS13C;refGene_

tx=XM_023255276.1;refGene_gn=VPS13C;refGene_tx=XM_023255277.1;refGene_gn=VPS13C;refGene_tx=XM_023255278.1;refGene_gn=VPS13C;refGene_tx=XM_023255279.1
B3 44976234 44976341 refGene_gn=VPS13C;refGene_tx=XM_023255275.1;refGene_gn=VPS13C;refGene_tx=XM_023255276.1;refGene_gn=VPS13C;refGene_tx=XM_023255277.1;refGene_gn=VPS13C;refGene_tx=XM_023255278.1;refGene_gn=VPS13C;refGene_tx=XM_023255279.1
B3 44976882 44977054 refGene_gn=VPS13C;refGene_tx=XM_023255275.1;refGene_gn=VPS13C;refGene_tx=XM_023255276.1;refGene_gn=VPS13C;refGene_tx=XM_023255277.1;refGene_gn=VPS13C;refGene_tx=XM_023255278.1;refGene_gn=VPS13C;refGene_tx=XM_023255279.1
B3 44978192 44978255 refGene_gn=VPS13C;refGene_tx=XM_023255275.1;refGene_gn=VPS13C;refGene_tx=XM_023255276.1;refGene_gn=VPS13C;refGene_tx=XM_023255277.1;refGene_gn=VPS13C;refGene_tx=XM_023255278.1;refGene_gn=VPS13C;refGene_tx=XM_023255279.1
B3 44989071 44989201 refGene_gn=VPS13C;refGene_tx=XM_023255275.1;refGene_gn=VPS13C;refGene_tx=XM_023255276.1;refGene_gn=VPS13C;refGene_tx=XM_023255277.1;refGene_gn=VPS13C;refGene_tx=XM_023255278.1;refGene_gn=VPS13C;refGene_tx=XM_023255279.1
B3 44989764 44989859 refGene_gn=VPS13C;refGene_tx=XM_023255275.1;refGene_gn=VPS13C;refGene_tx=XM_023255276.1;refGene_gn=VPS13C;refGene_tx=XM_023255277.1;refGene_gn=VPS13C;refGene_tx=XM_023255278.1;refGene_gn=VPS13C;refGene_tx=XM_023255279.1
B3 44995826 44995969 refGene_gn=VPS13C;refGene_tx=XM_023255275.1;refGene_gn=VPS13C;refGene_tx=XM_023255276.1;refGene_gn=VPS13C;refGene_tx=XM_023255277.1;refGene_gn=VPS13C;refGene_tx=XM_023255278.1;refGene_gn=VPS13C;refGene_tx=XM_023255279.1
B3 44996805 44996998 refGene_gn=VPS13C;refGene_tx=XM_023255275.1;refGene_gn=VPS13C;refGene_tx=XM_023255276.1;refGene_gn=VPS13C;refGene_tx=XM_023255277.1;refGene_gn=VPS13C;refGene_tx=XM_023255278.1;refGene_gn=VPS13C;refGene_tx=XM_023255279.1
B3 44997973 44998088 refGene_gn=VPS13C;refGene_tx=XM_023255275.1;refGene_gn=VPS13C;refGene_tx=XM_023255276.1;refGene_gn=VPS13C;refGene_tx=XM_023255277.1;refGene_gn=VPS13C;refGene_tx=XM_023255278.1;refGene_gn=VPS13C;refGene_tx=XM_023255279.1
B3 44998674 44998811 refGene_gn=VPS13C;refGene_tx=XM_023255275.1;refGene_gn=VPS13C;refGene_tx=XM_023255276.1;refGene_gn=VPS13C;refGene_tx=XM_023255277.1;refGene_gn=VPS13C;refGene_tx=XM_023255278.1;refGene_gn=VPS13C;refGene_tx=XM_023255279.1
B3 44999750 44999874 refGene_gn=VPS13C;refGene_tx=XM_023255275.1;refGene_gn=VPS13C;refGene_

tx=XM_023255276.1;refGene_gn=VPS13C;refGene_tx=XM_023255277.1;refGene_gn=VPS13C;refGene_tx=XM_023255278.1;refGene_gn=VPS13C;refGene_tx=XM_023255279.1
B3 45001753 45001871
refGene_gn=VPS13C;refGene_tx=XM_023255275.1;refGene_gn=VPS13C;refGene_tx=XM_023255276.1;refGene_gn=VPS13C;refGene_tx=XM_023255277.1;refGene_gn=VPS13C;refGene_tx=XM_023255278.1;refGene_gn=VPS13C;refGene_tx=XM_023255279.1
B3 45005651 45005781
refGene_gn=VPS13C;refGene_tx=XM_023255275.1;refGene_gn=VPS13C;refGene_tx=XM_023255276.1;refGene_gn=VPS13C;refGene_tx=XM_023255277.1;refGene_gn=VPS13C;refGene_tx=XM_023255278.1;refGene_gn=VPS13C;refGene_tx=XM_023255279.1
B3 45006824 45006903
refGene_gn=VPS13C;refGene_tx=XM_023255275.1;refGene_gn=VPS13C;refGene_tx=XM_023255276.1;refGene_gn=VPS13C;refGene_tx=XM_023255277.1;refGene_gn=VPS13C;refGene_tx=XM_023255278.1;refGene_gn=VPS13C;refGene_tx=XM_023255279.1
B3 45007579 45007719
refGene_gn=VPS13C;refGene_tx=XM_023255275.1;refGene_gn=VPS13C;refGene_tx=XM_023255276.1;refGene_gn=VPS13C;refGene_tx=XM_023255277.1;refGene_gn=VPS13C;refGene_tx=XM_023255278.1;refGene_gn=VPS13C;refGene_tx=XM_023255279.1
B3 45011170 45011324
refGene_gn=VPS13C;refGene_tx=XM_023255275.1;refGene_gn=VPS13C;refGene_tx=XM_023255276.1;refGene_gn=VPS13C;refGene_tx=XM_023255277.1;refGene_gn=VPS13C;refGene_tx=XM_023255278.1;refGene_gn=VPS13C;refGene_tx=XM_023255279.1
B3 45013166 45013246
refGene_gn=VPS13C;refGene_tx=XM_023255275.1;refGene_gn=VPS13C;refGene_tx=XM_023255276.1;refGene_gn=VPS13C;refGene_tx=XM_023255277.1;refGene_gn=VPS13C;refGene_tx=XM_023255278.1;refGene_gn=VPS13C;refGene_tx=XM_023255279.1
B3 45014672 45014732
refGene_gn=VPS13C;refGene_tx=XM_023255275.1;refGene_gn=VPS13C;refGene_tx=XM_023255276.1;refGene_gn=VPS13C;refGene_tx=XM_023255277.1;refGene_gn=VPS13C;refGene_tx=XM_023255278.1;refGene_gn=VPS13C;refGene_tx=XM_023255279.1
B3 45016835 45016995
refGene_gn=VPS13C;refGene_tx=XM_023255275.1;refGene_gn=VPS13C;refGene_tx=XM_023255276.1;refGene_gn=VPS13C;refGene_tx=XM_023255277.1;refGene_gn=VPS13C;refGene_tx=XM_023255278.1;refGene_gn=VPS13C;refGene_tx=XM_023255279.1
B3 45017732 45017849
refGene_gn=VPS13C;refGene_tx=XM_023255275.1;refGene_gn=VPS13C;refGene_tx=XM_023255276.1;refGene_gn=VPS13C;refGene_tx=XM_023255277.1;refGene_gn=VPS13C;refGene_tx=XM_023255278.1;refGene_gn=VPS13C;refGene_tx=XM_023255279.1
B3 45018951 45019055
refGene_gn=VPS13C;refGene_tx=XM_023255275.1;refGene_gn=VPS13C;refGene_

tx=XM_023255276.1;refGene_gn=VPS13C;refGene_tx=XM_023255277.1;refGene_gn=VPS13C;refGene_tx=XM_023255278.1;refGene_gn=VPS13C;refGene_tx=XM_023255279.1
B3 45019415 45019583
refGene_gn=VPS13C;refGene_tx=XM_023255275.1;refGene_gn=VPS13C;refGene_tx=XM_023255276.1;refGene_gn=VPS13C;refGene_tx=XM_023255277.1;refGene_gn=VPS13C;refGene_tx=XM_023255278.1;refGene_gn=VPS13C;refGene_tx=XM_023255279.1
B3 45020073 45020378
refGene_gn=VPS13C;refGene_tx=XM_023255275.1;refGene_gn=VPS13C;refGene_tx=XM_023255276.1;refGene_gn=VPS13C;refGene_tx=XM_023255277.1;refGene_gn=VPS13C;refGene_tx=XM_023255278.1;refGene_gn=VPS13C;refGene_tx=XM_023255279.1
B3 45022413 45022561
refGene_gn=VPS13C;refGene_tx=XM_023255275.1;refGene_gn=VPS13C;refGene_tx=XM_023255276.1;refGene_gn=VPS13C;refGene_tx=XM_023255277.1;refGene_gn=VPS13C;refGene_tx=XM_023255278.1;refGene_gn=VPS13C;refGene_tx=XM_023255279.1
B3 45031765 45031874
refGene_gn=VPS13C;refGene_tx=XM_023255275.1;refGene_gn=VPS13C;refGene_tx=XM_023255276.1;refGene_gn=VPS13C;refGene_tx=XM_023255277.1;refGene_gn=VPS13C;refGene_tx=XM_023255278.1;refGene_gn=VPS13C;refGene_tx=XM_023255279.1
B3 45036979 45037116
refGene_gn=VPS13C;refGene_tx=XM_023255275.1;refGene_gn=VPS13C;refGene_tx=XM_023255276.1;refGene_gn=VPS13C;refGene_tx=XM_023255277.1;refGene_gn=VPS13C;refGene_tx=XM_023255278.1;refGene_gn=VPS13C;refGene_tx=XM_023255279.1
B3 45038958 45039115
refGene_gn=VPS13C;refGene_tx=XM_023255275.1;refGene_gn=VPS13C;refGene_tx=XM_023255276.1;refGene_gn=VPS13C;refGene_tx=XM_023255277.1;refGene_gn=VPS13C;refGene_tx=XM_023255278.1;refGene_gn=VPS13C;refGene_tx=XM_023255279.1
B3 45040183 45040263
refGene_gn=VPS13C;refGene_tx=XM_023255275.1;refGene_gn=VPS13C;refGene_tx=XM_023255276.1;refGene_gn=VPS13C;refGene_tx=XM_023255277.1;refGene_gn=VPS13C;refGene_tx=XM_023255278.1;refGene_gn=VPS13C;refGene_tx=XM_023255279.1
B3 45040839 45040899
refGene_gn=VPS13C;refGene_tx=XM_023255275.1;refGene_gn=VPS13C;refGene_tx=XM_023255276.1;refGene_gn=VPS13C;refGene_tx=XM_023255277.1;refGene_gn=VPS13C;refGene_tx=XM_023255278.1;refGene_gn=VPS13C;refGene_tx=XM_023255279.1
B3 45042583 45042746
refGene_gn=VPS13C;refGene_tx=XM_023255275.1;refGene_gn=VPS13C;refGene_tx=XM_023255276.1;refGene_gn=VPS13C;refGene_tx=XM_023255277.1;refGene_gn=VPS13C;refGene_tx=XM_023255278.1;refGene_gn=VPS13C;refGene_tx=XM_023255279.1
B3 45044508 45044625
refGene_gn=VPS13C;refGene_tx=XM_023255275.1;refGene_gn=VPS13C;refGene_

tx=XM_023255276.1;refGene_gn=VPS13C;refGene_tx=XM_023255277.1;refGene_gn=VPS13C;refGene_tx=XM_023255278.1;refGene_gn=VPS13C;refGene_tx=XM_023255279.1
B3 45045937 45046041
refGene_gn=VPS13C;refGene_tx=XM_023255275.1;refGene_gn=VPS13C;refGene_tx=XM_023255276.1;refGene_gn=VPS13C;refGene_tx=XM_023255277.1;refGene_gn=VPS13C;refGene_tx=XM_023255278.1;refGene_gn=VPS13C;refGene_tx=XM_023255279.1
B3 45050519 45050687
refGene_gn=VPS13C;refGene_tx=XM_023255275.1;refGene_gn=VPS13C;refGene_tx=XM_023255276.1;refGene_gn=VPS13C;refGene_tx=XM_023255277.1;refGene_gn=VPS13C;refGene_tx=XM_023255278.1;refGene_gn=VPS13C;refGene_tx=XM_023255279.1
B3 45052091 45052396
refGene_gn=VPS13C;refGene_tx=XM_023255275.1;refGene_gn=VPS13C;refGene_tx=XM_023255276.1;refGene_gn=VPS13C;refGene_tx=XM_023255277.1;refGene_gn=VPS13C;refGene_tx=XM_023255278.1;refGene_gn=VPS13C;refGene_tx=XM_023255279.1
B3 45053324 45053472
refGene_gn=VPS13C;refGene_tx=XM_023255275.1;refGene_gn=VPS13C;refGene_tx=XM_023255276.1;refGene_gn=VPS13C;refGene_tx=XM_023255277.1;refGene_gn=VPS13C;refGene_tx=XM_023255278.1;refGene_gn=VPS13C;refGene_tx=XM_023255279.1
B3 45057457 45057608
refGene_gn=VPS13C;refGene_tx=XM_023255275.1;refGene_gn=VPS13C;refGene_tx=XM_023255276.1;refGene_gn=VPS13C;refGene_tx=XM_023255277.1;refGene_gn=VPS13C;refGene_tx=XM_023255278.1;refGene_gn=VPS13C;refGene_tx=XM_023255279.1
B3 45059631 45059741
refGene_gn=VPS13C;refGene_tx=XM_023255275.1;refGene_gn=VPS13C;refGene_tx=XM_023255276.1;refGene_gn=VPS13C;refGene_tx=XM_023255277.1;refGene_gn=VPS13C;refGene_tx=XM_023255278.1;refGene_gn=VPS13C;refGene_tx=XM_023255279.1
B3 45063226 45063396
refGene_gn=VPS13C;refGene_tx=XM_023255275.1;refGene_gn=VPS13C;refGene_tx=XM_023255276.1;refGene_gn=VPS13C;refGene_tx=XM_023255277.1;refGene_gn=VPS13C;refGene_tx=XM_023255278.1;refGene_gn=VPS13C;refGene_tx=XM_023255279.1
B3 45064669 45064917
refGene_gn=VPS13C;refGene_tx=XM_023255275.1;refGene_gn=VPS13C;refGene_tx=XM_023255276.1;refGene_gn=VPS13C;refGene_tx=XM_023255277.1;refGene_gn=VPS13C;refGene_tx=XM_023255278.1;refGene_gn=VPS13C;refGene_tx=XM_023255279.1
B3 45066033 45066263
refGene_gn=VPS13C;refGene_tx=XM_023255275.1;refGene_gn=VPS13C;refGene_tx=XM_023255276.1;refGene_gn=VPS13C;refGene_tx=XM_023255277.1;refGene_gn=VPS13C;refGene_tx=XM_023255278.1;refGene_gn=VPS13C;refGene_tx=XM_023255279.1
B3 45068720 45068813
refGene_gn=VPS13C;refGene_tx=XM_023255275.1;refGene_gn=VPS13C;refGene_

tx=XM_023255276.1;refGene_gn=VPS13C;refGene_tx=XM_023255277.1;refGene_gn=VPS13C;refGene_tx=XM_023255278.1;refGene_gn=VPS13C;refGene_tx=XM_023255279.1
B3 45071278 45071644
refGene_gn=VPS13C;refGene_tx=XM_023255275.1;refGene_gn=VPS13C;refGene_tx=XM_023255276.1;refGene_gn=VPS13C;refGene_tx=XM_023255277.1;refGene_gn=VPS13C;refGene_tx=XM_023255278.1;refGene_gn=VPS13C;refGene_tx=XM_023255279.1
B3 45072082 45072169
refGene_gn=VPS13C;refGene_tx=XM_023255275.1;refGene_gn=VPS13C;refGene_tx=XM_023255276.1;refGene_gn=VPS13C;refGene_tx=XM_023255277.1;refGene_gn=VPS13C;refGene_tx=XM_023255278.1;refGene_gn=VPS13C;refGene_tx=XM_023255279.1
B3 45073448 45073604
refGene_gn=VPS13C;refGene_tx=XM_023255275.1;refGene_gn=VPS13C;refGene_tx=XM_023255276.1;refGene_gn=VPS13C;refGene_tx=XM_023255277.1;refGene_gn=VPS13C;refGene_tx=XM_023255278.1;refGene_gn=VPS13C;refGene_tx=XM_023255279.1
B3 45073766 45074031
refGene_gn=VPS13C;refGene_tx=XM_023255275.1;refGene_gn=VPS13C;refGene_tx=XM_023255276.1;refGene_gn=VPS13C;refGene_tx=XM_023255277.1;refGene_gn=VPS13C;refGene_tx=XM_023255278.1;refGene_gn=VPS13C;refGene_tx=XM_023255279.1
B3 45074897 45075058
refGene_gn=VPS13C;refGene_tx=XM_023255275.1;refGene_gn=VPS13C;refGene_tx=XM_023255276.1;refGene_gn=VPS13C;refGene_tx=XM_023255277.1;refGene_gn=VPS13C;refGene_tx=XM_023255278.1;refGene_gn=VPS13C;refGene_tx=XM_023255279.1
B3 45076037 45076159
refGene_gn=VPS13C;refGene_tx=XM_023255275.1;refGene_gn=VPS13C;refGene_tx=XM_023255276.1;refGene_gn=VPS13C;refGene_tx=XM_023255277.1;refGene_gn=VPS13C;refGene_tx=XM_023255278.1;refGene_gn=VPS13C;refGene_tx=XM_023255279.1
B3 45076691 45076986
refGene_gn=VPS13C;refGene_tx=XM_023255275.1;refGene_gn=VPS13C;refGene_tx=XM_023255276.1;refGene_gn=VPS13C;refGene_tx=XM_023255277.1;refGene_gn=VPS13C;refGene_tx=XM_023255278.1;refGene_gn=VPS13C;refGene_tx=XM_023255279.1
B3 45078251 45078641
refGene_gn=VPS13C;refGene_tx=XM_023255275.1;refGene_gn=VPS13C;refGene_tx=XM_023255276.1;refGene_gn=VPS13C;refGene_tx=XM_023255277.1;refGene_gn=VPS13C;refGene_tx=XM_023255278.1;refGene_gn=VPS13C;refGene_tx=XM_023255279.1
B3 45080461 45080566
refGene_gn=VPS13C;refGene_tx=XM_023255275.1;refGene_gn=VPS13C;refGene_tx=XM_023255276.1;refGene_gn=VPS13C;refGene_tx=XM_023255277.1;refGene_gn=VPS13C;refGene_tx=XM_023255278.1;refGene_gn=VPS13C;refGene_tx=XM_023255279.1
B3 45081257 45081422
refGene_gn=VPS13C;refGene_tx=XM_023255275.1;refGene_gn=VPS13C;refGene_

tx=XM_023255276.1;refGene_gn=VPS13C;refGene_tx=XM_023255277.1;refGene_gn=VPS13C;refGene_tx=XM_023255278.1;refGene_gn=VPS13C;refGene_tx=XM_023255279.1
B3 45082963 45083092
refGene_gn=VPS13C;refGene_tx=XM_023255275.1;refGene_gn=VPS13C;refGene_tx=XM_023255276.1;refGene_gn=VPS13C;refGene_tx=XM_023255277.1;refGene_gn=VPS13C;refGene_tx=XM_023255278.1;refGene_gn=VPS13C;refGene_tx=XM_023255279.1
B3 45083602 45083736
refGene_gn=VPS13C;refGene_tx=XM_023255275.1;refGene_gn=VPS13C;refGene_tx=XM_023255276.1;refGene_gn=VPS13C;refGene_tx=XM_023255277.1;refGene_gn=VPS13C;refGene_tx=XM_023255278.1;refGene_gn=VPS13C;refGene_tx=XM_023255279.1
B3 45087705 45087832
refGene_gn=VPS13C;refGene_tx=XM_023255275.1;refGene_gn=VPS13C;refGene_tx=XM_023255276.1;refGene_gn=VPS13C;refGene_tx=XM_023255277.1;refGene_gn=VPS13C;refGene_tx=XM_023255278.1;refGene_gn=VPS13C;refGene_tx=XM_023255279.1
B3 45093446 45093682
refGene_gn=VPS13C;refGene_tx=XM_023255275.1;refGene_gn=VPS13C;refGene_tx=XM_023255276.1;refGene_gn=VPS13C;refGene_tx=XM_023255277.1;refGene_gn=VPS13C;refGene_tx=XM_023255278.1;refGene_gn=VPS13C;refGene_tx=XM_023255279.1
B3 45099673 45099815
refGene_gn=VPS13C;refGene_tx=XM_023255275.1;refGene_gn=VPS13C;refGene_tx=XM_023255276.1;refGene_gn=VPS13C;refGene_tx=XM_023255277.1;refGene_gn=VPS13C;refGene_tx=XM_023255278.1;refGene_gn=VPS13C;refGene_tx=XM_023255279.1
B3 45101603 45101744
refGene_gn=VPS13C;refGene_tx=XM_023255275.1;refGene_gn=VPS13C;refGene_tx=XM_023255276.1;refGene_gn=VPS13C;refGene_tx=XM_023255277.1;refGene_gn=VPS13C;refGene_tx=XM_023255278.1;refGene_gn=VPS13C;refGene_tx=XM_023255279.1
B3 45102545 45102627
refGene_gn=VPS13C;refGene_tx=XM_023255275.1;refGene_gn=VPS13C;refGene_tx=XM_023255276.1;refGene_gn=VPS13C;refGene_tx=XM_023255277.1;refGene_gn=VPS13C;refGene_tx=XM_023255278.1;refGene_gn=VPS13C;refGene_tx=XM_023255279.1
B3 45103202 45103272
refGene_gn=VPS13C;refGene_tx=XM_023255275.1;refGene_gn=VPS13C;refGene_tx=XM_023255276.1;refGene_gn=VPS13C;refGene_tx=XM_023255277.1;refGene_gn=VPS13C;refGene_tx=XM_023255278.1;refGene_gn=VPS13C;refGene_tx=XM_023255279.1
B3 45103857 45103969
refGene_gn=VPS13C;refGene_tx=XM_023255275.1;refGene_gn=VPS13C;refGene_tx=XM_023255276.1;refGene_gn=VPS13C;refGene_tx=XM_023255277.1;refGene_gn=VPS13C;refGene_tx=XM_023255278.1;refGene_gn=VPS13C;refGene_tx=XM_023255279.1
B3 45104098 45104212
refGene_gn=VPS13C;refGene_tx=XM_023255275.1;refGene_gn=VPS13C;refGene_

tx=XM_023255276.1;refGene_gn=VPS13C;refGene_tx=XM_023255277.1;refGene_gn=VPS13C;refGene_tx=XM_023255278.1;refGene_gn=VPS13C;refGene_tx=XM_023255279.1
B3 45104724 45104733 refGene_gn=VPS13C;refGene_tx=XM_023255279.1
B3 45106354 45106494
refGene_gn=VPS13C;refGene_tx=XM_023255275.1;refGene_gn=VPS13C;refGene_tx=XM_023255276.1;refGene_gn=VPS13C;refGene_tx=XM_023255277.1;refGene_gn=VPS13C;refGene_tx=XM_023255278.1
B3 45108173 45108255
refGene_gn=VPS13C;refGene_tx=XM_023255275.1;refGene_gn=VPS13C;refGene_tx=XM_023255276.1;refGene_gn=VPS13C;refGene_tx=XM_023255277.1;refGene_gn=VPS13C;refGene_tx=XM_023255278.1
B3 45109209 45109323
refGene_gn=VPS13C;refGene_tx=XM_023255275.1;refGene_gn=VPS13C;refGene_tx=XM_023255276.1;refGene_gn=VPS13C;refGene_tx=XM_023255277.1;refGene_gn=VPS13C;refGene_tx=XM_023255278.1
B3 45110067 45110143
refGene_gn=VPS13C;refGene_tx=XM_023255275.1;refGene_gn=VPS13C;refGene_tx=XM_023255276.1;refGene_gn=VPS13C;refGene_tx=XM_023255277.1;refGene_gn=VPS13C;refGene_tx=XM_023255278.1
B3 45110934 45111098
refGene_gn=VPS13C;refGene_tx=XM_023255275.1;refGene_gn=VPS13C;refGene_tx=XM_023255276.1;refGene_gn=VPS13C;refGene_tx=XM_023255277.1;refGene_gn=VPS13C;refGene_tx=XM_023255278.1
B3 45112314 45112360
refGene_gn=VPS13C;refGene_tx=XM_023255275.1;refGene_gn=VPS13C;refGene_tx=XM_023255276.1;refGene_gn=VPS13C;refGene_tx=XM_023255277.1;refGene_gn=VPS13C;refGene_tx=XM_023255278.1
B3 45114252 45114376
refGene_gn=VPS13C;refGene_tx=XM_023255275.1;refGene_gn=VPS13C;refGene_tx=XM_023255276.1;refGene_gn=VPS13C;refGene_tx=XM_023255277.1;refGene_gn=VPS13C;refGene_tx=XM_023255278.1
B3 45114884 45114999
refGene_gn=VPS13C;refGene_tx=XM_023255275.1;refGene_gn=VPS13C;refGene_tx=XM_023255276.1;refGene_gn=VPS13C;refGene_tx=XM_023255277.1;refGene_gn=VPS13C;refGene_tx=XM_023255278.1
B3 45115744 45115768
refGene_gn=VPS13C;refGene_tx=XM_023255277.1;refGene_gn=VPS13C;refGene_tx=XM_023255278.1
B3 45119874 45119963
refGene_gn=VPS13C;refGene_tx=XM_023255275.1;refGene_gn=VPS13C;refGene_tx=XM_023255276.1
B3 45126325 45126449
refGene_gn=VPS13C;refGene_tx=XM_023255275.1;refGene_gn=VPS13C;refGene_tx=XM_023255276.1
B3 45127839 45127920
refGene_gn=VPS13C;refGene_tx=XM_023255275.1;refGene_gn=VPS13C;refGene_tx=XM_023255276.1
B3 45128235 45128337
refGene_gn=VPS13C;refGene_tx=XM_023255275.1;refGene_gn=VPS13C;refGene_

tx=XM_023255276.1
B3 45727694 45727860 refGene_gn=RORA;refGene_tx=XM_023255280.1
B3 46260314 46260344 refGene_gn=RORA;refGene_tx=XM_023255280.1
B3 46341312 46341316 refGene_gn=RORA;refGene_tx=XM_019832321.2
B3 46343018 46343049 refGene_gn=RORA;refGene_tx=XM_023255282.1
B3 46400859 46400945
refGene_gn=RORA;refGene_tx=XM_019832321.2;refGene_gn=RORA;refGene_tx=XM_023255280.1;refGene_gn=RORA;refGene_tx=XM_023255282.1
B3 46419382 46419524
refGene_gn=RORA;refGene_tx=XM_019832321.2;refGene_gn=RORA;refGene_tx=XM_023255280.1;refGene_gn=RORA;refGene_tx=XM_023255282.1
B3 46423239 46423635
refGene_gn=RORA;refGene_tx=XM_019832321.2;refGene_gn=RORA;refGene_tx=XM_023255280.1;refGene_gn=RORA;refGene_tx=XM_023255282.1
B3 46429069 46429191
refGene_gn=RORA;refGene_tx=XM_019832321.2;refGene_gn=RORA;refGene_tx=XM_023255280.1;refGene_gn=RORA;refGene_tx=XM_023255282.1
B3 46431192 46431325
refGene_gn=RORA;refGene_tx=XM_019832321.2;refGene_gn=RORA;refGene_tx=XM_023255280.1;refGene_gn=RORA;refGene_tx=XM_023255282.1
B3 46431998 46432106
refGene_gn=RORA;refGene_tx=XM_019832321.2;refGene_gn=RORA;refGene_tx=XM_023255280.1;refGene_gn=RORA;refGene_tx=XM_023255282.1
B3 46433610 46433721
refGene_gn=RORA;refGene_tx=XM_019832321.2;refGene_gn=RORA;refGene_tx=XM_023255280.1;refGene_gn=RORA;refGene_tx=XM_023255282.1
B3 46435268 46435381
refGene_gn=RORA;refGene_tx=XM_019832321.2;refGene_gn=RORA;refGene_tx=XM_023255280.1;refGene_gn=RORA;refGene_tx=XM_023255282.1
B3 46437053 46437218
refGene_gn=RORA;refGene_tx=XM_019832321.2;refGene_gn=RORA;refGene_tx=XM_023255280.1;refGene_gn=RORA;refGene_tx=XM_023255282.1
B3 46454825 46454866
refGene_gn=ICE2;refGene_tx=XM_019832323.2;refGene_gn=ICE2;refGene_tx=XM_019832324.2;refGene_gn=ICE2;refGene_tx=XM_019832325.2;refGene_gn=ICE2;refGene_tx=XM_019832326.2;refGene_gn=ICE2;refGene_tx=XM_019832327.2;
refGene_gn=ICE2;refGene_tx=XM_019832328.2
B3 46456791 46456896
refGene_gn=ICE2;refGene_tx=XM_019832323.2;refGene_gn=ICE2;refGene_tx=XM_019832324.2;refGene_gn=ICE2;refGene_tx=XM_019832325.2;refGene_gn=ICE2;refGene_tx=XM_019832326.2;refGene_gn=ICE2;refGene_tx=XM_019832327.2;
refGene_gn=ICE2;refGene_tx=XM_019832328.2
B3 46462991 46463253
refGene_gn=ICE2;refGene_tx=XM_019832323.2;refGene_gn=ICE2;refGene_tx=XM_019832324.2;refGene_gn=ICE2;refGene_tx=XM_019832325.2;refGene_gn=ICE2;refGene_tx=XM_019832326.2;refGene_gn=ICE2;refGene_tx=XM_019832327.2;
refGene_gn=ICE2;refGene_tx=XM_019832328.2
B3 46464305 46464425
refGene_gn=ICE2;refGene_tx=XM_019832323.2;refGene_gn=ICE2;refGene_tx=XM_019832324.2;refGene_gn=ICE2;refGene_tx=XM_019832325.2;refGene_gn=ICE

2;refGene_tx=XM_019832326.2;refGene_gn=ICE2;refGene_tx=XM_019832327.2;
refGene_gn=ICE2;refGene_tx=XM_019832328.2;refGene_gn=ICE2;refGene_tx=X
M_019832329.2
B3 46476275 46476413
refGene_gn=ICE2;refGene_tx=XM_019832323.2;refGene_gn=ICE2;refGene_tx=X
M_019832324.2;refGene_gn=ICE2;refGene_tx=XM_019832325.2;refGene_gn=ICE
2;refGene_tx=XM_019832326.2;refGene_gn=ICE2;refGene_tx=XM_019832327.2;
refGene_gn=ICE2;refGene_tx=XM_019832328.2;refGene_gn=ICE2;refGene_tx=X
M_019832329.2
B3 46477000 46477117
refGene_gn=ICE2;refGene_tx=XM_019832323.2;refGene_gn=ICE2;refGene_tx=X
M_019832324.2;refGene_gn=ICE2;refGene_tx=XM_019832325.2;refGene_gn=ICE
2;refGene_tx=XM_019832326.2;refGene_gn=ICE2;refGene_tx=XM_019832327.2;
refGene_gn=ICE2;refGene_tx=XM_019832328.2;refGene_gn=ICE2;refGene_tx=X
M_019832329.2
B3 46477292 46477452
refGene_gn=ICE2;refGene_tx=XM_019832323.2;refGene_gn=ICE2;refGene_tx=X
M_019832324.2;refGene_gn=ICE2;refGene_tx=XM_019832325.2;refGene_gn=ICE
2;refGene_tx=XM_019832326.2;refGene_gn=ICE2;refGene_tx=XM_019832327.2;
refGene_gn=ICE2;refGene_tx=XM_019832328.2;refGene_gn=ICE2;refGene_tx=X
M_019832329.2
B3 46478328 46478510
refGene_gn=ICE2;refGene_tx=XM_019832323.2;refGene_gn=ICE2;refGene_tx=X
M_019832324.2;refGene_gn=ICE2;refGene_tx=XM_019832325.2;refGene_gn=ICE
2;refGene_tx=XM_019832326.2;refGene_gn=ICE2;refGene_tx=XM_019832327.2;
refGene_gn=ICE2;refGene_tx=XM_019832328.2;refGene_gn=ICE2;refGene_tx=X
M_019832329.2
B3 46484956 46484998
refGene_gn=ICE2;refGene_tx=XM_019832323.2;refGene_gn=ICE2;refGene_tx=X
M_019832324.2;refGene_gn=ICE2;refGene_tx=XM_019832325.2;refGene_gn=ICE
2;refGene_tx=XM_019832326.2;refGene_gn=ICE2;refGene_tx=XM_019832327.2;
refGene_gn=ICE2;refGene_tx=XM_019832329.2
B3 46486358 46487352
refGene_gn=ICE2;refGene_tx=XM_019832323.2;refGene_gn=ICE2;refGene_tx=X
M_019832324.2;refGene_gn=ICE2;refGene_tx=XM_019832325.2;refGene_gn=ICE
2;refGene_tx=XM_019832326.2;refGene_gn=ICE2;refGene_tx=XM_019832327.2;
refGene_gn=ICE2;refGene_tx=XM_019832328.2;refGene_gn=ICE2;refGene_tx=X
M_019832329.2
B3 46488362 46488538
refGene_gn=ICE2;refGene_tx=XM_019832323.2;refGene_gn=ICE2;refGene_tx=X
M_019832324.2;refGene_gn=ICE2;refGene_tx=XM_019832325.2;refGene_gn=ICE
2;refGene_tx=XM_019832326.2;refGene_gn=ICE2;refGene_tx=XM_019832327.2;
refGene_gn=ICE2;refGene_tx=XM_019832328.2;refGene_gn=ICE2;refGene_tx=X
M_019832329.2
B3 46492813 46492943
refGene_gn=ICE2;refGene_tx=XM_019832323.2;refGene_gn=ICE2;refGene_tx=X
M_019832324.2;refGene_gn=ICE2;refGene_tx=XM_019832325.2;refGene_gn=ICE
2;refGene_tx=XM_019832326.2;refGene_gn=ICE2;refGene_tx=XM_019832327.2;
refGene_gn=ICE2;refGene_tx=XM_019832328.2;refGene_gn=ICE2;refGene_tx=X
M_019832329.2

B3 46509622 46509707
refGene_gn=ICE2;refGene_tx=XM_019832323.2;refGene_gn=ICE2;refGene_tx=XM_019832324.2;refGene_gn=ICE2;refGene_tx=XM_019832325.2;refGene_gn=ICE2;refGene_tx=XM_019832326.2;refGene_gn=ICE2;refGene_tx=XM_019832327.2;refGene_gn=ICE2;refGene_tx=XM_019832328.2;refGene_gn=ICE2;refGene_tx=XM_019832329.2
B3 46511560 46511611
refGene_gn=ICE2;refGene_tx=XM_019832323.2;refGene_gn=ICE2;refGene_tx=XM_019832324.2;refGene_gn=ICE2;refGene_tx=XM_019832325.2;refGene_gn=ICE2;refGene_tx=XM_019832326.2;refGene_gn=ICE2;refGene_tx=XM_019832327.2;refGene_gn=ICE2;refGene_tx=XM_019832328.2;refGene_gn=ICE2;refGene_tx=XM_019832329.2
B3 46514597 46514856
refGene_gn=ICE2;refGene_tx=XM_019832323.2;refGene_gn=ICE2;refGene_tx=XM_019832324.2;refGene_gn=ICE2;refGene_tx=XM_019832325.2;refGene_gn=ICE2;refGene_tx=XM_019832326.2;refGene_gn=ICE2;refGene_tx=XM_019832327.2;refGene_gn=ICE2;refGene_tx=XM_019832328.2;refGene_gn=ICE2;refGene_tx=XM_019832329.2
B3 46518413 46518536
refGene_gn=ICE2;refGene_tx=XM_019832323.2;refGene_gn=ICE2;refGene_tx=XM_019832324.2;refGene_gn=ICE2;refGene_tx=XM_019832325.2;refGene_gn=ICE2;refGene_tx=XM_019832326.2;refGene_gn=ICE2;refGene_tx=XM_019832327.2;refGene_gn=ICE2;refGene_tx=XM_019832328.2;refGene_gn=ICE2;refGene_tx=XM_019832329.2
B3 46555584 46555632
refGene_gn=ANXA2;refGene_tx=XM_003987068.5;refGene_gn=ANXA2;refGene_tx=XM_006932480.3
B3 46558533 46558633
refGene_gn=ANXA2;refGene_tx=XM_003987068.5;refGene_gn=ANXA2;refGene_tx=XM_006932480.3
B3 46572184 46572279
refGene_gn=ANXA2;refGene_tx=XM_003987068.5;refGene_gn=ANXA2;refGene_tx=XM_006932480.3
B3 46575242 46575356
refGene_gn=ANXA2;refGene_tx=XM_003987068.5;refGene_gn=ANXA2;refGene_tx=XM_006932480.3
B3 46578388 46578479
refGene_gn=ANXA2;refGene_tx=XM_003987068.5;refGene_gn=ANXA2;refGene_tx=XM_006932480.3
B3 46579434 46579514
refGene_gn=ANXA2;refGene_tx=XM_003987068.5;refGene_gn=ANXA2;refGene_tx=XM_006932480.3
B3 46581320 46581380
refGene_gn=ANXA2;refGene_tx=XM_003987068.5;refGene_gn=ANXA2;refGene_tx=XM_006932480.3
B3 46583032 46583126
refGene_gn=ANXA2;refGene_tx=XM_003987068.5;refGene_gn=ANXA2;refGene_tx=XM_006932480.3
B3 46583702 46583798
refGene_gn=ANXA2;refGene_tx=XM_003987068.5;refGene_gn=ANXA2;refGene_tx

=XM_006932480.3
B3 46584256 46584315
refGene_gn=ANXA2;refGene_tx=XM_003987068.5;refGene_gn=ANXA2;refGene_tx
=XM_006932480.3
B3 46585400 46585523
refGene_gn=ANXA2;refGene_tx=XM_003987068.5;refGene_gn=ANXA2;refGene_tx
=XM_006932480.3
B3 46586935 46586995
refGene_gn=ANXA2;refGene_tx=XM_003987068.5;refGene_gn=ANXA2;refGene_tx
=XM_006932480.3
B3 46932000 46932978 refGene_gn=FOXB1;refGene_tx=XM_019832332.2
B3 47177245 47177309 refGene_gn=BNIP2;refGene_tx=XM_023256164.1
B3 47178625 47178716 refGene_gn=BNIP2;refGene_tx=XM_023256164.1
B3 47181219 47181272 refGene_gn=BNIP2;refGene_tx=XM_023256164.1
B3 47223467 47223642 refGene_gn=BNIP2;refGene_tx=XM_023256164.1
B3 47223643 47223785 refGene_gn=BNIP2;refGene_tx=XM_023256164.1
B3 47231600 47231707 refGene_gn=BNIP2;refGene_tx=XM_023256164.1
B3 47232999 47233067 refGene_gn=BNIP2;refGene_tx=XM_023256164.1
B3 47233587 47233755 refGene_gn=BNIP2;refGene_tx=XM_023256164.1
B3 47235394 47235571 refGene_gn=BNIP2;refGene_tx=XM_023256164.1
B3 47238315 47238418 refGene_gn=BNIP2;refGene_tx=XM_023256164.1
B3 47239595 47239727 refGene_gn=BNIP2;refGene_tx=XM_023256164.1
B3 47240868 47240955 refGene_gn=BNIP2;refGene_tx=XM_023256164.1
B3 47241084 47241183 refGene_gn=BNIP2;refGene_tx=XM_023256164.1
B3 47241929 47241965 refGene_gn=BNIP2;refGene_tx=XM_023256164.1
B3 47244869 47244921 refGene_gn=BNIP2;refGene_tx=XM_023256164.1
B3 47264858 47264930
refGene_gn=GTF2A2;refGene_tx=XM_003987091.4;refGene_gn=GTF2A2;refGene_
tx=XM_023255284.1
B3 47265697 47265802
refGene_gn=GTF2A2;refGene_tx=XM_003987091.4;refGene_gn=GTF2A2;refGene_
tx=XM_023255284.1
B3 47270538 47270665
refGene_gn=GTF2A2;refGene_tx=XM_003987091.4;refGene_gn=GTF2A2;refGene_
tx=XM_023255284.1
B3 47273007 47273033
refGene_gn=GTF2A2;refGene_tx=XM_003987091.4;refGene_gn=GTF2A2;refGene_
tx=XM_023255284.1
B3 47291976 47293293
refGene_gn=GCNT3;refGene_tx=XM_003987092.4;refGene_gn=GCNT3;refGene_tx
=XM_019832335.2
B3 47389135 47389260
refGene_gn=FAM81A;refGene_tx=XM_023255286.1;refGene_gn=FAM81A;refGene_
tx=XM_023255287.1
B3 47392786 47392982
refGene_gn=FAM81A;refGene_tx=XM_023255286.1;refGene_gn=FAM81A;refGene_
tx=XM_023255287.1
B3 47394565 47394701
refGene_gn=FAM81A;refGene_tx=XM_023255286.1;refGene_gn=FAM81A;refGene_
tx=XM_023255287.1

B3 47401786 47401893
refGene_gn=FAM81A;refGene_tx=XM_023255286.1;refGene_gn=FAM81A;refGene_tx=XM_023255287.1
B3 47403395 47403525
refGene_gn=FAM81A;refGene_tx=XM_023255286.1;refGene_gn=FAM81A;refGene_tx=XM_023255287.1
B3 47419657 47419776
refGene_gn=FAM81A;refGene_tx=XM_023255286.1;refGene_gn=FAM81A;refGene_tx=XM_023255287.1
B3 47443128 47443402
refGene_gn=FAM81A;refGene_tx=XM_023255286.1;refGene_gn=FAM81A;refGene_tx=XM_023255287.1
B3 47444406 47444426 refGene_gn=FAM81A;refGene_tx=XM_023255287.1
B3 47444435 47444482 refGene_gn=FAM81A;refGene_tx=XM_023255286.1
B3 47511047 47511050 refGene_gn=MYO1E;refGene_tx=XM_011282905.3
B3 47596551 47596695 refGene_gn=MYO1E;refGene_tx=XM_011282905.3
B3 47603198 47603288 refGene_gn=MYO1E;refGene_tx=XM_011282905.3
B3 47605454 47605549 refGene_gn=MYO1E;refGene_tx=XM_011282905.3
B3 47626340 47626428 refGene_gn=MYO1E;refGene_tx=XM_011282905.3
B3 47630116 47630206 refGene_gn=MYO1E;refGene_tx=XM_011282905.3
B3 47634328 47634460 refGene_gn=MYO1E;refGene_tx=XM_011282905.3
B3 47636428 47636563 refGene_gn=MYO1E;refGene_tx=XM_011282905.3
B3 47637967 47638100 refGene_gn=MYO1E;refGene_tx=XM_011282905.3
B3 47644446 47644643 refGene_gn=MYO1E;refGene_tx=XM_011282905.3
B3 47647279 47647360 refGene_gn=MYO1E;refGene_tx=XM_011282905.3
B3 47647668 47647755 refGene_gn=MYO1E;refGene_tx=XM_011282905.3
B3 47650335 47650422 refGene_gn=MYO1E;refGene_tx=XM_011282905.3
B3 47651392 47651560 refGene_gn=MYO1E;refGene_tx=XM_011282905.3
B3 47652548 47652634 refGene_gn=MYO1E;refGene_tx=XM_011282905.3
B3 47655040 47655122 refGene_gn=MYO1E;refGene_tx=XM_011282905.3
B3 47661004 47661111 refGene_gn=MYO1E;refGene_tx=XM_011282905.3
B3 47668961 47669060 refGene_gn=MYO1E;refGene_tx=XM_011282905.3
B3 47677944 47678089 refGene_gn=MYO1E;refGene_tx=XM_011282905.3
B3 47681863 47681978 refGene_gn=MYO1E;refGene_tx=XM_011282905.3
B3 47682184 47682354 refGene_gn=MYO1E;refGene_tx=XM_011282905.3
B3 47682437 47682583 refGene_gn=MYO1E;refGene_tx=XM_011282905.3
B3 47696692 47696839 refGene_gn=MYO1E;refGene_tx=XM_011282905.3
B3 47697729 47697887 refGene_gn=MYO1E;refGene_tx=XM_011282905.3
B3 47701680 47701773 refGene_gn=MYO1E;refGene_tx=XM_011282905.3
B3 47704529 47704731 refGene_gn=MYO1E;refGene_tx=XM_011282905.3
B3 47719684 47719854 refGene_gn=MYO1E;refGene_tx=XM_011282905.3
B3 47724851 47724928 refGene_gn=MYO1E;refGene_tx=XM_011282905.3
B3 47732433 47732565 refGene_gn=CCNB2;refGene_tx=XM_019832341.2
B3 47733031 47733142 refGene_gn=CCNB2;refGene_tx=XM_003987096.4
B3 47734227 47734338
refGene_gn=CCNB2;refGene_tx=XM_003987096.4;refGene_gn=CCNB2;refGene_tx=XM_019832341.2
B3 47739322 47739463
refGene_gn=CCNB2;refGene_tx=XM_003987096.4;refGene_gn=CCNB2;refGene_tx=XM_019832341.2

B3 47739766 47740003
refGene_gn=CCNB2;refGene_tx=XM_003987096.4;refGene_gn=CCNB2;refGene_tx=XM_019832341.2
B3 47742102 47742261
refGene_gn=CCNB2;refGene_tx=XM_003987096.4;refGene_gn=CCNB2;refGene_tx=XM_019832341.2
B3 47742364 47742535
refGene_gn=CCNB2;refGene_tx=XM_003987096.4;refGene_gn=CCNB2;refGene_tx=XM_019832341.2
B3 47756896 47757007
refGene_gn=CCNB2;refGene_tx=XM_003987096.4;refGene_gn=CCNB2;refGene_tx=XM_019832341.2
B3 47757109 47757238
refGene_gn=CCNB2;refGene_tx=XM_003987096.4;refGene_gn=CCNB2;refGene_tx=XM_019832341.2
B3 47759430 47759454
refGene_gn=CCNB2;refGene_tx=XM_003987096.4;refGene_gn=CCNB2;refGene_tx=XM_019832341.2
B3 47773118 47773236
refGene_gn=RNF111;refGene_tx=XM_003987100.5;refGene_gn=RNF111;refGene_tx=XM_019832343.1;refGene_gn=RNF111;refGene_tx=XM_019832344.1;refGene_gn=RNF111;refGene_tx=XM_019832345.1;refGene_gn=RNF111;refGene_tx=XM_019832346.1;refGene_gn=RNF111;refGene_tx=XM_019832347.2;refGene_gn=RNF111;refGene_tx=XM_019832348.2
B3 47775020 47775148
refGene_gn=RNF111;refGene_tx=XM_003987100.5;refGene_gn=RNF111;refGene_tx=XM_019832343.1;refGene_gn=RNF111;refGene_tx=XM_019832344.1;refGene_gn=RNF111;refGene_tx=XM_019832345.1;refGene_gn=RNF111;refGene_tx=XM_019832346.1;refGene_gn=RNF111;refGene_tx=XM_019832347.2;refGene_gn=RNF111;refGene_tx=XM_019832348.2
B3 47776481 47776577
refGene_gn=RNF111;refGene_tx=XM_003987100.5;refGene_gn=RNF111;refGene_tx=XM_019832343.1;refGene_gn=RNF111;refGene_tx=XM_019832344.1;refGene_gn=RNF111;refGene_tx=XM_019832345.1;refGene_gn=RNF111;refGene_tx=XM_019832346.1;refGene_gn=RNF111;refGene_tx=XM_019832347.2;refGene_gn=RNF111;refGene_tx=XM_019832348.2
B3 47777388 47777481
refGene_gn=RNF111;refGene_tx=XM_003987100.5;refGene_gn=RNF111;refGene_tx=XM_019832343.1;refGene_gn=RNF111;refGene_tx=XM_019832344.1;refGene_gn=RNF111;refGene_tx=XM_019832345.1;refGene_gn=RNF111;refGene_tx=XM_019832346.1;refGene_gn=RNF111;refGene_tx=XM_019832347.2;refGene_gn=RNF111;refGene_tx=XM_019832348.2
B3 47780468 47780595
refGene_gn=RNF111;refGene_tx=XM_003987100.5;refGene_gn=RNF111;refGene_tx=XM_019832343.1;refGene_gn=RNF111;refGene_tx=XM_019832344.1;refGene_gn=RNF111;refGene_tx=XM_019832345.1;refGene_gn=RNF111;refGene_tx=XM_019832346.1;refGene_gn=RNF111;refGene_tx=XM_019832347.2;refGene_gn=RNF111;refGene_tx=XM_019832348.2
B3 47781616 47781769
refGene_gn=RNF111;refGene_tx=XM_003987100.5;refGene_gn=RNF111;refGene_

tx=XM_019832343.1;refGene_gn=RNF111;refGene_tx=XM_019832344.1;refGene_gn=RNF111;refGene_tx=XM_019832345.1;refGene_gn=RNF111;refGene_tx=XM_019832346.1;refGene_gn=RNF111;refGene_tx=XM_019832347.2;refGene_gn=RNF111;refGene_tx=XM_019832348.2
B3 47783411 47783760
refGene_gn=RNF111;refGene_tx=XM_003987100.5;refGene_gn=RNF111;refGene_tx=XM_019832343.1;refGene_gn=RNF111;refGene_tx=XM_019832344.1;refGene_gn=RNF111;refGene_tx=XM_019832345.1;refGene_gn=RNF111;refGene_tx=XM_019832346.1;refGene_gn=RNF111;refGene_tx=XM_019832347.2;refGene_gn=RNF111;refGene_tx=XM_019832348.2
B3 47787657 47787916
refGene_gn=RNF111;refGene_tx=XM_003987100.5;refGene_gn=RNF111;refGene_tx=XM_019832343.1;refGene_gn=RNF111;refGene_tx=XM_019832344.1;refGene_gn=RNF111;refGene_tx=XM_019832345.1;refGene_gn=RNF111;refGene_tx=XM_019832346.1;refGene_gn=RNF111;refGene_tx=XM_019832347.2;refGene_gn=RNF111;refGene_tx=XM_019832348.2
B3 47792064 47792384
refGene_gn=RNF111;refGene_tx=XM_003987100.5;refGene_gn=RNF111;refGene_tx=XM_019832343.1;refGene_gn=RNF111;refGene_tx=XM_019832344.1;refGene_gn=RNF111;refGene_tx=XM_019832345.1;refGene_gn=RNF111;refGene_tx=XM_019832346.1;refGene_gn=RNF111;refGene_tx=XM_019832347.2;refGene_gn=RNF111;refGene_tx=XM_019832348.2
B3 47802908 47803103
refGene_gn=RNF111;refGene_tx=XM_003987100.5;refGene_gn=RNF111;refGene_tx=XM_019832343.1;refGene_gn=RNF111;refGene_tx=XM_019832344.1;refGene_gn=RNF111;refGene_tx=XM_019832345.1;refGene_gn=RNF111;refGene_tx=XM_019832346.1;refGene_gn=RNF111;refGene_tx=XM_019832347.2;refGene_gn=RNF111;refGene_tx=XM_019832348.2
B3 47806075 47806239
refGene_gn=RNF111;refGene_tx=XM_003987100.5;refGene_gn=RNF111;refGene_tx=XM_019832343.1;refGene_gn=RNF111;refGene_tx=XM_019832344.1;refGene_gn=RNF111;refGene_tx=XM_019832345.1;refGene_gn=RNF111;refGene_tx=XM_019832346.1;refGene_gn=RNF111;refGene_tx=XM_019832347.2;refGene_gn=RNF111;refGene_tx=XM_019832348.2
B3 47809691 47809818
refGene_gn=RNF111;refGene_tx=XM_003987100.5;refGene_gn=RNF111;refGene_tx=XM_019832343.1;refGene_gn=RNF111;refGene_tx=XM_019832344.1;refGene_gn=RNF111;refGene_tx=XM_019832345.1;refGene_gn=RNF111;refGene_tx=XM_019832346.1;refGene_gn=RNF111;refGene_tx=XM_019832347.2;refGene_gn=RNF111;refGene_tx=XM_019832348.2
B3 47827198 47828078
refGene_gn=RNF111;refGene_tx=XM_003987100.5;refGene_gn=RNF111;refGene_tx=XM_019832343.1;refGene_gn=RNF111;refGene_tx=XM_019832344.1;refGene_gn=RNF111;refGene_tx=XM_019832345.1;refGene_gn=RNF111;refGene_tx=XM_019832346.1;refGene_gn=RNF111;refGene_tx=XM_019832347.2;refGene_gn=RNF111;refGene_tx=XM_019832348.2
B3 47905931 47906093
refGene_gn=SLTM;refGene_tx=XM_023255288.1;refGene_gn=SLTM;refGene_tx=XM_023255289.1;refGene_gn=SLTM;refGene_tx=XM_023255290.1;refGene_gn=SLTM;refGene_tx=XM_023255292.1;refGene_gn=SLTM;refGene_tx=XM_023255293.1;

refGene_gn=SLTM;refGene_tx=XM_023255294.1;refGene_gn=SLTM;refGene_tx=X
M_023255295.1
B3 47907061 47907149
refGene_gn=SLTM;refGene_tx=XM_023255288.1;refGene_gn=SLTM;refGene_tx=X
M_023255289.1;refGene_gn=SLTM;refGene_tx=XM_023255290.1;refGene_gn=SLT
M;refGene_tx=XM_023255292.1;refGene_gn=SLTM;refGene_tx=XM_023255293.1;
refGene_gn=SLTM;refGene_tx=XM_023255294.1;refGene_gn=SLTM;refGene_tx=X
M_023255295.1
B3 47918605 47918670
refGene_gn=SLTM;refGene_tx=XM_023255288.1;refGene_gn=SLTM;refGene_tx=X
M_023255289.1;refGene_gn=SLTM;refGene_tx=XM_023255290.1;refGene_gn=SLT
M;refGene_tx=XM_023255292.1;refGene_gn=SLTM;refGene_tx=XM_023255293.1;
refGene_gn=SLTM;refGene_tx=XM_023255294.1;refGene_gn=SLTM;refGene_tx=X
M_023255295.1
B3 47921433 47921631
refGene_gn=SLTM;refGene_tx=XM_023255288.1;refGene_gn=SLTM;refGene_tx=X
M_023255289.1;refGene_gn=SLTM;refGene_tx=XM_023255290.1
B3 47922514 47922562
refGene_gn=SLTM;refGene_tx=XM_023255288.1;refGene_gn=SLTM;refGene_tx=X
M_023255289.1;refGene_gn=SLTM;refGene_tx=XM_023255290.1;refGene_gn=SLT
M;refGene_tx=XM_023255292.1;refGene_gn=SLTM;refGene_tx=XM_023255294.1
B3 47930970 47930998
refGene_gn=SLTM;refGene_tx=XM_023255288.1;refGene_gn=SLTM;refGene_tx=X
M_023255289.1;refGene_gn=SLTM;refGene_tx=XM_023255290.1;refGene_gn=SLT
M;refGene_tx=XM_023255292.1;refGene_gn=SLTM;refGene_tx=XM_023255293.1;
refGene_gn=SLTM;refGene_tx=XM_023255294.1;refGene_gn=SLTM;refGene_tx=X
M_023255295.1
B3 47932331 47932800
refGene_gn=SLTM;refGene_tx=XM_023255288.1;refGene_gn=SLTM;refGene_tx=X
M_023255289.1;refGene_gn=SLTM;refGene_tx=XM_023255290.1;refGene_gn=SLT
M;refGene_tx=XM_023255292.1;refGene_gn=SLTM;refGene_tx=XM_023255293.1;
refGene_gn=SLTM;refGene_tx=XM_023255294.1;refGene_gn=SLTM;refGene_tx=X
M_023255295.1
B3 47933416 47933466
refGene_gn=SLTM;refGene_tx=XM_023255288.1;refGene_gn=SLTM;refGene_tx=X
M_023255289.1;refGene_gn=SLTM;refGene_tx=XM_023255290.1;refGene_gn=SLT
M;refGene_tx=XM_023255292.1;refGene_gn=SLTM;refGene_tx=XM_023255293.1;
refGene_gn=SLTM;refGene_tx=XM_023255294.1;refGene_gn=SLTM;refGene_tx=X
M_023255295.1
B3 47935641 47935760
refGene_gn=SLTM;refGene_tx=XM_023255288.1;refGene_gn=SLTM;refGene_tx=X
M_023255289.1;refGene_gn=SLTM;refGene_tx=XM_023255290.1;refGene_gn=SLT
M;refGene_tx=XM_023255292.1;refGene_gn=SLTM;refGene_tx=XM_023255293.1;
refGene_gn=SLTM;refGene_tx=XM_023255294.1;refGene_gn=SLTM;refGene_tx=X
M_023255295.1
B3 47938552 47938702
refGene_gn=SLTM;refGene_tx=XM_023255288.1;refGene_gn=SLTM;refGene_tx=X
M_023255289.1;refGene_gn=SLTM;refGene_tx=XM_023255290.1;refGene_gn=SLT
M;refGene_tx=XM_023255292.1;refGene_gn=SLTM;refGene_tx=XM_023255293.1;
refGene_gn=SLTM;refGene_tx=XM_023255294.1;refGene_gn=SLTM;refGene_tx=X

M_023255295.1
B3 47938932 47939036
refGene_gn=SLTM;refGene_tx=XM_023255288.1;refGene_gn=SLTM;refGene_tx=XM_023255289.1;refGene_gn=SLTM;refGene_tx=XM_023255290.1;refGene_gn=SLTM;refGene_tx=XM_023255292.1;refGene_gn=SLTM;refGene_tx=XM_023255293.1;refGene_gn=SLTM;refGene_tx=XM_023255294.1;refGene_gn=SLTM;refGene_tx=XM_023255295.1
B3 47939128 47939295
refGene_gn=SLTM;refGene_tx=XM_023255288.1;refGene_gn=SLTM;refGene_tx=XM_023255289.1;refGene_gn=SLTM;refGene_tx=XM_023255290.1;refGene_gn=SLTM;refGene_tx=XM_023255292.1;refGene_gn=SLTM;refGene_tx=XM_023255293.1;refGene_gn=SLTM;refGene_tx=XM_023255294.1;refGene_gn=SLTM;refGene_tx=XM_023255295.1
B3 47939816 47939902
refGene_gn=SLTM;refGene_tx=XM_023255288.1;refGene_gn=SLTM;refGene_tx=XM_023255289.1;refGene_gn=SLTM;refGene_tx=XM_023255290.1;refGene_gn=SLTM;refGene_tx=XM_023255292.1;refGene_gn=SLTM;refGene_tx=XM_023255293.1;refGene_gn=SLTM;refGene_tx=XM_023255294.1;refGene_gn=SLTM;refGene_tx=XM_023255295.1
B3 47940098 47940262
refGene_gn=SLTM;refGene_tx=XM_023255288.1;refGene_gn=SLTM;refGene_tx=XM_023255289.1;refGene_gn=SLTM;refGene_tx=XM_023255290.1;refGene_gn=SLTM;refGene_tx=XM_023255292.1;refGene_gn=SLTM;refGene_tx=XM_023255293.1;refGene_gn=SLTM;refGene_tx=XM_023255294.1;refGene_gn=SLTM;refGene_tx=XM_023255295.1
B3 47942700 47942881
refGene_gn=SLTM;refGene_tx=XM_023255288.1;refGene_gn=SLTM;refGene_tx=XM_023255289.1;refGene_gn=SLTM;refGene_tx=XM_023255290.1;refGene_gn=SLTM;refGene_tx=XM_023255292.1;refGene_gn=SLTM;refGene_tx=XM_023255293.1;refGene_gn=SLTM;refGene_tx=XM_023255294.1;refGene_gn=SLTM;refGene_tx=XM_023255295.1
B3 47943624 47943749
refGene_gn=SLTM;refGene_tx=XM_023255288.1;refGene_gn=SLTM;refGene_tx=XM_023255289.1;refGene_gn=SLTM;refGene_tx=XM_023255290.1;refGene_gn=SLTM;refGene_tx=XM_023255292.1;refGene_gn=SLTM;refGene_tx=XM_023255293.1;refGene_gn=SLTM;refGene_tx=XM_023255294.1;refGene_gn=SLTM;refGene_tx=XM_023255295.1
B3 47944545 47944716
refGene_gn=SLTM;refGene_tx=XM_023255288.1;refGene_gn=SLTM;refGene_tx=XM_023255289.1;refGene_gn=SLTM;refGene_tx=XM_023255290.1;refGene_gn=SLTM;refGene_tx=XM_023255292.1;refGene_gn=SLTM;refGene_tx=XM_023255293.1;refGene_gn=SLTM;refGene_tx=XM_023255294.1;refGene_gn=SLTM;refGene_tx=XM_023255295.1
B3 47945418 47945733
refGene_gn=SLTM;refGene_tx=XM_023255288.1;refGene_gn=SLTM;refGene_tx=XM_023255289.1;refGene_gn=SLTM;refGene_tx=XM_023255290.1;refGene_gn=SLTM;refGene_tx=XM_023255292.1;refGene_gn=SLTM;refGene_tx=XM_023255293.1;refGene_gn=SLTM;refGene_tx=XM_023255294.1;refGene_gn=SLTM;refGene_tx=XM_023255295.1
B3 47945978 47946123

refGene_gn=SLTM;refGene_tx=XM_023255288.1;refGene_gn=SLTM;refGene_tx=XM_023255289.1;refGene_gn=SLTM;refGene_tx=XM_023255290.1;refGene_gn=SLTM;refGene_tx=XM_023255292.1;refGene_gn=SLTM;refGene_tx=XM_023255293.1;refGene_gn=SLTM;refGene_tx=XM_023255294.1;refGene_gn=SLTM;refGene_tx=XM_023255295.1
B3 47948058 47948219
refGene_gn=SLTM;refGene_tx=XM_023255288.1;refGene_gn=SLTM;refGene_tx=XM_023255289.1;refGene_gn=SLTM;refGene_tx=XM_023255290.1;refGene_gn=SLTM;refGene_tx=XM_023255292.1;refGene_gn=SLTM;refGene_tx=XM_023255293.1;refGene_gn=SLTM;refGene_tx=XM_023255294.1;refGene_gn=SLTM;refGene_tx=XM_023255295.1
B3 47950275 47950288 refGene_gn=SLTM;refGene_tx=XM_023255290.1
B3 47950299 47950408
refGene_gn=SLTM;refGene_tx=XM_023255288.1;refGene_gn=SLTM;refGene_tx=XM_023255289.1;refGene_gn=SLTM;refGene_tx=XM_023255292.1;refGene_gn=SLTM;refGene_tx=XM_023255293.1;refGene_gn=SLTM;refGene_tx=XM_023255294.1;refGene_gn=SLTM;refGene_tx=XM_023255295.1
B3 47989007 47989136
refGene_gn=MINDY2;refGene_tx=XM_003987101.4;refGene_gn=MINDY2;refGene_tx=XM_011282904.3;refGene_gn=MINDY2;refGene_tx=XM_019832349.2;refGene_gn=MINDY2;refGene_tx=XM_019832350.2
B3 47992994 47993189
refGene_gn=MINDY2;refGene_tx=XM_003987101.4;refGene_gn=MINDY2;refGene_tx=XM_011282904.3;refGene_gn=MINDY2;refGene_tx=XM_019832349.2;refGene_gn=MINDY2;refGene_tx=XM_019832350.2
B3 47996071 47996245
refGene_gn=MINDY2;refGene_tx=XM_003987101.4;refGene_gn=MINDY2;refGene_tx=XM_011282904.3;refGene_gn=MINDY2;refGene_tx=XM_019832349.2;refGene_gn=MINDY2;refGene_tx=XM_019832350.2
B3 48005229 48005325
refGene_gn=MINDY2;refGene_tx=XM_019832349.2;refGene_gn=MINDY2;refGene_tx=XM_019832350.2
B3 48012083 48012226
refGene_gn=MINDY2;refGene_tx=XM_003987101.4;refGene_gn=MINDY2;refGene_tx=XM_019832349.2;refGene_gn=MINDY2;refGene_tx=XM_019832350.2
B3 48020907 48021010
refGene_gn=MINDY2;refGene_tx=XM_003987101.4;refGene_gn=MINDY2;refGene_tx=XM_019832349.2;refGene_gn=MINDY2;refGene_tx=XM_019832350.2
B3 48030995 48031154
refGene_gn=MINDY2;refGene_tx=XM_003987101.4;refGene_gn=MINDY2;refGene_tx=XM_011282904.3;refGene_gn=MINDY2;refGene_tx=XM_019832349.2;refGene_gn=MINDY2;refGene_tx=XM_019832350.2
B3 48036462 48036527
refGene_gn=MINDY2;refGene_tx=XM_003987101.4;refGene_gn=MINDY2;refGene_tx=XM_011282904.3;refGene_gn=MINDY2;refGene_tx=XM_019832349.2;refGene_gn=MINDY2;refGene_tx=XM_019832350.2
B3 48049003 48049061
refGene_gn=MINDY2;refGene_tx=XM_003987101.4;refGene_gn=MINDY2;refGene_tx=XM_011282904.3;refGene_gn=MINDY2;refGene_tx=XM_019832349.2;refGene_gn=MINDY2;refGene_tx=XM_019832350.2

B3 48064337 48065195
refGene_gn=MINDY2;refGene_tx=XM_003987101.4;refGene_gn=MINDY2;refGene_tx=XM_011282904.3;refGene_gn=MINDY2;refGene_tx=XM_019832349.2;refGene_gn=MINDY2;refGene_tx=XM_019832350.2
B3 48089021 48089076 refGene_gn=ADAM10;refGene_tx=XM_011282902.3
B3 48122847 48122998 refGene_gn=ADAM10;refGene_tx=XM_011282902.3
B3 48150961 48151080 refGene_gn=ADAM10;refGene_tx=XM_011282902.3
B3 48153062 48153221 refGene_gn=ADAM10;refGene_tx=XM_011282902.3
B3 48169645 48169746 refGene_gn=ADAM10;refGene_tx=XM_011282902.3
B3 48181950 48182100 refGene_gn=ADAM10;refGene_tx=XM_011282902.3
B3 48185338 48185431 refGene_gn=ADAM10;refGene_tx=XM_011282902.3
B3 48188627 48188811 refGene_gn=ADAM10;refGene_tx=XM_011282902.3
B3 48196947 48197111 refGene_gn=ADAM10;refGene_tx=XM_011282902.3
B3 48203144 48203328 refGene_gn=ADAM10;refGene_tx=XM_011282902.3
B3 48208292 48208443 refGene_gn=ADAM10;refGene_tx=XM_011282902.3
B3 48209258 48209442 refGene_gn=ADAM10;refGene_tx=XM_011282902.3
B3 48210155 48210264 refGene_gn=ADAM10;refGene_tx=XM_011282902.3
B3 48211060 48211281 refGene_gn=ADAM10;refGene_tx=XM_011282902.3
B3 48222243 48222370 refGene_gn=ADAM10;refGene_tx=XM_011282902.3
B3 48224152 48224247 refGene_gn=ADAM10;refGene_tx=XM_011282902.3
B3 48251631 48251743 refGene_gn=LIPC;refGene_tx=XM_003987102.5
B3 48256844 48257063 refGene_gn=LIPC;refGene_tx=XM_003987102.5
B3 48258743 48258861 refGene_gn=LIPC;refGene_tx=XM_003987102.5
B3 48270563 48270809 refGene_gn=LIPC;refGene_tx=XM_003987102.5
B3 48273633 48273867 refGene_gn=LIPC;refGene_tx=XM_003987102.5
B3 48276119 48276237 refGene_gn=LIPC;refGene_tx=XM_003987102.5
B3 48277004 48277187 refGene_gn=LIPC;refGene_tx=XM_003987102.5
B3 48283570 48283678 refGene_gn=LIPC;refGene_tx=XM_003987102.5
B3 48283680 48283757 refGene_gn=LIPC;refGene_tx=XM_003987102.5
B3 48391227 48391312 refGene_gn=LIPC;refGene_tx=XM_003987102.5
B3 48592027 48592202 refGene_gn=AQP9;refGene_tx=XM_003987104.5
B3 48596843 48597061 refGene_gn=AQP9;refGene_tx=XM_003987104.5
B3 48602257 48602376 refGene_gn=AQP9;refGene_tx=XM_003987104.5
B3 48603951 48604089 refGene_gn=AQP9;refGene_tx=XM_003987104.5
B3 48607568 48607695 refGene_gn=AQP9;refGene_tx=XM_003987104.5
B3 48636153 48636264 refGene_gn=AQP9;refGene_tx=XM_003987104.5
B3 48696172 48696289
refGene_gn=ALDH1A2;refGene_tx=XM_006932501.4;refGene_gn=ALDH1A2;refGene_tx=XM_011282900.2;refGene_gn=ALDH1A2;refGene_tx=XM_011282901.2;refGene_gn=ALDH1A2;refGene_tx=XM_019832352.2
B3 48732949 48733054
refGene_gn=ALDH1A2;refGene_tx=XM_006932501.4;refGene_gn=ALDH1A2;refGene_tx=XM_011282900.2;refGene_gn=ALDH1A2;refGene_tx=XM_011282901.2;refGene_gn=ALDH1A2;refGene_tx=XM_019832352.2
B3 48733236 48733377
refGene_gn=ALDH1A2;refGene_tx=XM_006932501.4;refGene_gn=ALDH1A2;refGene_tx=XM_011282900.2;refGene_gn=ALDH1A2;refGene_tx=XM_011282901.2;refGene_gn=ALDH1A2;refGene_tx=XM_019832352.2;refGene_gn=ALDH1A2;refGene_tx=XM_019832353.2;refGene_gn=ALDH1A2;refGene_tx=XM_019832354.2
B3 48736561 48736691

refGene_gn=ALDH1A2;refGene_tx=XM_006932501.4;refGene_gn=ALDH1A2;refGene_tx=XM_011282900.2;refGene_gn=ALDH1A2;refGene_tx=XM_011282901.2;refGene_gn=ALDH1A2;refGene_tx=XM_019832352.2;refGene_gn=ALDH1A2;refGene_tx=XM_019832353.2;refGene_gn=ALDH1A2;refGene_tx=XM_019832354.2;refGene_gn=ALDH1A2;refGene_tx=XM_023255297.1
B3 48749096 48749158
refGene_gn=ALDH1A2;refGene_tx=XM_006932501.4;refGene_gn=ALDH1A2;refGene_tx=XM_011282900.2;refGene_gn=ALDH1A2;refGene_tx=XM_011282901.2;refGene_gn=ALDH1A2;refGene_tx=XM_019832352.2;refGene_gn=ALDH1A2;refGene_tx=XM_019832353.2;refGene_gn=ALDH1A2;refGene_tx=XM_019832354.2;refGene_gn=ALDH1A2;refGene_tx=XM_023255297.1
B3 48751170 48751299
refGene_gn=ALDH1A2;refGene_tx=XM_006932501.4;refGene_gn=ALDH1A2;refGene_tx=XM_011282900.2;refGene_gn=ALDH1A2;refGene_tx=XM_011282901.2;refGene_gn=ALDH1A2;refGene_tx=XM_019832352.2;refGene_gn=ALDH1A2;refGene_tx=XM_019832353.2;refGene_gn=ALDH1A2;refGene_tx=XM_019832354.2;refGene_gn=ALDH1A2;refGene_tx=XM_023255297.1
B3 48751425 48751539
refGene_gn=ALDH1A2;refGene_tx=XM_006932501.4;refGene_gn=ALDH1A2;refGene_tx=XM_011282900.2;refGene_gn=ALDH1A2;refGene_tx=XM_011282901.2;refGene_gn=ALDH1A2;refGene_tx=XM_019832352.2;refGene_gn=ALDH1A2;refGene_tx=XM_019832353.2;refGene_gn=ALDH1A2;refGene_tx=XM_019832354.2;refGene_gn=ALDH1A2;refGene_tx=XM_023255297.1
B3 48782112 48782215
refGene_gn=ALDH1A2;refGene_tx=XM_006932501.4;refGene_gn=ALDH1A2;refGene_tx=XM_011282900.2;refGene_gn=ALDH1A2;refGene_tx=XM_011282901.2;refGene_gn=ALDH1A2;refGene_tx=XM_019832352.2;refGene_gn=ALDH1A2;refGene_tx=XM_019832353.2;refGene_gn=ALDH1A2;refGene_tx=XM_019832354.2;refGene_gn=ALDH1A2;refGene_tx=XM_023255297.1
B3 48783927 48784112
refGene_gn=ALDH1A2;refGene_tx=XM_006932501.4;refGene_gn=ALDH1A2;refGene_tx=XM_011282900.2;refGene_gn=ALDH1A2;refGene_tx=XM_011282901.2;refGene_gn=ALDH1A2;refGene_tx=XM_019832352.2;refGene_gn=ALDH1A2;refGene_tx=XM_019832353.2;refGene_gn=ALDH1A2;refGene_tx=XM_019832354.2;refGene_gn=ALDH1A2;refGene_tx=XM_023255297.1
B3 48785473 48785638
refGene_gn=ALDH1A2;refGene_tx=XM_006932501.4;refGene_gn=ALDH1A2;refGene_tx=XM_011282900.2;refGene_gn=ALDH1A2;refGene_tx=XM_011282901.2;refGene_gn=ALDH1A2;refGene_tx=XM_019832352.2;refGene_gn=ALDH1A2;refGene_tx=XM_019832353.2;refGene_gn=ALDH1A2;refGene_tx=XM_019832354.2;refGene_gn=ALDH1A2;refGene_tx=XM_023255297.1
B3 48786322 48786480
refGene_gn=ALDH1A2;refGene_tx=XM_006932501.4;refGene_gn=ALDH1A2;refGene_tx=XM_011282900.2;refGene_gn=ALDH1A2;refGene_tx=XM_011282901.2;refGene_gn=ALDH1A2;refGene_tx=XM_019832352.2;refGene_gn=ALDH1A2;refGene_tx=XM_019832353.2;refGene_gn=ALDH1A2;refGene_tx=XM_019832354.2;refGene_gn=ALDH1A2;refGene_tx=XM_023255297.1
B3 48786826 48786901
refGene_gn=ALDH1A2;refGene_tx=XM_006932501.4;refGene_gn=ALDH1A2;refGene_tx=XM_011282900.2;refGene_gn=ALDH1A2;refGene_tx=XM_011282901.2;refGe

ne_gn=ALDH1A2;refGene_tx=XM_019832352.2;refGene_gn=ALDH1A2;refGene_tx=XM_019832353.2;refGene_gn=ALDH1A2;refGene_tx=XM_019832354.2;refGene_gn=ALDH1A2;refGene_tx=XM_023255297.1
B3 48787795 48787817 refGene_gn=ALDH1A2;refGene_tx=XM_019832352.2
B3 48791519 48791581 refGene_gn=ALDH1A2;refGene_tx=XM_011282900.2
B3 48793636 48793802
refGene_gn=ALDH1A2;refGene_tx=XM_006932501.4;refGene_gn=ALDH1A2;refGene_tx=XM_019832353.2;refGene_gn=ALDH1A2;refGene_tx=XM_019832354.2;refGene_gn=ALDH1A2;refGene_tx=XM_023255297.1
B3 48793898 48793971
refGene_gn=ALDH1A2;refGene_tx=XM_011282900.2;refGene_gn=ALDH1A2;refGene_tx=XM_011282901.2
B3 49004186 49004333 refGene_gn=POLR2M;refGene_tx=XM_003987106.5
B3 49006549 49006754 refGene_gn=POLR2M;refGene_tx=XM_003987106.5
B3 49009162 49009798 refGene_gn=POLR2M;refGene_tx=XM_003987106.5
B3 49011509 49011574 refGene_gn=POLR2M;refGene_tx=XM_003987106.5
B3 49011575 49011623 refGene_gn=POLR2M;refGene_tx=XM_003987106.5
B3 49035304 49035401 refGene_gn=MYZAP;refGene_tx=XM_011282897.3
B3 49043902 49043909 refGene_gn=MYZAP;refGene_tx=XM_019832355.2
B3 49044025 49044126
refGene_gn=MYZAP;refGene_tx=XM_011282897.3;refGene_gn=MYZAP;refGene_tx=XM_019832355.2
B3 49057111 49057195
refGene_gn=MYZAP;refGene_tx=XM_011282897.3;refGene_gn=MYZAP;refGene_tx=XM_019832355.2
B3 49071023 49071077 refGene_gn=MYZAP;refGene_tx=XM_019832356.2
B3 49077842 49077948
refGene_gn=MYZAP;refGene_tx=XM_011282897.3;refGene_gn=MYZAP;refGene_tx=XM_019832355.2;refGene_gn=MYZAP;refGene_tx=XM_019832356.2
B3 49079632 49079712
refGene_gn=MYZAP;refGene_tx=XM_011282897.3;refGene_gn=MYZAP;refGene_tx=XM_019832355.2;refGene_gn=MYZAP;refGene_tx=XM_019832356.2
B3 49083880 49084009
refGene_gn=MYZAP;refGene_tx=XM_011282897.3;refGene_gn=MYZAP;refGene_tx=XM_019832355.2;refGene_gn=MYZAP;refGene_tx=XM_019832356.2
B3 49085264 49085390
refGene_gn=MYZAP;refGene_tx=XM_011282897.3;refGene_gn=MYZAP;refGene_tx=XM_019832355.2;refGene_gn=MYZAP;refGene_tx=XM_019832356.2
B3 49089248 49089401
refGene_gn=MYZAP;refGene_tx=XM_011282897.3;refGene_gn=MYZAP;refGene_tx=XM_019832355.2;refGene_gn=MYZAP;refGene_tx=XM_019832356.2
B3 49093459 49093573
refGene_gn=MYZAP;refGene_tx=XM_011282897.3;refGene_gn=MYZAP;refGene_tx=XM_019832355.2;refGene_gn=MYZAP;refGene_tx=XM_019832356.2
B3 49097412 49097505
refGene_gn=MYZAP;refGene_tx=XM_011282897.3;refGene_gn=MYZAP;refGene_tx=XM_019832355.2;refGene_gn=MYZAP;refGene_tx=XM_019832356.2
B3 49101294 49101450
refGene_gn=MYZAP;refGene_tx=XM_011282897.3;refGene_gn=MYZAP;refGene_tx=XM_019832355.2;refGene_gn=MYZAP;refGene_tx=XM_019832356.2

B3 49115350 49115437
refGene_gn=MYZAP;refGene_tx=XM_011282897.3;refGene_gn=MYZAP;refGene_tx=XM_019832355.2;refGene_gn=MYZAP;refGene_tx=XM_019832356.2
B3 49125109 49125175
refGene_gn=MYZAP;refGene_tx=XM_011282897.3;refGene_gn=MYZAP;refGene_tx=XM_019832355.2;refGene_gn=MYZAP;refGene_tx=XM_019832356.2
B3 49169870 49170006
refGene_gn=CGNL1;refGene_tx=XM_023255299.1;refGene_gn=CGNL1;refGene_tx=XM_023255300.1;refGene_gn=CGNL1;refGene_tx=XM_023255301.1;refGene_gn=CGNL1;refGene_tx=XM_023255302.1;refGene_gn=CGNL1;refGene_tx=XM_023255303.1
B3 49171315 49171479
refGene_gn=CGNL1;refGene_tx=XM_023255299.1;refGene_gn=CGNL1;refGene_tx=XM_023255300.1;refGene_gn=CGNL1;refGene_tx=XM_023255301.1;refGene_gn=CGNL1;refGene_tx=XM_023255302.1;refGene_gn=CGNL1;refGene_tx=XM_023255303.1
B3 49171787 49171896
refGene_gn=CGNL1;refGene_tx=XM_023255299.1;refGene_gn=CGNL1;refGene_tx=XM_023255300.1;refGene_gn=CGNL1;refGene_tx=XM_023255301.1;refGene_gn=CGNL1;refGene_tx=XM_023255302.1;refGene_gn=CGNL1;refGene_tx=XM_023255303.1
B3 49172853 49172978
refGene_gn=CGNL1;refGene_tx=XM_023255299.1;refGene_gn=CGNL1;refGene_tx=XM_023255300.1;refGene_gn=CGNL1;refGene_tx=XM_023255301.1;refGene_gn=CGNL1;refGene_tx=XM_023255302.1;refGene_gn=CGNL1;refGene_tx=XM_023255303.1
B3 49173645 49173729
refGene_gn=CGNL1;refGene_tx=XM_023255299.1;refGene_gn=CGNL1;refGene_tx=XM_023255300.1;refGene_gn=CGNL1;refGene_tx=XM_023255301.1;refGene_gn=CGNL1;refGene_tx=XM_023255302.1;refGene_gn=CGNL1;refGene_tx=XM_023255303.1
B3 49184744 49184834
refGene_gn=CGNL1;refGene_tx=XM_023255299.1;refGene_gn=CGNL1;refGene_tx=XM_023255300.1;refGene_gn=CGNL1;refGene_tx=XM_023255301.1;refGene_gn=CGNL1;refGene_tx=XM_023255302.1;refGene_gn=CGNL1;refGene_tx=XM_023255303.1
B3 49187626 49187788
refGene_gn=CGNL1;refGene_tx=XM_023255299.1;refGene_gn=CGNL1;refGene_tx=XM_023255300.1;refGene_gn=CGNL1;refGene_tx=XM_023255301.1;refGene_gn=CGNL1;refGene_tx=XM_023255303.1
B3 49192041 49192212
refGene_gn=CGNL1;refGene_tx=XM_023255299.1;refGene_gn=CGNL1;refGene_tx=XM_023255300.1;refGene_gn=CGNL1;refGene_tx=XM_023255301.1;refGene_gn=CGNL1;refGene_tx=XM_023255302.1;refGene_gn=CGNL1;refGene_tx=XM_023255303.1
B3 49193112 49193265
refGene_gn=CGNL1;refGene_tx=XM_023255299.1;refGene_gn=CGNL1;refGene_tx=XM_023255300.1;refGene_gn=CGNL1;refGene_tx=XM_023255301.1;refGene_gn=CGNL1;refGene_tx=XM_023255302.1;refGene_gn=CGNL1;refGene_tx=XM_023255303.1

B3	49198325 49198430
refGene_gn=CGNL1;refGene_tx=XM_023255299.1;refGene_gn=CGNL1;refGene_tx
=XM_023255300.1;refGene_gn=CGNL1;refGene_tx=XM_023255301.1;refGene_gn=
CGNL1;refGene_tx=XM_023255302.1;refGene_gn=CGNL1;refGene_tx=XM_0232553
03.1
B3	49199661 49199868
refGene_gn=CGNL1;refGene_tx=XM_023255299.1;refGene_gn=CGNL1;refGene_tx
=XM_023255300.1;refGene_gn=CGNL1;refGene_tx=XM_023255301.1;refGene_gn=
CGNL1;refGene_tx=XM_023255302.1;refGene_gn=CGNL1;refGene_tx=XM_0232553
03.1
B3	49248894 49249107
refGene_gn=CGNL1;refGene_tx=XM_023255299.1;refGene_gn=CGNL1;refGene_tx
=XM_023255300.1;refGene_gn=CGNL1;refGene_tx=XM_023255301.1;refGene_gn=
CGNL1;refGene_tx=XM_023255302.1
B3	49257974 49258110
refGene_gn=CGNL1;refGene_tx=XM_023255299.1;refGene_gn=CGNL1;refGene_tx
=XM_023255300.1;refGene_gn=CGNL1;refGene_tx=XM_023255301.1;refGene_gn=
CGNL1;refGene_tx=XM_023255302.1;refGene_gn=CGNL1;refGene_tx=XM_0232553
03.1
B3	49259485 49259634
refGene_gn=CGNL1;refGene_tx=XM_023255299.1;refGene_gn=CGNL1;refGene_tx
=XM_023255300.1;refGene_gn=CGNL1;refGene_tx=XM_023255301.1;refGene_gn=
CGNL1;refGene_tx=XM_023255302.1;refGene_gn=CGNL1;refGene_tx=XM_0232553
03.1
B3	49260173 49260275
refGene_gn=CGNL1;refGene_tx=XM_023255299.1;refGene_gn=CGNL1;refGene_tx
=XM_023255300.1;refGene_gn=CGNL1;refGene_tx=XM_023255301.1;refGene_gn=
CGNL1;refGene_tx=XM_023255302.1;refGene_gn=CGNL1;refGene_tx=XM_0232553
03.1
B3	49263708 49263814
refGene_gn=CGNL1;refGene_tx=XM_023255299.1;refGene_gn=CGNL1;refGene_tx
=XM_023255300.1;refGene_gn=CGNL1;refGene_tx=XM_023255301.1;refGene_gn=
CGNL1;refGene_tx=XM_023255302.1;refGene_gn=CGNL1;refGene_tx=XM_0232553
03.1
B3	49265436 49265531
refGene_gn=CGNL1;refGene_tx=XM_023255299.1;refGene_gn=CGNL1;refGene_tx
=XM_023255300.1;refGene_gn=CGNL1;refGene_tx=XM_023255301.1;refGene_gn=
CGNL1;refGene_tx=XM_023255302.1;refGene_gn=CGNL1;refGene_tx=XM_0232553
03.1
B3	49266329 49267907
refGene_gn=CGNL1;refGene_tx=XM_023255299.1;refGene_gn=CGNL1;refGene_tx
=XM_023255300.1;refGene_gn=CGNL1;refGene_tx=XM_023255301.1;refGene_gn=
CGNL1;refGene_tx=XM_023255302.1;refGene_gn=CGNL1;refGene_tx=XM_0232553
03.1
B3	49407453 49407596
refGene_gn=TCF12;refGene_tx=XM_003987109.5;refGene_gn=TCF12;refGene_tx
=XM_006932510.4;refGene_gn=TCF12;refGene_tx=XM_011282890.3;refGene_gn=
TCF12;refGene_tx=XM_011282891.3;refGene_gn=TCF12;refGene_tx=XM_0112828
92.3;refGene_gn=TCF12;refGene_tx=XM_011282893.2;refGene_gn=TCF12;refGe
ne_tx=XM_019832365.2;refGene_gn=TCF12;refGene_tx=XM_023255304.1;refGen

e_gn=TCF12;refGene_tx=XM_023255305.1;refGene_gn=TCF12;refGene_tx=XM_02
3255306.1;refGene_gn=TCF12;refGene_tx=XM_023255307.1;refGene_gn=TCF12;
refGene_tx=XM_023255308.1;refGene_gn=TCF12;refGene_tx=XM_023255309.1;r
efGene_gn=TCF12;refGene_tx=XM_023255310.1
B3 49414456 49414689
refGene_gn=TCF12;refGene_tx=XM_003987109.5;refGene_gn=TCF12;refGene_tx
=XM_006932510.4;refGene_gn=TCF12;refGene_tx=XM_011282890.3;refGene_gn=
TCF12;refGene_tx=XM_011282891.3;refGene_gn=TCF12;refGene_tx=XM_0112828
92.3;refGene_gn=TCF12;refGene_tx=XM_011282893.2;refGene_gn=TCF12;refGe
ne_tx=XM_019832365.2;refGene_gn=TCF12;refGene_tx=XM_023255304.1;refGen
e_gn=TCF12;refGene_tx=XM_023255305.1;refGene_gn=TCF12;refGene_tx=XM_02
3255306.1;refGene_gn=TCF12;refGene_tx=XM_023255307.1;refGene_gn=TCF12;
refGene_tx=XM_023255308.1;refGene_gn=TCF12;refGene_tx=XM_023255309.1;r
efGene_gn=TCF12;refGene_tx=XM_023255310.1
B3 49419261 49419442
refGene_gn=TCF12;refGene_tx=XM_003987109.5;refGene_gn=TCF12;refGene_tx
=XM_006932510.4;refGene_gn=TCF12;refGene_tx=XM_011282890.3;refGene_gn=
TCF12;refGene_tx=XM_011282891.3;refGene_gn=TCF12;refGene_tx=XM_0112828
92.3;refGene_gn=TCF12;refGene_tx=XM_011282893.2;refGene_gn=TCF12;refGe
ne_tx=XM_019832365.2;refGene_gn=TCF12;refGene_tx=XM_023255304.1;refGen
e_gn=TCF12;refGene_tx=XM_023255305.1;refGene_gn=TCF12;refGene_tx=XM_02
3255306.1;refGene_gn=TCF12;refGene_tx=XM_023255307.1;refGene_gn=TCF12;
refGene_tx=XM_023255308.1;refGene_gn=TCF12;refGene_tx=XM_023255309.1;r
efGene_gn=TCF12;refGene_tx=XM_023255310.1
B3 49420609 49420724
refGene_gn=TCF12;refGene_tx=XM_003987109.5;refGene_gn=TCF12;refGene_tx
=XM_006932510.4;refGene_gn=TCF12;refGene_tx=XM_011282890.3;refGene_gn=
TCF12;refGene_tx=XM_011282891.3;refGene_gn=TCF12;refGene_tx=XM_0112828
92.3;refGene_gn=TCF12;refGene_tx=XM_011282893.2;refGene_gn=TCF12;refGe
ne_tx=XM_019832365.2;refGene_gn=TCF12;refGene_tx=XM_023255304.1;refGen
e_gn=TCF12;refGene_tx=XM_023255305.1;refGene_gn=TCF12;refGene_tx=XM_02
3255306.1;refGene_gn=TCF12;refGene_tx=XM_023255307.1;refGene_gn=TCF12;
refGene_tx=XM_023255308.1;refGene_gn=TCF12;refGene_tx=XM_023255309.1;r
efGene_gn=TCF12;refGene_tx=XM_023255310.1
B3 49431908 49432115
refGene_gn=TCF12;refGene_tx=XM_003987109.5;refGene_gn=TCF12;refGene_tx
=XM_006932510.4;refGene_gn=TCF12;refGene_tx=XM_011282890.3;refGene_gn=
TCF12;refGene_tx=XM_011282891.3;refGene_gn=TCF12;refGene_tx=XM_0112828
92.3;refGene_gn=TCF12;refGene_tx=XM_011282893.2;refGene_gn=TCF12;refGe
ne_tx=XM_019832365.2;refGene_gn=TCF12;refGene_tx=XM_023255304.1;refGen
e_gn=TCF12;refGene_tx=XM_023255305.1;refGene_gn=TCF12;refGene_tx=XM_02
3255306.1;refGene_gn=TCF12;refGene_tx=XM_023255307.1;refGene_gn=TCF12;
refGene_tx=XM_023255308.1;refGene_gn=TCF12;refGene_tx=XM_023255309.1;r
efGene_gn=TCF12;refGene_tx=XM_023255310.1
B3 49432867 49432939
refGene_gn=TCF12;refGene_tx=XM_011282890.3;refGene_gn=TCF12;refGene_tx
=XM_011282891.3;refGene_gn=TCF12;refGene_tx=XM_019832365.2;refGene_gn=
TCF12;refGene_tx=XM_023255304.1;refGene_gn=TCF12;refGene_tx=XM_0232553
05.1;refGene_gn=TCF12;refGene_tx=XM_023255306.1;refGene_gn=TCF12;refGe
ne_tx=XM_023255308.1;refGene_gn=TCF12;refGene_tx=XM_023255309.1

B3 49433948 49434022
refGene_gn=TCF12;refGene_tx=XM_003987109.5;refGene_gn=TCF12;refGene_tx=XM_006932510.4;refGene_gn=TCF12;refGene_tx=XM_011282890.3;refGene_gn=TCF12;refGene_tx=XM_011282891.3;refGene_gn=TCF12;refGene_tx=XM_011282892.3;refGene_gn=TCF12;refGene_tx=XM_011282893.2;refGene_gn=TCF12;refGene_tx=XM_019832365.2;refGene_gn=TCF12;refGene_tx=XM_023255304.1;refGene_gn=TCF12;refGene_tx=XM_023255305.1;refGene_gn=TCF12;refGene_tx=XM_023255306.1;refGene_gn=TCF12;refGene_tx=XM_023255307.1;refGene_gn=TCF12;refGene_tx=XM_023255308.1;refGene_gn=TCF12;refGene_tx=XM_023255309.1;refGene_gn=TCF12;refGene_tx=XM_023255310.1
B3 49440973 49441052
refGene_gn=TCF12;refGene_tx=XM_003987109.5;refGene_gn=TCF12;refGene_tx=XM_006932510.4;refGene_gn=TCF12;refGene_tx=XM_011282890.3;refGene_gn=TCF12;refGene_tx=XM_011282891.3;refGene_gn=TCF12;refGene_tx=XM_011282892.3;refGene_gn=TCF12;refGene_tx=XM_011282893.2;refGene_gn=TCF12;refGene_tx=XM_019832365.2;refGene_gn=TCF12;refGene_tx=XM_023255304.1;refGene_gn=TCF12;refGene_tx=XM_023255305.1;refGene_gn=TCF12;refGene_tx=XM_023255306.1;refGene_gn=TCF12;refGene_tx=XM_023255307.1;refGene_gn=TCF12;refGene_tx=XM_023255308.1;refGene_gn=TCF12;refGene_tx=XM_023255309.1;refGene_gn=TCF12;refGene_tx=XM_023255310.1
B3 49449789 49449854
refGene_gn=TCF12;refGene_tx=XM_003987109.5;refGene_gn=TCF12;refGene_tx=XM_006932510.4;refGene_gn=TCF12;refGene_tx=XM_011282890.3;refGene_gn=TCF12;refGene_tx=XM_011282891.3;refGene_gn=TCF12;refGene_tx=XM_011282892.3;refGene_gn=TCF12;refGene_tx=XM_011282893.2;refGene_gn=TCF12;refGene_tx=XM_019832365.2;refGene_gn=TCF12;refGene_tx=XM_023255304.1;refGene_gn=TCF12;refGene_tx=XM_023255305.1;refGene_gn=TCF12;refGene_tx=XM_023255306.1;refGene_gn=TCF12;refGene_tx=XM_023255307.1;refGene_gn=TCF12;refGene_tx=XM_023255308.1;refGene_gn=TCF12;refGene_tx=XM_023255309.1;refGene_gn=TCF12;refGene_tx=XM_023255310.1
B3 49450180 49450325
refGene_gn=TCF12;refGene_tx=XM_003987109.5;refGene_gn=TCF12;refGene_tx=XM_006932510.4;refGene_gn=TCF12;refGene_tx=XM_011282890.3;refGene_gn=TCF12;refGene_tx=XM_011282891.3;refGene_gn=TCF12;refGene_tx=XM_011282892.3;refGene_gn=TCF12;refGene_tx=XM_011282893.2;refGene_gn=TCF12;refGene_tx=XM_019832365.2;refGene_gn=TCF12;refGene_tx=XM_023255304.1;refGene_gn=TCF12;refGene_tx=XM_023255305.1;refGene_gn=TCF12;refGene_tx=XM_023255306.1;refGene_gn=TCF12;refGene_tx=XM_023255307.1;refGene_gn=TCF12;refGene_tx=XM_023255308.1;refGene_gn=TCF12;refGene_tx=XM_023255309.1;refGene_gn=TCF12;refGene_tx=XM_023255310.1
B3 49450602 49450742
refGene_gn=TCF12;refGene_tx=XM_003987109.5;refGene_gn=TCF12;refGene_tx=XM_006932510.4;refGene_gn=TCF12;refGene_tx=XM_011282890.3;refGene_gn=TCF12;refGene_tx=XM_011282891.3;refGene_gn=TCF12;refGene_tx=XM_011282892.3;refGene_gn=TCF12;refGene_tx=XM_011282893.2;refGene_gn=TCF12;refGene_tx=XM_019832365.2;refGene_gn=TCF12;refGene_tx=XM_023255304.1;refGene_gn=TCF12;refGene_tx=XM_023255305.1;refGene_gn=TCF12;refGene_tx=XM_023255306.1;refGene_gn=TCF12;refGene_tx=XM_023255307.1;refGene_gn=TCF12;refGene_tx=XM_023255308.1;refGene_gn=TCF12;refGene_tx=XM_023255309.1;refGene_gn=TCF12;refGene_tx=XM_023255310.1

B3 49451806 49451912
refGene_gn=TCF12;refGene_tx=XM_003987109.5;refGene_gn=TCF12;refGene_tx=XM_006932510.4;refGene_gn=TCF12;refGene_tx=XM_011282890.3;refGene_gn=TCF12;refGene_tx=XM_011282891.3;refGene_gn=TCF12;refGene_tx=XM_011282892.3;refGene_gn=TCF12;refGene_tx=XM_011282893.2;refGene_gn=TCF12;refGene_tx=XM_019832365.2;refGene_gn=TCF12;refGene_tx=XM_023255304.1;refGene_gn=TCF12;refGene_tx=XM_023255305.1;refGene_gn=TCF12;refGene_tx=XM_023255306.1;refGene_gn=TCF12;refGene_tx=XM_023255307.1;refGene_gn=TCF12;refGene_tx=XM_023255308.1;refGene_gn=TCF12;refGene_tx=XM_023255309.1;refGene_gn=TCF12;refGene_tx=XM_023255310.1
B3 49463290 49463359
refGene_gn=TCF12;refGene_tx=XM_003987109.5;refGene_gn=TCF12;refGene_tx=XM_006932510.4;refGene_gn=TCF12;refGene_tx=XM_019832365.2;refGene_gn=TCF12;refGene_tx=XM_023255310.1
B3 49483589 49483642
refGene_gn=TCF12;refGene_tx=XM_011282890.3;refGene_gn=TCF12;refGene_tx=XM_011282891.3;refGene_gn=TCF12;refGene_tx=XM_011282892.3;refGene_gn=TCF12;refGene_tx=XM_011282893.2;refGene_gn=TCF12;refGene_tx=XM_023255304.1;refGene_gn=TCF12;refGene_tx=XM_023255305.1;refGene_gn=TCF12;refGene_tx=XM_023255306.1;refGene_gn=TCF12;refGene_tx=XM_023255307.1;refGene_gn=TCF12;refGene_tx=XM_023255308.1;refGene_gn=TCF12;refGene_tx=XM_023255309.1
B3 49487610 49487746
refGene_gn=TCF12;refGene_tx=XM_011282890.3;refGene_gn=TCF12;refGene_tx=XM_011282891.3;refGene_gn=TCF12;refGene_tx=XM_011282892.3;refGene_gn=TCF12;refGene_tx=XM_011282893.2;refGene_gn=TCF12;refGene_tx=XM_023255304.1;refGene_gn=TCF12;refGene_tx=XM_023255305.1;refGene_gn=TCF12;refGene_tx=XM_023255306.1;refGene_gn=TCF12;refGene_tx=XM_023255307.1;refGene_gn=TCF12;refGene_tx=XM_023255308.1;refGene_gn=TCF12;refGene_tx=XM_023255309.1
B3 49517786 49517851
refGene_gn=TCF12;refGene_tx=XM_011282890.3;refGene_gn=TCF12;refGene_tx=XM_011282891.3;refGene_gn=TCF12;refGene_tx=XM_011282892.3;refGene_gn=TCF12;refGene_tx=XM_011282893.2;refGene_gn=TCF12;refGene_tx=XM_023255304.1;refGene_gn=TCF12;refGene_tx=XM_023255305.1;refGene_gn=TCF12;refGene_tx=XM_023255306.1;refGene_gn=TCF12;refGene_tx=XM_023255307.1;refGene_gn=TCF12;refGene_tx=XM_023255308.1;refGene_gn=TCF12;refGene_tx=XM_023255309.1
B3 49588227 49588330
refGene_gn=TCF12;refGene_tx=XM_011282890.3;refGene_gn=TCF12;refGene_tx=XM_011282891.3;refGene_gn=TCF12;refGene_tx=XM_011282892.3;refGene_gn=TCF12;refGene_tx=XM_011282893.2;refGene_gn=TCF12;refGene_tx=XM_023255304.1;refGene_gn=TCF12;refGene_tx=XM_023255305.1;refGene_gn=TCF12;refGene_tx=XM_023255306.1;refGene_gn=TCF12;refGene_tx=XM_023255307.1;refGene_gn=TCF12;refGene_tx=XM_023255308.1;refGene_gn=TCF12;refGene_tx=XM_023255309.1
B3 49612996 49613044 refGene_gn=TCF12;refGene_tx=XM_023255308.1
B3 49622576 49622650
refGene_gn=TCF12;refGene_tx=XM_011282890.3;refGene_gn=TCF12;refGene_tx=XM_011282891.3;refGene_gn=TCF12;refGene_tx=XM_011282892.3;refGene_gn=

TCF12;refGene_tx=XM_011282893.2;refGene_gn=TCF12;refGene_tx=XM_0232553
04.1;refGene_gn=TCF12;refGene_tx=XM_023255305.1;refGene_gn=TCF12;refGe
ne_tx=XM_023255306.1;refGene_gn=TCF12;refGene_tx=XM_023255307.1
B3 49781628 49781701
refGene_gn=TCF12;refGene_tx=XM_011282890.3;refGene_gn=TCF12;refGene_tx
=XM_011282891.3;refGene_gn=TCF12;refGene_tx=XM_011282892.3;refGene_gn=
TCF12;refGene_tx=XM_011282893.2;refGene_gn=TCF12;refGene_tx=XM_0232553
04.1;refGene_gn=TCF12;refGene_tx=XM_023255305.1;refGene_gn=TCF12;refGe
ne_tx=XM_023255306.1;refGene_gn=TCF12;refGene_tx=XM_023255307.1
B3 49782782 49782857
refGene_gn=TCF12;refGene_tx=XM_011282890.3;refGene_gn=TCF12;refGene_tx
=XM_011282891.3;refGene_gn=TCF12;refGene_tx=XM_011282892.3;refGene_gn=
TCF12;refGene_tx=XM_011282893.2;refGene_gn=TCF12;refGene_tx=XM_0232553
04.1;refGene_gn=TCF12;refGene_tx=XM_023255305.1;refGene_gn=TCF12;refGe
ne_tx=XM_023255306.1;refGene_gn=TCF12;refGene_tx=XM_023255307.1
B3 50004773 50004801
refGene_gn=ZNF280D;refGene_tx=XM_011282887.3;refGene_gn=ZNF280D;refGen
e_tx=XM_011282888.3;refGene_gn=ZNF280D;refGene_tx=XM_023255311.1;refGe
ne_gn=ZNF280D;refGene_tx=XM_023255312.1;refGene_gn=ZNF280D;refGene_tx=
XM_023255313.1;refGene_gn=ZNF280D;refGene_tx=XM_023255315.1;refGene_gn
=ZNF280D;refGene_tx=XM_023255316.1;refGene_gn=ZNF280D;refGene_tx=XM_02
3255317.1;refGene_gn=ZNF280D;refGene_tx=XM_023255318.1
B3 50007874 50008021
refGene_gn=ZNF280D;refGene_tx=XM_011282887.3;refGene_gn=ZNF280D;refGen
e_tx=XM_011282888.3;refGene_gn=ZNF280D;refGene_tx=XM_019832366.2;refGe
ne_gn=ZNF280D;refGene_tx=XM_023255311.1;refGene_gn=ZNF280D;refGene_tx=
XM_023255312.1;refGene_gn=ZNF280D;refGene_tx=XM_023255313.1;refGene_gn
=ZNF280D;refGene_tx=XM_023255314.1;refGene_gn=ZNF280D;refGene_tx=XM_02
3255315.1;refGene_gn=ZNF280D;refGene_tx=XM_023255316.1;refGene_gn=ZNF2
80D;refGene_tx=XM_023255317.1;refGene_gn=ZNF280D;refGene_tx=XM_0232553
18.1
B3 50010085 50010151
refGene_gn=ZNF280D;refGene_tx=XM_011282887.3;refGene_gn=ZNF280D;refGen
e_tx=XM_011282888.3;refGene_gn=ZNF280D;refGene_tx=XM_019832366.2;refGe
ne_gn=ZNF280D;refGene_tx=XM_023255311.1;refGene_gn=ZNF280D;refGene_tx=
XM_023255312.1;refGene_gn=ZNF280D;refGene_tx=XM_023255313.1;refGene_gn
=ZNF280D;refGene_tx=XM_023255314.1;refGene_gn=ZNF280D;refGene_tx=XM_02
3255315.1;refGene_gn=ZNF280D;refGene_tx=XM_023255316.1;refGene_gn=ZNF2
80D;refGene_tx=XM_023255317.1;refGene_gn=ZNF280D;refGene_tx=XM_0232553
18.1
B3 50010250 50010390
refGene_gn=ZNF280D;refGene_tx=XM_011282887.3;refGene_gn=ZNF280D;refGen
e_tx=XM_011282888.3;refGene_gn=ZNF280D;refGene_tx=XM_019832366.2;refGe
ne_gn=ZNF280D;refGene_tx=XM_023255311.1;refGene_gn=ZNF280D;refGene_tx=
XM_023255312.1;refGene_gn=ZNF280D;refGene_tx=XM_023255313.1;refGene_gn
=ZNF280D;refGene_tx=XM_023255314.1;refGene_gn=ZNF280D;refGene_tx=XM_02
3255315.1;refGene_gn=ZNF280D;refGene_tx=XM_023255316.1;refGene_gn=ZNF2
80D;refGene_tx=XM_023255317.1;refGene_gn=ZNF280D;refGene_tx=XM_0232553
18.1
B3 50015977 50016095

refGene_gn=ZNF280D;refGene_tx=XM_011282887.3;refGene_gn=ZNF280D;refGene_tx=XM_011282888.3;refGene_gn=ZNF280D;refGene_tx=XM_019832366.2;refGene_gn=ZNF280D;refGene_tx=XM_023255311.1;refGene_gn=ZNF280D;refGene_tx=XM_023255312.1;refGene_gn=ZNF280D;refGene_tx=XM_023255313.1;refGene_gn=ZNF280D;refGene_tx=XM_023255314.1;refGene_gn=ZNF280D;refGene_tx=XM_023255315.1;refGene_gn=ZNF280D;refGene_tx=XM_023255316.1;refGene_gn=ZNF280D;refGene_tx=XM_023255317.1;refGene_gn=ZNF280D;refGene_tx=XM_023255318.1
B3 50029024 50029195
refGene_gn=ZNF280D;refGene_tx=XM_011282887.3;refGene_gn=ZNF280D;refGene_tx=XM_011282888.3;refGene_gn=ZNF280D;refGene_tx=XM_019832366.2;refGene_gn=ZNF280D;refGene_tx=XM_023255311.1;refGene_gn=ZNF280D;refGene_tx=XM_023255312.1;refGene_gn=ZNF280D;refGene_tx=XM_023255313.1;refGene_gn=ZNF280D;refGene_tx=XM_023255314.1;refGene_gn=ZNF280D;refGene_tx=XM_023255315.1;refGene_gn=ZNF280D;refGene_tx=XM_023255316.1;refGene_gn=ZNF280D;refGene_tx=XM_023255317.1;refGene_gn=ZNF280D;refGene_tx=XM_023255318.1
B3 50029333 50029443
refGene_gn=ZNF280D;refGene_tx=XM_011282887.3;refGene_gn=ZNF280D;refGene_tx=XM_011282888.3;refGene_gn=ZNF280D;refGene_tx=XM_019832366.2;refGene_gn=ZNF280D;refGene_tx=XM_023255311.1;refGene_gn=ZNF280D;refGene_tx=XM_023255312.1;refGene_gn=ZNF280D;refGene_tx=XM_023255313.1;refGene_gn=ZNF280D;refGene_tx=XM_023255314.1;refGene_gn=ZNF280D;refGene_tx=XM_023255315.1;refGene_gn=ZNF280D;refGene_tx=XM_023255316.1;refGene_gn=ZNF280D;refGene_tx=XM_023255317.1;refGene_gn=ZNF280D;refGene_tx=XM_023255318.1
B3 50050904 50051128
refGene_gn=ZNF280D;refGene_tx=XM_011282887.3;refGene_gn=ZNF280D;refGene_tx=XM_011282888.3;refGene_gn=ZNF280D;refGene_tx=XM_019832366.2;refGene_gn=ZNF280D;refGene_tx=XM_023255311.1;refGene_gn=ZNF280D;refGene_tx=XM_023255312.1;refGene_gn=ZNF280D;refGene_tx=XM_023255313.1;refGene_gn=ZNF280D;refGene_tx=XM_023255314.1;refGene_gn=ZNF280D;refGene_tx=XM_023255315.1;refGene_gn=ZNF280D;refGene_tx=XM_023255316.1;refGene_gn=ZNF280D;refGene_tx=XM_023255317.1;refGene_gn=ZNF280D;refGene_tx=XM_023255318.1
B3 50054514 50054672
refGene_gn=ZNF280D;refGene_tx=XM_011282887.3;refGene_gn=ZNF280D;refGene_tx=XM_011282888.3;refGene_gn=ZNF280D;refGene_tx=XM_019832366.2;refGene_gn=ZNF280D;refGene_tx=XM_023255311.1;refGene_gn=ZNF280D;refGene_tx=XM_023255312.1;refGene_gn=ZNF280D;refGene_tx=XM_023255313.1;refGene_gn=ZNF280D;refGene_tx=XM_023255314.1;refGene_gn=ZNF280D;refGene_tx=XM_023255315.1;refGene_gn=ZNF280D;refGene_tx=XM_023255316.1;refGene_gn=ZNF280D;refGene_tx=XM_023255317.1;refGene_gn=ZNF280D;refGene_tx=XM_023255318.1
B3 50055365 50055466
refGene_gn=ZNF280D;refGene_tx=XM_011282887.3;refGene_gn=ZNF280D;refGene_tx=XM_011282888.3;refGene_gn=ZNF280D;refGene_tx=XM_019832366.2;refGene_gn=ZNF280D;refGene_tx=XM_023255311.1;refGene_gn=ZNF280D;refGene_tx=XM_023255312.1;refGene_gn=ZNF280D;refGene_tx=XM_023255313.1;refGene_gn=ZNF280D;refGene_tx=XM_023255314.1;refGene_gn=ZNF280D;refGene_tx=XM_02

3255315.1;refGene_gn=ZNF280D;refGene_tx=XM_023255316.1;refGene_gn=ZNF2
80D;refGene_tx=XM_023255317.1;refGene_gn=ZNF280D;refGene_tx=XM_0232553
18.1
B3 50056443 50056590
refGene_gn=ZNF280D;refGene_tx=XM_011282887.3;refGene_gn=ZNF280D;refGen
e_tx=XM_011282888.3;refGene_gn=ZNF280D;refGene_tx=XM_019832366.2;refGe
ne_gn=ZNF280D;refGene_tx=XM_023255311.1;refGene_gn=ZNF280D;refGene_tx=
XM_023255312.1;refGene_gn=ZNF280D;refGene_tx=XM_023255313.1;refGene_gn
=ZNF280D;refGene_tx=XM_023255314.1;refGene_gn=ZNF280D;refGene_tx=XM_02
3255315.1;refGene_gn=ZNF280D;refGene_tx=XM_023255316.1;refGene_gn=ZNF2
80D;refGene_tx=XM_023255317.1;refGene_gn=ZNF280D;refGene_tx=XM_0232553
18.1
B3 50062594 50062729
refGene_gn=ZNF280D;refGene_tx=XM_011282887.3;refGene_gn=ZNF280D;refGen
e_tx=XM_011282888.3;refGene_gn=ZNF280D;refGene_tx=XM_019832366.2;refGe
ne_gn=ZNF280D;refGene_tx=XM_023255311.1;refGene_gn=ZNF280D;refGene_tx=
XM_023255312.1;refGene_gn=ZNF280D;refGene_tx=XM_023255313.1;refGene_gn
=ZNF280D;refGene_tx=XM_023255314.1;refGene_gn=ZNF280D;refGene_tx=XM_02
3255315.1;refGene_gn=ZNF280D;refGene_tx=XM_023255316.1;refGene_gn=ZNF2
80D;refGene_tx=XM_023255317.1;refGene_gn=ZNF280D;refGene_tx=XM_0232553
18.1
B3 50064552 50064860
refGene_gn=ZNF280D;refGene_tx=XM_011282887.3;refGene_gn=ZNF280D;refGen
e_tx=XM_011282888.3;refGene_gn=ZNF280D;refGene_tx=XM_019832366.2;refGe
ne_gn=ZNF280D;refGene_tx=XM_023255311.1;refGene_gn=ZNF280D;refGene_tx=
XM_023255312.1;refGene_gn=ZNF280D;refGene_tx=XM_023255313.1;refGene_gn
=ZNF280D;refGene_tx=XM_023255314.1;refGene_gn=ZNF280D;refGene_tx=XM_02
3255315.1;refGene_gn=ZNF280D;refGene_tx=XM_023255316.1;refGene_gn=ZNF2
80D;refGene_tx=XM_023255317.1;refGene_gn=ZNF280D;refGene_tx=XM_0232553
18.1
B3 50065326 50065467
refGene_gn=ZNF280D;refGene_tx=XM_011282887.3;refGene_gn=ZNF280D;refGen
e_tx=XM_011282888.3;refGene_gn=ZNF280D;refGene_tx=XM_019832366.2;refGe
ne_gn=ZNF280D;refGene_tx=XM_023255311.1;refGene_gn=ZNF280D;refGene_tx=
XM_023255312.1;refGene_gn=ZNF280D;refGene_tx=XM_023255314.1;refGene_gn
=ZNF280D;refGene_tx=XM_023255315.1;refGene_gn=ZNF280D;refGene_tx=XM_02
3255316.1;refGene_gn=ZNF280D;refGene_tx=XM_023255317.1;refGene_gn=ZNF2
80D;refGene_tx=XM_023255318.1
B3 50075936 50075999
refGene_gn=ZNF280D;refGene_tx=XM_011282887.3;refGene_gn=ZNF280D;refGen
e_tx=XM_011282888.3;refGene_gn=ZNF280D;refGene_tx=XM_019832366.2;refGe
ne_gn=ZNF280D;refGene_tx=XM_023255311.1;refGene_gn=ZNF280D;refGene_tx=
XM_023255312.1;refGene_gn=ZNF280D;refGene_tx=XM_023255313.1;refGene_gn
=ZNF280D;refGene_tx=XM_023255314.1;refGene_gn=ZNF280D;refGene_tx=XM_02
3255315.1;refGene_gn=ZNF280D;refGene_tx=XM_023255316.1;refGene_gn=ZNF2
80D;refGene_tx=XM_023255317.1;refGene_gn=ZNF280D;refGene_tx=XM_0232553
18.1
B3 50082120 50082239
refGene_gn=ZNF280D;refGene_tx=XM_011282887.3;refGene_gn=ZNF280D;refGen
e_tx=XM_011282888.3;refGene_gn=ZNF280D;refGene_tx=XM_019832366.2;refGe

ne_gn=ZNF280D;refGene_tx=XM_023255311.1;refGene_gn=ZNF280D;refGene_tx=XM_023255312.1;refGene_gn=ZNF280D;refGene_tx=XM_023255313.1;refGene_gn=ZNF280D;refGene_tx=XM_023255314.1;refGene_gn=ZNF280D;refGene_tx=XM_023255315.1;refGene_gn=ZNF280D;refGene_tx=XM_023255316.1;refGene_gn=ZNF280D;refGene_tx=XM_023255317.1;refGene_gn=ZNF280D;refGene_tx=XM_023255318.1
B3 50082386 50082423
refGene_gn=ZNF280D;refGene_tx=XM_011282887.3;refGene_gn=ZNF280D;refGene_tx=XM_011282888.3;refGene_gn=ZNF280D;refGene_tx=XM_019832366.2;refGene_gn=ZNF280D;refGene_tx=XM_023255311.1;refGene_gn=ZNF280D;refGene_tx=XM_023255312.1;refGene_gn=ZNF280D;refGene_tx=XM_023255313.1;refGene_gn=ZNF280D;refGene_tx=XM_023255314.1;refGene_gn=ZNF280D;refGene_tx=XM_023255315.1;refGene_gn=ZNF280D;refGene_tx=XM_023255316.1;refGene_gn=ZNF280D;refGene_tx=XM_023255317.1;refGene_gn=ZNF280D;refGene_tx=XM_023255318.1
B3 50082551 50082567 refGene_gn=ZNF280D;refGene_tx=XM_023255318.1
B3 50083046 50083260 refGene_gn=ZNF280D;refGene_tx=XM_023255315.1
B3 50101385 50101431
refGene_gn=ZNF280D;refGene_tx=XM_011282887.3;refGene_gn=ZNF280D;refGene_tx=XM_011282888.3;refGene_gn=ZNF280D;refGene_tx=XM_019832366.2;refGene_gn=ZNF280D;refGene_tx=XM_023255311.1;refGene_gn=ZNF280D;refGene_tx=XM_023255312.1;refGene_gn=ZNF280D;refGene_tx=XM_023255313.1;refGene_gn=ZNF280D;refGene_tx=XM_023255314.1;refGene_gn=ZNF280D;refGene_tx=XM_023255316.1;refGene_gn=ZNF280D;refGene_tx=XM_023255317.1
B3 50119413 50119452 refGene_gn=ZNF280D;refGene_tx=XM_023255316.1
B3 50124815 50124871
refGene_gn=ZNF280D;refGene_tx=XM_011282887.3;refGene_gn=ZNF280D;refGene_tx=XM_011282888.3;refGene_gn=ZNF280D;refGene_tx=XM_019832366.2;refGene_gn=ZNF280D;refGene_tx=XM_023255311.1;refGene_gn=ZNF280D;refGene_tx=XM_023255312.1;refGene_gn=ZNF280D;refGene_tx=XM_023255313.1;refGene_gn=ZNF280D;refGene_tx=XM_023255314.1
B3 50127504 50128129
refGene_gn=ZNF280D;refGene_tx=XM_011282887.3;refGene_gn=ZNF280D;refGene_tx=XM_011282888.3;refGene_gn=ZNF280D;refGene_tx=XM_019832366.2;refGene_gn=ZNF280D;refGene_tx=XM_023255311.1;refGene_gn=ZNF280D;refGene_tx=XM_023255312.1;refGene_gn=ZNF280D;refGene_tx=XM_023255313.1;refGene_gn=ZNF280D;refGene_tx=XM_023255314.1
B3 50253657 50253660 refGene_gn=MNS1;refGene_tx=XM_023255319.1
B3 50254650 50254872
refGene_gn=MNS1;refGene_tx=XM_023255319.1;refGene_gn=MNS1;refGene_tx=XM_023255320.1
B3 50271852 50271980
refGene_gn=MNS1;refGene_tx=XM_023255319.1;refGene_gn=MNS1;refGene_tx=XM_023255320.1
B3 50280071 50280174
refGene_gn=MNS1;refGene_tx=XM_023255319.1;refGene_gn=MNS1;refGene_tx=XM_023255320.1
B3 50289065 50289295
refGene_gn=MNS1;refGene_tx=XM_023255319.1;refGene_gn=MNS1;refGene_tx=XM_023255320.1

B3 50289883 50290100
refGene_gn=MNS1;refGene_tx=XM_023255319.1;refGene_gn=MNS1;refGene_tx=XM_023255320.1
B3 50290198 50290306
refGene_gn=MNS1;refGene_tx=XM_023255319.1;refGene_gn=MNS1;refGene_tx=XM_023255320.1
B3 50299043 50299301
refGene_gn=MNS1;refGene_tx=XM_023255319.1;refGene_gn=MNS1;refGene_tx=XM_023255320.1
B3 50305419 50305545
refGene_gn=MNS1;refGene_tx=XM_023255319.1;refGene_gn=MNS1;refGene_tx=XM_023255320.1
B3 50307173 50307266
refGene_gn=MNS1;refGene_tx=XM_023255319.1;refGene_gn=MNS1;refGene_tx=XM_023255320.1
B3 50308007 50308082
refGene_gn=TEX9;refGene_tx=XM_003987127.5;refGene_gn=TEX9;refGene_tx=XM_019832374.2;refGene_gn=TEX9;refGene_tx=XM_023255321.1
B3 50308705 50308840
refGene_gn=TEX9;refGene_tx=XM_003987127.5;refGene_gn=TEX9;refGene_tx=XM_019832374.2;refGene_gn=TEX9;refGene_tx=XM_023255321.1
B3 50326963 50327098
refGene_gn=TEX9;refGene_tx=XM_003987127.5;refGene_gn=TEX9;refGene_tx=XM_019832374.2
B3 50346018 50346192
refGene_gn=TEX9;refGene_tx=XM_003987127.5;refGene_gn=TEX9;refGene_tx=XM_019832374.2;refGene_gn=TEX9;refGene_tx=XM_023255321.1
B3 50346615 50346698
refGene_gn=TEX9;refGene_tx=XM_003987127.5;refGene_gn=TEX9;refGene_tx=XM_019832374.2;refGene_gn=TEX9;refGene_tx=XM_023255321.1
B3 50349405 50349584
refGene_gn=TEX9;refGene_tx=XM_003987127.5;refGene_gn=TEX9;refGene_tx=XM_019832374.2;refGene_gn=TEX9;refGene_tx=XM_023255321.1
B3 50351417 50351500
refGene_gn=TEX9;refGene_tx=XM_003987127.5;refGene_gn=TEX9;refGene_tx=XM_019832374.2;refGene_gn=TEX9;refGene_tx=XM_023255321.1
B3 50352275 50352321
refGene_gn=TEX9;refGene_tx=XM_003987127.5;refGene_gn=TEX9;refGene_tx=XM_019832374.2;refGene_gn=TEX9;refGene_tx=XM_023255321.1
B3 50357735 50357815
refGene_gn=TEX9;refGene_tx=XM_003987127.5;refGene_gn=TEX9;refGene_tx=XM_019832374.2;refGene_gn=TEX9;refGene_tx=XM_023255321.1
B3 50359475 50359539
refGene_gn=TEX9;refGene_tx=XM_003987127.5;refGene_gn=TEX9;refGene_tx=XM_019832374.2;refGene_gn=TEX9;refGene_tx=XM_023255321.1
B3 50359699 50359713 refGene_gn=TEX9;refGene_tx=XM_019832374.2
B3 50369164 50369256
refGene_gn=TEX9;refGene_tx=XM_003987127.5;refGene_gn=TEX9;refGene_tx=XM_023255321.1
B3 50369360 50369387

refGene_gn=TEX9;refGene_tx=XM_003987127.5;refGene_gn=TEX9;refGene_tx=XM_023255321.1
B3	50533161	50533322	refGene_gn=RFX7;refGene_tx=XM_023255322.1
B3	50589064	50589130	refGene_gn=RFX7;refGene_tx=XM_023255322.1
B3	50597166	50597175	refGene_gn=RFX7;refGene_tx=XM_023255322.1
B3	50623172	50623323	refGene_gn=RFX7;refGene_tx=XM_023255322.1
B3	50655531	50655648	refGene_gn=RFX7;refGene_tx=XM_023255322.1
B3	50656920	50657005	refGene_gn=RFX7;refGene_tx=XM_023255322.1
B3	50657742	50657950	refGene_gn=RFX7;refGene_tx=XM_023255322.1
B3	50660922	50661218	refGene_gn=RFX7;refGene_tx=XM_023255322.1
B3	50662916	50666189	refGene_gn=RFX7;refGene_tx=XM_023255322.1
B3	50741970	50742015
refGene_gn=NEDD4;refGene_tx=XM_004001640.5;refGene_gn=NEDD4;refGene_tx=XM_019832378.2
B3	50757635	50757709
refGene_gn=NEDD4;refGene_tx=XM_004001640.5;refGene_gn=NEDD4;refGene_tx=XM_019832378.2
B3	50759475	50759522	refGene_gn=NEDD4;refGene_tx=XM_019832379.2
B3	50768387	50768466
refGene_gn=NEDD4;refGene_tx=XM_004001640.5;refGene_gn=NEDD4;refGene_tx=XM_019832378.2;refGene_gn=NEDD4;refGene_tx=XM_019832379.2
B3	50768568	50768607
refGene_gn=NEDD4;refGene_tx=XM_004001640.5;refGene_gn=NEDD4;refGene_tx=XM_019832378.2;refGene_gn=NEDD4;refGene_tx=XM_019832379.2
B3	50792932	50792986
refGene_gn=NEDD4;refGene_tx=XM_004001640.5;refGene_gn=NEDD4;refGene_tx=XM_019832379.2
B3	50799676	50801236	refGene_gn=NEDD4;refGene_tx=XM_011282881.3
B3	50837083	50837134
refGene_gn=NEDD4;refGene_tx=XM_004001640.5;refGene_gn=NEDD4;refGene_tx=XM_011282881.3;refGene_gn=NEDD4;refGene_tx=XM_019832378.2;refGene_gn=NEDD4;refGene_tx=XM_019832379.2
B3	50838636	50838698
refGene_gn=NEDD4;refGene_tx=XM_004001640.5;refGene_gn=NEDD4;refGene_tx=XM_011282881.3;refGene_gn=NEDD4;refGene_tx=XM_019832378.2;refGene_gn=NEDD4;refGene_tx=XM_019832379.2;refGene_gn=NEDD4;refGene_tx=XM_019832380.2
B3	50841085	50841188
refGene_gn=NEDD4;refGene_tx=XM_004001640.5;refGene_gn=NEDD4;refGene_tx=XM_011282881.3;refGene_gn=NEDD4;refGene_tx=XM_019832378.2;refGene_gn=NEDD4;refGene_tx=XM_019832379.2;refGene_gn=NEDD4;refGene_tx=XM_019832380.2
B3	50846991	50847158
refGene_gn=NEDD4;refGene_tx=XM_004001640.5;refGene_gn=NEDD4;refGene_tx=XM_011282881.3;refGene_gn=NEDD4;refGene_tx=XM_019832378.2;refGene_gn=NEDD4;refGene_tx=XM_019832379.2;refGene_gn=NEDD4;refGene_tx=XM_019832380.2
B3	50849039	50849157
refGene_gn=NEDD4;refGene_tx=XM_004001640.5;refGene_gn=NEDD4;refGene_tx=XM_011282881.3;refGene_gn=NEDD4;refGene_tx=XM_019832378.2;refGene_gn=

NEDD4;refGene_tx=XM_019832379.2;refGene_gn=NEDD4;refGene_tx=XM_0198323
80.2
B3 50849248 50849416
refGene_gn=NEDD4;refGene_tx=XM_004001640.5;refGene_gn=NEDD4;refGene_tx
=XM_011282881.3;refGene_gn=NEDD4;refGene_tx=XM_019832378.2;refGene_gn=
NEDD4;refGene_tx=XM_019832379.2;refGene_gn=NEDD4;refGene_tx=XM_0198323
80.2
B3 50853102 50853168
refGene_gn=NEDD4;refGene_tx=XM_004001640.5;refGene_gn=NEDD4;refGene_tx
=XM_011282881.3;refGene_gn=NEDD4;refGene_tx=XM_019832378.2;refGene_gn=
NEDD4;refGene_tx=XM_019832379.2;refGene_gn=NEDD4;refGene_tx=XM_0198323
80.2
B3 50857066 50857186
refGene_gn=NEDD4;refGene_tx=XM_004001640.5;refGene_gn=NEDD4;refGene_tx
=XM_011282881.3;refGene_gn=NEDD4;refGene_tx=XM_019832378.2;refGene_gn=
NEDD4;refGene_tx=XM_019832379.2;refGene_gn=NEDD4;refGene_tx=XM_0198323
80.2
B3 50858639 50858840
refGene_gn=NEDD4;refGene_tx=XM_004001640.5;refGene_gn=NEDD4;refGene_tx
=XM_011282881.3;refGene_gn=NEDD4;refGene_tx=XM_019832378.2;refGene_gn=
NEDD4;refGene_tx=XM_019832379.2;refGene_gn=NEDD4;refGene_tx=XM_0198323
80.2
B3 50861465 50861546
refGene_gn=NEDD4;refGene_tx=XM_004001640.5;refGene_gn=NEDD4;refGene_tx
=XM_011282881.3;refGene_gn=NEDD4;refGene_tx=XM_019832378.2;refGene_gn=
NEDD4;refGene_tx=XM_019832379.2;refGene_gn=NEDD4;refGene_tx=XM_0198323
80.2
B3 50861767 50861822
refGene_gn=NEDD4;refGene_tx=XM_004001640.5;refGene_gn=NEDD4;refGene_tx
=XM_011282881.3;refGene_gn=NEDD4;refGene_tx=XM_019832378.2;refGene_gn=
NEDD4;refGene_tx=XM_019832379.2;refGene_gn=NEDD4;refGene_tx=XM_0198323
80.2
B3 50861914 50861973
refGene_gn=NEDD4;refGene_tx=XM_004001640.5;refGene_gn=NEDD4;refGene_tx
=XM_011282881.3;refGene_gn=NEDD4;refGene_tx=XM_019832378.2;refGene_gn=
NEDD4;refGene_tx=XM_019832379.2;refGene_gn=NEDD4;refGene_tx=XM_0198323
80.2
B3 50863016 50863082
refGene_gn=NEDD4;refGene_tx=XM_004001640.5;refGene_gn=NEDD4;refGene_tx
=XM_011282881.3;refGene_gn=NEDD4;refGene_tx=XM_019832378.2;refGene_gn=
NEDD4;refGene_tx=XM_019832379.2;refGene_gn=NEDD4;refGene_tx=XM_0198323
80.2
B3 50866900 50867130
refGene_gn=NEDD4;refGene_tx=XM_004001640.5;refGene_gn=NEDD4;refGene_tx
=XM_011282881.3;refGene_gn=NEDD4;refGene_tx=XM_019832378.2;refGene_gn=
NEDD4;refGene_tx=XM_019832379.2;refGene_gn=NEDD4;refGene_tx=XM_0198323
80.2
B3 50868075 50868197
refGene_gn=NEDD4;refGene_tx=XM_004001640.5;refGene_gn=NEDD4;refGene_tx
=XM_011282881.3;refGene_gn=NEDD4;refGene_tx=XM_019832378.2;refGene_gn=

NEDD4;refGene_tx=XM_019832379.2;refGene_gn=NEDD4;refGene_tx=XM_0198323
80.2
B3 50868290 50868361
refGene_gn=NEDD4;refGene_tx=XM_004001640.5;refGene_gn=NEDD4;refGene_tx
=XM_011282881.3;refGene_gn=NEDD4;refGene_tx=XM_019832378.2;refGene_gn=
NEDD4;refGene_tx=XM_019832379.2;refGene_gn=NEDD4;refGene_tx=XM_0198323
80.2
B3 50869838 50869934
refGene_gn=NEDD4;refGene_tx=XM_004001640.5;refGene_gn=NEDD4;refGene_tx
=XM_011282881.3;refGene_gn=NEDD4;refGene_tx=XM_019832378.2;refGene_gn=
NEDD4;refGene_tx=XM_019832379.2;refGene_gn=NEDD4;refGene_tx=XM_0198323
80.2
B3 50870266 50870340
refGene_gn=NEDD4;refGene_tx=XM_004001640.5;refGene_gn=NEDD4;refGene_tx
=XM_011282881.3;refGene_gn=NEDD4;refGene_tx=XM_019832378.2;refGene_gn=
NEDD4;refGene_tx=XM_019832379.2;refGene_gn=NEDD4;refGene_tx=XM_0198323
80.2
B3 50870647 50870708
refGene_gn=NEDD4;refGene_tx=XM_004001640.5;refGene_gn=NEDD4;refGene_tx
=XM_011282881.3;refGene_gn=NEDD4;refGene_tx=XM_019832378.2;refGene_gn=
NEDD4;refGene_tx=XM_019832379.2;refGene_gn=NEDD4;refGene_tx=XM_0198323
80.2
B3 50873721 50873781
refGene_gn=NEDD4;refGene_tx=XM_004001640.5;refGene_gn=NEDD4;refGene_tx
=XM_011282881.3;refGene_gn=NEDD4;refGene_tx=XM_019832378.2;refGene_gn=
NEDD4;refGene_tx=XM_019832379.2;refGene_gn=NEDD4;refGene_tx=XM_0198323
80.2
B3 50873863 50873971
refGene_gn=NEDD4;refGene_tx=XM_004001640.5;refGene_gn=NEDD4;refGene_tx
=XM_011282881.3;refGene_gn=NEDD4;refGene_tx=XM_019832378.2;refGene_gn=
NEDD4;refGene_tx=XM_019832379.2;refGene_gn=NEDD4;refGene_tx=XM_0198323
80.2
B3 50874587 50874684
refGene_gn=NEDD4;refGene_tx=XM_004001640.5;refGene_gn=NEDD4;refGene_tx
=XM_011282881.3;refGene_gn=NEDD4;refGene_tx=XM_019832378.2;refGene_gn=
NEDD4;refGene_tx=XM_019832379.2;refGene_gn=NEDD4;refGene_tx=XM_0198323
80.2
B3 50878070 50878143
refGene_gn=NEDD4;refGene_tx=XM_004001640.5;refGene_gn=NEDD4;refGene_tx
=XM_011282881.3;refGene_gn=NEDD4;refGene_tx=XM_019832378.2;refGene_gn=
NEDD4;refGene_tx=XM_019832379.2;refGene_gn=NEDD4;refGene_tx=XM_0198323
80.2
B3 50878652 50878755
refGene_gn=NEDD4;refGene_tx=XM_004001640.5;refGene_gn=NEDD4;refGene_tx
=XM_011282881.3;refGene_gn=NEDD4;refGene_tx=XM_019832378.2;refGene_gn=
NEDD4;refGene_tx=XM_019832379.2;refGene_gn=NEDD4;refGene_tx=XM_0198323
80.2
B3 50938297 50938391
refGene_gn=PRTG;refGene_tx=XM_023255323.1;refGene_gn=PRTG;refGene_tx=X
M_023255324.1

B3 50940566 50940869
refGene_gn=PRTG;refGene_tx=XM_023255323.1;refGene_gn=PRTG;refGene_tx=XM_023255324.1
B3 50999722 50999867
refGene_gn=PRTG;refGene_tx=XM_023255323.1;refGene_gn=PRTG;refGene_tx=XM_023255324.1
B3 51000862 51000996
refGene_gn=PRTG;refGene_tx=XM_023255323.1;refGene_gn=PRTG;refGene_tx=XM_023255324.1
B3 51002735 51002873
refGene_gn=PRTG;refGene_tx=XM_023255323.1;refGene_gn=PRTG;refGene_tx=XM_023255324.1
B3 51003125 51003284
refGene_gn=PRTG;refGene_tx=XM_023255323.1;refGene_gn=PRTG;refGene_tx=XM_023255324.1
B3 51003879 51004039
refGene_gn=PRTG;refGene_tx=XM_023255323.1;refGene_gn=PRTG;refGene_tx=XM_023255324.1
B3 51004846 51005094
refGene_gn=PRTG;refGene_tx=XM_023255323.1;refGene_gn=PRTG;refGene_tx=XM_023255324.1
B3 51007094 51007259
refGene_gn=PRTG;refGene_tx=XM_023255323.1;refGene_gn=PRTG;refGene_tx=XM_023255324.1
B3 51009985 51010291
refGene_gn=PRTG;refGene_tx=XM_023255323.1;refGene_gn=PRTG;refGene_tx=XM_023255324.1
B3 51011392 51011581
refGene_gn=PRTG;refGene_tx=XM_023255323.1;refGene_gn=PRTG;refGene_tx=XM_023255324.1
B3 51045526 51045622
refGene_gn=PRTG;refGene_tx=XM_023255323.1;refGene_gn=PRTG;refGene_tx=XM_023255324.1
B3 51046271 51046458
refGene_gn=PRTG;refGene_tx=XM_023255323.1;refGene_gn=PRTG;refGene_tx=XM_023255324.1
B3 51047389 51047517
refGene_gn=PRTG;refGene_tx=XM_023255323.1;refGene_gn=PRTG;refGene_tx=XM_023255324.1
B3 51048744 51048915
refGene_gn=PRTG;refGene_tx=XM_023255323.1;refGene_gn=PRTG;refGene_tx=XM_023255324.1
B3 51056640 51056823
refGene_gn=PRTG;refGene_tx=XM_023255323.1;refGene_gn=PRTG;refGene_tx=XM_023255324.1
B3 51058178 51058299
refGene_gn=PRTG;refGene_tx=XM_023255323.1;refGene_gn=PRTG;refGene_tx=XM_023255324.1
B3 51059818 51059984
refGene_gn=PRTG;refGene_tx=XM_023255323.1;refGene_gn=PRTG;refGene_tx=X

M_023255324.1
B3	51062272	51062377
refGene_gn=PRTG;refGene_tx=XM_023255323.1;refGene_gn=PRTG;refGene_tx=XM_023255324.1
B3	51062841	51063246
refGene_gn=PRTG;refGene_tx=XM_023255323.1;refGene_gn=PRTG;refGene_tx=XM_023255324.1
B3	51080216	51080281	refGene_gn=PRTG;refGene_tx=XM_023255324.1
B3	51089161	51090193
refGene_gn=PYGO1;refGene_tx=XM_003987114.5;refGene_gn=PYGO1;refGene_tx=XM_023255325.1;refGene_gn=PYGO1;refGene_tx=XM_023255326.1;refGene_gn=PYGO1;refGene_tx=XM_023255327.1
B3	51112477	51112563	refGene_gn=PYGO1;refGene_tx=XM_003987114.5
B3	51113629	51114754	refGene_gn=PYGO1;refGene_tx=XM_003987114.5
B3	51150247	51150370
refGene_gn=DNAAF4;refGene_tx=XM_003987115.5;refGene_gn=DNAAF4;refGene_tx=XM_011282879.3;refGene_gn=DNAAF4;refGene_tx=XM_019832383.2;refGene_gn=DNAAF4;refGene_tx=XM_023255328.1;refGene_gn=DNAAF4;refGene_tx=XM_023255329.1
B3	51150711	51150859
refGene_gn=DNAAF4;refGene_tx=XM_003987115.5;refGene_gn=DNAAF4;refGene_tx=XM_011282879.3;refGene_gn=DNAAF4;refGene_tx=XM_019832383.2;refGene_gn=DNAAF4;refGene_tx=XM_023255328.1;refGene_gn=DNAAF4;refGene_tx=XM_023255329.1;refGene_gn=DNAAF4;refGene_tx=XM_023255330.1
B3	51154263	51154397
refGene_gn=DNAAF4;refGene_tx=XM_003987115.5;refGene_gn=DNAAF4;refGene_tx=XM_011282879.3;refGene_gn=DNAAF4;refGene_tx=XM_019832383.2;refGene_gn=DNAAF4;refGene_tx=XM_023255328.1;refGene_gn=DNAAF4;refGene_tx=XM_023255329.1;refGene_gn=DNAAF4;refGene_tx=XM_023255330.1
B3	51176514	51176746
refGene_gn=DNAAF4;refGene_tx=XM_003987115.5;refGene_gn=DNAAF4;refGene_tx=XM_011282879.3;refGene_gn=DNAAF4;refGene_tx=XM_019832383.2;refGene_gn=DNAAF4;refGene_tx=XM_023255328.1;refGene_gn=DNAAF4;refGene_tx=XM_023255329.1;refGene_gn=DNAAF4;refGene_tx=XM_023255330.1
B3	51191248	51191394
refGene_gn=DNAAF4;refGene_tx=XM_003987115.5;refGene_gn=DNAAF4;refGene_tx=XM_011282879.3;refGene_gn=DNAAF4;refGene_tx=XM_019832383.2;refGene_gn=DNAAF4;refGene_tx=XM_023255328.1;refGene_gn=DNAAF4;refGene_tx=XM_023255329.1;refGene_gn=DNAAF4;refGene_tx=XM_023255330.1
B3	51193105	51193215
refGene_gn=DNAAF4;refGene_tx=XM_003987115.5;refGene_gn=DNAAF4;refGene_tx=XM_011282879.3;refGene_gn=DNAAF4;refGene_tx=XM_019832383.2;refGene_gn=DNAAF4;refGene_tx=XM_023255328.1;refGene_gn=DNAAF4;refGene_tx=XM_023255329.1;refGene_gn=DNAAF4;refGene_tx=XM_023255330.1
B3	51208830	51208984
refGene_gn=DNAAF4;refGene_tx=XM_003987115.5;refGene_gn=DNAAF4;refGene_tx=XM_011282879.3;refGene_gn=DNAAF4;refGene_tx=XM_019832383.2;refGene_gn=DNAAF4;refGene_tx=XM_023255328.1;refGene_gn=DNAAF4;refGene_tx=XM_023255329.1;refGene_gn=DNAAF4;refGene_tx=XM_023255330.1
B3	51210057	51210163

refGene_gn=DNAAF4;refGene_tx=XM_011282879.3;refGene_gn=DNAAF4;refGene_tx=XM_019832383.2;refGene_gn=DNAAF4;refGene_tx=XM_023255328.1;refGene_gn=DNAAF4;refGene_tx=XM_023255329.1;refGene_gn=DNAAF4;refGene_tx=XM_023255330.1
B3 51212481 51212591
refGene_gn=DNAAF4;refGene_tx=XM_003987115.5;refGene_gn=DNAAF4;refGene_tx=XM_011282879.3;refGene_gn=DNAAF4;refGene_tx=XM_019832383.2;refGene_gn=DNAAF4;refGene_tx=XM_023255328.1;refGene_gn=DNAAF4;refGene_tx=XM_023255329.1;refGene_gn=DNAAF4;refGene_tx=XM_023255330.1
B3 51223309 51223657
refGene_gn=CB3H15orf65;refGene_tx=XM_006932517.4
B3 51229989 51230007
refGene_gn=CB3H15orf65;refGene_tx=XM_006932517.4
B3 51251480 51251540
refGene_gn=CCPG1;refGene_tx=XM_003987117.5;refGene_gn=CCPG1;refGene_tx=XM_006932514.4;refGene_gn=CCPG1;refGene_tx=XM_006932515.4;refGene_gn=CCPG1;refGene_tx=XM_006932516.4;refGene_gn=CCPG1;refGene_tx=XM_019832385.2;refGene_gn=CCPG1;refGene_tx=XM_023255331.1
B3 51254079 51254200
refGene_gn=CCPG1;refGene_tx=XM_003987117.5;refGene_gn=CCPG1;refGene_tx=XM_006932514.4;refGene_gn=CCPG1;refGene_tx=XM_006932515.4;refGene_gn=CCPG1;refGene_tx=XM_006932516.4;refGene_gn=CCPG1;refGene_tx=XM_019832385.2;refGene_gn=CCPG1;refGene_tx=XM_023255331.1
B3 51255839 51255916
refGene_gn=CCPG1;refGene_tx=XM_003987117.5;refGene_gn=CCPG1;refGene_tx=XM_006932514.4;refGene_gn=CCPG1;refGene_tx=XM_006932515.4;refGene_gn=CCPG1;refGene_tx=XM_006932516.4;refGene_gn=CCPG1;refGene_tx=XM_019832385.2;refGene_gn=CCPG1;refGene_tx=XM_023255331.1
B3 51256770 51256969
refGene_gn=CCPG1;refGene_tx=XM_003987117.5;refGene_gn=CCPG1;refGene_tx=XM_006932514.4;refGene_gn=CCPG1;refGene_tx=XM_006932515.4;refGene_gn=CCPG1;refGene_tx=XM_006932516.4;refGene_gn=CCPG1;refGene_tx=XM_019832385.2;refGene_gn=CCPG1;refGene_tx=XM_023255331.1
B3 51258947 51259043
refGene_gn=CCPG1;refGene_tx=XM_006932514.4;refGene_gn=CCPG1;refGene_tx=XM_006932515.4;refGene_gn=CCPG1;refGene_tx=XM_019832385.2;refGene_gn=CCPG1;refGene_tx=XM_023255331.1
B3 51260872 51261124
refGene_gn=CCPG1;refGene_tx=XM_003987117.5;refGene_gn=CCPG1;refGene_tx=XM_006932514.4;refGene_gn=CCPG1;refGene_tx=XM_006932515.4;refGene_gn=CCPG1;refGene_tx=XM_006932516.4;refGene_gn=CCPG1;refGene_tx=XM_019832385.2;refGene_gn=CCPG1;refGene_tx=XM_023255331.1
B3 51262333 51262345
refGene_gn=CCPG1;refGene_tx=XM_006932515.4;refGene_gn=CCPG1;refGene_tx=XM_006932516.4;refGene_gn=CCPG1;refGene_tx=XM_019832385.2;refGene_gn=CCPG1;refGene_tx=XM_023255331.1
B3 51264752 51264874
refGene_gn=CCPG1;refGene_tx=XM_003987117.5;refGene_gn=CCPG1;refGene_tx=XM_006932514.4;refGene_gn=CCPG1;refGene_tx=XM_006932515.4;refGene_gn=CCPG1;refGene_tx=XM_006932516.4;refGene_gn=CCPG1;refGene_tx=XM_0198323

85.2;refGene_gn=CCPG1;refGene_tx=XM_023255331.1
B3 51268358 51269767
refGene_gn=CCPG1;refGene_tx=XM_003987117.5;refGene_gn=CCPG1;refGene_tx=XM_006932514.4;refGene_gn=CCPG1;refGene_tx=XM_006932515.4;refGene_gn=CCPG1;refGene_tx=XM_006932516.4;refGene_gn=CCPG1;refGene_tx=XM_019832385.2;refGene_gn=CCPG1;refGene_tx=XM_023255331.1
B3 51272030 51272220
refGene_gn=CCPG1;refGene_tx=XM_003987117.5;refGene_gn=CCPG1;refGene_tx=XM_006932514.4;refGene_gn=CCPG1;refGene_tx=XM_006932515.4;refGene_gn=CCPG1;refGene_tx=XM_006932516.4;refGene_gn=CCPG1;refGene_tx=XM_019832385.2;refGene_gn=CCPG1;refGene_tx=XM_023255331.1
B3 51272674 51272691 refGene_gn=CCPG1;refGene_tx=XM_006932515.4
B3 51273022 51273160
refGene_gn=PIGB;refGene_tx=XM_006932511.3;refGene_gn=PIGB;refGene_tx=XM_006932512.3;refGene_gn=PIGB;refGene_tx=XM_019832386.2
B3 51273670 51273851
refGene_gn=PIGB;refGene_tx=XM_006932511.3;refGene_gn=PIGB;refGene_tx=XM_006932512.3;refGene_gn=PIGB;refGene_tx=XM_019832386.2
B3 51275477 51275691
refGene_gn=PIGB;refGene_tx=XM_006932511.3;refGene_gn=PIGB;refGene_tx=XM_006932512.3;refGene_gn=PIGB;refGene_tx=XM_019832386.2
B3 51281991 51282056
refGene_gn=PIGB;refGene_tx=XM_006932511.3;refGene_gn=PIGB;refGene_tx=XM_006932512.3;refGene_gn=PIGB;refGene_tx=XM_019832386.2
B3 51282664 51282876
refGene_gn=PIGB;refGene_tx=XM_006932511.3;refGene_gn=PIGB;refGene_tx=XM_006932512.3
B3 51283998 51284050
refGene_gn=PIGB;refGene_tx=XM_006932511.3;refGene_gn=PIGB;refGene_tx=XM_006932512.3;refGene_gn=PIGB;refGene_tx=XM_019832386.2
B3 51291397 51291538
refGene_gn=PIGB;refGene_tx=XM_006932511.3;refGene_gn=PIGB;refGene_tx=XM_006932512.3;refGene_gn=PIGB;refGene_tx=XM_019832386.2
B3 51303204 51303335
refGene_gn=PIGB;refGene_tx=XM_006932511.3;refGene_gn=PIGB;refGene_tx=XM_006932512.3;refGene_gn=PIGB;refGene_tx=XM_019832386.2
B3 51305254 51305359
refGene_gn=PIGB;refGene_tx=XM_006932511.3;refGene_gn=PIGB;refGene_tx=XM_019832386.2
B3 51310230 51310348
refGene_gn=PIGB;refGene_tx=XM_006932511.3;refGene_gn=PIGB;refGene_tx=XM_006932512.3;refGene_gn=PIGB;refGene_tx=XM_019832386.2
B3 51310778 51310914
refGene_gn=PIGB;refGene_tx=XM_006932511.3;refGene_gn=PIGB;refGene_tx=XM_006932512.3;refGene_gn=PIGB;refGene_tx=XM_019832386.2
B3 51312935 51313068
refGene_gn=PIGB;refGene_tx=XM_006932511.3;refGene_gn=PIGB;refGene_tx=XM_006932512.3;refGene_gn=PIGB;refGene_tx=XM_019832386.2
B3 51314668 51314833
refGene_gn=PIGBOS1;refGene_tx=XM_019832387.2;refGene_gn=PIGBOS1;refGen

e_tx=XM_019832388.1
B3 51376031 51376184
refGene_gn=RAB27A;refGene_tx=XM_011282873.3;refGene_gn=RAB27A;refGene_tx=XM_011282874.3;refGene_gn=RAB27A;refGene_tx=XM_011282876.3;refGene_gn=RAB27A;refGene_tx=XM_011282877.3;refGene_gn=RAB27A;refGene_tx=XM_019832389.2
B3 51384345 51384431
refGene_gn=RAB27A;refGene_tx=XM_011282873.3;refGene_gn=RAB27A;refGene_tx=XM_011282874.3;refGene_gn=RAB27A;refGene_tx=XM_011282876.3;refGene_gn=RAB27A;refGene_tx=XM_011282877.3;refGene_gn=RAB27A;refGene_tx=XM_019832389.2
B3 51386629 51386733
refGene_gn=RAB27A;refGene_tx=XM_011282873.3;refGene_gn=RAB27A;refGene_tx=XM_011282874.3;refGene_gn=RAB27A;refGene_tx=XM_011282876.3;refGene_gn=RAB27A;refGene_tx=XM_011282877.3;refGene_gn=RAB27A;refGene_tx=XM_019832389.2
B3 51393018 51393142
refGene_gn=RAB27A;refGene_tx=XM_011282873.3;refGene_gn=RAB27A;refGene_tx=XM_011282874.3;refGene_gn=RAB27A;refGene_tx=XM_011282876.3;refGene_gn=RAB27A;refGene_tx=XM_011282877.3;refGene_gn=RAB27A;refGene_tx=XM_019832389.2
B3 51404612 51404811
refGene_gn=RAB27A;refGene_tx=XM_011282873.3;refGene_gn=RAB27A;refGene_tx=XM_011282874.3;refGene_gn=RAB27A;refGene_tx=XM_011282876.3;refGene_gn=RAB27A;refGene_tx=XM_011282877.3;refGene_gn=RAB27A;refGene_tx=XM_019832389.2
B3 51412385 51412466 refGene_gn=RSL24D1;refGene_tx=XM_003987121.5
B3 51416995 51417109 refGene_gn=RSL24D1;refGene_tx=XM_003987121.5
B3 51418843 51418916 refGene_gn=RSL24D1;refGene_tx=XM_003987121.5
B3 51424269 51424333 refGene_gn=RSL24D1;refGene_tx=XM_003987121.5
B3 51426106 51426192 refGene_gn=RSL24D1;refGene_tx=XM_003987121.5
B3 51427378 51427452 refGene_gn=RSL24D1;refGene_tx=XM_003987121.5
B3 51967355 51967641
refGene_gn=UNC13C;refGene_tx=XM_011282872.3;refGene_gn=UNC13C;refGene_tx=XM_019832390.2;refGene_gn=UNC13C;refGene_tx=XM_019832391.2;refGene_gn=UNC13C;refGene_tx=XM_019832392.2;refGene_gn=UNC13C;refGene_tx=XM_019832393.2;refGene_gn=UNC13C;refGene_tx=XM_019832394.2
B3 51970765 51970925
refGene_gn=UNC13C;refGene_tx=XM_011282872.3;refGene_gn=UNC13C;refGene_tx=XM_019832390.2;refGene_gn=UNC13C;refGene_tx=XM_019832391.2;refGene_gn=UNC13C;refGene_tx=XM_019832392.2;refGene_gn=UNC13C;refGene_tx=XM_019832393.2;refGene_gn=UNC13C;refGene_tx=XM_019832394.2
B3 51973127 51973220
refGene_gn=UNC13C;refGene_tx=XM_011282872.3;refGene_gn=UNC13C;refGene_tx=XM_019832390.2;refGene_gn=UNC13C;refGene_tx=XM_019832391.2;refGene_gn=UNC13C;refGene_tx=XM_019832392.2;refGene_gn=UNC13C;refGene_tx=XM_019832393.2;refGene_gn=UNC13C;refGene_tx=XM_019832394.2
B3 52024195 52024343
refGene_gn=UNC13C;refGene_tx=XM_011282872.3;refGene_gn=UNC13C;refGene_tx=XM_019832390.2;refGene_gn=UNC13C;refGene_tx=XM_019832391.2;refGene_

gn=UNC13C;refGene_tx=XM_019832392.2;refGene_gn=UNC13C;refGene_tx=XM_01
9832393.2;refGene_gn=UNC13C;refGene_tx=XM_019832394.2
B3 52037938 52038019
refGene_gn=UNC13C;refGene_tx=XM_011282872.3;refGene_gn=UNC13C;refGene_
tx=XM_019832390.2;refGene_gn=UNC13C;refGene_tx=XM_019832391.2;refGene_
gn=UNC13C;refGene_tx=XM_019832392.2;refGene_gn=UNC13C;refGene_tx=XM_01
9832393.2;refGene_gn=UNC13C;refGene_tx=XM_019832394.2
B3 52043657 52043714
refGene_gn=UNC13C;refGene_tx=XM_011282872.3;refGene_gn=UNC13C;refGene_
tx=XM_019832390.2;refGene_gn=UNC13C;refGene_tx=XM_019832391.2;refGene_
gn=UNC13C;refGene_tx=XM_019832392.2;refGene_gn=UNC13C;refGene_tx=XM_01
9832393.2;refGene_gn=UNC13C;refGene_tx=XM_019832394.2
B3 52052896 52053020
refGene_gn=UNC13C;refGene_tx=XM_011282872.3;refGene_gn=UNC13C;refGene_
tx=XM_019832390.2;refGene_gn=UNC13C;refGene_tx=XM_019832391.2;refGene_
gn=UNC13C;refGene_tx=XM_019832392.2;refGene_gn=UNC13C;refGene_tx=XM_01
9832393.2;refGene_gn=UNC13C;refGene_tx=XM_019832394.2
B3 52057906 52058056
refGene_gn=UNC13C;refGene_tx=XM_011282872.3;refGene_gn=UNC13C;refGene_
tx=XM_019832390.2;refGene_gn=UNC13C;refGene_tx=XM_019832391.2;refGene_
gn=UNC13C;refGene_tx=XM_019832392.2;refGene_gn=UNC13C;refGene_tx=XM_01
9832393.2;refGene_gn=UNC13C;refGene_tx=XM_019832394.2
B3 52065547 52065636
refGene_gn=UNC13C;refGene_tx=XM_011282872.3;refGene_gn=UNC13C;refGene_
tx=XM_019832390.2;refGene_gn=UNC13C;refGene_tx=XM_019832391.2;refGene_
gn=UNC13C;refGene_tx=XM_019832392.2;refGene_gn=UNC13C;refGene_tx=XM_01
9832393.2;refGene_gn=UNC13C;refGene_tx=XM_019832394.2
B3 52078856 52078886
refGene_gn=UNC13C;refGene_tx=XM_019832390.2;refGene_gn=UNC13C;refGene_
tx=XM_019832391.2;refGene_gn=UNC13C;refGene_tx=XM_019832392.2
B3 52079392 52079470
refGene_gn=UNC13C;refGene_tx=XM_011282872.3;refGene_gn=UNC13C;refGene_
tx=XM_019832390.2;refGene_gn=UNC13C;refGene_tx=XM_019832391.2;refGene_
gn=UNC13C;refGene_tx=XM_019832392.2;refGene_gn=UNC13C;refGene_tx=XM_01
9832393.2;refGene_gn=UNC13C;refGene_tx=XM_019832394.2
B3 52084726 52084804
refGene_gn=UNC13C;refGene_tx=XM_011282872.3;refGene_gn=UNC13C;refGene_
tx=XM_019832390.2;refGene_gn=UNC13C;refGene_tx=XM_019832391.2;refGene_
gn=UNC13C;refGene_tx=XM_019832392.2;refGene_gn=UNC13C;refGene_tx=XM_01
9832393.2;refGene_gn=UNC13C;refGene_tx=XM_019832394.2
B3 52091137 52091281
refGene_gn=UNC13C;refGene_tx=XM_011282872.3;refGene_gn=UNC13C;refGene_
tx=XM_019832390.2;refGene_gn=UNC13C;refGene_tx=XM_019832391.2;refGene_
gn=UNC13C;refGene_tx=XM_019832392.2;refGene_gn=UNC13C;refGene_tx=XM_01
9832393.2;refGene_gn=UNC13C;refGene_tx=XM_019832394.2
B3 52092054 52092151
refGene_gn=UNC13C;refGene_tx=XM_011282872.3;refGene_gn=UNC13C;refGene_
tx=XM_019832390.2;refGene_gn=UNC13C;refGene_tx=XM_019832391.2;refGene_
gn=UNC13C;refGene_tx=XM_019832392.2;refGene_gn=UNC13C;refGene_tx=XM_01
9832393.2;refGene_gn=UNC13C;refGene_tx=XM_019832394.2

B3 52100509 52100636
refGene_gn=UNC13C;refGene_tx=XM_011282872.3;refGene_gn=UNC13C;refGene_tx=XM_019832390.2;refGene_gn=UNC13C;refGene_tx=XM_019832391.2;refGene_gn=UNC13C;refGene_tx=XM_019832392.2;refGene_gn=UNC13C;refGene_tx=XM_019832393.2;refGene_gn=UNC13C;refGene_tx=XM_019832394.2
B3 52146825 52146911
refGene_gn=UNC13C;refGene_tx=XM_011282872.3;refGene_gn=UNC13C;refGene_tx=XM_019832390.2;refGene_gn=UNC13C;refGene_tx=XM_019832391.2;refGene_gn=UNC13C;refGene_tx=XM_019832392.2;refGene_gn=UNC13C;refGene_tx=XM_019832393.2;refGene_gn=UNC13C;refGene_tx=XM_019832394.2
B3 52176151 52176285
refGene_gn=UNC13C;refGene_tx=XM_011282872.3;refGene_gn=UNC13C;refGene_tx=XM_019832390.2;refGene_gn=UNC13C;refGene_tx=XM_019832391.2;refGene_gn=UNC13C;refGene_tx=XM_019832392.2;refGene_gn=UNC13C;refGene_tx=XM_019832393.2;refGene_gn=UNC13C;refGene_tx=XM_019832394.2
B3 52223244 52223373
refGene_gn=UNC13C;refGene_tx=XM_011282872.3;refGene_gn=UNC13C;refGene_tx=XM_019832390.2;refGene_gn=UNC13C;refGene_tx=XM_019832391.2;refGene_gn=UNC13C;refGene_tx=XM_019832392.2;refGene_gn=UNC13C;refGene_tx=XM_019832393.2;refGene_gn=UNC13C;refGene_tx=XM_019832394.2
B3 52227311 52227401
refGene_gn=UNC13C;refGene_tx=XM_011282872.3;refGene_gn=UNC13C;refGene_tx=XM_019832390.2;refGene_gn=UNC13C;refGene_tx=XM_019832391.2;refGene_gn=UNC13C;refGene_tx=XM_019832392.2;refGene_gn=UNC13C;refGene_tx=XM_019832393.2;refGene_gn=UNC13C;refGene_tx=XM_019832394.2
B3 52229059 52229128
refGene_gn=UNC13C;refGene_tx=XM_011282872.3;refGene_gn=UNC13C;refGene_tx=XM_019832390.2;refGene_gn=UNC13C;refGene_tx=XM_019832391.2;refGene_gn=UNC13C;refGene_tx=XM_019832392.2;refGene_gn=UNC13C;refGene_tx=XM_019832393.2;refGene_gn=UNC13C;refGene_tx=XM_019832394.2
B3 52240603 52240760
refGene_gn=UNC13C;refGene_tx=XM_011282872.3;refGene_gn=UNC13C;refGene_tx=XM_019832390.2;refGene_gn=UNC13C;refGene_tx=XM_019832391.2;refGene_gn=UNC13C;refGene_tx=XM_019832392.2;refGene_gn=UNC13C;refGene_tx=XM_019832393.2;refGene_gn=UNC13C;refGene_tx=XM_019832394.2
B3 52256529 52256693
refGene_gn=UNC13C;refGene_tx=XM_011282872.3;refGene_gn=UNC13C;refGene_tx=XM_019832390.2;refGene_gn=UNC13C;refGene_tx=XM_019832391.2;refGene_gn=UNC13C;refGene_tx=XM_019832392.2;refGene_gn=UNC13C;refGene_tx=XM_019832393.2;refGene_gn=UNC13C;refGene_tx=XM_019832394.2
B3 52258530 52258646
refGene_gn=UNC13C;refGene_tx=XM_011282872.3;refGene_gn=UNC13C;refGene_tx=XM_019832390.2;refGene_gn=UNC13C;refGene_tx=XM_019832391.2;refGene_gn=UNC13C;refGene_tx=XM_019832392.2;refGene_gn=UNC13C;refGene_tx=XM_019832393.2;refGene_gn=UNC13C;refGene_tx=XM_019832394.2
B3 52262433 52262603
refGene_gn=UNC13C;refGene_tx=XM_011282872.3;refGene_gn=UNC13C;refGene_tx=XM_019832390.2;refGene_gn=UNC13C;refGene_tx=XM_019832391.2;refGene_gn=UNC13C;refGene_tx=XM_019832392.2;refGene_gn=UNC13C;refGene_tx=XM_019832393.2;refGene_gn=UNC13C;refGene_tx=XM_019832394.2

B3 52295221 52295363
refGene_gn=UNC13C;refGene_tx=XM_011282872.3;refGene_gn=UNC13C;refGene_tx=XM_019832390.2;refGene_gn=UNC13C;refGene_tx=XM_019832391.2;refGene_gn=UNC13C;refGene_tx=XM_019832392.2;refGene_gn=UNC13C;refGene_tx=XM_019832393.2;refGene_gn=UNC13C;refGene_tx=XM_019832394.2
B3 52296532 52296760
refGene_gn=UNC13C;refGene_tx=XM_011282872.3;refGene_gn=UNC13C;refGene_tx=XM_019832390.2;refGene_gn=UNC13C;refGene_tx=XM_019832391.2;refGene_gn=UNC13C;refGene_tx=XM_019832392.2;refGene_gn=UNC13C;refGene_tx=XM_019832393.2;refGene_gn=UNC13C;refGene_tx=XM_019832394.2
B3 52308839 52309059
refGene_gn=UNC13C;refGene_tx=XM_011282872.3;refGene_gn=UNC13C;refGene_tx=XM_019832390.2;refGene_gn=UNC13C;refGene_tx=XM_019832391.2;refGene_gn=UNC13C;refGene_tx=XM_019832392.2;refGene_gn=UNC13C;refGene_tx=XM_019832393.2;refGene_gn=UNC13C;refGene_tx=XM_019832394.2
B3 52326847 52326919
refGene_gn=UNC13C;refGene_tx=XM_011282872.3;refGene_gn=UNC13C;refGene_tx=XM_019832390.2;refGene_gn=UNC13C;refGene_tx=XM_019832391.2;refGene_gn=UNC13C;refGene_tx=XM_019832392.2;refGene_gn=UNC13C;refGene_tx=XM_019832393.2;refGene_gn=UNC13C;refGene_tx=XM_019832394.2
B3 52328044 52328050
refGene_gn=UNC13C;refGene_tx=XM_011282872.3;refGene_gn=UNC13C;refGene_tx=XM_019832390.2;refGene_gn=UNC13C;refGene_tx=XM_019832392.2;refGene_gn=UNC13C;refGene_tx=XM_019832393.2
B3 52329189 52329268
refGene_gn=UNC13C;refGene_tx=XM_011282872.3;refGene_gn=UNC13C;refGene_tx=XM_019832390.2;refGene_gn=UNC13C;refGene_tx=XM_019832391.2;refGene_gn=UNC13C;refGene_tx=XM_019832392.2;refGene_gn=UNC13C;refGene_tx=XM_019832393.2;refGene_gn=UNC13C;refGene_tx=XM_019832394.2
B3 52422357 52422422
refGene_gn=UNC13C;refGene_tx=XM_011282872.3;refGene_gn=UNC13C;refGene_tx=XM_019832390.2;refGene_gn=UNC13C;refGene_tx=XM_019832391.2;refGene_gn=UNC13C;refGene_tx=XM_019832392.2;refGene_gn=UNC13C;refGene_tx=XM_019832393.2;refGene_gn=UNC13C;refGene_tx=XM_019832394.2
B3 52422996 52423019
refGene_gn=UNC13C;refGene_tx=XM_011282872.3;refGene_gn=UNC13C;refGene_tx=XM_019832390.2;refGene_gn=UNC13C;refGene_tx=XM_019832391.2;refGene_gn=UNC13C;refGene_tx=XM_019832392.2;refGene_gn=UNC13C;refGene_tx=XM_019832393.2;refGene_gn=UNC13C;refGene_tx=XM_019832394.2
B3 52536942 52539934
refGene_gn=UNC13C;refGene_tx=XM_011282872.3;refGene_gn=UNC13C;refGene_tx=XM_019832390.2;refGene_gn=UNC13C;refGene_tx=XM_019832391.2;refGene_gn=UNC13C;refGene_tx=XM_019832392.2;refGene_gn=UNC13C;refGene_tx=XM_019832393.2;refGene_gn=UNC13C;refGene_tx=XM_019832394.2
B3 52789046 52789199
refGene_gn=WDR72;refGene_tx=XM_023255332.1;refGene_gn=WDR72;refGene_tx=XM_023255333.1;refGene_gn=WDR72;refGene_tx=XM_023255334.1;refGene_gn=WDR72;refGene_tx=XM_023255335.1
B3 52793297 52793404
refGene_gn=WDR72;refGene_tx=XM_023255332.1;refGene_gn=WDR72;refGene_tx

=XM_023255333.1;refGene_gn=WDR72;refGene_tx=XM_023255334.1;refGene_gn=WDR72;refGene_tx=XM_023255335.1
B3 52799257 52799336
refGene_gn=WDR72;refGene_tx=XM_023255332.1;refGene_gn=WDR72;refGene_tx=XM_023255333.1;refGene_gn=WDR72;refGene_tx=XM_023255334.1;refGene_gn=WDR72;refGene_tx=XM_023255335.1
B3 52800475 52800650
refGene_gn=WDR72;refGene_tx=XM_023255332.1;refGene_gn=WDR72;refGene_tx=XM_023255333.1;refGene_gn=WDR72;refGene_tx=XM_023255334.1;refGene_gn=WDR72;refGene_tx=XM_023255335.1
B3 52801308 52801385
refGene_gn=WDR72;refGene_tx=XM_023255332.1;refGene_gn=WDR72;refGene_tx=XM_023255333.1;refGene_gn=WDR72;refGene_tx=XM_023255334.1;refGene_gn=WDR72;refGene_tx=XM_023255335.1
B3 52802463 52802583
refGene_gn=WDR72;refGene_tx=XM_023255332.1;refGene_gn=WDR72;refGene_tx=XM_023255333.1;refGene_gn=WDR72;refGene_tx=XM_023255334.1;refGene_gn=WDR72;refGene_tx=XM_023255335.1
B3 52803435 52803581
refGene_gn=WDR72;refGene_tx=XM_023255332.1;refGene_gn=WDR72;refGene_tx=XM_023255333.1;refGene_gn=WDR72;refGene_tx=XM_023255334.1;refGene_gn=WDR72;refGene_tx=XM_023255335.1
B3 52803963 52804060
refGene_gn=WDR72;refGene_tx=XM_023255332.1;refGene_gn=WDR72;refGene_tx=XM_023255333.1;refGene_gn=WDR72;refGene_tx=XM_023255334.1;refGene_gn=WDR72;refGene_tx=XM_023255335.1
B3 52808891 52809033
refGene_gn=WDR72;refGene_tx=XM_023255332.1;refGene_gn=WDR72;refGene_tx=XM_023255333.1;refGene_gn=WDR72;refGene_tx=XM_023255334.1;refGene_gn=WDR72;refGene_tx=XM_023255335.1
B3 52817597 52817825
refGene_gn=WDR72;refGene_tx=XM_023255332.1;refGene_gn=WDR72;refGene_tx=XM_023255333.1;refGene_gn=WDR72;refGene_tx=XM_023255334.1;refGene_gn=WDR72;refGene_tx=XM_023255335.1
B3 52820126 52820347
refGene_gn=WDR72;refGene_tx=XM_023255332.1;refGene_gn=WDR72;refGene_tx=XM_023255333.1;refGene_gn=WDR72;refGene_tx=XM_023255334.1;refGene_gn=WDR72;refGene_tx=XM_023255335.1
B3 52823743 52823939
refGene_gn=WDR72;refGene_tx=XM_023255332.1;refGene_gn=WDR72;refGene_tx=XM_023255333.1;refGene_gn=WDR72;refGene_tx=XM_023255334.1;refGene_gn=WDR72;refGene_tx=XM_023255335.1
B3 52853917 52854120
refGene_gn=WDR72;refGene_tx=XM_023255332.1;refGene_gn=WDR72;refGene_tx=XM_023255333.1;refGene_gn=WDR72;refGene_tx=XM_023255334.1;refGene_gn=WDR72;refGene_tx=XM_023255335.1
B3 52900704 52901531
refGene_gn=WDR72;refGene_tx=XM_023255332.1;refGene_gn=WDR72;refGene_tx=XM_023255333.1;refGene_gn=WDR72;refGene_tx=XM_023255334.1;refGene_gn=WDR72;refGene_tx=XM_023255335.1

B3 52904168 52904260
refGene_gn=WDR72;refGene_tx=XM_023255332.1;refGene_gn=WDR72;refGene_tx=XM_023255333.1;refGene_gn=WDR72;refGene_tx=XM_023255334.1;refGene_gn=WDR72;refGene_tx=XM_023255335.1
B3 52908664 52908744
refGene_gn=WDR72;refGene_tx=XM_023255332.1;refGene_gn=WDR72;refGene_tx=XM_023255333.1;refGene_gn=WDR72;refGene_tx=XM_023255334.1;refGene_gn=WDR72;refGene_tx=XM_023255335.1
B3 52914196 52914395
refGene_gn=WDR72;refGene_tx=XM_023255332.1;refGene_gn=WDR72;refGene_tx=XM_023255333.1;refGene_gn=WDR72;refGene_tx=XM_023255334.1;refGene_gn=WDR72;refGene_tx=XM_023255335.1
B3 52986120 52986225
refGene_gn=WDR72;refGene_tx=XM_023255332.1;refGene_gn=WDR72;refGene_tx=XM_023255333.1;refGene_gn=WDR72;refGene_tx=XM_023255334.1;refGene_gn=WDR72;refGene_tx=XM_023255335.1
B3 52989346 52989398 refGene_gn=WDR72;refGene_tx=XM_023255334.1
B3 52991787 52991843
refGene_gn=WDR72;refGene_tx=XM_023255332.1;refGene_gn=WDR72;refGene_tx=XM_023255333.1;refGene_gn=WDR72;refGene_tx=XM_023255334.1
B3 52992881 52992919 refGene_gn=WDR72;refGene_tx=XM_023255335.1
B3 53675066 53676171 refGene_gn=ONECUT1;refGene_tx=XM_003987131.4
B3 53713732 53714025 refGene_gn=ONECUT1;refGene_tx=XM_003987131.4
B3 53881967 53882012
refGene_gn=ARPP19;refGene_tx=XM_023255336.1;refGene_gn=ARPP19;refGene_tx=XM_023255337.1
B3 53895267 53895282 refGene_gn=ARPP19;refGene_tx=XM_023255338.1
B3 53895380 53895503
refGene_gn=ARPP19;refGene_tx=XM_023255336.1;refGene_gn=ARPP19;refGene_tx=XM_023255337.1;refGene_gn=ARPP19;refGene_tx=XM_023255338.1
B3 53899923 53900094
refGene_gn=ARPP19;refGene_tx=XM_023255336.1;refGene_gn=ARPP19;refGene_tx=XM_023255337.1;refGene_gn=ARPP19;refGene_tx=XM_023255338.1
B3 53917029 53917056
refGene_gn=MYO5A;refGene_tx=XM_023255339.1;refGene_gn=MYO5A;refGene_tx=XM_023255340.1;refGene_gn=MYO5A;refGene_tx=XM_023255341.1;refGene_gn=MYO5A;refGene_tx=XM_023255342.1;refGene_gn=MYO5A;refGene_tx=XM_023255343.1;refGene_gn=MYO5A;refGene_tx=XM_023255344.1;refGene_gn=MYO5A;refGene_tx=XM_023255345.1
B3 53980075 53980186
refGene_gn=MYO5A;refGene_tx=XM_023255339.1;refGene_gn=MYO5A;refGene_tx=XM_023255340.1;refGene_gn=MYO5A;refGene_tx=XM_023255341.1;refGene_gn=MYO5A;refGene_tx=XM_023255342.1;refGene_gn=MYO5A;refGene_tx=XM_023255343.1;refGene_gn=MYO5A;refGene_tx=XM_023255344.1;refGene_gn=MYO5A;refGene_tx=XM_023255345.1
B3 53983430 53983602
refGene_gn=MYO5A;refGene_tx=XM_023255339.1;refGene_gn=MYO5A;refGene_tx=XM_023255340.1;refGene_gn=MYO5A;refGene_tx=XM_023255341.1;refGene_gn=MYO5A;refGene_tx=XM_023255342.1;refGene_gn=MYO5A;refGene_tx=XM_023255343.1;refGene_gn=MYO5A;refGene_tx=XM_023255344.1;refGene_gn=MYO5A;refGe

ne_tx=XM_023255345.1
B3 53985499 53985644
refGene_gn=MYO5A;refGene_tx=XM_023255339.1;refGene_gn=MYO5A;refGene_tx=XM_023255340.1;refGene_gn=MYO5A;refGene_tx=XM_023255341.1;refGene_gn=MYO5A;refGene_tx=XM_023255342.1;refGene_gn=MYO5A;refGene_tx=XM_023255343.1;refGene_gn=MYO5A;refGene_tx=XM_023255344.1;refGene_gn=MYO5A;refGene_tx=XM_023255345.1
B3 54005384 54005541
refGene_gn=MYO5A;refGene_tx=XM_023255339.1;refGene_gn=MYO5A;refGene_tx=XM_023255340.1;refGene_gn=MYO5A;refGene_tx=XM_023255341.1;refGene_gn=MYO5A;refGene_tx=XM_023255342.1;refGene_gn=MYO5A;refGene_tx=XM_023255343.1;refGene_gn=MYO5A;refGene_tx=XM_023255344.1;refGene_gn=MYO5A;refGene_tx=XM_023255345.1
B3 54010161 54010305
refGene_gn=MYO5A;refGene_tx=XM_023255339.1;refGene_gn=MYO5A;refGene_tx=XM_023255340.1;refGene_gn=MYO5A;refGene_tx=XM_023255341.1;refGene_gn=MYO5A;refGene_tx=XM_023255342.1;refGene_gn=MYO5A;refGene_tx=XM_023255343.1;refGene_gn=MYO5A;refGene_tx=XM_023255344.1;refGene_gn=MYO5A;refGene_tx=XM_023255345.1
B3 54018391 54018473
refGene_gn=MYO5A;refGene_tx=XM_023255339.1;refGene_gn=MYO5A;refGene_tx=XM_023255340.1;refGene_gn=MYO5A;refGene_tx=XM_023255341.1;refGene_gn=MYO5A;refGene_tx=XM_023255342.1;refGene_gn=MYO5A;refGene_tx=XM_023255343.1;refGene_gn=MYO5A;refGene_tx=XM_023255344.1;refGene_gn=MYO5A;refGene_tx=XM_023255345.1
B3 54018823 54018931
refGene_gn=MYO5A;refGene_tx=XM_023255339.1;refGene_gn=MYO5A;refGene_tx=XM_023255340.1;refGene_gn=MYO5A;refGene_tx=XM_023255341.1;refGene_gn=MYO5A;refGene_tx=XM_023255342.1;refGene_gn=MYO5A;refGene_tx=XM_023255343.1;refGene_gn=MYO5A;refGene_tx=XM_023255344.1;refGene_gn=MYO5A;refGene_tx=XM_023255345.1
B3 54020669 54020776
refGene_gn=MYO5A;refGene_tx=XM_023255339.1;refGene_gn=MYO5A;refGene_tx=XM_023255340.1;refGene_gn=MYO5A;refGene_tx=XM_023255341.1;refGene_gn=MYO5A;refGene_tx=XM_023255342.1;refGene_gn=MYO5A;refGene_tx=XM_023255343.1;refGene_gn=MYO5A;refGene_tx=XM_023255344.1;refGene_gn=MYO5A;refGene_tx=XM_023255345.1
B3 54027944 54028210
refGene_gn=MYO5A;refGene_tx=XM_023255339.1;refGene_gn=MYO5A;refGene_tx=XM_023255340.1;refGene_gn=MYO5A;refGene_tx=XM_023255341.1;refGene_gn=MYO5A;refGene_tx=XM_023255342.1;refGene_gn=MYO5A;refGene_tx=XM_023255343.1;refGene_gn=MYO5A;refGene_tx=XM_023255344.1;refGene_gn=MYO5A;refGene_tx=XM_023255345.1
B3 54028839 54028921
refGene_gn=MYO5A;refGene_tx=XM_023255339.1;refGene_gn=MYO5A;refGene_tx=XM_023255340.1;refGene_gn=MYO5A;refGene_tx=XM_023255341.1;refGene_gn=MYO5A;refGene_tx=XM_023255342.1;refGene_gn=MYO5A;refGene_tx=XM_023255343.1;refGene_gn=MYO5A;refGene_tx=XM_023255344.1;refGene_gn=MYO5A;refGene_tx=XM_023255345.1
B3 54033590 54033731

refGene_gn=MYO5A;refGene_tx=XM_023255339.1;refGene_gn=MYO5A;refGene_tx=XM_023255340.1;refGene_gn=MYO5A;refGene_tx=XM_023255341.1;refGene_gn=MYO5A;refGene_tx=XM_023255342.1;refGene_gn=MYO5A;refGene_tx=XM_023255343.1;refGene_gn=MYO5A;refGene_tx=XM_023255344.1;refGene_gn=MYO5A;refGene_tx=XM_023255345.1
B3 54035820 54035946
refGene_gn=MYO5A;refGene_tx=XM_023255339.1;refGene_gn=MYO5A;refGene_tx=XM_023255340.1;refGene_gn=MYO5A;refGene_tx=XM_023255341.1;refGene_gn=MYO5A;refGene_tx=XM_023255342.1;refGene_gn=MYO5A;refGene_tx=XM_023255343.1;refGene_gn=MYO5A;refGene_tx=XM_023255344.1;refGene_gn=MYO5A;refGene_tx=XM_023255345.1
B3 54037191 54037275
refGene_gn=MYO5A;refGene_tx=XM_023255339.1;refGene_gn=MYO5A;refGene_tx=XM_023255340.1;refGene_gn=MYO5A;refGene_tx=XM_023255341.1;refGene_gn=MYO5A;refGene_tx=XM_023255342.1;refGene_gn=MYO5A;refGene_tx=XM_023255343.1;refGene_gn=MYO5A;refGene_tx=XM_023255344.1;refGene_gn=MYO5A;refGene_tx=XM_023255345.1
B3 54040518 54040680
refGene_gn=MYO5A;refGene_tx=XM_023255339.1;refGene_gn=MYO5A;refGene_tx=XM_023255340.1;refGene_gn=MYO5A;refGene_tx=XM_023255341.1;refGene_gn=MYO5A;refGene_tx=XM_023255342.1;refGene_gn=MYO5A;refGene_tx=XM_023255343.1;refGene_gn=MYO5A;refGene_tx=XM_023255344.1;refGene_gn=MYO5A;refGene_tx=XM_023255345.1
B3 54042087 54042185
refGene_gn=MYO5A;refGene_tx=XM_023255339.1;refGene_gn=MYO5A;refGene_tx=XM_023255340.1;refGene_gn=MYO5A;refGene_tx=XM_023255341.1;refGene_gn=MYO5A;refGene_tx=XM_023255342.1;refGene_gn=MYO5A;refGene_tx=XM_023255343.1;refGene_gn=MYO5A;refGene_tx=XM_023255344.1;refGene_gn=MYO5A;refGene_tx=XM_023255345.1
B3 54044126 54044213
refGene_gn=MYO5A;refGene_tx=XM_023255339.1;refGene_gn=MYO5A;refGene_tx=XM_023255340.1;refGene_gn=MYO5A;refGene_tx=XM_023255341.1;refGene_gn=MYO5A;refGene_tx=XM_023255342.1;refGene_gn=MYO5A;refGene_tx=XM_023255343.1;refGene_gn=MYO5A;refGene_tx=XM_023255344.1;refGene_gn=MYO5A;refGene_tx=XM_023255345.1
B3 54044300 54044409
refGene_gn=MYO5A;refGene_tx=XM_023255339.1;refGene_gn=MYO5A;refGene_tx=XM_023255340.1;refGene_gn=MYO5A;refGene_tx=XM_023255341.1;refGene_gn=MYO5A;refGene_tx=XM_023255342.1;refGene_gn=MYO5A;refGene_tx=XM_023255343.1;refGene_gn=MYO5A;refGene_tx=XM_023255344.1;refGene_gn=MYO5A;refGene_tx=XM_023255345.1
B3 54046698 54046910
refGene_gn=MYO5A;refGene_tx=XM_023255339.1;refGene_gn=MYO5A;refGene_tx=XM_023255340.1;refGene_gn=MYO5A;refGene_tx=XM_023255341.1;refGene_gn=MYO5A;refGene_tx=XM_023255342.1;refGene_gn=MYO5A;refGene_tx=XM_023255343.1;refGene_gn=MYO5A;refGene_tx=XM_023255344.1;refGene_gn=MYO5A;refGene_tx=XM_023255345.1
B3 54047797 54047954
refGene_gn=MYO5A;refGene_tx=XM_023255339.1;refGene_gn=MYO5A;refGene_tx=XM_023255340.1;refGene_gn=MYO5A;refGene_tx=XM_023255341.1;refGene_gn=

MYO5A;refGene_tx=XM_023255342.1;refGene_gn=MYO5A;refGene_tx=XM_0232553
43.1;refGene_gn=MYO5A;refGene_tx=XM_023255344.1;refGene_gn=MYO5A;refGe
ne_tx=XM_023255345.1
B3 54051418 54051658
refGene_gn=MYO5A;refGene_tx=XM_023255339.1;refGene_gn=MYO5A;refGene_tx
=XM_023255340.1;refGene_gn=MYO5A;refGene_tx=XM_023255341.1;refGene_gn=
MYO5A;refGene_tx=XM_023255342.1;refGene_gn=MYO5A;refGene_tx=XM_0232553
43.1;refGene_gn=MYO5A;refGene_tx=XM_023255344.1;refGene_gn=MYO5A;refGe
ne_tx=XM_023255345.1
B3 54052869 54053118
refGene_gn=MYO5A;refGene_tx=XM_023255339.1;refGene_gn=MYO5A;refGene_tx
=XM_023255340.1;refGene_gn=MYO5A;refGene_tx=XM_023255341.1;refGene_gn=
MYO5A;refGene_tx=XM_023255342.1;refGene_gn=MYO5A;refGene_tx=XM_0232553
43.1;refGene_gn=MYO5A;refGene_tx=XM_023255344.1;refGene_gn=MYO5A;refGe
ne_tx=XM_023255345.1
B3 54056113 54056207
refGene_gn=MYO5A;refGene_tx=XM_023255339.1;refGene_gn=MYO5A;refGene_tx
=XM_023255340.1;refGene_gn=MYO5A;refGene_tx=XM_023255341.1;refGene_gn=
MYO5A;refGene_tx=XM_023255342.1;refGene_gn=MYO5A;refGene_tx=XM_0232553
43.1;refGene_gn=MYO5A;refGene_tx=XM_023255344.1;refGene_gn=MYO5A;refGe
ne_tx=XM_023255345.1
B3 54058525 54058674
refGene_gn=MYO5A;refGene_tx=XM_023255339.1;refGene_gn=MYO5A;refGene_tx
=XM_023255340.1;refGene_gn=MYO5A;refGene_tx=XM_023255341.1;refGene_gn=
MYO5A;refGene_tx=XM_023255342.1;refGene_gn=MYO5A;refGene_tx=XM_0232553
43.1;refGene_gn=MYO5A;refGene_tx=XM_023255344.1;refGene_gn=MYO5A;refGe
ne_tx=XM_023255345.1
B3 54063275 54063389
refGene_gn=MYO5A;refGene_tx=XM_023255339.1;refGene_gn=MYO5A;refGene_tx
=XM_023255340.1;refGene_gn=MYO5A;refGene_tx=XM_023255341.1;refGene_gn=
MYO5A;refGene_tx=XM_023255342.1;refGene_gn=MYO5A;refGene_tx=XM_0232553
43.1;refGene_gn=MYO5A;refGene_tx=XM_023255344.1;refGene_gn=MYO5A;refGe
ne_tx=XM_023255345.1
B3 54070060 54070204
refGene_gn=MYO5A;refGene_tx=XM_023255339.1;refGene_gn=MYO5A;refGene_tx
=XM_023255340.1;refGene_gn=MYO5A;refGene_tx=XM_023255341.1;refGene_gn=
MYO5A;refGene_tx=XM_023255342.1;refGene_gn=MYO5A;refGene_tx=XM_0232553
43.1;refGene_gn=MYO5A;refGene_tx=XM_023255344.1;refGene_gn=MYO5A;refGe
ne_tx=XM_023255345.1
B3 54070460 54070514
refGene_gn=MYO5A;refGene_tx=XM_023255339.1;refGene_gn=MYO5A;refGene_tx
=XM_023255340.1;refGene_gn=MYO5A;refGene_tx=XM_023255341.1;refGene_gn=
MYO5A;refGene_tx=XM_023255342.1;refGene_gn=MYO5A;refGene_tx=XM_0232553
43.1;refGene_gn=MYO5A;refGene_tx=XM_023255344.1;refGene_gn=MYO5A;refGe
ne_tx=XM_023255345.1
B3 54074653 54074881
refGene_gn=MYO5A;refGene_tx=XM_023255339.1;refGene_gn=MYO5A;refGene_tx
=XM_023255340.1;refGene_gn=MYO5A;refGene_tx=XM_023255341.1;refGene_gn=
MYO5A;refGene_tx=XM_023255342.1;refGene_gn=MYO5A;refGene_tx=XM_0232553
43.1;refGene_gn=MYO5A;refGene_tx=XM_023255344.1;refGene_gn=MYO5A;refGe

ne_tx=XM_023255345.1
B3	54077088 54077097
refGene_gn=MYO5A;refGene_tx=XM_023255339.1;refGene_gn=MYO5A;refGene_tx=XM_023255340.1;refGene_gn=MYO5A;refGene_tx=XM_023255342.1;refGene_gn=MYO5A;refGene_tx=XM_023255343.1;refGene_gn=MYO5A;refGene_tx=XM_023255345.1
B3	54079414 54079515
refGene_gn=MYO5A;refGene_tx=XM_023255339.1;refGene_gn=MYO5A;refGene_tx=XM_023255340.1;refGene_gn=MYO5A;refGene_tx=XM_023255341.1;refGene_gn=MYO5A;refGene_tx=XM_023255342.1;refGene_gn=MYO5A;refGene_tx=XM_023255343.1;refGene_gn=MYO5A;refGene_tx=XM_023255344.1;refGene_gn=MYO5A;refGene_tx=XM_023255345.1
B3	54082383 54082464
refGene_gn=MYO5A;refGene_tx=XM_023255339.1;refGene_gn=MYO5A;refGene_tx=XM_023255340.1;refGene_gn=MYO5A;refGene_tx=XM_023255341.1;refGene_gn=MYO5A;refGene_tx=XM_023255342.1;refGene_gn=MYO5A;refGene_tx=XM_023255344.1
B3	54084256 54084455
refGene_gn=MYO5A;refGene_tx=XM_023255339.1;refGene_gn=MYO5A;refGene_tx=XM_023255340.1;refGene_gn=MYO5A;refGene_tx=XM_023255341.1;refGene_gn=MYO5A;refGene_tx=XM_023255342.1;refGene_gn=MYO5A;refGene_tx=XM_023255343.1;refGene_gn=MYO5A;refGene_tx=XM_023255344.1;refGene_gn=MYO5A;refGene_tx=XM_023255345.1
B3	54086876 54086951
refGene_gn=MYO5A;refGene_tx=XM_023255339.1;refGene_gn=MYO5A;refGene_tx=XM_023255340.1;refGene_gn=MYO5A;refGene_tx=XM_023255341.1;refGene_gn=MYO5A;refGene_tx=XM_023255343.1
B3	54088201 54088295
refGene_gn=MYO5A;refGene_tx=XM_023255339.1;refGene_gn=MYO5A;refGene_tx=XM_023255340.1;refGene_gn=MYO5A;refGene_tx=XM_023255341.1;refGene_gn=MYO5A;refGene_tx=XM_023255342.1;refGene_gn=MYO5A;refGene_tx=XM_023255343.1;refGene_gn=MYO5A;refGene_tx=XM_023255344.1;refGene_gn=MYO5A;refGene_tx=XM_023255345.1
B3	54093396 54093543
refGene_gn=MYO5A;refGene_tx=XM_023255339.1;refGene_gn=MYO5A;refGene_tx=XM_023255340.1;refGene_gn=MYO5A;refGene_tx=XM_023255341.1;refGene_gn=MYO5A;refGene_tx=XM_023255342.1;refGene_gn=MYO5A;refGene_tx=XM_023255343.1;refGene_gn=MYO5A;refGene_tx=XM_023255344.1;refGene_gn=MYO5A;refGene_tx=XM_023255345.1
B3	54095602 54095757
refGene_gn=MYO5A;refGene_tx=XM_023255339.1;refGene_gn=MYO5A;refGene_tx=XM_023255340.1;refGene_gn=MYO5A;refGene_tx=XM_023255341.1;refGene_gn=MYO5A;refGene_tx=XM_023255342.1;refGene_gn=MYO5A;refGene_tx=XM_023255343.1;refGene_gn=MYO5A;refGene_tx=XM_023255344.1;refGene_gn=MYO5A;refGene_tx=XM_023255345.1
B3	54098209 54098299
refGene_gn=MYO5A;refGene_tx=XM_023255339.1;refGene_gn=MYO5A;refGene_tx=XM_023255340.1;refGene_gn=MYO5A;refGene_tx=XM_023255341.1;refGene_gn=MYO5A;refGene_tx=XM_023255342.1;refGene_gn=MYO5A;refGene_tx=XM_023255343.1;refGene_gn=MYO5A;refGene_tx=XM_023255344.1;refGene_gn=MYO5A;refGe

ne_tx=XM_023255345.1
B3 54099860 54100011
refGene_gn=MYO5A;refGene_tx=XM_023255339.1;refGene_gn=MYO5A;refGene_tx=XM_023255340.1;refGene_gn=MYO5A;refGene_tx=XM_023255341.1;refGene_gn=MYO5A;refGene_tx=XM_023255342.1;refGene_gn=MYO5A;refGene_tx=XM_023255343.1;refGene_gn=MYO5A;refGene_tx=XM_023255344.1;refGene_gn=MYO5A;refGene_tx=XM_023255345.1
B3 54101917 54102200
refGene_gn=MYO5A;refGene_tx=XM_023255339.1;refGene_gn=MYO5A;refGene_tx=XM_023255340.1;refGene_gn=MYO5A;refGene_tx=XM_023255341.1;refGene_gn=MYO5A;refGene_tx=XM_023255342.1;refGene_gn=MYO5A;refGene_tx=XM_023255343.1;refGene_gn=MYO5A;refGene_tx=XM_023255344.1;refGene_gn=MYO5A;refGene_tx=XM_023255345.1
B3 54103661 54103836
refGene_gn=MYO5A;refGene_tx=XM_023255339.1;refGene_gn=MYO5A;refGene_tx=XM_023255340.1;refGene_gn=MYO5A;refGene_tx=XM_023255341.1;refGene_gn=MYO5A;refGene_tx=XM_023255342.1;refGene_gn=MYO5A;refGene_tx=XM_023255343.1;refGene_gn=MYO5A;refGene_tx=XM_023255344.1;refGene_gn=MYO5A;refGene_tx=XM_023255345.1
B3 54105199 54105280
refGene_gn=MYO5A;refGene_tx=XM_023255339.1;refGene_gn=MYO5A;refGene_tx=XM_023255340.1;refGene_gn=MYO5A;refGene_tx=XM_023255341.1;refGene_gn=MYO5A;refGene_tx=XM_023255342.1;refGene_gn=MYO5A;refGene_tx=XM_023255343.1;refGene_gn=MYO5A;refGene_tx=XM_023255344.1;refGene_gn=MYO5A;refGene_tx=XM_023255345.1
B3 54105558 54105711
refGene_gn=MYO5A;refGene_tx=XM_023255339.1;refGene_gn=MYO5A;refGene_tx=XM_023255340.1;refGene_gn=MYO5A;refGene_tx=XM_023255341.1;refGene_gn=MYO5A;refGene_tx=XM_023255342.1;refGene_gn=MYO5A;refGene_tx=XM_023255343.1;refGene_gn=MYO5A;refGene_tx=XM_023255344.1;refGene_gn=MYO5A;refGene_tx=XM_023255345.1
B3 54118328 54118355 refGene_gn=MYO5C;refGene_tx=XM_019832406.2
B3 54128931 54129042 refGene_gn=MYO5C;refGene_tx=XM_019832406.2
B3 54134921 54135087 refGene_gn=MYO5C;refGene_tx=XM_019832406.2
B3 54135673 54135818 refGene_gn=MYO5C;refGene_tx=XM_019832406.2
B3 54138867 54139024 refGene_gn=MYO5C;refGene_tx=XM_019832406.2
B3 54140986 54141010 refGene_gn=MYO5C;refGene_tx=XM_019832407.2
B3 54141166 54141310
refGene_gn=MYO5C;refGene_tx=XM_019832406.2;refGene_gn=MYO5C;refGene_tx=XM_019832407.2
B3 54142313 54142395
refGene_gn=MYO5C;refGene_tx=XM_019832406.2;refGene_gn=MYO5C;refGene_tx=XM_019832407.2
B3 54143433 54143541
refGene_gn=MYO5C;refGene_tx=XM_019832406.2;refGene_gn=MYO5C;refGene_tx=XM_019832407.2
B3 54150048 54150155
refGene_gn=MYO5C;refGene_tx=XM_019832406.2;refGene_gn=MYO5C;refGene_tx=XM_019832407.2
B3 54153500 54153766

refGene_gn=MYO5C;refGene_tx=XM_019832406.2;refGene_gn=MYO5C;refGene_tx=XM_019832407.2
B3 54157843 54157925
refGene_gn=MYO5C;refGene_tx=XM_019832406.2;refGene_gn=MYO5C;refGene_tx=XM_019832407.2
B3 54159180 54159321
refGene_gn=MYO5C;refGene_tx=XM_019832406.2;refGene_gn=MYO5C;refGene_tx=XM_019832407.2
B3 54161340 54161466
refGene_gn=MYO5C;refGene_tx=XM_019832406.2;refGene_gn=MYO5C;refGene_tx=XM_019832407.2
B3 54167525 54167609
refGene_gn=MYO5C;refGene_tx=XM_019832406.2;refGene_gn=MYO5C;refGene_tx=XM_019832407.2
B3 54169358 54169493
refGene_gn=MYO5C;refGene_tx=XM_019832406.2;refGene_gn=MYO5C;refGene_tx=XM_019832407.2
B3 54169913 54170011
refGene_gn=MYO5C;refGene_tx=XM_019832406.2;refGene_gn=MYO5C;refGene_tx=XM_019832407.2
B3 54172432 54172519
refGene_gn=MYO5C;refGene_tx=XM_019832406.2;refGene_gn=MYO5C;refGene_tx=XM_019832407.2
B3 54172978 54173090
refGene_gn=MYO5C;refGene_tx=XM_019832406.2;refGene_gn=MYO5C;refGene_tx=XM_019832407.2
B3 54174134 54174346
refGene_gn=MYO5C;refGene_tx=XM_019832406.2;refGene_gn=MYO5C;refGene_tx=XM_019832407.2
B3 54176951 54177117
refGene_gn=MYO5C;refGene_tx=XM_019832406.2;refGene_gn=MYO5C;refGene_tx=XM_019832407.2
B3 54177451 54177598
refGene_gn=MYO5C;refGene_tx=XM_019832406.2;refGene_gn=MYO5C;refGene_tx=XM_019832407.2
B3 54178988 54179153
refGene_gn=MYO5C;refGene_tx=XM_019832406.2;refGene_gn=MYO5C;refGene_tx=XM_019832407.2
B3 54180439 54180533
refGene_gn=MYO5C;refGene_tx=XM_019832406.2;refGene_gn=MYO5C;refGene_tx=XM_019832407.2
B3 54183111 54183175
refGene_gn=MYO5C;refGene_tx=XM_019832406.2;refGene_gn=MYO5C;refGene_tx=XM_019832407.2
B3 54185249 54185430
refGene_gn=MYO5C;refGene_tx=XM_019832406.2;refGene_gn=MYO5C;refGene_tx=XM_019832407.2
B3 54187327 54187421
refGene_gn=MYO5C;refGene_tx=XM_019832406.2;refGene_gn=MYO5C;refGene_tx=XM_019832407.2

B3 54187655 54187719
refGene_gn=MYO5C;refGene_tx=XM_019832406.2;refGene_gn=MYO5C;refGene_tx=XM_019832407.2
B3 54187805 54187886
refGene_gn=MYO5C;refGene_tx=XM_019832406.2;refGene_gn=MYO5C;refGene_tx=XM_019832407.2
B3 54188741 54188922
refGene_gn=MYO5C;refGene_tx=XM_019832406.2;refGene_gn=MYO5C;refGene_tx=XM_019832407.2
B3 54190817 54190911
refGene_gn=MYO5C;refGene_tx=XM_019832406.2;refGene_gn=MYO5C;refGene_tx=XM_019832407.2
B3 54191792 54191856
refGene_gn=MYO5C;refGene_tx=XM_019832406.2;refGene_gn=MYO5C;refGene_tx=XM_019832407.2
B3 54193191 54193360
refGene_gn=MYO5C;refGene_tx=XM_019832406.2;refGene_gn=MYO5C;refGene_tx=XM_019832407.2
B3 54197247 54197335
refGene_gn=MYO5C;refGene_tx=XM_019832406.2;refGene_gn=MYO5C;refGene_tx=XM_019832407.2
B3 54200496 54200595
refGene_gn=MYO5C;refGene_tx=XM_019832406.2;refGene_gn=MYO5C;refGene_tx=XM_019832407.2
B3 54201906 54202061
refGene_gn=MYO5C;refGene_tx=XM_019832406.2;refGene_gn=MYO5C;refGene_tx=XM_019832407.2
B3 54203603 54203693
refGene_gn=MYO5C;refGene_tx=XM_019832406.2;refGene_gn=MYO5C;refGene_tx=XM_019832407.2
B3 54205997 54206148
refGene_gn=MYO5C;refGene_tx=XM_019832406.2;refGene_gn=MYO5C;refGene_tx=XM_019832407.2
B3 54206882 54207165
refGene_gn=MYO5C;refGene_tx=XM_019832406.2;refGene_gn=MYO5C;refGene_tx=XM_019832407.2
B3 54218264 54218439
refGene_gn=MYO5C;refGene_tx=XM_019832406.2;refGene_gn=MYO5C;refGene_tx=XM_019832407.2
B3 54218843 54218924
refGene_gn=MYO5C;refGene_tx=XM_019832406.2;refGene_gn=MYO5C;refGene_tx=XM_019832407.2
B3 54222055 54222208
refGene_gn=MYO5C;refGene_tx=XM_019832406.2;refGene_gn=MYO5C;refGene_tx=XM_019832407.2
B3 54230213 54230339
refGene_gn=GNB5;refGene_tx=XM_023255346.1;refGene_gn=GNB5;refGene_tx=XM_023255349.1
B3 54234676 54234788
refGene_gn=GNB5;refGene_tx=XM_023255346.1;refGene_gn=GNB5;refGene_tx=X

M_023255347.1;refGene_gn=GNB5;refGene_tx=XM_023255348.1
B3 54246632 54246769
refGene_gn=GNB5;refGene_tx=XM_023255346.1;refGene_gn=GNB5;refGene_tx=XM_023255347.1;refGene_gn=GNB5;refGene_tx=XM_023255348.1;refGene_gn=GNB5;refGene_tx=XM_023255349.1
B3 54250810 54250852
refGene_gn=GNB5;refGene_tx=XM_023255346.1;refGene_gn=GNB5;refGene_tx=XM_023255347.1;refGene_gn=GNB5;refGene_tx=XM_023255348.1;refGene_gn=GNB5;refGene_tx=XM_023255349.1
B3 54253078 54253155
refGene_gn=GNB5;refGene_tx=XM_023255346.1;refGene_gn=GNB5;refGene_tx=XM_023255347.1;refGene_gn=GNB5;refGene_tx=XM_023255348.1;refGene_gn=GNB5;refGene_tx=XM_023255349.1
B3 54259066 54259199
refGene_gn=GNB5;refGene_tx=XM_023255346.1;refGene_gn=GNB5;refGene_tx=XM_023255347.1;refGene_gn=GNB5;refGene_tx=XM_023255348.1;refGene_gn=GNB5;refGene_tx=XM_023255349.1
B3 54274421 54274565
refGene_gn=GNB5;refGene_tx=XM_023255346.1;refGene_gn=GNB5;refGene_tx=XM_023255347.1;refGene_gn=GNB5;refGene_tx=XM_023255348.1;refGene_gn=GNB5;refGene_tx=XM_023255349.1
B3 54276095 54276187
refGene_gn=GNB5;refGene_tx=XM_023255346.1;refGene_gn=GNB5;refGene_tx=XM_023255347.1;refGene_gn=GNB5;refGene_tx=XM_023255348.1;refGene_gn=GNB5;refGene_tx=XM_023255349.1
B3 54278720 54278769
refGene_gn=GNB5;refGene_tx=XM_023255346.1;refGene_gn=GNB5;refGene_tx=XM_023255347.1;refGene_gn=GNB5;refGene_tx=XM_023255348.1;refGene_gn=GNB5;refGene_tx=XM_023255349.1
B3 54280805 54280902
refGene_gn=GNB5;refGene_tx=XM_023255346.1;refGene_gn=GNB5;refGene_tx=XM_023255347.1;refGene_gn=GNB5;refGene_tx=XM_023255348.1;refGene_gn=GNB5;refGene_tx=XM_023255349.1
B3 54281824 54281991
refGene_gn=GNB5;refGene_tx=XM_023255346.1;refGene_gn=GNB5;refGene_tx=XM_023255347.1;refGene_gn=GNB5;refGene_tx=XM_023255348.1;refGene_gn=GNB5;refGene_tx=XM_023255349.1
B3 54283478 54283490
refGene_gn=GNB5;refGene_tx=XM_023255346.1;refGene_gn=GNB5;refGene_tx=XM_023255347.1;refGene_gn=GNB5;refGene_tx=XM_023255348.1;refGene_gn=GNB5;refGene_tx=XM_023255349.1
B3 54291736 54292192 refGene_gn=BCL2L10;refGene_tx=XM_023256228.1
B3 54295329 54295452 refGene_gn=BCL2L10;refGene_tx=XM_023256228.1
B3 54363478 54364580
refGene_gn=MAPK6;refGene_tx=XM_011282866.3;refGene_gn=MAPK6;refGene_tx=XM_019832410.2;refGene_gn=MAPK6;refGene_tx=XM_019832411.1;refGene_gn=MAPK6;refGene_tx=XM_023255350.1
B3 54366483 54366685
refGene_gn=MAPK6;refGene_tx=XM_011282866.3;refGene_gn=MAPK6;refGene_tx=XM_019832410.2;refGene_gn=MAPK6;refGene_tx=XM_019832411.1;refGene_gn=

MAPK6;refGene_tx=XM_023255350.1
B3 54370770 54370935
refGene_gn=MAPK6;refGene_tx=XM_011282866.3;refGene_gn=MAPK6;refGene_tx=XM_019832410.2;refGene_gn=MAPK6;refGene_tx=XM_019832411.1;refGene_gn=MAPK6;refGene_tx=XM_023255350.1
B3 54377069 54377219
refGene_gn=MAPK6;refGene_tx=XM_011282866.3;refGene_gn=MAPK6;refGene_tx=XM_019832410.2;refGene_gn=MAPK6;refGene_tx=XM_019832411.1;refGene_gn=MAPK6;refGene_tx=XM_023255350.1;refGene_gn=MAPK6;refGene_tx=XM_023255351.1
B3 54378402 54378957
refGene_gn=MAPK6;refGene_tx=XM_011282866.3;refGene_gn=MAPK6;refGene_tx=XM_019832410.2;refGene_gn=MAPK6;refGene_tx=XM_019832411.1;refGene_gn=MAPK6;refGene_tx=XM_023255350.1;refGene_gn=MAPK6;refGene_tx=XM_023255351.1
B3 54422022 54422092
refGene_gn=LEO1;refGene_tx=XM_019832412.2;refGene_gn=LEO1;refGene_tx=XM_023255352.1;refGene_gn=LEO1;refGene_tx=XM_023255353.1;refGene_gn=LEO1;refGene_tx=XM_023255354.1;refGene_gn=LEO1;refGene_tx=XM_023255355.1
B3 54438033 54438091 refGene_gn=LEO1;refGene_tx=XM_023255357.1
B3 54444868 54444908 refGene_gn=LEO1;refGene_tx=XM_023255356.1
B3 54446993 54447746
refGene_gn=LEO1;refGene_tx=XM_019832412.2;refGene_gn=LEO1;refGene_tx=XM_023255352.1;refGene_gn=LEO1;refGene_tx=XM_023255353.1;refGene_gn=LEO1;refGene_tx=XM_023255354.1;refGene_gn=LEO1;refGene_tx=XM_023255355.1;refGene_gn=LEO1;refGene_tx=XM_023255356.1;refGene_gn=LEO1;refGene_tx=XM_023255357.1
B3 54450533 54450638
refGene_gn=LEO1;refGene_tx=XM_019832412.2;refGene_gn=LEO1;refGene_tx=XM_023255352.1;refGene_gn=LEO1;refGene_tx=XM_023255353.1;refGene_gn=LEO1;refGene_tx=XM_023255354.1;refGene_gn=LEO1;refGene_tx=XM_023255355.1;refGene_gn=LEO1;refGene_tx=XM_023255356.1;refGene_gn=LEO1;refGene_tx=XM_023255357.1
B3 54453452 54453547
refGene_gn=LEO1;refGene_tx=XM_019832412.2;refGene_gn=LEO1;refGene_tx=XM_023255352.1;refGene_gn=LEO1;refGene_tx=XM_023255353.1;refGene_gn=LEO1;refGene_tx=XM_023255354.1;refGene_gn=LEO1;refGene_tx=XM_023255355.1;refGene_gn=LEO1;refGene_tx=XM_023255356.1;refGene_gn=LEO1;refGene_tx=XM_023255357.1
B3 54454542 54454688
refGene_gn=LEO1;refGene_tx=XM_019832412.2;refGene_gn=LEO1;refGene_tx=XM_023255352.1;refGene_gn=LEO1;refGene_tx=XM_023255353.1;refGene_gn=LEO1;refGene_tx=XM_023255354.1;refGene_gn=LEO1;refGene_tx=XM_023255355.1;refGene_gn=LEO1;refGene_tx=XM_023255356.1;refGene_gn=LEO1;refGene_tx=XM_023255357.1
B3 54455515 54455600
refGene_gn=LEO1;refGene_tx=XM_019832412.2;refGene_gn=LEO1;refGene_tx=XM_023255352.1;refGene_gn=LEO1;refGene_tx=XM_023255353.1;refGene_gn=LEO1;refGene_tx=XM_023255354.1;refGene_gn=LEO1;refGene_tx=XM_023255355.1;refGene_gn=LEO1;refGene_tx=XM_023255356.1;refGene_gn=LEO1;refGene_tx=X

M_023255357.1
B3 54463081 54463176
refGene_gn=LEO1;refGene_tx=XM_019832412.2;refGene_gn=LEO1;refGene_tx=XM_023255352.1;refGene_gn=LEO1;refGene_tx=XM_023255353.1;refGene_gn=LEO1;refGene_tx=XM_023255354.1;refGene_gn=LEO1;refGene_tx=XM_023255355.1;refGene_gn=LEO1;refGene_tx=XM_023255356.1;refGene_gn=LEO1;refGene_tx=XM_023255357.1
B3 54463708 54463843
refGene_gn=LEO1;refGene_tx=XM_019832412.2;refGene_gn=LEO1;refGene_tx=XM_023255352.1;refGene_gn=LEO1;refGene_tx=XM_023255353.1;refGene_gn=LEO1;refGene_tx=XM_023255354.1;refGene_gn=LEO1;refGene_tx=XM_023255355.1;refGene_gn=LEO1;refGene_tx=XM_023255356.1;refGene_gn=LEO1;refGene_tx=XM_023255357.1
B3 54464960 54465096
refGene_gn=LEO1;refGene_tx=XM_019832412.2;refGene_gn=LEO1;refGene_tx=XM_023255352.1;refGene_gn=LEO1;refGene_tx=XM_023255353.1;refGene_gn=LEO1;refGene_tx=XM_023255354.1;refGene_gn=LEO1;refGene_tx=XM_023255355.1;refGene_gn=LEO1;refGene_tx=XM_023255356.1;refGene_gn=LEO1;refGene_tx=XM_023255357.1
B3 54467541 54467727
refGene_gn=LEO1;refGene_tx=XM_019832412.2;refGene_gn=LEO1;refGene_tx=XM_023255352.1;refGene_gn=LEO1;refGene_tx=XM_023255353.1;refGene_gn=LEO1;refGene_tx=XM_023255354.1;refGene_gn=LEO1;refGene_tx=XM_023255355.1;refGene_gn=LEO1;refGene_tx=XM_023255356.1;refGene_gn=LEO1;refGene_tx=XM_023255357.1
B3 54469365 54469463
refGene_gn=LEO1;refGene_tx=XM_019832412.2;refGene_gn=LEO1;refGene_tx=XM_023255352.1;refGene_gn=LEO1;refGene_tx=XM_023255353.1;refGene_gn=LEO1;refGene_tx=XM_023255354.1;refGene_gn=LEO1;refGene_tx=XM_023255355.1;refGene_gn=LEO1;refGene_tx=XM_023255356.1;refGene_gn=LEO1;refGene_tx=XM_023255357.1
B3 54477964 54478069 refGene_gn=LEO1;refGene_tx=XM_023255357.1
B3 54484473 54484488
refGene_gn=LOC109500670;refGene_tx=XM_023255358.1
B3 54486047 54486142
refGene_gn=LOC109500670;refGene_tx=XM_023255358.1
B3 54486904 54487039
refGene_gn=LOC109500670;refGene_tx=XM_023255358.1
B3 54489145 54489281
refGene_gn=LOC109500670;refGene_tx=XM_023255358.1
B3 54490177 54490336
refGene_gn=LOC109500670;refGene_tx=XM_023255358.1
B3 54502544 54502579 refGene_gn=TMOD3;refGene_tx=XM_003987135.4
B3 54506410 54506555 refGene_gn=TMOD3;refGene_tx=XM_003987135.4
B3 54507503 54507647 refGene_gn=TMOD3;refGene_tx=XM_003987135.4
B3 54510414 54510522 refGene_gn=TMOD3;refGene_tx=XM_003987135.4
B3 54512151 54512282 refGene_gn=TMOD3;refGene_tx=XM_003987135.4
B3 54522465 54522555 refGene_gn=TMOD3;refGene_tx=XM_003987135.4
B3 54524009 54524132 refGene_gn=TMOD3;refGene_tx=XM_003987135.4
B3 54540818 54540975 refGene_gn=TMOD3;refGene_tx=XM_003987135.4

B3 54547154 54547280 refGene_gn=TMOD3;refGene_tx=XM_003987135.4
B3 54598856 54598891
refGene_gn=TMOD2;refGene_tx=XM_023255359.1;refGene_gn=TMOD2;refGene_tx=XM_023255360.1;refGene_gn=TMOD2;refGene_tx=XM_023255361.1
B3 54600844 54600989
refGene_gn=TMOD2;refGene_tx=XM_023255359.1;refGene_gn=TMOD2;refGene_tx=XM_023255360.1;refGene_gn=TMOD2;refGene_tx=XM_023255361.1
B3 54602124 54602196 refGene_gn=TMOD2;refGene_tx=XM_023255362.1
B3 54607718 54607862
refGene_gn=TMOD2;refGene_tx=XM_023255359.1;refGene_gn=TMOD2;refGene_tx=XM_023255360.1;refGene_gn=TMOD2;refGene_tx=XM_023255361.1;refGene_gn=TMOD2;refGene_tx=XM_023255362.1
B3 54614538 54614646
refGene_gn=TMOD2;refGene_tx=XM_023255359.1;refGene_gn=TMOD2;refGene_tx=XM_023255360.1;refGene_gn=TMOD2;refGene_tx=XM_023255361.1;refGene_gn=TMOD2;refGene_tx=XM_023255362.1
B3 54616941 54617072
refGene_gn=TMOD2;refGene_tx=XM_023255359.1;refGene_gn=TMOD2;refGene_tx=XM_023255360.1;refGene_gn=TMOD2;refGene_tx=XM_023255361.1;refGene_gn=TMOD2;refGene_tx=XM_023255362.1
B3 54620273 54620360
refGene_gn=TMOD2;refGene_tx=XM_023255359.1;refGene_gn=TMOD2;refGene_tx=XM_023255360.1;refGene_gn=TMOD2;refGene_tx=XM_023255361.1;refGene_gn=TMOD2;refGene_tx=XM_023255362.1
B3 54624665 54624788
refGene_gn=TMOD2;refGene_tx=XM_023255359.1;refGene_gn=TMOD2;refGene_tx=XM_023255360.1;refGene_gn=TMOD2;refGene_tx=XM_023255361.1;refGene_gn=TMOD2;refGene_tx=XM_023255362.1
B3 54629857 54630014
refGene_gn=TMOD2;refGene_tx=XM_023255359.1;refGene_gn=TMOD2;refGene_tx=XM_023255360.1;refGene_gn=TMOD2;refGene_tx=XM_023255361.1;refGene_gn=TMOD2;refGene_tx=XM_023255362.1
B3 54631616 54631742
refGene_gn=TMOD2;refGene_tx=XM_023255359.1;refGene_gn=TMOD2;refGene_tx=XM_023255360.1;refGene_gn=TMOD2;refGene_tx=XM_023255361.1;refGene_gn=TMOD2;refGene_tx=XM_023255362.1
B3 54665477 54665750 refGene_gn=LYSMD2;refGene_tx=XM_023255363.1
B3 54678785 54679117
refGene_gn=LYSMD2;refGene_tx=XM_011282864.3;refGene_gn=LYSMD2;refGene_tx=XM_019832418.2;refGene_gn=LYSMD2;refGene_tx=XM_023255363.1;refGene_gn=LYSMD2;refGene_tx=XM_023255364.1
B3 54680402 54680445
refGene_gn=LYSMD2;refGene_tx=XM_011282864.3;refGene_gn=LYSMD2;refGene_tx=XM_019832418.2;refGene_gn=LYSMD2;refGene_tx=XM_023255363.1;refGene_gn=LYSMD2;refGene_tx=XM_023255364.1
B3 54684377 54684496
refGene_gn=SCG3;refGene_tx=XM_003987138.5;refGene_gn=SCG3;refGene_tx=XM_019832419.2
B3 54689700 54689781
refGene_gn=SCG3;refGene_tx=XM_003987138.5;refGene_gn=SCG3;refGene_tx=X

M_019832419.2
B3	54701142	54701274	refGene_gn=SCG3;refGene_tx=XM_003987138.5;refGene_gn=SCG3;refGene_tx=XM_019832419.2
B3	54703156	54703240	refGene_gn=SCG3;refGene_tx=XM_003987138.5;refGene_gn=SCG3;refGene_tx=XM_019832419.2
B3	54705655	54705772	refGene_gn=SCG3;refGene_tx=XM_003987138.5;refGene_gn=SCG3;refGene_tx=XM_019832419.2
B3	54708589	54708767	refGene_gn=SCG3;refGene_tx=XM_003987138.5;refGene_gn=SCG3;refGene_tx=XM_019832419.2
B3	54711209	54711359	refGene_gn=SCG3;refGene_tx=XM_003987138.5
B3	54712353	54712496	refGene_gn=SCG3;refGene_tx=XM_003987138.5
B3	54717057	54717273	refGene_gn=SCG3;refGene_tx=XM_003987138.5
B3	54717368	54717414	refGene_gn=SCG3;refGene_tx=XM_003987138.5
B3	54717910	54717963	refGene_gn=SCG3;refGene_tx=XM_003987138.5
B3	54718662	54718753	refGene_gn=SCG3;refGene_tx=XM_003987138.5
B3	54781743	54781830	refGene_gn=DMXL2;refGene_tx=XM_011282863.3;refGene_gn=DMXL2;refGene_tx=XM_019832420.2;refGene_gn=DMXL2;refGene_tx=XM_019832421.2;refGene_gn=DMXL2;refGene_tx=XM_019832423.2
B3	54811038	54811164	refGene_gn=DMXL2;refGene_tx=XM_011282863.3;refGene_gn=DMXL2;refGene_tx=XM_019832420.2;refGene_gn=DMXL2;refGene_tx=XM_019832421.2;refGene_gn=DMXL2;refGene_tx=XM_019832423.2
B3	54814256	54814328	refGene_gn=DMXL2;refGene_tx=XM_011282863.3;refGene_gn=DMXL2;refGene_tx=XM_019832420.2;refGene_gn=DMXL2;refGene_tx=XM_019832421.2;refGene_gn=DMXL2;refGene_tx=XM_019832423.2
B3	54816578	54816657	refGene_gn=DMXL2;refGene_tx=XM_011282863.3;refGene_gn=DMXL2;refGene_tx=XM_019832420.2;refGene_gn=DMXL2;refGene_tx=XM_019832421.2;refGene_gn=DMXL2;refGene_tx=XM_019832423.2
B3	54817443	54817579	refGene_gn=DMXL2;refGene_tx=XM_011282863.3;refGene_gn=DMXL2;refGene_tx=XM_019832420.2;refGene_gn=DMXL2;refGene_tx=XM_019832421.2;refGene_gn=DMXL2;refGene_tx=XM_019832423.2
B3	54818212	54818279	refGene_gn=DMXL2;refGene_tx=XM_011282863.3;refGene_gn=DMXL2;refGene_tx=XM_019832420.2;refGene_gn=DMXL2;refGene_tx=XM_019832421.2;refGene_gn=DMXL2;refGene_tx=XM_019832423.2
B3	54841424	54841603	refGene_gn=DMXL2;refGene_tx=XM_011282863.3;refGene_gn=DMXL2;refGene_tx=XM_019832420.2;refGene_gn=DMXL2;refGene_tx=XM_019832421.2;refGene_gn=DMXL2;refGene_tx=XM_019832423.2
B3	54843488	54843672	refGene_gn=DMXL2;refGene_tx=XM_011282863.3;refGene_gn=DMXL2;refGene_tx=XM_019832420.2;refGene_gn=DMXL2;refGene_tx=XM_019832421.2;refGene_gn=

DMXL2;refGene_tx=XM_019832423.2
B3 54847031 54847206
refGene_gn=DMXL2;refGene_tx=XM_011282863.3;refGene_gn=DMXL2;refGene_tx=XM_019832420.2;refGene_gn=DMXL2;refGene_tx=XM_019832421.2;refGene_gn=DMXL2;refGene_tx=XM_019832423.2
B3 54852020 54852260
refGene_gn=DMXL2;refGene_tx=XM_011282863.3;refGene_gn=DMXL2;refGene_tx=XM_019832420.2;refGene_gn=DMXL2;refGene_tx=XM_019832421.2;refGene_gn=DMXL2;refGene_tx=XM_019832423.2
B3 54852767 54853039
refGene_gn=DMXL2;refGene_tx=XM_011282863.3;refGene_gn=DMXL2;refGene_tx=XM_019832420.2;refGene_gn=DMXL2;refGene_tx=XM_019832421.2;refGene_gn=DMXL2;refGene_tx=XM_019832423.2
B3 54853680 54854377
refGene_gn=DMXL2;refGene_tx=XM_011282863.3;refGene_gn=DMXL2;refGene_tx=XM_019832420.2;refGene_gn=DMXL2;refGene_tx=XM_019832421.2;refGene_gn=DMXL2;refGene_tx=XM_019832423.2
B3 54854718 54854840
refGene_gn=DMXL2;refGene_tx=XM_011282863.3;refGene_gn=DMXL2;refGene_tx=XM_019832420.2;refGene_gn=DMXL2;refGene_tx=XM_019832421.2;refGene_gn=DMXL2;refGene_tx=XM_019832423.2
B3 54872364 54872454
refGene_gn=DMXL2;refGene_tx=XM_011282863.3;refGene_gn=DMXL2;refGene_tx=XM_019832420.2;refGene_gn=DMXL2;refGene_tx=XM_019832421.2;refGene_gn=DMXL2;refGene_tx=XM_019832423.2
B3 54874591 54874709
refGene_gn=DMXL2;refGene_tx=XM_011282863.3;refGene_gn=DMXL2;refGene_tx=XM_019832420.2;refGene_gn=DMXL2;refGene_tx=XM_019832421.2;refGene_gn=DMXL2;refGene_tx=XM_019832423.2
B3 54880209 54880329
refGene_gn=DMXL2;refGene_tx=XM_011282863.3;refGene_gn=DMXL2;refGene_tx=XM_019832420.2;refGene_gn=DMXL2;refGene_tx=XM_019832421.2;refGene_gn=DMXL2;refGene_tx=XM_019832423.2
B3 54883629 54883857
refGene_gn=DMXL2;refGene_tx=XM_011282863.3;refGene_gn=DMXL2;refGene_tx=XM_019832420.2;refGene_gn=DMXL2;refGene_tx=XM_019832421.2;refGene_gn=DMXL2;refGene_tx=XM_019832423.2
B3 54886018 54887695
refGene_gn=DMXL2;refGene_tx=XM_011282863.3;refGene_gn=DMXL2;refGene_tx=XM_019832420.2;refGene_gn=DMXL2;refGene_tx=XM_019832421.2;refGene_gn=DMXL2;refGene_tx=XM_019832423.2
B3 54890625 54890736
refGene_gn=DMXL2;refGene_tx=XM_011282863.3;refGene_gn=DMXL2;refGene_tx=XM_019832420.2;refGene_gn=DMXL2;refGene_tx=XM_019832421.2;refGene_gn=DMXL2;refGene_tx=XM_019832423.2
B3 54894770 54894940
refGene_gn=DMXL2;refGene_tx=XM_011282863.3;refGene_gn=DMXL2;refGene_tx=XM_019832420.2;refGene_gn=DMXL2;refGene_tx=XM_019832421.2;refGene_gn=DMXL2;refGene_tx=XM_019832423.2
B3 54898120 54898218

refGene_gn=DMXL2;refGene_tx=XM_011282863.3;refGene_gn=DMXL2;refGene_tx=XM_019832420.2;refGene_gn=DMXL2;refGene_tx=XM_019832421.2;refGene_gn=DMXL2;refGene_tx=XM_019832423.2
B3 54898658 54898824
refGene_gn=DMXL2;refGene_tx=XM_011282863.3;refGene_gn=DMXL2;refGene_tx=XM_019832420.2;refGene_gn=DMXL2;refGene_tx=XM_019832421.2;refGene_gn=DMXL2;refGene_tx=XM_019832423.2
B3 54900160 54900425
refGene_gn=DMXL2;refGene_tx=XM_011282863.3;refGene_gn=DMXL2;refGene_tx=XM_019832420.2;refGene_gn=DMXL2;refGene_tx=XM_019832421.2;refGene_gn=DMXL2;refGene_tx=XM_019832423.2
B3 54904255 54905334
refGene_gn=DMXL2;refGene_tx=XM_011282863.3;refGene_gn=DMXL2;refGene_tx=XM_019832420.2;refGene_gn=DMXL2;refGene_tx=XM_019832421.2;refGene_gn=DMXL2;refGene_tx=XM_019832423.2
B3 54906108 54906300
refGene_gn=DMXL2;refGene_tx=XM_011282863.3;refGene_gn=DMXL2;refGene_tx=XM_019832420.2;refGene_gn=DMXL2;refGene_tx=XM_019832421.2;refGene_gn=DMXL2;refGene_tx=XM_019832423.2
B3 54907852 54907929
refGene_gn=DMXL2;refGene_tx=XM_011282863.3;refGene_gn=DMXL2;refGene_tx=XM_019832420.2;refGene_gn=DMXL2;refGene_tx=XM_019832421.2;refGene_gn=DMXL2;refGene_tx=XM_019832423.2
B3 54909281 54909412
refGene_gn=DMXL2;refGene_tx=XM_011282863.3;refGene_gn=DMXL2;refGene_tx=XM_019832420.2;refGene_gn=DMXL2;refGene_tx=XM_019832421.2;refGene_gn=DMXL2;refGene_tx=XM_019832423.2
B3 54911179 54911428
refGene_gn=DMXL2;refGene_tx=XM_011282863.3;refGene_gn=DMXL2;refGene_tx=XM_019832420.2;refGene_gn=DMXL2;refGene_tx=XM_019832421.2;refGene_gn=DMXL2;refGene_tx=XM_019832423.2
B3 54914828 54915007
refGene_gn=DMXL2;refGene_tx=XM_011282863.3;refGene_gn=DMXL2;refGene_tx=XM_019832420.2;refGene_gn=DMXL2;refGene_tx=XM_019832421.2;refGene_gn=DMXL2;refGene_tx=XM_019832423.2
B3 54917212 54917343
refGene_gn=DMXL2;refGene_tx=XM_011282863.3;refGene_gn=DMXL2;refGene_tx=XM_019832420.2;refGene_gn=DMXL2;refGene_tx=XM_019832421.2;refGene_gn=DMXL2;refGene_tx=XM_019832423.2
B3 54917935 54918021
refGene_gn=DMXL2;refGene_tx=XM_011282863.3;refGene_gn=DMXL2;refGene_tx=XM_019832420.2;refGene_gn=DMXL2;refGene_tx=XM_019832421.2;refGene_gn=DMXL2;refGene_tx=XM_019832423.2
B3 54918423 54918625
refGene_gn=DMXL2;refGene_tx=XM_011282863.3;refGene_gn=DMXL2;refGene_tx=XM_019832420.2;refGene_gn=DMXL2;refGene_tx=XM_019832421.2;refGene_gn=DMXL2;refGene_tx=XM_019832423.2
B3 54919465 54919583
refGene_gn=DMXL2;refGene_tx=XM_011282863.3;refGene_gn=DMXL2;refGene_tx=XM_019832420.2;refGene_gn=DMXL2;refGene_tx=XM_019832421.2;refGene_gn=

DMXL2;refGene_tx=XM_019832423.2
B3	54922972	54923035
refGene_gn=DMXL2;refGene_tx=XM_011282863.3;refGene_gn=DMXL2;refGene_tx=XM_019832420.2
B3	54923836	54923923
refGene_gn=DMXL2;refGene_tx=XM_011282863.3;refGene_gn=DMXL2;refGene_tx=XM_019832420.2;refGene_gn=DMXL2;refGene_tx=XM_019832421.2;refGene_gn=DMXL2;refGene_tx=XM_019832423.2
B3	54924005	54924127
refGene_gn=DMXL2;refGene_tx=XM_011282863.3;refGene_gn=DMXL2;refGene_tx=XM_019832420.2;refGene_gn=DMXL2;refGene_tx=XM_019832421.2;refGene_gn=DMXL2;refGene_tx=XM_019832423.2
B3	54925506	54925645
refGene_gn=DMXL2;refGene_tx=XM_011282863.3;refGene_gn=DMXL2;refGene_tx=XM_019832420.2;refGene_gn=DMXL2;refGene_tx=XM_019832421.2;refGene_gn=DMXL2;refGene_tx=XM_019832423.2
B3	54926306	54926367
refGene_gn=DMXL2;refGene_tx=XM_011282863.3;refGene_gn=DMXL2;refGene_tx=XM_019832420.2;refGene_gn=DMXL2;refGene_tx=XM_019832421.2;refGene_gn=DMXL2;refGene_tx=XM_019832423.2
B3	54926500	54926628
refGene_gn=DMXL2;refGene_tx=XM_011282863.3;refGene_gn=DMXL2;refGene_tx=XM_019832420.2;refGene_gn=DMXL2;refGene_tx=XM_019832421.2;refGene_gn=DMXL2;refGene_tx=XM_019832423.2
B3	54927135	54927213
refGene_gn=DMXL2;refGene_tx=XM_011282863.3;refGene_gn=DMXL2;refGene_tx=XM_019832420.2;refGene_gn=DMXL2;refGene_tx=XM_019832421.2;refGene_gn=DMXL2;refGene_tx=XM_019832423.2
B3	54928871	54928963
refGene_gn=DMXL2;refGene_tx=XM_011282863.3;refGene_gn=DMXL2;refGene_tx=XM_019832420.2;refGene_gn=DMXL2;refGene_tx=XM_019832421.2;refGene_gn=DMXL2;refGene_tx=XM_019832423.2
B3	54929612	54929665
refGene_gn=DMXL2;refGene_tx=XM_011282863.3;refGene_gn=DMXL2;refGene_tx=XM_019832420.2;refGene_gn=DMXL2;refGene_tx=XM_019832421.2;refGene_gn=DMXL2;refGene_tx=XM_019832423.2
B3	54930887	54931105
refGene_gn=DMXL2;refGene_tx=XM_011282863.3;refGene_gn=DMXL2;refGene_tx=XM_019832420.2;refGene_gn=DMXL2;refGene_tx=XM_019832421.2;refGene_gn=DMXL2;refGene_tx=XM_019832423.2
B3	54931777	54931987
refGene_gn=DMXL2;refGene_tx=XM_011282863.3;refGene_gn=DMXL2;refGene_tx=XM_019832420.2;refGene_gn=DMXL2;refGene_tx=XM_019832421.2;refGene_gn=DMXL2;refGene_tx=XM_019832423.2
B3	54946027	54946505	refGene_gn=GLDN;refGene_tx=XM_003987139.5
B3	54948862	54949013	refGene_gn=GLDN;refGene_tx=XM_003987139.5
B3	54950022	54950148	refGene_gn=GLDN;refGene_tx=XM_003987139.5
B3	54950245	54950329	refGene_gn=GLDN;refGene_tx=XM_003987139.5
B3	54953081	54953219	refGene_gn=GLDN;refGene_tx=XM_003987139.5
B3	54955233	54955371	refGene_gn=GLDN;refGene_tx=XM_003987139.5

B3 54963978 54964086 refGene_gn=GLDN;refGene_tx=XM_003987139.5
B3 54964454 54964472 refGene_gn=GLDN;refGene_tx=XM_003987139.5
B3 54972691 54972743 refGene_gn=GLDN;refGene_tx=XM_003987139.5
B3 55005095 55005458 refGene_gn=GLDN;refGene_tx=XM_003987139.5
B3 55104888 55105033
refGene_gn=LOC101094932;refGene_tx=XM_006932525.3
B3 55111019 55111170
refGene_gn=LOC101094932;refGene_tx=XM_006932525.3
B3 55122742 55122897
refGene_gn=LOC101094932;refGene_tx=XM_006932525.3
B3 55129622 55129799
refGene_gn=LOC101094932;refGene_tx=XM_006932525.3
B3 55132753 55132868
refGene_gn=LOC101094932;refGene_tx=XM_006932525.3
B3 55136885 55137000
refGene_gn=LOC101094932;refGene_tx=XM_006932525.3
B3 55137439 55137602
refGene_gn=LOC101094932;refGene_tx=XM_006932525.3
B3 55140693 55140935
refGene_gn=LOC101094932;refGene_tx=XM_006932525.3
B3 55142138 55142396
refGene_gn=LOC101094932;refGene_tx=XM_006932525.3
B3 55247467 55247519
refGene_gn=TNFAIP8L3;refGene_tx=XM_019832430.2
B3 55269600 55269649
refGene_gn=TNFAIP8L3;refGene_tx=XM_023255365.1
B3 55281850 55282413
refGene_gn=TNFAIP8L3;refGene_tx=XM_019832430.2;refGene_gn=TNFAIP8L3;re
fGene_tx=XM_023255365.1
B3 55304228 55304243 refGene_gn=AP4E1;refGene_tx=XM_023255367.1
B3 55307513 55307674
refGene_gn=AP4E1;refGene_tx=XM_023255366.1;refGene_gn=AP4E1;refGene_tx
=XM_023255367.1;refGene_gn=AP4E1;refGene_tx=XM_023255368.1
B3 55308726 55308884
refGene_gn=AP4E1;refGene_tx=XM_023255366.1;refGene_gn=AP4E1;refGene_tx
=XM_023255367.1;refGene_gn=AP4E1;refGene_tx=XM_023255368.1
B3 55310394 55310585
refGene_gn=AP4E1;refGene_tx=XM_023255366.1;refGene_gn=AP4E1;refGene_tx
=XM_023255367.1;refGene_gn=AP4E1;refGene_tx=XM_023255368.1
B3 55312210 55312765
refGene_gn=AP4E1;refGene_tx=XM_023255366.1;refGene_gn=AP4E1;refGene_tx
=XM_023255367.1;refGene_gn=AP4E1;refGene_tx=XM_023255368.1
B3 55316073 55316332
refGene_gn=AP4E1;refGene_tx=XM_023255366.1;refGene_gn=AP4E1;refGene_tx
=XM_023255367.1;refGene_gn=AP4E1;refGene_tx=XM_023255368.1
B3 55321261 55321385
refGene_gn=AP4E1;refGene_tx=XM_023255366.1;refGene_gn=AP4E1;refGene_tx
=XM_023255367.1;refGene_gn=AP4E1;refGene_tx=XM_023255368.1
B3 55325686 55325801
refGene_gn=AP4E1;refGene_tx=XM_023255366.1;refGene_gn=AP4E1;refGene_tx

=XM_023255367.1;refGene_gn=AP4E1;refGene_tx=XM_023255368.1
B3 55328473 55328776
refGene_gn=AP4E1;refGene_tx=XM_023255366.1;refGene_gn=AP4E1;refGene_tx=XM_023255367.1;refGene_gn=AP4E1;refGene_tx=XM_023255368.1
B3 55333175 55333294
refGene_gn=AP4E1;refGene_tx=XM_023255366.1;refGene_gn=AP4E1;refGene_tx=XM_023255367.1;refGene_gn=AP4E1;refGene_tx=XM_023255368.1
B3 55333402 55333515
refGene_gn=AP4E1;refGene_tx=XM_023255366.1;refGene_gn=AP4E1;refGene_tx=XM_023255367.1;refGene_gn=AP4E1;refGene_tx=XM_023255368.1
B3 55335239 55335379
refGene_gn=AP4E1;refGene_tx=XM_023255366.1;refGene_gn=AP4E1;refGene_tx=XM_023255367.1;refGene_gn=AP4E1;refGene_tx=XM_023255368.1
B3 55337858 55337968
refGene_gn=AP4E1;refGene_tx=XM_023255366.1;refGene_gn=AP4E1;refGene_tx=XM_023255367.1;refGene_gn=AP4E1;refGene_tx=XM_023255368.1
B3 55338071 55338194
refGene_gn=AP4E1;refGene_tx=XM_023255366.1;refGene_gn=AP4E1;refGene_tx=XM_023255367.1;refGene_gn=AP4E1;refGene_tx=XM_023255368.1
B3 55346215 55346289
refGene_gn=AP4E1;refGene_tx=XM_023255366.1;refGene_gn=AP4E1;refGene_tx=XM_023255367.1;refGene_gn=AP4E1;refGene_tx=XM_023255368.1
B3 55349538 55349705
refGene_gn=AP4E1;refGene_tx=XM_023255366.1;refGene_gn=AP4E1;refGene_tx=XM_023255367.1;refGene_gn=AP4E1;refGene_tx=XM_023255368.1
B3 55353544 55353704
refGene_gn=AP4E1;refGene_tx=XM_023255366.1;refGene_gn=AP4E1;refGene_tx=XM_023255367.1;refGene_gn=AP4E1;refGene_tx=XM_023255368.1
B3 55357095 55357217
refGene_gn=AP4E1;refGene_tx=XM_023255366.1;refGene_gn=AP4E1;refGene_tx=XM_023255367.1;refGene_gn=AP4E1;refGene_tx=XM_023255368.1
B3 55358472 55358546
refGene_gn=AP4E1;refGene_tx=XM_023255366.1;refGene_gn=AP4E1;refGene_tx=XM_023255367.1
B3 55362108 55362232
refGene_gn=AP4E1;refGene_tx=XM_023255366.1;refGene_gn=AP4E1;refGene_tx=XM_023255367.1
B3 55365231 55365303
refGene_gn=AP4E1;refGene_tx=XM_023255366.1;refGene_gn=AP4E1;refGene_tx=XM_023255367.1
B3 55368140 55368290
refGene_gn=AP4E1;refGene_tx=XM_023255366.1;refGene_gn=AP4E1;refGene_tx=XM_023255367.1
B3 55489741 55489768
refGene_gn=SPPL2A;refGene_tx=XM_019832433.2;refGene_gn=SPPL2A;refGene_tx=XM_019832434.2;refGene_gn=SPPL2A;refGene_tx=XM_019832435.2
B3 55498230 55498350
refGene_gn=SPPL2A;refGene_tx=XM_019832433.2;refGene_gn=SPPL2A;refGene_tx=XM_019832434.2;refGene_gn=SPPL2A;refGene_tx=XM_019832435.2
B3 55498658 55498841

refGene_gn=SPPL2A;refGene_tx=XM_019832433.2;refGene_gn=SPPL2A;refGene_tx=XM_019832434.2
B3 55499559 55499649
refGene_gn=SPPL2A;refGene_tx=XM_019832433.2;refGene_gn=SPPL2A;refGene_tx=XM_019832434.2;refGene_gn=SPPL2A;refGene_tx=XM_019832435.2
B3 55500132 55500266
refGene_gn=SPPL2A;refGene_tx=XM_019832433.2;refGene_gn=SPPL2A;refGene_tx=XM_019832434.2;refGene_gn=SPPL2A;refGene_tx=XM_019832435.2
B3 55506870 55507022
refGene_gn=SPPL2A;refGene_tx=XM_019832433.2;refGene_gn=SPPL2A;refGene_tx=XM_019832434.2;refGene_gn=SPPL2A;refGene_tx=XM_019832435.2
B3 55510294 55510391
refGene_gn=SPPL2A;refGene_tx=XM_019832433.2;refGene_gn=SPPL2A;refGene_tx=XM_019832434.2;refGene_gn=SPPL2A;refGene_tx=XM_019832435.2
B3 55510812 55510914
refGene_gn=SPPL2A;refGene_tx=XM_019832433.2;refGene_gn=SPPL2A;refGene_tx=XM_019832434.2;refGene_gn=SPPL2A;refGene_tx=XM_019832435.2
B3 55512092 55512174
refGene_gn=SPPL2A;refGene_tx=XM_019832433.2;refGene_gn=SPPL2A;refGene_tx=XM_019832434.2;refGene_gn=SPPL2A;refGene_tx=XM_019832435.2
B3 55512715 55512790
refGene_gn=SPPL2A;refGene_tx=XM_019832433.2;refGene_gn=SPPL2A;refGene_tx=XM_019832434.2;refGene_gn=SPPL2A;refGene_tx=XM_019832435.2
B3 55516155 55516212
refGene_gn=SPPL2A;refGene_tx=XM_019832433.2;refGene_gn=SPPL2A;refGene_tx=XM_019832434.2;refGene_gn=SPPL2A;refGene_tx=XM_019832435.2
B3 55516407 55516461
refGene_gn=SPPL2A;refGene_tx=XM_019832433.2;refGene_gn=SPPL2A;refGene_tx=XM_019832435.2
B3 55516899 55517002
refGene_gn=SPPL2A;refGene_tx=XM_019832433.2;refGene_gn=SPPL2A;refGene_tx=XM_019832434.2;refGene_gn=SPPL2A;refGene_tx=XM_019832435.2
B3 55518607 55518685
refGene_gn=SPPL2A;refGene_tx=XM_019832433.2;refGene_gn=SPPL2A;refGene_tx=XM_019832434.2;refGene_gn=SPPL2A;refGene_tx=XM_019832435.2
B3 55520073 55520234
refGene_gn=SPPL2A;refGene_tx=XM_019832433.2;refGene_gn=SPPL2A;refGene_tx=XM_019832434.2;refGene_gn=SPPL2A;refGene_tx=XM_019832435.2
B3 55530747 55530822
refGene_gn=SPPL2A;refGene_tx=XM_019832433.2;refGene_gn=SPPL2A;refGene_tx=XM_019832434.2;refGene_gn=SPPL2A;refGene_tx=XM_019832435.2
B3 55552978 55552981
refGene_gn=TRPM7;refGene_tx=XM_011282861.3;refGene_gn=TRPM7;refGene_tx=XM_019832437.2;refGene_gn=TRPM7;refGene_tx=XM_019832438.2;refGene_gn=TRPM7;refGene_tx=XM_023255369.1;refGene_gn=TRPM7;refGene_tx=XM_023255370.1
B3 55574193 55574273
refGene_gn=TRPM7;refGene_tx=XM_011282861.3;refGene_gn=TRPM7;refGene_tx=XM_019832437.2;refGene_gn=TRPM7;refGene_tx=XM_019832438.2;refGene_gn=TRPM7;refGene_tx=XM_023255369.1;refGene_gn=TRPM7;refGene_tx=XM_0232553

70.1
B3	55580981 55581020
refGene_gn=TRPM7;refGene_tx=XM_011282861.3;refGene_gn=TRPM7;refGene_tx=XM_019832437.2;refGene_gn=TRPM7;refGene_tx=XM_019832438.2;refGene_gn=TRPM7;refGene_tx=XM_023255369.1;refGene_gn=TRPM7;refGene_tx=XM_023255370.1
B3	55588811 55589010
refGene_gn=TRPM7;refGene_tx=XM_011282861.3;refGene_gn=TRPM7;refGene_tx=XM_019832437.2;refGene_gn=TRPM7;refGene_tx=XM_019832438.2;refGene_gn=TRPM7;refGene_tx=XM_023255369.1;refGene_gn=TRPM7;refGene_tx=XM_023255370.1
B3	55592297 55592511
refGene_gn=TRPM7;refGene_tx=XM_011282861.3;refGene_gn=TRPM7;refGene_tx=XM_019832437.2;refGene_gn=TRPM7;refGene_tx=XM_019832438.2;refGene_gn=TRPM7;refGene_tx=XM_023255369.1;refGene_gn=TRPM7;refGene_tx=XM_023255370.1
B3	55594756 55594881
refGene_gn=TRPM7;refGene_tx=XM_011282861.3;refGene_gn=TRPM7;refGene_tx=XM_019832437.2;refGene_gn=TRPM7;refGene_tx=XM_019832438.2;refGene_gn=TRPM7;refGene_tx=XM_023255369.1;refGene_gn=TRPM7;refGene_tx=XM_023255370.1
B3	55596563 55596735
refGene_gn=TRPM7;refGene_tx=XM_011282861.3;refGene_gn=TRPM7;refGene_tx=XM_019832437.2;refGene_gn=TRPM7;refGene_tx=XM_019832438.2;refGene_gn=TRPM7;refGene_tx=XM_023255369.1;refGene_gn=TRPM7;refGene_tx=XM_023255370.1
B3	55598341 55598516
refGene_gn=TRPM7;refGene_tx=XM_011282861.3;refGene_gn=TRPM7;refGene_tx=XM_019832437.2;refGene_gn=TRPM7;refGene_tx=XM_019832438.2;refGene_gn=TRPM7;refGene_tx=XM_023255369.1;refGene_gn=TRPM7;refGene_tx=XM_023255370.1
B3	55599470 55599594
refGene_gn=TRPM7;refGene_tx=XM_011282861.3;refGene_gn=TRPM7;refGene_tx=XM_019832437.2;refGene_gn=TRPM7;refGene_tx=XM_019832438.2;refGene_gn=TRPM7;refGene_tx=XM_023255369.1;refGene_gn=TRPM7;refGene_tx=XM_023255370.1
B3	55600945 55601018
refGene_gn=TRPM7;refGene_tx=XM_011282861.3;refGene_gn=TRPM7;refGene_tx=XM_019832437.2;refGene_gn=TRPM7;refGene_tx=XM_019832438.2;refGene_gn=TRPM7;refGene_tx=XM_023255369.1;refGene_gn=TRPM7;refGene_tx=XM_023255370.1
B3	55602818 55602919
refGene_gn=TRPM7;refGene_tx=XM_011282861.3;refGene_gn=TRPM7;refGene_tx=XM_019832437.2;refGene_gn=TRPM7;refGene_tx=XM_019832438.2;refGene_gn=TRPM7;refGene_tx=XM_023255369.1;refGene_gn=TRPM7;refGene_tx=XM_023255370.1
B3	55609635 55609770
refGene_gn=TRPM7;refGene_tx=XM_011282861.3;refGene_gn=TRPM7;refGene_tx=XM_019832437.2;refGene_gn=TRPM7;refGene_tx=XM_019832438.2;refGene_gn=TRPM7;refGene_tx=XM_023255369.1;refGene_gn=TRPM7;refGene_tx=XM_0232553

70.1
B3	55620237 55620291
refGene_gn=TRPM7;refGene_tx=XM_011282861.3;refGene_gn=TRPM7;refGene_tx=XM_019832437.2;refGene_gn=TRPM7;refGene_tx=XM_019832438.2;refGene_gn=TRPM7;refGene_tx=XM_023255369.1
B3	55624489 55624630
refGene_gn=TRPM7;refGene_tx=XM_011282861.3;refGene_gn=TRPM7;refGene_tx=XM_019832437.2;refGene_gn=TRPM7;refGene_tx=XM_019832438.2;refGene_gn=TRPM7;refGene_tx=XM_023255369.1;refGene_gn=TRPM7;refGene_tx=XM_023255370.1
B3	55625020 55625155
refGene_gn=TRPM7;refGene_tx=XM_011282861.3;refGene_gn=TRPM7;refGene_tx=XM_019832437.2;refGene_gn=TRPM7;refGene_tx=XM_019832438.2;refGene_gn=TRPM7;refGene_tx=XM_023255369.1;refGene_gn=TRPM7;refGene_tx=XM_023255370.1
B3	55626609 55626890
refGene_gn=TRPM7;refGene_tx=XM_011282861.3;refGene_gn=TRPM7;refGene_tx=XM_019832437.2;refGene_gn=TRPM7;refGene_tx=XM_019832438.2;refGene_gn=TRPM7;refGene_tx=XM_023255369.1;refGene_gn=TRPM7;refGene_tx=XM_023255370.1
B3	55627902 55628131
refGene_gn=TRPM7;refGene_tx=XM_011282861.3;refGene_gn=TRPM7;refGene_tx=XM_019832437.2;refGene_gn=TRPM7;refGene_tx=XM_019832438.2;refGene_gn=TRPM7;refGene_tx=XM_023255369.1;refGene_gn=TRPM7;refGene_tx=XM_023255370.1
B3	55629501 55629657
refGene_gn=TRPM7;refGene_tx=XM_011282861.3;refGene_gn=TRPM7;refGene_tx=XM_019832437.2;refGene_gn=TRPM7;refGene_tx=XM_019832438.2;refGene_gn=TRPM7;refGene_tx=XM_023255369.1;refGene_gn=TRPM7;refGene_tx=XM_023255370.1
B3	55629734 55629878
refGene_gn=TRPM7;refGene_tx=XM_011282861.3;refGene_gn=TRPM7;refGene_tx=XM_019832437.2;refGene_gn=TRPM7;refGene_tx=XM_019832438.2;refGene_gn=TRPM7;refGene_tx=XM_023255369.1;refGene_gn=TRPM7;refGene_tx=XM_023255370.1
B3	55632101 55632230
refGene_gn=TRPM7;refGene_tx=XM_011282861.3;refGene_gn=TRPM7;refGene_tx=XM_019832437.2;refGene_gn=TRPM7;refGene_tx=XM_019832438.2;refGene_gn=TRPM7;refGene_tx=XM_023255369.1;refGene_gn=TRPM7;refGene_tx=XM_023255370.1
B3	55634369 55634648
refGene_gn=TRPM7;refGene_tx=XM_011282861.3;refGene_gn=TRPM7;refGene_tx=XM_019832437.2;refGene_gn=TRPM7;refGene_tx=XM_019832438.2;refGene_gn=TRPM7;refGene_tx=XM_023255369.1;refGene_gn=TRPM7;refGene_tx=XM_023255370.1
B3	55636845 55637020
refGene_gn=TRPM7;refGene_tx=XM_011282861.3;refGene_gn=TRPM7;refGene_tx=XM_019832437.2;refGene_gn=TRPM7;refGene_tx=XM_019832438.2;refGene_gn=TRPM7;refGene_tx=XM_023255369.1;refGene_gn=TRPM7;refGene_tx=XM_023255370.1

B3 55639271 55639398
refGene_gn=TRPM7;refGene_tx=XM_011282861.3;refGene_gn=TRPM7;refGene_tx=XM_019832437.2;refGene_gn=TRPM7;refGene_tx=XM_019832438.2;refGene_gn=TRPM7;refGene_tx=XM_023255369.1;refGene_gn=TRPM7;refGene_tx=XM_023255370.1
B3 55640631 55640816
refGene_gn=TRPM7;refGene_tx=XM_011282861.3;refGene_gn=TRPM7;refGene_tx=XM_019832437.2;refGene_gn=TRPM7;refGene_tx=XM_019832438.2;refGene_gn=TRPM7;refGene_tx=XM_023255369.1;refGene_gn=TRPM7;refGene_tx=XM_023255370.1
B3 55641486 55641619
refGene_gn=TRPM7;refGene_tx=XM_011282861.3;refGene_gn=TRPM7;refGene_tx=XM_019832437.2;refGene_gn=TRPM7;refGene_tx=XM_019832438.2;refGene_gn=TRPM7;refGene_tx=XM_023255369.1;refGene_gn=TRPM7;refGene_tx=XM_023255370.1
B3 55642228 55642944
refGene_gn=TRPM7;refGene_tx=XM_011282861.3;refGene_gn=TRPM7;refGene_tx=XM_019832437.2;refGene_gn=TRPM7;refGene_tx=XM_019832438.2;refGene_gn=TRPM7;refGene_tx=XM_023255369.1;refGene_gn=TRPM7;refGene_tx=XM_023255370.1
B3 55645625 55645696
refGene_gn=TRPM7;refGene_tx=XM_011282861.3;refGene_gn=TRPM7;refGene_tx=XM_019832437.2;refGene_gn=TRPM7;refGene_tx=XM_019832438.2;refGene_gn=TRPM7;refGene_tx=XM_023255369.1;refGene_gn=TRPM7;refGene_tx=XM_023255370.1
B3 55648976 55649073
refGene_gn=TRPM7;refGene_tx=XM_011282861.3;refGene_gn=TRPM7;refGene_tx=XM_019832437.2;refGene_gn=TRPM7;refGene_tx=XM_019832438.2;refGene_gn=TRPM7;refGene_tx=XM_023255369.1;refGene_gn=TRPM7;refGene_tx=XM_023255370.1
B3 55652343 55652414
refGene_gn=TRPM7;refGene_tx=XM_011282861.3;refGene_gn=TRPM7;refGene_tx=XM_019832437.2;refGene_gn=TRPM7;refGene_tx=XM_019832438.2;refGene_gn=TRPM7;refGene_tx=XM_023255369.1;refGene_gn=TRPM7;refGene_tx=XM_023255370.1
B3 55655876 55655911
refGene_gn=TRPM7;refGene_tx=XM_011282861.3;refGene_gn=TRPM7;refGene_tx=XM_019832437.2;refGene_gn=TRPM7;refGene_tx=XM_019832438.2;refGene_gn=TRPM7;refGene_tx=XM_023255369.1;refGene_gn=TRPM7;refGene_tx=XM_023255370.1
B3 55656993 55657019
refGene_gn=TRPM7;refGene_tx=XM_011282861.3;refGene_gn=TRPM7;refGene_tx=XM_019832437.2;refGene_gn=TRPM7;refGene_tx=XM_019832438.2;refGene_gn=TRPM7;refGene_tx=XM_023255369.1;refGene_gn=TRPM7;refGene_tx=XM_023255370.1
B3 55658207 55658308 refGene_gn=TRPM7;refGene_tx=XM_019832438.2
B3 55659269 55659320
refGene_gn=TRPM7;refGene_tx=XM_011282861.3;refGene_gn=TRPM7;refGene_tx=XM_019832437.2;refGene_gn=TRPM7;refGene_tx=XM_023255369.1;refGene_gn=TRPM7;refGene_tx=XM_023255370.1

B3	55659402	55659468
refGene_gn=TRPM7;refGene_tx=XM_011282861.3;refGene_gn=TRPM7;refGene_tx=XM_019832437.2;refGene_gn=TRPM7;refGene_tx=XM_023255369.1;refGene_gn=TRPM7;refGene_tx=XM_023255370.1
B3	55660040	55660324
refGene_gn=TRPM7;refGene_tx=XM_011282861.3;refGene_gn=TRPM7;refGene_tx=XM_019832437.2;refGene_gn=TRPM7;refGene_tx=XM_023255369.1;refGene_gn=TRPM7;refGene_tx=XM_023255370.1
B3	55660468	55660551
refGene_gn=TRPM7;refGene_tx=XM_011282861.3;refGene_gn=TRPM7;refGene_tx=XM_019832437.2;refGene_gn=TRPM7;refGene_tx=XM_023255369.1;refGene_gn=TRPM7;refGene_tx=XM_023255370.1
B3	55660714	55660920
refGene_gn=TRPM7;refGene_tx=XM_011282861.3;refGene_gn=TRPM7;refGene_tx=XM_019832437.2;refGene_gn=TRPM7;refGene_tx=XM_023255369.1;refGene_gn=TRPM7;refGene_tx=XM_023255370.1
B3	55664491	55664543
refGene_gn=TRPM7;refGene_tx=XM_023255369.1;refGene_gn=TRPM7;refGene_tx=XM_023255370.1
B3	55665461	55665513	refGene_gn=TRPM7;refGene_tx=XM_011282861.3
B3	55665634	55665741
refGene_gn=TRPM7;refGene_tx=XM_011282861.3;refGene_gn=TRPM7;refGene_tx=XM_023255369.1;refGene_gn=TRPM7;refGene_tx=XM_023255370.1
B3	55674507	55674638
refGene_gn=TRPM7;refGene_tx=XM_011282861.3;refGene_gn=TRPM7;refGene_tx=XM_019832437.2;refGene_gn=TRPM7;refGene_tx=XM_023255369.1;refGene_gn=TRPM7;refGene_tx=XM_023255370.1
B3	55687286	55687339	refGene_gn=USP50;refGene_tx=XM_003987182.5
B3	55689109	55689304
refGene_gn=USP50;refGene_tx=XM_003987182.5;refGene_gn=USP50;refGene_tx=XM_019832439.2
B3	55689544	55689740
refGene_gn=USP50;refGene_tx=XM_003987182.5;refGene_gn=USP50;refGene_tx=XM_019832439.2
B3	55691359	55691566
refGene_gn=USP50;refGene_tx=XM_003987182.5;refGene_gn=USP50;refGene_tx=XM_019832439.2
B3	55692821	55692964
refGene_gn=USP50;refGene_tx=XM_003987182.5;refGene_gn=USP50;refGene_tx=XM_019832439.2
B3	55699750	55699883
refGene_gn=USP50;refGene_tx=XM_003987182.5;refGene_gn=USP50;refGene_tx=XM_019832439.2
B3	55727806	55727998
refGene_gn=USP50;refGene_tx=XM_003987182.5;refGene_gn=USP50;refGene_tx=XM_019832439.2
B3	55729607	55729793	refGene_gn=USP8;refGene_tx=XM_023255371.1
B3	55729961	55730094	refGene_gn=USP8;refGene_tx=XM_023255371.1
B3	55732053	55732196	refGene_gn=USP8;refGene_tx=XM_023255371.1
B3	55733327	55733564	refGene_gn=USP8;refGene_tx=XM_023255371.1

B3 55735612 55735823 refGene_gn=USP8;refGene_tx=XM_023255371.1
B3 55736318 55736531 refGene_gn=USP8;refGene_tx=XM_023255371.1
B3 55738027 55738290 refGene_gn=USP8;refGene_tx=XM_023255371.1
B3 55738709 55738787 refGene_gn=USP8;refGene_tx=XM_023255371.1
B3 55740682 55740769 refGene_gn=USP8;refGene_tx=XM_023255371.1
B3 55743375 55743960 refGene_gn=USP8;refGene_tx=XM_023255371.1
B3 55759529 55759753 refGene_gn=USP8;refGene_tx=XM_023255371.1
B3 55760039 55760184 refGene_gn=USP8;refGene_tx=XM_023255371.1
B3 55763650 55763813 refGene_gn=USP8;refGene_tx=XM_023255371.1
B3 55769058 55769203 refGene_gn=USP8;refGene_tx=XM_023255371.1
B3 55771036 55771079 refGene_gn=USP8;refGene_tx=XM_023255371.1
B3 55773323 55773483 refGene_gn=USP8;refGene_tx=XM_023255371.1
B3 55774752 55774838 refGene_gn=USP8;refGene_tx=XM_023255371.1
B3 55781127 55781272 refGene_gn=USP8;refGene_tx=XM_023255371.1
B3 55783951 55784055 refGene_gn=USP8;refGene_tx=XM_023255371.1
B3 55883217 55883325
refGene_gn=GABPB1;refGene_tx=XM_003987146.5;refGene_gn=GABPB1;refGene_tx=XM_006932529.4;refGene_gn=GABPB1;refGene_tx=XM_023255372.1;refGene_gn=GABPB1;refGene_tx=XM_023255373.1;refGene_gn=GABPB1;refGene_tx=XM_023255374.1;refGene_gn=GABPB1;refGene_tx=XM_023255375.1
B3 55887637 55887805
refGene_gn=GABPB1;refGene_tx=XM_003987146.5;refGene_gn=GABPB1;refGene_tx=XM_006932529.4;refGene_gn=GABPB1;refGene_tx=XM_023255372.1;refGene_gn=GABPB1;refGene_tx=XM_023255373.1;refGene_gn=GABPB1;refGene_tx=XM_023255374.1;refGene_gn=GABPB1;refGene_tx=XM_023255375.1
B3 55889008 55889203
refGene_gn=GABPB1;refGene_tx=XM_003987146.5;refGene_gn=GABPB1;refGene_tx=XM_006932529.4;refGene_gn=GABPB1;refGene_tx=XM_023255372.1;refGene_gn=GABPB1;refGene_tx=XM_023255373.1;refGene_gn=GABPB1;refGene_tx=XM_023255374.1;refGene_gn=GABPB1;refGene_tx=XM_023255375.1
B3 55890695 55890843
refGene_gn=GABPB1;refGene_tx=XM_003987146.5;refGene_gn=GABPB1;refGene_tx=XM_006932529.4;refGene_gn=GABPB1;refGene_tx=XM_023255372.1;refGene_gn=GABPB1;refGene_tx=XM_023255373.1;refGene_gn=GABPB1;refGene_tx=XM_023255374.1;refGene_gn=GABPB1;refGene_tx=XM_023255375.1
B3 55891195 55891309
refGene_gn=GABPB1;refGene_tx=XM_003987146.5;refGene_gn=GABPB1;refGene_tx=XM_006932529.4;refGene_gn=GABPB1;refGene_tx=XM_023255372.1;refGene_gn=GABPB1;refGene_tx=XM_023255373.1;refGene_gn=GABPB1;refGene_tx=XM_023255374.1;refGene_gn=GABPB1;refGene_tx=XM_023255375.1
B3 55900161 55900347
refGene_gn=GABPB1;refGene_tx=XM_003987146.5;refGene_gn=GABPB1;refGene_tx=XM_006932529.4;refGene_gn=GABPB1;refGene_tx=XM_023255372.1;refGene_gn=GABPB1;refGene_tx=XM_023255373.1;refGene_gn=GABPB1;refGene_tx=XM_023255374.1;refGene_gn=GABPB1;refGene_tx=XM_023255375.1
B3 55903410 55903545
refGene_gn=GABPB1;refGene_tx=XM_003987146.5;refGene_gn=GABPB1;refGene_tx=XM_006932529.4;refGene_gn=GABPB1;refGene_tx=XM_023255372.1;refGene_gn=GABPB1;refGene_tx=XM_023255373.1;refGene_gn=GABPB1;refGene_tx=XM_023255374.1;refGene_gn=GABPB1;refGene_tx=XM_023255375.1

B3 55911182 55911335
refGene_gn=GABPB1;refGene_tx=XM_003987146.5;refGene_gn=GABPB1;refGene_tx=XM_006932529.4;refGene_gn=GABPB1;refGene_tx=XM_023255372.1;refGene_gn=GABPB1;refGene_tx=XM_023255373.1;refGene_gn=GABPB1;refGene_tx=XM_023255374.1;refGene_gn=GABPB1;refGene_tx=XM_023255375.1
B3 55933930 55933958 refGene_gn=HDC;refGene_tx=XM_023255376.1
B3 55936155 55936328
refGene_gn=HDC;refGene_tx=XM_023255376.1;refGene_gn=HDC;refGene_tx=XM_023255377.1
B3 55940472 55940586
refGene_gn=HDC;refGene_tx=XM_023255376.1;refGene_gn=HDC;refGene_tx=XM_023255377.1
B3 55941520 55941643
refGene_gn=HDC;refGene_tx=XM_023255376.1;refGene_gn=HDC;refGene_tx=XM_023255377.1
B3 55944213 55944348
refGene_gn=HDC;refGene_tx=XM_023255376.1;refGene_gn=HDC;refGene_tx=XM_023255377.1
B3 55944612 55944756
refGene_gn=HDC;refGene_tx=XM_023255376.1;refGene_gn=HDC;refGene_tx=XM_023255377.1
B3 55945214 55945281
refGene_gn=HDC;refGene_tx=XM_023255376.1;refGene_gn=HDC;refGene_tx=XM_023255377.1
B3 55946110 55946273
refGene_gn=HDC;refGene_tx=XM_023255376.1;refGene_gn=HDC;refGene_tx=XM_023255377.1
B3 55946363 55946454
refGene_gn=HDC;refGene_tx=XM_023255376.1;refGene_gn=HDC;refGene_tx=XM_023255377.1
B3 55950492 55950591
refGene_gn=HDC;refGene_tx=XM_023255376.1;refGene_gn=HDC;refGene_tx=XM_023255377.1
B3 55954559 55954661
refGene_gn=HDC;refGene_tx=XM_023255376.1;refGene_gn=HDC;refGene_tx=XM_023255377.1
B3 55954792 55955536
refGene_gn=HDC;refGene_tx=XM_023255376.1;refGene_gn=HDC;refGene_tx=XM_023255377.1
B3 55959684 55959861 refGene_gn=SLC27A2;refGene_tx=XM_003987149.4
B3 55961034 55961165 refGene_gn=SLC27A2;refGene_tx=XM_003987149.4
B3 55965337 55965435 refGene_gn=SLC27A2;refGene_tx=XM_003987149.4
B3 55966935 55967134 refGene_gn=SLC27A2;refGene_tx=XM_003987149.4
B3 55969287 55969378 refGene_gn=SLC27A2;refGene_tx=XM_003987149.4
B3 55973206 55973401 refGene_gn=SLC27A2;refGene_tx=XM_003987149.4
B3 55985652 55985777 refGene_gn=SLC27A2;refGene_tx=XM_003987149.4
B3 55988057 55988216 refGene_gn=SLC27A2;refGene_tx=XM_003987149.4
B3 55993291 55993501 refGene_gn=SLC27A2;refGene_tx=XM_003987149.4
B3 56002345 56002823 refGene_gn=SLC27A2;refGene_tx=XM_003987149.4
B3 56054587 56054615

refGene_gn=ATP8B4;refGene_tx=XM_019832446.2;refGene_gn=ATP8B4;refGene_tx=XM_019832447.2;refGene_gn=ATP8B4;refGene_tx=XM_019832448.2;refGene_gn=ATP8B4;refGene_tx=XM_023255378.1;refGene_gn=ATP8B4;refGene_tx=XM_023255380.1
B3 56085169 56085228
refGene_gn=ATP8B4;refGene_tx=XM_019832446.2;refGene_gn=ATP8B4;refGene_tx=XM_019832447.2;refGene_gn=ATP8B4;refGene_tx=XM_019832448.2;refGene_gn=ATP8B4;refGene_tx=XM_023255378.1;refGene_gn=ATP8B4;refGene_tx=XM_023255380.1
B3 56112224 56112338
refGene_gn=ATP8B4;refGene_tx=XM_019832446.2;refGene_gn=ATP8B4;refGene_tx=XM_019832447.2;refGene_gn=ATP8B4;refGene_tx=XM_019832448.2;refGene_gn=ATP8B4;refGene_tx=XM_023255378.1;refGene_gn=ATP8B4;refGene_tx=XM_023255380.1
B3 56115158 56115257
refGene_gn=ATP8B4;refGene_tx=XM_019832446.2;refGene_gn=ATP8B4;refGene_tx=XM_019832447.2;refGene_gn=ATP8B4;refGene_tx=XM_019832448.2;refGene_gn=ATP8B4;refGene_tx=XM_023255378.1;refGene_gn=ATP8B4;refGene_tx=XM_023255380.1
B3 56119126 56119188
refGene_gn=ATP8B4;refGene_tx=XM_019832446.2;refGene_gn=ATP8B4;refGene_tx=XM_019832447.2;refGene_gn=ATP8B4;refGene_tx=XM_019832448.2;refGene_gn=ATP8B4;refGene_tx=XM_023255378.1;refGene_gn=ATP8B4;refGene_tx=XM_023255380.1
B3 56168412 56168485
refGene_gn=ATP8B4;refGene_tx=XM_019832446.2;refGene_gn=ATP8B4;refGene_tx=XM_019832447.2;refGene_gn=ATP8B4;refGene_tx=XM_019832448.2;refGene_gn=ATP8B4;refGene_tx=XM_023255378.1;refGene_gn=ATP8B4;refGene_tx=XM_023255380.1
B3 56179970 56180041
refGene_gn=ATP8B4;refGene_tx=XM_019832446.2;refGene_gn=ATP8B4;refGene_tx=XM_019832447.2;refGene_gn=ATP8B4;refGene_tx=XM_019832448.2;refGene_gn=ATP8B4;refGene_tx=XM_023255378.1;refGene_gn=ATP8B4;refGene_tx=XM_023255380.1
B3 56183717 56183800
refGene_gn=ATP8B4;refGene_tx=XM_019832446.2;refGene_gn=ATP8B4;refGene_tx=XM_019832447.2;refGene_gn=ATP8B4;refGene_tx=XM_019832448.2;refGene_gn=ATP8B4;refGene_tx=XM_023255378.1;refGene_gn=ATP8B4;refGene_tx=XM_023255380.1
B3 56192857 56193016
refGene_gn=ATP8B4;refGene_tx=XM_019832446.2;refGene_gn=ATP8B4;refGene_tx=XM_019832447.2;refGene_gn=ATP8B4;refGene_tx=XM_019832448.2;refGene_gn=ATP8B4;refGene_tx=XM_023255378.1;refGene_gn=ATP8B4;refGene_tx=XM_023255380.1
B3 56199826 56199915
refGene_gn=ATP8B4;refGene_tx=XM_019832446.2;refGene_gn=ATP8B4;refGene_tx=XM_019832447.2;refGene_gn=ATP8B4;refGene_tx=XM_019832448.2;refGene_gn=ATP8B4;refGene_tx=XM_023255378.1;refGene_gn=ATP8B4;refGene_tx=XM_023255380.1
B3 56202493 56202690

refGene_gn=ATP8B4;refGene_tx=XM_019832446.2;refGene_gn=ATP8B4;refGene_tx=XM_019832447.2;refGene_gn=ATP8B4;refGene_tx=XM_019832448.2;refGene_gn=ATP8B4;refGene_tx=XM_023255378.1;refGene_gn=ATP8B4;refGene_tx=XM_023255380.1
B3	56208709	56208918
refGene_gn=ATP8B4;refGene_tx=XM_019832446.2;refGene_gn=ATP8B4;refGene_tx=XM_019832447.2;refGene_gn=ATP8B4;refGene_tx=XM_019832448.2;refGene_gn=ATP8B4;refGene_tx=XM_023255378.1;refGene_gn=ATP8B4;refGene_tx=XM_023255380.1
B3	56212388	56212432
refGene_gn=ATP8B4;refGene_tx=XM_019832446.2;refGene_gn=ATP8B4;refGene_tx=XM_019832447.2;refGene_gn=ATP8B4;refGene_tx=XM_019832448.2;refGene_gn=ATP8B4;refGene_tx=XM_023255378.1;refGene_gn=ATP8B4;refGene_tx=XM_023255380.1
B3	56217556	56217722
refGene_gn=ATP8B4;refGene_tx=XM_019832446.2;refGene_gn=ATP8B4;refGene_tx=XM_019832447.2;refGene_gn=ATP8B4;refGene_tx=XM_019832448.2;refGene_gn=ATP8B4;refGene_tx=XM_023255378.1;refGene_gn=ATP8B4;refGene_tx=XM_023255380.1
B3	56224605	56224794
refGene_gn=ATP8B4;refGene_tx=XM_019832446.2;refGene_gn=ATP8B4;refGene_tx=XM_019832447.2;refGene_gn=ATP8B4;refGene_tx=XM_019832448.2;refGene_gn=ATP8B4;refGene_tx=XM_023255378.1;refGene_gn=ATP8B4;refGene_tx=XM_023255380.1
B3	56230468	56230584
refGene_gn=ATP8B4;refGene_tx=XM_019832446.2;refGene_gn=ATP8B4;refGene_tx=XM_019832447.2;refGene_gn=ATP8B4;refGene_tx=XM_019832448.2;refGene_gn=ATP8B4;refGene_tx=XM_023255378.1;refGene_gn=ATP8B4;refGene_tx=XM_023255380.1
B3	56233596	56233761
refGene_gn=ATP8B4;refGene_tx=XM_019832446.2;refGene_gn=ATP8B4;refGene_tx=XM_019832447.2;refGene_gn=ATP8B4;refGene_tx=XM_019832448.2;refGene_gn=ATP8B4;refGene_tx=XM_023255378.1;refGene_gn=ATP8B4;refGene_tx=XM_023255380.1
B3	56234726	56234838
refGene_gn=ATP8B4;refGene_tx=XM_019832446.2;refGene_gn=ATP8B4;refGene_tx=XM_019832447.2;refGene_gn=ATP8B4;refGene_tx=XM_019832448.2;refGene_gn=ATP8B4;refGene_tx=XM_023255378.1;refGene_gn=ATP8B4;refGene_tx=XM_023255380.1
B3	56237081	56237187
refGene_gn=ATP8B4;refGene_tx=XM_019832446.2;refGene_gn=ATP8B4;refGene_tx=XM_019832447.2;refGene_gn=ATP8B4;refGene_tx=XM_019832448.2;refGene_gn=ATP8B4;refGene_tx=XM_023255378.1;refGene_gn=ATP8B4;refGene_tx=XM_023255380.1
B3	56253504	56253649
refGene_gn=ATP8B4;refGene_tx=XM_019832446.2;refGene_gn=ATP8B4;refGene_tx=XM_019832447.2;refGene_gn=ATP8B4;refGene_tx=XM_019832448.2;refGene_gn=ATP8B4;refGene_tx=XM_023255378.1;refGene_gn=ATP8B4;refGene_tx=XM_023255380.1
B3	56256444	56256628

refGene_gn=ATP8B4;refGene_tx=XM_019832446.2;refGene_gn=ATP8B4;refGene_tx=XM_019832447.2;refGene_gn=ATP8B4;refGene_tx=XM_019832448.2;refGene_gn=ATP8B4;refGene_tx=XM_023255380.1
B3 56257216 56257440
refGene_gn=ATP8B4;refGene_tx=XM_019832446.2;refGene_gn=ATP8B4;refGene_tx=XM_019832447.2;refGene_gn=ATP8B4;refGene_tx=XM_019832448.2;refGene_gn=ATP8B4;refGene_tx=XM_023255380.1
B3 56277953 56278037
refGene_gn=ATP8B4;refGene_tx=XM_019832446.2;refGene_gn=ATP8B4;refGene_tx=XM_019832447.2;refGene_gn=ATP8B4;refGene_tx=XM_019832448.2;refGene_gn=ATP8B4;refGene_tx=XM_023255378.1;refGene_gn=ATP8B4;refGene_tx=XM_023255380.1
B3 56280756 56281002
refGene_gn=ATP8B4;refGene_tx=XM_019832446.2;refGene_gn=ATP8B4;refGene_tx=XM_019832447.2;refGene_gn=ATP8B4;refGene_tx=XM_019832448.2;refGene_gn=ATP8B4;refGene_tx=XM_023255378.1;refGene_gn=ATP8B4;refGene_tx=XM_023255380.1
B3 56290431 56290437 refGene_gn=ATP8B4;refGene_tx=XM_023255380.1
B3 56290651 56290790
refGene_gn=ATP8B4;refGene_tx=XM_019832446.2;refGene_gn=ATP8B4;refGene_tx=XM_019832447.2;refGene_gn=ATP8B4;refGene_tx=XM_019832448.2;refGene_gn=ATP8B4;refGene_tx=XM_023255378.1
B3 56293674 56293805
refGene_gn=ATP8B4;refGene_tx=XM_019832446.2;refGene_gn=ATP8B4;refGene_tx=XM_019832447.2;refGene_gn=ATP8B4;refGene_tx=XM_019832448.2;refGene_gn=ATP8B4;refGene_tx=XM_023255378.1
B3 56295413 56295695
refGene_gn=ATP8B4;refGene_tx=XM_019832446.2;refGene_gn=ATP8B4;refGene_tx=XM_019832447.2;refGene_gn=ATP8B4;refGene_tx=XM_019832448.2;refGene_gn=ATP8B4;refGene_tx=XM_023255378.1
B3 56472964 56473212 refGene_gn=DTWD1;refGene_tx=XM_006932532.4
B3 56482111 56482358 refGene_gn=DTWD1;refGene_tx=XM_006932532.4
B3 56483056 56483200 refGene_gn=DTWD1;refGene_tx=XM_006932532.4
B3 56487595 56487859 refGene_gn=DTWD1;refGene_tx=XM_006932532.4
B3 56495884 56495932
refGene_gn=FAM227B;refGene_tx=XM_019832452.2;refGene_gn=FAM227B;refGene_tx=XM_019832453.2;refGene_gn=FAM227B;refGene_tx=XM_019832454.2;refGene_gn=FAM227B;refGene_tx=XM_019832455.2;refGene_gn=FAM227B;refGene_tx=XM_019832456.2;refGene_gn=FAM227B;refGene_tx=XM_019832458.2;refGene_gn=FAM227B;refGene_tx=XM_019832461.2;refGene_gn=FAM227B;refGene_tx=XM_023255381.1
B3 56498782 56498836
refGene_gn=FAM227B;refGene_tx=XM_019832452.2;refGene_gn=FAM227B;refGene_tx=XM_019832453.2;refGene_gn=FAM227B;refGene_tx=XM_019832454.2;refGene_gn=FAM227B;refGene_tx=XM_019832455.2;refGene_gn=FAM227B;refGene_tx=XM_019832456.2;refGene_gn=FAM227B;refGene_tx=XM_019832458.2;refGene_gn=FAM227B;refGene_tx=XM_019832461.2;refGene_gn=FAM227B;refGene_tx=XM_023255381.1
B3 56506867 56507099
refGene_gn=FAM227B;refGene_tx=XM_019832452.2;refGene_gn=FAM227B;refGen

e_tx=XM_019832453.2;refGene_gn=FAM227B;refGene_tx=XM_019832454.2;refGe
ne_gn=FAM227B;refGene_tx=XM_019832455.2;refGene_gn=FAM227B;refGene_tx=
XM_019832456.2;refGene_gn=FAM227B;refGene_tx=XM_019832458.2;refGene_gn
=FAM227B;refGene_tx=XM_019832461.2;refGene_gn=FAM227B;refGene_tx=XM_02
3255381.1
B3 56508454 56508522
refGene_gn=FAM227B;refGene_tx=XM_019832452.2;refGene_gn=FAM227B;refGen
e_tx=XM_019832453.2;refGene_gn=FAM227B;refGene_tx=XM_019832454.2;refGe
ne_gn=FAM227B;refGene_tx=XM_019832455.2;refGene_gn=FAM227B;refGene_tx=
XM_019832456.2;refGene_gn=FAM227B;refGene_tx=XM_019832458.2;refGene_gn
=FAM227B;refGene_tx=XM_019832461.2;refGene_gn=FAM227B;refGene_tx=XM_02
3255381.1
B3 56514299 56514335
refGene_gn=FAM227B;refGene_tx=XM_019832452.2;refGene_gn=FAM227B;refGen
e_tx=XM_019832453.2;refGene_gn=FAM227B;refGene_tx=XM_019832454.2;refGe
ne_gn=FAM227B;refGene_tx=XM_019832455.2;refGene_gn=FAM227B;refGene_tx=
XM_019832456.2;refGene_gn=FAM227B;refGene_tx=XM_019832458.2;refGene_gn
=FAM227B;refGene_tx=XM_019832461.2
B3 56515013 56515118
refGene_gn=FAM227B;refGene_tx=XM_019832452.2;refGene_gn=FAM227B;refGen
e_tx=XM_019832453.2;refGene_gn=FAM227B;refGene_tx=XM_019832455.2;refGe
ne_gn=FAM227B;refGene_tx=XM_019832456.2;refGene_gn=FAM227B;refGene_tx=
XM_019832458.2;refGene_gn=FAM227B;refGene_tx=XM_019832461.2
B3 56524997 56525096
refGene_gn=FAM227B;refGene_tx=XM_019832452.2;refGene_gn=FAM227B;refGen
e_tx=XM_019832453.2;refGene_gn=FAM227B;refGene_tx=XM_019832454.2;refGe
ne_gn=FAM227B;refGene_tx=XM_019832455.2;refGene_gn=FAM227B;refGene_tx=
XM_019832456.2;refGene_gn=FAM227B;refGene_tx=XM_019832458.2;refGene_gn
=FAM227B;refGene_tx=XM_019832461.2;refGene_gn=FAM227B;refGene_tx=XM_02
3255381.1
B3 56531561 56531663
refGene_gn=FAM227B;refGene_tx=XM_019832452.2;refGene_gn=FAM227B;refGen
e_tx=XM_019832453.2;refGene_gn=FAM227B;refGene_tx=XM_019832454.2;refGe
ne_gn=FAM227B;refGene_tx=XM_019832455.2;refGene_gn=FAM227B;refGene_tx=
XM_019832456.2;refGene_gn=FAM227B;refGene_tx=XM_019832458.2;refGene_gn
=FAM227B;refGene_tx=XM_019832461.2;refGene_gn=FAM227B;refGene_tx=XM_02
3255381.1
B3 56536229 56536356
refGene_gn=FAM227B;refGene_tx=XM_019832452.2;refGene_gn=FAM227B;refGen
e_tx=XM_019832453.2;refGene_gn=FAM227B;refGene_tx=XM_019832454.2;refGe
ne_gn=FAM227B;refGene_tx=XM_019832455.2;refGene_gn=FAM227B;refGene_tx=
XM_019832456.2;refGene_gn=FAM227B;refGene_tx=XM_019832458.2;refGene_gn
=FAM227B;refGene_tx=XM_019832461.2;refGene_gn=FAM227B;refGene_tx=XM_02
3255381.1
B3 56556143 56556281
refGene_gn=FAM227B;refGene_tx=XM_019832452.2;refGene_gn=FAM227B;refGen
e_tx=XM_019832453.2;refGene_gn=FAM227B;refGene_tx=XM_019832454.2;refGe
ne_gn=FAM227B;refGene_tx=XM_019832455.2;refGene_gn=FAM227B;refGene_tx=
XM_019832456.2;refGene_gn=FAM227B;refGene_tx=XM_019832458.2;refGene_gn
=FAM227B;refGene_tx=XM_019832461.2;refGene_gn=FAM227B;refGene_tx=XM_02

3255381.1
B3 56575462 56575657
refGene_gn=FGF7;refGene_tx=XM_006932534.4;refGene_gn=FGF7;refGene_tx=XM_006932535.4
B3 56576737 56576841
refGene_gn=FGF7;refGene_tx=XM_006932534.4;refGene_gn=FGF7;refGene_tx=XM_006932535.4
B3 56630899 56631185
refGene_gn=FGF7;refGene_tx=XM_006932534.4;refGene_gn=FGF7;refGene_tx=XM_006932535.4
B3 56666851 56666949
refGene_gn=FAM227B;refGene_tx=XM_019832452.2;refGene_gn=FAM227B;refGene_tx=XM_019832453.2;refGene_gn=FAM227B;refGene_tx=XM_019832454.2;refGene_gn=FAM227B;refGene_tx=XM_019832455.2;refGene_gn=FAM227B;refGene_tx=XM_019832456.2;refGene_gn=FAM227B;refGene_tx=XM_019832458.2;refGene_gn=FAM227B;refGene_tx=XM_019832461.2;refGene_gn=FAM227B;refGene_tx=XM_023255381.1
B3 56668390 56668579 refGene_gn=FAM227B;refGene_tx=XM_019832456.2
B3 56671154 56671315
refGene_gn=FAM227B;refGene_tx=XM_019832452.2;refGene_gn=FAM227B;refGene_tx=XM_019832453.2;refGene_gn=FAM227B;refGene_tx=XM_019832454.2;refGene_gn=FAM227B;refGene_tx=XM_023255381.1
B3 56689888 56690024
refGene_gn=FAM227B;refGene_tx=XM_019832455.2;refGene_gn=FAM227B;refGene_tx=XM_019832458.2
B3 56694351 56694429
refGene_gn=FAM227B;refGene_tx=XM_019832452.2;refGene_gn=FAM227B;refGene_tx=XM_019832453.2;refGene_gn=FAM227B;refGene_tx=XM_019832454.2;refGene_gn=FAM227B;refGene_tx=XM_023255381.1
B3 56695264 56695339
refGene_gn=FAM227B;refGene_tx=XM_019832452.2;refGene_gn=FAM227B;refGene_tx=XM_019832453.2;refGene_gn=FAM227B;refGene_tx=XM_019832454.2;refGene_gn=FAM227B;refGene_tx=XM_019832458.2;refGene_gn=FAM227B;refGene_tx=XM_023255381.1
B3 56696959 56697080 refGene_gn=GALK2;refGene_tx=XM_023255384.1
B3 56698852 56698922
refGene_gn=FAM227B;refGene_tx=XM_019832452.2;refGene_gn=FAM227B;refGene_tx=XM_019832453.2;refGene_gn=FAM227B;refGene_tx=XM_019832454.2;refGene_gn=FAM227B;refGene_tx=XM_019832455.2;refGene_gn=FAM227B;refGene_tx=XM_019832461.2;refGene_gn=FAM227B;refGene_tx=XM_023255381.1
B3 56701732 56701895
refGene_gn=FAM227B;refGene_tx=XM_019832452.2;refGene_gn=FAM227B;refGene_tx=XM_019832453.2;refGene_gn=FAM227B;refGene_tx=XM_019832454.2;refGene_gn=FAM227B;refGene_tx=XM_019832455.2;refGene_gn=FAM227B;refGene_tx=XM_023255381.1
B3 56702233 56702441
refGene_gn=GALK2;refGene_tx=XM_003987152.4;refGene_gn=GALK2;refGene_tx=XM_023255382.1;refGene_gn=GALK2;refGene_tx=XM_023255383.1
B3 56712080 56712282
refGene_gn=GALK2;refGene_tx=XM_003987152.4;refGene_gn=GALK2;refGene_tx

=XM_023255382.1;refGene_gn=GALK2;refGene_tx=XM_023255383.1;refGene_gn=GALK2;refGene_tx=XM_023255384.1
B3 56731834 56731891
refGene_gn=GALK2;refGene_tx=XM_023255382.1;refGene_gn=GALK2;refGene_tx=XM_023255383.1;refGene_gn=GALK2;refGene_tx=XM_023255384.1
B3 56741582 56741793
refGene_gn=GALK2;refGene_tx=XM_003987152.4;refGene_gn=GALK2;refGene_tx=XM_023255382.1;refGene_gn=GALK2;refGene_tx=XM_023255383.1;refGene_gn=GALK2;refGene_tx=XM_023255384.1
B3 56751143 56751296
refGene_gn=GALK2;refGene_tx=XM_003987152.4;refGene_gn=GALK2;refGene_tx=XM_023255382.1;refGene_gn=GALK2;refGene_tx=XM_023255383.1;refGene_gn=GALK2;refGene_tx=XM_023255384.1
B3 56752763 56752862
refGene_gn=GALK2;refGene_tx=XM_003987152.4;refGene_gn=GALK2;refGene_tx=XM_023255382.1;refGene_gn=GALK2;refGene_tx=XM_023255383.1;refGene_gn=GALK2;refGene_tx=XM_023255384.1
B3 56796678 56796825
refGene_gn=GALK2;refGene_tx=XM_003987152.4;refGene_gn=GALK2;refGene_tx=XM_023255382.1;refGene_gn=GALK2;refGene_tx=XM_023255383.1;refGene_gn=GALK2;refGene_tx=XM_023255384.1
B3 56799803 56799894
refGene_gn=GALK2;refGene_tx=XM_003987152.4;refGene_gn=GALK2;refGene_tx=XM_023255382.1;refGene_gn=GALK2;refGene_tx=XM_023255383.1;refGene_gn=GALK2;refGene_tx=XM_023255384.1
B3 56808090 56808214
refGene_gn=GALK2;refGene_tx=XM_003987152.4;refGene_gn=GALK2;refGene_tx=XM_023255382.1;refGene_gn=GALK2;refGene_tx=XM_023255383.1;refGene_gn=GALK2;refGene_tx=XM_023255384.1
B3 56813120 56813209
refGene_gn=GALK2;refGene_tx=XM_003987152.4;refGene_gn=GALK2;refGene_tx=XM_023255382.1;refGene_gn=GALK2;refGene_tx=XM_023255383.1;refGene_gn=GALK2;refGene_tx=XM_023255384.1
B3 56828933 56828986
refGene_gn=GALK2;refGene_tx=XM_003987152.4;refGene_gn=GALK2;refGene_tx=XM_023255382.1;refGene_gn=GALK2;refGene_tx=XM_023255384.1
B3 56838700 56838720 refGene_gn=GALK2;refGene_tx=XM_023255383.1
B3 56839151 56839205
refGene_gn=COPS2;refGene_tx=XM_003987155.5;refGene_gn=COPS2;refGene_tx=XM_003987156.5
B3 56851113 56851227
refGene_gn=COPS2;refGene_tx=XM_003987155.5;refGene_gn=COPS2;refGene_tx=XM_003987156.5
B3 56851896 56851974
refGene_gn=COPS2;refGene_tx=XM_003987155.5;refGene_gn=COPS2;refGene_tx=XM_003987156.5
B3 56856541 56856667
refGene_gn=COPS2;refGene_tx=XM_003987155.5;refGene_gn=COPS2;refGene_tx=XM_003987156.5
B3 56859122 56859233

refGene_gn=COPS2;refGene_tx=XM_003987155.5;refGene_gn=COPS2;refGene_tx=XM_003987156.5
B3 56859364 56859442
refGene_gn=COPS2;refGene_tx=XM_003987155.5;refGene_gn=COPS2;refGene_tx=XM_003987156.5
B3 56861710 56861885
refGene_gn=COPS2;refGene_tx=XM_003987155.5;refGene_gn=COPS2;refGene_tx=XM_003987156.5
B3 56862567 56862746
refGene_gn=COPS2;refGene_tx=XM_003987155.5;refGene_gn=COPS2;refGene_tx=XM_003987156.5
B3 56862855 56862908
refGene_gn=COPS2;refGene_tx=XM_003987155.5;refGene_gn=COPS2;refGene_tx=XM_003987156.5
B3 56865501 56865599
refGene_gn=COPS2;refGene_tx=XM_003987155.5;refGene_gn=COPS2;refGene_tx=XM_003987156.5
B3 56866947 56867030
refGene_gn=COPS2;refGene_tx=XM_003987155.5;refGene_gn=COPS2;refGene_tx=XM_003987156.5
B3 56867554 56867613
refGene_gn=COPS2;refGene_tx=XM_003987155.5;refGene_gn=COPS2;refGene_tx=XM_003987156.5
B3 56868502 56868647
refGene_gn=COPS2;refGene_tx=XM_003987155.5;refGene_gn=COPS2;refGene_tx=XM_003987156.5
B3 56933294 56933318
refGene_gn=SECISBP2L;refGene_tx=XM_003987157.5;refGene_gn=SECISBP2L;refGene_tx=XM_019832463.2
B3 56941937 56942116
refGene_gn=SECISBP2L;refGene_tx=XM_003987157.5;refGene_gn=SECISBP2L;refGene_tx=XM_019832463.2
B3 56944150 56944475
refGene_gn=SECISBP2L;refGene_tx=XM_003987157.5;refGene_gn=SECISBP2L;refGene_tx=XM_019832463.2
B3 56946102 56946238
refGene_gn=SECISBP2L;refGene_tx=XM_003987157.5;refGene_gn=SECISBP2L;refGene_tx=XM_019832463.2
B3 56951068 56951298
refGene_gn=SECISBP2L;refGene_tx=XM_003987157.5;refGene_gn=SECISBP2L;refGene_tx=XM_019832463.2
B3 56951577 56951602
refGene_gn=SECISBP2L;refGene_tx=XM_003987157.5;refGene_gn=SECISBP2L;refGene_tx=XM_019832463.2
B3 56952028 56952144
refGene_gn=SECISBP2L;refGene_tx=XM_003987157.5;refGene_gn=SECISBP2L;refGene_tx=XM_019832463.2
B3 56958304 56958439
refGene_gn=SECISBP2L;refGene_tx=XM_003987157.5
B3 56960170 56960251

refGene_gn=SECISBP2L;refGene_tx=XM_003987157.5;refGene_gn=SECISBP2L;refGene_tx=XM_019832463.2
B3 56960756 56960924
refGene_gn=SECISBP2L;refGene_tx=XM_003987157.5;refGene_gn=SECISBP2L;refGene_tx=XM_019832463.2
B3 56961048 56961190
refGene_gn=SECISBP2L;refGene_tx=XM_003987157.5;refGene_gn=SECISBP2L;refGene_tx=XM_019832463.2
B3 56968580 56968750
refGene_gn=SECISBP2L;refGene_tx=XM_003987157.5;refGene_gn=SECISBP2L;refGene_tx=XM_019832463.2
B3 56969669 56969802
refGene_gn=SECISBP2L;refGene_tx=XM_003987157.5;refGene_gn=SECISBP2L;refGene_tx=XM_019832463.2
B3 56971399 56971562
refGene_gn=SECISBP2L;refGene_tx=XM_003987157.5;refGene_gn=SECISBP2L;refGene_tx=XM_019832463.2
B3 56983730 56983951
refGene_gn=SECISBP2L;refGene_tx=XM_003987157.5;refGene_gn=SECISBP2L;refGene_tx=XM_019832463.2
B3 56985021 56985176
refGene_gn=SECISBP2L;refGene_tx=XM_003987157.5;refGene_gn=SECISBP2L;refGene_tx=XM_019832463.2
B3 56986917 56987137
refGene_gn=SECISBP2L;refGene_tx=XM_003987157.5;refGene_gn=SECISBP2L;refGene_tx=XM_019832463.2
B3 56994345 56995031
refGene_gn=SECISBP2L;refGene_tx=XM_003987157.5;refGene_gn=SECISBP2L;refGene_tx=XM_019832463.2
B3 57027388 57027973 refGene_gn=SHC4;refGene_tx=XM_023255385.1
B3 57067314 57067385 refGene_gn=SHC4;refGene_tx=XM_023255385.1
B3 57098038 57098102 refGene_gn=SHC4;refGene_tx=XM_023255385.1
B3 57101208 57101328 refGene_gn=SHC4;refGene_tx=XM_023255385.1
B3 57106889 57107465 refGene_gn=EID1;refGene_tx=XM_019832464.2
B3 57111215 57111269 refGene_gn=SHC4;refGene_tx=XM_023255385.1
B3 57115549 57115601 refGene_gn=SHC4;refGene_tx=XM_023255385.1
B3 57127538 57127662 refGene_gn=SHC4;refGene_tx=XM_023255385.1
B3 57129702 57129874 refGene_gn=SHC4;refGene_tx=XM_023255385.1
B3 57131993 57132054 refGene_gn=SHC4;refGene_tx=XM_023255385.1
B3 57140622 57140802 refGene_gn=SHC4;refGene_tx=XM_023255385.1
B3 57148766 57149020 refGene_gn=SHC4;refGene_tx=XM_023255385.1
B3 57155959 57156115 refGene_gn=SHC4;refGene_tx=XM_023255385.1
B3 57167132 57167389 refGene_gn=CEP152;refGene_tx=XM_019832470.2
B3 57173316 57173413 refGene_gn=CEP152;refGene_tx=XM_019832470.2
B3 57181155 57181259 refGene_gn=CEP152;refGene_tx=XM_019832470.2
B3 57181462 57181535 refGene_gn=CEP152;refGene_tx=XM_019832470.2
B3 57181615 57181894 refGene_gn=CEP152;refGene_tx=XM_019832470.2
B3 57183249 57183307 refGene_gn=CEP152;refGene_tx=XM_019832470.2
B3 57183309 57183414 refGene_gn=CEP152;refGene_tx=XM_019832470.2
B3 57185717 57185858 refGene_gn=CEP152;refGene_tx=XM_019832470.2

B3	57187168	57187308	refGene_gn=CEP152;refGene_tx=XM_019832470.2
B3	57190885	57191086	refGene_gn=CEP152;refGene_tx=XM_019832470.2
B3	57195100	57195248	refGene_gn=CEP152;refGene_tx=XM_019832470.2
B3	57196871	57196963	refGene_gn=CEP152;refGene_tx=XM_019832470.2
B3	57197800	57197964	refGene_gn=CEP152;refGene_tx=XM_019832470.2
B3	57218986	57219191	refGene_gn=CEP152;refGene_tx=XM_019832470.2
B3	57227232	57227361	refGene_gn=CEP152;refGene_tx=XM_019832470.2
B3	57227949	57228059	refGene_gn=CEP152;refGene_tx=XM_019832470.2
B3	57228803	57228932	refGene_gn=CEP152;refGene_tx=XM_019832470.2
B3	57229075	57229211	refGene_gn=CEP152;refGene_tx=XM_019832470.2
B3	57235464	57235746	refGene_gn=CEP152;refGene_tx=XM_019832470.2
B3	57238398	57238530	refGene_gn=CEP152;refGene_tx=XM_019832470.2
B3	57243166	57243709	refGene_gn=CEP152;refGene_tx=XM_019832470.2
B3	57243710	57243818	refGene_gn=CEP152;refGene_tx=XM_019832470.2
B3	57246688	57246830	refGene_gn=CEP152;refGene_tx=XM_019832470.2
B3	57251160	57251328	refGene_gn=CEP152;refGene_tx=XM_019832470.2
B3	57254719	57254816	refGene_gn=CEP152;refGene_tx=XM_019832470.2
B3	57255903	57256007	refGene_gn=CEP152;refGene_tx=XM_019832470.2
B3	57258423	57258580	refGene_gn=CEP152;refGene_tx=XM_019832470.2
B3	57258829	57258933	refGene_gn=CEP152;refGene_tx=XM_019832470.2
B3	57265065	57266099	refGene_gn=CEP152;refGene_tx=XM_019832470.2
B3	57355501	57356064
refGene_gn=LOC111560945;refGene_tx=XM_023256165.1
B3	57356696	57356805
refGene_gn=LOC111560945;refGene_tx=XM_023256165.1
B3	57356877	57357041	refGene_gn=FBN1;refGene_tx=XM_023255387.1
B3	57385051	57385134	refGene_gn=FBN1;refGene_tx=XM_023255387.1
B3	57387418	57387517	refGene_gn=FBN1;refGene_tx=XM_023255387.1
B3	57396782	57396878	refGene_gn=FBN1;refGene_tx=XM_023255387.1
B3	57400235	57400331	refGene_gn=FBN1;refGene_tx=XM_023255387.1
B3	57462540	57462738	refGene_gn=FBN1;refGene_tx=XM_023255387.1
B3	57466157	57466283	refGene_gn=FBN1;refGene_tx=XM_023255387.1
B3	57473368	57473494	refGene_gn=FBN1;refGene_tx=XM_023255387.1
B3	57478952	57479111	refGene_gn=FBN1;refGene_tx=XM_023255387.1
B3	57483554	57483734	refGene_gn=FBN1;refGene_tx=XM_023255387.1
B3	57484363	57484504	refGene_gn=FBN1;refGene_tx=XM_023255387.1
B3	57486193	57486313	refGene_gn=FBN1;refGene_tx=XM_023255387.1
B3	57489163	57489289	refGene_gn=FBN1;refGene_tx=XM_023255387.1
B3	57490985	57491108	refGene_gn=FBN1;refGene_tx=XM_023255387.1
B3	57494323	57494446	refGene_gn=FBN1;refGene_tx=XM_023255387.1
B3	57495519	57495672	refGene_gn=FBN1;refGene_tx=XM_023255387.1
B3	57499304	57499358	refGene_gn=FBN1;refGene_tx=XM_023255387.1
B3	57500924	57501050	refGene_gn=FBN1;refGene_tx=XM_023255387.1
B3	57504652	57504778	refGene_gn=FBN1;refGene_tx=XM_023255387.1
B3	57505704	57505824	refGene_gn=FBN1;refGene_tx=XM_023255387.1
B3	57506040	57506178	refGene_gn=FBN1;refGene_tx=XM_023255387.1
B3	57507021	57507072	refGene_gn=FBN1;refGene_tx=XM_023255387.1
B3	57508939	57509065	refGene_gn=FBN1;refGene_tx=XM_023255387.1
B3	57511147	57511375	refGene_gn=FBN1;refGene_tx=XM_023255387.1
B3	57513840	57513966	refGene_gn=FBN1;refGene_tx=XM_023255387.1

B3 57514085 57514214 refGene_gn=FBN1;refGene_tx=XM_023255387.1
B3 57514995 57515121 refGene_gn=FBN1;refGene_tx=XM_023255387.1
B3 57515232 57515358 refGene_gn=FBN1;refGene_tx=XM_023255387.1
B3 57516526 57516649 refGene_gn=FBN1;refGene_tx=XM_023255387.1
B3 57517730 57517856 refGene_gn=FBN1;refGene_tx=XM_023255387.1
B3 57520188 57520314 refGene_gn=FBN1;refGene_tx=XM_023255387.1
B3 57529208 57529331 refGene_gn=FBN1;refGene_tx=XM_023255387.1
B3 57529474 57529597 refGene_gn=FBN1;refGene_tx=XM_023255387.1
B3 57530980 57531106 refGene_gn=FBN1;refGene_tx=XM_023255387.1
B3 57533122 57533245 refGene_gn=FBN1;refGene_tx=XM_023255387.1
B3 57535016 57535139 refGene_gn=FBN1;refGene_tx=XM_023255387.1
B3 57535439 57535604 refGene_gn=FBN1;refGene_tx=XM_023255387.1
B3 57538177 57538246 refGene_gn=FBN1;refGene_tx=XM_023255387.1
B3 57538342 57538468 refGene_gn=FBN1;refGene_tx=XM_023255387.1
B3 57539659 57539782 refGene_gn=FBN1;refGene_tx=XM_023255387.1
B3 57540433 57540592 refGene_gn=FBN1;refGene_tx=XM_023255387.1
B3 57543556 57543628 refGene_gn=FBN1;refGene_tx=XM_023255387.1
B3 57547072 57547198 refGene_gn=FBN1;refGene_tx=XM_023255387.1
B3 57549976 57550099 refGene_gn=FBN1;refGene_tx=XM_023255387.1
B3 57553521 57553647 refGene_gn=FBN1;refGene_tx=XM_023255387.1
B3 57555602 57555719 refGene_gn=FBN1;refGene_tx=XM_023255387.1
B3 57556679 57556808 refGene_gn=FBN1;refGene_tx=XM_023255387.1
B3 57557442 57557562 refGene_gn=FBN1;refGene_tx=XM_023255387.1
B3 57559929 57560055 refGene_gn=FBN1;refGene_tx=XM_023255387.1
B3 57563776 57563926 refGene_gn=FBN1;refGene_tx=XM_023255387.1
B3 57564294 57564360 refGene_gn=FBN1;refGene_tx=XM_023255387.1
B3 57564872 57564989 refGene_gn=FBN1;refGene_tx=XM_023255387.1
B3 57566784 57566904 refGene_gn=FBN1;refGene_tx=XM_023255387.1
B3 57568406 57568529 refGene_gn=FBN1;refGene_tx=XM_023255387.1
B3 57569866 57569998 refGene_gn=FBN1;refGene_tx=XM_023255387.1
B3 57572724 57572850 refGene_gn=FBN1;refGene_tx=XM_023255387.1
B3 57573416 57573623 refGene_gn=FBN1;refGene_tx=XM_023255387.1
B3 57575514 57575640 refGene_gn=FBN1;refGene_tx=XM_023255387.1
B3 57575890 57576013 refGene_gn=FBN1;refGene_tx=XM_023255387.1
B3 57578964 57579081 refGene_gn=FBN1;refGene_tx=XM_023255387.1
B3 57579290 57579419 refGene_gn=FBN1;refGene_tx=XM_023255387.1
B3 57580408 57580528 refGene_gn=FBN1;refGene_tx=XM_023255387.1
B3 57584929 57585161 refGene_gn=FBN1;refGene_tx=XM_023255387.1
B3 57587649 57587824 refGene_gn=FBN1;refGene_tx=XM_023255387.1
B3 57589033 57589423 refGene_gn=FBN1;refGene_tx=XM_023255387.1
B3 57665360 57665417
refGene_gn=DUT;refGene_tx=XM_003987163.5;refGene_gn=DUT;refGene_tx=XM_006932538.4;refGene_gn=DUT;refGene_tx=XM_006932540.4;refGene_gn=DUT;refGene_tx=XM_019832479.2
B3 57665820 57665891
refGene_gn=DUT;refGene_tx=XM_003987163.5;refGene_gn=DUT;refGene_tx=XM_006932538.4;refGene_gn=DUT;refGene_tx=XM_006932540.4;refGene_gn=DUT;refGene_tx=XM_019832479.2
B3 57666030 57666105
refGene_gn=DUT;refGene_tx=XM_003987163.5;refGene_gn=DUT;refGene_tx=XM_

006932538.4;refGene_gn=DUT;refGene_tx=XM_006932540.4;refGene_gn=DUT;refGene_tx=XM_019832479.2
B3 57672098 57672143
refGene_gn=DUT;refGene_tx=XM_003987163.5;refGene_gn=DUT;refGene_tx=XM_006932538.4;refGene_gn=DUT;refGene_tx=XM_006932540.4;refGene_gn=DUT;refGene_tx=XM_019832479.2
B3 57673781 57673873
refGene_gn=DUT;refGene_tx=XM_003987163.5;refGene_gn=DUT;refGene_tx=XM_006932538.4;refGene_gn=DUT;refGene_tx=XM_006932540.4;refGene_gn=DUT;refGene_tx=XM_019832479.2
B3 57676300 57676502
refGene_gn=DUT;refGene_tx=XM_003987163.5;refGene_gn=DUT;refGene_tx=XM_006932538.4;refGene_gn=DUT;refGene_tx=XM_006932540.4;refGene_gn=DUT;refGene_tx=XM_019832479.2
B3 57676935 57677101
refGene_gn=DUT;refGene_tx=XM_003987163.5;refGene_gn=DUT;refGene_tx=XM_006932538.4
B3 57677392 57677417
refGene_gn=DUT;refGene_tx=XM_006932540.4;refGene_gn=DUT;refGene_tx=XM_019832479.2
B3 57697788 57697924 refGene_gn=SLC12A1;refGene_tx=XM_003987165.2
B3 57698963 57699031 refGene_gn=SLC12A1;refGene_tx=XM_003987165.2
B3 57700869 57701005 refGene_gn=SLC12A1;refGene_tx=XM_003987165.2
B3 57709644 57709731 refGene_gn=SLC12A1;refGene_tx=XM_003987165.2
B3 57711594 57711706 refGene_gn=SLC12A1;refGene_tx=XM_003987165.2
B3 57711944 57711999 refGene_gn=SLC12A1;refGene_tx=XM_003987165.2
B3 57712000 57712077 refGene_gn=SLC12A1;refGene_tx=XM_003987165.2
B3 57714871 57715015 refGene_gn=SLC12A1;refGene_tx=XM_003987165.2
B3 57724569 57724652 refGene_gn=SLC12A1;refGene_tx=XM_003987165.2
B3 57728577 57728684 refGene_gn=SLC12A1;refGene_tx=XM_003987165.2
B3 57731210 57731351 refGene_gn=SLC12A1;refGene_tx=XM_003987165.2
B3 57739549 57739661 refGene_gn=SLC12A1;refGene_tx=XM_003987165.2
B3 57740443 57740543 refGene_gn=SLC12A1;refGene_tx=XM_003987165.2
B3 57743925 57744081 refGene_gn=SLC12A1;refGene_tx=XM_003987165.2
B3 57746231 57746333 refGene_gn=SLC12A1;refGene_tx=XM_003987165.2
B3 57748118 57748242 refGene_gn=SLC12A1;refGene_tx=XM_003987165.2
B3 57748561 57748669 refGene_gn=SLC12A1;refGene_tx=XM_003987165.2
B3 57749818 57749970 refGene_gn=SLC12A1;refGene_tx=XM_003987165.2
B3 57752409 57752494 refGene_gn=SLC12A1;refGene_tx=XM_003987165.2
B3 57771484 57771612 refGene_gn=SLC12A1;refGene_tx=XM_003987165.2
B3 57773561 57773673 refGene_gn=SLC12A1;refGene_tx=XM_003987165.2
B3 57775596 57775707 refGene_gn=SLC12A1;refGene_tx=XM_003987165.2
B3 57776759 57776899 refGene_gn=SLC12A1;refGene_tx=XM_003987165.2
B3 57778859 57778955 refGene_gn=SLC12A1;refGene_tx=XM_003987165.2
B3 57785480 57785556 refGene_gn=SLC12A1;refGene_tx=XM_003987165.2
B3 57785711 57785846 refGene_gn=SLC12A1;refGene_tx=XM_003987165.2
B3 57798547 57798967 refGene_gn=SLC12A1;refGene_tx=XM_003987165.2
B3 57804210 57804456
refGene_gn=CTXN2;refGene_tx=XM_006932543.4;refGene_gn=CTXN2;refGene_tx=XM_006932544.4;refGene_gn=CTXN2;refGene_tx=XM_006932546.4;refGene_gn=

CTXN2;refGene_tx=XM_019832485.2;refGene_gn=CTXN2;refGene_tx=XM_0198324
88.2;refGene_gn=CTXN2;refGene_tx=XM_023255396.1;refGene_gn=CTXN2;refGe
ne_tx=XM_023255397.1
B3 57830488 57830649
refGene_gn=MYEF2;refGene_tx=XM_023255389.1;refGene_gn=MYEF2;refGene_tx
=XM_023255390.1;refGene_gn=MYEF2;refGene_tx=XM_023255391.1;refGene_gn=
MYEF2;refGene_tx=XM_023255392.1;refGene_gn=MYEF2;refGene_tx=XM_0232553
93.1
B3 57835665 57835874
refGene_gn=MYEF2;refGene_tx=XM_023255389.1;refGene_gn=MYEF2;refGene_tx
=XM_023255390.1;refGene_gn=MYEF2;refGene_tx=XM_023255391.1;refGene_gn=
MYEF2;refGene_tx=XM_023255392.1;refGene_gn=MYEF2;refGene_tx=XM_0232553
93.1;refGene_gn=MYEF2;refGene_tx=XM_023255394.1
B3 57837103 57837156
refGene_gn=MYEF2;refGene_tx=XM_023255389.1;refGene_gn=MYEF2;refGene_tx
=XM_023255390.1;refGene_gn=MYEF2;refGene_tx=XM_023255391.1;refGene_gn=
MYEF2;refGene_tx=XM_023255392.1;refGene_gn=MYEF2;refGene_tx=XM_0232553
93.1;refGene_gn=MYEF2;refGene_tx=XM_023255394.1
B3 57838384 57838392
refGene_gn=MYEF2;refGene_tx=XM_023255389.1;refGene_gn=MYEF2;refGene_tx
=XM_023255390.1;refGene_gn=MYEF2;refGene_tx=XM_023255391.1;refGene_gn=
MYEF2;refGene_tx=XM_023255392.1;refGene_gn=MYEF2;refGene_tx=XM_0232553
93.1;refGene_gn=MYEF2;refGene_tx=XM_023255394.1
B3 57838495 57838589
refGene_gn=MYEF2;refGene_tx=XM_023255389.1;refGene_gn=MYEF2;refGene_tx
=XM_023255390.1;refGene_gn=MYEF2;refGene_tx=XM_023255391.1;refGene_gn=
MYEF2;refGene_tx=XM_023255392.1;refGene_gn=MYEF2;refGene_tx=XM_0232553
93.1;refGene_gn=MYEF2;refGene_tx=XM_023255394.1
B3 57842273 57842465
refGene_gn=MYEF2;refGene_tx=XM_023255389.1;refGene_gn=MYEF2;refGene_tx
=XM_023255390.1;refGene_gn=MYEF2;refGene_tx=XM_023255391.1;refGene_gn=
MYEF2;refGene_tx=XM_023255393.1;refGene_gn=MYEF2;refGene_tx=XM_0232553
94.1
B3 57843212 57843366
refGene_gn=MYEF2;refGene_tx=XM_023255389.1;refGene_gn=MYEF2;refGene_tx
=XM_023255390.1;refGene_gn=MYEF2;refGene_tx=XM_023255391.1;refGene_gn=
MYEF2;refGene_tx=XM_023255392.1;refGene_gn=MYEF2;refGene_tx=XM_0232553
93.1;refGene_gn=MYEF2;refGene_tx=XM_023255394.1
B3 57843891 57843941
refGene_gn=MYEF2;refGene_tx=XM_023255389.1;refGene_gn=MYEF2;refGene_tx
=XM_023255390.1;refGene_gn=MYEF2;refGene_tx=XM_023255391.1;refGene_gn=
MYEF2;refGene_tx=XM_023255392.1;refGene_gn=MYEF2;refGene_tx=XM_0232553
93.1;refGene_gn=MYEF2;refGene_tx=XM_023255394.1
B3 57844059 57844123
refGene_gn=MYEF2;refGene_tx=XM_023255389.1;refGene_gn=MYEF2;refGene_tx
=XM_023255390.1;refGene_gn=MYEF2;refGene_tx=XM_023255391.1;refGene_gn=
MYEF2;refGene_tx=XM_023255392.1;refGene_gn=MYEF2;refGene_tx=XM_0232553
93.1;refGene_gn=MYEF2;refGene_tx=XM_023255394.1
B3 57848731 57848833
refGene_gn=MYEF2;refGene_tx=XM_023255389.1;refGene_gn=MYEF2;refGene_tx

=XM_023255390.1;refGene_gn=MYEF2;refGene_tx=XM_023255391.1;refGene_gn=MYEF2;refGene_tx=XM_023255392.1;refGene_gn=MYEF2;refGene_tx=XM_023255393.1;refGene_gn=MYEF2;refGene_tx=XM_023255394.1
B3 57850335 57850386
refGene_gn=MYEF2;refGene_tx=XM_023255389.1;refGene_gn=MYEF2;refGene_tx=XM_023255390.1;refGene_gn=MYEF2;refGene_tx=XM_023255391.1;refGene_gn=MYEF2;refGene_tx=XM_023255392.1;refGene_gn=MYEF2;refGene_tx=XM_023255393.1;refGene_gn=MYEF2;refGene_tx=XM_023255394.1
B3 57850678 57850747
refGene_gn=MYEF2;refGene_tx=XM_023255389.1;refGene_gn=MYEF2;refGene_tx=XM_023255390.1;refGene_gn=MYEF2;refGene_tx=XM_023255391.1;refGene_gn=MYEF2;refGene_tx=XM_023255392.1;refGene_gn=MYEF2;refGene_tx=XM_023255393.1;refGene_gn=MYEF2;refGene_tx=XM_023255394.1
B3 57850990 57851089
refGene_gn=MYEF2;refGene_tx=XM_023255389.1;refGene_gn=MYEF2;refGene_tx=XM_023255390.1;refGene_gn=MYEF2;refGene_tx=XM_023255392.1;refGene_gn=MYEF2;refGene_tx=XM_023255393.1;refGene_gn=MYEF2;refGene_tx=XM_023255394.1
B3 57851378 57851441
refGene_gn=MYEF2;refGene_tx=XM_023255389.1;refGene_gn=MYEF2;refGene_tx=XM_023255392.1;refGene_gn=MYEF2;refGene_tx=XM_023255393.1;refGene_gn=MYEF2;refGene_tx=XM_023255394.1
B3 57853153 57853362
refGene_gn=MYEF2;refGene_tx=XM_023255389.1;refGene_gn=MYEF2;refGene_tx=XM_023255390.1;refGene_gn=MYEF2;refGene_tx=XM_023255391.1;refGene_gn=MYEF2;refGene_tx=XM_023255392.1;refGene_gn=MYEF2;refGene_tx=XM_023255393.1;refGene_gn=MYEF2;refGene_tx=XM_023255394.1
B3 57853441 57853493
refGene_gn=MYEF2;refGene_tx=XM_023255389.1;refGene_gn=MYEF2;refGene_tx=XM_023255390.1;refGene_gn=MYEF2;refGene_tx=XM_023255391.1;refGene_gn=MYEF2;refGene_tx=XM_023255392.1;refGene_gn=MYEF2;refGene_tx=XM_023255394.1
B3 57860293 57860457
refGene_gn=MYEF2;refGene_tx=XM_023255389.1;refGene_gn=MYEF2;refGene_tx=XM_023255390.1;refGene_gn=MYEF2;refGene_tx=XM_023255391.1;refGene_gn=MYEF2;refGene_tx=XM_023255392.1;refGene_gn=MYEF2;refGene_tx=XM_023255393.1;refGene_gn=MYEF2;refGene_tx=XM_023255394.1
B3 57861033 57861356 refGene_gn=SLC24A5;refGene_tx=XM_011282858.3
B3 57862071 57862173 refGene_gn=SLC24A5;refGene_tx=XM_011282858.3
B3 57864065 57864272 refGene_gn=SLC24A5;refGene_tx=XM_011282858.3
B3 57866344 57866628 refGene_gn=SLC24A5;refGene_tx=XM_011282858.3
B3 57868422 57868523 refGene_gn=SLC24A5;refGene_tx=XM_011282858.3
B3 57868858 57868962 refGene_gn=SLC24A5;refGene_tx=XM_011282858.3
B3 57869055 57869139 refGene_gn=SLC24A5;refGene_tx=XM_011282858.3
B3 57886162 57886342 refGene_gn=SLC24A5;refGene_tx=XM_011282858.3
B3 57887052 57887173 refGene_gn=SLC24A5;refGene_tx=XM_011282858.3
B3 58229964 58231253
refGene_gn=SEMA6D;refGene_tx=XM_011282855.3;refGene_gn=SEMA6D;refGene_tx=XM_019832489.2;refGene_gn=SEMA6D;refGene_tx=XM_019832491.2;refGene_gn=SEMA6D;refGene_tx=XM_019832492.2;refGene_gn=SEMA6D;refGene_tx=XM_02

3255398.1;refGene_gn=SEMA6D;refGene_tx=XM_023255399.1;refGene_gn=SEMA6D;refGene_tx=XM_023255400.1;refGene_gn=SEMA6D;refGene_tx=XM_023255401.1;refGene_gn=SEMA6D;refGene_tx=XM_023255402.1;refGene_gn=SEMA6D;refGene_tx=XM_023255403.1;refGene_gn=SEMA6D;refGene_tx=XM_023255404.1;refGene_gn=SEMA6D;refGene_tx=XM_023255405.1;refGene_gn=SEMA6D;refGene_tx=XM_023255406.1;refGene_gn=SEMA6D;refGene_tx=XM_023255407.1
B3 58233044 58233212
refGene_gn=SEMA6D;refGene_tx=XM_011282855.3;refGene_gn=SEMA6D;refGene_tx=XM_019832489.2;refGene_gn=SEMA6D;refGene_tx=XM_019832491.2;refGene_gn=SEMA6D;refGene_tx=XM_019832492.2;refGene_gn=SEMA6D;refGene_tx=XM_023255398.1;refGene_gn=SEMA6D;refGene_tx=XM_023255399.1;refGene_gn=SEMA6D;refGene_tx=XM_023255400.1;refGene_gn=SEMA6D;refGene_tx=XM_023255401.1;refGene_gn=SEMA6D;refGene_tx=XM_023255402.1;refGene_gn=SEMA6D;refGene_tx=XM_023255403.1
B3 58234676 58234733
refGene_gn=SEMA6D;refGene_tx=XM_011282855.3;refGene_gn=SEMA6D;refGene_tx=XM_019832489.2;refGene_gn=SEMA6D;refGene_tx=XM_019832491.2;refGene_gn=SEMA6D;refGene_tx=XM_019832492.2;refGene_gn=SEMA6D;refGene_tx=XM_023255398.1;refGene_gn=SEMA6D;refGene_tx=XM_023255399.1;refGene_gn=SEMA6D;refGene_tx=XM_023255400.1;refGene_gn=SEMA6D;refGene_tx=XM_023255401.1;refGene_gn=SEMA6D;refGene_tx=XM_023255404.1;refGene_gn=SEMA6D;refGene_tx=XM_023255405.1
B3 58235094 58235195
refGene_gn=SEMA6D;refGene_tx=XM_011282855.3;refGene_gn=SEMA6D;refGene_tx=XM_019832489.2;refGene_gn=SEMA6D;refGene_tx=XM_019832491.2;refGene_gn=SEMA6D;refGene_tx=XM_019832492.2;refGene_gn=SEMA6D;refGene_tx=XM_023255398.1;refGene_gn=SEMA6D;refGene_tx=XM_023255399.1;refGene_gn=SEMA6D;refGene_tx=XM_023255400.1;refGene_gn=SEMA6D;refGene_tx=XM_023255401.1;refGene_gn=SEMA6D;refGene_tx=XM_023255402.1;refGene_gn=SEMA6D;refGene_tx=XM_023255403.1;refGene_gn=SEMA6D;refGene_tx=XM_023255404.1;refGene_gn=SEMA6D;refGene_tx=XM_023255405.1;refGene_gn=SEMA6D;refGene_tx=XM_023255406.1;refGene_gn=SEMA6D;refGene_tx=XM_023255407.1
B3 58235626 58235704
refGene_gn=SEMA6D;refGene_tx=XM_011282855.3;refGene_gn=SEMA6D;refGene_tx=XM_019832489.2;refGene_gn=SEMA6D;refGene_tx=XM_019832491.2;refGene_gn=SEMA6D;refGene_tx=XM_019832492.2;refGene_gn=SEMA6D;refGene_tx=XM_023255398.1;refGene_gn=SEMA6D;refGene_tx=XM_023255399.1;refGene_gn=SEMA6D;refGene_tx=XM_023255400.1;refGene_gn=SEMA6D;refGene_tx=XM_023255401.1;refGene_gn=SEMA6D;refGene_tx=XM_023255402.1;refGene_gn=SEMA6D;refGene_tx=XM_023255403.1;refGene_gn=SEMA6D;refGene_tx=XM_023255404.1;refGene_gn=SEMA6D;refGene_tx=XM_023255405.1;refGene_gn=SEMA6D;refGene_tx=XM_023255406.1;refGene_gn=SEMA6D;refGene_tx=XM_023255407.1
B3 58235799 58235940
refGene_gn=SEMA6D;refGene_tx=XM_011282855.3;refGene_gn=SEMA6D;refGene_tx=XM_019832489.2;refGene_gn=SEMA6D;refGene_tx=XM_019832491.2;refGene_gn=SEMA6D;refGene_tx=XM_019832492.2;refGene_gn=SEMA6D;refGene_tx=XM_023255398.1;refGene_gn=SEMA6D;refGene_tx=XM_023255399.1;refGene_gn=SEMA6D;refGene_tx=XM_023255400.1;refGene_gn=SEMA6D;refGene_tx=XM_023255401.1;refGene_gn=SEMA6D;refGene_tx=XM_023255402.1;refGene_gn=SEMA6D;refGene_tx=XM_023255403.1;refGene_gn=SEMA6D;refGene_tx=XM_023255404.1;refGen

e_gn=SEMA6D;refGene_tx=XM_023255405.1;refGene_gn=SEMA6D;refGene_tx=XM_023255406.1;refGene_gn=SEMA6D;refGene_tx=XM_023255407.1
B3 58236723 58236897
refGene_gn=SEMA6D;refGene_tx=XM_011282855.3;refGene_gn=SEMA6D;refGene_tx=XM_019832489.2;refGene_gn=SEMA6D;refGene_tx=XM_019832491.2;refGene_gn=SEMA6D;refGene_tx=XM_019832492.2;refGene_gn=SEMA6D;refGene_tx=XM_023255398.1;refGene_gn=SEMA6D;refGene_tx=XM_023255399.1;refGene_gn=SEMA6D;refGene_tx=XM_023255400.1;refGene_gn=SEMA6D;refGene_tx=XM_023255401.1;refGene_gn=SEMA6D;refGene_tx=XM_023255402.1;refGene_gn=SEMA6D;refGene_tx=XM_023255403.1;refGene_gn=SEMA6D;refGene_tx=XM_023255404.1;refGene_gn=SEMA6D;refGene_tx=XM_023255405.1;refGene_gn=SEMA6D;refGene_tx=XM_023255406.1;refGene_gn=SEMA6D;refGene_tx=XM_023255407.1
B3 58236987 58237143
refGene_gn=SEMA6D;refGene_tx=XM_011282855.3;refGene_gn=SEMA6D;refGene_tx=XM_019832489.2;refGene_gn=SEMA6D;refGene_tx=XM_019832491.2;refGene_gn=SEMA6D;refGene_tx=XM_019832492.2;refGene_gn=SEMA6D;refGene_tx=XM_023255398.1;refGene_gn=SEMA6D;refGene_tx=XM_023255399.1;refGene_gn=SEMA6D;refGene_tx=XM_023255400.1;refGene_gn=SEMA6D;refGene_tx=XM_023255401.1;refGene_gn=SEMA6D;refGene_tx=XM_023255402.1;refGene_gn=SEMA6D;refGene_tx=XM_023255403.1;refGene_gn=SEMA6D;refGene_tx=XM_023255404.1;refGene_gn=SEMA6D;refGene_tx=XM_023255405.1;refGene_gn=SEMA6D;refGene_tx=XM_023255406.1;refGene_gn=SEMA6D;refGene_tx=XM_023255407.1
B3 58237449 58237581
refGene_gn=SEMA6D;refGene_tx=XM_011282855.3;refGene_gn=SEMA6D;refGene_tx=XM_019832489.2;refGene_gn=SEMA6D;refGene_tx=XM_019832491.2;refGene_gn=SEMA6D;refGene_tx=XM_019832492.2;refGene_gn=SEMA6D;refGene_tx=XM_023255398.1;refGene_gn=SEMA6D;refGene_tx=XM_023255399.1;refGene_gn=SEMA6D;refGene_tx=XM_023255400.1;refGene_gn=SEMA6D;refGene_tx=XM_023255401.1;refGene_gn=SEMA6D;refGene_tx=XM_023255402.1;refGene_gn=SEMA6D;refGene_tx=XM_023255403.1;refGene_gn=SEMA6D;refGene_tx=XM_023255404.1;refGene_gn=SEMA6D;refGene_tx=XM_023255405.1;refGene_gn=SEMA6D;refGene_tx=XM_023255406.1;refGene_gn=SEMA6D;refGene_tx=XM_023255407.1
B3 58237690 58237908
refGene_gn=SEMA6D;refGene_tx=XM_011282855.3;refGene_gn=SEMA6D;refGene_tx=XM_019832489.2;refGene_gn=SEMA6D;refGene_tx=XM_019832491.2;refGene_gn=SEMA6D;refGene_tx=XM_019832492.2;refGene_gn=SEMA6D;refGene_tx=XM_023255398.1;refGene_gn=SEMA6D;refGene_tx=XM_023255399.1;refGene_gn=SEMA6D;refGene_tx=XM_023255400.1;refGene_gn=SEMA6D;refGene_tx=XM_023255401.1;refGene_gn=SEMA6D;refGene_tx=XM_023255402.1;refGene_gn=SEMA6D;refGene_tx=XM_023255403.1;refGene_gn=SEMA6D;refGene_tx=XM_023255404.1;refGene_gn=SEMA6D;refGene_tx=XM_023255405.1;refGene_gn=SEMA6D;refGene_tx=XM_023255406.1;refGene_gn=SEMA6D;refGene_tx=XM_023255407.1
B3 58238644 58238733
refGene_gn=SEMA6D;refGene_tx=XM_011282855.3;refGene_gn=SEMA6D;refGene_tx=XM_019832489.2;refGene_gn=SEMA6D;refGene_tx=XM_019832491.2;refGene_gn=SEMA6D;refGene_tx=XM_019832492.2;refGene_gn=SEMA6D;refGene_tx=XM_023255398.1;refGene_gn=SEMA6D;refGene_tx=XM_023255399.1;refGene_gn=SEMA6D;refGene_tx=XM_023255400.1;refGene_gn=SEMA6D;refGene_tx=XM_023255401.1;refGene_gn=SEMA6D;refGene_tx=XM_023255402.1;refGene_gn=SEMA6D;refGene_tx=XM_023255403.1;refGene_gn=SEMA6D;refGene_tx=XM_023255404.1;refGen

e_gn=SEMA6D;refGene_tx=XM_023255405.1;refGene_gn=SEMA6D;refGene_tx=XM_023255406.1;refGene_gn=SEMA6D;refGene_tx=XM_023255407.1
B3 58239426 58239546
refGene_gn=SEMA6D;refGene_tx=XM_011282855.3;refGene_gn=SEMA6D;refGene_tx=XM_019832489.2;refGene_gn=SEMA6D;refGene_tx=XM_019832491.2;refGene_gn=SEMA6D;refGene_tx=XM_019832492.2;refGene_gn=SEMA6D;refGene_tx=XM_023255398.1;refGene_gn=SEMA6D;refGene_tx=XM_023255399.1;refGene_gn=SEMA6D;refGene_tx=XM_023255400.1;refGene_gn=SEMA6D;refGene_tx=XM_023255401.1;refGene_gn=SEMA6D;refGene_tx=XM_023255402.1;refGene_gn=SEMA6D;refGene_tx=XM_023255403.1;refGene_gn=SEMA6D;refGene_tx=XM_023255404.1;refGene_gn=SEMA6D;refGene_tx=XM_023255405.1;refGene_gn=SEMA6D;refGene_tx=XM_023255406.1;refGene_gn=SEMA6D;refGene_tx=XM_023255407.1
B3 58240217 58240308
refGene_gn=SEMA6D;refGene_tx=XM_011282855.3;refGene_gn=SEMA6D;refGene_tx=XM_019832489.2;refGene_gn=SEMA6D;refGene_tx=XM_019832491.2;refGene_gn=SEMA6D;refGene_tx=XM_019832492.2;refGene_gn=SEMA6D;refGene_tx=XM_023255398.1;refGene_gn=SEMA6D;refGene_tx=XM_023255399.1;refGene_gn=SEMA6D;refGene_tx=XM_023255400.1;refGene_gn=SEMA6D;refGene_tx=XM_023255401.1;refGene_gn=SEMA6D;refGene_tx=XM_023255402.1;refGene_gn=SEMA6D;refGene_tx=XM_023255403.1;refGene_gn=SEMA6D;refGene_tx=XM_023255404.1;refGene_gn=SEMA6D;refGene_tx=XM_023255405.1;refGene_gn=SEMA6D;refGene_tx=XM_023255406.1;refGene_gn=SEMA6D;refGene_tx=XM_023255407.1
B3 58240536 58240638
refGene_gn=SEMA6D;refGene_tx=XM_011282855.3;refGene_gn=SEMA6D;refGene_tx=XM_019832489.2;refGene_gn=SEMA6D;refGene_tx=XM_019832491.2;refGene_gn=SEMA6D;refGene_tx=XM_019832492.2;refGene_gn=SEMA6D;refGene_tx=XM_023255398.1;refGene_gn=SEMA6D;refGene_tx=XM_023255399.1;refGene_gn=SEMA6D;refGene_tx=XM_023255400.1;refGene_gn=SEMA6D;refGene_tx=XM_023255401.1;refGene_gn=SEMA6D;refGene_tx=XM_023255402.1;refGene_gn=SEMA6D;refGene_tx=XM_023255403.1;refGene_gn=SEMA6D;refGene_tx=XM_023255404.1;refGene_gn=SEMA6D;refGene_tx=XM_023255405.1;refGene_gn=SEMA6D;refGene_tx=XM_023255406.1;refGene_gn=SEMA6D;refGene_tx=XM_023255407.1
B3 58240746 58240809
refGene_gn=SEMA6D;refGene_tx=XM_011282855.3;refGene_gn=SEMA6D;refGene_tx=XM_019832489.2;refGene_gn=SEMA6D;refGene_tx=XM_019832491.2;refGene_gn=SEMA6D;refGene_tx=XM_019832492.2;refGene_gn=SEMA6D;refGene_tx=XM_023255398.1;refGene_gn=SEMA6D;refGene_tx=XM_023255399.1;refGene_gn=SEMA6D;refGene_tx=XM_023255400.1;refGene_gn=SEMA6D;refGene_tx=XM_023255401.1;refGene_gn=SEMA6D;refGene_tx=XM_023255402.1;refGene_gn=SEMA6D;refGene_tx=XM_023255403.1;refGene_gn=SEMA6D;refGene_tx=XM_023255404.1;refGene_gn=SEMA6D;refGene_tx=XM_023255405.1;refGene_gn=SEMA6D;refGene_tx=XM_023255406.1;refGene_gn=SEMA6D;refGene_tx=XM_023255407.1
B3 58240908 58240969
refGene_gn=SEMA6D;refGene_tx=XM_011282855.3;refGene_gn=SEMA6D;refGene_tx=XM_019832489.2;refGene_gn=SEMA6D;refGene_tx=XM_019832491.2;refGene_gn=SEMA6D;refGene_tx=XM_019832492.2;refGene_gn=SEMA6D;refGene_tx=XM_023255398.1;refGene_gn=SEMA6D;refGene_tx=XM_023255399.1;refGene_gn=SEMA6D;refGene_tx=XM_023255400.1;refGene_gn=SEMA6D;refGene_tx=XM_023255401.1;refGene_gn=SEMA6D;refGene_tx=XM_023255402.1;refGene_gn=SEMA6D;refGene_tx=XM_023255403.1;refGene_gn=SEMA6D;refGene_tx=XM_023255404.1;refGen

e_gn=SEMA6D;refGene_tx=XM_023255405.1;refGene_gn=SEMA6D;refGene_tx=XM_023255406.1;refGene_gn=SEMA6D;refGene_tx=XM_023255407.1
B3 58241529 58241641
refGene_gn=SEMA6D;refGene_tx=XM_011282855.3;refGene_gn=SEMA6D;refGene_tx=XM_019832489.2;refGene_gn=SEMA6D;refGene_tx=XM_019832491.2;refGene_gn=SEMA6D;refGene_tx=XM_019832492.2;refGene_gn=SEMA6D;refGene_tx=XM_023255398.1;refGene_gn=SEMA6D;refGene_tx=XM_023255399.1;refGene_gn=SEMA6D;refGene_tx=XM_023255400.1;refGene_gn=SEMA6D;refGene_tx=XM_023255401.1;refGene_gn=SEMA6D;refGene_tx=XM_023255402.1;refGene_gn=SEMA6D;refGene_tx=XM_023255403.1;refGene_gn=SEMA6D;refGene_tx=XM_023255404.1;refGene_gn=SEMA6D;refGene_tx=XM_023255405.1;refGene_gn=SEMA6D;refGene_tx=XM_023255406.1;refGene_gn=SEMA6D;refGene_tx=XM_023255407.1
B3 58242054 58242163
refGene_gn=SEMA6D;refGene_tx=XM_011282855.3;refGene_gn=SEMA6D;refGene_tx=XM_019832489.2;refGene_gn=SEMA6D;refGene_tx=XM_019832491.2;refGene_gn=SEMA6D;refGene_tx=XM_019832492.2;refGene_gn=SEMA6D;refGene_tx=XM_023255398.1;refGene_gn=SEMA6D;refGene_tx=XM_023255399.1;refGene_gn=SEMA6D;refGene_tx=XM_023255400.1;refGene_gn=SEMA6D;refGene_tx=XM_023255401.1;refGene_gn=SEMA6D;refGene_tx=XM_023255402.1;refGene_gn=SEMA6D;refGene_tx=XM_023255403.1;refGene_gn=SEMA6D;refGene_tx=XM_023255404.1;refGene_gn=SEMA6D;refGene_tx=XM_023255405.1;refGene_gn=SEMA6D;refGene_tx=XM_023255406.1;refGene_gn=SEMA6D;refGene_tx=XM_023255407.1
B3 60280382 60280440 refGene_gn=SQOR;refGene_tx=XM_011282854.3
B3 60281590 60281769 refGene_gn=SQOR;refGene_tx=XM_011282854.3
B3 60282053 60282121 refGene_gn=SQOR;refGene_tx=XM_011282854.3
B3 60287447 60287631 refGene_gn=SQOR;refGene_tx=XM_011282854.3
B3 60293453 60293663 refGene_gn=SQOR;refGene_tx=XM_011282854.3
B3 60295670 60295865 refGene_gn=SQOR;refGene_tx=XM_011282854.3
B3 60298223 60298277 refGene_gn=SQOR;refGene_tx=XM_011282854.3
B3 60306015 60306186 refGene_gn=SQOR;refGene_tx=XM_011282854.3
B3 60309124 60309358 refGene_gn=SQOR;refGene_tx=XM_011282854.3
B3 60348863 60348983
refGene_gn=BLOC1S6;refGene_tx=XM_011282853.3;refGene_gn=BLOC1S6;refGene_tx=XM_019832499.2
B3 60349400 60349487
refGene_gn=BLOC1S6;refGene_tx=XM_011282853.3;refGene_gn=BLOC1S6;refGene_tx=XM_019832499.2
B3 60351452 60351540
refGene_gn=BLOC1S6;refGene_tx=XM_011282853.3;refGene_gn=BLOC1S6;refGene_tx=XM_019832499.2
B3 60359895 60360037
refGene_gn=BLOC1S6;refGene_tx=XM_011282853.3;refGene_gn=BLOC1S6;refGene_tx=XM_019832499.2
B3 60364800 60364807 refGene_gn=BLOC1S6;refGene_tx=XM_019832499.2
B3 60364979 60365061 refGene_gn=BLOC1S6;refGene_tx=XM_011282853.3
B3 60432780 60433171
refGene_gn=SLC30A4;refGene_tx=XM_003987200.5;refGene_gn=SLC30A4;refGene_tx=XM_023255408.1;refGene_gn=SLC30A4;refGene_tx=XM_023255409.1;refGene_gn=SLC30A4;refGene_tx=XM_023255410.1;refGene_gn=SLC30A4;refGene_tx=XM_023255411.1;refGene_gn=SLC30A4;refGene_tx=XM_023255412.1;refGene_gn

=SLC30A4;refGene_tx=XM_023255413.1
B3 60440321 60440468
refGene_gn=SLC30A4;refGene_tx=XM_003987200.5;refGene_gn=SLC30A4;refGene_tx=XM_023255408.1;refGene_gn=SLC30A4;refGene_tx=XM_023255409.1;refGene_gn=SLC30A4;refGene_tx=XM_023255410.1;refGene_gn=SLC30A4;refGene_tx=XM_023255411.1;refGene_gn=SLC30A4;refGene_tx=XM_023255412.1;refGene_gn=SLC30A4;refGene_tx=XM_023255413.1
B3 60440813 60440917 refGene_gn=SLC30A4;refGene_tx=XM_023255413.1
B3 60452710 60452864
refGene_gn=SLC30A4;refGene_tx=XM_003987200.5;refGene_gn=SLC30A4;refGene_tx=XM_023255408.1;refGene_gn=SLC30A4;refGene_tx=XM_023255409.1;refGene_gn=SLC30A4;refGene_tx=XM_023255410.1;refGene_gn=SLC30A4;refGene_tx=XM_023255411.1;refGene_gn=SLC30A4;refGene_tx=XM_023255412.1
B3 60454319 60454518
refGene_gn=SLC30A4;refGene_tx=XM_003987200.5;refGene_gn=SLC30A4;refGene_tx=XM_023255408.1;refGene_gn=SLC30A4;refGene_tx=XM_023255409.1;refGene_gn=SLC30A4;refGene_tx=XM_023255410.1;refGene_gn=SLC30A4;refGene_tx=XM_023255411.1;refGene_gn=SLC30A4;refGene_tx=XM_023255412.1
B3 60455416 60455522
refGene_gn=SLC30A4;refGene_tx=XM_003987200.5;refGene_gn=SLC30A4;refGene_tx=XM_023255408.1;refGene_gn=SLC30A4;refGene_tx=XM_023255409.1;refGene_gn=SLC30A4;refGene_tx=XM_023255410.1;refGene_gn=SLC30A4;refGene_tx=XM_023255411.1;refGene_gn=SLC30A4;refGene_tx=XM_023255412.1
B3 60456639 60456774
refGene_gn=SLC30A4;refGene_tx=XM_003987200.5;refGene_gn=SLC30A4;refGene_tx=XM_023255408.1;refGene_gn=SLC30A4;refGene_tx=XM_023255409.1;refGene_gn=SLC30A4;refGene_tx=XM_023255411.1
B3 60457376 60457531
refGene_gn=SLC30A4;refGene_tx=XM_003987200.5;refGene_gn=SLC30A4;refGene_tx=XM_023255408.1;refGene_gn=SLC30A4;refGene_tx=XM_023255409.1;refGene_gn=SLC30A4;refGene_tx=XM_023255410.1;refGene_gn=SLC30A4;refGene_tx=XM_023255411.1;refGene_gn=SLC30A4;refGene_tx=XM_023255412.1
B3 60496544 60496613
refGene_gn=CB3H15orf48;refGene_tx=XM_011282851.3;refGene_gn=CB3H15orf48;refGene_tx=XM_019832503.1
B3 60498535 60498596
refGene_gn=CB3H15orf48;refGene_tx=XM_011282851.3;refGene_gn=CB3H15orf48;refGene_tx=XM_019832503.1
B3 60499434 60499520
refGene_gn=CB3H15orf48;refGene_tx=XM_011282851.3;refGene_gn=CB3H15orf48;refGene_tx=XM_019832503.1
B3 60499692 60499728
refGene_gn=CB3H15orf48;refGene_tx=XM_011282851.3;refGene_gn=CB3H15orf48;refGene_tx=XM_019832503.1
B3 60506315 60506483 refGene_gn=SPATA5L1;refGene_tx=XM_006932564.3
B3 60508008 60508134 refGene_gn=SPATA5L1;refGene_tx=XM_006932564.3
B3 60509046 60509181
refGene_gn=SPATA5L1;refGene_tx=XM_006932564.3;refGene_gn=SPATA5L1;refGene_tx=XM_019832505.1
B3 60513004 60513229

refGene_gn=SPATA5L1;refGene_tx=XM_006932564.3;refGene_gn=SPATA5L1;refGene_tx=XM_019832505.1
B3 60513744 60513954
refGene_gn=SPATA5L1;refGene_tx=XM_006932564.3;refGene_gn=SPATA5L1;refGene_tx=XM_019832505.1
B3 60516854 60517013
refGene_gn=SPATA5L1;refGene_tx=XM_006932564.3;refGene_gn=SPATA5L1;refGene_tx=XM_019832505.1
B3 60522397 60522583
refGene_gn=SPATA5L1;refGene_tx=XM_006932564.3;refGene_gn=SPATA5L1;refGene_tx=XM_019832505.1
B3 60524057 60525152
refGene_gn=SPATA5L1;refGene_tx=XM_006932564.3;refGene_gn=SPATA5L1;refGene_tx=XM_019832505.1
B3 60553379 60553448 refGene_gn=GATM;refGene_tx=XM_003987192.4
B3 60554974 60555193 refGene_gn=GATM;refGene_tx=XM_003987192.4
B3 60561091 60561287 refGene_gn=GATM;refGene_tx=XM_003987192.4
B3 60562310 60562501 refGene_gn=GATM;refGene_tx=XM_003987192.4
B3 60563465 60563603 refGene_gn=GATM;refGene_tx=XM_003987192.4
B3 60563958 60564123 refGene_gn=GATM;refGene_tx=XM_003987192.4
B3 60565537 60565601 refGene_gn=GATM;refGene_tx=XM_003987192.4
B3 60566063 60566180 refGene_gn=GATM;refGene_tx=XM_003987192.4
B3 60568219 60568332 refGene_gn=GATM;refGene_tx=XM_003987192.4
B3 60653418 60653516 refGene_gn=SLC28A2;refGene_tx=XM_023255423.1
B3 60653518 60653533 refGene_gn=SLC28A2;refGene_tx=XM_023255423.1
B3 60653739 60653806 refGene_gn=SLC28A2;refGene_tx=XM_023255423.1
B3 60653808 60653840 refGene_gn=SLC28A2;refGene_tx=XM_023255423.1
B3 60655176 60655254 refGene_gn=SLC28A2;refGene_tx=XM_023255423.1
B3 60655846 60655980 refGene_gn=SLC28A2;refGene_tx=XM_023255423.1
B3 60655981 60656044 refGene_gn=SLC28A2;refGene_tx=XM_023255423.1
B3 60656237 60656314 refGene_gn=SLC28A2;refGene_tx=XM_023255423.1
B3 60656316 60656372 refGene_gn=SLC28A2;refGene_tx=XM_023255423.1
B3 60656374 60656402 refGene_gn=SLC28A2;refGene_tx=XM_023255423.1
B3 60656814 60656875 refGene_gn=SLC28A2;refGene_tx=XM_023255423.1
B3 60656876 60656973 refGene_gn=SLC28A2;refGene_tx=XM_023255423.1
B3 60657019 60657107 refGene_gn=SLC28A2;refGene_tx=XM_023255423.1
B3 60689570 60690071
refGene_gn=SHF;refGene_tx=XM_011282848.2;refGene_gn=SHF;refGene_tx=XM_011282849.2;refGene_gn=SHF;refGene_tx=XM_023255419.1;refGene_gn=SHF;refGene_tx=XM_023255420.1;refGene_gn=SHF;refGene_tx=XM_023255421.1;refGene_gn=SHF;refGene_tx=XM_023255422.1
B3 60699424 60699566
refGene_gn=SHF;refGene_tx=XM_011282848.2;refGene_gn=SHF;refGene_tx=XM_011282849.2;refGene_gn=SHF;refGene_tx=XM_023255419.1;refGene_gn=SHF;refGene_tx=XM_023255420.1;refGene_gn=SHF;refGene_tx=XM_023255421.1;refGene_gn=SHF;refGene_tx=XM_023255422.1
B3 60701796 60702003
refGene_gn=SHF;refGene_tx=XM_011282848.2;refGene_gn=SHF;refGene_tx=XM_011282849.2;refGene_gn=SHF;refGene_tx=XM_023255419.1;refGene_gn=SHF;refGene_tx=XM_023255420.1;refGene_gn=SHF;refGene_tx=XM_023255421.1;refGe

ne_gn=SHF;refGene_tx=XM_023255422.1
B3 60703846 60703987
refGene_gn=SHF;refGene_tx=XM_011282848.2;refGene_gn=SHF;refGene_tx=XM_023255419.1
B3 60705237 60705409
refGene_gn=SHF;refGene_tx=XM_011282848.2;refGene_gn=SHF;refGene_tx=XM_011282849.2;refGene_gn=SHF;refGene_tx=XM_023255419.1
B3 60705467 60705674
refGene_gn=SHF;refGene_tx=XM_011282848.2;refGene_gn=SHF;refGene_tx=XM_011282849.2;refGene_gn=SHF;refGene_tx=XM_023255419.1;refGene_gn=SHF;refGene_tx=XM_023255420.1;refGene_gn=SHF;refGene_tx=XM_023255421.1;refGene_gn=SHF;refGene_tx=XM_023255422.1
B3 60706450 60706587
refGene_gn=SHF;refGene_tx=XM_023255419.1;refGene_gn=SHF;refGene_tx=XM_023255420.1;refGene_gn=SHF;refGene_tx=XM_023255421.1
B3 60709197 60709384
refGene_gn=SHF;refGene_tx=XM_011282848.2;refGene_gn=SHF;refGene_tx=XM_011282849.2;refGene_gn=SHF;refGene_tx=XM_023255422.1
B3 60712193 60712316
refGene_gn=DUOX1;refGene_tx=XM_023255414.1;refGene_gn=DUOX1;refGene_tx=XM_023255415.1;refGene_gn=DUOX1;refGene_tx=XM_023255416.1;refGene_gn=DUOX1;refGene_tx=XM_023255417.1
B3 60712970 60713099
refGene_gn=DUOX1;refGene_tx=XM_023255414.1;refGene_gn=DUOX1;refGene_tx=XM_023255415.1;refGene_gn=DUOX1;refGene_tx=XM_023255416.1;refGene_gn=DUOX1;refGene_tx=XM_023255417.1
B3 60713342 60713498
refGene_gn=DUOX1;refGene_tx=XM_023255414.1;refGene_gn=DUOX1;refGene_tx=XM_023255415.1;refGene_gn=DUOX1;refGene_tx=XM_023255416.1;refGene_gn=DUOX1;refGene_tx=XM_023255417.1
B3 60714755 60714914
refGene_gn=DUOX1;refGene_tx=XM_023255414.1;refGene_gn=DUOX1;refGene_tx=XM_023255415.1;refGene_gn=DUOX1;refGene_tx=XM_023255416.1;refGene_gn=DUOX1;refGene_tx=XM_023255417.1
B3 60715141 60715374
refGene_gn=DUOX1;refGene_tx=XM_023255414.1;refGene_gn=DUOX1;refGene_tx=XM_023255415.1;refGene_gn=DUOX1;refGene_tx=XM_023255416.1;refGene_gn=DUOX1;refGene_tx=XM_023255417.1;refGene_gn=DUOX1;refGene_tx=XM_023255418.1
B3 60715815 60715969
refGene_gn=DUOX1;refGene_tx=XM_023255414.1;refGene_gn=DUOX1;refGene_tx=XM_023255415.1;refGene_gn=DUOX1;refGene_tx=XM_023255416.1;refGene_gn=DUOX1;refGene_tx=XM_023255417.1;refGene_gn=DUOX1;refGene_tx=XM_023255418.1
B3 60719843 60719971
refGene_gn=DUOX1;refGene_tx=XM_023255414.1;refGene_gn=DUOX1;refGene_tx=XM_023255415.1;refGene_gn=DUOX1;refGene_tx=XM_023255416.1;refGene_gn=DUOX1;refGene_tx=XM_023255417.1
B3 60721364 60721414
refGene_gn=DUOX1;refGene_tx=XM_023255414.1;refGene_gn=DUOX1;refGene_tx

=XM_023255415.1;refGene_gn=DUOX1;refGene_tx=XM_023255416.1;refGene_gn=DUOX1;refGene_tx=XM_023255417.1;refGene_gn=DUOX1;refGene_tx=XM_023255418.1
B3 60721717 60721817
refGene_gn=DUOX1;refGene_tx=XM_023255414.1;refGene_gn=DUOX1;refGene_tx=XM_023255415.1;refGene_gn=DUOX1;refGene_tx=XM_023255416.1;refGene_gn=DUOX1;refGene_tx=XM_023255417.1;refGene_gn=DUOX1;refGene_tx=XM_023255418.1
B3 60722370 60722601
refGene_gn=DUOX1;refGene_tx=XM_023255414.1;refGene_gn=DUOX1;refGene_tx=XM_023255415.1;refGene_gn=DUOX1;refGene_tx=XM_023255416.1;refGene_gn=DUOX1;refGene_tx=XM_023255417.1;refGene_gn=DUOX1;refGene_tx=XM_023255418.1
B3 60722746 60722925
refGene_gn=DUOX1;refGene_tx=XM_023255414.1;refGene_gn=DUOX1;refGene_tx=XM_023255415.1;refGene_gn=DUOX1;refGene_tx=XM_023255416.1;refGene_gn=DUOX1;refGene_tx=XM_023255417.1;refGene_gn=DUOX1;refGene_tx=XM_023255418.1
B3 60723513 60723639
refGene_gn=DUOX1;refGene_tx=XM_023255414.1;refGene_gn=DUOX1;refGene_tx=XM_023255415.1;refGene_gn=DUOX1;refGene_tx=XM_023255416.1;refGene_gn=DUOX1;refGene_tx=XM_023255417.1;refGene_gn=DUOX1;refGene_tx=XM_023255418.1
B3 60724051 60724121
refGene_gn=DUOX1;refGene_tx=XM_023255414.1;refGene_gn=DUOX1;refGene_tx=XM_023255415.1;refGene_gn=DUOX1;refGene_tx=XM_023255416.1;refGene_gn=DUOX1;refGene_tx=XM_023255417.1;refGene_gn=DUOX1;refGene_tx=XM_023255418.1
B3 60726328 60726504
refGene_gn=DUOX1;refGene_tx=XM_023255414.1;refGene_gn=DUOX1;refGene_tx=XM_023255415.1;refGene_gn=DUOX1;refGene_tx=XM_023255416.1;refGene_gn=DUOX1;refGene_tx=XM_023255417.1;refGene_gn=DUOX1;refGene_tx=XM_023255418.1
B3 60726790 60726884
refGene_gn=DUOX1;refGene_tx=XM_023255414.1;refGene_gn=DUOX1;refGene_tx=XM_023255415.1;refGene_gn=DUOX1;refGene_tx=XM_023255416.1;refGene_gn=DUOX1;refGene_tx=XM_023255417.1;refGene_gn=DUOX1;refGene_tx=XM_023255418.1
B3 60727145 60727371
refGene_gn=DUOX1;refGene_tx=XM_023255414.1;refGene_gn=DUOX1;refGene_tx=XM_023255415.1;refGene_gn=DUOX1;refGene_tx=XM_023255416.1;refGene_gn=DUOX1;refGene_tx=XM_023255417.1;refGene_gn=DUOX1;refGene_tx=XM_023255418.1
B3 60729455 60729641
refGene_gn=DUOX1;refGene_tx=XM_023255414.1;refGene_gn=DUOX1;refGene_tx=XM_023255415.1;refGene_gn=DUOX1;refGene_tx=XM_023255416.1;refGene_gn=DUOX1;refGene_tx=XM_023255417.1;refGene_gn=DUOX1;refGene_tx=XM_023255418.1
B3 60730267 60730467
refGene_gn=DUOX1;refGene_tx=XM_023255414.1;refGene_gn=DUOX1;refGene_tx

=XM_023255415.1;refGene_gn=DUOX1;refGene_tx=XM_023255416.1;refGene_gn=
DUOX1;refGene_tx=XM_023255417.1;refGene_gn=DUOX1;refGene_tx=XM_0232554
18.1
B3	60731195	60731309
refGene_gn=DUOX1;refGene_tx=XM_023255414.1;refGene_gn=DUOX1;refGene_tx
=XM_023255415.1;refGene_gn=DUOX1;refGene_tx=XM_023255416.1;refGene_gn=
DUOX1;refGene_tx=XM_023255417.1;refGene_gn=DUOX1;refGene_tx=XM_0232554
18.1
B3	60731954	60732110
refGene_gn=DUOX1;refGene_tx=XM_023255414.1;refGene_gn=DUOX1;refGene_tx
=XM_023255415.1;refGene_gn=DUOX1;refGene_tx=XM_023255416.1;refGene_gn=
DUOX1;refGene_tx=XM_023255418.1
B3	60732706	60732825
refGene_gn=DUOX1;refGene_tx=XM_023255414.1;refGene_gn=DUOX1;refGene_tx
=XM_023255415.1;refGene_gn=DUOX1;refGene_tx=XM_023255416.1;refGene_gn=
DUOX1;refGene_tx=XM_023255417.1;refGene_gn=DUOX1;refGene_tx=XM_0232554
18.1
B3	60733015	60733191
refGene_gn=DUOX1;refGene_tx=XM_023255414.1;refGene_gn=DUOX1;refGene_tx
=XM_023255415.1;refGene_gn=DUOX1;refGene_tx=XM_023255416.1;refGene_gn=
DUOX1;refGene_tx=XM_023255417.1;refGene_gn=DUOX1;refGene_tx=XM_0232554
18.1
B3	60736136	60736309
refGene_gn=DUOX1;refGene_tx=XM_023255414.1;refGene_gn=DUOX1;refGene_tx
=XM_023255415.1;refGene_gn=DUOX1;refGene_tx=XM_023255416.1;refGene_gn=
DUOX1;refGene_tx=XM_023255417.1;refGene_gn=DUOX1;refGene_tx=XM_0232554
18.1
B3	60736562	60736665
refGene_gn=DUOX1;refGene_tx=XM_023255414.1;refGene_gn=DUOX1;refGene_tx
=XM_023255415.1;refGene_gn=DUOX1;refGene_tx=XM_023255416.1;refGene_gn=
DUOX1;refGene_tx=XM_023255417.1;refGene_gn=DUOX1;refGene_tx=XM_0232554
18.1
B3	60737640	60737731
refGene_gn=DUOX1;refGene_tx=XM_023255414.1;refGene_gn=DUOX1;refGene_tx
=XM_023255415.1;refGene_gn=DUOX1;refGene_tx=XM_023255416.1;refGene_gn=
DUOX1;refGene_tx=XM_023255417.1;refGene_gn=DUOX1;refGene_tx=XM_0232554
18.1
B3	60738590	60738687
refGene_gn=DUOX1;refGene_tx=XM_023255414.1;refGene_gn=DUOX1;refGene_tx
=XM_023255415.1;refGene_gn=DUOX1;refGene_tx=XM_023255416.1;refGene_gn=
DUOX1;refGene_tx=XM_023255417.1;refGene_gn=DUOX1;refGene_tx=XM_0232554
18.1
B3	60738806	60738867
refGene_gn=DUOX1;refGene_tx=XM_023255414.1;refGene_gn=DUOX1;refGene_tx
=XM_023255415.1;refGene_gn=DUOX1;refGene_tx=XM_023255416.1;refGene_gn=
DUOX1;refGene_tx=XM_023255417.1;refGene_gn=DUOX1;refGene_tx=XM_0232554
18.1
B3	60739264	60739431
refGene_gn=DUOX1;refGene_tx=XM_023255414.1;refGene_gn=DUOX1;refGene_tx
=XM_023255415.1;refGene_gn=DUOX1;refGene_tx=XM_023255416.1;refGene_gn=

DUOX1;refGene_tx=XM_023255417.1;refGene_gn=DUOX1;refGene_tx=XM_0232554
18.1
B3 60739561 60739763
refGene_gn=DUOX1;refGene_tx=XM_023255414.1;refGene_gn=DUOX1;refGene_tx
=XM_023255415.1;refGene_gn=DUOX1;refGene_tx=XM_023255416.1;refGene_gn=
DUOX1;refGene_tx=XM_023255417.1;refGene_gn=DUOX1;refGene_tx=XM_0232554
18.1
B3 60739945 60740133
refGene_gn=DUOX1;refGene_tx=XM_023255414.1;refGene_gn=DUOX1;refGene_tx
=XM_023255415.1;refGene_gn=DUOX1;refGene_tx=XM_023255416.1;refGene_gn=
DUOX1;refGene_tx=XM_023255417.1;refGene_gn=DUOX1;refGene_tx=XM_0232554
18.1
B3 60741420 60741585
refGene_gn=DUOX1;refGene_tx=XM_023255414.1;refGene_gn=DUOX1;refGene_tx
=XM_023255415.1;refGene_gn=DUOX1;refGene_tx=XM_023255416.1;refGene_gn=
DUOX1;refGene_tx=XM_023255417.1;refGene_gn=DUOX1;refGene_tx=XM_0232554
18.1
B3 60741762 60741846
refGene_gn=DUOX1;refGene_tx=XM_023255414.1;refGene_gn=DUOX1;refGene_tx
=XM_023255415.1;refGene_gn=DUOX1;refGene_tx=XM_023255416.1;refGene_gn=
DUOX1;refGene_tx=XM_023255417.1;refGene_gn=DUOX1;refGene_tx=XM_0232554
18.1
B3 60743703 60743761
refGene_gn=DUOX1;refGene_tx=XM_023255414.1;refGene_gn=DUOX1;refGene_tx
=XM_023255415.1;refGene_gn=DUOX1;refGene_tx=XM_023255416.1;refGene_gn=
DUOX1;refGene_tx=XM_023255417.1;refGene_gn=DUOX1;refGene_tx=XM_0232554
18.1
B3 60751721 60751868 refGene_gn=DUOXA1;refGene_tx=XM_019832517.2
B3 60752196 60752254 refGene_gn=DUOXA1;refGene_tx=XM_019832517.2
B3 60752594 60752729 refGene_gn=DUOXA1;refGene_tx=XM_019832517.2
B3 60752935 60753149 refGene_gn=DUOXA1;refGene_tx=XM_019832517.2
B3 60753392 60753610 refGene_gn=DUOXA1;refGene_tx=XM_019832517.2
B3 60754306 60754563 refGene_gn=DUOXA1;refGene_tx=XM_019832517.2
B3 60755813 60756007
refGene_gn=DUOXA2;refGene_tx=XM_003987204.5;refGene_gn=DUOXA2;refGene_
tx=XM_006932565.4
B3 60756365 60756580
refGene_gn=DUOXA2;refGene_tx=XM_003987204.5;refGene_gn=DUOXA2;refGene_
tx=XM_006932565.4
B3 60756886 60757100
refGene_gn=DUOXA2;refGene_tx=XM_003987204.5;refGene_gn=DUOXA2;refGene_
tx=XM_006932565.4
B3 60757349 60757484 refGene_gn=DUOXA2;refGene_tx=XM_003987204.5
B3 60757757 60757815
refGene_gn=DUOXA2;refGene_tx=XM_003987204.5;refGene_gn=DUOXA2;refGene_
tx=XM_006932565.4
B3 60758952 60759099
refGene_gn=DUOXA2;refGene_tx=XM_003987204.5;refGene_gn=DUOXA2;refGene_
tx=XM_006932565.4
B3 60760277 60760347 refGene_gn=DUOX2;refGene_tx=XM_023255425.1

B3	60760601 60760691 refGene_gn=DUOX2;refGene_tx=XM_023255425.1
B3	60760947 60761112 refGene_gn=DUOX2;refGene_tx=XM_023255425.1
B3	60761650 60761838 refGene_gn=DUOX2;refGene_tx=XM_023255425.1
B3	60762020 60762222 refGene_gn=DUOX2;refGene_tx=XM_023255425.1
B3	60762352 60762519 refGene_gn=DUOX2;refGene_tx=XM_023255425.1
B3	60762916 60762977 refGene_gn=DUOX2;refGene_tx=XM_023255425.1
B3	60763096 60763193 refGene_gn=DUOX2;refGene_tx=XM_023255425.1
B3	60763622 60763713 refGene_gn=DUOX2;refGene_tx=XM_023255425.1
B3	60764259 60764362 refGene_gn=DUOX2;refGene_tx=XM_023255425.1
B3	60764915 60765079 refGene_gn=DUOX2;refGene_tx=XM_023255425.1
B3	60765677 60765853 refGene_gn=DUOX2;refGene_tx=XM_023255425.1
B3	60766070 60766189 refGene_gn=DUOX2;refGene_tx=XM_023255425.1
B3	60766530 60766668 refGene_gn=DUOX2;refGene_tx=XM_023255425.1
B3	60766886 60767000 refGene_gn=DUOX2;refGene_tx=XM_023255425.1
B3	60767192 60767395 refGene_gn=DUOX2;refGene_tx=XM_023255425.1
B3	60767704 60767890 refGene_gn=DUOX2;refGene_tx=XM_023255425.1
B3	60768828 60769054 refGene_gn=DUOX2;refGene_tx=XM_023255425.1
B3	60769133 60769227 refGene_gn=DUOX2;refGene_tx=XM_023255425.1
B3	60770874 60771062 refGene_gn=DUOX2;refGene_tx=XM_023255425.1
B3	60771805 60771875 refGene_gn=DUOX2;refGene_tx=XM_023255425.1
B3	60772292 60772376 refGene_gn=DUOX2;refGene_tx=XM_023255425.1
B3	60772925 60773104 refGene_gn=DUOX2;refGene_tx=XM_023255425.1
B3	60773245 60773476 refGene_gn=DUOX2;refGene_tx=XM_023255425.1
B3	60773655 60773755 refGene_gn=DUOX2;refGene_tx=XM_023255425.1
B3	60774447 60774497 refGene_gn=DUOX2;refGene_tx=XM_023255425.1
B3	60774763 60774891 refGene_gn=DUOX2;refGene_tx=XM_023255425.1
B3	60775123 60775256 refGene_gn=DUOX2;refGene_tx=XM_023255425.1
B3	60775258 60775276 refGene_gn=DUOX2;refGene_tx=XM_023255425.1
B3	60776672 60776905 refGene_gn=DUOX2;refGene_tx=XM_023255425.1
B3	60777128 60777287 refGene_gn=DUOX2;refGene_tx=XM_023255425.1
B3	60777636 60777792 refGene_gn=DUOX2;refGene_tx=XM_023255425.1
B3	60777900 60778029 refGene_gn=DUOX2;refGene_tx=XM_023255425.1
B3	60778343 60778466 refGene_gn=DUOX2;refGene_tx=XM_023255425.1
B3	60801754 60801920 refGene_gn=SORD;refGene_tx=XM_003987197.5
B3	60802814 60802936 refGene_gn=SORD;refGene_tx=XM_003987197.5
B3	60806069 60806245 refGene_gn=SORD;refGene_tx=XM_003987197.5
B3	60807279 60807345 refGene_gn=SORD;refGene_tx=XM_003987197.5
B3	60810665 60810784 refGene_gn=SORD;refGene_tx=XM_003987197.5
B3	60814483 60814643 refGene_gn=SORD;refGene_tx=XM_003987197.5
B3	60818349 60818514 refGene_gn=SORD;refGene_tx=XM_003987197.5
B3	60821064 60821098 refGene_gn=SORD;refGene_tx=XM_003987197.5
B3	60834794 60834857 refGene_gn=SORD;refGene_tx=XM_003987197.5
B3	60856498 60856554 refGene_gn=TERB2;refGene_tx=XM_023255426.1
B3	60867271 60867411 refGene_gn=TERB2;refGene_tx=XM_003987198.3
B3	60876260 60876349
refGene_gn=TERB2;refGene_tx=XM_003987198.3;refGene_gn=TERB2;refGene_tx
=XM_023255426.1
B3	60884141 60884227
refGene_gn=TERB2;refGene_tx=XM_003987198.3;refGene_gn=TERB2;refGene_tx
=XM_023255426.1

B3 60888302 60888364
refGene_gn=TERB2;refGene_tx=XM_003987198.3;refGene_gn=TERB2;refGene_tx
=XM_023255426.1
B3 60890712 60890852
refGene_gn=TERB2;refGene_tx=XM_003987198.3;refGene_gn=TERB2;refGene_tx
=XM_023255426.1
B3 60891851 60891933
refGene_gn=TERB2;refGene_tx=XM_003987198.3;refGene_gn=TERB2;refGene_tx
=XM_023255426.1
B3 60892047 60892111
refGene_gn=TERB2;refGene_tx=XM_003987198.3;refGene_gn=TERB2;refGene_tx
=XM_023255426.1
B3 60900151 60900693
refGene_gn=TRIM69;refGene_tx=XM_019832522.2;refGene_gn=TRIM69;refGene_
tx=XM_019832523.2;refGene_gn=TRIM69;refGene_tx=XM_019832524.2;refGene_
gn=TRIM69;refGene_tx=XM_019832525.2;refGene_gn=TRIM69;refGene_tx=XM_01
9832526.2;refGene_gn=TRIM69;refGene_tx=XM_019832528.2
B3 60908590 60908715
refGene_gn=TRIM69;refGene_tx=XM_019832522.2;refGene_gn=TRIM69;refGene_
tx=XM_019832523.2;refGene_gn=TRIM69;refGene_tx=XM_019832524.2;refGene_
gn=TRIM69;refGene_tx=XM_019832525.2;refGene_gn=TRIM69;refGene_tx=XM_01
9832526.2;refGene_gn=TRIM69;refGene_tx=XM_019832528.2
B3 60908797 60908820
refGene_gn=TRIM69;refGene_tx=XM_019832522.2;refGene_gn=TRIM69;refGene_
tx=XM_019832523.2;refGene_gn=TRIM69;refGene_tx=XM_019832524.2;refGene_
gn=TRIM69;refGene_tx=XM_019832525.2;refGene_gn=TRIM69;refGene_tx=XM_01
9832526.2;refGene_gn=TRIM69;refGene_tx=XM_019832528.2
B3 60909527 60909769
refGene_gn=TRIM69;refGene_tx=XM_019832522.2;refGene_gn=TRIM69;refGene_
tx=XM_019832523.2;refGene_gn=TRIM69;refGene_tx=XM_019832524.2;refGene_
gn=TRIM69;refGene_tx=XM_019832525.2;refGene_gn=TRIM69;refGene_tx=XM_01
9832526.2;refGene_gn=TRIM69;refGene_tx=XM_019832528.2
B3 60912815 60912977
refGene_gn=TRIM69;refGene_tx=XM_019832522.2;refGene_gn=TRIM69;refGene_
tx=XM_019832523.2;refGene_gn=TRIM69;refGene_tx=XM_019832524.2;refGene_
gn=TRIM69;refGene_tx=XM_019832525.2;refGene_gn=TRIM69;refGene_tx=XM_01
9832526.2;refGene_gn=TRIM69;refGene_tx=XM_019832528.2
B3 60913753 60914080
refGene_gn=TRIM69;refGene_tx=XM_019832522.2;refGene_gn=TRIM69;refGene_
tx=XM_019832523.2;refGene_gn=TRIM69;refGene_tx=XM_019832524.2;refGene_
gn=TRIM69;refGene_tx=XM_019832525.2;refGene_gn=TRIM69;refGene_tx=XM_01
9832526.2
B3 60972389 60972665 refGene_gn=B2M;refGene_tx=NM_001009876.1
B3 60975849 60975916 refGene_gn=B2M;refGene_tx=NM_001009876.1
B3 60995716 60995854 refGene_gn=PATL2;refGene_tx=XM_023256166.1
B3 60996500 60996581 refGene_gn=PATL2;refGene_tx=XM_023256166.1
B3 60997272 60997415 refGene_gn=PATL2;refGene_tx=XM_023256166.1
B3 60997934 60998003 refGene_gn=PATL2;refGene_tx=XM_023256166.1
B3 60998235 60998377 refGene_gn=PATL2;refGene_tx=XM_023256166.1
B3 60999658 60999779 refGene_gn=PATL2;refGene_tx=XM_023256166.1

| | | | |
|---|---|---|---|
| B3 | 60999879 | 60999977 | refGene_gn=PATL2;refGene_tx=XM_023256166.1 |
| B3 | 61000189 | 61000243 | refGene_gn=PATL2;refGene_tx=XM_023256166.1 |
| B3 | 61000344 | 61000478 | refGene_gn=PATL2;refGene_tx=XM_023256166.1 |
| B3 | 61000620 | 61000780 | refGene_gn=PATL2;refGene_tx=XM_023256166.1 |
| B3 | 61001262 | 61001403 | refGene_gn=PATL2;refGene_tx=XM_023256166.1 |
| B3 | 61002526 | 61002624 | refGene_gn=PATL2;refGene_tx=XM_023256166.1 |
| B3 | 61004484 | 61004634 | refGene_gn=PATL2;refGene_tx=XM_023256166.1 |
| B3 | 61007203 | 61007384 | refGene_gn=PATL2;refGene_tx=XM_023256166.1 |
| B3 | 61007407 | 61007661 | refGene_gn=SPG11;refGene_tx=XM_003987209.5 |
| B3 | 61010129 | 61010314 | refGene_gn=SPG11;refGene_tx=XM_003987209.5 |
| B3 | 61011296 | 61011521 | refGene_gn=SPG11;refGene_tx=XM_003987209.5 |
| B3 | 61012940 | 61013142 | refGene_gn=SPG11;refGene_tx=XM_003987209.5 |
| B3 | 61020737 | 61020875 | refGene_gn=SPG11;refGene_tx=XM_003987209.5 |
| B3 | 61021071 | 61021520 | refGene_gn=SPG11;refGene_tx=XM_003987209.5 |
| B3 | 61022840 | 61022986 | refGene_gn=SPG11;refGene_tx=XM_003987209.5 |
| B3 | 61034090 | 61034223 | refGene_gn=SPG11;refGene_tx=XM_003987209.5 |
| B3 | 61039934 | 61040090 | refGene_gn=SPG11;refGene_tx=XM_003987209.5 |
| B3 | 61040506 | 61040682 | refGene_gn=SPG11;refGene_tx=XM_003987209.5 |
| B3 | 61043195 | 61043372 | refGene_gn=SPG11;refGene_tx=XM_003987209.5 |
| B3 | 61044214 | 61044286 | refGene_gn=SPG11;refGene_tx=XM_003987209.5 |
| B3 | 61044571 | 61044699 | refGene_gn=SPG11;refGene_tx=XM_003987209.5 |
| B3 | 61045002 | 61045178 | refGene_gn=SPG11;refGene_tx=XM_003987209.5 |
| B3 | 61046061 | 61046275 | refGene_gn=SPG11;refGene_tx=XM_003987209.5 |
| B3 | 61051213 | 61051417 | refGene_gn=SPG11;refGene_tx=XM_003987209.5 |
| B3 | 61052714 | 61052821 | refGene_gn=SPG11;refGene_tx=XM_003987209.5 |
| B3 | 61054273 | 61054419 | refGene_gn=SPG11;refGene_tx=XM_003987209.5 |
| B3 | 61056417 | 61056579 | refGene_gn=SPG11;refGene_tx=XM_003987209.5 |
| B3 | 61058758 | 61058825 | refGene_gn=SPG11;refGene_tx=XM_003987209.5 |
| B3 | 61061850 | 61062016 | refGene_gn=SPG11;refGene_tx=XM_003987209.5 |
| B3 | 61063197 | 61063403 | refGene_gn=SPG11;refGene_tx=XM_003987209.5 |
| B3 | 61063666 | 61063775 | refGene_gn=SPG11;refGene_tx=XM_003987209.5 |
| B3 | 61064798 | 61064958 | refGene_gn=SPG11;refGene_tx=XM_003987209.5 |
| B3 | 61065566 | 61065839 | refGene_gn=SPG11;refGene_tx=XM_003987209.5 |
| B3 | 61066463 | 61066664 | refGene_gn=SPG11;refGene_tx=XM_003987209.5 |
| B3 | 61067398 | 61067506 | refGene_gn=SPG11;refGene_tx=XM_003987209.5 |
| B3 | 61077544 | 61077707 | refGene_gn=SPG11;refGene_tx=XM_003987209.5 |
| B3 | 61079901 | 61080116 | refGene_gn=SPG11;refGene_tx=XM_003987209.5 |
| B3 | 61080565 | 61081316 | refGene_gn=SPG11;refGene_tx=XM_003987209.5 |
| B3 | 61087156 | 61087296 | refGene_gn=SPG11;refGene_tx=XM_003987209.5 |
| B3 | 61088192 | 61088391 | refGene_gn=SPG11;refGene_tx=XM_003987209.5 |
| B3 | 61088792 | 61088930 | refGene_gn=SPG11;refGene_tx=XM_003987209.5 |
| B3 | 61089846 | 61089980 | refGene_gn=SPG11;refGene_tx=XM_003987209.5 |
| B3 | 61090696 | 61090804 | refGene_gn=SPG11;refGene_tx=XM_003987209.5 |
| B3 | 61092329 | 61092498 | refGene_gn=SPG11;refGene_tx=XM_003987209.5 |
| B3 | 61092934 | 61093023 | refGene_gn=SPG11;refGene_tx=XM_003987209.5 |
| B3 | 61093220 | 61093376 | refGene_gn=SPG11;refGene_tx=XM_003987209.5 |
| B3 | 61094052 | 61094204 | refGene_gn=SPG11;refGene_tx=XM_003987209.5 |
| B3 | 61094837 | 61095018 | refGene_gn=SPG11;refGene_tx=XM_003987209.5 |
| B3 | 61097010 | 61097142 | refGene_gn=EIF3J;refGene_tx=XM_003987207.5 |
| B3 | 61097455 | 61097529 | refGene_gn=EIF3J;refGene_tx=XM_003987207.5 |

B3 61098985 61099147 refGene_gn=EIF3J;refGene_tx=XM_003987207.5
B3 61101501 61101616 refGene_gn=EIF3J;refGene_tx=XM_003987207.5
B3 61106726 61106818 refGene_gn=EIF3J;refGene_tx=XM_003987207.5
B3 61107319 61107374 refGene_gn=EIF3J;refGene_tx=XM_003987207.5
B3 61116515 61116619 refGene_gn=EIF3J;refGene_tx=XM_003987207.5
B3 61116706 61116743 refGene_gn=EIF3J;refGene_tx=XM_003987207.5
B3 61131853 61131919
refGene_gn=CTDSPL2;refGene_tx=XM_003987210.5;refGene_gn=CTDSPL2;refGene_tx=XM_011282844.3;refGene_gn=CTDSPL2;refGene_tx=XM_019832530.2;refGene_gn=CTDSPL2;refGene_tx=XM_019832531.2
B3 61132536 61132632
refGene_gn=CTDSPL2;refGene_tx=XM_003987210.5;refGene_gn=CTDSPL2;refGene_tx=XM_011282844.3;refGene_gn=CTDSPL2;refGene_tx=XM_019832530.2;refGene_gn=CTDSPL2;refGene_tx=XM_019832531.2
B3 61134087 61134214
refGene_gn=CTDSPL2;refGene_tx=XM_003987210.5;refGene_gn=CTDSPL2;refGene_tx=XM_011282844.3;refGene_gn=CTDSPL2;refGene_tx=XM_019832530.2;refGene_gn=CTDSPL2;refGene_tx=XM_019832531.2
B3 61137179 61137259
refGene_gn=CTDSPL2;refGene_tx=XM_003987210.5;refGene_gn=CTDSPL2;refGene_tx=XM_011282844.3;refGene_gn=CTDSPL2;refGene_tx=XM_019832530.2;refGene_gn=CTDSPL2;refGene_tx=XM_019832531.2
B3 61137363 61137426
refGene_gn=CTDSPL2;refGene_tx=XM_003987210.5;refGene_gn=CTDSPL2;refGene_tx=XM_011282844.3;refGene_gn=CTDSPL2;refGene_tx=XM_019832530.2;refGene_gn=CTDSPL2;refGene_tx=XM_019832531.2
B3 61147651 61147738
refGene_gn=CTDSPL2;refGene_tx=XM_003987210.5;refGene_gn=CTDSPL2;refGene_tx=XM_011282844.3;refGene_gn=CTDSPL2;refGene_tx=XM_019832530.2;refGene_gn=CTDSPL2;refGene_tx=XM_019832531.2
B3 61149906 61150018
refGene_gn=CTDSPL2;refGene_tx=XM_003987210.5;refGene_gn=CTDSPL2;refGene_tx=XM_011282844.3;refGene_gn=CTDSPL2;refGene_tx=XM_019832530.2;refGene_gn=CTDSPL2;refGene_tx=XM_019832531.2
B3 61150355 61150434
refGene_gn=CTDSPL2;refGene_tx=XM_003987210.5;refGene_gn=CTDSPL2;refGene_tx=XM_011282844.3;refGene_gn=CTDSPL2;refGene_tx=XM_019832530.2;refGene_gn=CTDSPL2;refGene_tx=XM_019832531.2
B3 61154496 61154712
refGene_gn=CTDSPL2;refGene_tx=XM_003987210.5;refGene_gn=CTDSPL2;refGene_tx=XM_011282844.3;refGene_gn=CTDSPL2;refGene_tx=XM_019832530.2;refGene_gn=CTDSPL2;refGene_tx=XM_019832531.2
B3 61160975 61161125
refGene_gn=CTDSPL2;refGene_tx=XM_003987210.5;refGene_gn=CTDSPL2;refGene_tx=XM_011282844.3;refGene_gn=CTDSPL2;refGene_tx=XM_019832530.2;refGene_gn=CTDSPL2;refGene_tx=XM_019832531.2
B3 61162683 61162822
refGene_gn=CTDSPL2;refGene_tx=XM_003987210.5;refGene_gn=CTDSPL2;refGene_tx=XM_011282844.3;refGene_gn=CTDSPL2;refGene_tx=XM_019832530.2;refGene_gn=CTDSPL2;refGene_tx=XM_019832531.2

B3	61182451	61182637
refGene_gn=CTDSPL2;refGene_tx=XM_003987210.5;refGene_gn=CTDSPL2;refGene_tx=XM_011282844.3;refGene_gn=CTDSPL2;refGene_tx=XM_019832530.2;refGene_gn=CTDSPL2;refGene_tx=XM_019832531.2
B3	61220863	61220934
refGene_gn=CASC4;refGene_tx=XM_011282841.3;refGene_gn=CASC4;refGene_tx=XM_011282842.3;refGene_gn=CASC4;refGene_tx=XM_011282843.3
B3	61227737	61227905
refGene_gn=CASC4;refGene_tx=XM_011282841.3;refGene_gn=CASC4;refGene_tx=XM_011282842.3
B3	61258994	61259165
refGene_gn=CASC4;refGene_tx=XM_011282841.3;refGene_gn=CASC4;refGene_tx=XM_011282842.3;refGene_gn=CASC4;refGene_tx=XM_011282843.3
B3	61260893	61260992
refGene_gn=CASC4;refGene_tx=XM_011282841.3;refGene_gn=CASC4;refGene_tx=XM_011282843.3
B3	61277235	61277316
refGene_gn=CASC4;refGene_tx=XM_011282841.3;refGene_gn=CASC4;refGene_tx=XM_011282842.3;refGene_gn=CASC4;refGene_tx=XM_011282843.3
B3	61277640	61277785
refGene_gn=CASC4;refGene_tx=XM_011282841.3;refGene_gn=CASC4;refGene_tx=XM_011282842.3;refGene_gn=CASC4;refGene_tx=XM_011282843.3
B3	61279867	61279958
refGene_gn=CASC4;refGene_tx=XM_011282841.3;refGene_gn=CASC4;refGene_tx=XM_011282842.3;refGene_gn=CASC4;refGene_tx=XM_011282843.3
B3	61281662	61281765
refGene_gn=CASC4;refGene_tx=XM_011282841.3;refGene_gn=CASC4;refGene_tx=XM_011282842.3;refGene_gn=CASC4;refGene_tx=XM_011282843.3
B3	61288142	61288197
refGene_gn=CASC4;refGene_tx=XM_011282841.3;refGene_gn=CASC4;refGene_tx=XM_011282842.3;refGene_gn=CASC4;refGene_tx=XM_011282843.3
B3	61325081	61325408
refGene_gn=CASC4;refGene_tx=XM_011282841.3;refGene_gn=CASC4;refGene_tx=XM_011282842.3;refGene_gn=CASC4;refGene_tx=XM_011282843.3
B3	61404862	61404964
refGene_gn=FRMD5;refGene_tx=XM_023255429.1;refGene_gn=FRMD5;refGene_tx=XM_023255430.1;refGene_gn=FRMD5;refGene_tx=XM_023255431.1;refGene_gn=FRMD5;refGene_tx=XM_023255432.1;refGene_gn=FRMD5;refGene_tx=XM_023255434.1;refGene_gn=FRMD5;refGene_tx=XM_023255435.1
B3	61677072	61677177
refGene_gn=FRMD5;refGene_tx=XM_023255429.1;refGene_gn=FRMD5;refGene_tx=XM_023255430.1;refGene_gn=FRMD5;refGene_tx=XM_023255431.1;refGene_gn=FRMD5;refGene_tx=XM_023255432.1;refGene_gn=FRMD5;refGene_tx=XM_023255434.1;refGene_gn=FRMD5;refGene_tx=XM_023255435.1
B3	61681821	61681864
refGene_gn=FRMD5;refGene_tx=XM_023255429.1;refGene_gn=FRMD5;refGene_tx=XM_023255430.1;refGene_gn=FRMD5;refGene_tx=XM_023255431.1;refGene_gn=FRMD5;refGene_tx=XM_023255432.1;refGene_gn=FRMD5;refGene_tx=XM_023255434.1;refGene_gn=FRMD5;refGene_tx=XM_023255435.1
B3	61682092	61682171

refGene_gn=FRMD5;refGene_tx=XM_019832537.2;refGene_gn=FRMD5;refGene_tx
=XM_023255429.1;refGene_gn=FRMD5;refGene_tx=XM_023255430.1;refGene_gn=
FRMD5;refGene_tx=XM_023255431.1;refGene_gn=FRMD5;refGene_tx=XM_0232554
32.1;refGene_gn=FRMD5;refGene_tx=XM_023255434.1;refGene_gn=FRMD5;refGe
ne_tx=XM_023255435.1
B3 61689958 61690056
refGene_gn=FRMD5;refGene_tx=XM_019832537.2;refGene_gn=FRMD5;refGene_tx
=XM_023255429.1;refGene_gn=FRMD5;refGene_tx=XM_023255430.1;refGene_gn=
FRMD5;refGene_tx=XM_023255431.1;refGene_gn=FRMD5;refGene_tx=XM_0232554
32.1;refGene_gn=FRMD5;refGene_tx=XM_023255434.1;refGene_gn=FRMD5;refGe
ne_tx=XM_023255435.1
B3 61692528 61692652
refGene_gn=FRMD5;refGene_tx=XM_019832537.2;refGene_gn=FRMD5;refGene_tx
=XM_023255429.1;refGene_gn=FRMD5;refGene_tx=XM_023255430.1;refGene_gn=
FRMD5;refGene_tx=XM_023255431.1;refGene_gn=FRMD5;refGene_tx=XM_0232554
32.1;refGene_gn=FRMD5;refGene_tx=XM_023255434.1;refGene_gn=FRMD5;refGe
ne_tx=XM_023255435.1
B3 61695630 61695718
refGene_gn=FRMD5;refGene_tx=XM_019832537.2;refGene_gn=FRMD5;refGene_tx
=XM_023255429.1;refGene_gn=FRMD5;refGene_tx=XM_023255430.1;refGene_gn=
FRMD5;refGene_tx=XM_023255431.1;refGene_gn=FRMD5;refGene_tx=XM_0232554
32.1;refGene_gn=FRMD5;refGene_tx=XM_023255434.1;refGene_gn=FRMD5;refGe
ne_tx=XM_023255435.1
B3 61703737 61703826
refGene_gn=FRMD5;refGene_tx=XM_019832537.2;refGene_gn=FRMD5;refGene_tx
=XM_023255429.1;refGene_gn=FRMD5;refGene_tx=XM_023255430.1;refGene_gn=
FRMD5;refGene_tx=XM_023255431.1;refGene_gn=FRMD5;refGene_tx=XM_0232554
32.1;refGene_gn=FRMD5;refGene_tx=XM_023255434.1;refGene_gn=FRMD5;refGe
ne_tx=XM_023255435.1
B3 61706050 61706137
refGene_gn=FRMD5;refGene_tx=XM_019832537.2;refGene_gn=FRMD5;refGene_tx
=XM_023255429.1;refGene_gn=FRMD5;refGene_tx=XM_023255430.1;refGene_gn=
FRMD5;refGene_tx=XM_023255432.1;refGene_gn=FRMD5;refGene_tx=XM_0232554
34.1
B3 61706727 61706791
refGene_gn=FRMD5;refGene_tx=XM_019832537.2;refGene_gn=FRMD5;refGene_tx
=XM_023255429.1;refGene_gn=FRMD5;refGene_tx=XM_023255430.1;refGene_gn=
FRMD5;refGene_tx=XM_023255431.1;refGene_gn=FRMD5;refGene_tx=XM_0232554
32.1;refGene_gn=FRMD5;refGene_tx=XM_023255434.1;refGene_gn=FRMD5;refGe
ne_tx=XM_023255435.1
B3 61707383 61707475
refGene_gn=FRMD5;refGene_tx=XM_019832537.2;refGene_gn=FRMD5;refGene_tx
=XM_023255429.1;refGene_gn=FRMD5;refGene_tx=XM_023255430.1;refGene_gn=
FRMD5;refGene_tx=XM_023255431.1;refGene_gn=FRMD5;refGene_tx=XM_0232554
32.1;refGene_gn=FRMD5;refGene_tx=XM_023255434.1;refGene_gn=FRMD5;refGe
ne_tx=XM_023255435.1
B3 61710090 61710165
refGene_gn=FRMD5;refGene_tx=XM_019832537.2;refGene_gn=FRMD5;refGene_tx
=XM_023255429.1;refGene_gn=FRMD5;refGene_tx=XM_023255431.1;refGene_gn=
FRMD5;refGene_tx=XM_023255432.1;refGene_gn=FRMD5;refGene_tx=XM_0232554

34.1;refGene_gn=FRMD5;refGene_tx=XM_023255435.1
B3 61710734 61710803
refGene_gn=FRMD5;refGene_tx=XM_019832537.2;refGene_gn=FRMD5;refGene_tx=XM_023255429.1;refGene_gn=FRMD5;refGene_tx=XM_023255430.1;refGene_gn=FRMD5;refGene_tx=XM_023255431.1;refGene_gn=FRMD5;refGene_tx=XM_023255432.1;refGene_gn=FRMD5;refGene_tx=XM_023255434.1;refGene_gn=FRMD5;refGene_tx=XM_023255435.1
B3 61711906 61712013
refGene_gn=FRMD5;refGene_tx=XM_019832537.2;refGene_gn=FRMD5;refGene_tx=XM_023255429.1;refGene_gn=FRMD5;refGene_tx=XM_023255430.1;refGene_gn=FRMD5;refGene_tx=XM_023255431.1;refGene_gn=FRMD5;refGene_tx=XM_023255432.1;refGene_gn=FRMD5;refGene_tx=XM_023255434.1;refGene_gn=FRMD5;refGene_tx=XM_023255435.1
B3 61719434 61720012
refGene_gn=FRMD5;refGene_tx=XM_019832537.2;refGene_gn=FRMD5;refGene_tx=XM_023255429.1;refGene_gn=FRMD5;refGene_tx=XM_023255430.1;refGene_gn=FRMD5;refGene_tx=XM_023255431.1;refGene_gn=FRMD5;refGene_tx=XM_023255432.1;refGene_gn=FRMD5;refGene_tx=XM_023255434.1;refGene_gn=FRMD5;refGene_tx=XM_023255435.1
B3 61720654 61720729
refGene_gn=FRMD5;refGene_tx=XM_023255432.1;refGene_gn=FRMD5;refGene_tx=XM_023255434.1;refGene_gn=FRMD5;refGene_tx=XM_023255435.1
B3 61721049 61721063 refGene_gn=FRMD5;refGene_tx=XM_023255434.1
B3 61727793 61728049
refGene_gn=WDR76;refGene_tx=XM_019832540.2;refGene_gn=WDR76;refGene_tx=XM_019832541.2;refGene_gn=WDR76;refGene_tx=XM_019832542.1;refGene_gn=WDR76;refGene_tx=XM_019832543.2;refGene_gn=WDR76;refGene_tx=XM_019832545.2;refGene_gn=WDR76;refGene_tx=XM_023255436.1
B3 61733875 61733929
refGene_gn=WDR76;refGene_tx=XM_019832540.2;refGene_gn=WDR76;refGene_tx=XM_019832541.2;refGene_gn=WDR76;refGene_tx=XM_019832542.1;refGene_gn=WDR76;refGene_tx=XM_019832543.2;refGene_gn=WDR76;refGene_tx=XM_019832545.2;refGene_gn=WDR76;refGene_tx=XM_023255436.1
B3 61735555 61735708
refGene_gn=WDR76;refGene_tx=XM_019832540.2;refGene_gn=WDR76;refGene_tx=XM_019832541.2;refGene_gn=WDR76;refGene_tx=XM_019832542.1;refGene_gn=WDR76;refGene_tx=XM_019832543.2;refGene_gn=WDR76;refGene_tx=XM_019832545.2;refGene_gn=WDR76;refGene_tx=XM_023255436.1
B3 61736271 61736489
refGene_gn=WDR76;refGene_tx=XM_019832540.2;refGene_gn=WDR76;refGene_tx=XM_019832541.2;refGene_gn=WDR76;refGene_tx=XM_019832542.1;refGene_gn=WDR76;refGene_tx=XM_019832543.2;refGene_gn=WDR76;refGene_tx=XM_019832545.2;refGene_gn=WDR76;refGene_tx=XM_023255436.1
B3 61746764 61746923
refGene_gn=WDR76;refGene_tx=XM_019832540.2;refGene_gn=WDR76;refGene_tx=XM_019832541.2;refGene_gn=WDR76;refGene_tx=XM_019832542.1;refGene_gn=WDR76;refGene_tx=XM_019832543.2;refGene_gn=WDR76;refGene_tx=XM_019832545.2;refGene_gn=WDR76;refGene_tx=XM_023255436.1
B3 61749139 61749166
refGene_gn=WDR76;refGene_tx=XM_019832540.2;refGene_gn=WDR76;refGene_tx

=XM_019832542.1;refGene_gn=WDR76;refGene_tx=XM_019832543.2;refGene_gn=WDR76;refGene_tx=XM_019832545.2;refGene_gn=WDR76;refGene_tx=XM_023255436.1
B3 61760190 61760344 refGene_gn=WDR76;refGene_tx=XM_019832540.2;refGene_gn=WDR76;refGene_tx=XM_019832541.2;refGene_gn=WDR76;refGene_tx=XM_019832542.1;refGene_gn=WDR76;refGene_tx=XM_019832543.2;refGene_gn=WDR76;refGene_tx=XM_019832545.2;refGene_gn=WDR76;refGene_tx=XM_023255436.1
B3 61761388 61761432 refGene_gn=WDR76;refGene_tx=XM_019832540.2;refGene_gn=WDR76;refGene_tx=XM_019832541.2;refGene_gn=WDR76;refGene_tx=XM_019832542.1;refGene_gn=WDR76;refGene_tx=XM_019832543.2;refGene_gn=WDR76;refGene_tx=XM_019832545.2;refGene_gn=WDR76;refGene_tx=XM_023255436.1
B3 61761529 61761631 refGene_gn=WDR76;refGene_tx=XM_019832540.2;refGene_gn=WDR76;refGene_tx=XM_019832541.2;refGene_gn=WDR76;refGene_tx=XM_019832542.1;refGene_gn=WDR76;refGene_tx=XM_019832543.2;refGene_gn=WDR76;refGene_tx=XM_019832545.2;refGene_gn=WDR76;refGene_tx=XM_023255436.1
B3 61763545 61763672 refGene_gn=WDR76;refGene_tx=XM_019832540.2;refGene_gn=WDR76;refGene_tx=XM_019832541.2;refGene_gn=WDR76;refGene_tx=XM_019832542.1;refGene_gn=WDR76;refGene_tx=XM_019832543.2;refGene_gn=WDR76;refGene_tx=XM_019832545.2;refGene_gn=WDR76;refGene_tx=XM_023255436.1
B3 61779005 61779061 refGene_gn=WDR76;refGene_tx=XM_019832540.2;refGene_gn=WDR76;refGene_tx=XM_019832541.2;refGene_gn=WDR76;refGene_tx=XM_019832542.1;refGene_gn=WDR76;refGene_tx=XM_019832543.2;refGene_gn=WDR76;refGene_tx=XM_019832545.2;refGene_gn=WDR76;refGene_tx=XM_023255436.1
B3 61781223 61781316 refGene_gn=WDR76;refGene_tx=XM_019832540.2;refGene_gn=WDR76;refGene_tx=XM_019832541.2;refGene_gn=WDR76;refGene_tx=XM_019832542.1;refGene_gn=WDR76;refGene_tx=XM_019832545.2;refGene_gn=WDR76;refGene_tx=XM_023255436.1
B3 61789144 61789555 refGene_gn=WDR76;refGene_tx=XM_019832540.2;refGene_gn=WDR76;refGene_tx=XM_019832541.2;refGene_gn=WDR76;refGene_tx=XM_019832542.1;refGene_gn=WDR76;refGene_tx=XM_019832543.2;refGene_gn=WDR76;refGene_tx=XM_019832545.2
B3 61790192 61790291 refGene_gn=WDR76;refGene_tx=XM_019832540.2;refGene_gn=WDR76;refGene_tx=XM_019832541.2;refGene_gn=WDR76;refGene_tx=XM_019832543.2;refGene_gn=WDR76;refGene_tx=XM_023255436.1
B3 61790515 61790621 refGene_gn=WDR76;refGene_tx=XM_019832542.1
B3 61792052 61792161 refGene_gn=WDR76;refGene_tx=XM_019832542.1
B3 61792385 61792464 refGene_gn=MFAP1;refGene_tx=XM_003987216.5
B3 61798376 61798596 refGene_gn=MFAP1;refGene_tx=XM_003987216.5
B3 61801005 61801135 refGene_gn=MFAP1;refGene_tx=XM_003987216.5
B3 61801678 61801866 refGene_gn=MFAP1;refGene_tx=XM_003987216.5
B3 61802351 61802460 refGene_gn=MFAP1;refGene_tx=XM_003987216.5
B3 61802562 61802723 refGene_gn=MFAP1;refGene_tx=XM_003987216.5

B3 61804011 61804171 refGene_gn=MFAP1;refGene_tx=XM_003987216.5
B3 61807892 61807982 refGene_gn=MFAP1;refGene_tx=XM_003987216.5
B3 61808080 61808263 refGene_gn=MFAP1;refGene_tx=XM_003987216.5
B3 61810495 61810695
refGene_gn=HYPK;refGene_tx=XM_003987217.5;refGene_gn=HYPK;refGene_tx=XM_006932578.4
B3 61810846 61810898 refGene_gn=HYPK;refGene_tx=XM_003987217.5
B3 61810941 61811009 refGene_gn=HYPK;refGene_tx=XM_006932578.4
B3 61811274 61811330
refGene_gn=HYPK;refGene_tx=XM_003987217.5;refGene_gn=HYPK;refGene_tx=XM_006932578.4
B3 61811652 61811814
refGene_gn=HYPK;refGene_tx=XM_003987217.5;refGene_gn=HYPK;refGene_tx=XM_006932578.4
B3 61812429 61812543
refGene_gn=LOC109500696;refGene_tx=XM_019832546.2
B3 61813027 61813204
refGene_gn=LOC109500696;refGene_tx=XM_019832546.2
B3 61813374 61813638
refGene_gn=LOC109500696;refGene_tx=XM_019832546.2
B3 61814081 61814099
refGene_gn=LOC101099886;refGene_tx=XM_019832547.2
B3 61814229 61814323
refGene_gn=LOC101099886;refGene_tx=XM_019832547.2
B3 61814970 61815066
refGene_gn=LOC101099886;refGene_tx=XM_019832547.2
B3 61815165 61815292
refGene_gn=LOC101099886;refGene_tx=XM_019832547.2
B3 61815691 61815764
refGene_gn=LOC101099886;refGene_tx=XM_019832547.2
B3 61816198 61816247
refGene_gn=LOC101099886;refGene_tx=XM_019832547.2
B3 61816393 61816547
refGene_gn=LOC101099886;refGene_tx=XM_019832547.2
B3 61816730 61816926
refGene_gn=LOC101099886;refGene_tx=XM_019832547.2
B3 61817937 61818223
refGene_gn=SERF2;refGene_tx=XM_003987218.4;refGene_gn=SERF2;refGene_tx=XM_011283027.2
B3 61818817 61818926
refGene_gn=SERF2;refGene_tx=XM_003987218.4;refGene_gn=SERF2;refGene_tx=XM_011283027.2
B3 61819538 61819545
refGene_gn=SERF2;refGene_tx=XM_003987218.4;refGene_gn=SERF2;refGene_tx=XM_011283027.2
B3 61832859 61832865 refGene_gn=ELL3;refGene_tx=XM_006932581.3
B3 61840652 61840784
refGene_gn=ELL3;refGene_tx=XM_019832549.1;refGene_gn=ELL3;refGene_tx=XM_019832550.1
B3 61841044 61841080

refGene_gn=ELL3;refGene_tx=XM_006932581.3;refGene_gn=ELL3;refGene_tx=XM_019832549.1;refGene_gn=ELL3;refGene_tx=XM_019832550.1
B3 61841436 61841546
refGene_gn=ELL3;refGene_tx=XM_006932581.3;refGene_gn=ELL3;refGene_tx=XM_019832549.1;refGene_gn=ELL3;refGene_tx=XM_019832550.1
B3 61841659 61841861
refGene_gn=ELL3;refGene_tx=XM_006932581.3;refGene_gn=ELL3;refGene_tx=XM_019832549.1;refGene_gn=ELL3;refGene_tx=XM_019832550.1
B3 61841950 61842030
refGene_gn=ELL3;refGene_tx=XM_006932581.3;refGene_gn=ELL3;refGene_tx=XM_019832549.1;refGene_gn=ELL3;refGene_tx=XM_019832550.1
B3 61842167 61842243
refGene_gn=ELL3;refGene_tx=XM_006932581.3;refGene_gn=ELL3;refGene_tx=XM_019832549.1;refGene_gn=ELL3;refGene_tx=XM_019832550.1
B3 61842417 61842598
refGene_gn=ELL3;refGene_tx=XM_006932581.3;refGene_gn=ELL3;refGene_tx=XM_019832549.1;refGene_gn=ELL3;refGene_tx=XM_019832550.1
B3 61842680 61842723
refGene_gn=ELL3;refGene_tx=XM_006932581.3;refGene_gn=ELL3;refGene_tx=XM_019832549.1;refGene_gn=ELL3;refGene_tx=XM_019832550.1
B3 61842847 61843019
refGene_gn=ELL3;refGene_tx=XM_006932581.3;refGene_gn=ELL3;refGene_tx=XM_019832549.1;refGene_gn=ELL3;refGene_tx=XM_019832550.1
B3 61843436 61843481
refGene_gn=ELL3;refGene_tx=XM_006932581.3;refGene_gn=ELL3;refGene_tx=XM_019832549.1;refGene_gn=ELL3;refGene_tx=XM_019832550.1
B3 61843573 61843684
refGene_gn=ELL3;refGene_tx=XM_006932581.3;refGene_gn=ELL3;refGene_tx=XM_019832549.1;refGene_gn=ELL3;refGene_tx=XM_019832550.1
B3 61844634 61844636 refGene_gn=ELL3;refGene_tx=XM_019832550.1
B3 61845591 61845705 refGene_gn=PDIA3;refGene_tx=XM_003987221.4
B3 61845914 61845972 refGene_gn=PDIA3;refGene_tx=XM_003987221.4
B3 61846290 61846370 refGene_gn=PDIA3;refGene_tx=XM_003987221.4
B3 61846905 61847034 refGene_gn=PDIA3;refGene_tx=XM_003987221.4
B3 61847868 61847977 refGene_gn=PDIA3;refGene_tx=XM_003987221.4
B3 61848759 61848942 refGene_gn=PDIA3;refGene_tx=XM_003987221.4
B3 61849551 61849677 refGene_gn=PDIA3;refGene_tx=XM_003987221.4
B3 61849967 61850084 refGene_gn=PDIA3;refGene_tx=XM_003987221.4
B3 61853575 61853705 refGene_gn=PDIA3;refGene_tx=XM_003987221.4
B3 61854887 61854995 refGene_gn=PDIA3;refGene_tx=XM_003987221.4
B3 61863328 61863446 refGene_gn=PDIA3;refGene_tx=XM_003987221.4
B3 61865146 61865225 refGene_gn=PDIA3;refGene_tx=XM_003987221.4
B3 61870739 61870906 refGene_gn=PDIA3;refGene_tx=XM_003987221.4
B3 61871458 61871705
refGene_gn=CATSPER2;refGene_tx=XM_003987222.3;refGene_gn=CATSPER2;refGene_tx=XM_006932582.2;refGene_gn=CATSPER2;refGene_tx=XM_019832551.1
B3 61871903 61872050
refGene_gn=CATSPER2;refGene_tx=XM_003987222.3;refGene_gn=CATSPER2;refGene_tx=XM_006932582.2;refGene_gn=CATSPER2;refGene_tx=XM_006932583.4;refGene_gn=CATSPER2;refGene_tx=XM_011283028.3;refGene_gn=CATSPER2;refGen

e_tx=XM_019832551.1
B3 61872554 61872728
refGene_gn=CATSPER2;refGene_tx=XM_003987222.3;refGene_gn=CATSPER2;refGene_tx=XM_006932583.4;refGene_gn=CATSPER2;refGene_tx=XM_011283028.3;refGene_gn=CATSPER2;refGene_tx=XM_019832551.1
B3 61872919 61872988
refGene_gn=CATSPER2;refGene_tx=XM_003987222.3;refGene_gn=CATSPER2;refGene_tx=XM_006932583.4;refGene_gn=CATSPER2;refGene_tx=XM_011283028.3;refGene_gn=CATSPER2;refGene_tx=XM_019832551.1
B3 61877857 61878030
refGene_gn=CATSPER2;refGene_tx=XM_003987222.3;refGene_gn=CATSPER2;refGene_tx=XM_006932582.2;refGene_gn=CATSPER2;refGene_tx=XM_006932583.4;refGene_gn=CATSPER2;refGene_tx=XM_011283028.3;refGene_gn=CATSPER2;refGene_tx=XM_019832551.1
B3 61878539 61878695
refGene_gn=CATSPER2;refGene_tx=XM_003987222.3;refGene_gn=CATSPER2;refGene_tx=XM_006932582.2;refGene_gn=CATSPER2;refGene_tx=XM_006932583.4;refGene_gn=CATSPER2;refGene_tx=XM_011283028.3;refGene_gn=CATSPER2;refGene_tx=XM_019832551.1
B3 61878973 61879098
refGene_gn=CATSPER2;refGene_tx=XM_003987222.3;refGene_gn=CATSPER2;refGene_tx=XM_006932582.2;refGene_gn=CATSPER2;refGene_tx=XM_006932583.4;refGene_gn=CATSPER2;refGene_tx=XM_011283028.3;refGene_gn=CATSPER2;refGene_tx=XM_019832551.1
B3 61880341 61880520
refGene_gn=CATSPER2;refGene_tx=XM_003987222.3;refGene_gn=CATSPER2;refGene_tx=XM_006932582.2;refGene_gn=CATSPER2;refGene_tx=XM_006932583.4;refGene_gn=CATSPER2;refGene_tx=XM_011283028.3
B3 61880734 61880834
refGene_gn=CATSPER2;refGene_tx=XM_003987222.3;refGene_gn=CATSPER2;refGene_tx=XM_006932582.2;refGene_gn=CATSPER2;refGene_tx=XM_006932583.4;refGene_gn=CATSPER2;refGene_tx=XM_011283028.3
B3 61881156 61881213
refGene_gn=CATSPER2;refGene_tx=XM_003987222.3;refGene_gn=CATSPER2;refGene_tx=XM_006932582.2;refGene_gn=CATSPER2;refGene_tx=XM_006932583.4;refGene_gn=CATSPER2;refGene_tx=XM_011283028.3
B3 61883654 61883896
refGene_gn=CATSPER2;refGene_tx=XM_003987222.3;refGene_gn=CATSPER2;refGene_tx=XM_006932582.2;refGene_gn=CATSPER2;refGene_tx=XM_006932583.4;refGene_gn=CATSPER2;refGene_tx=XM_011283028.3;refGene_gn=CATSPER2;refGene_tx=XM_019832551.1
B3 61884255 61884423
refGene_gn=CATSPER2;refGene_tx=XM_003987222.3;refGene_gn=CATSPER2;refGene_tx=XM_006932582.2;refGene_gn=CATSPER2;refGene_tx=XM_006932583.4;refGene_gn=CATSPER2;refGene_tx=XM_011283028.3;refGene_gn=CATSPER2;refGene_tx=XM_019832551.1
B3 61885571 61885603
refGene_gn=CATSPER2;refGene_tx=XM_003987222.3;refGene_gn=CATSPER2;refGene_tx=XM_006932582.2;refGene_gn=CATSPER2;refGene_tx=XM_006932583.4;refGene_gn=CATSPER2;refGene_tx=XM_011283028.3;refGene_gn=CATSPER2;refGen

e_tx=XM_019832551.1
B3 61903335 61903381
refGene_gn=STRC;refGene_tx=XM_003987223.3;refGene_gn=STRC;refGene_tx=XM_019832552.1;refGene_gn=STRC;refGene_tx=XM_019832553.1;refGene_gn=STRC;refGene_tx=XM_019832554.1;refGene_gn=STRC;refGene_tx=XM_019832560.1
B3 61903695 61904481
refGene_gn=STRC;refGene_tx=XM_003987223.3;refGene_gn=STRC;refGene_tx=XM_019832552.1;refGene_gn=STRC;refGene_tx=XM_019832553.1;refGene_gn=STRC;refGene_tx=XM_019832554.1;refGene_gn=STRC;refGene_tx=XM_019832560.1
B3 61904572 61904597
refGene_gn=STRC;refGene_tx=XM_003987223.3;refGene_gn=STRC;refGene_tx=XM_019832552.1;refGene_gn=STRC;refGene_tx=XM_019832553.1;refGene_gn=STRC;refGene_tx=XM_019832554.1;refGene_gn=STRC;refGene_tx=XM_019832560.1
B3 61905347 61906602
refGene_gn=STRC;refGene_tx=XM_003987223.3;refGene_gn=STRC;refGene_tx=XM_019832552.1;refGene_gn=STRC;refGene_tx=XM_019832553.1;refGene_gn=STRC;refGene_tx=XM_019832554.1;refGene_gn=STRC;refGene_tx=XM_019832560.1
B3 61907569 61907632
refGene_gn=STRC;refGene_tx=XM_003987223.3;refGene_gn=STRC;refGene_tx=XM_019832552.1;refGene_gn=STRC;refGene_tx=XM_019832553.1;refGene_gn=STRC;refGene_tx=XM_019832554.1;refGene_gn=STRC;refGene_tx=XM_019832560.1
B3 61907729 61907840
refGene_gn=STRC;refGene_tx=XM_003987223.3;refGene_gn=STRC;refGene_tx=XM_019832552.1;refGene_gn=STRC;refGene_tx=XM_019832553.1;refGene_gn=STRC;refGene_tx=XM_019832554.1;refGene_gn=STRC;refGene_tx=XM_019832560.1
B3 61908014 61908181
refGene_gn=STRC;refGene_tx=XM_003987223.3;refGene_gn=STRC;refGene_tx=XM_019832552.1;refGene_gn=STRC;refGene_tx=XM_019832553.1;refGene_gn=STRC;refGene_tx=XM_019832554.1;refGene_gn=STRC;refGene_tx=XM_019832560.1
B3 61908368 61908555
refGene_gn=STRC;refGene_tx=XM_003987223.3;refGene_gn=STRC;refGene_tx=XM_019832552.1;refGene_gn=STRC;refGene_tx=XM_019832553.1;refGene_gn=STRC;refGene_tx=XM_019832554.1;refGene_gn=STRC;refGene_tx=XM_019832560.1
B3 61908686 61908802
refGene_gn=STRC;refGene_tx=XM_003987223.3;refGene_gn=STRC;refGene_tx=XM_019832552.1;refGene_gn=STRC;refGene_tx=XM_019832553.1;refGene_gn=STRC;refGene_tx=XM_019832554.1;refGene_gn=STRC;refGene_tx=XM_019832560.1
B3 61909062 61909195
refGene_gn=STRC;refGene_tx=XM_003987223.3;refGene_gn=STRC;refGene_tx=XM_019832552.1;refGene_gn=STRC;refGene_tx=XM_019832553.1;refGene_gn=STRC;refGene_tx=XM_019832554.1;refGene_gn=STRC;refGene_tx=XM_019832560.1
B3 61909553 61909736
refGene_gn=STRC;refGene_tx=XM_003987223.3;refGene_gn=STRC;refGene_tx=XM_019832552.1;refGene_gn=STRC;refGene_tx=XM_019832553.1;refGene_gn=STRC;refGene_tx=XM_019832554.1;refGene_gn=STRC;refGene_tx=XM_019832560.1
B3 61910019 61910058
refGene_gn=STRC;refGene_tx=XM_003987223.3;refGene_gn=STRC;refGene_tx=XM_019832552.1;refGene_gn=STRC;refGene_tx=XM_019832553.1;refGene_gn=STRC;refGene_tx=XM_019832554.1;refGene_gn=STRC;refGene_tx=XM_019832560.1
B3 61910238 61910406

refGene_gn=STRC;refGene_tx=XM_003987223.3;refGene_gn=STRC;refGene_tx=XM_019832552.1;refGene_gn=STRC;refGene_tx=XM_019832560.1
B3 61910578 61910644
refGene_gn=STRC;refGene_tx=XM_003987223.3;refGene_gn=STRC;refGene_tx=XM_019832552.1;refGene_gn=STRC;refGene_tx=XM_019832553.1;refGene_gn=STRC;refGene_tx=XM_019832554.1;refGene_gn=STRC;refGene_tx=XM_019832560.1
B3 61911100 61911229
refGene_gn=STRC;refGene_tx=XM_003987223.3;refGene_gn=STRC;refGene_tx=XM_019832552.1;refGene_gn=STRC;refGene_tx=XM_019832553.1;refGene_gn=STRC;refGene_tx=XM_019832554.1;refGene_gn=STRC;refGene_tx=XM_019832560.1
B3 61911525 61911584
refGene_gn=STRC;refGene_tx=XM_003987223.3;refGene_gn=STRC;refGene_tx=XM_019832552.1;refGene_gn=STRC;refGene_tx=XM_019832553.1;refGene_gn=STRC;refGene_tx=XM_019832554.1;refGene_gn=STRC;refGene_tx=XM_019832560.1
B3 61912369 61912493
refGene_gn=STRC;refGene_tx=XM_003987223.3;refGene_gn=STRC;refGene_tx=XM_019832552.1;refGene_gn=STRC;refGene_tx=XM_019832553.1;refGene_gn=STRC;refGene_tx=XM_019832554.1;refGene_gn=STRC;refGene_tx=XM_019832560.1
B3 61912604 61912717
refGene_gn=STRC;refGene_tx=XM_003987223.3;refGene_gn=STRC;refGene_tx=XM_019832552.1;refGene_gn=STRC;refGene_tx=XM_019832553.1;refGene_gn=STRC;refGene_tx=XM_019832554.1;refGene_gn=STRC;refGene_tx=XM_019832560.1
B3 61914492 61914628
refGene_gn=STRC;refGene_tx=XM_003987223.3;refGene_gn=STRC;refGene_tx=XM_019832552.1;refGene_gn=STRC;refGene_tx=XM_019832553.1;refGene_gn=STRC;refGene_tx=XM_019832554.1;refGene_gn=STRC;refGene_tx=XM_019832560.1
B3 61915042 61915239
refGene_gn=STRC;refGene_tx=XM_003987223.3;refGene_gn=STRC;refGene_tx=XM_019832552.1;refGene_gn=STRC;refGene_tx=XM_019832553.1;refGene_gn=STRC;refGene_tx=XM_019832554.1;refGene_gn=STRC;refGene_tx=XM_019832560.1
B3 61915433 61915524
refGene_gn=STRC;refGene_tx=XM_003987223.3;refGene_gn=STRC;refGene_tx=XM_019832552.1;refGene_gn=STRC;refGene_tx=XM_019832553.1;refGene_gn=STRC;refGene_tx=XM_019832554.1;refGene_gn=STRC;refGene_tx=XM_019832560.1
B3 61915736 61915893
refGene_gn=STRC;refGene_tx=XM_003987223.3;refGene_gn=STRC;refGene_tx=XM_019832552.1;refGene_gn=STRC;refGene_tx=XM_019832553.1;refGene_gn=STRC;refGene_tx=XM_019832554.1;refGene_gn=STRC;refGene_tx=XM_019832560.1
B3 61916525 61916695
refGene_gn=STRC;refGene_tx=XM_003987223.3;refGene_gn=STRC;refGene_tx=XM_019832552.1;refGene_gn=STRC;refGene_tx=XM_019832553.1;refGene_gn=STRC;refGene_tx=XM_019832554.1
B3 61917298 61917454
refGene_gn=STRC;refGene_tx=XM_003987223.3;refGene_gn=STRC;refGene_tx=XM_019832552.1;refGene_gn=STRC;refGene_tx=XM_019832553.1;refGene_gn=STRC;refGene_tx=XM_019832554.1;refGene_gn=STRC;refGene_tx=XM_019832560.1
B3 61917833 61917976
refGene_gn=STRC;refGene_tx=XM_003987223.3;refGene_gn=STRC;refGene_tx=XM_019832552.1;refGene_gn=STRC;refGene_tx=XM_019832553.1;refGene_gn=STRC;refGene_tx=XM_019832554.1

B3	61918166 61918315
refGene_gn=STRC;refGene_tx=XM_003987223.3;refGene_gn=STRC;refGene_tx=XM_019832552.1;refGene_gn=STRC;refGene_tx=XM_019832553.1;refGene_gn=STRC;refGene_tx=XM_019832554.1
B3	61918578 61918676
refGene_gn=STRC;refGene_tx=XM_003987223.3;refGene_gn=STRC;refGene_tx=XM_019832552.1;refGene_gn=STRC;refGene_tx=XM_019832553.1;refGene_gn=STRC;refGene_tx=XM_019832554.1
B3	61918771 61918919
refGene_gn=STRC;refGene_tx=XM_003987223.3;refGene_gn=STRC;refGene_tx=XM_019832552.1;refGene_gn=STRC;refGene_tx=XM_019832553.1;refGene_gn=STRC;refGene_tx=XM_019832554.1
B3	61919114 61919203
refGene_gn=STRC;refGene_tx=XM_003987223.3;refGene_gn=STRC;refGene_tx=XM_019832552.1;refGene_gn=STRC;refGene_tx=XM_019832553.1;refGene_gn=STRC;refGene_tx=XM_019832554.1
B3	61919591 61919708
refGene_gn=CKMT1A;refGene_tx=XM_011283029.3;refGene_gn=CKMT1A;refGene_tx=XM_023255438.1
B3	61919920 61920046
refGene_gn=CKMT1A;refGene_tx=XM_011283029.3;refGene_gn=CKMT1A;refGene_tx=XM_023255438.1
B3	61920238 61920373
refGene_gn=CKMT1A;refGene_tx=XM_011283029.3;refGene_gn=CKMT1A;refGene_tx=XM_023255438.1
B3	61922009 61922133
refGene_gn=CKMT1A;refGene_tx=XM_011283029.3;refGene_gn=CKMT1A;refGene_tx=XM_023255438.1
B3	61922252 61922338
refGene_gn=CKMT1A;refGene_tx=XM_011283029.3;refGene_gn=CKMT1A;refGene_tx=XM_023255438.1
B3	61922473 61922695
refGene_gn=CKMT1A;refGene_tx=XM_011283029.3;refGene_gn=CKMT1A;refGene_tx=XM_023255438.1
B3	61923099 61923195
refGene_gn=CKMT1A;refGene_tx=XM_011283029.3;refGene_gn=CKMT1A;refGene_tx=XM_023255438.1
B3	61923398 61923597
refGene_gn=CKMT1A;refGene_tx=XM_011283029.3;refGene_gn=CKMT1A;refGene_tx=XM_023255438.1
B3	61924262 61924411
refGene_gn=CKMT1A;refGene_tx=XM_011283029.3;refGene_gn=CKMT1A;refGene_tx=XM_023255438.1
B3	61932276 61932423
refGene_gn=PPIP5K1;refGene_tx=XM_006932584.4;refGene_gn=PPIP5K1;refGene_tx=XM_011283034.3;refGene_gn=PPIP5K1;refGene_tx=XM_011283035.3;refGene_gn=PPIP5K1;refGene_tx=XM_011283037.3;refGene_gn=PPIP5K1;refGene_tx=XM_011283038.3;refGene_gn=PPIP5K1;refGene_tx=XM_011283039.3;refGene_gn=PPIP5K1;refGene_tx=XM_011283040.3;refGene_gn=PPIP5K1;refGene_tx=XM_019832561.2;refGene_gn=PPIP5K1;refGene_tx=XM_019832562.2;refGene_gn=PPIP

5K1;refGene_tx=XM_019832563.2;refGene_gn=PPIP5K1;refGene_tx=XM_019832564.2;refGene_gn=PPIP5K1;refGene_tx=XM_019832565.2;refGene_gn=PPIP5K1;refGene_tx=XM_019832566.2;refGene_gn=PPIP5K1;refGene_tx=XM_023255439.1;refGene_gn=PPIP5K1;refGene_tx=XM_023255440.1;refGene_gn=PPIP5K1;refGene_tx=XM_023255441.1
B3 61932884 61933080
refGene_gn=PPIP5K1;refGene_tx=XM_006932584.4;refGene_gn=PPIP5K1;refGene_tx=XM_011283034.3;refGene_gn=PPIP5K1;refGene_tx=XM_011283035.3;refGene_gn=PPIP5K1;refGene_tx=XM_011283037.3;refGene_gn=PPIP5K1;refGene_tx=XM_011283038.3;refGene_gn=PPIP5K1;refGene_tx=XM_011283039.3;refGene_gn=PPIP5K1;refGene_tx=XM_011283040.3;refGene_gn=PPIP5K1;refGene_tx=XM_019832561.2;refGene_gn=PPIP5K1;refGene_tx=XM_019832562.2;refGene_gn=PPIP5K1;refGene_tx=XM_019832563.2;refGene_gn=PPIP5K1;refGene_tx=XM_019832564.2;refGene_gn=PPIP5K1;refGene_tx=XM_019832565.2;refGene_gn=PPIP5K1;refGene_tx=XM_019832566.2;refGene_gn=PPIP5K1;refGene_tx=XM_023255439.1;refGene_gn=PPIP5K1;refGene_tx=XM_023255440.1;refGene_gn=PPIP5K1;refGene_tx=XM_023255441.1
B3 61933523 61933614
refGene_gn=PPIP5K1;refGene_tx=XM_006932584.4;refGene_gn=PPIP5K1;refGene_tx=XM_011283034.3;refGene_gn=PPIP5K1;refGene_tx=XM_011283035.3;refGene_gn=PPIP5K1;refGene_tx=XM_011283037.3;refGene_gn=PPIP5K1;refGene_tx=XM_011283038.3;refGene_gn=PPIP5K1;refGene_tx=XM_011283039.3;refGene_gn=PPIP5K1;refGene_tx=XM_011283040.3;refGene_gn=PPIP5K1;refGene_tx=XM_019832561.2;refGene_gn=PPIP5K1;refGene_tx=XM_019832562.2;refGene_gn=PPIP5K1;refGene_tx=XM_019832563.2;refGene_gn=PPIP5K1;refGene_tx=XM_019832564.2;refGene_gn=PPIP5K1;refGene_tx=XM_019832565.2;refGene_gn=PPIP5K1;refGene_tx=XM_019832566.2;refGene_gn=PPIP5K1;refGene_tx=XM_023255439.1;refGene_gn=PPIP5K1;refGene_tx=XM_023255440.1;refGene_gn=PPIP5K1;refGene_tx=XM_023255441.1
B3 61933840 61933926
refGene_gn=PPIP5K1;refGene_tx=XM_006932584.4;refGene_gn=PPIP5K1;refGene_tx=XM_011283034.3;refGene_gn=PPIP5K1;refGene_tx=XM_011283035.3;refGene_gn=PPIP5K1;refGene_tx=XM_011283037.3;refGene_gn=PPIP5K1;refGene_tx=XM_011283038.3;refGene_gn=PPIP5K1;refGene_tx=XM_011283039.3;refGene_gn=PPIP5K1;refGene_tx=XM_011283040.3;refGene_gn=PPIP5K1;refGene_tx=XM_019832561.2;refGene_gn=PPIP5K1;refGene_tx=XM_019832562.2;refGene_gn=PPIP5K1;refGene_tx=XM_019832563.2;refGene_gn=PPIP5K1;refGene_tx=XM_019832564.2;refGene_gn=PPIP5K1;refGene_tx=XM_019832565.2;refGene_gn=PPIP5K1;refGene_tx=XM_019832566.2;refGene_gn=PPIP5K1;refGene_tx=XM_023255439.1;refGene_gn=PPIP5K1;refGene_tx=XM_023255440.1;refGene_gn=PPIP5K1;refGene_tx=XM_023255441.1
B3 61934255 61934410
refGene_gn=PPIP5K1;refGene_tx=XM_006932584.4;refGene_gn=PPIP5K1;refGene_tx=XM_011283034.3;refGene_gn=PPIP5K1;refGene_tx=XM_011283035.3;refGene_gn=PPIP5K1;refGene_tx=XM_011283037.3;refGene_gn=PPIP5K1;refGene_tx=XM_011283038.3;refGene_gn=PPIP5K1;refGene_tx=XM_011283039.3;refGene_gn=PPIP5K1;refGene_tx=XM_011283040.3;refGene_gn=PPIP5K1;refGene_tx=XM_019832561.2;refGene_gn=PPIP5K1;refGene_tx=XM_019832562.2;refGene_gn=PPIP5K1;refGene_tx=XM_019832563.2;refGene_gn=PPIP5K1;refGene_tx=XM_019832564.2;refGene_gn=PPIP5K1;refGene_tx=XM_019832565.2;refGene_gn=PPIP5K1;r

efGene_tx=XM_019832566.2;refGene_gn=PPIP5K1;refGene_tx=XM_023255439.1;
refGene_gn=PPIP5K1;refGene_tx=XM_023255440.1;refGene_gn=PPIP5K1;refGen
e_tx=XM_023255441.1
B3 61934521 61934623
refGene_gn=PPIP5K1;refGene_tx=XM_006932584.4;refGene_gn=PPIP5K1;refGen
e_tx=XM_011283034.3;refGene_gn=PPIP5K1;refGene_tx=XM_011283035.3;refGe
ne_gn=PPIP5K1;refGene_tx=XM_011283037.3;refGene_gn=PPIP5K1;refGene_tx=
XM_011283038.3;refGene_gn=PPIP5K1;refGene_tx=XM_011283039.3;refGene_gn
=PPIP5K1;refGene_tx=XM_011283040.3;refGene_gn=PPIP5K1;refGene_tx=XM_01
9832561.2;refGene_gn=PPIP5K1;refGene_tx=XM_019832562.2;refGene_gn=PPIP
5K1;refGene_tx=XM_019832563.2;refGene_gn=PPIP5K1;refGene_tx=XM_0198325
64.2;refGene_gn=PPIP5K1;refGene_tx=XM_019832565.2;refGene_gn=PPIP5K1;r
efGene_tx=XM_019832566.2;refGene_gn=PPIP5K1;refGene_tx=XM_023255439.1;
refGene_gn=PPIP5K1;refGene_tx=XM_023255440.1;refGene_gn=PPIP5K1;refGen
e_tx=XM_023255441.1
B3 61935069 61935243
refGene_gn=PPIP5K1;refGene_tx=XM_006932584.4;refGene_gn=PPIP5K1;refGen
e_tx=XM_011283034.3;refGene_gn=PPIP5K1;refGene_tx=XM_011283035.3;refGe
ne_gn=PPIP5K1;refGene_tx=XM_011283037.3;refGene_gn=PPIP5K1;refGene_tx=
XM_011283038.3;refGene_gn=PPIP5K1;refGene_tx=XM_011283039.3;refGene_gn
=PPIP5K1;refGene_tx=XM_011283040.3;refGene_gn=PPIP5K1;refGene_tx=XM_01
9832561.2;refGene_gn=PPIP5K1;refGene_tx=XM_019832562.2;refGene_gn=PPIP
5K1;refGene_tx=XM_019832563.2;refGene_gn=PPIP5K1;refGene_tx=XM_0198325
64.2;refGene_gn=PPIP5K1;refGene_tx=XM_019832565.2;refGene_gn=PPIP5K1;r
efGene_tx=XM_019832566.2;refGene_gn=PPIP5K1;refGene_tx=XM_019832567.1;
refGene_gn=PPIP5K1;refGene_tx=XM_023255439.1;refGene_gn=PPIP5K1;refGen
e_tx=XM_023255440.1;refGene_gn=PPIP5K1;refGene_tx=XM_023255441.1
B3 61935355 61935477
refGene_gn=PPIP5K1;refGene_tx=XM_006932584.4;refGene_gn=PPIP5K1;refGen
e_tx=XM_011283034.3;refGene_gn=PPIP5K1;refGene_tx=XM_011283035.3;refGe
ne_gn=PPIP5K1;refGene_tx=XM_011283037.3;refGene_gn=PPIP5K1;refGene_tx=
XM_011283038.3;refGene_gn=PPIP5K1;refGene_tx=XM_011283039.3;refGene_gn
=PPIP5K1;refGene_tx=XM_011283040.3;refGene_gn=PPIP5K1;refGene_tx=XM_01
9832561.2;refGene_gn=PPIP5K1;refGene_tx=XM_019832562.2;refGene_gn=PPIP
5K1;refGene_tx=XM_019832563.2;refGene_gn=PPIP5K1;refGene_tx=XM_0198325
64.2;refGene_gn=PPIP5K1;refGene_tx=XM_019832565.2;refGene_gn=PPIP5K1;r
efGene_tx=XM_019832566.2;refGene_gn=PPIP5K1;refGene_tx=XM_019832567.1;
refGene_gn=PPIP5K1;refGene_tx=XM_023255439.1;refGene_gn=PPIP5K1;refGen
e_tx=XM_023255440.1;refGene_gn=PPIP5K1;refGene_tx=XM_023255441.1
B3 61937481 61937583
refGene_gn=PPIP5K1;refGene_tx=XM_006932584.4;refGene_gn=PPIP5K1;refGen
e_tx=XM_011283034.3;refGene_gn=PPIP5K1;refGene_tx=XM_011283035.3;refGe
ne_gn=PPIP5K1;refGene_tx=XM_011283037.3;refGene_gn=PPIP5K1;refGene_tx=
XM_011283038.3;refGene_gn=PPIP5K1;refGene_tx=XM_011283039.3;refGene_gn
=PPIP5K1;refGene_tx=XM_011283040.3;refGene_gn=PPIP5K1;refGene_tx=XM_01
9832561.2;refGene_gn=PPIP5K1;refGene_tx=XM_019832562.2;refGene_gn=PPIP
5K1;refGene_tx=XM_019832563.2;refGene_gn=PPIP5K1;refGene_tx=XM_0198325
64.2;refGene_gn=PPIP5K1;refGene_tx=XM_019832565.2;refGene_gn=PPIP5K1;r
efGene_tx=XM_019832566.2;refGene_gn=PPIP5K1;refGene_tx=XM_019832567.1;
refGene_gn=PPIP5K1;refGene_tx=XM_023255439.1;refGene_gn=PPIP5K1;refGen

e_tx=XM_023255440.1;refGene_gn=PPIP5K1;refGene_tx=XM_023255441.1
B3 61937828 61937915
refGene_gn=PPIP5K1;refGene_tx=XM_006932584.4;refGene_gn=PPIP5K1;refGene_tx=XM_011283034.3;refGene_gn=PPIP5K1;refGene_tx=XM_011283035.3;refGene_gn=PPIP5K1;refGene_tx=XM_011283037.3;refGene_gn=PPIP5K1;refGene_tx=XM_011283038.3;refGene_gn=PPIP5K1;refGene_tx=XM_011283039.3;refGene_gn=PPIP5K1;refGene_tx=XM_011283040.3;refGene_gn=PPIP5K1;refGene_tx=XM_019832561.2;refGene_gn=PPIP5K1;refGene_tx=XM_019832562.2;refGene_gn=PPIP5K1;refGene_tx=XM_019832563.2;refGene_gn=PPIP5K1;refGene_tx=XM_019832564.2;refGene_gn=PPIP5K1;refGene_tx=XM_019832565.2;refGene_gn=PPIP5K1;refGene_tx=XM_019832566.2;refGene_gn=PPIP5K1;refGene_tx=XM_019832567.1;refGene_gn=PPIP5K1;refGene_tx=XM_023255439.1;refGene_gn=PPIP5K1;refGene_tx=XM_023255440.1;refGene_gn=PPIP5K1;refGene_tx=XM_023255441.1
B3 61938032 61938108
refGene_gn=PPIP5K1;refGene_tx=XM_006932584.4;refGene_gn=PPIP5K1;refGene_tx=XM_011283034.3;refGene_gn=PPIP5K1;refGene_tx=XM_011283035.3;refGene_gn=PPIP5K1;refGene_tx=XM_011283037.3;refGene_gn=PPIP5K1;refGene_tx=XM_011283038.3;refGene_gn=PPIP5K1;refGene_tx=XM_011283039.3;refGene_gn=PPIP5K1;refGene_tx=XM_011283040.3;refGene_gn=PPIP5K1;refGene_tx=XM_019832561.2;refGene_gn=PPIP5K1;refGene_tx=XM_019832562.2;refGene_gn=PPIP5K1;refGene_tx=XM_019832563.2;refGene_gn=PPIP5K1;refGene_tx=XM_019832564.2;refGene_gn=PPIP5K1;refGene_tx=XM_019832565.2;refGene_gn=PPIP5K1;refGene_tx=XM_019832566.2;refGene_gn=PPIP5K1;refGene_tx=XM_019832567.1;refGene_gn=PPIP5K1;refGene_tx=XM_023255439.1;refGene_gn=PPIP5K1;refGene_tx=XM_023255440.1;refGene_gn=PPIP5K1;refGene_tx=XM_023255441.1
B3 61938237 61938347
refGene_gn=PPIP5K1;refGene_tx=XM_006932584.4;refGene_gn=PPIP5K1;refGene_tx=XM_011283034.3;refGene_gn=PPIP5K1;refGene_tx=XM_011283035.3;refGene_gn=PPIP5K1;refGene_tx=XM_011283037.3;refGene_gn=PPIP5K1;refGene_tx=XM_011283038.3;refGene_gn=PPIP5K1;refGene_tx=XM_011283039.3;refGene_gn=PPIP5K1;refGene_tx=XM_011283040.3;refGene_gn=PPIP5K1;refGene_tx=XM_019832561.2;refGene_gn=PPIP5K1;refGene_tx=XM_019832562.2;refGene_gn=PPIP5K1;refGene_tx=XM_019832563.2;refGene_gn=PPIP5K1;refGene_tx=XM_019832564.2;refGene_gn=PPIP5K1;refGene_tx=XM_019832565.2;refGene_gn=PPIP5K1;refGene_tx=XM_019832566.2;refGene_gn=PPIP5K1;refGene_tx=XM_019832567.1;refGene_gn=PPIP5K1;refGene_tx=XM_023255439.1;refGene_gn=PPIP5K1;refGene_tx=XM_023255440.1;refGene_gn=PPIP5K1;refGene_tx=XM_023255441.1
B3 61938430 61938513
refGene_gn=PPIP5K1;refGene_tx=XM_006932584.4;refGene_gn=PPIP5K1;refGene_tx=XM_011283034.3;refGene_gn=PPIP5K1;refGene_tx=XM_011283035.3;refGene_gn=PPIP5K1;refGene_tx=XM_011283037.3;refGene_gn=PPIP5K1;refGene_tx=XM_011283038.3;refGene_gn=PPIP5K1;refGene_tx=XM_011283039.3;refGene_gn=PPIP5K1;refGene_tx=XM_011283040.3;refGene_gn=PPIP5K1;refGene_tx=XM_019832561.2;refGene_gn=PPIP5K1;refGene_tx=XM_019832562.2;refGene_gn=PPIP5K1;refGene_tx=XM_019832563.2;refGene_gn=PPIP5K1;refGene_tx=XM_019832564.2;refGene_gn=PPIP5K1;refGene_tx=XM_019832565.2;refGene_gn=PPIP5K1;refGene_tx=XM_019832566.2;refGene_gn=PPIP5K1;refGene_tx=XM_019832567.1;refGene_gn=PPIP5K1;refGene_tx=XM_023255439.1;refGene_gn=PPIP5K1;refGene_tx=XM_023255440.1;refGene_gn=PPIP5K1;refGene_tx=XM_023255441.1
B3 61939244 61939376

refGene_gn=PPIP5K1;refGene_tx=XM_006932584.4;refGene_gn=PPIP5K1;refGen
e_tx=XM_011283034.3;refGene_gn=PPIP5K1;refGene_tx=XM_011283035.3;refGe
ne_gn=PPIP5K1;refGene_tx=XM_011283037.3;refGene_gn=PPIP5K1;refGene_tx=
XM_011283038.3;refGene_gn=PPIP5K1;refGene_tx=XM_011283039.3;refGene_gn
=PPIP5K1;refGene_tx=XM_011283040.3;refGene_gn=PPIP5K1;refGene_tx=XM_01
9832561.2;refGene_gn=PPIP5K1;refGene_tx=XM_019832562.2;refGene_gn=PPIP
5K1;refGene_tx=XM_019832563.2;refGene_gn=PPIP5K1;refGene_tx=XM_0198325
64.2;refGene_gn=PPIP5K1;refGene_tx=XM_019832565.2;refGene_gn=PPIP5K1;r
efGene_tx=XM_019832566.2;refGene_gn=PPIP5K1;refGene_tx=XM_019832567.1;
refGene_gn=PPIP5K1;refGene_tx=XM_023255439.1;refGene_gn=PPIP5K1;refGen
e_tx=XM_023255440.1;refGene_gn=PPIP5K1;refGene_tx=XM_023255441.1
B3 61939544 61939666
refGene_gn=PPIP5K1;refGene_tx=XM_006932584.4;refGene_gn=PPIP5K1;refGen
e_tx=XM_011283034.3;refGene_gn=PPIP5K1;refGene_tx=XM_011283035.3;refGe
ne_gn=PPIP5K1;refGene_tx=XM_011283037.3;refGene_gn=PPIP5K1;refGene_tx=
XM_011283038.3;refGene_gn=PPIP5K1;refGene_tx=XM_011283039.3;refGene_gn
=PPIP5K1;refGene_tx=XM_011283040.3;refGene_gn=PPIP5K1;refGene_tx=XM_01
9832561.2;refGene_gn=PPIP5K1;refGene_tx=XM_019832562.2;refGene_gn=PPIP
5K1;refGene_tx=XM_019832563.2;refGene_gn=PPIP5K1;refGene_tx=XM_0198325
64.2;refGene_gn=PPIP5K1;refGene_tx=XM_019832565.2;refGene_gn=PPIP5K1;r
efGene_tx=XM_019832566.2;refGene_gn=PPIP5K1;refGene_tx=XM_019832567.1;
refGene_gn=PPIP5K1;refGene_tx=XM_023255439.1;refGene_gn=PPIP5K1;refGen
e_tx=XM_023255440.1;refGene_gn=PPIP5K1;refGene_tx=XM_023255441.1
B3 61939958 61940141
refGene_gn=PPIP5K1;refGene_tx=XM_006932584.4;refGene_gn=PPIP5K1;refGen
e_tx=XM_011283034.3;refGene_gn=PPIP5K1;refGene_tx=XM_011283035.3;refGe
ne_gn=PPIP5K1;refGene_tx=XM_011283037.3;refGene_gn=PPIP5K1;refGene_tx=
XM_011283038.3;refGene_gn=PPIP5K1;refGene_tx=XM_011283039.3;refGene_gn
=PPIP5K1;refGene_tx=XM_011283040.3;refGene_gn=PPIP5K1;refGene_tx=XM_01
9832561.2;refGene_gn=PPIP5K1;refGene_tx=XM_019832562.2;refGene_gn=PPIP
5K1;refGene_tx=XM_019832563.2;refGene_gn=PPIP5K1;refGene_tx=XM_0198325
64.2;refGene_gn=PPIP5K1;refGene_tx=XM_019832565.2;refGene_gn=PPIP5K1;r
efGene_tx=XM_019832566.2;refGene_gn=PPIP5K1;refGene_tx=XM_019832567.1;
refGene_gn=PPIP5K1;refGene_tx=XM_023255439.1;refGene_gn=PPIP5K1;refGen
e_tx=XM_023255440.1;refGene_gn=PPIP5K1;refGene_tx=XM_023255441.1
B3 61943028 61943170
refGene_gn=PPIP5K1;refGene_tx=XM_006932584.4;refGene_gn=PPIP5K1;refGen
e_tx=XM_011283034.3;refGene_gn=PPIP5K1;refGene_tx=XM_011283035.3;refGe
ne_gn=PPIP5K1;refGene_tx=XM_011283037.3;refGene_gn=PPIP5K1;refGene_tx=
XM_011283038.3;refGene_gn=PPIP5K1;refGene_tx=XM_011283039.3;refGene_gn
=PPIP5K1;refGene_tx=XM_011283040.3;refGene_gn=PPIP5K1;refGene_tx=XM_01
9832561.2;refGene_gn=PPIP5K1;refGene_tx=XM_019832562.2;refGene_gn=PPIP
5K1;refGene_tx=XM_019832563.2;refGene_gn=PPIP5K1;refGene_tx=XM_0198325
64.2;refGene_gn=PPIP5K1;refGene_tx=XM_019832565.2;refGene_gn=PPIP5K1;r
efGene_tx=XM_019832566.2;refGene_gn=PPIP5K1;refGene_tx=XM_019832567.1;
refGene_gn=PPIP5K1;refGene_tx=XM_023255439.1;refGene_gn=PPIP5K1;refGen
e_tx=XM_023255440.1;refGene_gn=PPIP5K1;refGene_tx=XM_023255441.1
B3 61943359 61943583
refGene_gn=PPIP5K1;refGene_tx=XM_006932584.4;refGene_gn=PPIP5K1;refGen
e_tx=XM_011283034.3;refGene_gn=PPIP5K1;refGene_tx=XM_011283035.3;refGe

ne_gn=PPIP5K1;refGene_tx=XM_011283037.3;refGene_gn=PPIP5K1;refGene_tx=XM_011283038.3;refGene_gn=PPIP5K1;refGene_tx=XM_011283039.3;refGene_gn=PPIP5K1;refGene_tx=XM_011283040.3;refGene_gn=PPIP5K1;refGene_tx=XM_019832561.2;refGene_gn=PPIP5K1;refGene_tx=XM_019832562.2;refGene_gn=PPIP5K1;refGene_tx=XM_019832563.2;refGene_gn=PPIP5K1;refGene_tx=XM_019832564.2;refGene_gn=PPIP5K1;refGene_tx=XM_019832565.2;refGene_gn=PPIP5K1;refGene_tx=XM_019832566.2;refGene_gn=PPIP5K1;refGene_tx=XM_019832567.1;refGene_gn=PPIP5K1;refGene_tx=XM_023255439.1;refGene_gn=PPIP5K1;refGene_tx=XM_023255440.1;refGene_gn=PPIP5K1;refGene_tx=XM_023255441.1
B3 61943890 61944018
refGene_gn=PPIP5K1;refGene_tx=XM_006932584.4;refGene_gn=PPIP5K1;refGene_tx=XM_011283034.3;refGene_gn=PPIP5K1;refGene_tx=XM_011283035.3;refGene_gn=PPIP5K1;refGene_tx=XM_011283037.3;refGene_gn=PPIP5K1;refGene_tx=XM_011283038.3;refGene_gn=PPIP5K1;refGene_tx=XM_011283039.3;refGene_gn=PPIP5K1;refGene_tx=XM_011283040.3;refGene_gn=PPIP5K1;refGene_tx=XM_019832561.2;refGene_gn=PPIP5K1;refGene_tx=XM_019832562.2;refGene_gn=PPIP5K1;refGene_tx=XM_019832563.2;refGene_gn=PPIP5K1;refGene_tx=XM_019832564.2;refGene_gn=PPIP5K1;refGene_tx=XM_019832565.2;refGene_gn=PPIP5K1;refGene_tx=XM_019832566.2;refGene_gn=PPIP5K1;refGene_tx=XM_019832567.1;refGene_gn=PPIP5K1;refGene_tx=XM_023255439.1;refGene_gn=PPIP5K1;refGene_tx=XM_023255440.1;refGene_gn=PPIP5K1;refGene_tx=XM_023255441.1
B3 61944227 61944342
refGene_gn=PPIP5K1;refGene_tx=XM_006932584.4;refGene_gn=PPIP5K1;refGene_tx=XM_011283034.3;refGene_gn=PPIP5K1;refGene_tx=XM_011283035.3;refGene_gn=PPIP5K1;refGene_tx=XM_011283037.3;refGene_gn=PPIP5K1;refGene_tx=XM_011283038.3;refGene_gn=PPIP5K1;refGene_tx=XM_011283039.3;refGene_gn=PPIP5K1;refGene_tx=XM_011283040.3;refGene_gn=PPIP5K1;refGene_tx=XM_019832561.2;refGene_gn=PPIP5K1;refGene_tx=XM_019832562.2;refGene_gn=PPIP5K1;refGene_tx=XM_019832563.2;refGene_gn=PPIP5K1;refGene_tx=XM_019832564.2;refGene_gn=PPIP5K1;refGene_tx=XM_019832565.2;refGene_gn=PPIP5K1;refGene_tx=XM_019832566.2;refGene_gn=PPIP5K1;refGene_tx=XM_019832567.1;refGene_gn=PPIP5K1;refGene_tx=XM_023255439.1;refGene_gn=PPIP5K1;refGene_tx=XM_023255440.1;refGene_gn=PPIP5K1;refGene_tx=XM_023255441.1
B3 61944507 61944615
refGene_gn=PPIP5K1;refGene_tx=XM_006932584.4;refGene_gn=PPIP5K1;refGene_tx=XM_011283034.3;refGene_gn=PPIP5K1;refGene_tx=XM_011283035.3;refGene_gn=PPIP5K1;refGene_tx=XM_011283037.3;refGene_gn=PPIP5K1;refGene_tx=XM_011283038.3;refGene_gn=PPIP5K1;refGene_tx=XM_011283039.3;refGene_gn=PPIP5K1;refGene_tx=XM_011283040.3;refGene_gn=PPIP5K1;refGene_tx=XM_019832561.2;refGene_gn=PPIP5K1;refGene_tx=XM_019832562.2;refGene_gn=PPIP5K1;refGene_tx=XM_019832563.2;refGene_gn=PPIP5K1;refGene_tx=XM_019832564.2;refGene_gn=PPIP5K1;refGene_tx=XM_019832565.2;refGene_gn=PPIP5K1;refGene_tx=XM_019832566.2;refGene_gn=PPIP5K1;refGene_tx=XM_019832567.1;refGene_gn=PPIP5K1;refGene_tx=XM_023255439.1;refGene_gn=PPIP5K1;refGene_tx=XM_023255440.1;refGene_gn=PPIP5K1;refGene_tx=XM_023255441.1
B3 61945261 61945378
refGene_gn=PPIP5K1;refGene_tx=XM_006932584.4;refGene_gn=PPIP5K1;refGene_tx=XM_011283034.3;refGene_gn=PPIP5K1;refGene_tx=XM_011283035.3;refGene_gn=PPIP5K1;refGene_tx=XM_011283037.3;refGene_gn=PPIP5K1;refGene_tx=XM_011283038.3;refGene_gn=PPIP5K1;refGene_tx=XM_011283039.3;refGene_gn

=PPIP5K1;refGene_tx=XM_011283040.3;refGene_gn=PPIP5K1;refGene_tx=XM_019832561.2;refGene_gn=PPIP5K1;refGene_tx=XM_019832562.2;refGene_gn=PPIP5K1;refGene_tx=XM_019832563.2;refGene_gn=PPIP5K1;refGene_tx=XM_019832564.2;refGene_gn=PPIP5K1;refGene_tx=XM_019832565.2;refGene_gn=PPIP5K1;refGene_tx=XM_019832566.2;refGene_gn=PPIP5K1;refGene_tx=XM_019832567.1;refGene_gn=PPIP5K1;refGene_tx=XM_023255439.1;refGene_gn=PPIP5K1;refGene_tx=XM_023255440.1;refGene_gn=PPIP5K1;refGene_tx=XM_023255441.1
B3 61945532 61945697
refGene_gn=PPIP5K1;refGene_tx=XM_006932584.4;refGene_gn=PPIP5K1;refGene_tx=XM_011283034.3;refGene_gn=PPIP5K1;refGene_tx=XM_011283035.3;refGene_gn=PPIP5K1;refGene_tx=XM_011283037.3;refGene_gn=PPIP5K1;refGene_tx=XM_011283038.3;refGene_gn=PPIP5K1;refGene_tx=XM_011283039.3;refGene_gn=PPIP5K1;refGene_tx=XM_011283040.3;refGene_gn=PPIP5K1;refGene_tx=XM_019832561.2;refGene_gn=PPIP5K1;refGene_tx=XM_019832562.2;refGene_gn=PPIP5K1;refGene_tx=XM_019832563.2;refGene_gn=PPIP5K1;refGene_tx=XM_019832564.2;refGene_gn=PPIP5K1;refGene_tx=XM_019832565.2;refGene_gn=PPIP5K1;refGene_tx=XM_019832566.2;refGene_gn=PPIP5K1;refGene_tx=XM_019832567.1;refGene_gn=PPIP5K1;refGene_tx=XM_023255439.1;refGene_gn=PPIP5K1;refGene_tx=XM_023255440.1;refGene_gn=PPIP5K1;refGene_tx=XM_023255441.1
B3 61947725 61947875
refGene_gn=PPIP5K1;refGene_tx=XM_011283034.3;refGene_gn=PPIP5K1;refGene_tx=XM_011283035.3;refGene_gn=PPIP5K1;refGene_tx=XM_011283037.3;refGene_gn=PPIP5K1;refGene_tx=XM_011283038.3;refGene_gn=PPIP5K1;refGene_tx=XM_019832561.2;refGene_gn=PPIP5K1;refGene_tx=XM_019832562.2;refGene_gn=PPIP5K1;refGene_tx=XM_019832563.2;refGene_gn=PPIP5K1;refGene_tx=XM_019832564.2;refGene_gn=PPIP5K1;refGene_tx=XM_019832565.2;refGene_gn=PPIP5K1;refGene_tx=XM_019832567.1;refGene_gn=PPIP5K1;refGene_tx=XM_023255439.1
B3 61948498 61948589
refGene_gn=PPIP5K1;refGene_tx=XM_006932584.4;refGene_gn=PPIP5K1;refGene_tx=XM_011283034.3;refGene_gn=PPIP5K1;refGene_tx=XM_011283035.3;refGene_gn=PPIP5K1;refGene_tx=XM_011283037.3;refGene_gn=PPIP5K1;refGene_tx=XM_011283038.3;refGene_gn=PPIP5K1;refGene_tx=XM_011283039.3;refGene_gn=PPIP5K1;refGene_tx=XM_011283040.3;refGene_gn=PPIP5K1;refGene_tx=XM_019832561.2;refGene_gn=PPIP5K1;refGene_tx=XM_019832562.2;refGene_gn=PPIP5K1;refGene_tx=XM_019832563.2;refGene_gn=PPIP5K1;refGene_tx=XM_019832564.2;refGene_gn=PPIP5K1;refGene_tx=XM_019832565.2;refGene_gn=PPIP5K1;refGene_tx=XM_019832566.2;refGene_gn=PPIP5K1;refGene_tx=XM_019832567.1;refGene_gn=PPIP5K1;refGene_tx=XM_023255439.1;refGene_gn=PPIP5K1;refGene_tx=XM_023255440.1;refGene_gn=PPIP5K1;refGene_tx=XM_023255441.1
B3 61951407 61951533
refGene_gn=PPIP5K1;refGene_tx=XM_006932584.4;refGene_gn=PPIP5K1;refGene_tx=XM_011283034.3;refGene_gn=PPIP5K1;refGene_tx=XM_011283037.3;refGene_gn=PPIP5K1;refGene_tx=XM_011283039.3;refGene_gn=PPIP5K1;refGene_tx=XM_019832561.2;refGene_gn=PPIP5K1;refGene_tx=XM_019832562.2;refGene_gn=PPIP5K1;refGene_tx=XM_019832563.2;refGene_gn=PPIP5K1;refGene_tx=XM_019832564.2;refGene_gn=PPIP5K1;refGene_tx=XM_019832565.2;refGene_gn=PPIP5K1;refGene_tx=XM_019832566.2;refGene_gn=PPIP5K1;refGene_tx=XM_019832567.1;refGene_gn=PPIP5K1;refGene_tx=XM_023255439.1;refGene_gn=PPIP5K1;refGene_tx=XM_023255440.1;refGene_gn=PPIP5K1;refGene_tx=XM_023255441.1

B3	61952244	61952307
refGene_gn=PPIP5K1;refGene_tx=XM_006932584.4;refGene_gn=PPIP5K1;refGene_tx=XM_011283034.3;refGene_gn=PPIP5K1;refGene_tx=XM_019832561.2;refGene_gn=PPIP5K1;refGene_tx=XM_019832562.2;refGene_gn=PPIP5K1;refGene_tx=XM_019832563.2;refGene_gn=PPIP5K1;refGene_tx=XM_019832564.2;refGene_gn=PPIP5K1;refGene_tx=XM_019832565.2;refGene_gn=PPIP5K1;refGene_tx=XM_019832566.2;refGene_gn=PPIP5K1;refGene_tx=XM_019832567.1;refGene_gn=PPIP5K1;refGene_tx=XM_023255439.1;refGene_gn=PPIP5K1;refGene_tx=XM_023255440.1
B3	61955766	61955949
refGene_gn=PPIP5K1;refGene_tx=XM_011283035.3;refGene_gn=PPIP5K1;refGene_tx=XM_019832561.2;refGene_gn=PPIP5K1;refGene_tx=XM_019832562.2;refGene_gn=PPIP5K1;refGene_tx=XM_019832563.2;refGene_gn=PPIP5K1;refGene_tx=XM_019832564.2;refGene_gn=PPIP5K1;refGene_tx=XM_019832565.2;refGene_gn=PPIP5K1;refGene_tx=XM_019832566.2;refGene_gn=PPIP5K1;refGene_tx=XM_019832567.1;refGene_gn=PPIP5K1;refGene_tx=XM_023255439.1
B3	61957445	61957583
refGene_gn=PPIP5K1;refGene_tx=XM_006932584.4;refGene_gn=PPIP5K1;refGene_tx=XM_011283034.3;refGene_gn=PPIP5K1;refGene_tx=XM_011283035.3;refGene_gn=PPIP5K1;refGene_tx=XM_011283037.3;refGene_gn=PPIP5K1;refGene_tx=XM_011283038.3;refGene_gn=PPIP5K1;refGene_tx=XM_011283039.3;refGene_gn=PPIP5K1;refGene_tx=XM_011283040.3;refGene_gn=PPIP5K1;refGene_tx=XM_019832561.2;refGene_gn=PPIP5K1;refGene_tx=XM_019832562.2;refGene_gn=PPIP5K1;refGene_tx=XM_019832563.2;refGene_gn=PPIP5K1;refGene_tx=XM_019832564.2;refGene_gn=PPIP5K1;refGene_tx=XM_019832565.2;refGene_gn=PPIP5K1;refGene_tx=XM_019832566.2;refGene_gn=PPIP5K1;refGene_tx=XM_019832567.1;refGene_gn=PPIP5K1;refGene_tx=XM_023255439.1;refGene_gn=PPIP5K1;refGene_tx=XM_023255440.1;refGene_gn=PPIP5K1;refGene_tx=XM_023255441.1
B3	61965738	61965852
refGene_gn=PPIP5K1;refGene_tx=XM_006932584.4;refGene_gn=PPIP5K1;refGene_tx=XM_011283034.3;refGene_gn=PPIP5K1;refGene_tx=XM_011283035.3;refGene_gn=PPIP5K1;refGene_tx=XM_011283037.3;refGene_gn=PPIP5K1;refGene_tx=XM_011283038.3;refGene_gn=PPIP5K1;refGene_tx=XM_011283039.3;refGene_gn=PPIP5K1;refGene_tx=XM_011283040.3;refGene_gn=PPIP5K1;refGene_tx=XM_019832561.2;refGene_gn=PPIP5K1;refGene_tx=XM_019832562.2;refGene_gn=PPIP5K1;refGene_tx=XM_019832563.2;refGene_gn=PPIP5K1;refGene_tx=XM_019832564.2;refGene_gn=PPIP5K1;refGene_tx=XM_019832565.2;refGene_gn=PPIP5K1;refGene_tx=XM_019832566.2;refGene_gn=PPIP5K1;refGene_tx=XM_019832567.1;refGene_gn=PPIP5K1;refGene_tx=XM_023255439.1;refGene_gn=PPIP5K1;refGene_tx=XM_023255440.1;refGene_gn=PPIP5K1;refGene_tx=XM_023255441.1
B3	61969857	61970807
refGene_gn=PPIP5K1;refGene_tx=XM_006932584.4;refGene_gn=PPIP5K1;refGene_tx=XM_011283034.3;refGene_gn=PPIP5K1;refGene_tx=XM_011283035.3;refGene_gn=PPIP5K1;refGene_tx=XM_011283037.3;refGene_gn=PPIP5K1;refGene_tx=XM_011283038.3;refGene_gn=PPIP5K1;refGene_tx=XM_011283039.3;refGene_gn=PPIP5K1;refGene_tx=XM_011283040.3;refGene_gn=PPIP5K1;refGene_tx=XM_019832561.2;refGene_gn=PPIP5K1;refGene_tx=XM_019832562.2;refGene_gn=PPIP5K1;refGene_tx=XM_019832563.2;refGene_gn=PPIP5K1;refGene_tx=XM_019832564.2;refGene_gn=PPIP5K1;refGene_tx=XM_019832565.2;refGene_gn=PPIP5K1;refGene_tx=XM_019832566.2;refGene_gn=PPIP5K1;refGene_tx=XM_019832567.1;

refGene_gn=PPIP5K1;refGene_tx=XM_023255439.1;refGene_gn=PPIP5K1;refGene_tx=XM_023255440.1;refGene_gn=PPIP5K1;refGene_tx=XM_023255441.1
B3 61974472 61974628
refGene_gn=MAP1A;refGene_tx=XM_003987227.4;refGene_gn=MAP1A;refGene_tx=XM_019832571.1;refGene_gn=MAP1A;refGene_tx=XM_019832572.1
B3 61974823 61975044
refGene_gn=MAP1A;refGene_tx=XM_003987227.4;refGene_gn=MAP1A;refGene_tx=XM_019832572.1
B3 61975185 61983349
refGene_gn=MAP1A;refGene_tx=XM_003987227.4;refGene_gn=MAP1A;refGene_tx=XM_019832571.1;refGene_gn=MAP1A;refGene_tx=XM_019832572.1
B3 61983583 61983724
refGene_gn=MAP1A;refGene_tx=XM_003987227.4;refGene_gn=MAP1A;refGene_tx=XM_019832571.1
B3 61983989 61984072
refGene_gn=MAP1A;refGene_tx=XM_003987227.4;refGene_gn=MAP1A;refGene_tx=XM_019832571.1
B3 61991862 61991964
refGene_gn=MAP1A;refGene_tx=XM_003987227.4;refGene_gn=MAP1A;refGene_tx=XM_019832571.1
B3 61992917 61993155
refGene_gn=MAP1A;refGene_tx=XM_003987227.4;refGene_gn=MAP1A;refGene_tx=XM_019832571.1
B3 62018090 62018097
refGene_gn=TP53BP1;refGene_tx=XM_006932585.4;refGene_gn=TP53BP1;refGene_tx=XM_019832573.2;refGene_gn=TP53BP1;refGene_tx=XM_019832574.2;refGene_gn=TP53BP1;refGene_tx=XM_019832576.2;refGene_gn=TP53BP1;refGene_tx=XM_019832577.2
B3 62018673 62018858
refGene_gn=TP53BP1;refGene_tx=XM_006932585.4;refGene_gn=TP53BP1;refGene_tx=XM_019832573.2;refGene_gn=TP53BP1;refGene_tx=XM_019832574.2;refGene_gn=TP53BP1;refGene_tx=XM_019832576.2;refGene_gn=TP53BP1;refGene_tx=XM_019832577.2
B3 62019064 62019155
refGene_gn=TP53BP1;refGene_tx=XM_006932585.4;refGene_gn=TP53BP1;refGene_tx=XM_019832573.2;refGene_gn=TP53BP1;refGene_tx=XM_019832574.2;refGene_gn=TP53BP1;refGene_tx=XM_019832576.2;refGene_gn=TP53BP1;refGene_tx=XM_019832577.2
B3 62019593 62019678
refGene_gn=TP53BP1;refGene_tx=XM_006932585.4;refGene_gn=TP53BP1;refGene_tx=XM_019832573.2;refGene_gn=TP53BP1;refGene_tx=XM_019832574.2;refGene_gn=TP53BP1;refGene_tx=XM_019832576.2;refGene_gn=TP53BP1;refGene_tx=XM_019832577.2
B3 62027105 62027239
refGene_gn=TP53BP1;refGene_tx=XM_006932585.4;refGene_gn=TP53BP1;refGene_tx=XM_019832573.2;refGene_gn=TP53BP1;refGene_tx=XM_019832574.2;refGene_gn=TP53BP1;refGene_tx=XM_019832576.2;refGene_gn=TP53BP1;refGene_tx=XM_019832577.2;refGene_gn=TP53BP1;refGene_tx=XM_019832578.1
B3 62027812 62027971
refGene_gn=TP53BP1;refGene_tx=XM_006932585.4;refGene_gn=TP53BP1;refGen

e_tx=XM_019832573.2;refGene_gn=TP53BP1;refGene_tx=XM_019832574.2;refGene_gn=TP53BP1;refGene_tx=XM_019832576.2;refGene_gn=TP53BP1;refGene_tx=XM_019832577.2;refGene_gn=TP53BP1;refGene_tx=XM_019832578.1
B3 62028348 62028478
refGene_gn=TP53BP1;refGene_tx=XM_006932585.4;refGene_gn=TP53BP1;refGene_tx=XM_019832573.2;refGene_gn=TP53BP1;refGene_tx=XM_019832574.2;refGene_gn=TP53BP1;refGene_tx=XM_019832576.2;refGene_gn=TP53BP1;refGene_tx=XM_019832577.2;refGene_gn=TP53BP1;refGene_tx=XM_019832578.1
B3 62030244 62030411
refGene_gn=TP53BP1;refGene_tx=XM_006932585.4;refGene_gn=TP53BP1;refGene_tx=XM_019832573.2;refGene_gn=TP53BP1;refGene_tx=XM_019832574.2;refGene_gn=TP53BP1;refGene_tx=XM_019832576.2;refGene_gn=TP53BP1;refGene_tx=XM_019832577.2;refGene_gn=TP53BP1;refGene_tx=XM_019832578.1
B3 62031971 62032098
refGene_gn=TP53BP1;refGene_tx=XM_006932585.4;refGene_gn=TP53BP1;refGene_tx=XM_019832573.2;refGene_gn=TP53BP1;refGene_tx=XM_019832574.2;refGene_gn=TP53BP1;refGene_tx=XM_019832576.2;refGene_gn=TP53BP1;refGene_tx=XM_019832577.2;refGene_gn=TP53BP1;refGene_tx=XM_019832578.1
B3 62033003 62033098
refGene_gn=TP53BP1;refGene_tx=XM_006932585.4;refGene_gn=TP53BP1;refGene_tx=XM_019832573.2;refGene_gn=TP53BP1;refGene_tx=XM_019832574.2;refGene_gn=TP53BP1;refGene_tx=XM_019832576.2;refGene_gn=TP53BP1;refGene_tx=XM_019832577.2;refGene_gn=TP53BP1;refGene_tx=XM_019832578.1
B3 62035992 62036201
refGene_gn=TP53BP1;refGene_tx=XM_006932585.4;refGene_gn=TP53BP1;refGene_tx=XM_019832573.2;refGene_gn=TP53BP1;refGene_tx=XM_019832574.2;refGene_gn=TP53BP1;refGene_tx=XM_019832576.2;refGene_gn=TP53BP1;refGene_tx=XM_019832577.2;refGene_gn=TP53BP1;refGene_tx=XM_019832578.1
B3 62045708 62047032
refGene_gn=TP53BP1;refGene_tx=XM_006932585.4;refGene_gn=TP53BP1;refGene_tx=XM_019832573.2;refGene_gn=TP53BP1;refGene_tx=XM_019832574.2;refGene_gn=TP53BP1;refGene_tx=XM_019832576.2;refGene_gn=TP53BP1;refGene_tx=XM_019832577.2;refGene_gn=TP53BP1;refGene_tx=XM_019832578.1
B3 62053068 62053188
refGene_gn=TP53BP1;refGene_tx=XM_006932585.4;refGene_gn=TP53BP1;refGene_tx=XM_019832573.2;refGene_gn=TP53BP1;refGene_tx=XM_019832574.2;refGene_gn=TP53BP1;refGene_tx=XM_019832576.2;refGene_gn=TP53BP1;refGene_tx=XM_019832577.2;refGene_gn=TP53BP1;refGene_tx=XM_019832578.1
B3 62053946 62054150
refGene_gn=TP53BP1;refGene_tx=XM_006932585.4;refGene_gn=TP53BP1;refGene_tx=XM_019832573.2;refGene_gn=TP53BP1;refGene_tx=XM_019832574.2;refGene_gn=TP53BP1;refGene_tx=XM_019832576.2;refGene_gn=TP53BP1;refGene_tx=XM_019832577.2;refGene_gn=TP53BP1;refGene_tx=XM_019832578.1;refGene_gn=TP53BP1;refGene_tx=XM_023255443.1
B3 62057612 62057670
refGene_gn=TP53BP1;refGene_tx=XM_006932585.4;refGene_gn=TP53BP1;refGene_tx=XM_019832573.2;refGene_gn=TP53BP1;refGene_tx=XM_019832574.2;refGene_gn=TP53BP1;refGene_tx=XM_019832576.2;refGene_gn=TP53BP1;refGene_tx=XM_019832577.2;refGene_gn=TP53BP1;refGene_tx=XM_019832578.1;refGene_gn=TP53BP1;refGene_tx=XM_023255443.1

B3 62058653 62058743
refGene_gn=TP53BP1;refGene_tx=XM_006932585.4;refGene_gn=TP53BP1;refGene_tx=XM_019832573.2;refGene_gn=TP53BP1;refGene_tx=XM_019832574.2;refGene_gn=TP53BP1;refGene_tx=XM_019832577.2;refGene_gn=TP53BP1;refGene_tx=XM_019832578.1;refGene_gn=TP53BP1;refGene_tx=XM_023255443.1
B3 62062982 62063466
refGene_gn=TP53BP1;refGene_tx=XM_006932585.4;refGene_gn=TP53BP1;refGene_tx=XM_019832573.2;refGene_gn=TP53BP1;refGene_tx=XM_019832574.2;refGene_gn=TP53BP1;refGene_tx=XM_019832576.2;refGene_gn=TP53BP1;refGene_tx=XM_019832577.2;refGene_gn=TP53BP1;refGene_tx=XM_019832578.1;refGene_gn=TP53BP1;refGene_tx=XM_023255443.1
B3 62067144 62067297
refGene_gn=TP53BP1;refGene_tx=XM_006932585.4;refGene_gn=TP53BP1;refGene_tx=XM_019832573.2;refGene_gn=TP53BP1;refGene_tx=XM_019832574.2;refGene_gn=TP53BP1;refGene_tx=XM_019832576.2;refGene_gn=TP53BP1;refGene_tx=XM_019832577.2;refGene_gn=TP53BP1;refGene_tx=XM_019832578.1;refGene_gn=TP53BP1;refGene_tx=XM_023255443.1
B3 62073247 62073519
refGene_gn=TP53BP1;refGene_tx=XM_006932585.4;refGene_gn=TP53BP1;refGene_tx=XM_019832573.2;refGene_gn=TP53BP1;refGene_tx=XM_019832574.2;refGene_gn=TP53BP1;refGene_tx=XM_019832576.2;refGene_gn=TP53BP1;refGene_tx=XM_019832577.2;refGene_gn=TP53BP1;refGene_tx=XM_019832578.1;refGene_gn=TP53BP1;refGene_tx=XM_023255443.1
B3 62074195 62074345
refGene_gn=TP53BP1;refGene_tx=XM_006932585.4;refGene_gn=TP53BP1;refGene_tx=XM_019832573.2;refGene_gn=TP53BP1;refGene_tx=XM_019832574.2;refGene_gn=TP53BP1;refGene_tx=XM_019832576.2;refGene_gn=TP53BP1;refGene_tx=XM_019832578.1;refGene_gn=TP53BP1;refGene_tx=XM_023255443.1
B3 62074637 62075068
refGene_gn=TP53BP1;refGene_tx=XM_006932585.4;refGene_gn=TP53BP1;refGene_tx=XM_019832573.2;refGene_gn=TP53BP1;refGene_tx=XM_019832574.2;refGene_gn=TP53BP1;refGene_tx=XM_019832576.2;refGene_gn=TP53BP1;refGene_tx=XM_019832577.2;refGene_gn=TP53BP1;refGene_tx=XM_019832578.1;refGene_gn=TP53BP1;refGene_tx=XM_023255443.1
B3 62077888 62078080
refGene_gn=TP53BP1;refGene_tx=XM_006932585.4;refGene_gn=TP53BP1;refGene_tx=XM_019832573.2;refGene_gn=TP53BP1;refGene_tx=XM_019832574.2;refGene_gn=TP53BP1;refGene_tx=XM_019832576.2;refGene_gn=TP53BP1;refGene_tx=XM_019832577.2;refGene_gn=TP53BP1;refGene_tx=XM_019832578.1;refGene_gn=TP53BP1;refGene_tx=XM_023255443.1
B3 62078483 62078702
refGene_gn=TP53BP1;refGene_tx=XM_006932585.4;refGene_gn=TP53BP1;refGene_tx=XM_019832573.2;refGene_gn=TP53BP1;refGene_tx=XM_019832574.2;refGene_gn=TP53BP1;refGene_tx=XM_019832576.2;refGene_gn=TP53BP1;refGene_tx=XM_019832577.2;refGene_gn=TP53BP1;refGene_tx=XM_019832578.1;refGene_gn=TP53BP1;refGene_tx=XM_023255443.1
B3 62080345 62080561
refGene_gn=TP53BP1;refGene_tx=XM_006932585.4;refGene_gn=TP53BP1;refGene_tx=XM_019832573.2;refGene_gn=TP53BP1;refGene_tx=XM_019832574.2;refGene_gn=TP53BP1;refGene_tx=XM_019832576.2;refGene_gn=TP53BP1;refGene_tx=

XM_019832577.2;refGene_gn=TP53BP1;refGene_tx=XM_019832578.1;refGene_gn=TP53BP1;refGene_tx=XM_023255443.1
B3 62084376 62084471
refGene_gn=TP53BP1;refGene_tx=XM_006932585.4;refGene_gn=TP53BP1;refGene_tx=XM_019832573.2;refGene_gn=TP53BP1;refGene_tx=XM_019832574.2;refGene_gn=TP53BP1;refGene_tx=XM_019832576.2;refGene_gn=TP53BP1;refGene_tx=XM_019832577.2;refGene_gn=TP53BP1;refGene_tx=XM_019832578.1;refGene_gn=TP53BP1;refGene_tx=XM_023255443.1
B3 62085205 62085405
refGene_gn=TP53BP1;refGene_tx=XM_006932585.4;refGene_gn=TP53BP1;refGene_tx=XM_019832573.2;refGene_gn=TP53BP1;refGene_tx=XM_019832574.2;refGene_gn=TP53BP1;refGene_tx=XM_019832576.2;refGene_gn=TP53BP1;refGene_tx=XM_019832577.2;refGene_gn=TP53BP1;refGene_tx=XM_019832578.1;refGene_gn=TP53BP1;refGene_tx=XM_023255443.1
B3 62085874 62086020
refGene_gn=TP53BP1;refGene_tx=XM_006932585.4;refGene_gn=TP53BP1;refGene_tx=XM_019832573.2;refGene_gn=TP53BP1;refGene_tx=XM_019832574.2;refGene_gn=TP53BP1;refGene_tx=XM_019832576.2;refGene_gn=TP53BP1;refGene_tx=XM_019832577.2;refGene_gn=TP53BP1;refGene_tx=XM_019832578.1;refGene_gn=TP53BP1;refGene_tx=XM_023255443.1
B3 62086368 62086556
refGene_gn=TP53BP1;refGene_tx=XM_006932585.4;refGene_gn=TP53BP1;refGene_tx=XM_019832573.2;refGene_gn=TP53BP1;refGene_tx=XM_019832574.2;refGene_gn=TP53BP1;refGene_tx=XM_019832576.2;refGene_gn=TP53BP1;refGene_tx=XM_019832577.2;refGene_gn=TP53BP1;refGene_tx=XM_019832578.1;refGene_gn=TP53BP1;refGene_tx=XM_023255443.1
B3 62088246 62088259 refGene_gn=TUBGCP4;refGene_tx=XM_003987229.3
B3 62088932 62089072 refGene_gn=TUBGCP4;refGene_tx=XM_003987229.3
B3 62089694 62089811 refGene_gn=TUBGCP4;refGene_tx=XM_003987229.3
B3 62091291 62091426 refGene_gn=TUBGCP4;refGene_tx=XM_003987229.3
B3 62091974 62092152 refGene_gn=TUBGCP4;refGene_tx=XM_003987229.3
B3 62095148 62095287 refGene_gn=TUBGCP4;refGene_tx=XM_003987229.3
B3 62095708 62095816 refGene_gn=TUBGCP4;refGene_tx=XM_003987229.3
B3 62097288 62097394 refGene_gn=TUBGCP4;refGene_tx=XM_003987229.3
B3 62097809 62097860 refGene_gn=TUBGCP4;refGene_tx=XM_003987229.3
B3 62102834 62102959 refGene_gn=TUBGCP4;refGene_tx=XM_003987229.3
B3 62103179 62103345 refGene_gn=TUBGCP4;refGene_tx=XM_003987229.3
B3 62106324 62106526 refGene_gn=TUBGCP4;refGene_tx=XM_003987229.3
B3 62109059 62109139 refGene_gn=TUBGCP4;refGene_tx=XM_003987229.3
B3 62112303 62112360 refGene_gn=TUBGCP4;refGene_tx=XM_003987229.3
B3 62113395 62113449 refGene_gn=TUBGCP4;refGene_tx=XM_003987229.3
B3 62113814 62113937 refGene_gn=TUBGCP4;refGene_tx=XM_003987229.3
B3 62116718 62116847 refGene_gn=TUBGCP4;refGene_tx=XM_003987229.3
B3 62122652 62122730 refGene_gn=TUBGCP4;refGene_tx=XM_003987229.3
B3 62124170 62124488
refGene_gn=ZSCAN29;refGene_tx=XM_006932590.2;refGene_gn=ZSCAN29;refGene_tx=XM_019832583.1;refGene_gn=ZSCAN29;refGene_tx=XM_023255444.1;refGene_gn=ZSCAN29;refGene_tx=XM_023255445.1;refGene_gn=ZSCAN29;refGene_tx=XM_023255446.1
B3 62124973 62125178

refGene_gn=ZSCAN29;refGene_tx=XM_023255444.1;refGene_gn=ZSCAN29;refGene_tx=XM_023255445.1;refGene_gn=ZSCAN29;refGene_tx=XM_023255446.1
B3 62126985 62127684
refGene_gn=ZSCAN29;refGene_tx=XM_006932590.2;refGene_gn=ZSCAN29;refGene_tx=XM_019832583.1;refGene_gn=ZSCAN29;refGene_tx=XM_023255444.1;refGene_gn=ZSCAN29;refGene_tx=XM_023255445.1;refGene_gn=ZSCAN29;refGene_tx=XM_023255446.1
B3 62131014 62131485
refGene_gn=ZSCAN29;refGene_tx=XM_006932590.2;refGene_gn=ZSCAN29;refGene_tx=XM_019832583.1;refGene_gn=ZSCAN29;refGene_tx=XM_023255444.1;refGene_gn=ZSCAN29;refGene_tx=XM_023255445.1;refGene_gn=ZSCAN29;refGene_tx=XM_023255446.1
B3 62133719 62134588
refGene_gn=ZSCAN29;refGene_tx=XM_006932590.2;refGene_gn=ZSCAN29;refGene_tx=XM_019832583.1;refGene_gn=ZSCAN29;refGene_tx=XM_023255444.1;refGene_gn=ZSCAN29;refGene_tx=XM_023255445.1;refGene_gn=ZSCAN29;refGene_tx=XM_023255446.1
B3 62139090 62139285
refGene_gn=ADAL;refGene_tx=XM_011283042.3;refGene_gn=ADAL;refGene_tx=XM_011283043.2
B3 62139838 62139939
refGene_gn=ADAL;refGene_tx=XM_011283042.3;refGene_gn=ADAL;refGene_tx=XM_011283043.2
B3 62142456 62142595
refGene_gn=ADAL;refGene_tx=XM_011283042.3;refGene_gn=ADAL;refGene_tx=XM_011283043.2
B3 62144527 62144608
refGene_gn=ADAL;refGene_tx=XM_011283042.3;refGene_gn=ADAL;refGene_tx=XM_011283043.2
B3 62145695 62145822
refGene_gn=ADAL;refGene_tx=XM_011283042.3;refGene_gn=ADAL;refGene_tx=XM_011283043.2
B3 62148204 62148286
refGene_gn=ADAL;refGene_tx=XM_011283042.3;refGene_gn=ADAL;refGene_tx=XM_011283043.2
B3 62150801 62150889
refGene_gn=ADAL;refGene_tx=XM_011283042.3;refGene_gn=ADAL;refGene_tx=XM_011283043.2
B3 62155696 62155757
refGene_gn=ADAL;refGene_tx=XM_011283042.3;refGene_gn=ADAL;refGene_tx=XM_011283043.2
B3 62156470 62156601
refGene_gn=ADAL;refGene_tx=XM_011283042.3;refGene_gn=ADAL;refGene_tx=XM_011283043.2
B3 62156875 62156941
refGene_gn=ADAL;refGene_tx=XM_011283042.3;refGene_gn=ADAL;refGene_tx=XM_011283043.2
B3 62163973 62166031 refGene_gn=LCMT2;refGene_tx=XM_003987233.3
B3 62204356 62204561 refGene_gn=TGM7;refGene_tx=XM_019833781.2
B3 62205227 62205473 refGene_gn=TGM7;refGene_tx=XM_019833781.2

B3 62206121 62206240 refGene_gn=TGM7;refGene_tx=XM_019833781.2
B3 62207870 62207999 refGene_gn=TGM7;refGene_tx=XM_019833781.2
B3 62208087 62208265 refGene_gn=TGM7;refGene_tx=XM_019833781.2
B3 62210741 62210880 refGene_gn=TGM7;refGene_tx=XM_019833781.2
B3 62215215 62215319 refGene_gn=TGM7;refGene_tx=XM_019833781.2
B3 62215667 62215910 refGene_gn=TGM7;refGene_tx=XM_019833781.2
B3 62217350 62217677 refGene_gn=TGM7;refGene_tx=XM_019833781.2
B3 62218079 62218240 refGene_gn=TGM7;refGene_tx=XM_019833781.2
B3 62220491 62220625 refGene_gn=TGM7;refGene_tx=XM_019833781.2
B3 62220856 62221118 refGene_gn=TGM7;refGene_tx=XM_019833781.2
B3 62227333 62227343
refGene_gn=TGM5;refGene_tx=XM_006932597.4;refGene_gn=TGM5;refGene_tx=XM_019832585.2
B3 62233231 62233411
refGene_gn=TGM5;refGene_tx=XM_006932597.4;refGene_gn=TGM5;refGene_tx=XM_019832585.2
B3 62233527 62233773 refGene_gn=TGM5;refGene_tx=XM_006932597.4
B3 62237093 62237212
refGene_gn=TGM5;refGene_tx=XM_006932597.4;refGene_gn=TGM5;refGene_tx=XM_019832585.2
B3 62240170 62240299
refGene_gn=TGM5;refGene_tx=XM_006932597.4;refGene_gn=TGM5;refGene_tx=XM_019832585.2
B3 62240895 62241073
refGene_gn=TGM5;refGene_tx=XM_006932597.4;refGene_gn=TGM5;refGene_tx=XM_019832585.2
B3 62249848 62249987
refGene_gn=TGM5;refGene_tx=XM_006932597.4;refGene_gn=TGM5;refGene_tx=XM_019832585.2
B3 62250632 62250736
refGene_gn=TGM5;refGene_tx=XM_006932597.4;refGene_gn=TGM5;refGene_tx=XM_019832585.2
B3 62250841 62251081
refGene_gn=TGM5;refGene_tx=XM_006932597.4;refGene_gn=TGM5;refGene_tx=XM_019832585.2
B3 62254378 62254744
refGene_gn=TGM5;refGene_tx=XM_006932597.4;refGene_gn=TGM5;refGene_tx=XM_019832585.2
B3 62255296 62255457
refGene_gn=TGM5;refGene_tx=XM_006932597.4;refGene_gn=TGM5;refGene_tx=XM_019832585.2
B3 62256906 62257040
refGene_gn=TGM5;refGene_tx=XM_006932597.4;refGene_gn=TGM5;refGene_tx=XM_019832585.2
B3 62257232 62257386
refGene_gn=TGM5;refGene_tx=XM_006932597.4;refGene_gn=TGM5;refGene_tx=XM_019832585.2
B3 62261153 62261199 refGene_gn=CCNDBP1;refGene_tx=XM_023255449.1
B3 62267482 62267492
refGene_gn=EPB42;refGene_tx=XM_019832587.2;refGene_gn=EPB42;refGene_tx

=XM_019832588.2
B3	62268838	62269027
refGene_gn=EPB42;refGene_tx=XM_019832587.2;refGene_gn=EPB42;refGene_tx=XM_019832588.2
B3	62269654	62269888
refGene_gn=EPB42;refGene_tx=XM_019832587.2;refGene_gn=EPB42;refGene_tx=XM_019832588.2
B3	62273978	62274097
refGene_gn=EPB42;refGene_tx=XM_019832587.2;refGene_gn=EPB42;refGene_tx=XM_019832588.2
B3	62275979	62276084
refGene_gn=EPB42;refGene_tx=XM_019832587.2;refGene_gn=EPB42;refGene_tx=XM_019832588.2
B3	62276693	62276871
refGene_gn=EPB42;refGene_tx=XM_019832587.2;refGene_gn=EPB42;refGene_tx=XM_019832588.2
B3	62277258	62277397
refGene_gn=EPB42;refGene_tx=XM_019832587.2;refGene_gn=EPB42;refGene_tx=XM_019832588.2
B3	62277685	62277789
refGene_gn=EPB42;refGene_tx=XM_019832587.2;refGene_gn=EPB42;refGene_tx=XM_019832588.2
B3	62278578	62278821
refGene_gn=EPB42;refGene_tx=XM_019832587.2;refGene_gn=EPB42;refGene_tx=XM_019832588.2
B3	62278838	62278845	refGene_gn=CCNDBP1;refGene_tx=XM_019832592.2
B3	62281463	62281763
refGene_gn=EPB42;refGene_tx=XM_019832587.2;refGene_gn=EPB42;refGene_tx=XM_019832588.2
B3	62284115	62284272
refGene_gn=CCNDBP1;refGene_tx=XM_019832589.1;refGene_gn=CCNDBP1;refGene_tx=XM_019832590.2;refGene_gn=CCNDBP1;refGene_tx=XM_019832591.1;refGene_gn=CCNDBP1;refGene_tx=XM_019832595.1
B3	62284676	62284846
refGene_gn=EPB42;refGene_tx=XM_019832587.2;refGene_gn=EPB42;refGene_tx=XM_019832588.2
B3	62286893	62287027
refGene_gn=EPB42;refGene_tx=XM_019832587.2;refGene_gn=EPB42;refGene_tx=XM_019832588.2
B3	62289563	62289726
refGene_gn=EPB42;refGene_tx=XM_019832587.2;refGene_gn=EPB42;refGene_tx=XM_019832588.2
B3	62290623	62290642	refGene_gn=CCNDBP1;refGene_tx=XM_023255450.1
B3	62291083	62291198
refGene_gn=CCNDBP1;refGene_tx=XM_003987234.5;refGene_gn=CCNDBP1;refGene_tx=XM_019832589.1;refGene_gn=CCNDBP1;refGene_tx=XM_019832590.2;refGene_gn=CCNDBP1;refGene_tx=XM_019832591.1;refGene_gn=CCNDBP1;refGene_tx=XM_019832592.2;refGene_gn=CCNDBP1;refGene_tx=XM_019832594.2;refGene_gn=CCNDBP1;refGene_tx=XM_019832595.1;refGene_gn=CCNDBP1;refGene_tx=XM_023255449.1;refGene_gn=CCNDBP1;refGene_tx=XM_023255450.1

B3 62291454 62291501
refGene_gn=CCNDBP1;refGene_tx=XM_003987234.5;refGene_gn=CCNDBP1;refGene_tx=XM_019832589.1;refGene_gn=CCNDBP1;refGene_tx=XM_019832590.2;refGene_gn=CCNDBP1;refGene_tx=XM_019832591.1;refGene_gn=CCNDBP1;refGene_tx=XM_019832592.2;refGene_gn=CCNDBP1;refGene_tx=XM_019832595.1;refGene_gn=CCNDBP1;refGene_tx=XM_023255449.1;refGene_gn=CCNDBP1;refGene_tx=XM_023255450.1
B3 62292717 62292778
refGene_gn=CCNDBP1;refGene_tx=XM_003987234.5;refGene_gn=CCNDBP1;refGene_tx=XM_019832589.1;refGene_gn=CCNDBP1;refGene_tx=XM_019832590.2;refGene_gn=CCNDBP1;refGene_tx=XM_019832591.1;refGene_gn=CCNDBP1;refGene_tx=XM_019832592.2;refGene_gn=CCNDBP1;refGene_tx=XM_019832594.2;refGene_gn=CCNDBP1;refGene_tx=XM_019832595.1;refGene_gn=CCNDBP1;refGene_tx=XM_023255449.1;refGene_gn=CCNDBP1;refGene_tx=XM_023255450.1
B3 62293837 62294118
refGene_gn=CCNDBP1;refGene_tx=XM_003987234.5;refGene_gn=CCNDBP1;refGene_tx=XM_019832589.1;refGene_gn=CCNDBP1;refGene_tx=XM_019832590.2;refGene_gn=CCNDBP1;refGene_tx=XM_019832591.1;refGene_gn=CCNDBP1;refGene_tx=XM_019832592.2;refGene_gn=CCNDBP1;refGene_tx=XM_019832594.2;refGene_gn=CCNDBP1;refGene_tx=XM_019832595.1;refGene_gn=CCNDBP1;refGene_tx=XM_023255449.1;refGene_gn=CCNDBP1;refGene_tx=XM_023255450.1
B3 62294438 62294515
refGene_gn=CCNDBP1;refGene_tx=XM_003987234.5;refGene_gn=CCNDBP1;refGene_tx=XM_019832589.1;refGene_gn=CCNDBP1;refGene_tx=XM_019832590.2;refGene_gn=CCNDBP1;refGene_tx=XM_019832591.1;refGene_gn=CCNDBP1;refGene_tx=XM_019832592.2;refGene_gn=CCNDBP1;refGene_tx=XM_019832594.2;refGene_gn=CCNDBP1;refGene_tx=XM_019832595.1;refGene_gn=CCNDBP1;refGene_tx=XM_023255449.1;refGene_gn=CCNDBP1;refGene_tx=XM_023255450.1
B3 62295313 62295387
refGene_gn=CCNDBP1;refGene_tx=XM_003987234.5;refGene_gn=CCNDBP1;refGene_tx=XM_019832589.1;refGene_gn=CCNDBP1;refGene_tx=XM_019832590.2;refGene_gn=CCNDBP1;refGene_tx=XM_019832591.1;refGene_gn=CCNDBP1;refGene_tx=XM_019832592.2;refGene_gn=CCNDBP1;refGene_tx=XM_019832594.2;refGene_gn=CCNDBP1;refGene_tx=XM_019832595.1;refGene_gn=CCNDBP1;refGene_tx=XM_023255449.1;refGene_gn=CCNDBP1;refGene_tx=XM_023255450.1
B3 62295566 62295663
refGene_gn=CCNDBP1;refGene_tx=XM_003987234.5;refGene_gn=CCNDBP1;refGene_tx=XM_019832589.1;refGene_gn=CCNDBP1;refGene_tx=XM_019832590.2;refGene_gn=CCNDBP1;refGene_tx=XM_019832591.1;refGene_gn=CCNDBP1;refGene_tx=XM_019832592.2;refGene_gn=CCNDBP1;refGene_tx=XM_019832594.2;refGene_gn=CCNDBP1;refGene_tx=XM_019832595.1;refGene_gn=CCNDBP1;refGene_tx=XM_023255449.1;refGene_gn=CCNDBP1;refGene_tx=XM_023255450.1
B3 62296406 62296488
refGene_gn=CCNDBP1;refGene_tx=XM_003987234.5;refGene_gn=CCNDBP1;refGene_tx=XM_019832589.1;refGene_gn=CCNDBP1;refGene_tx=XM_019832590.2;refGene_gn=CCNDBP1;refGene_tx=XM_019832591.1;refGene_gn=CCNDBP1;refGene_tx=XM_019832592.2;refGene_gn=CCNDBP1;refGene_tx=XM_019832594.2;refGene_gn=CCNDBP1;refGene_tx=XM_019832595.1;refGene_gn=CCNDBP1;refGene_tx=XM_023255449.1;refGene_gn=CCNDBP1;refGene_tx=XM_023255450.1
B3 62299083 62299163

refGene_gn=CCNDBP1;refGene_tx=XM_003987234.5;refGene_gn=CCNDBP1;refGene_tx=XM_019832589.1;refGene_gn=CCNDBP1;refGene_tx=XM_019832590.2;refGene_gn=CCNDBP1;refGene_tx=XM_019832591.1;refGene_gn=CCNDBP1;refGene_tx=XM_019832592.2;refGene_gn=CCNDBP1;refGene_tx=XM_019832594.2;refGene_gn=CCNDBP1;refGene_tx=XM_023255449.1;refGene_gn=CCNDBP1;refGene_tx=XM_023255450.1
B3 62299455 62299515
refGene_gn=CCNDBP1;refGene_tx=XM_003987234.5;refGene_gn=CCNDBP1;refGene_tx=XM_019832589.1;refGene_gn=CCNDBP1;refGene_tx=XM_019832590.2;refGene_gn=CCNDBP1;refGene_tx=XM_019832591.1;refGene_gn=CCNDBP1;refGene_tx=XM_019832592.2;refGene_gn=CCNDBP1;refGene_tx=XM_019832594.2;refGene_gn=CCNDBP1;refGene_tx=XM_023255449.1;refGene_gn=CCNDBP1;refGene_tx=XM_023255450.1
B3 62299691 62299800
refGene_gn=CCNDBP1;refGene_tx=XM_003987234.5;refGene_gn=CCNDBP1;refGene_tx=XM_019832589.1;refGene_gn=CCNDBP1;refGene_tx=XM_019832590.2;refGene_gn=CCNDBP1;refGene_tx=XM_019832591.1;refGene_gn=CCNDBP1;refGene_tx=XM_019832592.2;refGene_gn=CCNDBP1;refGene_tx=XM_019832594.2;refGene_gn=CCNDBP1;refGene_tx=XM_023255449.1;refGene_gn=CCNDBP1;refGene_tx=XM_023255450.1
B3 62300570 62300897
refGene_gn=TMEM62;refGene_tx=XM_011283044.3;refGene_gn=TMEM62;refGene_tx=XM_023255451.1;refGene_gn=TMEM62;refGene_tx=XM_023255452.1;refGene_gn=TMEM62;refGene_tx=XM_023255453.1;refGene_gn=TMEM62;refGene_tx=XM_023255454.1
B3 62304387 62304506
refGene_gn=TMEM62;refGene_tx=XM_011283044.3;refGene_gn=TMEM62;refGene_tx=XM_023255451.1;refGene_gn=TMEM62;refGene_tx=XM_023255452.1;refGene_gn=TMEM62;refGene_tx=XM_023255453.1;refGene_gn=TMEM62;refGene_tx=XM_023255454.1
B3 62306482 62306587
refGene_gn=TMEM62;refGene_tx=XM_011283044.3;refGene_gn=TMEM62;refGene_tx=XM_023255451.1;refGene_gn=TMEM62;refGene_tx=XM_023255452.1;refGene_gn=TMEM62;refGene_tx=XM_023255453.1;refGene_gn=TMEM62;refGene_tx=XM_023255454.1
B3 62308107 62308192
refGene_gn=TMEM62;refGene_tx=XM_011283044.3;refGene_gn=TMEM62;refGene_tx=XM_023255451.1;refGene_gn=TMEM62;refGene_tx=XM_023255452.1;refGene_gn=TMEM62;refGene_tx=XM_023255453.1;refGene_gn=TMEM62;refGene_tx=XM_023255454.1
B3 62310052 62310166
refGene_gn=TMEM62;refGene_tx=XM_011283044.3;refGene_gn=TMEM62;refGene_tx=XM_023255451.1;refGene_gn=TMEM62;refGene_tx=XM_023255452.1;refGene_gn=TMEM62;refGene_tx=XM_023255453.1;refGene_gn=TMEM62;refGene_tx=XM_023255454.1
B3 62315353 62315513
refGene_gn=TMEM62;refGene_tx=XM_011283044.3;refGene_gn=TMEM62;refGene_tx=XM_023255451.1;refGene_gn=TMEM62;refGene_tx=XM_023255452.1;refGene_gn=TMEM62;refGene_tx=XM_023255453.1;refGene_gn=TMEM62;refGene_tx=XM_023255454.1

B3	62318843	62318999
refGene_gn=TMEM62;refGene_tx=XM_011283044.3;refGene_gn=TMEM62;refGene_tx=XM_023255451.1;refGene_gn=TMEM62;refGene_tx=XM_023255452.1;refGene_gn=TMEM62;refGene_tx=XM_023255453.1;refGene_gn=TMEM62;refGene_tx=XM_023255454.1
B3	62321353	62321476
refGene_gn=TMEM62;refGene_tx=XM_011283044.3;refGene_gn=TMEM62;refGene_tx=XM_023255451.1;refGene_gn=TMEM62;refGene_tx=XM_023255452.1;refGene_gn=TMEM62;refGene_tx=XM_023255453.1;refGene_gn=TMEM62;refGene_tx=XM_023255454.1
B3	62321621	62321746
refGene_gn=TMEM62;refGene_tx=XM_011283044.3;refGene_gn=TMEM62;refGene_tx=XM_023255451.1;refGene_gn=TMEM62;refGene_tx=XM_023255452.1;refGene_gn=TMEM62;refGene_tx=XM_023255453.1;refGene_gn=TMEM62;refGene_tx=XM_023255454.1
B3	62324518	62324660
refGene_gn=TMEM62;refGene_tx=XM_011283044.3;refGene_gn=TMEM62;refGene_tx=XM_023255451.1
B3	62328759	62328767	refGene_gn=TMEM62;refGene_tx=XM_023255451.1
B3	62328822	62328868	refGene_gn=TMEM62;refGene_tx=XM_011283044.3
B3	62331869	62332007	refGene_gn=TMEM62;refGene_tx=XM_011283044.3
B3	62333133	62333245	refGene_gn=TMEM62;refGene_tx=XM_011283044.3
B3	62333499	62333679	refGene_gn=TMEM62;refGene_tx=XM_011283044.3
B3	62348144	62348225	refGene_gn=UBR1;refGene_tx=XM_023255455.1
B3	62368302	62368559	refGene_gn=UBR1;refGene_tx=XM_023255455.1
B3	62371510	62371589	refGene_gn=UBR1;refGene_tx=XM_023255455.1
B3	62385759	62385870	refGene_gn=UBR1;refGene_tx=XM_023255455.1
B3	62400188	62400319	refGene_gn=UBR1;refGene_tx=XM_023255455.1
B3	62403236	62403375	refGene_gn=UBR1;refGene_tx=XM_023255455.1
B3	62404331	62404394	refGene_gn=UBR1;refGene_tx=XM_023255455.1
B3	62404641	62404765	refGene_gn=UBR1;refGene_tx=XM_023255455.1
B3	62405526	62405634	refGene_gn=UBR1;refGene_tx=XM_023255455.1
B3	62406278	62406367	refGene_gn=UBR1;refGene_tx=XM_023255455.1
B3	62408834	62408933	refGene_gn=UBR1;refGene_tx=XM_023255455.1
B3	62409910	62410068	refGene_gn=UBR1;refGene_tx=XM_023255455.1
B3	62412512	62412612	refGene_gn=UBR1;refGene_tx=XM_023255455.1
B3	62413223	62413352	refGene_gn=UBR1;refGene_tx=XM_023255455.1
B3	62416753	62416934	refGene_gn=UBR1;refGene_tx=XM_023255455.1
B3	62419292	62419354	refGene_gn=UBR1;refGene_tx=XM_023255455.1
B3	62419634	62419745	refGene_gn=UBR1;refGene_tx=XM_023255455.1
B3	62420711	62420777	refGene_gn=UBR1;refGene_tx=XM_023255455.1
B3	62421043	62421145	refGene_gn=UBR1;refGene_tx=XM_023255455.1
B3	62422623	62422688	refGene_gn=UBR1;refGene_tx=XM_023255455.1
B3	62424842	62424966	refGene_gn=UBR1;refGene_tx=XM_023255455.1
B3	62429542	62429595	refGene_gn=UBR1;refGene_tx=XM_023255455.1
B3	62431607	62431710	refGene_gn=UBR1;refGene_tx=XM_023255455.1
B3	62435027	62435076	refGene_gn=UBR1;refGene_tx=XM_023255455.1
B3	62435480	62435635	refGene_gn=UBR1;refGene_tx=XM_023255455.1
B3	62440715	62440815	refGene_gn=UBR1;refGene_tx=XM_023255455.1
B3	62441767	62441877	refGene_gn=UBR1;refGene_tx=XM_023255455.1

B3 62444419 62444506 refGene_gn=UBR1;refGene_tx=XM_023255455.1
B3 62445450 62445632 refGene_gn=UBR1;refGene_tx=XM_023255455.1
B3 62449758 62449964 refGene_gn=UBR1;refGene_tx=XM_023255455.1
B3 62451369 62451463 refGene_gn=UBR1;refGene_tx=XM_023255455.1
B3 62452681 62452831 refGene_gn=UBR1;refGene_tx=XM_023255455.1
B3 62455205 62455303 refGene_gn=UBR1;refGene_tx=XM_023255455.1
B3 62458001 62458095 refGene_gn=UBR1;refGene_tx=XM_023255455.1
B3 62459395 62459544 refGene_gn=UBR1;refGene_tx=XM_023255455.1
B3 62461585 62461641 refGene_gn=UBR1;refGene_tx=XM_023255455.1
B3 62462647 62462744 refGene_gn=UBR1;refGene_tx=XM_023255455.1
B3 62463524 62463592 refGene_gn=UBR1;refGene_tx=XM_023255455.1
B3 62465202 62465353 refGene_gn=UBR1;refGene_tx=XM_023255455.1
B3 62468440 62468528 refGene_gn=UBR1;refGene_tx=XM_023255455.1
B3 62470341 62470475 refGene_gn=UBR1;refGene_tx=XM_023255455.1
B3 62471771 62471880 refGene_gn=UBR1;refGene_tx=XM_023255455.1
B3 62474312 62474369 refGene_gn=UBR1;refGene_tx=XM_023255455.1
B3 62476865 62476943 refGene_gn=UBR1;refGene_tx=XM_023255455.1
B3 62485667 62485838 refGene_gn=UBR1;refGene_tx=XM_023255455.1
B3 62487129 62487231 refGene_gn=UBR1;refGene_tx=XM_023255455.1
B3 62489417 62489559 refGene_gn=UBR1;refGene_tx=XM_023255455.1
B3 62552439 62552508
refGene_gn=TTBK2;refGene_tx=XM_011283046.3;refGene_gn=TTBK2;refGene_tx=XM_019832596.2
B3 62557362 62557510
refGene_gn=TTBK2;refGene_tx=XM_011283046.3;refGene_gn=TTBK2;refGene_tx=XM_019832596.2
B3 62587112 62587186
refGene_gn=TTBK2;refGene_tx=XM_011283046.3;refGene_gn=TTBK2;refGene_tx=XM_019832596.2;refGene_gn=TTBK2;refGene_tx=XM_019832597.2;refGene_gn=TTBK2;refGene_tx=XM_019832599.2;refGene_gn=TTBK2;refGene_tx=XM_019832600.2;refGene_gn=TTBK2;refGene_tx=XM_023255457.1
B3 62593456 62593597
refGene_gn=TTBK2;refGene_tx=XM_011283046.3;refGene_gn=TTBK2;refGene_tx=XM_019832596.2;refGene_gn=TTBK2;refGene_tx=XM_019832597.2;refGene_gn=TTBK2;refGene_tx=XM_019832599.2;refGene_gn=TTBK2;refGene_tx=XM_019832600.2;refGene_gn=TTBK2;refGene_tx=XM_023255457.1
B3 62595077 62595182
refGene_gn=TTBK2;refGene_tx=XM_011283046.3;refGene_gn=TTBK2;refGene_tx=XM_019832596.2;refGene_gn=TTBK2;refGene_tx=XM_019832597.2;refGene_gn=TTBK2;refGene_tx=XM_019832599.2;refGene_gn=TTBK2;refGene_tx=XM_019832600.2;refGene_gn=TTBK2;refGene_tx=XM_023255457.1;refGene_gn=TTBK2;refGene_tx=XM_023255458.1
B3 62618489 62618555
refGene_gn=TTBK2;refGene_tx=XM_011283046.3;refGene_gn=TTBK2;refGene_tx=XM_019832596.2;refGene_gn=TTBK2;refGene_tx=XM_019832597.2;refGene_gn=TTBK2;refGene_tx=XM_019832599.2;refGene_gn=TTBK2;refGene_tx=XM_019832600.2;refGene_gn=TTBK2;refGene_tx=XM_023255457.1;refGene_gn=TTBK2;refGene_tx=XM_023255458.1
B3 62622769 62622862
refGene_gn=TTBK2;refGene_tx=XM_011283046.3;refGene_gn=TTBK2;refGene_tx

=XM_019832596.2;refGene_gn=TTBK2;refGene_tx=XM_019832597.2;refGene_gn=TTBK2;refGene_tx=XM_019832599.2;refGene_gn=TTBK2;refGene_tx=XM_019832600.2;refGene_gn=TTBK2;refGene_tx=XM_023255457.1;refGene_gn=TTBK2;refGene_tx=XM_023255458.1
B3 62623475 62623601 refGene_gn=TTBK2;refGene_tx=XM_011283046.3;refGene_gn=TTBK2;refGene_tx=XM_019832596.2;refGene_gn=TTBK2;refGene_tx=XM_019832597.2;refGene_gn=TTBK2;refGene_tx=XM_019832599.2;refGene_gn=TTBK2;refGene_tx=XM_019832600.2;refGene_gn=TTBK2;refGene_tx=XM_023255457.1;refGene_gn=TTBK2;refGene_tx=XM_023255458.1
B3 62634474 62634632 refGene_gn=TTBK2;refGene_tx=XM_011283046.3;refGene_gn=TTBK2;refGene_tx=XM_019832596.2;refGene_gn=TTBK2;refGene_tx=XM_019832597.2;refGene_gn=TTBK2;refGene_tx=XM_019832599.2;refGene_gn=TTBK2;refGene_tx=XM_019832600.2;refGene_gn=TTBK2;refGene_tx=XM_023255457.1;refGene_gn=TTBK2;refGene_tx=XM_023255458.1
B3 62642565 62642782 refGene_gn=TTBK2;refGene_tx=XM_011283046.3;refGene_gn=TTBK2;refGene_tx=XM_019832596.2;refGene_gn=TTBK2;refGene_tx=XM_019832597.2;refGene_gn=TTBK2;refGene_tx=XM_019832599.2;refGene_gn=TTBK2;refGene_tx=XM_019832600.2;refGene_gn=TTBK2;refGene_tx=XM_023255457.1;refGene_gn=TTBK2;refGene_tx=XM_023255458.1
B3 62653033 62653245 refGene_gn=TTBK2;refGene_tx=XM_011283046.3;refGene_gn=TTBK2;refGene_tx=XM_019832596.2;refGene_gn=TTBK2;refGene_tx=XM_019832597.2;refGene_gn=TTBK2;refGene_tx=XM_019832599.2;refGene_gn=TTBK2;refGene_tx=XM_019832600.2;refGene_gn=TTBK2;refGene_tx=XM_023255457.1;refGene_gn=TTBK2;refGene_tx=XM_023255458.1
B3 62655165 62655754 refGene_gn=TTBK2;refGene_tx=XM_011283046.3;refGene_gn=TTBK2;refGene_tx=XM_019832596.2;refGene_gn=TTBK2;refGene_tx=XM_019832597.2;refGene_gn=TTBK2;refGene_tx=XM_019832599.2;refGene_gn=TTBK2;refGene_tx=XM_019832600.2;refGene_gn=TTBK2;refGene_tx=XM_023255457.1;refGene_gn=TTBK2;refGene_tx=XM_023255458.1
B3 62668219 62669493 refGene_gn=TTBK2;refGene_tx=XM_011283046.3;refGene_gn=TTBK2;refGene_tx=XM_019832596.2;refGene_gn=TTBK2;refGene_tx=XM_019832597.2;refGene_gn=TTBK2;refGene_tx=XM_019832599.2;refGene_gn=TTBK2;refGene_tx=XM_019832600.2;refGene_gn=TTBK2;refGene_tx=XM_023255457.1;refGene_gn=TTBK2;refGene_tx=XM_023255458.1
B3 62673053 62673516 refGene_gn=TTBK2;refGene_tx=XM_011283046.3;refGene_gn=TTBK2;refGene_tx=XM_019832596.2;refGene_gn=TTBK2;refGene_tx=XM_019832597.2;refGene_gn=TTBK2;refGene_tx=XM_019832599.2;refGene_gn=TTBK2;refGene_tx=XM_019832600.2;refGene_gn=TTBK2;refGene_tx=XM_023255457.1;refGene_gn=TTBK2;refGene_tx=XM_023255458.1
B3 62681505 62681595 refGene_gn=CDAN1;refGene_tx=XM_023255459.1
B3 62681823 62682314 refGene_gn=CDAN1;refGene_tx=XM_023255459.1
B3 62682555 62682759 refGene_gn=CDAN1;refGene_tx=XM_023255459.1
B3 62682958 62683128 refGene_gn=CDAN1;refGene_tx=XM_023255459.1

| | | | |
|---|---|---|---|
| B3 | 62683260 | 62683374 | refGene_gn=CDAN1;refGene_tx=XM_023255459.1 |
| B3 | 62683456 | 62683535 | refGene_gn=CDAN1;refGene_tx=XM_023255459.1 |
| B3 | 62683940 | 62684061 | refGene_gn=CDAN1;refGene_tx=XM_023255459.1 |
| B3 | 62684218 | 62684328 | refGene_gn=CDAN1;refGene_tx=XM_023255459.1 |
| B3 | 62684794 | 62684884 | refGene_gn=CDAN1;refGene_tx=XM_023255459.1 |
| B3 | 62685523 | 62685599 | refGene_gn=CDAN1;refGene_tx=XM_023255459.1 |
| B3 | 62686130 | 62686289 | refGene_gn=CDAN1;refGene_tx=XM_023255459.1 |
| B3 | 62686290 | 62686337 | refGene_gn=CDAN1;refGene_tx=XM_023255459.1 |
| B3 | 62686647 | 62686768 | refGene_gn=CDAN1;refGene_tx=XM_023255459.1 |
| B3 | 62686903 | 62687050 | refGene_gn=CDAN1;refGene_tx=XM_023255459.1 |
| B3 | 62687160 | 62687327 | refGene_gn=CDAN1;refGene_tx=XM_023255459.1 |
| B3 | 62687671 | 62687759 | refGene_gn=CDAN1;refGene_tx=XM_023255459.1 |
| B3 | 62688012 | 62688102 | refGene_gn=CDAN1;refGene_tx=XM_023255459.1 |
| B3 | 62688278 | 62688333 | refGene_gn=CDAN1;refGene_tx=XM_023255459.1 |
| B3 | 62688538 | 62688672 | refGene_gn=CDAN1;refGene_tx=XM_023255459.1 |
| B3 | 62688753 | 62688857 | refGene_gn=CDAN1;refGene_tx=XM_023255459.1 |
| B3 | 62689106 | 62689265 | refGene_gn=CDAN1;refGene_tx=XM_023255459.1 |
| B3 | 62689634 | 62689698 | refGene_gn=CDAN1;refGene_tx=XM_023255459.1 |
| B3 | 62689891 | 62689970 | refGene_gn=CDAN1;refGene_tx=XM_023255459.1 |
| B3 | 62690129 | 62690278 | refGene_gn=CDAN1;refGene_tx=XM_023255459.1 |
| B3 | 62691418 | 62691526 | refGene_gn=CDAN1;refGene_tx=XM_023255459.1 |
| B3 | 62691642 | 62691706 | refGene_gn=CDAN1;refGene_tx=XM_023255459.1 |
| B3 | 62691873 | 62692055 | refGene_gn=CDAN1;refGene_tx=XM_023255459.1 |
| B3 | 62692304 | 62692412 | refGene_gn=CDAN1;refGene_tx=XM_023255459.1 |
| B3 | 62693032 | 62693158 | refGene_gn=CDAN1;refGene_tx=XM_023255459.1 |
| B3 | 62696860 | 62696962 | refGene_gn=STARD9;refGene_tx=XM_023256167.1 |
| B3 | 62697517 | 62697676 | refGene_gn=STARD9;refGene_tx=XM_023256167.1 |
| B3 | 62697865 | 62697945 | refGene_gn=STARD9;refGene_tx=XM_023256167.1 |
| B3 | 62698180 | 62698383 | refGene_gn=STARD9;refGene_tx=XM_023256167.1 |
| B3 | 62698551 | 62698616 | refGene_gn=STARD9;refGene_tx=XM_023256167.1 |
| B3 | 62698869 | 62698991 | refGene_gn=STARD9;refGene_tx=XM_023256167.1 |
| B3 | 62699180 | 62699298 | refGene_gn=STARD9;refGene_tx=XM_023256167.1 |
| B3 | 62710229 | 62710370 | refGene_gn=STARD9;refGene_tx=XM_023256167.1 |
| B3 | 62710716 | 62710900 | refGene_gn=STARD9;refGene_tx=XM_023256167.1 |
| B3 | 62711337 | 62711535 | refGene_gn=STARD9;refGene_tx=XM_023256167.1 |
| B3 | 62711721 | 62721891 | refGene_gn=STARD9;refGene_tx=XM_023256167.1 |
| B3 | 62724098 | 62724579 | refGene_gn=STARD9;refGene_tx=XM_023256167.1 |
| B3 | 62725033 | 62725221 | refGene_gn=STARD9;refGene_tx=XM_023256167.1 |
| B3 | 62730652 | 62730756 | refGene_gn=STARD9;refGene_tx=XM_023256167.1 |
| B3 | 62730889 | 62730972 | refGene_gn=STARD9;refGene_tx=XM_023256167.1 |
| B3 | 62731665 | 62731803 | refGene_gn=STARD9;refGene_tx=XM_023256167.1 |
| B3 | 62732123 | 62732175 | refGene_gn=STARD9;refGene_tx=XM_023256167.1 |
| B3 | 62736206 | 62736257 | refGene_gn=STARD9;refGene_tx=XM_023256167.1 |
| B3 | 62738208 | 62738263 | refGene_gn=STARD9;refGene_tx=XM_023256167.1 |
| B3 | 62738624 | 62738791 | refGene_gn=STARD9;refGene_tx=XM_023256167.1 |
| B3 | 62739230 | 62739308 | refGene_gn=STARD9;refGene_tx=XM_023256167.1 |
| B3 | 62740465 | 62740563 | refGene_gn=STARD9;refGene_tx=XM_023256167.1 |
| B3 | 62740745 | 62740955 | refGene_gn=STARD9;refGene_tx=XM_023256167.1 |
| B3 | 62741607 | 62741705 | refGene_gn=STARD9;refGene_tx=XM_023256167.1 |
| B3 | 62742884 | 62742952 | refGene_gn=STARD9;refGene_tx=XM_023256167.1 |

B3 62750289 62750362 refGene_gn=STARD9;refGene_tx=XM_023256167.1
B3 62752223 62752293 refGene_gn=STARD9;refGene_tx=XM_023256167.1
B3 62761356 62761469 refGene_gn=STARD9;refGene_tx=XM_023256167.1
B3 62763251 62763313 refGene_gn=STARD9;refGene_tx=XM_023256167.1
B3 62763401 62763434 refGene_gn=STARD9;refGene_tx=XM_023256167.1
B3 62767373 62767490 refGene_gn=STARD9;refGene_tx=XM_023256167.1
B3 62810784 62810901 refGene_gn=STARD9;refGene_tx=XM_023256167.1
B3 62812533 62812603 refGene_gn=STARD9;refGene_tx=XM_023256167.1
B3 62818185 62818232 refGene_gn=STARD9;refGene_tx=XM_023256167.1
B3 62836396 62836434 refGene_gn=HAUS2;refGene_tx=XM_023255460.1
B3 62838314 62838524
refGene_gn=HAUS2;refGene_tx=XM_003987235.5;refGene_gn=HAUS2;refGene_tx=XM_006932608.4;refGene_gn=HAUS2;refGene_tx=XM_019832614.2;refGene_gn=HAUS2;refGene_tx=XM_019832615.2;refGene_gn=HAUS2;refGene_tx=XM_019832616.2;refGene_gn=HAUS2;refGene_tx=XM_023255460.1
B3 62840774 62840883
refGene_gn=HAUS2;refGene_tx=XM_003987235.5;refGene_gn=HAUS2;refGene_tx=XM_006932608.4;refGene_gn=HAUS2;refGene_tx=XM_019832614.2;refGene_gn=HAUS2;refGene_tx=XM_019832615.2;refGene_gn=HAUS2;refGene_tx=XM_019832616.2;refGene_gn=HAUS2;refGene_tx=XM_023255460.1
B3 62842351 62842484
refGene_gn=HAUS2;refGene_tx=XM_003987235.5;refGene_gn=HAUS2;refGene_tx=XM_006932608.4;refGene_gn=HAUS2;refGene_tx=XM_019832614.2;refGene_gn=HAUS2;refGene_tx=XM_023255460.1
B3 62845123 62845193
refGene_gn=HAUS2;refGene_tx=XM_003987235.5;refGene_gn=HAUS2;refGene_tx=XM_006932608.4;refGene_gn=HAUS2;refGene_tx=XM_019832614.2;refGene_gn=HAUS2;refGene_tx=XM_023255460.1
B3 62845517 62845610
refGene_gn=HAUS2;refGene_tx=XM_003987235.5;refGene_gn=HAUS2;refGene_tx=XM_006932608.4;refGene_gn=HAUS2;refGene_tx=XM_019832614.2;refGene_gn=HAUS2;refGene_tx=XM_023255460.1
B3 62852901 62853012
refGene_gn=HAUS2;refGene_tx=XM_006932608.4;refGene_gn=HAUS2;refGene_tx=XM_023255460.1
B3 62853143 62853236 refGene_gn=HAUS2;refGene_tx=XM_003987235.5
B3 62853685 62853769
refGene_gn=LRRC57;refGene_tx=XM_011283048.3;refGene_gn=LRRC57;refGene_tx=XM_011283049.3;refGene_gn=LRRC57;refGene_tx=XM_011283050.3;refGene_gn=LRRC57;refGene_tx=XM_011283051.3;refGene_gn=LRRC57;refGene_tx=XM_011283052.3
B3 62853875 62854014
refGene_gn=LRRC57;refGene_tx=XM_011283048.3;refGene_gn=LRRC57;refGene_tx=XM_011283049.3;refGene_gn=LRRC57;refGene_tx=XM_011283050.3;refGene_gn=LRRC57;refGene_tx=XM_011283051.3;refGene_gn=LRRC57;refGene_tx=XM_011283052.3
B3 62854588 62854857
refGene_gn=LRRC57;refGene_tx=XM_011283048.3;refGene_gn=LRRC57;refGene_tx=XM_011283049.3;refGene_gn=LRRC57;refGene_tx=XM_011283050.3;refGene_gn=LRRC57;refGene_tx=XM_01

1283052.3
B3	62857376 62857562
refGene_gn=LRRC57;refGene_tx=XM_011283048.3;refGene_gn=LRRC57;refGene_tx=XM_011283049.3;refGene_gn=LRRC57;refGene_tx=XM_011283050.3;refGene_gn=LRRC57;refGene_tx=XM_011283051.3;refGene_gn=LRRC57;refGene_tx=XM_011283052.3
B3	62858204 62858246
refGene_gn=LRRC57;refGene_tx=XM_011283048.3;refGene_gn=LRRC57;refGene_tx=XM_011283049.3;refGene_gn=LRRC57;refGene_tx=XM_011283050.3;refGene_gn=LRRC57;refGene_tx=XM_011283051.3;refGene_gn=LRRC57;refGene_tx=XM_011283052.3
B3	62863898 62863964
refGene_gn=SNAP23;refGene_tx=XM_023255463.1;refGene_gn=SNAP23;refGene_tx=XM_023255464.1;refGene_gn=SNAP23;refGene_tx=XM_023255465.1;refGene_gn=SNAP23;refGene_tx=XM_023255466.1
B3	62866880 62867025
refGene_gn=SNAP23;refGene_tx=XM_023255463.1;refGene_gn=SNAP23;refGene_tx=XM_023255464.1;refGene_gn=SNAP23;refGene_tx=XM_023255465.1;refGene_gn=SNAP23;refGene_tx=XM_023255466.1
B3	62868386 62868545
refGene_gn=SNAP23;refGene_tx=XM_023255463.1;refGene_gn=SNAP23;refGene_tx=XM_023255464.1;refGene_gn=SNAP23;refGene_tx=XM_023255465.1;refGene_gn=SNAP23;refGene_tx=XM_023255466.1
B3	62874547 62874580
refGene_gn=SNAP23;refGene_tx=XM_023255463.1;refGene_gn=SNAP23;refGene_tx=XM_023255464.1;refGene_gn=SNAP23;refGene_tx=XM_023255465.1
B3	62879130 62879248
refGene_gn=SNAP23;refGene_tx=XM_023255463.1;refGene_gn=SNAP23;refGene_tx=XM_023255464.1;refGene_gn=SNAP23;refGene_tx=XM_023255465.1;refGene_gn=SNAP23;refGene_tx=XM_023255466.1
B3	62880196 62880245
refGene_gn=SNAP23;refGene_tx=XM_023255463.1;refGene_gn=SNAP23;refGene_tx=XM_023255464.1;refGene_gn=SNAP23;refGene_tx=XM_023255465.1;refGene_gn=SNAP23;refGene_tx=XM_023255466.1
B3	62880659 62880701
refGene_gn=SNAP23;refGene_tx=XM_023255463.1;refGene_gn=SNAP23;refGene_tx=XM_023255464.1;refGene_gn=SNAP23;refGene_tx=XM_023255465.1;refGene_gn=SNAP23;refGene_tx=XM_023255466.1
B3	62881496 62881553
refGene_gn=SNAP23;refGene_tx=XM_023255463.1;refGene_gn=SNAP23;refGene_tx=XM_023255464.1;refGene_gn=SNAP23;refGene_tx=XM_023255465.1;refGene_gn=SNAP23;refGene_tx=XM_023255466.1
B3	62926963 62927017
refGene_gn=ZNF106;refGene_tx=XM_011283053.2;refGene_gn=ZNF106;refGene_tx=XM_011283054.2;refGene_gn=ZNF106;refGene_tx=XM_019832620.2;refGene_gn=ZNF106;refGene_tx=XM_023255467.1
B3	62932135 62932197
refGene_gn=ZNF106;refGene_tx=XM_011283053.2;refGene_gn=ZNF106;refGene_tx=XM_011283054.2;refGene_gn=ZNF106;refGene_tx=XM_019832619.2;refGene_gn=ZNF106;refGene_tx=XM_019832620.2;refGene_gn=ZNF106;refGene_tx=XM_02

3255467.1
B3 62939060 62939261
refGene_gn=ZNF106;refGene_tx=XM_011283053.2;refGene_gn=ZNF106;refGene_tx=XM_011283054.2;refGene_gn=ZNF106;refGene_tx=XM_019832619.2;refGene_gn=ZNF106;refGene_tx=XM_023255467.1
B3 62942094 62944275
refGene_gn=ZNF106;refGene_tx=XM_011283053.2;refGene_gn=ZNF106;refGene_tx=XM_011283054.2;refGene_gn=ZNF106;refGene_tx=XM_019832619.2;refGene_gn=ZNF106;refGene_tx=XM_023255467.1
B3 62945333 62945961
refGene_gn=ZNF106;refGene_tx=XM_011283053.2;refGene_gn=ZNF106;refGene_tx=XM_011283054.2;refGene_gn=ZNF106;refGene_tx=XM_019832619.2;refGene_gn=ZNF106;refGene_tx=XM_019832620.2;refGene_gn=ZNF106;refGene_tx=XM_023255467.1
B3 62947221 62947291
refGene_gn=ZNF106;refGene_tx=XM_011283053.2;refGene_gn=ZNF106;refGene_tx=XM_011283054.2;refGene_gn=ZNF106;refGene_tx=XM_019832619.2;refGene_gn=ZNF106;refGene_tx=XM_019832620.2;refGene_gn=ZNF106;refGene_tx=XM_023255467.1
B3 62948502 62948657
refGene_gn=ZNF106;refGene_tx=XM_011283053.2;refGene_gn=ZNF106;refGene_tx=XM_011283054.2;refGene_gn=ZNF106;refGene_tx=XM_019832619.2;refGene_gn=ZNF106;refGene_tx=XM_019832620.2;refGene_gn=ZNF106;refGene_tx=XM_023255467.1
B3 62948741 62948802
refGene_gn=ZNF106;refGene_tx=XM_011283053.2;refGene_gn=ZNF106;refGene_tx=XM_011283054.2;refGene_gn=ZNF106;refGene_tx=XM_019832619.2;refGene_gn=ZNF106;refGene_tx=XM_019832620.2;refGene_gn=ZNF106;refGene_tx=XM_023255467.1
B3 62949279 62949621
refGene_gn=ZNF106;refGene_tx=XM_011283053.2;refGene_gn=ZNF106;refGene_tx=XM_011283054.2;refGene_gn=ZNF106;refGene_tx=XM_019832619.2;refGene_gn=ZNF106;refGene_tx=XM_019832620.2
B3 62951546 62952324
refGene_gn=ZNF106;refGene_tx=XM_011283053.2;refGene_gn=ZNF106;refGene_tx=XM_011283054.2;refGene_gn=ZNF106;refGene_tx=XM_019832619.2;refGene_gn=ZNF106;refGene_tx=XM_019832620.2;refGene_gn=ZNF106;refGene_tx=XM_023255467.1
B3 62952680 62952736
refGene_gn=ZNF106;refGene_tx=XM_011283053.2;refGene_gn=ZNF106;refGene_tx=XM_011283054.2;refGene_gn=ZNF106;refGene_tx=XM_019832619.2;refGene_gn=ZNF106;refGene_tx=XM_019832620.2;refGene_gn=ZNF106;refGene_tx=XM_023255467.1
B3 62953547 62953693
refGene_gn=ZNF106;refGene_tx=XM_011283053.2;refGene_gn=ZNF106;refGene_tx=XM_011283054.2;refGene_gn=ZNF106;refGene_tx=XM_019832619.2;refGene_gn=ZNF106;refGene_tx=XM_019832620.2;refGene_gn=ZNF106;refGene_tx=XM_023255467.1
B3 62955073 62955208
refGene_gn=ZNF106;refGene_tx=XM_011283053.2;refGene_gn=ZNF106;refGene_

tx=XM_011283054.2;refGene_gn=ZNF106;refGene_tx=XM_019832619.2;refGene_gn=ZNF106;refGene_tx=XM_019832620.2;refGene_gn=ZNF106;refGene_tx=XM_023255467.1
B3 62955919 62956036
refGene_gn=ZNF106;refGene_tx=XM_011283053.2;refGene_gn=ZNF106;refGene_tx=XM_011283054.2;refGene_gn=ZNF106;refGene_tx=XM_019832619.2;refGene_gn=ZNF106;refGene_tx=XM_019832620.2;refGene_gn=ZNF106;refGene_tx=XM_023255467.1
B3 62958836 62959028
refGene_gn=ZNF106;refGene_tx=XM_011283053.2;refGene_gn=ZNF106;refGene_tx=XM_011283054.2;refGene_gn=ZNF106;refGene_tx=XM_019832619.2;refGene_gn=ZNF106;refGene_tx=XM_019832620.2;refGene_gn=ZNF106;refGene_tx=XM_023255467.1
B3 62959545 62959608
refGene_gn=ZNF106;refGene_tx=XM_011283053.2;refGene_gn=ZNF106;refGene_tx=XM_011283054.2;refGene_gn=ZNF106;refGene_tx=XM_019832619.2;refGene_gn=ZNF106;refGene_tx=XM_019832620.2;refGene_gn=ZNF106;refGene_tx=XM_023255467.1
B3 62960445 62960565
refGene_gn=ZNF106;refGene_tx=XM_011283053.2;refGene_gn=ZNF106;refGene_tx=XM_011283054.2;refGene_gn=ZNF106;refGene_tx=XM_019832619.2;refGene_gn=ZNF106;refGene_tx=XM_019832620.2;refGene_gn=ZNF106;refGene_tx=XM_023255467.1
B3 62961082 62961154
refGene_gn=ZNF106;refGene_tx=XM_011283053.2;refGene_gn=ZNF106;refGene_tx=XM_011283054.2;refGene_gn=ZNF106;refGene_tx=XM_019832619.2;refGene_gn=ZNF106;refGene_tx=XM_019832620.2;refGene_gn=ZNF106;refGene_tx=XM_023255467.1
B3 62961969 62962041
refGene_gn=ZNF106;refGene_tx=XM_011283053.2;refGene_gn=ZNF106;refGene_tx=XM_011283054.2;refGene_gn=ZNF106;refGene_tx=XM_019832619.2;refGene_gn=ZNF106;refGene_tx=XM_019832620.2;refGene_gn=ZNF106;refGene_tx=XM_023255467.1
B3 62964734 62964881
refGene_gn=ZNF106;refGene_tx=XM_011283053.2;refGene_gn=ZNF106;refGene_tx=XM_011283054.2;refGene_gn=ZNF106;refGene_tx=XM_019832619.2;refGene_gn=ZNF106;refGene_tx=XM_019832620.2;refGene_gn=ZNF106;refGene_tx=XM_023255467.1
B3 62965563 62965620
refGene_gn=ZNF106;refGene_tx=XM_011283053.2;refGene_gn=ZNF106;refGene_tx=XM_011283054.2;refGene_gn=ZNF106;refGene_tx=XM_019832619.2;refGene_gn=ZNF106;refGene_tx=XM_019832620.2;refGene_gn=ZNF106;refGene_tx=XM_023255467.1
B3 62975491 62975518
refGene_gn=CAPN3;refGene_tx=XM_003987244.5;refGene_gn=CAPN3;refGene_tx=XM_019832621.2;refGene_gn=CAPN3;refGene_tx=XM_019832622.2;refGene_gn=CAPN3;refGene_tx=XM_019832623.2
B3 62975881 62975940
refGene_gn=CAPN3;refGene_tx=XM_003987244.5;refGene_gn=CAPN3;refGene_tx=XM_019832621.2;refGene_gn=CAPN3;refGene_tx=XM_019832622.2;refGene_gn=

CAPN3;refGene_tx=XM_019832623.2
B3 62976243 62976360
refGene_gn=CAPN3;refGene_tx=XM_003987244.5;refGene_gn=CAPN3;refGene_tx=XM_019832621.2;refGene_gn=CAPN3;refGene_tx=XM_019832622.2;refGene_gn=CAPN3;refGene_tx=XM_019832623.2
B3 62976574 62976653
refGene_gn=CAPN3;refGene_tx=XM_003987244.5;refGene_gn=CAPN3;refGene_tx=XM_019832621.2;refGene_gn=CAPN3;refGene_tx=XM_019832622.2;refGene_gn=CAPN3;refGene_tx=XM_019832623.2
B3 62976742 62976811
refGene_gn=CAPN3;refGene_tx=XM_003987244.5;refGene_gn=CAPN3;refGene_tx=XM_019832621.2;refGene_gn=CAPN3;refGene_tx=XM_019832622.2;refGene_gn=CAPN3;refGene_tx=XM_019832623.2
B3 62977226 62977291
refGene_gn=CAPN3;refGene_tx=XM_003987244.5;refGene_gn=CAPN3;refGene_tx=XM_019832621.2;refGene_gn=CAPN3;refGene_tx=XM_019832622.2;refGene_gn=CAPN3;refGene_tx=XM_019832623.2
B3 62977386 62977444
refGene_gn=CAPN3;refGene_tx=XM_003987244.5;refGene_gn=CAPN3;refGene_tx=XM_019832621.2;refGene_gn=CAPN3;refGene_tx=XM_019832622.2;refGene_gn=CAPN3;refGene_tx=XM_019832623.2
B3 62977844 62977922
refGene_gn=CAPN3;refGene_tx=XM_003987244.5;refGene_gn=CAPN3;refGene_tx=XM_019832621.2;refGene_gn=CAPN3;refGene_tx=XM_019832622.2;refGene_gn=CAPN3;refGene_tx=XM_019832623.2
B3 62978889 62979003
refGene_gn=CAPN3;refGene_tx=XM_003987244.5;refGene_gn=CAPN3;refGene_tx=XM_019832621.2
B3 62981922 62981940
refGene_gn=CAPN3;refGene_tx=XM_003987244.5;refGene_gn=CAPN3;refGene_tx=XM_019832621.2
B3 62983888 62983925
refGene_gn=CAPN3;refGene_tx=XM_003987244.5;refGene_gn=CAPN3;refGene_tx=XM_019832621.2;refGene_gn=CAPN3;refGene_tx=XM_019832622.2;refGene_gn=CAPN3;refGene_tx=XM_019832623.2
B3 62984709 62984918
refGene_gn=CAPN3;refGene_tx=XM_003987244.5;refGene_gn=CAPN3;refGene_tx=XM_019832621.2;refGene_gn=CAPN3;refGene_tx=XM_019832622.2;refGene_gn=CAPN3;refGene_tx=XM_019832623.2
B3 62985634 62985646
refGene_gn=CAPN3;refGene_tx=XM_003987244.5;refGene_gn=CAPN3;refGene_tx=XM_019832621.2;refGene_gn=CAPN3;refGene_tx=XM_019832622.2;refGene_gn=CAPN3;refGene_tx=XM_019832623.2
B3 62985994 62986164
refGene_gn=CAPN3;refGene_tx=XM_003987244.5;refGene_gn=CAPN3;refGene_tx=XM_019832621.2;refGene_gn=CAPN3;refGene_tx=XM_019832622.2;refGene_gn=CAPN3;refGene_tx=XM_019832623.2
B3 62987406 62987567
refGene_gn=CAPN3;refGene_tx=XM_003987244.5;refGene_gn=CAPN3;refGene_tx=XM_019832621.2;refGene_gn=CAPN3;refGene_tx=XM_019832622.2;refGene_gn=

CAPN3;refGene_tx=XM_019832623.2
B3 62988352 62988430
refGene_gn=CAPN3;refGene_tx=XM_003987244.5;refGene_gn=CAPN3;refGene_tx=XM_019832621.2;refGene_gn=CAPN3;refGene_tx=XM_019832622.2;refGene_gn=CAPN3;refGene_tx=XM_019832623.2
B3 62991044 62991130
refGene_gn=CAPN3;refGene_tx=XM_003987244.5;refGene_gn=CAPN3;refGene_tx=XM_019832621.2;refGene_gn=CAPN3;refGene_tx=XM_019832622.2;refGene_gn=CAPN3;refGene_tx=XM_019832623.2
B3 62992231 62992315
refGene_gn=CAPN3;refGene_tx=XM_003987244.5;refGene_gn=CAPN3;refGene_tx=XM_019832621.2;refGene_gn=CAPN3;refGene_tx=XM_019832622.2;refGene_gn=CAPN3;refGene_tx=XM_019832623.2
B3 62994368 62994512
refGene_gn=CAPN3;refGene_tx=XM_003987244.5;refGene_gn=CAPN3;refGene_tx=XM_019832621.2;refGene_gn=CAPN3;refGene_tx=XM_019832622.2;refGene_gn=CAPN3;refGene_tx=XM_019832623.2
B3 62995375 62995544
refGene_gn=CAPN3;refGene_tx=XM_003987244.5;refGene_gn=CAPN3;refGene_tx=XM_019832621.2;refGene_gn=CAPN3;refGene_tx=XM_019832622.2;refGene_gn=CAPN3;refGene_tx=XM_019832623.2
B3 62996263 62996397
refGene_gn=CAPN3;refGene_tx=XM_003987244.5;refGene_gn=CAPN3;refGene_tx=XM_019832621.2;refGene_gn=CAPN3;refGene_tx=XM_019832622.2;refGene_gn=CAPN3;refGene_tx=XM_019832623.2
B3 62998226 62998345
refGene_gn=CAPN3;refGene_tx=XM_003987244.5;refGene_gn=CAPN3;refGene_tx=XM_019832621.2;refGene_gn=CAPN3;refGene_tx=XM_019832622.2;refGene_gn=CAPN3;refGene_tx=XM_019832623.2
B3 62999973 63000043
refGene_gn=CAPN3;refGene_tx=XM_003987244.5;refGene_gn=CAPN3;refGene_tx=XM_019832621.2;refGene_gn=CAPN3;refGene_tx=XM_019832622.2;refGene_gn=CAPN3;refGene_tx=XM_019832623.2
B3 63007050 63007299
refGene_gn=CAPN3;refGene_tx=XM_019832621.2;refGene_gn=CAPN3;refGene_tx=XM_019832623.2
B3 63024050 63024359
refGene_gn=CAPN3;refGene_tx=XM_003987244.5;refGene_gn=CAPN3;refGene_tx=XM_019832622.2
B3 63034872 63034982
refGene_gn=GANC;refGene_tx=XM_003987247.5;refGene_gn=GANC;refGene_tx=XM_019832624.2;refGene_gn=GANC;refGene_tx=XM_019832625.2;refGene_gn=GANC;refGene_tx=XM_019832626.2;refGene_gn=GANC;refGene_tx=XM_019832627.2
B3 63035852 63035956
refGene_gn=GANC;refGene_tx=XM_003987247.5;refGene_gn=GANC;refGene_tx=XM_019832624.2;refGene_gn=GANC;refGene_tx=XM_019832625.2;refGene_gn=GANC;refGene_tx=XM_019832626.2;refGene_gn=GANC;refGene_tx=XM_019832627.2
B3 63039783 63039896
refGene_gn=GANC;refGene_tx=XM_003987247.5;refGene_gn=GANC;refGene_tx=XM_019832624.2;refGene_gn=GANC;refGene_tx=XM_019832625.2;refGene_gn=GAN

C;refGene_tx=XM_019832626.2;refGene_gn=GANC;refGene_tx=XM_019832627.2
B3 63041103 63041217
refGene_gn=GANC;refGene_tx=XM_003987247.5;refGene_gn=GANC;refGene_tx=XM_019832624.2;refGene_gn=GANC;refGene_tx=XM_019832625.2;refGene_gn=GANC;refGene_tx=XM_019832626.2;refGene_gn=GANC;refGene_tx=XM_019832627.2
B3 63043903 63043978
refGene_gn=GANC;refGene_tx=XM_003987247.5;refGene_gn=GANC;refGene_tx=XM_019832624.2;refGene_gn=GANC;refGene_tx=XM_019832625.2;refGene_gn=GANC;refGene_tx=XM_019832626.2;refGene_gn=GANC;refGene_tx=XM_019832627.2
B3 63047825 63047902
refGene_gn=GANC;refGene_tx=XM_003987247.5;refGene_gn=GANC;refGene_tx=XM_019832624.2;refGene_gn=GANC;refGene_tx=XM_019832625.2;refGene_gn=GANC;refGene_tx=XM_019832626.2;refGene_gn=GANC;refGene_tx=XM_019832627.2
B3 63049783 63049848
refGene_gn=GANC;refGene_tx=XM_003987247.5;refGene_gn=GANC;refGene_tx=XM_019832624.2;refGene_gn=GANC;refGene_tx=XM_019832625.2;refGene_gn=GANC;refGene_tx=XM_019832626.2;refGene_gn=GANC;refGene_tx=XM_019832627.2
B3 63051517 63051761
refGene_gn=GANC;refGene_tx=XM_003987247.5;refGene_gn=GANC;refGene_tx=XM_019832624.2;refGene_gn=GANC;refGene_tx=XM_019832625.2;refGene_gn=GANC;refGene_tx=XM_019832626.2;refGene_gn=GANC;refGene_tx=XM_019832627.2
B3 63053058 63053160
refGene_gn=GANC;refGene_tx=XM_003987247.5;refGene_gn=GANC;refGene_tx=XM_019832624.2;refGene_gn=GANC;refGene_tx=XM_019832625.2;refGene_gn=GANC;refGene_tx=XM_019832626.2;refGene_gn=GANC;refGene_tx=XM_019832627.2
B3 63057722 63057819
refGene_gn=GANC;refGene_tx=XM_003987247.5;refGene_gn=GANC;refGene_tx=XM_019832624.2;refGene_gn=GANC;refGene_tx=XM_019832625.2;refGene_gn=GANC;refGene_tx=XM_019832626.2;refGene_gn=GANC;refGene_tx=XM_019832627.2
B3 63058876 63059020
refGene_gn=GANC;refGene_tx=XM_003987247.5;refGene_gn=GANC;refGene_tx=XM_019832624.2;refGene_gn=GANC;refGene_tx=XM_019832625.2;refGene_gn=GANC;refGene_tx=XM_019832626.2;refGene_gn=GANC;refGene_tx=XM_019832627.2
B3 63061009 63061089
refGene_gn=GANC;refGene_tx=XM_003987247.5;refGene_gn=GANC;refGene_tx=XM_019832624.2;refGene_gn=GANC;refGene_tx=XM_019832625.2;refGene_gn=GANC;refGene_tx=XM_019832626.2;refGene_gn=GANC;refGene_tx=XM_019832627.2
B3 63062031 63062158
refGene_gn=GANC;refGene_tx=XM_003987247.5;refGene_gn=GANC;refGene_tx=XM_019832624.2;refGene_gn=GANC;refGene_tx=XM_019832625.2;refGene_gn=GANC;refGene_tx=XM_019832626.2;refGene_gn=GANC;refGene_tx=XM_019832627.2
B3 63065328 63065564
refGene_gn=GANC;refGene_tx=XM_003987247.5;refGene_gn=GANC;refGene_tx=XM_019832624.2;refGene_gn=GANC;refGene_tx=XM_019832625.2;refGene_gn=GANC;refGene_tx=XM_019832626.2;refGene_gn=GANC;refGene_tx=XM_019832627.2
B3 63072770 63072924
refGene_gn=GANC;refGene_tx=XM_003987247.5;refGene_gn=GANC;refGene_tx=XM_019832624.2;refGene_gn=GANC;refGene_tx=XM_019832625.2;refGene_gn=GANC;refGene_tx=XM_019832626.2;refGene_gn=GANC;refGene_tx=XM_019832627.2
B3 63073145 63073326

refGene_gn=GANC;refGene_tx=XM_003987247.5;refGene_gn=GANC;refGene_tx=XM_019832624.2;refGene_gn=GANC;refGene_tx=XM_019832625.2;refGene_gn=GANC;refGene_tx=XM_019832626.2;refGene_gn=GANC;refGene_tx=XM_019832627.2
B3 63075078 63075175
refGene_gn=GANC;refGene_tx=XM_003987247.5;refGene_gn=GANC;refGene_tx=XM_019832624.2;refGene_gn=GANC;refGene_tx=XM_019832625.2;refGene_gn=GANC;refGene_tx=XM_019832626.2;refGene_gn=GANC;refGene_tx=XM_019832627.2
B3 63076572 63076639
refGene_gn=GANC;refGene_tx=XM_003987247.5;refGene_gn=GANC;refGene_tx=XM_019832624.2;refGene_gn=GANC;refGene_tx=XM_019832625.2;refGene_gn=GANC;refGene_tx=XM_019832626.2;refGene_gn=GANC;refGene_tx=XM_019832627.2
B3 63082232 63082278
refGene_gn=GANC;refGene_tx=XM_003987247.5;refGene_gn=GANC;refGene_tx=XM_019832624.2;refGene_gn=GANC;refGene_tx=XM_019832625.2;refGene_gn=GANC;refGene_tx=XM_019832626.2;refGene_gn=GANC;refGene_tx=XM_019832627.2
B3 63092403 63092586
refGene_gn=GANC;refGene_tx=XM_003987247.5;refGene_gn=GANC;refGene_tx=XM_019832624.2;refGene_gn=GANC;refGene_tx=XM_019832625.2;refGene_gn=GANC;refGene_tx=XM_019832626.2;refGene_gn=GANC;refGene_tx=XM_019832627.2
B3 63097083 63097211
refGene_gn=GANC;refGene_tx=XM_003987247.5;refGene_gn=GANC;refGene_tx=XM_019832624.2;refGene_gn=GANC;refGene_tx=XM_019832625.2;refGene_gn=GANC;refGene_tx=XM_019832626.2;refGene_gn=GANC;refGene_tx=XM_019832627.2
B3 63106527 63106661
refGene_gn=GANC;refGene_tx=XM_003987247.5;refGene_gn=GANC;refGene_tx=XM_019832624.2;refGene_gn=GANC;refGene_tx=XM_019832625.2;refGene_gn=GANC;refGene_tx=XM_019832626.2;refGene_gn=GANC;refGene_tx=XM_019832627.2
B3 63107991 63108054
refGene_gn=GANC;refGene_tx=XM_003987247.5;refGene_gn=GANC;refGene_tx=XM_019832624.2;refGene_gn=GANC;refGene_tx=XM_019832625.2;refGene_gn=GANC;refGene_tx=XM_019832626.2;refGene_gn=GANC;refGene_tx=XM_019832627.2
B3 63109701 63109730 refGene_gn=GANC;refGene_tx=XM_003987247.5
B3 63110819 63110975
refGene_gn=TMEM87A;refGene_tx=XM_003987248.4;refGene_gn=TMEM87A;refGene_tx=XM_006932617.3
B3 63112247 63112308
refGene_gn=TMEM87A;refGene_tx=XM_003987248.4;refGene_gn=TMEM87A;refGene_tx=XM_006932617.3
B3 63115223 63115309
refGene_gn=TMEM87A;refGene_tx=XM_003987248.4;refGene_gn=TMEM87A;refGene_tx=XM_006932617.3
B3 63118160 63118274
refGene_gn=TMEM87A;refGene_tx=XM_003987248.4;refGene_gn=TMEM87A;refGene_tx=XM_006932617.3
B3 63120453 63120507
refGene_gn=TMEM87A;refGene_tx=XM_003987248.4;refGene_gn=TMEM87A;refGene_tx=XM_006932617.3
B3 63120583 63120628
refGene_gn=TMEM87A;refGene_tx=XM_003987248.4;refGene_gn=TMEM87A;refGene_tx=XM_006932617.3

B3 63126278 63126396
refGene_gn=TMEM87A;refGene_tx=XM_003987248.4;refGene_gn=TMEM87A;refGen
e_tx=XM_006932617.3
B3 63130622 63130684
refGene_gn=TMEM87A;refGene_tx=XM_003987248.4;refGene_gn=TMEM87A;refGen
e_tx=XM_006932617.3
B3 63131455 63131639
refGene_gn=TMEM87A;refGene_tx=XM_003987248.4;refGene_gn=TMEM87A;refGen
e_tx=XM_006932617.3
B3 63132635 63132735
refGene_gn=TMEM87A;refGene_tx=XM_003987248.4;refGene_gn=TMEM87A;refGen
e_tx=XM_006932617.3
B3 63135241 63135335
refGene_gn=TMEM87A;refGene_tx=XM_003987248.4;refGene_gn=TMEM87A;refGen
e_tx=XM_006932617.3
B3 63136097 63136169 refGene_gn=TMEM87A;refGene_tx=XM_006932617.3
B3 63136780 63136849 refGene_gn=TMEM87A;refGene_tx=XM_003987248.4
B3 63139096 63139205
refGene_gn=TMEM87A;refGene_tx=XM_003987248.4;refGene_gn=TMEM87A;refGen
e_tx=XM_006932617.3
B3 63140196 63140255
refGene_gn=TMEM87A;refGene_tx=XM_003987248.4;refGene_gn=TMEM87A;refGen
e_tx=XM_006932617.3
B3 63141021 63141125
refGene_gn=TMEM87A;refGene_tx=XM_003987248.4;refGene_gn=TMEM87A;refGen
e_tx=XM_006932617.3
B3 63146094 63146168
refGene_gn=TMEM87A;refGene_tx=XM_003987248.4;refGene_gn=TMEM87A;refGen
e_tx=XM_006932617.3
B3 63146582 63146644
refGene_gn=TMEM87A;refGene_tx=XM_003987248.4;refGene_gn=TMEM87A;refGen
e_tx=XM_006932617.3
B3 63147302 63147358
refGene_gn=TMEM87A;refGene_tx=XM_003987248.4;refGene_gn=TMEM87A;refGen
e_tx=XM_006932617.3
B3 63147850 63147881
refGene_gn=TMEM87A;refGene_tx=XM_003987248.4;refGene_gn=TMEM87A;refGen
e_tx=XM_006932617.3
B3 63150349 63150391
refGene_gn=TMEM87A;refGene_tx=XM_003987248.4;refGene_gn=TMEM87A;refGen
e_tx=XM_006932617.3
B3 63153399 63153472
refGene_gn=VPS39;refGene_tx=XM_003987249.5;refGene_gn=VPS39;refGene_tx
=XM_019832628.2
B3 63161196 63161262
refGene_gn=VPS39;refGene_tx=XM_003987249.5;refGene_gn=VPS39;refGene_tx
=XM_019832628.2
B3 63163982 63164018 refGene_gn=VPS39;refGene_tx=XM_019832628.2
B3 63164501 63164566
refGene_gn=VPS39;refGene_tx=XM_003987249.5;refGene_gn=VPS39;refGene_tx

=XM_019832628.2
B3	63165102	63165145
refGene_gn=VPS39;refGene_tx=XM_003987249.5;refGene_gn=VPS39;refGene_tx
=XM_019832628.2
B3	63166325	63166420
refGene_gn=VPS39;refGene_tx=XM_003987249.5;refGene_gn=VPS39;refGene_tx
=XM_019832628.2
B3	63167598	63167697
refGene_gn=VPS39;refGene_tx=XM_003987249.5;refGene_gn=VPS39;refGene_tx
=XM_019832628.2
B3	63168090	63168183
refGene_gn=VPS39;refGene_tx=XM_003987249.5;refGene_gn=VPS39;refGene_tx
=XM_019832628.2
B3	63170146	63170330
refGene_gn=VPS39;refGene_tx=XM_003987249.5;refGene_gn=VPS39;refGene_tx
=XM_019832628.2
B3	63173450	63173571
refGene_gn=VPS39;refGene_tx=XM_003987249.5;refGene_gn=VPS39;refGene_tx
=XM_019832628.2
B3	63173680	63173801
refGene_gn=VPS39;refGene_tx=XM_003987249.5;refGene_gn=VPS39;refGene_tx
=XM_019832628.2
B3	63180480	63180610
refGene_gn=VPS39;refGene_tx=XM_003987249.5;refGene_gn=VPS39;refGene_tx
=XM_019832628.2
B3	63187550	63187693
refGene_gn=VPS39;refGene_tx=XM_003987249.5;refGene_gn=VPS39;refGene_tx
=XM_019832628.2
B3	63190625	63190769
refGene_gn=VPS39;refGene_tx=XM_003987249.5;refGene_gn=VPS39;refGene_tx
=XM_019832628.2
B3	63191246	63191388
refGene_gn=VPS39;refGene_tx=XM_003987249.5;refGene_gn=VPS39;refGene_tx
=XM_019832628.2
B3	63191528	63191615
refGene_gn=VPS39;refGene_tx=XM_003987249.5;refGene_gn=VPS39;refGene_tx
=XM_019832628.2
B3	63191915	63191989
refGene_gn=VPS39;refGene_tx=XM_003987249.5;refGene_gn=VPS39;refGene_tx
=XM_019832628.2
B3	63192361	63192460
refGene_gn=VPS39;refGene_tx=XM_003987249.5;refGene_gn=VPS39;refGene_tx
=XM_019832628.2
B3	63193054	63193172
refGene_gn=VPS39;refGene_tx=XM_003987249.5;refGene_gn=VPS39;refGene_tx
=XM_019832628.2
B3	63193653	63193782
refGene_gn=VPS39;refGene_tx=XM_003987249.5;refGene_gn=VPS39;refGene_tx
=XM_019832628.2
B3	63194426	63194529

refGene_gn=VPS39;refGene_tx=XM_003987249.5;refGene_gn=VPS39;refGene_tx=XM_019832628.2
B3 63194726 63194772
refGene_gn=VPS39;refGene_tx=XM_003987249.5;refGene_gn=VPS39;refGene_tx=XM_019832628.2
B3 63195677 63195827
refGene_gn=VPS39;refGene_tx=XM_003987249.5;refGene_gn=VPS39;refGene_tx=XM_019832628.2
B3 63195992 63196127
refGene_gn=VPS39;refGene_tx=XM_003987249.5;refGene_gn=VPS39;refGene_tx=XM_019832628.2
B3 63196390 63196482
refGene_gn=VPS39;refGene_tx=XM_003987249.5;refGene_gn=VPS39;refGene_tx=XM_019832628.2
B3 63197268 63197344
refGene_gn=VPS39;refGene_tx=XM_003987249.5;refGene_gn=VPS39;refGene_tx=XM_019832628.2
B3 63201303 63201414
refGene_gn=PLA2G4F;refGene_tx=XM_011283056.3;refGene_gn=PLA2G4F;refGene_tx=XM_019832632.2
B3 63202421 63202494 refGene_gn=PLA2G4F;refGene_tx=XM_011283056.3
B3 63203091 63203228 refGene_gn=PLA2G4F;refGene_tx=XM_011283056.3
B3 63203319 63203448
refGene_gn=PLA2G4F;refGene_tx=XM_011283056.3;refGene_gn=PLA2G4F;refGene_tx=XM_019832632.2
B3 63203899 63203940
refGene_gn=PLA2G4F;refGene_tx=XM_011283056.3;refGene_gn=PLA2G4F;refGene_tx=XM_019832632.2
B3 63204174 63204246
refGene_gn=PLA2G4F;refGene_tx=XM_011283056.3;refGene_gn=PLA2G4F;refGene_tx=XM_019832632.2;refGene_gn=PLA2G4F;refGene_tx=XM_019832633.1
B3 63204724 63204791
refGene_gn=PLA2G4F;refGene_tx=XM_011283056.3;refGene_gn=PLA2G4F;refGene_tx=XM_019832632.2;refGene_gn=PLA2G4F;refGene_tx=XM_019832633.1
B3 63206879 63207049
refGene_gn=PLA2G4F;refGene_tx=XM_011283056.3;refGene_gn=PLA2G4F;refGene_tx=XM_019832632.2;refGene_gn=PLA2G4F;refGene_tx=XM_019832633.1
B3 63207180 63207294
refGene_gn=PLA2G4F;refGene_tx=XM_011283056.3;refGene_gn=PLA2G4F;refGene_tx=XM_019832632.2;refGene_gn=PLA2G4F;refGene_tx=XM_019832633.1
B3 63207507 63207545
refGene_gn=PLA2G4F;refGene_tx=XM_011283056.3;refGene_gn=PLA2G4F;refGene_tx=XM_019832632.2;refGene_gn=PLA2G4F;refGene_tx=XM_019832633.1
B3 63207802 63207938
refGene_gn=PLA2G4F;refGene_tx=XM_011283056.3;refGene_gn=PLA2G4F;refGene_tx=XM_019832632.2;refGene_gn=PLA2G4F;refGene_tx=XM_019832633.1
B3 63209623 63209760
refGene_gn=PLA2G4F;refGene_tx=XM_011283056.3;refGene_gn=PLA2G4F;refGene_tx=XM_019832632.2;refGene_gn=PLA2G4F;refGene_tx=XM_019832633.1
B3 63210030 63210253

refGene_gn=PLA2G4F;refGene_tx=XM_011283056.3;refGene_gn=PLA2G4F;refGene_tx=XM_019832632.2;refGene_gn=PLA2G4F;refGene_tx=XM_019832633.1
B3 63211604 63211719
refGene_gn=PLA2G4F;refGene_tx=XM_011283056.3;refGene_gn=PLA2G4F;refGene_tx=XM_019832632.2;refGene_gn=PLA2G4F;refGene_tx=XM_019832633.1
B3 63211944 63212082
refGene_gn=PLA2G4F;refGene_tx=XM_011283056.3;refGene_gn=PLA2G4F;refGene_tx=XM_019832632.2;refGene_gn=PLA2G4F;refGene_tx=XM_019832633.1
B3 63212184 63212292
refGene_gn=PLA2G4F;refGene_tx=XM_011283056.3;refGene_gn=PLA2G4F;refGene_tx=XM_019832632.2;refGene_gn=PLA2G4F;refGene_tx=XM_019832633.1
B3 63213502 63213697
refGene_gn=PLA2G4F;refGene_tx=XM_011283056.3;refGene_gn=PLA2G4F;refGene_tx=XM_019832632.2;refGene_gn=PLA2G4F;refGene_tx=XM_019832633.1
B3 63213948 63214115
refGene_gn=PLA2G4F;refGene_tx=XM_011283056.3;refGene_gn=PLA2G4F;refGene_tx=XM_019832632.2;refGene_gn=PLA2G4F;refGene_tx=XM_019832633.1
B3 63214917 63215104
refGene_gn=PLA2G4F;refGene_tx=XM_011283056.3;refGene_gn=PLA2G4F;refGene_tx=XM_019832632.2;refGene_gn=PLA2G4F;refGene_tx=XM_019832633.1
B3 63215396 63215617
refGene_gn=PLA2G4F;refGene_tx=XM_011283056.3;refGene_gn=PLA2G4F;refGene_tx=XM_019832632.2;refGene_gn=PLA2G4F;refGene_tx=XM_019832633.1
B3 63249248 63249293 refGene_gn=PLA2G4D;refGene_tx=XM_011283072.3
B3 63253626 63253699 refGene_gn=PLA2G4D;refGene_tx=XM_011283072.3
B3 63253865 63254002 refGene_gn=PLA2G4D;refGene_tx=XM_011283072.3
B3 63254589 63254721 refGene_gn=PLA2G4D;refGene_tx=XM_011283072.3
B3 63255436 63255477 refGene_gn=PLA2G4D;refGene_tx=XM_011283072.3
B3 63255820 63255863 refGene_gn=PLA2G4D;refGene_tx=XM_011283072.3
B3 63257198 63257262 refGene_gn=PLA2G4D;refGene_tx=XM_011283072.3
B3 63257661 63257789 refGene_gn=PLA2G4D;refGene_tx=XM_011283072.3
B3 63258570 63258681 refGene_gn=PLA2G4D;refGene_tx=XM_011283072.3
B3 63258808 63258846 refGene_gn=PLA2G4D;refGene_tx=XM_011283072.3
B3 63259026 63259162 refGene_gn=PLA2G4D;refGene_tx=XM_011283072.3
B3 63259481 63259618 refGene_gn=PLA2G4D;refGene_tx=XM_011283072.3
B3 63260806 63261029 refGene_gn=PLA2G4D;refGene_tx=XM_011283072.3
B3 63265341 63265459 refGene_gn=PLA2G4D;refGene_tx=XM_011283072.3
B3 63265864 63266002 refGene_gn=PLA2G4D;refGene_tx=XM_011283072.3
B3 63266243 63266351 refGene_gn=PLA2G4D;refGene_tx=XM_011283072.3
B3 63266491 63266683 refGene_gn=PLA2G4D;refGene_tx=XM_011283072.3
B3 63266960 63267127 refGene_gn=PLA2G4D;refGene_tx=XM_011283072.3
B3 63268002 63268189 refGene_gn=PLA2G4D;refGene_tx=XM_011283072.3
B3 63269291 63269518 refGene_gn=PLA2G4D;refGene_tx=XM_011283072.3
B3 63282373 63282553
refGene_gn=PLA2G4E;refGene_tx=XM_003987251.4;refGene_gn=PLA2G4E;refGene_tx=XM_019832635.2
B3 63303952 63304048 refGene_gn=PLA2G4E;refGene_tx=XM_019832634.1
B3 63310829 63310902
refGene_gn=PLA2G4E;refGene_tx=XM_003987251.4;refGene_gn=PLA2G4E;refGene_tx=XM_019832634.1;refGene_gn=PLA2G4E;refGene_tx=XM_019832635.2

B3 63316213 63316350
refGene_gn=PLA2G4E;refGene_tx=XM_003987251.4;refGene_gn=PLA2G4E;refGene_tx=XM_019832634.1;refGene_gn=PLA2G4E;refGene_tx=XM_019832635.2
B3 63317986 63318118
refGene_gn=PLA2G4E;refGene_tx=XM_003987251.4;refGene_gn=PLA2G4E;refGene_tx=XM_019832634.1;refGene_gn=PLA2G4E;refGene_tx=XM_019832635.2
B3 63318915 63318956
refGene_gn=PLA2G4E;refGene_tx=XM_003987251.4;refGene_gn=PLA2G4E;refGene_tx=XM_019832634.1;refGene_gn=PLA2G4E;refGene_tx=XM_019832635.2
B3 63320586 63320629
refGene_gn=PLA2G4E;refGene_tx=XM_003987251.4;refGene_gn=PLA2G4E;refGene_tx=XM_019832634.1;refGene_gn=PLA2G4E;refGene_tx=XM_019832635.2
B3 63321489 63321553
refGene_gn=PLA2G4E;refGene_tx=XM_003987251.4;refGene_gn=PLA2G4E;refGene_tx=XM_019832634.1;refGene_gn=PLA2G4E;refGene_tx=XM_019832635.2
B3 63322388 63322549
refGene_gn=PLA2G4E;refGene_tx=XM_003987251.4;refGene_gn=PLA2G4E;refGene_tx=XM_019832634.1;refGene_gn=PLA2G4E;refGene_tx=XM_019832635.2
B3 63322649 63322736
refGene_gn=PLA2G4E;refGene_tx=XM_003987251.4;refGene_gn=PLA2G4E;refGene_tx=XM_019832634.1;refGene_gn=PLA2G4E;refGene_tx=XM_019832635.2
B3 63323118 63323156
refGene_gn=PLA2G4E;refGene_tx=XM_003987251.4;refGene_gn=PLA2G4E;refGene_tx=XM_019832634.1;refGene_gn=PLA2G4E;refGene_tx=XM_019832635.2
B3 63325287 63325423
refGene_gn=PLA2G4E;refGene_tx=XM_003987251.4;refGene_gn=PLA2G4E;refGene_tx=XM_019832634.1;refGene_gn=PLA2G4E;refGene_tx=XM_019832635.2
B3 63326664 63326801
refGene_gn=PLA2G4E;refGene_tx=XM_003987251.4;refGene_gn=PLA2G4E;refGene_tx=XM_019832634.1;refGene_gn=PLA2G4E;refGene_tx=XM_019832635.2
B3 63329352 63329575
refGene_gn=PLA2G4E;refGene_tx=XM_003987251.4;refGene_gn=PLA2G4E;refGene_tx=XM_019832634.1;refGene_gn=PLA2G4E;refGene_tx=XM_019832635.2
B3 63332219 63332334
refGene_gn=PLA2G4E;refGene_tx=XM_003987251.4;refGene_gn=PLA2G4E;refGene_tx=XM_019832634.1;refGene_gn=PLA2G4E;refGene_tx=XM_019832635.2
B3 63332853 63332991
refGene_gn=PLA2G4E;refGene_tx=XM_003987251.4;refGene_gn=PLA2G4E;refGene_tx=XM_019832634.1;refGene_gn=PLA2G4E;refGene_tx=XM_019832635.2
B3 63333383 63333491
refGene_gn=PLA2G4E;refGene_tx=XM_003987251.4;refGene_gn=PLA2G4E;refGene_tx=XM_019832634.1;refGene_gn=PLA2G4E;refGene_tx=XM_019832635.2
B3 63334032 63334236
refGene_gn=PLA2G4E;refGene_tx=XM_003987251.4;refGene_gn=PLA2G4E;refGene_tx=XM_019832634.1;refGene_gn=PLA2G4E;refGene_tx=XM_019832635.2
B3 63335108 63335275
refGene_gn=PLA2G4E;refGene_tx=XM_003987251.4;refGene_gn=PLA2G4E;refGene_tx=XM_019832634.1;refGene_gn=PLA2G4E;refGene_tx=XM_019832635.2
B3 63336514 63336698
refGene_gn=PLA2G4E;refGene_tx=XM_003987251.4;refGene_gn=PLA2G4E;refGen

e_tx=XM_019832634.1;refGene_gn=PLA2G4E;refGene_tx=XM_019832635.2
B3	63337150	63337371
refGene_gn=PLA2G4E;refGene_tx=XM_003987251.4;refGene_gn=PLA2G4E;refGene_tx=XM_019832634.1
B3	63342087	63342098	refGene_gn=PLA2G4E;refGene_tx=XM_019832635.2
B3	63350913	63351149	refGene_gn=EHD4;refGene_tx=XM_003987252.5
B3	63372093	63372270	refGene_gn=EHD4;refGene_tx=XM_003987252.5
B3	63381755	63381853	refGene_gn=EHD4;refGene_tx=XM_003987252.5
B3	63415782	63416195	refGene_gn=EHD4;refGene_tx=XM_003987252.5
B3	63426068	63426233	refGene_gn=EHD4;refGene_tx=XM_003987252.5
B3	63434819	63435356	refGene_gn=EHD4;refGene_tx=XM_003987252.5
B3	63442636	63442747	refGene_gn=SPTBN5;refGene_tx=XM_023256168.1
B3	63442986	63443154	refGene_gn=SPTBN5;refGene_tx=XM_023256168.1
B3	63445877	63445994	refGene_gn=SPTBN5;refGene_tx=XM_023256168.1
B3	63448035	63448193	refGene_gn=SPTBN5;refGene_tx=XM_023256168.1
B3	63448727	63448758	refGene_gn=SPTBN5;refGene_tx=XM_023256168.1
B3	63448759	63448957	refGene_gn=SPTBN5;refGene_tx=XM_023256168.1
B3	63449766	63450404	refGene_gn=SPTBN5;refGene_tx=XM_023256168.1
B3	63451676	63451815	refGene_gn=SPTBN5;refGene_tx=XM_023256168.1
B3	63451915	63452148	refGene_gn=SPTBN5;refGene_tx=XM_023256168.1
B3	63452386	63452540	refGene_gn=SPTBN5;refGene_tx=XM_023256168.1
B3	63452632	63452833	refGene_gn=SPTBN5;refGene_tx=XM_023256168.1
B3	63452910	63453117	refGene_gn=SPTBN5;refGene_tx=XM_023256168.1
B3	63453521	63453722	refGene_gn=SPTBN5;refGene_tx=XM_023256168.1
B3	63454416	63454569	refGene_gn=SPTBN5;refGene_tx=XM_023256168.1
B3	63457867	63457998	refGene_gn=SPTBN5;refGene_tx=XM_023256168.1
B3	63458236	63458476	refGene_gn=SPTBN5;refGene_tx=XM_023256168.1
B3	63458763	63459051	refGene_gn=SPTBN5;refGene_tx=XM_023256168.1
B3	63460047	63460288	refGene_gn=SPTBN5;refGene_tx=XM_023256168.1
B3	63460456	63460596	refGene_gn=SPTBN5;refGene_tx=XM_023256168.1
B3	63460764	63460864	refGene_gn=SPTBN5;refGene_tx=XM_023256168.1
B3	63461598	63461769	refGene_gn=SPTBN5;refGene_tx=XM_023256168.1
B3	63462283	63462448	refGene_gn=SPTBN5;refGene_tx=XM_023256168.1
B3	63462893	63463108	refGene_gn=SPTBN5;refGene_tx=XM_023256168.1
B3	63463498	63463685	refGene_gn=SPTBN5;refGene_tx=XM_023256168.1
B3	63463913	63464114	refGene_gn=SPTBN5;refGene_tx=XM_023256168.1
B3	63464369	63464486	refGene_gn=SPTBN5;refGene_tx=XM_023256168.1
B3	63465598	63465756	refGene_gn=SPTBN5;refGene_tx=XM_023256168.1
B3	63466149	63466285	refGene_gn=SPTBN5;refGene_tx=XM_023256168.1
B3	63466543	63466689	refGene_gn=SPTBN5;refGene_tx=XM_023256168.1
B3	63467465	63467580	refGene_gn=SPTBN5;refGene_tx=XM_023256168.1
B3	63467686	63467797	refGene_gn=SPTBN5;refGene_tx=XM_023256168.1
B3	63468013	63468193	refGene_gn=SPTBN5;refGene_tx=XM_023256168.1
B3	63468910	63469114	refGene_gn=SPTBN5;refGene_tx=XM_023256168.1
B3	63469290	63469440	refGene_gn=SPTBN5;refGene_tx=XM_023256168.1
B3	63469664	63469769	refGene_gn=SPTBN5;refGene_tx=XM_023256168.1
B3	63470166	63470334	refGene_gn=SPTBN5;refGene_tx=XM_023256168.1
B3	63470873	63471023	refGene_gn=SPTBN5;refGene_tx=XM_023256168.1
B3	63472015	63472207	refGene_gn=SPTBN5;refGene_tx=XM_023256168.1
B3	63472472	63472568	refGene_gn=SPTBN5;refGene_tx=XM_023256168.1

B3 63473766 63473882 refGene_gn=SPTBN5;refGene_tx=XM_023256168.1
B3 63473971 63474086 refGene_gn=SPTBN5;refGene_tx=XM_023256168.1
B3 63474881 63474995 refGene_gn=SPTBN5;refGene_tx=XM_023256168.1
B3 63475204 63475326 refGene_gn=SPTBN5;refGene_tx=XM_023256168.1
B3 63475589 63475752 refGene_gn=SPTBN5;refGene_tx=XM_023256168.1
B3 63475970 63476159 refGene_gn=SPTBN5;refGene_tx=XM_023256168.1
B3 63476432 63476510 refGene_gn=SPTBN5;refGene_tx=XM_023256168.1
B3 63477143 63477316 refGene_gn=SPTBN5;refGene_tx=XM_023256168.1
B3 63478338 63478429 refGene_gn=SPTBN5;refGene_tx=XM_023256168.1
B3 63478561 63478708 refGene_gn=SPTBN5;refGene_tx=XM_023256168.1
B3 63479329 63479467 refGene_gn=SPTBN5;refGene_tx=XM_023256168.1
B3 63479546 63479803 refGene_gn=SPTBN5;refGene_tx=XM_023256168.1
B3 63479989 63480176 refGene_gn=SPTBN5;refGene_tx=XM_023256168.1
B3 63480431 63480641 refGene_gn=SPTBN5;refGene_tx=XM_023256168.1
B3 63481125 63481322 refGene_gn=SPTBN5;refGene_tx=XM_023256168.1
B3 63481425 63481630 refGene_gn=SPTBN5;refGene_tx=XM_023256168.1
B3 63481922 63482117 refGene_gn=SPTBN5;refGene_tx=XM_023256168.1
B3 63483217 63483373 refGene_gn=SPTBN5;refGene_tx=XM_023256168.1
B3 63483613 63483831 refGene_gn=SPTBN5;refGene_tx=XM_023256168.1
B3 63483995 63484185 refGene_gn=SPTBN5;refGene_tx=XM_023256168.1
B3 63484390 63484567 refGene_gn=SPTBN5;refGene_tx=XM_023256168.1
B3 63484734 63484839 refGene_gn=SPTBN5;refGene_tx=XM_023256168.1
B3 63485174 63485405 refGene_gn=SPTBN5;refGene_tx=XM_023256168.1
B3 63485582 63485651 refGene_gn=SPTBN5;refGene_tx=XM_023256168.1
B3 63486138 63486377 refGene_gn=SPTBN5;refGene_tx=XM_023256168.1
B3 63486555 63486644 refGene_gn=SPTBN5;refGene_tx=XM_023256168.1
B3 63487458 63487509 refGene_gn=SPTBN5;refGene_tx=XM_023256168.1
B3 63487732 63487826 refGene_gn=SPTBN5;refGene_tx=XM_023256168.1
B3 63488063 63488316 refGene_gn=SPTBN5;refGene_tx=XM_023256168.1
B3 63489893 63490105
refGene_gn=LOC101083342;refGene_tx=XM_006932620.4;refGene_gn=LOC101083
342;refGene_tx=XM_006932622.4
B3 63490194 63490381
refGene_gn=LOC101083342;refGene_tx=XM_006932620.4;refGene_gn=LOC101083
342;refGene_tx=XM_006932622.4
B3 63490845 63491012
refGene_gn=LOC101083342;refGene_tx=XM_006932620.4;refGene_gn=LOC101083
342;refGene_tx=XM_006932622.4
B3 63491361 63491541
refGene_gn=LOC101083342;refGene_tx=XM_006932620.4;refGene_gn=LOC101083
342;refGene_tx=XM_006932622.4
B3 63491712 63491817
refGene_gn=LOC101083342;refGene_tx=XM_006932620.4;refGene_gn=LOC101083
342;refGene_tx=XM_006932622.4
B3 63492039 63492177
refGene_gn=LOC101083342;refGene_tx=XM_006932620.4;refGene_gn=LOC101083
342;refGene_tx=XM_006932622.4
B3 63492472 63492590
refGene_gn=LOC101083342;refGene_tx=XM_006932620.4;refGene_gn=LOC101083
342;refGene_tx=XM_006932622.4

B3 63492718 63492941
refGene_gn=LOC101083342;refGene_tx=XM_006932620.4;refGene_gn=LOC101083
342;refGene_tx=XM_006932622.4
B3 63493151 63493288
refGene_gn=LOC101083342;refGene_tx=XM_006932620.4;refGene_gn=LOC101083
342;refGene_tx=XM_006932622.4
B3 63493941 63494077
refGene_gn=LOC101083342;refGene_tx=XM_006932620.4;refGene_gn=LOC101083
342;refGene_tx=XM_006932622.4
B3 63494660 63494698
refGene_gn=LOC101083342;refGene_tx=XM_006932620.4;refGene_gn=LOC101083
342;refGene_tx=XM_006932622.4
B3 63494997 63495081
refGene_gn=LOC101083342;refGene_tx=XM_006932620.4;refGene_gn=LOC101083
342;refGene_tx=XM_006932622.4
B3 63495326 63495457
refGene_gn=LOC101083342;refGene_tx=XM_006932620.4;refGene_gn=LOC101083
342;refGene_tx=XM_006932622.4
B3 63496034 63496089
refGene_gn=LOC101083342;refGene_tx=XM_006932620.4;refGene_gn=LOC101083
342;refGene_tx=XM_006932622.4
B3 63496349 63496392
refGene_gn=LOC101083342;refGene_tx=XM_006932620.4;refGene_gn=LOC101083
342;refGene_tx=XM_006932622.4
B3 63496594 63496635
refGene_gn=LOC101083342;refGene_tx=XM_006932620.4;refGene_gn=LOC101083
342;refGene_tx=XM_006932622.4
B3 63496992 63497124
refGene_gn=LOC101083342;refGene_tx=XM_006932620.4;refGene_gn=LOC101083
342;refGene_tx=XM_006932622.4
B3 63497248 63497385
refGene_gn=LOC101083342;refGene_tx=XM_006932620.4;refGene_gn=LOC101083
342;refGene_tx=XM_006932622.4
B3 63497688 63497761
refGene_gn=LOC101083342;refGene_tx=XM_006932620.4;refGene_gn=LOC101083
342;refGene_tx=XM_006932622.4
B3 63499108 63499117
refGene_gn=LOC101083342;refGene_tx=XM_006932620.4
B3 63500576 63500665
refGene_gn=LOC101083342;refGene_tx=XM_006932621.3
B3 63500761 63500921
refGene_gn=LOC101083342;refGene_tx=XM_006932621.3
B3 63501145 63501222
refGene_gn=LOC101083342;refGene_tx=XM_006932621.3;refGene_gn=LOC101083
342;refGene_tx=XM_006932622.4
B3 63501419 63501515
refGene_gn=LOC101083342;refGene_tx=XM_006932621.3;refGene_gn=LOC101083
342;refGene_tx=XM_006932622.4
B3 63501986 63502043
refGene_gn=LOC101083342;refGene_tx=XM_006932621.3;refGene_gn=LOC101083

342;refGene_tx=XM_006932622.4
B3 63502662 63502916
refGene_gn=LOC101083342;refGene_tx=XM_006932621.3;refGene_gn=LOC101083342;refGene_tx=XM_006932622.4
B3 63502992 63503146
refGene_gn=LOC101083342;refGene_tx=XM_006932621.3;refGene_gn=LOC101083342;refGene_tx=XM_006932622.4
B3 63506749 63506813
refGene_gn=LOC101083342;refGene_tx=XM_006932621.3;refGene_gn=LOC101083342;refGene_tx=XM_006932622.4
B3 63509274 63509502
refGene_gn=MAPKBP1;refGene_tx=XM_006932624.4;refGene_gn=MAPKBP1;refGene_tx=XM_019832645.2;refGene_gn=MAPKBP1;refGene_tx=XM_019832646.2;refGene_gn=MAPKBP1;refGene_tx=XM_019832647.2;refGene_gn=MAPKBP1;refGene_tx=XM_019832648.2;refGene_gn=MAPKBP1;refGene_tx=XM_023255469.1
B3 63509941 63510027
refGene_gn=MAPKBP1;refGene_tx=XM_006932624.4;refGene_gn=MAPKBP1;refGene_tx=XM_019832645.2;refGene_gn=MAPKBP1;refGene_tx=XM_019832646.2;refGene_gn=MAPKBP1;refGene_tx=XM_019832647.2;refGene_gn=MAPKBP1;refGene_tx=XM_019832648.2;refGene_gn=MAPKBP1;refGene_tx=XM_023255469.1
B3 63510459 63511056
refGene_gn=MAPKBP1;refGene_tx=XM_006932624.4;refGene_gn=MAPKBP1;refGene_tx=XM_019832645.2;refGene_gn=MAPKBP1;refGene_tx=XM_019832646.2;refGene_gn=MAPKBP1;refGene_tx=XM_019832647.2;refGene_gn=MAPKBP1;refGene_tx=XM_019832648.2;refGene_gn=MAPKBP1;refGene_tx=XM_023255469.1
B3 63511278 63511562
refGene_gn=MAPKBP1;refGene_tx=XM_006932624.4;refGene_gn=MAPKBP1;refGene_tx=XM_019832645.2;refGene_gn=MAPKBP1;refGene_tx=XM_019832646.2;refGene_gn=MAPKBP1;refGene_tx=XM_019832647.2;refGene_gn=MAPKBP1;refGene_tx=XM_019832648.2;refGene_gn=MAPKBP1;refGene_tx=XM_023255469.1
B3 63511844 63511929
refGene_gn=MAPKBP1;refGene_tx=XM_006932624.4;refGene_gn=MAPKBP1;refGene_tx=XM_019832645.2;refGene_gn=MAPKBP1;refGene_tx=XM_019832646.2;refGene_gn=MAPKBP1;refGene_tx=XM_019832647.2;refGene_gn=MAPKBP1;refGene_tx=XM_019832648.2;refGene_gn=MAPKBP1;refGene_tx=XM_023255469.1
B3 63512103 63512301
refGene_gn=MAPKBP1;refGene_tx=XM_006932624.4;refGene_gn=MAPKBP1;refGene_tx=XM_019832645.2;refGene_gn=MAPKBP1;refGene_tx=XM_019832646.2;refGene_gn=MAPKBP1;refGene_tx=XM_019832647.2;refGene_gn=MAPKBP1;refGene_tx=XM_019832648.2;refGene_gn=MAPKBP1;refGene_tx=XM_023255469.1
B3 63512404 63512550
refGene_gn=MAPKBP1;refGene_tx=XM_006932624.4;refGene_gn=MAPKBP1;refGene_tx=XM_019832645.2;refGene_gn=MAPKBP1;refGene_tx=XM_019832646.2;refGene_gn=MAPKBP1;refGene_tx=XM_019832647.2;refGene_gn=MAPKBP1;refGene_tx=XM_019832648.2;refGene_gn=MAPKBP1;refGene_tx=XM_023255469.1
B3 63512775 63512942
refGene_gn=MAPKBP1;refGene_tx=XM_006932624.4;refGene_gn=MAPKBP1;refGene_tx=XM_019832645.2;refGene_gn=MAPKBP1;refGene_tx=XM_019832646.2;refGene_gn=MAPKBP1;refGene_tx=XM_019832647.2;refGene_gn=MAPKBP1;refGene_tx=XM_019832648.2;refGene_gn=MAPKBP1;refGene_tx=XM_023255469.1

B3 63513372 63513612
refGene_gn=MAPKBP1;refGene_tx=XM_006932624.4;refGene_gn=MAPKBP1;refGene_tx=XM_019832645.2;refGene_gn=MAPKBP1;refGene_tx=XM_019832646.2;refGene_gn=MAPKBP1;refGene_tx=XM_019832647.2;refGene_gn=MAPKBP1;refGene_tx=XM_019832648.2;refGene_gn=MAPKBP1;refGene_tx=XM_023255469.1
B3 63515512 63515568
refGene_gn=MAPKBP1;refGene_tx=XM_006932624.4;refGene_gn=MAPKBP1;refGene_tx=XM_019832645.2;refGene_gn=MAPKBP1;refGene_tx=XM_019832646.2;refGene_gn=MAPKBP1;refGene_tx=XM_019832647.2;refGene_gn=MAPKBP1;refGene_tx=XM_019832648.2;refGene_gn=MAPKBP1;refGene_tx=XM_023255469.1
B3 63515775 63515912
refGene_gn=MAPKBP1;refGene_tx=XM_006932624.4;refGene_gn=MAPKBP1;refGene_tx=XM_019832645.2;refGene_gn=MAPKBP1;refGene_tx=XM_019832646.2;refGene_gn=MAPKBP1;refGene_tx=XM_019832647.2;refGene_gn=MAPKBP1;refGene_tx=XM_019832648.2;refGene_gn=MAPKBP1;refGene_tx=XM_023255469.1
B3 63516131 63516266
refGene_gn=MAPKBP1;refGene_tx=XM_006932624.4;refGene_gn=MAPKBP1;refGene_tx=XM_019832645.2;refGene_gn=MAPKBP1;refGene_tx=XM_019832646.2;refGene_gn=MAPKBP1;refGene_tx=XM_019832647.2;refGene_gn=MAPKBP1;refGene_tx=XM_019832648.2;refGene_gn=MAPKBP1;refGene_tx=XM_023255469.1
B3 63516525 63516589
refGene_gn=MAPKBP1;refGene_tx=XM_006932624.4;refGene_gn=MAPKBP1;refGene_tx=XM_019832645.2;refGene_gn=MAPKBP1;refGene_tx=XM_019832646.2;refGene_gn=MAPKBP1;refGene_tx=XM_019832647.2;refGene_gn=MAPKBP1;refGene_tx=XM_019832648.2;refGene_gn=MAPKBP1;refGene_tx=XM_023255469.1
B3 63516767 63516879
refGene_gn=MAPKBP1;refGene_tx=XM_006932624.4;refGene_gn=MAPKBP1;refGene_tx=XM_019832645.2;refGene_gn=MAPKBP1;refGene_tx=XM_019832646.2;refGene_gn=MAPKBP1;refGene_tx=XM_019832647.2;refGene_gn=MAPKBP1;refGene_tx=XM_019832648.2;refGene_gn=MAPKBP1;refGene_tx=XM_023255469.1
B3 63516992 63517068
refGene_gn=MAPKBP1;refGene_tx=XM_006932624.4;refGene_gn=MAPKBP1;refGene_tx=XM_019832645.2;refGene_gn=MAPKBP1;refGene_tx=XM_019832646.2;refGene_gn=MAPKBP1;refGene_tx=XM_019832647.2;refGene_gn=MAPKBP1;refGene_tx=XM_019832648.2;refGene_gn=MAPKBP1;refGene_tx=XM_023255469.1
B3 63517363 63517485
refGene_gn=MAPKBP1;refGene_tx=XM_006932624.4;refGene_gn=MAPKBP1;refGene_tx=XM_019832645.2;refGene_gn=MAPKBP1;refGene_tx=XM_019832646.2;refGene_gn=MAPKBP1;refGene_tx=XM_019832647.2;refGene_gn=MAPKBP1;refGene_tx=XM_019832648.2;refGene_gn=MAPKBP1;refGene_tx=XM_023255469.1
B3 63517623 63517694
refGene_gn=MAPKBP1;refGene_tx=XM_006932624.4;refGene_gn=MAPKBP1;refGene_tx=XM_019832645.2;refGene_gn=MAPKBP1;refGene_tx=XM_019832646.2;refGene_gn=MAPKBP1;refGene_tx=XM_019832647.2;refGene_gn=MAPKBP1;refGene_tx=XM_019832648.2;refGene_gn=MAPKBP1;refGene_tx=XM_023255469.1
B3 63518059 63518185
refGene_gn=MAPKBP1;refGene_tx=XM_006932624.4;refGene_gn=MAPKBP1;refGene_tx=XM_019832645.2;refGene_gn=MAPKBP1;refGene_tx=XM_019832646.2;refGene_gn=MAPKBP1;refGene_tx=XM_019832647.2;refGene_gn=MAPKBP1;refGene_tx=XM_019832648.2;refGene_gn=MAPKBP1;refGene_tx=XM_023255469.1

B3 63518455 63518547
refGene_gn=MAPKBP1;refGene_tx=XM_006932624.4;refGene_gn=MAPKBP1;refGene_tx=XM_019832645.2;refGene_gn=MAPKBP1;refGene_tx=XM_019832646.2;refGene_gn=MAPKBP1;refGene_tx=XM_019832647.2;refGene_gn=MAPKBP1;refGene_tx=XM_019832648.2;refGene_gn=MAPKBP1;refGene_tx=XM_023255469.1
B3 63519327 63519503
refGene_gn=MAPKBP1;refGene_tx=XM_006932624.4;refGene_gn=MAPKBP1;refGene_tx=XM_019832645.2;refGene_gn=MAPKBP1;refGene_tx=XM_019832646.2;refGene_gn=MAPKBP1;refGene_tx=XM_019832647.2;refGene_gn=MAPKBP1;refGene_tx=XM_019832648.2;refGene_gn=MAPKBP1;refGene_tx=XM_023255469.1
B3 63519716 63519863
refGene_gn=MAPKBP1;refGene_tx=XM_006932624.4;refGene_gn=MAPKBP1;refGene_tx=XM_019832645.2;refGene_gn=MAPKBP1;refGene_tx=XM_019832646.2;refGene_gn=MAPKBP1;refGene_tx=XM_019832647.2;refGene_gn=MAPKBP1;refGene_tx=XM_019832648.2;refGene_gn=MAPKBP1;refGene_tx=XM_023255469.1
B3 63520442 63520632
refGene_gn=MAPKBP1;refGene_tx=XM_006932624.4;refGene_gn=MAPKBP1;refGene_tx=XM_019832645.2;refGene_gn=MAPKBP1;refGene_tx=XM_019832646.2;refGene_gn=MAPKBP1;refGene_tx=XM_019832647.2;refGene_gn=MAPKBP1;refGene_tx=XM_019832648.2;refGene_gn=MAPKBP1;refGene_tx=XM_023255469.1
B3 63521443 63521604
refGene_gn=MAPKBP1;refGene_tx=XM_006932624.4;refGene_gn=MAPKBP1;refGene_tx=XM_019832645.2;refGene_gn=MAPKBP1;refGene_tx=XM_019832646.2;refGene_gn=MAPKBP1;refGene_tx=XM_019832647.2;refGene_gn=MAPKBP1;refGene_tx=XM_019832648.2;refGene_gn=MAPKBP1;refGene_tx=XM_023255469.1
B3 63521930 63521948
refGene_gn=MAPKBP1;refGene_tx=XM_019832645.2;refGene_gn=MAPKBP1;refGene_tx=XM_019832647.2;refGene_gn=MAPKBP1;refGene_tx=XM_019832648.2;refGene_gn=MAPKBP1;refGene_tx=XM_023255469.1
B3 63522182 63522365
refGene_gn=MAPKBP1;refGene_tx=XM_006932624.4;refGene_gn=MAPKBP1;refGene_tx=XM_019832645.2;refGene_gn=MAPKBP1;refGene_tx=XM_019832646.2;refGene_gn=MAPKBP1;refGene_tx=XM_019832647.2;refGene_gn=MAPKBP1;refGene_tx=XM_019832648.2;refGene_gn=MAPKBP1;refGene_tx=XM_023255469.1
B3 63522619 63522757
refGene_gn=MAPKBP1;refGene_tx=XM_006932624.4;refGene_gn=MAPKBP1;refGene_tx=XM_019832645.2;refGene_gn=MAPKBP1;refGene_tx=XM_019832646.2;refGene_gn=MAPKBP1;refGene_tx=XM_019832647.2;refGene_gn=MAPKBP1;refGene_tx=XM_019832648.2;refGene_gn=MAPKBP1;refGene_tx=XM_023255469.1
B3 63523153 63523324
refGene_gn=MAPKBP1;refGene_tx=XM_006932624.4;refGene_gn=MAPKBP1;refGene_tx=XM_019832645.2;refGene_gn=MAPKBP1;refGene_tx=XM_019832646.2;refGene_gn=MAPKBP1;refGene_tx=XM_019832647.2;refGene_gn=MAPKBP1;refGene_tx=XM_019832648.2;refGene_gn=MAPKBP1;refGene_tx=XM_023255469.1
B3 63524086 63524144
refGene_gn=MAPKBP1;refGene_tx=XM_006932624.4;refGene_gn=MAPKBP1;refGene_tx=XM_019832645.2;refGene_gn=MAPKBP1;refGene_tx=XM_019832646.2;refGene_gn=MAPKBP1;refGene_tx=XM_019832647.2;refGene_gn=MAPKBP1;refGene_tx=XM_019832648.2;refGene_gn=MAPKBP1;refGene_tx=XM_023255469.1
B3 63524381 63524444

refGene_gn=MAPKBP1;refGene_tx=XM_006932624.4;refGene_gn=MAPKBP1;refGene_tx=XM_019832645.2;refGene_gn=MAPKBP1;refGene_tx=XM_019832646.2;refGene_gn=MAPKBP1;refGene_tx=XM_019832647.2;refGene_gn=MAPKBP1;refGene_tx=XM_019832648.2;refGene_gn=MAPKBP1;refGene_tx=XM_023255469.1
B3 63535466 63535558
refGene_gn=MAPKBP1;refGene_tx=XM_006932624.4;refGene_gn=MAPKBP1;refGene_tx=XM_019832645.2;refGene_gn=MAPKBP1;refGene_tx=XM_019832646.2;refGene_gn=MAPKBP1;refGene_tx=XM_019832647.2;refGene_gn=MAPKBP1;refGene_tx=XM_019832648.2;refGene_gn=MAPKBP1;refGene_tx=XM_023255469.1
B3 63556086 63556383
refGene_gn=MAPKBP1;refGene_tx=XM_019832645.2;refGene_gn=MAPKBP1;refGene_tx=XM_019832646.2;refGene_gn=MAPKBP1;refGene_tx=XM_019832647.2;refGene_gn=MAPKBP1;refGene_tx=XM_023255469.1
B3 63559804 63559921
refGene_gn=MAPKBP1;refGene_tx=XM_006932624.4;refGene_gn=MAPKBP1;refGene_tx=XM_019832648.2
B3 63566703 63567971
refGene_gn=MGA;refGene_tx=XM_011283057.3;refGene_gn=MGA;refGene_tx=XM_019832637.2;refGene_gn=MGA;refGene_tx=XM_019832638.2;refGene_gn=MGA;refGene_tx=XM_019832639.2;refGene_gn=MGA;refGene_tx=XM_019832640.2;refGene_gn=MGA;refGene_tx=XM_019832641.2;refGene_gn=MGA;refGene_tx=XM_019832642.2;refGene_gn=MGA;refGene_tx=XM_019832643.2;refGene_gn=MGA;refGene_tx=XM_019832644.2
B3 63569329 63569506
refGene_gn=MGA;refGene_tx=XM_011283057.3;refGene_gn=MGA;refGene_tx=XM_019832637.2;refGene_gn=MGA;refGene_tx=XM_019832638.2;refGene_gn=MGA;refGene_tx=XM_019832639.2;refGene_gn=MGA;refGene_tx=XM_019832640.2;refGene_gn=MGA;refGene_tx=XM_019832641.2;refGene_gn=MGA;refGene_tx=XM_019832642.2;refGene_gn=MGA;refGene_tx=XM_019832643.2;refGene_gn=MGA;refGene_tx=XM_019832644.2
B3 63570604 63570838
refGene_gn=MGA;refGene_tx=XM_011283057.3;refGene_gn=MGA;refGene_tx=XM_019832637.2;refGene_gn=MGA;refGene_tx=XM_019832638.2;refGene_gn=MGA;refGene_tx=XM_019832639.2;refGene_gn=MGA;refGene_tx=XM_019832640.2;refGene_gn=MGA;refGene_tx=XM_019832641.2;refGene_gn=MGA;refGene_tx=XM_019832642.2;refGene_gn=MGA;refGene_tx=XM_019832643.2;refGene_gn=MGA;refGene_tx=XM_019832644.2
B3 63571126 63571238
refGene_gn=MGA;refGene_tx=XM_011283057.3;refGene_gn=MGA;refGene_tx=XM_019832637.2;refGene_gn=MGA;refGene_tx=XM_019832638.2;refGene_gn=MGA;refGene_tx=XM_019832639.2;refGene_gn=MGA;refGene_tx=XM_019832640.2;refGene_gn=MGA;refGene_tx=XM_019832641.2;refGene_gn=MGA;refGene_tx=XM_019832642.2;refGene_gn=MGA;refGene_tx=XM_019832643.2;refGene_gn=MGA;refGene_tx=XM_019832644.2
B3 63572832 63573039
refGene_gn=MGA;refGene_tx=XM_011283057.3;refGene_gn=MGA;refGene_tx=XM_019832637.2;refGene_gn=MGA;refGene_tx=XM_019832638.2;refGene_gn=MGA;refGene_tx=XM_019832639.2;refGene_gn=MGA;refGene_tx=XM_019832640.2;refGene_gn=MGA;refGene_tx=XM_019832641.2;refGene_gn=MGA;refGene_tx=XM_019832642.2;refGene_gn=MGA;refGene_tx=XM_019832643.2;refGene_gn=MGA;refGene

_tx=XM_019832644.2
B3 63575557 63575609
refGene_gn=MGA;refGene_tx=XM_011283057.3;refGene_gn=MGA;refGene_tx=XM_019832637.2;refGene_gn=MGA;refGene_tx=XM_019832638.2;refGene_gn=MGA;refGene_tx=XM_019832639.2;refGene_gn=MGA;refGene_tx=XM_019832640.2;refGene_gn=MGA;refGene_tx=XM_019832641.2;refGene_gn=MGA;refGene_tx=XM_019832642.2;refGene_gn=MGA;refGene_tx=XM_019832643.2;refGene_gn=MGA;refGene_tx=XM_019832644.2
B3 63578119 63578250
refGene_gn=MGA;refGene_tx=XM_011283057.3;refGene_gn=MGA;refGene_tx=XM_019832637.2;refGene_gn=MGA;refGene_tx=XM_019832638.2;refGene_gn=MGA;refGene_tx=XM_019832639.2;refGene_gn=MGA;refGene_tx=XM_019832640.2;refGene_gn=MGA;refGene_tx=XM_019832641.2;refGene_gn=MGA;refGene_tx=XM_019832642.2;refGene_gn=MGA;refGene_tx=XM_019832643.2;refGene_gn=MGA;refGene_tx=XM_019832644.2
B3 63581236 63582750
refGene_gn=MGA;refGene_tx=XM_011283057.3;refGene_gn=MGA;refGene_tx=XM_019832637.2;refGene_gn=MGA;refGene_tx=XM_019832638.2;refGene_gn=MGA;refGene_tx=XM_019832639.2;refGene_gn=MGA;refGene_tx=XM_019832640.2;refGene_gn=MGA;refGene_tx=XM_019832641.2;refGene_gn=MGA;refGene_tx=XM_019832642.2;refGene_gn=MGA;refGene_tx=XM_019832643.2;refGene_gn=MGA;refGene_tx=XM_019832644.2
B3 63582938 63583229
refGene_gn=MGA;refGene_tx=XM_011283057.3;refGene_gn=MGA;refGene_tx=XM_019832637.2;refGene_gn=MGA;refGene_tx=XM_019832638.2;refGene_gn=MGA;refGene_tx=XM_019832639.2;refGene_gn=MGA;refGene_tx=XM_019832640.2;refGene_gn=MGA;refGene_tx=XM_019832641.2;refGene_gn=MGA;refGene_tx=XM_019832642.2;refGene_gn=MGA;refGene_tx=XM_019832643.2;refGene_gn=MGA;refGene_tx=XM_019832644.2
B3 63586517 63587144
refGene_gn=MGA;refGene_tx=XM_011283057.3;refGene_gn=MGA;refGene_tx=XM_019832637.2;refGene_gn=MGA;refGene_tx=XM_019832638.2;refGene_gn=MGA;refGene_tx=XM_019832639.2;refGene_gn=MGA;refGene_tx=XM_019832640.2;refGene_gn=MGA;refGene_tx=XM_019832641.2;refGene_gn=MGA;refGene_tx=XM_019832642.2;refGene_gn=MGA;refGene_tx=XM_019832643.2;refGene_gn=MGA;refGene_tx=XM_019832644.2
B3 63588420 63588718
refGene_gn=MGA;refGene_tx=XM_011283057.3;refGene_gn=MGA;refGene_tx=XM_019832637.2;refGene_gn=MGA;refGene_tx=XM_019832638.2;refGene_gn=MGA;refGene_tx=XM_019832639.2;refGene_gn=MGA;refGene_tx=XM_019832640.2;refGene_gn=MGA;refGene_tx=XM_019832641.2;refGene_gn=MGA;refGene_tx=XM_019832642.2;refGene_gn=MGA;refGene_tx=XM_019832643.2;refGene_gn=MGA;refGene_tx=XM_019832644.2
B3 63591154 63591669
refGene_gn=MGA;refGene_tx=XM_011283057.3;refGene_gn=MGA;refGene_tx=XM_019832637.2;refGene_gn=MGA;refGene_tx=XM_019832638.2;refGene_gn=MGA;refGene_tx=XM_019832639.2;refGene_gn=MGA;refGene_tx=XM_019832640.2;refGene_gn=MGA;refGene_tx=XM_019832641.2;refGene_gn=MGA;refGene_tx=XM_019832642.2;refGene_gn=MGA;refGene_tx=XM_019832643.2;refGene_gn=MGA;refGene_tx=XM_019832644.2

B3 63593629 63593702
refGene_gn=MGA;refGene_tx=XM_011283057.3;refGene_gn=MGA;refGene_tx=XM_019832637.2;refGene_gn=MGA;refGene_tx=XM_019832638.2;refGene_gn=MGA;refGene_tx=XM_019832639.2;refGene_gn=MGA;refGene_tx=XM_019832640.2;refGene_gn=MGA;refGene_tx=XM_019832641.2;refGene_gn=MGA;refGene_tx=XM_019832642.2;refGene_gn=MGA;refGene_tx=XM_019832643.2;refGene_gn=MGA;refGene_tx=XM_019832644.2
B3 63594301 63594478
refGene_gn=MGA;refGene_tx=XM_011283057.3;refGene_gn=MGA;refGene_tx=XM_019832637.2;refGene_gn=MGA;refGene_tx=XM_019832638.2;refGene_gn=MGA;refGene_tx=XM_019832639.2;refGene_gn=MGA;refGene_tx=XM_019832640.2;refGene_gn=MGA;refGene_tx=XM_019832641.2;refGene_gn=MGA;refGene_tx=XM_019832642.2;refGene_gn=MGA;refGene_tx=XM_019832643.2;refGene_gn=MGA;refGene_tx=XM_019832644.2
B3 63595579 63595806
refGene_gn=MGA;refGene_tx=XM_011283057.3;refGene_gn=MGA;refGene_tx=XM_019832637.2;refGene_gn=MGA;refGene_tx=XM_019832638.2;refGene_gn=MGA;refGene_tx=XM_019832639.2;refGene_gn=MGA;refGene_tx=XM_019832640.2;refGene_gn=MGA;refGene_tx=XM_019832641.2;refGene_gn=MGA;refGene_tx=XM_019832642.2;refGene_gn=MGA;refGene_tx=XM_019832643.2;refGene_gn=MGA;refGene_tx=XM_019832644.2
B3 63618232 63618581
refGene_gn=MGA;refGene_tx=XM_011283057.3;refGene_gn=MGA;refGene_tx=XM_019832637.2;refGene_gn=MGA;refGene_tx=XM_019832638.2;refGene_gn=MGA;refGene_tx=XM_019832639.2;refGene_gn=MGA;refGene_tx=XM_019832640.2;refGene_gn=MGA;refGene_tx=XM_019832641.2;refGene_gn=MGA;refGene_tx=XM_019832642.2;refGene_gn=MGA;refGene_tx=XM_019832643.2;refGene_gn=MGA;refGene_tx=XM_019832644.2
B3 63620405 63621067
refGene_gn=MGA;refGene_tx=XM_011283057.3;refGene_gn=MGA;refGene_tx=XM_019832637.2;refGene_gn=MGA;refGene_tx=XM_019832638.2;refGene_gn=MGA;refGene_tx=XM_019832639.2;refGene_gn=MGA;refGene_tx=XM_019832640.2;refGene_gn=MGA;refGene_tx=XM_019832641.2;refGene_gn=MGA;refGene_tx=XM_019832642.2;refGene_gn=MGA;refGene_tx=XM_019832643.2;refGene_gn=MGA;refGene_tx=XM_019832644.2
B3 63622225 63622330
refGene_gn=MGA;refGene_tx=XM_011283057.3;refGene_gn=MGA;refGene_tx=XM_019832637.2;refGene_gn=MGA;refGene_tx=XM_019832638.2;refGene_gn=MGA;refGene_tx=XM_019832639.2;refGene_gn=MGA;refGene_tx=XM_019832640.2;refGene_gn=MGA;refGene_tx=XM_019832641.2;refGene_gn=MGA;refGene_tx=XM_019832642.2;refGene_gn=MGA;refGene_tx=XM_019832643.2;refGene_gn=MGA;refGene_tx=XM_019832644.2
B3 63622797 63622929
refGene_gn=MGA;refGene_tx=XM_011283057.3;refGene_gn=MGA;refGene_tx=XM_019832637.2;refGene_gn=MGA;refGene_tx=XM_019832638.2;refGene_gn=MGA;refGene_tx=XM_019832639.2;refGene_gn=MGA;refGene_tx=XM_019832640.2;refGene_gn=MGA;refGene_tx=XM_019832641.2;refGene_gn=MGA;refGene_tx=XM_019832642.2;refGene_gn=MGA;refGene_tx=XM_019832643.2;refGene_gn=MGA;refGene_tx=XM_019832644.2
B3 63634400 63634496

refGene_gn=MGA;refGene_tx=XM_011283057.3;refGene_gn=MGA;refGene_tx=XM_019832637.2;refGene_gn=MGA;refGene_tx=XM_019832638.2;refGene_gn=MGA;refGene_tx=XM_019832639.2;refGene_gn=MGA;refGene_tx=XM_019832640.2;refGene_gn=MGA;refGene_tx=XM_019832641.2;refGene_gn=MGA;refGene_tx=XM_019832642.2;refGene_gn=MGA;refGene_tx=XM_019832643.2;refGene_gn=MGA;refGene_tx=XM_019832644.2
B3 63634614 63634693
refGene_gn=MGA;refGene_tx=XM_011283057.3;refGene_gn=MGA;refGene_tx=XM_019832637.2;refGene_gn=MGA;refGene_tx=XM_019832638.2;refGene_gn=MGA;refGene_tx=XM_019832639.2;refGene_gn=MGA;refGene_tx=XM_019832640.2;refGene_gn=MGA;refGene_tx=XM_019832641.2;refGene_gn=MGA;refGene_tx=XM_019832642.2;refGene_gn=MGA;refGene_tx=XM_019832643.2;refGene_gn=MGA;refGene_tx=XM_019832644.2
B3 63635490 63636439
refGene_gn=MGA;refGene_tx=XM_011283057.3;refGene_gn=MGA;refGene_tx=XM_019832637.2;refGene_gn=MGA;refGene_tx=XM_019832638.2;refGene_gn=MGA;refGene_tx=XM_019832639.2;refGene_gn=MGA;refGene_tx=XM_019832640.2;refGene_gn=MGA;refGene_tx=XM_019832641.2;refGene_gn=MGA;refGene_tx=XM_019832642.2;refGene_gn=MGA;refGene_tx=XM_019832643.2;refGene_gn=MGA;refGene_tx=XM_019832644.2
B3 63661183 63662247
refGene_gn=MGA;refGene_tx=XM_011283057.3;refGene_gn=MGA;refGene_tx=XM_019832637.2;refGene_gn=MGA;refGene_tx=XM_019832638.2;refGene_gn=MGA;refGene_tx=XM_019832639.2;refGene_gn=MGA;refGene_tx=XM_019832640.2;refGene_gn=MGA;refGene_tx=XM_019832641.2;refGene_gn=MGA;refGene_tx=XM_019832642.2;refGene_gn=MGA;refGene_tx=XM_019832643.2;refGene_gn=MGA;refGene_tx=XM_019832644.2
B3 63778403 63778794
refGene_gn=TYRO3;refGene_tx=XM_023255470.1;refGene_gn=TYRO3;refGene_tx=XM_023255471.1
B3 63780956 63781093
refGene_gn=TYRO3;refGene_tx=XM_023255470.1;refGene_gn=TYRO3;refGene_tx=XM_023255471.1
B3 63781337 63781497
refGene_gn=TYRO3;refGene_tx=XM_023255470.1;refGene_gn=TYRO3;refGene_tx=XM_023255471.1
B3 63781680 63781790
refGene_gn=TYRO3;refGene_tx=XM_023255470.1;refGene_gn=TYRO3;refGene_tx=XM_023255471.1
B3 63782201 63782323
refGene_gn=TYRO3;refGene_tx=XM_023255470.1;refGene_gn=TYRO3;refGene_tx=XM_023255471.1
B3 63782751 63782844
refGene_gn=TYRO3;refGene_tx=XM_023255470.1;refGene_gn=TYRO3;refGene_tx=XM_023255471.1
B3 63783290 63783371
refGene_gn=TYRO3;refGene_tx=XM_023255470.1;refGene_gn=TYRO3;refGene_tx=XM_023255471.1
B3 63783709 63783805
refGene_gn=TYRO3;refGene_tx=XM_023255470.1;refGene_gn=TYRO3;refGene_tx

=XM_023255471.1
B3 63784344 63784445
refGene_gn=TYRO3;refGene_tx=XM_023255470.1;refGene_gn=TYRO3;refGene_tx
=XM_023255471.1
B3 63784528 63784658
refGene_gn=TYRO3;refGene_tx=XM_023255470.1;refGene_gn=TYRO3;refGene_tx
=XM_023255471.1
B3 63785039 63785184
refGene_gn=TYRO3;refGene_tx=XM_023255470.1;refGene_gn=TYRO3;refGene_tx
=XM_023255471.1
B3 63785752 63785898
refGene_gn=TYRO3;refGene_tx=XM_023255470.1;refGene_gn=TYRO3;refGene_tx
=XM_023255471.1
B3 63786586 63786764
refGene_gn=TYRO3;refGene_tx=XM_023255470.1;refGene_gn=TYRO3;refGene_tx
=XM_023255471.1
B3 63788310 63788426
refGene_gn=TYRO3;refGene_tx=XM_023255470.1;refGene_gn=TYRO3;refGene_tx
=XM_023255471.1
B3 63789224 63789311
refGene_gn=TYRO3;refGene_tx=XM_023255470.1;refGene_gn=TYRO3;refGene_tx
=XM_023255471.1
B3 63790788 63790959
refGene_gn=TYRO3;refGene_tx=XM_023255470.1;refGene_gn=TYRO3;refGene_tx
=XM_023255471.1
B3 63791730 63791831
refGene_gn=TYRO3;refGene_tx=XM_023255470.1;refGene_gn=TYRO3;refGene_tx
=XM_023255471.1
B3 63792068 63792252
refGene_gn=TYRO3;refGene_tx=XM_023255470.1;refGene_gn=TYRO3;refGene_tx
=XM_023255471.1
B3 63794076 63794197
refGene_gn=TYRO3;refGene_tx=XM_023255470.1;refGene_gn=TYRO3;refGene_tx
=XM_023255471.1
B3 63802996 63803177
refGene_gn=RPAP1;refGene_tx=XM_019832649.2;refGene_gn=RPAP1;refGene_tx
=XM_023255472.1;refGene_gn=RPAP1;refGene_tx=XM_023255473.1;refGene_gn=
RPAP1;refGene_tx=XM_023255474.1
B3 63803483 63803632
refGene_gn=RPAP1;refGene_tx=XM_019832649.2;refGene_gn=RPAP1;refGene_tx
=XM_023255472.1;refGene_gn=RPAP1;refGene_tx=XM_023255473.1;refGene_gn=
RPAP1;refGene_tx=XM_023255474.1
B3 63803915 63804005
refGene_gn=RPAP1;refGene_tx=XM_019832649.2;refGene_gn=RPAP1;refGene_tx
=XM_023255472.1;refGene_gn=RPAP1;refGene_tx=XM_023255473.1;refGene_gn=
RPAP1;refGene_tx=XM_023255474.1
B3 63804692 63804813
refGene_gn=RPAP1;refGene_tx=XM_019832649.2;refGene_gn=RPAP1;refGene_tx
=XM_023255472.1;refGene_gn=RPAP1;refGene_tx=XM_023255473.1;refGene_gn=
RPAP1;refGene_tx=XM_023255474.1

B3 63805584 63805806
refGene_gn=RPAP1;refGene_tx=XM_019832649.2;refGene_gn=RPAP1;refGene_tx=XM_023255472.1;refGene_gn=RPAP1;refGene_tx=XM_023255473.1;refGene_gn=RPAP1;refGene_tx=XM_023255474.1
B3 63806684 63806864
refGene_gn=RPAP1;refGene_tx=XM_019832649.2;refGene_gn=RPAP1;refGene_tx=XM_023255472.1;refGene_gn=RPAP1;refGene_tx=XM_023255473.1;refGene_gn=RPAP1;refGene_tx=XM_023255474.1
B3 63807897 63808013
refGene_gn=RPAP1;refGene_tx=XM_019832649.2;refGene_gn=RPAP1;refGene_tx=XM_023255472.1;refGene_gn=RPAP1;refGene_tx=XM_023255473.1;refGene_gn=RPAP1;refGene_tx=XM_023255474.1
B3 63808294 63808393
refGene_gn=RPAP1;refGene_tx=XM_019832649.2;refGene_gn=RPAP1;refGene_tx=XM_023255472.1;refGene_gn=RPAP1;refGene_tx=XM_023255473.1;refGene_gn=RPAP1;refGene_tx=XM_023255474.1
B3 63809240 63809342
refGene_gn=RPAP1;refGene_tx=XM_019832649.2;refGene_gn=RPAP1;refGene_tx=XM_023255472.1;refGene_gn=RPAP1;refGene_tx=XM_023255473.1;refGene_gn=RPAP1;refGene_tx=XM_023255474.1
B3 63809564 63809732
refGene_gn=RPAP1;refGene_tx=XM_019832649.2;refGene_gn=RPAP1;refGene_tx=XM_023255472.1;refGene_gn=RPAP1;refGene_tx=XM_023255473.1;refGene_gn=RPAP1;refGene_tx=XM_023255474.1
B3 63809978 63810164
refGene_gn=RPAP1;refGene_tx=XM_019832649.2;refGene_gn=RPAP1;refGene_tx=XM_023255472.1;refGene_gn=RPAP1;refGene_tx=XM_023255473.1;refGene_gn=RPAP1;refGene_tx=XM_023255474.1
B3 63810288 63810423
refGene_gn=RPAP1;refGene_tx=XM_019832649.2;refGene_gn=RPAP1;refGene_tx=XM_023255472.1;refGene_gn=RPAP1;refGene_tx=XM_023255473.1;refGene_gn=RPAP1;refGene_tx=XM_023255474.1
B3 63810530 63810701
refGene_gn=RPAP1;refGene_tx=XM_019832649.2;refGene_gn=RPAP1;refGene_tx=XM_023255472.1;refGene_gn=RPAP1;refGene_tx=XM_023255473.1;refGene_gn=RPAP1;refGene_tx=XM_023255474.1
B3 63811221 63811379
refGene_gn=RPAP1;refGene_tx=XM_019832649.2;refGene_gn=RPAP1;refGene_tx=XM_023255472.1;refGene_gn=RPAP1;refGene_tx=XM_023255473.1;refGene_gn=RPAP1;refGene_tx=XM_023255474.1
B3 63811843 63812002
refGene_gn=RPAP1;refGene_tx=XM_019832649.2;refGene_gn=RPAP1;refGene_tx=XM_023255472.1;refGene_gn=RPAP1;refGene_tx=XM_023255473.1;refGene_gn=RPAP1;refGene_tx=XM_023255474.1
B3 63812126 63812328
refGene_gn=RPAP1;refGene_tx=XM_019832649.2;refGene_gn=RPAP1;refGene_tx=XM_023255472.1;refGene_gn=RPAP1;refGene_tx=XM_023255473.1;refGene_gn=RPAP1;refGene_tx=XM_023255474.1
B3 63812759 63812869
refGene_gn=RPAP1;refGene_tx=XM_019832649.2;refGene_gn=RPAP1;refGene_tx

=XM_023255472.1;refGene_gn=RPAP1;refGene_tx=XM_023255473.1;refGene_gn=RPAP1;refGene_tx=XM_023255474.1
B3	63813152	63813348	refGene_gn=RPAP1;refGene_tx=XM_019832649.2;refGene_gn=RPAP1;refGene_tx=XM_023255472.1;refGene_gn=RPAP1;refGene_tx=XM_023255473.1;refGene_gn=RPAP1;refGene_tx=XM_023255474.1
B3	63813562	63813715	refGene_gn=RPAP1;refGene_tx=XM_019832649.2;refGene_gn=RPAP1;refGene_tx=XM_023255472.1;refGene_gn=RPAP1;refGene_tx=XM_023255473.1;refGene_gn=RPAP1;refGene_tx=XM_023255474.1
B3	63813913	63814056	refGene_gn=RPAP1;refGene_tx=XM_019832649.2;refGene_gn=RPAP1;refGene_tx=XM_023255472.1;refGene_gn=RPAP1;refGene_tx=XM_023255473.1;refGene_gn=RPAP1;refGene_tx=XM_023255474.1
B3	63814644	63815401	refGene_gn=RPAP1;refGene_tx=XM_019832649.2;refGene_gn=RPAP1;refGene_tx=XM_023255472.1;refGene_gn=RPAP1;refGene_tx=XM_023255473.1;refGene_gn=RPAP1;refGene_tx=XM_023255474.1
B3	63818645	63818822	refGene_gn=RPAP1;refGene_tx=XM_019832649.2;refGene_gn=RPAP1;refGene_tx=XM_023255472.1;refGene_gn=RPAP1;refGene_tx=XM_023255473.1;refGene_gn=RPAP1;refGene_tx=XM_023255474.1
B3	63818960	63819020	refGene_gn=RPAP1;refGene_tx=XM_019832649.2;refGene_gn=RPAP1;refGene_tx=XM_023255472.1;refGene_gn=RPAP1;refGene_tx=XM_023255473.1;refGene_gn=RPAP1;refGene_tx=XM_023255474.1
B3	63819127	63819277	refGene_gn=RPAP1;refGene_tx=XM_019832649.2;refGene_gn=RPAP1;refGene_tx=XM_023255472.1;refGene_gn=RPAP1;refGene_tx=XM_023255473.1;refGene_gn=RPAP1;refGene_tx=XM_023255474.1
B3	63830722	63830926	refGene_gn=ITPKA;refGene_tx=XM_023255475.1
B3	63831039	63831111	refGene_gn=ITPKA;refGene_tx=XM_023255475.1
B3	63831385	63831487	refGene_gn=ITPKA;refGene_tx=XM_023255475.1
B3	63831705	63831910	refGene_gn=ITPKA;refGene_tx=XM_023255475.1
B3	63832041	63832258	refGene_gn=ITPKA;refGene_tx=XM_023255475.1
B3	63832343	63832440	refGene_gn=ITPKA;refGene_tx=XM_023255475.1
B3	63838371	63838860	refGene_gn=ITPKA;refGene_tx=XM_023255475.1
B3	63849598	63849705	refGene_gn=RTF1;refGene_tx=XM_023255476.1
B3	63849961	63850073	refGene_gn=RTF1;refGene_tx=XM_023255476.1
B3	63851008	63851104	refGene_gn=RTF1;refGene_tx=XM_023255476.1
B3	63851583	63851661	refGene_gn=RTF1;refGene_tx=XM_023255476.1
B3	63852092	63852150	refGene_gn=RTF1;refGene_tx=XM_023255476.1
B3	63852344	63852466	refGene_gn=RTF1;refGene_tx=XM_023255476.1
B3	63853088	63853166	refGene_gn=RTF1;refGene_tx=XM_023255476.1
B3	63853748	63853856	refGene_gn=RTF1;refGene_tx=XM_023255476.1
B3	63853949	63854037	refGene_gn=RTF1;refGene_tx=XM_023255476.1
B3	63855358	63855441	refGene_gn=RTF1;refGene_tx=XM_023255476.1
B3	63856407	63856585	refGene_gn=RTF1;refGene_tx=XM_023255476.1
B3	63857073	63857209	refGene_gn=RTF1;refGene_tx=XM_023255476.1
B3	63864801	63864913	refGene_gn=RTF1;refGene_tx=XM_023255476.1

B3 63865599 63865714 refGene_gn=RTF1;refGene_tx=XM_023255476.1
B3 63872811 63873016 refGene_gn=RTF1;refGene_tx=XM_023255476.1
B3 63875426 63875574 refGene_gn=RTF1;refGene_tx=XM_023255476.1
B3 63890723 63890834 refGene_gn=RTF1;refGene_tx=XM_023255476.1
B3 63902569 63902767 refGene_gn=RTF1;refGene_tx=XM_023255476.1
B3 63919000 63919576
refGene_gn=NDUFAF1;refGene_tx=XM_006932629.4;refGene_gn=NDUFAF1;refGene_tx=XM_011283058.3;refGene_gn=NDUFAF1;refGene_tx=XM_023255478.1
B3 63923560 63923746
refGene_gn=NDUFAF1;refGene_tx=XM_006932629.4;refGene_gn=NDUFAF1;refGene_tx=XM_011283058.3
B3 63938034 63938109
refGene_gn=NDUFAF1;refGene_tx=XM_006932629.4;refGene_gn=NDUFAF1;refGene_tx=XM_011283058.3;refGene_gn=NDUFAF1;refGene_tx=XM_023255478.1
B3 63938864 63939014
refGene_gn=NDUFAF1;refGene_tx=XM_006932629.4;refGene_gn=NDUFAF1;refGene_tx=XM_011283058.3;refGene_gn=NDUFAF1;refGene_tx=XM_023255478.1
B3 63940998 63941092
refGene_gn=NUSAP1;refGene_tx=XM_003987262.5;refGene_gn=NUSAP1;refGene_tx=XM_006932632.4;refGene_gn=NUSAP1;refGene_tx=XM_006932633.4;refGene_gn=NUSAP1;refGene_tx=XM_006932634.4
B3 63942240 63942349
refGene_gn=NUSAP1;refGene_tx=XM_003987262.5;refGene_gn=NUSAP1;refGene_tx=XM_006932632.4;refGene_gn=NUSAP1;refGene_tx=XM_006932633.4;refGene_gn=NUSAP1;refGene_tx=XM_006932634.4
B3 63944420 63944537
refGene_gn=NUSAP1;refGene_tx=XM_003987262.5;refGene_gn=NUSAP1;refGene_tx=XM_006932632.4;refGene_gn=NUSAP1;refGene_tx=XM_006932633.4;refGene_gn=NUSAP1;refGene_tx=XM_006932634.4
B3 63947363 63947521
refGene_gn=NUSAP1;refGene_tx=XM_003987262.5;refGene_gn=NUSAP1;refGene_tx=XM_006932632.4;refGene_gn=NUSAP1;refGene_tx=XM_006932633.4;refGene_gn=NUSAP1;refGene_tx=XM_006932634.4
B3 63951387 63951563
refGene_gn=NUSAP1;refGene_tx=XM_003987262.5;refGene_gn=NUSAP1;refGene_tx=XM_006932632.4;refGene_gn=NUSAP1;refGene_tx=XM_006932633.4;refGene_gn=NUSAP1;refGene_tx=XM_006932634.4
B3 63955979 63956089
refGene_gn=NUSAP1;refGene_tx=XM_003987262.5;refGene_gn=NUSAP1;refGene_tx=XM_006932632.4;refGene_gn=NUSAP1;refGene_tx=XM_006932633.4;refGene_gn=NUSAP1;refGene_tx=XM_006932634.4
B3 63963133 63963235
refGene_gn=NUSAP1;refGene_tx=XM_003987262.5;refGene_gn=NUSAP1;refGene_tx=XM_006932632.4;refGene_gn=NUSAP1;refGene_tx=XM_006932633.4;refGene_gn=NUSAP1;refGene_tx=XM_006932634.4
B3 63965423 63965580
refGene_gn=NUSAP1;refGene_tx=XM_003987262.5;refGene_gn=NUSAP1;refGene_tx=XM_006932632.4;refGene_gn=NUSAP1;refGene_tx=XM_006932633.4;refGene_gn=NUSAP1;refGene_tx=XM_006932634.4
B3 63966340 63966484

refGene_gn=NUSAP1;refGene_tx=XM_003987262.5;refGene_gn=NUSAP1;refGene_tx=XM_006932632.4;refGene_gn=NUSAP1;refGene_tx=XM_006932633.4;refGene_gn=NUSAP1;refGene_tx=XM_006932634.4
B3 63972034 63972103
refGene_gn=NUSAP1;refGene_tx=XM_003987262.5;refGene_gn=NUSAP1;refGene_tx=XM_006932632.4;refGene_gn=NUSAP1;refGene_tx=XM_006932633.4;refGene_gn=NUSAP1;refGene_tx=XM_006932634.4
B3 63981224 63981329
refGene_gn=NUSAP1;refGene_tx=XM_003987262.5;refGene_gn=NUSAP1;refGene_tx=XM_006932632.4;refGene_gn=NUSAP1;refGene_tx=XM_006932633.4;refGene_gn=NUSAP1;refGene_tx=XM_006932634.4
B3 63981691 63982016 refGene_gn=OIP5;refGene_tx=XM_003987333.5
B3 63982265 63982332 refGene_gn=OIP5;refGene_tx=XM_003987333.5
B3 63986531 63986654 refGene_gn=OIP5;refGene_tx=XM_003987333.5
B3 63990267 63990349 refGene_gn=OIP5;refGene_tx=XM_003987333.5
B3 63991557 63991653 refGene_gn=OIP5;refGene_tx=XM_003987333.5
B3 64028292 64028346 refGene_gn=CHP1;refGene_tx=XM_003987264.5
B3 64028502 64028625 refGene_gn=CHP1;refGene_tx=XM_003987264.5
B3 64033833 64033895 refGene_gn=CHP1;refGene_tx=XM_003987264.5
B3 64041815 64041943 refGene_gn=CHP1;refGene_tx=XM_003987264.5
B3 64047375 64047456 refGene_gn=CHP1;refGene_tx=XM_003987264.5
B3 64061760 64061833 refGene_gn=CHP1;refGene_tx=XM_003987264.5
B3 64071110 64071177 refGene_gn=CHP1;refGene_tx=XM_003987264.5
B3 64073094 64073227
refGene_gn=EXD1;refGene_tx=XM_006932635.3;refGene_gn=EXD1;refGene_tx=XM_023255479.1
B3 64082498 64082567
refGene_gn=EXD1;refGene_tx=XM_006932635.3;refGene_gn=EXD1;refGene_tx=XM_023255479.1
B3 64086416 64086474
refGene_gn=EXD1;refGene_tx=XM_006932635.3;refGene_gn=EXD1;refGene_tx=XM_023255479.1
B3 64086716 64086844
refGene_gn=EXD1;refGene_tx=XM_006932635.3;refGene_gn=EXD1;refGene_tx=XM_023255479.1
B3 64091663 64091722
refGene_gn=EXD1;refGene_tx=XM_006932635.3;refGene_gn=EXD1;refGene_tx=XM_023255479.1
B3 64095301 64095388
refGene_gn=EXD1;refGene_tx=XM_006932635.3;refGene_gn=EXD1;refGene_tx=XM_023255479.1
B3 64098783 64098888
refGene_gn=EXD1;refGene_tx=XM_006932635.3;refGene_gn=EXD1;refGene_tx=XM_023255479.1
B3 64098970 64099051
refGene_gn=EXD1;refGene_tx=XM_006932635.3;refGene_gn=EXD1;refGene_tx=XM_023255479.1
B3 64103030 64103120 refGene_gn=EXD1;refGene_tx=XM_023255479.1
B3 64104114 64104258
refGene_gn=EXD1;refGene_tx=XM_006932635.3;refGene_gn=EXD1;refGene_tx=X

M_023255479.1
B3	64104922	64105114
refGene_gn=EXD1;refGene_tx=XM_006932635.3;refGene_gn=EXD1;refGene_tx=XM_023255479.1
B3	64109462	64110101
refGene_gn=EXD1;refGene_tx=XM_006932635.3;refGene_gn=EXD1;refGene_tx=XM_023255479.1
B3	64188362	64188505
refGene_gn=INO80;refGene_tx=XM_023255480.1;refGene_gn=INO80;refGene_tx=XM_023255481.1;refGene_gn=INO80;refGene_tx=XM_023255482.1
B3	64188740	64188910
refGene_gn=INO80;refGene_tx=XM_023255480.1;refGene_gn=INO80;refGene_tx=XM_023255481.1;refGene_gn=INO80;refGene_tx=XM_023255482.1
B3	64189003	64189071
refGene_gn=INO80;refGene_tx=XM_023255480.1;refGene_gn=INO80;refGene_tx=XM_023255481.1;refGene_gn=INO80;refGene_tx=XM_023255482.1
B3	64191006	64191174
refGene_gn=INO80;refGene_tx=XM_023255480.1;refGene_gn=INO80;refGene_tx=XM_023255481.1;refGene_gn=INO80;refGene_tx=XM_023255482.1
B3	64196557	64196678
refGene_gn=INO80;refGene_tx=XM_023255480.1;refGene_gn=INO80;refGene_tx=XM_023255481.1;refGene_gn=INO80;refGene_tx=XM_023255482.1
B3	64197986	64198201
refGene_gn=INO80;refGene_tx=XM_023255480.1;refGene_gn=INO80;refGene_tx=XM_023255481.1;refGene_gn=INO80;refGene_tx=XM_023255482.1
B3	64205555	64205609
refGene_gn=INO80;refGene_tx=XM_023255480.1;refGene_gn=INO80;refGene_tx=XM_023255481.1;refGene_gn=INO80;refGene_tx=XM_023255482.1
B3	64206425	64206629
refGene_gn=INO80;refGene_tx=XM_023255480.1;refGene_gn=INO80;refGene_tx=XM_023255481.1;refGene_gn=INO80;refGene_tx=XM_023255482.1
B3	64208670	64208866
refGene_gn=INO80;refGene_tx=XM_023255480.1;refGene_gn=INO80;refGene_tx=XM_023255481.1;refGene_gn=INO80;refGene_tx=XM_023255482.1
B3	64209742	64209810
refGene_gn=INO80;refGene_tx=XM_023255480.1;refGene_gn=INO80;refGene_tx=XM_023255481.1;refGene_gn=INO80;refGene_tx=XM_023255482.1
B3	64210660	64210870
refGene_gn=INO80;refGene_tx=XM_023255480.1;refGene_gn=INO80;refGene_tx=XM_023255481.1;refGene_gn=INO80;refGene_tx=XM_023255482.1
B3	64211381	64211462
refGene_gn=INO80;refGene_tx=XM_023255480.1;refGene_gn=INO80;refGene_tx=XM_023255481.1;refGene_gn=INO80;refGene_tx=XM_023255482.1
B3	64212265	64212361
refGene_gn=INO80;refGene_tx=XM_023255480.1;refGene_gn=INO80;refGene_tx=XM_023255481.1;refGene_gn=INO80;refGene_tx=XM_023255482.1
B3	64221775	64221835
refGene_gn=INO80;refGene_tx=XM_023255480.1;refGene_gn=INO80;refGene_tx=XM_023255481.1;refGene_gn=INO80;refGene_tx=XM_023255482.1
B3	64222259	64222402

refGene_gn=INO80;refGene_tx=XM_023255480.1;refGene_gn=INO80;refGene_tx
=XM_023255481.1;refGene_gn=INO80;refGene_tx=XM_023255482.1
B3 64223946 64224031
refGene_gn=INO80;refGene_tx=XM_023255480.1;refGene_gn=INO80;refGene_tx
=XM_023255481.1;refGene_gn=INO80;refGene_tx=XM_023255482.1
B3 64225025 64225143
refGene_gn=INO80;refGene_tx=XM_023255480.1;refGene_gn=INO80;refGene_tx
=XM_023255481.1;refGene_gn=INO80;refGene_tx=XM_023255482.1
B3 64226077 64226163
refGene_gn=INO80;refGene_tx=XM_023255480.1;refGene_gn=INO80;refGene_tx
=XM_023255481.1;refGene_gn=INO80;refGene_tx=XM_023255482.1
B3 64229402 64229570
refGene_gn=INO80;refGene_tx=XM_023255480.1;refGene_gn=INO80;refGene_tx
=XM_023255481.1;refGene_gn=INO80;refGene_tx=XM_023255482.1
B3 64229788 64229922
refGene_gn=INO80;refGene_tx=XM_023255480.1;refGene_gn=INO80;refGene_tx
=XM_023255481.1;refGene_gn=INO80;refGene_tx=XM_023255482.1
B3 64231205 64231273
refGene_gn=INO80;refGene_tx=XM_023255480.1;refGene_gn=INO80;refGene_tx
=XM_023255481.1;refGene_gn=INO80;refGene_tx=XM_023255482.1
B3 64231970 64232064
refGene_gn=INO80;refGene_tx=XM_023255480.1;refGene_gn=INO80;refGene_tx
=XM_023255481.1;refGene_gn=INO80;refGene_tx=XM_023255482.1
B3 64232835 64233007
refGene_gn=INO80;refGene_tx=XM_023255480.1;refGene_gn=INO80;refGene_tx
=XM_023255481.1;refGene_gn=INO80;refGene_tx=XM_023255482.1
B3 64250237 64250378
refGene_gn=INO80;refGene_tx=XM_023255480.1;refGene_gn=INO80;refGene_tx
=XM_023255481.1;refGene_gn=INO80;refGene_tx=XM_023255482.1
B3 64256663 64256889
refGene_gn=INO80;refGene_tx=XM_023255480.1;refGene_gn=INO80;refGene_tx
=XM_023255481.1;refGene_gn=INO80;refGene_tx=XM_023255482.1
B3 64260967 64261095
refGene_gn=INO80;refGene_tx=XM_023255480.1;refGene_gn=INO80;refGene_tx
=XM_023255481.1;refGene_gn=INO80;refGene_tx=XM_023255482.1
B3 64272103 64272198
refGene_gn=INO80;refGene_tx=XM_023255480.1;refGene_gn=INO80;refGene_tx
=XM_023255481.1;refGene_gn=INO80;refGene_tx=XM_023255482.1
B3 64280030 64280103
refGene_gn=INO80;refGene_tx=XM_023255480.1;refGene_gn=INO80;refGene_tx
=XM_023255481.1;refGene_gn=INO80;refGene_tx=XM_023255482.1
B3 64288102 64288261
refGene_gn=INO80;refGene_tx=XM_023255480.1;refGene_gn=INO80;refGene_tx
=XM_023255481.1;refGene_gn=INO80;refGene_tx=XM_023255482.1
B3 64288878 64288981
refGene_gn=INO80;refGene_tx=XM_023255480.1;refGene_gn=INO80;refGene_tx
=XM_023255481.1;refGene_gn=INO80;refGene_tx=XM_023255482.1
B3 64290476 64290565
refGene_gn=INO80;refGene_tx=XM_023255480.1;refGene_gn=INO80;refGene_tx
=XM_023255481.1;refGene_gn=INO80;refGene_tx=XM_023255482.1

B3 64292994 64293150
refGene_gn=INO80;refGene_tx=XM_023255480.1;refGene_gn=INO80;refGene_tx
=XM_023255481.1;refGene_gn=INO80;refGene_tx=XM_023255482.1
B3 64293442 64293602
refGene_gn=INO80;refGene_tx=XM_023255480.1;refGene_gn=INO80;refGene_tx
=XM_023255481.1;refGene_gn=INO80;refGene_tx=XM_023255482.1
B3 64294244 64294460
refGene_gn=INO80;refGene_tx=XM_023255480.1;refGene_gn=INO80;refGene_tx
=XM_023255481.1;refGene_gn=INO80;refGene_tx=XM_023255482.1
B3 64296385 64296603
refGene_gn=INO80;refGene_tx=XM_023255480.1;refGene_gn=INO80;refGene_tx
=XM_023255481.1;refGene_gn=INO80;refGene_tx=XM_023255482.1
B3 64301208 64301389
refGene_gn=INO80;refGene_tx=XM_023255480.1;refGene_gn=INO80;refGene_tx
=XM_023255481.1
B3 64318559 64318961 refGene_gn=CHAC1;refGene_tx=XM_006932672.4
B3 64319415 64319451 refGene_gn=CHAC1;refGene_tx=XM_006932672.4
B3 64319857 64320108 refGene_gn=CHAC1;refGene_tx=XM_006932672.4
B3 64320110 64320471 refGene_gn=CHAC1;refGene_tx=XM_006932672.4
B3 64332134 64332140 refGene_gn=DLL4;refGene_tx=XM_003987266.5
B3 64332637 64332746 refGene_gn=DLL4;refGene_tx=XM_003987266.5
B3 64333261 64333964 refGene_gn=DLL4;refGene_tx=XM_003987266.5
B3 64335112 64335332 refGene_gn=DLL4;refGene_tx=XM_003987266.5
B3 64335500 64335670 refGene_gn=DLL4;refGene_tx=XM_003987266.5
B3 64337239 64337370 refGene_gn=DLL4;refGene_tx=XM_003987266.5
B3 64337457 64337518 refGene_gn=DLL4;refGene_tx=XM_003987266.5
B3 64337942 64338206 refGene_gn=DLL4;refGene_tx=XM_003987266.5
B3 64339035 64339093 refGene_gn=DLL4;refGene_tx=XM_003987266.5
B3 64339591 64339861 refGene_gn=DLL4;refGene_tx=XM_003987266.5
B3 64339972 64340038 refGene_gn=DLL4;refGene_tx=XM_003987266.5
B3 64365558 64366284
refGene_gn=VPS18;refGene_tx=XM_003987267.5;refGene_gn=VPS18;refGene_tx
=XM_019832662.2
B3 64367313 64369184
refGene_gn=VPS18;refGene_tx=XM_003987267.5;refGene_gn=VPS18;refGene_tx
=XM_019832662.2
B3 64369355 64369447
refGene_gn=VPS18;refGene_tx=XM_003987267.5;refGene_gn=VPS18;refGene_tx
=XM_019832662.2
B3 64371564 64371706 refGene_gn=VPS18;refGene_tx=XM_003987267.5
B3 64372732 64372823 refGene_gn=VPS18;refGene_tx=XM_003987267.5
B3 64393763 64393971 refGene_gn=RHOV;refGene_tx=XM_023255483.1
B3 64394167 64394226 refGene_gn=RHOV;refGene_tx=XM_023255483.1
B3 64394403 64394847 refGene_gn=RHOV;refGene_tx=XM_023255483.1
B3 64408735 64408959 refGene_gn=SPINT1;refGene_tx=XM_023256169.1
B3 64409321 64409369 refGene_gn=SPINT1;refGene_tx=XM_023256169.1
B3 64409538 64409709 refGene_gn=SPINT1;refGene_tx=XM_023256169.1
B3 64410633 64410684 refGene_gn=SPINT1;refGene_tx=XM_023256169.1
B3 64410800 64410926 refGene_gn=SPINT1;refGene_tx=XM_023256169.1
B3 64411249 64411276 refGene_gn=SPINT1;refGene_tx=XM_023256169.1

B3 64411459 64411630 refGene_gn=SPINT1;refGene_tx=XM_023256169.1
B3 64411726 64411865 refGene_gn=SPINT1;refGene_tx=XM_023256169.1
B3 64412097 64412225 refGene_gn=SPINT1;refGene_tx=XM_023256169.1
B3 64419993 64420533 refGene_gn=SPINT1;refGene_tx=XM_023256169.1
B3 64420746 64421199 refGene_gn=SPINT1;refGene_tx=XM_023256169.1
B3 64421245 64421351 refGene_gn=SPINT1;refGene_tx=XM_023256169.1
B3 64421719 64421872 refGene_gn=SPINT1;refGene_tx=XM_023256169.1
B3 64434427 64434682
refGene_gn=PPP1R14D;refGene_tx=XM_023255484.1;refGene_gn=PPP1R14D;refGene_tx=XM_023255485.1
B3 64445992 64446299
refGene_gn=PPP1R14D;refGene_tx=XM_023255484.1;refGene_gn=PPP1R14D;refGene_tx=XM_023255485.1
B3 64446504 64446570
refGene_gn=PPP1R14D;refGene_tx=XM_023255484.1;refGene_gn=PPP1R14D;refGene_tx=XM_023255485.1
B3 64448047 64448126 refGene_gn=ZFYVE19;refGene_tx=XM_019832663.2
B3 64448210 64448341 refGene_gn=ZFYVE19;refGene_tx=XM_019832663.2
B3 64448448 64448547 refGene_gn=ZFYVE19;refGene_tx=XM_019832663.2
B3 64448868 64448948 refGene_gn=ZFYVE19;refGene_tx=XM_019832663.2
B3 64449390 64449600 refGene_gn=ZFYVE19;refGene_tx=XM_019832663.2
B3 64451300 64451409 refGene_gn=ZFYVE19;refGene_tx=XM_019832663.2
B3 64451738 64451884 refGene_gn=ZFYVE19;refGene_tx=XM_019832663.2
B3 64451983 64452102 refGene_gn=ZFYVE19;refGene_tx=XM_019832663.2
B3 64452484 64452538 refGene_gn=ZFYVE19;refGene_tx=XM_019832663.2
B3 64452704 64452826 refGene_gn=ZFYVE19;refGene_tx=XM_019832663.2
B3 64454081 64454135 refGene_gn=ZFYVE19;refGene_tx=XM_019832663.2
B3 64454516 64454594 refGene_gn=DNAJC17;refGene_tx=XM_003987269.5
B3 64485300 64485370 refGene_gn=DNAJC17;refGene_tx=XM_003987269.5
B3 64485671 64485730 refGene_gn=DNAJC17;refGene_tx=XM_003987269.5
B3 64485965 64486053 refGene_gn=DNAJC17;refGene_tx=XM_003987269.5
B3 64487926 64488012 refGene_gn=DNAJC17;refGene_tx=XM_003987269.5
B3 64488238 64488335 refGene_gn=DNAJC17;refGene_tx=XM_003987269.5
B3 64488928 64488972 refGene_gn=DNAJC17;refGene_tx=XM_003987269.5
B3 64489427 64489505 refGene_gn=DNAJC17;refGene_tx=XM_003987269.5
B3 64490021 64490102 refGene_gn=DNAJC17;refGene_tx=XM_003987269.5
B3 64490614 64490725 refGene_gn=DNAJC17;refGene_tx=XM_003987269.5
B3 64493100 64493628
refGene_gn=CB3H15orf62;refGene_tx=XM_006932637.4
B3 64496057 64496180 refGene_gn=DNAJC17;refGene_tx=XM_003987269.5
B3 64496833 64496957 refGene_gn=GCHFR;refGene_tx=XM_003987271.5
B3 64498081 64498176 refGene_gn=GCHFR;refGene_tx=XM_003987271.5
B3 64500101 64500137 refGene_gn=GCHFR;refGene_tx=XM_003987271.5
B3 64507179 64507366
refGene_gn=RMDN3;refGene_tx=XM_006932641.4;refGene_gn=RMDN3;refGene_tx=XM_006932642.3
B3 64509616 64509809
refGene_gn=RMDN3;refGene_tx=XM_006932641.4;refGene_gn=RMDN3;refGene_tx=XM_006932642.3
B3 64510240 64510384

refGene_gn=RMDN3;refGene_tx=XM_006932641.4;refGene_gn=RMDN3;refGene_tx=XM_006932642.3;refGene_gn=RMDN3;refGene_tx=XM_019832665.2;refGene_gn=RMDN3;refGene_tx=XM_023255486.1
B3 64515657 64515949
refGene_gn=RMDN3;refGene_tx=XM_006932641.4;refGene_gn=RMDN3;refGene_tx=XM_006932642.3;refGene_gn=RMDN3;refGene_tx=XM_019832665.2;refGene_gn=RMDN3;refGene_tx=XM_023255486.1
B3 64516721 64516824
refGene_gn=RMDN3;refGene_tx=XM_006932641.4;refGene_gn=RMDN3;refGene_tx=XM_006932642.3;refGene_gn=RMDN3;refGene_tx=XM_019832665.2;refGene_gn=RMDN3;refGene_tx=XM_023255486.1
B3 64520029 64520090
refGene_gn=RMDN3;refGene_tx=XM_006932641.4;refGene_gn=RMDN3;refGene_tx=XM_006932642.3;refGene_gn=RMDN3;refGene_tx=XM_019832665.2;refGene_gn=RMDN3;refGene_tx=XM_023255486.1
B3 64521506 64521582
refGene_gn=RMDN3;refGene_tx=XM_006932641.4;refGene_gn=RMDN3;refGene_tx=XM_006932642.3;refGene_gn=RMDN3;refGene_tx=XM_019832665.2;refGene_gn=RMDN3;refGene_tx=XM_023255486.1
B3 64521736 64521814
refGene_gn=RMDN3;refGene_tx=XM_006932641.4;refGene_gn=RMDN3;refGene_tx=XM_006932642.3;refGene_gn=RMDN3;refGene_tx=XM_019832665.2;refGene_gn=RMDN3;refGene_tx=XM_023255486.1
B3 64522056 64522155
refGene_gn=RMDN3;refGene_tx=XM_006932641.4;refGene_gn=RMDN3;refGene_tx=XM_006932642.3;refGene_gn=RMDN3;refGene_tx=XM_019832665.2;refGene_gn=RMDN3;refGene_tx=XM_023255486.1
B3 64522360 64522414
refGene_gn=RMDN3;refGene_tx=XM_006932641.4;refGene_gn=RMDN3;refGene_tx=XM_006932642.3;refGene_gn=RMDN3;refGene_tx=XM_019832665.2;refGene_gn=RMDN3;refGene_tx=XM_023255486.1
B3 64522506 64522587
refGene_gn=RMDN3;refGene_tx=XM_006932641.4;refGene_gn=RMDN3;refGene_tx=XM_006932642.3;refGene_gn=RMDN3;refGene_tx=XM_019832665.2;refGene_gn=RMDN3;refGene_tx=XM_023255486.1
B3 64522827 64522881
refGene_gn=RMDN3;refGene_tx=XM_006932641.4;refGene_gn=RMDN3;refGene_tx=XM_019832665.2;refGene_gn=RMDN3;refGene_tx=XM_023255486.1
B3 64524893 64525017
refGene_gn=RAD51;refGene_tx=XM_003987273.5;refGene_gn=RAD51;refGene_tx=XM_006932640.4;refGene_gn=RAD51;refGene_tx=XM_019832666.2
B3 64525465 64525587
refGene_gn=RAD51;refGene_tx=XM_003987273.5;refGene_gn=RAD51;refGene_tx=XM_006932640.4;refGene_gn=RAD51;refGene_tx=XM_019832666.2
B3 64525723 64525738 refGene_gn=RMDN3;refGene_tx=XM_006932642.3
B3 64525794 64525924
refGene_gn=RAD51;refGene_tx=XM_003987273.5;refGene_gn=RAD51;refGene_tx=XM_006932640.4;refGene_gn=RAD51;refGene_tx=XM_019832666.2
B3 64526409 64526523
refGene_gn=RAD51;refGene_tx=XM_003987273.5;refGene_gn=RAD51;refGene_tx

=XM_006932640.4;refGene_gn=RAD51;refGene_tx=XM_019832666.2
B3 64533976 64534071
refGene_gn=RAD51;refGene_tx=XM_003987273.5;refGene_gn=RAD51;refGene_tx=XM_006932640.4;refGene_gn=RAD51;refGene_tx=XM_019832666.2
B3 64541252 64541344
refGene_gn=RAD51;refGene_tx=XM_003987273.5;refGene_gn=RAD51;refGene_tx=XM_006932640.4;refGene_gn=RAD51;refGene_tx=XM_019832666.2
B3 64542739 64542857
refGene_gn=RAD51;refGene_tx=XM_003987273.5;refGene_gn=RAD51;refGene_tx=XM_006932640.4;refGene_gn=RAD51;refGene_tx=XM_019832666.2
B3 64550211 64550349
refGene_gn=RAD51;refGene_tx=XM_003987273.5;refGene_gn=RAD51;refGene_tx=XM_006932640.4;refGene_gn=RAD51;refGene_tx=XM_019832666.2
B3 64551825 64551912
refGene_gn=RAD51;refGene_tx=XM_003987273.5;refGene_gn=RAD51;refGene_tx=XM_006932640.4;refGene_gn=RAD51;refGene_tx=XM_019832666.2
B3 64573487 64573602 refGene_gn=KNL1;refGene_tx=XM_011283064.3
B3 64574349 64574472 refGene_gn=KNL1;refGene_tx=XM_011283064.3
B3 64575158 64575277 refGene_gn=KNL1;refGene_tx=XM_011283064.3
B3 64576399 64576509 refGene_gn=KNL1;refGene_tx=XM_011283064.3
B3 64577171 64577240 refGene_gn=KNL1;refGene_tx=XM_011283064.3
B3 64581777 64581878 refGene_gn=KNL1;refGene_tx=XM_011283064.3
B3 64582603 64582705 refGene_gn=KNL1;refGene_tx=XM_011283064.3
B3 64583479 64583519 refGene_gn=KNL1;refGene_tx=XM_011283064.3
B3 64583680 64583758 refGene_gn=KNL1;refGene_tx=XM_011283064.3
B3 64587178 64587266 refGene_gn=KNL1;refGene_tx=XM_011283064.3
B3 64588785 64588902 refGene_gn=KNL1;refGene_tx=XM_011283064.3
B3 64589888 64589979 refGene_gn=KNL1;refGene_tx=XM_011283064.3
B3 64592260 64592376 refGene_gn=KNL1;refGene_tx=XM_011283064.3
B3 64600854 64600953 refGene_gn=KNL1;refGene_tx=XM_011283064.3
B3 64601722 64601790 refGene_gn=KNL1;refGene_tx=XM_011283064.3
B3 64602172 64602311 refGene_gn=KNL1;refGene_tx=XM_011283064.3
B3 64604116 64606450 refGene_gn=KNL1;refGene_tx=XM_011283064.3
B3 64606452 64609194 refGene_gn=KNL1;refGene_tx=XM_011283064.3
B3 64609950 64610003 refGene_gn=KNL1;refGene_tx=XM_011283064.3
B3 64614078 64614116 refGene_gn=KNL1;refGene_tx=XM_011283064.3
B3 64615444 64615478 refGene_gn=KNL1;refGene_tx=XM_011283064.3
B3 64616428 64616478 refGene_gn=KNL1;refGene_tx=XM_011283064.3
B3 64628060 64628122 refGene_gn=KNL1;refGene_tx=XM_011283064.3
B3 64632842 64632902 refGene_gn=KNL1;refGene_tx=XM_011283064.3
B3 64633807 64633847 refGene_gn=KNL1;refGene_tx=XM_011283064.3
B3 64634945 64634980 refGene_gn=KNL1;refGene_tx=XM_011283064.3
B3 64659331 64660066 refGene_gn=RPUSD2;refGene_tx=XM_003987340.4
B3 64661247 64661544 refGene_gn=RPUSD2;refGene_tx=XM_003987340.4
B3 64663280 64663907 refGene_gn=RPUSD2;refGene_tx=XM_003987340.4
B3 64672016 64672112 refGene_gn=CCDC32;refGene_tx=XM_019832670.2
B3 64673958 64674214
refGene_gn=CCDC32;refGene_tx=XM_006932643.4;refGene_gn=CCDC32;refGene_tx=XM_006932644.4;refGene_gn=CCDC32;refGene_tx=XM_019832670.2
B3 64678884 64679041

refGene_gn=CCDC32;refGene_tx=XM_006932643.4;refGene_gn=CCDC32;refGene_tx=XM_006932644.4;refGene_gn=CCDC32;refGene_tx=XM_019832670.2
B3 64682021 64682160
refGene_gn=CCDC32;refGene_tx=XM_006932643.4;refGene_gn=CCDC32;refGene_tx=XM_006932644.4;refGene_gn=CCDC32;refGene_tx=XM_019832670.2
B3 64756206 64757337 refGene_gn=CHST14;refGene_tx=XM_003987275.4
B3 64762062 64762252
refGene_gn=BAHD1;refGene_tx=XM_006932647.4;refGene_gn=BAHD1;refGene_tx=XM_019832672.2;refGene_gn=BAHD1;refGene_tx=XM_023255487.1;refGene_gn=BAHD1;refGene_tx=XM_023255488.1;refGene_gn=BAHD1;refGene_tx=XM_023255489.1;refGene_gn=BAHD1;refGene_tx=XM_023255490.1;refGene_gn=BAHD1;refGene_tx=XM_023255491.1;refGene_gn=BAHD1;refGene_tx=XM_023255492.1
B3 64762725 64762826
refGene_gn=BAHD1;refGene_tx=XM_006932647.4;refGene_gn=BAHD1;refGene_tx=XM_019832672.2;refGene_gn=BAHD1;refGene_tx=XM_023255487.1;refGene_gn=BAHD1;refGene_tx=XM_023255488.1;refGene_gn=BAHD1;refGene_tx=XM_023255489.1;refGene_gn=BAHD1;refGene_tx=XM_023255490.1;refGene_gn=BAHD1;refGene_tx=XM_023255491.1;refGene_gn=BAHD1;refGene_tx=XM_023255492.1
B3 64763612 64763689
refGene_gn=BAHD1;refGene_tx=XM_006932647.4;refGene_gn=BAHD1;refGene_tx=XM_019832672.2;refGene_gn=BAHD1;refGene_tx=XM_023255487.1;refGene_gn=BAHD1;refGene_tx=XM_023255488.1;refGene_gn=BAHD1;refGene_tx=XM_023255489.1;refGene_gn=BAHD1;refGene_tx=XM_023255490.1;refGene_gn=BAHD1;refGene_tx=XM_023255491.1;refGene_gn=BAHD1;refGene_tx=XM_023255492.1
B3 64764108 64764268
refGene_gn=BAHD1;refGene_tx=XM_006932647.4;refGene_gn=BAHD1;refGene_tx=XM_019832672.2;refGene_gn=BAHD1;refGene_tx=XM_023255487.1;refGene_gn=BAHD1;refGene_tx=XM_023255488.1;refGene_gn=BAHD1;refGene_tx=XM_023255489.1;refGene_gn=BAHD1;refGene_tx=XM_023255490.1;refGene_gn=BAHD1;refGene_tx=XM_023255491.1;refGene_gn=BAHD1;refGene_tx=XM_023255492.1
B3 64765237 64765620
refGene_gn=BAHD1;refGene_tx=XM_006932647.4;refGene_gn=BAHD1;refGene_tx=XM_019832672.2;refGene_gn=BAHD1;refGene_tx=XM_023255487.1;refGene_gn=BAHD1;refGene_tx=XM_023255488.1;refGene_gn=BAHD1;refGene_tx=XM_023255489.1;refGene_gn=BAHD1;refGene_tx=XM_023255490.1;refGene_gn=BAHD1;refGene_tx=XM_023255491.1;refGene_gn=BAHD1;refGene_tx=XM_023255492.1
B3 64767437 64768877
refGene_gn=BAHD1;refGene_tx=XM_006932647.4;refGene_gn=BAHD1;refGene_tx=XM_019832672.2;refGene_gn=BAHD1;refGene_tx=XM_023255487.1;refGene_gn=BAHD1;refGene_tx=XM_023255488.1;refGene_gn=BAHD1;refGene_tx=XM_023255489.1;refGene_gn=BAHD1;refGene_tx=XM_023255490.1;refGene_gn=BAHD1;refGene_tx=XM_023255491.1;refGene_gn=BAHD1;refGene_tx=XM_023255492.1
B3 64784542 64784573 refGene_gn=BAHD1;refGene_tx=XM_006932647.4
B3 64784757 64784953 refGene_gn=BAHD1;refGene_tx=XM_023255487.1
B3 64804966 64805100 refGene_gn=IVD;refGene_tx=XM_003987277.4
B3 64805799 64805872 refGene_gn=IVD;refGene_tx=XM_003987277.4
B3 64805947 64806052 refGene_gn=IVD;refGene_tx=XM_003987277.4
B3 64806598 64806680 refGene_gn=IVD;refGene_tx=XM_003987277.4
B3 64807221 64807315 refGene_gn=IVD;refGene_tx=XM_003987277.4
B3 64808973 64809070 refGene_gn=IVD;refGene_tx=XM_003987277.4

B3 64810321 64810458 refGene_gn=IVD;refGene_tx=XM_003987277.4
B3 64810652 64810746 refGene_gn=IVD;refGene_tx=XM_003987277.4
B3 64811196 64811366 refGene_gn=IVD;refGene_tx=XM_003987277.4
B3 64813420 64813472 refGene_gn=IVD;refGene_tx=XM_003987277.4
B3 64813729 64813819 refGene_gn=IVD;refGene_tx=XM_003987277.4
B3 64815223 64815376 refGene_gn=IVD;refGene_tx=XM_003987277.4
B3 64823275 64823407
refGene_gn=KNSTRN;refGene_tx=XM_019832674.2;refGene_gn=KNSTRN;refGene_tx=XM_019832675.2;refGene_gn=KNSTRN;refGene_tx=XM_019832676.2
B3 64824131 64824206
refGene_gn=KNSTRN;refGene_tx=XM_019832674.2;refGene_gn=KNSTRN;refGene_tx=XM_019832675.2;refGene_gn=KNSTRN;refGene_tx=XM_019832676.2
B3 64824985 64825047
refGene_gn=KNSTRN;refGene_tx=XM_019832674.2;refGene_gn=KNSTRN;refGene_tx=XM_019832675.2;refGene_gn=KNSTRN;refGene_tx=XM_019832676.2
B3 64825231 64825303
refGene_gn=KNSTRN;refGene_tx=XM_019832674.2;refGene_gn=KNSTRN;refGene_tx=XM_019832675.2;refGene_gn=KNSTRN;refGene_tx=XM_019832676.2
B3 64828043 64828137
refGene_gn=KNSTRN;refGene_tx=XM_019832674.2;refGene_gn=KNSTRN;refGene_tx=XM_019832675.2;refGene_gn=KNSTRN;refGene_tx=XM_019832676.2
B3 64828354 64828460
refGene_gn=KNSTRN;refGene_tx=XM_019832674.2;refGene_gn=KNSTRN;refGene_tx=XM_019832675.2;refGene_gn=KNSTRN;refGene_tx=XM_019832676.2
B3 64829859 64829907
refGene_gn=KNSTRN;refGene_tx=XM_019832674.2;refGene_gn=KNSTRN;refGene_tx=XM_019832675.2;refGene_gn=KNSTRN;refGene_tx=XM_019832676.2
B3 64830649 64830779
refGene_gn=KNSTRN;refGene_tx=XM_019832674.2;refGene_gn=KNSTRN;refGene_tx=XM_019832675.2;refGene_gn=KNSTRN;refGene_tx=XM_019832676.2
B3 64833876 64833971
refGene_gn=KNSTRN;refGene_tx=XM_019832674.2;refGene_gn=KNSTRN;refGene_tx=XM_019832675.2;refGene_gn=KNSTRN;refGene_tx=XM_019832676.2
B3 64834186 64834551 refGene_gn=KNSTRN;refGene_tx=XM_019832674.2
B3 64844327 64847203 refGene_gn=DISP2;refGene_tx=XM_003987278.5
B3 64847205 64847587 refGene_gn=DISP2;refGene_tx=XM_003987278.5
B3 64849120 64849218 refGene_gn=DISP2;refGene_tx=XM_003987278.5
B3 64849578 64849706 refGene_gn=DISP2;refGene_tx=XM_003987278.5
B3 64849884 64850000 refGene_gn=DISP2;refGene_tx=XM_003987278.5
B3 64850263 64850387 refGene_gn=DISP2;refGene_tx=XM_003987278.5
B3 64850535 64850565 refGene_gn=DISP2;refGene_tx=XM_003987278.5
B3 64850834 64851164 refGene_gn=DISP2;refGene_tx=XM_003987278.5
B3 64856392 64856505 refGene_gn=DISP2;refGene_tx=XM_003987278.5
B3 64888883 64889342
refGene_gn=INAFM2;refGene_tx=XM_023255493.1;refGene_gn=INAFM2;refGene_tx=XM_023255494.1
B3 64903090 64903174
refGene_gn=PLCB2;refGene_tx=XM_023255495.1;refGene_gn=PLCB2;refGene_tx=XM_023255496.1;refGene_gn=PLCB2;refGene_tx=XM_023255497.1
B3 64906235 64906313

refGene_gn=PLCB2;refGene_tx=XM_023255495.1;refGene_gn=PLCB2;refGene_tx=XM_023255496.1;refGene_gn=PLCB2;refGene_tx=XM_023255497.1
B3 64906970 64907039
refGene_gn=PLCB2;refGene_tx=XM_023255495.1;refGene_gn=PLCB2;refGene_tx=XM_023255496.1;refGene_gn=PLCB2;refGene_tx=XM_023255497.1
B3 64907674 64907815
refGene_gn=PLCB2;refGene_tx=XM_023255495.1;refGene_gn=PLCB2;refGene_tx=XM_023255496.1;refGene_gn=PLCB2;refGene_tx=XM_023255497.1
B3 64907952 64908032
refGene_gn=PLCB2;refGene_tx=XM_023255495.1;refGene_gn=PLCB2;refGene_tx=XM_023255496.1;refGene_gn=PLCB2;refGene_tx=XM_023255497.1
B3 64908117 64908171
refGene_gn=PLCB2;refGene_tx=XM_023255495.1;refGene_gn=PLCB2;refGene_tx=XM_023255496.1;refGene_gn=PLCB2;refGene_tx=XM_023255497.1
B3 64908277 64908353
refGene_gn=PLCB2;refGene_tx=XM_023255495.1;refGene_gn=PLCB2;refGene_tx=XM_023255496.1;refGene_gn=PLCB2;refGene_tx=XM_023255497.1
B3 64910815 64910916
refGene_gn=PLCB2;refGene_tx=XM_023255495.1;refGene_gn=PLCB2;refGene_tx=XM_023255496.1;refGene_gn=PLCB2;refGene_tx=XM_023255497.1
B3 64911060 64911227
refGene_gn=PLCB2;refGene_tx=XM_023255495.1;refGene_gn=PLCB2;refGene_tx=XM_023255496.1;refGene_gn=PLCB2;refGene_tx=XM_023255497.1
B3 64911316 64911463
refGene_gn=PLCB2;refGene_tx=XM_023255495.1;refGene_gn=PLCB2;refGene_tx=XM_023255496.1;refGene_gn=PLCB2;refGene_tx=XM_023255497.1
B3 64911642 64911800
refGene_gn=PLCB2;refGene_tx=XM_023255495.1;refGene_gn=PLCB2;refGene_tx=XM_023255496.1;refGene_gn=PLCB2;refGene_tx=XM_023255497.1
B3 64912065 64912148
refGene_gn=PLCB2;refGene_tx=XM_023255495.1;refGene_gn=PLCB2;refGene_tx=XM_023255496.1;refGene_gn=PLCB2;refGene_tx=XM_023255497.1
B3 64912410 64912495
refGene_gn=PLCB2;refGene_tx=XM_023255495.1;refGene_gn=PLCB2;refGene_tx=XM_023255496.1;refGene_gn=PLCB2;refGene_tx=XM_023255497.1
B3 64913071 64913234
refGene_gn=PLCB2;refGene_tx=XM_023255495.1;refGene_gn=PLCB2;refGene_tx=XM_023255496.1;refGene_gn=PLCB2;refGene_tx=XM_023255497.1
B3 64913356 64913463
refGene_gn=PLCB2;refGene_tx=XM_023255495.1;refGene_gn=PLCB2;refGene_tx=XM_023255496.1;refGene_gn=PLCB2;refGene_tx=XM_023255497.1
B3 64913592 64913689
refGene_gn=PLCB2;refGene_tx=XM_023255495.1;refGene_gn=PLCB2;refGene_tx=XM_023255496.1;refGene_gn=PLCB2;refGene_tx=XM_023255497.1
B3 64914757 64914842
refGene_gn=PLCB2;refGene_tx=XM_023255495.1;refGene_gn=PLCB2;refGene_tx=XM_023255496.1;refGene_gn=PLCB2;refGene_tx=XM_023255497.1
B3 64914971 64915096
refGene_gn=PLCB2;refGene_tx=XM_023255495.1;refGene_gn=PLCB2;refGene_tx=XM_023255496.1;refGene_gn=PLCB2;refGene_tx=XM_023255497.1

B3 64915642 64915797
refGene_gn=PLCB2;refGene_tx=XM_023255495.1;refGene_gn=PLCB2;refGene_tx=XM_023255496.1;refGene_gn=PLCB2;refGene_tx=XM_023255497.1
B3 64916351 64916516
refGene_gn=PLCB2;refGene_tx=XM_023255495.1;refGene_gn=PLCB2;refGene_tx=XM_023255496.1;refGene_gn=PLCB2;refGene_tx=XM_023255497.1
B3 64917094 64917194
refGene_gn=PLCB2;refGene_tx=XM_023255495.1;refGene_gn=PLCB2;refGene_tx=XM_023255496.1;refGene_gn=PLCB2;refGene_tx=XM_023255497.1
B3 64917719 64917824
refGene_gn=PLCB2;refGene_tx=XM_023255495.1;refGene_gn=PLCB2;refGene_tx=XM_023255496.1;refGene_gn=PLCB2;refGene_tx=XM_023255497.1
B3 64917991 64918086
refGene_gn=PLCB2;refGene_tx=XM_023255495.1;refGene_gn=PLCB2;refGene_tx=XM_023255496.1;refGene_gn=PLCB2;refGene_tx=XM_023255497.1
B3 64918209 64918285
refGene_gn=PLCB2;refGene_tx=XM_023255495.1;refGene_gn=PLCB2;refGene_tx=XM_023255496.1;refGene_gn=PLCB2;refGene_tx=XM_023255497.1
B3 64918498 64918570
refGene_gn=PLCB2;refGene_tx=XM_023255495.1;refGene_gn=PLCB2;refGene_tx=XM_023255496.1;refGene_gn=PLCB2;refGene_tx=XM_023255497.1
B3 64918685 64918908
refGene_gn=PLCB2;refGene_tx=XM_023255495.1;refGene_gn=PLCB2;refGene_tx=XM_023255496.1;refGene_gn=PLCB2;refGene_tx=XM_023255497.1
B3 64918989 64919164
refGene_gn=PLCB2;refGene_tx=XM_023255495.1;refGene_gn=PLCB2;refGene_tx=XM_023255496.1;refGene_gn=PLCB2;refGene_tx=XM_023255497.1
B3 64919300 64919377
refGene_gn=PLCB2;refGene_tx=XM_023255495.1;refGene_gn=PLCB2;refGene_tx=XM_023255496.1;refGene_gn=PLCB2;refGene_tx=XM_023255497.1
B3 64919467 64919563
refGene_gn=PLCB2;refGene_tx=XM_023255495.1;refGene_gn=PLCB2;refGene_tx=XM_023255496.1;refGene_gn=PLCB2;refGene_tx=XM_023255497.1
B3 64920074 64920132
refGene_gn=PLCB2;refGene_tx=XM_023255495.1;refGene_gn=PLCB2;refGene_tx=XM_023255496.1;refGene_gn=PLCB2;refGene_tx=XM_023255497.1
B3 64920794 64920881
refGene_gn=PLCB2;refGene_tx=XM_023255495.1;refGene_gn=PLCB2;refGene_tx=XM_023255496.1;refGene_gn=PLCB2;refGene_tx=XM_023255497.1
B3 64921226 64921430
refGene_gn=PLCB2;refGene_tx=XM_023255495.1;refGene_gn=PLCB2;refGene_tx=XM_023255496.1;refGene_gn=PLCB2;refGene_tx=XM_023255497.1
B3 64927462 64928611 refGene_gn=ANKRD63;refGene_tx=XM_011283066.3
B3 64934039 64934207
refGene_gn=LOC101091201;refGene_tx=XM_003987280.5;refGene_gn=LOC101091201;refGene_tx=XM_019832679.2;refGene_gn=LOC101091201;refGene_tx=XM_019832680.2;refGene_gn=LOC101091201;refGene_tx=XM_019832681.2;refGene_gn=LOC101091201;refGene_tx=XM_019832682.2;refGene_gn=LOC101091201;refGene_tx=XM_019832684.2;refGene_gn=LOC101091201;refGene_tx=XM_023255498.1;refGene_gn=LOC101091201;refGene_tx=XM_023255499.1

B3 64935579 64935714
refGene_gn=LOC101091201;refGene_tx=XM_003987280.5;refGene_gn=LOC101091201;refGene_tx=XM_019832679.2;refGene_gn=LOC101091201;refGene_tx=XM_019832680.2;refGene_gn=LOC101091201;refGene_tx=XM_019832681.2;refGene_gn=LOC101091201;refGene_tx=XM_019832682.2;refGene_gn=LOC101091201;refGene_tx=XM_019832684.2;refGene_gn=LOC101091201;refGene_tx=XM_023255498.1;refGene_gn=LOC101091201;refGene_tx=XM_023255499.1
B3 64936181 64936307
refGene_gn=LOC101091201;refGene_tx=XM_003987280.5;refGene_gn=LOC101091201;refGene_tx=XM_019832679.2;refGene_gn=LOC101091201;refGene_tx=XM_019832680.2;refGene_gn=LOC101091201;refGene_tx=XM_019832681.2;refGene_gn=LOC101091201;refGene_tx=XM_019832682.2;refGene_gn=LOC101091201;refGene_tx=XM_019832684.2;refGene_gn=LOC101091201;refGene_tx=XM_023255498.1;refGene_gn=LOC101091201;refGene_tx=XM_023255499.1
B3 64936419 64936546
refGene_gn=LOC101091201;refGene_tx=XM_003987280.5;refGene_gn=LOC101091201;refGene_tx=XM_019832679.2;refGene_gn=LOC101091201;refGene_tx=XM_019832680.2;refGene_gn=LOC101091201;refGene_tx=XM_019832681.2;refGene_gn=LOC101091201;refGene_tx=XM_019832682.2;refGene_gn=LOC101091201;refGene_tx=XM_019832684.2;refGene_gn=LOC101091201;refGene_tx=XM_023255498.1;refGene_gn=LOC101091201;refGene_tx=XM_023255499.1
B3 64936903 64937037
refGene_gn=LOC101091201;refGene_tx=XM_003987280.5;refGene_gn=LOC101091201;refGene_tx=XM_019832679.2;refGene_gn=LOC101091201;refGene_tx=XM_019832680.2;refGene_gn=LOC101091201;refGene_tx=XM_019832681.2;refGene_gn=LOC101091201;refGene_tx=XM_019832682.2;refGene_gn=LOC101091201;refGene_tx=XM_019832684.2;refGene_gn=LOC101091201;refGene_tx=XM_023255498.1;refGene_gn=LOC101091201;refGene_tx=XM_023255499.1
B3 64937145 64937643
refGene_gn=LOC101091201;refGene_tx=XM_003987280.5;refGene_gn=LOC101091201;refGene_tx=XM_019832679.2;refGene_gn=LOC101091201;refGene_tx=XM_019832680.2;refGene_gn=LOC101091201;refGene_tx=XM_019832681.2;refGene_gn=LOC101091201;refGene_tx=XM_019832682.2;refGene_gn=LOC101091201;refGene_tx=XM_019832684.2;refGene_gn=LOC101091201;refGene_tx=XM_023255498.1;refGene_gn=LOC101091201;refGene_tx=XM_023255499.1
B3 64943135 64943789
refGene_gn=LOC101091201;refGene_tx=XM_003987280.5;refGene_gn=LOC101091201;refGene_tx=XM_019832679.2;refGene_gn=LOC101091201;refGene_tx=XM_019832680.2;refGene_gn=LOC101091201;refGene_tx=XM_019832681.2;refGene_gn=LOC101091201;refGene_tx=XM_019832682.2;refGene_gn=LOC101091201;refGene_tx=XM_019832684.2;refGene_gn=LOC101091201;refGene_tx=XM_023255498.1;refGene_gn=LOC101091201;refGene_tx=XM_023255499.1
B3 64944664 64944916
refGene_gn=LOC101091201;refGene_tx=XM_003987280.5;refGene_gn=LOC101091201;refGene_tx=XM_019832679.2;refGene_gn=LOC101091201;refGene_tx=XM_019832680.2;refGene_gn=LOC101091201;refGene_tx=XM_019832681.2;refGene_gn=LOC101091201;refGene_tx=XM_019832682.2;refGene_gn=LOC101091201;refGene_tx=XM_019832684.2;refGene_gn=LOC101091201;refGene_tx=XM_023255498.1;refGene_gn=LOC101091201;refGene_tx=XM_023255499.1
B3 64956723 64957436

refGene_gn=LOC109500712;refGene_tx=XM_023255501.1
B3 64958846 64958934
refGene_gn=LOC109500712;refGene_tx=XM_023255501.1
B3 65001526 65001602 refGene_gn=BUB1B;refGene_tx=XM_023255500.1
B3 65002155 65002271 refGene_gn=BUB1B;refGene_tx=XM_023255500.1
B3 65002797 65002910 refGene_gn=BUB1B;refGene_tx=XM_023255500.1
B3 65004478 65004708 refGene_gn=BUB1B;refGene_tx=XM_023255500.1
B3 65008210 65008302 refGene_gn=BUB1B;refGene_tx=XM_023255500.1
B3 65008474 65008689 refGene_gn=BUB1B;refGene_tx=XM_023255500.1
B3 65009803 65009973 refGene_gn=BUB1B;refGene_tx=XM_023255500.1
B3 65017596 65017793 refGene_gn=BUB1B;refGene_tx=XM_023255500.1
B3 65022675 65022820 refGene_gn=BUB1B;refGene_tx=XM_023255500.1
B3 65023246 65023306 refGene_gn=BUB1B;refGene_tx=XM_023255500.1
B3 65030003 65030150 refGene_gn=BUB1B;refGene_tx=XM_023255500.1
B3 65032704 65032739 refGene_gn=BUB1B;refGene_tx=XM_023255500.1
B3 65068703 65068768 refGene_gn=BMF;refGene_tx=XM_006932651.3
B3 65068769 65068825 refGene_gn=BMF;refGene_tx=XM_006932651.3
B3 65069210 65069249 refGene_gn=BMF;refGene_tx=XM_006932651.3
B3 65069251 65069338 refGene_gn=BMF;refGene_tx=XM_006932651.3
B3 65071501 65071798 refGene_gn=BMF;refGene_tx=XM_006932651.3
B3 65073247 65073408 refGene_gn=BMF;refGene_tx=XM_006932651.3
B3 65085441 65085543 refGene_gn=BMF;refGene_tx=XM_006932651.3
B3 65136326 65136350 refGene_gn=SRP14;refGene_tx=XM_003987282.5
B3 65136490 65136563 refGene_gn=SRP14;refGene_tx=XM_003987282.5
B3 65137080 65137193 refGene_gn=SRP14;refGene_tx=XM_003987282.5
B3 65138574 65138607 refGene_gn=SRP14;refGene_tx=XM_003987282.5
B3 65139133 65139223 refGene_gn=SRP14;refGene_tx=XM_003987282.5
B3 65145389 65145447
refGene_gn=EIF2AK4;refGene_tx=XM_023255502.1;refGene_gn=EIF2AK4;refGen
e_tx=XM_023255503.1;refGene_gn=EIF2AK4;refGene_tx=XM_023255504.1;refGe
ne_gn=EIF2AK4;refGene_tx=XM_023255505.1;refGene_gn=EIF2AK4;refGene_tx=
XM_023255506.1;refGene_gn=EIF2AK4;refGene_tx=XM_023255507.1
B3 65145955 65146074
refGene_gn=EIF2AK4;refGene_tx=XM_023255502.1;refGene_gn=EIF2AK4;refGen
e_tx=XM_023255503.1;refGene_gn=EIF2AK4;refGene_tx=XM_023255504.1;refGe
ne_gn=EIF2AK4;refGene_tx=XM_023255505.1;refGene_gn=EIF2AK4;refGene_tx=
XM_023255506.1;refGene_gn=EIF2AK4;refGene_tx=XM_023255507.1
B3 65148843 65148888
refGene_gn=EIF2AK4;refGene_tx=XM_023255502.1;refGene_gn=EIF2AK4;refGen
e_tx=XM_023255503.1;refGene_gn=EIF2AK4;refGene_tx=XM_023255504.1;refGe
ne_gn=EIF2AK4;refGene_tx=XM_023255505.1;refGene_gn=EIF2AK4;refGene_tx=
XM_023255506.1;refGene_gn=EIF2AK4;refGene_tx=XM_023255507.1
B3 65149411 65149480
refGene_gn=EIF2AK4;refGene_tx=XM_023255502.1;refGene_gn=EIF2AK4;refGen
e_tx=XM_023255503.1;refGene_gn=EIF2AK4;refGene_tx=XM_023255504.1;refGe
ne_gn=EIF2AK4;refGene_tx=XM_023255505.1;refGene_gn=EIF2AK4;refGene_tx=
XM_023255506.1;refGene_gn=EIF2AK4;refGene_tx=XM_023255507.1
B3 65150675 65150773
refGene_gn=EIF2AK4;refGene_tx=XM_023255502.1;refGene_gn=EIF2AK4;refGen
e_tx=XM_023255503.1;refGene_gn=EIF2AK4;refGene_tx=XM_023255504.1;refGe

ne_gn=EIF2AK4;refGene_tx=XM_023255505.1;refGene_gn=EIF2AK4;refGene_tx=XM_023255506.1;refGene_gn=EIF2AK4;refGene_tx=XM_023255507.1
B3 65151663 65151722
refGene_gn=EIF2AK4;refGene_tx=XM_023255502.1;refGene_gn=EIF2AK4;refGene_tx=XM_023255503.1;refGene_gn=EIF2AK4;refGene_tx=XM_023255504.1;refGene_gn=EIF2AK4;refGene_tx=XM_023255505.1;refGene_gn=EIF2AK4;refGene_tx=XM_023255506.1;refGene_gn=EIF2AK4;refGene_tx=XM_023255507.1;refGene_gn=EIF2AK4;refGene_tx=XM_023255508.1
B3 65154169 65154282
refGene_gn=EIF2AK4;refGene_tx=XM_023255502.1;refGene_gn=EIF2AK4;refGene_tx=XM_023255503.1;refGene_gn=EIF2AK4;refGene_tx=XM_023255504.1;refGene_gn=EIF2AK4;refGene_tx=XM_023255505.1;refGene_gn=EIF2AK4;refGene_tx=XM_023255506.1;refGene_gn=EIF2AK4;refGene_tx=XM_023255507.1
B3 65161109 65161196
refGene_gn=EIF2AK4;refGene_tx=XM_023255502.1;refGene_gn=EIF2AK4;refGene_tx=XM_023255503.1;refGene_gn=EIF2AK4;refGene_tx=XM_023255504.1;refGene_gn=EIF2AK4;refGene_tx=XM_023255505.1;refGene_gn=EIF2AK4;refGene_tx=XM_023255506.1;refGene_gn=EIF2AK4;refGene_tx=XM_023255507.1;refGene_gn=EIF2AK4;refGene_tx=XM_023255508.1
B3 65162690 65162819
refGene_gn=EIF2AK4;refGene_tx=XM_023255502.1;refGene_gn=EIF2AK4;refGene_tx=XM_023255503.1;refGene_gn=EIF2AK4;refGene_tx=XM_023255504.1;refGene_gn=EIF2AK4;refGene_tx=XM_023255505.1;refGene_gn=EIF2AK4;refGene_tx=XM_023255506.1;refGene_gn=EIF2AK4;refGene_tx=XM_023255507.1;refGene_gn=EIF2AK4;refGene_tx=XM_023255508.1
B3 65163433 65163541
refGene_gn=EIF2AK4;refGene_tx=XM_023255502.1;refGene_gn=EIF2AK4;refGene_tx=XM_023255503.1;refGene_gn=EIF2AK4;refGene_tx=XM_023255504.1;refGene_gn=EIF2AK4;refGene_tx=XM_023255505.1;refGene_gn=EIF2AK4;refGene_tx=XM_023255506.1;refGene_gn=EIF2AK4;refGene_tx=XM_023255507.1;refGene_gn=EIF2AK4;refGene_tx=XM_023255508.1
B3 65168368 65168503
refGene_gn=EIF2AK4;refGene_tx=XM_023255502.1;refGene_gn=EIF2AK4;refGene_tx=XM_023255503.1;refGene_gn=EIF2AK4;refGene_tx=XM_023255504.1;refGene_gn=EIF2AK4;refGene_tx=XM_023255505.1;refGene_gn=EIF2AK4;refGene_tx=XM_023255506.1;refGene_gn=EIF2AK4;refGene_tx=XM_023255507.1;refGene_gn=EIF2AK4;refGene_tx=XM_023255508.1
B3 65168848 65169019
refGene_gn=EIF2AK4;refGene_tx=XM_023255502.1;refGene_gn=EIF2AK4;refGene_tx=XM_023255503.1;refGene_gn=EIF2AK4;refGene_tx=XM_023255504.1;refGene_gn=EIF2AK4;refGene_tx=XM_023255505.1;refGene_gn=EIF2AK4;refGene_tx=XM_023255506.1;refGene_gn=EIF2AK4;refGene_tx=XM_023255507.1;refGene_gn=EIF2AK4;refGene_tx=XM_023255508.1
B3 65171497 65171563
refGene_gn=EIF2AK4;refGene_tx=XM_023255502.1;refGene_gn=EIF2AK4;refGene_tx=XM_023255503.1;refGene_gn=EIF2AK4;refGene_tx=XM_023255504.1;refGene_gn=EIF2AK4;refGene_tx=XM_023255505.1;refGene_gn=EIF2AK4;refGene_tx=XM_023255506.1;refGene_gn=EIF2AK4;refGene_tx=XM_023255507.1;refGene_gn=EIF2AK4;refGene_tx=XM_023255508.1
B3 65172970 65173087

refGene_gn=EIF2AK4;refGene_tx=XM_023255502.1;refGene_gn=EIF2AK4;refGene_tx=XM_023255503.1;refGene_gn=EIF2AK4;refGene_tx=XM_023255504.1;refGene_gn=EIF2AK4;refGene_tx=XM_023255505.1;refGene_gn=EIF2AK4;refGene_tx=XM_023255506.1;refGene_gn=EIF2AK4;refGene_tx=XM_023255507.1;refGene_gn=EIF2AK4;refGene_tx=XM_023255508.1
B3 65174560 65174729
refGene_gn=EIF2AK4;refGene_tx=XM_023255502.1;refGene_gn=EIF2AK4;refGene_tx=XM_023255503.1;refGene_gn=EIF2AK4;refGene_tx=XM_023255504.1;refGene_gn=EIF2AK4;refGene_tx=XM_023255505.1;refGene_gn=EIF2AK4;refGene_tx=XM_023255506.1;refGene_gn=EIF2AK4;refGene_tx=XM_023255507.1;refGene_gn=EIF2AK4;refGene_tx=XM_023255508.1
B3 65175677 65175727
refGene_gn=EIF2AK4;refGene_tx=XM_023255502.1;refGene_gn=EIF2AK4;refGene_tx=XM_023255503.1;refGene_gn=EIF2AK4;refGene_tx=XM_023255504.1;refGene_gn=EIF2AK4;refGene_tx=XM_023255505.1;refGene_gn=EIF2AK4;refGene_tx=XM_023255506.1;refGene_gn=EIF2AK4;refGene_tx=XM_023255507.1;refGene_gn=EIF2AK4;refGene_tx=XM_023255508.1
B3 65179085 65179207
refGene_gn=EIF2AK4;refGene_tx=XM_023255502.1;refGene_gn=EIF2AK4;refGene_tx=XM_023255503.1;refGene_gn=EIF2AK4;refGene_tx=XM_023255504.1;refGene_gn=EIF2AK4;refGene_tx=XM_023255505.1;refGene_gn=EIF2AK4;refGene_tx=XM_023255506.1;refGene_gn=EIF2AK4;refGene_tx=XM_023255507.1;refGene_gn=EIF2AK4;refGene_tx=XM_023255508.1
B3 65179705 65179781
refGene_gn=EIF2AK4;refGene_tx=XM_023255502.1;refGene_gn=EIF2AK4;refGene_tx=XM_023255503.1;refGene_gn=EIF2AK4;refGene_tx=XM_023255504.1;refGene_gn=EIF2AK4;refGene_tx=XM_023255505.1;refGene_gn=EIF2AK4;refGene_tx=XM_023255506.1;refGene_gn=EIF2AK4;refGene_tx=XM_023255507.1;refGene_gn=EIF2AK4;refGene_tx=XM_023255508.1
B3 65181757 65181994
refGene_gn=EIF2AK4;refGene_tx=XM_023255502.1;refGene_gn=EIF2AK4;refGene_tx=XM_023255503.1;refGene_gn=EIF2AK4;refGene_tx=XM_023255504.1;refGene_gn=EIF2AK4;refGene_tx=XM_023255505.1;refGene_gn=EIF2AK4;refGene_tx=XM_023255506.1;refGene_gn=EIF2AK4;refGene_tx=XM_023255507.1;refGene_gn=EIF2AK4;refGene_tx=XM_023255508.1
B3 65184496 65184550
refGene_gn=EIF2AK4;refGene_tx=XM_023255502.1;refGene_gn=EIF2AK4;refGene_tx=XM_023255504.1;refGene_gn=EIF2AK4;refGene_tx=XM_023255505.1;refGene_gn=EIF2AK4;refGene_tx=XM_023255506.1;refGene_gn=EIF2AK4;refGene_tx=XM_023255507.1;refGene_gn=EIF2AK4;refGene_tx=XM_023255508.1
B3 65186199 65186301
refGene_gn=EIF2AK4;refGene_tx=XM_023255502.1;refGene_gn=EIF2AK4;refGene_tx=XM_023255503.1;refGene_gn=EIF2AK4;refGene_tx=XM_023255504.1;refGene_gn=EIF2AK4;refGene_tx=XM_023255505.1;refGene_gn=EIF2AK4;refGene_tx=XM_023255506.1;refGene_gn=EIF2AK4;refGene_tx=XM_023255507.1;refGene_gn=EIF2AK4;refGene_tx=XM_023255508.1
B3 65189431 65189511
refGene_gn=EIF2AK4;refGene_tx=XM_023255502.1;refGene_gn=EIF2AK4;refGene_tx=XM_023255503.1;refGene_gn=EIF2AK4;refGene_tx=XM_023255504.1;refGene_gn=EIF2AK4;refGene_tx=XM_023255505.1;refGene_gn=EIF2AK4;refGene_tx=

XM_023255506.1;refGene_gn=EIF2AK4;refGene_tx=XM_023255507.1;refGene_gn
=EIF2AK4;refGene_tx=XM_023255508.1
B3 65191229 65191284
refGene_gn=EIF2AK4;refGene_tx=XM_023255502.1;refGene_gn=EIF2AK4;refGen
e_tx=XM_023255503.1;refGene_gn=EIF2AK4;refGene_tx=XM_023255504.1;refGe
ne_gn=EIF2AK4;refGene_tx=XM_023255505.1;refGene_gn=EIF2AK4;refGene_tx=
XM_023255506.1;refGene_gn=EIF2AK4;refGene_tx=XM_023255507.1;refGene_gn
=EIF2AK4;refGene_tx=XM_023255508.1
B3 65193947 65194052
refGene_gn=EIF2AK4;refGene_tx=XM_023255502.1;refGene_gn=EIF2AK4;refGen
e_tx=XM_023255503.1;refGene_gn=EIF2AK4;refGene_tx=XM_023255504.1;refGe
ne_gn=EIF2AK4;refGene_tx=XM_023255505.1;refGene_gn=EIF2AK4;refGene_tx=
XM_023255506.1;refGene_gn=EIF2AK4;refGene_tx=XM_023255507.1;refGene_gn
=EIF2AK4;refGene_tx=XM_023255508.1
B3 65196518 65196641
refGene_gn=EIF2AK4;refGene_tx=XM_023255502.1;refGene_gn=EIF2AK4;refGen
e_tx=XM_023255503.1;refGene_gn=EIF2AK4;refGene_tx=XM_023255504.1;refGe
ne_gn=EIF2AK4;refGene_tx=XM_023255505.1;refGene_gn=EIF2AK4;refGene_tx=
XM_023255506.1;refGene_gn=EIF2AK4;refGene_tx=XM_023255507.1;refGene_gn
=EIF2AK4;refGene_tx=XM_023255508.1
B3 65198534 65198618
refGene_gn=EIF2AK4;refGene_tx=XM_023255502.1;refGene_gn=EIF2AK4;refGen
e_tx=XM_023255503.1;refGene_gn=EIF2AK4;refGene_tx=XM_023255504.1;refGe
ne_gn=EIF2AK4;refGene_tx=XM_023255505.1;refGene_gn=EIF2AK4;refGene_tx=
XM_023255506.1;refGene_gn=EIF2AK4;refGene_tx=XM_023255507.1;refGene_gn
=EIF2AK4;refGene_tx=XM_023255508.1
B3 65200328 65200392
refGene_gn=EIF2AK4;refGene_tx=XM_023255502.1;refGene_gn=EIF2AK4;refGen
e_tx=XM_023255503.1;refGene_gn=EIF2AK4;refGene_tx=XM_023255504.1;refGe
ne_gn=EIF2AK4;refGene_tx=XM_023255505.1;refGene_gn=EIF2AK4;refGene_tx=
XM_023255506.1;refGene_gn=EIF2AK4;refGene_tx=XM_023255507.1;refGene_gn
=EIF2AK4;refGene_tx=XM_023255508.1
B3 65201141 65201593
refGene_gn=EIF2AK4;refGene_tx=XM_023255502.1;refGene_gn=EIF2AK4;refGen
e_tx=XM_023255503.1;refGene_gn=EIF2AK4;refGene_tx=XM_023255504.1;refGe
ne_gn=EIF2AK4;refGene_tx=XM_023255505.1;refGene_gn=EIF2AK4;refGene_tx=
XM_023255506.1;refGene_gn=EIF2AK4;refGene_tx=XM_023255507.1;refGene_gn
=EIF2AK4;refGene_tx=XM_023255508.1
B3 65205079 65205237
refGene_gn=EIF2AK4;refGene_tx=XM_023255502.1;refGene_gn=EIF2AK4;refGen
e_tx=XM_023255503.1;refGene_gn=EIF2AK4;refGene_tx=XM_023255504.1;refGe
ne_gn=EIF2AK4;refGene_tx=XM_023255505.1;refGene_gn=EIF2AK4;refGene_tx=
XM_023255506.1;refGene_gn=EIF2AK4;refGene_tx=XM_023255507.1;refGene_gn
=EIF2AK4;refGene_tx=XM_023255508.1
B3 65205818 65205925
refGene_gn=EIF2AK4;refGene_tx=XM_023255502.1;refGene_gn=EIF2AK4;refGen
e_tx=XM_023255503.1;refGene_gn=EIF2AK4;refGene_tx=XM_023255504.1;refGe
ne_gn=EIF2AK4;refGene_tx=XM_023255505.1;refGene_gn=EIF2AK4;refGene_tx=
XM_023255506.1;refGene_gn=EIF2AK4;refGene_tx=XM_023255507.1;refGene_gn
=EIF2AK4;refGene_tx=XM_023255508.1

B3 65211452 65211988
refGene_gn=EIF2AK4;refGene_tx=XM_023255502.1;refGene_gn=EIF2AK4;refGene_tx=XM_023255503.1;refGene_gn=EIF2AK4;refGene_tx=XM_023255504.1;refGene_gn=EIF2AK4;refGene_tx=XM_023255505.1;refGene_gn=EIF2AK4;refGene_tx=XM_023255506.1;refGene_gn=EIF2AK4;refGene_tx=XM_023255507.1;refGene_gn=EIF2AK4;refGene_tx=XM_023255508.1
B3 65213622 65213780
refGene_gn=EIF2AK4;refGene_tx=XM_023255502.1;refGene_gn=EIF2AK4;refGene_tx=XM_023255503.1;refGene_gn=EIF2AK4;refGene_tx=XM_023255504.1;refGene_gn=EIF2AK4;refGene_tx=XM_023255505.1;refGene_gn=EIF2AK4;refGene_tx=XM_023255506.1;refGene_gn=EIF2AK4;refGene_tx=XM_023255507.1;refGene_gn=EIF2AK4;refGene_tx=XM_023255508.1
B3 65216759 65216875
refGene_gn=EIF2AK4;refGene_tx=XM_023255502.1;refGene_gn=EIF2AK4;refGene_tx=XM_023255503.1;refGene_gn=EIF2AK4;refGene_tx=XM_023255504.1;refGene_gn=EIF2AK4;refGene_tx=XM_023255505.1;refGene_gn=EIF2AK4;refGene_tx=XM_023255506.1;refGene_gn=EIF2AK4;refGene_tx=XM_023255507.1;refGene_gn=EIF2AK4;refGene_tx=XM_023255508.1
B3 65219754 65219903
refGene_gn=EIF2AK4;refGene_tx=XM_023255502.1;refGene_gn=EIF2AK4;refGene_tx=XM_023255503.1;refGene_gn=EIF2AK4;refGene_tx=XM_023255504.1;refGene_gn=EIF2AK4;refGene_tx=XM_023255505.1;refGene_gn=EIF2AK4;refGene_tx=XM_023255506.1;refGene_gn=EIF2AK4;refGene_tx=XM_023255507.1;refGene_gn=EIF2AK4;refGene_tx=XM_023255508.1
B3 65221058 65221136
refGene_gn=EIF2AK4;refGene_tx=XM_023255502.1;refGene_gn=EIF2AK4;refGene_tx=XM_023255503.1;refGene_gn=EIF2AK4;refGene_tx=XM_023255504.1;refGene_gn=EIF2AK4;refGene_tx=XM_023255505.1;refGene_gn=EIF2AK4;refGene_tx=XM_023255506.1;refGene_gn=EIF2AK4;refGene_tx=XM_023255507.1;refGene_gn=EIF2AK4;refGene_tx=XM_023255508.1
B3 65223860 65224013
refGene_gn=EIF2AK4;refGene_tx=XM_023255502.1;refGene_gn=EIF2AK4;refGene_tx=XM_023255503.1;refGene_gn=EIF2AK4;refGene_tx=XM_023255504.1;refGene_gn=EIF2AK4;refGene_tx=XM_023255505.1;refGene_gn=EIF2AK4;refGene_tx=XM_023255506.1;refGene_gn=EIF2AK4;refGene_tx=XM_023255507.1;refGene_gn=EIF2AK4;refGene_tx=XM_023255508.1
B3 65227489 65227592
refGene_gn=EIF2AK4;refGene_tx=XM_023255502.1;refGene_gn=EIF2AK4;refGene_tx=XM_023255503.1;refGene_gn=EIF2AK4;refGene_tx=XM_023255504.1;refGene_gn=EIF2AK4;refGene_tx=XM_023255508.1
B3 65231617 65231730
refGene_gn=EIF2AK4;refGene_tx=XM_023255502.1;refGene_gn=EIF2AK4;refGene_tx=XM_023255503.1;refGene_gn=EIF2AK4;refGene_tx=XM_023255504.1;refGene_gn=EIF2AK4;refGene_tx=XM_023255505.1;refGene_gn=EIF2AK4;refGene_tx=XM_023255506.1;refGene_gn=EIF2AK4;refGene_tx=XM_023255507.1;refGene_gn=EIF2AK4;refGene_tx=XM_023255508.1
B3 65234472 65234499 refGene_gn=EIF2AK4;refGene_tx=XM_023255504.1
B3 65243265 65243409
refGene_gn=EIF2AK4;refGene_tx=XM_023255502.1;refGene_gn=EIF2AK4;refGene_tx=XM_023255503.1;refGene_gn=EIF2AK4;refGene_tx=XM_023255508.1

B3 65267399 65267571 refGene_gn=GPR176;refGene_tx=XM_003987283.5
B3 65386531 65386784 refGene_gn=GPR176;refGene_tx=XM_003987283.5
B3 65392451 65393574 refGene_gn=GPR176;refGene_tx=XM_003987283.5
B3 65413090 65413216 refGene_gn=FSIP1;refGene_tx=XM_006932677.4
B3 65420456 65420640 refGene_gn=FSIP1;refGene_tx=XM_006932677.4
B3 65425093 65425248 refGene_gn=FSIP1;refGene_tx=XM_006932677.4
B3 65426602 65426696 refGene_gn=FSIP1;refGene_tx=XM_006932677.4
B3 65452350 65452449 refGene_gn=FSIP1;refGene_tx=XM_006932677.4
B3 65454764 65454889 refGene_gn=FSIP1;refGene_tx=XM_006932677.4
B3 65457611 65457722 refGene_gn=FSIP1;refGene_tx=XM_006932677.4
B3 65464458 65464617 refGene_gn=FSIP1;refGene_tx=XM_006932677.4
B3 65480754 65480895 refGene_gn=FSIP1;refGene_tx=XM_006932677.4
B3 65598938 65599449 refGene_gn=FSIP1;refGene_tx=XM_006932677.4
B3 65608619 65608666 refGene_gn=FSIP1;refGene_tx=XM_006932677.4
B3 65614882 65614890 refGene_gn=THBS1;refGene_tx=XM_011283077.3
B3 65615725 65615865 refGene_gn=THBS1;refGene_tx=XM_011283077.3
B3 65615969 65616067 refGene_gn=THBS1;refGene_tx=XM_011283077.3
B3 65616500 65616772 refGene_gn=THBS1;refGene_tx=XM_011283077.3
B3 65616942 65617170 refGene_gn=THBS1;refGene_tx=XM_011283077.3
B3 65617351 65617586 refGene_gn=THBS1;refGene_tx=XM_011283077.3
B3 65618790 65618909 refGene_gn=THBS1;refGene_tx=XM_011283077.3
B3 65619022 65619182 refGene_gn=THBS1;refGene_tx=XM_011283077.3
B3 65619697 65619805 refGene_gn=THBS1;refGene_tx=XM_011283077.3
B3 65620258 65620477 refGene_gn=THBS1;refGene_tx=XM_011283077.3
B3 65620850 65621003 refGene_gn=THBS1;refGene_tx=XM_011283077.3
B3 65621114 65621242 refGene_gn=THBS1;refGene_tx=XM_011283077.3
B3 65621496 65621670 refGene_gn=THBS1;refGene_tx=XM_011283077.3
B3 65621968 65622145 refGene_gn=THBS1;refGene_tx=XM_011283077.3
B3 65622440 65622614 refGene_gn=THBS1;refGene_tx=XM_011283077.3
B3 65624073 65624167 refGene_gn=THBS1;refGene_tx=XM_011283077.3
B3 65625079 65625202 refGene_gn=THBS1;refGene_tx=XM_011283077.3
B3 65625314 65625514 refGene_gn=THBS1;refGene_tx=XM_011283077.3
B3 65625811 65625887 refGene_gn=THBS1;refGene_tx=XM_011283077.3
B3 65626811 65627371 refGene_gn=THBS1;refGene_tx=XM_011283077.3
B3 65627624 65627691 refGene_gn=THBS1;refGene_tx=XM_011283077.3
B3 66505578 66505613 refGene_gn=RASGRP1;refGene_tx=XM_023255509.1
B3 66510042 66510227 refGene_gn=RASGRP1;refGene_tx=XM_023255509.1
B3 66550129 66550235 refGene_gn=RASGRP1;refGene_tx=XM_023255509.1
B3 66555369 66555704 refGene_gn=RASGRP1;refGene_tx=XM_019832692.2
B3 66555862 66555931
refGene_gn=RASGRP1;refGene_tx=XM_019832692.2;refGene_gn=RASGRP1;refGen
e_tx=XM_023255509.1
B3 66556894 66557026
refGene_gn=RASGRP1;refGene_tx=XM_019832692.2;refGene_gn=RASGRP1;refGen
e_tx=XM_023255509.1
B3 66559127 66559281
refGene_gn=RASGRP1;refGene_tx=XM_019832692.2;refGene_gn=RASGRP1;refGen
e_tx=XM_023255509.1
B3 66563149 66563323
refGene_gn=RASGRP1;refGene_tx=XM_019832692.2;refGene_gn=RASGRP1;refGen

e_tx=XM_023255509.1
B3 66564258 66564375
refGene_gn=RASGRP1;refGene_tx=XM_019832692.2;refGene_gn=RASGRP1;refGen
e_tx=XM_023255509.1
B3 66566548 66566824
refGene_gn=RASGRP1;refGene_tx=XM_019832692.2;refGene_gn=RASGRP1;refGen
e_tx=XM_023255509.1
B3 66568149 66568230
refGene_gn=RASGRP1;refGene_tx=XM_019832692.2;refGene_gn=RASGRP1;refGen
e_tx=XM_023255509.1
B3 66569496 66569601
refGene_gn=RASGRP1;refGene_tx=XM_019832692.2;refGene_gn=RASGRP1;refGen
e_tx=XM_023255509.1
B3 66570317 66570427
refGene_gn=RASGRP1;refGene_tx=XM_019832692.2;refGene_gn=RASGRP1;refGen
e_tx=XM_023255509.1
B3 66571070 66571215
refGene_gn=RASGRP1;refGene_tx=XM_019832692.2;refGene_gn=RASGRP1;refGen
e_tx=XM_023255509.1
B3 66572178 66572215
refGene_gn=RASGRP1;refGene_tx=XM_019832692.2;refGene_gn=RASGRP1;refGen
e_tx=XM_023255509.1
B3 66572971 66573124
refGene_gn=RASGRP1;refGene_tx=XM_019832692.2;refGene_gn=RASGRP1;refGen
e_tx=XM_023255509.1
B3 66574276 66574662
refGene_gn=RASGRP1;refGene_tx=XM_019832692.2;refGene_gn=RASGRP1;refGen
e_tx=XM_023255509.1
B3 66583300 66583435
refGene_gn=RASGRP1;refGene_tx=XM_019832692.2;refGene_gn=RASGRP1;refGen
e_tx=XM_023255509.1
B3 66589351 66589756 refGene_gn=FAM98B;refGene_tx=XM_003987287.5
B3 66592046 66592214 refGene_gn=FAM98B;refGene_tx=XM_003987287.5
B3 66605673 66605790 refGene_gn=FAM98B;refGene_tx=XM_003987287.5
B3 66606353 66606434 refGene_gn=FAM98B;refGene_tx=XM_003987287.5
B3 66609259 66609438 refGene_gn=FAM98B;refGene_tx=XM_003987287.5
B3 66616318 66616453 refGene_gn=FAM98B;refGene_tx=XM_003987287.5
B3 66617565 66617711 refGene_gn=FAM98B;refGene_tx=XM_003987287.5
B3 66625928 66625999 refGene_gn=FAM98B;refGene_tx=XM_003987287.5
B3 66746571 66747222
refGene_gn=SPRED1;refGene_tx=XM_011283078.3;refGene_gn=SPRED1;refGene_
tx=XM_019832694.2
B3 66749925 66750027 refGene_gn=SPRED1;refGene_tx=XM_011283078.3
B3 66763570 66763723
refGene_gn=SPRED1;refGene_tx=XM_011283078.3;refGene_gn=SPRED1;refGene_
tx=XM_019832694.2
B3 66782141 66782191
refGene_gn=SPRED1;refGene_tx=XM_011283078.3;refGene_gn=SPRED1;refGene_
tx=XM_019832694.2
B3 66786621 66786790

refGene_gn=SPRED1;refGene_tx=XM_011283078.3;refGene_gn=SPRED1;refGene_tx=XM_019832694.2
B3 66814931 66815106
refGene_gn=SPRED1;refGene_tx=XM_011283078.3;refGene_gn=SPRED1;refGene_tx=XM_019832694.2
B3 66883488 66883520
refGene_gn=SPRED1;refGene_tx=XM_011283078.3;refGene_gn=SPRED1;refGene_tx=XM_019832694.2
B3 67141421 67141620
refGene_gn=TMCO5A;refGene_tx=XM_006932652.3;refGene_gn=TMCO5A;refGene_tx=XM_019832695.2;refGene_gn=TMCO5A;refGene_tx=XM_019832696.2;refGene_gn=TMCO5A;refGene_tx=XM_019832697.2;refGene_gn=TMCO5A;refGene_tx=XM_019832698.2;refGene_gn=TMCO5A;refGene_tx=XM_019832700.2;refGene_gn=TMCO5A;refGene_tx=XM_023255510.1;refGene_gn=TMCO5A;refGene_tx=XM_023255511.1
B3 67145709 67145750
refGene_gn=TMCO5A;refGene_tx=XM_006932652.3;refGene_gn=TMCO5A;refGene_tx=XM_019832695.2;refGene_gn=TMCO5A;refGene_tx=XM_019832696.2;refGene_gn=TMCO5A;refGene_tx=XM_019832697.2;refGene_gn=TMCO5A;refGene_tx=XM_019832698.2;refGene_gn=TMCO5A;refGene_tx=XM_019832700.2;refGene_gn=TMCO5A;refGene_tx=XM_023255510.1;refGene_gn=TMCO5A;refGene_tx=XM_023255511.1
B3 67147470 67147528
refGene_gn=TMCO5A;refGene_tx=XM_006932652.3;refGene_gn=TMCO5A;refGene_tx=XM_019832695.2;refGene_gn=TMCO5A;refGene_tx=XM_019832696.2;refGene_gn=TMCO5A;refGene_tx=XM_019832697.2;refGene_gn=TMCO5A;refGene_tx=XM_019832698.2;refGene_gn=TMCO5A;refGene_tx=XM_019832700.2;refGene_gn=TMCO5A;refGene_tx=XM_023255510.1;refGene_gn=TMCO5A;refGene_tx=XM_023255511.1
B3 67148898 67148963
refGene_gn=TMCO5A;refGene_tx=XM_006932652.3;refGene_gn=TMCO5A;refGene_tx=XM_019832695.2;refGene_gn=TMCO5A;refGene_tx=XM_019832696.2;refGene_gn=TMCO5A;refGene_tx=XM_019832697.2;refGene_gn=TMCO5A;refGene_tx=XM_019832698.2;refGene_gn=TMCO5A;refGene_tx=XM_019832700.2;refGene_gn=TMCO5A;refGene_tx=XM_023255510.1;refGene_gn=TMCO5A;refGene_tx=XM_023255511.1
B3 67149391 67149451
refGene_gn=TMCO5A;refGene_tx=XM_006932652.3;refGene_gn=TMCO5A;refGene_tx=XM_019832695.2;refGene_gn=TMCO5A;refGene_tx=XM_019832696.2;refGene_gn=TMCO5A;refGene_tx=XM_019832697.2;refGene_gn=TMCO5A;refGene_tx=XM_019832698.2;refGene_gn=TMCO5A;refGene_tx=XM_019832700.2;refGene_gn=TMCO5A;refGene_tx=XM_023255510.1;refGene_gn=TMCO5A;refGene_tx=XM_023255511.1
B3 67149859 67149916
refGene_gn=TMCO5A;refGene_tx=XM_006932652.3;refGene_gn=TMCO5A;refGene_tx=XM_019832695.2;refGene_gn=TMCO5A;refGene_tx=XM_019832696.2;refGene_gn=TMCO5A;refGene_tx=XM_019832697.2;refGene_gn=TMCO5A;refGene_tx=XM_019832698.2;refGene_gn=TMCO5A;refGene_tx=XM_019832700.2;refGene_gn=TMCO5A;refGene_tx=XM_023255510.1;refGene_gn=TMCO5A;refGene_tx=XM_023255511.1

B3 67153042 67153114
refGene_gn=TMCO5A;refGene_tx=XM_006932652.3;refGene_gn=TMCO5A;refGene_tx=XM_019832695.2;refGene_gn=TMCO5A;refGene_tx=XM_019832696.2;refGene_gn=TMCO5A;refGene_tx=XM_019832697.2;refGene_gn=TMCO5A;refGene_tx=XM_019832698.2;refGene_gn=TMCO5A;refGene_tx=XM_019832700.2;refGene_gn=TMCO5A;refGene_tx=XM_023255510.1;refGene_gn=TMCO5A;refGene_tx=XM_023255511.1
B3 67153776 67153827
refGene_gn=TMCO5A;refGene_tx=XM_006932652.3;refGene_gn=TMCO5A;refGene_tx=XM_019832695.2;refGene_gn=TMCO5A;refGene_tx=XM_019832696.2;refGene_gn=TMCO5A;refGene_tx=XM_019832697.2;refGene_gn=TMCO5A;refGene_tx=XM_019832698.2;refGene_gn=TMCO5A;refGene_tx=XM_019832700.2;refGene_gn=TMCO5A;refGene_tx=XM_023255510.1
B3 67154470 67154594
refGene_gn=TMCO5A;refGene_tx=XM_006932652.3;refGene_gn=TMCO5A;refGene_tx=XM_019832695.2;refGene_gn=TMCO5A;refGene_tx=XM_019832696.2;refGene_gn=TMCO5A;refGene_tx=XM_019832697.2;refGene_gn=TMCO5A;refGene_tx=XM_019832698.2;refGene_gn=TMCO5A;refGene_tx=XM_019832700.2;refGene_gn=TMCO5A;refGene_tx=XM_023255510.1;refGene_gn=TMCO5A;refGene_tx=XM_023255511.1
B3 67155252 67155402
refGene_gn=TMCO5A;refGene_tx=XM_006932652.3;refGene_gn=TMCO5A;refGene_tx=XM_019832695.2;refGene_gn=TMCO5A;refGene_tx=XM_019832696.2;refGene_gn=TMCO5A;refGene_tx=XM_019832697.2;refGene_gn=TMCO5A;refGene_tx=XM_019832698.2;refGene_gn=TMCO5A;refGene_tx=XM_019832700.2;refGene_gn=TMCO5A;refGene_tx=XM_023255510.1;refGene_gn=TMCO5A;refGene_tx=XM_023255511.1
B3 67156370 67156396 refGene_gn=TMCO5A;refGene_tx=XM_019832696.2
B3 67156412 67156457
refGene_gn=TMCO5A;refGene_tx=XM_006932652.3;refGene_gn=TMCO5A;refGene_tx=XM_019832695.2;refGene_gn=TMCO5A;refGene_tx=XM_023255510.1;refGene_gn=TMCO5A;refGene_tx=XM_023255511.1
B3 67156701 67156703
refGene_gn=TMCO5A;refGene_tx=XM_019832695.2;refGene_gn=TMCO5A;refGene_tx=XM_023255511.1
B3 67882651 67882663
refGene_gn=MEIS2;refGene_tx=XM_023255512.1;refGene_gn=MEIS2;refGene_tx=XM_023255514.1;refGene_gn=MEIS2;refGene_tx=XM_023255515.1;refGene_gn=MEIS2;refGene_tx=XM_023255517.1
B3 67883217 67883226
refGene_gn=MEIS2;refGene_tx=XM_023255513.1;refGene_gn=MEIS2;refGene_tx=XM_023255516.1;refGene_gn=MEIS2;refGene_tx=XM_023255518.1
B3 67883879 67884112
refGene_gn=MEIS2;refGene_tx=XM_006932654.4;refGene_gn=MEIS2;refGene_tx=XM_023255512.1;refGene_gn=MEIS2;refGene_tx=XM_023255513.1;refGene_gn=MEIS2;refGene_tx=XM_023255514.1;refGene_gn=MEIS2;refGene_tx=XM_023255515.1;refGene_gn=MEIS2;refGene_tx=XM_023255516.1;refGene_gn=MEIS2;refGene_tx=XM_023255517.1;refGene_gn=MEIS2;refGene_tx=XM_023255518.1
B3 67885666 67885808
refGene_gn=MEIS2;refGene_tx=XM_006932654.4;refGene_gn=MEIS2;refGene_tx

=XM_023255512.1;refGene_gn=MEIS2;refGene_tx=XM_023255513.1;refGene_gn=MEIS2;refGene_tx=XM_023255514.1;refGene_gn=MEIS2;refGene_tx=XM_023255515.1;refGene_gn=MEIS2;refGene_tx=XM_023255516.1;refGene_gn=MEIS2;refGene_tx=XM_023255517.1;refGene_gn=MEIS2;refGene_tx=XM_023255518.1
B3 67886519 67886570
refGene_gn=MEIS2;refGene_tx=XM_006932654.4;refGene_gn=MEIS2;refGene_tx=XM_023255512.1;refGene_gn=MEIS2;refGene_tx=XM_023255513.1;refGene_gn=MEIS2;refGene_tx=XM_023255514.1;refGene_gn=MEIS2;refGene_tx=XM_023255515.1;refGene_gn=MEIS2;refGene_tx=XM_023255516.1;refGene_gn=MEIS2;refGene_tx=XM_023255517.1;refGene_gn=MEIS2;refGene_tx=XM_023255518.1
B3 67887539 67887590
refGene_gn=MEIS2;refGene_tx=XM_006932654.4;refGene_gn=MEIS2;refGene_tx=XM_023255512.1;refGene_gn=MEIS2;refGene_tx=XM_023255513.1;refGene_gn=MEIS2;refGene_tx=XM_023255514.1;refGene_gn=MEIS2;refGene_tx=XM_023255515.1;refGene_gn=MEIS2;refGene_tx=XM_023255516.1;refGene_gn=MEIS2;refGene_tx=XM_023255517.1;refGene_gn=MEIS2;refGene_tx=XM_023255518.1
B3 67888409 67888559
refGene_gn=MEIS2;refGene_tx=XM_006932654.4;refGene_gn=MEIS2;refGene_tx=XM_023255512.1;refGene_gn=MEIS2;refGene_tx=XM_023255513.1;refGene_gn=MEIS2;refGene_tx=XM_023255514.1;refGene_gn=MEIS2;refGene_tx=XM_023255515.1;refGene_gn=MEIS2;refGene_tx=XM_023255516.1;refGene_gn=MEIS2;refGene_tx=XM_023255517.1;refGene_gn=MEIS2;refGene_tx=XM_023255518.1
B3 67898370 67898485
refGene_gn=MEIS2;refGene_tx=XM_006932654.4;refGene_gn=MEIS2;refGene_tx=XM_023255512.1;refGene_gn=MEIS2;refGene_tx=XM_023255513.1;refGene_gn=MEIS2;refGene_tx=XM_023255514.1;refGene_gn=MEIS2;refGene_tx=XM_023255515.1;refGene_gn=MEIS2;refGene_tx=XM_023255516.1;refGene_gn=MEIS2;refGene_tx=XM_023255517.1;refGene_gn=MEIS2;refGene_tx=XM_023255518.1
B3 67947351 67947497
refGene_gn=MEIS2;refGene_tx=XM_006932654.4;refGene_gn=MEIS2;refGene_tx=XM_023255512.1;refGene_gn=MEIS2;refGene_tx=XM_023255513.1;refGene_gn=MEIS2;refGene_tx=XM_023255514.1;refGene_gn=MEIS2;refGene_tx=XM_023255515.1;refGene_gn=MEIS2;refGene_tx=XM_023255516.1;refGene_gn=MEIS2;refGene_tx=XM_023255517.1;refGene_gn=MEIS2;refGene_tx=XM_023255518.1
B3 68032008 68032085
refGene_gn=MEIS2;refGene_tx=XM_006932654.4;refGene_gn=MEIS2;refGene_tx=XM_023255512.1;refGene_gn=MEIS2;refGene_tx=XM_023255513.1;refGene_gn=MEIS2;refGene_tx=XM_023255514.1;refGene_gn=MEIS2;refGene_tx=XM_023255515.1;refGene_gn=MEIS2;refGene_tx=XM_023255516.1;refGene_gn=MEIS2;refGene_tx=XM_023255517.1;refGene_gn=MEIS2;refGene_tx=XM_023255518.1
B3 68087102 68087161
refGene_gn=MEIS2;refGene_tx=XM_006932654.4;refGene_gn=MEIS2;refGene_tx=XM_023255512.1;refGene_gn=MEIS2;refGene_tx=XM_023255513.1;refGene_gn=MEIS2;refGene_tx=XM_023255514.1;refGene_gn=MEIS2;refGene_tx=XM_023255515.1;refGene_gn=MEIS2;refGene_tx=XM_023255516.1;refGene_gn=MEIS2;refGene_tx=XM_023255517.1;refGene_gn=MEIS2;refGene_tx=XM_023255518.1
B3 68088542 68088653
refGene_gn=MEIS2;refGene_tx=XM_006932654.4;refGene_gn=MEIS2;refGene_tx=XM_023255512.1;refGene_gn=MEIS2;refGene_tx=XM_023255513.1;refGene_gn=MEIS2;refGene_tx=XM_023255514.1;refGene_gn=MEIS2;refGene_tx=XM_0232555

15.1;refGene_gn=MEIS2;refGene_tx=XM_023255516.1;refGene_gn=MEIS2;refGene_tx=XM_023255517.1;refGene_gn=MEIS2;refGene_tx=XM_023255518.1
B3 68089000 68089059
refGene_gn=MEIS2;refGene_tx=XM_023255515.1;refGene_gn=MEIS2;refGene_tx=XM_023255516.1;refGene_gn=MEIS2;refGene_tx=XM_023255517.1;refGene_gn=MEIS2;refGene_tx=XM_023255518.1
B3 68091342 68091629
refGene_gn=MEIS2;refGene_tx=XM_006932654.4;refGene_gn=MEIS2;refGene_tx=XM_023255512.1;refGene_gn=MEIS2;refGene_tx=XM_023255513.1;refGene_gn=MEIS2;refGene_tx=XM_023255514.1
B3 68116540 68116562
refGene_gn=CB3H15orf41;refGene_tx=XM_023255524.1
B3 68121693 68121811
refGene_gn=CB3H15orf41;refGene_tx=XM_023255520.1
B3 68175320 68175450
refGene_gn=CB3H15orf41;refGene_tx=XM_003987293.5;refGene_gn=CB3H15orf41;refGene_tx=XM_006932656.4;refGene_gn=CB3H15orf41;refGene_tx=XM_023255522.1;refGene_gn=CB3H15orf41;refGene_tx=XM_023255523.1;refGene_gn=CB3H15orf41;refGene_tx=XM_023255525.1;refGene_gn=CB3H15orf41;refGene_tx=XM_023255526.1
B3 68200412 68200500
refGene_gn=CB3H15orf41;refGene_tx=XM_023255521.1
B3 68276745 68276851
refGene_gn=CB3H15orf41;refGene_tx=XM_003987293.5;refGene_gn=CB3H15orf41;refGene_tx=XM_006932656.4;refGene_gn=CB3H15orf41;refGene_tx=XM_023255520.1;refGene_gn=CB3H15orf41;refGene_tx=XM_023255521.1;refGene_gn=CB3H15orf41;refGene_tx=XM_023255522.1;refGene_gn=CB3H15orf41;refGene_tx=XM_023255523.1;refGene_gn=CB3H15orf41;refGene_tx=XM_023255524.1;refGene_gn=CB3H15orf41;refGene_tx=XM_023255526.1
B3 68277461 68277527
refGene_gn=CB3H15orf41;refGene_tx=XM_003987293.5;refGene_gn=CB3H15orf41;refGene_tx=XM_006932656.4;refGene_gn=CB3H15orf41;refGene_tx=XM_023255520.1;refGene_gn=CB3H15orf41;refGene_tx=XM_023255521.1;refGene_gn=CB3H15orf41;refGene_tx=XM_023255522.1;refGene_gn=CB3H15orf41;refGene_tx=XM_023255523.1;refGene_gn=CB3H15orf41;refGene_tx=XM_023255524.1;refGene_gn=CB3H15orf41;refGene_tx=XM_023255525.1;refGene_gn=CB3H15orf41;refGene_tx=XM_023255526.1
B3 68290628 68290696
refGene_gn=CB3H15orf41;refGene_tx=XM_003987293.5;refGene_gn=CB3H15orf41;refGene_tx=XM_006932656.4;refGene_gn=CB3H15orf41;refGene_tx=XM_023255520.1;refGene_gn=CB3H15orf41;refGene_tx=XM_023255521.1;refGene_gn=CB3H15orf41;refGene_tx=XM_023255522.1;refGene_gn=CB3H15orf41;refGene_tx=XM_023255523.1;refGene_gn=CB3H15orf41;refGene_tx=XM_023255524.1;refGene_gn=CB3H15orf41;refGene_tx=XM_023255525.1;refGene_gn=CB3H15orf41;refGene_tx=XM_023255526.1
B3 68296519 68296569
refGene_gn=CB3H15orf41;refGene_tx=XM_003987293.5;refGene_gn=CB3H15orf41;refGene_tx=XM_006932656.4;refGene_gn=CB3H15orf41;refGene_tx=XM_023255520.1;refGene_gn=CB3H15orf41;refGene_tx=XM_023255521.1;refGene_gn=CB3H15orf41;refGene_tx=XM_023255522.1;refGene_gn=CB3H15orf41;refGene_tx=X

M_023255523.1;refGene_gn=CB3H15orf41;refGene_tx=XM_023255524.1;refGene_gn=CB3H15orf41;refGene_tx=XM_023255525.1;refGene_gn=CB3H15orf41;refGene_tx=XM_023255526.1
B3 68296923 68297003
refGene_gn=CB3H15orf41;refGene_tx=XM_003987293.5;refGene_gn=CB3H15orf41;refGene_tx=XM_006932656.4;refGene_gn=CB3H15orf41;refGene_tx=XM_023255520.1;refGene_gn=CB3H15orf41;refGene_tx=XM_023255521.1;refGene_gn=CB3H15orf41;refGene_tx=XM_023255522.1;refGene_gn=CB3H15orf41;refGene_tx=XM_023255523.1;refGene_gn=CB3H15orf41;refGene_tx=XM_023255524.1;refGene_gn=CB3H15orf41;refGene_tx=XM_023255525.1;refGene_gn=CB3H15orf41;refGene_tx=XM_023255526.1
B3 68311557 68311597
refGene_gn=CB3H15orf41;refGene_tx=XM_023255526.1
B3 68327952 68328025
refGene_gn=CB3H15orf41;refGene_tx=XM_003987293.5;refGene_gn=CB3H15orf41;refGene_tx=XM_006932656.4;refGene_gn=CB3H15orf41;refGene_tx=XM_023255520.1;refGene_gn=CB3H15orf41;refGene_tx=XM_023255521.1;refGene_gn=CB3H15orf41;refGene_tx=XM_023255522.1;refGene_gn=CB3H15orf41;refGene_tx=XM_023255523.1;refGene_gn=CB3H15orf41;refGene_tx=XM_023255524.1;refGene_gn=CB3H15orf41;refGene_tx=XM_023255525.1
B3 68331337 68331398
refGene_gn=CB3H15orf41;refGene_tx=XM_003987293.5;refGene_gn=CB3H15orf41;refGene_tx=XM_006932656.4;refGene_gn=CB3H15orf41;refGene_tx=XM_023255520.1;refGene_gn=CB3H15orf41;refGene_tx=XM_023255521.1;refGene_gn=CB3H15orf41;refGene_tx=XM_023255522.1;refGene_gn=CB3H15orf41;refGene_tx=XM_023255523.1;refGene_gn=CB3H15orf41;refGene_tx=XM_023255524.1;refGene_gn=CB3H15orf41;refGene_tx=XM_023255525.1
B3 68334637 68334765
refGene_gn=CB3H15orf41;refGene_tx=XM_023255522.1;refGene_gn=CB3H15orf41;refGene_tx=XM_023255523.1
B3 68338052 68338117
refGene_gn=CB3H15orf41;refGene_tx=XM_003987293.5;refGene_gn=CB3H15orf41;refGene_tx=XM_023255520.1;refGene_gn=CB3H15orf41;refGene_tx=XM_023255521.1;refGene_gn=CB3H15orf41;refGene_tx=XM_023255524.1;refGene_gn=CB3H15orf41;refGene_tx=XM_023255525.1
B3 68339005 68339051
refGene_gn=CB3H15orf41;refGene_tx=XM_003987293.5;refGene_gn=CB3H15orf41;refGene_tx=XM_023255520.1;refGene_gn=CB3H15orf41;refGene_tx=XM_023255521.1;refGene_gn=CB3H15orf41;refGene_tx=XM_023255524.1;refGene_gn=CB3H15orf41;refGene_tx=XM_023255525.1
B3 68388828 68388929
refGene_gn=CB3H15orf41;refGene_tx=XM_003987293.5;refGene_gn=CB3H15orf41;refGene_tx=XM_006932656.4;refGene_gn=CB3H15orf41;refGene_tx=XM_023255520.1;refGene_gn=CB3H15orf41;refGene_tx=XM_023255521.1;refGene_gn=CB3H15orf41;refGene_tx=XM_023255524.1;refGene_gn=CB3H15orf41;refGene_tx=XM_023255525.1
B3 69433758 69433781
refGene_gn=DPH6;refGene_tx=XM_003987294.5;refGene_gn=DPH6;refGene_tx=XM_006932657.4
B3 69438651 69438746

refGene_gn=DPH6;refGene_tx=XM_003987294.5;refGene_gn=DPH6;refGene_tx=XM_006932657.4
B3 69452232 69452426
refGene_gn=DPH6;refGene_tx=XM_003987294.5;refGene_gn=DPH6;refGene_tx=XM_006932657.4
B3 69471274 69471460 refGene_gn=DPH6;refGene_tx=XM_003987294.5
B3 69539540 69539614 refGene_gn=DPH6;refGene_tx=XM_006932657.4
B3 69543501 69543620 refGene_gn=DPH6;refGene_tx=XM_006932657.4
B3 69575217 69575279 refGene_gn=DPH6;refGene_tx=XM_006932657.4
B3 69604217 69604312 refGene_gn=DPH6;refGene_tx=XM_006932657.4
B3 69610658 69610746 refGene_gn=DPH6;refGene_tx=XM_006932657.4
B3 69612652 69612706 refGene_gn=DPH6;refGene_tx=XM_006932657.4
B3 69933938 69936020
refGene_gn=ZNF770;refGene_tx=XM_003987295.5;refGene_gn=ZNF770;refGene_tx=XM_006932658.3;refGene_gn=ZNF770;refGene_tx=XM_019832709.2
B3 69941391 69941466 refGene_gn=AQR;refGene_tx=XM_011283083.3
B3 69944883 69944940 refGene_gn=AQR;refGene_tx=XM_011283083.3
B3 69948410 69948451 refGene_gn=AQR;refGene_tx=XM_011283083.3
B3 69956086 69956122 refGene_gn=AQR;refGene_tx=XM_011283083.3
B3 69958798 69958919 refGene_gn=AQR;refGene_tx=XM_011283083.3
B3 69961554 69961695 refGene_gn=AQR;refGene_tx=XM_011283083.3
B3 69963391 69963460 refGene_gn=AQR;refGene_tx=XM_011283083.3
B3 69964554 69964655 refGene_gn=AQR;refGene_tx=XM_011283083.3
B3 69968264 69968341 refGene_gn=AQR;refGene_tx=XM_011283083.3
B3 69970186 69970251 refGene_gn=AQR;refGene_tx=XM_011283083.3
B3 69971390 69971507 refGene_gn=AQR;refGene_tx=XM_011283083.3
B3 69972831 69972945 refGene_gn=AQR;refGene_tx=XM_011283083.3
B3 69974691 69974795 refGene_gn=AQR;refGene_tx=XM_011283083.3
B3 69979873 69979976 refGene_gn=AQR;refGene_tx=XM_011283083.3
B3 69982468 69982589 refGene_gn=AQR;refGene_tx=XM_011283083.3
B3 69985752 69985894 refGene_gn=AQR;refGene_tx=XM_011283083.3
B3 69988958 69989137 refGene_gn=AQR;refGene_tx=XM_011283083.3
B3 69992464 69992632 refGene_gn=AQR;refGene_tx=XM_011283083.3
B3 69993912 69994082 refGene_gn=AQR;refGene_tx=XM_011283083.3
B3 69997360 69997602 refGene_gn=AQR;refGene_tx=XM_011283083.3
B3 69999069 69999216 refGene_gn=AQR;refGene_tx=XM_011283083.3
B3 70000038 70000108 refGene_gn=AQR;refGene_tx=XM_011283083.3
B3 70002340 70002451 refGene_gn=AQR;refGene_tx=XM_011283083.3
B3 70005141 70005251 refGene_gn=AQR;refGene_tx=XM_011283083.3
B3 70008224 70008360 refGene_gn=AQR;refGene_tx=XM_011283083.3
B3 70010328 70010538 refGene_gn=AQR;refGene_tx=XM_011283083.3
B3 70012764 70012902 refGene_gn=AQR;refGene_tx=XM_011283083.3
B3 70016677 70016749 refGene_gn=AQR;refGene_tx=XM_011283083.3
B3 70017837 70018025 refGene_gn=AQR;refGene_tx=XM_011283083.3
B3 70018734 70018906 refGene_gn=AQR;refGene_tx=XM_011283083.3
B3 70021076 70021247 refGene_gn=AQR;refGene_tx=XM_011283083.3
B3 70024926 70025012 refGene_gn=AQR;refGene_tx=XM_011283083.3
B3 70026299 70026474 refGene_gn=AQR;refGene_tx=XM_011283083.3
B3 70029565 70029679 refGene_gn=AQR;refGene_tx=XM_011283083.3
B3 70031128 70031440 refGene_gn=AQR;refGene_tx=XM_011283083.3

B3 70081099 70081228 refGene_gn=ACTC1;refGene_tx=XM_023255528.1
B3 70082387 70082712 refGene_gn=ACTC1;refGene_tx=XM_023255528.1
B3 70083367 70083529 refGene_gn=ACTC1;refGene_tx=XM_023255528.1
B3 70083651 70083843 refGene_gn=ACTC1;refGene_tx=XM_023255528.1
B3 70084653 70084835 refGene_gn=ACTC1;refGene_tx=XM_023255528.1
B3 70085338 70085482 refGene_gn=ACTC1;refGene_tx=XM_023255528.1
B3 70124445 70124516 refGene_gn=GJD2;refGene_tx=XM_003987299.4
B3 70125626 70126521 refGene_gn=GJD2;refGene_tx=XM_003987299.4
B3 70270563 70270717
refGene_gn=ARHGAP11A;refGene_tx=XM_023256170.1
B3 70275212 70275352
refGene_gn=ARHGAP11A;refGene_tx=XM_023256170.1
B3 70276054 70276308
refGene_gn=ARHGAP11A;refGene_tx=XM_023256170.1
B3 70276957 70277121
refGene_gn=ARHGAP11A;refGene_tx=XM_023256170.1
B3 70277285 70277432
refGene_gn=ARHGAP11A;refGene_tx=XM_023256170.1
B3 70277966 70278041
refGene_gn=ARHGAP11A;refGene_tx=XM_023256170.1
B3 70279749 70279914
refGene_gn=ARHGAP11A;refGene_tx=XM_023256170.1
B3 70282563 70282693
refGene_gn=ARHGAP11A;refGene_tx=XM_023256170.1
B3 70283456 70283565
refGene_gn=ARHGAP11A;refGene_tx=XM_023256170.1
B3 70285293 70285432
refGene_gn=ARHGAP11A;refGene_tx=XM_023256170.1
B3 70285818 70287377
refGene_gn=ARHGAP11A;refGene_tx=XM_023256170.1
B3 70291039 70291140
refGene_gn=SCG5;refGene_tx=XM_003987301.5;refGene_gn=SCG5;refGene_tx=XM_006932660.4;refGene_gn=SCG5;refGene_tx=XM_019832711.2
B3 70292644 70292877
refGene_gn=SCG5;refGene_tx=XM_003987301.5;refGene_gn=SCG5;refGene_tx=XM_006932660.4;refGene_gn=SCG5;refGene_tx=XM_019832711.2
B3 70334765 70334915
refGene_gn=SCG5;refGene_tx=XM_003987301.5;refGene_gn=SCG5;refGene_tx=XM_006932660.4;refGene_gn=SCG5;refGene_tx=XM_019832711.2
B3 70339189 70339302
refGene_gn=SCG5;refGene_tx=XM_003987301.5;refGene_gn=SCG5;refGene_tx=XM_006932660.4
B3 70346388 70346442
refGene_gn=SCG5;refGene_tx=XM_003987301.5;refGene_gn=SCG5;refGene_tx=XM_006932660.4;refGene_gn=SCG5;refGene_tx=XM_019832711.2
B3 70350333 70350429
refGene_gn=SCG5;refGene_tx=XM_003987301.5;refGene_gn=SCG5;refGene_tx=XM_006932660.4;refGene_gn=SCG5;refGene_tx=XM_019832711.2
B3 70384124 70384679 refGene_gn=GREM1;refGene_tx=XM_011283085.3
B3 70421505 70421550

refGene_gn=FMN1;refGene_tx=XM_023255529.1;refGene_gn=FMN1;refGene_tx=XM_023255530.1;refGene_gn=FMN1;refGene_tx=XM_023255531.1;refGene_gn=FMN1;refGene_tx=XM_023255532.1;refGene_gn=FMN1;refGene_tx=XM_023255533.1
B3 70424227 70424312
refGene_gn=FMN1;refGene_tx=XM_023255529.1;refGene_gn=FMN1;refGene_tx=XM_023255530.1;refGene_gn=FMN1;refGene_tx=XM_023255531.1;refGene_gn=FMN1;refGene_tx=XM_023255532.1;refGene_gn=FMN1;refGene_tx=XM_023255533.1
B3 70449774 70449924
refGene_gn=FMN1;refGene_tx=XM_023255529.1;refGene_gn=FMN1;refGene_tx=XM_023255530.1;refGene_gn=FMN1;refGene_tx=XM_023255531.1;refGene_gn=FMN1;refGene_tx=XM_023255532.1;refGene_gn=FMN1;refGene_tx=XM_023255533.1
B3 70456367 70456419
refGene_gn=FMN1;refGene_tx=XM_023255529.1;refGene_gn=FMN1;refGene_tx=XM_023255530.1;refGene_gn=FMN1;refGene_tx=XM_023255531.1;refGene_gn=FMN1;refGene_tx=XM_023255532.1;refGene_gn=FMN1;refGene_tx=XM_023255533.1
B3 70509727 70509820
refGene_gn=FMN1;refGene_tx=XM_023255529.1;refGene_gn=FMN1;refGene_tx=XM_023255530.1;refGene_gn=FMN1;refGene_tx=XM_023255531.1;refGene_gn=FMN1;refGene_tx=XM_023255532.1;refGene_gn=FMN1;refGene_tx=XM_023255533.1
B3 70540078 70540199
refGene_gn=FMN1;refGene_tx=XM_023255529.1;refGene_gn=FMN1;refGene_tx=XM_023255530.1;refGene_gn=FMN1;refGene_tx=XM_023255531.1;refGene_gn=FMN1;refGene_tx=XM_023255532.1;refGene_gn=FMN1;refGene_tx=XM_023255533.1
B3 70545462 70545522
refGene_gn=FMN1;refGene_tx=XM_023255529.1;refGene_gn=FMN1;refGene_tx=XM_023255530.1;refGene_gn=FMN1;refGene_tx=XM_023255531.1;refGene_gn=FMN1;refGene_tx=XM_023255532.1;refGene_gn=FMN1;refGene_tx=XM_023255533.1
B3 70546629 70546776
refGene_gn=FMN1;refGene_tx=XM_023255529.1;refGene_gn=FMN1;refGene_tx=XM_023255530.1;refGene_gn=FMN1;refGene_tx=XM_023255531.1;refGene_gn=FMN1;refGene_tx=XM_023255532.1;refGene_gn=FMN1;refGene_tx=XM_023255533.1
B3 70548456 70548586
refGene_gn=FMN1;refGene_tx=XM_023255529.1;refGene_gn=FMN1;refGene_tx=XM_023255530.1;refGene_gn=FMN1;refGene_tx=XM_023255531.1;refGene_gn=FMN1;refGene_tx=XM_023255532.1;refGene_gn=FMN1;refGene_tx=XM_023255533.1
B3 70555063 70555152
refGene_gn=FMN1;refGene_tx=XM_023255529.1;refGene_gn=FMN1;refGene_tx=XM_023255530.1;refGene_gn=FMN1;refGene_tx=XM_023255531.1;refGene_gn=FMN1;refGene_tx=XM_023255532.1;refGene_gn=FMN1;refGene_tx=XM_023255533.1
B3 70557314 70557376
refGene_gn=FMN1;refGene_tx=XM_023255529.1;refGene_gn=FMN1;refGene_tx=XM_023255530.1;refGene_gn=FMN1;refGene_tx=XM_023255531.1;refGene_gn=FMN1;refGene_tx=XM_023255532.1;refGene_gn=FMN1;refGene_tx=XM_023255533.1
B3 70573567 70573655
refGene_gn=FMN1;refGene_tx=XM_023255529.1;refGene_gn=FMN1;refGene_tx=XM_023255530.1;refGene_gn=FMN1;refGene_tx=XM_023255531.1;refGene_gn=FMN1;refGene_tx=XM_023255532.1;refGene_gn=FMN1;refGene_tx=XM_023255533.1
B3 70610865 70611016
refGene_gn=FMN1;refGene_tx=XM_023255529.1;refGene_gn=FMN1;refGene_tx=XM_023255530.1;refGene_gn=FMN1;refGene_tx=XM_023255531.1;refGene_gn=FMN

1;refGene_tx=XM_023255532.1;refGene_gn=FMN1;refGene_tx=XM_023255533.1
B3 70615637 70616404
refGene_gn=FMN1;refGene_tx=XM_023255529.1;refGene_gn=FMN1;refGene_tx=XM_023255530.1;refGene_gn=FMN1;refGene_tx=XM_023255531.1;refGene_gn=FMN1;refGene_tx=XM_023255532.1;refGene_gn=FMN1;refGene_tx=XM_023255533.1
B3 70660972 70661034
refGene_gn=FMN1;refGene_tx=XM_023255529.1;refGene_gn=FMN1;refGene_tx=XM_023255530.1;refGene_gn=FMN1;refGene_tx=XM_023255531.1;refGene_gn=FMN1;refGene_tx=XM_023255532.1;refGene_gn=FMN1;refGene_tx=XM_023255533.1
B3 70704315 70704433
refGene_gn=FMN1;refGene_tx=XM_023255529.1;refGene_gn=FMN1;refGene_tx=XM_023255530.1;refGene_gn=FMN1;refGene_tx=XM_023255531.1;refGene_gn=FMN1;refGene_tx=XM_023255533.1
B3 70705806 70707186 refGene_gn=FMN1;refGene_tx=XM_023255533.1
B3 70729149 70729328
refGene_gn=FMN1;refGene_tx=XM_023255529.1;refGene_gn=FMN1;refGene_tx=XM_023255530.1;refGene_gn=FMN1;refGene_tx=XM_023255531.1
B3 70807655 70809510
refGene_gn=FMN1;refGene_tx=XM_023255529.1;refGene_gn=FMN1;refGene_tx=XM_023255530.1;refGene_gn=FMN1;refGene_tx=XM_023255531.1;refGene_gn=FMN1;refGene_tx=XM_023255532.1
B3 70876880 70877079
refGene_gn=LOC101096989;refGene_tx=XM_006932661.4
B3 70878148 70878192
refGene_gn=LOC101096989;refGene_tx=XM_006932661.4
B3 70878474 70878550
refGene_gn=LOC101096989;refGene_tx=XM_006932661.4
B3 70879179 70879244
refGene_gn=LOC101096989;refGene_tx=XM_006932661.4
B3 70880031 70880091
refGene_gn=LOC101096989;refGene_tx=XM_006932661.4
B3 70880578 70880635
refGene_gn=LOC101096989;refGene_tx=XM_006932661.4
B3 70881987 70882056
refGene_gn=LOC101096989;refGene_tx=XM_006932661.4
B3 70882585 70882636
refGene_gn=LOC101096989;refGene_tx=XM_006932661.4
B3 70882961 70883079
refGene_gn=LOC101096989;refGene_tx=XM_006932661.4
B3 70883389 70883539
refGene_gn=LOC101096989;refGene_tx=XM_006932661.4
B3 70884274 70884309
refGene_gn=LOC101096989;refGene_tx=XM_006932661.4
B3 70938849 70938900
refGene_gn=RYR3;refGene_tx=XM_023255535.1;refGene_gn=RYR3;refGene_tx=XM_023255536.1;refGene_gn=RYR3;refGene_tx=XM_023255537.1;refGene_gn=RYR3;refGene_tx=XM_023255538.1
B3 71107992 71108112
refGene_gn=RYR3;refGene_tx=XM_023255535.1;refGene_gn=RYR3;refGene_tx=XM_023255536.1;refGene_gn=RYR3;refGene_tx=XM_023255537.1;refGene_gn=RYR

3;refGene_tx=XM_023255538.1
B3	71139264	71139372
refGene_gn=RYR3;refGene_tx=XM_023255535.1;refGene_gn=RYR3;refGene_tx=XM_023255536.1;refGene_gn=RYR3;refGene_tx=XM_023255537.1;refGene_gn=RYR3;refGene_tx=XM_023255538.1
B3	71169097	71169172
refGene_gn=RYR3;refGene_tx=XM_023255535.1;refGene_gn=RYR3;refGene_tx=XM_023255536.1;refGene_gn=RYR3;refGene_tx=XM_023255537.1;refGene_gn=RYR3;refGene_tx=XM_023255538.1
B3	71171454	71171533
refGene_gn=RYR3;refGene_tx=XM_023255535.1;refGene_gn=RYR3;refGene_tx=XM_023255536.1;refGene_gn=RYR3;refGene_tx=XM_023255537.1;refGene_gn=RYR3;refGene_tx=XM_023255538.1
B3	71175331	71175444
refGene_gn=RYR3;refGene_tx=XM_023255535.1;refGene_gn=RYR3;refGene_tx=XM_023255536.1;refGene_gn=RYR3;refGene_tx=XM_023255537.1;refGene_gn=RYR3;refGene_tx=XM_023255538.1
B3	71176318	71176418
refGene_gn=RYR3;refGene_tx=XM_023255535.1;refGene_gn=RYR3;refGene_tx=XM_023255536.1;refGene_gn=RYR3;refGene_tx=XM_023255537.1;refGene_gn=RYR3;refGene_tx=XM_023255538.1
B3	71180244	71180338
refGene_gn=RYR3;refGene_tx=XM_023255535.1;refGene_gn=RYR3;refGene_tx=XM_023255536.1;refGene_gn=RYR3;refGene_tx=XM_023255537.1;refGene_gn=RYR3;refGene_tx=XM_023255538.1
B3	71185450	71185525
refGene_gn=RYR3;refGene_tx=XM_023255535.1;refGene_gn=RYR3;refGene_tx=XM_023255536.1;refGene_gn=RYR3;refGene_tx=XM_023255537.1;refGene_gn=RYR3;refGene_tx=XM_023255538.1
B3	71188118	71188275
refGene_gn=RYR3;refGene_tx=XM_023255535.1;refGene_gn=RYR3;refGene_tx=XM_023255536.1;refGene_gn=RYR3;refGene_tx=XM_023255537.1;refGene_gn=RYR3;refGene_tx=XM_023255538.1
B3	71199255	71199429
refGene_gn=RYR3;refGene_tx=XM_023255535.1;refGene_gn=RYR3;refGene_tx=XM_023255536.1;refGene_gn=RYR3;refGene_tx=XM_023255537.1;refGene_gn=RYR3;refGene_tx=XM_023255538.1
B3	71203324	71203446
refGene_gn=RYR3;refGene_tx=XM_023255535.1;refGene_gn=RYR3;refGene_tx=XM_023255536.1;refGene_gn=RYR3;refGene_tx=XM_023255537.1;refGene_gn=RYR3;refGene_tx=XM_023255538.1
B3	71214063	71214232
refGene_gn=RYR3;refGene_tx=XM_023255535.1;refGene_gn=RYR3;refGene_tx=XM_023255536.1;refGene_gn=RYR3;refGene_tx=XM_023255537.1;refGene_gn=RYR3;refGene_tx=XM_023255538.1
B3	71215581	71215717
refGene_gn=RYR3;refGene_tx=XM_023255535.1;refGene_gn=RYR3;refGene_tx=XM_023255536.1;refGene_gn=RYR3;refGene_tx=XM_023255537.1;refGene_gn=RYR3;refGene_tx=XM_023255538.1
B3	71218618	71218714

refGene_gn=RYR3;refGene_tx=XM_023255535.1;refGene_gn=RYR3;refGene_tx=XM_023255536.1;refGene_gn=RYR3;refGene_tx=XM_023255537.1;refGene_gn=RYR3;refGene_tx=XM_023255538.1
B3 71220489 71220608
refGene_gn=RYR3;refGene_tx=XM_023255535.1;refGene_gn=RYR3;refGene_tx=XM_023255536.1;refGene_gn=RYR3;refGene_tx=XM_023255537.1;refGene_gn=RYR3;refGene_tx=XM_023255538.1
B3 71227800 71227934
refGene_gn=RYR3;refGene_tx=XM_023255535.1;refGene_gn=RYR3;refGene_tx=XM_023255536.1;refGene_gn=RYR3;refGene_tx=XM_023255537.1;refGene_gn=RYR3;refGene_tx=XM_023255538.1
B3 71230436 71230678
refGene_gn=RYR3;refGene_tx=XM_023255535.1;refGene_gn=RYR3;refGene_tx=XM_023255536.1;refGene_gn=RYR3;refGene_tx=XM_023255537.1;refGene_gn=RYR3;refGene_tx=XM_023255538.1
B3 71239724 71239917
refGene_gn=RYR3;refGene_tx=XM_023255535.1;refGene_gn=RYR3;refGene_tx=XM_023255536.1;refGene_gn=RYR3;refGene_tx=XM_023255537.1;refGene_gn=RYR3;refGene_tx=XM_023255538.1
B3 71251465 71251682
refGene_gn=RYR3;refGene_tx=XM_023255535.1;refGene_gn=RYR3;refGene_tx=XM_023255536.1;refGene_gn=RYR3;refGene_tx=XM_023255537.1;refGene_gn=RYR3;refGene_tx=XM_023255538.1
B3 71256868 71256973
refGene_gn=RYR3;refGene_tx=XM_023255535.1;refGene_gn=RYR3;refGene_tx=XM_023255536.1;refGene_gn=RYR3;refGene_tx=XM_023255537.1;refGene_gn=RYR3;refGene_tx=XM_023255538.1
B3 71258809 71258913
refGene_gn=RYR3;refGene_tx=XM_023255535.1;refGene_gn=RYR3;refGene_tx=XM_023255536.1;refGene_gn=RYR3;refGene_tx=XM_023255537.1;refGene_gn=RYR3;refGene_tx=XM_023255538.1
B3 71259179 71259263
refGene_gn=RYR3;refGene_tx=XM_023255535.1;refGene_gn=RYR3;refGene_tx=XM_023255536.1;refGene_gn=RYR3;refGene_tx=XM_023255537.1;refGene_gn=RYR3;refGene_tx=XM_023255538.1
B3 71260931 71261091
refGene_gn=RYR3;refGene_tx=XM_023255535.1;refGene_gn=RYR3;refGene_tx=XM_023255536.1;refGene_gn=RYR3;refGene_tx=XM_023255537.1;refGene_gn=RYR3;refGene_tx=XM_023255538.1
B3 71262450 71262598
refGene_gn=RYR3;refGene_tx=XM_023255535.1;refGene_gn=RYR3;refGene_tx=XM_023255536.1;refGene_gn=RYR3;refGene_tx=XM_023255537.1;refGene_gn=RYR3;refGene_tx=XM_023255538.1
B3 71263468 71263674
refGene_gn=RYR3;refGene_tx=XM_023255535.1;refGene_gn=RYR3;refGene_tx=XM_023255536.1;refGene_gn=RYR3;refGene_tx=XM_023255537.1;refGene_gn=RYR3;refGene_tx=XM_023255538.1
B3 71264257 71264432
refGene_gn=RYR3;refGene_tx=XM_023255535.1;refGene_gn=RYR3;refGene_tx=XM_023255536.1;refGene_gn=RYR3;refGene_tx=XM_023255537.1;refGene_gn=RYR

3;refGene_tx=XM_023255538.1
B3	71272927	71273136
refGene_gn=RYR3;refGene_tx=XM_023255535.1;refGene_gn=RYR3;refGene_tx=XM_023255536.1;refGene_gn=RYR3;refGene_tx=XM_023255537.1;refGene_gn=RYR3;refGene_tx=XM_023255538.1
B3	71275032	71275208
refGene_gn=RYR3;refGene_tx=XM_023255535.1;refGene_gn=RYR3;refGene_tx=XM_023255536.1;refGene_gn=RYR3;refGene_tx=XM_023255537.1;refGene_gn=RYR3;refGene_tx=XM_023255538.1
B3	71276533	71276570
refGene_gn=RYR3;refGene_tx=XM_023255535.1;refGene_gn=RYR3;refGene_tx=XM_023255536.1;refGene_gn=RYR3;refGene_tx=XM_023255537.1;refGene_gn=RYR3;refGene_tx=XM_023255538.1
B3	71278303	71278467
refGene_gn=RYR3;refGene_tx=XM_023255535.1;refGene_gn=RYR3;refGene_tx=XM_023255536.1;refGene_gn=RYR3;refGene_tx=XM_023255537.1;refGene_gn=RYR3;refGene_tx=XM_023255538.1
B3	71281528	71281694
refGene_gn=RYR3;refGene_tx=XM_023255535.1;refGene_gn=RYR3;refGene_tx=XM_023255536.1;refGene_gn=RYR3;refGene_tx=XM_023255537.1;refGene_gn=RYR3;refGene_tx=XM_023255538.1
B3	71288057	71288144
refGene_gn=RYR3;refGene_tx=XM_023255535.1;refGene_gn=RYR3;refGene_tx=XM_023255536.1;refGene_gn=RYR3;refGene_tx=XM_023255537.1;refGene_gn=RYR3;refGene_tx=XM_023255538.1
B3	71288476	71288703
refGene_gn=RYR3;refGene_tx=XM_023255535.1;refGene_gn=RYR3;refGene_tx=XM_023255536.1;refGene_gn=RYR3;refGene_tx=XM_023255537.1;refGene_gn=RYR3;refGene_tx=XM_023255538.1
B3	71290393	71291189
refGene_gn=RYR3;refGene_tx=XM_023255535.1;refGene_gn=RYR3;refGene_tx=XM_023255536.1;refGene_gn=RYR3;refGene_tx=XM_023255537.1;refGene_gn=RYR3;refGene_tx=XM_023255538.1
B3	71293305	71293506
refGene_gn=RYR3;refGene_tx=XM_023255535.1;refGene_gn=RYR3;refGene_tx=XM_023255536.1;refGene_gn=RYR3;refGene_tx=XM_023255537.1;refGene_gn=RYR3;refGene_tx=XM_023255538.1
B3	71298647	71298750
refGene_gn=RYR3;refGene_tx=XM_023255535.1;refGene_gn=RYR3;refGene_tx=XM_023255536.1;refGene_gn=RYR3;refGene_tx=XM_023255537.1;refGene_gn=RYR3;refGene_tx=XM_023255538.1
B3	71300321	71300459
refGene_gn=RYR3;refGene_tx=XM_023255535.1;refGene_gn=RYR3;refGene_tx=XM_023255536.1;refGene_gn=RYR3;refGene_tx=XM_023255537.1;refGene_gn=RYR3;refGene_tx=XM_023255538.1
B3	71318998	71319272
refGene_gn=RYR3;refGene_tx=XM_023255535.1;refGene_gn=RYR3;refGene_tx=XM_023255536.1;refGene_gn=RYR3;refGene_tx=XM_023255537.1;refGene_gn=RYR3;refGene_tx=XM_023255538.1
B3	71322301	71322416

refGene_gn=RYR3;refGene_tx=XM_023255535.1;refGene_gn=RYR3;refGene_tx=XM_023255536.1;refGene_gn=RYR3;refGene_tx=XM_023255537.1;refGene_gn=RYR3;refGene_tx=XM_023255538.1
B3 71324060 71324190
refGene_gn=RYR3;refGene_tx=XM_023255535.1;refGene_gn=RYR3;refGene_tx=XM_023255536.1;refGene_gn=RYR3;refGene_tx=XM_023255537.1;refGene_gn=RYR3;refGene_tx=XM_023255538.1
B3 71325296 71325400
refGene_gn=RYR3;refGene_tx=XM_023255535.1;refGene_gn=RYR3;refGene_tx=XM_023255536.1;refGene_gn=RYR3;refGene_tx=XM_023255537.1;refGene_gn=RYR3;refGene_tx=XM_023255538.1
B3 71328578 71328714
refGene_gn=RYR3;refGene_tx=XM_023255535.1;refGene_gn=RYR3;refGene_tx=XM_023255536.1;refGene_gn=RYR3;refGene_tx=XM_023255537.1;refGene_gn=RYR3;refGene_tx=XM_023255538.1
B3 71339125 71339306
refGene_gn=RYR3;refGene_tx=XM_023255535.1;refGene_gn=RYR3;refGene_tx=XM_023255536.1;refGene_gn=RYR3;refGene_tx=XM_023255537.1;refGene_gn=RYR3;refGene_tx=XM_023255538.1
B3 71340723 71340832
refGene_gn=RYR3;refGene_tx=XM_023255535.1;refGene_gn=RYR3;refGene_tx=XM_023255536.1;refGene_gn=RYR3;refGene_tx=XM_023255537.1;refGene_gn=RYR3;refGene_tx=XM_023255538.1
B3 71343265 71343386
refGene_gn=RYR3;refGene_tx=XM_023255535.1;refGene_gn=RYR3;refGene_tx=XM_023255536.1;refGene_gn=RYR3;refGene_tx=XM_023255537.1;refGene_gn=RYR3;refGene_tx=XM_023255538.1
B3 71351221 71351391
refGene_gn=RYR3;refGene_tx=XM_023255535.1;refGene_gn=RYR3;refGene_tx=XM_023255536.1;refGene_gn=RYR3;refGene_tx=XM_023255537.1;refGene_gn=RYR3;refGene_tx=XM_023255538.1
B3 71353775 71353996
refGene_gn=RYR3;refGene_tx=XM_023255535.1;refGene_gn=RYR3;refGene_tx=XM_023255536.1;refGene_gn=RYR3;refGene_tx=XM_023255537.1;refGene_gn=RYR3;refGene_tx=XM_023255538.1
B3 71356423 71356514
refGene_gn=RYR3;refGene_tx=XM_023255535.1;refGene_gn=RYR3;refGene_tx=XM_023255536.1;refGene_gn=RYR3;refGene_tx=XM_023255537.1;refGene_gn=RYR3;refGene_tx=XM_023255538.1
B3 71358134 71358275
refGene_gn=RYR3;refGene_tx=XM_023255535.1;refGene_gn=RYR3;refGene_tx=XM_023255536.1;refGene_gn=RYR3;refGene_tx=XM_023255537.1;refGene_gn=RYR3;refGene_tx=XM_023255538.1
B3 71359502 71359666
refGene_gn=RYR3;refGene_tx=XM_023255535.1;refGene_gn=RYR3;refGene_tx=XM_023255536.1;refGene_gn=RYR3;refGene_tx=XM_023255537.1;refGene_gn=RYR3;refGene_tx=XM_023255538.1
B3 71361749 71361828
refGene_gn=RYR3;refGene_tx=XM_023255535.1;refGene_gn=RYR3;refGene_tx=XM_023255536.1;refGene_gn=RYR3;refGene_tx=XM_023255537.1;refGene_gn=RYR

3;refGene_tx=XM_023255538.1
B3	71365778	71365868
refGene_gn=RYR3;refGene_tx=XM_023255535.1;refGene_gn=RYR3;refGene_tx=XM_023255536.1;refGene_gn=RYR3;refGene_tx=XM_023255537.1;refGene_gn=RYR3;refGene_tx=XM_023255538.1
B3	71366878	71367025
refGene_gn=RYR3;refGene_tx=XM_023255535.1;refGene_gn=RYR3;refGene_tx=XM_023255536.1;refGene_gn=RYR3;refGene_tx=XM_023255537.1;refGene_gn=RYR3;refGene_tx=XM_023255538.1
B3	71367235	71367298
refGene_gn=RYR3;refGene_tx=XM_023255535.1;refGene_gn=RYR3;refGene_tx=XM_023255536.1;refGene_gn=RYR3;refGene_tx=XM_023255537.1;refGene_gn=RYR3;refGene_tx=XM_023255538.1
B3	71368716	71368792
refGene_gn=RYR3;refGene_tx=XM_023255535.1;refGene_gn=RYR3;refGene_tx=XM_023255536.1;refGene_gn=RYR3;refGene_tx=XM_023255537.1;refGene_gn=RYR3;refGene_tx=XM_023255538.1
B3	71368911	71369035
refGene_gn=RYR3;refGene_tx=XM_023255535.1;refGene_gn=RYR3;refGene_tx=XM_023255536.1;refGene_gn=RYR3;refGene_tx=XM_023255537.1;refGene_gn=RYR3;refGene_tx=XM_023255538.1
B3	71371790	71371906
refGene_gn=RYR3;refGene_tx=XM_023255535.1;refGene_gn=RYR3;refGene_tx=XM_023255536.1;refGene_gn=RYR3;refGene_tx=XM_023255537.1;refGene_gn=RYR3;refGene_tx=XM_023255538.1
B3	71372454	71372522
refGene_gn=RYR3;refGene_tx=XM_023255535.1;refGene_gn=RYR3;refGene_tx=XM_023255536.1;refGene_gn=RYR3;refGene_tx=XM_023255537.1;refGene_gn=RYR3;refGene_tx=XM_023255538.1
B3	71373453	71373575
refGene_gn=RYR3;refGene_tx=XM_023255535.1;refGene_gn=RYR3;refGene_tx=XM_023255536.1;refGene_gn=RYR3;refGene_tx=XM_023255537.1;refGene_gn=RYR3;refGene_tx=XM_023255538.1
B3	71378522	71378572
refGene_gn=RYR3;refGene_tx=XM_023255535.1;refGene_gn=RYR3;refGene_tx=XM_023255536.1;refGene_gn=RYR3;refGene_tx=XM_023255537.1;refGene_gn=RYR3;refGene_tx=XM_023255538.1
B3	71379164	71379225
refGene_gn=RYR3;refGene_tx=XM_023255535.1;refGene_gn=RYR3;refGene_tx=XM_023255536.1;refGene_gn=RYR3;refGene_tx=XM_023255537.1;refGene_gn=RYR3;refGene_tx=XM_023255538.1
B3	71381924	71382163
refGene_gn=RYR3;refGene_tx=XM_023255535.1;refGene_gn=RYR3;refGene_tx=XM_023255536.1;refGene_gn=RYR3;refGene_tx=XM_023255537.1;refGene_gn=RYR3;refGene_tx=XM_023255538.1
B3	71383800	71383882
refGene_gn=RYR3;refGene_tx=XM_023255535.1;refGene_gn=RYR3;refGene_tx=XM_023255536.1;refGene_gn=RYR3;refGene_tx=XM_023255537.1;refGene_gn=RYR3;refGene_tx=XM_023255538.1
B3	71392753	71392884

refGene_gn=RYR3;refGene_tx=XM_023255535.1;refGene_gn=RYR3;refGene_tx=XM_023255536.1;refGene_gn=RYR3;refGene_tx=XM_023255537.1;refGene_gn=RYR3;refGene_tx=XM_023255538.1
B3 71399157 71399478
refGene_gn=RYR3;refGene_tx=XM_023255535.1;refGene_gn=RYR3;refGene_tx=XM_023255536.1;refGene_gn=RYR3;refGene_tx=XM_023255537.1;refGene_gn=RYR3;refGene_tx=XM_023255538.1
B3 71401797 71402038
refGene_gn=RYR3;refGene_tx=XM_023255535.1;refGene_gn=RYR3;refGene_tx=XM_023255536.1;refGene_gn=RYR3;refGene_tx=XM_023255537.1;refGene_gn=RYR3;refGene_tx=XM_023255538.1
B3 71408966 71409054
refGene_gn=RYR3;refGene_tx=XM_023255535.1;refGene_gn=RYR3;refGene_tx=XM_023255536.1;refGene_gn=RYR3;refGene_tx=XM_023255537.1;refGene_gn=RYR3;refGene_tx=XM_023255538.1
B3 71409162 71409255
refGene_gn=RYR3;refGene_tx=XM_023255535.1;refGene_gn=RYR3;refGene_tx=XM_023255536.1;refGene_gn=RYR3;refGene_tx=XM_023255537.1;refGene_gn=RYR3;refGene_tx=XM_023255538.1
B3 71413504 71413519
refGene_gn=RYR3;refGene_tx=XM_023255535.1;refGene_gn=RYR3;refGene_tx=XM_023255536.1;refGene_gn=RYR3;refGene_tx=XM_023255537.1;refGene_gn=RYR3;refGene_tx=XM_023255538.1
B3 71416238 71416409
refGene_gn=RYR3;refGene_tx=XM_023255535.1;refGene_gn=RYR3;refGene_tx=XM_023255536.1;refGene_gn=RYR3;refGene_tx=XM_023255537.1;refGene_gn=RYR3;refGene_tx=XM_023255538.1
B3 71417675 71417735
refGene_gn=RYR3;refGene_tx=XM_023255535.1;refGene_gn=RYR3;refGene_tx=XM_023255536.1;refGene_gn=RYR3;refGene_tx=XM_023255537.1;refGene_gn=RYR3;refGene_tx=XM_023255538.1
B3 71419262 71419394
refGene_gn=RYR3;refGene_tx=XM_023255535.1;refGene_gn=RYR3;refGene_tx=XM_023255536.1;refGene_gn=RYR3;refGene_tx=XM_023255537.1;refGene_gn=RYR3;refGene_tx=XM_023255538.1
B3 71420396 71420509
refGene_gn=RYR3;refGene_tx=XM_023255535.1;refGene_gn=RYR3;refGene_tx=XM_023255536.1;refGene_gn=RYR3;refGene_tx=XM_023255537.1;refGene_gn=RYR3;refGene_tx=XM_023255538.1
B3 71423304 71423401
refGene_gn=RYR3;refGene_tx=XM_023255535.1;refGene_gn=RYR3;refGene_tx=XM_023255536.1;refGene_gn=RYR3;refGene_tx=XM_023255537.1;refGene_gn=RYR3;refGene_tx=XM_023255538.1
B3 71424985 71425092
refGene_gn=RYR3;refGene_tx=XM_023255535.1;refGene_gn=RYR3;refGene_tx=XM_023255536.1;refGene_gn=RYR3;refGene_tx=XM_023255537.1;refGene_gn=RYR3;refGene_tx=XM_023255538.1
B3 71425512 71425564
refGene_gn=RYR3;refGene_tx=XM_023255535.1;refGene_gn=RYR3;refGene_tx=XM_023255536.1;refGene_gn=RYR3;refGene_tx=XM_023255537.1;refGene_gn=RYR

3;refGene_tx=XM_023255538.1
B3	71426547	71426604
refGene_gn=RYR3;refGene_tx=XM_023255535.1;refGene_gn=RYR3;refGene_tx=XM_023255536.1;refGene_gn=RYR3;refGene_tx=XM_023255537.1;refGene_gn=RYR3;refGene_tx=XM_023255538.1
B3	71427791	71427891
refGene_gn=RYR3;refGene_tx=XM_023255535.1;refGene_gn=RYR3;refGene_tx=XM_023255536.1;refGene_gn=RYR3;refGene_tx=XM_023255537.1;refGene_gn=RYR3;refGene_tx=XM_023255538.1
B3	71428042	71428122
refGene_gn=RYR3;refGene_tx=XM_023255535.1;refGene_gn=RYR3;refGene_tx=XM_023255536.1;refGene_gn=RYR3;refGene_tx=XM_023255537.1;refGene_gn=RYR3;refGene_tx=XM_023255538.1
B3	71429663	71429740
refGene_gn=RYR3;refGene_tx=XM_023255535.1;refGene_gn=RYR3;refGene_tx=XM_023255536.1;refGene_gn=RYR3;refGene_tx=XM_023255537.1;refGene_gn=RYR3;refGene_tx=XM_023255538.1
B3	71433011	71433085
refGene_gn=RYR3;refGene_tx=XM_023255535.1;refGene_gn=RYR3;refGene_tx=XM_023255536.1;refGene_gn=RYR3;refGene_tx=XM_023255537.1;refGene_gn=RYR3;refGene_tx=XM_023255538.1
B3	71433333	71433351
refGene_gn=RYR3;refGene_tx=XM_023255535.1;refGene_gn=RYR3;refGene_tx=XM_023255536.1;refGene_gn=RYR3;refGene_tx=XM_023255537.1;refGene_gn=RYR3;refGene_tx=XM_023255538.1
B3	71433788	71433869
refGene_gn=RYR3;refGene_tx=XM_023255535.1;refGene_gn=RYR3;refGene_tx=XM_023255536.1;refGene_gn=RYR3;refGene_tx=XM_023255537.1;refGene_gn=RYR3;refGene_tx=XM_023255538.1
B3	71434356	71434445
refGene_gn=RYR3;refGene_tx=XM_023255535.1;refGene_gn=RYR3;refGene_tx=XM_023255536.1;refGene_gn=RYR3;refGene_tx=XM_023255537.1;refGene_gn=RYR3;refGene_tx=XM_023255538.1
B3	71435874	71436003
refGene_gn=RYR3;refGene_tx=XM_023255535.1;refGene_gn=RYR3;refGene_tx=XM_023255536.1;refGene_gn=RYR3;refGene_tx=XM_023255537.1;refGene_gn=RYR3;refGene_tx=XM_023255538.1
B3	71439095	71439200
refGene_gn=RYR3;refGene_tx=XM_023255535.1;refGene_gn=RYR3;refGene_tx=XM_023255536.1;refGene_gn=RYR3;refGene_tx=XM_023255537.1;refGene_gn=RYR3;refGene_tx=XM_023255538.1
B3	71440937	71441019
refGene_gn=RYR3;refGene_tx=XM_023255535.1;refGene_gn=RYR3;refGene_tx=XM_023255536.1;refGene_gn=RYR3;refGene_tx=XM_023255537.1
B3	71441671	71442999
refGene_gn=RYR3;refGene_tx=XM_023255535.1;refGene_gn=RYR3;refGene_tx=XM_023255536.1;refGene_gn=RYR3;refGene_tx=XM_023255537.1
B3	71445257	71445316
refGene_gn=RYR3;refGene_tx=XM_023255535.1;refGene_gn=RYR3;refGene_tx=XM_023255536.1;refGene_gn=RYR3;refGene_tx=XM_023255537.1

B3 71446222 71446394
refGene_gn=RYR3;refGene_tx=XM_023255535.1;refGene_gn=RYR3;refGene_tx=XM_023255536.1;refGene_gn=RYR3;refGene_tx=XM_023255537.1
B3 71447677 71447764
refGene_gn=RYR3;refGene_tx=XM_023255535.1;refGene_gn=RYR3;refGene_tx=XM_023255536.1;refGene_gn=RYR3;refGene_tx=XM_023255537.1
B3 71449198 71449399
refGene_gn=RYR3;refGene_tx=XM_023255535.1;refGene_gn=RYR3;refGene_tx=XM_023255536.1;refGene_gn=RYR3;refGene_tx=XM_023255537.1
B3 71452472 71452603
refGene_gn=RYR3;refGene_tx=XM_023255535.1;refGene_gn=RYR3;refGene_tx=XM_023255536.1;refGene_gn=RYR3;refGene_tx=XM_023255537.1
B3 71456942 71456985
refGene_gn=RYR3;refGene_tx=XM_023255535.1;refGene_gn=RYR3;refGene_tx=XM_023255536.1;refGene_gn=RYR3;refGene_tx=XM_023255537.1
B3 71457729 71457857
refGene_gn=RYR3;refGene_tx=XM_023255535.1;refGene_gn=RYR3;refGene_tx=XM_023255536.1;refGene_gn=RYR3;refGene_tx=XM_023255537.1
B3 71458270 71458331
refGene_gn=RYR3;refGene_tx=XM_023255535.1;refGene_gn=RYR3;refGene_tx=XM_023255536.1;refGene_gn=RYR3;refGene_tx=XM_023255537.1
B3 71458612 71458759
refGene_gn=RYR3;refGene_tx=XM_023255535.1;refGene_gn=RYR3;refGene_tx=XM_023255536.1;refGene_gn=RYR3;refGene_tx=XM_023255537.1
B3 71462350 71462485
refGene_gn=RYR3;refGene_tx=XM_023255535.1;refGene_gn=RYR3;refGene_tx=XM_023255536.1;refGene_gn=RYR3;refGene_tx=XM_023255537.1
B3 71463782 71463939
refGene_gn=RYR3;refGene_tx=XM_023255535.1;refGene_gn=RYR3;refGene_tx=XM_023255536.1;refGene_gn=RYR3;refGene_tx=XM_023255537.1
B3 71465017 71465082
refGene_gn=RYR3;refGene_tx=XM_023255535.1;refGene_gn=RYR3;refGene_tx=XM_023255536.1;refGene_gn=RYR3;refGene_tx=XM_023255537.1
B3 71465732 71465833
refGene_gn=RYR3;refGene_tx=XM_023255535.1;refGene_gn=RYR3;refGene_tx=XM_023255536.1;refGene_gn=RYR3;refGene_tx=XM_023255537.1
B3 71467847 71467899
refGene_gn=RYR3;refGene_tx=XM_023255535.1;refGene_gn=RYR3;refGene_tx=XM_023255536.1;refGene_gn=RYR3;refGene_tx=XM_023255537.1
B3 71469192 71469288
refGene_gn=RYR3;refGene_tx=XM_023255535.1;refGene_gn=RYR3;refGene_tx=XM_023255536.1;refGene_gn=RYR3;refGene_tx=XM_023255537.1
B3 71470632 71470748 refGene_gn=AVEN;refGene_tx=XM_006932680.2
B3 71471524 71471876 refGene_gn=AVEN;refGene_tx=XM_006932680.2
B3 71474666 71474762 refGene_gn=AVEN;refGene_tx=XM_006932680.2
B3 71476572 71476643 refGene_gn=AVEN;refGene_tx=XM_006932680.2
B3 71620583 71620761 refGene_gn=AVEN;refGene_tx=XM_006932680.2
B3 71669502 71669983 refGene_gn=AVEN;refGene_tx=XM_006932680.2
B3 71670616 71670621 refGene_gn=AVEN;refGene_tx=XM_006932680.2
B3 71684578 71686177 refGene_gn=CHRM5;refGene_tx=XM_003987304.5

B3 71703921 71704074 refGene_gn=EMC7;refGene_tx=XM_003987305.5
B3 71707868 71707949 refGene_gn=EMC7;refGene_tx=XM_003987305.5
B3 71709222 71709361 refGene_gn=EMC7;refGene_tx=XM_003987305.5
B3 71713587 71713707 refGene_gn=EMC7;refGene_tx=XM_003987305.5
B3 71719071 71719304 refGene_gn=EMC7;refGene_tx=XM_003987305.5
B3 71758029 71758062 refGene_gn=KATNBL1;refGene_tx=XM_023255540.1
B3 71760941 71761035 refGene_gn=KATNBL1;refGene_tx=XM_023255540.1
B3 71761842 71761932 refGene_gn=KATNBL1;refGene_tx=XM_023255540.1
B3 71762313 71762403 refGene_gn=KATNBL1;refGene_tx=XM_023255540.1
B3 71762492 71762543 refGene_gn=KATNBL1;refGene_tx=XM_023255540.1
B3 71763664 71763783 refGene_gn=KATNBL1;refGene_tx=XM_023255540.1
B3 71769099 71769379 refGene_gn=KATNBL1;refGene_tx=XM_023255540.1
B3 71770832 71770873 refGene_gn=KATNBL1;refGene_tx=XM_023255540.1
B3 71771801 71771918 refGene_gn=KATNBL1;refGene_tx=XM_023255540.1
B3 71818612 71818698
refGene_gn=EMC4;refGene_tx=XM_003987307.3;refGene_gn=EMC4;refGene_tx=XM_006932662.2
B3 71819038 71819153
refGene_gn=EMC4;refGene_tx=XM_003987307.3;refGene_gn=EMC4;refGene_tx=XM_006932662.2
B3 71822349 71822503
refGene_gn=EMC4;refGene_tx=XM_003987307.3;refGene_gn=EMC4;refGene_tx=XM_006932662.2;refGene_gn=EMC4;refGene_tx=XM_023255541.1;refGene_gn=EMC4;refGene_tx=XM_023255542.1
B3 71822795 71822956
refGene_gn=EMC4;refGene_tx=XM_003987307.3;refGene_gn=EMC4;refGene_tx=XM_023255541.1;refGene_gn=EMC4;refGene_tx=XM_023255542.1
B3 71823184 71823303
refGene_gn=EMC4;refGene_tx=XM_003987307.3;refGene_gn=EMC4;refGene_tx=XM_006932662.2;refGene_gn=EMC4;refGene_tx=XM_023255541.1;refGene_gn=EMC4;refGene_tx=XM_023255542.1
B3 71826637 71826729
refGene_gn=SLC12A6;refGene_tx=XM_003987308.5;refGene_gn=SLC12A6;refGene_tx=XM_011283087.3;refGene_gn=SLC12A6;refGene_tx=XM_019832715.2
B3 71827660 71827794
refGene_gn=SLC12A6;refGene_tx=XM_003987308.5;refGene_gn=SLC12A6;refGene_tx=XM_011283087.3;refGene_gn=SLC12A6;refGene_tx=XM_019832715.2
B3 71828185 71828370
refGene_gn=SLC12A6;refGene_tx=XM_003987308.5;refGene_gn=SLC12A6;refGene_tx=XM_011283087.3;refGene_gn=SLC12A6;refGene_tx=XM_019832715.2
B3 71828583 71828691
refGene_gn=SLC12A6;refGene_tx=XM_003987308.5;refGene_gn=SLC12A6;refGene_tx=XM_011283087.3;refGene_gn=SLC12A6;refGene_tx=XM_019832715.2
B3 71829288 71829420
refGene_gn=SLC12A6;refGene_tx=XM_003987308.5;refGene_gn=SLC12A6;refGene_tx=XM_011283087.3;refGene_gn=SLC12A6;refGene_tx=XM_019832715.2
B3 71830291 71830461
refGene_gn=SLC12A6;refGene_tx=XM_003987308.5;refGene_gn=SLC12A6;refGene_tx=XM_011283087.3;refGene_gn=SLC12A6;refGene_tx=XM_019832715.2
B3 71831482 71831678

refGene_gn=SLC12A6;refGene_tx=XM_003987308.5;refGene_gn=SLC12A6;refGene_tx=XM_011283087.3;refGene_gn=SLC12A6;refGene_tx=XM_019832715.2
B3 71833206 71833375
refGene_gn=SLC12A6;refGene_tx=XM_003987308.5;refGene_gn=SLC12A6;refGene_tx=XM_011283087.3;refGene_gn=SLC12A6;refGene_tx=XM_019832715.2
B3 71833798 71833903
refGene_gn=SLC12A6;refGene_tx=XM_003987308.5;refGene_gn=SLC12A6;refGene_tx=XM_011283087.3;refGene_gn=SLC12A6;refGene_tx=XM_019832715.2
B3 71834683 71834803
refGene_gn=SLC12A6;refGene_tx=XM_003987308.5;refGene_gn=SLC12A6;refGene_tx=XM_011283087.3;refGene_gn=SLC12A6;refGene_tx=XM_019832715.2
B3 71836501 71836600
refGene_gn=SLC12A6;refGene_tx=XM_003987308.5;refGene_gn=SLC12A6;refGene_tx=XM_011283087.3;refGene_gn=SLC12A6;refGene_tx=XM_019832715.2
B3 71838004 71838123
refGene_gn=SLC12A6;refGene_tx=XM_003987308.5;refGene_gn=SLC12A6;refGene_tx=XM_011283087.3;refGene_gn=SLC12A6;refGene_tx=XM_019832715.2
B3 71838608 71838783
refGene_gn=SLC12A6;refGene_tx=XM_003987308.5;refGene_gn=SLC12A6;refGene_tx=XM_011283087.3;refGene_gn=SLC12A6;refGene_tx=XM_019832715.2
B3 71841598 71841656
refGene_gn=SLC12A6;refGene_tx=XM_003987308.5;refGene_gn=SLC12A6;refGene_tx=XM_011283087.3;refGene_gn=SLC12A6;refGene_tx=XM_019832715.2
B3 71841926 71842025
refGene_gn=SLC12A6;refGene_tx=XM_003987308.5;refGene_gn=SLC12A6;refGene_tx=XM_011283087.3;refGene_gn=SLC12A6;refGene_tx=XM_019832715.2
B3 71842434 71842593
refGene_gn=SLC12A6;refGene_tx=XM_003987308.5;refGene_gn=SLC12A6;refGene_tx=XM_011283087.3;refGene_gn=SLC12A6;refGene_tx=XM_019832715.2
B3 71843394 71843609
refGene_gn=SLC12A6;refGene_tx=XM_003987308.5;refGene_gn=SLC12A6;refGene_tx=XM_011283087.3;refGene_gn=SLC12A6;refGene_tx=XM_019832715.2
B3 71845558 71845800
refGene_gn=SLC12A6;refGene_tx=XM_003987308.5;refGene_gn=SLC12A6;refGene_tx=XM_011283087.3;refGene_gn=SLC12A6;refGene_tx=XM_019832715.2
B3 71846490 71846621
refGene_gn=SLC12A6;refGene_tx=XM_003987308.5;refGene_gn=SLC12A6;refGene_tx=XM_011283087.3;refGene_gn=SLC12A6;refGene_tx=XM_019832715.2
B3 71847886 71847941
refGene_gn=SLC12A6;refGene_tx=XM_003987308.5;refGene_gn=SLC12A6;refGene_tx=XM_011283087.3;refGene_gn=SLC12A6;refGene_tx=XM_019832715.2
B3 71849252 71849399
refGene_gn=SLC12A6;refGene_tx=XM_003987308.5;refGene_gn=SLC12A6;refGene_tx=XM_011283087.3;refGene_gn=SLC12A6;refGene_tx=XM_019832715.2
B3 71850504 71850636
refGene_gn=SLC12A6;refGene_tx=XM_003987308.5;refGene_gn=SLC12A6;refGene_tx=XM_011283087.3;refGene_gn=SLC12A6;refGene_tx=XM_019832715.2
B3 71851879 71851977
refGene_gn=SLC12A6;refGene_tx=XM_003987308.5;refGene_gn=SLC12A6;refGene_tx=XM_011283087.3;refGene_gn=SLC12A6;refGene_tx=XM_019832715.2

B3 71866946 71866991
refGene_gn=SLC12A6;refGene_tx=XM_003987308.5;refGene_gn=SLC12A6;refGene_tx=XM_011283087.3
B3 71892597 71892721 refGene_gn=SLC12A6;refGene_tx=XM_011283087.3
B3 71911836 71912107
refGene_gn=SLC12A6;refGene_tx=XM_003987308.5;refGene_gn=SLC12A6;refGene_tx=XM_019832715.2
B3 71917032 71917173 refGene_gn=NOP10;refGene_tx=XM_003987309.4
B3 71917477 71917531 refGene_gn=NOP10;refGene_tx=XM_003987309.4
B3 71919503 71919519
refGene_gn=NUTM1;refGene_tx=XM_003987352.4;refGene_gn=NUTM1;refGene_tx=XM_019832716.2
B3 71921144 71921844
refGene_gn=NUTM1;refGene_tx=XM_003987352.4;refGene_gn=NUTM1;refGene_tx=XM_019832716.2
B3 71923787 71923916
refGene_gn=NUTM1;refGene_tx=XM_003987352.4;refGene_gn=NUTM1;refGene_tx=XM_019832716.2
B3 71925273 71925413
refGene_gn=NUTM1;refGene_tx=XM_003987352.4;refGene_gn=NUTM1;refGene_tx=XM_019832716.2
B3 71925925 71926215
refGene_gn=NUTM1;refGene_tx=XM_003987352.4;refGene_gn=NUTM1;refGene_tx=XM_019832716.2
B3 71926484 71926601
refGene_gn=NUTM1;refGene_tx=XM_003987352.4;refGene_gn=NUTM1;refGene_tx=XM_019832716.2
B3 71926951 71928964
refGene_gn=NUTM1;refGene_tx=XM_003987352.4;refGene_gn=NUTM1;refGene_tx=XM_019832716.2
B3 71930274 71930429 refGene_gn=LPCAT4;refGene_tx=XM_011283088.3
B3 71930687 71930844 refGene_gn=LPCAT4;refGene_tx=XM_011283088.3
B3 71931171 71931270 refGene_gn=LPCAT4;refGene_tx=XM_011283088.3
B3 71932408 71932541 refGene_gn=LPCAT4;refGene_tx=XM_011283088.3
B3 71933093 71933219 refGene_gn=LPCAT4;refGene_tx=XM_011283088.3
B3 71933476 71933559 refGene_gn=LPCAT4;refGene_tx=XM_011283088.3
B3 71933699 71933754 refGene_gn=LPCAT4;refGene_tx=XM_011283088.3
B3 71934306 71934341 refGene_gn=LPCAT4;refGene_tx=XM_011283088.3
B3 71934531 71934590 refGene_gn=LPCAT4;refGene_tx=XM_011283088.3
B3 71934875 71934936 refGene_gn=LPCAT4;refGene_tx=XM_011283088.3
B3 71935052 71935165 refGene_gn=LPCAT4;refGene_tx=XM_011283088.3
B3 71935542 71935763 refGene_gn=LPCAT4;refGene_tx=XM_011283088.3
B3 71936098 71936241 refGene_gn=LPCAT4;refGene_tx=XM_011283088.3
B3 71937532 71937646 refGene_gn=LPCAT4;refGene_tx=XM_011283088.3
B3 71976967 71977906
refGene_gn=LOC101098718;refGene_tx=XM_003987310.3
B3 72007018 72007999
refGene_gn=LOC101091202;refGene_tx=XM_011283341.3
B3 72021450 72022389
refGene_gn=LOC101098959;refGene_tx=XM_023256229.1

B3 72051281 72052220
refGene_gn=LOC105260633;refGene_tx=XM_023256230.1
B3 72074353 72075292
refGene_gn=LOC101091702;refGene_tx=XM_011283091.2
B3 72097752 72098691
refGene_gn=LOC101099214;refGene_tx=XM_003987312.2
B3 72123529 72124468
refGene_gn=LOC101099470;refGene_tx=XM_003987313.2
B3 72159549 72160488
refGene_gn=LOC105260634;refGene_tx=XM_011283092.2
B3 72182045 72183008
refGene_gn=LOC101091946;refGene_tx=XM_003987355.3
B3 72201607 72202352
refGene_gn=LOC101092189;refGene_tx=XM_011283093.2
B3 72202354 72202545
refGene_gn=LOC101092189;refGene_tx=XM_011283093.2
B3 72243408 72244344
refGene_gn=LOC101092440;refGene_tx=XM_023256171.1
B3 72301884 72302802
refGene_gn=LOC101099972;refGene_tx=XM_003987314.5
B3 72309465 72310404
refGene_gn=LOC101100217;refGene_tx=XM_003987315.1
B3 72324522 72325458
refGene_gn=LOC101100475;refGene_tx=XM_003987316.3
B3 72346990 72347926
refGene_gn=LOC101093190;refGene_tx=XM_003987356.1
B3 72367882 72368818
refGene_gn=LOC101093442;refGene_tx=XM_011283342.2
B3 72382582 72383521
refGene_gn=LOC111560946;refGene_tx=XM_023256172.1
B3 72399108 72400047
refGene_gn=LOC101093693;refGene_tx=XM_003987357.2
B3 72412225 72412393
refGene_gn=LOC101093942;refGene_tx=XM_023256173.1
B3 72412394 72413138
refGene_gn=LOC101093942;refGene_tx=XM_023256173.1
B3 72436624 72437563
refGene_gn=LOC101094185;refGene_tx=XM_003987359.2
B3 72486881 72487790
refGene_gn=LOC111560844;refGene_tx=XM_023255544.1
B3 72511627 72512484
refGene_gn=LOC101094425;refGene_tx=XM_006932682.3
B3 72512486 72512565
refGene_gn=LOC101094425;refGene_tx=XM_006932682.3
B3 72521062 72522004
refGene_gn=LOC101094677;refGene_tx=XM_003987360.2
B3 72537176 72538103
refGene_gn=LOC101100726;refGene_tx=XM_003987317.2
B3 72557716 72558646
refGene_gn=LOC101094933;refGene_tx=XM_003987361.2

B3 72573065 72574022
refGene_gn=LOC101095178;refGene_tx=XM_023256231.1
B3 72592890 72592944
refGene_gn=LOC101095421;refGene_tx=XM_003987362.3
B3 72592945 72593845
refGene_gn=LOC101095421;refGene_tx=XM_003987362.3
B3 72611661 72612594
refGene_gn=LOC101100969;refGene_tx=XM_003987318.4
B3 72640670 72641609
refGene_gn=LOC101095903;refGene_tx=XM_003987363.4
B3 72675048 72675981
refGene_gn=LOC101096153;refGene_tx=XM_006932683.4
B3 72683568 72684510
refGene_gn=LOC101096397;refGene_tx=XM_023256174.1
B3 72703444 72704386
refGene_gn=LOC101096655;refGene_tx=XM_003987365.3
B3 72716828 72717755
refGene_gn=LOC101101223;refGene_tx=XM_003987319.2
B3 72758395 72758759
refGene_gn=LOC101097153;refGene_tx=XM_003987366.2
B3 72758761 72759348
refGene_gn=LOC101097153;refGene_tx=XM_003987366.2
B3 72816145 72817078
refGene_gn=LOC101101474;refGene_tx=XM_011283094.1
B3 72845188 72845237
refGene_gn=LOC101084884;refGene_tx=XM_003987402.2
B3 72846399 72847379
refGene_gn=LOC101084884;refGene_tx=XM_003987402.2
B3 72867589 72867601
refGene_gn=LOC101085131;refGene_tx=XM_023256176.1
B3 72872949 72873562
refGene_gn=LOC101085131;refGene_tx=XM_023256176.1
B3 72873564 72873701
refGene_gn=LOC101085131;refGene_tx=XM_023256176.1
B3 72883508 72884483
refGene_gn=LOC101097811;refGene_tx=XM_003987367.2
B3 72906944 72907909
refGene_gn=LOC101085382;refGene_tx=XM_003987403.1
B3 72917526 72917533
refGene_gn=LOC101085382;refGene_tx=XM_003987403.1
B3 72925539 72926475
refGene_gn=LOC101098057;refGene_tx=XM_003987368.3
B3 72940904 72941876
refGene_gn=LOC101098304;refGene_tx=XM_003987369.2
B3 72955580 72956507
refGene_gn=LOC101085637;refGene_tx=XM_003987404.4
B3 72973123 72974095
refGene_gn=LOC101085895;refGene_tx=XM_006932705.3
B3 73002358 73003309
refGene_gn=LOC101086151;refGene_tx=XM_023256232.1

B3 73020978 73021956
refGene_gn=LOC101098543;refGene_tx=XM_003987370.2
B3 73034295 73035255
refGene_gn=LOC101086399;refGene_tx=XM_023256233.1
B3 73062701 73063625
refGene_gn=LOC101098795;refGene_tx=XM_003987371.3
B3 73078111 73079038
refGene_gn=LOC101086656;refGene_tx=XM_011283095.1
B3 73094779 73095721
refGene_gn=LOC101099051;refGene_tx=XM_023256234.1
B3 73109754 73110693
refGene_gn=LOC101099303;refGene_tx=XM_003987373.1
B3 73123347 73124289
refGene_gn=LOC101099558;refGene_tx=XM_023256235.1
B3 73137132 73138059
refGene_gn=LOC111560988;refGene_tx=XM_023256236.1
B3 73152483 73153470
refGene_gn=LOC111560949;refGene_tx=XM_023256177.1
B3 73166707 73167248
refGene_gn=LOC111560951;refGene_tx=XM_023256178.1
B3 73167250 73167675
refGene_gn=LOC111560951;refGene_tx=XM_023256178.1
B3 73208561 73209530
refGene_gn=LOC111560953;refGene_tx=XM_023256179.1
B3 73230300 73231269
refGene_gn=LOC101088169;refGene_tx=XM_003987412.3
B3 73257396 73258338
refGene_gn=LOC101088672;refGene_tx=XM_003987413.2
B3 73326646 73327573
refGene_gn=LOC101089447;refGene_tx=XM_011283097.3
B3 73355649 73356612
refGene_gn=LOC101089699;refGene_tx=XM_006932685.4
B3 73369828 73370777
refGene_gn=LOC105260660;refGene_tx=XM_011283360.2
B3 73371775 73371864
refGene_gn=LOC105260660;refGene_tx=XM_011283360.2
B3 73378381 73378634
refGene_gn=LOC101099806;refGene_tx=XM_011283361.2
B3 73378636 73378663
refGene_gn=LOC101099806;refGene_tx=XM_011283361.2
B3 73378665 73378742
refGene_gn=LOC101099806;refGene_tx=XM_011283361.2
B3 73378743 73378875
refGene_gn=LOC101099806;refGene_tx=XM_011283361.2
B3 73378876 73379374
refGene_gn=LOC101099806;refGene_tx=XM_011283361.2
B3 73383306 73384251
refGene_gn=LOC111560846;refGene_tx=XM_023255546.1
B3 73458748 73458868 refGene_gn=TTC5;refGene_tx=XM_003987375.4
B3 73461819 73461964 refGene_gn=TTC5;refGene_tx=XM_003987375.4

B3	73465181 73465396 refGene_gn=TTC5;refGene_tx=XM_003987375.4
B3	73466254 73466401 refGene_gn=TTC5;refGene_tx=XM_003987375.4
B3	73467330 73467387 refGene_gn=TTC5;refGene_tx=XM_003987375.4
B3	73470877 73470969 refGene_gn=TTC5;refGene_tx=XM_003987375.4
B3	73472730 73472881 refGene_gn=TTC5;refGene_tx=XM_003987375.4
B3	73473264 73473476 refGene_gn=TTC5;refGene_tx=XM_003987375.4
B3	73475194 73475327 refGene_gn=TTC5;refGene_tx=XM_003987375.4
B3	73477505 73477556 refGene_gn=TTC5;refGene_tx=XM_003987375.4
B3	73479153 73479315
refGene_gn=LOC101090696;refGene_tx=XM_011283351.2
B3	73479316 73479418
refGene_gn=LOC101090696;refGene_tx=XM_011283351.2
B3	73482633 73482836
refGene_gn=CCNB1IP1;refGene_tx=XM_003987376.5;refGene_gn=CCNB1IP1;refGene_tx=XM_019832725.2;refGene_gn=CCNB1IP1;refGene_tx=XM_023255547.1;refGene_gn=CCNB1IP1;refGene_tx=XM_023255549.1
B3	73485104 73485438
refGene_gn=CCNB1IP1;refGene_tx=XM_003987376.5;refGene_gn=CCNB1IP1;refGene_tx=XM_019832725.2;refGene_gn=CCNB1IP1;refGene_tx=XM_023255547.1;refGene_gn=CCNB1IP1;refGene_tx=XM_023255549.1
B3	73492733 73493030
refGene_gn=CCNB1IP1;refGene_tx=XM_003987376.5;refGene_gn=CCNB1IP1;refGene_tx=XM_019832725.2;refGene_gn=CCNB1IP1;refGene_tx=XM_023255547.1;refGene_gn=CCNB1IP1;refGene_tx=XM_023255549.1
B3	73517869 73517912
refGene_gn=PARP2;refGene_tx=XM_003987418.4;refGene_gn=PARP2;refGene_tx=XM_006932687.3;refGene_gn=PARP2;refGene_tx=XM_019832730.2
B3	73518897 73519059
refGene_gn=PARP2;refGene_tx=XM_003987418.4;refGene_gn=PARP2;refGene_tx=XM_006932687.3;refGene_gn=PARP2;refGene_tx=XM_019832730.2
B3	73519350 73519421
refGene_gn=PARP2;refGene_tx=XM_003987418.4;refGene_gn=PARP2;refGene_tx=XM_006932687.3;refGene_gn=PARP2;refGene_tx=XM_019832730.2
B3	73520017 73520068
refGene_gn=PARP2;refGene_tx=XM_003987418.4;refGene_gn=PARP2;refGene_tx=XM_006932687.3;refGene_gn=PARP2;refGene_tx=XM_019832730.2
B3	73520280 73520377
refGene_gn=PARP2;refGene_tx=XM_003987418.4;refGene_gn=PARP2;refGene_tx=XM_006932687.3;refGene_gn=PARP2;refGene_tx=XM_019832730.2
B3	73520786 73520862
refGene_gn=PARP2;refGene_tx=XM_003987418.4;refGene_gn=PARP2;refGene_tx=XM_006932687.3;refGene_gn=PARP2;refGene_tx=XM_019832730.2
B3	73521494 73521597
refGene_gn=PARP2;refGene_tx=XM_003987418.4;refGene_gn=PARP2;refGene_tx=XM_006932687.3;refGene_gn=PARP2;refGene_tx=XM_019832730.2
B3	73522337 73522497
refGene_gn=PARP2;refGene_tx=XM_003987418.4;refGene_gn=PARP2;refGene_tx=XM_006932687.3;refGene_gn=PARP2;refGene_tx=XM_019832730.2
B3	73522756 73522895
refGene_gn=PARP2;refGene_tx=XM_003987418.4;refGene_gn=PARP2;refGene_tx

=XM_006932687.3;refGene_gn=PARP2;refGene_tx=XM_019832730.2
B3 73524448 73524512
refGene_gn=PARP2;refGene_tx=XM_003987418.4;refGene_gn=PARP2;refGene_tx=XM_006932687.3;refGene_gn=PARP2;refGene_tx=XM_019832730.2
B3 73524590 73524725
refGene_gn=PARP2;refGene_tx=XM_003987418.4;refGene_gn=PARP2;refGene_tx=XM_006932687.3;refGene_gn=PARP2;refGene_tx=XM_019832730.2
B3 73524995 73525123
refGene_gn=PARP2;refGene_tx=XM_003987418.4;refGene_gn=PARP2;refGene_tx=XM_006932687.3;refGene_gn=PARP2;refGene_tx=XM_019832730.2
B3 73525280 73525380
refGene_gn=PARP2;refGene_tx=XM_003987418.4;refGene_gn=PARP2;refGene_tx=XM_006932687.3;refGene_gn=PARP2;refGene_tx=XM_019832730.2
B3 73525746 73525845
refGene_gn=PARP2;refGene_tx=XM_003987418.4;refGene_gn=PARP2;refGene_tx=XM_006932687.3;refGene_gn=PARP2;refGene_tx=XM_019832730.2
B3 73526086 73526211
refGene_gn=PARP2;refGene_tx=XM_003987418.4;refGene_gn=PARP2;refGene_tx=XM_006932687.3;refGene_gn=PARP2;refGene_tx=XM_019832730.2
B3 73526329 73526489
refGene_gn=PARP2;refGene_tx=XM_003987418.4;refGene_gn=PARP2;refGene_tx=XM_006932687.3;refGene_gn=PARP2;refGene_tx=XM_019832730.2
B3 73526809 73527022
refGene_gn=TEP1;refGene_tx=XM_023255550.1;refGene_gn=TEP1;refGene_tx=XM_023255551.1;refGene_gn=TEP1;refGene_tx=XM_023255552.1;refGene_gn=TEP1;refGene_tx=XM_023255557.1
B3 73530504 73530528 refGene_gn=TEP1;refGene_tx=XM_023255554.1
B3 73531713 73531836 refGene_gn=TEP1;refGene_tx=XM_023255553.1
B3 73532289 73532394
refGene_gn=TEP1;refGene_tx=XM_023255550.1;refGene_gn=TEP1;refGene_tx=XM_023255551.1;refGene_gn=TEP1;refGene_tx=XM_023255552.1;refGene_gn=TEP1;refGene_tx=XM_023255553.1;refGene_gn=TEP1;refGene_tx=XM_023255554.1;refGene_gn=TEP1;refGene_tx=XM_023255557.1
B3 73532497 73532703
refGene_gn=TEP1;refGene_tx=XM_023255550.1;refGene_gn=TEP1;refGene_tx=XM_023255551.1;refGene_gn=TEP1;refGene_tx=XM_023255552.1;refGene_gn=TEP1;refGene_tx=XM_023255553.1;refGene_gn=TEP1;refGene_tx=XM_023255554.1;refGene_gn=TEP1;refGene_tx=XM_023255555.1;refGene_gn=TEP1;refGene_tx=XM_023255557.1;refGene_gn=TEP1;refGene_tx=XM_023255559.1
B3 73532808 73532914
refGene_gn=TEP1;refGene_tx=XM_023255550.1;refGene_gn=TEP1;refGene_tx=XM_023255551.1;refGene_gn=TEP1;refGene_tx=XM_023255552.1;refGene_gn=TEP1;refGene_tx=XM_023255553.1;refGene_gn=TEP1;refGene_tx=XM_023255554.1;refGene_gn=TEP1;refGene_tx=XM_023255557.1;refGene_gn=TEP1;refGene_tx=XM_023255558.1
B3 73533835 73533932
refGene_gn=TEP1;refGene_tx=XM_023255550.1;refGene_gn=TEP1;refGene_tx=XM_023255551.1;refGene_gn=TEP1;refGene_tx=XM_023255552.1;refGene_gn=TEP1;refGene_tx=XM_023255553.1;refGene_gn=TEP1;refGene_tx=XM_023255554.1;refGene_gn=TEP1;refGene_tx=XM_023255555.1;refGene_gn=TEP1;refGene_tx=X

M_023255557.1;refGene_gn=TEP1;refGene_tx=XM_023255558.1;refGene_gn=TEP1;refGene_tx=XM_023255559.1
B3 73534083 73534227
refGene_gn=TEP1;refGene_tx=XM_023255550.1;refGene_gn=TEP1;refGene_tx=XM_023255551.1;refGene_gn=TEP1;refGene_tx=XM_023255552.1;refGene_gn=TEP1;refGene_tx=XM_023255553.1;refGene_gn=TEP1;refGene_tx=XM_023255554.1;refGene_gn=TEP1;refGene_tx=XM_023255555.1;refGene_gn=TEP1;refGene_tx=XM_023255557.1;refGene_gn=TEP1;refGene_tx=XM_023255558.1;refGene_gn=TEP1;refGene_tx=XM_023255559.1
B3 73535540 73535665
refGene_gn=TEP1;refGene_tx=XM_023255550.1;refGene_gn=TEP1;refGene_tx=XM_023255551.1;refGene_gn=TEP1;refGene_tx=XM_023255552.1;refGene_gn=TEP1;refGene_tx=XM_023255553.1;refGene_gn=TEP1;refGene_tx=XM_023255554.1;refGene_gn=TEP1;refGene_tx=XM_023255555.1;refGene_gn=TEP1;refGene_tx=XM_023255557.1
B3 73535821 73535958
refGene_gn=TEP1;refGene_tx=XM_023255550.1;refGene_gn=TEP1;refGene_tx=XM_023255551.1;refGene_gn=TEP1;refGene_tx=XM_023255552.1;refGene_gn=TEP1;refGene_tx=XM_023255553.1;refGene_gn=TEP1;refGene_tx=XM_023255554.1;refGene_gn=TEP1;refGene_tx=XM_023255555.1;refGene_gn=TEP1;refGene_tx=XM_023255557.1;refGene_gn=TEP1;refGene_tx=XM_023255558.1;refGene_gn=TEP1;refGene_tx=XM_023255559.1;refGene_gn=TEP1;refGene_tx=XM_023255560.1;refGene_gn=TEP1;refGene_tx=XM_023255561.1
B3 73536096 73536229
refGene_gn=TEP1;refGene_tx=XM_023255550.1;refGene_gn=TEP1;refGene_tx=XM_023255551.1;refGene_gn=TEP1;refGene_tx=XM_023255552.1;refGene_gn=TEP1;refGene_tx=XM_023255553.1;refGene_gn=TEP1;refGene_tx=XM_023255554.1;refGene_gn=TEP1;refGene_tx=XM_023255555.1;refGene_gn=TEP1;refGene_tx=XM_023255557.1;refGene_gn=TEP1;refGene_tx=XM_023255558.1;refGene_gn=TEP1;refGene_tx=XM_023255559.1
B3 73536604 73536681
refGene_gn=TEP1;refGene_tx=XM_023255550.1;refGene_gn=TEP1;refGene_tx=XM_023255551.1;refGene_gn=TEP1;refGene_tx=XM_023255552.1;refGene_gn=TEP1;refGene_tx=XM_023255553.1;refGene_gn=TEP1;refGene_tx=XM_023255554.1;refGene_gn=TEP1;refGene_tx=XM_023255555.1;refGene_gn=TEP1;refGene_tx=XM_023255557.1;refGene_gn=TEP1;refGene_tx=XM_023255558.1;refGene_gn=TEP1;refGene_tx=XM_023255559.1;refGene_gn=TEP1;refGene_tx=XM_023255560.1;refGene_gn=TEP1;refGene_tx=XM_023255561.1
B3 73536761 73536925
refGene_gn=TEP1;refGene_tx=XM_023255550.1;refGene_gn=TEP1;refGene_tx=XM_023255551.1;refGene_gn=TEP1;refGene_tx=XM_023255552.1;refGene_gn=TEP1;refGene_tx=XM_023255553.1;refGene_gn=TEP1;refGene_tx=XM_023255554.1;refGene_gn=TEP1;refGene_tx=XM_023255555.1;refGene_gn=TEP1;refGene_tx=XM_023255557.1;refGene_gn=TEP1;refGene_tx=XM_023255558.1;refGene_gn=TEP1;refGene_tx=XM_023255559.1;refGene_gn=TEP1;refGene_tx=XM_023255560.1;refGene_gn=TEP1;refGene_tx=XM_023255561.1;refGene_gn=TEP1;refGene_tx=XM_023255562.1
B3 73537259 73537367
refGene_gn=TEP1;refGene_tx=XM_023255550.1;refGene_gn=TEP1;refGene_tx=XM_023255551.1;refGene_gn=TEP1;refGene_tx=XM_023255552.1;refGene_gn=TEP

1;refGene_tx=XM_023255553.1;refGene_gn=TEP1;refGene_tx=XM_023255554.1;
refGene_gn=TEP1;refGene_tx=XM_023255555.1;refGene_gn=TEP1;refGene_tx=X
M_023255557.1;refGene_gn=TEP1;refGene_tx=XM_023255558.1;refGene_gn=TEP
1;refGene_tx=XM_023255559.1;refGene_gn=TEP1;refGene_tx=XM_023255560.1;
refGene_gn=TEP1;refGene_tx=XM_023255561.1;refGene_gn=TEP1;refGene_tx=X
M_023255562.1
B3 73538570 73538684
refGene_gn=TEP1;refGene_tx=XM_023255550.1;refGene_gn=TEP1;refGene_tx=X
M_023255551.1;refGene_gn=TEP1;refGene_tx=XM_023255552.1;refGene_gn=TEP
1;refGene_tx=XM_023255553.1;refGene_gn=TEP1;refGene_tx=XM_023255554.1;
refGene_gn=TEP1;refGene_tx=XM_023255555.1;refGene_gn=TEP1;refGene_tx=X
M_023255557.1;refGene_gn=TEP1;refGene_tx=XM_023255558.1;refGene_gn=TEP
1;refGene_tx=XM_023255559.1;refGene_gn=TEP1;refGene_tx=XM_023255560.1;
refGene_gn=TEP1;refGene_tx=XM_023255561.1;refGene_gn=TEP1;refGene_tx=X
M_023255562.1
B3 73538925 73539086
refGene_gn=TEP1;refGene_tx=XM_023255550.1;refGene_gn=TEP1;refGene_tx=X
M_023255551.1;refGene_gn=TEP1;refGene_tx=XM_023255552.1;refGene_gn=TEP
1;refGene_tx=XM_023255553.1;refGene_gn=TEP1;refGene_tx=XM_023255554.1;
refGene_gn=TEP1;refGene_tx=XM_023255555.1;refGene_gn=TEP1;refGene_tx=X
M_023255557.1;refGene_gn=TEP1;refGene_tx=XM_023255558.1;refGene_gn=TEP
1;refGene_tx=XM_023255559.1;refGene_gn=TEP1;refGene_tx=XM_023255560.1;
refGene_gn=TEP1;refGene_tx=XM_023255561.1;refGene_gn=TEP1;refGene_tx=X
M_023255562.1
B3 73539713 73539926
refGene_gn=TEP1;refGene_tx=XM_023255550.1;refGene_gn=TEP1;refGene_tx=X
M_023255551.1;refGene_gn=TEP1;refGene_tx=XM_023255552.1;refGene_gn=TEP
1;refGene_tx=XM_023255553.1;refGene_gn=TEP1;refGene_tx=XM_023255554.1;
refGene_gn=TEP1;refGene_tx=XM_023255555.1;refGene_gn=TEP1;refGene_tx=X
M_023255557.1;refGene_gn=TEP1;refGene_tx=XM_023255558.1;refGene_gn=TEP
1;refGene_tx=XM_023255559.1;refGene_gn=TEP1;refGene_tx=XM_023255560.1;
refGene_gn=TEP1;refGene_tx=XM_023255561.1;refGene_gn=TEP1;refGene_tx=X
M_023255562.1
B3 73540032 73540186
refGene_gn=TEP1;refGene_tx=XM_023255550.1;refGene_gn=TEP1;refGene_tx=X
M_023255551.1;refGene_gn=TEP1;refGene_tx=XM_023255552.1;refGene_gn=TEP
1;refGene_tx=XM_023255553.1;refGene_gn=TEP1;refGene_tx=XM_023255554.1;
refGene_gn=TEP1;refGene_tx=XM_023255555.1;refGene_gn=TEP1;refGene_tx=X
M_023255557.1;refGene_gn=TEP1;refGene_tx=XM_023255558.1;refGene_gn=TEP
1;refGene_tx=XM_023255559.1;refGene_gn=TEP1;refGene_tx=XM_023255560.1;
refGene_gn=TEP1;refGene_tx=XM_023255561.1;refGene_gn=TEP1;refGene_tx=X
M_023255562.1
B3 73540439 73540652
refGene_gn=TEP1;refGene_tx=XM_023255550.1;refGene_gn=TEP1;refGene_tx=X
M_023255551.1;refGene_gn=TEP1;refGene_tx=XM_023255552.1;refGene_gn=TEP
1;refGene_tx=XM_023255553.1;refGene_gn=TEP1;refGene_tx=XM_023255554.1;
refGene_gn=TEP1;refGene_tx=XM_023255555.1;refGene_gn=TEP1;refGene_tx=X
M_023255557.1;refGene_gn=TEP1;refGene_tx=XM_023255558.1;refGene_gn=TEP
1;refGene_tx=XM_023255559.1;refGene_gn=TEP1;refGene_tx=XM_023255560.1;
refGene_gn=TEP1;refGene_tx=XM_023255561.1;refGene_gn=TEP1;refGene_tx=X

M_023255562.1
B3 73540798 73540954
refGene_gn=TEP1;refGene_tx=XM_023255550.1;refGene_gn=TEP1;refGene_tx=XM_023255551.1;refGene_gn=TEP1;refGene_tx=XM_023255552.1;refGene_gn=TEP1;refGene_tx=XM_023255553.1;refGene_gn=TEP1;refGene_tx=XM_023255554.1;refGene_gn=TEP1;refGene_tx=XM_023255555.1;refGene_gn=TEP1;refGene_tx=XM_023255557.1;refGene_gn=TEP1;refGene_tx=XM_023255558.1;refGene_gn=TEP1;refGene_tx=XM_023255559.1;refGene_gn=TEP1;refGene_tx=XM_023255560.1;refGene_gn=TEP1;refGene_tx=XM_023255561.1;refGene_gn=TEP1;refGene_tx=XM_023255562.1
B3 73541203 73541303
refGene_gn=TEP1;refGene_tx=XM_023255550.1;refGene_gn=TEP1;refGene_tx=XM_023255551.1;refGene_gn=TEP1;refGene_tx=XM_023255552.1;refGene_gn=TEP1;refGene_tx=XM_023255553.1;refGene_gn=TEP1;refGene_tx=XM_023255554.1;refGene_gn=TEP1;refGene_tx=XM_023255555.1;refGene_gn=TEP1;refGene_tx=XM_023255557.1;refGene_gn=TEP1;refGene_tx=XM_023255558.1;refGene_gn=TEP1;refGene_tx=XM_023255559.1;refGene_gn=TEP1;refGene_tx=XM_023255560.1;refGene_gn=TEP1;refGene_tx=XM_023255561.1;refGene_gn=TEP1;refGene_tx=XM_023255562.1
B3 73541427 73541552
refGene_gn=TEP1;refGene_tx=XM_023255550.1;refGene_gn=TEP1;refGene_tx=XM_023255551.1;refGene_gn=TEP1;refGene_tx=XM_023255552.1;refGene_gn=TEP1;refGene_tx=XM_023255553.1;refGene_gn=TEP1;refGene_tx=XM_023255554.1;refGene_gn=TEP1;refGene_tx=XM_023255555.1;refGene_gn=TEP1;refGene_tx=XM_023255557.1;refGene_gn=TEP1;refGene_tx=XM_023255558.1;refGene_gn=TEP1;refGene_tx=XM_023255559.1;refGene_gn=TEP1;refGene_tx=XM_023255560.1;refGene_gn=TEP1;refGene_tx=XM_023255561.1;refGene_gn=TEP1;refGene_tx=XM_023255562.1
B3 73542395 73542519
refGene_gn=TEP1;refGene_tx=XM_023255550.1;refGene_gn=TEP1;refGene_tx=XM_023255551.1;refGene_gn=TEP1;refGene_tx=XM_023255552.1;refGene_gn=TEP1;refGene_tx=XM_023255553.1;refGene_gn=TEP1;refGene_tx=XM_023255554.1;refGene_gn=TEP1;refGene_tx=XM_023255555.1;refGene_gn=TEP1;refGene_tx=XM_023255557.1;refGene_gn=TEP1;refGene_tx=XM_023255558.1;refGene_gn=TEP1;refGene_tx=XM_023255559.1;refGene_gn=TEP1;refGene_tx=XM_023255560.1;refGene_gn=TEP1;refGene_tx=XM_023255561.1;refGene_gn=TEP1;refGene_tx=XM_023255562.1
B3 73542711 73542955
refGene_gn=TEP1;refGene_tx=XM_023255550.1;refGene_gn=TEP1;refGene_tx=XM_023255551.1;refGene_gn=TEP1;refGene_tx=XM_023255552.1;refGene_gn=TEP1;refGene_tx=XM_023255553.1;refGene_gn=TEP1;refGene_tx=XM_023255554.1;refGene_gn=TEP1;refGene_tx=XM_023255555.1;refGene_gn=TEP1;refGene_tx=XM_023255557.1;refGene_gn=TEP1;refGene_tx=XM_023255558.1;refGene_gn=TEP1;refGene_tx=XM_023255559.1;refGene_gn=TEP1;refGene_tx=XM_023255560.1;refGene_gn=TEP1;refGene_tx=XM_023255561.1;refGene_gn=TEP1;refGene_tx=XM_023255562.1
B3 73543591 73543706
refGene_gn=TEP1;refGene_tx=XM_023255550.1;refGene_gn=TEP1;refGene_tx=XM_023255551.1;refGene_gn=TEP1;refGene_tx=XM_023255552.1;refGene_gn=TEP1;refGene_tx=XM_023255553.1;refGene_gn=TEP1;refGene_tx=XM_023255554.1;

refGene_gn=TEP1;refGene_tx=XM_023255555.1;refGene_gn=TEP1;refGene_tx=XM_023255557.1;refGene_gn=TEP1;refGene_tx=XM_023255558.1;refGene_gn=TEP1;refGene_tx=XM_023255559.1;refGene_gn=TEP1;refGene_tx=XM_023255560.1;refGene_gn=TEP1;refGene_tx=XM_023255561.1;refGene_gn=TEP1;refGene_tx=XM_023255562.1
B3 73543965 73544054
refGene_gn=TEP1;refGene_tx=XM_023255550.1;refGene_gn=TEP1;refGene_tx=XM_023255551.1;refGene_gn=TEP1;refGene_tx=XM_023255552.1;refGene_gn=TEP1;refGene_tx=XM_023255553.1;refGene_gn=TEP1;refGene_tx=XM_023255554.1;refGene_gn=TEP1;refGene_tx=XM_023255555.1;refGene_gn=TEP1;refGene_tx=XM_023255557.1;refGene_gn=TEP1;refGene_tx=XM_023255558.1;refGene_gn=TEP1;refGene_tx=XM_023255559.1;refGene_gn=TEP1;refGene_tx=XM_023255560.1;refGene_gn=TEP1;refGene_tx=XM_023255561.1;refGene_gn=TEP1;refGene_tx=XM_023255562.1
B3 73544257 73544391
refGene_gn=TEP1;refGene_tx=XM_023255550.1;refGene_gn=TEP1;refGene_tx=XM_023255551.1;refGene_gn=TEP1;refGene_tx=XM_023255552.1;refGene_gn=TEP1;refGene_tx=XM_023255553.1;refGene_gn=TEP1;refGene_tx=XM_023255554.1;refGene_gn=TEP1;refGene_tx=XM_023255555.1;refGene_gn=TEP1;refGene_tx=XM_023255557.1;refGene_gn=TEP1;refGene_tx=XM_023255558.1;refGene_gn=TEP1;refGene_tx=XM_023255559.1;refGene_gn=TEP1;refGene_tx=XM_023255560.1;refGene_gn=TEP1;refGene_tx=XM_023255561.1;refGene_gn=TEP1;refGene_tx=XM_023255562.1
B3 73544877 73545028
refGene_gn=TEP1;refGene_tx=XM_023255550.1;refGene_gn=TEP1;refGene_tx=XM_023255551.1;refGene_gn=TEP1;refGene_tx=XM_023255552.1;refGene_gn=TEP1;refGene_tx=XM_023255553.1;refGene_gn=TEP1;refGene_tx=XM_023255554.1;refGene_gn=TEP1;refGene_tx=XM_023255555.1;refGene_gn=TEP1;refGene_tx=XM_023255557.1;refGene_gn=TEP1;refGene_tx=XM_023255558.1;refGene_gn=TEP1;refGene_tx=XM_023255559.1;refGene_gn=TEP1;refGene_tx=XM_023255560.1;refGene_gn=TEP1;refGene_tx=XM_023255561.1;refGene_gn=TEP1;refGene_tx=XM_023255562.1
B3 73545184 73545317
refGene_gn=TEP1;refGene_tx=XM_023255550.1;refGene_gn=TEP1;refGene_tx=XM_023255551.1;refGene_gn=TEP1;refGene_tx=XM_023255552.1;refGene_gn=TEP1;refGene_tx=XM_023255553.1;refGene_gn=TEP1;refGene_tx=XM_023255554.1;refGene_gn=TEP1;refGene_tx=XM_023255555.1;refGene_gn=TEP1;refGene_tx=XM_023255557.1;refGene_gn=TEP1;refGene_tx=XM_023255558.1;refGene_gn=TEP1;refGene_tx=XM_023255559.1;refGene_gn=TEP1;refGene_tx=XM_023255560.1;refGene_gn=TEP1;refGene_tx=XM_023255561.1;refGene_gn=TEP1;refGene_tx=XM_023255562.1
B3 73545611 73545704
refGene_gn=TEP1;refGene_tx=XM_023255550.1;refGene_gn=TEP1;refGene_tx=XM_023255551.1;refGene_gn=TEP1;refGene_tx=XM_023255552.1;refGene_gn=TEP1;refGene_tx=XM_023255553.1;refGene_gn=TEP1;refGene_tx=XM_023255554.1;refGene_gn=TEP1;refGene_tx=XM_023255555.1;refGene_gn=TEP1;refGene_tx=XM_023255557.1;refGene_gn=TEP1;refGene_tx=XM_023255558.1;refGene_gn=TEP1;refGene_tx=XM_023255559.1;refGene_gn=TEP1;refGene_tx=XM_023255560.1;refGene_gn=TEP1;refGene_tx=XM_023255561.1;refGene_gn=TEP1;refGene_tx=XM_023255562.1

B3	73546153 73546333
refGene_gn=TEP1;refGene_tx=XM_023255550.1;refGene_gn=TEP1;refGene_tx=XM_023255551.1;refGene_gn=TEP1;refGene_tx=XM_023255552.1;refGene_gn=TEP1;refGene_tx=XM_023255553.1;refGene_gn=TEP1;refGene_tx=XM_023255554.1;refGene_gn=TEP1;refGene_tx=XM_023255555.1;refGene_gn=TEP1;refGene_tx=XM_023255557.1;refGene_gn=TEP1;refGene_tx=XM_023255558.1;refGene_gn=TEP1;refGene_tx=XM_023255559.1;refGene_gn=TEP1;refGene_tx=XM_023255560.1;refGene_gn=TEP1;refGene_tx=XM_023255561.1;refGene_gn=TEP1;refGene_tx=XM_023255562.1
B3	73546446 73546603
refGene_gn=TEP1;refGene_tx=XM_023255550.1;refGene_gn=TEP1;refGene_tx=XM_023255551.1;refGene_gn=TEP1;refGene_tx=XM_023255552.1;refGene_gn=TEP1;refGene_tx=XM_023255553.1;refGene_gn=TEP1;refGene_tx=XM_023255554.1;refGene_gn=TEP1;refGene_tx=XM_023255555.1;refGene_gn=TEP1;refGene_tx=XM_023255557.1;refGene_gn=TEP1;refGene_tx=XM_023255558.1;refGene_gn=TEP1;refGene_tx=XM_023255559.1;refGene_gn=TEP1;refGene_tx=XM_023255560.1;refGene_gn=TEP1;refGene_tx=XM_023255561.1;refGene_gn=TEP1;refGene_tx=XM_023255562.1
B3	73546689 73546865
refGene_gn=TEP1;refGene_tx=XM_023255550.1;refGene_gn=TEP1;refGene_tx=XM_023255551.1;refGene_gn=TEP1;refGene_tx=XM_023255552.1;refGene_gn=TEP1;refGene_tx=XM_023255553.1;refGene_gn=TEP1;refGene_tx=XM_023255554.1;refGene_gn=TEP1;refGene_tx=XM_023255555.1;refGene_gn=TEP1;refGene_tx=XM_023255557.1;refGene_gn=TEP1;refGene_tx=XM_023255558.1;refGene_gn=TEP1;refGene_tx=XM_023255559.1;refGene_gn=TEP1;refGene_tx=XM_023255560.1;refGene_gn=TEP1;refGene_tx=XM_023255561.1;refGene_gn=TEP1;refGene_tx=XM_023255562.1
B3	73546984 73547179
refGene_gn=TEP1;refGene_tx=XM_023255550.1;refGene_gn=TEP1;refGene_tx=XM_023255551.1;refGene_gn=TEP1;refGene_tx=XM_023255552.1;refGene_gn=TEP1;refGene_tx=XM_023255553.1;refGene_gn=TEP1;refGene_tx=XM_023255554.1;refGene_gn=TEP1;refGene_tx=XM_023255555.1;refGene_gn=TEP1;refGene_tx=XM_023255557.1;refGene_gn=TEP1;refGene_tx=XM_023255558.1;refGene_gn=TEP1;refGene_tx=XM_023255559.1;refGene_gn=TEP1;refGene_tx=XM_023255560.1;refGene_gn=TEP1;refGene_tx=XM_023255561.1;refGene_gn=TEP1;refGene_tx=XM_023255562.1
B3	73547338 73547456
refGene_gn=TEP1;refGene_tx=XM_023255550.1;refGene_gn=TEP1;refGene_tx=XM_023255551.1;refGene_gn=TEP1;refGene_tx=XM_023255552.1;refGene_gn=TEP1;refGene_tx=XM_023255553.1;refGene_gn=TEP1;refGene_tx=XM_023255554.1;refGene_gn=TEP1;refGene_tx=XM_023255555.1;refGene_gn=TEP1;refGene_tx=XM_023255557.1;refGene_gn=TEP1;refGene_tx=XM_023255558.1;refGene_gn=TEP1;refGene_tx=XM_023255559.1;refGene_gn=TEP1;refGene_tx=XM_023255560.1;refGene_gn=TEP1;refGene_tx=XM_023255561.1;refGene_gn=TEP1;refGene_tx=XM_023255562.1
B3	73547561 73547675
refGene_gn=TEP1;refGene_tx=XM_023255550.1;refGene_gn=TEP1;refGene_tx=XM_023255551.1;refGene_gn=TEP1;refGene_tx=XM_023255552.1;refGene_gn=TEP1;refGene_tx=XM_023255553.1;refGene_gn=TEP1;refGene_tx=XM_023255554.1;refGene_gn=TEP1;refGene_tx=XM_023255555.1;refGene_gn=TEP1;refGene_tx=X

M_023255557.1;refGene_gn=TEP1;refGene_tx=XM_023255558.1;refGene_gn=TEP1;refGene_tx=XM_023255559.1;refGene_gn=TEP1;refGene_tx=XM_023255560.1;refGene_gn=TEP1;refGene_tx=XM_023255561.1;refGene_gn=TEP1;refGene_tx=XM_023255562.1
B3 73547937 73548062
refGene_gn=TEP1;refGene_tx=XM_023255550.1;refGene_gn=TEP1;refGene_tx=XM_023255551.1;refGene_gn=TEP1;refGene_tx=XM_023255552.1;refGene_gn=TEP1;refGene_tx=XM_023255553.1;refGene_gn=TEP1;refGene_tx=XM_023255554.1;refGene_gn=TEP1;refGene_tx=XM_023255555.1;refGene_gn=TEP1;refGene_tx=XM_023255557.1;refGene_gn=TEP1;refGene_tx=XM_023255558.1;refGene_gn=TEP1;refGene_tx=XM_023255559.1;refGene_gn=TEP1;refGene_tx=XM_023255560.1;refGene_gn=TEP1;refGene_tx=XM_023255561.1;refGene_gn=TEP1;refGene_tx=XM_023255562.1
B3 73548645 73548766
refGene_gn=TEP1;refGene_tx=XM_023255550.1;refGene_gn=TEP1;refGene_tx=XM_023255551.1;refGene_gn=TEP1;refGene_tx=XM_023255552.1;refGene_gn=TEP1;refGene_tx=XM_023255553.1;refGene_gn=TEP1;refGene_tx=XM_023255554.1;refGene_gn=TEP1;refGene_tx=XM_023255555.1;refGene_gn=TEP1;refGene_tx=XM_023255557.1;refGene_gn=TEP1;refGene_tx=XM_023255558.1;refGene_gn=TEP1;refGene_tx=XM_023255559.1;refGene_gn=TEP1;refGene_tx=XM_023255560.1;refGene_gn=TEP1;refGene_tx=XM_023255561.1;refGene_gn=TEP1;refGene_tx=XM_023255562.1
B3 73548995 73549172
refGene_gn=TEP1;refGene_tx=XM_023255550.1;refGene_gn=TEP1;refGene_tx=XM_023255551.1;refGene_gn=TEP1;refGene_tx=XM_023255552.1;refGene_gn=TEP1;refGene_tx=XM_023255553.1;refGene_gn=TEP1;refGene_tx=XM_023255554.1;refGene_gn=TEP1;refGene_tx=XM_023255555.1;refGene_gn=TEP1;refGene_tx=XM_023255557.1;refGene_gn=TEP1;refGene_tx=XM_023255558.1;refGene_gn=TEP1;refGene_tx=XM_023255559.1;refGene_gn=TEP1;refGene_tx=XM_023255560.1;refGene_gn=TEP1;refGene_tx=XM_023255561.1;refGene_gn=TEP1;refGene_tx=XM_023255562.1
B3 73549844 73550003
refGene_gn=TEP1;refGene_tx=XM_023255550.1;refGene_gn=TEP1;refGene_tx=XM_023255551.1;refGene_gn=TEP1;refGene_tx=XM_023255552.1;refGene_gn=TEP1;refGene_tx=XM_023255553.1;refGene_gn=TEP1;refGene_tx=XM_023255554.1;refGene_gn=TEP1;refGene_tx=XM_023255555.1;refGene_gn=TEP1;refGene_tx=XM_023255557.1;refGene_gn=TEP1;refGene_tx=XM_023255558.1;refGene_gn=TEP1;refGene_tx=XM_023255559.1;refGene_gn=TEP1;refGene_tx=XM_023255560.1;refGene_gn=TEP1;refGene_tx=XM_023255561.1;refGene_gn=TEP1;refGene_tx=XM_023255562.1
B3 73550884 73550944
refGene_gn=TEP1;refGene_tx=XM_023255550.1;refGene_gn=TEP1;refGene_tx=XM_023255551.1;refGene_gn=TEP1;refGene_tx=XM_023255552.1;refGene_gn=TEP1;refGene_tx=XM_023255553.1;refGene_gn=TEP1;refGene_tx=XM_023255554.1;refGene_gn=TEP1;refGene_tx=XM_023255555.1;refGene_gn=TEP1;refGene_tx=XM_023255557.1;refGene_gn=TEP1;refGene_tx=XM_023255558.1;refGene_gn=TEP1;refGene_tx=XM_023255559.1;refGene_gn=TEP1;refGene_tx=XM_023255560.1;refGene_gn=TEP1;refGene_tx=XM_023255561.1;refGene_gn=TEP1;refGene_tx=XM_023255562.1
B3 73551257 73551388

refGene_gn=TEP1;refGene_tx=XM_023255550.1;refGene_gn=TEP1;refGene_tx=X
M_023255551.1;refGene_gn=TEP1;refGene_tx=XM_023255552.1;refGene_gn=TEP
1;refGene_tx=XM_023255553.1;refGene_gn=TEP1;refGene_tx=XM_023255554.1;
refGene_gn=TEP1;refGene_tx=XM_023255555.1;refGene_gn=TEP1;refGene_tx=X
M_023255557.1;refGene_gn=TEP1;refGene_tx=XM_023255558.1;refGene_gn=TEP
1;refGene_tx=XM_023255559.1;refGene_gn=TEP1;refGene_tx=XM_023255560.1;
refGene_gn=TEP1;refGene_tx=XM_023255561.1;refGene_gn=TEP1;refGene_tx=X
M_023255562.1
B3 73552157 73552235
refGene_gn=TEP1;refGene_tx=XM_023255550.1;refGene_gn=TEP1;refGene_tx=X
M_023255551.1;refGene_gn=TEP1;refGene_tx=XM_023255552.1;refGene_gn=TEP
1;refGene_tx=XM_023255553.1;refGene_gn=TEP1;refGene_tx=XM_023255554.1;
refGene_gn=TEP1;refGene_tx=XM_023255555.1;refGene_gn=TEP1;refGene_tx=X
M_023255557.1;refGene_gn=TEP1;refGene_tx=XM_023255558.1;refGene_gn=TEP
1;refGene_tx=XM_023255559.1;refGene_gn=TEP1;refGene_tx=XM_023255560.1;
refGene_gn=TEP1;refGene_tx=XM_023255561.1;refGene_gn=TEP1;refGene_tx=X
M_023255562.1
B3 73552414 73552573
refGene_gn=TEP1;refGene_tx=XM_023255550.1;refGene_gn=TEP1;refGene_tx=X
M_023255551.1;refGene_gn=TEP1;refGene_tx=XM_023255552.1;refGene_gn=TEP
1;refGene_tx=XM_023255553.1;refGene_gn=TEP1;refGene_tx=XM_023255554.1;
refGene_gn=TEP1;refGene_tx=XM_023255555.1;refGene_gn=TEP1;refGene_tx=X
M_023255557.1;refGene_gn=TEP1;refGene_tx=XM_023255558.1;refGene_gn=TEP
1;refGene_tx=XM_023255559.1;refGene_gn=TEP1;refGene_tx=XM_023255560.1;
refGene_gn=TEP1;refGene_tx=XM_023255561.1;refGene_gn=TEP1;refGene_tx=X
M_023255562.1
B3 73553156 73553325
refGene_gn=TEP1;refGene_tx=XM_023255550.1;refGene_gn=TEP1;refGene_tx=X
M_023255551.1;refGene_gn=TEP1;refGene_tx=XM_023255552.1;refGene_gn=TEP
1;refGene_tx=XM_023255553.1;refGene_gn=TEP1;refGene_tx=XM_023255554.1;
refGene_gn=TEP1;refGene_tx=XM_023255555.1;refGene_gn=TEP1;refGene_tx=X
M_023255557.1;refGene_gn=TEP1;refGene_tx=XM_023255558.1;refGene_gn=TEP
1;refGene_tx=XM_023255559.1;refGene_gn=TEP1;refGene_tx=XM_023255560.1;
refGene_gn=TEP1;refGene_tx=XM_023255561.1;refGene_gn=TEP1;refGene_tx=X
M_023255562.1
B3 73557252 73557439
refGene_gn=TEP1;refGene_tx=XM_023255550.1;refGene_gn=TEP1;refGene_tx=X
M_023255551.1;refGene_gn=TEP1;refGene_tx=XM_023255552.1;refGene_gn=TEP
1;refGene_tx=XM_023255553.1;refGene_gn=TEP1;refGene_tx=XM_023255554.1;
refGene_gn=TEP1;refGene_tx=XM_023255555.1;refGene_gn=TEP1;refGene_tx=X
M_023255557.1;refGene_gn=TEP1;refGene_tx=XM_023255558.1;refGene_gn=TEP
1;refGene_tx=XM_023255559.1;refGene_gn=TEP1;refGene_tx=XM_023255560.1;
refGene_gn=TEP1;refGene_tx=XM_023255561.1;refGene_gn=TEP1;refGene_tx=X
M_023255562.1
B3 73557670 73557761
refGene_gn=TEP1;refGene_tx=XM_023255550.1;refGene_gn=TEP1;refGene_tx=X
M_023255551.1;refGene_gn=TEP1;refGene_tx=XM_023255552.1;refGene_gn=TEP
1;refGene_tx=XM_023255553.1;refGene_gn=TEP1;refGene_tx=XM_023255554.1;
refGene_gn=TEP1;refGene_tx=XM_023255555.1;refGene_gn=TEP1;refGene_tx=X
M_023255557.1;refGene_gn=TEP1;refGene_tx=XM_023255558.1;refGene_gn=TEP

1;refGene_tx=XM_023255559.1;refGene_gn=TEP1;refGene_tx=XM_023255560.1;
refGene_gn=TEP1;refGene_tx=XM_023255561.1;refGene_gn=TEP1;refGene_tx=X
M_023255562.1
B3 73558435 73558545
refGene_gn=TEP1;refGene_tx=XM_023255550.1;refGene_gn=TEP1;refGene_tx=X
M_023255551.1;refGene_gn=TEP1;refGene_tx=XM_023255552.1;refGene_gn=TEP
1;refGene_tx=XM_023255553.1;refGene_gn=TEP1;refGene_tx=XM_023255554.1;
refGene_gn=TEP1;refGene_tx=XM_023255555.1;refGene_gn=TEP1;refGene_tx=X
M_023255557.1;refGene_gn=TEP1;refGene_tx=XM_023255558.1;refGene_gn=TEP
1;refGene_tx=XM_023255559.1;refGene_gn=TEP1;refGene_tx=XM_023255560.1;
refGene_gn=TEP1;refGene_tx=XM_023255561.1;refGene_gn=TEP1;refGene_tx=X
M_023255562.1
B3 73562075 73562233
refGene_gn=TEP1;refGene_tx=XM_023255550.1;refGene_gn=TEP1;refGene_tx=X
M_023255551.1;refGene_gn=TEP1;refGene_tx=XM_023255552.1;refGene_gn=TEP
1;refGene_tx=XM_023255553.1;refGene_gn=TEP1;refGene_tx=XM_023255554.1;
refGene_gn=TEP1;refGene_tx=XM_023255555.1;refGene_gn=TEP1;refGene_tx=X
M_023255557.1;refGene_gn=TEP1;refGene_tx=XM_023255558.1;refGene_gn=TEP
1;refGene_tx=XM_023255559.1;refGene_gn=TEP1;refGene_tx=XM_023255560.1;
refGene_gn=TEP1;refGene_tx=XM_023255561.1;refGene_gn=TEP1;refGene_tx=X
M_023255562.1
B3 73562442 73562564
refGene_gn=TEP1;refGene_tx=XM_023255550.1;refGene_gn=TEP1;refGene_tx=X
M_023255551.1;refGene_gn=TEP1;refGene_tx=XM_023255552.1;refGene_gn=TEP
1;refGene_tx=XM_023255553.1;refGene_gn=TEP1;refGene_tx=XM_023255554.1;
refGene_gn=TEP1;refGene_tx=XM_023255555.1;refGene_gn=TEP1;refGene_tx=X
M_023255557.1;refGene_gn=TEP1;refGene_tx=XM_023255558.1;refGene_gn=TEP
1;refGene_tx=XM_023255559.1;refGene_gn=TEP1;refGene_tx=XM_023255560.1;
refGene_gn=TEP1;refGene_tx=XM_023255561.1;refGene_gn=TEP1;refGene_tx=X
M_023255562.1
B3 73563941 73564016
refGene_gn=TEP1;refGene_tx=XM_023255550.1;refGene_gn=TEP1;refGene_tx=X
M_023255551.1;refGene_gn=TEP1;refGene_tx=XM_023255552.1;refGene_gn=TEP
1;refGene_tx=XM_023255553.1;refGene_gn=TEP1;refGene_tx=XM_023255554.1;
refGene_gn=TEP1;refGene_tx=XM_023255555.1;refGene_gn=TEP1;refGene_tx=X
M_023255557.1;refGene_gn=TEP1;refGene_tx=XM_023255558.1;refGene_gn=TEP
1;refGene_tx=XM_023255559.1;refGene_gn=TEP1;refGene_tx=XM_023255560.1;
refGene_gn=TEP1;refGene_tx=XM_023255561.1;refGene_gn=TEP1;refGene_tx=X
M_023255562.1
B3 73564795 73564957
refGene_gn=TEP1;refGene_tx=XM_023255550.1;refGene_gn=TEP1;refGene_tx=X
M_023255551.1;refGene_gn=TEP1;refGene_tx=XM_023255552.1;refGene_gn=TEP
1;refGene_tx=XM_023255553.1;refGene_gn=TEP1;refGene_tx=XM_023255554.1;
refGene_gn=TEP1;refGene_tx=XM_023255555.1;refGene_gn=TEP1;refGene_tx=X
M_023255557.1;refGene_gn=TEP1;refGene_tx=XM_023255558.1;refGene_gn=TEP
1;refGene_tx=XM_023255559.1;refGene_gn=TEP1;refGene_tx=XM_023255560.1;
refGene_gn=TEP1;refGene_tx=XM_023255561.1;refGene_gn=TEP1;refGene_tx=X
M_023255562.1
B3 73565351 73565513
refGene_gn=TEP1;refGene_tx=XM_023255550.1;refGene_gn=TEP1;refGene_tx=X

M_023255551.1;refGene_gn=TEP1;refGene_tx=XM_023255552.1;refGene_gn=TEP1;refGene_tx=XM_023255553.1;refGene_gn=TEP1;refGene_tx=XM_023255554.1;refGene_gn=TEP1;refGene_tx=XM_023255555.1;refGene_gn=TEP1;refGene_tx=XM_023255557.1;refGene_gn=TEP1;refGene_tx=XM_023255558.1;refGene_gn=TEP1;refGene_tx=XM_023255559.1;refGene_gn=TEP1;refGene_tx=XM_023255560.1;refGene_gn=TEP1;refGene_tx=XM_023255561.1;refGene_gn=TEP1;refGene_tx=XM_023255562.1
B3 73566268 73566403
refGene_gn=TEP1;refGene_tx=XM_023255550.1;refGene_gn=TEP1;refGene_tx=XM_023255551.1;refGene_gn=TEP1;refGene_tx=XM_023255552.1;refGene_gn=TEP1;refGene_tx=XM_023255553.1;refGene_gn=TEP1;refGene_tx=XM_023255554.1;refGene_gn=TEP1;refGene_tx=XM_023255555.1;refGene_gn=TEP1;refGene_tx=XM_023255557.1;refGene_gn=TEP1;refGene_tx=XM_023255558.1;refGene_gn=TEP1;refGene_tx=XM_023255559.1;refGene_gn=TEP1;refGene_tx=XM_023255560.1;refGene_gn=TEP1;refGene_tx=XM_023255561.1;refGene_gn=TEP1;refGene_tx=XM_023255562.1
B3 73566713 73566896
refGene_gn=TEP1;refGene_tx=XM_023255550.1;refGene_gn=TEP1;refGene_tx=XM_023255551.1;refGene_gn=TEP1;refGene_tx=XM_023255552.1;refGene_gn=TEP1;refGene_tx=XM_023255553.1;refGene_gn=TEP1;refGene_tx=XM_023255554.1;refGene_gn=TEP1;refGene_tx=XM_023255555.1;refGene_gn=TEP1;refGene_tx=XM_023255557.1;refGene_gn=TEP1;refGene_tx=XM_023255558.1;refGene_gn=TEP1;refGene_tx=XM_023255559.1;refGene_gn=TEP1;refGene_tx=XM_023255560.1;refGene_gn=TEP1;refGene_tx=XM_023255561.1;refGene_gn=TEP1;refGene_tx=XM_023255562.1
B3 73568226 73568880
refGene_gn=TEP1;refGene_tx=XM_023255550.1;refGene_gn=TEP1;refGene_tx=XM_023255551.1;refGene_gn=TEP1;refGene_tx=XM_023255552.1;refGene_gn=TEP1;refGene_tx=XM_023255553.1;refGene_gn=TEP1;refGene_tx=XM_023255554.1;refGene_gn=TEP1;refGene_tx=XM_023255555.1;refGene_gn=TEP1;refGene_tx=XM_023255558.1;refGene_gn=TEP1;refGene_tx=XM_023255559.1;refGene_gn=TEP1;refGene_tx=XM_023255560.1;refGene_gn=TEP1;refGene_tx=XM_023255561.1;refGene_gn=TEP1;refGene_tx=XM_023255562.1
B3 73586367 73586920
refGene_gn=KLHL33;refGene_tx=XM_003987377.5;refGene_gn=KLHL33;refGene_tx=XM_011283100.3;refGene_gn=KLHL33;refGene_tx=XM_011283101.3
B3 73587027 73587195
refGene_gn=KLHL33;refGene_tx=XM_003987377.5;refGene_gn=KLHL33;refGene_tx=XM_011283100.3;refGene_gn=KLHL33;refGene_tx=XM_011283101.3
B3 73587306 73588231
refGene_gn=KLHL33;refGene_tx=XM_003987377.5;refGene_gn=KLHL33;refGene_tx=XM_011283100.3;refGene_gn=KLHL33;refGene_tx=XM_011283101.3
B3 73592916 73593670
refGene_gn=KLHL33;refGene_tx=XM_003987377.5;refGene_gn=KLHL33;refGene_tx=XM_011283100.3;refGene_gn=KLHL33;refGene_tx=XM_011283101.3
B3 73611455 73611495
refGene_gn=OSGEP;refGene_tx=XM_003987378.5;refGene_gn=OSGEP;refGene_tx=XM_019832732.2
B3 73611640 73611739
refGene_gn=OSGEP;refGene_tx=XM_003987378.5;refGene_gn=OSGEP;refGene_tx

=XM_019832732.2
B3	73611835	73611911
refGene_gn=OSGEP;refGene_tx=XM_003987378.5;refGene_gn=OSGEP;refGene_tx
=XM_019832732.2
B3	73612137	73612228
refGene_gn=OSGEP;refGene_tx=XM_003987378.5;refGene_gn=OSGEP;refGene_tx
=XM_019832732.2
B3	73612341	73612407
refGene_gn=OSGEP;refGene_tx=XM_003987378.5;refGene_gn=OSGEP;refGene_tx
=XM_019832732.2
B3	73612978	73613057
refGene_gn=OSGEP;refGene_tx=XM_003987378.5;refGene_gn=OSGEP;refGene_tx
=XM_019832732.2
B3	73613204	73613254
refGene_gn=OSGEP;refGene_tx=XM_003987378.5;refGene_gn=OSGEP;refGene_tx
=XM_019832732.2
B3	73613377	73613473
refGene_gn=OSGEP;refGene_tx=XM_003987378.5;refGene_gn=OSGEP;refGene_tx
=XM_019832732.2
B3	73615054	73615230
refGene_gn=OSGEP;refGene_tx=XM_003987378.5;refGene_gn=OSGEP;refGene_tx
=XM_019832732.2
B3	73615413	73615533	refGene_gn=OSGEP;refGene_tx=XM_003987378.5
B3	73617299	73617414
refGene_gn=OSGEP;refGene_tx=XM_003987378.5;refGene_gn=OSGEP;refGene_tx
=XM_019832732.2
B3	73617943	73618034	refGene_gn=APEX1;refGene_tx=XM_003987382.4
B3	73618369	73618495
refGene_gn=APEX1;refGene_tx=XM_003987381.5;refGene_gn=APEX1;refGene_tx
=XM_003987382.4;refGene_gn=APEX1;refGene_tx=XM_003987383.4
B3	73618696	73618884
refGene_gn=APEX1;refGene_tx=XM_003987381.5;refGene_gn=APEX1;refGene_tx
=XM_003987382.4;refGene_gn=APEX1;refGene_tx=XM_003987383.4
B3	73619508	73619701
refGene_gn=APEX1;refGene_tx=XM_003987381.5;refGene_gn=APEX1;refGene_tx
=XM_003987382.4;refGene_gn=APEX1;refGene_tx=XM_003987383.4
B3	73619842	73620360
refGene_gn=APEX1;refGene_tx=XM_003987381.5;refGene_gn=APEX1;refGene_tx
=XM_003987382.4;refGene_gn=APEX1;refGene_tx=XM_003987383.4
B3	73621424	73621568
refGene_gn=PIP4P1;refGene_tx=XM_003987384.5;refGene_gn=PIP4P1;refGene_
tx=XM_003987385.5
B3	73622057	73622148
refGene_gn=PIP4P1;refGene_tx=XM_003987384.5;refGene_gn=PIP4P1;refGene_
tx=XM_003987385.5
B3	73622250	73622303
refGene_gn=PIP4P1;refGene_tx=XM_003987384.5;refGene_gn=PIP4P1;refGene_
tx=XM_003987385.5
B3	73622509	73622615
refGene_gn=PIP4P1;refGene_tx=XM_003987384.5;refGene_gn=PIP4P1;refGene_

tx=XM_003987385.5
B3 73623053 73623160
refGene_gn=PIP4P1;refGene_tx=XM_003987384.5;refGene_gn=PIP4P1;refGene_tx=XM_003987385.5
B3 73623550 73623741
refGene_gn=PIP4P1;refGene_tx=XM_003987384.5;refGene_gn=PIP4P1;refGene_tx=XM_003987385.5
B3 73624064 73624227
refGene_gn=PIP4P1;refGene_tx=XM_003987384.5;refGene_gn=PIP4P1;refGene_tx=XM_003987385.5
B3 73644841 73644852 refGene_gn=PNP;refGene_tx=XM_003987386.4
B3 73648506 73648676 refGene_gn=PNP;refGene_tx=XM_003987386.4
B3 73652043 73652147 refGene_gn=PNP;refGene_tx=XM_003987386.4
B3 73652337 73652513 refGene_gn=PNP;refGene_tx=XM_003987386.4
B3 73652619 73652810 refGene_gn=PNP;refGene_tx=XM_003987386.4
B3 73653375 73653593 refGene_gn=PNP;refGene_tx=XM_003987386.4
B3 73680751 73681399 refGene_gn=RNASE10;refGene_tx=XM_019832735.2
B3 73717483 73718101
refGene_gn=RNASE9;refGene_tx=XM_019832737.2;refGene_gn=RNASE9;refGene_tx=XM_019832738.1;refGene_gn=RNASE9;refGene_tx=XM_019832740.1;refGene_gn=RNASE9;refGene_tx=XM_023255565.1;refGene_gn=RNASE9;refGene_tx=XM_023255567.1
B3 73723351 73723375
refGene_gn=RNASE9;refGene_tx=XM_019832737.2;refGene_gn=RNASE9;refGene_tx=XM_019832738.1;refGene_gn=RNASE9;refGene_tx=XM_023255565.1
B3 73741217 73741817 refGene_gn=RNASE11;refGene_tx=XM_003987388.4
B3 73746131 73746604 refGene_gn=RNASE12;refGene_tx=XM_023255568.1
B3 73747082 73747116 refGene_gn=RNASE12;refGene_tx=XM_023255568.1
B3 73790037 73791081
refGene_gn=LOC101091703;refGene_tx=XM_003987421.3
B3 73854377 73854818
refGene_gn=ANG;refGene_tx=XM_019832745.1;refGene_gn=ANG;refGene_tx=XM_019832746.1;refGene_gn=ANG;refGene_tx=XM_019832747.1;refGene_gn=ANG;refGene_tx=XM_019832748.1;refGene_gn=ANG;refGene_tx=XM_019832749.1
B3 73860145 73860583
refGene_gn=RNASE4;refGene_tx=XM_006932693.4;refGene_gn=RNASE4;refGene_tx=XM_019832750.2;refGene_gn=RNASE4;refGene_tx=XM_019832752.2;refGene_gn=RNASE4;refGene_tx=XM_019832753.2;refGene_gn=RNASE4;refGene_tx=XM_019832754.2
B3 73872234 73872672
refGene_gn=EDDM3B;refGene_tx=XM_019832755.2;refGene_gn=EDDM3B;refGene_tx=XM_023256237.1
B3 73884910 73885375
refGene_gn=LOC101092190;refGene_tx=XM_006932694.3
B3 73905165 73905621 refGene_gn=RNASE1;refGene_tx=XM_023255569.1
B3 73996282 73996363 refGene_gn=METTL17;refGene_tx=XM_003987424.4
B3 73996513 73996667 refGene_gn=METTL17;refGene_tx=XM_003987424.4
B3 73996755 73996890 refGene_gn=METTL17;refGene_tx=XM_003987424.4
B3 73997917 73997999 refGene_gn=METTL17;refGene_tx=XM_003987424.4
B3 73998262 73998344 refGene_gn=METTL17;refGene_tx=XM_003987424.4

B3 73998669 73998743 refGene_gn=METTL17;refGene_tx=XM_003987424.4
B3 73999221 73999316 refGene_gn=METTL17;refGene_tx=XM_003987424.4
B3 73999657 73999728 refGene_gn=METTL17;refGene_tx=XM_003987424.4
B3 73999928 74000036 refGene_gn=METTL17;refGene_tx=XM_003987424.4
B3 74000534 74000603 refGene_gn=METTL17;refGene_tx=XM_003987424.4
B3 74001406 74001457 refGene_gn=METTL17;refGene_tx=XM_003987424.4
B3 74001722 74001815 refGene_gn=METTL17;refGene_tx=XM_003987424.4
B3 74002042 74002230 refGene_gn=METTL17;refGene_tx=XM_003987424.4
B3 74002304 74002416 refGene_gn=METTL17;refGene_tx=XM_003987424.4
B3 74004269 74004384
refGene_gn=SLC39A2;refGene_tx=XM_003987393.4;refGene_gn=SLC39A2;refGene_tx=XM_019832758.2
B3 74004565 74004696
refGene_gn=SLC39A2;refGene_tx=XM_003987393.4;refGene_gn=SLC39A2;refGene_tx=XM_019832758.2
B3 74004931 74004982
refGene_gn=SLC39A2;refGene_tx=XM_003987393.4;refGene_gn=SLC39A2;refGene_tx=XM_019832758.2
B3 74005541 74006171 refGene_gn=SLC39A2;refGene_tx=XM_003987393.4
B3 74013551 74013718
refGene_gn=NDRG2;refGene_tx=XM_003987394.5;refGene_gn=NDRG2;refGene_tx=XM_006932698.4;refGene_gn=NDRG2;refGene_tx=XM_006932699.4;refGene_gn=NDRG2;refGene_tx=XM_011283104.3;refGene_gn=NDRG2;refGene_tx=XM_011283105.3
B3 74013991 74014043
refGene_gn=NDRG2;refGene_tx=XM_003987394.5;refGene_gn=NDRG2;refGene_tx=XM_006932698.4;refGene_gn=NDRG2;refGene_tx=XM_006932699.4;refGene_gn=NDRG2;refGene_tx=XM_011283104.3;refGene_gn=NDRG2;refGene_tx=XM_011283105.3
B3 74014429 74014465
refGene_gn=NDRG2;refGene_tx=XM_003987394.5;refGene_gn=NDRG2;refGene_tx=XM_006932698.4;refGene_gn=NDRG2;refGene_tx=XM_006932699.4;refGene_gn=NDRG2;refGene_tx=XM_011283104.3;refGene_gn=NDRG2;refGene_tx=XM_011283105.3
B3 74014704 74014752
refGene_gn=NDRG2;refGene_tx=XM_003987394.5;refGene_gn=NDRG2;refGene_tx=XM_006932698.4;refGene_gn=NDRG2;refGene_tx=XM_006932699.4;refGene_gn=NDRG2;refGene_tx=XM_011283104.3;refGene_gn=NDRG2;refGene_tx=XM_011283105.3
B3 74015007 74015059
refGene_gn=NDRG2;refGene_tx=XM_003987394.5;refGene_gn=NDRG2;refGene_tx=XM_006932698.4;refGene_gn=NDRG2;refGene_tx=XM_006932699.4;refGene_gn=NDRG2;refGene_tx=XM_011283104.3;refGene_gn=NDRG2;refGene_tx=XM_011283105.3
B3 74015342 74015387
refGene_gn=NDRG2;refGene_tx=XM_003987394.5;refGene_gn=NDRG2;refGene_tx=XM_006932698.4;refGene_gn=NDRG2;refGene_tx=XM_006932699.4;refGene_gn=NDRG2;refGene_tx=XM_011283104.3;refGene_gn=NDRG2;refGene_tx=XM_011283105.3
B3 74016052 74016156

refGene_gn=NDRG2;refGene_tx=XM_003987394.5;refGene_gn=NDRG2;refGene_tx=XM_006932698.4;refGene_gn=NDRG2;refGene_tx=XM_006932699.4;refGene_gn=NDRG2;refGene_tx=XM_011283104.3;refGene_gn=NDRG2;refGene_tx=XM_011283105.3
B3 74016331 74016388
refGene_gn=NDRG2;refGene_tx=XM_003987394.5;refGene_gn=NDRG2;refGene_tx=XM_006932698.4;refGene_gn=NDRG2;refGene_tx=XM_006932699.4;refGene_gn=NDRG2;refGene_tx=XM_011283104.3;refGene_gn=NDRG2;refGene_tx=XM_011283105.3
B3 74016595 74016682
refGene_gn=NDRG2;refGene_tx=XM_003987394.5;refGene_gn=NDRG2;refGene_tx=XM_006932698.4;refGene_gn=NDRG2;refGene_tx=XM_006932699.4;refGene_gn=NDRG2;refGene_tx=XM_011283104.3;refGene_gn=NDRG2;refGene_tx=XM_011283105.3
B3 74016897 74016958
refGene_gn=NDRG2;refGene_tx=XM_003987394.5;refGene_gn=NDRG2;refGene_tx=XM_006932698.4;refGene_gn=NDRG2;refGene_tx=XM_006932699.4;refGene_gn=NDRG2;refGene_tx=XM_011283104.3;refGene_gn=NDRG2;refGene_tx=XM_011283105.3
B3 74017917 74017980
refGene_gn=NDRG2;refGene_tx=XM_003987394.5;refGene_gn=NDRG2;refGene_tx=XM_006932698.4;refGene_gn=NDRG2;refGene_tx=XM_006932699.4;refGene_gn=NDRG2;refGene_tx=XM_011283104.3;refGene_gn=NDRG2;refGene_tx=XM_011283105.3
B3 74018156 74018277
refGene_gn=NDRG2;refGene_tx=XM_003987394.5;refGene_gn=NDRG2;refGene_tx=XM_006932698.4;refGene_gn=NDRG2;refGene_tx=XM_006932699.4;refGene_gn=NDRG2;refGene_tx=XM_011283104.3;refGene_gn=NDRG2;refGene_tx=XM_011283105.3
B3 74018491 74018597
refGene_gn=NDRG2;refGene_tx=XM_003987394.5;refGene_gn=NDRG2;refGene_tx=XM_006932698.4;refGene_gn=NDRG2;refGene_tx=XM_006932699.4;refGene_gn=NDRG2;refGene_tx=XM_011283104.3;refGene_gn=NDRG2;refGene_tx=XM_011283105.3
B3 74019006 74019048
refGene_gn=NDRG2;refGene_tx=XM_003987394.5;refGene_gn=NDRG2;refGene_tx=XM_006932698.4;refGene_gn=NDRG2;refGene_tx=XM_006932699.4
B3 74019382 74019457
refGene_gn=NDRG2;refGene_tx=XM_003987394.5;refGene_gn=NDRG2;refGene_tx=XM_006932698.4;refGene_gn=NDRG2;refGene_tx=XM_006932699.4;refGene_gn=NDRG2;refGene_tx=XM_011283104.3
B3 74025584 74025757
refGene_gn=TPPP2;refGene_tx=XM_003987398.5;refGene_gn=TPPP2;refGene_tx=XM_019832769.2;refGene_gn=TPPP2;refGene_tx=XM_019832770.2
B3 74025990 74026144
refGene_gn=TPPP2;refGene_tx=XM_003987398.5;refGene_gn=TPPP2;refGene_tx=XM_019832769.2;refGene_gn=TPPP2;refGene_tx=XM_019832770.2
B3 74026777 74027014
refGene_gn=TPPP2;refGene_tx=XM_019832769.2;refGene_gn=TPPP2;refGene_tx=XM_019832770.2

B3	74027114	74027303	refGene_gn=TPPP2;refGene_tx=XM_003987398.5
B3	74028939	74029404	refGene_gn=RNASE13;refGene_tx=XM_003987399.3
B3	74039674	74040136	refGene_gn=RNASE7;refGene_tx=XM_006932702.4
B3	74053992	74054562
refGene_gn=LOC111560955;refGene_tx=XM_023256180.1
B3	74055055	74055373
refGene_gn=LOC111560955;refGene_tx=XM_023256180.1
B3	74068250	74068253	refGene_gn=ARHGEF40;refGene_tx=XM_023255570.1
B3	74070816	74071014	refGene_gn=ARHGEF40;refGene_tx=XM_023255570.1
B3	74071602	74072848	refGene_gn=ARHGEF40;refGene_tx=XM_023255570.1
B3	74072985	74073153	refGene_gn=ARHGEF40;refGene_tx=XM_023255570.1
B3	74073281	74073402	refGene_gn=ARHGEF40;refGene_tx=XM_023255570.1
B3	74073985	74074082	refGene_gn=ARHGEF40;refGene_tx=XM_023255570.1
B3	74074193	74074274	refGene_gn=ARHGEF40;refGene_tx=XM_023255570.1
B3	74074391	74074508	refGene_gn=ARHGEF40;refGene_tx=XM_023255570.1
B3	74075844	74075940	refGene_gn=ARHGEF40;refGene_tx=XM_023255570.1
B3	74076021	74076137	refGene_gn=ARHGEF40;refGene_tx=XM_023255570.1
B3	74076512	74076639	refGene_gn=ARHGEF40;refGene_tx=XM_023255570.1
B3	74078262	74078385	refGene_gn=ARHGEF40;refGene_tx=XM_023255570.1
B3	74078487	74078505	refGene_gn=ARHGEF40;refGene_tx=XM_023255570.1
B3	74078506	74078632	refGene_gn=ARHGEF40;refGene_tx=XM_023255570.1
B3	74079129	74079740	refGene_gn=ARHGEF40;refGene_tx=XM_023255570.1
B3	74079875	74080107	refGene_gn=ARHGEF40;refGene_tx=XM_023255570.1
B3	74080446	74080533	refGene_gn=ARHGEF40;refGene_tx=XM_023255570.1
B3	74081125	74081341	refGene_gn=ARHGEF40;refGene_tx=XM_023255570.1
B3	74082055	74082226	refGene_gn=ARHGEF40;refGene_tx=XM_023255570.1
B3	74083439	74083617	refGene_gn=ARHGEF40;refGene_tx=XM_023255570.1
B3	74084382	74084487	refGene_gn=ARHGEF40;refGene_tx=XM_023255570.1
B3	74084715	74084859	refGene_gn=ARHGEF40;refGene_tx=XM_023255570.1
B3	74085363	74085494	refGene_gn=ARHGEF40;refGene_tx=XM_023255570.1
B3	74086152	74086194	refGene_gn=ARHGEF40;refGene_tx=XM_023255570.1
B3	74087816	74088418
refGene_gn=ZNF219;refGene_tx=XM_011283108.3;refGene_gn=ZNF219;refGene_tx=XM_019832766.2;refGene_gn=ZNF219;refGene_tx=XM_019832767.2;refGene_gn=ZNF219;refGene_tx=XM_023255571.1
B3	74088645	74088777
refGene_gn=ZNF219;refGene_tx=XM_011283108.3;refGene_gn=ZNF219;refGene_tx=XM_019832766.2;refGene_gn=ZNF219;refGene_tx=XM_019832767.2;refGene_gn=ZNF219;refGene_tx=XM_023255571.1
B3	74089086	74090494
refGene_gn=ZNF219;refGene_tx=XM_011283108.3;refGene_gn=ZNF219;refGene_tx=XM_019832766.2;refGene_gn=ZNF219;refGene_tx=XM_019832767.2;refGene_gn=ZNF219;refGene_tx=XM_023255571.1
B3	74090786	74090792
refGene_gn=ZNF219;refGene_tx=XM_011283108.3;refGene_gn=ZNF219;refGene_tx=XM_019832766.2;refGene_gn=ZNF219;refGene_tx=XM_019832767.2;refGene_gn=ZNF219;refGene_tx=XM_023255571.1
B3	74098641	74098749	refGene_gn=TMEM253;refGene_tx=XM_003987427.3
B3	74099098	74099209	refGene_gn=TMEM253;refGene_tx=XM_003987427.3
B3	74099301	74099358	refGene_gn=TMEM253;refGene_tx=XM_003987427.3

B3 74099683 74099794 refGene_gn=TMEM253;refGene_tx=XM_003987427.3
B3 74099912 74100059 refGene_gn=TMEM253;refGene_tx=XM_003987427.3
B3 74100425 74100506 refGene_gn=TMEM253;refGene_tx=XM_003987427.3
B3 74129460 74130418
refGene_gn=LOC101096398;refGene_tx=XM_011283109.3
B3 74130419 74130550
refGene_gn=LOC101096398;refGene_tx=XM_011283109.3
B3 74157790 74157874
refGene_gn=HNRNPC;refGene_tx=XM_003987428.5;refGene_gn=HNRNPC;refGene_tx=XM_006932709.4;refGene_gn=HNRNPC;refGene_tx=XM_011283110.3;refGene_gn=HNRNPC;refGene_tx=XM_011283111.3;refGene_gn=HNRNPC;refGene_tx=XM_011283112.3;refGene_gn=HNRNPC;refGene_tx=XM_011283113.3;refGene_gn=HNRNPC;refGene_tx=XM_011283114.3;refGene_gn=HNRNPC;refGene_tx=XM_011283115.3;refGene_gn=HNRNPC;refGene_tx=XM_011283116.3;refGene_gn=HNRNPC;refGene_tx=XM_019832772.2
B3 74157973 74158134
refGene_gn=HNRNPC;refGene_tx=XM_003987428.5;refGene_gn=HNRNPC;refGene_tx=XM_006932709.4;refGene_gn=HNRNPC;refGene_tx=XM_011283110.3;refGene_gn=HNRNPC;refGene_tx=XM_011283111.3;refGene_gn=HNRNPC;refGene_tx=XM_011283112.3;refGene_gn=HNRNPC;refGene_tx=XM_011283113.3;refGene_gn=HNRNPC;refGene_tx=XM_011283114.3;refGene_gn=HNRNPC;refGene_tx=XM_011283115.3;refGene_gn=HNRNPC;refGene_tx=XM_011283116.3;refGene_gn=HNRNPC;refGene_tx=XM_019832772.2
B3 74158376 74158490
refGene_gn=HNRNPC;refGene_tx=XM_003987428.5;refGene_gn=HNRNPC;refGene_tx=XM_006932709.4;refGene_gn=HNRNPC;refGene_tx=XM_011283110.3;refGene_gn=HNRNPC;refGene_tx=XM_011283111.3;refGene_gn=HNRNPC;refGene_tx=XM_011283112.3;refGene_gn=HNRNPC;refGene_tx=XM_011283113.3;refGene_gn=HNRNPC;refGene_tx=XM_011283114.3;refGene_gn=HNRNPC;refGene_tx=XM_011283115.3;refGene_gn=HNRNPC;refGene_tx=XM_011283116.3;refGene_gn=HNRNPC;refGene_tx=XM_019832772.2
B3 74159030 74159191
refGene_gn=HNRNPC;refGene_tx=XM_003987428.5;refGene_gn=HNRNPC;refGene_tx=XM_006932709.4;refGene_gn=HNRNPC;refGene_tx=XM_011283110.3;refGene_gn=HNRNPC;refGene_tx=XM_011283111.3;refGene_gn=HNRNPC;refGene_tx=XM_011283112.3;refGene_gn=HNRNPC;refGene_tx=XM_011283113.3;refGene_gn=HNRNPC;refGene_tx=XM_011283114.3;refGene_gn=HNRNPC;refGene_tx=XM_011283115.3;refGene_gn=HNRNPC;refGene_tx=XM_011283116.3;refGene_gn=HNRNPC;refGene_tx=XM_019832772.2
B3 74179557 74179605
refGene_gn=HNRNPC;refGene_tx=XM_003987428.5;refGene_gn=HNRNPC;refGene_tx=XM_006932709.4;refGene_gn=HNRNPC;refGene_tx=XM_011283110.3;refGene_gn=HNRNPC;refGene_tx=XM_011283111.3;refGene_gn=HNRNPC;refGene_tx=XM_011283112.3;refGene_gn=HNRNPC;refGene_tx=XM_011283113.3;refGene_gn=HNRNPC;refGene_tx=XM_011283114.3;refGene_gn=HNRNPC;refGene_tx=XM_011283115.3;refGene_gn=HNRNPC;refGene_tx=XM_011283116.3;refGene_gn=HNRNPC;refGene_tx=XM_019832772.2
B3 74180201 74180316
refGene_gn=HNRNPC;refGene_tx=XM_003987428.5;refGene_gn=HNRNPC;refGene_tx=XM_006932709.4;refGene_gn=HNRNPC;refGene_tx=XM_011283110.3;refGene_

gn=HNRNPC;refGene_tx=XM_011283111.3;refGene_gn=HNRNPC;refGene_tx=XM_011283112.3;refGene_gn=HNRNPC;refGene_tx=XM_011283113.3;refGene_gn=HNRNPC;refGene_tx=XM_011283114.3;refGene_gn=HNRNPC;refGene_tx=XM_011283115.3;refGene_gn=HNRNPC;refGene_tx=XM_011283116.3;refGene_gn=HNRNPC;refGene_tx=XM_019832772.2
B3 74185771 74186012
refGene_gn=HNRNPC;refGene_tx=XM_003987428.5;refGene_gn=HNRNPC;refGene_tx=XM_006932709.4;refGene_gn=HNRNPC;refGene_tx=XM_011283110.3;refGene_gn=HNRNPC;refGene_tx=XM_011283111.3;refGene_gn=HNRNPC;refGene_tx=XM_011283112.3;refGene_gn=HNRNPC;refGene_tx=XM_011283113.3;refGene_gn=HNRNPC;refGene_tx=XM_011283114.3;refGene_gn=HNRNPC;refGene_tx=XM_011283115.3;refGene_gn=HNRNPC;refGene_tx=XM_011283116.3;refGene_gn=HNRNPC;refGene_tx=XM_019832772.2
B3 74241525 74241544 refGene_gn=RPGRIP1;refGene_tx=XM_019832776.2
B3 74261632 74261765 refGene_gn=RPGRIP1;refGene_tx=XM_019832776.2
B3 74264219 74264485 refGene_gn=RPGRIP1;refGene_tx=XM_019832776.2
B3 74265754 74265854 refGene_gn=RPGRIP1;refGene_tx=XM_019832776.2
B3 74266369 74266501 refGene_gn=RPGRIP1;refGene_tx=XM_019832776.2
B3 74268883 74268989 refGene_gn=RPGRIP1;refGene_tx=XM_019832776.2
B3 74271975 74271999 refGene_gn=RPGRIP1;refGene_tx=XM_019832776.2
B3 74274130 74274277 refGene_gn=RPGRIP1;refGene_tx=XM_019832776.2
B3 74274751 74274825 refGene_gn=RPGRIP1;refGene_tx=XM_019832776.2
B3 74276686 74276832 refGene_gn=RPGRIP1;refGene_tx=XM_019832776.2
B3 74278928 74279089 refGene_gn=RPGRIP1;refGene_tx=XM_019832776.2
B3 74279544 74279688 refGene_gn=RPGRIP1;refGene_tx=XM_019832776.2
B3 74280152 74280303 refGene_gn=RPGRIP1;refGene_tx=XM_019832776.2
B3 74282327 74282780 refGene_gn=RPGRIP1;refGene_tx=XM_019832776.2
B3 74282941 74283093 refGene_gn=RPGRIP1;refGene_tx=XM_019832776.2
B3 74283570 74283913 refGene_gn=RPGRIP1;refGene_tx=XM_019832776.2
B3 74285425 74285610 refGene_gn=RPGRIP1;refGene_tx=XM_019832776.2
B3 74285917 74286121 refGene_gn=RPGRIP1;refGene_tx=XM_019832776.2
B3 74286938 74287077 refGene_gn=RPGRIP1;refGene_tx=XM_019832776.2
B3 74300135 74300233 refGene_gn=RPGRIP1;refGene_tx=XM_019832776.2
B3 74309381 74309574 refGene_gn=RPGRIP1;refGene_tx=XM_019832776.2
B3 74318019 74318104 refGene_gn=RPGRIP1;refGene_tx=XM_019832776.2
B3 74320991 74321122 refGene_gn=RPGRIP1;refGene_tx=XM_019832776.2
B3 74324533 74324646 refGene_gn=RPGRIP1;refGene_tx=XM_019832776.2
B3 74326110 74326256 refGene_gn=SUPT16H;refGene_tx=XM_003987429.5
B3 74326915 74326993 refGene_gn=SUPT16H;refGene_tx=XM_003987429.5
B3 74327120 74327250 refGene_gn=SUPT16H;refGene_tx=XM_003987429.5
B3 74327901 74328031 refGene_gn=SUPT16H;refGene_tx=XM_003987429.5
B3 74335129 74335299 refGene_gn=SUPT16H;refGene_tx=XM_003987429.5
B3 74335639 74335715 refGene_gn=SUPT16H;refGene_tx=XM_003987429.5
B3 74336030 74336143 refGene_gn=SUPT16H;refGene_tx=XM_003987429.5
B3 74336472 74336598 refGene_gn=SUPT16H;refGene_tx=XM_003987429.5
B3 74337445 74337564 refGene_gn=SUPT16H;refGene_tx=XM_003987429.5
B3 74337822 74337949 refGene_gn=SUPT16H;refGene_tx=XM_003987429.5
B3 74338054 74338190 refGene_gn=SUPT16H;refGene_tx=XM_003987429.5
B3 74338651 74338779 refGene_gn=SUPT16H;refGene_tx=XM_003987429.5
B3 74339480 74339634 refGene_gn=SUPT16H;refGene_tx=XM_003987429.5

B3 74339723 74339839 refGene_gn=SUPT16H;refGene_tx=XM_003987429.5
B3 74339946 74340042 refGene_gn=SUPT16H;refGene_tx=XM_003987429.5
B3 74340156 74340222 refGene_gn=SUPT16H;refGene_tx=XM_003987429.5
B3 74342240 74342353 refGene_gn=SUPT16H;refGene_tx=XM_003987429.5
B3 74342488 74342562 refGene_gn=SUPT16H;refGene_tx=XM_003987429.5
B3 74343740 74343831 refGene_gn=SUPT16H;refGene_tx=XM_003987429.5
B3 74345635 74345808 refGene_gn=SUPT16H;refGene_tx=XM_003987429.5
B3 74347016 74347168 refGene_gn=SUPT16H;refGene_tx=XM_003987429.5
B3 74347595 74347742 refGene_gn=SUPT16H;refGene_tx=XM_003987429.5
B3 74348520 74348673 refGene_gn=SUPT16H;refGene_tx=XM_003987429.5
B3 74349455 74349626 refGene_gn=SUPT16H;refGene_tx=XM_003987429.5
B3 74350744 74350837 refGene_gn=SUPT16H;refGene_tx=XM_003987429.5
B3 74365112 74365178 refGene_gn=SUPT16H;refGene_tx=XM_003987429.5
B3 74366838 74367405
refGene_gn=CHD8;refGene_tx=XM_011283119.3;refGene_gn=CHD8;refGene_tx=XM_011283120.3;refGene_gn=CHD8;refGene_tx=XM_011283122.3;refGene_gn=CHD8;refGene_tx=XM_011283123.3;refGene_gn=CHD8;refGene_tx=XM_019832774.2;refGene_gn=CHD8;refGene_tx=XM_019832775.2
B3 74370030 74370147
refGene_gn=CHD8;refGene_tx=XM_011283119.3;refGene_gn=CHD8;refGene_tx=XM_011283120.3;refGene_gn=CHD8;refGene_tx=XM_011283122.3;refGene_gn=CHD8;refGene_tx=XM_011283123.3;refGene_gn=CHD8;refGene_tx=XM_019832774.2;refGene_gn=CHD8;refGene_tx=XM_019832775.2
B3 74370540 74370720
refGene_gn=CHD8;refGene_tx=XM_011283119.3;refGene_gn=CHD8;refGene_tx=XM_011283120.3;refGene_gn=CHD8;refGene_tx=XM_011283122.3;refGene_gn=CHD8;refGene_tx=XM_011283123.3;refGene_gn=CHD8;refGene_tx=XM_019832774.2
B3 74370910 74371024
refGene_gn=CHD8;refGene_tx=XM_011283119.3;refGene_gn=CHD8;refGene_tx=XM_011283120.3;refGene_gn=CHD8;refGene_tx=XM_011283122.3;refGene_gn=CHD8;refGene_tx=XM_011283123.3;refGene_gn=CHD8;refGene_tx=XM_019832774.2;refGene_gn=CHD8;refGene_tx=XM_019832775.2
B3 74371830 74372133
refGene_gn=CHD8;refGene_tx=XM_011283119.3;refGene_gn=CHD8;refGene_tx=XM_011283120.3;refGene_gn=CHD8;refGene_tx=XM_011283122.3;refGene_gn=CHD8;refGene_tx=XM_011283123.3;refGene_gn=CHD8;refGene_tx=XM_019832774.2;refGene_gn=CHD8;refGene_tx=XM_019832775.2
B3 74372415 74372567
refGene_gn=CHD8;refGene_tx=XM_011283119.3;refGene_gn=CHD8;refGene_tx=XM_011283120.3;refGene_gn=CHD8;refGene_tx=XM_011283122.3;refGene_gn=CHD8;refGene_tx=XM_011283123.3;refGene_gn=CHD8;refGene_tx=XM_019832774.2;refGene_gn=CHD8;refGene_tx=XM_019832775.2
B3 74372802 74373522
refGene_gn=CHD8;refGene_tx=XM_011283119.3;refGene_gn=CHD8;refGene_tx=XM_011283120.3;refGene_gn=CHD8;refGene_tx=XM_011283122.3;refGene_gn=CHD8;refGene_tx=XM_011283123.3;refGene_gn=CHD8;refGene_tx=XM_019832774.2;refGene_gn=CHD8;refGene_tx=XM_019832775.2
B3 74373613 74373822
refGene_gn=CHD8;refGene_tx=XM_011283119.3;refGene_gn=CHD8;refGene_tx=XM_011283120.3;refGene_gn=CHD8;refGene_tx=XM_011283122.3;refGene_gn=CHD

8;refGene_tx=XM_011283123.3;refGene_gn=CHD8;refGene_tx=XM_019832774.2;
refGene_gn=CHD8;refGene_tx=XM_019832775.2
B3 74374734 74374942
refGene_gn=CHD8;refGene_tx=XM_011283119.3;refGene_gn=CHD8;refGene_tx=XM_011283120.3;refGene_gn=CHD8;refGene_tx=XM_011283122.3;refGene_gn=CHD8;refGene_tx=XM_011283123.3;refGene_gn=CHD8;refGene_tx=XM_019832774.2;refGene_gn=CHD8;refGene_tx=XM_019832775.2
B3 74375110 74375165
refGene_gn=CHD8;refGene_tx=XM_011283119.3;refGene_gn=CHD8;refGene_tx=XM_011283120.3;refGene_gn=CHD8;refGene_tx=XM_011283122.3;refGene_gn=CHD8;refGene_tx=XM_011283123.3;refGene_gn=CHD8;refGene_tx=XM_019832774.2;refGene_gn=CHD8;refGene_tx=XM_019832775.2
B3 74375633 74375709
refGene_gn=CHD8;refGene_tx=XM_011283119.3;refGene_gn=CHD8;refGene_tx=XM_011283120.3;refGene_gn=CHD8;refGene_tx=XM_011283122.3;refGene_gn=CHD8;refGene_tx=XM_011283123.3;refGene_gn=CHD8;refGene_tx=XM_019832774.2;refGene_gn=CHD8;refGene_tx=XM_019832775.2
B3 74376810 74376940
refGene_gn=CHD8;refGene_tx=XM_011283119.3;refGene_gn=CHD8;refGene_tx=XM_011283120.3;refGene_gn=CHD8;refGene_tx=XM_011283122.3;refGene_gn=CHD8;refGene_tx=XM_011283123.3;refGene_gn=CHD8;refGene_tx=XM_019832774.2;refGene_gn=CHD8;refGene_tx=XM_019832775.2
B3 74378353 74378457
refGene_gn=CHD8;refGene_tx=XM_011283119.3;refGene_gn=CHD8;refGene_tx=XM_011283120.3;refGene_gn=CHD8;refGene_tx=XM_011283122.3;refGene_gn=CHD8;refGene_tx=XM_011283123.3;refGene_gn=CHD8;refGene_tx=XM_019832774.2;refGene_gn=CHD8;refGene_tx=XM_019832775.2
B3 74378726 74378816
refGene_gn=CHD8;refGene_tx=XM_011283119.3;refGene_gn=CHD8;refGene_tx=XM_011283120.3;refGene_gn=CHD8;refGene_tx=XM_011283122.3;refGene_gn=CHD8;refGene_tx=XM_011283123.3;refGene_gn=CHD8;refGene_tx=XM_019832774.2;refGene_gn=CHD8;refGene_tx=XM_019832775.2
B3 74378906 74379087
refGene_gn=CHD8;refGene_tx=XM_011283119.3;refGene_gn=CHD8;refGene_tx=XM_011283120.3;refGene_gn=CHD8;refGene_tx=XM_011283122.3;refGene_gn=CHD8;refGene_tx=XM_011283123.3;refGene_gn=CHD8;refGene_tx=XM_019832774.2;refGene_gn=CHD8;refGene_tx=XM_019832775.2
B3 74379167 74379367
refGene_gn=CHD8;refGene_tx=XM_011283119.3;refGene_gn=CHD8;refGene_tx=XM_011283120.3;refGene_gn=CHD8;refGene_tx=XM_011283122.3;refGene_gn=CHD8;refGene_tx=XM_011283123.3;refGene_gn=CHD8;refGene_tx=XM_019832774.2;refGene_gn=CHD8;refGene_tx=XM_019832775.2
B3 74379567 74379764
refGene_gn=CHD8;refGene_tx=XM_011283119.3;refGene_gn=CHD8;refGene_tx=XM_011283120.3;refGene_gn=CHD8;refGene_tx=XM_011283122.3;refGene_gn=CHD8;refGene_tx=XM_011283123.3;refGene_gn=CHD8;refGene_tx=XM_019832774.2;refGene_gn=CHD8;refGene_tx=XM_019832775.2
B3 74380103 74380214
refGene_gn=CHD8;refGene_tx=XM_011283119.3;refGene_gn=CHD8;refGene_tx=XM_011283120.3;refGene_gn=CHD8;refGene_tx=XM_011283122.3;refGene_gn=CHD

8;refGene_tx=XM_011283123.3;refGene_gn=CHD8;refGene_tx=XM_019832774.2;
refGene_gn=CHD8;refGene_tx=XM_019832775.2
B3 74380355 74380535
refGene_gn=CHD8;refGene_tx=XM_011283119.3;refGene_gn=CHD8;refGene_tx=X
M_011283120.3;refGene_gn=CHD8;refGene_tx=XM_011283122.3;refGene_gn=CHD
8;refGene_tx=XM_011283123.3;refGene_gn=CHD8;refGene_tx=XM_019832774.2;
refGene_gn=CHD8;refGene_tx=XM_019832775.2
B3 74380732 74380900
refGene_gn=CHD8;refGene_tx=XM_011283119.3;refGene_gn=CHD8;refGene_tx=X
M_011283120.3;refGene_gn=CHD8;refGene_tx=XM_011283122.3;refGene_gn=CHD
8;refGene_tx=XM_011283123.3;refGene_gn=CHD8;refGene_tx=XM_019832774.2;
refGene_gn=CHD8;refGene_tx=XM_019832775.2
B3 74381374 74381570
refGene_gn=CHD8;refGene_tx=XM_011283119.3;refGene_gn=CHD8;refGene_tx=X
M_011283120.3;refGene_gn=CHD8;refGene_tx=XM_011283122.3;refGene_gn=CHD
8;refGene_tx=XM_011283123.3;refGene_gn=CHD8;refGene_tx=XM_019832774.2;
refGene_gn=CHD8;refGene_tx=XM_019832775.2
B3 74381819 74382030
refGene_gn=CHD8;refGene_tx=XM_011283119.3;refGene_gn=CHD8;refGene_tx=X
M_011283120.3;refGene_gn=CHD8;refGene_tx=XM_011283122.3;refGene_gn=CHD
8;refGene_tx=XM_011283123.3;refGene_gn=CHD8;refGene_tx=XM_019832774.2;
refGene_gn=CHD8;refGene_tx=XM_019832775.2
B3 74382612 74382868
refGene_gn=CHD8;refGene_tx=XM_011283119.3;refGene_gn=CHD8;refGene_tx=X
M_011283120.3;refGene_gn=CHD8;refGene_tx=XM_011283122.3;refGene_gn=CHD
8;refGene_tx=XM_011283123.3;refGene_gn=CHD8;refGene_tx=XM_019832774.2;
refGene_gn=CHD8;refGene_tx=XM_019832775.2
B3 74383099 74383243
refGene_gn=CHD8;refGene_tx=XM_011283119.3;refGene_gn=CHD8;refGene_tx=X
M_011283120.3;refGene_gn=CHD8;refGene_tx=XM_011283122.3;refGene_gn=CHD
8;refGene_tx=XM_011283123.3;refGene_gn=CHD8;refGene_tx=XM_019832774.2;
refGene_gn=CHD8;refGene_tx=XM_019832775.2
B3 74384251 74384428
refGene_gn=CHD8;refGene_tx=XM_011283119.3;refGene_gn=CHD8;refGene_tx=X
M_011283120.3;refGene_gn=CHD8;refGene_tx=XM_011283122.3;refGene_gn=CHD
8;refGene_tx=XM_011283123.3;refGene_gn=CHD8;refGene_tx=XM_019832774.2;
refGene_gn=CHD8;refGene_tx=XM_019832775.2
B3 74385264 74385508
refGene_gn=CHD8;refGene_tx=XM_011283119.3;refGene_gn=CHD8;refGene_tx=X
M_011283120.3;refGene_gn=CHD8;refGene_tx=XM_011283122.3;refGene_gn=CHD
8;refGene_tx=XM_011283123.3;refGene_gn=CHD8;refGene_tx=XM_019832774.2;
refGene_gn=CHD8;refGene_tx=XM_019832775.2
B3 74385663 74385785
refGene_gn=CHD8;refGene_tx=XM_011283119.3;refGene_gn=CHD8;refGene_tx=X
M_011283120.3;refGene_gn=CHD8;refGene_tx=XM_011283122.3;refGene_gn=CHD
8;refGene_tx=XM_011283123.3;refGene_gn=CHD8;refGene_tx=XM_019832774.2;
refGene_gn=CHD8;refGene_tx=XM_019832775.2
B3 74387329 74387467
refGene_gn=CHD8;refGene_tx=XM_011283119.3;refGene_gn=CHD8;refGene_tx=X
M_011283120.3;refGene_gn=CHD8;refGene_tx=XM_011283122.3;refGene_gn=CHD

8;refGene_tx=XM_011283123.3;refGene_gn=CHD8;refGene_tx=XM_019832774.2;
refGene_gn=CHD8;refGene_tx=XM_019832775.2
B3 74391106 74391190
refGene_gn=CHD8;refGene_tx=XM_011283119.3;refGene_gn=CHD8;refGene_tx=X
M_011283120.3;refGene_gn=CHD8;refGene_tx=XM_011283122.3;refGene_gn=CHD
8;refGene_tx=XM_011283123.3;refGene_gn=CHD8;refGene_tx=XM_019832774.2;
refGene_gn=CHD8;refGene_tx=XM_019832775.2
B3 74395006 74395124
refGene_gn=CHD8;refGene_tx=XM_011283119.3;refGene_gn=CHD8;refGene_tx=X
M_011283120.3;refGene_gn=CHD8;refGene_tx=XM_011283122.3;refGene_gn=CHD
8;refGene_tx=XM_011283123.3;refGene_gn=CHD8;refGene_tx=XM_019832774.2;
refGene_gn=CHD8;refGene_tx=XM_019832775.2
B3 74395636 74395692
refGene_gn=CHD8;refGene_tx=XM_011283119.3;refGene_gn=CHD8;refGene_tx=X
M_011283120.3;refGene_gn=CHD8;refGene_tx=XM_011283122.3;refGene_gn=CHD
8;refGene_tx=XM_011283123.3;refGene_gn=CHD8;refGene_tx=XM_019832774.2;
refGene_gn=CHD8;refGene_tx=XM_019832775.2
B3 74396052 74396121
refGene_gn=CHD8;refGene_tx=XM_011283119.3;refGene_gn=CHD8;refGene_tx=X
M_011283120.3;refGene_gn=CHD8;refGene_tx=XM_011283122.3;refGene_gn=CHD
8;refGene_tx=XM_011283123.3;refGene_gn=CHD8;refGene_tx=XM_019832774.2;
refGene_gn=CHD8;refGene_tx=XM_019832775.2
B3 74396201 74396384
refGene_gn=CHD8;refGene_tx=XM_011283119.3;refGene_gn=CHD8;refGene_tx=X
M_011283120.3;refGene_gn=CHD8;refGene_tx=XM_011283122.3;refGene_gn=CHD
8;refGene_tx=XM_011283123.3;refGene_gn=CHD8;refGene_tx=XM_019832774.2;
refGene_gn=CHD8;refGene_tx=XM_019832775.2
B3 74404665 74404780
refGene_gn=CHD8;refGene_tx=XM_011283119.3;refGene_gn=CHD8;refGene_tx=X
M_011283120.3;refGene_gn=CHD8;refGene_tx=XM_011283122.3;refGene_gn=CHD
8;refGene_tx=XM_011283123.3;refGene_gn=CHD8;refGene_tx=XM_019832774.2;
refGene_gn=CHD8;refGene_tx=XM_019832775.2
B3 74406176 74406562
refGene_gn=CHD8;refGene_tx=XM_011283119.3;refGene_gn=CHD8;refGene_tx=X
M_011283120.3;refGene_gn=CHD8;refGene_tx=XM_011283122.3;refGene_gn=CHD
8;refGene_tx=XM_011283123.3;refGene_gn=CHD8;refGene_tx=XM_019832774.2;
refGene_gn=CHD8;refGene_tx=XM_019832775.2
B3 74407289 74407661
refGene_gn=CHD8;refGene_tx=XM_011283119.3;refGene_gn=CHD8;refGene_tx=X
M_011283120.3;refGene_gn=CHD8;refGene_tx=XM_011283122.3;refGene_gn=CHD
8;refGene_tx=XM_011283123.3;refGene_gn=CHD8;refGene_tx=XM_019832774.2;
refGene_gn=CHD8;refGene_tx=XM_019832775.2
B3 74408520 74409363
refGene_gn=CHD8;refGene_tx=XM_011283119.3;refGene_gn=CHD8;refGene_tx=X
M_011283120.3;refGene_gn=CHD8;refGene_tx=XM_019832774.2;refGene_gn=CHD
8;refGene_tx=XM_019832775.2
B3 74409532 74409538
refGene_gn=CHD8;refGene_tx=XM_011283122.3;refGene_gn=CHD8;refGene_tx=X
M_011283123.3
B3 74434485 74434593

refGene_gn=RAB2B;refGene_tx=XM_003987431.4;refGene_gn=RAB2B;refGene_tx=XM_006932710.3;refGene_gn=RAB2B;refGene_tx=XM_019832785.1
B3 74435438 74435507 refGene_gn=RAB2B;refGene_tx=XM_003987431.4;refGene_gn=RAB2B;refGene_tx=XM_006932710.3;refGene_gn=RAB2B;refGene_tx=XM_019832785.1
B3 74436069 74436181 refGene_gn=RAB2B;refGene_tx=XM_003987431.4;refGene_gn=RAB2B;refGene_tx=XM_006932710.3;refGene_gn=RAB2B;refGene_tx=XM_019832785.1
B3 74440322 74440415 refGene_gn=RAB2B;refGene_tx=XM_003987431.4;refGene_gn=RAB2B;refGene_tx=XM_006932710.3;refGene_gn=RAB2B;refGene_tx=XM_019832785.1
B3 74440627 74440710 refGene_gn=RAB2B;refGene_tx=XM_003987431.4;refGene_gn=RAB2B;refGene_tx=XM_006932710.3;refGene_gn=RAB2B;refGene_tx=XM_019832785.1
B3 74445838 74445906 refGene_gn=RAB2B;refGene_tx=XM_003987431.4;refGene_gn=RAB2B;refGene_tx=XM_006932710.3;refGene_gn=RAB2B;refGene_tx=XM_019832785.1
B3 74448687 74448759 refGene_gn=RAB2B;refGene_tx=XM_003987431.4
B3 74448985 74449031 refGene_gn=RAB2B;refGene_tx=XM_003987431.4
B3 74449435 74449441 refGene_gn=TOX4;refGene_tx=XM_011283124.3
B3 74449652 74449721 refGene_gn=TOX4;refGene_tx=XM_011283124.3
B3 74455452 74455695 refGene_gn=TOX4;refGene_tx=XM_011283124.3
B3 74456535 74456796 refGene_gn=TOX4;refGene_tx=XM_011283124.3
B3 74457148 74457379 refGene_gn=TOX4;refGene_tx=XM_011283124.3
B3 74458912 74458993 refGene_gn=TOX4;refGene_tx=XM_011283124.3
B3 74459393 74460137 refGene_gn=TOX4;refGene_tx=XM_011283124.3
B3 74461113 74461277 refGene_gn=TOX4;refGene_tx=XM_011283124.3
B3 74462029 74462090 refGene_gn=TOX4;refGene_tx=XM_011283124.3
B3 74463375 74463487 refGene_gn=METTL3;refGene_tx=XM_011283125.3
B3 74464378 74464491 refGene_gn=METTL3;refGene_tx=XM_011283125.3
B3 74464627 74464693 refGene_gn=METTL3;refGene_tx=XM_011283125.3
B3 74464816 74464925 refGene_gn=METTL3;refGene_tx=XM_011283125.3
B3 74465109 74465148 refGene_gn=METTL3;refGene_tx=XM_011283125.3
B3 74465521 74465709 refGene_gn=METTL3;refGene_tx=XM_011283125.3
B3 74465934 74466151 refGene_gn=METTL3;refGene_tx=XM_011283125.3
B3 74467132 74467308 refGene_gn=METTL3;refGene_tx=XM_011283125.3
B3 74468439 74468844 refGene_gn=METTL3;refGene_tx=XM_011283125.3
B3 74468938 74469156 refGene_gn=METTL3;refGene_tx=XM_011283125.3
B3 74477759 74477859 refGene_gn=METTL3;refGene_tx=XM_011283125.3
B3 74487637 74487878 refGene_gn=SALL2;refGene_tx=XM_019832788.2;refGene_gn=SALL2;refGene_tx=XM_023255574.1
B3 74488375 74491335 refGene_gn=SALL2;refGene_tx=XM_019832786.2;refGene_gn=SALL2;refGene_tx=XM_019832787.2;refGene_gn=SALL2;refGene_tx=XM_019832788.2;refGene_gn=SALL2;refGene_tx=XM_023255573.1;refGene_gn=SALL2;refGene_tx=XM_023255574.1
B3 74491723 74491790 refGene_gn=SALL2;refGene_tx=XM_023255573.1;refGene_gn=SALL2;refGene_tx=XM_023255574.1

B3 74501842 74501915
refGene_gn=SALL2;refGene_tx=XM_019832786.2;refGene_gn=SALL2;refGene_tx=XM_019832787.2;refGene_gn=SALL2;refGene_tx=XM_019832788.2
B3 74538140 74539082
refGene_gn=LOC101098305;refGene_tx=XM_003987434.2
B3 74557852 74558890
refGene_gn=LOC109500755;refGene_tx=XM_023255575.1
B3 74579601 74580639
refGene_gn=LOC101099559;refGene_tx=XM_003987439.5
B3 74607139 74608078
refGene_gn=LOC101099807;refGene_tx=XM_003987440.4
B3 74613014 74613956
refGene_gn=LOC101100058;refGene_tx=XM_023255576.1
B3 74662764 74664299 refGene_gn=LOC101100310
B3 74671592 74672091 refGene_gn=LOC109501106
B3 74673653 74675568 refGene_gn=LOC109500742
B3 74678628 74680963 refGene_gn=LOC101101063
B3 74680980 74681771 refGene_gn=LOC101101319
B3 74693622 74694713 refGene_gn=LOC105260657
B3 74707493 74708402 refGene_gn=LOC101080387
B3 74732847 74733705 refGene_gn=LOC101081143
B3 74738698 74739655 refGene_gn=LOC101081388
B3 74765773 74766866 refGene_gn=LOC101081900
B3 74782702 74783218 refGene_gn=LOC101082160
B3 74792037 74793566 refGene_gn=LOC105260658
B3 74809664 74810177 refGene_gn=LOC101082412
B3 74814846 74815831 refGene_gn=LOC101082664
B3 74864345 74866345 refGene_gn=LOC101083943
B3 74868586 74870727 refGene_gn=LOC102899903
B3 74901608 74902762 refGene_gn=LOC111560851
B3 74906265 74906874 refGene_gn=LOC111560958
B3 74911652 74913035 refGene_gn=LOC111560959
B3 74917813 74918793 refGene_gn=LOC111560852
B3 74924372 74925878 refGene_gn=LOC105260635
B3 74939289 74942474 refGene_gn=LOC111560960
B3 74968713 74970428 refGene_gn=LOC109500744
B3 74983023 74985807 refGene_gn=LOC101086484
B3 74998155 74999412 refGene_gn=LOC111560849
B3 75012822 75014525 refGene_gn=LOC101086979
B3 75020069 75022106 refGene_gn=LOC109500743
B3 75022509 75023902 refGene_gn=LOC109500748
B3 75045448 75051353 refGene_gn=LOC111560961
B3 75063511 75065307 refGene_gn=LOC111560850
B3 75066426 75067781 refGene_gn=LOC109500746
B3 75090987 75092360 refGene_gn=LOC109500750
B3 75110118 75111999 refGene_gn=LOC109500749
B3 75117759 75118453 refGene_gn=LOC111560853
B3 75148130 75148769 refGene_gn=LOC111560962
B3 75189984 75190684 refGene_gn=LOC109500745
B3 75216765 75218942 refGene_gn=LOC109500747

B3	75386044 75387989 refGene_gn=LOC111556216
B3	75452715 75471329 refGene_gn=LOC101094024
B3	75483824 75483967 refGene_gn=DAD1;refGene_tx=XM_019832791.2
B3	75494461 75494592 refGene_gn=DAD1;refGene_tx=XM_003987450.4
B3	75504036 75504247
refGene_gn=DAD1;refGene_tx=XM_003987450.4;refGene_gn=DAD1;refGene_tx=XM_019832791.2
B3	75515396 75515458 refGene_gn=ABHD4;refGene_tx=XM_011283126.3
B3	75518880 75518969
refGene_gn=ABHD4;refGene_tx=XM_011283126.3;refGene_gn=ABHD4;refGene_tx=XM_011283127.3;refGene_gn=ABHD4;refGene_tx=XM_011283128.3
B3	75520011 75520384
refGene_gn=ABHD4;refGene_tx=XM_011283126.3;refGene_gn=ABHD4;refGene_tx=XM_011283127.3;refGene_gn=ABHD4;refGene_tx=XM_011283128.3
B3	75520541 75520696
refGene_gn=ABHD4;refGene_tx=XM_011283126.3;refGene_gn=ABHD4;refGene_tx=XM_011283127.3;refGene_gn=ABHD4;refGene_tx=XM_011283128.3
B3	75522646 75522758
refGene_gn=ABHD4;refGene_tx=XM_011283126.3;refGene_gn=ABHD4;refGene_tx=XM_011283127.3;refGene_gn=ABHD4;refGene_tx=XM_011283128.3
B3	75525812 75525999
refGene_gn=ABHD4;refGene_tx=XM_011283126.3;refGene_gn=ABHD4;refGene_tx=XM_011283127.3;refGene_gn=ABHD4;refGene_tx=XM_011283128.3
B3	75526734 75526824
refGene_gn=ABHD4;refGene_tx=XM_011283126.3;refGene_gn=ABHD4;refGene_tx=XM_011283127.3;refGene_gn=ABHD4;refGene_tx=XM_011283128.3
B3	75588202 75589180
refGene_gn=LOC111560858;refGene_tx=XM_023255577.1
B3	75639394 75639515 refGene_gn=OXA1L;refGene_tx=XM_019832796.2
B3	75639517 75639675 refGene_gn=OXA1L;refGene_tx=XM_019832796.2
B3	75640106 75640268 refGene_gn=OXA1L;refGene_tx=XM_019832796.2
B3	75640983 75641197 refGene_gn=OXA1L;refGene_tx=XM_019832796.2
B3	75658797 75658941 refGene_gn=OXA1L;refGene_tx=XM_019832796.2
B3	75659194 75659250 refGene_gn=OXA1L;refGene_tx=XM_019832796.2
B3	75659251 75659281 refGene_gn=OXA1L;refGene_tx=XM_019832796.2
B3	75659436 75659601 refGene_gn=OXA1L;refGene_tx=XM_019832796.2
B3	75659771 75659876 refGene_gn=OXA1L;refGene_tx=XM_019832796.2
B3	75659988 75660151 refGene_gn=OXA1L;refGene_tx=XM_019832796.2
B3	75660236 75660317 refGene_gn=OXA1L;refGene_tx=XM_019832796.2
B3	75660399 75660527 refGene_gn=OXA1L;refGene_tx=XM_019832796.2
B3	75661577 75661684
refGene_gn=SLC7A7;refGene_tx=XM_003987452.5;refGene_gn=SLC7A7;refGene_tx=XM_006932719.4;refGene_gn=SLC7A7;refGene_tx=XM_006932720.3;refGene_gn=SLC7A7;refGene_tx=XM_011283129.3
B3	75662044 75662228
refGene_gn=SLC7A7;refGene_tx=XM_003987452.5;refGene_gn=SLC7A7;refGene_tx=XM_006932719.4;refGene_gn=SLC7A7;refGene_tx=XM_006932720.3;refGene_gn=SLC7A7;refGene_tx=XM_011283129.3
B3	75662411 75662561
refGene_gn=SLC7A7;refGene_tx=XM_003987452.5;refGene_gn=SLC7A7;refGene_

tx=XM_006932719.4;refGene_gn=SLC7A7;refGene_tx=XM_006932720.3;refGene_gn=SLC7A7;refGene_tx=XM_011283129.3
B3 75663089 75663186
refGene_gn=SLC7A7;refGene_tx=XM_003987452.5;refGene_gn=SLC7A7;refGene_tx=XM_006932719.4;refGene_gn=SLC7A7;refGene_tx=XM_006932720.3;refGene_gn=SLC7A7;refGene_tx=XM_011283129.3
B3 75663493 75663597
refGene_gn=SLC7A7;refGene_tx=XM_003987452.5;refGene_gn=SLC7A7;refGene_tx=XM_006932719.4;refGene_gn=SLC7A7;refGene_tx=XM_006932720.3;refGene_gn=SLC7A7;refGene_tx=XM_011283129.3
B3 75664157 75664281
refGene_gn=SLC7A7;refGene_tx=XM_003987452.5;refGene_gn=SLC7A7;refGene_tx=XM_006932719.4;refGene_gn=SLC7A7;refGene_tx=XM_006932720.3;refGene_gn=SLC7A7;refGene_tx=XM_011283129.3
B3 75665642 75665787
refGene_gn=SLC7A7;refGene_tx=XM_003987452.5;refGene_gn=SLC7A7;refGene_tx=XM_006932719.4;refGene_gn=SLC7A7;refGene_tx=XM_006932720.3;refGene_gn=SLC7A7;refGene_tx=XM_011283129.3
B3 75666756 75666882
refGene_gn=SLC7A7;refGene_tx=XM_003987452.5;refGene_gn=SLC7A7;refGene_tx=XM_006932719.4;refGene_gn=SLC7A7;refGene_tx=XM_006932720.3;refGene_gn=SLC7A7;refGene_tx=XM_011283129.3
B3 75688764 75689263
refGene_gn=SLC7A7;refGene_tx=XM_003987452.5;refGene_gn=SLC7A7;refGene_tx=XM_006932719.4;refGene_gn=SLC7A7;refGene_tx=XM_006932720.3;refGene_gn=SLC7A7;refGene_tx=XM_011283129.3
B3 75702474 75702502
refGene_gn=MRPL52;refGene_tx=XM_003987453.4;refGene_gn=MRPL52;refGene_tx=XM_006932721.3;refGene_gn=MRPL52;refGene_tx=XM_006932722.3;refGene_gn=MRPL52;refGene_tx=XM_006932723.3;refGene_gn=MRPL52;refGene_tx=XM_019832797.2;refGene_gn=MRPL52;refGene_tx=XM_023255578.1;refGene_gn=MRPL52;refGene_tx=XM_023255579.1
B3 75702606 75702667
refGene_gn=MRPL52;refGene_tx=XM_003987453.4;refGene_gn=MRPL52;refGene_tx=XM_006932721.3;refGene_gn=MRPL52;refGene_tx=XM_006932722.3;refGene_gn=MRPL52;refGene_tx=XM_019832797.2;refGene_gn=MRPL52;refGene_tx=XM_023255578.1;refGene_gn=MRPL52;refGene_tx=XM_023255579.1
B3 75702745 75702813
refGene_gn=MRPL52;refGene_tx=XM_003987453.4;refGene_gn=MRPL52;refGene_tx=XM_006932721.3;refGene_gn=MRPL52;refGene_tx=XM_006932722.3;refGene_gn=MRPL52;refGene_tx=XM_019832797.2;refGene_gn=MRPL52;refGene_tx=XM_023255578.1;refGene_gn=MRPL52;refGene_tx=XM_023255579.1
B3 75704611 75704683
refGene_gn=MRPL52;refGene_tx=XM_003987453.4;refGene_gn=MRPL52;refGene_tx=XM_006932721.3;refGene_gn=MRPL52;refGene_tx=XM_006932722.3;refGene_gn=MRPL52;refGene_tx=XM_006932723.3;refGene_gn=MRPL52;refGene_tx=XM_019832797.2;refGene_gn=MRPL52;refGene_tx=XM_019832798.2;refGene_gn=MRPL52;refGene_tx=XM_023255578.1
B3 75705052 75705202
refGene_gn=MRPL52;refGene_tx=XM_003987453.4;refGene_gn=MRPL52;refGene_

tx=XM_006932721.3;refGene_gn=MRPL52;refGene_tx=XM_006932722.3;refGene_gn=MRPL52;refGene_tx=XM_006932723.3;refGene_gn=MRPL52;refGene_tx=XM_019832797.2;refGene_gn=MRPL52;refGene_tx=XM_019832798.2;refGene_gn=MRPL52;refGene_tx=XM_023255578.1;refGene_gn=MRPL52;refGene_tx=XM_023255579.1
B3	75707394	75707499	refGene_gn=MMP14;refGene_tx=XM_003987454.4
B3	75711490	75711639	refGene_gn=MMP14;refGene_tx=XM_003987454.4
B3	75712018	75712141	refGene_gn=MMP14;refGene_tx=XM_003987454.4
B3	75712520	75712828	refGene_gn=MMP14;refGene_tx=XM_003987454.4
B3	75713324	75713486	refGene_gn=MMP14;refGene_tx=XM_003987454.4
B3	75714052	75714213	refGene_gn=MMP14;refGene_tx=XM_003987454.4
B3	75714390	75714529	refGene_gn=MMP14;refGene_tx=XM_003987454.4
B3	75714661	75714812	refGene_gn=MMP14;refGene_tx=XM_003987454.4
B3	75715288	75715404	refGene_gn=MMP14;refGene_tx=XM_003987454.4
B3	75715781	75716116	refGene_gn=MMP14;refGene_tx=XM_003987454.4
B3	75731079	75731113	refGene_gn=LRP10;refGene_tx=XM_003987455.5
B3	75731508	75731553	refGene_gn=LRP10;refGene_tx=XM_003987455.5
B3	75732073	75732206	refGene_gn=LRP10;refGene_tx=XM_003987455.5
B3	75732939	75733130	refGene_gn=LRP10;refGene_tx=XM_003987455.5
B3	75733246	75734264	refGene_gn=LRP10;refGene_tx=XM_003987455.5
B3	75734597	75734727	refGene_gn=LRP10;refGene_tx=XM_003987455.5
B3	75734856	75735444	refGene_gn=LRP10;refGene_tx=XM_003987455.5
B3	75740233	75740336	refGene_gn=REM2;refGene_tx=XM_003987456.5
B3	75741573	75741915	refGene_gn=REM2;refGene_tx=XM_003987456.5
B3	75742156	75742230	refGene_gn=REM2;refGene_tx=XM_003987456.5
B3	75742957	75743165	refGene_gn=REM2;refGene_tx=XM_003987456.5
B3	75743523	75743819	refGene_gn=REM2;refGene_tx=XM_003987456.5
B3	75755522	75755526	refGene_gn=RBM23;refGene_tx=XM_011283130.3;refGene_gn=RBM23;refGene_tx=XM_019832800.2
B3	75755618	75755707	refGene_gn=RBM23;refGene_tx=XM_003987457.5;refGene_gn=RBM23;refGene_tx=XM_011283130.3;refGene_gn=RBM23;refGene_tx=XM_019832800.2;refGene_gn=RBM23;refGene_tx=XM_019832801.2
B3	75755802	75755922	refGene_gn=RBM23;refGene_tx=XM_003987457.5;refGene_gn=RBM23;refGene_tx=XM_011283130.3;refGene_gn=RBM23;refGene_tx=XM_019832800.2;refGene_gn=RBM23;refGene_tx=XM_019832801.2
B3	75756013	75756209	refGene_gn=RBM23;refGene_tx=XM_003987457.5;refGene_gn=RBM23;refGene_tx=XM_011283130.3;refGene_gn=RBM23;refGene_tx=XM_019832800.2;refGene_gn=RBM23;refGene_tx=XM_019832801.2
B3	75757726	75757792	refGene_gn=RBM23;refGene_tx=XM_003987457.5;refGene_gn=RBM23;refGene_tx=XM_011283130.3;refGene_gn=RBM23;refGene_tx=XM_019832800.2;refGene_gn=RBM23;refGene_tx=XM_019832801.2
B3	75758393	75758531	refGene_gn=RBM23;refGene_tx=XM_003987457.5;refGene_gn=RBM23;refGene_tx=XM_011283130.3;refGene_gn=RBM23;refGene_tx=XM_019832800.2;refGene_gn=RBM23;refGene_tx=XM_019832801.2

B3 75758616 75758769
refGene_gn=RBM23;refGene_tx=XM_003987457.5;refGene_gn=RBM23;refGene_tx=XM_011283130.3;refGene_gn=RBM23;refGene_tx=XM_019832800.2;refGene_gn=RBM23;refGene_tx=XM_019832801.2
B3 75758863 75758981
refGene_gn=RBM23;refGene_tx=XM_003987457.5;refGene_gn=RBM23;refGene_tx=XM_011283130.3;refGene_gn=RBM23;refGene_tx=XM_019832800.2;refGene_gn=RBM23;refGene_tx=XM_019832801.2
B3 75759133 75759187
refGene_gn=RBM23;refGene_tx=XM_003987457.5;refGene_gn=RBM23;refGene_tx=XM_011283130.3;refGene_gn=RBM23;refGene_tx=XM_019832800.2;refGene_gn=RBM23;refGene_tx=XM_019832801.2
B3 75759440 75759614
refGene_gn=RBM23;refGene_tx=XM_003987457.5;refGene_gn=RBM23;refGene_tx=XM_011283130.3;refGene_gn=RBM23;refGene_tx=XM_019832800.2;refGene_gn=RBM23;refGene_tx=XM_019832801.2
B3 75759748 75759796
refGene_gn=RBM23;refGene_tx=XM_003987457.5;refGene_gn=RBM23;refGene_tx=XM_011283130.3
B3 75760276 75760392
refGene_gn=RBM23;refGene_tx=XM_003987457.5;refGene_gn=RBM23;refGene_tx=XM_011283130.3;refGene_gn=RBM23;refGene_tx=XM_019832800.2;refGene_gn=RBM23;refGene_tx=XM_019832801.2
B3 75760972 75761038
refGene_gn=RBM23;refGene_tx=XM_003987457.5;refGene_gn=RBM23;refGene_tx=XM_011283130.3;refGene_gn=RBM23;refGene_tx=XM_019832800.2;refGene_gn=RBM23;refGene_tx=XM_019832801.2
B3 75767911 75768064
refGene_gn=PRMT5;refGene_tx=XM_003987460.5;refGene_gn=PRMT5;refGene_tx=XM_019832799.2
B3 75768984 75769049
refGene_gn=PRMT5;refGene_tx=XM_003987460.5;refGene_gn=PRMT5;refGene_tx=XM_019832799.2
B3 75769257 75769374
refGene_gn=PRMT5;refGene_tx=XM_003987460.5;refGene_gn=PRMT5;refGene_tx=XM_019832799.2
B3 75769552 75769646
refGene_gn=PRMT5;refGene_tx=XM_003987460.5;refGene_gn=PRMT5;refGene_tx=XM_019832799.2
B3 75769890 75770000
refGene_gn=PRMT5;refGene_tx=XM_003987460.5;refGene_gn=PRMT5;refGene_tx=XM_019832799.2
B3 75770556 75770732
refGene_gn=PRMT5;refGene_tx=XM_003987460.5;refGene_gn=PRMT5;refGene_tx=XM_019832799.2
B3 75770818 75770941
refGene_gn=PRMT5;refGene_tx=XM_003987460.5;refGene_gn=PRMT5;refGene_tx=XM_019832799.2
B3 75771024 75771083
refGene_gn=PRMT5;refGene_tx=XM_003987460.5;refGene_gn=PRMT5;refGene_tx

=XM_019832799.2
B3	75771175	75771253
refGene_gn=PRMT5;refGene_tx=XM_003987460.5;refGene_gn=PRMT5;refGene_tx
=XM_019832799.2
B3	75771426	75771588
refGene_gn=PRMT5;refGene_tx=XM_003987460.5;refGene_gn=PRMT5;refGene_tx
=XM_019832799.2
B3	75772062	75772226
refGene_gn=PRMT5;refGene_tx=XM_003987460.5;refGene_gn=PRMT5;refGene_tx
=XM_019832799.2
B3	75774161	75774211
refGene_gn=PRMT5;refGene_tx=XM_003987460.5;refGene_gn=PRMT5;refGene_tx
=XM_019832799.2
B3	75774356	75774469
refGene_gn=PRMT5;refGene_tx=XM_003987460.5;refGene_gn=PRMT5;refGene_tx
=XM_019832799.2
B3	75775046	75775181
refGene_gn=PRMT5;refGene_tx=XM_003987460.5;refGene_gn=PRMT5;refGene_tx
=XM_019832799.2
B3	75775355	75775441	refGene_gn=PRMT5;refGene_tx=XM_003987460.5
B3	75775749	75775868	refGene_gn=PRMT5;refGene_tx=XM_003987460.5
B3	75776534	75776644	refGene_gn=PRMT5;refGene_tx=XM_003987460.5
B3	75781207	75781214	refGene_gn=HAUS4;refGene_tx=XM_019832802.2
B3	75792284	75792390
refGene_gn=HAUS4;refGene_tx=XM_006932725.4;refGene_gn=HAUS4;refGene_tx
=XM_006932726.4;refGene_gn=HAUS4;refGene_tx=XM_006932727.4;refGene_gn=
HAUS4;refGene_tx=XM_006932729.4;refGene_gn=HAUS4;refGene_tx=XM_0232555
80.1
B3	75793591	75793660
refGene_gn=HAUS4;refGene_tx=XM_006932725.4;refGene_gn=HAUS4;refGene_tx
=XM_006932726.4;refGene_gn=HAUS4;refGene_tx=XM_006932727.4;refGene_gn=
HAUS4;refGene_tx=XM_006932729.4;refGene_gn=HAUS4;refGene_tx=XM_0198328
02.2;refGene_gn=HAUS4;refGene_tx=XM_023255580.1
B3	75794004	75794135
refGene_gn=HAUS4;refGene_tx=XM_006932725.4;refGene_gn=HAUS4;refGene_tx
=XM_006932726.4;refGene_gn=HAUS4;refGene_tx=XM_006932727.4;refGene_gn=
HAUS4;refGene_tx=XM_006932729.4;refGene_gn=HAUS4;refGene_tx=XM_0198328
02.2;refGene_gn=HAUS4;refGene_tx=XM_023255580.1
B3	75794498	75794644
refGene_gn=HAUS4;refGene_tx=XM_006932725.4;refGene_gn=HAUS4;refGene_tx
=XM_006932726.4;refGene_gn=HAUS4;refGene_tx=XM_006932727.4;refGene_gn=
HAUS4;refGene_tx=XM_006932729.4;refGene_gn=HAUS4;refGene_tx=XM_0198328
02.2;refGene_gn=HAUS4;refGene_tx=XM_023255580.1
B3	75795260	75795357
refGene_gn=HAUS4;refGene_tx=XM_006932725.4;refGene_gn=HAUS4;refGene_tx
=XM_006932726.4;refGene_gn=HAUS4;refGene_tx=XM_006932727.4;refGene_gn=
HAUS4;refGene_tx=XM_006932729.4;refGene_gn=HAUS4;refGene_tx=XM_0198328
02.2;refGene_gn=HAUS4;refGene_tx=XM_023255580.1
B3	75796324	75796459
refGene_gn=HAUS4;refGene_tx=XM_006932725.4;refGene_gn=HAUS4;refGene_tx

=XM_006932726.4;refGene_gn=HAUS4;refGene_tx=XM_006932727.4;refGene_gn=HAUS4;refGene_tx=XM_006932729.4;refGene_gn=HAUS4;refGene_tx=XM_019832802.2;refGene_gn=HAUS4;refGene_tx=XM_023255580.1
B3 75796922 75797054
refGene_gn=HAUS4;refGene_tx=XM_006932725.4;refGene_gn=HAUS4;refGene_tx=XM_006932726.4;refGene_gn=HAUS4;refGene_tx=XM_006932727.4;refGene_gn=HAUS4;refGene_tx=XM_006932729.4;refGene_gn=HAUS4;refGene_tx=XM_019832802.2;refGene_gn=HAUS4;refGene_tx=XM_023255580.1
B3 75797143 75797286
refGene_gn=HAUS4;refGene_tx=XM_006932725.4;refGene_gn=HAUS4;refGene_tx=XM_006932726.4;refGene_gn=HAUS4;refGene_tx=XM_006932727.4;refGene_gn=HAUS4;refGene_tx=XM_006932729.4;refGene_gn=HAUS4;refGene_tx=XM_019832802.2;refGene_gn=HAUS4;refGene_tx=XM_023255580.1
B3 75799423 75799478
refGene_gn=HAUS4;refGene_tx=XM_006932725.4;refGene_gn=HAUS4;refGene_tx=XM_006932726.4;refGene_gn=HAUS4;refGene_tx=XM_006932727.4;refGene_gn=HAUS4;refGene_tx=XM_006932729.4;refGene_gn=HAUS4;refGene_tx=XM_019832802.2;refGene_gn=HAUS4;refGene_tx=XM_023255580.1
B3 75812890 75813016 refGene_gn=AJUBA;refGene_tx=XM_003987465.5
B3 75813512 75813581 refGene_gn=AJUBA;refGene_tx=XM_003987465.5
B3 75814338 75814390 refGene_gn=AJUBA;refGene_tx=XM_003987465.5
B3 75814473 75814604 refGene_gn=AJUBA;refGene_tx=XM_003987465.5
B3 75815353 75815416 refGene_gn=AJUBA;refGene_tx=XM_003987465.5
B3 75815549 75815617 refGene_gn=AJUBA;refGene_tx=XM_003987465.5
B3 75817297 75817399 refGene_gn=AJUBA;refGene_tx=XM_003987465.5
B3 75820176 75821212 refGene_gn=AJUBA;refGene_tx=XM_003987465.5
B3 75825307 75825727
refGene_gn=CB3H14orf93;refGene_tx=XM_011283132.3;refGene_gn=CB3H14orf93;refGene_tx=XM_011283133.3;refGene_gn=CB3H14orf93;refGene_tx=XM_011283134.3;refGene_gn=CB3H14orf93;refGene_tx=XM_023255581.1
B3 75825978 75826091
refGene_gn=CB3H14orf93;refGene_tx=XM_011283132.3;refGene_gn=CB3H14orf93;refGene_tx=XM_011283133.3;refGene_gn=CB3H14orf93;refGene_tx=XM_011283134.3;refGene_gn=CB3H14orf93;refGene_tx=XM_023255581.1
B3 75827272 75827376
refGene_gn=CB3H14orf93;refGene_tx=XM_011283132.3;refGene_gn=CB3H14orf93;refGene_tx=XM_011283133.3;refGene_gn=CB3H14orf93;refGene_tx=XM_011283134.3;refGene_gn=CB3H14orf93;refGene_tx=XM_023255581.1
B3 75827601 75827663
refGene_gn=CB3H14orf93;refGene_tx=XM_011283132.3;refGene_gn=CB3H14orf93;refGene_tx=XM_011283133.3;refGene_gn=CB3H14orf93;refGene_tx=XM_011283134.3;refGene_gn=CB3H14orf93;refGene_tx=XM_023255581.1
B3 75831063 75831384
refGene_gn=CB3H14orf93;refGene_tx=XM_011283132.3;refGene_gn=CB3H14orf93;refGene_tx=XM_011283133.3;refGene_gn=CB3H14orf93;refGene_tx=XM_023255581.1
B3 75833573 75834173
refGene_gn=CB3H14orf93;refGene_tx=XM_011283132.3;refGene_gn=CB3H14orf93;refGene_tx=XM_011283133.3;refGene_gn=CB3H14orf93;refGene_tx=XM_011283134.3;refGene_gn=CB3H14orf93;refGene_tx=XM_023255581.1

B3 75861336 75861623 refGene_gn=PSMB5;refGene_tx=XM_003987466.5
B3 75866154 75866461 refGene_gn=PSMB5;refGene_tx=XM_003987466.5
B3 75867182 75867380 refGene_gn=PSMB5;refGene_tx=XM_003987466.5
B3 75873077 75873932 refGene_gn=PSMB11;refGene_tx=XM_003987557.3
B3 75878569 75879069
refGene_gn=CDH24;refGene_tx=XM_011283135.3;refGene_gn=CDH24;refGene_tx=XM_023255582.1;refGene_gn=CDH24;refGene_tx=XM_023255583.1
B3 75879635 75879884
refGene_gn=CDH24;refGene_tx=XM_011283135.3;refGene_gn=CDH24;refGene_tx=XM_023255582.1;refGene_gn=CDH24;refGene_tx=XM_023255583.1
B3 75880131 75880243
refGene_gn=CDH24;refGene_tx=XM_011283135.3;refGene_gn=CDH24;refGene_tx=XM_023255582.1;refGene_gn=CDH24;refGene_tx=XM_023255583.1
B3 75880324 75880446
refGene_gn=CDH24;refGene_tx=XM_011283135.3;refGene_gn=CDH24;refGene_tx=XM_023255582.1;refGene_gn=CDH24;refGene_tx=XM_023255583.1
B3 75882773 75882910
refGene_gn=CDH24;refGene_tx=XM_011283135.3;refGene_gn=CDH24;refGene_tx=XM_023255582.1;refGene_gn=CDH24;refGene_tx=XM_023255583.1
B3 75883858 75884112
refGene_gn=CDH24;refGene_tx=XM_011283135.3;refGene_gn=CDH24;refGene_tx=XM_023255582.1;refGene_gn=CDH24;refGene_tx=XM_023255583.1
B3 75884523 75884711
refGene_gn=CDH24;refGene_tx=XM_011283135.3;refGene_gn=CDH24;refGene_tx=XM_023255582.1;refGene_gn=CDH24;refGene_tx=XM_023255583.1
B3 75884886 75885054
refGene_gn=CDH24;refGene_tx=XM_011283135.3;refGene_gn=CDH24;refGene_tx=XM_023255582.1;refGene_gn=CDH24;refGene_tx=XM_023255583.1
B3 75885128 75885248
refGene_gn=CDH24;refGene_tx=XM_011283135.3;refGene_gn=CDH24;refGene_tx=XM_023255582.1;refGene_gn=CDH24;refGene_tx=XM_023255583.1
B3 75885403 75885698
refGene_gn=CDH24;refGene_tx=XM_011283135.3;refGene_gn=CDH24;refGene_tx=XM_023255582.1;refGene_gn=CDH24;refGene_tx=XM_023255583.1
B3 75885888 75886089
refGene_gn=CDH24;refGene_tx=XM_011283135.3;refGene_gn=CDH24;refGene_tx=XM_023255582.1;refGene_gn=CDH24;refGene_tx=XM_023255583.1
B3 75889469 75889796
refGene_gn=ACIN1;refGene_tx=XM_003987468.5;refGene_gn=ACIN1;refGene_tx=XM_003987469.5;refGene_gn=ACIN1;refGene_tx=XM_003987470.5;refGene_gn=ACIN1;refGene_tx=XM_006932734.4;refGene_gn=ACIN1;refGene_tx=XM_019832805.2;refGene_gn=ACIN1;refGene_tx=XM_019832806.1;refGene_gn=ACIN1;refGene_tx=XM_023255584.1
B3 75891122 75891223
refGene_gn=ACIN1;refGene_tx=XM_003987468.5;refGene_gn=ACIN1;refGene_tx=XM_003987469.5;refGene_gn=ACIN1;refGene_tx=XM_003987470.5;refGene_gn=ACIN1;refGene_tx=XM_006932734.4;refGene_gn=ACIN1;refGene_tx=XM_019832805.2;refGene_gn=ACIN1;refGene_tx=XM_019832806.1;refGene_gn=ACIN1;refGene_tx=XM_023255584.1
B3 75891324 75891649

refGene_gn=ACIN1;refGene_tx=XM_003987468.5;refGene_gn=ACIN1;refGene_tx=XM_003987469.5;refGene_gn=ACIN1;refGene_tx=XM_003987470.5;refGene_gn=ACIN1;refGene_tx=XM_006932734.4;refGene_gn=ACIN1;refGene_tx=XM_019832805.2;refGene_gn=ACIN1;refGene_tx=XM_019832806.1;refGene_gn=ACIN1;refGene_tx=XM_023255584.1
B3 75891885 75891993
refGene_gn=ACIN1;refGene_tx=XM_003987468.5;refGene_gn=ACIN1;refGene_tx=XM_003987469.5;refGene_gn=ACIN1;refGene_tx=XM_003987470.5;refGene_gn=ACIN1;refGene_tx=XM_006932734.4;refGene_gn=ACIN1;refGene_tx=XM_019832805.2;refGene_gn=ACIN1;refGene_tx=XM_019832806.1;refGene_gn=ACIN1;refGene_tx=XM_023255584.1
B3 75892132 75892240
refGene_gn=ACIN1;refGene_tx=XM_003987468.5;refGene_gn=ACIN1;refGene_tx=XM_003987469.5;refGene_gn=ACIN1;refGene_tx=XM_003987470.5;refGene_gn=ACIN1;refGene_tx=XM_006932734.4;refGene_gn=ACIN1;refGene_tx=XM_019832805.2;refGene_gn=ACIN1;refGene_tx=XM_019832806.1;refGene_gn=ACIN1;refGene_tx=XM_023255584.1
B3 75892625 75892771
refGene_gn=ACIN1;refGene_tx=XM_003987468.5;refGene_gn=ACIN1;refGene_tx=XM_003987469.5;refGene_gn=ACIN1;refGene_tx=XM_003987470.5;refGene_gn=ACIN1;refGene_tx=XM_006932734.4;refGene_gn=ACIN1;refGene_tx=XM_019832805.2;refGene_gn=ACIN1;refGene_tx=XM_019832806.1;refGene_gn=ACIN1;refGene_tx=XM_023255584.1
B3 75893348 75893490
refGene_gn=ACIN1;refGene_tx=XM_003987468.5;refGene_gn=ACIN1;refGene_tx=XM_003987469.5;refGene_gn=ACIN1;refGene_tx=XM_003987470.5;refGene_gn=ACIN1;refGene_tx=XM_006932734.4;refGene_gn=ACIN1;refGene_tx=XM_019832805.2;refGene_gn=ACIN1;refGene_tx=XM_019832806.1;refGene_gn=ACIN1;refGene_tx=XM_023255584.1
B3 75893979 75894132
refGene_gn=ACIN1;refGene_tx=XM_003987468.5;refGene_gn=ACIN1;refGene_tx=XM_003987469.5;refGene_gn=ACIN1;refGene_tx=XM_003987470.5;refGene_gn=ACIN1;refGene_tx=XM_006932734.4;refGene_gn=ACIN1;refGene_tx=XM_019832805.2;refGene_gn=ACIN1;refGene_tx=XM_019832806.1;refGene_gn=ACIN1;refGene_tx=XM_023255584.1
B3 75894229 75894363
refGene_gn=ACIN1;refGene_tx=XM_003987468.5;refGene_gn=ACIN1;refGene_tx=XM_003987469.5;refGene_gn=ACIN1;refGene_tx=XM_003987470.5;refGene_gn=ACIN1;refGene_tx=XM_006932734.4;refGene_gn=ACIN1;refGene_tx=XM_019832805.2;refGene_gn=ACIN1;refGene_tx=XM_019832806.1;refGene_gn=ACIN1;refGene_tx=XM_023255584.1
B3 75895231 75895274
refGene_gn=ACIN1;refGene_tx=XM_003987468.5;refGene_gn=ACIN1;refGene_tx=XM_003987469.5;refGene_gn=ACIN1;refGene_tx=XM_003987470.5;refGene_gn=ACIN1;refGene_tx=XM_006932734.4;refGene_gn=ACIN1;refGene_tx=XM_019832805.2;refGene_gn=ACIN1;refGene_tx=XM_019832806.1;refGene_gn=ACIN1;refGene_tx=XM_023255584.1
B3 75898697 75898839
refGene_gn=ACIN1;refGene_tx=XM_003987468.5;refGene_gn=ACIN1;refGene_tx=XM_003987469.5;refGene_gn=ACIN1;refGene_tx=XM_003987470.5;refGene_gn=

ACIN1;refGene_tx=XM_006932734.4;refGene_gn=ACIN1;refGene_tx=XM_0198328
05.2;refGene_gn=ACIN1;refGene_tx=XM_019832806.1;refGene_gn=ACIN1;refGe
ne_tx=XM_023255584.1
B3 75900639 75900758 refGene_gn=ACIN1;refGene_tx=XM_019832806.1
B3 75906464 75906580
refGene_gn=ACIN1;refGene_tx=XM_003987468.5;refGene_gn=ACIN1;refGene_tx
=XM_003987469.5;refGene_gn=ACIN1;refGene_tx=XM_003987470.5;refGene_gn=
ACIN1;refGene_tx=XM_006932734.4;refGene_gn=ACIN1;refGene_tx=XM_0198328
05.2;refGene_gn=ACIN1;refGene_tx=XM_023255584.1
B3 75907080 75907296
refGene_gn=ACIN1;refGene_tx=XM_003987468.5;refGene_gn=ACIN1;refGene_tx
=XM_003987469.5;refGene_gn=ACIN1;refGene_tx=XM_003987470.5;refGene_gn=
ACIN1;refGene_tx=XM_006932734.4;refGene_gn=ACIN1;refGene_tx=XM_0232555
84.1
B3 75907797 75909057
refGene_gn=ACIN1;refGene_tx=XM_003987468.5;refGene_gn=ACIN1;refGene_tx
=XM_003987469.5;refGene_gn=ACIN1;refGene_tx=XM_003987470.5;refGene_gn=
ACIN1;refGene_tx=XM_006932734.4;refGene_gn=ACIN1;refGene_tx=XM_0198328
05.2;refGene_gn=ACIN1;refGene_tx=XM_023255584.1
B3 75910543 75910632
refGene_gn=ACIN1;refGene_tx=XM_003987468.5;refGene_gn=ACIN1;refGene_tx
=XM_003987469.5;refGene_gn=ACIN1;refGene_tx=XM_003987470.5;refGene_gn=
ACIN1;refGene_tx=XM_006932734.4;refGene_gn=ACIN1;refGene_tx=XM_0198328
05.2;refGene_gn=ACIN1;refGene_tx=XM_023255584.1
B3 75918710 75918830
refGene_gn=ACIN1;refGene_tx=XM_003987468.5;refGene_gn=ACIN1;refGene_tx
=XM_003987470.5;refGene_gn=ACIN1;refGene_tx=XM_006932734.4;refGene_gn=
ACIN1;refGene_tx=XM_019832805.2;refGene_gn=ACIN1;refGene_tx=XM_0232555
84.1
B3 75919243 75919355
refGene_gn=ACIN1;refGene_tx=XM_003987468.5;refGene_gn=ACIN1;refGene_tx
=XM_003987469.5;refGene_gn=ACIN1;refGene_tx=XM_003987470.5;refGene_gn=
ACIN1;refGene_tx=XM_006932734.4;refGene_gn=ACIN1;refGene_tx=XM_0198328
05.2;refGene_gn=ACIN1;refGene_tx=XM_023255584.1
B3 75921540 75921606
refGene_gn=ACIN1;refGene_tx=XM_003987468.5;refGene_gn=ACIN1;refGene_tx
=XM_003987469.5;refGene_gn=ACIN1;refGene_tx=XM_003987470.5;refGene_gn=
ACIN1;refGene_tx=XM_006932734.4;refGene_gn=ACIN1;refGene_tx=XM_0198328
05.2;refGene_gn=ACIN1;refGene_tx=XM_023255584.1
B3 75923016 75923328
refGene_gn=ACIN1;refGene_tx=XM_003987468.5;refGene_gn=ACIN1;refGene_tx
=XM_003987469.5;refGene_gn=ACIN1;refGene_tx=XM_003987470.5;refGene_gn=
ACIN1;refGene_tx=XM_019832805.2
B3 75925044 75925473
refGene_gn=CB3H14orf119;refGene_tx=XM_003987472.5;refGene_gn=CB3H14orf
119;refGene_tx=XM_019832807.1;refGene_gn=CB3H14orf119;refGene_tx=XM_01
9832808.2
B3 75926324 75926532
refGene_gn=LOC111560964;refGene_tx=XM_023256181.1
B3 75927008 75927188

refGene_gn=LOC111560964;refGene_tx=XM_023256181.1
B3 75927363 75927593
refGene_gn=LOC111560964;refGene_tx=XM_023256181.1
B3 75927802 75928075
refGene_gn=LOC111560964;refGene_tx=XM_023256181.1
B3 75928179 75928276
refGene_gn=LOC111560964;refGene_tx=XM_023256181.1
B3 75928351 75928421
refGene_gn=LOC111560964;refGene_tx=XM_023256181.1
B3 75928604 75928776
refGene_gn=LOC111560964;refGene_tx=XM_023256181.1
B3 75928851 75928997
refGene_gn=LOC111560964;refGene_tx=XM_023256181.1
B3 75929143 75929228
refGene_gn=LOC111560964;refGene_tx=XM_023256181.1
B3 75929438 75929564
refGene_gn=LOC111560964;refGene_tx=XM_023256181.1
B3 75929618 75929627
refGene_gn=LOC111560964;refGene_tx=XM_023256181.1
B3 75929681 75929987
refGene_gn=LOC111560964;refGene_tx=XM_023256181.1
B3 75930284 75930336
refGene_gn=LOC111560964;refGene_tx=XM_023256181.1
B3 75930427 75930512
refGene_gn=LOC111560964;refGene_tx=XM_023256181.1
B3 75930642 75930937
refGene_gn=LOC111560964;refGene_tx=XM_023256181.1
B3 75939953 75940289 refGene_gn=CEBPE;refGene_tx=XM_003987473.3
B3 75940915 75941425 refGene_gn=CEBPE;refGene_tx=XM_003987473.3
B3 75948502 75948666 refGene_gn=SLC7A8;refGene_tx=XM_006932735.4
B3 75949169 75949347 refGene_gn=SLC7A8;refGene_tx=XM_006932735.4
B3 75950950 75951100 refGene_gn=SLC7A8;refGene_tx=XM_006932735.4
B3 75952826 75952923 refGene_gn=SLC7A8;refGene_tx=XM_006932735.4
B3 75956665 75956769 refGene_gn=SLC7A8;refGene_tx=XM_006932735.4
B3 75958231 75958355 refGene_gn=SLC7A8;refGene_tx=XM_006932735.4
B3 75958984 75959025 refGene_gn=SLC7A8;refGene_tx=XM_006932735.4
B3 75959026 75959139 refGene_gn=SLC7A8;refGene_tx=XM_006932735.4
B3 75961212 75961338 refGene_gn=SLC7A8;refGene_tx=XM_006932735.4
B3 75981244 75981396 refGene_gn=SLC7A8;refGene_tx=XM_006932735.4
B3 75982305 75982510 refGene_gn=SLC7A8;refGene_tx=XM_006932735.4
B3 75998055 75998215 refGene_gn=SLC7A8;refGene_tx=XM_006932735.4
B3 76027985 76028085 refGene_gn=RNF212B;refGene_tx=XM_006932743.3
B3 76029846 76029899 refGene_gn=RNF212B;refGene_tx=XM_006932743.3
B3 76030409 76030484 refGene_gn=RNF212B;refGene_tx=XM_006932743.3
B3 76040740 76040856 refGene_gn=RNF212B;refGene_tx=XM_006932743.3
B3 76041485 76041546 refGene_gn=RNF212B;refGene_tx=XM_006932743.3
B3 76042501 76042530 refGene_gn=RNF212B;refGene_tx=XM_006932743.3
B3 76043514 76043561 refGene_gn=RNF212B;refGene_tx=XM_006932743.3
B3 76043784 76043827 refGene_gn=RNF212B;refGene_tx=XM_006932743.3
B3 76043987 76044048 refGene_gn=RNF212B;refGene_tx=XM_006932743.3

B3 76044437 76044486 refGene_gn=RNF212B;refGene_tx=XM_006932743.3
B3 76049787 76049827 refGene_gn=RNF212B;refGene_tx=XM_006932743.3
B3 76050116 76050217 refGene_gn=RNF212B;refGene_tx=XM_006932743.3
B3 76050473 76050535 refGene_gn=RNF212B;refGene_tx=XM_006932743.3
B3 76052261 76052327 refGene_gn=RNF212B;refGene_tx=XM_006932743.3
B3 76054546 76056147
refGene_gn=HOMEZ;refGene_tx=XM_003987478.5;refGene_gn=HOMEZ;refGene_tx=XM_006932737.4;refGene_gn=HOMEZ;refGene_tx=XM_006932738.4
B3 76061339 76061379 refGene_gn=HOMEZ;refGene_tx=XM_003987478.5
B3 76061946 76062066 refGene_gn=HOMEZ;refGene_tx=XM_006932737.4
B3 76065346 76065434
refGene_gn=HOMEZ;refGene_tx=XM_006932737.4;refGene_gn=HOMEZ;refGene_tx=XM_006932738.4
B3 76070330 76070753 refGene_gn=PPP1R3E;refGene_tx=XM_023255588.1
B3 76071000 76071417 refGene_gn=PPP1R3E;refGene_tx=XM_023255588.1
B3 76076945 76077377 refGene_gn=BCL2L2;refGene_tx=XM_003987479.4
B3 76077939 76078089 refGene_gn=BCL2L2;refGene_tx=XM_003987479.4
B3 76087189 76087540 refGene_gn=PABPN1;refGene_tx=XM_023255587.1
B3 76087898 76088013 refGene_gn=PABPN1;refGene_tx=XM_023255587.1
B3 76088720 76088788 refGene_gn=PABPN1;refGene_tx=XM_023255587.1
B3 76089104 76089211 refGene_gn=PABPN1;refGene_tx=XM_023255587.1
B3 76089676 76089764 refGene_gn=PABPN1;refGene_tx=XM_023255587.1
B3 76089852 76090004 refGene_gn=PABPN1;refGene_tx=XM_023255587.1
B3 76090943 76090983 refGene_gn=PABPN1;refGene_tx=XM_023255587.1
B3 76099762 76100051
refGene_gn=SLC22A17;refGene_tx=XM_011283140.2;refGene_gn=SLC22A17;refGene_tx=XM_019832815.2;refGene_gn=SLC22A17;refGene_tx=XM_023255589.1
B3 76100168 76100280
refGene_gn=SLC22A17;refGene_tx=XM_011283140.2;refGene_gn=SLC22A17;refGene_tx=XM_019832815.2;refGene_gn=SLC22A17;refGene_tx=XM_023255589.1
B3 76100518 76100790
refGene_gn=SLC22A17;refGene_tx=XM_011283140.2;refGene_gn=SLC22A17;refGene_tx=XM_019832815.2;refGene_gn=SLC22A17;refGene_tx=XM_023255589.1
B3 76100949 76101055
refGene_gn=SLC22A17;refGene_tx=XM_011283140.2;refGene_gn=SLC22A17;refGene_tx=XM_019832815.2;refGene_gn=SLC22A17;refGene_tx=XM_023255589.1
B3 76101217 76101363
refGene_gn=SLC22A17;refGene_tx=XM_011283140.2;refGene_gn=SLC22A17;refGene_tx=XM_019832815.2;refGene_gn=SLC22A17;refGene_tx=XM_023255589.1
B3 76101552 76101718
refGene_gn=SLC22A17;refGene_tx=XM_011283140.2;refGene_gn=SLC22A17;refGene_tx=XM_019832815.2;refGene_gn=SLC22A17;refGene_tx=XM_023255589.1
B3 76102311 76102466
refGene_gn=SLC22A17;refGene_tx=XM_011283140.2;refGene_gn=SLC22A17;refGene_tx=XM_019832815.2;refGene_gn=SLC22A17;refGene_tx=XM_023255589.1
B3 76104607 76104711
refGene_gn=SLC22A17;refGene_tx=XM_011283140.2;refGene_gn=SLC22A17;refGene_tx=XM_019832815.2;refGene_gn=SLC22A17;refGene_tx=XM_023255589.1
B3 76104803 76105211
refGene_gn=SLC22A17;refGene_tx=XM_011283140.2;refGene_gn=SLC22A17;refG

ene_tx=XM_019832815.2;refGene_gn=SLC22A17;refGene_tx=XM_023255589.1
B3	76107496	76107555
refGene_gn=EFS;refGene_tx=XM_019832812.2;refGene_gn=EFS;refGene_tx=XM_019832813.2
B3	76107852	76107960
refGene_gn=EFS;refGene_tx=XM_006932745.4;refGene_gn=EFS;refGene_tx=XM_019832811.2;refGene_gn=EFS;refGene_tx=XM_019832812.2
B3	76109592	76110027
refGene_gn=EFS;refGene_tx=XM_003987484.5;refGene_gn=EFS;refGene_tx=XM_006932745.4;refGene_gn=EFS;refGene_tx=XM_019832811.2;refGene_gn=EFS;refGene_tx=XM_019832812.2;refGene_gn=EFS;refGene_tx=XM_019832813.2;refGene_gn=EFS;refGene_tx=XM_019832814.2
B3	76111158	76111248
refGene_gn=EFS;refGene_tx=XM_003987484.5;refGene_gn=EFS;refGene_tx=XM_006932745.4;refGene_gn=EFS;refGene_tx=XM_019832811.2;refGene_gn=EFS;refGene_tx=XM_019832812.2;refGene_gn=EFS;refGene_tx=XM_019832813.2;refGene_gn=EFS;refGene_tx=XM_019832814.2
B3	76111612	76112335
refGene_gn=EFS;refGene_tx=XM_003987484.5;refGene_gn=EFS;refGene_tx=XM_006932745.4;refGene_gn=EFS;refGene_tx=XM_019832811.2;refGene_gn=EFS;refGene_tx=XM_019832812.2;refGene_gn=EFS;refGene_tx=XM_019832813.2;refGene_gn=EFS;refGene_tx=XM_019832814.2
B3	76112437	76112578
refGene_gn=EFS;refGene_tx=XM_003987484.5;refGene_gn=EFS;refGene_tx=XM_006932745.4;refGene_gn=EFS;refGene_tx=XM_019832811.2;refGene_gn=EFS;refGene_tx=XM_019832812.2;refGene_gn=EFS;refGene_tx=XM_019832813.2;refGene_gn=EFS;refGene_tx=XM_019832814.2
B3	76112838	76113117
refGene_gn=EFS;refGene_tx=XM_003987484.5;refGene_gn=EFS;refGene_tx=XM_006932745.4;refGene_gn=EFS;refGene_tx=XM_019832811.2;refGene_gn=EFS;refGene_tx=XM_019832812.2;refGene_gn=EFS;refGene_tx=XM_019832813.2;refGene_gn=EFS;refGene_tx=XM_019832814.2
B3	76117394	76117412
refGene_gn=EFS;refGene_tx=XM_003987484.5;refGene_gn=EFS;refGene_tx=XM_006932745.4;refGene_gn=EFS;refGene_tx=XM_019832811.2;refGene_gn=EFS;refGene_tx=XM_019832812.2;refGene_gn=EFS;refGene_tx=XM_019832813.2
B3	76125375	76125549
refGene_gn=IL25;refGene_tx=XM_003987559.4;refGene_gn=IL25;refGene_tx=XM_019832817.2
B3	76125727	76125972
refGene_gn=IL25;refGene_tx=XM_003987559.4;refGene_gn=IL25;refGene_tx=XM_019832817.2
B3	76127833	76128089
refGene_gn=IL25;refGene_tx=XM_003987559.4;refGene_gn=IL25;refGene_tx=XM_019832817.2
B3	76129495	76129621
refGene_gn=CMTM5;refGene_tx=XM_003987486.5;refGene_gn=CMTM5;refGene_tx=XM_019832821.2
B3	76130502	76130655
refGene_gn=CMTM5;refGene_tx=XM_003987486.5;refGene_gn=CMTM5;refGene_tx

=XM_019832821.2
B3	76131163 76131256 refGene_gn=CMTM5;refGene_tx=XM_003987486.5
B3	76131453 76131538
refGene_gn=CMTM5;refGene_tx=XM_003987486.5;refGene_gn=CMTM5;refGene_tx
=XM_019832821.2
B3	76131638 76131652
refGene_gn=CMTM5;refGene_tx=XM_003987486.5;refGene_gn=CMTM5;refGene_tx
=XM_019832821.2
B3	76134157 76134181 refGene_gn=MYH6;refGene_tx=XM_019832816.2
B3	76134539 76134674 refGene_gn=MYH6;refGene_tx=XM_019832816.2
B3	76135382 76135478 refGene_gn=MYH6;refGene_tx=XM_019832816.2
B3	76136642 76136918 refGene_gn=MYH6;refGene_tx=XM_019832816.2
B3	76137121 76137247 refGene_gn=MYH6;refGene_tx=XM_019832816.2
B3	76138157 76138361 refGene_gn=MYH6;refGene_tx=XM_019832816.2
B3	76138665 76138974 refGene_gn=MYH6;refGene_tx=XM_019832816.2
B3	76140104 76140229 refGene_gn=MYH6;refGene_tx=XM_019832816.2
B3	76140339 76140505 refGene_gn=MYH6;refGene_tx=XM_019832816.2
B3	76140696 76140880 refGene_gn=MYH6;refGene_tx=XM_019832816.2
B3	76141420 76141617 refGene_gn=MYH6;refGene_tx=XM_019832816.2
B3	76141994 76142113 refGene_gn=MYH6;refGene_tx=XM_019832816.2
B3	76142211 76142338 refGene_gn=MYH6;refGene_tx=XM_019832816.2
B3	76142654 76143044 refGene_gn=MYH6;refGene_tx=XM_019832816.2
B3	76145299 76145390 refGene_gn=MYH6;refGene_tx=XM_019832816.2
B3	76145634 76145780 refGene_gn=MYH6;refGene_tx=XM_019832816.2
B3	76146067 76146244 refGene_gn=MYH6;refGene_tx=XM_019832816.2
B3	76146393 76146636 refGene_gn=MYH6;refGene_tx=XM_019832816.2
B3	76146772 76147028 refGene_gn=MYH6;refGene_tx=XM_019832816.2
B3	76149093 76149230 refGene_gn=MYH6;refGene_tx=XM_019832816.2
B3	76149502 76149626 refGene_gn=MYH6;refGene_tx=XM_019832816.2
B3	76149771 76149889 refGene_gn=MYH6;refGene_tx=XM_019832816.2
B3	76149978 76150066 refGene_gn=MYH6;refGene_tx=XM_019832816.2
B3	76150341 76150412 refGene_gn=MYH6;refGene_tx=XM_019832816.2
B3	76151082 76151392 refGene_gn=MYH6;refGene_tx=XM_019832816.2
B3	76152562 76152733 refGene_gn=MYH6;refGene_tx=XM_019832816.2
B3	76152997 76153266 refGene_gn=MYH6;refGene_tx=XM_019832816.2
B3	76154975 76155114 refGene_gn=MYH6;refGene_tx=XM_019832816.2
B3	76155201 76155305 refGene_gn=MYH6;refGene_tx=XM_019832816.2
B3	76155776 76155875 refGene_gn=MYH6;refGene_tx=XM_019832816.2
B3	76156097 76156161 refGene_gn=MYH6;refGene_tx=XM_019832816.2
B3	76157048 76157141 refGene_gn=MYH6;refGene_tx=XM_019832816.2
B3	76157411 76157523 refGene_gn=MYH6;refGene_tx=XM_019832816.2
B3	76157803 76157831 refGene_gn=MYH6;refGene_tx=XM_019832816.2
B3	76157916 76158073 refGene_gn=MYH6;refGene_tx=XM_019832816.2
B3	76158314 76158458 refGene_gn=MYH6;refGene_tx=XM_019832816.2
B3	76159915 76160116 refGene_gn=MYH6;refGene_tx=XM_019832816.2
B3	76165417 76165435
refGene_gn=LOC101096736;refGene_tx=XM_006932746.4;refGene_gn=LOC101096
736;refGene_tx=XM_011283143.3
B3	76166029 76166164
refGene_gn=LOC101096736;refGene_tx=XM_006932746.4;refGene_gn=LOC101096

736;refGene_tx=XM_011283143.3
B3 76166287 76166383
refGene_gn=LOC101096736;refGene_tx=XM_006932746.4;refGene_gn=LOC101096736;refGene_tx=XM_011283143.3
B3 76167355 76167631
refGene_gn=LOC101096736;refGene_tx=XM_006932746.4;refGene_gn=LOC101096736;refGene_tx=XM_011283143.3
B3 76167834 76167960
refGene_gn=LOC101096736;refGene_tx=XM_006932746.4;refGene_gn=LOC101096736;refGene_tx=XM_011283143.3
B3 76168083 76168287
refGene_gn=LOC101096736;refGene_tx=XM_006932746.4;refGene_gn=LOC101096736;refGene_tx=XM_011283143.3
B3 76168508 76168817
refGene_gn=LOC101096736;refGene_tx=XM_006932746.4;refGene_gn=LOC101096736;refGene_tx=XM_011283143.3
B3 76169406 76169531
refGene_gn=LOC101096736;refGene_tx=XM_006932746.4;refGene_gn=LOC101096736;refGene_tx=XM_011283143.3
B3 76169667 76169833
refGene_gn=LOC101096736;refGene_tx=XM_006932746.4;refGene_gn=LOC101096736;refGene_tx=XM_011283143.3
B3 76170013 76170197
refGene_gn=LOC101096736;refGene_tx=XM_006932746.4;refGene_gn=LOC101096736;refGene_tx=XM_011283143.3
B3 76170779 76170976
refGene_gn=LOC101096736;refGene_tx=XM_006932746.4;refGene_gn=LOC101096736;refGene_tx=XM_011283143.3
B3 76171902 76172021
refGene_gn=LOC101096736;refGene_tx=XM_006932746.4;refGene_gn=LOC101096736;refGene_tx=XM_011283143.3
B3 76172210 76172337
refGene_gn=LOC101096736;refGene_tx=XM_006932746.4;refGene_gn=LOC101096736;refGene_tx=XM_011283143.3
B3 76172551 76172941
refGene_gn=LOC101096736;refGene_tx=XM_006932746.4;refGene_gn=LOC101096736;refGene_tx=XM_011283143.3
B3 76173642 76173733
refGene_gn=LOC101096736;refGene_tx=XM_006932746.4;refGene_gn=LOC101096736;refGene_tx=XM_011283143.3
B3 76174844 76174990
refGene_gn=LOC101096736;refGene_tx=XM_006932746.4;refGene_gn=LOC101096736;refGene_tx=XM_011283143.3
B3 76175779 76175956
refGene_gn=LOC101096736;refGene_tx=XM_006932746.4;refGene_gn=LOC101096736;refGene_tx=XM_011283143.3
B3 76176105 76176348
refGene_gn=LOC101096736;refGene_tx=XM_006932746.4;refGene_gn=LOC101096736;refGene_tx=XM_011283143.3
B3 76176969 76177225

refGene_gn=LOC101096736;refGene_tx=XM_006932746.4;refGene_gn=LOC101096736;refGene_tx=XM_011283143.3
B3 76177466 76177603
refGene_gn=LOC101096736;refGene_tx=XM_006932746.4;refGene_gn=LOC101096736;refGene_tx=XM_011283143.3
B3 76177868 76177992
refGene_gn=LOC101096736;refGene_tx=XM_006932746.4;refGene_gn=LOC101096736;refGene_tx=XM_011283143.3
B3 76178137 76178255
refGene_gn=LOC101096736;refGene_tx=XM_006932746.4;refGene_gn=LOC101096736;refGene_tx=XM_011283143.3
B3 76179157 76179245
refGene_gn=LOC101096736;refGene_tx=XM_006932746.4;refGene_gn=LOC101096736;refGene_tx=XM_011283143.3
B3 76179599 76179667
refGene_gn=LOC101096736;refGene_tx=XM_006932746.4;refGene_gn=LOC101096736;refGene_tx=XM_011283143.3
B3 76179951 76180261
refGene_gn=LOC101096736;refGene_tx=XM_006932746.4;refGene_gn=LOC101096736;refGene_tx=XM_011283143.3
B3 76180843 76181014
refGene_gn=LOC101096736;refGene_tx=XM_006932746.4;refGene_gn=LOC101096736;refGene_tx=XM_011283143.3
B3 76181278 76181428
refGene_gn=LOC101096736;refGene_tx=XM_006932746.4;refGene_gn=LOC101096736;refGene_tx=XM_011283143.3
B3 76181541 76181660
refGene_gn=LOC101096736;refGene_tx=XM_006932746.4;refGene_gn=LOC101096736;refGene_tx=XM_011283143.3
B3 76181762 76181901
refGene_gn=LOC101096736;refGene_tx=XM_006932746.4;refGene_gn=LOC101096736;refGene_tx=XM_011283143.3
B3 76182544 76182648
refGene_gn=LOC101096736;refGene_tx=XM_006932746.4;refGene_gn=LOC101096736;refGene_tx=XM_011283143.3
B3 76182882 76182981
refGene_gn=LOC101096736;refGene_tx=XM_006932746.4;refGene_gn=LOC101096736;refGene_tx=XM_011283143.3
B3 76183306 76183370
refGene_gn=LOC101096736;refGene_tx=XM_006932746.4;refGene_gn=LOC101096736;refGene_tx=XM_011283143.3
B3 76183473 76183566
refGene_gn=LOC101096736;refGene_tx=XM_006932746.4;refGene_gn=LOC101096736;refGene_tx=XM_011283143.3
B3 76183649 76183758
refGene_gn=LOC101096736;refGene_tx=XM_006932746.4;refGene_gn=LOC101096736;refGene_tx=XM_011283143.3
B3 76184358 76184386
refGene_gn=LOC101096736;refGene_tx=XM_006932746.4;refGene_gn=LOC101096736;refGene_tx=XM_011283143.3

B3 76184566 76184723
refGene_gn=LOC101096736;refGene_tx=XM_006932746.4;refGene_gn=LOC101096736;refGene_tx=XM_011283143.3
B3 76185022 76185166
refGene_gn=LOC101096736;refGene_tx=XM_006932746.4;refGene_gn=LOC101096736;refGene_tx=XM_011283143.3
B3 76185459 76185660
refGene_gn=LOC101096736;refGene_tx=XM_006932746.4;refGene_gn=LOC101096736;refGene_tx=XM_011283143.3
B3 76206586 76206607
refGene_gn=NGDN;refGene_tx=XM_003987561.5;refGene_gn=NGDN;refGene_tx=XM_006932749.4
B3 76206919 76206979
refGene_gn=NGDN;refGene_tx=XM_003987561.5;refGene_gn=NGDN;refGene_tx=XM_006932749.4
B3 76207808 76207880 refGene_gn=NGDN;refGene_tx=XM_003987561.5
B3 76211257 76211395
refGene_gn=NGDN;refGene_tx=XM_003987561.5;refGene_gn=NGDN;refGene_tx=XM_006932749.4
B3 76211641 76211726
refGene_gn=NGDN;refGene_tx=XM_003987561.5;refGene_gn=NGDN;refGene_tx=XM_006932749.4
B3 76211816 76211869
refGene_gn=NGDN;refGene_tx=XM_003987561.5;refGene_gn=NGDN;refGene_tx=XM_006932749.4
B3 76212112 76212233
refGene_gn=NGDN;refGene_tx=XM_003987561.5;refGene_gn=NGDN;refGene_tx=XM_006932749.4
B3 76212328 76212497
refGene_gn=NGDN;refGene_tx=XM_003987561.5;refGene_gn=NGDN;refGene_tx=XM_006932749.4
B3 76213211 76213368
refGene_gn=NGDN;refGene_tx=XM_003987561.5;refGene_gn=NGDN;refGene_tx=XM_006932749.4
B3 76213511 76213569
refGene_gn=NGDN;refGene_tx=XM_003987561.5;refGene_gn=NGDN;refGene_tx=XM_006932749.4
B3 76214023 76214043
refGene_gn=NGDN;refGene_tx=XM_003987561.5;refGene_gn=NGDN;refGene_tx=XM_006932749.4
B3 76249489 76250469
refGene_gn=ZFHX2;refGene_tx=XM_019832824.2;refGene_gn=ZFHX2;refGene_tx=XM_019832825.2;refGene_gn=ZFHX2;refGene_tx=XM_019832826.2;refGene_gn=ZFHX2;refGene_tx=XM_019832827.2;refGene_gn=ZFHX2;refGene_tx=XM_019832828.2;refGene_gn=ZFHX2;refGene_tx=XM_019832831.2;refGene_gn=ZFHX2;refGene_tx=XM_019832832.2;refGene_gn=ZFHX2;refGene_tx=XM_023255590.1;refGene_gn=ZFHX2;refGene_tx=XM_023255591.1;refGene_gn=ZFHX2;refGene_tx=XM_023255592.1;refGene_gn=ZFHX2;refGene_tx=XM_023255593.1
B3 76250719 76254190
refGene_gn=ZFHX2;refGene_tx=XM_019832824.2;refGene_gn=ZFHX2;refGene_tx

=XM_019832825.2;refGene_gn=ZFHX2;refGene_tx=XM_019832826.2;refGene_gn=
ZFHX2;refGene_tx=XM_019832827.2;refGene_gn=ZFHX2;refGene_tx=XM_0198328
28.2;refGene_gn=ZFHX2;refGene_tx=XM_019832831.2;refGene_gn=ZFHX2;refGe
ne_tx=XM_019832832.2;refGene_gn=ZFHX2;refGene_tx=XM_023255590.1;refGen
e_gn=ZFHX2;refGene_tx=XM_023255591.1;refGene_gn=ZFHX2;refGene_tx=XM_02
3255592.1;refGene_gn=ZFHX2;refGene_tx=XM_023255593.1
B3 76254329 76254456
refGene_gn=ZFHX2;refGene_tx=XM_019832824.2;refGene_gn=ZFHX2;refGene_tx
=XM_019832825.2;refGene_gn=ZFHX2;refGene_tx=XM_019832826.2;refGene_gn=
ZFHX2;refGene_tx=XM_019832827.2;refGene_gn=ZFHX2;refGene_tx=XM_0198328
28.2;refGene_gn=ZFHX2;refGene_tx=XM_019832831.2;refGene_gn=ZFHX2;refGe
ne_tx=XM_019832832.2;refGene_gn=ZFHX2;refGene_tx=XM_023255590.1;refGen
e_gn=ZFHX2;refGene_tx=XM_023255591.1;refGene_gn=ZFHX2;refGene_tx=XM_02
3255592.1;refGene_gn=ZFHX2;refGene_tx=XM_023255593.1
B3 76254820 76255021
refGene_gn=ZFHX2;refGene_tx=XM_019832824.2;refGene_gn=ZFHX2;refGene_tx
=XM_019832825.2;refGene_gn=ZFHX2;refGene_tx=XM_019832826.2;refGene_gn=
ZFHX2;refGene_tx=XM_019832827.2;refGene_gn=ZFHX2;refGene_tx=XM_0198328
28.2;refGene_gn=ZFHX2;refGene_tx=XM_019832831.2;refGene_gn=ZFHX2;refGe
ne_tx=XM_019832832.2;refGene_gn=ZFHX2;refGene_tx=XM_023255590.1;refGen
e_gn=ZFHX2;refGene_tx=XM_023255591.1;refGene_gn=ZFHX2;refGene_tx=XM_02
3255592.1;refGene_gn=ZFHX2;refGene_tx=XM_023255593.1
B3 76256687 76256758
refGene_gn=ZFHX2;refGene_tx=XM_019832824.2;refGene_gn=ZFHX2;refGene_tx
=XM_019832825.2;refGene_gn=ZFHX2;refGene_tx=XM_019832826.2;refGene_gn=
ZFHX2;refGene_tx=XM_019832827.2;refGene_gn=ZFHX2;refGene_tx=XM_0198328
28.2;refGene_gn=ZFHX2;refGene_tx=XM_019832831.2;refGene_gn=ZFHX2;refGe
ne_tx=XM_019832832.2;refGene_gn=ZFHX2;refGene_tx=XM_023255590.1;refGen
e_gn=ZFHX2;refGene_tx=XM_023255591.1;refGene_gn=ZFHX2;refGene_tx=XM_02
3255592.1;refGene_gn=ZFHX2;refGene_tx=XM_023255593.1
B3 76257165 76257240
refGene_gn=ZFHX2;refGene_tx=XM_019832824.2;refGene_gn=ZFHX2;refGene_tx
=XM_019832825.2;refGene_gn=ZFHX2;refGene_tx=XM_019832826.2;refGene_gn=
ZFHX2;refGene_tx=XM_019832827.2;refGene_gn=ZFHX2;refGene_tx=XM_0198328
28.2;refGene_gn=ZFHX2;refGene_tx=XM_019832831.2;refGene_gn=ZFHX2;refGe
ne_tx=XM_019832832.2;refGene_gn=ZFHX2;refGene_tx=XM_023255590.1;refGen
e_gn=ZFHX2;refGene_tx=XM_023255591.1;refGene_gn=ZFHX2;refGene_tx=XM_02
3255592.1;refGene_gn=ZFHX2;refGene_tx=XM_023255593.1
B3 76258510 76258751
refGene_gn=ZFHX2;refGene_tx=XM_019832824.2;refGene_gn=ZFHX2;refGene_tx
=XM_019832825.2;refGene_gn=ZFHX2;refGene_tx=XM_019832826.2;refGene_gn=
ZFHX2;refGene_tx=XM_019832827.2;refGene_gn=ZFHX2;refGene_tx=XM_0198328
28.2;refGene_gn=ZFHX2;refGene_tx=XM_019832831.2;refGene_gn=ZFHX2;refGe
ne_tx=XM_019832832.2;refGene_gn=ZFHX2;refGene_tx=XM_023255590.1;refGen
e_gn=ZFHX2;refGene_tx=XM_023255591.1;refGene_gn=ZFHX2;refGene_tx=XM_02
3255592.1;refGene_gn=ZFHX2;refGene_tx=XM_023255593.1
B3 76259296 76259814
refGene_gn=ZFHX2;refGene_tx=XM_019832824.2;refGene_gn=ZFHX2;refGene_tx
=XM_019832825.2;refGene_gn=ZFHX2;refGene_tx=XM_019832826.2;refGene_gn=
ZFHX2;refGene_tx=XM_019832827.2;refGene_gn=ZFHX2;refGene_tx=XM_0198328

28.2;refGene_gn=ZFHX2;refGene_tx=XM_019832831.2;refGene_gn=ZFHX2;refGene_tx=XM_019832832.2;refGene_gn=ZFHX2;refGene_tx=XM_023255590.1;refGene_gn=ZFHX2;refGene_tx=XM_023255591.1;refGene_gn=ZFHX2;refGene_tx=XM_023255592.1;refGene_gn=ZFHX2;refGene_tx=XM_023255593.1
B3 76259994 76262035
refGene_gn=ZFHX2;refGene_tx=XM_019832824.2;refGene_gn=ZFHX2;refGene_tx=XM_019832825.2;refGene_gn=ZFHX2;refGene_tx=XM_019832826.2;refGene_gn=ZFHX2;refGene_tx=XM_019832827.2;refGene_gn=ZFHX2;refGene_tx=XM_019832828.2;refGene_gn=ZFHX2;refGene_tx=XM_019832831.2;refGene_gn=ZFHX2;refGene_tx=XM_019832832.2;refGene_gn=ZFHX2;refGene_tx=XM_023255590.1;refGene_gn=ZFHX2;refGene_tx=XM_023255591.1;refGene_gn=ZFHX2;refGene_tx=XM_023255592.1;refGene_gn=ZFHX2;refGene_tx=XM_023255593.1
B3 76280322 76280383 refGene_gn=THTPA;refGene_tx=XM_006932754.4
B3 76280870 76281422
refGene_gn=THTPA;refGene_tx=XM_003987489.3;refGene_gn=THTPA;refGene_tx=XM_006932754.4
B3 76282519 76282665
refGene_gn=THTPA;refGene_tx=XM_003987489.3;refGene_gn=THTPA;refGene_tx=XM_006932754.4
B3 76283548 76283650
refGene_gn=AP1G2;refGene_tx=XM_019832835.2;refGene_gn=AP1G2;refGene_tx=XM_019832836.2
B3 76283752 76283851
refGene_gn=AP1G2;refGene_tx=XM_019832835.2;refGene_gn=AP1G2;refGene_tx=XM_019832836.2
B3 76284066 76284230
refGene_gn=AP1G2;refGene_tx=XM_019832835.2;refGene_gn=AP1G2;refGene_tx=XM_019832836.2
B3 76285306 76285442
refGene_gn=AP1G2;refGene_tx=XM_019832835.2;refGene_gn=AP1G2;refGene_tx=XM_019832836.2
B3 76285525 76285646
refGene_gn=AP1G2;refGene_tx=XM_019832835.2;refGene_gn=AP1G2;refGene_tx=XM_019832836.2
B3 76285978 76286083
refGene_gn=AP1G2;refGene_tx=XM_019832835.2;refGene_gn=AP1G2;refGene_tx=XM_019832836.2
B3 76286340 76286468
refGene_gn=AP1G2;refGene_tx=XM_019832835.2;refGene_gn=AP1G2;refGene_tx=XM_019832836.2
B3 76286555 76286645
refGene_gn=AP1G2;refGene_tx=XM_019832835.2;refGene_gn=AP1G2;refGene_tx=XM_019832836.2
B3 76287329 76287452
refGene_gn=AP1G2;refGene_tx=XM_019832835.2;refGene_gn=AP1G2;refGene_tx=XM_019832836.2
B3 76287535 76287590
refGene_gn=AP1G2;refGene_tx=XM_019832835.2;refGene_gn=AP1G2;refGene_tx=XM_019832836.2
B3 76287688 76287829

refGene_gn=AP1G2;refGene_tx=XM_019832835.2;refGene_gn=AP1G2;refGene_tx=XM_019832836.2
B3 76288025 76288139
refGene_gn=AP1G2;refGene_tx=XM_019832835.2;refGene_gn=AP1G2;refGene_tx=XM_019832836.2
B3 76288310 76288366
refGene_gn=AP1G2;refGene_tx=XM_019832835.2;refGene_gn=AP1G2;refGene_tx=XM_019832836.2
B3 76288541 76288640
refGene_gn=AP1G2;refGene_tx=XM_019832835.2;refGene_gn=AP1G2;refGene_tx=XM_019832836.2
B3 76288939 76289020
refGene_gn=AP1G2;refGene_tx=XM_019832835.2;refGene_gn=AP1G2;refGene_tx=XM_019832836.2
B3 76289399 76289495
refGene_gn=AP1G2;refGene_tx=XM_019832835.2;refGene_gn=AP1G2;refGene_tx=XM_019832836.2
B3 76289613 76289690 refGene_gn=AP1G2;refGene_tx=XM_019832835.2
B3 76289864 76289961
refGene_gn=AP1G2;refGene_tx=XM_019832835.2;refGene_gn=AP1G2;refGene_tx=XM_019832836.2
B3 76290082 76290224 refGene_gn=AP1G2;refGene_tx=XM_019832835.2
B3 76290365 76290490 refGene_gn=AP1G2;refGene_tx=XM_019832835.2
B3 76290949 76291153 refGene_gn=AP1G2;refGene_tx=XM_019832835.2
B3 76293521 76293605 refGene_gn=JPH4;refGene_tx=XM_019833789.2
B3 76294540 76295073 refGene_gn=JPH4;refGene_tx=XM_019833789.2
B3 76295397 76295516 refGene_gn=JPH4;refGene_tx=XM_019833789.2
B3 76298543 76299315 refGene_gn=JPH4;refGene_tx=XM_019833789.2
B3 76300094 76300473 refGene_gn=JPH4;refGene_tx=XM_019833789.2
B3 76333774 76333914
refGene_gn=LOC101097899;refGene_tx=XM_019832853.2;refGene_gn=LOC101097899;refGene_tx=XM_023255594.1;refGene_gn=LOC101097899;refGene_tx=XM_023255595.1
B3 76334082 76334260
refGene_gn=LOC101097899;refGene_tx=XM_019832853.2;refGene_gn=LOC101097899;refGene_tx=XM_023255594.1;refGene_gn=LOC101097899;refGene_tx=XM_023255595.1
B3 76334793 76334895
refGene_gn=LOC101097899;refGene_tx=XM_019832853.2;refGene_gn=LOC101097899;refGene_tx=XM_023255594.1;refGene_gn=LOC101097899;refGene_tx=XM_023255595.1
B3 76337769 76337840
refGene_gn=LOC101097899;refGene_tx=XM_019832853.2;refGene_gn=LOC101097899;refGene_tx=XM_023255594.1
B3 76338901 76338953
refGene_gn=LOC101097899;refGene_tx=XM_019832853.2;refGene_gn=LOC101097899;refGene_tx=XM_023255594.1
B3 76339460 76339595
refGene_gn=LOC101097899;refGene_tx=XM_019832853.2;refGene_gn=LOC101097899;refGene_tx=XM_023255594.1;refGene_gn=LOC101097899;refGene_tx=XM_02

3255595.1
B3	76339847 76339903
refGene_gn=LOC101097899;refGene_tx=XM_019832853.2;refGene_gn=LOC101097899;refGene_tx=XM_023255594.1;refGene_gn=LOC101097899;refGene_tx=XM_023255595.1
B3	76340149 76340261
refGene_gn=LOC101097899;refGene_tx=XM_019832853.2;refGene_gn=LOC101097899;refGene_tx=XM_023255594.1;refGene_gn=LOC101097899;refGene_tx=XM_023255595.1
B3	76553070 76553201
refGene_gn=LOC101098145;refGene_tx=XM_003987493.4
B3	76554313 76554491
refGene_gn=LOC101098145;refGene_tx=XM_003987493.4
B3	76558208 76558310
refGene_gn=LOC101098145;refGene_tx=XM_003987493.4
B3	76560327 76560398
refGene_gn=LOC101098145;refGene_tx=XM_003987493.4
B3	76560495 76560547
refGene_gn=LOC101098145;refGene_tx=XM_003987493.4
B3	76560845 76560980
refGene_gn=LOC101098145;refGene_tx=XM_003987493.4
B3	76561403 76561459
refGene_gn=LOC101098145;refGene_tx=XM_003987493.4
B3	76563000 76563115
refGene_gn=LOC101098145;refGene_tx=XM_003987493.4
B3	76564343 76564383
refGene_gn=CARMIL3;refGene_tx=XM_011283152.3;refGene_gn=CARMIL3;refGene_tx=XM_019832837.2;refGene_gn=CARMIL3;refGene_tx=XM_019832838.2;refGene_gn=CARMIL3;refGene_tx=XM_019832839.2;refGene_gn=CARMIL3;refGene_tx=XM_019832840.2;refGene_gn=CARMIL3;refGene_tx=XM_019832841.2;refGene_gn=CARMIL3;refGene_tx=XM_019832842.2;refGene_gn=CARMIL3;refGene_tx=XM_019832844.2
B3	76565893 76565988
refGene_gn=CARMIL3;refGene_tx=XM_011283152.3;refGene_gn=CARMIL3;refGene_tx=XM_019832837.2;refGene_gn=CARMIL3;refGene_tx=XM_019832838.2;refGene_gn=CARMIL3;refGene_tx=XM_019832839.2;refGene_gn=CARMIL3;refGene_tx=XM_019832840.2;refGene_gn=CARMIL3;refGene_tx=XM_019832841.2;refGene_gn=CARMIL3;refGene_tx=XM_019832842.2;refGene_gn=CARMIL3;refGene_tx=XM_019832844.2
B3	76566327 76566378
refGene_gn=CARMIL3;refGene_tx=XM_011283152.3;refGene_gn=CARMIL3;refGene_tx=XM_019832837.2;refGene_gn=CARMIL3;refGene_tx=XM_019832838.2;refGene_gn=CARMIL3;refGene_tx=XM_019832839.2;refGene_gn=CARMIL3;refGene_tx=XM_019832840.2;refGene_gn=CARMIL3;refGene_tx=XM_019832841.2;refGene_gn=CARMIL3;refGene_tx=XM_019832842.2;refGene_gn=CARMIL3;refGene_tx=XM_019832844.2
B3	76566480 76566540
refGene_gn=CARMIL3;refGene_tx=XM_011283152.3;refGene_gn=CARMIL3;refGene_tx=XM_019832837.2;refGene_gn=CARMIL3;refGene_tx=XM_019832838.2;refGene_gn=CARMIL3;refGene_tx=XM_019832839.2;refGene_gn=CARMIL3;refGene_tx=

XM_019832840.2;refGene_gn=CARMIL3;refGene_tx=XM_019832841.2;refGene_gn=CARMIL3;refGene_tx=XM_019832842.2;refGene_gn=CARMIL3;refGene_tx=XM_019832844.2
B3 76566625 76566741
refGene_gn=CARMIL3;refGene_tx=XM_011283152.3;refGene_gn=CARMIL3;refGene_tx=XM_019832837.2;refGene_gn=CARMIL3;refGene_tx=XM_019832838.2;refGene_gn=CARMIL3;refGene_tx=XM_019832839.2;refGene_gn=CARMIL3;refGene_tx=XM_019832840.2;refGene_gn=CARMIL3;refGene_tx=XM_019832841.2;refGene_gn=CARMIL3;refGene_tx=XM_019832842.2;refGene_gn=CARMIL3;refGene_tx=XM_019832844.2
B3 76566956 76567054
refGene_gn=CARMIL3;refGene_tx=XM_011283152.3;refGene_gn=CARMIL3;refGene_tx=XM_019832837.2;refGene_gn=CARMIL3;refGene_tx=XM_019832838.2;refGene_gn=CARMIL3;refGene_tx=XM_019832839.2;refGene_gn=CARMIL3;refGene_tx=XM_019832840.2;refGene_gn=CARMIL3;refGene_tx=XM_019832841.2;refGene_gn=CARMIL3;refGene_tx=XM_019832842.2;refGene_gn=CARMIL3;refGene_tx=XM_019832844.2;refGene_gn=CARMIL3;refGene_tx=XM_019832845.2
B3 76567294 76567365
refGene_gn=CARMIL3;refGene_tx=XM_011283152.3;refGene_gn=CARMIL3;refGene_tx=XM_019832837.2;refGene_gn=CARMIL3;refGene_tx=XM_019832838.2;refGene_gn=CARMIL3;refGene_tx=XM_019832839.2;refGene_gn=CARMIL3;refGene_tx=XM_019832840.2;refGene_gn=CARMIL3;refGene_tx=XM_019832841.2;refGene_gn=CARMIL3;refGene_tx=XM_019832842.2;refGene_gn=CARMIL3;refGene_tx=XM_019832844.2;refGene_gn=CARMIL3;refGene_tx=XM_019832845.2
B3 76567463 76567537
refGene_gn=CARMIL3;refGene_tx=XM_011283152.3;refGene_gn=CARMIL3;refGene_tx=XM_019832837.2;refGene_gn=CARMIL3;refGene_tx=XM_019832838.2;refGene_gn=CARMIL3;refGene_tx=XM_019832839.2;refGene_gn=CARMIL3;refGene_tx=XM_019832840.2;refGene_gn=CARMIL3;refGene_tx=XM_019832841.2;refGene_gn=CARMIL3;refGene_tx=XM_019832842.2;refGene_gn=CARMIL3;refGene_tx=XM_019832844.2;refGene_gn=CARMIL3;refGene_tx=XM_019832845.2
B3 76567758 76567834
refGene_gn=CARMIL3;refGene_tx=XM_011283152.3;refGene_gn=CARMIL3;refGene_tx=XM_019832837.2;refGene_gn=CARMIL3;refGene_tx=XM_019832838.2;refGene_gn=CARMIL3;refGene_tx=XM_019832839.2;refGene_gn=CARMIL3;refGene_tx=XM_019832840.2;refGene_gn=CARMIL3;refGene_tx=XM_019832841.2;refGene_gn=CARMIL3;refGene_tx=XM_019832842.2;refGene_gn=CARMIL3;refGene_tx=XM_019832844.2;refGene_gn=CARMIL3;refGene_tx=XM_019832845.2
B3 76567917 76568006
refGene_gn=CARMIL3;refGene_tx=XM_011283152.3;refGene_gn=CARMIL3;refGene_tx=XM_019832837.2;refGene_gn=CARMIL3;refGene_tx=XM_019832838.2;refGene_gn=CARMIL3;refGene_tx=XM_019832839.2;refGene_gn=CARMIL3;refGene_tx=XM_019832840.2;refGene_gn=CARMIL3;refGene_tx=XM_019832841.2;refGene_gn=CARMIL3;refGene_tx=XM_019832842.2;refGene_gn=CARMIL3;refGene_tx=XM_019832844.2;refGene_gn=CARMIL3;refGene_tx=XM_019832845.2
B3 76568474 76568569
refGene_gn=CARMIL3;refGene_tx=XM_011283152.3;refGene_gn=CARMIL3;refGene_tx=XM_019832837.2;refGene_gn=CARMIL3;refGene_tx=XM_019832838.2;refGene_gn=CARMIL3;refGene_tx=XM_019832839.2;refGene_gn=CARMIL3;refGene_tx=XM_019832840.2;refGene_gn=CARMIL3;refGene_tx=XM_019832841.2;refGene_gn

=CARMIL3;refGene_tx=XM_019832842.2;refGene_gn=CARMIL3;refGene_tx=XM_019832844.2;refGene_gn=CARMIL3;refGene_tx=XM_019832845.2
B3 76568796 76568883
refGene_gn=CARMIL3;refGene_tx=XM_011283152.3;refGene_gn=CARMIL3;refGene_tx=XM_019832837.2;refGene_gn=CARMIL3;refGene_tx=XM_019832838.2;refGene_gn=CARMIL3;refGene_tx=XM_019832839.2;refGene_gn=CARMIL3;refGene_tx=XM_019832840.2;refGene_gn=CARMIL3;refGene_tx=XM_019832841.2;refGene_gn=CARMIL3;refGene_tx=XM_019832842.2;refGene_gn=CARMIL3;refGene_tx=XM_019832844.2;refGene_gn=CARMIL3;refGene_tx=XM_019832845.2
B3 76569096 76569206
refGene_gn=CARMIL3;refGene_tx=XM_011283152.3;refGene_gn=CARMIL3;refGene_tx=XM_019832837.2;refGene_gn=CARMIL3;refGene_tx=XM_019832838.2;refGene_gn=CARMIL3;refGene_tx=XM_019832839.2;refGene_gn=CARMIL3;refGene_tx=XM_019832840.2;refGene_gn=CARMIL3;refGene_tx=XM_019832841.2;refGene_gn=CARMIL3;refGene_tx=XM_019832842.2;refGene_gn=CARMIL3;refGene_tx=XM_019832844.2;refGene_gn=CARMIL3;refGene_tx=XM_019832845.2
B3 76569342 76569420
refGene_gn=CARMIL3;refGene_tx=XM_011283152.3;refGene_gn=CARMIL3;refGene_tx=XM_019832837.2;refGene_gn=CARMIL3;refGene_tx=XM_019832838.2;refGene_gn=CARMIL3;refGene_tx=XM_019832839.2;refGene_gn=CARMIL3;refGene_tx=XM_019832840.2;refGene_gn=CARMIL3;refGene_tx=XM_019832841.2;refGene_gn=CARMIL3;refGene_tx=XM_019832842.2;refGene_gn=CARMIL3;refGene_tx=XM_019832844.2;refGene_gn=CARMIL3;refGene_tx=XM_019832845.2
B3 76569964 76570041
refGene_gn=CARMIL3;refGene_tx=XM_011283152.3;refGene_gn=CARMIL3;refGene_tx=XM_019832837.2;refGene_gn=CARMIL3;refGene_tx=XM_019832838.2;refGene_gn=CARMIL3;refGene_tx=XM_019832839.2;refGene_gn=CARMIL3;refGene_tx=XM_019832840.2;refGene_gn=CARMIL3;refGene_tx=XM_019832841.2;refGene_gn=CARMIL3;refGene_tx=XM_019832842.2;refGene_gn=CARMIL3;refGene_tx=XM_019832844.2;refGene_gn=CARMIL3;refGene_tx=XM_019832845.2
B3 76570122 76570227
refGene_gn=CARMIL3;refGene_tx=XM_011283152.3;refGene_gn=CARMIL3;refGene_tx=XM_019832837.2;refGene_gn=CARMIL3;refGene_tx=XM_019832838.2;refGene_gn=CARMIL3;refGene_tx=XM_019832839.2;refGene_gn=CARMIL3;refGene_tx=XM_019832840.2;refGene_gn=CARMIL3;refGene_tx=XM_019832841.2;refGene_gn=CARMIL3;refGene_tx=XM_019832842.2;refGene_gn=CARMIL3;refGene_tx=XM_019832844.2;refGene_gn=CARMIL3;refGene_tx=XM_019832845.2
B3 76570321 76570391
refGene_gn=CARMIL3;refGene_tx=XM_011283152.3;refGene_gn=CARMIL3;refGene_tx=XM_019832837.2;refGene_gn=CARMIL3;refGene_tx=XM_019832838.2;refGene_gn=CARMIL3;refGene_tx=XM_019832839.2;refGene_gn=CARMIL3;refGene_tx=XM_019832840.2;refGene_gn=CARMIL3;refGene_tx=XM_019832841.2;refGene_gn=CARMIL3;refGene_tx=XM_019832842.2;refGene_gn=CARMIL3;refGene_tx=XM_019832844.2;refGene_gn=CARMIL3;refGene_tx=XM_019832845.2
B3 76570848 76570930
refGene_gn=CARMIL3;refGene_tx=XM_011283152.3;refGene_gn=CARMIL3;refGene_tx=XM_019832837.2;refGene_gn=CARMIL3;refGene_tx=XM_019832838.2;refGene_gn=CARMIL3;refGene_tx=XM_019832839.2;refGene_gn=CARMIL3;refGene_tx=XM_019832840.2;refGene_gn=CARMIL3;refGene_tx=XM_019832841.2;refGene_gn=CARMIL3;refGene_tx=XM_019832842.2;refGene_gn=CARMIL3;refGene_tx=XM_01

9832844.2;refGene_gn=CARMIL3;refGene_tx=XM_019832845.2
B3 76571047 76571144
refGene_gn=CARMIL3;refGene_tx=XM_011283152.3;refGene_gn=CARMIL3;refGene_tx=XM_019832837.2;refGene_gn=CARMIL3;refGene_tx=XM_019832838.2;refGene_gn=CARMIL3;refGene_tx=XM_019832839.2;refGene_gn=CARMIL3;refGene_tx=XM_019832840.2;refGene_gn=CARMIL3;refGene_tx=XM_019832841.2;refGene_gn=CARMIL3;refGene_tx=XM_019832842.2;refGene_gn=CARMIL3;refGene_tx=XM_019832844.2;refGene_gn=CARMIL3;refGene_tx=XM_019832845.2
B3 76571306 76571361
refGene_gn=CARMIL3;refGene_tx=XM_011283152.3;refGene_gn=CARMIL3;refGene_tx=XM_019832837.2;refGene_gn=CARMIL3;refGene_tx=XM_019832838.2;refGene_gn=CARMIL3;refGene_tx=XM_019832839.2;refGene_gn=CARMIL3;refGene_tx=XM_019832840.2;refGene_gn=CARMIL3;refGene_tx=XM_019832841.2;refGene_gn=CARMIL3;refGene_tx=XM_019832842.2;refGene_gn=CARMIL3;refGene_tx=XM_019832844.2;refGene_gn=CARMIL3;refGene_tx=XM_019832845.2
B3 76571441 76571614
refGene_gn=CARMIL3;refGene_tx=XM_011283152.3;refGene_gn=CARMIL3;refGene_tx=XM_019832837.2;refGene_gn=CARMIL3;refGene_tx=XM_019832838.2;refGene_gn=CARMIL3;refGene_tx=XM_019832839.2;refGene_gn=CARMIL3;refGene_tx=XM_019832840.2;refGene_gn=CARMIL3;refGene_tx=XM_019832841.2;refGene_gn=CARMIL3;refGene_tx=XM_019832842.2;refGene_gn=CARMIL3;refGene_tx=XM_019832844.2;refGene_gn=CARMIL3;refGene_tx=XM_019832845.2
B3 76571879 76571948
refGene_gn=CARMIL3;refGene_tx=XM_011283152.3;refGene_gn=CARMIL3;refGene_tx=XM_019832837.2;refGene_gn=CARMIL3;refGene_tx=XM_019832838.2;refGene_gn=CARMIL3;refGene_tx=XM_019832839.2;refGene_gn=CARMIL3;refGene_tx=XM_019832840.2;refGene_gn=CARMIL3;refGene_tx=XM_019832841.2;refGene_gn=CARMIL3;refGene_tx=XM_019832842.2;refGene_gn=CARMIL3;refGene_tx=XM_019832844.2;refGene_gn=CARMIL3;refGene_tx=XM_019832845.2
B3 76572161 76572255
refGene_gn=CARMIL3;refGene_tx=XM_011283152.3;refGene_gn=CARMIL3;refGene_tx=XM_019832837.2;refGene_gn=CARMIL3;refGene_tx=XM_019832838.2;refGene_gn=CARMIL3;refGene_tx=XM_019832839.2;refGene_gn=CARMIL3;refGene_tx=XM_019832840.2;refGene_gn=CARMIL3;refGene_tx=XM_019832841.2;refGene_gn=CARMIL3;refGene_tx=XM_019832842.2;refGene_gn=CARMIL3;refGene_tx=XM_019832844.2;refGene_gn=CARMIL3;refGene_tx=XM_019832845.2
B3 76572341 76572440
refGene_gn=CARMIL3;refGene_tx=XM_011283152.3;refGene_gn=CARMIL3;refGene_tx=XM_019832837.2;refGene_gn=CARMIL3;refGene_tx=XM_019832838.2;refGene_gn=CARMIL3;refGene_tx=XM_019832839.2;refGene_gn=CARMIL3;refGene_tx=XM_019832840.2;refGene_gn=CARMIL3;refGene_tx=XM_019832841.2;refGene_gn=CARMIL3;refGene_tx=XM_019832842.2;refGene_gn=CARMIL3;refGene_tx=XM_019832844.2;refGene_gn=CARMIL3;refGene_tx=XM_019832845.2
B3 76572862 76572991
refGene_gn=CARMIL3;refGene_tx=XM_011283152.3;refGene_gn=CARMIL3;refGene_tx=XM_019832837.2;refGene_gn=CARMIL3;refGene_tx=XM_019832838.2;refGene_gn=CARMIL3;refGene_tx=XM_019832839.2;refGene_gn=CARMIL3;refGene_tx=XM_019832840.2;refGene_gn=CARMIL3;refGene_tx=XM_019832841.2;refGene_gn=CARMIL3;refGene_tx=XM_019832842.2;refGene_gn=CARMIL3;refGene_tx=XM_019832844.2;refGene_gn=CARMIL3;refGene_tx=XM_019832845.2

B3 76573166 76573280
refGene_gn=CARMIL3;refGene_tx=XM_011283152.3;refGene_gn=CARMIL3;refGene_tx=XM_019832837.2;refGene_gn=CARMIL3;refGene_tx=XM_019832838.2;refGene_gn=CARMIL3;refGene_tx=XM_019832839.2;refGene_gn=CARMIL3;refGene_tx=XM_019832840.2;refGene_gn=CARMIL3;refGene_tx=XM_019832841.2;refGene_gn=CARMIL3;refGene_tx=XM_019832842.2;refGene_gn=CARMIL3;refGene_tx=XM_019832844.2;refGene_gn=CARMIL3;refGene_tx=XM_019832845.2
B3 76573701 76573877
refGene_gn=CARMIL3;refGene_tx=XM_011283152.3;refGene_gn=CARMIL3;refGene_tx=XM_019832837.2;refGene_gn=CARMIL3;refGene_tx=XM_019832838.2;refGene_gn=CARMIL3;refGene_tx=XM_019832839.2;refGene_gn=CARMIL3;refGene_tx=XM_019832840.2;refGene_gn=CARMIL3;refGene_tx=XM_019832841.2;refGene_gn=CARMIL3;refGene_tx=XM_019832842.2;refGene_gn=CARMIL3;refGene_tx=XM_019832844.2;refGene_gn=CARMIL3;refGene_tx=XM_019832845.2
B3 76574655 76574743
refGene_gn=CARMIL3;refGene_tx=XM_011283152.3;refGene_gn=CARMIL3;refGene_tx=XM_019832837.2;refGene_gn=CARMIL3;refGene_tx=XM_019832838.2;refGene_gn=CARMIL3;refGene_tx=XM_019832839.2;refGene_gn=CARMIL3;refGene_tx=XM_019832840.2;refGene_gn=CARMIL3;refGene_tx=XM_019832841.2;refGene_gn=CARMIL3;refGene_tx=XM_019832842.2;refGene_gn=CARMIL3;refGene_tx=XM_019832844.2;refGene_gn=CARMIL3;refGene_tx=XM_019832845.2
B3 76574881 76575019
refGene_gn=CARMIL3;refGene_tx=XM_011283152.3;refGene_gn=CARMIL3;refGene_tx=XM_019832837.2;refGene_gn=CARMIL3;refGene_tx=XM_019832838.2;refGene_gn=CARMIL3;refGene_tx=XM_019832839.2;refGene_gn=CARMIL3;refGene_tx=XM_019832840.2;refGene_gn=CARMIL3;refGene_tx=XM_019832841.2;refGene_gn=CARMIL3;refGene_tx=XM_019832842.2;refGene_gn=CARMIL3;refGene_tx=XM_019832844.2;refGene_gn=CARMIL3;refGene_tx=XM_019832845.2
B3 76575308 76575372
refGene_gn=CARMIL3;refGene_tx=XM_011283152.3;refGene_gn=CARMIL3;refGene_tx=XM_019832837.2;refGene_gn=CARMIL3;refGene_tx=XM_019832838.2;refGene_gn=CARMIL3;refGene_tx=XM_019832839.2;refGene_gn=CARMIL3;refGene_tx=XM_019832840.2;refGene_gn=CARMIL3;refGene_tx=XM_019832841.2;refGene_gn=CARMIL3;refGene_tx=XM_019832842.2;refGene_gn=CARMIL3;refGene_tx=XM_019832844.2;refGene_gn=CARMIL3;refGene_tx=XM_019832845.2
B3 76575513 76575722
refGene_gn=CARMIL3;refGene_tx=XM_011283152.3;refGene_gn=CARMIL3;refGene_tx=XM_019832837.2;refGene_gn=CARMIL3;refGene_tx=XM_019832838.2;refGene_gn=CARMIL3;refGene_tx=XM_019832839.2;refGene_gn=CARMIL3;refGene_tx=XM_019832840.2;refGene_gn=CARMIL3;refGene_tx=XM_019832841.2;refGene_gn=CARMIL3;refGene_tx=XM_019832842.2;refGene_gn=CARMIL3;refGene_tx=XM_019832844.2;refGene_gn=CARMIL3;refGene_tx=XM_019832845.2
B3 76576104 76576208
refGene_gn=CARMIL3;refGene_tx=XM_011283152.3;refGene_gn=CARMIL3;refGene_tx=XM_019832837.2;refGene_gn=CARMIL3;refGene_tx=XM_019832838.2;refGene_gn=CARMIL3;refGene_tx=XM_019832839.2;refGene_gn=CARMIL3;refGene_tx=XM_019832840.2;refGene_gn=CARMIL3;refGene_tx=XM_019832841.2;refGene_gn=CARMIL3;refGene_tx=XM_019832842.2;refGene_gn=CARMIL3;refGene_tx=XM_019832844.2;refGene_gn=CARMIL3;refGene_tx=XM_019832845.2
B3 76576859 76577175

refGene_gn=CARMIL3;refGene_tx=XM_011283152.3;refGene_gn=CARMIL3;refGen
e_tx=XM_019832837.2;refGene_gn=CARMIL3;refGene_tx=XM_019832838.2;refGe
ne_gn=CARMIL3;refGene_tx=XM_019832839.2;refGene_gn=CARMIL3;refGene_tx=
XM_019832840.2;refGene_gn=CARMIL3;refGene_tx=XM_019832841.2;refGene_gn
=CARMIL3;refGene_tx=XM_019832842.2;refGene_gn=CARMIL3;refGene_tx=XM_01
9832844.2;refGene_gn=CARMIL3;refGene_tx=XM_019832845.2
B3 76577539 76577668
refGene_gn=CARMIL3;refGene_tx=XM_011283152.3;refGene_gn=CARMIL3;refGen
e_tx=XM_019832837.2;refGene_gn=CARMIL3;refGene_tx=XM_019832838.2;refGe
ne_gn=CARMIL3;refGene_tx=XM_019832839.2;refGene_gn=CARMIL3;refGene_tx=
XM_019832840.2;refGene_gn=CARMIL3;refGene_tx=XM_019832841.2;refGene_gn
=CARMIL3;refGene_tx=XM_019832844.2;refGene_gn=CARMIL3;refGene_tx=XM_01
9832845.2
B3 76578325 76578392
refGene_gn=CARMIL3;refGene_tx=XM_011283152.3;refGene_gn=CARMIL3;refGen
e_tx=XM_019832837.2;refGene_gn=CARMIL3;refGene_tx=XM_019832838.2;refGe
ne_gn=CARMIL3;refGene_tx=XM_019832839.2;refGene_gn=CARMIL3;refGene_tx=
XM_019832840.2;refGene_gn=CARMIL3;refGene_tx=XM_019832841.2;refGene_gn
=CARMIL3;refGene_tx=XM_019832842.2;refGene_gn=CARMIL3;refGene_tx=XM_01
9832844.2;refGene_gn=CARMIL3;refGene_tx=XM_019832845.2
B3 76578494 76578584
refGene_gn=CARMIL3;refGene_tx=XM_011283152.3;refGene_gn=CARMIL3;refGen
e_tx=XM_019832837.2;refGene_gn=CARMIL3;refGene_tx=XM_019832838.2;refGe
ne_gn=CARMIL3;refGene_tx=XM_019832839.2;refGene_gn=CARMIL3;refGene_tx=
XM_019832840.2;refGene_gn=CARMIL3;refGene_tx=XM_019832841.2;refGene_gn
=CARMIL3;refGene_tx=XM_019832842.2;refGene_gn=CARMIL3;refGene_tx=XM_01
9832844.2;refGene_gn=CARMIL3;refGene_tx=XM_019832845.2
B3 76580056 76580200
refGene_gn=CARMIL3;refGene_tx=XM_011283152.3;refGene_gn=CARMIL3;refGen
e_tx=XM_019832837.2;refGene_gn=CARMIL3;refGene_tx=XM_019832838.2;refGe
ne_gn=CARMIL3;refGene_tx=XM_019832839.2;refGene_gn=CARMIL3;refGene_tx=
XM_019832840.2;refGene_gn=CARMIL3;refGene_tx=XM_019832841.2;refGene_gn
=CARMIL3;refGene_tx=XM_019832842.2;refGene_gn=CARMIL3;refGene_tx=XM_01
9832844.2;refGene_gn=CARMIL3;refGene_tx=XM_019832845.2
B3 76580283 76580464
refGene_gn=CARMIL3;refGene_tx=XM_011283152.3;refGene_gn=CARMIL3;refGen
e_tx=XM_019832837.2;refGene_gn=CARMIL3;refGene_tx=XM_019832838.2;refGe
ne_gn=CARMIL3;refGene_tx=XM_019832839.2;refGene_gn=CARMIL3;refGene_tx=
XM_019832840.2;refGene_gn=CARMIL3;refGene_tx=XM_019832841.2;refGene_gn
=CARMIL3;refGene_tx=XM_019832842.2;refGene_gn=CARMIL3;refGene_tx=XM_01
9832845.2
B3 76580622 76580733
refGene_gn=CARMIL3;refGene_tx=XM_011283152.3;refGene_gn=CARMIL3;refGen
e_tx=XM_019832837.2;refGene_gn=CARMIL3;refGene_tx=XM_019832838.2;refGe
ne_gn=CARMIL3;refGene_tx=XM_019832839.2;refGene_gn=CARMIL3;refGene_tx=
XM_019832840.2;refGene_gn=CARMIL3;refGene_tx=XM_019832841.2;refGene_gn
=CARMIL3;refGene_tx=XM_019832842.2;refGene_gn=CARMIL3;refGene_tx=XM_01
9832845.2
B3 76580846 76580875
refGene_gn=CARMIL3;refGene_tx=XM_011283152.3;refGene_gn=CARMIL3;refGen

e_tx=XM_019832837.2;refGene_gn=CARMIL3;refGene_tx=XM_019832838.2;refGene_gn=CARMIL3;refGene_tx=XM_019832839.2;refGene_gn=CARMIL3;refGene_tx=XM_019832840.2;refGene_gn=CARMIL3;refGene_tx=XM_019832841.2;refGene_gn=CARMIL3;refGene_tx=XM_019832842.2;refGene_gn=CARMIL3;refGene_tx=XM_019832845.2
B3 76584436 76584604 refGene_gn=CPNE6;refGene_tx=XM_006932755.4
B3 76585021 76585201 refGene_gn=CPNE6;refGene_tx=XM_006932755.4
B3 76585538 76585613 refGene_gn=CPNE6;refGene_tx=XM_006932755.4
B3 76585777 76585852 refGene_gn=CPNE6;refGene_tx=XM_006932755.4
B3 76585998 76586082 refGene_gn=CPNE6;refGene_tx=XM_006932755.4
B3 76586173 76586263 refGene_gn=CPNE6;refGene_tx=XM_006932755.4
B3 76586620 76586725 refGene_gn=CPNE6;refGene_tx=XM_006932755.4
B3 76586951 76587038 refGene_gn=CPNE6;refGene_tx=XM_006932755.4
B3 76587258 76587318 refGene_gn=CPNE6;refGene_tx=XM_006932755.4
B3 76587582 76587716 refGene_gn=CPNE6;refGene_tx=XM_006932755.4
B3 76587802 76587857 refGene_gn=CPNE6;refGene_tx=XM_006932755.4
B3 76587950 76588002 refGene_gn=CPNE6;refGene_tx=XM_006932755.4
B3 76588330 76588464 refGene_gn=CPNE6;refGene_tx=XM_006932755.4
B3 76588608 76588845 refGene_gn=CPNE6;refGene_tx=XM_006932755.4
B3 76589051 76589189 refGene_gn=CPNE6;refGene_tx=XM_006932755.4
B3 76592353 76592686
refGene_gn=NRL;refGene_tx=XM_019832855.2;refGene_gn=NRL;refGene_tx=XM_023255599.1;refGene_gn=NRL;refGene_tx=XM_023255600.1
B3 76593596 76593977
refGene_gn=NRL;refGene_tx=XM_019832855.2;refGene_gn=NRL;refGene_tx=XM_023255599.1;refGene_gn=NRL;refGene_tx=XM_023255600.1
B3 76603406 76603435
refGene_gn=PCK2;refGene_tx=XM_003987496.5;refGene_gn=PCK2;refGene_tx=XM_019832846.2
B3 76605444 76605690
refGene_gn=PCK2;refGene_tx=XM_003987496.5;refGene_gn=PCK2;refGene_tx=XM_019832846.2
B3 76606145 76606330
refGene_gn=PCK2;refGene_tx=XM_003987496.5;refGene_gn=PCK2;refGene_tx=XM_019832846.2
B3 76606417 76606621
refGene_gn=PCK2;refGene_tx=XM_003987496.5;refGene_gn=PCK2;refGene_tx=XM_019832846.2
B3 76607000 76607188
refGene_gn=PCK2;refGene_tx=XM_003987496.5;refGene_gn=PCK2;refGene_tx=XM_019832846.2
B3 76607495 76607658
refGene_gn=PCK2;refGene_tx=XM_003987496.5;refGene_gn=PCK2;refGene_tx=XM_019832846.2
B3 76607905 76608124
refGene_gn=PCK2;refGene_tx=XM_003987496.5;refGene_gn=PCK2;refGene_tx=XM_019832846.2
B3 76610534 76610672
refGene_gn=PCK2;refGene_tx=XM_003987496.5;refGene_gn=PCK2;refGene_tx=XM_019832846.2

B3 76610950 76611046
refGene_gn=PCK2;refGene_tx=XM_003987496.5;refGene_gn=PCK2;refGene_tx=XM_019832846.2
B3 76611245 76611741
refGene_gn=PCK2;refGene_tx=XM_003987496.5;refGene_gn=PCK2;refGene_tx=XM_019832846.2
B3 76620217 76620323 refGene_gn=DCAF11;refGene_tx=XM_006932756.4
B3 76620958 76621165
refGene_gn=DCAF11;refGene_tx=XM_003987497.5;refGene_gn=DCAF11;refGene_tx=XM_003987498.5;refGene_gn=DCAF11;refGene_tx=XM_006932756.4;refGene_gn=DCAF11;refGene_tx=XM_006932757.4;refGene_gn=DCAF11;refGene_tx=XM_006932758.4;refGene_gn=DCAF11;refGene_tx=XM_019832847.2;refGene_gn=DCAF11;refGene_tx=XM_019832848.2
B3 76622050 76622178
refGene_gn=DCAF11;refGene_tx=XM_003987497.5;refGene_gn=DCAF11;refGene_tx=XM_003987498.5;refGene_gn=DCAF11;refGene_tx=XM_006932756.4;refGene_gn=DCAF11;refGene_tx=XM_006932757.4;refGene_gn=DCAF11;refGene_tx=XM_006932758.4;refGene_gn=DCAF11;refGene_tx=XM_019832847.2;refGene_gn=DCAF11;refGene_tx=XM_019832848.2
B3 76622436 76622564
refGene_gn=DCAF11;refGene_tx=XM_003987497.5;refGene_gn=DCAF11;refGene_tx=XM_003987498.5;refGene_gn=DCAF11;refGene_tx=XM_006932756.4;refGene_gn=DCAF11;refGene_tx=XM_006932757.4;refGene_gn=DCAF11;refGene_tx=XM_006932758.4;refGene_gn=DCAF11;refGene_tx=XM_019832847.2;refGene_gn=DCAF11;refGene_tx=XM_019832848.2
B3 76622841 76622906
refGene_gn=DCAF11;refGene_tx=XM_003987497.5;refGene_gn=DCAF11;refGene_tx=XM_003987498.5;refGene_gn=DCAF11;refGene_tx=XM_006932756.4;refGene_gn=DCAF11;refGene_tx=XM_006932757.4;refGene_gn=DCAF11;refGene_tx=XM_006932758.4;refGene_gn=DCAF11;refGene_tx=XM_019832847.2;refGene_gn=DCAF11;refGene_tx=XM_019832848.2
B3 76623214 76623315
refGene_gn=DCAF11;refGene_tx=XM_003987497.5;refGene_gn=DCAF11;refGene_tx=XM_003987498.5;refGene_gn=DCAF11;refGene_tx=XM_006932756.4;refGene_gn=DCAF11;refGene_tx=XM_006932757.4;refGene_gn=DCAF11;refGene_tx=XM_006932758.4;refGene_gn=DCAF11;refGene_tx=XM_019832847.2;refGene_gn=DCAF11;refGene_tx=XM_019832848.2
B3 76623544 76623691
refGene_gn=DCAF11;refGene_tx=XM_003987497.5;refGene_gn=DCAF11;refGene_tx=XM_003987498.5;refGene_gn=DCAF11;refGene_tx=XM_006932756.4;refGene_gn=DCAF11;refGene_tx=XM_006932757.4;refGene_gn=DCAF11;refGene_tx=XM_006932758.4;refGene_gn=DCAF11;refGene_tx=XM_019832847.2;refGene_gn=DCAF11;refGene_tx=XM_019832848.2
B3 76623904 76623959
refGene_gn=DCAF11;refGene_tx=XM_003987497.5;refGene_gn=DCAF11;refGene_tx=XM_003987498.5;refGene_gn=DCAF11;refGene_tx=XM_006932756.4;refGene_gn=DCAF11;refGene_tx=XM_006932757.4;refGene_gn=DCAF11;refGene_tx=XM_006932758.4;refGene_gn=DCAF11;refGene_tx=XM_019832847.2;refGene_gn=DCAF11;refGene_tx=XM_019832848.2
B3 76624318 76624387

refGene_gn=DCAF11;refGene_tx=XM_003987497.5;refGene_gn=DCAF11;refGene_tx=XM_003987498.5;refGene_gn=DCAF11;refGene_tx=XM_006932756.4;refGene_gn=DCAF11;refGene_tx=XM_006932757.4;refGene_gn=DCAF11;refGene_tx=XM_006932758.4;refGene_gn=DCAF11;refGene_tx=XM_019832847.2;refGene_gn=DCAF11;refGene_tx=XM_019832848.2
B3 76624718 76624776
refGene_gn=DCAF11;refGene_tx=XM_003987497.5;refGene_gn=DCAF11;refGene_tx=XM_003987498.5;refGene_gn=DCAF11;refGene_tx=XM_006932756.4;refGene_gn=DCAF11;refGene_tx=XM_006932757.4;refGene_gn=DCAF11;refGene_tx=XM_006932758.4;refGene_gn=DCAF11;refGene_tx=XM_019832847.2;refGene_gn=DCAF11;refGene_tx=XM_019832848.2
B3 76624874 76625060
refGene_gn=DCAF11;refGene_tx=XM_003987497.5;refGene_gn=DCAF11;refGene_tx=XM_003987498.5;refGene_gn=DCAF11;refGene_tx=XM_006932756.4;refGene_gn=DCAF11;refGene_tx=XM_006932757.4;refGene_gn=DCAF11;refGene_tx=XM_006932758.4;refGene_gn=DCAF11;refGene_tx=XM_019832847.2;refGene_gn=DCAF11;refGene_tx=XM_019832848.2
B3 76625492 76625646
refGene_gn=DCAF11;refGene_tx=XM_003987497.5;refGene_gn=DCAF11;refGene_tx=XM_003987498.5;refGene_gn=DCAF11;refGene_tx=XM_006932756.4;refGene_gn=DCAF11;refGene_tx=XM_006932757.4;refGene_gn=DCAF11;refGene_tx=XM_006932758.4;refGene_gn=DCAF11;refGene_tx=XM_019832847.2;refGene_gn=DCAF11;refGene_tx=XM_019832848.2
B3 76626049 76626202
refGene_gn=DCAF11;refGene_tx=XM_003987497.5;refGene_gn=DCAF11;refGene_tx=XM_003987498.5;refGene_gn=DCAF11;refGene_tx=XM_006932756.4;refGene_gn=DCAF11;refGene_tx=XM_006932757.4;refGene_gn=DCAF11;refGene_tx=XM_006932758.4;refGene_gn=DCAF11;refGene_tx=XM_019832847.2;refGene_gn=DCAF11;refGene_tx=XM_019832848.2
B3 76627484 76627591
refGene_gn=DCAF11;refGene_tx=XM_003987497.5;refGene_gn=DCAF11;refGene_tx=XM_003987498.5;refGene_gn=DCAF11;refGene_tx=XM_006932756.4;refGene_gn=DCAF11;refGene_tx=XM_006932757.4;refGene_gn=DCAF11;refGene_tx=XM_006932758.4;refGene_gn=DCAF11;refGene_tx=XM_019832847.2;refGene_gn=DCAF11;refGene_tx=XM_019832848.2
B3 76627686 76627821
refGene_gn=DCAF11;refGene_tx=XM_003987497.5;refGene_gn=DCAF11;refGene_tx=XM_003987498.5;refGene_gn=DCAF11;refGene_tx=XM_006932756.4;refGene_gn=DCAF11;refGene_tx=XM_006932757.4;refGene_gn=DCAF11;refGene_tx=XM_006932758.4;refGene_gn=DCAF11;refGene_tx=XM_019832847.2;refGene_gn=DCAF11;refGene_tx=XM_019832848.2
B3 76633349 76633615 refGene_gn=FITM1;refGene_tx=XM_003987500.3
B3 76633994 76634607 refGene_gn=FITM1;refGene_tx=XM_003987500.3
B3 76637438 76637477 refGene_gn=PSME1;refGene_tx=XM_003987501.4
B3 76638103 76638136 refGene_gn=PSME1;refGene_tx=XM_003987501.4
B3 76638280 76638343 refGene_gn=PSME1;refGene_tx=XM_003987501.4
B3 76638458 76638569 refGene_gn=PSME1;refGene_tx=XM_003987501.4
B3 76638657 76638703 refGene_gn=PSME1;refGene_tx=XM_003987501.4
B3 76638818 76638916 refGene_gn=PSME1;refGene_tx=XM_003987501.4
B3 76639160 76639229 refGene_gn=PSME1;refGene_tx=XM_003987501.4

B3 76639314 76639382 refGene_gn=PSME1;refGene_tx=XM_003987501.4
B3 76639484 76639539 refGene_gn=PSME1;refGene_tx=XM_003987501.4
B3 76639616 76639703 refGene_gn=PSME1;refGene_tx=XM_003987501.4
B3 76639876 76639957 refGene_gn=PSME1;refGene_tx=XM_003987501.4
B3 76640157 76640338 refGene_gn=EMC9;refGene_tx=XM_003987502.5
B3 76640491 76640586 refGene_gn=EMC9;refGene_tx=XM_003987502.5
B3 76640675 76640745 refGene_gn=EMC9;refGene_tx=XM_003987502.5
B3 76641643 76641720 refGene_gn=EMC9;refGene_tx=XM_003987502.5
B3 76641862 76642060 refGene_gn=EMC9;refGene_tx=XM_003987502.5
B3 76643419 76643500 refGene_gn=PSME2;refGene_tx=XM_003987503.5
B3 76643591 76643678 refGene_gn=PSME2;refGene_tx=XM_003987503.5
B3 76643975 76644030 refGene_gn=PSME2;refGene_tx=XM_003987503.5
B3 76644192 76644260 refGene_gn=PSME2;refGene_tx=XM_003987503.5
B3 76644416 76644485 refGene_gn=PSME2;refGene_tx=XM_003987503.5
B3 76645520 76645618 refGene_gn=PSME2;refGene_tx=XM_003987503.5
B3 76645704 76645735 refGene_gn=PSME2;refGene_tx=XM_003987503.5
B3 76645839 76645926 refGene_gn=PSME2;refGene_tx=XM_003987503.5
B3 76646162 76646225 refGene_gn=PSME2;refGene_tx=XM_003987503.5
B3 76646640 76646673 refGene_gn=PSME2;refGene_tx=XM_003987503.5
B3 76646983 76647031 refGene_gn=PSME2;refGene_tx=XM_003987503.5
B3 76648150 76648342
refGene_gn=RNF31;refGene_tx=XM_006932761.4;refGene_gn=RNF31;refGene_tx
=XM_006932762.3;refGene_gn=RNF31;refGene_tx=XM_006932763.4
B3 76648429 76648576
refGene_gn=RNF31;refGene_tx=XM_006932761.4;refGene_gn=RNF31;refGene_tx
=XM_006932762.3;refGene_gn=RNF31;refGene_tx=XM_006932763.4
B3 76648727 76648883
refGene_gn=RNF31;refGene_tx=XM_006932761.4;refGene_gn=RNF31;refGene_tx
=XM_006932762.3;refGene_gn=RNF31;refGene_tx=XM_006932763.4
B3 76649117 76649186
refGene_gn=RNF31;refGene_tx=XM_006932761.4;refGene_gn=RNF31;refGene_tx
=XM_006932762.3;refGene_gn=RNF31;refGene_tx=XM_006932763.4
B3 76649286 76649362
refGene_gn=RNF31;refGene_tx=XM_006932761.4;refGene_gn=RNF31;refGene_tx
=XM_006932762.3;refGene_gn=RNF31;refGene_tx=XM_006932763.4
B3 76649591 76649769
refGene_gn=RNF31;refGene_tx=XM_006932761.4;refGene_gn=RNF31;refGene_tx
=XM_006932762.3;refGene_gn=RNF31;refGene_tx=XM_006932763.4
B3 76650434 76650837
refGene_gn=RNF31;refGene_tx=XM_006932761.4;refGene_gn=RNF31;refGene_tx
=XM_006932762.3;refGene_gn=RNF31;refGene_tx=XM_006932763.4
B3 76651033 76651324
refGene_gn=RNF31;refGene_tx=XM_006932761.4;refGene_gn=RNF31;refGene_tx
=XM_006932762.3;refGene_gn=RNF31;refGene_tx=XM_006932763.4
B3 76651580 76651829
refGene_gn=RNF31;refGene_tx=XM_006932761.4;refGene_gn=RNF31;refGene_tx
=XM_006932762.3;refGene_gn=RNF31;refGene_tx=XM_006932763.4
B3 76651937 76652123
refGene_gn=RNF31;refGene_tx=XM_006932761.4;refGene_gn=RNF31;refGene_tx
=XM_006932762.3;refGene_gn=RNF31;refGene_tx=XM_006932763.4

B3	76652234	76652441
refGene_gn=RNF31;refGene_tx=XM_006932761.4;refGene_gn=RNF31;refGene_tx=XM_006932762.3;refGene_gn=RNF31;refGene_tx=XM_006932763.4
B3	76655354	76655528
refGene_gn=RNF31;refGene_tx=XM_006932761.4;refGene_gn=RNF31;refGene_tx=XM_006932762.3;refGene_gn=RNF31;refGene_tx=XM_006932763.4
B3	76655611	76655710
refGene_gn=RNF31;refGene_tx=XM_006932761.4;refGene_gn=RNF31;refGene_tx=XM_006932762.3;refGene_gn=RNF31;refGene_tx=XM_006932763.4
B3	76655806	76655896
refGene_gn=RNF31;refGene_tx=XM_006932761.4;refGene_gn=RNF31;refGene_tx=XM_006932762.3;refGene_gn=RNF31;refGene_tx=XM_006932763.4
B3	76657058	76657173
refGene_gn=RNF31;refGene_tx=XM_006932761.4;refGene_gn=RNF31;refGene_tx=XM_006932762.3;refGene_gn=RNF31;refGene_tx=XM_006932763.4
B3	76657288	76657407
refGene_gn=RNF31;refGene_tx=XM_006932761.4;refGene_gn=RNF31;refGene_tx=XM_006932762.3;refGene_gn=RNF31;refGene_tx=XM_006932763.4
B3	76657679	76657793
refGene_gn=RNF31;refGene_tx=XM_006932761.4;refGene_gn=RNF31;refGene_tx=XM_006932762.3;refGene_gn=RNF31;refGene_tx=XM_006932763.4
B3	76657942	76658000
refGene_gn=RNF31;refGene_tx=XM_006932761.4;refGene_gn=RNF31;refGene_tx=XM_006932762.3;refGene_gn=RNF31;refGene_tx=XM_006932763.4
B3	76658718	76658815
refGene_gn=RNF31;refGene_tx=XM_006932761.4;refGene_gn=RNF31;refGene_tx=XM_006932762.3;refGene_gn=RNF31;refGene_tx=XM_006932763.4
B3	76659144	76659313
refGene_gn=RNF31;refGene_tx=XM_006932761.4;refGene_gn=RNF31;refGene_tx=XM_006932762.3;refGene_gn=RNF31;refGene_tx=XM_006932763.4
B3	76659422	76659476
refGene_gn=RNF31;refGene_tx=XM_006932761.4;refGene_gn=RNF31;refGene_tx=XM_006932762.3;refGene_gn=RNF31;refGene_tx=XM_006932763.4
B3	76660604	76660717
refGene_gn=IRF9;refGene_tx=XM_006932767.4;refGene_gn=IRF9;refGene_tx=XM_019832851.2
B3	76660780	76660788	refGene_gn=IRF9;refGene_tx=XM_019832849.1
B3	76660985	76661166
refGene_gn=IRF9;refGene_tx=XM_006932767.4;refGene_gn=IRF9;refGene_tx=XM_019832849.1;refGene_gn=IRF9;refGene_tx=XM_019832850.1;refGene_gn=IRF9;refGene_tx=XM_019832851.2
B3	76661480	76661664
refGene_gn=IRF9;refGene_tx=XM_006932767.4;refGene_gn=IRF9;refGene_tx=XM_019832849.1;refGene_gn=IRF9;refGene_tx=XM_019832850.1;refGene_gn=IRF9;refGene_tx=XM_019832851.2
B3	76661916	76662047
refGene_gn=IRF9;refGene_tx=XM_006932767.4;refGene_gn=IRF9;refGene_tx=XM_019832849.1;refGene_gn=IRF9;refGene_tx=XM_019832850.1
B3	76662137	76662219
refGene_gn=IRF9;refGene_tx=XM_006932767.4;refGene_gn=IRF9;refGene_tx=X

M_019832849.1;refGene_gn=IRF9;refGene_tx=XM_019832850.1
B3	76662322	76662394
refGene_gn=IRF9;refGene_tx=XM_006932767.4;refGene_gn=IRF9;refGene_tx=XM_019832849.1;refGene_gn=IRF9;refGene_tx=XM_019832850.1;refGene_gn=IRF9;refGene_tx=XM_019832851.2
B3	76663268	76663610
refGene_gn=IRF9;refGene_tx=XM_006932767.4;refGene_gn=IRF9;refGene_tx=XM_019832849.1;refGene_gn=IRF9;refGene_tx=XM_019832850.1;refGene_gn=IRF9;refGene_tx=XM_019832851.2
B3	76664530	76664646
refGene_gn=IRF9;refGene_tx=XM_006932767.4;refGene_gn=IRF9;refGene_tx=XM_019832849.1;refGene_gn=IRF9;refGene_tx=XM_019832850.1;refGene_gn=IRF9;refGene_tx=XM_019832851.2
B3	76664801	76664930
refGene_gn=IRF9;refGene_tx=XM_006932767.4;refGene_gn=IRF9;refGene_tx=XM_019832849.1;refGene_gn=IRF9;refGene_tx=XM_019832850.1;refGene_gn=IRF9;refGene_tx=XM_019832851.2
B3	76671552	76671608	refGene_gn=REC8;refGene_tx=XM_003987564.4
B3	76671702	76671771	refGene_gn=REC8;refGene_tx=XM_003987564.4
B3	76671902	76672045
refGene_gn=REC8;refGene_tx=XM_003987564.4;refGene_gn=REC8;refGene_tx=XM_023255603.1
B3	76672130	76672206	refGene_gn=REC8;refGene_tx=XM_003987564.4
B3	76672529	76672650
refGene_gn=REC8;refGene_tx=XM_003987564.4;refGene_gn=REC8;refGene_tx=XM_023255603.1
B3	76673569	76673654
refGene_gn=REC8;refGene_tx=XM_003987564.4;refGene_gn=REC8;refGene_tx=XM_023255603.1
B3	76673994	76674074
refGene_gn=REC8;refGene_tx=XM_003987564.4;refGene_gn=REC8;refGene_tx=XM_023255603.1
B3	76674326	76674417
refGene_gn=REC8;refGene_tx=XM_003987564.4;refGene_gn=REC8;refGene_tx=XM_023255603.1
B3	76674546	76674619
refGene_gn=REC8;refGene_tx=XM_003987564.4;refGene_gn=REC8;refGene_tx=XM_023255603.1
B3	76674698	76674784
refGene_gn=REC8;refGene_tx=XM_003987564.4;refGene_gn=REC8;refGene_tx=XM_023255603.1
B3	76674928	76674978
refGene_gn=REC8;refGene_tx=XM_003987564.4;refGene_gn=REC8;refGene_tx=XM_023255603.1
B3	76675297	76675444
refGene_gn=REC8;refGene_tx=XM_003987564.4;refGene_gn=REC8;refGene_tx=XM_023255603.1
B3	76675789	76675856
refGene_gn=REC8;refGene_tx=XM_003987564.4;refGene_gn=REC8;refGene_tx=XM_023255603.1

B3 76675961 76676107
refGene_gn=REC8;refGene_tx=XM_003987564.4;refGene_gn=REC8;refGene_tx=XM_023255603.1
B3 76676289 76676338
refGene_gn=REC8;refGene_tx=XM_003987564.4;refGene_gn=REC8;refGene_tx=XM_023255603.1
B3 76676580 76676647
refGene_gn=REC8;refGene_tx=XM_003987564.4;refGene_gn=REC8;refGene_tx=XM_023255603.1
B3 76676741 76676873
refGene_gn=REC8;refGene_tx=XM_003987564.4;refGene_gn=REC8;refGene_tx=XM_023255603.1
B3 76676950 76677057
refGene_gn=REC8;refGene_tx=XM_003987564.4;refGene_gn=REC8;refGene_tx=XM_023255603.1
B3 76677151 76677243
refGene_gn=REC8;refGene_tx=XM_003987564.4;refGene_gn=REC8;refGene_tx=XM_023255603.1
B3 76677577 76677708 refGene_gn=IPO4;refGene_tx=XM_003987507.5
B3 76677807 76677877 refGene_gn=IPO4;refGene_tx=XM_003987507.5
B3 76678564 76678669 refGene_gn=IPO4;refGene_tx=XM_003987507.5
B3 76678757 76678890 refGene_gn=IPO4;refGene_tx=XM_003987507.5
B3 76678995 76679204 refGene_gn=IPO4;refGene_tx=XM_003987507.5
B3 76679304 76679430 refGene_gn=IPO4;refGene_tx=XM_003987507.5
B3 76679770 76679815 refGene_gn=IPO4;refGene_tx=XM_003987507.5
B3 76679968 76680162 refGene_gn=IPO4;refGene_tx=XM_003987507.5
B3 76680253 76680359 refGene_gn=IPO4;refGene_tx=XM_003987507.5
B3 76680455 76680513 refGene_gn=IPO4;refGene_tx=XM_003987507.5
B3 76680589 76680667 refGene_gn=IPO4;refGene_tx=XM_003987507.5
B3 76680776 76680889 refGene_gn=IPO4;refGene_tx=XM_003987507.5
B3 76680999 76681111 refGene_gn=IPO4;refGene_tx=XM_003987507.5
B3 76681266 76681387 refGene_gn=IPO4;refGene_tx=XM_003987507.5
B3 76681637 76681751 refGene_gn=IPO4;refGene_tx=XM_003987507.5
B3 76681859 76681973 refGene_gn=IPO4;refGene_tx=XM_003987507.5
B3 76682197 76682327 refGene_gn=IPO4;refGene_tx=XM_003987507.5
B3 76682490 76682599 refGene_gn=IPO4;refGene_tx=XM_003987507.5
B3 76682903 76683013 refGene_gn=IPO4;refGene_tx=XM_003987507.5
B3 76683170 76683224 refGene_gn=IPO4;refGene_tx=XM_003987507.5
B3 76683311 76683476 refGene_gn=IPO4;refGene_tx=XM_003987507.5
B3 76683718 76683802 refGene_gn=IPO4;refGene_tx=XM_003987507.5
B3 76683888 76683990 refGene_gn=IPO4;refGene_tx=XM_003987507.5
B3 76684095 76684161 refGene_gn=IPO4;refGene_tx=XM_003987507.5
B3 76684388 76684568 refGene_gn=IPO4;refGene_tx=XM_003987507.5
B3 76684662 76684792 refGene_gn=IPO4;refGene_tx=XM_003987507.5
B3 76685241 76685283 refGene_gn=IPO4;refGene_tx=XM_003987507.5
B3 76685368 76685448 refGene_gn=IPO4;refGene_tx=XM_003987507.5
B3 76685575 76685662 refGene_gn=IPO4;refGene_tx=XM_003987507.5
B3 76685742 76685811 refGene_gn=IPO4;refGene_tx=XM_003987507.5
B3 76686435 76686829
refGene_gn=TM9SF1;refGene_tx=XM_011283155.3;refGene_gn=TM9SF1;refGene_

tx=XM_011283156.3;refGene_gn=TM9SF1;refGene_tx=XM_011283157.3
B3 76687404 76687678
refGene_gn=TM9SF1;refGene_tx=XM_011283155.3;refGene_gn=TM9SF1;refGene_tx=XM_011283156.3;refGene_gn=TM9SF1;refGene_tx=XM_011283157.3
B3 76688962 76689148
refGene_gn=TM9SF1;refGene_tx=XM_011283155.3;refGene_gn=TM9SF1;refGene_tx=XM_011283156.3;refGene_gn=TM9SF1;refGene_tx=XM_011283157.3
B3 76689434 76690056
refGene_gn=TM9SF1;refGene_tx=XM_011283155.3;refGene_gn=TM9SF1;refGene_tx=XM_011283156.3;refGene_gn=TM9SF1;refGene_tx=XM_011283157.3
B3 76690856 76691201
refGene_gn=TM9SF1;refGene_tx=XM_011283155.3;refGene_gn=TM9SF1;refGene_tx=XM_011283156.3;refGene_gn=TM9SF1;refGene_tx=XM_011283157.3
B3 76696517 76696742
refGene_gn=TSSK4;refGene_tx=XM_003987508.4;refGene_gn=TSSK4;refGene_tx=XM_003987509.4;refGene_gn=TSSK4;refGene_tx=XM_006932770.4;refGene_gn=TSSK4;refGene_tx=XM_006932772.4;refGene_gn=TSSK4;refGene_tx=XM_019832859.2;refGene_gn=TSSK4;refGene_tx=XM_023255606.1
B3 76697065 76697308
refGene_gn=TSSK4;refGene_tx=XM_003987508.4;refGene_gn=TSSK4;refGene_tx=XM_003987509.4;refGene_gn=TSSK4;refGene_tx=XM_006932770.4;refGene_gn=TSSK4;refGene_tx=XM_006932772.4;refGene_gn=TSSK4;refGene_tx=XM_019832859.2;refGene_gn=TSSK4;refGene_tx=XM_023255606.1
B3 76697693 76698087
refGene_gn=TSSK4;refGene_tx=XM_003987508.4;refGene_gn=TSSK4;refGene_tx=XM_003987509.4;refGene_gn=TSSK4;refGene_tx=XM_006932770.4;refGene_gn=TSSK4;refGene_tx=XM_006932772.4;refGene_gn=TSSK4;refGene_tx=XM_019832859.2;refGene_gn=TSSK4;refGene_tx=XM_023255606.1
B3 76698439 76698695
refGene_gn=TSSK4;refGene_tx=XM_003987508.4;refGene_gn=TSSK4;refGene_tx=XM_003987509.4;refGene_gn=TSSK4;refGene_tx=XM_006932770.4;refGene_gn=TSSK4;refGene_tx=XM_006932772.4;refGene_gn=TSSK4;refGene_tx=XM_019832859.2;refGene_gn=TSSK4;refGene_tx=XM_023255606.1
B3 76699945 76700004
refGene_gn=CHMP4A;refGene_tx=XM_003987510.5;refGene_gn=CHMP4A;refGene_tx=XM_011283159.2;refGene_gn=CHMP4A;refGene_tx=XM_011283160.3;refGene_gn=CHMP4A;refGene_tx=XM_019832860.2;refGene_gn=CHMP4A;refGene_tx=XM_019832861.2;refGene_gn=CHMP4A;refGene_tx=XM_019832862.2
B3 76700434 76700570
refGene_gn=CHMP4A;refGene_tx=XM_003987510.5;refGene_gn=CHMP4A;refGene_tx=XM_011283159.2;refGene_gn=CHMP4A;refGene_tx=XM_011283160.3;refGene_gn=CHMP4A;refGene_tx=XM_019832860.2;refGene_gn=CHMP4A;refGene_tx=XM_019832861.2;refGene_gn=CHMP4A;refGene_tx=XM_019832862.2
B3 76700740 76700849
refGene_gn=CHMP4A;refGene_tx=XM_003987510.5;refGene_gn=CHMP4A;refGene_tx=XM_011283159.2;refGene_gn=CHMP4A;refGene_tx=XM_011283160.3;refGene_gn=CHMP4A;refGene_tx=XM_019832860.2;refGene_gn=CHMP4A;refGene_tx=XM_019832861.2;refGene_gn=CHMP4A;refGene_tx=XM_019832862.2
B3 76701590 76701768
refGene_gn=CHMP4A;refGene_tx=XM_003987510.5;refGene_gn=CHMP4A;refGene_

tx=XM_011283159.2;refGene_gn=CHMP4A;refGene_tx=XM_011283160.3;refGene_gn=CHMP4A;refGene_tx=XM_019832860.2;refGene_gn=CHMP4A;refGene_tx=XM_019832861.2;refGene_gn=CHMP4A;refGene_tx=XM_019832862.2
B3 76701856 76702006 refGene_gn=CHMP4A;refGene_tx=XM_003987510.5;refGene_gn=CHMP4A;refGene_tx=XM_011283159.2;refGene_gn=CHMP4A;refGene_tx=XM_011283160.3;refGene_gn=CHMP4A;refGene_tx=XM_019832860.2;refGene_gn=CHMP4A;refGene_tx=XM_019832861.2;refGene_gn=CHMP4A;refGene_tx=XM_019832862.2
B3 76703319 76703344 refGene_gn=CHMP4A;refGene_tx=XM_011283160.3;refGene_gn=CHMP4A;refGene_tx=XM_019832861.2;refGene_gn=CHMP4A;refGene_tx=XM_019832862.2
B3 76703633 76703745 refGene_gn=CHMP4A;refGene_tx=XM_003987510.5;refGene_gn=CHMP4A;refGene_tx=XM_019832860.2
B3 76703975 76704213 refGene_gn=CHMP4A;refGene_tx=XM_011283159.2
B3 76704248 76704396 refGene_gn=LOC101081306;refGene_tx=XM_003987512.5;refGene_gn=LOC101081306;refGene_tx=XM_019832863.2
B3 76704550 76704646 refGene_gn=LOC101081306;refGene_tx=XM_003987512.5;refGene_gn=LOC101081306;refGene_tx=XM_019832863.2
B3 76704736 76704844 refGene_gn=LOC101081306;refGene_tx=XM_003987512.5;refGene_gn=LOC101081306;refGene_tx=XM_019832863.2
B3 76705184 76705295 refGene_gn=LOC101081306;refGene_tx=XM_003987512.5;refGene_gn=LOC101081306;refGene_tx=XM_019832863.2
B3 76705364 76705425 refGene_gn=LOC101081306;refGene_tx=XM_003987512.5;refGene_gn=LOC101081306;refGene_tx=XM_019832863.2
B3 76705538 76705575 refGene_gn=LOC101081306;refGene_tx=XM_003987512.5;refGene_gn=LOC101081306;refGene_tx=XM_019832863.2
B3 76706682 76706779 refGene_gn=NEDD8;refGene_tx=XM_003987511.5
B3 76707525 76707608 refGene_gn=NEDD8;refGene_tx=XM_003987511.5
B3 76707775 76707823 refGene_gn=NEDD8;refGene_tx=XM_003987511.5
B3 76715847 76715865 refGene_gn=NEDD8;refGene_tx=XM_003987511.5
B3 76716859 76716946 refGene_gn=GMPR2;refGene_tx=XM_003987513.5;refGene_gn=GMPR2;refGene_tx=XM_023255604.1
B3 76717083 76717203 refGene_gn=GMPR2;refGene_tx=XM_003987513.5
B3 76717820 76717871 refGene_gn=GMPR2;refGene_tx=XM_023255605.1
B3 76717939 76718011 refGene_gn=GMPR2;refGene_tx=XM_011283161.3
B3 76719364 76719448 refGene_gn=GMPR2;refGene_tx=XM_003987513.5;refGene_gn=GMPR2;refGene_tx=XM_011283161.3;refGene_gn=GMPR2;refGene_tx=XM_023255604.1;refGene_gn=GMPR2;refGene_tx=XM_023255605.1
B3 76719822 76719996 refGene_gn=GMPR2;refGene_tx=XM_003987513.5;refGene_gn=GMPR2;refGene_tx=XM_011283161.3;refGene_gn=GMPR2;refGene_tx=XM_023255604.1;refGene_gn=

GMPR2;refGene_tx=XM_023255605.1
B3 76720746 76720828
refGene_gn=GMPR2;refGene_tx=XM_003987513.5;refGene_gn=GMPR2;refGene_tx
=XM_011283161.3;refGene_gn=GMPR2;refGene_tx=XM_023255604.1;refGene_gn=
GMPR2;refGene_tx=XM_023255605.1
B3 76720916 76721023
refGene_gn=GMPR2;refGene_tx=XM_003987513.5;refGene_gn=GMPR2;refGene_tx
=XM_011283161.3;refGene_gn=GMPR2;refGene_tx=XM_023255604.1;refGene_gn=
GMPR2;refGene_tx=XM_023255605.1
B3 76721176 76721283
refGene_gn=GMPR2;refGene_tx=XM_003987513.5;refGene_gn=GMPR2;refGene_tx
=XM_011283161.3;refGene_gn=GMPR2;refGene_tx=XM_023255604.1;refGene_gn=
GMPR2;refGene_tx=XM_023255605.1;refGene_gn=TINF2;refGene_tx=XM_0039875
14.5;refGene_gn=TINF2;refGene_tx=XM_006932775.4
B3 76721852 76722012
refGene_gn=GMPR2;refGene_tx=XM_003987513.5;refGene_gn=GMPR2;refGene_tx
=XM_011283161.3;refGene_gn=GMPR2;refGene_tx=XM_023255604.1;refGene_gn=
GMPR2;refGene_tx=XM_023255605.1
B3 76722214 76722404
refGene_gn=GMPR2;refGene_tx=XM_003987513.5;refGene_gn=GMPR2;refGene_tx
=XM_011283161.3;refGene_gn=GMPR2;refGene_tx=XM_023255604.1;refGene_gn=
GMPR2;refGene_tx=XM_023255605.1
B3 76722629 76722656 refGene_gn=TINF2;refGene_tx=XM_019832858.2
B3 76723345 76723519 refGene_gn=TINF2;refGene_tx=XM_006932774.4
B3 76723648 76723740
refGene_gn=TINF2;refGene_tx=XM_006932774.4;refGene_gn=TINF2;refGene_tx
=XM_006932775.4;refGene_gn=TINF2;refGene_tx=XM_019832858.2
B3 76723837 76723905
refGene_gn=TINF2;refGene_tx=XM_003987514.5;refGene_gn=TINF2;refGene_tx
=XM_006932774.4;refGene_gn=TINF2;refGene_tx=XM_006932775.4;refGene_gn=
TINF2;refGene_tx=XM_019832858.2
B3 76723994 76724457
refGene_gn=TINF2;refGene_tx=XM_003987514.5;refGene_gn=TINF2;refGene_tx
=XM_006932774.4;refGene_gn=TINF2;refGene_tx=XM_006932775.4;refGene_gn=
TINF2;refGene_tx=XM_019832858.2
B3 76724617 76724714
refGene_gn=TINF2;refGene_tx=XM_003987514.5;refGene_gn=TINF2;refGene_tx
=XM_006932774.4;refGene_gn=TINF2;refGene_tx=XM_006932775.4;refGene_gn=
TINF2;refGene_tx=XM_019832858.2
B3 76724799 76724907
refGene_gn=TINF2;refGene_tx=XM_003987514.5;refGene_gn=TINF2;refGene_tx
=XM_006932774.4;refGene_gn=TINF2;refGene_tx=XM_006932775.4;refGene_gn=
TINF2;refGene_tx=XM_019832858.2
B3 76724986 76725088
refGene_gn=TINF2;refGene_tx=XM_003987514.5;refGene_gn=TINF2;refGene_tx
=XM_006932774.4;refGene_gn=TINF2;refGene_tx=XM_006932775.4;refGene_gn=
TINF2;refGene_tx=XM_019832858.2
B3 76725202 76725307
refGene_gn=TINF2;refGene_tx=XM_003987514.5;refGene_gn=TINF2;refGene_tx
=XM_006932774.4;refGene_gn=TINF2;refGene_tx=XM_006932775.4;refGene_gn=

TINF2;refGene_tx=XM_019832858.2
B3 76725444 76725636
refGene_gn=TINF2;refGene_tx=XM_003987514.5;refGene_gn=TINF2;refGene_tx=XM_006932774.4;refGene_gn=TINF2;refGene_tx=XM_006932775.4;refGene_gn=TINF2;refGene_tx=XM_019832858.2
B3 76734890 76735119
refGene_gn=TGM1;refGene_tx=XM_003987515.4;refGene_gn=TGM1;refGene_tx=XM_019832869.2
B3 76739453 76739590
refGene_gn=TGM1;refGene_tx=XM_003987515.4;refGene_gn=TGM1;refGene_tx=XM_019832869.2
B3 76739963 76740124
refGene_gn=TGM1;refGene_tx=XM_003987515.4;refGene_gn=TGM1;refGene_tx=XM_019832869.2
B3 76740247 76740529
refGene_gn=TGM1;refGene_tx=XM_003987515.4;refGene_gn=TGM1;refGene_tx=XM_019832869.2
B3 76740636 76740790
refGene_gn=TGM1;refGene_tx=XM_003987515.4;refGene_gn=TGM1;refGene_tx=XM_019832869.2
B3 76741335 76741424
refGene_gn=TGM1;refGene_tx=XM_003987515.4;refGene_gn=TGM1;refGene_tx=XM_019832869.2
B3 76744319 76744423
refGene_gn=TGM1;refGene_tx=XM_003987515.4;refGene_gn=TGM1;refGene_tx=XM_019832869.2
B3 76744555 76744694
refGene_gn=TGM1;refGene_tx=XM_003987515.4;refGene_gn=TGM1;refGene_tx=XM_019832869.2
B3 76745075 76745250
refGene_gn=TGM1;refGene_tx=XM_003987515.4;refGene_gn=TGM1;refGene_tx=XM_019832869.2
B3 76745702 76745810
refGene_gn=TGM1;refGene_tx=XM_003987515.4;refGene_gn=TGM1;refGene_tx=XM_019832869.2
B3 76745917 76746036
refGene_gn=TGM1;refGene_tx=XM_003987515.4;refGene_gn=TGM1;refGene_tx=XM_019832869.2
B3 76746415 76746664
refGene_gn=TGM1;refGene_tx=XM_003987515.4;refGene_gn=TGM1;refGene_tx=XM_019832869.2
B3 76747652 76747841
refGene_gn=TGM1;refGene_tx=XM_003987515.4;refGene_gn=TGM1;refGene_tx=XM_019832869.2
B3 76747968 76748296
refGene_gn=TGM1;refGene_tx=XM_003987515.4;refGene_gn=TGM1;refGene_tx=XM_019832869.2
B3 76751536 76751685
refGene_gn=RABGGTA;refGene_tx=XM_003987517.5;refGene_gn=RABGGTA;refGene_tx=XM_011283162.3;refGene_gn=RABGGTA;refGene_tx=XM_011283163.3;refGe

ne_gn=RABGGTA;refGene_tx=XM_011283165.3
B3	76752439	76752527
refGene_gn=RABGGTA;refGene_tx=XM_003987517.5;refGene_gn=RABGGTA;refGene_tx=XM_011283162.3;refGene_gn=RABGGTA;refGene_tx=XM_011283163.3;refGene_gn=RABGGTA;refGene_tx=XM_011283165.3
B3	76752792	76752906
refGene_gn=RABGGTA;refGene_tx=XM_003987517.5;refGene_gn=RABGGTA;refGene_tx=XM_011283162.3;refGene_gn=RABGGTA;refGene_tx=XM_011283163.3;refGene_gn=RABGGTA;refGene_tx=XM_011283165.3
B3	76754016	76754133
refGene_gn=RABGGTA;refGene_tx=XM_003987517.5;refGene_gn=RABGGTA;refGene_tx=XM_011283162.3;refGene_gn=RABGGTA;refGene_tx=XM_011283163.3;refGene_gn=RABGGTA;refGene_tx=XM_011283165.3
B3	76754227	76754316
refGene_gn=RABGGTA;refGene_tx=XM_003987517.5;refGene_gn=RABGGTA;refGene_tx=XM_011283162.3;refGene_gn=RABGGTA;refGene_tx=XM_011283163.3;refGene_gn=RABGGTA;refGene_tx=XM_011283165.3
B3	76754454	76754543
refGene_gn=RABGGTA;refGene_tx=XM_003987517.5;refGene_gn=RABGGTA;refGene_tx=XM_011283162.3;refGene_gn=RABGGTA;refGene_tx=XM_011283163.3;refGene_gn=RABGGTA;refGene_tx=XM_011283165.3
B3	76754694	76754746
refGene_gn=RABGGTA;refGene_tx=XM_003987517.5;refGene_gn=RABGGTA;refGene_tx=XM_011283162.3;refGene_gn=RABGGTA;refGene_tx=XM_011283163.3;refGene_gn=RABGGTA;refGene_tx=XM_011283165.3
B3	76754844	76754950
refGene_gn=RABGGTA;refGene_tx=XM_003987517.5;refGene_gn=RABGGTA;refGene_tx=XM_011283162.3;refGene_gn=RABGGTA;refGene_tx=XM_011283163.3;refGene_gn=RABGGTA;refGene_tx=XM_011283165.3
B3	76755055	76755157
refGene_gn=RABGGTA;refGene_tx=XM_003987517.5;refGene_gn=RABGGTA;refGene_tx=XM_011283162.3;refGene_gn=RABGGTA;refGene_tx=XM_011283163.3;refGene_gn=RABGGTA;refGene_tx=XM_011283165.3
B3	76755246	76755329
refGene_gn=RABGGTA;refGene_tx=XM_003987517.5;refGene_gn=RABGGTA;refGene_tx=XM_011283162.3;refGene_gn=RABGGTA;refGene_tx=XM_011283163.3;refGene_gn=RABGGTA;refGene_tx=XM_011283165.3
B3	76755418	76755502
refGene_gn=RABGGTA;refGene_tx=XM_003987517.5;refGene_gn=RABGGTA;refGene_tx=XM_011283162.3;refGene_gn=RABGGTA;refGene_tx=XM_011283163.3;refGene_gn=RABGGTA;refGene_tx=XM_011283165.3
B3	76755799	76756003
refGene_gn=RABGGTA;refGene_tx=XM_003987517.5;refGene_gn=RABGGTA;refGene_tx=XM_011283162.3;refGene_gn=RABGGTA;refGene_tx=XM_011283163.3;refGene_gn=RABGGTA;refGene_tx=XM_011283165.3
B3	76756240	76756428
refGene_gn=RABGGTA;refGene_tx=XM_003987517.5;refGene_gn=RABGGTA;refGene_tx=XM_011283162.3;refGene_gn=RABGGTA;refGene_tx=XM_011283163.3;refGene_gn=RABGGTA;refGene_tx=XM_011283165.3
B3	76756613	76756738

refGene_gn=RABGGTA;refGene_tx=XM_003987517.5;refGene_gn=RABGGTA;refGene_tx=XM_011283162.3;refGene_gn=RABGGTA;refGene_tx=XM_011283163.3;refGene_gn=RABGGTA;refGene_tx=XM_011283165.3
B3	76757083	76757194
refGene_gn=RABGGTA;refGene_tx=XM_003987517.5;refGene_gn=RABGGTA;refGene_tx=XM_011283162.3;refGene_gn=RABGGTA;refGene_tx=XM_011283163.3;refGene_gn=RABGGTA;refGene_tx=XM_011283165.3
B3	76757343	76757355
refGene_gn=RABGGTA;refGene_tx=XM_003987517.5;refGene_gn=RABGGTA;refGene_tx=XM_011283162.3;refGene_gn=RABGGTA;refGene_tx=XM_011283163.3;refGene_gn=RABGGTA;refGene_tx=XM_011283165.3
B3	76770255	76770392
refGene_gn=DHRS1;refGene_tx=XM_003987518.5;refGene_gn=DHRS1;refGene_tx=XM_006932779.4
B3	76770576	76770657
refGene_gn=DHRS1;refGene_tx=XM_003987518.5;refGene_gn=DHRS1;refGene_tx=XM_006932779.4
B3	76770972	76771042
refGene_gn=DHRS1;refGene_tx=XM_003987518.5;refGene_gn=DHRS1;refGene_tx=XM_006932779.4
B3	76771583	76771730
refGene_gn=DHRS1;refGene_tx=XM_003987518.5;refGene_gn=DHRS1;refGene_tx=XM_006932779.4
B3	76772039	76772172
refGene_gn=DHRS1;refGene_tx=XM_003987518.5;refGene_gn=DHRS1;refGene_tx=XM_006932779.4
B3	76773486	76773566
refGene_gn=DHRS1;refGene_tx=XM_003987518.5;refGene_gn=DHRS1;refGene_tx=XM_006932779.4
B3	76773726	76773870
refGene_gn=DHRS1;refGene_tx=XM_003987518.5;refGene_gn=DHRS1;refGene_tx=XM_006932779.4
B3	76775020	76775170
refGene_gn=DHRS1;refGene_tx=XM_003987518.5;refGene_gn=DHRS1;refGene_tx=XM_006932779.4
B3	76776021	76776256	refGene_gn=NOP9;refGene_tx=XM_003987519.4
B3	76776469	76776928
refGene_gn=NOP9;refGene_tx=XM_003987519.4;refGene_gn=NOP9;refGene_tx=XM_006932780.3
B3	76777683	76777794
refGene_gn=NOP9;refGene_tx=XM_003987519.4;refGene_gn=NOP9;refGene_tx=XM_006932780.3
B3	76778037	76778179
refGene_gn=NOP9;refGene_tx=XM_003987519.4;refGene_gn=NOP9;refGene_tx=XM_006932780.3
B3	76778295	76778488
refGene_gn=NOP9;refGene_tx=XM_003987519.4;refGene_gn=NOP9;refGene_tx=XM_006932780.3
B3	76779177	76779318
refGene_gn=NOP9;refGene_tx=XM_003987519.4;refGene_gn=NOP9;refGene_tx=X

M_006932780.3
B3 76779706 76779832
refGene_gn=NOP9;refGene_tx=XM_003987519.4;refGene_gn=NOP9;refGene_tx=XM_006932780.3
B3 76779987 76780224
refGene_gn=NOP9;refGene_tx=XM_003987519.4;refGene_gn=NOP9;refGene_tx=XM_006932780.3
B3 76780436 76780542
refGene_gn=NOP9;refGene_tx=XM_003987519.4;refGene_gn=NOP9;refGene_tx=XM_006932780.3
B3 76780874 76781032
refGene_gn=NOP9;refGene_tx=XM_003987519.4;refGene_gn=NOP9;refGene_tx=XM_006932780.3
B3 76781517 76781542 refGene_gn=CIDEB;refGene_tx=XM_019832874.2
B3 76781573 76781712
refGene_gn=CIDEB;refGene_tx=XM_003987520.4;refGene_gn=CIDEB;refGene_tx=XM_019832873.2
B3 76781879 76782070
refGene_gn=CIDEB;refGene_tx=XM_003987520.4;refGene_gn=CIDEB;refGene_tx=XM_019832873.2;refGene_gn=CIDEB;refGene_tx=XM_019832874.2
B3 76782322 76782472
refGene_gn=CIDEB;refGene_tx=XM_003987520.4;refGene_gn=CIDEB;refGene_tx=XM_019832873.2;refGene_gn=CIDEB;refGene_tx=XM_019832874.2
B3 76783316 76783461
refGene_gn=CIDEB;refGene_tx=XM_003987520.4;refGene_gn=CIDEB;refGene_tx=XM_019832873.2;refGene_gn=CIDEB;refGene_tx=XM_019832874.2
B3 76783757 76783798
refGene_gn=CIDEB;refGene_tx=XM_003987520.4;refGene_gn=CIDEB;refGene_tx=XM_019832874.2
B3 76786885 76787962
refGene_gn=LTB4R2;refGene_tx=XM_023255608.1;refGene_gn=LTB4R2;refGene_tx=XM_023255609.1
B3 76791802 76792801
refGene_gn=LTB4R;refGene_tx=XM_011283166.3;refGene_gn=LTB4R;refGene_tx=XM_011283167.3;refGene_gn=LTB4R;refGene_tx=XM_019832870.2;refGene_gn=LTB4R;refGene_tx=XM_019832871.2
B3 76795175 76795328 refGene_gn=ADCY4;refGene_tx=XM_023256182.1
B3 76795414 76795539 refGene_gn=ADCY4;refGene_tx=XM_023256182.1
B3 76795851 76795966 refGene_gn=ADCY4;refGene_tx=XM_023256182.1
B3 76796059 76796167 refGene_gn=ADCY4;refGene_tx=XM_023256182.1
B3 76796461 76796608 refGene_gn=ADCY4;refGene_tx=XM_023256182.1
B3 76797916 76798075 refGene_gn=ADCY4;refGene_tx=XM_023256182.1
B3 76798547 76798632 refGene_gn=ADCY4;refGene_tx=XM_023256182.1
B3 76798825 76799010 refGene_gn=ADCY4;refGene_tx=XM_023256182.1
B3 76799268 76799467 refGene_gn=ADCY4;refGene_tx=XM_023256182.1
B3 76800198 76800210 refGene_gn=ADCY4;refGene_tx=XM_023256182.1
B3 76800880 76800978 refGene_gn=ADCY4;refGene_tx=XM_023256182.1
B3 76801341 76801411 refGene_gn=ADCY4;refGene_tx=XM_023256182.1
B3 76801611 76801698 refGene_gn=ADCY4;refGene_tx=XM_023256182.1
B3 76801816 76801860 refGene_gn=ADCY4;refGene_tx=XM_023256182.1

B3	76804570 76804744 refGene_gn=ADCY4;refGene_tx=XM_023256182.1
B3	76804946 76805079 refGene_gn=ADCY4;refGene_tx=XM_023256182.1
B3	76805337 76805496 refGene_gn=ADCY4;refGene_tx=XM_023256182.1
B3	76805649 76805777 refGene_gn=ADCY4;refGene_tx=XM_023256182.1
B3	76806466 76806578 refGene_gn=ADCY4;refGene_tx=XM_023256182.1
B3	76806656 76806805 refGene_gn=ADCY4;refGene_tx=XM_023256182.1
B3	76807246 76807396 refGene_gn=ADCY4;refGene_tx=XM_023256182.1
B3	76808048 76808210 refGene_gn=ADCY4;refGene_tx=XM_023256182.1
B3	76808326 76808524 refGene_gn=ADCY4;refGene_tx=XM_023256182.1
B3	76809811 76809970 refGene_gn=ADCY4;refGene_tx=XM_023256182.1
B3	76811816 76811965
refGene_gn=RIPK3;refGene_tx=XM_003987566.5;refGene_gn=RIPK3;refGene_tx=XM_006932785.4
B3	76812518 76812579
refGene_gn=RIPK3;refGene_tx=XM_003987566.5;refGene_gn=RIPK3;refGene_tx=XM_006932785.4
B3	76812722 76813085
refGene_gn=RIPK3;refGene_tx=XM_003987566.5;refGene_gn=RIPK3;refGene_tx=XM_006932785.4
B3	76813327 76813395
refGene_gn=RIPK3;refGene_tx=XM_003987566.5;refGene_gn=RIPK3;refGene_tx=XM_006932785.4
B3	76813503 76813671
refGene_gn=RIPK3;refGene_tx=XM_003987566.5;refGene_gn=RIPK3;refGene_tx=XM_006932785.4
B3	76813881 76813928
refGene_gn=RIPK3;refGene_tx=XM_003987566.5;refGene_gn=RIPK3;refGene_tx=XM_006932785.4
B3	76814054 76814194
refGene_gn=RIPK3;refGene_tx=XM_003987566.5;refGene_gn=RIPK3;refGene_tx=XM_006932785.4
B3	76814649 76814959
refGene_gn=RIPK3;refGene_tx=XM_003987566.5;refGene_gn=RIPK3;refGene_tx=XM_006932785.4
B3	76815097 76815238
refGene_gn=RIPK3;refGene_tx=XM_003987566.5;refGene_gn=RIPK3;refGene_tx=XM_006932785.4
B3	76815491 76815517
refGene_gn=RIPK3;refGene_tx=XM_003987566.5;refGene_gn=RIPK3;refGene_tx=XM_006932785.4
B3	76841556 76841656 refGene_gn=NFATC4;refGene_tx=XM_023255611.1
B3	76842755 76843851 refGene_gn=NFATC4;refGene_tx=XM_023255611.1
B3	76845272 76845435 refGene_gn=NFATC4;refGene_tx=XM_023255611.1
B3	76846022 76846222 refGene_gn=NFATC4;refGene_tx=XM_023255611.1
B3	76846552 76846725 refGene_gn=NFATC4;refGene_tx=XM_023255611.1
B3	76847187 76847328 refGene_gn=NFATC4;refGene_tx=XM_023255611.1
B3	76848229 76848285 refGene_gn=NFATC4;refGene_tx=XM_023255611.1
B3	76848562 76848689 refGene_gn=NFATC4;refGene_tx=XM_023255611.1
B3	76848881 76849463 refGene_gn=NFATC4;refGene_tx=XM_023255611.1
B3	76850213 76850281 refGene_gn=NFATC4;refGene_tx=XM_023255611.1

B3	76871618	76871816	refGene_gn=NYNRIN;refGene_tx=XM_011283171.3
B3	76882316	76882984	refGene_gn=NYNRIN;refGene_tx=XM_011283171.3
B3	76883104	76884667	refGene_gn=NYNRIN;refGene_tx=XM_011283171.3
B3	76885631	76885762	refGene_gn=NYNRIN;refGene_tx=XM_011283171.3
B3	76885904	76886001	refGene_gn=NYNRIN;refGene_tx=XM_011283171.3
B3	76887673	76887775	refGene_gn=NYNRIN;refGene_tx=XM_011283171.3
B3	76887986	76888088	refGene_gn=NYNRIN;refGene_tx=XM_011283171.3
B3	76889409	76892260	refGene_gn=NYNRIN;refGene_tx=XM_011283171.3
B3	76905756	76905954	refGene_gn=CBLN3;refGene_tx=XM_003987522.4
B3	76906313	76906433	refGene_gn=CBLN3;refGene_tx=XM_003987522.4
B3	76906740	76907040	refGene_gn=CBLN3;refGene_tx=XM_003987522.4
B3	76908676	76908686
refGene_gn=KHNYN;refGene_tx=XM_006932788.4;refGene_gn=KHNYN;refGene_tx
=XM_006932789.4
B3	76908796	76909017
refGene_gn=KHNYN;refGene_tx=XM_003987523.5;refGene_gn=KHNYN;refGene_tx
=XM_006932788.4;refGene_gn=KHNYN;refGene_tx=XM_006932789.4;refGene_gn=
KHNYN;refGene_tx=XM_023255613.1
B3	76910119	76911231
refGene_gn=KHNYN;refGene_tx=XM_003987523.5;refGene_gn=KHNYN;refGene_tx
=XM_006932788.4;refGene_gn=KHNYN;refGene_tx=XM_006932789.4;refGene_gn=
KHNYN;refGene_tx=XM_023255613.1
B3	76911358	76911489
refGene_gn=KHNYN;refGene_tx=XM_003987523.5;refGene_gn=KHNYN;refGene_tx
=XM_006932788.4;refGene_gn=KHNYN;refGene_tx=XM_006932789.4;refGene_gn=
KHNYN;refGene_tx=XM_023255613.1
B3	76911574	76911671
refGene_gn=KHNYN;refGene_tx=XM_003987523.5;refGene_gn=KHNYN;refGene_tx
=XM_006932788.4;refGene_gn=KHNYN;refGene_tx=XM_006932789.4;refGene_gn=
KHNYN;refGene_tx=XM_023255613.1
B3	76914483	76914591
refGene_gn=KHNYN;refGene_tx=XM_003987523.5;refGene_gn=KHNYN;refGene_tx
=XM_006932788.4;refGene_gn=KHNYN;refGene_tx=XM_023255613.1
B3	76914800	76914902
refGene_gn=KHNYN;refGene_tx=XM_003987523.5;refGene_gn=KHNYN;refGene_tx
=XM_006932788.4;refGene_gn=KHNYN;refGene_tx=XM_006932789.4;refGene_gn=
KHNYN;refGene_tx=XM_023255613.1
B3	76915454	76915707
refGene_gn=KHNYN;refGene_tx=XM_003987523.5;refGene_gn=KHNYN;refGene_tx
=XM_006932788.4;refGene_gn=KHNYN;refGene_tx=XM_006932789.4;refGene_gn=
KHNYN;refGene_tx=XM_023255613.1
B3	76918334	76918747
refGene_gn=SDR39U1;refGene_tx=XM_003987524.4;refGene_gn=SDR39U1;refGen
e_tx=XM_011283172.3
B3	76918994	76919138
refGene_gn=SDR39U1;refGene_tx=XM_003987524.4;refGene_gn=SDR39U1;refGen
e_tx=XM_011283172.3
B3	76919918	76920040	refGene_gn=SDR39U1;refGene_tx=XM_003987524.4
B3	76920431	76920514
refGene_gn=SDR39U1;refGene_tx=XM_003987524.4;refGene_gn=SDR39U1;refGen

e_tx=XM_011283172.3
B3	76920608 76920715 refGene_gn=SDR39U1;refGene_tx=XM_003987524.4
B3	76921026 76921042 refGene_gn=SDR39U1;refGene_tx=XM_003987524.4
B3	76949166 76949310
refGene_gn=LOC101084796;refGene_tx=XM_003987525.3;refGene_gn=LOC101084796;refGene_tx=XM_006932790.3;refGene_gn=LOC101084796;refGene_tx=XM_019832880.2
B3	76949718 76949973
refGene_gn=LOC101084796;refGene_tx=XM_003987525.3;refGene_gn=LOC101084796;refGene_tx=XM_006932790.3;refGene_gn=LOC101084796;refGene_tx=XM_019832880.2
B3	76950199 76950335
refGene_gn=LOC101084796;refGene_tx=XM_003987525.3;refGene_gn=LOC101084796;refGene_tx=XM_006932790.3;refGene_gn=LOC101084796;refGene_tx=XM_019832880.2
B3	76951048 76951199
refGene_gn=LOC101084796;refGene_tx=XM_003987525.3;refGene_gn=LOC101084796;refGene_tx=XM_006932790.3
B3	76951424 76951523
refGene_gn=LOC101084796;refGene_tx=XM_006932790.3
B3	76951869 76951927
refGene_gn=LOC101084796;refGene_tx=XM_003987525.3;refGene_gn=LOC101084796;refGene_tx=XM_006932790.3
B3	76965693 76965837
refGene_gn=LOC101098468;refGene_tx=XM_003987568.3
B3	76966277 76966477
refGene_gn=LOC101098468;refGene_tx=XM_003987568.3
B3	76966479 76966531
refGene_gn=LOC101098468;refGene_tx=XM_003987568.3
B3	76966766 76966902
refGene_gn=LOC101098468;refGene_tx=XM_003987568.3
B3	76967352 76967503
refGene_gn=LOC101098468;refGene_tx=XM_003987568.3
B3	76968028 76968083
refGene_gn=LOC101098468;refGene_tx=XM_003987568.3
B3	77003501 77003675
refGene_gn=LOC101098719;refGene_tx=XM_003987569.3
B3	77004074 77004329
refGene_gn=LOC101098719;refGene_tx=XM_003987569.3
B3	77004496 77004632
refGene_gn=LOC101098719;refGene_tx=XM_003987569.3
B3	77005072 77005220
refGene_gn=LOC101098719;refGene_tx=XM_003987569.3
B3	77005989 77006044
refGene_gn=LOC101098719;refGene_tx=XM_003987569.3
B3	77019128 77019284 refGene_gn=GZMH;refGene_tx=XM_006932792.3
B3	77019712 77019970 refGene_gn=GZMH;refGene_tx=XM_006932792.3
B3	77020173 77020293 refGene_gn=GZMH;refGene_tx=XM_006932792.3
B3	77020294 77020310 refGene_gn=GZMH;refGene_tx=XM_006932792.3
B3	77020815 77021002 refGene_gn=GZMH;refGene_tx=XM_006932792.3

B3 77022048 77022103 refGene_gn=GZMH;refGene_tx=XM_006932792.3
B3 77023945 77024218
refGene_gn=LOC101099215;refGene_tx=XM_006932791.3
B3 77024575 77024839
refGene_gn=LOC101099215;refGene_tx=XM_006932791.3
B3 77025033 77025169
refGene_gn=LOC101099215;refGene_tx=XM_006932791.3
B3 77025398 77025549
refGene_gn=LOC101099215;refGene_tx=XM_006932791.3
B3 77026682 77026737
refGene_gn=LOC101099215;refGene_tx=XM_006932791.3
B3 77035672 77035843 refGene_gn=GZMB;refGene_tx=XM_006932823.3
B3 77036325 77036583 refGene_gn=GZMB;refGene_tx=XM_006932823.3
B3 77036786 77036922 refGene_gn=GZMB;refGene_tx=XM_006932823.3
B3 77037379 77037527 refGene_gn=GZMB;refGene_tx=XM_006932823.3
B3 77039102 77039157 refGene_gn=GZMB;refGene_tx=XM_006932823.3
B3 77182727 77182751
refGene_gn=STXBP6;refGene_tx=XM_006932794.3;refGene_gn=STXBP6;refGene_tx=XM_019832882.2;refGene_gn=STXBP6;refGene_tx=XM_023255615.1
B3 77188479 77188637
refGene_gn=STXBP6;refGene_tx=XM_006932794.3;refGene_gn=STXBP6;refGene_tx=XM_019832882.2;refGene_gn=STXBP6;refGene_tx=XM_023255615.1
B3 77230717 77230883
refGene_gn=STXBP6;refGene_tx=XM_006932794.3;refGene_gn=STXBP6;refGene_tx=XM_019832882.2;refGene_gn=STXBP6;refGene_tx=XM_023255615.1
B3 77231817 77231948
refGene_gn=STXBP6;refGene_tx=XM_006932794.3;refGene_gn=STXBP6;refGene_tx=XM_019832882.2;refGene_gn=STXBP6;refGene_tx=XM_023255615.1
B3 77350617 77350771
refGene_gn=STXBP6;refGene_tx=XM_006932794.3;refGene_gn=STXBP6;refGene_tx=XM_019832882.2;refGene_gn=STXBP6;refGene_tx=XM_023255615.1
B3 78740500 78741505
refGene_gn=NOVA1;refGene_tx=XM_003987526.5;refGene_gn=NOVA1;refGene_tx=XM_019832884.2;refGene_gn=NOVA1;refGene_tx=XM_019832885.2
B3 78765818 78765890
refGene_gn=NOVA1;refGene_tx=XM_003987526.5;refGene_gn=NOVA1;refGene_tx=XM_019832885.2
B3 78775956 78776123
refGene_gn=NOVA1;refGene_tx=XM_003987526.5;refGene_gn=NOVA1;refGene_tx=XM_019832884.2;refGene_gn=NOVA1;refGene_tx=XM_019832885.2
B3 78885810 78885954
refGene_gn=NOVA1;refGene_tx=XM_003987526.5;refGene_gn=NOVA1;refGene_tx=XM_019832884.2;refGene_gn=NOVA1;refGene_tx=XM_019832885.2
B3 78886988 78887013 refGene_gn=NOVA1;refGene_tx=XM_019832885.2
B3 78887699 78887835
refGene_gn=NOVA1;refGene_tx=XM_003987526.5;refGene_gn=NOVA1;refGene_tx=XM_019832884.2
B3 80940412 80941777 refGene_gn=FOXG1;refGene_tx=XM_019833790.2
B3 80954843 80955006
refGene_gn=LOC109501108;refGene_tx=XM_019833791.2

B3	80956070	80956159
refGene_gn=LOC109501108;refGene_tx=XM_019833791.2
B3	80958597	80958890
refGene_gn=LOC109501108;refGene_tx=XM_019833791.2
B3	80964383	80964612
refGene_gn=LOC109501108;refGene_tx=XM_019833791.2
B3	81663676	81663895
refGene_gn=PRKD1;refGene_tx=XM_023255617.1;refGene_gn=PRKD1;refGene_tx=XM_023255618.1;refGene_gn=PRKD1;refGene_tx=XM_023255619.1;refGene_gn=PRKD1;refGene_tx=XM_023255620.1;refGene_gn=PRKD1;refGene_tx=XM_023255621.1
B3	81664684	81664770
refGene_gn=PRKD1;refGene_tx=XM_023255617.1;refGene_gn=PRKD1;refGene_tx=XM_023255618.1;refGene_gn=PRKD1;refGene_tx=XM_023255619.1;refGene_gn=PRKD1;refGene_tx=XM_023255620.1;refGene_gn=PRKD1;refGene_tx=XM_023255621.1
B3	81683345	81683613
refGene_gn=PRKD1;refGene_tx=XM_023255617.1;refGene_gn=PRKD1;refGene_tx=XM_023255618.1;refGene_gn=PRKD1;refGene_tx=XM_023255619.1;refGene_gn=PRKD1;refGene_tx=XM_023255620.1;refGene_gn=PRKD1;refGene_tx=XM_023255621.1
B3	81685229	81685328
refGene_gn=PRKD1;refGene_tx=XM_023255617.1;refGene_gn=PRKD1;refGene_tx=XM_023255618.1;refGene_gn=PRKD1;refGene_tx=XM_023255619.1;refGene_gn=PRKD1;refGene_tx=XM_023255620.1;refGene_gn=PRKD1;refGene_tx=XM_023255621.1
B3	81685885	81686047
refGene_gn=PRKD1;refGene_tx=XM_023255617.1;refGene_gn=PRKD1;refGene_tx=XM_023255618.1;refGene_gn=PRKD1;refGene_tx=XM_023255619.1;refGene_gn=PRKD1;refGene_tx=XM_023255620.1;refGene_gn=PRKD1;refGene_tx=XM_023255621.1
B3	81705068	81705175
refGene_gn=PRKD1;refGene_tx=XM_023255617.1;refGene_gn=PRKD1;refGene_tx=XM_023255618.1;refGene_gn=PRKD1;refGene_tx=XM_023255619.1;refGene_gn=PRKD1;refGene_tx=XM_023255620.1;refGene_gn=PRKD1;refGene_tx=XM_023255621.1
B3	81707528	81707601
refGene_gn=PRKD1;refGene_tx=XM_023255617.1;refGene_gn=PRKD1;refGene_tx=XM_023255618.1;refGene_gn=PRKD1;refGene_tx=XM_023255619.1;refGene_gn=PRKD1;refGene_tx=XM_023255620.1;refGene_gn=PRKD1;refGene_tx=XM_023255621.1
B3	81709848	81709901
refGene_gn=PRKD1;refGene_tx=XM_023255617.1;refGene_gn=PRKD1;refGene_tx=XM_023255618.1;refGene_gn=PRKD1;refGene_tx=XM_023255619.1;refGene_gn=PRKD1;refGene_tx=XM_023255620.1;refGene_gn=PRKD1;refGene_tx=XM_023255621.1
B3	81710947	81711227
refGene_gn=PRKD1;refGene_tx=XM_023255617.1;refGene_gn=PRKD1;refGene_tx=XM_023255618.1;refGene_gn=PRKD1;refGene_tx=XM_023255619.1;refGene_gn=PRKD1;refGene_tx=XM_023255620.1;refGene_gn=PRKD1;refGene_tx=XM_0232556

21.1
B3	81712938	81713016
refGene_gn=PRKD1;refGene_tx=XM_023255617.1;refGene_gn=PRKD1;refGene_tx=XM_023255618.1;refGene_gn=PRKD1;refGene_tx=XM_023255619.1;refGene_gn=PRKD1;refGene_tx=XM_023255620.1;refGene_gn=PRKD1;refGene_tx=XM_023255621.1
B3	81714668	81714792
refGene_gn=PRKD1;refGene_tx=XM_023255617.1;refGene_gn=PRKD1;refGene_tx=XM_023255618.1;refGene_gn=PRKD1;refGene_tx=XM_023255619.1;refGene_gn=PRKD1;refGene_tx=XM_023255620.1;refGene_gn=PRKD1;refGene_tx=XM_023255621.1
B3	81716250	81716497
refGene_gn=PRKD1;refGene_tx=XM_023255617.1;refGene_gn=PRKD1;refGene_tx=XM_023255618.1;refGene_gn=PRKD1;refGene_tx=XM_023255619.1;refGene_gn=PRKD1;refGene_tx=XM_023255620.1;refGene_gn=PRKD1;refGene_tx=XM_023255621.1
B3	81718577	81718655
refGene_gn=PRKD1;refGene_tx=XM_023255617.1;refGene_gn=PRKD1;refGene_tx=XM_023255618.1;refGene_gn=PRKD1;refGene_tx=XM_023255619.1;refGene_gn=PRKD1;refGene_tx=XM_023255620.1;refGene_gn=PRKD1;refGene_tx=XM_023255621.1
B3	81718834	81719045
refGene_gn=PRKD1;refGene_tx=XM_023255617.1;refGene_gn=PRKD1;refGene_tx=XM_023255618.1;refGene_gn=PRKD1;refGene_tx=XM_023255619.1;refGene_gn=PRKD1;refGene_tx=XM_023255620.1;refGene_gn=PRKD1;refGene_tx=XM_023255621.1
B3	81730747	81730771
refGene_gn=PRKD1;refGene_tx=XM_023255617.1;refGene_gn=PRKD1;refGene_tx=XM_023255619.1;refGene_gn=PRKD1;refGene_tx=XM_023255621.1
B3	81741818	81741979
refGene_gn=PRKD1;refGene_tx=XM_023255617.1;refGene_gn=PRKD1;refGene_tx=XM_023255618.1;refGene_gn=PRKD1;refGene_tx=XM_023255619.1;refGene_gn=PRKD1;refGene_tx=XM_023255620.1;refGene_gn=PRKD1;refGene_tx=XM_023255621.1
B3	81743847	81743979
refGene_gn=PRKD1;refGene_tx=XM_023255617.1;refGene_gn=PRKD1;refGene_tx=XM_023255618.1;refGene_gn=PRKD1;refGene_tx=XM_023255619.1;refGene_gn=PRKD1;refGene_tx=XM_023255620.1;refGene_gn=PRKD1;refGene_tx=XM_023255621.1
B3	81807886	81808025
refGene_gn=PRKD1;refGene_tx=XM_023255617.1;refGene_gn=PRKD1;refGene_tx=XM_023255618.1;refGene_gn=PRKD1;refGene_tx=XM_023255619.1;refGene_gn=PRKD1;refGene_tx=XM_023255620.1;refGene_gn=PRKD1;refGene_tx=XM_023255621.1
B3	81998433	81998691
refGene_gn=PRKD1;refGene_tx=XM_023255617.1;refGene_gn=PRKD1;refGene_tx=XM_023255618.1;refGene_gn=PRKD1;refGene_tx=XM_023255619.1;refGene_gn=PRKD1;refGene_tx=XM_023255620.1
B3	82599388	82599425
refGene_gn=G2E3;refGene_tx=XM_003987528.5;refGene_gn=G2E3;refGene_tx=X

M_006932797.4;refGene_gn=G2E3;refGene_tx=XM_019832887.2;refGene_gn=G2E3;refGene_tx=XM_019832888.2
B3 82604800 82604898
refGene_gn=G2E3;refGene_tx=XM_003987528.5;refGene_gn=G2E3;refGene_tx=XM_019832887.2
B3 82607913 82608015
refGene_gn=G2E3;refGene_tx=XM_003987528.5;refGene_gn=G2E3;refGene_tx=XM_006932797.4;refGene_gn=G2E3;refGene_tx=XM_019832887.2;refGene_gn=G2E3;refGene_tx=XM_019832888.2
B3 82611961 82612086
refGene_gn=G2E3;refGene_tx=XM_003987528.5;refGene_gn=G2E3;refGene_tx=XM_006932797.4;refGene_gn=G2E3;refGene_tx=XM_019832887.2;refGene_gn=G2E3;refGene_tx=XM_019832888.2
B3 82613714 82613880
refGene_gn=G2E3;refGene_tx=XM_003987528.5;refGene_gn=G2E3;refGene_tx=XM_006932797.4;refGene_gn=G2E3;refGene_tx=XM_019832887.2;refGene_gn=G2E3;refGene_tx=XM_019832888.2
B3 82621767 82621874
refGene_gn=G2E3;refGene_tx=XM_003987528.5;refGene_gn=G2E3;refGene_tx=XM_006932797.4;refGene_gn=G2E3;refGene_tx=XM_019832887.2;refGene_gn=G2E3;refGene_tx=XM_019832888.2
B3 82622564 82622681
refGene_gn=G2E3;refGene_tx=XM_003987528.5;refGene_gn=G2E3;refGene_tx=XM_006932797.4;refGene_gn=G2E3;refGene_tx=XM_019832887.2;refGene_gn=G2E3;refGene_tx=XM_019832888.2
B3 82626031 82626156
refGene_gn=G2E3;refGene_tx=XM_003987528.5;refGene_gn=G2E3;refGene_tx=XM_006932797.4;refGene_gn=G2E3;refGene_tx=XM_019832887.2;refGene_gn=G2E3;refGene_tx=XM_019832888.2
B3 82626261 82626394
refGene_gn=G2E3;refGene_tx=XM_003987528.5;refGene_gn=G2E3;refGene_tx=XM_006932797.4;refGene_gn=G2E3;refGene_tx=XM_019832887.2;refGene_gn=G2E3;refGene_tx=XM_019832888.2
B3 82629280 82629588
refGene_gn=G2E3;refGene_tx=XM_003987528.5;refGene_gn=G2E3;refGene_tx=XM_006932797.4;refGene_gn=G2E3;refGene_tx=XM_019832887.2;refGene_gn=G2E3;refGene_tx=XM_019832888.2
B3 82632472 82632654
refGene_gn=G2E3;refGene_tx=XM_003987528.5;refGene_gn=G2E3;refGene_tx=XM_006932797.4;refGene_gn=G2E3;refGene_tx=XM_019832887.2;refGene_gn=G2E3;refGene_tx=XM_019832888.2
B3 82636343 82636516
refGene_gn=G2E3;refGene_tx=XM_003987528.5;refGene_gn=G2E3;refGene_tx=XM_006932797.4;refGene_gn=G2E3;refGene_tx=XM_019832887.2;refGene_gn=G2E3;refGene_tx=XM_019832888.2
B3 82641216 82641407
refGene_gn=G2E3;refGene_tx=XM_003987528.5;refGene_gn=G2E3;refGene_tx=XM_006932797.4;refGene_gn=G2E3;refGene_tx=XM_019832887.2;refGene_gn=G2E3;refGene_tx=XM_019832888.2
B3 82642530 82642787

refGene_gn=G2E3;refGene_tx=XM_003987528.5;refGene_gn=G2E3;refGene_tx=XM_006932797.4;refGene_gn=G2E3;refGene_tx=XM_019832887.2;refGene_gn=G2E3;refGene_tx=XM_019832888.2
B3 82647956 82648014
refGene_gn=SCFD1;refGene_tx=XM_023255623.1;refGene_gn=SCFD1;refGene_tx=XM_023255624.1
B3 82654482 82654553
refGene_gn=SCFD1;refGene_tx=XM_023255623.1;refGene_gn=SCFD1;refGene_tx=XM_023255624.1
B3 82656823 82656912
refGene_gn=SCFD1;refGene_tx=XM_023255623.1;refGene_gn=SCFD1;refGene_tx=XM_023255624.1
B3 82658168 82658259
refGene_gn=SCFD1;refGene_tx=XM_023255623.1;refGene_gn=SCFD1;refGene_tx=XM_023255624.1
B3 82660457 82660580
refGene_gn=SCFD1;refGene_tx=XM_023255623.1;refGene_gn=SCFD1;refGene_tx=XM_023255624.1
B3 82662016 82662104
refGene_gn=SCFD1;refGene_tx=XM_023255623.1;refGene_gn=SCFD1;refGene_tx=XM_023255624.1
B3 82666164 82666254
refGene_gn=SCFD1;refGene_tx=XM_023255623.1;refGene_gn=SCFD1;refGene_tx=XM_023255624.1
B3 82669483 82669539
refGene_gn=SCFD1;refGene_tx=XM_023255623.1;refGene_gn=SCFD1;refGene_tx=XM_023255624.1
B3 82670483 82670569
refGene_gn=SCFD1;refGene_tx=XM_023255623.1;refGene_gn=SCFD1;refGene_tx=XM_023255624.1
B3 82672662 82672762
refGene_gn=SCFD1;refGene_tx=XM_023255623.1;refGene_gn=SCFD1;refGene_tx=XM_023255624.1
B3 82691260 82691400
refGene_gn=SCFD1;refGene_tx=XM_023255623.1;refGene_gn=SCFD1;refGene_tx=XM_023255624.1
B3 82694722 82694813
refGene_gn=SCFD1;refGene_tx=XM_023255623.1;refGene_gn=SCFD1;refGene_tx=XM_023255624.1
B3 82695376 82695450
refGene_gn=SCFD1;refGene_tx=XM_023255623.1;refGene_gn=SCFD1;refGene_tx=XM_023255624.1
B3 82696126 82696208
refGene_gn=SCFD1;refGene_tx=XM_023255623.1;refGene_gn=SCFD1;refGene_tx=XM_023255624.1
B3 82715699 82715796
refGene_gn=SCFD1;refGene_tx=XM_023255623.1;refGene_gn=SCFD1;refGene_tx=XM_023255624.1
B3 82719530 82719601
refGene_gn=SCFD1;refGene_tx=XM_023255623.1;refGene_gn=SCFD1;refGene_tx

=XM_023255624.1
B3 82722320 82722400
refGene_gn=SCFD1;refGene_tx=XM_023255623.1;refGene_gn=SCFD1;refGene_tx
=XM_023255624.1
B3 82727594 82727657
refGene_gn=SCFD1;refGene_tx=XM_023255623.1;refGene_gn=SCFD1;refGene_tx
=XM_023255624.1
B3 82733878 82733954
refGene_gn=SCFD1;refGene_tx=XM_023255623.1;refGene_gn=SCFD1;refGene_tx
=XM_023255624.1
B3 82743053 82743107 refGene_gn=SCFD1;refGene_tx=XM_023255623.1
B3 82747400 82747453
refGene_gn=SCFD1;refGene_tx=XM_023255623.1;refGene_gn=SCFD1;refGene_tx
=XM_023255624.1
B3 82749916 82749950
refGene_gn=SCFD1;refGene_tx=XM_023255623.1;refGene_gn=SCFD1;refGene_tx
=XM_023255624.1
B3 82751499 82751565
refGene_gn=SCFD1;refGene_tx=XM_023255623.1;refGene_gn=SCFD1;refGene_tx
=XM_023255624.1
B3 82761257 82761326
refGene_gn=SCFD1;refGene_tx=XM_023255623.1;refGene_gn=SCFD1;refGene_tx
=XM_023255624.1
B3 82764934 82764958
refGene_gn=SCFD1;refGene_tx=XM_023255623.1;refGene_gn=SCFD1;refGene_tx
=XM_023255624.1
B3 82895125 82895159 refGene_gn=COCH;refGene_tx=XM_011283175.2
B3 82895248 82895296 refGene_gn=COCH;refGene_tx=XM_011283175.2
B3 82897354 82897511 refGene_gn=COCH;refGene_tx=XM_011283175.2
B3 82898585 82898719 refGene_gn=COCH;refGene_tx=XM_011283175.2
B3 82899225 82899288 refGene_gn=COCH;refGene_tx=XM_011283175.2
B3 82900361 82900406 refGene_gn=COCH;refGene_tx=XM_011283175.2
B3 82900492 82900640 refGene_gn=COCH;refGene_tx=XM_011283175.2
B3 82903611 82903715 refGene_gn=COCH;refGene_tx=XM_011283175.2
B3 82904430 82904657 refGene_gn=COCH;refGene_tx=XM_011283175.2
B3 82904826 82905343 refGene_gn=COCH;refGene_tx=XM_011283175.2
B3 82907955 82908131 refGene_gn=COCH;refGene_tx=XM_011283175.2
B3 82913163 82913333
refGene_gn=STRN3;refGene_tx=XM_023255631.1;refGene_gn=STRN3;refGene_tx
=XM_023255632.1;refGene_gn=STRN3;refGene_tx=XM_023255633.1;refGene_gn=
STRN3;refGene_tx=XM_023255634.1
B3 82913417 82913504
refGene_gn=STRN3;refGene_tx=XM_023255631.1;refGene_gn=STRN3;refGene_tx
=XM_023255632.1;refGene_gn=STRN3;refGene_tx=XM_023255633.1;refGene_gn=
STRN3;refGene_tx=XM_023255634.1
B3 82919605 82919713
refGene_gn=STRN3;refGene_tx=XM_023255631.1;refGene_gn=STRN3;refGene_tx
=XM_023255632.1;refGene_gn=STRN3;refGene_tx=XM_023255633.1;refGene_gn=
STRN3;refGene_tx=XM_023255634.1
B3 82924152 82924293

refGene_gn=STRN3;refGene_tx=XM_023255631.1;refGene_gn=STRN3;refGene_tx=XM_023255632.1;refGene_gn=STRN3;refGene_tx=XM_023255633.1;refGene_gn=STRN3;refGene_tx=XM_023255634.1
B3 82925579 82925747
refGene_gn=STRN3;refGene_tx=XM_023255631.1;refGene_gn=STRN3;refGene_tx=XM_023255632.1;refGene_gn=STRN3;refGene_tx=XM_023255633.1;refGene_gn=STRN3;refGene_tx=XM_023255634.1
B3 82929455 82929577
refGene_gn=STRN3;refGene_tx=XM_023255631.1;refGene_gn=STRN3;refGene_tx=XM_023255632.1;refGene_gn=STRN3;refGene_tx=XM_023255633.1;refGene_gn=STRN3;refGene_tx=XM_023255634.1
B3 82929873 82929921
refGene_gn=STRN3;refGene_tx=XM_023255631.1;refGene_gn=STRN3;refGene_tx=XM_023255632.1;refGene_gn=STRN3;refGene_tx=XM_023255633.1;refGene_gn=STRN3;refGene_tx=XM_023255634.1
B3 82930115 82930291
refGene_gn=STRN3;refGene_tx=XM_023255631.1;refGene_gn=STRN3;refGene_tx=XM_023255632.1;refGene_gn=STRN3;refGene_tx=XM_023255633.1;refGene_gn=STRN3;refGene_tx=XM_023255634.1
B3 82931655 82931789
refGene_gn=STRN3;refGene_tx=XM_023255631.1;refGene_gn=STRN3;refGene_tx=XM_023255632.1;refGene_gn=STRN3;refGene_tx=XM_023255633.1;refGene_gn=STRN3;refGene_tx=XM_023255634.1
B3 82936546 82936683 refGene_gn=STRN3;refGene_tx=XM_023255634.1
B3 82946043 82946158
refGene_gn=STRN3;refGene_tx=XM_023255631.1;refGene_gn=STRN3;refGene_tx=XM_023255633.1;refGene_gn=STRN3;refGene_tx=XM_023255634.1
B3 82951215 82951357
refGene_gn=STRN3;refGene_tx=XM_023255631.1;refGene_gn=STRN3;refGene_tx=XM_023255632.1;refGene_gn=STRN3;refGene_tx=XM_023255633.1;refGene_gn=STRN3;refGene_tx=XM_023255634.1
B3 82953791 82953921
refGene_gn=STRN3;refGene_tx=XM_023255631.1;refGene_gn=STRN3;refGene_tx=XM_023255632.1;refGene_gn=STRN3;refGene_tx=XM_023255633.1;refGene_gn=STRN3;refGene_tx=XM_023255634.1
B3 82958182 82958356
refGene_gn=STRN3;refGene_tx=XM_023255631.1;refGene_gn=STRN3;refGene_tx=XM_023255632.1;refGene_gn=STRN3;refGene_tx=XM_023255633.1;refGene_gn=STRN3;refGene_tx=XM_023255634.1
B3 82960678 82960760
refGene_gn=STRN3;refGene_tx=XM_023255631.1;refGene_gn=STRN3;refGene_tx=XM_023255632.1;refGene_gn=STRN3;refGene_tx=XM_023255633.1;refGene_gn=STRN3;refGene_tx=XM_023255634.1
B3 82965692 82965766
refGene_gn=STRN3;refGene_tx=XM_023255631.1;refGene_gn=STRN3;refGene_tx=XM_023255632.1;refGene_gn=STRN3;refGene_tx=XM_023255633.1;refGene_gn=STRN3;refGene_tx=XM_023255634.1
B3 82966398 82966502
refGene_gn=STRN3;refGene_tx=XM_023255631.1;refGene_gn=STRN3;refGene_tx=XM_023255632.1;refGene_gn=STRN3;refGene_tx=XM_023255633.1;refGene_gn=

STRN3;refGene_tx=XM_023255634.1
B3 83018118 83018400
refGene_gn=STRN3;refGene_tx=XM_023255631.1;refGene_gn=STRN3;refGene_tx=XM_023255632.1;refGene_gn=STRN3;refGene_tx=XM_023255634.1
B3 83047841 83047979
refGene_gn=AP4S1;refGene_tx=XM_023255635.1;refGene_gn=AP4S1;refGene_tx=XM_023255636.1;refGene_gn=AP4S1;refGene_tx=XM_023255637.1;refGene_gn=AP4S1;refGene_tx=XM_023255638.1;refGene_gn=AP4S1;refGene_tx=XM_023255639.1;refGene_gn=AP4S1;refGene_tx=XM_023255640.1;refGene_gn=AP4S1;refGene_tx=XM_023255641.1;refGene_gn=AP4S1;refGene_tx=XM_023255642.1
B3 83051167 83051254
refGene_gn=AP4S1;refGene_tx=XM_023255635.1;refGene_gn=AP4S1;refGene_tx=XM_023255636.1;refGene_gn=AP4S1;refGene_tx=XM_023255637.1;refGene_gn=AP4S1;refGene_tx=XM_023255638.1;refGene_gn=AP4S1;refGene_tx=XM_023255639.1;refGene_gn=AP4S1;refGene_tx=XM_023255640.1;refGene_gn=AP4S1;refGene_tx=XM_023255641.1;refGene_gn=AP4S1;refGene_tx=XM_023255642.1
B3 83052097 83052166
refGene_gn=AP4S1;refGene_tx=XM_023255635.1;refGene_gn=AP4S1;refGene_tx=XM_023255636.1;refGene_gn=AP4S1;refGene_tx=XM_023255637.1;refGene_gn=AP4S1;refGene_tx=XM_023255638.1;refGene_gn=AP4S1;refGene_tx=XM_023255639.1;refGene_gn=AP4S1;refGene_tx=XM_023255640.1;refGene_gn=AP4S1;refGene_tx=XM_023255641.1;refGene_gn=AP4S1;refGene_tx=XM_023255642.1
B3 83058515 83058587
refGene_gn=AP4S1;refGene_tx=XM_023255635.1;refGene_gn=AP4S1;refGene_tx=XM_023255636.1;refGene_gn=AP4S1;refGene_tx=XM_023255637.1;refGene_gn=AP4S1;refGene_tx=XM_023255638.1;refGene_gn=AP4S1;refGene_tx=XM_023255639.1;refGene_gn=AP4S1;refGene_tx=XM_023255640.1;refGene_gn=AP4S1;refGene_tx=XM_023255641.1;refGene_gn=AP4S1;refGene_tx=XM_023255642.1
B3 83063310 83063466
refGene_gn=AP4S1;refGene_tx=XM_023255635.1;refGene_gn=AP4S1;refGene_tx=XM_023255636.1;refGene_gn=AP4S1;refGene_tx=XM_023255637.1;refGene_gn=AP4S1;refGene_tx=XM_023255638.1;refGene_gn=AP4S1;refGene_tx=XM_023255639.1
B3 83067336 83067465 refGene_gn=AP4S1;refGene_tx=XM_023255640.1
B3 83071394 83071532
refGene_gn=HECTD1;refGene_tx=XM_023255625.1;refGene_gn=HECTD1;refGene_tx=XM_023255626.1;refGene_gn=HECTD1;refGene_tx=XM_023255627.1;refGene_gn=HECTD1;refGene_tx=XM_023255628.1;refGene_gn=HECTD1;refGene_tx=XM_023255629.1;refGene_gn=HECTD1;refGene_tx=XM_023255630.1
B3 83071641 83071786
refGene_gn=HECTD1;refGene_tx=XM_023255625.1;refGene_gn=HECTD1;refGene_tx=XM_023255626.1;refGene_gn=HECTD1;refGene_tx=XM_023255627.1;refGene_gn=HECTD1;refGene_tx=XM_023255628.1;refGene_gn=HECTD1;refGene_tx=XM_023255629.1;refGene_gn=HECTD1;refGene_tx=XM_023255630.1
B3 83074141 83074292
refGene_gn=HECTD1;refGene_tx=XM_023255625.1;refGene_gn=HECTD1;refGene_tx=XM_023255626.1;refGene_gn=HECTD1;refGene_tx=XM_023255627.1;refGene_gn=HECTD1;refGene_tx=XM_023255628.1;refGene_gn=HECTD1;refGene_tx=XM_023255629.1;refGene_gn=HECTD1;refGene_tx=XM_023255630.1
B3 83074996 83075090

refGene_gn=HECTD1;refGene_tx=XM_023255625.1;refGene_gn=HECTD1;refGene_tx=XM_023255626.1;refGene_gn=HECTD1;refGene_tx=XM_023255627.1;refGene_gn=HECTD1;refGene_tx=XM_023255628.1;refGene_gn=HECTD1;refGene_tx=XM_023255629.1;refGene_gn=HECTD1;refGene_tx=XM_023255630.1
B3 83075217 83075296
refGene_gn=HECTD1;refGene_tx=XM_023255625.1;refGene_gn=HECTD1;refGene_tx=XM_023255626.1;refGene_gn=HECTD1;refGene_tx=XM_023255627.1;refGene_gn=HECTD1;refGene_tx=XM_023255628.1;refGene_gn=HECTD1;refGene_tx=XM_023255629.1;refGene_gn=HECTD1;refGene_tx=XM_023255630.1
B3 83076303 83076911
refGene_gn=HECTD1;refGene_tx=XM_023255625.1;refGene_gn=HECTD1;refGene_tx=XM_023255626.1;refGene_gn=HECTD1;refGene_tx=XM_023255627.1;refGene_gn=HECTD1;refGene_tx=XM_023255628.1;refGene_gn=HECTD1;refGene_tx=XM_023255629.1;refGene_gn=HECTD1;refGene_tx=XM_023255630.1
B3 83078468 83078606
refGene_gn=HECTD1;refGene_tx=XM_023255625.1;refGene_gn=HECTD1;refGene_tx=XM_023255626.1;refGene_gn=HECTD1;refGene_tx=XM_023255627.1;refGene_gn=HECTD1;refGene_tx=XM_023255628.1;refGene_gn=HECTD1;refGene_tx=XM_023255629.1;refGene_gn=HECTD1;refGene_tx=XM_023255630.1
B3 83080087 83080289
refGene_gn=HECTD1;refGene_tx=XM_023255625.1;refGene_gn=HECTD1;refGene_tx=XM_023255626.1;refGene_gn=HECTD1;refGene_tx=XM_023255627.1;refGene_gn=HECTD1;refGene_tx=XM_023255628.1;refGene_gn=HECTD1;refGene_tx=XM_023255629.1;refGene_gn=HECTD1;refGene_tx=XM_023255630.1
B3 83082540 83082668
refGene_gn=HECTD1;refGene_tx=XM_023255625.1;refGene_gn=HECTD1;refGene_tx=XM_023255626.1;refGene_gn=HECTD1;refGene_tx=XM_023255627.1;refGene_gn=HECTD1;refGene_tx=XM_023255628.1;refGene_gn=HECTD1;refGene_tx=XM_023255629.1;refGene_gn=HECTD1;refGene_tx=XM_023255630.1
B3 83082882 83082968
refGene_gn=HECTD1;refGene_tx=XM_023255625.1;refGene_gn=HECTD1;refGene_tx=XM_023255626.1;refGene_gn=HECTD1;refGene_tx=XM_023255627.1;refGene_gn=HECTD1;refGene_tx=XM_023255628.1;refGene_gn=HECTD1;refGene_tx=XM_023255629.1;refGene_gn=HECTD1;refGene_tx=XM_023255630.1
B3 83083052 83083260
refGene_gn=HECTD1;refGene_tx=XM_023255625.1;refGene_gn=HECTD1;refGene_tx=XM_023255626.1;refGene_gn=HECTD1;refGene_tx=XM_023255627.1;refGene_gn=HECTD1;refGene_tx=XM_023255628.1;refGene_gn=HECTD1;refGene_tx=XM_023255629.1;refGene_gn=HECTD1;refGene_tx=XM_023255630.1
B3 83083648 83083831
refGene_gn=HECTD1;refGene_tx=XM_023255625.1;refGene_gn=HECTD1;refGene_tx=XM_023255626.1;refGene_gn=HECTD1;refGene_tx=XM_023255627.1;refGene_gn=HECTD1;refGene_tx=XM_023255628.1;refGene_gn=HECTD1;refGene_tx=XM_023255630.1
B3 83084052 83084107
refGene_gn=HECTD1;refGene_tx=XM_023255625.1;refGene_gn=HECTD1;refGene_tx=XM_023255626.1;refGene_gn=HECTD1;refGene_tx=XM_023255627.1;refGene_gn=HECTD1;refGene_tx=XM_023255628.1;refGene_gn=HECTD1;refGene_tx=XM_023255629.1;refGene_gn=HECTD1;refGene_tx=XM_023255630.1
B3 83085208 83085431

refGene_gn=HECTD1;refGene_tx=XM_023255625.1;refGene_gn=HECTD1;refGene_tx=XM_023255626.1;refGene_gn=HECTD1;refGene_tx=XM_023255627.1;refGene_gn=HECTD1;refGene_tx=XM_023255628.1;refGene_gn=HECTD1;refGene_tx=XM_023255629.1;refGene_gn=HECTD1;refGene_tx=XM_023255630.1
B3 83088035 83088192
refGene_gn=HECTD1;refGene_tx=XM_023255625.1;refGene_gn=HECTD1;refGene_tx=XM_023255626.1;refGene_gn=HECTD1;refGene_tx=XM_023255627.1;refGene_gn=HECTD1;refGene_tx=XM_023255628.1;refGene_gn=HECTD1;refGene_tx=XM_023255629.1;refGene_gn=HECTD1;refGene_tx=XM_023255630.1
B3 83089995 83090118
refGene_gn=HECTD1;refGene_tx=XM_023255625.1;refGene_gn=HECTD1;refGene_tx=XM_023255626.1;refGene_gn=HECTD1;refGene_tx=XM_023255627.1;refGene_gn=HECTD1;refGene_tx=XM_023255628.1;refGene_gn=HECTD1;refGene_tx=XM_023255629.1;refGene_gn=HECTD1;refGene_tx=XM_023255630.1
B3 83092193 83092327
refGene_gn=HECTD1;refGene_tx=XM_023255625.1;refGene_gn=HECTD1;refGene_tx=XM_023255626.1;refGene_gn=HECTD1;refGene_tx=XM_023255627.1;refGene_gn=HECTD1;refGene_tx=XM_023255628.1;refGene_gn=HECTD1;refGene_tx=XM_023255629.1;refGene_gn=HECTD1;refGene_tx=XM_023255630.1
B3 83096613 83096835
refGene_gn=HECTD1;refGene_tx=XM_023255625.1;refGene_gn=HECTD1;refGene_tx=XM_023255626.1;refGene_gn=HECTD1;refGene_tx=XM_023255627.1;refGene_gn=HECTD1;refGene_tx=XM_023255628.1;refGene_gn=HECTD1;refGene_tx=XM_023255629.1;refGene_gn=HECTD1;refGene_tx=XM_023255630.1
B3 83097528 83098365
refGene_gn=HECTD1;refGene_tx=XM_023255625.1;refGene_gn=HECTD1;refGene_tx=XM_023255626.1;refGene_gn=HECTD1;refGene_tx=XM_023255627.1;refGene_gn=HECTD1;refGene_tx=XM_023255628.1;refGene_gn=HECTD1;refGene_tx=XM_023255629.1
B3 83102044 83102221
refGene_gn=HECTD1;refGene_tx=XM_023255625.1;refGene_gn=HECTD1;refGene_tx=XM_023255626.1;refGene_gn=HECTD1;refGene_tx=XM_023255627.1;refGene_gn=HECTD1;refGene_tx=XM_023255628.1;refGene_gn=HECTD1;refGene_tx=XM_023255629.1;refGene_gn=HECTD1;refGene_tx=XM_023255630.1
B3 83102312 83102479
refGene_gn=HECTD1;refGene_tx=XM_023255625.1;refGene_gn=HECTD1;refGene_tx=XM_023255626.1;refGene_gn=HECTD1;refGene_tx=XM_023255627.1;refGene_gn=HECTD1;refGene_tx=XM_023255628.1;refGene_gn=HECTD1;refGene_tx=XM_023255629.1;refGene_gn=HECTD1;refGene_tx=XM_023255630.1
B3 83103668 83103992
refGene_gn=HECTD1;refGene_tx=XM_023255625.1;refGene_gn=HECTD1;refGene_tx=XM_023255626.1;refGene_gn=HECTD1;refGene_tx=XM_023255627.1;refGene_gn=HECTD1;refGene_tx=XM_023255628.1;refGene_gn=HECTD1;refGene_tx=XM_023255629.1;refGene_gn=HECTD1;refGene_tx=XM_023255630.1
B3 83104275 83104415
refGene_gn=HECTD1;refGene_tx=XM_023255625.1;refGene_gn=HECTD1;refGene_tx=XM_023255626.1;refGene_gn=HECTD1;refGene_tx=XM_023255627.1;refGene_gn=HECTD1;refGene_tx=XM_023255628.1;refGene_gn=HECTD1;refGene_tx=XM_023255629.1;refGene_gn=HECTD1;refGene_tx=XM_023255630.1
B3 83105174 83105309

refGene_gn=HECTD1;refGene_tx=XM_023255625.1;refGene_gn=HECTD1;refGene_tx=XM_023255626.1;refGene_gn=HECTD1;refGene_tx=XM_023255627.1;refGene_gn=HECTD1;refGene_tx=XM_023255628.1;refGene_gn=HECTD1;refGene_tx=XM_023255629.1;refGene_gn=HECTD1;refGene_tx=XM_023255630.1
B3 83107445 83107551
refGene_gn=HECTD1;refGene_tx=XM_023255625.1;refGene_gn=HECTD1;refGene_tx=XM_023255626.1;refGene_gn=HECTD1;refGene_tx=XM_023255627.1;refGene_gn=HECTD1;refGene_tx=XM_023255628.1;refGene_gn=HECTD1;refGene_tx=XM_023255629.1;refGene_gn=HECTD1;refGene_tx=XM_023255630.1
B3 83109265 83109360
refGene_gn=HECTD1;refGene_tx=XM_023255625.1;refGene_gn=HECTD1;refGene_tx=XM_023255626.1;refGene_gn=HECTD1;refGene_tx=XM_023255627.1;refGene_gn=HECTD1;refGene_tx=XM_023255628.1;refGene_gn=HECTD1;refGene_tx=XM_023255629.1;refGene_gn=HECTD1;refGene_tx=XM_023255630.1
B3 83110677 83110824
refGene_gn=HECTD1;refGene_tx=XM_023255625.1;refGene_gn=HECTD1;refGene_tx=XM_023255626.1;refGene_gn=HECTD1;refGene_tx=XM_023255627.1;refGene_gn=HECTD1;refGene_tx=XM_023255628.1;refGene_gn=HECTD1;refGene_tx=XM_023255629.1;refGene_gn=HECTD1;refGene_tx=XM_023255630.1
B3 83111376 83111511
refGene_gn=HECTD1;refGene_tx=XM_023255625.1;refGene_gn=HECTD1;refGene_tx=XM_023255626.1;refGene_gn=HECTD1;refGene_tx=XM_023255627.1;refGene_gn=HECTD1;refGene_tx=XM_023255628.1;refGene_gn=HECTD1;refGene_tx=XM_023255629.1;refGene_gn=HECTD1;refGene_tx=XM_023255630.1
B3 83115586 83115663
refGene_gn=HECTD1;refGene_tx=XM_023255625.1;refGene_gn=HECTD1;refGene_tx=XM_023255626.1;refGene_gn=HECTD1;refGene_tx=XM_023255627.1;refGene_gn=HECTD1;refGene_tx=XM_023255628.1;refGene_gn=HECTD1;refGene_tx=XM_023255629.1;refGene_gn=HECTD1;refGene_tx=XM_023255630.1
B3 83115761 83116025
refGene_gn=HECTD1;refGene_tx=XM_023255625.1;refGene_gn=HECTD1;refGene_tx=XM_023255626.1;refGene_gn=HECTD1;refGene_tx=XM_023255627.1;refGene_gn=HECTD1;refGene_tx=XM_023255628.1;refGene_gn=HECTD1;refGene_tx=XM_023255629.1;refGene_gn=HECTD1;refGene_tx=XM_023255630.1
B3 83116566 83116789
refGene_gn=HECTD1;refGene_tx=XM_023255625.1;refGene_gn=HECTD1;refGene_tx=XM_023255626.1;refGene_gn=HECTD1;refGene_tx=XM_023255627.1;refGene_gn=HECTD1;refGene_tx=XM_023255628.1;refGene_gn=HECTD1;refGene_tx=XM_023255629.1;refGene_gn=HECTD1;refGene_tx=XM_023255630.1
B3 83121919 83122090
refGene_gn=HECTD1;refGene_tx=XM_023255625.1;refGene_gn=HECTD1;refGene_tx=XM_023255626.1;refGene_gn=HECTD1;refGene_tx=XM_023255627.1;refGene_gn=HECTD1;refGene_tx=XM_023255628.1;refGene_gn=HECTD1;refGene_tx=XM_023255629.1;refGene_gn=HECTD1;refGene_tx=XM_023255630.1
B3 83122279 83122418
refGene_gn=HECTD1;refGene_tx=XM_023255625.1;refGene_gn=HECTD1;refGene_tx=XM_023255626.1;refGene_gn=HECTD1;refGene_tx=XM_023255627.1;refGene_gn=HECTD1;refGene_tx=XM_023255628.1;refGene_gn=HECTD1;refGene_tx=XM_023255629.1;refGene_gn=HECTD1;refGene_tx=XM_023255630.1
B3 83131389 83131564

refGene_gn=HECTD1;refGene_tx=XM_023255625.1;refGene_gn=HECTD1;refGene_tx=XM_023255626.1;refGene_gn=HECTD1;refGene_tx=XM_023255627.1;refGene_gn=HECTD1;refGene_tx=XM_023255628.1;refGene_gn=HECTD1;refGene_tx=XM_023255629.1;refGene_gn=HECTD1;refGene_tx=XM_023255630.1
B3 83132197 83132312
refGene_gn=HECTD1;refGene_tx=XM_023255625.1;refGene_gn=HECTD1;refGene_tx=XM_023255626.1;refGene_gn=HECTD1;refGene_tx=XM_023255627.1;refGene_gn=HECTD1;refGene_tx=XM_023255628.1;refGene_gn=HECTD1;refGene_tx=XM_023255629.1;refGene_gn=HECTD1;refGene_tx=XM_023255630.1
B3 83134959 83135058
refGene_gn=HECTD1;refGene_tx=XM_023255625.1;refGene_gn=HECTD1;refGene_tx=XM_023255626.1;refGene_gn=HECTD1;refGene_tx=XM_023255627.1;refGene_gn=HECTD1;refGene_tx=XM_023255628.1;refGene_gn=HECTD1;refGene_tx=XM_023255629.1;refGene_gn=HECTD1;refGene_tx=XM_023255630.1
B3 83135143 83135220
refGene_gn=HECTD1;refGene_tx=XM_023255625.1;refGene_gn=HECTD1;refGene_tx=XM_023255626.1;refGene_gn=HECTD1;refGene_tx=XM_023255627.1;refGene_gn=HECTD1;refGene_tx=XM_023255628.1;refGene_gn=HECTD1;refGene_tx=XM_023255629.1;refGene_gn=HECTD1;refGene_tx=XM_023255630.1
B3 83135743 83136014
refGene_gn=HECTD1;refGene_tx=XM_023255625.1;refGene_gn=HECTD1;refGene_tx=XM_023255626.1;refGene_gn=HECTD1;refGene_tx=XM_023255627.1;refGene_gn=HECTD1;refGene_tx=XM_023255628.1;refGene_gn=HECTD1;refGene_tx=XM_023255629.1;refGene_gn=HECTD1;refGene_tx=XM_023255630.1
B3 83136110 83136392
refGene_gn=HECTD1;refGene_tx=XM_023255625.1;refGene_gn=HECTD1;refGene_tx=XM_023255626.1;refGene_gn=HECTD1;refGene_tx=XM_023255627.1;refGene_gn=HECTD1;refGene_tx=XM_023255628.1;refGene_gn=HECTD1;refGene_tx=XM_023255629.1;refGene_gn=HECTD1;refGene_tx=XM_023255630.1
B3 83137348 83137576
refGene_gn=HECTD1;refGene_tx=XM_023255625.1;refGene_gn=HECTD1;refGene_tx=XM_023255626.1;refGene_gn=HECTD1;refGene_tx=XM_023255627.1;refGene_gn=HECTD1;refGene_tx=XM_023255628.1;refGene_gn=HECTD1;refGene_tx=XM_023255629.1;refGene_gn=HECTD1;refGene_tx=XM_023255630.1
B3 83141099 83141334
refGene_gn=HECTD1;refGene_tx=XM_023255625.1;refGene_gn=HECTD1;refGene_tx=XM_023255626.1;refGene_gn=HECTD1;refGene_tx=XM_023255627.1;refGene_gn=HECTD1;refGene_tx=XM_023255628.1;refGene_gn=HECTD1;refGene_tx=XM_023255629.1;refGene_gn=HECTD1;refGene_tx=XM_023255630.1
B3 83160445 83160585
refGene_gn=HECTD1;refGene_tx=XM_023255625.1;refGene_gn=HECTD1;refGene_tx=XM_023255626.1;refGene_gn=HECTD1;refGene_tx=XM_023255627.1;refGene_gn=HECTD1;refGene_tx=XM_023255628.1;refGene_gn=HECTD1;refGene_tx=XM_023255629.1;refGene_gn=HECTD1;refGene_tx=XM_023255630.1
B3 83161754 83162316
refGene_gn=LOC109500794;refGene_tx=XM_023256183.1
B3 83162378 83163064
refGene_gn=LOC109500794;refGene_tx=XM_023256183.1
B3 83167915 83167975
refGene_gn=LOC109500794;refGene_tx=XM_023256183.1

B3 83235304 83235612
refGene_gn=HEATR5A;refGene_tx=XM_011283176.3;refGene_gn=HEATR5A;refGene_tx=XM_019832900.2;refGene_gn=HEATR5A;refGene_tx=XM_019832901.2;refGene_gn=HEATR5A;refGene_tx=XM_019832902.2;refGene_gn=HEATR5A;refGene_tx=XM_019832903.2
B3 83235912 83236126
refGene_gn=HEATR5A;refGene_tx=XM_011283176.3;refGene_gn=HEATR5A;refGene_tx=XM_019832900.2;refGene_gn=HEATR5A;refGene_tx=XM_019832901.2;refGene_gn=HEATR5A;refGene_tx=XM_019832902.2;refGene_gn=HEATR5A;refGene_tx=XM_019832903.2
B3 83247211 83247366
refGene_gn=HEATR5A;refGene_tx=XM_011283176.3;refGene_gn=HEATR5A;refGene_tx=XM_019832900.2;refGene_gn=HEATR5A;refGene_tx=XM_019832901.2;refGene_gn=HEATR5A;refGene_tx=XM_019832902.2;refGene_gn=HEATR5A;refGene_tx=XM_019832903.2
B3 83252123 83252348
refGene_gn=HEATR5A;refGene_tx=XM_011283176.3;refGene_gn=HEATR5A;refGene_tx=XM_019832900.2;refGene_gn=HEATR5A;refGene_tx=XM_019832901.2;refGene_gn=HEATR5A;refGene_tx=XM_019832902.2;refGene_gn=HEATR5A;refGene_tx=XM_019832903.2
B3 83253874 83254147
refGene_gn=HEATR5A;refGene_tx=XM_011283176.3;refGene_gn=HEATR5A;refGene_tx=XM_019832900.2;refGene_gn=HEATR5A;refGene_tx=XM_019832901.2;refGene_gn=HEATR5A;refGene_tx=XM_019832902.2;refGene_gn=HEATR5A;refGene_tx=XM_019832903.2
B3 83255314 83255462
refGene_gn=HEATR5A;refGene_tx=XM_011283176.3;refGene_gn=HEATR5A;refGene_tx=XM_019832900.2;refGene_gn=HEATR5A;refGene_tx=XM_019832901.2;refGene_gn=HEATR5A;refGene_tx=XM_019832902.2;refGene_gn=HEATR5A;refGene_tx=XM_019832903.2
B3 83256765 83256893
refGene_gn=HEATR5A;refGene_tx=XM_011283176.3;refGene_gn=HEATR5A;refGene_tx=XM_019832900.2;refGene_gn=HEATR5A;refGene_tx=XM_019832901.2;refGene_gn=HEATR5A;refGene_tx=XM_019832902.2;refGene_gn=HEATR5A;refGene_tx=XM_019832903.2
B3 83259123 83259372
refGene_gn=HEATR5A;refGene_tx=XM_011283176.3;refGene_gn=HEATR5A;refGene_tx=XM_019832900.2;refGene_gn=HEATR5A;refGene_tx=XM_019832901.2;refGene_gn=HEATR5A;refGene_tx=XM_019832902.2;refGene_gn=HEATR5A;refGene_tx=XM_019832903.2
B3 83261240 83261463
refGene_gn=HEATR5A;refGene_tx=XM_011283176.3;refGene_gn=HEATR5A;refGene_tx=XM_019832900.2;refGene_gn=HEATR5A;refGene_tx=XM_019832901.2;refGene_gn=HEATR5A;refGene_tx=XM_019832902.2;refGene_gn=HEATR5A;refGene_tx=XM_019832903.2
B3 83263862 83264042
refGene_gn=HEATR5A;refGene_tx=XM_011283176.3;refGene_gn=HEATR5A;refGene_tx=XM_019832900.2;refGene_gn=HEATR5A;refGene_tx=XM_019832901.2;refGene_gn=HEATR5A;refGene_tx=XM_019832902.2;refGene_gn=HEATR5A;refGene_tx=XM_019832903.2

B3 83265242 83265311
refGene_gn=HEATR5A;refGene_tx=XM_011283176.3;refGene_gn=HEATR5A;refGene_tx=XM_019832900.2;refGene_gn=HEATR5A;refGene_tx=XM_019832901.2;refGene_gn=HEATR5A;refGene_tx=XM_019832902.2;refGene_gn=HEATR5A;refGene_tx=XM_019832903.2
B3 83266720 83266902
refGene_gn=HEATR5A;refGene_tx=XM_011283176.3;refGene_gn=HEATR5A;refGene_tx=XM_019832900.2;refGene_gn=HEATR5A;refGene_tx=XM_019832901.2;refGene_gn=HEATR5A;refGene_tx=XM_019832902.2;refGene_gn=HEATR5A;refGene_tx=XM_019832903.2
B3 83268391 83268631
refGene_gn=HEATR5A;refGene_tx=XM_011283176.3;refGene_gn=HEATR5A;refGene_tx=XM_019832900.2;refGene_gn=HEATR5A;refGene_tx=XM_019832901.2;refGene_gn=HEATR5A;refGene_tx=XM_019832902.2;refGene_gn=HEATR5A;refGene_tx=XM_019832903.2
B3 83272844 83273024
refGene_gn=HEATR5A;refGene_tx=XM_011283176.3;refGene_gn=HEATR5A;refGene_tx=XM_019832900.2;refGene_gn=HEATR5A;refGene_tx=XM_019832901.2;refGene_gn=HEATR5A;refGene_tx=XM_019832902.2;refGene_gn=HEATR5A;refGene_tx=XM_019832903.2
B3 83276151 83276290
refGene_gn=HEATR5A;refGene_tx=XM_011283176.3;refGene_gn=HEATR5A;refGene_tx=XM_019832900.2;refGene_gn=HEATR5A;refGene_tx=XM_019832901.2;refGene_gn=HEATR5A;refGene_tx=XM_019832902.2;refGene_gn=HEATR5A;refGene_tx=XM_019832903.2
B3 83278549 83278576
refGene_gn=HEATR5A;refGene_tx=XM_019832900.2;refGene_gn=HEATR5A;refGene_tx=XM_019832901.2;refGene_gn=HEATR5A;refGene_tx=XM_019832902.2;refGene_gn=HEATR5A;refGene_tx=XM_019832903.2
B3 83281218 83281388
refGene_gn=HEATR5A;refGene_tx=XM_011283176.3;refGene_gn=HEATR5A;refGene_tx=XM_019832900.2;refGene_gn=HEATR5A;refGene_tx=XM_019832901.2;refGene_gn=HEATR5A;refGene_tx=XM_019832902.2;refGene_gn=HEATR5A;refGene_tx=XM_019832903.2
B3 83282464 83282654
refGene_gn=HEATR5A;refGene_tx=XM_011283176.3;refGene_gn=HEATR5A;refGene_tx=XM_019832900.2;refGene_gn=HEATR5A;refGene_tx=XM_019832901.2;refGene_gn=HEATR5A;refGene_tx=XM_019832902.2;refGene_gn=HEATR5A;refGene_tx=XM_019832903.2
B3 83286004 83286164
refGene_gn=HEATR5A;refGene_tx=XM_011283176.3;refGene_gn=HEATR5A;refGene_tx=XM_019832900.2;refGene_gn=HEATR5A;refGene_tx=XM_019832901.2;refGene_gn=HEATR5A;refGene_tx=XM_019832902.2;refGene_gn=HEATR5A;refGene_tx=XM_019832903.2
B3 83297918 83298109
refGene_gn=HEATR5A;refGene_tx=XM_011283176.3;refGene_gn=HEATR5A;refGene_tx=XM_019832900.2;refGene_gn=HEATR5A;refGene_tx=XM_019832901.2;refGene_gn=HEATR5A;refGene_tx=XM_019832902.2;refGene_gn=HEATR5A;refGene_tx=XM_019832903.2
B3 83298952 83299058

refGene_gn=HEATR5A;refGene_tx=XM_011283176.3;refGene_gn=HEATR5A;refGene_tx=XM_019832900.2;refGene_gn=HEATR5A;refGene_tx=XM_019832901.2;refGene_gn=HEATR5A;refGene_tx=XM_019832902.2;refGene_gn=HEATR5A;refGene_tx=XM_019832903.2
B3 83307267 83307443
refGene_gn=HEATR5A;refGene_tx=XM_011283176.3;refGene_gn=HEATR5A;refGene_tx=XM_019832900.2;refGene_gn=HEATR5A;refGene_tx=XM_019832901.2;refGene_gn=HEATR5A;refGene_tx=XM_019832902.2;refGene_gn=HEATR5A;refGene_tx=XM_019832903.2
B3 83307534 83307698
refGene_gn=HEATR5A;refGene_tx=XM_011283176.3;refGene_gn=HEATR5A;refGene_tx=XM_019832900.2;refGene_gn=HEATR5A;refGene_tx=XM_019832901.2;refGene_gn=HEATR5A;refGene_tx=XM_019832902.2;refGene_gn=HEATR5A;refGene_tx=XM_019832903.2
B3 83311930 83312040
refGene_gn=HEATR5A;refGene_tx=XM_011283176.3;refGene_gn=HEATR5A;refGene_tx=XM_019832900.2;refGene_gn=HEATR5A;refGene_tx=XM_019832901.2;refGene_gn=HEATR5A;refGene_tx=XM_019832902.2;refGene_gn=HEATR5A;refGene_tx=XM_019832903.2
B3 83315519 83315619
refGene_gn=HEATR5A;refGene_tx=XM_011283176.3;refGene_gn=HEATR5A;refGene_tx=XM_019832900.2;refGene_gn=HEATR5A;refGene_tx=XM_019832901.2;refGene_gn=HEATR5A;refGene_tx=XM_019832902.2;refGene_gn=HEATR5A;refGene_tx=XM_019832903.2
B3 83316606 83316759
refGene_gn=HEATR5A;refGene_tx=XM_011283176.3;refGene_gn=HEATR5A;refGene_tx=XM_019832900.2;refGene_gn=HEATR5A;refGene_tx=XM_019832901.2;refGene_gn=HEATR5A;refGene_tx=XM_019832902.2;refGene_gn=HEATR5A;refGene_tx=XM_019832903.2
B3 83320926 83321038
refGene_gn=HEATR5A;refGene_tx=XM_011283176.3;refGene_gn=HEATR5A;refGene_tx=XM_019832900.2;refGene_gn=HEATR5A;refGene_tx=XM_019832901.2;refGene_gn=HEATR5A;refGene_tx=XM_019832902.2;refGene_gn=HEATR5A;refGene_tx=XM_019832903.2
B3 83323193 83323444
refGene_gn=HEATR5A;refGene_tx=XM_011283176.3;refGene_gn=HEATR5A;refGene_tx=XM_019832900.2;refGene_gn=HEATR5A;refGene_tx=XM_019832901.2;refGene_gn=HEATR5A;refGene_tx=XM_019832902.2;refGene_gn=HEATR5A;refGene_tx=XM_019832903.2
B3 83325040 83325196
refGene_gn=HEATR5A;refGene_tx=XM_011283176.3;refGene_gn=HEATR5A;refGene_tx=XM_019832900.2;refGene_gn=HEATR5A;refGene_tx=XM_019832901.2;refGene_gn=HEATR5A;refGene_tx=XM_019832902.2;refGene_gn=HEATR5A;refGene_tx=XM_019832903.2
B3 83325811 83326067
refGene_gn=HEATR5A;refGene_tx=XM_011283176.3;refGene_gn=HEATR5A;refGene_tx=XM_019832900.2;refGene_gn=HEATR5A;refGene_tx=XM_019832901.2;refGene_gn=HEATR5A;refGene_tx=XM_019832902.2;refGene_gn=HEATR5A;refGene_tx=XM_019832903.2
B3 83327218 83327379

refGene_gn=HEATR5A;refGene_tx=XM_011283176.3;refGene_gn=HEATR5A;refGene_tx=XM_019832900.2;refGene_gn=HEATR5A;refGene_tx=XM_019832901.2;refGene_gn=HEATR5A;refGene_tx=XM_019832902.2;refGene_gn=HEATR5A;refGene_tx=XM_019832903.2
B3 83333153 83333328
refGene_gn=HEATR5A;refGene_tx=XM_011283176.3;refGene_gn=HEATR5A;refGene_tx=XM_019832900.2;refGene_gn=HEATR5A;refGene_tx=XM_019832901.2;refGene_gn=HEATR5A;refGene_tx=XM_019832902.2;refGene_gn=HEATR5A;refGene_tx=XM_019832903.2
B3 83334152 83334302
refGene_gn=HEATR5A;refGene_tx=XM_011283176.3;refGene_gn=HEATR5A;refGene_tx=XM_019832900.2;refGene_gn=HEATR5A;refGene_tx=XM_019832901.2;refGene_gn=HEATR5A;refGene_tx=XM_019832902.2;refGene_gn=HEATR5A;refGene_tx=XM_019832903.2
B3 83336744 83336853
refGene_gn=HEATR5A;refGene_tx=XM_011283176.3;refGene_gn=HEATR5A;refGene_tx=XM_019832900.2;refGene_gn=HEATR5A;refGene_tx=XM_019832901.2;refGene_gn=HEATR5A;refGene_tx=XM_019832902.2;refGene_gn=HEATR5A;refGene_tx=XM_019832903.2
B3 83338312 83338524
refGene_gn=HEATR5A;refGene_tx=XM_011283176.3;refGene_gn=HEATR5A;refGene_tx=XM_019832900.2;refGene_gn=HEATR5A;refGene_tx=XM_019832901.2;refGene_gn=HEATR5A;refGene_tx=XM_019832902.2;refGene_gn=HEATR5A;refGene_tx=XM_019832903.2
B3 83339754 83339880
refGene_gn=HEATR5A;refGene_tx=XM_011283176.3;refGene_gn=HEATR5A;refGene_tx=XM_019832900.2;refGene_gn=HEATR5A;refGene_tx=XM_019832901.2;refGene_gn=HEATR5A;refGene_tx=XM_019832902.2;refGene_gn=HEATR5A;refGene_tx=XM_019832903.2
B3 83382177 83382503 refGene_gn=DTD2;refGene_tx=XM_003987531.5
B3 83386568 83386638 refGene_gn=DTD2;refGene_tx=XM_003987531.5
B3 83389873 83389984 refGene_gn=DTD2;refGene_tx=XM_003987531.5
B3 83404393 83405413 refGene_gn=GPR33;refGene_tx=XM_023256239.1
B3 83467848 83467956 refGene_gn=NUBPL;refGene_tx=XM_003987533.5
B3 83468511 83468659 refGene_gn=NUBPL;refGene_tx=XM_003987533.5
B3 83472412 83472447 refGene_gn=NUBPL;refGene_tx=XM_003987533.5
B3 83514914 83515005 refGene_gn=NUBPL;refGene_tx=XM_003987533.5
B3 83554100 83554140 refGene_gn=NUBPL;refGene_tx=XM_003987533.5
B3 83554230 83554321
refGene_gn=NUBPL;refGene_tx=XM_003987533.5;refGene_gn=NUBPL;refGene_tx=XM_019832904.2
B3 83631076 83631106
refGene_gn=NUBPL;refGene_tx=XM_011283179.3;refGene_gn=NUBPL;refGene_tx=XM_019832905.2;refGene_gn=NUBPL;refGene_tx=XM_019832907.2
B3 83645471 83645565
refGene_gn=NUBPL;refGene_tx=XM_003987533.5;refGene_gn=NUBPL;refGene_tx=XM_011283179.3;refGene_gn=NUBPL;refGene_tx=XM_019832904.2;refGene_gn=NUBPL;refGene_tx=XM_019832905.2;refGene_gn=NUBPL;refGene_tx=XM_019832907.2
B3 83661831 83661917

refGene_gn=NUBPL;refGene_tx=XM_003987533.5;refGene_gn=NUBPL;refGene_tx=XM_011283179.3;refGene_gn=NUBPL;refGene_tx=XM_019832904.2;refGene_gn=NUBPL;refGene_tx=XM_019832905.2;refGene_gn=NUBPL;refGene_tx=XM_019832907.2
B3 83677542 83677663
refGene_gn=NUBPL;refGene_tx=XM_003987533.5;refGene_gn=NUBPL;refGene_tx=XM_011283179.3;refGene_gn=NUBPL;refGene_tx=XM_019832904.2;refGene_gn=NUBPL;refGene_tx=XM_019832905.2;refGene_gn=NUBPL;refGene_tx=XM_019832907.2
B3 83680874 83680957
refGene_gn=NUBPL;refGene_tx=XM_003987533.5;refGene_gn=NUBPL;refGene_tx=XM_011283179.3;refGene_gn=NUBPL;refGene_tx=XM_019832904.2;refGene_gn=NUBPL;refGene_tx=XM_019832905.2;refGene_gn=NUBPL;refGene_tx=XM_019832907.2
B3 83688209 83688272
refGene_gn=NUBPL;refGene_tx=XM_003987533.5;refGene_gn=NUBPL;refGene_tx=XM_011283179.3;refGene_gn=NUBPL;refGene_tx=XM_019832904.2;refGene_gn=NUBPL;refGene_tx=XM_019832905.2;refGene_gn=NUBPL;refGene_tx=XM_019832907.2
B3 83896486 83900206
refGene_gn=ARHGAP5;refGene_tx=XM_019832909.2;refGene_gn=ARHGAP5;refGene_tx=XM_023255643.1;refGene_gn=ARHGAP5;refGene_tx=XM_023255644.1;refGene_gn=ARHGAP5;refGene_tx=XM_023255645.1;refGene_gn=ARHGAP5;refGene_tx=XM_023255646.1
B3 83925055 83925203
refGene_gn=ARHGAP5;refGene_tx=XM_019832909.2;refGene_gn=ARHGAP5;refGene_tx=XM_023255643.1;refGene_gn=ARHGAP5;refGene_tx=XM_023255644.1;refGene_gn=ARHGAP5;refGene_tx=XM_023255645.1;refGene_gn=ARHGAP5;refGene_tx=XM_023255646.1
B3 83949818 83949846 refGene_gn=ARHGAP5;refGene_tx=XM_023255646.1
B3 83952054 83952132
refGene_gn=ARHGAP5;refGene_tx=XM_019832909.2;refGene_gn=ARHGAP5;refGene_tx=XM_023255643.1;refGene_gn=ARHGAP5;refGene_tx=XM_023255644.1;refGene_gn=ARHGAP5;refGene_tx=XM_023255645.1;refGene_gn=ARHGAP5;refGene_tx=XM_023255646.1
B3 83954823 83954955
refGene_gn=ARHGAP5;refGene_tx=XM_019832909.2;refGene_gn=ARHGAP5;refGene_tx=XM_023255643.1;refGene_gn=ARHGAP5;refGene_tx=XM_023255644.1;refGene_gn=ARHGAP5;refGene_tx=XM_023255645.1
B3 83955679 83955785
refGene_gn=ARHGAP5;refGene_tx=XM_019832909.2;refGene_gn=ARHGAP5;refGene_tx=XM_023255643.1;refGene_gn=ARHGAP5;refGene_tx=XM_023255644.1;refGene_gn=ARHGAP5;refGene_tx=XM_023255645.1
B3 83960072 83960400
refGene_gn=ARHGAP5;refGene_tx=XM_019832909.2;refGene_gn=ARHGAP5;refGene_tx=XM_023255643.1;refGene_gn=ARHGAP5;refGene_tx=XM_023255644.1;refGene_gn=ARHGAP5;refGene_tx=XM_023255645.1
B3 84229184 84229508
refGene_gn=AKAP6;refGene_tx=XM_003987535.5;refGene_gn=AKAP6;refGene_tx=XM_011283183.3;refGene_gn=AKAP6;refGene_tx=XM_019832912.2;refGene_gn=

AKAP6;refGene_tx=XM_019832913.2;refGene_gn=AKAP6;refGene_tx=XM_0198329
14.2;refGene_gn=AKAP6;refGene_tx=XM_019832915.2
B3 84313752 84314004
refGene_gn=AKAP6;refGene_tx=XM_003987535.5;refGene_gn=AKAP6;refGene_tx
=XM_011283183.3;refGene_gn=AKAP6;refGene_tx=XM_019832912.2;refGene_gn=
AKAP6;refGene_tx=XM_019832913.2;refGene_gn=AKAP6;refGene_tx=XM_0198329
14.2;refGene_gn=AKAP6;refGene_tx=XM_019832915.2
B3 84323432 84325199
refGene_gn=AKAP6;refGene_tx=XM_003987535.5;refGene_gn=AKAP6;refGene_tx
=XM_011283183.3;refGene_gn=AKAP6;refGene_tx=XM_019832912.2;refGene_gn=
AKAP6;refGene_tx=XM_019832913.2;refGene_gn=AKAP6;refGene_tx=XM_0198329
14.2;refGene_gn=AKAP6;refGene_tx=XM_019832915.2
B3 84353538 84353661
refGene_gn=AKAP6;refGene_tx=XM_003987535.5;refGene_gn=AKAP6;refGene_tx
=XM_011283183.3;refGene_gn=AKAP6;refGene_tx=XM_019832912.2;refGene_gn=
AKAP6;refGene_tx=XM_019832913.2;refGene_gn=AKAP6;refGene_tx=XM_0198329
14.2;refGene_gn=AKAP6;refGene_tx=XM_019832915.2
B3 84372221 84372318
refGene_gn=AKAP6;refGene_tx=XM_003987535.5;refGene_gn=AKAP6;refGene_tx
=XM_011283183.3;refGene_gn=AKAP6;refGene_tx=XM_019832912.2;refGene_gn=
AKAP6;refGene_tx=XM_019832913.2;refGene_gn=AKAP6;refGene_tx=XM_0198329
14.2;refGene_gn=AKAP6;refGene_tx=XM_019832915.2
B3 84373455 84373622
refGene_gn=AKAP6;refGene_tx=XM_003987535.5;refGene_gn=AKAP6;refGene_tx
=XM_011283183.3;refGene_gn=AKAP6;refGene_tx=XM_019832912.2;refGene_gn=
AKAP6;refGene_tx=XM_019832913.2;refGene_gn=AKAP6;refGene_tx=XM_0198329
14.2;refGene_gn=AKAP6;refGene_tx=XM_019832915.2
B3 84452754 84452903
refGene_gn=AKAP6;refGene_tx=XM_003987535.5;refGene_gn=AKAP6;refGene_tx
=XM_011283183.3;refGene_gn=AKAP6;refGene_tx=XM_019832912.2;refGene_gn=
AKAP6;refGene_tx=XM_019832913.2;refGene_gn=AKAP6;refGene_tx=XM_0198329
14.2;refGene_gn=AKAP6;refGene_tx=XM_019832915.2
B3 84471493 84471614
refGene_gn=AKAP6;refGene_tx=XM_003987535.5;refGene_gn=AKAP6;refGene_tx
=XM_011283183.3;refGene_gn=AKAP6;refGene_tx=XM_019832912.2;refGene_gn=
AKAP6;refGene_tx=XM_019832913.2;refGene_gn=AKAP6;refGene_tx=XM_0198329
14.2;refGene_gn=AKAP6;refGene_tx=XM_019832915.2
B3 84505132 84505279
refGene_gn=AKAP6;refGene_tx=XM_003987535.5;refGene_gn=AKAP6;refGene_tx
=XM_011283183.3;refGene_gn=AKAP6;refGene_tx=XM_019832912.2;refGene_gn=
AKAP6;refGene_tx=XM_019832913.2;refGene_gn=AKAP6;refGene_tx=XM_0198329
14.2;refGene_gn=AKAP6;refGene_tx=XM_019832915.2
B3 84508326 84508551
refGene_gn=AKAP6;refGene_tx=XM_003987535.5;refGene_gn=AKAP6;refGene_tx
=XM_011283183.3;refGene_gn=AKAP6;refGene_tx=XM_019832912.2;refGene_gn=
AKAP6;refGene_tx=XM_019832913.2;refGene_gn=AKAP6;refGene_tx=XM_0198329
14.2;refGene_gn=AKAP6;refGene_tx=XM_019832915.2
B3 84539162 84539378
refGene_gn=AKAP6;refGene_tx=XM_003987535.5;refGene_gn=AKAP6;refGene_tx
=XM_011283183.3;refGene_gn=AKAP6;refGene_tx=XM_019832912.2;refGene_gn=

AKAP6;refGene_tx=XM_019832913.2;refGene_gn=AKAP6;refGene_tx=XM_0198329
14.2;refGene_gn=AKAP6;refGene_tx=XM_019832915.2
B3 84591013 84594385
refGene_gn=AKAP6;refGene_tx=XM_003987535.5;refGene_gn=AKAP6;refGene_tx
=XM_011283183.3;refGene_gn=AKAP6;refGene_tx=XM_019832912.2;refGene_gn=
AKAP6;refGene_tx=XM_019832913.2;refGene_gn=AKAP6;refGene_tx=XM_0198329
14.2;refGene_gn=AKAP6;refGene_tx=XM_019832915.2
B3 84682305 84682463
refGene_gn=NPAS3;refGene_tx=XM_023255647.1;refGene_gn=NPAS3;refGene_tx
=XM_023255648.1;refGene_gn=NPAS3;refGene_tx=XM_023255649.1;refGene_gn=
NPAS3;refGene_tx=XM_023255650.1;refGene_gn=NPAS3;refGene_tx=XM_0232556
51.1;refGene_gn=NPAS3;refGene_tx=XM_023255653.1
B3 84686751 84686801 refGene_gn=NPAS3;refGene_tx=XM_023255652.1
B3 84795121 84795211 refGene_gn=NPAS3;refGene_tx=XM_023255653.1
B3 84948253 84948498
refGene_gn=NPAS3;refGene_tx=XM_023255647.1;refGene_gn=NPAS3;refGene_tx
=XM_023255648.1;refGene_gn=NPAS3;refGene_tx=XM_023255649.1;refGene_gn=
NPAS3;refGene_tx=XM_023255650.1;refGene_gn=NPAS3;refGene_tx=XM_0232556
51.1;refGene_gn=NPAS3;refGene_tx=XM_023255652.1;refGene_gn=NPAS3;refGe
ne_tx=XM_023255653.1
B3 84948609 84948660
refGene_gn=NPAS3;refGene_tx=XM_023255647.1;refGene_gn=NPAS3;refGene_tx
=XM_023255648.1;refGene_gn=NPAS3;refGene_tx=XM_023255652.1
B3 85100329 85100412
refGene_gn=NPAS3;refGene_tx=XM_023255647.1;refGene_gn=NPAS3;refGene_tx
=XM_023255648.1;refGene_gn=NPAS3;refGene_tx=XM_023255649.1;refGene_gn=
NPAS3;refGene_tx=XM_023255650.1;refGene_gn=NPAS3;refGene_tx=XM_0232556
51.1;refGene_gn=NPAS3;refGene_tx=XM_023255652.1;refGene_gn=NPAS3;refGe
ne_tx=XM_023255653.1
B3 85300990 85301080
refGene_gn=NPAS3;refGene_tx=XM_023255647.1;refGene_gn=NPAS3;refGene_tx
=XM_023255648.1;refGene_gn=NPAS3;refGene_tx=XM_023255649.1;refGene_gn=
NPAS3;refGene_tx=XM_023255650.1;refGene_gn=NPAS3;refGene_tx=XM_0232556
51.1;refGene_gn=NPAS3;refGene_tx=XM_023255652.1;refGene_gn=NPAS3;refGe
ne_tx=XM_023255653.1
B3 85414719 85414894
refGene_gn=NPAS3;refGene_tx=XM_023255647.1;refGene_gn=NPAS3;refGene_tx
=XM_023255648.1;refGene_gn=NPAS3;refGene_tx=XM_023255649.1;refGene_gn=
NPAS3;refGene_tx=XM_023255650.1;refGene_gn=NPAS3;refGene_tx=XM_0232556
51.1;refGene_gn=NPAS3;refGene_tx=XM_023255652.1;refGene_gn=NPAS3;refGe
ne_tx=XM_023255653.1
B3 85475440 85475559
refGene_gn=NPAS3;refGene_tx=XM_023255647.1;refGene_gn=NPAS3;refGene_tx
=XM_023255648.1;refGene_gn=NPAS3;refGene_tx=XM_023255649.1;refGene_gn=
NPAS3;refGene_tx=XM_023255650.1;refGene_gn=NPAS3;refGene_tx=XM_0232556
51.1;refGene_gn=NPAS3;refGene_tx=XM_023255652.1;refGene_gn=NPAS3;refGe
ne_tx=XM_023255653.1
B3 85514116 85514310
refGene_gn=NPAS3;refGene_tx=XM_023255647.1;refGene_gn=NPAS3;refGene_tx
=XM_023255648.1;refGene_gn=NPAS3;refGene_tx=XM_023255649.1;refGene_gn=

NPAS3;refGene_tx=XM_023255650.1;refGene_gn=NPAS3;refGene_tx=XM_023255651.1;refGene_gn=NPAS3;refGene_tx=XM_023255652.1;refGene_gn=NPAS3;refGene_tx=XM_023255653.1
B3 85518963 85519070 refGene_gn=NPAS3;refGene_tx=XM_023255647.1;refGene_gn=NPAS3;refGene_tx=XM_023255648.1;refGene_gn=NPAS3;refGene_tx=XM_023255649.1;refGene_gn=NPAS3;refGene_tx=XM_023255650.1;refGene_gn=NPAS3;refGene_tx=XM_023255651.1;refGene_gn=NPAS3;refGene_tx=XM_023255652.1;refGene_gn=NPAS3;refGene_tx=XM_023255653.1
B3 85534055 85534203 refGene_gn=NPAS3;refGene_tx=XM_023255647.1;refGene_gn=NPAS3;refGene_tx=XM_023255648.1;refGene_gn=NPAS3;refGene_tx=XM_023255649.1;refGene_gn=NPAS3;refGene_tx=XM_023255650.1;refGene_gn=NPAS3;refGene_tx=XM_023255651.1;refGene_gn=NPAS3;refGene_tx=XM_023255652.1;refGene_gn=NPAS3;refGene_tx=XM_023255653.1
B3 85537776 85537901 refGene_gn=NPAS3;refGene_tx=XM_023255647.1;refGene_gn=NPAS3;refGene_tx=XM_023255648.1;refGene_gn=NPAS3;refGene_tx=XM_023255649.1;refGene_gn=NPAS3;refGene_tx=XM_023255650.1;refGene_gn=NPAS3;refGene_tx=XM_023255651.1;refGene_gn=NPAS3;refGene_tx=XM_023255652.1;refGene_gn=NPAS3;refGene_tx=XM_023255653.1
B3 85539682 85541043 refGene_gn=NPAS3;refGene_tx=XM_023255647.1;refGene_gn=NPAS3;refGene_tx=XM_023255648.1;refGene_gn=NPAS3;refGene_tx=XM_023255649.1;refGene_gn=NPAS3;refGene_tx=XM_023255650.1;refGene_gn=NPAS3;refGene_tx=XM_023255651.1;refGene_gn=NPAS3;refGene_tx=XM_023255652.1;refGene_gn=NPAS3;refGene_tx=XM_023255653.1
B3 85623251 85623283 refGene_gn=EGLN3;refGene_tx=XM_011283184.3
B3 85624746 85624820 refGene_gn=EGLN3;refGene_tx=XM_011283184.3
B3 85626799 85626936 refGene_gn=EGLN3;refGene_tx=XM_011283184.3
B3 85628743 85628863 refGene_gn=EGLN3;refGene_tx=XM_011283184.3
B3 85650519 85650876 refGene_gn=EGLN3;refGene_tx=XM_011283184.3
B3 86061196 86061670 refGene_gn=LOC101088757;refGene_tx=XM_011283185.3
B3 86092782 86092886 refGene_gn=SPTSSA;refGene_tx=XM_023255654.1
B3 86114182 86114285 refGene_gn=SPTSSA;refGene_tx=XM_023255654.1
B3 86150426 86150703 refGene_gn=EAPP;refGene_tx=XM_003987536.5
B3 86157904 86158018 refGene_gn=EAPP;refGene_tx=XM_003987536.5
B3 86160955 86161073 refGene_gn=EAPP;refGene_tx=XM_003987536.5
B3 86162233 86162302 refGene_gn=EAPP;refGene_tx=XM_003987536.5
B3 86162304 86162328 refGene_gn=EAPP;refGene_tx=XM_003987536.5
B3 86163877 86164059 refGene_gn=EAPP;refGene_tx=XM_003987536.5
B3 86168412 86168486 refGene_gn=EAPP;refGene_tx=XM_003987536.5
B3 86223444 86223498 refGene_gn=SNX6;refGene_tx=XM_023255655.1
B3 86231245 86231295 refGene_gn=SNX6;refGene_tx=XM_023255655.1
B3 86231297 86231330 refGene_gn=SNX6;refGene_tx=XM_023255655.1
B3 86231419 86231579 refGene_gn=SNX6;refGene_tx=XM_023255655.1
B3 86234423 86234510 refGene_gn=SNX6;refGene_tx=XM_023255655.1
B3 86239484 86239524 refGene_gn=SNX6;refGene_tx=XM_023255655.1
B3 86242207 86242283 refGene_gn=SNX6;refGene_tx=XM_023255655.1

B3 86251634 86251740 refGene_gn=SNX6;refGene_tx=XM_023255655.1
B3 86253450 86253546 refGene_gn=SNX6;refGene_tx=XM_023255655.1
B3 86261604 86261728 refGene_gn=SNX6;refGene_tx=XM_023255655.1
B3 86263720 86263842 refGene_gn=SNX6;refGene_tx=XM_023255655.1
B3 86266440 86266551 refGene_gn=SNX6;refGene_tx=XM_023255655.1
B3 86267422 86267527 refGene_gn=SNX6;refGene_tx=XM_023255655.1
B3 86283336 86283384 refGene_gn=SNX6;refGene_tx=XM_023255655.1
B3 86283540 86283546 refGene_gn=SNX6;refGene_tx=XM_023255655.1
B3 86343122 86343235 refGene_gn=CFL2;refGene_tx=XM_011283186.3
B3 86343317 86343394 refGene_gn=CFL2;refGene_tx=XM_011283186.3
B3 86343509 86343817 refGene_gn=CFL2;refGene_tx=XM_011283186.3
B3 86344891 86344894 refGene_gn=CFL2;refGene_tx=XM_011283186.3
B3 86371171 86371368 refGene_gn=BAZ1A;refGene_tx=XM_023255656.1
B3 86372162 86372250 refGene_gn=BAZ1A;refGene_tx=XM_023255656.1
B3 86373470 86373613 refGene_gn=BAZ1A;refGene_tx=XM_023255656.1
B3 86376089 86376559 refGene_gn=BAZ1A;refGene_tx=XM_023255656.1
B3 86378128 86378355 refGene_gn=BAZ1A;refGene_tx=XM_023255656.1
B3 86378446 86378694 refGene_gn=BAZ1A;refGene_tx=XM_023255656.1
B3 86386393 86386542 refGene_gn=BAZ1A;refGene_tx=XM_023255656.1
B3 86387623 86387778 refGene_gn=BAZ1A;refGene_tx=XM_023255656.1
B3 86388495 86388659 refGene_gn=BAZ1A;refGene_tx=XM_023255656.1
B3 86390256 86390853 refGene_gn=BAZ1A;refGene_tx=XM_023255656.1
B3 86394963 86395088 refGene_gn=BAZ1A;refGene_tx=XM_023255656.1
B3 86399507 86399578 refGene_gn=BAZ1A;refGene_tx=XM_023255656.1
B3 86399579 86399622 refGene_gn=BAZ1A;refGene_tx=XM_023255656.1
B3 86399881 86400047 refGene_gn=BAZ1A;refGene_tx=XM_023255656.1
B3 86401410 86401635 refGene_gn=BAZ1A;refGene_tx=XM_023255656.1
B3 86401760 86401856 refGene_gn=BAZ1A;refGene_tx=XM_023255656.1
B3 86407063 86407210 refGene_gn=BAZ1A;refGene_tx=XM_023255656.1
B3 86408531 86408670 refGene_gn=BAZ1A;refGene_tx=XM_023255656.1
B3 86409494 86409590 refGene_gn=BAZ1A;refGene_tx=XM_023255656.1
B3 86412921 86413088 refGene_gn=BAZ1A;refGene_tx=XM_023255656.1
B3 86413761 86413861 refGene_gn=BAZ1A;refGene_tx=XM_023255656.1
B3 86414876 86415011 refGene_gn=BAZ1A;refGene_tx=XM_023255656.1
B3 86420006 86420094 refGene_gn=BAZ1A;refGene_tx=XM_023255656.1
B3 86421888 86421990 refGene_gn=BAZ1A;refGene_tx=XM_023255656.1
B3 86431782 86431926 refGene_gn=BAZ1A;refGene_tx=XM_023255656.1
B3 86450792 86451071 refGene_gn=BAZ1A;refGene_tx=XM_023255656.1
B3 86464284 86464397 refGene_gn=BAZ1A;refGene_tx=XM_023255656.1
B3 86539404 86539590
refGene_gn=LOC101083598;refGene_tx=XM_019832923.2;refGene_gn=LOC101083
598;refGene_tx=XM_019832926.2
B3 86547302 86547419
refGene_gn=LOC101083598;refGene_tx=XM_003987585.5
B3 86549886 86549977
refGene_gn=LOC101083598;refGene_tx=XM_003987585.5;refGene_gn=LOC101083
598;refGene_tx=XM_019832923.2;refGene_gn=LOC101083598;refGene_tx=XM_01
9832924.2;refGene_gn=LOC101083598;refGene_tx=XM_019832926.2
B3 86553677 86553784
refGene_gn=LOC101083598;refGene_tx=XM_003987585.5;refGene_gn=LOC101083

598;refGene_tx=XM_019832923.2;refGene_gn=LOC101083598;refGene_tx=XM_019832924.2;refGene_gn=LOC101083598;refGene_tx=XM_019832926.2
B3 86561683 86561794
refGene_gn=LOC101083598;refGene_tx=XM_003987585.5;refGene_gn=LOC101083598;refGene_tx=XM_019832923.2;refGene_gn=LOC101083598;refGene_tx=XM_019832924.2;refGene_gn=LOC101083598;refGene_tx=XM_019832926.2
B3 86563490 86563612
refGene_gn=LOC101083598;refGene_tx=XM_003987585.5;refGene_gn=LOC101083598;refGene_tx=XM_019832923.2;refGene_gn=LOC101083598;refGene_tx=XM_019832924.2;refGene_gn=LOC101083598;refGene_tx=XM_019832926.2
B3 86567192 86567331
refGene_gn=LOC101083598;refGene_tx=XM_003987585.5;refGene_gn=LOC101083598;refGene_tx=XM_019832923.2;refGene_gn=LOC101083598;refGene_tx=XM_019832924.2;refGene_gn=LOC101083598;refGene_tx=XM_019832926.2
B3 86568904 86569026
refGene_gn=LOC101083598;refGene_tx=XM_003987585.5;refGene_gn=LOC101083598;refGene_tx=XM_019832923.2;refGene_gn=LOC101083598;refGene_tx=XM_019832924.2;refGene_gn=LOC101083598;refGene_tx=XM_019832926.2
B3 86589978 86590261
refGene_gn=LOC101083598;refGene_tx=XM_003987585.5;refGene_gn=LOC101083598;refGene_tx=XM_019832923.2;refGene_gn=LOC101083598;refGene_tx=XM_019832924.2;refGene_gn=LOC101083598;refGene_tx=XM_019832926.2
B3 86596739 86596770 refGene_gn=SRP54;refGene_tx=XM_003987538.5
B3 86601291 86601404
refGene_gn=SRP54;refGene_tx=XM_003987537.5;refGene_gn=SRP54;refGene_tx=XM_003987538.5;refGene_gn=SRP54;refGene_tx=XM_006932802.4
B3 86604207 86604299
refGene_gn=SRP54;refGene_tx=XM_003987537.5;refGene_gn=SRP54;refGene_tx=XM_003987538.5;refGene_gn=SRP54;refGene_tx=XM_006932802.4
B3 86605255 86605340
refGene_gn=SRP54;refGene_tx=XM_003987537.5;refGene_gn=SRP54;refGene_tx=XM_003987538.5;refGene_gn=SRP54;refGene_tx=XM_006932802.4
B3 86606895 86607000
refGene_gn=SRP54;refGene_tx=XM_003987537.5;refGene_gn=SRP54;refGene_tx=XM_003987538.5;refGene_gn=SRP54;refGene_tx=XM_006932802.4
B3 86609104 86609171
refGene_gn=SRP54;refGene_tx=XM_003987537.5;refGene_gn=SRP54;refGene_tx=XM_003987538.5;refGene_gn=SRP54;refGene_tx=XM_006932802.4
B3 86609251 86609309
refGene_gn=SRP54;refGene_tx=XM_003987537.5;refGene_gn=SRP54;refGene_tx=XM_003987538.5;refGene_gn=SRP54;refGene_tx=XM_006932802.4
B3 86613210 86613361
refGene_gn=SRP54;refGene_tx=XM_003987537.5;refGene_gn=SRP54;refGene_tx=XM_003987538.5;refGene_gn=SRP54;refGene_tx=XM_006932802.4
B3 86614460 86614609
refGene_gn=SRP54;refGene_tx=XM_003987537.5;refGene_gn=SRP54;refGene_tx=XM_003987538.5;refGene_gn=SRP54;refGene_tx=XM_006932802.4
B3 86615422 86615523
refGene_gn=SRP54;refGene_tx=XM_003987537.5;refGene_gn=SRP54;refGene_tx=XM_003987538.5;refGene_gn=SRP54;refGene_tx=XM_006932802.4

B3 86616014 86616101
refGene_gn=SRP54;refGene_tx=XM_003987537.5;refGene_gn=SRP54;refGene_tx=XM_003987538.5;refGene_gn=SRP54;refGene_tx=XM_006932802.4
B3 86619489 86619563
refGene_gn=SRP54;refGene_tx=XM_003987537.5;refGene_gn=SRP54;refGene_tx=XM_003987538.5;refGene_gn=SRP54;refGene_tx=XM_006932802.4
B3 86619781 86619890
refGene_gn=SRP54;refGene_tx=XM_003987537.5;refGene_gn=SRP54;refGene_tx=XM_003987538.5;refGene_gn=SRP54;refGene_tx=XM_006932802.4
B3 86625259 86625430
refGene_gn=SRP54;refGene_tx=XM_003987537.5;refGene_gn=SRP54;refGene_tx=XM_003987538.5;refGene_gn=SRP54;refGene_tx=XM_006932802.4
B3 86627838 86627934
refGene_gn=SRP54;refGene_tx=XM_003987537.5;refGene_gn=SRP54;refGene_tx=XM_003987538.5;refGene_gn=SRP54;refGene_tx=XM_006932802.4
B3 86629025 86629117
refGene_gn=SRP54;refGene_tx=XM_003987537.5;refGene_gn=SRP54;refGene_tx=XM_003987538.5;refGene_gn=SRP54;refGene_tx=XM_006932802.4
B3 86636827 86636884 refGene_gn=FAM177A1;refGene_tx=XM_011283187.3
B3 86636885 86636990 refGene_gn=FAM177A1;refGene_tx=XM_011283187.3
B3 86642632 86642806 refGene_gn=FAM177A1;refGene_tx=XM_011283187.3
B3 86656881 86656948 refGene_gn=FAM177A1;refGene_tx=XM_011283187.3
B3 86658016 86658114 refGene_gn=FAM177A1;refGene_tx=XM_011283187.3
B3 86660263 86660470 refGene_gn=FAM177A1;refGene_tx=XM_011283187.3
B3 86663069 86663258
refGene_gn=PPP2R3C;refGene_tx=XM_011283190.3;refGene_gn=PPP2R3C;refGene_tx=XM_019832920.2;refGene_gn=PPP2R3C;refGene_tx=XM_019832921.2;refGene_gn=PPP2R3C;refGene_tx=XM_019832922.2
B3 86666420 86666480
refGene_gn=PPP2R3C;refGene_tx=XM_011283190.3;refGene_gn=PPP2R3C;refGene_tx=XM_019832920.2;refGene_gn=PPP2R3C;refGene_tx=XM_019832921.2;refGene_gn=PPP2R3C;refGene_tx=XM_019832922.2
B3 86669512 86669650
refGene_gn=PPP2R3C;refGene_tx=XM_011283190.3;refGene_gn=PPP2R3C;refGene_tx=XM_019832920.2;refGene_gn=PPP2R3C;refGene_tx=XM_019832921.2;refGene_gn=PPP2R3C;refGene_tx=XM_019832922.2
B3 86671783 86671920
refGene_gn=PPP2R3C;refGene_tx=XM_011283190.3;refGene_gn=PPP2R3C;refGene_tx=XM_019832920.2;refGene_gn=PPP2R3C;refGene_tx=XM_019832921.2;refGene_gn=PPP2R3C;refGene_tx=XM_019832922.2
B3 86672641 86672717
refGene_gn=PPP2R3C;refGene_tx=XM_011283190.3;refGene_gn=PPP2R3C;refGene_tx=XM_019832920.2;refGene_gn=PPP2R3C;refGene_tx=XM_019832921.2;refGene_gn=PPP2R3C;refGene_tx=XM_019832922.2
B3 86672794 86672850
refGene_gn=PPP2R3C;refGene_tx=XM_011283190.3;refGene_gn=PPP2R3C;refGene_tx=XM_019832920.2;refGene_gn=PPP2R3C;refGene_tx=XM_019832921.2;refGene_gn=PPP2R3C;refGene_tx=XM_019832922.2
B3 86673251 86673384
refGene_gn=PPP2R3C;refGene_tx=XM_011283190.3;refGene_gn=PPP2R3C;refGen

e_tx=XM_019832920.2;refGene_gn=PPP2R3C;refGene_tx=XM_019832921.2;refGene_gn=PPP2R3C;refGene_tx=XM_019832922.2
B3 86676512 86676583
refGene_gn=PPP2R3C;refGene_tx=XM_011283190.3;refGene_gn=PPP2R3C;refGene_tx=XM_019832920.2;refGene_gn=PPP2R3C;refGene_tx=XM_019832921.2;refGene_gn=PPP2R3C;refGene_tx=XM_019832922.2
B3 86678209 86678307
refGene_gn=PPP2R3C;refGene_tx=XM_011283190.3;refGene_gn=PPP2R3C;refGene_tx=XM_019832920.2;refGene_gn=PPP2R3C;refGene_tx=XM_019832921.2;refGene_gn=PPP2R3C;refGene_tx=XM_019832922.2
B3 86678746 86678859
refGene_gn=PPP2R3C;refGene_tx=XM_011283190.3;refGene_gn=PPP2R3C;refGene_tx=XM_019832920.2;refGene_gn=PPP2R3C;refGene_tx=XM_019832921.2;refGene_gn=PPP2R3C;refGene_tx=XM_019832922.2
B3 86679502 86679607
refGene_gn=PPP2R3C;refGene_tx=XM_011283190.3;refGene_gn=PPP2R3C;refGene_tx=XM_019832920.2
B3 86682908 86683036
refGene_gn=PPP2R3C;refGene_tx=XM_011283190.3;refGene_gn=PPP2R3C;refGene_tx=XM_019832920.2
B3 86688473 86688531
refGene_gn=PPP2R3C;refGene_tx=XM_011283190.3;refGene_gn=PPP2R3C;refGene_tx=XM_019832920.2
B3 86689835 86690821 refGene_gn=KIAA0391;refGene_tx=XM_003987540.5
B3 86692273 86692284 refGene_gn=KIAA0391;refGene_tx=XM_011283193.3
B3 86692688 86692736
refGene_gn=KIAA0391;refGene_tx=XM_003987540.5;refGene_gn=KIAA0391;refGene_tx=XM_011283193.3
B3 86693280 86693413
refGene_gn=KIAA0391;refGene_tx=XM_003987540.5;refGene_gn=KIAA0391;refGene_tx=XM_011283193.3;refGene_gn=KIAA0391;refGene_tx=XM_011283194.2
B3 86736003 86736111
refGene_gn=KIAA0391;refGene_tx=XM_003987540.5;refGene_gn=KIAA0391;refGene_tx=XM_011283193.3;refGene_gn=KIAA0391;refGene_tx=XM_011283194.2
B3 86815249 86815398
refGene_gn=KIAA0391;refGene_tx=XM_003987540.5;refGene_gn=KIAA0391;refGene_tx=XM_011283193.3;refGene_gn=KIAA0391;refGene_tx=XM_011283194.2
B3 86818059 86818255
refGene_gn=KIAA0391;refGene_tx=XM_003987540.5;refGene_gn=KIAA0391;refGene_tx=XM_011283193.3;refGene_gn=KIAA0391;refGene_tx=XM_011283194.2
B3 86820027 86820159
refGene_gn=KIAA0391;refGene_tx=XM_003987540.5;refGene_gn=KIAA0391;refGene_tx=XM_011283193.3;refGene_gn=KIAA0391;refGene_tx=XM_011283194.2
B3 86826215 86826234 refGene_gn=PSMA6;refGene_tx=XM_019832930.2
B3 86838118 86838194
refGene_gn=PSMA6;refGene_tx=XM_003987541.5;refGene_gn=PSMA6;refGene_tx=XM_006932803.3
B3 86848683 86848778
refGene_gn=PSMA6;refGene_tx=XM_003987541.5;refGene_gn=PSMA6;refGene_tx=XM_006932803.3;refGene_gn=PSMA6;refGene_tx=XM_019832930.2

B3 86849253 86849335
refGene_gn=PSMA6;refGene_tx=XM_003987541.5;refGene_gn=PSMA6;refGene_tx=XM_006932803.3;refGene_gn=PSMA6;refGene_tx=XM_019832930.2
B3 86850055 86850211
refGene_gn=PSMA6;refGene_tx=XM_003987541.5;refGene_gn=PSMA6;refGene_tx=XM_006932803.3;refGene_gn=PSMA6;refGene_tx=XM_019832930.2
B3 86853075 86853254
refGene_gn=PSMA6;refGene_tx=XM_003987541.5;refGene_gn=PSMA6;refGene_tx=XM_006932803.3;refGene_gn=PSMA6;refGene_tx=XM_019832930.2
B3 86854556 86854651
refGene_gn=PSMA6;refGene_tx=XM_003987541.5;refGene_gn=PSMA6;refGene_tx=XM_019832930.2
B3 86856259 86856317
refGene_gn=PSMA6;refGene_tx=XM_003987541.5;refGene_gn=PSMA6;refGene_tx=XM_006932803.3;refGene_gn=PSMA6;refGene_tx=XM_019832930.2
B3 86921614 86921662 refGene_gn=NFKBIA;refGene_tx=XM_003987542.5
B3 86921982 86922243 refGene_gn=NFKBIA;refGene_tx=XM_003987542.5
B3 86922370 86922459 refGene_gn=NFKBIA;refGene_tx=XM_003987542.5
B3 86922752 86922963 refGene_gn=NFKBIA;refGene_tx=XM_003987542.5
B3 86923298 86923407 refGene_gn=NFKBIA;refGene_tx=XM_003987542.5
B3 86924005 86924232 refGene_gn=NFKBIA;refGene_tx=XM_003987542.5
B3 86963110 86963318
refGene_gn=LOC101090873;refGene_tx=XM_019832933.2
B3 86963319 86963990
refGene_gn=LOC101090873;refGene_tx=XM_019832933.2
B3 87042353 87044015 refGene_gn=INSM2;refGene_tx=XM_004001359.4
B3 87047362 87047511 refGene_gn=RALGAPA1;refGene_tx=XM_019832932.2
B3 87048962 87049016 refGene_gn=RALGAPA1;refGene_tx=XM_019832932.2
B3 87049445 87049448 refGene_gn=RALGAPA1;refGene_tx=XM_019832932.2
B3 87056095 87056220 refGene_gn=RALGAPA1;refGene_tx=XM_019832932.2
B3 87077648 87077776 refGene_gn=RALGAPA1;refGene_tx=XM_019832932.2
B3 87079441 87079600 refGene_gn=RALGAPA1;refGene_tx=XM_019832932.2
B3 87106051 87106207 refGene_gn=RALGAPA1;refGene_tx=XM_019832932.2
B3 87114293 87114417 refGene_gn=RALGAPA1;refGene_tx=XM_019832932.2
B3 87135563 87135635 refGene_gn=RALGAPA1;refGene_tx=XM_019832932.2
B3 87138199 87139064 refGene_gn=RALGAPA1;refGene_tx=XM_019832932.2
B3 87147522 87147706 refGene_gn=RALGAPA1;refGene_tx=XM_019832932.2
B3 87148112 87148247 refGene_gn=RALGAPA1;refGene_tx=XM_019832932.2
B3 87160772 87160841 refGene_gn=RALGAPA1;refGene_tx=XM_019832932.2
B3 87166896 87167007 refGene_gn=RALGAPA1;refGene_tx=XM_019832932.2
B3 87168258 87168367 refGene_gn=RALGAPA1;refGene_tx=XM_019832932.2
B3 87172172 87172231 refGene_gn=RALGAPA1;refGene_tx=XM_019832932.2
B3 87175251 87175377 refGene_gn=RALGAPA1;refGene_tx=XM_019832932.2
B3 87178129 87178258 refGene_gn=RALGAPA1;refGene_tx=XM_019832932.2
B3 87179663 87179819 refGene_gn=RALGAPA1;refGene_tx=XM_019832932.2
B3 87180578 87180677 refGene_gn=RALGAPA1;refGene_tx=XM_019832932.2
B3 87180942 87181091 refGene_gn=RALGAPA1;refGene_tx=XM_019832932.2
B3 87181093 87181132 refGene_gn=RALGAPA1;refGene_tx=XM_019832932.2
B3 87183704 87183857 refGene_gn=RALGAPA1;refGene_tx=XM_019832932.2
B3 87195108 87195285 refGene_gn=RALGAPA1;refGene_tx=XM_019832932.2

B3 87196735 87196952 refGene_gn=RALGAPA1;refGene_tx=XM_019832932.2
B3 87197948 87198073 refGene_gn=RALGAPA1;refGene_tx=XM_019832932.2
B3 87199835 87201383 refGene_gn=RALGAPA1;refGene_tx=XM_019832932.2
B3 87209619 87209760 refGene_gn=RALGAPA1;refGene_tx=XM_019832932.2
B3 87214926 87215088 refGene_gn=RALGAPA1;refGene_tx=XM_019832932.2
B3 87216117 87216355 refGene_gn=RALGAPA1;refGene_tx=XM_019832932.2
B3 87218330 87218460 refGene_gn=RALGAPA1;refGene_tx=XM_019832932.2
B3 87221239 87221388 refGene_gn=RALGAPA1;refGene_tx=XM_019832932.2
B3 87226201 87226339 refGene_gn=RALGAPA1;refGene_tx=XM_019832932.2
B3 87237280 87237478 refGene_gn=RALGAPA1;refGene_tx=XM_019832932.2
B3 87248741 87248981 refGene_gn=RALGAPA1;refGene_tx=XM_019832932.2
B3 87251681 87251890 refGene_gn=RALGAPA1;refGene_tx=XM_019832932.2
B3 87253167 87253306 refGene_gn=RALGAPA1;refGene_tx=XM_019832932.2
B3 87255465 87255581 refGene_gn=RALGAPA1;refGene_tx=XM_019832932.2
B3 87257037 87257215 refGene_gn=RALGAPA1;refGene_tx=XM_019832932.2
B3 87258762 87258806 refGene_gn=RALGAPA1;refGene_tx=XM_019832932.2
B3 87263266 87263324 refGene_gn=RALGAPA1;refGene_tx=XM_019832932.2
B3 87269074 87269124 refGene_gn=RALGAPA1;refGene_tx=XM_019832932.2
B3 87269667 87269703 refGene_gn=RALGAPA1;refGene_tx=XM_019832932.2
B3 87269705 87269777 refGene_gn=RALGAPA1;refGene_tx=XM_019832932.2
B3 87317279 87317385 refGene_gn=RALGAPA1;refGene_tx=XM_019832932.2
B3 87340825 87340967
refGene_gn=BRMS1L;refGene_tx=XM_003987543.4;refGene_gn=BRMS1L;refGene_tx=XM_006932804.3
B3 87343959 87344050
refGene_gn=BRMS1L;refGene_tx=XM_003987543.4;refGene_gn=BRMS1L;refGene_tx=XM_006932804.3
B3 87348620 87348748
refGene_gn=BRMS1L;refGene_tx=XM_003987543.4;refGene_gn=BRMS1L;refGene_tx=XM_006932804.3
B3 87350798 87350878
refGene_gn=BRMS1L;refGene_tx=XM_003987543.4;refGene_gn=BRMS1L;refGene_tx=XM_006932804.3
B3 87362995 87363092
refGene_gn=BRMS1L;refGene_tx=XM_003987543.4;refGene_gn=BRMS1L;refGene_tx=XM_006932804.3
B3 87364316 87364400
refGene_gn=BRMS1L;refGene_tx=XM_003987543.4;refGene_gn=BRMS1L;refGene_tx=XM_006932804.3
B3 87365529 87365594
refGene_gn=BRMS1L;refGene_tx=XM_003987543.4;refGene_gn=BRMS1L;refGene_tx=XM_006932804.3
B3 87366310 87366350
refGene_gn=BRMS1L;refGene_tx=XM_003987543.4;refGene_gn=BRMS1L;refGene_tx=XM_006932804.3
B3 87366795 87366943
refGene_gn=BRMS1L;refGene_tx=XM_003987543.4;refGene_gn=BRMS1L;refGene_tx=XM_006932804.3
B3 87368802 87368920
refGene_gn=BRMS1L;refGene_tx=XM_003987543.4;refGene_gn=BRMS1L;refGene_

tx=XM_006932804.3
B3	87759881 87759989
refGene_gn=MBIP;refGene_tx=XM_003987544.4;refGene_gn=MBIP;refGene_tx=XM_006932806.2;refGene_gn=MBIP;refGene_tx=XM_006932807.3;refGene_gn=MBIP;refGene_tx=XM_006932808.2;refGene_gn=MBIP;refGene_tx=XM_006932809.2;refGene_gn=MBIP;refGene_tx=XM_006932810.3;refGene_gn=MBIP;refGene_tx=XM_006932811.3
B3	87761107 87761157
refGene_gn=MBIP;refGene_tx=XM_019832939.1;refGene_gn=MBIP;refGene_tx=XM_019832940.1
B3	87761249 87761288
refGene_gn=MBIP;refGene_tx=XM_003987544.4;refGene_gn=MBIP;refGene_tx=XM_006932806.2;refGene_gn=MBIP;refGene_tx=XM_006932807.3;refGene_gn=MBIP;refGene_tx=XM_006932808.2;refGene_gn=MBIP;refGene_tx=XM_006932809.2;refGene_gn=MBIP;refGene_tx=XM_006932810.3;refGene_gn=MBIP;refGene_tx=XM_006932811.3;refGene_gn=MBIP;refGene_tx=XM_019832939.1;refGene_gn=MBIP;refGene_tx=XM_019832940.1
B3	87767905 87768003
refGene_gn=MBIP;refGene_tx=XM_003987544.4;refGene_gn=MBIP;refGene_tx=XM_006932806.2;refGene_gn=MBIP;refGene_tx=XM_006932807.3;refGene_gn=MBIP;refGene_tx=XM_006932808.2;refGene_gn=MBIP;refGene_tx=XM_006932809.2;refGene_gn=MBIP;refGene_tx=XM_019832939.1
B3	87771613 87771766
refGene_gn=MBIP;refGene_tx=XM_003987544.4;refGene_gn=MBIP;refGene_tx=XM_006932806.2;refGene_gn=MBIP;refGene_tx=XM_006932807.3;refGene_gn=MBIP;refGene_tx=XM_006932808.2;refGene_gn=MBIP;refGene_tx=XM_006932810.3;refGene_gn=MBIP;refGene_tx=XM_006932811.3;refGene_gn=MBIP;refGene_tx=XM_019832939.1;refGene_gn=MBIP;refGene_tx=XM_019832940.1
B3	87771994 87772060
refGene_gn=MBIP;refGene_tx=XM_003987544.4;refGene_gn=MBIP;refGene_tx=XM_006932806.2;refGene_gn=MBIP;refGene_tx=XM_006932807.3;refGene_gn=MBIP;refGene_tx=XM_006932808.2;refGene_gn=MBIP;refGene_tx=XM_006932809.2;refGene_gn=MBIP;refGene_tx=XM_006932810.3;refGene_gn=MBIP;refGene_tx=XM_006932811.3;refGene_gn=MBIP;refGene_tx=XM_019832939.1;refGene_gn=MBIP;refGene_tx=XM_019832940.1
B3	87774820 87774917
refGene_gn=MBIP;refGene_tx=XM_003987544.4;refGene_gn=MBIP;refGene_tx=XM_006932806.2;refGene_gn=MBIP;refGene_tx=XM_006932807.3;refGene_gn=MBIP;refGene_tx=XM_006932808.2;refGene_gn=MBIP;refGene_tx=XM_006932809.2;refGene_gn=MBIP;refGene_tx=XM_006932810.3;refGene_gn=MBIP;refGene_tx=XM_006932811.3;refGene_gn=MBIP;refGene_tx=XM_019832939.1;refGene_gn=MBIP;refGene_tx=XM_019832940.1
B3	87775006 87775231
refGene_gn=MBIP;refGene_tx=XM_003987544.4;refGene_gn=MBIP;refGene_tx=XM_006932806.2;refGene_gn=MBIP;refGene_tx=XM_006932807.3;refGene_gn=MBIP;refGene_tx=XM_006932808.2;refGene_gn=MBIP;refGene_tx=XM_006932809.2;refGene_gn=MBIP;refGene_tx=XM_006932810.3;refGene_gn=MBIP;refGene_tx=XM_006932811.3;refGene_gn=MBIP;refGene_tx=XM_019832939.1;refGene_gn=MBIP;refGene_tx=XM_019832940.1
B3	87777005 87777125

refGene_gn=MBIP;refGene_tx=XM_003987544.4;refGene_gn=MBIP;refGene_tx=X
M_006932806.2;refGene_gn=MBIP;refGene_tx=XM_006932807.3;refGene_gn=MBI
P;refGene_tx=XM_006932809.2;refGene_gn=MBIP;refGene_tx=XM_006932810.3;
refGene_gn=MBIP;refGene_tx=XM_006932811.3;refGene_gn=MBIP;refGene_tx=X
M_019832939.1;refGene_gn=MBIP;refGene_tx=XM_019832940.1
B3 87780903 87781032
refGene_gn=MBIP;refGene_tx=XM_003987544.4;refGene_gn=MBIP;refGene_tx=X
M_006932806.2;refGene_gn=MBIP;refGene_tx=XM_006932807.3;refGene_gn=MBI
P;refGene_tx=XM_006932808.2;refGene_gn=MBIP;refGene_tx=XM_006932809.2;
refGene_gn=MBIP;refGene_tx=XM_006932810.3;refGene_gn=MBIP;refGene_tx=X
M_006932811.3;refGene_gn=MBIP;refGene_tx=XM_019832939.1;refGene_gn=MBI
P;refGene_tx=XM_019832940.1
B3 87949378 87950115
refGene_gn=NKX2-1;refGene_tx=XM_019832936.2;refGene_gn=NKX2-1;refGene_
tx=XM_019832937.2
B3 87951084 87951457
refGene_gn=NKX2-1;refGene_tx=XM_019832936.2;refGene_gn=NKX2-1;refGene_
tx=XM_019832937.2
B3 88013525 88014091
refGene_gn=NKX2-8;refGene_tx=XM_011283196.3;refGene_gn=NKX2-8;refGene_
tx=XM_019832941.2
B3 88014755 88015020
refGene_gn=NKX2-8;refGene_tx=XM_011283196.3;refGene_gn=NKX2-8;refGene_
tx=XM_019832941.2
B3 88015220 88015319
refGene_gn=NKX2-8;refGene_tx=XM_011283196.3;refGene_gn=NKX2-8;refGene_
tx=XM_019832941.2
B3 88092327 88092331
refGene_gn=PAX9;refGene_tx=XM_023255659.1;refGene_gn=PAX9;refGene_tx=X
M_023255660.1
B3 88093099 88093726
refGene_gn=PAX9;refGene_tx=XM_023255659.1;refGene_gn=PAX9;refGene_tx=X
M_023255660.1
B3 88096712 88096855
refGene_gn=PAX9;refGene_tx=XM_023255659.1;refGene_gn=PAX9;refGene_tx=X
M_023255660.1
B3 88105916 88106171
refGene_gn=PAX9;refGene_tx=XM_023255659.1;refGene_gn=PAX9;refGene_tx=X
M_023255660.1
B3 88106340 88106354 refGene_gn=PAX9;refGene_tx=XM_023255660.1
B3 88108227 88108228 refGene_gn=SLC25A21;refGene_tx=XM_019832945.2
B3 88110264 88110326
refGene_gn=SLC25A21;refGene_tx=XM_019832945.2;refGene_gn=SLC25A21;refG
ene_tx=XM_023255661.1;refGene_gn=SLC25A21;refGene_tx=XM_023255662.1;re
fGene_gn=SLC25A21;refGene_tx=XM_023255663.1
B3 88113627 88113680
refGene_gn=SLC25A21;refGene_tx=XM_019832945.2;refGene_gn=SLC25A21;refG
ene_tx=XM_023255661.1;refGene_gn=SLC25A21;refGene_tx=XM_023255662.1;re
fGene_gn=SLC25A21;refGene_tx=XM_023255663.1
B3 88114568 88114750

refGene_gn=SLC25A21;refGene_tx=XM_019832945.2;refGene_gn=SLC25A21;refGene_tx=XM_023255661.1;refGene_gn=SLC25A21;refGene_tx=XM_023255662.1;refGene_gn=SLC25A21;refGene_tx=XM_023255663.1
B3 88143691 88143856
refGene_gn=SLC25A21;refGene_tx=XM_019832945.2;refGene_gn=SLC25A21;refGene_tx=XM_023255661.1;refGene_gn=SLC25A21;refGene_tx=XM_023255662.1;refGene_gn=SLC25A21;refGene_tx=XM_023255663.1
B3 88156086 88156194
refGene_gn=SLC25A21;refGene_tx=XM_019832945.2;refGene_gn=SLC25A21;refGene_tx=XM_023255661.1;refGene_gn=SLC25A21;refGene_tx=XM_023255662.1;refGene_gn=SLC25A21;refGene_tx=XM_023255663.1
B3 88159992 88160052
refGene_gn=SLC25A21;refGene_tx=XM_019832945.2;refGene_gn=SLC25A21;refGene_tx=XM_023255661.1;refGene_gn=SLC25A21;refGene_tx=XM_023255663.1
B3 88166974 88167041
refGene_gn=SLC25A21;refGene_tx=XM_019832945.2;refGene_gn=SLC25A21;refGene_tx=XM_023255661.1;refGene_gn=SLC25A21;refGene_tx=XM_023255662.1;refGene_gn=SLC25A21;refGene_tx=XM_023255663.1
B3 88242918 88243002
refGene_gn=SLC25A21;refGene_tx=XM_019832945.2;refGene_gn=SLC25A21;refGene_tx=XM_023255661.1;refGene_gn=SLC25A21;refGene_tx=XM_023255662.1;refGene_gn=SLC25A21;refGene_tx=XM_023255663.1
B3 88303227 88303276
refGene_gn=SLC25A21;refGene_tx=XM_019832945.2;refGene_gn=SLC25A21;refGene_tx=XM_023255661.1;refGene_gn=SLC25A21;refGene_tx=XM_023255662.1;refGene_gn=SLC25A21;refGene_tx=XM_023255663.1
B3 88602261 88602331
refGene_gn=SLC25A21;refGene_tx=XM_019832945.2;refGene_gn=SLC25A21;refGene_tx=XM_023255661.1;refGene_gn=SLC25A21;refGene_tx=XM_023255662.1
B3 88668686 88668705
refGene_gn=MIPOL1;refGene_tx=XM_011283197.3;refGene_gn=MIPOL1;refGene_tx=XM_011283198.3
B3 88687058 88687278
refGene_gn=MIPOL1;refGene_tx=XM_011283197.3;refGene_gn=MIPOL1;refGene_tx=XM_011283198.3
B3 88688692 88688828
refGene_gn=MIPOL1;refGene_tx=XM_011283197.3;refGene_gn=MIPOL1;refGene_tx=XM_011283198.3
B3 88690138 88690244
refGene_gn=MIPOL1;refGene_tx=XM_011283197.3;refGene_gn=MIPOL1;refGene_tx=XM_011283198.3
B3 88708257 88708387
refGene_gn=MIPOL1;refGene_tx=XM_011283197.3;refGene_gn=MIPOL1;refGene_tx=XM_011283198.3
B3 88720999 88721033
refGene_gn=MIPOL1;refGene_tx=XM_011283197.3;refGene_gn=MIPOL1;refGene_tx=XM_011283198.3
B3 88721292 88721463
refGene_gn=MIPOL1;refGene_tx=XM_011283197.3;refGene_gn=MIPOL1;refGene_tx=XM_011283198.3

B3 88761137 88761245
refGene_gn=MIPOL1;refGene_tx=XM_011283197.3;refGene_gn=MIPOL1;refGene_tx=XM_011283198.3
B3 88813067 88813162
refGene_gn=MIPOL1;refGene_tx=XM_011283197.3;refGene_gn=MIPOL1;refGene_tx=XM_011283198.3
B3 88882528 88882759
refGene_gn=MIPOL1;refGene_tx=XM_011283197.3;refGene_gn=MIPOL1;refGene_tx=XM_011283198.3
B3 88924359 88924426
refGene_gn=MIPOL1;refGene_tx=XM_011283197.3;refGene_gn=MIPOL1;refGene_tx=XM_011283198.3
B3 88966085 88967447 refGene_gn=FOXA1;refGene_tx=XM_011283199.3
B3 88969701 88969773 refGene_gn=FOXA1;refGene_tx=XM_011283199.3
B3 89048892 89048973 refGene_gn=TTC6;refGene_tx=XM_019832949.2
B3 89054612 89054816 refGene_gn=TTC6;refGene_tx=XM_019832949.2
B3 89061527 89061661 refGene_gn=TTC6;refGene_tx=XM_019832949.2
B3 89066765 89066960 refGene_gn=TTC6;refGene_tx=XM_019832949.2
B3 89079674 89079816 refGene_gn=TTC6;refGene_tx=XM_019832949.2
B3 89095959 89096064 refGene_gn=TTC6;refGene_tx=XM_019832949.2
B3 89105522 89105612 refGene_gn=TTC6;refGene_tx=XM_019832949.2
B3 89107956 89108031 refGene_gn=TTC6;refGene_tx=XM_019832949.2
B3 89122618 89122992 refGene_gn=TTC6;refGene_tx=XM_019832949.2
B3 89128718 89129178 refGene_gn=TTC6;refGene_tx=XM_019832949.2
B3 89129519 89129649 refGene_gn=TTC6;refGene_tx=XM_019832949.2
B3 89130500 89130673 refGene_gn=TTC6;refGene_tx=XM_019832949.2
B3 89132384 89132521 refGene_gn=TTC6;refGene_tx=XM_019832949.2
B3 89136777 89136947 refGene_gn=TTC6;refGene_tx=XM_019832949.2
B3 89139900 89140021 refGene_gn=TTC6;refGene_tx=XM_019832949.2
B3 89141420 89141571 refGene_gn=TTC6;refGene_tx=XM_019832949.2
B3 89143269 89143352 refGene_gn=TTC6;refGene_tx=XM_019832949.2
B3 89146979 89147056 refGene_gn=TTC6;refGene_tx=XM_019832949.2
B3 89147486 89147647 refGene_gn=TTC6;refGene_tx=XM_019832949.2
B3 89158071 89158206 refGene_gn=TTC6;refGene_tx=XM_019832949.2
B3 89159469 89159619 refGene_gn=TTC6;refGene_tx=XM_019832949.2
B3 89160414 89160555 refGene_gn=TTC6;refGene_tx=XM_019832949.2
B3 89162065 89162179 refGene_gn=TTC6;refGene_tx=XM_019832949.2
B3 89165519 89165639 refGene_gn=TTC6;refGene_tx=XM_019832949.2
B3 89170981 89171055 refGene_gn=TTC6;refGene_tx=XM_019832949.2
B3 89177445 89177656 refGene_gn=TTC6;refGene_tx=XM_019832949.2
B3 89180443 89180596 refGene_gn=TTC6;refGene_tx=XM_019832949.2
B3 89181466 89181637 refGene_gn=TTC6;refGene_tx=XM_019832949.2
B3 89189534 89189760 refGene_gn=TTC6;refGene_tx=XM_019832949.2
B3 89190215 89190352 refGene_gn=TTC6;refGene_tx=XM_019832949.2
B3 89506702 89507852
refGene_gn=SSTR1;refGene_tx=XM_019832950.2;refGene_gn=SSTR1;refGene_tx=XM_019832951.2
B3 89508209 89508250
refGene_gn=SSTR1;refGene_tx=XM_019832950.2;refGene_gn=SSTR1;refGene_tx=XM_019832951.2

B3 89545335 89546805 refGene_gn=CLEC14A;refGene_tx=XM_003987593.5
B3 89546976 89547100
refGene_gn=LOC109496334;refGene_tx=XM_023256184.1
B3 90194217 90194307 refGene_gn=SEC23A;refGene_tx=XM_011283203.3
B3 90205268 90205334 refGene_gn=SEC23A;refGene_tx=XM_011283203.3
B3 90207616 90207772 refGene_gn=SEC23A;refGene_tx=XM_011283203.3
B3 90209292 90209379 refGene_gn=SEC23A;refGene_tx=XM_011283203.3
B3 90211320 90211482 refGene_gn=SEC23A;refGene_tx=XM_011283203.3
B3 90214205 90214283 refGene_gn=SEC23A;refGene_tx=XM_011283203.3
B3 90222484 90222622 refGene_gn=SEC23A;refGene_tx=XM_011283203.3
B3 90222623 90222639 refGene_gn=SEC23A;refGene_tx=XM_011283203.3
B3 90229415 90229522 refGene_gn=SEC23A;refGene_tx=XM_011283203.3
B3 90230730 90230820 refGene_gn=SEC23A;refGene_tx=XM_011283203.3
B3 90231942 90232023 refGene_gn=SEC23A;refGene_tx=XM_011283203.3
B3 90235014 90235138 refGene_gn=SEC23A;refGene_tx=XM_011283203.3
B3 90240831 90240947 refGene_gn=SEC23A;refGene_tx=XM_011283203.3
B3 90242318 90242477 refGene_gn=SEC23A;refGene_tx=XM_011283203.3
B3 90244435 90244580 refGene_gn=SEC23A;refGene_tx=XM_011283203.3
B3 90245339 90245419 refGene_gn=SEC23A;refGene_tx=XM_011283203.3
B3 90249347 90249584 refGene_gn=SEC23A;refGene_tx=XM_011283203.3
B3 90250481 90250568 refGene_gn=SEC23A;refGene_tx=XM_011283203.3
B3 90250772 90250830 refGene_gn=SEC23A;refGene_tx=XM_011283203.3
B3 90252653 90252874 refGene_gn=SEC23A;refGene_tx=XM_011283203.3
B3 90272052 90272189 refGene_gn=GEMIN2;refGene_tx=XM_006932840.2
B3 90272545 90272630 refGene_gn=GEMIN2;refGene_tx=XM_006932840.2
B3 90274632 90274722 refGene_gn=GEMIN2;refGene_tx=XM_006932840.2
B3 90275235 90275295 refGene_gn=GEMIN2;refGene_tx=XM_006932840.2
B3 90277865 90277979 refGene_gn=GEMIN2;refGene_tx=XM_006932840.2
B3 90279781 90279826 refGene_gn=GEMIN2;refGene_tx=XM_006932840.2
B3 90281134 90281203 refGene_gn=GEMIN2;refGene_tx=XM_006932840.2
B3 90285741 90285852 refGene_gn=GEMIN2;refGene_tx=XM_006932840.2
B3 90289077 90289136 refGene_gn=GEMIN2;refGene_tx=XM_006932840.2
B3 90290791 90290831 refGene_gn=GEMIN2;refGene_tx=XM_006932840.2
B3 90302296 90302328
refGene_gn=TRAPPC6B;refGene_tx=XM_003987596.5;refGene_gn=TRAPPC6B;refG
ene_tx=XM_019832952.2
B3 90303291 90303385
refGene_gn=TRAPPC6B;refGene_tx=XM_003987596.5;refGene_gn=TRAPPC6B;refG
ene_tx=XM_019832952.2
B3 90305187 90305271
refGene_gn=TRAPPC6B;refGene_tx=XM_003987596.5;refGene_gn=TRAPPC6B;refG
ene_tx=XM_019832952.2
B3 90306572 90306690
refGene_gn=TRAPPC6B;refGene_tx=XM_003987596.5;refGene_gn=TRAPPC6B;refG
ene_tx=XM_019832952.2
B3 90307669 90307737
refGene_gn=TRAPPC6B;refGene_tx=XM_003987596.5;refGene_gn=TRAPPC6B;refG
ene_tx=XM_019832952.2
B3 90312655 90312736 refGene_gn=TRAPPC6B;refGene_tx=XM_003987596.5
B3 90316211 90316217 refGene_gn=TRAPPC6B;refGene_tx=XM_019832952.2

B3 90316616 90316729 refGene_gn=PNN;refGene_tx=XM_011283207.3
B3 90317404 90317476 refGene_gn=PNN;refGene_tx=XM_011283207.3
B3 90317869 90317938 refGene_gn=PNN;refGene_tx=XM_011283207.3
B3 90318429 90318502 refGene_gn=PNN;refGene_tx=XM_011283207.3
B3 90318602 90318697 refGene_gn=PNN;refGene_tx=XM_011283207.3
B3 90318861 90318937 refGene_gn=PNN;refGene_tx=XM_011283207.3
B3 90319933 90320089 refGene_gn=PNN;refGene_tx=XM_011283207.3
B3 90320171 90320310 refGene_gn=PNN;refGene_tx=XM_011283207.3
B3 90321003 90322562 refGene_gn=PNN;refGene_tx=XM_011283207.3
B3 90355637 90355752 refGene_gn=MIA2;refGene_tx=XM_019832953.2
B3 90358251 90358385 refGene_gn=MIA2;refGene_tx=XM_019832953.2
B3 90360703 90360790 refGene_gn=MIA2;refGene_tx=XM_019832953.2
B3 90366329 90367533 refGene_gn=MIA2;refGene_tx=XM_019832953.2
B3 90376669 90376888 refGene_gn=MIA2;refGene_tx=XM_019832953.2
B3 90376994 90377095 refGene_gn=MIA2;refGene_tx=XM_019832953.2
B3 90391541 90391673 refGene_gn=MIA2;refGene_tx=XM_019832953.2
B3 90399314 90399336 refGene_gn=MIA2;refGene_tx=XM_019832953.2
B3 90399428 90399475 refGene_gn=MIA2;refGene_tx=XM_019832953.2
B3 90399476 90399518 refGene_gn=MIA2;refGene_tx=XM_019832953.2
B3 90405772 90405844 refGene_gn=MIA2;refGene_tx=XM_019832953.2
B3 90407960 90407987 refGene_gn=MIA2;refGene_tx=XM_019832953.2
B3 90410239 90410350 refGene_gn=MIA2;refGene_tx=XM_019832953.2
B3 90410992 90411064 refGene_gn=MIA2;refGene_tx=XM_019832953.2
B3 90418351 90418456 refGene_gn=MIA2;refGene_tx=XM_019832953.2
B3 90423081 90423129 refGene_gn=MIA2;refGene_tx=XM_019832953.2
B3 90423130 90423205 refGene_gn=MIA2;refGene_tx=XM_019832953.2
B3 90424703 90424824 refGene_gn=MIA2;refGene_tx=XM_019832953.2
B3 90426290 90426337 refGene_gn=MIA2;refGene_tx=XM_019832953.2
B3 90427098 90427189 refGene_gn=MIA2;refGene_tx=XM_019832953.2
B3 90429311 90429450 refGene_gn=MIA2;refGene_tx=XM_019832953.2
B3 90433687 90433789 refGene_gn=MIA2;refGene_tx=XM_019832953.2
B3 90434717 90434778 refGene_gn=MIA2;refGene_tx=XM_019832953.2
B3 90435607 90435643 refGene_gn=MIA2;refGene_tx=XM_019832953.2
B3 90439861 90439929 refGene_gn=MIA2;refGene_tx=XM_019832953.2
B3 90441484 90441567 refGene_gn=MIA2;refGene_tx=XM_019832953.2
B3 90443205 90443334 refGene_gn=MIA2;refGene_tx=XM_019832953.2
B3 90446957 90447116 refGene_gn=MIA2;refGene_tx=XM_019832953.2
B3 90459544 90459667 refGene_gn=MIA2;refGene_tx=XM_019832953.2
B3 90464627 90464686 refGene_gn=MIA2;refGene_tx=XM_019832953.2
B3 90465553 90465788 refGene_gn=MIA2;refGene_tx=XM_019832953.2
B3 90466862 90467029 refGene_gn=MIA2;refGene_tx=XM_019832953.2
B3 90486166 90486438
refGene_gn=FBXO33;refGene_tx=XM_011283375.3;refGene_gn=FBXO33;refGene_
tx=XM_023255664.1
B3 90487616 90488302
refGene_gn=FBXO33;refGene_tx=XM_011283375.3;refGene_gn=FBXO33;refGene_
tx=XM_023255664.1
B3 90488832 90488943 refGene_gn=FBXO33;refGene_tx=XM_011283375.3
B3 90516541 90517149
refGene_gn=FBXO33;refGene_tx=XM_011283375.3;refGene_gn=FBXO33;refGene_

tx=XM_023255664.1
B3	92655454 92656839
refGene_gn=LRFN5;refGene_tx=XM_023255665.1;refGene_gn=LRFN5;refGene_tx=XM_023255666.1;refGene_gn=LRFN5;refGene_tx=XM_023255667.1
B3	92657188 92657225 refGene_gn=LRFN5;refGene_tx=XM_023255666.1
B3	92659628 92660341 refGene_gn=LRFN5;refGene_tx=XM_023255665.1
B3	92666770 92666814
refGene_gn=LRFN5;refGene_tx=XM_023255665.1;refGene_gn=LRFN5;refGene_tx=XM_023255667.1
B3	92672958 92672976 refGene_gn=LRFN5;refGene_tx=XM_023255665.1
B3	95005564 95005770
refGene_gn=LOC111560966;refGene_tx=XM_023256185.1
B3	95005772 95005966
refGene_gn=LOC111560966;refGene_tx=XM_023256185.1
B3	95013417 95013520
refGene_gn=LOC111560966;refGene_tx=XM_023256185.1
B3	95014435 95014564
refGene_gn=LOC111560966;refGene_tx=XM_023256185.1
B3	95018898 95019025
refGene_gn=LOC111560966;refGene_tx=XM_023256185.1
B3	95152361 95152811
refGene_gn=LOC101086065;refGene_tx=XM_023255668.1
B3	95153781 95154876
refGene_gn=LOC101086065;refGene_tx=XM_023255668.1
B3	95519864 95520388
refGene_gn=CB3H14orf28;refGene_tx=XM_003987602.4;refGene_gn=CB3H14orf28;refGene_tx=XM_019832966.1
B3	95522474 95522564
refGene_gn=CB3H14orf28;refGene_tx=XM_003987602.4;refGene_gn=CB3H14orf28;refGene_tx=XM_019832966.1
B3	95523743 95523878
refGene_gn=CB3H14orf28;refGene_tx=XM_003987602.4
B3	95524931 95525115
refGene_gn=CB3H14orf28;refGene_tx=XM_003987602.4;refGene_gn=CB3H14orf28;refGene_tx=XM_019832966.1
B3	95542192 95542356
refGene_gn=KLHL28;refGene_tx=XM_003987603.5;refGene_gn=KLHL28;refGene_tx=XM_011283217.3;refGene_gn=KLHL28;refGene_tx=XM_019832967.2;refGene_gn=KLHL28;refGene_tx=XM_019832968.2;refGene_gn=KLHL28;refGene_tx=XM_019832969.2;refGene_gn=KLHL28;refGene_tx=XM_019832971.2;refGene_gn=KLHL28;refGene_tx=XM_023255669.1
B3	95543625 95543834
refGene_gn=KLHL28;refGene_tx=XM_003987603.5;refGene_gn=KLHL28;refGene_tx=XM_011283217.3;refGene_gn=KLHL28;refGene_tx=XM_019832967.2;refGene_gn=KLHL28;refGene_tx=XM_019832968.2;refGene_gn=KLHL28;refGene_tx=XM_019832969.2;refGene_gn=KLHL28;refGene_tx=XM_019832971.2;refGene_gn=KLHL28;refGene_tx=XM_023255669.1
B3	95545761 95546205
refGene_gn=KLHL28;refGene_tx=XM_003987603.5;refGene_gn=KLHL28;refGene_tx=XM_011283217.3;refGene_gn=KLHL28;refGene_tx=XM_019832967.2;refGene_

gn=KLHL28;refGene_tx=XM_019832968.2;refGene_gn=KLHL28;refGene_tx=XM_01
9832969.2;refGene_gn=KLHL28;refGene_tx=XM_019832971.2;refGene_gn=KLHL2
8;refGene_tx=XM_023255669.1
B3 95554578 95555477
refGene_gn=KLHL28;refGene_tx=XM_003987603.5;refGene_gn=KLHL28;refGene_
tx=XM_011283217.3;refGene_gn=KLHL28;refGene_tx=XM_019832967.2;refGene_
gn=KLHL28;refGene_tx=XM_019832968.2;refGene_gn=KLHL28;refGene_tx=XM_01
9832969.2;refGene_gn=KLHL28;refGene_tx=XM_019832971.2;refGene_gn=KLHL2
8;refGene_tx=XM_023255669.1
B3 95558659 95558662
refGene_gn=KLHL28;refGene_tx=XM_019832968.2;refGene_gn=KLHL28;refGene_
tx=XM_023255669.1
B3 95560966 95561221 refGene_gn=KLHL28;refGene_tx=XM_019832967.2
B3 95571345 95571348 refGene_gn=KLHL28;refGene_tx=XM_019832969.2
B3 95678223 95678541 refGene_gn=PRPF39;refGene_tx=XM_011283218.3
B3 95679008 95679134 refGene_gn=PRPF39;refGene_tx=XM_011283218.3
B3 95679568 95679655
refGene_gn=PRPF39;refGene_tx=XM_011283219.3;refGene_gn=PRPF39;refGene_
tx=XM_011283220.2;refGene_gn=PRPF39;refGene_tx=XM_011283222.2;refGene_
gn=PRPF39;refGene_tx=XM_019832972.2
B3 95679783 95679902
refGene_gn=PRPF39;refGene_tx=XM_011283218.3;refGene_gn=PRPF39;refGene_
tx=XM_011283219.3;refGene_gn=PRPF39;refGene_tx=XM_011283220.2;refGene_
gn=PRPF39;refGene_tx=XM_011283222.2;refGene_gn=PRPF39;refGene_tx=XM_01
9832972.2
B3 95684569 95684737
refGene_gn=PRPF39;refGene_tx=XM_011283218.3;refGene_gn=PRPF39;refGene_
tx=XM_011283219.3;refGene_gn=PRPF39;refGene_tx=XM_011283220.2;refGene_
gn=PRPF39;refGene_tx=XM_011283222.2;refGene_gn=PRPF39;refGene_tx=XM_01
9832972.2
B3 95689105 95689271
refGene_gn=PRPF39;refGene_tx=XM_011283218.3;refGene_gn=PRPF39;refGene_
tx=XM_011283219.3;refGene_gn=PRPF39;refGene_tx=XM_011283220.2;refGene_
gn=PRPF39;refGene_tx=XM_011283222.2;refGene_gn=PRPF39;refGene_tx=XM_01
9832972.2
B3 95689728 95689836
refGene_gn=PRPF39;refGene_tx=XM_011283218.3;refGene_gn=PRPF39;refGene_
tx=XM_011283219.3;refGene_gn=PRPF39;refGene_tx=XM_011283220.2;refGene_
gn=PRPF39;refGene_tx=XM_011283222.2;refGene_gn=PRPF39;refGene_tx=XM_01
9832972.2
B3 95691403 95691568
refGene_gn=PRPF39;refGene_tx=XM_011283218.3;refGene_gn=PRPF39;refGene_
tx=XM_011283219.3;refGene_gn=PRPF39;refGene_tx=XM_011283220.2;refGene_
gn=PRPF39;refGene_tx=XM_011283222.2;refGene_gn=PRPF39;refGene_tx=XM_01
9832972.2
B3 95691882 95692009
refGene_gn=PRPF39;refGene_tx=XM_011283218.3;refGene_gn=PRPF39;refGene_
tx=XM_011283219.3;refGene_gn=PRPF39;refGene_tx=XM_011283220.2;refGene_
gn=PRPF39;refGene_tx=XM_011283222.2;refGene_gn=PRPF39;refGene_tx=XM_01
9832972.2

B3 95692416 95692685
refGene_gn=PRPF39;refGene_tx=XM_011283218.3;refGene_gn=PRPF39;refGene_tx=XM_011283219.3;refGene_gn=PRPF39;refGene_tx=XM_011283220.2;refGene_gn=PRPF39;refGene_tx=XM_011283222.2;refGene_gn=PRPF39;refGene_tx=XM_019832972.2
B3 95693487 95693672
refGene_gn=PRPF39;refGene_tx=XM_011283218.3;refGene_gn=PRPF39;refGene_tx=XM_011283219.3;refGene_gn=PRPF39;refGene_tx=XM_011283220.2;refGene_gn=PRPF39;refGene_tx=XM_011283222.2;refGene_gn=PRPF39;refGene_tx=XM_019832972.2
B3 95696580 95696655
refGene_gn=PRPF39;refGene_tx=XM_011283218.3;refGene_gn=PRPF39;refGene_tx=XM_011283219.3;refGene_gn=PRPF39;refGene_tx=XM_011283220.2;refGene_gn=PRPF39;refGene_tx=XM_011283222.2;refGene_gn=PRPF39;refGene_tx=XM_019832972.2
B3 95697161 95697282
refGene_gn=PRPF39;refGene_tx=XM_011283218.3;refGene_gn=PRPF39;refGene_tx=XM_011283219.3;refGene_gn=PRPF39;refGene_tx=XM_011283220.2;refGene_gn=PRPF39;refGene_tx=XM_011283222.2;refGene_gn=PRPF39;refGene_tx=XM_019832972.2
B3 95697595 95697652
refGene_gn=PRPF39;refGene_tx=XM_011283218.3;refGene_gn=PRPF39;refGene_tx=XM_011283219.3;refGene_gn=PRPF39;refGene_tx=XM_011283220.2;refGene_gn=PRPF39;refGene_tx=XM_011283222.2;refGene_gn=PRPF39;refGene_tx=XM_019832972.2
B3 95698925 95698980
refGene_gn=FKBP3;refGene_tx=XM_004001428.5;refGene_gn=FKBP3;refGene_tx=XM_019832973.1
B3 95700268 95700366
refGene_gn=FKBP3;refGene_tx=XM_004001428.5;refGene_gn=FKBP3;refGene_tx=XM_019832973.1
B3 95701944 95702012
refGene_gn=FKBP3;refGene_tx=XM_004001428.5;refGene_gn=FKBP3;refGene_tx=XM_019832973.1
B3 95702498 95702634
refGene_gn=FKBP3;refGene_tx=XM_004001428.5;refGene_gn=FKBP3;refGene_tx=XM_019832973.1
B3 95707987 95708095
refGene_gn=FKBP3;refGene_tx=XM_004001428.5;refGene_gn=FKBP3;refGene_tx=XM_019832973.1
B3 95708749 95708851
refGene_gn=FKBP3;refGene_tx=XM_004001428.5;refGene_gn=FKBP3;refGene_tx=XM_019832973.1
B3 95710928 95710961 refGene_gn=FKBP3;refGene_tx=XM_019832973.1
B3 95711348 95711456 refGene_gn=FKBP3;refGene_tx=XM_004001428.5
B3 95712960 95713477 refGene_gn=FANCM;refGene_tx=XM_023256186.1
B3 95714041 95714214 refGene_gn=FANCM;refGene_tx=XM_023256186.1
B3 95716704 95716782 refGene_gn=FANCM;refGene_tx=XM_023256186.1
B3 95722015 95722174 refGene_gn=FANCM;refGene_tx=XM_023256186.1
B3 95728681 95728813 refGene_gn=FANCM;refGene_tx=XM_023256186.1

B3 95730458 95730591 refGene_gn=FANCM;refGene_tx=XM_023256186.1
B3 95731703 95731826 refGene_gn=FANCM;refGene_tx=XM_023256186.1
B3 95732318 95732405 refGene_gn=FANCM;refGene_tx=XM_023256186.1
B3 95733320 95733505 refGene_gn=FANCM;refGene_tx=XM_023256186.1
B3 95744429 95744636 refGene_gn=FANCM;refGene_tx=XM_023256186.1
B3 95746073 95746287 refGene_gn=FANCM;refGene_tx=XM_023256186.1
B3 95751155 95751313 refGene_gn=FANCM;refGene_tx=XM_023256186.1
B3 95752049 95752205 refGene_gn=FANCM;refGene_tx=XM_023256186.1
B3 95753733 95755639 refGene_gn=FANCM;refGene_tx=XM_023256186.1
B3 95759239 95759256 refGene_gn=FANCM;refGene_tx=XM_023256186.1
B3 95759801 95759870 refGene_gn=FANCM;refGene_tx=XM_023256186.1
B3 95761499 95761628 refGene_gn=FANCM;refGene_tx=XM_023256186.1
B3 95762603 95762760 refGene_gn=FANCM;refGene_tx=XM_023256186.1
B3 95765683 95765790 refGene_gn=FANCM;refGene_tx=XM_023256186.1
B3 95766540 95767107 refGene_gn=FANCM;refGene_tx=XM_023256186.1
B3 95771273 95771649 refGene_gn=FANCM;refGene_tx=XM_023256186.1
B3 95773538 95773830 refGene_gn=FANCM;refGene_tx=XM_023256186.1
B3 95774411 95774550 refGene_gn=FANCM;refGene_tx=XM_023256186.1
B3 95777892 95777996
refGene_gn=MIS18BP1;refGene_tx=XM_006932849.4;refGene_gn=MIS18BP1;refGene_tx=XM_019832975.2;refGene_gn=MIS18BP1;refGene_tx=XM_019832976.2;refGene_gn=MIS18BP1;refGene_tx=XM_019832978.2;refGene_gn=MIS18BP1;refGene_tx=XM_023255670.1
B3 95778178 95778233
refGene_gn=MIS18BP1;refGene_tx=XM_006932849.4;refGene_gn=MIS18BP1;refGene_tx=XM_019832975.2;refGene_gn=MIS18BP1;refGene_tx=XM_019832976.2;refGene_gn=MIS18BP1;refGene_tx=XM_019832978.2;refGene_gn=MIS18BP1;refGene_tx=XM_023255670.1
B3 95780207 95780295
refGene_gn=MIS18BP1;refGene_tx=XM_006932849.4;refGene_gn=MIS18BP1;refGene_tx=XM_019832975.2;refGene_gn=MIS18BP1;refGene_tx=XM_019832976.2;refGene_gn=MIS18BP1;refGene_tx=XM_019832978.2;refGene_gn=MIS18BP1;refGene_tx=XM_023255670.1
B3 95783587 95783603 refGene_gn=MIS18BP1;refGene_tx=XM_019832977.2
B3 95784958 95785107
refGene_gn=MIS18BP1;refGene_tx=XM_006932849.4;refGene_gn=MIS18BP1;refGene_tx=XM_019832975.2;refGene_gn=MIS18BP1;refGene_tx=XM_019832976.2;refGene_gn=MIS18BP1;refGene_tx=XM_019832977.2;refGene_gn=MIS18BP1;refGene_tx=XM_023255670.1
B3 95787342 95787503
refGene_gn=MIS18BP1;refGene_tx=XM_006932849.4;refGene_gn=MIS18BP1;refGene_tx=XM_019832975.2;refGene_gn=MIS18BP1;refGene_tx=XM_019832976.2;refGene_gn=MIS18BP1;refGene_tx=XM_019832977.2;refGene_gn=MIS18BP1;refGene_tx=XM_023255670.1
B3 95788274 95788447
refGene_gn=MIS18BP1;refGene_tx=XM_006932849.4;refGene_gn=MIS18BP1;refGene_tx=XM_019832975.2;refGene_gn=MIS18BP1;refGene_tx=XM_019832976.2;refGene_gn=MIS18BP1;refGene_tx=XM_019832977.2;refGene_gn=MIS18BP1;refGene_tx=XM_023255670.1
B3 95797303 95798171

refGene_gn=MIS18BP1;refGene_tx=XM_006932849.4;refGene_gn=MIS18BP1;refGene_tx=XM_019832975.2;refGene_gn=MIS18BP1;refGene_tx=XM_019832976.2;refGene_gn=MIS18BP1;refGene_tx=XM_019832977.2;refGene_gn=MIS18BP1;refGene_tx=XM_019832978.2;refGene_gn=MIS18BP1;refGene_tx=XM_023255670.1
B3 95800440 95800537
refGene_gn=MIS18BP1;refGene_tx=XM_006932849.4;refGene_gn=MIS18BP1;refGene_tx=XM_019832975.2;refGene_gn=MIS18BP1;refGene_tx=XM_019832976.2;refGene_gn=MIS18BP1;refGene_tx=XM_019832977.2;refGene_gn=MIS18BP1;refGene_tx=XM_019832978.2;refGene_gn=MIS18BP1;refGene_tx=XM_023255670.1
B3 95801014 95801166
refGene_gn=MIS18BP1;refGene_tx=XM_006932849.4;refGene_gn=MIS18BP1;refGene_tx=XM_019832975.2;refGene_gn=MIS18BP1;refGene_tx=XM_019832976.2;refGene_gn=MIS18BP1;refGene_tx=XM_019832977.2;refGene_gn=MIS18BP1;refGene_tx=XM_019832978.2;refGene_gn=MIS18BP1;refGene_tx=XM_023255670.1
B3 95801791 95801949
refGene_gn=MIS18BP1;refGene_tx=XM_006932849.4;refGene_gn=MIS18BP1;refGene_tx=XM_019832975.2;refGene_gn=MIS18BP1;refGene_tx=XM_019832976.2;refGene_gn=MIS18BP1;refGene_tx=XM_019832977.2;refGene_gn=MIS18BP1;refGene_tx=XM_019832978.2;refGene_gn=MIS18BP1;refGene_tx=XM_023255670.1
B3 95802903 95802991
refGene_gn=MIS18BP1;refGene_tx=XM_006932849.4;refGene_gn=MIS18BP1;refGene_tx=XM_019832975.2;refGene_gn=MIS18BP1;refGene_tx=XM_019832976.2;refGene_gn=MIS18BP1;refGene_tx=XM_019832977.2;refGene_gn=MIS18BP1;refGene_tx=XM_019832978.2;refGene_gn=MIS18BP1;refGene_tx=XM_023255670.1
B3 95805271 95805402
refGene_gn=MIS18BP1;refGene_tx=XM_006932849.4;refGene_gn=MIS18BP1;refGene_tx=XM_019832975.2;refGene_gn=MIS18BP1;refGene_tx=XM_019832976.2;refGene_gn=MIS18BP1;refGene_tx=XM_019832977.2;refGene_gn=MIS18BP1;refGene_tx=XM_019832978.2;refGene_gn=MIS18BP1;refGene_tx=XM_023255670.1
B3 95807084 95807158
refGene_gn=MIS18BP1;refGene_tx=XM_006932849.4;refGene_gn=MIS18BP1;refGene_tx=XM_019832975.2;refGene_gn=MIS18BP1;refGene_tx=XM_019832976.2;refGene_gn=MIS18BP1;refGene_tx=XM_019832977.2;refGene_gn=MIS18BP1;refGene_tx=XM_019832978.2;refGene_gn=MIS18BP1;refGene_tx=XM_023255670.1
B3 95826228 95826713
refGene_gn=MIS18BP1;refGene_tx=XM_006932849.4;refGene_gn=MIS18BP1;refGene_tx=XM_019832975.2;refGene_gn=MIS18BP1;refGene_tx=XM_019832976.2;refGene_gn=MIS18BP1;refGene_tx=XM_019832977.2;refGene_gn=MIS18BP1;refGene_tx=XM_019832978.2;refGene_gn=MIS18BP1;refGene_tx=XM_023255670.1
B3 95826934 95827039
refGene_gn=MIS18BP1;refGene_tx=XM_006932849.4;refGene_gn=MIS18BP1;refGene_tx=XM_019832975.2;refGene_gn=MIS18BP1;refGene_tx=XM_019832977.2;refGene_gn=MIS18BP1;refGene_tx=XM_019832978.2;refGene_gn=MIS18BP1;refGene_tx=XM_023255670.1
B3 95830888 95831429
refGene_gn=MIS18BP1;refGene_tx=XM_006932849.4;refGene_gn=MIS18BP1;refGene_tx=XM_019832975.2;refGene_gn=MIS18BP1;refGene_tx=XM_019832976.2;refGene_gn=MIS18BP1;refGene_tx=XM_019832977.2;refGene_gn=MIS18BP1;refGene_tx=XM_019832978.2;refGene_gn=MIS18BP1;refGene_tx=XM_023255670.1
B3 96961852 96962560 refGene_gn=RPL10L;refGene_tx=XM_003987606.4

B3 97113385 97113474 refGene_gn=MDGA2;refGene_tx=XM_023255671.1
B3 97116686 97116792 refGene_gn=MDGA2;refGene_tx=XM_023255671.1
B3 97127920 97128051 refGene_gn=MDGA2;refGene_tx=XM_023255671.1
B3 97141266 97141425 refGene_gn=MDGA2;refGene_tx=XM_023255671.1
B3 97141879 97142035 refGene_gn=MDGA2;refGene_tx=XM_023255671.1
B3 97152733 97152754 refGene_gn=MDGA2;refGene_tx=XM_023255671.1
B3 97156790 97156968 refGene_gn=MDGA2;refGene_tx=XM_023255671.1
B3 97205525 97205674 refGene_gn=MDGA2;refGene_tx=XM_023255671.1
B3 97232135 97232405 refGene_gn=MDGA2;refGene_tx=XM_023255671.1
B3 97309864 97310158 refGene_gn=MDGA2;refGene_tx=XM_023255671.1
B3 97336500 97336830 refGene_gn=MDGA2;refGene_tx=XM_023255671.1
B3 97384354 97384624 refGene_gn=MDGA2;refGene_tx=XM_023255671.1
B3 97406707 97406840 refGene_gn=MDGA2;refGene_tx=XM_023255671.1
B3 97430730 97430927 refGene_gn=MDGA2;refGene_tx=XM_023255671.1
B3 97509249 97509424 refGene_gn=MDGA2;refGene_tx=XM_023255671.1
B3 97600753 97600893 refGene_gn=MDGA2;refGene_tx=XM_023255671.1
B3 97965284 97965564 refGene_gn=MDGA2;refGene_tx=XM_023255671.1
B3 99661208 99661217 refGene_gn=RPS29;refGene_tx=XM_003987610.3
B3 99662440 99662540 refGene_gn=RPS29;refGene_tx=XM_003987610.3
B3 99662774 99662836 refGene_gn=RPS29;refGene_tx=XM_003987610.3
B3 99671441 99671744
refGene_gn=LRR1;refGene_tx=XM_003987611.4;refGene_gn=LRR1;refGene_tx=XM_006932850.3;refGene_gn=LRR1;refGene_tx=XM_019832980.2
B3 99675057 99675159
refGene_gn=LRR1;refGene_tx=XM_003987611.4;refGene_gn=LRR1;refGene_tx=XM_019832980.2
B3 99677837 99678559
refGene_gn=LRR1;refGene_tx=XM_003987611.4;refGene_gn=LRR1;refGene_tx=XM_006932850.3;refGene_gn=LRR1;refGene_tx=XM_019832980.2
B3 99680317 99680558
refGene_gn=LRR1;refGene_tx=XM_003987611.4;refGene_gn=LRR1;refGene_tx=XM_006932850.3;refGene_gn=LRR1;refGene_tx=XM_019832980.2
B3 99683771 99684092 refGene_gn=RPL36AL;refGene_tx=XM_003987612.4
B3 99685564 99686905 refGene_gn=MGAT2;refGene_tx=XM_006932851.4
B3 99689577 99690081 refGene_gn=DNAAF2;refGene_tx=XM_003987614.4
B3 99691313 99691457 refGene_gn=DNAAF2;refGene_tx=XM_003987614.4
B3 99694388 99696218 refGene_gn=DNAAF2;refGene_tx=XM_003987614.4
B3 99702178 99702197
refGene_gn=POLE2;refGene_tx=XM_003987616.4;refGene_gn=POLE2;refGene_tx=XM_006932853.3;refGene_gn=POLE2;refGene_tx=XM_019832981.2
B3 99705927 99705995
refGene_gn=POLE2;refGene_tx=XM_003987616.4;refGene_gn=POLE2;refGene_tx=XM_006932853.3;refGene_gn=POLE2;refGene_tx=XM_019832981.2
B3 99707702 99707879
refGene_gn=POLE2;refGene_tx=XM_003987616.4;refGene_gn=POLE2;refGene_tx=XM_006932853.3;refGene_gn=POLE2;refGene_tx=XM_019832981.2
B3 99709165 99709274
refGene_gn=POLE2;refGene_tx=XM_003987616.4;refGene_gn=POLE2;refGene_tx=XM_006932853.3;refGene_gn=POLE2;refGene_tx=XM_019832981.2
B3 99711366 99711444

refGene_gn=POLE2;refGene_tx=XM_003987616.4;refGene_gn=POLE2;refGene_tx=XM_006932853.3;refGene_gn=POLE2;refGene_tx=XM_019832981.2
B3	99711531	99711591	refGene_gn=POLE2;refGene_tx=XM_003987616.4;refGene_gn=POLE2;refGene_tx=XM_006932853.3;refGene_gn=POLE2;refGene_tx=XM_019832981.2
B3	99712377	99712432	refGene_gn=POLE2;refGene_tx=XM_003987616.4;refGene_gn=POLE2;refGene_tx=XM_006932853.3;refGene_gn=POLE2;refGene_tx=XM_019832981.2
B3	99712609	99712699	refGene_gn=POLE2;refGene_tx=XM_003987616.4;refGene_gn=POLE2;refGene_tx=XM_006932853.3;refGene_gn=POLE2;refGene_tx=XM_019832981.2
B3	99713799	99713972	refGene_gn=POLE2;refGene_tx=XM_003987616.4;refGene_gn=POLE2;refGene_tx=XM_006932853.3;refGene_gn=POLE2;refGene_tx=XM_019832981.2
B3	99716913	99716986	refGene_gn=POLE2;refGene_tx=XM_003987616.4;refGene_gn=POLE2;refGene_tx=XM_006932853.3;refGene_gn=POLE2;refGene_tx=XM_019832981.2
B3	99717617	99717666	refGene_gn=POLE2;refGene_tx=XM_003987616.4;refGene_gn=POLE2;refGene_tx=XM_006932853.3;refGene_gn=POLE2;refGene_tx=XM_019832981.2
B3	99718088	99718145	refGene_gn=POLE2;refGene_tx=XM_003987616.4;refGene_gn=POLE2;refGene_tx=XM_006932853.3;refGene_gn=POLE2;refGene_tx=XM_019832981.2
B3	99718753	99718837	refGene_gn=POLE2;refGene_tx=XM_003987616.4;refGene_gn=POLE2;refGene_tx=XM_006932853.3;refGene_gn=POLE2;refGene_tx=XM_019832981.2
B3	99721702	99721777	refGene_gn=POLE2;refGene_tx=XM_003987616.4;refGene_gn=POLE2;refGene_tx=XM_006932853.3;refGene_gn=POLE2;refGene_tx=XM_019832981.2
B3	99725266	99725360	refGene_gn=POLE2;refGene_tx=XM_003987616.4;refGene_gn=POLE2;refGene_tx=XM_006932853.3;refGene_gn=POLE2;refGene_tx=XM_019832981.2
B3	99725490	99725568	refGene_gn=POLE2;refGene_tx=XM_003987616.4;refGene_gn=POLE2;refGene_tx=XM_006932853.3;refGene_gn=POLE2;refGene_tx=XM_019832981.2
B3	99728425	99728501	refGene_gn=POLE2;refGene_tx=XM_003987616.4;refGene_gn=POLE2;refGene_tx=XM_006932853.3;refGene_gn=POLE2;refGene_tx=XM_019832981.2
B3	99729406	99729507	refGene_gn=POLE2;refGene_tx=XM_003987616.4;refGene_gn=POLE2;refGene_tx=XM_006932853.3;refGene_gn=POLE2;refGene_tx=XM_019832981.2
B3	99732791	99732832	refGene_gn=POLE2;refGene_tx=XM_019832981.2
B3	99733275	99733343	refGene_gn=POLE2;refGene_tx=XM_003987616.4;refGene_gn=POLE2;refGene_tx=XM_006932853.3
B3	99746333	99746429	refGene_gn=KLHDC1;refGene_tx=XM_003987618.4;refGene_gn=KLHDC1;refGene_tx=XM_019832983.2
B3	99760712	99760783	refGene_gn=KLHDC1;refGene_tx=XM_003987618.4;refGene_gn=KLHDC1;refGene_

tx=XM_019832983.2
B3	99761258 99761412
refGene_gn=KLHDC1;refGene_tx=XM_003987618.4;refGene_gn=KLHDC1;refGene_tx=XM_019832983.2;refGene_gn=KLHDC1;refGene_tx=XM_023255672.1
B3	99761805 99761924
refGene_gn=KLHDC1;refGene_tx=XM_003987618.4;refGene_gn=KLHDC1;refGene_tx=XM_019832983.2;refGene_gn=KLHDC1;refGene_tx=XM_023255672.1
B3	99769811 99769890
refGene_gn=KLHDC1;refGene_tx=XM_003987618.4;refGene_gn=KLHDC1;refGene_tx=XM_019832983.2;refGene_gn=KLHDC1;refGene_tx=XM_023255672.1
B3	99771786 99771870
refGene_gn=KLHDC1;refGene_tx=XM_003987618.4;refGene_gn=KLHDC1;refGene_tx=XM_019832983.2;refGene_gn=KLHDC1;refGene_tx=XM_023255672.1
B3	99776970 99777054
refGene_gn=KLHDC1;refGene_tx=XM_003987618.4;refGene_gn=KLHDC1;refGene_tx=XM_019832983.2;refGene_gn=KLHDC1;refGene_tx=XM_023255672.1
B3	99777554 99777613
refGene_gn=KLHDC1;refGene_tx=XM_003987618.4;refGene_gn=KLHDC1;refGene_tx=XM_019832983.2;refGene_gn=KLHDC1;refGene_tx=XM_023255672.1
B3	99782292 99782405
refGene_gn=KLHDC1;refGene_tx=XM_003987618.4;refGene_gn=KLHDC1;refGene_tx=XM_019832983.2;refGene_gn=KLHDC1;refGene_tx=XM_023255672.1
B3	99783827 99783900
refGene_gn=KLHDC1;refGene_tx=XM_003987618.4;refGene_gn=KLHDC1;refGene_tx=XM_019832983.2;refGene_gn=KLHDC1;refGene_tx=XM_023255672.1
B3	99789157 99789242
refGene_gn=KLHDC1;refGene_tx=XM_003987618.4;refGene_gn=KLHDC1;refGene_tx=XM_019832983.2;refGene_gn=KLHDC1;refGene_tx=XM_023255672.1
B3	99791400 99791453
refGene_gn=KLHDC1;refGene_tx=XM_003987618.4;refGene_gn=KLHDC1;refGene_tx=XM_019832983.2;refGene_gn=KLHDC1;refGene_tx=XM_023255672.1
B3	99797964 99798151
refGene_gn=KLHDC1;refGene_tx=XM_003987618.4;refGene_gn=KLHDC1;refGene_tx=XM_019832983.2;refGene_gn=KLHDC1;refGene_tx=XM_023255672.1
B3	99804265 99804418 refGene_gn=KLHDC2;refGene_tx=XM_023255674.1
B3	99807456 99807536 refGene_gn=KLHDC2;refGene_tx=XM_023255674.1
B3	99809324 99809442 refGene_gn=KLHDC2;refGene_tx=XM_023255674.1
B3	99811448 99811564 refGene_gn=KLHDC2;refGene_tx=XM_023255674.1
B3	99811875 99811957 refGene_gn=KLHDC2;refGene_tx=XM_023255674.1
B3	99812123 99812207 refGene_gn=KLHDC2;refGene_tx=XM_023255674.1
B3	99812728 99812809 refGene_gn=KLHDC2;refGene_tx=XM_023255674.1
B3	99812884 99812943 refGene_gn=KLHDC2;refGene_tx=XM_023255674.1
B3	99813559 99813669 refGene_gn=KLHDC2;refGene_tx=XM_023255674.1
B3	99814055 99814128 refGene_gn=KLHDC2;refGene_tx=XM_023255674.1
B3	99815484 99815572 refGene_gn=KLHDC2;refGene_tx=XM_023255674.1
B3	99815678 99815731 refGene_gn=KLHDC2;refGene_tx=XM_023255674.1
B3	99815917 99816041 refGene_gn=KLHDC2;refGene_tx=XM_023255674.1
B3	99817451 99817529 refGene_gn=NEMF;refGene_tx=XM_023255673.1
B3	99817804 99817884 refGene_gn=NEMF;refGene_tx=XM_023255673.1
B3	99817963 99818007 refGene_gn=NEMF;refGene_tx=XM_023255673.1

B3 99818102 99818203 refGene_gn=NEMF;refGene_tx=XM_023255673.1
B3 99819118 99819151 refGene_gn=NEMF;refGene_tx=XM_023255673.1
B3 99820760 99820958 refGene_gn=NEMF;refGene_tx=XM_023255673.1
B3 99822486 99822540 refGene_gn=NEMF;refGene_tx=XM_023255673.1
B3 99822541 99822565 refGene_gn=NEMF;refGene_tx=XM_023255673.1
B3 99828700 99828854 refGene_gn=NEMF;refGene_tx=XM_023255673.1
B3 99830500 99830550 refGene_gn=NEMF;refGene_tx=XM_023255673.1
B3 99830696 99830739 refGene_gn=NEMF;refGene_tx=XM_023255673.1
B3 99831471 99831751 refGene_gn=NEMF;refGene_tx=XM_023255673.1
B3 99833380 99833501 refGene_gn=NEMF;refGene_tx=XM_023255673.1
B3 99833589 99833648 refGene_gn=NEMF;refGene_tx=XM_023255673.1
B3 99834061 99834119 refGene_gn=NEMF;refGene_tx=XM_023255673.1
B3 99835566 99835679 refGene_gn=NEMF;refGene_tx=XM_023255673.1
B3 99841889 99841952 refGene_gn=NEMF;refGene_tx=XM_023255673.1
B3 99843340 99843444 refGene_gn=NEMF;refGene_tx=XM_023255673.1
B3 99850280 99850369 refGene_gn=NEMF;refGene_tx=XM_023255673.1
B3 99852965 99853029 refGene_gn=NEMF;refGene_tx=XM_023255673.1
B3 99853341 99853533 refGene_gn=NEMF;refGene_tx=XM_023255673.1
B3 99853789 99853998 refGene_gn=NEMF;refGene_tx=XM_023255673.1
B3 99854137 99854215 refGene_gn=NEMF;refGene_tx=XM_023255673.1
B3 99855572 99855635 refGene_gn=NEMF;refGene_tx=XM_023255673.1
B3 99856023 99856099 refGene_gn=NEMF;refGene_tx=XM_023255673.1
B3 99856177 99856248 refGene_gn=NEMF;refGene_tx=XM_023255673.1
B3 99857359 99857433 refGene_gn=NEMF;refGene_tx=XM_023255673.1
B3 99858823 99858910 refGene_gn=NEMF;refGene_tx=XM_023255673.1
B3 99859884 99859952 refGene_gn=NEMF;refGene_tx=XM_023255673.1
B3 99860920 99861069 refGene_gn=NEMF;refGene_tx=XM_023255673.1
B3 99867971 99868097 refGene_gn=NEMF;refGene_tx=XM_023255673.1
B3 99870974 99871077 refGene_gn=NEMF;refGene_tx=XM_023255673.1
B3 99871223 99871292 refGene_gn=NEMF;refGene_tx=XM_023255673.1
B3 99872120 99872179 refGene_gn=NEMF;refGene_tx=XM_023255673.1
B3 99909268 99909796 refGene_gn=ARF6;refGene_tx=XM_003987620.4
B3 100102635 100102650
refGene_gn=VCPKMT;refGene_tx=XM_023255677.1
B3 100105707 100105875
refGene_gn=VCPKMT;refGene_tx=XM_023255676.1;refGene_gn=VCPKMT;refGene_tx=XM_023255677.1
B3 100108024 100108144
refGene_gn=VCPKMT;refGene_tx=XM_023255676.1;refGene_gn=VCPKMT;refGene_tx=XM_023255677.1
B3 100108960 100109033
refGene_gn=VCPKMT;refGene_tx=XM_023255676.1;refGene_gn=VCPKMT;refGene_tx=XM_023255677.1
B3 100109186 100109297
refGene_gn=VCPKMT;refGene_tx=XM_023255676.1;refGene_gn=VCPKMT;refGene_tx=XM_023255677.1
B3 100109394 100109660
refGene_gn=VCPKMT;refGene_tx=XM_023255676.1;refGene_gn=VCPKMT;refGene_tx=XM_023255677.1
B3 100111539 100112049

refGene_gn=SOS2;refGene_tx=XM_023255675.1
B3 100112887 100112997
refGene_gn=SOS2;refGene_tx=XM_023255675.1
B3 100121310 100121352
refGene_gn=SOS2;refGene_tx=XM_023255675.1
B3 100121882 100122144
refGene_gn=SOS2;refGene_tx=XM_023255675.1
B3 100123422 100123539
refGene_gn=SOS2;refGene_tx=XM_023255675.1
B3 100126667 100126840
refGene_gn=SOS2;refGene_tx=XM_023255675.1
B3 100127826 100127944
refGene_gn=SOS2;refGene_tx=XM_023255675.1
B3 100131665 100131828
refGene_gn=SOS2;refGene_tx=XM_023255675.1
B3 100131996 100132058
refGene_gn=SOS2;refGene_tx=XM_023255675.1
B3 100132059 100132117
refGene_gn=SOS2;refGene_tx=XM_023255675.1
B3 100135287 100135510
refGene_gn=SOS2;refGene_tx=XM_023255675.1
B3 100137326 100137430
refGene_gn=SOS2;refGene_tx=XM_023255675.1
B3 100141323 100141446
refGene_gn=SOS2;refGene_tx=XM_023255675.1
B3 100142849 100142931
refGene_gn=SOS2;refGene_tx=XM_023255675.1
B3 100144388 100145044
refGene_gn=SOS2;refGene_tx=XM_023255675.1
B3 100146103 100146231
refGene_gn=SOS2;refGene_tx=XM_023255675.1
B3 100159028 100159127
refGene_gn=SOS2;refGene_tx=XM_023255675.1
B3 100162956 100163067
refGene_gn=SOS2;refGene_tx=XM_023255675.1
B3 100165950 100166094
refGene_gn=SOS2;refGene_tx=XM_023255675.1
B3 100169286 100169490
refGene_gn=SOS2;refGene_tx=XM_023255675.1
B3 100172993 100173158
refGene_gn=SOS2;refGene_tx=XM_023255675.1
B3 100174057 100174189
refGene_gn=SOS2;refGene_tx=XM_023255675.1
B3 100176813 100176939
refGene_gn=SOS2;refGene_tx=XM_023255675.1
B3 100200893 100200980
refGene_gn=SOS2;refGene_tx=XM_023255675.1
B3 100207108 100207304
refGene_gn=L2HGDH;refGene_tx=XM_023255678.1;refGene_gn=L2HGDH;refGene_tx=XM_023255679.1;refGene_gn=L2HGDH;refGene_tx=XM_023255680.1

B3 100218630 100218762
refGene_gn=L2HGDH;refGene_tx=XM_023255678.1;refGene_gn=L2HGDH;refGene_tx=XM_023255679.1;refGene_gn=L2HGDH;refGene_tx=XM_023255680.1
B3 100220243 100220401
refGene_gn=L2HGDH;refGene_tx=XM_023255678.1;refGene_gn=L2HGDH;refGene_tx=XM_023255679.1;refGene_gn=L2HGDH;refGene_tx=XM_023255680.1
B3 100221354 100221522
refGene_gn=L2HGDH;refGene_tx=XM_023255678.1;refGene_gn=L2HGDH;refGene_tx=XM_023255679.1;refGene_gn=L2HGDH;refGene_tx=XM_023255680.1
B3 100227682 100227717
refGene_gn=L2HGDH;refGene_tx=XM_023255678.1;refGene_gn=L2HGDH;refGene_tx=XM_023255679.1
B3 100229692 100229855
refGene_gn=L2HGDH;refGene_tx=XM_023255678.1;refGene_gn=L2HGDH;refGene_tx=XM_023255679.1
B3 100236594 100236726
refGene_gn=L2HGDH;refGene_tx=XM_023255678.1;refGene_gn=L2HGDH;refGene_tx=XM_023255680.1
B3 100243479 100243631
refGene_gn=L2HGDH;refGene_tx=XM_023255678.1;refGene_gn=L2HGDH;refGene_tx=XM_023255679.1;refGene_gn=L2HGDH;refGene_tx=XM_023255680.1
B3 100244361 100244477
refGene_gn=L2HGDH;refGene_tx=XM_023255678.1;refGene_gn=L2HGDH;refGene_tx=XM_023255679.1;refGene_gn=L2HGDH;refGene_tx=XM_023255680.1
B3 100247971 100248111
refGene_gn=L2HGDH;refGene_tx=XM_023255678.1;refGene_gn=L2HGDH;refGene_tx=XM_023255679.1;refGene_gn=L2HGDH;refGene_tx=XM_023255680.1
B3 100261132 100261240
refGene_gn=CDKL1;refGene_tx=XM_003987625.4;refGene_gn=CDKL1;refGene_tx=XM_023255681.1
B3 100263334 100263490
refGene_gn=CDKL1;refGene_tx=XM_003987625.4;refGene_gn=CDKL1;refGene_tx=XM_023255681.1
B3 100264701 100264758
refGene_gn=CDKL1;refGene_tx=XM_003987625.4;refGene_gn=CDKL1;refGene_tx=XM_023255681.1
B3 100267587 100267670
refGene_gn=CDKL1;refGene_tx=XM_003987625.4;refGene_gn=CDKL1;refGene_tx=XM_023255681.1
B3 100269881 100270082
refGene_gn=CDKL1;refGene_tx=XM_003987625.4;refGene_gn=CDKL1;refGene_tx=XM_023255681.1
B3 100270698 100270789
refGene_gn=CDKL1;refGene_tx=XM_003987625.4;refGene_gn=CDKL1;refGene_tx=XM_023255681.1
B3 100273203 100273276
refGene_gn=CDKL1;refGene_tx=XM_003987625.4;refGene_gn=CDKL1;refGene_tx=XM_023255681.1
B3 100284057 100284179
refGene_gn=CDKL1;refGene_tx=XM_003987625.4;refGene_gn=CDKL1;refGene_tx

=XM_023255681.1
B3	100317146	100317317
refGene_gn=CDKL1;refGene_tx=XM_003987625.4;refGene_gn=CDKL1;refGene_tx
=XM_023255681.1
B3	100343879	100343967
refGene_gn=MAP4K5;refGene_tx=XM_003987627.5
B3	100345545	100345601
refGene_gn=MAP4K5;refGene_tx=XM_003987627.5
B3	100347852	100347923
refGene_gn=MAP4K5;refGene_tx=XM_003987627.5
B3	100350703	100350796
refGene_gn=MAP4K5;refGene_tx=XM_003987627.5
B3	100352750	100352819
refGene_gn=MAP4K5;refGene_tx=XM_003987627.5
B3	100355766	100355944
refGene_gn=MAP4K5;refGene_tx=XM_003987627.5
B3	100356331	100356435
refGene_gn=MAP4K5;refGene_tx=XM_003987627.5
B3	100358623	100358682
refGene_gn=MAP4K5;refGene_tx=XM_003987627.5
B3	100359048	100359163
refGene_gn=MAP4K5;refGene_tx=XM_003987627.5
B3	100359247	100359250
refGene_gn=MAP4K5;refGene_tx=XM_003987627.5
B3	100362591	100362652
refGene_gn=MAP4K5;refGene_tx=XM_003987627.5
B3	100362953	100363033
refGene_gn=MAP4K5;refGene_tx=XM_003987627.5
B3	100365814	100365899
refGene_gn=MAP4K5;refGene_tx=XM_003987627.5
B3	100366768	100366810
refGene_gn=MAP4K5;refGene_tx=XM_003987627.5
B3	100367001	100367099
refGene_gn=MAP4K5;refGene_tx=XM_003987627.5
B3	100368016	100368170
refGene_gn=MAP4K5;refGene_tx=XM_003987627.5
B3	100368793	100368836
refGene_gn=MAP4K5;refGene_tx=XM_003987627.5
B3	100370184	100370252
refGene_gn=MAP4K5;refGene_tx=XM_003987627.5
B3	100371272	100371331
refGene_gn=MAP4K5;refGene_tx=XM_003987627.5
B3	100376420	100376499
refGene_gn=MAP4K5;refGene_tx=XM_003987627.5
B3	100381376	100381493
refGene_gn=MAP4K5;refGene_tx=XM_003987627.5
B3	100382529	100382611
refGene_gn=MAP4K5;refGene_tx=XM_003987627.5
B3	100384660	100384723
refGene_gn=MAP4K5;refGene_tx=XM_003987627.5

B3	100387363	100387495
refGene_gn=MAP4K5;refGene_tx=XM_003987627.5
B3	100393463	100393483
refGene_gn=MAP4K5;refGene_tx=XM_003987627.5
B3	100393484	100393537
refGene_gn=MAP4K5;refGene_tx=XM_003987627.5
B3	100394342	100394385
refGene_gn=MAP4K5;refGene_tx=XM_003987627.5
B3	100394470	100394490
refGene_gn=MAP4K5;refGene_tx=XM_003987627.5
B3	100394492	100394517
refGene_gn=MAP4K5;refGene_tx=XM_003987627.5
B3	100399694	100399750
refGene_gn=MAP4K5;refGene_tx=XM_003987627.5
B3	100405686	100405751
refGene_gn=MAP4K5;refGene_tx=XM_003987627.5
B3	100406250	100406341
refGene_gn=MAP4K5;refGene_tx=XM_003987627.5
B3	100432776	100432834
refGene_gn=MAP4K5;refGene_tx=XM_003987627.5
B3	100455779	100455887
refGene_gn=MAP4K5;refGene_tx=XM_003987627.5
B3	100468812	100468846
refGene_gn=ATL1;refGene_tx=XM_003987628.5;refGene_gn=ATL1;refGene_tx=XM_003987629.5;refGene_gn=ATL1;refGene_tx=XM_006932855.4
B3	100507239	100507487
refGene_gn=ATL1;refGene_tx=XM_003987628.5;refGene_gn=ATL1;refGene_tx=XM_003987629.5;refGene_gn=ATL1;refGene_tx=XM_006932855.4
B3	100509769	100509904
refGene_gn=ATL1;refGene_tx=XM_003987628.5;refGene_gn=ATL1;refGene_tx=XM_003987629.5;refGene_gn=ATL1;refGene_tx=XM_006932855.4
B3	100510344	100510449
refGene_gn=ATL1;refGene_tx=XM_003987628.5;refGene_gn=ATL1;refGene_tx=XM_003987629.5;refGene_gn=ATL1;refGene_tx=XM_006932855.4
B3	100512518	100512569
refGene_gn=ATL1;refGene_tx=XM_003987628.5;refGene_gn=ATL1;refGene_tx=XM_003987629.5;refGene_gn=ATL1;refGene_tx=XM_006932855.4
B3	100514980	100515037
refGene_gn=ATL1;refGene_tx=XM_003987628.5;refGene_gn=ATL1;refGene_tx=XM_003987629.5
B3	100529921	100530014
refGene_gn=ATL1;refGene_tx=XM_003987628.5;refGene_gn=ATL1;refGene_tx=XM_003987629.5;refGene_gn=ATL1;refGene_tx=XM_006932855.4
B3	100531031	100531170
refGene_gn=ATL1;refGene_tx=XM_003987628.5;refGene_gn=ATL1;refGene_tx=XM_003987629.5;refGene_gn=ATL1;refGene_tx=XM_006932855.4
B3	100535930	100536058
refGene_gn=ATL1;refGene_tx=XM_003987628.5;refGene_gn=ATL1;refGene_tx=XM_003987629.5;refGene_gn=ATL1;refGene_tx=XM_006932855.4
B3	100537141	100537198

refGene_gn=ATL1;refGene_tx=XM_003987628.5;refGene_gn=ATL1;refGene_tx=XM_003987629.5;refGene_gn=ATL1;refGene_tx=XM_006932855.4
B3 100538270 100538342
refGene_gn=ATL1;refGene_tx=XM_003987628.5;refGene_gn=ATL1;refGene_tx=XM_003987629.5;refGene_gn=ATL1;refGene_tx=XM_006932855.4
B3 100539662 100540094
refGene_gn=ATL1;refGene_tx=XM_003987628.5;refGene_gn=ATL1;refGene_tx=XM_003987629.5;refGene_gn=ATL1;refGene_tx=XM_006932855.4
B3 100541296 100541311
refGene_gn=ATL1;refGene_tx=XM_003987628.5;refGene_gn=ATL1;refGene_tx=XM_006932855.4
B3 100543612 100543723
refGene_gn=ATL1;refGene_tx=XM_003987628.5;refGene_gn=ATL1;refGene_tx=XM_003987629.5;refGene_gn=ATL1;refGene_tx=XM_006932855.4
B3 100546154 100546356
refGene_gn=SAV1;refGene_tx=XM_003987630.4
B3 100552284 100552428
refGene_gn=SAV1;refGene_tx=XM_003987630.4
B3 100556098 100556369
refGene_gn=SAV1;refGene_tx=XM_003987630.4
B3 100566020 100566464
refGene_gn=SAV1;refGene_tx=XM_003987630.4
B3 100569587 100569681
refGene_gn=SAV1;refGene_tx=XM_003987630.4
B3 100607903 100608113
refGene_gn=NIN;refGene_tx=XM_023255682.1
B3 100610254 100610368
refGene_gn=NIN;refGene_tx=XM_023255682.1
B3 100613917 100614118
refGene_gn=NIN;refGene_tx=XM_023255682.1
B3 100617555 100617657
refGene_gn=NIN;refGene_tx=XM_023255682.1
B3 100619812 100619962
refGene_gn=NIN;refGene_tx=XM_023255682.1
B3 100621387 100621555
refGene_gn=NIN;refGene_tx=XM_023255682.1
B3 100623285 100623432
refGene_gn=NIN;refGene_tx=XM_023255682.1
B3 100624168 100624282
refGene_gn=NIN;refGene_tx=XM_023255682.1
B3 100624948 100625071
refGene_gn=NIN;refGene_tx=XM_023255682.1
B3 100627798 100627912
refGene_gn=NIN;refGene_tx=XM_023255682.1
B3 100630971 100631115
refGene_gn=NIN;refGene_tx=XM_023255682.1
B3 100631116 100631188
refGene_gn=NIN;refGene_tx=XM_023255682.1
B3 100632714 100632784
refGene_gn=NIN;refGene_tx=XM_023255682.1

B3 100632888 100633014
refGene_gn=NIN;refGene_tx=XM_023255682.1
B3 100634280 100636422
refGene_gn=NIN;refGene_tx=XM_023255682.1
B3 100637469 100637975
refGene_gn=NIN;refGene_tx=XM_023255682.1
B3 100639249 100639371
refGene_gn=NIN;refGene_tx=XM_023255682.1
B3 100641221 100641360
refGene_gn=NIN;refGene_tx=XM_023255682.1
B3 100644815 100644905
refGene_gn=NIN;refGene_tx=XM_023255682.1
B3 100645293 100645404
refGene_gn=NIN;refGene_tx=XM_023255682.1
B3 100648115 100648290
refGene_gn=NIN;refGene_tx=XM_023255682.1
B3 100648528 100648669
refGene_gn=NIN;refGene_tx=XM_023255682.1
B3 100648976 100649113
refGene_gn=NIN;refGene_tx=XM_023255682.1
B3 100649605 100649773
refGene_gn=NIN;refGene_tx=XM_023255682.1
B3 100650250 100650397
refGene_gn=NIN;refGene_tx=XM_023255682.1
B3 100654519 100654710
refGene_gn=NIN;refGene_tx=XM_023255682.1
B3 100655971 100656011
refGene_gn=NIN;refGene_tx=XM_023255682.1
B3 100671256 100671417
refGene_gn=NIN;refGene_tx=XM_023255682.1
B3 100683977 100684059
refGene_gn=NIN;refGene_tx=XM_023255682.1
B3 100694643 100694826
refGene_gn=NIN;refGene_tx=XM_023255682.1
B3 100739816 100739872
refGene_gn=ABHD12B;refGene_tx=XM_003987669.4
B3 100744887 100745015
refGene_gn=ABHD12B;refGene_tx=XM_003987669.4
B3 100746144 100746247
refGene_gn=ABHD12B;refGene_tx=XM_003987669.4
B3 100747874 100747994
refGene_gn=ABHD12B;refGene_tx=XM_003987669.4
B3 100749933 100749964
refGene_gn=ABHD12B;refGene_tx=XM_003987669.4
B3 100753165 100753211
refGene_gn=ABHD12B;refGene_tx=XM_003987669.4
B3 100753317 100753447
refGene_gn=ABHD12B;refGene_tx=XM_003987669.4
B3 100753867 100753905
refGene_gn=ABHD12B;refGene_tx=XM_003987669.4

B3 100758025 100758105
refGene_gn=ABHD12B;refGene_tx=XM_003987669.4
B3 100766905 100766988
refGene_gn=ABHD12B;refGene_tx=XM_003987669.4
B3 100767986 100768065
refGene_gn=ABHD12B;refGene_tx=XM_003987669.4
B3 100768649 100768768
refGene_gn=ABHD12B;refGene_tx=XM_003987669.4
B3 100768891 100768919
refGene_gn=ABHD12B;refGene_tx=XM_003987669.4
B3 100769591 100769768
refGene_gn=PYGL;refGene_tx=XM_023255684.1
B3 100772310 100772377
refGene_gn=PYGL;refGene_tx=XM_023255684.1
B3 100772811 100772946
refGene_gn=PYGL;refGene_tx=XM_023255684.1
B3 100781725 100781933
refGene_gn=PYGL;refGene_tx=XM_023255684.1
B3 100783354 100783496
refGene_gn=PYGL;refGene_tx=XM_023255684.1
B3 100783601 100783660
refGene_gn=PYGL;refGene_tx=XM_023255684.1
B3 100783782 100783930
refGene_gn=PYGL;refGene_tx=XM_023255684.1
B3 100784075 100784177
refGene_gn=PYGL;refGene_tx=XM_023255684.1
B3 100784706 100784821
refGene_gn=PYGL;refGene_tx=XM_023255684.1
B3 100785292 100785456
refGene_gn=PYGL;refGene_tx=XM_023255684.1
B3 100785799 100785946
refGene_gn=PYGL;refGene_tx=XM_023255684.1
B3 100786571 100786664
refGene_gn=PYGL;refGene_tx=XM_023255684.1
B3 100786892 100787036
refGene_gn=PYGL;refGene_tx=XM_023255684.1
B3 100789718 100789801
refGene_gn=PYGL;refGene_tx=XM_023255684.1
B3 100790273 100790385
refGene_gn=PYGL;refGene_tx=XM_023255684.1
B3 100794186 100794318
refGene_gn=PYGL;refGene_tx=XM_023255684.1
B3 100803813 100803917
refGene_gn=PYGL;refGene_tx=XM_023255684.1
B3 100807579 100807658
refGene_gn=PYGL;refGene_tx=XM_023255684.1
B3 100810353 100810455
refGene_gn=PYGL;refGene_tx=XM_023255684.1
B3 100815321 100815564
refGene_gn=PYGL;refGene_tx=XM_023255684.1

B3 100843398 100843461
refGene_gn=TRIM9;refGene_tx=XM_011283229.3;refGene_gn=TRIM9;refGene_tx=XM_011283230.3;refGene_gn=TRIM9;refGene_tx=XM_019833002.2;refGene_gn=TRIM9;refGene_tx=XM_019833003.2;refGene_gn=TRIM9;refGene_tx=XM_019833004.2;refGene_gn=TRIM9;refGene_tx=XM_019833006.2;refGene_gn=TRIM9;refGene_tx=XM_019833007.2;refGene_gn=TRIM9;refGene_tx=XM_019833008.2
B3 100845496 100845659
refGene_gn=TRIM9;refGene_tx=XM_011283229.3;refGene_gn=TRIM9;refGene_tx=XM_011283230.3;refGene_gn=TRIM9;refGene_tx=XM_019833002.2;refGene_gn=TRIM9;refGene_tx=XM_019833003.2;refGene_gn=TRIM9;refGene_tx=XM_019833004.2;refGene_gn=TRIM9;refGene_tx=XM_019833006.2;refGene_gn=TRIM9;refGene_tx=XM_019833007.2;refGene_gn=TRIM9;refGene_tx=XM_019833008.2
B3 100848231 100848535
refGene_gn=TRIM9;refGene_tx=XM_011283229.3;refGene_gn=TRIM9;refGene_tx=XM_011283230.3;refGene_gn=TRIM9;refGene_tx=XM_019833002.2;refGene_gn=TRIM9;refGene_tx=XM_019833003.2;refGene_gn=TRIM9;refGene_tx=XM_019833004.2;refGene_gn=TRIM9;refGene_tx=XM_019833006.2;refGene_gn=TRIM9;refGene_tx=XM_019833007.2;refGene_gn=TRIM9;refGene_tx=XM_019833008.2
B3 100849389 100849413
refGene_gn=TRIM9;refGene_tx=XM_019833002.2;refGene_gn=TRIM9;refGene_tx=XM_019833003.2;refGene_gn=TRIM9;refGene_tx=XM_019833004.2;refGene_gn=TRIM9;refGene_tx=XM_019833006.2;refGene_gn=TRIM9;refGene_tx=XM_019833007.2;refGene_gn=TRIM9;refGene_tx=XM_019833008.2
B3 100849818 100849860
refGene_gn=TRIM9;refGene_tx=XM_019833002.2;refGene_gn=TRIM9;refGene_tx=XM_019833003.2;refGene_gn=TRIM9;refGene_tx=XM_019833004.2;refGene_gn=TRIM9;refGene_tx=XM_019833006.2;refGene_gn=TRIM9;refGene_tx=XM_019833008.2
B3 100852493 100852682
refGene_gn=TRIM9;refGene_tx=XM_019833002.2;refGene_gn=TRIM9;refGene_tx=XM_019833003.2;refGene_gn=TRIM9;refGene_tx=XM_019833004.2
B3 100865106 100865245
refGene_gn=TRIM9;refGene_tx=XM_011283229.3;refGene_gn=TRIM9;refGene_tx=XM_011283230.3;refGene_gn=TRIM9;refGene_tx=XM_019833002.2;refGene_gn=TRIM9;refGene_tx=XM_019833003.2;refGene_gn=TRIM9;refGene_tx=XM_019833004.2;refGene_gn=TRIM9;refGene_tx=XM_019833006.2;refGene_gn=TRIM9;refGene_tx=XM_019833007.2;refGene_gn=TRIM9;refGene_tx=XM_019833008.2
B3 100866438 100866596
refGene_gn=TRIM9;refGene_tx=XM_011283229.3;refGene_gn=TRIM9;refGene_tx=XM_011283230.3;refGene_gn=TRIM9;refGene_tx=XM_019833002.2;refGene_gn=TRIM9;refGene_tx=XM_019833003.2;refGene_gn=TRIM9;refGene_tx=XM_019833004.2;refGene_gn=TRIM9;refGene_tx=XM_019833006.2;refGene_gn=TRIM9;refGene_tx=XM_019833007.2;refGene_gn=TRIM9;refGene_tx=XM_019833008.2
B3 100874470 100874624
refGene_gn=TRIM9;refGene_tx=XM_011283229.3;refGene_gn=TRIM9;refGene_tx=XM_011283230.3;refGene_gn=TRIM9;refGene_tx=XM_019833002.2;refGene_gn=TRIM9;refGene_tx=XM_019833003.2;refGene_gn=TRIM9;refGene_tx=XM_019833004.2;refGene_gn=TRIM9;refGene_tx=XM_019833006.2;refGene_gn=TRIM9;refGene_tx=XM_019833007.2;refGene_gn=TRIM9;refGene_tx=XM_019833008.2
B3 100876206 100876317

refGene_gn=TRIM9;refGene_tx=XM_011283229.3;refGene_gn=TRIM9;refGene_tx=XM_011283230.3;refGene_gn=TRIM9;refGene_tx=XM_019833002.2;refGene_gn=TRIM9;refGene_tx=XM_019833003.2;refGene_gn=TRIM9;refGene_tx=XM_019833004.2;refGene_gn=TRIM9;refGene_tx=XM_019833006.2;refGene_gn=TRIM9;refGene_tx=XM_019833007.2;refGene_gn=TRIM9;refGene_tx=XM_019833008.2
B3 100888732 100888855
refGene_gn=TRIM9;refGene_tx=XM_011283229.3;refGene_gn=TRIM9;refGene_tx=XM_011283230.3;refGene_gn=TRIM9;refGene_tx=XM_019833002.2;refGene_gn=TRIM9;refGene_tx=XM_019833003.2;refGene_gn=TRIM9;refGene_tx=XM_019833004.2;refGene_gn=TRIM9;refGene_tx=XM_019833006.2;refGene_gn=TRIM9;refGene_tx=XM_019833007.2;refGene_gn=TRIM9;refGene_tx=XM_019833008.2
B3 100891174 100891270
refGene_gn=TRIM9;refGene_tx=XM_011283229.3;refGene_gn=TRIM9;refGene_tx=XM_011283230.3;refGene_gn=TRIM9;refGene_tx=XM_019833002.2;refGene_gn=TRIM9;refGene_tx=XM_019833003.2;refGene_gn=TRIM9;refGene_tx=XM_019833004.2;refGene_gn=TRIM9;refGene_tx=XM_019833006.2;refGene_gn=TRIM9;refGene_tx=XM_019833007.2;refGene_gn=TRIM9;refGene_tx=XM_019833008.2
B3 100952922 100953744
refGene_gn=TRIM9;refGene_tx=XM_011283229.3;refGene_gn=TRIM9;refGene_tx=XM_011283230.3;refGene_gn=TRIM9;refGene_tx=XM_019833002.2;refGene_gn=TRIM9;refGene_tx=XM_019833003.2;refGene_gn=TRIM9;refGene_tx=XM_019833004.2;refGene_gn=TRIM9;refGene_tx=XM_019833006.2;refGene_gn=TRIM9;refGene_tx=XM_019833007.2;refGene_gn=TRIM9;refGene_tx=XM_019833008.2
B3 101061499 101061651
refGene_gn=TMX1;refGene_tx=XM_003987633.4
B3 101065416 101065532
refGene_gn=TMX1;refGene_tx=XM_003987633.4
B3 101067335 101067381
refGene_gn=TMX1;refGene_tx=XM_003987633.4
B3 101070224 101070353
refGene_gn=TMX1;refGene_tx=XM_003987633.4
B3 101072324 101072370
refGene_gn=TMX1;refGene_tx=XM_003987633.4
B3 101072471 101072573
refGene_gn=TMX1;refGene_tx=XM_003987633.4
B3 101072689 101072756
refGene_gn=TMX1;refGene_tx=XM_003987633.4
B3 101075596 101075775
refGene_gn=TMX1;refGene_tx=XM_003987633.4
B3 101506891 101506990
refGene_gn=FRMD6;refGene_tx=XM_011283232.2;refGene_gn=FRMD6;refGene_tx=XM_019833010.2;refGene_gn=FRMD6;refGene_tx=XM_019833011.2;refGene_gn=FRMD6;refGene_tx=XM_019833012.2;refGene_gn=FRMD6;refGene_tx=XM_023255685.1;refGene_gn=FRMD6;refGene_tx=XM_023255686.1
B3 101512696 101512787
refGene_gn=FRMD6;refGene_tx=XM_011283232.2;refGene_gn=FRMD6;refGene_tx=XM_019833010.2;refGene_gn=FRMD6;refGene_tx=XM_019833011.2;refGene_gn=FRMD6;refGene_tx=XM_019833012.2;refGene_gn=FRMD6;refGene_tx=XM_023255685.1;refGene_gn=FRMD6;refGene_tx=XM_023255686.1
B3 101515679 101515783

refGene_gn=FRMD6;refGene_tx=XM_006932860.4;refGene_gn=FRMD6;refGene_tx
=XM_011283232.2;refGene_gn=FRMD6;refGene_tx=XM_019833010.2;refGene_gn=
FRMD6;refGene_tx=XM_019833011.2;refGene_gn=FRMD6;refGene_tx=XM_0198330
12.2;refGene_gn=FRMD6;refGene_tx=XM_019833013.2;refGene_gn=FRMD6;refGe
ne_tx=XM_023255685.1;refGene_gn=FRMD6;refGene_tx=XM_023255686.1
B3 101516797 101516874
refGene_gn=FRMD6;refGene_tx=XM_006932860.4;refGene_gn=FRMD6;refGene_tx
=XM_011283232.2;refGene_gn=FRMD6;refGene_tx=XM_019833010.2;refGene_gn=
FRMD6;refGene_tx=XM_019833011.2;refGene_gn=FRMD6;refGene_tx=XM_0198330
12.2;refGene_gn=FRMD6;refGene_tx=XM_019833013.2;refGene_gn=FRMD6;refGe
ne_tx=XM_023255685.1;refGene_gn=FRMD6;refGene_tx=XM_023255686.1
B3 101519144 101519331
refGene_gn=FRMD6;refGene_tx=XM_006932860.4;refGene_gn=FRMD6;refGene_tx
=XM_011283232.2;refGene_gn=FRMD6;refGene_tx=XM_019833010.2;refGene_gn=
FRMD6;refGene_tx=XM_019833011.2;refGene_gn=FRMD6;refGene_tx=XM_0198330
12.2;refGene_gn=FRMD6;refGene_tx=XM_019833013.2;refGene_gn=FRMD6;refGe
ne_tx=XM_023255685.1;refGene_gn=FRMD6;refGene_tx=XM_023255686.1
B3 101522559 101522715
refGene_gn=FRMD6;refGene_tx=XM_006932860.4;refGene_gn=FRMD6;refGene_tx
=XM_011283232.2;refGene_gn=FRMD6;refGene_tx=XM_019833010.2;refGene_gn=
FRMD6;refGene_tx=XM_019833011.2;refGene_gn=FRMD6;refGene_tx=XM_0198330
12.2;refGene_gn=FRMD6;refGene_tx=XM_019833013.2;refGene_gn=FRMD6;refGe
ne_tx=XM_023255685.1;refGene_gn=FRMD6;refGene_tx=XM_023255686.1
B3 101526115 101526181
refGene_gn=FRMD6;refGene_tx=XM_006932860.4;refGene_gn=FRMD6;refGene_tx
=XM_011283232.2;refGene_gn=FRMD6;refGene_tx=XM_019833010.2;refGene_gn=
FRMD6;refGene_tx=XM_019833011.2;refGene_gn=FRMD6;refGene_tx=XM_0198330
12.2;refGene_gn=FRMD6;refGene_tx=XM_019833013.2;refGene_gn=FRMD6;refGe
ne_tx=XM_023255685.1;refGene_gn=FRMD6;refGene_tx=XM_023255686.1
B3 101527047 101527116
refGene_gn=FRMD6;refGene_tx=XM_006932860.4;refGene_gn=FRMD6;refGene_tx
=XM_011283232.2;refGene_gn=FRMD6;refGene_tx=XM_019833010.2;refGene_gn=
FRMD6;refGene_tx=XM_019833011.2;refGene_gn=FRMD6;refGene_tx=XM_0198330
13.2;refGene_gn=FRMD6;refGene_tx=XM_023255685.1;refGene_gn=FRMD6;refGe
ne_tx=XM_023255686.1
B3 101529993 101530168
refGene_gn=FRMD6;refGene_tx=XM_006932860.4;refGene_gn=FRMD6;refGene_tx
=XM_011283232.2;refGene_gn=FRMD6;refGene_tx=XM_019833010.2;refGene_gn=
FRMD6;refGene_tx=XM_019833011.2;refGene_gn=FRMD6;refGene_tx=XM_0198330
12.2;refGene_gn=FRMD6;refGene_tx=XM_019833013.2;refGene_gn=FRMD6;refGe
ne_tx=XM_023255685.1;refGene_gn=FRMD6;refGene_tx=XM_023255686.1
B3 101534617 101534953
refGene_gn=FRMD6;refGene_tx=XM_006932860.4;refGene_gn=FRMD6;refGene_tx
=XM_011283232.2;refGene_gn=FRMD6;refGene_tx=XM_019833010.2;refGene_gn=
FRMD6;refGene_tx=XM_019833011.2;refGene_gn=FRMD6;refGene_tx=XM_0198330
12.2;refGene_gn=FRMD6;refGene_tx=XM_019833013.2;refGene_gn=FRMD6;refGe
ne_tx=XM_023255685.1
B3 101536072 101536204
refGene_gn=FRMD6;refGene_tx=XM_006932860.4;refGene_gn=FRMD6;refGene_tx
=XM_011283232.2;refGene_gn=FRMD6;refGene_tx=XM_019833010.2;refGene_gn=

FRMD6;refGene_tx=XM_019833011.2;refGene_gn=FRMD6;refGene_tx=XM_019833012.2;refGene_gn=FRMD6;refGene_tx=XM_019833013.2;refGene_gn=FRMD6;refGene_tx=XM_023255685.1;refGene_gn=FRMD6;refGene_tx=XM_023255686.1
B3 101540061 101540153
refGene_gn=FRMD6;refGene_tx=XM_006932860.4;refGene_gn=FRMD6;refGene_tx=XM_011283232.2;refGene_gn=FRMD6;refGene_tx=XM_019833010.2;refGene_gn=FRMD6;refGene_tx=XM_019833011.2;refGene_gn=FRMD6;refGene_tx=XM_019833012.2;refGene_gn=FRMD6;refGene_tx=XM_019833013.2;refGene_gn=FRMD6;refGene_tx=XM_023255685.1;refGene_gn=FRMD6;refGene_tx=XM_023255686.1
B3 101542074 101542359
refGene_gn=FRMD6;refGene_tx=XM_006932860.4;refGene_gn=FRMD6;refGene_tx=XM_011283232.2;refGene_gn=FRMD6;refGene_tx=XM_019833010.2;refGene_gn=FRMD6;refGene_tx=XM_019833011.2;refGene_gn=FRMD6;refGene_tx=XM_019833012.2;refGene_gn=FRMD6;refGene_tx=XM_019833013.2;refGene_gn=FRMD6;refGene_tx=XM_023255685.1;refGene_gn=FRMD6;refGene_tx=XM_023255686.1
B3 101738126 101738213
refGene_gn=GNG2;refGene_tx=XM_011283234.3;refGene_gn=GNG2;refGene_tx=XM_019833015.2;refGene_gn=GNG2;refGene_tx=XM_023255687.1
B3 101761391 101761520
refGene_gn=GNG2;refGene_tx=XM_011283234.3;refGene_gn=GNG2;refGene_tx=XM_019833015.2;refGene_gn=GNG2;refGene_tx=XM_023255687.1
B3 101795476 101795537
refGene_gn=RTRAF;refGene_tx=XM_003987637.3
B3 101797337 101797462
refGene_gn=RTRAF;refGene_tx=XM_003987637.3
B3 101799431 101799531
refGene_gn=RTRAF;refGene_tx=XM_003987637.3
B3 101806392 101806479
refGene_gn=RTRAF;refGene_tx=XM_003987637.3
B3 101807355 101807444
refGene_gn=RTRAF;refGene_tx=XM_003987637.3
B3 101810062 101810131
refGene_gn=RTRAF;refGene_tx=XM_003987637.3
B3 101812377 101812426
refGene_gn=RTRAF;refGene_tx=XM_003987637.3
B3 101812545 101812700
refGene_gn=RTRAF;refGene_tx=XM_003987637.3
B3 101813635 101813646
refGene_gn=NID2;refGene_tx=XM_023255688.1
B3 101813936 101814049
refGene_gn=NID2;refGene_tx=XM_023255688.1
B3 101814909 101815033
refGene_gn=NID2;refGene_tx=XM_023255688.1
B3 101816502 101816660
refGene_gn=NID2;refGene_tx=XM_023255688.1
B3 101817174 101817346
refGene_gn=NID2;refGene_tx=XM_023255688.1
B3 101817821 101817951
refGene_gn=NID2;refGene_tx=XM_023255688.1
B3 101820391 101820561

refGene_gn=NID2;refGene_tx=XM_023255688.1
B3 101821173 101821395
refGene_gn=NID2;refGene_tx=XM_023255688.1
B3 101825120 101825351
refGene_gn=NID2;refGene_tx=XM_023255688.1
B3 101826165 101826285
refGene_gn=NID2;refGene_tx=XM_023255688.1
B3 101833040 101833184
refGene_gn=NID2;refGene_tx=XM_023255688.1
B3 101834082 101834211
refGene_gn=NID2;refGene_tx=XM_023255688.1
B3 101835150 101835294
refGene_gn=NID2;refGene_tx=XM_023255688.1
B3 101846950 101847181
refGene_gn=NID2;refGene_tx=XM_023255688.1
B3 101848725 101848926
refGene_gn=NID2;refGene_tx=XM_023255688.1
B3 101850079 101850325
refGene_gn=NID2;refGene_tx=XM_023255688.1
B3 101850756 101850906
refGene_gn=NID2;refGene_tx=XM_023255688.1
B3 101861843 101862598
refGene_gn=NID2;refGene_tx=XM_023255688.1
B3 101868025 101868258
refGene_gn=NID2;refGene_tx=XM_023255688.1
B3 101879118 101879427
refGene_gn=NID2;refGene_tx=XM_023255688.1
B3 101880082 101880295
refGene_gn=NID2;refGene_tx=XM_023255688.1
B3 102100681 102101518
refGene_gn=PTGDR;refGene_tx=XM_003987638.5
B3 102113882 102114122
refGene_gn=PTGDR;refGene_tx=XM_003987638.5
B3 102148285 102149128
refGene_gn=PTGER2;refGene_tx=XM_023255689.1
B3 102160457 102160679
refGene_gn=PTGER2;refGene_tx=XM_023255689.1
B3 102279774 102280043
refGene_gn=TXNDC16;refGene_tx=XM_006932861.4;refGene_gn=TXNDC16;refGene_tx=XM_006932862.4;refGene_gn=TXNDC16;refGene_tx=XM_019833019.2
B3 102300771 102300962
refGene_gn=TXNDC16;refGene_tx=XM_006932861.4;refGene_gn=TXNDC16;refGene_tx=XM_006932862.4
B3 102313675 102313836
refGene_gn=TXNDC16;refGene_tx=XM_006932861.4;refGene_gn=TXNDC16;refGene_tx=XM_006932862.4;refGene_gn=TXNDC16;refGene_tx=XM_019833019.2
B3 102318137 102318276
refGene_gn=TXNDC16;refGene_tx=XM_006932861.4;refGene_gn=TXNDC16;refGene_tx=XM_006932862.4;refGene_gn=TXNDC16;refGene_tx=XM_019833019.2
B3 102319944 102320029

refGene_gn=TXNDC16;refGene_tx=XM_006932861.4;refGene_gn=TXNDC16;refGene_tx=XM_006932862.4;refGene_gn=TXNDC16;refGene_tx=XM_019833019.2
B3 102327224 102327361
refGene_gn=TXNDC16;refGene_tx=XM_006932861.4;refGene_gn=TXNDC16;refGene_tx=XM_006932862.4;refGene_gn=TXNDC16;refGene_tx=XM_019833019.2
B3 102327708 102327877
refGene_gn=TXNDC16;refGene_tx=XM_006932861.4;refGene_gn=TXNDC16;refGene_tx=XM_006932862.4;refGene_gn=TXNDC16;refGene_tx=XM_019833019.2
B3 102334438 102334498
refGene_gn=TXNDC16;refGene_tx=XM_006932861.4;refGene_gn=TXNDC16;refGene_tx=XM_006932862.4;refGene_gn=TXNDC16;refGene_tx=XM_019833019.2
B3 102334965 102335109
refGene_gn=TXNDC16;refGene_tx=XM_006932861.4;refGene_gn=TXNDC16;refGene_tx=XM_006932862.4;refGene_gn=TXNDC16;refGene_tx=XM_019833019.2
B3 102338658 102338782
refGene_gn=TXNDC16;refGene_tx=XM_006932861.4;refGene_gn=TXNDC16;refGene_tx=XM_006932862.4;refGene_gn=TXNDC16;refGene_tx=XM_019833019.2
B3 102343201 102343262
refGene_gn=TXNDC16;refGene_tx=XM_006932861.4;refGene_gn=TXNDC16;refGene_tx=XM_006932862.4;refGene_gn=TXNDC16;refGene_tx=XM_019833019.2
B3 102343655 102343822
refGene_gn=TXNDC16;refGene_tx=XM_006932861.4;refGene_gn=TXNDC16;refGene_tx=XM_006932862.4;refGene_gn=TXNDC16;refGene_tx=XM_019833019.2
B3 102357578 102357729
refGene_gn=TXNDC16;refGene_tx=XM_006932861.4;refGene_gn=TXNDC16;refGene_tx=XM_006932862.4;refGene_gn=TXNDC16;refGene_tx=XM_019833019.2
B3 102361866 102361957
refGene_gn=TXNDC16;refGene_tx=XM_006932861.4;refGene_gn=TXNDC16;refGene_tx=XM_006932862.4;refGene_gn=TXNDC16;refGene_tx=XM_019833019.2
B3 102369506 102369628
refGene_gn=TXNDC16;refGene_tx=XM_006932861.4;refGene_gn=TXNDC16;refGene_tx=XM_006932862.4;refGene_gn=TXNDC16;refGene_tx=XM_019833019.2
B3 102399181 102399256
refGene_gn=TXNDC16;refGene_tx=XM_006932861.4;refGene_gn=TXNDC16;refGene_tx=XM_006932862.4;refGene_gn=TXNDC16;refGene_tx=XM_019833019.2
B3 102400364 102400438
refGene_gn=TXNDC16;refGene_tx=XM_006932861.4;refGene_gn=TXNDC16;refGene_tx=XM_006932862.4;refGene_gn=TXNDC16;refGene_tx=XM_019833019.2
B3 102405784 102405867
refGene_gn=TXNDC16;refGene_tx=XM_006932861.4;refGene_gn=TXNDC16;refGene_tx=XM_006932862.4;refGene_gn=TXNDC16;refGene_tx=XM_019833019.2
B3 102407138 102407301
refGene_gn=TXNDC16;refGene_tx=XM_006932861.4;refGene_gn=TXNDC16;refGene_tx=XM_006932862.4;refGene_gn=TXNDC16;refGene_tx=XM_019833019.2
B3 102418127 102418541
refGene_gn=GPR137C;refGene_tx=XM_011283235.3;refGene_gn=GPR137C;refGene_tx=XM_019833020.2
B3 102456934 102456978
refGene_gn=GPR137C;refGene_tx=XM_011283235.3;refGene_gn=GPR137C;refGene_tx=XM_019833020.2

B3	102458722	102458951
refGene_gn=GPR137C;refGene_tx=XM_011283235.3;refGene_gn=GPR137C;refGene_tx=XM_019833020.2
B3	102478156	102478306
refGene_gn=GPR137C;refGene_tx=XM_011283235.3;refGene_gn=GPR137C;refGene_tx=XM_019833020.2
B3	102479498	102479624
refGene_gn=GPR137C;refGene_tx=XM_011283235.3
B3	102479795	102479914
refGene_gn=GPR137C;refGene_tx=XM_011283235.3;refGene_gn=GPR137C;refGene_tx=XM_019833020.2
B3	102481175	102481353
refGene_gn=GPR137C;refGene_tx=XM_011283235.3;refGene_gn=GPR137C;refGene_tx=XM_019833020.2
B3	102487835	102487896
refGene_gn=ERO1A;refGene_tx=XM_023255690.1
B3	102489387	102489521
refGene_gn=ERO1A;refGene_tx=XM_023255690.1
B3	102489611	102489698
refGene_gn=ERO1A;refGene_tx=XM_023255690.1
B3	102495046	102495116
refGene_gn=ERO1A;refGene_tx=XM_023255690.1
B3	102495830	102496077
refGene_gn=ERO1A;refGene_tx=XM_023255690.1
B3	102497376	102497469
refGene_gn=ERO1A;refGene_tx=XM_023255690.1
B3	102497597	102497624
refGene_gn=ERO1A;refGene_tx=XM_023255690.1
B3	102499822	102499834
refGene_gn=ERO1A;refGene_tx=XM_023255690.1
B3	102502121	102502168
refGene_gn=ERO1A;refGene_tx=XM_023255690.1
B3	102504625	102504746
refGene_gn=ERO1A;refGene_tx=XM_023255690.1
B3	102509131	102509205
refGene_gn=ERO1A;refGene_tx=XM_023255690.1
B3	102509307	102509384
refGene_gn=ERO1A;refGene_tx=XM_023255690.1
B3	102517482	102517521
refGene_gn=ERO1A;refGene_tx=XM_023255690.1
B3	102521044	102521128
refGene_gn=ERO1A;refGene_tx=XM_023255690.1
B3	102521877	102521997
refGene_gn=ERO1A;refGene_tx=XM_023255690.1
B3	102532238	102532352
refGene_gn=ERO1A;refGene_tx=XM_023255690.1
B3	102543647	102543774
refGene_gn=PSMC6;refGene_tx=XM_003987641.4
B3	102545011	102545091
refGene_gn=PSMC6;refGene_tx=XM_003987641.4

B3 102545178 102545218
refGene_gn=PSMC6;refGene_tx=XM_003987641.4
B3 102545469 102545522
refGene_gn=PSMC6;refGene_tx=XM_003987641.4
B3 102547671 102547739
refGene_gn=PSMC6;refGene_tx=XM_003987641.4
B3 102547967 102548082
refGene_gn=PSMC6;refGene_tx=XM_003987641.4
B3 102551459 102551547
refGene_gn=PSMC6;refGene_tx=XM_003987641.4
B3 102553356 102553418
refGene_gn=PSMC6;refGene_tx=XM_003987641.4
B3 102553505 102553629
refGene_gn=PSMC6;refGene_tx=XM_003987641.4
B3 102554219 102554281
refGene_gn=PSMC6;refGene_tx=XM_003987641.4
B3 102556854 102556975
refGene_gn=PSMC6;refGene_tx=XM_003987641.4
B3 102557105 102557186
refGene_gn=PSMC6;refGene_tx=XM_003987641.4
B3 102562004 102562076
refGene_gn=PSMC6;refGene_tx=XM_003987641.4
B3 102565663 102565782
refGene_gn=PSMC6;refGene_tx=XM_003987641.4
B3 102568372 102568429
refGene_gn=STYX;refGene_tx=XM_003987642.5;refGene_gn=STYX;refGene_tx=XM_019833022.2
B3 102580952 102580985
refGene_gn=STYX;refGene_tx=XM_003987642.5;refGene_gn=STYX;refGene_tx=XM_019833022.2
B3 102582666 102582720
refGene_gn=STYX;refGene_tx=XM_003987642.5;refGene_gn=STYX;refGene_tx=XM_019833022.2
B3 102586793 102586891
refGene_gn=STYX;refGene_tx=XM_003987642.5;refGene_gn=STYX;refGene_tx=XM_019833022.2
B3 102590285 102590346
refGene_gn=STYX;refGene_tx=XM_003987642.5;refGene_gn=STYX;refGene_tx=XM_019833022.2
B3 102590755 102590792
refGene_gn=STYX;refGene_tx=XM_003987642.5;refGene_gn=STYX;refGene_tx=XM_019833022.2
B3 102591224 102591264
refGene_gn=STYX;refGene_tx=XM_003987642.5;refGene_gn=STYX;refGene_tx=XM_019833022.2
B3 102591355 102591406
refGene_gn=STYX;refGene_tx=XM_003987642.5
B3 102593578 102593651
refGene_gn=STYX;refGene_tx=XM_003987642.5;refGene_gn=STYX;refGene_tx=XM_019833022.2

B3 102599834 102599928
refGene_gn=STYX;refGene_tx=XM_003987642.5;refGene_gn=STYX;refGene_tx=XM_019833022.2
B3 102602107 102602181
refGene_gn=STYX;refGene_tx=XM_003987642.5;refGene_gn=STYX;refGene_tx=XM_019833022.2
B3 102608948 102609096
refGene_gn=GNPNAT1;refGene_tx=XM_003987643.5;refGene_gn=GNPNAT1;refGene_tx=XM_019833023.2;refGene_gn=GNPNAT1;refGene_tx=XM_023255691.1
B3 102610772 102610834
refGene_gn=GNPNAT1;refGene_tx=XM_003987643.5;refGene_gn=GNPNAT1;refGene_tx=XM_019833023.2;refGene_gn=GNPNAT1;refGene_tx=XM_023255691.1
B3 102612206 102612334
refGene_gn=GNPNAT1;refGene_tx=XM_003987643.5;refGene_gn=GNPNAT1;refGene_tx=XM_019833023.2;refGene_gn=GNPNAT1;refGene_tx=XM_023255691.1
B3 102615046 102615109
refGene_gn=GNPNAT1;refGene_tx=XM_003987643.5;refGene_gn=GNPNAT1;refGene_tx=XM_019833023.2;refGene_gn=GNPNAT1;refGene_tx=XM_023255691.1
B3 102616463 102616617
refGene_gn=GNPNAT1;refGene_tx=XM_003987643.5;refGene_gn=GNPNAT1;refGene_tx=XM_019833023.2;refGene_gn=GNPNAT1;refGene_tx=XM_023255691.1
B3 102712163 102712337
refGene_gn=FERMT2;refGene_tx=XM_023255692.1;refGene_gn=FERMT2;refGene_tx=XM_023255693.1;refGene_gn=FERMT2;refGene_tx=XM_023255694.1;refGene_gn=FERMT2;refGene_tx=XM_023255695.1;refGene_gn=FERMT2;refGene_tx=XM_023255696.1;refGene_gn=FERMT2;refGene_tx=XM_023255697.1
B3 102714055 102714197
refGene_gn=FERMT2;refGene_tx=XM_023255692.1;refGene_gn=FERMT2;refGene_tx=XM_023255693.1;refGene_gn=FERMT2;refGene_tx=XM_023255694.1;refGene_gn=FERMT2;refGene_tx=XM_023255695.1;refGene_gn=FERMT2;refGene_tx=XM_023255696.1;refGene_gn=FERMT2;refGene_tx=XM_023255697.1
B3 102714843 102714968
refGene_gn=FERMT2;refGene_tx=XM_023255692.1;refGene_gn=FERMT2;refGene_tx=XM_023255693.1;refGene_gn=FERMT2;refGene_tx=XM_023255694.1;refGene_gn=FERMT2;refGene_tx=XM_023255695.1;refGene_gn=FERMT2;refGene_tx=XM_023255696.1;refGene_gn=FERMT2;refGene_tx=XM_023255697.1
B3 102715600 102715621
refGene_gn=FERMT2;refGene_tx=XM_023255692.1;refGene_gn=FERMT2;refGene_tx=XM_023255693.1;refGene_gn=FERMT2;refGene_tx=XM_023255694.1;refGene_gn=FERMT2;refGene_tx=XM_023255696.1
B3 102716126 102716150
refGene_gn=FERMT2;refGene_tx=XM_023255692.1;refGene_gn=FERMT2;refGene_tx=XM_023255694.1
B3 102718192 102718414
refGene_gn=FERMT2;refGene_tx=XM_023255692.1;refGene_gn=FERMT2;refGene_tx=XM_023255693.1;refGene_gn=FERMT2;refGene_tx=XM_023255694.1;refGene_gn=FERMT2;refGene_tx=XM_023255695.1;refGene_gn=FERMT2;refGene_tx=XM_023255696.1;refGene_gn=FERMT2;refGene_tx=XM_023255697.1
B3 102718548 102718655
refGene_gn=FERMT2;refGene_tx=XM_023255692.1;refGene_gn=FERMT2;refGene_

tx=XM_023255693.1;refGene_gn=FERMT2;refGene_tx=XM_023255694.1;refGene_gn=FERMT2;refGene_tx=XM_023255695.1;refGene_gn=FERMT2;refGene_tx=XM_023255696.1;refGene_gn=FERMT2;refGene_tx=XM_023255697.1
B3 102724889 102725014
refGene_gn=FERMT2;refGene_tx=XM_023255692.1;refGene_gn=FERMT2;refGene_tx=XM_023255693.1;refGene_gn=FERMT2;refGene_tx=XM_023255694.1;refGene_gn=FERMT2;refGene_tx=XM_023255695.1;refGene_gn=FERMT2;refGene_tx=XM_023255696.1;refGene_gn=FERMT2;refGene_tx=XM_023255697.1
B3 102726143 102726193
refGene_gn=FERMT2;refGene_tx=XM_023255692.1;refGene_gn=FERMT2;refGene_tx=XM_023255693.1;refGene_gn=FERMT2;refGene_tx=XM_023255694.1;refGene_gn=FERMT2;refGene_tx=XM_023255695.1;refGene_gn=FERMT2;refGene_tx=XM_023255696.1;refGene_gn=FERMT2;refGene_tx=XM_023255697.1
B3 102727056 102727191
refGene_gn=FERMT2;refGene_tx=XM_023255692.1;refGene_gn=FERMT2;refGene_tx=XM_023255693.1;refGene_gn=FERMT2;refGene_tx=XM_023255694.1;refGene_gn=FERMT2;refGene_tx=XM_023255695.1;refGene_gn=FERMT2;refGene_tx=XM_023255696.1;refGene_gn=FERMT2;refGene_tx=XM_023255697.1
B3 102732683 102732791
refGene_gn=FERMT2;refGene_tx=XM_023255692.1;refGene_gn=FERMT2;refGene_tx=XM_023255693.1;refGene_gn=FERMT2;refGene_tx=XM_023255694.1;refGene_gn=FERMT2;refGene_tx=XM_023255695.1;refGene_gn=FERMT2;refGene_tx=XM_023255696.1;refGene_gn=FERMT2;refGene_tx=XM_023255697.1
B3 102734261 102734364
refGene_gn=FERMT2;refGene_tx=XM_023255692.1;refGene_gn=FERMT2;refGene_tx=XM_023255693.1;refGene_gn=FERMT2;refGene_tx=XM_023255694.1;refGene_gn=FERMT2;refGene_tx=XM_023255695.1;refGene_gn=FERMT2;refGene_tx=XM_023255696.1;refGene_gn=FERMT2;refGene_tx=XM_023255697.1
B3 102734474 102734700
refGene_gn=FERMT2;refGene_tx=XM_023255692.1;refGene_gn=FERMT2;refGene_tx=XM_023255693.1;refGene_gn=FERMT2;refGene_tx=XM_023255694.1;refGene_gn=FERMT2;refGene_tx=XM_023255695.1;refGene_gn=FERMT2;refGene_tx=XM_023255696.1;refGene_gn=FERMT2;refGene_tx=XM_023255697.1
B3 102735040 102735073
refGene_gn=FERMT2;refGene_tx=XM_023255692.1;refGene_gn=FERMT2;refGene_tx=XM_023255693.1;refGene_gn=FERMT2;refGene_tx=XM_023255695.1
B3 102744100 102744235
refGene_gn=FERMT2;refGene_tx=XM_023255692.1;refGene_gn=FERMT2;refGene_tx=XM_023255693.1;refGene_gn=FERMT2;refGene_tx=XM_023255694.1;refGene_gn=FERMT2;refGene_tx=XM_023255695.1;refGene_gn=FERMT2;refGene_tx=XM_023255696.1;refGene_gn=FERMT2;refGene_tx=XM_023255697.1
B3 102763824 102764058
refGene_gn=FERMT2;refGene_tx=XM_023255692.1;refGene_gn=FERMT2;refGene_tx=XM_023255693.1;refGene_gn=FERMT2;refGene_tx=XM_023255694.1;refGene_gn=FERMT2;refGene_tx=XM_023255695.1;refGene_gn=FERMT2;refGene_tx=XM_023255696.1;refGene_gn=FERMT2;refGene_tx=XM_023255697.1
B3 102796113 102796270
refGene_gn=FERMT2;refGene_tx=XM_023255692.1;refGene_gn=FERMT2;refGene_tx=XM_023255693.1;refGene_gn=FERMT2;refGene_tx=XM_023255694.1;refGene_gn=FERMT2;refGene_tx=XM_023255695.1;refGene_gn=FERMT2;refGene_tx=XM_02

3255696.1;refGene_gn=FERMT2;refGene_tx=XM_023255697.1
B3	102865539	102865721
refGene_gn=DDHD1;refGene_tx=XM_023255698.1
B3	102870549	102870633
refGene_gn=DDHD1;refGene_tx=XM_023255698.1
B3	102873433	102873625
refGene_gn=DDHD1;refGene_tx=XM_023255698.1
B3	102874635	102874888
refGene_gn=DDHD1;refGene_tx=XM_023255698.1
B3	102876793	102876943
refGene_gn=DDHD1;refGene_tx=XM_023255698.1
B3	102883187	102883263
refGene_gn=DDHD1;refGene_tx=XM_023255698.1
B3	102886465	102886728
refGene_gn=DDHD1;refGene_tx=XM_023255698.1
B3	102894523	102894630
refGene_gn=DDHD1;refGene_tx=XM_023255698.1
B3	102901772	102901879
refGene_gn=DDHD1;refGene_tx=XM_023255698.1
B3	102909615	102909763
refGene_gn=DDHD1;refGene_tx=XM_023255698.1
B3	102910821	102910950
refGene_gn=DDHD1;refGene_tx=XM_023255698.1
B3	102920238	102920412
refGene_gn=DDHD1;refGene_tx=XM_023255698.1
B3	102971143	102971803
refGene_gn=DDHD1;refGene_tx=XM_023255698.1
B3	102971805	102971971
refGene_gn=DDHD1;refGene_tx=XM_023255698.1
B3	103657101	103657958
refGene_gn=BMP4;refGene_tx=XM_006932864.3;refGene_gn=BMP4;refGene_tx=XM_019833027.1
B3	103658969	103659342
refGene_gn=BMP4;refGene_tx=XM_006932864.3;refGene_gn=BMP4;refGene_tx=XM_019833027.1
B3	103660437	103660562
refGene_gn=BMP4;refGene_tx=XM_019833027.1
B3	103663885	103663947
refGene_gn=BMP4;refGene_tx=XM_019833027.1
B3	104022616	104022625
refGene_gn=CDKN3;refGene_tx=XM_003987646.4;refGene_gn=CDKN3;refGene_tx=XM_006932866.3;refGene_gn=CDKN3;refGene_tx=XM_006932867.3;refGene_gn=CDKN3;refGene_tx=XM_006932868.3;refGene_gn=CDKN3;refGene_tx=XM_019833030.2
B3	104024794	104024877
refGene_gn=CDKN3;refGene_tx=XM_003987646.4;refGene_gn=CDKN3;refGene_tx=XM_006932866.3;refGene_gn=CDKN3;refGene_tx=XM_006932867.3;refGene_gn=CDKN3;refGene_tx=XM_006932868.3;refGene_gn=CDKN3;refGene_tx=XM_019833030.2
B3	104027516	104027572

refGene_gn=CDKN3;refGene_tx=XM_003987646.4;refGene_gn=CDKN3;refGene_tx=XM_006932866.3;refGene_gn=CDKN3;refGene_tx=XM_006932867.3;refGene_gn=CDKN3;refGene_tx=XM_006932868.3;refGene_gn=CDKN3;refGene_tx=XM_019833030.2
B3 104028087 104028132
refGene_gn=CDKN3;refGene_tx=XM_003987646.4;refGene_gn=CDKN3;refGene_tx=XM_006932866.3;refGene_gn=CDKN3;refGene_tx=XM_006932867.3
B3 104030803 104031026
refGene_gn=CDKN3;refGene_tx=XM_003987646.4;refGene_gn=CDKN3;refGene_tx=XM_006932866.3;refGene_gn=CDKN3;refGene_tx=XM_006932867.3;refGene_gn=CDKN3;refGene_tx=XM_006932868.3;refGene_gn=CDKN3;refGene_tx=XM_019833030.2
B3 104035197 104035229
refGene_gn=CDKN3;refGene_tx=XM_003987646.4;refGene_gn=CDKN3;refGene_tx=XM_006932866.3;refGene_gn=CDKN3;refGene_tx=XM_006932867.3;refGene_gn=CDKN3;refGene_tx=XM_006932868.3;refGene_gn=CDKN3;refGene_tx=XM_019833030.2
B3 104037514 104037618
refGene_gn=CDKN3;refGene_tx=XM_003987646.4;refGene_gn=CDKN3;refGene_tx=XM_006932866.3;refGene_gn=CDKN3;refGene_tx=XM_006932867.3;refGene_gn=CDKN3;refGene_tx=XM_006932868.3;refGene_gn=CDKN3;refGene_tx=XM_019833030.2
B3 104039512 104039599
refGene_gn=CDKN3;refGene_tx=XM_003987646.4;refGene_gn=CDKN3;refGene_tx=XM_006932866.3;refGene_gn=CDKN3;refGene_tx=XM_006932867.3;refGene_gn=CDKN3;refGene_tx=XM_006932868.3;refGene_gn=CDKN3;refGene_tx=XM_019833030.2
B3 104046396 104046424
refGene_gn=CNIH1;refGene_tx=XM_003987648.5
B3 104048515 104048659
refGene_gn=CNIH1;refGene_tx=XM_003987648.5
B3 104050073 104050186
refGene_gn=CNIH1;refGene_tx=XM_003987648.5
B3 104054114 104054183
refGene_gn=CNIH1;refGene_tx=XM_003987648.5
B3 104059380 104059461
refGene_gn=CNIH1;refGene_tx=XM_003987648.5
B3 104072256 104072295
refGene_gn=GMFB;refGene_tx=XM_019833032.2
B3 104077941 104078013
refGene_gn=GMFB;refGene_tx=XM_003987649.5;refGene_gn=GMFB;refGene_tx=XM_006932869.4
B3 104079700 104079774
refGene_gn=GMFB;refGene_tx=XM_003987649.5;refGene_gn=GMFB;refGene_tx=XM_006932869.4;refGene_gn=GMFB;refGene_tx=XM_019833032.2
B3 104080694 104080780
refGene_gn=GMFB;refGene_tx=XM_003987649.5;refGene_gn=GMFB;refGene_tx=XM_006932869.4;refGene_gn=GMFB;refGene_tx=XM_019833032.2
B3 104081484 104081534
refGene_gn=GMFB;refGene_tx=XM_003987649.5;refGene_gn=GMFB;refGene_tx=X

M_006932869.4;refGene_gn=GMFB;refGene_tx=XM_019833032.2
B3	104082227	104082277
refGene_gn=GMFB;refGene_tx=XM_003987649.5;refGene_gn=GMFB;refGene_tx=XM_006932869.4;refGene_gn=GMFB;refGene_tx=XM_019833032.2
B3	104083919	104084016
refGene_gn=GMFB;refGene_tx=XM_003987649.5;refGene_gn=GMFB;refGene_tx=XM_006932869.4;refGene_gn=GMFB;refGene_tx=XM_019833032.2
B3	104088669	104088672
refGene_gn=GMFB;refGene_tx=XM_003987649.5;refGene_gn=GMFB;refGene_tx=XM_006932869.4;refGene_gn=GMFB;refGene_tx=XM_019833032.2
B3	104099007	104099111
refGene_gn=CGRRF1;refGene_tx=XM_003987650.5
B3	104112767	104112907
refGene_gn=CGRRF1;refGene_tx=XM_003987650.5;refGene_gn=CGRRF1;refGene_tx=XM_019833031.2
B3	104120002	104120180
refGene_gn=CGRRF1;refGene_tx=XM_003987650.5;refGene_gn=CGRRF1;refGene_tx=XM_019833031.2
B3	104120585	104120733
refGene_gn=CGRRF1;refGene_tx=XM_003987650.5;refGene_gn=CGRRF1;refGene_tx=XM_019833031.2
B3	104126695	104126803
refGene_gn=CGRRF1;refGene_tx=XM_003987650.5;refGene_gn=CGRRF1;refGene_tx=XM_019833031.2
B3	104127020	104127341
refGene_gn=CGRRF1;refGene_tx=XM_003987650.5;refGene_gn=CGRRF1;refGene_tx=XM_019833031.2
B3	104147257	104147929
refGene_gn=LOC111560967;refGene_tx=XM_023256187.1
B3	104148489	104148829
refGene_gn=LOC111560967;refGene_tx=XM_023256187.1
B3	104149016	104149356
refGene_gn=LOC111560967;refGene_tx=XM_023256187.1;refGene_gn=SAMD4A;refGene_tx=XM_023255700.1;refGene_gn=SAMD4A;refGene_tx=XM_023255701.1;refGene_gn=SAMD4A;refGene_tx=XM_023255702.1
B3	104273438	104273957
refGene_gn=SAMD4A;refGene_tx=XM_019833034.2;refGene_gn=SAMD4A;refGene_tx=XM_023255700.1;refGene_gn=SAMD4A;refGene_tx=XM_023255701.1;refGene_gn=SAMD4A;refGene_tx=XM_023255702.1
B3	104308225	104308489
refGene_gn=SAMD4A;refGene_tx=XM_019833034.2;refGene_gn=SAMD4A;refGene_tx=XM_023255700.1;refGene_gn=SAMD4A;refGene_tx=XM_023255701.1
B3	104319802	104319912
refGene_gn=SAMD4A;refGene_tx=XM_019833034.2;refGene_gn=SAMD4A;refGene_tx=XM_023255700.1;refGene_gn=SAMD4A;refGene_tx=XM_023255701.1;refGene_gn=SAMD4A;refGene_tx=XM_023255702.1
B3	104322109	104322196
refGene_gn=SAMD4A;refGene_tx=XM_019833034.2;refGene_gn=SAMD4A;refGene_tx=XM_023255700.1;refGene_gn=SAMD4A;refGene_tx=XM_023255701.1;refGene_gn=SAMD4A;refGene_tx=XM_023255702.1

B3	104329922	104330256
refGene_gn=SAMD4A;refGene_tx=XM_006932873.4;refGene_gn=SAMD4A;refGene_tx=XM_019833034.2;refGene_gn=SAMD4A;refGene_tx=XM_023255700.1;refGene_gn=SAMD4A;refGene_tx=XM_023255701.1;refGene_gn=SAMD4A;refGene_tx=XM_023255702.1
B3	104332769	104332855
refGene_gn=SAMD4A;refGene_tx=XM_006932873.4;refGene_gn=SAMD4A;refGene_tx=XM_019833034.2;refGene_gn=SAMD4A;refGene_tx=XM_023255700.1;refGene_gn=SAMD4A;refGene_tx=XM_023255701.1;refGene_gn=SAMD4A;refGene_tx=XM_023255702.1
B3	104338090	104338209
refGene_gn=SAMD4A;refGene_tx=XM_006932873.4;refGene_gn=SAMD4A;refGene_tx=XM_019833034.2;refGene_gn=SAMD4A;refGene_tx=XM_023255700.1;refGene_gn=SAMD4A;refGene_tx=XM_023255701.1;refGene_gn=SAMD4A;refGene_tx=XM_023255702.1
B3	104342761	104342963
refGene_gn=SAMD4A;refGene_tx=XM_006932873.4;refGene_gn=SAMD4A;refGene_tx=XM_019833034.2;refGene_gn=SAMD4A;refGene_tx=XM_023255700.1;refGene_gn=SAMD4A;refGene_tx=XM_023255701.1;refGene_gn=SAMD4A;refGene_tx=XM_023255702.1
B3	104344299	104344426
refGene_gn=SAMD4A;refGene_tx=XM_006932873.4;refGene_gn=SAMD4A;refGene_tx=XM_019833034.2;refGene_gn=SAMD4A;refGene_tx=XM_023255700.1;refGene_gn=SAMD4A;refGene_tx=XM_023255701.1;refGene_gn=SAMD4A;refGene_tx=XM_023255702.1
B3	104352540	104352624
refGene_gn=SAMD4A;refGene_tx=XM_006932873.4;refGene_gn=SAMD4A;refGene_tx=XM_019833034.2;refGene_gn=SAMD4A;refGene_tx=XM_023255700.1;refGene_gn=SAMD4A;refGene_tx=XM_023255701.1;refGene_gn=SAMD4A;refGene_tx=XM_023255702.1
B3	104358167	104358196
refGene_gn=SAMD4A;refGene_tx=XM_006932873.4;refGene_gn=SAMD4A;refGene_tx=XM_019833034.2;refGene_gn=SAMD4A;refGene_tx=XM_023255700.1;refGene_gn=SAMD4A;refGene_tx=XM_023255701.1;refGene_gn=SAMD4A;refGene_tx=XM_023255702.1
B3	104407565	104407575
refGene_gn=GCH1;refGene_tx=XM_023255703.1
B3	104408282	104408373
refGene_gn=GCH1;refGene_tx=XM_023255705.1
B3	104408518	104408645
refGene_gn=GCH1;refGene_tx=XM_011283385.2;refGene_gn=GCH1;refGene_tx=XM_023255704.1
B3	104409551	104409636
refGene_gn=GCH1;refGene_tx=XM_011283385.2;refGene_gn=GCH1;refGene_tx=XM_023255703.1;refGene_gn=GCH1;refGene_tx=XM_023255705.1
B3	104410597	104410629
refGene_gn=GCH1;refGene_tx=XM_011283385.2;refGene_gn=GCH1;refGene_tx=XM_023255703.1;refGene_gn=GCH1;refGene_tx=XM_023255705.1
B3	104424631	104424687
refGene_gn=GCH1;refGene_tx=XM_011283385.2;refGene_gn=GCH1;refGene_tx=X

M_023255703.1;refGene_gn=GCH1;refGene_tx=XM_023255704.1;refGene_gn=GCH
1;refGene_tx=XM_023255705.1
B3 104427322 104427432
refGene_gn=GCH1;refGene_tx=XM_011283385.2;refGene_gn=GCH1;refGene_tx=X
M_023255703.1;refGene_gn=GCH1;refGene_tx=XM_023255704.1;refGene_gn=GCH
1;refGene_tx=XM_023255705.1
B3 104454145 104454485
refGene_gn=GCH1;refGene_tx=XM_011283385.2;refGene_gn=GCH1;refGene_tx=X
M_023255703.1;refGene_gn=GCH1;refGene_tx=XM_023255704.1;refGene_gn=GCH
1;refGene_tx=XM_023255705.1
B3 104494076 104494277
refGene_gn=WDHD1;refGene_tx=XM_011283237.3
B3 104497016 104497155
refGene_gn=WDHD1;refGene_tx=XM_011283237.3
B3 104500803 104500934
refGene_gn=WDHD1;refGene_tx=XM_011283237.3
B3 104502395 104502563
refGene_gn=WDHD1;refGene_tx=XM_011283237.3
B3 104503178 104503222
refGene_gn=WDHD1;refGene_tx=XM_011283237.3
B3 104505448 104505505
refGene_gn=WDHD1;refGene_tx=XM_011283237.3
B3 104505691 104505810
refGene_gn=WDHD1;refGene_tx=XM_011283237.3
B3 104505907 104506125
refGene_gn=WDHD1;refGene_tx=XM_011283237.3
B3 104511755 104511887
refGene_gn=WDHD1;refGene_tx=XM_011283237.3
B3 104512547 104512662
refGene_gn=WDHD1;refGene_tx=XM_011283237.3
B3 104519905 104520062
refGene_gn=WDHD1;refGene_tx=XM_011283237.3
B3 104521746 104521884
refGene_gn=WDHD1;refGene_tx=XM_011283237.3
B3 104524477 104524719
refGene_gn=WDHD1;refGene_tx=XM_011283237.3
B3 104527201 104527386
refGene_gn=WDHD1;refGene_tx=XM_011283237.3
B3 104529214 104529402
refGene_gn=WDHD1;refGene_tx=XM_011283237.3
B3 104532105 104532316
refGene_gn=WDHD1;refGene_tx=XM_011283237.3
B3 104538070 104538212
refGene_gn=WDHD1;refGene_tx=XM_011283237.3
B3 104538568 104538675
refGene_gn=WDHD1;refGene_tx=XM_011283237.3
B3 104539714 104539807
refGene_gn=WDHD1;refGene_tx=XM_011283237.3
B3 104541549 104541645
refGene_gn=WDHD1;refGene_tx=XM_011283237.3

B3 104542551 104542602
refGene_gn=WDHD1;refGene_tx=XM_011283237.3
B3 104542784 104542896
refGene_gn=WDHD1;refGene_tx=XM_011283237.3
B3 104544036 104544188
refGene_gn=WDHD1;refGene_tx=XM_011283237.3
B3 104550433 104550545
refGene_gn=WDHD1;refGene_tx=XM_011283237.3
B3 104560523 104560600
refGene_gn=WDHD1;refGene_tx=XM_011283237.3
B3 104574387 104575713
refGene_gn=SOCS4;refGene_tx=XM_004001462.5;refGene_gn=SOCS4;refGene_tx=XM_011283240.3
B3 104589970 104589988
refGene_gn=MAPK1IP1L;refGene_tx=XM_011283238.3;refGene_gn=MAPK1IP1L;refGene_tx=XM_011283239.3
B3 104590684 104591389
refGene_gn=MAPK1IP1L;refGene_tx=XM_011283238.3;refGene_gn=MAPK1IP1L;refGene_tx=XM_011283239.3
B3 104592615 104592627
refGene_gn=MAPK1IP1L;refGene_tx=XM_011283238.3;refGene_gn=MAPK1IP1L;refGene_tx=XM_011283239.3
B3 104656127 104656145
refGene_gn=LGALS3;refGene_tx=XM_003987655.3
B3 104657828 104658236
refGene_gn=LGALS3;refGene_tx=XM_003987655.3
B3 104660292 104660381
refGene_gn=LGALS3;refGene_tx=XM_003987655.3
B3 104663628 104663794
refGene_gn=LGALS3;refGene_tx=XM_003987655.3
B3 104666786 104666942
refGene_gn=LGALS3;refGene_tx=XM_003987655.3
B3 104670914 104671055
refGene_gn=DLGAP5;refGene_tx=XM_023255706.1;refGene_gn=DLGAP5;refGene_tx=XM_023255707.1;refGene_gn=DLGAP5;refGene_tx=XM_023255708.1
B3 104673222 104673272
refGene_gn=DLGAP5;refGene_tx=XM_023255706.1;refGene_gn=DLGAP5;refGene_tx=XM_023255707.1;refGene_gn=DLGAP5;refGene_tx=XM_023255708.1
B3 104674504 104674751
refGene_gn=DLGAP5;refGene_tx=XM_023255706.1;refGene_gn=DLGAP5;refGene_tx=XM_023255707.1;refGene_gn=DLGAP5;refGene_tx=XM_023255708.1
B3 104675339 104675391
refGene_gn=DLGAP5;refGene_tx=XM_023255706.1;refGene_gn=DLGAP5;refGene_tx=XM_023255707.1;refGene_gn=DLGAP5;refGene_tx=XM_023255708.1
B3 104677123 104677316
refGene_gn=DLGAP5;refGene_tx=XM_023255706.1;refGene_gn=DLGAP5;refGene_tx=XM_023255707.1;refGene_gn=DLGAP5;refGene_tx=XM_023255708.1
B3 104681481 104681698
refGene_gn=DLGAP5;refGene_tx=XM_023255706.1;refGene_gn=DLGAP5;refGene_tx=XM_023255707.1;refGene_gn=DLGAP5;refGene_tx=XM_023255708.1

B3 104685729 104685822
refGene_gn=DLGAP5;refGene_tx=XM_023255706.1;refGene_gn=DLGAP5;refGene_tx=XM_023255707.1;refGene_gn=DLGAP5;refGene_tx=XM_023255708.1
B3 104691064 104691225
refGene_gn=DLGAP5;refGene_tx=XM_023255706.1;refGene_gn=DLGAP5;refGene_tx=XM_023255707.1;refGene_gn=DLGAP5;refGene_tx=XM_023255708.1
B3 104691997 104692083
refGene_gn=DLGAP5;refGene_tx=XM_023255706.1;refGene_gn=DLGAP5;refGene_tx=XM_023255707.1;refGene_gn=DLGAP5;refGene_tx=XM_023255708.1
B3 104694971 104695095
refGene_gn=DLGAP5;refGene_tx=XM_023255706.1;refGene_gn=DLGAP5;refGene_tx=XM_023255707.1;refGene_gn=DLGAP5;refGene_tx=XM_023255708.1
B3 104695444 104695572
refGene_gn=DLGAP5;refGene_tx=XM_023255706.1;refGene_gn=DLGAP5;refGene_tx=XM_023255707.1;refGene_gn=DLGAP5;refGene_tx=XM_023255708.1
B3 104696744 104697019
refGene_gn=DLGAP5;refGene_tx=XM_023255706.1;refGene_gn=DLGAP5;refGene_tx=XM_023255707.1;refGene_gn=DLGAP5;refGene_tx=XM_023255708.1
B3 104698584 104698655
refGene_gn=DLGAP5;refGene_tx=XM_023255706.1;refGene_gn=DLGAP5;refGene_tx=XM_023255707.1;refGene_gn=DLGAP5;refGene_tx=XM_023255708.1
B3 104699632 104699752
refGene_gn=DLGAP5;refGene_tx=XM_023255706.1;refGene_gn=DLGAP5;refGene_tx=XM_023255707.1;refGene_gn=DLGAP5;refGene_tx=XM_023255708.1
B3 104699846 104699931
refGene_gn=DLGAP5;refGene_tx=XM_023255706.1;refGene_gn=DLGAP5;refGene_tx=XM_023255707.1;refGene_gn=DLGAP5;refGene_tx=XM_023255708.1
B3 104701673 104701736
refGene_gn=DLGAP5;refGene_tx=XM_023255706.1;refGene_gn=DLGAP5;refGene_tx=XM_023255707.1;refGene_gn=DLGAP5;refGene_tx=XM_023255708.1
B3 104703805 104703990
refGene_gn=DLGAP5;refGene_tx=XM_023255706.1;refGene_gn=DLGAP5;refGene_tx=XM_023255707.1;refGene_gn=DLGAP5;refGene_tx=XM_023255708.1
B3 104709617 104709855
refGene_gn=DLGAP5;refGene_tx=XM_023255706.1;refGene_gn=DLGAP5;refGene_tx=XM_023255707.1;refGene_gn=DLGAP5;refGene_tx=XM_023255708.1
B3 104784190 104784336
refGene_gn=FBXO34;refGene_tx=XM_023255709.1
B3 104868593 104870742
refGene_gn=FBXO34;refGene_tx=XM_003987657.5;refGene_gn=FBXO34;refGene_tx=XM_019833040.2;refGene_gn=FBXO34;refGene_tx=XM_019833041.2;refGene_gn=FBXO34;refGene_tx=XM_019833042.2;refGene_gn=FBXO34;refGene_tx=XM_019833043.2;refGene_gn=FBXO34;refGene_tx=XM_019833045.2;refGene_gn=FBXO34;refGene_tx=XM_019833046.2;refGene_gn=FBXO34;refGene_tx=XM_019833048.2;refGene_gn=FBXO34;refGene_tx=XM_023255709.1;refGene_gn=FBXO34;refGene_tx=XM_023255710.1
B3 104881566 104881873
refGene_gn=ATG14;refGene_tx=XM_023255712.1
B3 104888869 104888955
refGene_gn=ATG14;refGene_tx=XM_023255712.1

B3 104889032 104889123
refGene_gn=ATG14;refGene_tx=XM_023255712.1
B3 104890492 104890610
refGene_gn=ATG14;refGene_tx=XM_023255712.1
B3 104892274 104892504
refGene_gn=ATG14;refGene_tx=XM_023255712.1
B3 104896066 104896304
refGene_gn=ATG14;refGene_tx=XM_023255712.1
B3 104900587 104900669
refGene_gn=ATG14;refGene_tx=XM_023255712.1
B3 104903062 104903105
refGene_gn=ATG14;refGene_tx=XM_023255712.1
B3 104904679 104904742
refGene_gn=ATG14;refGene_tx=XM_023255712.1
B3 104919699 104919920
refGene_gn=ATG14;refGene_tx=XM_023255712.1
B3 104921733 104921810
refGene_gn=TBPL2;refGene_tx=XM_023256188.1
B3 104929672 104929767
refGene_gn=TBPL2;refGene_tx=XM_023256188.1
B3 104934093 104934261
refGene_gn=TBPL2;refGene_tx=XM_023256188.1
B3 104934431 104934439
refGene_gn=TBPL2;refGene_tx=XM_023256188.1
B3 104949265 104949353
refGene_gn=TBPL2;refGene_tx=XM_023256188.1
B3 104950445 104950906
refGene_gn=TBPL2;refGene_tx=XM_023256188.1
B3 104953811 104954114
refGene_gn=TBPL2;refGene_tx=XM_023256188.1
B3 105081852 105082375
refGene_gn=KTN1;refGene_tx=XM_023255713.1
B3 105085814 105085952
refGene_gn=KTN1;refGene_tx=XM_023255713.1
B3 105087696 105087867
refGene_gn=KTN1;refGene_tx=XM_023255713.1
B3 105088919 105089050
refGene_gn=KTN1;refGene_tx=XM_023255713.1
B3 105094233 105094350
refGene_gn=KTN1;refGene_tx=XM_023255713.1
B3 105097174 105097315
refGene_gn=KTN1;refGene_tx=XM_023255713.1
B3 105100812 105100919
refGene_gn=KTN1;refGene_tx=XM_023255713.1
B3 105101732 105101865
refGene_gn=KTN1;refGene_tx=XM_023255713.1
B3 105103213 105103301
refGene_gn=KTN1;refGene_tx=XM_023255713.1
B3 105106699 105106866
refGene_gn=KTN1;refGene_tx=XM_023255713.1

B3 105107498 105107567
refGene_gn=KTN1;refGene_tx=XM_023255713.1
B3 105109012 105109050
refGene_gn=KTN1;refGene_tx=XM_023255713.1
B3 105109739 105109830
refGene_gn=KTN1;refGene_tx=XM_023255713.1
B3 105110168 105110237
refGene_gn=KTN1;refGene_tx=XM_023255713.1
B3 105110759 105110797
refGene_gn=KTN1;refGene_tx=XM_023255713.1
B3 105110944 105111026
refGene_gn=KTN1;refGene_tx=XM_023255713.1
B3 105113085 105113154
refGene_gn=KTN1;refGene_tx=XM_023255713.1
B3 105117931 105117966
refGene_gn=KTN1;refGene_tx=XM_023255713.1
B3 105118380 105118471
refGene_gn=KTN1;refGene_tx=XM_023255713.1
B3 105119160 105119229
refGene_gn=KTN1;refGene_tx=XM_023255713.1
B3 105119859 105119897
refGene_gn=KTN1;refGene_tx=XM_023255713.1
B3 105126599 105126690
refGene_gn=KTN1;refGene_tx=XM_023255713.1
B3 105126839 105126881
refGene_gn=KTN1;refGene_tx=XM_023255713.1
B3 105126882 105126909
refGene_gn=KTN1;refGene_tx=XM_023255713.1
B3 105128191 105128229
refGene_gn=KTN1;refGene_tx=XM_023255713.1
B3 105128885 105128976
refGene_gn=KTN1;refGene_tx=XM_023255713.1
B3 105129054 105129123
refGene_gn=KTN1;refGene_tx=XM_023255713.1
B3 105129615 105129653
refGene_gn=KTN1;refGene_tx=XM_023255713.1
B3 105132037 105132128
refGene_gn=KTN1;refGene_tx=XM_023255713.1
B3 105134339 105134408
refGene_gn=KTN1;refGene_tx=XM_023255713.1
B3 105135676 105135714
refGene_gn=KTN1;refGene_tx=XM_023255713.1
B3 105137315 105137406
refGene_gn=KTN1;refGene_tx=XM_023255713.1
B3 105139393 105139480
refGene_gn=KTN1;refGene_tx=XM_023255713.1
B3 105142727 105142817
refGene_gn=KTN1;refGene_tx=XM_023255713.1
B3 105146652 105146733
refGene_gn=KTN1;refGene_tx=XM_023255713.1

B3 105147514 105147604
refGene_gn=KTN1;refGene_tx=XM_023255713.1
B3 105147732 105147825
refGene_gn=KTN1;refGene_tx=XM_023255713.1
B3 105149181 105149253
refGene_gn=KTN1;refGene_tx=XM_023255713.1
B3 105149491 105149575
refGene_gn=KTN1;refGene_tx=XM_023255713.1
B3 105149733 105149817
refGene_gn=KTN1;refGene_tx=XM_023255713.1
B3 105152450 105152534
refGene_gn=KTN1;refGene_tx=XM_023255713.1
B3 105154150 105154243
refGene_gn=KTN1;refGene_tx=XM_023255713.1
B3 105155553 105155667
refGene_gn=KTN1;refGene_tx=XM_023255713.1
B3 105543513 105543590
refGene_gn=PELI2;refGene_tx=XM_023255714.1
B3 105599512 105599642
refGene_gn=PELI2;refGene_tx=XM_023255714.1
B3 105709111 105709213
refGene_gn=PELI2;refGene_tx=XM_023255714.1
B3 105724402 105724600
refGene_gn=PELI2;refGene_tx=XM_023255714.1
B3 105731837 105732026
refGene_gn=PELI2;refGene_tx=XM_023255714.1
B3 105739528 105740095
refGene_gn=PELI2;refGene_tx=XM_023255714.1
B3 106203783 106204404
refGene_gn=OTX2;refGene_tx=XM_003987680.5;refGene_gn=OTX2;refGene_tx=XM_006932891.4;refGene_gn=OTX2;refGene_tx=XM_019833051.2;refGene_gn=OTX2;refGene_tx=XM_019833052.2;refGene_gn=OTX2;refGene_tx=XM_019833053.2;refGene_gn=OTX2;refGene_tx=XM_019833054.2;refGene_gn=OTX2;refGene_tx=XM_019833055.1;refGene_gn=OTX2;refGene_tx=XM_019833056.2;refGene_gn=OTX2;refGene_tx=XM_023255716.1
B3 106206303 106206479
refGene_gn=OTX2;refGene_tx=XM_003987680.5;refGene_gn=OTX2;refGene_tx=XM_006932891.4;refGene_gn=OTX2;refGene_tx=XM_019833051.2;refGene_gn=OTX2;refGene_tx=XM_019833052.2;refGene_gn=OTX2;refGene_tx=XM_019833053.2;refGene_gn=OTX2;refGene_tx=XM_019833054.2;refGene_gn=OTX2;refGene_tx=XM_019833055.1;refGene_gn=OTX2;refGene_tx=XM_019833056.2;refGene_gn=OTX2;refGene_tx=XM_023255716.1
B3 106207446 106207543
refGene_gn=OTX2;refGene_tx=XM_003987680.5;refGene_gn=OTX2;refGene_tx=XM_006932891.4;refGene_gn=OTX2;refGene_tx=XM_019833051.2;refGene_gn=OTX2;refGene_tx=XM_019833052.2;refGene_gn=OTX2;refGene_tx=XM_019833053.2;refGene_gn=OTX2;refGene_tx=XM_019833054.2;refGene_gn=OTX2;refGene_tx=XM_019833055.1;refGene_gn=OTX2;refGene_tx=XM_019833056.2;refGene_gn=OTX2;refGene_tx=XM_023255716.1
B3 106566270 106566459

refGene_gn=EXOC5;refGene_tx=XM_023255717.1
B3 106567110 106567326
refGene_gn=EXOC5;refGene_tx=XM_023255717.1
B3 106567523 106567632
refGene_gn=EXOC5;refGene_tx=XM_023255717.1
B3 106571053 106571140
refGene_gn=EXOC5;refGene_tx=XM_023255717.1
B3 106572366 106572487
refGene_gn=EXOC5;refGene_tx=XM_023255717.1
B3 106576868 106576977
refGene_gn=EXOC5;refGene_tx=XM_023255717.1
B3 106584730 106584878
refGene_gn=EXOC5;refGene_tx=XM_023255717.1
B3 106586718 106586928
refGene_gn=EXOC5;refGene_tx=XM_023255717.1
B3 106587948 106588031
refGene_gn=EXOC5;refGene_tx=XM_023255717.1
B3 106589864 106590005
refGene_gn=EXOC5;refGene_tx=XM_023255717.1
B3 106590107 106590152
refGene_gn=EXOC5;refGene_tx=XM_023255717.1
B3 106591620 106591730
refGene_gn=EXOC5;refGene_tx=XM_023255717.1
B3 106592902 106592931
refGene_gn=EXOC5;refGene_tx=XM_023255717.1
B3 106595717 106595782
refGene_gn=EXOC5;refGene_tx=XM_023255717.1
B3 106599306 106599501
refGene_gn=EXOC5;refGene_tx=XM_023255717.1
B3 106601476 106601624
refGene_gn=EXOC5;refGene_tx=XM_023255717.1
B3 106602462 106602557
refGene_gn=EXOC5;refGene_tx=XM_023255717.1
B3 106622731 106622758
refGene_gn=EXOC5;refGene_tx=XM_023255717.1
B3 106623427 106623501
refGene_gn=AP5M1;refGene_tx=XM_003987682.5;refGene_gn=AP5M1;refGene_tx
=XM_019833058.2;refGene_gn=AP5M1;refGene_tx=XM_023255718.1;refGene_gn=
AP5M1;refGene_tx=XM_023255719.1
B3 106627582 106628228
refGene_gn=AP5M1;refGene_tx=XM_003987682.5;refGene_gn=AP5M1;refGene_tx
=XM_019833058.2;refGene_gn=AP5M1;refGene_tx=XM_023255718.1;refGene_gn=
AP5M1;refGene_tx=XM_023255719.1
B3 106631692 106631920
refGene_gn=AP5M1;refGene_tx=XM_003987682.5;refGene_gn=AP5M1;refGene_tx
=XM_019833058.2;refGene_gn=AP5M1;refGene_tx=XM_023255718.1;refGene_gn=
AP5M1;refGene_tx=XM_023255719.1
B3 106633958 106634098
refGene_gn=AP5M1;refGene_tx=XM_003987682.5;refGene_gn=AP5M1;refGene_tx
=XM_019833058.2;refGene_gn=AP5M1;refGene_tx=XM_023255718.1;refGene_gn=

AP5M1;refGene_tx=XM_023255719.1
B3 106634839 106634925
refGene_gn=AP5M1;refGene_tx=XM_003987682.5;refGene_gn=AP5M1;refGene_tx=XM_019833058.2;refGene_gn=AP5M1;refGene_tx=XM_023255718.1;refGene_gn=AP5M1;refGene_tx=XM_023255719.1
B3 106635295 106635414
refGene_gn=AP5M1;refGene_tx=XM_003987682.5;refGene_gn=AP5M1;refGene_tx=XM_019833058.2;refGene_gn=AP5M1;refGene_tx=XM_023255718.1;refGene_gn=AP5M1;refGene_tx=XM_023255719.1
B3 106638539 106638636
refGene_gn=AP5M1;refGene_tx=XM_003987682.5;refGene_gn=AP5M1;refGene_tx=XM_019833058.2;refGene_gn=AP5M1;refGene_tx=XM_023255718.1
B3 106641473 106641556
refGene_gn=AP5M1;refGene_tx=XM_003987682.5;refGene_gn=AP5M1;refGene_tx=XM_019833058.2;refGene_gn=AP5M1;refGene_tx=XM_023255718.1;refGene_gn=AP5M1;refGene_tx=XM_023255719.1
B3 106643703 106643722
refGene_gn=AP5M1;refGene_tx=XM_019833058.2
B3 106652833 106652891
refGene_gn=AP5M1;refGene_tx=XM_023255718.1
B3 106724131 106724419
refGene_gn=NAA30;refGene_tx=XM_023255720.1
B3 106724420 106724900
refGene_gn=NAA30;refGene_tx=XM_023255720.1
B3 106730587 106730711
refGene_gn=NAA30;refGene_tx=XM_023255720.1
B3 106732499 106732555
refGene_gn=NAA30;refGene_tx=XM_023255720.1
B3 106742182 106742320
refGene_gn=NAA30;refGene_tx=XM_023255720.1
B3 106793345 106793581
refGene_gn=CCDC198;refGene_tx=XM_003987683.4;refGene_gn=CCDC198;refGene_tx=XM_006932892.3;refGene_gn=CCDC198;refGene_tx=XM_019833060.2
B3 106801543 106801703
refGene_gn=CCDC198;refGene_tx=XM_003987683.4;refGene_gn=CCDC198;refGene_tx=XM_006932892.3;refGene_gn=CCDC198;refGene_tx=XM_019833060.2
B3 106802514 106802616
refGene_gn=CCDC198;refGene_tx=XM_003987683.4;refGene_gn=CCDC198;refGene_tx=XM_006932892.3;refGene_gn=CCDC198;refGene_tx=XM_019833060.2
B3 106804201 106804288
refGene_gn=CCDC198;refGene_tx=XM_003987683.4;refGene_gn=CCDC198;refGene_tx=XM_006932892.3;refGene_gn=CCDC198;refGene_tx=XM_019833060.2
B3 106818041 106818083
refGene_gn=CCDC198;refGene_tx=XM_019833060.2
B3 106818088 106818171
refGene_gn=CCDC198;refGene_tx=XM_003987683.4;refGene_gn=CCDC198;refGene_tx=XM_006932892.3
B3 106820576 106820799
refGene_gn=CCDC198;refGene_tx=XM_003987683.4;refGene_gn=CCDC198;refGene_tx=XM_006932892.3

B3 106867856 106868098
refGene_gn=SLC35F4;refGene_tx=XM_003987684.4
B3 106871091 106871181
refGene_gn=SLC35F4;refGene_tx=XM_003987684.4
B3 106874808 106875001
refGene_gn=SLC35F4;refGene_tx=XM_003987684.4
B3 106877254 106877380
refGene_gn=SLC35F4;refGene_tx=XM_003987684.4
B3 106887937 106888157
refGene_gn=SLC35F4;refGene_tx=XM_003987684.4
B3 106893970 106894268
refGene_gn=SLC35F4;refGene_tx=XM_003987684.4
B3 106899066 106899252
refGene_gn=SLC35F4;refGene_tx=XM_003987684.4
B3 107116482 107116585
refGene_gn=SLC35F4;refGene_tx=XM_003987684.4
B3 107244572 107244641
refGene_gn=CB3H14orf37;refGene_tx=XM_023255721.1
B3 107244877 107245012
refGene_gn=CB3H14orf37;refGene_tx=XM_023255721.1
B3 107252224 107252256
refGene_gn=CB3H14orf37;refGene_tx=XM_023255721.1
B3 107305927 107305947
refGene_gn=CB3H14orf37;refGene_tx=XM_023255722.1
B3 107331112 107331370
refGene_gn=CB3H14orf37;refGene_tx=XM_023255721.1;refGene_gn=CB3H14orf3
7;refGene_tx=XM_023255722.1
B3 107354203 107354416
refGene_gn=CB3H14orf37;refGene_tx=XM_023255721.1;refGene_gn=CB3H14orf3
7;refGene_tx=XM_023255722.1
B3 107357247 107357502
refGene_gn=CB3H14orf37;refGene_tx=XM_023255721.1;refGene_gn=CB3H14orf3
7;refGene_tx=XM_023255722.1
B3 107360006 107361441
refGene_gn=CB3H14orf37;refGene_tx=XM_023255721.1;refGene_gn=CB3H14orf3
7;refGene_tx=XM_023255722.1
B3 107370729 107370873
refGene_gn=CB3H14orf37;refGene_tx=XM_023255721.1;refGene_gn=CB3H14orf3
7;refGene_tx=XM_023255722.1
B3 107411144 107411221
refGene_gn=ACTR10;refGene_tx=XM_003987686.5
B3 107411627 107411671
refGene_gn=ACTR10;refGene_tx=XM_023255723.1
B3 107414649 107414722
refGene_gn=ACTR10;refGene_tx=XM_003987686.5;refGene_gn=ACTR10;refGene_
tx=XM_023255723.1
B3 107417205 107417288
refGene_gn=ACTR10;refGene_tx=XM_003987686.5;refGene_gn=ACTR10;refGene_
tx=XM_023255723.1
B3 107418057 107418166

refGene_gn=ACTR10;refGene_tx=XM_003987686.5;refGene_gn=ACTR10;refGene_tx=XM_023255723.1
B3 107422207 107422315
refGene_gn=ACTR10;refGene_tx=XM_003987686.5;refGene_gn=ACTR10;refGene_tx=XM_023255723.1
B3 107424325 107424393
refGene_gn=ACTR10;refGene_tx=XM_003987686.5;refGene_gn=ACTR10;refGene_tx=XM_023255723.1
B3 107424770 107424850
refGene_gn=ACTR10;refGene_tx=XM_003987686.5;refGene_gn=ACTR10;refGene_tx=XM_023255723.1
B3 107428843 107428879
refGene_gn=ACTR10;refGene_tx=XM_003987686.5;refGene_gn=ACTR10;refGene_tx=XM_023255723.1
B3 107430989 107431069
refGene_gn=ACTR10;refGene_tx=XM_003987686.5;refGene_gn=ACTR10;refGene_tx=XM_023255723.1
B3 107431148 107431222
refGene_gn=ACTR10;refGene_tx=XM_003987686.5;refGene_gn=ACTR10;refGene_tx=XM_023255723.1
B3 107435495 107435577
refGene_gn=ACTR10;refGene_tx=XM_003987686.5;refGene_gn=ACTR10;refGene_tx=XM_023255723.1
B3 107436603 107436805
refGene_gn=ACTR10;refGene_tx=XM_003987686.5;refGene_gn=ACTR10;refGene_tx=XM_023255723.1
B3 107438333 107438515
refGene_gn=ACTR10;refGene_tx=XM_003987686.5;refGene_gn=ACTR10;refGene_tx=XM_023255723.1
B3 107448436 107448457
refGene_gn=PSMA3;refGene_tx=XM_023255726.1
B3 107452702 107452785
refGene_gn=PSMA3;refGene_tx=XM_023255726.1
B3 107455357 107455481
refGene_gn=PSMA3;refGene_tx=XM_023255726.1
B3 107458412 107458514
refGene_gn=PSMA3;refGene_tx=XM_023255726.1
B3 107459247 107459457
refGene_gn=PSMA3;refGene_tx=XM_023255727.1
B3 107459775 107459849
refGene_gn=PSMA3;refGene_tx=XM_023255726.1;refGene_gn=PSMA3;refGene_tx=XM_023255727.1
B3 107462376 107462449
refGene_gn=PSMA3;refGene_tx=XM_023255726.1;refGene_gn=PSMA3;refGene_tx=XM_023255727.1
B3 107464090 107464156
refGene_gn=PSMA3;refGene_tx=XM_023255726.1;refGene_gn=PSMA3;refGene_tx=XM_023255727.1
B3 107466505 107466552
refGene_gn=PSMA3;refGene_tx=XM_023255726.1;refGene_gn=PSMA3;refGene_tx

=XM_023255727.1
B3	107468462	107468530
refGene_gn=PSMA3;refGene_tx=XM_023255726.1;refGene_gn=PSMA3;refGene_tx
=XM_023255727.1
B3	107469280	107469345
refGene_gn=PSMA3;refGene_tx=XM_023255726.1;refGene_gn=PSMA3;refGene_tx
=XM_023255727.1
B3	107469745	107469790
refGene_gn=PSMA3;refGene_tx=XM_023255726.1;refGene_gn=PSMA3;refGene_tx
=XM_023255727.1
B3	107490881	107490887
refGene_gn=ARID4A;refGene_tx=XM_019833062.2;refGene_gn=ARID4A;refGene_
tx=XM_023255724.1;refGene_gn=ARID4A;refGene_tx=XM_023255725.1
B3	107492837	107492948
refGene_gn=ARID4A;refGene_tx=XM_019833062.2;refGene_gn=ARID4A;refGene_
tx=XM_023255724.1;refGene_gn=ARID4A;refGene_tx=XM_023255725.1
B3	107494902	107494968
refGene_gn=ARID4A;refGene_tx=XM_019833062.2;refGene_gn=ARID4A;refGene_
tx=XM_023255724.1;refGene_gn=ARID4A;refGene_tx=XM_023255725.1
B3	107495962	107496053
refGene_gn=ARID4A;refGene_tx=XM_019833062.2;refGene_gn=ARID4A;refGene_
tx=XM_023255724.1;refGene_gn=ARID4A;refGene_tx=XM_023255725.1
B3	107504127	107504207
refGene_gn=ARID4A;refGene_tx=XM_019833062.2;refGene_gn=ARID4A;refGene_
tx=XM_023255724.1;refGene_gn=ARID4A;refGene_tx=XM_023255725.1
B3	107504296	107504391
refGene_gn=ARID4A;refGene_tx=XM_019833062.2;refGene_gn=ARID4A;refGene_
tx=XM_023255724.1;refGene_gn=ARID4A;refGene_tx=XM_023255725.1
B3	107505972	107506105
refGene_gn=ARID4A;refGene_tx=XM_019833062.2;refGene_gn=ARID4A;refGene_
tx=XM_023255724.1;refGene_gn=ARID4A;refGene_tx=XM_023255725.1
B3	107509191	107509271
refGene_gn=ARID4A;refGene_tx=XM_019833062.2;refGene_gn=ARID4A;refGene_
tx=XM_023255724.1;refGene_gn=ARID4A;refGene_tx=XM_023255725.1
B3	107511096	107511173
refGene_gn=ARID4A;refGene_tx=XM_019833062.2;refGene_gn=ARID4A;refGene_
tx=XM_023255724.1;refGene_gn=ARID4A;refGene_tx=XM_023255725.1
B3	107511574	107511741
refGene_gn=ARID4A;refGene_tx=XM_019833062.2;refGene_gn=ARID4A;refGene_
tx=XM_023255724.1;refGene_gn=ARID4A;refGene_tx=XM_023255725.1
B3	107524611	107524684
refGene_gn=ARID4A;refGene_tx=XM_019833062.2;refGene_gn=ARID4A;refGene_
tx=XM_023255724.1;refGene_gn=ARID4A;refGene_tx=XM_023255725.1
B3	107525773	107525868
refGene_gn=ARID4A;refGene_tx=XM_019833062.2;refGene_gn=ARID4A;refGene_
tx=XM_023255724.1;refGene_gn=ARID4A;refGene_tx=XM_023255725.1
B3	107526016	107526114
refGene_gn=ARID4A;refGene_tx=XM_019833062.2;refGene_gn=ARID4A;refGene_
tx=XM_023255724.1;refGene_gn=ARID4A;refGene_tx=XM_023255725.1
B3	107526652	107526884

refGene_gn=ARID4A;refGene_tx=XM_019833062.2;refGene_gn=ARID4A;refGene_tx=XM_023255724.1;refGene_gn=ARID4A;refGene_tx=XM_023255725.1
B3 107531387 107531638
refGene_gn=ARID4A;refGene_tx=XM_019833062.2;refGene_gn=ARID4A;refGene_tx=XM_023255724.1;refGene_gn=ARID4A;refGene_tx=XM_023255725.1
B3 107534145 107534343
refGene_gn=ARID4A;refGene_tx=XM_019833062.2;refGene_gn=ARID4A;refGene_tx=XM_023255724.1;refGene_gn=ARID4A;refGene_tx=XM_023255725.1
B3 107539246 107539331
refGene_gn=ARID4A;refGene_tx=XM_019833062.2;refGene_gn=ARID4A;refGene_tx=XM_023255724.1;refGene_gn=ARID4A;refGene_tx=XM_023255725.1
B3 107540162 107540364
refGene_gn=ARID4A;refGene_tx=XM_019833062.2;refGene_gn=ARID4A;refGene_tx=XM_023255724.1;refGene_gn=ARID4A;refGene_tx=XM_023255725.1
B3 107541940 107543074
refGene_gn=ARID4A;refGene_tx=XM_019833062.2;refGene_gn=ARID4A;refGene_tx=XM_023255724.1;refGene_gn=ARID4A;refGene_tx=XM_023255725.1
B3 107543801 107543906
refGene_gn=ARID4A;refGene_tx=XM_019833062.2;refGene_gn=ARID4A;refGene_tx=XM_023255724.1;refGene_gn=ARID4A;refGene_tx=XM_023255725.1
B3 107548043 107548250
refGene_gn=ARID4A;refGene_tx=XM_019833062.2;refGene_gn=ARID4A;refGene_tx=XM_023255724.1;refGene_gn=ARID4A;refGene_tx=XM_023255725.1
B3 107548863 107549010
refGene_gn=ARID4A;refGene_tx=XM_019833062.2;refGene_gn=ARID4A;refGene_tx=XM_023255724.1;refGene_gn=ARID4A;refGene_tx=XM_023255725.1
B3 107552528 107552632
refGene_gn=ARID4A;refGene_tx=XM_019833062.2;refGene_gn=ARID4A;refGene_tx=XM_023255724.1;refGene_gn=ARID4A;refGene_tx=XM_023255725.1
B3 107569310 107569446
refGene_gn=TOMM20L;refGene_tx=XM_023255729.1
B3 107569576 107569620
refGene_gn=TOMM20L;refGene_tx=XM_023255729.1
B3 107570717 107570799
refGene_gn=TOMM20L;refGene_tx=XM_023255729.1
B3 107575904 107576047
refGene_gn=TOMM20L;refGene_tx=XM_023255729.1
B3 107576971 107577025
refGene_gn=TOMM20L;refGene_tx=XM_023255729.1
B3 107577480 107577615
refGene_gn=TIMM9;refGene_tx=XM_003987688.5;refGene_gn=TIMM9;refGene_tx=XM_019833063.2;refGene_gn=TIMM9;refGene_tx=XM_023255730.1
B3 107578519 107578615
refGene_gn=TIMM9;refGene_tx=XM_003987688.5;refGene_gn=TIMM9;refGene_tx=XM_019833063.2;refGene_gn=TIMM9;refGene_tx=XM_023255730.1
B3 107578991 107579030
refGene_gn=TIMM9;refGene_tx=XM_003987688.5;refGene_gn=TIMM9;refGene_tx=XM_019833063.2;refGene_gn=TIMM9;refGene_tx=XM_023255730.1
B3 107593718 107593917
refGene_gn=KIAA0586;refGene_tx=XM_023255728.1

B3 107594834 107594905
refGene_gn=KIAA0586;refGene_tx=XM_023255728.1
B3 107597679 107597749
refGene_gn=KIAA0586;refGene_tx=XM_023255728.1
B3 107599601 107599671
refGene_gn=KIAA0586;refGene_tx=XM_023255728.1
B3 107605440 107605615
refGene_gn=KIAA0586;refGene_tx=XM_023255728.1
B3 107606370 107606592
refGene_gn=KIAA0586;refGene_tx=XM_023255728.1
B3 107608582 107608736
refGene_gn=KIAA0586;refGene_tx=XM_023255728.1
B3 107611136 107611304
refGene_gn=KIAA0586;refGene_tx=XM_023255728.1
B3 107612733 107612825
refGene_gn=KIAA0586;refGene_tx=XM_023255728.1
B3 107612826 107612858
refGene_gn=KIAA0586;refGene_tx=XM_023255728.1
B3 107618342 107618451
refGene_gn=KIAA0586;refGene_tx=XM_023255728.1
B3 107619201 107619419
refGene_gn=KIAA0586;refGene_tx=XM_023255728.1
B3 107619841 107619914
refGene_gn=KIAA0586;refGene_tx=XM_023255728.1
B3 107621177 107621405
refGene_gn=KIAA0586;refGene_tx=XM_023255728.1
B3 107622361 107622536
refGene_gn=KIAA0586;refGene_tx=XM_023255728.1
B3 107626806 107627004
refGene_gn=KIAA0586;refGene_tx=XM_023255728.1
B3 107628465 107628653
refGene_gn=KIAA0586;refGene_tx=XM_023255728.1
B3 107631692 107631800
refGene_gn=KIAA0586;refGene_tx=XM_023255728.1
B3 107632972 107633053
refGene_gn=KIAA0586;refGene_tx=XM_023255728.1
B3 107637621 107637815
refGene_gn=KIAA0586;refGene_tx=XM_023255728.1
B3 107639256 107639375
refGene_gn=KIAA0586;refGene_tx=XM_023255728.1
B3 107643441 107643641
refGene_gn=KIAA0586;refGene_tx=XM_023255728.1
B3 107647304 107647464
refGene_gn=KIAA0586;refGene_tx=XM_023255728.1
B3 107648025 107648245
refGene_gn=KIAA0586;refGene_tx=XM_023255728.1
B3 107648779 107649036
refGene_gn=KIAA0586;refGene_tx=XM_023255728.1
B3 107651117 107651194
refGene_gn=KIAA0586;refGene_tx=XM_023255728.1

B3	107655995	107656130
refGene_gn=KIAA0586;refGene_tx=XM_023255728.1
B3	107658600	107658778
refGene_gn=KIAA0586;refGene_tx=XM_023255728.1
B3	107666606	107666761
refGene_gn=KIAA0586;refGene_tx=XM_023255728.1
B3	107668899	107669008
refGene_gn=KIAA0586;refGene_tx=XM_023255728.1
B3	107696761	107696827
refGene_gn=KIAA0586;refGene_tx=XM_023255728.1
B3	107704728	107704880
refGene_gn=KIAA0586;refGene_tx=XM_023255728.1
B3	107782257	107782602
refGene_gn=DACT1;refGene_tx=XM_023255731.1;refGene_gn=DACT1;refGene_tx=XM_023255732.1
B3	107784830	107784963
refGene_gn=DACT1;refGene_tx=XM_023255731.1;refGene_gn=DACT1;refGene_tx=XM_023255732.1
B3	107785716	107785872
refGene_gn=DACT1;refGene_tx=XM_023255731.1;refGene_gn=DACT1;refGene_tx=XM_023255732.1
B3	107789336	107791225
refGene_gn=DACT1;refGene_tx=XM_023255731.1;refGene_gn=DACT1;refGene_tx=XM_023255732.1
B3	108401106	108401289
refGene_gn=DAAM1;refGene_tx=XM_023255733.1;refGene_gn=DAAM1;refGene_tx=XM_023255734.1;refGene_gn=DAAM1;refGene_tx=XM_023255735.1
B3	108422459	108422549
refGene_gn=DAAM1;refGene_tx=XM_023255733.1;refGene_gn=DAAM1;refGene_tx=XM_023255734.1;refGene_gn=DAAM1;refGene_tx=XM_023255735.1
B3	108450102	108450174
refGene_gn=DAAM1;refGene_tx=XM_023255733.1;refGene_gn=DAAM1;refGene_tx=XM_023255734.1;refGene_gn=DAAM1;refGene_tx=XM_023255735.1
B3	108455803	108455898
refGene_gn=DAAM1;refGene_tx=XM_023255733.1;refGene_gn=DAAM1;refGene_tx=XM_023255734.1;refGene_gn=DAAM1;refGene_tx=XM_023255735.1
B3	108458761	108459095
refGene_gn=DAAM1;refGene_tx=XM_023255733.1;refGene_gn=DAAM1;refGene_tx=XM_023255734.1;refGene_gn=DAAM1;refGene_tx=XM_023255735.1
B3	108459756	108459867
refGene_gn=DAAM1;refGene_tx=XM_023255733.1;refGene_gn=DAAM1;refGene_tx=XM_023255734.1;refGene_gn=DAAM1;refGene_tx=XM_023255735.1
B3	108459991	108460095
refGene_gn=DAAM1;refGene_tx=XM_023255733.1;refGene_gn=DAAM1;refGene_tx=XM_023255734.1;refGene_gn=DAAM1;refGene_tx=XM_023255735.1
B3	108461249	108461316
refGene_gn=DAAM1;refGene_tx=XM_023255733.1;refGene_gn=DAAM1;refGene_tx=XM_023255734.1;refGene_gn=DAAM1;refGene_tx=XM_023255735.1
B3	108461544	108461662
refGene_gn=DAAM1;refGene_tx=XM_023255733.1;refGene_gn=DAAM1;refGene_tx

=XM_023255734.1;refGene_gn=DAAM1;refGene_tx=XM_023255735.1
B3 108462094 108462233
refGene_gn=DAAM1;refGene_tx=XM_023255733.1;refGene_gn=DAAM1;refGene_tx=XM_023255734.1;refGene_gn=DAAM1;refGene_tx=XM_023255735.1
B3 108462517 108462576
refGene_gn=DAAM1;refGene_tx=XM_023255733.1;refGene_gn=DAAM1;refGene_tx=XM_023255734.1;refGene_gn=DAAM1;refGene_tx=XM_023255735.1
B3 108465734 108465922
refGene_gn=DAAM1;refGene_tx=XM_023255733.1;refGene_gn=DAAM1;refGene_tx=XM_023255734.1;refGene_gn=DAAM1;refGene_tx=XM_023255735.1
B3 108466631 108466931
refGene_gn=DAAM1;refGene_tx=XM_023255733.1;refGene_gn=DAAM1;refGene_tx=XM_023255734.1;refGene_gn=DAAM1;refGene_tx=XM_023255735.1
B3 108467245 108467353
refGene_gn=DAAM1;refGene_tx=XM_023255733.1;refGene_gn=DAAM1;refGene_tx=XM_023255734.1;refGene_gn=DAAM1;refGene_tx=XM_023255735.1
B3 108474214 108474244
refGene_gn=DAAM1;refGene_tx=XM_023255733.1;refGene_gn=DAAM1;refGene_tx=XM_023255734.1
B3 108475379 108475486
refGene_gn=DAAM1;refGene_tx=XM_023255733.1;refGene_gn=DAAM1;refGene_tx=XM_023255734.1;refGene_gn=DAAM1;refGene_tx=XM_023255735.1
B3 108483046 108483131
refGene_gn=DAAM1;refGene_tx=XM_023255733.1;refGene_gn=DAAM1;refGene_tx=XM_023255734.1;refGene_gn=DAAM1;refGene_tx=XM_023255735.1
B3 108488193 108488300
refGene_gn=DAAM1;refGene_tx=XM_023255733.1;refGene_gn=DAAM1;refGene_tx=XM_023255734.1;refGene_gn=DAAM1;refGene_tx=XM_023255735.1
B3 108490107 108490196
refGene_gn=DAAM1;refGene_tx=XM_023255733.1;refGene_gn=DAAM1;refGene_tx=XM_023255734.1;refGene_gn=DAAM1;refGene_tx=XM_023255735.1
B3 108491058 108491227
refGene_gn=DAAM1;refGene_tx=XM_023255733.1;refGene_gn=DAAM1;refGene_tx=XM_023255734.1;refGene_gn=DAAM1;refGene_tx=XM_023255735.1
B3 108495112 108495220
refGene_gn=DAAM1;refGene_tx=XM_023255733.1;refGene_gn=DAAM1;refGene_tx=XM_023255734.1;refGene_gn=DAAM1;refGene_tx=XM_023255735.1
B3 108497109 108497170
refGene_gn=DAAM1;refGene_tx=XM_023255733.1;refGene_gn=DAAM1;refGene_tx=XM_023255734.1;refGene_gn=DAAM1;refGene_tx=XM_023255735.1
B3 108500922 108501054
refGene_gn=DAAM1;refGene_tx=XM_023255733.1;refGene_gn=DAAM1;refGene_tx=XM_023255734.1;refGene_gn=DAAM1;refGene_tx=XM_023255735.1
B3 108504484 108504655
refGene_gn=DAAM1;refGene_tx=XM_023255733.1;refGene_gn=DAAM1;refGene_tx=XM_023255734.1;refGene_gn=DAAM1;refGene_tx=XM_023255735.1
B3 108505946 108506156
refGene_gn=DAAM1;refGene_tx=XM_023255733.1;refGene_gn=DAAM1;refGene_tx=XM_023255734.1;refGene_gn=DAAM1;refGene_tx=XM_023255735.1
B3 108556992 108558519

refGene_gn=GPR135;refGene_tx=XM_023255736.1
B3	108566777	108566903
refGene_gn=L3HYPDH;refGene_tx=XM_023255737.1
B3	108568960	108569098
refGene_gn=L3HYPDH;refGene_tx=XM_023255737.1
B3	108569187	108569310
refGene_gn=L3HYPDH;refGene_tx=XM_023255737.1
B3	108571927	108572007
refGene_gn=L3HYPDH;refGene_tx=XM_023255737.1
B3	108572009	108572096
refGene_gn=L3HYPDH;refGene_tx=XM_023255737.1
B3	108576057	108576565
refGene_gn=L3HYPDH;refGene_tx=XM_023255737.1
B3	108576889	108576893
refGene_gn=JKAMP;refGene_tx=XM_003987698.5;refGene_gn=JKAMP;refGene_tx=XM_006932897.4
B3	108579930	108580022
refGene_gn=JKAMP;refGene_tx=XM_003987698.5;refGene_gn=JKAMP;refGene_tx=XM_006932897.4
B3	108580572	108580727
refGene_gn=JKAMP;refGene_tx=XM_003987698.5;refGene_gn=JKAMP;refGene_tx=XM_006932897.4
B3	108588230	108588437
refGene_gn=JKAMP;refGene_tx=XM_003987698.5;refGene_gn=JKAMP;refGene_tx=XM_006932897.4
B3	108591706	108591888
refGene_gn=JKAMP;refGene_tx=XM_003987698.5;refGene_gn=JKAMP;refGene_tx=XM_006932897.4
B3	108596430	108596507
refGene_gn=JKAMP;refGene_tx=XM_003987698.5;refGene_gn=JKAMP;refGene_tx=XM_006932897.4
B3	108605730	108605949
refGene_gn=JKAMP;refGene_tx=XM_003987698.5;refGene_gn=JKAMP;refGene_tx=XM_006932897.4
B3	108606978	108607064
refGene_gn=CCDC175;refGene_tx=XM_011283246.3;refGene_gn=CCDC175;refGene_tx=XM_019833067.2;refGene_gn=CCDC175;refGene_tx=XM_023255738.1
B3	108611252	108611415
refGene_gn=CCDC175;refGene_tx=XM_011283246.3;refGene_gn=CCDC175;refGene_tx=XM_019833067.2;refGene_gn=CCDC175;refGene_tx=XM_023255738.1
B3	108612703	108612747
refGene_gn=CCDC175;refGene_tx=XM_011283246.3;refGene_gn=CCDC175;refGene_tx=XM_019833067.2;refGene_gn=CCDC175;refGene_tx=XM_023255738.1
B3	108615968	108616071
refGene_gn=CCDC175;refGene_tx=XM_011283246.3;refGene_gn=CCDC175;refGene_tx=XM_019833067.2;refGene_gn=CCDC175;refGene_tx=XM_023255738.1
B3	108619458	108619611
refGene_gn=CCDC175;refGene_tx=XM_011283246.3;refGene_gn=CCDC175;refGene_tx=XM_019833067.2;refGene_gn=CCDC175;refGene_tx=XM_023255738.1
B3	108633853	108633933

refGene_gn=CCDC175;refGene_tx=XM_011283246.3;refGene_gn=CCDC175;refGene_tx=XM_019833067.2;refGene_gn=CCDC175;refGene_tx=XM_023255738.1
B3 108639287 108639426
refGene_gn=CCDC175;refGene_tx=XM_011283246.3;refGene_gn=CCDC175;refGene_tx=XM_019833067.2;refGene_gn=CCDC175;refGene_tx=XM_023255738.1
B3 108646186 108646318
refGene_gn=CCDC175;refGene_tx=XM_011283246.3;refGene_gn=CCDC175;refGene_tx=XM_019833067.2;refGene_gn=CCDC175;refGene_tx=XM_023255738.1
B3 108646920 108647056
refGene_gn=CCDC175;refGene_tx=XM_011283246.3;refGene_gn=CCDC175;refGene_tx=XM_019833067.2;refGene_gn=CCDC175;refGene_tx=XM_023255738.1
B3 108648091 108648163
refGene_gn=CCDC175;refGene_tx=XM_011283246.3;refGene_gn=CCDC175;refGene_tx=XM_019833067.2;refGene_gn=CCDC175;refGene_tx=XM_023255738.1
B3 108651121 108651232
refGene_gn=CCDC175;refGene_tx=XM_011283246.3;refGene_gn=CCDC175;refGene_tx=XM_019833067.2;refGene_gn=CCDC175;refGene_tx=XM_023255738.1
B3 108653899 108654036
refGene_gn=CCDC175;refGene_tx=XM_011283246.3;refGene_gn=CCDC175;refGene_tx=XM_019833067.2;refGene_gn=CCDC175;refGene_tx=XM_023255738.1
B3 108658083 108658165
refGene_gn=CCDC175;refGene_tx=XM_011283246.3;refGene_gn=CCDC175;refGene_tx=XM_019833067.2;refGene_gn=CCDC175;refGene_tx=XM_023255738.1
B3 108661703 108661813
refGene_gn=CCDC175;refGene_tx=XM_011283246.3;refGene_gn=CCDC175;refGene_tx=XM_019833067.2;refGene_gn=CCDC175;refGene_tx=XM_023255738.1
B3 108664673 108664796
refGene_gn=CCDC175;refGene_tx=XM_011283246.3;refGene_gn=CCDC175;refGene_tx=XM_023255738.1
B3 108669436 108669665
refGene_gn=CCDC175;refGene_tx=XM_011283246.3;refGene_gn=CCDC175;refGene_tx=XM_019833067.2;refGene_gn=CCDC175;refGene_tx=XM_023255738.1
B3 108672912 108673048
refGene_gn=CCDC175;refGene_tx=XM_011283246.3;refGene_gn=CCDC175;refGene_tx=XM_019833067.2;refGene_gn=CCDC175;refGene_tx=XM_023255738.1
B3 108678541 108678653
refGene_gn=CCDC175;refGene_tx=XM_011283246.3;refGene_gn=CCDC175;refGene_tx=XM_019833067.2;refGene_gn=CCDC175;refGene_tx=XM_023255738.1
B3 108681137 108681223
refGene_gn=CCDC175;refGene_tx=XM_011283246.3;refGene_gn=CCDC175;refGene_tx=XM_019833067.2;refGene_gn=CCDC175;refGene_tx=XM_023255738.1
B3 108683540 108683700
refGene_gn=CCDC175;refGene_tx=XM_011283246.3;refGene_gn=CCDC175;refGene_tx=XM_019833067.2;refGene_gn=CCDC175;refGene_tx=XM_023255738.1
B3 108703971 108704014
refGene_gn=RTN1;refGene_tx=XM_011283247.3;refGene_gn=RTN1;refGene_tx=XM_011283248.3
B3 108708141 108708200
refGene_gn=RTN1;refGene_tx=XM_011283247.3;refGene_gn=RTN1;refGene_tx=XM_011283248.3

B3	108708285	108708332
refGene_gn=RTN1;refGene_tx=XM_011283247.3;refGene_gn=RTN1;refGene_tx=XM_011283248.3
B3	108709100	108709170
refGene_gn=RTN1;refGene_tx=XM_011283247.3;refGene_gn=RTN1;refGene_tx=XM_011283248.3
B3	108710615	108710754
refGene_gn=RTN1;refGene_tx=XM_011283247.3;refGene_gn=RTN1;refGene_tx=XM_011283248.3
B3	108713156	108713364
refGene_gn=RTN1;refGene_tx=XM_011283247.3;refGene_gn=RTN1;refGene_tx=XM_011283248.3
B3	108728621	108728682
refGene_gn=RTN1;refGene_tx=XM_011283248.3
B3	108800483	108801212
refGene_gn=RTN1;refGene_tx=XM_011283247.3
B3	108809139	108809913
refGene_gn=RTN1;refGene_tx=XM_011283247.3
B3	108931309	108931550
refGene_gn=RTN1;refGene_tx=XM_011283247.3
B3	108975584	108975632
refGene_gn=LOC101100478;refGene_tx=XM_006932901.4;refGene_gn=LOC101100478;refGene_tx=XM_019833069.2;refGene_gn=LOC101100478;refGene_tx=XM_019833073.2;refGene_gn=LOC101100478;refGene_tx=XM_019833074.2;refGene_gn=LOC101100478;refGene_tx=XM_019833075.2
B3	108975888	108976107
refGene_gn=LOC101100478;refGene_tx=XM_006932901.4;refGene_gn=LOC101100478;refGene_tx=XM_019833069.2;refGene_gn=LOC101100478;refGene_tx=XM_019833073.2;refGene_gn=LOC101100478;refGene_tx=XM_019833074.2;refGene_gn=LOC101100478;refGene_tx=XM_019833075.2
B3	108977041	108977182
refGene_gn=LOC101100478;refGene_tx=XM_006932901.4;refGene_gn=LOC101100478;refGene_tx=XM_019833069.2;refGene_gn=LOC101100478;refGene_tx=XM_019833073.2;refGene_gn=LOC101100478;refGene_tx=XM_019833074.2;refGene_gn=LOC101100478;refGene_tx=XM_019833075.2
B3	108977737	108977801
refGene_gn=LOC101100478;refGene_tx=XM_006932901.4;refGene_gn=LOC101100478;refGene_tx=XM_019833069.2;refGene_gn=LOC101100478;refGene_tx=XM_019833073.2;refGene_gn=LOC101100478;refGene_tx=XM_019833074.2;refGene_gn=LOC101100478;refGene_tx=XM_019833075.2
B3	108978109	108978180
refGene_gn=LOC101100478;refGene_tx=XM_006932901.4;refGene_gn=LOC101100478;refGene_tx=XM_019833069.2;refGene_gn=LOC101100478;refGene_tx=XM_019833073.2;refGene_gn=LOC101100478;refGene_tx=XM_019833074.2;refGene_gn=LOC101100478;refGene_tx=XM_019833075.2
B3	108982632	108982815
refGene_gn=LOC101100478;refGene_tx=XM_006932901.4;refGene_gn=LOC101100478;refGene_tx=XM_019833069.2;refGene_gn=LOC101100478;refGene_tx=XM_019833074.2;refGene_gn=LOC101100478;refGene_tx=XM_019833075.2
B3	108989927	108990083

refGene_gn=LOC101100478;refGene_tx=XM_006932901.4;refGene_gn=LOC101100478;refGene_tx=XM_019833069.2;refGene_gn=LOC101100478;refGene_tx=XM_019833073.2;refGene_gn=LOC101100478;refGene_tx=XM_019833074.2;refGene_gn=LOC101100478;refGene_tx=XM_019833075.2
B3 108992601 108992798
refGene_gn=LOC101100478;refGene_tx=XM_006932901.4;refGene_gn=LOC101100478;refGene_tx=XM_019833069.2;refGene_gn=LOC101100478;refGene_tx=XM_019833073.2;refGene_gn=LOC101100478;refGene_tx=XM_019833074.2;refGene_gn=LOC101100478;refGene_tx=XM_019833075.2
B3 108993678 108993810
refGene_gn=LOC101100478;refGene_tx=XM_006932901.4;refGene_gn=LOC101100478;refGene_tx=XM_019833069.2;refGene_gn=LOC101100478;refGene_tx=XM_019833073.2;refGene_gn=LOC101100478;refGene_tx=XM_019833074.2;refGene_gn=LOC101100478;refGene_tx=XM_019833075.2
B3 108999934 109000111
refGene_gn=LOC101100478;refGene_tx=XM_006932901.4;refGene_gn=LOC101100478;refGene_tx=XM_019833069.2;refGene_gn=LOC101100478;refGene_tx=XM_019833073.2;refGene_gn=LOC101100478;refGene_tx=XM_019833074.2;refGene_gn=LOC101100478;refGene_tx=XM_019833075.2
B3 109000879 109000997
refGene_gn=LOC101100478;refGene_tx=XM_006932901.4;refGene_gn=LOC101100478;refGene_tx=XM_019833069.2;refGene_gn=LOC101100478;refGene_tx=XM_019833073.2;refGene_gn=LOC101100478;refGene_tx=XM_019833074.2;refGene_gn=LOC101100478;refGene_tx=XM_019833075.2
B3 109009256 109009386
refGene_gn=LOC101100478;refGene_tx=XM_006932901.4;refGene_gn=LOC101100478;refGene_tx=XM_019833069.2;refGene_gn=LOC101100478;refGene_tx=XM_019833073.2;refGene_gn=LOC101100478;refGene_tx=XM_019833074.2;refGene_gn=LOC101100478;refGene_tx=XM_019833075.2
B3 109013056 109013179
refGene_gn=LOC101100478;refGene_tx=XM_006932901.4;refGene_gn=LOC101100478;refGene_tx=XM_019833069.2;refGene_gn=LOC101100478;refGene_tx=XM_019833073.2;refGene_gn=LOC101100478;refGene_tx=XM_019833074.2;refGene_gn=LOC101100478;refGene_tx=XM_019833075.2
B3 109015099 109015215
refGene_gn=LOC101100478;refGene_tx=XM_006932901.4;refGene_gn=LOC101100478;refGene_tx=XM_019833069.2;refGene_gn=LOC101100478;refGene_tx=XM_019833073.2;refGene_gn=LOC101100478;refGene_tx=XM_019833074.2;refGene_gn=LOC101100478;refGene_tx=XM_019833075.2
B3 109019100 109019313
refGene_gn=LOC101100478;refGene_tx=XM_006932901.4;refGene_gn=LOC101100478;refGene_tx=XM_019833069.2;refGene_gn=LOC101100478;refGene_tx=XM_019833073.2;refGene_gn=LOC101100478;refGene_tx=XM_019833075.2
B3 109026642 109026762
refGene_gn=LOC101100478;refGene_tx=XM_006932901.4;refGene_gn=LOC101100478;refGene_tx=XM_019833069.2;refGene_gn=LOC101100478;refGene_tx=XM_019833073.2;refGene_gn=LOC101100478;refGene_tx=XM_019833074.2;refGene_gn=LOC101100478;refGene_tx=XM_019833075.2
B3 109035059 109035251
refGene_gn=LOC101100478;refGene_tx=XM_006932901.4;refGene_gn=LOC101100

478;refGene_tx=XM_019833069.2;refGene_gn=LOC101100478;refGene_tx=XM_019833073.2;refGene_gn=LOC101100478;refGene_tx=XM_019833074.2;refGene_gn=LOC101100478;refGene_tx=XM_019833075.2
B3 109036354 109036480
refGene_gn=LOC101100478;refGene_tx=XM_006932901.4;refGene_gn=LOC101100478;refGene_tx=XM_019833069.2;refGene_gn=LOC101100478;refGene_tx=XM_019833073.2;refGene_gn=LOC101100478;refGene_tx=XM_019833074.2;refGene_gn=LOC101100478;refGene_tx=XM_019833075.2
B3 109038841 109038976
refGene_gn=LOC101100478;refGene_tx=XM_006932901.4;refGene_gn=LOC101100478;refGene_tx=XM_019833069.2;refGene_gn=LOC101100478;refGene_tx=XM_019833073.2;refGene_gn=LOC101100478;refGene_tx=XM_019833074.2;refGene_gn=LOC101100478;refGene_tx=XM_019833075.2
B3 109040420 109040598
refGene_gn=LOC101100478;refGene_tx=XM_006932901.4;refGene_gn=LOC101100478;refGene_tx=XM_019833069.2;refGene_gn=LOC101100478;refGene_tx=XM_019833073.2;refGene_gn=LOC101100478;refGene_tx=XM_019833074.2;refGene_gn=LOC101100478;refGene_tx=XM_019833075.2
B3 109042859 109043080
refGene_gn=LOC101100478;refGene_tx=XM_006932901.4;refGene_gn=LOC101100478;refGene_tx=XM_019833069.2;refGene_gn=LOC101100478;refGene_tx=XM_019833073.2;refGene_gn=LOC101100478;refGene_tx=XM_019833074.2;refGene_gn=LOC101100478;refGene_tx=XM_019833075.2
B3 109044508 109044631
refGene_gn=LOC101100478;refGene_tx=XM_006932901.4;refGene_gn=LOC101100478;refGene_tx=XM_019833069.2;refGene_gn=LOC101100478;refGene_tx=XM_019833073.2;refGene_gn=LOC101100478;refGene_tx=XM_019833074.2;refGene_gn=LOC101100478;refGene_tx=XM_019833075.2
B3 109048795 109048926
refGene_gn=LOC101100478;refGene_tx=XM_006932901.4;refGene_gn=LOC101100478;refGene_tx=XM_019833069.2;refGene_gn=LOC101100478;refGene_tx=XM_019833073.2;refGene_gn=LOC101100478;refGene_tx=XM_019833074.2;refGene_gn=LOC101100478;refGene_tx=XM_019833075.2
B3 109050878 109050987
refGene_gn=LOC101100478;refGene_tx=XM_006932901.4;refGene_gn=LOC101100478;refGene_tx=XM_019833069.2;refGene_gn=LOC101100478;refGene_tx=XM_019833073.2;refGene_gn=LOC101100478;refGene_tx=XM_019833074.2;refGene_gn=LOC101100478;refGene_tx=XM_019833075.2
B3 109051670 109051810
refGene_gn=LOC101100478;refGene_tx=XM_006932901.4;refGene_gn=LOC101100478;refGene_tx=XM_019833069.2;refGene_gn=LOC101100478;refGene_tx=XM_019833073.2;refGene_gn=LOC101100478;refGene_tx=XM_019833074.2;refGene_gn=LOC101100478;refGene_tx=XM_019833075.2
B3 109055323 109055460
refGene_gn=LOC101100478;refGene_tx=XM_006932901.4;refGene_gn=LOC101100478;refGene_tx=XM_019833069.2;refGene_gn=LOC101100478;refGene_tx=XM_019833073.2;refGene_gn=LOC101100478;refGene_tx=XM_019833074.2;refGene_gn=LOC101100478;refGene_tx=XM_019833075.2
B3 109060038 109060256
refGene_gn=LOC101100478;refGene_tx=XM_006932901.4;refGene_gn=LOC101100

478;refGene_tx=XM_019833069.2;refGene_gn=LOC101100478;refGene_tx=XM_01
9833073.2;refGene_gn=LOC101100478;refGene_tx=XM_019833074.2;refGene_gn
=LOC101100478;refGene_tx=XM_019833075.2
B3 109064173 109064242
refGene_gn=LOC101100478;refGene_tx=XM_006932901.4;refGene_gn=LOC101100
478;refGene_tx=XM_019833069.2;refGene_gn=LOC101100478;refGene_tx=XM_01
9833073.2;refGene_gn=LOC101100478;refGene_tx=XM_019833074.2;refGene_gn
=LOC101100478;refGene_tx=XM_019833075.2
B3 109072129 109072259
refGene_gn=LOC101100478;refGene_tx=XM_019833075.2
B3 109072890 109073031
refGene_gn=LOC101100478;refGene_tx=XM_006932901.4;refGene_gn=LOC101100
478;refGene_tx=XM_019833069.2;refGene_gn=LOC101100478;refGene_tx=XM_01
9833073.2;refGene_gn=LOC101100478;refGene_tx=XM_019833074.2;refGene_gn
=LOC101100478;refGene_tx=XM_019833075.2
B3 109078010 109078155
refGene_gn=LOC101100478;refGene_tx=XM_006932901.4;refGene_gn=LOC101100
478;refGene_tx=XM_019833069.2;refGene_gn=LOC101100478;refGene_tx=XM_01
9833073.2;refGene_gn=LOC101100478;refGene_tx=XM_019833074.2
B3 109079595 109079764
refGene_gn=LOC101100478;refGene_tx=XM_006932901.4;refGene_gn=LOC101100
478;refGene_tx=XM_019833069.2;refGene_gn=LOC101100478;refGene_tx=XM_01
9833073.2;refGene_gn=LOC101100478;refGene_tx=XM_019833074.2
B3 109080606 109080646
refGene_gn=LOC101100478;refGene_tx=XM_019833069.2
B3 109087678 109087736
refGene_gn=LOC101100478;refGene_tx=XM_006932901.4;refGene_gn=LOC101100
478;refGene_tx=XM_019833073.2;refGene_gn=LOC101100478;refGene_tx=XM_01
9833074.2
B3 109120267 109120965
refGene_gn=PCNX4;refGene_tx=XM_011283249.3;refGene_gn=PCNX4;refGene_tx
=XM_023255740.1
B3 109121925 109121963
refGene_gn=PCNX4;refGene_tx=XM_019833078.2;refGene_gn=PCNX4;refGene_tx
=XM_019833079.2
B3 109123834 109123836
refGene_gn=PCNX4;refGene_tx=XM_023255741.1
B3 109126488 109126668
refGene_gn=PCNX4;refGene_tx=XM_011283249.3;refGene_gn=PCNX4;refGene_tx
=XM_019833078.2;refGene_gn=PCNX4;refGene_tx=XM_019833079.2;refGene_gn=
PCNX4;refGene_tx=XM_023255740.1;refGene_gn=PCNX4;refGene_tx=XM_0232557
41.1
B3 109126751 109127266
refGene_gn=PCNX4;refGene_tx=XM_011283249.3;refGene_gn=PCNX4;refGene_tx
=XM_019833078.2;refGene_gn=PCNX4;refGene_tx=XM_019833079.2;refGene_gn=
PCNX4;refGene_tx=XM_023255740.1;refGene_gn=PCNX4;refGene_tx=XM_0232557
41.1
B3 109127504 109127605
refGene_gn=PCNX4;refGene_tx=XM_011283249.3;refGene_gn=PCNX4;refGene_tx
=XM_019833078.2;refGene_gn=PCNX4;refGene_tx=XM_019833079.2;refGene_gn=

PCNX4;refGene_tx=XM_023255740.1;refGene_gn=PCNX4;refGene_tx=XM_0232557
41.1
B3 109127726 109127849
refGene_gn=PCNX4;refGene_tx=XM_011283249.3;refGene_gn=PCNX4;refGene_tx
=XM_019833078.2;refGene_gn=PCNX4;refGene_tx=XM_019833079.2;refGene_gn=
PCNX4;refGene_tx=XM_023255740.1;refGene_gn=PCNX4;refGene_tx=XM_0232557
41.1
B3 109132589 109132950
refGene_gn=PCNX4;refGene_tx=XM_011283249.3;refGene_gn=PCNX4;refGene_tx
=XM_019833078.2;refGene_gn=PCNX4;refGene_tx=XM_019833079.2;refGene_gn=
PCNX4;refGene_tx=XM_023255740.1;refGene_gn=PCNX4;refGene_tx=XM_0232557
41.1
B3 109136186 109136290
refGene_gn=PCNX4;refGene_tx=XM_011283249.3;refGene_gn=PCNX4;refGene_tx
=XM_019833078.2;refGene_gn=PCNX4;refGene_tx=XM_019833079.2;refGene_gn=
PCNX4;refGene_tx=XM_023255740.1;refGene_gn=PCNX4;refGene_tx=XM_0232557
41.1
B3 109139351 109140388
refGene_gn=PCNX4;refGene_tx=XM_011283249.3;refGene_gn=PCNX4;refGene_tx
=XM_019833078.2;refGene_gn=PCNX4;refGene_tx=XM_019833079.2;refGene_gn=
PCNX4;refGene_tx=XM_023255740.1;refGene_gn=PCNX4;refGene_tx=XM_0232557
41.1
B3 109140763 109140950
refGene_gn=PCNX4;refGene_tx=XM_011283249.3;refGene_gn=PCNX4;refGene_tx
=XM_019833078.2;refGene_gn=PCNX4;refGene_tx=XM_019833079.2;refGene_gn=
PCNX4;refGene_tx=XM_023255740.1;refGene_gn=PCNX4;refGene_tx=XM_0232557
41.1
B3 109144304 109144556
refGene_gn=PCNX4;refGene_tx=XM_011283249.3;refGene_gn=PCNX4;refGene_tx
=XM_019833078.2;refGene_gn=PCNX4;refGene_tx=XM_019833079.2;refGene_gn=
PCNX4;refGene_tx=XM_023255740.1;refGene_gn=PCNX4;refGene_tx=XM_0232557
41.1
B3 109151103 109151151
refGene_gn=DHRS7;refGene_tx=XM_003987704.5
B3 109154644 109154860
refGene_gn=DHRS7;refGene_tx=XM_003987704.5
B3 109156069 109156192
refGene_gn=DHRS7;refGene_tx=XM_003987704.5
B3 109158948 109159188
refGene_gn=DHRS7;refGene_tx=XM_003987704.5
B3 109159626 109159733
refGene_gn=DHRS7;refGene_tx=XM_003987704.5
B3 109162798 109162951
refGene_gn=DHRS7;refGene_tx=XM_003987704.5
B3 109169993 109170126
refGene_gn=DHRS7;refGene_tx=XM_003987704.5
B3 109249342 109249634
refGene_gn=PPM1A;refGene_tx=XM_006932902.4
B3 109284065 109284919
refGene_gn=PPM1A;refGene_tx=XM_006932902.4;refGene_gn=PPM1A;refGene_tx

=XM_023255742.1
B3 109287117 109287235
refGene_gn=PPM1A;refGene_tx=XM_006932902.4;refGene_gn=PPM1A;refGene_tx
=XM_023255742.1
B3 109291583 109291692
refGene_gn=PPM1A;refGene_tx=XM_006932902.4;refGene_gn=PPM1A;refGene_tx
=XM_023255742.1
B3 109293078 109293136
refGene_gn=PPM1A;refGene_tx=XM_006932902.4;refGene_gn=PPM1A;refGene_tx
=XM_023255742.1
B3 109294089 109294119
refGene_gn=PPM1A;refGene_tx=XM_006932902.4;refGene_gn=PPM1A;refGene_tx
=XM_023255742.1
B3 109384794 109384994
refGene_gn=CB3H14orf39;refGene_tx=XM_019833081.2;refGene_gn=CB3H14orf3
9;refGene_tx=XM_019833082.2;refGene_gn=CB3H14orf39;refGene_tx=XM_01983
3084.2;refGene_gn=CB3H14orf39;refGene_tx=XM_019833085.2;refGene_gn=CB3
H14orf39;refGene_tx=XM_019833087.2;refGene_gn=CB3H14orf39;refGene_tx=X
M_023255743.1
B3 109386208 109386268
refGene_gn=CB3H14orf39;refGene_tx=XM_019833081.2;refGene_gn=CB3H14orf3
9;refGene_tx=XM_019833082.2;refGene_gn=CB3H14orf39;refGene_tx=XM_01983
3084.2;refGene_gn=CB3H14orf39;refGene_tx=XM_019833085.2;refGene_gn=CB3
H14orf39;refGene_tx=XM_019833087.2
B3 109391147 109391205
refGene_gn=CB3H14orf39;refGene_tx=XM_019833081.2;refGene_gn=CB3H14orf3
9;refGene_tx=XM_019833082.2;refGene_gn=CB3H14orf39;refGene_tx=XM_01983
3084.2;refGene_gn=CB3H14orf39;refGene_tx=XM_019833085.2;refGene_gn=CB3
H14orf39;refGene_tx=XM_019833087.2;refGene_gn=CB3H14orf39;refGene_tx=X
M_023255743.1
B3 109398185 109398321
refGene_gn=CB3H14orf39;refGene_tx=XM_019833081.2;refGene_gn=CB3H14orf3
9;refGene_tx=XM_019833082.2;refGene_gn=CB3H14orf39;refGene_tx=XM_01983
3084.2;refGene_gn=CB3H14orf39;refGene_tx=XM_019833085.2;refGene_gn=CB3
H14orf39;refGene_tx=XM_019833087.2;refGene_gn=CB3H14orf39;refGene_tx=X
M_023255743.1
B3 109399999 109400178
refGene_gn=CB3H14orf39;refGene_tx=XM_019833081.2;refGene_gn=CB3H14orf3
9;refGene_tx=XM_019833082.2;refGene_gn=CB3H14orf39;refGene_tx=XM_01983
3084.2;refGene_gn=CB3H14orf39;refGene_tx=XM_019833085.2;refGene_gn=CB3
H14orf39;refGene_tx=XM_019833087.2;refGene_gn=CB3H14orf39;refGene_tx=X
M_023255743.1
B3 109402485 109402547
refGene_gn=CB3H14orf39;refGene_tx=XM_019833081.2;refGene_gn=CB3H14orf3
9;refGene_tx=XM_019833082.2;refGene_gn=CB3H14orf39;refGene_tx=XM_01983
3084.2;refGene_gn=CB3H14orf39;refGene_tx=XM_019833085.2;refGene_gn=CB3
H14orf39;refGene_tx=XM_019833087.2;refGene_gn=CB3H14orf39;refGene_tx=X
M_023255743.1
B3 109408700 109408759
refGene_gn=CB3H14orf39;refGene_tx=XM_019833081.2;refGene_gn=CB3H14orf3

9;refGene_tx=XM_019833082.2;refGene_gn=CB3H14orf39;refGene_tx=XM_01983
3084.2;refGene_gn=CB3H14orf39;refGene_tx=XM_019833085.2;refGene_gn=CB3
H14orf39;refGene_tx=XM_019833087.2;refGene_gn=CB3H14orf39;refGene_tx=X
M_023255743.1
B3 109409066 109409152
refGene_gn=CB3H14orf39;refGene_tx=XM_019833081.2;refGene_gn=CB3H14orf3
9;refGene_tx=XM_019833082.2;refGene_gn=CB3H14orf39;refGene_tx=XM_01983
3084.2;refGene_gn=CB3H14orf39;refGene_tx=XM_019833085.2;refGene_gn=CB3
H14orf39;refGene_tx=XM_019833087.2;refGene_gn=CB3H14orf39;refGene_tx=X
M_023255743.1
B3 109415262 109415339
refGene_gn=CB3H14orf39;refGene_tx=XM_019833081.2;refGene_gn=CB3H14orf3
9;refGene_tx=XM_019833082.2;refGene_gn=CB3H14orf39;refGene_tx=XM_01983
3084.2;refGene_gn=CB3H14orf39;refGene_tx=XM_019833085.2;refGene_gn=CB3
H14orf39;refGene_tx=XM_019833087.2;refGene_gn=CB3H14orf39;refGene_tx=X
M_023255743.1
B3 109416039 109416170
refGene_gn=CB3H14orf39;refGene_tx=XM_019833081.2;refGene_gn=CB3H14orf3
9;refGene_tx=XM_019833082.2;refGene_gn=CB3H14orf39;refGene_tx=XM_01983
3084.2;refGene_gn=CB3H14orf39;refGene_tx=XM_019833085.2;refGene_gn=CB3
H14orf39;refGene_tx=XM_019833087.2;refGene_gn=CB3H14orf39;refGene_tx=X
M_023255743.1
B3 109418612 109418698
refGene_gn=CB3H14orf39;refGene_tx=XM_019833081.2;refGene_gn=CB3H14orf3
9;refGene_tx=XM_019833082.2;refGene_gn=CB3H14orf39;refGene_tx=XM_01983
3084.2;refGene_gn=CB3H14orf39;refGene_tx=XM_019833085.2;refGene_gn=CB3
H14orf39;refGene_tx=XM_019833087.2;refGene_gn=CB3H14orf39;refGene_tx=X
M_023255743.1
B3 109419623 109419744
refGene_gn=CB3H14orf39;refGene_tx=XM_019833081.2;refGene_gn=CB3H14orf3
9;refGene_tx=XM_019833082.2;refGene_gn=CB3H14orf39;refGene_tx=XM_01983
3084.2;refGene_gn=CB3H14orf39;refGene_tx=XM_019833085.2;refGene_gn=CB3
H14orf39;refGene_tx=XM_019833087.2;refGene_gn=CB3H14orf39;refGene_tx=X
M_023255743.1
B3 109421482 109421525
refGene_gn=CB3H14orf39;refGene_tx=XM_019833081.2;refGene_gn=CB3H14orf3
9;refGene_tx=XM_019833082.2;refGene_gn=CB3H14orf39;refGene_tx=XM_01983
3084.2;refGene_gn=CB3H14orf39;refGene_tx=XM_019833085.2;refGene_gn=CB3
H14orf39;refGene_tx=XM_019833087.2;refGene_gn=CB3H14orf39;refGene_tx=X
M_023255743.1
B3 109421610 109421798
refGene_gn=CB3H14orf39;refGene_tx=XM_019833081.2;refGene_gn=CB3H14orf3
9;refGene_tx=XM_019833082.2;refGene_gn=CB3H14orf39;refGene_tx=XM_01983
3084.2;refGene_gn=CB3H14orf39;refGene_tx=XM_019833085.2;refGene_gn=CB3
H14orf39;refGene_tx=XM_019833087.2;refGene_gn=CB3H14orf39;refGene_tx=X
M_023255743.1
B3 109430325 109430415
refGene_gn=CB3H14orf39;refGene_tx=XM_019833081.2;refGene_gn=CB3H14orf3
9;refGene_tx=XM_019833082.2;refGene_gn=CB3H14orf39;refGene_tx=XM_01983
3084.2;refGene_gn=CB3H14orf39;refGene_tx=XM_019833085.2;refGene_gn=CB3

H14orf39;refGene_tx=XM_019833087.2;refGene_gn=CB3H14orf39;refGene_tx=XM_023255743.1
B3 109434726 109434853
refGene_gn=CB3H14orf39;refGene_tx=XM_019833081.2;refGene_gn=CB3H14orf39;refGene_tx=XM_019833082.2;refGene_gn=CB3H14orf39;refGene_tx=XM_019833084.2;refGene_gn=CB3H14orf39;refGene_tx=XM_019833085.2;refGene_gn=CB3H14orf39;refGene_tx=XM_019833087.2;refGene_gn=CB3H14orf39;refGene_tx=XM_023255743.1
B3 109437508 109437565
refGene_gn=CB3H14orf39;refGene_tx=XM_019833081.2;refGene_gn=CB3H14orf39;refGene_tx=XM_019833082.2;refGene_gn=CB3H14orf39;refGene_tx=XM_019833084.2;refGene_gn=CB3H14orf39;refGene_tx=XM_019833085.2;refGene_gn=CB3H14orf39;refGene_tx=XM_019833087.2;refGene_gn=CB3H14orf39;refGene_tx=XM_023255743.1
B3 109437659 109437708
refGene_gn=CB3H14orf39;refGene_tx=XM_019833081.2;refGene_gn=CB3H14orf39;refGene_tx=XM_019833082.2;refGene_gn=CB3H14orf39;refGene_tx=XM_019833084.2;refGene_gn=CB3H14orf39;refGene_tx=XM_019833085.2;refGene_gn=CB3H14orf39;refGene_tx=XM_023255743.1
B3 109468049 109468621
refGene_gn=SIX6;refGene_tx=XM_003987707.4;refGene_gn=SIX6;refGene_tx=XM_019833089.2
B3 109469434 109469619
refGene_gn=SIX6;refGene_tx=XM_019833089.2
B3 109469739 109469908
refGene_gn=SIX6;refGene_tx=XM_003987707.4;refGene_gn=SIX6;refGene_tx=XM_019833089.2
B3 109591737 109592032
refGene_gn=SIX1;refGene_tx=XM_003987708.5
B3 109594069 109594629
refGene_gn=SIX1;refGene_tx=XM_003987708.5
B3 109648756 109649553
refGene_gn=SIX4;refGene_tx=XM_023255745.1;refGene_gn=SIX4;refGene_tx=XM_023255746.1;refGene_gn=SIX4;refGene_tx=XM_023255747.1
B3 109653199 109653885
refGene_gn=SIX4;refGene_tx=XM_023255745.1;refGene_gn=SIX4;refGene_tx=XM_023255746.1;refGene_gn=SIX4;refGene_tx=XM_023255747.1
B3 109656612 109657484
refGene_gn=SIX4;refGene_tx=XM_023255745.1;refGene_gn=SIX4;refGene_tx=XM_023255746.1;refGene_gn=SIX4;refGene_tx=XM_023255747.1
B3 109657665 109657668
refGene_gn=SIX4;refGene_tx=XM_023255746.1
B3 109665355 109665444
refGene_gn=MNAT1;refGene_tx=XM_019833090.2
B3 109727016 109727169
refGene_gn=MNAT1;refGene_tx=XM_019833090.2
B3 109728868 109728942
refGene_gn=MNAT1;refGene_tx=XM_019833090.2
B3 109745415 109745519
refGene_gn=MNAT1;refGene_tx=XM_019833090.2

B3	109752000	109752141
refGene_gn=MNAT1;refGene_tx=XM_019833090.2
B3	109757315	109757441
refGene_gn=MNAT1;refGene_tx=XM_019833090.2
B3	109807709	109807831
refGene_gn=MNAT1;refGene_tx=XM_019833090.2
B3	109890151	109890272
refGene_gn=MNAT1;refGene_tx=XM_019833090.2
B3	109897209	109897325
refGene_gn=TRMT5;refGene_tx=XM_006932905.3;refGene_gn=TRMT5;refGene_tx=XM_019833091.2
B3	109897679	109898334
refGene_gn=TRMT5;refGene_tx=XM_006932905.3;refGene_gn=TRMT5;refGene_tx=XM_019833091.2
B3	109902558	109902683
refGene_gn=TRMT5;refGene_tx=XM_006932905.3;refGene_gn=TRMT5;refGene_tx=XM_019833091.2
B3	109905755	109906411
refGene_gn=TRMT5;refGene_tx=XM_006932905.3;refGene_gn=TRMT5;refGene_tx=XM_019833091.2
B3	109907611	109907616
refGene_gn=TRMT5;refGene_tx=XM_006932905.3
B3	109907857	109907962
refGene_gn=SLC38A6;refGene_tx=XM_003987711.5;refGene_gn=SLC38A6;refGene_tx=XM_019833092.2;refGene_gn=SLC38A6;refGene_tx=XM_019833093.2
B3	109909093	109909224
refGene_gn=SLC38A6;refGene_tx=XM_003987711.5;refGene_gn=SLC38A6;refGene_tx=XM_019833092.2;refGene_gn=SLC38A6;refGene_tx=XM_019833093.2
B3	109911820	109911894
refGene_gn=SLC38A6;refGene_tx=XM_003987711.5;refGene_gn=SLC38A6;refGene_tx=XM_019833092.2;refGene_gn=SLC38A6;refGene_tx=XM_019833093.2;refGene_gn=SLC38A6;refGene_tx=XM_019833095.2
B3	109924620	109924673
refGene_gn=SLC38A6;refGene_tx=XM_003987711.5;refGene_gn=SLC38A6;refGene_tx=XM_019833092.2;refGene_gn=SLC38A6;refGene_tx=XM_019833093.2;refGene_gn=SLC38A6;refGene_tx=XM_019833095.2
B3	109927807	109927847
refGene_gn=SLC38A6;refGene_tx=XM_003987711.5;refGene_gn=SLC38A6;refGene_tx=XM_019833092.2;refGene_gn=SLC38A6;refGene_tx=XM_019833093.2;refGene_gn=SLC38A6;refGene_tx=XM_019833095.2
B3	109939346	109939425
refGene_gn=SLC38A6;refGene_tx=XM_003987711.5;refGene_gn=SLC38A6;refGene_tx=XM_019833092.2;refGene_gn=SLC38A6;refGene_tx=XM_019833093.2;refGene_gn=SLC38A6;refGene_tx=XM_019833095.2
B3	109949794	109949877
refGene_gn=SLC38A6;refGene_tx=XM_003987711.5;refGene_gn=SLC38A6;refGene_tx=XM_019833092.2;refGene_gn=SLC38A6;refGene_tx=XM_019833093.2;refGene_gn=SLC38A6;refGene_tx=XM_019833095.2
B3	109950343	109950402
refGene_gn=SLC38A6;refGene_tx=XM_003987711.5;refGene_gn=SLC38A6;refGen

e_tx=XM_019833092.2;refGene_gn=SLC38A6;refGene_tx=XM_019833093.2;refGene_gn=SLC38A6;refGene_tx=XM_019833095.2
B3	109958125	109958191
refGene_gn=SLC38A6;refGene_tx=XM_003987711.5;refGene_gn=SLC38A6;refGene_tx=XM_019833092.2;refGene_gn=SLC38A6;refGene_tx=XM_019833093.2;refGene_gn=SLC38A6;refGene_tx=XM_019833095.2
B3	109958431	109958485
refGene_gn=SLC38A6;refGene_tx=XM_003987711.5;refGene_gn=SLC38A6;refGene_tx=XM_019833092.2;refGene_gn=SLC38A6;refGene_tx=XM_019833093.2;refGene_gn=SLC38A6;refGene_tx=XM_019833095.2
B3	109960112	109960192
refGene_gn=SLC38A6;refGene_tx=XM_003987711.5;refGene_gn=SLC38A6;refGene_tx=XM_019833092.2;refGene_gn=SLC38A6;refGene_tx=XM_019833093.2;refGene_gn=SLC38A6;refGene_tx=XM_019833095.2
B3	109962562	109962663
refGene_gn=SLC38A6;refGene_tx=XM_003987711.5;refGene_gn=SLC38A6;refGene_tx=XM_019833092.2;refGene_gn=SLC38A6;refGene_tx=XM_019833093.2;refGene_gn=SLC38A6;refGene_tx=XM_019833095.2
B3	109968054	109968179
refGene_gn=SLC38A6;refGene_tx=XM_003987711.5;refGene_gn=SLC38A6;refGene_tx=XM_019833092.2;refGene_gn=SLC38A6;refGene_tx=XM_019833093.2;refGene_gn=SLC38A6;refGene_tx=XM_019833095.2
B3	109969579	109969724
refGene_gn=SLC38A6;refGene_tx=XM_003987711.5;refGene_gn=SLC38A6;refGene_tx=XM_019833092.2;refGene_gn=SLC38A6;refGene_tx=XM_019833093.2;refGene_gn=SLC38A6;refGene_tx=XM_019833095.2
B3	109969843	109969938
refGene_gn=SLC38A6;refGene_tx=XM_003987711.5;refGene_gn=SLC38A6;refGene_tx=XM_019833092.2;refGene_gn=SLC38A6;refGene_tx=XM_019833093.2;refGene_gn=SLC38A6;refGene_tx=XM_019833095.2
B3	109970115	109970196
refGene_gn=SLC38A6;refGene_tx=XM_003987711.5
B3	109983503	109983632
refGene_gn=SLC38A6;refGene_tx=XM_019833092.2;refGene_gn=SLC38A6;refGene_tx=XM_019833093.2;refGene_gn=SLC38A6;refGene_tx=XM_019833095.2
B3	110141424	110142486
refGene_gn=TMEM30B;refGene_tx=XM_003987714.5
B3	110178745	110179108
refGene_gn=PRKCH;refGene_tx=XM_003987712.5
B3	110246680	110246744
refGene_gn=PRKCH;refGene_tx=XM_003987712.5
B3	110300837	110300988
refGene_gn=PRKCH;refGene_tx=XM_003987712.5
B3	110303478	110303513
refGene_gn=PRKCH;refGene_tx=XM_003987712.5
B3	110307286	110307375
refGene_gn=PRKCH;refGene_tx=XM_003987712.5
B3	110308721	110308851
refGene_gn=PRKCH;refGene_tx=XM_003987712.5
B3	110311020	110311148

refGene_gn=PRKCH;refGene_tx=XM_003987712.5
B3 110313831 110313975
refGene_gn=PRKCH;refGene_tx=XM_003987712.5
B3 110314136 110314310
refGene_gn=PRKCH;refGene_tx=XM_003987712.5
B3 110338376 110338531
refGene_gn=PRKCH;refGene_tx=XM_003987712.5
B3 110381502 110381641
refGene_gn=PRKCH;refGene_tx=XM_003987712.5
B3 110382268 110382457
refGene_gn=PRKCH;refGene_tx=XM_003987712.5
B3 110408940 110409084
refGene_gn=PRKCH;refGene_tx=XM_003987712.5
B3 110410389 110410536
refGene_gn=PRKCH;refGene_tx=XM_003987712.5
B3 110570835 110570870
refGene_gn=HIF1A;refGene_tx=XM_003987716.5
B3 110589269 110589460
refGene_gn=HIF1A;refGene_tx=XM_003987716.5
B3 110590059 110590205
refGene_gn=HIF1A;refGene_tx=XM_003987716.5
B3 110590287 110590372
refGene_gn=HIF1A;refGene_tx=XM_003987716.5
B3 110594103 110594216
refGene_gn=HIF1A;refGene_tx=XM_003987716.5
B3 110594810 110595013
refGene_gn=HIF1A;refGene_tx=XM_003987716.5
B3 110599453 110599560
refGene_gn=HIF1A;refGene_tx=XM_003987716.5
B3 110600692 110600840
refGene_gn=HIF1A;refGene_tx=XM_003987716.5
B3 110603072 110603293
refGene_gn=HIF1A;refGene_tx=XM_003987716.5
B3 110604186 110604473
refGene_gn=HIF1A;refGene_tx=XM_003987716.5
B3 110605721 110605844
refGene_gn=HIF1A;refGene_tx=XM_003987716.5
B3 110605976 110606404
refGene_gn=HIF1A;refGene_tx=XM_003987716.5
B3 110609070 110609179
refGene_gn=HIF1A;refGene_tx=XM_003987716.5
B3 110609967 110610094
refGene_gn=HIF1A;refGene_tx=XM_003987716.5
B3 110610570 110610722
refGene_gn=HIF1A;refGene_tx=XM_003987716.5
B3 110625115 110625243
refGene_gn=SNAPC1;refGene_tx=XM_011283254.3
B3 110627327 110627487
refGene_gn=SNAPC1;refGene_tx=XM_011283254.3;refGene_gn=SNAPC1;refGene_
tx=XM_019833098.1;refGene_gn=SNAPC1;refGene_tx=XM_019833099.2

B3 110627640 110627781
refGene_gn=SNAPC1;refGene_tx=XM_011283254.3;refGene_gn=SNAPC1;refGene_tx=XM_019833098.1;refGene_gn=SNAPC1;refGene_tx=XM_019833099.2
B3 110628635 110628740
refGene_gn=SNAPC1;refGene_tx=XM_011283254.3;refGene_gn=SNAPC1;refGene_tx=XM_019833098.1;refGene_gn=SNAPC1;refGene_tx=XM_019833099.2
B3 110631724 110631883
refGene_gn=SNAPC1;refGene_tx=XM_011283254.3;refGene_gn=SNAPC1;refGene_tx=XM_019833098.1;refGene_gn=SNAPC1;refGene_tx=XM_019833099.2
B3 110633704 110633773
refGene_gn=SNAPC1;refGene_tx=XM_011283254.3;refGene_gn=SNAPC1;refGene_tx=XM_019833098.1;refGene_gn=SNAPC1;refGene_tx=XM_019833099.2
B3 110634529 110634592
refGene_gn=SNAPC1;refGene_tx=XM_011283254.3;refGene_gn=SNAPC1;refGene_tx=XM_019833098.1;refGene_gn=SNAPC1;refGene_tx=XM_019833099.2
B3 110643532 110643683
refGene_gn=SNAPC1;refGene_tx=XM_011283254.3;refGene_gn=SNAPC1;refGene_tx=XM_019833098.1;refGene_gn=SNAPC1;refGene_tx=XM_019833099.2
B3 110655770 110655869
refGene_gn=SNAPC1;refGene_tx=XM_011283254.3;refGene_gn=SNAPC1;refGene_tx=XM_019833098.1;refGene_gn=SNAPC1;refGene_tx=XM_019833099.2
B3 110658741 110658794
refGene_gn=SNAPC1;refGene_tx=XM_011283254.3;refGene_gn=SNAPC1;refGene_tx=XM_019833098.1;refGene_gn=SNAPC1;refGene_tx=XM_019833099.2
B3 110676792 110676805
refGene_gn=SYT16;refGene_tx=XM_023255753.1
B3 110805178 110805355
refGene_gn=SYT16;refGene_tx=XM_023255753.1
B3 110831902 110832425
refGene_gn=SYT16;refGene_tx=XM_023255750.1;refGene_gn=SYT16;refGene_tx=XM_023255751.1;refGene_gn=SYT16;refGene_tx=XM_023255752.1
B3 110896110 110896323
refGene_gn=SYT16;refGene_tx=XM_023255750.1;refGene_gn=SYT16;refGene_tx=XM_023255751.1;refGene_gn=SYT16;refGene_tx=XM_023255752.1;refGene_gn=SYT16;refGene_tx=XM_023255753.1
B3 110901896 110902153
refGene_gn=SYT16;refGene_tx=XM_023255750.1;refGene_gn=SYT16;refGene_tx=XM_023255751.1;refGene_gn=SYT16;refGene_tx=XM_023255752.1;refGene_gn=SYT16;refGene_tx=XM_023255753.1
B3 110907443 110907884
refGene_gn=SYT16;refGene_tx=XM_023255750.1;refGene_gn=SYT16;refGene_tx=XM_023255751.1;refGene_gn=SYT16;refGene_tx=XM_023255752.1;refGene_gn=SYT16;refGene_tx=XM_023255753.1
B3 110910485 110910668
refGene_gn=SYT16;refGene_tx=XM_023255752.1
B3 110910719 110910909
refGene_gn=SYT16;refGene_tx=XM_023255750.1;refGene_gn=SYT16;refGene_tx=XM_023255751.1;refGene_gn=SYT16;refGene_tx=XM_023255753.1
B3 110917601 110917735
refGene_gn=SYT16;refGene_tx=XM_023255751.1

B3	110927652	110927966
refGene_gn=SYT16;refGene_tx=XM_023255750.1;refGene_gn=SYT16;refGene_tx=XM_023255753.1
B3	111060449	111060659
refGene_gn=LOC109500869;refGene_tx=XM_019833103.2
B3	111465567	111466515
refGene_gn=KCNH5;refGene_tx=XM_019833104.2
B3	111519188	111519385
refGene_gn=KCNH5;refGene_tx=XM_019833104.2
B3	111532974	111533227
refGene_gn=KCNH5;refGene_tx=XM_019833104.2
B3	111571219	111571419
refGene_gn=KCNH5;refGene_tx=XM_019833104.2
B3	111668750	111669177
refGene_gn=KCNH5;refGene_tx=XM_019833104.2
B3	111692688	111693081
refGene_gn=KCNH5;refGene_tx=XM_019833104.2
B3	111699315	111699431
refGene_gn=KCNH5;refGene_tx=XM_019833104.2
B3	111709184	111709313
refGene_gn=KCNH5;refGene_tx=XM_019833104.2
B3	111713723	111713830
refGene_gn=KCNH5;refGene_tx=XM_019833104.2
B3	111734565	111734689
refGene_gn=KCNH5;refGene_tx=XM_019833104.2
B3	111761909	111761982
refGene_gn=KCNH5;refGene_tx=XM_019833104.2
B3	111856582	111856826
refGene_gn=RHOJ;refGene_tx=XM_011283255.3
B3	111917154	111917213
refGene_gn=RHOJ;refGene_tx=XM_011283255.3
B3	111928064	111928229
refGene_gn=RHOJ;refGene_tx=XM_011283255.3
B3	111930552	111930648
refGene_gn=RHOJ;refGene_tx=XM_011283255.3
B3	111940514	111940661
refGene_gn=RHOJ;refGene_tx=XM_011283255.3
B3	111956825	111957014
refGene_gn=GPHB5;refGene_tx=XM_003987721.3
B3	111962087	111962291
refGene_gn=GPHB5;refGene_tx=XM_003987721.3
B3	112012201	112012301
refGene_gn=PPP2R5E;refGene_tx=XM_003987722.5;refGene_gn=PPP2R5E;refGene_tx=XM_011283256.3;refGene_gn=PPP2R5E;refGene_tx=XM_011283257.3
B3	112017238	112017340
refGene_gn=PPP2R5E;refGene_tx=XM_003987722.5;refGene_gn=PPP2R5E;refGene_tx=XM_011283256.3;refGene_gn=PPP2R5E;refGene_tx=XM_011283257.3
B3	112019392	112019520
refGene_gn=PPP2R5E;refGene_tx=XM_003987722.5;refGene_gn=PPP2R5E;refGene_tx=XM_011283256.3;refGene_gn=PPP2R5E;refGene_tx=XM_011283257.3

B3 112024195 112024315
refGene_gn=PPP2R5E;refGene_tx=XM_003987722.5;refGene_gn=PPP2R5E;refGene_tx=XM_011283256.3;refGene_gn=PPP2R5E;refGene_tx=XM_011283257.3
B3 112026098 112026149
refGene_gn=PPP2R5E;refGene_tx=XM_003987722.5;refGene_gn=PPP2R5E;refGene_tx=XM_011283256.3;refGene_gn=PPP2R5E;refGene_tx=XM_011283257.3
B3 112026251 112026305
refGene_gn=PPP2R5E;refGene_tx=XM_003987722.5;refGene_gn=PPP2R5E;refGene_tx=XM_011283256.3;refGene_gn=PPP2R5E;refGene_tx=XM_011283257.3
B3 112027733 112027842
refGene_gn=PPP2R5E;refGene_tx=XM_003987722.5;refGene_gn=PPP2R5E;refGene_tx=XM_011283256.3;refGene_gn=PPP2R5E;refGene_tx=XM_011283257.3
B3 112028822 112028882
refGene_gn=PPP2R5E;refGene_tx=XM_003987722.5;refGene_gn=PPP2R5E;refGene_tx=XM_011283256.3;refGene_gn=PPP2R5E;refGene_tx=XM_011283257.3
B3 112029001 112029132
refGene_gn=PPP2R5E;refGene_tx=XM_003987722.5;refGene_gn=PPP2R5E;refGene_tx=XM_011283256.3;refGene_gn=PPP2R5E;refGene_tx=XM_011283257.3
B3 112044519 112044612
refGene_gn=PPP2R5E;refGene_tx=XM_003987722.5;refGene_gn=PPP2R5E;refGene_tx=XM_011283256.3;refGene_gn=PPP2R5E;refGene_tx=XM_011283257.3
B3 112051183 112051285
refGene_gn=PPP2R5E;refGene_tx=XM_003987722.5;refGene_gn=PPP2R5E;refGene_tx=XM_011283256.3;refGene_gn=PPP2R5E;refGene_tx=XM_011283257.3
B3 112074964 112075161
refGene_gn=PPP2R5E;refGene_tx=XM_003987722.5;refGene_gn=PPP2R5E;refGene_tx=XM_011283256.3;refGene_gn=PPP2R5E;refGene_tx=XM_011283257.3
B3 112153928 112154085
refGene_gn=PPP2R5E;refGene_tx=XM_003987722.5;refGene_gn=PPP2R5E;refGene_tx=XM_011283256.3;refGene_gn=PPP2R5E;refGene_tx=XM_011283257.3
B3 112187287 112188451
refGene_gn=WDR89;refGene_tx=XM_003987723.5;refGene_gn=WDR89;refGene_tx=XM_006932906.4;refGene_gn=WDR89;refGene_tx=XM_019833112.2;refGene_gn=WDR89;refGene_tx=XM_019833113.2;refGene_gn=WDR89;refGene_tx=XM_019833116.2
B3 112269505 112270057
refGene_gn=SGPP1;refGene_tx=XM_011283398.3
B3 112282388 112282478
refGene_gn=SGPP1;refGene_tx=XM_011283398.3
B3 112302335 112303046
refGene_gn=SGPP1;refGene_tx=XM_011283398.3
B3 112495617 112495765
refGene_gn=SYNE2;refGene_tx=XM_023255756.1
B3 112501664 112501765
refGene_gn=SYNE2;refGene_tx=XM_023255756.1
B3 112504627 112504729
refGene_gn=SYNE2;refGene_tx=XM_023255756.1
B3 112507105 112507240
refGene_gn=SYNE2;refGene_tx=XM_023255756.1
B3 112515474 112515642

refGene_gn=SYNE2;refGene_tx=XM_023255756.1
B3 112516252 112516365
refGene_gn=SYNE2;refGene_tx=XM_023255756.1
B3 112517346 112517509
refGene_gn=SYNE2;refGene_tx=XM_023255756.1
B3 112518864 112518943
refGene_gn=SYNE2;refGene_tx=XM_023255756.1
B3 112519423 112519611
refGene_gn=SYNE2;refGene_tx=XM_023255756.1
B3 112521333 112521498
refGene_gn=SYNE2;refGene_tx=XM_023255756.1
B3 112522833 112522983
refGene_gn=SYNE2;refGene_tx=XM_023255756.1
B3 112524839 112525010
refGene_gn=SYNE2;refGene_tx=XM_023255756.1
B3 112527758 112527917
refGene_gn=SYNE2;refGene_tx=XM_023255756.1
B3 112528215 112528389
refGene_gn=SYNE2;refGene_tx=XM_023255756.1
B3 112533130 112533265
refGene_gn=SYNE2;refGene_tx=XM_023255756.1
B3 112534963 112535122
refGene_gn=SYNE2;refGene_tx=XM_023255756.1
B3 112537442 112537654
refGene_gn=SYNE2;refGene_tx=XM_023255756.1
B3 112537800 112537891
refGene_gn=SYNE2;refGene_tx=XM_023255756.1
B3 112538716 112538835
refGene_gn=SYNE2;refGene_tx=XM_023255756.1
B3 112539055 112539182
refGene_gn=SYNE2;refGene_tx=XM_023255756.1
B3 112540932 112541090
refGene_gn=SYNE2;refGene_tx=XM_023255756.1
B3 112541963 112542111
refGene_gn=SYNE2;refGene_tx=XM_023255756.1
B3 112542746 112543351
refGene_gn=SYNE2;refGene_tx=XM_023255756.1
B3 112546442 112546622
refGene_gn=SYNE2;refGene_tx=XM_023255756.1
B3 112548521 112548672
refGene_gn=SYNE2;refGene_tx=XM_023255756.1
B3 112558540 112558699
refGene_gn=SYNE2;refGene_tx=XM_023255756.1
B3 112559343 112559505
refGene_gn=SYNE2;refGene_tx=XM_023255756.1
B3 112561730 112561832
refGene_gn=SYNE2;refGene_tx=XM_023255756.1
B3 112563411 112563612
refGene_gn=SYNE2;refGene_tx=XM_023255756.1
B3 112563956 112564128

refGene_gn=SYNE2;refGene_tx=XM_023255756.1
B3 112564716 112564829
refGene_gn=SYNE2;refGene_tx=XM_023255756.1
B3 112566618 112566821
refGene_gn=SYNE2;refGene_tx=XM_023255756.1
B3 112567499 112567619
refGene_gn=SYNE2;refGene_tx=XM_023255756.1
B3 112567716 112568008
refGene_gn=SYNE2;refGene_tx=XM_023255756.1
B3 112569182 112569334
refGene_gn=SYNE2;refGene_tx=XM_023255756.1
B3 112570902 112571212
refGene_gn=SYNE2;refGene_tx=XM_023255756.1
B3 112572733 112572898
refGene_gn=SYNE2;refGene_tx=XM_023255756.1
B3 112575845 112576187
refGene_gn=SYNE2;refGene_tx=XM_023255756.1
B3 112585139 112585295
refGene_gn=SYNE2;refGene_tx=XM_023255756.1
B3 112586190 112586456
refGene_gn=SYNE2;refGene_tx=XM_023255756.1
B3 112588347 112590439
refGene_gn=SYNE2;refGene_tx=XM_023255756.1
B3 112592351 112592674
refGene_gn=SYNE2;refGene_tx=XM_023255756.1
B3 112595334 112595479
refGene_gn=SYNE2;refGene_tx=XM_023255756.1
B3 112596299 112596518
refGene_gn=SYNE2;refGene_tx=XM_023255756.1
B3 112602660 112602926
refGene_gn=SYNE2;refGene_tx=XM_023255756.1
B3 112605199 112605368
refGene_gn=SYNE2;refGene_tx=XM_023255756.1
B3 112608287 112608443
refGene_gn=SYNE2;refGene_tx=XM_023255756.1
B3 112610831 112610972
refGene_gn=SYNE2;refGene_tx=XM_023255756.1
B3 112612781 112612964
refGene_gn=SYNE2;refGene_tx=XM_023255756.1
B3 112613982 112614120
refGene_gn=SYNE2;refGene_tx=XM_023255756.1
B3 112621276 112621462
refGene_gn=SYNE2;refGene_tx=XM_023255756.1
B3 112625730 112625853
refGene_gn=SYNE2;refGene_tx=XM_023255756.1
B3 112627805 112627988
refGene_gn=SYNE2;refGene_tx=XM_023255756.1
B3 112630850 112630982
refGene_gn=SYNE2;refGene_tx=XM_023255756.1
B3 112634696 112634894

refGene_gn=SYNE2;refGene_tx=XM_023255756.1
B3 112635495 112635567
refGene_gn=SYNE2;refGene_tx=XM_023255756.1
B3 112636796 112636907
refGene_gn=SYNE2;refGene_tx=XM_023255756.1
B3 112640151 112640268
refGene_gn=SYNE2;refGene_tx=XM_023255756.1
B3 112643885 112644116
refGene_gn=SYNE2;refGene_tx=XM_023255756.1
B3 112647821 112648004
refGene_gn=SYNE2;refGene_tx=XM_023255756.1
B3 112649241 112649370
refGene_gn=SYNE2;refGene_tx=XM_023255756.1
B3 112649884 112650006
refGene_gn=SYNE2;refGene_tx=XM_023255756.1
B3 112650149 112650291
refGene_gn=SYNE2;refGene_tx=XM_023255756.1
B3 112651713 112651845
refGene_gn=SYNE2;refGene_tx=XM_023255756.1
B3 112653207 112653360
refGene_gn=SYNE2;refGene_tx=XM_023255756.1
B3 112653522 112653732
refGene_gn=SYNE2;refGene_tx=XM_023255756.1
B3 112655328 112655430
refGene_gn=SYNE2;refGene_tx=XM_023255756.1
B3 112656308 112656428
refGene_gn=SYNE2;refGene_tx=XM_023255756.1
B3 112656597 112656798
refGene_gn=SYNE2;refGene_tx=XM_023255756.1
B3 112659393 112659525
refGene_gn=SYNE2;refGene_tx=XM_023255756.1
B3 112659526 112659568
refGene_gn=SYNE2;refGene_tx=XM_023255756.1
B3 112660888 112661020
refGene_gn=SYNE2;refGene_tx=XM_023255756.1
B3 112664549 112664746
refGene_gn=SYNE2;refGene_tx=XM_023255756.1
B3 112665895 112666028
refGene_gn=SYNE2;refGene_tx=XM_023255756.1
B3 112667503 112667686
refGene_gn=SYNE2;refGene_tx=XM_023255756.1
B3 112668101 112668248
refGene_gn=SYNE2;refGene_tx=XM_023255756.1
B3 112670326 112670503
refGene_gn=SYNE2;refGene_tx=XM_023255756.1
B3 112672269 112672425
refGene_gn=SYNE2;refGene_tx=XM_023255756.1
B3 112673948 112674101
refGene_gn=SYNE2;refGene_tx=XM_023255756.1
B3 112680213 112680384

refGene_gn=SYNE2;refGene_tx=XM_023255756.1
B3	112681067	112681198
refGene_gn=SYNE2;refGene_tx=XM_023255756.1
B3	112683545	112683750
refGene_gn=SYNE2;refGene_tx=XM_023255756.1
B3	112684849	112685029
refGene_gn=SYNE2;refGene_tx=XM_023255756.1
B3	112685734	112685863
refGene_gn=SYNE2;refGene_tx=XM_023255756.1
B3	112687586	112687741
refGene_gn=SYNE2;refGene_tx=XM_023255756.1
B3	112687854	112687999
refGene_gn=SYNE2;refGene_tx=XM_023255756.1
B3	112688944	112689039
refGene_gn=SYNE2;refGene_tx=XM_023255756.1
B3	112690765	112691000
refGene_gn=SYNE2;refGene_tx=XM_023255756.1
B3	112694822	112695017
refGene_gn=SYNE2;refGene_tx=XM_023255756.1
B3	112697652	112697778
refGene_gn=SYNE2;refGene_tx=XM_023255756.1
B3	112702976	112703132
refGene_gn=SYNE2;refGene_tx=XM_023255756.1
B3	112705845	112706004
refGene_gn=SYNE2;refGene_tx=XM_023255756.1
B3	112707296	112707463
refGene_gn=SYNE2;refGene_tx=XM_023255756.1
B3	112720388	112720551
refGene_gn=SYNE2;refGene_tx=XM_023255756.1
B3	112728077	112728265
refGene_gn=SYNE2;refGene_tx=XM_023255756.1
B3	112728784	112728935
refGene_gn=SYNE2;refGene_tx=XM_023255756.1
B3	112729265	112729448
refGene_gn=SYNE2;refGene_tx=XM_023255756.1
B3	112730846	112730984
refGene_gn=SYNE2;refGene_tx=XM_023255756.1
B3	112732307	112732502
refGene_gn=SYNE2;refGene_tx=XM_023255756.1
B3	112734409	112734692
refGene_gn=SYNE2;refGene_tx=XM_023255756.1
B3	112735227	112735296
refGene_gn=SYNE2;refGene_tx=XM_023255756.1
B3	112736220	112736360
refGene_gn=SYNE2;refGene_tx=XM_023255756.1
B3	112738261	112738376
refGene_gn=SYNE2;refGene_tx=XM_023255756.1
B3	112739829	112740032
refGene_gn=SYNE2;refGene_tx=XM_023255756.1
B3	112741066	112741267

refGene_gn=SYNE2;refGene_tx=XM_023255756.1
B3	112742178	112742307
refGene_gn=SYNE2;refGene_tx=XM_023255756.1
B3	112743850	112744042
refGene_gn=SYNE2;refGene_tx=XM_023255756.1
B3	112744816	112744903
refGene_gn=SYNE2;refGene_tx=XM_023255756.1
B3	112745272	112745319
refGene_gn=SYNE2;refGene_tx=XM_023255756.1
B3	112745611	112745825
refGene_gn=SYNE2;refGene_tx=XM_023255756.1
B3	112755124	112755314
refGene_gn=ESR2;refGene_tx=XM_023255757.1
B3	112757160	112757341
refGene_gn=ESR2;refGene_tx=XM_023255757.1
B3	112773833	112773967
refGene_gn=ESR2;refGene_tx=XM_023255757.1
B3	112782331	112782470
refGene_gn=ESR2;refGene_tx=XM_023255757.1
B3	112782858	112783158
refGene_gn=ESR2;refGene_tx=XM_023255757.1
B3	112794839	112794956
refGene_gn=ESR2;refGene_tx=XM_023255757.1
B3	112807830	112808003
refGene_gn=ESR2;refGene_tx=XM_023255757.1
B3	112810334	112810753
refGene_gn=ESR2;refGene_tx=XM_023255757.1
B3	112863380	112863542
refGene_gn=LOC101086571;refGene_tx=XM_019833119.2;refGene_gn=LOC101086571;refGene_tx=XM_019833120.2;refGene_gn=LOC101086571;refGene_tx=XM_019833121.2;refGene_gn=LOC101086571;refGene_tx=XM_019833124.2;refGene_gn=LOC101086571;refGene_tx=XM_023255758.1;refGene_gn=LOC101086571;refGene_tx=XM_023255759.1;refGene_gn=LOC101086571;refGene_tx=XM_023255760.1
B3	112866592	112866808
refGene_gn=LOC101086571;refGene_tx=XM_019833119.2;refGene_gn=LOC101086571;refGene_tx=XM_019833120.2;refGene_gn=LOC101086571;refGene_tx=XM_019833121.2;refGene_gn=LOC101086571;refGene_tx=XM_019833124.2;refGene_gn=LOC101086571;refGene_tx=XM_023255758.1;refGene_gn=LOC101086571;refGene_tx=XM_023255759.1;refGene_gn=LOC101086571;refGene_tx=XM_023255760.1
B3	112868096	112868336
refGene_gn=LOC101086571;refGene_tx=XM_019833119.2;refGene_gn=LOC101086571;refGene_tx=XM_019833120.2;refGene_gn=LOC101086571;refGene_tx=XM_019833121.2;refGene_gn=LOC101086571;refGene_tx=XM_019833124.2;refGene_gn=LOC101086571;refGene_tx=XM_023255758.1;refGene_gn=LOC101086571;refGene_tx=XM_023255759.1;refGene_gn=LOC101086571;refGene_tx=XM_023255760.1
B3	112874836	112875037
refGene_gn=LOC101086571;refGene_tx=XM_019833119.2;refGene_gn=LOC101086571;refGene_tx=XM_019833120.2;refGene_gn=LOC101086571;refGene_tx=XM_019833121.2;refGene_gn=LOC101086571;refGene_tx=XM_019833124.2;refGene_gn=LOC101086571;refGene_tx=XM_023255758.1;refGene_gn=LOC101086571;refGen

e_tx=XM_023255759.1;refGene_gn=LOC101086571;refGene_tx=XM_023255760.1
B3 112878026 112878162
refGene_gn=LOC101086571;refGene_tx=XM_019833119.2;refGene_gn=LOC101086571;refGene_tx=XM_019833120.2;refGene_gn=LOC101086571;refGene_tx=XM_019833121.2;refGene_gn=LOC101086571;refGene_tx=XM_019833124.2;refGene_gn=LOC101086571;refGene_tx=XM_023255758.1;refGene_gn=LOC101086571;refGene_tx=XM_023255759.1;refGene_gn=LOC101086571;refGene_tx=XM_023255760.1
B3 112888944 112889201
refGene_gn=LOC101086571;refGene_tx=XM_019833119.2;refGene_gn=LOC101086571;refGene_tx=XM_019833120.2;refGene_gn=LOC101086571;refGene_tx=XM_019833121.2;refGene_gn=LOC101086571;refGene_tx=XM_019833124.2;refGene_gn=LOC101086571;refGene_tx=XM_023255758.1;refGene_gn=LOC101086571;refGene_tx=XM_023255759.1;refGene_gn=LOC101086571;refGene_tx=XM_023255760.1
B3 112896779 112896962
refGene_gn=LOC101086571;refGene_tx=XM_019833119.2;refGene_gn=LOC101086571;refGene_tx=XM_019833120.2;refGene_gn=LOC101086571;refGene_tx=XM_019833121.2;refGene_gn=LOC101086571;refGene_tx=XM_019833124.2;refGene_gn=LOC101086571;refGene_tx=XM_023255758.1;refGene_gn=LOC101086571;refGene_tx=XM_023255759.1;refGene_gn=LOC101086571;refGene_tx=XM_023255760.1
B3 112903605 112903743
refGene_gn=LOC101086571;refGene_tx=XM_019833120.2;refGene_gn=LOC101086571;refGene_tx=XM_019833124.2
B3 112912331 112912628
refGene_gn=LOC101086571;refGene_tx=XM_019833119.2;refGene_gn=LOC101086571;refGene_tx=XM_019833121.2;refGene_gn=LOC101086571;refGene_tx=XM_023255758.1;refGene_gn=MTHFD1;refGene_tx=XM_003987726.5
B3 112928885 112928970
refGene_gn=MTHFD1;refGene_tx=XM_003987726.5
B3 112935938 112936004
refGene_gn=MTHFD1;refGene_tx=XM_006932908.4
B3 112936309 112936369
refGene_gn=MTHFD1;refGene_tx=XM_003987726.5;refGene_gn=MTHFD1;refGene_tx=XM_006932908.4
B3 112937038 112937092
refGene_gn=MTHFD1;refGene_tx=XM_003987726.5;refGene_gn=MTHFD1;refGene_tx=XM_006932908.4
B3 112938178 112938315
refGene_gn=MTHFD1;refGene_tx=XM_003987726.5;refGene_gn=MTHFD1;refGene_tx=XM_006932908.4
B3 112938454 112938555
refGene_gn=MTHFD1;refGene_tx=XM_003987726.5;refGene_gn=MTHFD1;refGene_tx=XM_006932908.4
B3 112940611 112940748
refGene_gn=MTHFD1;refGene_tx=XM_003987726.5;refGene_gn=MTHFD1;refGene_tx=XM_006932908.4
B3 112941598 112941710
refGene_gn=MTHFD1;refGene_tx=XM_003987726.5;refGene_gn=MTHFD1;refGene_tx=XM_006932908.4
B3 112946069 112946197
refGene_gn=MTHFD1;refGene_tx=XM_003987726.5;refGene_gn=MTHFD1;refGene_

tx=XM_006932908.4
B3	112946598	112946696
refGene_gn=MTHFD1;refGene_tx=XM_003987726.5;refGene_gn=MTHFD1;refGene_tx=XM_006932908.4
B3	112947330	112947504
refGene_gn=MTHFD1;refGene_tx=XM_003987726.5;refGene_gn=MTHFD1;refGene_tx=XM_006932908.4
B3	112948135	112948272
refGene_gn=MTHFD1;refGene_tx=XM_003987726.5;refGene_gn=MTHFD1;refGene_tx=XM_006932908.4
B3	112949799	112949846
refGene_gn=MTHFD1;refGene_tx=XM_003987726.5;refGene_gn=MTHFD1;refGene_tx=XM_006932908.4
B3	112950486	112950594
refGene_gn=MTHFD1;refGene_tx=XM_003987726.5;refGene_gn=MTHFD1;refGene_tx=XM_006932908.4
B3	112950736	112950811
refGene_gn=MTHFD1;refGene_tx=XM_003987726.5;refGene_gn=MTHFD1;refGene_tx=XM_006932908.4
B3	112954054	112954157
refGene_gn=MTHFD1;refGene_tx=XM_003987726.5;refGene_gn=MTHFD1;refGene_tx=XM_006932908.4
B3	112957565	112957642
refGene_gn=MTHFD1;refGene_tx=XM_003987726.5;refGene_gn=MTHFD1;refGene_tx=XM_006932908.4
B3	112958261	112958402
refGene_gn=MTHFD1;refGene_tx=XM_003987726.5;refGene_gn=MTHFD1;refGene_tx=XM_006932908.4
B3	112960280	112960349
refGene_gn=MTHFD1;refGene_tx=XM_003987726.5;refGene_gn=MTHFD1;refGene_tx=XM_006932908.4
B3	112960622	112960734
refGene_gn=MTHFD1;refGene_tx=XM_003987726.5;refGene_gn=MTHFD1;refGene_tx=XM_006932908.4
B3	112960832	112960972
refGene_gn=MTHFD1;refGene_tx=XM_003987726.5;refGene_gn=MTHFD1;refGene_tx=XM_006932908.4
B3	112963468	112963510
refGene_gn=MTHFD1;refGene_tx=XM_003987726.5;refGene_gn=MTHFD1;refGene_tx=XM_006932908.4
B3	112965822	112965923
refGene_gn=MTHFD1;refGene_tx=XM_003987726.5;refGene_gn=MTHFD1;refGene_tx=XM_006932908.4
B3	112966753	112966931
refGene_gn=MTHFD1;refGene_tx=XM_003987726.5;refGene_gn=MTHFD1;refGene_tx=XM_006932908.4
B3	112969241	112969349
refGene_gn=MTHFD1;refGene_tx=XM_003987726.5;refGene_gn=MTHFD1;refGene_tx=XM_006932908.4
B3	112970117	112970270

refGene_gn=MTHFD1;refGene_tx=XM_003987726.5;refGene_gn=MTHFD1;refGene_tx=XM_006932908.4
B3 112972601 112972691
refGene_gn=MTHFD1;refGene_tx=XM_003987726.5;refGene_gn=MTHFD1;refGene_tx=XM_006932908.4
B3 112982271 112983558
refGene_gn=AKAP5;refGene_tx=XM_006932909.4;refGene_gn=AKAP5;refGene_tx=XM_019833128.2;refGene_gn=AKAP5;refGene_tx=XM_023255763.1
B3 112999387 113000522
refGene_gn=ZBTB25;refGene_tx=XM_019833125.2;refGene_gn=ZBTB25;refGene_tx=XM_019833126.2;refGene_gn=ZBTB25;refGene_tx=XM_023255762.1
B3 113003905 113004078
refGene_gn=ZBTB25;refGene_tx=XM_019833125.2;refGene_gn=ZBTB25;refGene_tx=XM_023255762.1
B3 113038542 113040684
refGene_gn=ZBTB1;refGene_tx=XM_003987729.5;refGene_gn=ZBTB1;refGene_tx=XM_023255764.1;refGene_gn=ZBTB1;refGene_tx=XM_023255765.1;refGene_gn=ZBTB1;refGene_tx=XM_023255766.1
B3 113045917 113045978
refGene_gn=ZBTB1;refGene_tx=XM_023255765.1
B3 113047564 113047601
refGene_gn=ZBTB1;refGene_tx=XM_023255766.1
B3 113055620 113057540
refGene_gn=HSPA2;refGene_tx=XM_006932910.4;refGene_gn=HSPA2;refGene_tx=XM_006932911.4;refGene_gn=HSPA2;refGene_tx=XM_019833131.2
B3 113063336 113063411
refGene_gn=PPP1R36;refGene_tx=XM_011283263.3;refGene_gn=PPP1R36;refGene_tx=XM_019833132.2;refGene_gn=PPP1R36;refGene_tx=XM_019833133.2;refGene_gn=PPP1R36;refGene_tx=XM_019833136.2;refGene_gn=PPP1R36;refGene_tx=XM_019833137.2;refGene_gn=PPP1R36;refGene_tx=XM_023255768.1
B3 113064215 113064280
refGene_gn=PPP1R36;refGene_tx=XM_011283263.3;refGene_gn=PPP1R36;refGene_tx=XM_019833132.2;refGene_gn=PPP1R36;refGene_tx=XM_019833133.2;refGene_gn=PPP1R36;refGene_tx=XM_019833136.2;refGene_gn=PPP1R36;refGene_tx=XM_019833137.2;refGene_gn=PPP1R36;refGene_tx=XM_023255768.1
B3 113074400 113074487
refGene_gn=PPP1R36;refGene_tx=XM_011283263.3;refGene_gn=PPP1R36;refGene_tx=XM_019833132.2;refGene_gn=PPP1R36;refGene_tx=XM_019833133.2;refGene_gn=PPP1R36;refGene_tx=XM_019833136.2;refGene_gn=PPP1R36;refGene_tx=XM_019833137.2;refGene_gn=PPP1R36;refGene_tx=XM_023255768.1
B3 113075251 113075349
refGene_gn=PPP1R36;refGene_tx=XM_011283263.3;refGene_gn=PPP1R36;refGene_tx=XM_019833132.2;refGene_gn=PPP1R36;refGene_tx=XM_019833133.2;refGene_gn=PPP1R36;refGene_tx=XM_019833136.2;refGene_gn=PPP1R36;refGene_tx=XM_023255768.1
B3 113075456 113075523
refGene_gn=PPP1R36;refGene_tx=XM_011283263.3;refGene_gn=PPP1R36;refGene_tx=XM_019833132.2;refGene_gn=PPP1R36;refGene_tx=XM_019833133.2;refGene_gn=PPP1R36;refGene_tx=XM_019833136.2;refGene_gn=PPP1R36;refGene_tx=XM_023255768.1

B3	113078109	113078208
refGene_gn=PPP1R36;refGene_tx=XM_011283263.3;refGene_gn=PPP1R36;refGen
e_tx=XM_019833132.2;refGene_gn=PPP1R36;refGene_tx=XM_019833136.2;refGe
ne_gn=PPP1R36;refGene_tx=XM_023255768.1
B3	113083261	113083396
refGene_gn=PPP1R36;refGene_tx=XM_011283263.3;refGene_gn=PPP1R36;refGen
e_tx=XM_019833132.2;refGene_gn=PPP1R36;refGene_tx=XM_019833133.2;refGe
ne_gn=PPP1R36;refGene_tx=XM_019833136.2;refGene_gn=PPP1R36;refGene_tx=
XM_019833137.2;refGene_gn=PPP1R36;refGene_tx=XM_023255768.1
B3	113092224	113092267
refGene_gn=PPP1R36;refGene_tx=XM_011283263.3;refGene_gn=PPP1R36;refGen
e_tx=XM_019833132.2;refGene_gn=PPP1R36;refGene_tx=XM_019833133.2;refGe
ne_gn=PPP1R36;refGene_tx=XM_019833136.2;refGene_gn=PPP1R36;refGene_tx=
XM_019833137.2;refGene_gn=PPP1R36;refGene_tx=XM_023255768.1
B3	113092577	113092756
refGene_gn=PPP1R36;refGene_tx=XM_011283263.3;refGene_gn=PPP1R36;refGen
e_tx=XM_019833132.2;refGene_gn=PPP1R36;refGene_tx=XM_019833133.2;refGe
ne_gn=PPP1R36;refGene_tx=XM_019833136.2;refGene_gn=PPP1R36;refGene_tx=
XM_019833137.2;refGene_gn=PPP1R36;refGene_tx=XM_023255768.1
B3	113092840	113093032
refGene_gn=PPP1R36;refGene_tx=XM_011283263.3;refGene_gn=PPP1R36;refGen
e_tx=XM_019833132.2;refGene_gn=PPP1R36;refGene_tx=XM_019833133.2;refGe
ne_gn=PPP1R36;refGene_tx=XM_019833136.2;refGene_gn=PPP1R36;refGene_tx=
XM_019833137.2;refGene_gn=PPP1R36;refGene_tx=XM_023255768.1
B3	113093196	113093302
refGene_gn=PPP1R36;refGene_tx=XM_023255768.1
B3	113093555	113093742
refGene_gn=PPP1R36;refGene_tx=XM_011283263.3
B3	113102433	113102563
refGene_gn=PPP1R36;refGene_tx=XM_019833132.2;refGene_gn=PPP1R36;refGen
e_tx=XM_019833133.2;refGene_gn=PPP1R36;refGene_tx=XM_019833136.2;refGe
ne_gn=PPP1R36;refGene_tx=XM_019833137.2
B3	113106974	113107095
refGene_gn=PPP1R36;refGene_tx=XM_019833132.2;refGene_gn=PPP1R36;refGen
e_tx=XM_019833133.2;refGene_gn=PPP1R36;refGene_tx=XM_019833137.2
B3	113225218	113225566
refGene_gn=PLEKHG3;refGene_tx=XM_011283266.2;refGene_gn=PLEKHG3;refGen
e_tx=XM_011283267.2;refGene_gn=PLEKHG3;refGene_tx=XM_011283268.2;refGe
ne_gn=PLEKHG3;refGene_tx=XM_011283269.2;refGene_gn=PLEKHG3;refGene_tx=
XM_023255771.1
B3	113226643	113226741
refGene_gn=PLEKHG3;refGene_tx=XM_011283266.2;refGene_gn=PLEKHG3;refGen
e_tx=XM_011283267.2;refGene_gn=PLEKHG3;refGene_tx=XM_011283268.2;refGe
ne_gn=PLEKHG3;refGene_tx=XM_011283269.2;refGene_gn=PLEKHG3;refGene_tx=
XM_023255771.1
B3	113228005	113228075
refGene_gn=PLEKHG3;refGene_tx=XM_011283266.2;refGene_gn=PLEKHG3;refGen
e_tx=XM_011283267.2;refGene_gn=PLEKHG3;refGene_tx=XM_011283268.2;refGe
ne_gn=PLEKHG3;refGene_tx=XM_011283269.2;refGene_gn=PLEKHG3;refGene_tx=
XM_023255771.1

B3	113228382	113228429
refGene_gn=PLEKHG3;refGene_tx=XM_011283266.2;refGene_gn=PLEKHG3;refGen
e_tx=XM_011283267.2;refGene_gn=PLEKHG3;refGene_tx=XM_011283268.2;refGe
ne_gn=PLEKHG3;refGene_tx=XM_011283269.2;refGene_gn=PLEKHG3;refGene_tx=
XM_023255771.1
B3	113228539	113228690
refGene_gn=PLEKHG3;refGene_tx=XM_011283266.2;refGene_gn=PLEKHG3;refGen
e_tx=XM_011283267.2;refGene_gn=PLEKHG3;refGene_tx=XM_011283268.2;refGe
ne_gn=PLEKHG3;refGene_tx=XM_011283269.2;refGene_gn=PLEKHG3;refGene_tx=
XM_023255771.1
B3	113228778	113228910
refGene_gn=PLEKHG3;refGene_tx=XM_011283266.2;refGene_gn=PLEKHG3;refGen
e_tx=XM_011283267.2;refGene_gn=PLEKHG3;refGene_tx=XM_011283268.2;refGe
ne_gn=PLEKHG3;refGene_tx=XM_011283269.2;refGene_gn=PLEKHG3;refGene_tx=
XM_023255771.1
B3	113229168	113229351
refGene_gn=PLEKHG3;refGene_tx=XM_011283266.2;refGene_gn=PLEKHG3;refGen
e_tx=XM_011283267.2;refGene_gn=PLEKHG3;refGene_tx=XM_011283268.2;refGe
ne_gn=PLEKHG3;refGene_tx=XM_011283269.2;refGene_gn=PLEKHG3;refGene_tx=
XM_023255771.1
B3	113229927	113230020
refGene_gn=PLEKHG3;refGene_tx=XM_011283266.2;refGene_gn=PLEKHG3;refGen
e_tx=XM_011283267.2;refGene_gn=PLEKHG3;refGene_tx=XM_011283268.2;refGe
ne_gn=PLEKHG3;refGene_tx=XM_011283269.2;refGene_gn=PLEKHG3;refGene_tx=
XM_023255771.1
B3	113230270	113230357
refGene_gn=PLEKHG3;refGene_tx=XM_011283266.2;refGene_gn=PLEKHG3;refGen
e_tx=XM_011283267.2;refGene_gn=PLEKHG3;refGene_tx=XM_011283268.2;refGe
ne_gn=PLEKHG3;refGene_tx=XM_011283269.2;refGene_gn=PLEKHG3;refGene_tx=
XM_023255771.1
B3	113230607	113230641
refGene_gn=PLEKHG3;refGene_tx=XM_011283266.2;refGene_gn=PLEKHG3;refGen
e_tx=XM_011283267.2;refGene_gn=PLEKHG3;refGene_tx=XM_011283268.2;refGe
ne_gn=PLEKHG3;refGene_tx=XM_011283269.2;refGene_gn=PLEKHG3;refGene_tx=
XM_023255771.1
B3	113230974	113231073
refGene_gn=PLEKHG3;refGene_tx=XM_011283266.2;refGene_gn=PLEKHG3;refGen
e_tx=XM_011283267.2;refGene_gn=PLEKHG3;refGene_tx=XM_011283268.2;refGe
ne_gn=PLEKHG3;refGene_tx=XM_011283269.2;refGene_gn=PLEKHG3;refGene_tx=
XM_023255771.1
B3	113233766	113233856
refGene_gn=PLEKHG3;refGene_tx=XM_011283266.2;refGene_gn=PLEKHG3;refGen
e_tx=XM_011283267.2;refGene_gn=PLEKHG3;refGene_tx=XM_023255771.1
B3	113234268	113234307
refGene_gn=PLEKHG3;refGene_tx=XM_011283266.2;refGene_gn=PLEKHG3;refGen
e_tx=XM_011283267.2;refGene_gn=PLEKHG3;refGene_tx=XM_011283268.2;refGe
ne_gn=PLEKHG3;refGene_tx=XM_011283269.2;refGene_gn=PLEKHG3;refGene_tx=
XM_023255771.1
B3	113234747	113234770
refGene_gn=PLEKHG3;refGene_tx=XM_011283266.2;refGene_gn=PLEKHG3;refGen

e_tx=XM_011283267.2;refGene_gn=PLEKHG3;refGene_tx=XM_011283268.2;refGene_gn=PLEKHG3;refGene_tx=XM_011283269.2;refGene_gn=PLEKHG3;refGene_tx=XM_023255771.1
B3	113236123	113236237
refGene_gn=PLEKHG3;refGene_tx=XM_011283266.2;refGene_gn=PLEKHG3;refGene_tx=XM_011283267.2;refGene_gn=PLEKHG3;refGene_tx=XM_011283268.2;refGene_gn=PLEKHG3;refGene_tx=XM_011283269.2;refGene_gn=PLEKHG3;refGene_tx=XM_023255771.1
B3	113237506	113238908
refGene_gn=PLEKHG3;refGene_tx=XM_011283266.2;refGene_gn=PLEKHG3;refGene_tx=XM_011283267.2;refGene_gn=PLEKHG3;refGene_tx=XM_011283268.2;refGene_gn=PLEKHG3;refGene_tx=XM_011283269.2;refGene_gn=PLEKHG3;refGene_tx=XM_023255771.1
B3	113239382	113240116
refGene_gn=PLEKHG3;refGene_tx=XM_011283266.2;refGene_gn=PLEKHG3;refGene_tx=XM_011283267.2;refGene_gn=PLEKHG3;refGene_tx=XM_011283268.2;refGene_gn=PLEKHG3;refGene_tx=XM_011283269.2;refGene_gn=PLEKHG3;refGene_tx=XM_023255771.1
B3	113245102	113245270
refGene_gn=SPTB;refGene_tx=XM_023255769.1
B3	113245382	113245425
refGene_gn=SPTB;refGene_tx=XM_023255769.1
B3	113245666	113245840
refGene_gn=SPTB;refGene_tx=XM_023255769.1
B3	113248154	113248402
refGene_gn=SPTB;refGene_tx=XM_023255769.1
B3	113257609	113257688
refGene_gn=SPTB;refGene_tx=XM_023255769.1
B3	113258176	113258226
refGene_gn=SPTB;refGene_tx=XM_023255769.1
B3	113258532	113258729
refGene_gn=SPTB;refGene_tx=XM_023255769.1
B3	113259562	113259647
refGene_gn=SPTB;refGene_tx=XM_023255769.1
B3	113260118	113260257
refGene_gn=SPTB;refGene_tx=XM_023255769.1
B3	113261323	113261568
refGene_gn=SPTB;refGene_tx=XM_023255769.1
B3	113263045	113263420
refGene_gn=SPTB;refGene_tx=XM_023255769.1
B3	113263679	113263884
refGene_gn=SPTB;refGene_tx=XM_023255769.1
B3	113264772	113264903
refGene_gn=SPTB;refGene_tx=XM_023255769.1
B3	113265364	113265643
refGene_gn=SPTB;refGene_tx=XM_023255769.1
B3	113268960	113269050
refGene_gn=SPTB;refGene_tx=XM_023255769.1
B3	113269676	113269883
refGene_gn=SPTB;refGene_tx=XM_023255769.1

B3 113270247 113270511
refGene_gn=SPTB;refGene_tx=XM_023255769.1
B3 113271720 113271867
refGene_gn=SPTB;refGene_tx=XM_023255769.1
B3 113273247 113273338
refGene_gn=SPTB;refGene_tx=XM_023255769.1
B3 113273450 113273653
refGene_gn=SPTB;refGene_tx=XM_023255769.1
B3 113274080 113274837
refGene_gn=SPTB;refGene_tx=XM_023255769.1
B3 113280377 113280515
refGene_gn=SPTB;refGene_tx=XM_023255769.1
B3 113281610 113282481
refGene_gn=SPTB;refGene_tx=XM_023255769.1
B3 113283082 113283233
refGene_gn=SPTB;refGene_tx=XM_023255769.1
B3 113283960 113284263
refGene_gn=SPTB;refGene_tx=XM_023255769.1
B3 113285004 113285163
refGene_gn=SPTB;refGene_tx=XM_023255769.1
B3 113285786 113285904
refGene_gn=SPTB;refGene_tx=XM_023255769.1
B3 113287562 113287750
refGene_gn=SPTB;refGene_tx=XM_023255769.1
B3 113288564 113288677
refGene_gn=SPTB;refGene_tx=XM_023255769.1
B3 113289127 113289243
refGene_gn=SPTB;refGene_tx=XM_023255769.1
B3 113289563 113289644
refGene_gn=SPTB;refGene_tx=XM_023255769.1
B3 113290050 113290142
refGene_gn=SPTB;refGene_tx=XM_023255769.1
B3 113291132 113291306
refGene_gn=SPTB;refGene_tx=XM_023255769.1
B3 113292542 113292694
refGene_gn=SPTB;refGene_tx=XM_023255769.1
B3 113310339 113310487
refGene_gn=SPTB;refGene_tx=XM_023255769.1
B3 113400009 113400048
refGene_gn=CHURC1;refGene_tx=XM_019833144.2
B3 113405082 113405218
refGene_gn=CHURC1;refGene_tx=XM_019833144.2
B3 113406742 113406813
refGene_gn=CHURC1;refGene_tx=XM_019833144.2
B3 113419242 113419593
refGene_gn=GPX2;refGene_tx=XM_003987735.4
B3 113421918 113422140
refGene_gn=GPX2;refGene_tx=XM_003987735.4
B3 113422922 113422978
refGene_gn=CHURC1;refGene_tx=XM_019833144.2

B3 113423379 113423452
refGene_gn=CHURC1;refGene_tx=XM_019833144.2
B3 113427537 113427696
refGene_gn=RAB15;refGene_tx=XM_011283399.3;refGene_gn=RAB15;refGene_tx=XM_019833139.2;refGene_gn=RAB15;refGene_tx=XM_019833142.2;refGene_gn=RAB15;refGene_tx=XM_019833143.2;refGene_gn=RAB15;refGene_tx=XM_023255772.1
B3 113427880 113427946
refGene_gn=RAB15;refGene_tx=XM_011283399.3;refGene_gn=RAB15;refGene_tx=XM_019833139.2;refGene_gn=RAB15;refGene_tx=XM_019833142.2;refGene_gn=RAB15;refGene_tx=XM_019833143.2;refGene_gn=RAB15;refGene_tx=XM_023255772.1
B3 113429138 113429228
refGene_gn=RAB15;refGene_tx=XM_011283399.3;refGene_gn=RAB15;refGene_tx=XM_019833139.2;refGene_gn=RAB15;refGene_tx=XM_019833142.2;refGene_gn=RAB15;refGene_tx=XM_019833143.2;refGene_gn=RAB15;refGene_tx=XM_023255772.1
B3 113429885 113429963
refGene_gn=RAB15;refGene_tx=XM_011283399.3;refGene_gn=RAB15;refGene_tx=XM_019833139.2;refGene_gn=RAB15;refGene_tx=XM_019833142.2;refGene_gn=RAB15;refGene_tx=XM_019833143.2
B3 113430468 113430529
refGene_gn=RAB15;refGene_tx=XM_011283399.3;refGene_gn=RAB15;refGene_tx=XM_019833139.2;refGene_gn=RAB15;refGene_tx=XM_019833142.2;refGene_gn=RAB15;refGene_tx=XM_019833143.2;refGene_gn=RAB15;refGene_tx=XM_023255772.1
B3 113431291 113431352
refGene_gn=RAB15;refGene_tx=XM_011283399.3;refGene_gn=RAB15;refGene_tx=XM_019833139.2;refGene_gn=RAB15;refGene_tx=XM_019833142.2;refGene_gn=RAB15;refGene_tx=XM_019833143.2;refGene_gn=RAB15;refGene_tx=XM_023255772.1
B3 113435550 113435671
refGene_gn=RAB15;refGene_tx=XM_019833139.2
B3 113447997 113448121
refGene_gn=RAB15;refGene_tx=XM_011283399.3;refGene_gn=RAB15;refGene_tx=XM_023255772.1
B3 113459938 113460082
refGene_gn=FNTB;refGene_tx=XM_003987734.5
B3 113474926 113474991
refGene_gn=FNTB;refGene_tx=XM_003987734.5
B3 113487566 113487639
refGene_gn=FNTB;refGene_tx=XM_003987734.5
B3 113489918 113490010
refGene_gn=FNTB;refGene_tx=XM_003987734.5
B3 113497707 113497854
refGene_gn=FNTB;refGene_tx=XM_003987734.5
B3 113497955 113498039
refGene_gn=FNTB;refGene_tx=XM_003987734.5
B3 113503531 113503618
refGene_gn=FNTB;refGene_tx=XM_003987734.5

B3	113509648	113509778
refGene_gn=FNTB;refGene_tx=XM_003987734.5
B3	113512665	113512798
refGene_gn=FNTB;refGene_tx=XM_003987734.5
B3	113518876	113518988
refGene_gn=FNTB;refGene_tx=XM_003987734.5
B3	113519861	113519976
refGene_gn=FNTB;refGene_tx=XM_003987734.5
B3	113525370	113525502
refGene_gn=FNTB;refGene_tx=XM_003987734.5
B3	113539669	113539857
refGene_gn=MAX;refGene_tx=NM_001009866.1;refGene_gn=MAX;refGene_tx=XM_023254759.1;refGene_gn=MAX;refGene_tx=XM_023254760.1;refGene_gn=MAX;refGene_tx=XM_023254761.1;refGene_gn=MAX;refGene_tx=XM_023254762.1
B3	113540561	113540646
refGene_gn=MAX;refGene_tx=XM_023254762.1
B3	113541125	113541255
refGene_gn=MAX;refGene_tx=NM_001009866.1;refGene_gn=MAX;refGene_tx=XM_023254759.1;refGene_gn=MAX;refGene_tx=XM_023254760.1;refGene_gn=MAX;refGene_tx=XM_023254761.1;refGene_gn=MAX;refGene_tx=XM_023254762.1
B3	113553905	113554013
refGene_gn=MAX;refGene_tx=NM_001009866.1;refGene_gn=MAX;refGene_tx=XM_023254759.1;refGene_gn=MAX;refGene_tx=XM_023254760.1;refGene_gn=MAX;refGene_tx=XM_023254761.1
B3	113561681	113561708
refGene_gn=MAX;refGene_tx=NM_001009866.1;refGene_gn=MAX;refGene_tx=XM_023254759.1
B3	113562461	113562497
refGene_gn=MAX;refGene_tx=NM_001009866.1;refGene_gn=MAX;refGene_tx=XM_023254759.1;refGene_gn=MAX;refGene_tx=XM_023254760.1;refGene_gn=MAX;refGene_tx=XM_023254761.1
B3	113937308	113937511
refGene_gn=FUT8;refGene_tx=XM_011283271.3;refGene_gn=FUT8;refGene_tx=XM_011283272.3;refGene_gn=FUT8;refGene_tx=XM_011283273.3;refGene_gn=FUT8;refGene_tx=XM_011283274.3;refGene_gn=FUT8;refGene_tx=XM_011283275.2;refGene_gn=FUT8;refGene_tx=XM_011283276.3;refGene_gn=FUT8;refGene_tx=XM_011283277.3;refGene_gn=FUT8;refGene_tx=XM_019833146.2;refGene_gn=FUT8;refGene_tx=XM_019833147.2
B3	113993312	113993428
refGene_gn=FUT8;refGene_tx=XM_011283271.3;refGene_gn=FUT8;refGene_tx=XM_011283272.3;refGene_gn=FUT8;refGene_tx=XM_011283273.3;refGene_gn=FUT8;refGene_tx=XM_011283274.3;refGene_gn=FUT8;refGene_tx=XM_011283275.2;refGene_gn=FUT8;refGene_tx=XM_011283276.3;refGene_gn=FUT8;refGene_tx=XM_011283277.3;refGene_gn=FUT8;refGene_tx=XM_019833146.2;refGene_gn=FUT8;refGene_tx=XM_019833147.2
B3	113993543	113993706
refGene_gn=FUT8;refGene_tx=XM_011283271.3;refGene_gn=FUT8;refGene_tx=XM_011283272.3;refGene_gn=FUT8;refGene_tx=XM_011283273.3;refGene_gn=FUT8;refGene_tx=XM_011283274.3;refGene_gn=FUT8;refGene_tx=XM_011283275.2;refGene_gn=FUT8;refGene_tx=XM_011283276.3;refGene_gn=FUT8;refGene_tx=X

M_011283277.3;refGene_gn=FUT8;refGene_tx=XM_019833146.2;refGene_gn=FUT8;refGene_tx=XM_019833147.2
B3 114001972 114002087
refGene_gn=FUT8;refGene_tx=XM_011283271.3;refGene_gn=FUT8;refGene_tx=XM_011283272.3;refGene_gn=FUT8;refGene_tx=XM_011283273.3;refGene_gn=FUT8;refGene_tx=XM_011283274.3;refGene_gn=FUT8;refGene_tx=XM_011283275.2;refGene_gn=FUT8;refGene_tx=XM_011283276.3;refGene_gn=FUT8;refGene_tx=XM_011283277.3;refGene_gn=FUT8;refGene_tx=XM_019833146.2;refGene_gn=FUT8;refGene_tx=XM_019833147.2
B3 114036483 114036721
refGene_gn=FUT8;refGene_tx=XM_011283271.3;refGene_gn=FUT8;refGene_tx=XM_011283272.3;refGene_gn=FUT8;refGene_tx=XM_011283273.3;refGene_gn=FUT8;refGene_tx=XM_011283274.3;refGene_gn=FUT8;refGene_tx=XM_011283275.2;refGene_gn=FUT8;refGene_tx=XM_011283276.3;refGene_gn=FUT8;refGene_tx=XM_011283277.3;refGene_gn=FUT8;refGene_tx=XM_019833146.2;refGene_gn=FUT8;refGene_tx=XM_019833147.2
B3 114077468 114077715
refGene_gn=FUT8;refGene_tx=XM_011283271.3;refGene_gn=FUT8;refGene_tx=XM_011283272.3;refGene_gn=FUT8;refGene_tx=XM_011283273.3;refGene_gn=FUT8;refGene_tx=XM_011283274.3;refGene_gn=FUT8;refGene_tx=XM_011283275.2;refGene_gn=FUT8;refGene_tx=XM_011283276.3;refGene_gn=FUT8;refGene_tx=XM_011283277.3;refGene_gn=FUT8;refGene_tx=XM_019833146.2;refGene_gn=FUT8;refGene_tx=XM_019833147.2
B3 114081731 114081908
refGene_gn=FUT8;refGene_tx=XM_011283271.3;refGene_gn=FUT8;refGene_tx=XM_011283272.3;refGene_gn=FUT8;refGene_tx=XM_011283273.3;refGene_gn=FUT8;refGene_tx=XM_011283274.3;refGene_gn=FUT8;refGene_tx=XM_011283275.2;refGene_gn=FUT8;refGene_tx=XM_011283276.3;refGene_gn=FUT8;refGene_tx=XM_011283277.3;refGene_gn=FUT8;refGene_tx=XM_019833146.2;refGene_gn=FUT8;refGene_tx=XM_019833147.2
B3 114087904 114088055
refGene_gn=FUT8;refGene_tx=XM_011283271.3;refGene_gn=FUT8;refGene_tx=XM_011283272.3;refGene_gn=FUT8;refGene_tx=XM_011283273.3;refGene_gn=FUT8;refGene_tx=XM_011283274.3;refGene_gn=FUT8;refGene_tx=XM_011283275.2;refGene_gn=FUT8;refGene_tx=XM_011283276.3;refGene_gn=FUT8;refGene_tx=XM_011283277.3;refGene_gn=FUT8;refGene_tx=XM_019833146.2;refGene_gn=FUT8;refGene_tx=XM_019833147.2
B3 114095952 114096270
refGene_gn=FUT8;refGene_tx=XM_011283271.3;refGene_gn=FUT8;refGene_tx=XM_011283272.3;refGene_gn=FUT8;refGene_tx=XM_011283273.3;refGene_gn=FUT8;refGene_tx=XM_011283274.3;refGene_gn=FUT8;refGene_tx=XM_011283275.2;refGene_gn=FUT8;refGene_tx=XM_011283276.3;refGene_gn=FUT8;refGene_tx=XM_011283277.3;refGene_gn=FUT8;refGene_tx=XM_019833146.2;refGene_gn=FUT8;refGene_tx=XM_019833147.2
B3 114699452 114699502
refGene_gn=LOC101096574;refGene_tx=XM_023255777.1;refGene_gn=LOC101096574;refGene_tx=XM_023255778.1;refGene_gn=LOC101096574;refGene_tx=XM_023255779.1;refGene_gn=LOC101096574;refGene_tx=XM_023255780.1
B3 114699600 114699753
refGene_gn=LOC101096574;refGene_tx=XM_023255777.1;refGene_gn=LOC101096

574;refGene_tx=XM_023255778.1;refGene_gn=LOC101096574;refGene_tx=XM_023255779.1;refGene_gn=LOC101096574;refGene_tx=XM_023255780.1
B3 114701027 114701124
refGene_gn=LOC101096574;refGene_tx=XM_023255777.1;refGene_gn=LOC101096574;refGene_tx=XM_023255778.1;refGene_gn=LOC101096574;refGene_tx=XM_023255779.1;refGene_gn=LOC101096574;refGene_tx=XM_023255780.1
B3 114701813 114701864
refGene_gn=LOC101096574;refGene_tx=XM_023255777.1;refGene_gn=LOC101096574;refGene_tx=XM_023255779.1;refGene_gn=LOC101096574;refGene_tx=XM_023255780.1
B3 114703817 114703895
refGene_gn=LOC101096574;refGene_tx=XM_023255777.1;refGene_gn=LOC101096574;refGene_tx=XM_023255778.1;refGene_gn=LOC101096574;refGene_tx=XM_023255779.1;refGene_gn=LOC101096574;refGene_tx=XM_023255780.1
B3 114704101 114704185
refGene_gn=LOC101096574;refGene_tx=XM_023255777.1;refGene_gn=LOC101096574;refGene_tx=XM_023255778.1;refGene_gn=LOC101096574;refGene_tx=XM_023255779.1;refGene_gn=LOC101096574;refGene_tx=XM_023255780.1
B3 114704685 114704764
refGene_gn=LOC101096574;refGene_tx=XM_023255777.1;refGene_gn=LOC101096574;refGene_tx=XM_023255778.1;refGene_gn=LOC101096574;refGene_tx=XM_023255779.1;refGene_gn=LOC101096574;refGene_tx=XM_023255780.1
B3 114705148 114705290
refGene_gn=LOC101096574;refGene_tx=XM_023255777.1;refGene_gn=LOC101096574;refGene_tx=XM_023255778.1
B3 114712153 114712212
refGene_gn=LOC101096574;refGene_tx=XM_023255777.1;refGene_gn=LOC101096574;refGene_tx=XM_023255778.1;refGene_gn=LOC101096574;refGene_tx=XM_023255779.1;refGene_gn=LOC101096574;refGene_tx=XM_023255780.1
B3 114712401 114712521
refGene_gn=LOC101096574;refGene_tx=XM_023255777.1;refGene_gn=LOC101096574;refGene_tx=XM_023255778.1;refGene_gn=LOC101096574;refGene_tx=XM_023255779.1;refGene_gn=LOC101096574;refGene_tx=XM_023255780.1
B3 114732390 114732454
refGene_gn=GPHN;refGene_tx=XM_011283278.3;refGene_gn=GPHN;refGene_tx=XM_011283279.3;refGene_gn=GPHN;refGene_tx=XM_011283280.3;refGene_gn=GPHN;refGene_tx=XM_011283281.2;refGene_gn=GPHN;refGene_tx=XM_011283282.2;refGene_gn=GPHN;refGene_tx=XM_011283283.2;refGene_gn=GPHN;refGene_tx=XM_019833151.2;refGene_gn=GPHN;refGene_tx=XM_019833152.2;refGene_gn=GPHN;refGene_tx=XM_019833154.2;refGene_gn=GPHN;refGene_tx=XM_019833155.2
B3 114742363 114742364
refGene_gn=GPHN;refGene_tx=XM_019833153.2
B3 114896646 114896725
refGene_gn=GPHN;refGene_tx=XM_011283278.3;refGene_gn=GPHN;refGene_tx=XM_011283279.3;refGene_gn=GPHN;refGene_tx=XM_011283280.3;refGene_gn=GPHN;refGene_tx=XM_011283281.2;refGene_gn=GPHN;refGene_tx=XM_011283282.2;refGene_gn=GPHN;refGene_tx=XM_011283283.2;refGene_gn=GPHN;refGene_tx=XM_019833151.2;refGene_gn=GPHN;refGene_tx=XM_019833152.2;refGene_gn=GPHN;refGene_tx=XM_019833153.2;refGene_gn=GPHN;refGene_tx=XM_019833154.2;refGene_gn=GPHN;refGene_tx=XM_019833155.2

B3 115007907 115007965
refGene_gn=GPHN;refGene_tx=XM_011283278.3;refGene_gn=GPHN;refGene_tx=XM_011283279.3;refGene_gn=GPHN;refGene_tx=XM_011283280.3;refGene_gn=GPHN;refGene_tx=XM_011283281.2;refGene_gn=GPHN;refGene_tx=XM_011283282.2;refGene_gn=GPHN;refGene_tx=XM_011283283.2;refGene_gn=GPHN;refGene_tx=XM_019833151.2;refGene_gn=GPHN;refGene_tx=XM_019833152.2;refGene_gn=GPHN;refGene_tx=XM_019833153.2;refGene_gn=GPHN;refGene_tx=XM_019833154.2;refGene_gn=GPHN;refGene_tx=XM_019833155.2
B3 115040922 115041015
refGene_gn=GPHN;refGene_tx=XM_011283278.3;refGene_gn=GPHN;refGene_tx=XM_011283279.3;refGene_gn=GPHN;refGene_tx=XM_011283280.3;refGene_gn=GPHN;refGene_tx=XM_011283281.2;refGene_gn=GPHN;refGene_tx=XM_011283282.2;refGene_gn=GPHN;refGene_tx=XM_011283283.2;refGene_gn=GPHN;refGene_tx=XM_019833151.2;refGene_gn=GPHN;refGene_tx=XM_019833152.2;refGene_gn=GPHN;refGene_tx=XM_019833153.2;refGene_gn=GPHN;refGene_tx=XM_019833154.2;refGene_gn=GPHN;refGene_tx=XM_019833155.2
B3 115053945 115053984
refGene_gn=GPHN;refGene_tx=XM_011283278.3;refGene_gn=GPHN;refGene_tx=XM_011283279.3;refGene_gn=GPHN;refGene_tx=XM_011283280.3;refGene_gn=GPHN;refGene_tx=XM_011283282.2;refGene_gn=GPHN;refGene_tx=XM_019833151.2;refGene_gn=GPHN;refGene_tx=XM_019833153.2;refGene_gn=GPHN;refGene_tx=XM_019833154.2;refGene_gn=GPHN;refGene_tx=XM_019833155.2
B3 115079271 115079366
refGene_gn=GPHN;refGene_tx=XM_011283278.3;refGene_gn=GPHN;refGene_tx=XM_011283279.3;refGene_gn=GPHN;refGene_tx=XM_011283280.3;refGene_gn=GPHN;refGene_tx=XM_011283281.2;refGene_gn=GPHN;refGene_tx=XM_011283282.2;refGene_gn=GPHN;refGene_tx=XM_011283283.2;refGene_gn=GPHN;refGene_tx=XM_019833151.2;refGene_gn=GPHN;refGene_tx=XM_019833152.2;refGene_gn=GPHN;refGene_tx=XM_019833153.2;refGene_gn=GPHN;refGene_tx=XM_019833154.2;refGene_gn=GPHN;refGene_tx=XM_019833155.2
B3 115122765 115122832
refGene_gn=GPHN;refGene_tx=XM_011283278.3;refGene_gn=GPHN;refGene_tx=XM_011283279.3;refGene_gn=GPHN;refGene_tx=XM_011283280.3;refGene_gn=GPHN;refGene_tx=XM_011283281.2;refGene_gn=GPHN;refGene_tx=XM_011283282.2;refGene_gn=GPHN;refGene_tx=XM_011283283.2;refGene_gn=GPHN;refGene_tx=XM_019833151.2;refGene_gn=GPHN;refGene_tx=XM_019833152.2;refGene_gn=GPHN;refGene_tx=XM_019833153.2;refGene_gn=GPHN;refGene_tx=XM_019833154.2;refGene_gn=GPHN;refGene_tx=XM_019833155.2
B3 115128738 115129011
refGene_gn=GPHN;refGene_tx=XM_011283278.3;refGene_gn=GPHN;refGene_tx=XM_011283279.3;refGene_gn=GPHN;refGene_tx=XM_011283280.3;refGene_gn=GPHN;refGene_tx=XM_011283281.2;refGene_gn=GPHN;refGene_tx=XM_011283282.2;refGene_gn=GPHN;refGene_tx=XM_011283283.2;refGene_gn=GPHN;refGene_tx=XM_019833151.2;refGene_gn=GPHN;refGene_tx=XM_019833152.2;refGene_gn=GPHN;refGene_tx=XM_019833153.2;refGene_gn=GPHN;refGene_tx=XM_019833154.2;refGene_gn=GPHN;refGene_tx=XM_019833155.2
B3 115130339 115130447
refGene_gn=GPHN;refGene_tx=XM_011283278.3;refGene_gn=GPHN;refGene_tx=XM_011283279.3;refGene_gn=GPHN;refGene_tx=XM_011283280.3;refGene_gn=GPHN;refGene_tx=XM_011283281.2;refGene_gn=GPHN;refGene_tx=XM_019833151.2;

refGene_gn=GPHN;refGene_tx=XM_019833152.2;refGene_gn=GPHN;refGene_tx=XM_019833153.2;refGene_gn=GPHN;refGene_tx=XM_019833154.2
B3 115152378 115152513
refGene_gn=GPHN;refGene_tx=XM_011283278.3;refGene_gn=GPHN;refGene_tx=XM_011283279.3;refGene_gn=GPHN;refGene_tx=XM_011283280.3;refGene_gn=GPHN;refGene_tx=XM_011283281.2;refGene_gn=GPHN;refGene_tx=XM_011283282.2;refGene_gn=GPHN;refGene_tx=XM_011283283.2;refGene_gn=GPHN;refGene_tx=XM_019833151.2;refGene_gn=GPHN;refGene_tx=XM_019833152.2;refGene_gn=GPHN;refGene_tx=XM_019833153.2;refGene_gn=GPHN;refGene_tx=XM_019833154.2;refGene_gn=GPHN;refGene_tx=XM_019833155.2
B3 115167426 115167483
refGene_gn=GPHN;refGene_tx=XM_011283278.3;refGene_gn=GPHN;refGene_tx=XM_011283282.2;refGene_gn=GPHN;refGene_tx=XM_019833151.2;refGene_gn=GPHN;refGene_tx=XM_019833152.2;refGene_gn=GPHN;refGene_tx=XM_019833153.2;refGene_gn=GPHN;refGene_tx=XM_019833154.2;refGene_gn=GPHN;refGene_tx=XM_019833155.2
B3 115169706 115169778
refGene_gn=GPHN;refGene_tx=XM_011283278.3;refGene_gn=GPHN;refGene_tx=XM_019833151.2;refGene_gn=GPHN;refGene_tx=XM_019833152.2;refGene_gn=GPHN;refGene_tx=XM_019833153.2;refGene_gn=GPHN;refGene_tx=XM_019833155.2
B3 115206563 115206606
refGene_gn=GPHN;refGene_tx=XM_011283278.3;refGene_gn=GPHN;refGene_tx=XM_011283279.3;refGene_gn=GPHN;refGene_tx=XM_011283280.3;refGene_gn=GPHN;refGene_tx=XM_011283281.2;refGene_gn=GPHN;refGene_tx=XM_011283282.2;refGene_gn=GPHN;refGene_tx=XM_011283283.2;refGene_gn=GPHN;refGene_tx=XM_019833151.2;refGene_gn=GPHN;refGene_tx=XM_019833152.2;refGene_gn=GPHN;refGene_tx=XM_019833153.2;refGene_gn=GPHN;refGene_tx=XM_019833154.2;refGene_gn=GPHN;refGene_tx=XM_019833155.2
B3 115248578 115248716
refGene_gn=GPHN;refGene_tx=XM_011283278.3;refGene_gn=GPHN;refGene_tx=XM_011283279.3;refGene_gn=GPHN;refGene_tx=XM_011283280.3;refGene_gn=GPHN;refGene_tx=XM_011283281.2;refGene_gn=GPHN;refGene_tx=XM_011283282.2;refGene_gn=GPHN;refGene_tx=XM_011283283.2;refGene_gn=GPHN;refGene_tx=XM_019833151.2;refGene_gn=GPHN;refGene_tx=XM_019833152.2;refGene_gn=GPHN;refGene_tx=XM_019833153.2;refGene_gn=GPHN;refGene_tx=XM_019833154.2;refGene_gn=GPHN;refGene_tx=XM_019833155.2
B3 115269321 115269414
refGene_gn=GPHN;refGene_tx=XM_011283278.3;refGene_gn=GPHN;refGene_tx=XM_011283279.3;refGene_gn=GPHN;refGene_tx=XM_011283280.3;refGene_gn=GPHN;refGene_tx=XM_011283281.2;refGene_gn=GPHN;refGene_tx=XM_011283282.2;refGene_gn=GPHN;refGene_tx=XM_011283283.2;refGene_gn=GPHN;refGene_tx=XM_019833151.2;refGene_gn=GPHN;refGene_tx=XM_019833152.2;refGene_gn=GPHN;refGene_tx=XM_019833153.2;refGene_gn=GPHN;refGene_tx=XM_019833154.2;refGene_gn=GPHN;refGene_tx=XM_019833155.2
B3 115279222 115279278
refGene_gn=GPHN;refGene_tx=XM_011283278.3;refGene_gn=GPHN;refGene_tx=XM_011283279.3;refGene_gn=GPHN;refGene_tx=XM_011283280.3;refGene_gn=GPHN;refGene_tx=XM_011283281.2;refGene_gn=GPHN;refGene_tx=XM_011283282.2;refGene_gn=GPHN;refGene_tx=XM_011283283.2;refGene_gn=GPHN;refGene_tx=XM_019833151.2;refGene_gn=GPHN;refGene_tx=XM_019833152.2;refGene_gn=GPH

N;refGene_tx=XM_019833153.2;refGene_gn=GPHN;refGene_tx=XM_019833154.2;
refGene_gn=GPHN;refGene_tx=XM_019833155.2
B3 115284510 115284645
refGene_gn=GPHN;refGene_tx=XM_011283278.3;refGene_gn=GPHN;refGene_tx=X
M_011283279.3;refGene_gn=GPHN;refGene_tx=XM_011283280.3;refGene_gn=GPH
N;refGene_tx=XM_011283281.2;refGene_gn=GPHN;refGene_tx=XM_011283282.2;
refGene_gn=GPHN;refGene_tx=XM_011283283.2;refGene_gn=GPHN;refGene_tx=X
M_019833151.2;refGene_gn=GPHN;refGene_tx=XM_019833152.2;refGene_gn=GPH
N;refGene_tx=XM_019833153.2;refGene_gn=GPHN;refGene_tx=XM_019833154.2;
refGene_gn=GPHN;refGene_tx=XM_019833155.2
B3 115286544 115286603
refGene_gn=GPHN;refGene_tx=XM_011283278.3;refGene_gn=GPHN;refGene_tx=X
M_011283279.3;refGene_gn=GPHN;refGene_tx=XM_011283280.3;refGene_gn=GPH
N;refGene_tx=XM_011283281.2;refGene_gn=GPHN;refGene_tx=XM_011283282.2;
refGene_gn=GPHN;refGene_tx=XM_011283283.2;refGene_gn=GPHN;refGene_tx=X
M_019833151.2;refGene_gn=GPHN;refGene_tx=XM_019833152.2;refGene_gn=GPH
N;refGene_tx=XM_019833153.2;refGene_gn=GPHN;refGene_tx=XM_019833154.2;
refGene_gn=GPHN;refGene_tx=XM_019833155.2
B3 115287662 115287816
refGene_gn=GPHN;refGene_tx=XM_011283278.3;refGene_gn=GPHN;refGene_tx=X
M_011283279.3;refGene_gn=GPHN;refGene_tx=XM_011283280.3;refGene_gn=GPH
N;refGene_tx=XM_011283281.2;refGene_gn=GPHN;refGene_tx=XM_011283282.2;
refGene_gn=GPHN;refGene_tx=XM_011283283.2;refGene_gn=GPHN;refGene_tx=X
M_019833151.2;refGene_gn=GPHN;refGene_tx=XM_019833152.2;refGene_gn=GPH
N;refGene_tx=XM_019833153.2;refGene_gn=GPHN;refGene_tx=XM_019833154.2;
refGene_gn=GPHN;refGene_tx=XM_019833155.2
B3 115303949 115304071
refGene_gn=GPHN;refGene_tx=XM_011283278.3;refGene_gn=GPHN;refGene_tx=X
M_011283279.3;refGene_gn=GPHN;refGene_tx=XM_011283280.3;refGene_gn=GPH
N;refGene_tx=XM_011283281.2;refGene_gn=GPHN;refGene_tx=XM_011283282.2;
refGene_gn=GPHN;refGene_tx=XM_011283283.2;refGene_gn=GPHN;refGene_tx=X
M_019833151.2;refGene_gn=GPHN;refGene_tx=XM_019833152.2;refGene_gn=GPH
N;refGene_tx=XM_019833153.2;refGene_gn=GPHN;refGene_tx=XM_019833154.2;
refGene_gn=GPHN;refGene_tx=XM_019833155.2
B3 115327854 115327942
refGene_gn=GPHN;refGene_tx=XM_011283278.3;refGene_gn=GPHN;refGene_tx=X
M_011283279.3;refGene_gn=GPHN;refGene_tx=XM_011283280.3;refGene_gn=GPH
N;refGene_tx=XM_011283281.2;refGene_gn=GPHN;refGene_tx=XM_011283282.2;
refGene_gn=GPHN;refGene_tx=XM_011283283.2;refGene_gn=GPHN;refGene_tx=X
M_019833151.2;refGene_gn=GPHN;refGene_tx=XM_019833152.2;refGene_gn=GPH
N;refGene_tx=XM_019833153.2;refGene_gn=GPHN;refGene_tx=XM_019833154.2;
refGene_gn=GPHN;refGene_tx=XM_019833155.2
B3 115340823 115340897
refGene_gn=GPHN;refGene_tx=XM_011283278.3;refGene_gn=GPHN;refGene_tx=X
M_011283279.3;refGene_gn=GPHN;refGene_tx=XM_011283280.3;refGene_gn=GPH
N;refGene_tx=XM_011283281.2;refGene_gn=GPHN;refGene_tx=XM_011283282.2;
refGene_gn=GPHN;refGene_tx=XM_011283283.2;refGene_gn=GPHN;refGene_tx=X
M_019833151.2;refGene_gn=GPHN;refGene_tx=XM_019833152.2;refGene_gn=GPH
N;refGene_tx=XM_019833153.2;refGene_gn=GPHN;refGene_tx=XM_019833154.2;
refGene_gn=GPHN;refGene_tx=XM_019833155.2

B3 115345374 115345439
refGene_gn=GPHN;refGene_tx=XM_011283278.3;refGene_gn=GPHN;refGene_tx=XM_011283279.3;refGene_gn=GPHN;refGene_tx=XM_011283280.3;refGene_gn=GPHN;refGene_tx=XM_011283281.2;refGene_gn=GPHN;refGene_tx=XM_011283282.2;refGene_gn=GPHN;refGene_tx=XM_011283283.2;refGene_gn=GPHN;refGene_tx=XM_019833151.2;refGene_gn=GPHN;refGene_tx=XM_019833152.2;refGene_gn=GPHN;refGene_tx=XM_019833153.2;refGene_gn=GPHN;refGene_tx=XM_019833154.2;refGene_gn=GPHN;refGene_tx=XM_019833155.2
B3 115349253 115349357
refGene_gn=GPHN;refGene_tx=XM_011283278.3;refGene_gn=GPHN;refGene_tx=XM_011283279.3;refGene_gn=GPHN;refGene_tx=XM_011283280.3;refGene_gn=GPHN;refGene_tx=XM_011283281.2;refGene_gn=GPHN;refGene_tx=XM_011283282.2;refGene_gn=GPHN;refGene_tx=XM_011283283.2;refGene_gn=GPHN;refGene_tx=XM_019833151.2;refGene_gn=GPHN;refGene_tx=XM_019833152.2;refGene_gn=GPHN;refGene_tx=XM_019833153.2;refGene_gn=GPHN;refGene_tx=XM_019833154.2;refGene_gn=GPHN;refGene_tx=XM_019833155.2
B3 115360385 115360482
refGene_gn=GPHN;refGene_tx=XM_011283278.3;refGene_gn=GPHN;refGene_tx=XM_011283279.3;refGene_gn=GPHN;refGene_tx=XM_011283280.3;refGene_gn=GPHN;refGene_tx=XM_011283281.2;refGene_gn=GPHN;refGene_tx=XM_011283282.2;refGene_gn=GPHN;refGene_tx=XM_011283283.2;refGene_gn=GPHN;refGene_tx=XM_019833151.2;refGene_gn=GPHN;refGene_tx=XM_019833152.2;refGene_gn=GPHN;refGene_tx=XM_019833153.2;refGene_gn=GPHN;refGene_tx=XM_019833154.2;refGene_gn=GPHN;refGene_tx=XM_019833155.2
B3 115361582 115361716
refGene_gn=GPHN;refGene_tx=XM_011283278.3;refGene_gn=GPHN;refGene_tx=XM_011283279.3;refGene_gn=GPHN;refGene_tx=XM_011283280.3;refGene_gn=GPHN;refGene_tx=XM_011283281.2;refGene_gn=GPHN;refGene_tx=XM_011283282.2;refGene_gn=GPHN;refGene_tx=XM_011283283.2;refGene_gn=GPHN;refGene_tx=XM_019833151.2;refGene_gn=GPHN;refGene_tx=XM_019833152.2;refGene_gn=GPHN;refGene_tx=XM_019833153.2;refGene_gn=GPHN;refGene_tx=XM_019833154.2;refGene_gn=GPHN;refGene_tx=XM_019833155.2
B3 115382675 115382834
refGene_gn=FAM71D;refGene_tx=XM_019833156.2;refGene_gn=FAM71D;refGene_tx=XM_019833157.2
B3 115385145 115385345
refGene_gn=FAM71D;refGene_tx=XM_019833156.2;refGene_gn=FAM71D;refGene_tx=XM_019833157.2
B3 115386269 115386804
refGene_gn=FAM71D;refGene_tx=XM_019833156.2;refGene_gn=FAM71D;refGene_tx=XM_019833157.2
B3 115389607 115389858
refGene_gn=FAM71D;refGene_tx=XM_019833156.2;refGene_gn=FAM71D;refGene_tx=XM_019833157.2
B3 115399113 115399142
refGene_gn=FAM71D;refGene_tx=XM_019833156.2;refGene_gn=FAM71D;refGene_tx=XM_019833157.2
B3 115400372 115400392
refGene_gn=FAM71D;refGene_tx=XM_019833157.2
B3 115404545 115404568

refGene_gn=FAM71D;refGene_tx=XM_019833156.2
B3 115449214 115449581
refGene_gn=MPP5;refGene_tx=XM_011283287.2;refGene_gn=MPP5;refGene_tx=XM_011283288.2
B3 115462778 115462987
refGene_gn=MPP5;refGene_tx=XM_011283287.2;refGene_gn=MPP5;refGene_tx=XM_011283288.2
B3 115471073 115471151
refGene_gn=MPP5;refGene_tx=XM_011283287.2;refGene_gn=MPP5;refGene_tx=XM_011283288.2
B3 115472465 115472612
refGene_gn=MPP5;refGene_tx=XM_011283287.2;refGene_gn=MPP5;refGene_tx=XM_011283288.2
B3 115474110 115474272
refGene_gn=MPP5;refGene_tx=XM_011283287.2;refGene_gn=MPP5;refGene_tx=XM_011283288.2
B3 115476182 115476260
refGene_gn=MPP5;refGene_tx=XM_011283287.2;refGene_gn=MPP5;refGene_tx=XM_011283288.2
B3 115487075 115487259
refGene_gn=MPP5;refGene_tx=XM_011283287.2;refGene_gn=MPP5;refGene_tx=XM_011283288.2
B3 115491199 115491271
refGene_gn=MPP5;refGene_tx=XM_011283287.2;refGene_gn=MPP5;refGene_tx=XM_011283288.2
B3 115491622 115491694
refGene_gn=MPP5;refGene_tx=XM_011283287.2;refGene_gn=MPP5;refGene_tx=XM_011283288.2
B3 115493901 115494069
refGene_gn=MPP5;refGene_tx=XM_011283287.2;refGene_gn=MPP5;refGene_tx=XM_011283288.2
B3 115494991 115495194
refGene_gn=MPP5;refGene_tx=XM_011283287.2;refGene_gn=MPP5;refGene_tx=XM_011283288.2
B3 115496455 115496566
refGene_gn=MPP5;refGene_tx=XM_011283287.2;refGene_gn=MPP5;refGene_tx=XM_011283288.2
B3 115504282 115504459
refGene_gn=MPP5;refGene_tx=XM_011283287.2;refGene_gn=MPP5;refGene_tx=XM_011283288.2
B3 115509462 115509604
refGene_gn=ATP6V1D;refGene_tx=XM_003987744.5;refGene_gn=ATP6V1D;refGene_tx=XM_006932927.4;refGene_gn=ATP6V1D;refGene_tx=XM_006932928.3
B3 115510545 115510624
refGene_gn=ATP6V1D;refGene_tx=XM_003987744.5;refGene_gn=ATP6V1D;refGene_tx=XM_006932927.4;refGene_gn=ATP6V1D;refGene_tx=XM_006932928.3
B3 115511941 115512008
refGene_gn=ATP6V1D;refGene_tx=XM_003987744.5;refGene_gn=ATP6V1D;refGene_tx=XM_006932927.4;refGene_gn=ATP6V1D;refGene_tx=XM_006932928.3
B3 115512593 115512697

refGene_gn=ATP6V1D;refGene_tx=XM_003987744.5;refGene_gn=ATP6V1D;refGene_tx=XM_006932927.4;refGene_gn=ATP6V1D;refGene_tx=XM_006932928.3
B3	115513828	115513873
refGene_gn=ATP6V1D;refGene_tx=XM_003987744.5;refGene_gn=ATP6V1D;refGene_tx=XM_006932927.4;refGene_gn=ATP6V1D;refGene_tx=XM_006932928.3
B3	115514816	115514884
refGene_gn=ATP6V1D;refGene_tx=XM_003987744.5;refGene_gn=ATP6V1D;refGene_tx=XM_006932927.4;refGene_gn=ATP6V1D;refGene_tx=XM_006932928.3
B3	115516638	115516718
refGene_gn=ATP6V1D;refGene_tx=XM_003987744.5;refGene_gn=ATP6V1D;refGene_tx=XM_006932927.4;refGene_gn=ATP6V1D;refGene_tx=XM_006932928.3
B3	115519813	115519966
refGene_gn=ATP6V1D;refGene_tx=XM_006932927.4;refGene_gn=ATP6V1D;refGene_tx=XM_006932928.3
B3	115526229	115526347
refGene_gn=ATP6V1D;refGene_tx=XM_003987744.5
B3	115531495	115531536
refGene_gn=ATP6V1D;refGene_tx=XM_003987744.5
B3	115536264	115536505
refGene_gn=EIF2S1;refGene_tx=XM_011283290.3
B3	115540322	115540402
refGene_gn=EIF2S1;refGene_tx=XM_011283290.3
B3	115543553	115543705
refGene_gn=EIF2S1;refGene_tx=XM_011283290.3
B3	115547326	115547433
refGene_gn=EIF2S1;refGene_tx=XM_011283290.3
B3	115548775	115548873
refGene_gn=EIF2S1;refGene_tx=XM_011283290.3
B3	115549788	115549932
refGene_gn=EIF2S1;refGene_tx=XM_011283290.3
B3	115550844	115550970
refGene_gn=EIF2S1;refGene_tx=XM_011283290.3
B3	115554666	115554794
refGene_gn=PLEK2;refGene_tx=XM_003987746.3
B3	115555500	115555579
refGene_gn=PLEK2;refGene_tx=XM_003987746.3
B3	115556739	115556823
refGene_gn=PLEK2;refGene_tx=XM_003987746.3
B3	115558257	115558359
refGene_gn=PLEK2;refGene_tx=XM_003987746.3
B3	115558592	115558780
refGene_gn=PLEK2;refGene_tx=XM_003987746.3
B3	115559038	115559130
refGene_gn=PLEK2;refGene_tx=XM_003987746.3
B3	115560385	115560567
refGene_gn=PLEK2;refGene_tx=XM_003987746.3
B3	115563984	115564149
refGene_gn=PLEK2;refGene_tx=XM_003987746.3
B3	115573424	115573466
refGene_gn=PLEK2;refGene_tx=XM_003987746.3

B3	115630649	115631171
refGene_gn=TMEM229B;refGene_tx=XM_006932930.4;refGene_gn=TMEM229B;refGene_tx=XM_006932931.4;refGene_gn=TMEM229B;refGene_tx=XM_019833159.2;refGene_gn=TMEM229B;refGene_tx=XM_019833160.2;refGene_gn=TMEM229B;refGene_tx=XM_019833161.2;refGene_gn=TMEM229B;refGene_tx=XM_019833162.2;refGene_gn=TMEM229B;refGene_tx=XM_019833163.2;refGene_gn=TMEM229B;refGene_tx=XM_019833164.2;refGene_gn=TMEM229B;refGene_tx=XM_019833165.2;refGene_gn=TMEM229B;refGene_tx=XM_019833166.2;refGene_gn=TMEM229B;refGene_tx=XM_019833167.2;refGene_gn=TMEM229B;refGene_tx=XM_019833168.2;refGene_gn=TMEM229B;refGene_tx=XM_023255781.1
B3	115637659	115637731
refGene_gn=TMEM229B;refGene_tx=XM_019833163.2;refGene_gn=TMEM229B;refGene_tx=XM_019833164.2
B3	115642435	115642528
refGene_gn=TMEM229B;refGene_tx=XM_006932930.4;refGene_gn=TMEM229B;refGene_tx=XM_006932931.4;refGene_gn=TMEM229B;refGene_tx=XM_019833159.2;refGene_gn=TMEM229B;refGene_tx=XM_019833160.2;refGene_gn=TMEM229B;refGene_tx=XM_019833161.2;refGene_gn=TMEM229B;refGene_tx=XM_019833162.2;refGene_gn=TMEM229B;refGene_tx=XM_023255781.1
B3	115699761	115699887
refGene_gn=PLEKHH1;refGene_tx=XM_023255782.1;refGene_gn=PLEKHH1;refGene_tx=XM_023255783.1;refGene_gn=PLEKHH1;refGene_tx=XM_023255784.1;refGene_gn=PLEKHH1;refGene_tx=XM_023255785.1;refGene_gn=PLEKHH1;refGene_tx=XM_023255786.1;refGene_gn=PLEKHH1;refGene_tx=XM_023255787.1
B3	115714648	115714711
refGene_gn=PLEKHH1;refGene_tx=XM_023255782.1;refGene_gn=PLEKHH1;refGene_tx=XM_023255783.1;refGene_gn=PLEKHH1;refGene_tx=XM_023255784.1;refGene_gn=PLEKHH1;refGene_tx=XM_023255785.1;refGene_gn=PLEKHH1;refGene_tx=XM_023255787.1
B3	115715581	115715731
refGene_gn=PLEKHH1;refGene_tx=XM_023255782.1;refGene_gn=PLEKHH1;refGene_tx=XM_023255783.1;refGene_gn=PLEKHH1;refGene_tx=XM_023255784.1;refGene_gn=PLEKHH1;refGene_tx=XM_023255785.1;refGene_gn=PLEKHH1;refGene_tx=XM_023255787.1
B3	115718733	115718817
refGene_gn=PLEKHH1;refGene_tx=XM_023255782.1;refGene_gn=PLEKHH1;refGene_tx=XM_023255783.1;refGene_gn=PLEKHH1;refGene_tx=XM_023255784.1;refGene_gn=PLEKHH1;refGene_tx=XM_023255785.1;refGene_gn=PLEKHH1;refGene_tx=XM_023255786.1;refGene_gn=PLEKHH1;refGene_tx=XM_023255787.1
B3	115720873	115720955
refGene_gn=PLEKHH1;refGene_tx=XM_023255782.1;refGene_gn=PLEKHH1;refGene_tx=XM_023255783.1;refGene_gn=PLEKHH1;refGene_tx=XM_023255784.1;refGene_gn=PLEKHH1;refGene_tx=XM_023255785.1;refGene_gn=PLEKHH1;refGene_tx=XM_023255786.1;refGene_gn=PLEKHH1;refGene_tx=XM_023255787.1
B3	115721075	115721836
refGene_gn=PLEKHH1;refGene_tx=XM_023255782.1;refGene_gn=PLEKHH1;refGene_tx=XM_023255783.1;refGene_gn=PLEKHH1;refGene_tx=XM_023255784.1;refGene_gn=PLEKHH1;refGene_tx=XM_023255785.1;refGene_gn=PLEKHH1;refGene_tx=XM_023255786.1;refGene_gn=PLEKHH1;refGene_tx=XM_023255787.1
B3	115725114	115725193

refGene_gn=PLEKHH1;refGene_tx=XM_023255782.1;refGene_gn=PLEKHH1;refGene_tx=XM_023255783.1;refGene_gn=PLEKHH1;refGene_tx=XM_023255784.1;refGene_gn=PLEKHH1;refGene_tx=XM_023255785.1;refGene_gn=PLEKHH1;refGene_tx=XM_023255786.1;refGene_gn=PLEKHH1;refGene_tx=XM_023255787.1
B3 115725838 115725930
refGene_gn=PLEKHH1;refGene_tx=XM_023255782.1;refGene_gn=PLEKHH1;refGene_tx=XM_023255783.1;refGene_gn=PLEKHH1;refGene_tx=XM_023255784.1;refGene_gn=PLEKHH1;refGene_tx=XM_023255785.1;refGene_gn=PLEKHH1;refGene_tx=XM_023255786.1;refGene_gn=PLEKHH1;refGene_tx=XM_023255787.1
B3 115727560 115727711
refGene_gn=PLEKHH1;refGene_tx=XM_023255782.1;refGene_gn=PLEKHH1;refGene_tx=XM_023255783.1;refGene_gn=PLEKHH1;refGene_tx=XM_023255784.1;refGene_gn=PLEKHH1;refGene_tx=XM_023255785.1;refGene_gn=PLEKHH1;refGene_tx=XM_023255786.1;refGene_gn=PLEKHH1;refGene_tx=XM_023255787.1
B3 115727922 115728068
refGene_gn=PLEKHH1;refGene_tx=XM_023255782.1;refGene_gn=PLEKHH1;refGene_tx=XM_023255783.1;refGene_gn=PLEKHH1;refGene_tx=XM_023255784.1;refGene_gn=PLEKHH1;refGene_tx=XM_023255785.1;refGene_gn=PLEKHH1;refGene_tx=XM_023255786.1;refGene_gn=PLEKHH1;refGene_tx=XM_023255787.1
B3 115729028 115729139
refGene_gn=PLEKHH1;refGene_tx=XM_023255782.1;refGene_gn=PLEKHH1;refGene_tx=XM_023255783.1;refGene_gn=PLEKHH1;refGene_tx=XM_023255784.1;refGene_gn=PLEKHH1;refGene_tx=XM_023255785.1;refGene_gn=PLEKHH1;refGene_tx=XM_023255786.1;refGene_gn=PLEKHH1;refGene_tx=XM_023255787.1
B3 115730070 115730157
refGene_gn=PLEKHH1;refGene_tx=XM_023255782.1;refGene_gn=PLEKHH1;refGene_tx=XM_023255783.1;refGene_gn=PLEKHH1;refGene_tx=XM_023255784.1;refGene_gn=PLEKHH1;refGene_tx=XM_023255785.1;refGene_gn=PLEKHH1;refGene_tx=XM_023255786.1;refGene_gn=PLEKHH1;refGene_tx=XM_023255787.1
B3 115730492 115730654
refGene_gn=PLEKHH1;refGene_tx=XM_023255782.1;refGene_gn=PLEKHH1;refGene_tx=XM_023255783.1;refGene_gn=PLEKHH1;refGene_tx=XM_023255784.1;refGene_gn=PLEKHH1;refGene_tx=XM_023255785.1;refGene_gn=PLEKHH1;refGene_tx=XM_023255786.1;refGene_gn=PLEKHH1;refGene_tx=XM_023255787.1
B3 115731851 115731932
refGene_gn=PLEKHH1;refGene_tx=XM_023255782.1;refGene_gn=PLEKHH1;refGene_tx=XM_023255783.1;refGene_gn=PLEKHH1;refGene_tx=XM_023255784.1;refGene_gn=PLEKHH1;refGene_tx=XM_023255785.1;refGene_gn=PLEKHH1;refGene_tx=XM_023255786.1;refGene_gn=PLEKHH1;refGene_tx=XM_023255787.1
B3 115732269 115732452
refGene_gn=PLEKHH1;refGene_tx=XM_023255782.1;refGene_gn=PLEKHH1;refGene_tx=XM_023255783.1;refGene_gn=PLEKHH1;refGene_tx=XM_023255784.1;refGene_gn=PLEKHH1;refGene_tx=XM_023255785.1;refGene_gn=PLEKHH1;refGene_tx=XM_023255786.1;refGene_gn=PLEKHH1;refGene_tx=XM_023255787.1
B3 115732850 115732959
refGene_gn=PLEKHH1;refGene_tx=XM_023255782.1;refGene_gn=PLEKHH1;refGene_tx=XM_023255783.1;refGene_gn=PLEKHH1;refGene_tx=XM_023255784.1;refGene_gn=PLEKHH1;refGene_tx=XM_023255785.1;refGene_gn=PLEKHH1;refGene_tx=XM_023255786.1;refGene_gn=PLEKHH1;refGene_tx=XM_023255787.1
B3 115733734 115733847

refGene_gn=PLEKHH1;refGene_tx=XM_023255782.1;refGene_gn=PLEKHH1;refGene_tx=XM_023255783.1;refGene_gn=PLEKHH1;refGene_tx=XM_023255784.1;refGene_gn=PLEKHH1;refGene_tx=XM_023255785.1;refGene_gn=PLEKHH1;refGene_tx=XM_023255786.1;refGene_gn=PLEKHH1;refGene_tx=XM_023255787.1
B3 115734479 115734656
refGene_gn=PLEKHH1;refGene_tx=XM_023255782.1;refGene_gn=PLEKHH1;refGene_tx=XM_023255783.1;refGene_gn=PLEKHH1;refGene_tx=XM_023255784.1;refGene_gn=PLEKHH1;refGene_tx=XM_023255785.1;refGene_gn=PLEKHH1;refGene_tx=XM_023255786.1;refGene_gn=PLEKHH1;refGene_tx=XM_023255787.1
B3 115735015 115735113
refGene_gn=PLEKHH1;refGene_tx=XM_023255782.1;refGene_gn=PLEKHH1;refGene_tx=XM_023255783.1;refGene_gn=PLEKHH1;refGene_tx=XM_023255784.1;refGene_gn=PLEKHH1;refGene_tx=XM_023255785.1;refGene_gn=PLEKHH1;refGene_tx=XM_023255786.1;refGene_gn=PLEKHH1;refGene_tx=XM_023255787.1
B3 115735603 115735781
refGene_gn=PLEKHH1;refGene_tx=XM_023255782.1;refGene_gn=PLEKHH1;refGene_tx=XM_023255783.1;refGene_gn=PLEKHH1;refGene_tx=XM_023255784.1;refGene_gn=PLEKHH1;refGene_tx=XM_023255785.1;refGene_gn=PLEKHH1;refGene_tx=XM_023255786.1;refGene_gn=PLEKHH1;refGene_tx=XM_023255787.1
B3 115736326 115736482
refGene_gn=PLEKHH1;refGene_tx=XM_023255782.1;refGene_gn=PLEKHH1;refGene_tx=XM_023255783.1;refGene_gn=PLEKHH1;refGene_tx=XM_023255784.1;refGene_gn=PLEKHH1;refGene_tx=XM_023255785.1;refGene_gn=PLEKHH1;refGene_tx=XM_023255786.1;refGene_gn=PLEKHH1;refGene_tx=XM_023255787.1
B3 115737567 115737668
refGene_gn=PLEKHH1;refGene_tx=XM_023255782.1;refGene_gn=PLEKHH1;refGene_tx=XM_023255783.1;refGene_gn=PLEKHH1;refGene_tx=XM_023255784.1;refGene_gn=PLEKHH1;refGene_tx=XM_023255785.1;refGene_gn=PLEKHH1;refGene_tx=XM_023255786.1;refGene_gn=PLEKHH1;refGene_tx=XM_023255787.1
B3 115738450 115738592
refGene_gn=PLEKHH1;refGene_tx=XM_023255782.1;refGene_gn=PLEKHH1;refGene_tx=XM_023255783.1;refGene_gn=PLEKHH1;refGene_tx=XM_023255784.1;refGene_gn=PLEKHH1;refGene_tx=XM_023255785.1;refGene_gn=PLEKHH1;refGene_tx=XM_023255786.1;refGene_gn=PLEKHH1;refGene_tx=XM_023255787.1
B3 115739735 115739878
refGene_gn=PLEKHH1;refGene_tx=XM_023255782.1;refGene_gn=PLEKHH1;refGene_tx=XM_023255783.1;refGene_gn=PLEKHH1;refGene_tx=XM_023255784.1;refGene_gn=PLEKHH1;refGene_tx=XM_023255785.1;refGene_gn=PLEKHH1;refGene_tx=XM_023255786.1;refGene_gn=PLEKHH1;refGene_tx=XM_023255787.1
B3 115740008 115740138
refGene_gn=PLEKHH1;refGene_tx=XM_023255782.1;refGene_gn=PLEKHH1;refGene_tx=XM_023255783.1;refGene_gn=PLEKHH1;refGene_tx=XM_023255784.1;refGene_gn=PLEKHH1;refGene_tx=XM_023255785.1;refGene_gn=PLEKHH1;refGene_tx=XM_023255786.1;refGene_gn=PLEKHH1;refGene_tx=XM_023255787.1
B3 115742066 115742153
refGene_gn=PLEKHH1;refGene_tx=XM_023255782.1;refGene_gn=PLEKHH1;refGene_tx=XM_023255783.1;refGene_gn=PLEKHH1;refGene_tx=XM_023255784.1;refGene_gn=PLEKHH1;refGene_tx=XM_023255785.1;refGene_gn=PLEKHH1;refGene_tx=XM_023255786.1;refGene_gn=PLEKHH1;refGene_tx=XM_023255787.1
B3 115742442 115742589

refGene_gn=PLEKHH1;refGene_tx=XM_023255782.1;refGene_gn=PLEKHH1;refGene_tx=XM_023255783.1;refGene_gn=PLEKHH1;refGene_tx=XM_023255784.1;refGene_gn=PLEKHH1;refGene_tx=XM_023255785.1;refGene_gn=PLEKHH1;refGene_tx=XM_023255786.1;refGene_gn=PLEKHH1;refGene_tx=XM_023255787.1
B3 115743533 115743695
refGene_gn=PLEKHH1;refGene_tx=XM_023255782.1;refGene_gn=PLEKHH1;refGene_tx=XM_023255783.1;refGene_gn=PLEKHH1;refGene_tx=XM_023255784.1;refGene_gn=PLEKHH1;refGene_tx=XM_023255785.1;refGene_gn=PLEKHH1;refGene_tx=XM_023255786.1;refGene_gn=PLEKHH1;refGene_tx=XM_023255787.1
B3 115746289 115746382
refGene_gn=PIGH;refGene_tx=XM_003987749.5
B3 115746519 115746543
refGene_gn=PIGH;refGene_tx=XM_019833176.2
B3 115747729 115747813
refGene_gn=PIGH;refGene_tx=XM_003987749.5;refGene_gn=PIGH;refGene_tx=XM_019833176.2
B3 115750384 115750594
refGene_gn=PIGH;refGene_tx=XM_003987749.5;refGene_gn=PIGH;refGene_tx=XM_019833176.2
B3 115755952 115756132
refGene_gn=PIGH;refGene_tx=XM_003987749.5;refGene_gn=PIGH;refGene_tx=XM_019833176.2
B3 115770000 115770111
refGene_gn=ARG2;refGene_tx=XM_003987751.5;refGene_gn=ARG2;refGene_tx=XM_019833179.2
B3 115770293 115770377
refGene_gn=ARG2;refGene_tx=XM_019833180.2
B3 115771009 115771082
refGene_gn=ARG2;refGene_tx=XM_003987751.5;refGene_gn=ARG2;refGene_tx=XM_019833179.2;refGene_gn=ARG2;refGene_tx=XM_019833180.2
B3 115793386 115793564
refGene_gn=ARG2;refGene_tx=XM_003987751.5;refGene_gn=ARG2;refGene_tx=XM_019833179.2;refGene_gn=ARG2;refGene_tx=XM_019833180.2
B3 115795958 115796118
refGene_gn=ARG2;refGene_tx=XM_003987751.5;refGene_gn=ARG2;refGene_tx=XM_019833179.2;refGene_gn=ARG2;refGene_tx=XM_019833180.2
B3 115797025 115797120
refGene_gn=ARG2;refGene_tx=XM_003987751.5;refGene_gn=ARG2;refGene_tx=XM_019833179.2;refGene_gn=ARG2;refGene_tx=XM_019833180.2
B3 115797309 115797414
refGene_gn=ARG2;refGene_tx=XM_003987751.5;refGene_gn=ARG2;refGene_tx=XM_019833179.2;refGene_gn=ARG2;refGene_tx=XM_019833180.2
B3 115798390 115798527
refGene_gn=ARG2;refGene_tx=XM_003987751.5;refGene_gn=ARG2;refGene_tx=XM_019833179.2;refGene_gn=ARG2;refGene_tx=XM_019833180.2
B3 115800631 115800837
refGene_gn=ARG2;refGene_tx=XM_003987751.5;refGene_gn=ARG2;refGene_tx=XM_019833179.2;refGene_gn=ARG2;refGene_tx=XM_019833180.2
B3 115801309 115801406
refGene_gn=VTI1B;refGene_tx=XM_003987750.4

B3	115802665	115802727
refGene_gn=VTI1B;refGene_tx=XM_003987750.4
B3	115804276	115804450
refGene_gn=VTI1B;refGene_tx=XM_003987750.4
B3	115807215	115807407
refGene_gn=VTI1B;refGene_tx=XM_003987750.4
B3	115809549	115809608
refGene_gn=VTI1B;refGene_tx=XM_003987750.4
B3	115831781	115831896
refGene_gn=VTI1B;refGene_tx=XM_003987750.4
B3	115832519	115832574
refGene_gn=RDH11;refGene_tx=XM_019833177.2
B3	115835504	115835607
refGene_gn=RDH11;refGene_tx=XM_011283292.3;refGene_gn=RDH11;refGene_tx
=XM_011283293.3
B3	115842558	115842565
refGene_gn=RDH11;refGene_tx=XM_019833178.2
B3	115844836	115845026
refGene_gn=RDH11;refGene_tx=XM_011283292.3;refGene_gn=RDH11;refGene_tx
=XM_011283293.3;refGene_gn=RDH11;refGene_tx=XM_019833177.2;refGene_gn=
RDH11;refGene_tx=XM_019833178.2
B3	115848159	115848369
refGene_gn=RDH11;refGene_tx=XM_011283292.3;refGene_gn=RDH11;refGene_tx
=XM_011283293.3;refGene_gn=RDH11;refGene_tx=XM_019833177.2;refGene_gn=
RDH11;refGene_tx=XM_019833178.2
B3	115848926	115849031
refGene_gn=RDH11;refGene_tx=XM_011283292.3;refGene_gn=RDH11;refGene_tx
=XM_011283293.3;refGene_gn=RDH11;refGene_tx=XM_019833177.2;refGene_gn=
RDH11;refGene_tx=XM_019833178.2
B3	115850393	115850549
refGene_gn=RDH11;refGene_tx=XM_011283292.3;refGene_gn=RDH11;refGene_tx
=XM_011283293.3;refGene_gn=RDH11;refGene_tx=XM_019833177.2;refGene_gn=
RDH11;refGene_tx=XM_019833178.2
B3	115850899	115851018
refGene_gn=RDH11;refGene_tx=XM_011283292.3;refGene_gn=RDH11;refGene_tx
=XM_011283293.3;refGene_gn=RDH11;refGene_tx=XM_019833177.2;refGene_gn=
RDH11;refGene_tx=XM_019833178.2
B3	115851278	115851387
refGene_gn=RDH11;refGene_tx=XM_011283292.3;refGene_gn=RDH11;refGene_tx
=XM_019833177.2;refGene_gn=RDH11;refGene_tx=XM_019833178.2
B3	115853741	115854030
refGene_gn=RDH11;refGene_tx=XM_011283292.3;refGene_gn=RDH11;refGene_tx
=XM_011283293.3;refGene_gn=RDH11;refGene_tx=XM_019833177.2;refGene_gn=
RDH11;refGene_tx=XM_019833178.2
B3	115865791	115865859
refGene_gn=RDH12;refGene_tx=XM_003987754.5;refGene_gn=RDH12;refGene_tx
=XM_019833181.1;refGene_gn=RDH12;refGene_tx=XM_019833182.1
B3	115867519	115867638
refGene_gn=RDH12;refGene_tx=XM_003987754.5;refGene_gn=RDH12;refGene_tx
=XM_019833181.1;refGene_gn=RDH12;refGene_tx=XM_019833182.1

B3	115868126	115868282
refGene_gn=RDH12;refGene_tx=XM_003987754.5;refGene_gn=RDH12;refGene_tx=XM_019833181.1;refGene_gn=RDH12;refGene_tx=XM_019833182.1
B3	115868943	115869048
refGene_gn=RDH12;refGene_tx=XM_003987754.5;refGene_gn=RDH12;refGene_tx=XM_019833181.1;refGene_gn=RDH12;refGene_tx=XM_019833182.1
B3	115869857	115870067
refGene_gn=RDH12;refGene_tx=XM_003987754.5;refGene_gn=RDH12;refGene_tx=XM_019833181.1;refGene_gn=RDH12;refGene_tx=XM_019833182.1
B3	115871820	115872053
refGene_gn=RDH12;refGene_tx=XM_003987754.5;refGene_gn=RDH12;refGene_tx=XM_019833181.1;refGene_gn=RDH12;refGene_tx=XM_019833182.1
B3	115874179	115874282
refGene_gn=RDH12;refGene_tx=XM_003987754.5;refGene_gn=RDH12;refGene_tx=XM_019833181.1;refGene_gn=RDH12;refGene_tx=XM_019833182.1
B3	115891884	115892088
refGene_gn=ZFYVE26;refGene_tx=XM_019833183.2
B3	115894803	115894848
refGene_gn=ZFYVE26;refGene_tx=XM_019833183.2
B3	115895627	115895810
refGene_gn=ZFYVE26;refGene_tx=XM_019833183.2
B3	115897009	115897069
refGene_gn=ZFYVE26;refGene_tx=XM_019833183.2
B3	115897361	115897503
refGene_gn=ZFYVE26;refGene_tx=XM_019833183.2
B3	115898338	115898538
refGene_gn=ZFYVE26;refGene_tx=XM_019833183.2
B3	115899226	115899424
refGene_gn=ZFYVE26;refGene_tx=XM_019833183.2
B3	115903336	115903555
refGene_gn=ZFYVE26;refGene_tx=XM_019833183.2
B3	115904034	115904244
refGene_gn=ZFYVE26;refGene_tx=XM_019833183.2
B3	115904501	115904649
refGene_gn=ZFYVE26;refGene_tx=XM_019833183.2
B3	115908739	115908960
refGene_gn=ZFYVE26;refGene_tx=XM_019833183.2
B3	115909866	115910003
refGene_gn=ZFYVE26;refGene_tx=XM_019833183.2
B3	115910681	115910713
refGene_gn=ZFYVE26;refGene_tx=XM_019833183.2
B3	115912520	115912657
refGene_gn=ZFYVE26;refGene_tx=XM_019833183.2
B3	115913958	115914066
refGene_gn=ZFYVE26;refGene_tx=XM_019833183.2
B3	115914067	115914123
refGene_gn=ZFYVE26;refGene_tx=XM_019833183.2
B3	115916297	115916393
refGene_gn=ZFYVE26;refGene_tx=XM_019833183.2
B3	115917178	115917425

refGene_gn=ZFYVE26;refGene_tx=XM_019833183.2
B3 115918878 115919055
refGene_gn=ZFYVE26;refGene_tx=XM_019833183.2
B3 115919454 115919577
refGene_gn=ZFYVE26;refGene_tx=XM_019833183.2
B3 115923441 115923546
refGene_gn=ZFYVE26;refGene_tx=XM_019833183.2
B3 115924570 115924767
refGene_gn=ZFYVE26;refGene_tx=XM_019833183.2
B3 115926062 115926808
refGene_gn=ZFYVE26;refGene_tx=XM_019833183.2
B3 115927664 115927767
refGene_gn=ZFYVE26;refGene_tx=XM_019833183.2
B3 115928410 115928629
refGene_gn=ZFYVE26;refGene_tx=XM_019833183.2
B3 115929182 115929347
refGene_gn=ZFYVE26;refGene_tx=XM_019833183.2
B3 115929454 115929574
refGene_gn=ZFYVE26;refGene_tx=XM_019833183.2
B3 115931618 115931882
refGene_gn=ZFYVE26;refGene_tx=XM_019833183.2
B3 115933084 115933286
refGene_gn=ZFYVE26;refGene_tx=XM_019833183.2
B3 115935393 115935545
refGene_gn=ZFYVE26;refGene_tx=XM_019833183.2
B3 115936260 115936329
refGene_gn=ZFYVE26;refGene_tx=XM_019833183.2
B3 115939323 115939410
refGene_gn=ZFYVE26;refGene_tx=XM_019833183.2
B3 115939649 115940261
refGene_gn=ZFYVE26;refGene_tx=XM_019833183.2
B3 115941513 115941717
refGene_gn=ZFYVE26;refGene_tx=XM_019833183.2
B3 115946724 115946888
refGene_gn=ZFYVE26;refGene_tx=XM_019833183.2
B3 115947844 115947933
refGene_gn=ZFYVE26;refGene_tx=XM_019833183.2
B3 115948080 115948245
refGene_gn=ZFYVE26;refGene_tx=XM_019833183.2
B3 115948830 115948961
refGene_gn=ZFYVE26;refGene_tx=XM_019833183.2
B3 115949332 115949855
refGene_gn=ZFYVE26;refGene_tx=XM_019833183.2
B3 115951710 115951800
refGene_gn=ZFYVE26;refGene_tx=XM_019833183.2
B3 115956883 115956962
refGene_gn=ZFYVE26;refGene_tx=XM_019833183.2
B3 115958607 115958801
refGene_gn=ZFYVE26;refGene_tx=XM_019833183.2
B3 115964230 115964314

refGene_gn=RAD51B;refGene_tx=XM_011283295.3;refGene_gn=RAD51B;refGene_tx=XM_019833185.1;refGene_gn=RAD51B;refGene_tx=XM_019833191.2;refGene_gn=RAD51B;refGene_tx=XM_023255788.1;refGene_gn=RAD51B;refGene_tx=XM_023255789.1
B3	115966915	115967029
refGene_gn=RAD51B;refGene_tx=XM_011283295.3;refGene_gn=RAD51B;refGene_tx=XM_019833185.1;refGene_gn=RAD51B;refGene_tx=XM_019833191.2;refGene_gn=RAD51B;refGene_tx=XM_023255788.1;refGene_gn=RAD51B;refGene_tx=XM_023255789.1
B3	115971783	115971900
refGene_gn=RAD51B;refGene_tx=XM_011283295.3;refGene_gn=RAD51B;refGene_tx=XM_019833185.1;refGene_gn=RAD51B;refGene_tx=XM_019833191.2;refGene_gn=RAD51B;refGene_tx=XM_023255788.1;refGene_gn=RAD51B;refGene_tx=XM_023255789.1
B3	115990196	115990333
refGene_gn=RAD51B;refGene_tx=XM_011283295.3;refGene_gn=RAD51B;refGene_tx=XM_019833185.1;refGene_gn=RAD51B;refGene_tx=XM_019833191.2;refGene_gn=RAD51B;refGene_tx=XM_023255788.1;refGene_gn=RAD51B;refGene_tx=XM_023255789.1
B3	116000290	116000410
refGene_gn=RAD51B;refGene_tx=XM_011283295.3;refGene_gn=RAD51B;refGene_tx=XM_019833185.1;refGene_gn=RAD51B;refGene_tx=XM_019833191.2;refGene_gn=RAD51B;refGene_tx=XM_023255788.1;refGene_gn=RAD51B;refGene_tx=XM_023255789.1
B3	116001464	116001648
refGene_gn=RAD51B;refGene_tx=XM_011283295.3;refGene_gn=RAD51B;refGene_tx=XM_019833185.1;refGene_gn=RAD51B;refGene_tx=XM_019833191.2;refGene_gn=RAD51B;refGene_tx=XM_023255788.1;refGene_gn=RAD51B;refGene_tx=XM_023255789.1
B3	116376525	116376622
refGene_gn=RAD51B;refGene_tx=XM_011283295.3;refGene_gn=RAD51B;refGene_tx=XM_019833185.1;refGene_gn=RAD51B;refGene_tx=XM_019833191.2;refGene_gn=RAD51B;refGene_tx=XM_023255788.1;refGene_gn=RAD51B;refGene_tx=XM_023255789.1
B3	116484769	116484873
refGene_gn=RAD51B;refGene_tx=XM_011283295.3;refGene_gn=RAD51B;refGene_tx=XM_019833185.1;refGene_gn=RAD51B;refGene_tx=XM_019833191.2;refGene_gn=RAD51B;refGene_tx=XM_023255788.1;refGene_gn=RAD51B;refGene_tx=XM_023255789.1
B3	116489096	116489114
refGene_gn=RAD51B;refGene_tx=XM_023255789.1
B3	116527834	116527913
refGene_gn=RAD51B;refGene_tx=XM_011283295.3;refGene_gn=RAD51B;refGene_tx=XM_019833185.1;refGene_gn=RAD51B;refGene_tx=XM_019833191.2
B3	116537667	116537684
refGene_gn=RAD51B;refGene_tx=XM_019833191.2
B3	116607081	116607195
refGene_gn=RAD51B;refGene_tx=XM_011283295.3;refGene_gn=RAD51B;refGene_tx=XM_019833185.1;refGene_gn=RAD51B;refGene_tx=XM_023255788.1
B3	116832273	116833233

refGene_gn=ZFP36L1;refGene_tx=XM_006932939.4
B3 116835634 116835691
refGene_gn=ZFP36L1;refGene_tx=XM_006932939.4
B3 116882035 116882194
refGene_gn=ACTN1;refGene_tx=XM_003987767.5;refGene_gn=ACTN1;refGene_tx=XM_003987768.4;refGene_gn=ACTN1;refGene_tx=XM_003987769.5;refGene_gn=ACTN1;refGene_tx=XM_019833192.2;refGene_gn=ACTN1;refGene_tx=XM_019833194.2;refGene_gn=ACTN1;refGene_tx=XM_019833195.1;refGene_gn=ACTN1;refGene_tx=XM_019833196.2;refGene_gn=ACTN1;refGene_tx=XM_019833197.2
B3 116883586 116883745
refGene_gn=ACTN1;refGene_tx=XM_003987767.5;refGene_gn=ACTN1;refGene_tx=XM_003987768.4;refGene_gn=ACTN1;refGene_tx=XM_003987769.5;refGene_gn=ACTN1;refGene_tx=XM_019833192.2;refGene_gn=ACTN1;refGene_tx=XM_019833194.2;refGene_gn=ACTN1;refGene_tx=XM_019833195.1;refGene_gn=ACTN1;refGene_tx=XM_019833196.2;refGene_gn=ACTN1;refGene_tx=XM_019833197.2
B3 116884946 116885012
refGene_gn=ACTN1;refGene_tx=XM_003987768.4;refGene_gn=ACTN1;refGene_tx=XM_003987769.5;refGene_gn=ACTN1;refGene_tx=XM_019833192.2;refGene_gn=ACTN1;refGene_tx=XM_019833195.1;refGene_gn=ACTN1;refGene_tx=XM_019833196.2;refGene_gn=ACTN1;refGene_tx=XM_019833197.2
B3 116885451 116885532
refGene_gn=ACTN1;refGene_tx=XM_003987767.5;refGene_gn=ACTN1;refGene_tx=XM_003987769.5;refGene_gn=ACTN1;refGene_tx=XM_019833192.2;refGene_gn=ACTN1;refGene_tx=XM_019833194.2;refGene_gn=ACTN1;refGene_tx=XM_019833196.2;refGene_gn=ACTN1;refGene_tx=XM_019833197.2
B3 116886437 116886584
refGene_gn=ACTN1;refGene_tx=XM_003987767.5;refGene_gn=ACTN1;refGene_tx=XM_003987768.4;refGene_gn=ACTN1;refGene_tx=XM_003987769.5;refGene_gn=ACTN1;refGene_tx=XM_019833192.2;refGene_gn=ACTN1;refGene_tx=XM_019833194.2;refGene_gn=ACTN1;refGene_tx=XM_019833195.1;refGene_gn=ACTN1;refGene_tx=XM_019833196.2;refGene_gn=ACTN1;refGene_tx=XM_019833197.2
B3 116887330 116887510
refGene_gn=ACTN1;refGene_tx=XM_003987767.5;refGene_gn=ACTN1;refGene_tx=XM_003987768.4;refGene_gn=ACTN1;refGene_tx=XM_003987769.5;refGene_gn=ACTN1;refGene_tx=XM_019833192.2;refGene_gn=ACTN1;refGene_tx=XM_019833194.2;refGene_gn=ACTN1;refGene_tx=XM_019833195.1;refGene_gn=ACTN1;refGene_tx=XM_019833196.2;refGene_gn=ACTN1;refGene_tx=XM_019833197.2
B3 116888661 116888796
refGene_gn=ACTN1;refGene_tx=XM_003987767.5;refGene_gn=ACTN1;refGene_tx=XM_003987768.4;refGene_gn=ACTN1;refGene_tx=XM_003987769.5;refGene_gn=ACTN1;refGene_tx=XM_019833192.2;refGene_gn=ACTN1;refGene_tx=XM_019833194.2;refGene_gn=ACTN1;refGene_tx=XM_019833195.1;refGene_gn=ACTN1;refGene_tx=XM_019833196.2;refGene_gn=ACTN1;refGene_tx=XM_019833197.2
B3 116889099 116889282
refGene_gn=ACTN1;refGene_tx=XM_003987767.5;refGene_gn=ACTN1;refGene_tx=XM_003987768.4;refGene_gn=ACTN1;refGene_tx=XM_003987769.5;refGene_gn=ACTN1;refGene_tx=XM_019833192.2;refGene_gn=ACTN1;refGene_tx=XM_019833194.2;refGene_gn=ACTN1;refGene_tx=XM_019833195.1;refGene_gn=ACTN1;refGene_tx=XM_019833196.2;refGene_gn=ACTN1;refGene_tx=XM_019833197.2
B3 116890081 116890222

refGene_gn=ACTN1;refGene_tx=XM_003987767.5;refGene_gn=ACTN1;refGene_tx
=XM_003987768.4;refGene_gn=ACTN1;refGene_tx=XM_003987769.5;refGene_gn=
ACTN1;refGene_tx=XM_019833192.2;refGene_gn=ACTN1;refGene_tx=XM_0198331
94.2;refGene_gn=ACTN1;refGene_tx=XM_019833195.1;refGene_gn=ACTN1;refGe
ne_tx=XM_019833196.2;refGene_gn=ACTN1;refGene_tx=XM_019833197.2
B3 116890700 116890809
refGene_gn=ACTN1;refGene_tx=XM_003987767.5;refGene_gn=ACTN1;refGene_tx
=XM_003987768.4;refGene_gn=ACTN1;refGene_tx=XM_003987769.5;refGene_gn=
ACTN1;refGene_tx=XM_019833192.2;refGene_gn=ACTN1;refGene_tx=XM_0198331
94.2;refGene_gn=ACTN1;refGene_tx=XM_019833195.1;refGene_gn=ACTN1;refGe
ne_tx=XM_019833196.2;refGene_gn=ACTN1;refGene_tx=XM_019833197.2
B3 116891289 116891440
refGene_gn=ACTN1;refGene_tx=XM_003987767.5;refGene_gn=ACTN1;refGene_tx
=XM_003987768.4;refGene_gn=ACTN1;refGene_tx=XM_003987769.5;refGene_gn=
ACTN1;refGene_tx=XM_019833192.2;refGene_gn=ACTN1;refGene_tx=XM_0198331
94.2;refGene_gn=ACTN1;refGene_tx=XM_019833195.1;refGene_gn=ACTN1;refGe
ne_tx=XM_019833196.2;refGene_gn=ACTN1;refGene_tx=XM_019833197.2
B3 116894687 116894835
refGene_gn=ACTN1;refGene_tx=XM_003987767.5;refGene_gn=ACTN1;refGene_tx
=XM_003987768.4;refGene_gn=ACTN1;refGene_tx=XM_003987769.5;refGene_gn=
ACTN1;refGene_tx=XM_019833192.2;refGene_gn=ACTN1;refGene_tx=XM_0198331
94.2;refGene_gn=ACTN1;refGene_tx=XM_019833195.1;refGene_gn=ACTN1;refGe
ne_tx=XM_019833196.2;refGene_gn=ACTN1;refGene_tx=XM_019833197.2
B3 116897393 116897624
refGene_gn=ACTN1;refGene_tx=XM_003987767.5;refGene_gn=ACTN1;refGene_tx
=XM_003987768.4;refGene_gn=ACTN1;refGene_tx=XM_003987769.5;refGene_gn=
ACTN1;refGene_tx=XM_019833192.2;refGene_gn=ACTN1;refGene_tx=XM_0198331
94.2;refGene_gn=ACTN1;refGene_tx=XM_019833195.1;refGene_gn=ACTN1;refGe
ne_tx=XM_019833196.2;refGene_gn=ACTN1;refGene_tx=XM_019833197.2
B3 116898942 116899035
refGene_gn=ACTN1;refGene_tx=XM_003987767.5;refGene_gn=ACTN1;refGene_tx
=XM_003987768.4;refGene_gn=ACTN1;refGene_tx=XM_003987769.5;refGene_gn=
ACTN1;refGene_tx=XM_019833192.2;refGene_gn=ACTN1;refGene_tx=XM_0198331
94.2;refGene_gn=ACTN1;refGene_tx=XM_019833195.1;refGene_gn=ACTN1;refGe
ne_tx=XM_019833196.2;refGene_gn=ACTN1;refGene_tx=XM_019833197.2
B3 116906162 116906248
refGene_gn=ACTN1;refGene_tx=XM_003987767.5;refGene_gn=ACTN1;refGene_tx
=XM_003987768.4;refGene_gn=ACTN1;refGene_tx=XM_003987769.5;refGene_gn=
ACTN1;refGene_tx=XM_019833192.2;refGene_gn=ACTN1;refGene_tx=XM_0198331
94.2;refGene_gn=ACTN1;refGene_tx=XM_019833195.1;refGene_gn=ACTN1;refGe
ne_tx=XM_019833196.2;refGene_gn=ACTN1;refGene_tx=XM_019833197.2
B3 116908086 116908168
refGene_gn=ACTN1;refGene_tx=XM_003987767.5;refGene_gn=ACTN1;refGene_tx
=XM_003987768.4;refGene_gn=ACTN1;refGene_tx=XM_003987769.5;refGene_gn=
ACTN1;refGene_tx=XM_019833192.2;refGene_gn=ACTN1;refGene_tx=XM_0198331
94.2;refGene_gn=ACTN1;refGene_tx=XM_019833195.1;refGene_gn=ACTN1;refGe
ne_tx=XM_019833196.2;refGene_gn=ACTN1;refGene_tx=XM_019833197.2
B3 116912127 116912206
refGene_gn=ACTN1;refGene_tx=XM_003987767.5;refGene_gn=ACTN1;refGene_tx
=XM_003987768.4;refGene_gn=ACTN1;refGene_tx=XM_003987769.5;refGene_gn=

ACTN1;refGene_tx=XM_019833192.2;refGene_gn=ACTN1;refGene_tx=XM_019833194.2;refGene_gn=ACTN1;refGene_tx=XM_019833195.1;refGene_gn=ACTN1;refGene_tx=XM_019833196.2;refGene_gn=ACTN1;refGene_tx=XM_019833197.2
B3 116912721 116912809
refGene_gn=ACTN1;refGene_tx=XM_003987767.5;refGene_gn=ACTN1;refGene_tx=XM_003987768.4;refGene_gn=ACTN1;refGene_tx=XM_003987769.5;refGene_gn=ACTN1;refGene_tx=XM_019833192.2;refGene_gn=ACTN1;refGene_tx=XM_019833194.2;refGene_gn=ACTN1;refGene_tx=XM_019833195.1;refGene_gn=ACTN1;refGene_tx=XM_019833196.2;refGene_gn=ACTN1;refGene_tx=XM_019833197.2
B3 116915125 116915212
refGene_gn=ACTN1;refGene_tx=XM_003987767.5;refGene_gn=ACTN1;refGene_tx=XM_003987768.4;refGene_gn=ACTN1;refGene_tx=XM_003987769.5;refGene_gn=ACTN1;refGene_tx=XM_019833192.2;refGene_gn=ACTN1;refGene_tx=XM_019833194.2;refGene_gn=ACTN1;refGene_tx=XM_019833195.1;refGene_gn=ACTN1;refGene_tx=XM_019833196.2;refGene_gn=ACTN1;refGene_tx=XM_019833197.2
B3 116922006 116922126
refGene_gn=ACTN1;refGene_tx=XM_003987767.5;refGene_gn=ACTN1;refGene_tx=XM_003987768.4;refGene_gn=ACTN1;refGene_tx=XM_003987769.5;refGene_gn=ACTN1;refGene_tx=XM_019833192.2;refGene_gn=ACTN1;refGene_tx=XM_019833194.2;refGene_gn=ACTN1;refGene_tx=XM_019833195.1;refGene_gn=ACTN1;refGene_tx=XM_019833196.2;refGene_gn=ACTN1;refGene_tx=XM_019833197.2
B3 116926533 116926648
refGene_gn=ACTN1;refGene_tx=XM_003987767.5;refGene_gn=ACTN1;refGene_tx=XM_003987768.4;refGene_gn=ACTN1;refGene_tx=XM_003987769.5;refGene_gn=ACTN1;refGene_tx=XM_019833192.2;refGene_gn=ACTN1;refGene_tx=XM_019833194.2;refGene_gn=ACTN1;refGene_tx=XM_019833195.1;refGene_gn=ACTN1;refGene_tx=XM_019833196.2;refGene_gn=ACTN1;refGene_tx=XM_019833197.2
B3 116934232 116934451
refGene_gn=ACTN1;refGene_tx=XM_019833192.2;refGene_gn=ACTN1;refGene_tx=XM_019833194.2;refGene_gn=ACTN1;refGene_tx=XM_019833195.1
B3 116944974 116945019
refGene_gn=ACTN1;refGene_tx=XM_019833196.2
B3 116976703 116976808
refGene_gn=ACTN1;refGene_tx=XM_003987767.5;refGene_gn=ACTN1;refGene_tx=XM_003987768.4;refGene_gn=ACTN1;refGene_tx=XM_003987769.5
B3 117047122 117048892
refGene_gn=DCAF5;refGene_tx=XM_003987770.5;refGene_gn=DCAF5;refGene_tx=XM_006932940.4;refGene_gn=DCAF5;refGene_tx=XM_019833199.2
B3 117055688 117055816
refGene_gn=DCAF5;refGene_tx=XM_003987770.5;refGene_gn=DCAF5;refGene_tx=XM_006932940.4;refGene_gn=DCAF5;refGene_tx=XM_019833199.2
B3 117072639 117072706
refGene_gn=DCAF5;refGene_tx=XM_003987770.5;refGene_gn=DCAF5;refGene_tx=XM_006932940.4;refGene_gn=DCAF5;refGene_tx=XM_019833199.2
B3 117092540 117092754
refGene_gn=DCAF5;refGene_tx=XM_003987770.5;refGene_gn=DCAF5;refGene_tx=XM_006932940.4;refGene_gn=DCAF5;refGene_tx=XM_019833199.2
B3 117115786 117115916
refGene_gn=DCAF5;refGene_tx=XM_003987770.5;refGene_gn=DCAF5;refGene_tx=XM_006932940.4;refGene_gn=DCAF5;refGene_tx=XM_019833199.2

B3 117117692 117117832
refGene_gn=DCAF5;refGene_tx=XM_003987770.5;refGene_gn=DCAF5;refGene_tx=XM_006932940.4;refGene_gn=DCAF5;refGene_tx=XM_019833199.2
B3 117118899 117118936
refGene_gn=DCAF5;refGene_tx=XM_003987770.5;refGene_gn=DCAF5;refGene_tx=XM_006932940.4;refGene_gn=DCAF5;refGene_tx=XM_019833199.2
B3 117122435 117122579
refGene_gn=DCAF5;refGene_tx=XM_003987770.5;refGene_gn=DCAF5;refGene_tx=XM_006932940.4;refGene_gn=DCAF5;refGene_tx=XM_019833199.2
B3 117151118 117151332
refGene_gn=DCAF5;refGene_tx=XM_003987770.5;refGene_gn=DCAF5;refGene_tx=XM_019833199.2
B3 117203859 117204192
refGene_gn=EXD2;refGene_tx=XM_003987771.5;refGene_gn=EXD2;refGene_tx=XM_011283296.3
B3 117218516 117218773
refGene_gn=EXD2;refGene_tx=XM_003987771.5;refGene_gn=EXD2;refGene_tx=XM_011283296.3
B3 117223261 117223388
refGene_gn=EXD2;refGene_tx=XM_003987771.5;refGene_gn=EXD2;refGene_tx=XM_011283296.3
B3 117230048 117230380
refGene_gn=EXD2;refGene_tx=XM_003987771.5;refGene_gn=EXD2;refGene_tx=XM_011283296.3
B3 117231412 117231519
refGene_gn=EXD2;refGene_tx=XM_003987771.5;refGene_gn=EXD2;refGene_tx=XM_011283296.3
B3 117232057 117232193
refGene_gn=EXD2;refGene_tx=XM_003987771.5;refGene_gn=EXD2;refGene_tx=XM_011283296.3
B3 117236574 117236931
refGene_gn=EXD2;refGene_tx=XM_003987771.5;refGene_gn=EXD2;refGene_tx=XM_011283296.3
B3 117239990 117240207
refGene_gn=EXD2;refGene_tx=XM_003987771.5;refGene_gn=EXD2;refGene_tx=XM_011283296.3
B3 117259126 117259303
refGene_gn=GALNT16;refGene_tx=XM_003987773.5;refGene_gn=GALNT16;refGene_tx=XM_019833202.2
B3 117320258 117320416
refGene_gn=GALNT16;refGene_tx=XM_003987773.5;refGene_gn=GALNT16;refGene_tx=XM_019833202.2
B3 117324140 117324239
refGene_gn=GALNT16;refGene_tx=XM_003987773.5;refGene_gn=GALNT16;refGene_tx=XM_019833202.2
B3 117325167 117325235
refGene_gn=GALNT16;refGene_tx=XM_003987773.5;refGene_gn=GALNT16;refGene_tx=XM_019833202.2
B3 117325944 117326010
refGene_gn=GALNT16;refGene_tx=XM_003987773.5;refGene_gn=GALNT16;refGen

e_tx=XM_019833202.2
B3 117328117 117328239
refGene_gn=GALNT16;refGene_tx=XM_003987773.5;refGene_gn=GALNT16;refGen
e_tx=XM_019833202.2
B3 117330481 117330569
refGene_gn=GALNT16;refGene_tx=XM_003987773.5;refGene_gn=GALNT16;refGen
e_tx=XM_019833202.2
B3 117332310 117332395
refGene_gn=GALNT16;refGene_tx=XM_003987773.5;refGene_gn=GALNT16;refGen
e_tx=XM_019833202.2
B3 117332692 117332796
refGene_gn=GALNT16;refGene_tx=XM_003987773.5;refGene_gn=GALNT16;refGen
e_tx=XM_019833202.2
B3 117338338 117338465
refGene_gn=GALNT16;refGene_tx=XM_003987773.5;refGene_gn=GALNT16;refGen
e_tx=XM_019833202.2
B3 117339736 117339829
refGene_gn=GALNT16;refGene_tx=XM_003987773.5;refGene_gn=GALNT16;refGen
e_tx=XM_019833202.2
B3 117341462 117341546
refGene_gn=GALNT16;refGene_tx=XM_003987773.5;refGene_gn=GALNT16;refGen
e_tx=XM_019833202.2
B3 117342660 117342811
refGene_gn=GALNT16;refGene_tx=XM_019833202.2
B3 117346783 117346925
refGene_gn=GALNT16;refGene_tx=XM_003987773.5
B3 117347625 117347748
refGene_gn=GALNT16;refGene_tx=XM_003987773.5
B3 117351247 117351385
refGene_gn=GALNT16;refGene_tx=XM_003987773.5
B3 117370584 117370687
refGene_gn=ERH;refGene_tx=XM_011283297.1
B3 117376523 117376644
refGene_gn=ERH;refGene_tx=XM_011283297.1
B3 117383489 117383577
refGene_gn=ERH;refGene_tx=XM_011283297.1
B3 117387986 117388121
refGene_gn=ERH;refGene_tx=XM_011283297.1
B3 117389109 117389205
refGene_gn=SLC39A9;refGene_tx=XM_003987775.5
B3 117408875 117408984
refGene_gn=SLC39A9;refGene_tx=XM_003987775.5
B3 117425479 117425674
refGene_gn=SLC39A9;refGene_tx=XM_003987775.5
B3 117432419 117432488
refGene_gn=SLC39A9;refGene_tx=XM_003987775.5
B3 117433670 117433756
refGene_gn=SLC39A9;refGene_tx=XM_003987775.5
B3 117434231 117434366
refGene_gn=SLC39A9;refGene_tx=XM_003987775.5

B3 117436369 117436600
refGene_gn=SLC39A9;refGene_tx=XM_003987775.5
B3 117449577 117449726
refGene_gn=PLEKHD1;refGene_tx=XM_003987777.4
B3 117461126 117461220
refGene_gn=PLEKHD1;refGene_tx=XM_003987777.4
B3 117461552 117461642
refGene_gn=PLEKHD1;refGene_tx=XM_003987777.4
B3 117461884 117461961
refGene_gn=PLEKHD1;refGene_tx=XM_003987777.4
B3 117462778 117462870
refGene_gn=PLEKHD1;refGene_tx=XM_003987777.4
B3 117463830 117463883
refGene_gn=PLEKHD1;refGene_tx=XM_003987777.4
B3 117472090 117472185
refGene_gn=PLEKHD1;refGene_tx=XM_003987777.4
B3 117473397 117473491
refGene_gn=PLEKHD1;refGene_tx=XM_003987777.4
B3 117476058 117476237
refGene_gn=PLEKHD1;refGene_tx=XM_003987777.4
B3 117476756 117476889
refGene_gn=PLEKHD1;refGene_tx=XM_003987777.4
B3 117477256 117477401
refGene_gn=PLEKHD1;refGene_tx=XM_003987777.4
B3 117477729 117477879
refGene_gn=PLEKHD1;refGene_tx=XM_003987777.4
B3 117478098 117478268
refGene_gn=PLEKHD1;refGene_tx=XM_003987777.4
B3 117507515 117509618
refGene_gn=CCDC177;refGene_tx=XM_023255791.1
B3 117589557 117589678
refGene_gn=SUSD6;refGene_tx=XM_003987778.5;refGene_gn=SUSD6;refGene_tx=XM_006932941.4;refGene_gn=SUSD6;refGene_tx=XM_019833204.2;refGene_gn=SUSD6;refGene_tx=XM_019833206.2
B3 117629882 117630080
refGene_gn=SUSD6;refGene_tx=XM_003987778.5;refGene_gn=SUSD6;refGene_tx=XM_006932941.4;refGene_gn=SUSD6;refGene_tx=XM_019833204.2;refGene_gn=SUSD6;refGene_tx=XM_019833206.2
B3 117631065 117631204
refGene_gn=SUSD6;refGene_tx=XM_003987778.5;refGene_gn=SUSD6;refGene_tx=XM_006932941.4;refGene_gn=SUSD6;refGene_tx=XM_019833204.2;refGene_gn=SUSD6;refGene_tx=XM_019833206.2
B3 117635493 117635921
refGene_gn=SUSD6;refGene_tx=XM_003987778.5;refGene_gn=SUSD6;refGene_tx=XM_006932941.4;refGene_gn=SUSD6;refGene_tx=XM_019833204.2;refGene_gn=SUSD6;refGene_tx=XM_019833206.2
B3 117638292 117638318
refGene_gn=SUSD6;refGene_tx=XM_003987778.5;refGene_gn=SUSD6;refGene_tx=XM_006932941.4;refGene_gn=SUSD6;refGene_tx=XM_019833204.2;refGene_gn=SUSD6;refGene_tx=XM_019833206.2

B3 117688958 117689084
refGene_gn=SRSF5;refGene_tx=XM_003987779.3;refGene_gn=SRSF5;refGene_tx=XM_003987780.4;refGene_gn=SRSF5;refGene_tx=XM_006932944.3;refGene_gn=SRSF5;refGene_tx=XM_019833208.2;refGene_gn=SRSF5;refGene_tx=XM_019833209.2
B3 117689382 117689453
refGene_gn=SRSF5;refGene_tx=XM_003987779.3;refGene_gn=SRSF5;refGene_tx=XM_003987780.4;refGene_gn=SRSF5;refGene_tx=XM_006932944.3;refGene_gn=SRSF5;refGene_tx=XM_019833208.2;refGene_gn=SRSF5;refGene_tx=XM_019833209.2
B3 117689580 117689679
refGene_gn=SRSF5;refGene_tx=XM_003987779.3;refGene_gn=SRSF5;refGene_tx=XM_003987780.4;refGene_gn=SRSF5;refGene_tx=XM_006932944.3;refGene_gn=SRSF5;refGene_tx=XM_019833208.2;refGene_gn=SRSF5;refGene_tx=XM_019833209.2
B3 117689982 117690052
refGene_gn=SRSF5;refGene_tx=XM_003987779.3;refGene_gn=SRSF5;refGene_tx=XM_003987780.4;refGene_gn=SRSF5;refGene_tx=XM_006932944.3;refGene_gn=SRSF5;refGene_tx=XM_019833208.2;refGene_gn=SRSF5;refGene_tx=XM_019833209.2
B3 117691296 117691370
refGene_gn=SRSF5;refGene_tx=XM_003987779.3;refGene_gn=SRSF5;refGene_tx=XM_003987780.4;refGene_gn=SRSF5;refGene_tx=XM_006932944.3;refGene_gn=SRSF5;refGene_tx=XM_019833208.2;refGene_gn=SRSF5;refGene_tx=XM_019833209.2
B3 117691819 117691930
refGene_gn=SRSF5;refGene_tx=XM_003987779.3;refGene_gn=SRSF5;refGene_tx=XM_003987780.4;refGene_gn=SRSF5;refGene_tx=XM_006932944.3;refGene_gn=SRSF5;refGene_tx=XM_019833208.2;refGene_gn=SRSF5;refGene_tx=XM_019833209.2
B3 117692017 117692285
refGene_gn=SRSF5;refGene_tx=XM_003987779.3;refGene_gn=SRSF5;refGene_tx=XM_003987780.4;refGene_gn=SRSF5;refGene_tx=XM_006932944.3;refGene_gn=SRSF5;refGene_tx=XM_019833208.2;refGene_gn=SRSF5;refGene_tx=XM_019833209.2
B3 117695924 117696031
refGene_gn=SLC10A1;refGene_tx=XM_003987782.4;refGene_gn=SLC10A1;refGene_tx=XM_019833207.2
B3 117697964 117698161
refGene_gn=SLC10A1;refGene_tx=XM_003987782.4;refGene_gn=SLC10A1;refGene_tx=XM_019833207.2
B3 117699723 117699902
refGene_gn=SLC10A1;refGene_tx=XM_003987782.4;refGene_gn=SLC10A1;refGene_tx=XM_019833207.2
B3 117705776 117705987
refGene_gn=SLC10A1;refGene_tx=XM_003987782.4;refGene_gn=SLC10A1;refGene_tx=XM_019833207.2
B3 117714955 117715311
refGene_gn=SLC10A1;refGene_tx=XM_003987782.4;refGene_gn=SLC10A1;refGene_tx=XM_019833207.2

B3 117774514 117774613
refGene_gn=SMOC1;refGene_tx=XM_006932946.4;refGene_gn=SMOC1;refGene_tx=XM_006932947.4;refGene_gn=SMOC1;refGene_tx=XM_006932948.4
B3 117851340 117851506
refGene_gn=SMOC1;refGene_tx=XM_006932946.4;refGene_gn=SMOC1;refGene_tx=XM_006932947.4;refGene_gn=SMOC1;refGene_tx=XM_006932948.4
B3 117852665 117852778
refGene_gn=SMOC1;refGene_tx=XM_006932946.4;refGene_gn=SMOC1;refGene_tx=XM_006932947.4;refGene_gn=SMOC1;refGene_tx=XM_006932948.4
B3 117882091 117882191
refGene_gn=SMOC1;refGene_tx=XM_006932946.4;refGene_gn=SMOC1;refGene_tx=XM_006932947.4;refGene_gn=SMOC1;refGene_tx=XM_006932948.4
B3 117884342 117884390
refGene_gn=SMOC1;refGene_tx=XM_006932946.4;refGene_gn=SMOC1;refGene_tx=XM_006932947.4;refGene_gn=SMOC1;refGene_tx=XM_006932948.4
B3 117898593 117898683
refGene_gn=SMOC1;refGene_tx=XM_006932946.4;refGene_gn=SMOC1;refGene_tx=XM_006932947.4;refGene_gn=SMOC1;refGene_tx=XM_006932948.4
B3 117900665 117900746
refGene_gn=SMOC1;refGene_tx=XM_006932946.4;refGene_gn=SMOC1;refGene_tx=XM_006932947.4;refGene_gn=SMOC1;refGene_tx=XM_006932948.4
B3 117916417 117916610
refGene_gn=SMOC1;refGene_tx=XM_006932946.4;refGene_gn=SMOC1;refGene_tx=XM_006932947.4;refGene_gn=SMOC1;refGene_tx=XM_006932948.4
B3 117917185 117917268
refGene_gn=SMOC1;refGene_tx=XM_006932946.4;refGene_gn=SMOC1;refGene_tx=XM_006932947.4;refGene_gn=SMOC1;refGene_tx=XM_006932948.4
B3 117917451 117917557
refGene_gn=SMOC1;refGene_tx=XM_006932946.4;refGene_gn=SMOC1;refGene_tx=XM_006932947.4;refGene_gn=SMOC1;refGene_tx=XM_006932948.4
B3 117926625 117926870
refGene_gn=SMOC1;refGene_tx=XM_006932946.4;refGene_gn=SMOC1;refGene_tx=XM_006932947.4;refGene_gn=SMOC1;refGene_tx=XM_006932948.4
B3 117932724 117932741
refGene_gn=SMOC1;refGene_tx=XM_006932946.4;refGene_gn=SMOC1;refGene_tx=XM_006932947.4;refGene_gn=SMOC1;refGene_tx=XM_006932948.4
B3 117952101 117952112
refGene_gn=SLC8A3;refGene_tx=XM_011283306.3
B3 117954225 117954602
refGene_gn=SLC8A3;refGene_tx=XM_011283300.3;refGene_gn=SLC8A3;refGene_tx=XM_011283301.3;refGene_gn=SLC8A3;refGene_tx=XM_011283302.3;refGene_gn=SLC8A3;refGene_tx=XM_011283303.3;refGene_gn=SLC8A3;refGene_tx=XM_011283305.3;refGene_gn=SLC8A3;refGene_tx=XM_011283308.3;refGene_gn=SLC8A3;refGene_tx=XM_019833212.2
B3 117957211 117957487
refGene_gn=SLC8A3;refGene_tx=XM_011283300.3;refGene_gn=SLC8A3;refGene_tx=XM_011283301.3;refGene_gn=SLC8A3;refGene_tx=XM_011283302.3;refGene_gn=SLC8A3;refGene_tx=XM_011283303.3;refGene_gn=SLC8A3;refGene_tx=XM_011283305.3;refGene_gn=SLC8A3;refGene_tx=XM_011283306.3;refGene_gn=SLC8A3;refGene_tx=XM_011283308.3;refGene_gn=SLC8A3;refGene_tx=XM_019833212.

2
B3 117959134 117959234
refGene_gn=SLC8A3;refGene_tx=XM_011283300.3;refGene_gn=SLC8A3;refGene_tx=XM_011283301.3;refGene_gn=SLC8A3;refGene_tx=XM_011283302.3;refGene_gn=SLC8A3;refGene_tx=XM_011283303.3;refGene_gn=SLC8A3;refGene_tx=XM_011283305.3;refGene_gn=SLC8A3;refGene_tx=XM_011283306.3;refGene_gn=SLC8A3;refGene_tx=XM_011283308.3;refGene_gn=SLC8A3;refGene_tx=XM_019833212.2
B3 117959775 117959900
refGene_gn=SLC8A3;refGene_tx=XM_011283300.3;refGene_gn=SLC8A3;refGene_tx=XM_011283301.3;refGene_gn=SLC8A3;refGene_tx=XM_011283302.3;refGene_gn=SLC8A3;refGene_tx=XM_011283303.3;refGene_gn=SLC8A3;refGene_tx=XM_011283305.3;refGene_gn=SLC8A3;refGene_tx=XM_011283306.3;refGene_gn=SLC8A3;refGene_tx=XM_011283308.3;refGene_gn=SLC8A3;refGene_tx=XM_019833212.2
B3 117963844 117963862
refGene_gn=SLC8A3;refGene_tx=XM_011283300.3;refGene_gn=SLC8A3;refGene_tx=XM_011283301.3;refGene_gn=SLC8A3;refGene_tx=XM_011283302.3;refGene_gn=SLC8A3;refGene_tx=XM_011283306.3;refGene_gn=SLC8A3;refGene_tx=XM_011283308.3;refGene_gn=SLC8A3;refGene_tx=XM_019833212.2
B3 117968850 117968954
refGene_gn=SLC8A3;refGene_tx=XM_011283301.3;refGene_gn=SLC8A3;refGene_tx=XM_011283305.3;refGene_gn=SLC8A3;refGene_tx=XM_011283308.3;refGene_gn=SLC8A3;refGene_tx=XM_019833212.2
B3 117971518 117971625
refGene_gn=SLC8A3;refGene_tx=XM_011283300.3;refGene_gn=SLC8A3;refGene_tx=XM_011283302.3;refGene_gn=SLC8A3;refGene_tx=XM_011283303.3;refGene_gn=SLC8A3;refGene_tx=XM_011283306.3
B3 118077037 118078821
refGene_gn=SLC8A3;refGene_tx=XM_011283300.3;refGene_gn=SLC8A3;refGene_tx=XM_011283301.3;refGene_gn=SLC8A3;refGene_tx=XM_011283302.3;refGene_gn=SLC8A3;refGene_tx=XM_011283303.3;refGene_gn=SLC8A3;refGene_tx=XM_011283305.3;refGene_gn=SLC8A3;refGene_tx=XM_011283306.3;refGene_gn=SLC8A3;refGene_tx=XM_011283308.3
B3 118179349 118179472
refGene_gn=LOC101081902;refGene_tx=XM_003987788.5
B3 118182593 118182656
refGene_gn=LOC101081902;refGene_tx=XM_003987788.5
B3 118187438 118187510
refGene_gn=LOC101081902;refGene_tx=XM_003987788.5
B3 118189264 118189333
refGene_gn=LOC101081902;refGene_tx=XM_003987788.5
B3 118209466 118209607
refGene_gn=SYNJ2BP;refGene_tx=XM_003987787.5
B3 118211591 118211687
refGene_gn=SYNJ2BP;refGene_tx=XM_003987787.5
B3 118221369 118221506
refGene_gn=SYNJ2BP;refGene_tx=XM_003987787.5
B3 118242672 118242736
refGene_gn=SYNJ2BP;refGene_tx=XM_003987787.5

B3 118274726 118276736
refGene_gn=LOC101088229;refGene_tx=XM_011283403.2
B3 118290891 118293027
refGene_gn=LOC101082414;refGene_tx=XM_003987789.5
B3 118317222 118319412
refGene_gn=LOC101084205;refGene_tx=XM_006932956.4;refGene_gn=LOC101084205;refGene_tx=XM_019833214.2
B3 118356198 118356329
refGene_gn=MED6;refGene_tx=XM_003987790.5;refGene_gn=MED6;refGene_tx=XM_006932957.4
B3 118356911 118356939
refGene_gn=MED6;refGene_tx=XM_003987790.5;refGene_gn=MED6;refGene_tx=XM_006932957.4
B3 118367810 118367926
refGene_gn=MED6;refGene_tx=XM_003987790.5;refGene_gn=MED6;refGene_tx=XM_006932957.4
B3 118369813 118369922
refGene_gn=MED6;refGene_tx=XM_003987790.5;refGene_gn=MED6;refGene_tx=XM_006932957.4
B3 118370278 118370361
refGene_gn=MED6;refGene_tx=XM_003987790.5;refGene_gn=MED6;refGene_tx=XM_006932957.4
B3 118373636 118373728
refGene_gn=MED6;refGene_tx=XM_003987790.5;refGene_gn=MED6;refGene_tx=XM_006932957.4
B3 118374990 118375150
refGene_gn=MED6;refGene_tx=XM_003987790.5;refGene_gn=MED6;refGene_tx=XM_006932957.4
B3 118376243 118376262
refGene_gn=MED6;refGene_tx=XM_006932957.4
B3 118379890 118379912
refGene_gn=MED6;refGene_tx=XM_003987790.5
B3 118419158 118419576
refGene_gn=TTC9;refGene_tx=XM_023255792.1
B3 118445924 118446107
refGene_gn=TTC9;refGene_tx=XM_023255792.1
B3 118448039 118448119
refGene_gn=TTC9;refGene_tx=XM_023255792.1
B3 118502832 118503317
refGene_gn=MAP3K9;refGene_tx=XM_023255793.1
B3 118504611 118505415
refGene_gn=MAP3K9;refGene_tx=XM_023255793.1
B3 118506391 118506504
refGene_gn=MAP3K9;refGene_tx=XM_023255793.1
B3 118507917 118507986
refGene_gn=MAP3K9;refGene_tx=XM_023255793.1
B3 118510305 118510459
refGene_gn=MAP3K9;refGene_tx=XM_023255793.1
B3 118511835 118511958
refGene_gn=MAP3K9;refGene_tx=XM_023255793.1

B3	118513678	118513919
refGene_gn=MAP3K9;refGene_tx=XM_023255793.1
B3	118518717	118518893
refGene_gn=MAP3K9;refGene_tx=XM_023255793.1
B3	118519770	118519919
refGene_gn=MAP3K9;refGene_tx=XM_023255793.1
B3	118530708	118530889
refGene_gn=MAP3K9;refGene_tx=XM_023255793.1
B3	118570547	118570961
refGene_gn=MAP3K9;refGene_tx=XM_023255793.1
B3	118578201	118578601
refGene_gn=MAP3K9;refGene_tx=XM_023255793.1
B3	118633163	118633316
refGene_gn=PCNX1;refGene_tx=XM_023255794.1;refGene_gn=PCNX1;refGene_tx=XM_023255795.1;refGene_gn=PCNX1;refGene_tx=XM_023255796.1;refGene_gn=PCNX1;refGene_tx=XM_023255797.1;refGene_gn=PCNX1;refGene_tx=XM_023255798.1;refGene_gn=PCNX1;refGene_tx=XM_023255799.1;refGene_gn=PCNX1;refGene_tx=XM_023255801.1
B3	118671883	118672092
refGene_gn=PCNX1;refGene_tx=XM_023255794.1;refGene_gn=PCNX1;refGene_tx=XM_023255795.1;refGene_gn=PCNX1;refGene_tx=XM_023255796.1;refGene_gn=PCNX1;refGene_tx=XM_023255797.1;refGene_gn=PCNX1;refGene_tx=XM_023255798.1;refGene_gn=PCNX1;refGene_tx=XM_023255799.1;refGene_gn=PCNX1;refGene_tx=XM_023255801.1
B3	118682905	118683011
refGene_gn=PCNX1;refGene_tx=XM_023255794.1;refGene_gn=PCNX1;refGene_tx=XM_023255795.1;refGene_gn=PCNX1;refGene_tx=XM_023255796.1;refGene_gn=PCNX1;refGene_tx=XM_023255797.1;refGene_gn=PCNX1;refGene_tx=XM_023255798.1;refGene_gn=PCNX1;refGene_tx=XM_023255799.1;refGene_gn=PCNX1;refGene_tx=XM_023255800.1;refGene_gn=PCNX1;refGene_tx=XM_023255801.1
B3	118688708	118688754
refGene_gn=PCNX1;refGene_tx=XM_023255794.1;refGene_gn=PCNX1;refGene_tx=XM_023255795.1;refGene_gn=PCNX1;refGene_tx=XM_023255796.1;refGene_gn=PCNX1;refGene_tx=XM_023255797.1;refGene_gn=PCNX1;refGene_tx=XM_023255798.1;refGene_gn=PCNX1;refGene_tx=XM_023255799.1;refGene_gn=PCNX1;refGene_tx=XM_023255800.1;refGene_gn=PCNX1;refGene_tx=XM_023255801.1
B3	118689795	118689885
refGene_gn=PCNX1;refGene_tx=XM_023255794.1;refGene_gn=PCNX1;refGene_tx=XM_023255795.1;refGene_gn=PCNX1;refGene_tx=XM_023255796.1;refGene_gn=PCNX1;refGene_tx=XM_023255797.1;refGene_gn=PCNX1;refGene_tx=XM_023255798.1;refGene_gn=PCNX1;refGene_tx=XM_023255799.1;refGene_gn=PCNX1;refGene_tx=XM_023255800.1;refGene_gn=PCNX1;refGene_tx=XM_023255801.1
B3	118697483	118699193
refGene_gn=PCNX1;refGene_tx=XM_023255794.1;refGene_gn=PCNX1;refGene_tx=XM_023255795.1;refGene_gn=PCNX1;refGene_tx=XM_023255796.1;refGene_gn=PCNX1;refGene_tx=XM_023255797.1;refGene_gn=PCNX1;refGene_tx=XM_023255798.1;refGene_gn=PCNX1;refGene_tx=XM_023255799.1;refGene_gn=PCNX1;refGene_tx=XM_023255800.1
B3	118709273	118709406
refGene_gn=PCNX1;refGene_tx=XM_023255794.1;refGene_gn=PCNX1;refGene_tx

=XM_023255795.1;refGene_gn=PCNX1;refGene_tx=XM_023255796.1;refGene_gn=PCNX1;refGene_tx=XM_023255797.1;refGene_gn=PCNX1;refGene_tx=XM_023255800.1;refGene_gn=PCNX1;refGene_tx=XM_023255801.1
B3 118714932 118715117
refGene_gn=PCNX1;refGene_tx=XM_023255794.1;refGene_gn=PCNX1;refGene_tx=XM_023255795.1;refGene_gn=PCNX1;refGene_tx=XM_023255796.1;refGene_gn=PCNX1;refGene_tx=XM_023255797.1;refGene_gn=PCNX1;refGene_tx=XM_023255800.1;refGene_gn=PCNX1;refGene_tx=XM_023255801.1
B3 118728219 118728310
refGene_gn=PCNX1;refGene_tx=XM_023255794.1;refGene_gn=PCNX1;refGene_tx=XM_023255795.1;refGene_gn=PCNX1;refGene_tx=XM_023255796.1;refGene_gn=PCNX1;refGene_tx=XM_023255797.1;refGene_gn=PCNX1;refGene_tx=XM_023255798.1;refGene_gn=PCNX1;refGene_tx=XM_023255799.1;refGene_gn=PCNX1;refGene_tx=XM_023255800.1;refGene_gn=PCNX1;refGene_tx=XM_023255801.1
B3 118730047 118730105
refGene_gn=PCNX1;refGene_tx=XM_023255794.1;refGene_gn=PCNX1;refGene_tx=XM_023255795.1;refGene_gn=PCNX1;refGene_tx=XM_023255796.1;refGene_gn=PCNX1;refGene_tx=XM_023255797.1;refGene_gn=PCNX1;refGene_tx=XM_023255798.1;refGene_gn=PCNX1;refGene_tx=XM_023255799.1;refGene_gn=PCNX1;refGene_tx=XM_023255800.1;refGene_gn=PCNX1;refGene_tx=XM_023255801.1
B3 118732211 118732429
refGene_gn=PCNX1;refGene_tx=XM_023255794.1;refGene_gn=PCNX1;refGene_tx=XM_023255795.1;refGene_gn=PCNX1;refGene_tx=XM_023255796.1;refGene_gn=PCNX1;refGene_tx=XM_023255797.1;refGene_gn=PCNX1;refGene_tx=XM_023255798.1;refGene_gn=PCNX1;refGene_tx=XM_023255799.1;refGene_gn=PCNX1;refGene_tx=XM_023255800.1;refGene_gn=PCNX1;refGene_tx=XM_023255801.1
B3 118739090 118739244
refGene_gn=PCNX1;refGene_tx=XM_023255794.1;refGene_gn=PCNX1;refGene_tx=XM_023255795.1;refGene_gn=PCNX1;refGene_tx=XM_023255796.1;refGene_gn=PCNX1;refGene_tx=XM_023255797.1;refGene_gn=PCNX1;refGene_tx=XM_023255798.1;refGene_gn=PCNX1;refGene_tx=XM_023255799.1;refGene_gn=PCNX1;refGene_tx=XM_023255800.1;refGene_gn=PCNX1;refGene_tx=XM_023255801.1
B3 118741825 118741858
refGene_gn=PCNX1;refGene_tx=XM_023255794.1;refGene_gn=PCNX1;refGene_tx=XM_023255795.1;refGene_gn=PCNX1;refGene_tx=XM_023255796.1;refGene_gn=PCNX1;refGene_tx=XM_023255797.1;refGene_gn=PCNX1;refGene_tx=XM_023255798.1;refGene_gn=PCNX1;refGene_tx=XM_023255799.1;refGene_gn=PCNX1;refGene_tx=XM_023255800.1;refGene_gn=PCNX1;refGene_tx=XM_023255801.1
B3 118745000 118745172
refGene_gn=PCNX1;refGene_tx=XM_023255794.1;refGene_gn=PCNX1;refGene_tx=XM_023255795.1;refGene_gn=PCNX1;refGene_tx=XM_023255796.1;refGene_gn=PCNX1;refGene_tx=XM_023255797.1;refGene_gn=PCNX1;refGene_tx=XM_023255798.1;refGene_gn=PCNX1;refGene_tx=XM_023255799.1;refGene_gn=PCNX1;refGene_tx=XM_023255800.1;refGene_gn=PCNX1;refGene_tx=XM_023255801.1
B3 118745863 118745974
refGene_gn=PCNX1;refGene_tx=XM_023255794.1;refGene_gn=PCNX1;refGene_tx=XM_023255795.1;refGene_gn=PCNX1;refGene_tx=XM_023255796.1;refGene_gn=PCNX1;refGene_tx=XM_023255797.1;refGene_gn=PCNX1;refGene_tx=XM_023255798.1;refGene_gn=PCNX1;refGene_tx=XM_023255799.1;refGene_gn=PCNX1;refGene_tx=XM_023255800.1;refGene_gn=PCNX1;refGene_tx=XM_023255801.1

B3	118749074	118749166
refGene_gn=PCNX1;refGene_tx=XM_023255794.1;refGene_gn=PCNX1;refGene_tx=XM_023255795.1;refGene_gn=PCNX1;refGene_tx=XM_023255796.1;refGene_gn=PCNX1;refGene_tx=XM_023255797.1;refGene_gn=PCNX1;refGene_tx=XM_023255798.1;refGene_gn=PCNX1;refGene_tx=XM_023255799.1;refGene_gn=PCNX1;refGene_tx=XM_023255800.1;refGene_gn=PCNX1;refGene_tx=XM_023255801.1
B3	118756112	118756222
refGene_gn=PCNX1;refGene_tx=XM_023255794.1;refGene_gn=PCNX1;refGene_tx=XM_023255795.1;refGene_gn=PCNX1;refGene_tx=XM_023255796.1;refGene_gn=PCNX1;refGene_tx=XM_023255797.1;refGene_gn=PCNX1;refGene_tx=XM_023255798.1;refGene_gn=PCNX1;refGene_tx=XM_023255799.1;refGene_gn=PCNX1;refGene_tx=XM_023255800.1;refGene_gn=PCNX1;refGene_tx=XM_023255801.1
B3	118756609	118756715
refGene_gn=PCNX1;refGene_tx=XM_023255794.1;refGene_gn=PCNX1;refGene_tx=XM_023255795.1;refGene_gn=PCNX1;refGene_tx=XM_023255796.1;refGene_gn=PCNX1;refGene_tx=XM_023255797.1;refGene_gn=PCNX1;refGene_tx=XM_023255798.1;refGene_gn=PCNX1;refGene_tx=XM_023255799.1;refGene_gn=PCNX1;refGene_tx=XM_023255800.1;refGene_gn=PCNX1;refGene_tx=XM_023255801.1
B3	118759593	118759686
refGene_gn=PCNX1;refGene_tx=XM_023255794.1;refGene_gn=PCNX1;refGene_tx=XM_023255795.1;refGene_gn=PCNX1;refGene_tx=XM_023255796.1;refGene_gn=PCNX1;refGene_tx=XM_023255797.1;refGene_gn=PCNX1;refGene_tx=XM_023255798.1;refGene_gn=PCNX1;refGene_tx=XM_023255799.1;refGene_gn=PCNX1;refGene_tx=XM_023255800.1;refGene_gn=PCNX1;refGene_tx=XM_023255801.1
B3	118760333	118760484
refGene_gn=PCNX1;refGene_tx=XM_023255794.1;refGene_gn=PCNX1;refGene_tx=XM_023255795.1;refGene_gn=PCNX1;refGene_tx=XM_023255796.1;refGene_gn=PCNX1;refGene_tx=XM_023255797.1;refGene_gn=PCNX1;refGene_tx=XM_023255798.1;refGene_gn=PCNX1;refGene_tx=XM_023255799.1;refGene_gn=PCNX1;refGene_tx=XM_023255800.1;refGene_gn=PCNX1;refGene_tx=XM_023255801.1
B3	118765899	118766041
refGene_gn=PCNX1;refGene_tx=XM_023255794.1;refGene_gn=PCNX1;refGene_tx=XM_023255795.1;refGene_gn=PCNX1;refGene_tx=XM_023255796.1;refGene_gn=PCNX1;refGene_tx=XM_023255797.1;refGene_gn=PCNX1;refGene_tx=XM_023255798.1;refGene_gn=PCNX1;refGene_tx=XM_023255799.1;refGene_gn=PCNX1;refGene_tx=XM_023255800.1;refGene_gn=PCNX1;refGene_tx=XM_023255801.1
B3	118766750	118766928
refGene_gn=PCNX1;refGene_tx=XM_023255794.1;refGene_gn=PCNX1;refGene_tx=XM_023255795.1;refGene_gn=PCNX1;refGene_tx=XM_023255796.1;refGene_gn=PCNX1;refGene_tx=XM_023255797.1;refGene_gn=PCNX1;refGene_tx=XM_023255798.1;refGene_gn=PCNX1;refGene_tx=XM_023255799.1;refGene_gn=PCNX1;refGene_tx=XM_023255800.1;refGene_gn=PCNX1;refGene_tx=XM_023255801.1
B3	118769538	118769647
refGene_gn=PCNX1;refGene_tx=XM_023255794.1;refGene_gn=PCNX1;refGene_tx=XM_023255795.1;refGene_gn=PCNX1;refGene_tx=XM_023255796.1;refGene_gn=PCNX1;refGene_tx=XM_023255797.1;refGene_gn=PCNX1;refGene_tx=XM_023255798.1;refGene_gn=PCNX1;refGene_tx=XM_023255799.1;refGene_gn=PCNX1;refGene_tx=XM_023255800.1;refGene_gn=PCNX1;refGene_tx=XM_023255801.1
B3	118770532	118770662
refGene_gn=PCNX1;refGene_tx=XM_023255794.1;refGene_gn=PCNX1;refGene_tx

=XM_023255795.1;refGene_gn=PCNX1;refGene_tx=XM_023255796.1;refGene_gn=PCNX1;refGene_tx=XM_023255797.1;refGene_gn=PCNX1;refGene_tx=XM_023255798.1;refGene_gn=PCNX1;refGene_tx=XM_023255799.1;refGene_gn=PCNX1;refGene_tx=XM_023255800.1;refGene_gn=PCNX1;refGene_tx=XM_023255801.1
B3 118773481 118773540
refGene_gn=PCNX1;refGene_tx=XM_023255794.1;refGene_gn=PCNX1;refGene_tx=XM_023255795.1;refGene_gn=PCNX1;refGene_tx=XM_023255796.1;refGene_gn=PCNX1;refGene_tx=XM_023255797.1;refGene_gn=PCNX1;refGene_tx=XM_023255798.1;refGene_gn=PCNX1;refGene_tx=XM_023255799.1;refGene_gn=PCNX1;refGene_tx=XM_023255800.1;refGene_gn=PCNX1;refGene_tx=XM_023255801.1
B3 118775378 118775594
refGene_gn=PCNX1;refGene_tx=XM_023255794.1;refGene_gn=PCNX1;refGene_tx=XM_023255795.1;refGene_gn=PCNX1;refGene_tx=XM_023255796.1;refGene_gn=PCNX1;refGene_tx=XM_023255797.1;refGene_gn=PCNX1;refGene_tx=XM_023255798.1;refGene_gn=PCNX1;refGene_tx=XM_023255799.1;refGene_gn=PCNX1;refGene_tx=XM_023255800.1;refGene_gn=PCNX1;refGene_tx=XM_023255801.1
B3 118781914 118782168
refGene_gn=PCNX1;refGene_tx=XM_023255794.1;refGene_gn=PCNX1;refGene_tx=XM_023255795.1;refGene_gn=PCNX1;refGene_tx=XM_023255796.1;refGene_gn=PCNX1;refGene_tx=XM_023255797.1;refGene_gn=PCNX1;refGene_tx=XM_023255798.1;refGene_gn=PCNX1;refGene_tx=XM_023255799.1;refGene_gn=PCNX1;refGene_tx=XM_023255800.1;refGene_gn=PCNX1;refGene_tx=XM_023255801.1
B3 118783882 118784113
refGene_gn=PCNX1;refGene_tx=XM_023255794.1;refGene_gn=PCNX1;refGene_tx=XM_023255795.1;refGene_gn=PCNX1;refGene_tx=XM_023255796.1;refGene_gn=PCNX1;refGene_tx=XM_023255797.1;refGene_gn=PCNX1;refGene_tx=XM_023255798.1;refGene_gn=PCNX1;refGene_tx=XM_023255799.1;refGene_gn=PCNX1;refGene_tx=XM_023255800.1;refGene_gn=PCNX1;refGene_tx=XM_023255801.1
B3 118784744 118784845
refGene_gn=PCNX1;refGene_tx=XM_023255794.1;refGene_gn=PCNX1;refGene_tx=XM_023255795.1;refGene_gn=PCNX1;refGene_tx=XM_023255796.1;refGene_gn=PCNX1;refGene_tx=XM_023255797.1;refGene_gn=PCNX1;refGene_tx=XM_023255798.1;refGene_gn=PCNX1;refGene_tx=XM_023255799.1;refGene_gn=PCNX1;refGene_tx=XM_023255800.1;refGene_gn=PCNX1;refGene_tx=XM_023255801.1
B3 118785619 118785770
refGene_gn=PCNX1;refGene_tx=XM_023255794.1;refGene_gn=PCNX1;refGene_tx=XM_023255795.1;refGene_gn=PCNX1;refGene_tx=XM_023255796.1;refGene_gn=PCNX1;refGene_tx=XM_023255797.1;refGene_gn=PCNX1;refGene_tx=XM_023255798.1;refGene_gn=PCNX1;refGene_tx=XM_023255799.1;refGene_gn=PCNX1;refGene_tx=XM_023255800.1;refGene_gn=PCNX1;refGene_tx=XM_023255801.1
B3 118790880 118791111
refGene_gn=PCNX1;refGene_tx=XM_023255794.1;refGene_gn=PCNX1;refGene_tx=XM_023255795.1;refGene_gn=PCNX1;refGene_tx=XM_023255796.1;refGene_gn=PCNX1;refGene_tx=XM_023255797.1;refGene_gn=PCNX1;refGene_tx=XM_023255798.1;refGene_gn=PCNX1;refGene_tx=XM_023255799.1;refGene_gn=PCNX1;refGene_tx=XM_023255800.1;refGene_gn=PCNX1;refGene_tx=XM_023255801.1
B3 118791931 118792206
refGene_gn=PCNX1;refGene_tx=XM_023255794.1;refGene_gn=PCNX1;refGene_tx=XM_023255795.1;refGene_gn=PCNX1;refGene_tx=XM_023255796.1;refGene_gn=PCNX1;refGene_tx=XM_023255797.1;refGene_gn=PCNX1;refGene_tx=XM_0232557

98.1;refGene_gn=PCNX1;refGene_tx=XM_023255799.1;refGene_gn=PCNX1;refGene_tx=XM_023255800.1;refGene_gn=PCNX1;refGene_tx=XM_023255801.1
B3 118793851 118794057
refGene_gn=PCNX1;refGene_tx=XM_023255794.1;refGene_gn=PCNX1;refGene_tx=XM_023255795.1;refGene_gn=PCNX1;refGene_tx=XM_023255796.1;refGene_gn=PCNX1;refGene_tx=XM_023255797.1;refGene_gn=PCNX1;refGene_tx=XM_023255798.1;refGene_gn=PCNX1;refGene_tx=XM_023255799.1;refGene_gn=PCNX1;refGene_tx=XM_023255800.1;refGene_gn=PCNX1;refGene_tx=XM_023255801.1
B3 118796511 118796954
refGene_gn=PCNX1;refGene_tx=XM_023255794.1;refGene_gn=PCNX1;refGene_tx=XM_023255795.1;refGene_gn=PCNX1;refGene_tx=XM_023255796.1;refGene_gn=PCNX1;refGene_tx=XM_023255797.1;refGene_gn=PCNX1;refGene_tx=XM_023255798.1;refGene_gn=PCNX1;refGene_tx=XM_023255799.1;refGene_gn=PCNX1;refGene_tx=XM_023255800.1;refGene_gn=PCNX1;refGene_tx=XM_023255801.1
B3 118797360 118797501
refGene_gn=PCNX1;refGene_tx=XM_023255794.1;refGene_gn=PCNX1;refGene_tx=XM_023255795.1;refGene_gn=PCNX1;refGene_tx=XM_023255796.1;refGene_gn=PCNX1;refGene_tx=XM_023255797.1;refGene_gn=PCNX1;refGene_tx=XM_023255798.1;refGene_gn=PCNX1;refGene_tx=XM_023255799.1;refGene_gn=PCNX1;refGene_tx=XM_023255800.1;refGene_gn=PCNX1;refGene_tx=XM_023255801.1
B3 118797710 118797851
refGene_gn=PCNX1;refGene_tx=XM_023255794.1;refGene_gn=PCNX1;refGene_tx=XM_023255795.1;refGene_gn=PCNX1;refGene_tx=XM_023255796.1;refGene_gn=PCNX1;refGene_tx=XM_023255797.1;refGene_gn=PCNX1;refGene_tx=XM_023255798.1;refGene_gn=PCNX1;refGene_tx=XM_023255799.1;refGene_gn=PCNX1;refGene_tx=XM_023255800.1;refGene_gn=PCNX1;refGene_tx=XM_023255801.1
B3 118797895 118798023
refGene_gn=PCNX1;refGene_tx=XM_023255794.1;refGene_gn=PCNX1;refGene_tx=XM_023255795.1;refGene_gn=PCNX1;refGene_tx=XM_023255796.1;refGene_gn=PCNX1;refGene_tx=XM_023255798.1;refGene_gn=PCNX1;refGene_tx=XM_023255800.1;refGene_gn=PCNX1;refGene_tx=XM_023255801.1
B3 119205875 119206634
refGene_gn=SIPA1L1;refGene_tx=XM_019833223.2
B3 119206635 119207374
refGene_gn=SIPA1L1;refGene_tx=XM_019833223.2
B3 119225668 119225799
refGene_gn=SIPA1L1;refGene_tx=XM_019833223.2
B3 119231303 119231492
refGene_gn=SIPA1L1;refGene_tx=XM_019833223.2
B3 119247417 119247592
refGene_gn=SIPA1L1;refGene_tx=XM_019833223.2
B3 119257092 119257196
refGene_gn=SIPA1L1;refGene_tx=XM_019833223.2
B3 119260050 119260208
refGene_gn=SIPA1L1;refGene_tx=XM_019833223.2
B3 119269198 119269772
refGene_gn=SIPA1L1;refGene_tx=XM_019833223.2
B3 119270409 119270684
refGene_gn=SIPA1L1;refGene_tx=XM_019833223.2
B3 119278351 119278621

refGene_gn=SIPA1L1;refGene_tx=XM_019833223.2
B3 119289213 119289363
refGene_gn=SIPA1L1;refGene_tx=XM_019833223.2
B3 119293829 119293954
refGene_gn=SIPA1L1;refGene_tx=XM_019833223.2
B3 119296205 119296268
refGene_gn=SIPA1L1;refGene_tx=XM_019833223.2
B3 119296705 119296824
refGene_gn=SIPA1L1;refGene_tx=XM_019833223.2
B3 119300674 119301117
refGene_gn=SIPA1L1;refGene_tx=XM_019833223.2
B3 119311760 119311994
refGene_gn=SIPA1L1;refGene_tx=XM_019833223.2
B3 119312761 119312927
refGene_gn=SIPA1L1;refGene_tx=XM_019833223.2
B3 119317790 119318037
refGene_gn=SIPA1L1;refGene_tx=XM_019833223.2
B3 119329894 119330041
refGene_gn=SIPA1L1;refGene_tx=XM_019833223.2
B3 119331370 119331485
refGene_gn=SIPA1L1;refGene_tx=XM_019833223.2
B3 119333986 119334071
refGene_gn=SIPA1L1;refGene_tx=XM_019833223.2
B3 119334710 119334854
refGene_gn=SIPA1L1;refGene_tx=XM_019833223.2
B3 119542721 119542805
refGene_gn=RGS6;refGene_tx=XM_006932961.4;refGene_gn=RGS6;refGene_tx=XM_011283315.2;refGene_gn=RGS6;refGene_tx=XM_011283317.2;refGene_gn=RGS6;refGene_tx=XM_011283318.2;refGene_gn=RGS6;refGene_tx=XM_019833233.1;refGene_gn=RGS6;refGene_tx=XM_019833234.2;refGene_gn=RGS6;refGene_tx=XM_019833235.1;refGene_gn=RGS6;refGene_tx=XM_019833236.2;refGene_gn=RGS6;refGene_tx=XM_019833237.2;refGene_gn=RGS6;refGene_tx=XM_019833238.2;refGene_gn=RGS6;refGene_tx=XM_019833239.2;refGene_gn=RGS6;refGene_tx=XM_019833242.2
B3 119905568 119905668
refGene_gn=RGS6;refGene_tx=XM_006932961.4;refGene_gn=RGS6;refGene_tx=XM_011283315.2;refGene_gn=RGS6;refGene_tx=XM_011283317.2;refGene_gn=RGS6;refGene_tx=XM_011283318.2;refGene_gn=RGS6;refGene_tx=XM_019833233.1;refGene_gn=RGS6;refGene_tx=XM_019833234.2;refGene_gn=RGS6;refGene_tx=XM_019833235.1;refGene_gn=RGS6;refGene_tx=XM_019833236.2;refGene_gn=RGS6;refGene_tx=XM_019833237.2;refGene_gn=RGS6;refGene_tx=XM_019833238.2;refGene_gn=RGS6;refGene_tx=XM_019833239.2;refGene_gn=RGS6;refGene_tx=XM_019833242.2
B3 119987068 119987119
refGene_gn=RGS6;refGene_tx=XM_006932961.4;refGene_gn=RGS6;refGene_tx=XM_011283315.2;refGene_gn=RGS6;refGene_tx=XM_011283317.2;refGene_gn=RGS6;refGene_tx=XM_011283318.2;refGene_gn=RGS6;refGene_tx=XM_019833233.1;refGene_gn=RGS6;refGene_tx=XM_019833234.2;refGene_gn=RGS6;refGene_tx=XM_019833235.1;refGene_gn=RGS6;refGene_tx=XM_019833236.2;refGene_gn=RGS6;refGene_tx=XM_019833237.2;refGene_gn=RGS6;refGene_tx=XM_019833238.2;

refGene_gn=RGS6;refGene_tx=XM_019833239.2;refGene_gn=RGS6;refGene_tx=XM_019833242.2
B3 119989551 119989658
refGene_gn=RGS6;refGene_tx=XM_006932961.4;refGene_gn=RGS6;refGene_tx=XM_011283315.2;refGene_gn=RGS6;refGene_tx=XM_011283317.2;refGene_gn=RGS6;refGene_tx=XM_011283318.2;refGene_gn=RGS6;refGene_tx=XM_019833233.1;refGene_gn=RGS6;refGene_tx=XM_019833234.2;refGene_gn=RGS6;refGene_tx=XM_019833235.1;refGene_gn=RGS6;refGene_tx=XM_019833236.2;refGene_gn=RGS6;refGene_tx=XM_019833237.2;refGene_gn=RGS6;refGene_tx=XM_019833238.2;refGene_gn=RGS6;refGene_tx=XM_019833239.2;refGene_gn=RGS6;refGene_tx=XM_019833242.2
B3 119990865 119990917
refGene_gn=RGS6;refGene_tx=XM_006932961.4;refGene_gn=RGS6;refGene_tx=XM_011283315.2;refGene_gn=RGS6;refGene_tx=XM_011283317.2;refGene_gn=RGS6;refGene_tx=XM_011283318.2;refGene_gn=RGS6;refGene_tx=XM_019833233.1;refGene_gn=RGS6;refGene_tx=XM_019833234.2;refGene_gn=RGS6;refGene_tx=XM_019833235.1;refGene_gn=RGS6;refGene_tx=XM_019833236.2;refGene_gn=RGS6;refGene_tx=XM_019833237.2;refGene_gn=RGS6;refGene_tx=XM_019833238.2;refGene_gn=RGS6;refGene_tx=XM_019833239.2;refGene_gn=RGS6;refGene_tx=XM_019833242.2
B3 119996319 119996384
refGene_gn=RGS6;refGene_tx=XM_006932961.4;refGene_gn=RGS6;refGene_tx=XM_011283315.2;refGene_gn=RGS6;refGene_tx=XM_011283317.2;refGene_gn=RGS6;refGene_tx=XM_011283318.2;refGene_gn=RGS6;refGene_tx=XM_019833233.1;refGene_gn=RGS6;refGene_tx=XM_019833234.2;refGene_gn=RGS6;refGene_tx=XM_019833235.1;refGene_gn=RGS6;refGene_tx=XM_019833236.2;refGene_gn=RGS6;refGene_tx=XM_019833237.2;refGene_gn=RGS6;refGene_tx=XM_019833238.2;refGene_gn=RGS6;refGene_tx=XM_019833239.2;refGene_gn=RGS6;refGene_tx=XM_019833242.2
B3 120000796 120000873
refGene_gn=RGS6;refGene_tx=XM_006932961.4;refGene_gn=RGS6;refGene_tx=XM_011283315.2;refGene_gn=RGS6;refGene_tx=XM_011283317.2;refGene_gn=RGS6;refGene_tx=XM_011283318.2;refGene_gn=RGS6;refGene_tx=XM_019833233.1;refGene_gn=RGS6;refGene_tx=XM_019833234.2;refGene_gn=RGS6;refGene_tx=XM_019833235.1;refGene_gn=RGS6;refGene_tx=XM_019833236.2;refGene_gn=RGS6;refGene_tx=XM_019833237.2;refGene_gn=RGS6;refGene_tx=XM_019833238.2;refGene_gn=RGS6;refGene_tx=XM_019833239.2;refGene_gn=RGS6;refGene_tx=XM_019833242.2
B3 120003273 120003355
refGene_gn=RGS6;refGene_tx=XM_006932961.4;refGene_gn=RGS6;refGene_tx=XM_011283315.2;refGene_gn=RGS6;refGene_tx=XM_011283317.2;refGene_gn=RGS6;refGene_tx=XM_011283318.2;refGene_gn=RGS6;refGene_tx=XM_019833233.1;refGene_gn=RGS6;refGene_tx=XM_019833234.2;refGene_gn=RGS6;refGene_tx=XM_019833235.1;refGene_gn=RGS6;refGene_tx=XM_019833236.2;refGene_gn=RGS6;refGene_tx=XM_019833237.2;refGene_gn=RGS6;refGene_tx=XM_019833238.2;refGene_gn=RGS6;refGene_tx=XM_019833239.2;refGene_gn=RGS6;refGene_tx=XM_019833242.2
B3 120005139 120005214
refGene_gn=RGS6;refGene_tx=XM_006932961.4;refGene_gn=RGS6;refGene_tx=XM_011283315.2;refGene_gn=RGS6;refGene_tx=XM_011283317.2;refGene_gn=RGS

6;refGene_tx=XM_011283318.2;refGene_gn=RGS6;refGene_tx=XM_019833233.1;
refGene_gn=RGS6;refGene_tx=XM_019833234.2;refGene_gn=RGS6;refGene_tx=X
M_019833235.1;refGene_gn=RGS6;refGene_tx=XM_019833236.2;refGene_gn=RGS
6;refGene_tx=XM_019833237.2;refGene_gn=RGS6;refGene_tx=XM_019833238.2;
refGene_gn=RGS6;refGene_tx=XM_019833239.2;refGene_gn=RGS6;refGene_tx=X
M_019833242.2
B3 120006880 120006979
refGene_gn=RGS6;refGene_tx=XM_006932961.4;refGene_gn=RGS6;refGene_tx=X
M_011283315.2;refGene_gn=RGS6;refGene_tx=XM_011283317.2;refGene_gn=RGS
6;refGene_tx=XM_011283318.2;refGene_gn=RGS6;refGene_tx=XM_019833233.1;
refGene_gn=RGS6;refGene_tx=XM_019833234.2;refGene_gn=RGS6;refGene_tx=X
M_019833235.1;refGene_gn=RGS6;refGene_tx=XM_019833236.2;refGene_gn=RGS
6;refGene_tx=XM_019833237.2;refGene_gn=RGS6;refGene_tx=XM_019833238.2;
refGene_gn=RGS6;refGene_tx=XM_019833239.2;refGene_gn=RGS6;refGene_tx=X
M_019833242.2
B3 120008113 120008175
refGene_gn=RGS6;refGene_tx=XM_006932961.4;refGene_gn=RGS6;refGene_tx=X
M_011283315.2;refGene_gn=RGS6;refGene_tx=XM_011283317.2;refGene_gn=RGS
6;refGene_tx=XM_011283318.2;refGene_gn=RGS6;refGene_tx=XM_019833233.1;
refGene_gn=RGS6;refGene_tx=XM_019833234.2;refGene_gn=RGS6;refGene_tx=X
M_019833235.1;refGene_gn=RGS6;refGene_tx=XM_019833236.2;refGene_gn=RGS
6;refGene_tx=XM_019833237.2;refGene_gn=RGS6;refGene_tx=XM_019833238.2;
refGene_gn=RGS6;refGene_tx=XM_019833239.2;refGene_gn=RGS6;refGene_tx=X
M_019833242.2
B3 120024656 120024767
refGene_gn=RGS6;refGene_tx=XM_006932961.4;refGene_gn=RGS6;refGene_tx=X
M_011283315.2;refGene_gn=RGS6;refGene_tx=XM_011283317.2;refGene_gn=RGS
6;refGene_tx=XM_011283318.2;refGene_gn=RGS6;refGene_tx=XM_019833233.1;
refGene_gn=RGS6;refGene_tx=XM_019833234.2;refGene_gn=RGS6;refGene_tx=X
M_019833235.1;refGene_gn=RGS6;refGene_tx=XM_019833236.2;refGene_gn=RGS
6;refGene_tx=XM_019833237.2;refGene_gn=RGS6;refGene_tx=XM_019833238.2;
refGene_gn=RGS6;refGene_tx=XM_019833239.2
B3 120036625 120036751
refGene_gn=RGS6;refGene_tx=XM_006932961.4;refGene_gn=RGS6;refGene_tx=X
M_011283315.2;refGene_gn=RGS6;refGene_tx=XM_011283317.2;refGene_gn=RGS
6;refGene_tx=XM_011283318.2;refGene_gn=RGS6;refGene_tx=XM_019833233.1;
refGene_gn=RGS6;refGene_tx=XM_019833234.2;refGene_gn=RGS6;refGene_tx=X
M_019833235.1;refGene_gn=RGS6;refGene_tx=XM_019833236.2;refGene_gn=RGS
6;refGene_tx=XM_019833237.2;refGene_gn=RGS6;refGene_tx=XM_019833238.2;
refGene_gn=RGS6;refGene_tx=XM_019833239.2;refGene_gn=RGS6;refGene_tx=X
M_019833242.2
B3 120044722 120044909
refGene_gn=RGS6;refGene_tx=XM_006932961.4;refGene_gn=RGS6;refGene_tx=X
M_011283315.2;refGene_gn=RGS6;refGene_tx=XM_011283317.2;refGene_gn=RGS
6;refGene_tx=XM_011283318.2;refGene_gn=RGS6;refGene_tx=XM_019833233.1;
refGene_gn=RGS6;refGene_tx=XM_019833234.2;refGene_gn=RGS6;refGene_tx=X
M_019833235.1;refGene_gn=RGS6;refGene_tx=XM_019833236.2;refGene_gn=RGS
6;refGene_tx=XM_019833237.2;refGene_gn=RGS6;refGene_tx=XM_019833238.2;
refGene_gn=RGS6;refGene_tx=XM_019833239.2;refGene_gn=RGS6;refGene_tx=X
M_019833242.2

B3	120059756	120059846
refGene_gn=RGS6;refGene_tx=XM_006932961.4;refGene_gn=RGS6;refGene_tx=XM_011283315.2;refGene_gn=RGS6;refGene_tx=XM_011283317.2;refGene_gn=RGS6;refGene_tx=XM_011283318.2;refGene_gn=RGS6;refGene_tx=XM_019833233.1;refGene_gn=RGS6;refGene_tx=XM_019833234.2;refGene_gn=RGS6;refGene_tx=XM_019833235.1;refGene_gn=RGS6;refGene_tx=XM_019833236.2;refGene_gn=RGS6;refGene_tx=XM_019833237.2;refGene_gn=RGS6;refGene_tx=XM_019833238.2;refGene_gn=RGS6;refGene_tx=XM_019833239.2;refGene_gn=RGS6;refGene_tx=XM_019833242.2
B3	120063874	120063928
refGene_gn=RGS6;refGene_tx=XM_006932961.4;refGene_gn=RGS6;refGene_tx=XM_011283315.2;refGene_gn=RGS6;refGene_tx=XM_011283317.2;refGene_gn=RGS6;refGene_tx=XM_019833233.1;refGene_gn=RGS6;refGene_tx=XM_019833234.2;refGene_gn=RGS6;refGene_tx=XM_019833235.1;refGene_gn=RGS6;refGene_tx=XM_019833236.2;refGene_gn=RGS6;refGene_tx=XM_019833237.2;refGene_gn=RGS6;refGene_tx=XM_019833238.2;refGene_gn=RGS6;refGene_tx=XM_019833239.2;refGene_gn=RGS6;refGene_tx=XM_019833242.2
B3	120076026	120076093
refGene_gn=RGS6;refGene_tx=XM_011283315.2;refGene_gn=RGS6;refGene_tx=XM_019833233.1;refGene_gn=RGS6;refGene_tx=XM_019833234.2;refGene_gn=RGS6;refGene_tx=XM_019833235.1;refGene_gn=RGS6;refGene_tx=XM_019833236.2;refGene_gn=RGS6;refGene_tx=XM_019833237.2;refGene_gn=RGS6;refGene_tx=XM_019833238.2;refGene_gn=RGS6;refGene_tx=XM_019833239.2;refGene_gn=RGS6;refGene_tx=XM_019833242.2
B3	120084953	120085120
refGene_gn=RGS6;refGene_tx=XM_011283315.2;refGene_gn=RGS6;refGene_tx=XM_011283317.2;refGene_gn=RGS6;refGene_tx=XM_011283318.2;refGene_gn=RGS6;refGene_tx=XM_019833233.1;refGene_gn=RGS6;refGene_tx=XM_019833234.2;refGene_gn=RGS6;refGene_tx=XM_019833235.1;refGene_gn=RGS6;refGene_tx=XM_019833236.2;refGene_gn=RGS6;refGene_tx=XM_019833237.2;refGene_gn=RGS6;refGene_tx=XM_019833238.2;refGene_gn=RGS6;refGene_tx=XM_019833239.2;refGene_gn=RGS6;refGene_tx=XM_019833242.2
B3	120097636	120097669
refGene_gn=RGS6;refGene_tx=XM_006932961.4
B3	120143124	120143195
refGene_gn=DPF3;refGene_tx=XM_011283320.2
B3	120143714	120143796
refGene_gn=DPF3;refGene_tx=XM_011283320.2
B3	120153922	120154035
refGene_gn=DPF3;refGene_tx=XM_011283320.2
B3	120194720	120194827
refGene_gn=DPF3;refGene_tx=XM_023255803.1;refGene_gn=DPF3;refGene_tx=XM_023255804.1
B3	120195902	120196102
refGene_gn=DPF3;refGene_tx=XM_023255802.1
B3	120199004	120199133
refGene_gn=DPF3;refGene_tx=XM_011283320.2;refGene_gn=DPF3;refGene_tx=XM_023255802.1;refGene_gn=DPF3;refGene_tx=XM_023255803.1;refGene_gn=DPF3;refGene_tx=XM_023255804.1
B3	120216442	120216580

refGene_gn=DPF3;refGene_tx=XM_011283320.2;refGene_gn=DPF3;refGene_tx=XM_023255802.1;refGene_gn=DPF3;refGene_tx=XM_023255803.1;refGene_gn=DPF3;refGene_tx=XM_023255804.1
B3 120236844 120236923
refGene_gn=DPF3;refGene_tx=XM_011283320.2;refGene_gn=DPF3;refGene_tx=XM_023255802.1;refGene_gn=DPF3;refGene_tx=XM_023255803.1;refGene_gn=DPF3;refGene_tx=XM_023255804.1
B3 120245263 120245359
refGene_gn=DPF3;refGene_tx=XM_011283320.2;refGene_gn=DPF3;refGene_tx=XM_023255802.1;refGene_gn=DPF3;refGene_tx=XM_023255803.1;refGene_gn=DPF3;refGene_tx=XM_023255804.1
B3 120252332 120252460
refGene_gn=DPF3;refGene_tx=XM_011283320.2;refGene_gn=DPF3;refGene_tx=XM_023255802.1;refGene_gn=DPF3;refGene_tx=XM_023255803.1;refGene_gn=DPF3;refGene_tx=XM_023255804.1
B3 120270240 120270348
refGene_gn=DPF3;refGene_tx=XM_011283320.2;refGene_gn=DPF3;refGene_tx=XM_023255802.1;refGene_gn=DPF3;refGene_tx=XM_023255803.1;refGene_gn=DPF3;refGene_tx=XM_023255804.1
B3 120290601 120290762
refGene_gn=DPF3;refGene_tx=XM_011283320.2;refGene_gn=DPF3;refGene_tx=XM_023255802.1;refGene_gn=DPF3;refGene_tx=XM_023255803.1;refGene_gn=DPF3;refGene_tx=XM_023255804.1
B3 120402509 120402541
refGene_gn=DPF3;refGene_tx=XM_011283320.2;refGene_gn=DPF3;refGene_tx=XM_023255802.1;refGene_gn=DPF3;refGene_tx=XM_023255803.1;refGene_gn=DPF3;refGene_tx=XM_023255804.1
B3 120441297 120441352
refGene_gn=DCAF4;refGene_tx=XM_023255805.1
B3 120453629 120453726
refGene_gn=DCAF4;refGene_tx=XM_006932965.3;refGene_gn=DCAF4;refGene_tx=XM_006932967.3;refGene_gn=DCAF4;refGene_tx=XM_019833251.2;refGene_gn=DCAF4;refGene_tx=XM_023255805.1;refGene_gn=DCAF4;refGene_tx=XM_023255806.1
B3 120455885 120455989
refGene_gn=DCAF4;refGene_tx=XM_006932965.3;refGene_gn=DCAF4;refGene_tx=XM_006932967.3;refGene_gn=DCAF4;refGene_tx=XM_019833251.2;refGene_gn=DCAF4;refGene_tx=XM_023255805.1;refGene_gn=DCAF4;refGene_tx=XM_023255806.1
B3 120456414 120456572
refGene_gn=DCAF4;refGene_tx=XM_006932965.3;refGene_gn=DCAF4;refGene_tx=XM_006932967.3;refGene_gn=DCAF4;refGene_tx=XM_019833251.2;refGene_gn=DCAF4;refGene_tx=XM_023255805.1;refGene_gn=DCAF4;refGene_tx=XM_023255806.1
B3 120457269 120457352
refGene_gn=DCAF4;refGene_tx=XM_006932965.3;refGene_gn=DCAF4;refGene_tx=XM_006932967.3;refGene_gn=DCAF4;refGene_tx=XM_019833251.2;refGene_gn=DCAF4;refGene_tx=XM_023255805.1;refGene_gn=DCAF4;refGene_tx=XM_023255806.1
B3 120458603 120458706

refGene_gn=DCAF4;refGene_tx=XM_006932965.3;refGene_gn=DCAF4;refGene_tx=XM_006932967.3;refGene_gn=DCAF4;refGene_tx=XM_019833251.2;refGene_gn=DCAF4;refGene_tx=XM_023255805.1;refGene_gn=DCAF4;refGene_tx=XM_023255806.1
B3 120461234 120461378
refGene_gn=DCAF4;refGene_tx=XM_006932965.3;refGene_gn=DCAF4;refGene_tx=XM_006932967.3;refGene_gn=DCAF4;refGene_tx=XM_019833251.2;refGene_gn=DCAF4;refGene_tx=XM_023255805.1;refGene_gn=DCAF4;refGene_tx=XM_023255806.1
B3 120462338 120462388
refGene_gn=DCAF4;refGene_tx=XM_006932965.3;refGene_gn=DCAF4;refGene_tx=XM_006932967.3;refGene_gn=DCAF4;refGene_tx=XM_019833251.2;refGene_gn=DCAF4;refGene_tx=XM_023255805.1;refGene_gn=DCAF4;refGene_tx=XM_023255806.1
B3 120466579 120466659
refGene_gn=DCAF4;refGene_tx=XM_006932965.3;refGene_gn=DCAF4;refGene_tx=XM_006932967.3;refGene_gn=DCAF4;refGene_tx=XM_019833251.2;refGene_gn=DCAF4;refGene_tx=XM_023255805.1;refGene_gn=DCAF4;refGene_tx=XM_023255806.1
B3 120467933 120468029
refGene_gn=DCAF4;refGene_tx=XM_006932965.3;refGene_gn=DCAF4;refGene_tx=XM_006932967.3;refGene_gn=DCAF4;refGene_tx=XM_019833251.2;refGene_gn=DCAF4;refGene_tx=XM_023255805.1;refGene_gn=DCAF4;refGene_tx=XM_023255806.1
B3 120468141 120468239
refGene_gn=DCAF4;refGene_tx=XM_006932965.3;refGene_gn=DCAF4;refGene_tx=XM_006932967.3;refGene_gn=DCAF4;refGene_tx=XM_019833251.2;refGene_gn=DCAF4;refGene_tx=XM_023255805.1;refGene_gn=DCAF4;refGene_tx=XM_023255806.1
B3 120469230 120469404
refGene_gn=DCAF4;refGene_tx=XM_006932965.3;refGene_gn=DCAF4;refGene_tx=XM_006932967.3;refGene_gn=DCAF4;refGene_tx=XM_023255805.1;refGene_gn=DCAF4;refGene_tx=XM_023255806.1
B3 120469709 120469824
refGene_gn=DCAF4;refGene_tx=XM_006932965.3;refGene_gn=DCAF4;refGene_tx=XM_006932967.3;refGene_gn=DCAF4;refGene_tx=XM_019833251.2;refGene_gn=DCAF4;refGene_tx=XM_023255805.1;refGene_gn=DCAF4;refGene_tx=XM_023255806.1
B3 120471572 120471766
refGene_gn=DCAF4;refGene_tx=XM_006932965.3;refGene_gn=DCAF4;refGene_tx=XM_006932967.3;refGene_gn=DCAF4;refGene_tx=XM_019833251.2;refGene_gn=DCAF4;refGene_tx=XM_023255805.1;refGene_gn=DCAF4;refGene_tx=XM_023255806.1
B3 120479454 120479687
refGene_gn=ZFYVE1;refGene_tx=XM_003987802.5;refGene_gn=ZFYVE1;refGene_tx=XM_003987803.5;refGene_gn=ZFYVE1;refGene_tx=XM_023255807.1;refGene_gn=ZFYVE1;refGene_tx=XM_023255808.1
B3 120481129 120481243
refGene_gn=ZFYVE1;refGene_tx=XM_003987802.5;refGene_gn=ZFYVE1;refGene_tx=XM_003987803.5;refGene_gn=ZFYVE1;refGene_tx=XM_023255807.1;refGene_

gn=ZFYVE1;refGene_tx=XM_023255808.1
B3 120481918 120482099
refGene_gn=ZFYVE1;refGene_tx=XM_003987802.5;refGene_gn=ZFYVE1;refGene_tx=XM_003987803.5;refGene_gn=ZFYVE1;refGene_tx=XM_023255807.1;refGene_gn=ZFYVE1;refGene_tx=XM_023255808.1
B3 120482835 120483006
refGene_gn=ZFYVE1;refGene_tx=XM_003987802.5;refGene_gn=ZFYVE1;refGene_tx=XM_003987803.5;refGene_gn=ZFYVE1;refGene_tx=XM_023255807.1;refGene_gn=ZFYVE1;refGene_tx=XM_023255808.1
B3 120485369 120485487
refGene_gn=ZFYVE1;refGene_tx=XM_003987802.5;refGene_gn=ZFYVE1;refGene_tx=XM_003987803.5;refGene_gn=ZFYVE1;refGene_tx=XM_023255807.1;refGene_gn=ZFYVE1;refGene_tx=XM_023255808.1
B3 120485583 120485681
refGene_gn=ZFYVE1;refGene_tx=XM_003987802.5;refGene_gn=ZFYVE1;refGene_tx=XM_003987803.5;refGene_gn=ZFYVE1;refGene_tx=XM_023255807.1;refGene_gn=ZFYVE1;refGene_tx=XM_023255808.1
B3 120486237 120486346
refGene_gn=ZFYVE1;refGene_tx=XM_003987802.5;refGene_gn=ZFYVE1;refGene_tx=XM_003987803.5;refGene_gn=ZFYVE1;refGene_tx=XM_023255807.1;refGene_gn=ZFYVE1;refGene_tx=XM_023255808.1
B3 120488315 120488422
refGene_gn=ZFYVE1;refGene_tx=XM_003987802.5;refGene_gn=ZFYVE1;refGene_tx=XM_003987803.5;refGene_gn=ZFYVE1;refGene_tx=XM_023255807.1;refGene_gn=ZFYVE1;refGene_tx=XM_023255808.1
B3 120503103 120503318
refGene_gn=ZFYVE1;refGene_tx=XM_003987802.5
B3 120507792 120508297
refGene_gn=ZFYVE1;refGene_tx=XM_003987802.5
B3 120529828 120530311
refGene_gn=ZFYVE1;refGene_tx=XM_003987802.5
B3 120567151 120567193
refGene_gn=RBM25;refGene_tx=XM_006932968.4;refGene_gn=RBM25;refGene_tx=XM_023255810.1
B3 120567305 120567426
refGene_gn=RBM25;refGene_tx=XM_003987804.5;refGene_gn=RBM25;refGene_tx=XM_006932968.4;refGene_gn=RBM25;refGene_tx=XM_023255809.1;refGene_gn=RBM25;refGene_tx=XM_023255810.1
B3 120573936 120573986
refGene_gn=RBM25;refGene_tx=XM_003987804.5;refGene_gn=RBM25;refGene_tx=XM_006932968.4;refGene_gn=RBM25;refGene_tx=XM_023255809.1;refGene_gn=RBM25;refGene_tx=XM_023255810.1
B3 120575565 120575733
refGene_gn=RBM25;refGene_tx=XM_003987804.5;refGene_gn=RBM25;refGene_tx=XM_006932968.4;refGene_gn=RBM25;refGene_tx=XM_023255809.1;refGene_gn=RBM25;refGene_tx=XM_023255810.1
B3 120581602 120581660
refGene_gn=RBM25;refGene_tx=XM_003987804.5;refGene_gn=RBM25;refGene_tx=XM_006932968.4;refGene_gn=RBM25;refGene_tx=XM_023255809.1;refGene_gn=RBM25;refGene_tx=XM_023255810.1

B3 120586875 120587036
refGene_gn=RBM25;refGene_tx=XM_003987804.5;refGene_gn=RBM25;refGene_tx=XM_006932968.4;refGene_gn=RBM25;refGene_tx=XM_023255809.1;refGene_gn=RBM25;refGene_tx=XM_023255810.1
B3 120595901 120596087
refGene_gn=RBM25;refGene_tx=XM_003987804.5;refGene_gn=RBM25;refGene_tx=XM_006932968.4;refGene_gn=RBM25;refGene_tx=XM_023255809.1;refGene_gn=RBM25;refGene_tx=XM_023255810.1
B3 120597459 120597513
refGene_gn=RBM25;refGene_tx=XM_003987804.5;refGene_gn=RBM25;refGene_tx=XM_006932968.4;refGene_gn=RBM25;refGene_tx=XM_023255809.1;refGene_gn=RBM25;refGene_tx=XM_023255810.1
B3 120597751 120597835
refGene_gn=RBM25;refGene_tx=XM_003987804.5;refGene_gn=RBM25;refGene_tx=XM_006932968.4;refGene_gn=RBM25;refGene_tx=XM_023255809.1;refGene_gn=RBM25;refGene_tx=XM_023255810.1
B3 120601367 120601654
refGene_gn=RBM25;refGene_tx=XM_003987804.5;refGene_gn=RBM25;refGene_tx=XM_006932968.4;refGene_gn=RBM25;refGene_tx=XM_023255809.1;refGene_gn=RBM25;refGene_tx=XM_023255810.1
B3 120603002 120603225
refGene_gn=RBM25;refGene_tx=XM_003987804.5;refGene_gn=RBM25;refGene_tx=XM_006932968.4;refGene_gn=RBM25;refGene_tx=XM_023255809.1;refGene_gn=RBM25;refGene_tx=XM_023255810.1
B3 120603528 120603618
refGene_gn=RBM25;refGene_tx=XM_003987804.5;refGene_gn=RBM25;refGene_tx=XM_006932968.4;refGene_gn=RBM25;refGene_tx=XM_023255809.1;refGene_gn=RBM25;refGene_tx=XM_023255810.1
B3 120606134 120606208
refGene_gn=RBM25;refGene_tx=XM_003987804.5;refGene_gn=RBM25;refGene_tx=XM_006932968.4;refGene_gn=RBM25;refGene_tx=XM_023255809.1;refGene_gn=RBM25;refGene_tx=XM_023255810.1
B3 120608577 120608728
refGene_gn=RBM25;refGene_tx=XM_003987804.5;refGene_gn=RBM25;refGene_tx=XM_006932968.4;refGene_gn=RBM25;refGene_tx=XM_023255809.1;refGene_gn=RBM25;refGene_tx=XM_023255810.1
B3 120613201 120613526
refGene_gn=RBM25;refGene_tx=XM_003987804.5;refGene_gn=RBM25;refGene_tx=XM_006932968.4;refGene_gn=RBM25;refGene_tx=XM_023255809.1;refGene_gn=RBM25;refGene_tx=XM_023255810.1
B3 120613679 120613795
refGene_gn=RBM25;refGene_tx=XM_023255810.1
B3 120613898 120614173
refGene_gn=RBM25;refGene_tx=XM_003987804.5;refGene_gn=RBM25;refGene_tx=XM_006932968.4;refGene_gn=RBM25;refGene_tx=XM_023255809.1
B3 120614514 120614613
refGene_gn=RBM25;refGene_tx=XM_003987804.5;refGene_gn=RBM25;refGene_tx=XM_006932968.4;refGene_gn=RBM25;refGene_tx=XM_023255809.1
B3 120616133 120616181
refGene_gn=RBM25;refGene_tx=XM_003987804.5;refGene_gn=RBM25;refGene_tx

=XM_006932968.4;refGene_gn=RBM25;refGene_tx=XM_023255809.1
B3 120619960 120620053
refGene_gn=RBM25;refGene_tx=XM_003987804.5;refGene_gn=RBM25;refGene_tx=XM_006932968.4;refGene_gn=RBM25;refGene_tx=XM_023255809.1
B3 120640631 120640718
refGene_gn=PSEN1;refGene_tx=XM_003987805.4;refGene_gn=PSEN1;refGene_tx=XM_006932969.3;refGene_gn=PSEN1;refGene_tx=XM_019833255.2
B3 120664461 120664712
refGene_gn=PSEN1;refGene_tx=XM_003987805.4;refGene_gn=PSEN1;refGene_tx=XM_006932969.3;refGene_gn=PSEN1;refGene_tx=XM_019833255.2
B3 120667811 120667953
refGene_gn=PSEN1;refGene_tx=XM_003987805.4;refGene_gn=PSEN1;refGene_tx=XM_006932969.3;refGene_gn=PSEN1;refGene_tx=XM_019833255.2
B3 120673800 120673868
refGene_gn=PSEN1;refGene_tx=XM_003987805.4;refGene_gn=PSEN1;refGene_tx=XM_006932969.3;refGene_gn=PSEN1;refGene_tx=XM_019833255.2
B3 120675160 120675381
refGene_gn=PSEN1;refGene_tx=XM_003987805.4;refGene_gn=PSEN1;refGene_tx=XM_006932969.3;refGene_gn=PSEN1;refGene_tx=XM_019833255.2
B3 120680473 120680572
refGene_gn=PSEN1;refGene_tx=XM_003987805.4;refGene_gn=PSEN1;refGene_tx=XM_006932969.3;refGene_gn=PSEN1;refGene_tx=XM_019833255.2
B3 120690269 120690356
refGene_gn=PSEN1;refGene_tx=XM_003987805.4;refGene_gn=PSEN1;refGene_tx=XM_006932969.3;refGene_gn=PSEN1;refGene_tx=XM_019833255.2
B3 120693090 120693264
refGene_gn=PSEN1;refGene_tx=XM_003987805.4;refGene_gn=PSEN1;refGene_tx=XM_006932969.3;refGene_gn=PSEN1;refGene_tx=XM_019833255.2
B3 120700188 120700307
refGene_gn=PSEN1;refGene_tx=XM_003987805.4;refGene_gn=PSEN1;refGene_tx=XM_006932969.3;refGene_gn=PSEN1;refGene_tx=XM_019833255.2
B3 120702289 120702445
refGene_gn=PSEN1;refGene_tx=XM_003987805.4;refGene_gn=PSEN1;refGene_tx=XM_006932969.3;refGene_gn=PSEN1;refGene_tx=XM_019833255.2
B3 120722931 120722997
refGene_gn=PAPLN;refGene_tx=XM_019833252.2;refGene_gn=PAPLN;refGene_tx=XM_023255811.1;refGene_gn=PAPLN;refGene_tx=XM_023255812.1
B3 120725295 120725411
refGene_gn=PAPLN;refGene_tx=XM_019833252.2;refGene_gn=PAPLN;refGene_tx=XM_023255811.1;refGene_gn=PAPLN;refGene_tx=XM_023255812.1
B3 120726253 120726314
refGene_gn=PAPLN;refGene_tx=XM_019833252.2;refGene_gn=PAPLN;refGene_tx=XM_023255811.1;refGene_gn=PAPLN;refGene_tx=XM_023255812.1
B3 120726568 120726671
refGene_gn=PAPLN;refGene_tx=XM_019833252.2;refGene_gn=PAPLN;refGene_tx=XM_023255811.1;refGene_gn=PAPLN;refGene_tx=XM_023255812.1
B3 120728712 120728843
refGene_gn=PAPLN;refGene_tx=XM_019833252.2;refGene_gn=PAPLN;refGene_tx=XM_023255811.1;refGene_gn=PAPLN;refGene_tx=XM_023255812.1
B3 120729657 120729781

refGene_gn=PAPLN;refGene_tx=XM_019833252.2;refGene_gn=PAPLN;refGene_tx=XM_023255811.1;refGene_gn=PAPLN;refGene_tx=XM_023255812.1
B3 120730147 120730228
refGene_gn=PAPLN;refGene_tx=XM_019833252.2;refGene_gn=PAPLN;refGene_tx=XM_023255811.1;refGene_gn=PAPLN;refGene_tx=XM_023255812.1
B3 120730312 120730485
refGene_gn=PAPLN;refGene_tx=XM_019833252.2;refGene_gn=PAPLN;refGene_tx=XM_023255811.1;refGene_gn=PAPLN;refGene_tx=XM_023255812.1
B3 120730605 120730729
refGene_gn=PAPLN;refGene_tx=XM_019833252.2;refGene_gn=PAPLN;refGene_tx=XM_023255811.1;refGene_gn=PAPLN;refGene_tx=XM_023255812.1
B3 120731124 120731251
refGene_gn=PAPLN;refGene_tx=XM_019833252.2;refGene_gn=PAPLN;refGene_tx=XM_023255811.1;refGene_gn=PAPLN;refGene_tx=XM_023255812.1
B3 120731619 120731685
refGene_gn=PAPLN;refGene_tx=XM_019833252.2;refGene_gn=PAPLN;refGene_tx=XM_023255811.1
B3 120732230 120732438
refGene_gn=PAPLN;refGene_tx=XM_019833252.2;refGene_gn=PAPLN;refGene_tx=XM_023255811.1;refGene_gn=PAPLN;refGene_tx=XM_023255812.1
B3 120733020 120733203
refGene_gn=PAPLN;refGene_tx=XM_019833252.2;refGene_gn=PAPLN;refGene_tx=XM_023255811.1;refGene_gn=PAPLN;refGene_tx=XM_023255812.1
B3 120733398 120733540
refGene_gn=PAPLN;refGene_tx=XM_019833252.2;refGene_gn=PAPLN;refGene_tx=XM_023255811.1;refGene_gn=PAPLN;refGene_tx=XM_023255812.1
B3 120741474 120741555
refGene_gn=PAPLN;refGene_tx=XM_019833252.2;refGene_gn=PAPLN;refGene_tx=XM_023255811.1;refGene_gn=PAPLN;refGene_tx=XM_023255812.1
B3 120741720 120741961
refGene_gn=PAPLN;refGene_tx=XM_019833252.2;refGene_gn=PAPLN;refGene_tx=XM_023255811.1;refGene_gn=PAPLN;refGene_tx=XM_023255812.1
B3 120742961 120743082
refGene_gn=PAPLN;refGene_tx=XM_019833252.2;refGene_gn=PAPLN;refGene_tx=XM_023255811.1;refGene_gn=PAPLN;refGene_tx=XM_023255812.1
B3 120743350 120743489
refGene_gn=PAPLN;refGene_tx=XM_019833252.2;refGene_gn=PAPLN;refGene_tx=XM_023255811.1;refGene_gn=PAPLN;refGene_tx=XM_023255812.1
B3 120744550 120745028
refGene_gn=PAPLN;refGene_tx=XM_019833252.2;refGene_gn=PAPLN;refGene_tx=XM_023255811.1;refGene_gn=PAPLN;refGene_tx=XM_023255812.1
B3 120745735 120745873
refGene_gn=PAPLN;refGene_tx=XM_019833252.2;refGene_gn=PAPLN;refGene_tx=XM_023255811.1;refGene_gn=PAPLN;refGene_tx=XM_023255812.1
B3 120745979 120746104
refGene_gn=PAPLN;refGene_tx=XM_019833252.2;refGene_gn=PAPLN;refGene_tx=XM_023255811.1;refGene_gn=PAPLN;refGene_tx=XM_023255812.1
B3 120746397 120746551
refGene_gn=PAPLN;refGene_tx=XM_019833252.2;refGene_gn=PAPLN;refGene_tx=XM_023255811.1;refGene_gn=PAPLN;refGene_tx=XM_023255812.1

B3 120747117 120747255
refGene_gn=PAPLN;refGene_tx=XM_019833252.2;refGene_gn=PAPLN;refGene_tx=XM_023255811.1;refGene_gn=PAPLN;refGene_tx=XM_023255812.1
B3 120747966 120748094
refGene_gn=PAPLN;refGene_tx=XM_019833252.2;refGene_gn=PAPLN;refGene_tx=XM_023255811.1;refGene_gn=PAPLN;refGene_tx=XM_023255812.1
B3 120748177 120748286
refGene_gn=PAPLN;refGene_tx=XM_019833252.2;refGene_gn=PAPLN;refGene_tx=XM_023255811.1;refGene_gn=PAPLN;refGene_tx=XM_023255812.1
B3 120750596 120750763
refGene_gn=PAPLN;refGene_tx=XM_019833252.2;refGene_gn=PAPLN;refGene_tx=XM_023255811.1;refGene_gn=PAPLN;refGene_tx=XM_023255812.1
B3 120754448 120754624
refGene_gn=PAPLN;refGene_tx=XM_019833252.2;refGene_gn=PAPLN;refGene_tx=XM_023255811.1;refGene_gn=PAPLN;refGene_tx=XM_023255812.1
B3 120758385 120759101
refGene_gn=NUMB;refGene_tx=XM_003987807.4;refGene_gn=NUMB;refGene_tx=XM_019833256.2;refGene_gn=NUMB;refGene_tx=XM_019833257.2;refGene_gn=NUMB;refGene_tx=XM_019833258.2;refGene_gn=NUMB;refGene_tx=XM_019833259.2;refGene_gn=NUMB;refGene_tx=XM_019833260.2;refGene_gn=NUMB;refGene_tx=XM_019833261.1;refGene_gn=NUMB;refGene_tx=XM_019833262.2;refGene_gn=NUMB;refGene_tx=XM_019833263.2;refGene_gn=NUMB;refGene_tx=XM_019833264.2
B3 120760003 120760150
refGene_gn=NUMB;refGene_tx=XM_003987807.4;refGene_gn=NUMB;refGene_tx=XM_019833256.2;refGene_gn=NUMB;refGene_tx=XM_019833257.2;refGene_gn=NUMB;refGene_tx=XM_019833258.2;refGene_gn=NUMB;refGene_tx=XM_019833259.2;refGene_gn=NUMB;refGene_tx=XM_019833260.2;refGene_gn=NUMB;refGene_tx=XM_019833261.1;refGene_gn=NUMB;refGene_tx=XM_019833262.2
B3 120762628 120762778
refGene_gn=NUMB;refGene_tx=XM_003987807.4;refGene_gn=NUMB;refGene_tx=XM_019833256.2;refGene_gn=NUMB;refGene_tx=XM_019833257.2;refGene_gn=NUMB;refGene_tx=XM_019833258.2;refGene_gn=NUMB;refGene_tx=XM_019833259.2;refGene_gn=NUMB;refGene_tx=XM_019833260.2;refGene_gn=NUMB;refGene_tx=XM_019833261.1;refGene_gn=NUMB;refGene_tx=XM_019833262.2;refGene_gn=NUMB;refGene_tx=XM_019833263.2;refGene_gn=NUMB;refGene_tx=XM_019833264.2
B3 120764569 120764863
refGene_gn=NUMB;refGene_tx=XM_003987807.4;refGene_gn=NUMB;refGene_tx=XM_019833256.2;refGene_gn=NUMB;refGene_tx=XM_019833257.2;refGene_gn=NUMB;refGene_tx=XM_019833258.2;refGene_gn=NUMB;refGene_tx=XM_019833259.2;refGene_gn=NUMB;refGene_tx=XM_019833260.2;refGene_gn=NUMB;refGene_tx=XM_019833261.1;refGene_gn=NUMB;refGene_tx=XM_019833262.2;refGene_gn=NUMB;refGene_tx=XM_019833263.2;refGene_gn=NUMB;refGene_tx=XM_019833264.2
B3 120766754 120766959
refGene_gn=NUMB;refGene_tx=XM_003987807.4;refGene_gn=NUMB;refGene_tx=XM_019833256.2;refGene_gn=NUMB;refGene_tx=XM_019833257.2;refGene_gn=NUMB;refGene_tx=XM_019833258.2;refGene_gn=NUMB;refGene_tx=XM_019833259.2;refGene_gn=NUMB;refGene_tx=XM_019833260.2;refGene_gn=NUMB;refGene_tx=XM_019833261.1;refGene_gn=NUMB;refGene_tx=XM_019833262.2;refGene_gn=NUMB;refGene_tx=XM_019833263.2;refGene_gn=NUMB;refGene_tx=XM_019833264.2
B3 120776268 120776409

refGene_gn=NUMB;refGene_tx=XM_003987807.4;refGene_gn=NUMB;refGene_tx=XM_019833256.2;refGene_gn=NUMB;refGene_tx=XM_019833257.2;refGene_gn=NUMB;refGene_tx=XM_019833258.2;refGene_gn=NUMB;refGene_tx=XM_019833259.2;refGene_gn=NUMB;refGene_tx=XM_019833260.2;refGene_gn=NUMB;refGene_tx=XM_019833261.1;refGene_gn=NUMB;refGene_tx=XM_019833262.2;refGene_gn=NUMB;refGene_tx=XM_019833263.2;refGene_gn=NUMB;refGene_tx=XM_019833264.2
B3 120778723 120778798
refGene_gn=NUMB;refGene_tx=XM_003987807.4;refGene_gn=NUMB;refGene_tx=XM_019833256.2;refGene_gn=NUMB;refGene_tx=XM_019833257.2;refGene_gn=NUMB;refGene_tx=XM_019833258.2;refGene_gn=NUMB;refGene_tx=XM_019833259.2;refGene_gn=NUMB;refGene_tx=XM_019833260.2;refGene_gn=NUMB;refGene_tx=XM_019833261.1;refGene_gn=NUMB;refGene_tx=XM_019833262.2;refGene_gn=NUMB;refGene_tx=XM_019833263.2;refGene_gn=NUMB;refGene_tx=XM_019833264.2
B3 120796443 120796476
refGene_gn=NUMB;refGene_tx=XM_003987807.4;refGene_gn=NUMB;refGene_tx=XM_019833256.2;refGene_gn=NUMB;refGene_tx=XM_019833257.2;refGene_gn=NUMB;refGene_tx=XM_019833260.2;refGene_gn=NUMB;refGene_tx=XM_019833261.1;refGene_gn=NUMB;refGene_tx=XM_019833262.2;refGene_gn=NUMB;refGene_tx=XM_019833263.2
B3 120802129 120802204
refGene_gn=NUMB;refGene_tx=XM_003987807.4;refGene_gn=NUMB;refGene_tx=XM_019833256.2;refGene_gn=NUMB;refGene_tx=XM_019833257.2;refGene_gn=NUMB;refGene_tx=XM_019833258.2;refGene_gn=NUMB;refGene_tx=XM_019833259.2;refGene_gn=NUMB;refGene_tx=XM_019833260.2;refGene_gn=NUMB;refGene_tx=XM_019833261.1;refGene_gn=NUMB;refGene_tx=XM_019833262.2;refGene_gn=NUMB;refGene_tx=XM_019833263.2;refGene_gn=NUMB;refGene_tx=XM_019833264.2
B3 120833607 120833748
refGene_gn=NUMB;refGene_tx=XM_003987807.4;refGene_gn=NUMB;refGene_tx=XM_019833256.2;refGene_gn=NUMB;refGene_tx=XM_019833257.2;refGene_gn=NUMB;refGene_tx=XM_019833258.2;refGene_gn=NUMB;refGene_tx=XM_019833259.2;refGene_gn=NUMB;refGene_tx=XM_019833260.2;refGene_gn=NUMB;refGene_tx=XM_019833261.1;refGene_gn=NUMB;refGene_tx=XM_019833262.2;refGene_gn=NUMB;refGene_tx=XM_019833263.2;refGene_gn=NUMB;refGene_tx=XM_019833264.2
B3 120963248 120963308
refGene_gn=NUMB;refGene_tx=XM_019833256.2;refGene_gn=NUMB;refGene_tx=XM_019833257.2;refGene_gn=NUMB;refGene_tx=XM_019833258.2;refGene_gn=NUMB;refGene_tx=XM_019833259.2;refGene_gn=NUMB;refGene_tx=XM_019833263.2;refGene_gn=NUMB;refGene_tx=XM_019833264.2
B3 120982201 120982360
refGene_gn=HEATR4;refGene_tx=XM_003987825.4;refGene_gn=HEATR4;refGene_tx=XM_011283325.3;refGene_gn=HEATR4;refGene_tx=XM_019833266.2;refGene_gn=HEATR4;refGene_tx=XM_019833267.2
B3 120994004 120996188
refGene_gn=RIOX1;refGene_tx=XM_019833271.2
B3 120996333 120996392
refGene_gn=HEATR4;refGene_tx=XM_003987825.4;refGene_gn=HEATR4;refGene_tx=XM_011283325.3;refGene_gn=HEATR4;refGene_tx=XM_019833266.2;refGene_gn=HEATR4;refGene_tx=XM_019833267.2
B3 120997823 120997983
refGene_gn=HEATR4;refGene_tx=XM_003987825.4;refGene_gn=HEATR4;refGene_

tx=XM_011283325.3;refGene_gn=HEATR4;refGene_tx=XM_019833266.2;refGene_gn=HEATR4;refGene_tx=XM_019833267.2
B3 120998276 120998285
refGene_gn=HEATR4;refGene_tx=XM_023255815.1
B3 120998701 120998780
refGene_gn=HEATR4;refGene_tx=XM_003987825.4;refGene_gn=HEATR4;refGene_tx=XM_011283325.3;refGene_gn=HEATR4;refGene_tx=XM_019833266.2;refGene_gn=HEATR4;refGene_tx=XM_019833267.2;refGene_gn=HEATR4;refGene_tx=XM_023255815.1
B3 120999441 120999631
refGene_gn=HEATR4;refGene_tx=XM_003987825.4;refGene_gn=HEATR4;refGene_tx=XM_011283325.3;refGene_gn=HEATR4;refGene_tx=XM_019833266.2;refGene_gn=HEATR4;refGene_tx=XM_019833267.2;refGene_gn=HEATR4;refGene_tx=XM_023255815.1
B3 121000441 121000511
refGene_gn=HEATR4;refGene_tx=XM_003987825.4;refGene_gn=HEATR4;refGene_tx=XM_011283325.3;refGene_gn=HEATR4;refGene_tx=XM_019833266.2;refGene_gn=HEATR4;refGene_tx=XM_019833267.2;refGene_gn=HEATR4;refGene_tx=XM_023255815.1
B3 121001258 121001439
refGene_gn=HEATR4;refGene_tx=XM_003987825.4;refGene_gn=HEATR4;refGene_tx=XM_011283325.3;refGene_gn=HEATR4;refGene_tx=XM_019833266.2;refGene_gn=HEATR4;refGene_tx=XM_019833267.2;refGene_gn=HEATR4;refGene_tx=XM_023255815.1
B3 121002371 121002490
refGene_gn=HEATR4;refGene_tx=XM_003987825.4;refGene_gn=HEATR4;refGene_tx=XM_011283325.3;refGene_gn=HEATR4;refGene_tx=XM_019833266.2;refGene_gn=HEATR4;refGene_tx=XM_019833267.2;refGene_gn=HEATR4;refGene_tx=XM_023255815.1
B3 121004800 121004905
refGene_gn=HEATR4;refGene_tx=XM_003987825.4;refGene_gn=HEATR4;refGene_tx=XM_011283325.3;refGene_gn=HEATR4;refGene_tx=XM_019833266.2;refGene_gn=HEATR4;refGene_tx=XM_019833267.2;refGene_gn=HEATR4;refGene_tx=XM_023255815.1
B3 121005551 121005712
refGene_gn=HEATR4;refGene_tx=XM_003987825.4;refGene_gn=HEATR4;refGene_tx=XM_011283325.3;refGene_gn=HEATR4;refGene_tx=XM_019833266.2;refGene_gn=HEATR4;refGene_tx=XM_019833267.2;refGene_gn=HEATR4;refGene_tx=XM_023255815.1
B3 121007633 121007813
refGene_gn=HEATR4;refGene_tx=XM_003987825.4;refGene_gn=HEATR4;refGene_tx=XM_011283325.3;refGene_gn=HEATR4;refGene_tx=XM_019833266.2;refGene_gn=HEATR4;refGene_tx=XM_019833267.2;refGene_gn=HEATR4;refGene_tx=XM_023255815.1
B3 121011243 121011387
refGene_gn=HEATR4;refGene_tx=XM_003987825.4;refGene_gn=HEATR4;refGene_tx=XM_011283325.3;refGene_gn=HEATR4;refGene_tx=XM_019833266.2;refGene_gn=HEATR4;refGene_tx=XM_019833267.2;refGene_gn=HEATR4;refGene_tx=XM_023255815.1
B3 121012606 121012810

refGene_gn=HEATR4;refGene_tx=XM_003987825.4;refGene_gn=HEATR4;refGene_tx=XM_011283325.3;refGene_gn=HEATR4;refGene_tx=XM_019833266.2;refGene_gn=HEATR4;refGene_tx=XM_019833267.2;refGene_gn=HEATR4;refGene_tx=XM_023255815.1
B3 121017382 121017523
refGene_gn=HEATR4;refGene_tx=XM_003987825.4;refGene_gn=HEATR4;refGene_tx=XM_011283325.3;refGene_gn=HEATR4;refGene_tx=XM_019833266.2;refGene_gn=HEATR4;refGene_tx=XM_019833267.2;refGene_gn=HEATR4;refGene_tx=XM_023255815.1
B3 121018554 121018742
refGene_gn=HEATR4;refGene_tx=XM_003987825.4;refGene_gn=HEATR4;refGene_tx=XM_011283325.3;refGene_gn=HEATR4;refGene_tx=XM_019833266.2;refGene_gn=HEATR4;refGene_tx=XM_019833267.2;refGene_gn=HEATR4;refGene_tx=XM_023255815.1
B3 121019918 121020829
refGene_gn=HEATR4;refGene_tx=XM_003987825.4;refGene_gn=HEATR4;refGene_tx=XM_011283325.3;refGene_gn=HEATR4;refGene_tx=XM_019833266.2;refGene_gn=HEATR4;refGene_tx=XM_019833267.2;refGene_gn=HEATR4;refGene_tx=XM_023255815.1
B3 121030035 121030606
refGene_gn=LOC101092274;refGene_tx=XM_003987826.4
B3 121032434 121032637
refGene_gn=LOC101092274;refGene_tx=XM_003987826.4
B3 121034991 121035591
refGene_gn=LOC101092274;refGene_tx=XM_003987826.4
B3 121063163 121063617
refGene_gn=ACOT4;refGene_tx=XM_006932977.4
B3 121064572 121064775
refGene_gn=ACOT4;refGene_tx=XM_006932977.4
B3 121067377 121068097
refGene_gn=ACOT4;refGene_tx=XM_006932977.4
B3 121119782 121120469
refGene_gn=LOC105259864;refGene_tx=XM_023255817.1
B3 121135592 121136049
refGene_gn=ACOT6;refGene_tx=XM_023255818.1
B3 121138187 121138390
refGene_gn=ACOT6;refGene_tx=XM_023255818.1
B3 121142292 121142898
refGene_gn=ACOT6;refGene_tx=XM_023255818.1
B3 121146427 121146992
refGene_gn=LOC101092771;refGene_tx=XM_006932978.4;refGene_gn=LOC101092771;refGene_tx=XM_019833273.2;refGene_gn=LOC101092771;refGene_tx=XM_019833274.2;refGene_gn=LOC101092771;refGene_tx=XM_019833275.2
B3 121152111 121152311
refGene_gn=LOC101092771;refGene_tx=XM_006932978.4;refGene_gn=LOC101092771;refGene_tx=XM_019833273.2;refGene_gn=LOC101092771;refGene_tx=XM_019833274.2;refGene_gn=LOC101092771;refGene_tx=XM_019833275.2
B3 121152595 121153171
refGene_gn=LOC101092771;refGene_tx=XM_006932978.4
B3 121155902 121155920

refGene_gn=LOC101092771;refGene_tx=XM_019833273.2;refGene_gn=LOC101092771;refGene_tx=XM_019833274.2;refGene_gn=LOC101092771;refGene_tx=XM_019833275.2
B3 121158322 121158325
refGene_gn=DNAL1;refGene_tx=XM_003987810.5
B3 121170381 121170420
refGene_gn=DNAL1;refGene_tx=XM_003987810.5
B3 121173566 121173676
refGene_gn=DNAL1;refGene_tx=XM_003987810.5
B3 121176866 121176922
refGene_gn=DNAL1;refGene_tx=XM_003987810.5
B3 121187706 121187762
refGene_gn=DNAL1;refGene_tx=XM_003987810.5
B3 121195760 121195887
refGene_gn=DNAL1;refGene_tx=XM_003987810.5
B3 121196886 121197027
refGene_gn=DNAL1;refGene_tx=XM_003987810.5
B3 121201838 121201879
refGene_gn=DNAL1;refGene_tx=XM_003987810.5
B3 121209347 121210409
refGene_gn=PNMA1;refGene_tx=XM_003987811.4
B3 121215359 121215540
refGene_gn=ELMSAN1;refGene_tx=XM_023255820.1;refGene_gn=ELMSAN1;refGene_tx=XM_023255821.1;refGene_gn=ELMSAN1;refGene_tx=XM_023255822.1;refGene_gn=ELMSAN1;refGene_tx=XM_023255823.1;refGene_gn=ELMSAN1;refGene_tx=XM_023255824.1;refGene_gn=ELMSAN1;refGene_tx=XM_023255826.1
B3 121215934 121216135
refGene_gn=ELMSAN1;refGene_tx=XM_023255820.1;refGene_gn=ELMSAN1;refGene_tx=XM_023255821.1;refGene_gn=ELMSAN1;refGene_tx=XM_023255822.1;refGene_gn=ELMSAN1;refGene_tx=XM_023255823.1;refGene_gn=ELMSAN1;refGene_tx=XM_023255824.1;refGene_gn=ELMSAN1;refGene_tx=XM_023255825.1;refGene_gn=ELMSAN1;refGene_tx=XM_023255826.1
B3 121218065 121218278
refGene_gn=ELMSAN1;refGene_tx=XM_023255820.1;refGene_gn=ELMSAN1;refGene_tx=XM_023255821.1;refGene_gn=ELMSAN1;refGene_tx=XM_023255822.1;refGene_gn=ELMSAN1;refGene_tx=XM_023255823.1;refGene_gn=ELMSAN1;refGene_tx=XM_023255824.1;refGene_gn=ELMSAN1;refGene_tx=XM_023255825.1
B3 121219855 121220005
refGene_gn=ELMSAN1;refGene_tx=XM_023255820.1;refGene_gn=ELMSAN1;refGene_tx=XM_023255821.1;refGene_gn=ELMSAN1;refGene_tx=XM_023255822.1;refGene_gn=ELMSAN1;refGene_tx=XM_023255823.1;refGene_gn=ELMSAN1;refGene_tx=XM_023255824.1;refGene_gn=ELMSAN1;refGene_tx=XM_023255825.1;refGene_gn=ELMSAN1;refGene_tx=XM_023255826.1
B3 121222590 121222679
refGene_gn=ELMSAN1;refGene_tx=XM_023255820.1;refGene_gn=ELMSAN1;refGene_tx=XM_023255821.1;refGene_gn=ELMSAN1;refGene_tx=XM_023255822.1;refGene_gn=ELMSAN1;refGene_tx=XM_023255823.1;refGene_gn=ELMSAN1;refGene_tx=XM_023255824.1;refGene_gn=ELMSAN1;refGene_tx=XM_023255825.1;refGene_gn=ELMSAN1;refGene_tx=XM_023255826.1
B3 121223345 121223421

refGene_gn=ELMSAN1;refGene_tx=XM_023255820.1;refGene_gn=ELMSAN1;refGene_tx=XM_023255821.1;refGene_gn=ELMSAN1;refGene_tx=XM_023255822.1;refGene_gn=ELMSAN1;refGene_tx=XM_023255823.1;refGene_gn=ELMSAN1;refGene_tx=XM_023255824.1;refGene_gn=ELMSAN1;refGene_tx=XM_023255825.1;refGene_gn=ELMSAN1;refGene_tx=XM_023255826.1
B3 121223948 121224052
refGene_gn=ELMSAN1;refGene_tx=XM_023255820.1;refGene_gn=ELMSAN1;refGene_tx=XM_023255821.1;refGene_gn=ELMSAN1;refGene_tx=XM_023255822.1;refGene_gn=ELMSAN1;refGene_tx=XM_023255823.1;refGene_gn=ELMSAN1;refGene_tx=XM_023255824.1;refGene_gn=ELMSAN1;refGene_tx=XM_023255825.1;refGene_gn=ELMSAN1;refGene_tx=XM_023255826.1
B3 121224156 121224299
refGene_gn=ELMSAN1;refGene_tx=XM_023255820.1;refGene_gn=ELMSAN1;refGene_tx=XM_023255821.1;refGene_gn=ELMSAN1;refGene_tx=XM_023255822.1;refGene_gn=ELMSAN1;refGene_tx=XM_023255823.1;refGene_gn=ELMSAN1;refGene_tx=XM_023255824.1;refGene_gn=ELMSAN1;refGene_tx=XM_023255825.1;refGene_gn=ELMSAN1;refGene_tx=XM_023255826.1
B3 121224762 121224829
refGene_gn=ELMSAN1;refGene_tx=XM_023255820.1;refGene_gn=ELMSAN1;refGene_tx=XM_023255821.1;refGene_gn=ELMSAN1;refGene_tx=XM_023255822.1;refGene_gn=ELMSAN1;refGene_tx=XM_023255823.1;refGene_gn=ELMSAN1;refGene_tx=XM_023255824.1;refGene_gn=ELMSAN1;refGene_tx=XM_023255825.1;refGene_gn=ELMSAN1;refGene_tx=XM_023255826.1
B3 121226816 121227162
refGene_gn=ELMSAN1;refGene_tx=XM_023255820.1;refGene_gn=ELMSAN1;refGene_tx=XM_023255821.1;refGene_gn=ELMSAN1;refGene_tx=XM_023255822.1;refGene_gn=ELMSAN1;refGene_tx=XM_023255823.1;refGene_gn=ELMSAN1;refGene_tx=XM_023255824.1;refGene_gn=ELMSAN1;refGene_tx=XM_023255825.1;refGene_gn=ELMSAN1;refGene_tx=XM_023255826.1
B3 121233263 121233563
refGene_gn=ELMSAN1;refGene_tx=XM_023255820.1;refGene_gn=ELMSAN1;refGene_tx=XM_023255821.1;refGene_gn=ELMSAN1;refGene_tx=XM_023255822.1;refGene_gn=ELMSAN1;refGene_tx=XM_023255823.1;refGene_gn=ELMSAN1;refGene_tx=XM_023255824.1;refGene_gn=ELMSAN1;refGene_tx=XM_023255825.1;refGene_gn=ELMSAN1;refGene_tx=XM_023255826.1
B3 121234897 121236370
refGene_gn=ELMSAN1;refGene_tx=XM_023255820.1;refGene_gn=ELMSAN1;refGene_tx=XM_023255821.1;refGene_gn=ELMSAN1;refGene_tx=XM_023255822.1;refGene_gn=ELMSAN1;refGene_tx=XM_023255823.1;refGene_gn=ELMSAN1;refGene_tx=XM_023255824.1;refGene_gn=ELMSAN1;refGene_tx=XM_023255825.1;refGene_gn=ELMSAN1;refGene_tx=XM_023255826.1
B3 121339393 121339430
refGene_gn=PTGR2;refGene_tx=XM_003987813.4;refGene_gn=PTGR2;refGene_tx=XM_011283326.3;refGene_gn=PTGR2;refGene_tx=XM_019833281.2;refGene_gn=PTGR2;refGene_tx=XM_019833282.1
B3 121342000 121342119
refGene_gn=PTGR2;refGene_tx=XM_003987813.4;refGene_gn=PTGR2;refGene_tx=XM_011283326.3;refGene_gn=PTGR2;refGene_tx=XM_019833281.2;refGene_gn=PTGR2;refGene_tx=XM_019833282.1
B3 121346430 121346523

refGene_gn=PTGR2;refGene_tx=XM_019833283.1;refGene_gn=PTGR2;refGene_tx=XM_019833284.1
B3 121349287 121349479
refGene_gn=PTGR2;refGene_tx=XM_003987813.4;refGene_gn=PTGR2;refGene_tx=XM_011283326.3;refGene_gn=PTGR2;refGene_tx=XM_019833281.2;refGene_gn=PTGR2;refGene_tx=XM_019833282.1;refGene_gn=PTGR2;refGene_tx=XM_019833283.1;refGene_gn=PTGR2;refGene_tx=XM_019833284.1
B3 121352146 121352317
refGene_gn=PTGR2;refGene_tx=XM_003987813.4;refGene_gn=PTGR2;refGene_tx=XM_011283326.3;refGene_gn=PTGR2;refGene_tx=XM_019833281.2;refGene_gn=PTGR2;refGene_tx=XM_019833282.1;refGene_gn=PTGR2;refGene_tx=XM_019833283.1;refGene_gn=PTGR2;refGene_tx=XM_019833284.1
B3 121353064 121353274
refGene_gn=PTGR2;refGene_tx=XM_003987813.4;refGene_gn=PTGR2;refGene_tx=XM_011283326.3;refGene_gn=PTGR2;refGene_tx=XM_019833281.2;refGene_gn=PTGR2;refGene_tx=XM_019833282.1;refGene_gn=PTGR2;refGene_tx=XM_019833283.1;refGene_gn=PTGR2;refGene_tx=XM_019833284.1
B3 121354454 121354576
refGene_gn=PTGR2;refGene_tx=XM_003987813.4;refGene_gn=PTGR2;refGene_tx=XM_011283326.3;refGene_gn=PTGR2;refGene_tx=XM_019833281.2;refGene_gn=PTGR2;refGene_tx=XM_019833282.1;refGene_gn=PTGR2;refGene_tx=XM_019833283.1;refGene_gn=PTGR2;refGene_tx=XM_019833284.1
B3 121355599 121355687
refGene_gn=PTGR2;refGene_tx=XM_003987813.4;refGene_gn=PTGR2;refGene_tx=XM_011283326.3;refGene_gn=PTGR2;refGene_tx=XM_019833281.2;refGene_gn=PTGR2;refGene_tx=XM_019833282.1;refGene_gn=PTGR2;refGene_tx=XM_019833283.1;refGene_gn=PTGR2;refGene_tx=XM_019833284.1
B3 121357068 121357108
refGene_gn=PTGR2;refGene_tx=XM_003987813.4;refGene_gn=PTGR2;refGene_tx=XM_011283326.3;refGene_gn=PTGR2;refGene_tx=XM_019833281.2;refGene_gn=PTGR2;refGene_tx=XM_019833282.1;refGene_gn=PTGR2;refGene_tx=XM_019833283.1;refGene_gn=PTGR2;refGene_tx=XM_019833284.1
B3 121358040 121358117
refGene_gn=PTGR2;refGene_tx=XM_003987813.4;refGene_gn=PTGR2;refGene_tx=XM_011283326.3;refGene_gn=PTGR2;refGene_tx=XM_019833281.2;refGene_gn=PTGR2;refGene_tx=XM_019833282.1;refGene_gn=PTGR2;refGene_tx=XM_019833283.1;refGene_gn=PTGR2;refGene_tx=XM_019833284.1
B3 121373645 121373678
refGene_gn=ZNF410;refGene_tx=XM_003987814.4;refGene_gn=ZNF410;refGene_tx=XM_003987815.4;refGene_gn=ZNF410;refGene_tx=XM_019833277.1;refGene_gn=ZNF410;refGene_tx=XM_019833278.2;refGene_gn=ZNF410;refGene_tx=XM_019833279.2;refGene_gn=ZNF410;refGene_tx=XM_019833280.1
B3 121374639 121374775
refGene_gn=ZNF410;refGene_tx=XM_003987814.4;refGene_gn=ZNF410;refGene_tx=XM_003987815.4;refGene_gn=ZNF410;refGene_tx=XM_019833277.1;refGene_gn=ZNF410;refGene_tx=XM_019833278.2;refGene_gn=ZNF410;refGene_tx=XM_019833279.2;refGene_gn=ZNF410;refGene_tx=XM_019833280.1
B3 121378892 121379111
refGene_gn=ZNF410;refGene_tx=XM_003987814.4;refGene_gn=ZNF410;refGene_tx=XM_003987815.4;refGene_gn=ZNF410;refGene_tx=XM_019833277.1;refGene_

gn=ZNF410;refGene_tx=XM_019833278.2;refGene_gn=ZNF410;refGene_tx=XM_019833280.1
B3 121380296 121380488
refGene_gn=ZNF410;refGene_tx=XM_003987814.4;refGene_gn=ZNF410;refGene_tx=XM_003987815.4;refGene_gn=ZNF410;refGene_tx=XM_019833277.1;refGene_gn=ZNF410;refGene_tx=XM_019833278.2;refGene_gn=ZNF410;refGene_tx=XM_019833279.2;refGene_gn=ZNF410;refGene_tx=XM_019833280.1
B3 121392930 121393081
refGene_gn=ZNF410;refGene_tx=XM_003987814.4;refGene_gn=ZNF410;refGene_tx=XM_003987815.4;refGene_gn=ZNF410;refGene_tx=XM_019833277.1;refGene_gn=ZNF410;refGene_tx=XM_019833278.2;refGene_gn=ZNF410;refGene_tx=XM_019833279.2;refGene_gn=ZNF410;refGene_tx=XM_019833280.1
B3 121393688 121393870
refGene_gn=ZNF410;refGene_tx=XM_003987814.4;refGene_gn=ZNF410;refGene_tx=XM_003987815.4;refGene_gn=ZNF410;refGene_tx=XM_019833277.1;refGene_gn=ZNF410;refGene_tx=XM_019833278.2;refGene_gn=ZNF410;refGene_tx=XM_019833279.2;refGene_gn=ZNF410;refGene_tx=XM_019833280.1
B3 121398208 121398298
refGene_gn=ZNF410;refGene_tx=XM_003987814.4;refGene_gn=ZNF410;refGene_tx=XM_003987815.4;refGene_gn=ZNF410;refGene_tx=XM_019833277.1;refGene_gn=ZNF410;refGene_tx=XM_019833278.2;refGene_gn=ZNF410;refGene_tx=XM_019833279.2;refGene_gn=ZNF410;refGene_tx=XM_019833280.1
B3 121401284 121401410
refGene_gn=ZNF410;refGene_tx=XM_003987814.4;refGene_gn=ZNF410;refGene_tx=XM_003987815.4;refGene_gn=ZNF410;refGene_tx=XM_019833277.1;refGene_gn=ZNF410;refGene_tx=XM_019833278.2;refGene_gn=ZNF410;refGene_tx=XM_019833279.2;refGene_gn=ZNF410;refGene_tx=XM_019833280.1
B3 121402148 121402289
refGene_gn=ZNF410;refGene_tx=XM_003987814.4;refGene_gn=ZNF410;refGene_tx=XM_019833277.1;refGene_gn=ZNF410;refGene_tx=XM_019833278.2;refGene_gn=ZNF410;refGene_tx=XM_019833279.2
B3 121402584 121402607
refGene_gn=ZNF410;refGene_tx=XM_019833278.2
B3 121403733 121403861
refGene_gn=ZNF410;refGene_tx=XM_003987814.4;refGene_gn=ZNF410;refGene_tx=XM_003987815.4;refGene_gn=ZNF410;refGene_tx=XM_019833279.2
B3 121409364 121409403
refGene_gn=ZNF410;refGene_tx=XM_003987814.4;refGene_gn=ZNF410;refGene_tx=XM_003987815.4;refGene_gn=ZNF410;refGene_tx=XM_019833277.1;refGene_gn=ZNF410;refGene_tx=XM_019833279.2;refGene_gn=ZNF410;refGene_tx=XM_019833280.1
B3 121411532 121411671
refGene_gn=FAM161B;refGene_tx=XM_006932979.3
B3 121412435 121412578
refGene_gn=FAM161B;refGene_tx=XM_006932979.3
B3 121414477 121414577
refGene_gn=FAM161B;refGene_tx=XM_006932979.3
B3 121414782 121414947
refGene_gn=FAM161B;refGene_tx=XM_006932979.3
B3 121416865 121416993

refGene_gn=FAM161B;refGene_tx=XM_006932979.3
B3	121417300	121417647
refGene_gn=FAM161B;refGene_tx=XM_006932979.3
B3	121419233	121419784
refGene_gn=FAM161B;refGene_tx=XM_006932979.3
B3	121420825	121421148
refGene_gn=FAM161B;refGene_tx=XM_006932979.3
B3	121423281	121423335
refGene_gn=FAM161B;refGene_tx=XM_006932979.3
B3	121423651	121423817
refGene_gn=COQ6;refGene_tx=XM_023255828.1
B3	121428761	121428896
refGene_gn=COQ6;refGene_tx=XM_023255828.1
B3	121430246	121430305
refGene_gn=COQ6;refGene_tx=XM_023255828.1
B3	121430596	121430720
refGene_gn=COQ6;refGene_tx=XM_023255828.1
B3	121433479	121433610
refGene_gn=COQ6;refGene_tx=XM_023255828.1
B3	121434304	121434412
refGene_gn=COQ6;refGene_tx=XM_023255828.1
B3	121434505	121434568
refGene_gn=COQ6;refGene_tx=XM_023255828.1
B3	121434751	121434859
refGene_gn=COQ6;refGene_tx=XM_023255828.1
B3	121437357	121437560
refGene_gn=COQ6;refGene_tx=XM_023255828.1
B3	121437638	121437754
refGene_gn=COQ6;refGene_tx=XM_023255828.1
B3	121437917	121438084
refGene_gn=COQ6;refGene_tx=XM_023255828.1
B3	121438518	121438548
refGene_gn=COQ6;refGene_tx=XM_023255828.1
B3	121446122	121446209
refGene_gn=ENTPD5;refGene_tx=XM_023255829.1;refGene_gn=ENTPD5;refGene_tx=XM_023255830.1;refGene_gn=ENTPD5;refGene_tx=XM_023255831.1;refGene_gn=ENTPD5;refGene_tx=XM_023255832.1;refGene_gn=ENTPD5;refGene_tx=XM_023255833.1;refGene_gn=ENTPD5;refGene_tx=XM_023255834.1;refGene_gn=ENTPD5;refGene_tx=XM_023255835.1;refGene_gn=ENTPD5;refGene_tx=XM_023255836.1;refGene_gn=ENTPD5;refGene_tx=XM_023255837.1;refGene_gn=ENTPD5;refGene_tx=XM_023255838.1;refGene_gn=ENTPD5;refGene_tx=XM_023255839.1;refGene_gn=ENTPD5;refGene_tx=XM_023255840.1;refGene_gn=ENTPD5;refGene_tx=XM_023255841.1;refGene_gn=ENTPD5;refGene_tx=XM_023255842.1;refGene_gn=ENTPD5;refGene_tx=XM_023255843.1;refGene_gn=ENTPD5;refGene_tx=XM_023255844.1;refGene_gn=ENTPD5;refGene_tx=XM_023255845.1;refGene_gn=ENTPD5;refGene_tx=XM_023255846.1;refGene_gn=ENTPD5;refGene_tx=XM_023255847.1
B3	121448670	121448786
refGene_gn=ENTPD5;refGene_tx=XM_023255829.1;refGene_gn=ENTPD5;refGene_tx=XM_023255830.1;refGene_gn=ENTPD5;refGene_tx=XM_023255831.1;refGene_gn=ENTPD5;refGene_tx=XM_023255832.1;refGene_gn=ENTPD5;refGene_tx=XM_02

3255833.1;refGene_gn=ENTPD5;refGene_tx=XM_023255834.1;refGene_gn=ENTPD5;refGene_tx=XM_023255835.1;refGene_gn=ENTPD5;refGene_tx=XM_023255836.1;refGene_gn=ENTPD5;refGene_tx=XM_023255837.1;refGene_gn=ENTPD5;refGene_tx=XM_023255838.1;refGene_gn=ENTPD5;refGene_tx=XM_023255839.1;refGene_gn=ENTPD5;refGene_tx=XM_023255840.1;refGene_gn=ENTPD5;refGene_tx=XM_023255841.1;refGene_gn=ENTPD5;refGene_tx=XM_023255842.1;refGene_gn=ENTPD5;refGene_tx=XM_023255843.1;refGene_gn=ENTPD5;refGene_tx=XM_023255844.1;refGene_gn=ENTPD5;refGene_tx=XM_023255845.1;refGene_gn=ENTPD5;refGene_tx=XM_023255846.1;refGene_gn=ENTPD5;refGene_tx=XM_023255847.1
B3 121452656 121452713
refGene_gn=ENTPD5;refGene_tx=XM_023255829.1;refGene_gn=ENTPD5;refGene_tx=XM_023255830.1;refGene_gn=ENTPD5;refGene_tx=XM_023255831.1;refGene_gn=ENTPD5;refGene_tx=XM_023255832.1;refGene_gn=ENTPD5;refGene_tx=XM_023255833.1;refGene_gn=ENTPD5;refGene_tx=XM_023255834.1;refGene_gn=ENTPD5;refGene_tx=XM_023255835.1;refGene_gn=ENTPD5;refGene_tx=XM_023255836.1;refGene_gn=ENTPD5;refGene_tx=XM_023255837.1;refGene_gn=ENTPD5;refGene_tx=XM_023255838.1;refGene_gn=ENTPD5;refGene_tx=XM_023255839.1;refGene_gn=ENTPD5;refGene_tx=XM_023255840.1;refGene_gn=ENTPD5;refGene_tx=XM_023255841.1;refGene_gn=ENTPD5;refGene_tx=XM_023255842.1;refGene_gn=ENTPD5;refGene_tx=XM_023255843.1;refGene_gn=ENTPD5;refGene_tx=XM_023255844.1;refGene_gn=ENTPD5;refGene_tx=XM_023255845.1;refGene_gn=ENTPD5;refGene_tx=XM_023255846.1;refGene_gn=ENTPD5;refGene_tx=XM_023255847.1
B3 121453448 121453589
refGene_gn=ENTPD5;refGene_tx=XM_023255829.1;refGene_gn=ENTPD5;refGene_tx=XM_023255830.1;refGene_gn=ENTPD5;refGene_tx=XM_023255831.1;refGene_gn=ENTPD5;refGene_tx=XM_023255832.1;refGene_gn=ENTPD5;refGene_tx=XM_023255833.1;refGene_gn=ENTPD5;refGene_tx=XM_023255834.1;refGene_gn=ENTPD5;refGene_tx=XM_023255835.1;refGene_gn=ENTPD5;refGene_tx=XM_023255836.1;refGene_gn=ENTPD5;refGene_tx=XM_023255837.1;refGene_gn=ENTPD5;refGene_tx=XM_023255838.1;refGene_gn=ENTPD5;refGene_tx=XM_023255839.1;refGene_gn=ENTPD5;refGene_tx=XM_023255840.1;refGene_gn=ENTPD5;refGene_tx=XM_023255841.1;refGene_gn=ENTPD5;refGene_tx=XM_023255842.1;refGene_gn=ENTPD5;refGene_tx=XM_023255843.1;refGene_gn=ENTPD5;refGene_tx=XM_023255844.1;refGene_gn=ENTPD5;refGene_tx=XM_023255845.1;refGene_gn=ENTPD5;refGene_tx=XM_023255846.1;refGene_gn=ENTPD5;refGene_tx=XM_023255847.1
B3 121454760 121454862
refGene_gn=ENTPD5;refGene_tx=XM_023255829.1;refGene_gn=ENTPD5;refGene_tx=XM_023255830.1;refGene_gn=ENTPD5;refGene_tx=XM_023255831.1;refGene_gn=ENTPD5;refGene_tx=XM_023255832.1;refGene_gn=ENTPD5;refGene_tx=XM_023255833.1;refGene_gn=ENTPD5;refGene_tx=XM_023255834.1;refGene_gn=ENTPD5;refGene_tx=XM_023255835.1;refGene_gn=ENTPD5;refGene_tx=XM_023255836.1;refGene_gn=ENTPD5;refGene_tx=XM_023255837.1;refGene_gn=ENTPD5;refGene_tx=XM_023255838.1;refGene_gn=ENTPD5;refGene_tx=XM_023255839.1;refGene_gn=ENTPD5;refGene_tx=XM_023255840.1;refGene_gn=ENTPD5;refGene_tx=XM_023255841.1;refGene_gn=ENTPD5;refGene_tx=XM_023255842.1;refGene_gn=ENTPD5;refGene_tx=XM_023255843.1;refGene_gn=ENTPD5;refGene_tx=XM_023255844.1;refGene_gn=ENTPD5;refGene_tx=XM_023255845.1;refGene_gn=ENTPD5;refGene_tx=XM_023255846.1;refGene_gn=ENTPD5;refGene_tx=XM_023255847.1
B3 121456059 121456121
refGene_gn=ENTPD5;refGene_tx=XM_023255829.1;refGene_gn=ENTPD5;refGene_

tx=XM_023255830.1;refGene_gn=ENTPD5;refGene_tx=XM_023255831.1;refGene_gn=ENTPD5;refGene_tx=XM_023255832.1;refGene_gn=ENTPD5;refGene_tx=XM_023255833.1;refGene_gn=ENTPD5;refGene_tx=XM_023255834.1;refGene_gn=ENTPD5;refGene_tx=XM_023255835.1;refGene_gn=ENTPD5;refGene_tx=XM_023255836.1;refGene_gn=ENTPD5;refGene_tx=XM_023255837.1;refGene_gn=ENTPD5;refGene_tx=XM_023255838.1;refGene_gn=ENTPD5;refGene_tx=XM_023255839.1;refGene_gn=ENTPD5;refGene_tx=XM_023255840.1;refGene_gn=ENTPD5;refGene_tx=XM_023255841.1;refGene_gn=ENTPD5;refGene_tx=XM_023255842.1;refGene_gn=ENTPD5;refGene_tx=XM_023255843.1;refGene_gn=ENTPD5;refGene_tx=XM_023255844.1;refGene_gn=ENTPD5;refGene_tx=XM_023255845.1;refGene_gn=ENTPD5;refGene_tx=XM_023255846.1;refGene_gn=ENTPD5;refGene_tx=XM_023255847.1
B3 121456723 121456803
refGene_gn=ENTPD5;refGene_tx=XM_023255829.1;refGene_gn=ENTPD5;refGene_tx=XM_023255830.1;refGene_gn=ENTPD5;refGene_tx=XM_023255831.1;refGene_gn=ENTPD5;refGene_tx=XM_023255832.1;refGene_gn=ENTPD5;refGene_tx=XM_023255833.1;refGene_gn=ENTPD5;refGene_tx=XM_023255834.1;refGene_gn=ENTPD5;refGene_tx=XM_023255835.1;refGene_gn=ENTPD5;refGene_tx=XM_023255836.1;refGene_gn=ENTPD5;refGene_tx=XM_023255837.1;refGene_gn=ENTPD5;refGene_tx=XM_023255838.1;refGene_gn=ENTPD5;refGene_tx=XM_023255839.1;refGene_gn=ENTPD5;refGene_tx=XM_023255840.1;refGene_gn=ENTPD5;refGene_tx=XM_023255841.1;refGene_gn=ENTPD5;refGene_tx=XM_023255842.1;refGene_gn=ENTPD5;refGene_tx=XM_023255843.1;refGene_gn=ENTPD5;refGene_tx=XM_023255844.1;refGene_gn=ENTPD5;refGene_tx=XM_023255845.1;refGene_gn=ENTPD5;refGene_tx=XM_023255846.1;refGene_gn=ENTPD5;refGene_tx=XM_023255847.1
B3 121457114 121457203
refGene_gn=ENTPD5;refGene_tx=XM_023255829.1;refGene_gn=ENTPD5;refGene_tx=XM_023255830.1;refGene_gn=ENTPD5;refGene_tx=XM_023255831.1;refGene_gn=ENTPD5;refGene_tx=XM_023255832.1;refGene_gn=ENTPD5;refGene_tx=XM_023255833.1;refGene_gn=ENTPD5;refGene_tx=XM_023255834.1;refGene_gn=ENTPD5;refGene_tx=XM_023255835.1;refGene_gn=ENTPD5;refGene_tx=XM_023255836.1;refGene_gn=ENTPD5;refGene_tx=XM_023255837.1;refGene_gn=ENTPD5;refGene_tx=XM_023255838.1;refGene_gn=ENTPD5;refGene_tx=XM_023255839.1;refGene_gn=ENTPD5;refGene_tx=XM_023255840.1;refGene_gn=ENTPD5;refGene_tx=XM_023255841.1;refGene_gn=ENTPD5;refGene_tx=XM_023255842.1;refGene_gn=ENTPD5;refGene_tx=XM_023255843.1;refGene_gn=ENTPD5;refGene_tx=XM_023255844.1;refGene_gn=ENTPD5;refGene_tx=XM_023255845.1;refGene_gn=ENTPD5;refGene_tx=XM_023255846.1;refGene_gn=ENTPD5;refGene_tx=XM_023255847.1
B3 121457535 121457571
refGene_gn=ENTPD5;refGene_tx=XM_023255829.1;refGene_gn=ENTPD5;refGene_tx=XM_023255830.1;refGene_gn=ENTPD5;refGene_tx=XM_023255831.1;refGene_gn=ENTPD5;refGene_tx=XM_023255832.1;refGene_gn=ENTPD5;refGene_tx=XM_023255833.1;refGene_gn=ENTPD5;refGene_tx=XM_023255834.1;refGene_gn=ENTPD5;refGene_tx=XM_023255835.1;refGene_gn=ENTPD5;refGene_tx=XM_023255836.1;refGene_gn=ENTPD5;refGene_tx=XM_023255837.1;refGene_gn=ENTPD5;refGene_tx=XM_023255838.1;refGene_gn=ENTPD5;refGene_tx=XM_023255839.1;refGene_gn=ENTPD5;refGene_tx=XM_023255840.1;refGene_gn=ENTPD5;refGene_tx=XM_023255841.1;refGene_gn=ENTPD5;refGene_tx=XM_023255842.1;refGene_gn=ENTPD5;refGene_tx=XM_023255843.1;refGene_gn=ENTPD5;refGene_tx=XM_023255844.1;refGene_gn=ENTPD5;refGene_tx=XM_023255846.1
B3 121457846 121457922

refGene_gn=ENTPD5;refGene_tx=XM_023255829.1;refGene_gn=ENTPD5;refGene_
tx=XM_023255830.1;refGene_gn=ENTPD5;refGene_tx=XM_023255831.1;refGene_
gn=ENTPD5;refGene_tx=XM_023255832.1;refGene_gn=ENTPD5;refGene_tx=XM_02
3255833.1;refGene_gn=ENTPD5;refGene_tx=XM_023255834.1;refGene_gn=ENTPD
5;refGene_tx=XM_023255835.1;refGene_gn=ENTPD5;refGene_tx=XM_023255836.
1;refGene_gn=ENTPD5;refGene_tx=XM_023255837.1;refGene_gn=ENTPD5;refGen
e_tx=XM_023255838.1;refGene_gn=ENTPD5;refGene_tx=XM_023255839.1;refGen
e_gn=ENTPD5;refGene_tx=XM_023255840.1;refGene_gn=ENTPD5;refGene_tx=XM_
023255841.1;refGene_gn=ENTPD5;refGene_tx=XM_023255842.1;refGene_gn=ENT
PD5;refGene_tx=XM_023255843.1;refGene_gn=ENTPD5;refGene_tx=XM_02325584
4.1;refGene_gn=ENTPD5;refGene_tx=XM_023255845.1;refGene_gn=ENTPD5;refG
ene_tx=XM_023255846.1;refGene_gn=ENTPD5;refGene_tx=XM_023255847.1
B3 121466298 121466442
refGene_gn=ENTPD5;refGene_tx=XM_023255829.1;refGene_gn=ENTPD5;refGene_
tx=XM_023255830.1;refGene_gn=ENTPD5;refGene_tx=XM_023255831.1;refGene_
gn=ENTPD5;refGene_tx=XM_023255832.1;refGene_gn=ENTPD5;refGene_tx=XM_02
3255833.1;refGene_gn=ENTPD5;refGene_tx=XM_023255834.1;refGene_gn=ENTPD
5;refGene_tx=XM_023255835.1;refGene_gn=ENTPD5;refGene_tx=XM_023255836.
1;refGene_gn=ENTPD5;refGene_tx=XM_023255837.1;refGene_gn=ENTPD5;refGen
e_tx=XM_023255838.1;refGene_gn=ENTPD5;refGene_tx=XM_023255839.1;refGen
e_gn=ENTPD5;refGene_tx=XM_023255840.1;refGene_gn=ENTPD5;refGene_tx=XM_
023255841.1;refGene_gn=ENTPD5;refGene_tx=XM_023255842.1;refGene_gn=ENT
PD5;refGene_tx=XM_023255843.1;refGene_gn=ENTPD5;refGene_tx=XM_02325584
4.1;refGene_gn=ENTPD5;refGene_tx=XM_023255845.1
B3 121468195 121468275
refGene_gn=ENTPD5;refGene_tx=XM_023255829.1;refGene_gn=ENTPD5;refGene_
tx=XM_023255830.1;refGene_gn=ENTPD5;refGene_tx=XM_023255831.1;refGene_
gn=ENTPD5;refGene_tx=XM_023255832.1;refGene_gn=ENTPD5;refGene_tx=XM_02
3255833.1;refGene_gn=ENTPD5;refGene_tx=XM_023255834.1;refGene_gn=ENTPD
5;refGene_tx=XM_023255835.1;refGene_gn=ENTPD5;refGene_tx=XM_023255836.
1;refGene_gn=ENTPD5;refGene_tx=XM_023255837.1;refGene_gn=ENTPD5;refGen
e_tx=XM_023255838.1;refGene_gn=ENTPD5;refGene_tx=XM_023255839.1;refGen
e_gn=ENTPD5;refGene_tx=XM_023255840.1;refGene_gn=ENTPD5;refGene_tx=XM_
023255841.1;refGene_gn=ENTPD5;refGene_tx=XM_023255842.1;refGene_gn=ENT
PD5;refGene_tx=XM_023255843.1;refGene_gn=ENTPD5;refGene_tx=XM_02325584
4.1;refGene_gn=ENTPD5;refGene_tx=XM_023255845.1;refGene_gn=ENTPD5;refG
ene_tx=XM_023255846.1;refGene_gn=ENTPD5;refGene_tx=XM_023255847.1
B3 121468921 121469138
refGene_gn=ENTPD5;refGene_tx=XM_023255829.1;refGene_gn=ENTPD5;refGene_
tx=XM_023255830.1;refGene_gn=ENTPD5;refGene_tx=XM_023255831.1;refGene_
gn=ENTPD5;refGene_tx=XM_023255832.1;refGene_gn=ENTPD5;refGene_tx=XM_02
3255833.1;refGene_gn=ENTPD5;refGene_tx=XM_023255834.1;refGene_gn=ENTPD
5;refGene_tx=XM_023255835.1;refGene_gn=ENTPD5;refGene_tx=XM_023255836.
1;refGene_gn=ENTPD5;refGene_tx=XM_023255837.1;refGene_gn=ENTPD5;refGen
e_tx=XM_023255838.1;refGene_gn=ENTPD5;refGene_tx=XM_023255839.1;refGen
e_gn=ENTPD5;refGene_tx=XM_023255840.1;refGene_gn=ENTPD5;refGene_tx=XM_
023255841.1;refGene_gn=ENTPD5;refGene_tx=XM_023255842.1;refGene_gn=ENT
PD5;refGene_tx=XM_023255843.1;refGene_gn=ENTPD5;refGene_tx=XM_02325584
4.1;refGene_gn=ENTPD5;refGene_tx=XM_023255845.1;refGene_gn=ENTPD5;refG
ene_tx=XM_023255846.1;refGene_gn=ENTPD5;refGene_tx=XM_023255847.1

B3 121491764 121491820
refGene_gn=BBOF1;refGene_tx=XM_003987820.5;refGene_gn=BBOF1;refGene_tx=XM_006932987.4;refGene_gn=BBOF1;refGene_tx=XM_006932988.4;refGene_gn=BBOF1;refGene_tx=XM_019833285.2;refGene_gn=BBOF1;refGene_tx=XM_019833287.2;refGene_gn=BBOF1;refGene_tx=XM_019833288.2
B3 121495829 121496058
refGene_gn=BBOF1;refGene_tx=XM_003987820.5;refGene_gn=BBOF1;refGene_tx=XM_006932987.4;refGene_gn=BBOF1;refGene_tx=XM_006932988.4;refGene_gn=BBOF1;refGene_tx=XM_019833285.2;refGene_gn=BBOF1;refGene_tx=XM_019833287.2;refGene_gn=BBOF1;refGene_tx=XM_019833288.2
B3 121503428 121503494
refGene_gn=BBOF1;refGene_tx=XM_003987820.5;refGene_gn=BBOF1;refGene_tx=XM_006932987.4;refGene_gn=BBOF1;refGene_tx=XM_006932988.4;refGene_gn=BBOF1;refGene_tx=XM_019833285.2;refGene_gn=BBOF1;refGene_tx=XM_019833287.2
B3 121504875 121505019
refGene_gn=BBOF1;refGene_tx=XM_003987820.5;refGene_gn=BBOF1;refGene_tx=XM_006932987.4;refGene_gn=BBOF1;refGene_tx=XM_006932988.4;refGene_gn=BBOF1;refGene_tx=XM_019833285.2
B3 121514392 121514473
refGene_gn=BBOF1;refGene_tx=XM_003987820.5;refGene_gn=BBOF1;refGene_tx=XM_006932987.4;refGene_gn=BBOF1;refGene_tx=XM_006932988.4;refGene_gn=BBOF1;refGene_tx=XM_019833285.2;refGene_gn=BBOF1;refGene_tx=XM_019833287.2;refGene_gn=BBOF1;refGene_tx=XM_019833288.2
B3 121517242 121517313
refGene_gn=BBOF1;refGene_tx=XM_003987820.5;refGene_gn=BBOF1;refGene_tx=XM_006932987.4;refGene_gn=BBOF1;refGene_tx=XM_006932988.4;refGene_gn=BBOF1;refGene_tx=XM_019833285.2;refGene_gn=BBOF1;refGene_tx=XM_019833287.2;refGene_gn=BBOF1;refGene_tx=XM_019833288.2
B3 121518074 121518219
refGene_gn=BBOF1;refGene_tx=XM_003987820.5;refGene_gn=BBOF1;refGene_tx=XM_006932987.4;refGene_gn=BBOF1;refGene_tx=XM_006932988.4;refGene_gn=BBOF1;refGene_tx=XM_019833285.2;refGene_gn=BBOF1;refGene_tx=XM_019833287.2;refGene_gn=BBOF1;refGene_tx=XM_019833288.2
B3 121519558 121520052
refGene_gn=BBOF1;refGene_tx=XM_003987820.5;refGene_gn=BBOF1;refGene_tx=XM_006932987.4;refGene_gn=BBOF1;refGene_tx=XM_006932988.4;refGene_gn=BBOF1;refGene_tx=XM_019833285.2;refGene_gn=BBOF1;refGene_tx=XM_019833287.2;refGene_gn=BBOF1;refGene_tx=XM_019833288.2
B3 121521888 121521990
refGene_gn=BBOF1;refGene_tx=XM_003987820.5;refGene_gn=BBOF1;refGene_tx=XM_006932987.4;refGene_gn=BBOF1;refGene_tx=XM_006932988.4;refGene_gn=BBOF1;refGene_tx=XM_019833285.2;refGene_gn=BBOF1;refGene_tx=XM_019833287.2;refGene_gn=BBOF1;refGene_tx=XM_019833288.2
B3 121525931 121526006
refGene_gn=BBOF1;refGene_tx=XM_003987820.5;refGene_gn=BBOF1;refGene_tx=XM_006932987.4;refGene_gn=BBOF1;refGene_tx=XM_006932988.4;refGene_gn=BBOF1;refGene_tx=XM_019833285.2;refGene_gn=BBOF1;refGene_tx=XM_019833287.2;refGene_gn=BBOF1;refGene_tx=XM_019833288.2
B3 121526176 121526291

refGene_gn=BBOF1;refGene_tx=XM_003987820.5;refGene_gn=BBOF1;refGene_tx=XM_006932987.4;refGene_gn=BBOF1;refGene_tx=XM_006932988.4;refGene_gn=BBOF1;refGene_tx=XM_019833285.2;refGene_gn=BBOF1;refGene_tx=XM_019833287.2;refGene_gn=BBOF1;refGene_tx=XM_019833288.2
B3 121528829 121528934
refGene_gn=ALDH6A1;refGene_tx=XM_003987821.5
B3 121529406 121529424
refGene_gn=ALDH6A1;refGene_tx=XM_019833289.2
B3 121532822 121532834
refGene_gn=BBOF1;refGene_tx=XM_003987820.5
B3 121532958 121533057
refGene_gn=ALDH6A1;refGene_tx=XM_003987821.5;refGene_gn=ALDH6A1;refGene_tx=XM_019833289.2
B3 121533324 121533504
refGene_gn=ALDH6A1;refGene_tx=XM_003987821.5;refGene_gn=ALDH6A1;refGene_tx=XM_019833289.2
B3 121535359 121535541
refGene_gn=ALDH6A1;refGene_tx=XM_003987821.5;refGene_gn=ALDH6A1;refGene_tx=XM_019833289.2
B3 121535736 121535926
refGene_gn=ALDH6A1;refGene_tx=XM_003987821.5;refGene_gn=ALDH6A1;refGene_tx=XM_019833289.2
B3 121538147 121538269
refGene_gn=ALDH6A1;refGene_tx=XM_003987821.5;refGene_gn=ALDH6A1;refGene_tx=XM_019833289.2
B3 121539253 121539556
refGene_gn=ALDH6A1;refGene_tx=XM_003987821.5;refGene_gn=ALDH6A1;refGene_tx=XM_019833289.2
B3 121539978 121540057
refGene_gn=ALDH6A1;refGene_tx=XM_003987821.5;refGene_gn=ALDH6A1;refGene_tx=XM_019833289.2
B3 121540263 121540425
refGene_gn=ALDH6A1;refGene_tx=XM_003987821.5;refGene_gn=ALDH6A1;refGene_tx=XM_019833289.2
B3 121540599 121540674
refGene_gn=ALDH6A1;refGene_tx=XM_003987821.5;refGene_gn=ALDH6A1;refGene_tx=XM_019833289.2
B3 121541228 121541252
refGene_gn=BBOF1;refGene_tx=XM_006932988.4
B3 121542262 121542319
refGene_gn=ALDH6A1;refGene_tx=XM_003987821.5;refGene_gn=ALDH6A1;refGene_tx=XM_019833289.2
B3 121546470 121546518
refGene_gn=BBOF1;refGene_tx=XM_006932987.4;refGene_gn=BBOF1;refGene_tx=XM_019833287.2;refGene_gn=BBOF1;refGene_tx=XM_019833288.2
B3 121547110 121547128
refGene_gn=BBOF1;refGene_tx=XM_019833285.2
B3 121550625 121550676
refGene_gn=ALDH6A1;refGene_tx=XM_003987821.5;refGene_gn=ALDH6A1;refGene_tx=XM_019833289.2

B3 121551257 121551288
refGene_gn=LIN52;refGene_tx=XM_003987822.4;refGene_gn=LIN52;refGene_tx=XM_006932985.4;refGene_gn=LIN52;refGene_tx=XM_006932986.3;refGene_gn=LIN52;refGene_tx=XM_019833292.2;refGene_gn=LIN52;refGene_tx=XM_023255849.1
B3 121556286 121556361
refGene_gn=LIN52;refGene_tx=XM_003987822.4;refGene_gn=LIN52;refGene_tx=XM_006932985.4;refGene_gn=LIN52;refGene_tx=XM_006932986.3;refGene_gn=LIN52;refGene_tx=XM_019833292.2;refGene_gn=LIN52;refGene_tx=XM_023255849.1
B3 121558118 121558156
refGene_gn=LIN52;refGene_tx=XM_003987822.4;refGene_gn=LIN52;refGene_tx=XM_006932985.4;refGene_gn=LIN52;refGene_tx=XM_006932986.3;refGene_gn=LIN52;refGene_tx=XM_019833292.2;refGene_gn=LIN52;refGene_tx=XM_023255849.1
B3 121559061 121559128
refGene_gn=LIN52;refGene_tx=XM_003987822.4;refGene_gn=LIN52;refGene_tx=XM_006932985.4;refGene_gn=LIN52;refGene_tx=XM_006932986.3;refGene_gn=LIN52;refGene_tx=XM_019833292.2;refGene_gn=LIN52;refGene_tx=XM_023255849.1
B3 121560816 121560900
refGene_gn=LIN52;refGene_tx=XM_003987822.4;refGene_gn=LIN52;refGene_tx=XM_006932985.4;refGene_gn=LIN52;refGene_tx=XM_006932986.3;refGene_gn=LIN52;refGene_tx=XM_019833292.2;refGene_gn=LIN52;refGene_tx=XM_023255849.1
B3 121564666 121564710
refGene_gn=LIN52;refGene_tx=XM_006932985.4
B3 121580420 121580428
refGene_gn=LIN52;refGene_tx=XM_006932986.3
B3 121588535 121588555
refGene_gn=LIN52;refGene_tx=XM_019833292.2
B3 121610224 121610268
refGene_gn=LIN52;refGene_tx=XM_023255849.1
B3 121657234 121657290
refGene_gn=LIN52;refGene_tx=XM_003987822.4
B3 121696628 121696998
refGene_gn=VSX2;refGene_tx=XM_011283328.3;refGene_gn=VSX2;refGene_tx=XM_023255850.1
B3 121698285 121698370
refGene_gn=VSX2;refGene_tx=XM_011283328.3;refGene_gn=VSX2;refGene_tx=XM_023255850.1
B3 121702031 121702155
refGene_gn=VSX2;refGene_tx=XM_011283328.3;refGene_gn=VSX2;refGene_tx=XM_023255850.1
B3 121712406 121712587
refGene_gn=VSX2;refGene_tx=XM_011283328.3;refGene_gn=VSX2;refGene_tx=XM_023255850.1
B3 121713368 121713751
refGene_gn=VSX2;refGene_tx=XM_011283328.3;refGene_gn=VSX2;refGene_tx=XM_023255850.1

B3	121727467	121727536
refGene_gn=ABCD4;refGene_tx=XM_003987823.5;refGene_gn=ABCD4;refGene_tx=XM_023255851.1;refGene_gn=ABCD4;refGene_tx=XM_023255852.1;refGene_gn=ABCD4;refGene_tx=XM_023255853.1;refGene_gn=ABCD4;refGene_tx=XM_023255854.1;refGene_gn=ABCD4;refGene_tx=XM_023255855.1;refGene_gn=ABCD4;refGene_tx=XM_023255856.1
B3	121727707	121727823
refGene_gn=ABCD4;refGene_tx=XM_003987823.5;refGene_gn=ABCD4;refGene_tx=XM_006932991.3;refGene_gn=ABCD4;refGene_tx=XM_023255851.1;refGene_gn=ABCD4;refGene_tx=XM_023255852.1;refGene_gn=ABCD4;refGene_tx=XM_023255853.1;refGene_gn=ABCD4;refGene_tx=XM_023255854.1;refGene_gn=ABCD4;refGene_tx=XM_023255855.1;refGene_gn=ABCD4;refGene_tx=XM_023255856.1
B3	121728478	121728555
refGene_gn=ABCD4;refGene_tx=XM_003987823.5;refGene_gn=ABCD4;refGene_tx=XM_006932991.3;refGene_gn=ABCD4;refGene_tx=XM_023255851.1;refGene_gn=ABCD4;refGene_tx=XM_023255852.1;refGene_gn=ABCD4;refGene_tx=XM_023255853.1;refGene_gn=ABCD4;refGene_tx=XM_023255854.1;refGene_gn=ABCD4;refGene_tx=XM_023255855.1;refGene_gn=ABCD4;refGene_tx=XM_023255856.1
B3	121728832	121728885
refGene_gn=ABCD4;refGene_tx=XM_003987823.5;refGene_gn=ABCD4;refGene_tx=XM_006932991.3;refGene_gn=ABCD4;refGene_tx=XM_023255851.1;refGene_gn=ABCD4;refGene_tx=XM_023255852.1;refGene_gn=ABCD4;refGene_tx=XM_023255853.1;refGene_gn=ABCD4;refGene_tx=XM_023255854.1;refGene_gn=ABCD4;refGene_tx=XM_023255855.1;refGene_gn=ABCD4;refGene_tx=XM_023255856.1
B3	121729317	121729367
refGene_gn=ABCD4;refGene_tx=XM_003987823.5;refGene_gn=ABCD4;refGene_tx=XM_006932991.3;refGene_gn=ABCD4;refGene_tx=XM_023255851.1;refGene_gn=ABCD4;refGene_tx=XM_023255852.1;refGene_gn=ABCD4;refGene_tx=XM_023255853.1;refGene_gn=ABCD4;refGene_tx=XM_023255854.1;refGene_gn=ABCD4;refGene_tx=XM_023255855.1;refGene_gn=ABCD4;refGene_tx=XM_023255856.1
B3	121729782	121729819
refGene_gn=ABCD4;refGene_tx=XM_003987823.5;refGene_gn=ABCD4;refGene_tx=XM_006932991.3;refGene_gn=ABCD4;refGene_tx=XM_023255851.1;refGene_gn=ABCD4;refGene_tx=XM_023255852.1;refGene_gn=ABCD4;refGene_tx=XM_023255853.1;refGene_gn=ABCD4;refGene_tx=XM_023255854.1;refGene_gn=ABCD4;refGene_tx=XM_023255855.1;refGene_gn=ABCD4;refGene_tx=XM_023255856.1
B3	121730211	121730303
refGene_gn=ABCD4;refGene_tx=XM_003987823.5;refGene_gn=ABCD4;refGene_tx=XM_006932991.3;refGene_gn=ABCD4;refGene_tx=XM_023255851.1;refGene_gn=ABCD4;refGene_tx=XM_023255852.1;refGene_gn=ABCD4;refGene_tx=XM_023255853.1;refGene_gn=ABCD4;refGene_tx=XM_023255854.1;refGene_gn=ABCD4;refGene_tx=XM_023255855.1;refGene_gn=ABCD4;refGene_tx=XM_023255856.1
B3	121730476	121730685
refGene_gn=ABCD4;refGene_tx=XM_003987823.5;refGene_gn=ABCD4;refGene_tx=XM_006932991.3;refGene_gn=ABCD4;refGene_tx=XM_023255851.1;refGene_gn=ABCD4;refGene_tx=XM_023255852.1;refGene_gn=ABCD4;refGene_tx=XM_023255853.1;refGene_gn=ABCD4;refGene_tx=XM_023255854.1;refGene_gn=ABCD4;refGene_tx=XM_023255855.1;refGene_gn=ABCD4;refGene_tx=XM_023255856.1
B3	121732384	121732474
refGene_gn=ABCD4;refGene_tx=XM_003987823.5;refGene_gn=ABCD4;refGene_tx

=XM_006932991.3;refGene_gn=ABCD4;refGene_tx=XM_023255851.1;refGene_gn=ABCD4;refGene_tx=XM_023255852.1;refGene_gn=ABCD4;refGene_tx=XM_023255853.1;refGene_gn=ABCD4;refGene_tx=XM_023255854.1;refGene_gn=ABCD4;refGene_tx=XM_023255855.1;refGene_gn=ABCD4;refGene_tx=XM_023255856.1
B3 121732647 121732739
refGene_gn=ABCD4;refGene_tx=XM_003987823.5;refGene_gn=ABCD4;refGene_tx=XM_006932991.3;refGene_gn=ABCD4;refGene_tx=XM_023255851.1;refGene_gn=ABCD4;refGene_tx=XM_023255852.1;refGene_gn=ABCD4;refGene_tx=XM_023255853.1;refGene_gn=ABCD4;refGene_tx=XM_023255854.1;refGene_gn=ABCD4;refGene_tx=XM_023255855.1;refGene_gn=ABCD4;refGene_tx=XM_023255856.1
B3 121732854 121732976
refGene_gn=ABCD4;refGene_tx=XM_003987823.5;refGene_gn=ABCD4;refGene_tx=XM_006932991.3;refGene_gn=ABCD4;refGene_tx=XM_023255851.1;refGene_gn=ABCD4;refGene_tx=XM_023255852.1;refGene_gn=ABCD4;refGene_tx=XM_023255853.1;refGene_gn=ABCD4;refGene_tx=XM_023255854.1;refGene_gn=ABCD4;refGene_tx=XM_023255855.1;refGene_gn=ABCD4;refGene_tx=XM_023255856.1
B3 121733234 121733329
refGene_gn=ABCD4;refGene_tx=XM_003987823.5;refGene_gn=ABCD4;refGene_tx=XM_006932991.3;refGene_gn=ABCD4;refGene_tx=XM_023255851.1;refGene_gn=ABCD4;refGene_tx=XM_023255852.1;refGene_gn=ABCD4;refGene_tx=XM_023255853.1;refGene_gn=ABCD4;refGene_tx=XM_023255854.1;refGene_gn=ABCD4;refGene_tx=XM_023255855.1;refGene_gn=ABCD4;refGene_tx=XM_023255856.1
B3 121734971 121735022
refGene_gn=ABCD4;refGene_tx=XM_003987823.5;refGene_gn=ABCD4;refGene_tx=XM_006932991.3;refGene_gn=ABCD4;refGene_tx=XM_023255851.1;refGene_gn=ABCD4;refGene_tx=XM_023255852.1;refGene_gn=ABCD4;refGene_tx=XM_023255853.1;refGene_gn=ABCD4;refGene_tx=XM_023255854.1;refGene_gn=ABCD4;refGene_tx=XM_023255855.1;refGene_gn=ABCD4;refGene_tx=XM_023255856.1
B3 121735552 121735678
refGene_gn=ABCD4;refGene_tx=XM_003987823.5;refGene_gn=ABCD4;refGene_tx=XM_006932991.3;refGene_gn=ABCD4;refGene_tx=XM_023255851.1;refGene_gn=ABCD4;refGene_tx=XM_023255852.1;refGene_gn=ABCD4;refGene_tx=XM_023255853.1;refGene_gn=ABCD4;refGene_tx=XM_023255854.1;refGene_gn=ABCD4;refGene_tx=XM_023255855.1;refGene_gn=ABCD4;refGene_tx=XM_023255856.1
B3 121736054 121736171
refGene_gn=ABCD4;refGene_tx=XM_003987823.5;refGene_gn=ABCD4;refGene_tx=XM_006932991.3;refGene_gn=ABCD4;refGene_tx=XM_023255851.1;refGene_gn=ABCD4;refGene_tx=XM_023255852.1;refGene_gn=ABCD4;refGene_tx=XM_023255853.1;refGene_gn=ABCD4;refGene_tx=XM_023255854.1;refGene_gn=ABCD4;refGene_tx=XM_023255855.1
B3 121736839 121736898
refGene_gn=ABCD4;refGene_tx=XM_023255851.1
B3 121737366 121737506
refGene_gn=ABCD4;refGene_tx=XM_003987823.5;refGene_gn=ABCD4;refGene_tx=XM_006932991.3
B3 121738657 121738785
refGene_gn=ABCD4;refGene_tx=XM_003987823.5;refGene_gn=ABCD4;refGene_tx=XM_006932991.3
B3 121738926 121739045
refGene_gn=ABCD4;refGene_tx=XM_003987823.5;refGene_gn=ABCD4;refGene_tx

=XM_006932991.3
B3	121742802	121742840
refGene_gn=ABCD4;refGene_tx=XM_003987823.5;refGene_gn=ABCD4;refGene_tx=XM_006932991.3;refGene_gn=ABCD4;refGene_tx=XM_023255852.1
B3	121776947	121779059
refGene_gn=VRTN;refGene_tx=XM_023255857.1
B3	121805116	121805275
refGene_gn=SYNDIG1L;refGene_tx=XM_019833298.2;refGene_gn=SYNDIG1L;refGene_tx=XM_019833300.2;refGene_gn=SYNDIG1L;refGene_tx=XM_019833302.2
B3	121805433	121805574
refGene_gn=SYNDIG1L;refGene_tx=XM_019833298.2;refGene_gn=SYNDIG1L;refGene_tx=XM_019833300.2;refGene_gn=SYNDIG1L;refGene_tx=XM_019833302.2
B3	121807007	121807427
refGene_gn=SYNDIG1L;refGene_tx=XM_019833298.2;refGene_gn=SYNDIG1L;refGene_tx=XM_019833300.2;refGene_gn=SYNDIG1L;refGene_tx=XM_019833302.2
B3	121865135	121865222
refGene_gn=NPC2;refGene_tx=XM_003987833.5
B3	121867425	121867598
refGene_gn=NPC2;refGene_tx=XM_003987833.5
B3	121869130	121869238
refGene_gn=NPC2;refGene_tx=XM_003987833.5
B3	121872992	121873074
refGene_gn=NPC2;refGene_tx=XM_003987833.5
B3	121873533	121873604
refGene_gn=ISCA2;refGene_tx=XM_003987834.3
B3	121873807	121873910
refGene_gn=ISCA2;refGene_tx=XM_003987834.3
B3	121874028	121874144
refGene_gn=ISCA2;refGene_tx=XM_003987834.3
B3	121874782	121874957
refGene_gn=ISCA2;refGene_tx=XM_003987834.3
B3	121881352	121881498
refGene_gn=LTBP2;refGene_tx=XM_023255858.1;refGene_gn=LTBP2;refGene_tx=XM_023255859.1;refGene_gn=LTBP2;refGene_tx=XM_023255860.1
B3	121881950	121882100
refGene_gn=LTBP2;refGene_tx=XM_023255858.1;refGene_gn=LTBP2;refGene_tx=XM_023255859.1;refGene_gn=LTBP2;refGene_tx=XM_023255860.1
B3	121883082	121883364
refGene_gn=LTBP2;refGene_tx=XM_023255858.1;refGene_gn=LTBP2;refGene_tx=XM_023255859.1;refGene_gn=LTBP2;refGene_tx=XM_023255860.1
B3	121883668	121883836
refGene_gn=LTBP2;refGene_tx=XM_023255858.1;refGene_gn=LTBP2;refGene_tx=XM_023255859.1;refGene_gn=LTBP2;refGene_tx=XM_023255860.1
B3	121883924	121884062
refGene_gn=LTBP2;refGene_tx=XM_023255858.1;refGene_gn=LTBP2;refGene_tx=XM_023255859.1;refGene_gn=LTBP2;refGene_tx=XM_023255860.1;refGene_gn=LTBP2;refGene_tx=XM_023255861.1
B3	121884368	121884497
refGene_gn=LTBP2;refGene_tx=XM_023255858.1;refGene_gn=LTBP2;refGene_tx=XM_023255859.1;refGene_gn=LTBP2;refGene_tx=XM_023255860.1;refGene_gn=

LTBP2;refGene_tx=XM_023255861.1
B3 121885193 121885277
refGene_gn=LTBP2;refGene_tx=XM_023255858.1;refGene_gn=LTBP2;refGene_tx=XM_023255859.1;refGene_gn=LTBP2;refGene_tx=XM_023255860.1;refGene_gn=LTBP2;refGene_tx=XM_023255861.1
B3 121885397 121885589
refGene_gn=LTBP2;refGene_tx=XM_023255858.1;refGene_gn=LTBP2;refGene_tx=XM_023255859.1;refGene_gn=LTBP2;refGene_tx=XM_023255860.1;refGene_gn=LTBP2;refGene_tx=XM_023255861.1
B3 121886666 121886810
refGene_gn=LTBP2;refGene_tx=XM_023255858.1;refGene_gn=LTBP2;refGene_tx=XM_023255859.1;refGene_gn=LTBP2;refGene_tx=XM_023255860.1;refGene_gn=LTBP2;refGene_tx=XM_023255861.1
B3 121887380 121887506
refGene_gn=LTBP2;refGene_tx=XM_023255858.1;refGene_gn=LTBP2;refGene_tx=XM_023255859.1;refGene_gn=LTBP2;refGene_tx=XM_023255860.1;refGene_gn=LTBP2;refGene_tx=XM_023255861.1
B3 121887793 121887925
refGene_gn=LTBP2;refGene_tx=XM_023255858.1;refGene_gn=LTBP2;refGene_tx=XM_023255859.1;refGene_gn=LTBP2;refGene_tx=XM_023255860.1;refGene_gn=LTBP2;refGene_tx=XM_023255861.1
B3 121888623 121888746
refGene_gn=LTBP2;refGene_tx=XM_023255858.1;refGene_gn=LTBP2;refGene_tx=XM_023255859.1;refGene_gn=LTBP2;refGene_tx=XM_023255860.1;refGene_gn=LTBP2;refGene_tx=XM_023255861.1
B3 121889149 121889275
refGene_gn=LTBP2;refGene_tx=XM_023255858.1;refGene_gn=LTBP2;refGene_tx=XM_023255859.1;refGene_gn=LTBP2;refGene_tx=XM_023255860.1;refGene_gn=LTBP2;refGene_tx=XM_023255861.1
B3 121889376 121889499
refGene_gn=LTBP2;refGene_tx=XM_023255858.1;refGene_gn=LTBP2;refGene_tx=XM_023255859.1;refGene_gn=LTBP2;refGene_tx=XM_023255860.1;refGene_gn=LTBP2;refGene_tx=XM_023255861.1
B3 121890270 121890396
refGene_gn=LTBP2;refGene_tx=XM_023255858.1;refGene_gn=LTBP2;refGene_tx=XM_023255859.1;refGene_gn=LTBP2;refGene_tx=XM_023255860.1;refGene_gn=LTBP2;refGene_tx=XM_023255861.1
B3 121890840 121890966
refGene_gn=LTBP2;refGene_tx=XM_023255858.1;refGene_gn=LTBP2;refGene_tx=XM_023255859.1;refGene_gn=LTBP2;refGene_tx=XM_023255860.1;refGene_gn=LTBP2;refGene_tx=XM_023255861.1
B3 121891198 121891321
refGene_gn=LTBP2;refGene_tx=XM_023255858.1;refGene_gn=LTBP2;refGene_tx=XM_023255859.1;refGene_gn=LTBP2;refGene_tx=XM_023255860.1;refGene_gn=LTBP2;refGene_tx=XM_023255861.1
B3 121892400 121892520
refGene_gn=LTBP2;refGene_tx=XM_023255858.1;refGene_gn=LTBP2;refGene_tx=XM_023255860.1;refGene_gn=LTBP2;refGene_tx=XM_023255861.1
B3 121897773 121897893
refGene_gn=LTBP2;refGene_tx=XM_023255858.1;refGene_gn=LTBP2;refGene_tx

=XM_023255859.1;refGene_gn=LTBP2;refGene_tx=XM_023255860.1;refGene_gn=LTBP2;refGene_tx=XM_023255861.1
B3 121902237 121902366
refGene_gn=LTBP2;refGene_tx=XM_023255858.1;refGene_gn=LTBP2;refGene_tx=XM_023255859.1;refGene_gn=LTBP2;refGene_tx=XM_023255860.1;refGene_gn=LTBP2;refGene_tx=XM_023255861.1
B3 121903022 121903151
refGene_gn=LTBP2;refGene_tx=XM_023255858.1;refGene_gn=LTBP2;refGene_tx=XM_023255859.1;refGene_gn=LTBP2;refGene_tx=XM_023255860.1;refGene_gn=LTBP2;refGene_tx=XM_023255861.1
B3 121905307 121905421
refGene_gn=LTBP2;refGene_tx=XM_023255858.1;refGene_gn=LTBP2;refGene_tx=XM_023255859.1;refGene_gn=LTBP2;refGene_tx=XM_023255860.1;refGene_gn=LTBP2;refGene_tx=XM_023255861.1
B3 121906130 121906170
refGene_gn=LTBP2;refGene_tx=XM_023255858.1;refGene_gn=LTBP2;refGene_tx=XM_023255859.1;refGene_gn=LTBP2;refGene_tx=XM_023255860.1;refGene_gn=LTBP2;refGene_tx=XM_023255861.1
B3 121907319 121907339
refGene_gn=LTBP2;refGene_tx=XM_023255858.1;refGene_gn=LTBP2;refGene_tx=XM_023255859.1;refGene_gn=LTBP2;refGene_tx=XM_023255860.1;refGene_gn=LTBP2;refGene_tx=XM_023255861.1
B3 121908435 121908651
refGene_gn=LTBP2;refGene_tx=XM_023255858.1;refGene_gn=LTBP2;refGene_tx=XM_023255859.1;refGene_gn=LTBP2;refGene_tx=XM_023255860.1;refGene_gn=LTBP2;refGene_tx=XM_023255861.1
B3 121908911 121909076
refGene_gn=LTBP2;refGene_tx=XM_023255858.1;refGene_gn=LTBP2;refGene_tx=XM_023255859.1;refGene_gn=LTBP2;refGene_tx=XM_023255860.1;refGene_gn=LTBP2;refGene_tx=XM_023255861.1
B3 121912913 121913036
refGene_gn=LTBP2;refGene_tx=XM_023255858.1;refGene_gn=LTBP2;refGene_tx=XM_023255859.1;refGene_gn=LTBP2;refGene_tx=XM_023255860.1;refGene_gn=LTBP2;refGene_tx=XM_023255861.1
B3 121916462 121916537
refGene_gn=LTBP2;refGene_tx=XM_023255858.1;refGene_gn=LTBP2;refGene_tx=XM_023255859.1;refGene_gn=LTBP2;refGene_tx=XM_023255860.1;refGene_gn=LTBP2;refGene_tx=XM_023255861.1
B3 121928203 121928306
refGene_gn=LTBP2;refGene_tx=XM_023255858.1;refGene_gn=LTBP2;refGene_tx=XM_023255859.1;refGene_gn=LTBP2;refGene_tx=XM_023255860.1;refGene_gn=LTBP2;refGene_tx=XM_023255861.1
B3 121929375 121929653
refGene_gn=LTBP2;refGene_tx=XM_023255858.1;refGene_gn=LTBP2;refGene_tx=XM_023255859.1;refGene_gn=LTBP2;refGene_tx=XM_023255860.1;refGene_gn=LTBP2;refGene_tx=XM_023255861.1
B3 121930456 121930663
refGene_gn=LTBP2;refGene_tx=XM_023255858.1;refGene_gn=LTBP2;refGene_tx=XM_023255859.1;refGene_gn=LTBP2;refGene_tx=XM_023255860.1;refGene_gn=LTBP2;refGene_tx=XM_023255861.1

B3 121931224 121931395
refGene_gn=LTBP2;refGene_tx=XM_023255858.1;refGene_gn=LTBP2;refGene_tx=XM_023255859.1;refGene_gn=LTBP2;refGene_tx=XM_023255860.1;refGene_gn=LTBP2;refGene_tx=XM_023255861.1
B3 121933350 121933541
refGene_gn=LTBP2;refGene_tx=XM_023255858.1;refGene_gn=LTBP2;refGene_tx=XM_023255859.1;refGene_gn=LTBP2;refGene_tx=XM_023255860.1;refGene_gn=LTBP2;refGene_tx=XM_023255861.1
B3 121961300 121961550
refGene_gn=LTBP2;refGene_tx=XM_023255858.1;refGene_gn=LTBP2;refGene_tx=XM_023255859.1;refGene_gn=LTBP2;refGene_tx=XM_023255860.1;refGene_gn=LTBP2;refGene_tx=XM_023255861.1
B3 121962678 121962745
refGene_gn=LTBP2;refGene_tx=XM_023255860.1
B3 121976299 121976370
refGene_gn=LTBP2;refGene_tx=XM_023255858.1;refGene_gn=LTBP2;refGene_tx=XM_023255859.1;refGene_gn=LTBP2;refGene_tx=XM_023255861.1
B3 121983295 121983789
refGene_gn=LTBP2;refGene_tx=XM_023255858.1;refGene_gn=LTBP2;refGene_tx=XM_023255859.1;refGene_gn=LTBP2;refGene_tx=XM_023255861.1
B3 122028252 122028355
refGene_gn=AREL1;refGene_tx=XM_006932993.4;refGene_gn=AREL1;refGene_tx=XM_006932994.4;refGene_gn=AREL1;refGene_tx=XM_011283333.3;refGene_gn=AREL1;refGene_tx=XM_023255862.1;refGene_gn=AREL1;refGene_tx=XM_023255863.1
B3 122028460 122028636
refGene_gn=AREL1;refGene_tx=XM_006932993.4;refGene_gn=AREL1;refGene_tx=XM_006932994.4;refGene_gn=AREL1;refGene_tx=XM_011283333.3;refGene_gn=AREL1;refGene_tx=XM_023255862.1;refGene_gn=AREL1;refGene_tx=XM_023255863.1
B3 122029040 122029130
refGene_gn=AREL1;refGene_tx=XM_006932993.4;refGene_gn=AREL1;refGene_tx=XM_006932994.4;refGene_gn=AREL1;refGene_tx=XM_011283333.3;refGene_gn=AREL1;refGene_tx=XM_023255862.1;refGene_gn=AREL1;refGene_tx=XM_023255863.1
B3 122031769 122031828
refGene_gn=AREL1;refGene_tx=XM_006932993.4;refGene_gn=AREL1;refGene_tx=XM_006932994.4;refGene_gn=AREL1;refGene_tx=XM_011283333.3;refGene_gn=AREL1;refGene_tx=XM_023255862.1;refGene_gn=AREL1;refGene_tx=XM_023255863.1
B3 122031915 122032045
refGene_gn=AREL1;refGene_tx=XM_006932993.4;refGene_gn=AREL1;refGene_tx=XM_006932994.4;refGene_gn=AREL1;refGene_tx=XM_011283333.3;refGene_gn=AREL1;refGene_tx=XM_023255862.1;refGene_gn=AREL1;refGene_tx=XM_023255863.1
B3 122032741 122032867
refGene_gn=AREL1;refGene_tx=XM_006932993.4;refGene_gn=AREL1;refGene_tx=XM_006932994.4;refGene_gn=AREL1;refGene_tx=XM_011283333.3;refGene_gn=AREL1;refGene_tx=XM_023255862.1;refGene_gn=AREL1;refGene_tx=XM_023255863.1

B3	122033036	122033216
refGene_gn=AREL1;refGene_tx=XM_006932993.4;refGene_gn=AREL1;refGene_tx=XM_006932994.4;refGene_gn=AREL1;refGene_tx=XM_011283333.3;refGene_gn=AREL1;refGene_tx=XM_023255862.1;refGene_gn=AREL1;refGene_tx=XM_023255863.1
B3	122034127	122034237
refGene_gn=AREL1;refGene_tx=XM_006932993.4;refGene_gn=AREL1;refGene_tx=XM_006932994.4;refGene_gn=AREL1;refGene_tx=XM_011283333.3;refGene_gn=AREL1;refGene_tx=XM_023255862.1;refGene_gn=AREL1;refGene_tx=XM_023255863.1
B3	122034991	122035067
refGene_gn=AREL1;refGene_tx=XM_006932993.4;refGene_gn=AREL1;refGene_tx=XM_006932994.4;refGene_gn=AREL1;refGene_tx=XM_011283333.3;refGene_gn=AREL1;refGene_tx=XM_023255862.1;refGene_gn=AREL1;refGene_tx=XM_023255863.1
B3	122036042	122036164
refGene_gn=AREL1;refGene_tx=XM_006932993.4;refGene_gn=AREL1;refGene_tx=XM_006932994.4;refGene_gn=AREL1;refGene_tx=XM_011283333.3;refGene_gn=AREL1;refGene_tx=XM_023255862.1;refGene_gn=AREL1;refGene_tx=XM_023255863.1
B3	122036295	122036437
refGene_gn=AREL1;refGene_tx=XM_006932993.4;refGene_gn=AREL1;refGene_tx=XM_006932994.4;refGene_gn=AREL1;refGene_tx=XM_011283333.3;refGene_gn=AREL1;refGene_tx=XM_023255862.1;refGene_gn=AREL1;refGene_tx=XM_023255863.1
B3	122037313	122037391
refGene_gn=AREL1;refGene_tx=XM_006932993.4;refGene_gn=AREL1;refGene_tx=XM_006932994.4;refGene_gn=AREL1;refGene_tx=XM_011283333.3;refGene_gn=AREL1;refGene_tx=XM_023255862.1;refGene_gn=AREL1;refGene_tx=XM_023255863.1
B3	122039124	122039372
refGene_gn=AREL1;refGene_tx=XM_006932993.4;refGene_gn=AREL1;refGene_tx=XM_006932994.4;refGene_gn=AREL1;refGene_tx=XM_011283333.3;refGene_gn=AREL1;refGene_tx=XM_023255862.1;refGene_gn=AREL1;refGene_tx=XM_023255863.1
B3	122039579	122039760
refGene_gn=AREL1;refGene_tx=XM_006932993.4;refGene_gn=AREL1;refGene_tx=XM_006932994.4;refGene_gn=AREL1;refGene_tx=XM_011283333.3;refGene_gn=AREL1;refGene_tx=XM_023255862.1;refGene_gn=AREL1;refGene_tx=XM_023255863.1
B3	122039999	122040169
refGene_gn=AREL1;refGene_tx=XM_006932993.4;refGene_gn=AREL1;refGene_tx=XM_006932994.4;refGene_gn=AREL1;refGene_tx=XM_011283333.3;refGene_gn=AREL1;refGene_tx=XM_023255862.1;refGene_gn=AREL1;refGene_tx=XM_023255863.1
B3	122049471	122049709
refGene_gn=AREL1;refGene_tx=XM_006932993.4;refGene_gn=AREL1;refGene_tx=XM_006932994.4;refGene_gn=AREL1;refGene_tx=XM_011283333.3;refGene_gn=AREL1;refGene_tx=XM_023255862.1;refGene_gn=AREL1;refGene_tx=XM_023255863.1

B3 122050913 122051140
refGene_gn=AREL1;refGene_tx=XM_006932993.4;refGene_gn=AREL1;refGene_tx=XM_006932994.4;refGene_gn=AREL1;refGene_tx=XM_011283333.3;refGene_gn=AREL1;refGene_tx=XM_023255862.1;refGene_gn=AREL1;refGene_tx=XM_023255863.1
B3 122052022 122052038
refGene_gn=AREL1;refGene_tx=XM_006932993.4;refGene_gn=AREL1;refGene_tx=XM_006932994.4;refGene_gn=AREL1;refGene_tx=XM_011283333.3;refGene_gn=AREL1;refGene_tx=XM_023255862.1;refGene_gn=AREL1;refGene_tx=XM_023255863.1
B3 122074983 122074986
refGene_gn=FCF1;refGene_tx=XM_011283334.2
B3 122075330 122075398
refGene_gn=FCF1;refGene_tx=XM_011283334.2;refGene_gn=FCF1;refGene_tx=XM_011283335.3;refGene_gn=FCF1;refGene_tx=XM_019833306.2
B3 122077188 122077260
refGene_gn=FCF1;refGene_tx=XM_011283334.2;refGene_gn=FCF1;refGene_tx=XM_011283335.3;refGene_gn=FCF1;refGene_tx=XM_019833306.2
B3 122077857 122078006
refGene_gn=FCF1;refGene_tx=XM_011283334.2;refGene_gn=FCF1;refGene_tx=XM_011283335.3;refGene_gn=FCF1;refGene_tx=XM_019833306.2
B3 122082273 122082346
refGene_gn=FCF1;refGene_tx=XM_011283334.2;refGene_gn=FCF1;refGene_tx=XM_011283335.3;refGene_gn=FCF1;refGene_tx=XM_019833306.2
B3 122086922 122087010
refGene_gn=FCF1;refGene_tx=XM_011283334.2;refGene_gn=FCF1;refGene_tx=XM_011283335.3;refGene_gn=FCF1;refGene_tx=XM_019833306.2
B3 122088207 122088302
refGene_gn=FCF1;refGene_tx=XM_011283334.2;refGene_gn=FCF1;refGene_tx=XM_011283335.3;refGene_gn=FCF1;refGene_tx=XM_019833306.2
B3 122089017 122089066
refGene_gn=FCF1;refGene_tx=XM_011283334.2;refGene_gn=FCF1;refGene_tx=XM_011283335.3;refGene_gn=FCF1;refGene_tx=XM_019833306.2
B3 122112019 122112895
refGene_gn=YLPM1;refGene_tx=XM_011283407.3;refGene_gn=YLPM1;refGene_tx=XM_023255864.1;refGene_gn=YLPM1;refGene_tx=XM_023255865.1
B3 122121987 122122224
refGene_gn=YLPM1;refGene_tx=XM_011283407.3;refGene_gn=YLPM1;refGene_tx=XM_023255864.1;refGene_gn=YLPM1;refGene_tx=XM_023255865.1
B3 122123147 122123327
refGene_gn=YLPM1;refGene_tx=XM_011283407.3;refGene_gn=YLPM1;refGene_tx=XM_023255864.1;refGene_gn=YLPM1;refGene_tx=XM_023255865.1
B3 122124093 122125088
refGene_gn=YLPM1;refGene_tx=XM_011283407.3;refGene_gn=YLPM1;refGene_tx=XM_023255864.1;refGene_gn=YLPM1;refGene_tx=XM_023255865.1
B3 122141904 122144025
refGene_gn=YLPM1;refGene_tx=XM_011283407.3;refGene_gn=YLPM1;refGene_tx=XM_023255864.1;refGene_gn=YLPM1;refGene_tx=XM_023255865.1
B3 122146651 122146772
refGene_gn=YLPM1;refGene_tx=XM_011283407.3;refGene_gn=YLPM1;refGene_tx

=XM_023255864.1;refGene_gn=YLPM1;refGene_tx=XM_023255865.1
B3 122153832 122154253
refGene_gn=YLPM1;refGene_tx=XM_011283407.3;refGene_gn=YLPM1;refGene_tx
=XM_023255864.1;refGene_gn=YLPM1;refGene_tx=XM_023255865.1
B3 122154365 122154458
refGene_gn=YLPM1;refGene_tx=XM_011283407.3;refGene_gn=YLPM1;refGene_tx
=XM_023255864.1;refGene_gn=YLPM1;refGene_tx=XM_023255865.1
B3 122154673 122154869
refGene_gn=YLPM1;refGene_tx=XM_011283407.3;refGene_gn=YLPM1;refGene_tx
=XM_023255864.1;refGene_gn=YLPM1;refGene_tx=XM_023255865.1
B3 122155527 122155646
refGene_gn=YLPM1;refGene_tx=XM_011283407.3;refGene_gn=YLPM1;refGene_tx
=XM_023255864.1;refGene_gn=YLPM1;refGene_tx=XM_023255865.1
B3 122156550 122156705
refGene_gn=YLPM1;refGene_tx=XM_011283407.3;refGene_gn=YLPM1;refGene_tx
=XM_023255864.1;refGene_gn=YLPM1;refGene_tx=XM_023255865.1
B3 122160132 122160195
refGene_gn=YLPM1;refGene_tx=XM_011283407.3;refGene_gn=YLPM1;refGene_tx
=XM_023255864.1;refGene_gn=YLPM1;refGene_tx=XM_023255865.1
B3 122160482 122160602
refGene_gn=YLPM1;refGene_tx=XM_011283407.3;refGene_gn=YLPM1;refGene_tx
=XM_023255864.1;refGene_gn=YLPM1;refGene_tx=XM_023255865.1
B3 122161074 122161251
refGene_gn=YLPM1;refGene_tx=XM_011283407.3;refGene_gn=YLPM1;refGene_tx
=XM_023255864.1;refGene_gn=YLPM1;refGene_tx=XM_023255865.1
B3 122161330 122161414
refGene_gn=YLPM1;refGene_tx=XM_011283407.3;refGene_gn=YLPM1;refGene_tx
=XM_023255864.1;refGene_gn=YLPM1;refGene_tx=XM_023255865.1
B3 122161702 122161786
refGene_gn=YLPM1;refGene_tx=XM_011283407.3;refGene_gn=YLPM1;refGene_tx
=XM_023255864.1;refGene_gn=YLPM1;refGene_tx=XM_023255865.1
B3 122163964 122164045
refGene_gn=YLPM1;refGene_tx=XM_011283407.3;refGene_gn=YLPM1;refGene_tx
=XM_023255864.1;refGene_gn=YLPM1;refGene_tx=XM_023255865.1
B3 122164803 122164836
refGene_gn=YLPM1;refGene_tx=XM_023255864.1
B3 122167186 122167238
refGene_gn=YLPM1;refGene_tx=XM_011283407.3
B3 122172302 122172433
refGene_gn=YLPM1;refGene_tx=XM_011283407.3;refGene_gn=YLPM1;refGene_tx
=XM_023255865.1
B3 122185667 122185814
refGene_gn=YLPM1;refGene_tx=XM_011283407.3
B3 122207325 122207511
refGene_gn=PROX2;refGene_tx=XM_003987877.3
B3 122208971 122209166
refGene_gn=PROX2;refGene_tx=XM_003987877.3
B3 122210192 122210300
refGene_gn=PROX2;refGene_tx=XM_003987877.3
B3 122214258 122215581

refGene_gn=PROX2;refGene_tx=XM_003987877.3
B3 122217456 122217477
refGene_gn=DLST;refGene_tx=XM_023255870.1
B3 122234646 122234709
refGene_gn=DLST;refGene_tx=XM_003987837.5
B3 122235278 122235312
refGene_gn=DLST;refGene_tx=XM_003987837.5;refGene_gn=DLST;refGene_tx=XM_023255870.1
B3 122238139 122238188
refGene_gn=DLST;refGene_tx=XM_003987837.5;refGene_gn=DLST;refGene_tx=XM_023255870.1
B3 122241024 122241080
refGene_gn=DLST;refGene_tx=XM_003987837.5;refGene_gn=DLST;refGene_tx=XM_023255870.1
B3 122241208 122241283
refGene_gn=DLST;refGene_tx=XM_003987837.5;refGene_gn=DLST;refGene_tx=XM_023255870.1
B3 122241510 122241566
refGene_gn=DLST;refGene_tx=XM_003987837.5;refGene_gn=DLST;refGene_tx=XM_023255870.1
B3 122242139 122242251
refGene_gn=DLST;refGene_tx=XM_003987837.5;refGene_gn=DLST;refGene_tx=XM_023255870.1
B3 122243774 122243930
refGene_gn=DLST;refGene_tx=XM_003987837.5;refGene_gn=DLST;refGene_tx=XM_023255870.1
B3 122244299 122244376
refGene_gn=DLST;refGene_tx=XM_003987837.5;refGene_gn=DLST;refGene_tx=XM_023255870.1
B3 122245254 122245352
refGene_gn=DLST;refGene_tx=XM_003987837.5;refGene_gn=DLST;refGene_tx=XM_023255870.1
B3 122249163 122249294
refGene_gn=DLST;refGene_tx=XM_003987837.5;refGene_gn=DLST;refGene_tx=XM_023255870.1
B3 122250708 122250782
refGene_gn=DLST;refGene_tx=XM_003987837.5;refGene_gn=DLST;refGene_tx=XM_023255870.1
B3 122251080 122251164
refGene_gn=DLST;refGene_tx=XM_003987837.5;refGene_gn=DLST;refGene_tx=XM_023255870.1
B3 122251580 122251748
refGene_gn=DLST;refGene_tx=XM_003987837.5;refGene_gn=DLST;refGene_tx=XM_023255870.1
B3 122252719 122252854
refGene_gn=DLST;refGene_tx=XM_003987837.5;refGene_gn=DLST;refGene_tx=XM_023255870.1
B3 122256834 122256945
refGene_gn=RPS6KL1;refGene_tx=XM_023255869.1
B3 122257046 122257328

refGene_gn=RPS6KL1;refGene_tx=XM_023255866.1;refGene_gn=RPS6KL1;refGene_tx=XM_023255867.1;refGene_gn=RPS6KL1;refGene_tx=XM_023255868.1;refGene_gn=RPS6KL1;refGene_tx=XM_023255869.1
B3 122258657 122258740
refGene_gn=RPS6KL1;refGene_tx=XM_023255866.1;refGene_gn=RPS6KL1;refGene_tx=XM_023255867.1;refGene_gn=RPS6KL1;refGene_tx=XM_023255868.1;refGene_gn=RPS6KL1;refGene_tx=XM_023255869.1
B3 122258907 122258997
refGene_gn=RPS6KL1;refGene_tx=XM_023255866.1;refGene_gn=RPS6KL1;refGene_tx=XM_023255867.1;refGene_gn=RPS6KL1;refGene_tx=XM_023255868.1;refGene_gn=RPS6KL1;refGene_tx=XM_023255869.1
B3 122259407 122260016
refGene_gn=RPS6KL1;refGene_tx=XM_023255866.1;refGene_gn=RPS6KL1;refGene_tx=XM_023255867.1;refGene_gn=RPS6KL1;refGene_tx=XM_023255868.1;refGene_gn=RPS6KL1;refGene_tx=XM_023255869.1
B3 122260612 122260745
refGene_gn=RPS6KL1;refGene_tx=XM_023255866.1;refGene_gn=RPS6KL1;refGene_tx=XM_023255867.1;refGene_gn=RPS6KL1;refGene_tx=XM_023255868.1;refGene_gn=RPS6KL1;refGene_tx=XM_023255869.1
B3 122261126 122261174
refGene_gn=RPS6KL1;refGene_tx=XM_023255866.1;refGene_gn=RPS6KL1;refGene_tx=XM_023255867.1;refGene_gn=RPS6KL1;refGene_tx=XM_023255869.1
B3 122267697 122267790
refGene_gn=RPS6KL1;refGene_tx=XM_023255866.1;refGene_gn=RPS6KL1;refGene_tx=XM_023255867.1;refGene_gn=RPS6KL1;refGene_tx=XM_023255868.1;refGene_gn=RPS6KL1;refGene_tx=XM_023255869.1
B3 122268770 122268895
refGene_gn=RPS6KL1;refGene_tx=XM_023255866.1;refGene_gn=RPS6KL1;refGene_tx=XM_023255867.1;refGene_gn=RPS6KL1;refGene_tx=XM_023255868.1;refGene_gn=RPS6KL1;refGene_tx=XM_023255869.1
B3 122271545 122271810
refGene_gn=RPS6KL1;refGene_tx=XM_023255866.1;refGene_gn=RPS6KL1;refGene_tx=XM_023255867.1;refGene_gn=RPS6KL1;refGene_tx=XM_023255868.1;refGene_gn=RPS6KL1;refGene_tx=XM_023255869.1
B3 122291455 122291483
refGene_gn=PGF;refGene_tx=XM_003987840.4;refGene_gn=PGF;refGene_tx=XM_003987841.4;refGene_gn=PGF;refGene_tx=XM_019833310.1
B3 122294028 122294124
refGene_gn=PGF;refGene_tx=XM_003987840.4;refGene_gn=PGF;refGene_tx=XM_019833310.1
B3 122294196 122294226
refGene_gn=PGF;refGene_tx=XM_003987840.4;refGene_gn=PGF;refGene_tx=XM_003987841.4;refGene_gn=PGF;refGene_tx=XM_019833310.1
B3 122296318 122296395
refGene_gn=PGF;refGene_tx=XM_003987840.4;refGene_gn=PGF;refGene_tx=XM_003987841.4;refGene_gn=PGF;refGene_tx=XM_019833310.1
B3 122297162 122297359
refGene_gn=PGF;refGene_tx=XM_003987840.4;refGene_gn=PGF;refGene_tx=XM_003987841.4;refGene_gn=PGF;refGene_tx=XM_019833310.1
B3 122301684 122301727

refGene_gn=PGF;refGene_tx=XM_003987840.4;refGene_gn=PGF;refGene_tx=XM_003987841.4;refGene_gn=PGF;refGene_tx=XM_019833310.1
B3 122302962 122303037
refGene_gn=PGF;refGene_tx=XM_003987840.4;refGene_gn=PGF;refGene_tx=XM_003987841.4;refGene_gn=PGF;refGene_tx=XM_019833310.1
B3 122344399 122344562
refGene_gn=EIF2B2;refGene_tx=XM_003987842.4
B3 122344680 122344801
refGene_gn=EIF2B2;refGene_tx=XM_003987842.4
B3 122344928 122345077
refGene_gn=EIF2B2;refGene_tx=XM_003987842.4
B3 122345940 122346104
refGene_gn=EIF2B2;refGene_tx=XM_003987842.4
B3 122346754 122346850
refGene_gn=EIF2B2;refGene_tx=XM_003987842.4
B3 122347830 122347968
refGene_gn=EIF2B2;refGene_tx=XM_003987842.4
B3 122349197 122349264
refGene_gn=EIF2B2;refGene_tx=XM_003987842.4
B3 122350510 122350668
refGene_gn=EIF2B2;refGene_tx=XM_003987842.4
B3 122358938 122359061
refGene_gn=MLH3;refGene_tx=XM_003987843.5;refGene_gn=MLH3;refGene_tx=XM_019833311.2;refGene_gn=MLH3;refGene_tx=XM_019833313.2;refGene_gn=MLH3;refGene_tx=XM_023255871.1
B3 122360037 122360189
refGene_gn=MLH3;refGene_tx=XM_003987843.5;refGene_gn=MLH3;refGene_tx=XM_019833311.2;refGene_gn=MLH3;refGene_tx=XM_019833313.2;refGene_gn=MLH3;refGene_tx=XM_023255871.1
B3 122361413 122361492
refGene_gn=MLH3;refGene_tx=XM_003987843.5;refGene_gn=MLH3;refGene_tx=XM_019833311.2;refGene_gn=MLH3;refGene_tx=XM_019833313.2;refGene_gn=MLH3;refGene_tx=XM_023255871.1
B3 122361602 122361626
refGene_gn=MLH3;refGene_tx=XM_003987843.5;refGene_gn=MLH3;refGene_tx=XM_019833311.2;refGene_gn=MLH3;refGene_tx=XM_019833313.2;refGene_gn=MLH3;refGene_tx=XM_023255871.1
B3 122366872 122367032
refGene_gn=MLH3;refGene_tx=XM_003987843.5;refGene_gn=MLH3;refGene_tx=XM_019833311.2;refGene_gn=MLH3;refGene_tx=XM_019833313.2;refGene_gn=MLH3;refGene_tx=XM_023255871.1
B3 122368030 122368142
refGene_gn=MLH3;refGene_tx=XM_003987843.5;refGene_gn=MLH3;refGene_tx=XM_019833311.2;refGene_gn=MLH3;refGene_tx=XM_019833313.2;refGene_gn=MLH3;refGene_tx=XM_023255871.1
B3 122369452 122369524
refGene_gn=MLH3;refGene_tx=XM_003987843.5;refGene_gn=MLH3;refGene_tx=XM_019833311.2;refGene_gn=MLH3;refGene_tx=XM_019833313.2;refGene_gn=MLH3;refGene_tx=XM_023255871.1
B3 122374920 122374993

refGene_gn=MLH3;refGene_tx=XM_003987843.5;refGene_gn=MLH3;refGene_tx=XM_019833311.2;refGene_gn=MLH3;refGene_tx=XM_019833313.2;refGene_gn=MLH3;refGene_tx=XM_023255871.1
B3 122376369 122376474
refGene_gn=MLH3;refGene_tx=XM_003987843.5;refGene_gn=MLH3;refGene_tx=XM_019833311.2;refGene_gn=MLH3;refGene_tx=XM_019833313.2;refGene_gn=MLH3;refGene_tx=XM_023255871.1
B3 122377234 122377320
refGene_gn=MLH3;refGene_tx=XM_003987843.5;refGene_gn=MLH3;refGene_tx=XM_019833311.2;refGene_gn=MLH3;refGene_tx=XM_019833313.2;refGene_gn=MLH3;refGene_tx=XM_023255871.1
B3 122377986 122378085
refGene_gn=MLH3;refGene_tx=XM_003987843.5;refGene_gn=MLH3;refGene_tx=XM_019833311.2;refGene_gn=MLH3;refGene_tx=XM_019833313.2
B3 122381386 122384681
refGene_gn=MLH3;refGene_tx=XM_003987843.5;refGene_gn=MLH3;refGene_tx=XM_019833311.2;refGene_gn=MLH3;refGene_tx=XM_019833313.2;refGene_gn=MLH3;refGene_tx=XM_023255871.1
B3 122387983 122388199
refGene_gn=ACYP1;refGene_tx=XM_003987845.3;refGene_gn=ACYP1;refGene_tx=XM_011283412.3
B3 122394954 122395052
refGene_gn=ACYP1;refGene_tx=XM_003987845.3;refGene_gn=ACYP1;refGene_tx=XM_011283412.3
B3 122400661 122400824
refGene_gn=ACYP1;refGene_tx=XM_011283412.3
B3 122402092 122403418
refGene_gn=ZC2HC1C;refGene_tx=XM_006933003.4;refGene_gn=ZC2HC1C;refGene_tx=XM_019833316.2;refGene_gn=ZC2HC1C;refGene_tx=XM_019833317.2
B3 122408175 122408439
refGene_gn=ZC2HC1C;refGene_tx=XM_006933003.4;refGene_gn=ZC2HC1C;refGene_tx=XM_019833316.2;refGene_gn=ZC2HC1C;refGene_tx=XM_019833317.2
B3 122414643 122414766
refGene_gn=NEK9;refGene_tx=XM_023255872.1
B3 122416801 122417011
refGene_gn=NEK9;refGene_tx=XM_023255872.1
B3 122418214 122418376
refGene_gn=NEK9;refGene_tx=XM_023255872.1
B3 122419449 122419652
refGene_gn=NEK9;refGene_tx=XM_023255872.1
B3 122422783 122422843
refGene_gn=NEK9;refGene_tx=XM_023255872.1
B3 122423603 122423774
refGene_gn=NEK9;refGene_tx=XM_023255872.1
B3 122427618 122427780
refGene_gn=NEK9;refGene_tx=XM_023255872.1
B3 122428535 122428644
refGene_gn=NEK9;refGene_tx=XM_023255872.1
B3 122430238 122430394
refGene_gn=NEK9;refGene_tx=XM_023255872.1

B3 122431613 122431660
refGene_gn=NEK9;refGene_tx=XM_023255872.1
B3 122431845 122432046
refGene_gn=NEK9;refGene_tx=XM_023255872.1
B3 122432635 122432780
refGene_gn=NEK9;refGene_tx=XM_023255872.1
B3 122434620 122434674
refGene_gn=NEK9;refGene_tx=XM_023255872.1
B3 122434675 122434814
refGene_gn=NEK9;refGene_tx=XM_023255872.1
B3 122435279 122435330
refGene_gn=NEK9;refGene_tx=XM_023255872.1
B3 122437729 122437794
refGene_gn=NEK9;refGene_tx=XM_023255872.1
B3 122439091 122439202
refGene_gn=NEK9;refGene_tx=XM_023255872.1
B3 122440997 122441129
refGene_gn=NEK9;refGene_tx=XM_023255872.1
B3 122443591 122443697
refGene_gn=NEK9;refGene_tx=XM_023255872.1
B3 122445494 122445565
refGene_gn=NEK9;refGene_tx=XM_023255872.1
B3 122446099 122446155
refGene_gn=NEK9;refGene_tx=XM_023255872.1
B3 122448068 122448246
refGene_gn=NEK9;refGene_tx=XM_023255872.1
B3 122450553 122450772
refGene_gn=NEK9;refGene_tx=XM_023255872.1
B3 122459330 122459452
refGene_gn=TMED10;refGene_tx=XM_003987847.5
B3 122459849 122459976
refGene_gn=TMED10;refGene_tx=XM_003987847.5
B3 122473379 122473453
refGene_gn=TMED10;refGene_tx=XM_003987847.5
B3 122478410 122478522
refGene_gn=TMED10;refGene_tx=XM_003987847.5
B3 122497661 122497886
refGene_gn=TMED10;refGene_tx=XM_003987847.5
B3 122573870 122574011
refGene_gn=FOS;refGene_tx=NM_001009341.1
B3 122574779 122575034
refGene_gn=FOS;refGene_tx=NM_001009341.1
B3 122575437 122575545
refGene_gn=FOS;refGene_tx=NM_001009341.1
B3 122575668 122576310
refGene_gn=FOS;refGene_tx=NM_001009341.1
B3 122688201 122688208
refGene_gn=JDP2;refGene_tx=XM_019833320.2
B3 122692771 122692880
refGene_gn=JDP2;refGene_tx=XM_019833321.2

B3 122698486 122698710
refGene_gn=JDP2;refGene_tx=XM_011283415.3;refGene_gn=JDP2;refGene_tx=XM_019833320.2;refGene_gn=JDP2;refGene_tx=XM_019833321.2;refGene_gn=JDP2;refGene_tx=XM_023255873.1
B3 122719467 122719572
refGene_gn=JDP2;refGene_tx=XM_011283415.3;refGene_gn=JDP2;refGene_tx=XM_019833320.2;refGene_gn=JDP2;refGene_tx=XM_023255873.1
B3 122725866 122726052
refGene_gn=JDP2;refGene_tx=XM_011283415.3;refGene_gn=JDP2;refGene_tx=XM_019833320.2;refGene_gn=JDP2;refGene_tx=XM_019833321.2;refGene_gn=JDP2;refGene_tx=XM_023255873.1
B3 122779213 122779276
refGene_gn=BATF;refGene_tx=XM_003987849.3
B3 122782429 122782534
refGene_gn=BATF;refGene_tx=XM_003987849.3
B3 122802554 122802764
refGene_gn=BATF;refGene_tx=XM_003987849.3
B3 122844070 122844775
refGene_gn=FLVCR2;refGene_tx=XM_003987879.4;refGene_gn=FLVCR2;refGene_tx=XM_019833323.1
B3 122872821 122872963
refGene_gn=FLVCR2;refGene_tx=XM_003987879.4;refGene_gn=FLVCR2;refGene_tx=XM_019833323.1
B3 122876334 122876475
refGene_gn=FLVCR2;refGene_tx=XM_003987879.4;refGene_gn=FLVCR2;refGene_tx=XM_019833323.1
B3 122886820 122886888
refGene_gn=FLVCR2;refGene_tx=XM_003987879.4;refGene_gn=FLVCR2;refGene_tx=XM_019833323.1
B3 122888045 122888149
refGene_gn=FLVCR2;refGene_tx=XM_003987879.4;refGene_gn=FLVCR2;refGene_tx=XM_019833323.1
B3 122891500 122891611
refGene_gn=FLVCR2;refGene_tx=XM_003987879.4;refGene_gn=FLVCR2;refGene_tx=XM_019833323.1
B3 122892773 122892879
refGene_gn=FLVCR2;refGene_tx=XM_003987879.4;refGene_gn=FLVCR2;refGene_tx=XM_019833323.1
B3 122893029 122893141
refGene_gn=FLVCR2;refGene_tx=XM_003987879.4;refGene_gn=FLVCR2;refGene_tx=XM_019833323.1
B3 122893720 122893776
refGene_gn=FLVCR2;refGene_tx=XM_003987879.4;refGene_gn=FLVCR2;refGene_tx=XM_019833323.1
B3 122894029 122894083
refGene_gn=FLVCR2;refGene_tx=XM_019833323.1
B3 122897633 122897714
refGene_gn=FLVCR2;refGene_tx=XM_003987879.4
B3 122899936 122899950
refGene_gn=ERG28;refGene_tx=XM_006933008.4;refGene_gn=ERG28;refGene_tx

=XM_011283419.2
B3 122900223 122900351
refGene_gn=ERG28;refGene_tx=XM_006933007.4;refGene_gn=ERG28;refGene_tx=XM_011283417.3
B3 122902242 122902441
refGene_gn=ERG28;refGene_tx=XM_003987850.5;refGene_gn=ERG28;refGene_tx=XM_011283416.2;refGene_gn=ERG28;refGene_tx=XM_011283418.2;refGene_gn=ERG28;refGene_tx=XM_023255890.1
B3 122902569 122902688
refGene_gn=ERG28;refGene_tx=XM_003987850.5;refGene_gn=ERG28;refGene_tx=XM_006933007.4;refGene_gn=ERG28;refGene_tx=XM_006933008.4;refGene_gn=ERG28;refGene_tx=XM_011283416.2;refGene_gn=ERG28;refGene_tx=XM_011283417.3;refGene_gn=ERG28;refGene_tx=XM_011283418.2;refGene_gn=ERG28;refGene_tx=XM_011283419.2;refGene_gn=ERG28;refGene_tx=XM_023255890.1
B3 122903041 122903185
refGene_gn=ERG28;refGene_tx=XM_011283416.2;refGene_gn=ERG28;refGene_tx=XM_011283417.3;refGene_gn=ERG28;refGene_tx=XM_011283419.2;refGene_gn=ERG28;refGene_tx=XM_023255890.1
B3 122905645 122905740
refGene_gn=ERG28;refGene_tx=XM_003987850.5;refGene_gn=ERG28;refGene_tx=XM_006933007.4;refGene_gn=ERG28;refGene_tx=XM_006933008.4;refGene_gn=ERG28;refGene_tx=XM_011283416.2;refGene_gn=ERG28;refGene_tx=XM_011283417.3;refGene_gn=ERG28;refGene_tx=XM_011283418.2;refGene_gn=ERG28;refGene_tx=XM_011283419.2;refGene_gn=ERG28;refGene_tx=XM_023255890.1
B3 122908575 122908708
refGene_gn=ERG28;refGene_tx=XM_003987850.5;refGene_gn=ERG28;refGene_tx=XM_006933007.4;refGene_gn=ERG28;refGene_tx=XM_006933008.4;refGene_gn=ERG28;refGene_tx=XM_011283416.2;refGene_gn=ERG28;refGene_tx=XM_011283417.3;refGene_gn=ERG28;refGene_tx=XM_011283418.2;refGene_gn=ERG28;refGene_tx=XM_011283419.2;refGene_gn=ERG28;refGene_tx=XM_023255890.1
B3 122913375 122913449
refGene_gn=TTLL5;refGene_tx=XM_023255874.1;refGene_gn=TTLL5;refGene_tx=XM_023255875.1;refGene_gn=TTLL5;refGene_tx=XM_023255876.1;refGene_gn=TTLL5;refGene_tx=XM_023255877.1;refGene_gn=TTLL5;refGene_tx=XM_023255878.1;refGene_gn=TTLL5;refGene_tx=XM_023255879.1;refGene_gn=TTLL5;refGene_tx=XM_023255880.1;refGene_gn=TTLL5;refGene_tx=XM_023255881.1;refGene_gn=TTLL5;refGene_tx=XM_023255882.1;refGene_gn=TTLL5;refGene_tx=XM_023255883.1;refGene_gn=TTLL5;refGene_tx=XM_023255884.1;refGene_gn=TTLL5;refGene_tx=XM_023255885.1;refGene_gn=TTLL5;refGene_tx=XM_023255886.1;refGene_gn=TTLL5;refGene_tx=XM_023255887.1;refGene_gn=TTLL5;refGene_tx=XM_023255888.1;refGene_gn=TTLL5;refGene_tx=XM_023255889.1
B3 122919166 122919273
refGene_gn=TTLL5;refGene_tx=XM_023255874.1;refGene_gn=TTLL5;refGene_tx=XM_023255875.1;refGene_gn=TTLL5;refGene_tx=XM_023255876.1;refGene_gn=TTLL5;refGene_tx=XM_023255877.1;refGene_gn=TTLL5;refGene_tx=XM_023255878.1;refGene_gn=TTLL5;refGene_tx=XM_023255879.1;refGene_gn=TTLL5;refGene_tx=XM_023255880.1;refGene_gn=TTLL5;refGene_tx=XM_023255881.1;refGene_gn=TTLL5;refGene_tx=XM_023255882.1;refGene_gn=TTLL5;refGene_tx=XM_023255883.1;refGene_gn=TTLL5;refGene_tx=XM_023255884.1;refGene_gn=TTLL5;refGene_tx=XM_023255885.1;refGene_gn=TTLL5;refGene_tx=XM_023255886.1;r

efGene_gn=TTLL5;refGene_tx=XM_023255887.1;refGene_gn=TTLL5;refGene_tx=XM_023255888.1;refGene_gn=TTLL5;refGene_tx=XM_023255889.1
B3 122931156 122931239
refGene_gn=TTLL5;refGene_tx=XM_023255874.1;refGene_gn=TTLL5;refGene_tx=XM_023255875.1;refGene_gn=TTLL5;refGene_tx=XM_023255876.1;refGene_gn=TTLL5;refGene_tx=XM_023255877.1;refGene_gn=TTLL5;refGene_tx=XM_023255878.1;refGene_gn=TTLL5;refGene_tx=XM_023255879.1;refGene_gn=TTLL5;refGene_tx=XM_023255880.1;refGene_gn=TTLL5;refGene_tx=XM_023255881.1;refGene_gn=TTLL5;refGene_tx=XM_023255882.1;refGene_gn=TTLL5;refGene_tx=XM_023255883.1;refGene_gn=TTLL5;refGene_tx=XM_023255884.1;refGene_gn=TTLL5;refGene_tx=XM_023255885.1;refGene_gn=TTLL5;refGene_tx=XM_023255886.1;refGene_gn=TTLL5;refGene_tx=XM_023255887.1;refGene_gn=TTLL5;refGene_tx=XM_023255888.1;refGene_gn=TTLL5;refGene_tx=XM_023255889.1
B3 122932829 122932936
refGene_gn=TTLL5;refGene_tx=XM_023255874.1;refGene_gn=TTLL5;refGene_tx=XM_023255875.1;refGene_gn=TTLL5;refGene_tx=XM_023255876.1;refGene_gn=TTLL5;refGene_tx=XM_023255877.1;refGene_gn=TTLL5;refGene_tx=XM_023255878.1;refGene_gn=TTLL5;refGene_tx=XM_023255879.1;refGene_gn=TTLL5;refGene_tx=XM_023255880.1;refGene_gn=TTLL5;refGene_tx=XM_023255881.1;refGene_gn=TTLL5;refGene_tx=XM_023255882.1;refGene_gn=TTLL5;refGene_tx=XM_023255883.1;refGene_gn=TTLL5;refGene_tx=XM_023255884.1;refGene_gn=TTLL5;refGene_tx=XM_023255885.1;refGene_gn=TTLL5;refGene_tx=XM_023255886.1;refGene_gn=TTLL5;refGene_tx=XM_023255887.1;refGene_gn=TTLL5;refGene_tx=XM_023255888.1;refGene_gn=TTLL5;refGene_tx=XM_023255889.1
B3 122940381 122940512
refGene_gn=TTLL5;refGene_tx=XM_023255874.1;refGene_gn=TTLL5;refGene_tx=XM_023255875.1;refGene_gn=TTLL5;refGene_tx=XM_023255876.1;refGene_gn=TTLL5;refGene_tx=XM_023255877.1;refGene_gn=TTLL5;refGene_tx=XM_023255878.1;refGene_gn=TTLL5;refGene_tx=XM_023255879.1;refGene_gn=TTLL5;refGene_tx=XM_023255880.1;refGene_gn=TTLL5;refGene_tx=XM_023255881.1;refGene_gn=TTLL5;refGene_tx=XM_023255882.1;refGene_gn=TTLL5;refGene_tx=XM_023255883.1;refGene_gn=TTLL5;refGene_tx=XM_023255884.1;refGene_gn=TTLL5;refGene_tx=XM_023255885.1;refGene_gn=TTLL5;refGene_tx=XM_023255886.1;refGene_gn=TTLL5;refGene_tx=XM_023255887.1;refGene_gn=TTLL5;refGene_tx=XM_023255888.1;refGene_gn=TTLL5;refGene_tx=XM_023255889.1
B3 122945262 122945345
refGene_gn=TTLL5;refGene_tx=XM_023255874.1;refGene_gn=TTLL5;refGene_tx=XM_023255875.1;refGene_gn=TTLL5;refGene_tx=XM_023255876.1;refGene_gn=TTLL5;refGene_tx=XM_023255877.1;refGene_gn=TTLL5;refGene_tx=XM_023255878.1;refGene_gn=TTLL5;refGene_tx=XM_023255879.1;refGene_gn=TTLL5;refGene_tx=XM_023255880.1;refGene_gn=TTLL5;refGene_tx=XM_023255881.1;refGene_gn=TTLL5;refGene_tx=XM_023255882.1;refGene_gn=TTLL5;refGene_tx=XM_023255883.1;refGene_gn=TTLL5;refGene_tx=XM_023255884.1;refGene_gn=TTLL5;refGene_tx=XM_023255885.1;refGene_gn=TTLL5;refGene_tx=XM_023255886.1;refGene_gn=TTLL5;refGene_tx=XM_023255887.1;refGene_gn=TTLL5;refGene_tx=XM_023255888.1;refGene_gn=TTLL5;refGene_tx=XM_023255889.1
B3 122953492 122953562
refGene_gn=TTLL5;refGene_tx=XM_023255874.1;refGene_gn=TTLL5;refGene_tx=XM_023255875.1;refGene_gn=TTLL5;refGene_tx=XM_023255876.1;refGene_gn=TTLL5;refGene_tx=XM_023255877.1;refGene_gn=TTLL5;refGene_tx=XM_0232558

78.1;refGene_gn=TTLL5;refGene_tx=XM_023255879.1;refGene_gn=TTLL5;refGene_tx=XM_023255880.1;refGene_gn=TTLL5;refGene_tx=XM_023255881.1;refGene_gn=TTLL5;refGene_tx=XM_023255882.1;refGene_gn=TTLL5;refGene_tx=XM_023255883.1;refGene_gn=TTLL5;refGene_tx=XM_023255884.1;refGene_gn=TTLL5;refGene_tx=XM_023255885.1;refGene_gn=TTLL5;refGene_tx=XM_023255886.1;refGene_gn=TTLL5;refGene_tx=XM_023255887.1;refGene_gn=TTLL5;refGene_tx=XM_023255888.1;refGene_gn=TTLL5;refGene_tx=XM_023255889.1
B3 122954094 122954179
refGene_gn=TTLL5;refGene_tx=XM_023255874.1;refGene_gn=TTLL5;refGene_tx=XM_023255875.1;refGene_gn=TTLL5;refGene_tx=XM_023255876.1;refGene_gn=TTLL5;refGene_tx=XM_023255877.1;refGene_gn=TTLL5;refGene_tx=XM_023255878.1;refGene_gn=TTLL5;refGene_tx=XM_023255879.1;refGene_gn=TTLL5;refGene_tx=XM_023255880.1;refGene_gn=TTLL5;refGene_tx=XM_023255881.1;refGene_gn=TTLL5;refGene_tx=XM_023255882.1;refGene_gn=TTLL5;refGene_tx=XM_023255883.1;refGene_gn=TTLL5;refGene_tx=XM_023255884.1;refGene_gn=TTLL5;refGene_tx=XM_023255885.1;refGene_gn=TTLL5;refGene_tx=XM_023255886.1;refGene_gn=TTLL5;refGene_tx=XM_023255887.1;refGene_gn=TTLL5;refGene_tx=XM_023255888.1;refGene_gn=TTLL5;refGene_tx=XM_023255889.1
B3 122966919 122967021
refGene_gn=TTLL5;refGene_tx=XM_023255874.1;refGene_gn=TTLL5;refGene_tx=XM_023255875.1;refGene_gn=TTLL5;refGene_tx=XM_023255876.1;refGene_gn=TTLL5;refGene_tx=XM_023255877.1;refGene_gn=TTLL5;refGene_tx=XM_023255878.1;refGene_gn=TTLL5;refGene_tx=XM_023255879.1;refGene_gn=TTLL5;refGene_tx=XM_023255880.1;refGene_gn=TTLL5;refGene_tx=XM_023255881.1;refGene_gn=TTLL5;refGene_tx=XM_023255882.1;refGene_gn=TTLL5;refGene_tx=XM_023255883.1;refGene_gn=TTLL5;refGene_tx=XM_023255884.1;refGene_gn=TTLL5;refGene_tx=XM_023255885.1;refGene_gn=TTLL5;refGene_tx=XM_023255886.1;refGene_gn=TTLL5;refGene_tx=XM_023255887.1;refGene_gn=TTLL5;refGene_tx=XM_023255888.1;refGene_gn=TTLL5;refGene_tx=XM_023255889.1
B3 122968813 122968905
refGene_gn=TTLL5;refGene_tx=XM_023255874.1;refGene_gn=TTLL5;refGene_tx=XM_023255875.1;refGene_gn=TTLL5;refGene_tx=XM_023255876.1;refGene_gn=TTLL5;refGene_tx=XM_023255877.1;refGene_gn=TTLL5;refGene_tx=XM_023255878.1;refGene_gn=TTLL5;refGene_tx=XM_023255879.1;refGene_gn=TTLL5;refGene_tx=XM_023255880.1;refGene_gn=TTLL5;refGene_tx=XM_023255881.1;refGene_gn=TTLL5;refGene_tx=XM_023255882.1;refGene_gn=TTLL5;refGene_tx=XM_023255883.1;refGene_gn=TTLL5;refGene_tx=XM_023255884.1;refGene_gn=TTLL5;refGene_tx=XM_023255885.1;refGene_gn=TTLL5;refGene_tx=XM_023255886.1;refGene_gn=TTLL5;refGene_tx=XM_023255887.1;refGene_gn=TTLL5;refGene_tx=XM_023255888.1;refGene_gn=TTLL5;refGene_tx=XM_023255889.1
B3 122969937 122970045
refGene_gn=TTLL5;refGene_tx=XM_023255874.1;refGene_gn=TTLL5;refGene_tx=XM_023255875.1;refGene_gn=TTLL5;refGene_tx=XM_023255876.1;refGene_gn=TTLL5;refGene_tx=XM_023255877.1;refGene_gn=TTLL5;refGene_tx=XM_023255878.1;refGene_gn=TTLL5;refGene_tx=XM_023255879.1;refGene_gn=TTLL5;refGene_tx=XM_023255880.1;refGene_gn=TTLL5;refGene_tx=XM_023255881.1;refGene_gn=TTLL5;refGene_tx=XM_023255882.1;refGene_gn=TTLL5;refGene_tx=XM_023255883.1;refGene_gn=TTLL5;refGene_tx=XM_023255884.1;refGene_gn=TTLL5;refGene_tx=XM_023255885.1;refGene_gn=TTLL5;refGene_tx=XM_023255886.1;refGene_gn=TTLL5;refGene_tx=XM_023255887.1;refGene_gn=TTLL5;refGene_tx=

XM_023255888.1;refGene_gn=TTLL5;refGene_tx=XM_023255889.1
B3 122980678 122980760
refGene_gn=TTLL5;refGene_tx=XM_023255874.1;refGene_gn=TTLL5;refGene_tx=XM_023255875.1;refGene_gn=TTLL5;refGene_tx=XM_023255876.1;refGene_gn=TTLL5;refGene_tx=XM_023255877.1;refGene_gn=TTLL5;refGene_tx=XM_023255878.1;refGene_gn=TTLL5;refGene_tx=XM_023255879.1;refGene_gn=TTLL5;refGene_tx=XM_023255880.1;refGene_gn=TTLL5;refGene_tx=XM_023255881.1;refGene_gn=TTLL5;refGene_tx=XM_023255882.1;refGene_gn=TTLL5;refGene_tx=XM_023255883.1;refGene_gn=TTLL5;refGene_tx=XM_023255884.1;refGene_gn=TTLL5;refGene_tx=XM_023255885.1;refGene_gn=TTLL5;refGene_tx=XM_023255886.1;refGene_gn=TTLL5;refGene_tx=XM_023255887.1;refGene_gn=TTLL5;refGene_tx=XM_023255888.1;refGene_gn=TTLL5;refGene_tx=XM_023255889.1
B3 122982630 122982692
refGene_gn=TTLL5;refGene_tx=XM_023255874.1;refGene_gn=TTLL5;refGene_tx=XM_023255875.1;refGene_gn=TTLL5;refGene_tx=XM_023255876.1;refGene_gn=TTLL5;refGene_tx=XM_023255877.1;refGene_gn=TTLL5;refGene_tx=XM_023255878.1;refGene_gn=TTLL5;refGene_tx=XM_023255879.1;refGene_gn=TTLL5;refGene_tx=XM_023255880.1;refGene_gn=TTLL5;refGene_tx=XM_023255881.1;refGene_gn=TTLL5;refGene_tx=XM_023255882.1;refGene_gn=TTLL5;refGene_tx=XM_023255883.1;refGene_gn=TTLL5;refGene_tx=XM_023255884.1;refGene_gn=TTLL5;refGene_tx=XM_023255885.1;refGene_gn=TTLL5;refGene_tx=XM_023255886.1;refGene_gn=TTLL5;refGene_tx=XM_023255887.1;refGene_gn=TTLL5;refGene_tx=XM_023255888.1;refGene_gn=TTLL5;refGene_tx=XM_023255889.1
B3 122983840 122983935
refGene_gn=TTLL5;refGene_tx=XM_023255874.1;refGene_gn=TTLL5;refGene_tx=XM_023255875.1;refGene_gn=TTLL5;refGene_tx=XM_023255876.1;refGene_gn=TTLL5;refGene_tx=XM_023255877.1;refGene_gn=TTLL5;refGene_tx=XM_023255878.1;refGene_gn=TTLL5;refGene_tx=XM_023255879.1;refGene_gn=TTLL5;refGene_tx=XM_023255880.1;refGene_gn=TTLL5;refGene_tx=XM_023255881.1;refGene_gn=TTLL5;refGene_tx=XM_023255882.1;refGene_gn=TTLL5;refGene_tx=XM_023255883.1;refGene_gn=TTLL5;refGene_tx=XM_023255884.1;refGene_gn=TTLL5;refGene_tx=XM_023255885.1;refGene_gn=TTLL5;refGene_tx=XM_023255886.1;refGene_gn=TTLL5;refGene_tx=XM_023255887.1;refGene_gn=TTLL5;refGene_tx=XM_023255888.1;refGene_gn=TTLL5;refGene_tx=XM_023255889.1
B3 122986401 122986443
refGene_gn=TTLL5;refGene_tx=XM_023255874.1;refGene_gn=TTLL5;refGene_tx=XM_023255875.1;refGene_gn=TTLL5;refGene_tx=XM_023255876.1;refGene_gn=TTLL5;refGene_tx=XM_023255877.1;refGene_gn=TTLL5;refGene_tx=XM_023255879.1;refGene_gn=TTLL5;refGene_tx=XM_023255880.1;refGene_gn=TTLL5;refGene_tx=XM_023255881.1;refGene_gn=TTLL5;refGene_tx=XM_023255883.1;refGene_gn=TTLL5;refGene_tx=XM_023255885.1;refGene_gn=TTLL5;refGene_tx=XM_023255887.1;refGene_gn=TTLL5;refGene_tx=XM_023255888.1;refGene_gn=TTLL5;refGene_tx=XM_023255889.1
B3 122994374 122994488
refGene_gn=TTLL5;refGene_tx=XM_023255874.1;refGene_gn=TTLL5;refGene_tx=XM_023255875.1;refGene_gn=TTLL5;refGene_tx=XM_023255876.1;refGene_gn=TTLL5;refGene_tx=XM_023255877.1;refGene_gn=TTLL5;refGene_tx=XM_023255878.1;refGene_gn=TTLL5;refGene_tx=XM_023255879.1;refGene_gn=TTLL5;refGene_tx=XM_023255880.1;refGene_gn=TTLL5;refGene_tx=XM_023255881.1;refGene_gn=TTLL5;refGene_tx=XM_023255882.1;refGene_gn=TTLL5;refGene_tx=XM_02

3255883.1;refGene_gn=TTLL5;refGene_tx=XM_023255884.1;refGene_gn=TTLL5;
refGene_tx=XM_023255885.1;refGene_gn=TTLL5;refGene_tx=XM_023255886.1;r
efGene_gn=TTLL5;refGene_tx=XM_023255887.1;refGene_gn=TTLL5;refGene_tx=
XM_023255888.1;refGene_gn=TTLL5;refGene_tx=XM_023255889.1
B3 122994782 122994874
refGene_gn=TTLL5;refGene_tx=XM_023255874.1;refGene_gn=TTLL5;refGene_tx
=XM_023255875.1;refGene_gn=TTLL5;refGene_tx=XM_023255876.1;refGene_gn=
TTLL5;refGene_tx=XM_023255877.1;refGene_gn=TTLL5;refGene_tx=XM_0232558
78.1;refGene_gn=TTLL5;refGene_tx=XM_023255879.1;refGene_gn=TTLL5;refGe
ne_tx=XM_023255880.1;refGene_gn=TTLL5;refGene_tx=XM_023255881.1;refGen
e_gn=TTLL5;refGene_tx=XM_023255882.1;refGene_gn=TTLL5;refGene_tx=XM_02
3255883.1;refGene_gn=TTLL5;refGene_tx=XM_023255884.1;refGene_gn=TTLL5;
refGene_tx=XM_023255885.1;refGene_gn=TTLL5;refGene_tx=XM_023255886.1;r
efGene_gn=TTLL5;refGene_tx=XM_023255887.1;refGene_gn=TTLL5;refGene_tx=
XM_023255888.1;refGene_gn=TTLL5;refGene_tx=XM_023255889.1
B3 123002085 123002148
refGene_gn=TTLL5;refGene_tx=XM_023255874.1;refGene_gn=TTLL5;refGene_tx
=XM_023255875.1;refGene_gn=TTLL5;refGene_tx=XM_023255876.1;refGene_gn=
TTLL5;refGene_tx=XM_023255877.1;refGene_gn=TTLL5;refGene_tx=XM_0232558
78.1;refGene_gn=TTLL5;refGene_tx=XM_023255879.1;refGene_gn=TTLL5;refGe
ne_tx=XM_023255880.1;refGene_gn=TTLL5;refGene_tx=XM_023255881.1;refGen
e_gn=TTLL5;refGene_tx=XM_023255882.1;refGene_gn=TTLL5;refGene_tx=XM_02
3255883.1;refGene_gn=TTLL5;refGene_tx=XM_023255884.1;refGene_gn=TTLL5;
refGene_tx=XM_023255885.1;refGene_gn=TTLL5;refGene_tx=XM_023255886.1;r
efGene_gn=TTLL5;refGene_tx=XM_023255887.1;refGene_gn=TTLL5;refGene_tx=
XM_023255888.1;refGene_gn=TTLL5;refGene_tx=XM_023255889.1
B3 123007517 123007556
refGene_gn=TTLL5;refGene_tx=XM_023255874.1;refGene_gn=TTLL5;refGene_tx
=XM_023255875.1;refGene_gn=TTLL5;refGene_tx=XM_023255876.1;refGene_gn=
TTLL5;refGene_tx=XM_023255878.1;refGene_gn=TTLL5;refGene_tx=XM_0232558
79.1;refGene_gn=TTLL5;refGene_tx=XM_023255880.1;refGene_gn=TTLL5;refGe
ne_tx=XM_023255881.1;refGene_gn=TTLL5;refGene_tx=XM_023255882.1;refGen
e_gn=TTLL5;refGene_tx=XM_023255885.1;refGene_gn=TTLL5;refGene_tx=XM_02
3255886.1;refGene_gn=TTLL5;refGene_tx=XM_023255887.1;refGene_gn=TTLL5;
refGene_tx=XM_023255888.1;refGene_gn=TTLL5;refGene_tx=XM_023255889.1
B3 123014462 123014630
refGene_gn=TTLL5;refGene_tx=XM_023255874.1;refGene_gn=TTLL5;refGene_tx
=XM_023255875.1;refGene_gn=TTLL5;refGene_tx=XM_023255876.1;refGene_gn=
TTLL5;refGene_tx=XM_023255877.1;refGene_gn=TTLL5;refGene_tx=XM_0232558
78.1;refGene_gn=TTLL5;refGene_tx=XM_023255879.1;refGene_gn=TTLL5;refGe
ne_tx=XM_023255880.1;refGene_gn=TTLL5;refGene_tx=XM_023255881.1;refGen
e_gn=TTLL5;refGene_tx=XM_023255882.1;refGene_gn=TTLL5;refGene_tx=XM_02
3255883.1;refGene_gn=TTLL5;refGene_tx=XM_023255884.1;refGene_gn=TTLL5;
refGene_tx=XM_023255885.1;refGene_gn=TTLL5;refGene_tx=XM_023255886.1;r
efGene_gn=TTLL5;refGene_tx=XM_023255887.1;refGene_gn=TTLL5;refGene_tx=
XM_023255888.1;refGene_gn=TTLL5;refGene_tx=XM_023255889.1
B3 123015543 123015876
refGene_gn=TTLL5;refGene_tx=XM_023255874.1;refGene_gn=TTLL5;refGene_tx
=XM_023255875.1;refGene_gn=TTLL5;refGene_tx=XM_023255876.1;refGene_gn=
TTLL5;refGene_tx=XM_023255877.1;refGene_gn=TTLL5;refGene_tx=XM_0232558

78.1;refGene_gn=TTLL5;refGene_tx=XM_023255879.1;refGene_gn=TTLL5;refGene_tx=XM_023255880.1;refGene_gn=TTLL5;refGene_tx=XM_023255881.1;refGene_gn=TTLL5;refGene_tx=XM_023255882.1;refGene_gn=TTLL5;refGene_tx=XM_023255883.1;refGene_gn=TTLL5;refGene_tx=XM_023255884.1;refGene_gn=TTLL5;refGene_tx=XM_023255885.1;refGene_gn=TTLL5;refGene_tx=XM_023255886.1;refGene_gn=TTLL5;refGene_tx=XM_023255887.1;refGene_gn=TTLL5;refGene_tx=XM_023255888.1;refGene_gn=TTLL5;refGene_tx=XM_023255889.1
B3 123020408 123020529
refGene_gn=TTLL5;refGene_tx=XM_023255874.1;refGene_gn=TTLL5;refGene_tx=XM_023255875.1;refGene_gn=TTLL5;refGene_tx=XM_023255876.1;refGene_gn=TTLL5;refGene_tx=XM_023255877.1;refGene_gn=TTLL5;refGene_tx=XM_023255878.1;refGene_gn=TTLL5;refGene_tx=XM_023255879.1;refGene_gn=TTLL5;refGene_tx=XM_023255880.1;refGene_gn=TTLL5;refGene_tx=XM_023255881.1;refGene_gn=TTLL5;refGene_tx=XM_023255882.1;refGene_gn=TTLL5;refGene_tx=XM_023255883.1;refGene_gn=TTLL5;refGene_tx=XM_023255884.1;refGene_gn=TTLL5;refGene_tx=XM_023255885.1;refGene_gn=TTLL5;refGene_tx=XM_023255886.1;refGene_gn=TTLL5;refGene_tx=XM_023255887.1;refGene_gn=TTLL5;refGene_tx=XM_023255888.1;refGene_gn=TTLL5;refGene_tx=XM_023255889.1
B3 123023455 123023602
refGene_gn=TTLL5;refGene_tx=XM_023255874.1;refGene_gn=TTLL5;refGene_tx=XM_023255875.1;refGene_gn=TTLL5;refGene_tx=XM_023255876.1;refGene_gn=TTLL5;refGene_tx=XM_023255877.1;refGene_gn=TTLL5;refGene_tx=XM_023255878.1;refGene_gn=TTLL5;refGene_tx=XM_023255879.1;refGene_gn=TTLL5;refGene_tx=XM_023255880.1;refGene_gn=TTLL5;refGene_tx=XM_023255881.1;refGene_gn=TTLL5;refGene_tx=XM_023255882.1;refGene_gn=TTLL5;refGene_tx=XM_023255883.1;refGene_gn=TTLL5;refGene_tx=XM_023255884.1;refGene_gn=TTLL5;refGene_tx=XM_023255885.1;refGene_gn=TTLL5;refGene_tx=XM_023255886.1;refGene_gn=TTLL5;refGene_tx=XM_023255887.1;refGene_gn=TTLL5;refGene_tx=XM_023255888.1;refGene_gn=TTLL5;refGene_tx=XM_023255889.1
B3 123024952 123025056
refGene_gn=TTLL5;refGene_tx=XM_023255874.1;refGene_gn=TTLL5;refGene_tx=XM_023255875.1;refGene_gn=TTLL5;refGene_tx=XM_023255876.1;refGene_gn=TTLL5;refGene_tx=XM_023255877.1;refGene_gn=TTLL5;refGene_tx=XM_023255878.1;refGene_gn=TTLL5;refGene_tx=XM_023255879.1;refGene_gn=TTLL5;refGene_tx=XM_023255880.1;refGene_gn=TTLL5;refGene_tx=XM_023255881.1;refGene_gn=TTLL5;refGene_tx=XM_023255882.1;refGene_gn=TTLL5;refGene_tx=XM_023255883.1;refGene_gn=TTLL5;refGene_tx=XM_023255884.1;refGene_gn=TTLL5;refGene_tx=XM_023255885.1;refGene_gn=TTLL5;refGene_tx=XM_023255886.1;refGene_gn=TTLL5;refGene_tx=XM_023255887.1;refGene_gn=TTLL5;refGene_tx=XM_023255888.1;refGene_gn=TTLL5;refGene_tx=XM_023255889.1
B3 123027946 123028074
refGene_gn=TTLL5;refGene_tx=XM_023255874.1;refGene_gn=TTLL5;refGene_tx=XM_023255875.1;refGene_gn=TTLL5;refGene_tx=XM_023255876.1;refGene_gn=TTLL5;refGene_tx=XM_023255877.1;refGene_gn=TTLL5;refGene_tx=XM_023255878.1;refGene_gn=TTLL5;refGene_tx=XM_023255879.1;refGene_gn=TTLL5;refGene_tx=XM_023255880.1;refGene_gn=TTLL5;refGene_tx=XM_023255881.1;refGene_gn=TTLL5;refGene_tx=XM_023255882.1;refGene_gn=TTLL5;refGene_tx=XM_023255883.1;refGene_gn=TTLL5;refGene_tx=XM_023255884.1;refGene_gn=TTLL5;refGene_tx=XM_023255885.1;refGene_gn=TTLL5;refGene_tx=XM_023255886.1;refGene_gn=TTLL5;refGene_tx=XM_023255887.1;refGene_gn=TTLL5;refGene_tx=

XM_023255888.1;refGene_gn=TTLL5;refGene_tx=XM_023255889.1
B3 123030605 123030689
refGene_gn=TTLL5;refGene_tx=XM_023255874.1;refGene_gn=TTLL5;refGene_tx=XM_023255875.1;refGene_gn=TTLL5;refGene_tx=XM_023255876.1;refGene_gn=TTLL5;refGene_tx=XM_023255877.1;refGene_gn=TTLL5;refGene_tx=XM_023255878.1;refGene_gn=TTLL5;refGene_tx=XM_023255879.1;refGene_gn=TTLL5;refGene_tx=XM_023255880.1;refGene_gn=TTLL5;refGene_tx=XM_023255881.1;refGene_gn=TTLL5;refGene_tx=XM_023255882.1;refGene_gn=TTLL5;refGene_tx=XM_023255883.1;refGene_gn=TTLL5;refGene_tx=XM_023255884.1;refGene_gn=TTLL5;refGene_tx=XM_023255885.1;refGene_gn=TTLL5;refGene_tx=XM_023255886.1;refGene_gn=TTLL5;refGene_tx=XM_023255887.1;refGene_gn=TTLL5;refGene_tx=XM_023255888.1;refGene_gn=TTLL5;refGene_tx=XM_023255889.1
B3 123031278 123031680
refGene_gn=TTLL5;refGene_tx=XM_023255874.1;refGene_gn=TTLL5;refGene_tx=XM_023255875.1;refGene_gn=TTLL5;refGene_tx=XM_023255876.1;refGene_gn=TTLL5;refGene_tx=XM_023255877.1;refGene_gn=TTLL5;refGene_tx=XM_023255878.1;refGene_gn=TTLL5;refGene_tx=XM_023255879.1;refGene_gn=TTLL5;refGene_tx=XM_023255880.1;refGene_gn=TTLL5;refGene_tx=XM_023255881.1;refGene_gn=TTLL5;refGene_tx=XM_023255882.1;refGene_gn=TTLL5;refGene_tx=XM_023255883.1;refGene_gn=TTLL5;refGene_tx=XM_023255884.1;refGene_gn=TTLL5;refGene_tx=XM_023255885.1;refGene_gn=TTLL5;refGene_tx=XM_023255886.1;refGene_gn=TTLL5;refGene_tx=XM_023255887.1;refGene_gn=TTLL5;refGene_tx=XM_023255888.1;refGene_gn=TTLL5;refGene_tx=XM_023255889.1
B3 123040601 123040789
refGene_gn=TTLL5;refGene_tx=XM_023255874.1;refGene_gn=TTLL5;refGene_tx=XM_023255875.1;refGene_gn=TTLL5;refGene_tx=XM_023255876.1;refGene_gn=TTLL5;refGene_tx=XM_023255877.1;refGene_gn=TTLL5;refGene_tx=XM_023255878.1;refGene_gn=TTLL5;refGene_tx=XM_023255879.1;refGene_gn=TTLL5;refGene_tx=XM_023255880.1;refGene_gn=TTLL5;refGene_tx=XM_023255881.1;refGene_gn=TTLL5;refGene_tx=XM_023255882.1;refGene_gn=TTLL5;refGene_tx=XM_023255883.1;refGene_gn=TTLL5;refGene_tx=XM_023255884.1;refGene_gn=TTLL5;refGene_tx=XM_023255885.1;refGene_gn=TTLL5;refGene_tx=XM_023255886.1;refGene_gn=TTLL5;refGene_tx=XM_023255887.1;refGene_gn=TTLL5;refGene_tx=XM_023255888.1;refGene_gn=TTLL5;refGene_tx=XM_023255889.1
B3 123060519 123060674
refGene_gn=TTLL5;refGene_tx=XM_023255874.1;refGene_gn=TTLL5;refGene_tx=XM_023255875.1;refGene_gn=TTLL5;refGene_tx=XM_023255876.1;refGene_gn=TTLL5;refGene_tx=XM_023255877.1;refGene_gn=TTLL5;refGene_tx=XM_023255878.1;refGene_gn=TTLL5;refGene_tx=XM_023255879.1;refGene_gn=TTLL5;refGene_tx=XM_023255880.1;refGene_gn=TTLL5;refGene_tx=XM_023255881.1;refGene_gn=TTLL5;refGene_tx=XM_023255882.1;refGene_gn=TTLL5;refGene_tx=XM_023255883.1;refGene_gn=TTLL5;refGene_tx=XM_023255884.1;refGene_gn=TTLL5;refGene_tx=XM_023255885.1;refGene_gn=TTLL5;refGene_tx=XM_023255886.1;refGene_gn=TTLL5;refGene_tx=XM_023255887.1;refGene_gn=TTLL5;refGene_tx=XM_023255888.1;refGene_gn=TTLL5;refGene_tx=XM_023255889.1
B3 123103268 123103341
refGene_gn=TTLL5;refGene_tx=XM_023255887.1;refGene_gn=TTLL5;refGene_tx=XM_023255888.1
B3 123104075 123104096
refGene_gn=TTLL5;refGene_tx=XM_023255888.1

B3 123107523 123107719
refGene_gn=TTLL5;refGene_tx=XM_023255874.1;refGene_gn=TTLL5;refGene_tx=XM_023255875.1;refGene_gn=TTLL5;refGene_tx=XM_023255876.1;refGene_gn=TTLL5;refGene_tx=XM_023255877.1;refGene_gn=TTLL5;refGene_tx=XM_023255878.1;refGene_gn=TTLL5;refGene_tx=XM_023255879.1;refGene_gn=TTLL5;refGene_tx=XM_023255880.1;refGene_gn=TTLL5;refGene_tx=XM_023255881.1;refGene_gn=TTLL5;refGene_tx=XM_023255882.1;refGene_gn=TTLL5;refGene_tx=XM_023255883.1;refGene_gn=TTLL5;refGene_tx=XM_023255884.1;refGene_gn=TTLL5;refGene_tx=XM_023255885.1;refGene_gn=TTLL5;refGene_tx=XM_023255886.1;refGene_gn=TTLL5;refGene_tx=XM_023255887.1
B3 123126096 123126219
refGene_gn=TTLL5;refGene_tx=XM_023255874.1;refGene_gn=TTLL5;refGene_tx=XM_023255875.1;refGene_gn=TTLL5;refGene_tx=XM_023255876.1;refGene_gn=TTLL5;refGene_tx=XM_023255877.1;refGene_gn=TTLL5;refGene_tx=XM_023255878.1;refGene_gn=TTLL5;refGene_tx=XM_023255879.1;refGene_gn=TTLL5;refGene_tx=XM_023255881.1;refGene_gn=TTLL5;refGene_tx=XM_023255885.1
B3 123129159 123129377
refGene_gn=TTLL5;refGene_tx=XM_023255874.1;refGene_gn=TTLL5;refGene_tx=XM_023255875.1;refGene_gn=TTLL5;refGene_tx=XM_023255876.1;refGene_gn=TTLL5;refGene_tx=XM_023255877.1;refGene_gn=TTLL5;refGene_tx=XM_023255878.1;refGene_gn=TTLL5;refGene_tx=XM_023255879.1;refGene_gn=TTLL5;refGene_tx=XM_023255880.1;refGene_gn=TTLL5;refGene_tx=XM_023255881.1;refGene_gn=TTLL5;refGene_tx=XM_023255882.1;refGene_gn=TTLL5;refGene_tx=XM_023255883.1;refGene_gn=TTLL5;refGene_tx=XM_023255884.1;refGene_gn=TTLL5;refGene_tx=XM_023255885.1;refGene_gn=TTLL5;refGene_tx=XM_023255886.1;refGene_gn=TTLL5;refGene_tx=XM_023255889.1
B3 123147964 123148044
refGene_gn=TTLL5;refGene_tx=XM_023255874.1;refGene_gn=TTLL5;refGene_tx=XM_023255875.1;refGene_gn=TTLL5;refGene_tx=XM_023255876.1;refGene_gn=TTLL5;refGene_tx=XM_023255877.1;refGene_gn=TTLL5;refGene_tx=XM_023255878.1;refGene_gn=TTLL5;refGene_tx=XM_023255879.1;refGene_gn=TTLL5;refGene_tx=XM_023255880.1;refGene_gn=TTLL5;refGene_tx=XM_023255881.1;refGene_gn=TTLL5;refGene_tx=XM_023255882.1;refGene_gn=TTLL5;refGene_tx=XM_023255883.1;refGene_gn=TTLL5;refGene_tx=XM_023255884.1;refGene_gn=TTLL5;refGene_tx=XM_023255886.1
B3 123152217 123152339
refGene_gn=TTLL5;refGene_tx=XM_023255879.1;refGene_gn=TTLL5;refGene_tx=XM_023255886.1
B3 123158501 123158710
refGene_gn=TTLL5;refGene_tx=XM_023255874.1;refGene_gn=TTLL5;refGene_tx=XM_023255875.1;refGene_gn=TTLL5;refGene_tx=XM_023255876.1;refGene_gn=TTLL5;refGene_tx=XM_023255877.1;refGene_gn=TTLL5;refGene_tx=XM_023255878.1;refGene_gn=TTLL5;refGene_tx=XM_023255880.1;refGene_gn=TTLL5;refGene_tx=XM_023255882.1;refGene_gn=TTLL5;refGene_tx=XM_023255883.1;refGene_gn=TTLL5;refGene_tx=XM_023255884.1
B3 123197408 123197431
refGene_gn=TTLL5;refGene_tx=XM_023255881.1;refGene_gn=TTLL5;refGene_tx=XM_023255885.1
B3 123202075 123202234
refGene_gn=TGFB3;refGene_tx=XM_003987851.3;refGene_gn=TGFB3;refGene_tx

=XM_006933010.3
B3 123203115 123203269
refGene_gn=TGFB3;refGene_tx=XM_003987851.3;refGene_gn=TGFB3;refGene_tx=XM_006933010.3
B3 123205365 123205537
refGene_gn=TGFB3;refGene_tx=XM_003987851.3;refGene_gn=TGFB3;refGene_tx=XM_006933010.3
B3 123208163 123208271
refGene_gn=TGFB3;refGene_tx=XM_003987851.3;refGene_gn=TGFB3;refGene_tx=XM_006933010.3
B3 123214033 123214163
refGene_gn=TGFB3;refGene_tx=XM_003987851.3;refGene_gn=TGFB3;refGene_tx=XM_006933010.3
B3 123214468 123214632
refGene_gn=TGFB3;refGene_tx=XM_003987851.3;refGene_gn=TGFB3;refGene_tx=XM_006933010.3
B3 123222346 123222698
refGene_gn=TGFB3;refGene_tx=XM_003987851.3;refGene_gn=TGFB3;refGene_tx=XM_006933010.3
B3 123228111 123228165
refGene_gn=IFT43;refGene_tx=XM_003987852.5;refGene_gn=IFT43;refGene_tx=XM_006933011.2
B3 123228924 123229017
refGene_gn=IFT43;refGene_tx=XM_019833336.2
B3 123229278 123229371
refGene_gn=IFT43;refGene_tx=XM_003987852.5;refGene_gn=IFT43;refGene_tx=XM_006933011.2;refGene_gn=IFT43;refGene_tx=XM_019833336.2
B3 123256255 123256270
refGene_gn=IFT43;refGene_tx=XM_019833338.2
B3 123257298 123257363
refGene_gn=IFT43;refGene_tx=XM_003987852.5;refGene_gn=IFT43;refGene_tx=XM_006933011.2;refGene_gn=IFT43;refGene_tx=XM_019833336.2;refGene_gn=IFT43;refGene_tx=XM_019833338.2;refGene_gn=IFT43;refGene_tx=XM_019833339.1
B3 123286141 123286174
refGene_gn=IFT43;refGene_tx=XM_003987852.5;refGene_gn=IFT43;refGene_tx=XM_006933011.2;refGene_gn=IFT43;refGene_tx=XM_019833336.2;refGene_gn=IFT43;refGene_tx=XM_019833338.2
B3 123286829 123286876
refGene_gn=IFT43;refGene_tx=XM_003987852.5;refGene_gn=IFT43;refGene_tx=XM_006933011.2;refGene_gn=IFT43;refGene_tx=XM_019833336.2;refGene_gn=IFT43;refGene_tx=XM_019833338.2
B3 123308966 123309039
refGene_gn=IFT43;refGene_tx=XM_003987852.5;refGene_gn=IFT43;refGene_tx=XM_006933011.2;refGene_gn=IFT43;refGene_tx=XM_019833336.2;refGene_gn=IFT43;refGene_tx=XM_019833338.2;refGene_gn=IFT43;refGene_tx=XM_019833339.1
B3 123309316 123309392
refGene_gn=IFT43;refGene_tx=XM_003987852.5;refGene_gn=IFT43;refGene_tx=XM_006933011.2;refGene_gn=IFT43;refGene_tx=XM_019833336.2;refGene_gn=

IFT43;refGene_tx=XM_019833338.2;refGene_gn=IFT43;refGene_tx=XM_0198333
39.1
B3	123309911	123310039
refGene_gn=IFT43;refGene_tx=XM_003987852.5;refGene_gn=IFT43;refGene_tx
=XM_006933011.2;refGene_gn=IFT43;refGene_tx=XM_019833336.2;refGene_gn=
IFT43;refGene_tx=XM_019833338.2;refGene_gn=IFT43;refGene_tx=XM_0198333
39.1
B3	123310141	123310361
refGene_gn=IFT43;refGene_tx=XM_003987852.5;refGene_gn=IFT43;refGene_tx
=XM_006933011.2;refGene_gn=IFT43;refGene_tx=XM_019833336.2;refGene_gn=
IFT43;refGene_tx=XM_019833338.2;refGene_gn=IFT43;refGene_tx=XM_0198333
39.1
B3	123393531	123394193
refGene_gn=GPATCH2L;refGene_tx=XM_019833325.1;refGene_gn=GPATCH2L;refG
ene_tx=XM_019833327.2;refGene_gn=GPATCH2L;refGene_tx=XM_019833328.2;re
fGene_gn=GPATCH2L;refGene_tx=XM_019833333.2;refGene_gn=GPATCH2L;refGen
e_tx=XM_019833334.1
B3	123410766	123410831
refGene_gn=GPATCH2L;refGene_tx=XM_019833325.1;refGene_gn=GPATCH2L;refG
ene_tx=XM_019833327.2;refGene_gn=GPATCH2L;refGene_tx=XM_019833328.2;re
fGene_gn=GPATCH2L;refGene_tx=XM_019833333.2;refGene_gn=GPATCH2L;refGen
e_tx=XM_019833334.1
B3	123415537	123415714
refGene_gn=GPATCH2L;refGene_tx=XM_019833325.1;refGene_gn=GPATCH2L;refG
ene_tx=XM_019833327.2;refGene_gn=GPATCH2L;refGene_tx=XM_019833328.2;re
fGene_gn=GPATCH2L;refGene_tx=XM_019833333.2;refGene_gn=GPATCH2L;refGen
e_tx=XM_019833334.1
B3	123417232	123417312
refGene_gn=GPATCH2L;refGene_tx=XM_019833325.1;refGene_gn=GPATCH2L;refG
ene_tx=XM_019833327.2;refGene_gn=GPATCH2L;refGene_tx=XM_019833328.2;re
fGene_gn=GPATCH2L;refGene_tx=XM_019833333.2;refGene_gn=GPATCH2L;refGen
e_tx=XM_019833334.1
B3	123420028	123420096
refGene_gn=GPATCH2L;refGene_tx=XM_019833325.1;refGene_gn=GPATCH2L;refG
ene_tx=XM_019833327.2;refGene_gn=GPATCH2L;refGene_tx=XM_019833328.2;re
fGene_gn=GPATCH2L;refGene_tx=XM_019833333.2;refGene_gn=GPATCH2L;refGen
e_tx=XM_019833334.1
B3	123423462	123423502
refGene_gn=GPATCH2L;refGene_tx=XM_019833328.2
B3	123423558	123423613
refGene_gn=GPATCH2L;refGene_tx=XM_019833325.1;refGene_gn=GPATCH2L;refG
ene_tx=XM_019833327.2;refGene_gn=GPATCH2L;refGene_tx=XM_019833334.1
B3	123425465	123425551
refGene_gn=GPATCH2L;refGene_tx=XM_019833325.1;refGene_gn=GPATCH2L;refG
ene_tx=XM_019833327.2;refGene_gn=GPATCH2L;refGene_tx=XM_019833328.2;re
fGene_gn=GPATCH2L;refGene_tx=XM_019833334.1
B3	123437434	123437529
refGene_gn=GPATCH2L;refGene_tx=XM_019833325.1;refGene_gn=GPATCH2L;refG
ene_tx=XM_019833327.2;refGene_gn=GPATCH2L;refGene_tx=XM_019833328.2;re
fGene_gn=GPATCH2L;refGene_tx=XM_019833333.2;refGene_gn=GPATCH2L;refGen

e_tx=XM_019833334.1
B3 123439579 123439701
refGene_gn=GPATCH2L;refGene_tx=XM_019833334.1
B3 123443697 123443858
refGene_gn=GPATCH2L;refGene_tx=XM_019833327.2;refGene_gn=GPATCH2L;refGene_tx=XM_019833328.2;refGene_gn=GPATCH2L;refGene_tx=XM_019833333.2
B3 123466869 123467081
refGene_gn=GPATCH2L;refGene_tx=XM_019833325.1
B3 123542108 123542110
refGene_gn=ESRRB;refGene_tx=XM_011283421.2
B3 123603567 123603617
refGene_gn=ESRRB;refGene_tx=XM_003987856.4
B3 123658484 123658894
refGene_gn=ESRRB;refGene_tx=XM_003987856.4;refGene_gn=ESRRB;refGene_tx=XM_011283421.2;refGene_gn=ESRRB;refGene_tx=XM_019833335.2
B3 123676533 123676650
refGene_gn=ESRRB;refGene_tx=XM_003987856.4;refGene_gn=ESRRB;refGene_tx=XM_011283421.2;refGene_gn=ESRRB;refGene_tx=XM_019833335.2
B3 123697187 123697298
refGene_gn=ESRRB;refGene_tx=XM_003987856.4;refGene_gn=ESRRB;refGene_tx=XM_011283421.2;refGene_gn=ESRRB;refGene_tx=XM_019833335.2
B3 123697787 123697949
refGene_gn=ESRRB;refGene_tx=XM_003987856.4;refGene_gn=ESRRB;refGene_tx=XM_011283421.2;refGene_gn=ESRRB;refGene_tx=XM_019833335.2
B3 123706575 123706845
refGene_gn=ESRRB;refGene_tx=XM_003987856.4;refGene_gn=ESRRB;refGene_tx=XM_011283421.2;refGene_gn=ESRRB;refGene_tx=XM_019833335.2
B3 123712697 123712942
refGene_gn=ESRRB;refGene_tx=XM_003987856.4;refGene_gn=ESRRB;refGene_tx=XM_011283421.2;refGene_gn=ESRRB;refGene_tx=XM_019833335.2
B3 123917864 123918167
refGene_gn=VASH1;refGene_tx=XM_003987859.4;refGene_gn=VASH1;refGene_tx=XM_006933012.3;refGene_gn=VASH1;refGene_tx=XM_006933013.3;refGene_gn=VASH1;refGene_tx=XM_006933014.3;refGene_gn=VASH1;refGene_tx=XM_019833353.1;refGene_gn=VASH1;refGene_tx=XM_019833354.2;refGene_gn=VASH1;refGene_tx=XM_019833356.2
B3 123921141 123921264
refGene_gn=VASH1;refGene_tx=XM_006933012.3;refGene_gn=VASH1;refGene_tx=XM_006933013.3;refGene_gn=VASH1;refGene_tx=XM_019833353.1;refGene_gn=VASH1;refGene_tx=XM_019833354.2
B3 123923914 123923992
refGene_gn=VASH1;refGene_tx=XM_019833353.1;refGene_gn=VASH1;refGene_tx=XM_019833354.2;refGene_gn=VASH1;refGene_tx=XM_019833356.2
B3 123924171 123924260
refGene_gn=VASH1;refGene_tx=XM_003987859.4;refGene_gn=VASH1;refGene_tx=XM_006933012.3;refGene_gn=VASH1;refGene_tx=XM_006933013.3;refGene_gn=VASH1;refGene_tx=XM_006933014.3;refGene_gn=VASH1;refGene_tx=XM_019833353.1;refGene_gn=VASH1;refGene_tx=XM_019833354.2;refGene_gn=VASH1;refGene_tx=XM_019833356.2
B3 123925366 123925426

refGene_gn=VASH1;refGene_tx=XM_003987859.4;refGene_gn=VASH1;refGene_tx=XM_006933012.3;refGene_gn=VASH1;refGene_tx=XM_006933013.3;refGene_gn=VASH1;refGene_tx=XM_006933014.3;refGene_gn=VASH1;refGene_tx=XM_019833353.1;refGene_gn=VASH1;refGene_tx=XM_019833354.2;refGene_gn=VASH1;refGene_tx=XM_019833356.2
B3	123927039	123927114
refGene_gn=VASH1;refGene_tx=XM_003987859.4;refGene_gn=VASH1;refGene_tx=XM_006933012.3;refGene_gn=VASH1;refGene_tx=XM_006933013.3;refGene_gn=VASH1;refGene_tx=XM_006933014.3;refGene_gn=VASH1;refGene_tx=XM_019833353.1;refGene_gn=VASH1;refGene_tx=XM_019833354.2;refGene_gn=VASH1;refGene_tx=XM_019833356.2
B3	123929784	123930166
refGene_gn=VASH1;refGene_tx=XM_003987859.4;refGene_gn=VASH1;refGene_tx=XM_006933012.3;refGene_gn=VASH1;refGene_tx=XM_006933013.3;refGene_gn=VASH1;refGene_tx=XM_006933014.3;refGene_gn=VASH1;refGene_tx=XM_019833353.1;refGene_gn=VASH1;refGene_tx=XM_019833354.2;refGene_gn=VASH1;refGene_tx=XM_019833356.2
B3	123931732	123931845
refGene_gn=VASH1;refGene_tx=XM_003987859.4;refGene_gn=VASH1;refGene_tx=XM_006933012.3;refGene_gn=VASH1;refGene_tx=XM_006933013.3;refGene_gn=VASH1;refGene_tx=XM_006933014.3;refGene_gn=VASH1;refGene_tx=XM_019833353.1;refGene_gn=VASH1;refGene_tx=XM_019833354.2;refGene_gn=VASH1;refGene_tx=XM_019833356.2
B3	123932736	123932809
refGene_gn=VASH1;refGene_tx=XM_003987859.4;refGene_gn=VASH1;refGene_tx=XM_006933012.3;refGene_gn=VASH1;refGene_tx=XM_006933013.3;refGene_gn=VASH1;refGene_tx=XM_006933014.3;refGene_gn=VASH1;refGene_tx=XM_019833353.1;refGene_gn=VASH1;refGene_tx=XM_019833354.2;refGene_gn=VASH1;refGene_tx=XM_019833356.2
B3	123942094	123942255
refGene_gn=ANGEL1;refGene_tx=XM_006933019.3;refGene_gn=ANGEL1;refGene_tx=XM_011283424.3;refGene_gn=ANGEL1;refGene_tx=XM_019833340.2;refGene_gn=ANGEL1;refGene_tx=XM_023255891.1
B3	123942999	123943163
refGene_gn=ANGEL1;refGene_tx=XM_006933019.3;refGene_gn=ANGEL1;refGene_tx=XM_011283424.3;refGene_gn=ANGEL1;refGene_tx=XM_019833340.2;refGene_gn=ANGEL1;refGene_tx=XM_023255891.1
B3	123943711	123943781
refGene_gn=ANGEL1;refGene_tx=XM_006933019.3;refGene_gn=ANGEL1;refGene_tx=XM_011283424.3;refGene_gn=ANGEL1;refGene_tx=XM_019833340.2;refGene_gn=ANGEL1;refGene_tx=XM_023255891.1
B3	123956364	123956475
refGene_gn=ANGEL1;refGene_tx=XM_006933019.3;refGene_gn=ANGEL1;refGene_tx=XM_011283424.3;refGene_gn=ANGEL1;refGene_tx=XM_019833340.2;refGene_gn=ANGEL1;refGene_tx=XM_023255891.1
B3	123956995	123957122
refGene_gn=ANGEL1;refGene_tx=XM_006933019.3;refGene_gn=ANGEL1;refGene_tx=XM_011283424.3;refGene_gn=ANGEL1;refGene_tx=XM_019833340.2;refGene_gn=ANGEL1;refGene_tx=XM_023255891.1
B3	123959637	123960071

refGene_gn=ANGEL1;refGene_tx=XM_006933019.3;refGene_gn=ANGEL1;refGene_tx=XM_011283424.3;refGene_gn=ANGEL1;refGene_tx=XM_019833340.2;refGene_gn=ANGEL1;refGene_tx=XM_023255891.1
B3 123960690 123960760
refGene_gn=ANGEL1;refGene_tx=XM_006933019.3;refGene_gn=ANGEL1;refGene_tx=XM_011283424.3;refGene_gn=ANGEL1;refGene_tx=XM_019833340.2;refGene_gn=ANGEL1;refGene_tx=XM_023255891.1
B3 123961223 123961450
refGene_gn=ANGEL1;refGene_tx=XM_006933019.3;refGene_gn=ANGEL1;refGene_tx=XM_011283424.3;refGene_gn=ANGEL1;refGene_tx=XM_019833340.2;refGene_gn=ANGEL1;refGene_tx=XM_023255891.1
B3 123962339 123962915
refGene_gn=ANGEL1;refGene_tx=XM_006933019.3;refGene_gn=ANGEL1;refGene_tx=XM_011283424.3;refGene_gn=ANGEL1;refGene_tx=XM_019833340.2;refGene_gn=ANGEL1;refGene_tx=XM_023255891.1
B3 123965624 123965688
refGene_gn=ANGEL1;refGene_tx=XM_023255891.1
B3 123966609 123966712
refGene_gn=ANGEL1;refGene_tx=XM_006933019.3
B3 123979987 123980024
refGene_gn=LRRC74A;refGene_tx=XM_006933015.3;refGene_gn=LRRC74A;refGene_tx=XM_006933016.4;refGene_gn=LRRC74A;refGene_tx=XM_019833341.1;refGene_gn=LRRC74A;refGene_tx=XM_019833343.2;refGene_gn=LRRC74A;refGene_tx=XM_019833344.1;refGene_gn=LRRC74A;refGene_tx=XM_019833345.1;refGene_gn=LRRC74A;refGene_tx=XM_019833346.1
B3 123981197 123981225
refGene_gn=LRRC74A;refGene_tx=XM_019833342.2
B3 123981765 123981894
refGene_gn=LRRC74A;refGene_tx=XM_006933015.3;refGene_gn=LRRC74A;refGene_tx=XM_006933016.4;refGene_gn=LRRC74A;refGene_tx=XM_019833341.1;refGene_gn=LRRC74A;refGene_tx=XM_019833342.2;refGene_gn=LRRC74A;refGene_tx=XM_019833343.2;refGene_gn=LRRC74A;refGene_tx=XM_019833344.1;refGene_gn=LRRC74A;refGene_tx=XM_019833345.1;refGene_gn=LRRC74A;refGene_tx=XM_019833346.1
B3 123984582 123984755
refGene_gn=LRRC74A;refGene_tx=XM_006933015.3;refGene_gn=LRRC74A;refGene_tx=XM_006933016.4;refGene_gn=LRRC74A;refGene_tx=XM_019833341.1;refGene_gn=LRRC74A;refGene_tx=XM_019833342.2;refGene_gn=LRRC74A;refGene_tx=XM_019833343.2;refGene_gn=LRRC74A;refGene_tx=XM_019833344.1;refGene_gn=LRRC74A;refGene_tx=XM_019833345.1;refGene_gn=LRRC74A;refGene_tx=XM_019833346.1;refGene_gn=LRRC74A;refGene_tx=XM_019833347.1;refGene_gn=LRRC74A;refGene_tx=XM_019833348.1;refGene_gn=LRRC74A;refGene_tx=XM_019833349.2
B3 123987539 123987647
refGene_gn=LRRC74A;refGene_tx=XM_006933015.3;refGene_gn=LRRC74A;refGene_tx=XM_006933016.4;refGene_gn=LRRC74A;refGene_tx=XM_019833341.1;refGene_gn=LRRC74A;refGene_tx=XM_019833342.2;refGene_gn=LRRC74A;refGene_tx=XM_019833343.2;refGene_gn=LRRC74A;refGene_tx=XM_019833345.1;refGene_gn=LRRC74A;refGene_tx=XM_019833347.1;refGene_gn=LRRC74A;refGene_tx=XM_019833348.1;refGene_gn=LRRC74A;refGene_tx=XM_019833349.2

B3	123988639	123988736
refGene_gn=LRRC74A;refGene_tx=XM_006933015.3;refGene_gn=LRRC74A;refGene_tx=XM_006933016.4;refGene_gn=LRRC74A;refGene_tx=XM_019833341.1;refGene_gn=LRRC74A;refGene_tx=XM_019833342.2;refGene_gn=LRRC74A;refGene_tx=XM_019833343.2;refGene_gn=LRRC74A;refGene_tx=XM_019833344.1;refGene_gn=LRRC74A;refGene_tx=XM_019833345.1;refGene_gn=LRRC74A;refGene_tx=XM_019833346.1;refGene_gn=LRRC74A;refGene_tx=XM_019833347.1;refGene_gn=LRRC74A;refGene_tx=XM_019833348.1;refGene_gn=LRRC74A;refGene_tx=XM_019833349.2
B3	123993236	123993286
refGene_gn=LRRC74A;refGene_tx=XM_006933015.3;refGene_gn=LRRC74A;refGene_tx=XM_006933016.4;refGene_gn=LRRC74A;refGene_tx=XM_019833341.1;refGene_gn=LRRC74A;refGene_tx=XM_019833342.2;refGene_gn=LRRC74A;refGene_tx=XM_019833343.2;refGene_gn=LRRC74A;refGene_tx=XM_019833344.1;refGene_gn=LRRC74A;refGene_tx=XM_019833345.1;refGene_gn=LRRC74A;refGene_tx=XM_019833346.1;refGene_gn=LRRC74A;refGene_tx=XM_019833347.1;refGene_gn=LRRC74A;refGene_tx=XM_019833348.1;refGene_gn=LRRC74A;refGene_tx=XM_019833349.2
B3	123993662	123993744
refGene_gn=LRRC74A;refGene_tx=XM_006933015.3;refGene_gn=LRRC74A;refGene_tx=XM_006933016.4;refGene_gn=LRRC74A;refGene_tx=XM_019833341.1;refGene_gn=LRRC74A;refGene_tx=XM_019833342.2;refGene_gn=LRRC74A;refGene_tx=XM_019833343.2;refGene_gn=LRRC74A;refGene_tx=XM_019833344.1;refGene_gn=LRRC74A;refGene_tx=XM_019833345.1;refGene_gn=LRRC74A;refGene_tx=XM_019833346.1;refGene_gn=LRRC74A;refGene_tx=XM_019833347.1;refGene_gn=LRRC74A;refGene_tx=XM_019833348.1;refGene_gn=LRRC74A;refGene_tx=XM_019833349.2
B3	123998584	123998670
refGene_gn=LRRC74A;refGene_tx=XM_006933015.3;refGene_gn=LRRC74A;refGene_tx=XM_006933016.4;refGene_gn=LRRC74A;refGene_tx=XM_019833341.1;refGene_gn=LRRC74A;refGene_tx=XM_019833342.2;refGene_gn=LRRC74A;refGene_tx=XM_019833343.2;refGene_gn=LRRC74A;refGene_tx=XM_019833344.1;refGene_gn=LRRC74A;refGene_tx=XM_019833345.1;refGene_gn=LRRC74A;refGene_tx=XM_019833346.1;refGene_gn=LRRC74A;refGene_tx=XM_019833347.1;refGene_gn=LRRC74A;refGene_tx=XM_019833348.1;refGene_gn=LRRC74A;refGene_tx=XM_019833349.2
B3	123999108	123999303
refGene_gn=LRRC74A;refGene_tx=XM_006933015.3;refGene_gn=LRRC74A;refGene_tx=XM_006933016.4;refGene_gn=LRRC74A;refGene_tx=XM_019833341.1;refGene_gn=LRRC74A;refGene_tx=XM_019833342.2;refGene_gn=LRRC74A;refGene_tx=XM_019833343.2;refGene_gn=LRRC74A;refGene_tx=XM_019833344.1;refGene_gn=LRRC74A;refGene_tx=XM_019833345.1;refGene_gn=LRRC74A;refGene_tx=XM_019833346.1;refGene_gn=LRRC74A;refGene_tx=XM_019833347.1;refGene_gn=LRRC74A;refGene_tx=XM_019833348.1;refGene_gn=LRRC74A;refGene_tx=XM_019833349.2
B3	124002506	124002602
refGene_gn=LRRC74A;refGene_tx=XM_006933015.3;refGene_gn=LRRC74A;refGene_tx=XM_006933016.4;refGene_gn=LRRC74A;refGene_tx=XM_019833341.1;refGene_gn=LRRC74A;refGene_tx=XM_019833342.2;refGene_gn=LRRC74A;refGene_tx=XM_019833343.2;refGene_gn=LRRC74A;refGene_tx=XM_019833344.1;refGene_gn

=LRRC74A;refGene_tx=XM_019833345.1;refGene_gn=LRRC74A;refGene_tx=XM_019833346.1;refGene_gn=LRRC74A;refGene_tx=XM_019833347.1;refGene_gn=LRRC74A;refGene_tx=XM_019833348.1;refGene_gn=LRRC74A;refGene_tx=XM_019833349.2
B3 124005844 124005991
refGene_gn=LRRC74A;refGene_tx=XM_006933015.3;refGene_gn=LRRC74A;refGene_tx=XM_006933016.4;refGene_gn=LRRC74A;refGene_tx=XM_019833341.1;refGene_gn=LRRC74A;refGene_tx=XM_019833342.2;refGene_gn=LRRC74A;refGene_tx=XM_019833343.2;refGene_gn=LRRC74A;refGene_tx=XM_019833344.1;refGene_gn=LRRC74A;refGene_tx=XM_019833345.1;refGene_gn=LRRC74A;refGene_tx=XM_019833346.1;refGene_gn=LRRC74A;refGene_tx=XM_019833347.1;refGene_gn=LRRC74A;refGene_tx=XM_019833348.1;refGene_gn=LRRC74A;refGene_tx=XM_019833349.2
B3 124008292 124008400
refGene_gn=LRRC74A;refGene_tx=XM_006933015.3;refGene_gn=LRRC74A;refGene_tx=XM_006933016.4;refGene_gn=LRRC74A;refGene_tx=XM_019833341.1;refGene_gn=LRRC74A;refGene_tx=XM_019833342.2;refGene_gn=LRRC74A;refGene_tx=XM_019833343.2;refGene_gn=LRRC74A;refGene_tx=XM_019833344.1;refGene_gn=LRRC74A;refGene_tx=XM_019833345.1;refGene_gn=LRRC74A;refGene_tx=XM_019833346.1;refGene_gn=LRRC74A;refGene_tx=XM_019833347.1;refGene_gn=LRRC74A;refGene_tx=XM_019833348.1;refGene_gn=LRRC74A;refGene_tx=XM_019833349.2
B3 124008529 124008612
refGene_gn=LRRC74A;refGene_tx=XM_006933015.3;refGene_gn=LRRC74A;refGene_tx=XM_006933016.4;refGene_gn=LRRC74A;refGene_tx=XM_019833341.1;refGene_gn=LRRC74A;refGene_tx=XM_019833342.2;refGene_gn=LRRC74A;refGene_tx=XM_019833343.2;refGene_gn=LRRC74A;refGene_tx=XM_019833344.1;refGene_gn=LRRC74A;refGene_tx=XM_019833345.1;refGene_gn=LRRC74A;refGene_tx=XM_019833346.1;refGene_gn=LRRC74A;refGene_tx=XM_019833347.1;refGene_gn=LRRC74A;refGene_tx=XM_019833348.1;refGene_gn=LRRC74A;refGene_tx=XM_019833349.2
B3 124010965 124010990
refGene_gn=LRRC74A;refGene_tx=XM_006933016.4;refGene_gn=LRRC74A;refGene_tx=XM_019833343.2
B3 124012411 124012538
refGene_gn=LRRC74A;refGene_tx=XM_006933015.3;refGene_gn=LRRC74A;refGene_tx=XM_019833341.1;refGene_gn=LRRC74A;refGene_tx=XM_019833342.2;refGene_gn=LRRC74A;refGene_tx=XM_019833344.1;refGene_gn=LRRC74A;refGene_tx=XM_019833345.1;refGene_gn=LRRC74A;refGene_tx=XM_019833346.1;refGene_gn=LRRC74A;refGene_tx=XM_019833347.1;refGene_gn=LRRC74A;refGene_tx=XM_019833348.1;refGene_gn=LRRC74A;refGene_tx=XM_019833349.2
B3 124137121 124139491
refGene_gn=IRF2BPL;refGene_tx=XM_011279729.2
B3 124195923 124195975
refGene_gn=CIPC;refGene_tx=XM_006933023.4
B3 124197614 124197621
refGene_gn=CIPC;refGene_tx=XM_019833363.2
B3 124202622 124202758
refGene_gn=CIPC;refGene_tx=XM_003987860.4;refGene_gn=CIPC;refGene_tx=XM_006933021.4;refGene_gn=CIPC;refGene_tx=XM_006933022.4;refGene_gn=CIP

C;refGene_tx=XM_019833362.2
B3 124209925 124210095
refGene_gn=CIPC;refGene_tx=XM_003987860.4;refGene_gn=CIPC;refGene_tx=XM_006933021.4;refGene_gn=CIPC;refGene_tx=XM_006933022.4;refGene_gn=CIPC;refGene_tx=XM_006933023.4;refGene_gn=CIPC;refGene_tx=XM_019833362.2;refGene_gn=CIPC;refGene_tx=XM_019833363.2
B3 124213651 124214545
refGene_gn=CIPC;refGene_tx=XM_003987860.4;refGene_gn=CIPC;refGene_tx=XM_006933021.4;refGene_gn=CIPC;refGene_tx=XM_006933022.4;refGene_gn=CIPC;refGene_tx=XM_006933023.4;refGene_gn=CIPC;refGene_tx=XM_019833362.2;refGene_gn=CIPC;refGene_tx=XM_019833363.2
B3 124234458 124234724
refGene_gn=ZDHHC22;refGene_tx=XM_003987861.4;refGene_gn=ZDHHC22;refGene_tx=XM_019833364.2;refGene_gn=ZDHHC22;refGene_tx=XM_019833365.2;refGene_gn=ZDHHC22;refGene_tx=XM_023255896.1
B3 124239898 124240424
refGene_gn=ZDHHC22;refGene_tx=XM_003987861.4;refGene_gn=ZDHHC22;refGene_tx=XM_019833364.2;refGene_gn=ZDHHC22;refGene_tx=XM_019833365.2;refGene_gn=ZDHHC22;refGene_tx=XM_023255896.1
B3 124311298 124311448
refGene_gn=TMEM63C;refGene_tx=XM_003987862.5;refGene_gn=TMEM63C;refGene_tx=XM_019833358.2;refGene_gn=TMEM63C;refGene_tx=XM_019833359.1;refGene_gn=TMEM63C;refGene_tx=XM_019833360.2;refGene_gn=TMEM63C;refGene_tx=XM_019833361.2;refGene_gn=TMEM63C;refGene_tx=XM_023255892.1;refGene_gn=TMEM63C;refGene_tx=XM_023255893.1;refGene_gn=TMEM63C;refGene_tx=XM_023255894.1;refGene_gn=TMEM63C;refGene_tx=XM_023255895.1
B3 124311981 124312061
refGene_gn=TMEM63C;refGene_tx=XM_003987862.5;refGene_gn=TMEM63C;refGene_tx=XM_019833358.2;refGene_gn=TMEM63C;refGene_tx=XM_019833359.1;refGene_gn=TMEM63C;refGene_tx=XM_019833360.2;refGene_gn=TMEM63C;refGene_tx=XM_019833361.2;refGene_gn=TMEM63C;refGene_tx=XM_023255892.1;refGene_gn=TMEM63C;refGene_tx=XM_023255893.1;refGene_gn=TMEM63C;refGene_tx=XM_023255894.1;refGene_gn=TMEM63C;refGene_tx=XM_023255895.1
B3 124312477 124312559
refGene_gn=TMEM63C;refGene_tx=XM_003987862.5;refGene_gn=TMEM63C;refGene_tx=XM_019833358.2;refGene_gn=TMEM63C;refGene_tx=XM_019833359.1;refGene_gn=TMEM63C;refGene_tx=XM_019833360.2;refGene_gn=TMEM63C;refGene_tx=XM_019833361.2;refGene_gn=TMEM63C;refGene_tx=XM_023255892.1;refGene_gn=TMEM63C;refGene_tx=XM_023255893.1;refGene_gn=TMEM63C;refGene_tx=XM_023255894.1;refGene_gn=TMEM63C;refGene_tx=XM_023255895.1
B3 124317294 124317332
refGene_gn=TMEM63C;refGene_tx=XM_003987862.5;refGene_gn=TMEM63C;refGene_tx=XM_019833358.2;refGene_gn=TMEM63C;refGene_tx=XM_019833359.1;refGene_gn=TMEM63C;refGene_tx=XM_019833360.2;refGene_gn=TMEM63C;refGene_tx=XM_019833361.2;refGene_gn=TMEM63C;refGene_tx=XM_023255892.1;refGene_gn=TMEM63C;refGene_tx=XM_023255893.1;refGene_gn=TMEM63C;refGene_tx=XM_023255894.1;refGene_gn=TMEM63C;refGene_tx=XM_023255895.1
B3 124321476 124321619
refGene_gn=TMEM63C;refGene_tx=XM_003987862.5;refGene_gn=TMEM63C;refGene_tx=XM_019833358.2;refGene_gn=TMEM63C;refGene_tx=XM_019833359.1;refGe

ne_gn=TMEM63C;refGene_tx=XM_019833360.2;refGene_gn=TMEM63C;refGene_tx=XM_019833361.2;refGene_gn=TMEM63C;refGene_tx=XM_023255892.1;refGene_gn=TMEM63C;refGene_tx=XM_023255893.1;refGene_gn=TMEM63C;refGene_tx=XM_023255894.1;refGene_gn=TMEM63C;refGene_tx=XM_023255895.1
B3 124323007 124323056
refGene_gn=TMEM63C;refGene_tx=XM_003987862.5;refGene_gn=TMEM63C;refGene_tx=XM_019833358.2;refGene_gn=TMEM63C;refGene_tx=XM_019833359.1;refGene_gn=TMEM63C;refGene_tx=XM_019833360.2;refGene_gn=TMEM63C;refGene_tx=XM_019833361.2;refGene_gn=TMEM63C;refGene_tx=XM_023255892.1;refGene_gn=TMEM63C;refGene_tx=XM_023255893.1;refGene_gn=TMEM63C;refGene_tx=XM_023255894.1;refGene_gn=TMEM63C;refGene_tx=XM_023255895.1
B3 124325030 124325139
refGene_gn=TMEM63C;refGene_tx=XM_003987862.5;refGene_gn=TMEM63C;refGene_tx=XM_019833358.2;refGene_gn=TMEM63C;refGene_tx=XM_019833359.1;refGene_gn=TMEM63C;refGene_tx=XM_019833360.2;refGene_gn=TMEM63C;refGene_tx=XM_019833361.2;refGene_gn=TMEM63C;refGene_tx=XM_023255892.1;refGene_gn=TMEM63C;refGene_tx=XM_023255893.1;refGene_gn=TMEM63C;refGene_tx=XM_023255894.1;refGene_gn=TMEM63C;refGene_tx=XM_023255895.1
B3 124328106 124328180
refGene_gn=TMEM63C;refGene_tx=XM_003987862.5;refGene_gn=TMEM63C;refGene_tx=XM_019833358.2;refGene_gn=TMEM63C;refGene_tx=XM_019833359.1;refGene_gn=TMEM63C;refGene_tx=XM_019833360.2;refGene_gn=TMEM63C;refGene_tx=XM_019833361.2;refGene_gn=TMEM63C;refGene_tx=XM_023255892.1;refGene_gn=TMEM63C;refGene_tx=XM_023255893.1;refGene_gn=TMEM63C;refGene_tx=XM_023255894.1;refGene_gn=TMEM63C;refGene_tx=XM_023255895.1
B3 124328803 124328884
refGene_gn=TMEM63C;refGene_tx=XM_003987862.5;refGene_gn=TMEM63C;refGene_tx=XM_019833358.2;refGene_gn=TMEM63C;refGene_tx=XM_019833359.1;refGene_gn=TMEM63C;refGene_tx=XM_019833360.2;refGene_gn=TMEM63C;refGene_tx=XM_019833361.2;refGene_gn=TMEM63C;refGene_tx=XM_023255892.1;refGene_gn=TMEM63C;refGene_tx=XM_023255893.1;refGene_gn=TMEM63C;refGene_tx=XM_023255894.1;refGene_gn=TMEM63C;refGene_tx=XM_023255895.1
B3 124328997 124329121
refGene_gn=TMEM63C;refGene_tx=XM_003987862.5;refGene_gn=TMEM63C;refGene_tx=XM_019833358.2;refGene_gn=TMEM63C;refGene_tx=XM_019833359.1;refGene_gn=TMEM63C;refGene_tx=XM_019833360.2;refGene_gn=TMEM63C;refGene_tx=XM_019833361.2;refGene_gn=TMEM63C;refGene_tx=XM_023255892.1;refGene_gn=TMEM63C;refGene_tx=XM_023255893.1;refGene_gn=TMEM63C;refGene_tx=XM_023255894.1;refGene_gn=TMEM63C;refGene_tx=XM_023255895.1
B3 124329862 124329996
refGene_gn=TMEM63C;refGene_tx=XM_003987862.5;refGene_gn=TMEM63C;refGene_tx=XM_019833358.2;refGene_gn=TMEM63C;refGene_tx=XM_019833359.1;refGene_gn=TMEM63C;refGene_tx=XM_019833360.2;refGene_gn=TMEM63C;refGene_tx=XM_019833361.2;refGene_gn=TMEM63C;refGene_tx=XM_023255892.1;refGene_gn=TMEM63C;refGene_tx=XM_023255893.1;refGene_gn=TMEM63C;refGene_tx=XM_023255894.1;refGene_gn=TMEM63C;refGene_tx=XM_023255895.1
B3 124331433 124331556
refGene_gn=TMEM63C;refGene_tx=XM_003987862.5;refGene_gn=TMEM63C;refGene_tx=XM_019833358.2;refGene_gn=TMEM63C;refGene_tx=XM_019833359.1;refGene_gn=TMEM63C;refGene_tx=XM_019833360.2;refGene_gn=TMEM63C;refGene_tx=

XM_019833361.2;refGene_gn=TMEM63C;refGene_tx=XM_023255892.1;refGene_gn=TMEM63C;refGene_tx=XM_023255893.1;refGene_gn=TMEM63C;refGene_tx=XM_023255894.1;refGene_gn=TMEM63C;refGene_tx=XM_023255895.1
B3 124331940 124332094
refGene_gn=TMEM63C;refGene_tx=XM_003987862.5;refGene_gn=TMEM63C;refGene_tx=XM_019833358.2;refGene_gn=TMEM63C;refGene_tx=XM_019833359.1;refGene_gn=TMEM63C;refGene_tx=XM_019833360.2;refGene_gn=TMEM63C;refGene_tx=XM_019833361.2;refGene_gn=TMEM63C;refGene_tx=XM_023255892.1;refGene_gn=TMEM63C;refGene_tx=XM_023255893.1;refGene_gn=TMEM63C;refGene_tx=XM_023255894.1;refGene_gn=TMEM63C;refGene_tx=XM_023255895.1
B3 124333293 124333400
refGene_gn=TMEM63C;refGene_tx=XM_003987862.5;refGene_gn=TMEM63C;refGene_tx=XM_019833358.2;refGene_gn=TMEM63C;refGene_tx=XM_019833359.1;refGene_gn=TMEM63C;refGene_tx=XM_019833360.2;refGene_gn=TMEM63C;refGene_tx=XM_019833361.2;refGene_gn=TMEM63C;refGene_tx=XM_023255892.1;refGene_gn=TMEM63C;refGene_tx=XM_023255893.1;refGene_gn=TMEM63C;refGene_tx=XM_023255894.1;refGene_gn=TMEM63C;refGene_tx=XM_023255895.1
B3 124334368 124334455
refGene_gn=TMEM63C;refGene_tx=XM_003987862.5;refGene_gn=TMEM63C;refGene_tx=XM_019833358.2;refGene_gn=TMEM63C;refGene_tx=XM_019833359.1;refGene_gn=TMEM63C;refGene_tx=XM_019833360.2;refGene_gn=TMEM63C;refGene_tx=XM_019833361.2;refGene_gn=TMEM63C;refGene_tx=XM_023255892.1;refGene_gn=TMEM63C;refGene_tx=XM_023255893.1;refGene_gn=TMEM63C;refGene_tx=XM_023255894.1;refGene_gn=TMEM63C;refGene_tx=XM_023255895.1
B3 124335020 124335086
refGene_gn=TMEM63C;refGene_tx=XM_003987862.5;refGene_gn=TMEM63C;refGene_tx=XM_019833358.2;refGene_gn=TMEM63C;refGene_tx=XM_019833359.1;refGene_gn=TMEM63C;refGene_tx=XM_019833360.2;refGene_gn=TMEM63C;refGene_tx=XM_019833361.2;refGene_gn=TMEM63C;refGene_tx=XM_023255892.1;refGene_gn=TMEM63C;refGene_tx=XM_023255893.1;refGene_gn=TMEM63C;refGene_tx=XM_023255894.1;refGene_gn=TMEM63C;refGene_tx=XM_023255895.1
B3 124337010 124337173
refGene_gn=TMEM63C;refGene_tx=XM_003987862.5;refGene_gn=TMEM63C;refGene_tx=XM_019833358.2;refGene_gn=TMEM63C;refGene_tx=XM_019833359.1;refGene_gn=TMEM63C;refGene_tx=XM_019833360.2;refGene_gn=TMEM63C;refGene_tx=XM_019833361.2;refGene_gn=TMEM63C;refGene_tx=XM_023255892.1;refGene_gn=TMEM63C;refGene_tx=XM_023255893.1;refGene_gn=TMEM63C;refGene_tx=XM_023255894.1;refGene_gn=TMEM63C;refGene_tx=XM_023255895.1
B3 124337415 124337521
refGene_gn=TMEM63C;refGene_tx=XM_003987862.5;refGene_gn=TMEM63C;refGene_tx=XM_019833358.2;refGene_gn=TMEM63C;refGene_tx=XM_019833359.1;refGene_gn=TMEM63C;refGene_tx=XM_019833360.2;refGene_gn=TMEM63C;refGene_tx=XM_019833361.2;refGene_gn=TMEM63C;refGene_tx=XM_023255892.1;refGene_gn=TMEM63C;refGene_tx=XM_023255893.1;refGene_gn=TMEM63C;refGene_tx=XM_023255894.1;refGene_gn=TMEM63C;refGene_tx=XM_023255895.1
B3 124337926 124338094
refGene_gn=TMEM63C;refGene_tx=XM_003987862.5;refGene_gn=TMEM63C;refGene_tx=XM_019833358.2;refGene_gn=TMEM63C;refGene_tx=XM_019833359.1;refGene_gn=TMEM63C;refGene_tx=XM_019833360.2;refGene_gn=TMEM63C;refGene_tx=XM_019833361.2;refGene_gn=TMEM63C;refGene_tx=XM_023255892.1;refGene_gn

=TMEM63C;refGene_tx=XM_023255893.1;refGene_gn=TMEM63C;refGene_tx=XM_02
3255894.1;refGene_gn=TMEM63C;refGene_tx=XM_023255895.1
B3	124340104	124340214
refGene_gn=TMEM63C;refGene_tx=XM_003987862.5;refGene_gn=TMEM63C;refGen
e_tx=XM_019833358.2;refGene_gn=TMEM63C;refGene_tx=XM_019833359.1;refGe
ne_gn=TMEM63C;refGene_tx=XM_019833360.2;refGene_gn=TMEM63C;refGene_tx=
XM_019833361.2;refGene_gn=TMEM63C;refGene_tx=XM_023255892.1;refGene_gn
=TMEM63C;refGene_tx=XM_023255893.1;refGene_gn=TMEM63C;refGene_tx=XM_02
3255894.1;refGene_gn=TMEM63C;refGene_tx=XM_023255895.1
B3	124341610	124341679
refGene_gn=TMEM63C;refGene_tx=XM_003987862.5;refGene_gn=TMEM63C;refGen
e_tx=XM_019833358.2;refGene_gn=TMEM63C;refGene_tx=XM_019833359.1;refGe
ne_gn=TMEM63C;refGene_tx=XM_019833360.2;refGene_gn=TMEM63C;refGene_tx=
XM_019833361.2;refGene_gn=TMEM63C;refGene_tx=XM_023255892.1;refGene_gn
=TMEM63C;refGene_tx=XM_023255893.1;refGene_gn=TMEM63C;refGene_tx=XM_02
3255894.1;refGene_gn=TMEM63C;refGene_tx=XM_023255895.1
B3	124344500	124344704
refGene_gn=TMEM63C;refGene_tx=XM_003987862.5;refGene_gn=TMEM63C;refGen
e_tx=XM_019833358.2;refGene_gn=TMEM63C;refGene_tx=XM_019833359.1;refGe
ne_gn=TMEM63C;refGene_tx=XM_019833360.2;refGene_gn=TMEM63C;refGene_tx=
XM_019833361.2;refGene_gn=TMEM63C;refGene_tx=XM_023255892.1;refGene_gn
=TMEM63C;refGene_tx=XM_023255893.1;refGene_gn=TMEM63C;refGene_tx=XM_02
3255894.1;refGene_gn=TMEM63C;refGene_tx=XM_023255895.1
B3	124355489	124355624
refGene_gn=NGB;refGene_tx=XM_023255897.1
B3	124357026	124357146
refGene_gn=NGB;refGene_tx=XM_023255897.1
B3	124358696	124358808
refGene_gn=NGB;refGene_tx=XM_023255897.1
B3	124360153	124360242
refGene_gn=NGB;refGene_tx=XM_023255897.1
B3	124366200	124366306
refGene_gn=POMT2;refGene_tx=XM_003987863.3;refGene_gn=POMT2;refGene_tx
=XM_006933027.2;refGene_gn=POMT2;refGene_tx=XM_019833367.1;refGene_gn=
POMT2;refGene_tx=XM_019833368.2;refGene_gn=POMT2;refGene_tx=XM_0198333
69.1
B3	124367133	124367248
refGene_gn=POMT2;refGene_tx=XM_003987863.3;refGene_gn=POMT2;refGene_tx
=XM_006933027.2;refGene_gn=POMT2;refGene_tx=XM_019833367.1;refGene_gn=
POMT2;refGene_tx=XM_019833368.2;refGene_gn=POMT2;refGene_tx=XM_0198333
69.1
B3	124367399	124367540
refGene_gn=POMT2;refGene_tx=XM_003987863.3;refGene_gn=POMT2;refGene_tx
=XM_006933027.2;refGene_gn=POMT2;refGene_tx=XM_019833367.1;refGene_gn=
POMT2;refGene_tx=XM_019833368.2;refGene_gn=POMT2;refGene_tx=XM_0198333
69.1
B3	124368674	124368780
refGene_gn=POMT2;refGene_tx=XM_003987863.3;refGene_gn=POMT2;refGene_tx
=XM_006933027.2;refGene_gn=POMT2;refGene_tx=XM_019833367.1;refGene_gn=
POMT2;refGene_tx=XM_019833368.2;refGene_gn=POMT2;refGene_tx=XM_0198333

69.1
B3 124368866 124368926
refGene_gn=POMT2;refGene_tx=XM_003987863.3;refGene_gn=POMT2;refGene_tx=XM_006933027.2;refGene_gn=POMT2;refGene_tx=XM_019833367.1;refGene_gn=POMT2;refGene_tx=XM_019833368.2;refGene_gn=POMT2;refGene_tx=XM_019833369.1
B3 124369254 124369326
refGene_gn=POMT2;refGene_tx=XM_003987863.3;refGene_gn=POMT2;refGene_tx=XM_006933027.2;refGene_gn=POMT2;refGene_tx=XM_019833367.1;refGene_gn=POMT2;refGene_tx=XM_019833368.2;refGene_gn=POMT2;refGene_tx=XM_019833369.1
B3 124372220 124372297
refGene_gn=POMT2;refGene_tx=XM_003987863.3;refGene_gn=POMT2;refGene_tx=XM_006933027.2;refGene_gn=POMT2;refGene_tx=XM_019833367.1;refGene_gn=POMT2;refGene_tx=XM_019833368.2;refGene_gn=POMT2;refGene_tx=XM_019833369.1
B3 124373396 124373488
refGene_gn=POMT2;refGene_tx=XM_003987863.3;refGene_gn=POMT2;refGene_tx=XM_006933027.2;refGene_gn=POMT2;refGene_tx=XM_019833367.1;refGene_gn=POMT2;refGene_tx=XM_019833368.2;refGene_gn=POMT2;refGene_tx=XM_019833369.1
B3 124374165 124374317
refGene_gn=POMT2;refGene_tx=XM_003987863.3;refGene_gn=POMT2;refGene_tx=XM_006933027.2;refGene_gn=POMT2;refGene_tx=XM_019833367.1;refGene_gn=POMT2;refGene_tx=XM_019833368.2;refGene_gn=POMT2;refGene_tx=XM_019833369.1
B3 124375663 124375742
refGene_gn=POMT2;refGene_tx=XM_003987863.3;refGene_gn=POMT2;refGene_tx=XM_006933027.2;refGene_gn=POMT2;refGene_tx=XM_019833367.1;refGene_gn=POMT2;refGene_tx=XM_019833368.2;refGene_gn=POMT2;refGene_tx=XM_019833369.1
B3 124377471 124377541
refGene_gn=POMT2;refGene_tx=XM_003987863.3;refGene_gn=POMT2;refGene_tx=XM_006933027.2;refGene_gn=POMT2;refGene_tx=XM_019833367.1;refGene_gn=POMT2;refGene_tx=XM_019833368.2;refGene_gn=POMT2;refGene_tx=XM_019833369.1
B3 124378435 124378483
refGene_gn=POMT2;refGene_tx=XM_006933027.2;refGene_gn=POMT2;refGene_tx=XM_019833367.1;refGene_gn=POMT2;refGene_tx=XM_019833368.2;refGene_gn=POMT2;refGene_tx=XM_019833369.1
B3 124380080 124380147
refGene_gn=POMT2;refGene_tx=XM_003987863.3;refGene_gn=POMT2;refGene_tx=XM_006933027.2;refGene_gn=POMT2;refGene_tx=XM_019833367.1;refGene_gn=POMT2;refGene_tx=XM_019833368.2;refGene_gn=POMT2;refGene_tx=XM_019833369.1
B3 124384496 124384606
refGene_gn=POMT2;refGene_tx=XM_003987863.3;refGene_gn=POMT2;refGene_tx=XM_006933027.2;refGene_gn=POMT2;refGene_tx=XM_019833367.1;refGene_gn=POMT2;refGene_tx=XM_019833368.2;refGene_gn=POMT2;refGene_tx=XM_019833369.1

B3 124387360 124387443
refGene_gn=POMT2;refGene_tx=XM_003987863.3;refGene_gn=POMT2;refGene_tx=XM_006933027.2;refGene_gn=POMT2;refGene_tx=XM_019833367.1;refGene_gn=POMT2;refGene_tx=XM_019833368.2;refGene_gn=POMT2;refGene_tx=XM_019833369.1
B3 124388113 124388220
refGene_gn=POMT2;refGene_tx=XM_003987863.3;refGene_gn=POMT2;refGene_tx=XM_006933027.2;refGene_gn=POMT2;refGene_tx=XM_019833367.1;refGene_gn=POMT2;refGene_tx=XM_019833368.2;refGene_gn=POMT2;refGene_tx=XM_019833369.1
B3 124389526 124389686
refGene_gn=POMT2;refGene_tx=XM_003987863.3;refGene_gn=POMT2;refGene_tx=XM_006933027.2;refGene_gn=POMT2;refGene_tx=XM_019833367.1;refGene_gn=POMT2;refGene_tx=XM_019833368.2
B3 124391583 124391692
refGene_gn=POMT2;refGene_tx=XM_003987863.3;refGene_gn=POMT2;refGene_tx=XM_006933027.2;refGene_gn=POMT2;refGene_tx=XM_019833367.1;refGene_gn=POMT2;refGene_tx=XM_019833368.2
B3 124393415 124393524
refGene_gn=POMT2;refGene_tx=XM_003987863.3;refGene_gn=POMT2;refGene_tx=XM_006933027.2;refGene_gn=POMT2;refGene_tx=XM_019833367.1;refGene_gn=POMT2;refGene_tx=XM_019833368.2
B3 124395495 124395600
refGene_gn=POMT2;refGene_tx=XM_003987863.3;refGene_gn=POMT2;refGene_tx=XM_006933027.2;refGene_gn=POMT2;refGene_tx=XM_019833368.2
B3 124400877 124400962
refGene_gn=POMT2;refGene_tx=XM_003987863.3;refGene_gn=POMT2;refGene_tx=XM_006933027.2;refGene_gn=POMT2;refGene_tx=XM_019833367.1;refGene_gn=POMT2;refGene_tx=XM_019833368.2;refGene_gn=POMT2;refGene_tx=XM_019833369.1
B3 124407099 124407557
refGene_gn=POMT2;refGene_tx=XM_003987863.3;refGene_gn=POMT2;refGene_tx=XM_006933027.2;refGene_gn=POMT2;refGene_tx=XM_019833367.1;refGene_gn=POMT2;refGene_tx=XM_019833369.1
B3 124407828 124407843
refGene_gn=GSTZ1;refGene_tx=XM_003987864.3
B3 124410522 124410540
refGene_gn=GSTZ1;refGene_tx=XM_006933026.3
B3 124410818 124410870
refGene_gn=GSTZ1;refGene_tx=XM_003987864.3;refGene_gn=GSTZ1;refGene_tx=XM_006933026.3
B3 124412772 124412840
refGene_gn=GSTZ1;refGene_tx=XM_003987864.3;refGene_gn=GSTZ1;refGene_tx=XM_006933026.3
B3 124413313 124413394
refGene_gn=GSTZ1;refGene_tx=XM_003987864.3;refGene_gn=GSTZ1;refGene_tx=XM_006933026.3
B3 124413744 124413870
refGene_gn=GSTZ1;refGene_tx=XM_003987864.3;refGene_gn=GSTZ1;refGene_tx=XM_006933026.3

B3 124414972 124415051
refGene_gn=GSTZ1;refGene_tx=XM_003987864.3;refGene_gn=GSTZ1;refGene_tx=XM_006933026.3
B3 124415603 124415656
refGene_gn=GSTZ1;refGene_tx=XM_003987864.3;refGene_gn=GSTZ1;refGene_tx=XM_006933026.3
B3 124416085 124416135
refGene_gn=GSTZ1;refGene_tx=XM_003987864.3;refGene_gn=GSTZ1;refGene_tx=XM_006933026.3
B3 124416921 124417048
refGene_gn=GSTZ1;refGene_tx=XM_003987864.3;refGene_gn=GSTZ1;refGene_tx=XM_006933026.3
B3 124429593 124429811
refGene_gn=TMED8;refGene_tx=XM_023255898.1
B3 124431140 124431446
refGene_gn=TMED8;refGene_tx=XM_023255898.1
B3 124431674 124431801
refGene_gn=TMED8;refGene_tx=XM_023255898.1
B3 124433364 124433494
refGene_gn=TMED8;refGene_tx=XM_023255898.1
B3 124439405 124439487
refGene_gn=TMED8;refGene_tx=XM_023255898.1
B3 124457820 124457905
refGene_gn=TMED8;refGene_tx=XM_023255898.1
B3 124458253 124460287
refGene_gn=SAMD15;refGene_tx=XM_006933029.4;refGene_gn=SAMD15;refGene_tx=XM_019833370.2
B3 124461157 124461329
refGene_gn=SAMD15;refGene_tx=XM_006933029.4;refGene_gn=SAMD15;refGene_tx=XM_019833370.2
B3 124468188 124468425
refGene_gn=SAMD15;refGene_tx=XM_006933029.4
B3 124471118 124471296
refGene_gn=NOXRED1;refGene_tx=XM_003987867.4;refGene_gn=NOXRED1;refGene_tx=XM_006933030.2;refGene_gn=NOXRED1;refGene_tx=XM_019833377.1
B3 124479206 124479429
refGene_gn=NOXRED1;refGene_tx=XM_003987867.4;refGene_gn=NOXRED1;refGene_tx=XM_006933030.2;refGene_gn=NOXRED1;refGene_tx=XM_019833377.1
B3 124480085 124480237
refGene_gn=NOXRED1;refGene_tx=XM_003987867.4;refGene_gn=NOXRED1;refGene_tx=XM_006933030.2;refGene_gn=NOXRED1;refGene_tx=XM_019833377.1
B3 124480856 124481037
refGene_gn=NOXRED1;refGene_tx=XM_003987867.4;refGene_gn=NOXRED1;refGene_tx=XM_006933030.2;refGene_gn=NOXRED1;refGene_tx=XM_019833377.1
B3 124485175 124485378
refGene_gn=NOXRED1;refGene_tx=XM_003987867.4;refGene_gn=NOXRED1;refGene_tx=XM_006933030.2;refGene_gn=NOXRED1;refGene_tx=XM_019833377.1
B3 124487059 124487231
refGene_gn=NOXRED1;refGene_tx=XM_003987867.4;refGene_gn=NOXRED1;refGene_tx=XM_006933030.2;refGene_gn=NOXRED1;refGene_tx=XM_019833377.1

B3 124491270 124491307
refGene_gn=NOXRED1;refGene_tx=XM_019833377.1
B3 124491532 124491553
refGene_gn=VIPAS39;refGene_tx=XM_003987868.3;refGene_gn=VIPAS39;refGene_tx=XM_019833375.1
B3 124492301 124492406
refGene_gn=VIPAS39;refGene_tx=XM_003987868.3;refGene_gn=VIPAS39;refGene_tx=XM_019833375.1
B3 124492807 124492897
refGene_gn=VIPAS39;refGene_tx=XM_003987868.3
B3 124493484 124493571
refGene_gn=VIPAS39;refGene_tx=XM_003987868.3;refGene_gn=VIPAS39;refGene_tx=XM_019833375.1
B3 124494635 124494725
refGene_gn=VIPAS39;refGene_tx=XM_003987868.3;refGene_gn=VIPAS39;refGene_tx=XM_019833375.1
B3 124495050 124495092
refGene_gn=VIPAS39;refGene_tx=XM_003987868.3;refGene_gn=VIPAS39;refGene_tx=XM_019833375.1
B3 124495761 124495896
refGene_gn=VIPAS39;refGene_tx=XM_003987868.3;refGene_gn=VIPAS39;refGene_tx=XM_019833375.1
B3 124496337 124496413
refGene_gn=VIPAS39;refGene_tx=XM_003987868.3;refGene_gn=VIPAS39;refGene_tx=XM_019833375.1
B3 124498112 124498186
refGene_gn=VIPAS39;refGene_tx=XM_003987868.3;refGene_gn=VIPAS39;refGene_tx=XM_019833375.1
B3 124501682 124501710
refGene_gn=VIPAS39;refGene_tx=XM_003987868.3;refGene_gn=VIPAS39;refGene_tx=XM_019833375.1
B3 124503282 124503385
refGene_gn=VIPAS39;refGene_tx=XM_003987868.3;refGene_gn=VIPAS39;refGene_tx=XM_019833375.1
B3 124503871 124503905
refGene_gn=VIPAS39;refGene_tx=XM_003987868.3;refGene_gn=VIPAS39;refGene_tx=XM_019833375.1
B3 124504537 124504630
refGene_gn=VIPAS39;refGene_tx=XM_003987868.3;refGene_gn=VIPAS39;refGene_tx=XM_019833375.1
B3 124505810 124505867
refGene_gn=VIPAS39;refGene_tx=XM_003987868.3;refGene_gn=VIPAS39;refGene_tx=XM_019833375.1
B3 124506594 124506659
refGene_gn=VIPAS39;refGene_tx=XM_003987868.3;refGene_gn=VIPAS39;refGene_tx=XM_019833375.1
B3 124507009 124507048
refGene_gn=VIPAS39;refGene_tx=XM_003987868.3;refGene_gn=VIPAS39;refGene_tx=XM_019833375.1
B3 124507887 124508034

refGene_gn=VIPAS39;refGene_tx=XM_003987868.3;refGene_gn=VIPAS39;refGene_tx=XM_019833375.1
B3	124509705	124509808
refGene_gn=VIPAS39;refGene_tx=XM_003987868.3;refGene_gn=VIPAS39;refGene_tx=XM_019833375.1
B3	124511144	124511237
refGene_gn=VIPAS39;refGene_tx=XM_003987868.3;refGene_gn=VIPAS39;refGene_tx=XM_019833375.1
B3	124516101	124516181
refGene_gn=AHSA1;refGene_tx=XM_011283425.2
B3	124517531	124517722
refGene_gn=AHSA1;refGene_tx=XM_011283425.2
B3	124519452	124519535
refGene_gn=AHSA1;refGene_tx=XM_011283425.2
B3	124519931	124520049
refGene_gn=AHSA1;refGene_tx=XM_011283425.2
B3	124521140	124521229
refGene_gn=AHSA1;refGene_tx=XM_011283425.2
B3	124522040	124522169
refGene_gn=AHSA1;refGene_tx=XM_011283425.2
B3	124522649	124522751
refGene_gn=AHSA1;refGene_tx=XM_011283425.2
B3	124523232	124523284
refGene_gn=AHSA1;refGene_tx=XM_011283425.2
B3	124523401	124523574
refGene_gn=AHSA1;refGene_tx=XM_011283425.2
B3	124528725	124529243
refGene_gn=ISM2;refGene_tx=XM_023255899.1;refGene_gn=ISM2;refGene_tx=XM_023255900.1;refGene_gn=ISM2;refGene_tx=XM_023255901.1;refGene_gn=ISM2;refGene_tx=XM_023255902.1;refGene_gn=ISM2;refGene_tx=XM_023255903.1
B3	124530424	124530508
refGene_gn=ISM2;refGene_tx=XM_023255899.1;refGene_gn=ISM2;refGene_tx=XM_023255902.1;refGene_gn=ISM2;refGene_tx=XM_023255903.1
B3	124530760	124530901
refGene_gn=ISM2;refGene_tx=XM_023255899.1;refGene_gn=ISM2;refGene_tx=XM_023255900.1;refGene_gn=ISM2;refGene_tx=XM_023255902.1;refGene_gn=ISM2;refGene_tx=XM_023255903.1
B3	124532543	124532919
refGene_gn=ISM2;refGene_tx=XM_023255899.1;refGene_gn=ISM2;refGene_tx=XM_023255900.1;refGene_gn=ISM2;refGene_tx=XM_023255901.1;refGene_gn=ISM2;refGene_tx=XM_023255902.1;refGene_gn=ISM2;refGene_tx=XM_023255903.1
B3	124534193	124534436
refGene_gn=ISM2;refGene_tx=XM_023255899.1;refGene_gn=ISM2;refGene_tx=XM_023255900.1;refGene_gn=ISM2;refGene_tx=XM_023255901.1;refGene_gn=ISM2;refGene_tx=XM_023255902.1;refGene_gn=ISM2;refGene_tx=XM_023255903.1
B3	124541013	124541139
refGene_gn=ISM2;refGene_tx=XM_023255899.1;refGene_gn=ISM2;refGene_tx=XM_023255900.1;refGene_gn=ISM2;refGene_tx=XM_023255901.1
B3	124552783	124552903
refGene_gn=SPTLC2;refGene_tx=XM_003987869.4

B3 124557815 124557945
refGene_gn=SPTLC2;refGene_tx=XM_003987869.4
B3 124561055 124561191
refGene_gn=SPTLC2;refGene_tx=XM_003987869.4
B3 124604682 124604809
refGene_gn=SPTLC2;refGene_tx=XM_003987869.4
B3 124607509 124607729
refGene_gn=SPTLC2;refGene_tx=XM_003987869.4
B3 124608717 124608823
refGene_gn=SPTLC2;refGene_tx=XM_003987869.4
B3 124610417 124610511
refGene_gn=SPTLC2;refGene_tx=XM_003987869.4
B3 124615847 124615972
refGene_gn=SPTLC2;refGene_tx=XM_003987869.4
B3 124617667 124617816
refGene_gn=SPTLC2;refGene_tx=XM_003987869.4
B3 124619907 124620062
refGene_gn=SPTLC2;refGene_tx=XM_003987869.4
B3 124628030 124628225
refGene_gn=SPTLC2;refGene_tx=XM_003987869.4
B3 124655986 124656124
refGene_gn=SPTLC2;refGene_tx=XM_003987869.4
B3 124696009 124696439
refGene_gn=ALKBH1;refGene_tx=XM_023256189.1
B3 124697481 124697675
refGene_gn=ALKBH1;refGene_tx=XM_023256189.1
B3 124702644 124702770
refGene_gn=ALKBH1;refGene_tx=XM_023256189.1
B3 124719540 124719810
refGene_gn=ALKBH1;refGene_tx=XM_023256189.1
B3 124722265 124722448
refGene_gn=ALKBH1;refGene_tx=XM_023256189.1
B3 124722553 124722650
refGene_gn=SLIRP;refGene_tx=XM_003987871.4
B3 124726607 124726666
refGene_gn=SLIRP;refGene_tx=XM_003987871.4
B3 124730695 124730803
refGene_gn=SLIRP;refGene_tx=XM_003987871.4
B3 124731251 124731323
refGene_gn=SLIRP;refGene_tx=XM_003987871.4
B3 124731851 124732050
refGene_gn=SNW1;refGene_tx=XM_003987872.4
B3 124732130 124732294
refGene_gn=SNW1;refGene_tx=XM_003987872.4
B3 124734617 124734735
refGene_gn=SNW1;refGene_tx=XM_003987872.4
B3 124736994 124737091
refGene_gn=SNW1;refGene_tx=XM_003987872.4
B3 124741127 124741269
refGene_gn=SNW1;refGene_tx=XM_003987872.4

B3 124742324 124742441
refGene_gn=SNW1;refGene_tx=XM_003987872.4
B3 124744105 124744171
refGene_gn=SNW1;refGene_tx=XM_003987872.4
B3 124744861 124744931
refGene_gn=SNW1;refGene_tx=XM_003987872.4
B3 124746089 124746194
refGene_gn=SNW1;refGene_tx=XM_003987872.4
B3 124747133 124747240
refGene_gn=SNW1;refGene_tx=XM_003987872.4
B3 124747334 124747430
refGene_gn=SNW1;refGene_tx=XM_003987872.4
B3 124755805 124755967
refGene_gn=SNW1;refGene_tx=XM_003987872.4
B3 124759515 124759669
refGene_gn=SNW1;refGene_tx=XM_003987872.4
B3 124764245 124764259
refGene_gn=SNW1;refGene_tx=XM_003987872.4
B3 124809413 124809548
refGene_gn=ADCK1;refGene_tx=XM_003987889.3;refGene_gn=ADCK1;refGene_tx=XM_006933032.2;refGene_gn=ADCK1;refGene_tx=XM_019833382.1
B3 124812324 124812408
refGene_gn=ADCK1;refGene_tx=XM_003987889.3;refGene_gn=ADCK1;refGene_tx=XM_006933032.2;refGene_gn=ADCK1;refGene_tx=XM_019833382.1
B3 124851487 124851691
refGene_gn=ADCK1;refGene_tx=XM_003987889.3;refGene_gn=ADCK1;refGene_tx=XM_006933032.2;refGene_gn=ADCK1;refGene_tx=XM_019833382.1
B3 124877294 124877453
refGene_gn=ADCK1;refGene_tx=XM_003987889.3;refGene_gn=ADCK1;refGene_tx=XM_006933032.2
B3 124888206 124888365
refGene_gn=ADCK1;refGene_tx=XM_003987889.3;refGene_gn=ADCK1;refGene_tx=XM_006933032.2;refGene_gn=ADCK1;refGene_tx=XM_019833382.1
B3 124896926 124897094
refGene_gn=ADCK1;refGene_tx=XM_003987889.3;refGene_gn=ADCK1;refGene_tx=XM_006933032.2;refGene_gn=ADCK1;refGene_tx=XM_019833382.1
B3 124910203 124910353
refGene_gn=ADCK1;refGene_tx=XM_003987889.3;refGene_gn=ADCK1;refGene_tx=XM_006933032.2;refGene_gn=ADCK1;refGene_tx=XM_019833382.1
B3 124911479 124911677
refGene_gn=ADCK1;refGene_tx=XM_003987889.3;refGene_gn=ADCK1;refGene_tx=XM_006933032.2;refGene_gn=ADCK1;refGene_tx=XM_019833382.1
B3 124916915 124917109
refGene_gn=ADCK1;refGene_tx=XM_003987889.3;refGene_gn=ADCK1;refGene_tx=XM_006933032.2;refGene_gn=ADCK1;refGene_tx=XM_019833382.1
B3 124918691 124918863
refGene_gn=ADCK1;refGene_tx=XM_003987889.3;refGene_gn=ADCK1;refGene_tx=XM_006933032.2;refGene_gn=ADCK1;refGene_tx=XM_019833382.1
B3 125193945 125194654
refGene_gn=NRXN3;refGene_tx=XM_011283426.3;refGene_gn=NRXN3;refGene_tx

=XM_019833384.2;refGene_gn=NRXN3;refGene_tx=XM_019833385.2;refGene_gn=NRXN3;refGene_tx=XM_019833386.2;refGene_gn=NRXN3;refGene_tx=XM_019833387.2;refGene_gn=NRXN3;refGene_tx=XM_019833388.2;refGene_gn=NRXN3;refGene_tx=XM_019833389.2;refGene_gn=NRXN3;refGene_tx=XM_019833390.2;refGene_gn=NRXN3;refGene_tx=XM_019833391.2;refGene_gn=NRXN3;refGene_tx=XM_019833392.2;refGene_gn=NRXN3;refGene_tx=XM_019833393.2;refGene_gn=NRXN3;refGene_tx=XM_019833394.2;refGene_gn=NRXN3;refGene_tx=XM_019833395.2;refGene_gn=NRXN3;refGene_tx=XM_019833396.2;refGene_gn=NRXN3;refGene_tx=XM_019833397.2;refGene_gn=NRXN3;refGene_tx=XM_019833398.2;refGene_gn=NRXN3;refGene_tx=XM_019833399.2;refGene_gn=NRXN3;refGene_tx=XM_019833400.2;refGene_gn=NRXN3;refGene_tx=XM_019833401.2;refGene_gn=NRXN3;refGene_tx=XM_019833402.2;refGene_gn=NRXN3;refGene_tx=XM_019833403.2;refGene_gn=NRXN3;refGene_tx=XM_019833404.2;refGene_gn=NRXN3;refGene_tx=XM_019833405.2;refGene_gn=NRXN3;refGene_tx=XM_019833407.2;refGene_gn=NRXN3;refGene_tx=XM_019833412.2;refGene_gn=NRXN3;refGene_tx=XM_019833413.2;refGene_gn=NRXN3;refGene_tx=XM_019833414.2;refGene_gn=NRXN3;refGene_tx=XM_019833415.2;refGene_gn=NRXN3;refGene_tx=XM_019833416.2;refGene_gn=NRXN3;refGene_tx=XM_019833417.2
B3 125227737 125227755
refGene_gn=NRXN3;refGene_tx=XM_011283426.3;refGene_gn=NRXN3;refGene_tx=XM_019833384.2;refGene_gn=NRXN3;refGene_tx=XM_019833385.2;refGene_gn=NRXN3;refGene_tx=XM_019833386.2;refGene_gn=NRXN3;refGene_tx=XM_019833387.2;refGene_gn=NRXN3;refGene_tx=XM_019833388.2;refGene_gn=NRXN3;refGene_tx=XM_019833389.2;refGene_gn=NRXN3;refGene_tx=XM_019833390.2;refGene_gn=NRXN3;refGene_tx=XM_019833391.2;refGene_gn=NRXN3;refGene_tx=XM_019833392.2;refGene_gn=NRXN3;refGene_tx=XM_019833394.2;refGene_gn=NRXN3;refGene_tx=XM_019833395.2;refGene_gn=NRXN3;refGene_tx=XM_019833396.2;refGene_gn=NRXN3;refGene_tx=XM_019833398.2;refGene_gn=NRXN3;refGene_tx=XM_019833399.2;refGene_gn=NRXN3;refGene_tx=XM_019833400.2;refGene_gn=NRXN3;refGene_tx=XM_019833401.2;refGene_gn=NRXN3;refGene_tx=XM_019833402.2;refGene_gn=NRXN3;refGene_tx=XM_019833403.2;refGene_gn=NRXN3;refGene_tx=XM_019833404.2;refGene_gn=NRXN3;refGene_tx=XM_019833405.2;refGene_gn=NRXN3;refGene_tx=XM_019833407.2;refGene_gn=NRXN3;refGene_tx=XM_019833412.2;refGene_gn=NRXN3;refGene_tx=XM_019833417.2
B3 125246524 125246554
refGene_gn=NRXN3;refGene_tx=XM_011283426.3;refGene_gn=NRXN3;refGene_tx=XM_019833384.2;refGene_gn=NRXN3;refGene_tx=XM_019833385.2;refGene_gn=NRXN3;refGene_tx=XM_019833386.2;refGene_gn=NRXN3;refGene_tx=XM_019833387.2;refGene_gn=NRXN3;refGene_tx=XM_019833388.2;refGene_gn=NRXN3;refGene_tx=XM_019833389.2;refGene_gn=NRXN3;refGene_tx=XM_019833390.2;refGene_gn=NRXN3;refGene_tx=XM_019833391.2;refGene_gn=NRXN3;refGene_tx=XM_019833392.2;refGene_gn=NRXN3;refGene_tx=XM_019833394.2;refGene_gn=NRXN3;refGene_tx=XM_019833395.2;refGene_gn=NRXN3;refGene_tx=XM_019833396.2;refGene_gn=NRXN3;refGene_tx=XM_019833398.2;refGene_gn=NRXN3;refGene_tx=XM_019833399.2;refGene_gn=NRXN3;refGene_tx=XM_019833400.2;refGene_gn=NRXN3;refGene_tx=XM_019833401.2;refGene_gn=NRXN3;refGene_tx=XM_019833402.2;refGene_gn=NRXN3;refGene_tx=XM_019833403.2;refGene_gn=NRXN3;refGene_tx=XM_019833404.2;refGene_gn=NRXN3;refGene_tx=XM_019833405.2;refGene_gn=NRXN3;refGene_tx=XM_019833407.2;refGene_gn=NRXN3;refGene_tx=XM_019833412.2;refGene_gn=NRXN3;refGene_tx=XM_019833417.2

B3	125249513	125249525
refGene_gn=NRXN3;refGene_tx=XM_019833384.2;refGene_gn=NRXN3;refGene_tx=XM_019833385.2;refGene_gn=NRXN3;refGene_tx=XM_019833387.2;refGene_gn=NRXN3;refGene_tx=XM_019833388.2;refGene_gn=NRXN3;refGene_tx=XM_019833389.2;refGene_gn=NRXN3;refGene_tx=XM_019833391.2;refGene_gn=NRXN3;refGene_tx=XM_019833392.2;refGene_gn=NRXN3;refGene_tx=XM_019833394.2;refGene_gn=NRXN3;refGene_tx=XM_019833395.2;refGene_gn=NRXN3;refGene_tx=XM_019833398.2;refGene_gn=NRXN3;refGene_tx=XM_019833399.2;refGene_gn=NRXN3;refGene_tx=XM_019833400.2;refGene_gn=NRXN3;refGene_tx=XM_019833401.2;refGene_gn=NRXN3;refGene_tx=XM_019833403.2;refGene_gn=NRXN3;refGene_tx=XM_019833404.2;refGene_gn=NRXN3;refGene_tx=XM_019833405.2;refGene_gn=NRXN3;refGene_tx=XM_019833407.2
B3	125581448	125581750
refGene_gn=NRXN3;refGene_tx=XM_011283426.3;refGene_gn=NRXN3;refGene_tx=XM_019833384.2;refGene_gn=NRXN3;refGene_tx=XM_019833385.2;refGene_gn=NRXN3;refGene_tx=XM_019833386.2;refGene_gn=NRXN3;refGene_tx=XM_019833387.2;refGene_gn=NRXN3;refGene_tx=XM_019833388.2;refGene_gn=NRXN3;refGene_tx=XM_019833389.2;refGene_gn=NRXN3;refGene_tx=XM_019833390.2;refGene_gn=NRXN3;refGene_tx=XM_019833391.2;refGene_gn=NRXN3;refGene_tx=XM_019833392.2;refGene_gn=NRXN3;refGene_tx=XM_019833393.2;refGene_gn=NRXN3;refGene_tx=XM_019833394.2;refGene_gn=NRXN3;refGene_tx=XM_019833395.2;refGene_gn=NRXN3;refGene_tx=XM_019833396.2;refGene_gn=NRXN3;refGene_tx=XM_019833397.2;refGene_gn=NRXN3;refGene_tx=XM_019833398.2;refGene_gn=NRXN3;refGene_tx=XM_019833399.2;refGene_gn=NRXN3;refGene_tx=XM_019833400.2;refGene_gn=NRXN3;refGene_tx=XM_019833401.2;refGene_gn=NRXN3;refGene_tx=XM_019833402.2;refGene_gn=NRXN3;refGene_tx=XM_019833403.2;refGene_gn=NRXN3;refGene_tx=XM_019833404.2;refGene_gn=NRXN3;refGene_tx=XM_019833405.2;refGene_gn=NRXN3;refGene_tx=XM_019833407.2;refGene_gn=NRXN3;refGene_tx=XM_019833412.2;refGene_gn=NRXN3;refGene_tx=XM_019833413.2;refGene_gn=NRXN3;refGene_tx=XM_019833414.2;refGene_gn=NRXN3;refGene_tx=XM_019833415.2;refGene_gn=NRXN3;refGene_tx=XM_019833416.2;refGene_gn=NRXN3;refGene_tx=XM_019833417.2
B3	125585698	125585722
refGene_gn=NRXN3;refGene_tx=XM_019833384.2;refGene_gn=NRXN3;refGene_tx=XM_019833385.2;refGene_gn=NRXN3;refGene_tx=XM_019833386.2;refGene_gn=NRXN3;refGene_tx=XM_019833388.2;refGene_gn=NRXN3;refGene_tx=XM_019833389.2;refGene_gn=NRXN3;refGene_tx=XM_019833390.2;refGene_gn=NRXN3;refGene_tx=XM_019833392.2;refGene_gn=NRXN3;refGene_tx=XM_019833393.2;refGene_gn=NRXN3;refGene_tx=XM_019833394.2;refGene_gn=NRXN3;refGene_tx=XM_019833395.2;refGene_gn=NRXN3;refGene_tx=XM_019833398.2;refGene_gn=NRXN3;refGene_tx=XM_019833399.2;refGene_gn=NRXN3;refGene_tx=XM_019833403.2;refGene_gn=NRXN3;refGene_tx=XM_019833404.2;refGene_gn=NRXN3;refGene_tx=XM_019833407.2;refGene_gn=NRXN3;refGene_tx=XM_019833412.2;refGene_gn=NRXN3;refGene_tx=XM_019833413.2;refGene_gn=NRXN3;refGene_tx=XM_019833416.2
B3	125587609	125587771
refGene_gn=NRXN3;refGene_tx=XM_011283426.3;refGene_gn=NRXN3;refGene_tx=XM_019833384.2;refGene_gn=NRXN3;refGene_tx=XM_019833385.2;refGene_gn=NRXN3;refGene_tx=XM_019833386.2;refGene_gn=NRXN3;refGene_tx=XM_019833387.2;refGene_gn=NRXN3;refGene_tx=XM_019833388.2;refGene_gn=NRXN3;refGe

ne_tx=XM_019833389.2;refGene_gn=NRXN3;refGene_tx=XM_019833390.2;refGen
e_gn=NRXN3;refGene_tx=XM_019833391.2;refGene_gn=NRXN3;refGene_tx=XM_01
9833392.2;refGene_gn=NRXN3;refGene_tx=XM_019833393.2;refGene_gn=NRXN3;
refGene_tx=XM_019833394.2;refGene_gn=NRXN3;refGene_tx=XM_019833395.2;r
efGene_gn=NRXN3;refGene_tx=XM_019833396.2;refGene_gn=NRXN3;refGene_tx=
XM_019833397.2;refGene_gn=NRXN3;refGene_tx=XM_019833398.2;refGene_gn=N
RXN3;refGene_tx=XM_019833399.2;refGene_gn=NRXN3;refGene_tx=XM_01983340
0.2;refGene_gn=NRXN3;refGene_tx=XM_019833401.2;refGene_gn=NRXN3;refGen
e_tx=XM_019833402.2;refGene_gn=NRXN3;refGene_tx=XM_019833403.2;refGene
_gn=NRXN3;refGene_tx=XM_019833404.2;refGene_gn=NRXN3;refGene_tx=XM_019
833405.2;refGene_gn=NRXN3;refGene_tx=XM_019833407.2;refGene_gn=NRXN3;r
efGene_tx=XM_019833412.2;refGene_gn=NRXN3;refGene_tx=XM_019833413.2;re
fGene_gn=NRXN3;refGene_tx=XM_019833414.2;refGene_gn=NRXN3;refGene_tx=X
M_019833415.2;refGene_gn=NRXN3;refGene_tx=XM_019833416.2;refGene_gn=NR
XN3;refGene_tx=XM_019833417.2
B3 125644687 125645126
refGene_gn=NRXN3;refGene_tx=XM_011283426.3;refGene_gn=NRXN3;refGene_tx
=XM_019833384.2;refGene_gn=NRXN3;refGene_tx=XM_019833385.2;refGene_gn=
NRXN3;refGene_tx=XM_019833386.2;refGene_gn=NRXN3;refGene_tx=XM_0198333
87.2;refGene_gn=NRXN3;refGene_tx=XM_019833388.2;refGene_gn=NRXN3;refGe
ne_tx=XM_019833389.2;refGene_gn=NRXN3;refGene_tx=XM_019833390.2;refGen
e_gn=NRXN3;refGene_tx=XM_019833391.2;refGene_gn=NRXN3;refGene_tx=XM_01
9833392.2;refGene_gn=NRXN3;refGene_tx=XM_019833393.2;refGene_gn=NRXN3;
refGene_tx=XM_019833394.2;refGene_gn=NRXN3;refGene_tx=XM_019833395.2;r
efGene_gn=NRXN3;refGene_tx=XM_019833396.2;refGene_gn=NRXN3;refGene_tx=
XM_019833397.2;refGene_gn=NRXN3;refGene_tx=XM_019833398.2;refGene_gn=N
RXN3;refGene_tx=XM_019833399.2;refGene_gn=NRXN3;refGene_tx=XM_01983340
0.2;refGene_gn=NRXN3;refGene_tx=XM_019833401.2;refGene_gn=NRXN3;refGen
e_tx=XM_019833402.2;refGene_gn=NRXN3;refGene_tx=XM_019833403.2;refGene
_gn=NRXN3;refGene_tx=XM_019833404.2;refGene_gn=NRXN3;refGene_tx=XM_019
833405.2;refGene_gn=NRXN3;refGene_tx=XM_019833407.2;refGene_gn=NRXN3;r
efGene_tx=XM_019833412.2;refGene_gn=NRXN3;refGene_tx=XM_019833413.2;re
fGene_gn=NRXN3;refGene_tx=XM_019833414.2;refGene_gn=NRXN3;refGene_tx=X
M_019833415.2;refGene_gn=NRXN3;refGene_tx=XM_019833416.2;refGene_gn=NR
XN3;refGene_tx=XM_019833417.2
B3 125649594 125649978
refGene_gn=NRXN3;refGene_tx=XM_011283426.3;refGene_gn=NRXN3;refGene_tx
=XM_019833384.2;refGene_gn=NRXN3;refGene_tx=XM_019833385.2;refGene_gn=
NRXN3;refGene_tx=XM_019833386.2;refGene_gn=NRXN3;refGene_tx=XM_0198333
87.2;refGene_gn=NRXN3;refGene_tx=XM_019833388.2;refGene_gn=NRXN3;refGe
ne_tx=XM_019833389.2;refGene_gn=NRXN3;refGene_tx=XM_019833390.2;refGen
e_gn=NRXN3;refGene_tx=XM_019833391.2;refGene_gn=NRXN3;refGene_tx=XM_01
9833392.2;refGene_gn=NRXN3;refGene_tx=XM_019833393.2;refGene_gn=NRXN3;
refGene_tx=XM_019833394.2;refGene_gn=NRXN3;refGene_tx=XM_019833395.2;r
efGene_gn=NRXN3;refGene_tx=XM_019833396.2;refGene_gn=NRXN3;refGene_tx=
XM_019833397.2;refGene_gn=NRXN3;refGene_tx=XM_019833398.2;refGene_gn=N
RXN3;refGene_tx=XM_019833399.2;refGene_gn=NRXN3;refGene_tx=XM_01983340
0.2;refGene_gn=NRXN3;refGene_tx=XM_019833401.2;refGene_gn=NRXN3;refGen
e_tx=XM_019833402.2;refGene_gn=NRXN3;refGene_tx=XM_019833403.2;refGene
_gn=NRXN3;refGene_tx=XM_019833404.2;refGene_gn=NRXN3;refGene_tx=XM_019

833405.2;refGene_gn=NRXN3;refGene_tx=XM_019833407.2;refGene_gn=NRXN3;refGene_tx=XM_019833412.2;refGene_gn=NRXN3;refGene_tx=XM_019833413.2;refGene_gn=NRXN3;refGene_tx=XM_019833414.2;refGene_gn=NRXN3;refGene_tx=XM_019833415.2;refGene_gn=NRXN3;refGene_tx=XM_019833416.2;refGene_gn=NRXN3;refGene_tx=XM_019833417.2
B3 125731663 125731867
refGene_gn=NRXN3;refGene_tx=XM_011283426.3;refGene_gn=NRXN3;refGene_tx=XM_019833384.2;refGene_gn=NRXN3;refGene_tx=XM_019833385.2;refGene_gn=NRXN3;refGene_tx=XM_019833386.2;refGene_gn=NRXN3;refGene_tx=XM_019833387.2;refGene_gn=NRXN3;refGene_tx=XM_019833388.2;refGene_gn=NRXN3;refGene_tx=XM_019833389.2;refGene_gn=NRXN3;refGene_tx=XM_019833390.2;refGene_gn=NRXN3;refGene_tx=XM_019833391.2;refGene_gn=NRXN3;refGene_tx=XM_019833392.2;refGene_gn=NRXN3;refGene_tx=XM_019833393.2;refGene_gn=NRXN3;refGene_tx=XM_019833394.2;refGene_gn=NRXN3;refGene_tx=XM_019833395.2;refGene_gn=NRXN3;refGene_tx=XM_019833396.2;refGene_gn=NRXN3;refGene_tx=XM_019833397.2;refGene_gn=NRXN3;refGene_tx=XM_019833398.2;refGene_gn=NRXN3;refGene_tx=XM_019833399.2;refGene_gn=NRXN3;refGene_tx=XM_019833400.2;refGene_gn=NRXN3;refGene_tx=XM_019833401.2;refGene_gn=NRXN3;refGene_tx=XM_019833402.2;refGene_gn=NRXN3;refGene_tx=XM_019833403.2;refGene_gn=NRXN3;refGene_tx=XM_019833404.2;refGene_gn=NRXN3;refGene_tx=XM_019833405.2;refGene_gn=NRXN3;refGene_tx=XM_019833407.2;refGene_gn=NRXN3;refGene_tx=XM_019833412.2;refGene_gn=NRXN3;refGene_tx=XM_019833413.2;refGene_gn=NRXN3;refGene_tx=XM_019833414.2;refGene_gn=NRXN3;refGene_tx=XM_019833415.2;refGene_gn=NRXN3;refGene_tx=XM_019833416.2;refGene_gn=NRXN3;refGene_tx=XM_019833417.2
B3 125737836 125737863
refGene_gn=NRXN3;refGene_tx=XM_011283426.3;refGene_gn=NRXN3;refGene_tx=XM_019833384.2;refGene_gn=NRXN3;refGene_tx=XM_019833385.2;refGene_gn=NRXN3;refGene_tx=XM_019833386.2;refGene_gn=NRXN3;refGene_tx=XM_019833387.2;refGene_gn=NRXN3;refGene_tx=XM_019833388.2;refGene_gn=NRXN3;refGene_tx=XM_019833390.2;refGene_gn=NRXN3;refGene_tx=XM_019833391.2;refGene_gn=NRXN3;refGene_tx=XM_019833393.2;refGene_gn=NRXN3;refGene_tx=XM_019833394.2;refGene_gn=NRXN3;refGene_tx=XM_019833398.2;refGene_gn=NRXN3;refGene_tx=XM_019833399.2;refGene_gn=NRXN3;refGene_tx=XM_019833400.2;refGene_gn=NRXN3;refGene_tx=XM_019833401.2;refGene_gn=NRXN3;refGene_tx=XM_019833402.2;refGene_gn=NRXN3;refGene_tx=XM_019833403.2;refGene_gn=NRXN3;refGene_tx=XM_019833404.2;refGene_gn=NRXN3;refGene_tx=XM_019833405.2;refGene_gn=NRXN3;refGene_tx=XM_019833407.2;refGene_gn=NRXN3;refGene_tx=XM_019833412.2
B3 125879323 125879443
refGene_gn=NRXN3;refGene_tx=XM_011283426.3;refGene_gn=NRXN3;refGene_tx=XM_019833384.2;refGene_gn=NRXN3;refGene_tx=XM_019833385.2;refGene_gn=NRXN3;refGene_tx=XM_019833386.2;refGene_gn=NRXN3;refGene_tx=XM_019833387.2;refGene_gn=NRXN3;refGene_tx=XM_019833388.2;refGene_gn=NRXN3;refGene_tx=XM_019833389.2;refGene_gn=NRXN3;refGene_tx=XM_019833390.2;refGene_gn=NRXN3;refGene_tx=XM_019833391.2;refGene_gn=NRXN3;refGene_tx=XM_019833392.2;refGene_gn=NRXN3;refGene_tx=XM_019833393.2;refGene_gn=NRXN3;refGene_tx=XM_019833394.2;refGene_gn=NRXN3;refGene_tx=XM_019833395.2;refGene_gn=NRXN3;refGene_tx=XM_019833396.2;refGene_gn=NRXN3;refGene_tx=XM_019833397.2;refGene_gn=NRXN3;refGene_tx=XM_019833398.2;refGene_gn=N

RXN3;refGene_tx=XM_019833399.2;refGene_gn=NRXN3;refGene_tx=XM_01983340
0.2;refGene_gn=NRXN3;refGene_tx=XM_019833401.2;refGene_gn=NRXN3;refGen
e_tx=XM_019833402.2;refGene_gn=NRXN3;refGene_tx=XM_019833403.2;refGene
_gn=NRXN3;refGene_tx=XM_019833404.2;refGene_gn=NRXN3;refGene_tx=XM_019
833405.2;refGene_gn=NRXN3;refGene_tx=XM_019833407.2;refGene_gn=NRXN3;r
efGene_tx=XM_019833412.2;refGene_gn=NRXN3;refGene_tx=XM_019833413.2;re
fGene_gn=NRXN3;refGene_tx=XM_019833414.2;refGene_gn=NRXN3;refGene_tx=X
M_019833415.2;refGene_gn=NRXN3;refGene_tx=XM_019833416.2;refGene_gn=NR
XN3;refGene_tx=XM_019833417.2
B3 125888996 125889378
refGene_gn=NRXN3;refGene_tx=XM_011283426.3;refGene_gn=NRXN3;refGene_tx
=XM_019833384.2;refGene_gn=NRXN3;refGene_tx=XM_019833385.2;refGene_gn=
NRXN3;refGene_tx=XM_019833386.2;refGene_gn=NRXN3;refGene_tx=XM_0198333
87.2;refGene_gn=NRXN3;refGene_tx=XM_019833388.2;refGene_gn=NRXN3;refGe
ne_tx=XM_019833389.2;refGene_gn=NRXN3;refGene_tx=XM_019833390.2;refGen
e_gn=NRXN3;refGene_tx=XM_019833391.2;refGene_gn=NRXN3;refGene_tx=XM_01
9833392.2;refGene_gn=NRXN3;refGene_tx=XM_019833393.2;refGene_gn=NRXN3;
refGene_tx=XM_019833394.2;refGene_gn=NRXN3;refGene_tx=XM_019833395.2;r
efGene_gn=NRXN3;refGene_tx=XM_019833396.2;refGene_gn=NRXN3;refGene_tx=
XM_019833397.2;refGene_gn=NRXN3;refGene_tx=XM_019833398.2;refGene_gn=N
RXN3;refGene_tx=XM_019833399.2;refGene_gn=NRXN3;refGene_tx=XM_01983340
0.2;refGene_gn=NRXN3;refGene_tx=XM_019833401.2;refGene_gn=NRXN3;refGen
e_tx=XM_019833402.2;refGene_gn=NRXN3;refGene_tx=XM_019833403.2;refGene
_gn=NRXN3;refGene_tx=XM_019833404.2;refGene_gn=NRXN3;refGene_tx=XM_019
833405.2;refGene_gn=NRXN3;refGene_tx=XM_019833407.2;refGene_gn=NRXN3;r
efGene_tx=XM_019833412.2;refGene_gn=NRXN3;refGene_tx=XM_019833413.2;re
fGene_gn=NRXN3;refGene_tx=XM_019833414.2;refGene_gn=NRXN3;refGene_tx=X
M_019833415.2;refGene_gn=NRXN3;refGene_tx=XM_019833416.2;refGene_gn=NR
XN3;refGene_tx=XM_019833417.2
B3 125890180 125890371
refGene_gn=NRXN3;refGene_tx=XM_011283426.3;refGene_gn=NRXN3;refGene_tx
=XM_019833384.2;refGene_gn=NRXN3;refGene_tx=XM_019833385.2;refGene_gn=
NRXN3;refGene_tx=XM_019833386.2;refGene_gn=NRXN3;refGene_tx=XM_0198333
87.2;refGene_gn=NRXN3;refGene_tx=XM_019833388.2;refGene_gn=NRXN3;refGe
ne_tx=XM_019833389.2;refGene_gn=NRXN3;refGene_tx=XM_019833390.2;refGen
e_gn=NRXN3;refGene_tx=XM_019833391.2;refGene_gn=NRXN3;refGene_tx=XM_01
9833392.2;refGene_gn=NRXN3;refGene_tx=XM_019833393.2;refGene_gn=NRXN3;
refGene_tx=XM_019833394.2;refGene_gn=NRXN3;refGene_tx=XM_019833395.2;r
efGene_gn=NRXN3;refGene_tx=XM_019833396.2;refGene_gn=NRXN3;refGene_tx=
XM_019833397.2;refGene_gn=NRXN3;refGene_tx=XM_019833398.2;refGene_gn=N
RXN3;refGene_tx=XM_019833399.2;refGene_gn=NRXN3;refGene_tx=XM_01983340
0.2;refGene_gn=NRXN3;refGene_tx=XM_019833401.2;refGene_gn=NRXN3;refGen
e_tx=XM_019833402.2;refGene_gn=NRXN3;refGene_tx=XM_019833403.2;refGene
_gn=NRXN3;refGene_tx=XM_019833404.2;refGene_gn=NRXN3;refGene_tx=XM_019
833405.2;refGene_gn=NRXN3;refGene_tx=XM_019833407.2;refGene_gn=NRXN3;r
efGene_tx=XM_019833412.2;refGene_gn=NRXN3;refGene_tx=XM_019833413.2;re
fGene_gn=NRXN3;refGene_tx=XM_019833414.2;refGene_gn=NRXN3;refGene_tx=X
M_019833415.2;refGene_gn=NRXN3;refGene_tx=XM_019833416.2;refGene_gn=NR
XN3;refGene_tx=XM_019833417.2
B3 125891170 125891344

refGene_gn=NRXN3;refGene_tx=XM_011283426.3;refGene_gn=NRXN3;refGene_tx
=XM_019833384.2;refGene_gn=NRXN3;refGene_tx=XM_019833385.2;refGene_gn=
NRXN3;refGene_tx=XM_019833386.2;refGene_gn=NRXN3;refGene_tx=XM_0198333
87.2;refGene_gn=NRXN3;refGene_tx=XM_019833388.2;refGene_gn=NRXN3;refGe
ne_tx=XM_019833389.2;refGene_gn=NRXN3;refGene_tx=XM_019833390.2;refGen
e_gn=NRXN3;refGene_tx=XM_019833391.2;refGene_gn=NRXN3;refGene_tx=XM_01
9833392.2;refGene_gn=NRXN3;refGene_tx=XM_019833393.2;refGene_gn=NRXN3;
refGene_tx=XM_019833394.2;refGene_gn=NRXN3;refGene_tx=XM_019833395.2;r
efGene_gn=NRXN3;refGene_tx=XM_019833396.2;refGene_gn=NRXN3;refGene_tx=
XM_019833397.2;refGene_gn=NRXN3;refGene_tx=XM_019833398.2;refGene_gn=N
RXN3;refGene_tx=XM_019833399.2;refGene_gn=NRXN3;refGene_tx=XM_01983340
0.2;refGene_gn=NRXN3;refGene_tx=XM_019833401.2;refGene_gn=NRXN3;refGen
e_tx=XM_019833402.2;refGene_gn=NRXN3;refGene_tx=XM_019833403.2;refGene
_gn=NRXN3;refGene_tx=XM_019833404.2;refGene_gn=NRXN3;refGene_tx=XM_019
833405.2;refGene_gn=NRXN3;refGene_tx=XM_019833407.2;refGene_gn=NRXN3;r
efGene_tx=XM_019833412.2;refGene_gn=NRXN3;refGene_tx=XM_019833413.2;re
fGene_gn=NRXN3;refGene_tx=XM_019833414.2;refGene_gn=NRXN3;refGene_tx=X
M_019833415.2;refGene_gn=NRXN3;refGene_tx=XM_019833416.2;refGene_gn=NR
XN3;refGene_tx=XM_019833417.2
B3 125895532 125895559
refGene_gn=NRXN3;refGene_tx=XM_019833384.2;refGene_gn=NRXN3;refGene_tx
=XM_019833385.2;refGene_gn=NRXN3;refGene_tx=XM_019833386.2;refGene_gn=
NRXN3;refGene_tx=XM_019833387.2;refGene_gn=NRXN3;refGene_tx=XM_0198333
89.2;refGene_gn=NRXN3;refGene_tx=XM_019833393.2;refGene_gn=NRXN3;refGe
ne_tx=XM_019833394.2;refGene_gn=NRXN3;refGene_tx=XM_019833398.2;refGen
e_gn=NRXN3;refGene_tx=XM_019833399.2;refGene_gn=NRXN3;refGene_tx=XM_01
9833403.2;refGene_gn=NRXN3;refGene_tx=XM_019833404.2
B3 125907195 125907315
refGene_gn=NRXN3;refGene_tx=XM_011283426.3;refGene_gn=NRXN3;refGene_tx
=XM_019833384.2;refGene_gn=NRXN3;refGene_tx=XM_019833385.2;refGene_gn=
NRXN3;refGene_tx=XM_019833386.2;refGene_gn=NRXN3;refGene_tx=XM_0198333
87.2;refGene_gn=NRXN3;refGene_tx=XM_019833388.2;refGene_gn=NRXN3;refGe
ne_tx=XM_019833389.2;refGene_gn=NRXN3;refGene_tx=XM_019833390.2;refGen
e_gn=NRXN3;refGene_tx=XM_019833391.2;refGene_gn=NRXN3;refGene_tx=XM_01
9833392.2;refGene_gn=NRXN3;refGene_tx=XM_019833393.2;refGene_gn=NRXN3;
refGene_tx=XM_019833394.2;refGene_gn=NRXN3;refGene_tx=XM_019833395.2;r
efGene_gn=NRXN3;refGene_tx=XM_019833396.2;refGene_gn=NRXN3;refGene_tx=
XM_019833397.2;refGene_gn=NRXN3;refGene_tx=XM_019833398.2;refGene_gn=N
RXN3;refGene_tx=XM_019833399.2;refGene_gn=NRXN3;refGene_tx=XM_01983340
0.2;refGene_gn=NRXN3;refGene_tx=XM_019833401.2;refGene_gn=NRXN3;refGen
e_tx=XM_019833402.2;refGene_gn=NRXN3;refGene_tx=XM_019833403.2;refGene
_gn=NRXN3;refGene_tx=XM_019833404.2;refGene_gn=NRXN3;refGene_tx=XM_019
833405.2;refGene_gn=NRXN3;refGene_tx=XM_019833407.2;refGene_gn=NRXN3;r
efGene_tx=XM_019833412.2;refGene_gn=NRXN3;refGene_tx=XM_019833413.2;re
fGene_gn=NRXN3;refGene_tx=XM_019833414.2;refGene_gn=NRXN3;refGene_tx=X
M_019833415.2;refGene_gn=NRXN3;refGene_tx=XM_019833416.2;refGene_gn=NR
XN3;refGene_tx=XM_019833417.2
B3 126174021 126174274
refGene_gn=NRXN3;refGene_tx=XM_019833408.2;refGene_gn=NRXN3;refGene_tx
=XM_019833409.2;refGene_gn=NRXN3;refGene_tx=XM_019833410.2;refGene_gn=

NRXN3;refGene_tx=XM_019833411.2;refGene_gn=NRXN3;refGene_tx=XM_0198334
18.2
B3 126350190 126350372
refGene_gn=NRXN3;refGene_tx=XM_011283426.3;refGene_gn=NRXN3;refGene_tx
=XM_019833384.2;refGene_gn=NRXN3;refGene_tx=XM_019833385.2;refGene_gn=
NRXN3;refGene_tx=XM_019833386.2;refGene_gn=NRXN3;refGene_tx=XM_0198333
87.2;refGene_gn=NRXN3;refGene_tx=XM_019833388.2;refGene_gn=NRXN3;refGe
ne_tx=XM_019833389.2;refGene_gn=NRXN3;refGene_tx=XM_019833390.2;refGen
e_gn=NRXN3;refGene_tx=XM_019833391.2;refGene_gn=NRXN3;refGene_tx=XM_01
9833392.2;refGene_gn=NRXN3;refGene_tx=XM_019833393.2;refGene_gn=NRXN3;
refGene_tx=XM_019833394.2;refGene_gn=NRXN3;refGene_tx=XM_019833395.2;r
efGene_gn=NRXN3;refGene_tx=XM_019833396.2;refGene_gn=NRXN3;refGene_tx=
XM_019833397.2;refGene_gn=NRXN3;refGene_tx=XM_019833398.2;refGene_gn=N
RXN3;refGene_tx=XM_019833399.2;refGene_gn=NRXN3;refGene_tx=XM_01983340
0.2;refGene_gn=NRXN3;refGene_tx=XM_019833401.2;refGene_gn=NRXN3;refGen
e_tx=XM_019833402.2;refGene_gn=NRXN3;refGene_tx=XM_019833403.2;refGene
_gn=NRXN3;refGene_tx=XM_019833404.2;refGene_gn=NRXN3;refGene_tx=XM_019
833405.2;refGene_gn=NRXN3;refGene_tx=XM_019833407.2;refGene_gn=NRXN3;r
efGene_tx=XM_019833408.2;refGene_gn=NRXN3;refGene_tx=XM_019833409.2;re
fGene_gn=NRXN3;refGene_tx=XM_019833410.2;refGene_gn=NRXN3;refGene_tx=X
M_019833411.2;refGene_gn=NRXN3;refGene_tx=XM_019833412.2;refGene_gn=NR
XN3;refGene_tx=XM_019833413.2;refGene_gn=NRXN3;refGene_tx=XM_019833414
.
2;refGene_gn=NRXN3;refGene_tx=XM_019833415.2;refGene_gn=NRXN3;refGene_
tx=XM_019833416.2;refGene_gn=NRXN3;refGene_tx=XM_019833417.2;refGene_g
n=NRXN3;refGene_tx=XM_019833418.2
B3 126524143 126524315
refGene_gn=NRXN3;refGene_tx=XM_011283426.3;refGene_gn=NRXN3;refGene_tx
=XM_019833384.2;refGene_gn=NRXN3;refGene_tx=XM_019833385.2;refGene_gn=
NRXN3;refGene_tx=XM_019833386.2;refGene_gn=NRXN3;refGene_tx=XM_0198333
87.2;refGene_gn=NRXN3;refGene_tx=XM_019833388.2;refGene_gn=NRXN3;refGe
ne_tx=XM_019833389.2;refGene_gn=NRXN3;refGene_tx=XM_019833390.2;refGen
e_gn=NRXN3;refGene_tx=XM_019833391.2;refGene_gn=NRXN3;refGene_tx=XM_01
9833392.2;refGene_gn=NRXN3;refGene_tx=XM_019833393.2;refGene_gn=NRXN3;
refGene_tx=XM_019833394.2;refGene_gn=NRXN3;refGene_tx=XM_019833395.2;r
efGene_gn=NRXN3;refGene_tx=XM_019833396.2;refGene_gn=NRXN3;refGene_tx=
XM_019833397.2;refGene_gn=NRXN3;refGene_tx=XM_019833398.2;refGene_gn=N
RXN3;refGene_tx=XM_019833399.2;refGene_gn=NRXN3;refGene_tx=XM_01983340
0.2;refGene_gn=NRXN3;refGene_tx=XM_019833401.2;refGene_gn=NRXN3;refGen
e_tx=XM_019833402.2;refGene_gn=NRXN3;refGene_tx=XM_019833403.2;refGene
_gn=NRXN3;refGene_tx=XM_019833404.2;refGene_gn=NRXN3;refGene_tx=XM_019
833405.2;refGene_gn=NRXN3;refGene_tx=XM_019833407.2;refGene_gn=NRXN3;r
efGene_tx=XM_019833408.2;refGene_gn=NRXN3;refGene_tx=XM_019833409.2;re
fGene_gn=NRXN3;refGene_tx=XM_019833410.2;refGene_gn=NRXN3;refGene_tx=X
M_019833411.2;refGene_gn=NRXN3;refGene_tx=XM_019833412.2;refGene_gn=NR
XN3;refGene_tx=XM_019833413.2;refGene_gn=NRXN3;refGene_tx=XM_019833414
.
2;refGene_gn=NRXN3;refGene_tx=XM_019833415.2;refGene_gn=NRXN3;refGene_
tx=XM_019833416.2;refGene_gn=NRXN3;refGene_tx=XM_019833417.2;refGene_g
n=NRXN3;refGene_tx=XM_019833418.2

B3	126551858	126551948
refGene_gn=NRXN3;refGene_tx=XM_019833384.2;refGene_gn=NRXN3;refGene_tx=XM_019833385.2;refGene_gn=NRXN3;refGene_tx=XM_019833386.2;refGene_gn=NRXN3;refGene_tx=XM_019833387.2;refGene_gn=NRXN3;refGene_tx=XM_019833388.2;refGene_gn=NRXN3;refGene_tx=XM_019833389.2;refGene_gn=NRXN3;refGene_tx=XM_019833390.2;refGene_gn=NRXN3;refGene_tx=XM_019833391.2;refGene_gn=NRXN3;refGene_tx=XM_019833392.2;refGene_gn=NRXN3;refGene_tx=XM_019833393.2;refGene_gn=NRXN3;refGene_tx=XM_019833398.2;refGene_gn=NRXN3;refGene_tx=XM_019833399.2;refGene_gn=NRXN3;refGene_tx=XM_019833400.2;refGene_gn=NRXN3;refGene_tx=XM_019833401.2;refGene_gn=NRXN3;refGene_tx=XM_019833402.2;refGene_gn=NRXN3;refGene_tx=XM_019833403.2;refGene_gn=NRXN3;refGene_tx=XM_019833404.2;refGene_gn=NRXN3;refGene_tx=XM_019833405.2;refGene_gn=NRXN3;refGene_tx=XM_019833409.2;refGene_gn=NRXN3;refGene_tx=XM_019833410.2;refGene_gn=NRXN3;refGene_tx=XM_019833412.2;refGene_gn=NRXN3;refGene_tx=XM_019833413.2
B3	126557179	126557487
refGene_gn=NRXN3;refGene_tx=XM_011283426.3;refGene_gn=NRXN3;refGene_tx=XM_019833384.2;refGene_gn=NRXN3;refGene_tx=XM_019833385.2;refGene_gn=NRXN3;refGene_tx=XM_019833386.2;refGene_gn=NRXN3;refGene_tx=XM_019833387.2;refGene_gn=NRXN3;refGene_tx=XM_019833388.2;refGene_gn=NRXN3;refGene_tx=XM_019833389.2;refGene_gn=NRXN3;refGene_tx=XM_019833390.2;refGene_gn=NRXN3;refGene_tx=XM_019833391.2;refGene_gn=NRXN3;refGene_tx=XM_019833392.2;refGene_gn=NRXN3;refGene_tx=XM_019833393.2;refGene_gn=NRXN3;refGene_tx=XM_019833394.2;refGene_gn=NRXN3;refGene_tx=XM_019833395.2;refGene_gn=NRXN3;refGene_tx=XM_019833396.2;refGene_gn=NRXN3;refGene_tx=XM_019833397.2;refGene_gn=NRXN3;refGene_tx=XM_019833398.2;refGene_gn=NRXN3;refGene_tx=XM_019833399.2;refGene_gn=NRXN3;refGene_tx=XM_019833400.2;refGene_gn=NRXN3;refGene_tx=XM_019833401.2;refGene_gn=NRXN3;refGene_tx=XM_019833402.2;refGene_gn=NRXN3;refGene_tx=XM_019833403.2;refGene_gn=NRXN3;refGene_tx=XM_019833404.2;refGene_gn=NRXN3;refGene_tx=XM_019833405.2;refGene_gn=NRXN3;refGene_tx=XM_019833407.2;refGene_gn=NRXN3;refGene_tx=XM_019833408.2;refGene_gn=NRXN3;refGene_tx=XM_019833409.2;refGene_gn=NRXN3;refGene_tx=XM_019833410.2;refGene_gn=NRXN3;refGene_tx=XM_019833411.2;refGene_gn=NRXN3;refGene_tx=XM_019833412.2;refGene_gn=NRXN3;refGene_tx=XM_019833413.2;refGene_gn=NRXN3;refGene_tx=XM_019833414.
2;refGene_gn=NRXN3;refGene_tx=XM_019833415.2;refGene_gn=NRXN3;refGene_tx=XM_019833416.2;refGene_gn=NRXN3;refGene_tx=XM_019833417.2;refGene_gn=NRXN3;refGene_tx=XM_019833418.2
B3	126652445	126652533
refGene_gn=NRXN3;refGene_tx=XM_011283426.3;refGene_gn=NRXN3;refGene_tx=XM_019833384.2;refGene_gn=NRXN3;refGene_tx=XM_019833385.2;refGene_gn=NRXN3;refGene_tx=XM_019833386.2;refGene_gn=NRXN3;refGene_tx=XM_019833387.2;refGene_gn=NRXN3;refGene_tx=XM_019833388.2;refGene_gn=NRXN3;refGene_tx=XM_019833389.2;refGene_gn=NRXN3;refGene_tx=XM_019833390.2;refGene_gn=NRXN3;refGene_tx=XM_019833391.2;refGene_gn=NRXN3;refGene_tx=XM_019833392.2;refGene_gn=NRXN3;refGene_tx=XM_019833393.2;refGene_gn=NRXN3;refGene_tx=XM_019833394.2;refGene_gn=NRXN3;refGene_tx=XM_019833395.2;refGene_gn=NRXN3;refGene_tx=XM_019833396.2;refGene_gn=NRXN3;refGene_tx=XM_019833397.2;refGene_gn=NRXN3;refGene_tx=XM_019833398.2;refGene_gn=N

RXN3;refGene_tx=XM_019833399.2;refGene_gn=NRXN3;refGene_tx=XM_01983340
0.2;refGene_gn=NRXN3;refGene_tx=XM_019833401.2;refGene_gn=NRXN3;refGen
e_tx=XM_019833402.2;refGene_gn=NRXN3;refGene_tx=XM_019833403.2;refGene
_gn=NRXN3;refGene_tx=XM_019833404.2;refGene_gn=NRXN3;refGene_tx=XM_019
833405.2;refGene_gn=NRXN3;refGene_tx=XM_019833407.2;refGene_gn=NRXN3;r
efGene_tx=XM_019833408.2;refGene_gn=NRXN3;refGene_tx=XM_019833409.2;re
fGene_gn=NRXN3;refGene_tx=XM_019833410.2;refGene_gn=NRXN3;refGene_tx=X
M_019833411.2;refGene_gn=NRXN3;refGene_tx=XM_019833412.2;refGene_gn=NR
XN3;refGene_tx=XM_019833413.2;refGene_gn=NRXN3;refGene_tx=XM_019833414
.
2;refGene_gn=NRXN3;refGene_tx=XM_019833415.2;refGene_gn=NRXN3;refGene_
tx=XM_019833416.2;refGene_gn=NRXN3;refGene_tx=XM_019833417.2;refGene_g
n=NRXN3;refGene_tx=XM_019833418.2
B3 126702490 126703416
refGene_gn=NRXN3;refGene_tx=XM_011283426.3;refGene_gn=NRXN3;refGene_tx
=XM_019833384.2;refGene_gn=NRXN3;refGene_tx=XM_019833385.2;refGene_gn=
NRXN3;refGene_tx=XM_019833386.2;refGene_gn=NRXN3;refGene_tx=XM_0198333
87.2;refGene_gn=NRXN3;refGene_tx=XM_019833388.2;refGene_gn=NRXN3;refGe
ne_tx=XM_019833389.2;refGene_gn=NRXN3;refGene_tx=XM_019833390.2;refGen
e_gn=NRXN3;refGene_tx=XM_019833391.2;refGene_gn=NRXN3;refGene_tx=XM_01
9833392.2;refGene_gn=NRXN3;refGene_tx=XM_019833393.2;refGene_gn=NRXN3;
refGene_tx=XM_019833394.2;refGene_gn=NRXN3;refGene_tx=XM_019833395.2;r
efGene_gn=NRXN3;refGene_tx=XM_019833396.2;refGene_gn=NRXN3;refGene_tx=
XM_019833397.2;refGene_gn=NRXN3;refGene_tx=XM_019833398.2;refGene_gn=N
RXN3;refGene_tx=XM_019833399.2;refGene_gn=NRXN3;refGene_tx=XM_01983340
0.2;refGene_gn=NRXN3;refGene_tx=XM_019833401.2;refGene_gn=NRXN3;refGen
e_tx=XM_019833402.2;refGene_gn=NRXN3;refGene_tx=XM_019833403.2;refGene
_gn=NRXN3;refGene_tx=XM_019833404.2;refGene_gn=NRXN3;refGene_tx=XM_019
833405.2;refGene_gn=NRXN3;refGene_tx=XM_019833407.2;refGene_gn=NRXN3;r
efGene_tx=XM_019833408.2;refGene_gn=NRXN3;refGene_tx=XM_019833409.2;re
fGene_gn=NRXN3;refGene_tx=XM_019833410.2;refGene_gn=NRXN3;refGene_tx=X
M_019833411.2;refGene_gn=NRXN3;refGene_tx=XM_019833412.2;refGene_gn=NR
XN3;refGene_tx=XM_019833413.2;refGene_gn=NRXN3;refGene_tx=XM_019833414
.
2;refGene_gn=NRXN3;refGene_tx=XM_019833415.2;refGene_gn=NRXN3;refGene_
tx=XM_019833416.2;refGene_gn=NRXN3;refGene_tx=XM_019833417.2;refGene_g
n=NRXN3;refGene_tx=XM_019833418.2
B3 127052242 127052265
refGene_gn=DIO2;refGene_tx=XM_006933041.4
B3 127052267 127052838
refGene_gn=DIO2;refGene_tx=XM_006933041.4
B3 127060877 127061099
refGene_gn=DIO2;refGene_tx=XM_006933041.4
B3 127291599 127291759
refGene_gn=CEP128;refGene_tx=XM_019833419.2;refGene_gn=CEP128;refGene_
tx=XM_019833420.2;refGene_gn=CEP128;refGene_tx=XM_019833423.2
B3 127298656 127298784
refGene_gn=CEP128;refGene_tx=XM_019833419.2;refGene_gn=CEP128;refGene_
tx=XM_019833420.2;refGene_gn=CEP128;refGene_tx=XM_019833423.2;refGene_
gn=CEP128;refGene_tx=XM_019833425.1

B3	127304556	127304660
refGene_gn=CEP128;refGene_tx=XM_023255904.1
B3	127306251	127306360
refGene_gn=CEP128;refGene_tx=XM_019833419.2;refGene_gn=CEP128;refGene_tx=XM_019833420.2;refGene_gn=CEP128;refGene_tx=XM_019833423.2;refGene_gn=CEP128;refGene_tx=XM_019833425.1;refGene_gn=CEP128;refGene_tx=XM_023255904.1
B3	127318270	127318384
refGene_gn=CEP128;refGene_tx=XM_019833419.2;refGene_gn=CEP128;refGene_tx=XM_019833420.2;refGene_gn=CEP128;refGene_tx=XM_019833423.2;refGene_gn=CEP128;refGene_tx=XM_019833425.1;refGene_gn=CEP128;refGene_tx=XM_023255904.1
B3	127322849	127322927
refGene_gn=CEP128;refGene_tx=XM_019833419.2;refGene_gn=CEP128;refGene_tx=XM_019833420.2;refGene_gn=CEP128;refGene_tx=XM_019833423.2;refGene_gn=CEP128;refGene_tx=XM_019833425.1;refGene_gn=CEP128;refGene_tx=XM_023255904.1
B3	127352290	127352314
refGene_gn=CEP128;refGene_tx=XM_019833419.2;refGene_gn=CEP128;refGene_tx=XM_019833420.2;refGene_gn=CEP128;refGene_tx=XM_019833423.2;refGene_gn=CEP128;refGene_tx=XM_019833425.1;refGene_gn=CEP128;refGene_tx=XM_023255904.1
B3	127360263	127360353
refGene_gn=CEP128;refGene_tx=XM_019833419.2;refGene_gn=CEP128;refGene_tx=XM_019833420.2;refGene_gn=CEP128;refGene_tx=XM_019833425.1;refGene_gn=CEP128;refGene_tx=XM_023255904.1
B3	127373478	127373528
refGene_gn=CEP128;refGene_tx=XM_019833419.2;refGene_gn=CEP128;refGene_tx=XM_019833420.2;refGene_gn=CEP128;refGene_tx=XM_019833423.2;refGene_gn=CEP128;refGene_tx=XM_019833425.1;refGene_gn=CEP128;refGene_tx=XM_023255904.1
B3	127515377	127515570
refGene_gn=CEP128;refGene_tx=XM_019833419.2;refGene_gn=CEP128;refGene_tx=XM_019833420.2;refGene_gn=CEP128;refGene_tx=XM_019833423.2;refGene_gn=CEP128;refGene_tx=XM_019833425.1;refGene_gn=CEP128;refGene_tx=XM_023255904.1
B3	127529288	127529348
refGene_gn=CEP128;refGene_tx=XM_019833419.2;refGene_gn=CEP128;refGene_tx=XM_019833420.2;refGene_gn=CEP128;refGene_tx=XM_019833423.2;refGene_gn=CEP128;refGene_tx=XM_019833425.1;refGene_gn=CEP128;refGene_tx=XM_023255904.1
B3	127534022	127534199
refGene_gn=CEP128;refGene_tx=XM_019833419.2;refGene_gn=CEP128;refGene_tx=XM_019833420.2;refGene_gn=CEP128;refGene_tx=XM_019833423.2;refGene_gn=CEP128;refGene_tx=XM_019833425.1;refGene_gn=CEP128;refGene_tx=XM_023255904.1
B3	127550775	127550940
refGene_gn=CEP128;refGene_tx=XM_019833419.2;refGene_gn=CEP128;refGene_tx=XM_019833420.2;refGene_gn=CEP128;refGene_tx=XM_019833423.2;refGene_gn=CEP128;refGene_tx=XM_019833425.1;refGene_gn=CEP128;refGene_tx=XM_02

3255904.1
B3	127556150	127556801
refGene_gn=CEP128;refGene_tx=XM_019833419.2;refGene_gn=CEP128;refGene_tx=XM_019833420.2;refGene_gn=CEP128;refGene_tx=XM_019833423.2;refGene_gn=CEP128;refGene_tx=XM_019833425.1;refGene_gn=CEP128;refGene_tx=XM_023255904.1
B3	127564169	127564520
refGene_gn=CEP128;refGene_tx=XM_019833419.2;refGene_gn=CEP128;refGene_tx=XM_019833420.2;refGene_gn=CEP128;refGene_tx=XM_019833423.2;refGene_gn=CEP128;refGene_tx=XM_019833425.1;refGene_gn=CEP128;refGene_tx=XM_023255904.1
B3	127598760	127598912
refGene_gn=CEP128;refGene_tx=XM_019833419.2;refGene_gn=CEP128;refGene_tx=XM_019833420.2;refGene_gn=CEP128;refGene_tx=XM_019833423.2;refGene_gn=CEP128;refGene_tx=XM_019833425.1;refGene_gn=CEP128;refGene_tx=XM_023255904.1
B3	127605637	127605770
refGene_gn=CEP128;refGene_tx=XM_019833419.2;refGene_gn=CEP128;refGene_tx=XM_019833420.2;refGene_gn=CEP128;refGene_tx=XM_019833423.2;refGene_gn=CEP128;refGene_tx=XM_019833425.1;refGene_gn=CEP128;refGene_tx=XM_023255904.1
B3	127607646	127607721
refGene_gn=CEP128;refGene_tx=XM_019833419.2;refGene_gn=CEP128;refGene_tx=XM_019833420.2;refGene_gn=CEP128;refGene_tx=XM_019833423.2;refGene_gn=CEP128;refGene_tx=XM_019833425.1;refGene_gn=CEP128;refGene_tx=XM_023255904.1
B3	127610520	127610607
refGene_gn=CEP128;refGene_tx=XM_019833419.2;refGene_gn=CEP128;refGene_tx=XM_019833420.2;refGene_gn=CEP128;refGene_tx=XM_019833423.2;refGene_gn=CEP128;refGene_tx=XM_019833425.1;refGene_gn=CEP128;refGene_tx=XM_023255904.1
B3	127629999	127630116
refGene_gn=CEP128;refGene_tx=XM_019833419.2;refGene_gn=CEP128;refGene_tx=XM_019833420.2;refGene_gn=CEP128;refGene_tx=XM_019833423.2;refGene_gn=CEP128;refGene_tx=XM_019833425.1;refGene_gn=CEP128;refGene_tx=XM_023255904.1
B3	127650450	127650523
refGene_gn=CEP128;refGene_tx=XM_019833419.2;refGene_gn=CEP128;refGene_tx=XM_019833420.2;refGene_gn=CEP128;refGene_tx=XM_019833423.2;refGene_gn=CEP128;refGene_tx=XM_019833425.1;refGene_gn=CEP128;refGene_tx=XM_023255904.1
B3	127654262	127654354
refGene_gn=CEP128;refGene_tx=XM_019833419.2;refGene_gn=CEP128;refGene_tx=XM_019833420.2;refGene_gn=CEP128;refGene_tx=XM_019833423.2;refGene_gn=CEP128;refGene_tx=XM_019833425.1;refGene_gn=CEP128;refGene_tx=XM_023255904.1
B3	127657271	127657390
refGene_gn=CEP128;refGene_tx=XM_019833419.2;refGene_gn=CEP128;refGene_tx=XM_019833420.2;refGene_gn=CEP128;refGene_tx=XM_019833423.2;refGene_gn=CEP128;refGene_tx=XM_019833425.1;refGene_gn=CEP128;refGene_tx=XM_02

3255904.1
B3	127658428	127658555
refGene_gn=CEP128;refGene_tx=XM_019833419.2;refGene_gn=CEP128;refGene_tx=XM_019833420.2;refGene_gn=CEP128;refGene_tx=XM_019833423.2;refGene_gn=CEP128;refGene_tx=XM_019833425.1;refGene_gn=CEP128;refGene_tx=XM_023255904.1
B3	127671532	127671619
refGene_gn=CEP128;refGene_tx=XM_019833419.2;refGene_gn=CEP128;refGene_tx=XM_019833420.2;refGene_gn=CEP128;refGene_tx=XM_019833423.2;refGene_gn=CEP128;refGene_tx=XM_019833425.1;refGene_gn=CEP128;refGene_tx=XM_023255904.1
B3	127673637	127673784
refGene_gn=CEP128;refGene_tx=XM_019833419.2;refGene_gn=CEP128;refGene_tx=XM_019833420.2;refGene_gn=CEP128;refGene_tx=XM_019833423.2;refGene_gn=CEP128;refGene_tx=XM_019833425.1;refGene_gn=CEP128;refGene_tx=XM_023255904.1
B3	127702691	127702861
refGene_gn=TSHR;refGene_tx=NM_001009288.1;refGene_gn=TSHR;refGene_tx=XM_019831721.1;refGene_gn=TSHR;refGene_tx=XM_019831722.2;refGene_gn=TSHR;refGene_tx=XM_019831724.1;refGene_gn=TSHR;refGene_tx=XM_019831725.1;refGene_gn=TSHR;refGene_tx=XM_019831726.1;refGene_gn=TSHR;refGene_tx=XM_019831727.1;refGene_gn=TSHR;refGene_tx=XM_019831728.1
B3	127793699	127793771
refGene_gn=TSHR;refGene_tx=NM_001009288.1;refGene_gn=TSHR;refGene_tx=XM_019831721.1;refGene_gn=TSHR;refGene_tx=XM_019831722.2;refGene_gn=TSHR;refGene_tx=XM_019831724.1;refGene_gn=TSHR;refGene_tx=XM_019831725.1;refGene_gn=TSHR;refGene_tx=XM_019831726.1;refGene_gn=TSHR;refGene_tx=XM_019831727.1
B3	127798952	127799027
refGene_gn=TSHR;refGene_tx=NM_001009288.1;refGene_gn=TSHR;refGene_tx=XM_019831721.1;refGene_gn=TSHR;refGene_tx=XM_019831722.2;refGene_gn=TSHR;refGene_tx=XM_019831725.1;refGene_gn=TSHR;refGene_tx=XM_019831727.1;refGene_gn=TSHR;refGene_tx=XM_019831730.1
B3	127818682	127818757
refGene_gn=TSHR;refGene_tx=NM_001009288.1;refGene_gn=TSHR;refGene_tx=XM_019831721.1;refGene_gn=TSHR;refGene_tx=XM_019831722.2;refGene_gn=TSHR;refGene_tx=XM_019831724.1;refGene_gn=TSHR;refGene_tx=XM_019831726.1;refGene_gn=TSHR;refGene_tx=XM_019831728.1;refGene_gn=TSHR;refGene_tx=XM_019831730.1
B3	127822075	127822150
refGene_gn=TSHR;refGene_tx=NM_001009288.1;refGene_gn=TSHR;refGene_tx=XM_019831721.1;refGene_gn=TSHR;refGene_tx=XM_019831722.2;refGene_gn=TSHR;refGene_tx=XM_019831724.1;refGene_gn=TSHR;refGene_tx=XM_019831725.1;refGene_gn=TSHR;refGene_tx=XM_019831726.1;refGene_gn=TSHR;refGene_tx=XM_019831727.1;refGene_gn=TSHR;refGene_tx=XM_019831728.1;refGene_gn=TSHR;refGene_tx=XM_019831730.1
B3	127823564	127823642
refGene_gn=TSHR;refGene_tx=NM_001009288.1;refGene_gn=TSHR;refGene_tx=XM_019831721.1;refGene_gn=TSHR;refGene_tx=XM_019831722.2;refGene_gn=TSHR;refGene_tx=XM_019831724.1;refGene_gn=TSHR;refGene_tx=XM_019831725.1;

refGene_gn=TSHR;refGene_tx=XM_019831726.1;refGene_gn=TSHR;refGene_tx=XM_019831727.1;refGene_gn=TSHR;refGene_tx=XM_019831728.1;refGene_gn=TSHR;refGene_tx=XM_019831730.1
B3 127826909 127826978
refGene_gn=TSHR;refGene_tx=NM_001009288.1;refGene_gn=TSHR;refGene_tx=XM_019831721.1;refGene_gn=TSHR;refGene_tx=XM_019831722.2;refGene_gn=TSHR;refGene_tx=XM_019831724.1;refGene_gn=TSHR;refGene_tx=XM_019831725.1;refGene_gn=TSHR;refGene_tx=XM_019831726.1;refGene_gn=TSHR;refGene_tx=XM_019831727.1;refGene_gn=TSHR;refGene_tx=XM_019831728.1;refGene_gn=TSHR;refGene_tx=XM_019831730.1
B3 127833881 127833902
refGene_gn=TSHR;refGene_tx=XM_019831721.1;refGene_gn=TSHR;refGene_tx=XM_019831724.1;refGene_gn=TSHR;refGene_tx=XM_019831725.1;refGene_gn=TSHR;refGene_tx=XM_019831728.1;refGene_gn=TSHR;refGene_tx=XM_019831730.1
B3 127839356 127839434
refGene_gn=TSHR;refGene_tx=NM_001009288.1;refGene_gn=TSHR;refGene_tx=XM_019831721.1;refGene_gn=TSHR;refGene_tx=XM_019831722.2;refGene_gn=TSHR;refGene_tx=XM_019831724.1;refGene_gn=TSHR;refGene_tx=XM_019831725.1;refGene_gn=TSHR;refGene_tx=XM_019831726.1;refGene_gn=TSHR;refGene_tx=XM_019831727.1;refGene_gn=TSHR;refGene_tx=XM_019831728.1;refGene_gn=TSHR;refGene_tx=XM_019831730.1
B3 127860799 127860988
refGene_gn=TSHR;refGene_tx=NM_001009288.1;refGene_gn=TSHR;refGene_tx=XM_019831721.1;refGene_gn=TSHR;refGene_tx=XM_019831722.2;refGene_gn=TSHR;refGene_tx=XM_019831724.1;refGene_gn=TSHR;refGene_tx=XM_019831725.1;refGene_gn=TSHR;refGene_tx=XM_019831726.1;refGene_gn=TSHR;refGene_tx=XM_019831727.1;refGene_gn=TSHR;refGene_tx=XM_019831728.1;refGene_gn=TSHR;refGene_tx=XM_019831730.1
B3 127863350 127864761
refGene_gn=TSHR;refGene_tx=NM_001009288.1;refGene_gn=TSHR;refGene_tx=XM_019831721.1;refGene_gn=TSHR;refGene_tx=XM_019831722.2;refGene_gn=TSHR;refGene_tx=XM_019831724.1;refGene_gn=TSHR;refGene_tx=XM_019831725.1;refGene_gn=TSHR;refGene_tx=XM_019831726.1;refGene_gn=TSHR;refGene_tx=XM_019831727.1;refGene_gn=TSHR;refGene_tx=XM_019831728.1;refGene_gn=TSHR;refGene_tx=XM_019831730.1
B3 127915829 127915937
refGene_gn=GTF2A1;refGene_tx=XM_023255911.1
B3 127924397 127924487
refGene_gn=GTF2A1;refGene_tx=XM_023255911.1
B3 127930035 127930356
refGene_gn=GTF2A1;refGene_tx=XM_023255911.1
B3 127933462 127933596
refGene_gn=GTF2A1;refGene_tx=XM_023255911.1
B3 127934981 127935057
refGene_gn=GTF2A1;refGene_tx=XM_023255911.1
B3 127939246 127939311
refGene_gn=GTF2A1;refGene_tx=XM_023255911.1
B3 127941086 127941291
refGene_gn=GTF2A1;refGene_tx=XM_023255911.1
B3 127951670 127951772

refGene_gn=GTF2A1;refGene_tx=XM_023255911.1
B3	127955696	127955726
refGene_gn=GTF2A1;refGene_tx=XM_023255911.1
B3	127985429	127985513
refGene_gn=STON2;refGene_tx=XM_023255906.1;refGene_gn=STON2;refGene_tx=XM_023255907.1;refGene_gn=STON2;refGene_tx=XM_023255908.1;refGene_gn=STON2;refGene_tx=XM_023255909.1;refGene_gn=STON2;refGene_tx=XM_023255910.1
B3	127987069	127987272
refGene_gn=STON2;refGene_tx=XM_023255906.1;refGene_gn=STON2;refGene_tx=XM_023255907.1;refGene_gn=STON2;refGene_tx=XM_023255908.1;refGene_gn=STON2;refGene_tx=XM_023255909.1;refGene_gn=STON2;refGene_tx=XM_023255910.1
B3	127992838	127994677
refGene_gn=STON2;refGene_tx=XM_023255906.1;refGene_gn=STON2;refGene_tx=XM_023255907.1;refGene_gn=STON2;refGene_tx=XM_023255908.1;refGene_gn=STON2;refGene_tx=XM_023255909.1;refGene_gn=STON2;refGene_tx=XM_023255910.1
B3	128035640	128035811
refGene_gn=STON2;refGene_tx=XM_023255906.1;refGene_gn=STON2;refGene_tx=XM_023255907.1;refGene_gn=STON2;refGene_tx=XM_023255908.1;refGene_gn=STON2;refGene_tx=XM_023255909.1
B3	128068259	128068457
refGene_gn=STON2;refGene_tx=XM_023255906.1;refGene_gn=STON2;refGene_tx=XM_023255907.1;refGene_gn=STON2;refGene_tx=XM_023255908.1;refGene_gn=STON2;refGene_tx=XM_023255909.1;refGene_gn=STON2;refGene_tx=XM_023255910.1
B3	128090582	128090867
refGene_gn=STON2;refGene_tx=XM_023255906.1;refGene_gn=STON2;refGene_tx=XM_023255907.1;refGene_gn=STON2;refGene_tx=XM_023255908.1;refGene_gn=STON2;refGene_tx=XM_023255909.1;refGene_gn=STON2;refGene_tx=XM_023255910.1
B3	128092679	128092767
refGene_gn=STON2;refGene_tx=XM_023255906.1;refGene_gn=STON2;refGene_tx=XM_023255907.1;refGene_gn=STON2;refGene_tx=XM_023255908.1;refGene_gn=STON2;refGene_tx=XM_023255909.1;refGene_gn=STON2;refGene_tx=XM_023255910.1
B3	128161911	128162121
refGene_gn=SEL1L;refGene_tx=XM_003987895.5
B3	128164187	128164316
refGene_gn=SEL1L;refGene_tx=XM_003987895.5
B3	128168472	128168645
refGene_gn=SEL1L;refGene_tx=XM_003987895.5
B3	128169816	128169891
refGene_gn=SEL1L;refGene_tx=XM_003987895.5
B3	128170456	128170622
refGene_gn=SEL1L;refGene_tx=XM_003987895.5
B3	128171390	128171539
refGene_gn=SEL1L;refGene_tx=XM_003987895.5
B3	128171864	128171952

refGene_gn=SEL1L;refGene_tx=XM_003987895.5
B3 128173238 128173301
refGene_gn=SEL1L;refGene_tx=XM_003987895.5
B3 128174616 128174694
refGene_gn=SEL1L;refGene_tx=XM_003987895.5
B3 128175636 128175705
refGene_gn=SEL1L;refGene_tx=XM_003987895.5
B3 128177718 128177775
refGene_gn=SEL1L;refGene_tx=XM_003987895.5
B3 128181327 128181482
refGene_gn=SEL1L;refGene_tx=XM_003987895.5
B3 128181847 128181929
refGene_gn=SEL1L;refGene_tx=XM_003987895.5
B3 128183509 128183569
refGene_gn=SEL1L;refGene_tx=XM_003987895.5
B3 128183659 128183713
refGene_gn=SEL1L;refGene_tx=XM_003987895.5
B3 128186096 128186259
refGene_gn=SEL1L;refGene_tx=XM_003987895.5
B3 128187304 128187410
refGene_gn=SEL1L;refGene_tx=XM_003987895.5
B3 128188782 128188950
refGene_gn=SEL1L;refGene_tx=XM_003987895.5
B3 128203979 128204214
refGene_gn=SEL1L;refGene_tx=XM_003987895.5
B3 128205243 128205281
refGene_gn=SEL1L;refGene_tx=XM_003987895.5
B3 128210871 128210941
refGene_gn=SEL1L;refGene_tx=XM_003987895.5
B3 131903626 131905609
refGene_gn=FLRT2;refGene_tx=XM_006933044.4;refGene_gn=FLRT2;refGene_tx
=XM_023255912.1
B3 133866231 133866378
refGene_gn=GALC;refGene_tx=XM_023255914.1
B3 133916239 133916248
refGene_gn=GALC;refGene_tx=XM_023255917.1
B3 133924139 133924214
refGene_gn=GALC;refGene_tx=XM_023255916.1
B3 133955774 133955921
refGene_gn=GALC;refGene_tx=XM_019833437.2;refGene_gn=GALC;refGene_tx=X
M_019833439.1;refGene_gn=GALC;refGene_tx=XM_019833440.2;refGene_gn=GAL
C;refGene_tx=XM_019833441.2;refGene_gn=GALC;refGene_tx=XM_023255913.1
B3 133959361 133959469
refGene_gn=GALC;refGene_tx=XM_023255915.1
B3 133961839 133961916
refGene_gn=GALC;refGene_tx=XM_019833437.2;refGene_gn=GALC;refGene_tx=X
M_019833439.1;refGene_gn=GALC;refGene_tx=XM_019833440.2;refGene_gn=GAL
C;refGene_tx=XM_019833441.2;refGene_gn=GALC;refGene_tx=XM_023255913.1;
refGene_gn=GALC;refGene_tx=XM_023255914.1;refGene_gn=GALC;refGene_tx=X
M_023255915.1;refGene_gn=GALC;refGene_tx=XM_023255916.1;refGene_gn=GAL

C;refGene_tx=XM_023255917.1
B3	133963540	133963704
refGene_gn=GALC;refGene_tx=XM_019833437.2;refGene_gn=GALC;refGene_tx=XM_019833439.1;refGene_gn=GALC;refGene_tx=XM_019833440.2;refGene_gn=GALC;refGene_tx=XM_019833441.2;refGene_gn=GALC;refGene_tx=XM_023255913.1;refGene_gn=GALC;refGene_tx=XM_023255914.1;refGene_gn=GALC;refGene_tx=XM_023255915.1;refGene_gn=GALC;refGene_tx=XM_023255916.1;refGene_gn=GALC;refGene_tx=XM_023255917.1
B3	133967142	133967323
refGene_gn=GALC;refGene_tx=XM_019833437.2;refGene_gn=GALC;refGene_tx=XM_019833439.1;refGene_gn=GALC;refGene_tx=XM_019833440.2;refGene_gn=GALC;refGene_tx=XM_019833441.2;refGene_gn=GALC;refGene_tx=XM_023255913.1;refGene_gn=GALC;refGene_tx=XM_023255914.1;refGene_gn=GALC;refGene_tx=XM_023255915.1;refGene_gn=GALC;refGene_tx=XM_023255916.1;refGene_gn=GALC;refGene_tx=XM_023255917.1
B3	133969357	133969508
refGene_gn=GALC;refGene_tx=XM_019833437.2;refGene_gn=GALC;refGene_tx=XM_019833439.1;refGene_gn=GALC;refGene_tx=XM_019833440.2;refGene_gn=GALC;refGene_tx=XM_019833441.2;refGene_gn=GALC;refGene_tx=XM_023255913.1;refGene_gn=GALC;refGene_tx=XM_023255914.1;refGene_gn=GALC;refGene_tx=XM_023255915.1;refGene_gn=GALC;refGene_tx=XM_023255916.1;refGene_gn=GALC;refGene_tx=XM_023255917.1
B3	133971597	133971684
refGene_gn=GALC;refGene_tx=XM_019833437.2;refGene_gn=GALC;refGene_tx=XM_019833439.1;refGene_gn=GALC;refGene_tx=XM_019833440.2;refGene_gn=GALC;refGene_tx=XM_019833441.2;refGene_gn=GALC;refGene_tx=XM_023255913.1;refGene_gn=GALC;refGene_tx=XM_023255914.1;refGene_gn=GALC;refGene_tx=XM_023255915.1;refGene_gn=GALC;refGene_tx=XM_023255916.1;refGene_gn=GALC;refGene_tx=XM_023255917.1
B3	133972445	133972535
refGene_gn=GALC;refGene_tx=XM_019833437.2;refGene_gn=GALC;refGene_tx=XM_019833439.1;refGene_gn=GALC;refGene_tx=XM_019833440.2;refGene_gn=GALC;refGene_tx=XM_019833441.2;refGene_gn=GALC;refGene_tx=XM_023255913.1;refGene_gn=GALC;refGene_tx=XM_023255914.1;refGene_gn=GALC;refGene_tx=XM_023255915.1;refGene_gn=GALC;refGene_tx=XM_023255916.1;refGene_gn=GALC;refGene_tx=XM_023255917.1
B3	133986003	133986131
refGene_gn=GALC;refGene_tx=XM_019833437.2;refGene_gn=GALC;refGene_tx=XM_019833439.1;refGene_gn=GALC;refGene_tx=XM_019833440.2;refGene_gn=GALC;refGene_tx=XM_019833441.2;refGene_gn=GALC;refGene_tx=XM_023255913.1;refGene_gn=GALC;refGene_tx=XM_023255914.1;refGene_gn=GALC;refGene_tx=XM_023255915.1;refGene_gn=GALC;refGene_tx=XM_023255916.1;refGene_gn=GALC;refGene_tx=XM_023255917.1
B3	133986956	133987081
refGene_gn=GALC;refGene_tx=XM_019833437.2;refGene_gn=GALC;refGene_tx=XM_019833439.1;refGene_gn=GALC;refGene_tx=XM_019833440.2;refGene_gn=GALC;refGene_tx=XM_019833441.2;refGene_gn=GALC;refGene_tx=XM_023255913.1;refGene_gn=GALC;refGene_tx=XM_023255914.1;refGene_gn=GALC;refGene_tx=XM_023255915.1;refGene_gn=GALC;refGene_tx=XM_023255916.1;refGene_gn=GALC;refGene_tx=XM_023255917.1

B3	133988185	133988341
refGene_gn=GALC;refGene_tx=XM_019833437.2;refGene_gn=GALC;refGene_tx=XM_019833439.1;refGene_gn=GALC;refGene_tx=XM_019833440.2;refGene_gn=GALC;refGene_tx=XM_019833441.2;refGene_gn=GALC;refGene_tx=XM_023255913.1;refGene_gn=GALC;refGene_tx=XM_023255914.1;refGene_gn=GALC;refGene_tx=XM_023255915.1;refGene_gn=GALC;refGene_tx=XM_023255916.1;refGene_gn=GALC;refGene_tx=XM_023255917.1
B3	133996393	133996524
refGene_gn=GALC;refGene_tx=XM_019833437.2;refGene_gn=GALC;refGene_tx=XM_019833439.1;refGene_gn=GALC;refGene_tx=XM_019833440.2;refGene_gn=GALC;refGene_tx=XM_019833441.2;refGene_gn=GALC;refGene_tx=XM_023255913.1;refGene_gn=GALC;refGene_tx=XM_023255914.1;refGene_gn=GALC;refGene_tx=XM_023255915.1;refGene_gn=GALC;refGene_tx=XM_023255916.1;refGene_gn=GALC;refGene_tx=XM_023255917.1
B3	133999203	133999242
refGene_gn=GALC;refGene_tx=XM_019833437.2;refGene_gn=GALC;refGene_tx=XM_019833439.1;refGene_gn=GALC;refGene_tx=XM_019833440.2;refGene_gn=GALC;refGene_tx=XM_019833441.2;refGene_gn=GALC;refGene_tx=XM_023255913.1;refGene_gn=GALC;refGene_tx=XM_023255914.1;refGene_gn=GALC;refGene_tx=XM_023255915.1;refGene_gn=GALC;refGene_tx=XM_023255916.1;refGene_gn=GALC;refGene_tx=XM_023255917.1
B3	134001607	134001747
refGene_gn=GALC;refGene_tx=XM_019833437.2;refGene_gn=GALC;refGene_tx=XM_019833439.1;refGene_gn=GALC;refGene_tx=XM_019833440.2;refGene_gn=GALC;refGene_tx=XM_019833441.2;refGene_gn=GALC;refGene_tx=XM_023255913.1;refGene_gn=GALC;refGene_tx=XM_023255914.1;refGene_gn=GALC;refGene_tx=XM_023255915.1;refGene_gn=GALC;refGene_tx=XM_023255916.1;refGene_gn=GALC;refGene_tx=XM_023255917.1
B3	134003722	134003836
refGene_gn=GALC;refGene_tx=XM_019833437.2;refGene_gn=GALC;refGene_tx=XM_019833439.1;refGene_gn=GALC;refGene_tx=XM_019833440.2;refGene_gn=GALC;refGene_tx=XM_019833441.2;refGene_gn=GALC;refGene_tx=XM_023255913.1;refGene_gn=GALC;refGene_tx=XM_023255914.1;refGene_gn=GALC;refGene_tx=XM_023255915.1;refGene_gn=GALC;refGene_tx=XM_023255916.1;refGene_gn=GALC;refGene_tx=XM_023255917.1
B3	134005625	134005689
refGene_gn=GALC;refGene_tx=XM_023255913.1;refGene_gn=GALC;refGene_tx=XM_023255914.1;refGene_gn=GALC;refGene_tx=XM_023255915.1;refGene_gn=GALC;refGene_tx=XM_023255916.1;refGene_gn=GALC;refGene_tx=XM_023255917.1
B3	134005937	134006006
refGene_gn=GALC;refGene_tx=XM_023255913.1;refGene_gn=GALC;refGene_tx=XM_023255914.1;refGene_gn=GALC;refGene_tx=XM_023255915.1;refGene_gn=GALC;refGene_tx=XM_023255916.1;refGene_gn=GALC;refGene_tx=XM_023255917.1
B3	134013620	134013815
refGene_gn=GALC;refGene_tx=XM_023255913.1;refGene_gn=GALC;refGene_tx=XM_023255914.1;refGene_gn=GALC;refGene_tx=XM_023255915.1;refGene_gn=GALC;refGene_tx=XM_023255916.1;refGene_gn=GALC;refGene_tx=XM_023255917.1
B3	134035706	134036732
refGene_gn=GPR65;refGene_tx=XM_003987900.5
B3	134165262	134165883

refGene_gn=KCNK10;refGene_tx=XM_003987901.5;refGene_gn=KCNK10;refGene_tx=XM_019833446.2;refGene_gn=KCNK10;refGene_tx=XM_023255918.1
B3	134167748	134167891
refGene_gn=KCNK10;refGene_tx=XM_003987901.5;refGene_gn=KCNK10;refGene_tx=XM_019833446.2;refGene_gn=KCNK10;refGene_tx=XM_023255918.1
B3	134173640	134173827
refGene_gn=KCNK10;refGene_tx=XM_003987901.5;refGene_gn=KCNK10;refGene_tx=XM_019833446.2;refGene_gn=KCNK10;refGene_tx=XM_023255918.1
B3	134202825	134202986
refGene_gn=KCNK10;refGene_tx=XM_003987901.5;refGene_gn=KCNK10;refGene_tx=XM_019833446.2;refGene_gn=KCNK10;refGene_tx=XM_023255918.1
B3	134215324	134215442
refGene_gn=KCNK10;refGene_tx=XM_003987901.5;refGene_gn=KCNK10;refGene_tx=XM_019833446.2;refGene_gn=KCNK10;refGene_tx=XM_023255918.1
B3	134235691	134236026
refGene_gn=KCNK10;refGene_tx=XM_003987901.5;refGene_gn=KCNK10;refGene_tx=XM_019833446.2;refGene_gn=KCNK10;refGene_tx=XM_023255918.1
B3	134291051	134291103
refGene_gn=KCNK10;refGene_tx=XM_003987901.5
B3	134338665	134338684
refGene_gn=SPATA7;refGene_tx=XM_003987902.4;refGene_gn=SPATA7;refGene_tx=XM_011283428.2;refGene_gn=SPATA7;refGene_tx=XM_023255920.1
B3	134345380	134345455
refGene_gn=SPATA7;refGene_tx=XM_003987902.4;refGene_gn=SPATA7;refGene_tx=XM_011283428.2
B3	134347293	134347389
refGene_gn=SPATA7;refGene_tx=XM_003987902.4
B3	134351485	134351533
refGene_gn=SPATA7;refGene_tx=XM_003987902.4;refGene_gn=SPATA7;refGene_tx=XM_011283428.2;refGene_gn=SPATA7;refGene_tx=XM_023255920.1
B3	134360891	134361025
refGene_gn=SPATA7;refGene_tx=XM_003987902.4;refGene_gn=SPATA7;refGene_tx=XM_011283428.2;refGene_gn=SPATA7;refGene_tx=XM_023255920.1
B3	134367502	134367975
refGene_gn=SPATA7;refGene_tx=XM_003987902.4;refGene_gn=SPATA7;refGene_tx=XM_011283428.2;refGene_gn=SPATA7;refGene_tx=XM_023255920.1
B3	134369098	134369165
refGene_gn=SPATA7;refGene_tx=XM_003987902.4;refGene_gn=SPATA7;refGene_tx=XM_011283428.2;refGene_gn=SPATA7;refGene_tx=XM_023255920.1
B3	134371577	134371693
refGene_gn=SPATA7;refGene_tx=XM_003987902.4;refGene_gn=SPATA7;refGene_tx=XM_011283428.2;refGene_gn=SPATA7;refGene_tx=XM_023255920.1
B3	134373112	134373166
refGene_gn=SPATA7;refGene_tx=XM_003987902.4;refGene_gn=SPATA7;refGene_tx=XM_011283428.2;refGene_gn=SPATA7;refGene_tx=XM_023255920.1
B3	134373959	134374037
refGene_gn=SPATA7;refGene_tx=XM_003987902.4;refGene_gn=SPATA7;refGene_tx=XM_011283428.2;refGene_gn=SPATA7;refGene_tx=XM_023255920.1
B3	134377063	134377121
refGene_gn=SPATA7;refGene_tx=XM_003987902.4;refGene_gn=SPATA7;refGene_

tx=XM_011283428.2;refGene_gn=SPATA7;refGene_tx=XM_023255920.1
B3 134377367 134377952
refGene_gn=SPATA7;refGene_tx=XM_003987902.4;refGene_gn=SPATA7;refGene_tx=XM_011283428.2;refGene_gn=SPATA7;refGene_tx=XM_023255920.1
B3 134392137 134392266
refGene_gn=PTPN21;refGene_tx=XM_023255921.1;refGene_gn=PTPN21;refGene_tx=XM_023255922.1;refGene_gn=PTPN21;refGene_tx=XM_023255923.1
B3 134392505 134392666
refGene_gn=PTPN21;refGene_tx=XM_023255921.1;refGene_gn=PTPN21;refGene_tx=XM_023255922.1;refGene_gn=PTPN21;refGene_tx=XM_023255923.1
B3 134393178 134393413
refGene_gn=PTPN21;refGene_tx=XM_023255921.1;refGene_gn=PTPN21;refGene_tx=XM_023255922.1;refGene_gn=PTPN21;refGene_tx=XM_023255923.1
B3 134393808 134393937
refGene_gn=PTPN21;refGene_tx=XM_023255921.1;refGene_gn=PTPN21;refGene_tx=XM_023255922.1;refGene_gn=PTPN21;refGene_tx=XM_023255923.1
B3 134395913 134396135
refGene_gn=PTPN21;refGene_tx=XM_023255921.1;refGene_gn=PTPN21;refGene_tx=XM_023255922.1;refGene_gn=PTPN21;refGene_tx=XM_023255923.1
B3 134397413 134397551
refGene_gn=PTPN21;refGene_tx=XM_023255921.1;refGene_gn=PTPN21;refGene_tx=XM_023255922.1;refGene_gn=PTPN21;refGene_tx=XM_023255923.1
B3 134402113 134403525
refGene_gn=PTPN21;refGene_tx=XM_023255921.1;refGene_gn=PTPN21;refGene_tx=XM_023255922.1;refGene_gn=PTPN21;refGene_tx=XM_023255923.1
B3 134407532 134407617
refGene_gn=PTPN21;refGene_tx=XM_023255921.1;refGene_gn=PTPN21;refGene_tx=XM_023255922.1;refGene_gn=PTPN21;refGene_tx=XM_023255923.1
B3 134408252 134408313
refGene_gn=PTPN21;refGene_tx=XM_023255921.1;refGene_gn=PTPN21;refGene_tx=XM_023255922.1;refGene_gn=PTPN21;refGene_tx=XM_023255923.1
B3 134418905 134418985
refGene_gn=PTPN21;refGene_tx=XM_023255921.1;refGene_gn=PTPN21;refGene_tx=XM_023255922.1;refGene_gn=PTPN21;refGene_tx=XM_023255923.1
B3 134419900 134419988
refGene_gn=PTPN21;refGene_tx=XM_023255921.1;refGene_gn=PTPN21;refGene_tx=XM_023255922.1;refGene_gn=PTPN21;refGene_tx=XM_023255923.1
B3 134423344 134423433
refGene_gn=PTPN21;refGene_tx=XM_023255921.1;refGene_gn=PTPN21;refGene_tx=XM_023255922.1;refGene_gn=PTPN21;refGene_tx=XM_023255923.1
B3 134423816 134423904
refGene_gn=PTPN21;refGene_tx=XM_023255921.1;refGene_gn=PTPN21;refGene_tx=XM_023255922.1;refGene_gn=PTPN21;refGene_tx=XM_023255923.1
B3 134426593 134426664
refGene_gn=PTPN21;refGene_tx=XM_023255921.1;refGene_gn=PTPN21;refGene_tx=XM_023255922.1;refGene_gn=PTPN21;refGene_tx=XM_023255923.1
B3 134427466 134427534
refGene_gn=PTPN21;refGene_tx=XM_023255921.1;refGene_gn=PTPN21;refGene_tx=XM_023255922.1;refGene_gn=PTPN21;refGene_tx=XM_023255923.1
B3 134429448 134429546

refGene_gn=PTPN21;refGene_tx=XM_023255921.1;refGene_gn=PTPN21;refGene_tx=XM_023255922.1;refGene_gn=PTPN21;refGene_tx=XM_023255923.1
B3 134439274 134439444
refGene_gn=PTPN21;refGene_tx=XM_023255921.1;refGene_gn=PTPN21;refGene_tx=XM_023255922.1;refGene_gn=PTPN21;refGene_tx=XM_023255923.1
B3 134468053 134468233
refGene_gn=PTPN21;refGene_tx=XM_023255921.1;refGene_gn=PTPN21;refGene_tx=XM_023255922.1;refGene_gn=PTPN21;refGene_tx=XM_023255923.1
B3 134480965 134481001
refGene_gn=ZC3H14;refGene_tx=XM_023255935.1;refGene_gn=ZC3H14;refGene_tx=XM_023255936.1;refGene_gn=ZC3H14;refGene_tx=XM_023255937.1;refGene_gn=ZC3H14;refGene_tx=XM_023255938.1
B3 134481466 134481509
refGene_gn=ZC3H14;refGene_tx=XM_023255935.1;refGene_gn=ZC3H14;refGene_tx=XM_023255936.1;refGene_gn=ZC3H14;refGene_tx=XM_023255937.1;refGene_gn=ZC3H14;refGene_tx=XM_023255938.1
B3 134490018 134490133
refGene_gn=ZC3H14;refGene_tx=XM_023255935.1;refGene_gn=ZC3H14;refGene_tx=XM_023255936.1;refGene_gn=ZC3H14;refGene_tx=XM_023255937.1;refGene_gn=ZC3H14;refGene_tx=XM_023255938.1
B3 134492177 134492218
refGene_gn=ZC3H14;refGene_tx=XM_023255935.1;refGene_gn=ZC3H14;refGene_tx=XM_023255936.1;refGene_gn=ZC3H14;refGene_tx=XM_023255937.1;refGene_gn=ZC3H14;refGene_tx=XM_023255938.1
B3 134493091 134493287
refGene_gn=ZC3H14;refGene_tx=XM_023255935.1;refGene_gn=ZC3H14;refGene_tx=XM_023255936.1;refGene_gn=ZC3H14;refGene_tx=XM_023255937.1;refGene_gn=ZC3H14;refGene_tx=XM_023255938.1
B3 134493628 134494058
refGene_gn=ZC3H14;refGene_tx=XM_023255935.1;refGene_gn=ZC3H14;refGene_tx=XM_023255936.1;refGene_gn=ZC3H14;refGene_tx=XM_023255937.1;refGene_gn=ZC3H14;refGene_tx=XM_023255938.1
B3 134494749 134494910
refGene_gn=ZC3H14;refGene_tx=XM_023255935.1;refGene_gn=ZC3H14;refGene_tx=XM_023255936.1;refGene_gn=ZC3H14;refGene_tx=XM_023255937.1;refGene_gn=ZC3H14;refGene_tx=XM_023255938.1
B3 134495276 134495377
refGene_gn=ZC3H14;refGene_tx=XM_023255935.1;refGene_gn=ZC3H14;refGene_tx=XM_023255936.1;refGene_gn=ZC3H14;refGene_tx=XM_023255937.1;refGene_gn=ZC3H14;refGene_tx=XM_023255938.1
B3 134498439 134498595
refGene_gn=ZC3H14;refGene_tx=XM_023255935.1;refGene_gn=ZC3H14;refGene_tx=XM_023255936.1;refGene_gn=ZC3H14;refGene_tx=XM_023255937.1;refGene_gn=ZC3H14;refGene_tx=XM_023255938.1
B3 134510187 134510581
refGene_gn=ZC3H14;refGene_tx=XM_006933048.3
B3 134514952 134515027
refGene_gn=ZC3H14;refGene_tx=XM_006933048.3;refGene_gn=ZC3H14;refGene_tx=XM_023255935.1;refGene_gn=ZC3H14;refGene_tx=XM_023255936.1;refGene_gn=ZC3H14;refGene_tx=XM_023255937.1;refGene_gn=ZC3H14;refGene_tx=XM_02

3255938.1
B3 134519590 134519750
refGene_gn=ZC3H14;refGene_tx=XM_023255935.1;refGene_gn=ZC3H14;refGene_tx=XM_023255936.1
B3 134520513 134520746
refGene_gn=ZC3H14;refGene_tx=XM_023255935.1;refGene_gn=ZC3H14;refGene_tx=XM_023255936.1
B3 134524156 134524277
refGene_gn=ZC3H14;refGene_tx=XM_006933048.3;refGene_gn=ZC3H14;refGene_tx=XM_023255935.1;refGene_gn=ZC3H14;refGene_tx=XM_023255936.1;refGene_gn=ZC3H14;refGene_tx=XM_023255937.1;refGene_gn=ZC3H14;refGene_tx=XM_023255938.1
B3 134525435 134525572
refGene_gn=ZC3H14;refGene_tx=XM_006933048.3;refGene_gn=ZC3H14;refGene_tx=XM_023255935.1;refGene_gn=ZC3H14;refGene_tx=XM_023255936.1;refGene_gn=ZC3H14;refGene_tx=XM_023255937.1;refGene_gn=ZC3H14;refGene_tx=XM_023255938.1
B3 134526208 134526300
refGene_gn=ZC3H14;refGene_tx=XM_006933048.3;refGene_gn=ZC3H14;refGene_tx=XM_023255935.1;refGene_gn=ZC3H14;refGene_tx=XM_023255936.1;refGene_gn=ZC3H14;refGene_tx=XM_023255937.1;refGene_gn=ZC3H14;refGene_tx=XM_023255938.1
B3 134527034 134527141
refGene_gn=ZC3H14;refGene_tx=XM_006933048.3;refGene_gn=ZC3H14;refGene_tx=XM_023255935.1;refGene_gn=ZC3H14;refGene_tx=XM_023255936.1;refGene_gn=ZC3H14;refGene_tx=XM_023255937.1;refGene_gn=ZC3H14;refGene_tx=XM_023255938.1
B3 134527525 134527532
refGene_gn=ZC3H14;refGene_tx=XM_006933048.3;refGene_gn=ZC3H14;refGene_tx=XM_023255935.1;refGene_gn=ZC3H14;refGene_tx=XM_023255936.1;refGene_gn=ZC3H14;refGene_tx=XM_023255937.1;refGene_gn=ZC3H14;refGene_tx=XM_023255938.1
B3 134531425 134531462
refGene_gn=EML5;refGene_tx=XM_023255924.1;refGene_gn=EML5;refGene_tx=XM_023255925.1;refGene_gn=EML5;refGene_tx=XM_023255926.1;refGene_gn=EML5;refGene_tx=XM_023255927.1;refGene_gn=EML5;refGene_tx=XM_023255928.1;refGene_gn=EML5;refGene_tx=XM_023255930.1;refGene_gn=EML5;refGene_tx=XM_023255931.1;refGene_gn=EML5;refGene_tx=XM_023255933.1
B3 134531682 134531783
refGene_gn=EML5;refGene_tx=XM_023255924.1;refGene_gn=EML5;refGene_tx=XM_023255925.1;refGene_gn=EML5;refGene_tx=XM_023255926.1;refGene_gn=EML5;refGene_tx=XM_023255927.1;refGene_gn=EML5;refGene_tx=XM_023255928.1;refGene_gn=EML5;refGene_tx=XM_023255930.1;refGene_gn=EML5;refGene_tx=XM_023255931.1;refGene_gn=EML5;refGene_tx=XM_023255933.1
B3 134532495 134532649
refGene_gn=EML5;refGene_tx=XM_023255924.1;refGene_gn=EML5;refGene_tx=XM_023255925.1;refGene_gn=EML5;refGene_tx=XM_023255926.1;refGene_gn=EML5;refGene_tx=XM_023255927.1;refGene_gn=EML5;refGene_tx=XM_023255928.1;refGene_gn=EML5;refGene_tx=XM_023255930.1;refGene_gn=EML5;refGene_tx=XM_023255931.1;refGene_gn=EML5;refGene_tx=XM_023255933.1

B3	134533137	134533241
refGene_gn=EML5;refGene_tx=XM_023255924.1;refGene_gn=EML5;refGene_tx=XM_023255925.1;refGene_gn=EML5;refGene_tx=XM_023255926.1;refGene_gn=EML5;refGene_tx=XM_023255927.1;refGene_gn=EML5;refGene_tx=XM_023255928.1;refGene_gn=EML5;refGene_tx=XM_023255930.1;refGene_gn=EML5;refGene_tx=XM_023255931.1;refGene_gn=EML5;refGene_tx=XM_023255933.1
B3	134533664	134533827
refGene_gn=EML5;refGene_tx=XM_023255924.1;refGene_gn=EML5;refGene_tx=XM_023255925.1;refGene_gn=EML5;refGene_tx=XM_023255926.1;refGene_gn=EML5;refGene_tx=XM_023255927.1;refGene_gn=EML5;refGene_tx=XM_023255928.1;refGene_gn=EML5;refGene_tx=XM_023255930.1;refGene_gn=EML5;refGene_tx=XM_023255931.1;refGene_gn=EML5;refGene_tx=XM_023255933.1
B3	134535123	134535296
refGene_gn=EML5;refGene_tx=XM_023255924.1;refGene_gn=EML5;refGene_tx=XM_023255925.1;refGene_gn=EML5;refGene_tx=XM_023255926.1;refGene_gn=EML5;refGene_tx=XM_023255927.1;refGene_gn=EML5;refGene_tx=XM_023255928.1;refGene_gn=EML5;refGene_tx=XM_023255930.1;refGene_gn=EML5;refGene_tx=XM_023255931.1;refGene_gn=EML5;refGene_tx=XM_023255932.1;refGene_gn=EML5;refGene_tx=XM_023255933.1
B3	134535470	134535663
refGene_gn=EML5;refGene_tx=XM_023255924.1;refGene_gn=EML5;refGene_tx=XM_023255925.1;refGene_gn=EML5;refGene_tx=XM_023255926.1;refGene_gn=EML5;refGene_tx=XM_023255927.1;refGene_gn=EML5;refGene_tx=XM_023255928.1;refGene_gn=EML5;refGene_tx=XM_023255930.1;refGene_gn=EML5;refGene_tx=XM_023255931.1;refGene_gn=EML5;refGene_tx=XM_023255932.1;refGene_gn=EML5;refGene_tx=XM_023255933.1
B3	134537150	134537265
refGene_gn=EML5;refGene_tx=XM_023255924.1;refGene_gn=EML5;refGene_tx=XM_023255925.1;refGene_gn=EML5;refGene_tx=XM_023255926.1;refGene_gn=EML5;refGene_tx=XM_023255927.1;refGene_gn=EML5;refGene_tx=XM_023255928.1;refGene_gn=EML5;refGene_tx=XM_023255930.1;refGene_gn=EML5;refGene_tx=XM_023255931.1;refGene_gn=EML5;refGene_tx=XM_023255932.1;refGene_gn=EML5;refGene_tx=XM_023255933.1
B3	134538672	134538830
refGene_gn=EML5;refGene_tx=XM_023255924.1;refGene_gn=EML5;refGene_tx=XM_023255925.1;refGene_gn=EML5;refGene_tx=XM_023255926.1;refGene_gn=EML5;refGene_tx=XM_023255927.1;refGene_gn=EML5;refGene_tx=XM_023255928.1;refGene_gn=EML5;refGene_tx=XM_023255930.1;refGene_gn=EML5;refGene_tx=XM_023255931.1;refGene_gn=EML5;refGene_tx=XM_023255932.1;refGene_gn=EML5;refGene_tx=XM_023255933.1;refGene_gn=EML5;refGene_tx=XM_023255934.1
B3	134541054	134541263
refGene_gn=EML5;refGene_tx=XM_023255924.1;refGene_gn=EML5;refGene_tx=XM_023255925.1;refGene_gn=EML5;refGene_tx=XM_023255926.1;refGene_gn=EML5;refGene_tx=XM_023255927.1;refGene_gn=EML5;refGene_tx=XM_023255928.1;refGene_gn=EML5;refGene_tx=XM_023255930.1;refGene_gn=EML5;refGene_tx=XM_023255931.1;refGene_gn=EML5;refGene_tx=XM_023255932.1;refGene_gn=EML5;refGene_tx=XM_023255933.1;refGene_gn=EML5;refGene_tx=XM_023255934.1
B3	134542197	134542371
refGene_gn=EML5;refGene_tx=XM_023255924.1;refGene_gn=EML5;refGene_tx=XM_023255925.1;refGene_gn=EML5;refGene_tx=XM_023255926.1;refGene_gn=EML

5;refGene_tx=XM_023255927.1;refGene_gn=EML5;refGene_tx=XM_023255928.1;
refGene_gn=EML5;refGene_tx=XM_023255930.1;refGene_gn=EML5;refGene_tx=X
M_023255931.1;refGene_gn=EML5;refGene_tx=XM_023255932.1;refGene_gn=EML
5;refGene_tx=XM_023255933.1;refGene_gn=EML5;refGene_tx=XM_023255934.1
B3 134552292 134552313
refGene_gn=EML5;refGene_tx=XM_023255924.1;refGene_gn=EML5;refGene_tx=X
M_023255925.1;refGene_gn=EML5;refGene_tx=XM_023255927.1;refGene_gn=EML
5;refGene_tx=XM_023255930.1;refGene_gn=EML5;refGene_tx=XM_023255931.1;
refGene_gn=EML5;refGene_tx=XM_023255932.1;refGene_gn=EML5;refGene_tx=X
M_023255933.1;refGene_gn=EML5;refGene_tx=XM_023255934.1
B3 134554825 134554924
refGene_gn=EML5;refGene_tx=XM_023255924.1;refGene_gn=EML5;refGene_tx=X
M_023255925.1;refGene_gn=EML5;refGene_tx=XM_023255926.1;refGene_gn=EML
5;refGene_tx=XM_023255927.1;refGene_gn=EML5;refGene_tx=XM_023255928.1;
refGene_gn=EML5;refGene_tx=XM_023255930.1;refGene_gn=EML5;refGene_tx=X
M_023255931.1;refGene_gn=EML5;refGene_tx=XM_023255932.1;refGene_gn=EML
5;refGene_tx=XM_023255933.1;refGene_gn=EML5;refGene_tx=XM_023255934.1
B3 134557247 134557377
refGene_gn=EML5;refGene_tx=XM_023255924.1;refGene_gn=EML5;refGene_tx=X
M_023255925.1;refGene_gn=EML5;refGene_tx=XM_023255926.1;refGene_gn=EML
5;refGene_tx=XM_023255927.1;refGene_gn=EML5;refGene_tx=XM_023255928.1;
refGene_gn=EML5;refGene_tx=XM_023255930.1;refGene_gn=EML5;refGene_tx=X
M_023255931.1;refGene_gn=EML5;refGene_tx=XM_023255932.1;refGene_gn=EML
5;refGene_tx=XM_023255933.1;refGene_gn=EML5;refGene_tx=XM_023255934.1
B3 134558564 134558643
refGene_gn=EML5;refGene_tx=XM_023255924.1;refGene_gn=EML5;refGene_tx=X
M_023255925.1;refGene_gn=EML5;refGene_tx=XM_023255926.1;refGene_gn=EML
5;refGene_tx=XM_023255927.1;refGene_gn=EML5;refGene_tx=XM_023255928.1;
refGene_gn=EML5;refGene_tx=XM_023255930.1;refGene_gn=EML5;refGene_tx=X
M_023255931.1;refGene_gn=EML5;refGene_tx=XM_023255932.1;refGene_gn=EML
5;refGene_tx=XM_023255933.1;refGene_gn=EML5;refGene_tx=XM_023255934.1
B3 134562264 134562273
refGene_gn=EML5;refGene_tx=XM_023255924.1;refGene_gn=EML5;refGene_tx=X
M_023255926.1;refGene_gn=EML5;refGene_tx=XM_023255930.1;refGene_gn=EML
5;refGene_tx=XM_023255931.1;refGene_gn=EML5;refGene_tx=XM_023255932.1;
refGene_gn=EML5;refGene_tx=XM_023255933.1;refGene_gn=EML5;refGene_tx=X
M_023255934.1
B3 134564189 134564204
refGene_gn=EML5;refGene_tx=XM_023255924.1;refGene_gn=EML5;refGene_tx=X
M_023255925.1;refGene_gn=EML5;refGene_tx=XM_023255926.1;refGene_gn=EML
5;refGene_tx=XM_023255930.1;refGene_gn=EML5;refGene_tx=XM_023255931.1;
refGene_gn=EML5;refGene_tx=XM_023255932.1;refGene_gn=EML5;refGene_tx=X
M_023255933.1;refGene_gn=EML5;refGene_tx=XM_023255934.1
B3 134568252 134568379
refGene_gn=EML5;refGene_tx=XM_023255924.1;refGene_gn=EML5;refGene_tx=X
M_023255925.1;refGene_gn=EML5;refGene_tx=XM_023255926.1;refGene_gn=EML
5;refGene_tx=XM_023255927.1;refGene_gn=EML5;refGene_tx=XM_023255928.1;
refGene_gn=EML5;refGene_tx=XM_023255930.1;refGene_gn=EML5;refGene_tx=X
M_023255931.1;refGene_gn=EML5;refGene_tx=XM_023255932.1;refGene_gn=EML
5;refGene_tx=XM_023255933.1;refGene_gn=EML5;refGene_tx=XM_023255934.1

B3	134569057	134569259
refGene_gn=EML5;refGene_tx=XM_023255924.1;refGene_gn=EML5;refGene_tx=XM_023255925.1;refGene_gn=EML5;refGene_tx=XM_023255926.1;refGene_gn=EML5;refGene_tx=XM_023255927.1;refGene_gn=EML5;refGene_tx=XM_023255928.1;refGene_gn=EML5;refGene_tx=XM_023255930.1;refGene_gn=EML5;refGene_tx=XM_023255931.1;refGene_gn=EML5;refGene_tx=XM_023255932.1;refGene_gn=EML5;refGene_tx=XM_023255933.1;refGene_gn=EML5;refGene_tx=XM_023255934.1
B3	134572042	134572219
refGene_gn=EML5;refGene_tx=XM_023255924.1;refGene_gn=EML5;refGene_tx=XM_023255925.1;refGene_gn=EML5;refGene_tx=XM_023255926.1;refGene_gn=EML5;refGene_tx=XM_023255927.1;refGene_gn=EML5;refGene_tx=XM_023255928.1;refGene_gn=EML5;refGene_tx=XM_023255930.1;refGene_gn=EML5;refGene_tx=XM_023255931.1;refGene_gn=EML5;refGene_tx=XM_023255932.1;refGene_gn=EML5;refGene_tx=XM_023255933.1;refGene_gn=EML5;refGene_tx=XM_023255934.1
B3	134573290	134573379
refGene_gn=EML5;refGene_tx=XM_023255924.1;refGene_gn=EML5;refGene_tx=XM_023255925.1;refGene_gn=EML5;refGene_tx=XM_023255926.1;refGene_gn=EML5;refGene_tx=XM_023255927.1;refGene_gn=EML5;refGene_tx=XM_023255928.1;refGene_gn=EML5;refGene_tx=XM_023255930.1;refGene_gn=EML5;refGene_tx=XM_023255931.1;refGene_gn=EML5;refGene_tx=XM_023255932.1;refGene_gn=EML5;refGene_tx=XM_023255933.1;refGene_gn=EML5;refGene_tx=XM_023255934.1
B3	134588023	134588155
refGene_gn=EML5;refGene_tx=XM_023255924.1;refGene_gn=EML5;refGene_tx=XM_023255925.1;refGene_gn=EML5;refGene_tx=XM_023255926.1;refGene_gn=EML5;refGene_tx=XM_023255927.1;refGene_gn=EML5;refGene_tx=XM_023255928.1;refGene_gn=EML5;refGene_tx=XM_023255930.1;refGene_gn=EML5;refGene_tx=XM_023255931.1;refGene_gn=EML5;refGene_tx=XM_023255932.1;refGene_gn=EML5;refGene_tx=XM_023255933.1;refGene_gn=EML5;refGene_tx=XM_023255934.1
B3	134588882	134589035
refGene_gn=EML5;refGene_tx=XM_023255924.1;refGene_gn=EML5;refGene_tx=XM_023255925.1;refGene_gn=EML5;refGene_tx=XM_023255926.1;refGene_gn=EML5;refGene_tx=XM_023255927.1;refGene_gn=EML5;refGene_tx=XM_023255928.1;refGene_gn=EML5;refGene_tx=XM_023255930.1;refGene_gn=EML5;refGene_tx=XM_023255931.1;refGene_gn=EML5;refGene_tx=XM_023255932.1;refGene_gn=EML5;refGene_tx=XM_023255933.1;refGene_gn=EML5;refGene_tx=XM_023255934.1
B3	134607241	134607383
refGene_gn=EML5;refGene_tx=XM_023255924.1;refGene_gn=EML5;refGene_tx=XM_023255925.1;refGene_gn=EML5;refGene_tx=XM_023255926.1;refGene_gn=EML5;refGene_tx=XM_023255927.1;refGene_gn=EML5;refGene_tx=XM_023255928.1;refGene_gn=EML5;refGene_tx=XM_023255930.1;refGene_gn=EML5;refGene_tx=XM_023255931.1;refGene_gn=EML5;refGene_tx=XM_023255932.1;refGene_gn=EML5;refGene_tx=XM_023255933.1;refGene_gn=EML5;refGene_tx=XM_023255934.1
B3	134612917	134613045
refGene_gn=EML5;refGene_tx=XM_023255924.1;refGene_gn=EML5;refGene_tx=XM_023255925.1;refGene_gn=EML5;refGene_tx=XM_023255926.1;refGene_gn=EML5;refGene_tx=XM_023255927.1;refGene_gn=EML5;refGene_tx=XM_023255928.1;refGene_gn=EML5;refGene_tx=XM_023255930.1;refGene_gn=EML5;refGene_tx=XM_023255931.1;refGene_gn=EML5;refGene_tx=XM_023255932.1;refGene_gn=EML5;refGene_tx=XM_023255933.1;refGene_gn=EML5;refGene_tx=XM_023255934.1
B3	134614330	134614442

refGene_gn=EML5;refGene_tx=XM_023255924.1;refGene_gn=EML5;refGene_tx=XM_023255925.1;refGene_gn=EML5;refGene_tx=XM_023255926.1;refGene_gn=EML5;refGene_tx=XM_023255927.1;refGene_gn=EML5;refGene_tx=XM_023255928.1;refGene_gn=EML5;refGene_tx=XM_023255930.1;refGene_gn=EML5;refGene_tx=XM_023255931.1;refGene_gn=EML5;refGene_tx=XM_023255932.1;refGene_gn=EML5;refGene_tx=XM_023255933.1;refGene_gn=EML5;refGene_tx=XM_023255934.1
B3 134615057 134615260
refGene_gn=EML5;refGene_tx=XM_023255924.1;refGene_gn=EML5;refGene_tx=XM_023255925.1;refGene_gn=EML5;refGene_tx=XM_023255926.1;refGene_gn=EML5;refGene_tx=XM_023255927.1;refGene_gn=EML5;refGene_tx=XM_023255928.1;refGene_gn=EML5;refGene_tx=XM_023255930.1;refGene_gn=EML5;refGene_tx=XM_023255931.1;refGene_gn=EML5;refGene_tx=XM_023255932.1;refGene_gn=EML5;refGene_tx=XM_023255933.1;refGene_gn=EML5;refGene_tx=XM_023255934.1
B3 134620582 134620683
refGene_gn=EML5;refGene_tx=XM_023255924.1;refGene_gn=EML5;refGene_tx=XM_023255925.1;refGene_gn=EML5;refGene_tx=XM_023255926.1;refGene_gn=EML5;refGene_tx=XM_023255927.1;refGene_gn=EML5;refGene_tx=XM_023255928.1;refGene_gn=EML5;refGene_tx=XM_023255930.1;refGene_gn=EML5;refGene_tx=XM_023255931.1;refGene_gn=EML5;refGene_tx=XM_023255932.1;refGene_gn=EML5;refGene_tx=XM_023255933.1;refGene_gn=EML5;refGene_tx=XM_023255934.1
B3 134621626 134621720
refGene_gn=EML5;refGene_tx=XM_023255924.1;refGene_gn=EML5;refGene_tx=XM_023255925.1;refGene_gn=EML5;refGene_tx=XM_023255926.1;refGene_gn=EML5;refGene_tx=XM_023255927.1;refGene_gn=EML5;refGene_tx=XM_023255928.1;refGene_gn=EML5;refGene_tx=XM_023255930.1;refGene_gn=EML5;refGene_tx=XM_023255931.1;refGene_gn=EML5;refGene_tx=XM_023255932.1;refGene_gn=EML5;refGene_tx=XM_023255933.1;refGene_gn=EML5;refGene_tx=XM_023255934.1
B3 134623261 134623367
refGene_gn=EML5;refGene_tx=XM_023255924.1;refGene_gn=EML5;refGene_tx=XM_023255925.1;refGene_gn=EML5;refGene_tx=XM_023255926.1;refGene_gn=EML5;refGene_tx=XM_023255927.1;refGene_gn=EML5;refGene_tx=XM_023255928.1;refGene_gn=EML5;refGene_tx=XM_023255930.1;refGene_gn=EML5;refGene_tx=XM_023255931.1;refGene_gn=EML5;refGene_tx=XM_023255932.1;refGene_gn=EML5;refGene_tx=XM_023255933.1;refGene_gn=EML5;refGene_tx=XM_023255934.1
B3 134629597 134629784
refGene_gn=EML5;refGene_tx=XM_023255924.1;refGene_gn=EML5;refGene_tx=XM_023255925.1;refGene_gn=EML5;refGene_tx=XM_023255926.1;refGene_gn=EML5;refGene_tx=XM_023255927.1;refGene_gn=EML5;refGene_tx=XM_023255928.1;refGene_gn=EML5;refGene_tx=XM_023255930.1;refGene_gn=EML5;refGene_tx=XM_023255931.1;refGene_gn=EML5;refGene_tx=XM_023255932.1;refGene_gn=EML5;refGene_tx=XM_023255933.1;refGene_gn=EML5;refGene_tx=XM_023255934.1
B3 134630965 134631084
refGene_gn=EML5;refGene_tx=XM_023255924.1;refGene_gn=EML5;refGene_tx=XM_023255925.1;refGene_gn=EML5;refGene_tx=XM_023255926.1;refGene_gn=EML5;refGene_tx=XM_023255927.1;refGene_gn=EML5;refGene_tx=XM_023255928.1;refGene_gn=EML5;refGene_tx=XM_023255930.1;refGene_gn=EML5;refGene_tx=XM_023255931.1;refGene_gn=EML5;refGene_tx=XM_023255932.1;refGene_gn=EML5;refGene_tx=XM_023255933.1;refGene_gn=EML5;refGene_tx=XM_023255934.1
B3 134634218 134634325
refGene_gn=EML5;refGene_tx=XM_023255924.1;refGene_gn=EML5;refGene_tx=X

M_023255925.1;refGene_gn=EML5;refGene_tx=XM_023255926.1;refGene_gn=EML5;refGene_tx=XM_023255927.1;refGene_gn=EML5;refGene_tx=XM_023255928.1;refGene_gn=EML5;refGene_tx=XM_023255930.1;refGene_gn=EML5;refGene_tx=XM_023255931.1;refGene_gn=EML5;refGene_tx=XM_023255932.1;refGene_gn=EML5;refGene_tx=XM_023255933.1;refGene_gn=EML5;refGene_tx=XM_023255934.1
B3 134635181 134635349
refGene_gn=EML5;refGene_tx=XM_023255924.1;refGene_gn=EML5;refGene_tx=XM_023255925.1;refGene_gn=EML5;refGene_tx=XM_023255926.1;refGene_gn=EML5;refGene_tx=XM_023255927.1;refGene_gn=EML5;refGene_tx=XM_023255928.1;refGene_gn=EML5;refGene_tx=XM_023255930.1;refGene_gn=EML5;refGene_tx=XM_023255931.1;refGene_gn=EML5;refGene_tx=XM_023255932.1;refGene_gn=EML5;refGene_tx=XM_023255933.1;refGene_gn=EML5;refGene_tx=XM_023255934.1
B3 134643678 134643891
refGene_gn=EML5;refGene_tx=XM_023255924.1;refGene_gn=EML5;refGene_tx=XM_023255925.1;refGene_gn=EML5;refGene_tx=XM_023255926.1;refGene_gn=EML5;refGene_tx=XM_023255927.1;refGene_gn=EML5;refGene_tx=XM_023255928.1;refGene_gn=EML5;refGene_tx=XM_023255930.1;refGene_gn=EML5;refGene_tx=XM_023255931.1;refGene_gn=EML5;refGene_tx=XM_023255932.1;refGene_gn=EML5;refGene_tx=XM_023255933.1;refGene_gn=EML5;refGene_tx=XM_023255934.1
B3 134644988 134645245
refGene_gn=EML5;refGene_tx=XM_023255924.1;refGene_gn=EML5;refGene_tx=XM_023255925.1;refGene_gn=EML5;refGene_tx=XM_023255926.1;refGene_gn=EML5;refGene_tx=XM_023255927.1;refGene_gn=EML5;refGene_tx=XM_023255928.1;refGene_gn=EML5;refGene_tx=XM_023255930.1;refGene_gn=EML5;refGene_tx=XM_023255931.1;refGene_gn=EML5;refGene_tx=XM_023255932.1;refGene_gn=EML5;refGene_tx=XM_023255933.1;refGene_gn=EML5;refGene_tx=XM_023255934.1
B3 134649785 134649923
refGene_gn=EML5;refGene_tx=XM_023255924.1;refGene_gn=EML5;refGene_tx=XM_023255925.1;refGene_gn=EML5;refGene_tx=XM_023255926.1;refGene_gn=EML5;refGene_tx=XM_023255927.1;refGene_gn=EML5;refGene_tx=XM_023255928.1;refGene_gn=EML5;refGene_tx=XM_023255931.1;refGene_gn=EML5;refGene_tx=XM_023255932.1;refGene_gn=EML5;refGene_tx=XM_023255933.1;refGene_gn=EML5;refGene_tx=XM_023255934.1
B3 134668392 134668594
refGene_gn=EML5;refGene_tx=XM_023255924.1;refGene_gn=EML5;refGene_tx=XM_023255925.1;refGene_gn=EML5;refGene_tx=XM_023255926.1;refGene_gn=EML5;refGene_tx=XM_023255927.1;refGene_gn=EML5;refGene_tx=XM_023255928.1;refGene_gn=EML5;refGene_tx=XM_023255930.1;refGene_gn=EML5;refGene_tx=XM_023255931.1;refGene_gn=EML5;refGene_tx=XM_023255932.1;refGene_gn=EML5;refGene_tx=XM_023255933.1;refGene_gn=EML5;refGene_tx=XM_023255934.1
B3 134669869 134670005
refGene_gn=EML5;refGene_tx=XM_023255924.1;refGene_gn=EML5;refGene_tx=XM_023255925.1;refGene_gn=EML5;refGene_tx=XM_023255926.1;refGene_gn=EML5;refGene_tx=XM_023255927.1;refGene_gn=EML5;refGene_tx=XM_023255928.1;refGene_gn=EML5;refGene_tx=XM_023255930.1;refGene_gn=EML5;refGene_tx=XM_023255931.1;refGene_gn=EML5;refGene_tx=XM_023255932.1;refGene_gn=EML5;refGene_tx=XM_023255933.1;refGene_gn=EML5;refGene_tx=XM_023255934.1
B3 134671457 134671643
refGene_gn=EML5;refGene_tx=XM_023255924.1;refGene_gn=EML5;refGene_tx=XM_023255925.1;refGene_gn=EML5;refGene_tx=XM_023255926.1;refGene_gn=EML

5;refGene_tx=XM_023255927.1;refGene_gn=EML5;refGene_tx=XM_023255928.1;
refGene_gn=EML5;refGene_tx=XM_023255930.1;refGene_gn=EML5;refGene_tx=X
M_023255931.1;refGene_gn=EML5;refGene_tx=XM_023255932.1;refGene_gn=EML
5;refGene_tx=XM_023255934.1
B3 134674646 134674715
refGene_gn=EML5;refGene_tx=XM_023255924.1;refGene_gn=EML5;refGene_tx=X
M_023255925.1;refGene_gn=EML5;refGene_tx=XM_023255926.1;refGene_gn=EML
5;refGene_tx=XM_023255927.1;refGene_gn=EML5;refGene_tx=XM_023255928.1;
refGene_gn=EML5;refGene_tx=XM_023255930.1;refGene_gn=EML5;refGene_tx=X
M_023255931.1;refGene_gn=EML5;refGene_tx=XM_023255932.1;refGene_gn=EML
5;refGene_tx=XM_023255934.1
B3 134677272 134677371
refGene_gn=EML5;refGene_tx=XM_023255924.1;refGene_gn=EML5;refGene_tx=X
M_023255925.1;refGene_gn=EML5;refGene_tx=XM_023255926.1;refGene_gn=EML
5;refGene_tx=XM_023255927.1;refGene_gn=EML5;refGene_tx=XM_023255928.1;
refGene_gn=EML5;refGene_tx=XM_023255930.1;refGene_gn=EML5;refGene_tx=X
M_023255931.1;refGene_gn=EML5;refGene_tx=XM_023255932.1;refGene_gn=EML
5;refGene_tx=XM_023255934.1
B3 134686144 134686304
refGene_gn=EML5;refGene_tx=XM_023255924.1;refGene_gn=EML5;refGene_tx=X
M_023255925.1;refGene_gn=EML5;refGene_tx=XM_023255926.1;refGene_gn=EML
5;refGene_tx=XM_023255927.1;refGene_gn=EML5;refGene_tx=XM_023255928.1;
refGene_gn=EML5;refGene_tx=XM_023255930.1;refGene_gn=EML5;refGene_tx=X
M_023255932.1;refGene_gn=EML5;refGene_tx=XM_023255934.1
B3 134712956 134713153
refGene_gn=EML5;refGene_tx=XM_023255924.1;refGene_gn=EML5;refGene_tx=X
M_023255925.1;refGene_gn=EML5;refGene_tx=XM_023255926.1;refGene_gn=EML
5;refGene_tx=XM_023255927.1;refGene_gn=EML5;refGene_tx=XM_023255928.1;
refGene_gn=EML5;refGene_tx=XM_023255930.1;refGene_gn=EML5;refGene_tx=X
M_023255932.1;refGene_gn=EML5;refGene_tx=XM_023255934.1
B3 134743655 134743769
refGene_gn=TTC8;refGene_tx=XM_011283430.3;refGene_gn=TTC8;refGene_tx=X
M_011283431.3;refGene_gn=TTC8;refGene_tx=XM_011283432.3;refGene_gn=TTC
8;refGene_tx=XM_011283433.3;refGene_gn=TTC8;refGene_tx=XM_011283436.3
B3 134755136 134755166
refGene_gn=TTC8;refGene_tx=XM_011283430.3;refGene_gn=TTC8;refGene_tx=X
M_011283432.3;refGene_gn=TTC8;refGene_tx=XM_011283436.3
B3 134760504 134760625
refGene_gn=TTC8;refGene_tx=XM_011283430.3;refGene_gn=TTC8;refGene_tx=X
M_011283431.3;refGene_gn=TTC8;refGene_tx=XM_011283432.3;refGene_gn=TTC
8;refGene_tx=XM_011283433.3;refGene_gn=TTC8;refGene_tx=XM_011283434.2;
refGene_gn=TTC8;refGene_tx=XM_011283436.3
B3 134761968 134762032
refGene_gn=TTC8;refGene_tx=XM_011283430.3;refGene_gn=TTC8;refGene_tx=X
M_011283431.3;refGene_gn=TTC8;refGene_tx=XM_011283432.3;refGene_gn=TTC
8;refGene_tx=XM_011283433.3;refGene_gn=TTC8;refGene_tx=XM_011283434.2;
refGene_gn=TTC8;refGene_tx=XM_011283436.3
B3 134762139 134762299
refGene_gn=TTC8;refGene_tx=XM_011283430.3;refGene_gn=TTC8;refGene_tx=X
M_011283431.3;refGene_gn=TTC8;refGene_tx=XM_011283432.3;refGene_gn=TTC

8;refGene_tx=XM_011283433.3;refGene_gn=TTC8;refGene_tx=XM_011283434.2;
refGene_gn=TTC8;refGene_tx=XM_011283436.3
B3 134762495 134762585
refGene_gn=TTC8;refGene_tx=XM_011283430.3;refGene_gn=TTC8;refGene_tx=X
M_011283431.3;refGene_gn=TTC8;refGene_tx=XM_011283434.2;refGene_gn=TTC
8;refGene_tx=XM_011283436.3
B3 134764701 134764746
refGene_gn=TTC8;refGene_tx=XM_011283430.3;refGene_gn=TTC8;refGene_tx=X
M_011283431.3;refGene_gn=TTC8;refGene_tx=XM_011283432.3;refGene_gn=TTC
8;refGene_tx=XM_011283433.3;refGene_gn=TTC8;refGene_tx=XM_011283434.2;
refGene_gn=TTC8;refGene_tx=XM_011283436.3
B3 134773362 134773448
refGene_gn=TTC8;refGene_tx=XM_011283430.3;refGene_gn=TTC8;refGene_tx=X
M_011283431.3;refGene_gn=TTC8;refGene_tx=XM_011283432.3;refGene_gn=TTC
8;refGene_tx=XM_011283433.3;refGene_gn=TTC8;refGene_tx=XM_011283434.2
B3 134777671 134777759
refGene_gn=TTC8;refGene_tx=XM_011283430.3;refGene_gn=TTC8;refGene_tx=X
M_011283431.3;refGene_gn=TTC8;refGene_tx=XM_011283432.3;refGene_gn=TTC
8;refGene_tx=XM_011283433.3;refGene_gn=TTC8;refGene_tx=XM_011283434.2
B3 134780115 134780226
refGene_gn=TTC8;refGene_tx=XM_011283430.3;refGene_gn=TTC8;refGene_tx=X
M_011283431.3;refGene_gn=TTC8;refGene_tx=XM_011283432.3;refGene_gn=TTC
8;refGene_tx=XM_011283433.3;refGene_gn=TTC8;refGene_tx=XM_011283434.2
B3 134794159 134794299
refGene_gn=TTC8;refGene_tx=XM_011283430.3;refGene_gn=TTC8;refGene_tx=X
M_011283431.3;refGene_gn=TTC8;refGene_tx=XM_011283432.3;refGene_gn=TTC
8;refGene_tx=XM_011283433.3;refGene_gn=TTC8;refGene_tx=XM_011283434.2;
refGene_gn=TTC8;refGene_tx=XM_011283436.3
B3 134795576 134795751
refGene_gn=TTC8;refGene_tx=XM_011283430.3;refGene_gn=TTC8;refGene_tx=X
M_011283431.3;refGene_gn=TTC8;refGene_tx=XM_011283432.3;refGene_gn=TTC
8;refGene_tx=XM_011283433.3;refGene_gn=TTC8;refGene_tx=XM_011283434.2;
refGene_gn=TTC8;refGene_tx=XM_011283436.3
B3 134796138 134796261
refGene_gn=TTC8;refGene_tx=XM_011283430.3;refGene_gn=TTC8;refGene_tx=X
M_011283431.3;refGene_gn=TTC8;refGene_tx=XM_011283432.3;refGene_gn=TTC
8;refGene_tx=XM_011283433.3;refGene_gn=TTC8;refGene_tx=XM_011283434.2;
refGene_gn=TTC8;refGene_tx=XM_011283436.3
B3 134799378 134799462
refGene_gn=TTC8;refGene_tx=XM_011283430.3;refGene_gn=TTC8;refGene_tx=X
M_011283431.3;refGene_gn=TTC8;refGene_tx=XM_011283432.3;refGene_gn=TTC
8;refGene_tx=XM_011283433.3;refGene_gn=TTC8;refGene_tx=XM_011283434.2;
refGene_gn=TTC8;refGene_tx=XM_011283436.3
B3 134802023 134802140
refGene_gn=TTC8;refGene_tx=XM_011283430.3;refGene_gn=TTC8;refGene_tx=X
M_011283431.3;refGene_gn=TTC8;refGene_tx=XM_011283432.3;refGene_gn=TTC
8;refGene_tx=XM_011283433.3;refGene_gn=TTC8;refGene_tx=XM_011283434.2;
refGene_gn=TTC8;refGene_tx=XM_011283436.3
B3 135033165 135033715
refGene_gn=FOXN3;refGene_tx=XM_011283437.3;refGene_gn=FOXN3;refGene_tx

=XM_019833461.2;refGene_gn=FOXN3;refGene_tx=XM_019833462.2;refGene_gn=FOXN3;refGene_tx=XM_023255940.1;refGene_gn=FOXN3;refGene_tx=XM_023255941.1;refGene_gn=FOXN3;refGene_tx=XM_023255942.1
B3 135048022 135048128
refGene_gn=FOXN3;refGene_tx=XM_011283437.3;refGene_gn=FOXN3;refGene_tx=XM_019833461.2;refGene_gn=FOXN3;refGene_tx=XM_019833462.2;refGene_gn=FOXN3;refGene_tx=XM_023255940.1;refGene_gn=FOXN3;refGene_tx=XM_023255941.1;refGene_gn=FOXN3;refGene_tx=XM_023255942.1
B3 135057164 135057230
refGene_gn=FOXN3;refGene_tx=XM_011283437.3;refGene_gn=FOXN3;refGene_tx=XM_019833461.2;refGene_gn=FOXN3;refGene_tx=XM_019833462.2;refGene_gn=FOXN3;refGene_tx=XM_023255940.1;refGene_gn=FOXN3;refGene_tx=XM_023255941.1
B3 135143475 135143540
refGene_gn=FOXN3;refGene_tx=XM_011283437.3;refGene_gn=FOXN3;refGene_tx=XM_019833461.2;refGene_gn=FOXN3;refGene_tx=XM_019833462.2;refGene_gn=FOXN3;refGene_tx=XM_023255940.1;refGene_gn=FOXN3;refGene_tx=XM_023255941.1;refGene_gn=FOXN3;refGene_tx=XM_023255942.1
B3 135200315 135200452
refGene_gn=FOXN3;refGene_tx=XM_011283437.3;refGene_gn=FOXN3;refGene_tx=XM_019833461.2;refGene_gn=FOXN3;refGene_tx=XM_019833462.2;refGene_gn=FOXN3;refGene_tx=XM_023255940.1;refGene_gn=FOXN3;refGene_tx=XM_023255941.1;refGene_gn=FOXN3;refGene_tx=XM_023255942.1
B3 135252579 135253122
refGene_gn=FOXN3;refGene_tx=XM_011283437.3;refGene_gn=FOXN3;refGene_tx=XM_019833461.2;refGene_gn=FOXN3;refGene_tx=XM_019833462.2;refGene_gn=FOXN3;refGene_tx=XM_023255940.1;refGene_gn=FOXN3;refGene_tx=XM_023255941.1;refGene_gn=FOXN3;refGene_tx=XM_023255942.1
B3 135598788 135598870
refGene_gn=EFCAB11;refGene_tx=XM_006933051.4
B3 135657907 135657911
refGene_gn=EFCAB11;refGene_tx=XM_011283441.3
B3 135680430 135680485
refGene_gn=EFCAB11;refGene_tx=XM_023255947.1
B3 135717734 135717831
refGene_gn=EFCAB11;refGene_tx=XM_019833472.2
B3 135732270 135732361
refGene_gn=EFCAB11;refGene_tx=XM_006933051.4;refGene_gn=EFCAB11;refGene_tx=XM_011283441.3;refGene_gn=EFCAB11;refGene_tx=XM_019833472.2;refGene_gn=EFCAB11;refGene_tx=XM_023255947.1
B3 135733249 135733351
refGene_gn=EFCAB11;refGene_tx=XM_006933051.4;refGene_gn=EFCAB11;refGene_tx=XM_011283441.3;refGene_gn=EFCAB11;refGene_tx=XM_019833472.2;refGene_gn=EFCAB11;refGene_tx=XM_023255947.1
B3 135752223 135752269
refGene_gn=EFCAB11;refGene_tx=XM_006933051.4;refGene_gn=EFCAB11;refGene_tx=XM_011283441.3;refGene_gn=EFCAB11;refGene_tx=XM_019833472.2;refGene_gn=EFCAB11;refGene_tx=XM_023255947.1
B3 135756758 135756854
refGene_gn=EFCAB11;refGene_tx=XM_006933051.4;refGene_gn=EFCAB11;refGen

e_tx=XM_011283441.3;refGene_gn=EFCAB11;refGene_tx=XM_019833472.2;refGene_gn=EFCAB11;refGene_tx=XM_023255947.1
B3 135757386 135757461
refGene_gn=EFCAB11;refGene_tx=XM_006933051.4;refGene_gn=EFCAB11;refGene_tx=XM_011283441.3;refGene_gn=EFCAB11;refGene_tx=XM_019833472.2;refGene_gn=EFCAB11;refGene_tx=XM_023255947.1
B3 135765266 135765825
refGene_gn=TDP1;refGene_tx=XM_003987908.4;refGene_gn=TDP1;refGene_tx=XM_011283442.3;refGene_gn=TDP1;refGene_tx=XM_019833466.2;refGene_gn=TDP1;refGene_tx=XM_019833467.1;refGene_gn=TDP1;refGene_tx=XM_019833468.1;refGene_gn=TDP1;refGene_tx=XM_023255944.1;refGene_gn=TDP1;refGene_tx=XM_023255946.1
B3 135769325 135769369
refGene_gn=TDP1;refGene_tx=XM_003987908.4;refGene_gn=TDP1;refGene_tx=XM_011283442.3;refGene_gn=TDP1;refGene_tx=XM_019833466.2;refGene_gn=TDP1;refGene_tx=XM_019833467.1;refGene_gn=TDP1;refGene_tx=XM_019833468.1;refGene_gn=TDP1;refGene_tx=XM_023255944.1;refGene_gn=TDP1;refGene_tx=XM_023255946.1
B3 135770550 135770606
refGene_gn=TDP1;refGene_tx=XM_003987908.4;refGene_gn=TDP1;refGene_tx=XM_011283442.3;refGene_gn=TDP1;refGene_tx=XM_019833466.2;refGene_gn=TDP1;refGene_tx=XM_019833467.1;refGene_gn=TDP1;refGene_tx=XM_019833468.1;refGene_gn=TDP1;refGene_tx=XM_023255944.1;refGene_gn=TDP1;refGene_tx=XM_023255946.1
B3 135772293 135772390
refGene_gn=TDP1;refGene_tx=XM_003987908.4;refGene_gn=TDP1;refGene_tx=XM_011283442.3;refGene_gn=TDP1;refGene_tx=XM_019833466.2;refGene_gn=TDP1;refGene_tx=XM_019833467.1;refGene_gn=TDP1;refGene_tx=XM_019833468.1;refGene_gn=TDP1;refGene_tx=XM_023255944.1;refGene_gn=TDP1;refGene_tx=XM_023255946.1
B3 135776520 135776555
refGene_gn=TDP1;refGene_tx=XM_003987908.4;refGene_gn=TDP1;refGene_tx=XM_011283442.3;refGene_gn=TDP1;refGene_tx=XM_019833466.2;refGene_gn=TDP1;refGene_tx=XM_019833467.1;refGene_gn=TDP1;refGene_tx=XM_019833468.1;refGene_gn=TDP1;refGene_tx=XM_023255944.1;refGene_gn=TDP1;refGene_tx=XM_023255946.1
B3 135781251 135781344
refGene_gn=TDP1;refGene_tx=XM_003987908.4;refGene_gn=TDP1;refGene_tx=XM_011283442.3;refGene_gn=TDP1;refGene_tx=XM_019833466.2;refGene_gn=TDP1;refGene_tx=XM_019833467.1;refGene_gn=TDP1;refGene_tx=XM_019833468.1;refGene_gn=TDP1;refGene_tx=XM_023255944.1;refGene_gn=TDP1;refGene_tx=XM_023255946.1
B3 135785190 135785358
refGene_gn=TDP1;refGene_tx=XM_003987908.4;refGene_gn=TDP1;refGene_tx=XM_011283442.3;refGene_gn=TDP1;refGene_tx=XM_019833466.2;refGene_gn=TDP1;refGene_tx=XM_019833467.1;refGene_gn=TDP1;refGene_tx=XM_019833468.1;refGene_gn=TDP1;refGene_tx=XM_023255944.1;refGene_gn=TDP1;refGene_tx=XM_023255946.1
B3 135785807 135785886
refGene_gn=TDP1;refGene_tx=XM_003987908.4;refGene_gn=TDP1;refGene_tx=X

M_011283442.3;refGene_gn=TDP1;refGene_tx=XM_019833466.2;refGene_gn=TDP
1;refGene_tx=XM_019833467.1;refGene_gn=TDP1;refGene_tx=XM_019833468.1;
refGene_gn=TDP1;refGene_tx=XM_023255944.1;refGene_gn=TDP1;refGene_tx=X
M_023255946.1
B3 135787697 135787883
refGene_gn=TDP1;refGene_tx=XM_003987908.4;refGene_gn=TDP1;refGene_tx=X
M_011283442.3;refGene_gn=TDP1;refGene_tx=XM_019833466.2;refGene_gn=TDP
1;refGene_tx=XM_019833467.1;refGene_gn=TDP1;refGene_tx=XM_019833468.1;
refGene_gn=TDP1;refGene_tx=XM_023255944.1;refGene_gn=TDP1;refGene_tx=X
M_023255946.1
B3 135788501 135788550
refGene_gn=TDP1;refGene_tx=XM_003987908.4;refGene_gn=TDP1;refGene_tx=X
M_011283442.3;refGene_gn=TDP1;refGene_tx=XM_019833466.2;refGene_gn=TDP
1;refGene_tx=XM_019833467.1;refGene_gn=TDP1;refGene_tx=XM_019833468.1;
refGene_gn=TDP1;refGene_tx=XM_023255944.1;refGene_gn=TDP1;refGene_tx=X
M_023255946.1
B3 135791131 135791198
refGene_gn=TDP1;refGene_tx=XM_003987908.4;refGene_gn=TDP1;refGene_tx=X
M_011283442.3;refGene_gn=TDP1;refGene_tx=XM_019833466.2;refGene_gn=TDP
1;refGene_tx=XM_019833467.1;refGene_gn=TDP1;refGene_tx=XM_019833468.1;
refGene_gn=TDP1;refGene_tx=XM_023255944.1;refGene_gn=TDP1;refGene_tx=X
M_023255946.1
B3 135792849 135792957
refGene_gn=TDP1;refGene_tx=XM_003987908.4;refGene_gn=TDP1;refGene_tx=X
M_011283442.3;refGene_gn=TDP1;refGene_tx=XM_019833466.2;refGene_gn=TDP
1;refGene_tx=XM_019833467.1;refGene_gn=TDP1;refGene_tx=XM_019833468.1;
refGene_gn=TDP1;refGene_tx=XM_023255944.1;refGene_gn=TDP1;refGene_tx=X
M_023255946.1
B3 135816654 135816679
refGene_gn=TDP1;refGene_tx=XM_023255946.1
B3 135830842 135830945
refGene_gn=TDP1;refGene_tx=XM_003987908.4;refGene_gn=TDP1;refGene_tx=X
M_011283442.3;refGene_gn=TDP1;refGene_tx=XM_019833466.2;refGene_gn=TDP
1;refGene_tx=XM_019833467.1;refGene_gn=TDP1;refGene_tx=XM_019833468.1;
refGene_gn=TDP1;refGene_tx=XM_023255944.1
B3 135843519 135843628
refGene_gn=TDP1;refGene_tx=XM_003987908.4;refGene_gn=TDP1;refGene_tx=X
M_011283442.3;refGene_gn=TDP1;refGene_tx=XM_019833466.2;refGene_gn=TDP
1;refGene_tx=XM_019833467.1;refGene_gn=TDP1;refGene_tx=XM_019833468.1;
refGene_gn=TDP1;refGene_tx=XM_023255944.1
B3 135850388 135850462
refGene_gn=TDP1;refGene_tx=XM_003987908.4;refGene_gn=TDP1;refGene_tx=X
M_011283442.3;refGene_gn=TDP1;refGene_tx=XM_019833466.2;refGene_gn=TDP
1;refGene_tx=XM_019833467.1;refGene_gn=TDP1;refGene_tx=XM_019833468.1;
refGene_gn=TDP1;refGene_tx=XM_023255944.1
B3 135870769 135871109
refGene_gn=KCNK13;refGene_tx=XM_011283476.3
B3 135975849 135976742
refGene_gn=KCNK13;refGene_tx=XM_011283476.3
B3 136031544 136031547

refGene_gn=PSMC1;refGene_tx=XM_003987910.5
B3 136035478 136035532
refGene_gn=PSMC1;refGene_tx=XM_003987910.5
B3 136036770 136036867
refGene_gn=PSMC1;refGene_tx=XM_003987910.5
B3 136038339 136038464
refGene_gn=PSMC1;refGene_tx=XM_003987910.5
B3 136038713 136038899
refGene_gn=PSMC1;refGene_tx=XM_003987910.5
B3 136039052 136039181
refGene_gn=PSMC1;refGene_tx=XM_003987910.5
B3 136040004 136040101
refGene_gn=PSMC1;refGene_tx=XM_003987910.5
B3 136042288 136042478
refGene_gn=PSMC1;refGene_tx=XM_003987910.5
B3 136042972 136043124
refGene_gn=PSMC1;refGene_tx=XM_003987910.5
B3 136043673 136043828
refGene_gn=PSMC1;refGene_tx=XM_003987910.5
B3 136044749 136044884
refGene_gn=PSMC1;refGene_tx=XM_003987910.5
B3 136046752 136046878
refGene_gn=NRDE2;refGene_tx=XM_019833479.2
B3 136047035 136047401
refGene_gn=NRDE2;refGene_tx=XM_019833478.2;refGene_gn=NRDE2;refGene_tx=XM_019833479.2
B3 136050242 136050381
refGene_gn=NRDE2;refGene_tx=XM_019833478.2;refGene_gn=NRDE2;refGene_tx=XM_019833479.2;refGene_gn=NRDE2;refGene_tx=XM_019833480.2
B3 136051494 136052423
refGene_gn=NRDE2;refGene_tx=XM_019833478.2;refGene_gn=NRDE2;refGene_tx=XM_019833479.2;refGene_gn=NRDE2;refGene_tx=XM_019833480.2
B3 136053121 136053508
refGene_gn=NRDE2;refGene_tx=XM_019833478.2;refGene_gn=NRDE2;refGene_tx=XM_019833479.2;refGene_gn=NRDE2;refGene_tx=XM_019833480.2
B3 136055460 136055636
refGene_gn=NRDE2;refGene_tx=XM_019833478.2;refGene_gn=NRDE2;refGene_tx=XM_019833479.2;refGene_gn=NRDE2;refGene_tx=XM_019833480.2
B3 136060901 136061022
refGene_gn=NRDE2;refGene_tx=XM_019833478.2;refGene_gn=NRDE2;refGene_tx=XM_019833479.2;refGene_gn=NRDE2;refGene_tx=XM_019833480.2
B3 136063398 136063532
refGene_gn=NRDE2;refGene_tx=XM_019833478.2;refGene_gn=NRDE2;refGene_tx=XM_019833479.2;refGene_gn=NRDE2;refGene_tx=XM_019833480.2
B3 136065671 136066077
refGene_gn=NRDE2;refGene_tx=XM_019833478.2;refGene_gn=NRDE2;refGene_tx=XM_019833479.2;refGene_gn=NRDE2;refGene_tx=XM_019833480.2
B3 136066917 136067368
refGene_gn=NRDE2;refGene_tx=XM_019833478.2;refGene_gn=NRDE2;refGene_tx=XM_019833479.2;refGene_gn=NRDE2;refGene_tx=XM_019833480.2

B3	136073189	136073339
refGene_gn=NRDE2;refGene_tx=XM_019833478.2;refGene_gn=NRDE2;refGene_tx=XM_019833479.2;refGene_gn=NRDE2;refGene_tx=XM_019833480.2
B3	136076340	136076571
refGene_gn=NRDE2;refGene_tx=XM_019833478.2;refGene_gn=NRDE2;refGene_tx=XM_019833479.2;refGene_gn=NRDE2;refGene_tx=XM_019833480.2
B3	136079092	136079201
refGene_gn=NRDE2;refGene_tx=XM_019833478.2;refGene_gn=NRDE2;refGene_tx=XM_019833479.2;refGene_gn=NRDE2;refGene_tx=XM_019833480.2
B3	136093147	136093211
refGene_gn=NRDE2;refGene_tx=XM_019833478.2;refGene_gn=NRDE2;refGene_tx=XM_019833479.2;refGene_gn=NRDE2;refGene_tx=XM_019833480.2
B3	136161900	136161903
refGene_gn=CALM1;refGene_tx=XM_023255948.1
B3	136164678	136164709
refGene_gn=CALM1;refGene_tx=XM_023255948.1
B3	136165627	136165771
refGene_gn=CALM1;refGene_tx=XM_023255948.1
B3	136167950	136168057
refGene_gn=CALM1;refGene_tx=XM_023255948.1
B3	136168483	136168619
refGene_gn=CALM1;refGene_tx=XM_023255948.1
B3	136168801	136168830
refGene_gn=CALM1;refGene_tx=XM_023255948.1
B3	136275106	136275328
refGene_gn=TTC7B;refGene_tx=XM_019833487.1;refGene_gn=TTC7B;refGene_tx=XM_023255949.1;refGene_gn=TTC7B;refGene_tx=XM_023255950.1;refGene_gn=TTC7B;refGene_tx=XM_023255951.1;refGene_gn=TTC7B;refGene_tx=XM_023255952.1;refGene_gn=TTC7B;refGene_tx=XM_023255953.1;refGene_gn=TTC7B;refGene_tx=XM_023255954.1;refGene_gn=TTC7B;refGene_tx=XM_023255955.1
B3	136307986	136308189
refGene_gn=TTC7B;refGene_tx=XM_019833487.1;refGene_gn=TTC7B;refGene_tx=XM_023255949.1;refGene_gn=TTC7B;refGene_tx=XM_023255950.1;refGene_gn=TTC7B;refGene_tx=XM_023255951.1;refGene_gn=TTC7B;refGene_tx=XM_023255952.1;refGene_gn=TTC7B;refGene_tx=XM_023255953.1;refGene_gn=TTC7B;refGene_tx=XM_023255954.1;refGene_gn=TTC7B;refGene_tx=XM_023255955.1
B3	136322281	136322422
refGene_gn=TTC7B;refGene_tx=XM_019833487.1;refGene_gn=TTC7B;refGene_tx=XM_023255949.1;refGene_gn=TTC7B;refGene_tx=XM_023255950.1;refGene_gn=TTC7B;refGene_tx=XM_023255951.1;refGene_gn=TTC7B;refGene_tx=XM_023255952.1;refGene_gn=TTC7B;refGene_tx=XM_023255953.1;refGene_gn=TTC7B;refGene_tx=XM_023255954.1;refGene_gn=TTC7B;refGene_tx=XM_023255955.1
B3	136331340	136331391
refGene_gn=TTC7B;refGene_tx=XM_019833487.1;refGene_gn=TTC7B;refGene_tx=XM_023255949.1;refGene_gn=TTC7B;refGene_tx=XM_023255950.1;refGene_gn=TTC7B;refGene_tx=XM_023255952.1;refGene_gn=TTC7B;refGene_tx=XM_023255953.1
B3	136333736	136333901
refGene_gn=TTC7B;refGene_tx=XM_019833487.1;refGene_gn=TTC7B;refGene_tx=XM_023255949.1;refGene_gn=TTC7B;refGene_tx=XM_023255950.1;refGene_gn=

TTC7B;refGene_tx=XM_023255951.1
B3 136338320 136338418
refGene_gn=TTC7B;refGene_tx=XM_019833487.1;refGene_gn=TTC7B;refGene_tx=XM_023255949.1;refGene_gn=TTC7B;refGene_tx=XM_023255950.1;refGene_gn=TTC7B;refGene_tx=XM_023255951.1;refGene_gn=TTC7B;refGene_tx=XM_023255952.1;refGene_gn=TTC7B;refGene_tx=XM_023255953.1;refGene_gn=TTC7B;refGene_tx=XM_023255954.1;refGene_gn=TTC7B;refGene_tx=XM_023255955.1
B3 136345244 136345361
refGene_gn=TTC7B;refGene_tx=XM_019833487.1;refGene_gn=TTC7B;refGene_tx=XM_023255949.1;refGene_gn=TTC7B;refGene_tx=XM_023255950.1;refGene_gn=TTC7B;refGene_tx=XM_023255951.1;refGene_gn=TTC7B;refGene_tx=XM_023255952.1;refGene_gn=TTC7B;refGene_tx=XM_023255953.1;refGene_gn=TTC7B;refGene_tx=XM_023255954.1;refGene_gn=TTC7B;refGene_tx=XM_023255955.1
B3 136368626 136368787
refGene_gn=TTC7B;refGene_tx=XM_019833487.1;refGene_gn=TTC7B;refGene_tx=XM_023255949.1;refGene_gn=TTC7B;refGene_tx=XM_023255950.1;refGene_gn=TTC7B;refGene_tx=XM_023255951.1;refGene_gn=TTC7B;refGene_tx=XM_023255952.1;refGene_gn=TTC7B;refGene_tx=XM_023255953.1;refGene_gn=TTC7B;refGene_tx=XM_023255954.1;refGene_gn=TTC7B;refGene_tx=XM_023255955.1
B3 136371544 136371617
refGene_gn=TTC7B;refGene_tx=XM_019833487.1;refGene_gn=TTC7B;refGene_tx=XM_023255949.1;refGene_gn=TTC7B;refGene_tx=XM_023255950.1;refGene_gn=TTC7B;refGene_tx=XM_023255951.1;refGene_gn=TTC7B;refGene_tx=XM_023255952.1;refGene_gn=TTC7B;refGene_tx=XM_023255953.1;refGene_gn=TTC7B;refGene_tx=XM_023255954.1;refGene_gn=TTC7B;refGene_tx=XM_023255955.1
B3 136377991 136378049
refGene_gn=TTC7B;refGene_tx=XM_019833487.1;refGene_gn=TTC7B;refGene_tx=XM_023255949.1;refGene_gn=TTC7B;refGene_tx=XM_023255950.1;refGene_gn=TTC7B;refGene_tx=XM_023255951.1;refGene_gn=TTC7B;refGene_tx=XM_023255952.1;refGene_gn=TTC7B;refGene_tx=XM_023255953.1;refGene_gn=TTC7B;refGene_tx=XM_023255954.1;refGene_gn=TTC7B;refGene_tx=XM_023255955.1
B3 136380159 136380277
refGene_gn=TTC7B;refGene_tx=XM_019833487.1;refGene_gn=TTC7B;refGene_tx=XM_023255949.1;refGene_gn=TTC7B;refGene_tx=XM_023255950.1;refGene_gn=TTC7B;refGene_tx=XM_023255951.1;refGene_gn=TTC7B;refGene_tx=XM_023255952.1;refGene_gn=TTC7B;refGene_tx=XM_023255953.1;refGene_gn=TTC7B;refGene_tx=XM_023255954.1;refGene_gn=TTC7B;refGene_tx=XM_023255955.1
B3 136382223 136382328
refGene_gn=TTC7B;refGene_tx=XM_019833487.1;refGene_gn=TTC7B;refGene_tx=XM_023255949.1;refGene_gn=TTC7B;refGene_tx=XM_023255950.1;refGene_gn=TTC7B;refGene_tx=XM_023255951.1;refGene_gn=TTC7B;refGene_tx=XM_023255952.1;refGene_gn=TTC7B;refGene_tx=XM_023255953.1;refGene_gn=TTC7B;refGene_tx=XM_023255954.1;refGene_gn=TTC7B;refGene_tx=XM_023255955.1
B3 136383372 136383456
refGene_gn=TTC7B;refGene_tx=XM_019833487.1;refGene_gn=TTC7B;refGene_tx=XM_023255949.1;refGene_gn=TTC7B;refGene_tx=XM_023255950.1;refGene_gn=TTC7B;refGene_tx=XM_023255951.1;refGene_gn=TTC7B;refGene_tx=XM_023255952.1;refGene_gn=TTC7B;refGene_tx=XM_023255953.1;refGene_gn=TTC7B;refGene_tx=XM_023255954.1;refGene_gn=TTC7B;refGene_tx=XM_023255955.1
B3 136403693 136403831

refGene_gn=TTC7B;refGene_tx=XM_019833487.1;refGene_gn=TTC7B;refGene_tx=XM_023255949.1;refGene_gn=TTC7B;refGene_tx=XM_023255950.1;refGene_gn=TTC7B;refGene_tx=XM_023255951.1;refGene_gn=TTC7B;refGene_tx=XM_023255952.1;refGene_gn=TTC7B;refGene_tx=XM_023255953.1;refGene_gn=TTC7B;refGene_tx=XM_023255954.1;refGene_gn=TTC7B;refGene_tx=XM_023255955.1
B3 136408167 136408231
refGene_gn=TTC7B;refGene_tx=XM_019833487.1;refGene_gn=TTC7B;refGene_tx=XM_023255949.1;refGene_gn=TTC7B;refGene_tx=XM_023255950.1;refGene_gn=TTC7B;refGene_tx=XM_023255951.1;refGene_gn=TTC7B;refGene_tx=XM_023255952.1;refGene_gn=TTC7B;refGene_tx=XM_023255953.1;refGene_gn=TTC7B;refGene_tx=XM_023255954.1;refGene_gn=TTC7B;refGene_tx=XM_023255955.1
B3 136416138 136416311
refGene_gn=TTC7B;refGene_tx=XM_019833487.1;refGene_gn=TTC7B;refGene_tx=XM_023255949.1;refGene_gn=TTC7B;refGene_tx=XM_023255950.1;refGene_gn=TTC7B;refGene_tx=XM_023255951.1;refGene_gn=TTC7B;refGene_tx=XM_023255952.1;refGene_gn=TTC7B;refGene_tx=XM_023255953.1;refGene_gn=TTC7B;refGene_tx=XM_023255954.1;refGene_gn=TTC7B;refGene_tx=XM_023255955.1
B3 136422086 136422165
refGene_gn=TTC7B;refGene_tx=XM_023255949.1;refGene_gn=TTC7B;refGene_tx=XM_023255950.1;refGene_gn=TTC7B;refGene_tx=XM_023255951.1;refGene_gn=TTC7B;refGene_tx=XM_023255952.1;refGene_gn=TTC7B;refGene_tx=XM_023255953.1;refGene_gn=TTC7B;refGene_tx=XM_023255954.1;refGene_gn=TTC7B;refGene_tx=XM_023255955.1
B3 136449506 136449530
refGene_gn=TTC7B;refGene_tx=XM_023255949.1;refGene_gn=TTC7B;refGene_tx=XM_023255951.1;refGene_gn=TTC7B;refGene_tx=XM_023255952.1;refGene_gn=TTC7B;refGene_tx=XM_023255954.1
B3 136450187 136450309
refGene_gn=TTC7B;refGene_tx=XM_023255949.1;refGene_gn=TTC7B;refGene_tx=XM_023255950.1;refGene_gn=TTC7B;refGene_tx=XM_023255951.1;refGene_gn=TTC7B;refGene_tx=XM_023255952.1;refGene_gn=TTC7B;refGene_tx=XM_023255953.1;refGene_gn=TTC7B;refGene_tx=XM_023255954.1;refGene_gn=TTC7B;refGene_tx=XM_023255955.1
B3 136466418 136466549
refGene_gn=TTC7B;refGene_tx=XM_023255949.1;refGene_gn=TTC7B;refGene_tx=XM_023255950.1;refGene_gn=TTC7B;refGene_tx=XM_023255951.1;refGene_gn=TTC7B;refGene_tx=XM_023255952.1;refGene_gn=TTC7B;refGene_tx=XM_023255953.1;refGene_gn=TTC7B;refGene_tx=XM_023255954.1;refGene_gn=TTC7B;refGene_tx=XM_023255955.1
B3 136500751 136500920
refGene_gn=TTC7B;refGene_tx=XM_023255949.1;refGene_gn=TTC7B;refGene_tx=XM_023255950.1;refGene_gn=TTC7B;refGene_tx=XM_023255951.1;refGene_gn=TTC7B;refGene_tx=XM_023255952.1;refGene_gn=TTC7B;refGene_tx=XM_023255953.1;refGene_gn=TTC7B;refGene_tx=XM_023255954.1;refGene_gn=TTC7B;refGene_tx=XM_023255955.1
B3 136506923 136507078
refGene_gn=TTC7B;refGene_tx=XM_023255949.1;refGene_gn=TTC7B;refGene_tx=XM_023255950.1;refGene_gn=TTC7B;refGene_tx=XM_023255951.1;refGene_gn=TTC7B;refGene_tx=XM_023255952.1;refGene_gn=TTC7B;refGene_tx=XM_023255953.1;refGene_gn=TTC7B;refGene_tx=XM_023255954.1;refGene_gn=TTC7B;refGe

ne_tx=XM_023255955.1
B3 136530483 136530604
refGene_gn=TTC7B;refGene_tx=XM_023255949.1;refGene_gn=TTC7B;refGene_tx=XM_023255950.1;refGene_gn=TTC7B;refGene_tx=XM_023255951.1;refGene_gn=TTC7B;refGene_tx=XM_023255952.1;refGene_gn=TTC7B;refGene_tx=XM_023255953.1;refGene_gn=TTC7B;refGene_tx=XM_023255954.1;refGene_gn=TTC7B;refGene_tx=XM_023255955.1
B3 136555697 136556276
refGene_gn=LOC111560915;refGene_tx=XM_023255956.1
B3 136557773 136557923
refGene_gn=LOC111560915;refGene_tx=XM_023255956.1
B3 136570826 136570829
refGene_gn=LOC111560915;refGene_tx=XM_023255956.1
B3 136573160 136573198
refGene_gn=RPS6KA5;refGene_tx=XM_019833489.2;refGene_gn=RPS6KA5;refGene_tx=XM_019833492.2
B3 136577254 136577494
refGene_gn=RPS6KA5;refGene_tx=XM_003987915.5;refGene_gn=RPS6KA5;refGene_tx=XM_006933057.4;refGene_gn=RPS6KA5;refGene_tx=XM_019833490.1;refGene_gn=RPS6KA5;refGene_tx=XM_019833491.2;refGene_gn=RPS6KA5;refGene_tx=XM_019833493.1;refGene_gn=RPS6KA5;refGene_tx=XM_019833495.1;refGene_gn=RPS6KA5;refGene_tx=XM_019833496.1;refGene_gn=RPS6KA5;refGene_tx=XM_023255958.1
B3 136578994 136579158
refGene_gn=RPS6KA5;refGene_tx=XM_003987915.5;refGene_gn=RPS6KA5;refGene_tx=XM_006933057.4;refGene_gn=RPS6KA5;refGene_tx=XM_019833489.2;refGene_gn=RPS6KA5;refGene_tx=XM_019833490.1;refGene_gn=RPS6KA5;refGene_tx=XM_019833493.1;refGene_gn=RPS6KA5;refGene_tx=XM_019833495.1;refGene_gn=RPS6KA5;refGene_tx=XM_019833496.1;refGene_gn=RPS6KA5;refGene_tx=XM_023255958.1
B3 136580963 136581123
refGene_gn=RPS6KA5;refGene_tx=XM_003987915.5;refGene_gn=RPS6KA5;refGene_tx=XM_006933057.4;refGene_gn=RPS6KA5;refGene_tx=XM_019833489.2;refGene_gn=RPS6KA5;refGene_tx=XM_019833490.1;refGene_gn=RPS6KA5;refGene_tx=XM_019833492.2;refGene_gn=RPS6KA5;refGene_tx=XM_019833493.1;refGene_gn=RPS6KA5;refGene_tx=XM_019833495.1;refGene_gn=RPS6KA5;refGene_tx=XM_019833496.1;refGene_gn=RPS6KA5;refGene_tx=XM_023255958.1
B3 136589876 136590068
refGene_gn=RPS6KA5;refGene_tx=XM_003987915.5;refGene_gn=RPS6KA5;refGene_tx=XM_006933057.4;refGene_gn=RPS6KA5;refGene_tx=XM_019833489.2;refGene_gn=RPS6KA5;refGene_tx=XM_019833490.1;refGene_gn=RPS6KA5;refGene_tx=XM_019833491.2;refGene_gn=RPS6KA5;refGene_tx=XM_019833492.2;refGene_gn=RPS6KA5;refGene_tx=XM_019833493.1;refGene_gn=RPS6KA5;refGene_tx=XM_019833495.1;refGene_gn=RPS6KA5;refGene_tx=XM_019833496.1;refGene_gn=RPS6KA5;refGene_tx=XM_023255958.1
B3 136593616 136593787
refGene_gn=RPS6KA5;refGene_tx=XM_003987915.5;refGene_gn=RPS6KA5;refGene_tx=XM_006933057.4;refGene_gn=RPS6KA5;refGene_tx=XM_019833489.2;refGene_gn=RPS6KA5;refGene_tx=XM_019833490.1;refGene_gn=RPS6KA5;refGene_tx=XM_019833491.2;refGene_gn=RPS6KA5;refGene_tx=XM_019833492.2;refGene_gn

=RPS6KA5;refGene_tx=XM_019833493.1;refGene_gn=RPS6KA5;refGene_tx=XM_01
9833495.1;refGene_gn=RPS6KA5;refGene_tx=XM_019833496.1;refGene_gn=RPS6
KA5;refGene_tx=XM_023255958.1
B3	136597331	136597425
refGene_gn=RPS6KA5;refGene_tx=XM_003987915.5;refGene_gn=RPS6KA5;refGen
e_tx=XM_006933057.4;refGene_gn=RPS6KA5;refGene_tx=XM_019833489.2;refGe
ne_gn=RPS6KA5;refGene_tx=XM_019833490.1;refGene_gn=RPS6KA5;refGene_tx=
XM_019833491.2;refGene_gn=RPS6KA5;refGene_tx=XM_019833492.2;refGene_gn
=RPS6KA5;refGene_tx=XM_019833493.1;refGene_gn=RPS6KA5;refGene_tx=XM_01
9833495.1;refGene_gn=RPS6KA5;refGene_tx=XM_019833496.1;refGene_gn=RPS6
KA5;refGene_tx=XM_023255958.1
B3	136598111	136598245
refGene_gn=RPS6KA5;refGene_tx=XM_003987915.5;refGene_gn=RPS6KA5;refGen
e_tx=XM_006933057.4;refGene_gn=RPS6KA5;refGene_tx=XM_019833489.2;refGe
ne_gn=RPS6KA5;refGene_tx=XM_019833490.1;refGene_gn=RPS6KA5;refGene_tx=
XM_019833491.2;refGene_gn=RPS6KA5;refGene_tx=XM_019833492.2;refGene_gn
=RPS6KA5;refGene_tx=XM_019833493.1;refGene_gn=RPS6KA5;refGene_tx=XM_01
9833495.1;refGene_gn=RPS6KA5;refGene_tx=XM_019833496.1;refGene_gn=RPS6
KA5;refGene_tx=XM_023255958.1
B3	136599107	136599233
refGene_gn=RPS6KA5;refGene_tx=XM_003987915.5;refGene_gn=RPS6KA5;refGen
e_tx=XM_006933057.4;refGene_gn=RPS6KA5;refGene_tx=XM_019833489.2;refGe
ne_gn=RPS6KA5;refGene_tx=XM_019833490.1;refGene_gn=RPS6KA5;refGene_tx=
XM_019833491.2;refGene_gn=RPS6KA5;refGene_tx=XM_019833492.2;refGene_gn
=RPS6KA5;refGene_tx=XM_019833493.1;refGene_gn=RPS6KA5;refGene_tx=XM_01
9833495.1;refGene_gn=RPS6KA5;refGene_tx=XM_019833496.1;refGene_gn=RPS6
KA5;refGene_tx=XM_023255958.1
B3	136601153	136601315
refGene_gn=RPS6KA5;refGene_tx=XM_003987915.5;refGene_gn=RPS6KA5;refGen
e_tx=XM_006933057.4;refGene_gn=RPS6KA5;refGene_tx=XM_019833489.2;refGe
ne_gn=RPS6KA5;refGene_tx=XM_019833490.1;refGene_gn=RPS6KA5;refGene_tx=
XM_019833491.2;refGene_gn=RPS6KA5;refGene_tx=XM_019833492.2;refGene_gn
=RPS6KA5;refGene_tx=XM_019833493.1;refGene_gn=RPS6KA5;refGene_tx=XM_01
9833495.1;refGene_gn=RPS6KA5;refGene_tx=XM_019833496.1;refGene_gn=RPS6
KA5;refGene_tx=XM_023255958.1
B3	136604752	136604903
refGene_gn=RPS6KA5;refGene_tx=XM_003987915.5;refGene_gn=RPS6KA5;refGen
e_tx=XM_006933057.4;refGene_gn=RPS6KA5;refGene_tx=XM_019833489.2;refGe
ne_gn=RPS6KA5;refGene_tx=XM_019833490.1;refGene_gn=RPS6KA5;refGene_tx=
XM_019833491.2;refGene_gn=RPS6KA5;refGene_tx=XM_019833492.2;refGene_gn
=RPS6KA5;refGene_tx=XM_019833493.1;refGene_gn=RPS6KA5;refGene_tx=XM_01
9833495.1;refGene_gn=RPS6KA5;refGene_tx=XM_019833496.1;refGene_gn=RPS6
KA5;refGene_tx=XM_023255958.1
B3	136623810	136623914
refGene_gn=RPS6KA5;refGene_tx=XM_003987915.5;refGene_gn=RPS6KA5;refGen
e_tx=XM_006933057.4;refGene_gn=RPS6KA5;refGene_tx=XM_019833489.2;refGe
ne_gn=RPS6KA5;refGene_tx=XM_019833490.1;refGene_gn=RPS6KA5;refGene_tx=
XM_019833491.2;refGene_gn=RPS6KA5;refGene_tx=XM_019833492.2;refGene_gn
=RPS6KA5;refGene_tx=XM_019833493.1;refGene_gn=RPS6KA5;refGene_tx=XM_01
9833495.1;refGene_gn=RPS6KA5;refGene_tx=XM_023255958.1

B3 136625858 136625942
refGene_gn=RPS6KA5;refGene_tx=XM_003987915.5;refGene_gn=RPS6KA5;refGene_tx=XM_006933057.4;refGene_gn=RPS6KA5;refGene_tx=XM_019833489.2;refGene_gn=RPS6KA5;refGene_tx=XM_019833490.1;refGene_gn=RPS6KA5;refGene_tx=XM_019833491.2;refGene_gn=RPS6KA5;refGene_tx=XM_019833492.2;refGene_gn=RPS6KA5;refGene_tx=XM_019833493.1;refGene_gn=RPS6KA5;refGene_tx=XM_019833495.1;refGene_gn=RPS6KA5;refGene_tx=XM_023255958.1
B3 136650271 136650379
refGene_gn=RPS6KA5;refGene_tx=XM_003987915.5;refGene_gn=RPS6KA5;refGene_tx=XM_006933057.4;refGene_gn=RPS6KA5;refGene_tx=XM_019833489.2;refGene_gn=RPS6KA5;refGene_tx=XM_019833490.1;refGene_gn=RPS6KA5;refGene_tx=XM_019833491.2;refGene_gn=RPS6KA5;refGene_tx=XM_019833492.2;refGene_gn=RPS6KA5;refGene_tx=XM_023255958.1
B3 136653310 136653426
refGene_gn=RPS6KA5;refGene_tx=XM_003987915.5;refGene_gn=RPS6KA5;refGene_tx=XM_006933057.4;refGene_gn=RPS6KA5;refGene_tx=XM_019833489.2;refGene_gn=RPS6KA5;refGene_tx=XM_019833490.1;refGene_gn=RPS6KA5;refGene_tx=XM_019833491.2;refGene_gn=RPS6KA5;refGene_tx=XM_019833492.2;refGene_gn=RPS6KA5;refGene_tx=XM_023255958.1
B3 136682882 136683101
refGene_gn=RPS6KA5;refGene_tx=XM_003987915.5;refGene_gn=RPS6KA5;refGene_tx=XM_006933057.4;refGene_gn=RPS6KA5;refGene_tx=XM_019833489.2;refGene_gn=RPS6KA5;refGene_tx=XM_019833490.1;refGene_gn=RPS6KA5;refGene_tx=XM_019833491.2;refGene_gn=RPS6KA5;refGene_tx=XM_019833492.2;refGene_gn=RPS6KA5;refGene_tx=XM_023255958.1
B3 136703891 136703963
refGene_gn=RPS6KA5;refGene_tx=XM_003987915.5;refGene_gn=RPS6KA5;refGene_tx=XM_019833489.2;refGene_gn=RPS6KA5;refGene_tx=XM_019833491.2;refGene_gn=RPS6KA5;refGene_tx=XM_019833492.2
B3 136754186 136754286
refGene_gn=RPS6KA5;refGene_tx=XM_003987915.5;refGene_gn=RPS6KA5;refGene_tx=XM_019833489.2;refGene_gn=RPS6KA5;refGene_tx=XM_019833491.2;refGene_gn=RPS6KA5;refGene_tx=XM_019833492.2
B3 136882259 136883041
refGene_gn=GPR68;refGene_tx=XM_023255960.1
B3 136883043 136883344
refGene_gn=GPR68;refGene_tx=XM_023255960.1
B3 136914692 136915712
refGene_gn=CCDC88C;refGene_tx=XM_003987962.4;refGene_gn=CCDC88C;refGene_tx=XM_023255961.1;refGene_gn=CCDC88C;refGene_tx=XM_023255962.1
B3 136919267 136919557
refGene_gn=CCDC88C;refGene_tx=XM_003987962.4;refGene_gn=CCDC88C;refGene_tx=XM_023255961.1;refGene_gn=CCDC88C;refGene_tx=XM_023255962.1
B3 136920423 136920492
refGene_gn=CCDC88C;refGene_tx=XM_003987962.4;refGene_gn=CCDC88C;refGene_tx=XM_023255961.1;refGene_gn=CCDC88C;refGene_tx=XM_023255962.1
B3 136923179 136923245
refGene_gn=CCDC88C;refGene_tx=XM_003987962.4;refGene_gn=CCDC88C;refGene_tx=XM_023255961.1;refGene_gn=CCDC88C;refGene_tx=XM_023255962.1
B3 136925897 136926086

refGene_gn=CCDC88C;refGene_tx=XM_003987962.4;refGene_gn=CCDC88C;refGene_tx=XM_023255961.1;refGene_gn=CCDC88C;refGene_tx=XM_023255962.1
B3 136930929 136931168
refGene_gn=CCDC88C;refGene_tx=XM_003987962.4;refGene_gn=CCDC88C;refGene_tx=XM_023255961.1;refGene_gn=CCDC88C;refGene_tx=XM_023255962.1
B3 136932484 136932574
refGene_gn=CCDC88C;refGene_tx=XM_003987962.4;refGene_gn=CCDC88C;refGene_tx=XM_023255961.1;refGene_gn=CCDC88C;refGene_tx=XM_023255962.1
B3 136935123 136935269
refGene_gn=CCDC88C;refGene_tx=XM_003987962.4;refGene_gn=CCDC88C;refGene_tx=XM_023255961.1;refGene_gn=CCDC88C;refGene_tx=XM_023255962.1
B3 136937221 136937408
refGene_gn=CCDC88C;refGene_tx=XM_003987962.4;refGene_gn=CCDC88C;refGene_tx=XM_023255961.1;refGene_gn=CCDC88C;refGene_tx=XM_023255962.1
B3 136939460 136939604
refGene_gn=CCDC88C;refGene_tx=XM_003987962.4;refGene_gn=CCDC88C;refGene_tx=XM_023255961.1;refGene_gn=CCDC88C;refGene_tx=XM_023255962.1
B3 136942537 136942815
refGene_gn=CCDC88C;refGene_tx=XM_003987962.4;refGene_gn=CCDC88C;refGene_tx=XM_023255961.1;refGene_gn=CCDC88C;refGene_tx=XM_023255962.1
B3 136943944 136944106
refGene_gn=CCDC88C;refGene_tx=XM_003987962.4;refGene_gn=CCDC88C;refGene_tx=XM_023255961.1;refGene_gn=CCDC88C;refGene_tx=XM_023255962.1
B3 136945242 136945431
refGene_gn=CCDC88C;refGene_tx=XM_003987962.4;refGene_gn=CCDC88C;refGene_tx=XM_023255961.1;refGene_gn=CCDC88C;refGene_tx=XM_023255962.1
B3 136945596 136945713
refGene_gn=CCDC88C;refGene_tx=XM_023255961.1
B3 136946477 136946616
refGene_gn=CCDC88C;refGene_tx=XM_003987962.4;refGene_gn=CCDC88C;refGene_tx=XM_023255961.1;refGene_gn=CCDC88C;refGene_tx=XM_023255962.1
B3 136948142 136948270
refGene_gn=CCDC88C;refGene_tx=XM_003987962.4;refGene_gn=CCDC88C;refGene_tx=XM_023255961.1;refGene_gn=CCDC88C;refGene_tx=XM_023255962.1
B3 136950240 136951311
refGene_gn=CCDC88C;refGene_tx=XM_003987962.4;refGene_gn=CCDC88C;refGene_tx=XM_023255961.1;refGene_gn=CCDC88C;refGene_tx=XM_023255962.1
B3 136952346 136952484
refGene_gn=CCDC88C;refGene_tx=XM_003987962.4;refGene_gn=CCDC88C;refGene_tx=XM_023255961.1;refGene_gn=CCDC88C;refGene_tx=XM_023255962.1
B3 136956788 136956973
refGene_gn=CCDC88C;refGene_tx=XM_003987962.4;refGene_gn=CCDC88C;refGene_tx=XM_023255961.1;refGene_gn=CCDC88C;refGene_tx=XM_023255962.1
B3 136960116 136960261
refGene_gn=CCDC88C;refGene_tx=XM_003987962.4;refGene_gn=CCDC88C;refGene_tx=XM_023255961.1;refGene_gn=CCDC88C;refGene_tx=XM_023255962.1
B3 136960917 136961064
refGene_gn=CCDC88C;refGene_tx=XM_003987962.4;refGene_gn=CCDC88C;refGene_tx=XM_023255961.1;refGene_gn=CCDC88C;refGene_tx=XM_023255962.1
B3 136970342 136970501

refGene_gn=CCDC88C;refGene_tx=XM_003987962.4;refGene_gn=CCDC88C;refGene_tx=XM_023255961.1;refGene_gn=CCDC88C;refGene_tx=XM_023255962.1
B3 136970872 136970954
refGene_gn=CCDC88C;refGene_tx=XM_003987962.4;refGene_gn=CCDC88C;refGene_tx=XM_023255961.1;refGene_gn=CCDC88C;refGene_tx=XM_023255962.1
B3 136971687 136971872
refGene_gn=CCDC88C;refGene_tx=XM_003987962.4;refGene_gn=CCDC88C;refGene_tx=XM_023255961.1;refGene_gn=CCDC88C;refGene_tx=XM_023255962.1
B3 136972124 136972265
refGene_gn=CCDC88C;refGene_tx=XM_003987962.4;refGene_gn=CCDC88C;refGene_tx=XM_023255961.1;refGene_gn=CCDC88C;refGene_tx=XM_023255962.1
B3 136974846 136974930
refGene_gn=CCDC88C;refGene_tx=XM_003987962.4;refGene_gn=CCDC88C;refGene_tx=XM_023255961.1;refGene_gn=CCDC88C;refGene_tx=XM_023255962.1
B3 136975806 136975865
refGene_gn=CCDC88C;refGene_tx=XM_003987962.4;refGene_gn=CCDC88C;refGene_tx=XM_023255961.1;refGene_gn=CCDC88C;refGene_tx=XM_023255962.1
B3 136989773 136989843
refGene_gn=CCDC88C;refGene_tx=XM_003987962.4;refGene_gn=CCDC88C;refGene_tx=XM_023255961.1;refGene_gn=CCDC88C;refGene_tx=XM_023255962.1
B3 137034512 137034621
refGene_gn=CCDC88C;refGene_tx=XM_003987962.4;refGene_gn=CCDC88C;refGene_tx=XM_023255961.1;refGene_gn=CCDC88C;refGene_tx=XM_023255962.1
B3 137041632 137041733
refGene_gn=CCDC88C;refGene_tx=XM_003987962.4;refGene_gn=CCDC88C;refGene_tx=XM_023255961.1;refGene_gn=CCDC88C;refGene_tx=XM_023255962.1
B3 137042473 137042533
refGene_gn=CCDC88C;refGene_tx=XM_003987962.4;refGene_gn=CCDC88C;refGene_tx=XM_023255961.1;refGene_gn=CCDC88C;refGene_tx=XM_023255962.1
B3 137086137 137086248
refGene_gn=PPP4R3A;refGene_tx=XM_011283449.2;refGene_gn=PPP4R3A;refGene_tx=XM_011283450.2
B3 137087304 137087531
refGene_gn=PPP4R3A;refGene_tx=XM_011283449.2;refGene_gn=PPP4R3A;refGene_tx=XM_011283450.2
B3 137087911 137088102
refGene_gn=PPP4R3A;refGene_tx=XM_011283449.2;refGene_gn=PPP4R3A;refGene_tx=XM_011283450.2
B3 137088681 137088824
refGene_gn=PPP4R3A;refGene_tx=XM_011283449.2;refGene_gn=PPP4R3A;refGene_tx=XM_011283450.2
B3 137089951 137090121
refGene_gn=PPP4R3A;refGene_tx=XM_011283449.2;refGene_gn=PPP4R3A;refGene_tx=XM_011283450.2
B3 137095336 137095495
refGene_gn=PPP4R3A;refGene_tx=XM_011283449.2;refGene_gn=PPP4R3A;refGene_tx=XM_011283450.2
B3 137097184 137097287
refGene_gn=PPP4R3A;refGene_tx=XM_011283449.2;refGene_gn=PPP4R3A;refGene_tx=XM_011283450.2

B3 137097388 137097520
refGene_gn=PPP4R3A;refGene_tx=XM_011283449.2;refGene_gn=PPP4R3A;refGene_tx=XM_011283450.2
B3 137099349 137099505
refGene_gn=PPP4R3A;refGene_tx=XM_011283449.2;refGene_gn=PPP4R3A;refGene_tx=XM_011283450.2
B3 137099937 137100054
refGene_gn=PPP4R3A;refGene_tx=XM_011283449.2;refGene_gn=PPP4R3A;refGene_tx=XM_011283450.2
B3 137100134 137100212
refGene_gn=PPP4R3A;refGene_tx=XM_011283449.2;refGene_gn=PPP4R3A;refGene_tx=XM_011283450.2
B3 137103968 137104586
refGene_gn=PPP4R3A;refGene_tx=XM_011283449.2;refGene_gn=PPP4R3A;refGene_tx=XM_011283450.2
B3 137108004 137108103
refGene_gn=PPP4R3A;refGene_tx=XM_011283449.2;refGene_gn=PPP4R3A;refGene_tx=XM_011283450.2
B3 137111447 137111503
refGene_gn=PPP4R3A;refGene_tx=XM_011283449.2;refGene_gn=PPP4R3A;refGene_tx=XM_011283450.2
B3 137122248 137122390
refGene_gn=PPP4R3A;refGene_tx=XM_011283449.2;refGene_gn=PPP4R3A;refGene_tx=XM_011283450.2
B3 137122545 137122682
refGene_gn=LOC102902607;refGene_tx=XM_023256190.1
B3 137122772 137123663
refGene_gn=LOC102902607;refGene_tx=XM_023256190.1
B3 137125065 137125084
refGene_gn=LOC102902607;refGene_tx=XM_023256190.1
B3 137174168 137174372
refGene_gn=CATSPERB;refGene_tx=XM_023256191.1
B3 137184057 137184132
refGene_gn=CATSPERB;refGene_tx=XM_023256191.1
B3 137184814 137184915
refGene_gn=CATSPERB;refGene_tx=XM_023256191.1
B3 137186438 137186574
refGene_gn=CATSPERB;refGene_tx=XM_023256191.1
B3 137187902 137188013
refGene_gn=CATSPERB;refGene_tx=XM_023256191.1
B3 137192024 137192135
refGene_gn=CATSPERB;refGene_tx=XM_023256191.1
B3 137193447 137193645
refGene_gn=CATSPERB;refGene_tx=XM_023256191.1
B3 137196228 137196368
refGene_gn=CATSPERB;refGene_tx=XM_023256191.1
B3 137201624 137201963
refGene_gn=CATSPERB;refGene_tx=XM_023256191.1
B3 137203937 137204128
refGene_gn=CATSPERB;refGene_tx=XM_023256191.1

B3 137215775 137215930
refGene_gn=CATSPERB;refGene_tx=XM_023256191.1
B3 137218868 137219013
refGene_gn=CATSPERB;refGene_tx=XM_023256191.1
B3 137230832 137231050
refGene_gn=CATSPERB;refGene_tx=XM_023256191.1
B3 137238758 137238911
refGene_gn=CATSPERB;refGene_tx=XM_023256191.1
B3 137240447 137240600
refGene_gn=CATSPERB;refGene_tx=XM_023256191.1
B3 137241233 137241280
refGene_gn=CATSPERB;refGene_tx=XM_023256191.1
B3 137241554 137241621
refGene_gn=CATSPERB;refGene_tx=XM_023256191.1
B3 137246084 137246117
refGene_gn=CATSPERB;refGene_tx=XM_023256191.1
B3 137247783 137247902
refGene_gn=CATSPERB;refGene_tx=XM_023256191.1
B3 137248039 137248135
refGene_gn=CATSPERB;refGene_tx=XM_023256191.1
B3 137253326 137253476
refGene_gn=CATSPERB;refGene_tx=XM_023256191.1
B3 137257829 137257925
refGene_gn=CATSPERB;refGene_tx=XM_023256191.1
B3 137269362 137269423
refGene_gn=CATSPERB;refGene_tx=XM_023256191.1
B3 137273873 137274014
refGene_gn=CATSPERB;refGene_tx=XM_023256191.1
B3 137276439 137276528
refGene_gn=CATSPERB;refGene_tx=XM_023256191.1
B3 137279208 137279287
refGene_gn=CATSPERB;refGene_tx=XM_023256191.1
B3 137319367 137319478
refGene_gn=TC2N;refGene_tx=XM_019833517.2;refGene_gn=TC2N;refGene_tx=XM_023255963.1;refGene_gn=TC2N;refGene_tx=XM_023255964.1
B3 137321422 137321622
refGene_gn=TC2N;refGene_tx=XM_019833517.2;refGene_gn=TC2N;refGene_tx=XM_023255963.1;refGene_gn=TC2N;refGene_tx=XM_023255964.1
B3 137323894 137324009
refGene_gn=TC2N;refGene_tx=XM_019833517.2;refGene_gn=TC2N;refGene_tx=XM_023255963.1;refGene_gn=TC2N;refGene_tx=XM_023255964.1
B3 137331086 137331278
refGene_gn=TC2N;refGene_tx=XM_019833517.2;refGene_gn=TC2N;refGene_tx=XM_023255963.1;refGene_gn=TC2N;refGene_tx=XM_023255964.1
B3 137335644 137335761
refGene_gn=TC2N;refGene_tx=XM_019833517.2;refGene_gn=TC2N;refGene_tx=XM_023255963.1;refGene_gn=TC2N;refGene_tx=XM_023255964.1
B3 137336142 137336243
refGene_gn=TC2N;refGene_tx=XM_019833517.2;refGene_gn=TC2N;refGene_tx=XM_023255963.1;refGene_gn=TC2N;refGene_tx=XM_023255964.1

B3 137337008 137337084
refGene_gn=TC2N;refGene_tx=XM_019833517.2;refGene_gn=TC2N;refGene_tx=XM_023255963.1;refGene_gn=TC2N;refGene_tx=XM_023255964.1
B3 137338285 137338377
refGene_gn=TC2N;refGene_tx=XM_019833517.2;refGene_gn=TC2N;refGene_tx=XM_023255963.1;refGene_gn=TC2N;refGene_tx=XM_023255964.1
B3 137339972 137340140
refGene_gn=TC2N;refGene_tx=XM_019833517.2;refGene_gn=TC2N;refGene_tx=XM_023255963.1;refGene_gn=TC2N;refGene_tx=XM_023255964.1
B3 137347683 137347917
refGene_gn=TC2N;refGene_tx=XM_019833517.2;refGene_gn=TC2N;refGene_tx=XM_023255963.1;refGene_gn=TC2N;refGene_tx=XM_023255964.1
B3 137349281 137349348
refGene_gn=TC2N;refGene_tx=XM_019833517.2;refGene_gn=TC2N;refGene_tx=XM_023255963.1;refGene_gn=TC2N;refGene_tx=XM_023255964.1
B3 137394012 137394284
refGene_gn=FBLN5;refGene_tx=XM_003987921.4;refGene_gn=FBLN5;refGene_tx=XM_011283455.3;refGene_gn=FBLN5;refGene_tx=XM_019833518.1;refGene_gn=FBLN5;refGene_tx=XM_019833519.2;refGene_gn=FBLN5;refGene_tx=XM_019833520.2;refGene_gn=FBLN5;refGene_tx=XM_019833521.2
B3 137401700 137401806
refGene_gn=FBLN5;refGene_tx=XM_019833518.1;refGene_gn=FBLN5;refGene_tx=XM_019833519.2;refGene_gn=FBLN5;refGene_tx=XM_019833520.2;refGene_gn=FBLN5;refGene_tx=XM_019833521.2
B3 137402008 137402204
refGene_gn=FBLN5;refGene_tx=XM_003987921.4;refGene_gn=FBLN5;refGene_tx=XM_011283455.3;refGene_gn=FBLN5;refGene_tx=XM_019833518.1;refGene_gn=FBLN5;refGene_tx=XM_019833519.2;refGene_gn=FBLN5;refGene_tx=XM_019833520.2;refGene_gn=FBLN5;refGene_tx=XM_019833521.2
B3 137405453 137405580
refGene_gn=FBLN5;refGene_tx=XM_003987921.4;refGene_gn=FBLN5;refGene_tx=XM_011283455.3;refGene_gn=FBLN5;refGene_tx=XM_019833518.1;refGene_gn=FBLN5;refGene_tx=XM_019833519.2;refGene_gn=FBLN5;refGene_tx=XM_019833520.2;refGene_gn=FBLN5;refGene_tx=XM_019833521.2
B3 137406502 137406625
refGene_gn=FBLN5;refGene_tx=XM_003987921.4;refGene_gn=FBLN5;refGene_tx=XM_011283455.3;refGene_gn=FBLN5;refGene_tx=XM_019833518.1;refGene_gn=FBLN5;refGene_tx=XM_019833519.2;refGene_gn=FBLN5;refGene_tx=XM_019833520.2;refGene_gn=FBLN5;refGene_tx=XM_019833521.2
B3 137410995 137411115
refGene_gn=FBLN5;refGene_tx=XM_003987921.4;refGene_gn=FBLN5;refGene_tx=XM_011283455.3;refGene_gn=FBLN5;refGene_tx=XM_019833518.1;refGene_gn=FBLN5;refGene_tx=XM_019833519.2;refGene_gn=FBLN5;refGene_tx=XM_019833520.2;refGene_gn=FBLN5;refGene_tx=XM_019833521.2
B3 137414220 137414337
refGene_gn=FBLN5;refGene_tx=XM_003987921.4;refGene_gn=FBLN5;refGene_tx=XM_011283455.3;refGene_gn=FBLN5;refGene_tx=XM_019833518.1;refGene_gn=FBLN5;refGene_tx=XM_019833519.2;refGene_gn=FBLN5;refGene_tx=XM_019833520.2;refGene_gn=FBLN5;refGene_tx=XM_019833521.2
B3 137417416 137417539

refGene_gn=FBLN5;refGene_tx=XM_003987921.4;refGene_gn=FBLN5;refGene_tx=XM_011283455.3;refGene_gn=FBLN5;refGene_tx=XM_019833518.1;refGene_gn=FBLN5;refGene_tx=XM_019833519.2;refGene_gn=FBLN5;refGene_tx=XM_019833520.2;refGene_gn=FBLN5;refGene_tx=XM_019833521.2
B3 137463564 137463819
refGene_gn=FBLN5;refGene_tx=XM_003987921.4;refGene_gn=FBLN5;refGene_tx=XM_011283455.3;refGene_gn=FBLN5;refGene_tx=XM_019833518.1;refGene_gn=FBLN5;refGene_tx=XM_019833519.2;refGene_gn=FBLN5;refGene_tx=XM_019833520.2;refGene_gn=FBLN5;refGene_tx=XM_019833521.2
B3 137467691 137467743
refGene_gn=FBLN5;refGene_tx=XM_003987921.4;refGene_gn=FBLN5;refGene_tx=XM_011283455.3;refGene_gn=FBLN5;refGene_tx=XM_019833518.1;refGene_gn=FBLN5;refGene_tx=XM_019833519.2;refGene_gn=FBLN5;refGene_tx=XM_019833520.2;refGene_gn=FBLN5;refGene_tx=XM_019833521.2
B3 137469698 137469753
refGene_gn=FBLN5;refGene_tx=XM_003987921.4;refGene_gn=FBLN5;refGene_tx=XM_011283455.3;refGene_gn=FBLN5;refGene_tx=XM_019833518.1;refGene_gn=FBLN5;refGene_tx=XM_019833519.2;refGene_gn=FBLN5;refGene_tx=XM_019833520.2;refGene_gn=FBLN5;refGene_tx=XM_019833521.2
B3 137470669 137470680
refGene_gn=FBLN5;refGene_tx=XM_019833521.2
B3 137473002 137473058
refGene_gn=FBLN5;refGene_tx=XM_003987921.4;refGene_gn=FBLN5;refGene_tx=XM_019833518.1
B3 137473520 137473537
refGene_gn=FBLN5;refGene_tx=XM_011283455.3;refGene_gn=FBLN5;refGene_tx=XM_019833519.2;refGene_gn=FBLN5;refGene_tx=XM_019833520.2
B3 137491030 137491251
refGene_gn=TRIP11;refGene_tx=XM_011283456.3;refGene_gn=TRIP11;refGene_tx=XM_011283457.3
B3 137493348 137493493
refGene_gn=TRIP11;refGene_tx=XM_011283456.3;refGene_gn=TRIP11;refGene_tx=XM_011283457.3
B3 137494481 137494508
refGene_gn=TRIP11;refGene_tx=XM_019833523.1
B3 137494791 137494908
refGene_gn=TRIP11;refGene_tx=XM_011283456.3;refGene_gn=TRIP11;refGene_tx=XM_011283457.3;refGene_gn=TRIP11;refGene_tx=XM_019833523.1
B3 137495558 137495673
refGene_gn=TRIP11;refGene_tx=XM_011283456.3;refGene_gn=TRIP11;refGene_tx=XM_011283457.3;refGene_gn=TRIP11;refGene_tx=XM_019833523.1
B3 137496485 137496567
refGene_gn=TRIP11;refGene_tx=XM_011283456.3;refGene_gn=TRIP11;refGene_tx=XM_011283457.3;refGene_gn=TRIP11;refGene_tx=XM_019833523.1
B3 137507946 137508046
refGene_gn=TRIP11;refGene_tx=XM_011283456.3;refGene_gn=TRIP11;refGene_tx=XM_011283457.3;refGene_gn=TRIP11;refGene_tx=XM_019833523.1
B3 137512576 137512680
refGene_gn=TRIP11;refGene_tx=XM_011283456.3;refGene_gn=TRIP11;refGene_tx=XM_011283457.3;refGene_gn=TRIP11;refGene_tx=XM_019833523.1;refGene_

gn=TRIP11;refGene_tx=XM_019833524.1
B3	137514173	137514337
refGene_gn=TRIP11;refGene_tx=XM_011283456.3;refGene_gn=TRIP11;refGene_tx=XM_011283457.3;refGene_gn=TRIP11;refGene_tx=XM_019833523.1;refGene_gn=TRIP11;refGene_tx=XM_019833524.1
B3	137518530	137518724
refGene_gn=TRIP11;refGene_tx=XM_011283456.3;refGene_gn=TRIP11;refGene_tx=XM_011283457.3;refGene_gn=TRIP11;refGene_tx=XM_019833523.1
B3	137519963	137520104
refGene_gn=TRIP11;refGene_tx=XM_011283456.3;refGene_gn=TRIP11;refGene_tx=XM_011283457.3;refGene_gn=TRIP11;refGene_tx=XM_019833523.1;refGene_gn=TRIP11;refGene_tx=XM_019833524.1
B3	137520983	137524004
refGene_gn=TRIP11;refGene_tx=XM_011283456.3;refGene_gn=TRIP11;refGene_tx=XM_011283457.3;refGene_gn=TRIP11;refGene_tx=XM_019833523.1;refGene_gn=TRIP11;refGene_tx=XM_019833524.1
B3	137524539	137524752
refGene_gn=TRIP11;refGene_tx=XM_011283456.3;refGene_gn=TRIP11;refGene_tx=XM_011283457.3;refGene_gn=TRIP11;refGene_tx=XM_019833523.1;refGene_gn=TRIP11;refGene_tx=XM_019833524.1
B3	137528072	137528159
refGene_gn=TRIP11;refGene_tx=XM_011283456.3;refGene_gn=TRIP11;refGene_tx=XM_011283457.3;refGene_gn=TRIP11;refGene_tx=XM_019833523.1;refGene_gn=TRIP11;refGene_tx=XM_019833524.1
B3	137528739	137528780
refGene_gn=TRIP11;refGene_tx=XM_011283456.3;refGene_gn=TRIP11;refGene_tx=XM_011283457.3;refGene_gn=TRIP11;refGene_tx=XM_019833523.1;refGene_gn=TRIP11;refGene_tx=XM_019833524.1
B3	137532132	137532498
refGene_gn=TRIP11;refGene_tx=XM_011283456.3;refGene_gn=TRIP11;refGene_tx=XM_011283457.3;refGene_gn=TRIP11;refGene_tx=XM_019833523.1;refGene_gn=TRIP11;refGene_tx=XM_019833524.1
B3	137534834	137535000
refGene_gn=TRIP11;refGene_tx=XM_011283456.3;refGene_gn=TRIP11;refGene_tx=XM_011283457.3;refGene_gn=TRIP11;refGene_tx=XM_019833523.1;refGene_gn=TRIP11;refGene_tx=XM_019833524.1
B3	137536467	137536536
refGene_gn=TRIP11;refGene_tx=XM_011283456.3;refGene_gn=TRIP11;refGene_tx=XM_011283457.3;refGene_gn=TRIP11;refGene_tx=XM_019833523.1;refGene_gn=TRIP11;refGene_tx=XM_019833524.1
B3	137538701	137538980
refGene_gn=TRIP11;refGene_tx=XM_011283456.3;refGene_gn=TRIP11;refGene_tx=XM_011283457.3;refGene_gn=TRIP11;refGene_tx=XM_019833523.1;refGene_gn=TRIP11;refGene_tx=XM_019833524.1
B3	137541657	137541768
refGene_gn=TRIP11;refGene_tx=XM_011283456.3;refGene_gn=TRIP11;refGene_tx=XM_011283457.3;refGene_gn=TRIP11;refGene_tx=XM_019833523.1;refGene_gn=TRIP11;refGene_tx=XM_019833524.1
B3	137547891	137547953
refGene_gn=TRIP11;refGene_tx=XM_011283456.3;refGene_gn=TRIP11;refGene_

tx=XM_011283457.3;refGene_gn=TRIP11;refGene_tx=XM_019833523.1;refGene_gn=TRIP11;refGene_tx=XM_019833524.1
B3 137552097 137552236
refGene_gn=TRIP11;refGene_tx=XM_011283456.3;refGene_gn=TRIP11;refGene_tx=XM_011283457.3;refGene_gn=TRIP11;refGene_tx=XM_019833523.1;refGene_gn=TRIP11;refGene_tx=XM_019833524.1
B3 137574417 137574515
refGene_gn=ATXN3;refGene_tx=XM_006933060.3;refGene_gn=ATXN3;refGene_tx=XM_006933061.3;refGene_gn=ATXN3;refGene_tx=XM_006933062.3;refGene_gn=ATXN3;refGene_tx=XM_006933066.3;refGene_gn=ATXN3;refGene_tx=XM_019833525.2;refGene_gn=ATXN3;refGene_tx=XM_019833526.2;refGene_gn=ATXN3;refGene_tx=XM_019833529.2;refGene_gn=ATXN3;refGene_tx=XM_019833532.2;refGene_gn=ATXN3;refGene_tx=XM_023255966.1
B3 137579439 137579528
refGene_gn=ATXN3;refGene_tx=XM_006933060.3;refGene_gn=ATXN3;refGene_tx=XM_006933061.3;refGene_gn=ATXN3;refGene_tx=XM_006933062.3;refGene_gn=ATXN3;refGene_tx=XM_006933066.3;refGene_gn=ATXN3;refGene_tx=XM_019833525.2;refGene_gn=ATXN3;refGene_tx=XM_019833526.2;refGene_gn=ATXN3;refGene_tx=XM_019833529.2;refGene_gn=ATXN3;refGene_tx=XM_019833532.2;refGene_gn=ATXN3;refGene_tx=XM_023255966.1
B3 137581703 137581706
refGene_gn=ATXN3;refGene_tx=XM_006933060.3;refGene_gn=ATXN3;refGene_tx=XM_006933061.3;refGene_gn=ATXN3;refGene_tx=XM_006933066.3;refGene_gn=ATXN3;refGene_tx=XM_019833529.2;refGene_gn=ATXN3;refGene_tx=XM_019833532.2;refGene_gn=ATXN3;refGene_tx=XM_023255966.1
B3 137584265 137584299
refGene_gn=ATXN3;refGene_tx=XM_019833530.2;refGene_gn=ATXN3;refGene_tx=XM_019833531.2
B3 137584847 137584938
refGene_gn=ATXN3;refGene_tx=XM_019833527.2
B3 137585191 137585288
refGene_gn=ATXN3;refGene_tx=XM_006933060.3;refGene_gn=ATXN3;refGene_tx=XM_006933061.3;refGene_gn=ATXN3;refGene_tx=XM_006933062.3;refGene_gn=ATXN3;refGene_tx=XM_006933066.3;refGene_gn=ATXN3;refGene_tx=XM_019833525.2;refGene_gn=ATXN3;refGene_tx=XM_019833526.2;refGene_gn=ATXN3;refGene_tx=XM_019833527.2;refGene_gn=ATXN3;refGene_tx=XM_019833529.2;refGene_gn=ATXN3;refGene_tx=XM_019833530.2;refGene_gn=ATXN3;refGene_tx=XM_019833531.2;refGene_gn=ATXN3;refGene_tx=XM_019833532.2;refGene_gn=ATXN3;refGene_tx=XM_023255966.1
B3 137586839 137587006
refGene_gn=ATXN3;refGene_tx=XM_006933060.3;refGene_gn=ATXN3;refGene_tx=XM_006933061.3;refGene_gn=ATXN3;refGene_tx=XM_006933062.3;refGene_gn=ATXN3;refGene_tx=XM_006933066.3;refGene_gn=ATXN3;refGene_tx=XM_019833525.2;refGene_gn=ATXN3;refGene_tx=XM_019833526.2;refGene_gn=ATXN3;refGene_tx=XM_019833527.2;refGene_gn=ATXN3;refGene_tx=XM_019833529.2;refGene_gn=ATXN3;refGene_tx=XM_019833530.2;refGene_gn=ATXN3;refGene_tx=XM_019833531.2;refGene_gn=ATXN3;refGene_tx=XM_019833532.2;refGene_gn=ATXN3;refGene_tx=XM_023255966.1
B3 137587652 137587785
refGene_gn=ATXN3;refGene_tx=XM_006933060.3;refGene_gn=ATXN3;refGene_tx

=XM_006933061.3;refGene_gn=ATXN3;refGene_tx=XM_006933062.3;refGene_gn=
ATXN3;refGene_tx=XM_006933066.3;refGene_gn=ATXN3;refGene_tx=XM_0198335
25.2;refGene_gn=ATXN3;refGene_tx=XM_019833526.2;refGene_gn=ATXN3;refGe
ne_tx=XM_019833527.2;refGene_gn=ATXN3;refGene_tx=XM_019833529.2;refGen
e_gn=ATXN3;refGene_tx=XM_019833530.2;refGene_gn=ATXN3;refGene_tx=XM_01
9833531.2;refGene_gn=ATXN3;refGene_tx=XM_019833532.2;refGene_gn=ATXN3;
refGene_tx=XM_023255966.1
B3 137593341 137593429
refGene_gn=ATXN3;refGene_tx=XM_006933060.3;refGene_gn=ATXN3;refGene_tx
=XM_006933061.3;refGene_gn=ATXN3;refGene_tx=XM_006933062.3;refGene_gn=
ATXN3;refGene_tx=XM_006933066.3;refGene_gn=ATXN3;refGene_tx=XM_0198335
25.2;refGene_gn=ATXN3;refGene_tx=XM_019833526.2;refGene_gn=ATXN3;refGe
ne_tx=XM_019833527.2;refGene_gn=ATXN3;refGene_tx=XM_019833529.2;refGen
e_gn=ATXN3;refGene_tx=XM_019833530.2;refGene_gn=ATXN3;refGene_tx=XM_01
9833531.2;refGene_gn=ATXN3;refGene_tx=XM_019833532.2;refGene_gn=ATXN3;
refGene_tx=XM_023255966.1
B3 137598137 137598204
refGene_gn=ATXN3;refGene_tx=XM_006933060.3;refGene_gn=ATXN3;refGene_tx
=XM_006933061.3;refGene_gn=ATXN3;refGene_tx=XM_006933062.3;refGene_gn=
ATXN3;refGene_tx=XM_006933066.3;refGene_gn=ATXN3;refGene_tx=XM_0198335
25.2;refGene_gn=ATXN3;refGene_tx=XM_019833526.2;refGene_gn=ATXN3;refGe
ne_tx=XM_019833527.2;refGene_gn=ATXN3;refGene_tx=XM_019833529.2;refGen
e_gn=ATXN3;refGene_tx=XM_019833530.2;refGene_gn=ATXN3;refGene_tx=XM_01
9833531.2;refGene_gn=ATXN3;refGene_tx=XM_019833532.2;refGene_gn=ATXN3;
refGene_tx=XM_023255966.1
B3 137598677 137598763
refGene_gn=ATXN3;refGene_tx=XM_006933060.3;refGene_gn=ATXN3;refGene_tx
=XM_006933061.3;refGene_gn=ATXN3;refGene_tx=XM_006933062.3;refGene_gn=
ATXN3;refGene_tx=XM_006933066.3;refGene_gn=ATXN3;refGene_tx=XM_0198335
25.2;refGene_gn=ATXN3;refGene_tx=XM_019833526.2;refGene_gn=ATXN3;refGe
ne_tx=XM_019833527.2;refGene_gn=ATXN3;refGene_tx=XM_019833529.2;refGen
e_gn=ATXN3;refGene_tx=XM_019833530.2;refGene_gn=ATXN3;refGene_tx=XM_01
9833531.2;refGene_gn=ATXN3;refGene_tx=XM_019833532.2;refGene_gn=ATXN3;
refGene_tx=XM_023255966.1
B3 137600455 137600500
refGene_gn=ATXN3;refGene_tx=XM_006933060.3;refGene_gn=ATXN3;refGene_tx
=XM_006933061.3;refGene_gn=ATXN3;refGene_tx=XM_006933062.3;refGene_gn=
ATXN3;refGene_tx=XM_006933066.3;refGene_gn=ATXN3;refGene_tx=XM_0198335
25.2;refGene_gn=ATXN3;refGene_tx=XM_019833526.2;refGene_gn=ATXN3;refGe
ne_tx=XM_019833527.2;refGene_gn=ATXN3;refGene_tx=XM_019833530.2;refGen
e_gn=ATXN3;refGene_tx=XM_019833531.2;refGene_gn=ATXN3;refGene_tx=XM_02
3255966.1
B3 137600729 137600894
refGene_gn=ATXN3;refGene_tx=XM_006933060.3;refGene_gn=ATXN3;refGene_tx
=XM_006933061.3;refGene_gn=ATXN3;refGene_tx=XM_006933062.3;refGene_gn=
ATXN3;refGene_tx=XM_006933066.3;refGene_gn=ATXN3;refGene_tx=XM_0198335
25.2;refGene_gn=ATXN3;refGene_tx=XM_019833526.2;refGene_gn=ATXN3;refGe
ne_tx=XM_019833527.2;refGene_gn=ATXN3;refGene_tx=XM_019833529.2;refGen
e_gn=ATXN3;refGene_tx=XM_019833530.2;refGene_gn=ATXN3;refGene_tx=XM_01
9833531.2;refGene_gn=ATXN3;refGene_tx=XM_019833532.2;refGene_gn=ATXN3;

refGene_tx=XM_023255966.1
B3 137602240 137602277
refGene_gn=ATXN3;refGene_tx=XM_019833525.2;refGene_gn=ATXN3;refGene_tx=XM_019833526.2;refGene_gn=ATXN3;refGene_tx=XM_019833527.2;refGene_gn=ATXN3;refGene_tx=XM_019833530.2
B3 137602386 137602466
refGene_gn=ATXN3;refGene_tx=XM_019833525.2;refGene_gn=ATXN3;refGene_tx=XM_019833526.2;refGene_gn=ATXN3;refGene_tx=XM_019833527.2;refGene_gn=ATXN3;refGene_tx=XM_019833530.2
B3 137607831 137607855
refGene_gn=ATXN3;refGene_tx=XM_006933060.3;refGene_gn=ATXN3;refGene_tx=XM_006933061.3;refGene_gn=ATXN3;refGene_tx=XM_006933062.3;refGene_gn=ATXN3;refGene_tx=XM_019833529.2;refGene_gn=ATXN3;refGene_tx=XM_019833531.2;refGene_gn=ATXN3;refGene_tx=XM_019833532.2
B3 137614467 137614504
refGene_gn=NDUFB1;refGene_tx=XM_003987965.2
B3 137615443 137615588
refGene_gn=NDUFB1;refGene_tx=XM_003987965.2
B3 137620829 137620986
refGene_gn=NDUFB1;refGene_tx=XM_003987965.2
B3 137629070 137629219
refGene_gn=CPSF2;refGene_tx=XM_003987928.5;refGene_gn=CPSF2;refGene_tx=XM_019833533.2
B3 137630424 137630584
refGene_gn=CPSF2;refGene_tx=XM_003987928.5;refGene_gn=CPSF2;refGene_tx=XM_019833533.2
B3 137630663 137630769
refGene_gn=CPSF2;refGene_tx=XM_003987928.5;refGene_gn=CPSF2;refGene_tx=XM_019833533.2
B3 137632073 137632203
refGene_gn=CPSF2;refGene_tx=XM_003987928.5;refGene_gn=CPSF2;refGene_tx=XM_019833533.2
B3 137633913 137634029
refGene_gn=CPSF2;refGene_tx=XM_003987928.5;refGene_gn=CPSF2;refGene_tx=XM_019833533.2
B3 137635690 137635878
refGene_gn=CPSF2;refGene_tx=XM_003987928.5;refGene_gn=CPSF2;refGene_tx=XM_019833533.2
B3 137636550 137636841
refGene_gn=CPSF2;refGene_tx=XM_003987928.5;refGene_gn=CPSF2;refGene_tx=XM_019833533.2
B3 137642373 137642474
refGene_gn=CPSF2;refGene_tx=XM_003987928.5;refGene_gn=CPSF2;refGene_tx=XM_019833533.2
B3 137643917 137644118
refGene_gn=CPSF2;refGene_tx=XM_003987928.5;refGene_gn=CPSF2;refGene_tx=XM_019833533.2
B3 137646488 137646641
refGene_gn=CPSF2;refGene_tx=XM_003987928.5;refGene_gn=CPSF2;refGene_tx=XM_019833533.2

B3 137648045 137648271
refGene_gn=CPSF2;refGene_tx=XM_003987928.5;refGene_gn=CPSF2;refGene_tx
=XM_019833533.2
B3 137649414 137649714
refGene_gn=CPSF2;refGene_tx=XM_003987928.5;refGene_gn=CPSF2;refGene_tx
=XM_019833533.2
B3 137651118 137651253
refGene_gn=CPSF2;refGene_tx=XM_003987928.5;refGene_gn=CPSF2;refGene_tx
=XM_019833533.2
B3 137651642 137651735
refGene_gn=CPSF2;refGene_tx=XM_003987928.5;refGene_gn=CPSF2;refGene_tx
=XM_019833533.2
B3 137796324 137796454
refGene_gn=SLC24A4;refGene_tx=XM_003987930.5;refGene_gn=SLC24A4;refGen
e_tx=XM_011283458.3
B3 137798342 137798453
refGene_gn=SLC24A4;refGene_tx=XM_003987930.5;refGene_gn=SLC24A4;refGen
e_tx=XM_011283458.3
B3 137909075 137909152
refGene_gn=SLC24A4;refGene_tx=XM_003987930.5;refGene_gn=SLC24A4;refGen
e_tx=XM_011283458.3
B3 137911915 137911990
refGene_gn=SLC24A4;refGene_tx=XM_003987930.5;refGene_gn=SLC24A4;refGen
e_tx=XM_011283458.3
B3 137914785 137914870
refGene_gn=SLC24A4;refGene_tx=XM_003987930.5;refGene_gn=SLC24A4;refGen
e_tx=XM_011283458.3
B3 137915410 137915514
refGene_gn=SLC24A4;refGene_tx=XM_003987930.5;refGene_gn=SLC24A4;refGen
e_tx=XM_011283458.3
B3 137916339 137916414
refGene_gn=SLC24A4;refGene_tx=XM_003987930.5;refGene_gn=SLC24A4;refGen
e_tx=XM_011283458.3
B3 137919169 137919195
refGene_gn=SLC24A4;refGene_tx=XM_003987930.5;refGene_gn=SLC24A4;refGen
e_tx=XM_011283458.3
B3 137920793 137920847
refGene_gn=SLC24A4;refGene_tx=XM_003987930.5;refGene_gn=SLC24A4;refGen
e_tx=XM_011283458.3
B3 137922481 137922624
refGene_gn=SLC24A4;refGene_tx=XM_003987930.5;refGene_gn=SLC24A4;refGen
e_tx=XM_011283458.3
B3 137926874 137927044
refGene_gn=SLC24A4;refGene_tx=XM_003987930.5;refGene_gn=SLC24A4;refGen
e_tx=XM_011283458.3
B3 137931690 137931895
refGene_gn=SLC24A4;refGene_tx=XM_003987930.5;refGene_gn=SLC24A4;refGen
e_tx=XM_011283458.3
B3 137956840 137957007
refGene_gn=SLC24A4;refGene_tx=XM_003987930.5;refGene_gn=SLC24A4;refGen

e_tx=XM_011283458.3
B3 137961230 137961345
refGene_gn=SLC24A4;refGene_tx=XM_003987930.5;refGene_gn=SLC24A4;refGen
e_tx=XM_011283458.3
B3 137965459 137965572
refGene_gn=SLC24A4;refGene_tx=XM_003987930.5;refGene_gn=SLC24A4;refGen
e_tx=XM_011283458.3
B3 137966035 137966101
refGene_gn=SLC24A4;refGene_tx=XM_003987930.5;refGene_gn=SLC24A4;refGen
e_tx=XM_011283458.3
B3 137967312 137967465
refGene_gn=SLC24A4;refGene_tx=XM_003987930.5;refGene_gn=SLC24A4;refGen
e_tx=XM_011283458.3
B3 137994069 137994113
refGene_gn=RIN3;refGene_tx=XM_023255968.1
B3 138034170 138034375
refGene_gn=RIN3;refGene_tx=XM_023255968.1
B3 138049672 138049790
refGene_gn=RIN3;refGene_tx=XM_023255968.1
B3 138074588 138074661
refGene_gn=RIN3;refGene_tx=XM_023255968.1
B3 138082847 138082939
refGene_gn=RIN3;refGene_tx=XM_023255968.1
B3 138090402 138090803
refGene_gn=RIN3;refGene_tx=XM_023255968.1
B3 138090846 138091963
refGene_gn=RIN3;refGene_tx=XM_023255968.1
B3 138094133 138094442
refGene_gn=RIN3;refGene_tx=XM_023255968.1
B3 138109479 138109611
refGene_gn=RIN3;refGene_tx=XM_023255968.1
B3 138114850 138115014
refGene_gn=RIN3;refGene_tx=XM_023255968.1
B3 138117742 138118063
refGene_gn=RIN3;refGene_tx=XM_023255968.1
B3 138128518 138128561
refGene_gn=LGMN;refGene_tx=XM_003987931.4
B3 138128829 138128897
refGene_gn=LGMN;refGene_tx=XM_003987931.4
B3 138131806 138131977
refGene_gn=LGMN;refGene_tx=XM_003987931.4
B3 138133069 138133270
refGene_gn=LGMN;refGene_tx=XM_003987931.4
B3 138135185 138135275
refGene_gn=LGMN;refGene_tx=XM_003987931.4
B3 138135361 138135480
refGene_gn=LGMN;refGene_tx=XM_003987931.4
B3 138136557 138136624
refGene_gn=LGMN;refGene_tx=XM_003987931.4
B3 138137411 138137474

refGene_gn=LGMN;refGene_tx=XM_003987931.4
B3	138138183	138138259
refGene_gn=LGMN;refGene_tx=XM_003987931.4
B3	138139572	138139658
refGene_gn=LGMN;refGene_tx=XM_003987931.4
B3	138141346	138141428
refGene_gn=LGMN;refGene_tx=XM_003987931.4
B3	138142663	138142761
refGene_gn=LGMN;refGene_tx=XM_003987931.4
B3	138153870	138154008
refGene_gn=LGMN;refGene_tx=XM_003987931.4
B3	138199940	138200484
refGene_gn=GOLGA5;refGene_tx=XM_023255969.1;refGene_gn=GOLGA5;refGene_tx=XM_023255970.1
B3	138207563	138207791
refGene_gn=GOLGA5;refGene_tx=XM_023255969.1;refGene_gn=GOLGA5;refGene_tx=XM_023255970.1
B3	138209530	138209750
refGene_gn=GOLGA5;refGene_tx=XM_023255969.1;refGene_gn=GOLGA5;refGene_tx=XM_023255970.1
B3	138210205	138210329
refGene_gn=GOLGA5;refGene_tx=XM_023255969.1;refGene_gn=GOLGA5;refGene_tx=XM_023255970.1
B3	138213046	138213250
refGene_gn=GOLGA5;refGene_tx=XM_023255969.1;refGene_gn=GOLGA5;refGene_tx=XM_023255970.1
B3	138216908	138217079
refGene_gn=GOLGA5;refGene_tx=XM_023255969.1;refGene_gn=GOLGA5;refGene_tx=XM_023255970.1
B3	138219507	138219636
refGene_gn=GOLGA5;refGene_tx=XM_023255969.1;refGene_gn=GOLGA5;refGene_tx=XM_023255970.1
B3	138222116	138222215
refGene_gn=GOLGA5;refGene_tx=XM_023255969.1;refGene_gn=GOLGA5;refGene_tx=XM_023255970.1
B3	138224143	138224369
refGene_gn=GOLGA5;refGene_tx=XM_023255969.1;refGene_gn=GOLGA5;refGene_tx=XM_023255970.1
B3	138227421	138227527
refGene_gn=GOLGA5;refGene_tx=XM_023255969.1;refGene_gn=GOLGA5;refGene_tx=XM_023255970.1
B3	138228907	138228971
refGene_gn=GOLGA5;refGene_tx=XM_023255969.1;refGene_gn=GOLGA5;refGene_tx=XM_023255970.1
B3	138230844	138230925
refGene_gn=GOLGA5;refGene_tx=XM_023255969.1;refGene_gn=GOLGA5;refGene_tx=XM_023255970.1
B3	138293501	138293547
refGene_gn=CHGA;refGene_tx=XM_023255975.1
B3	138294369	138294416

refGene_gn=CHGA;refGene_tx=XM_023255975.1
B3 138296738 138296832
refGene_gn=CHGA;refGene_tx=XM_023255975.1
B3 138297616 138297685
refGene_gn=CHGA;refGene_tx=XM_023255975.1
B3 138299778 138299874
refGene_gn=CHGA;refGene_tx=XM_023255975.1
B3 138301751 138302189
refGene_gn=CHGA;refGene_tx=XM_023255975.1
B3 138302903 138303394
refGene_gn=CHGA;refGene_tx=XM_023255975.1
B3 138305011 138305095
refGene_gn=CHGA;refGene_tx=XM_023255975.1
B3 138310379 138310738
refGene_gn=ITPK1;refGene_tx=XM_019833537.2;refGene_gn=ITPK1;refGene_tx=XM_019833538.2;refGene_gn=ITPK1;refGene_tx=XM_019833539.2;refGene_gn=ITPK1;refGene_tx=XM_023255971.1;refGene_gn=ITPK1;refGene_tx=XM_023255972.1;refGene_gn=ITPK1;refGene_tx=XM_023255973.1;refGene_gn=ITPK1;refGene_tx=XM_023255974.1
B3 138315271 138315434
refGene_gn=ITPK1;refGene_tx=XM_019833537.2;refGene_gn=ITPK1;refGene_tx=XM_019833538.2;refGene_gn=ITPK1;refGene_tx=XM_019833539.2;refGene_gn=ITPK1;refGene_tx=XM_023255971.1;refGene_gn=ITPK1;refGene_tx=XM_023255972.1;refGene_gn=ITPK1;refGene_tx=XM_023255973.1;refGene_gn=ITPK1;refGene_tx=XM_023255974.1
B3 138319844 138319912
refGene_gn=ITPK1;refGene_tx=XM_019833537.2;refGene_gn=ITPK1;refGene_tx=XM_019833538.2;refGene_gn=ITPK1;refGene_tx=XM_019833539.2;refGene_gn=ITPK1;refGene_tx=XM_023255971.1;refGene_gn=ITPK1;refGene_tx=XM_023255972.1;refGene_gn=ITPK1;refGene_tx=XM_023255973.1;refGene_gn=ITPK1;refGene_tx=XM_023255974.1
B3 138321190 138321316
refGene_gn=ITPK1;refGene_tx=XM_019833537.2;refGene_gn=ITPK1;refGene_tx=XM_019833538.2;refGene_gn=ITPK1;refGene_tx=XM_019833539.2;refGene_gn=ITPK1;refGene_tx=XM_023255971.1;refGene_gn=ITPK1;refGene_tx=XM_023255972.1;refGene_gn=ITPK1;refGene_tx=XM_023255974.1
B3 138325966 138326132
refGene_gn=ITPK1;refGene_tx=XM_019833537.2;refGene_gn=ITPK1;refGene_tx=XM_019833538.2;refGene_gn=ITPK1;refGene_tx=XM_019833539.2;refGene_gn=ITPK1;refGene_tx=XM_023255971.1;refGene_gn=ITPK1;refGene_tx=XM_023255972.1;refGene_gn=ITPK1;refGene_tx=XM_023255973.1;refGene_gn=ITPK1;refGene_tx=XM_023255974.1
B3 138330037 138330078
refGene_gn=ITPK1;refGene_tx=XM_019833537.2;refGene_gn=ITPK1;refGene_tx=XM_019833538.2;refGene_gn=ITPK1;refGene_tx=XM_019833539.2;refGene_gn=ITPK1;refGene_tx=XM_023255971.1;refGene_gn=ITPK1;refGene_tx=XM_023255972.1;refGene_gn=ITPK1;refGene_tx=XM_023255973.1;refGene_gn=ITPK1;refGene_tx=XM_023255974.1
B3 138330442 138330541
refGene_gn=ITPK1;refGene_tx=XM_019833537.2;refGene_gn=ITPK1;refGene_tx

=XM_019833538.2;refGene_gn=ITPK1;refGene_tx=XM_019833539.2;refGene_gn=ITPK1;refGene_tx=XM_023255971.1;refGene_gn=ITPK1;refGene_tx=XM_023255972.1;refGene_gn=ITPK1;refGene_tx=XM_023255973.1;refGene_gn=ITPK1;refGene_tx=XM_023255974.1
B3 138355948 138356066
refGene_gn=ITPK1;refGene_tx=XM_019833537.2;refGene_gn=ITPK1;refGene_tx=XM_019833538.2;refGene_gn=ITPK1;refGene_tx=XM_019833539.2;refGene_gn=ITPK1;refGene_tx=XM_023255971.1;refGene_gn=ITPK1;refGene_tx=XM_023255972.1;refGene_gn=ITPK1;refGene_tx=XM_023255973.1;refGene_gn=ITPK1;refGene_tx=XM_023255974.1
B3 138380997 138381123
refGene_gn=ITPK1;refGene_tx=XM_023255971.1;refGene_gn=ITPK1;refGene_tx=XM_023255972.1;refGene_gn=ITPK1;refGene_tx=XM_023255973.1
B3 138432496 138432535
refGene_gn=ITPK1;refGene_tx=XM_023255972.1
B3 138444539 138444564
refGene_gn=ITPK1;refGene_tx=XM_023255971.1;refGene_gn=ITPK1;refGene_tx=XM_023255973.1;refGene_gn=ITPK1;refGene_tx=XM_023255974.1
B3 138486370 138486465
refGene_gn=ITPK1;refGene_tx=XM_023255971.1;refGene_gn=ITPK1;refGene_tx=XM_023255973.1;refGene_gn=ITPK1;refGene_tx=XM_023255974.1
B3 138542250 138543309
refGene_gn=MOAP1;refGene_tx=XM_023255976.1
B3 138544238 138544336
refGene_gn=TMEM251;refGene_tx=XM_003987934.4
B3 138544761 138545173
refGene_gn=TMEM251;refGene_tx=XM_003987934.4
B3 138559103 138559198
refGene_gn=GON7;refGene_tx=XM_003987936.5
B3 138561865 138562070
refGene_gn=GON7;refGene_tx=XM_003987936.5
B3 138562332 138562482
refGene_gn=UBR7;refGene_tx=XM_003987935.5
B3 138564300 138564434
refGene_gn=UBR7;refGene_tx=XM_003987935.5
B3 138564958 138565019
refGene_gn=UBR7;refGene_tx=XM_003987935.5
B3 138566485 138566581
refGene_gn=UBR7;refGene_tx=XM_003987935.5
B3 138568906 138568960
refGene_gn=UBR7;refGene_tx=XM_003987935.5
B3 138569434 138569540
refGene_gn=UBR7;refGene_tx=XM_003987935.5
B3 138571832 138572041
refGene_gn=UBR7;refGene_tx=XM_003987935.5
B3 138572205 138572355
refGene_gn=UBR7;refGene_tx=XM_003987935.5
B3 138573164 138573327
refGene_gn=UBR7;refGene_tx=XM_003987935.5
B3 138574353 138574415

refGene_gn=UBR7;refGene_tx=XM_003987935.5
B3	138579824	138579917
refGene_gn=UBR7;refGene_tx=XM_003987935.5
B3	138595615	138596431
refGene_gn=BTBD7;refGene_tx=XM_023255977.1;refGene_gn=BTBD7;refGene_tx=XM_023255978.1
B3	138598897	138599344
refGene_gn=BTBD7;refGene_tx=XM_023255977.1;refGene_gn=BTBD7;refGene_tx=XM_023255978.1
B3	138600842	138601021
refGene_gn=BTBD7;refGene_tx=XM_023255977.1;refGene_gn=BTBD7;refGene_tx=XM_023255978.1;refGene_gn=BTBD7;refGene_tx=XM_023255979.1
B3	138602319	138602383
refGene_gn=BTBD7;refGene_tx=XM_023255979.1
B3	138603426	138603473
refGene_gn=BTBD7;refGene_tx=XM_023255980.1
B3	138604034	138604224
refGene_gn=BTBD7;refGene_tx=XM_023255977.1;refGene_gn=BTBD7;refGene_tx=XM_023255978.1;refGene_gn=BTBD7;refGene_tx=XM_023255979.1;refGene_gn=BTBD7;refGene_tx=XM_023255980.1
B3	138606386	138606530
refGene_gn=BTBD7;refGene_tx=XM_023255977.1;refGene_gn=BTBD7;refGene_tx=XM_023255978.1;refGene_gn=BTBD7;refGene_tx=XM_023255979.1;refGene_gn=BTBD7;refGene_tx=XM_023255980.1
B3	138610077	138610238
refGene_gn=BTBD7;refGene_tx=XM_023255977.1;refGene_gn=BTBD7;refGene_tx=XM_023255978.1;refGene_gn=BTBD7;refGene_tx=XM_023255979.1;refGene_gn=BTBD7;refGene_tx=XM_023255980.1
B3	138614790	138614866
refGene_gn=BTBD7;refGene_tx=XM_023255977.1;refGene_gn=BTBD7;refGene_tx=XM_023255979.1;refGene_gn=BTBD7;refGene_tx=XM_023255980.1
B3	138616375	138616584
refGene_gn=BTBD7;refGene_tx=XM_023255977.1;refGene_gn=BTBD7;refGene_tx=XM_023255979.1;refGene_gn=BTBD7;refGene_tx=XM_023255980.1
B3	138642797	138642823
refGene_gn=BTBD7;refGene_tx=XM_023255982.1
B3	138646425	138646463
refGene_gn=BTBD7;refGene_tx=XM_023255981.1
B3	138647010	138648090
refGene_gn=BTBD7;refGene_tx=XM_023255977.1;refGene_gn=BTBD7;refGene_tx=XM_023255978.1;refGene_gn=BTBD7;refGene_tx=XM_023255979.1;refGene_gn=BTBD7;refGene_tx=XM_023255980.1;refGene_gn=BTBD7;refGene_tx=XM_023255981.1;refGene_gn=BTBD7;refGene_tx=XM_023255982.1
B3	138648809	138648891
refGene_gn=BTBD7;refGene_tx=XM_023255977.1;refGene_gn=BTBD7;refGene_tx=XM_023255978.1;refGene_gn=BTBD7;refGene_tx=XM_023255979.1;refGene_gn=BTBD7;refGene_tx=XM_023255980.1;refGene_gn=BTBD7;refGene_tx=XM_023255981.1;refGene_gn=BTBD7;refGene_tx=XM_023255982.1
B3	138698646	138698679
refGene_gn=LOC111560969;refGene_tx=XM_023256192.1

B3 138698936 138699237
refGene_gn=LOC111560969;refGene_tx=XM_023256192.1
B3 138699746 138699977
refGene_gn=LOC111560969;refGene_tx=XM_023256192.1
B3 138700104 138700223
refGene_gn=LOC111560969;refGene_tx=XM_023256192.1
B3 138794030 138794133
refGene_gn=UNC79;refGene_tx=XM_019833555.2
B3 138794891 138795012
refGene_gn=UNC79;refGene_tx=XM_019833555.2
B3 138802182 138802487
refGene_gn=UNC79;refGene_tx=XM_019833555.2
B3 138805824 138805995
refGene_gn=UNC79;refGene_tx=XM_019833555.2
B3 138817702 138817795
refGene_gn=UNC79;refGene_tx=XM_019833555.2
B3 138826866 138826922
refGene_gn=UNC79;refGene_tx=XM_019833555.2
B3 138827602 138827732
refGene_gn=UNC79;refGene_tx=XM_019833555.2
B3 138853930 138853995
refGene_gn=UNC79;refGene_tx=XM_019833555.2
B3 138858810 138858899
refGene_gn=UNC79;refGene_tx=XM_019833555.2
B3 138859529 138859570
refGene_gn=UNC79;refGene_tx=XM_019833555.2
B3 138862252 138862281
refGene_gn=UNC79;refGene_tx=XM_019833555.2
B3 138867301 138867531
refGene_gn=UNC79;refGene_tx=XM_019833555.2
B3 138869843 138870015
refGene_gn=UNC79;refGene_tx=XM_019833555.2
B3 138871380 138871611
refGene_gn=UNC79;refGene_tx=XM_019833555.2
B3 138884686 138884877
refGene_gn=UNC79;refGene_tx=XM_019833555.2
B3 138885447 138885571
refGene_gn=UNC79;refGene_tx=XM_019833555.2
B3 138887532 138887673
refGene_gn=UNC79;refGene_tx=XM_019833555.2
B3 138889784 138890006
refGene_gn=UNC79;refGene_tx=XM_019833555.2
B3 138892140 138892368
refGene_gn=UNC79;refGene_tx=XM_019833555.2
B3 138896521 138896620
refGene_gn=UNC79;refGene_tx=XM_019833555.2
B3 138896621 138896664
refGene_gn=UNC79;refGene_tx=XM_019833555.2
B3 138898711 138898791
refGene_gn=UNC79;refGene_tx=XM_019833555.2

B3	138898875	138899024
refGene_gn=UNC79;refGene_tx=XM_019833555.2
B3	138903786	138903944
refGene_gn=UNC79;refGene_tx=XM_019833555.2
B3	138907407	138907589
refGene_gn=UNC79;refGene_tx=XM_019833555.2
B3	138909744	138909946
refGene_gn=UNC79;refGene_tx=XM_019833555.2
B3	138911021	138911201
refGene_gn=UNC79;refGene_tx=XM_019833555.2
B3	138921018	138921305
refGene_gn=UNC79;refGene_tx=XM_019833555.2
B3	138923587	138923770
refGene_gn=UNC79;refGene_tx=XM_019833555.2
B3	138924357	138924520
refGene_gn=UNC79;refGene_tx=XM_019833555.2
B3	138927032	138927182
refGene_gn=UNC79;refGene_tx=XM_019833555.2
B3	138927765	138928983
refGene_gn=UNC79;refGene_tx=XM_019833555.2
B3	138936624	138936732
refGene_gn=UNC79;refGene_tx=XM_019833555.2
B3	138940019	138940136
refGene_gn=UNC79;refGene_tx=XM_019833555.2
B3	138942937	138943021
refGene_gn=UNC79;refGene_tx=XM_019833555.2
B3	138947566	138947669
refGene_gn=UNC79;refGene_tx=XM_019833555.2
B3	138949909	138950050
refGene_gn=UNC79;refGene_tx=XM_019833555.2
B3	138952889	138952928
refGene_gn=UNC79;refGene_tx=XM_019833555.2
B3	138962006	138962119
refGene_gn=UNC79;refGene_tx=XM_019833555.2
B3	138962207	138962293
refGene_gn=UNC79;refGene_tx=XM_019833555.2
B3	138963181	138963355
refGene_gn=UNC79;refGene_tx=XM_019833555.2
B3	138966717	138966786
refGene_gn=UNC79;refGene_tx=XM_019833555.2
B3	138969736	138969847
refGene_gn=UNC79;refGene_tx=XM_019833555.2
B3	138975391	138975496
refGene_gn=UNC79;refGene_tx=XM_019833555.2
B3	138983889	138983967
refGene_gn=UNC79;refGene_tx=XM_019833555.2
B3	138991092	138991182
refGene_gn=UNC79;refGene_tx=XM_019833555.2
B3	138993232	138993408
refGene_gn=UNC79;refGene_tx=XM_019833555.2

B3	138994366	138994553
refGene_gn=UNC79;refGene_tx=XM_019833555.2
B3	138996037	138996235
refGene_gn=UNC79;refGene_tx=XM_019833555.2
B3	138998286	138998364
refGene_gn=UNC79;refGene_tx=XM_019833555.2
B3	139016445	139016487
refGene_gn=UNC79;refGene_tx=XM_019833555.2
B3	139018689	139018890
refGene_gn=UNC79;refGene_tx=XM_019833555.2
B3	139031944	139032047
refGene_gn=PRIMA1;refGene_tx=XM_003987939.5;refGene_gn=PRIMA1;refGene_tx=XM_023255983.1
B3	139033184	139033230
refGene_gn=PRIMA1;refGene_tx=XM_006933070.4
B3	139046439	139046569
refGene_gn=PRIMA1;refGene_tx=XM_003987939.5;refGene_gn=PRIMA1;refGene_tx=XM_006933070.4;refGene_gn=PRIMA1;refGene_tx=XM_023255983.1
B3	139082751	139082887
refGene_gn=PRIMA1;refGene_tx=XM_003987939.5;refGene_gn=PRIMA1;refGene_tx=XM_006933070.4;refGene_gn=PRIMA1;refGene_tx=XM_023255983.1
B3	139089983	139090076
refGene_gn=PRIMA1;refGene_tx=XM_003987939.5;refGene_gn=PRIMA1;refGene_tx=XM_006933070.4;refGene_gn=PRIMA1;refGene_tx=XM_023255983.1
B3	139194333	139194543
refGene_gn=FAM181A;refGene_tx=XM_019833575.2;refGene_gn=FAM181A;refGene_tx=XM_019833576.2
B3	139195283	139196249
refGene_gn=FAM181A;refGene_tx=XM_006933071.3;refGene_gn=FAM181A;refGene_tx=XM_019833575.2;refGene_gn=FAM181A;refGene_tx=XM_019833576.2;refGene_gn=FAM181A;refGene_tx=XM_019833577.2;refGene_gn=FAM181A;refGene_tx=XM_019833578.2
B3	139200083	139200220
refGene_gn=ASB2;refGene_tx=XM_023255984.1;refGene_gn=ASB2;refGene_tx=XM_023255985.1;refGene_gn=ASB2;refGene_tx=XM_023255987.1
B3	139202472	139202626
refGene_gn=ASB2;refGene_tx=XM_023255984.1;refGene_gn=ASB2;refGene_tx=XM_023255985.1;refGene_gn=ASB2;refGene_tx=XM_023255987.1
B3	139203023	139203170
refGene_gn=ASB2;refGene_tx=XM_023255986.1
B3	139203458	139204023
refGene_gn=ASB2;refGene_tx=XM_023255984.1;refGene_gn=ASB2;refGene_tx=XM_023255985.1;refGene_gn=ASB2;refGene_tx=XM_023255986.1;refGene_gn=ASB2;refGene_tx=XM_023255987.1
B3	139211519	139211691
refGene_gn=ASB2;refGene_tx=XM_023255984.1;refGene_gn=ASB2;refGene_tx=XM_023255985.1;refGene_gn=ASB2;refGene_tx=XM_023255986.1;refGene_gn=ASB2;refGene_tx=XM_023255987.1
B3	139214609	139214855
refGene_gn=ASB2;refGene_tx=XM_023255984.1;refGene_gn=ASB2;refGene_tx=X

M_023255985.1;refGene_gn=ASB2;refGene_tx=XM_023255986.1;refGene_gn=ASB2;refGene_tx=XM_023255987.1
B3	139217182	139217338
refGene_gn=ASB2;refGene_tx=XM_023255984.1;refGene_gn=ASB2;refGene_tx=XM_023255985.1;refGene_gn=ASB2;refGene_tx=XM_023255986.1;refGene_gn=ASB2;refGene_tx=XM_023255987.1
B3	139218145	139218312
refGene_gn=ASB2;refGene_tx=XM_023255984.1;refGene_gn=ASB2;refGene_tx=XM_023255985.1;refGene_gn=ASB2;refGene_tx=XM_023255986.1;refGene_gn=ASB2;refGene_tx=XM_023255987.1
B3	139221362	139221452
refGene_gn=ASB2;refGene_tx=XM_023255984.1;refGene_gn=ASB2;refGene_tx=XM_023255985.1;refGene_gn=ASB2;refGene_tx=XM_023255986.1;refGene_gn=ASB2;refGene_tx=XM_023255987.1
B3	139228091	139228297
refGene_gn=ASB2;refGene_tx=XM_023255984.1;refGene_gn=ASB2;refGene_tx=XM_023255985.1;refGene_gn=ASB2;refGene_tx=XM_023255986.1
B3	139263667	139263847
refGene_gn=LOC111560970;refGene_tx=XM_023256193.1
B3	139263848	139263891
refGene_gn=LOC111560970;refGene_tx=XM_023256193.1
B3	139265400	139265579
refGene_gn=LOC111560970;refGene_tx=XM_023256193.1
B3	139267191	139267302
refGene_gn=LOC111560970;refGene_tx=XM_023256193.1
B3	139268111	139268183
refGene_gn=LOC111560970;refGene_tx=XM_023256193.1
B3	139268554	139268671
refGene_gn=LOC111560970;refGene_tx=XM_023256193.1
B3	139271861	139272341
refGene_gn=LOC111560970;refGene_tx=XM_023256193.1
B3	139287952	139287955
refGene_gn=OTUB2;refGene_tx=XM_011283467.3
B3	139299122	139299218
refGene_gn=OTUB2;refGene_tx=XM_011283467.3
B3	139300594	139300713
refGene_gn=OTUB2;refGene_tx=XM_011283467.3
B3	139304511	139304596
refGene_gn=OTUB2;refGene_tx=XM_011283467.3
B3	139305180	139305375
refGene_gn=OTUB2;refGene_tx=XM_011283467.3
B3	139306328	139306571
refGene_gn=OTUB2;refGene_tx=XM_011283467.3
B3	139311443	139311724
refGene_gn=DDX24;refGene_tx=XM_006933075.4;refGene_gn=DDX24;refGene_tx=XM_023255988.1;refGene_gn=DDX24;refGene_tx=XM_023255989.1
B3	139313403	139313533
refGene_gn=DDX24;refGene_tx=XM_006933075.4;refGene_gn=DDX24;refGene_tx=XM_023255988.1;refGene_gn=DDX24;refGene_tx=XM_023255989.1
B3	139315127	139315316

refGene_gn=DDX24;refGene_tx=XM_006933075.4;refGene_gn=DDX24;refGene_tx=XM_023255988.1;refGene_gn=DDX24;refGene_tx=XM_023255989.1
B3 139318086 139318162
refGene_gn=DDX24;refGene_tx=XM_006933075.4;refGene_gn=DDX24;refGene_tx=XM_023255988.1;refGene_gn=DDX24;refGene_tx=XM_023255989.1
B3 139320263 139320779
refGene_gn=DDX24;refGene_tx=XM_006933075.4;refGene_gn=DDX24;refGene_tx=XM_023255988.1;refGene_gn=DDX24;refGene_tx=XM_023255989.1
B3 139321062 139321216
refGene_gn=DDX24;refGene_tx=XM_006933075.4;refGene_gn=DDX24;refGene_tx=XM_023255988.1;refGene_gn=DDX24;refGene_tx=XM_023255989.1
B3 139322265 139322778
refGene_gn=DDX24;refGene_tx=XM_006933075.4;refGene_gn=DDX24;refGene_tx=XM_023255988.1;refGene_gn=DDX24;refGene_tx=XM_023255989.1
B3 139329712 139330424
refGene_gn=DDX24;refGene_tx=XM_006933075.4;refGene_gn=DDX24;refGene_tx=XM_023255988.1;refGene_gn=DDX24;refGene_tx=XM_023255989.1
B3 139360367 139360570
refGene_gn=IFI27L2;refGene_tx=XM_011283468.3
B3 139361150 139361312
refGene_gn=IFI27L2;refGene_tx=XM_011283468.3
B3 139361805 139361835
refGene_gn=IFI27L2;refGene_tx=XM_011283468.3
B3 139362001 139362008
refGene_gn=IFI27L2;refGene_tx=XM_011283468.3
B3 139397770 139397890
refGene_gn=PPP4R4;refGene_tx=XM_011283478.3;refGene_gn=PPP4R4;refGene_tx=XM_023255990.1;refGene_gn=PPP4R4;refGene_tx=XM_023255992.1;refGene_gn=PPP4R4;refGene_tx=XM_023255993.1
B3 139399284 139399358
refGene_gn=PPP4R4;refGene_tx=XM_011283478.3;refGene_gn=PPP4R4;refGene_tx=XM_023255990.1;refGene_gn=PPP4R4;refGene_tx=XM_023255992.1;refGene_gn=PPP4R4;refGene_tx=XM_023255993.1
B3 139434940 139435043
refGene_gn=PPP4R4;refGene_tx=XM_011283478.3;refGene_gn=PPP4R4;refGene_tx=XM_023255990.1;refGene_gn=PPP4R4;refGene_tx=XM_023255992.1;refGene_gn=PPP4R4;refGene_tx=XM_023255993.1
B3 139454349 139454497
refGene_gn=PPP4R4;refGene_tx=XM_011283478.3;refGene_gn=PPP4R4;refGene_tx=XM_023255990.1;refGene_gn=PPP4R4;refGene_tx=XM_023255992.1;refGene_gn=PPP4R4;refGene_tx=XM_023255993.1
B3 139455001 139455075
refGene_gn=PPP4R4;refGene_tx=XM_011283478.3;refGene_gn=PPP4R4;refGene_tx=XM_023255990.1;refGene_gn=PPP4R4;refGene_tx=XM_023255992.1;refGene_gn=PPP4R4;refGene_tx=XM_023255993.1
B3 139457621 139457728
refGene_gn=PPP4R4;refGene_tx=XM_011283478.3;refGene_gn=PPP4R4;refGene_tx=XM_023255990.1;refGene_gn=PPP4R4;refGene_tx=XM_023255992.1;refGene_gn=PPP4R4;refGene_tx=XM_023255993.1
B3 139458544 139458652

refGene_gn=PPP4R4;refGene_tx=XM_011283478.3;refGene_gn=PPP4R4;refGene_tx=XM_023255990.1;refGene_gn=PPP4R4;refGene_tx=XM_023255992.1;refGene_gn=PPP4R4;refGene_tx=XM_023255993.1
B3 139465588 139465710
refGene_gn=PPP4R4;refGene_tx=XM_011283478.3;refGene_gn=PPP4R4;refGene_tx=XM_023255990.1;refGene_gn=PPP4R4;refGene_tx=XM_023255992.1;refGene_gn=PPP4R4;refGene_tx=XM_023255993.1
B3 139468136 139468259
refGene_gn=PPP4R4;refGene_tx=XM_011283478.3;refGene_gn=PPP4R4;refGene_tx=XM_023255990.1;refGene_gn=PPP4R4;refGene_tx=XM_023255992.1;refGene_gn=PPP4R4;refGene_tx=XM_023255993.1
B3 139469307 139469477
refGene_gn=PPP4R4;refGene_tx=XM_011283478.3;refGene_gn=PPP4R4;refGene_tx=XM_023255990.1;refGene_gn=PPP4R4;refGene_tx=XM_023255992.1;refGene_gn=PPP4R4;refGene_tx=XM_023255993.1
B3 139469813 139469933
refGene_gn=PPP4R4;refGene_tx=XM_011283478.3;refGene_gn=PPP4R4;refGene_tx=XM_023255990.1;refGene_gn=PPP4R4;refGene_tx=XM_023255992.1;refGene_gn=PPP4R4;refGene_tx=XM_023255993.1
B3 139472242 139472320
refGene_gn=PPP4R4;refGene_tx=XM_011283478.3;refGene_gn=PPP4R4;refGene_tx=XM_023255990.1;refGene_gn=PPP4R4;refGene_tx=XM_023255992.1;refGene_gn=PPP4R4;refGene_tx=XM_023255993.1
B3 139473032 139473116
refGene_gn=PPP4R4;refGene_tx=XM_011283478.3;refGene_gn=PPP4R4;refGene_tx=XM_023255990.1;refGene_gn=PPP4R4;refGene_tx=XM_023255992.1;refGene_gn=PPP4R4;refGene_tx=XM_023255993.1
B3 139473710 139473893
refGene_gn=PPP4R4;refGene_tx=XM_011283478.3;refGene_gn=PPP4R4;refGene_tx=XM_023255990.1;refGene_gn=PPP4R4;refGene_tx=XM_023255992.1;refGene_gn=PPP4R4;refGene_tx=XM_023255993.1
B3 139477816 139477922
refGene_gn=PPP4R4;refGene_tx=XM_011283478.3;refGene_gn=PPP4R4;refGene_tx=XM_023255990.1;refGene_gn=PPP4R4;refGene_tx=XM_023255992.1;refGene_gn=PPP4R4;refGene_tx=XM_023255993.1
B3 139479446 139479594
refGene_gn=PPP4R4;refGene_tx=XM_011283478.3;refGene_gn=PPP4R4;refGene_tx=XM_023255990.1;refGene_gn=PPP4R4;refGene_tx=XM_023255992.1;refGene_gn=PPP4R4;refGene_tx=XM_023255993.1
B3 139483878 139484023
refGene_gn=PPP4R4;refGene_tx=XM_011283478.3;refGene_gn=PPP4R4;refGene_tx=XM_023255990.1;refGene_gn=PPP4R4;refGene_tx=XM_023255992.1;refGene_gn=PPP4R4;refGene_tx=XM_023255993.1
B3 139486182 139486224
refGene_gn=PPP4R4;refGene_tx=XM_011283478.3;refGene_gn=PPP4R4;refGene_tx=XM_023255990.1;refGene_gn=PPP4R4;refGene_tx=XM_023255992.1
B3 139486311 139486386
refGene_gn=PPP4R4;refGene_tx=XM_011283478.3;refGene_gn=PPP4R4;refGene_tx=XM_023255990.1
B3 139493239 139493309

refGene_gn=PPP4R4;refGene_tx=XM_011283478.3;refGene_gn=PPP4R4;refGene_tx=XM_023255990.1;refGene_gn=PPP4R4;refGene_tx=XM_023255992.1;refGene_gn=PPP4R4;refGene_tx=XM_023255993.1
B3 139493741 139493828
refGene_gn=PPP4R4;refGene_tx=XM_011283478.3;refGene_gn=PPP4R4;refGene_tx=XM_023255990.1;refGene_gn=PPP4R4;refGene_tx=XM_023255992.1;refGene_gn=PPP4R4;refGene_tx=XM_023255993.1
B3 139494140 139494234
refGene_gn=PPP4R4;refGene_tx=XM_011283478.3;refGene_gn=PPP4R4;refGene_tx=XM_023255990.1;refGene_gn=PPP4R4;refGene_tx=XM_023255992.1;refGene_gn=PPP4R4;refGene_tx=XM_023255993.1
B3 139495307 139495378
refGene_gn=PPP4R4;refGene_tx=XM_011283478.3;refGene_gn=PPP4R4;refGene_tx=XM_023255990.1;refGene_gn=PPP4R4;refGene_tx=XM_023255992.1;refGene_gn=PPP4R4;refGene_tx=XM_023255993.1
B3 139499455 139499603
refGene_gn=PPP4R4;refGene_tx=XM_011283478.3;refGene_gn=PPP4R4;refGene_tx=XM_023255990.1;refGene_gn=PPP4R4;refGene_tx=XM_023255992.1;refGene_gn=PPP4R4;refGene_tx=XM_023255993.1
B3 139502750 139502775
refGene_gn=PPP4R4;refGene_tx=XM_011283478.3
B3 139508126 139508285
refGene_gn=LOC109500998;refGene_tx=XM_019833583.1
B3 139508286 139508376
refGene_gn=LOC109500998;refGene_tx=XM_019833583.1
B3 139510320 139510422
refGene_gn=LOC109500998;refGene_tx=XM_019833583.1
B3 139511007 139511071
refGene_gn=PPP4R4;refGene_tx=XM_023255990.1;refGene_gn=PPP4R4;refGene_tx=XM_023255992.1;refGene_gn=PPP4R4;refGene_tx=XM_023255993.1
B3 139524205 139524391
refGene_gn=SERPINA6;refGene_tx=XM_003987945.2
B3 139525832 139525980
refGene_gn=SERPINA6;refGene_tx=XM_003987945.2
B3 139530891 139531162
refGene_gn=SERPINA6;refGene_tx=XM_003987945.2
B3 139537615 139538225
refGene_gn=SERPINA6;refGene_tx=XM_003987945.2
B3 139599172 139599409
refGene_gn=SERPINA1;refGene_tx=XM_006933076.3
B3 139600037 139600185
refGene_gn=SERPINA1;refGene_tx=XM_006933076.3
B3 139601478 139601749
refGene_gn=SERPINA1;refGene_tx=XM_006933076.3
B3 139603126 139603781
refGene_gn=SERPINA1;refGene_tx=XM_006933076.3
B3 139648984 139649188
refGene_gn=SERPINA11;refGene_tx=XM_006933077.2;refGene_gn=SERPINA11;refGene_tx=XM_019833585.1
B3 139651191 139651339

refGene_gn=SERPINA11;refGene_tx=XM_006933077.2;refGene_gn=SERPINA11;refGene_tx=XM_019833585.1
B3 139654312 139654586
refGene_gn=SERPINA11;refGene_tx=XM_006933077.2;refGene_gn=SERPINA11;refGene_tx=XM_019833585.1
B3 139659324 139659967
refGene_gn=SERPINA11;refGene_tx=XM_006933077.2;refGene_gn=SERPINA11;refGene_tx=XM_019833585.1
B3 139682558 139682762
refGene_gn=SERPINA9;refGene_tx=XM_003987948.2;refGene_gn=SERPINA9;refGene_tx=XM_019833586.1;refGene_gn=SERPINA9;refGene_tx=XM_019833587.2;refGene_gn=SERPINA9;refGene_tx=XM_023255995.1
B3 139684156 139684304
refGene_gn=SERPINA9;refGene_tx=XM_003987948.2;refGene_gn=SERPINA9;refGene_tx=XM_019833586.1;refGene_gn=SERPINA9;refGene_tx=XM_019833587.2;refGene_gn=SERPINA9;refGene_tx=XM_023255995.1
B3 139686468 139686742
refGene_gn=SERPINA9;refGene_tx=XM_003987948.2;refGene_gn=SERPINA9;refGene_tx=XM_019833586.1;refGene_gn=SERPINA9;refGene_tx=XM_019833587.2;refGene_gn=SERPINA9;refGene_tx=XM_023255995.1
B3 139692614 139693247
refGene_gn=SERPINA9;refGene_tx=XM_003987948.2;refGene_gn=SERPINA9;refGene_tx=XM_019833586.1;refGene_gn=SERPINA9;refGene_tx=XM_019833587.2;refGene_gn=SERPINA9;refGene_tx=XM_023255995.1
B3 139699115 139699185
refGene_gn=SERPINA9;refGene_tx=XM_019833586.1
B3 139705759 139705951
refGene_gn=SERPINA12;refGene_tx=XM_003987972.2;refGene_gn=SERPINA12;refGene_tx=XM_019833591.1;refGene_gn=SERPINA12;refGene_tx=XM_019833592.1
B3 139707519 139707667
refGene_gn=SERPINA12;refGene_tx=XM_003987972.2;refGene_gn=SERPINA12;refGene_tx=XM_019833591.1;refGene_gn=SERPINA12;refGene_tx=XM_019833592.1
B3 139711332 139711603
refGene_gn=SERPINA12;refGene_tx=XM_003987972.2;refGene_gn=SERPINA12;refGene_tx=XM_019833591.1;refGene_gn=SERPINA12;refGene_tx=XM_019833592.1
B3 139713331 139713982
refGene_gn=SERPINA12;refGene_tx=XM_003987972.2;refGene_gn=SERPINA12;refGene_tx=XM_019833591.1;refGene_gn=SERPINA12;refGene_tx=XM_019833592.1
B3 139714537 139714547
refGene_gn=SERPINA12;refGene_tx=XM_019833591.1
B3 139756503 139756564
refGene_gn=SERPINA4;refGene_tx=XM_006933078.2
B3 139758604 139759270
refGene_gn=SERPINA4;refGene_tx=XM_006933078.2
B3 139768481 139768755
refGene_gn=SERPINA4;refGene_tx=XM_006933078.2
B3 139769592 139769752
refGene_gn=SERPINA4;refGene_tx=XM_006933078.2
B3 139770519 139770720
refGene_gn=SERPINA4;refGene_tx=XM_006933078.2

B3 139777800 139777810
refGene_gn=SERPINA5;refGene_tx=XM_003987950.4;refGene_gn=SERPINA5;refGene_tx=XM_006933079.3
B3 139782393 139782492
refGene_gn=SERPINA5;refGene_tx=XM_006933079.3
B3 139783893 139784526
refGene_gn=SERPINA5;refGene_tx=XM_003987950.4;refGene_gn=SERPINA5;refGene_tx=XM_006933079.3
B3 139785858 139786129
refGene_gn=SERPINA5;refGene_tx=XM_003987950.4;refGene_gn=SERPINA5;refGene_tx=XM_006933079.3
B3 139786490 139786638
refGene_gn=SERPINA5;refGene_tx=XM_003987950.4;refGene_gn=SERPINA5;refGene_tx=XM_006933079.3
B3 139787168 139787357
refGene_gn=SERPINA5;refGene_tx=XM_003987950.4;refGene_gn=SERPINA5;refGene_tx=XM_006933079.3
B3 139799721 139800358
refGene_gn=SERPINA3;refGene_tx=XM_023255996.1
B3 139802265 139802298
refGene_gn=SERPINA3;refGene_tx=XM_023255996.1
B3 139802300 139802538
refGene_gn=SERPINA3;refGene_tx=XM_023255996.1
B3 139804735 139804886
refGene_gn=SERPINA3;refGene_tx=XM_023255996.1
B3 139806918 139807122
refGene_gn=SERPINA3;refGene_tx=XM_023255996.1
B3 139929917 139930073
refGene_gn=GSC;refGene_tx=XM_023255997.1
B3 139930392 139930652
refGene_gn=GSC;refGene_tx=XM_023255997.1
B3 139931127 139931482
refGene_gn=GSC;refGene_tx=XM_023255997.1
B3 140203506 140203672
refGene_gn=DICER1;refGene_tx=XM_023255998.1;refGene_gn=DICER1;refGene_tx=XM_023255999.1;refGene_gn=DICER1;refGene_tx=XM_023256000.1
B3 140204035 140204111
refGene_gn=DICER1;refGene_tx=XM_023255998.1;refGene_gn=DICER1;refGene_tx=XM_023255999.1;refGene_gn=DICER1;refGene_tx=XM_023256000.1
B3 140204203 140204366
refGene_gn=DICER1;refGene_tx=XM_023255998.1;refGene_gn=DICER1;refGene_tx=XM_023255999.1;refGene_gn=DICER1;refGene_tx=XM_023256000.1
B3 140206792 140207061
refGene_gn=DICER1;refGene_tx=XM_023255998.1;refGene_gn=DICER1;refGene_tx=XM_023255999.1;refGene_gn=DICER1;refGene_tx=XM_023256000.1
B3 140208887 140209776
refGene_gn=DICER1;refGene_tx=XM_023255998.1;refGene_gn=DICER1;refGene_tx=XM_023255999.1;refGene_gn=DICER1;refGene_tx=XM_023256000.1
B3 140212852 140213008
refGene_gn=DICER1;refGene_tx=XM_023255998.1;refGene_gn=DICER1;refGene_

tx=XM_023255999.1
B3 140216429 140217213
refGene_gn=DICER1;refGene_tx=XM_023255998.1;refGene_gn=DICER1;refGene_tx=XM_023255999.1;refGene_gn=DICER1;refGene_tx=XM_023256000.1
B3 140218168 140218344
refGene_gn=DICER1;refGene_tx=XM_023255998.1;refGene_gn=DICER1;refGene_tx=XM_023255999.1;refGene_gn=DICER1;refGene_tx=XM_023256000.1
B3 140218759 140218865
refGene_gn=DICER1;refGene_tx=XM_023255998.1;refGene_gn=DICER1;refGene_tx=XM_023255999.1;refGene_gn=DICER1;refGene_tx=XM_023256000.1
B3 140219146 140219329
refGene_gn=DICER1;refGene_tx=XM_023255998.1;refGene_gn=DICER1;refGene_tx=XM_023255999.1;refGene_gn=DICER1;refGene_tx=XM_023256000.1
B3 140220972 140221126
refGene_gn=DICER1;refGene_tx=XM_023255998.1;refGene_gn=DICER1;refGene_tx=XM_023255999.1;refGene_gn=DICER1;refGene_tx=XM_023256000.1
B3 140221328 140221542
refGene_gn=DICER1;refGene_tx=XM_023255998.1;refGene_gn=DICER1;refGene_tx=XM_023255999.1;refGene_gn=DICER1;refGene_tx=XM_023256000.1
B3 140221746 140221926
refGene_gn=DICER1;refGene_tx=XM_023255998.1;refGene_gn=DICER1;refGene_tx=XM_023255999.1;refGene_gn=DICER1;refGene_tx=XM_023256000.1
B3 140223893 140224033
refGene_gn=DICER1;refGene_tx=XM_023255998.1;refGene_gn=DICER1;refGene_tx=XM_023255999.1;refGene_gn=DICER1;refGene_tx=XM_023256000.1
B3 140224853 140224929
refGene_gn=DICER1;refGene_tx=XM_023255998.1;refGene_gn=DICER1;refGene_tx=XM_023255999.1;refGene_gn=DICER1;refGene_tx=XM_023256000.1
B3 140225894 140226027
refGene_gn=DICER1;refGene_tx=XM_023255998.1;refGene_gn=DICER1;refGene_tx=XM_023255999.1;refGene_gn=DICER1;refGene_tx=XM_023256000.1
B3 140228858 140229013
refGene_gn=DICER1;refGene_tx=XM_023255998.1;refGene_gn=DICER1;refGene_tx=XM_023255999.1;refGene_gn=DICER1;refGene_tx=XM_023256000.1
B3 140229621 140229864
refGene_gn=DICER1;refGene_tx=XM_023255998.1;refGene_gn=DICER1;refGene_tx=XM_023255999.1;refGene_gn=DICER1;refGene_tx=XM_023256000.1
B3 140231199 140231332
refGene_gn=DICER1;refGene_tx=XM_023255998.1;refGene_gn=DICER1;refGene_tx=XM_023255999.1;refGene_gn=DICER1;refGene_tx=XM_023256000.1
B3 140238393 140238866
refGene_gn=DICER1;refGene_tx=XM_023255998.1;refGene_gn=DICER1;refGene_tx=XM_023255999.1;refGene_gn=DICER1;refGene_tx=XM_023256000.1
B3 140240330 140240499
refGene_gn=DICER1;refGene_tx=XM_023255998.1;refGene_gn=DICER1;refGene_tx=XM_023255999.1;refGene_gn=DICER1;refGene_tx=XM_023256000.1
B3 140242716 140242877
refGene_gn=DICER1;refGene_tx=XM_023255998.1;refGene_gn=DICER1;refGene_tx=XM_023255999.1;refGene_gn=DICER1;refGene_tx=XM_023256000.1
B3 140243322 140243457

refGene_gn=DICER1;refGene_tx=XM_023255998.1;refGene_gn=DICER1;refGene_tx=XM_023255999.1;refGene_gn=DICER1;refGene_tx=XM_023256000.1
B3 140244440 140244571
refGene_gn=DICER1;refGene_tx=XM_023255998.1;refGene_gn=DICER1;refGene_tx=XM_023255999.1;refGene_gn=DICER1;refGene_tx=XM_023256000.1
B3 140245766 140245929
refGene_gn=DICER1;refGene_tx=XM_023255998.1;refGene_gn=DICER1;refGene_tx=XM_023255999.1;refGene_gn=DICER1;refGene_tx=XM_023256000.1
B3 140246561 140246708
refGene_gn=DICER1;refGene_tx=XM_023255998.1;refGene_gn=DICER1;refGene_tx=XM_023255999.1;refGene_gn=DICER1;refGene_tx=XM_023256000.1
B3 140312289 140312458
refGene_gn=CLMN;refGene_tx=XM_023256001.1;refGene_gn=CLMN;refGene_tx=XM_023256002.1;refGene_gn=CLMN;refGene_tx=XM_023256003.1;refGene_gn=CLMN;refGene_tx=XM_023256004.1
B3 140313882 140313979
refGene_gn=CLMN;refGene_tx=XM_023256001.1;refGene_gn=CLMN;refGene_tx=XM_023256002.1
B3 140314360 140314431
refGene_gn=CLMN;refGene_tx=XM_023256001.1;refGene_gn=CLMN;refGene_tx=XM_023256002.1;refGene_gn=CLMN;refGene_tx=XM_023256003.1;refGene_gn=CLMN;refGene_tx=XM_023256004.1
B3 140314469 140314487
refGene_gn=CLMN;refGene_tx=XM_023256005.1
B3 140315050 140315111
refGene_gn=CLMN;refGene_tx=XM_023256001.1;refGene_gn=CLMN;refGene_tx=XM_023256002.1;refGene_gn=CLMN;refGene_tx=XM_023256003.1;refGene_gn=CLMN;refGene_tx=XM_023256004.1;refGene_gn=CLMN;refGene_tx=XM_023256005.1
B3 140319353 140319550
refGene_gn=CLMN;refGene_tx=XM_023256001.1;refGene_gn=CLMN;refGene_tx=XM_023256002.1;refGene_gn=CLMN;refGene_tx=XM_023256003.1;refGene_gn=CLMN;refGene_tx=XM_023256004.1;refGene_gn=CLMN;refGene_tx=XM_023256005.1
B3 140322802 140324464
refGene_gn=CLMN;refGene_tx=XM_023256001.1;refGene_gn=CLMN;refGene_tx=XM_023256002.1;refGene_gn=CLMN;refGene_tx=XM_023256003.1;refGene_gn=CLMN;refGene_tx=XM_023256004.1;refGene_gn=CLMN;refGene_tx=XM_023256005.1
B3 140329676 140329759
refGene_gn=CLMN;refGene_tx=XM_023256001.1;refGene_gn=CLMN;refGene_tx=XM_023256002.1;refGene_gn=CLMN;refGene_tx=XM_023256003.1;refGene_gn=CLMN;refGene_tx=XM_023256004.1;refGene_gn=CLMN;refGene_tx=XM_023256005.1
B3 140331657 140331851
refGene_gn=CLMN;refGene_tx=XM_023256001.1;refGene_gn=CLMN;refGene_tx=XM_023256002.1;refGene_gn=CLMN;refGene_tx=XM_023256003.1;refGene_gn=CLMN;refGene_tx=XM_023256004.1;refGene_gn=CLMN;refGene_tx=XM_023256005.1
B3 140333763 140333954
refGene_gn=CLMN;refGene_tx=XM_023256001.1;refGene_gn=CLMN;refGene_tx=XM_023256002.1;refGene_gn=CLMN;refGene_tx=XM_023256003.1;refGene_gn=CLMN;refGene_tx=XM_023256004.1;refGene_gn=CLMN;refGene_tx=XM_023256005.1
B3 140336030 140336123
refGene_gn=CLMN;refGene_tx=XM_023256001.1;refGene_gn=CLMN;refGene_tx=X

M_023256002.1;refGene_gn=CLMN;refGene_tx=XM_023256003.1;refGene_gn=CLMN;refGene_tx=XM_023256004.1;refGene_gn=CLMN;refGene_tx=XM_023256005.1
B3 140341557 140341641
refGene_gn=CLMN;refGene_tx=XM_023256001.1;refGene_gn=CLMN;refGene_tx=XM_023256002.1;refGene_gn=CLMN;refGene_tx=XM_023256003.1;refGene_gn=CLMN;refGene_tx=XM_023256004.1;refGene_gn=CLMN;refGene_tx=XM_023256005.1
B3 140344635 140344731
refGene_gn=CLMN;refGene_tx=XM_023256001.1;refGene_gn=CLMN;refGene_tx=XM_023256002.1;refGene_gn=CLMN;refGene_tx=XM_023256003.1;refGene_gn=CLMN;refGene_tx=XM_023256004.1;refGene_gn=CLMN;refGene_tx=XM_023256005.1
B3 140350449 140350511
refGene_gn=CLMN;refGene_tx=XM_023256001.1;refGene_gn=CLMN;refGene_tx=XM_023256002.1;refGene_gn=CLMN;refGene_tx=XM_023256003.1;refGene_gn=CLMN;refGene_tx=XM_023256004.1;refGene_gn=CLMN;refGene_tx=XM_023256005.1
B3 140374615 140374671
refGene_gn=CLMN;refGene_tx=XM_023256001.1;refGene_gn=CLMN;refGene_tx=XM_023256003.1;refGene_gn=CLMN;refGene_tx=XM_023256005.1
B3 140425921 140426003
refGene_gn=CLMN;refGene_tx=XM_023256001.1;refGene_gn=CLMN;refGene_tx=XM_023256002.1;refGene_gn=CLMN;refGene_tx=XM_023256003.1;refGene_gn=CLMN;refGene_tx=XM_023256004.1;refGene_gn=CLMN;refGene_tx=XM_023256005.1
B3 140514562 140514763
refGene_gn=SYNE3;refGene_tx=XM_003987951.5;refGene_gn=SYNE3;refGene_tx=XM_006933080.4;refGene_gn=SYNE3;refGene_tx=XM_019833599.2;refGene_gn=SYNE3;refGene_tx=XM_019833600.2;refGene_gn=SYNE3;refGene_tx=XM_023256006.1;refGene_gn=SYNE3;refGene_tx=XM_023256007.1;refGene_gn=SYNE3;refGene_tx=XM_023256008.1;refGene_gn=SYNE3;refGene_tx=XM_023256009.1
B3 140524641 140524680
refGene_gn=SYNE3;refGene_tx=XM_003987951.5;refGene_gn=SYNE3;refGene_tx=XM_006933080.4;refGene_gn=SYNE3;refGene_tx=XM_019833599.2;refGene_gn=SYNE3;refGene_tx=XM_019833600.2;refGene_gn=SYNE3;refGene_tx=XM_023256006.1;refGene_gn=SYNE3;refGene_tx=XM_023256007.1;refGene_gn=SYNE3;refGene_tx=XM_023256008.1
B3 140525691 140525841
refGene_gn=SYNE3;refGene_tx=XM_003987951.5;refGene_gn=SYNE3;refGene_tx=XM_006933080.4;refGene_gn=SYNE3;refGene_tx=XM_019833599.2;refGene_gn=SYNE3;refGene_tx=XM_019833600.2;refGene_gn=SYNE3;refGene_tx=XM_023256006.1;refGene_gn=SYNE3;refGene_tx=XM_023256007.1;refGene_gn=SYNE3;refGene_tx=XM_023256008.1;refGene_gn=SYNE3;refGene_tx=XM_023256009.1
B3 140529029 140529188
refGene_gn=SYNE3;refGene_tx=XM_003987951.5;refGene_gn=SYNE3;refGene_tx=XM_006933080.4;refGene_gn=SYNE3;refGene_tx=XM_019833599.2;refGene_gn=SYNE3;refGene_tx=XM_019833600.2;refGene_gn=SYNE3;refGene_tx=XM_023256006.1;refGene_gn=SYNE3;refGene_tx=XM_023256007.1;refGene_gn=SYNE3;refGene_tx=XM_023256008.1;refGene_gn=SYNE3;refGene_tx=XM_023256009.1
B3 140530404 140530534
refGene_gn=SYNE3;refGene_tx=XM_003987951.5;refGene_gn=SYNE3;refGene_tx=XM_006933080.4;refGene_gn=SYNE3;refGene_tx=XM_019833599.2;refGene_gn=SYNE3;refGene_tx=XM_019833600.2;refGene_gn=SYNE3;refGene_tx=XM_023256006.1;refGene_gn=SYNE3;refGene_tx=XM_023256007.1;refGene_gn=SYNE3;refGe

ne_tx=XM_023256008.1;refGene_gn=SYNE3;refGene_tx=XM_023256009.1
B3 140530961 140531134
refGene_gn=SYNE3;refGene_tx=XM_003987951.5;refGene_gn=SYNE3;refGene_tx=XM_006933080.4;refGene_gn=SYNE3;refGene_tx=XM_019833599.2;refGene_gn=SYNE3;refGene_tx=XM_019833600.2;refGene_gn=SYNE3;refGene_tx=XM_023256006.1;refGene_gn=SYNE3;refGene_tx=XM_023256007.1;refGene_gn=SYNE3;refGene_tx=XM_023256008.1;refGene_gn=SYNE3;refGene_tx=XM_023256009.1
B3 140531242 140531404
refGene_gn=SYNE3;refGene_tx=XM_003987951.5;refGene_gn=SYNE3;refGene_tx=XM_006933080.4;refGene_gn=SYNE3;refGene_tx=XM_019833599.2;refGene_gn=SYNE3;refGene_tx=XM_019833600.2;refGene_gn=SYNE3;refGene_tx=XM_023256006.1;refGene_gn=SYNE3;refGene_tx=XM_023256007.1;refGene_gn=SYNE3;refGene_tx=XM_023256008.1;refGene_gn=SYNE3;refGene_tx=XM_023256009.1
B3 140534361 140534496
refGene_gn=SYNE3;refGene_tx=XM_003987951.5;refGene_gn=SYNE3;refGene_tx=XM_006933080.4;refGene_gn=SYNE3;refGene_tx=XM_019833599.2;refGene_gn=SYNE3;refGene_tx=XM_019833600.2;refGene_gn=SYNE3;refGene_tx=XM_023256006.1;refGene_gn=SYNE3;refGene_tx=XM_023256007.1;refGene_gn=SYNE3;refGene_tx=XM_023256008.1;refGene_gn=SYNE3;refGene_tx=XM_023256009.1
B3 140535617 140535761
refGene_gn=SYNE3;refGene_tx=XM_003987951.5;refGene_gn=SYNE3;refGene_tx=XM_006933080.4;refGene_gn=SYNE3;refGene_tx=XM_019833599.2;refGene_gn=SYNE3;refGene_tx=XM_019833600.2;refGene_gn=SYNE3;refGene_tx=XM_023256006.1;refGene_gn=SYNE3;refGene_tx=XM_023256007.1;refGene_gn=SYNE3;refGene_tx=XM_023256008.1;refGene_gn=SYNE3;refGene_tx=XM_023256009.1
B3 140536709 140536892
refGene_gn=SYNE3;refGene_tx=XM_003987951.5;refGene_gn=SYNE3;refGene_tx=XM_006933080.4;refGene_gn=SYNE3;refGene_tx=XM_019833599.2;refGene_gn=SYNE3;refGene_tx=XM_019833600.2;refGene_gn=SYNE3;refGene_tx=XM_023256006.1;refGene_gn=SYNE3;refGene_tx=XM_023256007.1;refGene_gn=SYNE3;refGene_tx=XM_023256008.1;refGene_gn=SYNE3;refGene_tx=XM_023256009.1
B3 140544288 140544463
refGene_gn=SYNE3;refGene_tx=XM_003987951.5;refGene_gn=SYNE3;refGene_tx=XM_006933080.4;refGene_gn=SYNE3;refGene_tx=XM_019833599.2;refGene_gn=SYNE3;refGene_tx=XM_019833600.2;refGene_gn=SYNE3;refGene_tx=XM_023256006.1;refGene_gn=SYNE3;refGene_tx=XM_023256007.1;refGene_gn=SYNE3;refGene_tx=XM_023256008.1;refGene_gn=SYNE3;refGene_tx=XM_023256009.1
B3 140545609 140545746
refGene_gn=SYNE3;refGene_tx=XM_003987951.5;refGene_gn=SYNE3;refGene_tx=XM_006933080.4;refGene_gn=SYNE3;refGene_tx=XM_019833599.2;refGene_gn=SYNE3;refGene_tx=XM_019833600.2;refGene_gn=SYNE3;refGene_tx=XM_023256006.1;refGene_gn=SYNE3;refGene_tx=XM_023256007.1;refGene_gn=SYNE3;refGene_tx=XM_023256008.1;refGene_gn=SYNE3;refGene_tx=XM_023256009.1
B3 140549667 140550015
refGene_gn=SYNE3;refGene_tx=XM_003987951.5;refGene_gn=SYNE3;refGene_tx=XM_006933080.4;refGene_gn=SYNE3;refGene_tx=XM_019833599.2;refGene_gn=SYNE3;refGene_tx=XM_019833600.2;refGene_gn=SYNE3;refGene_tx=XM_023256006.1;refGene_gn=SYNE3;refGene_tx=XM_023256007.1;refGene_gn=SYNE3;refGene_tx=XM_023256008.1;refGene_gn=SYNE3;refGene_tx=XM_023256009.1
B3 140551714 140551876

refGene_gn=SYNE3;refGene_tx=XM_003987951.5;refGene_gn=SYNE3;refGene_tx=XM_006933080.4;refGene_gn=SYNE3;refGene_tx=XM_019833599.2;refGene_gn=SYNE3;refGene_tx=XM_019833600.2;refGene_gn=SYNE3;refGene_tx=XM_023256006.1;refGene_gn=SYNE3;refGene_tx=XM_023256007.1;refGene_gn=SYNE3;refGene_tx=XM_023256008.1;refGene_gn=SYNE3;refGene_tx=XM_023256009.1
B3 140559096 140559406
refGene_gn=SYNE3;refGene_tx=XM_003987951.5;refGene_gn=SYNE3;refGene_tx=XM_006933080.4;refGene_gn=SYNE3;refGene_tx=XM_019833599.2;refGene_gn=SYNE3;refGene_tx=XM_019833600.2;refGene_gn=SYNE3;refGene_tx=XM_023256006.1;refGene_gn=SYNE3;refGene_tx=XM_023256007.1;refGene_gn=SYNE3;refGene_tx=XM_023256008.1;refGene_gn=SYNE3;refGene_tx=XM_023256009.1
B3 140560430 140560603
refGene_gn=SYNE3;refGene_tx=XM_003987951.5;refGene_gn=SYNE3;refGene_tx=XM_006933080.4;refGene_gn=SYNE3;refGene_tx=XM_019833599.2;refGene_gn=SYNE3;refGene_tx=XM_019833600.2;refGene_gn=SYNE3;refGene_tx=XM_023256006.1;refGene_gn=SYNE3;refGene_tx=XM_023256007.1;refGene_gn=SYNE3;refGene_tx=XM_023256008.1;refGene_gn=SYNE3;refGene_tx=XM_023256009.1
B3 140568978 140569122
refGene_gn=SYNE3;refGene_tx=XM_003987951.5;refGene_gn=SYNE3;refGene_tx=XM_006933080.4;refGene_gn=SYNE3;refGene_tx=XM_019833599.2;refGene_gn=SYNE3;refGene_tx=XM_019833600.2;refGene_gn=SYNE3;refGene_tx=XM_023256006.1;refGene_gn=SYNE3;refGene_tx=XM_023256007.1;refGene_gn=SYNE3;refGene_tx=XM_023256008.1;refGene_gn=SYNE3;refGene_tx=XM_023256009.1
B3 140619433 140619728
refGene_gn=GLRX5;refGene_tx=XM_003987952.3
B3 140629374 140629553
refGene_gn=GLRX5;refGene_tx=XM_003987952.3
B3 140778285 140778429
refGene_gn=TCL1A;refGene_tx=XM_023256194.1
B3 140778654 140778828
refGene_gn=TCL1A;refGene_tx=XM_023256194.1
B3 140782490 140782610
refGene_gn=TCL1A;refGene_tx=XM_023256194.1
B3 141059324 141059351
refGene_gn=CB3H14orf132;refGene_tx=XM_023256010.1
B3 141078077 141078095
refGene_gn=CB3H14orf132;refGene_tx=XM_023256011.1
B3 141111958 141112183
refGene_gn=CB3H14orf132;refGene_tx=XM_023256010.1;refGene_gn=CB3H14orf132;refGene_tx=XM_023256011.1
B3 141262522 141262596
refGene_gn=LOC101082667;refGene_tx=XM_006933085.4;refGene_gn=LOC101082667;refGene_tx=XM_006933086.4;refGene_gn=LOC101082667;refGene_tx=XM_006933087.4
B3 141266240 141267345
refGene_gn=LOC101082667;refGene_tx=XM_006933085.4;refGene_gn=LOC101082667;refGene_tx=XM_006933086.4
B3 141288042 141289114
refGene_gn=LOC101082667;refGene_tx=XM_006933087.4;refGene_gn=LOC101082667;refGene_tx=XM_006933088.2;refGene_gn=LOC101082667;refGene_tx=XM_00

6933090.1
B3 141302462 141302600
refGene_gn=ATG2B;refGene_tx=XM_019833610.2
B3 141306803 141307034
refGene_gn=ATG2B;refGene_tx=XM_011283531.3;refGene_gn=ATG2B;refGene_tx=XM_019833608.2;refGene_gn=ATG2B;refGene_tx=XM_019833609.2
B3 141310490 141310640
refGene_gn=ATG2B;refGene_tx=XM_011283531.3;refGene_gn=ATG2B;refGene_tx=XM_019833608.2;refGene_gn=ATG2B;refGene_tx=XM_019833609.2;refGene_gn=ATG2B;refGene_tx=XM_019833610.2
B3 141311517 141311672
refGene_gn=ATG2B;refGene_tx=XM_011283531.3;refGene_gn=ATG2B;refGene_tx=XM_019833608.2;refGene_gn=ATG2B;refGene_tx=XM_019833609.2;refGene_gn=ATG2B;refGene_tx=XM_019833610.2
B3 141311924 141312059
refGene_gn=ATG2B;refGene_tx=XM_011283531.3;refGene_gn=ATG2B;refGene_tx=XM_019833608.2;refGene_gn=ATG2B;refGene_tx=XM_019833609.2;refGene_gn=ATG2B;refGene_tx=XM_019833610.2;refGene_gn=ATG2B;refGene_tx=XM_019833611.2
B3 141312602 141312685
refGene_gn=ATG2B;refGene_tx=XM_011283531.3;refGene_gn=ATG2B;refGene_tx=XM_019833608.2;refGene_gn=ATG2B;refGene_tx=XM_019833609.2;refGene_gn=ATG2B;refGene_tx=XM_019833610.2;refGene_gn=ATG2B;refGene_tx=XM_019833611.2
B3 141313054 141313124
refGene_gn=ATG2B;refGene_tx=XM_011283531.3;refGene_gn=ATG2B;refGene_tx=XM_019833608.2;refGene_gn=ATG2B;refGene_tx=XM_019833609.2;refGene_gn=ATG2B;refGene_tx=XM_019833610.2;refGene_gn=ATG2B;refGene_tx=XM_019833611.2
B3 141316068 141316276
refGene_gn=ATG2B;refGene_tx=XM_011283531.3;refGene_gn=ATG2B;refGene_tx=XM_019833608.2;refGene_gn=ATG2B;refGene_tx=XM_019833609.2;refGene_gn=ATG2B;refGene_tx=XM_019833610.2;refGene_gn=ATG2B;refGene_tx=XM_019833611.2
B3 141316584 141316663
refGene_gn=ATG2B;refGene_tx=XM_011283531.3;refGene_gn=ATG2B;refGene_tx=XM_019833608.2;refGene_gn=ATG2B;refGene_tx=XM_019833609.2;refGene_gn=ATG2B;refGene_tx=XM_019833610.2;refGene_gn=ATG2B;refGene_tx=XM_019833611.2
B3 141321981 141322083
refGene_gn=ATG2B;refGene_tx=XM_011283531.3;refGene_gn=ATG2B;refGene_tx=XM_019833608.2;refGene_gn=ATG2B;refGene_tx=XM_019833609.2;refGene_gn=ATG2B;refGene_tx=XM_019833610.2;refGene_gn=ATG2B;refGene_tx=XM_019833611.2
B3 141322767 141322962
refGene_gn=ATG2B;refGene_tx=XM_011283531.3;refGene_gn=ATG2B;refGene_tx=XM_019833608.2;refGene_gn=ATG2B;refGene_tx=XM_019833609.2;refGene_gn=ATG2B;refGene_tx=XM_019833610.2;refGene_gn=ATG2B;refGene_tx=XM_019833611.2
B3 141323887 141323996

refGene_gn=ATG2B;refGene_tx=XM_011283531.3;refGene_gn=ATG2B;refGene_tx=XM_019833608.2;refGene_gn=ATG2B;refGene_tx=XM_019833609.2;refGene_gn=ATG2B;refGene_tx=XM_019833610.2;refGene_gn=ATG2B;refGene_tx=XM_019833611.2
B3 141324686 141324913
refGene_gn=ATG2B;refGene_tx=XM_011283531.3;refGene_gn=ATG2B;refGene_tx=XM_019833608.2;refGene_gn=ATG2B;refGene_tx=XM_019833609.2;refGene_gn=ATG2B;refGene_tx=XM_019833610.2;refGene_gn=ATG2B;refGene_tx=XM_019833611.2
B3 141326129 141326332
refGene_gn=ATG2B;refGene_tx=XM_011283531.3;refGene_gn=ATG2B;refGene_tx=XM_019833608.2;refGene_gn=ATG2B;refGene_tx=XM_019833609.2;refGene_gn=ATG2B;refGene_tx=XM_019833610.2;refGene_gn=ATG2B;refGene_tx=XM_019833611.2
B3 141328041 141328183
refGene_gn=ATG2B;refGene_tx=XM_011283531.3;refGene_gn=ATG2B;refGene_tx=XM_019833608.2;refGene_gn=ATG2B;refGene_tx=XM_019833609.2;refGene_gn=ATG2B;refGene_tx=XM_019833610.2;refGene_gn=ATG2B;refGene_tx=XM_019833611.2
B3 141329735 141329906
refGene_gn=ATG2B;refGene_tx=XM_011283531.3;refGene_gn=ATG2B;refGene_tx=XM_019833608.2;refGene_gn=ATG2B;refGene_tx=XM_019833609.2;refGene_gn=ATG2B;refGene_tx=XM_019833610.2;refGene_gn=ATG2B;refGene_tx=XM_019833611.2
B3 141330378 141330455
refGene_gn=ATG2B;refGene_tx=XM_011283531.3;refGene_gn=ATG2B;refGene_tx=XM_019833608.2;refGene_gn=ATG2B;refGene_tx=XM_019833609.2;refGene_gn=ATG2B;refGene_tx=XM_019833610.2;refGene_gn=ATG2B;refGene_tx=XM_019833611.2
B3 141330841 141330912
refGene_gn=ATG2B;refGene_tx=XM_011283531.3;refGene_gn=ATG2B;refGene_tx=XM_019833608.2;refGene_gn=ATG2B;refGene_tx=XM_019833609.2;refGene_gn=ATG2B;refGene_tx=XM_019833610.2;refGene_gn=ATG2B;refGene_tx=XM_019833611.2
B3 141331476 141331569
refGene_gn=ATG2B;refGene_tx=XM_011283531.3;refGene_gn=ATG2B;refGene_tx=XM_019833608.2;refGene_gn=ATG2B;refGene_tx=XM_019833609.2;refGene_gn=ATG2B;refGene_tx=XM_019833610.2;refGene_gn=ATG2B;refGene_tx=XM_019833611.2
B3 141331721 141331828
refGene_gn=ATG2B;refGene_tx=XM_011283531.3;refGene_gn=ATG2B;refGene_tx=XM_019833608.2;refGene_gn=ATG2B;refGene_tx=XM_019833609.2;refGene_gn=ATG2B;refGene_tx=XM_019833610.2;refGene_gn=ATG2B;refGene_tx=XM_019833611.2
B3 141332997 141333078
refGene_gn=ATG2B;refGene_tx=XM_011283531.3;refGene_gn=ATG2B;refGene_tx=XM_019833608.2;refGene_gn=ATG2B;refGene_tx=XM_019833609.2;refGene_gn=ATG2B;refGene_tx=XM_019833610.2;refGene_gn=ATG2B;refGene_tx=XM_019833611.2
B3 141333247 141333447

refGene_gn=ATG2B;refGene_tx=XM_011283531.3;refGene_gn=ATG2B;refGene_tx=XM_019833608.2;refGene_gn=ATG2B;refGene_tx=XM_019833609.2;refGene_gn=ATG2B;refGene_tx=XM_019833610.2;refGene_gn=ATG2B;refGene_tx=XM_019833611.2
B3 141334430 141334581
refGene_gn=ATG2B;refGene_tx=XM_011283531.3;refGene_gn=ATG2B;refGene_tx=XM_019833608.2;refGene_gn=ATG2B;refGene_tx=XM_019833609.2;refGene_gn=ATG2B;refGene_tx=XM_019833610.2;refGene_gn=ATG2B;refGene_tx=XM_019833611.2
B3 141335078 141335251
refGene_gn=ATG2B;refGene_tx=XM_011283531.3;refGene_gn=ATG2B;refGene_tx=XM_019833608.2;refGene_gn=ATG2B;refGene_tx=XM_019833609.2;refGene_gn=ATG2B;refGene_tx=XM_019833610.2;refGene_gn=ATG2B;refGene_tx=XM_019833611.2
B3 141335652 141335810
refGene_gn=ATG2B;refGene_tx=XM_011283531.3;refGene_gn=ATG2B;refGene_tx=XM_019833608.2;refGene_gn=ATG2B;refGene_tx=XM_019833609.2;refGene_gn=ATG2B;refGene_tx=XM_019833610.2;refGene_gn=ATG2B;refGene_tx=XM_019833611.2
B3 141340654 141340797
refGene_gn=ATG2B;refGene_tx=XM_011283531.3;refGene_gn=ATG2B;refGene_tx=XM_019833608.2;refGene_gn=ATG2B;refGene_tx=XM_019833609.2;refGene_gn=ATG2B;refGene_tx=XM_019833610.2;refGene_gn=ATG2B;refGene_tx=XM_019833611.2
B3 141341075 141341268
refGene_gn=ATG2B;refGene_tx=XM_011283531.3;refGene_gn=ATG2B;refGene_tx=XM_019833608.2;refGene_gn=ATG2B;refGene_tx=XM_019833609.2;refGene_gn=ATG2B;refGene_tx=XM_019833610.2;refGene_gn=ATG2B;refGene_tx=XM_019833611.2
B3 141343061 141343164
refGene_gn=ATG2B;refGene_tx=XM_011283531.3;refGene_gn=ATG2B;refGene_tx=XM_019833608.2;refGene_gn=ATG2B;refGene_tx=XM_019833609.2;refGene_gn=ATG2B;refGene_tx=XM_019833610.2;refGene_gn=ATG2B;refGene_tx=XM_019833611.2
B3 141343989 141344263
refGene_gn=ATG2B;refGene_tx=XM_011283531.3;refGene_gn=ATG2B;refGene_tx=XM_019833608.2;refGene_gn=ATG2B;refGene_tx=XM_019833609.2;refGene_gn=ATG2B;refGene_tx=XM_019833610.2;refGene_gn=ATG2B;refGene_tx=XM_019833611.2
B3 141346536 141346725
refGene_gn=ATG2B;refGene_tx=XM_011283531.3;refGene_gn=ATG2B;refGene_tx=XM_019833608.2;refGene_gn=ATG2B;refGene_tx=XM_019833609.2;refGene_gn=ATG2B;refGene_tx=XM_019833610.2;refGene_gn=ATG2B;refGene_tx=XM_019833611.2
B3 141346863 141346956
refGene_gn=ATG2B;refGene_tx=XM_011283531.3;refGene_gn=ATG2B;refGene_tx=XM_019833608.2;refGene_gn=ATG2B;refGene_tx=XM_019833609.2;refGene_gn=ATG2B;refGene_tx=XM_019833610.2;refGene_gn=ATG2B;refGene_tx=XM_019833611.2
B3 141347786 141347937

refGene_gn=ATG2B;refGene_tx=XM_011283531.3;refGene_gn=ATG2B;refGene_tx=XM_019833608.2;refGene_gn=ATG2B;refGene_tx=XM_019833609.2;refGene_gn=ATG2B;refGene_tx=XM_019833610.2;refGene_gn=ATG2B;refGene_tx=XM_019833611.2
B3 141349163 141349425
refGene_gn=ATG2B;refGene_tx=XM_011283531.3;refGene_gn=ATG2B;refGene_tx=XM_019833608.2;refGene_gn=ATG2B;refGene_tx=XM_019833609.2;refGene_gn=ATG2B;refGene_tx=XM_019833610.2;refGene_gn=ATG2B;refGene_tx=XM_019833611.2
B3 141350575 141350681
refGene_gn=ATG2B;refGene_tx=XM_011283531.3;refGene_gn=ATG2B;refGene_tx=XM_019833608.2;refGene_gn=ATG2B;refGene_tx=XM_019833609.2;refGene_gn=ATG2B;refGene_tx=XM_019833610.2;refGene_gn=ATG2B;refGene_tx=XM_019833611.2
B3 141350774 141350929
refGene_gn=ATG2B;refGene_tx=XM_011283531.3;refGene_gn=ATG2B;refGene_tx=XM_019833608.2;refGene_gn=ATG2B;refGene_tx=XM_019833609.2;refGene_gn=ATG2B;refGene_tx=XM_019833610.2;refGene_gn=ATG2B;refGene_tx=XM_019833611.2
B3 141351937 141352123
refGene_gn=ATG2B;refGene_tx=XM_011283531.3;refGene_gn=ATG2B;refGene_tx=XM_019833608.2;refGene_gn=ATG2B;refGene_tx=XM_019833609.2;refGene_gn=ATG2B;refGene_tx=XM_019833610.2;refGene_gn=ATG2B;refGene_tx=XM_019833611.2
B3 141352370 141352467
refGene_gn=ATG2B;refGene_tx=XM_011283531.3;refGene_gn=ATG2B;refGene_tx=XM_019833608.2;refGene_gn=ATG2B;refGene_tx=XM_019833609.2;refGene_gn=ATG2B;refGene_tx=XM_019833610.2;refGene_gn=ATG2B;refGene_tx=XM_019833611.2
B3 141355490 141355670
refGene_gn=ATG2B;refGene_tx=XM_011283531.3;refGene_gn=ATG2B;refGene_tx=XM_019833608.2;refGene_gn=ATG2B;refGene_tx=XM_019833609.2;refGene_gn=ATG2B;refGene_tx=XM_019833610.2;refGene_gn=ATG2B;refGene_tx=XM_019833611.2
B3 141357006 141357169
refGene_gn=ATG2B;refGene_tx=XM_011283531.3;refGene_gn=ATG2B;refGene_tx=XM_019833608.2;refGene_gn=ATG2B;refGene_tx=XM_019833609.2;refGene_gn=ATG2B;refGene_tx=XM_019833610.2;refGene_gn=ATG2B;refGene_tx=XM_019833611.2
B3 141358678 141358781
refGene_gn=ATG2B;refGene_tx=XM_011283531.3;refGene_gn=ATG2B;refGene_tx=XM_019833608.2;refGene_gn=ATG2B;refGene_tx=XM_019833609.2;refGene_gn=ATG2B;refGene_tx=XM_019833610.2;refGene_gn=ATG2B;refGene_tx=XM_019833611.2
B3 141359477 141359630
refGene_gn=ATG2B;refGene_tx=XM_011283531.3;refGene_gn=ATG2B;refGene_tx=XM_019833608.2;refGene_gn=ATG2B;refGene_tx=XM_019833609.2;refGene_gn=ATG2B;refGene_tx=XM_019833610.2;refGene_gn=ATG2B;refGene_tx=XM_019833611.2
B3 141362639 141362802

refGene_gn=ATG2B;refGene_tx=XM_011283531.3;refGene_gn=ATG2B;refGene_tx
=XM_019833608.2;refGene_gn=ATG2B;refGene_tx=XM_019833609.2;refGene_gn=
ATG2B;refGene_tx=XM_019833610.2;refGene_gn=ATG2B;refGene_tx=XM_0198336
11.2
B3	141376022	141376184
refGene_gn=ATG2B;refGene_tx=XM_011283531.3;refGene_gn=ATG2B;refGene_tx
=XM_019833608.2;refGene_gn=ATG2B;refGene_tx=XM_019833609.2;refGene_gn=
ATG2B;refGene_tx=XM_019833610.2;refGene_gn=ATG2B;refGene_tx=XM_0198336
11.2
B3	141398033	141398291
refGene_gn=GSKIP;refGene_tx=XM_006933091.4;refGene_gn=GSKIP;refGene_tx
=XM_006933092.4;refGene_gn=GSKIP;refGene_tx=XM_006933093.4;refGene_gn=
GSKIP;refGene_tx=XM_006933094.4;refGene_gn=GSKIP;refGene_tx=XM_0198336
12.2;refGene_gn=GSKIP;refGene_tx=XM_023256013.1;refGene_gn=GSKIP;refGe
ne_tx=XM_023256014.1
B3	141400433	141400595
refGene_gn=GSKIP;refGene_tx=XM_006933091.4;refGene_gn=GSKIP;refGene_tx
=XM_006933092.4;refGene_gn=GSKIP;refGene_tx=XM_006933093.4;refGene_gn=
GSKIP;refGene_tx=XM_006933094.4;refGene_gn=GSKIP;refGene_tx=XM_0198336
12.2;refGene_gn=GSKIP;refGene_tx=XM_023256013.1;refGene_gn=GSKIP;refGe
ne_tx=XM_023256014.1
B3	141407780	141407879
refGene_gn=AK7;refGene_tx=XM_023256195.1
B3	141408295	141408487
refGene_gn=AK7;refGene_tx=XM_023256195.1
B3	141411091	141411271
refGene_gn=AK7;refGene_tx=XM_023256195.1
B3	141417910	141418019
refGene_gn=AK7;refGene_tx=XM_023256195.1
B3	141424556	141424651
refGene_gn=AK7;refGene_tx=XM_023256195.1
B3	141431905	141432016
refGene_gn=AK7;refGene_tx=XM_023256195.1
B3	141441992	141442073
refGene_gn=AK7;refGene_tx=XM_023256195.1
B3	141444424	141444513
refGene_gn=AK7;refGene_tx=XM_023256195.1
B3	141446861	141446952
refGene_gn=AK7;refGene_tx=XM_023256195.1
B3	141447864	141447942
refGene_gn=AK7;refGene_tx=XM_023256195.1
B3	141449008	141449158
refGene_gn=AK7;refGene_tx=XM_023256195.1
B3	141451941	141452070
refGene_gn=AK7;refGene_tx=XM_023256195.1
B3	141453316	141453455
refGene_gn=AK7;refGene_tx=XM_023256195.1
B3	141459173	141459302
refGene_gn=AK7;refGene_tx=XM_023256195.1
B3	141459672	141459741

refGene_gn=AK7;refGene_tx=XM_023256195.1
B3 141465930 141466128
refGene_gn=AK7;refGene_tx=XM_023256195.1
B3 141469702 141469923
refGene_gn=AK7;refGene_tx=XM_023256195.1
B3 141478090 141478249
refGene_gn=AK7;refGene_tx=XM_023256195.1
B3 141478993 141479032
refGene_gn=AK7;refGene_tx=XM_023256195.1
B3 141495192 141495200
refGene_gn=PAPOLA;refGene_tx=XM_003987956.5
B3 141495539 141495598
refGene_gn=PAPOLA;refGene_tx=XM_019833613.2;refGene_gn=PAPOLA;refGene_tx=XM_019833614.2;refGene_gn=PAPOLA;refGene_tx=XM_019833615.2;refGene_gn=PAPOLA;refGene_tx=XM_019833616.2;refGene_gn=PAPOLA;refGene_tx=XM_019833617.2
B3 141511816 141511990
refGene_gn=PAPOLA;refGene_tx=XM_003987956.5;refGene_gn=PAPOLA;refGene_tx=XM_019833613.2;refGene_gn=PAPOLA;refGene_tx=XM_019833614.2;refGene_gn=PAPOLA;refGene_tx=XM_019833615.2;refGene_gn=PAPOLA;refGene_tx=XM_019833616.2;refGene_gn=PAPOLA;refGene_tx=XM_019833617.2
B3 141512418 141512485
refGene_gn=PAPOLA;refGene_tx=XM_003987956.5;refGene_gn=PAPOLA;refGene_tx=XM_019833613.2;refGene_gn=PAPOLA;refGene_tx=XM_019833614.2;refGene_gn=PAPOLA;refGene_tx=XM_019833615.2;refGene_gn=PAPOLA;refGene_tx=XM_019833616.2;refGene_gn=PAPOLA;refGene_tx=XM_019833617.2
B3 141516093 141516175
refGene_gn=PAPOLA;refGene_tx=XM_003987956.5;refGene_gn=PAPOLA;refGene_tx=XM_019833613.2;refGene_gn=PAPOLA;refGene_tx=XM_019833614.2;refGene_gn=PAPOLA;refGene_tx=XM_019833615.2;refGene_gn=PAPOLA;refGene_tx=XM_019833616.2;refGene_gn=PAPOLA;refGene_tx=XM_019833617.2
B3 141517938 141518048
refGene_gn=PAPOLA;refGene_tx=XM_003987956.5;refGene_gn=PAPOLA;refGene_tx=XM_019833613.2;refGene_gn=PAPOLA;refGene_tx=XM_019833614.2;refGene_gn=PAPOLA;refGene_tx=XM_019833615.2;refGene_gn=PAPOLA;refGene_tx=XM_019833616.2;refGene_gn=PAPOLA;refGene_tx=XM_019833617.2
B3 141518503 141518557
refGene_gn=PAPOLA;refGene_tx=XM_003987956.5;refGene_gn=PAPOLA;refGene_tx=XM_019833613.2;refGene_gn=PAPOLA;refGene_tx=XM_019833614.2;refGene_gn=PAPOLA;refGene_tx=XM_019833615.2;refGene_gn=PAPOLA;refGene_tx=XM_019833616.2;refGene_gn=PAPOLA;refGene_tx=XM_019833617.2
B3 141521408 141521520
refGene_gn=PAPOLA;refGene_tx=XM_003987956.5;refGene_gn=PAPOLA;refGene_tx=XM_019833613.2;refGene_gn=PAPOLA;refGene_tx=XM_019833614.2;refGene_gn=PAPOLA;refGene_tx=XM_019833615.2;refGene_gn=PAPOLA;refGene_tx=XM_019833616.2;refGene_gn=PAPOLA;refGene_tx=XM_019833617.2
B3 141522207 141522297
refGene_gn=PAPOLA;refGene_tx=XM_003987956.5;refGene_gn=PAPOLA;refGene_tx=XM_019833613.2;refGene_gn=PAPOLA;refGene_tx=XM_019833614.2;refGene_gn=PAPOLA;refGene_tx=XM_019833615.2;refGene_gn=PAPOLA;refGene_tx=XM_01

9833616.2;refGene_gn=PAPOLA;refGene_tx=XM_019833617.2
B3	141522387	141522526
refGene_gn=PAPOLA;refGene_tx=XM_003987956.5;refGene_gn=PAPOLA;refGene_tx=XM_019833613.2;refGene_gn=PAPOLA;refGene_tx=XM_019833614.2;refGene_gn=PAPOLA;refGene_tx=XM_019833615.2;refGene_gn=PAPOLA;refGene_tx=XM_019833616.2;refGene_gn=PAPOLA;refGene_tx=XM_019833617.2
B3	141524073	141524146
refGene_gn=PAPOLA;refGene_tx=XM_003987956.5;refGene_gn=PAPOLA;refGene_tx=XM_019833613.2;refGene_gn=PAPOLA;refGene_tx=XM_019833614.2;refGene_gn=PAPOLA;refGene_tx=XM_019833615.2;refGene_gn=PAPOLA;refGene_tx=XM_019833616.2;refGene_gn=PAPOLA;refGene_tx=XM_019833617.2
B3	141525468	141525589
refGene_gn=PAPOLA;refGene_tx=XM_003987956.5;refGene_gn=PAPOLA;refGene_tx=XM_019833613.2;refGene_gn=PAPOLA;refGene_tx=XM_019833614.2;refGene_gn=PAPOLA;refGene_tx=XM_019833615.2;refGene_gn=PAPOLA;refGene_tx=XM_019833616.2;refGene_gn=PAPOLA;refGene_tx=XM_019833617.2
B3	141526544	141526629
refGene_gn=PAPOLA;refGene_tx=XM_003987956.5;refGene_gn=PAPOLA;refGene_tx=XM_019833613.2;refGene_gn=PAPOLA;refGene_tx=XM_019833614.2;refGene_gn=PAPOLA;refGene_tx=XM_019833615.2;refGene_gn=PAPOLA;refGene_tx=XM_019833616.2;refGene_gn=PAPOLA;refGene_tx=XM_019833617.2
B3	141531926	141531980
refGene_gn=PAPOLA;refGene_tx=XM_003987956.5;refGene_gn=PAPOLA;refGene_tx=XM_019833613.2;refGene_gn=PAPOLA;refGene_tx=XM_019833614.2;refGene_gn=PAPOLA;refGene_tx=XM_019833615.2;refGene_gn=PAPOLA;refGene_tx=XM_019833616.2;refGene_gn=PAPOLA;refGene_tx=XM_019833617.2
B3	141532500	141532620
refGene_gn=PAPOLA;refGene_tx=XM_003987956.5;refGene_gn=PAPOLA;refGene_tx=XM_019833613.2;refGene_gn=PAPOLA;refGene_tx=XM_019833614.2;refGene_gn=PAPOLA;refGene_tx=XM_019833615.2;refGene_gn=PAPOLA;refGene_tx=XM_019833616.2;refGene_gn=PAPOLA;refGene_tx=XM_019833617.2
B3	141533969	141534079
refGene_gn=PAPOLA;refGene_tx=XM_003987956.5;refGene_gn=PAPOLA;refGene_tx=XM_019833613.2;refGene_gn=PAPOLA;refGene_tx=XM_019833614.2;refGene_gn=PAPOLA;refGene_tx=XM_019833615.2;refGene_gn=PAPOLA;refGene_tx=XM_019833616.2;refGene_gn=PAPOLA;refGene_tx=XM_019833617.2
B3	141537930	141538052
refGene_gn=PAPOLA;refGene_tx=XM_003987956.5;refGene_gn=PAPOLA;refGene_tx=XM_019833613.2;refGene_gn=PAPOLA;refGene_tx=XM_019833614.2;refGene_gn=PAPOLA;refGene_tx=XM_019833615.2;refGene_gn=PAPOLA;refGene_tx=XM_019833616.2;refGene_gn=PAPOLA;refGene_tx=XM_019833617.2
B3	141542547	141542690
refGene_gn=PAPOLA;refGene_tx=XM_003987956.5;refGene_gn=PAPOLA;refGene_tx=XM_019833613.2;refGene_gn=PAPOLA;refGene_tx=XM_019833614.2;refGene_gn=PAPOLA;refGene_tx=XM_019833615.2;refGene_gn=PAPOLA;refGene_tx=XM_019833616.2;refGene_gn=PAPOLA;refGene_tx=XM_019833617.2
B3	141546236	141546337
refGene_gn=PAPOLA;refGene_tx=XM_003987956.5;refGene_gn=PAPOLA;refGene_tx=XM_019833613.2;refGene_gn=PAPOLA;refGene_tx=XM_019833614.2;refGene_gn=PAPOLA;refGene_tx=XM_019833615.2;refGene_gn=PAPOLA;refGene_tx=XM_01

9833616.2;refGene_gn=PAPOLA;refGene_tx=XM_019833617.2
B3 141546565 141546786
refGene_gn=PAPOLA;refGene_tx=XM_003987956.5;refGene_gn=PAPOLA;refGene_tx=XM_019833613.2;refGene_gn=PAPOLA;refGene_tx=XM_019833614.2;refGene_gn=PAPOLA;refGene_tx=XM_019833615.2;refGene_gn=PAPOLA;refGene_tx=XM_019833616.2;refGene_gn=PAPOLA;refGene_tx=XM_019833617.2
B3 141550599 141550662
refGene_gn=PAPOLA;refGene_tx=XM_003987956.5;refGene_gn=PAPOLA;refGene_tx=XM_019833613.2;refGene_gn=PAPOLA;refGene_tx=XM_019833614.2;refGene_gn=PAPOLA;refGene_tx=XM_019833616.2
B3 141552531 141552606
refGene_gn=PAPOLA;refGene_tx=XM_003987956.5;refGene_gn=PAPOLA;refGene_tx=XM_019833613.2;refGene_gn=PAPOLA;refGene_tx=XM_019833614.2;refGene_gn=PAPOLA;refGene_tx=XM_019833615.2
B3 141554635 141554731
refGene_gn=PAPOLA;refGene_tx=XM_003987956.5;refGene_gn=PAPOLA;refGene_tx=XM_019833613.2;refGene_gn=PAPOLA;refGene_tx=XM_019833614.2;refGene_gn=PAPOLA;refGene_tx=XM_019833615.2;refGene_gn=PAPOLA;refGene_tx=XM_019833616.2;refGene_gn=PAPOLA;refGene_tx=XM_019833617.2
B3 141785906 141786066
refGene_gn=VRK1;refGene_tx=XM_011283524.3;refGene_gn=VRK1;refGene_tx=XM_011283526.3;refGene_gn=VRK1;refGene_tx=XM_011283527.3;refGene_gn=VRK1;refGene_tx=XM_011283528.3;refGene_gn=VRK1;refGene_tx=XM_019833618.2;refGene_gn=VRK1;refGene_tx=XM_023256015.1
B3 141791192 141791248
refGene_gn=VRK1;refGene_tx=XM_011283524.3;refGene_gn=VRK1;refGene_tx=XM_011283526.3;refGene_gn=VRK1;refGene_tx=XM_011283527.3;refGene_gn=VRK1;refGene_tx=XM_019833618.2;refGene_gn=VRK1;refGene_tx=XM_023256015.1
B3 141798574 141798644
refGene_gn=VRK1;refGene_tx=XM_011283524.3;refGene_gn=VRK1;refGene_tx=XM_011283526.3;refGene_gn=VRK1;refGene_tx=XM_011283527.3;refGene_gn=VRK1;refGene_tx=XM_019833618.2;refGene_gn=VRK1;refGene_tx=XM_023256015.1
B3 141799723 141799811
refGene_gn=VRK1;refGene_tx=XM_011283524.3;refGene_gn=VRK1;refGene_tx=XM_011283526.3;refGene_gn=VRK1;refGene_tx=XM_011283527.3;refGene_gn=VRK1;refGene_tx=XM_011283528.3;refGene_gn=VRK1;refGene_tx=XM_019833618.2;refGene_gn=VRK1;refGene_tx=XM_023256015.1
B3 141803638 141803747
refGene_gn=VRK1;refGene_tx=XM_011283524.3;refGene_gn=VRK1;refGene_tx=XM_011283526.3;refGene_gn=VRK1;refGene_tx=XM_011283527.3;refGene_gn=VRK1;refGene_tx=XM_011283528.3;refGene_gn=VRK1;refGene_tx=XM_019833618.2;refGene_gn=VRK1;refGene_tx=XM_023256015.1
B3 141803862 141803955
refGene_gn=VRK1;refGene_tx=XM_011283524.3;refGene_gn=VRK1;refGene_tx=XM_011283526.3;refGene_gn=VRK1;refGene_tx=XM_011283527.3;refGene_gn=VRK1;refGene_tx=XM_011283528.3;refGene_gn=VRK1;refGene_tx=XM_019833618.2;refGene_gn=VRK1;refGene_tx=XM_023256015.1
B3 141805989 141806122
refGene_gn=VRK1;refGene_tx=XM_011283524.3;refGene_gn=VRK1;refGene_tx=XM_011283526.3;refGene_gn=VRK1;refGene_tx=XM_011283527.3;refGene_gn=VRK

1;refGene_tx=XM_011283528.3;refGene_gn=VRK1;refGene_tx=XM_019833618.2;
refGene_gn=VRK1;refGene_tx=XM_023256015.1
B3 141807160 141807281
refGene_gn=VRK1;refGene_tx=XM_011283524.3;refGene_gn=VRK1;refGene_tx=X
M_011283526.3;refGene_gn=VRK1;refGene_tx=XM_011283527.3;refGene_gn=VRK
1;refGene_tx=XM_011283528.3;refGene_gn=VRK1;refGene_tx=XM_019833618.2;
refGene_gn=VRK1;refGene_tx=XM_023256015.1
B3 141807574 141807633
refGene_gn=VRK1;refGene_tx=XM_011283524.3;refGene_gn=VRK1;refGene_tx=X
M_011283526.3;refGene_gn=VRK1;refGene_tx=XM_011283527.3;refGene_gn=VRK
1;refGene_tx=XM_011283528.3;refGene_gn=VRK1;refGene_tx=XM_019833618.2;
refGene_gn=VRK1;refGene_tx=XM_023256015.1
B3 141812085 141812264
refGene_gn=VRK1;refGene_tx=XM_011283524.3;refGene_gn=VRK1;refGene_tx=X
M_011283526.3;refGene_gn=VRK1;refGene_tx=XM_011283527.3;refGene_gn=VRK
1;refGene_tx=XM_011283528.3;refGene_gn=VRK1;refGene_tx=XM_019833618.2;
refGene_gn=VRK1;refGene_tx=XM_023256015.1
B3 141823599 141823687
refGene_gn=VRK1;refGene_tx=XM_011283524.3;refGene_gn=VRK1;refGene_tx=X
M_011283526.3;refGene_gn=VRK1;refGene_tx=XM_011283527.3;refGene_gn=VRK
1;refGene_tx=XM_011283528.3;refGene_gn=VRK1;refGene_tx=XM_019833618.2;
refGene_gn=VRK1;refGene_tx=XM_023256015.1
B3 141824900 141824969
refGene_gn=VRK1;refGene_tx=XM_011283524.3;refGene_gn=VRK1;refGene_tx=X
M_011283526.3;refGene_gn=VRK1;refGene_tx=XM_011283528.3;refGene_gn=VRK
1;refGene_tx=XM_023256015.1
B3 141828252 141828284
refGene_gn=VRK1;refGene_tx=XM_011283524.3;refGene_gn=VRK1;refGene_tx=X
M_011283526.3;refGene_gn=VRK1;refGene_tx=XM_011283527.3;refGene_gn=VRK
1;refGene_tx=XM_011283528.3;refGene_gn=VRK1;refGene_tx=XM_019833618.2;
refGene_gn=VRK1;refGene_tx=XM_023256015.1
B3 143962201 143964252
refGene_gn=BCL11B;refGene_tx=XM_023256017.1;refGene_gn=BCL11B;refGene_
tx=XM_023256018.1;refGene_gn=BCL11B;refGene_tx=XM_023256019.1;refGene_
gn=BCL11B;refGene_tx=XM_023256020.1;refGene_gn=BCL11B;refGene_tx=XM_02
3256021.1
B3 143981878 143982230
refGene_gn=BCL11B;refGene_tx=XM_023256021.1
B3 144013371 144013584
refGene_gn=BCL11B;refGene_tx=XM_023256017.1;refGene_gn=BCL11B;refGene_
tx=XM_023256018.1
B3 144040291 144040660
refGene_gn=BCL11B;refGene_tx=XM_023256017.1;refGene_gn=BCL11B;refGene_
tx=XM_023256018.1;refGene_gn=BCL11B;refGene_tx=XM_023256019.1;refGene_
gn=BCL11B;refGene_tx=XM_023256020.1
B3 144055417 144055475
refGene_gn=BCL11B;refGene_tx=XM_023256017.1;refGene_gn=BCL11B;refGene_
tx=XM_023256018.1;refGene_gn=BCL11B;refGene_tx=XM_023256019.1;refGene_
gn=BCL11B;refGene_tx=XM_023256020.1
B3 144164042 144164510

refGene_gn=SETD3;refGene_tx=XM_023256210.1
B3 144165414 144165575
refGene_gn=SETD3;refGene_tx=XM_023256210.1
B3 144171995 144172081
refGene_gn=SETD3;refGene_tx=XM_023256210.1
B3 144172947 144173114
refGene_gn=SETD3;refGene_tx=XM_023256210.1
B3 144174094 144174169
refGene_gn=SETD3;refGene_tx=XM_023256210.1
B3 144179139 144179254
refGene_gn=SETD3;refGene_tx=XM_023256210.1
B3 144180263 144180322
refGene_gn=SETD3;refGene_tx=XM_023256210.1
B3 144220398 144220655
refGene_gn=SETD3;refGene_tx=XM_023256210.1
B3 144220940 144221013
refGene_gn=SETD3;refGene_tx=XM_023256210.1
B3 144223280 144223429
refGene_gn=SETD3;refGene_tx=XM_023256210.1
B3 144225923 144226016
refGene_gn=SETD3;refGene_tx=XM_023256210.1
B3 144228048 144228151
refGene_gn=SETD3;refGene_tx=XM_023256210.1
B3 144255673 144255870
refGene_gn=CCNK;refGene_tx=XM_023256211.1
B3 144256321 144256403
refGene_gn=CCNK;refGene_tx=XM_023256211.1
B3 144257925 144258057
refGene_gn=CCNK;refGene_tx=XM_023256211.1
B3 144262917 144263023
refGene_gn=CCNK;refGene_tx=XM_023256211.1
B3 144263725 144263783
refGene_gn=CCNK;refGene_tx=XM_023256211.1
B3 144264498 144264668
refGene_gn=CCNK;refGene_tx=XM_023256211.1
B3 144264990 144265262
refGene_gn=CCNK;refGene_tx=XM_023256211.1
B3 144265862 144265896
refGene_gn=CCNK;refGene_tx=XM_023256211.1
B3 144269049 144269121
refGene_gn=CCNK;refGene_tx=XM_023256211.1
B3 144271886 144272512
refGene_gn=CCNK;refGene_tx=XM_023256211.1
B3 144276902 144276992
refGene_gn=CCDC85C;refGene_tx=XM_023256222.1
B3 144277782 144277881
refGene_gn=CCDC85C;refGene_tx=XM_023256222.1
B3 144278763 144278859
refGene_gn=CCDC85C;refGene_tx=XM_023256222.1
B3 144283313 144283421

refGene_gn=CCDC85C;refGene_tx=XM_023256222.1
B3 144292857 144292907
refGene_gn=CCDC85C;refGene_tx=XM_023256222.1
B3 144351966 144352282
refGene_gn=CCDC85C;refGene_tx=XM_023256222.1
B3 144352283 144352778
refGene_gn=CCDC85C;refGene_tx=XM_023256222.1
B3 144380167 144380437
refGene_gn=HHIPL1;refGene_tx=XM_023256207.1
B3 144388252 144388899
refGene_gn=HHIPL1;refGene_tx=XM_023256207.1;refGene_gn=HHIPL1;refGene_tx=XM_023256208.1
B3 144391449 144391593
refGene_gn=HHIPL1;refGene_tx=XM_023256207.1;refGene_gn=HHIPL1;refGene_tx=XM_023256208.1
B3 144393414 144393743
refGene_gn=HHIPL1;refGene_tx=XM_023256207.1;refGene_gn=HHIPL1;refGene_tx=XM_023256208.1
B3 144394235 144394362
refGene_gn=HHIPL1;refGene_tx=XM_023256207.1;refGene_gn=HHIPL1;refGene_tx=XM_023256208.1
B3 144397576 144397722
refGene_gn=HHIPL1;refGene_tx=XM_023256207.1;refGene_gn=HHIPL1;refGene_tx=XM_023256208.1
B3 144400768 144400850
refGene_gn=HHIPL1;refGene_tx=XM_023256207.1;refGene_gn=HHIPL1;refGene_tx=XM_023256208.1
B3 144404083 144404166
refGene_gn=HHIPL1;refGene_tx=XM_023256207.1;refGene_gn=HHIPL1;refGene_tx=XM_023256208.1
B3 144406401 144406937
refGene_gn=HHIPL1;refGene_tx=XM_023256207.1;refGene_gn=HHIPL1;refGene_tx=XM_023256208.1
B3 144413035 144413157
refGene_gn=LOC101093112;refGene_tx=XM_023256212.1
B3 144420355 144420436
refGene_gn=LOC101093112;refGene_tx=XM_023256212.1
B3 144420978 144421060
refGene_gn=LOC101093112;refGene_tx=XM_023256212.1
B3 144426885 144426959
refGene_gn=LOC101093112;refGene_tx=XM_023256212.1
B3 144427436 144427523
refGene_gn=LOC101093112;refGene_tx=XM_023256212.1
B3 144432635 144432774
refGene_gn=LOC101093112;refGene_tx=XM_023256212.1
B3 144433460 144433571
refGene_gn=LOC101093112;refGene_tx=XM_023256212.1
B3 144434424 144434575
refGene_gn=LOC101093112;refGene_tx=XM_023256212.1
B3 144434758 144434821

refGene_gn=LOC101093112;refGene_tx=XM_023256212.1
B3 144436771 144436844
refGene_gn=LOC101093112;refGene_tx=XM_023256212.1
B3 144439378 144439463
refGene_gn=LOC101093112;refGene_tx=XM_023256212.1
B3 144439904 144440015
refGene_gn=LOC101093112;refGene_tx=XM_023256212.1
B3 144443230 144443319
refGene_gn=LOC101093112;refGene_tx=XM_023256212.1
B3 144444142 144444209
refGene_gn=LOC101093112;refGene_tx=XM_023256212.1
B3 144444383 144444551
refGene_gn=LOC101093112;refGene_tx=XM_023256212.1
B3 144455214 144455242
refGene_gn=EML1;refGene_tx=XM_006933114.4
B3 144510788 144510855
refGene_gn=EML1;refGene_tx=XM_023256205.1;refGene_gn=EML1;refGene_tx=XM_023256206.1
B3 144564324 144564507
refGene_gn=EML1;refGene_tx=XM_006933114.4;refGene_gn=EML1;refGene_tx=XM_019833627.2;refGene_gn=EML1;refGene_tx=XM_023256205.1;refGene_gn=EML1;refGene_tx=XM_023256206.1
B3 144577005 144577141
refGene_gn=EML1;refGene_tx=XM_006933114.4;refGene_gn=EML1;refGene_tx=XM_019833627.2;refGene_gn=EML1;refGene_tx=XM_023256205.1;refGene_gn=EML1;refGene_tx=XM_023256206.1
B3 144591011 144591146
refGene_gn=EML1;refGene_tx=XM_006933114.4;refGene_gn=EML1;refGene_tx=XM_019833627.2;refGene_gn=EML1;refGene_tx=XM_023256205.1;refGene_gn=EML1;refGene_tx=XM_023256206.1
B3 144603575 144603604
refGene_gn=EML1;refGene_tx=XM_006933114.4;refGene_gn=EML1;refGene_tx=XM_019833627.2;refGene_gn=EML1;refGene_tx=XM_023256205.1;refGene_gn=EML1;refGene_tx=XM_023256206.1
B3 144604499 144604556
refGene_gn=EML1;refGene_tx=XM_006933114.4;refGene_gn=EML1;refGene_tx=XM_019833627.2;refGene_gn=EML1;refGene_tx=XM_023256205.1
B3 144606959 144607089
refGene_gn=EML1;refGene_tx=XM_006933114.4;refGene_gn=EML1;refGene_tx=XM_019833627.2;refGene_gn=EML1;refGene_tx=XM_023256205.1;refGene_gn=EML1;refGene_tx=XM_023256206.1
B3 144609796 144609946
refGene_gn=EML1;refGene_tx=XM_006933114.4;refGene_gn=EML1;refGene_tx=XM_019833627.2;refGene_gn=EML1;refGene_tx=XM_023256205.1;refGene_gn=EML1;refGene_tx=XM_023256206.1
B3 144611173 144611243
refGene_gn=EML1;refGene_tx=XM_006933114.4;refGene_gn=EML1;refGene_tx=XM_019833627.2;refGene_gn=EML1;refGene_tx=XM_023256205.1;refGene_gn=EML1;refGene_tx=XM_023256206.1
B3 144613012 144613123

refGene_gn=EML1;refGene_tx=XM_006933114.4;refGene_gn=EML1;refGene_tx=XM_019833627.2;refGene_gn=EML1;refGene_tx=XM_023256205.1;refGene_gn=EML1;refGene_tx=XM_023256206.1
B3 144613956 144614052
refGene_gn=EML1;refGene_tx=XM_006933114.4;refGene_gn=EML1;refGene_tx=XM_019833627.2;refGene_gn=EML1;refGene_tx=XM_023256205.1;refGene_gn=EML1;refGene_tx=XM_023256206.1
B3 144615666 144615801
refGene_gn=EML1;refGene_tx=XM_006933114.4;refGene_gn=EML1;refGene_tx=XM_019833627.2;refGene_gn=EML1;refGene_tx=XM_023256205.1;refGene_gn=EML1;refGene_tx=XM_023256206.1
B3 144616485 144616585
refGene_gn=EML1;refGene_tx=XM_006933114.4;refGene_gn=EML1;refGene_tx=XM_019833627.2;refGene_gn=EML1;refGene_tx=XM_023256205.1;refGene_gn=EML1;refGene_tx=XM_023256206.1
B3 144617499 144617654
refGene_gn=EML1;refGene_tx=XM_006933114.4;refGene_gn=EML1;refGene_tx=XM_019833627.2;refGene_gn=EML1;refGene_tx=XM_023256205.1;refGene_gn=EML1;refGene_tx=XM_023256206.1
B3 144619706 144619832
refGene_gn=EML1;refGene_tx=XM_006933114.4;refGene_gn=EML1;refGene_tx=XM_019833627.2;refGene_gn=EML1;refGene_tx=XM_023256205.1;refGene_gn=EML1;refGene_tx=XM_023256206.1
B3 144620072 144620204
refGene_gn=EML1;refGene_tx=XM_006933114.4;refGene_gn=EML1;refGene_tx=XM_019833627.2;refGene_gn=EML1;refGene_tx=XM_023256205.1;refGene_gn=EML1;refGene_tx=XM_023256206.1
B3 144623118 144623186
refGene_gn=EML1;refGene_tx=XM_006933114.4;refGene_gn=EML1;refGene_tx=XM_019833627.2;refGene_gn=EML1;refGene_tx=XM_023256205.1;refGene_gn=EML1;refGene_tx=XM_023256206.1
B3 144625400 144625489
refGene_gn=EML1;refGene_tx=XM_006933114.4;refGene_gn=EML1;refGene_tx=XM_019833627.2;refGene_gn=EML1;refGene_tx=XM_023256205.1;refGene_gn=EML1;refGene_tx=XM_023256206.1
B3 144636641 144636739
refGene_gn=EML1;refGene_tx=XM_006933114.4;refGene_gn=EML1;refGene_tx=XM_019833627.2;refGene_gn=EML1;refGene_tx=XM_023256205.1;refGene_gn=EML1;refGene_tx=XM_023256206.1
B3 144636844 144636932
refGene_gn=EML1;refGene_tx=XM_006933114.4;refGene_gn=EML1;refGene_tx=XM_019833627.2;refGene_gn=EML1;refGene_tx=XM_023256205.1;refGene_gn=EML1;refGene_tx=XM_023256206.1
B3 144638346 144638442
refGene_gn=EML1;refGene_tx=XM_006933114.4;refGene_gn=EML1;refGene_tx=XM_019833627.2;refGene_gn=EML1;refGene_tx=XM_023256205.1;refGene_gn=EML1;refGene_tx=XM_023256206.1
B3 144639649 144639780
refGene_gn=EML1;refGene_tx=XM_006933114.4;refGene_gn=EML1;refGene_tx=XM_019833627.2;refGene_gn=EML1;refGene_tx=XM_023256205.1;refGene_gn=EML

1;refGene_tx=XM_023256206.1
B3	144640431	144640557
refGene_gn=EML1;refGene_tx=XM_006933114.4;refGene_gn=EML1;refGene_tx=XM_019833627.2;refGene_gn=EML1;refGene_tx=XM_023256205.1;refGene_gn=EML1;refGene_tx=XM_023256206.1
B3	144662827	144662961
refGene_gn=EVL;refGene_tx=XM_023256220.1
B3	144663282	144663371
refGene_gn=EVL;refGene_tx=XM_019833629.1
B3	144700645	144700668
refGene_gn=EVL;refGene_tx=XM_011283523.3
B3	144723533	144723583
refGene_gn=EVL;refGene_tx=XM_023256219.1;refGene_gn=EVL;refGene_tx=XM_023256221.1
B3	144737313	144737324
refGene_gn=EVL;refGene_tx=XM_003988023.4
B3	144755558	144755727
refGene_gn=EVL;refGene_tx=XM_003988023.4;refGene_gn=EVL;refGene_tx=XM_011283523.3;refGene_gn=EVL;refGene_tx=XM_019833629.1;refGene_gn=EVL;refGene_tx=XM_023256219.1;refGene_gn=EVL;refGene_tx=XM_023256220.1;refGene_gn=EVL;refGene_tx=XM_023256221.1
B3	144761790	144761968
refGene_gn=EVL;refGene_tx=XM_003988023.4;refGene_gn=EVL;refGene_tx=XM_011283523.3;refGene_gn=EVL;refGene_tx=XM_019833629.1;refGene_gn=EVL;refGene_tx=XM_023256219.1;refGene_gn=EVL;refGene_tx=XM_023256220.1;refGene_gn=EVL;refGene_tx=XM_023256221.1
B3	144791593	144791657
refGene_gn=EVL;refGene_tx=XM_003988023.4;refGene_gn=EVL;refGene_tx=XM_011283523.3;refGene_gn=EVL;refGene_tx=XM_019833629.1;refGene_gn=EVL;refGene_tx=XM_023256219.1;refGene_gn=EVL;refGene_tx=XM_023256220.1;refGene_gn=EVL;refGene_tx=XM_023256221.1
B3	144794676	144794747
refGene_gn=EVL;refGene_tx=XM_003988023.4;refGene_gn=EVL;refGene_tx=XM_011283523.3;refGene_gn=EVL;refGene_tx=XM_019833629.1;refGene_gn=EVL;refGene_tx=XM_023256219.1;refGene_gn=EVL;refGene_tx=XM_023256220.1;refGene_gn=EVL;refGene_tx=XM_023256221.1
B3	144796349	144796579
refGene_gn=EVL;refGene_tx=XM_003988023.4;refGene_gn=EVL;refGene_tx=XM_011283523.3;refGene_gn=EVL;refGene_tx=XM_019833629.1;refGene_gn=EVL;refGene_tx=XM_023256219.1;refGene_gn=EVL;refGene_tx=XM_023256220.1;refGene_gn=EVL;refGene_tx=XM_023256221.1
B3	144797306	144797428
refGene_gn=EVL;refGene_tx=XM_003988023.4;refGene_gn=EVL;refGene_tx=XM_011283523.3;refGene_gn=EVL;refGene_tx=XM_019833629.1;refGene_gn=EVL;refGene_tx=XM_023256219.1;refGene_gn=EVL;refGene_tx=XM_023256220.1;refGene_gn=EVL;refGene_tx=XM_023256221.1
B3	144800430	144800491
refGene_gn=EVL;refGene_tx=XM_003988023.4;refGene_gn=EVL;refGene_tx=XM_011283523.3;refGene_gn=EVL;refGene_tx=XM_019833629.1;refGene_gn=EVL;refGene_tx=XM_023256219.1;refGene_gn=EVL;refGene_tx=XM_023256220.1;refGe

ne_gn=EVL;refGene_tx=XM_023256221.1
B3 144804287 144804351
refGene_gn=EVL;refGene_tx=XM_003988023.4;refGene_gn=EVL;refGene_tx=XM_011283523.3;refGene_gn=EVL;refGene_tx=XM_019833629.1;refGene_gn=EVL;refGene_tx=XM_023256219.1;refGene_gn=EVL;refGene_tx=XM_023256220.1;refGene_gn=EVL;refGene_tx=XM_023256221.1
B3 144805947 144806014
refGene_gn=EVL;refGene_tx=XM_003988023.4;refGene_gn=EVL;refGene_tx=XM_011283523.3;refGene_gn=EVL;refGene_tx=XM_019833629.1;refGene_gn=EVL;refGene_tx=XM_023256219.1;refGene_gn=EVL;refGene_tx=XM_023256220.1;refGene_gn=EVL;refGene_tx=XM_023256221.1
B3 144806106 144806169
refGene_gn=EVL;refGene_tx=XM_003988023.4;refGene_gn=EVL;refGene_tx=XM_011283523.3;refGene_gn=EVL;refGene_tx=XM_019833629.1;refGene_gn=EVL;refGene_tx=XM_023256219.1;refGene_gn=EVL;refGene_tx=XM_023256220.1
B3 144809369 144809436
refGene_gn=EVL;refGene_tx=XM_003988023.4;refGene_gn=EVL;refGene_tx=XM_011283523.3;refGene_gn=EVL;refGene_tx=XM_019833629.1;refGene_gn=EVL;refGene_tx=XM_023256219.1;refGene_gn=EVL;refGene_tx=XM_023256220.1;refGene_gn=EVL;refGene_tx=XM_023256221.1
B3 144809745 144809803
refGene_gn=EVL;refGene_tx=XM_003988023.4;refGene_gn=EVL;refGene_tx=XM_011283523.3;refGene_gn=EVL;refGene_tx=XM_019833629.1;refGene_gn=EVL;refGene_tx=XM_023256219.1;refGene_gn=EVL;refGene_tx=XM_023256220.1;refGene_gn=EVL;refGene_tx=XM_023256221.1
B3 144811751 144811789
refGene_gn=EVL;refGene_tx=XM_003988023.4;refGene_gn=EVL;refGene_tx=XM_011283523.3;refGene_gn=EVL;refGene_tx=XM_019833629.1;refGene_gn=EVL;refGene_tx=XM_023256219.1;refGene_gn=EVL;refGene_tx=XM_023256220.1;refGene_gn=EVL;refGene_tx=XM_023256221.1
B3 144814487 144814634
refGene_gn=DEGS2;refGene_tx=XM_023256213.1;refGene_gn=DEGS2;refGene_tx=XM_023256214.1;refGene_gn=DEGS2;refGene_tx=XM_023256215.1;refGene_gn=DEGS2;refGene_tx=XM_023256216.1;refGene_gn=DEGS2;refGene_tx=XM_023256218.1
B3 144816347 144817090
refGene_gn=DEGS2;refGene_tx=XM_023256213.1;refGene_gn=DEGS2;refGene_tx=XM_023256214.1;refGene_gn=DEGS2;refGene_tx=XM_023256215.1;refGene_gn=DEGS2;refGene_tx=XM_023256216.1;refGene_gn=DEGS2;refGene_tx=XM_023256218.1
B3 144832340 144832436
refGene_gn=DEGS2;refGene_tx=XM_023256214.1
B3 144833644 144833726
refGene_gn=DEGS2;refGene_tx=XM_023256214.1;refGene_gn=DEGS2;refGene_tx=XM_023256218.1
B3 144844327 144844372
refGene_gn=DEGS2;refGene_tx=XM_023256216.1
B3 144845628 144845689
refGene_gn=DEGS2;refGene_tx=XM_023256216.1
B3 144876521 144876660

refGene_gn=DEGS2;refGene_tx=XM_023256215.1
B3	144883255	144883264
refGene_gn=DEGS2;refGene_tx=XM_023256215.1
B3	144884244	144884911
refGene_gn=YY1;refGene_tx=XM_023256223.1
B3	144900662	144900825
refGene_gn=YY1;refGene_tx=XM_023256223.1
B3	144908611	144908672
refGene_gn=YY1;refGene_tx=XM_023256223.1
B3	144910360	144910519
refGene_gn=YY1;refGene_tx=XM_023256223.1
B3	144911856	144912039
refGene_gn=YY1;refGene_tx=XM_023256223.1
B3	144924038	144925097
refGene_gn=SLC25A29;refGene_tx=XM_011283522.2;refGene_gn=SLC25A29;refGene_tx=XM_019833631.2;refGene_gn=SLC25A29;refGene_tx=XM_023256224.1
B3	144925428	144925512
refGene_gn=SLC25A29;refGene_tx=XM_011283522.2;refGene_gn=SLC25A29;refGene_tx=XM_019833631.2;refGene_gn=SLC25A29;refGene_tx=XM_023256224.1
B3	144931419	144931463
refGene_gn=SLC25A29;refGene_tx=XM_011283522.2
B3	144937015	144937049
refGene_gn=SLC25A29;refGene_tx=XM_011283522.2
B3	144953747	144953775
refGene_gn=SLC25A47;refGene_tx=XM_023256225.1;refGene_gn=SLC25A47;refGene_tx=XM_023256226.1;refGene_gn=SLC25A47;refGene_tx=XM_023256227.1
B3	144955690	144955734
refGene_gn=SLC25A47;refGene_tx=XM_023256225.1;refGene_gn=SLC25A47;refGene_tx=XM_023256226.1;refGene_gn=SLC25A47;refGene_tx=XM_023256227.1
B3	144956023	144956095
refGene_gn=SLC25A47;refGene_tx=XM_023256225.1;refGene_gn=SLC25A47;refGene_tx=XM_023256226.1;refGene_gn=SLC25A47;refGene_tx=XM_023256227.1
B3	144956801	144956984
refGene_gn=SLC25A47;refGene_tx=XM_023256225.1;refGene_gn=SLC25A47;refGene_tx=XM_023256226.1;refGene_gn=SLC25A47;refGene_tx=XM_023256227.1
B3	144958075	144958388
refGene_gn=SLC25A47;refGene_tx=XM_023256225.1;refGene_gn=SLC25A47;refGene_tx=XM_023256226.1;refGene_gn=SLC25A47;refGene_tx=XM_023256227.1
B3	144958673	144958954
refGene_gn=SLC25A47;refGene_tx=XM_023256225.1;refGene_gn=SLC25A47;refGene_tx=XM_023256226.1;refGene_gn=SLC25A47;refGene_tx=XM_023256227.1
B3	144963310	144963472
refGene_gn=WARS;refGene_tx=XM_003987989.5;refGene_gn=WARS;refGene_tx=XM_006933117.4
B3	144964604	144964745
refGene_gn=WARS;refGene_tx=XM_003987989.5;refGene_gn=WARS;refGene_tx=XM_006933117.4
B3	144968941	144969115
refGene_gn=WARS;refGene_tx=XM_003987989.5;refGene_gn=WARS;refGene_tx=XM_006933117.4

B3 144969321 144969434
refGene_gn=WARS;refGene_tx=XM_003987989.5;refGene_gn=WARS;refGene_tx=XM_006933117.4
B3 144971348 144971449
refGene_gn=WARS;refGene_tx=XM_003987989.5;refGene_gn=WARS;refGene_tx=XM_006933117.4
B3 144975749 144975932
refGene_gn=WARS;refGene_tx=XM_003987989.5;refGene_gn=WARS;refGene_tx=XM_006933117.4
B3 144976446 144976566
refGene_gn=WARS;refGene_tx=XM_003987989.5;refGene_gn=WARS;refGene_tx=XM_006933117.4
B3 144980594 144980703
refGene_gn=WARS;refGene_tx=XM_003987989.5;refGene_gn=WARS;refGene_tx=XM_006933117.4
B3 144981704 144981918
refGene_gn=WARS;refGene_tx=XM_003987989.5;refGene_gn=WARS;refGene_tx=XM_006933117.4
B3 144986888 144987002
refGene_gn=WARS;refGene_tx=XM_003987989.5;refGene_gn=WARS;refGene_tx=XM_006933117.4
B3 144997070 144997877
refGene_gn=WDR25;refGene_tx=XM_006933118.4;refGene_gn=WDR25;refGene_tx=XM_006933121.4;refGene_gn=WDR25;refGene_tx=XM_006933123.4;refGene_gn=WDR25;refGene_tx=XM_006933124.4;refGene_gn=WDR25;refGene_tx=XM_006933125.3;refGene_gn=WDR25;refGene_tx=XM_006933126.4;refGene_gn=WDR25;refGene_tx=XM_019833632.2;refGene_gn=WDR25;refGene_tx=XM_019833633.2;refGene_gn=WDR25;refGene_tx=XM_023256022.1
B3 145077611 145077759
refGene_gn=WDR25;refGene_tx=XM_006933118.4;refGene_gn=WDR25;refGene_tx=XM_006933121.4;refGene_gn=WDR25;refGene_tx=XM_006933123.4;refGene_gn=WDR25;refGene_tx=XM_006933124.4;refGene_gn=WDR25;refGene_tx=XM_006933125.3;refGene_gn=WDR25;refGene_tx=XM_006933126.4;refGene_gn=WDR25;refGene_tx=XM_019833632.2;refGene_gn=WDR25;refGene_tx=XM_019833633.2;refGene_gn=WDR25;refGene_tx=XM_023256022.1
B3 145090116 145090247
refGene_gn=WDR25;refGene_tx=XM_006933118.4;refGene_gn=WDR25;refGene_tx=XM_006933121.4;refGene_gn=WDR25;refGene_tx=XM_006933123.4;refGene_gn=WDR25;refGene_tx=XM_006933124.4;refGene_gn=WDR25;refGene_tx=XM_006933125.3;refGene_gn=WDR25;refGene_tx=XM_006933126.4;refGene_gn=WDR25;refGene_tx=XM_019833632.2;refGene_gn=WDR25;refGene_tx=XM_019833633.2;refGene_gn=WDR25;refGene_tx=XM_023256022.1
B3 145129680 145129851
refGene_gn=WDR25;refGene_tx=XM_006933118.4;refGene_gn=WDR25;refGene_tx=XM_006933121.4;refGene_gn=WDR25;refGene_tx=XM_006933123.4;refGene_gn=WDR25;refGene_tx=XM_006933124.4;refGene_gn=WDR25;refGene_tx=XM_006933125.3;refGene_gn=WDR25;refGene_tx=XM_006933126.4;refGene_gn=WDR25;refGene_tx=XM_019833632.2;refGene_gn=WDR25;refGene_tx=XM_019833633.2;refGene_gn=WDR25;refGene_tx=XM_023256022.1
B3 145131852 145131975

refGene_gn=WDR25;refGene_tx=XM_006933118.4
B3 145132488 145132629
refGene_gn=WDR25;refGene_tx=XM_006933121.4;refGene_gn=WDR25;refGene_tx=XM_006933123.4;refGene_gn=WDR25;refGene_tx=XM_006933124.4;refGene_gn=WDR25;refGene_tx=XM_006933125.3;refGene_gn=WDR25;refGene_tx=XM_006933126.4;refGene_gn=WDR25;refGene_tx=XM_019833632.2;refGene_gn=WDR25;refGene_tx=XM_019833633.2;refGene_gn=WDR25;refGene_tx=XM_023256022.1
B3 145133270 145133492
refGene_gn=WDR25;refGene_tx=XM_006933121.4;refGene_gn=WDR25;refGene_tx=XM_006933123.4;refGene_gn=WDR25;refGene_tx=XM_006933124.4;refGene_gn=WDR25;refGene_tx=XM_006933125.3;refGene_gn=WDR25;refGene_tx=XM_006933126.4;refGene_gn=WDR25;refGene_tx=XM_019833632.2;refGene_gn=WDR25;refGene_tx=XM_019833633.2;refGene_gn=WDR25;refGene_tx=XM_023256022.1
B3 145141547 145142897
refGene_gn=BEGAIN;refGene_tx=XM_023256024.1;refGene_gn=BEGAIN;refGene_tx=XM_023256025.1;refGene_gn=BEGAIN;refGene_tx=XM_023256026.1;refGene_gn=BEGAIN;refGene_tx=XM_023256027.1;refGene_gn=BEGAIN;refGene_tx=XM_023256028.1;refGene_gn=BEGAIN;refGene_tx=XM_023256029.1;refGene_gn=BEGAIN;refGene_tx=XM_023256030.1;refGene_gn=BEGAIN;refGene_tx=XM_023256031.1;refGene_gn=BEGAIN;refGene_tx=XM_023256032.1;refGene_gn=BEGAIN;refGene_tx=XM_023256033.1;refGene_gn=BEGAIN;refGene_tx=XM_023256034.1;refGene_gn=BEGAIN;refGene_tx=XM_023256036.1;refGene_gn=BEGAIN;refGene_tx=XM_023256037.1;refGene_gn=BEGAIN;refGene_tx=XM_023256038.1
B3 145143950 145144034
refGene_gn=BEGAIN;refGene_tx=XM_023256024.1;refGene_gn=BEGAIN;refGene_tx=XM_023256026.1;refGene_gn=BEGAIN;refGene_tx=XM_023256027.1;refGene_gn=BEGAIN;refGene_tx=XM_023256028.1;refGene_gn=BEGAIN;refGene_tx=XM_023256029.1;refGene_gn=BEGAIN;refGene_tx=XM_023256030.1;refGene_gn=BEGAIN;refGene_tx=XM_023256031.1;refGene_gn=BEGAIN;refGene_tx=XM_023256032.1;refGene_gn=BEGAIN;refGene_tx=XM_023256033.1;refGene_gn=BEGAIN;refGene_tx=XM_023256034.1;refGene_gn=BEGAIN;refGene_tx=XM_023256036.1;refGene_gn=BEGAIN;refGene_tx=XM_023256037.1;refGene_gn=BEGAIN;refGene_tx=XM_023256038.1
B3 145145996 145146104
refGene_gn=BEGAIN;refGene_tx=XM_023256024.1;refGene_gn=BEGAIN;refGene_tx=XM_023256025.1;refGene_gn=BEGAIN;refGene_tx=XM_023256026.1;refGene_gn=BEGAIN;refGene_tx=XM_023256027.1;refGene_gn=BEGAIN;refGene_tx=XM_023256028.1;refGene_gn=BEGAIN;refGene_tx=XM_023256029.1;refGene_gn=BEGAIN;refGene_tx=XM_023256030.1;refGene_gn=BEGAIN;refGene_tx=XM_023256031.1;refGene_gn=BEGAIN;refGene_tx=XM_023256032.1;refGene_gn=BEGAIN;refGene_tx=XM_023256033.1;refGene_gn=BEGAIN;refGene_tx=XM_023256034.1;refGene_gn=BEGAIN;refGene_tx=XM_023256036.1;refGene_gn=BEGAIN;refGene_tx=XM_023256037.1;refGene_gn=BEGAIN;refGene_tx=XM_023256038.1
B3 145147272 145147656
refGene_gn=BEGAIN;refGene_tx=XM_023256024.1;refGene_gn=BEGAIN;refGene_tx=XM_023256025.1;refGene_gn=BEGAIN;refGene_tx=XM_023256026.1;refGene_gn=BEGAIN;refGene_tx=XM_023256027.1;refGene_gn=BEGAIN;refGene_tx=XM_023256028.1;refGene_gn=BEGAIN;refGene_tx=XM_023256029.1;refGene_gn=BEGAIN;refGene_tx=XM_023256030.1;refGene_gn=BEGAIN;refGene_tx=XM_023256031.1;refGene_gn=BEGAIN;refGene_tx=XM_023256032.1;refGene_gn=BEGAIN;refGen

e_tx=XM_023256033.1;refGene_gn=BEGAIN;refGene_tx=XM_023256034.1;refGene_gn=BEGAIN;refGene_tx=XM_023256036.1;refGene_gn=BEGAIN;refGene_tx=XM_023256037.1;refGene_gn=BEGAIN;refGene_tx=XM_023256038.1
B3 145148207 145148369
refGene_gn=BEGAIN;refGene_tx=XM_023256024.1;refGene_gn=BEGAIN;refGene_tx=XM_023256025.1;refGene_gn=BEGAIN;refGene_tx=XM_023256026.1;refGene_gn=BEGAIN;refGene_tx=XM_023256027.1;refGene_gn=BEGAIN;refGene_tx=XM_023256028.1;refGene_gn=BEGAIN;refGene_tx=XM_023256029.1;refGene_gn=BEGAIN;refGene_tx=XM_023256030.1;refGene_gn=BEGAIN;refGene_tx=XM_023256031.1;refGene_gn=BEGAIN;refGene_tx=XM_023256032.1;refGene_gn=BEGAIN;refGene_tx=XM_023256033.1;refGene_gn=BEGAIN;refGene_tx=XM_023256034.1;refGene_gn=BEGAIN;refGene_tx=XM_023256036.1;refGene_gn=BEGAIN;refGene_tx=XM_023256037.1;refGene_gn=BEGAIN;refGene_tx=XM_023256038.1
B3 145152408 145152498
refGene_gn=BEGAIN;refGene_tx=XM_023256024.1;refGene_gn=BEGAIN;refGene_tx=XM_023256025.1;refGene_gn=BEGAIN;refGene_tx=XM_023256028.1;refGene_gn=BEGAIN;refGene_tx=XM_023256029.1;refGene_gn=BEGAIN;refGene_tx=XM_023256030.1;refGene_gn=BEGAIN;refGene_tx=XM_023256031.1;refGene_gn=BEGAIN;refGene_tx=XM_023256032.1;refGene_gn=BEGAIN;refGene_tx=XM_023256038.1
B3 145167712 145167721
refGene_gn=BEGAIN;refGene_tx=XM_023256027.1
B3 145168700 145168729
refGene_gn=BEGAIN;refGene_tx=XM_023256024.1;refGene_gn=BEGAIN;refGene_tx=XM_023256025.1;refGene_gn=BEGAIN;refGene_tx=XM_023256026.1;refGene_gn=BEGAIN;refGene_tx=XM_023256028.1;refGene_gn=BEGAIN;refGene_tx=XM_023256029.1;refGene_gn=BEGAIN;refGene_tx=XM_023256030.1;refGene_gn=BEGAIN;refGene_tx=XM_023256031.1;refGene_gn=BEGAIN;refGene_tx=XM_023256032.1;refGene_gn=BEGAIN;refGene_tx=XM_023256033.1;refGene_gn=BEGAIN;refGene_tx=XM_023256034.1;refGene_gn=BEGAIN;refGene_tx=XM_023256036.1;refGene_gn=BEGAIN;refGene_tx=XM_023256037.1;refGene_gn=BEGAIN;refGene_tx=XM_023256038.1
B3 145169274 145169295
refGene_gn=BEGAIN;refGene_tx=XM_023256031.1;refGene_gn=BEGAIN;refGene_tx=XM_023256036.1
B3 145169732 145169756
refGene_gn=BEGAIN;refGene_tx=XM_023256030.1;refGene_gn=BEGAIN;refGene_tx=XM_023256034.1
B3 145170258 145170338
refGene_gn=BEGAIN;refGene_tx=XM_023256028.1
B3 145177836 145178475
refGene_gn=BEGAIN;refGene_tx=XM_023256024.1;refGene_gn=BEGAIN;refGene_tx=XM_023256025.1;refGene_gn=BEGAIN;refGene_tx=XM_023256026.1
B3 145185475 145185517
refGene_gn=BEGAIN;refGene_tx=XM_023256029.1;refGene_gn=BEGAIN;refGene_tx=XM_023256033.1
B3 145312712 145312779
refGene_gn=DLK1;refGene_tx=XM_023256039.1;refGene_gn=DLK1;refGene_tx=XM_023256040.1;refGene_gn=DLK1;refGene_tx=XM_023256041.1
B3 145314074 145314138

refGene_gn=DLK1;refGene_tx=XM_023256039.1;refGene_gn=DLK1;refGene_tx=XM_023256040.1;refGene_gn=DLK1;refGene_tx=XM_023256041.1
B3 145314645 145314776
refGene_gn=DLK1;refGene_tx=XM_023256039.1;refGene_gn=DLK1;refGene_tx=XM_023256040.1;refGene_gn=DLK1;refGene_tx=XM_023256041.1
B3 145317787 145317929
refGene_gn=DLK1;refGene_tx=XM_023256039.1;refGene_gn=DLK1;refGene_tx=XM_023256040.1;refGene_gn=DLK1;refGene_tx=XM_023256041.1
B3 145319528 145320276
refGene_gn=DLK1;refGene_tx=XM_023256039.1;refGene_gn=DLK1;refGene_tx=XM_023256040.1;refGene_gn=DLK1;refGene_tx=XM_023256041.1
B3 145401520 145401590
refGene_gn=LOC102900156;refGene_tx=XM_019833824.1
B3 145402359 145402485
refGene_gn=LOC102900156;refGene_tx=XM_019833824.1
B3 145406950 145407131
refGene_gn=LOC102900156;refGene_tx=XM_019833824.1
B3 145408832 145408946
refGene_gn=LOC102900156;refGene_tx=XM_019833824.1
B3 145413619 145413630
refGene_gn=LOC102900156;refGene_tx=XM_019833824.1
B3 145413711 145413886
refGene_gn=LOC102900156;refGene_tx=XM_019833824.1
B3 145422976 145423049
refGene_gn=LOC102900156;refGene_tx=XM_019833824.1
B3 145423393 145423606
refGene_gn=LOC102900156;refGene_tx=XM_019833824.1
B3 145454754 145458759
refGene_gn=RTL1;refGene_tx=XM_023256042.1
B3 145878790 145878809
refGene_gn=LOC102899591;refGene_tx=XM_023256197.1
B3 145879000 145879793
refGene_gn=LOC102899591;refGene_tx=XM_023256197.1
B3 145880576 145880598
refGene_gn=LOC102899591;refGene_tx=XM_023256197.1
B3 146049961 146049983
refGene_gn=LOC111560971;refGene_tx=XM_023256198.1
B3 146050653 146050799
refGene_gn=LOC111560971;refGene_tx=XM_023256198.1
B3 146050892 146050988
refGene_gn=LOC111560971;refGene_tx=XM_023256198.1
B3 146051649 146052447
refGene_gn=LOC111560971;refGene_tx=XM_023256198.1
B3 146052490 146053414
refGene_gn=DIO3;refGene_tx=XM_004001454.5
B3 146205001 146205028
refGene_gn=PPP2R5C;refGene_tx=XM_023256045.1
B3 146205468 146205474
refGene_gn=PPP2R5C;refGene_tx=XM_006933131.4
B3 146205911 146205977

refGene_gn=PPP2R5C;refGene_tx=XM_006933131.4;refGene_gn=PPP2R5C;refGene_tx=XM_023256045.1
B3 146219695 146219858
refGene_gn=PPP2R5C;refGene_tx=XM_006933131.4;refGene_gn=PPP2R5C;refGene_tx=XM_023256045.1
B3 146236234 146236328
refGene_gn=PPP2R5C;refGene_tx=XM_011283519.3;refGene_gn=PPP2R5C;refGene_tx=XM_011283520.3
B3 146285800 146286000
refGene_gn=PPP2R5C;refGene_tx=XM_006933131.4;refGene_gn=PPP2R5C;refGene_tx=XM_011283519.3;refGene_gn=PPP2R5C;refGene_tx=XM_011283520.3;refGene_gn=PPP2R5C;refGene_tx=XM_023256045.1
B3 146302106 146302217
refGene_gn=PPP2R5C;refGene_tx=XM_006933131.4;refGene_gn=PPP2R5C;refGene_tx=XM_011283519.3;refGene_gn=PPP2R5C;refGene_tx=XM_011283520.3;refGene_gn=PPP2R5C;refGene_tx=XM_023256045.1
B3 146303274 146303367
refGene_gn=PPP2R5C;refGene_tx=XM_006933131.4;refGene_gn=PPP2R5C;refGene_tx=XM_011283519.3;refGene_gn=PPP2R5C;refGene_tx=XM_011283520.3;refGene_gn=PPP2R5C;refGene_tx=XM_023256045.1
B3 146303452 146303583
refGene_gn=PPP2R5C;refGene_tx=XM_006933131.4;refGene_gn=PPP2R5C;refGene_tx=XM_011283519.3;refGene_gn=PPP2R5C;refGene_tx=XM_011283520.3;refGene_gn=PPP2R5C;refGene_tx=XM_023256045.1
B3 146308312 146308372
refGene_gn=PPP2R5C;refGene_tx=XM_006933131.4;refGene_gn=PPP2R5C;refGene_tx=XM_011283519.3;refGene_gn=PPP2R5C;refGene_tx=XM_011283520.3;refGene_gn=PPP2R5C;refGene_tx=XM_023256045.1
B3 146310349 146310458
refGene_gn=PPP2R5C;refGene_tx=XM_006933131.4;refGene_gn=PPP2R5C;refGene_tx=XM_011283519.3;refGene_gn=PPP2R5C;refGene_tx=XM_011283520.3;refGene_gn=PPP2R5C;refGene_tx=XM_023256045.1
B3 146311877 146311931
refGene_gn=PPP2R5C;refGene_tx=XM_006933131.4;refGene_gn=PPP2R5C;refGene_tx=XM_011283519.3;refGene_gn=PPP2R5C;refGene_tx=XM_011283520.3;refGene_gn=PPP2R5C;refGene_tx=XM_023256045.1
B3 146317165 146317336
refGene_gn=PPP2R5C;refGene_tx=XM_006933131.4;refGene_gn=PPP2R5C;refGene_tx=XM_011283519.3;refGene_gn=PPP2R5C;refGene_tx=XM_011283520.3;refGene_gn=PPP2R5C;refGene_tx=XM_023256045.1
B3 146318475 146318603
refGene_gn=PPP2R5C;refGene_tx=XM_006933131.4;refGene_gn=PPP2R5C;refGene_tx=XM_011283519.3;refGene_gn=PPP2R5C;refGene_tx=XM_011283520.3;refGene_gn=PPP2R5C;refGene_tx=XM_023256045.1
B3 146321299 146321401
refGene_gn=PPP2R5C;refGene_tx=XM_006933131.4;refGene_gn=PPP2R5C;refGene_tx=XM_011283519.3;refGene_gn=PPP2R5C;refGene_tx=XM_011283520.3;refGene_gn=PPP2R5C;refGene_tx=XM_023256045.1
B3 146322829 146322902
refGene_gn=PPP2R5C;refGene_tx=XM_006933131.4;refGene_gn=PPP2R5C;refGen

e_tx=XM_011283519.3;refGene_gn=PPP2R5C;refGene_tx=XM_011283520.3;refGene_gn=PPP2R5C;refGene_tx=XM_023256045.1
B3 146327196 146327313
refGene_gn=PPP2R5C;refGene_tx=XM_011283520.3
B3 146332859 146332991
refGene_gn=PPP2R5C;refGene_tx=XM_006933131.4;refGene_gn=PPP2R5C;refGene_tx=XM_011283519.3;refGene_gn=PPP2R5C;refGene_tx=XM_011283520.3;refGene_gn=PPP2R5C;refGene_tx=XM_023256045.1
B3 146367634 146367890
refGene_gn=DYNC1H1;refGene_tx=XM_023256046.1
B3 146376454 146376542
refGene_gn=DYNC1H1;refGene_tx=XM_023256046.1
B3 146378699 146378873
refGene_gn=DYNC1H1;refGene_tx=XM_023256046.1
B3 146379110 146379366
refGene_gn=DYNC1H1;refGene_tx=XM_023256046.1
B3 146379764 146379951
refGene_gn=DYNC1H1;refGene_tx=XM_023256046.1
B3 146381306 146381578
refGene_gn=DYNC1H1;refGene_tx=XM_023256046.1
B3 146381669 146381897
refGene_gn=DYNC1H1;refGene_tx=XM_023256046.1
B3 146391077 146392154
refGene_gn=DYNC1H1;refGene_tx=XM_023256046.1
B3 146392808 146392988
refGene_gn=DYNC1H1;refGene_tx=XM_023256046.1
B3 146394428 146394578
refGene_gn=DYNC1H1;refGene_tx=XM_023256046.1
B3 146397334 146397481
refGene_gn=DYNC1H1;refGene_tx=XM_023256046.1
B3 146399977 146400118
refGene_gn=DYNC1H1;refGene_tx=XM_023256046.1
B3 146400454 146400631
refGene_gn=DYNC1H1;refGene_tx=XM_023256046.1
B3 146400768 146400879
refGene_gn=DYNC1H1;refGene_tx=XM_023256046.1
B3 146400961 146401081
refGene_gn=DYNC1H1;refGene_tx=XM_023256046.1
B3 146401572 146401812
refGene_gn=DYNC1H1;refGene_tx=XM_023256046.1
B3 146408373 146408529
refGene_gn=DYNC1H1;refGene_tx=XM_023256046.1
B3 146408656 146408770
refGene_gn=DYNC1H1;refGene_tx=XM_023256046.1
B3 146409217 146409328
refGene_gn=DYNC1H1;refGene_tx=XM_023256046.1
B3 146409405 146409615
refGene_gn=DYNC1H1;refGene_tx=XM_023256046.1
B3 146409792 146409939
refGene_gn=DYNC1H1;refGene_tx=XM_023256046.1

B3 146410146 146410313
refGene_gn=DYNC1H1;refGene_tx=XM_023256046.1
B3 146410412 146410586
refGene_gn=DYNC1H1;refGene_tx=XM_023256046.1
B3 146411747 146411913
refGene_gn=DYNC1H1;refGene_tx=XM_023256046.1
B3 146411994 146412183
refGene_gn=DYNC1H1;refGene_tx=XM_023256046.1
B3 146412275 146412470
refGene_gn=DYNC1H1;refGene_tx=XM_023256046.1
B3 146413096 146413379
refGene_gn=DYNC1H1;refGene_tx=XM_023256046.1
B3 146413603 146413704
refGene_gn=DYNC1H1;refGene_tx=XM_023256046.1
B3 146414233 146414393
refGene_gn=DYNC1H1;refGene_tx=XM_023256046.1
B3 146415443 146415687
refGene_gn=DYNC1H1;refGene_tx=XM_023256046.1
B3 146415861 146416045
refGene_gn=DYNC1H1;refGene_tx=XM_023256046.1
B3 146416329 146416542
refGene_gn=DYNC1H1;refGene_tx=XM_023256046.1
B3 146417283 146417522
refGene_gn=DYNC1H1;refGene_tx=XM_023256046.1
B3 146417693 146417850
refGene_gn=DYNC1H1;refGene_tx=XM_023256046.1
B3 146419578 146419806
refGene_gn=DYNC1H1;refGene_tx=XM_023256046.1
B3 146420282 146420513
refGene_gn=DYNC1H1;refGene_tx=XM_023256046.1
B3 146420787 146420928
refGene_gn=DYNC1H1;refGene_tx=XM_023256046.1
B3 146421352 146421586
refGene_gn=DYNC1H1;refGene_tx=XM_023256046.1
B3 146421676 146421883
refGene_gn=DYNC1H1;refGene_tx=XM_023256046.1
B3 146421961 146422083
refGene_gn=DYNC1H1;refGene_tx=XM_023256046.1
B3 146422569 146422735
refGene_gn=DYNC1H1;refGene_tx=XM_023256046.1
B3 146423683 146423847
refGene_gn=DYNC1H1;refGene_tx=XM_023256046.1
B3 146426720 146426850
refGene_gn=DYNC1H1;refGene_tx=XM_023256046.1
B3 146428424 146428558
refGene_gn=DYNC1H1;refGene_tx=XM_023256046.1
B3 146428992 146429107
refGene_gn=DYNC1H1;refGene_tx=XM_023256046.1
B3 146429208 146429370
refGene_gn=DYNC1H1;refGene_tx=XM_023256046.1

B3 146429451 146429666
refGene_gn=DYNC1H1;refGene_tx=XM_023256046.1
B3 146429778 146429983
refGene_gn=DYNC1H1;refGene_tx=XM_023256046.1
B3 146431242 146431416
refGene_gn=DYNC1H1;refGene_tx=XM_023256046.1
B3 146431529 146431649
refGene_gn=DYNC1H1;refGene_tx=XM_023256046.1
B3 146431792 146431913
refGene_gn=DYNC1H1;refGene_tx=XM_023256046.1
B3 146433023 146433219
refGene_gn=DYNC1H1;refGene_tx=XM_023256046.1
B3 146433547 146433665
refGene_gn=DYNC1H1;refGene_tx=XM_023256046.1
B3 146433753 146433969
refGene_gn=DYNC1H1;refGene_tx=XM_023256046.1
B3 146434571 146434784
refGene_gn=DYNC1H1;refGene_tx=XM_023256046.1
B3 146434894 146435022
refGene_gn=DYNC1H1;refGene_tx=XM_023256046.1
B3 146436644 146436798
refGene_gn=DYNC1H1;refGene_tx=XM_023256046.1
B3 146437467 146437614
refGene_gn=DYNC1H1;refGene_tx=XM_023256046.1
B3 146437708 146437859
refGene_gn=DYNC1H1;refGene_tx=XM_023256046.1
B3 146438358 146438612
refGene_gn=DYNC1H1;refGene_tx=XM_023256046.1
B3 146438696 146438831
refGene_gn=DYNC1H1;refGene_tx=XM_023256046.1
B3 146438920 146439015
refGene_gn=DYNC1H1;refGene_tx=XM_023256046.1
B3 146439186 146439361
refGene_gn=DYNC1H1;refGene_tx=XM_023256046.1
B3 146439523 146439599
refGene_gn=DYNC1H1;refGene_tx=XM_023256046.1
B3 146440229 146440390
refGene_gn=DYNC1H1;refGene_tx=XM_023256046.1
B3 146440600 146440712
refGene_gn=DYNC1H1;refGene_tx=XM_023256046.1
B3 146440814 146440875
refGene_gn=DYNC1H1;refGene_tx=XM_023256046.1
B3 146440962 146441086
refGene_gn=DYNC1H1;refGene_tx=XM_023256046.1
B3 146441211 146441325
refGene_gn=DYNC1H1;refGene_tx=XM_023256046.1
B3 146442217 146442388
refGene_gn=DYNC1H1;refGene_tx=XM_023256046.1
B3 146442569 146442787
refGene_gn=DYNC1H1;refGene_tx=XM_023256046.1

B3 146442881 146442985
refGene_gn=DYNC1H1;refGene_tx=XM_023256046.1
B3 146444628 146444840
refGene_gn=DYNC1H1;refGene_tx=XM_023256046.1
B3 146445216 146445370
refGene_gn=DYNC1H1;refGene_tx=XM_023256046.1
B3 146445973 146446116
refGene_gn=DYNC1H1;refGene_tx=XM_023256046.1
B3 146446232 146446401
refGene_gn=DYNC1H1;refGene_tx=XM_023256046.1
B3 146446544 146446672
refGene_gn=DYNC1H1;refGene_tx=XM_023256046.1
B3 146446893 146447022
refGene_gn=DYNC1H1;refGene_tx=XM_023256046.1
B3 146476628 146476738
refGene_gn=HSP90AA1;refGene_tx=XM_023256047.1
B3 146476887 146477221
refGene_gn=HSP90AA1;refGene_tx=XM_023256047.1
B3 146477582 146477851
refGene_gn=HSP90AA1;refGene_tx=XM_023256047.1
B3 146478085 146478233
refGene_gn=HSP90AA1;refGene_tx=XM_023256047.1
B3 146478348 146478539
refGene_gn=HSP90AA1;refGene_tx=XM_023256047.1
B3 146478658 146478824
refGene_gn=HSP90AA1;refGene_tx=XM_023256047.1
B3 146479034 146479355
refGene_gn=HSP90AA1;refGene_tx=XM_023256047.1
B3 146479645 146479779
refGene_gn=HSP90AA1;refGene_tx=XM_023256047.1
B3 146480159 146480526
refGene_gn=HSP90AA1;refGene_tx=XM_023256047.1
B3 146480620 146480782
refGene_gn=HSP90AA1;refGene_tx=XM_023256047.1
B3 146525120 146525390
refGene_gn=WDR20;refGene_tx=XM_003987998.4;refGene_gn=WDR20;refGene_tx
=XM_003988000.4;refGene_gn=WDR20;refGene_tx=XM_006933135.3;refGene_gn=
WDR20;refGene_tx=XM_006933136.3;refGene_gn=WDR20;refGene_tx=XM_0069331
37.3;refGene_gn=WDR20;refGene_tx=XM_019833639.2
B3 146565826 146565878
refGene_gn=WDR20;refGene_tx=XM_006933135.3
B3 146567292 146567475
refGene_gn=WDR20;refGene_tx=XM_003987998.4;refGene_gn=WDR20;refGene_tx
=XM_006933135.3;refGene_gn=WDR20;refGene_tx=XM_006933136.3;refGene_gn=
WDR20;refGene_tx=XM_019833639.2
B3 146571683 146571728
refGene_gn=WDR20;refGene_tx=XM_023256048.1
B3 146578832 146580092
refGene_gn=WDR20;refGene_tx=XM_003987998.4;refGene_gn=WDR20;refGene_tx
=XM_003988000.4;refGene_gn=WDR20;refGene_tx=XM_006933135.3;refGene_gn=

WDR20;refGene_tx=XM_019833639.2;refGene_gn=WDR20;refGene_tx=XM_0232560
48.1;refGene_gn=WDR20;refGene_tx=XM_023256050.1;refGene_gn=WDR20;refGe
ne_tx=XM_023256051.1;refGene_gn=WDR20;refGene_tx=XM_023256052.1;refGen
e_gn=WDR20;refGene_tx=XM_023256053.1;refGene_gn=WDR20;refGene_tx=XM_02
3256054.1
B3 146589782 146590010
refGene_gn=MOK;refGene_tx=XM_023256059.1;refGene_gn=WDR20;refGene_tx=X
M_003987998.4;refGene_gn=WDR20;refGene_tx=XM_003988000.4;refGene_gn=WD
R20;refGene_tx=XM_006933135.3;refGene_gn=WDR20;refGene_tx=XM_006933136
.
3;refGene_gn=WDR20;refGene_tx=XM_006933137.3;refGene_gn=WDR20;refGene_
tx=XM_023256048.1;refGene_gn=WDR20;refGene_tx=XM_023256050.1;refGene_g
n=WDR20;refGene_tx=XM_023256051.1;refGene_gn=WDR20;refGene_tx=XM_02325
6052.1;refGene_gn=WDR20;refGene_tx=XM_023256053.1;refGene_gn=WDR20;ref
Gene_tx=XM_023256054.1
B3 146594599 146594680
refGene_gn=MOK;refGene_tx=XM_003988032.5;refGene_gn=MOK;refGene_tx=XM_
023256055.1;refGene_gn=MOK;refGene_tx=XM_023256056.1;refGene_gn=MOK;re
fGene_tx=XM_023256057.1;refGene_gn=MOK;refGene_tx=XM_023256058.1;refGe
ne_gn=MOK;refGene_tx=XM_023256061.1;refGene_gn=MOK;refGene_tx=XM_02325
6062.1;refGene_gn=MOK;refGene_tx=XM_023256063.1;refGene_gn=MOK;refGene
_tx=XM_023256064.1;refGene_gn=MOK;refGene_tx=XM_023256065.1;refGene_gn
=MOK;refGene_tx=XM_023256066.1;refGene_gn=MOK;refGene_tx=XM_023256067.
1;refGene_gn=MOK;refGene_tx=XM_023256068.1
B3 146594764 146594965
refGene_gn=MOK;refGene_tx=XM_003988032.5;refGene_gn=MOK;refGene_tx=XM_
023256055.1;refGene_gn=MOK;refGene_tx=XM_023256056.1;refGene_gn=MOK;re
fGene_tx=XM_023256057.1;refGene_gn=MOK;refGene_tx=XM_023256058.1;refGe
ne_gn=MOK;refGene_tx=XM_023256061.1;refGene_gn=MOK;refGene_tx=XM_02325
6062.1;refGene_gn=MOK;refGene_tx=XM_023256063.1;refGene_gn=MOK;refGene
_tx=XM_023256064.1;refGene_gn=MOK;refGene_tx=XM_023256065.1;refGene_gn
=MOK;refGene_tx=XM_023256066.1;refGene_gn=MOK;refGene_tx=XM_023256067.
1;refGene_gn=MOK;refGene_tx=XM_023256068.1
B3 146596383 146596498
refGene_gn=MOK;refGene_tx=XM_003988032.5;refGene_gn=MOK;refGene_tx=XM_
023256055.1;refGene_gn=MOK;refGene_tx=XM_023256056.1;refGene_gn=MOK;re
fGene_tx=XM_023256057.1;refGene_gn=MOK;refGene_tx=XM_023256058.1;refGe
ne_gn=MOK;refGene_tx=XM_023256059.1;refGene_gn=MOK;refGene_tx=XM_02325
6061.1;refGene_gn=MOK;refGene_tx=XM_023256062.1;refGene_gn=MOK;refGene
_tx=XM_023256063.1;refGene_gn=MOK;refGene_tx=XM_023256064.1;refGene_gn
=MOK;refGene_tx=XM_023256065.1;refGene_gn=MOK;refGene_tx=XM_023256066.
1;refGene_gn=MOK;refGene_tx=XM_023256067.1;refGene_gn=MOK;refGene_tx=X
M_023256068.1
B3 146597054 146597228
refGene_gn=MOK;refGene_tx=XM_003988032.5;refGene_gn=MOK;refGene_tx=XM_
023256055.1;refGene_gn=MOK;refGene_tx=XM_023256056.1;refGene_gn=MOK;re
fGene_tx=XM_023256057.1;refGene_gn=MOK;refGene_tx=XM_023256058.1;refGe
ne_gn=MOK;refGene_tx=XM_023256059.1;refGene_gn=MOK;refGene_tx=XM_02325
6061.1;refGene_gn=MOK;refGene_tx=XM_023256062.1;refGene_gn=MOK;refGene
_tx=XM_023256063.1;refGene_gn=MOK;refGene_tx=XM_023256064.1;refGene_gn

=MOK;refGene_tx=XM_023256065.1;refGene_gn=MOK;refGene_tx=XM_023256066.1;refGene_gn=MOK;refGene_tx=XM_023256067.1;refGene_gn=MOK;refGene_tx=XM_023256068.1
B3 146598213 146598315
refGene_gn=MOK;refGene_tx=XM_003988032.5;refGene_gn=MOK;refGene_tx=XM_023256055.1;refGene_gn=MOK;refGene_tx=XM_023256056.1;refGene_gn=MOK;refGene_tx=XM_023256057.1;refGene_gn=MOK;refGene_tx=XM_023256058.1;refGene_gn=MOK;refGene_tx=XM_023256059.1;refGene_gn=MOK;refGene_tx=XM_023256061.1;refGene_gn=MOK;refGene_tx=XM_023256062.1;refGene_gn=MOK;refGene_tx=XM_023256063.1;refGene_gn=MOK;refGene_tx=XM_023256064.1;refGene_gn=MOK;refGene_tx=XM_023256065.1;refGene_gn=MOK;refGene_tx=XM_023256066.1;refGene_gn=MOK;refGene_tx=XM_023256067.1;refGene_gn=MOK;refGene_tx=XM_023256068.1
B3 146599825 146600004
refGene_gn=MOK;refGene_tx=XM_003988032.5;refGene_gn=MOK;refGene_tx=XM_023256055.1;refGene_gn=MOK;refGene_tx=XM_023256056.1;refGene_gn=MOK;refGene_tx=XM_023256057.1;refGene_gn=MOK;refGene_tx=XM_023256058.1;refGene_gn=MOK;refGene_tx=XM_023256059.1;refGene_gn=MOK;refGene_tx=XM_023256061.1;refGene_gn=MOK;refGene_tx=XM_023256062.1;refGene_gn=MOK;refGene_tx=XM_023256063.1;refGene_gn=MOK;refGene_tx=XM_023256064.1;refGene_gn=MOK;refGene_tx=XM_023256065.1;refGene_gn=MOK;refGene_tx=XM_023256066.1;refGene_gn=MOK;refGene_tx=XM_023256067.1;refGene_gn=MOK;refGene_tx=XM_023256068.1
B3 146606599 146606648
refGene_gn=MOK;refGene_tx=XM_003988032.5;refGene_gn=MOK;refGene_tx=XM_023256055.1;refGene_gn=MOK;refGene_tx=XM_023256056.1;refGene_gn=MOK;refGene_tx=XM_023256057.1;refGene_gn=MOK;refGene_tx=XM_023256058.1;refGene_gn=MOK;refGene_tx=XM_023256059.1;refGene_gn=MOK;refGene_tx=XM_023256061.1;refGene_gn=MOK;refGene_tx=XM_023256062.1;refGene_gn=MOK;refGene_tx=XM_023256063.1;refGene_gn=MOK;refGene_tx=XM_023256064.1;refGene_gn=MOK;refGene_tx=XM_023256065.1;refGene_gn=MOK;refGene_tx=XM_023256066.1
B3 146606770 146606849
refGene_gn=MOK;refGene_tx=XM_003988032.5;refGene_gn=MOK;refGene_tx=XM_023256055.1;refGene_gn=MOK;refGene_tx=XM_023256056.1;refGene_gn=MOK;refGene_tx=XM_023256057.1;refGene_gn=MOK;refGene_tx=XM_023256058.1;refGene_gn=MOK;refGene_tx=XM_023256059.1;refGene_gn=MOK;refGene_tx=XM_023256061.1;refGene_gn=MOK;refGene_tx=XM_023256062.1;refGene_gn=MOK;refGene_tx=XM_023256063.1;refGene_gn=MOK;refGene_tx=XM_023256064.1;refGene_gn=MOK;refGene_tx=XM_023256065.1;refGene_gn=MOK;refGene_tx=XM_023256066.1
B3 146616620 146616691
refGene_gn=MOK;refGene_tx=XM_003988032.5;refGene_gn=MOK;refGene_tx=XM_023256055.1;refGene_gn=MOK;refGene_tx=XM_023256056.1;refGene_gn=MOK;refGene_tx=XM_023256057.1;refGene_gn=MOK;refGene_tx=XM_023256058.1;refGene_gn=MOK;refGene_tx=XM_023256059.1;refGene_gn=MOK;refGene_tx=XM_023256061.1;refGene_gn=MOK;refGene_tx=XM_023256062.1
B3 146618233 146618323
refGene_gn=MOK;refGene_tx=XM_003988032.5;refGene_gn=MOK;refGene_tx=XM_023256055.1;refGene_gn=MOK;refGene_tx=XM_023256056.1;refGene_gn=MOK;re

fGene_tx=XM_023256057.1;refGene_gn=MOK;refGene_tx=XM_023256058.1;refGene_gn=MOK;refGene_tx=XM_023256059.1;refGene_gn=MOK;refGene_tx=XM_023256061.1
B3 146631427 146631542
refGene_gn=MOK;refGene_tx=XM_003988032.5;refGene_gn=MOK;refGene_tx=XM_023256061.1
B3 146654545 146654739
refGene_gn=MOK;refGene_tx=XM_003988032.5;refGene_gn=MOK;refGene_tx=XM_023256055.1;refGene_gn=MOK;refGene_tx=XM_023256056.1;refGene_gn=MOK;refGene_tx=XM_023256057.1;refGene_gn=MOK;refGene_tx=XM_023256058.1;refGene_gn=MOK;refGene_tx=XM_023256059.1
B3 146668252 146668546
refGene_gn=ZNF839;refGene_tx=XM_023256069.1
B3 146673533 146674337
refGene_gn=ZNF839;refGene_tx=XM_019833645.2;refGene_gn=ZNF839;refGene_tx=XM_023256069.1
B3 146677007 146677247
refGene_gn=ZNF839;refGene_tx=XM_019833645.2;refGene_gn=ZNF839;refGene_tx=XM_023256069.1
B3 146678910 146679003
refGene_gn=ZNF839;refGene_tx=XM_019833645.2;refGene_gn=ZNF839;refGene_tx=XM_023256069.1
B3 146679653 146679803
refGene_gn=ZNF839;refGene_tx=XM_019833645.2;refGene_gn=ZNF839;refGene_tx=XM_023256069.1
B3 146682720 146682867
refGene_gn=ZNF839;refGene_tx=XM_019833645.2;refGene_gn=ZNF839;refGene_tx=XM_023256069.1
B3 146683003 146683097
refGene_gn=ZNF839;refGene_tx=XM_019833645.2;refGene_gn=ZNF839;refGene_tx=XM_023256069.1
B3 146683788 146684657
refGene_gn=ZNF839;refGene_tx=XM_019833645.2;refGene_gn=ZNF839;refGene_tx=XM_023256069.1
B3 146688714 146688917
refGene_gn=CINP;refGene_tx=XM_023256078.1;refGene_gn=CINP;refGene_tx=XM_023256079.1
B3 146690577 146690707
refGene_gn=CINP;refGene_tx=XM_023256078.1;refGene_gn=CINP;refGene_tx=XM_023256079.1
B3 146697768 146697898
refGene_gn=CINP;refGene_tx=XM_023256078.1;refGene_gn=CINP;refGene_tx=XM_023256079.1
B3 146700505 146700674
refGene_gn=CINP;refGene_tx=XM_023256078.1;refGene_gn=CINP;refGene_tx=XM_023256079.1
B3 146708117 146708139
refGene_gn=CINP;refGene_tx=XM_023256078.1
B3 146710967 146710974
refGene_gn=CINP;refGene_tx=XM_023256079.1

B3	146725118	146725337
refGene_gn=TECPR2;refGene_tx=XM_023256070.1;refGene_gn=TECPR2;refGene_tx=XM_023256071.1;refGene_gn=TECPR2;refGene_tx=XM_023256072.1;refGene_gn=TECPR2;refGene_tx=XM_023256073.1;refGene_gn=TECPR2;refGene_tx=XM_023256074.1;refGene_gn=TECPR2;refGene_tx=XM_023256075.1
B3	146740526	146740655
refGene_gn=TECPR2;refGene_tx=XM_023256070.1;refGene_gn=TECPR2;refGene_tx=XM_023256071.1;refGene_gn=TECPR2;refGene_tx=XM_023256072.1;refGene_gn=TECPR2;refGene_tx=XM_023256073.1;refGene_gn=TECPR2;refGene_tx=XM_023256074.1;refGene_gn=TECPR2;refGene_tx=XM_023256075.1
B3	146741037	146741169
refGene_gn=TECPR2;refGene_tx=XM_023256070.1;refGene_gn=TECPR2;refGene_tx=XM_023256071.1;refGene_gn=TECPR2;refGene_tx=XM_023256072.1;refGene_gn=TECPR2;refGene_tx=XM_023256073.1;refGene_gn=TECPR2;refGene_tx=XM_023256074.1;refGene_gn=TECPR2;refGene_tx=XM_023256075.1
B3	146744596	146744754
refGene_gn=TECPR2;refGene_tx=XM_023256070.1;refGene_gn=TECPR2;refGene_tx=XM_023256071.1;refGene_gn=TECPR2;refGene_tx=XM_023256072.1;refGene_gn=TECPR2;refGene_tx=XM_023256073.1;refGene_gn=TECPR2;refGene_tx=XM_023256074.1;refGene_gn=TECPR2;refGene_tx=XM_023256075.1
B3	146754049	146754362
refGene_gn=TECPR2;refGene_tx=XM_023256070.1;refGene_gn=TECPR2;refGene_tx=XM_023256071.1;refGene_gn=TECPR2;refGene_tx=XM_023256072.1;refGene_gn=TECPR2;refGene_tx=XM_023256073.1;refGene_gn=TECPR2;refGene_tx=XM_023256074.1;refGene_gn=TECPR2;refGene_tx=XM_023256075.1
B3	146756153	146756286
refGene_gn=TECPR2;refGene_tx=XM_023256070.1;refGene_gn=TECPR2;refGene_tx=XM_023256071.1;refGene_gn=TECPR2;refGene_tx=XM_023256072.1;refGene_gn=TECPR2;refGene_tx=XM_023256073.1;refGene_gn=TECPR2;refGene_tx=XM_023256074.1;refGene_gn=TECPR2;refGene_tx=XM_023256075.1
B3	146759241	146759556
refGene_gn=TECPR2;refGene_tx=XM_023256070.1;refGene_gn=TECPR2;refGene_tx=XM_023256071.1;refGene_gn=TECPR2;refGene_tx=XM_023256072.1;refGene_gn=TECPR2;refGene_tx=XM_023256073.1;refGene_gn=TECPR2;refGene_tx=XM_023256074.1;refGene_gn=TECPR2;refGene_tx=XM_023256075.1
B3	146760323	146761306
refGene_gn=TECPR2;refGene_tx=XM_023256070.1;refGene_gn=TECPR2;refGene_tx=XM_023256071.1;refGene_gn=TECPR2;refGene_tx=XM_023256072.1;refGene_gn=TECPR2;refGene_tx=XM_023256073.1;refGene_gn=TECPR2;refGene_tx=XM_023256074.1;refGene_gn=TECPR2;refGene_tx=XM_023256075.1
B3	146762867	146763051
refGene_gn=TECPR2;refGene_tx=XM_023256070.1;refGene_gn=TECPR2;refGene_tx=XM_023256071.1;refGene_gn=TECPR2;refGene_tx=XM_023256072.1;refGene_gn=TECPR2;refGene_tx=XM_023256073.1;refGene_gn=TECPR2;refGene_tx=XM_023256074.1;refGene_gn=TECPR2;refGene_tx=XM_023256075.1
B3	146765789	146765963
refGene_gn=TECPR2;refGene_tx=XM_023256070.1;refGene_gn=TECPR2;refGene_tx=XM_023256071.1;refGene_gn=TECPR2;refGene_tx=XM_023256072.1;refGene_gn=TECPR2;refGene_tx=XM_023256073.1;refGene_gn=TECPR2;refGene_tx=XM_023256074.1;refGene_gn=TECPR2;refGene_tx=XM_023256075.1

B3 146768111 146768292
refGene_gn=TECPR2;refGene_tx=XM_023256070.1;refGene_gn=TECPR2;refGene_tx=XM_023256071.1;refGene_gn=TECPR2;refGene_tx=XM_023256072.1;refGene_gn=TECPR2;refGene_tx=XM_023256073.1;refGene_gn=TECPR2;refGene_tx=XM_023256074.1;refGene_gn=TECPR2;refGene_tx=XM_023256075.1
B3 146769432 146769574
refGene_gn=TECPR2;refGene_tx=XM_023256070.1;refGene_gn=TECPR2;refGene_tx=XM_023256071.1;refGene_gn=TECPR2;refGene_tx=XM_023256072.1;refGene_gn=TECPR2;refGene_tx=XM_023256073.1;refGene_gn=TECPR2;refGene_tx=XM_023256074.1;refGene_gn=TECPR2;refGene_tx=XM_023256075.1
B3 146772200 146772441
refGene_gn=TECPR2;refGene_tx=XM_023256070.1;refGene_gn=TECPR2;refGene_tx=XM_023256071.1;refGene_gn=TECPR2;refGene_tx=XM_023256072.1;refGene_gn=TECPR2;refGene_tx=XM_023256073.1;refGene_gn=TECPR2;refGene_tx=XM_023256074.1;refGene_gn=TECPR2;refGene_tx=XM_023256075.1
B3 146773164 146773254
refGene_gn=TECPR2;refGene_tx=XM_023256070.1;refGene_gn=TECPR2;refGene_tx=XM_023256071.1;refGene_gn=TECPR2;refGene_tx=XM_023256072.1;refGene_gn=TECPR2;refGene_tx=XM_023256073.1;refGene_gn=TECPR2;refGene_tx=XM_023256074.1;refGene_gn=TECPR2;refGene_tx=XM_023256075.1
B3 146774807 146775041
refGene_gn=TECPR2;refGene_tx=XM_023256070.1;refGene_gn=TECPR2;refGene_tx=XM_023256071.1;refGene_gn=TECPR2;refGene_tx=XM_023256072.1;refGene_gn=TECPR2;refGene_tx=XM_023256073.1;refGene_gn=TECPR2;refGene_tx=XM_023256074.1;refGene_gn=TECPR2;refGene_tx=XM_023256075.1
B3 146775738 146775833
refGene_gn=TECPR2;refGene_tx=XM_023256075.1
B3 146782988 146783137
refGene_gn=TECPR2;refGene_tx=XM_023256070.1;refGene_gn=TECPR2;refGene_tx=XM_023256071.1;refGene_gn=TECPR2;refGene_tx=XM_023256072.1;refGene_gn=TECPR2;refGene_tx=XM_023256073.1;refGene_gn=TECPR2;refGene_tx=XM_023256074.1
B3 146783311 146783353
refGene_gn=TECPR2;refGene_tx=XM_023256074.1
B3 146806819 146807236
refGene_gn=TECPR2;refGene_tx=XM_023256072.1
B3 146815481 146815775
refGene_gn=TECPR2;refGene_tx=XM_023256070.1;refGene_gn=TECPR2;refGene_tx=XM_023256071.1;refGene_gn=TECPR2;refGene_tx=XM_023256073.1
B3 146816046 146816196
refGene_gn=TECPR2;refGene_tx=XM_023256070.1;refGene_gn=TECPR2;refGene_tx=XM_023256071.1
B3 146816528 146816680
refGene_gn=TECPR2;refGene_tx=XM_023256070.1;refGene_gn=TECPR2;refGene_tx=XM_023256071.1
B3 146824490 146825453
refGene_gn=ANKRD9;refGene_tx=XM_023256076.1;refGene_gn=ANKRD9;refGene_tx=XM_023256077.1
B3 146889724 146890028
refGene_gn=RCOR1;refGene_tx=XM_023256081.1

B3	146890102	146890162
refGene_gn=RCOR1;refGene_tx=XM_023256081.1
B3	146971063	146971147
refGene_gn=RCOR1;refGene_tx=XM_023256081.1
B3	146993480	146993533
refGene_gn=RCOR1;refGene_tx=XM_023256081.1
B3	146996563	146996725
refGene_gn=RCOR1;refGene_tx=XM_023256081.1
B3	146997434	146997553
refGene_gn=RCOR1;refGene_tx=XM_023256081.1
B3	147000825	147000904
refGene_gn=RCOR1;refGene_tx=XM_023256081.1
B3	147003755	147003950
refGene_gn=RCOR1;refGene_tx=XM_023256081.1
B3	147010754	147010788
refGene_gn=RCOR1;refGene_tx=XM_023256081.1
B3	147010790	147010831
refGene_gn=RCOR1;refGene_tx=XM_023256081.1
B3	147011309	147011367
refGene_gn=RCOR1;refGene_tx=XM_023256081.1
B3	147011877	147012107
refGene_gn=RCOR1;refGene_tx=XM_023256081.1
B3	147015411	147015453
refGene_gn=RCOR1;refGene_tx=XM_023256081.1
B3	147145873	147146118
refGene_gn=TRAF3;refGene_tx=XM_003988003.5;refGene_gn=TRAF3;refGene_tx=XM_006933142.4;refGene_gn=TRAF3;refGene_tx=XM_011283518.3;refGene_gn=TRAF3;refGene_tx=XM_019833654.2;refGene_gn=TRAF3;refGene_tx=XM_019833655.2;refGene_gn=TRAF3;refGene_tx=XM_023256082.1;refGene_gn=TRAF3;refGene_tx=XM_023256083.1
B3	147148171	147148223
refGene_gn=TRAF3;refGene_tx=XM_003988003.5;refGene_gn=TRAF3;refGene_tx=XM_006933142.4;refGene_gn=TRAF3;refGene_tx=XM_011283518.3;refGene_gn=TRAF3;refGene_tx=XM_019833654.2;refGene_gn=TRAF3;refGene_tx=XM_019833655.2;refGene_gn=TRAF3;refGene_tx=XM_023256082.1;refGene_gn=TRAF3;refGene_tx=XM_023256083.1
B3	147151544	147151649
refGene_gn=TRAF3;refGene_tx=XM_003988003.5;refGene_gn=TRAF3;refGene_tx=XM_006933142.4;refGene_gn=TRAF3;refGene_tx=XM_011283518.3;refGene_gn=TRAF3;refGene_tx=XM_019833654.2;refGene_gn=TRAF3;refGene_tx=XM_019833655.2;refGene_gn=TRAF3;refGene_tx=XM_023256082.1;refGene_gn=TRAF3;refGene_tx=XM_023256083.1
B3	147152510	147152678
refGene_gn=TRAF3;refGene_tx=XM_003988003.5;refGene_gn=TRAF3;refGene_tx=XM_006933142.4;refGene_gn=TRAF3;refGene_tx=XM_011283518.3;refGene_gn=TRAF3;refGene_tx=XM_019833654.2;refGene_gn=TRAF3;refGene_tx=XM_019833655.2;refGene_gn=TRAF3;refGene_tx=XM_023256082.1;refGene_gn=TRAF3;refGene_tx=XM_023256083.1
B3	147159425	147159506
refGene_gn=TRAF3;refGene_tx=XM_003988003.5;refGene_gn=TRAF3;refGene_tx

=XM_006933142.4;refGene_gn=TRAF3;refGene_tx=XM_011283518.3;refGene_gn=TRAF3;refGene_tx=XM_019833654.2;refGene_gn=TRAF3;refGene_tx=XM_019833655.2;refGene_gn=TRAF3;refGene_tx=XM_023256082.1;refGene_gn=TRAF3;refGene_tx=XM_023256083.1
B3 147162217 147162292
refGene_gn=TRAF3;refGene_tx=XM_003988003.5;refGene_gn=TRAF3;refGene_tx=XM_006933142.4;refGene_gn=TRAF3;refGene_tx=XM_019833654.2;refGene_gn=TRAF3;refGene_tx=XM_019833655.2;refGene_gn=TRAF3;refGene_tx=XM_023256082.1;refGene_gn=TRAF3;refGene_tx=XM_023256083.1
B3 147163866 147163959
refGene_gn=TRAF3;refGene_tx=XM_003988003.5;refGene_gn=TRAF3;refGene_tx=XM_006933142.4;refGene_gn=TRAF3;refGene_tx=XM_011283518.3;refGene_gn=TRAF3;refGene_tx=XM_019833654.2;refGene_gn=TRAF3;refGene_tx=XM_019833655.2;refGene_gn=TRAF3;refGene_tx=XM_023256082.1;refGene_gn=TRAF3;refGene_tx=XM_023256083.1
B3 147168618 147168759
refGene_gn=TRAF3;refGene_tx=XM_003988003.5;refGene_gn=TRAF3;refGene_tx=XM_006933142.4;refGene_gn=TRAF3;refGene_tx=XM_011283518.3;refGene_gn=TRAF3;refGene_tx=XM_019833654.2;refGene_gn=TRAF3;refGene_tx=XM_019833655.2;refGene_gn=TRAF3;refGene_tx=XM_023256082.1;refGene_gn=TRAF3;refGene_tx=XM_023256083.1
B3 147173835 147174010
refGene_gn=TRAF3;refGene_tx=XM_003988003.5;refGene_gn=TRAF3;refGene_tx=XM_006933142.4;refGene_gn=TRAF3;refGene_tx=XM_011283518.3;refGene_gn=TRAF3;refGene_tx=XM_019833654.2;refGene_gn=TRAF3;refGene_tx=XM_019833655.2;refGene_gn=TRAF3;refGene_tx=XM_023256082.1;refGene_gn=TRAF3;refGene_tx=XM_023256083.1
B3 147174910 147175482
refGene_gn=TRAF3;refGene_tx=XM_003988003.5;refGene_gn=TRAF3;refGene_tx=XM_006933142.4;refGene_gn=TRAF3;refGene_tx=XM_011283518.3;refGene_gn=TRAF3;refGene_tx=XM_019833654.2;refGene_gn=TRAF3;refGene_tx=XM_019833655.2;refGene_gn=TRAF3;refGene_tx=XM_023256082.1;refGene_gn=TRAF3;refGene_tx=XM_023256083.1
B3 147187132 147187175
refGene_gn=AMN;refGene_tx=XM_023256086.1
B3 147188015 147188134
refGene_gn=AMN;refGene_tx=XM_023256086.1
B3 147188238 147188283
refGene_gn=AMN;refGene_tx=XM_023256086.1
B3 147194287 147194375
refGene_gn=AMN;refGene_tx=XM_023256086.1
B3 147194480 147194698
refGene_gn=AMN;refGene_tx=XM_023256086.1
B3 147194844 147194982
refGene_gn=AMN;refGene_tx=XM_023256086.1
B3 147195099 147195208
refGene_gn=AMN;refGene_tx=XM_023256086.1
B3 147195302 147195379
refGene_gn=AMN;refGene_tx=XM_023256086.1
B3 147195564 147195730

refGene_gn=AMN;refGene_tx=XM_023256086.1
B3 147195806 147195996
refGene_gn=AMN;refGene_tx=XM_023256086.1
B3 147196076 147196170
refGene_gn=AMN;refGene_tx=XM_023256086.1
B3 147196250 147196331
refGene_gn=AMN;refGene_tx=XM_023256086.1
B3 147219487 147219712
refGene_gn=CDC42BPB;refGene_tx=XM_023256085.1
B3 147219713 147219740
refGene_gn=CDC42BPB;refGene_tx=XM_023256085.1
B3 147226073 147226144
refGene_gn=CDC42BPB;refGene_tx=XM_023256085.1
B3 147226227 147226333
refGene_gn=CDC42BPB;refGene_tx=XM_023256085.1
B3 147226869 147226987
refGene_gn=CDC42BPB;refGene_tx=XM_023256085.1
B3 147227085 147227203
refGene_gn=CDC42BPB;refGene_tx=XM_023256085.1
B3 147227308 147227393
refGene_gn=CDC42BPB;refGene_tx=XM_023256085.1
B3 147227489 147227587
refGene_gn=CDC42BPB;refGene_tx=XM_023256085.1
B3 147230767 147231364
refGene_gn=CDC42BPB;refGene_tx=XM_023256085.1
B3 147232461 147232524
refGene_gn=CDC42BPB;refGene_tx=XM_023256085.1
B3 147232731 147232948
refGene_gn=CDC42BPB;refGene_tx=XM_023256085.1
B3 147234469 147234551
refGene_gn=CDC42BPB;refGene_tx=XM_023256085.1
B3 147235579 147235719
refGene_gn=CDC42BPB;refGene_tx=XM_023256085.1
B3 147236306 147236443
refGene_gn=CDC42BPB;refGene_tx=XM_023256085.1
B3 147237302 147237408
refGene_gn=CDC42BPB;refGene_tx=XM_023256085.1
B3 147238667 147238745
refGene_gn=CDC42BPB;refGene_tx=XM_023256085.1
B3 147239106 147239196
refGene_gn=CDC42BPB;refGene_tx=XM_023256085.1
B3 147242823 147242903
refGene_gn=CDC42BPB;refGene_tx=XM_023256085.1
B3 147244631 147244726
refGene_gn=CDC42BPB;refGene_tx=XM_023256085.1
B3 147246360 147246509
refGene_gn=CDC42BPB;refGene_tx=XM_023256085.1
B3 147248948 147249054
refGene_gn=CDC42BPB;refGene_tx=XM_023256085.1
B3 147249468 147249593

refGene_gn=CDC42BPB;refGene_tx=XM_023256085.1
B3	147249948	147250054
refGene_gn=CDC42BPB;refGene_tx=XM_023256085.1
B3	147250160	147250405
refGene_gn=CDC42BPB;refGene_tx=XM_023256085.1
B3	147251700	147251811
refGene_gn=CDC42BPB;refGene_tx=XM_023256085.1
B3	147253224	147253467
refGene_gn=CDC42BPB;refGene_tx=XM_023256085.1
B3	147254836	147254970
refGene_gn=CDC42BPB;refGene_tx=XM_023256085.1
B3	147255749	147255869
refGene_gn=CDC42BPB;refGene_tx=XM_023256085.1
B3	147256473	147256640
refGene_gn=CDC42BPB;refGene_tx=XM_023256085.1
B3	147258339	147258419
refGene_gn=CDC42BPB;refGene_tx=XM_023256085.1
B3	147260742	147260991
refGene_gn=CDC42BPB;refGene_tx=XM_023256085.1
B3	147262528	147262729
refGene_gn=CDC42BPB;refGene_tx=XM_023256085.1
B3	147264339	147264433
refGene_gn=CDC42BPB;refGene_tx=XM_023256085.1
B3	147268899	147269048
refGene_gn=CDC42BPB;refGene_tx=XM_023256085.1
B3	147271450	147271546
refGene_gn=CDC42BPB;refGene_tx=XM_023256085.1
B3	147274600	147274684
refGene_gn=CDC42BPB;refGene_tx=XM_023256085.1
B3	147280023	147280115
refGene_gn=CDC42BPB;refGene_tx=XM_023256085.1
B3	147320596	147320771
refGene_gn=CDC42BPB;refGene_tx=XM_023256085.1
B3	147340040	147340046
refGene_gn=LBHD2;refGene_tx=XM_019833830.1
B3	147340307	147340463
refGene_gn=LBHD2;refGene_tx=XM_019833830.1
B3	147342064	147342151
refGene_gn=LBHD2;refGene_tx=XM_019833830.1
B3	147343862	147344019
refGene_gn=LBHD2;refGene_tx=XM_019833830.1
B3	147344994	147345086
refGene_gn=LBHD2;refGene_tx=XM_019833830.1
B3	147353621	147354012
refGene_gn=EXOC3L4;refGene_tx=XM_019833657.2;refGene_gn=EXOC3L4;refGene_tx=XM_019833659.2;refGene_gn=EXOC3L4;refGene_tx=XM_019833662.2;refGene_gn=EXOC3L4;refGene_tx=XM_019833664.2;refGene_gn=EXOC3L4;refGene_tx=XM_023256087.1
B3	147354923	147355575
refGene_gn=EXOC3L4;refGene_tx=XM_019833657.2;refGene_gn=EXOC3L4;refGen

e_tx=XM_019833659.2;refGene_gn=EXOC3L4;refGene_tx=XM_019833662.2;refGene_gn=EXOC3L4;refGene_tx=XM_019833664.2;refGene_gn=EXOC3L4;refGene_tx=XM_023256087.1
B3 147356338 147356450
refGene_gn=EXOC3L4;refGene_tx=XM_019833657.2;refGene_gn=EXOC3L4;refGene_tx=XM_019833659.2;refGene_gn=EXOC3L4;refGene_tx=XM_019833662.2;refGene_gn=EXOC3L4;refGene_tx=XM_019833664.2;refGene_gn=EXOC3L4;refGene_tx=XM_023256087.1
B3 147356721 147356844
refGene_gn=EXOC3L4;refGene_tx=XM_019833657.2;refGene_gn=EXOC3L4;refGene_tx=XM_019833659.2;refGene_gn=EXOC3L4;refGene_tx=XM_019833662.2;refGene_gn=EXOC3L4;refGene_tx=XM_019833664.2;refGene_gn=EXOC3L4;refGene_tx=XM_023256087.1
B3 147357212 147357313
refGene_gn=EXOC3L4;refGene_tx=XM_019833657.2;refGene_gn=EXOC3L4;refGene_tx=XM_019833659.2;refGene_gn=EXOC3L4;refGene_tx=XM_019833662.2;refGene_gn=EXOC3L4;refGene_tx=XM_019833664.2;refGene_gn=EXOC3L4;refGene_tx=XM_023256087.1
B3 147357548 147357629
refGene_gn=EXOC3L4;refGene_tx=XM_019833657.2;refGene_gn=EXOC3L4;refGene_tx=XM_019833659.2;refGene_gn=EXOC3L4;refGene_tx=XM_019833662.2;refGene_gn=EXOC3L4;refGene_tx=XM_019833664.2;refGene_gn=EXOC3L4;refGene_tx=XM_023256087.1
B3 147359157 147359272
refGene_gn=EXOC3L4;refGene_tx=XM_019833657.2;refGene_gn=EXOC3L4;refGene_tx=XM_019833659.2;refGene_gn=EXOC3L4;refGene_tx=XM_019833662.2;refGene_gn=EXOC3L4;refGene_tx=XM_019833664.2;refGene_gn=EXOC3L4;refGene_tx=XM_023256087.1
B3 147359831 147359951
refGene_gn=EXOC3L4;refGene_tx=XM_019833657.2;refGene_gn=EXOC3L4;refGene_tx=XM_019833659.2;refGene_gn=EXOC3L4;refGene_tx=XM_019833662.2;refGene_gn=EXOC3L4;refGene_tx=XM_019833664.2;refGene_gn=EXOC3L4;refGene_tx=XM_023256087.1
B3 147360043 147360196
refGene_gn=EXOC3L4;refGene_tx=XM_019833657.2;refGene_gn=EXOC3L4;refGene_tx=XM_019833659.2;refGene_gn=EXOC3L4;refGene_tx=XM_019833662.2;refGene_gn=EXOC3L4;refGene_tx=XM_023256087.1
B3 147360779 147360901
refGene_gn=EXOC3L4;refGene_tx=XM_019833657.2;refGene_gn=EXOC3L4;refGene_tx=XM_019833659.2;refGene_gn=EXOC3L4;refGene_tx=XM_019833662.2;refGene_gn=EXOC3L4;refGene_tx=XM_019833664.2;refGene_gn=EXOC3L4;refGene_tx=XM_023256087.1
B3 147362051 147362292
refGene_gn=EXOC3L4;refGene_tx=XM_019833657.2;refGene_gn=EXOC3L4;refGene_tx=XM_019833659.2;refGene_gn=EXOC3L4;refGene_tx=XM_019833662.2;refGene_gn=EXOC3L4;refGene_tx=XM_019833664.2;refGene_gn=EXOC3L4;refGene_tx=XM_023256087.1
B3 147374535 147374887
refGene_gn=TNFAIP2;refGene_tx=XM_023256088.1;refGene_gn=TNFAIP2;refGene_tx=XM_023256090.1

B3	147375105	147375748
refGene_gn=TNFAIP2;refGene_tx=XM_023256088.1;refGene_gn=TNFAIP2;refGene_tx=XM_023256090.1
B3	147377688	147377803
refGene_gn=TNFAIP2;refGene_tx=XM_023256088.1;refGene_gn=TNFAIP2;refGene_tx=XM_023256090.1
B3	147377957	147378080
refGene_gn=TNFAIP2;refGene_tx=XM_023256088.1;refGene_gn=TNFAIP2;refGene_tx=XM_023256090.1
B3	147378276	147378377
refGene_gn=TNFAIP2;refGene_tx=XM_023256088.1;refGene_gn=TNFAIP2;refGene_tx=XM_023256090.1
B3	147378975	147379074
refGene_gn=TNFAIP2;refGene_tx=XM_023256088.1;refGene_gn=TNFAIP2;refGene_tx=XM_023256090.1
B3	147379601	147379722
refGene_gn=TNFAIP2;refGene_tx=XM_023256088.1;refGene_gn=TNFAIP2;refGene_tx=XM_023256090.1
B3	147380606	147380729
refGene_gn=TNFAIP2;refGene_tx=XM_023256088.1;refGene_gn=TNFAIP2;refGene_tx=XM_023256090.1
B3	147380967	147381123
refGene_gn=TNFAIP2;refGene_tx=XM_023256088.1;refGene_gn=TNFAIP2;refGene_tx=XM_023256090.1
B3	147381308	147381430
refGene_gn=TNFAIP2;refGene_tx=XM_023256088.1;refGene_gn=TNFAIP2;refGene_tx=XM_023256090.1
B3	147383711	147383853
refGene_gn=TNFAIP2;refGene_tx=XM_023256088.1;refGene_gn=TNFAIP2;refGene_tx=XM_023256090.1
B3	147439176	147439379
refGene_gn=LOC101085134;refGene_tx=XM_023256091.1
B3	147439381	147439778
refGene_gn=LOC101085134;refGene_tx=XM_023256091.1
B3	147463554	147463572
refGene_gn=LOC102901183;refGene_tx=XM_006933148.4
B3	147465666	147465739
refGene_gn=LOC102901183;refGene_tx=XM_006933145.4;refGene_gn=LOC102901183;refGene_tx=XM_006933146.4;refGene_gn=LOC102901183;refGene_tx=XM_019833666.2;refGene_gn=LOC102901183;refGene_tx=XM_019833667.2
B3	147466309	147466434
refGene_gn=LOC102901183;refGene_tx=XM_006933145.4;refGene_gn=LOC102901183;refGene_tx=XM_006933146.4;refGene_gn=LOC102901183;refGene_tx=XM_019833666.2;refGene_gn=LOC102901183;refGene_tx=XM_019833667.2
B3	147466867	147467023
refGene_gn=LOC102901183;refGene_tx=XM_006933145.4;refGene_gn=LOC102901183;refGene_tx=XM_006933146.4;refGene_gn=LOC102901183;refGene_tx=XM_006933148.4;refGene_gn=LOC102901183;refGene_tx=XM_019833666.2;refGene_gn=LOC102901183;refGene_tx=XM_019833667.2
B3	147467288	147467402

refGene_gn=LOC102901183;refGene_tx=XM_006933145.4;refGene_gn=LOC102901183;refGene_tx=XM_006933146.4;refGene_gn=LOC102901183;refGene_tx=XM_006933148.4;refGene_gn=LOC102901183;refGene_tx=XM_019833666.2;refGene_gn=LOC102901183;refGene_tx=XM_019833667.2
B3 147468919 147469052
refGene_gn=LOC102901183;refGene_tx=XM_006933145.4;refGene_gn=LOC102901183;refGene_tx=XM_006933146.4;refGene_gn=LOC102901183;refGene_tx=XM_006933148.4;refGene_gn=LOC102901183;refGene_tx=XM_019833666.2;refGene_gn=LOC102901183;refGene_tx=XM_019833667.2
B3 147469394 147469487
refGene_gn=LOC102901183;refGene_tx=XM_006933145.4;refGene_gn=LOC102901183;refGene_tx=XM_006933146.4;refGene_gn=LOC102901183;refGene_tx=XM_006933148.4;refGene_gn=LOC102901183;refGene_tx=XM_019833666.2;refGene_gn=LOC102901183;refGene_tx=XM_019833667.2
B3 147471208 147471309
refGene_gn=LOC102901183;refGene_tx=XM_006933145.4;refGene_gn=LOC102901183;refGene_tx=XM_006933146.4;refGene_gn=LOC102901183;refGene_tx=XM_006933148.4;refGene_gn=LOC102901183;refGene_tx=XM_019833666.2;refGene_gn=LOC102901183;refGene_tx=XM_019833667.2
B3 147472216 147472309
refGene_gn=LOC102901183;refGene_tx=XM_006933145.4;refGene_gn=LOC102901183;refGene_tx=XM_006933146.4;refGene_gn=LOC102901183;refGene_tx=XM_006933148.4;refGene_gn=LOC102901183;refGene_tx=XM_019833666.2
B3 147474245 147474369
refGene_gn=LOC102901183;refGene_tx=XM_006933145.4;refGene_gn=LOC102901183;refGene_tx=XM_006933146.4;refGene_gn=LOC102901183;refGene_tx=XM_006933148.4;refGene_gn=LOC102901183;refGene_tx=XM_019833666.2;refGene_gn=LOC102901183;refGene_tx=XM_019833667.2
B3 147474850 147475412
refGene_gn=LOC102901183;refGene_tx=XM_006933145.4;refGene_gn=LOC102901183;refGene_tx=XM_006933146.4;refGene_gn=LOC102901183;refGene_tx=XM_006933148.4;refGene_gn=LOC102901183;refGene_tx=XM_019833666.2;refGene_gn=LOC102901183;refGene_tx=XM_019833667.2
B3 147477534 147477763
refGene_gn=LOC102901183;refGene_tx=XM_006933145.4;refGene_gn=LOC102901183;refGene_tx=XM_006933146.4;refGene_gn=LOC102901183;refGene_tx=XM_006933148.4;refGene_gn=LOC102901183;refGene_tx=XM_019833666.2;refGene_gn=LOC102901183;refGene_tx=XM_019833667.2
B3 147500003 147500049
refGene_gn=LOC105260685;refGene_tx=XM_023256199.1
B3 147500342 147500471
refGene_gn=LOC105260685;refGene_tx=XM_023256199.1
B3 147500944 147501009
refGene_gn=LOC105260685;refGene_tx=XM_023256199.1
B3 147501135 147501172
refGene_gn=LOC105260685;refGene_tx=XM_023256199.1
B3 147506461 147506720
refGene_gn=LOC105260685;refGene_tx=XM_023256199.1
B3 147507153 147507256
refGene_gn=LOC105260685;refGene_tx=XM_023256199.1

B3 147508036 147508108
refGene_gn=EIF5;refGene_tx=XM_003988006.5;refGene_gn=EIF5;refGene_tx=XM_023256093.1
B3 147508212 147508294
refGene_gn=EIF5;refGene_tx=XM_003988006.5;refGene_gn=EIF5;refGene_tx=XM_023256093.1
B3 147508566 147508739
refGene_gn=EIF5;refGene_tx=XM_003988006.5;refGene_gn=EIF5;refGene_tx=XM_023256093.1
B3 147509049 147509161
refGene_gn=EIF5;refGene_tx=XM_003988006.5;refGene_gn=EIF5;refGene_tx=XM_023256093.1
B3 147510007 147510150
refGene_gn=EIF5;refGene_tx=XM_003988006.5;refGene_gn=EIF5;refGene_tx=XM_023256093.1
B3 147510401 147510560
refGene_gn=EIF5;refGene_tx=XM_003988006.5;refGene_gn=EIF5;refGene_tx=XM_023256093.1
B3 147511078 147511240
refGene_gn=EIF5;refGene_tx=XM_003988006.5;refGene_gn=EIF5;refGene_tx=XM_023256093.1
B3 147511523 147511688
refGene_gn=EIF5;refGene_tx=XM_003988006.5;refGene_gn=EIF5;refGene_tx=XM_023256093.1
B3 147511956 147512091
refGene_gn=EIF5;refGene_tx=XM_003988006.5;refGene_gn=EIF5;refGene_tx=XM_023256093.1
B3 147512437 147512524
refGene_gn=EIF5;refGene_tx=XM_003988006.5;refGene_gn=EIF5;refGene_tx=XM_023256093.1
B3 147556081 147556132
refGene_gn=MARK3;refGene_tx=XM_023256094.1;refGene_gn=MARK3;refGene_tx=XM_023256095.1;refGene_gn=MARK3;refGene_tx=XM_023256096.1;refGene_gn=MARK3;refGene_tx=XM_023256097.1;refGene_gn=MARK3;refGene_tx=XM_023256098.1;refGene_gn=MARK3;refGene_tx=XM_023256099.1;refGene_gn=MARK3;refGene_tx=XM_023256100.1;refGene_gn=MARK3;refGene_tx=XM_023256101.1
B3 147579187 147579379
refGene_gn=MARK3;refGene_tx=XM_023256094.1;refGene_gn=MARK3;refGene_tx=XM_023256095.1;refGene_gn=MARK3;refGene_tx=XM_023256096.1;refGene_gn=MARK3;refGene_tx=XM_023256097.1;refGene_gn=MARK3;refGene_tx=XM_023256098.1;refGene_gn=MARK3;refGene_tx=XM_023256099.1;refGene_gn=MARK3;refGene_tx=XM_023256100.1;refGene_gn=MARK3;refGene_tx=XM_023256101.1
B3 147596387 147596441
refGene_gn=MARK3;refGene_tx=XM_023256094.1;refGene_gn=MARK3;refGene_tx=XM_023256095.1;refGene_gn=MARK3;refGene_tx=XM_023256096.1;refGene_gn=MARK3;refGene_tx=XM_023256097.1;refGene_gn=MARK3;refGene_tx=XM_023256099.1;refGene_gn=MARK3;refGene_tx=XM_023256100.1;refGene_gn=MARK3;refGene_tx=XM_023256101.1
B3 147625511 147625559
refGene_gn=MARK3;refGene_tx=XM_019833668.1

B3	147627302	147627351
refGene_gn=MARK3;refGene_tx=XM_019833668.1;refGene_gn=MARK3;refGene_tx=XM_023256094.1;refGene_gn=MARK3;refGene_tx=XM_023256095.1;refGene_gn=MARK3;refGene_tx=XM_023256096.1;refGene_gn=MARK3;refGene_tx=XM_023256097.1;refGene_gn=MARK3;refGene_tx=XM_023256098.1;refGene_gn=MARK3;refGene_tx=XM_023256099.1;refGene_gn=MARK3;refGene_tx=XM_023256100.1;refGene_gn=MARK3;refGene_tx=XM_023256101.1
B3	147630263	147630329
refGene_gn=MARK3;refGene_tx=XM_019833668.1;refGene_gn=MARK3;refGene_tx=XM_019833673.1;refGene_gn=MARK3;refGene_tx=XM_023256094.1;refGene_gn=MARK3;refGene_tx=XM_023256095.1;refGene_gn=MARK3;refGene_tx=XM_023256096.1;refGene_gn=MARK3;refGene_tx=XM_023256097.1;refGene_gn=MARK3;refGene_tx=XM_023256098.1;refGene_gn=MARK3;refGene_tx=XM_023256099.1;refGene_gn=MARK3;refGene_tx=XM_023256100.1;refGene_gn=MARK3;refGene_tx=XM_023256101.1
B3	147634782	147634853
refGene_gn=MARK3;refGene_tx=XM_019833668.1;refGene_gn=MARK3;refGene_tx=XM_019833673.1;refGene_gn=MARK3;refGene_tx=XM_023256094.1;refGene_gn=MARK3;refGene_tx=XM_023256095.1;refGene_gn=MARK3;refGene_tx=XM_023256096.1;refGene_gn=MARK3;refGene_tx=XM_023256097.1;refGene_gn=MARK3;refGene_tx=XM_023256098.1;refGene_gn=MARK3;refGene_tx=XM_023256099.1;refGene_gn=MARK3;refGene_tx=XM_023256100.1;refGene_gn=MARK3;refGene_tx=XM_023256101.1
B3	147639951	147640008
refGene_gn=MARK3;refGene_tx=XM_019833668.1;refGene_gn=MARK3;refGene_tx=XM_019833673.1;refGene_gn=MARK3;refGene_tx=XM_023256094.1;refGene_gn=MARK3;refGene_tx=XM_023256095.1;refGene_gn=MARK3;refGene_tx=XM_023256096.1;refGene_gn=MARK3;refGene_tx=XM_023256097.1;refGene_gn=MARK3;refGene_tx=XM_023256098.1;refGene_gn=MARK3;refGene_tx=XM_023256099.1;refGene_gn=MARK3;refGene_tx=XM_023256100.1;refGene_gn=MARK3;refGene_tx=XM_023256101.1
B3	147642259	147642496
refGene_gn=MARK3;refGene_tx=XM_019833668.1;refGene_gn=MARK3;refGene_tx=XM_019833673.1;refGene_gn=MARK3;refGene_tx=XM_023256094.1;refGene_gn=MARK3;refGene_tx=XM_023256095.1;refGene_gn=MARK3;refGene_tx=XM_023256096.1;refGene_gn=MARK3;refGene_tx=XM_023256097.1;refGene_gn=MARK3;refGene_tx=XM_023256098.1;refGene_gn=MARK3;refGene_tx=XM_023256099.1;refGene_gn=MARK3;refGene_tx=XM_023256100.1
B3	147642658	147642778
refGene_gn=MARK3;refGene_tx=XM_019833668.1;refGene_gn=MARK3;refGene_tx=XM_019833673.1;refGene_gn=MARK3;refGene_tx=XM_023256094.1;refGene_gn=MARK3;refGene_tx=XM_023256095.1;refGene_gn=MARK3;refGene_tx=XM_023256096.1;refGene_gn=MARK3;refGene_tx=XM_023256097.1;refGene_gn=MARK3;refGene_tx=XM_023256098.1;refGene_gn=MARK3;refGene_tx=XM_023256099.1;refGene_gn=MARK3;refGene_tx=XM_023256100.1
B3	147643512	147643612
refGene_gn=MARK3;refGene_tx=XM_019833668.1;refGene_gn=MARK3;refGene_tx=XM_019833673.1;refGene_gn=MARK3;refGene_tx=XM_023256094.1;refGene_gn=MARK3;refGene_tx=XM_023256095.1;refGene_gn=MARK3;refGene_tx=XM_023256096.1;refGene_gn=MARK3;refGene_tx=XM_023256097.1;refGene_gn=MARK3;refGe

ne_tx=XM_023256098.1;refGene_gn=MARK3;refGene_tx=XM_023256099.1;refGene_gn=MARK3;refGene_tx=XM_023256100.1;refGene_gn=MARK3;refGene_tx=XM_023256101.1
B3 147643928 147644041
refGene_gn=MARK3;refGene_tx=XM_019833668.1;refGene_gn=MARK3;refGene_tx=XM_019833673.1;refGene_gn=MARK3;refGene_tx=XM_023256094.1;refGene_gn=MARK3;refGene_tx=XM_023256095.1;refGene_gn=MARK3;refGene_tx=XM_023256096.1;refGene_gn=MARK3;refGene_tx=XM_023256097.1;refGene_gn=MARK3;refGene_tx=XM_023256098.1;refGene_gn=MARK3;refGene_tx=XM_023256099.1;refGene_gn=MARK3;refGene_tx=XM_023256100.1;refGene_gn=MARK3;refGene_tx=XM_023256101.1
B3 147644565 147644719
refGene_gn=MARK3;refGene_tx=XM_019833668.1;refGene_gn=MARK3;refGene_tx=XM_019833673.1;refGene_gn=MARK3;refGene_tx=XM_023256094.1;refGene_gn=MARK3;refGene_tx=XM_023256095.1;refGene_gn=MARK3;refGene_tx=XM_023256096.1;refGene_gn=MARK3;refGene_tx=XM_023256097.1;refGene_gn=MARK3;refGene_tx=XM_023256098.1;refGene_gn=MARK3;refGene_tx=XM_023256099.1;refGene_gn=MARK3;refGene_tx=XM_023256100.1;refGene_gn=MARK3;refGene_tx=XM_023256101.1
B3 147649262 147649480
refGene_gn=MARK3;refGene_tx=XM_019833668.1;refGene_gn=MARK3;refGene_tx=XM_019833673.1;refGene_gn=MARK3;refGene_tx=XM_023256094.1;refGene_gn=MARK3;refGene_tx=XM_023256095.1;refGene_gn=MARK3;refGene_tx=XM_023256096.1;refGene_gn=MARK3;refGene_tx=XM_023256097.1;refGene_gn=MARK3;refGene_tx=XM_023256098.1;refGene_gn=MARK3;refGene_tx=XM_023256099.1;refGene_gn=MARK3;refGene_tx=XM_023256100.1;refGene_gn=MARK3;refGene_tx=XM_023256101.1
B3 147651133 147651234
refGene_gn=MARK3;refGene_tx=XM_019833668.1;refGene_gn=MARK3;refGene_tx=XM_019833673.1;refGene_gn=MARK3;refGene_tx=XM_023256094.1;refGene_gn=MARK3;refGene_tx=XM_023256095.1;refGene_gn=MARK3;refGene_tx=XM_023256096.1;refGene_gn=MARK3;refGene_tx=XM_023256097.1;refGene_gn=MARK3;refGene_tx=XM_023256098.1;refGene_gn=MARK3;refGene_tx=XM_023256099.1;refGene_gn=MARK3;refGene_tx=XM_023256100.1;refGene_gn=MARK3;refGene_tx=XM_023256101.1
B3 147659817 147660075
refGene_gn=MARK3;refGene_tx=XM_019833668.1;refGene_gn=MARK3;refGene_tx=XM_019833673.1;refGene_gn=MARK3;refGene_tx=XM_023256094.1;refGene_gn=MARK3;refGene_tx=XM_023256095.1;refGene_gn=MARK3;refGene_tx=XM_023256096.1;refGene_gn=MARK3;refGene_tx=XM_023256097.1;refGene_gn=MARK3;refGene_tx=XM_023256098.1;refGene_gn=MARK3;refGene_tx=XM_023256099.1;refGene_gn=MARK3;refGene_tx=XM_023256100.1;refGene_gn=MARK3;refGene_tx=XM_023256101.1
B3 147664897 147664924
refGene_gn=MARK3;refGene_tx=XM_019833668.1;refGene_gn=MARK3;refGene_tx=XM_019833673.1;refGene_gn=MARK3;refGene_tx=XM_023256094.1;refGene_gn=MARK3;refGene_tx=XM_023256096.1;refGene_gn=MARK3;refGene_tx=XM_023256097.1;refGene_gn=MARK3;refGene_tx=XM_023256098.1;refGene_gn=MARK3;refGene_tx=XM_023256101.1
B3 147666211 147666256

refGene_gn=MARK3;refGene_tx=XM_019833668.1;refGene_gn=MARK3;refGene_tx=XM_019833673.1;refGene_gn=MARK3;refGene_tx=XM_023256094.1;refGene_gn=MARK3;refGene_tx=XM_023256095.1;refGene_gn=MARK3;refGene_tx=XM_023256097.1;refGene_gn=MARK3;refGene_tx=XM_023256098.1;refGene_gn=MARK3;refGene_tx=XM_023256101.1
B3 147668678 147669024
refGene_gn=MARK3;refGene_tx=XM_019833668.1;refGene_gn=MARK3;refGene_tx=XM_019833673.1;refGene_gn=MARK3;refGene_tx=XM_023256094.1;refGene_gn=MARK3;refGene_tx=XM_023256095.1;refGene_gn=MARK3;refGene_tx=XM_023256096.1;refGene_gn=MARK3;refGene_tx=XM_023256097.1;refGene_gn=MARK3;refGene_tx=XM_023256098.1;refGene_gn=MARK3;refGene_tx=XM_023256099.1;refGene_gn=MARK3;refGene_tx=XM_023256100.1;refGene_gn=MARK3;refGene_tx=XM_023256101.1
B3 147682713 147682892
refGene_gn=CKB;refGene_tx=XM_023256102.1;refGene_gn=CKB;refGene_tx=XM_023256103.1
B3 147682967 147683157
refGene_gn=CKB;refGene_tx=XM_023256102.1;refGene_gn=CKB;refGene_tx=XM_023256103.1
B3 147683401 147683525
refGene_gn=CKB;refGene_tx=XM_023256102.1;refGene_gn=CKB;refGene_tx=XM_023256103.1
B3 147684277 147684449
refGene_gn=CKB;refGene_tx=XM_023256102.1;refGene_gn=CKB;refGene_tx=XM_023256103.1
B3 147684667 147684800
refGene_gn=CKB;refGene_tx=XM_023256102.1;refGene_gn=CKB;refGene_tx=XM_023256103.1
B3 147684873 147685028
refGene_gn=CKB;refGene_tx=XM_023256102.1;refGene_gn=CKB;refGene_tx=XM_023256103.1
B3 147685235 147685428
refGene_gn=CKB;refGene_tx=XM_023256102.1;refGene_gn=CKB;refGene_tx=XM_023256103.1
B3 147692757 147693145
refGene_gn=TRMT61A;refGene_tx=XM_003988010.5;refGene_gn=TRMT61A;refGene_tx=XM_023256104.1;refGene_gn=TRMT61A;refGene_tx=XM_023256105.1
B3 147695521 147695788
refGene_gn=TRMT61A;refGene_tx=XM_003988010.5;refGene_gn=TRMT61A;refGene_tx=XM_023256104.1;refGene_gn=TRMT61A;refGene_tx=XM_023256105.1
B3 147697699 147697971
refGene_gn=TRMT61A;refGene_tx=XM_003988010.5;refGene_gn=TRMT61A;refGene_tx=XM_023256104.1;refGene_gn=TRMT61A;refGene_tx=XM_023256105.1
B3 147719729 147721073
refGene_gn=BAG5;refGene_tx=XM_003988011.5;refGene_gn=BAG5;refGene_tx=XM_006933149.4;refGene_gn=BAG5;refGene_tx=XM_019833686.2;refGene_gn=BAG5;refGene_tx=XM_023256107.1
B3 147722353 147723137
refGene_gn=APOPT1;refGene_tx=XM_023256200.1
B3 147723139 147723264

refGene_gn=APOPT1;refGene_tx=XM_023256200.1
B3	147740299	147740497
refGene_gn=APOPT1;refGene_tx=XM_023256200.1
B3	147741492	147741556
refGene_gn=APOPT1;refGene_tx=XM_023256200.1
B3	147748039	147748130
refGene_gn=APOPT1;refGene_tx=XM_023256200.1
B3	147750973	147751079
refGene_gn=APOPT1;refGene_tx=XM_023256200.1
B3	147788885	147788899
refGene_gn=KLC1;refGene_tx=XM_019833679.2;refGene_gn=KLC1;refGene_tx=XM_019833680.2;refGene_gn=KLC1;refGene_tx=XM_019833681.2;refGene_gn=KLC1;refGene_tx=XM_019833682.2;refGene_gn=KLC1;refGene_tx=XM_019833683.2;refGene_gn=KLC1;refGene_tx=XM_019833684.2;refGene_gn=KLC1;refGene_tx=XM_019833685.2
B3	147799738	147800000
refGene_gn=KLC1;refGene_tx=XM_019833679.2;refGene_gn=KLC1;refGene_tx=XM_019833680.2;refGene_gn=KLC1;refGene_tx=XM_019833681.2;refGene_gn=KLC1;refGene_tx=XM_019833682.2;refGene_gn=KLC1;refGene_tx=XM_019833683.2;refGene_gn=KLC1;refGene_tx=XM_019833684.2;refGene_gn=KLC1;refGene_tx=XM_019833685.2
B3	147802251	147802482
refGene_gn=KLC1;refGene_tx=XM_019833679.2;refGene_gn=KLC1;refGene_tx=XM_019833680.2;refGene_gn=KLC1;refGene_tx=XM_019833681.2;refGene_gn=KLC1;refGene_tx=XM_019833682.2;refGene_gn=KLC1;refGene_tx=XM_019833683.2;refGene_gn=KLC1;refGene_tx=XM_019833684.2;refGene_gn=KLC1;refGene_tx=XM_019833685.2
B3	147805842	147805921
refGene_gn=KLC1;refGene_tx=XM_019833679.2;refGene_gn=KLC1;refGene_tx=XM_019833680.2;refGene_gn=KLC1;refGene_tx=XM_019833681.2;refGene_gn=KLC1;refGene_tx=XM_019833682.2;refGene_gn=KLC1;refGene_tx=XM_019833683.2;refGene_gn=KLC1;refGene_tx=XM_019833684.2;refGene_gn=KLC1;refGene_tx=XM_019833685.2
B3	147806427	147806653
refGene_gn=KLC1;refGene_tx=XM_019833679.2;refGene_gn=KLC1;refGene_tx=XM_019833680.2;refGene_gn=KLC1;refGene_tx=XM_019833681.2;refGene_gn=KLC1;refGene_tx=XM_019833682.2;refGene_gn=KLC1;refGene_tx=XM_019833683.2;refGene_gn=KLC1;refGene_tx=XM_019833684.2;refGene_gn=KLC1;refGene_tx=XM_019833685.2
B3	147809017	147809105
refGene_gn=KLC1;refGene_tx=XM_019833679.2;refGene_gn=KLC1;refGene_tx=XM_019833680.2;refGene_gn=KLC1;refGene_tx=XM_019833681.2;refGene_gn=KLC1;refGene_tx=XM_019833682.2;refGene_gn=KLC1;refGene_tx=XM_019833683.2;refGene_gn=KLC1;refGene_tx=XM_019833684.2;refGene_gn=KLC1;refGene_tx=XM_019833685.2
B3	147809742	147809844
refGene_gn=KLC1;refGene_tx=XM_019833679.2;refGene_gn=KLC1;refGene_tx=XM_019833680.2;refGene_gn=KLC1;refGene_tx=XM_019833681.2;refGene_gn=KLC1;refGene_tx=XM_019833682.2;refGene_gn=KLC1;refGene_tx=XM_019833683.2;refGene_gn=KLC1;refGene_tx=XM_019833684.2;refGene_gn=KLC1;refGene_tx=X

M_019833685.2
B3 147811850 147812024
refGene_gn=KLC1;refGene_tx=XM_019833679.2;refGene_gn=KLC1;refGene_tx=XM_019833680.2;refGene_gn=KLC1;refGene_tx=XM_019833681.2;refGene_gn=KLC1;refGene_tx=XM_019833682.2;refGene_gn=KLC1;refGene_tx=XM_019833683.2;refGene_gn=KLC1;refGene_tx=XM_019833684.2;refGene_gn=KLC1;refGene_tx=XM_019833685.2
B3 147812170 147812270
refGene_gn=KLC1;refGene_tx=XM_019833679.2;refGene_gn=KLC1;refGene_tx=XM_019833680.2;refGene_gn=KLC1;refGene_tx=XM_019833681.2;refGene_gn=KLC1;refGene_tx=XM_019833682.2;refGene_gn=KLC1;refGene_tx=XM_019833683.2;refGene_gn=KLC1;refGene_tx=XM_019833684.2;refGene_gn=KLC1;refGene_tx=XM_019833685.2
B3 147814809 147814859
refGene_gn=KLC1;refGene_tx=XM_019833679.2;refGene_gn=KLC1;refGene_tx=XM_019833680.2;refGene_gn=KLC1;refGene_tx=XM_019833681.2;refGene_gn=KLC1;refGene_tx=XM_019833682.2;refGene_gn=KLC1;refGene_tx=XM_019833683.2;refGene_gn=KLC1;refGene_tx=XM_019833684.2;refGene_gn=KLC1;refGene_tx=XM_019833685.2
B3 147814938 147815006
refGene_gn=KLC1;refGene_tx=XM_019833679.2;refGene_gn=KLC1;refGene_tx=XM_019833680.2;refGene_gn=KLC1;refGene_tx=XM_019833681.2;refGene_gn=KLC1;refGene_tx=XM_019833682.2;refGene_gn=KLC1;refGene_tx=XM_019833683.2;refGene_gn=KLC1;refGene_tx=XM_019833684.2;refGene_gn=KLC1;refGene_tx=XM_019833685.2
B3 147816032 147816141
refGene_gn=KLC1;refGene_tx=XM_019833679.2;refGene_gn=KLC1;refGene_tx=XM_019833680.2;refGene_gn=KLC1;refGene_tx=XM_019833681.2;refGene_gn=KLC1;refGene_tx=XM_019833682.2;refGene_gn=KLC1;refGene_tx=XM_019833683.2;refGene_gn=KLC1;refGene_tx=XM_019833684.2;refGene_gn=KLC1;refGene_tx=XM_019833685.2
B3 147819452 147819614
refGene_gn=KLC1;refGene_tx=XM_019833679.2;refGene_gn=KLC1;refGene_tx=XM_019833680.2;refGene_gn=KLC1;refGene_tx=XM_019833681.2;refGene_gn=KLC1;refGene_tx=XM_019833682.2;refGene_gn=KLC1;refGene_tx=XM_019833683.2;refGene_gn=KLC1;refGene_tx=XM_019833684.2;refGene_gn=KLC1;refGene_tx=XM_019833685.2
B3 147823658 147823691
refGene_gn=KLC1;refGene_tx=XM_019833684.2
B3 147824410 147824416
refGene_gn=KLC1;refGene_tx=XM_019833685.2
B3 147825994 147826125
refGene_gn=KLC1;refGene_tx=XM_019833679.2;refGene_gn=KLC1;refGene_tx=XM_019833680.2;refGene_gn=KLC1;refGene_tx=XM_019833681.2;refGene_gn=KLC1;refGene_tx=XM_019833682.2
B3 147830569 147830639
refGene_gn=KLC1;refGene_tx=XM_019833679.2;refGene_gn=KLC1;refGene_tx=XM_019833680.2;refGene_gn=KLC1;refGene_tx=XM_019833681.2
B3 147831665 147831695
refGene_gn=KLC1;refGene_tx=XM_019833680.2

B3 147835162 147835394
refGene_gn=XRCC3;refGene_tx=XM_006933151.4
B3 147835488 147835535
refGene_gn=XRCC3;refGene_tx=XM_006933151.4
B3 147835693 147835906
refGene_gn=XRCC3;refGene_tx=XM_006933151.4
B3 147836386 147836459
refGene_gn=KLC1;refGene_tx=XM_019833679.2;refGene_gn=KLC1;refGene_tx=XM_019833681.2;refGene_gn=KLC1;refGene_tx=XM_019833682.2;refGene_gn=KLC1;refGene_tx=XM_019833683.2
B3 147836777 147836780
refGene_gn=KLC1;refGene_tx=XM_019833682.2
B3 147838015 147838170
refGene_gn=XRCC3;refGene_tx=XM_006933151.4
B3 147839920 147840133
refGene_gn=XRCC3;refGene_tx=XM_006933151.4
B3 147840455 147840593
refGene_gn=XRCC3;refGene_tx=XM_006933151.4
B3 147842265 147842320
refGene_gn=XRCC3;refGene_tx=XM_006933151.4
B3 147846071 147846209
refGene_gn=ZFYVE21;refGene_tx=XM_003988013.5;refGene_gn=ZFYVE21;refGene_tx=XM_003988014.5
B3 147853534 147853585
refGene_gn=ZFYVE21;refGene_tx=XM_003988013.5;refGene_gn=ZFYVE21;refGene_tx=XM_003988014.5
B3 147853960 147854129
refGene_gn=ZFYVE21;refGene_tx=XM_003988013.5;refGene_gn=ZFYVE21;refGene_tx=XM_003988014.5
B3 147854833 147854909
refGene_gn=ZFYVE21;refGene_tx=XM_003988013.5;refGene_gn=ZFYVE21;refGene_tx=XM_003988014.5
B3 147854991 147855083
refGene_gn=ZFYVE21;refGene_tx=XM_003988013.5;refGene_gn=ZFYVE21;refGene_tx=XM_003988014.5
B3 147855660 147855714
refGene_gn=ZFYVE21;refGene_tx=XM_003988014.5
B3 147858617 147858760
refGene_gn=ZFYVE21;refGene_tx=XM_003988013.5;refGene_gn=ZFYVE21;refGene_tx=XM_003988014.5
B3 147858919 147858955
refGene_gn=ZFYVE21;refGene_tx=XM_003988013.5;refGene_gn=ZFYVE21;refGene_tx=XM_003988014.5
B3 147860774 147860816
refGene_gn=PPP1R13B;refGene_tx=XM_019833678.2;refGene_gn=PPP1R13B;refGene_tx=XM_023256108.1;refGene_gn=PPP1R13B;refGene_tx=XM_023256109.1;refGene_gn=PPP1R13B;refGene_tx=XM_023256110.1;refGene_gn=PPP1R13B;refGene_tx=XM_023256111.1;refGene_gn=PPP1R13B;refGene_tx=XM_023256112.1
B3 147861455 147861655
refGene_gn=PPP1R13B;refGene_tx=XM_019833678.2;refGene_gn=PPP1R13B;refG

ene_tx=XM_023256108.1;refGene_gn=PPP1R13B;refGene_tx=XM_023256109.1;refGene_gn=PPP1R13B;refGene_tx=XM_023256110.1;refGene_gn=PPP1R13B;refGene_tx=XM_023256111.1;refGene_gn=PPP1R13B;refGene_tx=XM_023256112.1
B3 147862017 147862184
refGene_gn=PPP1R13B;refGene_tx=XM_019833678.2;refGene_gn=PPP1R13B;refGene_tx=XM_023256108.1;refGene_gn=PPP1R13B;refGene_tx=XM_023256109.1;refGene_gn=PPP1R13B;refGene_tx=XM_023256110.1;refGene_gn=PPP1R13B;refGene_tx=XM_023256111.1;refGene_gn=PPP1R13B;refGene_tx=XM_023256112.1
B3 147862938 147863072
refGene_gn=PPP1R13B;refGene_tx=XM_019833678.2;refGene_gn=PPP1R13B;refGene_tx=XM_023256108.1;refGene_gn=PPP1R13B;refGene_tx=XM_023256109.1;refGene_gn=PPP1R13B;refGene_tx=XM_023256110.1;refGene_gn=PPP1R13B;refGene_tx=XM_023256111.1;refGene_gn=PPP1R13B;refGene_tx=XM_023256112.1
B3 147863148 147863286
refGene_gn=PPP1R13B;refGene_tx=XM_019833678.2;refGene_gn=PPP1R13B;refGene_tx=XM_023256108.1;refGene_gn=PPP1R13B;refGene_tx=XM_023256109.1;refGene_gn=PPP1R13B;refGene_tx=XM_023256110.1;refGene_gn=PPP1R13B;refGene_tx=XM_023256111.1;refGene_gn=PPP1R13B;refGene_tx=XM_023256112.1
B3 147863885 147864664
refGene_gn=PPP1R13B;refGene_tx=XM_019833678.2;refGene_gn=PPP1R13B;refGene_tx=XM_023256108.1;refGene_gn=PPP1R13B;refGene_tx=XM_023256109.1;refGene_gn=PPP1R13B;refGene_tx=XM_023256110.1;refGene_gn=PPP1R13B;refGene_tx=XM_023256111.1;refGene_gn=PPP1R13B;refGene_tx=XM_023256112.1
B3 147865972 147866471
refGene_gn=PPP1R13B;refGene_tx=XM_019833678.2;refGene_gn=PPP1R13B;refGene_tx=XM_023256108.1;refGene_gn=PPP1R13B;refGene_tx=XM_023256109.1;refGene_gn=PPP1R13B;refGene_tx=XM_023256110.1;refGene_gn=PPP1R13B;refGene_tx=XM_023256111.1;refGene_gn=PPP1R13B;refGene_tx=XM_023256112.1
B3 147866850 147867020
refGene_gn=PPP1R13B;refGene_tx=XM_019833678.2;refGene_gn=PPP1R13B;refGene_tx=XM_023256108.1;refGene_gn=PPP1R13B;refGene_tx=XM_023256109.1;refGene_gn=PPP1R13B;refGene_tx=XM_023256110.1;refGene_gn=PPP1R13B;refGene_tx=XM_023256111.1;refGene_gn=PPP1R13B;refGene_tx=XM_023256112.1
B3 147870662 147870843
refGene_gn=PPP1R13B;refGene_tx=XM_019833678.2;refGene_gn=PPP1R13B;refGene_tx=XM_023256108.1;refGene_gn=PPP1R13B;refGene_tx=XM_023256109.1;refGene_gn=PPP1R13B;refGene_tx=XM_023256110.1;refGene_gn=PPP1R13B;refGene_tx=XM_023256111.1;refGene_gn=PPP1R13B;refGene_tx=XM_023256112.1
B3 147873559 147873724
refGene_gn=PPP1R13B;refGene_tx=XM_019833678.2;refGene_gn=PPP1R13B;refGene_tx=XM_023256108.1;refGene_gn=PPP1R13B;refGene_tx=XM_023256109.1;refGene_gn=PPP1R13B;refGene_tx=XM_023256110.1;refGene_gn=PPP1R13B;refGene_tx=XM_023256111.1;refGene_gn=PPP1R13B;refGene_tx=XM_023256112.1
B3 147875890 147876087
refGene_gn=PPP1R13B;refGene_tx=XM_019833678.2;refGene_gn=PPP1R13B;refGene_tx=XM_023256108.1;refGene_gn=PPP1R13B;refGene_tx=XM_023256109.1;refGene_gn=PPP1R13B;refGene_tx=XM_023256110.1;refGene_gn=PPP1R13B;refGene_tx=XM_023256111.1;refGene_gn=PPP1R13B;refGene_tx=XM_023256112.1
B3 147876857 147877032
refGene_gn=PPP1R13B;refGene_tx=XM_019833678.2;refGene_gn=PPP1R13B;refG

ene_tx=XM_023256108.1;refGene_gn=PPP1R13B;refGene_tx=XM_023256109.1;refGene_gn=PPP1R13B;refGene_tx=XM_023256110.1;refGene_gn=PPP1R13B;refGene_tx=XM_023256111.1;refGene_gn=PPP1R13B;refGene_tx=XM_023256112.1
B3 147878768 147878870
refGene_gn=PPP1R13B;refGene_tx=XM_019833678.2;refGene_gn=PPP1R13B;refGene_tx=XM_023256108.1;refGene_gn=PPP1R13B;refGene_tx=XM_023256109.1;refGene_gn=PPP1R13B;refGene_tx=XM_023256110.1;refGene_gn=PPP1R13B;refGene_tx=XM_023256111.1;refGene_gn=PPP1R13B;refGene_tx=XM_023256112.1
B3 147902740 147902817
refGene_gn=PPP1R13B;refGene_tx=XM_019833678.2;refGene_gn=PPP1R13B;refGene_tx=XM_023256108.1;refGene_gn=PPP1R13B;refGene_tx=XM_023256109.1;refGene_gn=PPP1R13B;refGene_tx=XM_023256110.1;refGene_gn=PPP1R13B;refGene_tx=XM_023256111.1;refGene_gn=PPP1R13B;refGene_tx=XM_023256112.1
B3 147908307 147908427
refGene_gn=PPP1R13B;refGene_tx=XM_019833678.2;refGene_gn=PPP1R13B;refGene_tx=XM_023256108.1;refGene_gn=PPP1R13B;refGene_tx=XM_023256109.1;refGene_gn=PPP1R13B;refGene_tx=XM_023256110.1;refGene_gn=PPP1R13B;refGene_tx=XM_023256111.1;refGene_gn=PPP1R13B;refGene_tx=XM_023256112.1
B3 147918785 147918933
refGene_gn=PPP1R13B;refGene_tx=XM_019833678.2;refGene_gn=PPP1R13B;refGene_tx=XM_023256108.1;refGene_gn=PPP1R13B;refGene_tx=XM_023256109.1;refGene_gn=PPP1R13B;refGene_tx=XM_023256110.1;refGene_gn=PPP1R13B;refGene_tx=XM_023256111.1;refGene_gn=PPP1R13B;refGene_tx=XM_023256112.1
B3 147920333 147920336
refGene_gn=PPP1R13B;refGene_tx=XM_023256109.1
B3 147955704 147955713
refGene_gn=PPP1R13B;refGene_tx=XM_023256108.1;refGene_gn=PPP1R13B;refGene_tx=XM_023256112.1
B3 148006224 148006259
refGene_gn=ATP5MPL;refGene_tx=XM_003988015.5
B3 148006899 148006923
refGene_gn=ATP5MPL;refGene_tx=XM_003988015.5
B3 148008065 148008189
refGene_gn=ATP5MPL;refGene_tx=XM_003988015.5
B3 148018068 148018268
refGene_gn=TDRD9;refGene_tx=XM_019833693.2;refGene_gn=TDRD9;refGene_tx=XM_023256113.1;refGene_gn=TDRD9;refGene_tx=XM_023256114.1;refGene_gn=TDRD9;refGene_tx=XM_023256115.1;refGene_gn=TDRD9;refGene_tx=XM_023256116.1;refGene_gn=TDRD9;refGene_tx=XM_023256117.1
B3 148029047 148029345
refGene_gn=RD3L;refGene_tx=XM_003988017.5;refGene_gn=RD3L;refGene_tx=XM_019833703.2;refGene_gn=RD3L;refGene_tx=XM_019833704.2
B3 148029689 148029988
refGene_gn=RD3L;refGene_tx=XM_003988017.5;refGene_gn=RD3L;refGene_tx=XM_019833703.2;refGene_gn=RD3L;refGene_tx=XM_019833704.2
B3 148037173 148037280
refGene_gn=TDRD9;refGene_tx=XM_019833693.2;refGene_gn=TDRD9;refGene_tx=XM_023256113.1;refGene_gn=TDRD9;refGene_tx=XM_023256114.1;refGene_gn=TDRD9;refGene_tx=XM_023256115.1;refGene_gn=TDRD9;refGene_tx=XM_023256116.1;refGene_gn=TDRD9;refGene_tx=XM_023256117.1

B3	148042464	148042562
refGene_gn=TDRD9;refGene_tx=XM_019833693.2;refGene_gn=TDRD9;refGene_tx=XM_023256113.1;refGene_gn=TDRD9;refGene_tx=XM_023256114.1;refGene_gn=TDRD9;refGene_tx=XM_023256115.1;refGene_gn=TDRD9;refGene_tx=XM_023256116.1;refGene_gn=TDRD9;refGene_tx=XM_023256117.1
B3	148043762	148043984
refGene_gn=TDRD9;refGene_tx=XM_019833693.2;refGene_gn=TDRD9;refGene_tx=XM_023256113.1;refGene_gn=TDRD9;refGene_tx=XM_023256114.1;refGene_gn=TDRD9;refGene_tx=XM_023256115.1;refGene_gn=TDRD9;refGene_tx=XM_023256116.1;refGene_gn=TDRD9;refGene_tx=XM_023256117.1
B3	148044939	148045062
refGene_gn=TDRD9;refGene_tx=XM_019833693.2;refGene_gn=TDRD9;refGene_tx=XM_023256113.1;refGene_gn=TDRD9;refGene_tx=XM_023256114.1;refGene_gn=TDRD9;refGene_tx=XM_023256115.1;refGene_gn=TDRD9;refGene_tx=XM_023256116.1;refGene_gn=TDRD9;refGene_tx=XM_023256117.1
B3	148048194	148048275
refGene_gn=TDRD9;refGene_tx=XM_019833693.2;refGene_gn=TDRD9;refGene_tx=XM_023256113.1;refGene_gn=TDRD9;refGene_tx=XM_023256114.1;refGene_gn=TDRD9;refGene_tx=XM_023256115.1;refGene_gn=TDRD9;refGene_tx=XM_023256116.1;refGene_gn=TDRD9;refGene_tx=XM_023256117.1
B3	148051115	148051280
refGene_gn=TDRD9;refGene_tx=XM_019833693.2;refGene_gn=TDRD9;refGene_tx=XM_023256113.1;refGene_gn=TDRD9;refGene_tx=XM_023256114.1;refGene_gn=TDRD9;refGene_tx=XM_023256115.1;refGene_gn=TDRD9;refGene_tx=XM_023256116.1;refGene_gn=TDRD9;refGene_tx=XM_023256117.1;refGene_gn=TDRD9;refGene_tx=XM_023256118.1;refGene_gn=TDRD9;refGene_tx=XM_023256119.1
B3	148056999	148057103
refGene_gn=TDRD9;refGene_tx=XM_019833693.2;refGene_gn=TDRD9;refGene_tx=XM_023256113.1;refGene_gn=TDRD9;refGene_tx=XM_023256114.1;refGene_gn=TDRD9;refGene_tx=XM_023256115.1;refGene_gn=TDRD9;refGene_tx=XM_023256116.1;refGene_gn=TDRD9;refGene_tx=XM_023256117.1;refGene_gn=TDRD9;refGene_tx=XM_023256118.1;refGene_gn=TDRD9;refGene_tx=XM_023256119.1
B3	148065042	148065107
refGene_gn=TDRD9;refGene_tx=XM_019833693.2;refGene_gn=TDRD9;refGene_tx=XM_023256113.1;refGene_gn=TDRD9;refGene_tx=XM_023256114.1;refGene_gn=TDRD9;refGene_tx=XM_023256115.1;refGene_gn=TDRD9;refGene_tx=XM_023256116.1;refGene_gn=TDRD9;refGene_tx=XM_023256117.1;refGene_gn=TDRD9;refGene_tx=XM_023256118.1;refGene_gn=TDRD9;refGene_tx=XM_023256119.1
B3	148067473	148067528
refGene_gn=TDRD9;refGene_tx=XM_019833693.2;refGene_gn=TDRD9;refGene_tx=XM_023256113.1;refGene_gn=TDRD9;refGene_tx=XM_023256114.1;refGene_gn=TDRD9;refGene_tx=XM_023256115.1;refGene_gn=TDRD9;refGene_tx=XM_023256116.1;refGene_gn=TDRD9;refGene_tx=XM_023256117.1;refGene_gn=TDRD9;refGene_tx=XM_023256118.1;refGene_gn=TDRD9;refGene_tx=XM_023256119.1
B3	148067679	148067764
refGene_gn=TDRD9;refGene_tx=XM_019833693.2;refGene_gn=TDRD9;refGene_tx=XM_023256113.1;refGene_gn=TDRD9;refGene_tx=XM_023256114.1;refGene_gn=TDRD9;refGene_tx=XM_023256115.1;refGene_gn=TDRD9;refGene_tx=XM_023256116.1;refGene_gn=TDRD9;refGene_tx=XM_023256117.1;refGene_gn=TDRD9;refGene_tx=XM_023256118.1;refGene_gn=TDRD9;refGene_tx=XM_023256119.1

B3	148069129	148069187
refGene_gn=TDRD9;refGene_tx=XM_019833693.2;refGene_gn=TDRD9;refGene_tx=XM_023256113.1;refGene_gn=TDRD9;refGene_tx=XM_023256114.1;refGene_gn=TDRD9;refGene_tx=XM_023256115.1;refGene_gn=TDRD9;refGene_tx=XM_023256116.1;refGene_gn=TDRD9;refGene_tx=XM_023256117.1;refGene_gn=TDRD9;refGene_tx=XM_023256118.1;refGene_gn=TDRD9;refGene_tx=XM_023256119.1
B3	148072378	148072483
refGene_gn=TDRD9;refGene_tx=XM_019833693.2;refGene_gn=TDRD9;refGene_tx=XM_023256113.1;refGene_gn=TDRD9;refGene_tx=XM_023256114.1;refGene_gn=TDRD9;refGene_tx=XM_023256115.1;refGene_gn=TDRD9;refGene_tx=XM_023256116.1;refGene_gn=TDRD9;refGene_tx=XM_023256118.1;refGene_gn=TDRD9;refGene_tx=XM_023256119.1
B3	148077516	148077614
refGene_gn=TDRD9;refGene_tx=XM_019833693.2;refGene_gn=TDRD9;refGene_tx=XM_023256113.1;refGene_gn=TDRD9;refGene_tx=XM_023256114.1;refGene_gn=TDRD9;refGene_tx=XM_023256115.1;refGene_gn=TDRD9;refGene_tx=XM_023256116.1;refGene_gn=TDRD9;refGene_tx=XM_023256117.1;refGene_gn=TDRD9;refGene_tx=XM_023256118.1;refGene_gn=TDRD9;refGene_tx=XM_023256119.1
B3	148078235	148078367
refGene_gn=TDRD9;refGene_tx=XM_019833693.2;refGene_gn=TDRD9;refGene_tx=XM_023256113.1;refGene_gn=TDRD9;refGene_tx=XM_023256114.1;refGene_gn=TDRD9;refGene_tx=XM_023256115.1;refGene_gn=TDRD9;refGene_tx=XM_023256116.1;refGene_gn=TDRD9;refGene_tx=XM_023256117.1;refGene_gn=TDRD9;refGene_tx=XM_023256118.1;refGene_gn=TDRD9;refGene_tx=XM_023256119.1
B3	148079451	148079617
refGene_gn=TDRD9;refGene_tx=XM_019833693.2;refGene_gn=TDRD9;refGene_tx=XM_023256113.1;refGene_gn=TDRD9;refGene_tx=XM_023256114.1;refGene_gn=TDRD9;refGene_tx=XM_023256115.1;refGene_gn=TDRD9;refGene_tx=XM_023256116.1;refGene_gn=TDRD9;refGene_tx=XM_023256117.1;refGene_gn=TDRD9;refGene_tx=XM_023256118.1;refGene_gn=TDRD9;refGene_tx=XM_023256119.1
B3	148079700	148079764
refGene_gn=TDRD9;refGene_tx=XM_019833693.2;refGene_gn=TDRD9;refGene_tx=XM_023256113.1;refGene_gn=TDRD9;refGene_tx=XM_023256114.1;refGene_gn=TDRD9;refGene_tx=XM_023256115.1;refGene_gn=TDRD9;refGene_tx=XM_023256116.1;refGene_gn=TDRD9;refGene_tx=XM_023256117.1;refGene_gn=TDRD9;refGene_tx=XM_023256118.1;refGene_gn=TDRD9;refGene_tx=XM_023256119.1
B3	148080061	148080125
refGene_gn=TDRD9;refGene_tx=XM_019833693.2;refGene_gn=TDRD9;refGene_tx=XM_023256113.1;refGene_gn=TDRD9;refGene_tx=XM_023256114.1;refGene_gn=TDRD9;refGene_tx=XM_023256115.1;refGene_gn=TDRD9;refGene_tx=XM_023256116.1;refGene_gn=TDRD9;refGene_tx=XM_023256117.1;refGene_gn=TDRD9;refGene_tx=XM_023256118.1;refGene_gn=TDRD9;refGene_tx=XM_023256119.1
B3	148080424	148080469
refGene_gn=TDRD9;refGene_tx=XM_019833693.2;refGene_gn=TDRD9;refGene_tx=XM_023256113.1;refGene_gn=TDRD9;refGene_tx=XM_023256114.1;refGene_gn=TDRD9;refGene_tx=XM_023256115.1;refGene_gn=TDRD9;refGene_tx=XM_023256116.1;refGene_gn=TDRD9;refGene_tx=XM_023256117.1;refGene_gn=TDRD9;refGene_tx=XM_023256118.1;refGene_gn=TDRD9;refGene_tx=XM_023256119.1
B3	148082327	148082381
refGene_gn=TDRD9;refGene_tx=XM_023256113.1;refGene_gn=TDRD9;refGene_tx

=XM_023256114.1;refGene_gn=TDRD9;refGene_tx=XM_023256115.1;refGene_gn=TDRD9;refGene_tx=XM_023256116.1;refGene_gn=TDRD9;refGene_tx=XM_023256117.1;refGene_gn=TDRD9;refGene_tx=XM_023256118.1;refGene_gn=TDRD9;refGene_tx=XM_023256119.1
B3 148098910 148099027
refGene_gn=TDRD9;refGene_tx=XM_019833693.2;refGene_gn=TDRD9;refGene_tx=XM_023256113.1;refGene_gn=TDRD9;refGene_tx=XM_023256114.1;refGene_gn=TDRD9;refGene_tx=XM_023256115.1;refGene_gn=TDRD9;refGene_tx=XM_023256116.1;refGene_gn=TDRD9;refGene_tx=XM_023256117.1;refGene_gn=TDRD9;refGene_tx=XM_023256118.1;refGene_gn=TDRD9;refGene_tx=XM_023256119.1
B3 148100290 148100398
refGene_gn=TDRD9;refGene_tx=XM_019833693.2;refGene_gn=TDRD9;refGene_tx=XM_023256113.1;refGene_gn=TDRD9;refGene_tx=XM_023256114.1;refGene_gn=TDRD9;refGene_tx=XM_023256115.1;refGene_gn=TDRD9;refGene_tx=XM_023256116.1;refGene_gn=TDRD9;refGene_tx=XM_023256117.1;refGene_gn=TDRD9;refGene_tx=XM_023256118.1;refGene_gn=TDRD9;refGene_tx=XM_023256119.1
B3 148102272 148102373
refGene_gn=TDRD9;refGene_tx=XM_019833693.2;refGene_gn=TDRD9;refGene_tx=XM_023256113.1;refGene_gn=TDRD9;refGene_tx=XM_023256114.1;refGene_gn=TDRD9;refGene_tx=XM_023256115.1;refGene_gn=TDRD9;refGene_tx=XM_023256116.1;refGene_gn=TDRD9;refGene_tx=XM_023256117.1;refGene_gn=TDRD9;refGene_tx=XM_023256118.1;refGene_gn=TDRD9;refGene_tx=XM_023256119.1
B3 148105560 148105788
refGene_gn=TDRD9;refGene_tx=XM_019833693.2;refGene_gn=TDRD9;refGene_tx=XM_023256113.1;refGene_gn=TDRD9;refGene_tx=XM_023256114.1;refGene_gn=TDRD9;refGene_tx=XM_023256115.1;refGene_gn=TDRD9;refGene_tx=XM_023256116.1;refGene_gn=TDRD9;refGene_tx=XM_023256117.1;refGene_gn=TDRD9;refGene_tx=XM_023256118.1;refGene_gn=TDRD9;refGene_tx=XM_023256119.1
B3 148107431 148107543
refGene_gn=TDRD9;refGene_tx=XM_019833693.2;refGene_gn=TDRD9;refGene_tx=XM_023256113.1;refGene_gn=TDRD9;refGene_tx=XM_023256114.1;refGene_gn=TDRD9;refGene_tx=XM_023256115.1;refGene_gn=TDRD9;refGene_tx=XM_023256116.1;refGene_gn=TDRD9;refGene_tx=XM_023256117.1;refGene_gn=TDRD9;refGene_tx=XM_023256118.1;refGene_gn=TDRD9;refGene_tx=XM_023256119.1
B3 148108926 148109139
refGene_gn=TDRD9;refGene_tx=XM_019833693.2;refGene_gn=TDRD9;refGene_tx=XM_023256113.1;refGene_gn=TDRD9;refGene_tx=XM_023256114.1;refGene_gn=TDRD9;refGene_tx=XM_023256115.1;refGene_gn=TDRD9;refGene_tx=XM_023256116.1;refGene_gn=TDRD9;refGene_tx=XM_023256117.1;refGene_gn=TDRD9;refGene_tx=XM_023256118.1;refGene_gn=TDRD9;refGene_tx=XM_023256119.1
B3 148109392 148109482
refGene_gn=TDRD9;refGene_tx=XM_019833693.2;refGene_gn=TDRD9;refGene_tx=XM_023256113.1;refGene_gn=TDRD9;refGene_tx=XM_023256114.1;refGene_gn=TDRD9;refGene_tx=XM_023256115.1;refGene_gn=TDRD9;refGene_tx=XM_023256116.1;refGene_gn=TDRD9;refGene_tx=XM_023256117.1;refGene_gn=TDRD9;refGene_tx=XM_023256118.1;refGene_gn=TDRD9;refGene_tx=XM_023256119.1
B3 148110355 148110655
refGene_gn=TDRD9;refGene_tx=XM_019833693.2;refGene_gn=TDRD9;refGene_tx=XM_023256113.1;refGene_gn=TDRD9;refGene_tx=XM_023256114.1;refGene_gn=TDRD9;refGene_tx=XM_023256115.1;refGene_gn=TDRD9;refGene_tx=XM_0232561

16.1;refGene_gn=TDRD9;refGene_tx=XM_023256117.1;refGene_gn=TDRD9;refGene_tx=XM_023256118.1;refGene_gn=TDRD9;refGene_tx=XM_023256119.1
B3 148111564 148111720
refGene_gn=TDRD9;refGene_tx=XM_019833693.2;refGene_gn=TDRD9;refGene_tx=XM_023256113.1;refGene_gn=TDRD9;refGene_tx=XM_023256114.1;refGene_gn=TDRD9;refGene_tx=XM_023256115.1;refGene_gn=TDRD9;refGene_tx=XM_023256116.1;refGene_gn=TDRD9;refGene_tx=XM_023256117.1;refGene_gn=TDRD9;refGene_tx=XM_023256118.1;refGene_gn=TDRD9;refGene_tx=XM_023256119.1
B3 148112851 148112922
refGene_gn=TDRD9;refGene_tx=XM_019833693.2;refGene_gn=TDRD9;refGene_tx=XM_023256113.1;refGene_gn=TDRD9;refGene_tx=XM_023256114.1;refGene_gn=TDRD9;refGene_tx=XM_023256115.1;refGene_gn=TDRD9;refGene_tx=XM_023256116.1;refGene_gn=TDRD9;refGene_tx=XM_023256117.1;refGene_gn=TDRD9;refGene_tx=XM_023256118.1;refGene_gn=TDRD9;refGene_tx=XM_023256119.1
B3 148114363 148114473
refGene_gn=TDRD9;refGene_tx=XM_019833693.2;refGene_gn=TDRD9;refGene_tx=XM_023256113.1;refGene_gn=TDRD9;refGene_tx=XM_023256114.1;refGene_gn=TDRD9;refGene_tx=XM_023256115.1;refGene_gn=TDRD9;refGene_tx=XM_023256116.1;refGene_gn=TDRD9;refGene_tx=XM_023256117.1;refGene_gn=TDRD9;refGene_tx=XM_023256118.1;refGene_gn=TDRD9;refGene_tx=XM_023256119.1
B3 148114962 148115059
refGene_gn=TDRD9;refGene_tx=XM_019833693.2;refGene_gn=TDRD9;refGene_tx=XM_023256113.1;refGene_gn=TDRD9;refGene_tx=XM_023256114.1;refGene_gn=TDRD9;refGene_tx=XM_023256115.1;refGene_gn=TDRD9;refGene_tx=XM_023256116.1;refGene_gn=TDRD9;refGene_tx=XM_023256117.1;refGene_gn=TDRD9;refGene_tx=XM_023256118.1;refGene_gn=TDRD9;refGene_tx=XM_023256119.1
B3 148121231 148121370
refGene_gn=TDRD9;refGene_tx=XM_019833693.2;refGene_gn=TDRD9;refGene_tx=XM_023256113.1;refGene_gn=TDRD9;refGene_tx=XM_023256114.1;refGene_gn=TDRD9;refGene_tx=XM_023256115.1;refGene_gn=TDRD9;refGene_tx=XM_023256116.1;refGene_gn=TDRD9;refGene_tx=XM_023256117.1;refGene_gn=TDRD9;refGene_tx=XM_023256118.1;refGene_gn=TDRD9;refGene_tx=XM_023256119.1
B3 148125074 148125193
refGene_gn=TDRD9;refGene_tx=XM_019833693.2;refGene_gn=TDRD9;refGene_tx=XM_023256113.1;refGene_gn=TDRD9;refGene_tx=XM_023256114.1;refGene_gn=TDRD9;refGene_tx=XM_023256115.1;refGene_gn=TDRD9;refGene_tx=XM_023256116.1;refGene_gn=TDRD9;refGene_tx=XM_023256117.1;refGene_gn=TDRD9;refGene_tx=XM_023256118.1;refGene_gn=TDRD9;refGene_tx=XM_023256119.1
B3 148133591 148133664
refGene_gn=TDRD9;refGene_tx=XM_019833693.2;refGene_gn=TDRD9;refGene_tx=XM_023256113.1;refGene_gn=TDRD9;refGene_tx=XM_023256114.1;refGene_gn=TDRD9;refGene_tx=XM_023256115.1;refGene_gn=TDRD9;refGene_tx=XM_023256116.1;refGene_gn=TDRD9;refGene_tx=XM_023256117.1;refGene_gn=TDRD9;refGene_tx=XM_023256118.1;refGene_gn=TDRD9;refGene_tx=XM_023256119.1
B3 148134995 148135097
refGene_gn=TDRD9;refGene_tx=XM_019833693.2;refGene_gn=TDRD9;refGene_tx=XM_023256113.1;refGene_gn=TDRD9;refGene_tx=XM_023256114.1;refGene_gn=TDRD9;refGene_tx=XM_023256115.1;refGene_gn=TDRD9;refGene_tx=XM_023256116.1;refGene_gn=TDRD9;refGene_tx=XM_023256117.1;refGene_gn=TDRD9;refGene_tx=XM_023256118.1;refGene_gn=TDRD9;refGene_tx=XM_023256119.1

B3	148162780	148162862
refGene_gn=ASPG;refGene_tx=XM_019833705.2;refGene_gn=ASPG;refGene_tx=XM_019833708.2;refGene_gn=ASPG;refGene_tx=XM_023256120.1;refGene_gn=ASPG;refGene_tx=XM_023256121.1
B3	148169211	148169320
refGene_gn=ASPG;refGene_tx=XM_019833705.2;refGene_gn=ASPG;refGene_tx=XM_019833708.2;refGene_gn=ASPG;refGene_tx=XM_023256120.1;refGene_gn=ASPG;refGene_tx=XM_023256121.1
B3	148169995	148170107
refGene_gn=ASPG;refGene_tx=XM_019833705.2;refGene_gn=ASPG;refGene_tx=XM_019833708.2;refGene_gn=ASPG;refGene_tx=XM_023256120.1;refGene_gn=ASPG;refGene_tx=XM_023256121.1
B3	148170769	148170784
refGene_gn=ASPG;refGene_tx=XM_019833707.2
B3	148172439	148172565
refGene_gn=ASPG;refGene_tx=XM_019833705.2;refGene_gn=ASPG;refGene_tx=XM_019833707.2;refGene_gn=ASPG;refGene_tx=XM_019833708.2;refGene_gn=ASPG;refGene_tx=XM_023256120.1;refGene_gn=ASPG;refGene_tx=XM_023256121.1
B3	148172863	148172947
refGene_gn=ASPG;refGene_tx=XM_019833705.2;refGene_gn=ASPG;refGene_tx=XM_019833707.2;refGene_gn=ASPG;refGene_tx=XM_019833708.2;refGene_gn=ASPG;refGene_tx=XM_023256120.1;refGene_gn=ASPG;refGene_tx=XM_023256121.1
B3	148173767	148173894
refGene_gn=ASPG;refGene_tx=XM_019833705.2;refGene_gn=ASPG;refGene_tx=XM_019833707.2;refGene_gn=ASPG;refGene_tx=XM_019833708.2;refGene_gn=ASPG;refGene_tx=XM_023256120.1;refGene_gn=ASPG;refGene_tx=XM_023256121.1
B3	148177773	148177886
refGene_gn=ASPG;refGene_tx=XM_019833705.2;refGene_gn=ASPG;refGene_tx=XM_019833707.2;refGene_gn=ASPG;refGene_tx=XM_019833708.2;refGene_gn=ASPG;refGene_tx=XM_023256120.1;refGene_gn=ASPG;refGene_tx=XM_023256121.1
B3	148180114	148180297
refGene_gn=ASPG;refGene_tx=XM_019833705.2;refGene_gn=ASPG;refGene_tx=XM_019833707.2;refGene_gn=ASPG;refGene_tx=XM_019833708.2;refGene_gn=ASPG;refGene_tx=XM_023256120.1;refGene_gn=ASPG;refGene_tx=XM_023256121.1
B3	148180394	148180508
refGene_gn=ASPG;refGene_tx=XM_019833705.2;refGene_gn=ASPG;refGene_tx=XM_019833707.2;refGene_gn=ASPG;refGene_tx=XM_019833708.2;refGene_gn=ASPG;refGene_tx=XM_023256120.1;refGene_gn=ASPG;refGene_tx=XM_023256121.1
B3	148181063	148181180
refGene_gn=ASPG;refGene_tx=XM_019833705.2;refGene_gn=ASPG;refGene_tx=XM_019833707.2;refGene_gn=ASPG;refGene_tx=XM_019833708.2;refGene_gn=ASPG;refGene_tx=XM_023256120.1;refGene_gn=ASPG;refGene_tx=XM_023256121.1
B3	148182283	148182379
refGene_gn=ASPG;refGene_tx=XM_019833705.2;refGene_gn=ASPG;refGene_tx=XM_019833707.2;refGene_gn=ASPG;refGene_tx=XM_019833708.2;refGene_gn=ASPG;refGene_tx=XM_023256120.1;refGene_gn=ASPG;refGene_tx=XM_023256121.1
B3	148182607	148182771
refGene_gn=ASPG;refGene_tx=XM_019833705.2;refGene_gn=ASPG;refGene_tx=XM_019833707.2;refGene_gn=ASPG;refGene_tx=XM_023256120.1;refGene_gn=ASPG;refGene_tx=XM_023256121.1

B3 148184737 148184824
refGene_gn=ASPG;refGene_tx=XM_019833705.2;refGene_gn=ASPG;refGene_tx=XM_019833707.2;refGene_gn=ASPG;refGene_tx=XM_019833708.2;refGene_gn=ASPG;refGene_tx=XM_023256120.1;refGene_gn=ASPG;refGene_tx=XM_023256121.1
B3 148186718 148186818
refGene_gn=ASPG;refGene_tx=XM_019833705.2;refGene_gn=ASPG;refGene_tx=XM_019833707.2;refGene_gn=ASPG;refGene_tx=XM_019833708.2;refGene_gn=ASPG;refGene_tx=XM_023256120.1;refGene_gn=ASPG;refGene_tx=XM_023256121.1
B3 148187082 148187163
refGene_gn=ASPG;refGene_tx=XM_019833705.2;refGene_gn=ASPG;refGene_tx=XM_019833707.2;refGene_gn=ASPG;refGene_tx=XM_023256120.1;refGene_gn=ASPG;refGene_tx=XM_023256121.1
B3 148187415 148187427
refGene_gn=ASPG;refGene_tx=XM_019833705.2;refGene_gn=ASPG;refGene_tx=XM_019833707.2;refGene_gn=ASPG;refGene_tx=XM_023256120.1;refGene_gn=ASPG;refGene_tx=XM_023256121.1
B3 148209085 148209127
refGene_gn=KIF26A;refGene_tx=XM_023256122.1
B3 148209486 148209732
refGene_gn=KIF26A;refGene_tx=XM_023256122.1
B3 148222152 148222605
refGene_gn=KIF26A;refGene_tx=XM_023256122.1
B3 148229233 148229424
refGene_gn=KIF26A;refGene_tx=XM_023256122.1
B3 148239032 148239222
refGene_gn=KIF26A;refGene_tx=XM_023256122.1
B3 148243374 148243584
refGene_gn=KIF26A;refGene_tx=XM_023256122.1
B3 148244055 148244149
refGene_gn=KIF26A;refGene_tx=XM_023256122.1
B3 148244413 148244676
refGene_gn=KIF26A;refGene_tx=XM_023256122.1
B3 148244782 148244966
refGene_gn=KIF26A;refGene_tx=XM_023256122.1
B3 148245080 148245243
refGene_gn=KIF26A;refGene_tx=XM_023256122.1
B3 148245576 148245739
refGene_gn=KIF26A;refGene_tx=XM_023256122.1
B3 148246381 148246817
refGene_gn=KIF26A;refGene_tx=XM_023256122.1
B3 148246819 148247581
refGene_gn=KIF26A;refGene_tx=XM_023256122.1
B3 148247583 148249233
refGene_gn=KIF26A;refGene_tx=XM_023256122.1
B3 148249898 148250104
refGene_gn=KIF26A;refGene_tx=XM_023256122.1
B3 148250608 148250759
refGene_gn=KIF26A;refGene_tx=XM_023256122.1
B3 148251214 148251396
refGene_gn=KIF26A;refGene_tx=XM_023256122.1

B3 148411725 148411905
refGene_gn=LOC111560972;refGene_tx=XM_023256201.1
B3 148412700 148412856
refGene_gn=LOC111560972;refGene_tx=XM_023256201.1
B3 148412958 148413063
refGene_gn=LOC111560972;refGene_tx=XM_023256201.1
B3 148417507 148417563
refGene_gn=LOC111560972;refGene_tx=XM_023256201.1
B3 148417670 148417741
refGene_gn=LOC111560972;refGene_tx=XM_023256201.1
B3 148418579 148419080
refGene_gn=LOC111560972;refGene_tx=XM_023256201.1
B3 148419688 148419864
refGene_gn=LOC111560972;refGene_tx=XM_023256201.1
B3 148584015 148584126
refGene_gn=CB3H14orf180;refGene_tx=XM_003988046.5;refGene_gn=CB3H14orf180;refGene_tx=XM_019833710.2;refGene_gn=CB3H14orf180;refGene_tx=XM_019833711.2;refGene_gn=CB3H14orf180;refGene_tx=XM_019833712.2;refGene_gn=CB3H14orf180;refGene_tx=XM_019833713.2;refGene_gn=CB3H14orf180;refGene_tx=XM_019833714.2
B3 148585434 148585564
refGene_gn=CB3H14orf180;refGene_tx=XM_003988046.5;refGene_gn=CB3H14orf180;refGene_tx=XM_019833713.2;refGene_gn=CB3H14orf180;refGene_tx=XM_019833714.2
B3 148587091 148587127
refGene_gn=CB3H14orf180;refGene_tx=XM_003988046.5;refGene_gn=CB3H14orf180;refGene_tx=XM_019833710.2;refGene_gn=CB3H14orf180;refGene_tx=XM_019833711.2;refGene_gn=CB3H14orf180;refGene_tx=XM_019833712.2;refGene_gn=CB3H14orf180;refGene_tx=XM_019833713.2;refGene_gn=CB3H14orf180;refGene_tx=XM_019833714.2
B3 148587476 148588197
refGene_gn=CB3H14orf180;refGene_tx=XM_003988046.5;refGene_gn=CB3H14orf180;refGene_tx=XM_019833710.2;refGene_gn=CB3H14orf180;refGene_tx=XM_019833711.2;refGene_gn=CB3H14orf180;refGene_tx=XM_019833712.2;refGene_gn=CB3H14orf180;refGene_tx=XM_019833713.2;refGene_gn=CB3H14orf180;refGene_tx=XM_019833714.2
B3 148591885 148592065
refGene_gn=TMEM179;refGene_tx=XM_019833717.2
B3 148593771 148593850
refGene_gn=TMEM179;refGene_tx=XM_019833717.2
B3 148594416 148594554
refGene_gn=TMEM179;refGene_tx=XM_019833717.2
B3 148601095 148601400
refGene_gn=TMEM179;refGene_tx=XM_019833717.2
B3 148670625 148670712
refGene_gn=INF2;refGene_tx=XM_023256124.1
B3 148672348 148672748
refGene_gn=INF2;refGene_tx=XM_023256124.1
B3 148674306 148674422
refGene_gn=INF2;refGene_tx=XM_023256124.1

B3 148674490 148674650
refGene_gn=INF2;refGene_tx=XM_023256124.1
B3 148675000 148675034
refGene_gn=INF2;refGene_tx=XM_023256124.1
B3 148675628 148675770
refGene_gn=INF2;refGene_tx=XM_023256124.1
B3 148676459 148676601
refGene_gn=INF2;refGene_tx=XM_023256124.1
B3 148676814 148677192
refGene_gn=INF2;refGene_tx=XM_023256124.1
B3 148677266 148677563
refGene_gn=INF2;refGene_tx=XM_023256124.1
B3 148678149 148678295
refGene_gn=INF2;refGene_tx=XM_023256124.1
B3 148678372 148678434
refGene_gn=INF2;refGene_tx=XM_023256124.1
B3 148678864 148678967
refGene_gn=INF2;refGene_tx=XM_023256124.1
B3 148679180 148679266
refGene_gn=INF2;refGene_tx=XM_023256124.1
B3 148679594 148679695
refGene_gn=INF2;refGene_tx=XM_023256124.1
B3 148680279 148680350
refGene_gn=INF2;refGene_tx=XM_023256124.1
B3 148680420 148680528
refGene_gn=INF2;refGene_tx=XM_023256124.1
B3 148680599 148680670
refGene_gn=INF2;refGene_tx=XM_023256124.1
B3 148681349 148681470
refGene_gn=INF2;refGene_tx=XM_023256124.1
B3 148681810 148681975
refGene_gn=INF2;refGene_tx=XM_023256124.1
B3 148682057 148682160
refGene_gn=INF2;refGene_tx=XM_023256124.1
B3 148682331 148682493
refGene_gn=INF2;refGene_tx=XM_023256124.1
B3 148683025 148683628
refGene_gn=INF2;refGene_tx=XM_023256124.1
B3 148684065 148684121
refGene_gn=INF2;refGene_tx=XM_023256124.1
B3 148690823 148691018
refGene_gn=ADSSL1;refGene_tx=XM_023256125.1
B3 148698470 148698573
refGene_gn=ADSSL1;refGene_tx=XM_023256125.1
B3 148700750 148700813
refGene_gn=ADSSL1;refGene_tx=XM_023256125.1
B3 148701537 148701588
refGene_gn=ADSSL1;refGene_tx=XM_023256125.1
B3 148701904 148701971
refGene_gn=ADSSL1;refGene_tx=XM_023256125.1

B3 148702716 148702824
refGene_gn=ADSSL1;refGene_tx=XM_023256125.1
B3 148702925 148703007
refGene_gn=ADSSL1;refGene_tx=XM_019833720.1;refGene_gn=ADSSL1;refGene_tx=XM_023256125.1
B3 148703766 148703893
refGene_gn=ADSSL1;refGene_tx=XM_019833720.1;refGene_gn=ADSSL1;refGene_tx=XM_023256125.1
B3 148704465 148704620
refGene_gn=ADSSL1;refGene_tx=XM_019833720.1;refGene_gn=ADSSL1;refGene_tx=XM_023256125.1
B3 148706294 148706419
refGene_gn=ADSSL1;refGene_tx=XM_019833720.1;refGene_gn=ADSSL1;refGene_tx=XM_023256125.1
B3 148707207 148707305
refGene_gn=ADSSL1;refGene_tx=XM_019833720.1;refGene_gn=ADSSL1;refGene_tx=XM_023256125.1
B3 148708392 148708542
refGene_gn=ADSSL1;refGene_tx=XM_019833720.1;refGene_gn=ADSSL1;refGene_tx=XM_023256125.1
B3 148708623 148708676
refGene_gn=ADSSL1;refGene_tx=XM_019833720.1;refGene_gn=ADSSL1;refGene_tx=XM_023256125.1
B3 148713664 148713782
refGene_gn=SIVA1;refGene_tx=XM_003988018.5;refGene_gn=SIVA1;refGene_tx=XM_006933157.4;refGene_gn=SIVA1;refGene_tx=XM_019833721.2
B3 148715709 148715904
refGene_gn=SIVA1;refGene_tx=XM_003988018.5;refGene_gn=SIVA1;refGene_tx=XM_006933157.4;refGene_gn=SIVA1;refGene_tx=XM_019833721.2
B3 148716624 148716781
refGene_gn=SIVA1;refGene_tx=XM_003988018.5;refGene_gn=SIVA1;refGene_tx=XM_006933157.4
B3 148717035 148717098
refGene_gn=SIVA1;refGene_tx=XM_006933157.4
B3 148718704 148719084
refGene_gn=SIVA1;refGene_tx=XM_003988018.5;refGene_gn=SIVA1;refGene_tx=XM_006933157.4;refGene_gn=SIVA1;refGene_tx=XM_019833721.2
B3 148724334 148724414
refGene_gn=AKT1;refGene_tx=NM_001322435.1
B3 148724660 148724763
refGene_gn=AKT1;refGene_tx=NM_001322435.1
B3 148726251 148726339
refGene_gn=AKT1;refGene_tx=NM_001322435.1
B3 148726882 148727097
refGene_gn=AKT1;refGene_tx=NM_001322435.1
B3 148727182 148727311
refGene_gn=AKT1;refGene_tx=NM_001322435.1
B3 148727381 148727507
refGene_gn=AKT1;refGene_tx=NM_001322435.1
B3 148727699 148727768

refGene_gn=AKT1;refGene_tx=NM_001322435.1
B3	148729209	148729275
refGene_gn=AKT1;refGene_tx=NM_001322435.1
B3	148729344	148729476
refGene_gn=AKT1;refGene_tx=NM_001322435.1
B3	148729600	148729748
refGene_gn=AKT1;refGene_tx=NM_001322435.1
B3	148729914	148730026
refGene_gn=AKT1;refGene_tx=NM_001322435.1
B3	148732597	148732726
refGene_gn=AKT1;refGene_tx=NM_001322435.1
B3	148742488	148742534
refGene_gn=AKT1;refGene_tx=NM_001322435.1
B3	148751402	148752626
refGene_gn=ZBTB42;refGene_tx=XM_023256126.1
B3	148755453	148756179
refGene_gn=LOC111560935;refGene_tx=XM_023256127.1
B3	148756984	148757167
refGene_gn=LOC111560935;refGene_tx=XM_023256127.1
B3	148757396	148757435
refGene_gn=LOC111560935;refGene_tx=XM_023256127.1
B3	148784581	148784731
refGene_gn=CEP170B;refGene_tx=XM_023256129.1
B3	148788512	148788566
refGene_gn=LOC109501063;refGene_tx=XM_023256128.1
B3	148789296	148789719
refGene_gn=LOC109501063;refGene_tx=XM_019833722.2;refGene_gn=LOC109501063;refGene_tx=XM_023256128.1
B3	148790314	148790684
refGene_gn=LOC109501063;refGene_tx=XM_019833722.2;refGene_gn=LOC109501063;refGene_tx=XM_023256128.1
B3	148791288	148791420
refGene_gn=CEP170B;refGene_tx=XM_023256129.1;refGene_gn=LOC109501063;refGene_tx=XM_019833722.2;refGene_gn=LOC109501063;refGene_tx=XM_023256128.1
B3	148793501	148793580
refGene_gn=CEP170B;refGene_tx=XM_023256129.1
B3	148794418	148794477
refGene_gn=CEP170B;refGene_tx=XM_023256129.1
B3	148795537	148795606
refGene_gn=CEP170B;refGene_tx=XM_023256129.1
B3	148795607	148795677
refGene_gn=CEP170B;refGene_tx=XM_023256129.1
B3	148796310	148796415
refGene_gn=CEP170B;refGene_tx=XM_023256129.1
B3	148796538	148797021
refGene_gn=CEP170B;refGene_tx=XM_023256129.1
B3	148797258	148797980
refGene_gn=CEP170B;refGene_tx=XM_023256129.1
B3	148798072	148798243

refGene_gn=CEP170B;refGene_tx=XM_023256129.1
B3 148798757 148798852
refGene_gn=CEP170B;refGene_tx=XM_023256129.1
B3 148799048 148800727
refGene_gn=CEP170B;refGene_tx=XM_023256129.1
B3 148801794 148802026
refGene_gn=CEP170B;refGene_tx=XM_023256129.1
B3 148804186 148804346
refGene_gn=CEP170B;refGene_tx=XM_023256129.1
B3 148804647 148804791
refGene_gn=CEP170B;refGene_tx=XM_023256129.1
B3 148804935 148805024
refGene_gn=CEP170B;refGene_tx=XM_023256129.1
B3 148805371 148805465
refGene_gn=CEP170B;refGene_tx=XM_023256129.1
B3 148805589 148805641
refGene_gn=CEP170B;refGene_tx=XM_023256129.1
B3 148805815 148806051
refGene_gn=CEP170B;refGene_tx=XM_023256129.1
B3 148819709 148819736
refGene_gn=PLD4;refGene_tx=XM_003988052.5
B3 148821500 148821694
refGene_gn=PLD4;refGene_tx=XM_003988052.5
B3 148822066 148822250
refGene_gn=PLD4;refGene_tx=XM_003988052.5
B3 148822391 148822512
refGene_gn=PLD4;refGene_tx=XM_003988052.5
B3 148822897 148823025
refGene_gn=PLD4;refGene_tx=XM_003988052.5
B3 148823590 148823791
refGene_gn=PLD4;refGene_tx=XM_003988052.5
B3 148824350 148824490
refGene_gn=PLD4;refGene_tx=XM_003988052.5
B3 148824635 148824801
refGene_gn=PLD4;refGene_tx=XM_003988052.5
B3 148824889 148824986
refGene_gn=PLD4;refGene_tx=XM_003988052.5
B3 148825488 148825691
refGene_gn=PLD4;refGene_tx=XM_003988052.5
B3 148829140 148834483
refGene_gn=AHNAK2;refGene_tx=XM_023256202.1
B3 148835152 148836130
refGene_gn=AHNAK2;refGene_tx=XM_023256202.1
B3 148842058 148845242
refGene_gn=AHNAK2;refGene_tx=XM_023256202.1
B3 148845385 148845570
refGene_gn=AHNAK2;refGene_tx=XM_023256202.1
B3 148845843 148845994
refGene_gn=AHNAK2;refGene_tx=XM_023256202.1
B3 148846798 148846909

refGene_gn=AHNAK2;refGene_tx=XM_023256202.1
B3 148847515 148847614
refGene_gn=AHNAK2;refGene_tx=XM_023256202.1
B3 148847708 148847767
refGene_gn=AHNAK2;refGene_tx=XM_023256202.1
B3 148860469 148860524
refGene_gn=AHNAK2;refGene_tx=XM_023256202.1
B3 148866846 148867296
refGene_gn=CLBA1;refGene_tx=XM_006933159.4
B3 148868860 148869006
refGene_gn=CLBA1;refGene_tx=XM_006933159.4
B3 148870570 148870700
refGene_gn=CLBA1;refGene_tx=XM_006933159.4
B3 148871803 148871917
refGene_gn=CLBA1;refGene_tx=XM_006933159.4
B3 148872909 148873074
refGene_gn=CLBA1;refGene_tx=XM_006933159.4
B3 148880515 148881199
refGene_gn=CDCA4;refGene_tx=XM_011283492.3;refGene_gn=CDCA4;refGene_tx=XM_023256130.1
B3 148902992 148904116
refGene_gn=GPR132;refGene_tx=XM_019833728.2;refGene_gn=GPR132;refGene_tx=XM_023256131.1
B3 148909499 148909506
refGene_gn=GPR132;refGene_tx=XM_019833728.2;refGene_gn=GPR132;refGene_tx=XM_023256131.1
B3 148957925 148958410
refGene_gn=JAG2;refGene_tx=XM_023256132.1;refGene_gn=JAG2;refGene_tx=XM_023256133.1;refGene_gn=JAG2;refGene_tx=XM_023256134.1
B3 148958714 148958871
refGene_gn=JAG2;refGene_tx=XM_023256132.1;refGene_gn=JAG2;refGene_tx=XM_023256133.1;refGene_gn=JAG2;refGene_tx=XM_023256134.1
B3 148960350 148960482
refGene_gn=JAG2;refGene_tx=XM_023256132.1;refGene_gn=JAG2;refGene_tx=XM_023256133.1;refGene_gn=JAG2;refGene_tx=XM_023256134.1
B3 148960949 148961198
refGene_gn=JAG2;refGene_tx=XM_023256132.1;refGene_gn=JAG2;refGene_tx=XM_023256133.1;refGene_gn=JAG2;refGene_tx=XM_023256134.1
B3 148961657 148961773
refGene_gn=JAG2;refGene_tx=XM_023256132.1;refGene_gn=JAG2;refGene_tx=XM_023256133.1;refGene_gn=JAG2;refGene_tx=XM_023256134.1
B3 148961879 148961993
refGene_gn=JAG2;refGene_tx=XM_023256132.1;refGene_gn=JAG2;refGene_tx=XM_023256133.1;refGene_gn=JAG2;refGene_tx=XM_023256134.1
B3 148962492 148962578
refGene_gn=JAG2;refGene_tx=XM_023256132.1;refGene_gn=JAG2;refGene_tx=XM_023256133.1;refGene_gn=JAG2;refGene_tx=XM_023256134.1
B3 148962664 148962692
refGene_gn=JAG2;refGene_tx=XM_023256132.1;refGene_gn=JAG2;refGene_tx=XM_023256133.1;refGene_gn=JAG2;refGene_tx=XM_023256134.1

B3	148963030	148963147
refGene_gn=JAG2;refGene_tx=XM_023256132.1;refGene_gn=JAG2;refGene_tx=XM_023256133.1;refGene_gn=JAG2;refGene_tx=XM_023256134.1
B3	148963323	148963437
refGene_gn=JAG2;refGene_tx=XM_023256132.1;refGene_gn=JAG2;refGene_tx=XM_023256133.1;refGene_gn=JAG2;refGene_tx=XM_023256134.1
B3	148963519	148963633
refGene_gn=JAG2;refGene_tx=XM_023256132.1;refGene_gn=JAG2;refGene_tx=XM_023256133.1;refGene_gn=JAG2;refGene_tx=XM_023256134.1
B3	148963851	148963965
refGene_gn=JAG2;refGene_tx=XM_023256132.1;refGene_gn=JAG2;refGene_tx=XM_023256133.1;refGene_gn=JAG2;refGene_tx=XM_023256134.1
B3	148964046	148964199
refGene_gn=JAG2;refGene_tx=XM_023256132.1;refGene_gn=JAG2;refGene_tx=XM_023256133.1;refGene_gn=JAG2;refGene_tx=XM_023256134.1
B3	148964313	148964464
refGene_gn=JAG2;refGene_tx=XM_023256132.1;refGene_gn=JAG2;refGene_tx=XM_023256133.1;refGene_gn=JAG2;refGene_tx=XM_023256134.1
B3	148965310	148965481
refGene_gn=JAG2;refGene_tx=XM_023256132.1;refGene_gn=JAG2;refGene_tx=XM_023256133.1;refGene_gn=JAG2;refGene_tx=XM_023256134.1
B3	148965593	148965640
refGene_gn=JAG2;refGene_tx=XM_023256132.1;refGene_gn=JAG2;refGene_tx=XM_023256133.1;refGene_gn=JAG2;refGene_tx=XM_023256134.1
B3	148965721	148965838
refGene_gn=JAG2;refGene_tx=XM_023256134.1
B3	148965979	148966093
refGene_gn=JAG2;refGene_tx=XM_023256132.1;refGene_gn=JAG2;refGene_tx=XM_023256133.1;refGene_gn=JAG2;refGene_tx=XM_023256134.1
B3	148966315	148966429
refGene_gn=JAG2;refGene_tx=XM_023256132.1;refGene_gn=JAG2;refGene_tx=XM_023256133.1;refGene_gn=JAG2;refGene_tx=XM_023256134.1
B3	148966619	148966739
refGene_gn=JAG2;refGene_tx=XM_023256132.1;refGene_gn=JAG2;refGene_tx=XM_023256133.1;refGene_gn=JAG2;refGene_tx=XM_023256134.1
B3	148966826	148966957
refGene_gn=JAG2;refGene_tx=XM_023256132.1;refGene_gn=JAG2;refGene_tx=XM_023256133.1;refGene_gn=JAG2;refGene_tx=XM_023256134.1
B3	148970118	148970179
refGene_gn=JAG2;refGene_tx=XM_023256132.1;refGene_gn=JAG2;refGene_tx=XM_023256133.1;refGene_gn=JAG2;refGene_tx=XM_023256134.1
B3	148970341	148970593
refGene_gn=JAG2;refGene_tx=XM_023256132.1;refGene_gn=JAG2;refGene_tx=XM_023256133.1;refGene_gn=JAG2;refGene_tx=XM_023256134.1
B3	148971986	148972044
refGene_gn=JAG2;refGene_tx=XM_023256132.1;refGene_gn=JAG2;refGene_tx=XM_023256133.1;refGene_gn=JAG2;refGene_tx=XM_023256134.1
B3	148979093	148979444
refGene_gn=JAG2;refGene_tx=XM_023256132.1;refGene_gn=JAG2;refGene_tx=XM_023256133.1;refGene_gn=JAG2;refGene_tx=XM_023256134.1

B3 148979646 148979712
refGene_gn=JAG2;refGene_tx=XM_023256132.1;refGene_gn=JAG2;refGene_tx=XM_023256133.1;refGene_gn=JAG2;refGene_tx=XM_023256134.1
B3 148980560 148980747
refGene_gn=LOC109501120;refGene_tx=XM_023256203.1
B3 148982185 148982310
refGene_gn=LOC109501120;refGene_tx=XM_023256203.1
B3 148982928 148983247
refGene_gn=LOC109501120;refGene_tx=XM_023256203.1
B3 148983329 148983428
refGene_gn=LOC109501120;refGene_tx=XM_023256203.1
B3 148983600 148983840
refGene_gn=LOC109501120;refGene_tx=XM_023256203.1
B3 148984089 148984301
refGene_gn=LOC109501120;refGene_tx=XM_023256203.1
B3 148985007 148985248
refGene_gn=NUDT14;refGene_tx=XM_023256135.1;refGene_gn=NUDT14;refGene_tx=XM_023256136.1;refGene_gn=NUDT14;refGene_tx=XM_023256137.1
B3 148987978 148988216
refGene_gn=NUDT14;refGene_tx=XM_023256135.1;refGene_gn=NUDT14;refGene_tx=XM_023256136.1;refGene_gn=NUDT14;refGene_tx=XM_023256137.1
B3 148988432 148988497
refGene_gn=NUDT14;refGene_tx=XM_023256135.1;refGene_gn=NUDT14;refGene_tx=XM_023256136.1;refGene_gn=NUDT14;refGene_tx=XM_023256137.1
B3 148989010 148989054
refGene_gn=NUDT14;refGene_tx=XM_023256135.1;refGene_gn=NUDT14;refGene_tx=XM_023256136.1;refGene_gn=NUDT14;refGene_tx=XM_023256137.1
B3 148989301 148989358
refGene_gn=NUDT14;refGene_tx=XM_023256135.1;refGene_gn=NUDT14;refGene_tx=XM_023256136.1
B3 148992034 148992115
refGene_gn=NUDT14;refGene_tx=XM_023256135.1;refGene_gn=NUDT14;refGene_tx=XM_023256136.1;refGene_gn=NUDT14;refGene_tx=XM_023256137.1
B3 149015030 149015068
refGene_gn=BRF1;refGene_tx=XM_023256138.1;refGene_gn=BRF1;refGene_tx=XM_023256139.1;refGene_gn=BRF1;refGene_tx=XM_023256140.1
B3 149015547 149015719
refGene_gn=BRF1;refGene_tx=XM_023256138.1;refGene_gn=BRF1;refGene_tx=XM_023256139.1;refGene_gn=BRF1;refGene_tx=XM_023256140.1
B3 149016066 149016118
refGene_gn=BRF1;refGene_tx=XM_023256138.1;refGene_gn=BRF1;refGene_tx=XM_023256139.1;refGene_gn=BRF1;refGene_tx=XM_023256140.1
B3 149018888 149019145
refGene_gn=BRF1;refGene_tx=XM_023256138.1;refGene_gn=BRF1;refGene_tx=XM_023256139.1;refGene_gn=BRF1;refGene_tx=XM_023256140.1
B3 149019988 149020044
refGene_gn=BRF1;refGene_tx=XM_023256138.1;refGene_gn=BRF1;refGene_tx=XM_023256139.1;refGene_gn=BRF1;refGene_tx=XM_023256140.1
B3 149020123 149020205
refGene_gn=BRF1;refGene_tx=XM_023256138.1;refGene_gn=BRF1;refGene_tx=X

M_023256139.1;refGene_gn=BRF1;refGene_tx=XM_023256140.1
B3 149020576 149020638
refGene_gn=BRF1;refGene_tx=XM_023256138.1;refGene_gn=BRF1;refGene_tx=XM_023256139.1;refGene_gn=BRF1;refGene_tx=XM_023256140.1
B3 149021894 149022164
refGene_gn=BRF1;refGene_tx=XM_023256138.1;refGene_gn=BRF1;refGene_tx=XM_023256139.1;refGene_gn=BRF1;refGene_tx=XM_023256140.1
B3 149025402 149025495
refGene_gn=BRF1;refGene_tx=XM_023256138.1;refGene_gn=BRF1;refGene_tx=XM_023256139.1;refGene_gn=BRF1;refGene_tx=XM_023256140.1
B3 149025586 149025647
refGene_gn=BRF1;refGene_tx=XM_023256138.1;refGene_gn=BRF1;refGene_tx=XM_023256139.1;refGene_gn=BRF1;refGene_tx=XM_023256140.1
B3 149026089 149026216
refGene_gn=BRF1;refGene_tx=XM_023256138.1;refGene_gn=BRF1;refGene_tx=XM_023256139.1;refGene_gn=BRF1;refGene_tx=XM_023256140.1
B3 149027756 149027850
refGene_gn=BRF1;refGene_tx=XM_023256138.1;refGene_gn=BRF1;refGene_tx=XM_023256139.1;refGene_gn=BRF1;refGene_tx=XM_023256140.1
B3 149028788 149028938
refGene_gn=BRF1;refGene_tx=XM_023256138.1;refGene_gn=BRF1;refGene_tx=XM_023256139.1
B3 149031830 149032207
refGene_gn=BTBD6;refGene_tx=XM_006933161.3
B3 149032298 149032389
refGene_gn=BTBD6;refGene_tx=XM_006933161.3
B3 149032528 149032647
refGene_gn=BTBD6;refGene_tx=XM_006933161.3
B3 149032774 149033807
refGene_gn=BTBD6;refGene_tx=XM_006933161.3
B3 149035359 149035432
refGene_gn=BRF1;refGene_tx=XM_023256138.1;refGene_gn=BRF1;refGene_tx=XM_023256139.1;refGene_gn=BRF1;refGene_tx=XM_023256140.1
B3 149038013 149038045
refGene_gn=BRF1;refGene_tx=XM_023256138.1;refGene_gn=BRF1;refGene_tx=XM_023256139.1;refGene_gn=BRF1;refGene_tx=XM_023256140.1
B3 149049010 149049184
refGene_gn=BRF1;refGene_tx=XM_023256138.1;refGene_gn=BRF1;refGene_tx=XM_023256139.1;refGene_gn=BRF1;refGene_tx=XM_023256140.1
B3 149061120 149061201
refGene_gn=BRF1;refGene_tx=XM_023256138.1;refGene_gn=BRF1;refGene_tx=XM_023256139.1;refGene_gn=BRF1;refGene_tx=XM_023256140.1
B3 149068040 149068224
refGene_gn=BRF1;refGene_tx=XM_023256138.1;refGene_gn=BRF1;refGene_tx=XM_023256139.1;refGene_gn=BRF1;refGene_tx=XM_023256140.1
B3 149091775 149091894
refGene_gn=PACS2;refGene_tx=XM_023256141.1;refGene_gn=PACS2;refGene_tx=XM_023256142.1;refGene_gn=PACS2;refGene_tx=XM_023256143.1;refGene_gn=PACS2;refGene_tx=XM_023256144.1;refGene_gn=PACS2;refGene_tx=XM_023256145.1;refGene_gn=PACS2;refGene_tx=XM_023256146.1;refGene_gn=PACS2;refGe

ne_tx=XM_023256147.1
B3 149108099 149109080
refGene_gn=LOC109501066;refGene_tx=XM_019833734.2
B3 149120507 149120595
refGene_gn=PACS2;refGene_tx=XM_023256141.1;refGene_gn=PACS2;refGene_tx=XM_023256142.1;refGene_gn=PACS2;refGene_tx=XM_023256143.1;refGene_gn=PACS2;refGene_tx=XM_023256144.1;refGene_gn=PACS2;refGene_tx=XM_023256145.1;refGene_gn=PACS2;refGene_tx=XM_023256146.1;refGene_gn=PACS2;refGene_tx=XM_023256147.1
B3 149122864 149122954
refGene_gn=PACS2;refGene_tx=XM_023256141.1;refGene_gn=PACS2;refGene_tx=XM_023256142.1;refGene_gn=PACS2;refGene_tx=XM_023256143.1;refGene_gn=PACS2;refGene_tx=XM_023256144.1;refGene_gn=PACS2;refGene_tx=XM_023256145.1;refGene_gn=PACS2;refGene_tx=XM_023256146.1;refGene_gn=PACS2;refGene_tx=XM_023256147.1
B3 149124003 149124129
refGene_gn=PACS2;refGene_tx=XM_023256141.1;refGene_gn=PACS2;refGene_tx=XM_023256142.1;refGene_gn=PACS2;refGene_tx=XM_023256143.1;refGene_gn=PACS2;refGene_tx=XM_023256144.1;refGene_gn=PACS2;refGene_tx=XM_023256145.1;refGene_gn=PACS2;refGene_tx=XM_023256146.1;refGene_gn=PACS2;refGene_tx=XM_023256147.1
B3 149125282 149125445
refGene_gn=PACS2;refGene_tx=XM_023256141.1;refGene_gn=PACS2;refGene_tx=XM_023256142.1;refGene_gn=PACS2;refGene_tx=XM_023256143.1;refGene_gn=PACS2;refGene_tx=XM_023256144.1;refGene_gn=PACS2;refGene_tx=XM_023256145.1;refGene_gn=PACS2;refGene_tx=XM_023256146.1;refGene_gn=PACS2;refGene_tx=XM_023256147.1
B3 149126136 149126210
refGene_gn=PACS2;refGene_tx=XM_023256141.1;refGene_gn=PACS2;refGene_tx=XM_023256142.1;refGene_gn=PACS2;refGene_tx=XM_023256143.1;refGene_gn=PACS2;refGene_tx=XM_023256144.1;refGene_gn=PACS2;refGene_tx=XM_023256145.1;refGene_gn=PACS2;refGene_tx=XM_023256146.1;refGene_gn=PACS2;refGene_tx=XM_023256147.1
B3 149126398 149126479
refGene_gn=PACS2;refGene_tx=XM_023256141.1;refGene_gn=PACS2;refGene_tx=XM_023256142.1;refGene_gn=PACS2;refGene_tx=XM_023256143.1;refGene_gn=PACS2;refGene_tx=XM_023256144.1;refGene_gn=PACS2;refGene_tx=XM_023256145.1;refGene_gn=PACS2;refGene_tx=XM_023256146.1;refGene_gn=PACS2;refGene_tx=XM_023256147.1
B3 149127586 149127646
refGene_gn=PACS2;refGene_tx=XM_023256141.1;refGene_gn=PACS2;refGene_tx=XM_023256142.1;refGene_gn=PACS2;refGene_tx=XM_023256143.1;refGene_gn=PACS2;refGene_tx=XM_023256144.1;refGene_gn=PACS2;refGene_tx=XM_023256145.1;refGene_gn=PACS2;refGene_tx=XM_023256146.1;refGene_gn=PACS2;refGene_tx=XM_023256147.1
B3 149130011 149130169
refGene_gn=PACS2;refGene_tx=XM_023256141.1;refGene_gn=PACS2;refGene_tx=XM_023256142.1;refGene_gn=PACS2;refGene_tx=XM_023256143.1;refGene_gn=PACS2;refGene_tx=XM_023256144.1;refGene_gn=PACS2;refGene_tx=XM_023256145.1;refGene_gn=PACS2;refGene_tx=XM_023256146.1;refGene_gn=PACS2;refGe

ne_tx=XM_023256147.1
B3 149131792 149131883
refGene_gn=PACS2;refGene_tx=XM_023256141.1;refGene_gn=PACS2;refGene_tx=XM_023256142.1;refGene_gn=PACS2;refGene_tx=XM_023256143.1;refGene_gn=PACS2;refGene_tx=XM_023256144.1;refGene_gn=PACS2;refGene_tx=XM_023256145.1;refGene_gn=PACS2;refGene_tx=XM_023256146.1;refGene_gn=PACS2;refGene_tx=XM_023256147.1
B3 149132216 149132291
refGene_gn=PACS2;refGene_tx=XM_023256141.1;refGene_gn=PACS2;refGene_tx=XM_023256142.1;refGene_gn=PACS2;refGene_tx=XM_023256143.1;refGene_gn=PACS2;refGene_tx=XM_023256144.1;refGene_gn=PACS2;refGene_tx=XM_023256145.1;refGene_gn=PACS2;refGene_tx=XM_023256146.1;refGene_gn=PACS2;refGene_tx=XM_023256147.1
B3 149133234 149133377
refGene_gn=PACS2;refGene_tx=XM_023256141.1;refGene_gn=PACS2;refGene_tx=XM_023256142.1;refGene_gn=PACS2;refGene_tx=XM_023256143.1;refGene_gn=PACS2;refGene_tx=XM_023256144.1;refGene_gn=PACS2;refGene_tx=XM_023256145.1;refGene_gn=PACS2;refGene_tx=XM_023256146.1;refGene_gn=PACS2;refGene_tx=XM_023256147.1
B3 149134225 149134358
refGene_gn=PACS2;refGene_tx=XM_023256141.1;refGene_gn=PACS2;refGene_tx=XM_023256142.1;refGene_gn=PACS2;refGene_tx=XM_023256143.1;refGene_gn=PACS2;refGene_tx=XM_023256144.1;refGene_gn=PACS2;refGene_tx=XM_023256145.1;refGene_gn=PACS2;refGene_tx=XM_023256146.1;refGene_gn=PACS2;refGene_tx=XM_023256147.1
B3 149134609 149134714
refGene_gn=PACS2;refGene_tx=XM_023256141.1;refGene_gn=PACS2;refGene_tx=XM_023256142.1;refGene_gn=PACS2;refGene_tx=XM_023256143.1;refGene_gn=PACS2;refGene_tx=XM_023256144.1;refGene_gn=PACS2;refGene_tx=XM_023256145.1;refGene_gn=PACS2;refGene_tx=XM_023256146.1;refGene_gn=PACS2;refGene_tx=XM_023256147.1
B3 149134859 149134966
refGene_gn=PACS2;refGene_tx=XM_023256141.1;refGene_gn=PACS2;refGene_tx=XM_023256142.1;refGene_gn=PACS2;refGene_tx=XM_023256143.1;refGene_gn=PACS2;refGene_tx=XM_023256144.1;refGene_gn=PACS2;refGene_tx=XM_023256145.1;refGene_gn=PACS2;refGene_tx=XM_023256146.1;refGene_gn=PACS2;refGene_tx=XM_023256147.1
B3 149135414 149135569
refGene_gn=PACS2;refGene_tx=XM_023256141.1;refGene_gn=PACS2;refGene_tx=XM_023256142.1;refGene_gn=PACS2;refGene_tx=XM_023256143.1;refGene_gn=PACS2;refGene_tx=XM_023256144.1;refGene_gn=PACS2;refGene_tx=XM_023256145.1;refGene_gn=PACS2;refGene_tx=XM_023256146.1;refGene_gn=PACS2;refGene_tx=XM_023256147.1
B3 149135999 149136122
refGene_gn=PACS2;refGene_tx=XM_023256141.1;refGene_gn=PACS2;refGene_tx=XM_023256142.1;refGene_gn=PACS2;refGene_tx=XM_023256143.1;refGene_gn=PACS2;refGene_tx=XM_023256144.1;refGene_gn=PACS2;refGene_tx=XM_023256145.1;refGene_gn=PACS2;refGene_tx=XM_023256146.1;refGene_gn=PACS2;refGene_tx=XM_023256147.1
B3 149136425 149136534

refGene_gn=PACS2;refGene_tx=XM_023256141.1;refGene_gn=PACS2;refGene_tx=XM_023256142.1;refGene_gn=PACS2;refGene_tx=XM_023256143.1;refGene_gn=PACS2;refGene_tx=XM_023256144.1;refGene_gn=PACS2;refGene_tx=XM_023256145.1;refGene_gn=PACS2;refGene_tx=XM_023256146.1;refGene_gn=PACS2;refGene_tx=XM_023256147.1
B3 149137244 149137277
refGene_gn=PACS2;refGene_tx=XM_023256141.1;refGene_gn=PACS2;refGene_tx=XM_023256142.1;refGene_gn=PACS2;refGene_tx=XM_023256143.1;refGene_gn=PACS2;refGene_tx=XM_023256147.1
B3 149140521 149140564
refGene_gn=PACS2;refGene_tx=XM_023256141.1;refGene_gn=PACS2;refGene_tx=XM_023256142.1;refGene_gn=PACS2;refGene_tx=XM_023256143.1;refGene_gn=PACS2;refGene_tx=XM_023256144.1;refGene_gn=PACS2;refGene_tx=XM_023256145.1;refGene_gn=PACS2;refGene_tx=XM_023256146.1;refGene_gn=PACS2;refGene_tx=XM_023256147.1
B3 149141691 149141734
refGene_gn=PACS2;refGene_tx=XM_023256141.1;refGene_gn=PACS2;refGene_tx=XM_023256142.1;refGene_gn=PACS2;refGene_tx=XM_023256143.1;refGene_gn=PACS2;refGene_tx=XM_023256144.1;refGene_gn=PACS2;refGene_tx=XM_023256145.1;refGene_gn=PACS2;refGene_tx=XM_023256146.1;refGene_gn=PACS2;refGene_tx=XM_023256147.1
B3 149141916 149142052
refGene_gn=PACS2;refGene_tx=XM_023256141.1;refGene_gn=PACS2;refGene_tx=XM_023256142.1;refGene_gn=PACS2;refGene_tx=XM_023256143.1;refGene_gn=PACS2;refGene_tx=XM_023256144.1;refGene_gn=PACS2;refGene_tx=XM_023256145.1;refGene_gn=PACS2;refGene_tx=XM_023256146.1;refGene_gn=PACS2;refGene_tx=XM_023256147.1
B3 149142879 149143106
refGene_gn=PACS2;refGene_tx=XM_023256141.1;refGene_gn=PACS2;refGene_tx=XM_023256142.1;refGene_gn=PACS2;refGene_tx=XM_023256143.1;refGene_gn=PACS2;refGene_tx=XM_023256144.1;refGene_gn=PACS2;refGene_tx=XM_023256145.1;refGene_gn=PACS2;refGene_tx=XM_023256146.1;refGene_gn=PACS2;refGene_tx=XM_023256147.1
B3 149143310 149143424
refGene_gn=PACS2;refGene_tx=XM_023256141.1;refGene_gn=PACS2;refGene_tx=XM_023256142.1;refGene_gn=PACS2;refGene_tx=XM_023256143.1;refGene_gn=PACS2;refGene_tx=XM_023256144.1;refGene_gn=PACS2;refGene_tx=XM_023256145.1;refGene_gn=PACS2;refGene_tx=XM_023256146.1
B3 149144088 149144207
refGene_gn=PACS2;refGene_tx=XM_023256141.1;refGene_gn=PACS2;refGene_tx=XM_023256142.1;refGene_gn=PACS2;refGene_tx=XM_023256143.1;refGene_gn=PACS2;refGene_tx=XM_023256144.1;refGene_gn=PACS2;refGene_tx=XM_023256145.1;refGene_gn=PACS2;refGene_tx=XM_023256146.1
B3 149148468 149148633
refGene_gn=TEX22;refGene_tx=XM_023256148.1
B3 149160605 149160734
refGene_gn=TEX22;refGene_tx=XM_023256148.1
B3 149160923 149161079
refGene_gn=TEX22;refGene_tx=XM_023256148.1
B3 149162825 149162993

refGene_gn=TEX22;refGene_tx=XM_023256148.1
B3 149167669 149167697
refGene_gn=MTA1;refGene_tx=XM_023256149.1;refGene_gn=MTA1;refGene_tx=XM_023256150.1;refGene_gn=MTA1;refGene_tx=XM_023256151.1
B3 149180583 149180651
refGene_gn=MTA1;refGene_tx=XM_023256149.1;refGene_gn=MTA1;refGene_tx=XM_023256150.1;refGene_gn=MTA1;refGene_tx=XM_023256151.1
B3 149184810 149184904
refGene_gn=MTA1;refGene_tx=XM_023256149.1;refGene_gn=MTA1;refGene_tx=XM_023256150.1;refGene_gn=MTA1;refGene_tx=XM_023256151.1
B3 149188533 149188584
refGene_gn=MTA1;refGene_tx=XM_023256149.1;refGene_gn=MTA1;refGene_tx=XM_023256150.1;refGene_gn=MTA1;refGene_tx=XM_023256151.1
B3 149188685 149188812
refGene_gn=MTA1;refGene_tx=XM_023256149.1;refGene_gn=MTA1;refGene_tx=XM_023256150.1;refGene_gn=MTA1;refGene_tx=XM_023256151.1
B3 149188879 149188943
refGene_gn=MTA1;refGene_tx=XM_023256149.1;refGene_gn=MTA1;refGene_tx=XM_023256150.1;refGene_gn=MTA1;refGene_tx=XM_023256151.1
B3 149190254 149190372
refGene_gn=MTA1;refGene_tx=XM_023256149.1;refGene_gn=MTA1;refGene_tx=XM_023256150.1;refGene_gn=MTA1;refGene_tx=XM_023256151.1
B3 149192987 149193090
refGene_gn=MTA1;refGene_tx=XM_023256149.1;refGene_gn=MTA1;refGene_tx=XM_023256150.1;refGene_gn=MTA1;refGene_tx=XM_023256151.1
B3 149193862 149193962
refGene_gn=MTA1;refGene_tx=XM_023256149.1;refGene_gn=MTA1;refGene_tx=XM_023256150.1;refGene_gn=MTA1;refGene_tx=XM_023256151.1
B3 149194273 149194462
refGene_gn=MTA1;refGene_tx=XM_023256149.1;refGene_gn=MTA1;refGene_tx=XM_023256150.1;refGene_gn=MTA1;refGene_tx=XM_023256151.1
B3 149195512 149195587
refGene_gn=MTA1;refGene_tx=XM_023256149.1;refGene_gn=MTA1;refGene_tx=XM_023256150.1;refGene_gn=MTA1;refGene_tx=XM_023256151.1
B3 149195682 149195741
refGene_gn=MTA1;refGene_tx=XM_023256149.1;refGene_gn=MTA1;refGene_tx=XM_023256150.1;refGene_gn=MTA1;refGene_tx=XM_023256151.1
B3 149196055 149196171
refGene_gn=MTA1;refGene_tx=XM_023256149.1;refGene_gn=MTA1;refGene_tx=XM_023256150.1;refGene_gn=MTA1;refGene_tx=XM_023256151.1
B3 149196397 149196549
refGene_gn=MTA1;refGene_tx=XM_023256149.1;refGene_gn=MTA1;refGene_tx=XM_023256150.1;refGene_gn=MTA1;refGene_tx=XM_023256151.1
B3 149196641 149196831
refGene_gn=MTA1;refGene_tx=XM_023256149.1;refGene_gn=MTA1;refGene_tx=XM_023256150.1;refGene_gn=MTA1;refGene_tx=XM_023256151.1
B3 149196981 149197071
refGene_gn=MTA1;refGene_tx=XM_023256149.1;refGene_gn=MTA1;refGene_tx=XM_023256150.1;refGene_gn=MTA1;refGene_tx=XM_023256151.1
B3 149198061 149198214

refGene_gn=MTA1;refGene_tx=XM_023256149.1;refGene_gn=MTA1;refGene_tx=XM_023256150.1;refGene_gn=MTA1;refGene_tx=XM_023256151.1
B3 149198340 149198376
refGene_gn=MTA1;refGene_tx=XM_023256149.1;refGene_gn=MTA1;refGene_tx=XM_023256150.1
B3 149200080 149200092
refGene_gn=MTA1;refGene_tx=XM_023256149.1
B3 149200983 149201015
refGene_gn=MTA1;refGene_tx=XM_023256149.1;refGene_gn=MTA1;refGene_tx=XM_023256150.1;refGene_gn=MTA1;refGene_tx=XM_023256151.1
B3 149201368 149201520
refGene_gn=MTA1;refGene_tx=XM_023256149.1;refGene_gn=MTA1;refGene_tx=XM_023256150.1;refGene_gn=MTA1;refGene_tx=XM_023256151.1
B3 149201672 149201823
refGene_gn=MTA1;refGene_tx=XM_023256149.1;refGene_gn=MTA1;refGene_tx=XM_023256150.1;refGene_gn=MTA1;refGene_tx=XM_023256151.1
B3 149205728 149205771
refGene_gn=CRIP2;refGene_tx=XM_023256153.1
B3 149206195 149206388
refGene_gn=CRIP2;refGene_tx=XM_023256152.1
B3 149208539 149208634
refGene_gn=CRIP2;refGene_tx=XM_023256152.1;refGene_gn=CRIP2;refGene_tx=XM_023256153.1
B3 149208724 149208782
refGene_gn=CRIP2;refGene_tx=XM_023256152.1;refGene_gn=CRIP2;refGene_tx=XM_023256153.1
B3 149209138 149209279
refGene_gn=CRIP2;refGene_tx=XM_023256152.1;refGene_gn=CRIP2;refGene_tx=XM_023256153.1
B3 149209358 149209427
refGene_gn=CRIP2;refGene_tx=XM_023256152.1;refGene_gn=CRIP2;refGene_tx=XM_023256153.1
B3 149209506 149209601
refGene_gn=CRIP2;refGene_tx=XM_023256152.1;refGene_gn=CRIP2;refGene_tx=XM_023256153.1
B3 149210063 149210121
refGene_gn=CRIP2;refGene_tx=XM_023256152.1;refGene_gn=CRIP2;refGene_tx=XM_023256153.1
B3 149210200 149210268
refGene_gn=CRIP2;refGene_tx=XM_023256152.1;refGene_gn=CRIP2;refGene_tx=XM_023256153.1
B3 149217979 149218019
refGene_gn=CRIP1;refGene_tx=XM_003988022.3;refGene_gn=CRIP1;refGene_tx=XM_006933168.2
B3 149218919 149219014
refGene_gn=CRIP1;refGene_tx=XM_003988022.3;refGene_gn=CRIP1;refGene_tx=XM_006933168.2
B3 149219088 149219146
refGene_gn=CRIP1;refGene_tx=XM_003988022.3;refGene_gn=CRIP1;refGene_tx=XM_006933168.2

B3 149219253 149219294
refGene_gn=CRIP1;refGene_tx=XM_003988022.3;refGene_gn=CRIP1;refGene_tx=XM_006933168.2
B3 149221762 149222095
refGene_gn=TEDC1;refGene_tx=XM_023256154.1;refGene_gn=TEDC1;refGene_tx=XM_023256155.1;refGene_gn=TEDC1;refGene_tx=XM_023256157.1;refGene_gn=TEDC1;refGene_tx=XM_023256158.1
B3 149222191 149222270
refGene_gn=TEDC1;refGene_tx=XM_023256154.1;refGene_gn=TEDC1;refGene_tx=XM_023256155.1;refGene_gn=TEDC1;refGene_tx=XM_023256157.1;refGene_gn=TEDC1;refGene_tx=XM_023256158.1
B3 149222821 149223042
refGene_gn=TEDC1;refGene_tx=XM_023256154.1;refGene_gn=TEDC1;refGene_tx=XM_023256155.1;refGene_gn=TEDC1;refGene_tx=XM_023256157.1;refGene_gn=TEDC1;refGene_tx=XM_023256158.1
B3 149223306 149223462
refGene_gn=TEDC1;refGene_tx=XM_023256154.1;refGene_gn=TEDC1;refGene_tx=XM_023256155.1;refGene_gn=TEDC1;refGene_tx=XM_023256157.1
B3 149224569 149224668
refGene_gn=TEDC1;refGene_tx=XM_023256154.1;refGene_gn=TEDC1;refGene_tx=XM_023256155.1;refGene_gn=TEDC1;refGene_tx=XM_023256157.1;refGene_gn=TEDC1;refGene_tx=XM_023256158.1
B3 149227100 149227187
refGene_gn=TEDC1;refGene_tx=XM_023256154.1;refGene_gn=TEDC1;refGene_tx=XM_023256155.1;refGene_gn=TEDC1;refGene_tx=XM_023256158.1
B3 149227582 149227750
refGene_gn=TEDC1;refGene_tx=XM_023256154.1;refGene_gn=TEDC1;refGene_tx=XM_023256155.1;refGene_gn=TEDC1;refGene_tx=XM_023256157.1;refGene_gn=TEDC1;refGene_tx=XM_023256158.1
B3 149227855 149228179
refGene_gn=TEDC1;refGene_tx=XM_023256154.1;refGene_gn=TEDC1;refGene_tx=XM_023256155.1;refGene_gn=TEDC1;refGene_tx=XM_023256157.1;refGene_gn=TEDC1;refGene_tx=XM_023256158.1
B3 149245497 149246454
refGene_gn=TMEM121;refGene_tx=XM_023256159.1
B3 149291626 149293449 refGene_gn=LOC101090949
B3 149308293 149310699 refGene_gn=LOC101091208
B3 149330391 149343374 refGene_gn=LOC105259932
B3 149344052 149345123
refGene_gn=LOC111560937;refGene_tx=XM_023256160.1
B3 149365157 149383040 refGene_gn=LOC105260702
B3 149383229 149417446 refGene_gn=LOC101091707
B3 149456218 149457050 refGene_gn=LOC111560938
B3 149485636 149487196
refGene_gn=LOC105260149;refGene_tx=XM_011279963.3
B3 149487242 149556969 refGene_gn=LOC101093530
B4 300888 301004
refGene_gn=ZMYND11;refGene_tx=XM_011283590.3;refGene_gn=ZMYND11;refGene_tx=XM_023256260.1;refGene_gn=ZMYND11;refGene_tx=XM_023256261.1;refGene_gn=ZMYND11;refGene_tx=XM_023256262.1;refGene_gn=ZMYND11;refGene_tx=

XM_023256263.1;refGene_gn=ZMYND11;refGene_tx=XM_023256264.1;refGene_gn=ZMYND11;refGene_tx=XM_023256265.1;refGene_gn=ZMYND11;refGene_tx=XM_023256266.1;refGene_gn=ZMYND11;refGene_tx=XM_023256267.1;refGene_gn=ZMYND11;refGene_tx=XM_023256268.1
B4 326959 327119
refGene_gn=ZMYND11;refGene_tx=XM_011283590.3;refGene_gn=ZMYND11;refGene_tx=XM_023256260.1;refGene_gn=ZMYND11;refGene_tx=XM_023256261.1;refGene_gn=ZMYND11;refGene_tx=XM_023256262.1;refGene_gn=ZMYND11;refGene_tx=XM_023256263.1;refGene_gn=ZMYND11;refGene_tx=XM_023256264.1;refGene_gn=ZMYND11;refGene_tx=XM_023256265.1;refGene_gn=ZMYND11;refGene_tx=XM_023256266.1;refGene_gn=ZMYND11;refGene_tx=XM_023256267.1;refGene_gn=ZMYND11;refGene_tx=XM_023256268.1
B4 334444 334489
refGene_gn=ZMYND11;refGene_tx=XM_011283590.3;refGene_gn=ZMYND11;refGene_tx=XM_023256260.1;refGene_gn=ZMYND11;refGene_tx=XM_023256261.1;refGene_gn=ZMYND11;refGene_tx=XM_023256262.1;refGene_gn=ZMYND11;refGene_tx=XM_023256263.1;refGene_gn=ZMYND11;refGene_tx=XM_023256264.1
B4 336537 336603
refGene_gn=ZMYND11;refGene_tx=XM_011283597.3;refGene_gn=ZMYND11;refGene_tx=XM_011283598.3
B4 337068 337230
refGene_gn=ZMYND11;refGene_tx=XM_011283590.3;refGene_gn=ZMYND11;refGene_tx=XM_011283597.3;refGene_gn=ZMYND11;refGene_tx=XM_011283598.3;refGene_gn=ZMYND11;refGene_tx=XM_023256260.1;refGene_gn=ZMYND11;refGene_tx=XM_023256261.1;refGene_gn=ZMYND11;refGene_tx=XM_023256262.1;refGene_gn=ZMYND11;refGene_tx=XM_023256263.1;refGene_gn=ZMYND11;refGene_tx=XM_023256264.1;refGene_gn=ZMYND11;refGene_tx=XM_023256265.1;refGene_gn=ZMYND11;refGene_tx=XM_023256266.1
B4 349025 349103
refGene_gn=ZMYND11;refGene_tx=XM_011283590.3;refGene_gn=ZMYND11;refGene_tx=XM_011283597.3;refGene_gn=ZMYND11;refGene_tx=XM_011283598.3;refGene_gn=ZMYND11;refGene_tx=XM_023256260.1;refGene_gn=ZMYND11;refGene_tx=XM_023256261.1;refGene_gn=ZMYND11;refGene_tx=XM_023256262.1;refGene_gn=ZMYND11;refGene_tx=XM_023256263.1;refGene_gn=ZMYND11;refGene_tx=XM_023256264.1;refGene_gn=ZMYND11;refGene_tx=XM_023256265.1;refGene_gn=ZMYND11;refGene_tx=XM_023256266.1;refGene_gn=ZMYND11;refGene_tx=XM_023256267.1;refGene_gn=ZMYND11;refGene_tx=XM_023256268.1
B4 349970 350063
refGene_gn=ZMYND11;refGene_tx=XM_011283590.3;refGene_gn=ZMYND11;refGene_tx=XM_011283597.3;refGene_gn=ZMYND11;refGene_tx=XM_011283598.3;refGene_gn=ZMYND11;refGene_tx=XM_023256260.1;refGene_gn=ZMYND11;refGene_tx=XM_023256261.1;refGene_gn=ZMYND11;refGene_tx=XM_023256262.1;refGene_gn=ZMYND11;refGene_tx=XM_023256263.1;refGene_gn=ZMYND11;refGene_tx=XM_023256264.1;refGene_gn=ZMYND11;refGene_tx=XM_023256265.1;refGene_gn=ZMYND11;refGene_tx=XM_023256266.1;refGene_gn=ZMYND11;refGene_tx=XM_023256267.1;refGene_gn=ZMYND11;refGene_tx=XM_023256268.1
B4 352087 352175
refGene_gn=ZMYND11;refGene_tx=XM_011283590.3;refGene_gn=ZMYND11;refGene_tx=XM_011283597.3;refGene_gn=ZMYND11;refGene_tx=XM_011283598.3;refGene_gn=ZMYND11;refGene_tx=XM_023256260.1;refGene_gn=ZMYND11;refGene_tx=

XM_023256261.1;refGene_gn=ZMYND11;refGene_tx=XM_023256262.1;refGene_gn=ZMYND11;refGene_tx=XM_023256263.1;refGene_gn=ZMYND11;refGene_tx=XM_023256264.1;refGene_gn=ZMYND11;refGene_tx=XM_023256265.1;refGene_gn=ZMYND11;refGene_tx=XM_023256266.1;refGene_gn=ZMYND11;refGene_tx=XM_023256267.1;refGene_gn=ZMYND11;refGene_tx=XM_023256268.1
B4 352928 352984
refGene_gn=ZMYND11;refGene_tx=XM_011283590.3;refGene_gn=ZMYND11;refGene_tx=XM_011283597.3;refGene_gn=ZMYND11;refGene_tx=XM_011283598.3;refGene_gn=ZMYND11;refGene_tx=XM_023256260.1;refGene_gn=ZMYND11;refGene_tx=XM_023256261.1;refGene_gn=ZMYND11;refGene_tx=XM_023256262.1;refGene_gn=ZMYND11;refGene_tx=XM_023256263.1;refGene_gn=ZMYND11;refGene_tx=XM_023256264.1;refGene_gn=ZMYND11;refGene_tx=XM_023256265.1;refGene_gn=ZMYND11;refGene_tx=XM_023256266.1;refGene_gn=ZMYND11;refGene_tx=XM_023256267.1;refGene_gn=ZMYND11;refGene_tx=XM_023256268.1
B4 357743 357821
refGene_gn=ZMYND11;refGene_tx=XM_011283590.3;refGene_gn=ZMYND11;refGene_tx=XM_011283597.3;refGene_gn=ZMYND11;refGene_tx=XM_011283598.3;refGene_gn=ZMYND11;refGene_tx=XM_023256260.1;refGene_gn=ZMYND11;refGene_tx=XM_023256261.1;refGene_gn=ZMYND11;refGene_tx=XM_023256262.1;refGene_gn=ZMYND11;refGene_tx=XM_023256263.1;refGene_gn=ZMYND11;refGene_tx=XM_023256264.1;refGene_gn=ZMYND11;refGene_tx=XM_023256265.1;refGene_gn=ZMYND11;refGene_tx=XM_023256266.1;refGene_gn=ZMYND11;refGene_tx=XM_023256267.1;refGene_gn=ZMYND11;refGene_tx=XM_023256268.1
B4 358663 358782
refGene_gn=ZMYND11;refGene_tx=XM_011283590.3;refGene_gn=ZMYND11;refGene_tx=XM_011283597.3;refGene_gn=ZMYND11;refGene_tx=XM_011283598.3;refGene_gn=ZMYND11;refGene_tx=XM_023256260.1;refGene_gn=ZMYND11;refGene_tx=XM_023256261.1;refGene_gn=ZMYND11;refGene_tx=XM_023256262.1;refGene_gn=ZMYND11;refGene_tx=XM_023256263.1;refGene_gn=ZMYND11;refGene_tx=XM_023256264.1;refGene_gn=ZMYND11;refGene_tx=XM_023256265.1;refGene_gn=ZMYND11;refGene_tx=XM_023256266.1;refGene_gn=ZMYND11;refGene_tx=XM_023256267.1;refGene_gn=ZMYND11;refGene_tx=XM_023256268.1
B4 363712 363920
refGene_gn=ZMYND11;refGene_tx=XM_011283590.3;refGene_gn=ZMYND11;refGene_tx=XM_011283597.3;refGene_gn=ZMYND11;refGene_tx=XM_011283598.3;refGene_gn=ZMYND11;refGene_tx=XM_023256260.1;refGene_gn=ZMYND11;refGene_tx=XM_023256261.1;refGene_gn=ZMYND11;refGene_tx=XM_023256262.1;refGene_gn=ZMYND11;refGene_tx=XM_023256263.1;refGene_gn=ZMYND11;refGene_tx=XM_023256264.1;refGene_gn=ZMYND11;refGene_tx=XM_023256265.1;refGene_gn=ZMYND11;refGene_tx=XM_023256266.1;refGene_gn=ZMYND11;refGene_tx=XM_023256267.1;refGene_gn=ZMYND11;refGene_tx=XM_023256268.1
B4 364268 364337
refGene_gn=ZMYND11;refGene_tx=XM_011283590.3;refGene_gn=ZMYND11;refGene_tx=XM_011283597.3;refGene_gn=ZMYND11;refGene_tx=XM_011283598.3;refGene_gn=ZMYND11;refGene_tx=XM_023256260.1;refGene_gn=ZMYND11;refGene_tx=XM_023256261.1;refGene_gn=ZMYND11;refGene_tx=XM_023256262.1;refGene_gn=ZMYND11;refGene_tx=XM_023256263.1;refGene_gn=ZMYND11;refGene_tx=XM_023256264.1;refGene_gn=ZMYND11;refGene_tx=XM_023256265.1;refGene_gn=ZMYND11;refGene_tx=XM_023256266.1;refGene_gn=ZMYND11;refGene_tx=XM_023256267.1;refGene_gn=ZMYND11;refGene_tx=XM_023256268.1

B4	365127	365400
refGene_gn=ZMYND11;refGene_tx=XM_011283590.3;refGene_gn=ZMYND11;refGene_tx=XM_011283597.3;refGene_gn=ZMYND11;refGene_tx=XM_011283598.3;refGene_gn=ZMYND11;refGene_tx=XM_023256260.1;refGene_gn=ZMYND11;refGene_tx=XM_023256261.1;refGene_gn=ZMYND11;refGene_tx=XM_023256262.1;refGene_gn=ZMYND11;refGene_tx=XM_023256263.1;refGene_gn=ZMYND11;refGene_tx=XM_023256264.1;refGene_gn=ZMYND11;refGene_tx=XM_023256265.1;refGene_gn=ZMYND11;refGene_tx=XM_023256266.1;refGene_gn=ZMYND11;refGene_tx=XM_023256267.1;refGene_gn=ZMYND11;refGene_tx=XM_023256268.1
B4	365710	365896
refGene_gn=ZMYND11;refGene_tx=XM_011283590.3;refGene_gn=ZMYND11;refGene_tx=XM_011283597.3;refGene_gn=ZMYND11;refGene_tx=XM_011283598.3;refGene_gn=ZMYND11;refGene_tx=XM_023256260.1;refGene_gn=ZMYND11;refGene_tx=XM_023256261.1;refGene_gn=ZMYND11;refGene_tx=XM_023256262.1;refGene_gn=ZMYND11;refGene_tx=XM_023256263.1;refGene_gn=ZMYND11;refGene_tx=XM_023256264.1;refGene_gn=ZMYND11;refGene_tx=XM_023256265.1;refGene_gn=ZMYND11;refGene_tx=XM_023256266.1;refGene_gn=ZMYND11;refGene_tx=XM_023256267.1;refGene_gn=ZMYND11;refGene_tx=XM_023256268.1
B4	375135	375258
refGene_gn=ZMYND11;refGene_tx=XM_011283590.3;refGene_gn=ZMYND11;refGene_tx=XM_011283597.3;refGene_gn=ZMYND11;refGene_tx=XM_011283598.3;refGene_gn=ZMYND11;refGene_tx=XM_023256260.1;refGene_gn=ZMYND11;refGene_tx=XM_023256261.1;refGene_gn=ZMYND11;refGene_tx=XM_023256262.1;refGene_gn=ZMYND11;refGene_tx=XM_023256263.1;refGene_gn=ZMYND11;refGene_tx=XM_023256264.1;refGene_gn=ZMYND11;refGene_tx=XM_023256265.1;refGene_gn=ZMYND11;refGene_tx=XM_023256266.1;refGene_gn=ZMYND11;refGene_tx=XM_023256267.1;refGene_gn=ZMYND11;refGene_tx=XM_023256268.1
B4	412690	412943
refGene_gn=DIP2C;refGene_tx=XM_019833868.2;refGene_gn=DIP2C;refGene_tx=XM_023256253.1;refGene_gn=DIP2C;refGene_tx=XM_023256254.1;refGene_gn=DIP2C;refGene_tx=XM_023256255.1;refGene_gn=DIP2C;refGene_tx=XM_023256256.1;refGene_gn=DIP2C;refGene_tx=XM_023256257.1;refGene_gn=DIP2C;refGene_tx=XM_023256258.1;refGene_gn=DIP2C;refGene_tx=XM_023256259.1
B4	415828	415952
refGene_gn=DIP2C;refGene_tx=XM_019833868.2;refGene_gn=DIP2C;refGene_tx=XM_023256253.1;refGene_gn=DIP2C;refGene_tx=XM_023256254.1;refGene_gn=DIP2C;refGene_tx=XM_023256255.1;refGene_gn=DIP2C;refGene_tx=XM_023256256.1;refGene_gn=DIP2C;refGene_tx=XM_023256257.1;refGene_gn=DIP2C;refGene_tx=XM_023256258.1;refGene_gn=DIP2C;refGene_tx=XM_023256259.1
B4	417571	417746
refGene_gn=DIP2C;refGene_tx=XM_019833868.2;refGene_gn=DIP2C;refGene_tx=XM_023256253.1;refGene_gn=DIP2C;refGene_tx=XM_023256254.1;refGene_gn=DIP2C;refGene_tx=XM_023256255.1;refGene_gn=DIP2C;refGene_tx=XM_023256256.1;refGene_gn=DIP2C;refGene_tx=XM_023256257.1;refGene_gn=DIP2C;refGene_tx=XM_023256258.1;refGene_gn=DIP2C;refGene_tx=XM_023256259.1
B4	420444	420519
refGene_gn=DIP2C;refGene_tx=XM_019833868.2;refGene_gn=DIP2C;refGene_tx=XM_023256253.1;refGene_gn=DIP2C;refGene_tx=XM_023256254.1;refGene_gn=DIP2C;refGene_tx=XM_023256255.1;refGene_gn=DIP2C;refGene_tx=XM_023256256.1;refGene_gn=DIP2C;refGene_tx=XM_023256257.1;refGene_gn=DIP2C;refGe

ne_tx=XM_023256258.1;refGene_gn=DIP2C;refGene_tx=XM_023256259.1
B4 422626 422684
refGene_gn=DIP2C;refGene_tx=XM_019833868.2;refGene_gn=DIP2C;refGene_tx=XM_023256253.1;refGene_gn=DIP2C;refGene_tx=XM_023256254.1;refGene_gn=DIP2C;refGene_tx=XM_023256255.1;refGene_gn=DIP2C;refGene_tx=XM_023256256.1;refGene_gn=DIP2C;refGene_tx=XM_023256257.1;refGene_gn=DIP2C;refGene_tx=XM_023256258.1;refGene_gn=DIP2C;refGene_tx=XM_023256259.1
B4 437607 437669
refGene_gn=DIP2C;refGene_tx=XM_019833868.2;refGene_gn=DIP2C;refGene_tx=XM_023256253.1;refGene_gn=DIP2C;refGene_tx=XM_023256254.1;refGene_gn=DIP2C;refGene_tx=XM_023256255.1;refGene_gn=DIP2C;refGene_tx=XM_023256256.1;refGene_gn=DIP2C;refGene_tx=XM_023256257.1;refGene_gn=DIP2C;refGene_tx=XM_023256258.1;refGene_gn=DIP2C;refGene_tx=XM_023256259.1
B4 439037 439067
refGene_gn=DIP2C;refGene_tx=XM_023256253.1;refGene_gn=DIP2C;refGene_tx=XM_023256254.1;refGene_gn=DIP2C;refGene_tx=XM_023256255.1;refGene_gn=DIP2C;refGene_tx=XM_023256256.1;refGene_gn=DIP2C;refGene_tx=XM_023256257.1
B4 449029 449065 refGene_gn=DIP2C;refGene_tx=XM_019833872.2
B4 460676 460847
refGene_gn=DIP2C;refGene_tx=XM_019833868.2;refGene_gn=DIP2C;refGene_tx=XM_019833872.2;refGene_gn=DIP2C;refGene_tx=XM_023256253.1;refGene_gn=DIP2C;refGene_tx=XM_023256254.1;refGene_gn=DIP2C;refGene_tx=XM_023256255.1;refGene_gn=DIP2C;refGene_tx=XM_023256256.1;refGene_gn=DIP2C;refGene_tx=XM_023256257.1;refGene_gn=DIP2C;refGene_tx=XM_023256258.1;refGene_gn=DIP2C;refGene_tx=XM_023256259.1
B4 462840 463009
refGene_gn=DIP2C;refGene_tx=XM_019833868.2;refGene_gn=DIP2C;refGene_tx=XM_019833872.2;refGene_gn=DIP2C;refGene_tx=XM_023256253.1;refGene_gn=DIP2C;refGene_tx=XM_023256254.1;refGene_gn=DIP2C;refGene_tx=XM_023256255.1;refGene_gn=DIP2C;refGene_tx=XM_023256256.1;refGene_gn=DIP2C;refGene_tx=XM_023256257.1;refGene_gn=DIP2C;refGene_tx=XM_023256258.1;refGene_gn=DIP2C;refGene_tx=XM_023256259.1
B4 471810 471941
refGene_gn=DIP2C;refGene_tx=XM_019833868.2;refGene_gn=DIP2C;refGene_tx=XM_019833872.2;refGene_gn=DIP2C;refGene_tx=XM_023256253.1;refGene_gn=DIP2C;refGene_tx=XM_023256254.1;refGene_gn=DIP2C;refGene_tx=XM_023256255.1;refGene_gn=DIP2C;refGene_tx=XM_023256256.1;refGene_gn=DIP2C;refGene_tx=XM_023256257.1;refGene_gn=DIP2C;refGene_tx=XM_023256258.1;refGene_gn=DIP2C;refGene_tx=XM_023256259.1
B4 479973 480083
refGene_gn=DIP2C;refGene_tx=XM_019833868.2;refGene_gn=DIP2C;refGene_tx=XM_019833872.2;refGene_gn=DIP2C;refGene_tx=XM_023256253.1;refGene_gn=DIP2C;refGene_tx=XM_023256254.1;refGene_gn=DIP2C;refGene_tx=XM_023256255.1;refGene_gn=DIP2C;refGene_tx=XM_023256256.1;refGene_gn=DIP2C;refGene_tx=XM_023256257.1;refGene_gn=DIP2C;refGene_tx=XM_023256258.1;refGene_gn=DIP2C;refGene_tx=XM_023256259.1
B4 480161 480273
refGene_gn=DIP2C;refGene_tx=XM_019833868.2;refGene_gn=DIP2C;refGene_tx=XM_019833872.2;refGene_gn=DIP2C;refGene_tx=XM_023256253.1;refGene_gn=

DIP2C;refGene_tx=XM_023256254.1;refGene_gn=DIP2C;refGene_tx=XM_0232562
55.1;refGene_gn=DIP2C;refGene_tx=XM_023256256.1;refGene_gn=DIP2C;refGe
ne_tx=XM_023256257.1;refGene_gn=DIP2C;refGene_tx=XM_023256258.1;refGen
e_gn=DIP2C;refGene_tx=XM_023256259.1
B4 481286 481408
refGene_gn=DIP2C;refGene_tx=XM_019833868.2;refGene_gn=DIP2C;refGene_tx
=XM_019833872.2;refGene_gn=DIP2C;refGene_tx=XM_023256253.1;refGene_gn=
DIP2C;refGene_tx=XM_023256254.1;refGene_gn=DIP2C;refGene_tx=XM_0232562
55.1;refGene_gn=DIP2C;refGene_tx=XM_023256256.1;refGene_gn=DIP2C;refGe
ne_tx=XM_023256257.1;refGene_gn=DIP2C;refGene_tx=XM_023256258.1;refGen
e_gn=DIP2C;refGene_tx=XM_023256259.1
B4 481852 481976
refGene_gn=DIP2C;refGene_tx=XM_019833868.2;refGene_gn=DIP2C;refGene_tx
=XM_019833872.2;refGene_gn=DIP2C;refGene_tx=XM_023256253.1;refGene_gn=
DIP2C;refGene_tx=XM_023256254.1;refGene_gn=DIP2C;refGene_tx=XM_0232562
55.1;refGene_gn=DIP2C;refGene_tx=XM_023256256.1;refGene_gn=DIP2C;refGe
ne_tx=XM_023256257.1;refGene_gn=DIP2C;refGene_tx=XM_023256258.1;refGen
e_gn=DIP2C;refGene_tx=XM_023256259.1
B4 487330 487411
refGene_gn=DIP2C;refGene_tx=XM_019833868.2;refGene_gn=DIP2C;refGene_tx
=XM_019833872.2;refGene_gn=DIP2C;refGene_tx=XM_023256253.1;refGene_gn=
DIP2C;refGene_tx=XM_023256254.1;refGene_gn=DIP2C;refGene_tx=XM_0232562
55.1;refGene_gn=DIP2C;refGene_tx=XM_023256256.1;refGene_gn=DIP2C;refGe
ne_tx=XM_023256257.1;refGene_gn=DIP2C;refGene_tx=XM_023256258.1;refGen
e_gn=DIP2C;refGene_tx=XM_023256259.1
B4 488644 488754
refGene_gn=DIP2C;refGene_tx=XM_019833868.2;refGene_gn=DIP2C;refGene_tx
=XM_019833872.2;refGene_gn=DIP2C;refGene_tx=XM_023256253.1;refGene_gn=
DIP2C;refGene_tx=XM_023256254.1;refGene_gn=DIP2C;refGene_tx=XM_0232562
55.1;refGene_gn=DIP2C;refGene_tx=XM_023256256.1;refGene_gn=DIP2C;refGe
ne_tx=XM_023256257.1;refGene_gn=DIP2C;refGene_tx=XM_023256258.1;refGen
e_gn=DIP2C;refGene_tx=XM_023256259.1
B4 494205 494407
refGene_gn=DIP2C;refGene_tx=XM_019833868.2;refGene_gn=DIP2C;refGene_tx
=XM_019833872.2;refGene_gn=DIP2C;refGene_tx=XM_023256253.1;refGene_gn=
DIP2C;refGene_tx=XM_023256254.1;refGene_gn=DIP2C;refGene_tx=XM_0232562
55.1;refGene_gn=DIP2C;refGene_tx=XM_023256256.1;refGene_gn=DIP2C;refGe
ne_tx=XM_023256257.1;refGene_gn=DIP2C;refGene_tx=XM_023256258.1;refGen
e_gn=DIP2C;refGene_tx=XM_023256259.1
B4 494900 495015
refGene_gn=DIP2C;refGene_tx=XM_019833868.2;refGene_gn=DIP2C;refGene_tx
=XM_019833872.2;refGene_gn=DIP2C;refGene_tx=XM_023256253.1;refGene_gn=
DIP2C;refGene_tx=XM_023256254.1;refGene_gn=DIP2C;refGene_tx=XM_0232562
55.1;refGene_gn=DIP2C;refGene_tx=XM_023256256.1;refGene_gn=DIP2C;refGe
ne_tx=XM_023256257.1;refGene_gn=DIP2C;refGene_tx=XM_023256258.1;refGen
e_gn=DIP2C;refGene_tx=XM_023256259.1
B4 495871 496080
refGene_gn=DIP2C;refGene_tx=XM_019833868.2;refGene_gn=DIP2C;refGene_tx
=XM_019833872.2;refGene_gn=DIP2C;refGene_tx=XM_023256253.1;refGene_gn=
DIP2C;refGene_tx=XM_023256254.1;refGene_gn=DIP2C;refGene_tx=XM_0232562

55.1;refGene_gn=DIP2C;refGene_tx=XM_023256256.1;refGene_gn=DIP2C;refGene_tx=XM_023256257.1;refGene_gn=DIP2C;refGene_tx=XM_023256258.1;refGene_gn=DIP2C;refGene_tx=XM_023256259.1
B4	497714	497851
refGene_gn=DIP2C;refGene_tx=XM_019833868.2;refGene_gn=DIP2C;refGene_tx=XM_019833872.2;refGene_gn=DIP2C;refGene_tx=XM_023256253.1;refGene_gn=DIP2C;refGene_tx=XM_023256254.1;refGene_gn=DIP2C;refGene_tx=XM_023256255.1;refGene_gn=DIP2C;refGene_tx=XM_023256256.1;refGene_gn=DIP2C;refGene_tx=XM_023256257.1;refGene_gn=DIP2C;refGene_tx=XM_023256258.1;refGene_gn=DIP2C;refGene_tx=XM_023256259.1
B4	499066	499206
refGene_gn=DIP2C;refGene_tx=XM_019833868.2;refGene_gn=DIP2C;refGene_tx=XM_019833872.2;refGene_gn=DIP2C;refGene_tx=XM_023256253.1;refGene_gn=DIP2C;refGene_tx=XM_023256254.1;refGene_gn=DIP2C;refGene_tx=XM_023256255.1;refGene_gn=DIP2C;refGene_tx=XM_023256256.1;refGene_gn=DIP2C;refGene_tx=XM_023256257.1;refGene_gn=DIP2C;refGene_tx=XM_023256258.1;refGene_gn=DIP2C;refGene_tx=XM_023256259.1
B4	515550	515665
refGene_gn=DIP2C;refGene_tx=XM_019833868.2;refGene_gn=DIP2C;refGene_tx=XM_019833872.2;refGene_gn=DIP2C;refGene_tx=XM_023256253.1;refGene_gn=DIP2C;refGene_tx=XM_023256254.1;refGene_gn=DIP2C;refGene_tx=XM_023256255.1;refGene_gn=DIP2C;refGene_tx=XM_023256256.1;refGene_gn=DIP2C;refGene_tx=XM_023256257.1;refGene_gn=DIP2C;refGene_tx=XM_023256258.1;refGene_gn=DIP2C;refGene_tx=XM_023256259.1
B4	516850	516970
refGene_gn=DIP2C;refGene_tx=XM_019833868.2;refGene_gn=DIP2C;refGene_tx=XM_019833872.2;refGene_gn=DIP2C;refGene_tx=XM_023256253.1;refGene_gn=DIP2C;refGene_tx=XM_023256254.1;refGene_gn=DIP2C;refGene_tx=XM_023256255.1;refGene_gn=DIP2C;refGene_tx=XM_023256256.1;refGene_gn=DIP2C;refGene_tx=XM_023256257.1;refGene_gn=DIP2C;refGene_tx=XM_023256258.1;refGene_gn=DIP2C;refGene_tx=XM_023256259.1
B4	517067	517161
refGene_gn=DIP2C;refGene_tx=XM_019833868.2;refGene_gn=DIP2C;refGene_tx=XM_019833872.2;refGene_gn=DIP2C;refGene_tx=XM_023256253.1;refGene_gn=DIP2C;refGene_tx=XM_023256254.1;refGene_gn=DIP2C;refGene_tx=XM_023256255.1;refGene_gn=DIP2C;refGene_tx=XM_023256256.1;refGene_gn=DIP2C;refGene_tx=XM_023256257.1;refGene_gn=DIP2C;refGene_tx=XM_023256258.1;refGene_gn=DIP2C;refGene_tx=XM_023256259.1
B4	518575	518640
refGene_gn=DIP2C;refGene_tx=XM_019833868.2;refGene_gn=DIP2C;refGene_tx=XM_019833872.2;refGene_gn=DIP2C;refGene_tx=XM_023256253.1;refGene_gn=DIP2C;refGene_tx=XM_023256254.1;refGene_gn=DIP2C;refGene_tx=XM_023256255.1;refGene_gn=DIP2C;refGene_tx=XM_023256256.1;refGene_gn=DIP2C;refGene_tx=XM_023256257.1;refGene_gn=DIP2C;refGene_tx=XM_023256258.1;refGene_gn=DIP2C;refGene_tx=XM_023256259.1
B4	520405	520508
refGene_gn=DIP2C;refGene_tx=XM_019833868.2;refGene_gn=DIP2C;refGene_tx=XM_019833872.2;refGene_gn=DIP2C;refGene_tx=XM_023256253.1;refGene_gn=DIP2C;refGene_tx=XM_023256254.1;refGene_gn=DIP2C;refGene_tx=XM_023256255.1;refGene_gn=DIP2C;refGene_tx=XM_023256256.1;refGene_gn=DIP2C;refGe

ne_tx=XM_023256257.1;refGene_gn=DIP2C;refGene_tx=XM_023256258.1;refGene_gn=DIP2C;refGene_tx=XM_023256259.1
B4	520621	520731
refGene_gn=DIP2C;refGene_tx=XM_019833868.2;refGene_gn=DIP2C;refGene_tx=XM_019833872.2;refGene_gn=DIP2C;refGene_tx=XM_023256253.1;refGene_gn=DIP2C;refGene_tx=XM_023256254.1;refGene_gn=DIP2C;refGene_tx=XM_023256255.1;refGene_gn=DIP2C;refGene_tx=XM_023256256.1;refGene_gn=DIP2C;refGene_tx=XM_023256257.1;refGene_gn=DIP2C;refGene_tx=XM_023256258.1;refGene_gn=DIP2C;refGene_tx=XM_023256259.1
B4	521026	521150
refGene_gn=DIP2C;refGene_tx=XM_019833868.2;refGene_gn=DIP2C;refGene_tx=XM_019833872.2;refGene_gn=DIP2C;refGene_tx=XM_023256253.1;refGene_gn=DIP2C;refGene_tx=XM_023256254.1;refGene_gn=DIP2C;refGene_tx=XM_023256255.1;refGene_gn=DIP2C;refGene_tx=XM_023256256.1;refGene_gn=DIP2C;refGene_tx=XM_023256257.1;refGene_gn=DIP2C;refGene_tx=XM_023256258.1;refGene_gn=DIP2C;refGene_tx=XM_023256259.1
B4	533850	533961
refGene_gn=DIP2C;refGene_tx=XM_019833868.2;refGene_gn=DIP2C;refGene_tx=XM_019833872.2;refGene_gn=DIP2C;refGene_tx=XM_023256253.1;refGene_gn=DIP2C;refGene_tx=XM_023256254.1;refGene_gn=DIP2C;refGene_tx=XM_023256255.1;refGene_gn=DIP2C;refGene_tx=XM_023256256.1;refGene_gn=DIP2C;refGene_tx=XM_023256257.1;refGene_gn=DIP2C;refGene_tx=XM_023256258.1;refGene_gn=DIP2C;refGene_tx=XM_023256259.1
B4	540143	540235
refGene_gn=DIP2C;refGene_tx=XM_019833868.2;refGene_gn=DIP2C;refGene_tx=XM_019833872.2;refGene_gn=DIP2C;refGene_tx=XM_023256253.1;refGene_gn=DIP2C;refGene_tx=XM_023256254.1;refGene_gn=DIP2C;refGene_tx=XM_023256255.1;refGene_gn=DIP2C;refGene_tx=XM_023256256.1;refGene_gn=DIP2C;refGene_tx=XM_023256257.1;refGene_gn=DIP2C;refGene_tx=XM_023256258.1;refGene_gn=DIP2C;refGene_tx=XM_023256259.1
B4	548748	548946
refGene_gn=DIP2C;refGene_tx=XM_019833868.2;refGene_gn=DIP2C;refGene_tx=XM_019833872.2;refGene_gn=DIP2C;refGene_tx=XM_023256253.1;refGene_gn=DIP2C;refGene_tx=XM_023256254.1;refGene_gn=DIP2C;refGene_tx=XM_023256255.1;refGene_gn=DIP2C;refGene_tx=XM_023256256.1;refGene_gn=DIP2C;refGene_tx=XM_023256257.1;refGene_gn=DIP2C;refGene_tx=XM_023256258.1;refGene_gn=DIP2C;refGene_tx=XM_023256259.1
B4	550230	550350
refGene_gn=DIP2C;refGene_tx=XM_019833868.2;refGene_gn=DIP2C;refGene_tx=XM_019833872.2;refGene_gn=DIP2C;refGene_tx=XM_023256253.1;refGene_gn=DIP2C;refGene_tx=XM_023256254.1;refGene_gn=DIP2C;refGene_tx=XM_023256255.1;refGene_gn=DIP2C;refGene_tx=XM_023256256.1;refGene_gn=DIP2C;refGene_tx=XM_023256257.1;refGene_gn=DIP2C;refGene_tx=XM_023256258.1;refGene_gn=DIP2C;refGene_tx=XM_023256259.1
B4	552961	553096
refGene_gn=DIP2C;refGene_tx=XM_019833868.2;refGene_gn=DIP2C;refGene_tx=XM_019833872.2;refGene_gn=DIP2C;refGene_tx=XM_023256253.1;refGene_gn=DIP2C;refGene_tx=XM_023256254.1;refGene_gn=DIP2C;refGene_tx=XM_023256255.1;refGene_gn=DIP2C;refGene_tx=XM_023256256.1;refGene_gn=DIP2C;refGene_tx=XM_023256257.1;refGene_gn=DIP2C;refGene_tx=XM_023256258.1;refGen

e_gn=DIP2C;refGene_tx=XM_023256259.1
B4	565444	565654
refGene_gn=DIP2C;refGene_tx=XM_019833868.2;refGene_gn=DIP2C;refGene_tx=XM_019833872.2;refGene_gn=DIP2C;refGene_tx=XM_023256253.1;refGene_gn=DIP2C;refGene_tx=XM_023256254.1;refGene_gn=DIP2C;refGene_tx=XM_023256255.1;refGene_gn=DIP2C;refGene_tx=XM_023256256.1;refGene_gn=DIP2C;refGene_tx=XM_023256257.1;refGene_gn=DIP2C;refGene_tx=XM_023256258.1;refGene_gn=DIP2C;refGene_tx=XM_023256259.1
B4	575679	575805
refGene_gn=DIP2C;refGene_tx=XM_019833868.2;refGene_gn=DIP2C;refGene_tx=XM_019833872.2;refGene_gn=DIP2C;refGene_tx=XM_023256253.1;refGene_gn=DIP2C;refGene_tx=XM_023256254.1;refGene_gn=DIP2C;refGene_tx=XM_023256255.1;refGene_gn=DIP2C;refGene_tx=XM_023256256.1;refGene_gn=DIP2C;refGene_tx=XM_023256257.1;refGene_gn=DIP2C;refGene_tx=XM_023256258.1;refGene_gn=DIP2C;refGene_tx=XM_023256259.1
B4	606381	606492
refGene_gn=DIP2C;refGene_tx=XM_019833868.2;refGene_gn=DIP2C;refGene_tx=XM_019833872.2;refGene_gn=DIP2C;refGene_tx=XM_023256253.1;refGene_gn=DIP2C;refGene_tx=XM_023256254.1;refGene_gn=DIP2C;refGene_tx=XM_023256255.1;refGene_gn=DIP2C;refGene_tx=XM_023256256.1;refGene_gn=DIP2C;refGene_tx=XM_023256257.1;refGene_gn=DIP2C;refGene_tx=XM_023256258.1;refGene_gn=DIP2C;refGene_tx=XM_023256259.1
B4	616011	616179
refGene_gn=DIP2C;refGene_tx=XM_019833868.2;refGene_gn=DIP2C;refGene_tx=XM_019833872.2;refGene_gn=DIP2C;refGene_tx=XM_023256253.1;refGene_gn=DIP2C;refGene_tx=XM_023256254.1;refGene_gn=DIP2C;refGene_tx=XM_023256255.1;refGene_gn=DIP2C;refGene_tx=XM_023256259.1
B4	617733	617805
refGene_gn=DIP2C;refGene_tx=XM_019833868.2;refGene_gn=DIP2C;refGene_tx=XM_019833872.2;refGene_gn=DIP2C;refGene_tx=XM_023256253.1;refGene_gn=DIP2C;refGene_tx=XM_023256254.1;refGene_gn=DIP2C;refGene_tx=XM_023256255.1;refGene_gn=DIP2C;refGene_tx=XM_023256256.1;refGene_gn=DIP2C;refGene_tx=XM_023256257.1;refGene_gn=DIP2C;refGene_tx=XM_023256258.1;refGene_gn=DIP2C;refGene_tx=XM_023256259.1
B4	638705	638808
refGene_gn=DIP2C;refGene_tx=XM_023256253.1;refGene_gn=DIP2C;refGene_tx=XM_023256256.1
B4	702264	702370
refGene_gn=DIP2C;refGene_tx=XM_019833868.2;refGene_gn=DIP2C;refGene_tx=XM_019833872.2
B4	823830	823915
refGene_gn=DIP2C;refGene_tx=XM_023256254.1;refGene_gn=DIP2C;refGene_tx=XM_023256255.1;refGene_gn=DIP2C;refGene_tx=XM_023256257.1;refGene_gn=DIP2C;refGene_tx=XM_023256258.1;refGene_gn=DIP2C;refGene_tx=XM_023256259.1
B4	932393	933147
refGene_gn=LOC102901321;refGene_tx=XM_023256270.1;refGene_gn=LOC102901321;refGene_tx=XM_023256271.1;refGene_gn=LOC102901321;refGene_tx=XM_023256272.1
B4	940767	940955

refGene_gn=LOC102901321;refGene_tx=XM_023256270.1;refGene_gn=LOC102901
321;refGene_tx=XM_023256271.1;refGene_gn=LOC102901321;refGene_tx=XM_02
3256272.1
B4	942302	942575
refGene_gn=LARP4B;refGene_tx=XM_011283588.3;refGene_gn=LARP4B;refGene_
tx=XM_011283589.3;refGene_gn=LARP4B;refGene_tx=XM_023256269.1
B4	945050	945159
refGene_gn=LARP4B;refGene_tx=XM_011283588.3;refGene_gn=LARP4B;refGene_
tx=XM_011283589.3;refGene_gn=LARP4B;refGene_tx=XM_023256269.1
B4	945250	945381
refGene_gn=LARP4B;refGene_tx=XM_011283588.3;refGene_gn=LARP4B;refGene_
tx=XM_011283589.3;refGene_gn=LARP4B;refGene_tx=XM_023256269.1
B4	947505	947670
refGene_gn=LARP4B;refGene_tx=XM_011283588.3;refGene_gn=LARP4B;refGene_
tx=XM_011283589.3;refGene_gn=LARP4B;refGene_tx=XM_023256269.1
B4	955240	955286
refGene_gn=LARP4B;refGene_tx=XM_011283588.3;refGene_gn=LARP4B;refGene_
tx=XM_011283589.3;refGene_gn=LARP4B;refGene_tx=XM_023256269.1
B4	959395	959647
refGene_gn=LARP4B;refGene_tx=XM_011283588.3;refGene_gn=LARP4B;refGene_
tx=XM_011283589.3;refGene_gn=LARP4B;refGene_tx=XM_023256269.1
B4	960170	960277
refGene_gn=LARP4B;refGene_tx=XM_011283588.3;refGene_gn=LARP4B;refGene_
tx=XM_011283589.3;refGene_gn=LARP4B;refGene_tx=XM_023256269.1
B4	962791	963001
refGene_gn=LARP4B;refGene_tx=XM_011283588.3;refGene_gn=LARP4B;refGene_
tx=XM_011283589.3;refGene_gn=LARP4B;refGene_tx=XM_023256269.1
B4	963085	963139
refGene_gn=LARP4B;refGene_tx=XM_011283588.3;refGene_gn=LARP4B;refGene_
tx=XM_011283589.3;refGene_gn=LARP4B;refGene_tx=XM_023256269.1
B4	964602	964713
refGene_gn=LARP4B;refGene_tx=XM_011283588.3;refGene_gn=LARP4B;refGene_
tx=XM_011283589.3;refGene_gn=LARP4B;refGene_tx=XM_023256269.1
B4	968420	968524
refGene_gn=LARP4B;refGene_tx=XM_011283588.3;refGene_gn=LARP4B;refGene_
tx=XM_011283589.3;refGene_gn=LARP4B;refGene_tx=XM_023256269.1
B4	974220	974357
refGene_gn=LARP4B;refGene_tx=XM_011283588.3;refGene_gn=LARP4B;refGene_
tx=XM_011283589.3;refGene_gn=LARP4B;refGene_tx=XM_023256269.1
B4	975946	976025
refGene_gn=LARP4B;refGene_tx=XM_011283588.3;refGene_gn=LARP4B;refGene_
tx=XM_011283589.3;refGene_gn=LARP4B;refGene_tx=XM_023256269.1
B4	988753	988894
refGene_gn=LARP4B;refGene_tx=XM_011283588.3;refGene_gn=LARP4B;refGene_
tx=XM_011283589.3;refGene_gn=LARP4B;refGene_tx=XM_023256269.1
B4	989152	989303
refGene_gn=LARP4B;refGene_tx=XM_011283588.3;refGene_gn=LARP4B;refGene_
tx=XM_011283589.3;refGene_gn=LARP4B;refGene_tx=XM_023256269.1
B4	993123	994728	refGene_gn=LARP4B;refGene_tx=XM_011283588.3
B4	1005290	1005350

refGene_gn=LARP4B;refGene_tx=XM_011283589.3;refGene_gn=LARP4B;refGene_tx=XM_023256269.1
B4 1005992 1006073
refGene_gn=LARP4B;refGene_tx=XM_011283589.3;refGene_gn=LARP4B;refGene_tx=XM_023256269.1
B4 1095096 1095144 refGene_gn=GTPBP4;refGene_tx=XM_003988063.5
B4 1099230 1099401 refGene_gn=GTPBP4;refGene_tx=XM_003988063.5
B4 1101983 1102087 refGene_gn=GTPBP4;refGene_tx=XM_003988063.5
B4 1102166 1102303 refGene_gn=GTPBP4;refGene_tx=XM_003988063.5
B4 1104128 1104229 refGene_gn=GTPBP4;refGene_tx=XM_003988063.5
B4 1105500 1105593 refGene_gn=GTPBP4;refGene_tx=XM_003988063.5
B4 1106629 1106821 refGene_gn=GTPBP4;refGene_tx=XM_003988063.5
B4 1106901 1106967 refGene_gn=GTPBP4;refGene_tx=XM_003988063.5
B4 1108938 1109028 refGene_gn=GTPBP4;refGene_tx=XM_003988063.5
B4 1109783 1109894 refGene_gn=GTPBP4;refGene_tx=XM_003988063.5
B4 1111509 1111587 refGene_gn=GTPBP4;refGene_tx=XM_003988063.5
B4 1111899 1111951 refGene_gn=GTPBP4;refGene_tx=XM_003988063.5
B4 1112334 1112435 refGene_gn=GTPBP4;refGene_tx=XM_003988063.5
B4 1113829 1114027 refGene_gn=GTPBP4;refGene_tx=XM_003988063.5
B4 1115318 1115384 refGene_gn=GTPBP4;refGene_tx=XM_003988063.5
B4 1119573 1119717 refGene_gn=GTPBP4;refGene_tx=XM_003988063.5
B4 1120601 1120754 refGene_gn=GTPBP4;refGene_tx=XM_003988063.5
B4 1135403 1135721
refGene_gn=LOC102901899;refGene_tx=XM_023256273.1;refGene_gn=LOC102901899;refGene_tx=XM_023256274.1;refGene_gn=LOC102901899;refGene_tx=XM_023256275.1
B4 1136481 1136612
refGene_gn=LOC102901899;refGene_tx=XM_023256273.1;refGene_gn=LOC102901899;refGene_tx=XM_023256274.1;refGene_gn=LOC102901899;refGene_tx=XM_023256275.1
B4 1137492 1137585
refGene_gn=LOC102901899;refGene_tx=XM_023256273.1;refGene_gn=LOC102901899;refGene_tx=XM_023256274.1;refGene_gn=LOC102901899;refGene_tx=XM_023256275.1
B4 1138022 1138195
refGene_gn=LOC102901899;refGene_tx=XM_023256273.1;refGene_gn=LOC102901899;refGene_tx=XM_023256274.1;refGene_gn=LOC102901899;refGene_tx=XM_023256275.1
B4 1138565 1138696
refGene_gn=LOC102901899;refGene_tx=XM_023256273.1
B4 1147345 1147572
refGene_gn=WDR37;refGene_tx=XM_023256276.1;refGene_gn=WDR37;refGene_tx=XM_023256277.1;refGene_gn=WDR37;refGene_tx=XM_023256280.1;refGene_gn=WDR37;refGene_tx=XM_023256281.1
B4 1163078 1163256
refGene_gn=WDR37;refGene_tx=XM_023256276.1;refGene_gn=WDR37;refGene_tx=XM_023256277.1;refGene_gn=WDR37;refGene_tx=XM_023256278.1;refGene_gn=WDR37;refGene_tx=XM_023256280.1;refGene_gn=WDR37;refGene_tx=XM_023256281.1
B4 1163884 1163887 refGene_gn=WDR37;refGene_tx=XM_023256279.1

B4	1166218	1166315
refGene_gn=WDR37;refGene_tx=XM_023256276.1;refGene_gn=WDR37;refGene_tx=XM_023256277.1;refGene_gn=WDR37;refGene_tx=XM_023256278.1;refGene_gn=WDR37;refGene_tx=XM_023256279.1;refGene_gn=WDR37;refGene_tx=XM_023256280.1;refGene_gn=WDR37;refGene_tx=XM_023256281.1
B4	1168815	1168911
refGene_gn=WDR37;refGene_tx=XM_023256276.1;refGene_gn=WDR37;refGene_tx=XM_023256277.1;refGene_gn=WDR37;refGene_tx=XM_023256278.1;refGene_gn=WDR37;refGene_tx=XM_023256279.1;refGene_gn=WDR37;refGene_tx=XM_023256280.1;refGene_gn=WDR37;refGene_tx=XM_023256281.1
B4	1169240	1169305
refGene_gn=WDR37;refGene_tx=XM_023256276.1;refGene_gn=WDR37;refGene_tx=XM_023256277.1;refGene_gn=WDR37;refGene_tx=XM_023256278.1;refGene_gn=WDR37;refGene_tx=XM_023256279.1;refGene_gn=WDR37;refGene_tx=XM_023256280.1;refGene_gn=WDR37;refGene_tx=XM_023256281.1
B4	1172969	1173105
refGene_gn=WDR37;refGene_tx=XM_023256276.1;refGene_gn=WDR37;refGene_tx=XM_023256277.1;refGene_gn=WDR37;refGene_tx=XM_023256278.1;refGene_gn=WDR37;refGene_tx=XM_023256279.1;refGene_gn=WDR37;refGene_tx=XM_023256280.1;refGene_gn=WDR37;refGene_tx=XM_023256281.1
B4	1174672	1174744
refGene_gn=WDR37;refGene_tx=XM_023256276.1;refGene_gn=WDR37;refGene_tx=XM_023256277.1;refGene_gn=WDR37;refGene_tx=XM_023256278.1;refGene_gn=WDR37;refGene_tx=XM_023256279.1;refGene_gn=WDR37;refGene_tx=XM_023256280.1;refGene_gn=WDR37;refGene_tx=XM_023256281.1
B4	1178037	1178082
refGene_gn=WDR37;refGene_tx=XM_023256276.1;refGene_gn=WDR37;refGene_tx=XM_023256277.1;refGene_gn=WDR37;refGene_tx=XM_023256278.1;refGene_gn=WDR37;refGene_tx=XM_023256279.1;refGene_gn=WDR37;refGene_tx=XM_023256280.1;refGene_gn=WDR37;refGene_tx=XM_023256281.1
B4	1179724	1179801
refGene_gn=WDR37;refGene_tx=XM_023256276.1;refGene_gn=WDR37;refGene_tx=XM_023256277.1;refGene_gn=WDR37;refGene_tx=XM_023256278.1;refGene_gn=WDR37;refGene_tx=XM_023256279.1;refGene_gn=WDR37;refGene_tx=XM_023256280.1;refGene_gn=WDR37;refGene_tx=XM_023256281.1
B4	1187236	1187477
refGene_gn=WDR37;refGene_tx=XM_023256276.1;refGene_gn=WDR37;refGene_tx=XM_023256277.1;refGene_gn=WDR37;refGene_tx=XM_023256278.1;refGene_gn=WDR37;refGene_tx=XM_023256279.1;refGene_gn=WDR37;refGene_tx=XM_023256280.1;refGene_gn=WDR37;refGene_tx=XM_023256281.1
B4	1188276	1188350	refGene_gn=WDR37;refGene_tx=XM_023256280.1
B4	1188816	1188958
refGene_gn=WDR37;refGene_tx=XM_023256276.1;refGene_gn=WDR37;refGene_tx=XM_023256277.1;refGene_gn=WDR37;refGene_tx=XM_023256278.1;refGene_gn=WDR37;refGene_tx=XM_023256279.1
B4	1204456	1204591
refGene_gn=WDR37;refGene_tx=XM_023256276.1;refGene_gn=WDR37;refGene_tx=XM_023256277.1;refGene_gn=WDR37;refGene_tx=XM_023256278.1;refGene_gn=WDR37;refGene_tx=XM_023256279.1;refGene_gn=WDR37;refGene_tx=XM_023256281.1

B4 1205036 1205151
refGene_gn=WDR37;refGene_tx=XM_023256276.1;refGene_gn=WDR37;refGene_tx=XM_023256277.1;refGene_gn=WDR37;refGene_tx=XM_023256278.1;refGene_gn=WDR37;refGene_tx=XM_023256279.1
B4 1208628 1208760
refGene_gn=WDR37;refGene_tx=XM_023256276.1;refGene_gn=WDR37;refGene_tx=XM_023256277.1;refGene_gn=WDR37;refGene_tx=XM_023256278.1;refGene_gn=WDR37;refGene_tx=XM_023256279.1
B4 1247421 1247598 refGene_gn=ADARB2;refGene_tx=XM_011283586.3
B4 1249283 1249462 refGene_gn=ADARB2;refGene_tx=XM_011283586.3
B4 1264373 1264555 refGene_gn=ADARB2;refGene_tx=XM_011283586.3
B4 1278276 1278445 refGene_gn=ADARB2;refGene_tx=XM_011283586.3
B4 1286521 1286673 refGene_gn=ADARB2;refGene_tx=XM_011283586.3
B4 1287554 1287723 refGene_gn=ADARB2;refGene_tx=XM_011283586.3
B4 1303451 1303566 refGene_gn=ADARB2;refGene_tx=XM_011283586.3
B4 1372633 1373508 refGene_gn=ADARB2;refGene_tx=XM_011283586.3
B4 1381482 1381569 refGene_gn=ADARB2;refGene_tx=XM_011283586.3
B4 1665593 1665693 refGene_gn=ADARB2;refGene_tx=XM_011283586.3
B4 2990873 2990985 refGene_gn=PFKP;refGene_tx=XM_023256282.1
B4 3004041 3004115 refGene_gn=PFKP;refGene_tx=XM_023256282.1
B4 3014279 3014357 refGene_gn=PFKP;refGene_tx=XM_023256282.1
B4 3017349 3017542 refGene_gn=PFKP;refGene_tx=XM_023256282.1
B4 3017805 3017829 refGene_gn=PFKP;refGene_tx=XM_023256282.1
B4 3018489 3018628 refGene_gn=PFKP;refGene_tx=XM_023256282.1
B4 3025635 3025680 refGene_gn=PFKP;refGene_tx=XM_023256282.1
B4 3025791 3025900 refGene_gn=PFKP;refGene_tx=XM_023256282.1
B4 3028490 3028586 refGene_gn=PFKP;refGene_tx=XM_023256282.1
B4 3029717 3029809 refGene_gn=PFKP;refGene_tx=XM_023256282.1
B4 3030040 3030041 refGene_gn=PFKP;refGene_tx=XM_023256282.1
B4 3030382 3030508 refGene_gn=PFKP;refGene_tx=XM_023256282.1
B4 3036145 3036210 refGene_gn=PFKP;refGene_tx=XM_023256282.1
B4 3036668 3036738 refGene_gn=PFKP;refGene_tx=XM_023256282.1
B4 3036861 3037008 refGene_gn=PFKP;refGene_tx=XM_023256282.1
B4 3040022 3040093 refGene_gn=PFKP;refGene_tx=XM_023256282.1
B4 3041992 3042080 refGene_gn=PFKP;refGene_tx=XM_023256282.1
B4 3042653 3042806 refGene_gn=PFKP;refGene_tx=XM_023256282.1
B4 3043734 3043815 refGene_gn=PFKP;refGene_tx=XM_023256282.1
B4 3056482 3056557 refGene_gn=PITRM1;refGene_tx=XM_023256283.1
B4 3056652 3056745 refGene_gn=PITRM1;refGene_tx=XM_023256283.1
B4 3056746 3056775 refGene_gn=PITRM1;refGene_tx=XM_023256283.1
B4 3057071 3057217 refGene_gn=PITRM1;refGene_tx=XM_023256283.1
B4 3057884 3058010 refGene_gn=PITRM1;refGene_tx=XM_023256283.1
B4 3058916 3059029 refGene_gn=PITRM1;refGene_tx=XM_023256283.1
B4 3059460 3059533 refGene_gn=PITRM1;refGene_tx=XM_023256283.1
B4 3060187 3060325 refGene_gn=PITRM1;refGene_tx=XM_023256283.1
B4 3062164 3062265 refGene_gn=PITRM1;refGene_tx=XM_023256283.1
B4 3062497 3062663 refGene_gn=PITRM1;refGene_tx=XM_023256283.1
B4 3062915 3062992 refGene_gn=PITRM1;refGene_tx=XM_023256283.1
B4 3063094 3063215 refGene_gn=PITRM1;refGene_tx=XM_023256283.1
B4 3064217 3064350 refGene_gn=PITRM1;refGene_tx=XM_023256283.1

B4	3064987	3065104	refGene_gn=PITRM1;refGene_tx=XM_023256283.1
B4	3069230	3069369	refGene_gn=PITRM1;refGene_tx=XM_023256283.1
B4	3080879	3081014	refGene_gn=PITRM1;refGene_tx=XM_023256283.1
B4	3081543	3081640	refGene_gn=PITRM1;refGene_tx=XM_023256283.1
B4	3081822	3081936	refGene_gn=PITRM1;refGene_tx=XM_023256283.1
B4	3082566	3082695	refGene_gn=PITRM1;refGene_tx=XM_023256283.1
B4	3083421	3083510	refGene_gn=PITRM1;refGene_tx=XM_023256283.1
B4	3083775	3083902	refGene_gn=PITRM1;refGene_tx=XM_023256283.1
B4	3091901	3092062	refGene_gn=PITRM1;refGene_tx=XM_023256283.1
B4	3093270	3093367	refGene_gn=PITRM1;refGene_tx=XM_023256283.1
B4	3093441	3093556	refGene_gn=PITRM1;refGene_tx=XM_023256283.1
B4	3094063	3094215	refGene_gn=PITRM1;refGene_tx=XM_023256283.1
B4	3094551	3094658	refGene_gn=PITRM1;refGene_tx=XM_023256283.1
B4	3097386	3097486	refGene_gn=PITRM1;refGene_tx=XM_023256283.1
B4	3100163	3100216	refGene_gn=PITRM1;refGene_tx=XM_023256283.1
B4	3654757	3654809	refGene_gn=KLF6;refGene_tx=XM_003988067.5
B4	3655448	3655572	refGene_gn=KLF6;refGene_tx=XM_003988067.5
B4	3656935	3657509	refGene_gn=KLF6;refGene_tx=XM_003988067.5
B4	3660437	3660539	refGene_gn=KLF6;refGene_tx=XM_003988067.5
B4	4629414	4629960
refGene_gn=LOC101097564;refGene_tx=XM_003988072.5
B4	4647482	4647521	refGene_gn=AKR1E2;refGene_tx=XM_023256285.1
B4	4648112	4648154	refGene_gn=AKR1E2;refGene_tx=XM_023256284.1
B4	4651308	4651476
refGene_gn=AKR1E2;refGene_tx=XM_023256284.1;refGene_gn=AKR1E2;refGene_
tx=XM_023256285.1
B4	4653255	4653372
refGene_gn=AKR1E2;refGene_tx=XM_023256284.1;refGene_gn=AKR1E2;refGene_
tx=XM_023256285.1
B4	4653509	4653587
refGene_gn=AKR1E2;refGene_tx=XM_023256284.1;refGene_gn=AKR1E2;refGene_
tx=XM_023256285.1
B4	4655309	4655432
refGene_gn=AKR1E2;refGene_tx=XM_023256284.1;refGene_gn=AKR1E2;refGene_
tx=XM_023256285.1
B4	4657578	4657676
refGene_gn=AKR1E2;refGene_tx=XM_023256284.1;refGene_gn=AKR1E2;refGene_
tx=XM_023256285.1
B4	4659594	4659667
refGene_gn=AKR1E2;refGene_tx=XM_023256284.1;refGene_gn=AKR1E2;refGene_
tx=XM_023256285.1
B4	4660142	4660226
refGene_gn=AKR1E2;refGene_tx=XM_023256284.1;refGene_gn=AKR1E2;refGene_
tx=XM_023256285.1
B4	4661320	4661403
refGene_gn=AKR1E2;refGene_tx=XM_023256284.1;refGene_gn=AKR1E2;refGene_
tx=XM_023256285.1
B4	4661663	4661706
refGene_gn=AKR1E2;refGene_tx=XM_023256284.1;refGene_gn=AKR1E2;refGene_
tx=XM_023256285.1

B4	4667918	4667928	refGene_gn=AKR1C3;refGene_tx=XM_023256288.1
B4	4667952	4667995
refGene_gn=AKR1C3;refGene_tx=XM_023256287.1;refGene_gn=AKR1C3;refGene_tx=XM_023256289.1
B4	4670849	4670932
refGene_gn=AKR1C3;refGene_tx=XM_023256287.1;refGene_gn=AKR1C3;refGene_tx=XM_023256288.1;refGene_gn=AKR1C3;refGene_tx=XM_023256289.1
B4	4671486	4671652
refGene_gn=AKR1C3;refGene_tx=XM_023256287.1;refGene_gn=AKR1C3;refGene_tx=XM_023256288.1;refGene_gn=AKR1C3;refGene_tx=XM_023256289.1
B4	4672359	4672469
refGene_gn=AKR1C3;refGene_tx=XM_023256287.1;refGene_gn=AKR1C3;refGene_tx=XM_023256288.1;refGene_gn=AKR1C3;refGene_tx=XM_023256289.1
B4	4676273	4676396
refGene_gn=AKR1C3;refGene_tx=XM_023256287.1;refGene_gn=AKR1C3;refGene_tx=XM_023256288.1
B4	4677389	4677467
refGene_gn=AKR1C3;refGene_tx=XM_023256287.1;refGene_gn=AKR1C3;refGene_tx=XM_023256288.1;refGene_gn=AKR1C3;refGene_tx=XM_023256289.1
B4	4678406	4678523
refGene_gn=AKR1C3;refGene_tx=XM_023256287.1;refGene_gn=AKR1C3;refGene_tx=XM_023256288.1;refGene_gn=AKR1C3;refGene_tx=XM_023256289.1
B4	4682112	4682280
refGene_gn=AKR1C3;refGene_tx=XM_023256287.1;refGene_gn=AKR1C3;refGene_tx=XM_023256288.1;refGene_gn=AKR1C3;refGene_tx=XM_023256289.1
B4	4683971	4684058
refGene_gn=AKR1C3;refGene_tx=XM_023256287.1;refGene_gn=AKR1C3;refGene_tx=XM_023256288.1;refGene_gn=AKR1C3;refGene_tx=XM_023256289.1
B4	4761964	4762468	refGene_gn=UCN3;refGene_tx=XM_003988075.3
B4	4775446	4776391	refGene_gn=TUBAL3;refGene_tx=XM_023256291.1
B4	4778192	4778341	refGene_gn=TUBAL3;refGene_tx=XM_023256291.1
B4	4785681	4785784	refGene_gn=TUBAL3;refGene_tx=XM_023256291.1
B4	4801846	4801974	refGene_gn=NET1;refGene_tx=XM_023256290.1
B4	4819692	4819759	refGene_gn=NET1;refGene_tx=XM_023256290.1
B4	4823324	4823384	refGene_gn=NET1;refGene_tx=XM_023256290.1
B4	4848269	4848362	refGene_gn=NET1;refGene_tx=XM_003988076.5
B4	4854225	4854333
refGene_gn=NET1;refGene_tx=XM_003988076.5;refGene_gn=NET1;refGene_tx=XM_023256290.1
B4	4854771	4854939
refGene_gn=NET1;refGene_tx=XM_003988076.5;refGene_gn=NET1;refGene_tx=XM_023256290.1
B4	4855328	4855391
refGene_gn=NET1;refGene_tx=XM_003988076.5;refGene_gn=NET1;refGene_tx=XM_023256290.1
B4	4855670	4855767
refGene_gn=NET1;refGene_tx=XM_003988076.5;refGene_gn=NET1;refGene_tx=XM_023256290.1
B4	4855905	4855982
refGene_gn=NET1;refGene_tx=XM_003988076.5;refGene_gn=NET1;refGene_tx=X

M_023256290.1
B4 4856688 4856946
refGene_gn=NET1;refGene_tx=XM_003988076.5;refGene_gn=NET1;refGene_tx=XM_023256290.1
B4 4857375 4857546
refGene_gn=NET1;refGene_tx=XM_003988076.5;refGene_gn=NET1;refGene_tx=XM_023256290.1
B4 4858686 4858873
refGene_gn=NET1;refGene_tx=XM_003988076.5;refGene_gn=NET1;refGene_tx=XM_023256290.1
B4 4859200 4859592
refGene_gn=NET1;refGene_tx=XM_003988076.5;refGene_gn=NET1;refGene_tx=XM_023256290.1
B4 4882322 4882976
refGene_gn=LOC102902667;refGene_tx=XM_011283583.3
B4 4900868 4901318
refGene_gn=LOC101099307;refGene_tx=XM_006933209.3
B4 5013609 5013737 refGene_gn=ASB13;refGene_tx=XM_023256293.1
B4 5015429 5015621 refGene_gn=ASB13;refGene_tx=XM_023256293.1
B4 5021462 5021597 refGene_gn=ASB13;refGene_tx=XM_023256293.1
B4 5023663 5023814 refGene_gn=ASB13;refGene_tx=XM_023256293.1
B4 5024813 5025001 refGene_gn=ASB13;refGene_tx=XM_023256293.1
B4 5032769 5032812 refGene_gn=ASB13;refGene_tx=XM_023256293.1
B4 5119679 5119749 refGene_gn=FAM208B;refGene_tx=XM_019833889.1
B4 5130359 5130364
refGene_gn=FAM208B;refGene_tx=XM_023256295.1;refGene_gn=FAM208B;refGene_tx=XM_023256296.1;refGene_gn=FAM208B;refGene_tx=XM_023256297.1
B4 5140826 5140960
refGene_gn=FAM208B;refGene_tx=XM_023256295.1;refGene_gn=FAM208B;refGene_tx=XM_023256296.1;refGene_gn=FAM208B;refGene_tx=XM_023256297.1
B4 5146926 5147072
refGene_gn=FAM208B;refGene_tx=XM_019833889.1;refGene_gn=FAM208B;refGene_tx=XM_023256295.1;refGene_gn=FAM208B;refGene_tx=XM_023256296.1;refGene_gn=FAM208B;refGene_tx=XM_023256297.1
B4 5147353 5147449
refGene_gn=FAM208B;refGene_tx=XM_019833889.1;refGene_gn=FAM208B;refGene_tx=XM_023256295.1;refGene_gn=FAM208B;refGene_tx=XM_023256296.1;refGene_gn=FAM208B;refGene_tx=XM_023256297.1
B4 5148737 5148810
refGene_gn=FAM208B;refGene_tx=XM_019833889.1;refGene_gn=FAM208B;refGene_tx=XM_023256295.1;refGene_gn=FAM208B;refGene_tx=XM_023256296.1;refGene_gn=FAM208B;refGene_tx=XM_023256297.1
B4 5152968 5153060
refGene_gn=FAM208B;refGene_tx=XM_019833889.1;refGene_gn=FAM208B;refGene_tx=XM_023256295.1;refGene_gn=FAM208B;refGene_tx=XM_023256296.1;refGene_gn=FAM208B;refGene_tx=XM_023256297.1
B4 5155161 5155299
refGene_gn=FAM208B;refGene_tx=XM_019833889.1;refGene_gn=FAM208B;refGene_tx=XM_023256296.1;refGene_gn=FAM208B;refGene_tx=XM_023256297.1
B4 5155445 5155570

refGene_gn=FAM208B;refGene_tx=XM_006933211.2;refGene_gn=FAM208B;refGene_tx=XM_019833889.1;refGene_gn=FAM208B;refGene_tx=XM_019833890.2;refGene_gn=FAM208B;refGene_tx=XM_019833895.2;refGene_gn=FAM208B;refGene_tx=XM_019833896.1;refGene_gn=FAM208B;refGene_tx=XM_023256295.1;refGene_gn=FAM208B;refGene_tx=XM_023256296.1;refGene_gn=FAM208B;refGene_tx=XM_023256297.1;refGene_gn=FAM208B;refGene_tx=XM_023256298.1
B4 5155649 5155669
refGene_gn=FAM208B;refGene_tx=XM_006933211.2;refGene_gn=FAM208B;refGene_tx=XM_019833889.1;refGene_gn=FAM208B;refGene_tx=XM_019833890.2;refGene_gn=FAM208B;refGene_tx=XM_019833895.2;refGene_gn=FAM208B;refGene_tx=XM_019833896.1;refGene_gn=FAM208B;refGene_tx=XM_023256295.1;refGene_gn=FAM208B;refGene_tx=XM_023256296.1;refGene_gn=FAM208B;refGene_tx=XM_023256297.1;refGene_gn=FAM208B;refGene_tx=XM_023256298.1
B4 5155761 5155861
refGene_gn=FAM208B;refGene_tx=XM_006933211.2;refGene_gn=FAM208B;refGene_tx=XM_019833889.1;refGene_gn=FAM208B;refGene_tx=XM_019833890.2;refGene_gn=FAM208B;refGene_tx=XM_019833895.2;refGene_gn=FAM208B;refGene_tx=XM_019833896.1;refGene_gn=FAM208B;refGene_tx=XM_023256295.1;refGene_gn=FAM208B;refGene_tx=XM_023256296.1;refGene_gn=FAM208B;refGene_tx=XM_023256297.1;refGene_gn=FAM208B;refGene_tx=XM_023256298.1
B4 5157573 5157674
refGene_gn=FAM208B;refGene_tx=XM_006933211.2;refGene_gn=FAM208B;refGene_tx=XM_019833889.1;refGene_gn=FAM208B;refGene_tx=XM_019833890.2;refGene_gn=FAM208B;refGene_tx=XM_019833895.2;refGene_gn=FAM208B;refGene_tx=XM_019833896.1;refGene_gn=FAM208B;refGene_tx=XM_023256295.1;refGene_gn=FAM208B;refGene_tx=XM_023256296.1;refGene_gn=FAM208B;refGene_tx=XM_023256297.1;refGene_gn=FAM208B;refGene_tx=XM_023256298.1
B4 5158355 5158459
refGene_gn=FAM208B;refGene_tx=XM_006933211.2;refGene_gn=FAM208B;refGene_tx=XM_019833889.1;refGene_gn=FAM208B;refGene_tx=XM_019833890.2;refGene_gn=FAM208B;refGene_tx=XM_019833895.2;refGene_gn=FAM208B;refGene_tx=XM_019833896.1;refGene_gn=FAM208B;refGene_tx=XM_023256295.1;refGene_gn=FAM208B;refGene_tx=XM_023256296.1;refGene_gn=FAM208B;refGene_tx=XM_023256297.1;refGene_gn=FAM208B;refGene_tx=XM_023256298.1
B4 5165307 5165380
refGene_gn=FAM208B;refGene_tx=XM_006933211.2;refGene_gn=FAM208B;refGene_tx=XM_019833889.1;refGene_gn=FAM208B;refGene_tx=XM_019833890.2;refGene_gn=FAM208B;refGene_tx=XM_019833895.2;refGene_gn=FAM208B;refGene_tx=XM_019833896.1;refGene_gn=FAM208B;refGene_tx=XM_023256295.1;refGene_gn=FAM208B;refGene_tx=XM_023256296.1;refGene_gn=FAM208B;refGene_tx=XM_023256297.1;refGene_gn=FAM208B;refGene_tx=XM_023256298.1
B4 5165479 5165542
refGene_gn=FAM208B;refGene_tx=XM_006933211.2;refGene_gn=FAM208B;refGene_tx=XM_019833889.1;refGene_gn=FAM208B;refGene_tx=XM_019833890.2;refGene_gn=FAM208B;refGene_tx=XM_019833895.2;refGene_gn=FAM208B;refGene_tx=XM_019833896.1;refGene_gn=FAM208B;refGene_tx=XM_023256295.1;refGene_gn=FAM208B;refGene_tx=XM_023256296.1;refGene_gn=FAM208B;refGene_tx=XM_023256297.1;refGene_gn=FAM208B;refGene_tx=XM_023256298.1
B4 5171007 5171724
refGene_gn=FAM208B;refGene_tx=XM_006933211.2;refGene_gn=FAM208B;refGen

e_tx=XM_019833889.1;refGene_gn=FAM208B;refGene_tx=XM_019833890.2;refGene_gn=FAM208B;refGene_tx=XM_019833895.2;refGene_gn=FAM208B;refGene_tx=XM_019833896.1;refGene_gn=FAM208B;refGene_tx=XM_023256295.1;refGene_gn=FAM208B;refGene_tx=XM_023256296.1;refGene_gn=FAM208B;refGene_tx=XM_023256297.1;refGene_gn=FAM208B;refGene_tx=XM_023256298.1
B4 5174854 5175097
refGene_gn=FAM208B;refGene_tx=XM_006933211.2;refGene_gn=FAM208B;refGene_tx=XM_019833889.1;refGene_gn=FAM208B;refGene_tx=XM_019833890.2;refGene_gn=FAM208B;refGene_tx=XM_019833895.2;refGene_gn=FAM208B;refGene_tx=XM_019833896.1;refGene_gn=FAM208B;refGene_tx=XM_023256295.1;refGene_gn=FAM208B;refGene_tx=XM_023256297.1;refGene_gn=FAM208B;refGene_tx=XM_023256298.1
B4 5179257 5180131
refGene_gn=FAM208B;refGene_tx=XM_006933211.2;refGene_gn=FAM208B;refGene_tx=XM_019833889.1;refGene_gn=FAM208B;refGene_tx=XM_019833890.2;refGene_gn=FAM208B;refGene_tx=XM_019833895.2;refGene_gn=FAM208B;refGene_tx=XM_019833896.1;refGene_gn=FAM208B;refGene_tx=XM_023256295.1;refGene_gn=FAM208B;refGene_tx=XM_023256296.1;refGene_gn=FAM208B;refGene_tx=XM_023256297.1;refGene_gn=FAM208B;refGene_tx=XM_023256298.1
B4 5181776 5182206
refGene_gn=FAM208B;refGene_tx=XM_006933211.2;refGene_gn=FAM208B;refGene_tx=XM_019833889.1;refGene_gn=FAM208B;refGene_tx=XM_019833890.2;refGene_gn=FAM208B;refGene_tx=XM_019833895.2;refGene_gn=FAM208B;refGene_tx=XM_019833896.1;refGene_gn=FAM208B;refGene_tx=XM_023256295.1;refGene_gn=FAM208B;refGene_tx=XM_023256296.1;refGene_gn=FAM208B;refGene_tx=XM_023256297.1;refGene_gn=FAM208B;refGene_tx=XM_023256298.1
B4 5184723 5188575
refGene_gn=FAM208B;refGene_tx=XM_006933211.2;refGene_gn=FAM208B;refGene_tx=XM_019833889.1;refGene_gn=FAM208B;refGene_tx=XM_019833890.2;refGene_gn=FAM208B;refGene_tx=XM_019833895.2;refGene_gn=FAM208B;refGene_tx=XM_019833896.1;refGene_gn=FAM208B;refGene_tx=XM_023256295.1;refGene_gn=FAM208B;refGene_tx=XM_023256296.1;refGene_gn=FAM208B;refGene_tx=XM_023256297.1;refGene_gn=FAM208B;refGene_tx=XM_023256298.1
B4 5194608 5194734
refGene_gn=FAM208B;refGene_tx=XM_006933211.2;refGene_gn=FAM208B;refGene_tx=XM_019833889.1;refGene_gn=FAM208B;refGene_tx=XM_019833890.2;refGene_gn=FAM208B;refGene_tx=XM_019833895.2;refGene_gn=FAM208B;refGene_tx=XM_019833896.1;refGene_gn=FAM208B;refGene_tx=XM_023256295.1;refGene_gn=FAM208B;refGene_tx=XM_023256296.1;refGene_gn=FAM208B;refGene_tx=XM_023256297.1;refGene_gn=FAM208B;refGene_tx=XM_023256298.1
B4 5195490 5195644
refGene_gn=FAM208B;refGene_tx=XM_006933211.2;refGene_gn=FAM208B;refGene_tx=XM_019833889.1;refGene_gn=FAM208B;refGene_tx=XM_019833890.2;refGene_gn=FAM208B;refGene_tx=XM_019833895.2;refGene_gn=FAM208B;refGene_tx=XM_019833896.1;refGene_gn=FAM208B;refGene_tx=XM_023256295.1;refGene_gn=FAM208B;refGene_tx=XM_023256296.1;refGene_gn=FAM208B;refGene_tx=XM_023256297.1;refGene_gn=FAM208B;refGene_tx=XM_023256298.1
B4 5196499 5196605
refGene_gn=FAM208B;refGene_tx=XM_006933211.2;refGene_gn=FAM208B;refGene_tx=XM_019833889.1;refGene_gn=FAM208B;refGene_tx=XM_019833890.2;refGe

ne_gn=FAM208B;refGene_tx=XM_019833895.2;refGene_gn=FAM208B;refGene_tx=XM_019833896.1;refGene_gn=FAM208B;refGene_tx=XM_023256295.1;refGene_gn=FAM208B;refGene_tx=XM_023256296.1;refGene_gn=FAM208B;refGene_tx=XM_023256297.1;refGene_gn=FAM208B;refGene_tx=XM_023256298.1
B4 5198587 5198769
refGene_gn=FAM208B;refGene_tx=XM_006933211.2;refGene_gn=FAM208B;refGene_tx=XM_019833889.1;refGene_gn=FAM208B;refGene_tx=XM_019833890.2;refGene_gn=FAM208B;refGene_tx=XM_019833895.2;refGene_gn=FAM208B;refGene_tx=XM_019833896.1;refGene_gn=FAM208B;refGene_tx=XM_023256295.1;refGene_gn=FAM208B;refGene_tx=XM_023256296.1;refGene_gn=FAM208B;refGene_tx=XM_023256297.1;refGene_gn=FAM208B;refGene_tx=XM_023256298.1
B4 5199251 5199370
refGene_gn=FAM208B;refGene_tx=XM_006933211.2;refGene_gn=FAM208B;refGene_tx=XM_019833889.1;refGene_gn=FAM208B;refGene_tx=XM_019833890.2;refGene_gn=FAM208B;refGene_tx=XM_019833895.2;refGene_gn=FAM208B;refGene_tx=XM_019833896.1;refGene_gn=FAM208B;refGene_tx=XM_023256295.1;refGene_gn=FAM208B;refGene_tx=XM_023256296.1;refGene_gn=FAM208B;refGene_tx=XM_023256297.1;refGene_gn=FAM208B;refGene_tx=XM_023256298.1
B4 5202091 5202238 refGene_gn=GDI2;refGene_tx=XM_023256299.1
B4 5202331 5202386 refGene_gn=GDI2;refGene_tx=XM_023256299.1
B4 5202588 5202733 refGene_gn=GDI2;refGene_tx=XM_023256299.1
B4 5204007 5204179 refGene_gn=GDI2;refGene_tx=XM_023256299.1
B4 5205368 5205468 refGene_gn=GDI2;refGene_tx=XM_023256299.1
B4 5211305 5211437 refGene_gn=GDI2;refGene_tx=XM_023256299.1
B4 5211672 5211871 refGene_gn=GDI2;refGene_tx=XM_023256299.1
B4 5220124 5220259 refGene_gn=GDI2;refGene_tx=XM_023256299.1
B4 5223670 5223770 refGene_gn=GDI2;refGene_tx=XM_023256299.1
B4 5226800 5226908 refGene_gn=GDI2;refGene_tx=XM_023256299.1
B4 5235065 5235110 refGene_gn=GDI2;refGene_tx=XM_023256299.1
B4 5273555 5273713 refGene_gn=ANKRD16;refGene_tx=XM_011283601.3
B4 5274223 5274234 refGene_gn=ANKRD16;refGene_tx=XM_023256306.1
B4 5274701 5274780
refGene_gn=ANKRD16;refGene_tx=XM_011283601.3;refGene_gn=ANKRD16;refGene_tx=XM_023256306.1
B4 5275824 5275986
refGene_gn=ANKRD16;refGene_tx=XM_011283601.3;refGene_gn=ANKRD16;refGene_tx=XM_023256306.1
B4 5276424 5276533
refGene_gn=ANKRD16;refGene_tx=XM_011283601.3;refGene_gn=ANKRD16;refGene_tx=XM_023256306.1
B4 5279583 5279626
refGene_gn=ANKRD16;refGene_tx=XM_011283601.3;refGene_gn=ANKRD16;refGene_tx=XM_023256306.1
B4 5281469 5281690
refGene_gn=ANKRD16;refGene_tx=XM_011283601.3;refGene_gn=ANKRD16;refGene_tx=XM_023256306.1
B4 5283632 5283952
refGene_gn=ANKRD16;refGene_tx=XM_011283601.3;refGene_gn=ANKRD16;refGene_tx=XM_023256306.1
B4 5284764 5284765 refGene_gn=FBXO18;refGene_tx=XM_023256302.1

B4	5289951	5290009
refGene_gn=FBXO18;refGene_tx=XM_023256300.1;refGene_gn=FBXO18;refGene_tx=XM_023256301.1
B4	5296542	5296698
refGene_gn=FBXO18;refGene_tx=XM_023256300.1;refGene_gn=FBXO18;refGene_tx=XM_023256301.1;refGene_gn=FBXO18;refGene_tx=XM_023256302.1
B4	5298855	5299457
refGene_gn=FBXO18;refGene_tx=XM_023256300.1;refGene_gn=FBXO18;refGene_tx=XM_023256301.1;refGene_gn=FBXO18;refGene_tx=XM_023256302.1;refGene_gn=FBXO18;refGene_tx=XM_023256303.1;refGene_gn=FBXO18;refGene_tx=XM_023256304.1;refGene_gn=FBXO18;refGene_tx=XM_023256305.1
B4	5301800	5301931
refGene_gn=FBXO18;refGene_tx=XM_023256300.1;refGene_gn=FBXO18;refGene_tx=XM_023256301.1;refGene_gn=FBXO18;refGene_tx=XM_023256302.1;refGene_gn=FBXO18;refGene_tx=XM_023256303.1;refGene_gn=FBXO18;refGene_tx=XM_023256304.1;refGene_gn=FBXO18;refGene_tx=XM_023256305.1
B4	5302036	5302172
refGene_gn=FBXO18;refGene_tx=XM_023256300.1;refGene_gn=FBXO18;refGene_tx=XM_023256301.1;refGene_gn=FBXO18;refGene_tx=XM_023256302.1;refGene_gn=FBXO18;refGene_tx=XM_023256303.1;refGene_gn=FBXO18;refGene_tx=XM_023256304.1;refGene_gn=FBXO18;refGene_tx=XM_023256305.1
B4	5303444	5303635
refGene_gn=FBXO18;refGene_tx=XM_023256300.1;refGene_gn=FBXO18;refGene_tx=XM_023256301.1;refGene_gn=FBXO18;refGene_tx=XM_023256302.1;refGene_gn=FBXO18;refGene_tx=XM_023256303.1;refGene_gn=FBXO18;refGene_tx=XM_023256304.1;refGene_gn=FBXO18;refGene_tx=XM_023256305.1
B4	5306006	5306099
refGene_gn=FBXO18;refGene_tx=XM_023256300.1;refGene_gn=FBXO18;refGene_tx=XM_023256301.1;refGene_gn=FBXO18;refGene_tx=XM_023256302.1;refGene_gn=FBXO18;refGene_tx=XM_023256303.1;refGene_gn=FBXO18;refGene_tx=XM_023256304.1;refGene_gn=FBXO18;refGene_tx=XM_023256305.1
B4	5306431	5306526
refGene_gn=FBXO18;refGene_tx=XM_023256300.1;refGene_gn=FBXO18;refGene_tx=XM_023256301.1;refGene_gn=FBXO18;refGene_tx=XM_023256302.1;refGene_gn=FBXO18;refGene_tx=XM_023256303.1;refGene_gn=FBXO18;refGene_tx=XM_023256304.1;refGene_gn=FBXO18;refGene_tx=XM_023256305.1
B4	5308264	5308433
refGene_gn=FBXO18;refGene_tx=XM_023256300.1;refGene_gn=FBXO18;refGene_tx=XM_023256301.1;refGene_gn=FBXO18;refGene_tx=XM_023256302.1;refGene_gn=FBXO18;refGene_tx=XM_023256303.1;refGene_gn=FBXO18;refGene_tx=XM_023256304.1;refGene_gn=FBXO18;refGene_tx=XM_023256305.1
B4	5309149	5309372
refGene_gn=FBXO18;refGene_tx=XM_023256300.1;refGene_gn=FBXO18;refGene_tx=XM_023256301.1;refGene_gn=FBXO18;refGene_tx=XM_023256302.1;refGene_gn=FBXO18;refGene_tx=XM_023256303.1;refGene_gn=FBXO18;refGene_tx=XM_023256304.1;refGene_gn=FBXO18;refGene_tx=XM_023256305.1
B4	5310153	5310241
refGene_gn=FBXO18;refGene_tx=XM_023256300.1;refGene_gn=FBXO18;refGene_tx=XM_023256301.1;refGene_gn=FBXO18;refGene_tx=XM_023256302.1;refGene_gn=FBXO18;refGene_tx=XM_023256303.1;refGene_gn=FBXO18;refGene_tx=XM_02

3256304.1;refGene_gn=FBXO18;refGene_tx=XM_023256305.1
B4 5310325 5310412
refGene_gn=FBXO18;refGene_tx=XM_023256300.1;refGene_gn=FBXO18;refGene_tx=XM_023256301.1;refGene_gn=FBXO18;refGene_tx=XM_023256302.1;refGene_gn=FBXO18;refGene_tx=XM_023256303.1;refGene_gn=FBXO18;refGene_tx=XM_023256304.1;refGene_gn=FBXO18;refGene_tx=XM_023256305.1
B4 5311109 5311246
refGene_gn=FBXO18;refGene_tx=XM_023256300.1;refGene_gn=FBXO18;refGene_tx=XM_023256301.1;refGene_gn=FBXO18;refGene_tx=XM_023256302.1;refGene_gn=FBXO18;refGene_tx=XM_023256303.1;refGene_gn=FBXO18;refGene_tx=XM_023256304.1;refGene_gn=FBXO18;refGene_tx=XM_023256305.1
B4 5313256 5313356
refGene_gn=FBXO18;refGene_tx=XM_023256300.1;refGene_gn=FBXO18;refGene_tx=XM_023256301.1;refGene_gn=FBXO18;refGene_tx=XM_023256302.1;refGene_gn=FBXO18;refGene_tx=XM_023256303.1;refGene_gn=FBXO18;refGene_tx=XM_023256304.1;refGene_gn=FBXO18;refGene_tx=XM_023256305.1
B4 5313447 5313569
refGene_gn=FBXO18;refGene_tx=XM_023256300.1;refGene_gn=FBXO18;refGene_tx=XM_023256301.1;refGene_gn=FBXO18;refGene_tx=XM_023256302.1;refGene_gn=FBXO18;refGene_tx=XM_023256303.1;refGene_gn=FBXO18;refGene_tx=XM_023256304.1;refGene_gn=FBXO18;refGene_tx=XM_023256305.1
B4 5315583 5315659
refGene_gn=FBXO18;refGene_tx=XM_023256300.1;refGene_gn=FBXO18;refGene_tx=XM_023256301.1;refGene_gn=FBXO18;refGene_tx=XM_023256302.1;refGene_gn=FBXO18;refGene_tx=XM_023256303.1;refGene_gn=FBXO18;refGene_tx=XM_023256304.1;refGene_gn=FBXO18;refGene_tx=XM_023256305.1
B4 5316247 5316445
refGene_gn=FBXO18;refGene_tx=XM_023256300.1;refGene_gn=FBXO18;refGene_tx=XM_023256301.1;refGene_gn=FBXO18;refGene_tx=XM_023256302.1;refGene_gn=FBXO18;refGene_tx=XM_023256303.1;refGene_gn=FBXO18;refGene_tx=XM_023256304.1;refGene_gn=FBXO18;refGene_tx=XM_023256305.1
B4 5316948 5317074
refGene_gn=FBXO18;refGene_tx=XM_023256300.1;refGene_gn=FBXO18;refGene_tx=XM_023256301.1;refGene_gn=FBXO18;refGene_tx=XM_023256302.1;refGene_gn=FBXO18;refGene_tx=XM_023256303.1;refGene_gn=FBXO18;refGene_tx=XM_023256304.1;refGene_gn=FBXO18;refGene_tx=XM_023256305.1
B4 5318595 5318702
refGene_gn=FBXO18;refGene_tx=XM_023256300.1;refGene_gn=FBXO18;refGene_tx=XM_023256301.1;refGene_gn=FBXO18;refGene_tx=XM_023256302.1;refGene_gn=FBXO18;refGene_tx=XM_023256303.1;refGene_gn=FBXO18;refGene_tx=XM_023256304.1;refGene_gn=FBXO18;refGene_tx=XM_023256305.1
B4 5331830 5331962
refGene_gn=FBXO18;refGene_tx=XM_023256300.1;refGene_gn=FBXO18;refGene_tx=XM_023256301.1;refGene_gn=FBXO18;refGene_tx=XM_023256302.1;refGene_gn=FBXO18;refGene_tx=XM_023256303.1;refGene_gn=FBXO18;refGene_tx=XM_023256304.1;refGene_gn=FBXO18;refGene_tx=XM_023256305.1
B4 5332457 5332628
refGene_gn=FBXO18;refGene_tx=XM_023256300.1;refGene_gn=FBXO18;refGene_tx=XM_023256301.1;refGene_gn=FBXO18;refGene_tx=XM_023256302.1;refGene_gn=FBXO18;refGene_tx=XM_023256303.1;refGene_gn=FBXO18;refGene_tx=XM_02

3256304.1;refGene_gn=FBXO18;refGene_tx=XM_023256305.1
B4 5339374 5339414 refGene_gn=IL15RA;refGene_tx=XM_023256309.1
B4 5348953 5349125
refGene_gn=IL15RA;refGene_tx=XM_023256307.1;refGene_gn=IL15RA;refGene_tx=XM_023256308.1
B4 5351854 5351927
refGene_gn=IL15RA;refGene_tx=XM_023256307.1;refGene_gn=IL15RA;refGene_tx=XM_023256308.1;refGene_gn=IL15RA;refGene_tx=XM_023256309.1
B4 5357636 5357669
refGene_gn=IL15RA;refGene_tx=XM_023256307.1;refGene_gn=IL15RA;refGene_tx=XM_023256308.1;refGene_gn=IL15RA;refGene_tx=XM_023256309.1
B4 5359289 5359484
refGene_gn=IL15RA;refGene_tx=XM_023256307.1;refGene_gn=IL15RA;refGene_tx=XM_023256308.1;refGene_gn=IL15RA;refGene_tx=XM_023256309.1
B4 5361946 5362039
refGene_gn=IL15RA;refGene_tx=XM_023256307.1;refGene_gn=IL15RA;refGene_tx=XM_023256309.1
B4 5366760 5366955
refGene_gn=IL15RA;refGene_tx=XM_023256307.1;refGene_gn=IL15RA;refGene_tx=XM_023256308.1;refGene_gn=IL15RA;refGene_tx=XM_023256309.1
B4 5397206 5397309
refGene_gn=IL15RA;refGene_tx=XM_023256307.1;refGene_gn=IL15RA;refGene_tx=XM_023256308.1;refGene_gn=IL15RA;refGene_tx=XM_023256309.1
B4 5445888 5445913
refGene_gn=IL2RA;refGene_tx=NM_001009355.1;refGene_gn=IL2RA;refGene_tx=XM_019833862.2;refGene_gn=IL2RA;refGene_tx=XM_019833863.2
B4 5450924 5450991
refGene_gn=IL2RA;refGene_tx=NM_001009355.1;refGene_gn=IL2RA;refGene_tx=XM_019833862.2;refGene_gn=IL2RA;refGene_tx=XM_019833863.2
B4 5452522 5452594
refGene_gn=IL2RA;refGene_tx=NM_001009355.1;refGene_gn=IL2RA;refGene_tx=XM_019833862.2;refGene_gn=IL2RA;refGene_tx=XM_019833863.2
B4 5452929 5453010
refGene_gn=IL2RA;refGene_tx=NM_001009355.1;refGene_gn=IL2RA;refGene_tx=XM_019833862.2;refGene_gn=IL2RA;refGene_tx=XM_019833863.2
B4 5454645 5454861
refGene_gn=IL2RA;refGene_tx=NM_001009355.1;refGene_gn=IL2RA;refGene_tx=XM_019833862.2;refGene_gn=IL2RA;refGene_tx=XM_019833863.2
B4 5457247 5457358
refGene_gn=IL2RA;refGene_tx=NM_001009355.1;refGene_gn=IL2RA;refGene_tx=XM_019833862.2;refGene_gn=IL2RA;refGene_tx=XM_019833863.2
B4 5460373 5460565
refGene_gn=IL2RA;refGene_tx=NM_001009355.1;refGene_gn=IL2RA;refGene_tx=XM_019833862.2;refGene_gn=IL2RA;refGene_tx=XM_019833863.2
B4 5499514 5499578 refGene_gn=IL2RA;refGene_tx=NM_001009355.1
B4 5552965 5553088
refGene_gn=RBM17;refGene_tx=XM_003988079.4;refGene_gn=RBM17;refGene_tx=XM_023256310.1;refGene_gn=RBM17;refGene_tx=XM_023256311.1
B4 5558269 5558386
refGene_gn=RBM17;refGene_tx=XM_003988079.4;refGene_gn=RBM17;refGene_tx

=XM_023256310.1;refGene_gn=RBM17;refGene_tx=XM_023256311.1
B4 5561386 5561553
refGene_gn=RBM17;refGene_tx=XM_003988079.4;refGene_gn=RBM17;refGene_tx=XM_023256310.1;refGene_gn=RBM17;refGene_tx=XM_023256311.1
B4 5563445 5563543
refGene_gn=RBM17;refGene_tx=XM_003988079.4;refGene_gn=RBM17;refGene_tx=XM_023256310.1;refGene_gn=RBM17;refGene_tx=XM_023256311.1
B4 5566505 5566562
refGene_gn=RBM17;refGene_tx=XM_003988079.4;refGene_gn=RBM17;refGene_tx=XM_023256310.1
B4 5567752 5567894
refGene_gn=RBM17;refGene_tx=XM_003988079.4;refGene_gn=RBM17;refGene_tx=XM_023256310.1;refGene_gn=RBM17;refGene_tx=XM_023256311.1
B4 5570038 5570190
refGene_gn=RBM17;refGene_tx=XM_003988079.4;refGene_gn=RBM17;refGene_tx=XM_023256310.1;refGene_gn=RBM17;refGene_tx=XM_023256311.1
B4 5571276 5571350
refGene_gn=RBM17;refGene_tx=XM_003988079.4;refGene_gn=RBM17;refGene_tx=XM_023256310.1;refGene_gn=RBM17;refGene_tx=XM_023256311.1
B4 5572186 5572285
refGene_gn=RBM17;refGene_tx=XM_003988079.4;refGene_gn=RBM17;refGene_tx=XM_023256310.1;refGene_gn=RBM17;refGene_tx=XM_023256311.1
B4 5573361 5573434
refGene_gn=RBM17;refGene_tx=XM_003988079.4;refGene_gn=RBM17;refGene_tx=XM_023256310.1;refGene_gn=RBM17;refGene_tx=XM_023256311.1
B4 5573561 5573665
refGene_gn=RBM17;refGene_tx=XM_003988079.4;refGene_gn=RBM17;refGene_tx=XM_023256310.1;refGene_gn=RBM17;refGene_tx=XM_023256311.1
B4 5599963 5599979 refGene_gn=PFKFB3;refGene_tx=XM_023256312.1
B4 5648664 5648740
refGene_gn=PFKFB3;refGene_tx=XM_003988081.5;refGene_gn=PFKFB3;refGene_tx=XM_006933216.4;refGene_gn=PFKFB3;refGene_tx=XM_019833901.2
B4 5655792 5655918
refGene_gn=PFKFB3;refGene_tx=XM_003988081.5;refGene_gn=PFKFB3;refGene_tx=XM_006933216.4;refGene_gn=PFKFB3;refGene_tx=XM_019833901.2;refGene_gn=PFKFB3;refGene_tx=XM_023256312.1
B4 5657082 5657179
refGene_gn=PFKFB3;refGene_tx=XM_003988081.5;refGene_gn=PFKFB3;refGene_tx=XM_006933216.4;refGene_gn=PFKFB3;refGene_tx=XM_019833901.2;refGene_gn=PFKFB3;refGene_tx=XM_023256312.1
B4 5657868 5657935
refGene_gn=PFKFB3;refGene_tx=XM_003988081.5;refGene_gn=PFKFB3;refGene_tx=XM_006933216.4;refGene_gn=PFKFB3;refGene_tx=XM_019833901.2;refGene_gn=PFKFB3;refGene_tx=XM_023256312.1
B4 5658367 5658442
refGene_gn=PFKFB3;refGene_tx=XM_003988081.5;refGene_gn=PFKFB3;refGene_tx=XM_006933216.4;refGene_gn=PFKFB3;refGene_tx=XM_019833901.2;refGene_gn=PFKFB3;refGene_tx=XM_023256312.1
B4 5658803 5658860
refGene_gn=PFKFB3;refGene_tx=XM_003988081.5;refGene_gn=PFKFB3;refGene_

tx=XM_006933216.4;refGene_gn=PFKFB3;refGene_tx=XM_019833901.2;refGene_gn=PFKFB3;refGene_tx=XM_023256312.1
B4 5661499 5661624
refGene_gn=PFKFB3;refGene_tx=XM_003988081.5;refGene_gn=PFKFB3;refGene_tx=XM_006933216.4;refGene_gn=PFKFB3;refGene_tx=XM_019833901.2;refGene_gn=PFKFB3;refGene_tx=XM_023256312.1
B4 5662106 5662314
refGene_gn=PFKFB3;refGene_tx=XM_003988081.5;refGene_gn=PFKFB3;refGene_tx=XM_006933216.4;refGene_gn=PFKFB3;refGene_tx=XM_019833901.2;refGene_gn=PFKFB3;refGene_tx=XM_023256312.1
B4 5662514 5662661
refGene_gn=PFKFB3;refGene_tx=XM_003988081.5;refGene_gn=PFKFB3;refGene_tx=XM_006933216.4;refGene_gn=PFKFB3;refGene_tx=XM_019833901.2;refGene_gn=PFKFB3;refGene_tx=XM_023256312.1
B4 5662796 5662901
refGene_gn=PFKFB3;refGene_tx=XM_003988081.5;refGene_gn=PFKFB3;refGene_tx=XM_006933216.4;refGene_gn=PFKFB3;refGene_tx=XM_019833901.2;refGene_gn=PFKFB3;refGene_tx=XM_023256312.1
B4 5664248 5664378
refGene_gn=PFKFB3;refGene_tx=XM_003988081.5;refGene_gn=PFKFB3;refGene_tx=XM_006933216.4;refGene_gn=PFKFB3;refGene_tx=XM_019833901.2;refGene_gn=PFKFB3;refGene_tx=XM_023256312.1
B4 5665340 5665403
refGene_gn=PFKFB3;refGene_tx=XM_003988081.5;refGene_gn=PFKFB3;refGene_tx=XM_006933216.4;refGene_gn=PFKFB3;refGene_tx=XM_019833901.2;refGene_gn=PFKFB3;refGene_tx=XM_023256312.1
B4 5665526 5665591
refGene_gn=PFKFB3;refGene_tx=XM_003988081.5;refGene_gn=PFKFB3;refGene_tx=XM_006933216.4;refGene_gn=PFKFB3;refGene_tx=XM_019833901.2;refGene_gn=PFKFB3;refGene_tx=XM_023256312.1
B4 5667308 5667482
refGene_gn=PFKFB3;refGene_tx=XM_003988081.5;refGene_gn=PFKFB3;refGene_tx=XM_006933216.4;refGene_gn=PFKFB3;refGene_tx=XM_019833901.2;refGene_gn=PFKFB3;refGene_tx=XM_023256312.1
B4 5668908 5668931
refGene_gn=PFKFB3;refGene_tx=XM_006933216.4;refGene_gn=PFKFB3;refGene_tx=XM_019833901.2;refGene_gn=PFKFB3;refGene_tx=XM_023256312.1
B4 5671261 5671304
refGene_gn=PFKFB3;refGene_tx=XM_019833901.2;refGene_gn=PFKFB3;refGene_tx=XM_023256312.1
B4 5672920 5672965
refGene_gn=PFKFB3;refGene_tx=XM_003988081.5;refGene_gn=PFKFB3;refGene_tx=XM_006933216.4
B4 5938178 5938334
refGene_gn=PRKCQ;refGene_tx=XM_003988082.5;refGene_gn=PRKCQ;refGene_tx=XM_006933217.4;refGene_gn=PRKCQ;refGene_tx=XM_019833904.2
B4 5941076 5941205
refGene_gn=PRKCQ;refGene_tx=XM_003988082.5;refGene_gn=PRKCQ;refGene_tx=XM_006933217.4;refGene_gn=PRKCQ;refGene_tx=XM_019833904.2
B4 5949359 5949548

refGene_gn=PRKCQ;refGene_tx=XM_003988082.5;refGene_gn=PRKCQ;refGene_tx=XM_006933217.4;refGene_gn=PRKCQ;refGene_tx=XM_019833904.2
B4 5960954 5961093
refGene_gn=PRKCQ;refGene_tx=XM_003988082.5;refGene_gn=PRKCQ;refGene_tx=XM_006933217.4;refGene_gn=PRKCQ;refGene_tx=XM_019833904.2
B4 5975073 5975136
refGene_gn=PRKCQ;refGene_tx=XM_003988082.5;refGene_gn=PRKCQ;refGene_tx=XM_006933217.4;refGene_gn=PRKCQ;refGene_tx=XM_019833904.2
B4 5976667 5976759
refGene_gn=PRKCQ;refGene_tx=XM_003988082.5;refGene_gn=PRKCQ;refGene_tx=XM_006933217.4;refGene_gn=PRKCQ;refGene_tx=XM_019833904.2
B4 6031864 6032038
refGene_gn=PRKCQ;refGene_tx=XM_003988082.5;refGene_gn=PRKCQ;refGene_tx=XM_006933217.4;refGene_gn=PRKCQ;refGene_tx=XM_019833904.2
B4 6035274 6035435
refGene_gn=PRKCQ;refGene_tx=XM_003988082.5;refGene_gn=PRKCQ;refGene_tx=XM_006933217.4;refGene_gn=PRKCQ;refGene_tx=XM_019833904.2
B4 6044293 6044411
refGene_gn=PRKCQ;refGene_tx=XM_003988082.5;refGene_gn=PRKCQ;refGene_tx=XM_006933217.4;refGene_gn=PRKCQ;refGene_tx=XM_019833904.2
B4 6045139 6045249
refGene_gn=PRKCQ;refGene_tx=XM_003988082.5;refGene_gn=PRKCQ;refGene_tx=XM_006933217.4;refGene_gn=PRKCQ;refGene_tx=XM_019833904.2
B4 6049143 6049273
refGene_gn=PRKCQ;refGene_tx=XM_003988082.5;refGene_gn=PRKCQ;refGene_tx=XM_006933217.4;refGene_gn=PRKCQ;refGene_tx=XM_019833904.2
B4 6054438 6054524
refGene_gn=PRKCQ;refGene_tx=XM_003988082.5;refGene_gn=PRKCQ;refGene_tx=XM_006933217.4;refGene_gn=PRKCQ;refGene_tx=XM_019833904.2
B4 6054620 6054652
refGene_gn=PRKCQ;refGene_tx=XM_003988082.5;refGene_gn=PRKCQ;refGene_tx=XM_006933217.4;refGene_gn=PRKCQ;refGene_tx=XM_019833904.2
B4 6055789 6055952
refGene_gn=PRKCQ;refGene_tx=XM_003988082.5;refGene_gn=PRKCQ;refGene_tx=XM_006933217.4;refGene_gn=PRKCQ;refGene_tx=XM_019833904.2
B4 6062115 6062176
refGene_gn=PRKCQ;refGene_tx=XM_003988082.5;refGene_gn=PRKCQ;refGene_tx=XM_006933217.4
B4 6065540 6065740
refGene_gn=PRKCQ;refGene_tx=XM_003988082.5;refGene_gn=PRKCQ;refGene_tx=XM_006933217.4;refGene_gn=PRKCQ;refGene_tx=XM_019833904.2
B4 6069604 6069731
refGene_gn=PRKCQ;refGene_tx=XM_003988082.5;refGene_gn=PRKCQ;refGene_tx=XM_006933217.4
B4 6070591 6070609 refGene_gn=PRKCQ;refGene_tx=XM_003988082.5
B4 6581029 6581170
refGene_gn=SFMBT2;refGene_tx=XM_023256313.1;refGene_gn=SFMBT2;refGene_tx=XM_023256314.1;refGene_gn=SFMBT2;refGene_tx=XM_023256315.1;refGene_gn=SFMBT2;refGene_tx=XM_023256316.1;refGene_gn=SFMBT2;refGene_tx=XM_023256317.1;refGene_gn=SFMBT2;refGene_tx=XM_023256319.1

B4 6588669 6588702 refGene_gn=SFMBT2;refGene_tx=XM_023256318.1
B4 6589889 6590018
refGene_gn=SFMBT2;refGene_tx=XM_023256313.1;refGene_gn=SFMBT2;refGene_tx=XM_023256314.1;refGene_gn=SFMBT2;refGene_tx=XM_023256315.1;refGene_gn=SFMBT2;refGene_tx=XM_023256316.1;refGene_gn=SFMBT2;refGene_tx=XM_023256317.1;refGene_gn=SFMBT2;refGene_tx=XM_023256318.1;refGene_gn=SFMBT2;refGene_tx=XM_023256319.1
B4 6590997 6591261
refGene_gn=SFMBT2;refGene_tx=XM_023256313.1;refGene_gn=SFMBT2;refGene_tx=XM_023256314.1;refGene_gn=SFMBT2;refGene_tx=XM_023256315.1;refGene_gn=SFMBT2;refGene_tx=XM_023256316.1;refGene_gn=SFMBT2;refGene_tx=XM_023256317.1;refGene_gn=SFMBT2;refGene_tx=XM_023256318.1;refGene_gn=SFMBT2;refGene_tx=XM_023256319.1
B4 6591517 6591684
refGene_gn=SFMBT2;refGene_tx=XM_023256313.1;refGene_gn=SFMBT2;refGene_tx=XM_023256314.1;refGene_gn=SFMBT2;refGene_tx=XM_023256315.1;refGene_gn=SFMBT2;refGene_tx=XM_023256316.1;refGene_gn=SFMBT2;refGene_tx=XM_023256317.1;refGene_gn=SFMBT2;refGene_tx=XM_023256318.1;refGene_gn=SFMBT2;refGene_tx=XM_023256319.1
B4 6594191 6594367
refGene_gn=SFMBT2;refGene_tx=XM_023256313.1;refGene_gn=SFMBT2;refGene_tx=XM_023256314.1;refGene_gn=SFMBT2;refGene_tx=XM_023256315.1;refGene_gn=SFMBT2;refGene_tx=XM_023256316.1;refGene_gn=SFMBT2;refGene_tx=XM_023256317.1;refGene_gn=SFMBT2;refGene_tx=XM_023256318.1;refGene_gn=SFMBT2;refGene_tx=XM_023256319.1
B4 6604243 6604353
refGene_gn=SFMBT2;refGene_tx=XM_023256313.1;refGene_gn=SFMBT2;refGene_tx=XM_023256314.1;refGene_gn=SFMBT2;refGene_tx=XM_023256315.1;refGene_gn=SFMBT2;refGene_tx=XM_023256316.1;refGene_gn=SFMBT2;refGene_tx=XM_023256317.1;refGene_gn=SFMBT2;refGene_tx=XM_023256318.1;refGene_gn=SFMBT2;refGene_tx=XM_023256319.1
B4 6616244 6616384
refGene_gn=SFMBT2;refGene_tx=XM_023256313.1;refGene_gn=SFMBT2;refGene_tx=XM_023256314.1;refGene_gn=SFMBT2;refGene_tx=XM_023256315.1;refGene_gn=SFMBT2;refGene_tx=XM_023256316.1;refGene_gn=SFMBT2;refGene_tx=XM_023256317.1;refGene_gn=SFMBT2;refGene_tx=XM_023256318.1;refGene_gn=SFMBT2;refGene_tx=XM_023256319.1
B4 6621350 6621421
refGene_gn=SFMBT2;refGene_tx=XM_023256313.1;refGene_gn=SFMBT2;refGene_tx=XM_023256314.1;refGene_gn=SFMBT2;refGene_tx=XM_023256315.1;refGene_gn=SFMBT2;refGene_tx=XM_023256316.1;refGene_gn=SFMBT2;refGene_tx=XM_023256317.1;refGene_gn=SFMBT2;refGene_tx=XM_023256318.1;refGene_gn=SFMBT2;refGene_tx=XM_023256319.1
B4 6622889 6622932
refGene_gn=SFMBT2;refGene_tx=XM_023256313.1;refGene_gn=SFMBT2;refGene_tx=XM_023256314.1;refGene_gn=SFMBT2;refGene_tx=XM_023256315.1;refGene_gn=SFMBT2;refGene_tx=XM_023256316.1;refGene_gn=SFMBT2;refGene_tx=XM_023256317.1;refGene_gn=SFMBT2;refGene_tx=XM_023256318.1;refGene_gn=SFMBT2;refGene_tx=XM_023256319.1
B4 6626012 6626126

refGene_gn=SFMBT2;refGene_tx=XM_023256313.1;refGene_gn=SFMBT2;refGene_tx=XM_023256314.1;refGene_gn=SFMBT2;refGene_tx=XM_023256315.1;refGene_gn=SFMBT2;refGene_tx=XM_023256316.1;refGene_gn=SFMBT2;refGene_tx=XM_023256317.1;refGene_gn=SFMBT2;refGene_tx=XM_023256318.1;refGene_gn=SFMBT2;refGene_tx=XM_023256319.1
B4 6639608 6639735
refGene_gn=SFMBT2;refGene_tx=XM_023256313.1;refGene_gn=SFMBT2;refGene_tx=XM_023256314.1;refGene_gn=SFMBT2;refGene_tx=XM_023256315.1;refGene_gn=SFMBT2;refGene_tx=XM_023256316.1;refGene_gn=SFMBT2;refGene_tx=XM_023256317.1;refGene_gn=SFMBT2;refGene_tx=XM_023256318.1;refGene_gn=SFMBT2;refGene_tx=XM_023256319.1
B4 6645864 6645947
refGene_gn=SFMBT2;refGene_tx=XM_023256313.1;refGene_gn=SFMBT2;refGene_tx=XM_023256314.1;refGene_gn=SFMBT2;refGene_tx=XM_023256315.1;refGene_gn=SFMBT2;refGene_tx=XM_023256316.1;refGene_gn=SFMBT2;refGene_tx=XM_023256317.1;refGene_gn=SFMBT2;refGene_tx=XM_023256318.1;refGene_gn=SFMBT2;refGene_tx=XM_023256319.1
B4 6660041 6660189
refGene_gn=SFMBT2;refGene_tx=XM_023256313.1;refGene_gn=SFMBT2;refGene_tx=XM_023256314.1;refGene_gn=SFMBT2;refGene_tx=XM_023256315.1;refGene_gn=SFMBT2;refGene_tx=XM_023256316.1;refGene_gn=SFMBT2;refGene_tx=XM_023256317.1;refGene_gn=SFMBT2;refGene_tx=XM_023256318.1;refGene_gn=SFMBT2;refGene_tx=XM_023256319.1
B4 6663501 6663603
refGene_gn=SFMBT2;refGene_tx=XM_023256313.1;refGene_gn=SFMBT2;refGene_tx=XM_023256314.1;refGene_gn=SFMBT2;refGene_tx=XM_023256315.1;refGene_gn=SFMBT2;refGene_tx=XM_023256316.1;refGene_gn=SFMBT2;refGene_tx=XM_023256317.1;refGene_gn=SFMBT2;refGene_tx=XM_023256318.1;refGene_gn=SFMBT2;refGene_tx=XM_023256319.1
B4 6688605 6688703
refGene_gn=SFMBT2;refGene_tx=XM_023256313.1;refGene_gn=SFMBT2;refGene_tx=XM_023256314.1;refGene_gn=SFMBT2;refGene_tx=XM_023256315.1;refGene_gn=SFMBT2;refGene_tx=XM_023256316.1;refGene_gn=SFMBT2;refGene_tx=XM_023256317.1;refGene_gn=SFMBT2;refGene_tx=XM_023256318.1;refGene_gn=SFMBT2;refGene_tx=XM_023256319.1
B4 6697476 6697723
refGene_gn=SFMBT2;refGene_tx=XM_023256313.1;refGene_gn=SFMBT2;refGene_tx=XM_023256314.1;refGene_gn=SFMBT2;refGene_tx=XM_023256315.1;refGene_gn=SFMBT2;refGene_tx=XM_023256316.1;refGene_gn=SFMBT2;refGene_tx=XM_023256317.1;refGene_gn=SFMBT2;refGene_tx=XM_023256318.1
B4 6699409 6699498
refGene_gn=SFMBT2;refGene_tx=XM_023256313.1;refGene_gn=SFMBT2;refGene_tx=XM_023256314.1;refGene_gn=SFMBT2;refGene_tx=XM_023256315.1;refGene_gn=SFMBT2;refGene_tx=XM_023256316.1;refGene_gn=SFMBT2;refGene_tx=XM_023256317.1;refGene_gn=SFMBT2;refGene_tx=XM_023256318.1
B4 6769940 6770181
refGene_gn=SFMBT2;refGene_tx=XM_023256313.1;refGene_gn=SFMBT2;refGene_tx=XM_023256314.1;refGene_gn=SFMBT2;refGene_tx=XM_023256315.1;refGene_gn=SFMBT2;refGene_tx=XM_023256316.1;refGene_gn=SFMBT2;refGene_tx=XM_023256317.1;refGene_gn=SFMBT2;refGene_tx=XM_023256318.1;refGene_gn=SFMBT

2;refGene_tx=XM_023256319.1
B4 6772300 6772395
refGene_gn=SFMBT2;refGene_tx=XM_023256313.1;refGene_gn=SFMBT2;refGene_tx=XM_023256314.1;refGene_gn=SFMBT2;refGene_tx=XM_023256315.1;refGene_gn=SFMBT2;refGene_tx=XM_023256316.1;refGene_gn=SFMBT2;refGene_tx=XM_023256317.1;refGene_gn=SFMBT2;refGene_tx=XM_023256318.1;refGene_gn=SFMBT2;refGene_tx=XM_023256319.1
B4 6781232 6781332
refGene_gn=SFMBT2;refGene_tx=XM_023256313.1;refGene_gn=SFMBT2;refGene_tx=XM_023256314.1;refGene_gn=SFMBT2;refGene_tx=XM_023256315.1;refGene_gn=SFMBT2;refGene_tx=XM_023256316.1;refGene_gn=SFMBT2;refGene_tx=XM_023256317.1;refGene_gn=SFMBT2;refGene_tx=XM_023256318.1;refGene_gn=SFMBT2;refGene_tx=XM_023256319.1
B4 6934021 6934323 refGene_gn=ITIH5;refGene_tx=XM_011283573.2
B4 6935302 6935680 refGene_gn=ITIH5;refGene_tx=XM_011283573.2
B4 6937503 6937620 refGene_gn=ITIH5;refGene_tx=XM_011283573.2
B4 6939413 6939467 refGene_gn=ITIH5;refGene_tx=XM_011283573.2
B4 6941601 6942161 refGene_gn=ITIH5;refGene_tx=XM_011283573.2
B4 6944548 6944858 refGene_gn=ITIH5;refGene_tx=XM_011283573.2
B4 6948147 6948316 refGene_gn=ITIH5;refGene_tx=XM_011283573.2
B4 6966533 6966650 refGene_gn=ITIH5;refGene_tx=XM_011283573.2
B4 6967347 6967517 refGene_gn=ITIH5;refGene_tx=XM_011283573.2
B4 6984348 6984599 refGene_gn=ITIH5;refGene_tx=XM_011283573.2
B4 6988339 6988441 refGene_gn=ITIH5;refGene_tx=XM_011283573.2
B4 6989430 6989594 refGene_gn=ITIH5;refGene_tx=XM_011283573.2
B4 7003792 7003837 refGene_gn=ITIH5;refGene_tx=XM_011283573.2
B4 7012648 7012738 refGene_gn=ITIH5;refGene_tx=XM_011283573.2
B4 7055711 7055795 refGene_gn=ITIH2;refGene_tx=XM_003988085.4
B4 7057071 7057146 refGene_gn=ITIH2;refGene_tx=XM_003988085.4
B4 7058846 7058879 refGene_gn=ITIH2;refGene_tx=XM_003988085.4
B4 7060289 7060459 refGene_gn=ITIH2;refGene_tx=XM_003988085.4
B4 7063387 7063492 refGene_gn=ITIH2;refGene_tx=XM_003988085.4
B4 7064242 7064405 refGene_gn=ITIH2;refGene_tx=XM_003988085.4
B4 7068080 7068188 refGene_gn=ITIH2;refGene_tx=XM_003988085.4
B4 7068915 7069044 refGene_gn=ITIH2;refGene_tx=XM_003988085.4
B4 7070441 7070558 refGene_gn=ITIH2;refGene_tx=XM_003988085.4
B4 7073886 7074055 refGene_gn=ITIH2;refGene_tx=XM_003988085.4
B4 7074634 7074760 refGene_gn=ITIH2;refGene_tx=XM_003988085.4
B4 7076562 7076744 refGene_gn=ITIH2;refGene_tx=XM_003988085.4
B4 7077836 7078022 refGene_gn=ITIH2;refGene_tx=XM_003988085.4
B4 7078403 7078543 refGene_gn=ITIH2;refGene_tx=XM_003988085.4
B4 7080517 7080687 refGene_gn=ITIH2;refGene_tx=XM_003988085.4
B4 7082820 7082958 refGene_gn=ITIH2;refGene_tx=XM_003988085.4
B4 7087942 7088056 refGene_gn=ITIH2;refGene_tx=XM_003988085.4
B4 7088394 7088593 refGene_gn=ITIH2;refGene_tx=XM_003988085.4
B4 7089339 7089512 refGene_gn=ITIH2;refGene_tx=XM_003988085.4
B4 7090886 7090998 refGene_gn=ITIH2;refGene_tx=XM_003988085.4
B4 7093451 7093599 refGene_gn=ITIH2;refGene_tx=XM_003988085.4
B4 7099319 7099421 refGene_gn=KIN;refGene_tx=XM_023256320.1
B4 7101494 7101557 refGene_gn=KIN;refGene_tx=XM_003988087.4

B4 7105636 7105737
refGene_gn=KIN;refGene_tx=XM_003988087.4;refGene_gn=KIN;refGene_tx=XM_023256320.1
B4 7107397 7107497
refGene_gn=KIN;refGene_tx=XM_003988087.4;refGene_gn=KIN;refGene_tx=XM_023256320.1
B4 7108422 7108491
refGene_gn=KIN;refGene_tx=XM_003988087.4;refGene_gn=KIN;refGene_tx=XM_023256320.1
B4 7111759 7111810
refGene_gn=KIN;refGene_tx=XM_003988087.4;refGene_gn=KIN;refGene_tx=XM_023256320.1
B4 7114392 7114522
refGene_gn=KIN;refGene_tx=XM_003988087.4;refGene_gn=KIN;refGene_tx=XM_023256320.1
B4 7121889 7121947
refGene_gn=KIN;refGene_tx=XM_003988087.4;refGene_gn=KIN;refGene_tx=XM_023256320.1
B4 7122920 7122969
refGene_gn=KIN;refGene_tx=XM_003988087.4;refGene_gn=KIN;refGene_tx=XM_023256320.1
B4 7126291 7126473
refGene_gn=KIN;refGene_tx=XM_003988087.4;refGene_gn=KIN;refGene_tx=XM_023256320.1
B4 7127264 7127387
refGene_gn=KIN;refGene_tx=XM_003988087.4;refGene_gn=KIN;refGene_tx=XM_023256320.1
B4 7127470 7127514
refGene_gn=KIN;refGene_tx=XM_003988087.4;refGene_gn=KIN;refGene_tx=XM_023256320.1
B4 7131272 7131367
refGene_gn=KIN;refGene_tx=XM_003988087.4;refGene_gn=KIN;refGene_tx=XM_023256320.1
B4 7137409 7137523
refGene_gn=KIN;refGene_tx=XM_003988087.4;refGene_gn=KIN;refGene_tx=XM_023256320.1
B4 7137798 7137854
refGene_gn=ATP5F1C;refGene_tx=XM_003988088.5;refGene_gn=ATP5F1C;refGene_tx=XM_003988089.4;refGene_gn=ATP5F1C;refGene_tx=XM_006933219.3
B4 7146912 7146947
refGene_gn=ATP5F1C;refGene_tx=XM_003988088.5;refGene_gn=ATP5F1C;refGene_tx=XM_003988089.4;refGene_gn=ATP5F1C;refGene_tx=XM_006933219.3
B4 7147416 7147548
refGene_gn=ATP5F1C;refGene_tx=XM_003988088.5;refGene_gn=ATP5F1C;refGene_tx=XM_003988089.4;refGene_gn=ATP5F1C;refGene_tx=XM_006933219.3
B4 7149972 7150177
refGene_gn=ATP5F1C;refGene_tx=XM_003988088.5;refGene_gn=ATP5F1C;refGene_tx=XM_003988089.4;refGene_gn=ATP5F1C;refGene_tx=XM_006933219.3
B4 7150744 7150888
refGene_gn=ATP5F1C;refGene_tx=XM_003988088.5;refGene_gn=ATP5F1C;refGen

e_tx=XM_003988089.4;refGene_gn=ATP5F1C;refGene_tx=XM_006933219.3
B4	7151225	7151290
refGene_gn=ATP5F1C;refGene_tx=XM_003988088.5;refGene_gn=ATP5F1C;refGen
e_tx=XM_003988089.4;refGene_gn=ATP5F1C;refGene_tx=XM_006933219.3
B4	7152441	7152597
refGene_gn=ATP5F1C;refGene_tx=XM_003988088.5;refGene_gn=ATP5F1C;refGen
e_tx=XM_003988089.4;refGene_gn=ATP5F1C;refGene_tx=XM_006933219.3
B4	7153013	7153114
refGene_gn=ATP5F1C;refGene_tx=XM_003988088.5;refGene_gn=ATP5F1C;refGen
e_tx=XM_003988089.4;refGene_gn=ATP5F1C;refGene_tx=XM_006933219.3
B4	7156264	7156271	refGene_gn=ATP5F1C;refGene_tx=XM_003988088.5
B4	7156918	7156922	refGene_gn=ATP5F1C;refGene_tx=XM_003988089.4
B4	7161061	7161141
refGene_gn=LOC111561440;refGene_tx=XM_023257681.1
B4	7161828	7161971
refGene_gn=LOC111561440;refGene_tx=XM_023257681.1
B4	7164547	7165084
refGene_gn=LOC111561440;refGene_tx=XM_023257681.1;refGene_gn=TAF3;refG
ene_tx=XM_011283569.3
B4	7169941	7170184
refGene_gn=TAF3;refGene_tx=XM_011283569.3;refGene_gn=TAF3;refGene_tx=X
M_011283570.3;refGene_gn=TAF3;refGene_tx=XM_011283571.3
B4	7298159	7299985
refGene_gn=TAF3;refGene_tx=XM_011283569.3;refGene_gn=TAF3;refGene_tx=X
M_011283570.3;refGene_gn=TAF3;refGene_tx=XM_011283571.3
B4	7310474	7310557
refGene_gn=TAF3;refGene_tx=XM_011283569.3;refGene_gn=TAF3;refGene_tx=X
M_011283570.3;refGene_gn=TAF3;refGene_tx=XM_011283571.3
B4	7341510	7341766
refGene_gn=TAF3;refGene_tx=XM_011283569.3;refGene_gn=TAF3;refGene_tx=X
M_011283570.3;refGene_gn=TAF3;refGene_tx=XM_011283571.3
B4	7345181	7345288
refGene_gn=TAF3;refGene_tx=XM_011283569.3;refGene_gn=TAF3;refGene_tx=X
M_011283570.3;refGene_gn=TAF3;refGene_tx=XM_011283571.3
B4	7346142	7346257
refGene_gn=TAF3;refGene_tx=XM_011283569.3;refGene_gn=TAF3;refGene_tx=X
M_011283570.3;refGene_gn=TAF3;refGene_tx=XM_011283571.3
B4	7380626	7380867
refGene_gn=GATA3;refGene_tx=XM_023256321.1;refGene_gn=GATA3;refGene_tx
=XM_023256322.1;refGene_gn=GATA3;refGene_tx=XM_023256323.1
B4	7383315	7383852
refGene_gn=GATA3;refGene_tx=XM_023256321.1;refGene_gn=GATA3;refGene_tx
=XM_023256322.1;refGene_gn=GATA3;refGene_tx=XM_023256323.1
B4	7389217	7389363
refGene_gn=GATA3;refGene_tx=XM_023256321.1;refGene_gn=GATA3;refGene_tx
=XM_023256322.1;refGene_gn=GATA3;refGene_tx=XM_023256323.1
B4	7393990	7394116
refGene_gn=GATA3;refGene_tx=XM_023256321.1;refGene_gn=GATA3;refGene_tx
=XM_023256322.1;refGene_gn=GATA3;refGene_tx=XM_023256323.1
B4	7398190	7398475

refGene_gn=GATA3;refGene_tx=XM_023256321.1;refGene_gn=GATA3;refGene_tx=XM_023256322.1;refGene_gn=GATA3;refGene_tx=XM_023256323.1
B4 10052988 10053041 refGene_gn=CELF2;refGene_tx=XM_023256326.1
B4 10171943 10171979 refGene_gn=CELF2;refGene_tx=XM_023256326.1
B4 10255398 10255451 refGene_gn=CELF2;refGene_tx=XM_019833917.2
B4 10263440 10263592
refGene_gn=CELF2;refGene_tx=XM_023256329.1;refGene_gn=CELF2;refGene_tx=XM_023256330.1
B4 10268051 10268125
refGene_gn=CELF2;refGene_tx=XM_023256327.1;refGene_gn=CELF2;refGene_tx=XM_023256328.1
B4 10413623 10413820
refGene_gn=CELF2;refGene_tx=XM_006933225.4;refGene_gn=CELF2;refGene_tx=XM_011283568.3;refGene_gn=CELF2;refGene_tx=XM_019833917.2;refGene_gn=CELF2;refGene_tx=XM_019833918.2;refGene_gn=CELF2;refGene_tx=XM_023256324.1;refGene_gn=CELF2;refGene_tx=XM_023256325.1;refGene_gn=CELF2;refGene_tx=XM_023256326.1;refGene_gn=CELF2;refGene_tx=XM_023256327.1;refGene_gn=CELF2;refGene_tx=XM_023256328.1;refGene_gn=CELF2;refGene_tx=XM_023256329.1;refGene_gn=CELF2;refGene_tx=XM_023256330.1
B4 10465553 10465636
refGene_gn=CELF2;refGene_tx=XM_006933225.4;refGene_gn=CELF2;refGene_tx=XM_011283568.3;refGene_gn=CELF2;refGene_tx=XM_019833917.2;refGene_gn=CELF2;refGene_tx=XM_019833918.2;refGene_gn=CELF2;refGene_tx=XM_023256324.1;refGene_gn=CELF2;refGene_tx=XM_023256325.1;refGene_gn=CELF2;refGene_tx=XM_023256326.1;refGene_gn=CELF2;refGene_tx=XM_023256327.1;refGene_gn=CELF2;refGene_tx=XM_023256328.1;refGene_gn=CELF2;refGene_tx=XM_023256329.1;refGene_gn=CELF2;refGene_tx=XM_023256330.1
B4 10496654 10496703
refGene_gn=CELF2;refGene_tx=XM_006933225.4;refGene_gn=CELF2;refGene_tx=XM_011283568.3;refGene_gn=CELF2;refGene_tx=XM_019833917.2;refGene_gn=CELF2;refGene_tx=XM_019833918.2;refGene_gn=CELF2;refGene_tx=XM_023256324.1;refGene_gn=CELF2;refGene_tx=XM_023256325.1;refGene_gn=CELF2;refGene_tx=XM_023256326.1;refGene_gn=CELF2;refGene_tx=XM_023256327.1;refGene_gn=CELF2;refGene_tx=XM_023256328.1;refGene_gn=CELF2;refGene_tx=XM_023256329.1;refGene_gn=CELF2;refGene_tx=XM_023256330.1
B4 10504818 10504953
refGene_gn=CELF2;refGene_tx=XM_006933225.4;refGene_gn=CELF2;refGene_tx=XM_011283568.3;refGene_gn=CELF2;refGene_tx=XM_019833917.2;refGene_gn=CELF2;refGene_tx=XM_019833918.2;refGene_gn=CELF2;refGene_tx=XM_023256324.1;refGene_gn=CELF2;refGene_tx=XM_023256325.1;refGene_gn=CELF2;refGene_tx=XM_023256326.1;refGene_gn=CELF2;refGene_tx=XM_023256327.1;refGene_gn=CELF2;refGene_tx=XM_023256328.1;refGene_gn=CELF2;refGene_tx=XM_023256329.1;refGene_gn=CELF2;refGene_tx=XM_023256330.1
B4 10513171 10513251
refGene_gn=CELF2;refGene_tx=XM_006933225.4;refGene_gn=CELF2;refGene_tx=XM_011283568.3;refGene_gn=CELF2;refGene_tx=XM_019833917.2;refGene_gn=CELF2;refGene_tx=XM_019833918.2;refGene_gn=CELF2;refGene_tx=XM_023256324.1;refGene_gn=CELF2;refGene_tx=XM_023256325.1;refGene_gn=CELF2;refGene_tx=XM_023256326.1;refGene_gn=CELF2;refGene_tx=XM_023256327.1;refGene_gn=CELF2;refGene_tx=XM_023256328.1;refGene_gn=CELF2;refGene_tx=XM_02

3256329.1;refGene_gn=CELF2;refGene_tx=XM_023256330.1
B4	10517356 10517515
refGene_gn=CELF2;refGene_tx=XM_006933225.4;refGene_gn=CELF2;refGene_tx=XM_011283568.3;refGene_gn=CELF2;refGene_tx=XM_019833917.2;refGene_gn=CELF2;refGene_tx=XM_019833918.2;refGene_gn=CELF2;refGene_tx=XM_019833920.2;refGene_gn=CELF2;refGene_tx=XM_023256324.1;refGene_gn=CELF2;refGene_tx=XM_023256325.1;refGene_gn=CELF2;refGene_tx=XM_023256326.1;refGene_gn=CELF2;refGene_tx=XM_023256327.1;refGene_gn=CELF2;refGene_tx=XM_023256328.1;refGene_gn=CELF2;refGene_tx=XM_023256329.1;refGene_gn=CELF2;refGene_tx=XM_023256330.1
B4	10521605 10521669
refGene_gn=CELF2;refGene_tx=XM_006933225.4;refGene_gn=CELF2;refGene_tx=XM_011283568.3;refGene_gn=CELF2;refGene_tx=XM_019833917.2;refGene_gn=CELF2;refGene_tx=XM_019833918.2;refGene_gn=CELF2;refGene_tx=XM_019833920.2;refGene_gn=CELF2;refGene_tx=XM_023256324.1;refGene_gn=CELF2;refGene_tx=XM_023256325.1;refGene_gn=CELF2;refGene_tx=XM_023256326.1;refGene_gn=CELF2;refGene_tx=XM_023256327.1;refGene_gn=CELF2;refGene_tx=XM_023256328.1;refGene_gn=CELF2;refGene_tx=XM_023256329.1;refGene_gn=CELF2;refGene_tx=XM_023256330.1
B4	10533186 10533321
refGene_gn=CELF2;refGene_tx=XM_006933225.4;refGene_gn=CELF2;refGene_tx=XM_011283568.3;refGene_gn=CELF2;refGene_tx=XM_019833917.2;refGene_gn=CELF2;refGene_tx=XM_019833918.2;refGene_gn=CELF2;refGene_tx=XM_019833920.2;refGene_gn=CELF2;refGene_tx=XM_023256324.1;refGene_gn=CELF2;refGene_tx=XM_023256325.1;refGene_gn=CELF2;refGene_tx=XM_023256326.1;refGene_gn=CELF2;refGene_tx=XM_023256327.1;refGene_gn=CELF2;refGene_tx=XM_023256328.1;refGene_gn=CELF2;refGene_tx=XM_023256329.1;refGene_gn=CELF2;refGene_tx=XM_023256330.1
B4	10559683 10559815
refGene_gn=CELF2;refGene_tx=XM_006933225.4;refGene_gn=CELF2;refGene_tx=XM_011283568.3;refGene_gn=CELF2;refGene_tx=XM_019833917.2;refGene_gn=CELF2;refGene_tx=XM_019833918.2;refGene_gn=CELF2;refGene_tx=XM_019833920.2;refGene_gn=CELF2;refGene_tx=XM_023256324.1;refGene_gn=CELF2;refGene_tx=XM_023256325.1;refGene_gn=CELF2;refGene_tx=XM_023256326.1;refGene_gn=CELF2;refGene_tx=XM_023256327.1;refGene_gn=CELF2;refGene_tx=XM_023256328.1;refGene_gn=CELF2;refGene_tx=XM_023256329.1;refGene_gn=CELF2;refGene_tx=XM_023256330.1
B4	10566724 10566922
refGene_gn=CELF2;refGene_tx=XM_006933225.4;refGene_gn=CELF2;refGene_tx=XM_011283568.3;refGene_gn=CELF2;refGene_tx=XM_019833917.2;refGene_gn=CELF2;refGene_tx=XM_019833918.2;refGene_gn=CELF2;refGene_tx=XM_019833920.2;refGene_gn=CELF2;refGene_tx=XM_023256324.1;refGene_gn=CELF2;refGene_tx=XM_023256325.1;refGene_gn=CELF2;refGene_tx=XM_023256326.1;refGene_gn=CELF2;refGene_tx=XM_023256327.1;refGene_gn=CELF2;refGene_tx=XM_023256328.1;refGene_gn=CELF2;refGene_tx=XM_023256329.1;refGene_gn=CELF2;refGene_tx=XM_023256330.1
B4	10571178 10571322
refGene_gn=CELF2;refGene_tx=XM_006933225.4;refGene_gn=CELF2;refGene_tx=XM_011283568.3;refGene_gn=CELF2;refGene_tx=XM_019833917.2;refGene_gn=CELF2;refGene_tx=XM_019833918.2;refGene_gn=CELF2;refGene_tx=XM_0198339

20.2;refGene_gn=CELF2;refGene_tx=XM_023256324.1;refGene_gn=CELF2;refGene_tx=XM_023256325.1;refGene_gn=CELF2;refGene_tx=XM_023256326.1;refGene_gn=CELF2;refGene_tx=XM_023256327.1;refGene_gn=CELF2;refGene_tx=XM_023256328.1
B4 10574490 10574618
refGene_gn=CELF2;refGene_tx=XM_006933225.4;refGene_gn=CELF2;refGene_tx=XM_011283568.3;refGene_gn=CELF2;refGene_tx=XM_019833917.2;refGene_gn=CELF2;refGene_tx=XM_019833918.2;refGene_gn=CELF2;refGene_tx=XM_019833920.2;refGene_gn=CELF2;refGene_tx=XM_023256324.1;refGene_gn=CELF2;refGene_tx=XM_023256325.1;refGene_gn=CELF2;refGene_tx=XM_023256326.1;refGene_gn=CELF2;refGene_tx=XM_023256327.1;refGene_gn=CELF2;refGene_tx=XM_023256328.1;refGene_gn=CELF2;refGene_tx=XM_023256329.1;refGene_gn=CELF2;refGene_tx=XM_023256330.1
B4 10699570 10700952 refGene_gn=USP6NL;refGene_tx=XM_019833909.2
B4 10727033 10727127 refGene_gn=USP6NL;refGene_tx=XM_019833909.2
B4 10727128 10727187 refGene_gn=USP6NL;refGene_tx=XM_019833909.2
B4 10730102 10730202 refGene_gn=USP6NL;refGene_tx=XM_019833909.2
B4 10730296 10730362 refGene_gn=USP6NL;refGene_tx=XM_019833909.2
B4 10731253 10731348 refGene_gn=USP6NL;refGene_tx=XM_019833909.2
B4 10734505 10734626 refGene_gn=USP6NL;refGene_tx=XM_019833909.2
B4 10736150 10736199 refGene_gn=USP6NL;refGene_tx=XM_019833909.2
B4 10738379 10738489 refGene_gn=USP6NL;refGene_tx=XM_019833909.2
B4 10744118 10744226 refGene_gn=USP6NL;refGene_tx=XM_019833909.2
B4 10751659 10751740 refGene_gn=USP6NL;refGene_tx=XM_019833909.2
B4 10764142 10764182 refGene_gn=USP6NL;refGene_tx=XM_019833909.2
B4 10774854 10774937 refGene_gn=USP6NL;refGene_tx=XM_019833909.2
B4 10776928 10776996 refGene_gn=USP6NL;refGene_tx=XM_019833909.2
B4 10780718 10780773 refGene_gn=USP6NL;refGene_tx=XM_019833909.2
B4 10995202 10995345
refGene_gn=ECHDC3;refGene_tx=XM_023256331.1;refGene_gn=ECHDC3;refGene_tx=XM_023256332.1;refGene_gn=ECHDC3;refGene_tx=XM_023256333.1;refGene_gn=ECHDC3;refGene_tx=XM_023256334.1
B4 10999562 10999684
refGene_gn=ECHDC3;refGene_tx=XM_023256331.1;refGene_gn=ECHDC3;refGene_tx=XM_023256332.1;refGene_gn=ECHDC3;refGene_tx=XM_023256333.1;refGene_gn=ECHDC3;refGene_tx=XM_023256334.1
B4 11001090 11001188
refGene_gn=ECHDC3;refGene_tx=XM_023256331.1;refGene_gn=ECHDC3;refGene_tx=XM_023256332.1;refGene_gn=ECHDC3;refGene_tx=XM_023256333.1;refGene_gn=ECHDC3;refGene_tx=XM_023256334.1
B4 11006099 11006300
refGene_gn=ECHDC3;refGene_tx=XM_023256331.1;refGene_gn=ECHDC3;refGene_tx=XM_023256332.1;refGene_gn=ECHDC3;refGene_tx=XM_023256333.1;refGene_gn=ECHDC3;refGene_tx=XM_023256334.1
B4 11015820 11016128
refGene_gn=ECHDC3;refGene_tx=XM_023256331.1;refGene_gn=ECHDC3;refGene_tx=XM_023256332.1;refGene_gn=ECHDC3;refGene_tx=XM_023256333.1;refGene_gn=ECHDC3;refGene_tx=XM_023256334.1
B4 11022885 11022964 refGene_gn=ECHDC3;refGene_tx=XM_023256332.1
B4 11022966 11023009 refGene_gn=ECHDC3;refGene_tx=XM_023256334.1

B4 11024260 11024371 refGene_gn=ECHDC3;refGene_tx=XM_023256331.1
B4 11026586 11026632
refGene_gn=ECHDC3;refGene_tx=XM_023256331.1;refGene_gn=ECHDC3;refGene_tx=XM_023256333.1
B4 11096585 11096723 refGene_gn=PROSER2;refGene_tx=XM_023256335.1
B4 11116070 11116323 refGene_gn=PROSER2;refGene_tx=XM_023256335.1
B4 11118372 11119406 refGene_gn=PROSER2;refGene_tx=XM_023256335.1
B4 11159337 11159347
refGene_gn=UPF2;refGene_tx=XM_011283561.3;refGene_gn=UPF2;refGene_tx=XM_019833926.2
B4 11166011 11166132
refGene_gn=UPF2;refGene_tx=XM_011283561.3;refGene_gn=UPF2;refGene_tx=XM_019833926.2
B4 11167536 11167678
refGene_gn=UPF2;refGene_tx=XM_011283561.3;refGene_gn=UPF2;refGene_tx=XM_019833926.2
B4 11177518 11177686
refGene_gn=UPF2;refGene_tx=XM_011283561.3;refGene_gn=UPF2;refGene_tx=XM_019833926.2
B4 11180620 11180719
refGene_gn=UPF2;refGene_tx=XM_011283561.3;refGene_gn=UPF2;refGene_tx=XM_019833926.2
B4 11181019 11181124
refGene_gn=UPF2;refGene_tx=XM_011283561.3;refGene_gn=UPF2;refGene_tx=XM_019833926.2
B4 11184621 11184761
refGene_gn=UPF2;refGene_tx=XM_011283561.3;refGene_gn=UPF2;refGene_tx=XM_019833926.2
B4 11189975 11190159
refGene_gn=UPF2;refGene_tx=XM_011283561.3;refGene_gn=UPF2;refGene_tx=XM_019833926.2
B4 11192216 11192492
refGene_gn=UPF2;refGene_tx=XM_011283561.3;refGene_gn=UPF2;refGene_tx=XM_019833926.2
B4 11193208 11193412
refGene_gn=UPF2;refGene_tx=XM_011283561.3;refGene_gn=UPF2;refGene_tx=XM_019833926.2
B4 11197351 11197537
refGene_gn=UPF2;refGene_tx=XM_011283561.3;refGene_gn=UPF2;refGene_tx=XM_019833926.2
B4 11198614 11198731
refGene_gn=UPF2;refGene_tx=XM_011283561.3;refGene_gn=UPF2;refGene_tx=XM_019833926.2
B4 11201491 11201605
refGene_gn=UPF2;refGene_tx=XM_011283561.3;refGene_gn=UPF2;refGene_tx=XM_019833926.2
B4 11209661 11209770
refGene_gn=UPF2;refGene_tx=XM_011283561.3;refGene_gn=UPF2;refGene_tx=XM_019833926.2
B4 11226744 11226830

refGene_gn=UPF2;refGene_tx=XM_011283561.3;refGene_gn=UPF2;refGene_tx=XM_019833926.2
B4 11232378 11232482
refGene_gn=UPF2;refGene_tx=XM_011283561.3;refGene_gn=UPF2;refGene_tx=XM_019833926.2
B4 11234515 11234665
refGene_gn=UPF2;refGene_tx=XM_011283561.3;refGene_gn=UPF2;refGene_tx=XM_019833926.2
B4 11244014 11244212
refGene_gn=UPF2;refGene_tx=XM_011283561.3;refGene_gn=UPF2;refGene_tx=XM_019833926.2
B4 11250312 11250473
refGene_gn=UPF2;refGene_tx=XM_011283561.3;refGene_gn=UPF2;refGene_tx=XM_019833926.2
B4 11259641 11260421
refGene_gn=UPF2;refGene_tx=XM_011283561.3;refGene_gn=UPF2;refGene_tx=XM_019833926.2
B4 11264389 11264751
refGene_gn=UPF2;refGene_tx=XM_011283561.3;refGene_gn=UPF2;refGene_tx=XM_019833926.2
B4 11289512 11289669
refGene_gn=DHTKD1;refGene_tx=XM_011283559.3;refGene_gn=DHTKD1;refGene_tx=XM_011283560.3
B4 11304209 11304365
refGene_gn=DHTKD1;refGene_tx=XM_011283559.3;refGene_gn=DHTKD1;refGene_tx=XM_011283560.3
B4 11309241 11309453
refGene_gn=DHTKD1;refGene_tx=XM_011283559.3;refGene_gn=DHTKD1;refGene_tx=XM_011283560.3;refGene_gn=DHTKD1;refGene_tx=XM_019833927.2
B4 11310234 11310429
refGene_gn=DHTKD1;refGene_tx=XM_011283559.3;refGene_gn=DHTKD1;refGene_tx=XM_011283560.3;refGene_gn=DHTKD1;refGene_tx=XM_019833927.2
B4 11313089 11313359
refGene_gn=DHTKD1;refGene_tx=XM_011283559.3;refGene_gn=DHTKD1;refGene_tx=XM_011283560.3;refGene_gn=DHTKD1;refGene_tx=XM_019833927.2
B4 11318136 11318308
refGene_gn=DHTKD1;refGene_tx=XM_011283559.3;refGene_gn=DHTKD1;refGene_tx=XM_011283560.3;refGene_gn=DHTKD1;refGene_tx=XM_019833927.2
B4 11319332 11319531
refGene_gn=DHTKD1;refGene_tx=XM_011283559.3;refGene_gn=DHTKD1;refGene_tx=XM_011283560.3;refGene_gn=DHTKD1;refGene_tx=XM_019833927.2
B4 11321850 11322163
refGene_gn=DHTKD1;refGene_tx=XM_011283559.3;refGene_gn=DHTKD1;refGene_tx=XM_011283560.3;refGene_gn=DHTKD1;refGene_tx=XM_019833927.2
B4 11323427 11323512
refGene_gn=DHTKD1;refGene_tx=XM_011283559.3;refGene_gn=DHTKD1;refGene_tx=XM_019833927.2
B4 11324252 11324392
refGene_gn=DHTKD1;refGene_tx=XM_011283559.3;refGene_gn=DHTKD1;refGene_tx=XM_019833927.2

B4 11330266 11330417
refGene_gn=DHTKD1;refGene_tx=XM_011283559.3;refGene_gn=DHTKD1;refGene_tx=XM_011283560.3;refGene_gn=DHTKD1;refGene_tx=XM_019833927.2
B4 11331802 11331909
refGene_gn=DHTKD1;refGene_tx=XM_011283559.3;refGene_gn=DHTKD1;refGene_tx=XM_011283560.3;refGene_gn=DHTKD1;refGene_tx=XM_019833927.2
B4 11332966 11333131
refGene_gn=DHTKD1;refGene_tx=XM_011283559.3;refGene_gn=DHTKD1;refGene_tx=XM_011283560.3;refGene_gn=DHTKD1;refGene_tx=XM_019833927.2
B4 11335543 11335626
refGene_gn=DHTKD1;refGene_tx=XM_011283559.3;refGene_gn=DHTKD1;refGene_tx=XM_011283560.3;refGene_gn=DHTKD1;refGene_tx=XM_019833927.2
B4 11335961 11336131
refGene_gn=DHTKD1;refGene_tx=XM_011283559.3;refGene_gn=DHTKD1;refGene_tx=XM_011283560.3;refGene_gn=DHTKD1;refGene_tx=XM_019833927.2
B4 11337888 11337974
refGene_gn=DHTKD1;refGene_tx=XM_011283559.3;refGene_gn=DHTKD1;refGene_tx=XM_011283560.3;refGene_gn=DHTKD1;refGene_tx=XM_019833927.2
B4 11338894 11338996
refGene_gn=DHTKD1;refGene_tx=XM_011283559.3;refGene_gn=DHTKD1;refGene_tx=XM_011283560.3;refGene_gn=DHTKD1;refGene_tx=XM_019833927.2
B4 11348316 11348323
refGene_gn=SEC61A2;refGene_tx=XM_023256336.1;refGene_gn=SEC61A2;refGene_tx=XM_023256337.1
B4 11350391 11350459
refGene_gn=SEC61A2;refGene_tx=XM_023256336.1;refGene_gn=SEC61A2;refGene_tx=XM_023256337.1
B4 11351541 11351607 refGene_gn=SEC61A2;refGene_tx=XM_023256336.1
B4 11357442 11357521
refGene_gn=SEC61A2;refGene_tx=XM_023256336.1;refGene_gn=SEC61A2;refGene_tx=XM_023256337.1
B4 11362393 11362525
refGene_gn=SEC61A2;refGene_tx=XM_023256336.1;refGene_gn=SEC61A2;refGene_tx=XM_023256337.1
B4 11362647 11362757
refGene_gn=SEC61A2;refGene_tx=XM_023256336.1;refGene_gn=SEC61A2;refGene_tx=XM_023256337.1
B4 11367941 11368095
refGene_gn=SEC61A2;refGene_tx=XM_023256336.1;refGene_gn=SEC61A2;refGene_tx=XM_023256337.1
B4 11369077 11369238
refGene_gn=SEC61A2;refGene_tx=XM_023256336.1;refGene_gn=SEC61A2;refGene_tx=XM_023256337.1
B4 11369562 11369760
refGene_gn=SEC61A2;refGene_tx=XM_023256336.1;refGene_gn=SEC61A2;refGene_tx=XM_023256337.1
B4 11371797 11371989
refGene_gn=SEC61A2;refGene_tx=XM_023256336.1;refGene_gn=SEC61A2;refGene_tx=XM_023256337.1
B4 11379041 11379118

refGene_gn=SEC61A2;refGene_tx=XM_023256336.1;refGene_gn=SEC61A2;refGene_tx=XM_023256337.1
B4 11381828 11382015
refGene_gn=SEC61A2;refGene_tx=XM_023256336.1;refGene_gn=SEC61A2;refGene_tx=XM_023256337.1
B4 11385209 11385319
refGene_gn=NUDT5;refGene_tx=XM_003988104.5;refGene_gn=NUDT5;refGene_tx=XM_006933238.4;refGene_gn=NUDT5;refGene_tx=XM_006933239.4;refGene_gn=NUDT5;refGene_tx=XM_006933240.4;refGene_gn=NUDT5;refGene_tx=XM_019833937.2;refGene_gn=NUDT5;refGene_tx=XM_019833939.2
B4 11387768 11387822
refGene_gn=NUDT5;refGene_tx=XM_003988104.5;refGene_gn=NUDT5;refGene_tx=XM_006933238.4;refGene_gn=NUDT5;refGene_tx=XM_006933239.4;refGene_gn=NUDT5;refGene_tx=XM_006933240.4;refGene_gn=NUDT5;refGene_tx=XM_019833937.2;refGene_gn=NUDT5;refGene_tx=XM_019833939.2
B4 11387912 11387965
refGene_gn=NUDT5;refGene_tx=XM_003988104.5;refGene_gn=NUDT5;refGene_tx=XM_006933238.4;refGene_gn=NUDT5;refGene_tx=XM_006933239.4;refGene_gn=NUDT5;refGene_tx=XM_006933240.4;refGene_gn=NUDT5;refGene_tx=XM_019833938.2;refGene_gn=NUDT5;refGene_tx=XM_019833939.2
B4 11389657 11389759
refGene_gn=NUDT5;refGene_tx=XM_003988104.5;refGene_gn=NUDT5;refGene_tx=XM_006933238.4;refGene_gn=NUDT5;refGene_tx=XM_006933239.4;refGene_gn=NUDT5;refGene_tx=XM_006933240.4;refGene_gn=NUDT5;refGene_tx=XM_019833937.2;refGene_gn=NUDT5;refGene_tx=XM_019833938.2;refGene_gn=NUDT5;refGene_tx=XM_019833939.2
B4 11390316 11390412
refGene_gn=NUDT5;refGene_tx=XM_003988104.5;refGene_gn=NUDT5;refGene_tx=XM_006933238.4;refGene_gn=NUDT5;refGene_tx=XM_006933239.4;refGene_gn=NUDT5;refGene_tx=XM_006933240.4;refGene_gn=NUDT5;refGene_tx=XM_019833937.2;refGene_gn=NUDT5;refGene_tx=XM_019833938.2;refGene_gn=NUDT5;refGene_tx=XM_019833939.2
B4 11391315 11391423
refGene_gn=NUDT5;refGene_tx=XM_003988104.5;refGene_gn=NUDT5;refGene_tx=XM_006933238.4;refGene_gn=NUDT5;refGene_tx=XM_006933239.4;refGene_gn=NUDT5;refGene_tx=XM_006933240.4;refGene_gn=NUDT5;refGene_tx=XM_019833937.2;refGene_gn=NUDT5;refGene_tx=XM_019833938.2;refGene_gn=NUDT5;refGene_tx=XM_019833939.2
B4 11392394 11392444
refGene_gn=NUDT5;refGene_tx=XM_003988104.5;refGene_gn=NUDT5;refGene_tx=XM_006933238.4;refGene_gn=NUDT5;refGene_tx=XM_006933239.4;refGene_gn=NUDT5;refGene_tx=XM_006933240.4;refGene_gn=NUDT5;refGene_tx=XM_019833937.2;refGene_gn=NUDT5;refGene_tx=XM_019833938.2
B4 11396707 11396775
refGene_gn=NUDT5;refGene_tx=XM_003988104.5;refGene_gn=NUDT5;refGene_tx=XM_006933238.4;refGene_gn=NUDT5;refGene_tx=XM_006933239.4;refGene_gn=NUDT5;refGene_tx=XM_006933240.4;refGene_gn=NUDT5;refGene_tx=XM_019833937.2;refGene_gn=NUDT5;refGene_tx=XM_019833938.2
B4 11398005 11398068
refGene_gn=NUDT5;refGene_tx=XM_003988104.5;refGene_gn=NUDT5;refGene_tx

=XM_006933238.4;refGene_gn=NUDT5;refGene_tx=XM_006933239.4;refGene_gn=NUDT5;refGene_tx=XM_006933240.4;refGene_gn=NUDT5;refGene_tx=XM_019833937.2;refGene_gn=NUDT5;refGene_tx=XM_019833938.2
B4 11408310 11408384
refGene_gn=CDC123;refGene_tx=XM_003988105.5;refGene_gn=CDC123;refGene_tx=XM_006933242.4;refGene_gn=CDC123;refGene_tx=XM_006933243.4;refGene_gn=CDC123;refGene_tx=XM_019833934.1;refGene_gn=CDC123;refGene_tx=XM_019833936.2
B4 11411859 11411931
refGene_gn=CDC123;refGene_tx=XM_003988105.5;refGene_gn=CDC123;refGene_tx=XM_006933243.4;refGene_gn=CDC123;refGene_tx=XM_019833934.1
B4 11426485 11426543
refGene_gn=CDC123;refGene_tx=XM_003988105.5;refGene_gn=CDC123;refGene_tx=XM_006933242.4;refGene_gn=CDC123;refGene_tx=XM_006933243.4;refGene_gn=CDC123;refGene_tx=XM_019833934.1;refGene_gn=CDC123;refGene_tx=XM_019833936.2
B4 11426785 11426818
refGene_gn=CDC123;refGene_tx=XM_003988105.5;refGene_gn=CDC123;refGene_tx=XM_006933242.4;refGene_gn=CDC123;refGene_tx=XM_006933243.4;refGene_gn=CDC123;refGene_tx=XM_019833934.1;refGene_gn=CDC123;refGene_tx=XM_019833936.2
B4 11432176 11432272
refGene_gn=CDC123;refGene_tx=XM_003988105.5;refGene_gn=CDC123;refGene_tx=XM_006933242.4;refGene_gn=CDC123;refGene_tx=XM_019833934.1
B4 11434728 11434835
refGene_gn=CDC123;refGene_tx=XM_003988105.5;refGene_gn=CDC123;refGene_tx=XM_006933242.4;refGene_gn=CDC123;refGene_tx=XM_006933243.4;refGene_gn=CDC123;refGene_tx=XM_019833934.1;refGene_gn=CDC123;refGene_tx=XM_019833936.2
B4 11448816 11448865
refGene_gn=CDC123;refGene_tx=XM_003988105.5;refGene_gn=CDC123;refGene_tx=XM_006933242.4;refGene_gn=CDC123;refGene_tx=XM_006933243.4;refGene_gn=CDC123;refGene_tx=XM_019833934.1;refGene_gn=CDC123;refGene_tx=XM_019833936.2
B4 11452949 11453025
refGene_gn=CDC123;refGene_tx=XM_003988105.5;refGene_gn=CDC123;refGene_tx=XM_006933242.4;refGene_gn=CDC123;refGene_tx=XM_006933243.4;refGene_gn=CDC123;refGene_tx=XM_019833934.1;refGene_gn=CDC123;refGene_tx=XM_019833936.2
B4 11454660 11454783
refGene_gn=CDC123;refGene_tx=XM_003988105.5;refGene_gn=CDC123;refGene_tx=XM_006933242.4;refGene_gn=CDC123;refGene_tx=XM_006933243.4;refGene_gn=CDC123;refGene_tx=XM_019833936.2
B4 11456262 11456291
refGene_gn=CDC123;refGene_tx=XM_003988105.5;refGene_gn=CDC123;refGene_tx=XM_006933242.4;refGene_gn=CDC123;refGene_tx=XM_006933243.4;refGene_gn=CDC123;refGene_tx=XM_019833934.1;refGene_gn=CDC123;refGene_tx=XM_019833936.2
B4 11463360 11463489
refGene_gn=CDC123;refGene_tx=XM_003988105.5;refGene_gn=CDC123;refGene_

tx=XM_006933242.4;refGene_gn=CDC123;refGene_tx=XM_006933243.4;refGene_gn=CDC123;refGene_tx=XM_019833934.1;refGene_gn=CDC123;refGene_tx=XM_019833936.2
B4 11467467 11467605 refGene_gn=CDC123;refGene_tx=XM_003988105.5;refGene_gn=CDC123;refGene_tx=XM_006933242.4;refGene_gn=CDC123;refGene_tx=XM_006933243.4;refGene_gn=CDC123;refGene_tx=XM_019833934.1;refGene_gn=CDC123;refGene_tx=XM_019833936.2
B4 11468455 11468482 refGene_gn=CDC123;refGene_tx=XM_003988105.5;refGene_gn=CDC123;refGene_tx=XM_006933242.4;refGene_gn=CDC123;refGene_tx=XM_006933243.4;refGene_gn=CDC123;refGene_tx=XM_019833934.1;refGene_gn=CDC123;refGene_tx=XM_019833936.2
B4 11547541 11547633 refGene_gn=CAMK1D;refGene_tx=XM_019833928.2;refGene_gn=CAMK1D;refGene_tx=XM_019833933.2
B4 11709552 11709684 refGene_gn=CAMK1D;refGene_tx=XM_019833928.2;refGene_gn=CAMK1D;refGene_tx=XM_019833933.2
B4 11815577 11815652 refGene_gn=CAMK1D;refGene_tx=XM_019833928.2;refGene_gn=CAMK1D;refGene_tx=XM_019833933.2
B4 11916598 11916737 refGene_gn=CAMK1D;refGene_tx=XM_019833928.2;refGene_gn=CAMK1D;refGene_tx=XM_019833933.2
B4 11923868 11923995 refGene_gn=CAMK1D;refGene_tx=XM_019833928.2;refGene_gn=CAMK1D;refGene_tx=XM_019833933.2
B4 11940886 11940962 refGene_gn=CAMK1D;refGene_tx=XM_019833928.2;refGene_gn=CAMK1D;refGene_tx=XM_019833933.2
B4 11963057 11963170 refGene_gn=CAMK1D;refGene_tx=XM_019833928.2;refGene_gn=CAMK1D;refGene_tx=XM_019833933.2
B4 11964453 11964532 refGene_gn=CAMK1D;refGene_tx=XM_019833928.2;refGene_gn=CAMK1D;refGene_tx=XM_019833933.2
B4 11973784 11973872 refGene_gn=CAMK1D;refGene_tx=XM_019833928.2;refGene_gn=CAMK1D;refGene_tx=XM_019833933.2
B4 11975021 11975143 refGene_gn=CAMK1D;refGene_tx=XM_019833928.2;refGene_gn=CAMK1D;refGene_tx=XM_019833933.2
B4 11976641 11976760 refGene_gn=CAMK1D;refGene_tx=XM_019833928.2;refGene_gn=CAMK1D;refGene_tx=XM_019833933.2
B4 12087387 12087663 refGene_gn=CCDC3;refGene_tx=XM_019833940.2
B4 12198617 12198792 refGene_gn=CCDC3;refGene_tx=XM_019833940.2
B4 12201644 12202018 refGene_gn=CCDC3;refGene_tx=XM_019833940.2
B4 12284603 12284718

refGene_gn=OPTN;refGene_tx=XM_019833941.2;refGene_gn=OPTN;refGene_tx=XM_023256339.1;refGene_gn=OPTN;refGene_tx=XM_023256340.1;refGene_gn=OPTN;refGene_tx=XM_023256341.1;refGene_gn=OPTN;refGene_tx=XM_023256342.1
B4 12290168 12290345
refGene_gn=OPTN;refGene_tx=XM_003988112.5;refGene_gn=OPTN;refGene_tx=XM_019833941.2;refGene_gn=OPTN;refGene_tx=XM_019833942.2;refGene_gn=OPTN;refGene_tx=XM_023256339.1;refGene_gn=OPTN;refGene_tx=XM_023256340.1;refGene_gn=OPTN;refGene_tx=XM_023256341.1;refGene_gn=OPTN;refGene_tx=XM_023256342.1;refGene_gn=OPTN;refGene_tx=XM_023256344.1
B4 12290977 12291180
refGene_gn=OPTN;refGene_tx=XM_003988112.5;refGene_gn=OPTN;refGene_tx=XM_019833941.2;refGene_gn=OPTN;refGene_tx=XM_019833942.2;refGene_gn=OPTN;refGene_tx=XM_023256339.1;refGene_gn=OPTN;refGene_tx=XM_023256340.1;refGene_gn=OPTN;refGene_tx=XM_023256341.1;refGene_gn=OPTN;refGene_tx=XM_023256342.1;refGene_gn=OPTN;refGene_tx=XM_023256344.1
B4 12295671 12295854
refGene_gn=OPTN;refGene_tx=XM_003988112.5;refGene_gn=OPTN;refGene_tx=XM_019833941.2;refGene_gn=OPTN;refGene_tx=XM_019833942.2;refGene_gn=OPTN;refGene_tx=XM_023256339.1;refGene_gn=OPTN;refGene_tx=XM_023256340.1;refGene_gn=OPTN;refGene_tx=XM_023256341.1;refGene_gn=OPTN;refGene_tx=XM_023256342.1;refGene_gn=OPTN;refGene_tx=XM_023256344.1
B4 12298309 12298383
refGene_gn=OPTN;refGene_tx=XM_003988112.5;refGene_gn=OPTN;refGene_tx=XM_019833941.2;refGene_gn=OPTN;refGene_tx=XM_019833942.2;refGene_gn=OPTN;refGene_tx=XM_023256339.1;refGene_gn=OPTN;refGene_tx=XM_023256340.1;refGene_gn=OPTN;refGene_tx=XM_023256341.1;refGene_gn=OPTN;refGene_tx=XM_023256342.1;refGene_gn=OPTN;refGene_tx=XM_023256344.1
B4 12300774 12300909
refGene_gn=OPTN;refGene_tx=XM_003988112.5;refGene_gn=OPTN;refGene_tx=XM_019833941.2;refGene_gn=OPTN;refGene_tx=XM_019833942.2;refGene_gn=OPTN;refGene_tx=XM_023256339.1;refGene_gn=OPTN;refGene_tx=XM_023256340.1;refGene_gn=OPTN;refGene_tx=XM_023256341.1;refGene_gn=OPTN;refGene_tx=XM_023256342.1;refGene_gn=OPTN;refGene_tx=XM_023256344.1
B4 12308355 12308458
refGene_gn=OPTN;refGene_tx=XM_003988112.5;refGene_gn=OPTN;refGene_tx=XM_019833941.2;refGene_gn=OPTN;refGene_tx=XM_019833942.2;refGene_gn=OPTN;refGene_tx=XM_023256339.1;refGene_gn=OPTN;refGene_tx=XM_023256340.1;refGene_gn=OPTN;refGene_tx=XM_023256341.1;refGene_gn=OPTN;refGene_tx=XM_023256342.1;refGene_gn=OPTN;refGene_tx=XM_023256344.1
B4 12309998 12310114
refGene_gn=OPTN;refGene_tx=XM_003988112.5;refGene_gn=OPTN;refGene_tx=XM_019833941.2;refGene_gn=OPTN;refGene_tx=XM_019833942.2;refGene_gn=OPTN;refGene_tx=XM_023256339.1;refGene_gn=OPTN;refGene_tx=XM_023256340.1;refGene_gn=OPTN;refGene_tx=XM_023256341.1;refGene_gn=OPTN;refGene_tx=XM_023256342.1;refGene_gn=OPTN;refGene_tx=XM_023256344.1
B4 12311981 12312131
refGene_gn=OPTN;refGene_tx=XM_003988112.5;refGene_gn=OPTN;refGene_tx=XM_019833941.2;refGene_gn=OPTN;refGene_tx=XM_019833942.2;refGene_gn=OPTN;refGene_tx=XM_023256339.1;refGene_gn=OPTN;refGene_tx=XM_023256340.1;refGene_gn=OPTN;refGene_tx=XM_023256341.1;refGene_gn=OPTN;refGene_tx=X

M_023256342.1;refGene_gn=OPTN;refGene_tx=XM_023256344.1
B4 12312509 12312603
refGene_gn=OPTN;refGene_tx=XM_003988112.5;refGene_gn=OPTN;refGene_tx=XM_019833941.2;refGene_gn=OPTN;refGene_tx=XM_019833942.2;refGene_gn=OPTN;refGene_tx=XM_023256339.1;refGene_gn=OPTN;refGene_tx=XM_023256340.1;refGene_gn=OPTN;refGene_tx=XM_023256341.1;refGene_gn=OPTN;refGene_tx=XM_023256342.1;refGene_gn=OPTN;refGene_tx=XM_023256344.1
B4 12315902 12316061
refGene_gn=OPTN;refGene_tx=XM_003988112.5;refGene_gn=OPTN;refGene_tx=XM_019833941.2;refGene_gn=OPTN;refGene_tx=XM_019833942.2;refGene_gn=OPTN;refGene_tx=XM_023256339.1;refGene_gn=OPTN;refGene_tx=XM_023256340.1;refGene_gn=OPTN;refGene_tx=XM_023256341.1;refGene_gn=OPTN;refGene_tx=XM_023256342.1;refGene_gn=OPTN;refGene_tx=XM_023256344.1
B4 12317665 12317796
refGene_gn=OPTN;refGene_tx=XM_003988112.5;refGene_gn=OPTN;refGene_tx=XM_019833941.2;refGene_gn=OPTN;refGene_tx=XM_019833942.2;refGene_gn=OPTN;refGene_tx=XM_023256339.1;refGene_gn=OPTN;refGene_tx=XM_023256340.1;refGene_gn=OPTN;refGene_tx=XM_023256341.1;refGene_gn=OPTN;refGene_tx=XM_023256342.1;refGene_gn=OPTN;refGene_tx=XM_023256344.1
B4 12318703 12318786
refGene_gn=OPTN;refGene_tx=XM_003988112.5;refGene_gn=OPTN;refGene_tx=XM_019833941.2;refGene_gn=OPTN;refGene_tx=XM_019833942.2;refGene_gn=OPTN;refGene_tx=XM_023256339.1;refGene_gn=OPTN;refGene_tx=XM_023256340.1;refGene_gn=OPTN;refGene_tx=XM_023256341.1;refGene_gn=OPTN;refGene_tx=XM_023256342.1;refGene_gn=OPTN;refGene_tx=XM_023256344.1
B4 12319144 12319188
refGene_gn=OPTN;refGene_tx=XM_023256341.1;refGene_gn=OPTN;refGene_tx=XM_023256342.1
B4 12322216 12322344
refGene_gn=OPTN;refGene_tx=XM_003988112.5;refGene_gn=OPTN;refGene_tx=XM_019833941.2;refGene_gn=OPTN;refGene_tx=XM_019833942.2;refGene_gn=OPTN;refGene_tx=XM_023256339.1;refGene_gn=OPTN;refGene_tx=XM_023256340.1;refGene_gn=OPTN;refGene_tx=XM_023256344.1
B4 12338111 12338118
refGene_gn=MCM10;refGene_tx=XM_023256345.1;refGene_gn=MCM10;refGene_tx=XM_023256346.1;refGene_gn=MCM10;refGene_tx=XM_023256347.1;refGene_gn=MCM10;refGene_tx=XM_023256348.1;refGene_gn=MCM10;refGene_tx=XM_023256349.1;refGene_gn=MCM10;refGene_tx=XM_023256350.1
B4 12341528 12341870
refGene_gn=MCM10;refGene_tx=XM_023256345.1;refGene_gn=MCM10;refGene_tx=XM_023256346.1;refGene_gn=MCM10;refGene_tx=XM_023256347.1;refGene_gn=MCM10;refGene_tx=XM_023256348.1;refGene_gn=MCM10;refGene_tx=XM_023256349.1;refGene_gn=MCM10;refGene_tx=XM_023256350.1
B4 12342655 12342760
refGene_gn=MCM10;refGene_tx=XM_023256345.1;refGene_gn=MCM10;refGene_tx=XM_023256346.1;refGene_gn=MCM10;refGene_tx=XM_023256347.1;refGene_gn=MCM10;refGene_tx=XM_023256348.1;refGene_gn=MCM10;refGene_tx=XM_023256349.1;refGene_gn=MCM10;refGene_tx=XM_023256350.1
B4 12343272 12343410
refGene_gn=MCM10;refGene_tx=XM_023256345.1;refGene_gn=MCM10;refGene_tx

=XM_023256346.1;refGene_gn=MCM10;refGene_tx=XM_023256347.1;refGene_gn=MCM10;refGene_tx=XM_023256348.1;refGene_gn=MCM10;refGene_tx=XM_023256349.1;refGene_gn=MCM10;refGene_tx=XM_023256350.1
B4 12346307 12346479
refGene_gn=MCM10;refGene_tx=XM_023256345.1;refGene_gn=MCM10;refGene_tx=XM_023256346.1;refGene_gn=MCM10;refGene_tx=XM_023256347.1;refGene_gn=MCM10;refGene_tx=XM_023256348.1;refGene_gn=MCM10;refGene_tx=XM_023256349.1;refGene_gn=MCM10;refGene_tx=XM_023256350.1
B4 12348092 12348258
refGene_gn=MCM10;refGene_tx=XM_023256345.1;refGene_gn=MCM10;refGene_tx=XM_023256346.1;refGene_gn=MCM10;refGene_tx=XM_023256347.1;refGene_gn=MCM10;refGene_tx=XM_023256348.1;refGene_gn=MCM10;refGene_tx=XM_023256349.1;refGene_gn=MCM10;refGene_tx=XM_023256350.1
B4 12350263 12350431
refGene_gn=MCM10;refGene_tx=XM_023256345.1;refGene_gn=MCM10;refGene_tx=XM_023256346.1;refGene_gn=MCM10;refGene_tx=XM_023256347.1;refGene_gn=MCM10;refGene_tx=XM_023256348.1;refGene_gn=MCM10;refGene_tx=XM_023256349.1;refGene_gn=MCM10;refGene_tx=XM_023256350.1
B4 12351612 12351729
refGene_gn=MCM10;refGene_tx=XM_023256345.1;refGene_gn=MCM10;refGene_tx=XM_023256346.1;refGene_gn=MCM10;refGene_tx=XM_023256347.1;refGene_gn=MCM10;refGene_tx=XM_023256348.1;refGene_gn=MCM10;refGene_tx=XM_023256349.1;refGene_gn=MCM10;refGene_tx=XM_023256350.1
B4 12354248 12354448
refGene_gn=MCM10;refGene_tx=XM_023256345.1;refGene_gn=MCM10;refGene_tx=XM_023256346.1;refGene_gn=MCM10;refGene_tx=XM_023256347.1;refGene_gn=MCM10;refGene_tx=XM_023256348.1;refGene_gn=MCM10;refGene_tx=XM_023256349.1;refGene_gn=MCM10;refGene_tx=XM_023256350.1
B4 12355753 12355854
refGene_gn=MCM10;refGene_tx=XM_023256345.1;refGene_gn=MCM10;refGene_tx=XM_023256346.1;refGene_gn=MCM10;refGene_tx=XM_023256347.1;refGene_gn=MCM10;refGene_tx=XM_023256348.1;refGene_gn=MCM10;refGene_tx=XM_023256349.1;refGene_gn=MCM10;refGene_tx=XM_023256350.1
B4 12356898 12357009
refGene_gn=MCM10;refGene_tx=XM_023256345.1;refGene_gn=MCM10;refGene_tx=XM_023256346.1;refGene_gn=MCM10;refGene_tx=XM_023256347.1;refGene_gn=MCM10;refGene_tx=XM_023256348.1;refGene_gn=MCM10;refGene_tx=XM_023256349.1;refGene_gn=MCM10;refGene_tx=XM_023256350.1
B4 12357093 12357211
refGene_gn=MCM10;refGene_tx=XM_023256345.1;refGene_gn=MCM10;refGene_tx=XM_023256346.1;refGene_gn=MCM10;refGene_tx=XM_023256347.1;refGene_gn=MCM10;refGene_tx=XM_023256348.1;refGene_gn=MCM10;refGene_tx=XM_023256349.1;refGene_gn=MCM10;refGene_tx=XM_023256350.1
B4 12358428 12358657
refGene_gn=MCM10;refGene_tx=XM_023256345.1;refGene_gn=MCM10;refGene_tx=XM_023256346.1;refGene_gn=MCM10;refGene_tx=XM_023256347.1;refGene_gn=MCM10;refGene_tx=XM_023256348.1;refGene_gn=MCM10;refGene_tx=XM_023256349.1;refGene_gn=MCM10;refGene_tx=XM_023256350.1
B4 12360048 12360193
refGene_gn=MCM10;refGene_tx=XM_023256345.1;refGene_gn=MCM10;refGene_tx

=XM_023256346.1;refGene_gn=MCM10;refGene_tx=XM_023256347.1;refGene_gn=MCM10;refGene_tx=XM_023256348.1;refGene_gn=MCM10;refGene_tx=XM_023256349.1;refGene_gn=MCM10;refGene_tx=XM_023256350.1
B4 12360822 12360935
refGene_gn=MCM10;refGene_tx=XM_023256345.1;refGene_gn=MCM10;refGene_tx=XM_023256346.1;refGene_gn=MCM10;refGene_tx=XM_023256347.1;refGene_gn=MCM10;refGene_tx=XM_023256348.1;refGene_gn=MCM10;refGene_tx=XM_023256349.1;refGene_gn=MCM10;refGene_tx=XM_023256350.1
B4 12362834 12362948
refGene_gn=MCM10;refGene_tx=XM_023256345.1;refGene_gn=MCM10;refGene_tx=XM_023256346.1;refGene_gn=MCM10;refGene_tx=XM_023256347.1;refGene_gn=MCM10;refGene_tx=XM_023256348.1;refGene_gn=MCM10;refGene_tx=XM_023256349.1;refGene_gn=MCM10;refGene_tx=XM_023256350.1
B4 12365114 12365260
refGene_gn=MCM10;refGene_tx=XM_023256345.1;refGene_gn=MCM10;refGene_tx=XM_023256346.1;refGene_gn=MCM10;refGene_tx=XM_023256347.1;refGene_gn=MCM10;refGene_tx=XM_023256348.1;refGene_gn=MCM10;refGene_tx=XM_023256349.1;refGene_gn=MCM10;refGene_tx=XM_023256350.1
B4 12371660 12371703
refGene_gn=MCM10;refGene_tx=XM_023256345.1;refGene_gn=MCM10;refGene_tx=XM_023256346.1;refGene_gn=MCM10;refGene_tx=XM_023256347.1;refGene_gn=MCM10;refGene_tx=XM_023256348.1;refGene_gn=MCM10;refGene_tx=XM_023256349.1;refGene_gn=MCM10;refGene_tx=XM_023256350.1
B4 12371800 12371884
refGene_gn=MCM10;refGene_tx=XM_023256345.1;refGene_gn=MCM10;refGene_tx=XM_023256346.1;refGene_gn=MCM10;refGene_tx=XM_023256347.1;refGene_gn=MCM10;refGene_tx=XM_023256348.1;refGene_gn=MCM10;refGene_tx=XM_023256349.1;refGene_gn=MCM10;refGene_tx=XM_023256350.1
B4 12381213 12381311
refGene_gn=UCMA;refGene_tx=XM_003988155.4;refGene_gn=UCMA;refGene_tx=XM_019833946.2
B4 12385229 12385328 refGene_gn=UCMA;refGene_tx=XM_003988155.4
B4 12387027 12387311 refGene_gn=UCMA;refGene_tx=XM_023256352.1
B4 12388908 12389004
refGene_gn=UCMA;refGene_tx=XM_003988155.4;refGene_gn=UCMA;refGene_tx=XM_019833946.2;refGene_gn=UCMA;refGene_tx=XM_023256352.1
B4 12389093 12389159
refGene_gn=UCMA;refGene_tx=XM_003988155.4;refGene_gn=UCMA;refGene_tx=XM_019833946.2;refGene_gn=UCMA;refGene_tx=XM_023256352.1
B4 12389480 12389640
refGene_gn=UCMA;refGene_tx=XM_003988155.4;refGene_gn=UCMA;refGene_tx=XM_019833946.2;refGene_gn=UCMA;refGene_tx=XM_023256352.1
B4 12429751 12429805 refGene_gn=PHYH;refGene_tx=XM_006933252.4
B4 12432629 12432764 refGene_gn=PHYH;refGene_tx=XM_006933252.4
B4 12432917 12432956 refGene_gn=PHYH;refGene_tx=XM_019833945.2
B4 12434881 12435031
refGene_gn=PHYH;refGene_tx=XM_006933252.4;refGene_gn=PHYH;refGene_tx=XM_019833945.2
B4 12436974 12437156
refGene_gn=PHYH;refGene_tx=XM_006933252.4;refGene_gn=PHYH;refGene_tx=X

M_019833945.2
B4 12439254 12439336
refGene_gn=PHYH;refGene_tx=XM_006933252.4;refGene_gn=PHYH;refGene_tx=XM_019833945.2
B4 12442216 12442385
refGene_gn=PHYH;refGene_tx=XM_006933252.4;refGene_gn=PHYH;refGene_tx=XM_019833945.2
B4 12442729 12442840
refGene_gn=PHYH;refGene_tx=XM_006933252.4;refGene_gn=PHYH;refGene_tx=XM_019833945.2
B4 12444635 12444694
refGene_gn=PHYH;refGene_tx=XM_006933252.4;refGene_gn=PHYH;refGene_tx=XM_019833945.2
B4 12446118 12446193
refGene_gn=PHYH;refGene_tx=XM_006933252.4;refGene_gn=PHYH;refGene_tx=XM_019833945.2
B4 12460533 12460548 refGene_gn=SEPHS1;refGene_tx=XM_023256356.1
B4 12461988 12462203
refGene_gn=SEPHS1;refGene_tx=XM_006933254.4;refGene_gn=SEPHS1;refGene_tx=XM_023256353.1;refGene_gn=SEPHS1;refGene_tx=XM_023256354.1;refGene_gn=SEPHS1;refGene_tx=XM_023256356.1;refGene_gn=SEPHS1;refGene_tx=XM_023256357.1
B4 12464008 12464221
refGene_gn=SEPHS1;refGene_tx=XM_006933254.4;refGene_gn=SEPHS1;refGene_tx=XM_023256353.1;refGene_gn=SEPHS1;refGene_tx=XM_023256354.1;refGene_gn=SEPHS1;refGene_tx=XM_023256356.1;refGene_gn=SEPHS1;refGene_tx=XM_023256357.1
B4 12473419 12473519
refGene_gn=SEPHS1;refGene_tx=XM_006933254.4;refGene_gn=SEPHS1;refGene_tx=XM_023256353.1;refGene_gn=SEPHS1;refGene_tx=XM_023256354.1;refGene_gn=SEPHS1;refGene_tx=XM_023256356.1;refGene_gn=SEPHS1;refGene_tx=XM_023256357.1
B4 12475746 12475837
refGene_gn=SEPHS1;refGene_tx=XM_006933254.4;refGene_gn=SEPHS1;refGene_tx=XM_023256353.1;refGene_gn=SEPHS1;refGene_tx=XM_023256354.1;refGene_gn=SEPHS1;refGene_tx=XM_023256356.1;refGene_gn=SEPHS1;refGene_tx=XM_023256357.1
B4 12479326 12479481
refGene_gn=SEPHS1;refGene_tx=XM_006933254.4;refGene_gn=SEPHS1;refGene_tx=XM_023256353.1;refGene_gn=SEPHS1;refGene_tx=XM_023256354.1;refGene_gn=SEPHS1;refGene_tx=XM_023256356.1;refGene_gn=SEPHS1;refGene_tx=XM_023256357.1
B4 12480717 12480825
refGene_gn=SEPHS1;refGene_tx=XM_006933254.4;refGene_gn=SEPHS1;refGene_tx=XM_023256353.1;refGene_gn=SEPHS1;refGene_tx=XM_023256354.1;refGene_gn=SEPHS1;refGene_tx=XM_023256356.1;refGene_gn=SEPHS1;refGene_tx=XM_023256357.1
B4 12482840 12482944
refGene_gn=SEPHS1;refGene_tx=XM_006933254.4;refGene_gn=SEPHS1;refGene_tx=XM_023256353.1;refGene_gn=SEPHS1;refGene_tx=XM_023256354.1;refGene_

gn=SEPHS1;refGene_tx=XM_023256356.1;refGene_gn=SEPHS1;refGene_tx=XM_023256357.1
B4 12487721 12487914
refGene_gn=SEPHS1;refGene_tx=XM_006933254.4;refGene_gn=SEPHS1;refGene_tx=XM_023256353.1;refGene_gn=SEPHS1;refGene_tx=XM_023256354.1;refGene_gn=SEPHS1;refGene_tx=XM_023256356.1;refGene_gn=SEPHS1;refGene_tx=XM_023256357.1
B4 12587122 12587301
refGene_gn=BEND7;refGene_tx=XM_023256358.1;refGene_gn=BEND7;refGene_tx=XM_023256359.1;refGene_gn=BEND7;refGene_tx=XM_023256360.1;refGene_gn=BEND7;refGene_tx=XM_023256361.1;refGene_gn=BEND7;refGene_tx=XM_023256362.1;refGene_gn=BEND7;refGene_tx=XM_023256364.1;refGene_gn=BEND7;refGene_tx=XM_023256368.1
B4 12589829 12589929
refGene_gn=BEND7;refGene_tx=XM_023256360.1;refGene_gn=BEND7;refGene_tx=XM_023256361.1
B4 12590601 12590615
refGene_gn=BEND7;refGene_tx=XM_023256369.1;refGene_gn=BEND7;refGene_tx=XM_023256370.1
B4 12590722 12590772
refGene_gn=BEND7;refGene_tx=XM_023256365.1;refGene_gn=BEND7;refGene_tx=XM_023256366.1
B4 12592218 12592272
refGene_gn=BEND7;refGene_tx=XM_023256358.1;refGene_gn=BEND7;refGene_tx=XM_023256359.1;refGene_gn=BEND7;refGene_tx=XM_023256360.1;refGene_gn=BEND7;refGene_tx=XM_023256361.1;refGene_gn=BEND7;refGene_tx=XM_023256362.1;refGene_gn=BEND7;refGene_tx=XM_023256364.1;refGene_gn=BEND7;refGene_tx=XM_023256365.1;refGene_gn=BEND7;refGene_tx=XM_023256366.1;refGene_gn=BEND7;refGene_tx=XM_023256368.1;refGene_gn=BEND7;refGene_tx=XM_023256369.1;refGene_gn=BEND7;refGene_tx=XM_023256370.1
B4 12601212 12601332
refGene_gn=BEND7;refGene_tx=XM_023256358.1;refGene_gn=BEND7;refGene_tx=XM_023256359.1;refGene_gn=BEND7;refGene_tx=XM_023256360.1;refGene_gn=BEND7;refGene_tx=XM_023256361.1;refGene_gn=BEND7;refGene_tx=XM_023256362.1;refGene_gn=BEND7;refGene_tx=XM_023256364.1;refGene_gn=BEND7;refGene_tx=XM_023256365.1;refGene_gn=BEND7;refGene_tx=XM_023256366.1;refGene_gn=BEND7;refGene_tx=XM_023256368.1;refGene_gn=BEND7;refGene_tx=XM_023256369.1;refGene_gn=BEND7;refGene_tx=XM_023256370.1
B4 12623765 12623991
refGene_gn=BEND7;refGene_tx=XM_023256358.1;refGene_gn=BEND7;refGene_tx=XM_023256359.1;refGene_gn=BEND7;refGene_tx=XM_023256360.1;refGene_gn=BEND7;refGene_tx=XM_023256361.1;refGene_gn=BEND7;refGene_tx=XM_023256362.1;refGene_gn=BEND7;refGene_tx=XM_023256364.1;refGene_gn=BEND7;refGene_tx=XM_023256365.1;refGene_gn=BEND7;refGene_tx=XM_023256366.1;refGene_gn=BEND7;refGene_tx=XM_023256367.1;refGene_gn=BEND7;refGene_tx=XM_023256368.1;refGene_gn=BEND7;refGene_tx=XM_023256369.1;refGene_gn=BEND7;refGene_tx=XM_023256370.1
B4 12633631 12633719 refGene_gn=BEND7;refGene_tx=XM_023256367.1
B4 12634132 12634286 refGene_gn=BEND7;refGene_tx=XM_023256367.1
B4 12634844 12635107

refGene_gn=BEND7;refGene_tx=XM_023256358.1;refGene_gn=BEND7;refGene_tx=XM_023256359.1;refGene_gn=BEND7;refGene_tx=XM_023256360.1;refGene_gn=BEND7;refGene_tx=XM_023256361.1;refGene_gn=BEND7;refGene_tx=XM_023256362.1;refGene_gn=BEND7;refGene_tx=XM_023256364.1;refGene_gn=BEND7;refGene_tx=XM_023256365.1;refGene_gn=BEND7;refGene_tx=XM_023256366.1;refGene_gn=BEND7;refGene_tx=XM_023256367.1;refGene_gn=BEND7;refGene_tx=XM_023256368.1;refGene_gn=BEND7;refGene_tx=XM_023256369.1;refGene_gn=BEND7;refGene_tx=XM_023256370.1
B4 12638642 12638765
refGene_gn=BEND7;refGene_tx=XM_023256358.1;refGene_gn=BEND7;refGene_tx=XM_023256359.1;refGene_gn=BEND7;refGene_tx=XM_023256360.1;refGene_gn=BEND7;refGene_tx=XM_023256361.1;refGene_gn=BEND7;refGene_tx=XM_023256362.1;refGene_gn=BEND7;refGene_tx=XM_023256364.1;refGene_gn=BEND7;refGene_tx=XM_023256365.1;refGene_gn=BEND7;refGene_tx=XM_023256366.1;refGene_gn=BEND7;refGene_tx=XM_023256367.1;refGene_gn=BEND7;refGene_tx=XM_023256368.1;refGene_gn=BEND7;refGene_tx=XM_023256369.1;refGene_gn=BEND7;refGene_tx=XM_023256370.1
B4 12641820 12642123
refGene_gn=BEND7;refGene_tx=XM_023256358.1;refGene_gn=BEND7;refGene_tx=XM_023256359.1;refGene_gn=BEND7;refGene_tx=XM_023256360.1;refGene_gn=BEND7;refGene_tx=XM_023256361.1;refGene_gn=BEND7;refGene_tx=XM_023256362.1;refGene_gn=BEND7;refGene_tx=XM_023256364.1;refGene_gn=BEND7;refGene_tx=XM_023256365.1;refGene_gn=BEND7;refGene_tx=XM_023256366.1;refGene_gn=BEND7;refGene_tx=XM_023256367.1;refGene_gn=BEND7;refGene_tx=XM_023256368.1;refGene_gn=BEND7;refGene_tx=XM_023256369.1;refGene_gn=BEND7;refGene_tx=XM_023256370.1
B4 12665791 12665875
refGene_gn=BEND7;refGene_tx=XM_023256358.1;refGene_gn=BEND7;refGene_tx=XM_023256359.1;refGene_gn=BEND7;refGene_tx=XM_023256360.1;refGene_gn=BEND7;refGene_tx=XM_023256361.1;refGene_gn=BEND7;refGene_tx=XM_023256364.1;refGene_gn=BEND7;refGene_tx=XM_023256365.1;refGene_gn=BEND7;refGene_tx=XM_023256366.1;refGene_gn=BEND7;refGene_tx=XM_023256367.1;refGene_gn=BEND7;refGene_tx=XM_023256369.1;refGene_gn=BEND7;refGene_tx=XM_023256370.1
B4 12668204 12668265
refGene_gn=BEND7;refGene_tx=XM_023256358.1;refGene_gn=BEND7;refGene_tx=XM_023256359.1;refGene_gn=BEND7;refGene_tx=XM_023256360.1;refGene_gn=BEND7;refGene_tx=XM_023256361.1;refGene_gn=BEND7;refGene_tx=XM_023256362.1;refGene_gn=BEND7;refGene_tx=XM_023256364.1;refGene_gn=BEND7;refGene_tx=XM_023256365.1;refGene_gn=BEND7;refGene_tx=XM_023256366.1;refGene_gn=BEND7;refGene_tx=XM_023256367.1;refGene_gn=BEND7;refGene_tx=XM_023256369.1;refGene_gn=BEND7;refGene_tx=XM_023256370.1
B4 12721783 12721849
refGene_gn=PRPF18;refGene_tx=XM_003988115.5;refGene_gn=PRPF18;refGene_tx=XM_006933263.4
B4 12738235 12738313
refGene_gn=PRPF18;refGene_tx=XM_003988115.5;refGene_gn=PRPF18;refGene_tx=XM_006933263.4
B4 12738428 12738455 refGene_gn=PRPF18;refGene_tx=XM_006933263.4
B4 12741032 12741137

refGene_gn=PRPF18;refGene_tx=XM_003988115.5;refGene_gn=PRPF18;refGene_tx=XM_006933263.4
B4 12743240 12743354
refGene_gn=PRPF18;refGene_tx=XM_003988115.5;refGene_gn=PRPF18;refGene_tx=XM_006933263.4
B4 12749407 12749554
refGene_gn=PRPF18;refGene_tx=XM_003988115.5;refGene_gn=PRPF18;refGene_tx=XM_006933263.4
B4 12751067 12751136
refGene_gn=PRPF18;refGene_tx=XM_003988115.5;refGene_gn=PRPF18;refGene_tx=XM_006933263.4
B4 12753068 12753209
refGene_gn=PRPF18;refGene_tx=XM_003988115.5;refGene_gn=PRPF18;refGene_tx=XM_006933263.4
B4 12753334 12753406
refGene_gn=PRPF18;refGene_tx=XM_003988115.5;refGene_gn=PRPF18;refGene_tx=XM_006933263.4
B4 12755727 12755883
refGene_gn=PRPF18;refGene_tx=XM_003988115.5;refGene_gn=PRPF18;refGene_tx=XM_006933263.4
B4 12769182 12769263
refGene_gn=PRPF18;refGene_tx=XM_003988115.5;refGene_gn=PRPF18;refGene_tx=XM_006933263.4
B4 12785447 12785550
refGene_gn=FRMD4A;refGene_tx=XM_023256371.1;refGene_gn=FRMD4A;refGene_tx=XM_023256373.1;refGene_gn=FRMD4A;refGene_tx=XM_023256374.1;refGene_gn=FRMD4A;refGene_tx=XM_023256375.1;refGene_gn=FRMD4A;refGene_tx=XM_023256377.1;refGene_gn=FRMD4A;refGene_tx=XM_023256378.1;refGene_gn=FRMD4A;refGene_tx=XM_023256379.1;refGene_gn=FRMD4A;refGene_tx=XM_023256380.1
B4 12799379 12799449
refGene_gn=FRMD4A;refGene_tx=XM_023256372.1;refGene_gn=FRMD4A;refGene_tx=XM_023256376.1
B4 12801720 12801817
refGene_gn=FRMD4A;refGene_tx=XM_023256371.1;refGene_gn=FRMD4A;refGene_tx=XM_023256372.1;refGene_gn=FRMD4A;refGene_tx=XM_023256373.1;refGene_gn=FRMD4A;refGene_tx=XM_023256374.1;refGene_gn=FRMD4A;refGene_tx=XM_023256375.1;refGene_gn=FRMD4A;refGene_tx=XM_023256376.1;refGene_gn=FRMD4A;refGene_tx=XM_023256377.1;refGene_gn=FRMD4A;refGene_tx=XM_023256378.1;refGene_gn=FRMD4A;refGene_tx=XM_023256379.1;refGene_gn=FRMD4A;refGene_tx=XM_023256380.1
B4 12803488 12804351
refGene_gn=FRMD4A;refGene_tx=XM_023256371.1;refGene_gn=FRMD4A;refGene_tx=XM_023256372.1;refGene_gn=FRMD4A;refGene_tx=XM_023256373.1;refGene_gn=FRMD4A;refGene_tx=XM_023256374.1;refGene_gn=FRMD4A;refGene_tx=XM_023256375.1;refGene_gn=FRMD4A;refGene_tx=XM_023256376.1;refGene_gn=FRMD4A;refGene_tx=XM_023256377.1;refGene_gn=FRMD4A;refGene_tx=XM_023256378.1;refGene_gn=FRMD4A;refGene_tx=XM_023256379.1;refGene_gn=FRMD4A;refGene_tx=XM_023256380.1
B4 12806195 12806363

refGene_gn=FRMD4A;refGene_tx=XM_023256371.1;refGene_gn=FRMD4A;refGene_tx=XM_023256372.1;refGene_gn=FRMD4A;refGene_tx=XM_023256373.1;refGene_gn=FRMD4A;refGene_tx=XM_023256374.1;refGene_gn=FRMD4A;refGene_tx=XM_023256375.1;refGene_gn=FRMD4A;refGene_tx=XM_023256376.1;refGene_gn=FRMD4A;refGene_tx=XM_023256377.1;refGene_gn=FRMD4A;refGene_tx=XM_023256378.1;refGene_gn=FRMD4A;refGene_tx=XM_023256379.1;refGene_gn=FRMD4A;refGene_tx=XM_023256380.1

B4 12807319 12807557
refGene_gn=FRMD4A;refGene_tx=XM_023256371.1;refGene_gn=FRMD4A;refGene_tx=XM_023256372.1;refGene_gn=FRMD4A;refGene_tx=XM_023256373.1;refGene_gn=FRMD4A;refGene_tx=XM_023256374.1;refGene_gn=FRMD4A;refGene_tx=XM_023256375.1;refGene_gn=FRMD4A;refGene_tx=XM_023256376.1;refGene_gn=FRMD4A;refGene_tx=XM_023256377.1;refGene_gn=FRMD4A;refGene_tx=XM_023256378.1;refGene_gn=FRMD4A;refGene_tx=XM_023256379.1;refGene_gn=FRMD4A;refGene_tx=XM_023256380.1

B4 12810008 12810065
refGene_gn=FRMD4A;refGene_tx=XM_023256371.1;refGene_gn=FRMD4A;refGene_tx=XM_023256372.1;refGene_gn=FRMD4A;refGene_tx=XM_023256373.1;refGene_gn=FRMD4A;refGene_tx=XM_023256374.1;refGene_gn=FRMD4A;refGene_tx=XM_023256375.1;refGene_gn=FRMD4A;refGene_tx=XM_023256376.1;refGene_gn=FRMD4A;refGene_tx=XM_023256377.1;refGene_gn=FRMD4A;refGene_tx=XM_023256378.1;refGene_gn=FRMD4A;refGene_tx=XM_023256379.1;refGene_gn=FRMD4A;refGene_tx=XM_023256380.1

B4 12812588 12812817
refGene_gn=FRMD4A;refGene_tx=XM_023256371.1;refGene_gn=FRMD4A;refGene_tx=XM_023256372.1;refGene_gn=FRMD4A;refGene_tx=XM_023256373.1;refGene_gn=FRMD4A;refGene_tx=XM_023256374.1;refGene_gn=FRMD4A;refGene_tx=XM_023256375.1;refGene_gn=FRMD4A;refGene_tx=XM_023256376.1;refGene_gn=FRMD4A;refGene_tx=XM_023256377.1;refGene_gn=FRMD4A;refGene_tx=XM_023256378.1;refGene_gn=FRMD4A;refGene_tx=XM_023256379.1;refGene_gn=FRMD4A;refGene_tx=XM_023256380.1

B4 12816022 12816145
refGene_gn=FRMD4A;refGene_tx=XM_023256371.1;refGene_gn=FRMD4A;refGene_tx=XM_023256372.1;refGene_gn=FRMD4A;refGene_tx=XM_023256373.1;refGene_gn=FRMD4A;refGene_tx=XM_023256374.1;refGene_gn=FRMD4A;refGene_tx=XM_023256375.1;refGene_gn=FRMD4A;refGene_tx=XM_023256376.1;refGene_gn=FRMD4A;refGene_tx=XM_023256377.1;refGene_gn=FRMD4A;refGene_tx=XM_023256378.1;refGene_gn=FRMD4A;refGene_tx=XM_023256379.1;refGene_gn=FRMD4A;refGene_tx=XM_023256380.1

B4 12819839 12819973
refGene_gn=FRMD4A;refGene_tx=XM_023256371.1;refGene_gn=FRMD4A;refGene_tx=XM_023256372.1;refGene_gn=FRMD4A;refGene_tx=XM_023256373.1;refGene_gn=FRMD4A;refGene_tx=XM_023256374.1;refGene_gn=FRMD4A;refGene_tx=XM_023256375.1;refGene_gn=FRMD4A;refGene_tx=XM_023256376.1;refGene_gn=FRMD4A;refGene_tx=XM_023256377.1;refGene_gn=FRMD4A;refGene_tx=XM_023256378.1;refGene_gn=FRMD4A;refGene_tx=XM_023256379.1;refGene_gn=FRMD4A;refGene_tx=XM_023256380.1

B4 12836645 12836787
refGene_gn=FRMD4A;refGene_tx=XM_023256371.1;refGene_gn=FRMD4A;refGene_tx=XM_023256372.1;refGene_gn=FRMD4A;refGene_tx=XM_023256373.1;refGene_

gn=FRMD4A;refGene_tx=XM_023256374.1;refGene_gn=FRMD4A;refGene_tx=XM_023256375.1;refGene_gn=FRMD4A;refGene_tx=XM_023256376.1;refGene_gn=FRMD4A;refGene_tx=XM_023256377.1;refGene_gn=FRMD4A;refGene_tx=XM_023256378.1;refGene_gn=FRMD4A;refGene_tx=XM_023256379.1;refGene_gn=FRMD4A;refGene_tx=XM_023256380.1
B4 12843039 12843178
refGene_gn=FRMD4A;refGene_tx=XM_023256371.1;refGene_gn=FRMD4A;refGene_tx=XM_023256372.1;refGene_gn=FRMD4A;refGene_tx=XM_023256373.1;refGene_gn=FRMD4A;refGene_tx=XM_023256374.1;refGene_gn=FRMD4A;refGene_tx=XM_023256375.1;refGene_gn=FRMD4A;refGene_tx=XM_023256376.1;refGene_gn=FRMD4A;refGene_tx=XM_023256377.1;refGene_gn=FRMD4A;refGene_tx=XM_023256378.1;refGene_gn=FRMD4A;refGene_tx=XM_023256379.1;refGene_gn=FRMD4A;refGene_tx=XM_023256380.1
B4 12848310 12848387
refGene_gn=FRMD4A;refGene_tx=XM_023256371.1;refGene_gn=FRMD4A;refGene_tx=XM_023256372.1;refGene_gn=FRMD4A;refGene_tx=XM_023256373.1;refGene_gn=FRMD4A;refGene_tx=XM_023256374.1;refGene_gn=FRMD4A;refGene_tx=XM_023256375.1;refGene_gn=FRMD4A;refGene_tx=XM_023256376.1;refGene_gn=FRMD4A;refGene_tx=XM_023256377.1;refGene_gn=FRMD4A;refGene_tx=XM_023256378.1;refGene_gn=FRMD4A;refGene_tx=XM_023256379.1;refGene_gn=FRMD4A;refGene_tx=XM_023256380.1
B4 12872993 12873080
refGene_gn=FRMD4A;refGene_tx=XM_023256371.1;refGene_gn=FRMD4A;refGene_tx=XM_023256372.1;refGene_gn=FRMD4A;refGene_tx=XM_023256373.1;refGene_gn=FRMD4A;refGene_tx=XM_023256374.1;refGene_gn=FRMD4A;refGene_tx=XM_023256375.1;refGene_gn=FRMD4A;refGene_tx=XM_023256376.1;refGene_gn=FRMD4A;refGene_tx=XM_023256377.1;refGene_gn=FRMD4A;refGene_tx=XM_023256378.1;refGene_gn=FRMD4A;refGene_tx=XM_023256379.1;refGene_gn=FRMD4A;refGene_tx=XM_023256380.1
B4 12877330 12877388
refGene_gn=FRMD4A;refGene_tx=XM_023256371.1;refGene_gn=FRMD4A;refGene_tx=XM_023256372.1;refGene_gn=FRMD4A;refGene_tx=XM_023256373.1;refGene_gn=FRMD4A;refGene_tx=XM_023256374.1;refGene_gn=FRMD4A;refGene_tx=XM_023256375.1;refGene_gn=FRMD4A;refGene_tx=XM_023256376.1;refGene_gn=FRMD4A;refGene_tx=XM_023256377.1;refGene_gn=FRMD4A;refGene_tx=XM_023256378.1;refGene_gn=FRMD4A;refGene_tx=XM_023256379.1;refGene_gn=FRMD4A;refGene_tx=XM_023256380.1
B4 12877663 12877729
refGene_gn=FRMD4A;refGene_tx=XM_023256371.1;refGene_gn=FRMD4A;refGene_tx=XM_023256372.1;refGene_gn=FRMD4A;refGene_tx=XM_023256373.1;refGene_gn=FRMD4A;refGene_tx=XM_023256374.1;refGene_gn=FRMD4A;refGene_tx=XM_023256375.1;refGene_gn=FRMD4A;refGene_tx=XM_023256376.1;refGene_gn=FRMD4A;refGene_tx=XM_023256377.1;refGene_gn=FRMD4A;refGene_tx=XM_023256378.1;refGene_gn=FRMD4A;refGene_tx=XM_023256379.1;refGene_gn=FRMD4A;refGene_tx=XM_023256380.1
B4 12882693 12882777
refGene_gn=FRMD4A;refGene_tx=XM_023256371.1;refGene_gn=FRMD4A;refGene_tx=XM_023256372.1;refGene_gn=FRMD4A;refGene_tx=XM_023256373.1;refGene_gn=FRMD4A;refGene_tx=XM_023256374.1;refGene_gn=FRMD4A;refGene_tx=XM_023256375.1;refGene_gn=FRMD4A;refGene_tx=XM_023256376.1;refGene_gn=FRMD4

A;refGene_tx=XM_023256377.1;refGene_gn=FRMD4A;refGene_tx=XM_023256378.1;refGene_gn=FRMD4A;refGene_tx=XM_023256379.1;refGene_gn=FRMD4A;refGene_tx=XM_023256380.1
B4 12894330 12894353
refGene_gn=FRMD4A;refGene_tx=XM_023256371.1;refGene_gn=FRMD4A;refGene_tx=XM_023256372.1;refGene_gn=FRMD4A;refGene_tx=XM_023256373.1;refGene_gn=FRMD4A;refGene_tx=XM_023256374.1;refGene_gn=FRMD4A;refGene_tx=XM_023256375.1;refGene_gn=FRMD4A;refGene_tx=XM_023256376.1;refGene_gn=FRMD4A;refGene_tx=XM_023256377.1;refGene_gn=FRMD4A;refGene_tx=XM_023256378.1;refGene_gn=FRMD4A;refGene_tx=XM_023256379.1;refGene_gn=FRMD4A;refGene_tx=XM_023256380.1
B4 12895266 12895323
refGene_gn=FRMD4A;refGene_tx=XM_023256371.1;refGene_gn=FRMD4A;refGene_tx=XM_023256372.1;refGene_gn=FRMD4A;refGene_tx=XM_023256373.1;refGene_gn=FRMD4A;refGene_tx=XM_023256374.1;refGene_gn=FRMD4A;refGene_tx=XM_023256375.1;refGene_gn=FRMD4A;refGene_tx=XM_023256376.1;refGene_gn=FRMD4A;refGene_tx=XM_023256377.1;refGene_gn=FRMD4A;refGene_tx=XM_023256378.1;refGene_gn=FRMD4A;refGene_tx=XM_023256379.1;refGene_gn=FRMD4A;refGene_tx=XM_023256380.1
B4 12907951 12908036
refGene_gn=FRMD4A;refGene_tx=XM_023256371.1;refGene_gn=FRMD4A;refGene_tx=XM_023256372.1;refGene_gn=FRMD4A;refGene_tx=XM_023256373.1;refGene_gn=FRMD4A;refGene_tx=XM_023256374.1;refGene_gn=FRMD4A;refGene_tx=XM_023256375.1;refGene_gn=FRMD4A;refGene_tx=XM_023256376.1;refGene_gn=FRMD4A;refGene_tx=XM_023256377.1;refGene_gn=FRMD4A;refGene_tx=XM_023256378.1;refGene_gn=FRMD4A;refGene_tx=XM_023256379.1;refGene_gn=FRMD4A;refGene_tx=XM_023256380.1
B4 12919064 12919157
refGene_gn=FRMD4A;refGene_tx=XM_023256371.1;refGene_gn=FRMD4A;refGene_tx=XM_023256372.1;refGene_gn=FRMD4A;refGene_tx=XM_023256373.1;refGene_gn=FRMD4A;refGene_tx=XM_023256374.1;refGene_gn=FRMD4A;refGene_tx=XM_023256375.1;refGene_gn=FRMD4A;refGene_tx=XM_023256376.1;refGene_gn=FRMD4A;refGene_tx=XM_023256377.1;refGene_gn=FRMD4A;refGene_tx=XM_023256378.1;refGene_gn=FRMD4A;refGene_tx=XM_023256379.1;refGene_gn=FRMD4A;refGene_tx=XM_023256380.1
B4 12931671 12931766
refGene_gn=FRMD4A;refGene_tx=XM_023256371.1;refGene_gn=FRMD4A;refGene_tx=XM_023256372.1;refGene_gn=FRMD4A;refGene_tx=XM_023256373.1;refGene_gn=FRMD4A;refGene_tx=XM_023256374.1;refGene_gn=FRMD4A;refGene_tx=XM_023256375.1;refGene_gn=FRMD4A;refGene_tx=XM_023256376.1;refGene_gn=FRMD4A;refGene_tx=XM_023256377.1;refGene_gn=FRMD4A;refGene_tx=XM_023256378.1;refGene_gn=FRMD4A;refGene_tx=XM_023256379.1;refGene_gn=FRMD4A;refGene_tx=XM_023256380.1
B4 12973098 12973164
refGene_gn=FRMD4A;refGene_tx=XM_023256371.1;refGene_gn=FRMD4A;refGene_tx=XM_023256372.1;refGene_gn=FRMD4A;refGene_tx=XM_023256373.1;refGene_gn=FRMD4A;refGene_tx=XM_023256374.1;refGene_gn=FRMD4A;refGene_tx=XM_023256375.1;refGene_gn=FRMD4A;refGene_tx=XM_023256376.1;refGene_gn=FRMD4A;refGene_tx=XM_023256377.1;refGene_gn=FRMD4A;refGene_tx=XM_023256378.1;refGene_gn=FRMD4A;refGene_tx=XM_023256379.1;refGene_gn=FRMD4A;refGen

e_tx=XM_023256380.1
B4	12988148	12988157	refGene_gn=FRMD4A;refGene_tx=XM_023256378.1
B4	13066696	13066855
refGene_gn=FRMD4A;refGene_tx=XM_023256371.1;refGene_gn=FRMD4A;refGene_
tx=XM_023256372.1
B4	13098206	13098299
refGene_gn=FRMD4A;refGene_tx=XM_023256374.1;refGene_gn=FRMD4A;refGene_
tx=XM_023256376.1
B4	13115704	13115770	refGene_gn=FRMD4A;refGene_tx=XM_023256375.1
B4	13237024	13237057	refGene_gn=FRMD4A;refGene_tx=XM_023256377.1
B4	13396857	13396959	refGene_gn=FRMD4A;refGene_tx=XM_023256373.1
B4	13582206	13582323
refGene_gn=FAM107B;refGene_tx=XM_006933265.4;refGene_gn=FAM107B;refGen
e_tx=XM_006933266.4;refGene_gn=FAM107B;refGene_tx=XM_023256381.1;refGe
ne_gn=FAM107B;refGene_tx=XM_023256382.1
B4	13583303	13583454
refGene_gn=FAM107B;refGene_tx=XM_006933265.4;refGene_gn=FAM107B;refGen
e_tx=XM_006933266.4;refGene_gn=FAM107B;refGene_tx=XM_023256381.1;refGe
ne_gn=FAM107B;refGene_tx=XM_023256382.1
B4	13591987	13592171
refGene_gn=FAM107B;refGene_tx=XM_006933265.4;refGene_gn=FAM107B;refGen
e_tx=XM_006933266.4;refGene_gn=FAM107B;refGene_tx=XM_023256381.1;refGe
ne_gn=FAM107B;refGene_tx=XM_023256382.1
B4	13721877	13721887	refGene_gn=FAM107B;refGene_tx=XM_023256382.1
B4	13866739	13866918	refGene_gn=CDNF;refGene_tx=XM_006933269.3
B4	13872581	13872723	refGene_gn=CDNF;refGene_tx=XM_006933269.3
B4	13874026	13874154	refGene_gn=CDNF;refGene_tx=XM_006933269.3
B4	13886441	13886544	refGene_gn=CDNF;refGene_tx=XM_006933269.3
B4	13887032	13887089	refGene_gn=HSPA14;refGene_tx=XM_003988117.5
B4	13888350	13888431	refGene_gn=HSPA14;refGene_tx=XM_003988117.5
B4	13888504	13888587
refGene_gn=HSPA14;refGene_tx=XM_003988117.5;refGene_gn=HSPA14;refGene_
tx=XM_011283611.3;refGene_gn=HSPA14;refGene_tx=XM_023256389.1
B4	13896944	13896993
refGene_gn=HSPA14;refGene_tx=XM_003988117.5;refGene_gn=HSPA14;refGene_
tx=XM_011283611.3;refGene_gn=HSPA14;refGene_tx=XM_023256389.1
B4	13897127	13897233
refGene_gn=HSPA14;refGene_tx=XM_003988117.5;refGene_gn=HSPA14;refGene_
tx=XM_011283611.3;refGene_gn=HSPA14;refGene_tx=XM_023256389.1
B4	13898503	13898594
refGene_gn=HSPA14;refGene_tx=XM_003988117.5;refGene_gn=HSPA14;refGene_
tx=XM_011283611.3;refGene_gn=HSPA14;refGene_tx=XM_023256389.1
B4	13899342	13899447
refGene_gn=HSPA14;refGene_tx=XM_003988117.5;refGene_gn=HSPA14;refGene_
tx=XM_011283611.3;refGene_gn=HSPA14;refGene_tx=XM_023256389.1
B4	13900641	13900803
refGene_gn=HSPA14;refGene_tx=XM_003988117.5;refGene_gn=HSPA14;refGene_
tx=XM_011283611.3;refGene_gn=HSPA14;refGene_tx=XM_023256389.1
B4	13901973	13902129
refGene_gn=HSPA14;refGene_tx=XM_003988117.5;refGene_gn=HSPA14;refGene_

tx=XM_011283611.3;refGene_gn=HSPA14;refGene_tx=XM_023256389.1
B4 13908152 13908255
refGene_gn=HSPA14;refGene_tx=XM_003988117.5;refGene_gn=HSPA14;refGene_tx=XM_011283611.3;refGene_gn=HSPA14;refGene_tx=XM_023256389.1
B4 13924601 13924814
refGene_gn=HSPA14;refGene_tx=XM_003988117.5;refGene_gn=HSPA14;refGene_tx=XM_011283611.3;refGene_gn=HSPA14;refGene_tx=XM_023256389.1
B4 13925215 13925389
refGene_gn=HSPA14;refGene_tx=XM_003988117.5;refGene_gn=HSPA14;refGene_tx=XM_011283611.3;refGene_gn=HSPA14;refGene_tx=XM_023256389.1
B4 13928313 13928384
refGene_gn=HSPA14;refGene_tx=XM_003988117.5;refGene_gn=HSPA14;refGene_tx=XM_011283611.3;refGene_gn=HSPA14;refGene_tx=XM_023256389.1
B4 13929255 13929334
refGene_gn=HSPA14;refGene_tx=XM_003988117.5;refGene_gn=HSPA14;refGene_tx=XM_011283611.3;refGene_gn=HSPA14;refGene_tx=XM_023256389.1
B4 13936279 13936310
refGene_gn=SUV39H2;refGene_tx=XM_023256390.1;refGene_gn=SUV39H2;refGene_tx=XM_023256392.1;refGene_gn=SUV39H2;refGene_tx=XM_023256393.1
B4 13939026 13939172
refGene_gn=SUV39H2;refGene_tx=XM_023256390.1;refGene_gn=SUV39H2;refGene_tx=XM_023256392.1;refGene_gn=SUV39H2;refGene_tx=XM_023256393.1
B4 13951333 13951999
refGene_gn=SUV39H2;refGene_tx=XM_006933270.4;refGene_gn=SUV39H2;refGene_tx=XM_019833973.1;refGene_gn=SUV39H2;refGene_tx=XM_023256390.1;refGene_gn=SUV39H2;refGene_tx=XM_023256391.1;refGene_gn=SUV39H2;refGene_tx=XM_023256392.1;refGene_gn=SUV39H2;refGene_tx=XM_023256393.1
B4 13954366 13954527
refGene_gn=SUV39H2;refGene_tx=XM_006933270.4;refGene_gn=SUV39H2;refGene_tx=XM_019833973.1;refGene_gn=SUV39H2;refGene_tx=XM_023256390.1;refGene_gn=SUV39H2;refGene_tx=XM_023256391.1;refGene_gn=SUV39H2;refGene_tx=XM_023256392.1;refGene_gn=SUV39H2;refGene_tx=XM_023256393.1
B4 13955834 13955964
refGene_gn=SUV39H2;refGene_tx=XM_006933270.4;refGene_gn=SUV39H2;refGene_tx=XM_019833973.1;refGene_gn=SUV39H2;refGene_tx=XM_023256390.1;refGene_gn=SUV39H2;refGene_tx=XM_023256391.1;refGene_gn=SUV39H2;refGene_tx=XM_023256393.1
B4 13956955 13957062
refGene_gn=SUV39H2;refGene_tx=XM_006933270.4;refGene_gn=SUV39H2;refGene_tx=XM_019833973.1;refGene_gn=SUV39H2;refGene_tx=XM_023256390.1;refGene_gn=SUV39H2;refGene_tx=XM_023256391.1;refGene_gn=SUV39H2;refGene_tx=XM_023256393.1
B4 13961863 13962780
refGene_gn=LOC101089021;refGene_tx=XM_006933272.4;refGene_gn=LOC101089021;refGene_tx=XM_019833962.2;refGene_gn=LOC101089021;refGene_tx=XM_019833963.2;refGene_gn=LOC101089021;refGene_tx=XM_019833967.2;refGene_gn=LOC101089021;refGene_tx=XM_019833968.2;refGene_gn=LOC101089021;refGene_tx=XM_019833969.2;refGene_gn=LOC101089021;refGene_tx=XM_019833971.2;refGene_gn=LOC101089021;refGene_tx=XM_023256383.1;refGene_gn=LOC101089021;refGene_tx=XM_023256384.1;refGene_gn=LOC101089021;refGene_tx=XM_02

3256385.1;refGene_gn=LOC101089021;refGene_tx=XM_023256386.1;refGene_gn=LOC101089021;refGene_tx=XM_023256387.1;refGene_gn=LOC101089021;refGene_tx=XM_023256388.1
B4 13973735 13973830
refGene_gn=LOC101089021;refGene_tx=XM_006933272.4;refGene_gn=LOC101089021;refGene_tx=XM_019833962.2;refGene_gn=LOC101089021;refGene_tx=XM_019833963.2;refGene_gn=LOC101089021;refGene_tx=XM_019833967.2;refGene_gn=LOC101089021;refGene_tx=XM_019833968.2;refGene_gn=LOC101089021;refGene_tx=XM_019833969.2;refGene_gn=LOC101089021;refGene_tx=XM_019833971.2;refGene_gn=LOC101089021;refGene_tx=XM_023256383.1;refGene_gn=LOC101089021;refGene_tx=XM_023256384.1;refGene_gn=LOC101089021;refGene_tx=XM_023256385.1;refGene_gn=LOC101089021;refGene_tx=XM_023256386.1;refGene_gn=LOC101089021;refGene_tx=XM_023256387.1;refGene_gn=LOC101089021;refGene_tx=XM_023256388.1
B4 13974520 13974609
refGene_gn=LOC101089021;refGene_tx=XM_006933272.4;refGene_gn=LOC101089021;refGene_tx=XM_019833962.2;refGene_gn=LOC101089021;refGene_tx=XM_019833963.2;refGene_gn=LOC101089021;refGene_tx=XM_019833967.2;refGene_gn=LOC101089021;refGene_tx=XM_019833968.2;refGene_gn=LOC101089021;refGene_tx=XM_019833969.2;refGene_gn=LOC101089021;refGene_tx=XM_019833971.2;refGene_gn=LOC101089021;refGene_tx=XM_023256383.1;refGene_gn=LOC101089021;refGene_tx=XM_023256384.1;refGene_gn=LOC101089021;refGene_tx=XM_023256385.1;refGene_gn=LOC101089021;refGene_tx=XM_023256386.1;refGene_gn=LOC101089021;refGene_tx=XM_023256387.1;refGene_gn=LOC101089021;refGene_tx=XM_023256388.1
B4 13979112 13979167
refGene_gn=LOC101089021;refGene_tx=XM_006933272.4;refGene_gn=LOC101089021;refGene_tx=XM_019833962.2;refGene_gn=LOC101089021;refGene_tx=XM_019833963.2;refGene_gn=LOC101089021;refGene_tx=XM_019833967.2;refGene_gn=LOC101089021;refGene_tx=XM_019833968.2;refGene_gn=LOC101089021;refGene_tx=XM_019833969.2;refGene_gn=LOC101089021;refGene_tx=XM_019833971.2;refGene_gn=LOC101089021;refGene_tx=XM_023256383.1;refGene_gn=LOC101089021;refGene_tx=XM_023256384.1;refGene_gn=LOC101089021;refGene_tx=XM_023256385.1;refGene_gn=LOC101089021;refGene_tx=XM_023256386.1;refGene_gn=LOC101089021;refGene_tx=XM_023256387.1;refGene_gn=LOC101089021;refGene_tx=XM_023256388.1
B4 13979692 13979829
refGene_gn=LOC101089021;refGene_tx=XM_006933272.4;refGene_gn=LOC101089021;refGene_tx=XM_019833962.2;refGene_gn=LOC101089021;refGene_tx=XM_019833963.2;refGene_gn=LOC101089021;refGene_tx=XM_019833967.2;refGene_gn=LOC101089021;refGene_tx=XM_019833968.2;refGene_gn=LOC101089021;refGene_tx=XM_019833969.2;refGene_gn=LOC101089021;refGene_tx=XM_019833971.2;refGene_gn=LOC101089021;refGene_tx=XM_023256383.1;refGene_gn=LOC101089021;refGene_tx=XM_023256384.1;refGene_gn=LOC101089021;refGene_tx=XM_023256385.1;refGene_gn=LOC101089021;refGene_tx=XM_023256386.1;refGene_gn=LOC101089021;refGene_tx=XM_023256387.1;refGene_gn=LOC101089021;refGene_tx=XM_023256388.1
B4 13982869 13982971
refGene_gn=LOC101089021;refGene_tx=XM_006933272.4;refGene_gn=LOC101089021;refGene_tx=XM_019833962.2;refGene_gn=LOC101089021;refGene_tx=XM_01

9833963.2;refGene_gn=LOC101089021;refGene_tx=XM_019833967.2;refGene_gn=LOC101089021;refGene_tx=XM_019833968.2;refGene_gn=LOC101089021;refGene_tx=XM_019833969.2;refGene_gn=LOC101089021;refGene_tx=XM_019833971.2;refGene_gn=LOC101089021;refGene_tx=XM_023256383.1;refGene_gn=LOC101089021;refGene_tx=XM_023256384.1;refGene_gn=LOC101089021;refGene_tx=XM_023256385.1;refGene_gn=LOC101089021;refGene_tx=XM_023256386.1;refGene_gn=LOC101089021;refGene_tx=XM_023256387.1;refGene_gn=LOC101089021;refGene_tx=XM_023256388.1
B4 13986620 13986761
refGene_gn=LOC101089021;refGene_tx=XM_006933272.4;refGene_gn=LOC101089021;refGene_tx=XM_019833962.2;refGene_gn=LOC101089021;refGene_tx=XM_019833963.2;refGene_gn=LOC101089021;refGene_tx=XM_019833967.2;refGene_gn=LOC101089021;refGene_tx=XM_019833968.2;refGene_gn=LOC101089021;refGene_tx=XM_019833971.2;refGene_gn=LOC101089021;refGene_tx=XM_023256383.1;refGene_gn=LOC101089021;refGene_tx=XM_023256384.1;refGene_gn=LOC101089021;refGene_tx=XM_023256385.1;refGene_gn=LOC101089021;refGene_tx=XM_023256386.1;refGene_gn=LOC101089021;refGene_tx=XM_023256387.1;refGene_gn=LOC101089021;refGene_tx=XM_023256388.1
B4 13986943 13987016
refGene_gn=LOC101089021;refGene_tx=XM_006933272.4;refGene_gn=LOC101089021;refGene_tx=XM_019833962.2;refGene_gn=LOC101089021;refGene_tx=XM_019833963.2;refGene_gn=LOC101089021;refGene_tx=XM_019833967.2;refGene_gn=LOC101089021;refGene_tx=XM_019833968.2;refGene_gn=LOC101089021;refGene_tx=XM_019833971.2;refGene_gn=LOC101089021;refGene_tx=XM_023256383.1;refGene_gn=LOC101089021;refGene_tx=XM_023256384.1;refGene_gn=LOC101089021;refGene_tx=XM_023256385.1;refGene_gn=LOC101089021;refGene_tx=XM_023256386.1;refGene_gn=LOC101089021;refGene_tx=XM_023256387.1;refGene_gn=LOC101089021;refGene_tx=XM_023256388.1
B4 13987413 13987532
refGene_gn=LOC101089021;refGene_tx=XM_006933272.4;refGene_gn=LOC101089021;refGene_tx=XM_019833962.2;refGene_gn=LOC101089021;refGene_tx=XM_019833963.2;refGene_gn=LOC101089021;refGene_tx=XM_019833967.2;refGene_gn=LOC101089021;refGene_tx=XM_019833968.2;refGene_gn=LOC101089021;refGene_tx=XM_019833971.2;refGene_gn=LOC101089021;refGene_tx=XM_023256383.1;refGene_gn=LOC101089021;refGene_tx=XM_023256384.1;refGene_gn=LOC101089021;refGene_tx=XM_023256385.1;refGene_gn=LOC101089021;refGene_tx=XM_023256386.1;refGene_gn=LOC101089021;refGene_tx=XM_023256387.1;refGene_gn=LOC101089021;refGene_tx=XM_023256388.1
B4 13988062 13988118
refGene_gn=LOC101089021;refGene_tx=XM_019833962.2;refGene_gn=LOC101089021;refGene_tx=XM_019833963.2;refGene_gn=LOC101089021;refGene_tx=XM_019833971.2;refGene_gn=LOC101089021;refGene_tx=XM_023256383.1;refGene_gn=LOC101089021;refGene_tx=XM_023256384.1;refGene_gn=LOC101089021;refGene_tx=XM_023256388.1
B4 13990899 13990959
refGene_gn=LOC101089021;refGene_tx=XM_006933272.4;refGene_gn=LOC101089021;refGene_tx=XM_019833962.2;refGene_gn=LOC101089021;refGene_tx=XM_019833963.2;refGene_gn=LOC101089021;refGene_tx=XM_019833967.2;refGene_gn=LOC101089021;refGene_tx=XM_019833968.2;refGene_gn=LOC101089021;refGene_tx=XM_019833969.2;refGene_gn=LOC101089021;refGene_tx=XM_023256383.1;

refGene_gn=LOC101089021;refGene_tx=XM_023256384.1;refGene_gn=LOC101089021;refGene_tx=XM_023256385.1;refGene_gn=LOC101089021;refGene_tx=XM_023256387.1
B4 13995019 13995026
refGene_gn=LOC101089021;refGene_tx=XM_023256384.1
B4 14000774 14000826
refGene_gn=LOC101089021;refGene_tx=XM_019833962.2;refGene_gn=LOC101089021;refGene_tx=XM_019833963.2;refGene_gn=LOC101089021;refGene_tx=XM_019833969.2;refGene_gn=LOC101089021;refGene_tx=XM_023256383.1;refGene_gn=LOC101089021;refGene_tx=XM_023256384.1
B4 14005236 14005448
refGene_gn=LOC101089021;refGene_tx=XM_019833962.2;refGene_gn=LOC101089021;refGene_tx=XM_019833963.2;refGene_gn=LOC101089021;refGene_tx=XM_019833969.2;refGene_gn=LOC101089021;refGene_tx=XM_023256383.1;refGene_gn=LOC101089021;refGene_tx=XM_023256384.1
B4 14016725 14016863
refGene_gn=MEIG1;refGene_tx=XM_006933273.4;refGene_gn=MEIG1;refGene_tx=XM_011283613.3;refGene_gn=MEIG1;refGene_tx=XM_019833975.2
B4 14020039 14020168
refGene_gn=MEIG1;refGene_tx=XM_006933273.4;refGene_gn=MEIG1;refGene_tx=XM_011283613.3;refGene_gn=MEIG1;refGene_tx=XM_019833975.2
B4 14061631 14061705
refGene_gn=ACBD7;refGene_tx=XM_003988123.4;refGene_gn=ACBD7;refGene_tx=XM_019833976.2;refGene_gn=ACBD7;refGene_tx=XM_019833977.2
B4 14061785 14061848
refGene_gn=ACBD7;refGene_tx=XM_003988123.4;refGene_gn=ACBD7;refGene_tx=XM_019833976.2;refGene_gn=ACBD7;refGene_tx=XM_019833977.2
B4 14061994 14062112
refGene_gn=ACBD7;refGene_tx=XM_003988123.4;refGene_gn=ACBD7;refGene_tx=XM_019833976.2;refGene_gn=ACBD7;refGene_tx=XM_019833977.2
B4 14070518 14070530 refGene_gn=ACBD7;refGene_tx=XM_003988123.4
B4 14085625 14086050
refGene_gn=CB4H10orf111;refGene_tx=XM_019835798.2
B4 14086354 14086418
refGene_gn=CB4H10orf111;refGene_tx=XM_019835798.2
B4 14090098 14090944
refGene_gn=RPP38;refGene_tx=XM_003988161.5;refGene_gn=RPP38;refGene_tx=XM_006933274.4;refGene_gn=RPP38;refGene_tx=XM_006933275.4;refGene_gn=RPP38;refGene_tx=XM_006933276.4;refGene_gn=RPP38;refGene_tx=XM_019833979.2;refGene_gn=RPP38;refGene_tx=XM_019833980.2;refGene_gn=RPP38;refGene_tx=XM_019833982.2
B4 14094526 14094547
refGene_gn=NMT2;refGene_tx=XM_023256394.1;refGene_gn=NMT2;refGene_tx=XM_023256395.1
B4 14094718 14094856
refGene_gn=NMT2;refGene_tx=XM_023256394.1;refGene_gn=NMT2;refGene_tx=XM_023256395.1
B4 14096800 14096968
refGene_gn=NMT2;refGene_tx=XM_023256394.1;refGene_gn=NMT2;refGene_tx=XM_023256395.1;refGene_gn=NMT2;refGene_tx=XM_023256396.1

B4 14103543 14103714
refGene_gn=NMT2;refGene_tx=XM_023256394.1;refGene_gn=NMT2;refGene_tx=XM_023256395.1;refGene_gn=NMT2;refGene_tx=XM_023256396.1
B4 14112656 14112765
refGene_gn=NMT2;refGene_tx=XM_023256394.1;refGene_gn=NMT2;refGene_tx=XM_023256395.1;refGene_gn=NMT2;refGene_tx=XM_023256396.1
B4 14113933 14114104
refGene_gn=NMT2;refGene_tx=XM_023256394.1;refGene_gn=NMT2;refGene_tx=XM_023256395.1;refGene_gn=NMT2;refGene_tx=XM_023256396.1
B4 14115387 14115504
refGene_gn=NMT2;refGene_tx=XM_023256394.1;refGene_gn=NMT2;refGene_tx=XM_023256395.1;refGene_gn=NMT2;refGene_tx=XM_023256396.1
B4 14115629 14115721
refGene_gn=NMT2;refGene_tx=XM_023256394.1;refGene_gn=NMT2;refGene_tx=XM_023256395.1;refGene_gn=NMT2;refGene_tx=XM_023256396.1
B4 14115817 14115936
refGene_gn=NMT2;refGene_tx=XM_023256394.1;refGene_gn=NMT2;refGene_tx=XM_023256395.1;refGene_gn=NMT2;refGene_tx=XM_023256396.1
B4 14119058 14119203
refGene_gn=NMT2;refGene_tx=XM_023256394.1;refGene_gn=NMT2;refGene_tx=XM_023256395.1;refGene_gn=NMT2;refGene_tx=XM_023256396.1
B4 14123146 14123189 refGene_gn=NMT2;refGene_tx=XM_023256395.1
B4 14123579 14123715
refGene_gn=NMT2;refGene_tx=XM_023256394.1;refGene_gn=NMT2;refGene_tx=XM_023256395.1;refGene_gn=NMT2;refGene_tx=XM_023256396.1
B4 14168924 14169034
refGene_gn=NMT2;refGene_tx=XM_023256394.1;refGene_gn=NMT2;refGene_tx=XM_023256395.1;refGene_gn=NMT2;refGene_tx=XM_023256396.1
B4 14211250 14212952
refGene_gn=FAM171A1;refGene_tx=XM_023256397.1;refGene_gn=FAM171A1;refGene_tx=XM_023256398.1
B4 14213569 14213684
refGene_gn=FAM171A1;refGene_tx=XM_023256397.1;refGene_gn=FAM171A1;refGene_tx=XM_023256398.1
B4 14216896 14217013
refGene_gn=FAM171A1;refGene_tx=XM_023256397.1;refGene_gn=FAM171A1;refGene_tx=XM_023256398.1
B4 14242995 14243172 refGene_gn=FAM171A1;refGene_tx=XM_023256397.1
B4 14247680 14247839
refGene_gn=FAM171A1;refGene_tx=XM_023256397.1;refGene_gn=FAM171A1;refGene_tx=XM_023256398.1
B4 14263565 14263658
refGene_gn=FAM171A1;refGene_tx=XM_023256397.1;refGene_gn=FAM171A1;refGene_tx=XM_023256398.1
B4 14272967 14273195
refGene_gn=FAM171A1;refGene_tx=XM_023256397.1;refGene_gn=FAM171A1;refGene_tx=XM_023256398.1
B4 14344396 14344493
refGene_gn=FAM171A1;refGene_tx=XM_023256397.1;refGene_gn=FAM171A1;refGene_tx=XM_023256398.1

B4 14496320 14496407 refGene_gn=ITGA8;refGene_tx=XM_011283614.3
B4 14498095 14498218 refGene_gn=ITGA8;refGene_tx=XM_011283614.3
B4 14507545 14507647 refGene_gn=ITGA8;refGene_tx=XM_011283614.3
B4 14514272 14514386 refGene_gn=ITGA8;refGene_tx=XM_011283614.3
B4 14519136 14519265 refGene_gn=ITGA8;refGene_tx=XM_011283614.3
B4 14528666 14528825 refGene_gn=ITGA8;refGene_tx=XM_011283614.3
B4 14531551 14531657 refGene_gn=ITGA8;refGene_tx=XM_011283614.3
B4 14541845 14541926 refGene_gn=ITGA8;refGene_tx=XM_011283614.3
B4 14547443 14547523 refGene_gn=ITGA8;refGene_tx=XM_011283614.3
B4 14551980 14552073 refGene_gn=ITGA8;refGene_tx=XM_011283614.3
B4 14558355 14558503 refGene_gn=ITGA8;refGene_tx=XM_011283614.3
B4 14559874 14559942 refGene_gn=ITGA8;refGene_tx=XM_011283614.3
B4 14560433 14560571 refGene_gn=ITGA8;refGene_tx=XM_011283614.3
B4 14561677 14561832 refGene_gn=ITGA8;refGene_tx=XM_011283614.3
B4 14562247 14562303 refGene_gn=ITGA8;refGene_tx=XM_011283614.3
B4 14569196 14569304 refGene_gn=ITGA8;refGene_tx=XM_011283614.3
B4 14570945 14570991 refGene_gn=ITGA8;refGene_tx=XM_011283614.3
B4 14604264 14604456 refGene_gn=ITGA8;refGene_tx=XM_011283614.3
B4 14607453 14607659 refGene_gn=ITGA8;refGene_tx=XM_011283614.3
B4 14621940 14621993 refGene_gn=ITGA8;refGene_tx=XM_011283614.3
B4 14624699 14624756 refGene_gn=ITGA8;refGene_tx=XM_011283614.3
B4 14626441 14626485 refGene_gn=ITGA8;refGene_tx=XM_011283614.3
B4 14633274 14633319 refGene_gn=ITGA8;refGene_tx=XM_011283614.3
B4 14634432 14634558 refGene_gn=ITGA8;refGene_tx=XM_011283614.3
B4 14645124 14645170 refGene_gn=ITGA8;refGene_tx=XM_011283614.3
B4 14646284 14646346 refGene_gn=ITGA8;refGene_tx=XM_011283614.3
B4 14652980 14653104 refGene_gn=ITGA8;refGene_tx=XM_011283614.3
B4 14656214 14656315 refGene_gn=ITGA8;refGene_tx=XM_011283614.3
B4 14682575 14682709 refGene_gn=ITGA8;refGene_tx=XM_011283614.3
B4 14683429 14683635 refGene_gn=ITGA8;refGene_tx=XM_011283614.3
B4 14750653 14750803
refGene_gn=MINDY3;refGene_tx=XM_003988125.5;refGene_gn=MINDY3;refGene_
tx=XM_011283617.3;refGene_gn=MINDY3;refGene_tx=XM_019833986.2;refGene_
gn=MINDY3;refGene_tx=XM_019833988.1;refGene_gn=MINDY3;refGene_tx=XM_01
9833989.2
B4 14753205 14753277
refGene_gn=MINDY3;refGene_tx=XM_003988125.5;refGene_gn=MINDY3;refGene_
tx=XM_011283617.3;refGene_gn=MINDY3;refGene_tx=XM_019833986.2;refGene_
gn=MINDY3;refGene_tx=XM_019833988.1;refGene_gn=MINDY3;refGene_tx=XM_01
9833989.2
B4 14755302 14755390
refGene_gn=MINDY3;refGene_tx=XM_003988125.5;refGene_gn=MINDY3;refGene_
tx=XM_011283617.3;refGene_gn=MINDY3;refGene_tx=XM_019833986.2;refGene_
gn=MINDY3;refGene_tx=XM_019833988.1;refGene_gn=MINDY3;refGene_tx=XM_01
9833989.2
B4 14756732 14756805
refGene_gn=MINDY3;refGene_tx=XM_003988125.5;refGene_gn=MINDY3;refGene_
tx=XM_011283617.3;refGene_gn=MINDY3;refGene_tx=XM_019833986.2;refGene_
gn=MINDY3;refGene_tx=XM_019833988.1;refGene_gn=MINDY3;refGene_tx=XM_01
9833989.2

B4 14771999 14772172 refGene_gn=MINDY3;refGene_tx=XM_019833987.2
B4 14780316 14780389
refGene_gn=MINDY3;refGene_tx=XM_003988125.5;refGene_gn=MINDY3;refGene_tx=XM_011283617.3;refGene_gn=MINDY3;refGene_tx=XM_019833986.2;refGene_gn=MINDY3;refGene_tx=XM_019833987.2;refGene_gn=MINDY3;refGene_tx=XM_019833988.1;refGene_gn=MINDY3;refGene_tx=XM_019833989.2
B4 14800118 14800199
refGene_gn=MINDY3;refGene_tx=XM_003988125.5;refGene_gn=MINDY3;refGene_tx=XM_019833986.2;refGene_gn=MINDY3;refGene_tx=XM_019833987.2;refGene_gn=MINDY3;refGene_tx=XM_019833988.1;refGene_gn=MINDY3;refGene_tx=XM_019833989.2
B4 14805058 14805129
refGene_gn=MINDY3;refGene_tx=XM_003988125.5;refGene_gn=MINDY3;refGene_tx=XM_011283617.3;refGene_gn=MINDY3;refGene_tx=XM_019833986.2;refGene_gn=MINDY3;refGene_tx=XM_019833987.2;refGene_gn=MINDY3;refGene_tx=XM_019833988.1;refGene_gn=MINDY3;refGene_tx=XM_019833989.2
B4 14818085 14818165
refGene_gn=MINDY3;refGene_tx=XM_003988125.5;refGene_gn=MINDY3;refGene_tx=XM_011283617.3;refGene_gn=MINDY3;refGene_tx=XM_019833986.2;refGene_gn=MINDY3;refGene_tx=XM_019833987.2;refGene_gn=MINDY3;refGene_tx=XM_019833988.1;refGene_gn=MINDY3;refGene_tx=XM_019833989.2
B4 14819040 14819114
refGene_gn=MINDY3;refGene_tx=XM_003988125.5;refGene_gn=MINDY3;refGene_tx=XM_011283617.3;refGene_gn=MINDY3;refGene_tx=XM_019833986.2;refGene_gn=MINDY3;refGene_tx=XM_019833987.2;refGene_gn=MINDY3;refGene_tx=XM_019833988.1;refGene_gn=MINDY3;refGene_tx=XM_019833989.2
B4 14821732 14821847
refGene_gn=MINDY3;refGene_tx=XM_003988125.5;refGene_gn=MINDY3;refGene_tx=XM_011283617.3;refGene_gn=MINDY3;refGene_tx=XM_019833986.2;refGene_gn=MINDY3;refGene_tx=XM_019833987.2;refGene_gn=MINDY3;refGene_tx=XM_019833988.1;refGene_gn=MINDY3;refGene_tx=XM_019833989.2
B4 14822724 14822776
refGene_gn=MINDY3;refGene_tx=XM_003988125.5;refGene_gn=MINDY3;refGene_tx=XM_011283617.3;refGene_gn=MINDY3;refGene_tx=XM_019833986.2;refGene_gn=MINDY3;refGene_tx=XM_019833987.2
B4 14826059 14826236
refGene_gn=MINDY3;refGene_tx=XM_003988125.5;refGene_gn=MINDY3;refGene_tx=XM_011283617.3;refGene_gn=MINDY3;refGene_tx=XM_019833986.2;refGene_gn=MINDY3;refGene_tx=XM_019833987.2
B4 14827930 14827991
refGene_gn=MINDY3;refGene_tx=XM_003988125.5;refGene_gn=MINDY3;refGene_tx=XM_011283617.3;refGene_gn=MINDY3;refGene_tx=XM_019833987.2
B4 14831633 14831713
refGene_gn=MINDY3;refGene_tx=XM_003988125.5;refGene_gn=MINDY3;refGene_tx=XM_011283617.3;refGene_gn=MINDY3;refGene_tx=XM_019833987.2
B4 14843626 14843720
refGene_gn=MINDY3;refGene_tx=XM_003988125.5;refGene_gn=MINDY3;refGene_tx=XM_011283617.3;refGene_gn=MINDY3;refGene_tx=XM_019833986.2;refGene_gn=MINDY3;refGene_tx=XM_019833987.2
B4 15473141 15473573

refGene_gn=PTER;refGene_tx=XM_006933278.3;refGene_gn=PTER;refGene_tx=XM_019833990.1;refGene_gn=PTER;refGene_tx=XM_019833991.1
B4 15474914 15475180
refGene_gn=PTER;refGene_tx=XM_006933278.3;refGene_gn=PTER;refGene_tx=XM_019833990.1;refGene_gn=PTER;refGene_tx=XM_019833991.1
B4 15500558 15500699
refGene_gn=PTER;refGene_tx=XM_006933278.3;refGene_gn=PTER;refGene_tx=XM_019833990.1;refGene_gn=PTER;refGene_tx=XM_019833991.1
B4 15506350 15506561
refGene_gn=PTER;refGene_tx=XM_006933278.3;refGene_gn=PTER;refGene_tx=XM_019833990.1;refGene_gn=PTER;refGene_tx=XM_019833991.1
B4 15509857 15510037 refGene_gn=C1QL3;refGene_tx=XM_011283618.3
B4 15515846 15516434 refGene_gn=C1QL3;refGene_tx=XM_011283618.3
B4 15575129 15575232
refGene_gn=RSU1;refGene_tx=XM_003988128.3;refGene_gn=RSU1;refGene_tx=XM_019833992.2;refGene_gn=RSU1;refGene_tx=XM_019833993.1
B4 15663517 15663650
refGene_gn=RSU1;refGene_tx=XM_003988128.3;refGene_gn=RSU1;refGene_tx=XM_019833992.2
B4 15709758 15709873
refGene_gn=RSU1;refGene_tx=XM_003988128.3;refGene_gn=RSU1;refGene_tx=XM_019833992.2;refGene_gn=RSU1;refGene_tx=XM_019833993.1
B4 15710144 15710227
refGene_gn=RSU1;refGene_tx=XM_003988128.3;refGene_gn=RSU1;refGene_tx=XM_019833992.2;refGene_gn=RSU1;refGene_tx=XM_019833993.1
B4 15711833 15711952
refGene_gn=RSU1;refGene_tx=XM_003988128.3;refGene_gn=RSU1;refGene_tx=XM_019833992.2;refGene_gn=RSU1;refGene_tx=XM_019833993.1
B4 15719091 15719212
refGene_gn=RSU1;refGene_tx=XM_003988128.3;refGene_gn=RSU1;refGene_tx=XM_019833992.2;refGene_gn=RSU1;refGene_tx=XM_019833993.1
B4 15737800 15737851
refGene_gn=RSU1;refGene_tx=XM_003988128.3;refGene_gn=RSU1;refGene_tx=XM_019833993.1
B4 15776628 15776737
refGene_gn=RSU1;refGene_tx=XM_003988128.3;refGene_gn=RSU1;refGene_tx=XM_019833992.2;refGene_gn=RSU1;refGene_tx=XM_019833993.1
B4 15784335 15784443
refGene_gn=CUBN;refGene_tx=XM_003988129.4;refGene_gn=CUBN;refGene_tx=XM_023256399.1
B4 15787321 15787557
refGene_gn=CUBN;refGene_tx=XM_003988129.4;refGene_gn=CUBN;refGene_tx=XM_023256399.1
B4 15789281 15789447
refGene_gn=CUBN;refGene_tx=XM_003988129.4;refGene_gn=CUBN;refGene_tx=XM_023256399.1
B4 15792654 15792836
refGene_gn=CUBN;refGene_tx=XM_003988129.4;refGene_gn=CUBN;refGene_tx=XM_023256399.1
B4 15793919 15794067

refGene_gn=CUBN;refGene_tx=XM_003988129.4;refGene_gn=CUBN;refGene_tx=XM_023256399.1
B4 15797017 15797223
refGene_gn=CUBN;refGene_tx=XM_003988129.4;refGene_gn=CUBN;refGene_tx=XM_023256399.1
B4 15797584 15797747
refGene_gn=CUBN;refGene_tx=XM_003988129.4;refGene_gn=CUBN;refGene_tx=XM_023256399.1
B4 15807432 15807641
refGene_gn=CUBN;refGene_tx=XM_003988129.4;refGene_gn=CUBN;refGene_tx=XM_023256399.1
B4 15825014 15825232
refGene_gn=CUBN;refGene_tx=XM_003988129.4;refGene_gn=CUBN;refGene_tx=XM_023256399.1
B4 15828053 15828183
refGene_gn=CUBN;refGene_tx=XM_003988129.4;refGene_gn=CUBN;refGene_tx=XM_023256399.1
B4 15830935 15831136
refGene_gn=CUBN;refGene_tx=XM_003988129.4;refGene_gn=CUBN;refGene_tx=XM_023256399.1
B4 15841869 15842019
refGene_gn=CUBN;refGene_tx=XM_003988129.4;refGene_gn=CUBN;refGene_tx=XM_023256399.1
B4 15844561 15844718
refGene_gn=CUBN;refGene_tx=XM_003988129.4;refGene_gn=CUBN;refGene_tx=XM_023256399.1
B4 15851509 15851697
refGene_gn=CUBN;refGene_tx=XM_003988129.4;refGene_gn=CUBN;refGene_tx=XM_023256399.1
B4 15853421 15853647
refGene_gn=CUBN;refGene_tx=XM_003988129.4;refGene_gn=CUBN;refGene_tx=XM_023256399.1
B4 15854377 15854499
refGene_gn=CUBN;refGene_tx=XM_003988129.4;refGene_gn=CUBN;refGene_tx=XM_023256399.1
B4 15856687 15856837
refGene_gn=CUBN;refGene_tx=XM_003988129.4;refGene_gn=CUBN;refGene_tx=XM_023256399.1
B4 15859345 15859552
refGene_gn=CUBN;refGene_tx=XM_003988129.4;refGene_gn=CUBN;refGene_tx=XM_023256399.1
B4 15860638 15860810
refGene_gn=CUBN;refGene_tx=XM_003988129.4;refGene_gn=CUBN;refGene_tx=XM_023256399.1
B4 15867065 15867247
refGene_gn=CUBN;refGene_tx=XM_003988129.4;refGene_gn=CUBN;refGene_tx=XM_023256399.1
B4 15867762 15867903
refGene_gn=CUBN;refGene_tx=XM_003988129.4;refGene_gn=CUBN;refGene_tx=XM_023256399.1

B4 15868542 15868752
refGene_gn=CUBN;refGene_tx=XM_003988129.4;refGene_gn=CUBN;refGene_tx=XM_023256399.1
B4 15870216 15870395
refGene_gn=CUBN;refGene_tx=XM_003988129.4;refGene_gn=CUBN;refGene_tx=XM_023256399.1
B4 15871572 15871747
refGene_gn=CUBN;refGene_tx=XM_003988129.4;refGene_gn=CUBN;refGene_tx=XM_023256399.1
B4 15875539 15875723
refGene_gn=CUBN;refGene_tx=XM_003988129.4;refGene_gn=CUBN;refGene_tx=XM_023256399.1
B4 15875880 15876071
refGene_gn=CUBN;refGene_tx=XM_003988129.4;refGene_gn=CUBN;refGene_tx=XM_023256399.1
B4 15878271 15878418
refGene_gn=CUBN;refGene_tx=XM_003988129.4;refGene_gn=CUBN;refGene_tx=XM_023256399.1
B4 15882976 15883174
refGene_gn=CUBN;refGene_tx=XM_003988129.4;refGene_gn=CUBN;refGene_tx=XM_023256399.1
B4 15890362 15890555
refGene_gn=CUBN;refGene_tx=XM_003988129.4;refGene_gn=CUBN;refGene_tx=XM_023256399.1
B4 15892450 15892635
refGene_gn=CUBN;refGene_tx=XM_003988129.4;refGene_gn=CUBN;refGene_tx=XM_023256399.1
B4 15893538 15893744
refGene_gn=CUBN;refGene_tx=XM_003988129.4;refGene_gn=CUBN;refGene_tx=XM_023256399.1
B4 15902566 15902699
refGene_gn=CUBN;refGene_tx=XM_003988129.4;refGene_gn=CUBN;refGene_tx=XM_023256399.1
B4 15903693 15903822
refGene_gn=CUBN;refGene_tx=XM_003988129.4;refGene_gn=CUBN;refGene_tx=XM_023256399.1
B4 15905383 15905494
refGene_gn=CUBN;refGene_tx=XM_003988129.4;refGene_gn=CUBN;refGene_tx=XM_023256399.1
B4 15907163 15907277
refGene_gn=CUBN;refGene_tx=XM_003988129.4;refGene_gn=CUBN;refGene_tx=XM_023256399.1
B4 15909005 15909165
refGene_gn=CUBN;refGene_tx=XM_003988129.4;refGene_gn=CUBN;refGene_tx=XM_023256399.1
B4 15929052 15929222
refGene_gn=CUBN;refGene_tx=XM_003988129.4;refGene_gn=CUBN;refGene_tx=XM_023256399.1
B4 15930661 15930836
refGene_gn=CUBN;refGene_tx=XM_003988129.4;refGene_gn=CUBN;refGene_tx=X

M_023256399.1
B4 15935668 15935850
refGene_gn=CUBN;refGene_tx=XM_003988129.4;refGene_gn=CUBN;refGene_tx=XM_023256399.1
B4 15958526 15958677
refGene_gn=CUBN;refGene_tx=XM_003988129.4;refGene_gn=CUBN;refGene_tx=XM_023256399.1
B4 15977628 15977816
refGene_gn=CUBN;refGene_tx=XM_003988129.4;refGene_gn=CUBN;refGene_tx=XM_023256399.1
B4 15980566 15980723
refGene_gn=CUBN;refGene_tx=XM_003988129.4;refGene_gn=CUBN;refGene_tx=XM_023256399.1
B4 15981709 15981891
refGene_gn=CUBN;refGene_tx=XM_003988129.4;refGene_gn=CUBN;refGene_tx=XM_023256399.1
B4 15982301 15982462
refGene_gn=CUBN;refGene_tx=XM_003988129.4;refGene_gn=CUBN;refGene_tx=XM_023256399.1
B4 15984032 15984225
refGene_gn=CUBN;refGene_tx=XM_003988129.4;refGene_gn=CUBN;refGene_tx=XM_023256399.1
B4 16000029 16000160
refGene_gn=CUBN;refGene_tx=XM_003988129.4;refGene_gn=CUBN;refGene_tx=XM_023256399.1
B4 16003327 16003544
refGene_gn=CUBN;refGene_tx=XM_003988129.4;refGene_gn=CUBN;refGene_tx=XM_023256399.1
B4 16003892 16004058
refGene_gn=CUBN;refGene_tx=XM_003988129.4;refGene_gn=CUBN;refGene_tx=XM_023256399.1
B4 16006421 16006600
refGene_gn=CUBN;refGene_tx=XM_003988129.4;refGene_gn=CUBN;refGene_tx=XM_023256399.1
B4 16006816 16006961
refGene_gn=CUBN;refGene_tx=XM_003988129.4;refGene_gn=CUBN;refGene_tx=XM_023256399.1
B4 16026313 16026501
refGene_gn=CUBN;refGene_tx=XM_003988129.4;refGene_gn=CUBN;refGene_tx=XM_023256399.1
B4 16028162 16028325
refGene_gn=CUBN;refGene_tx=XM_003988129.4;refGene_gn=CUBN;refGene_tx=XM_023256399.1
B4 16030124 16030306
refGene_gn=CUBN;refGene_tx=XM_003988129.4;refGene_gn=CUBN;refGene_tx=XM_023256399.1
B4 16037022 16037257
refGene_gn=CUBN;refGene_tx=XM_003988129.4;refGene_gn=CUBN;refGene_tx=XM_023256399.1
B4 16039820 16039933

refGene_gn=CUBN;refGene_tx=XM_003988129.4;refGene_gn=CUBN;refGene_tx=XM_023256399.1
B4 16041088 16041275
refGene_gn=CUBN;refGene_tx=XM_003988129.4;refGene_gn=CUBN;refGene_tx=XM_023256399.1
B4 16041458 16041577
refGene_gn=CUBN;refGene_tx=XM_003988129.4;refGene_gn=CUBN;refGene_tx=XM_023256399.1
B4 16043589 16043685
refGene_gn=CUBN;refGene_tx=XM_003988129.4;refGene_gn=CUBN;refGene_tx=XM_023256399.1
B4 16044550 16044682
refGene_gn=CUBN;refGene_tx=XM_003988129.4;refGene_gn=CUBN;refGene_tx=XM_023256399.1
B4 16047471 16047634
refGene_gn=CUBN;refGene_tx=XM_003988129.4;refGene_gn=CUBN;refGene_tx=XM_023256399.1
B4 16048627 16048754
refGene_gn=CUBN;refGene_tx=XM_003988129.4;refGene_gn=CUBN;refGene_tx=XM_023256399.1
B4 16054458 16054562
refGene_gn=CUBN;refGene_tx=XM_003988129.4;refGene_gn=CUBN;refGene_tx=XM_023256399.1
B4 16055753 16055855
refGene_gn=CUBN;refGene_tx=XM_003988129.4;refGene_gn=CUBN;refGene_tx=XM_023256399.1
B4 16058378 16058417
refGene_gn=CUBN;refGene_tx=XM_003988129.4;refGene_gn=CUBN;refGene_tx=XM_023256399.1
B4 16058976 16059072
refGene_gn=CUBN;refGene_tx=XM_003988129.4;refGene_gn=CUBN;refGene_tx=XM_023256399.1
B4 16060218 16060348
refGene_gn=CUBN;refGene_tx=XM_003988129.4;refGene_gn=CUBN;refGene_tx=XM_023256399.1
B4 16060735 16060848 refGene_gn=CUBN;refGene_tx=XM_003988129.4
B4 16075960 16076073
refGene_gn=TRDMT1;refGene_tx=XM_006933280.4;refGene_gn=TRDMT1;refGene_tx=XM_011283620.3;refGene_gn=TRDMT1;refGene_tx=XM_011283621.3;refGene_gn=TRDMT1;refGene_tx=XM_019833998.2;refGene_gn=TRDMT1;refGene_tx=XM_019833999.2;refGene_gn=TRDMT1;refGene_tx=XM_023256400.1
B4 16078847 16078948
refGene_gn=TRDMT1;refGene_tx=XM_003988130.5;refGene_gn=TRDMT1;refGene_tx=XM_023256401.1
B4 16084034 16084054 refGene_gn=TRDMT1;refGene_tx=XM_023256402.1
B4 16084820 16084950
refGene_gn=TRDMT1;refGene_tx=XM_003988130.5;refGene_gn=TRDMT1;refGene_tx=XM_006933280.4;refGene_gn=TRDMT1;refGene_tx=XM_011283620.3;refGene_gn=TRDMT1;refGene_tx=XM_011283621.3;refGene_gn=TRDMT1;refGene_tx=XM_019833998.2;refGene_gn=TRDMT1;refGene_tx=XM_019833999.2;refGene_gn=TRDMT

1;refGene_tx=XM_023256400.1;refGene_gn=TRDMT1;refGene_tx=XM_023256401.1;refGene_gn=TRDMT1;refGene_tx=XM_023256402.1
B4 16086739 16086797
refGene_gn=TRDMT1;refGene_tx=XM_003988130.5;refGene_gn=TRDMT1;refGene_tx=XM_006933280.4;refGene_gn=TRDMT1;refGene_tx=XM_011283620.3;refGene_gn=TRDMT1;refGene_tx=XM_011283621.3;refGene_gn=TRDMT1;refGene_tx=XM_019833998.2;refGene_gn=TRDMT1;refGene_tx=XM_019833999.2;refGene_gn=TRDMT1;refGene_tx=XM_023256400.1;refGene_gn=TRDMT1;refGene_tx=XM_023256401.1;refGene_gn=TRDMT1;refGene_tx=XM_023256402.1
B4 16088832 16089176
refGene_gn=TRDMT1;refGene_tx=XM_003988130.5;refGene_gn=TRDMT1;refGene_tx=XM_006933280.4;refGene_gn=TRDMT1;refGene_tx=XM_011283620.3;refGene_gn=TRDMT1;refGene_tx=XM_011283621.3;refGene_gn=TRDMT1;refGene_tx=XM_019833998.2;refGene_gn=TRDMT1;refGene_tx=XM_019833999.2;refGene_gn=TRDMT1;refGene_tx=XM_023256400.1;refGene_gn=TRDMT1;refGene_tx=XM_023256401.1;refGene_gn=TRDMT1;refGene_tx=XM_023256402.1
B4 16090572 16090656
refGene_gn=TRDMT1;refGene_tx=XM_003988130.5;refGene_gn=TRDMT1;refGene_tx=XM_006933280.4;refGene_gn=TRDMT1;refGene_tx=XM_011283620.3;refGene_gn=TRDMT1;refGene_tx=XM_011283621.3;refGene_gn=TRDMT1;refGene_tx=XM_019833998.2;refGene_gn=TRDMT1;refGene_tx=XM_019833999.2;refGene_gn=TRDMT1;refGene_tx=XM_023256400.1;refGene_gn=TRDMT1;refGene_tx=XM_023256401.1;refGene_gn=TRDMT1;refGene_tx=XM_023256402.1
B4 16092070 16092140
refGene_gn=TRDMT1;refGene_tx=XM_003988130.5;refGene_gn=TRDMT1;refGene_tx=XM_006933280.4;refGene_gn=TRDMT1;refGene_tx=XM_011283620.3;refGene_gn=TRDMT1;refGene_tx=XM_011283621.3;refGene_gn=TRDMT1;refGene_tx=XM_019833998.2;refGene_gn=TRDMT1;refGene_tx=XM_019833999.2;refGene_gn=TRDMT1;refGene_tx=XM_023256400.1;refGene_gn=TRDMT1;refGene_tx=XM_023256401.1;refGene_gn=TRDMT1;refGene_tx=XM_023256402.1
B4 16092914 16092980
refGene_gn=TRDMT1;refGene_tx=XM_003988130.5;refGene_gn=TRDMT1;refGene_tx=XM_006933280.4;refGene_gn=TRDMT1;refGene_tx=XM_011283620.3;refGene_gn=TRDMT1;refGene_tx=XM_011283621.3;refGene_gn=TRDMT1;refGene_tx=XM_019833998.2;refGene_gn=TRDMT1;refGene_tx=XM_019833999.2;refGene_gn=TRDMT1;refGene_tx=XM_023256402.1
B4 16093626 16093698
refGene_gn=TRDMT1;refGene_tx=XM_003988130.5;refGene_gn=TRDMT1;refGene_tx=XM_006933280.4;refGene_gn=TRDMT1;refGene_tx=XM_011283620.3;refGene_gn=TRDMT1;refGene_tx=XM_011283621.3;refGene_gn=TRDMT1;refGene_tx=XM_019833998.2;refGene_gn=TRDMT1;refGene_tx=XM_019833999.2;refGene_gn=TRDMT1;refGene_tx=XM_023256400.1;refGene_gn=TRDMT1;refGene_tx=XM_023256401.1;refGene_gn=TRDMT1;refGene_tx=XM_023256402.1
B4 16096262 16096339
refGene_gn=TRDMT1;refGene_tx=XM_003988130.5;refGene_gn=TRDMT1;refGene_tx=XM_006933280.4;refGene_gn=TRDMT1;refGene_tx=XM_023256400.1;refGene_gn=TRDMT1;refGene_tx=XM_023256401.1;refGene_gn=TRDMT1;refGene_tx=XM_023256402.1
B4 16112946 16113056
refGene_gn=TRDMT1;refGene_tx=XM_003988130.5;refGene_gn=TRDMT1;refGene_

tx=XM_006933280.4;refGene_gn=TRDMT1;refGene_tx=XM_011283620.3;refGene_gn=TRDMT1;refGene_tx=XM_011283621.3;refGene_gn=TRDMT1;refGene_tx=XM_019833998.2;refGene_gn=TRDMT1;refGene_tx=XM_019833999.2;refGene_gn=TRDMT1;refGene_tx=XM_023256400.1;refGene_gn=TRDMT1;refGene_tx=XM_023256401.1;refGene_gn=TRDMT1;refGene_tx=XM_023256402.1
B4 16140178 16140242 refGene_gn=TRDMT1;refGene_tx=XM_003988130.5;refGene_gn=TRDMT1;refGene_tx=XM_006933280.4;refGene_gn=TRDMT1;refGene_tx=XM_023256400.1;refGene_gn=TRDMT1;refGene_tx=XM_023256401.1;refGene_gn=TRDMT1;refGene_tx=XM_023256402.1
B4 16166360 16166923 refGene_gn=VIM;refGene_tx=XM_003988131.3
B4 16167585 16167646 refGene_gn=VIM;refGene_tx=XM_003988131.3
B4 16169988 16170084 refGene_gn=VIM;refGene_tx=XM_003988131.3
B4 16170181 16170343 refGene_gn=VIM;refGene_tx=XM_003988131.3
B4 16170871 16170997 refGene_gn=VIM;refGene_tx=XM_003988131.3
B4 16171228 16171449 refGene_gn=VIM;refGene_tx=XM_003988131.3
B4 16171916 16171960 refGene_gn=VIM;refGene_tx=XM_003988131.3
B4 16172335 16172421 refGene_gn=VIM;refGene_tx=XM_003988131.3
B4 16173269 16173311 refGene_gn=VIM;refGene_tx=XM_003988131.3
B4 16233321 16233796 refGene_gn=ST8SIA6;refGene_tx=XM_023256403.1
B4 16235425 16235518 refGene_gn=ST8SIA6;refGene_tx=XM_023256403.1
B4 16239021 16239134 refGene_gn=ST8SIA6;refGene_tx=XM_023256403.1
B4 16243085 16243230 refGene_gn=ST8SIA6;refGene_tx=XM_023256403.1
B4 16276375 16276462 refGene_gn=ST8SIA6;refGene_tx=XM_023256403.1
B4 16320915 16321005 refGene_gn=ST8SIA6;refGene_tx=XM_023256403.1
B4 16386871 16386970 refGene_gn=ST8SIA6;refGene_tx=XM_023256403.1
B4 16387480 16387581 refGene_gn=ST8SIA6;refGene_tx=XM_023256403.1
B4 16498649 16498732 refGene_gn=HACD1;refGene_tx=XM_006944170.4;refGene_gn=HACD1;refGene_tx=XM_023256404.1
B4 16502346 16502525 refGene_gn=HACD1;refGene_tx=XM_006944170.4;refGene_gn=HACD1;refGene_tx=XM_023256404.1
B4 16507332 16507454 refGene_gn=HACD1;refGene_tx=XM_006944170.4;refGene_gn=HACD1;refGene_tx=XM_023256404.1
B4 16510698 16510787 refGene_gn=HACD1;refGene_tx=XM_006944170.4;refGene_gn=HACD1;refGene_tx=XM_023256404.1
B4 16511078 16511097 refGene_gn=HACD1;refGene_tx=XM_006944170.4;refGene_gn=HACD1;refGene_tx=XM_023256404.1
B4 16511284 16511402 refGene_gn=HACD1;refGene_tx=XM_006944170.4;refGene_gn=HACD1;refGene_tx=XM_023256404.1
B4 16520544 16520801 refGene_gn=HACD1;refGene_tx=XM_023256404.1
B4 16521791 16521838 refGene_gn=HACD1;refGene_tx=XM_006944170.4
B4 16540871 16540911 refGene_gn=STAM;refGene_tx=XM_004001111.5
B4 16562638 16562723 refGene_gn=STAM;refGene_tx=XM_004001111.5
B4 16579707 16579783 refGene_gn=STAM;refGene_tx=XM_004001111.5

B4	16579864	16579960	refGene_gn=STAM;refGene_tx=XM_004001111.5
B4	16581534	16581681	refGene_gn=STAM;refGene_tx=XM_004001111.5
B4	16594447	16594538	refGene_gn=STAM;refGene_tx=XM_004001111.5
B4	16596189	16596382	refGene_gn=STAM;refGene_tx=XM_004001111.5
B4	16597657	16597752	refGene_gn=STAM;refGene_tx=XM_004001111.5
B4	16601128	16601217	refGene_gn=STAM;refGene_tx=XM_004001111.5
B4	16603303	16603391	refGene_gn=STAM;refGene_tx=XM_004001111.5
B4	16603616	16603671	refGene_gn=STAM;refGene_tx=XM_004001111.5
B4	16604238	16604392	refGene_gn=STAM;refGene_tx=XM_004001111.5
B4	16606127	16606303	refGene_gn=STAM;refGene_tx=XM_004001111.5
B4	16621018	16621250	refGene_gn=STAM;refGene_tx=XM_004001111.5
B4	16662634	16662891	refGene_gn=TMEM236;refGene_tx=XM_019834003.2
B4	16675199	16675272	refGene_gn=TMEM236;refGene_tx=XM_019834003.2
B4	16678980	16679122	refGene_gn=TMEM236;refGene_tx=XM_019834003.2
B4	16697115	16697699	refGene_gn=TMEM236;refGene_tx=XM_019834003.2
B4	16722338	16722399	refGene_gn=MRC1;refGene_tx=XM_011283622.3
B4	16736839	16737241	refGene_gn=MRC1;refGene_tx=XM_011283622.3
B4	16740754	16740928	refGene_gn=MRC1;refGene_tx=XM_011283622.3
B4	16744966	16745131	refGene_gn=MRC1;refGene_tx=XM_011283622.3
B4	16752686	16752800	refGene_gn=MRC1;refGene_tx=XM_011283622.3
B4	16755223	16755370	refGene_gn=MRC1;refGene_tx=XM_011283622.3
B4	16759675	16759861	refGene_gn=MRC1;refGene_tx=XM_011283622.3
B4	16768883	16769041	refGene_gn=MRC1;refGene_tx=XM_011283622.3
B4	16770831	16770942	refGene_gn=MRC1;refGene_tx=XM_011283622.3
B4	16778669	16778785	refGene_gn=MRC1;refGene_tx=XM_011283622.3
B4	16780772	16780921	refGene_gn=MRC1;refGene_tx=XM_011283622.3
B4	16783300	16783500	refGene_gn=MRC1;refGene_tx=XM_011283622.3
B4	16785804	16785932	refGene_gn=MRC1;refGene_tx=XM_011283622.3
B4	16786447	16786535	refGene_gn=MRC1;refGene_tx=XM_011283622.3
B4	16788841	16788986	refGene_gn=MRC1;refGene_tx=XM_011283622.3
B4	16790561	16790603	refGene_gn=MRC1;refGene_tx=XM_011283622.3
B4	16791723	16791887	refGene_gn=MRC1;refGene_tx=XM_011283622.3
B4	16793759	16793827	refGene_gn=MRC1;refGene_tx=XM_011283622.3
B4	16795488	16795589	refGene_gn=MRC1;refGene_tx=XM_011283622.3
B4	16796516	16796659	refGene_gn=MRC1;refGene_tx=XM_011283622.3
B4	16797039	16797154	refGene_gn=MRC1;refGene_tx=XM_011283622.3
B4	16801257	16801424	refGene_gn=MRC1;refGene_tx=XM_011283622.3
B4	16804298	16804401	refGene_gn=MRC1;refGene_tx=XM_011283622.3
B4	16808401	16808634	refGene_gn=MRC1;refGene_tx=XM_011283622.3
B4	16811012	16811178	refGene_gn=MRC1;refGene_tx=XM_011283622.3
B4	16812140	16812290	refGene_gn=MRC1;refGene_tx=XM_011283622.3
B4	16817200	16817314	refGene_gn=MRC1;refGene_tx=XM_011283622.3
B4	16817998	16818163	refGene_gn=MRC1;refGene_tx=XM_011283622.3
B4	16819403	16819445	refGene_gn=MRC1;refGene_tx=XM_011283622.3
B4	16820587	16820838	refGene_gn=MRC1;refGene_tx=XM_011283622.3
B4	16849268	16849529
refGene_gn=SLC39A12;refGene_tx=XM_003988133.5;refGene_gn=SLC39A12;refGene_tx=XM_003988135.4;refGene_gn=SLC39A12;refGene_tx=XM_019834004.2;refGene_gn=SLC39A12;refGene_tx=XM_019834005.2
B4	16857234	16857516

refGene_gn=SLC39A12;refGene_tx=XM_003988133.5;refGene_gn=SLC39A12;refGene_tx=XM_003988135.4;refGene_gn=SLC39A12;refGene_tx=XM_019834004.2;refGene_gn=SLC39A12;refGene_tx=XM_019834005.2
B4 16861368 16861576
refGene_gn=SLC39A12;refGene_tx=XM_003988133.5;refGene_gn=SLC39A12;refGene_tx=XM_003988135.4;refGene_gn=SLC39A12;refGene_tx=XM_019834004.2;refGene_gn=SLC39A12;refGene_tx=XM_019834005.2
B4 16874588 16874761
refGene_gn=SLC39A12;refGene_tx=XM_003988133.5;refGene_gn=SLC39A12;refGene_tx=XM_003988135.4;refGene_gn=SLC39A12;refGene_tx=XM_019834004.2;refGene_gn=SLC39A12;refGene_tx=XM_019834005.2
B4 16880056 16880228
refGene_gn=SLC39A12;refGene_tx=XM_003988133.5;refGene_gn=SLC39A12;refGene_tx=XM_003988135.4;refGene_gn=SLC39A12;refGene_tx=XM_019834004.2;refGene_gn=SLC39A12;refGene_tx=XM_019834005.2
B4 16885446 16885619
refGene_gn=SLC39A12;refGene_tx=XM_003988133.5;refGene_gn=SLC39A12;refGene_tx=XM_003988135.4;refGene_gn=SLC39A12;refGene_tx=XM_019834004.2;refGene_gn=SLC39A12;refGene_tx=XM_019834005.2
B4 16888979 16889132
refGene_gn=SLC39A12;refGene_tx=XM_003988133.5;refGene_gn=SLC39A12;refGene_tx=XM_003988135.4;refGene_gn=SLC39A12;refGene_tx=XM_019834004.2;refGene_gn=SLC39A12;refGene_tx=XM_019834005.2
B4 16891015 16891126
refGene_gn=SLC39A12;refGene_tx=XM_003988133.5;refGene_gn=SLC39A12;refGene_tx=XM_019834004.2
B4 16894722 16894789
refGene_gn=SLC39A12;refGene_tx=XM_003988133.5;refGene_gn=SLC39A12;refGene_tx=XM_003988135.4;refGene_gn=SLC39A12;refGene_tx=XM_019834004.2;refGene_gn=SLC39A12;refGene_tx=XM_019834005.2
B4 16898831 16898990
refGene_gn=SLC39A12;refGene_tx=XM_003988133.5;refGene_gn=SLC39A12;refGene_tx=XM_003988135.4;refGene_gn=SLC39A12;refGene_tx=XM_019834004.2;refGene_gn=SLC39A12;refGene_tx=XM_019834005.2
B4 16901020 16901208
refGene_gn=SLC39A12;refGene_tx=XM_003988133.5;refGene_gn=SLC39A12;refGene_tx=XM_003988135.4;refGene_gn=SLC39A12;refGene_tx=XM_019834004.2;refGene_gn=SLC39A12;refGene_tx=XM_019834005.2
B4 16934743 16934872
refGene_gn=SLC39A12;refGene_tx=XM_003988133.5;refGene_gn=SLC39A12;refGene_tx=XM_003988135.4;refGene_gn=SLC39A12;refGene_tx=XM_019834004.2;refGene_gn=SLC39A12;refGene_tx=XM_019834005.2
B4 17030109 17030232
refGene_gn=CACNB2;refGene_tx=XM_023256405.1;refGene_gn=CACNB2;refGene_tx=XM_023256406.1;refGene_gn=CACNB2;refGene_tx=XM_023256408.1
B4 17030588 17030624 refGene_gn=CACNB2;refGene_tx=XM_019834006.2
B4 17039235 17039328
refGene_gn=CACNB2;refGene_tx=XM_019834006.2;refGene_gn=CACNB2;refGene_tx=XM_023256405.1;refGene_gn=CACNB2;refGene_tx=XM_023256406.1;refGene_gn=CACNB2;refGene_tx=XM_023256408.1

B4 17155866 17155914 refGene_gn=CACNB2;refGene_tx=XM_019834010.2
B4 17184321 17184402 refGene_gn=CACNB2;refGene_tx=XM_019834007.2
B4 17220432 17220483 refGene_gn=CACNB2;refGene_tx=XM_019834009.2
B4 17249281 17249314 refGene_gn=CACNB2;refGene_tx=XM_019834012.1
B4 17284870 17284939 refGene_gn=CACNB2;refGene_tx=XM_011283625.2
B4 17285777 17285897
refGene_gn=CACNB2;refGene_tx=XM_011283625.2;refGene_gn=CACNB2;refGene_tx=XM_019834006.2;refGene_gn=CACNB2;refGene_tx=XM_019834007.2;refGene_gn=CACNB2;refGene_tx=XM_019834009.2;refGene_gn=CACNB2;refGene_tx=XM_019834010.2;refGene_gn=CACNB2;refGene_tx=XM_019834012.1;refGene_gn=CACNB2;refGene_tx=XM_023256405.1;refGene_gn=CACNB2;refGene_tx=XM_023256406.1;refGene_gn=CACNB2;refGene_tx=XM_023256408.1
B4 17379758 17379881
refGene_gn=CACNB2;refGene_tx=XM_011283625.2;refGene_gn=CACNB2;refGene_tx=XM_019834006.2;refGene_gn=CACNB2;refGene_tx=XM_019834007.2;refGene_gn=CACNB2;refGene_tx=XM_019834009.2;refGene_gn=CACNB2;refGene_tx=XM_019834010.2;refGene_gn=CACNB2;refGene_tx=XM_019834012.1;refGene_gn=CACNB2;refGene_tx=XM_023256405.1;refGene_gn=CACNB2;refGene_tx=XM_023256406.1;refGene_gn=CACNB2;refGene_tx=XM_023256408.1
B4 17382407 17382544
refGene_gn=CACNB2;refGene_tx=XM_011283625.2;refGene_gn=CACNB2;refGene_tx=XM_019834006.2;refGene_gn=CACNB2;refGene_tx=XM_019834007.2;refGene_gn=CACNB2;refGene_tx=XM_019834009.2;refGene_gn=CACNB2;refGene_tx=XM_019834010.2;refGene_gn=CACNB2;refGene_tx=XM_019834012.1;refGene_gn=CACNB2;refGene_tx=XM_023256405.1;refGene_gn=CACNB2;refGene_tx=XM_023256406.1;refGene_gn=CACNB2;refGene_tx=XM_023256408.1
B4 17386268 17386345
refGene_gn=CACNB2;refGene_tx=XM_011283625.2;refGene_gn=CACNB2;refGene_tx=XM_019834006.2;refGene_gn=CACNB2;refGene_tx=XM_019834007.2;refGene_gn=CACNB2;refGene_tx=XM_019834009.2;refGene_gn=CACNB2;refGene_tx=XM_019834010.2;refGene_gn=CACNB2;refGene_tx=XM_019834012.1;refGene_gn=CACNB2;refGene_tx=XM_023256405.1;refGene_gn=CACNB2;refGene_tx=XM_023256406.1;refGene_gn=CACNB2;refGene_tx=XM_023256408.1
B4 17391661 17391795
refGene_gn=CACNB2;refGene_tx=XM_011283625.2;refGene_gn=CACNB2;refGene_tx=XM_019834006.2;refGene_gn=CACNB2;refGene_tx=XM_019834007.2;refGene_gn=CACNB2;refGene_tx=XM_019834009.2;refGene_gn=CACNB2;refGene_tx=XM_019834010.2;refGene_gn=CACNB2;refGene_tx=XM_019834012.1;refGene_gn=CACNB2;refGene_tx=XM_023256405.1
B4 17391940 17391960 refGene_gn=CACNB2;refGene_tx=XM_023256408.1
B4 17392430 17392492 refGene_gn=CACNB2;refGene_tx=XM_023256406.1
B4 17395575 17395656
refGene_gn=CACNB2;refGene_tx=XM_011283625.2;refGene_gn=CACNB2;refGene_tx=XM_019834006.2;refGene_gn=CACNB2;refGene_tx=XM_019834007.2;refGene_gn=CACNB2;refGene_tx=XM_019834009.2;refGene_gn=CACNB2;refGene_tx=XM_019834010.2;refGene_gn=CACNB2;refGene_tx=XM_019834012.1;refGene_gn=CACNB2;refGene_tx=XM_023256405.1;refGene_gn=CACNB2;refGene_tx=XM_023256406.1;refGene_gn=CACNB2;refGene_tx=XM_023256408.1
B4 17396138 17396197
refGene_gn=CACNB2;refGene_tx=XM_011283625.2;refGene_gn=CACNB2;refGene_

tx=XM_019834006.2;refGene_gn=CACNB2;refGene_tx=XM_019834007.2;refGene_gn=CACNB2;refGene_tx=XM_019834009.2;refGene_gn=CACNB2;refGene_tx=XM_019834010.2;refGene_gn=CACNB2;refGene_tx=XM_019834012.1;refGene_gn=CACNB2;refGene_tx=XM_023256405.1;refGene_gn=CACNB2;refGene_tx=XM_023256406.1;refGene_gn=CACNB2;refGene_tx=XM_023256408.1
B4 17406584 17406694
refGene_gn=CACNB2;refGene_tx=XM_011283625.2;refGene_gn=CACNB2;refGene_tx=XM_019834006.2;refGene_gn=CACNB2;refGene_tx=XM_019834007.2;refGene_gn=CACNB2;refGene_tx=XM_019834009.2;refGene_gn=CACNB2;refGene_tx=XM_019834010.2;refGene_gn=CACNB2;refGene_tx=XM_019834012.1;refGene_gn=CACNB2;refGene_tx=XM_023256405.1;refGene_gn=CACNB2;refGene_tx=XM_023256406.1;refGene_gn=CACNB2;refGene_tx=XM_023256408.1
B4 17412307 17412459
refGene_gn=CACNB2;refGene_tx=XM_011283625.2;refGene_gn=CACNB2;refGene_tx=XM_019834006.2;refGene_gn=CACNB2;refGene_tx=XM_019834007.2;refGene_gn=CACNB2;refGene_tx=XM_019834009.2;refGene_gn=CACNB2;refGene_tx=XM_019834010.2;refGene_gn=CACNB2;refGene_tx=XM_019834012.1;refGene_gn=CACNB2;refGene_tx=XM_023256405.1;refGene_gn=CACNB2;refGene_tx=XM_023256406.1;refGene_gn=CACNB2;refGene_tx=XM_023256408.1
B4 17413878 17413974
refGene_gn=CACNB2;refGene_tx=XM_011283625.2;refGene_gn=CACNB2;refGene_tx=XM_019834006.2;refGene_gn=CACNB2;refGene_tx=XM_019834007.2;refGene_gn=CACNB2;refGene_tx=XM_019834009.2;refGene_gn=CACNB2;refGene_tx=XM_019834010.2;refGene_gn=CACNB2;refGene_tx=XM_019834012.1;refGene_gn=CACNB2;refGene_tx=XM_023256405.1;refGene_gn=CACNB2;refGene_tx=XM_023256406.1;refGene_gn=CACNB2;refGene_tx=XM_023256408.1
B4 17414952 17415138
refGene_gn=CACNB2;refGene_tx=XM_011283625.2;refGene_gn=CACNB2;refGene_tx=XM_019834006.2;refGene_gn=CACNB2;refGene_tx=XM_019834007.2;refGene_gn=CACNB2;refGene_tx=XM_019834009.2;refGene_gn=CACNB2;refGene_tx=XM_019834010.2;refGene_gn=CACNB2;refGene_tx=XM_019834012.1;refGene_gn=CACNB2;refGene_tx=XM_023256405.1;refGene_gn=CACNB2;refGene_tx=XM_023256406.1;refGene_gn=CACNB2;refGene_tx=XM_023256408.1
B4 17416036 17416531
refGene_gn=CACNB2;refGene_tx=XM_011283625.2;refGene_gn=CACNB2;refGene_tx=XM_019834006.2;refGene_gn=CACNB2;refGene_tx=XM_019834007.2;refGene_gn=CACNB2;refGene_tx=XM_019834009.2;refGene_gn=CACNB2;refGene_tx=XM_019834010.2;refGene_gn=CACNB2;refGene_tx=XM_019834012.1;refGene_gn=CACNB2;refGene_tx=XM_023256405.1;refGene_gn=CACNB2;refGene_tx=XM_023256406.1;refGene_gn=CACNB2;refGene_tx=XM_023256408.1
B4 17422022 17422235
refGene_gn=NSUN6;refGene_tx=XM_023256409.1;refGene_gn=NSUN6;refGene_tx=XM_023256410.1;refGene_gn=NSUN6;refGene_tx=XM_023256411.1;refGene_gn=NSUN6;refGene_tx=XM_023256412.1
B4 17423947 17424073
refGene_gn=NSUN6;refGene_tx=XM_023256409.1;refGene_gn=NSUN6;refGene_tx=XM_023256410.1;refGene_gn=NSUN6;refGene_tx=XM_023256411.1
B4 17427428 17427577
refGene_gn=NSUN6;refGene_tx=XM_023256409.1;refGene_gn=NSUN6;refGene_tx=XM_023256410.1;refGene_gn=NSUN6;refGene_tx=XM_023256411.1;refGene_gn=

NSUN6;refGene_tx=XM_023256412.1
B4 17442674 17442819
refGene_gn=NSUN6;refGene_tx=XM_023256409.1;refGene_gn=NSUN6;refGene_tx=XM_023256410.1;refGene_gn=NSUN6;refGene_tx=XM_023256411.1;refGene_gn=NSUN6;refGene_tx=XM_023256412.1
B4 17448920 17449040
refGene_gn=NSUN6;refGene_tx=XM_023256409.1;refGene_gn=NSUN6;refGene_tx=XM_023256410.1;refGene_gn=NSUN6;refGene_tx=XM_023256411.1;refGene_gn=NSUN6;refGene_tx=XM_023256412.1
B4 17454031 17454113
refGene_gn=NSUN6;refGene_tx=XM_023256409.1;refGene_gn=NSUN6;refGene_tx=XM_023256410.1;refGene_gn=NSUN6;refGene_tx=XM_023256411.1;refGene_gn=NSUN6;refGene_tx=XM_023256412.1
B4 17464994 17465148
refGene_gn=NSUN6;refGene_tx=XM_023256409.1;refGene_gn=NSUN6;refGene_tx=XM_023256410.1;refGene_gn=NSUN6;refGene_tx=XM_023256411.1;refGene_gn=NSUN6;refGene_tx=XM_023256412.1
B4 17466960 17467070
refGene_gn=NSUN6;refGene_tx=XM_023256409.1;refGene_gn=NSUN6;refGene_tx=XM_023256410.1;refGene_gn=NSUN6;refGene_tx=XM_023256411.1;refGene_gn=NSUN6;refGene_tx=XM_023256412.1
B4 17480508 17480588
refGene_gn=NSUN6;refGene_tx=XM_023256409.1;refGene_gn=NSUN6;refGene_tx=XM_023256410.1;refGene_gn=NSUN6;refGene_tx=XM_023256411.1;refGene_gn=NSUN6;refGene_tx=XM_023256412.1
B4 17482667 17482823
refGene_gn=NSUN6;refGene_tx=XM_023256409.1;refGene_gn=NSUN6;refGene_tx=XM_023256410.1;refGene_gn=NSUN6;refGene_tx=XM_023256411.1;refGene_gn=NSUN6;refGene_tx=XM_023256412.1
B4 17483351 17483393 refGene_gn=NSUN6;refGene_tx=XM_023256411.1
B4 17484827 17484902
refGene_gn=NSUN6;refGene_tx=XM_023256409.1;refGene_gn=NSUN6;refGene_tx=XM_023256410.1;refGene_gn=NSUN6;refGene_tx=XM_023256412.1
B4 17490455 17490504 refGene_gn=ARL5B;refGene_tx=XM_006933289.4
B4 17504915 17504976 refGene_gn=ARL5B;refGene_tx=XM_006933289.4
B4 17506541 17506689
refGene_gn=ARL5B;refGene_tx=XM_006933289.4;refGene_gn=ARL5B;refGene_tx=XM_023256413.1
B4 17509563 17509647
refGene_gn=ARL5B;refGene_tx=XM_006933289.4;refGene_gn=ARL5B;refGene_tx=XM_023256413.1
B4 17511738 17511890
refGene_gn=ARL5B;refGene_tx=XM_006933289.4;refGene_gn=ARL5B;refGene_tx=XM_023256413.1
B4 17513011 17513060
refGene_gn=ARL5B;refGene_tx=XM_006933289.4;refGene_gn=ARL5B;refGene_tx=XM_023256413.1
B4 17852971 17853149 refGene_gn=MALRD1;refGene_tx=XM_023257733.1
B4 17872684 17872825 refGene_gn=MALRD1;refGene_tx=XM_023257733.1
B4 17888240 17888335 refGene_gn=MALRD1;refGene_tx=XM_023257733.1

B4 17888423 17888585 refGene_gn=MALRD1;refGene_tx=XM_023257733.1
B4 17898697 17898794 refGene_gn=MALRD1;refGene_tx=XM_023257733.1
B4 17917948 17918050 refGene_gn=MALRD1;refGene_tx=XM_023257733.1
B4 17918693 17918840 refGene_gn=MALRD1;refGene_tx=XM_023257733.1
B4 17927252 17927419 refGene_gn=MALRD1;refGene_tx=XM_023257733.1
B4 17931746 17931839 refGene_gn=MALRD1;refGene_tx=XM_023257733.1
B4 17935150 17935358 refGene_gn=MALRD1;refGene_tx=XM_023257733.1
B4 17946317 17946464 refGene_gn=MALRD1;refGene_tx=XM_023257733.1
B4 17952910 17953008 refGene_gn=MALRD1;refGene_tx=XM_023257733.1
B4 17961024 17961198 refGene_gn=MALRD1;refGene_tx=XM_023257733.1
B4 17971072 17971193 refGene_gn=MALRD1;refGene_tx=XM_023257733.1
B4 17973410 17973551 refGene_gn=MALRD1;refGene_tx=XM_023257733.1
B4 18013671 18013824 refGene_gn=MALRD1;refGene_tx=XM_023257733.1
B4 18014233 18014339 refGene_gn=MALRD1;refGene_tx=XM_023257733.1
B4 18015253 18015621 refGene_gn=MALRD1;refGene_tx=XM_023257733.1
B4 18019066 18019479 refGene_gn=MALRD1;refGene_tx=XM_023257733.1
B4 18098803 18098891 refGene_gn=MALRD1;refGene_tx=XM_023257733.1
B4 18108865 18109042 refGene_gn=MALRD1;refGene_tx=XM_023257733.1
B4 18112233 18112396 refGene_gn=MALRD1;refGene_tx=XM_023257733.1
B4 18172978 18173135 refGene_gn=MALRD1;refGene_tx=XM_023257733.1
B4 18176915 18177026 refGene_gn=MALRD1;refGene_tx=XM_023257733.1
B4 18181142 18181356 refGene_gn=MALRD1;refGene_tx=XM_023257733.1
B4 18213377 18213625 refGene_gn=MALRD1;refGene_tx=XM_023257733.1
B4 18217217 18217509 refGene_gn=MALRD1;refGene_tx=XM_023257733.1
B4 18294180 18294426 refGene_gn=MALRD1;refGene_tx=XM_023257733.1
B4 18295990 18296148 refGene_gn=MALRD1;refGene_tx=XM_023257733.1
B4 18368227 18368411 refGene_gn=MALRD1;refGene_tx=XM_023257733.1
B4 18398336 18398465 refGene_gn=MALRD1;refGene_tx=XM_023257733.1
B4 18407847 18408009 refGene_gn=MALRD1;refGene_tx=XM_023257733.1
B4 18440197 18440355 refGene_gn=MALRD1;refGene_tx=XM_023257733.1
B4 18466249 18466451 refGene_gn=MALRD1;refGene_tx=XM_023257733.1
B4 18481776 18482040 refGene_gn=MALRD1;refGene_tx=XM_023257733.1
B4 18492892 18493018 refGene_gn=MALRD1;refGene_tx=XM_023257733.1
B4 18500487 18500554 refGene_gn=MALRD1;refGene_tx=XM_023257733.1
B4 18595124 18595204 refGene_gn=MALRD1;refGene_tx=XM_023257733.1
B4 18595297 18595314 refGene_gn=MALRD1;refGene_tx=XM_023257733.1
B4 18595315 18595395 refGene_gn=MALRD1;refGene_tx=XM_023257733.1
B4 18630855 18630931 refGene_gn=MALRD1;refGene_tx=XM_023257733.1
B4 18633696 18633777 refGene_gn=MALRD1;refGene_tx=XM_023257733.1
B4 18726166 18726278
refGene_gn=PLXDC2;refGene_tx=XM_003988142.5;refGene_gn=PLXDC2;refGene_
tx=XM_019834014.1
B4 18896014 18896226
refGene_gn=PLXDC2;refGene_tx=XM_003988142.5;refGene_gn=PLXDC2;refGene_
tx=XM_019834014.1
B4 18955076 18955223
refGene_gn=PLXDC2;refGene_tx=XM_003988142.5;refGene_gn=PLXDC2;refGene_
tx=XM_019834014.1
B4 18972757 18972827
refGene_gn=PLXDC2;refGene_tx=XM_003988142.5;refGene_gn=PLXDC2;refGene_

tx=XM_019834014.1
B4 19039481 19039604
refGene_gn=PLXDC2;refGene_tx=XM_003988142.5;refGene_gn=PLXDC2;refGene_
tx=XM_019834014.1
B4 19046967 19047086
refGene_gn=PLXDC2;refGene_tx=XM_003988142.5;refGene_gn=PLXDC2;refGene_
tx=XM_019834014.1
B4 19062019 19062119
refGene_gn=PLXDC2;refGene_tx=XM_003988142.5;refGene_gn=PLXDC2;refGene_
tx=XM_019834014.1
B4 19074976 19075072
refGene_gn=PLXDC2;refGene_tx=XM_003988142.5;refGene_gn=PLXDC2;refGene_
tx=XM_019834014.1
B4 19075263 19075345
refGene_gn=PLXDC2;refGene_tx=XM_003988142.5;refGene_gn=PLXDC2;refGene_
tx=XM_019834014.1
B4 19101764 19101825
refGene_gn=PLXDC2;refGene_tx=XM_003988142.5;refGene_gn=PLXDC2;refGene_
tx=XM_019834014.1
B4 19110194 19110345
refGene_gn=PLXDC2;refGene_tx=XM_003988142.5;refGene_gn=PLXDC2;refGene_
tx=XM_019834014.1
B4 19111321 19111360
refGene_gn=PLXDC2;refGene_tx=XM_003988142.5;refGene_gn=PLXDC2;refGene_
tx=XM_019834014.1
B4 19133252 19133413
refGene_gn=PLXDC2;refGene_tx=XM_003988142.5;refGene_gn=PLXDC2;refGene_
tx=XM_019834014.1
B4 19137672 19137741 refGene_gn=PLXDC2;refGene_tx=XM_019834014.1
B4 19170462 19170579 refGene_gn=PLXDC2;refGene_tx=XM_003988142.5
B4 19551234 19551411
refGene_gn=NEBL;refGene_tx=XM_019834015.2;refGene_gn=NEBL;refGene_tx=X
M_019834016.2;refGene_gn=NEBL;refGene_tx=XM_019834017.2;refGene_gn=NEB
L;refGene_tx=XM_019834018.2;refGene_gn=NEBL;refGene_tx=XM_019834019.2;
refGene_gn=NEBL;refGene_tx=XM_019834020.2;refGene_gn=NEBL;refGene_tx=X
M_019834021.2;refGene_gn=NEBL;refGene_tx=XM_019834022.2;refGene_gn=NEB
L;refGene_tx=XM_019834024.2;refGene_gn=NEBL;refGene_tx=XM_019834025.2;
refGene_gn=NEBL;refGene_tx=XM_019834026.2;refGene_gn=NEBL;refGene_tx=X
M_019834027.2;refGene_gn=NEBL;refGene_tx=XM_019834028.2
B4 19553277 19553384
refGene_gn=NEBL;refGene_tx=XM_019834015.2;refGene_gn=NEBL;refGene_tx=X
M_019834016.2;refGene_gn=NEBL;refGene_tx=XM_019834017.2;refGene_gn=NEB
L;refGene_tx=XM_019834018.2;refGene_gn=NEBL;refGene_tx=XM_019834019.2;
refGene_gn=NEBL;refGene_tx=XM_019834020.2;refGene_gn=NEBL;refGene_tx=X
M_019834021.2;refGene_gn=NEBL;refGene_tx=XM_019834022.2;refGene_gn=NEB
L;refGene_tx=XM_019834024.2;refGene_gn=NEBL;refGene_tx=XM_019834025.2;
refGene_gn=NEBL;refGene_tx=XM_019834026.2;refGene_gn=NEBL;refGene_tx=X
M_019834027.2;refGene_gn=NEBL;refGene_tx=XM_019834028.2
B4 19575429 19575633
refGene_gn=NEBL;refGene_tx=XM_019834015.2;refGene_gn=NEBL;refGene_tx=X

M_019834016.2;refGene_gn=NEBL;refGene_tx=XM_019834017.2;refGene_gn=NEB
L;refGene_tx=XM_019834018.2;refGene_gn=NEBL;refGene_tx=XM_019834019.2;
refGene_gn=NEBL;refGene_tx=XM_019834020.2;refGene_gn=NEBL;refGene_tx=X
M_019834021.2;refGene_gn=NEBL;refGene_tx=XM_019834022.2;refGene_gn=NEB
L;refGene_tx=XM_019834024.2;refGene_gn=NEBL;refGene_tx=XM_019834025.2
B4 19576832 19576925
refGene_gn=NEBL;refGene_tx=XM_019834015.2;refGene_gn=NEBL;refGene_tx=X
M_019834016.2;refGene_gn=NEBL;refGene_tx=XM_019834017.2;refGene_gn=NEB
L;refGene_tx=XM_019834018.2;refGene_gn=NEBL;refGene_tx=XM_019834019.2;
refGene_gn=NEBL;refGene_tx=XM_019834020.2;refGene_gn=NEBL;refGene_tx=X
M_019834021.2;refGene_gn=NEBL;refGene_tx=XM_019834022.2;refGene_gn=NEB
L;refGene_tx=XM_019834024.2;refGene_gn=NEBL;refGene_tx=XM_019834025.2;
refGene_gn=NEBL;refGene_tx=XM_019834026.2
B4 19579099 19579271
refGene_gn=NEBL;refGene_tx=XM_019834015.2;refGene_gn=NEBL;refGene_tx=X
M_019834016.2;refGene_gn=NEBL;refGene_tx=XM_019834017.2;refGene_gn=NEB
L;refGene_tx=XM_019834018.2;refGene_gn=NEBL;refGene_tx=XM_019834019.2;
refGene_gn=NEBL;refGene_tx=XM_019834020.2;refGene_gn=NEBL;refGene_tx=X
M_019834021.2;refGene_gn=NEBL;refGene_tx=XM_019834022.2;refGene_gn=NEB
L;refGene_tx=XM_019834024.2;refGene_gn=NEBL;refGene_tx=XM_019834025.2;
refGene_gn=NEBL;refGene_tx=XM_019834026.2;refGene_gn=NEBL;refGene_tx=X
M_019834027.2;refGene_gn=NEBL;refGene_tx=XM_019834028.2
B4 19580294 19580399
refGene_gn=NEBL;refGene_tx=XM_019834015.2;refGene_gn=NEBL;refGene_tx=X
M_019834016.2;refGene_gn=NEBL;refGene_tx=XM_019834017.2;refGene_gn=NEB
L;refGene_tx=XM_019834018.2;refGene_gn=NEBL;refGene_tx=XM_019834019.2;
refGene_gn=NEBL;refGene_tx=XM_019834020.2;refGene_gn=NEBL;refGene_tx=X
M_019834021.2;refGene_gn=NEBL;refGene_tx=XM_019834022.2;refGene_gn=NEB
L;refGene_tx=XM_019834024.2;refGene_gn=NEBL;refGene_tx=XM_019834025.2;
refGene_gn=NEBL;refGene_tx=XM_019834026.2;refGene_gn=NEBL;refGene_tx=X
M_019834027.2
B4 19581683 19581776
refGene_gn=NEBL;refGene_tx=XM_019834015.2;refGene_gn=NEBL;refGene_tx=X
M_019834016.2;refGene_gn=NEBL;refGene_tx=XM_019834017.2;refGene_gn=NEB
L;refGene_tx=XM_019834018.2;refGene_gn=NEBL;refGene_tx=XM_019834019.2;
refGene_gn=NEBL;refGene_tx=XM_019834020.2;refGene_gn=NEBL;refGene_tx=X
M_019834021.2;refGene_gn=NEBL;refGene_tx=XM_019834022.2;refGene_gn=NEB
L;refGene_tx=XM_019834024.2;refGene_gn=NEBL;refGene_tx=XM_019834025.2;
refGene_gn=NEBL;refGene_tx=XM_019834026.2;refGene_gn=NEBL;refGene_tx=X
M_019834027.2
B4 19583295 19583388
refGene_gn=NEBL;refGene_tx=XM_019834015.2;refGene_gn=NEBL;refGene_tx=X
M_019834016.2;refGene_gn=NEBL;refGene_tx=XM_019834017.2;refGene_gn=NEB
L;refGene_tx=XM_019834018.2;refGene_gn=NEBL;refGene_tx=XM_019834019.2;
refGene_gn=NEBL;refGene_tx=XM_019834020.2;refGene_gn=NEBL;refGene_tx=X
M_019834021.2;refGene_gn=NEBL;refGene_tx=XM_019834022.2;refGene_gn=NEB
L;refGene_tx=XM_019834024.2;refGene_gn=NEBL;refGene_tx=XM_019834025.2;
refGene_gn=NEBL;refGene_tx=XM_019834026.2;refGene_gn=NEBL;refGene_tx=X
M_019834027.2
B4 19584946 19585039

refGene_gn=NEBL;refGene_tx=XM_019834015.2;refGene_gn=NEBL;refGene_tx=X
M_019834016.2;refGene_gn=NEBL;refGene_tx=XM_019834017.2;refGene_gn=NEB
L;refGene_tx=XM_019834019.2;refGene_gn=NEBL;refGene_tx=XM_019834020.2;
refGene_gn=NEBL;refGene_tx=XM_019834021.2;refGene_gn=NEBL;refGene_tx=X
M_019834022.2;refGene_gn=NEBL;refGene_tx=XM_019834024.2;refGene_gn=NEB
L;refGene_tx=XM_019834026.2;refGene_gn=NEBL;refGene_tx=XM_019834027.2
B4 19590529 19590622
refGene_gn=NEBL;refGene_tx=XM_019834015.2;refGene_gn=NEBL;refGene_tx=X
M_019834016.2;refGene_gn=NEBL;refGene_tx=XM_019834017.2;refGene_gn=NEB
L;refGene_tx=XM_019834018.2;refGene_gn=NEBL;refGene_tx=XM_019834020.2;
refGene_gn=NEBL;refGene_tx=XM_019834021.2;refGene_gn=NEBL;refGene_tx=X
M_019834022.2;refGene_gn=NEBL;refGene_tx=XM_019834024.2;refGene_gn=NEB
L;refGene_tx=XM_019834026.2;refGene_gn=NEBL;refGene_tx=XM_019834027.2
B4 19592982 19593087
refGene_gn=NEBL;refGene_tx=XM_019834015.2;refGene_gn=NEBL;refGene_tx=X
M_019834016.2;refGene_gn=NEBL;refGene_tx=XM_019834017.2;refGene_gn=NEB
L;refGene_tx=XM_019834018.2;refGene_gn=NEBL;refGene_tx=XM_019834019.2;
refGene_gn=NEBL;refGene_tx=XM_019834020.2;refGene_gn=NEBL;refGene_tx=X
M_019834021.2;refGene_gn=NEBL;refGene_tx=XM_019834022.2;refGene_gn=NEB
L;refGene_tx=XM_019834024.2;refGene_gn=NEBL;refGene_tx=XM_019834025.2;
refGene_gn=NEBL;refGene_tx=XM_019834026.2;refGene_gn=NEBL;refGene_tx=X
M_019834027.2
B4 19596007 19596118
refGene_gn=NEBL;refGene_tx=XM_019834015.2;refGene_gn=NEBL;refGene_tx=X
M_019834016.2;refGene_gn=NEBL;refGene_tx=XM_019834017.2;refGene_gn=NEB
L;refGene_tx=XM_019834018.2;refGene_gn=NEBL;refGene_tx=XM_019834019.2;
refGene_gn=NEBL;refGene_tx=XM_019834020.2;refGene_gn=NEBL;refGene_tx=X
M_019834021.2;refGene_gn=NEBL;refGene_tx=XM_019834022.2;refGene_gn=NEB
L;refGene_tx=XM_019834024.2;refGene_gn=NEBL;refGene_tx=XM_019834025.2;
refGene_gn=NEBL;refGene_tx=XM_019834026.2;refGene_gn=NEBL;refGene_tx=X
M_019834027.2
B4 19596286 19596397
refGene_gn=NEBL;refGene_tx=XM_019834015.2;refGene_gn=NEBL;refGene_tx=X
M_019834016.2;refGene_gn=NEBL;refGene_tx=XM_019834017.2;refGene_gn=NEB
L;refGene_tx=XM_019834018.2;refGene_gn=NEBL;refGene_tx=XM_019834019.2;
refGene_gn=NEBL;refGene_tx=XM_019834020.2;refGene_gn=NEBL;refGene_tx=X
M_019834021.2;refGene_gn=NEBL;refGene_tx=XM_019834022.2;refGene_gn=NEB
L;refGene_tx=XM_019834024.2;refGene_gn=NEBL;refGene_tx=XM_019834025.2;
refGene_gn=NEBL;refGene_tx=XM_019834026.2;refGene_gn=NEBL;refGene_tx=X
M_019834027.2
B4 19601591 19601702
refGene_gn=NEBL;refGene_tx=XM_019834015.2;refGene_gn=NEBL;refGene_tx=X
M_019834016.2;refGene_gn=NEBL;refGene_tx=XM_019834017.2;refGene_gn=NEB
L;refGene_tx=XM_019834018.2;refGene_gn=NEBL;refGene_tx=XM_019834019.2;
refGene_gn=NEBL;refGene_tx=XM_019834020.2;refGene_gn=NEBL;refGene_tx=X
M_019834021.2;refGene_gn=NEBL;refGene_tx=XM_019834022.2;refGene_gn=NEB
L;refGene_tx=XM_019834025.2;refGene_gn=NEBL;refGene_tx=XM_019834026.2;
refGene_gn=NEBL;refGene_tx=XM_019834027.2
B4 19603678 19603789
refGene_gn=NEBL;refGene_tx=XM_019834015.2;refGene_gn=NEBL;refGene_tx=X

M_019834016.2;refGene_gn=NEBL;refGene_tx=XM_019834017.2;refGene_gn=NEB
L;refGene_tx=XM_019834018.2;refGene_gn=NEBL;refGene_tx=XM_019834019.2;
refGene_gn=NEBL;refGene_tx=XM_019834020.2;refGene_gn=NEBL;refGene_tx=X
M_019834021.2;refGene_gn=NEBL;refGene_tx=XM_019834022.2;refGene_gn=NEB
L;refGene_tx=XM_019834024.2;refGene_gn=NEBL;refGene_tx=XM_019834025.2;
refGene_gn=NEBL;refGene_tx=XM_019834026.2;refGene_gn=NEBL;refGene_tx=X
M_019834027.2
B4 19606430 19606541
refGene_gn=NEBL;refGene_tx=XM_019834015.2;refGene_gn=NEBL;refGene_tx=X
M_019834016.2;refGene_gn=NEBL;refGene_tx=XM_019834017.2;refGene_gn=NEB
L;refGene_tx=XM_019834018.2;refGene_gn=NEBL;refGene_tx=XM_019834019.2;
refGene_gn=NEBL;refGene_tx=XM_019834020.2;refGene_gn=NEBL;refGene_tx=X
M_019834021.2;refGene_gn=NEBL;refGene_tx=XM_019834022.2;refGene_gn=NEB
L;refGene_tx=XM_019834024.2;refGene_gn=NEBL;refGene_tx=XM_019834025.2;
refGene_gn=NEBL;refGene_tx=XM_019834026.2;refGene_gn=NEBL;refGene_tx=X
M_019834027.2
B4 19611287 19611395
refGene_gn=NEBL;refGene_tx=XM_019834015.2;refGene_gn=NEBL;refGene_tx=X
M_019834016.2;refGene_gn=NEBL;refGene_tx=XM_019834017.2;refGene_gn=NEB
L;refGene_tx=XM_019834018.2;refGene_gn=NEBL;refGene_tx=XM_019834019.2;
refGene_gn=NEBL;refGene_tx=XM_019834020.2;refGene_gn=NEBL;refGene_tx=X
M_019834021.2;refGene_gn=NEBL;refGene_tx=XM_019834022.2;refGene_gn=NEB
L;refGene_tx=XM_019834024.2;refGene_gn=NEBL;refGene_tx=XM_019834025.2;
refGene_gn=NEBL;refGene_tx=XM_019834026.2;refGene_gn=NEBL;refGene_tx=X
M_019834027.2
B4 19613374 19613479
refGene_gn=NEBL;refGene_tx=XM_019834015.2;refGene_gn=NEBL;refGene_tx=X
M_019834016.2;refGene_gn=NEBL;refGene_tx=XM_019834017.2;refGene_gn=NEB
L;refGene_tx=XM_019834018.2;refGene_gn=NEBL;refGene_tx=XM_019834019.2;
refGene_gn=NEBL;refGene_tx=XM_019834020.2;refGene_gn=NEBL;refGene_tx=X
M_019834021.2;refGene_gn=NEBL;refGene_tx=XM_019834022.2;refGene_gn=NEB
L;refGene_tx=XM_019834024.2;refGene_gn=NEBL;refGene_tx=XM_019834025.2;
refGene_gn=NEBL;refGene_tx=XM_019834026.2;refGene_gn=NEBL;refGene_tx=X
M_019834027.2
B4 19617643 19617745
refGene_gn=NEBL;refGene_tx=XM_019834015.2;refGene_gn=NEBL;refGene_tx=X
M_019834016.2;refGene_gn=NEBL;refGene_tx=XM_019834017.2;refGene_gn=NEB
L;refGene_tx=XM_019834018.2;refGene_gn=NEBL;refGene_tx=XM_019834019.2;
refGene_gn=NEBL;refGene_tx=XM_019834021.2;refGene_gn=NEBL;refGene_tx=X
M_019834022.2;refGene_gn=NEBL;refGene_tx=XM_019834024.2;refGene_gn=NEB
L;refGene_tx=XM_019834025.2;refGene_gn=NEBL;refGene_tx=XM_019834026.2;
refGene_gn=NEBL;refGene_tx=XM_019834027.2
B4 19619332 19619446
refGene_gn=NEBL;refGene_tx=XM_019834015.2;refGene_gn=NEBL;refGene_tx=X
M_019834016.2;refGene_gn=NEBL;refGene_tx=XM_019834017.2;refGene_gn=NEB
L;refGene_tx=XM_019834018.2;refGene_gn=NEBL;refGene_tx=XM_019834019.2;
refGene_gn=NEBL;refGene_tx=XM_019834020.2;refGene_gn=NEBL;refGene_tx=X
M_019834021.2;refGene_gn=NEBL;refGene_tx=XM_019834022.2;refGene_gn=NEB
L;refGene_tx=XM_019834024.2;refGene_gn=NEBL;refGene_tx=XM_019834025.2;
refGene_gn=NEBL;refGene_tx=XM_019834026.2;refGene_gn=NEBL;refGene_tx=X

M_019834027.2
B4 19629897 19629999
refGene_gn=NEBL;refGene_tx=XM_019834015.2;refGene_gn=NEBL;refGene_tx=XM_019834016.2;refGene_gn=NEBL;refGene_tx=XM_019834017.2;refGene_gn=NEBL;refGene_tx=XM_019834018.2;refGene_gn=NEBL;refGene_tx=XM_019834019.2;refGene_gn=NEBL;refGene_tx=XM_019834020.2;refGene_gn=NEBL;refGene_tx=XM_019834021.2;refGene_gn=NEBL;refGene_tx=XM_019834022.2;refGene_gn=NEBL;refGene_tx=XM_019834024.2;refGene_gn=NEBL;refGene_tx=XM_019834025.2;refGene_gn=NEBL;refGene_tx=XM_019834026.2;refGene_gn=NEBL;refGene_tx=XM_019834027.2
B4 19630970 19631072
refGene_gn=NEBL;refGene_tx=XM_019834015.2;refGene_gn=NEBL;refGene_tx=XM_019834016.2;refGene_gn=NEBL;refGene_tx=XM_019834017.2;refGene_gn=NEBL;refGene_tx=XM_019834018.2;refGene_gn=NEBL;refGene_tx=XM_019834019.2;refGene_gn=NEBL;refGene_tx=XM_019834020.2;refGene_gn=NEBL;refGene_tx=XM_019834021.2;refGene_gn=NEBL;refGene_tx=XM_019834022.2;refGene_gn=NEBL;refGene_tx=XM_019834024.2;refGene_gn=NEBL;refGene_tx=XM_019834025.2;refGene_gn=NEBL;refGene_tx=XM_019834026.2;refGene_gn=NEBL;refGene_tx=XM_019834027.2
B4 19639479 19639590
refGene_gn=NEBL;refGene_tx=XM_019834015.2;refGene_gn=NEBL;refGene_tx=XM_019834016.2;refGene_gn=NEBL;refGene_tx=XM_019834017.2;refGene_gn=NEBL;refGene_tx=XM_019834018.2;refGene_gn=NEBL;refGene_tx=XM_019834019.2;refGene_gn=NEBL;refGene_tx=XM_019834020.2;refGene_gn=NEBL;refGene_tx=XM_019834021.2;refGene_gn=NEBL;refGene_tx=XM_019834022.2;refGene_gn=NEBL;refGene_tx=XM_019834024.2;refGene_gn=NEBL;refGene_tx=XM_019834025.2;refGene_gn=NEBL;refGene_tx=XM_019834026.2;refGene_gn=NEBL;refGene_tx=XM_019834027.2
B4 19646386 19646497
refGene_gn=NEBL;refGene_tx=XM_019834015.2;refGene_gn=NEBL;refGene_tx=XM_019834016.2;refGene_gn=NEBL;refGene_tx=XM_019834017.2;refGene_gn=NEBL;refGene_tx=XM_019834018.2;refGene_gn=NEBL;refGene_tx=XM_019834019.2;refGene_gn=NEBL;refGene_tx=XM_019834020.2;refGene_gn=NEBL;refGene_tx=XM_019834021.2;refGene_gn=NEBL;refGene_tx=XM_019834022.2;refGene_gn=NEBL;refGene_tx=XM_019834024.2;refGene_gn=NEBL;refGene_tx=XM_019834025.2;refGene_gn=NEBL;refGene_tx=XM_019834026.2;refGene_gn=NEBL;refGene_tx=XM_019834027.2
B4 19648384 19648489
refGene_gn=NEBL;refGene_tx=XM_019834015.2;refGene_gn=NEBL;refGene_tx=XM_019834016.2;refGene_gn=NEBL;refGene_tx=XM_019834017.2;refGene_gn=NEBL;refGene_tx=XM_019834018.2;refGene_gn=NEBL;refGene_tx=XM_019834019.2;refGene_gn=NEBL;refGene_tx=XM_019834020.2;refGene_gn=NEBL;refGene_tx=XM_019834021.2;refGene_gn=NEBL;refGene_tx=XM_019834022.2;refGene_gn=NEBL;refGene_tx=XM_019834024.2;refGene_gn=NEBL;refGene_tx=XM_019834025.2;refGene_gn=NEBL;refGene_tx=XM_019834026.2;refGene_gn=NEBL;refGene_tx=XM_019834027.2
B4 19655149 19655221
refGene_gn=NEBL;refGene_tx=XM_019834015.2;refGene_gn=NEBL;refGene_tx=XM_019834016.2;refGene_gn=NEBL;refGene_tx=XM_019834017.2;refGene_gn=NEBL;refGene_tx=XM_019834018.2;refGene_gn=NEBL;refGene_tx=XM_019834019.2;

refGene_gn=NEBL;refGene_tx=XM_019834020.2;refGene_gn=NEBL;refGene_tx=XM_019834021.2;refGene_gn=NEBL;refGene_tx=XM_019834022.2;refGene_gn=NEBL;refGene_tx=XM_019834024.2;refGene_gn=NEBL;refGene_tx=XM_019834025.2;refGene_gn=NEBL;refGene_tx=XM_019834026.2;refGene_gn=NEBL;refGene_tx=XM_019834027.2
B4 19655312 19655390
refGene_gn=NEBL;refGene_tx=XM_019834015.2;refGene_gn=NEBL;refGene_tx=XM_019834016.2;refGene_gn=NEBL;refGene_tx=XM_019834017.2;refGene_gn=NEBL;refGene_tx=XM_019834018.2;refGene_gn=NEBL;refGene_tx=XM_019834019.2;refGene_gn=NEBL;refGene_tx=XM_019834020.2;refGene_gn=NEBL;refGene_tx=XM_019834024.2;refGene_gn=NEBL;refGene_tx=XM_019834025.2;refGene_gn=NEBL;refGene_tx=XM_019834026.2;refGene_gn=NEBL;refGene_tx=XM_019834027.2
B4 19715667 19715775 refGene_gn=NEBL;refGene_tx=XM_019834028.2
B4 19781244 19781329 refGene_gn=NEBL;refGene_tx=XM_019834028.2
B4 19864696 19864894
refGene_gn=CB4H10orf113;refGene_tx=XM_023257679.1
B4 19864896 19865034
refGene_gn=CB4H10orf113;refGene_tx=XM_023257679.1
B4 19882626 19882845
refGene_gn=CB4H10orf113;refGene_tx=XM_023257679.1
B4 19883016 19883025
refGene_gn=CB4H10orf113;refGene_tx=XM_023257679.1
B4 19907683 19907778 refGene_gn=NEBL;refGene_tx=XM_019834028.2
B4 19909021 19909090 refGene_gn=NEBL;refGene_tx=XM_019834028.2
B4 20160537 20160826 refGene_gn=CASC10;refGene_tx=XM_023256415.1
B4 20161695 20161796 refGene_gn=CASC10;refGene_tx=XM_023256415.1
B4 20172023 20172043 refGene_gn=SKIDA1;refGene_tx=XM_023256420.1
B4 20179377 20182241
refGene_gn=SKIDA1;refGene_tx=XM_023256416.1;refGene_gn=SKIDA1;refGene_tx=XM_023256417.1;refGene_gn=SKIDA1;refGene_tx=XM_023256418.1;refGene_gn=SKIDA1;refGene_tx=XM_023256419.1;refGene_gn=SKIDA1;refGene_tx=XM_023256420.1
B4 20182352 20182410
refGene_gn=SKIDA1;refGene_tx=XM_023256416.1;refGene_gn=SKIDA1;refGene_tx=XM_023256417.1;refGene_gn=SKIDA1;refGene_tx=XM_023256419.1;refGene_gn=SKIDA1;refGene_tx=XM_023256420.1
B4 20199306 20199466
refGene_gn=MLLT10;refGene_tx=XM_006933291.3;refGene_gn=MLLT10;refGene_tx=XM_006933292.3;refGene_gn=MLLT10;refGene_tx=XM_006933294.3;refGene_gn=MLLT10;refGene_tx=XM_006933296.3;refGene_gn=MLLT10;refGene_tx=XM_006933297.3;refGene_gn=MLLT10;refGene_tx=XM_006933298.3;refGene_gn=MLLT10;refGene_tx=XM_019834031.2;refGene_gn=MLLT10;refGene_tx=XM_019834032.1;refGene_gn=MLLT10;refGene_tx=XM_019834033.1;refGene_gn=MLLT10;refGene_tx=XM_019834034.1;refGene_gn=MLLT10;refGene_tx=XM_019834035.1;refGene_gn=MLLT10;refGene_tx=XM_019834038.1;refGene_gn=MLLT10;refGene_tx=XM_019834039.1;refGene_gn=MLLT10;refGene_tx=XM_019834040.1;refGene_gn=MLLT10;refGene_tx=XM_019834042.1
B4 20202716 20202796
refGene_gn=MLLT10;refGene_tx=XM_006933291.3;refGene_gn=MLLT10;refGene_tx=XM_006933292.3;refGene_gn=MLLT10;refGene_tx=XM_006933294.3;refGene_

gn=MLLT10;refGene_tx=XM_006933296.3;refGene_gn=MLLT10;refGene_tx=XM_00
6933297.3;refGene_gn=MLLT10;refGene_tx=XM_006933298.3;refGene_gn=MLLT1
0;refGene_tx=XM_019834031.2;refGene_gn=MLLT10;refGene_tx=XM_019834032.
1;refGene_gn=MLLT10;refGene_tx=XM_019834033.1;refGene_gn=MLLT10;refGen
e_tx=XM_019834034.1;refGene_gn=MLLT10;refGene_tx=XM_019834035.1;refGen
e_gn=MLLT10;refGene_tx=XM_019834038.1;refGene_gn=MLLT10;refGene_tx=XM_
019834039.1;refGene_gn=MLLT10;refGene_tx=XM_019834040.1;refGene_gn=MLL
T10;refGene_tx=XM_019834042.1
B4 20242925 20242980
refGene_gn=MLLT10;refGene_tx=XM_006933291.3;refGene_gn=MLLT10;refGene_
tx=XM_006933292.3;refGene_gn=MLLT10;refGene_tx=XM_006933294.3;refGene_
gn=MLLT10;refGene_tx=XM_006933296.3;refGene_gn=MLLT10;refGene_tx=XM_00
6933297.3;refGene_gn=MLLT10;refGene_tx=XM_006933298.3;refGene_gn=MLLT1
0;refGene_tx=XM_019834031.2;refGene_gn=MLLT10;refGene_tx=XM_019834032.
1;refGene_gn=MLLT10;refGene_tx=XM_019834033.1;refGene_gn=MLLT10;refGen
e_tx=XM_019834034.1;refGene_gn=MLLT10;refGene_tx=XM_019834035.1;refGen
e_gn=MLLT10;refGene_tx=XM_019834038.1;refGene_gn=MLLT10;refGene_tx=XM_
019834039.1;refGene_gn=MLLT10;refGene_tx=XM_019834040.1;refGene_gn=MLL
T10;refGene_tx=XM_019834042.1
B4 20285313 20285423
refGene_gn=MLLT10;refGene_tx=XM_006933291.3;refGene_gn=MLLT10;refGene_
tx=XM_006933292.3;refGene_gn=MLLT10;refGene_tx=XM_006933294.3;refGene_
gn=MLLT10;refGene_tx=XM_006933296.3;refGene_gn=MLLT10;refGene_tx=XM_00
6933297.3;refGene_gn=MLLT10;refGene_tx=XM_006933298.3;refGene_gn=MLLT1
0;refGene_tx=XM_011283629.3;refGene_gn=MLLT10;refGene_tx=XM_019834031.
2;refGene_gn=MLLT10;refGene_tx=XM_019834032.1;refGene_gn=MLLT10;refGen
e_tx=XM_019834033.1;refGene_gn=MLLT10;refGene_tx=XM_019834034.1;refGen
e_gn=MLLT10;refGene_tx=XM_019834035.1;refGene_gn=MLLT10;refGene_tx=XM_
019834038.1;refGene_gn=MLLT10;refGene_tx=XM_019834039.1;refGene_gn=MLL
T10;refGene_tx=XM_019834040.1;refGene_gn=MLLT10;refGene_tx=XM_01983404
2.1;refGene_gn=MLLT10;refGene_tx=XM_019834043.2
B4 20311498 20311602
refGene_gn=MLLT10;refGene_tx=XM_006933291.3;refGene_gn=MLLT10;refGene_
tx=XM_006933292.3;refGene_gn=MLLT10;refGene_tx=XM_006933294.3;refGene_
gn=MLLT10;refGene_tx=XM_006933296.3;refGene_gn=MLLT10;refGene_tx=XM_00
6933297.3;refGene_gn=MLLT10;refGene_tx=XM_006933298.3;refGene_gn=MLLT1
0;refGene_tx=XM_011283629.3;refGene_gn=MLLT10;refGene_tx=XM_019834031.
2;refGene_gn=MLLT10;refGene_tx=XM_019834032.1;refGene_gn=MLLT10;refGen
e_tx=XM_019834033.1;refGene_gn=MLLT10;refGene_tx=XM_019834034.1;refGen
e_gn=MLLT10;refGene_tx=XM_019834035.1;refGene_gn=MLLT10;refGene_tx=XM_
019834038.1;refGene_gn=MLLT10;refGene_tx=XM_019834039.1;refGene_gn=MLL
T10;refGene_tx=XM_019834040.1;refGene_gn=MLLT10;refGene_tx=XM_01983404
2.1;refGene_gn=MLLT10;refGene_tx=XM_019834043.2
B4 20313010 20313104
refGene_gn=MLLT10;refGene_tx=XM_006933291.3;refGene_gn=MLLT10;refGene_
tx=XM_006933292.3;refGene_gn=MLLT10;refGene_tx=XM_006933294.3;refGene_
gn=MLLT10;refGene_tx=XM_006933296.3;refGene_gn=MLLT10;refGene_tx=XM_00
6933297.3;refGene_gn=MLLT10;refGene_tx=XM_006933298.3;refGene_gn=MLLT1
0;refGene_tx=XM_011283629.3;refGene_gn=MLLT10;refGene_tx=XM_019834031.
2;refGene_gn=MLLT10;refGene_tx=XM_019834032.1;refGene_gn=MLLT10;refGen

e_tx=XM_019834033.1;refGene_gn=MLLT10;refGene_tx=XM_019834034.1;refGen
e_gn=MLLT10;refGene_tx=XM_019834035.1;refGene_gn=MLLT10;refGene_tx=XM_
019834038.1;refGene_gn=MLLT10;refGene_tx=XM_019834039.1;refGene_gn=MLL
T10;refGene_tx=XM_019834040.1;refGene_gn=MLLT10;refGene_tx=XM_01983404
2.1;refGene_gn=MLLT10;refGene_tx=XM_019834043.2
B4 20314967 20315063
refGene_gn=MLLT10;refGene_tx=XM_006933291.3;refGene_gn=MLLT10;refGene_
tx=XM_006933292.3;refGene_gn=MLLT10;refGene_tx=XM_006933294.3;refGene_
gn=MLLT10;refGene_tx=XM_006933297.3;refGene_gn=MLLT10;refGene_tx=XM_00
6933298.3;refGene_gn=MLLT10;refGene_tx=XM_011283629.3;refGene_gn=MLLT1
0;refGene_tx=XM_019834031.2;refGene_gn=MLLT10;refGene_tx=XM_019834032.
1;refGene_gn=MLLT10;refGene_tx=XM_019834033.1;refGene_gn=MLLT10;refGen
e_tx=XM_019834034.1;refGene_gn=MLLT10;refGene_tx=XM_019834035.1;refGen
e_gn=MLLT10;refGene_tx=XM_019834038.1;refGene_gn=MLLT10;refGene_tx=XM_
019834039.1;refGene_gn=MLLT10;refGene_tx=XM_019834040.1;refGene_gn=MLL
T10;refGene_tx=XM_019834042.1;refGene_gn=MLLT10;refGene_tx=XM_01983404
3.2
B4 20342252 20342348
refGene_gn=MLLT10;refGene_tx=XM_006933291.3;refGene_gn=MLLT10;refGene_
tx=XM_006933292.3;refGene_gn=MLLT10;refGene_tx=XM_006933294.3;refGene_
gn=MLLT10;refGene_tx=XM_006933296.3;refGene_gn=MLLT10;refGene_tx=XM_00
6933297.3;refGene_gn=MLLT10;refGene_tx=XM_006933298.3;refGene_gn=MLLT1
0;refGene_tx=XM_011283629.3;refGene_gn=MLLT10;refGene_tx=XM_019834031.
2;refGene_gn=MLLT10;refGene_tx=XM_019834032.1;refGene_gn=MLLT10;refGen
e_tx=XM_019834033.1;refGene_gn=MLLT10;refGene_tx=XM_019834034.1;refGen
e_gn=MLLT10;refGene_tx=XM_019834035.1;refGene_gn=MLLT10;refGene_tx=XM_
019834038.1;refGene_gn=MLLT10;refGene_tx=XM_019834039.1;refGene_gn=MLL
T10;refGene_tx=XM_019834040.1;refGene_gn=MLLT10;refGene_tx=XM_01983404
2.1;refGene_gn=MLLT10;refGene_tx=XM_019834043.2
B4 20364347 20364600
refGene_gn=MLLT10;refGene_tx=XM_006933291.3;refGene_gn=MLLT10;refGene_
tx=XM_006933292.3;refGene_gn=MLLT10;refGene_tx=XM_006933294.3;refGene_
gn=MLLT10;refGene_tx=XM_006933296.3;refGene_gn=MLLT10;refGene_tx=XM_00
6933297.3;refGene_gn=MLLT10;refGene_tx=XM_006933298.3;refGene_gn=MLLT1
0;refGene_tx=XM_011283629.3;refGene_gn=MLLT10;refGene_tx=XM_019834031.
2;refGene_gn=MLLT10;refGene_tx=XM_019834032.1;refGene_gn=MLLT10;refGen
e_tx=XM_019834033.1;refGene_gn=MLLT10;refGene_tx=XM_019834034.1;refGen
e_gn=MLLT10;refGene_tx=XM_019834035.1;refGene_gn=MLLT10;refGene_tx=XM_
019834038.1;refGene_gn=MLLT10;refGene_tx=XM_019834039.1;refGene_gn=MLL
T10;refGene_tx=XM_019834040.1;refGene_gn=MLLT10;refGene_tx=XM_01983404
2.1;refGene_gn=MLLT10;refGene_tx=XM_019834043.2
B4 20367917 20368487
refGene_gn=MLLT10;refGene_tx=XM_006933291.3;refGene_gn=MLLT10;refGene_
tx=XM_006933292.3;refGene_gn=MLLT10;refGene_tx=XM_006933294.3;refGene_
gn=MLLT10;refGene_tx=XM_006933296.3;refGene_gn=MLLT10;refGene_tx=XM_00
6933297.3;refGene_gn=MLLT10;refGene_tx=XM_006933298.3;refGene_gn=MLLT1
0;refGene_tx=XM_011283629.3;refGene_gn=MLLT10;refGene_tx=XM_019834031.
2;refGene_gn=MLLT10;refGene_tx=XM_019834032.1;refGene_gn=MLLT10;refGen
e_tx=XM_019834033.1;refGene_gn=MLLT10;refGene_tx=XM_019834034.1;refGen
e_gn=MLLT10;refGene_tx=XM_019834035.1;refGene_gn=MLLT10;refGene_tx=XM_

019834043.2
B4 20376349 20376394
refGene_gn=MLLT10;refGene_tx=XM_006933291.3;refGene_gn=MLLT10;refGene_tx=XM_006933292.3;refGene_gn=MLLT10;refGene_tx=XM_006933294.3;refGene_gn=MLLT10;refGene_tx=XM_006933296.3;refGene_gn=MLLT10;refGene_tx=XM_006933298.3;refGene_gn=MLLT10;refGene_tx=XM_011283629.3;refGene_gn=MLLT10;refGene_tx=XM_019834031.2;refGene_gn=MLLT10;refGene_tx=XM_019834032.1;refGene_gn=MLLT10;refGene_tx=XM_019834033.1;refGene_gn=MLLT10;refGene_tx=XM_019834035.1;refGene_gn=MLLT10;refGene_tx=XM_019834038.1;refGene_gn=MLLT10;refGene_tx=XM_019834039.1;refGene_gn=MLLT10;refGene_tx=XM_019834040.1;refGene_gn=MLLT10;refGene_tx=XM_019834043.2
B4 20377239 20377272
refGene_gn=MLLT10;refGene_tx=XM_006933291.3;refGene_gn=MLLT10;refGene_tx=XM_006933292.3;refGene_gn=MLLT10;refGene_tx=XM_006933294.3;refGene_gn=MLLT10;refGene_tx=XM_006933296.3;refGene_gn=MLLT10;refGene_tx=XM_006933298.3;refGene_gn=MLLT10;refGene_tx=XM_011283629.3;refGene_gn=MLLT10;refGene_tx=XM_019834031.2;refGene_gn=MLLT10;refGene_tx=XM_019834032.1;refGene_gn=MLLT10;refGene_tx=XM_019834038.1;refGene_gn=MLLT10;refGene_tx=XM_019834040.1;refGene_gn=MLLT10;refGene_tx=XM_019834043.2
B4 20390540 20390588
refGene_gn=MLLT10;refGene_tx=XM_006933291.3;refGene_gn=MLLT10;refGene_tx=XM_006933292.3;refGene_gn=MLLT10;refGene_tx=XM_006933296.3;refGene_gn=MLLT10;refGene_tx=XM_006933298.3;refGene_gn=MLLT10;refGene_tx=XM_011283629.3;refGene_gn=MLLT10;refGene_tx=XM_019834031.2;refGene_gn=MLLT10;refGene_tx=XM_019834032.1;refGene_gn=MLLT10;refGene_tx=XM_019834033.1;refGene_gn=MLLT10;refGene_tx=XM_019834034.1;refGene_gn=MLLT10;refGene_tx=XM_019834038.1;refGene_gn=MLLT10;refGene_tx=XM_019834039.1;refGene_gn=MLLT10;refGene_tx=XM_019834043.2
B4 20409914 20410093
refGene_gn=MLLT10;refGene_tx=XM_006933291.3;refGene_gn=MLLT10;refGene_tx=XM_006933292.3;refGene_gn=MLLT10;refGene_tx=XM_006933294.3;refGene_gn=MLLT10;refGene_tx=XM_006933296.3;refGene_gn=MLLT10;refGene_tx=XM_006933297.3;refGene_gn=MLLT10;refGene_tx=XM_006933298.3;refGene_gn=MLLT10;refGene_tx=XM_011283629.3;refGene_gn=MLLT10;refGene_tx=XM_019834031.2;refGene_gn=MLLT10;refGene_tx=XM_019834032.1;refGene_gn=MLLT10;refGene_tx=XM_019834033.1;refGene_gn=MLLT10;refGene_tx=XM_019834034.1;refGene_gn=MLLT10;refGene_tx=XM_019834035.1;refGene_gn=MLLT10;refGene_tx=XM_019834038.1;refGene_gn=MLLT10;refGene_tx=XM_019834039.1;refGene_gn=MLLT10;refGene_tx=XM_019834040.1;refGene_gn=MLLT10;refGene_tx=XM_019834042.1;refGene_gn=MLLT10;refGene_tx=XM_019834043.2
B4 20421215 20421333
refGene_gn=MLLT10;refGene_tx=XM_006933291.3;refGene_gn=MLLT10;refGene_tx=XM_006933292.3;refGene_gn=MLLT10;refGene_tx=XM_006933294.3;refGene_gn=MLLT10;refGene_tx=XM_006933296.3;refGene_gn=MLLT10;refGene_tx=XM_006933297.3;refGene_gn=MLLT10;refGene_tx=XM_006933298.3;refGene_gn=MLLT10;refGene_tx=XM_011283629.3;refGene_gn=MLLT10;refGene_tx=XM_019834031.2;refGene_gn=MLLT10;refGene_tx=XM_019834032.1;refGene_gn=MLLT10;refGene_tx=XM_019834033.1;refGene_gn=MLLT10;refGene_tx=XM_019834034.1;refGene_gn=MLLT10;refGene_tx=XM_019834035.1;refGene_gn=MLLT10;refGene_tx=XM_019834038.1;refGene_gn=MLLT10;refGene_tx=XM_019834039.1;refGene_gn=MLL

T10;refGene_tx=XM_019834040.1;refGene_gn=MLLT10;refGene_tx=XM_01983404
2.1;refGene_gn=MLLT10;refGene_tx=XM_019834043.2
B4 20422688 20422761
refGene_gn=MLLT10;refGene_tx=XM_006933291.3;refGene_gn=MLLT10;refGene_
tx=XM_006933292.3;refGene_gn=MLLT10;refGene_tx=XM_006933294.3;refGene_
gn=MLLT10;refGene_tx=XM_006933296.3;refGene_gn=MLLT10;refGene_tx=XM_00
6933297.3;refGene_gn=MLLT10;refGene_tx=XM_006933298.3;refGene_gn=MLLT1
0;refGene_tx=XM_011283629.3;refGene_gn=MLLT10;refGene_tx=XM_019834031.
2;refGene_gn=MLLT10;refGene_tx=XM_019834032.1;refGene_gn=MLLT10;refGen
e_tx=XM_019834033.1;refGene_gn=MLLT10;refGene_tx=XM_019834034.1;refGen
e_gn=MLLT10;refGene_tx=XM_019834035.1;refGene_gn=MLLT10;refGene_tx=XM_
019834038.1;refGene_gn=MLLT10;refGene_tx=XM_019834039.1;refGene_gn=MLL
T10;refGene_tx=XM_019834040.1;refGene_gn=MLLT10;refGene_tx=XM_01983404
2.1;refGene_gn=MLLT10;refGene_tx=XM_019834043.2
B4 20425534 20425689
refGene_gn=MLLT10;refGene_tx=XM_006933291.3;refGene_gn=MLLT10;refGene_
tx=XM_006933292.3;refGene_gn=MLLT10;refGene_tx=XM_006933294.3;refGene_
gn=MLLT10;refGene_tx=XM_006933296.3;refGene_gn=MLLT10;refGene_tx=XM_00
6933297.3;refGene_gn=MLLT10;refGene_tx=XM_006933298.3;refGene_gn=MLLT1
0;refGene_tx=XM_011283629.3;refGene_gn=MLLT10;refGene_tx=XM_019834031.
2;refGene_gn=MLLT10;refGene_tx=XM_019834032.1;refGene_gn=MLLT10;refGen
e_tx=XM_019834033.1;refGene_gn=MLLT10;refGene_tx=XM_019834034.1;refGen
e_gn=MLLT10;refGene_tx=XM_019834035.1;refGene_gn=MLLT10;refGene_tx=XM_
019834038.1;refGene_gn=MLLT10;refGene_tx=XM_019834039.1;refGene_gn=MLL
T10;refGene_tx=XM_019834040.1;refGene_gn=MLLT10;refGene_tx=XM_01983404
2.1;refGene_gn=MLLT10;refGene_tx=XM_019834043.2
B4 20427674 20427863
refGene_gn=MLLT10;refGene_tx=XM_006933291.3;refGene_gn=MLLT10;refGene_
tx=XM_006933292.3;refGene_gn=MLLT10;refGene_tx=XM_006933294.3;refGene_
gn=MLLT10;refGene_tx=XM_006933296.3;refGene_gn=MLLT10;refGene_tx=XM_00
6933297.3;refGene_gn=MLLT10;refGene_tx=XM_006933298.3;refGene_gn=MLLT1
0;refGene_tx=XM_011283629.3;refGene_gn=MLLT10;refGene_tx=XM_019834031.
2;refGene_gn=MLLT10;refGene_tx=XM_019834032.1;refGene_gn=MLLT10;refGen
e_tx=XM_019834033.1;refGene_gn=MLLT10;refGene_tx=XM_019834034.1;refGen
e_gn=MLLT10;refGene_tx=XM_019834035.1;refGene_gn=MLLT10;refGene_tx=XM_
019834038.1;refGene_gn=MLLT10;refGene_tx=XM_019834039.1;refGene_gn=MLL
T10;refGene_tx=XM_019834040.1;refGene_gn=MLLT10;refGene_tx=XM_01983404
2.1;refGene_gn=MLLT10;refGene_tx=XM_019834043.2
B4 20428291 20428380
refGene_gn=MLLT10;refGene_tx=XM_006933291.3;refGene_gn=MLLT10;refGene_
tx=XM_006933292.3;refGene_gn=MLLT10;refGene_tx=XM_006933294.3;refGene_
gn=MLLT10;refGene_tx=XM_006933296.3;refGene_gn=MLLT10;refGene_tx=XM_00
6933297.3;refGene_gn=MLLT10;refGene_tx=XM_006933298.3;refGene_gn=MLLT1
0;refGene_tx=XM_011283629.3;refGene_gn=MLLT10;refGene_tx=XM_019834031.
2;refGene_gn=MLLT10;refGene_tx=XM_019834032.1;refGene_gn=MLLT10;refGen
e_tx=XM_019834033.1;refGene_gn=MLLT10;refGene_tx=XM_019834034.1;refGen
e_gn=MLLT10;refGene_tx=XM_019834035.1;refGene_gn=MLLT10;refGene_tx=XM_
019834038.1;refGene_gn=MLLT10;refGene_tx=XM_019834039.1;refGene_gn=MLL
T10;refGene_tx=XM_019834040.1;refGene_gn=MLLT10;refGene_tx=XM_01983404
2.1;refGene_gn=MLLT10;refGene_tx=XM_019834043.2

B4 20428516 20428878
refGene_gn=MLLT10;refGene_tx=XM_006933291.3;refGene_gn=MLLT10;refGene_tx=XM_006933292.3;refGene_gn=MLLT10;refGene_tx=XM_006933294.3;refGene_gn=MLLT10;refGene_tx=XM_006933296.3;refGene_gn=MLLT10;refGene_tx=XM_006933297.3;refGene_gn=MLLT10;refGene_tx=XM_006933298.3;refGene_gn=MLLT10;refGene_tx=XM_011283629.3;refGene_gn=MLLT10;refGene_tx=XM_019834031.2;refGene_gn=MLLT10;refGene_tx=XM_019834032.1;refGene_gn=MLLT10;refGene_tx=XM_019834033.1;refGene_gn=MLLT10;refGene_tx=XM_019834034.1;refGene_gn=MLLT10;refGene_tx=XM_019834035.1;refGene_gn=MLLT10;refGene_tx=XM_019834038.1;refGene_gn=MLLT10;refGene_tx=XM_019834039.1;refGene_gn=MLLT10;refGene_tx=XM_019834040.1;refGene_gn=MLLT10;refGene_tx=XM_019834042.1;refGene_gn=MLLT10;refGene_tx=XM_019834043.2
B4 20429919 20430016
refGene_gn=MLLT10;refGene_tx=XM_006933291.3;refGene_gn=MLLT10;refGene_tx=XM_006933292.3;refGene_gn=MLLT10;refGene_tx=XM_006933294.3;refGene_gn=MLLT10;refGene_tx=XM_006933296.3;refGene_gn=MLLT10;refGene_tx=XM_006933297.3;refGene_gn=MLLT10;refGene_tx=XM_006933298.3;refGene_gn=MLLT10;refGene_tx=XM_011283629.3;refGene_gn=MLLT10;refGene_tx=XM_019834031.2;refGene_gn=MLLT10;refGene_tx=XM_019834032.1;refGene_gn=MLLT10;refGene_tx=XM_019834033.1;refGene_gn=MLLT10;refGene_tx=XM_019834034.1;refGene_gn=MLLT10;refGene_tx=XM_019834035.1;refGene_gn=MLLT10;refGene_tx=XM_019834038.1;refGene_gn=MLLT10;refGene_tx=XM_019834039.1;refGene_gn=MLLT10;refGene_tx=XM_019834040.1;refGene_gn=MLLT10;refGene_tx=XM_019834042.1;refGene_gn=MLLT10;refGene_tx=XM_019834043.2
B4 20434228 20434438
refGene_gn=MLLT10;refGene_tx=XM_006933291.3;refGene_gn=MLLT10;refGene_tx=XM_006933292.3;refGene_gn=MLLT10;refGene_tx=XM_006933294.3;refGene_gn=MLLT10;refGene_tx=XM_006933296.3;refGene_gn=MLLT10;refGene_tx=XM_006933297.3;refGene_gn=MLLT10;refGene_tx=XM_006933298.3;refGene_gn=MLLT10;refGene_tx=XM_011283629.3;refGene_gn=MLLT10;refGene_tx=XM_019834031.2;refGene_gn=MLLT10;refGene_tx=XM_019834032.1;refGene_gn=MLLT10;refGene_tx=XM_019834033.1;refGene_gn=MLLT10;refGene_tx=XM_019834034.1;refGene_gn=MLLT10;refGene_tx=XM_019834035.1;refGene_gn=MLLT10;refGene_tx=XM_019834038.1;refGene_gn=MLLT10;refGene_tx=XM_019834039.1;refGene_gn=MLLT10;refGene_tx=XM_019834040.1;refGene_gn=MLLT10;refGene_tx=XM_019834042.1;refGene_gn=MLLT10;refGene_tx=XM_019834043.2
B4 20436734 20436779
refGene_gn=MLLT10;refGene_tx=XM_006933291.3;refGene_gn=MLLT10;refGene_tx=XM_006933292.3;refGene_gn=MLLT10;refGene_tx=XM_006933294.3;refGene_gn=MLLT10;refGene_tx=XM_006933296.3;refGene_gn=MLLT10;refGene_tx=XM_006933297.3;refGene_gn=MLLT10;refGene_tx=XM_006933298.3;refGene_gn=MLLT10;refGene_tx=XM_011283629.3;refGene_gn=MLLT10;refGene_tx=XM_019834031.2;refGene_gn=MLLT10;refGene_tx=XM_019834032.1;refGene_gn=MLLT10;refGene_tx=XM_019834033.1;refGene_gn=MLLT10;refGene_tx=XM_019834034.1;refGene_gn=MLLT10;refGene_tx=XM_019834035.1;refGene_gn=MLLT10;refGene_tx=XM_019834038.1;refGene_gn=MLLT10;refGene_tx=XM_019834039.1;refGene_gn=MLLT10;refGene_tx=XM_019834040.1;refGene_gn=MLLT10;refGene_tx=XM_019834042.1;refGene_gn=MLLT10;refGene_tx=XM_019834043.2
B4 20458486 20458555 refGene_gn=DNAJC1;refGene_tx=XM_003988145.5
B4 20461002 20461463 refGene_gn=DNAJC1;refGene_tx=XM_003988145.5

B4 20462734 20462783 refGene_gn=DNAJC1;refGene_tx=XM_003988145.5
B4 20491101 20491221 refGene_gn=DNAJC1;refGene_tx=XM_003988145.5
B4 20562371 20562535 refGene_gn=DNAJC1;refGene_tx=XM_003988145.5
B4 20583918 20584009 refGene_gn=DNAJC1;refGene_tx=XM_003988145.5
B4 20597418 20597512 refGene_gn=DNAJC1;refGene_tx=XM_003988145.5
B4 20598481 20598579 refGene_gn=DNAJC1;refGene_tx=XM_003988145.5
B4 20599472 20599638 refGene_gn=DNAJC1;refGene_tx=XM_003988145.5
B4 20606050 20606097 refGene_gn=DNAJC1;refGene_tx=XM_003988145.5
B4 20606586 20606688 refGene_gn=DNAJC1;refGene_tx=XM_003988145.5
B4 20682044 20682257 refGene_gn=DNAJC1;refGene_tx=XM_003988145.5
B4 20989556 20989695
refGene_gn=COMMD3;refGene_tx=XM_003988146.4;refGene_gn=COMMD3;refGene_tx=XM_019834045.2
B4 20991032 20991144
refGene_gn=COMMD3;refGene_tx=XM_003988146.4;refGene_gn=COMMD3;refGene_tx=XM_019834045.2
B4 20991244 20991433
refGene_gn=COMMD3;refGene_tx=XM_003988146.4;refGene_gn=COMMD3;refGene_tx=XM_019834045.2
B4 20991797 20991858
refGene_gn=COMMD3;refGene_tx=XM_003988146.4;refGene_gn=COMMD3;refGene_tx=XM_019834045.2
B4 20991946 20992003
refGene_gn=COMMD3;refGene_tx=XM_003988146.4;refGene_gn=COMMD3;refGene_tx=XM_019834045.2
B4 20992103 20992163
refGene_gn=COMMD3;refGene_tx=XM_003988146.4;refGene_gn=COMMD3;refGene_tx=XM_019834045.2
B4 20993087 20993147
refGene_gn=COMMD3;refGene_tx=XM_003988146.4;refGene_gn=COMMD3;refGene_tx=XM_019834045.2
B4 20999861 20999973
refGene_gn=BMI1;refGene_tx=NM_001024962.1;refGene_gn=BMI1;refGene_tx=XM_006933246.4;refGene_gn=BMI1;refGene_tx=XM_006933247.3;refGene_gn=BMI1;refGene_tx=XM_006933248.3;refGene_gn=BMI1;refGene_tx=XM_011283607.3;refGene_gn=BMI1;refGene_tx=XM_023256249.1
B4 21000321 21000418
refGene_gn=BMI1;refGene_tx=NM_001024962.1;refGene_gn=BMI1;refGene_tx=XM_006933246.4;refGene_gn=BMI1;refGene_tx=XM_006933247.3;refGene_gn=BMI1;refGene_tx=XM_006933248.3;refGene_gn=BMI1;refGene_tx=XM_011283607.3;refGene_gn=BMI1;refGene_tx=XM_023256249.1
B4 21001022 21001078
refGene_gn=BMI1;refGene_tx=NM_001024962.1;refGene_gn=BMI1;refGene_tx=XM_006933246.4;refGene_gn=BMI1;refGene_tx=XM_006933247.3;refGene_gn=BMI1;refGene_tx=XM_006933248.3;refGene_gn=BMI1;refGene_tx=XM_011283607.3;refGene_gn=BMI1;refGene_tx=XM_023256249.1
B4 21001171 21001222
refGene_gn=BMI1;refGene_tx=NM_001024962.1;refGene_gn=BMI1;refGene_tx=XM_006933246.4;refGene_gn=BMI1;refGene_tx=XM_006933247.3;refGene_gn=BMI1;refGene_tx=XM_006933248.3;refGene_gn=BMI1;refGene_tx=XM_011283607.3;

refGene_gn=BMI1;refGene_tx=XM_023256249.1
B4 21001386 21001495
refGene_gn=BMI1;refGene_tx=NM_001024962.1;refGene_gn=BMI1;refGene_tx=XM_006933246.4;refGene_gn=BMI1;refGene_tx=XM_006933247.3;refGene_gn=BMI1;refGene_tx=XM_006933248.3;refGene_gn=BMI1;refGene_tx=XM_011283607.3;refGene_gn=BMI1;refGene_tx=XM_023256249.1
B4 21001573 21001619
refGene_gn=BMI1;refGene_tx=NM_001024962.1;refGene_gn=BMI1;refGene_tx=XM_006933246.4;refGene_gn=BMI1;refGene_tx=XM_006933247.3;refGene_gn=BMI1;refGene_tx=XM_006933248.3;refGene_gn=BMI1;refGene_tx=XM_011283607.3;refGene_gn=BMI1;refGene_tx=XM_023256249.1
B4 21002052 21002151
refGene_gn=BMI1;refGene_tx=NM_001024962.1;refGene_gn=BMI1;refGene_tx=XM_006933246.4;refGene_gn=BMI1;refGene_tx=XM_006933247.3;refGene_gn=BMI1;refGene_tx=XM_006933248.3;refGene_gn=BMI1;refGene_tx=XM_011283607.3;refGene_gn=BMI1;refGene_tx=XM_023256249.1
B4 21002457 21002538
refGene_gn=BMI1;refGene_tx=NM_001024962.1;refGene_gn=BMI1;refGene_tx=XM_006933246.4;refGene_gn=BMI1;refGene_tx=XM_006933247.3;refGene_gn=BMI1;refGene_tx=XM_006933248.3;refGene_gn=BMI1;refGene_tx=XM_011283607.3;refGene_gn=BMI1;refGene_tx=XM_023256249.1
B4 21002630 21002960
refGene_gn=BMI1;refGene_tx=NM_001024962.1;refGene_gn=BMI1;refGene_tx=XM_006933246.4;refGene_gn=BMI1;refGene_tx=XM_006933247.3;refGene_gn=BMI1;refGene_tx=XM_006933248.3;refGene_gn=BMI1;refGene_tx=XM_011283607.3;refGene_gn=BMI1;refGene_tx=XM_023256249.1
B4 21019170 21019195
refGene_gn=SPAG6;refGene_tx=XM_003988147.4;refGene_gn=SPAG6;refGene_tx=XM_019834047.2;refGene_gn=SPAG6;refGene_tx=XM_023256422.1
B4 21019280 21019376
refGene_gn=SPAG6;refGene_tx=XM_003988147.4;refGene_gn=SPAG6;refGene_tx=XM_019834047.2;refGene_gn=SPAG6;refGene_tx=XM_023256422.1
B4 21035005 21035057 refGene_gn=SPAG6;refGene_tx=XM_019834046.1
B4 21038909 21039076
refGene_gn=SPAG6;refGene_tx=XM_003988147.4;refGene_gn=SPAG6;refGene_tx=XM_019834046.1;refGene_gn=SPAG6;refGene_tx=XM_019834047.2;refGene_gn=SPAG6;refGene_tx=XM_023256421.1;refGene_gn=SPAG6;refGene_tx=XM_023256422.1
B4 21042698 21042882
refGene_gn=SPAG6;refGene_tx=XM_003988147.4;refGene_gn=SPAG6;refGene_tx=XM_019834046.1;refGene_gn=SPAG6;refGene_tx=XM_019834047.2;refGene_gn=SPAG6;refGene_tx=XM_023256421.1;refGene_gn=SPAG6;refGene_tx=XM_023256422.1
B4 21061501 21061707
refGene_gn=SPAG6;refGene_tx=XM_003988147.4;refGene_gn=SPAG6;refGene_tx=XM_019834046.1;refGene_gn=SPAG6;refGene_tx=XM_019834047.2;refGene_gn=SPAG6;refGene_tx=XM_023256421.1;refGene_gn=SPAG6;refGene_tx=XM_023256422.1
B4 21062657 21062831
refGene_gn=SPAG6;refGene_tx=XM_003988147.4;refGene_gn=SPAG6;refGene_tx

=XM_019834046.1;refGene_gn=SPAG6;refGene_tx=XM_019834047.2;refGene_gn=SPAG6;refGene_tx=XM_023256421.1;refGene_gn=SPAG6;refGene_tx=XM_023256422.1
B4 21064421 21064574
refGene_gn=SPAG6;refGene_tx=XM_003988147.4;refGene_gn=SPAG6;refGene_tx=XM_019834046.1;refGene_gn=SPAG6;refGene_tx=XM_019834047.2;refGene_gn=SPAG6;refGene_tx=XM_023256421.1;refGene_gn=SPAG6;refGene_tx=XM_023256422.1
B4 21068641 21068833
refGene_gn=SPAG6;refGene_tx=XM_003988147.4;refGene_gn=SPAG6;refGene_tx=XM_019834046.1;refGene_gn=SPAG6;refGene_tx=XM_023256421.1;refGene_gn=SPAG6;refGene_tx=XM_023256422.1
B4 21077372 21077489
refGene_gn=SPAG6;refGene_tx=XM_003988147.4;refGene_gn=SPAG6;refGene_tx=XM_019834046.1;refGene_gn=SPAG6;refGene_tx=XM_019834047.2;refGene_gn=SPAG6;refGene_tx=XM_023256421.1
B4 21079892 21080038
refGene_gn=SPAG6;refGene_tx=XM_003988147.4;refGene_gn=SPAG6;refGene_tx=XM_019834046.1;refGene_gn=SPAG6;refGene_tx=XM_019834047.2;refGene_gn=SPAG6;refGene_tx=XM_023256421.1
B4 21084475 21084545
refGene_gn=SPAG6;refGene_tx=XM_003988147.4;refGene_gn=SPAG6;refGene_tx=XM_019834046.1;refGene_gn=SPAG6;refGene_tx=XM_019834047.2;refGene_gn=SPAG6;refGene_tx=XM_023256421.1;refGene_gn=SPAG6;refGene_tx=XM_023256422.1
B4 21105881 21106868
refGene_gn=LOC111561444;refGene_tx=XM_023257735.1
B4 21187677 21187758
refGene_gn=PIP4K2A;refGene_tx=XM_003988149.4;refGene_gn=PIP4K2A;refGene_tx=XM_019834048.2
B4 21190355 21190459
refGene_gn=PIP4K2A;refGene_tx=XM_003988149.4;refGene_gn=PIP4K2A;refGene_tx=XM_019834048.2
B4 21192535 21192779
refGene_gn=PIP4K2A;refGene_tx=XM_003988149.4;refGene_gn=PIP4K2A;refGene_tx=XM_019834048.2
B4 21201746 21201860
refGene_gn=PIP4K2A;refGene_tx=XM_003988149.4;refGene_gn=PIP4K2A;refGene_tx=XM_019834048.2
B4 21216475 21216514
refGene_gn=PIP4K2A;refGene_tx=XM_003988149.4;refGene_gn=PIP4K2A;refGene_tx=XM_019834048.2
B4 21222720 21222867
refGene_gn=PIP4K2A;refGene_tx=XM_003988149.4;refGene_gn=PIP4K2A;refGene_tx=XM_019834048.2
B4 21237033 21237186
refGene_gn=PIP4K2A;refGene_tx=XM_003988149.4;refGene_gn=PIP4K2A;refGene_tx=XM_019834048.2
B4 21254341 21254438
refGene_gn=PIP4K2A;refGene_tx=XM_003988149.4;refGene_gn=PIP4K2A;refGen

e_tx=XM_019834048.2
B4 21255938 21256036
refGene_gn=PIP4K2A;refGene_tx=XM_003988149.4;refGene_gn=PIP4K2A;refGen
e_tx=XM_019834048.2
B4 21384130 21384274 refGene_gn=PIP4K2A;refGene_tx=XM_003988149.4
B4 21564619 21564667
refGene_gn=ARMC3;refGene_tx=XM_011283631.3;refGene_gn=ARMC3;refGene_tx
=XM_011283633.3;refGene_gn=ARMC3;refGene_tx=XM_019834049.2;refGene_gn=
ARMC3;refGene_tx=XM_019834050.2;refGene_gn=ARMC3;refGene_tx=XM_0198340
51.2;refGene_gn=ARMC3;refGene_tx=XM_019834052.2;refGene_gn=ARMC3;refGe
ne_tx=XM_019834056.2;refGene_gn=ARMC3;refGene_tx=XM_019834059.2;refGen
e_gn=ARMC3;refGene_tx=XM_019834060.2;refGene_gn=ARMC3;refGene_tx=XM_01
9834065.2;refGene_gn=ARMC3;refGene_tx=XM_023256423.1
B4 21579610 21579728
refGene_gn=ARMC3;refGene_tx=XM_011283631.3;refGene_gn=ARMC3;refGene_tx
=XM_011283633.3;refGene_gn=ARMC3;refGene_tx=XM_019834049.2;refGene_gn=
ARMC3;refGene_tx=XM_019834050.2;refGene_gn=ARMC3;refGene_tx=XM_0198340
51.2;refGene_gn=ARMC3;refGene_tx=XM_019834052.2;refGene_gn=ARMC3;refGe
ne_tx=XM_019834056.2;refGene_gn=ARMC3;refGene_tx=XM_019834059.2;refGen
e_gn=ARMC3;refGene_tx=XM_019834060.2;refGene_gn=ARMC3;refGene_tx=XM_01
9834065.2;refGene_gn=ARMC3;refGene_tx=XM_023256423.1
B4 21591124 21591250
refGene_gn=ARMC3;refGene_tx=XM_011283631.3;refGene_gn=ARMC3;refGene_tx
=XM_011283633.3;refGene_gn=ARMC3;refGene_tx=XM_019834049.2;refGene_gn=
ARMC3;refGene_tx=XM_019834050.2;refGene_gn=ARMC3;refGene_tx=XM_0198340
51.2;refGene_gn=ARMC3;refGene_tx=XM_019834052.2;refGene_gn=ARMC3;refGe
ne_tx=XM_019834056.2;refGene_gn=ARMC3;refGene_tx=XM_019834058.1;refGen
e_gn=ARMC3;refGene_tx=XM_019834059.2;refGene_gn=ARMC3;refGene_tx=XM_01
9834060.2;refGene_gn=ARMC3;refGene_tx=XM_019834065.2;refGene_gn=ARMC3;
refGene_tx=XM_023256423.1
B4 21597717 21597786
refGene_gn=ARMC3;refGene_tx=XM_011283631.3;refGene_gn=ARMC3;refGene_tx
=XM_011283633.3;refGene_gn=ARMC3;refGene_tx=XM_019834049.2;refGene_gn=
ARMC3;refGene_tx=XM_019834050.2;refGene_gn=ARMC3;refGene_tx=XM_0198340
51.2;refGene_gn=ARMC3;refGene_tx=XM_019834052.2;refGene_gn=ARMC3;refGe
ne_tx=XM_019834056.2;refGene_gn=ARMC3;refGene_tx=XM_019834058.1;refGen
e_gn=ARMC3;refGene_tx=XM_019834059.2;refGene_gn=ARMC3;refGene_tx=XM_01
9834060.2;refGene_gn=ARMC3;refGene_tx=XM_019834065.2;refGene_gn=ARMC3;
refGene_tx=XM_023256423.1
B4 21598280 21598456
refGene_gn=ARMC3;refGene_tx=XM_011283631.3;refGene_gn=ARMC3;refGene_tx
=XM_011283633.3;refGene_gn=ARMC3;refGene_tx=XM_019834049.2;refGene_gn=
ARMC3;refGene_tx=XM_019834050.2;refGene_gn=ARMC3;refGene_tx=XM_0198340
51.2;refGene_gn=ARMC3;refGene_tx=XM_019834052.2;refGene_gn=ARMC3;refGe
ne_tx=XM_019834056.2;refGene_gn=ARMC3;refGene_tx=XM_019834058.1;refGen
e_gn=ARMC3;refGene_tx=XM_019834059.2;refGene_gn=ARMC3;refGene_tx=XM_01
9834060.2;refGene_gn=ARMC3;refGene_tx=XM_019834065.2;refGene_gn=ARMC3;
refGene_tx=XM_023256423.1
B4 21602830 21603025
refGene_gn=ARMC3;refGene_tx=XM_011283631.3;refGene_gn=ARMC3;refGene_tx

=XM_011283633.3;refGene_gn=ARMC3;refGene_tx=XM_019834049.2;refGene_gn=ARMC3;refGene_tx=XM_019834050.2;refGene_gn=ARMC3;refGene_tx=XM_019834051.2;refGene_gn=ARMC3;refGene_tx=XM_019834052.2;refGene_gn=ARMC3;refGene_tx=XM_019834056.2;refGene_gn=ARMC3;refGene_tx=XM_019834058.1;refGene_gn=ARMC3;refGene_tx=XM_019834059.2;refGene_gn=ARMC3;refGene_tx=XM_019834060.2;refGene_gn=ARMC3;refGene_tx=XM_019834065.2;refGene_gn=ARMC3;refGene_tx=XM_023256423.1
B4 21605686 21605870
refGene_gn=ARMC3;refGene_tx=XM_011283631.3;refGene_gn=ARMC3;refGene_tx=XM_011283633.3;refGene_gn=ARMC3;refGene_tx=XM_019834049.2;refGene_gn=ARMC3;refGene_tx=XM_019834050.2;refGene_gn=ARMC3;refGene_tx=XM_019834051.2;refGene_gn=ARMC3;refGene_tx=XM_019834052.2;refGene_gn=ARMC3;refGene_tx=XM_019834056.2;refGene_gn=ARMC3;refGene_tx=XM_019834058.1;refGene_gn=ARMC3;refGene_tx=XM_019834059.2;refGene_gn=ARMC3;refGene_tx=XM_019834060.2;refGene_gn=ARMC3;refGene_tx=XM_019834065.2;refGene_gn=ARMC3;refGene_tx=XM_023256423.1
B4 21616916 21617069
refGene_gn=ARMC3;refGene_tx=XM_011283631.3;refGene_gn=ARMC3;refGene_tx=XM_011283633.3;refGene_gn=ARMC3;refGene_tx=XM_019834049.2;refGene_gn=ARMC3;refGene_tx=XM_019834050.2;refGene_gn=ARMC3;refGene_tx=XM_019834051.2;refGene_gn=ARMC3;refGene_tx=XM_019834052.2;refGene_gn=ARMC3;refGene_tx=XM_019834056.2;refGene_gn=ARMC3;refGene_tx=XM_019834058.1;refGene_gn=ARMC3;refGene_tx=XM_019834059.2;refGene_gn=ARMC3;refGene_tx=XM_019834060.2;refGene_gn=ARMC3;refGene_tx=XM_019834065.2;refGene_gn=ARMC3;refGene_tx=XM_023256423.1
B4 21617171 21617277
refGene_gn=ARMC3;refGene_tx=XM_011283631.3;refGene_gn=ARMC3;refGene_tx=XM_011283633.3;refGene_gn=ARMC3;refGene_tx=XM_019834049.2;refGene_gn=ARMC3;refGene_tx=XM_019834050.2;refGene_gn=ARMC3;refGene_tx=XM_019834051.2;refGene_gn=ARMC3;refGene_tx=XM_019834052.2;refGene_gn=ARMC3;refGene_tx=XM_019834056.2;refGene_gn=ARMC3;refGene_tx=XM_019834058.1;refGene_gn=ARMC3;refGene_tx=XM_019834059.2;refGene_gn=ARMC3;refGene_tx=XM_019834060.2;refGene_gn=ARMC3;refGene_tx=XM_019834065.2;refGene_gn=ARMC3;refGene_tx=XM_023256423.1
B4 21635420 21635670
refGene_gn=ARMC3;refGene_tx=XM_011283631.3;refGene_gn=ARMC3;refGene_tx=XM_011283633.3;refGene_gn=ARMC3;refGene_tx=XM_019834049.2;refGene_gn=ARMC3;refGene_tx=XM_019834050.2;refGene_gn=ARMC3;refGene_tx=XM_019834051.2;refGene_gn=ARMC3;refGene_tx=XM_019834052.2;refGene_gn=ARMC3;refGene_tx=XM_019834056.2;refGene_gn=ARMC3;refGene_tx=XM_019834058.1;refGene_gn=ARMC3;refGene_tx=XM_019834059.2;refGene_gn=ARMC3;refGene_tx=XM_019834060.2;refGene_gn=ARMC3;refGene_tx=XM_019834065.2;refGene_gn=ARMC3;refGene_tx=XM_023256423.1
B4 21639125 21639262
refGene_gn=ARMC3;refGene_tx=XM_011283631.3;refGene_gn=ARMC3;refGene_tx=XM_011283633.3;refGene_gn=ARMC3;refGene_tx=XM_019834049.2;refGene_gn=ARMC3;refGene_tx=XM_019834050.2;refGene_gn=ARMC3;refGene_tx=XM_019834051.2;refGene_gn=ARMC3;refGene_tx=XM_019834052.2;refGene_gn=ARMC3;refGene_tx=XM_019834056.2;refGene_gn=ARMC3;refGene_tx=XM_019834058.1;refGene_gn=ARMC3;refGene_tx=XM_019834059.2;refGene_gn=ARMC3;refGene_tx=XM_01

9834060.2;refGene_gn=ARMC3;refGene_tx=XM_019834065.2;refGene_gn=ARMC3;refGene_tx=XM_023256423.1
B4 21640919 21641088
refGene_gn=ARMC3;refGene_tx=XM_011283631.3;refGene_gn=ARMC3;refGene_tx=XM_011283633.3;refGene_gn=ARMC3;refGene_tx=XM_019834049.2;refGene_gn=ARMC3;refGene_tx=XM_019834050.2;refGene_gn=ARMC3;refGene_tx=XM_019834051.2;refGene_gn=ARMC3;refGene_tx=XM_019834052.2;refGene_gn=ARMC3;refGene_tx=XM_019834056.2;refGene_gn=ARMC3;refGene_tx=XM_019834058.1;refGene_gn=ARMC3;refGene_tx=XM_019834059.2;refGene_gn=ARMC3;refGene_tx=XM_019834060.2;refGene_gn=ARMC3;refGene_tx=XM_019834065.2;refGene_gn=ARMC3;refGene_tx=XM_023256423.1
B4 21644710 21644811
refGene_gn=ARMC3;refGene_tx=XM_011283633.3;refGene_gn=ARMC3;refGene_tx=XM_019834049.2;refGene_gn=ARMC3;refGene_tx=XM_019834050.2;refGene_gn=ARMC3;refGene_tx=XM_019834051.2;refGene_gn=ARMC3;refGene_tx=XM_019834052.2;refGene_gn=ARMC3;refGene_tx=XM_019834056.2;refGene_gn=ARMC3;refGene_tx=XM_019834058.1;refGene_gn=ARMC3;refGene_tx=XM_019834059.2;refGene_gn=ARMC3;refGene_tx=XM_019834060.2;refGene_gn=ARMC3;refGene_tx=XM_019834065.2;refGene_gn=ARMC3;refGene_tx=XM_023256423.1
B4 21645517 21645619
refGene_gn=ARMC3;refGene_tx=XM_011283633.3;refGene_gn=ARMC3;refGene_tx=XM_019834049.2;refGene_gn=ARMC3;refGene_tx=XM_019834050.2;refGene_gn=ARMC3;refGene_tx=XM_019834051.2;refGene_gn=ARMC3;refGene_tx=XM_019834052.2;refGene_gn=ARMC3;refGene_tx=XM_019834056.2;refGene_gn=ARMC3;refGene_tx=XM_019834058.1;refGene_gn=ARMC3;refGene_tx=XM_019834059.2;refGene_gn=ARMC3;refGene_tx=XM_019834060.2;refGene_gn=ARMC3;refGene_tx=XM_019834065.2;refGene_gn=ARMC3;refGene_tx=XM_023256423.1
B4 21646308 21646425
refGene_gn=ARMC3;refGene_tx=XM_011283633.3;refGene_gn=ARMC3;refGene_tx=XM_019834049.2;refGene_gn=ARMC3;refGene_tx=XM_019834050.2;refGene_gn=ARMC3;refGene_tx=XM_019834051.2;refGene_gn=ARMC3;refGene_tx=XM_019834052.2;refGene_gn=ARMC3;refGene_tx=XM_019834058.1;refGene_gn=ARMC3;refGene_tx=XM_019834059.2;refGene_gn=ARMC3;refGene_tx=XM_019834060.2;refGene_gn=ARMC3;refGene_tx=XM_019834065.2;refGene_gn=ARMC3;refGene_tx=XM_023256423.1
B4 21654196 21654462
refGene_gn=ARMC3;refGene_tx=XM_011283631.3;refGene_gn=ARMC3;refGene_tx=XM_011283633.3;refGene_gn=ARMC3;refGene_tx=XM_019834049.2;refGene_gn=ARMC3;refGene_tx=XM_019834050.2;refGene_gn=ARMC3;refGene_tx=XM_019834051.2;refGene_gn=ARMC3;refGene_tx=XM_019834052.2;refGene_gn=ARMC3;refGene_tx=XM_019834056.2;refGene_gn=ARMC3;refGene_tx=XM_019834058.1;refGene_gn=ARMC3;refGene_tx=XM_019834059.2;refGene_gn=ARMC3;refGene_tx=XM_019834060.2;refGene_gn=ARMC3;refGene_tx=XM_019834065.2;refGene_gn=ARMC3;refGene_tx=XM_023256423.1
B4 21654545 21654591 refGene_gn=ARMC3;refGene_tx=XM_019834060.2
B4 21656861 21657024
refGene_gn=ARMC3;refGene_tx=XM_011283631.3;refGene_gn=ARMC3;refGene_tx=XM_011283633.3;refGene_gn=ARMC3;refGene_tx=XM_019834049.2;refGene_gn=ARMC3;refGene_tx=XM_019834050.2;refGene_gn=ARMC3;refGene_tx=XM_019834051.2;refGene_gn=ARMC3;refGene_tx=XM_019834052.2;refGene_gn=ARMC3;refGe

ne_tx=XM_019834056.2;refGene_gn=ARMC3;refGene_tx=XM_019834058.1;refGene_gn=ARMC3;refGene_tx=XM_023256423.1
B4 21659925 21660135
refGene_gn=ARMC3;refGene_tx=XM_011283631.3;refGene_gn=ARMC3;refGene_tx=XM_011283633.3;refGene_gn=ARMC3;refGene_tx=XM_019834049.2;refGene_gn=ARMC3;refGene_tx=XM_019834050.2;refGene_gn=ARMC3;refGene_tx=XM_019834051.2;refGene_gn=ARMC3;refGene_tx=XM_019834052.2;refGene_gn=ARMC3;refGene_tx=XM_019834056.2;refGene_gn=ARMC3;refGene_tx=XM_019834058.1;refGene_gn=ARMC3;refGene_tx=XM_019834059.2;refGene_gn=ARMC3;refGene_tx=XM_023256423.1
B4 21784222 21784663 refGene_gn=PTF1A;refGene_tx=XM_023257788.1
B4 21784664 21784916 refGene_gn=PTF1A;refGene_tx=XM_023257788.1
B4 21784917 21785011 refGene_gn=PTF1A;refGene_tx=XM_023257788.1
B4 21785405 21785608 refGene_gn=PTF1A;refGene_tx=XM_023257788.1
B4 21789521 21789586
refGene_gn=CB4H10orf67;refGene_tx=XM_019834072.2
B4 21795908 21796000
refGene_gn=CB4H10orf67;refGene_tx=XM_019834070.2;refGene_gn=CB4H10orf67;refGene_tx=XM_023256424.1
B4 21814659 21814720
refGene_gn=CB4H10orf67;refGene_tx=XM_019834070.2;refGene_gn=CB4H10orf67;refGene_tx=XM_019834072.2;refGene_gn=CB4H10orf67;refGene_tx=XM_023256424.1
B4 21814806 21814887
refGene_gn=CB4H10orf67;refGene_tx=XM_019834070.2;refGene_gn=CB4H10orf67;refGene_tx=XM_019834072.2;refGene_gn=CB4H10orf67;refGene_tx=XM_023256424.1
B4 21828385 21828473
refGene_gn=CB4H10orf67;refGene_tx=XM_019834070.2;refGene_gn=CB4H10orf67;refGene_tx=XM_019834072.2;refGene_gn=CB4H10orf67;refGene_tx=XM_023256424.1
B4 21841260 21841326
refGene_gn=CB4H10orf67;refGene_tx=XM_019834070.2;refGene_gn=CB4H10orf67;refGene_tx=XM_019834072.2;refGene_gn=CB4H10orf67;refGene_tx=XM_023256424.1
B4 21841410 21841490
refGene_gn=CB4H10orf67;refGene_tx=XM_019834070.2;refGene_gn=CB4H10orf67;refGene_tx=XM_019834072.2;refGene_gn=CB4H10orf67;refGene_tx=XM_023256424.1
B4 21842434 21842497
refGene_gn=CB4H10orf67;refGene_tx=XM_019834070.2;refGene_gn=CB4H10orf67;refGene_tx=XM_019834072.2
B4 21856965 21857142
refGene_gn=CB4H10orf67;refGene_tx=XM_019834070.2;refGene_gn=CB4H10orf67;refGene_tx=XM_019834072.2;refGene_gn=CB4H10orf67;refGene_tx=XM_023256424.1
B4 21857812 21857872
refGene_gn=CB4H10orf67;refGene_tx=XM_019834070.2;refGene_gn=CB4H10orf67;refGene_tx=XM_019834072.2;refGene_gn=CB4H10orf67;refGene_tx=XM_023256424.1

B4 21875912 21875978
refGene_gn=CB4H10orf67;refGene_tx=XM_019834070.2;refGene_gn=CB4H10orf67;refGene_tx=XM_019834072.2;refGene_gn=CB4H10orf67;refGene_tx=XM_023256424.1
B4 21884645 21884704
refGene_gn=CB4H10orf67;refGene_tx=XM_019834070.2;refGene_gn=CB4H10orf67;refGene_tx=XM_019834072.2;refGene_gn=CB4H10orf67;refGene_tx=XM_023256424.1
B4 21885743 21885876
refGene_gn=CB4H10orf67;refGene_tx=XM_019834070.2;refGene_gn=CB4H10orf67;refGene_tx=XM_019834072.2;refGene_gn=CB4H10orf67;refGene_tx=XM_023256424.1
B4 21892242 21892395
refGene_gn=CB4H10orf67;refGene_tx=XM_019834070.2;refGene_gn=CB4H10orf67;refGene_tx=XM_019834072.2;refGene_gn=CB4H10orf67;refGene_tx=XM_023256424.1
B4 21927813 21927888
refGene_gn=CB4H10orf67;refGene_tx=XM_019834070.2;refGene_gn=CB4H10orf67;refGene_tx=XM_019834072.2;refGene_gn=CB4H10orf67;refGene_tx=XM_023256424.1
B4 21929574 21929718
refGene_gn=CB4H10orf67;refGene_tx=XM_019834070.2;refGene_gn=CB4H10orf67;refGene_tx=XM_019834072.2;refGene_gn=CB4H10orf67;refGene_tx=XM_023256424.1
B4 21937191 21937312
refGene_gn=CB4H10orf67;refGene_tx=XM_019834070.2;refGene_gn=CB4H10orf67;refGene_tx=XM_019834072.2;refGene_gn=CB4H10orf67;refGene_tx=XM_023256424.1
B4 21953381 21953560
refGene_gn=CB4H10orf67;refGene_tx=XM_019834070.2;refGene_gn=CB4H10orf67;refGene_tx=XM_019834072.2;refGene_gn=CB4H10orf67;refGene_tx=XM_023256424.1
B4 22029765 22031220 refGene_gn=OTUD1;refGene_tx=XM_003988183.5
B4 22593574 22593611 refGene_gn=KIAA1217;refGene_tx=XM_023256429.1
B4 22687888 22687958
refGene_gn=KIAA1217;refGene_tx=XM_011283634.3;refGene_gn=KIAA1217;refGene_tx=XM_023256426.1;refGene_gn=KIAA1217;refGene_tx=XM_023256427.1;refGene_gn=KIAA1217;refGene_tx=XM_023256430.1;refGene_gn=KIAA1217;refGene_tx=XM_023256431.1;refGene_gn=KIAA1217;refGene_tx=XM_023256433.1;refGene_gn=KIAA1217;refGene_tx=XM_023256441.1;refGene_gn=KIAA1217;refGene_tx=XM_023256442.1;refGene_gn=KIAA1217;refGene_tx=XM_023256444.1;refGene_gn=KIAA1217;refGene_tx=XM_023256445.1;refGene_gn=KIAA1217;refGene_tx=XM_023256446.1;refGene_gn=KIAA1217;refGene_tx=XM_023256447.1;refGene_gn=KIAA1217;refGene_tx=XM_023256448.1;refGene_gn=KIAA1217;refGene_tx=XM_023256449.1;refGene_gn=KIAA1217;refGene_tx=XM_023256450.1
B4 22696136 22696420
refGene_gn=KIAA1217;refGene_tx=XM_011283634.3;refGene_gn=KIAA1217;refGene_tx=XM_023256426.1;refGene_gn=KIAA1217;refGene_tx=XM_023256427.1;refGene_gn=KIAA1217;refGene_tx=XM_023256428.1;refGene_gn=KIAA1217;refGene_tx=XM_023256429.1;refGene_gn=KIAA1217;refGene_tx=XM_023256430.1;refG

ene_gn=KIAA1217;refGene_tx=XM_023256431.1;refGene_gn=KIAA1217;refGene_
tx=XM_023256433.1;refGene_gn=KIAA1217;refGene_tx=XM_023256436.1;refGen
e_gn=KIAA1217;refGene_tx=XM_023256441.1;refGene_gn=KIAA1217;refGene_tx
=XM_023256442.1;refGene_gn=KIAA1217;refGene_tx=XM_023256444.1;refGene_
gn=KIAA1217;refGene_tx=XM_023256445.1;refGene_gn=KIAA1217;refGene_tx=X
M_023256446.1;refGene_gn=KIAA1217;refGene_tx=XM_023256447.1;refGene_gn
=KIAA1217;refGene_tx=XM_023256448.1;refGene_gn=KIAA1217;refGene_tx=XM_
023256449.1;refGene_gn=KIAA1217;refGene_tx=XM_023256450.1
B4 22831932 22832013
refGene_gn=KIAA1217;refGene_tx=XM_023256426.1;refGene_gn=KIAA1217;refG
ene_tx=XM_023256427.1;refGene_gn=KIAA1217;refGene_tx=XM_023256428.1;re
fGene_gn=KIAA1217;refGene_tx=XM_023256429.1;refGene_gn=KIAA1217;refGen
e_tx=XM_023256430.1;refGene_gn=KIAA1217;refGene_tx=XM_023256431.1;refG
ene_gn=KIAA1217;refGene_tx=XM_023256436.1;refGene_gn=KIAA1217;refGene_
tx=XM_023256441.1;refGene_gn=KIAA1217;refGene_tx=XM_023256442.1;refGen
e_gn=KIAA1217;refGene_tx=XM_023256444.1;refGene_gn=KIAA1217;refGene_tx
=XM_023256445.1;refGene_gn=KIAA1217;refGene_tx=XM_023256449.1;refGene_
gn=KIAA1217;refGene_tx=XM_023256450.1
B4 22846798 22846997
refGene_gn=KIAA1217;refGene_tx=XM_011283634.3;refGene_gn=KIAA1217;refG
ene_tx=XM_023256426.1;refGene_gn=KIAA1217;refGene_tx=XM_023256427.1;re
fGene_gn=KIAA1217;refGene_tx=XM_023256428.1;refGene_gn=KIAA1217;refGen
e_tx=XM_023256429.1;refGene_gn=KIAA1217;refGene_tx=XM_023256430.1;refG
ene_gn=KIAA1217;refGene_tx=XM_023256431.1;refGene_gn=KIAA1217;refGene_
tx=XM_023256433.1;refGene_gn=KIAA1217;refGene_tx=XM_023256435.1;refGen
e_gn=KIAA1217;refGene_tx=XM_023256436.1;refGene_gn=KIAA1217;refGene_tx
=XM_023256441.1;refGene_gn=KIAA1217;refGene_tx=XM_023256442.1;refGene_
gn=KIAA1217;refGene_tx=XM_023256444.1;refGene_gn=KIAA1217;refGene_tx=X
M_023256445.1;refGene_gn=KIAA1217;refGene_tx=XM_023256446.1;refGene_gn
=KIAA1217;refGene_tx=XM_023256447.1;refGene_gn=KIAA1217;refGene_tx=XM_
023256448.1;refGene_gn=KIAA1217;refGene_tx=XM_023256449.1;refGene_gn=K
IAA1217;refGene_tx=XM_023256450.1
B4 22890495 22890694
refGene_gn=KIAA1217;refGene_tx=XM_011283634.3;refGene_gn=KIAA1217;refG
ene_tx=XM_023256426.1;refGene_gn=KIAA1217;refGene_tx=XM_023256427.1;re
fGene_gn=KIAA1217;refGene_tx=XM_023256428.1;refGene_gn=KIAA1217;refGen
e_tx=XM_023256429.1;refGene_gn=KIAA1217;refGene_tx=XM_023256430.1;refG
ene_gn=KIAA1217;refGene_tx=XM_023256431.1;refGene_gn=KIAA1217;refGene_
tx=XM_023256433.1;refGene_gn=KIAA1217;refGene_tx=XM_023256434.1;refGen
e_gn=KIAA1217;refGene_tx=XM_023256435.1;refGene_gn=KIAA1217;refGene_tx
=XM_023256436.1;refGene_gn=KIAA1217;refGene_tx=XM_023256441.1;refGene_
gn=KIAA1217;refGene_tx=XM_023256442.1;refGene_gn=KIAA1217;refGene_tx=X
M_023256444.1;refGene_gn=KIAA1217;refGene_tx=XM_023256445.1;refGene_gn
=KIAA1217;refGene_tx=XM_023256446.1;refGene_gn=KIAA1217;refGene_tx=XM_
023256447.1;refGene_gn=KIAA1217;refGene_tx=XM_023256448.1;refGene_gn=K
IAA1217;refGene_tx=XM_023256449.1;refGene_gn=KIAA1217;refGene_tx=XM_02
3256450.1
B4 22894773 22894867
refGene_gn=KIAA1217;refGene_tx=XM_011283634.3;refGene_gn=KIAA1217;refG
ene_tx=XM_023256426.1;refGene_gn=KIAA1217;refGene_tx=XM_023256427.1;re

fGene_gn=KIAA1217;refGene_tx=XM_023256428.1;refGene_gn=KIAA1217;refGene_tx=XM_023256429.1;refGene_gn=KIAA1217;refGene_tx=XM_023256430.1;refGene_gn=KIAA1217;refGene_tx=XM_023256431.1;refGene_gn=KIAA1217;refGene_tx=XM_023256433.1;refGene_gn=KIAA1217;refGene_tx=XM_023256434.1;refGene_gn=KIAA1217;refGene_tx=XM_023256435.1;refGene_gn=KIAA1217;refGene_tx=XM_023256436.1;refGene_gn=KIAA1217;refGene_tx=XM_023256441.1;refGene_gn=KIAA1217;refGene_tx=XM_023256442.1;refGene_gn=KIAA1217;refGene_tx=XM_023256444.1;refGene_gn=KIAA1217;refGene_tx=XM_023256445.1;refGene_gn=KIAA1217;refGene_tx=XM_023256446.1;refGene_gn=KIAA1217;refGene_tx=XM_023256447.1;refGene_gn=KIAA1217;refGene_tx=XM_023256448.1;refGene_gn=KIAA1217;refGene_tx=XM_023256449.1;refGene_gn=KIAA1217;refGene_tx=XM_023256450.1
B4 22926102 22926935
refGene_gn=KIAA1217;refGene_tx=XM_011283634.3;refGene_gn=KIAA1217;refGene_tx=XM_023256426.1;refGene_gn=KIAA1217;refGene_tx=XM_023256427.1;refGene_gn=KIAA1217;refGene_tx=XM_023256428.1;refGene_gn=KIAA1217;refGene_tx=XM_023256429.1;refGene_gn=KIAA1217;refGene_tx=XM_023256430.1;refGene_gn=KIAA1217;refGene_tx=XM_023256431.1;refGene_gn=KIAA1217;refGene_tx=XM_023256433.1;refGene_gn=KIAA1217;refGene_tx=XM_023256434.1;refGene_gn=KIAA1217;refGene_tx=XM_023256435.1;refGene_gn=KIAA1217;refGene_tx=XM_023256436.1;refGene_gn=KIAA1217;refGene_tx=XM_023256437.1;refGene_gn=KIAA1217;refGene_tx=XM_023256438.1;refGene_gn=KIAA1217;refGene_tx=XM_023256439.1;refGene_gn=KIAA1217;refGene_tx=XM_023256440.1;refGene_gn=KIAA1217;refGene_tx=XM_023256441.1;refGene_gn=KIAA1217;refGene_tx=XM_023256442.1;refGene_gn=KIAA1217;refGene_tx=XM_023256444.1;refGene_gn=KIAA1217;refGene_tx=XM_023256445.1;refGene_gn=KIAA1217;refGene_tx=XM_023256446.1;refGene_gn=KIAA1217;refGene_tx=XM_023256447.1;refGene_gn=KIAA1217;refGene_tx=XM_023256448.1;refGene_gn=KIAA1217;refGene_tx=XM_023256449.1;refGene_gn=KIAA1217;refGene_tx=XM_023256450.1
B4 22941826 22941931
refGene_gn=KIAA1217;refGene_tx=XM_011283634.3;refGene_gn=KIAA1217;refGene_tx=XM_023256426.1;refGene_gn=KIAA1217;refGene_tx=XM_023256427.1;refGene_gn=KIAA1217;refGene_tx=XM_023256428.1;refGene_gn=KIAA1217;refGene_tx=XM_023256429.1;refGene_gn=KIAA1217;refGene_tx=XM_023256431.1;refGene_gn=KIAA1217;refGene_tx=XM_023256434.1;refGene_gn=KIAA1217;refGene_tx=XM_023256435.1;refGene_gn=KIAA1217;refGene_tx=XM_023256436.1;refGene_gn=KIAA1217;refGene_tx=XM_023256437.1;refGene_gn=KIAA1217;refGene_tx=XM_023256438.1;refGene_gn=KIAA1217;refGene_tx=XM_023256439.1;refGene_gn=KIAA1217;refGene_tx=XM_023256440.1;refGene_gn=KIAA1217;refGene_tx=XM_023256441.1;refGene_gn=KIAA1217;refGene_tx=XM_023256442.1;refGene_gn=KIAA1217;refGene_tx=XM_023256444.1;refGene_gn=KIAA1217;refGene_tx=XM_023256445.1;refGene_gn=KIAA1217;refGene_tx=XM_023256448.1
B4 22942446 22942496
refGene_gn=KIAA1217;refGene_tx=XM_011283634.3;refGene_gn=KIAA1217;refGene_tx=XM_023256426.1;refGene_gn=KIAA1217;refGene_tx=XM_023256427.1;refGene_gn=KIAA1217;refGene_tx=XM_023256428.1;refGene_gn=KIAA1217;refGene_tx=XM_023256429.1;refGene_gn=KIAA1217;refGene_tx=XM_023256430.1;refGene_gn=KIAA1217;refGene_tx=XM_023256431.1;refGene_gn=KIAA1217;refGene_tx=XM_023256433.1;refGene_gn=KIAA1217;refGene_tx=XM_023256434.1;refGene_gn=KIAA1217;refGene_tx=XM_023256435.1;refGene_gn=KIAA1217;refGene_tx

=XM_023256436.1;refGene_gn=KIAA1217;refGene_tx=XM_023256437.1;refGene_gn=KIAA1217;refGene_tx=XM_023256438.1;refGene_gn=KIAA1217;refGene_tx=XM_023256439.1;refGene_gn=KIAA1217;refGene_tx=XM_023256440.1;refGene_gn=KIAA1217;refGene_tx=XM_023256441.1;refGene_gn=KIAA1217;refGene_tx=XM_023256442.1;refGene_gn=KIAA1217;refGene_tx=XM_023256444.1;refGene_gn=KIAA1217;refGene_tx=XM_023256445.1;refGene_gn=KIAA1217;refGene_tx=XM_023256446.1;refGene_gn=KIAA1217;refGene_tx=XM_023256447.1;refGene_gn=KIAA1217;refGene_tx=XM_023256448.1;refGene_gn=KIAA1217;refGene_tx=XM_023256449.1;refGene_gn=KIAA1217;refGene_tx=XM_023256450.1
B4 22948919 22949071
refGene_gn=KIAA1217;refGene_tx=XM_011283634.3;refGene_gn=KIAA1217;refGene_tx=XM_023256426.1;refGene_gn=KIAA1217;refGene_tx=XM_023256427.1;refGene_gn=KIAA1217;refGene_tx=XM_023256428.1;refGene_gn=KIAA1217;refGene_tx=XM_023256429.1;refGene_gn=KIAA1217;refGene_tx=XM_023256430.1;refGene_gn=KIAA1217;refGene_tx=XM_023256431.1;refGene_gn=KIAA1217;refGene_tx=XM_023256433.1;refGene_gn=KIAA1217;refGene_tx=XM_023256434.1;refGene_gn=KIAA1217;refGene_tx=XM_023256435.1;refGene_gn=KIAA1217;refGene_tx=XM_023256436.1;refGene_gn=KIAA1217;refGene_tx=XM_023256437.1;refGene_gn=KIAA1217;refGene_tx=XM_023256438.1;refGene_gn=KIAA1217;refGene_tx=XM_023256439.1;refGene_gn=KIAA1217;refGene_tx=XM_023256440.1;refGene_gn=KIAA1217;refGene_tx=XM_023256441.1;refGene_gn=KIAA1217;refGene_tx=XM_023256442.1;refGene_gn=KIAA1217;refGene_tx=XM_023256444.1;refGene_gn=KIAA1217;refGene_tx=XM_023256445.1;refGene_gn=KIAA1217;refGene_tx=XM_023256446.1;refGene_gn=KIAA1217;refGene_tx=XM_023256447.1;refGene_gn=KIAA1217;refGene_tx=XM_023256448.1;refGene_gn=KIAA1217;refGene_tx=XM_023256449.1;refGene_gn=KIAA1217;refGene_tx=XM_023256450.1
B4 22961387 22961563
refGene_gn=KIAA1217;refGene_tx=XM_011283634.3;refGene_gn=KIAA1217;refGene_tx=XM_023256426.1;refGene_gn=KIAA1217;refGene_tx=XM_023256427.1;refGene_gn=KIAA1217;refGene_tx=XM_023256428.1;refGene_gn=KIAA1217;refGene_tx=XM_023256429.1;refGene_gn=KIAA1217;refGene_tx=XM_023256430.1;refGene_gn=KIAA1217;refGene_tx=XM_023256431.1;refGene_gn=KIAA1217;refGene_tx=XM_023256433.1;refGene_gn=KIAA1217;refGene_tx=XM_023256434.1;refGene_gn=KIAA1217;refGene_tx=XM_023256435.1;refGene_gn=KIAA1217;refGene_tx=XM_023256436.1;refGene_gn=KIAA1217;refGene_tx=XM_023256437.1;refGene_gn=KIAA1217;refGene_tx=XM_023256438.1;refGene_gn=KIAA1217;refGene_tx=XM_023256439.1;refGene_gn=KIAA1217;refGene_tx=XM_023256440.1;refGene_gn=KIAA1217;refGene_tx=XM_023256441.1;refGene_gn=KIAA1217;refGene_tx=XM_023256442.1;refGene_gn=KIAA1217;refGene_tx=XM_023256444.1;refGene_gn=KIAA1217;refGene_tx=XM_023256445.1;refGene_gn=KIAA1217;refGene_tx=XM_023256446.1;refGene_gn=KIAA1217;refGene_tx=XM_023256447.1;refGene_gn=KIAA1217;refGene_tx=XM_023256448.1;refGene_gn=KIAA1217;refGene_tx=XM_023256449.1;refGene_gn=KIAA1217;refGene_tx=XM_023256450.1
B4 22966956 22967087
refGene_gn=KIAA1217;refGene_tx=XM_011283634.3;refGene_gn=KIAA1217;refGene_tx=XM_023256426.1;refGene_gn=KIAA1217;refGene_tx=XM_023256427.1;refGene_gn=KIAA1217;refGene_tx=XM_023256428.1;refGene_gn=KIAA1217;refGene_tx=XM_023256429.1;refGene_gn=KIAA1217;refGene_tx=XM_023256430.1;refGene_gn=KIAA1217;refGene_tx=XM_023256431.1;refGene_gn=KIAA1217;refGene_tx=XM_023256433.1;refGene_gn=KIAA1217;refGene_tx=XM_023256434.1;refGen

e_gn=KIAA1217;refGene_tx=XM_023256435.1;refGene_gn=KIAA1217;refGene_tx=XM_023256436.1;refGene_gn=KIAA1217;refGene_tx=XM_023256437.1;refGene_gn=KIAA1217;refGene_tx=XM_023256438.1;refGene_gn=KIAA1217;refGene_tx=XM_023256439.1;refGene_gn=KIAA1217;refGene_tx=XM_023256440.1;refGene_gn=KIAA1217;refGene_tx=XM_023256441.1;refGene_gn=KIAA1217;refGene_tx=XM_023256442.1;refGene_gn=KIAA1217;refGene_tx=XM_023256444.1;refGene_gn=KIAA1217;refGene_tx=XM_023256445.1;refGene_gn=KIAA1217;refGene_tx=XM_023256446.1;refGene_gn=KIAA1217;refGene_tx=XM_023256447.1;refGene_gn=KIAA1217;refGene_tx=XM_023256448.1;refGene_gn=KIAA1217;refGene_tx=XM_023256449.1;refGene_gn=KIAA1217;refGene_tx=XM_023256450.1
B4 22967661 22967809
refGene_gn=KIAA1217;refGene_tx=XM_011283634.3;refGene_gn=KIAA1217;refGene_tx=XM_023256426.1;refGene_gn=KIAA1217;refGene_tx=XM_023256427.1;refGene_gn=KIAA1217;refGene_tx=XM_023256428.1;refGene_gn=KIAA1217;refGene_tx=XM_023256429.1;refGene_gn=KIAA1217;refGene_tx=XM_023256430.1;refGene_gn=KIAA1217;refGene_tx=XM_023256431.1;refGene_gn=KIAA1217;refGene_tx=XM_023256433.1;refGene_gn=KIAA1217;refGene_tx=XM_023256434.1;refGene_gn=KIAA1217;refGene_tx=XM_023256435.1;refGene_gn=KIAA1217;refGene_tx=XM_023256436.1;refGene_gn=KIAA1217;refGene_tx=XM_023256437.1;refGene_gn=KIAA1217;refGene_tx=XM_023256438.1;refGene_gn=KIAA1217;refGene_tx=XM_023256439.1;refGene_gn=KIAA1217;refGene_tx=XM_023256440.1;refGene_gn=KIAA1217;refGene_tx=XM_023256441.1;refGene_gn=KIAA1217;refGene_tx=XM_023256442.1;refGene_gn=KIAA1217;refGene_tx=XM_023256444.1;refGene_gn=KIAA1217;refGene_tx=XM_023256445.1;refGene_gn=KIAA1217;refGene_tx=XM_023256446.1;refGene_gn=KIAA1217;refGene_tx=XM_023256447.1;refGene_gn=KIAA1217;refGene_tx=XM_023256448.1;refGene_gn=KIAA1217;refGene_tx=XM_023256449.1;refGene_gn=KIAA1217;refGene_tx=XM_023256450.1
B4 22969810 22970258
refGene_gn=KIAA1217;refGene_tx=XM_011283634.3;refGene_gn=KIAA1217;refGene_tx=XM_023256426.1;refGene_gn=KIAA1217;refGene_tx=XM_023256427.1;refGene_gn=KIAA1217;refGene_tx=XM_023256428.1;refGene_gn=KIAA1217;refGene_tx=XM_023256429.1;refGene_gn=KIAA1217;refGene_tx=XM_023256430.1;refGene_gn=KIAA1217;refGene_tx=XM_023256431.1;refGene_gn=KIAA1217;refGene_tx=XM_023256433.1;refGene_gn=KIAA1217;refGene_tx=XM_023256434.1;refGene_gn=KIAA1217;refGene_tx=XM_023256435.1;refGene_gn=KIAA1217;refGene_tx=XM_023256436.1;refGene_gn=KIAA1217;refGene_tx=XM_023256437.1;refGene_gn=KIAA1217;refGene_tx=XM_023256438.1;refGene_gn=KIAA1217;refGene_tx=XM_023256439.1;refGene_gn=KIAA1217;refGene_tx=XM_023256440.1;refGene_gn=KIAA1217;refGene_tx=XM_023256441.1;refGene_gn=KIAA1217;refGene_tx=XM_023256442.1;refGene_gn=KIAA1217;refGene_tx=XM_023256444.1;refGene_gn=KIAA1217;refGene_tx=XM_023256445.1;refGene_gn=KIAA1217;refGene_tx=XM_023256446.1;refGene_gn=KIAA1217;refGene_tx=XM_023256447.1;refGene_gn=KIAA1217;refGene_tx=XM_023256448.1;refGene_gn=KIAA1217;refGene_tx=XM_023256449.1;refGene_gn=KIAA1217;refGene_tx=XM_023256450.1
B4 22972774 22972958
refGene_gn=KIAA1217;refGene_tx=XM_011283634.3;refGene_gn=KIAA1217;refGene_tx=XM_023256426.1;refGene_gn=KIAA1217;refGene_tx=XM_023256427.1;refGene_gn=KIAA1217;refGene_tx=XM_023256428.1;refGene_gn=KIAA1217;refGene_tx=XM_023256429.1;refGene_gn=KIAA1217;refGene_tx=XM_023256430.1;refGene_gn=KIAA1217;refGene_tx=XM_023256431.1;refGene_gn=KIAA1217;refGene_

tx=XM_023256433.1;refGene_gn=KIAA1217;refGene_tx=XM_023256434.1;refGen
e_gn=KIAA1217;refGene_tx=XM_023256435.1;refGene_gn=KIAA1217;refGene_tx
=XM_023256436.1;refGene_gn=KIAA1217;refGene_tx=XM_023256437.1;refGene_
gn=KIAA1217;refGene_tx=XM_023256438.1;refGene_gn=KIAA1217;refGene_tx=X
M_023256439.1;refGene_gn=KIAA1217;refGene_tx=XM_023256440.1;refGene_gn
=KIAA1217;refGene_tx=XM_023256441.1;refGene_gn=KIAA1217;refGene_tx=XM_
023256442.1;refGene_gn=KIAA1217;refGene_tx=XM_023256444.1;refGene_gn=K
IAA1217;refGene_tx=XM_023256445.1;refGene_gn=KIAA1217;refGene_tx=XM_02
3256446.1;refGene_gn=KIAA1217;refGene_tx=XM_023256447.1;refGene_gn=KIA
A1217;refGene_tx=XM_023256448.1;refGene_gn=KIAA1217;refGene_tx=XM_0232
56449.1;refGene_gn=KIAA1217;refGene_tx=XM_023256450.1
B4 22977039 22977203
refGene_gn=KIAA1217;refGene_tx=XM_011283634.3;refGene_gn=KIAA1217;refG
ene_tx=XM_023256426.1;refGene_gn=KIAA1217;refGene_tx=XM_023256427.1;re
fGene_gn=KIAA1217;refGene_tx=XM_023256428.1;refGene_gn=KIAA1217;refGen
e_tx=XM_023256429.1;refGene_gn=KIAA1217;refGene_tx=XM_023256430.1;refG
ene_gn=KIAA1217;refGene_tx=XM_023256431.1;refGene_gn=KIAA1217;refGene_
tx=XM_023256433.1;refGene_gn=KIAA1217;refGene_tx=XM_023256434.1;refGen
e_gn=KIAA1217;refGene_tx=XM_023256435.1;refGene_gn=KIAA1217;refGene_tx
=XM_023256436.1;refGene_gn=KIAA1217;refGene_tx=XM_023256437.1;refGene_
gn=KIAA1217;refGene_tx=XM_023256438.1;refGene_gn=KIAA1217;refGene_tx=X
M_023256439.1;refGene_gn=KIAA1217;refGene_tx=XM_023256440.1;refGene_gn
=KIAA1217;refGene_tx=XM_023256441.1;refGene_gn=KIAA1217;refGene_tx=XM_
023256442.1;refGene_gn=KIAA1217;refGene_tx=XM_023256444.1;refGene_gn=K
IAA1217;refGene_tx=XM_023256445.1;refGene_gn=KIAA1217;refGene_tx=XM_02
3256446.1;refGene_gn=KIAA1217;refGene_tx=XM_023256447.1;refGene_gn=KIA
A1217;refGene_tx=XM_023256448.1;refGene_gn=KIAA1217;refGene_tx=XM_0232
56449.1;refGene_gn=KIAA1217;refGene_tx=XM_023256450.1
B4 22977919 22978084
refGene_gn=KIAA1217;refGene_tx=XM_011283634.3;refGene_gn=KIAA1217;refG
ene_tx=XM_023256426.1;refGene_gn=KIAA1217;refGene_tx=XM_023256427.1;re
fGene_gn=KIAA1217;refGene_tx=XM_023256428.1;refGene_gn=KIAA1217;refGen
e_tx=XM_023256429.1;refGene_gn=KIAA1217;refGene_tx=XM_023256430.1;refG
ene_gn=KIAA1217;refGene_tx=XM_023256431.1;refGene_gn=KIAA1217;refGene_
tx=XM_023256433.1;refGene_gn=KIAA1217;refGene_tx=XM_023256434.1;refGen
e_gn=KIAA1217;refGene_tx=XM_023256435.1;refGene_gn=KIAA1217;refGene_tx
=XM_023256436.1;refGene_gn=KIAA1217;refGene_tx=XM_023256437.1;refGene_
gn=KIAA1217;refGene_tx=XM_023256438.1;refGene_gn=KIAA1217;refGene_tx=X
M_023256439.1;refGene_gn=KIAA1217;refGene_tx=XM_023256440.1;refGene_gn
=KIAA1217;refGene_tx=XM_023256441.1;refGene_gn=KIAA1217;refGene_tx=XM_
023256442.1;refGene_gn=KIAA1217;refGene_tx=XM_023256444.1;refGene_gn=K
IAA1217;refGene_tx=XM_023256445.1;refGene_gn=KIAA1217;refGene_tx=XM_02
3256446.1;refGene_gn=KIAA1217;refGene_tx=XM_023256447.1;refGene_gn=KIA
A1217;refGene_tx=XM_023256448.1;refGene_gn=KIAA1217;refGene_tx=XM_0232
56449.1;refGene_gn=KIAA1217;refGene_tx=XM_023256450.1
B4 22981600 22981720
refGene_gn=KIAA1217;refGene_tx=XM_011283634.3;refGene_gn=KIAA1217;refG
ene_tx=XM_023256426.1;refGene_gn=KIAA1217;refGene_tx=XM_023256427.1;re
fGene_gn=KIAA1217;refGene_tx=XM_023256428.1;refGene_gn=KIAA1217;refGen
e_tx=XM_023256429.1;refGene_gn=KIAA1217;refGene_tx=XM_023256430.1;refG

ene_gn=KIAA1217;refGene_tx=XM_023256433.1;refGene_gn=KIAA1217;refGene_tx=XM_023256434.1;refGene_gn=KIAA1217;refGene_tx=XM_023256435.1;refGene_gn=KIAA1217;refGene_tx=XM_023256436.1;refGene_gn=KIAA1217;refGene_tx=XM_023256437.1;refGene_gn=KIAA1217;refGene_tx=XM_023256438.1;refGene_gn=KIAA1217;refGene_tx=XM_023256439.1;refGene_gn=KIAA1217;refGene_tx=XM_023256440.1;refGene_gn=KIAA1217;refGene_tx=XM_023256441.1;refGene_gn=KIAA1217;refGene_tx=XM_023256442.1;refGene_gn=KIAA1217;refGene_tx=XM_023256445.1;refGene_gn=KIAA1217;refGene_tx=XM_023256446.1;refGene_gn=KIAA1217;refGene_tx=XM_023256447.1;refGene_gn=KIAA1217;refGene_tx=XM_023256449.1
B4 22985442 22985520
refGene_gn=KIAA1217;refGene_tx=XM_011283634.3;refGene_gn=KIAA1217;refGene_tx=XM_023256426.1;refGene_gn=KIAA1217;refGene_tx=XM_023256427.1;refGene_gn=KIAA1217;refGene_tx=XM_023256428.1;refGene_gn=KIAA1217;refGene_tx=XM_023256429.1;refGene_gn=KIAA1217;refGene_tx=XM_023256430.1;refGene_gn=KIAA1217;refGene_tx=XM_023256431.1;refGene_gn=KIAA1217;refGene_tx=XM_023256433.1;refGene_gn=KIAA1217;refGene_tx=XM_023256434.1;refGene_gn=KIAA1217;refGene_tx=XM_023256435.1;refGene_gn=KIAA1217;refGene_tx=XM_023256436.1;refGene_gn=KIAA1217;refGene_tx=XM_023256437.1;refGene_gn=KIAA1217;refGene_tx=XM_023256438.1;refGene_gn=KIAA1217;refGene_tx=XM_023256439.1;refGene_gn=KIAA1217;refGene_tx=XM_023256440.1;refGene_gn=KIAA1217;refGene_tx=XM_023256441.1;refGene_gn=KIAA1217;refGene_tx=XM_023256445.1;refGene_gn=KIAA1217;refGene_tx=XM_023256446.1;refGene_gn=KIAA1217;refGene_tx=XM_023256449.1
B4 22985631 22987227
refGene_gn=KIAA1217;refGene_tx=XM_011283634.3;refGene_gn=KIAA1217;refGene_tx=XM_023256426.1;refGene_gn=KIAA1217;refGene_tx=XM_023256427.1;refGene_gn=KIAA1217;refGene_tx=XM_023256428.1;refGene_gn=KIAA1217;refGene_tx=XM_023256429.1;refGene_gn=KIAA1217;refGene_tx=XM_023256430.1;refGene_gn=KIAA1217;refGene_tx=XM_023256431.1;refGene_gn=KIAA1217;refGene_tx=XM_023256433.1;refGene_gn=KIAA1217;refGene_tx=XM_023256434.1;refGene_gn=KIAA1217;refGene_tx=XM_023256435.1;refGene_gn=KIAA1217;refGene_tx=XM_023256436.1;refGene_gn=KIAA1217;refGene_tx=XM_023256437.1;refGene_gn=KIAA1217;refGene_tx=XM_023256438.1;refGene_gn=KIAA1217;refGene_tx=XM_023256439.1;refGene_gn=KIAA1217;refGene_tx=XM_023256440.1
B4 22987746 22987884
refGene_gn=KIAA1217;refGene_tx=XM_011283634.3;refGene_gn=KIAA1217;refGene_tx=XM_023256426.1;refGene_gn=KIAA1217;refGene_tx=XM_023256427.1;refGene_gn=KIAA1217;refGene_tx=XM_023256428.1;refGene_gn=KIAA1217;refGene_tx=XM_023256429.1;refGene_gn=KIAA1217;refGene_tx=XM_023256430.1;refGene_gn=KIAA1217;refGene_tx=XM_023256431.1;refGene_gn=KIAA1217;refGene_tx=XM_023256433.1;refGene_gn=KIAA1217;refGene_tx=XM_023256434.1;refGene_gn=KIAA1217;refGene_tx=XM_023256435.1;refGene_gn=KIAA1217;refGene_tx=XM_023256436.1;refGene_gn=KIAA1217;refGene_tx=XM_023256437.1;refGene_gn=KIAA1217;refGene_tx=XM_023256438.1;refGene_gn=KIAA1217;refGene_tx=XM_023256439.1;refGene_gn=KIAA1217;refGene_tx=XM_023256440.1;refGene_gn=KIAA1217;refGene_tx=XM_023256441.1;refGene_gn=KIAA1217;refGene_tx=XM_023256442.1;refGene_gn=KIAA1217;refGene_tx=XM_023256444.1;refGene_gn=KIAA1217;refGene_tx=XM_023256445.1;refGene_gn=KIAA1217;refGene_tx=XM_023256446.1;refGene_gn=KIAA1217;refGene_tx=XM_023256447.1;refGene_gn=KIA

A1217;refGene_tx=XM_023256448.1;refGene_gn=KIAA1217;refGene_tx=XM_0232
56449.1;refGene_gn=KIAA1217;refGene_tx=XM_023256450.1
B4 22988581 22989082
refGene_gn=KIAA1217;refGene_tx=XM_011283634.3;refGene_gn=KIAA1217;refG
ene_tx=XM_023256426.1;refGene_gn=KIAA1217;refGene_tx=XM_023256427.1;re
fGene_gn=KIAA1217;refGene_tx=XM_023256428.1;refGene_gn=KIAA1217;refGen
e_tx=XM_023256429.1;refGene_gn=KIAA1217;refGene_tx=XM_023256430.1;refG
ene_gn=KIAA1217;refGene_tx=XM_023256431.1;refGene_gn=KIAA1217;refGene_
tx=XM_023256433.1;refGene_gn=KIAA1217;refGene_tx=XM_023256434.1;refGen
e_gn=KIAA1217;refGene_tx=XM_023256435.1;refGene_gn=KIAA1217;refGene_tx
=XM_023256436.1;refGene_gn=KIAA1217;refGene_tx=XM_023256437.1;refGene_
gn=KIAA1217;refGene_tx=XM_023256438.1;refGene_gn=KIAA1217;refGene_tx=X
M_023256439.1;refGene_gn=KIAA1217;refGene_tx=XM_023256440.1;refGene_gn
=KIAA1217;refGene_tx=XM_023256441.1;refGene_gn=KIAA1217;refGene_tx=XM_
023256442.1;refGene_gn=KIAA1217;refGene_tx=XM_023256444.1;refGene_gn=K
IAA1217;refGene_tx=XM_023256446.1;refGene_gn=KIAA1217;refGene_tx=XM_02
3256447.1;refGene_gn=KIAA1217;refGene_tx=XM_023256448.1
B4 23018989 23019055 refGene_gn=ARHGAP21;refGene_tx=XM_006933312.4
B4 23019056 23020691 refGene_gn=ARHGAP21;refGene_tx=XM_006933312.4
B4 23023578 23023610 refGene_gn=ARHGAP21;refGene_tx=XM_006933312.4
B4 23025547 23025653 refGene_gn=ARHGAP21;refGene_tx=XM_006933312.4
B4 23025973 23026015 refGene_gn=ARHGAP21;refGene_tx=XM_006933312.4
B4 23026234 23026360 refGene_gn=ARHGAP21;refGene_tx=XM_006933312.4
B4 23029328 23029418 refGene_gn=ARHGAP21;refGene_tx=XM_006933312.4
B4 23029495 23029569 refGene_gn=ARHGAP21;refGene_tx=XM_006933312.4
B4 23030489 23030568 refGene_gn=ARHGAP21;refGene_tx=XM_006933312.4
B4 23030668 23030824 refGene_gn=ARHGAP21;refGene_tx=XM_006933312.4
B4 23031638 23031781 refGene_gn=ARHGAP21;refGene_tx=XM_006933312.4
B4 23032374 23032511 refGene_gn=ARHGAP21;refGene_tx=XM_006933312.4
B4 23032590 23032655 refGene_gn=ARHGAP21;refGene_tx=XM_006933312.4
B4 23035246 23035531 refGene_gn=ARHGAP21;refGene_tx=XM_006933312.4
B4 23036878 23037004 refGene_gn=ARHGAP21;refGene_tx=XM_006933312.4
B4 23039109 23039146 refGene_gn=ARHGAP21;refGene_tx=XM_006933312.4
B4 23043942 23044045 refGene_gn=ARHGAP21;refGene_tx=XM_006933312.4
B4 23044195 23044372 refGene_gn=ARHGAP21;refGene_tx=XM_006933312.4
B4 23055912 23057809 refGene_gn=ARHGAP21;refGene_tx=XM_006933312.4
B4 23059154 23059184 refGene_gn=ARHGAP21;refGene_tx=XM_006933312.4
B4 23066056 23066111 refGene_gn=ARHGAP21;refGene_tx=XM_006933312.4
B4 23068635 23068714 refGene_gn=ARHGAP21;refGene_tx=XM_006933312.4
B4 23070698 23070791 refGene_gn=ARHGAP21;refGene_tx=XM_006933312.4
B4 23095452 23095477 refGene_gn=ARHGAP21;refGene_tx=XM_006933312.4
B4 23095732 23095912 refGene_gn=ARHGAP21;refGene_tx=XM_006933312.4
B4 23153045 23153123 refGene_gn=ARHGAP21;refGene_tx=XM_006933312.4
B4 23246834 23246882
refGene_gn=PRTFDC1;refGene_tx=XM_006933313.4;refGene_gn=PRTFDC1;refGen
e_tx=XM_023256451.1
B4 23248025 23248102
refGene_gn=PRTFDC1;refGene_tx=XM_006933313.4;refGene_gn=PRTFDC1;refGen
e_tx=XM_023256451.1
B4 23251095 23251142

refGene_gn=PRTFDC1;refGene_tx=XM_006933313.4;refGene_gn=PRTFDC1;refGen
e_tx=XM_023256451.1
B4	23251602	23251685
refGene_gn=PRTFDC1;refGene_tx=XM_006933313.4;refGene_gn=PRTFDC1;refGen
e_tx=XM_023256451.1
B4	23253046	23253064
refGene_gn=PRTFDC1;refGene_tx=XM_006933313.4;refGene_gn=PRTFDC1;refGen
e_tx=XM_023256451.1
B4	23264695	23264761
refGene_gn=PRTFDC1;refGene_tx=XM_006933313.4;refGene_gn=PRTFDC1;refGen
e_tx=XM_023256451.1
B4	23328416	23328600
refGene_gn=PRTFDC1;refGene_tx=XM_006933313.4;refGene_gn=PRTFDC1;refGen
e_tx=XM_023256451.1
B4	23334406	23334513	refGene_gn=PRTFDC1;refGene_tx=XM_006933313.4
B4	23344251	23344299	refGene_gn=PRTFDC1;refGene_tx=XM_006933313.4
B4	23372768	23372942
refGene_gn=ENKUR;refGene_tx=XM_003988170.5;refGene_gn=ENKUR;refGene_tx
=XM_019834078.2;refGene_gn=ENKUR;refGene_tx=XM_019834079.2
B4	23380941	23381088
refGene_gn=ENKUR;refGene_tx=XM_003988170.5;refGene_gn=ENKUR;refGene_tx
=XM_019834078.2;refGene_gn=ENKUR;refGene_tx=XM_019834079.2
B4	23384230	23384454
refGene_gn=ENKUR;refGene_tx=XM_003988170.5;refGene_gn=ENKUR;refGene_tx
=XM_019834078.2;refGene_gn=ENKUR;refGene_tx=XM_019834079.2
B4	23388503	23388649
refGene_gn=ENKUR;refGene_tx=XM_003988170.5;refGene_gn=ENKUR;refGene_tx
=XM_019834078.2;refGene_gn=ENKUR;refGene_tx=XM_019834079.2
B4	23396279	23396356	refGene_gn=ENKUR;refGene_tx=XM_003988170.5
B4	23396744	23396773	refGene_gn=ENKUR;refGene_tx=XM_019834078.2
B4	23404584	23406816
refGene_gn=THNSL1;refGene_tx=XM_006933315.4;refGene_gn=THNSL1;refGene_
tx=XM_006933316.4;refGene_gn=THNSL1;refGene_tx=XM_006933317.3;refGene_
gn=THNSL1;refGene_tx=XM_019834075.2;refGene_gn=THNSL1;refGene_tx=XM_01
9834076.2;refGene_gn=THNSL1;refGene_tx=XM_019834077.2
B4	23424936	23424969	refGene_gn=THNSL1;refGene_tx=XM_023256452.1
B4	23425985	23426125	refGene_gn=THNSL1;refGene_tx=XM_023256452.1
B4	23435832	23435966	refGene_gn=THNSL1;refGene_tx=XM_023256452.1
B4	23447020	23447124	refGene_gn=THNSL1;refGene_tx=XM_023256452.1
B4	23521827	23522720	refGene_gn=GPR158;refGene_tx=XM_011283685.2
B4	23559009	23559115	refGene_gn=GPR158;refGene_tx=XM_011283685.2
B4	23720543	23720646	refGene_gn=GPR158;refGene_tx=XM_011283685.2
B4	23751411	23751635	refGene_gn=GPR158;refGene_tx=XM_011283685.2
B4	23809245	23809314	refGene_gn=GPR158;refGene_tx=XM_011283685.2
B4	23884684	23884794	refGene_gn=GPR158;refGene_tx=XM_011283685.2
B4	23914267	23914506	refGene_gn=GPR158;refGene_tx=XM_011283685.2
B4	23930612	23930751	refGene_gn=GPR158;refGene_tx=XM_011283685.2
B4	23935832	23935938	refGene_gn=GPR158;refGene_tx=XM_011283685.2
B4	23939069	23939216	refGene_gn=GPR158;refGene_tx=XM_011283685.2
B4	23939944	23941420	refGene_gn=GPR158;refGene_tx=XM_011283685.2

B4 24229919 24230087
refGene_gn=MYO3A;refGene_tx=XM_023256453.1;refGene_gn=MYO3A;refGene_tx=XM_023256454.1;refGene_gn=MYO3A;refGene_tx=XM_023256455.1;refGene_gn=MYO3A;refGene_tx=XM_023256456.1;refGene_gn=MYO3A;refGene_tx=XM_023256457.1
B4 24232671 24232806
refGene_gn=MYO3A;refGene_tx=XM_023256453.1;refGene_gn=MYO3A;refGene_tx=XM_023256454.1;refGene_gn=MYO3A;refGene_tx=XM_023256455.1;refGene_gn=MYO3A;refGene_tx=XM_023256456.1;refGene_gn=MYO3A;refGene_tx=XM_023256457.1
B4 24255924 24256029
refGene_gn=MYO3A;refGene_tx=XM_023256453.1;refGene_gn=MYO3A;refGene_tx=XM_023256454.1;refGene_gn=MYO3A;refGene_tx=XM_023256455.1;refGene_gn=MYO3A;refGene_tx=XM_023256456.1;refGene_gn=MYO3A;refGene_tx=XM_023256457.1
B4 24256587 24256687
refGene_gn=MYO3A;refGene_tx=XM_023256453.1;refGene_gn=MYO3A;refGene_tx=XM_023256454.1;refGene_gn=MYO3A;refGene_tx=XM_023256455.1;refGene_gn=MYO3A;refGene_tx=XM_023256456.1;refGene_gn=MYO3A;refGene_tx=XM_023256457.1
B4 24274809 24274886
refGene_gn=MYO3A;refGene_tx=XM_023256453.1;refGene_gn=MYO3A;refGene_tx=XM_023256454.1;refGene_gn=MYO3A;refGene_tx=XM_023256455.1;refGene_gn=MYO3A;refGene_tx=XM_023256456.1;refGene_gn=MYO3A;refGene_tx=XM_023256457.1
B4 24278283 24278429
refGene_gn=MYO3A;refGene_tx=XM_023256453.1;refGene_gn=MYO3A;refGene_tx=XM_023256454.1;refGene_gn=MYO3A;refGene_tx=XM_023256455.1;refGene_gn=MYO3A;refGene_tx=XM_023256456.1;refGene_gn=MYO3A;refGene_tx=XM_023256457.1
B4 24281018 24281084
refGene_gn=MYO3A;refGene_tx=XM_023256453.1;refGene_gn=MYO3A;refGene_tx=XM_023256455.1;refGene_gn=MYO3A;refGene_tx=XM_023256456.1;refGene_gn=MYO3A;refGene_tx=XM_023256457.1
B4 24285210 24285366
refGene_gn=MYO3A;refGene_tx=XM_023256453.1;refGene_gn=MYO3A;refGene_tx=XM_023256454.1;refGene_gn=MYO3A;refGene_tx=XM_023256455.1;refGene_gn=MYO3A;refGene_tx=XM_023256456.1;refGene_gn=MYO3A;refGene_tx=XM_023256457.1
B4 24316508 24316608
refGene_gn=MYO3A;refGene_tx=XM_023256453.1;refGene_gn=MYO3A;refGene_tx=XM_023256454.1;refGene_gn=MYO3A;refGene_tx=XM_023256455.1;refGene_gn=MYO3A;refGene_tx=XM_023256456.1;refGene_gn=MYO3A;refGene_tx=XM_023256457.1
B4 24318230 24318347
refGene_gn=MYO3A;refGene_tx=XM_023256453.1;refGene_gn=MYO3A;refGene_tx=XM_023256454.1;refGene_gn=MYO3A;refGene_tx=XM_023256455.1;refGene_gn=MYO3A;refGene_tx=XM_023256456.1;refGene_gn=MYO3A;refGene_tx=XM_023256457.1
B4 24319819 24319924

refGene_gn=MYO3A;refGene_tx=XM_023256453.1;refGene_gn=MYO3A;refGene_tx=XM_023256454.1;refGene_gn=MYO3A;refGene_tx=XM_023256455.1;refGene_gn=MYO3A;refGene_tx=XM_023256456.1;refGene_gn=MYO3A;refGene_tx=XM_023256457.1
B4 24320031 24320115
refGene_gn=MYO3A;refGene_tx=XM_023256453.1;refGene_gn=MYO3A;refGene_tx=XM_023256454.1;refGene_gn=MYO3A;refGene_tx=XM_023256455.1;refGene_gn=MYO3A;refGene_tx=XM_023256456.1;refGene_gn=MYO3A;refGene_tx=XM_023256457.1
B4 24325043 24325246
refGene_gn=MYO3A;refGene_tx=XM_023256453.1;refGene_gn=MYO3A;refGene_tx=XM_023256454.1;refGene_gn=MYO3A;refGene_tx=XM_023256455.1;refGene_gn=MYO3A;refGene_tx=XM_023256456.1;refGene_gn=MYO3A;refGene_tx=XM_023256457.1
B4 24342060 24342159
refGene_gn=MYO3A;refGene_tx=XM_023256453.1;refGene_gn=MYO3A;refGene_tx=XM_023256454.1;refGene_gn=MYO3A;refGene_tx=XM_023256455.1;refGene_gn=MYO3A;refGene_tx=XM_023256456.1;refGene_gn=MYO3A;refGene_tx=XM_023256457.1
B4 24342258 24342373
refGene_gn=MYO3A;refGene_tx=XM_023256453.1;refGene_gn=MYO3A;refGene_tx=XM_023256454.1;refGene_gn=MYO3A;refGene_tx=XM_023256455.1;refGene_gn=MYO3A;refGene_tx=XM_023256456.1;refGene_gn=MYO3A;refGene_tx=XM_023256457.1
B4 24360101 24360228
refGene_gn=MYO3A;refGene_tx=XM_023256453.1;refGene_gn=MYO3A;refGene_tx=XM_023256454.1;refGene_gn=MYO3A;refGene_tx=XM_023256455.1;refGene_gn=MYO3A;refGene_tx=XM_023256456.1;refGene_gn=MYO3A;refGene_tx=XM_023256457.1
B4 24364899 24365110
refGene_gn=MYO3A;refGene_tx=XM_023256453.1;refGene_gn=MYO3A;refGene_tx=XM_023256454.1;refGene_gn=MYO3A;refGene_tx=XM_023256455.1;refGene_gn=MYO3A;refGene_tx=XM_023256456.1;refGene_gn=MYO3A;refGene_tx=XM_023256457.1
B4 24370212 24370360
refGene_gn=MYO3A;refGene_tx=XM_023256453.1;refGene_gn=MYO3A;refGene_tx=XM_023256454.1;refGene_gn=MYO3A;refGene_tx=XM_023256455.1;refGene_gn=MYO3A;refGene_tx=XM_023256456.1;refGene_gn=MYO3A;refGene_tx=XM_023256457.1
B4 24387356 24387510
refGene_gn=MYO3A;refGene_tx=XM_023256453.1;refGene_gn=MYO3A;refGene_tx=XM_023256454.1;refGene_gn=MYO3A;refGene_tx=XM_023256455.1;refGene_gn=MYO3A;refGene_tx=XM_023256456.1;refGene_gn=MYO3A;refGene_tx=XM_023256457.1
B4 24389287 24389376
refGene_gn=MYO3A;refGene_tx=XM_023256453.1;refGene_gn=MYO3A;refGene_tx=XM_023256454.1;refGene_gn=MYO3A;refGene_tx=XM_023256455.1;refGene_gn=MYO3A;refGene_tx=XM_023256456.1;refGene_gn=MYO3A;refGene_tx=XM_023256457.1
B4 24390458 24390588

refGene_gn=MYO3A;refGene_tx=XM_023256453.1;refGene_gn=MYO3A;refGene_tx=XM_023256454.1;refGene_gn=MYO3A;refGene_tx=XM_023256455.1;refGene_gn=MYO3A;refGene_tx=XM_023256456.1;refGene_gn=MYO3A;refGene_tx=XM_023256457.1
B4 24397451 24397531
refGene_gn=MYO3A;refGene_tx=XM_023256453.1;refGene_gn=MYO3A;refGene_tx=XM_023256454.1;refGene_gn=MYO3A;refGene_tx=XM_023256455.1;refGene_gn=MYO3A;refGene_tx=XM_023256456.1;refGene_gn=MYO3A;refGene_tx=XM_023256457.1
B4 24397997 24398075
refGene_gn=MYO3A;refGene_tx=XM_023256453.1;refGene_gn=MYO3A;refGene_tx=XM_023256454.1;refGene_gn=MYO3A;refGene_tx=XM_023256455.1;refGene_gn=MYO3A;refGene_tx=XM_023256456.1;refGene_gn=MYO3A;refGene_tx=XM_023256457.1
B4 24400673 24400879
refGene_gn=MYO3A;refGene_tx=XM_023256453.1;refGene_gn=MYO3A;refGene_tx=XM_023256454.1;refGene_gn=MYO3A;refGene_tx=XM_023256455.1;refGene_gn=MYO3A;refGene_tx=XM_023256456.1;refGene_gn=MYO3A;refGene_tx=XM_023256457.1
B4 24408617 24408729
refGene_gn=MYO3A;refGene_tx=XM_023256453.1;refGene_gn=MYO3A;refGene_tx=XM_023256454.1;refGene_gn=MYO3A;refGene_tx=XM_023256455.1;refGene_gn=MYO3A;refGene_tx=XM_023256456.1;refGene_gn=MYO3A;refGene_tx=XM_023256457.1
B4 24411077 24411240
refGene_gn=MYO3A;refGene_tx=XM_023256453.1;refGene_gn=MYO3A;refGene_tx=XM_023256454.1;refGene_gn=MYO3A;refGene_tx=XM_023256455.1;refGene_gn=MYO3A;refGene_tx=XM_023256456.1;refGene_gn=MYO3A;refGene_tx=XM_023256457.1
B4 24412764 24412888
refGene_gn=MYO3A;refGene_tx=XM_023256453.1;refGene_gn=MYO3A;refGene_tx=XM_023256454.1;refGene_gn=MYO3A;refGene_tx=XM_023256455.1;refGene_gn=MYO3A;refGene_tx=XM_023256456.1;refGene_gn=MYO3A;refGene_tx=XM_023256457.1
B4 24416656 24417560
refGene_gn=MYO3A;refGene_tx=XM_023256453.1;refGene_gn=MYO3A;refGene_tx=XM_023256454.1;refGene_gn=MYO3A;refGene_tx=XM_023256455.1;refGene_gn=MYO3A;refGene_tx=XM_023256456.1;refGene_gn=MYO3A;refGene_tx=XM_023256457.1
B4 24419373 24419518
refGene_gn=MYO3A;refGene_tx=XM_023256453.1;refGene_gn=MYO3A;refGene_tx=XM_023256454.1;refGene_gn=MYO3A;refGene_tx=XM_023256455.1;refGene_gn=MYO3A;refGene_tx=XM_023256457.1
B4 24447834 24447941
refGene_gn=MYO3A;refGene_tx=XM_023256453.1;refGene_gn=MYO3A;refGene_tx=XM_023256454.1;refGene_gn=MYO3A;refGene_tx=XM_023256455.1
B4 24451172 24451955
refGene_gn=LOC102899465;refGene_tx=XM_019834087.2
B4 24454586 24454712 refGene_gn=MYO3A;refGene_tx=XM_023256455.1
B4 24455930 24455971

refGene_gn=MYO3A;refGene_tx=XM_023256453.1;refGene_gn=MYO3A;refGene_tx=XM_023256454.1;refGene_gn=MYO3A;refGene_tx=XM_023256456.1;refGene_gn=MYO3A;refGene_tx=XM_023256457.1
B4 24457325 24457469
refGene_gn=MYO3A;refGene_tx=XM_023256453.1;refGene_gn=MYO3A;refGene_tx=XM_023256454.1;refGene_gn=MYO3A;refGene_tx=XM_023256456.1
B4 24460443 24460561
refGene_gn=MYO3A;refGene_tx=XM_023256453.1;refGene_gn=MYO3A;refGene_tx=XM_023256454.1;refGene_gn=MYO3A;refGene_tx=XM_023256456.1
B4 24465346 24465422 refGene_gn=GAD2;refGene_tx=XM_003988173.5
B4 24466126 24466186 refGene_gn=GAD2;refGene_tx=XM_003988173.5
B4 24466362 24466512 refGene_gn=GAD2;refGene_tx=XM_003988173.5
B4 24467484 24467718 refGene_gn=GAD2;refGene_tx=XM_003988173.5
B4 24472380 24472471 refGene_gn=GAD2;refGene_tx=XM_003988173.5
B4 24473024 24473137 refGene_gn=GAD2;refGene_tx=XM_003988173.5
B4 24478319 24478435 refGene_gn=GAD2;refGene_tx=XM_003988173.5
B4 24490155 24490235 refGene_gn=GAD2;refGene_tx=XM_003988173.5
B4 24516711 24516766 refGene_gn=GAD2;refGene_tx=XM_003988173.5
B4 24518385 24518502 refGene_gn=GAD2;refGene_tx=XM_003988173.5
B4 24520519 24520584 refGene_gn=GAD2;refGene_tx=XM_003988173.5
B4 24526771 24526850 refGene_gn=GAD2;refGene_tx=XM_003988173.5
B4 24532738 24532888 refGene_gn=GAD2;refGene_tx=XM_003988173.5
B4 24539549 24539657 refGene_gn=GAD2;refGene_tx=XM_003988173.5
B4 24540436 24540526 refGene_gn=GAD2;refGene_tx=XM_003988173.5
B4 24549139 24549313 refGene_gn=GAD2;refGene_tx=XM_003988173.5
B4 24711148 24711220
refGene_gn=APBB1IP;refGene_tx=XM_003988174.4;refGene_gn=APBB1IP;refGene_tx=XM_023256458.1
B4 24712975 24713047 refGene_gn=APBB1IP;refGene_tx=XM_023256459.1
B4 24718312 24718400
refGene_gn=APBB1IP;refGene_tx=XM_003988174.4;refGene_gn=APBB1IP;refGene_tx=XM_023256458.1;refGene_gn=APBB1IP;refGene_tx=XM_023256459.1
B4 24722004 24722297
refGene_gn=APBB1IP;refGene_tx=XM_003988174.4;refGene_gn=APBB1IP;refGene_tx=XM_023256458.1;refGene_gn=APBB1IP;refGene_tx=XM_023256459.1
B4 24723634 24723712
refGene_gn=APBB1IP;refGene_tx=XM_003988174.4;refGene_gn=APBB1IP;refGene_tx=XM_023256458.1;refGene_gn=APBB1IP;refGene_tx=XM_023256459.1
B4 24729962 24730122
refGene_gn=APBB1IP;refGene_tx=XM_003988174.4;refGene_gn=APBB1IP;refGene_tx=XM_023256458.1;refGene_gn=APBB1IP;refGene_tx=XM_023256459.1
B4 24731290 24731412
refGene_gn=APBB1IP;refGene_tx=XM_003988174.4;refGene_gn=APBB1IP;refGene_tx=XM_023256458.1;refGene_gn=APBB1IP;refGene_tx=XM_023256459.1
B4 24749634 24749721
refGene_gn=APBB1IP;refGene_tx=XM_003988174.4;refGene_gn=APBB1IP;refGene_tx=XM_023256458.1;refGene_gn=APBB1IP;refGene_tx=XM_023256459.1
B4 24752045 24752189
refGene_gn=APBB1IP;refGene_tx=XM_003988174.4;refGene_gn=APBB1IP;refGene_tx=XM_023256458.1;refGene_gn=APBB1IP;refGene_tx=XM_023256459.1

B4 24756153 24756264
refGene_gn=APBB1IP;refGene_tx=XM_003988174.4;refGene_gn=APBB1IP;refGene_tx=XM_023256458.1;refGene_gn=APBB1IP;refGene_tx=XM_023256459.1
B4 24762572 24762671
refGene_gn=APBB1IP;refGene_tx=XM_003988174.4;refGene_gn=APBB1IP;refGene_tx=XM_023256458.1;refGene_gn=APBB1IP;refGene_tx=XM_023256459.1
B4 24763268 24763380
refGene_gn=APBB1IP;refGene_tx=XM_003988174.4;refGene_gn=APBB1IP;refGene_tx=XM_023256458.1;refGene_gn=APBB1IP;refGene_tx=XM_023256459.1
B4 24770725 24770829
refGene_gn=APBB1IP;refGene_tx=XM_003988174.4;refGene_gn=APBB1IP;refGene_tx=XM_023256458.1;refGene_gn=APBB1IP;refGene_tx=XM_023256459.1
B4 24773335 24773845
refGene_gn=APBB1IP;refGene_tx=XM_003988174.4;refGene_gn=APBB1IP;refGene_tx=XM_023256458.1;refGene_gn=APBB1IP;refGene_tx=XM_023256459.1
B4 24792613 24792736
refGene_gn=LOC101087240;refGene_tx=XM_011283638.3
B4 24796107 24796187
refGene_gn=LOC101087240;refGene_tx=XM_011283638.3
B4 24804464 24804508
refGene_gn=LOC101087240;refGene_tx=XM_011283638.3
B4 24805290 24805370
refGene_gn=LOC101087240;refGene_tx=XM_011283638.3
B4 24808036 24808129
refGene_gn=LOC101087240;refGene_tx=XM_011283638.3
B4 24808219 24808301
refGene_gn=LOC101087240;refGene_tx=XM_011283638.3
B4 24808397 24808442
refGene_gn=LOC101087240;refGene_tx=XM_011283638.3
B4 24812855 24812985
refGene_gn=LOC101087240;refGene_tx=XM_011283638.3
B4 24813682 24813766
refGene_gn=LOC101087240;refGene_tx=XM_011283638.3
B4 24815390 24815463
refGene_gn=LOC101087240;refGene_tx=XM_011283638.3
B4 24816694 24816764
refGene_gn=LOC101087240;refGene_tx=XM_011283638.3
B4 24818314 24818402
refGene_gn=LOC101087240;refGene_tx=XM_011283638.3
B4 24820179 24820247
refGene_gn=LOC101087240;refGene_tx=XM_011283638.3
B4 24820436 24820535
refGene_gn=LOC101087240;refGene_tx=XM_011283638.3
B4 24820627 24820684
refGene_gn=LOC101087240;refGene_tx=XM_011283638.3
B4 24820768 24820833
refGene_gn=LOC101087240;refGene_tx=XM_011283638.3
B4 24824127 24824256
refGene_gn=LOC101087240;refGene_tx=XM_011283638.3
B4 24824888 24825007

refGene_gn=LOC101087240;refGene_tx=XM_011283638.3
B4	24828440	24828687
refGene_gn=LOC101087240;refGene_tx=XM_011283638.3
B4	24829128	24829257
refGene_gn=PDSS1;refGene_tx=XM_023256462.1;refGene_gn=PDSS1;refGene_tx=XM_023256463.1;refGene_gn=PDSS1;refGene_tx=XM_023256464.1
B4	24833572	24833637
refGene_gn=PDSS1;refGene_tx=XM_023256462.1;refGene_gn=PDSS1;refGene_tx=XM_023256463.1;refGene_gn=PDSS1;refGene_tx=XM_023256464.1
B4	24834069	24834190
refGene_gn=PDSS1;refGene_tx=XM_019834093.1;refGene_gn=PDSS1;refGene_tx=XM_023256462.1;refGene_gn=PDSS1;refGene_tx=XM_023256463.1;refGene_gn=PDSS1;refGene_tx=XM_023256464.1
B4	24836913	24837044
refGene_gn=PDSS1;refGene_tx=XM_023256462.1;refGene_gn=PDSS1;refGene_tx=XM_023256463.1;refGene_gn=PDSS1;refGene_tx=XM_023256464.1
B4	24845489	24845527	refGene_gn=PDSS1;refGene_tx=XM_019834092.2
B4	24847097	24847239
refGene_gn=PDSS1;refGene_tx=XM_019834092.2;refGene_gn=PDSS1;refGene_tx=XM_019834093.1;refGene_gn=PDSS1;refGene_tx=XM_023256462.1;refGene_gn=PDSS1;refGene_tx=XM_023256463.1;refGene_gn=PDSS1;refGene_tx=XM_023256464.1
B4	24850079	24850191
refGene_gn=PDSS1;refGene_tx=XM_019834092.2;refGene_gn=PDSS1;refGene_tx=XM_019834093.1;refGene_gn=PDSS1;refGene_tx=XM_023256462.1;refGene_gn=PDSS1;refGene_tx=XM_023256463.1
B4	24850290	24850400
refGene_gn=PDSS1;refGene_tx=XM_019834092.2;refGene_gn=PDSS1;refGene_tx=XM_019834093.1;refGene_gn=PDSS1;refGene_tx=XM_023256462.1;refGene_gn=PDSS1;refGene_tx=XM_023256463.1
B4	24861722	24861803
refGene_gn=PDSS1;refGene_tx=XM_019834092.2;refGene_gn=PDSS1;refGene_tx=XM_019834093.1;refGene_gn=PDSS1;refGene_tx=XM_023256462.1;refGene_gn=PDSS1;refGene_tx=XM_023256463.1;refGene_gn=PDSS1;refGene_tx=XM_023256464.1
B4	24861906	24862020
refGene_gn=PDSS1;refGene_tx=XM_019834092.2;refGene_gn=PDSS1;refGene_tx=XM_019834093.1;refGene_gn=PDSS1;refGene_tx=XM_023256462.1;refGene_gn=PDSS1;refGene_tx=XM_023256463.1;refGene_gn=PDSS1;refGene_tx=XM_023256464.1
B4	24867602	24867683
refGene_gn=PDSS1;refGene_tx=XM_019834092.2;refGene_gn=PDSS1;refGene_tx=XM_019834093.1;refGene_gn=PDSS1;refGene_tx=XM_023256462.1;refGene_gn=PDSS1;refGene_tx=XM_023256463.1;refGene_gn=PDSS1;refGene_tx=XM_023256464.1
B4	24869720	24869810	refGene_gn=PDSS1;refGene_tx=XM_023256463.1
B4	24870799	24870940
refGene_gn=PDSS1;refGene_tx=XM_019834092.2;refGene_gn=PDSS1;refGene_tx=XM_019834093.1;refGene_gn=PDSS1;refGene_tx=XM_023256462.1;refGene_gn=PDSS1;refGene_tx=XM_023256464.1

B4 24873013 24873189
refGene_gn=ABI1;refGene_tx=XM_011283639.3;refGene_gn=ABI1;refGene_tx=XM_011283640.2;refGene_gn=ABI1;refGene_tx=XM_011283641.2;refGene_gn=ABI1;refGene_tx=XM_011283642.2;refGene_gn=ABI1;refGene_tx=XM_011283643.2;refGene_gn=ABI1;refGene_tx=XM_011283644.3;refGene_gn=ABI1;refGene_tx=XM_011283645.2;refGene_gn=ABI1;refGene_tx=XM_011283646.3;refGene_gn=ABI1;refGene_tx=XM_011283647.2;refGene_gn=ABI1;refGene_tx=XM_011283648.2;refGene_gn=ABI1;refGene_tx=XM_011283649.3;refGene_gn=ABI1;refGene_tx=XM_011283650.3;refGene_gn=ABI1;refGene_tx=XM_011283651.3;refGene_gn=ABI1;refGene_tx=XM_019834089.1;refGene_gn=ABI1;refGene_tx=XM_023256460.1;refGene_gn=ABI1;refGene_tx=XM_023256461.1
B4 24877928 24878114
refGene_gn=ABI1;refGene_tx=XM_011283639.3;refGene_gn=ABI1;refGene_tx=XM_011283640.2;refGene_gn=ABI1;refGene_tx=XM_011283641.2;refGene_gn=ABI1;refGene_tx=XM_011283642.2;refGene_gn=ABI1;refGene_tx=XM_011283643.2;refGene_gn=ABI1;refGene_tx=XM_011283644.3;refGene_gn=ABI1;refGene_tx=XM_011283645.2;refGene_gn=ABI1;refGene_tx=XM_011283646.3;refGene_gn=ABI1;refGene_tx=XM_011283647.2;refGene_gn=ABI1;refGene_tx=XM_011283648.2;refGene_gn=ABI1;refGene_tx=XM_011283649.3;refGene_gn=ABI1;refGene_tx=XM_011283650.3;refGene_gn=ABI1;refGene_tx=XM_011283651.3;refGene_gn=ABI1;refGene_tx=XM_019834089.1;refGene_gn=ABI1;refGene_tx=XM_023256460.1;refGene_gn=ABI1;refGene_tx=XM_023256461.1
B4 24881752 24881839
refGene_gn=ABI1;refGene_tx=XM_011283639.3;refGene_gn=ABI1;refGene_tx=XM_011283640.2;refGene_gn=ABI1;refGene_tx=XM_011283641.2;refGene_gn=ABI1;refGene_tx=XM_011283642.2;refGene_gn=ABI1;refGene_tx=XM_011283643.2;refGene_gn=ABI1;refGene_tx=XM_011283645.2;refGene_gn=ABI1;refGene_tx=XM_011283646.3;refGene_gn=ABI1;refGene_tx=XM_019834089.1
B4 24884964 24885141
refGene_gn=ABI1;refGene_tx=XM_011283639.3;refGene_gn=ABI1;refGene_tx=XM_011283640.2;refGene_gn=ABI1;refGene_tx=XM_011283641.2;refGene_gn=ABI1;refGene_tx=XM_011283642.2;refGene_gn=ABI1;refGene_tx=XM_011283643.2;refGene_gn=ABI1;refGene_tx=XM_011283644.3;refGene_gn=ABI1;refGene_tx=XM_011283645.2;refGene_gn=ABI1;refGene_tx=XM_011283646.3;refGene_gn=ABI1;refGene_tx=XM_011283647.2;refGene_gn=ABI1;refGene_tx=XM_011283648.2;refGene_gn=ABI1;refGene_tx=XM_011283649.3;refGene_gn=ABI1;refGene_tx=XM_019834089.1;refGene_gn=ABI1;refGene_tx=XM_023256460.1;refGene_gn=ABI1;refGene_tx=XM_023256461.1
B4 24889227 24889308
refGene_gn=ABI1;refGene_tx=XM_011283639.3;refGene_gn=ABI1;refGene_tx=XM_011283640.2;refGene_gn=ABI1;refGene_tx=XM_011283641.2;refGene_gn=ABI1;refGene_tx=XM_011283644.3;refGene_gn=ABI1;refGene_tx=XM_019834089.1;refGene_gn=ABI1;refGene_tx=XM_023256460.1
B4 24890177 24890278
refGene_gn=ABI1;refGene_tx=XM_011283639.3;refGene_gn=ABI1;refGene_tx=XM_011283640.2;refGene_gn=ABI1;refGene_tx=XM_011283641.2;refGene_gn=ABI1;refGene_tx=XM_011283642.2;refGene_gn=ABI1;refGene_tx=XM_011283643.2;refGene_gn=ABI1;refGene_tx=XM_011283644.3;refGene_gn=ABI1;refGene_tx=XM_011283645.2;refGene_gn=ABI1;refGene_tx=XM_011283646.3;refGene_gn=ABI1;refGene_tx=XM_011283647.2;refGene_gn=ABI1;refGene_tx=XM_011283648.2;

refGene_gn=ABI1;refGene_tx=XM_011283649.3;refGene_gn=ABI1;refGene_tx=XM_011283650.3;refGene_gn=ABI1;refGene_tx=XM_011283651.3;refGene_gn=ABI1;refGene_tx=XM_019834089.1;refGene_gn=ABI1;refGene_tx=XM_023256460.1;refGene_gn=ABI1;refGene_tx=XM_023256461.1
B4 24893606 24893747
refGene_gn=ABI1;refGene_tx=XM_011283639.3;refGene_gn=ABI1;refGene_tx=XM_011283640.2;refGene_gn=ABI1;refGene_tx=XM_011283641.2;refGene_gn=ABI1;refGene_tx=XM_011283642.2;refGene_gn=ABI1;refGene_tx=XM_011283643.2;refGene_gn=ABI1;refGene_tx=XM_011283644.3;refGene_gn=ABI1;refGene_tx=XM_011283645.2;refGene_gn=ABI1;refGene_tx=XM_011283646.3;refGene_gn=ABI1;refGene_tx=XM_011283647.2;refGene_gn=ABI1;refGene_tx=XM_011283648.2;refGene_gn=ABI1;refGene_tx=XM_011283649.3;refGene_gn=ABI1;refGene_tx=XM_011283650.3;refGene_gn=ABI1;refGene_tx=XM_011283651.3;refGene_gn=ABI1;refGene_tx=XM_019834089.1;refGene_gn=ABI1;refGene_tx=XM_023256460.1;refGene_gn=ABI1;refGene_tx=XM_023256461.1
B4 24895313 24895414
refGene_gn=ABI1;refGene_tx=XM_011283639.3;refGene_gn=ABI1;refGene_tx=XM_011283640.2;refGene_gn=ABI1;refGene_tx=XM_011283641.2;refGene_gn=ABI1;refGene_tx=XM_011283642.2;refGene_gn=ABI1;refGene_tx=XM_011283643.2;refGene_gn=ABI1;refGene_tx=XM_011283644.3;refGene_gn=ABI1;refGene_tx=XM_011283645.2;refGene_gn=ABI1;refGene_tx=XM_011283646.3;refGene_gn=ABI1;refGene_tx=XM_011283647.2;refGene_gn=ABI1;refGene_tx=XM_011283648.2;refGene_gn=ABI1;refGene_tx=XM_011283649.3;refGene_gn=ABI1;refGene_tx=XM_011283650.3;refGene_gn=ABI1;refGene_tx=XM_011283651.3;refGene_gn=ABI1;refGene_tx=XM_019834089.1;refGene_gn=ABI1;refGene_tx=XM_023256460.1;refGene_gn=ABI1;refGene_tx=XM_023256461.1
B4 24896143 24896158
refGene_gn=ABI1;refGene_tx=XM_011283639.3;refGene_gn=ABI1;refGene_tx=XM_011283640.2;refGene_gn=ABI1;refGene_tx=XM_011283642.2;refGene_gn=ABI1;refGene_tx=XM_011283643.2;refGene_gn=ABI1;refGene_tx=XM_011283644.3;refGene_gn=ABI1;refGene_tx=XM_011283647.2;refGene_gn=ABI1;refGene_tx=XM_011283648.2;refGene_gn=ABI1;refGene_tx=XM_011283650.3;refGene_gn=ABI1;refGene_tx=XM_019834089.1;refGene_gn=ABI1;refGene_tx=XM_023256460.1
B4 24902143 24902320
refGene_gn=ABI1;refGene_tx=XM_011283639.3;refGene_gn=ABI1;refGene_tx=XM_011283640.2;refGene_gn=ABI1;refGene_tx=XM_011283641.2;refGene_gn=ABI1;refGene_tx=XM_011283642.2;refGene_gn=ABI1;refGene_tx=XM_011283643.2;refGene_gn=ABI1;refGene_tx=XM_011283644.3;refGene_gn=ABI1;refGene_tx=XM_011283645.2;refGene_gn=ABI1;refGene_tx=XM_011283646.3;refGene_gn=ABI1;refGene_tx=XM_011283647.2;refGene_gn=ABI1;refGene_tx=XM_011283648.2;refGene_gn=ABI1;refGene_tx=XM_011283649.3;refGene_gn=ABI1;refGene_tx=XM_011283650.3;refGene_gn=ABI1;refGene_tx=XM_011283651.3;refGene_gn=ABI1;refGene_tx=XM_019834089.1;refGene_gn=ABI1;refGene_tx=XM_023256460.1;refGene_gn=ABI1;refGene_tx=XM_023256461.1
B4 24948496 24948664
refGene_gn=ABI1;refGene_tx=XM_011283639.3;refGene_gn=ABI1;refGene_tx=XM_011283640.2;refGene_gn=ABI1;refGene_tx=XM_011283641.2;refGene_gn=ABI1;refGene_tx=XM_011283642.2;refGene_gn=ABI1;refGene_tx=XM_011283643.2;refGene_gn=ABI1;refGene_tx=XM_011283644.3;refGene_gn=ABI1;refGene_tx=XM_011283645.2;refGene_gn=ABI1;refGene_tx=XM_011283646.3;refGene_gn=ABI

1;refGene_tx=XM_011283647.2;refGene_gn=ABI1;refGene_tx=XM_011283648.2;
refGene_gn=ABI1;refGene_tx=XM_011283649.3;refGene_gn=ABI1;refGene_tx=X
M_011283650.3;refGene_gn=ABI1;refGene_tx=XM_011283651.3;refGene_gn=ABI
1;refGene_tx=XM_019834089.1;refGene_gn=ABI1;refGene_tx=XM_023256460.1;
refGene_gn=ABI1;refGene_tx=XM_023256461.1
B4 25002727 25002844
refGene_gn=ABI1;refGene_tx=XM_011283639.3;refGene_gn=ABI1;refGene_tx=X
M_011283640.2;refGene_gn=ABI1;refGene_tx=XM_011283641.2;refGene_gn=ABI
1;refGene_tx=XM_011283642.2;refGene_gn=ABI1;refGene_tx=XM_011283643.2;
refGene_gn=ABI1;refGene_tx=XM_011283644.3;refGene_gn=ABI1;refGene_tx=X
M_011283645.2;refGene_gn=ABI1;refGene_tx=XM_011283646.3;refGene_gn=ABI
1;refGene_tx=XM_011283647.2;refGene_gn=ABI1;refGene_tx=XM_011283648.2;
refGene_gn=ABI1;refGene_tx=XM_011283649.3;refGene_gn=ABI1;refGene_tx=X
M_011283650.3;refGene_gn=ABI1;refGene_tx=XM_011283651.3;refGene_gn=ABI
1;refGene_tx=XM_023256460.1;refGene_gn=ABI1;refGene_tx=XM_023256461.1
B4 25020552 25020558 refGene_gn=ABI1;refGene_tx=XM_019834089.1
B4 25022648 25022663 refGene_gn=ACBD5;refGene_tx=XM_006933328.3
B4 25022847 25023019 refGene_gn=ACBD5;refGene_tx=XM_006933328.3
B4 25027848 25027969 refGene_gn=ACBD5;refGene_tx=XM_006933328.3
B4 25029860 25029933 refGene_gn=ACBD5;refGene_tx=XM_006933328.3
B4 25038929 25039044 refGene_gn=ACBD5;refGene_tx=XM_006933328.3
B4 25039141 25039174 refGene_gn=ACBD5;refGene_tx=XM_006933328.3
B4 25043144 25043279 refGene_gn=ACBD5;refGene_tx=XM_006933328.3
B4 25044496 25044700 refGene_gn=ACBD5;refGene_tx=XM_006933328.3
B4 25045544 25045651 refGene_gn=ACBD5;refGene_tx=XM_006933328.3
B4 25049080 25049348 refGene_gn=ACBD5;refGene_tx=XM_006933328.3
B4 25053877 25054077 refGene_gn=ACBD5;refGene_tx=XM_006933328.3
B4 25061578 25061623 refGene_gn=ACBD5;refGene_tx=XM_006933328.3
B4 25061942 25062052 refGene_gn=ACBD5;refGene_tx=XM_006933328.3
B4 25066871 25066884 refGene_gn=ACBD5;refGene_tx=XM_006933328.3
B4 25083743 25083901
refGene_gn=MASTL;refGene_tx=XM_003988178.4;refGene_gn=MASTL;refGene_tx
=XM_019834100.1
B4 25086566 25086668
refGene_gn=MASTL;refGene_tx=XM_003988178.4;refGene_gn=MASTL;refGene_tx
=XM_019834100.1
B4 25086750 25086864 refGene_gn=MASTL;refGene_tx=XM_003988178.4
B4 25092915 25093057
refGene_gn=MASTL;refGene_tx=XM_003988178.4;refGene_gn=MASTL;refGene_tx
=XM_019834100.1
B4 25094370 25095519
refGene_gn=MASTL;refGene_tx=XM_003988178.4;refGene_gn=MASTL;refGene_tx
=XM_019834100.1
B4 25098244 25098417
refGene_gn=MASTL;refGene_tx=XM_003988178.4;refGene_gn=MASTL;refGene_tx
=XM_019834100.1
B4 25100946 25101097
refGene_gn=MASTL;refGene_tx=XM_003988178.4;refGene_gn=MASTL;refGene_tx
=XM_019834100.1
B4 25101544 25101651

refGene_gn=MASTL;refGene_tx=XM_003988178.4;refGene_gn=MASTL;refGene_tx=XM_019834100.1
B4 25107141 25107230
refGene_gn=MASTL;refGene_tx=XM_003988178.4;refGene_gn=MASTL;refGene_tx=XM_019834100.1
B4 25107958 25108098
refGene_gn=MASTL;refGene_tx=XM_003988178.4;refGene_gn=MASTL;refGene_tx=XM_019834100.1
B4 25115958 25116096
refGene_gn=MASTL;refGene_tx=XM_003988178.4;refGene_gn=MASTL;refGene_tx=XM_019834100.1
B4 25119060 25119246
refGene_gn=MASTL;refGene_tx=XM_003988178.4;refGene_gn=MASTL;refGene_tx=XM_019834100.1
B4 25120492 25120525 refGene_gn=YME1L1;refGene_tx=XM_023256465.1
B4 25129529 25129661 refGene_gn=YME1L1;refGene_tx=XM_023256465.1
B4 25136310 25136473 refGene_gn=YME1L1;refGene_tx=XM_023256465.1
B4 25138908 25139007 refGene_gn=YME1L1;refGene_tx=XM_023256465.1
B4 25143378 25143488 refGene_gn=YME1L1;refGene_tx=XM_023256465.1
B4 25145245 25145396 refGene_gn=YME1L1;refGene_tx=XM_023256465.1
B4 25145789 25145873 refGene_gn=YME1L1;refGene_tx=XM_023256465.1
B4 25148798 25148881 refGene_gn=YME1L1;refGene_tx=XM_023256465.1
B4 25155655 25155746 refGene_gn=YME1L1;refGene_tx=XM_023256465.1
B4 25158989 25159142 refGene_gn=YME1L1;refGene_tx=XM_023256465.1
B4 25159705 25159838 refGene_gn=YME1L1;refGene_tx=XM_023256465.1
B4 25160596 25160659 refGene_gn=YME1L1;refGene_tx=XM_023256465.1
B4 25163171 25163284 refGene_gn=YME1L1;refGene_tx=XM_023256465.1
B4 25164172 25164328 refGene_gn=YME1L1;refGene_tx=XM_023256465.1
B4 25165083 25165235 refGene_gn=YME1L1;refGene_tx=XM_023256465.1
B4 25165816 25165943 refGene_gn=YME1L1;refGene_tx=XM_023256465.1
B4 25166038 25166112 refGene_gn=YME1L1;refGene_tx=XM_023256465.1
B4 25166968 25167055 refGene_gn=YME1L1;refGene_tx=XM_023256465.1
B4 25169551 25169695 refGene_gn=YME1L1;refGene_tx=XM_023256465.1
B4 25191717 25191944
refGene_gn=ANKRD26;refGene_tx=XM_023256466.1;refGene_gn=ANKRD26;refGene_tx=XM_023256467.1;refGene_gn=ANKRD26;refGene_tx=XM_023256468.1;refGene_gn=ANKRD26;refGene_tx=XM_023256469.1;refGene_gn=ANKRD26;refGene_tx=XM_023256470.1;refGene_gn=ANKRD26;refGene_tx=XM_023256471.1
B4 25196867 25196982
refGene_gn=ANKRD26;refGene_tx=XM_023256466.1;refGene_gn=ANKRD26;refGene_tx=XM_023256467.1;refGene_gn=ANKRD26;refGene_tx=XM_023256468.1;refGene_gn=ANKRD26;refGene_tx=XM_023256469.1;refGene_gn=ANKRD26;refGene_tx=XM_023256470.1;refGene_gn=ANKRD26;refGene_tx=XM_023256471.1
B4 25197143 25197317
refGene_gn=ANKRD26;refGene_tx=XM_023256466.1;refGene_gn=ANKRD26;refGene_tx=XM_023256467.1;refGene_gn=ANKRD26;refGene_tx=XM_023256468.1;refGene_gn=ANKRD26;refGene_tx=XM_023256469.1;refGene_gn=ANKRD26;refGene_tx=XM_023256470.1;refGene_gn=ANKRD26;refGene_tx=XM_023256471.1
B4 25204646 25204753
refGene_gn=ANKRD26;refGene_tx=XM_023256466.1;refGene_gn=ANKRD26;refGen

e_tx=XM_023256467.1;refGene_gn=ANKRD26;refGene_tx=XM_023256468.1;refGene_gn=ANKRD26;refGene_tx=XM_023256469.1;refGene_gn=ANKRD26;refGene_tx=XM_023256470.1;refGene_gn=ANKRD26;refGene_tx=XM_023256471.1
B4 25208640 25208711
refGene_gn=ANKRD26;refGene_tx=XM_023256466.1;refGene_gn=ANKRD26;refGene_tx=XM_023256467.1;refGene_gn=ANKRD26;refGene_tx=XM_023256468.1;refGene_gn=ANKRD26;refGene_tx=XM_023256469.1;refGene_gn=ANKRD26;refGene_tx=XM_023256470.1;refGene_gn=ANKRD26;refGene_tx=XM_023256471.1
B4 25217445 25217482
refGene_gn=ANKRD26;refGene_tx=XM_023256466.1;refGene_gn=ANKRD26;refGene_tx=XM_023256467.1;refGene_gn=ANKRD26;refGene_tx=XM_023256468.1;refGene_gn=ANKRD26;refGene_tx=XM_023256469.1;refGene_gn=ANKRD26;refGene_tx=XM_023256470.1;refGene_gn=ANKRD26;refGene_tx=XM_023256471.1
B4 25229204 25229277
refGene_gn=ANKRD26;refGene_tx=XM_023256466.1;refGene_gn=ANKRD26;refGene_tx=XM_023256467.1;refGene_gn=ANKRD26;refGene_tx=XM_023256468.1;refGene_gn=ANKRD26;refGene_tx=XM_023256469.1;refGene_gn=ANKRD26;refGene_tx=XM_023256470.1;refGene_gn=ANKRD26;refGene_tx=XM_023256471.1
B4 25232360 25232421
refGene_gn=ANKRD26;refGene_tx=XM_023256466.1;refGene_gn=ANKRD26;refGene_tx=XM_023256467.1;refGene_gn=ANKRD26;refGene_tx=XM_023256468.1;refGene_gn=ANKRD26;refGene_tx=XM_023256469.1;refGene_gn=ANKRD26;refGene_tx=XM_023256470.1;refGene_gn=ANKRD26;refGene_tx=XM_023256471.1
B4 25232529 25232732
refGene_gn=ANKRD26;refGene_tx=XM_023256466.1;refGene_gn=ANKRD26;refGene_tx=XM_023256467.1;refGene_gn=ANKRD26;refGene_tx=XM_023256468.1;refGene_gn=ANKRD26;refGene_tx=XM_023256469.1;refGene_gn=ANKRD26;refGene_tx=XM_023256470.1;refGene_gn=ANKRD26;refGene_tx=XM_023256471.1
B4 25239119 25239246
refGene_gn=ANKRD26;refGene_tx=XM_023256466.1;refGene_gn=ANKRD26;refGene_tx=XM_023256467.1;refGene_gn=ANKRD26;refGene_tx=XM_023256468.1;refGene_gn=ANKRD26;refGene_tx=XM_023256469.1;refGene_gn=ANKRD26;refGene_tx=XM_023256470.1;refGene_gn=ANKRD26;refGene_tx=XM_023256471.1
B4 25239811 25239873
refGene_gn=ANKRD26;refGene_tx=XM_023256466.1;refGene_gn=ANKRD26;refGene_tx=XM_023256467.1;refGene_gn=ANKRD26;refGene_tx=XM_023256468.1;refGene_gn=ANKRD26;refGene_tx=XM_023256469.1;refGene_gn=ANKRD26;refGene_tx=XM_023256470.1;refGene_gn=ANKRD26;refGene_tx=XM_023256471.1
B4 25244716 25244807
refGene_gn=ANKRD26;refGene_tx=XM_023256466.1;refGene_gn=ANKRD26;refGene_tx=XM_023256467.1;refGene_gn=ANKRD26;refGene_tx=XM_023256468.1;refGene_gn=ANKRD26;refGene_tx=XM_023256469.1;refGene_gn=ANKRD26;refGene_tx=XM_023256470.1;refGene_gn=ANKRD26;refGene_tx=XM_023256471.1
B4 25247131 25247230
refGene_gn=ANKRD26;refGene_tx=XM_023256466.1;refGene_gn=ANKRD26;refGene_tx=XM_023256467.1;refGene_gn=ANKRD26;refGene_tx=XM_023256468.1;refGene_gn=ANKRD26;refGene_tx=XM_023256470.1;refGene_gn=ANKRD26;refGene_tx=XM_023256471.1
B4 25247860 25247889
refGene_gn=ANKRD26;refGene_tx=XM_023256466.1;refGene_gn=ANKRD26;refGen

e_tx=XM_023256467.1;refGene_gn=ANKRD26;refGene_tx=XM_023256468.1;refGene_gn=ANKRD26;refGene_tx=XM_023256469.1;refGene_gn=ANKRD26;refGene_tx=XM_023256470.1;refGene_gn=ANKRD26;refGene_tx=XM_023256471.1
B4 25247982 25248055
refGene_gn=ANKRD26;refGene_tx=XM_023256466.1;refGene_gn=ANKRD26;refGene_tx=XM_023256467.1;refGene_gn=ANKRD26;refGene_tx=XM_023256468.1;refGene_gn=ANKRD26;refGene_tx=XM_023256469.1;refGene_gn=ANKRD26;refGene_tx=XM_023256470.1;refGene_gn=ANKRD26;refGene_tx=XM_023256471.1
B4 25249692 25249950
refGene_gn=ANKRD26;refGene_tx=XM_023256466.1;refGene_gn=ANKRD26;refGene_tx=XM_023256467.1;refGene_gn=ANKRD26;refGene_tx=XM_023256468.1;refGene_gn=ANKRD26;refGene_tx=XM_023256469.1;refGene_gn=ANKRD26;refGene_tx=XM_023256470.1;refGene_gn=ANKRD26;refGene_tx=XM_023256471.1
B4 25252867 25252941
refGene_gn=ANKRD26;refGene_tx=XM_023256466.1;refGene_gn=ANKRD26;refGene_tx=XM_023256467.1;refGene_gn=ANKRD26;refGene_tx=XM_023256468.1;refGene_gn=ANKRD26;refGene_tx=XM_023256469.1;refGene_gn=ANKRD26;refGene_tx=XM_023256470.1;refGene_gn=ANKRD26;refGene_tx=XM_023256471.1
B4 25258355 25258526
refGene_gn=ANKRD26;refGene_tx=XM_023256466.1;refGene_gn=ANKRD26;refGene_tx=XM_023256467.1;refGene_gn=ANKRD26;refGene_tx=XM_023256468.1;refGene_gn=ANKRD26;refGene_tx=XM_023256469.1;refGene_gn=ANKRD26;refGene_tx=XM_023256470.1;refGene_gn=ANKRD26;refGene_tx=XM_023256471.1
B4 25313715 25313861
refGene_gn=LOC109496862;refGene_tx=XM_023256472.1
B4 25316127 25316404
refGene_gn=LOC109496862;refGene_tx=XM_023256472.1
B4 25333741 25333809
refGene_gn=LOC105260720;refGene_tx=XM_019835807.2
B4 25335981 25336156
refGene_gn=LOC105260720;refGene_tx=XM_019835807.2
B4 25336158 25336478
refGene_gn=LOC105260720;refGene_tx=XM_019835807.2
B4 25373230 25373647 refGene_gn=PTCHD3;refGene_tx=XM_019835808.2
B4 25373648 25373728 refGene_gn=PTCHD3;refGene_tx=XM_019835808.2
B4 25373730 25374729 refGene_gn=PTCHD3;refGene_tx=XM_019835808.2
B4 25377337 25377483 refGene_gn=PTCHD3;refGene_tx=XM_019835808.2
B4 25392921 25393043 refGene_gn=PTCHD3;refGene_tx=XM_019835808.2
B4 25393957 25394253 refGene_gn=PTCHD3;refGene_tx=XM_019835808.2
B4 25394255 25394835 refGene_gn=PTCHD3;refGene_tx=XM_019835808.2
B4 25483113 25483181 refGene_gn=RAB18;refGene_tx=XM_003988192.5
B4 25487968 25488024 refGene_gn=RAB18;refGene_tx=XM_003988192.5
B4 25509937 25509999 refGene_gn=RAB18;refGene_tx=XM_003988192.5
B4 25514506 25514579 refGene_gn=RAB18;refGene_tx=XM_003988192.5
B4 25515761 25515880 refGene_gn=RAB18;refGene_tx=XM_003988192.5
B4 25515952 25516019 refGene_gn=RAB18;refGene_tx=XM_003988192.5
B4 25519940 25520116 refGene_gn=RAB18;refGene_tx=XM_003988192.5
B4 25640532 25640719
refGene_gn=MKX;refGene_tx=XM_006933336.4;refGene_gn=MKX;refGene_tx=XM_019834130.2;refGene_gn=MKX;refGene_tx=XM_019834132.2;refGene_gn=MKX;re

fGene_tx=XM_019834133.2
B4	25640824	25640858
refGene_gn=MKX;refGene_tx=XM_006933336.4;refGene_gn=MKX;refGene_tx=XM_019834130.2;refGene_gn=MKX;refGene_tx=XM_019834132.2;refGene_gn=MKX;refGene_tx=XM_019834133.2
B4	25673747	25673767	refGene_gn=MKX;refGene_tx=XM_023256474.1
B4	25692502	25692838
refGene_gn=MKX;refGene_tx=XM_006933336.4;refGene_gn=MKX;refGene_tx=XM_019834130.2;refGene_gn=MKX;refGene_tx=XM_019834132.2;refGene_gn=MKX;refGene_tx=XM_019834133.2;refGene_gn=MKX;refGene_tx=XM_023256474.1
B4	25693266	25693420
refGene_gn=MKX;refGene_tx=XM_006933336.4;refGene_gn=MKX;refGene_tx=XM_019834130.2;refGene_gn=MKX;refGene_tx=XM_019834132.2;refGene_gn=MKX;refGene_tx=XM_019834133.2;refGene_gn=MKX;refGene_tx=XM_023256474.1
B4	25702173	25702333
refGene_gn=MKX;refGene_tx=XM_006933336.4;refGene_gn=MKX;refGene_tx=XM_019834130.2;refGene_gn=MKX;refGene_tx=XM_019834132.2;refGene_gn=MKX;refGene_tx=XM_019834133.2;refGene_gn=MKX;refGene_tx=XM_023256474.1
B4	25704136	25704403
refGene_gn=MKX;refGene_tx=XM_006933336.4;refGene_gn=MKX;refGene_tx=XM_019834130.2;refGene_gn=MKX;refGene_tx=XM_019834132.2;refGene_gn=MKX;refGene_tx=XM_019834133.2;refGene_gn=MKX;refGene_tx=XM_023256474.1
B4	25705252	25705359
refGene_gn=MKX;refGene_tx=XM_019834130.2;refGene_gn=MKX;refGene_tx=XM_023256474.1
B4	25780288	25780402	refGene_gn=ARMC4;refGene_tx=XM_023256476.1
B4	25807568	25807766
refGene_gn=ARMC4;refGene_tx=XM_023256475.1;refGene_gn=ARMC4;refGene_tx=XM_023256478.1;refGene_gn=ARMC4;refGene_tx=XM_023256479.1
B4	25818270	25818273	refGene_gn=ARMC4;refGene_tx=XM_023256477.1
B4	25827712	25827934
refGene_gn=ARMC4;refGene_tx=XM_023256475.1;refGene_gn=ARMC4;refGene_tx=XM_023256476.1;refGene_gn=ARMC4;refGene_tx=XM_023256477.1;refGene_gn=ARMC4;refGene_tx=XM_023256478.1;refGene_gn=ARMC4;refGene_tx=XM_023256479.1
B4	25829224	25829413
refGene_gn=ARMC4;refGene_tx=XM_023256475.1;refGene_gn=ARMC4;refGene_tx=XM_023256476.1;refGene_gn=ARMC4;refGene_tx=XM_023256477.1;refGene_gn=ARMC4;refGene_tx=XM_023256478.1;refGene_gn=ARMC4;refGene_tx=XM_023256479.1
B4	25863460	25863575
refGene_gn=ARMC4;refGene_tx=XM_023256475.1;refGene_gn=ARMC4;refGene_tx=XM_023256476.1;refGene_gn=ARMC4;refGene_tx=XM_023256477.1;refGene_gn=ARMC4;refGene_tx=XM_023256478.1;refGene_gn=ARMC4;refGene_tx=XM_023256479.1
B4	25901624	25901867
refGene_gn=ARMC4;refGene_tx=XM_023256475.1;refGene_gn=ARMC4;refGene_tx=XM_023256476.1;refGene_gn=ARMC4;refGene_tx=XM_023256477.1;refGene_gn=ARMC4;refGene_tx=XM_023256478.1;refGene_gn=ARMC4;refGene_tx=XM_023256479.1

B4	25902966 25903121
refGene_gn=ARMC4;refGene_tx=XM_023256475.1;refGene_gn=ARMC4;refGene_tx
=XM_023256476.1;refGene_gn=ARMC4;refGene_tx=XM_023256477.1;refGene_gn=
ARMC4;refGene_tx=XM_023256478.1;refGene_gn=ARMC4;refGene_tx=XM_0232564
79.1
B4	25905328 25905439
refGene_gn=ARMC4;refGene_tx=XM_023256475.1;refGene_gn=ARMC4;refGene_tx
=XM_023256476.1;refGene_gn=ARMC4;refGene_tx=XM_023256477.1;refGene_gn=
ARMC4;refGene_tx=XM_023256478.1;refGene_gn=ARMC4;refGene_tx=XM_0232564
79.1
B4	25905770 25906013
refGene_gn=ARMC4;refGene_tx=XM_023256475.1;refGene_gn=ARMC4;refGene_tx
=XM_023256476.1;refGene_gn=ARMC4;refGene_tx=XM_023256477.1;refGene_gn=
ARMC4;refGene_tx=XM_023256478.1;refGene_gn=ARMC4;refGene_tx=XM_0232564
79.1
B4	25908214 25908424
refGene_gn=ARMC4;refGene_tx=XM_023256475.1;refGene_gn=ARMC4;refGene_tx
=XM_023256476.1;refGene_gn=ARMC4;refGene_tx=XM_023256477.1;refGene_gn=
ARMC4;refGene_tx=XM_023256478.1;refGene_gn=ARMC4;refGene_tx=XM_0232564
79.1
B4	25908815 25908962
refGene_gn=ARMC4;refGene_tx=XM_023256475.1;refGene_gn=ARMC4;refGene_tx
=XM_023256476.1;refGene_gn=ARMC4;refGene_tx=XM_023256477.1;refGene_gn=
ARMC4;refGene_tx=XM_023256478.1;refGene_gn=ARMC4;refGene_tx=XM_0232564
79.1
B4	25927617 25927762
refGene_gn=ARMC4;refGene_tx=XM_023256475.1;refGene_gn=ARMC4;refGene_tx
=XM_023256476.1;refGene_gn=ARMC4;refGene_tx=XM_023256477.1;refGene_gn=
ARMC4;refGene_tx=XM_023256478.1;refGene_gn=ARMC4;refGene_tx=XM_0232564
79.1
B4	25932204 25932303
refGene_gn=ARMC4;refGene_tx=XM_023256475.1;refGene_gn=ARMC4;refGene_tx
=XM_023256476.1;refGene_gn=ARMC4;refGene_tx=XM_023256477.1;refGene_gn=
ARMC4;refGene_tx=XM_023256478.1;refGene_gn=ARMC4;refGene_tx=XM_0232564
79.1
B4	25933638 25933841
refGene_gn=ARMC4;refGene_tx=XM_023256475.1;refGene_gn=ARMC4;refGene_tx
=XM_023256476.1;refGene_gn=ARMC4;refGene_tx=XM_023256477.1;refGene_gn=
ARMC4;refGene_tx=XM_023256478.1;refGene_gn=ARMC4;refGene_tx=XM_0232564
79.1
B4	25944354 25944471
refGene_gn=ARMC4;refGene_tx=XM_023256475.1;refGene_gn=ARMC4;refGene_tx
=XM_023256476.1;refGene_gn=ARMC4;refGene_tx=XM_023256477.1;refGene_gn=
ARMC4;refGene_tx=XM_023256478.1;refGene_gn=ARMC4;refGene_tx=XM_0232564
79.1
B4	25946757 25946894
refGene_gn=ARMC4;refGene_tx=XM_023256475.1;refGene_gn=ARMC4;refGene_tx
=XM_023256476.1;refGene_gn=ARMC4;refGene_tx=XM_023256477.1;refGene_gn=
ARMC4;refGene_tx=XM_023256478.1;refGene_gn=ARMC4;refGene_tx=XM_0232564
79.1

B4	25947078 25947185
refGene_gn=ARMC4;refGene_tx=XM_023256475.1;refGene_gn=ARMC4;refGene_tx=XM_023256476.1;refGene_gn=ARMC4;refGene_tx=XM_023256477.1;refGene_gn=ARMC4;refGene_tx=XM_023256478.1;refGene_gn=ARMC4;refGene_tx=XM_023256479.1
B4	25947818 25948011
refGene_gn=ARMC4;refGene_tx=XM_023256475.1;refGene_gn=ARMC4;refGene_tx=XM_023256476.1;refGene_gn=ARMC4;refGene_tx=XM_023256477.1;refGene_gn=ARMC4;refGene_tx=XM_023256478.1
B4	25950165 25950323
refGene_gn=ARMC4;refGene_tx=XM_023256475.1;refGene_gn=ARMC4;refGene_tx=XM_023256476.1;refGene_gn=ARMC4;refGene_tx=XM_023256477.1
B4	25956765 25956989
refGene_gn=ARMC4;refGene_tx=XM_023256475.1;refGene_gn=ARMC4;refGene_tx=XM_023256476.1;refGene_gn=ARMC4;refGene_tx=XM_023256477.1
B4	26041196 26041376
refGene_gn=MPP7;refGene_tx=XM_006933337.3;refGene_gn=MPP7;refGene_tx=XM_019834123.1;refGene_gn=MPP7;refGene_tx=XM_019834124.1;refGene_gn=MPP7;refGene_tx=XM_019834125.1;refGene_gn=MPP7;refGene_tx=XM_019834126.1;refGene_gn=MPP7;refGene_tx=XM_019834127.1;refGene_gn=MPP7;refGene_tx=XM_019834128.1;refGene_gn=MPP7;refGene_tx=XM_019834129.1
B4	26041884 26042028
refGene_gn=MPP7;refGene_tx=XM_006933337.3;refGene_gn=MPP7;refGene_tx=XM_019834123.1;refGene_gn=MPP7;refGene_tx=XM_019834124.1;refGene_gn=MPP7;refGene_tx=XM_019834125.1;refGene_gn=MPP7;refGene_tx=XM_019834126.1;refGene_gn=MPP7;refGene_tx=XM_019834127.1;refGene_gn=MPP7;refGene_tx=XM_019834128.1;refGene_gn=MPP7;refGene_tx=XM_019834129.1
B4	26043601 26043710
refGene_gn=MPP7;refGene_tx=XM_006933337.3;refGene_gn=MPP7;refGene_tx=XM_019834123.1;refGene_gn=MPP7;refGene_tx=XM_019834124.1;refGene_gn=MPP7;refGene_tx=XM_019834125.1;refGene_gn=MPP7;refGene_tx=XM_019834126.1;refGene_gn=MPP7;refGene_tx=XM_019834127.1;refGene_gn=MPP7;refGene_tx=XM_019834128.1;refGene_gn=MPP7;refGene_tx=XM_019834129.1
B4	26044724 26044818
refGene_gn=MPP7;refGene_tx=XM_006933337.3;refGene_gn=MPP7;refGene_tx=XM_019834123.1;refGene_gn=MPP7;refGene_tx=XM_019834124.1;refGene_gn=MPP7;refGene_tx=XM_019834125.1;refGene_gn=MPP7;refGene_tx=XM_019834126.1;refGene_gn=MPP7;refGene_tx=XM_019834127.1;refGene_gn=MPP7;refGene_tx=XM_019834128.1;refGene_gn=MPP7;refGene_tx=XM_019834129.1
B4	26059413 26059494
refGene_gn=MPP7;refGene_tx=XM_006933337.3;refGene_gn=MPP7;refGene_tx=XM_019834123.1;refGene_gn=MPP7;refGene_tx=XM_019834124.1;refGene_gn=MPP7;refGene_tx=XM_019834125.1;refGene_gn=MPP7;refGene_tx=XM_019834127.1;refGene_gn=MPP7;refGene_tx=XM_019834128.1;refGene_gn=MPP7;refGene_tx=XM_019834129.1
B4	26096478 26096649
refGene_gn=MPP7;refGene_tx=XM_006933337.3;refGene_gn=MPP7;refGene_tx=XM_019834123.1;refGene_gn=MPP7;refGene_tx=XM_019834124.1;refGene_gn=MPP7;refGene_tx=XM_019834125.1;refGene_gn=MPP7;refGene_tx=XM_019834126.1;refGene_gn=MPP7;refGene_tx=XM_019834127.1

B4 26115864 26115906
refGene_gn=MPP7;refGene_tx=XM_019834123.1;refGene_gn=MPP7;refGene_tx=XM_019834124.1;refGene_gn=MPP7;refGene_tx=XM_019834125.1;refGene_gn=MPP7;refGene_tx=XM_019834126.1;refGene_gn=MPP7;refGene_tx=XM_019834127.1;refGene_gn=MPP7;refGene_tx=XM_019834128.1
B4 26116910 26116922
refGene_gn=MPP7;refGene_tx=XM_019834123.1;refGene_gn=MPP7;refGene_tx=XM_019834124.1;refGene_gn=MPP7;refGene_tx=XM_019834125.1;refGene_gn=MPP7;refGene_tx=XM_019834126.1;refGene_gn=MPP7;refGene_tx=XM_019834127.1;refGene_gn=MPP7;refGene_tx=XM_019834128.1
B4 26117021 26117042
refGene_gn=MPP7;refGene_tx=XM_019834123.1;refGene_gn=MPP7;refGene_tx=XM_019834124.1;refGene_gn=MPP7;refGene_tx=XM_019834125.1;refGene_gn=MPP7;refGene_tx=XM_019834126.1;refGene_gn=MPP7;refGene_tx=XM_019834127.1;refGene_gn=MPP7;refGene_tx=XM_019834128.1
B4 26125258 26125323
refGene_gn=MPP7;refGene_tx=XM_006933337.3;refGene_gn=MPP7;refGene_tx=XM_019834123.1;refGene_gn=MPP7;refGene_tx=XM_019834124.1;refGene_gn=MPP7;refGene_tx=XM_019834125.1;refGene_gn=MPP7;refGene_tx=XM_019834126.1;refGene_gn=MPP7;refGene_tx=XM_019834127.1;refGene_gn=MPP7;refGene_tx=XM_019834128.1;refGene_gn=MPP7;refGene_tx=XM_019834129.1
B4 26125813 26126010
refGene_gn=MPP7;refGene_tx=XM_006933337.3;refGene_gn=MPP7;refGene_tx=XM_019834123.1;refGene_gn=MPP7;refGene_tx=XM_019834124.1;refGene_gn=MPP7;refGene_tx=XM_019834125.1;refGene_gn=MPP7;refGene_tx=XM_019834126.1;refGene_gn=MPP7;refGene_tx=XM_019834127.1;refGene_gn=MPP7;refGene_tx=XM_019834128.1;refGene_gn=MPP7;refGene_tx=XM_019834129.1
B4 26126224 26126299
refGene_gn=MPP7;refGene_tx=XM_006933337.3;refGene_gn=MPP7;refGene_tx=XM_019834123.1;refGene_gn=MPP7;refGene_tx=XM_019834124.1;refGene_gn=MPP7;refGene_tx=XM_019834125.1;refGene_gn=MPP7;refGene_tx=XM_019834126.1;refGene_gn=MPP7;refGene_tx=XM_019834127.1;refGene_gn=MPP7;refGene_tx=XM_019834128.1;refGene_gn=MPP7;refGene_tx=XM_019834129.1
B4 26130162 26130248
refGene_gn=MPP7;refGene_tx=XM_006933337.3;refGene_gn=MPP7;refGene_tx=XM_019834123.1;refGene_gn=MPP7;refGene_tx=XM_019834124.1;refGene_gn=MPP7;refGene_tx=XM_019834125.1;refGene_gn=MPP7;refGene_tx=XM_019834126.1;refGene_gn=MPP7;refGene_tx=XM_019834127.1;refGene_gn=MPP7;refGene_tx=XM_019834128.1;refGene_gn=MPP7;refGene_tx=XM_019834129.1
B4 26130783 26130865
refGene_gn=MPP7;refGene_tx=XM_006933337.3;refGene_gn=MPP7;refGene_tx=XM_019834123.1;refGene_gn=MPP7;refGene_tx=XM_019834124.1;refGene_gn=MPP7;refGene_tx=XM_019834125.1;refGene_gn=MPP7;refGene_tx=XM_019834126.1;refGene_gn=MPP7;refGene_tx=XM_019834127.1;refGene_gn=MPP7;refGene_tx=XM_019834128.1;refGene_gn=MPP7;refGene_tx=XM_019834129.1
B4 26145554 26145686
refGene_gn=MPP7;refGene_tx=XM_006933337.3;refGene_gn=MPP7;refGene_tx=XM_019834123.1;refGene_gn=MPP7;refGene_tx=XM_019834124.1;refGene_gn=MPP7;refGene_tx=XM_019834125.1;refGene_gn=MPP7;refGene_tx=XM_019834126.1;refGene_gn=MPP7;refGene_tx=XM_019834127.1;refGene_gn=MPP7;refGene_tx=X

M_019834128.1;refGene_gn=MPP7;refGene_tx=XM_019834129.1
B4 26156082 26156163
refGene_gn=MPP7;refGene_tx=XM_006933337.3;refGene_gn=MPP7;refGene_tx=XM_019834123.1;refGene_gn=MPP7;refGene_tx=XM_019834124.1;refGene_gn=MPP7;refGene_tx=XM_019834125.1;refGene_gn=MPP7;refGene_tx=XM_019834126.1;refGene_gn=MPP7;refGene_tx=XM_019834128.1;refGene_gn=MPP7;refGene_tx=XM_019834129.1
B4 26158786 26158864
refGene_gn=MPP7;refGene_tx=XM_006933337.3;refGene_gn=MPP7;refGene_tx=XM_019834123.1;refGene_gn=MPP7;refGene_tx=XM_019834124.1;refGene_gn=MPP7;refGene_tx=XM_019834125.1;refGene_gn=MPP7;refGene_tx=XM_019834126.1;refGene_gn=MPP7;refGene_tx=XM_019834128.1;refGene_gn=MPP7;refGene_tx=XM_019834129.1
B4 26216667 26216786
refGene_gn=MPP7;refGene_tx=XM_006933337.3;refGene_gn=MPP7;refGene_tx=XM_019834123.1;refGene_gn=MPP7;refGene_tx=XM_019834124.1;refGene_gn=MPP7;refGene_tx=XM_019834125.1;refGene_gn=MPP7;refGene_tx=XM_019834126.1;refGene_gn=MPP7;refGene_tx=XM_019834127.1;refGene_gn=MPP7;refGene_tx=XM_019834128.1;refGene_gn=MPP7;refGene_tx=XM_019834129.1
B4 26251917 26251954
refGene_gn=MPP7;refGene_tx=XM_006933337.3;refGene_gn=MPP7;refGene_tx=XM_019834123.1;refGene_gn=MPP7;refGene_tx=XM_019834124.1;refGene_gn=MPP7;refGene_tx=XM_019834125.1;refGene_gn=MPP7;refGene_tx=XM_019834126.1;refGene_gn=MPP7;refGene_tx=XM_019834127.1;refGene_gn=MPP7;refGene_tx=XM_019834128.1;refGene_gn=MPP7;refGene_tx=XM_019834129.1
B4 26534066 26534107
refGene_gn=WAC;refGene_tx=XM_023256480.1;refGene_gn=WAC;refGene_tx=XM_023256481.1;refGene_gn=WAC;refGene_tx=XM_023256482.1;refGene_gn=WAC;refGene_tx=XM_023256483.1;refGene_gn=WAC;refGene_tx=XM_023256487.1;refGene_gn=WAC;refGene_tx=XM_023256488.1;refGene_gn=WAC;refGene_tx=XM_023256489.1
B4 26534523 26534560
refGene_gn=WAC;refGene_tx=XM_023256480.1;refGene_gn=WAC;refGene_tx=XM_023256481.1;refGene_gn=WAC;refGene_tx=XM_023256482.1;refGene_gn=WAC;refGene_tx=XM_023256483.1;refGene_gn=WAC;refGene_tx=XM_023256487.1;refGene_gn=WAC;refGene_tx=XM_023256488.1;refGene_gn=WAC;refGene_tx=XM_023256489.1
B4 26536095 26536309
refGene_gn=WAC;refGene_tx=XM_019834134.2;refGene_gn=WAC;refGene_tx=XM_023256480.1;refGene_gn=WAC;refGene_tx=XM_023256481.1;refGene_gn=WAC;refGene_tx=XM_023256482.1;refGene_gn=WAC;refGene_tx=XM_023256483.1;refGene_gn=WAC;refGene_tx=XM_023256484.1;refGene_gn=WAC;refGene_tx=XM_023256485.1;refGene_gn=WAC;refGene_tx=XM_023256486.1;refGene_gn=WAC;refGene_tx=XM_023256487.1;refGene_gn=WAC;refGene_tx=XM_023256488.1;refGene_gn=WAC;refGene_tx=XM_023256489.1
B4 26583247 26583354
refGene_gn=WAC;refGene_tx=XM_019834134.2;refGene_gn=WAC;refGene_tx=XM_023256480.1;refGene_gn=WAC;refGene_tx=XM_023256481.1;refGene_gn=WAC;refGene_tx=XM_023256482.1;refGene_gn=WAC;refGene_tx=XM_023256483.1;refGene_gn=WAC;refGene_tx=XM_023256484.1;refGene_gn=WAC;refGene_tx=XM_02325

6485.1;refGene_gn=WAC;refGene_tx=XM_023256486.1;refGene_gn=WAC;refGene_tx=XM_023256487.1;refGene_gn=WAC;refGene_tx=XM_023256488.1;refGene_gn=WAC;refGene_tx=XM_023256489.1
B4 26589016 26589132
refGene_gn=WAC;refGene_tx=XM_019834134.2;refGene_gn=WAC;refGene_tx=XM_023256480.1;refGene_gn=WAC;refGene_tx=XM_023256481.1;refGene_gn=WAC;refGene_tx=XM_023256482.1;refGene_gn=WAC;refGene_tx=XM_023256483.1;refGene_gn=WAC;refGene_tx=XM_023256484.1;refGene_gn=WAC;refGene_tx=XM_023256485.1;refGene_gn=WAC;refGene_tx=XM_023256486.1;refGene_gn=WAC;refGene_tx=XM_023256487.1;refGene_gn=WAC;refGene_tx=XM_023256488.1;refGene_gn=WAC;refGene_tx=XM_023256489.1
B4 26589693 26589806
refGene_gn=WAC;refGene_tx=XM_019834134.2;refGene_gn=WAC;refGene_tx=XM_023256480.1;refGene_gn=WAC;refGene_tx=XM_023256481.1;refGene_gn=WAC;refGene_tx=XM_023256482.1;refGene_gn=WAC;refGene_tx=XM_023256483.1;refGene_gn=WAC;refGene_tx=XM_023256484.1;refGene_gn=WAC;refGene_tx=XM_023256485.1;refGene_gn=WAC;refGene_tx=XM_023256486.1;refGene_gn=WAC;refGene_tx=XM_023256487.1;refGene_gn=WAC;refGene_tx=XM_023256488.1;refGene_gn=WAC;refGene_tx=XM_023256489.1
B4 26594108 26594417
refGene_gn=WAC;refGene_tx=XM_019834134.2;refGene_gn=WAC;refGene_tx=XM_023256480.1;refGene_gn=WAC;refGene_tx=XM_023256481.1;refGene_gn=WAC;refGene_tx=XM_023256482.1;refGene_gn=WAC;refGene_tx=XM_023256483.1;refGene_gn=WAC;refGene_tx=XM_023256484.1;refGene_gn=WAC;refGene_tx=XM_023256485.1;refGene_gn=WAC;refGene_tx=XM_023256486.1
B4 26606288 26606558
refGene_gn=WAC;refGene_tx=XM_019834134.2;refGene_gn=WAC;refGene_tx=XM_023256480.1;refGene_gn=WAC;refGene_tx=XM_023256481.1;refGene_gn=WAC;refGene_tx=XM_023256482.1;refGene_gn=WAC;refGene_tx=XM_023256483.1;refGene_gn=WAC;refGene_tx=XM_023256484.1;refGene_gn=WAC;refGene_tx=XM_023256485.1;refGene_gn=WAC;refGene_tx=XM_023256486.1;refGene_gn=WAC;refGene_tx=XM_023256487.1;refGene_gn=WAC;refGene_tx=XM_023256488.1;refGene_gn=WAC;refGene_tx=XM_023256489.1
B4 26607970 26608093
refGene_gn=WAC;refGene_tx=XM_019834134.2;refGene_gn=WAC;refGene_tx=XM_023256480.1;refGene_gn=WAC;refGene_tx=XM_023256481.1;refGene_gn=WAC;refGene_tx=XM_023256482.1;refGene_gn=WAC;refGene_tx=XM_023256483.1;refGene_gn=WAC;refGene_tx=XM_023256484.1;refGene_gn=WAC;refGene_tx=XM_023256485.1;refGene_gn=WAC;refGene_tx=XM_023256486.1;refGene_gn=WAC;refGene_tx=XM_023256487.1;refGene_gn=WAC;refGene_tx=XM_023256488.1;refGene_gn=WAC;refGene_tx=XM_023256489.1
B4 26609017 26609166
refGene_gn=WAC;refGene_tx=XM_019834134.2;refGene_gn=WAC;refGene_tx=XM_023256480.1;refGene_gn=WAC;refGene_tx=XM_023256481.1;refGene_gn=WAC;refGene_tx=XM_023256482.1;refGene_gn=WAC;refGene_tx=XM_023256483.1;refGene_gn=WAC;refGene_tx=XM_023256484.1;refGene_gn=WAC;refGene_tx=XM_023256485.1;refGene_gn=WAC;refGene_tx=XM_023256486.1;refGene_gn=WAC;refGene_tx=XM_023256487.1;refGene_gn=WAC;refGene_tx=XM_023256488.1;refGene_gn=WAC;refGene_tx=XM_023256489.1
B4 26611905 26612024

refGene_gn=WAC;refGene_tx=XM_019834134.2;refGene_gn=WAC;refGene_tx=XM_023256480.1;refGene_gn=WAC;refGene_tx=XM_023256481.1;refGene_gn=WAC;refGene_tx=XM_023256482.1;refGene_gn=WAC;refGene_tx=XM_023256483.1;refGene_gn=WAC;refGene_tx=XM_023256484.1;refGene_gn=WAC;refGene_tx=XM_023256485.1;refGene_gn=WAC;refGene_tx=XM_023256486.1;refGene_gn=WAC;refGene_tx=XM_023256487.1;refGene_gn=WAC;refGene_tx=XM_023256488.1;refGene_gn=WAC;refGene_tx=XM_023256489.1
B4 26613509 26613696
refGene_gn=WAC;refGene_tx=XM_019834134.2;refGene_gn=WAC;refGene_tx=XM_023256480.1;refGene_gn=WAC;refGene_tx=XM_023256481.1;refGene_gn=WAC;refGene_tx=XM_023256482.1;refGene_gn=WAC;refGene_tx=XM_023256483.1;refGene_gn=WAC;refGene_tx=XM_023256484.1;refGene_gn=WAC;refGene_tx=XM_023256485.1;refGene_gn=WAC;refGene_tx=XM_023256486.1;refGene_gn=WAC;refGene_tx=XM_023256487.1;refGene_gn=WAC;refGene_tx=XM_023256488.1;refGene_gn=WAC;refGene_tx=XM_023256489.1
B4 26614961 26615089
refGene_gn=WAC;refGene_tx=XM_019834134.2;refGene_gn=WAC;refGene_tx=XM_023256480.1;refGene_gn=WAC;refGene_tx=XM_023256481.1;refGene_gn=WAC;refGene_tx=XM_023256482.1;refGene_gn=WAC;refGene_tx=XM_023256483.1;refGene_gn=WAC;refGene_tx=XM_023256484.1;refGene_gn=WAC;refGene_tx=XM_023256485.1;refGene_gn=WAC;refGene_tx=XM_023256486.1;refGene_gn=WAC;refGene_tx=XM_023256487.1;refGene_gn=WAC;refGene_tx=XM_023256488.1;refGene_gn=WAC;refGene_tx=XM_023256489.1
B4 26617070 26617140
refGene_gn=WAC;refGene_tx=XM_019834134.2;refGene_gn=WAC;refGene_tx=XM_023256480.1;refGene_gn=WAC;refGene_tx=XM_023256481.1;refGene_gn=WAC;refGene_tx=XM_023256482.1;refGene_gn=WAC;refGene_tx=XM_023256483.1;refGene_gn=WAC;refGene_tx=XM_023256484.1;refGene_gn=WAC;refGene_tx=XM_023256485.1;refGene_gn=WAC;refGene_tx=XM_023256486.1;refGene_gn=WAC;refGene_tx=XM_023256487.1;refGene_gn=WAC;refGene_tx=XM_023256488.1;refGene_gn=WAC;refGene_tx=XM_023256489.1
B4 26654693 26654769 refGene_gn=BAMBI;refGene_tx=XM_023256490.1
B4 26658000 26658288 refGene_gn=BAMBI;refGene_tx=XM_023256490.1
B4 26658737 26659156 refGene_gn=BAMBI;refGene_tx=XM_023256490.1
B4 27211560 27211699
refGene_gn=LOC101090367;refGene_tx=XM_003988199.4
B4 27212190 27212349
refGene_gn=LOC101090367;refGene_tx=XM_003988199.4
B4 27225747 27225826
refGene_gn=LOC101090367;refGene_tx=XM_003988199.4
B4 27226646 27226716
refGene_gn=LOC101090367;refGene_tx=XM_003988199.4
B4 27409180 27409311
refGene_gn=MAP3K8;refGene_tx=XM_006933341.4;refGene_gn=MAP3K8;refGene_tx=XM_006933342.4;refGene_gn=MAP3K8;refGene_tx=XM_011283656.3;refGene_gn=MAP3K8;refGene_tx=XM_011283657.3;refGene_gn=MAP3K8;refGene_tx=XM_011283659.3;refGene_gn=MAP3K8;refGene_tx=XM_019834137.2
B4 27410269 27410516
refGene_gn=MAP3K8;refGene_tx=XM_006933341.4;refGene_gn=MAP3K8;refGene_tx=XM_006933342.4;refGene_gn=MAP3K8;refGene_tx=XM_011283656.3;refGene_

gn=MAP3K8;refGene_tx=XM_011283657.3;refGene_gn=MAP3K8;refGene_tx=XM_011283659.3;refGene_gn=MAP3K8;refGene_tx=XM_019834137.2
B4 27411152 27411305
refGene_gn=MAP3K8;refGene_tx=XM_006933341.4;refGene_gn=MAP3K8;refGene_tx=XM_006933342.4;refGene_gn=MAP3K8;refGene_tx=XM_011283656.3;refGene_gn=MAP3K8;refGene_tx=XM_011283657.3;refGene_gn=MAP3K8;refGene_tx=XM_011283659.3;refGene_gn=MAP3K8;refGene_tx=XM_019834137.2
B4 27415654 27415761
refGene_gn=MAP3K8;refGene_tx=XM_006933341.4;refGene_gn=MAP3K8;refGene_tx=XM_006933342.4;refGene_gn=MAP3K8;refGene_tx=XM_011283656.3;refGene_gn=MAP3K8;refGene_tx=XM_011283657.3;refGene_gn=MAP3K8;refGene_tx=XM_011283659.3;refGene_gn=MAP3K8;refGene_tx=XM_019834137.2
B4 27417419 27417681
refGene_gn=MAP3K8;refGene_tx=XM_006933341.4;refGene_gn=MAP3K8;refGene_tx=XM_006933342.4;refGene_gn=MAP3K8;refGene_tx=XM_011283656.3;refGene_gn=MAP3K8;refGene_tx=XM_011283657.3;refGene_gn=MAP3K8;refGene_tx=XM_011283659.3;refGene_gn=MAP3K8;refGene_tx=XM_019834137.2
B4 27420226 27420394
refGene_gn=MAP3K8;refGene_tx=XM_006933341.4;refGene_gn=MAP3K8;refGene_tx=XM_006933342.4;refGene_gn=MAP3K8;refGene_tx=XM_011283656.3;refGene_gn=MAP3K8;refGene_tx=XM_011283657.3;refGene_gn=MAP3K8;refGene_tx=XM_011283659.3;refGene_gn=MAP3K8;refGene_tx=XM_019834137.2
B4 27427216 27427552
refGene_gn=MAP3K8;refGene_tx=XM_006933341.4;refGene_gn=MAP3K8;refGene_tx=XM_006933342.4;refGene_gn=MAP3K8;refGene_tx=XM_011283656.3;refGene_gn=MAP3K8;refGene_tx=XM_011283657.3;refGene_gn=MAP3K8;refGene_tx=XM_011283659.3;refGene_gn=MAP3K8;refGene_tx=XM_019834137.2
B4 27455946 27456100 refGene_gn=MTPAP;refGene_tx=XM_003988201.5
B4 27462235 27462408 refGene_gn=MTPAP;refGene_tx=XM_003988201.5
B4 27462927 27463161 refGene_gn=MTPAP;refGene_tx=XM_003988201.5
B4 27465016 27465241 refGene_gn=MTPAP;refGene_tx=XM_003988201.5
B4 27471557 27471769 refGene_gn=MTPAP;refGene_tx=XM_003988201.5
B4 27473603 27473830 refGene_gn=MTPAP;refGene_tx=XM_003988201.5
B4 27484780 27484873 refGene_gn=MTPAP;refGene_tx=XM_003988201.5
B4 27484956 27485030 refGene_gn=MTPAP;refGene_tx=XM_003988201.5
B4 27487091 27487454 refGene_gn=MTPAP;refGene_tx=XM_003988201.5
B4 27676393 27676674
refGene_gn=JCAD;refGene_tx=XM_003988202.5;refGene_gn=JCAD;refGene_tx=XM_023256492.1;refGene_gn=JCAD;refGene_tx=XM_023256493.1
B4 27692420 27696151
refGene_gn=JCAD;refGene_tx=XM_003988202.5;refGene_gn=JCAD;refGene_tx=XM_019834139.2;refGene_gn=JCAD;refGene_tx=XM_023256492.1;refGene_gn=JCAD;refGene_tx=XM_023256493.1;refGene_gn=JCAD;refGene_tx=XM_023256494.1
B4 27701657 27701692
refGene_gn=JCAD;refGene_tx=XM_003988202.5;refGene_gn=JCAD;refGene_tx=XM_019834139.2;refGene_gn=JCAD;refGene_tx=XM_023256492.1;refGene_gn=JCAD;refGene_tx=XM_023256494.1
B4 27701752 27701772 refGene_gn=JCAD;refGene_tx=XM_023256493.1
B4 28016868 28016884 refGene_gn=SVIL;refGene_tx=XM_006933343.4
B4 28031950 28032011 refGene_gn=SVIL;refGene_tx=XM_019834141.2

B4	28053028	28053089
refGene_gn=SVIL;refGene_tx=XM_019834140.2;refGene_gn=SVIL;refGene_tx=XM_019834142.2;refGene_gn=SVIL;refGene_tx=XM_019834143.2;refGene_gn=SVIL;refGene_tx=XM_019834144.2;refGene_gn=SVIL;refGene_tx=XM_019834146.2;refGene_gn=SVIL;refGene_tx=XM_019834147.2;refGene_gn=SVIL;refGene_tx=XM_019834148.2;refGene_gn=SVIL;refGene_tx=XM_019834149.2;refGene_gn=SVIL;refGene_tx=XM_019834152.2;refGene_gn=SVIL;refGene_tx=XM_019834153.2;refGene_gn=SVIL;refGene_tx=XM_019834154.2
B4	28054792	28054850
refGene_gn=SVIL;refGene_tx=XM_006933343.4;refGene_gn=SVIL;refGene_tx=XM_019834140.2;refGene_gn=SVIL;refGene_tx=XM_019834141.2;refGene_gn=SVIL;refGene_tx=XM_019834142.2;refGene_gn=SVIL;refGene_tx=XM_019834143.2;refGene_gn=SVIL;refGene_tx=XM_019834144.2;refGene_gn=SVIL;refGene_tx=XM_019834146.2;refGene_gn=SVIL;refGene_tx=XM_019834147.2;refGene_gn=SVIL;refGene_tx=XM_019834148.2;refGene_gn=SVIL;refGene_tx=XM_019834149.2;refGene_gn=SVIL;refGene_tx=XM_019834152.2;refGene_gn=SVIL;refGene_tx=XM_019834153.2;refGene_gn=SVIL;refGene_tx=XM_019834154.2
B4	28060553	28060645
refGene_gn=SVIL;refGene_tx=XM_006933343.4;refGene_gn=SVIL;refGene_tx=XM_019834140.2;refGene_gn=SVIL;refGene_tx=XM_019834141.2;refGene_gn=SVIL;refGene_tx=XM_019834142.2;refGene_gn=SVIL;refGene_tx=XM_019834143.2;refGene_gn=SVIL;refGene_tx=XM_019834144.2;refGene_gn=SVIL;refGene_tx=XM_019834146.2;refGene_gn=SVIL;refGene_tx=XM_019834147.2;refGene_gn=SVIL;refGene_tx=XM_019834148.2;refGene_gn=SVIL;refGene_tx=XM_019834149.2;refGene_gn=SVIL;refGene_tx=XM_019834152.2;refGene_gn=SVIL;refGene_tx=XM_019834153.2;refGene_gn=SVIL;refGene_tx=XM_019834154.2
B4	28069369	28069427
refGene_gn=SVIL;refGene_tx=XM_006933343.4;refGene_gn=SVIL;refGene_tx=XM_019834140.2;refGene_gn=SVIL;refGene_tx=XM_019834141.2;refGene_gn=SVIL;refGene_tx=XM_019834142.2;refGene_gn=SVIL;refGene_tx=XM_019834143.2;refGene_gn=SVIL;refGene_tx=XM_019834144.2;refGene_gn=SVIL;refGene_tx=XM_019834146.2;refGene_gn=SVIL;refGene_tx=XM_019834147.2;refGene_gn=SVIL;refGene_tx=XM_019834148.2;refGene_gn=SVIL;refGene_tx=XM_019834149.2;refGene_gn=SVIL;refGene_tx=XM_019834150.2;refGene_gn=SVIL;refGene_tx=XM_019834152.2;refGene_gn=SVIL;refGene_tx=XM_019834153.2;refGene_gn=SVIL;refGene_tx=XM_019834154.2;refGene_gn=SVIL;refGene_tx=XM_023256495.1
B4	28069542	28069694
refGene_gn=SVIL;refGene_tx=XM_006933343.4;refGene_gn=SVIL;refGene_tx=XM_019834140.2;refGene_gn=SVIL;refGene_tx=XM_019834141.2;refGene_gn=SVIL;refGene_tx=XM_019834142.2;refGene_gn=SVIL;refGene_tx=XM_019834143.2;refGene_gn=SVIL;refGene_tx=XM_019834144.2;refGene_gn=SVIL;refGene_tx=XM_019834146.2;refGene_gn=SVIL;refGene_tx=XM_019834147.2;refGene_gn=SVIL;refGene_tx=XM_019834148.2;refGene_gn=SVIL;refGene_tx=XM_019834149.2;refGene_gn=SVIL;refGene_tx=XM_019834150.2;refGene_gn=SVIL;refGene_tx=XM_019834152.2;refGene_gn=SVIL;refGene_tx=XM_019834153.2;refGene_gn=SVIL;refGene_tx=XM_019834154.2;refGene_gn=SVIL;refGene_tx=XM_023256495.1
B4	28072520	28073160
refGene_gn=SVIL;refGene_tx=XM_006933343.4;refGene_gn=SVIL;refGene_tx=XM_019834140.2;refGene_gn=SVIL;refGene_tx=XM_019834141.2;refGene_gn=SVIL;refGene_tx=XM_019834142.2;refGene_gn=SVIL;refGene_tx=XM_019834143.2;

refGene_gn=SVIL;refGene_tx=XM_019834144.2;refGene_gn=SVIL;refGene_tx=XM_019834146.2;refGene_gn=SVIL;refGene_tx=XM_019834147.2;refGene_gn=SVIL;refGene_tx=XM_019834148.2;refGene_gn=SVIL;refGene_tx=XM_019834149.2;refGene_gn=SVIL;refGene_tx=XM_019834150.2;refGene_gn=SVIL;refGene_tx=XM_019834152.2;refGene_gn=SVIL;refGene_tx=XM_019834153.2;refGene_gn=SVIL;refGene_tx=XM_019834154.2;refGene_gn=SVIL;refGene_tx=XM_023256495.1
B4 28089174 28089255
refGene_gn=SVIL;refGene_tx=XM_006933343.4;refGene_gn=SVIL;refGene_tx=XM_019834140.2;refGene_gn=SVIL;refGene_tx=XM_019834141.2;refGene_gn=SVIL;refGene_tx=XM_019834142.2;refGene_gn=SVIL;refGene_tx=XM_019834144.2;refGene_gn=SVIL;refGene_tx=XM_019834146.2;refGene_gn=SVIL;refGene_tx=XM_019834147.2;refGene_gn=SVIL;refGene_tx=XM_019834148.2;refGene_gn=SVIL;refGene_tx=XM_019834150.2;refGene_gn=SVIL;refGene_tx=XM_023256495.1
B4 28091272 28092169
refGene_gn=SVIL;refGene_tx=XM_006933343.4;refGene_gn=SVIL;refGene_tx=XM_019834140.2;refGene_gn=SVIL;refGene_tx=XM_019834141.2;refGene_gn=SVIL;refGene_tx=XM_019834142.2;refGene_gn=SVIL;refGene_tx=XM_019834143.2;refGene_gn=SVIL;refGene_tx=XM_019834144.2;refGene_gn=SVIL;refGene_tx=XM_019834146.2;refGene_gn=SVIL;refGene_tx=XM_019834147.2;refGene_gn=SVIL;refGene_tx=XM_019834148.2;refGene_gn=SVIL;refGene_tx=XM_019834149.2;refGene_gn=SVIL;refGene_tx=XM_019834150.2;refGene_gn=SVIL;refGene_tx=XM_023256495.1
B4 28092550 28092721
refGene_gn=SVIL;refGene_tx=XM_006933343.4;refGene_gn=SVIL;refGene_tx=XM_019834140.2;refGene_gn=SVIL;refGene_tx=XM_019834141.2;refGene_gn=SVIL;refGene_tx=XM_019834142.2;refGene_gn=SVIL;refGene_tx=XM_019834143.2;refGene_gn=SVIL;refGene_tx=XM_019834144.2;refGene_gn=SVIL;refGene_tx=XM_019834146.2;refGene_gn=SVIL;refGene_tx=XM_019834148.2;refGene_gn=SVIL;refGene_tx=XM_019834150.2;refGene_gn=SVIL;refGene_tx=XM_023256495.1
B4 28093419 28093454
refGene_gn=SVIL;refGene_tx=XM_006933343.4;refGene_gn=SVIL;refGene_tx=XM_019834140.2;refGene_gn=SVIL;refGene_tx=XM_019834141.2;refGene_gn=SVIL;refGene_tx=XM_019834142.2;refGene_gn=SVIL;refGene_tx=XM_019834143.2;refGene_gn=SVIL;refGene_tx=XM_019834144.2;refGene_gn=SVIL;refGene_tx=XM_019834146.2;refGene_gn=SVIL;refGene_tx=XM_019834147.2;refGene_gn=SVIL;refGene_tx=XM_019834148.2;refGene_gn=SVIL;refGene_tx=XM_019834149.2;refGene_gn=SVIL;refGene_tx=XM_019834150.2;refGene_gn=SVIL;refGene_tx=XM_019834152.2;refGene_gn=SVIL;refGene_tx=XM_019834153.2;refGene_gn=SVIL;refGene_tx=XM_019834154.2;refGene_gn=SVIL;refGene_tx=XM_023256495.1
B4 28094020 28094082
refGene_gn=SVIL;refGene_tx=XM_006933343.4;refGene_gn=SVIL;refGene_tx=XM_019834140.2;refGene_gn=SVIL;refGene_tx=XM_019834141.2;refGene_gn=SVIL;refGene_tx=XM_019834142.2;refGene_gn=SVIL;refGene_tx=XM_019834143.2;refGene_gn=SVIL;refGene_tx=XM_019834144.2;refGene_gn=SVIL;refGene_tx=XM_019834146.2;refGene_gn=SVIL;refGene_tx=XM_019834147.2;refGene_gn=SVIL;refGene_tx=XM_019834148.2;refGene_gn=SVIL;refGene_tx=XM_019834149.2;refGene_gn=SVIL;refGene_tx=XM_019834150.2;refGene_gn=SVIL;refGene_tx=XM_019834152.2;refGene_gn=SVIL;refGene_tx=XM_019834153.2;refGene_gn=SVIL;refGene_tx=XM_019834154.2;refGene_gn=SVIL;refGene_tx=XM_023256495.1
B4 28094381 28094521

refGene_gn=SVIL;refGene_tx=XM_006933343.4;refGene_gn=SVIL;refGene_tx=XM_019834140.2;refGene_gn=SVIL;refGene_tx=XM_019834141.2;refGene_gn=SVIL;refGene_tx=XM_019834142.2;refGene_gn=SVIL;refGene_tx=XM_019834143.2;refGene_gn=SVIL;refGene_tx=XM_019834144.2;refGene_gn=SVIL;refGene_tx=XM_019834146.2;refGene_gn=SVIL;refGene_tx=XM_019834147.2;refGene_gn=SVIL;refGene_tx=XM_019834148.2;refGene_gn=SVIL;refGene_tx=XM_019834149.2;refGene_gn=SVIL;refGene_tx=XM_019834150.2;refGene_gn=SVIL;refGene_tx=XM_019834152.2;refGene_gn=SVIL;refGene_tx=XM_019834153.2;refGene_gn=SVIL;refGene_tx=XM_019834154.2;refGene_gn=SVIL;refGene_tx=XM_023256495.1
B4 28096367 28096463
refGene_gn=SVIL;refGene_tx=XM_006933343.4;refGene_gn=SVIL;refGene_tx=XM_019834140.2;refGene_gn=SVIL;refGene_tx=XM_019834141.2;refGene_gn=SVIL;refGene_tx=XM_019834142.2;refGene_gn=SVIL;refGene_tx=XM_019834143.2;refGene_gn=SVIL;refGene_tx=XM_019834146.2;refGene_gn=SVIL;refGene_tx=XM_019834147.2;refGene_gn=SVIL;refGene_tx=XM_019834150.2;refGene_gn=SVIL;refGene_tx=XM_019834152.2;refGene_gn=SVIL;refGene_tx=XM_019834153.2;refGene_gn=SVIL;refGene_tx=XM_023256495.1
B4 28098187 28098431
refGene_gn=SVIL;refGene_tx=XM_006933343.4;refGene_gn=SVIL;refGene_tx=XM_019834140.2;refGene_gn=SVIL;refGene_tx=XM_019834141.2;refGene_gn=SVIL;refGene_tx=XM_019834142.2;refGene_gn=SVIL;refGene_tx=XM_019834143.2;refGene_gn=SVIL;refGene_tx=XM_019834144.2;refGene_gn=SVIL;refGene_tx=XM_019834146.2;refGene_gn=SVIL;refGene_tx=XM_019834147.2;refGene_gn=SVIL;refGene_tx=XM_019834148.2;refGene_gn=SVIL;refGene_tx=XM_019834149.2;refGene_gn=SVIL;refGene_tx=XM_019834150.2;refGene_gn=SVIL;refGene_tx=XM_019834152.2;refGene_gn=SVIL;refGene_tx=XM_019834153.2;refGene_gn=SVIL;refGene_tx=XM_019834154.2;refGene_gn=SVIL;refGene_tx=XM_023256495.1
B4 28098890 28099464
refGene_gn=SVIL;refGene_tx=XM_006933343.4;refGene_gn=SVIL;refGene_tx=XM_019834140.2;refGene_gn=SVIL;refGene_tx=XM_019834141.2;refGene_gn=SVIL;refGene_tx=XM_019834142.2;refGene_gn=SVIL;refGene_tx=XM_019834143.2;refGene_gn=SVIL;refGene_tx=XM_019834144.2;refGene_gn=SVIL;refGene_tx=XM_019834146.2;refGene_gn=SVIL;refGene_tx=XM_019834147.2;refGene_gn=SVIL;refGene_tx=XM_019834148.2;refGene_gn=SVIL;refGene_tx=XM_019834149.2;refGene_gn=SVIL;refGene_tx=XM_019834150.2;refGene_gn=SVIL;refGene_tx=XM_019834152.2;refGene_gn=SVIL;refGene_tx=XM_019834153.2;refGene_gn=SVIL;refGene_tx=XM_019834154.2;refGene_gn=SVIL;refGene_tx=XM_023256495.1
B4 28100303 28100529
refGene_gn=SVIL;refGene_tx=XM_006933343.4;refGene_gn=SVIL;refGene_tx=XM_019834140.2;refGene_gn=SVIL;refGene_tx=XM_019834141.2;refGene_gn=SVIL;refGene_tx=XM_019834142.2;refGene_gn=SVIL;refGene_tx=XM_019834143.2;refGene_gn=SVIL;refGene_tx=XM_019834144.2;refGene_gn=SVIL;refGene_tx=XM_019834146.2;refGene_gn=SVIL;refGene_tx=XM_019834147.2;refGene_gn=SVIL;refGene_tx=XM_019834148.2;refGene_gn=SVIL;refGene_tx=XM_019834149.2;refGene_gn=SVIL;refGene_tx=XM_019834150.2;refGene_gn=SVIL;refGene_tx=XM_019834152.2;refGene_gn=SVIL;refGene_tx=XM_019834153.2;refGene_gn=SVIL;refGene_tx=XM_019834154.2;refGene_gn=SVIL;refGene_tx=XM_023256495.1
B4 28109284 28109414
refGene_gn=SVIL;refGene_tx=XM_006933343.4;refGene_gn=SVIL;refGene_tx=XM_019834140.2;refGene_gn=SVIL;refGene_tx=XM_019834141.2;refGene_gn=SVI

L;refGene_tx=XM_019834142.2;refGene_gn=SVIL;refGene_tx=XM_019834143.2;
refGene_gn=SVIL;refGene_tx=XM_019834144.2;refGene_gn=SVIL;refGene_tx=X
M_019834146.2;refGene_gn=SVIL;refGene_tx=XM_019834147.2;refGene_gn=SVI
L;refGene_tx=XM_019834148.2;refGene_gn=SVIL;refGene_tx=XM_019834149.2;
refGene_gn=SVIL;refGene_tx=XM_019834150.2;refGene_gn=SVIL;refGene_tx=X
M_019834152.2;refGene_gn=SVIL;refGene_tx=XM_019834153.2;refGene_gn=SVI
L;refGene_tx=XM_019834154.2;refGene_gn=SVIL;refGene_tx=XM_023256495.1
B4 28113062 28113110
refGene_gn=SVIL;refGene_tx=XM_006933343.4;refGene_gn=SVIL;refGene_tx=X
M_019834140.2;refGene_gn=SVIL;refGene_tx=XM_019834141.2;refGene_gn=SVI
L;refGene_tx=XM_019834142.2;refGene_gn=SVIL;refGene_tx=XM_019834143.2;
refGene_gn=SVIL;refGene_tx=XM_019834144.2;refGene_gn=SVIL;refGene_tx=X
M_019834147.2;refGene_gn=SVIL;refGene_tx=XM_019834150.2;refGene_gn=SVI
L;refGene_tx=XM_019834152.2;refGene_gn=SVIL;refGene_tx=XM_023256495.1
B4 28113662 28113740
refGene_gn=SVIL;refGene_tx=XM_006933343.4;refGene_gn=SVIL;refGene_tx=X
M_019834140.2;refGene_gn=SVIL;refGene_tx=XM_019834141.2;refGene_gn=SVI
L;refGene_tx=XM_019834143.2;refGene_gn=SVIL;refGene_tx=XM_019834144.2;
refGene_gn=SVIL;refGene_tx=XM_019834147.2;refGene_gn=SVIL;refGene_tx=X
M_019834150.2;refGene_gn=SVIL;refGene_tx=XM_019834152.2;refGene_gn=SVI
L;refGene_tx=XM_023256495.1
B4 28122068 28122216
refGene_gn=SVIL;refGene_tx=XM_006933343.4;refGene_gn=SVIL;refGene_tx=X
M_019834140.2;refGene_gn=SVIL;refGene_tx=XM_019834141.2;refGene_gn=SVI
L;refGene_tx=XM_019834142.2;refGene_gn=SVIL;refGene_tx=XM_019834143.2;
refGene_gn=SVIL;refGene_tx=XM_019834144.2;refGene_gn=SVIL;refGene_tx=X
M_019834146.2;refGene_gn=SVIL;refGene_tx=XM_019834147.2;refGene_gn=SVI
L;refGene_tx=XM_019834148.2;refGene_gn=SVIL;refGene_tx=XM_019834149.2;
refGene_gn=SVIL;refGene_tx=XM_019834150.2;refGene_gn=SVIL;refGene_tx=X
M_019834152.2;refGene_gn=SVIL;refGene_tx=XM_019834153.2;refGene_gn=SVI
L;refGene_tx=XM_019834154.2;refGene_gn=SVIL;refGene_tx=XM_023256495.1
B4 28124265 28124355
refGene_gn=SVIL;refGene_tx=XM_006933343.4;refGene_gn=SVIL;refGene_tx=X
M_019834140.2;refGene_gn=SVIL;refGene_tx=XM_019834141.2;refGene_gn=SVI
L;refGene_tx=XM_019834142.2;refGene_gn=SVIL;refGene_tx=XM_019834143.2;
refGene_gn=SVIL;refGene_tx=XM_019834144.2;refGene_gn=SVIL;refGene_tx=X
M_019834146.2;refGene_gn=SVIL;refGene_tx=XM_019834147.2;refGene_gn=SVI
L;refGene_tx=XM_019834148.2;refGene_gn=SVIL;refGene_tx=XM_019834149.2;
refGene_gn=SVIL;refGene_tx=XM_019834150.2;refGene_gn=SVIL;refGene_tx=X
M_019834152.2;refGene_gn=SVIL;refGene_tx=XM_019834153.2;refGene_gn=SVI
L;refGene_tx=XM_019834154.2;refGene_gn=SVIL;refGene_tx=XM_023256495.1
B4 28124445 28124532
refGene_gn=SVIL;refGene_tx=XM_006933343.4;refGene_gn=SVIL;refGene_tx=X
M_019834140.2;refGene_gn=SVIL;refGene_tx=XM_019834141.2;refGene_gn=SVI
L;refGene_tx=XM_019834142.2;refGene_gn=SVIL;refGene_tx=XM_019834143.2;
refGene_gn=SVIL;refGene_tx=XM_019834144.2;refGene_gn=SVIL;refGene_tx=X
M_019834146.2;refGene_gn=SVIL;refGene_tx=XM_019834147.2;refGene_gn=SVI
L;refGene_tx=XM_019834148.2;refGene_gn=SVIL;refGene_tx=XM_019834149.2;
refGene_gn=SVIL;refGene_tx=XM_019834150.2;refGene_gn=SVIL;refGene_tx=X
M_019834152.2;refGene_gn=SVIL;refGene_tx=XM_019834153.2;refGene_gn=SVI

L;refGene_tx=XM_019834154.2;refGene_gn=SVIL;refGene_tx=XM_023256495.1
B4 28126019 28126197
refGene_gn=SVIL;refGene_tx=XM_006933343.4;refGene_gn=SVIL;refGene_tx=XM_019834140.2;refGene_gn=SVIL;refGene_tx=XM_019834141.2;refGene_gn=SVIL;refGene_tx=XM_019834142.2;refGene_gn=SVIL;refGene_tx=XM_019834143.2;refGene_gn=SVIL;refGene_tx=XM_019834144.2;refGene_gn=SVIL;refGene_tx=XM_019834146.2;refGene_gn=SVIL;refGene_tx=XM_019834147.2;refGene_gn=SVIL;refGene_tx=XM_019834148.2;refGene_gn=SVIL;refGene_tx=XM_019834149.2;refGene_gn=SVIL;refGene_tx=XM_019834150.2;refGene_gn=SVIL;refGene_tx=XM_019834152.2;refGene_gn=SVIL;refGene_tx=XM_019834153.2;refGene_gn=SVIL;refGene_tx=XM_019834154.2;refGene_gn=SVIL;refGene_tx=XM_023256495.1
B4 28128520 28128693
refGene_gn=SVIL;refGene_tx=XM_006933343.4;refGene_gn=SVIL;refGene_tx=XM_019834140.2;refGene_gn=SVIL;refGene_tx=XM_019834141.2;refGene_gn=SVIL;refGene_tx=XM_019834142.2;refGene_gn=SVIL;refGene_tx=XM_019834143.2;refGene_gn=SVIL;refGene_tx=XM_019834144.2;refGene_gn=SVIL;refGene_tx=XM_019834146.2;refGene_gn=SVIL;refGene_tx=XM_019834147.2;refGene_gn=SVIL;refGene_tx=XM_019834148.2;refGene_gn=SVIL;refGene_tx=XM_019834149.2;refGene_gn=SVIL;refGene_tx=XM_019834150.2;refGene_gn=SVIL;refGene_tx=XM_019834152.2;refGene_gn=SVIL;refGene_tx=XM_019834153.2;refGene_gn=SVIL;refGene_tx=XM_019834154.2;refGene_gn=SVIL;refGene_tx=XM_023256495.1
B4 28130229 28130385
refGene_gn=SVIL;refGene_tx=XM_006933343.4;refGene_gn=SVIL;refGene_tx=XM_019834140.2;refGene_gn=SVIL;refGene_tx=XM_019834141.2;refGene_gn=SVIL;refGene_tx=XM_019834142.2;refGene_gn=SVIL;refGene_tx=XM_019834143.2;refGene_gn=SVIL;refGene_tx=XM_019834144.2;refGene_gn=SVIL;refGene_tx=XM_019834146.2;refGene_gn=SVIL;refGene_tx=XM_019834147.2;refGene_gn=SVIL;refGene_tx=XM_019834148.2;refGene_gn=SVIL;refGene_tx=XM_019834149.2;refGene_gn=SVIL;refGene_tx=XM_019834150.2;refGene_gn=SVIL;refGene_tx=XM_019834152.2;refGene_gn=SVIL;refGene_tx=XM_019834153.2;refGene_gn=SVIL;refGene_tx=XM_019834154.2;refGene_gn=SVIL;refGene_tx=XM_023256495.1
B4 28131847 28131984
refGene_gn=SVIL;refGene_tx=XM_006933343.4;refGene_gn=SVIL;refGene_tx=XM_019834140.2;refGene_gn=SVIL;refGene_tx=XM_019834141.2;refGene_gn=SVIL;refGene_tx=XM_019834142.2;refGene_gn=SVIL;refGene_tx=XM_019834143.2;refGene_gn=SVIL;refGene_tx=XM_019834144.2;refGene_gn=SVIL;refGene_tx=XM_019834146.2;refGene_gn=SVIL;refGene_tx=XM_019834147.2;refGene_gn=SVIL;refGene_tx=XM_019834148.2;refGene_gn=SVIL;refGene_tx=XM_019834149.2;refGene_gn=SVIL;refGene_tx=XM_019834150.2;refGene_gn=SVIL;refGene_tx=XM_019834152.2;refGene_gn=SVIL;refGene_tx=XM_019834153.2;refGene_gn=SVIL;refGene_tx=XM_019834154.2;refGene_gn=SVIL;refGene_tx=XM_023256495.1
B4 28132557 28132705
refGene_gn=SVIL;refGene_tx=XM_006933343.4;refGene_gn=SVIL;refGene_tx=XM_019834140.2;refGene_gn=SVIL;refGene_tx=XM_019834141.2;refGene_gn=SVIL;refGene_tx=XM_019834142.2;refGene_gn=SVIL;refGene_tx=XM_019834143.2;refGene_gn=SVIL;refGene_tx=XM_019834144.2;refGene_gn=SVIL;refGene_tx=XM_019834146.2;refGene_gn=SVIL;refGene_tx=XM_019834147.2;refGene_gn=SVIL;refGene_tx=XM_019834148.2;refGene_gn=SVIL;refGene_tx=XM_019834149.2;refGene_gn=SVIL;refGene_tx=XM_019834150.2;refGene_gn=SVIL;refGene_tx=XM_019834152.2;refGene_gn=SVIL;refGene_tx=XM_019834153.2;refGene_gn=SVI

L;refGene_tx=XM_019834154.2;refGene_gn=SVIL;refGene_tx=XM_023256495.1
B4 28132824 28132970
refGene_gn=SVIL;refGene_tx=XM_006933343.4;refGene_gn=SVIL;refGene_tx=XM_019834140.2;refGene_gn=SVIL;refGene_tx=XM_019834141.2;refGene_gn=SVIL;refGene_tx=XM_019834142.2;refGene_gn=SVIL;refGene_tx=XM_019834143.2;refGene_gn=SVIL;refGene_tx=XM_019834144.2;refGene_gn=SVIL;refGene_tx=XM_019834146.2;refGene_gn=SVIL;refGene_tx=XM_019834147.2;refGene_gn=SVIL;refGene_tx=XM_019834148.2;refGene_gn=SVIL;refGene_tx=XM_019834149.2;refGene_gn=SVIL;refGene_tx=XM_019834150.2;refGene_gn=SVIL;refGene_tx=XM_019834152.2;refGene_gn=SVIL;refGene_tx=XM_019834153.2;refGene_gn=SVIL;refGene_tx=XM_019834154.2;refGene_gn=SVIL;refGene_tx=XM_023256495.1
B4 28134568 28134744
refGene_gn=SVIL;refGene_tx=XM_006933343.4;refGene_gn=SVIL;refGene_tx=XM_019834140.2;refGene_gn=SVIL;refGene_tx=XM_019834141.2;refGene_gn=SVIL;refGene_tx=XM_019834142.2;refGene_gn=SVIL;refGene_tx=XM_019834143.2;refGene_gn=SVIL;refGene_tx=XM_019834144.2;refGene_gn=SVIL;refGene_tx=XM_019834146.2;refGene_gn=SVIL;refGene_tx=XM_019834147.2;refGene_gn=SVIL;refGene_tx=XM_019834148.2;refGene_gn=SVIL;refGene_tx=XM_019834149.2;refGene_gn=SVIL;refGene_tx=XM_019834150.2;refGene_gn=SVIL;refGene_tx=XM_019834152.2;refGene_gn=SVIL;refGene_tx=XM_019834153.2;refGene_gn=SVIL;refGene_tx=XM_019834154.2;refGene_gn=SVIL;refGene_tx=XM_023256495.1
B4 28137421 28137566
refGene_gn=SVIL;refGene_tx=XM_006933343.4;refGene_gn=SVIL;refGene_tx=XM_019834140.2;refGene_gn=SVIL;refGene_tx=XM_019834141.2;refGene_gn=SVIL;refGene_tx=XM_019834142.2;refGene_gn=SVIL;refGene_tx=XM_019834143.2;refGene_gn=SVIL;refGene_tx=XM_019834144.2;refGene_gn=SVIL;refGene_tx=XM_019834146.2;refGene_gn=SVIL;refGene_tx=XM_019834147.2;refGene_gn=SVIL;refGene_tx=XM_019834148.2;refGene_gn=SVIL;refGene_tx=XM_019834149.2;refGene_gn=SVIL;refGene_tx=XM_019834150.2;refGene_gn=SVIL;refGene_tx=XM_019834152.2;refGene_gn=SVIL;refGene_tx=XM_019834153.2;refGene_gn=SVIL;refGene_tx=XM_019834154.2;refGene_gn=SVIL;refGene_tx=XM_023256495.1
B4 28137908 28138185
refGene_gn=SVIL;refGene_tx=XM_006933343.4;refGene_gn=SVIL;refGene_tx=XM_019834140.2;refGene_gn=SVIL;refGene_tx=XM_019834141.2;refGene_gn=SVIL;refGene_tx=XM_019834142.2;refGene_gn=SVIL;refGene_tx=XM_019834143.2;refGene_gn=SVIL;refGene_tx=XM_019834144.2;refGene_gn=SVIL;refGene_tx=XM_019834146.2;refGene_gn=SVIL;refGene_tx=XM_019834147.2;refGene_gn=SVIL;refGene_tx=XM_019834148.2;refGene_gn=SVIL;refGene_tx=XM_019834149.2;refGene_gn=SVIL;refGene_tx=XM_019834150.2;refGene_gn=SVIL;refGene_tx=XM_019834152.2;refGene_gn=SVIL;refGene_tx=XM_019834153.2;refGene_gn=SVIL;refGene_tx=XM_019834154.2;refGene_gn=SVIL;refGene_tx=XM_023256495.1
B4 28143362 28143514
refGene_gn=SVIL;refGene_tx=XM_006933343.4;refGene_gn=SVIL;refGene_tx=XM_019834140.2;refGene_gn=SVIL;refGene_tx=XM_019834141.2;refGene_gn=SVIL;refGene_tx=XM_019834142.2;refGene_gn=SVIL;refGene_tx=XM_019834143.2;refGene_gn=SVIL;refGene_tx=XM_019834144.2;refGene_gn=SVIL;refGene_tx=XM_019834146.2;refGene_gn=SVIL;refGene_tx=XM_019834147.2;refGene_gn=SVIL;refGene_tx=XM_019834148.2;refGene_gn=SVIL;refGene_tx=XM_019834149.2;refGene_gn=SVIL;refGene_tx=XM_019834150.2;refGene_gn=SVIL;refGene_tx=XM_019834152.2;refGene_gn=SVIL;refGene_tx=XM_019834153.2;refGene_gn=SVI

L;refGene_tx=XM_019834154.2;refGene_gn=SVIL;refGene_tx=XM_023256495.1
B4 28145814 28145920
refGene_gn=SVIL;refGene_tx=XM_006933343.4;refGene_gn=SVIL;refGene_tx=XM_019834140.2;refGene_gn=SVIL;refGene_tx=XM_019834141.2;refGene_gn=SVIL;refGene_tx=XM_019834142.2;refGene_gn=SVIL;refGene_tx=XM_019834143.2;refGene_gn=SVIL;refGene_tx=XM_019834144.2;refGene_gn=SVIL;refGene_tx=XM_019834146.2;refGene_gn=SVIL;refGene_tx=XM_019834147.2;refGene_gn=SVIL;refGene_tx=XM_019834148.2;refGene_gn=SVIL;refGene_tx=XM_019834149.2;refGene_gn=SVIL;refGene_tx=XM_019834150.2;refGene_gn=SVIL;refGene_tx=XM_019834152.2;refGene_gn=SVIL;refGene_tx=XM_019834153.2;refGene_gn=SVIL;refGene_tx=XM_019834154.2;refGene_gn=SVIL;refGene_tx=XM_023256495.1
B4 28146628 28146836
refGene_gn=SVIL;refGene_tx=XM_006933343.4;refGene_gn=SVIL;refGene_tx=XM_019834140.2;refGene_gn=SVIL;refGene_tx=XM_019834141.2;refGene_gn=SVIL;refGene_tx=XM_019834142.2;refGene_gn=SVIL;refGene_tx=XM_019834143.2;refGene_gn=SVIL;refGene_tx=XM_019834144.2;refGene_gn=SVIL;refGene_tx=XM_019834146.2;refGene_gn=SVIL;refGene_tx=XM_019834147.2;refGene_gn=SVIL;refGene_tx=XM_019834148.2;refGene_gn=SVIL;refGene_tx=XM_019834149.2;refGene_gn=SVIL;refGene_tx=XM_019834150.2;refGene_gn=SVIL;refGene_tx=XM_019834152.2;refGene_gn=SVIL;refGene_tx=XM_019834153.2;refGene_gn=SVIL;refGene_tx=XM_019834154.2;refGene_gn=SVIL;refGene_tx=XM_023256495.1
B4 28148190 28148324
refGene_gn=SVIL;refGene_tx=XM_006933343.4;refGene_gn=SVIL;refGene_tx=XM_019834140.2;refGene_gn=SVIL;refGene_tx=XM_019834141.2;refGene_gn=SVIL;refGene_tx=XM_019834142.2;refGene_gn=SVIL;refGene_tx=XM_019834143.2;refGene_gn=SVIL;refGene_tx=XM_019834144.2;refGene_gn=SVIL;refGene_tx=XM_019834146.2;refGene_gn=SVIL;refGene_tx=XM_019834147.2;refGene_gn=SVIL;refGene_tx=XM_019834148.2;refGene_gn=SVIL;refGene_tx=XM_019834149.2;refGene_gn=SVIL;refGene_tx=XM_019834150.2;refGene_gn=SVIL;refGene_tx=XM_019834152.2;refGene_gn=SVIL;refGene_tx=XM_019834153.2;refGene_gn=SVIL;refGene_tx=XM_019834154.2;refGene_gn=SVIL;refGene_tx=XM_023256495.1
B4 28150036 28150192
refGene_gn=SVIL;refGene_tx=XM_006933343.4;refGene_gn=SVIL;refGene_tx=XM_019834140.2;refGene_gn=SVIL;refGene_tx=XM_019834141.2;refGene_gn=SVIL;refGene_tx=XM_019834142.2;refGene_gn=SVIL;refGene_tx=XM_019834143.2;refGene_gn=SVIL;refGene_tx=XM_019834144.2;refGene_gn=SVIL;refGene_tx=XM_019834146.2;refGene_gn=SVIL;refGene_tx=XM_019834147.2;refGene_gn=SVIL;refGene_tx=XM_019834148.2;refGene_gn=SVIL;refGene_tx=XM_019834149.2;refGene_gn=SVIL;refGene_tx=XM_019834150.2;refGene_gn=SVIL;refGene_tx=XM_019834152.2;refGene_gn=SVIL;refGene_tx=XM_019834153.2;refGene_gn=SVIL;refGene_tx=XM_019834154.2;refGene_gn=SVIL;refGene_tx=XM_023256495.1
B4 28151780 28151924
refGene_gn=SVIL;refGene_tx=XM_006933343.4;refGene_gn=SVIL;refGene_tx=XM_019834140.2;refGene_gn=SVIL;refGene_tx=XM_019834141.2;refGene_gn=SVIL;refGene_tx=XM_019834142.2;refGene_gn=SVIL;refGene_tx=XM_019834143.2;refGene_gn=SVIL;refGene_tx=XM_019834144.2;refGene_gn=SVIL;refGene_tx=XM_019834146.2;refGene_gn=SVIL;refGene_tx=XM_019834147.2;refGene_gn=SVIL;refGene_tx=XM_019834148.2;refGene_gn=SVIL;refGene_tx=XM_019834149.2;refGene_gn=SVIL;refGene_tx=XM_019834150.2;refGene_gn=SVIL;refGene_tx=XM_019834152.2;refGene_gn=SVIL;refGene_tx=XM_019834153.2;refGene_gn=SVI

L;refGene_tx=XM_019834154.2;refGene_gn=SVIL;refGene_tx=XM_023256495.1
B4 28152954 28153079
refGene_gn=SVIL;refGene_tx=XM_006933343.4;refGene_gn=SVIL;refGene_tx=XM_019834140.2;refGene_gn=SVIL;refGene_tx=XM_019834141.2;refGene_gn=SVIL;refGene_tx=XM_019834142.2;refGene_gn=SVIL;refGene_tx=XM_019834143.2;refGene_gn=SVIL;refGene_tx=XM_019834144.2;refGene_gn=SVIL;refGene_tx=XM_019834146.2;refGene_gn=SVIL;refGene_tx=XM_019834147.2;refGene_gn=SVIL;refGene_tx=XM_019834148.2;refGene_gn=SVIL;refGene_tx=XM_019834149.2;refGene_gn=SVIL;refGene_tx=XM_019834150.2;refGene_gn=SVIL;refGene_tx=XM_019834152.2;refGene_gn=SVIL;refGene_tx=XM_019834153.2;refGene_gn=SVIL;refGene_tx=XM_019834154.2;refGene_gn=SVIL;refGene_tx=XM_023256495.1
B4 28155485 28155641
refGene_gn=SVIL;refGene_tx=XM_006933343.4;refGene_gn=SVIL;refGene_tx=XM_019834140.2;refGene_gn=SVIL;refGene_tx=XM_019834141.2;refGene_gn=SVIL;refGene_tx=XM_019834142.2;refGene_gn=SVIL;refGene_tx=XM_019834143.2;refGene_gn=SVIL;refGene_tx=XM_019834144.2;refGene_gn=SVIL;refGene_tx=XM_019834146.2;refGene_gn=SVIL;refGene_tx=XM_019834147.2;refGene_gn=SVIL;refGene_tx=XM_019834148.2;refGene_gn=SVIL;refGene_tx=XM_019834149.2;refGene_gn=SVIL;refGene_tx=XM_019834150.2;refGene_gn=SVIL;refGene_tx=XM_019834152.2;refGene_gn=SVIL;refGene_tx=XM_019834153.2;refGene_gn=SVIL;refGene_tx=XM_019834154.2;refGene_gn=SVIL;refGene_tx=XM_023256495.1
B4 28155738 28155825
refGene_gn=SVIL;refGene_tx=XM_006933343.4;refGene_gn=SVIL;refGene_tx=XM_019834140.2;refGene_gn=SVIL;refGene_tx=XM_019834141.2;refGene_gn=SVIL;refGene_tx=XM_019834142.2;refGene_gn=SVIL;refGene_tx=XM_019834143.2;refGene_gn=SVIL;refGene_tx=XM_019834144.2;refGene_gn=SVIL;refGene_tx=XM_019834146.2;refGene_gn=SVIL;refGene_tx=XM_019834147.2;refGene_gn=SVIL;refGene_tx=XM_019834148.2;refGene_gn=SVIL;refGene_tx=XM_019834149.2;refGene_gn=SVIL;refGene_tx=XM_019834150.2;refGene_gn=SVIL;refGene_tx=XM_019834152.2;refGene_gn=SVIL;refGene_tx=XM_019834153.2;refGene_gn=SVIL;refGene_tx=XM_019834154.2;refGene_gn=SVIL;refGene_tx=XM_023256495.1
B4 28276953 28277572
refGene_gn=ZNF438;refGene_tx=XM_011283660.3;refGene_gn=ZNF438;refGene_tx=XM_011283662.3;refGene_gn=ZNF438;refGene_tx=XM_011283663.3;refGene_gn=ZNF438;refGene_tx=XM_011283664.3;refGene_gn=ZNF438;refGene_tx=XM_011283665.3;refGene_gn=ZNF438;refGene_tx=XM_019834156.2
B4 28279183 28281002
refGene_gn=ZNF438;refGene_tx=XM_011283660.3;refGene_gn=ZNF438;refGene_tx=XM_011283662.3;refGene_gn=ZNF438;refGene_tx=XM_011283663.3;refGene_gn=ZNF438;refGene_tx=XM_011283664.3;refGene_gn=ZNF438;refGene_tx=XM_011283665.3;refGene_gn=ZNF438;refGene_tx=XM_019834156.2
B4 28303821 28303861
refGene_gn=ZNF438;refGene_tx=XM_011283660.3;refGene_gn=ZNF438;refGene_tx=XM_011283662.3;refGene_gn=ZNF438;refGene_tx=XM_011283663.3;refGene_gn=ZNF438;refGene_tx=XM_011283664.3;refGene_gn=ZNF438;refGene_tx=XM_011283665.3;refGene_gn=ZNF438;refGene_tx=XM_019834156.2
B4 28693727 28693785
refGene_gn=ZEB1;refGene_tx=XM_023256497.1;refGene_gn=ZEB1;refGene_tx=XM_023256498.1
B4 28829289 28829490

refGene_gn=ZEB1;refGene_tx=XM_023256497.1;refGene_gn=ZEB1;refGene_tx=XM_023256498.1
B4 28862196 28862259
refGene_gn=ZEB1;refGene_tx=XM_023256497.1;refGene_gn=ZEB1;refGene_tx=XM_023256498.1
B4 28869471 28869633
refGene_gn=ZEB1;refGene_tx=XM_023256497.1;refGene_gn=ZEB1;refGene_tx=XM_023256498.1
B4 28875535 28875738
refGene_gn=ZEB1;refGene_tx=XM_019834160.1;refGene_gn=ZEB1;refGene_tx=XM_019834161.1;refGene_gn=ZEB1;refGene_tx=XM_023256497.1;refGene_gn=ZEB1;refGene_tx=XM_023256498.1
B4 28879758 28879864
refGene_gn=ZEB1;refGene_tx=XM_019834160.1;refGene_gn=ZEB1;refGene_tx=XM_019834161.1;refGene_gn=ZEB1;refGene_tx=XM_023256497.1;refGene_gn=ZEB1;refGene_tx=XM_023256498.1
B4 28885860 28887668
refGene_gn=ZEB1;refGene_tx=XM_019834160.1;refGene_gn=ZEB1;refGene_tx=XM_019834161.1;refGene_gn=ZEB1;refGene_tx=XM_023256497.1;refGene_gn=ZEB1;refGene_tx=XM_023256498.1
B4 28890116 28890297
refGene_gn=ZEB1;refGene_tx=XM_019834160.1;refGene_gn=ZEB1;refGene_tx=XM_019834161.1;refGene_gn=ZEB1;refGene_tx=XM_023256497.1;refGene_gn=ZEB1;refGene_tx=XM_023256498.1
B4 28890410 28891006
refGene_gn=ZEB1;refGene_tx=XM_019834160.1;refGene_gn=ZEB1;refGene_tx=XM_019834161.1;refGene_gn=ZEB1;refGene_tx=XM_023256497.1;refGene_gn=ZEB1;refGene_tx=XM_023256498.1
B4 29110504 29110679
refGene_gn=ARHGAP12;refGene_tx=XM_023256499.1;refGene_gn=ARHGAP12;refGene_tx=XM_023256500.1;refGene_gn=ARHGAP12;refGene_tx=XM_023256501.1;refGene_gn=ARHGAP12;refGene_tx=XM_023256502.1;refGene_gn=ARHGAP12;refGene_tx=XM_023256503.1
B4 29111871 29111974
refGene_gn=ARHGAP12;refGene_tx=XM_023256499.1;refGene_gn=ARHGAP12;refGene_tx=XM_023256500.1;refGene_gn=ARHGAP12;refGene_tx=XM_023256501.1;refGene_gn=ARHGAP12;refGene_tx=XM_023256502.1;refGene_gn=ARHGAP12;refGene_tx=XM_023256503.1
B4 29112199 29112334
refGene_gn=ARHGAP12;refGene_tx=XM_023256499.1;refGene_gn=ARHGAP12;refGene_tx=XM_023256500.1;refGene_gn=ARHGAP12;refGene_tx=XM_023256501.1;refGene_gn=ARHGAP12;refGene_tx=XM_023256502.1;refGene_gn=ARHGAP12;refGene_tx=XM_023256503.1
B4 29112430 29112508
refGene_gn=ARHGAP12;refGene_tx=XM_023256499.1;refGene_gn=ARHGAP12;refGene_tx=XM_023256500.1;refGene_gn=ARHGAP12;refGene_tx=XM_023256501.1;refGene_gn=ARHGAP12;refGene_tx=XM_023256502.1;refGene_gn=ARHGAP12;refGene_tx=XM_023256503.1
B4 29113823 29113922
refGene_gn=ARHGAP12;refGene_tx=XM_023256499.1;refGene_gn=ARHGAP12;refG

ene_tx=XM_023256500.1;refGene_gn=ARHGAP12;refGene_tx=XM_023256501.1;re
fGene_gn=ARHGAP12;refGene_tx=XM_023256502.1;refGene_gn=ARHGAP12;refGen
e_tx=XM_023256503.1
B4	29115999	29116116
refGene_gn=ARHGAP12;refGene_tx=XM_023256499.1;refGene_gn=ARHGAP12;refG
ene_tx=XM_023256500.1;refGene_gn=ARHGAP12;refGene_tx=XM_023256501.1;re
fGene_gn=ARHGAP12;refGene_tx=XM_023256502.1;refGene_gn=ARHGAP12;refGen
e_tx=XM_023256503.1
B4	29116922	29117025
refGene_gn=ARHGAP12;refGene_tx=XM_023256499.1;refGene_gn=ARHGAP12;refG
ene_tx=XM_023256500.1;refGene_gn=ARHGAP12;refGene_tx=XM_023256501.1;re
fGene_gn=ARHGAP12;refGene_tx=XM_023256502.1;refGene_gn=ARHGAP12;refGen
e_tx=XM_023256503.1
B4	29120925	29121024
refGene_gn=ARHGAP12;refGene_tx=XM_023256499.1;refGene_gn=ARHGAP12;refG
ene_tx=XM_023256500.1;refGene_gn=ARHGAP12;refGene_tx=XM_023256501.1;re
fGene_gn=ARHGAP12;refGene_tx=XM_023256503.1
B4	29125716	29125818
refGene_gn=ARHGAP12;refGene_tx=XM_023256499.1;refGene_gn=ARHGAP12;refG
ene_tx=XM_023256500.1;refGene_gn=ARHGAP12;refGene_tx=XM_023256501.1;re
fGene_gn=ARHGAP12;refGene_tx=XM_023256502.1;refGene_gn=ARHGAP12;refGen
e_tx=XM_023256503.1
B4	29132353	29132435
refGene_gn=ARHGAP12;refGene_tx=XM_023256499.1;refGene_gn=ARHGAP12;refG
ene_tx=XM_023256500.1;refGene_gn=ARHGAP12;refGene_tx=XM_023256501.1;re
fGene_gn=ARHGAP12;refGene_tx=XM_023256502.1;refGene_gn=ARHGAP12;refGen
e_tx=XM_023256503.1
B4	29136469	29136531
refGene_gn=ARHGAP12;refGene_tx=XM_023256499.1;refGene_gn=ARHGAP12;refG
ene_tx=XM_023256500.1;refGene_gn=ARHGAP12;refGene_tx=XM_023256501.1;re
fGene_gn=ARHGAP12;refGene_tx=XM_023256502.1;refGene_gn=ARHGAP12;refGen
e_tx=XM_023256503.1
B4	29141034	29141049
refGene_gn=ARHGAP12;refGene_tx=XM_023256499.1;refGene_gn=ARHGAP12;refG
ene_tx=XM_023256502.1;refGene_gn=ARHGAP12;refGene_tx=XM_023256503.1
B4	29141379	29141454
refGene_gn=ARHGAP12;refGene_tx=XM_023256499.1;refGene_gn=ARHGAP12;refG
ene_tx=XM_023256500.1;refGene_gn=ARHGAP12;refGene_tx=XM_023256502.1;re
fGene_gn=ARHGAP12;refGene_tx=XM_023256503.1
B4	29145493	29145619
refGene_gn=ARHGAP12;refGene_tx=XM_023256499.1;refGene_gn=ARHGAP12;refG
ene_tx=XM_023256500.1;refGene_gn=ARHGAP12;refGene_tx=XM_023256501.1;re
fGene_gn=ARHGAP12;refGene_tx=XM_023256502.1;refGene_gn=ARHGAP12;refGen
e_tx=XM_023256503.1
B4	29156391	29156472
refGene_gn=ARHGAP12;refGene_tx=XM_023256499.1;refGene_gn=ARHGAP12;refG
ene_tx=XM_023256500.1;refGene_gn=ARHGAP12;refGene_tx=XM_023256501.1;re
fGene_gn=ARHGAP12;refGene_tx=XM_023256502.1;refGene_gn=ARHGAP12;refGen
e_tx=XM_023256503.1
B4	29157648	29157789

refGene_gn=ARHGAP12;refGene_tx=XM_023256499.1;refGene_gn=ARHGAP12;refGene_tx=XM_023256500.1;refGene_gn=ARHGAP12;refGene_tx=XM_023256501.1;refGene_gn=ARHGAP12;refGene_tx=XM_023256502.1
B4 29165646 29165910
refGene_gn=ARHGAP12;refGene_tx=XM_023256499.1;refGene_gn=ARHGAP12;refGene_tx=XM_023256500.1;refGene_gn=ARHGAP12;refGene_tx=XM_023256501.1;refGene_gn=ARHGAP12;refGene_tx=XM_023256502.1;refGene_gn=ARHGAP12;refGene_tx=XM_023256503.1
B4 29209927 29210605
refGene_gn=ARHGAP12;refGene_tx=XM_023256499.1;refGene_gn=ARHGAP12;refGene_tx=XM_023256500.1;refGene_gn=ARHGAP12;refGene_tx=XM_023256501.1;refGene_gn=ARHGAP12;refGene_tx=XM_023256502.1;refGene_gn=ARHGAP12;refGene_tx=XM_023256503.1
B4 29307919 29308050 refGene_gn=KIF5B;refGene_tx=XM_003988210.5
B4 29308776 29308993 refGene_gn=KIF5B;refGene_tx=XM_003988210.5
B4 29310512 29310617 refGene_gn=KIF5B;refGene_tx=XM_003988210.5
B4 29310788 29310860 refGene_gn=KIF5B;refGene_tx=XM_003988210.5
B4 29310971 29311032 refGene_gn=KIF5B;refGene_tx=XM_003988210.5
B4 29312392 29312494 refGene_gn=KIF5B;refGene_tx=XM_003988210.5
B4 29313256 29313366 refGene_gn=KIF5B;refGene_tx=XM_003988210.5
B4 29313462 29313524 refGene_gn=KIF5B;refGene_tx=XM_003988210.5
B4 29314058 29314176 refGene_gn=KIF5B;refGene_tx=XM_003988210.5
B4 29314709 29314898 refGene_gn=KIF5B;refGene_tx=XM_003988210.5
B4 29319214 29319358 refGene_gn=KIF5B;refGene_tx=XM_003988210.5
B4 29321873 29322080 refGene_gn=KIF5B;refGene_tx=XM_003988210.5
B4 29324210 29324279 refGene_gn=KIF5B;refGene_tx=XM_003988210.5
B4 29327918 29328112 refGene_gn=KIF5B;refGene_tx=XM_003988210.5
B4 29328557 29328706 refGene_gn=KIF5B;refGene_tx=XM_003988210.5
B4 29329445 29329591 refGene_gn=KIF5B;refGene_tx=XM_003988210.5
B4 29329903 29330008 refGene_gn=KIF5B;refGene_tx=XM_003988210.5
B4 29331111 29331236 refGene_gn=KIF5B;refGene_tx=XM_003988210.5
B4 29331398 29331486 refGene_gn=KIF5B;refGene_tx=XM_003988210.5
B4 29331778 29331834 refGene_gn=KIF5B;refGene_tx=XM_003988210.5
B4 29332388 29332437 refGene_gn=KIF5B;refGene_tx=XM_003988210.5
B4 29333035 29333140 refGene_gn=KIF5B;refGene_tx=XM_003988210.5
B4 29334041 29334115 refGene_gn=KIF5B;refGene_tx=XM_003988210.5
B4 29343116 29343204 refGene_gn=KIF5B;refGene_tx=XM_003988210.5
B4 29350570 29350696 refGene_gn=KIF5B;refGene_tx=XM_003988210.5
B4 29514354 29514427
refGene_gn=EPC1;refGene_tx=XM_023256504.1;refGene_gn=EPC1;refGene_tx=XM_023256505.1;refGene_gn=EPC1;refGene_tx=XM_023256506.1;refGene_gn=EPC1;refGene_tx=XM_023256507.1;refGene_gn=EPC1;refGene_tx=XM_023256508.1;refGene_gn=EPC1;refGene_tx=XM_023256509.1;refGene_gn=EPC1;refGene_tx=XM_023256510.1
B4 29516425 29516789
refGene_gn=EPC1;refGene_tx=XM_023256504.1;refGene_gn=EPC1;refGene_tx=XM_023256505.1;refGene_gn=EPC1;refGene_tx=XM_023256506.1;refGene_gn=EPC1;refGene_tx=XM_023256507.1;refGene_gn=EPC1;refGene_tx=XM_023256508.1;refGene_gn=EPC1;refGene_tx=XM_023256509.1;refGene_gn=EPC1;refGene_tx=XM_023256510.1

B4	29516890 29517032
refGene_gn=EPC1;refGene_tx=XM_023256504.1;refGene_gn=EPC1;refGene_tx=XM_023256505.1;refGene_gn=EPC1;refGene_tx=XM_023256506.1;refGene_gn=EPC1;refGene_tx=XM_023256507.1;refGene_gn=EPC1;refGene_tx=XM_023256508.1;refGene_gn=EPC1;refGene_tx=XM_023256509.1;refGene_gn=EPC1;refGene_tx=XM_023256510.1
B4	29517861 29517948
refGene_gn=EPC1;refGene_tx=XM_023256504.1;refGene_gn=EPC1;refGene_tx=XM_023256506.1;refGene_gn=EPC1;refGene_tx=XM_023256507.1;refGene_gn=EPC1;refGene_tx=XM_023256508.1;refGene_gn=EPC1;refGene_tx=XM_023256509.1
B4	29518028 29518147
refGene_gn=EPC1;refGene_tx=XM_023256504.1;refGene_gn=EPC1;refGene_tx=XM_023256505.1;refGene_gn=EPC1;refGene_tx=XM_023256506.1;refGene_gn=EPC1;refGene_tx=XM_023256507.1;refGene_gn=EPC1;refGene_tx=XM_023256508.1;refGene_gn=EPC1;refGene_tx=XM_023256509.1;refGene_gn=EPC1;refGene_tx=XM_023256510.1
B4	29525661 29526014
refGene_gn=EPC1;refGene_tx=XM_023256504.1;refGene_gn=EPC1;refGene_tx=XM_023256505.1;refGene_gn=EPC1;refGene_tx=XM_023256506.1;refGene_gn=EPC1;refGene_tx=XM_023256507.1;refGene_gn=EPC1;refGene_tx=XM_023256508.1;refGene_gn=EPC1;refGene_tx=XM_023256509.1;refGene_gn=EPC1;refGene_tx=XM_023256510.1
B4	29527859 29528008
refGene_gn=EPC1;refGene_tx=XM_023256504.1;refGene_gn=EPC1;refGene_tx=XM_023256505.1;refGene_gn=EPC1;refGene_tx=XM_023256506.1;refGene_gn=EPC1;refGene_tx=XM_023256507.1;refGene_gn=EPC1;refGene_tx=XM_023256508.1;refGene_gn=EPC1;refGene_tx=XM_023256509.1;refGene_gn=EPC1;refGene_tx=XM_023256510.1
B4	29528091 29528181
refGene_gn=EPC1;refGene_tx=XM_023256504.1;refGene_gn=EPC1;refGene_tx=XM_023256505.1;refGene_gn=EPC1;refGene_tx=XM_023256506.1;refGene_gn=EPC1;refGene_tx=XM_023256507.1;refGene_gn=EPC1;refGene_tx=XM_023256508.1;refGene_gn=EPC1;refGene_tx=XM_023256509.1;refGene_gn=EPC1;refGene_tx=XM_023256510.1
B4	29528262 29528439
refGene_gn=EPC1;refGene_tx=XM_023256504.1;refGene_gn=EPC1;refGene_tx=XM_023256505.1;refGene_gn=EPC1;refGene_tx=XM_023256506.1;refGene_gn=EPC1;refGene_tx=XM_023256507.1;refGene_gn=EPC1;refGene_tx=XM_023256508.1;refGene_gn=EPC1;refGene_tx=XM_023256509.1;refGene_gn=EPC1;refGene_tx=XM_023256510.1
B4	29529685 29529845
refGene_gn=EPC1;refGene_tx=XM_023256504.1;refGene_gn=EPC1;refGene_tx=XM_023256505.1;refGene_gn=EPC1;refGene_tx=XM_023256506.1;refGene_gn=EPC1;refGene_tx=XM_023256507.1;refGene_gn=EPC1;refGene_tx=XM_023256508.1;refGene_gn=EPC1;refGene_tx=XM_023256509.1;refGene_gn=EPC1;refGene_tx=XM_023256510.1
B4	29530740 29530889
refGene_gn=EPC1;refGene_tx=XM_023256504.1;refGene_gn=EPC1;refGene_tx=XM_023256505.1;refGene_gn=EPC1;refGene_tx=XM_023256506.1;refGene_gn=EPC1;refGene_tx=XM_023256507.1;refGene_gn=EPC1;refGene_tx=XM_023256508.1;

refGene_gn=EPC1;refGene_tx=XM_023256509.1;refGene_gn=EPC1;refGene_tx=XM_023256510.1
B4 29531267 29531474
refGene_gn=EPC1;refGene_tx=XM_023256504.1;refGene_gn=EPC1;refGene_tx=XM_023256505.1;refGene_gn=EPC1;refGene_tx=XM_023256506.1;refGene_gn=EPC1;refGene_tx=XM_023256507.1;refGene_gn=EPC1;refGene_tx=XM_023256508.1;refGene_gn=EPC1;refGene_tx=XM_023256509.1;refGene_gn=EPC1;refGene_tx=XM_023256510.1
B4 29531861 29532007
refGene_gn=EPC1;refGene_tx=XM_023256504.1;refGene_gn=EPC1;refGene_tx=XM_023256505.1;refGene_gn=EPC1;refGene_tx=XM_023256506.1;refGene_gn=EPC1;refGene_tx=XM_023256507.1;refGene_gn=EPC1;refGene_tx=XM_023256508.1;refGene_gn=EPC1;refGene_tx=XM_023256509.1;refGene_gn=EPC1;refGene_tx=XM_023256510.1
B4 29544866 29544966
refGene_gn=EPC1;refGene_tx=XM_023256509.1;refGene_gn=EPC1;refGene_tx=XM_023256510.1
B4 29545472 29545632
refGene_gn=EPC1;refGene_tx=XM_023256504.1;refGene_gn=EPC1;refGene_tx=XM_023256505.1;refGene_gn=EPC1;refGene_tx=XM_023256506.1;refGene_gn=EPC1;refGene_tx=XM_023256507.1;refGene_gn=EPC1;refGene_tx=XM_023256508.1
B4 29582317 29582470
refGene_gn=EPC1;refGene_tx=XM_023256504.1;refGene_gn=EPC1;refGene_tx=XM_023256505.1
B4 29607330 29607333
refGene_gn=EPC1;refGene_tx=XM_023256506.1;refGene_gn=EPC1;refGene_tx=XM_023256507.1;refGene_gn=EPC1;refGene_tx=XM_023256508.1
B4 29654260 29655136
refGene_gn=LOC101093782;refGene_tx=XM_011283669.3
B4 29680439 29680718
refGene_gn=CCDC7;refGene_tx=XM_011283670.3;refGene_gn=CCDC7;refGene_tx=XM_019834169.2;refGene_gn=CCDC7;refGene_tx=XM_019834170.2;refGene_gn=CCDC7;refGene_tx=XM_019834171.2
B4 29681898 29681991
refGene_gn=CCDC7;refGene_tx=XM_011283670.3;refGene_gn=CCDC7;refGene_tx=XM_019834169.2;refGene_gn=CCDC7;refGene_tx=XM_019834170.2;refGene_gn=CCDC7;refGene_tx=XM_019834171.2
B4 29686927 29687011
refGene_gn=CCDC7;refGene_tx=XM_011283670.3;refGene_gn=CCDC7;refGene_tx=XM_019834169.2;refGene_gn=CCDC7;refGene_tx=XM_019834170.2;refGene_gn=CCDC7;refGene_tx=XM_019834171.2
B4 29707659 29707680
refGene_gn=CCDC7;refGene_tx=XM_011283670.3;refGene_gn=CCDC7;refGene_tx=XM_019834169.2;refGene_gn=CCDC7;refGene_tx=XM_019834170.2;refGene_gn=CCDC7;refGene_tx=XM_019834171.2
B4 29708018 29708051
refGene_gn=CCDC7;refGene_tx=XM_011283670.3;refGene_gn=CCDC7;refGene_tx=XM_019834169.2;refGene_gn=CCDC7;refGene_tx=XM_019834170.2;refGene_gn=CCDC7;refGene_tx=XM_019834171.2
B4 29717660 29717827

refGene_gn=CCDC7;refGene_tx=XM_011283670.3;refGene_gn=CCDC7;refGene_tx=XM_019834169.2;refGene_gn=CCDC7;refGene_tx=XM_019834170.2;refGene_gn=CCDC7;refGene_tx=XM_019834171.2
B4 29719645 29719707
refGene_gn=CCDC7;refGene_tx=XM_011283670.3;refGene_gn=CCDC7;refGene_tx=XM_019834169.2;refGene_gn=CCDC7;refGene_tx=XM_019834170.2;refGene_gn=CCDC7;refGene_tx=XM_019834171.2
B4 29720786 29720843
refGene_gn=CCDC7;refGene_tx=XM_011283670.3;refGene_gn=CCDC7;refGene_tx=XM_019834169.2;refGene_gn=CCDC7;refGene_tx=XM_019834170.2;refGene_gn=CCDC7;refGene_tx=XM_019834171.2
B4 29734029 29734105
refGene_gn=CCDC7;refGene_tx=XM_011283670.3;refGene_gn=CCDC7;refGene_tx=XM_019834169.2;refGene_gn=CCDC7;refGene_tx=XM_019834170.2;refGene_gn=CCDC7;refGene_tx=XM_019834171.2
B4 29744490 29744521
refGene_gn=CCDC7;refGene_tx=XM_011283670.3;refGene_gn=CCDC7;refGene_tx=XM_019834169.2;refGene_gn=CCDC7;refGene_tx=XM_019834170.2;refGene_gn=CCDC7;refGene_tx=XM_019834171.2
B4 29745649 29745739
refGene_gn=CCDC7;refGene_tx=XM_011283670.3;refGene_gn=CCDC7;refGene_tx=XM_019834169.2;refGene_gn=CCDC7;refGene_tx=XM_019834170.2;refGene_gn=CCDC7;refGene_tx=XM_019834171.2
B4 29753753 29753839
refGene_gn=CCDC7;refGene_tx=XM_011283670.3;refGene_gn=CCDC7;refGene_tx=XM_019834169.2;refGene_gn=CCDC7;refGene_tx=XM_019834170.2;refGene_gn=CCDC7;refGene_tx=XM_019834171.2
B4 29757688 29757743
refGene_gn=CCDC7;refGene_tx=XM_011283670.3;refGene_gn=CCDC7;refGene_tx=XM_019834169.2;refGene_gn=CCDC7;refGene_tx=XM_019834170.2;refGene_gn=CCDC7;refGene_tx=XM_019834171.2
B4 29784609 29784675
refGene_gn=CCDC7;refGene_tx=XM_011283670.3;refGene_gn=CCDC7;refGene_tx=XM_019834169.2;refGene_gn=CCDC7;refGene_tx=XM_019834170.2
B4 29798218 29798437
refGene_gn=CCDC7;refGene_tx=XM_011283670.3;refGene_gn=CCDC7;refGene_tx=XM_019834169.2;refGene_gn=CCDC7;refGene_tx=XM_019834170.2;refGene_gn=CCDC7;refGene_tx=XM_019834171.2
B4 29801550 29801588
refGene_gn=CCDC7;refGene_tx=XM_011283670.3;refGene_gn=CCDC7;refGene_tx=XM_019834169.2;refGene_gn=CCDC7;refGene_tx=XM_019834170.2;refGene_gn=CCDC7;refGene_tx=XM_019834171.2
B4 29803713 29803720
refGene_gn=CCDC7;refGene_tx=XM_011283670.3;refGene_gn=CCDC7;refGene_tx=XM_019834169.2;refGene_gn=CCDC7;refGene_tx=XM_019834170.2;refGene_gn=CCDC7;refGene_tx=XM_019834171.2
B4 29927987 29928048
refGene_gn=LOC102901117;refGene_tx=XM_023256511.1
B4 29936972 29937083
refGene_gn=LOC102901117;refGene_tx=XM_023256511.1

B4 29946830 29946941
refGene_gn=LOC102901117;refGene_tx=XM_023256511.1
B4 29961826 29961937
refGene_gn=LOC102901117;refGene_tx=XM_023256511.1
B4 29963038 29963149
refGene_gn=LOC102901117;refGene_tx=XM_023256511.1
B4 29966209 29966329
refGene_gn=LOC102901117;refGene_tx=XM_023256511.1
B4 29967315 29967345
refGene_gn=LOC102901117;refGene_tx=XM_023256511.1
B4 29996685 29996796
refGene_gn=LOC102901117;refGene_tx=XM_023256511.1
B4 30025628 30025733
refGene_gn=LOC102901117;refGene_tx=XM_023256511.1
B4 30036286 30036370
refGene_gn=LOC102901117;refGene_tx=XM_023256511.1
B4 30060519 30060642
refGene_gn=LOC102901117;refGene_tx=XM_023256511.1
B4 30082628 30082751
refGene_gn=LOC102901117;refGene_tx=XM_023256511.1
B4 30085454 30085495
refGene_gn=LOC102901117;refGene_tx=XM_023256511.1
B4 30109163 30109229
refGene_gn=ITGB1;refGene_tx=NM_001048160.2;refGene_gn=ITGB1;refGene_tx=XM_023256252.1
B4 30115605 30115680
refGene_gn=ITGB1;refGene_tx=XM_023256250.1;refGene_gn=ITGB1;refGene_tx=XM_023256251.1
B4 30116623 30116790
refGene_gn=ITGB1;refGene_tx=NM_001048160.2;refGene_gn=ITGB1;refGene_tx=XM_023256250.1;refGene_gn=ITGB1;refGene_tx=XM_023256251.1;refGene_gn=ITGB1;refGene_tx=XM_023256252.1
B4 30119162 30119395
refGene_gn=ITGB1;refGene_tx=NM_001048160.2;refGene_gn=ITGB1;refGene_tx=XM_023256250.1;refGene_gn=ITGB1;refGene_tx=XM_023256251.1;refGene_gn=ITGB1;refGene_tx=XM_023256252.1
B4 30120008 30120231
refGene_gn=ITGB1;refGene_tx=NM_001048160.2;refGene_gn=ITGB1;refGene_tx=XM_023256250.1;refGene_gn=ITGB1;refGene_tx=XM_023256251.1;refGene_gn=ITGB1;refGene_tx=XM_023256252.1
B4 30120453 30120692
refGene_gn=ITGB1;refGene_tx=NM_001048160.2;refGene_gn=ITGB1;refGene_tx=XM_023256250.1;refGene_gn=ITGB1;refGene_tx=XM_023256251.1;refGene_gn=ITGB1;refGene_tx=XM_023256252.1
B4 30122396 30122596
refGene_gn=ITGB1;refGene_tx=NM_001048160.2;refGene_gn=ITGB1;refGene_tx=XM_023256250.1;refGene_gn=ITGB1;refGene_tx=XM_023256251.1;refGene_gn=ITGB1;refGene_tx=XM_023256252.1
B4 30122782 30122923
refGene_gn=ITGB1;refGene_tx=NM_001048160.2;refGene_gn=ITGB1;refGene_tx

=XM_023256250.1;refGene_gn=ITGB1;refGene_tx=XM_023256251.1;refGene_gn=ITGB1;refGene_tx=XM_023256252.1
B4 30124656 30124746
refGene_gn=ITGB1;refGene_tx=NM_001048160.2;refGene_gn=ITGB1;refGene_tx=XM_023256250.1;refGene_gn=ITGB1;refGene_tx=XM_023256251.1;refGene_gn=ITGB1;refGene_tx=XM_023256252.1
B4 30125126 30125222
refGene_gn=ITGB1;refGene_tx=NM_001048160.2;refGene_gn=ITGB1;refGene_tx=XM_023256250.1;refGene_gn=ITGB1;refGene_tx=XM_023256251.1;refGene_gn=ITGB1;refGene_tx=XM_023256252.1
B4 30125998 30126154
refGene_gn=ITGB1;refGene_tx=NM_001048160.2;refGene_gn=ITGB1;refGene_tx=XM_023256250.1;refGene_gn=ITGB1;refGene_tx=XM_023256251.1;refGene_gn=ITGB1;refGene_tx=XM_023256252.1
B4 30128885 30129124
refGene_gn=ITGB1;refGene_tx=NM_001048160.2;refGene_gn=ITGB1;refGene_tx=XM_023256250.1;refGene_gn=ITGB1;refGene_tx=XM_023256251.1;refGene_gn=ITGB1;refGene_tx=XM_023256252.1
B4 30129630 30129801
refGene_gn=ITGB1;refGene_tx=NM_001048160.2;refGene_gn=ITGB1;refGene_tx=XM_023256250.1;refGene_gn=ITGB1;refGene_tx=XM_023256251.1;refGene_gn=ITGB1;refGene_tx=XM_023256252.1
B4 30131535 30131758
refGene_gn=ITGB1;refGene_tx=NM_001048160.2;refGene_gn=ITGB1;refGene_tx=XM_023256250.1;refGene_gn=ITGB1;refGene_tx=XM_023256251.1;refGene_gn=ITGB1;refGene_tx=XM_023256252.1
B4 30134518 30134604
refGene_gn=ITGB1;refGene_tx=NM_001048160.2;refGene_gn=ITGB1;refGene_tx=XM_023256250.1;refGene_gn=ITGB1;refGene_tx=XM_023256251.1;refGene_gn=ITGB1;refGene_tx=XM_023256252.1
B4 30137513 30137580
refGene_gn=ITGB1;refGene_tx=NM_001048160.2;refGene_gn=ITGB1;refGene_tx=XM_023256250.1;refGene_gn=ITGB1;refGene_tx=XM_023256251.1;refGene_gn=ITGB1;refGene_tx=XM_023256252.1
B4 30337328 30337618
refGene_gn=NRP1;refGene_tx=XM_003988215.5;refGene_gn=NRP1;refGene_tx=XM_003988216.5;refGene_gn=NRP1;refGene_tx=XM_006933352.4;refGene_gn=NRP1;refGene_tx=XM_006933353.4
B4 30339751 30339802
refGene_gn=NRP1;refGene_tx=XM_003988215.5;refGene_gn=NRP1;refGene_tx=XM_003988216.5;refGene_gn=NRP1;refGene_tx=XM_006933352.4;refGene_gn=NRP1;refGene_tx=XM_006933353.4
B4 30342090 30342187
refGene_gn=NRP1;refGene_tx=XM_003988215.5;refGene_gn=NRP1;refGene_tx=XM_003988216.5;refGene_gn=NRP1;refGene_tx=XM_006933352.4;refGene_gn=NRP1;refGene_tx=XM_006933353.4
B4 30342665 30342940
refGene_gn=NRP1;refGene_tx=XM_003988215.5;refGene_gn=NRP1;refGene_tx=XM_003988216.5;refGene_gn=NRP1;refGene_tx=XM_006933352.4;refGene_gn=NRP1;refGene_tx=XM_006933353.4

B4 30346781 30346919
refGene_gn=NRP1;refGene_tx=XM_003988215.5;refGene_gn=NRP1;refGene_tx=XM_003988216.5;refGene_gn=NRP1;refGene_tx=XM_006933352.4;refGene_gn=NRP1;refGene_tx=XM_006933353.4
B4 30352419 30352479
refGene_gn=NRP1;refGene_tx=XM_003988215.5;refGene_gn=NRP1;refGene_tx=XM_003988216.5;refGene_gn=NRP1;refGene_tx=XM_006933352.4;refGene_gn=NRP1;refGene_tx=XM_006933353.4
B4 30356843 30356948
refGene_gn=NRP1;refGene_tx=XM_003988215.5;refGene_gn=NRP1;refGene_tx=XM_003988216.5;refGene_gn=NRP1;refGene_tx=XM_006933352.4;refGene_gn=NRP1;refGene_tx=XM_006933353.4
B4 30361021 30361166
refGene_gn=NRP1;refGene_tx=XM_003988215.5;refGene_gn=NRP1;refGene_tx=XM_003988216.5;refGene_gn=NRP1;refGene_tx=XM_006933352.4;refGene_gn=NRP1;refGene_tx=XM_006933353.4
B4 30365681 30366013
refGene_gn=NRP1;refGene_tx=XM_003988215.5;refGene_gn=NRP1;refGene_tx=XM_003988216.5;refGene_gn=NRP1;refGene_tx=XM_006933352.4;refGene_gn=NRP1;refGene_tx=XM_006933353.4
B4 30373079 30373224
refGene_gn=NRP1;refGene_tx=XM_003988215.5;refGene_gn=NRP1;refGene_tx=XM_003988216.5;refGene_gn=NRP1;refGene_tx=XM_006933352.4;refGene_gn=NRP1;refGene_tx=XM_006933353.4
B4 30376887 30377043
refGene_gn=NRP1;refGene_tx=XM_003988215.5;refGene_gn=NRP1;refGene_tx=XM_003988216.5;refGene_gn=NRP1;refGene_tx=XM_006933352.4;refGene_gn=NRP1;refGene_tx=XM_006933353.4
B4 30401627 30401794
refGene_gn=NRP1;refGene_tx=XM_003988215.5;refGene_gn=NRP1;refGene_tx=XM_003988216.5;refGene_gn=NRP1;refGene_tx=XM_006933352.4;refGene_gn=NRP1;refGene_tx=XM_006933353.4
B4 30404324 30404480
refGene_gn=NRP1;refGene_tx=XM_003988215.5;refGene_gn=NRP1;refGene_tx=XM_003988216.5;refGene_gn=NRP1;refGene_tx=XM_006933352.4;refGene_gn=NRP1;refGene_tx=XM_006933353.4
B4 30410664 30410892
refGene_gn=NRP1;refGene_tx=XM_003988215.5;refGene_gn=NRP1;refGene_tx=XM_003988216.5;refGene_gn=NRP1;refGene_tx=XM_006933352.4;refGene_gn=NRP1;refGene_tx=XM_006933353.4
B4 30416846 30417028
refGene_gn=NRP1;refGene_tx=XM_003988215.5;refGene_gn=NRP1;refGene_tx=XM_003988216.5;refGene_gn=NRP1;refGene_tx=XM_006933352.4;refGene_gn=NRP1;refGene_tx=XM_006933353.4
B4 30468611 30468786
refGene_gn=NRP1;refGene_tx=XM_003988215.5;refGene_gn=NRP1;refGene_tx=XM_003988216.5;refGene_gn=NRP1;refGene_tx=XM_006933352.4;refGene_gn=NRP1;refGene_tx=XM_006933353.4
B4 30472219 30472292
refGene_gn=NRP1;refGene_tx=XM_003988215.5;refGene_gn=NRP1;refGene_tx=X

M_003988216.5;refGene_gn=NRP1;refGene_tx=XM_006933352.4;refGene_gn=NRP1;refGene_tx=XM_006933353.4
B4 31143324 31143721
refGene_gn=PARD3;refGene_tx=XM_019834173.2;refGene_gn=PARD3;refGene_tx=XM_019834174.2;refGene_gn=PARD3;refGene_tx=XM_019834175.2;refGene_gn=PARD3;refGene_tx=XM_019834176.2;refGene_gn=PARD3;refGene_tx=XM_019834178.2;refGene_gn=PARD3;refGene_tx=XM_019834179.2;refGene_gn=PARD3;refGene_tx=XM_019834181.2;refGene_gn=PARD3;refGene_tx=XM_019834182.2;refGene_gn=PARD3;refGene_tx=XM_019834183.2;refGene_gn=PARD3;refGene_tx=XM_019834184.2;refGene_gn=PARD3;refGene_tx=XM_019834185.2;refGene_gn=PARD3;refGene_tx=XM_019834186.2;refGene_gn=PARD3;refGene_tx=XM_019834188.2;refGene_gn=PARD3;refGene_tx=XM_019834189.2;refGene_gn=PARD3;refGene_tx=XM_023256513.1;refGene_gn=PARD3;refGene_tx=XM_023256514.1;refGene_gn=PARD3;refGene_tx=XM_023256515.1;refGene_gn=PARD3;refGene_tx=XM_023256517.1;refGene_gn=PARD3;refGene_tx=XM_023256518.1;refGene_gn=PARD3;refGene_tx=XM_023256519.1;refGene_gn=PARD3;refGene_tx=XM_023256523.1
B4 31149973 31150101
refGene_gn=PARD3;refGene_tx=XM_019834173.2;refGene_gn=PARD3;refGene_tx=XM_019834174.2;refGene_gn=PARD3;refGene_tx=XM_019834175.2;refGene_gn=PARD3;refGene_tx=XM_019834176.2;refGene_gn=PARD3;refGene_tx=XM_019834178.2;refGene_gn=PARD3;refGene_tx=XM_019834179.2;refGene_gn=PARD3;refGene_tx=XM_019834181.2;refGene_gn=PARD3;refGene_tx=XM_019834182.2;refGene_gn=PARD3;refGene_tx=XM_019834183.2;refGene_gn=PARD3;refGene_tx=XM_019834184.2;refGene_gn=PARD3;refGene_tx=XM_019834185.2;refGene_gn=PARD3;refGene_tx=XM_019834186.2;refGene_gn=PARD3;refGene_tx=XM_019834188.2;refGene_gn=PARD3;refGene_tx=XM_019834189.2;refGene_gn=PARD3;refGene_tx=XM_023256513.1;refGene_gn=PARD3;refGene_tx=XM_023256514.1;refGene_gn=PARD3;refGene_tx=XM_023256515.1;refGene_gn=PARD3;refGene_tx=XM_023256517.1;refGene_gn=PARD3;refGene_tx=XM_023256518.1;refGene_gn=PARD3;refGene_tx=XM_023256519.1;refGene_gn=PARD3;refGene_tx=XM_023256523.1
B4 31163351 31163472
refGene_gn=PARD3;refGene_tx=XM_019834173.2;refGene_gn=PARD3;refGene_tx=XM_019834174.2;refGene_gn=PARD3;refGene_tx=XM_019834175.2;refGene_gn=PARD3;refGene_tx=XM_019834176.2;refGene_gn=PARD3;refGene_tx=XM_019834178.2;refGene_gn=PARD3;refGene_tx=XM_019834179.2;refGene_gn=PARD3;refGene_tx=XM_019834181.2;refGene_gn=PARD3;refGene_tx=XM_019834182.2;refGene_gn=PARD3;refGene_tx=XM_019834183.2;refGene_gn=PARD3;refGene_tx=XM_019834184.2;refGene_gn=PARD3;refGene_tx=XM_019834185.2;refGene_gn=PARD3;refGene_tx=XM_019834186.2;refGene_gn=PARD3;refGene_tx=XM_019834188.2;refGene_gn=PARD3;refGene_tx=XM_019834189.2;refGene_gn=PARD3;refGene_tx=XM_023256513.1;refGene_gn=PARD3;refGene_tx=XM_023256514.1;refGene_gn=PARD3;refGene_tx=XM_023256515.1;refGene_gn=PARD3;refGene_tx=XM_023256517.1;refGene_gn=PARD3;refGene_tx=XM_023256518.1;refGene_gn=PARD3;refGene_tx=XM_023256519.1;refGene_gn=PARD3;refGene_tx=XM_023256523.1
B4 31297918 31298161
refGene_gn=PARD3;refGene_tx=XM_019834173.2;refGene_gn=PARD3;refGene_tx=XM_019834174.2;refGene_gn=PARD3;refGene_tx=XM_019834175.2;refGene_gn=PARD3;refGene_tx=XM_019834176.2;refGene_gn=PARD3;refGene_tx=XM_019834178.2;refGene_gn=PARD3;refGene_tx=XM_019834179.2;refGene_gn=PARD3;refGene_tx=XM_019834181.2;refGene_gn=PARD3;refGene_tx=XM_019834182.2;refGen

e_gn=PARD3;refGene_tx=XM_019834183.2;refGene_gn=PARD3;refGene_tx=XM_01
9834184.2;refGene_gn=PARD3;refGene_tx=XM_019834185.2;refGene_gn=PARD3;
refGene_tx=XM_019834186.2;refGene_gn=PARD3;refGene_tx=XM_019834188.2;r
efGene_gn=PARD3;refGene_tx=XM_019834189.2;refGene_gn=PARD3;refGene_tx=
XM_023256513.1;refGene_gn=PARD3;refGene_tx=XM_023256514.1;refGene_gn=P
ARD3;refGene_tx=XM_023256515.1;refGene_gn=PARD3;refGene_tx=XM_02325651
7.1;refGene_gn=PARD3;refGene_tx=XM_023256518.1;refGene_gn=PARD3;refGen
e_tx=XM_023256519.1;refGene_gn=PARD3;refGene_tx=XM_023256523.1
B4 31312486 31312597
refGene_gn=PARD3;refGene_tx=XM_019834173.2;refGene_gn=PARD3;refGene_tx
=XM_019834174.2;refGene_gn=PARD3;refGene_tx=XM_019834175.2;refGene_gn=
PARD3;refGene_tx=XM_019834176.2;refGene_gn=PARD3;refGene_tx=XM_0198341
78.2;refGene_gn=PARD3;refGene_tx=XM_019834179.2;refGene_gn=PARD3;refGe
ne_tx=XM_019834181.2;refGene_gn=PARD3;refGene_tx=XM_019834182.2;refGen
e_gn=PARD3;refGene_tx=XM_019834183.2;refGene_gn=PARD3;refGene_tx=XM_01
9834185.2;refGene_gn=PARD3;refGene_tx=XM_019834188.2;refGene_gn=PARD3;
refGene_tx=XM_019834189.2;refGene_gn=PARD3;refGene_tx=XM_023256513.1;r
efGene_gn=PARD3;refGene_tx=XM_023256514.1;refGene_gn=PARD3;refGene_tx=
XM_023256515.1;refGene_gn=PARD3;refGene_tx=XM_023256518.1
B4 31337058 31337089
refGene_gn=PARD3;refGene_tx=XM_023256520.1;refGene_gn=PARD3;refGene_tx
=XM_023256521.1;refGene_gn=PARD3;refGene_tx=XM_023256522.1
B4 31341716 31341948
refGene_gn=PARD3;refGene_tx=XM_019834173.2;refGene_gn=PARD3;refGene_tx
=XM_019834174.2;refGene_gn=PARD3;refGene_tx=XM_019834175.2;refGene_gn=
PARD3;refGene_tx=XM_019834176.2;refGene_gn=PARD3;refGene_tx=XM_0198341
78.2;refGene_gn=PARD3;refGene_tx=XM_019834179.2;refGene_gn=PARD3;refGe
ne_tx=XM_019834181.2;refGene_gn=PARD3;refGene_tx=XM_019834182.2;refGen
e_gn=PARD3;refGene_tx=XM_019834183.2;refGene_gn=PARD3;refGene_tx=XM_01
9834184.2;refGene_gn=PARD3;refGene_tx=XM_019834185.2;refGene_gn=PARD3;
refGene_tx=XM_019834186.2;refGene_gn=PARD3;refGene_tx=XM_019834188.2;r
efGene_gn=PARD3;refGene_tx=XM_019834189.2;refGene_gn=PARD3;refGene_tx=
XM_023256513.1;refGene_gn=PARD3;refGene_tx=XM_023256514.1;refGene_gn=P
ARD3;refGene_tx=XM_023256515.1;refGene_gn=PARD3;refGene_tx=XM_02325651
7.1;refGene_gn=PARD3;refGene_tx=XM_023256518.1;refGene_gn=PARD3;refGen
e_tx=XM_023256519.1;refGene_gn=PARD3;refGene_tx=XM_023256520.1;refGene
_gn=PARD3;refGene_tx=XM_023256521.1;refGene_gn=PARD3;refGene_tx=XM_023
256522.1;refGene_gn=PARD3;refGene_tx=XM_023256523.1
B4 31355887 31356115
refGene_gn=PARD3;refGene_tx=XM_019834173.2;refGene_gn=PARD3;refGene_tx
=XM_019834174.2;refGene_gn=PARD3;refGene_tx=XM_019834175.2;refGene_gn=
PARD3;refGene_tx=XM_019834176.2;refGene_gn=PARD3;refGene_tx=XM_0198341
78.2;refGene_gn=PARD3;refGene_tx=XM_019834179.2;refGene_gn=PARD3;refGe
ne_tx=XM_019834181.2;refGene_gn=PARD3;refGene_tx=XM_019834182.2;refGen
e_gn=PARD3;refGene_tx=XM_019834183.2;refGene_gn=PARD3;refGene_tx=XM_01
9834184.2;refGene_gn=PARD3;refGene_tx=XM_019834185.2;refGene_gn=PARD3;
refGene_tx=XM_019834186.2;refGene_gn=PARD3;refGene_tx=XM_019834188.2;r
efGene_gn=PARD3;refGene_tx=XM_023256513.1;refGene_gn=PARD3;refGene_tx=
XM_023256514.1;refGene_gn=PARD3;refGene_tx=XM_023256515.1;refGene_gn=P
ARD3;refGene_tx=XM_023256517.1;refGene_gn=PARD3;refGene_tx=XM_02325651

8.1;refGene_gn=PARD3;refGene_tx=XM_023256519.1;refGene_gn=PARD3;refGene_tx=XM_023256520.1;refGene_gn=PARD3;refGene_tx=XM_023256521.1;refGene_gn=PARD3;refGene_tx=XM_023256522.1;refGene_gn=PARD3;refGene_tx=XM_023256523.1
B4 31361638 31361683
refGene_gn=PARD3;refGene_tx=XM_019834173.2;refGene_gn=PARD3;refGene_tx=XM_019834174.2;refGene_gn=PARD3;refGene_tx=XM_019834175.2;refGene_gn=PARD3;refGene_tx=XM_019834176.2;refGene_gn=PARD3;refGene_tx=XM_019834181.2;refGene_gn=PARD3;refGene_tx=XM_019834182.2;refGene_gn=PARD3;refGene_tx=XM_019834183.2;refGene_gn=PARD3;refGene_tx=XM_019834184.2;refGene_gn=PARD3;refGene_tx=XM_019834185.2;refGene_gn=PARD3;refGene_tx=XM_019834188.2;refGene_gn=PARD3;refGene_tx=XM_023256513.1;refGene_gn=PARD3;refGene_tx=XM_023256514.1;refGene_gn=PARD3;refGene_tx=XM_023256515.1;refGene_gn=PARD3;refGene_tx=XM_023256517.1;refGene_gn=PARD3;refGene_tx=XM_023256518.1;refGene_gn=PARD3;refGene_tx=XM_023256519.1;refGene_gn=PARD3;refGene_tx=XM_023256520.1;refGene_gn=PARD3;refGene_tx=XM_023256521.1;refGene_gn=PARD3;refGene_tx=XM_023256522.1
B4 31362734 31362886
refGene_gn=PARD3;refGene_tx=XM_019834173.2;refGene_gn=PARD3;refGene_tx=XM_019834174.2;refGene_gn=PARD3;refGene_tx=XM_019834175.2;refGene_gn=PARD3;refGene_tx=XM_019834176.2;refGene_gn=PARD3;refGene_tx=XM_019834178.2;refGene_gn=PARD3;refGene_tx=XM_019834179.2;refGene_gn=PARD3;refGene_tx=XM_019834181.2;refGene_gn=PARD3;refGene_tx=XM_019834182.2;refGene_gn=PARD3;refGene_tx=XM_019834183.2;refGene_gn=PARD3;refGene_tx=XM_019834184.2;refGene_gn=PARD3;refGene_tx=XM_019834185.2;refGene_gn=PARD3;refGene_tx=XM_019834186.2;refGene_gn=PARD3;refGene_tx=XM_019834188.2;refGene_gn=PARD3;refGene_tx=XM_019834189.2;refGene_gn=PARD3;refGene_tx=XM_023256513.1;refGene_gn=PARD3;refGene_tx=XM_023256514.1;refGene_gn=PARD3;refGene_tx=XM_023256515.1;refGene_gn=PARD3;refGene_tx=XM_023256517.1;refGene_gn=PARD3;refGene_tx=XM_023256518.1;refGene_gn=PARD3;refGene_tx=XM_023256519.1;refGene_gn=PARD3;refGene_tx=XM_023256520.1;refGene_gn=PARD3;refGene_tx=XM_023256521.1;refGene_gn=PARD3;refGene_tx=XM_023256522.1;refGene_gn=PARD3;refGene_tx=XM_023256523.1
B4 31368201 31368391
refGene_gn=PARD3;refGene_tx=XM_019834173.2;refGene_gn=PARD3;refGene_tx=XM_019834174.2;refGene_gn=PARD3;refGene_tx=XM_019834175.2;refGene_gn=PARD3;refGene_tx=XM_019834176.2;refGene_gn=PARD3;refGene_tx=XM_019834178.2;refGene_gn=PARD3;refGene_tx=XM_019834179.2;refGene_gn=PARD3;refGene_tx=XM_019834181.2;refGene_gn=PARD3;refGene_tx=XM_019834182.2;refGene_gn=PARD3;refGene_tx=XM_019834183.2;refGene_gn=PARD3;refGene_tx=XM_019834184.2;refGene_gn=PARD3;refGene_tx=XM_019834185.2;refGene_gn=PARD3;refGene_tx=XM_019834186.2;refGene_gn=PARD3;refGene_tx=XM_019834188.2;refGene_gn=PARD3;refGene_tx=XM_019834189.2;refGene_gn=PARD3;refGene_tx=XM_023256513.1;refGene_gn=PARD3;refGene_tx=XM_023256514.1;refGene_gn=PARD3;refGene_tx=XM_023256515.1;refGene_gn=PARD3;refGene_tx=XM_023256517.1;refGene_gn=PARD3;refGene_tx=XM_023256518.1;refGene_gn=PARD3;refGene_tx=XM_023256519.1;refGene_gn=PARD3;refGene_tx=XM_023256520.1;refGene_gn=PARD3;refGene_tx=XM_023256521.1;refGene_gn=PARD3;refGene_tx=XM_023256522.1;refGene_gn=PARD3;refGene_tx=XM_023256523.1
B4 31375090 31375250

refGene_gn=PARD3;refGene_tx=XM_019834173.2;refGene_gn=PARD3;refGene_tx
=XM_019834174.2;refGene_gn=PARD3;refGene_tx=XM_019834175.2;refGene_gn=
PARD3;refGene_tx=XM_019834176.2;refGene_gn=PARD3;refGene_tx=XM_0198341
78.2;refGene_gn=PARD3;refGene_tx=XM_019834179.2;refGene_gn=PARD3;refGe
ne_tx=XM_019834181.2;refGene_gn=PARD3;refGene_tx=XM_019834182.2;refGen
e_gn=PARD3;refGene_tx=XM_019834183.2;refGene_gn=PARD3;refGene_tx=XM_01
9834184.2;refGene_gn=PARD3;refGene_tx=XM_019834185.2;refGene_gn=PARD3;
refGene_tx=XM_019834186.2;refGene_gn=PARD3;refGene_tx=XM_019834188.2;r
efGene_gn=PARD3;refGene_tx=XM_019834189.2;refGene_gn=PARD3;refGene_tx=
XM_023256513.1;refGene_gn=PARD3;refGene_tx=XM_023256514.1;refGene_gn=P
ARD3;refGene_tx=XM_023256515.1;refGene_gn=PARD3;refGene_tx=XM_02325651
7.1;refGene_gn=PARD3;refGene_tx=XM_023256518.1;refGene_gn=PARD3;refGen
e_tx=XM_023256519.1;refGene_gn=PARD3;refGene_tx=XM_023256520.1;refGene
_gn=PARD3;refGene_tx=XM_023256521.1;refGene_gn=PARD3;refGene_tx=XM_023
256522.1;refGene_gn=PARD3;refGene_tx=XM_023256523.1
B4 31384125 31384296
refGene_gn=PARD3;refGene_tx=XM_019834173.2;refGene_gn=PARD3;refGene_tx
=XM_019834174.2;refGene_gn=PARD3;refGene_tx=XM_019834175.2;refGene_gn=
PARD3;refGene_tx=XM_019834176.2;refGene_gn=PARD3;refGene_tx=XM_0198341
78.2;refGene_gn=PARD3;refGene_tx=XM_019834179.2;refGene_gn=PARD3;refGe
ne_tx=XM_019834181.2;refGene_gn=PARD3;refGene_tx=XM_019834182.2;refGen
e_gn=PARD3;refGene_tx=XM_019834183.2;refGene_gn=PARD3;refGene_tx=XM_01
9834184.2;refGene_gn=PARD3;refGene_tx=XM_019834185.2;refGene_gn=PARD3;
refGene_tx=XM_019834186.2;refGene_gn=PARD3;refGene_tx=XM_019834188.2;r
efGene_gn=PARD3;refGene_tx=XM_019834189.2;refGene_gn=PARD3;refGene_tx=
XM_023256513.1;refGene_gn=PARD3;refGene_tx=XM_023256514.1;refGene_gn=P
ARD3;refGene_tx=XM_023256515.1;refGene_gn=PARD3;refGene_tx=XM_02325651
7.1;refGene_gn=PARD3;refGene_tx=XM_023256518.1;refGene_gn=PARD3;refGen
e_tx=XM_023256519.1;refGene_gn=PARD3;refGene_tx=XM_023256520.1;refGene
_gn=PARD3;refGene_tx=XM_023256521.1;refGene_gn=PARD3;refGene_tx=XM_023
256522.1;refGene_gn=PARD3;refGene_tx=XM_023256523.1
B4 31385655 31385844
refGene_gn=PARD3;refGene_tx=XM_019834173.2;refGene_gn=PARD3;refGene_tx
=XM_019834174.2;refGene_gn=PARD3;refGene_tx=XM_019834175.2;refGene_gn=
PARD3;refGene_tx=XM_019834176.2;refGene_gn=PARD3;refGene_tx=XM_0198341
78.2;refGene_gn=PARD3;refGene_tx=XM_019834179.2;refGene_gn=PARD3;refGe
ne_tx=XM_019834181.2;refGene_gn=PARD3;refGene_tx=XM_019834182.2;refGen
e_gn=PARD3;refGene_tx=XM_019834183.2;refGene_gn=PARD3;refGene_tx=XM_01
9834184.2;refGene_gn=PARD3;refGene_tx=XM_019834185.2;refGene_gn=PARD3;
refGene_tx=XM_019834186.2;refGene_gn=PARD3;refGene_tx=XM_019834188.2;r
efGene_gn=PARD3;refGene_tx=XM_019834189.2;refGene_gn=PARD3;refGene_tx=
XM_023256513.1;refGene_gn=PARD3;refGene_tx=XM_023256514.1;refGene_gn=P
ARD3;refGene_tx=XM_023256515.1;refGene_gn=PARD3;refGene_tx=XM_02325651
7.1;refGene_gn=PARD3;refGene_tx=XM_023256518.1;refGene_gn=PARD3;refGen
e_tx=XM_023256519.1;refGene_gn=PARD3;refGene_tx=XM_023256520.1;refGene
_gn=PARD3;refGene_tx=XM_023256521.1;refGene_gn=PARD3;refGene_tx=XM_023
256522.1;refGene_gn=PARD3;refGene_tx=XM_023256523.1
B4 31401086 31401125
refGene_gn=PARD3;refGene_tx=XM_019834173.2;refGene_gn=PARD3;refGene_tx
=XM_019834174.2;refGene_gn=PARD3;refGene_tx=XM_019834175.2;refGene_gn=

PARD3;refGene_tx=XM_019834178.2;refGene_gn=PARD3;refGene_tx=XM_019834179.2;refGene_gn=PARD3;refGene_tx=XM_019834181.2;refGene_gn=PARD3;refGene_tx=XM_019834182.2;refGene_gn=PARD3;refGene_tx=XM_019834183.2;refGene_gn=PARD3;refGene_tx=XM_019834184.2;refGene_gn=PARD3;refGene_tx=XM_019834188.2;refGene_gn=PARD3;refGene_tx=XM_019834189.2;refGene_gn=PARD3;refGene_tx=XM_023256513.1;refGene_gn=PARD3;refGene_tx=XM_023256514.1;refGene_gn=PARD3;refGene_tx=XM_023256517.1;refGene_gn=PARD3;refGene_tx=XM_023256520.1;refGene_gn=PARD3;refGene_tx=XM_023256521.1
B4 31403510 31403639
refGene_gn=PARD3;refGene_tx=XM_019834173.2;refGene_gn=PARD3;refGene_tx=XM_019834174.2;refGene_gn=PARD3;refGene_tx=XM_019834175.2;refGene_gn=PARD3;refGene_tx=XM_019834176.2;refGene_gn=PARD3;refGene_tx=XM_019834178.2;refGene_gn=PARD3;refGene_tx=XM_019834179.2;refGene_gn=PARD3;refGene_tx=XM_019834181.2;refGene_gn=PARD3;refGene_tx=XM_019834182.2;refGene_gn=PARD3;refGene_tx=XM_019834183.2;refGene_gn=PARD3;refGene_tx=XM_019834184.2;refGene_gn=PARD3;refGene_tx=XM_019834185.2;refGene_gn=PARD3;refGene_tx=XM_019834186.2;refGene_gn=PARD3;refGene_tx=XM_019834188.2;refGene_gn=PARD3;refGene_tx=XM_019834189.2;refGene_gn=PARD3;refGene_tx=XM_023256513.1;refGene_gn=PARD3;refGene_tx=XM_023256514.1;refGene_gn=PARD3;refGene_tx=XM_023256515.1;refGene_gn=PARD3;refGene_tx=XM_023256517.1;refGene_gn=PARD3;refGene_tx=XM_023256518.1;refGene_gn=PARD3;refGene_tx=XM_023256519.1;refGene_gn=PARD3;refGene_tx=XM_023256520.1;refGene_gn=PARD3;refGene_tx=XM_023256521.1;refGene_gn=PARD3;refGene_tx=XM_023256522.1;refGene_gn=PARD3;refGene_tx=XM_023256523.1
B4 31406264 31406404
refGene_gn=PARD3;refGene_tx=XM_019834173.2;refGene_gn=PARD3;refGene_tx=XM_019834174.2;refGene_gn=PARD3;refGene_tx=XM_019834175.2;refGene_gn=PARD3;refGene_tx=XM_019834176.2;refGene_gn=PARD3;refGene_tx=XM_019834178.2;refGene_gn=PARD3;refGene_tx=XM_019834179.2;refGene_gn=PARD3;refGene_tx=XM_019834181.2;refGene_gn=PARD3;refGene_tx=XM_019834182.2;refGene_gn=PARD3;refGene_tx=XM_019834183.2;refGene_gn=PARD3;refGene_tx=XM_019834184.2;refGene_gn=PARD3;refGene_tx=XM_019834185.2;refGene_gn=PARD3;refGene_tx=XM_019834186.2;refGene_gn=PARD3;refGene_tx=XM_019834188.2;refGene_gn=PARD3;refGene_tx=XM_019834189.2;refGene_gn=PARD3;refGene_tx=XM_023256513.1;refGene_gn=PARD3;refGene_tx=XM_023256514.1;refGene_gn=PARD3;refGene_tx=XM_023256515.1;refGene_gn=PARD3;refGene_tx=XM_023256517.1;refGene_gn=PARD3;refGene_tx=XM_023256518.1;refGene_gn=PARD3;refGene_tx=XM_023256519.1;refGene_gn=PARD3;refGene_tx=XM_023256520.1;refGene_gn=PARD3;refGene_tx=XM_023256521.1;refGene_gn=PARD3;refGene_tx=XM_023256522.1;refGene_gn=PARD3;refGene_tx=XM_023256523.1
B4 31411247 31411633
refGene_gn=PARD3;refGene_tx=XM_019834173.2;refGene_gn=PARD3;refGene_tx=XM_019834174.2;refGene_gn=PARD3;refGene_tx=XM_019834175.2;refGene_gn=PARD3;refGene_tx=XM_019834176.2;refGene_gn=PARD3;refGene_tx=XM_019834178.2;refGene_gn=PARD3;refGene_tx=XM_019834179.2;refGene_gn=PARD3;refGene_tx=XM_019834181.2;refGene_gn=PARD3;refGene_tx=XM_019834182.2;refGene_gn=PARD3;refGene_tx=XM_019834183.2;refGene_gn=PARD3;refGene_tx=XM_019834184.2;refGene_gn=PARD3;refGene_tx=XM_019834185.2;refGene_gn=PARD3;refGene_tx=XM_019834186.2;refGene_gn=PARD3;refGene_tx=XM_019834188.2;refGene_gn=PARD3;refGene_tx=XM_019834189.2;refGene_gn=PARD3;refGene_tx=

XM_023256513.1;refGene_gn=PARD3;refGene_tx=XM_023256514.1;refGene_gn=PARD3;refGene_tx=XM_023256515.1;refGene_gn=PARD3;refGene_tx=XM_023256517.1;refGene_gn=PARD3;refGene_tx=XM_023256518.1;refGene_gn=PARD3;refGene_tx=XM_023256519.1;refGene_gn=PARD3;refGene_tx=XM_023256520.1;refGene_gn=PARD3;refGene_tx=XM_023256521.1;refGene_gn=PARD3;refGene_tx=XM_023256522.1;refGene_gn=PARD3;refGene_tx=XM_023256523.1
B4 31412743 31412869
refGene_gn=PARD3;refGene_tx=XM_019834173.2;refGene_gn=PARD3;refGene_tx=XM_019834174.2;refGene_gn=PARD3;refGene_tx=XM_019834175.2;refGene_gn=PARD3;refGene_tx=XM_019834176.2;refGene_gn=PARD3;refGene_tx=XM_019834178.2;refGene_gn=PARD3;refGene_tx=XM_019834179.2;refGene_gn=PARD3;refGene_tx=XM_019834181.2;refGene_gn=PARD3;refGene_tx=XM_019834182.2;refGene_gn=PARD3;refGene_tx=XM_019834183.2;refGene_gn=PARD3;refGene_tx=XM_019834184.2;refGene_gn=PARD3;refGene_tx=XM_019834185.2;refGene_gn=PARD3;refGene_tx=XM_019834186.2;refGene_gn=PARD3;refGene_tx=XM_019834188.2;refGene_gn=PARD3;refGene_tx=XM_019834189.2;refGene_gn=PARD3;refGene_tx=XM_023256513.1;refGene_gn=PARD3;refGene_tx=XM_023256514.1;refGene_gn=PARD3;refGene_tx=XM_023256515.1;refGene_gn=PARD3;refGene_tx=XM_023256517.1;refGene_gn=PARD3;refGene_tx=XM_023256518.1;refGene_gn=PARD3;refGene_tx=XM_023256519.1;refGene_gn=PARD3;refGene_tx=XM_023256520.1;refGene_gn=PARD3;refGene_tx=XM_023256521.1;refGene_gn=PARD3;refGene_tx=XM_023256522.1;refGene_gn=PARD3;refGene_tx=XM_023256523.1
B4 31436897 31436981
refGene_gn=PARD3;refGene_tx=XM_019834173.2;refGene_gn=PARD3;refGene_tx=XM_019834174.2;refGene_gn=PARD3;refGene_tx=XM_019834175.2;refGene_gn=PARD3;refGene_tx=XM_019834176.2;refGene_gn=PARD3;refGene_tx=XM_019834178.2;refGene_gn=PARD3;refGene_tx=XM_019834179.2;refGene_gn=PARD3;refGene_tx=XM_019834182.2;refGene_gn=PARD3;refGene_tx=XM_019834183.2;refGene_gn=PARD3;refGene_tx=XM_019834184.2;refGene_gn=PARD3;refGene_tx=XM_019834185.2;refGene_gn=PARD3;refGene_tx=XM_019834186.2;refGene_gn=PARD3;refGene_tx=XM_019834188.2;refGene_gn=PARD3;refGene_tx=XM_019834189.2;refGene_gn=PARD3;refGene_tx=XM_023256513.1;refGene_gn=PARD3;refGene_tx=XM_023256514.1;refGene_gn=PARD3;refGene_tx=XM_023256515.1;refGene_gn=PARD3;refGene_tx=XM_023256517.1;refGene_gn=PARD3;refGene_tx=XM_023256518.1;refGene_gn=PARD3;refGene_tx=XM_023256519.1;refGene_gn=PARD3;refGene_tx=XM_023256520.1;refGene_gn=PARD3;refGene_tx=XM_023256521.1;refGene_gn=PARD3;refGene_tx=XM_023256522.1;refGene_gn=PARD3;refGene_tx=XM_023256523.1
B4 31439446 31439538
refGene_gn=PARD3;refGene_tx=XM_019834173.2;refGene_gn=PARD3;refGene_tx=XM_019834174.2;refGene_gn=PARD3;refGene_tx=XM_019834175.2;refGene_gn=PARD3;refGene_tx=XM_019834176.2;refGene_gn=PARD3;refGene_tx=XM_019834178.2;refGene_gn=PARD3;refGene_tx=XM_019834179.2;refGene_gn=PARD3;refGene_tx=XM_019834181.2;refGene_gn=PARD3;refGene_tx=XM_019834182.2;refGene_gn=PARD3;refGene_tx=XM_019834183.2;refGene_gn=PARD3;refGene_tx=XM_019834184.2;refGene_gn=PARD3;refGene_tx=XM_019834185.2;refGene_gn=PARD3;refGene_tx=XM_019834186.2;refGene_gn=PARD3;refGene_tx=XM_019834188.2;refGene_gn=PARD3;refGene_tx=XM_019834189.2;refGene_gn=PARD3;refGene_tx=XM_023256513.1;refGene_gn=PARD3;refGene_tx=XM_023256514.1;refGene_gn=PARD3;refGene_tx=XM_023256515.1;refGene_gn=PARD3;refGene_tx=XM_02325651

7.1;refGene_gn=PARD3;refGene_tx=XM_023256518.1;refGene_gn=PARD3;refGene_tx=XM_023256519.1;refGene_gn=PARD3;refGene_tx=XM_023256520.1;refGene_gn=PARD3;refGene_tx=XM_023256521.1;refGene_gn=PARD3;refGene_tx=XM_023256522.1;refGene_gn=PARD3;refGene_tx=XM_023256523.1
B4 31481543 31481675
refGene_gn=PARD3;refGene_tx=XM_019834173.2;refGene_gn=PARD3;refGene_tx=XM_019834174.2;refGene_gn=PARD3;refGene_tx=XM_019834175.2;refGene_gn=PARD3;refGene_tx=XM_019834176.2;refGene_gn=PARD3;refGene_tx=XM_019834178.2;refGene_gn=PARD3;refGene_tx=XM_019834179.2;refGene_gn=PARD3;refGene_tx=XM_019834181.2;refGene_gn=PARD3;refGene_tx=XM_019834182.2;refGene_gn=PARD3;refGene_tx=XM_019834183.2;refGene_gn=PARD3;refGene_tx=XM_019834184.2;refGene_gn=PARD3;refGene_tx=XM_019834185.2;refGene_gn=PARD3;refGene_tx=XM_019834186.2;refGene_gn=PARD3;refGene_tx=XM_019834188.2;refGene_gn=PARD3;refGene_tx=XM_019834189.2;refGene_gn=PARD3;refGene_tx=XM_023256513.1;refGene_gn=PARD3;refGene_tx=XM_023256514.1;refGene_gn=PARD3;refGene_tx=XM_023256515.1;refGene_gn=PARD3;refGene_tx=XM_023256517.1;refGene_gn=PARD3;refGene_tx=XM_023256518.1;refGene_gn=PARD3;refGene_tx=XM_023256519.1;refGene_gn=PARD3;refGene_tx=XM_023256520.1;refGene_gn=PARD3;refGene_tx=XM_023256521.1;refGene_gn=PARD3;refGene_tx=XM_023256522.1;refGene_gn=PARD3;refGene_tx=XM_023256523.1
B4 31501478 31501657
refGene_gn=PARD3;refGene_tx=XM_019834173.2;refGene_gn=PARD3;refGene_tx=XM_019834174.2;refGene_gn=PARD3;refGene_tx=XM_019834175.2;refGene_gn=PARD3;refGene_tx=XM_019834176.2;refGene_gn=PARD3;refGene_tx=XM_019834178.2;refGene_gn=PARD3;refGene_tx=XM_019834179.2;refGene_gn=PARD3;refGene_tx=XM_019834181.2;refGene_gn=PARD3;refGene_tx=XM_019834182.2;refGene_gn=PARD3;refGene_tx=XM_019834183.2;refGene_gn=PARD3;refGene_tx=XM_019834184.2;refGene_gn=PARD3;refGene_tx=XM_019834185.2;refGene_gn=PARD3;refGene_tx=XM_019834186.2;refGene_gn=PARD3;refGene_tx=XM_019834188.2;refGene_gn=PARD3;refGene_tx=XM_019834189.2;refGene_gn=PARD3;refGene_tx=XM_023256513.1;refGene_gn=PARD3;refGene_tx=XM_023256514.1;refGene_gn=PARD3;refGene_tx=XM_023256515.1;refGene_gn=PARD3;refGene_tx=XM_023256517.1;refGene_gn=PARD3;refGene_tx=XM_023256518.1;refGene_gn=PARD3;refGene_tx=XM_023256519.1;refGene_gn=PARD3;refGene_tx=XM_023256520.1;refGene_gn=PARD3;refGene_tx=XM_023256521.1;refGene_gn=PARD3;refGene_tx=XM_023256522.1;refGene_gn=PARD3;refGene_tx=XM_023256523.1
B4 31542869 31543050
refGene_gn=PARD3;refGene_tx=XM_019834173.2;refGene_gn=PARD3;refGene_tx=XM_019834174.2;refGene_gn=PARD3;refGene_tx=XM_019834175.2;refGene_gn=PARD3;refGene_tx=XM_019834176.2;refGene_gn=PARD3;refGene_tx=XM_019834178.2;refGene_gn=PARD3;refGene_tx=XM_019834179.2;refGene_gn=PARD3;refGene_tx=XM_019834181.2;refGene_gn=PARD3;refGene_tx=XM_019834182.2;refGene_gn=PARD3;refGene_tx=XM_019834183.2;refGene_gn=PARD3;refGene_tx=XM_019834184.2;refGene_gn=PARD3;refGene_tx=XM_019834185.2;refGene_gn=PARD3;refGene_tx=XM_019834186.2;refGene_gn=PARD3;refGene_tx=XM_019834188.2;refGene_gn=PARD3;refGene_tx=XM_019834189.2;refGene_gn=PARD3;refGene_tx=XM_023256513.1;refGene_gn=PARD3;refGene_tx=XM_023256514.1;refGene_gn=PARD3;refGene_tx=XM_023256515.1;refGene_gn=PARD3;refGene_tx=XM_023256517.1;refGene_gn=PARD3;refGene_tx=XM_023256518.1;refGene_gn=PARD3;refGene_tx=XM_023256519.1;refGene_gn=PARD3;refGene_tx=XM_023256520.1;refGene

_gn=PARD3;refGene_tx=XM_023256521.1;refGene_gn=PARD3;refGene_tx=XM_023256522.1;refGene_gn=PARD3;refGene_tx=XM_023256523.1
B4 31675926 31676067 refGene_gn=PARD3;refGene_tx=XM_019834188.2
B4 31692977 31693079
refGene_gn=PARD3;refGene_tx=XM_019834173.2;refGene_gn=PARD3;refGene_tx=XM_019834174.2;refGene_gn=PARD3;refGene_tx=XM_019834175.2;refGene_gn=PARD3;refGene_tx=XM_019834176.2;refGene_gn=PARD3;refGene_tx=XM_019834178.2;refGene_gn=PARD3;refGene_tx=XM_019834179.2;refGene_gn=PARD3;refGene_tx=XM_019834181.2;refGene_gn=PARD3;refGene_tx=XM_019834182.2;refGene_gn=PARD3;refGene_tx=XM_019834183.2;refGene_gn=PARD3;refGene_tx=XM_019834184.2;refGene_gn=PARD3;refGene_tx=XM_019834185.2;refGene_gn=PARD3;refGene_tx=XM_019834186.2;refGene_gn=PARD3;refGene_tx=XM_019834189.2;refGene_gn=PARD3;refGene_tx=XM_023256513.1;refGene_gn=PARD3;refGene_tx=XM_023256514.1;refGene_gn=PARD3;refGene_tx=XM_023256515.1;refGene_gn=PARD3;refGene_tx=XM_023256517.1;refGene_gn=PARD3;refGene_tx=XM_023256518.1;refGene_gn=PARD3;refGene_tx=XM_023256519.1;refGene_gn=PARD3;refGene_tx=XM_023256520.1;refGene_gn=PARD3;refGene_tx=XM_023256521.1;refGene_gn=PARD3;refGene_tx=XM_023256522.1;refGene_gn=PARD3;refGene_tx=XM_023256523.1
B4 31704869 31705175
refGene_gn=PARD3;refGene_tx=XM_019834173.2;refGene_gn=PARD3;refGene_tx=XM_019834174.2;refGene_gn=PARD3;refGene_tx=XM_019834175.2;refGene_gn=PARD3;refGene_tx=XM_019834176.2;refGene_gn=PARD3;refGene_tx=XM_019834178.2;refGene_gn=PARD3;refGene_tx=XM_019834179.2;refGene_gn=PARD3;refGene_tx=XM_019834181.2;refGene_gn=PARD3;refGene_tx=XM_019834182.2;refGene_gn=PARD3;refGene_tx=XM_019834183.2;refGene_gn=PARD3;refGene_tx=XM_019834184.2;refGene_gn=PARD3;refGene_tx=XM_019834185.2;refGene_gn=PARD3;refGene_tx=XM_019834186.2;refGene_gn=PARD3;refGene_tx=XM_019834189.2;refGene_gn=PARD3;refGene_tx=XM_023256520.1
B4 31813211 31813331
refGene_gn=PARD3;refGene_tx=XM_023256513.1;refGene_gn=PARD3;refGene_tx=XM_023256514.1;refGene_gn=PARD3;refGene_tx=XM_023256515.1;refGene_gn=PARD3;refGene_tx=XM_023256517.1;refGene_gn=PARD3;refGene_tx=XM_023256518.1;refGene_gn=PARD3;refGene_tx=XM_023256519.1;refGene_gn=PARD3;refGene_tx=XM_023256521.1;refGene_gn=PARD3;refGene_tx=XM_023256522.1;refGene_gn=PARD3;refGene_tx=XM_023256523.1
B4 31982505 31982637
refGene_gn=CUL2;refGene_tx=XM_006933355.4;refGene_gn=CUL2;refGene_tx=XM_023256524.1;refGene_gn=CUL2;refGene_tx=XM_023256525.1;refGene_gn=CUL2;refGene_tx=XM_023256526.1;refGene_gn=CUL2;refGene_tx=XM_023256527.1;refGene_gn=CUL2;refGene_tx=XM_023256528.1
B4 31983660 31983777
refGene_gn=CUL2;refGene_tx=XM_006933355.4;refGene_gn=CUL2;refGene_tx=XM_023256524.1;refGene_gn=CUL2;refGene_tx=XM_023256525.1;refGene_gn=CUL2;refGene_tx=XM_023256526.1;refGene_gn=CUL2;refGene_tx=XM_023256527.1;refGene_gn=CUL2;refGene_tx=XM_023256528.1
B4 31984891 31984993
refGene_gn=CUL2;refGene_tx=XM_006933355.4;refGene_gn=CUL2;refGene_tx=XM_023256524.1;refGene_gn=CUL2;refGene_tx=XM_023256525.1;refGene_gn=CUL2;refGene_tx=XM_023256526.1;refGene_gn=CUL2;refGene_tx=XM_023256527.1;

refGene_gn=CUL2;refGene_tx=XM_023256528.1
B4	31986738	31986941
refGene_gn=CUL2;refGene_tx=XM_006933355.4;refGene_gn=CUL2;refGene_tx=XM_023256524.1;refGene_gn=CUL2;refGene_tx=XM_023256525.1;refGene_gn=CUL2;refGene_tx=XM_023256526.1;refGene_gn=CUL2;refGene_tx=XM_023256527.1;refGene_gn=CUL2;refGene_tx=XM_023256528.1
B4	31992776	31992843
refGene_gn=CUL2;refGene_tx=XM_006933355.4;refGene_gn=CUL2;refGene_tx=XM_023256524.1;refGene_gn=CUL2;refGene_tx=XM_023256525.1;refGene_gn=CUL2;refGene_tx=XM_023256526.1;refGene_gn=CUL2;refGene_tx=XM_023256527.1;refGene_gn=CUL2;refGene_tx=XM_023256528.1
B4	31998251	31998329
refGene_gn=CUL2;refGene_tx=XM_006933355.4;refGene_gn=CUL2;refGene_tx=XM_023256524.1;refGene_gn=CUL2;refGene_tx=XM_023256525.1;refGene_gn=CUL2;refGene_tx=XM_023256526.1;refGene_gn=CUL2;refGene_tx=XM_023256527.1;refGene_gn=CUL2;refGene_tx=XM_023256528.1
B4	31998836	31998989
refGene_gn=CUL2;refGene_tx=XM_006933355.4;refGene_gn=CUL2;refGene_tx=XM_023256524.1;refGene_gn=CUL2;refGene_tx=XM_023256525.1;refGene_gn=CUL2;refGene_tx=XM_023256526.1;refGene_gn=CUL2;refGene_tx=XM_023256527.1;refGene_gn=CUL2;refGene_tx=XM_023256528.1
B4	31999975	32000062
refGene_gn=CUL2;refGene_tx=XM_006933355.4;refGene_gn=CUL2;refGene_tx=XM_023256524.1;refGene_gn=CUL2;refGene_tx=XM_023256525.1;refGene_gn=CUL2;refGene_tx=XM_023256526.1;refGene_gn=CUL2;refGene_tx=XM_023256527.1;refGene_gn=CUL2;refGene_tx=XM_023256528.1
B4	32000171	32000300
refGene_gn=CUL2;refGene_tx=XM_006933355.4;refGene_gn=CUL2;refGene_tx=XM_023256524.1;refGene_gn=CUL2;refGene_tx=XM_023256525.1;refGene_gn=CUL2;refGene_tx=XM_023256526.1;refGene_gn=CUL2;refGene_tx=XM_023256527.1;refGene_gn=CUL2;refGene_tx=XM_023256528.1
B4	32000990	32001050
refGene_gn=CUL2;refGene_tx=XM_006933355.4;refGene_gn=CUL2;refGene_tx=XM_023256524.1;refGene_gn=CUL2;refGene_tx=XM_023256525.1;refGene_gn=CUL2;refGene_tx=XM_023256526.1;refGene_gn=CUL2;refGene_tx=XM_023256527.1;refGene_gn=CUL2;refGene_tx=XM_023256528.1
B4	32001990	32002098
refGene_gn=CUL2;refGene_tx=XM_006933355.4;refGene_gn=CUL2;refGene_tx=XM_023256524.1;refGene_gn=CUL2;refGene_tx=XM_023256525.1;refGene_gn=CUL2;refGene_tx=XM_023256526.1;refGene_gn=CUL2;refGene_tx=XM_023256527.1;refGene_gn=CUL2;refGene_tx=XM_023256528.1
B4	32003616	32003741
refGene_gn=CUL2;refGene_tx=XM_006933355.4;refGene_gn=CUL2;refGene_tx=XM_023256524.1;refGene_gn=CUL2;refGene_tx=XM_023256525.1;refGene_gn=CUL2;refGene_tx=XM_023256526.1;refGene_gn=CUL2;refGene_tx=XM_023256527.1;refGene_gn=CUL2;refGene_tx=XM_023256528.1
B4	32006630	32006793
refGene_gn=CUL2;refGene_tx=XM_006933355.4;refGene_gn=CUL2;refGene_tx=XM_023256524.1;refGene_gn=CUL2;refGene_tx=XM_023256525.1;refGene_gn=CUL2;refGene_tx=XM_023256526.1;refGene_gn=CUL2;refGene_tx=XM_023256527.1;

refGene_gn=CUL2;refGene_tx=XM_023256528.1
B4 32010084 32010195
refGene_gn=CUL2;refGene_tx=XM_006933355.4;refGene_gn=CUL2;refGene_tx=XM_023256524.1;refGene_gn=CUL2;refGene_tx=XM_023256525.1;refGene_gn=CUL2;refGene_tx=XM_023256526.1;refGene_gn=CUL2;refGene_tx=XM_023256527.1;refGene_gn=CUL2;refGene_tx=XM_023256528.1
B4 32010287 32010384
refGene_gn=CUL2;refGene_tx=XM_006933355.4;refGene_gn=CUL2;refGene_tx=XM_023256524.1;refGene_gn=CUL2;refGene_tx=XM_023256525.1;refGene_gn=CUL2;refGene_tx=XM_023256526.1;refGene_gn=CUL2;refGene_tx=XM_023256527.1;refGene_gn=CUL2;refGene_tx=XM_023256528.1
B4 32034048 32034131
refGene_gn=CUL2;refGene_tx=XM_006933355.4;refGene_gn=CUL2;refGene_tx=XM_023256524.1;refGene_gn=CUL2;refGene_tx=XM_023256525.1;refGene_gn=CUL2;refGene_tx=XM_023256526.1;refGene_gn=CUL2;refGene_tx=XM_023256527.1;refGene_gn=CUL2;refGene_tx=XM_023256528.1
B4 32041194 32041300
refGene_gn=CUL2;refGene_tx=XM_006933355.4;refGene_gn=CUL2;refGene_tx=XM_023256524.1;refGene_gn=CUL2;refGene_tx=XM_023256525.1;refGene_gn=CUL2;refGene_tx=XM_023256526.1;refGene_gn=CUL2;refGene_tx=XM_023256527.1;refGene_gn=CUL2;refGene_tx=XM_023256528.1
B4 32048216 32048306
refGene_gn=CUL2;refGene_tx=XM_006933355.4;refGene_gn=CUL2;refGene_tx=XM_023256524.1;refGene_gn=CUL2;refGene_tx=XM_023256525.1;refGene_gn=CUL2;refGene_tx=XM_023256526.1;refGene_gn=CUL2;refGene_tx=XM_023256527.1;refGene_gn=CUL2;refGene_tx=XM_023256528.1
B4 32051595 32051600 refGene_gn=CUL2;refGene_tx=XM_023256528.1
B4 32052045 32052148 refGene_gn=CUL2;refGene_tx=XM_023256528.1
B4 32062446 32062565 refGene_gn=CUL2;refGene_tx=XM_023256528.1
B4 32149243 32149287
refGene_gn=CREM;refGene_tx=XM_006933356.4;refGene_gn=CREM;refGene_tx=XM_023256529.1;refGene_gn=CREM;refGene_tx=XM_023256530.1;refGene_gn=CREM;refGene_tx=XM_023256531.1;refGene_gn=CREM;refGene_tx=XM_023256532.1;refGene_gn=CREM;refGene_tx=XM_023256533.1;refGene_gn=CREM;refGene_tx=XM_023256536.1;refGene_gn=CREM;refGene_tx=XM_023256537.1;refGene_gn=CREM;refGene_tx=XM_023256538.1;refGene_gn=CREM;refGene_tx=XM_023256539.1;refGene_gn=CREM;refGene_tx=XM_023256540.1;refGene_gn=CREM;refGene_tx=XM_023256541.1;refGene_gn=CREM;refGene_tx=XM_023256542.1;refGene_gn=CREM;refGene_tx=XM_023256543.1;refGene_gn=CREM;refGene_tx=XM_023256545.1;refGene_gn=CREM;refGene_tx=XM_023256546.1;refGene_gn=CREM;refGene_tx=XM_023256547.1;refGene_gn=CREM;refGene_tx=XM_023256548.1
B4 32159591 32159715
refGene_gn=CREM;refGene_tx=XM_006933356.4;refGene_gn=CREM;refGene_tx=XM_023256529.1;refGene_gn=CREM;refGene_tx=XM_023256530.1;refGene_gn=CREM;refGene_tx=XM_023256531.1;refGene_gn=CREM;refGene_tx=XM_023256532.1;refGene_gn=CREM;refGene_tx=XM_023256533.1;refGene_gn=CREM;refGene_tx=XM_023256536.1;refGene_gn=CREM;refGene_tx=XM_023256537.1;refGene_gn=CREM;refGene_tx=XM_023256538.1;refGene_gn=CREM;refGene_tx=XM_023256539.1;refGene_gn=CREM;refGene_tx=XM_023256540.1;refGene_gn=CREM;refGene_tx=XM_023256541.1;refGene_gn=CREM;refGene_tx=XM_023256542.1;refGene_gn=CRE

M;refGene_tx=XM_023256543.1;refGene_gn=CREM;refGene_tx=XM_023256545.1;
refGene_gn=CREM;refGene_tx=XM_023256546.1;refGene_gn=CREM;refGene_tx=X
M_023256547.1;refGene_gn=CREM;refGene_tx=XM_023256548.1
B4 32171540 32171615
refGene_gn=CREM;refGene_tx=XM_003988230.5;refGene_gn=CREM;refGene_tx=X
M_003988233.5;refGene_gn=CREM;refGene_tx=XM_003988235.5;refGene_gn=CRE
M;refGene_tx=XM_023256534.1
B4 32176718 32176751
refGene_gn=CREM;refGene_tx=XM_023256535.1;refGene_gn=CREM;refGene_tx=X
M_023256544.1
B4 32176909 32177056
refGene_gn=CREM;refGene_tx=XM_006933356.4;refGene_gn=CREM;refGene_tx=X
M_023256529.1;refGene_gn=CREM;refGene_tx=XM_023256530.1;refGene_gn=CRE
M;refGene_tx=XM_023256531.1;refGene_gn=CREM;refGene_tx=XM_023256532.1;
refGene_gn=CREM;refGene_tx=XM_023256533.1;refGene_gn=CREM;refGene_tx=X
M_023256534.1;refGene_gn=CREM;refGene_tx=XM_023256535.1;refGene_gn=CRE
M;refGene_tx=XM_023256540.1;refGene_gn=CREM;refGene_tx=XM_023256541.1;
refGene_gn=CREM;refGene_tx=XM_023256542.1;refGene_gn=CREM;refGene_tx=X
M_023256543.1
B4 32180895 32180993
refGene_gn=CREM;refGene_tx=XM_003988230.5;refGene_gn=CREM;refGene_tx=X
M_003988233.5;refGene_gn=CREM;refGene_tx=XM_003988235.5;refGene_gn=CRE
M;refGene_tx=XM_006933356.4;refGene_gn=CREM;refGene_tx=XM_023256529.1;
refGene_gn=CREM;refGene_tx=XM_023256530.1;refGene_gn=CREM;refGene_tx=X
M_023256531.1;refGene_gn=CREM;refGene_tx=XM_023256532.1;refGene_gn=CRE
M;refGene_tx=XM_023256533.1;refGene_gn=CREM;refGene_tx=XM_023256534.1;
refGene_gn=CREM;refGene_tx=XM_023256535.1;refGene_gn=CREM;refGene_tx=X
M_023256536.1;refGene_gn=CREM;refGene_tx=XM_023256537.1;refGene_gn=CRE
M;refGene_tx=XM_023256538.1;refGene_gn=CREM;refGene_tx=XM_023256539.1;
refGene_gn=CREM;refGene_tx=XM_023256540.1;refGene_gn=CREM;refGene_tx=X
M_023256541.1;refGene_gn=CREM;refGene_tx=XM_023256542.1;refGene_gn=CRE
M;refGene_tx=XM_023256543.1;refGene_gn=CREM;refGene_tx=XM_023256544.1;
refGene_gn=CREM;refGene_tx=XM_023256545.1;refGene_gn=CREM;refGene_tx=X
M_023256546.1;refGene_gn=CREM;refGene_tx=XM_023256547.1;refGene_gn=CRE
M;refGene_tx=XM_023256548.1
B4 32181349 32181492
refGene_gn=CREM;refGene_tx=XM_003988230.5;refGene_gn=CREM;refGene_tx=X
M_003988233.5;refGene_gn=CREM;refGene_tx=XM_003988235.5;refGene_gn=CRE
M;refGene_tx=XM_006933356.4;refGene_gn=CREM;refGene_tx=XM_023256529.1;
refGene_gn=CREM;refGene_tx=XM_023256530.1;refGene_gn=CREM;refGene_tx=X
M_023256531.1;refGene_gn=CREM;refGene_tx=XM_023256532.1;refGene_gn=CRE
M;refGene_tx=XM_023256533.1;refGene_gn=CREM;refGene_tx=XM_023256534.1;
refGene_gn=CREM;refGene_tx=XM_023256535.1;refGene_gn=CREM;refGene_tx=X
M_023256536.1;refGene_gn=CREM;refGene_tx=XM_023256537.1;refGene_gn=CRE
M;refGene_tx=XM_023256538.1;refGene_gn=CREM;refGene_tx=XM_023256539.1;
refGene_gn=CREM;refGene_tx=XM_023256540.1;refGene_gn=CREM;refGene_tx=X
M_023256541.1;refGene_gn=CREM;refGene_tx=XM_023256542.1;refGene_gn=CRE
M;refGene_tx=XM_023256543.1;refGene_gn=CREM;refGene_tx=XM_023256544.1;
refGene_gn=CREM;refGene_tx=XM_023256545.1;refGene_gn=CREM;refGene_tx=X
M_023256546.1;refGene_gn=CREM;refGene_tx=XM_023256547.1;refGene_gn=CRE

M;refGene_tx=XM_023256548.1;refGene_gn=CREM;refGene_tx=XM_023256549.1
B4	32190665	32190854
refGene_gn=CREM;refGene_tx=XM_003988230.5;refGene_gn=CREM;refGene_tx=XM_003988233.5;refGene_gn=CREM;refGene_tx=XM_006933356.4;refGene_gn=CREM;refGene_tx=XM_023256529.1;refGene_gn=CREM;refGene_tx=XM_023256530.1;refGene_gn=CREM;refGene_tx=XM_023256531.1;refGene_gn=CREM;refGene_tx=XM_023256532.1;refGene_gn=CREM;refGene_tx=XM_023256533.1;refGene_gn=CREM;refGene_tx=XM_023256534.1;refGene_gn=CREM;refGene_tx=XM_023256535.1;refGene_gn=CREM;refGene_tx=XM_023256536.1;refGene_gn=CREM;refGene_tx=XM_023256537.1;refGene_gn=CREM;refGene_tx=XM_023256538.1;refGene_gn=CREM;refGene_tx=XM_023256539.1;refGene_gn=CREM;refGene_tx=XM_023256544.1;refGene_gn=CREM;refGene_tx=XM_023256549.1
B4	32196945	32196985
refGene_gn=CREM;refGene_tx=XM_003988223.5;refGene_gn=CREM;refGene_tx=XM_003988234.5;refGene_gn=CREM;refGene_tx=XM_019834202.2;refGene_gn=CREM;refGene_tx=XM_019834203.2
B4	32201942	32201978
refGene_gn=CREM;refGene_tx=XM_003988233.5;refGene_gn=CREM;refGene_tx=XM_006933356.4;refGene_gn=CREM;refGene_tx=XM_019834202.2;refGene_gn=CREM;refGene_tx=XM_019834203.2;refGene_gn=CREM;refGene_tx=XM_023256529.1;refGene_gn=CREM;refGene_tx=XM_023256530.1;refGene_gn=CREM;refGene_tx=XM_023256531.1;refGene_gn=CREM;refGene_tx=XM_023256534.1;refGene_gn=CREM;refGene_tx=XM_023256535.1;refGene_gn=CREM;refGene_tx=XM_023256536.1;refGene_gn=CREM;refGene_tx=XM_023256537.1;refGene_gn=CREM;refGene_tx=XM_023256540.1;refGene_gn=CREM;refGene_tx=XM_023256541.1;refGene_gn=CREM;refGene_tx=XM_023256544.1;refGene_gn=CREM;refGene_tx=XM_023256545.1;refGene_gn=CREM;refGene_tx=XM_023256546.1;refGene_gn=CREM;refGene_tx=XM_023256549.1;refGene_gn=CREM;refGene_tx=XM_023256551.1
B4	32205983	32206140
refGene_gn=CREM;refGene_tx=XM_003988223.5;refGene_gn=CREM;refGene_tx=XM_003988230.5;refGene_gn=CREM;refGene_tx=XM_003988233.5;refGene_gn=CREM;refGene_tx=XM_003988234.5;refGene_gn=CREM;refGene_tx=XM_003988235.5;refGene_gn=CREM;refGene_tx=XM_006933356.4;refGene_gn=CREM;refGene_tx=XM_019834202.2;refGene_gn=CREM;refGene_tx=XM_019834203.2;refGene_gn=CREM;refGene_tx=XM_023256529.1;refGene_gn=CREM;refGene_tx=XM_023256530.1;refGene_gn=CREM;refGene_tx=XM_023256531.1;refGene_gn=CREM;refGene_tx=XM_023256532.1;refGene_gn=CREM;refGene_tx=XM_023256533.1;refGene_gn=CREM;refGene_tx=XM_023256534.1;refGene_gn=CREM;refGene_tx=XM_023256535.1;refGene_gn=CREM;refGene_tx=XM_023256536.1;refGene_gn=CREM;refGene_tx=XM_023256537.1;refGene_gn=CREM;refGene_tx=XM_023256538.1;refGene_gn=CREM;refGene_tx=XM_023256539.1;refGene_gn=CREM;refGene_tx=XM_023256540.1;refGene_gn=CREM;refGene_tx=XM_023256541.1;refGene_gn=CREM;refGene_tx=XM_023256542.1;refGene_gn=CREM;refGene_tx=XM_023256543.1;refGene_gn=CREM;refGene_tx=XM_023256544.1;refGene_gn=CREM;refGene_tx=XM_023256545.1;refGene_gn=CREM;refGene_tx=XM_023256546.1;refGene_gn=CREM;refGene_tx=XM_023256547.1;refGene_gn=CREM;refGene_tx=XM_023256548.1;refGene_gn=CREM;refGene_tx=XM_023256549.1;refGene_gn=CREM;refGene_tx=XM_023256550.1;refGene_gn=CREM;refGene_tx=XM_023256551.1;refGene_gn=CREM;refGene_tx=XM_023256552.1;refGene_gn=CREM;refGene_tx=XM_023256553.1;refGene_gn=CREM;refGene_tx=XM_023256554.1

B4 32211529 32211674
refGene_gn=CREM;refGene_tx=XM_003988223.5;refGene_gn=CREM;refGene_tx=XM_019834202.2;refGene_gn=CREM;refGene_tx=XM_023256531.1;refGene_gn=CREM;refGene_tx=XM_023256533.1;refGene_gn=CREM;refGene_tx=XM_023256537.1;refGene_gn=CREM;refGene_tx=XM_023256539.1;refGene_gn=CREM;refGene_tx=XM_023256541.1;refGene_gn=CREM;refGene_tx=XM_023256543.1;refGene_gn=CREM;refGene_tx=XM_023256546.1;refGene_gn=CREM;refGene_tx=XM_023256548.1
B4 32211940 32212085
refGene_gn=CREM;refGene_tx=XM_003988230.5;refGene_gn=CREM;refGene_tx=XM_003988233.5;refGene_gn=CREM;refGene_tx=XM_003988234.5;refGene_gn=CREM;refGene_tx=XM_003988235.5;refGene_gn=CREM;refGene_tx=XM_006933356.4;refGene_gn=CREM;refGene_tx=XM_019834203.2;refGene_gn=CREM;refGene_tx=XM_023256529.1;refGene_gn=CREM;refGene_tx=XM_023256530.1;refGene_gn=CREM;refGene_tx=XM_023256532.1;refGene_gn=CREM;refGene_tx=XM_023256534.1;refGene_gn=CREM;refGene_tx=XM_023256535.1;refGene_gn=CREM;refGene_tx=XM_023256536.1;refGene_gn=CREM;refGene_tx=XM_023256538.1;refGene_gn=CREM;refGene_tx=XM_023256540.1;refGene_gn=CREM;refGene_tx=XM_023256542.1;refGene_gn=CREM;refGene_tx=XM_023256544.1;refGene_gn=CREM;refGene_tx=XM_023256545.1;refGene_gn=CREM;refGene_tx=XM_023256547.1;refGene_gn=CREM;refGene_tx=XM_023256549.1;refGene_gn=CREM;refGene_tx=XM_023256550.1;refGene_gn=CREM;refGene_tx=XM_023256551.1;refGene_gn=CREM;refGene_tx=XM_023256552.1;refGene_gn=CREM;refGene_tx=XM_023256553.1;refGene_gn=CREM;refGene_tx=XM_023256554.1
B4 32318425 32318959
refGene_gn=LOC105260723;refGene_tx=XM_023256557.1
B4 32319132 32319286 refGene_gn=CCNY;refGene_tx=XM_011283702.3
B4 32472485 32472560
refGene_gn=CCNY;refGene_tx=XM_011283702.3;refGene_gn=CCNY;refGene_tx=XM_019834205.2;refGene_gn=CCNY;refGene_tx=XM_019834206.2;refGene_gn=CCNY;refGene_tx=XM_019834207.2;refGene_gn=CCNY;refGene_tx=XM_023256555.1;refGene_gn=CCNY;refGene_tx=XM_023256556.1
B4 32493426 32493461
refGene_gn=CCNY;refGene_tx=XM_011283702.3;refGene_gn=CCNY;refGene_tx=XM_019834205.2;refGene_gn=CCNY;refGene_tx=XM_019834206.2;refGene_gn=CCNY;refGene_tx=XM_019834207.2;refGene_gn=CCNY;refGene_tx=XM_023256555.1;refGene_gn=CCNY;refGene_tx=XM_023256556.1
B4 32508767 32508868
refGene_gn=CCNY;refGene_tx=XM_011283702.3;refGene_gn=CCNY;refGene_tx=XM_019834205.2;refGene_gn=CCNY;refGene_tx=XM_019834206.2;refGene_gn=CCNY;refGene_tx=XM_019834207.2;refGene_gn=CCNY;refGene_tx=XM_023256555.1;refGene_gn=CCNY;refGene_tx=XM_023256556.1
B4 32515225 32515261
refGene_gn=CCNY;refGene_tx=XM_011283702.3;refGene_gn=CCNY;refGene_tx=XM_019834205.2;refGene_gn=CCNY;refGene_tx=XM_019834206.2;refGene_gn=CCNY;refGene_tx=XM_019834207.2;refGene_gn=CCNY;refGene_tx=XM_023256555.1;refGene_gn=CCNY;refGene_tx=XM_023256556.1
B4 32518677 32518735
refGene_gn=CCNY;refGene_tx=XM_011283702.3;refGene_gn=CCNY;refGene_tx=XM_019834205.2;refGene_gn=CCNY;refGene_tx=XM_019834206.2;refGene_gn=CCNY;refGene_tx=XM_019834207.2;refGene_gn=CCNY;refGene_tx=XM_023256555.1;

refGene_gn=CCNY;refGene_tx=XM_023256556.1
B4	32518821	32518941
refGene_gn=CCNY;refGene_tx=XM_011283702.3;refGene_gn=CCNY;refGene_tx=XM_019834205.2;refGene_gn=CCNY;refGene_tx=XM_019834206.2;refGene_gn=CCNY;refGene_tx=XM_019834207.2;refGene_gn=CCNY;refGene_tx=XM_023256555.1;refGene_gn=CCNY;refGene_tx=XM_023256556.1
B4	32536925	32537092
refGene_gn=CCNY;refGene_tx=XM_011283702.3;refGene_gn=CCNY;refGene_tx=XM_019834205.2;refGene_gn=CCNY;refGene_tx=XM_019834206.2;refGene_gn=CCNY;refGene_tx=XM_019834207.2;refGene_gn=CCNY;refGene_tx=XM_023256555.1;refGene_gn=CCNY;refGene_tx=XM_023256556.1
B4	32545952	32546115
refGene_gn=CCNY;refGene_tx=XM_011283702.3;refGene_gn=CCNY;refGene_tx=XM_019834205.2;refGene_gn=CCNY;refGene_tx=XM_019834206.2;refGene_gn=CCNY;refGene_tx=XM_019834207.2;refGene_gn=CCNY;refGene_tx=XM_023256555.1;refGene_gn=CCNY;refGene_tx=XM_023256556.1
B4	32549955	32550072
refGene_gn=CCNY;refGene_tx=XM_011283702.3;refGene_gn=CCNY;refGene_tx=XM_019834205.2;refGene_gn=CCNY;refGene_tx=XM_019834206.2;refGene_gn=CCNY;refGene_tx=XM_019834207.2;refGene_gn=CCNY;refGene_tx=XM_023256555.1;refGene_gn=CCNY;refGene_tx=XM_023256556.1
B4	32579610	32579674	refGene_gn=GJD4;refGene_tx=XM_023256558.1
B4	32581111	32581710	refGene_gn=GJD4;refGene_tx=XM_023256558.1
B4	32581712	32582156	refGene_gn=GJD4;refGene_tx=XM_023256558.1
B4	32612224	32614243	refGene_gn=FZD8;refGene_tx=XM_023256559.1
B4	34202329	34202375	refGene_gn=USP18;refGene_tx=XM_011283700.3
B4	34203653	34203703	refGene_gn=USP18;refGene_tx=XM_011283700.3
B4	34204526	34204658	refGene_gn=USP18;refGene_tx=XM_011283700.3
B4	34210108	34210276	refGene_gn=USP18;refGene_tx=XM_011283700.3
B4	34210726	34210822	refGene_gn=USP18;refGene_tx=XM_011283700.3
B4	34211208	34211355	refGene_gn=USP18;refGene_tx=XM_011283700.3
B4	34211649	34211729	refGene_gn=USP18;refGene_tx=XM_011283700.3
B4	34238308	34238454	refGene_gn=USP18;refGene_tx=XM_011283700.3
B4	34251253	34251350	refGene_gn=USP18;refGene_tx=XM_011283700.3
B4	34252431	34252688	refGene_gn=USP18;refGene_tx=XM_011283700.3
B4	34267637	34267780	refGene_gn=USP18;refGene_tx=XM_011283700.3
B4	34281220	34281514
refGene_gn=TUBA8;refGene_tx=XM_004001072.3;refGene_gn=TUBA8;refGene_tx=XM_023256560.1
B4	34284809	34285490
refGene_gn=TUBA8;refGene_tx=XM_004001072.3;refGene_gn=TUBA8;refGene_tx=XM_023256560.1
B4	34287875	34288024
refGene_gn=TUBA8;refGene_tx=XM_004001072.3;refGene_gn=TUBA8;refGene_tx=XM_023256560.1
B4	34289747	34289970
refGene_gn=TUBA8;refGene_tx=XM_004001072.3;refGene_gn=TUBA8;refGene_tx=XM_023256560.1
B4	34299935	34299938	refGene_gn=TUBA8;refGene_tx=XM_004001072.3
B4	34322457	34322558	refGene_gn=PEX26;refGene_tx=XM_006933372.4

B4	34326249	34326396
refGene_gn=PEX26;refGene_tx=XM_006933372.4;refGene_gn=PEX26;refGene_tx=XM_019834210.2
B4	34328033	34328329
refGene_gn=PEX26;refGene_tx=XM_006933372.4;refGene_gn=PEX26;refGene_tx=XM_019834210.2
B4	34328812	34328953
refGene_gn=PEX26;refGene_tx=XM_006933372.4;refGene_gn=PEX26;refGene_tx=XM_019834210.2
B4	34330277	34330492
refGene_gn=PEX26;refGene_tx=XM_006933372.4;refGene_gn=PEX26;refGene_tx=XM_019834210.2
B4	34502171	34502435
refGene_gn=MICAL3;refGene_tx=XM_019834211.2;refGene_gn=MICAL3;refGene_tx=XM_019834212.2;refGene_gn=MICAL3;refGene_tx=XM_019834214.2;refGene_gn=MICAL3;refGene_tx=XM_023256561.1;refGene_gn=MICAL3;refGene_tx=XM_023256562.1;refGene_gn=MICAL3;refGene_tx=XM_023256563.1;refGene_gn=MICAL3;refGene_tx=XM_023256564.1;refGene_gn=MICAL3;refGene_tx=XM_023256565.1;refGene_gn=MICAL3;refGene_tx=XM_023256566.1;refGene_gn=MICAL3;refGene_tx=XM_023256567.1;refGene_gn=MICAL3;refGene_tx=XM_023256568.1;refGene_gn=MICAL3;refGene_tx=XM_023256569.1;refGene_gn=MICAL3;refGene_tx=XM_023256570.1;refGene_gn=MICAL3;refGene_tx=XM_023256571.1;refGene_gn=MICAL3;refGene_tx=XM_023256572.1;refGene_gn=MICAL3;refGene_tx=XM_023256573.1;refGene_gn=MICAL3;refGene_tx=XM_023256574.1;refGene_gn=MICAL3;refGene_tx=XM_023256575.1;refGene_gn=MICAL3;refGene_tx=XM_023256576.1;refGene_gn=MICAL3;refGene_tx=XM_023256577.1;refGene_gn=MICAL3;refGene_tx=XM_023256578.1
B4	34504500	34504708
refGene_gn=MICAL3;refGene_tx=XM_019834211.2;refGene_gn=MICAL3;refGene_tx=XM_019834212.2;refGene_gn=MICAL3;refGene_tx=XM_019834214.2;refGene_gn=MICAL3;refGene_tx=XM_023256561.1;refGene_gn=MICAL3;refGene_tx=XM_023256562.1;refGene_gn=MICAL3;refGene_tx=XM_023256563.1;refGene_gn=MICAL3;refGene_tx=XM_023256564.1;refGene_gn=MICAL3;refGene_tx=XM_023256565.1;refGene_gn=MICAL3;refGene_tx=XM_023256566.1;refGene_gn=MICAL3;refGene_tx=XM_023256567.1;refGene_gn=MICAL3;refGene_tx=XM_023256568.1;refGene_gn=MICAL3;refGene_tx=XM_023256569.1;refGene_gn=MICAL3;refGene_tx=XM_023256570.1;refGene_gn=MICAL3;refGene_tx=XM_023256571.1;refGene_gn=MICAL3;refGene_tx=XM_023256572.1;refGene_gn=MICAL3;refGene_tx=XM_023256573.1;refGene_gn=MICAL3;refGene_tx=XM_023256574.1;refGene_gn=MICAL3;refGene_tx=XM_023256575.1;refGene_gn=MICAL3;refGene_tx=XM_023256576.1;refGene_gn=MICAL3;refGene_tx=XM_023256577.1;refGene_gn=MICAL3;refGene_tx=XM_023256578.1
B4	34506587	34506704
refGene_gn=MICAL3;refGene_tx=XM_019834211.2;refGene_gn=MICAL3;refGene_tx=XM_019834212.2;refGene_gn=MICAL3;refGene_tx=XM_019834214.2;refGene_gn=MICAL3;refGene_tx=XM_023256561.1;refGene_gn=MICAL3;refGene_tx=XM_023256562.1;refGene_gn=MICAL3;refGene_tx=XM_023256563.1;refGene_gn=MICAL3;refGene_tx=XM_023256564.1;refGene_gn=MICAL3;refGene_tx=XM_023256565.1;refGene_gn=MICAL3;refGene_tx=XM_023256566.1;refGene_gn=MICAL3;refGene_tx=XM_023256567.1;refGene_gn=MICAL3;refGene_tx=XM_023256568.1;refGen

e_gn=MICAL3;refGene_tx=XM_023256569.1;refGene_gn=MICAL3;refGene_tx=XM_023256570.1;refGene_gn=MICAL3;refGene_tx=XM_023256571.1;refGene_gn=MICAL3;refGene_tx=XM_023256572.1;refGene_gn=MICAL3;refGene_tx=XM_023256573.1;refGene_gn=MICAL3;refGene_tx=XM_023256574.1;refGene_gn=MICAL3;refGene_tx=XM_023256575.1;refGene_gn=MICAL3;refGene_tx=XM_023256576.1;refGene_gn=MICAL3;refGene_tx=XM_023256577.1;refGene_gn=MICAL3;refGene_tx=XM_023256578.1
B4 34507105 34507207
refGene_gn=MICAL3;refGene_tx=XM_019834211.2;refGene_gn=MICAL3;refGene_tx=XM_019834212.2;refGene_gn=MICAL3;refGene_tx=XM_019834214.2;refGene_gn=MICAL3;refGene_tx=XM_023256561.1;refGene_gn=MICAL3;refGene_tx=XM_023256562.1;refGene_gn=MICAL3;refGene_tx=XM_023256563.1;refGene_gn=MICAL3;refGene_tx=XM_023256564.1;refGene_gn=MICAL3;refGene_tx=XM_023256565.1;refGene_gn=MICAL3;refGene_tx=XM_023256566.1;refGene_gn=MICAL3;refGene_tx=XM_023256567.1;refGene_gn=MICAL3;refGene_tx=XM_023256568.1;refGene_gn=MICAL3;refGene_tx=XM_023256569.1;refGene_gn=MICAL3;refGene_tx=XM_023256570.1;refGene_gn=MICAL3;refGene_tx=XM_023256571.1;refGene_gn=MICAL3;refGene_tx=XM_023256572.1;refGene_gn=MICAL3;refGene_tx=XM_023256573.1;refGene_gn=MICAL3;refGene_tx=XM_023256574.1;refGene_gn=MICAL3;refGene_tx=XM_023256575.1;refGene_gn=MICAL3;refGene_tx=XM_023256576.1;refGene_gn=MICAL3;refGene_tx=XM_023256577.1;refGene_gn=MICAL3;refGene_tx=XM_023256578.1
B4 34507817 34507973
refGene_gn=MICAL3;refGene_tx=XM_019834211.2;refGene_gn=MICAL3;refGene_tx=XM_019834212.2;refGene_gn=MICAL3;refGene_tx=XM_019834214.2;refGene_gn=MICAL3;refGene_tx=XM_023256561.1;refGene_gn=MICAL3;refGene_tx=XM_023256562.1;refGene_gn=MICAL3;refGene_tx=XM_023256563.1;refGene_gn=MICAL3;refGene_tx=XM_023256564.1;refGene_gn=MICAL3;refGene_tx=XM_023256565.1;refGene_gn=MICAL3;refGene_tx=XM_023256566.1;refGene_gn=MICAL3;refGene_tx=XM_023256567.1;refGene_gn=MICAL3;refGene_tx=XM_023256568.1;refGene_gn=MICAL3;refGene_tx=XM_023256569.1;refGene_gn=MICAL3;refGene_tx=XM_023256570.1;refGene_gn=MICAL3;refGene_tx=XM_023256571.1;refGene_gn=MICAL3;refGene_tx=XM_023256572.1;refGene_gn=MICAL3;refGene_tx=XM_023256573.1;refGene_gn=MICAL3;refGene_tx=XM_023256574.1;refGene_gn=MICAL3;refGene_tx=XM_023256575.1;refGene_gn=MICAL3;refGene_tx=XM_023256576.1;refGene_gn=MICAL3;refGene_tx=XM_023256577.1;refGene_gn=MICAL3;refGene_tx=XM_023256578.1
B4 34508981 34509082
refGene_gn=MICAL3;refGene_tx=XM_019834211.2;refGene_gn=MICAL3;refGene_tx=XM_019834212.2;refGene_gn=MICAL3;refGene_tx=XM_019834214.2;refGene_gn=MICAL3;refGene_tx=XM_023256561.1;refGene_gn=MICAL3;refGene_tx=XM_023256562.1;refGene_gn=MICAL3;refGene_tx=XM_023256563.1;refGene_gn=MICAL3;refGene_tx=XM_023256564.1;refGene_gn=MICAL3;refGene_tx=XM_023256565.1;refGene_gn=MICAL3;refGene_tx=XM_023256566.1;refGene_gn=MICAL3;refGene_tx=XM_023256567.1;refGene_gn=MICAL3;refGene_tx=XM_023256568.1;refGene_gn=MICAL3;refGene_tx=XM_023256569.1;refGene_gn=MICAL3;refGene_tx=XM_023256570.1;refGene_gn=MICAL3;refGene_tx=XM_023256571.1;refGene_gn=MICAL3;refGene_tx=XM_023256572.1;refGene_gn=MICAL3;refGene_tx=XM_023256573.1;refGene_gn=MICAL3;refGene_tx=XM_023256574.1;refGene_gn=MICAL3;refGene_tx=XM_023256575.1;refGene_gn=MICAL3;refGene_tx=XM_023256576.1;refG

ene_gn=MICAL3;refGene_tx=XM_023256577.1;refGene_gn=MICAL3;refGene_tx=XM_023256578.1
B4 34512053 34512311
refGene_gn=MICAL3;refGene_tx=XM_019834211.2;refGene_gn=MICAL3;refGene_tx=XM_019834212.2;refGene_gn=MICAL3;refGene_tx=XM_019834214.2;refGene_gn=MICAL3;refGene_tx=XM_023256561.1;refGene_gn=MICAL3;refGene_tx=XM_023256562.1;refGene_gn=MICAL3;refGene_tx=XM_023256563.1;refGene_gn=MICAL3;refGene_tx=XM_023256564.1;refGene_gn=MICAL3;refGene_tx=XM_023256565.1;refGene_gn=MICAL3;refGene_tx=XM_023256566.1;refGene_gn=MICAL3;refGene_tx=XM_023256567.1;refGene_gn=MICAL3;refGene_tx=XM_023256568.1;refGene_gn=MICAL3;refGene_tx=XM_023256569.1;refGene_gn=MICAL3;refGene_tx=XM_023256570.1;refGene_gn=MICAL3;refGene_tx=XM_023256571.1;refGene_gn=MICAL3;refGene_tx=XM_023256572.1;refGene_gn=MICAL3;refGene_tx=XM_023256573.1;refGene_gn=MICAL3;refGene_tx=XM_023256574.1;refGene_gn=MICAL3;refGene_tx=XM_023256575.1;refGene_gn=MICAL3;refGene_tx=XM_023256576.1;refGene_gn=MICAL3;refGene_tx=XM_023256577.1;refGene_gn=MICAL3;refGene_tx=XM_023256578.1
B4 34512688 34512804
refGene_gn=MICAL3;refGene_tx=XM_019834211.2;refGene_gn=MICAL3;refGene_tx=XM_019834212.2;refGene_gn=MICAL3;refGene_tx=XM_019834214.2;refGene_gn=MICAL3;refGene_tx=XM_023256561.1;refGene_gn=MICAL3;refGene_tx=XM_023256562.1;refGene_gn=MICAL3;refGene_tx=XM_023256563.1;refGene_gn=MICAL3;refGene_tx=XM_023256564.1;refGene_gn=MICAL3;refGene_tx=XM_023256565.1;refGene_gn=MICAL3;refGene_tx=XM_023256566.1;refGene_gn=MICAL3;refGene_tx=XM_023256567.1;refGene_gn=MICAL3;refGene_tx=XM_023256568.1;refGene_gn=MICAL3;refGene_tx=XM_023256569.1;refGene_gn=MICAL3;refGene_tx=XM_023256570.1;refGene_gn=MICAL3;refGene_tx=XM_023256571.1;refGene_gn=MICAL3;refGene_tx=XM_023256572.1;refGene_gn=MICAL3;refGene_tx=XM_023256573.1;refGene_gn=MICAL3;refGene_tx=XM_023256574.1;refGene_gn=MICAL3;refGene_tx=XM_023256575.1;refGene_gn=MICAL3;refGene_tx=XM_023256576.1;refGene_gn=MICAL3;refGene_tx=XM_023256577.1;refGene_gn=MICAL3;refGene_tx=XM_023256578.1
B4 34513820 34513947
refGene_gn=MICAL3;refGene_tx=XM_019834211.2;refGene_gn=MICAL3;refGene_tx=XM_019834212.2;refGene_gn=MICAL3;refGene_tx=XM_019834214.2;refGene_gn=MICAL3;refGene_tx=XM_023256561.1;refGene_gn=MICAL3;refGene_tx=XM_023256562.1;refGene_gn=MICAL3;refGene_tx=XM_023256563.1;refGene_gn=MICAL3;refGene_tx=XM_023256564.1;refGene_gn=MICAL3;refGene_tx=XM_023256565.1;refGene_gn=MICAL3;refGene_tx=XM_023256566.1;refGene_gn=MICAL3;refGene_tx=XM_023256567.1;refGene_gn=MICAL3;refGene_tx=XM_023256568.1;refGene_gn=MICAL3;refGene_tx=XM_023256569.1;refGene_gn=MICAL3;refGene_tx=XM_023256570.1;refGene_gn=MICAL3;refGene_tx=XM_023256571.1;refGene_gn=MICAL3;refGene_tx=XM_023256572.1;refGene_gn=MICAL3;refGene_tx=XM_023256573.1;refGene_gn=MICAL3;refGene_tx=XM_023256574.1;refGene_gn=MICAL3;refGene_tx=XM_023256575.1;refGene_gn=MICAL3;refGene_tx=XM_023256576.1;refGene_gn=MICAL3;refGene_tx=XM_023256577.1;refGene_gn=MICAL3;refGene_tx=XM_023256578.1
B4 34515392 34515489
refGene_gn=MICAL3;refGene_tx=XM_019834211.2;refGene_gn=MICAL3;refGene_tx=XM_019834212.2;refGene_gn=MICAL3;refGene_tx=XM_019834214.2;refGene_

gn=MICAL3;refGene_tx=XM_023256561.1;refGene_gn=MICAL3;refGene_tx=XM_023256562.1;refGene_gn=MICAL3;refGene_tx=XM_023256563.1;refGene_gn=MICAL3;refGene_tx=XM_023256564.1;refGene_gn=MICAL3;refGene_tx=XM_023256565.1;refGene_gn=MICAL3;refGene_tx=XM_023256566.1;refGene_gn=MICAL3;refGene_tx=XM_023256567.1;refGene_gn=MICAL3;refGene_tx=XM_023256568.1;refGene_gn=MICAL3;refGene_tx=XM_023256569.1;refGene_gn=MICAL3;refGene_tx=XM_023256570.1;refGene_gn=MICAL3;refGene_tx=XM_023256571.1;refGene_gn=MICAL3;refGene_tx=XM_023256572.1;refGene_gn=MICAL3;refGene_tx=XM_023256573.1;refGene_gn=MICAL3;refGene_tx=XM_023256574.1;refGene_gn=MICAL3;refGene_tx=XM_023256575.1;refGene_gn=MICAL3;refGene_tx=XM_023256576.1;refGene_gn=MICAL3;refGene_tx=XM_023256577.1;refGene_gn=MICAL3;refGene_tx=XM_023256578.1
B4 34518389 34518537
refGene_gn=MICAL3;refGene_tx=XM_019834211.2;refGene_gn=MICAL3;refGene_tx=XM_019834212.2;refGene_gn=MICAL3;refGene_tx=XM_019834214.2;refGene_gn=MICAL3;refGene_tx=XM_023256561.1;refGene_gn=MICAL3;refGene_tx=XM_023256562.1;refGene_gn=MICAL3;refGene_tx=XM_023256563.1;refGene_gn=MICAL3;refGene_tx=XM_023256564.1;refGene_gn=MICAL3;refGene_tx=XM_023256565.1;refGene_gn=MICAL3;refGene_tx=XM_023256566.1;refGene_gn=MICAL3;refGene_tx=XM_023256567.1;refGene_gn=MICAL3;refGene_tx=XM_023256568.1;refGene_gn=MICAL3;refGene_tx=XM_023256569.1;refGene_gn=MICAL3;refGene_tx=XM_023256570.1;refGene_gn=MICAL3;refGene_tx=XM_023256571.1;refGene_gn=MICAL3;refGene_tx=XM_023256572.1;refGene_gn=MICAL3;refGene_tx=XM_023256573.1;refGene_gn=MICAL3;refGene_tx=XM_023256574.1;refGene_gn=MICAL3;refGene_tx=XM_023256575.1;refGene_gn=MICAL3;refGene_tx=XM_023256576.1;refGene_gn=MICAL3;refGene_tx=XM_023256577.1;refGene_gn=MICAL3;refGene_tx=XM_023256578.1
B4 34520586 34520783
refGene_gn=MICAL3;refGene_tx=XM_019834211.2;refGene_gn=MICAL3;refGene_tx=XM_019834212.2;refGene_gn=MICAL3;refGene_tx=XM_019834214.2;refGene_gn=MICAL3;refGene_tx=XM_023256561.1;refGene_gn=MICAL3;refGene_tx=XM_023256562.1;refGene_gn=MICAL3;refGene_tx=XM_023256563.1;refGene_gn=MICAL3;refGene_tx=XM_023256564.1;refGene_gn=MICAL3;refGene_tx=XM_023256565.1;refGene_gn=MICAL3;refGene_tx=XM_023256566.1;refGene_gn=MICAL3;refGene_tx=XM_023256567.1;refGene_gn=MICAL3;refGene_tx=XM_023256568.1;refGene_gn=MICAL3;refGene_tx=XM_023256569.1;refGene_gn=MICAL3;refGene_tx=XM_023256570.1;refGene_gn=MICAL3;refGene_tx=XM_023256571.1;refGene_gn=MICAL3;refGene_tx=XM_023256572.1;refGene_gn=MICAL3;refGene_tx=XM_023256573.1;refGene_gn=MICAL3;refGene_tx=XM_023256574.1;refGene_gn=MICAL3;refGene_tx=XM_023256575.1;refGene_gn=MICAL3;refGene_tx=XM_023256576.1;refGene_gn=MICAL3;refGene_tx=XM_023256577.1;refGene_gn=MICAL3;refGene_tx=XM_023256578.1
B4 34522353 34522466
refGene_gn=MICAL3;refGene_tx=XM_019834211.2;refGene_gn=MICAL3;refGene_tx=XM_019834212.2;refGene_gn=MICAL3;refGene_tx=XM_019834214.2;refGene_gn=MICAL3;refGene_tx=XM_023256561.1;refGene_gn=MICAL3;refGene_tx=XM_023256562.1;refGene_gn=MICAL3;refGene_tx=XM_023256563.1;refGene_gn=MICAL3;refGene_tx=XM_023256564.1;refGene_gn=MICAL3;refGene_tx=XM_023256565.1;refGene_gn=MICAL3;refGene_tx=XM_023256566.1;refGene_gn=MICAL3;refGene_tx=XM_023256567.1;refGene_gn=MICAL3;refGene_tx=XM_023256568.1;refGen

e_gn=MICAL3;refGene_tx=XM_023256569.1;refGene_gn=MICAL3;refGene_tx=XM_023256570.1;refGene_gn=MICAL3;refGene_tx=XM_023256571.1;refGene_gn=MICAL3;refGene_tx=XM_023256572.1;refGene_gn=MICAL3;refGene_tx=XM_023256573.1;refGene_gn=MICAL3;refGene_tx=XM_023256574.1;refGene_gn=MICAL3;refGene_tx=XM_023256575.1;refGene_gn=MICAL3;refGene_tx=XM_023256576.1;refGene_gn=MICAL3;refGene_tx=XM_023256577.1;refGene_gn=MICAL3;refGene_tx=XM_023256578.1
B4 34522556 34522619
refGene_gn=MICAL3;refGene_tx=XM_019834211.2;refGene_gn=MICAL3;refGene_tx=XM_019834212.2;refGene_gn=MICAL3;refGene_tx=XM_019834214.2;refGene_gn=MICAL3;refGene_tx=XM_023256561.1;refGene_gn=MICAL3;refGene_tx=XM_023256562.1;refGene_gn=MICAL3;refGene_tx=XM_023256563.1;refGene_gn=MICAL3;refGene_tx=XM_023256564.1;refGene_gn=MICAL3;refGene_tx=XM_023256565.1;refGene_gn=MICAL3;refGene_tx=XM_023256566.1;refGene_gn=MICAL3;refGene_tx=XM_023256567.1;refGene_gn=MICAL3;refGene_tx=XM_023256568.1;refGene_gn=MICAL3;refGene_tx=XM_023256569.1;refGene_gn=MICAL3;refGene_tx=XM_023256570.1;refGene_gn=MICAL3;refGene_tx=XM_023256571.1;refGene_gn=MICAL3;refGene_tx=XM_023256572.1;refGene_gn=MICAL3;refGene_tx=XM_023256573.1;refGene_gn=MICAL3;refGene_tx=XM_023256574.1;refGene_gn=MICAL3;refGene_tx=XM_023256575.1;refGene_gn=MICAL3;refGene_tx=XM_023256576.1;refGene_gn=MICAL3;refGene_tx=XM_023256577.1;refGene_gn=MICAL3;refGene_tx=XM_023256578.1
B4 34523305 34523503
refGene_gn=MICAL3;refGene_tx=XM_019834211.2;refGene_gn=MICAL3;refGene_tx=XM_019834212.2;refGene_gn=MICAL3;refGene_tx=XM_019834214.2;refGene_gn=MICAL3;refGene_tx=XM_023256561.1;refGene_gn=MICAL3;refGene_tx=XM_023256562.1;refGene_gn=MICAL3;refGene_tx=XM_023256563.1;refGene_gn=MICAL3;refGene_tx=XM_023256564.1;refGene_gn=MICAL3;refGene_tx=XM_023256565.1;refGene_gn=MICAL3;refGene_tx=XM_023256566.1;refGene_gn=MICAL3;refGene_tx=XM_023256567.1;refGene_gn=MICAL3;refGene_tx=XM_023256568.1;refGene_gn=MICAL3;refGene_tx=XM_023256569.1;refGene_gn=MICAL3;refGene_tx=XM_023256570.1;refGene_gn=MICAL3;refGene_tx=XM_023256571.1;refGene_gn=MICAL3;refGene_tx=XM_023256572.1;refGene_gn=MICAL3;refGene_tx=XM_023256573.1;refGene_gn=MICAL3;refGene_tx=XM_023256574.1;refGene_gn=MICAL3;refGene_tx=XM_023256575.1;refGene_gn=MICAL3;refGene_tx=XM_023256576.1;refGene_gn=MICAL3;refGene_tx=XM_023256577.1;refGene_gn=MICAL3;refGene_tx=XM_023256578.1
B4 34525279 34525363
refGene_gn=MICAL3;refGene_tx=XM_019834211.2;refGene_gn=MICAL3;refGene_tx=XM_019834212.2;refGene_gn=MICAL3;refGene_tx=XM_019834214.2;refGene_gn=MICAL3;refGene_tx=XM_023256561.1;refGene_gn=MICAL3;refGene_tx=XM_023256562.1;refGene_gn=MICAL3;refGene_tx=XM_023256563.1;refGene_gn=MICAL3;refGene_tx=XM_023256564.1;refGene_gn=MICAL3;refGene_tx=XM_023256565.1;refGene_gn=MICAL3;refGene_tx=XM_023256566.1;refGene_gn=MICAL3;refGene_tx=XM_023256567.1;refGene_gn=MICAL3;refGene_tx=XM_023256568.1;refGene_gn=MICAL3;refGene_tx=XM_023256569.1;refGene_gn=MICAL3;refGene_tx=XM_023256570.1;refGene_gn=MICAL3;refGene_tx=XM_023256571.1;refGene_gn=MICAL3;refGene_tx=XM_023256572.1;refGene_gn=MICAL3;refGene_tx=XM_023256573.1;refGene_gn=MICAL3;refGene_tx=XM_023256578.1
B4 34528491 34528590

refGene_gn=MICAL3;refGene_tx=XM_019834211.2;refGene_gn=MICAL3;refGene_
tx=XM_019834212.2;refGene_gn=MICAL3;refGene_tx=XM_019834214.2;refGene_
gn=MICAL3;refGene_tx=XM_023256561.1;refGene_gn=MICAL3;refGene_tx=XM_02
3256562.1;refGene_gn=MICAL3;refGene_tx=XM_023256563.1;refGene_gn=MICAL
3;refGene_tx=XM_023256564.1;refGene_gn=MICAL3;refGene_tx=XM_023256565.
1;refGene_gn=MICAL3;refGene_tx=XM_023256566.1;refGene_gn=MICAL3;refGen
e_tx=XM_023256567.1;refGene_gn=MICAL3;refGene_tx=XM_023256568.1;refGen
e_gn=MICAL3;refGene_tx=XM_023256569.1;refGene_gn=MICAL3;refGene_tx=XM_
023256570.1;refGene_gn=MICAL3;refGene_tx=XM_023256571.1;refGene_gn=MIC
AL3;refGene_tx=XM_023256573.1;refGene_gn=MICAL3;refGene_tx=XM_02325657
8.1
B4 34530386 34530443
refGene_gn=MICAL3;refGene_tx=XM_019834211.2;refGene_gn=MICAL3;refGene_
tx=XM_019834212.2;refGene_gn=MICAL3;refGene_tx=XM_019834214.2;refGene_
gn=MICAL3;refGene_tx=XM_023256561.1;refGene_gn=MICAL3;refGene_tx=XM_02
3256562.1;refGene_gn=MICAL3;refGene_tx=XM_023256563.1;refGene_gn=MICAL
3;refGene_tx=XM_023256564.1;refGene_gn=MICAL3;refGene_tx=XM_023256565.
1;refGene_gn=MICAL3;refGene_tx=XM_023256566.1;refGene_gn=MICAL3;refGen
e_tx=XM_023256567.1;refGene_gn=MICAL3;refGene_tx=XM_023256568.1;refGen
e_gn=MICAL3;refGene_tx=XM_023256569.1;refGene_gn=MICAL3;refGene_tx=XM_
023256570.1;refGene_gn=MICAL3;refGene_tx=XM_023256571.1;refGene_gn=MIC
AL3;refGene_tx=XM_023256573.1;refGene_gn=MICAL3;refGene_tx=XM_02325657
8.1
B4 34532860 34532968
refGene_gn=MICAL3;refGene_tx=XM_019834211.2;refGene_gn=MICAL3;refGene_
tx=XM_019834212.2;refGene_gn=MICAL3;refGene_tx=XM_019834214.2;refGene_
gn=MICAL3;refGene_tx=XM_023256561.1;refGene_gn=MICAL3;refGene_tx=XM_02
3256562.1;refGene_gn=MICAL3;refGene_tx=XM_023256563.1;refGene_gn=MICAL
3;refGene_tx=XM_023256564.1;refGene_gn=MICAL3;refGene_tx=XM_023256565.
1;refGene_gn=MICAL3;refGene_tx=XM_023256566.1;refGene_gn=MICAL3;refGen
e_tx=XM_023256567.1;refGene_gn=MICAL3;refGene_tx=XM_023256568.1;refGen
e_gn=MICAL3;refGene_tx=XM_023256570.1;refGene_gn=MICAL3;refGene_tx=XM_
023256571.1;refGene_gn=MICAL3;refGene_tx=XM_023256573.1;refGene_gn=MIC
AL3;refGene_tx=XM_023256578.1
B4 34535784 34535892
refGene_gn=MICAL3;refGene_tx=XM_019834211.2;refGene_gn=MICAL3;refGene_
tx=XM_019834212.2;refGene_gn=MICAL3;refGene_tx=XM_019834214.2;refGene_
gn=MICAL3;refGene_tx=XM_023256561.1;refGene_gn=MICAL3;refGene_tx=XM_02
3256562.1;refGene_gn=MICAL3;refGene_tx=XM_023256563.1;refGene_gn=MICAL
3;refGene_tx=XM_023256564.1;refGene_gn=MICAL3;refGene_tx=XM_023256565.
1;refGene_gn=MICAL3;refGene_tx=XM_023256566.1;refGene_gn=MICAL3;refGen
e_tx=XM_023256567.1;refGene_gn=MICAL3;refGene_tx=XM_023256568.1;refGen
e_gn=MICAL3;refGene_tx=XM_023256569.1;refGene_gn=MICAL3;refGene_tx=XM_
023256570.1;refGene_gn=MICAL3;refGene_tx=XM_023256571.1;refGene_gn=MIC
AL3;refGene_tx=XM_023256573.1;refGene_gn=MICAL3;refGene_tx=XM_02325657
8.1
B4 34536504 34536691
refGene_gn=MICAL3;refGene_tx=XM_019834211.2;refGene_gn=MICAL3;refGene_
tx=XM_019834212.2;refGene_gn=MICAL3;refGene_tx=XM_019834214.2;refGene_
gn=MICAL3;refGene_tx=XM_023256561.1;refGene_gn=MICAL3;refGene_tx=XM_02

3256562.1;refGene_gn=MICAL3;refGene_tx=XM_023256563.1;refGene_gn=MICAL3;refGene_tx=XM_023256564.1;refGene_gn=MICAL3;refGene_tx=XM_023256565.1;refGene_gn=MICAL3;refGene_tx=XM_023256566.1;refGene_gn=MICAL3;refGene_tx=XM_023256567.1;refGene_gn=MICAL3;refGene_tx=XM_023256568.1;refGene_gn=MICAL3;refGene_tx=XM_023256569.1;refGene_gn=MICAL3;refGene_tx=XM_023256570.1;refGene_gn=MICAL3;refGene_tx=XM_023256571.1;refGene_gn=MICAL3;refGene_tx=XM_023256572.1;refGene_gn=MICAL3;refGene_tx=XM_023256573.1;refGene_gn=MICAL3;refGene_tx=XM_023256574.1;refGene_gn=MICAL3;refGene_tx=XM_023256575.1;refGene_gn=MICAL3;refGene_tx=XM_023256576.1;refGene_gn=MICAL3;refGene_tx=XM_023256577.1;refGene_gn=MICAL3;refGene_tx=XM_023256578.1
B4 34548107 34548196
refGene_gn=MICAL3;refGene_tx=XM_019834211.2;refGene_gn=MICAL3;refGene_tx=XM_019834212.2;refGene_gn=MICAL3;refGene_tx=XM_019834214.2;refGene_gn=MICAL3;refGene_tx=XM_023256561.1;refGene_gn=MICAL3;refGene_tx=XM_023256562.1;refGene_gn=MICAL3;refGene_tx=XM_023256563.1;refGene_gn=MICAL3;refGene_tx=XM_023256564.1;refGene_gn=MICAL3;refGene_tx=XM_023256565.1;refGene_gn=MICAL3;refGene_tx=XM_023256566.1;refGene_gn=MICAL3;refGene_tx=XM_023256567.1;refGene_gn=MICAL3;refGene_tx=XM_023256568.1;refGene_gn=MICAL3;refGene_tx=XM_023256569.1;refGene_gn=MICAL3;refGene_tx=XM_023256570.1;refGene_gn=MICAL3;refGene_tx=XM_023256571.1;refGene_gn=MICAL3;refGene_tx=XM_023256572.1;refGene_gn=MICAL3;refGene_tx=XM_023256573.1;refGene_gn=MICAL3;refGene_tx=XM_023256574.1;refGene_gn=MICAL3;refGene_tx=XM_023256575.1;refGene_gn=MICAL3;refGene_tx=XM_023256576.1;refGene_gn=MICAL3;refGene_tx=XM_023256577.1;refGene_gn=MICAL3;refGene_tx=XM_023256578.1
B4 34548951 34549039
refGene_gn=MICAL3;refGene_tx=XM_019834211.2;refGene_gn=MICAL3;refGene_tx=XM_019834212.2;refGene_gn=MICAL3;refGene_tx=XM_019834214.2;refGene_gn=MICAL3;refGene_tx=XM_023256561.1;refGene_gn=MICAL3;refGene_tx=XM_023256562.1;refGene_gn=MICAL3;refGene_tx=XM_023256563.1;refGene_gn=MICAL3;refGene_tx=XM_023256564.1;refGene_gn=MICAL3;refGene_tx=XM_023256565.1;refGene_gn=MICAL3;refGene_tx=XM_023256566.1;refGene_gn=MICAL3;refGene_tx=XM_023256567.1;refGene_gn=MICAL3;refGene_tx=XM_023256568.1;refGene_gn=MICAL3;refGene_tx=XM_023256569.1;refGene_gn=MICAL3;refGene_tx=XM_023256570.1;refGene_gn=MICAL3;refGene_tx=XM_023256571.1;refGene_gn=MICAL3;refGene_tx=XM_023256572.1;refGene_gn=MICAL3;refGene_tx=XM_023256573.1;refGene_gn=MICAL3;refGene_tx=XM_023256574.1;refGene_gn=MICAL3;refGene_tx=XM_023256575.1;refGene_gn=MICAL3;refGene_tx=XM_023256576.1;refGene_gn=MICAL3;refGene_tx=XM_023256577.1;refGene_gn=MICAL3;refGene_tx=XM_023256578.1
B4 34570934 34571130
refGene_gn=MICAL3;refGene_tx=XM_019834211.2;refGene_gn=MICAL3;refGene_tx=XM_019834212.2;refGene_gn=MICAL3;refGene_tx=XM_019834214.2;refGene_gn=MICAL3;refGene_tx=XM_023256561.1;refGene_gn=MICAL3;refGene_tx=XM_023256562.1;refGene_gn=MICAL3;refGene_tx=XM_023256563.1;refGene_gn=MICAL3;refGene_tx=XM_023256564.1;refGene_gn=MICAL3;refGene_tx=XM_023256565.1;refGene_gn=MICAL3;refGene_tx=XM_023256566.1;refGene_gn=MICAL3;refGene_tx=XM_023256567.1;refGene_gn=MICAL3;refGene_tx=XM_023256568.1;refGene_gn=MICAL3;refGene_tx=XM_023256569.1;refGene_gn=MICAL3;refGene_tx=XM_

023256570.1;refGene_gn=MICAL3;refGene_tx=XM_023256571.1;refGene_gn=MICAL3;refGene_tx=XM_023256572.1;refGene_gn=MICAL3;refGene_tx=XM_023256573.1;refGene_gn=MICAL3;refGene_tx=XM_023256574.1;refGene_gn=MICAL3;refGene_tx=XM_023256575.1;refGene_gn=MICAL3;refGene_tx=XM_023256576.1;refGene_gn=MICAL3;refGene_tx=XM_023256577.1;refGene_gn=MICAL3;refGene_tx=XM_023256578.1
B4 34579621 34579869
refGene_gn=MICAL3;refGene_tx=XM_019834211.2;refGene_gn=MICAL3;refGene_tx=XM_019834212.2;refGene_gn=MICAL3;refGene_tx=XM_019834214.2;refGene_gn=MICAL3;refGene_tx=XM_023256561.1;refGene_gn=MICAL3;refGene_tx=XM_023256562.1;refGene_gn=MICAL3;refGene_tx=XM_023256563.1;refGene_gn=MICAL3;refGene_tx=XM_023256564.1;refGene_gn=MICAL3;refGene_tx=XM_023256565.1;refGene_gn=MICAL3;refGene_tx=XM_023256566.1;refGene_gn=MICAL3;refGene_tx=XM_023256567.1;refGene_gn=MICAL3;refGene_tx=XM_023256568.1;refGene_gn=MICAL3;refGene_tx=XM_023256569.1;refGene_gn=MICAL3;refGene_tx=XM_023256570.1;refGene_gn=MICAL3;refGene_tx=XM_023256571.1;refGene_gn=MICAL3;refGene_tx=XM_023256572.1;refGene_gn=MICAL3;refGene_tx=XM_023256573.1;refGene_gn=MICAL3;refGene_tx=XM_023256574.1;refGene_gn=MICAL3;refGene_tx=XM_023256575.1;refGene_gn=MICAL3;refGene_tx=XM_023256576.1;refGene_gn=MICAL3;refGene_tx=XM_023256577.1;refGene_gn=MICAL3;refGene_tx=XM_023256578.1
B4 34580666 34580770 refGene_gn=MICAL3;refGene_tx=XM_023256578.1
B4 34581745 34581835
refGene_gn=MICAL3;refGene_tx=XM_019834211.2;refGene_gn=MICAL3;refGene_tx=XM_019834212.2;refGene_gn=MICAL3;refGene_tx=XM_019834214.2;refGene_gn=MICAL3;refGene_tx=XM_023256561.1;refGene_gn=MICAL3;refGene_tx=XM_023256562.1;refGene_gn=MICAL3;refGene_tx=XM_023256563.1;refGene_gn=MICAL3;refGene_tx=XM_023256564.1;refGene_gn=MICAL3;refGene_tx=XM_023256565.1;refGene_gn=MICAL3;refGene_tx=XM_023256566.1;refGene_gn=MICAL3;refGene_tx=XM_023256567.1;refGene_gn=MICAL3;refGene_tx=XM_023256569.1;refGene_gn=MICAL3;refGene_tx=XM_023256570.1;refGene_gn=MICAL3;refGene_tx=XM_023256572.1;refGene_gn=MICAL3;refGene_tx=XM_023256574.1;refGene_gn=MICAL3;refGene_tx=XM_023256575.1
B4 34583887 34584025
refGene_gn=MICAL3;refGene_tx=XM_019834211.2;refGene_gn=MICAL3;refGene_tx=XM_019834212.2;refGene_gn=MICAL3;refGene_tx=XM_019834214.2;refGene_gn=MICAL3;refGene_tx=XM_023256561.1;refGene_gn=MICAL3;refGene_tx=XM_023256562.1;refGene_gn=MICAL3;refGene_tx=XM_023256563.1;refGene_gn=MICAL3;refGene_tx=XM_023256564.1;refGene_gn=MICAL3;refGene_tx=XM_023256565.1;refGene_gn=MICAL3;refGene_tx=XM_023256566.1;refGene_gn=MICAL3;refGene_tx=XM_023256567.1;refGene_gn=MICAL3;refGene_tx=XM_023256568.1;refGene_gn=MICAL3;refGene_tx=XM_023256569.1;refGene_gn=MICAL3;refGene_tx=XM_023256570.1;refGene_gn=MICAL3;refGene_tx=XM_023256572.1;refGene_gn=MICAL3;refGene_tx=XM_023256574.1;refGene_gn=MICAL3;refGene_tx=XM_023256575.1;refGene_gn=MICAL3;refGene_tx=XM_023256576.1;refGene_gn=MICAL3;refGene_tx=XM_023256577.1
B4 34585160 34585223
refGene_gn=MICAL3;refGene_tx=XM_019834211.2;refGene_gn=MICAL3;refGene_tx=XM_019834212.2;refGene_gn=MICAL3;refGene_tx=XM_019834214.2;refGene_gn=MICAL3;refGene_tx=XM_023256561.1;refGene_gn=MICAL3;refGene_tx=XM_02

3256562.1;refGene_gn=MICAL3;refGene_tx=XM_023256563.1;refGene_gn=MICAL3;refGene_tx=XM_023256564.1;refGene_gn=MICAL3;refGene_tx=XM_023256565.1;refGene_gn=MICAL3;refGene_tx=XM_023256567.1;refGene_gn=MICAL3;refGene_tx=XM_023256568.1;refGene_gn=MICAL3;refGene_tx=XM_023256569.1;refGene_gn=MICAL3;refGene_tx=XM_023256570.1;refGene_gn=MICAL3;refGene_tx=XM_023256571.1;refGene_gn=MICAL3;refGene_tx=XM_023256572.1;refGene_gn=MICAL3;refGene_tx=XM_023256574.1;refGene_gn=MICAL3;refGene_tx=XM_023256575.1;refGene_gn=MICAL3;refGene_tx=XM_023256576.1
B4 34589242 34589356
refGene_gn=MICAL3;refGene_tx=XM_019834211.2;refGene_gn=MICAL3;refGene_tx=XM_019834212.2;refGene_gn=MICAL3;refGene_tx=XM_019834214.2;refGene_gn=MICAL3;refGene_tx=XM_023256561.1;refGene_gn=MICAL3;refGene_tx=XM_023256562.1;refGene_gn=MICAL3;refGene_tx=XM_023256563.1;refGene_gn=MICAL3;refGene_tx=XM_023256564.1;refGene_gn=MICAL3;refGene_tx=XM_023256565.1;refGene_gn=MICAL3;refGene_tx=XM_023256566.1;refGene_gn=MICAL3;refGene_tx=XM_023256567.1;refGene_gn=MICAL3;refGene_tx=XM_023256568.1;refGene_gn=MICAL3;refGene_tx=XM_023256569.1;refGene_gn=MICAL3;refGene_tx=XM_023256570.1;refGene_gn=MICAL3;refGene_tx=XM_023256571.1;refGene_gn=MICAL3;refGene_tx=XM_023256572.1;refGene_gn=MICAL3;refGene_tx=XM_023256574.1;refGene_gn=MICAL3;refGene_tx=XM_023256575.1;refGene_gn=MICAL3;refGene_tx=XM_023256576.1;refGene_gn=MICAL3;refGene_tx=XM_023256577.1
B4 34590126 34590273
refGene_gn=MICAL3;refGene_tx=XM_019834211.2;refGene_gn=MICAL3;refGene_tx=XM_019834212.2;refGene_gn=MICAL3;refGene_tx=XM_019834214.2;refGene_gn=MICAL3;refGene_tx=XM_023256561.1;refGene_gn=MICAL3;refGene_tx=XM_023256562.1;refGene_gn=MICAL3;refGene_tx=XM_023256563.1;refGene_gn=MICAL3;refGene_tx=XM_023256564.1;refGene_gn=MICAL3;refGene_tx=XM_023256565.1;refGene_gn=MICAL3;refGene_tx=XM_023256566.1;refGene_gn=MICAL3;refGene_tx=XM_023256567.1;refGene_gn=MICAL3;refGene_tx=XM_023256568.1;refGene_gn=MICAL3;refGene_tx=XM_023256569.1;refGene_gn=MICAL3;refGene_tx=XM_023256570.1;refGene_gn=MICAL3;refGene_tx=XM_023256571.1;refGene_gn=MICAL3;refGene_tx=XM_023256572.1;refGene_gn=MICAL3;refGene_tx=XM_023256573.1;refGene_gn=MICAL3;refGene_tx=XM_023256574.1;refGene_gn=MICAL3;refGene_tx=XM_023256575.1;refGene_gn=MICAL3;refGene_tx=XM_023256576.1;refGene_gn=MICAL3;refGene_tx=XM_023256577.1
B4 34590737 34590820
refGene_gn=MICAL3;refGene_tx=XM_019834211.2;refGene_gn=MICAL3;refGene_tx=XM_019834212.2;refGene_gn=MICAL3;refGene_tx=XM_019834214.2;refGene_gn=MICAL3;refGene_tx=XM_023256561.1;refGene_gn=MICAL3;refGene_tx=XM_023256562.1;refGene_gn=MICAL3;refGene_tx=XM_023256563.1;refGene_gn=MICAL3;refGene_tx=XM_023256564.1;refGene_gn=MICAL3;refGene_tx=XM_023256565.1;refGene_gn=MICAL3;refGene_tx=XM_023256566.1;refGene_gn=MICAL3;refGene_tx=XM_023256567.1;refGene_gn=MICAL3;refGene_tx=XM_023256568.1;refGene_gn=MICAL3;refGene_tx=XM_023256569.1;refGene_gn=MICAL3;refGene_tx=XM_023256570.1;refGene_gn=MICAL3;refGene_tx=XM_023256571.1;refGene_gn=MICAL3;refGene_tx=XM_023256572.1;refGene_gn=MICAL3;refGene_tx=XM_023256573.1;refGene_gn=MICAL3;refGene_tx=XM_023256574.1;refGene_gn=MICAL3;refGene_tx=XM_023256575.1;refGene_gn=MICAL3;refGene_tx=XM_023256576.1;refGene_gn=MICAL3;refGene_tx=XM_023256577.1
B4 34592396 34594248

refGene_gn=MICAL3;refGene_tx=XM_019834211.2;refGene_gn=MICAL3;refGene_tx=XM_019834212.2;refGene_gn=MICAL3;refGene_tx=XM_019834214.2;refGene_gn=MICAL3;refGene_tx=XM_023256561.1;refGene_gn=MICAL3;refGene_tx=XM_023256562.1;refGene_gn=MICAL3;refGene_tx=XM_023256563.1;refGene_gn=MICAL3;refGene_tx=XM_023256564.1;refGene_gn=MICAL3;refGene_tx=XM_023256565.1;refGene_gn=MICAL3;refGene_tx=XM_023256566.1;refGene_gn=MICAL3;refGene_tx=XM_023256567.1;refGene_gn=MICAL3;refGene_tx=XM_023256568.1;refGene_gn=MICAL3;refGene_tx=XM_023256569.1;refGene_gn=MICAL3;refGene_tx=XM_023256570.1;refGene_gn=MICAL3;refGene_tx=XM_023256571.1;refGene_gn=MICAL3;refGene_tx=XM_023256572.1;refGene_gn=MICAL3;refGene_tx=XM_023256573.1;refGene_gn=MICAL3;refGene_tx=XM_023256574.1;refGene_gn=MICAL3;refGene_tx=XM_023256575.1;refGene_gn=MICAL3;refGene_tx=XM_023256576.1;refGene_gn=MICAL3;refGene_tx=XM_023256577.1
B4 34594763 34594858
refGene_gn=MICAL3;refGene_tx=XM_019834211.2;refGene_gn=MICAL3;refGene_tx=XM_019834212.2;refGene_gn=MICAL3;refGene_tx=XM_019834214.2;refGene_gn=MICAL3;refGene_tx=XM_023256561.1;refGene_gn=MICAL3;refGene_tx=XM_023256562.1;refGene_gn=MICAL3;refGene_tx=XM_023256563.1;refGene_gn=MICAL3;refGene_tx=XM_023256564.1;refGene_gn=MICAL3;refGene_tx=XM_023256565.1;refGene_gn=MICAL3;refGene_tx=XM_023256566.1;refGene_gn=MICAL3;refGene_tx=XM_023256567.1;refGene_gn=MICAL3;refGene_tx=XM_023256568.1;refGene_gn=MICAL3;refGene_tx=XM_023256569.1;refGene_gn=MICAL3;refGene_tx=XM_023256570.1;refGene_gn=MICAL3;refGene_tx=XM_023256571.1;refGene_gn=MICAL3;refGene_tx=XM_023256572.1;refGene_gn=MICAL3;refGene_tx=XM_023256573.1;refGene_gn=MICAL3;refGene_tx=XM_023256574.1;refGene_gn=MICAL3;refGene_tx=XM_023256575.1;refGene_gn=MICAL3;refGene_tx=XM_023256576.1;refGene_gn=MICAL3;refGene_tx=XM_023256577.1
B4 34599268 34599319
refGene_gn=MICAL3;refGene_tx=XM_019834211.2;refGene_gn=MICAL3;refGene_tx=XM_019834212.2;refGene_gn=MICAL3;refGene_tx=XM_019834214.2;refGene_gn=MICAL3;refGene_tx=XM_023256561.1;refGene_gn=MICAL3;refGene_tx=XM_023256562.1;refGene_gn=MICAL3;refGene_tx=XM_023256563.1;refGene_gn=MICAL3;refGene_tx=XM_023256564.1;refGene_gn=MICAL3;refGene_tx=XM_023256566.1;refGene_gn=MICAL3;refGene_tx=XM_023256567.1;refGene_gn=MICAL3;refGene_tx=XM_023256568.1;refGene_gn=MICAL3;refGene_tx=XM_023256569.1;refGene_gn=MICAL3;refGene_tx=XM_023256571.1;refGene_gn=MICAL3;refGene_tx=XM_023256572.1;refGene_gn=MICAL3;refGene_tx=XM_023256573.1;refGene_gn=MICAL3;refGene_tx=XM_023256574.1;refGene_gn=MICAL3;refGene_tx=XM_023256575.1;refGene_gn=MICAL3;refGene_tx=XM_023256576.1
B4 34600722 34600833
refGene_gn=MICAL3;refGene_tx=XM_019834211.2;refGene_gn=MICAL3;refGene_tx=XM_019834212.2;refGene_gn=MICAL3;refGene_tx=XM_019834214.2;refGene_gn=MICAL3;refGene_tx=XM_023256561.1;refGene_gn=MICAL3;refGene_tx=XM_023256562.1;refGene_gn=MICAL3;refGene_tx=XM_023256563.1;refGene_gn=MICAL3;refGene_tx=XM_023256564.1;refGene_gn=MICAL3;refGene_tx=XM_023256565.1;refGene_gn=MICAL3;refGene_tx=XM_023256566.1;refGene_gn=MICAL3;refGene_tx=XM_023256567.1;refGene_gn=MICAL3;refGene_tx=XM_023256568.1;refGene_gn=MICAL3;refGene_tx=XM_023256569.1;refGene_gn=MICAL3;refGene_tx=XM_023256570.1;refGene_gn=MICAL3;refGene_tx=XM_023256571.1;refGene_gn=MICAL3;refGene_tx=XM_023256572.1;refGene_gn=MICAL3;refGene_tx=XM_02325657

3.1;refGene_gn=MICAL3;refGene_tx=XM_023256574.1;refGene_gn=MICAL3;refGene_tx=XM_023256575.1;refGene_gn=MICAL3;refGene_tx=XM_023256576.1;refGene_gn=MICAL3;refGene_tx=XM_023256577.1
B4 34602385 34602479
refGene_gn=MICAL3;refGene_tx=XM_019834211.2;refGene_gn=MICAL3;refGene_tx=XM_019834212.2;refGene_gn=MICAL3;refGene_tx=XM_019834214.2;refGene_gn=MICAL3;refGene_tx=XM_023256561.1;refGene_gn=MICAL3;refGene_tx=XM_023256562.1;refGene_gn=MICAL3;refGene_tx=XM_023256563.1;refGene_gn=MICAL3;refGene_tx=XM_023256564.1;refGene_gn=MICAL3;refGene_tx=XM_023256565.1;refGene_gn=MICAL3;refGene_tx=XM_023256566.1;refGene_gn=MICAL3;refGene_tx=XM_023256567.1;refGene_gn=MICAL3;refGene_tx=XM_023256568.1;refGene_gn=MICAL3;refGene_tx=XM_023256569.1;refGene_gn=MICAL3;refGene_tx=XM_023256570.1;refGene_gn=MICAL3;refGene_tx=XM_023256571.1;refGene_gn=MICAL3;refGene_tx=XM_023256572.1;refGene_gn=MICAL3;refGene_tx=XM_023256573.1;refGene_gn=MICAL3;refGene_tx=XM_023256574.1;refGene_gn=MICAL3;refGene_tx=XM_023256575.1;refGene_gn=MICAL3;refGene_tx=XM_023256576.1;refGene_gn=MICAL3;refGene_tx=XM_023256577.1
B4 34607420 34607471 refGene_gn=MICAL3;refGene_tx=XM_023256563.1
B4 34616229 34616295
refGene_gn=MICAL3;refGene_tx=XM_019834211.2;refGene_gn=MICAL3;refGene_tx=XM_019834212.2;refGene_gn=MICAL3;refGene_tx=XM_019834214.2;refGene_gn=MICAL3;refGene_tx=XM_023256561.1;refGene_gn=MICAL3;refGene_tx=XM_023256562.1;refGene_gn=MICAL3;refGene_tx=XM_023256564.1;refGene_gn=MICAL3;refGene_tx=XM_023256565.1;refGene_gn=MICAL3;refGene_tx=XM_023256566.1;refGene_gn=MICAL3;refGene_tx=XM_023256568.1;refGene_gn=MICAL3;refGene_tx=XM_023256569.1;refGene_gn=MICAL3;refGene_tx=XM_023256571.1;refGene_gn=MICAL3;refGene_tx=XM_023256572.1;refGene_gn=MICAL3;refGene_tx=XM_023256573.1;refGene_gn=MICAL3;refGene_tx=XM_023256574.1;refGene_gn=MICAL3;refGene_tx=XM_023256575.1;refGene_gn=MICAL3;refGene_tx=XM_023256576.1
B4 34619010 34619110
refGene_gn=MICAL3;refGene_tx=XM_019834211.2;refGene_gn=MICAL3;refGene_tx=XM_019834212.2;refGene_gn=MICAL3;refGene_tx=XM_019834214.2;refGene_gn=MICAL3;refGene_tx=XM_023256561.1;refGene_gn=MICAL3;refGene_tx=XM_023256562.1;refGene_gn=MICAL3;refGene_tx=XM_023256563.1;refGene_gn=MICAL3;refGene_tx=XM_023256564.1;refGene_gn=MICAL3;refGene_tx=XM_023256565.1;refGene_gn=MICAL3;refGene_tx=XM_023256566.1;refGene_gn=MICAL3;refGene_tx=XM_023256567.1;refGene_gn=MICAL3;refGene_tx=XM_023256568.1;refGene_gn=MICAL3;refGene_tx=XM_023256569.1;refGene_gn=MICAL3;refGene_tx=XM_023256570.1;refGene_gn=MICAL3;refGene_tx=XM_023256571.1;refGene_gn=MICAL3;refGene_tx=XM_023256572.1;refGene_gn=MICAL3;refGene_tx=XM_023256573.1;refGene_gn=MICAL3;refGene_tx=XM_023256574.1;refGene_gn=MICAL3;refGene_tx=XM_023256575.1;refGene_gn=MICAL3;refGene_tx=XM_023256576.1;refGene_gn=MICAL3;refGene_tx=XM_023256577.1
B4 34619261 34619335
refGene_gn=MICAL3;refGene_tx=XM_019834211.2;refGene_gn=MICAL3;refGene_tx=XM_019834212.2;refGene_gn=MICAL3;refGene_tx=XM_019834214.2;refGene_gn=MICAL3;refGene_tx=XM_023256561.1;refGene_gn=MICAL3;refGene_tx=XM_023256562.1;refGene_gn=MICAL3;refGene_tx=XM_023256563.1;refGene_gn=MICAL3;refGene_tx=XM_023256564.1;refGene_gn=MICAL3;refGene_tx=XM_023256565.

1;refGene_gn=MICAL3;refGene_tx=XM_023256566.1;refGene_gn=MICAL3;refGene_tx=XM_023256567.1;refGene_gn=MICAL3;refGene_tx=XM_023256568.1;refGene_gn=MICAL3;refGene_tx=XM_023256569.1;refGene_gn=MICAL3;refGene_tx=XM_023256570.1;refGene_gn=MICAL3;refGene_tx=XM_023256571.1;refGene_gn=MICAL3;refGene_tx=XM_023256572.1;refGene_gn=MICAL3;refGene_tx=XM_023256573.1;refGene_gn=MICAL3;refGene_tx=XM_023256574.1;refGene_gn=MICAL3;refGene_tx=XM_023256575.1;refGene_gn=MICAL3;refGene_tx=XM_023256576.1;refGene_gn=MICAL3;refGene_tx=XM_023256577.1
B4 34619422 34619607
refGene_gn=MICAL3;refGene_tx=XM_019834211.2;refGene_gn=MICAL3;refGene_tx=XM_019834212.2;refGene_gn=MICAL3;refGene_tx=XM_019834214.2;refGene_gn=MICAL3;refGene_tx=XM_023256561.1;refGene_gn=MICAL3;refGene_tx=XM_023256562.1;refGene_gn=MICAL3;refGene_tx=XM_023256563.1;refGene_gn=MICAL3;refGene_tx=XM_023256564.1;refGene_gn=MICAL3;refGene_tx=XM_023256565.1;refGene_gn=MICAL3;refGene_tx=XM_023256566.1;refGene_gn=MICAL3;refGene_tx=XM_023256567.1;refGene_gn=MICAL3;refGene_tx=XM_023256568.1;refGene_gn=MICAL3;refGene_tx=XM_023256569.1;refGene_gn=MICAL3;refGene_tx=XM_023256570.1;refGene_gn=MICAL3;refGene_tx=XM_023256571.1;refGene_gn=MICAL3;refGene_tx=XM_023256572.1;refGene_gn=MICAL3;refGene_tx=XM_023256573.1;refGene_gn=MICAL3;refGene_tx=XM_023256574.1;refGene_gn=MICAL3;refGene_tx=XM_023256575.1;refGene_gn=MICAL3;refGene_tx=XM_023256576.1;refGene_gn=MICAL3;refGene_tx=XM_023256577.1
B4 34661050 34661062 refGene_gn=BID;refGene_tx=XM_003988294.5
B4 34664818 34665032 refGene_gn=BID;refGene_tx=XM_003988294.5
B4 34668159 34668299 refGene_gn=BID;refGene_tx=XM_003988294.5
B4 34670501 34670714 refGene_gn=BID;refGene_tx=XM_003988294.5
B4 34671900 34671912 refGene_gn=BID;refGene_tx=XM_003988294.5
B4 34680303 34681122
refGene_gn=BCL2L13;refGene_tx=XM_023256579.1;refGene_gn=BCL2L13;refGene_tx=XM_023256580.1;refGene_gn=BCL2L13;refGene_tx=XM_023256581.1;refGene_gn=BCL2L13;refGene_tx=XM_023256582.1;refGene_gn=BCL2L13;refGene_tx=XM_023256583.1;refGene_gn=BCL2L13;refGene_tx=XM_023256584.1;refGene_gn=BCL2L13;refGene_tx=XM_023256585.1;refGene_gn=BCL2L13;refGene_tx=XM_023256586.1
B4 34696794 34696876
refGene_gn=BCL2L13;refGene_tx=XM_023256583.1;refGene_gn=BCL2L13;refGene_tx=XM_023256586.1
B4 34714825 34714969
refGene_gn=BCL2L13;refGene_tx=XM_023256579.1;refGene_gn=BCL2L13;refGene_tx=XM_023256580.1;refGene_gn=BCL2L13;refGene_tx=XM_023256581.1;refGene_gn=BCL2L13;refGene_tx=XM_023256582.1;refGene_gn=BCL2L13;refGene_tx=XM_023256583.1;refGene_gn=BCL2L13;refGene_tx=XM_023256584.1;refGene_gn=BCL2L13;refGene_tx=XM_023256585.1
B4 34723352 34723446
refGene_gn=BCL2L13;refGene_tx=XM_023256579.1;refGene_gn=BCL2L13;refGene_tx=XM_023256580.1;refGene_gn=BCL2L13;refGene_tx=XM_023256581.1;refGene_gn=BCL2L13;refGene_tx=XM_023256582.1
B4 34727559 34727716
refGene_gn=BCL2L13;refGene_tx=XM_023256579.1;refGene_gn=BCL2L13;refGene_tx=XM_023256580.1;refGene_gn=BCL2L13;refGene_tx=XM_023256581.1;refGe

ne_gn=BCL2L13;refGene_tx=XM_023256582.1
B4 34732417 34732507
refGene_gn=BCL2L13;refGene_tx=XM_023256579.1;refGene_gn=BCL2L13;refGene_tx=XM_023256580.1;refGene_gn=BCL2L13;refGene_tx=XM_023256581.1
B4 34755728 34755849
refGene_gn=BCL2L13;refGene_tx=XM_023256579.1;refGene_gn=BCL2L13;refGene_tx=XM_023256580.1;refGene_gn=BCL2L13;refGene_tx=XM_023256581.1;refGene_gn=BCL2L13;refGene_tx=XM_023256582.1
B4 34781295 34781328 refGene_gn=ATP6V1E1;refGene_tx=XM_003988246.5
B4 34786852 34786918 refGene_gn=ATP6V1E1;refGene_tx=XM_003988246.5
B4 34789045 34789155 refGene_gn=ATP6V1E1;refGene_tx=XM_003988246.5
B4 34789530 34789597 refGene_gn=ATP6V1E1;refGene_tx=XM_003988246.5
B4 34803988 34804078 refGene_gn=ATP6V1E1;refGene_tx=XM_003988246.5
B4 34804659 34804728 refGene_gn=ATP6V1E1;refGene_tx=XM_003988246.5
B4 34807347 34807442 refGene_gn=ATP6V1E1;refGene_tx=XM_003988246.5
B4 34810789 34810877 refGene_gn=ATP6V1E1;refGene_tx=XM_003988246.5
B4 34812361 34812424 refGene_gn=ATP6V1E1;refGene_tx=XM_003988246.5
B4 34814564 34814706
refGene_gn=SLC25A18;refGene_tx=XM_011283695.3;refGene_gn=SLC25A18;refGene_tx=XM_019834229.2;refGene_gn=SLC25A18;refGene_tx=XM_019834230.2;refGene_gn=SLC25A18;refGene_tx=XM_019834231.2;refGene_gn=SLC25A18;refGene_tx=XM_019834232.2;refGene_gn=SLC25A18;refGene_tx=XM_019834233.2;refGene_gn=SLC25A18;refGene_tx=XM_019834234.2;refGene_gn=SLC25A18;refGene_tx=XM_019834235.2;refGene_gn=SLC25A18;refGene_tx=XM_019834236.2
B4 34815139 34815215
refGene_gn=SLC25A18;refGene_tx=XM_011283695.3;refGene_gn=SLC25A18;refGene_tx=XM_019834229.2;refGene_gn=SLC25A18;refGene_tx=XM_019834230.2;refGene_gn=SLC25A18;refGene_tx=XM_019834231.2;refGene_gn=SLC25A18;refGene_tx=XM_019834232.2;refGene_gn=SLC25A18;refGene_tx=XM_019834233.2;refGene_gn=SLC25A18;refGene_tx=XM_019834234.2;refGene_gn=SLC25A18;refGene_tx=XM_019834235.2;refGene_gn=SLC25A18;refGene_tx=XM_019834236.2
B4 34815802 34815957
refGene_gn=SLC25A18;refGene_tx=XM_011283695.3;refGene_gn=SLC25A18;refGene_tx=XM_019834229.2;refGene_gn=SLC25A18;refGene_tx=XM_019834230.2;refGene_gn=SLC25A18;refGene_tx=XM_019834231.2;refGene_gn=SLC25A18;refGene_tx=XM_019834232.2;refGene_gn=SLC25A18;refGene_tx=XM_019834233.2;refGene_gn=SLC25A18;refGene_tx=XM_019834234.2;refGene_gn=SLC25A18;refGene_tx=XM_019834235.2;refGene_gn=SLC25A18;refGene_tx=XM_019834236.2
B4 34816779 34816945
refGene_gn=SLC25A18;refGene_tx=XM_011283695.3;refGene_gn=SLC25A18;refGene_tx=XM_019834229.2;refGene_gn=SLC25A18;refGene_tx=XM_019834230.2;refGene_gn=SLC25A18;refGene_tx=XM_019834231.2;refGene_gn=SLC25A18;refGene_tx=XM_019834232.2;refGene_gn=SLC25A18;refGene_tx=XM_019834233.2;refGene_gn=SLC25A18;refGene_tx=XM_019834234.2;refGene_gn=SLC25A18;refGene_tx=XM_019834235.2;refGene_gn=SLC25A18;refGene_tx=XM_019834236.2
B4 34820078 34820197
refGene_gn=SLC25A18;refGene_tx=XM_011283695.3;refGene_gn=SLC25A18;refGene_tx=XM_019834229.2;refGene_gn=SLC25A18;refGene_tx=XM_019834230.2;refGene_gn=SLC25A18;refGene_tx=XM_019834231.2;refGene_gn=SLC25A18;refGene_tx=XM_019834232.2;refGene_gn=SLC25A18;refGene_tx=XM_019834233.2;refG

ene_gn=SLC25A18;refGene_tx=XM_019834234.2;refGene_gn=SLC25A18;refGene_
tx=XM_019834235.2;refGene_gn=SLC25A18;refGene_tx=XM_019834236.2
B4 34822769 34822860
refGene_gn=SLC25A18;refGene_tx=XM_011283695.3;refGene_gn=SLC25A18;refG
ene_tx=XM_019834229.2;refGene_gn=SLC25A18;refGene_tx=XM_019834230.2;re
fGene_gn=SLC25A18;refGene_tx=XM_019834231.2;refGene_gn=SLC25A18;refGen
e_tx=XM_019834232.2;refGene_gn=SLC25A18;refGene_tx=XM_019834233.2;refG
ene_gn=SLC25A18;refGene_tx=XM_019834234.2;refGene_gn=SLC25A18;refGene_
tx=XM_019834235.2;refGene_gn=SLC25A18;refGene_tx=XM_019834236.2
B4 34824002 34824058
refGene_gn=SLC25A18;refGene_tx=XM_011283695.3;refGene_gn=SLC25A18;refG
ene_tx=XM_019834229.2;refGene_gn=SLC25A18;refGene_tx=XM_019834230.2;re
fGene_gn=SLC25A18;refGene_tx=XM_019834231.2;refGene_gn=SLC25A18;refGen
e_tx=XM_019834232.2;refGene_gn=SLC25A18;refGene_tx=XM_019834233.2;refG
ene_gn=SLC25A18;refGene_tx=XM_019834234.2;refGene_gn=SLC25A18;refGene_
tx=XM_019834235.2;refGene_gn=SLC25A18;refGene_tx=XM_019834236.2
B4 34824302 34824425
refGene_gn=SLC25A18;refGene_tx=XM_011283695.3;refGene_gn=SLC25A18;refG
ene_tx=XM_019834229.2;refGene_gn=SLC25A18;refGene_tx=XM_019834230.2;re
fGene_gn=SLC25A18;refGene_tx=XM_019834231.2;refGene_gn=SLC25A18;refGen
e_tx=XM_019834232.2;refGene_gn=SLC25A18;refGene_tx=XM_019834233.2;refG
ene_gn=SLC25A18;refGene_tx=XM_019834234.2;refGene_gn=SLC25A18;refGene_
tx=XM_019834235.2;refGene_gn=SLC25A18;refGene_tx=XM_019834236.2
B4 34825754 34825955
refGene_gn=SLC25A18;refGene_tx=XM_011283695.3;refGene_gn=SLC25A18;refG
ene_tx=XM_019834229.2;refGene_gn=SLC25A18;refGene_tx=XM_019834230.2;re
fGene_gn=SLC25A18;refGene_tx=XM_019834231.2;refGene_gn=SLC25A18;refGen
e_tx=XM_019834232.2;refGene_gn=SLC25A18;refGene_tx=XM_019834233.2;refG
ene_gn=SLC25A18;refGene_tx=XM_019834234.2;refGene_gn=SLC25A18;refGene_
tx=XM_019834235.2;refGene_gn=SLC25A18;refGene_tx=XM_019834236.2
B4 34836890 34836988
refGene_gn=SLC25A18;refGene_tx=XM_019834229.2;refGene_gn=SLC25A18;refG
ene_tx=XM_019834230.2
B4 34851770 34851776
refGene_gn=LOC101094605;refGene_tx=XM_023256587.1;refGene_gn=LOC101094
605;refGene_tx=XM_023256588.1;refGene_gn=LOC101094605;refGene_tx=XM_02
3256589.1;refGene_gn=LOC101094605;refGene_tx=XM_023256590.1;refGene_gn
=LOC101094605;refGene_tx=XM_023256591.1
B4 34852523 34852635
refGene_gn=LOC101094605;refGene_tx=XM_023256587.1;refGene_gn=LOC101094
605;refGene_tx=XM_023256588.1;refGene_gn=LOC101094605;refGene_tx=XM_02
3256589.1;refGene_gn=LOC101094605;refGene_tx=XM_023256590.1;refGene_gn
=LOC101094605;refGene_tx=XM_023256591.1
B4 34854814 34856234
refGene_gn=LOC101094605;refGene_tx=XM_023256587.1;refGene_gn=LOC101094
605;refGene_tx=XM_023256588.1;refGene_gn=LOC101094605;refGene_tx=XM_02
3256589.1;refGene_gn=LOC101094605;refGene_tx=XM_023256590.1;refGene_gn
=LOC101094605;refGene_tx=XM_023256591.1
B4 34866865 34867712
refGene_gn=LOC101094605;refGene_tx=XM_023256587.1;refGene_gn=LOC101094

605;refGene_tx=XM_023256588.1;refGene_gn=LOC101094605;refGene_tx=XM_023256589.1;refGene_gn=LOC101094605;refGene_tx=XM_023256590.1;refGene_gn=LOC101094605;refGene_tx=XM_023256591.1
B4 34867901 34868030
refGene_gn=LOC101094605;refGene_tx=XM_023256587.1;refGene_gn=LOC101094605;refGene_tx=XM_023256588.1;refGene_gn=LOC101094605;refGene_tx=XM_023256589.1;refGene_gn=LOC101094605;refGene_tx=XM_023256590.1;refGene_gn=LOC101094605;refGene_tx=XM_023256591.1
B4 34868651 34869040
refGene_gn=LOC101094605;refGene_tx=XM_023256587.1;refGene_gn=LOC101094605;refGene_tx=XM_023256588.1;refGene_gn=LOC101094605;refGene_tx=XM_023256589.1;refGene_gn=LOC101094605;refGene_tx=XM_023256590.1;refGene_gn=LOC101094605;refGene_tx=XM_023256591.1
B4 34869782 34869909
refGene_gn=LOC101094605;refGene_tx=XM_023256587.1;refGene_gn=LOC101094605;refGene_tx=XM_023256588.1;refGene_gn=LOC101094605;refGene_tx=XM_023256589.1;refGene_gn=LOC101094605;refGene_tx=XM_023256590.1;refGene_gn=LOC101094605;refGene_tx=XM_023256591.1
B4 34870124 34870216
refGene_gn=LOC101094605;refGene_tx=XM_023256587.1;refGene_gn=LOC101094605;refGene_tx=XM_023256588.1;refGene_gn=LOC101094605;refGene_tx=XM_023256589.1;refGene_gn=LOC101094605;refGene_tx=XM_023256590.1;refGene_gn=LOC101094605;refGene_tx=XM_023256591.1
B4 34870304 34870342
refGene_gn=LOC101094605;refGene_tx=XM_023256587.1;refGene_gn=LOC101094605;refGene_tx=XM_023256588.1;refGene_gn=LOC101094605;refGene_tx=XM_023256589.1;refGene_gn=LOC101094605;refGene_tx=XM_023256590.1;refGene_gn=LOC101094605;refGene_tx=XM_023256591.1
B4 34870847 34870977
refGene_gn=LOC101094605;refGene_tx=XM_023256587.1;refGene_gn=LOC101094605;refGene_tx=XM_023256588.1;refGene_gn=LOC101094605;refGene_tx=XM_023256589.1;refGene_gn=LOC101094605;refGene_tx=XM_023256590.1;refGene_gn=LOC101094605;refGene_tx=XM_023256591.1
B4 34881677 34881831
refGene_gn=LOC101094605;refGene_tx=XM_023256587.1;refGene_gn=LOC101094605;refGene_tx=XM_023256588.1;refGene_gn=LOC101094605;refGene_tx=XM_023256589.1;refGene_gn=LOC101094605;refGene_tx=XM_023256590.1;refGene_gn=LOC101094605;refGene_tx=XM_023256591.1
B4 34892295 34892379
refGene_gn=LOC101094605;refGene_tx=XM_023256587.1;refGene_gn=LOC101094605;refGene_tx=XM_023256588.1;refGene_gn=LOC101094605;refGene_tx=XM_023256590.1;refGene_gn=LOC101094605;refGene_tx=XM_023256591.1
B4 34899900 34900070
refGene_gn=LOC101094605;refGene_tx=XM_023256587.1;refGene_gn=LOC101094605;refGene_tx=XM_023256588.1;refGene_gn=LOC101094605;refGene_tx=XM_023256589.1;refGene_gn=LOC101094605;refGene_tx=XM_023256590.1;refGene_gn=LOC101094605;refGene_tx=XM_023256591.1
B4 34901057 34901107
refGene_gn=LOC101094605;refGene_tx=XM_023256587.1;refGene_gn=LOC101094605;refGene_tx=XM_023256588.1;refGene_gn=LOC101094605;refGene_tx=XM_02

3256589.1;refGene_gn=LOC101094605;refGene_tx=XM_023256590.1;refGene_gn=LOC101094605;refGene_tx=XM_023256591.1
B4 34903158 34903263
refGene_gn=LOC101094605;refGene_tx=XM_023256587.1;refGene_gn=LOC101094605;refGene_tx=XM_023256588.1;refGene_gn=LOC101094605;refGene_tx=XM_023256589.1;refGene_gn=LOC101094605;refGene_tx=XM_023256590.1;refGene_gn=LOC101094605;refGene_tx=XM_023256591.1
B4 34904220 34904360
refGene_gn=LOC101094605;refGene_tx=XM_023256587.1;refGene_gn=LOC101094605;refGene_tx=XM_023256588.1;refGene_gn=LOC101094605;refGene_tx=XM_023256589.1;refGene_gn=LOC101094605;refGene_tx=XM_023256590.1;refGene_gn=LOC101094605;refGene_tx=XM_023256591.1
B4 34905556 34905740
refGene_gn=LOC101094605;refGene_tx=XM_023256587.1;refGene_gn=LOC101094605;refGene_tx=XM_023256588.1;refGene_gn=LOC101094605;refGene_tx=XM_023256589.1;refGene_gn=LOC101094605;refGene_tx=XM_023256591.1
B4 34928129 34928224
refGene_gn=LOC101094605;refGene_tx=XM_023256587.1;refGene_gn=LOC101094605;refGene_tx=XM_023256588.1;refGene_gn=LOC101094605;refGene_tx=XM_023256589.1;refGene_gn=LOC101094605;refGene_tx=XM_023256591.1
B4 35029534 35029846
refGene_gn=LOC101094605;refGene_tx=XM_023256587.1;refGene_gn=LOC101094605;refGene_tx=XM_023256588.1;refGene_gn=LOC101094605;refGene_tx=XM_023256589.1;refGene_gn=LOC101094605;refGene_tx=XM_023256591.1
B4 35143749 35144071
refGene_gn=ADA2;refGene_tx=XM_023256592.1;refGene_gn=ADA2;refGene_tx=XM_023256593.1
B4 35146618 35146844
refGene_gn=ADA2;refGene_tx=XM_023256592.1;refGene_gn=ADA2;refGene_tx=XM_023256593.1;refGene_gn=ADA2;refGene_tx=XM_023256594.1
B4 35148252 35148463
refGene_gn=ADA2;refGene_tx=XM_023256592.1;refGene_gn=ADA2;refGene_tx=XM_023256593.1;refGene_gn=ADA2;refGene_tx=XM_023256594.1
B4 35158573 35158701
refGene_gn=ADA2;refGene_tx=XM_023256592.1;refGene_gn=ADA2;refGene_tx=XM_023256593.1;refGene_gn=ADA2;refGene_tx=XM_023256594.1
B4 35160410 35160501
refGene_gn=ADA2;refGene_tx=XM_023256592.1;refGene_gn=ADA2;refGene_tx=XM_023256593.1;refGene_gn=ADA2;refGene_tx=XM_023256594.1
B4 35162243 35162352
refGene_gn=ADA2;refGene_tx=XM_023256592.1;refGene_gn=ADA2;refGene_tx=XM_023256593.1;refGene_gn=ADA2;refGene_tx=XM_023256594.1
B4 35165508 35165666
refGene_gn=ADA2;refGene_tx=XM_023256592.1;refGene_gn=ADA2;refGene_tx=XM_023256593.1;refGene_gn=ADA2;refGene_tx=XM_023256594.1
B4 35165992 35166082 refGene_gn=ADA2;refGene_tx=XM_023256593.1
B4 35166210 35166413
refGene_gn=ADA2;refGene_tx=XM_023256592.1;refGene_gn=ADA2;refGene_tx=XM_023256594.1
B4 35166653 35166753

refGene_gn=ADA2;refGene_tx=XM_023256592.1;refGene_gn=ADA2;refGene_tx=XM_023256594.1
B4 35169091 35169095 refGene_gn=ADA2;refGene_tx=XM_023256592.1
B4 35191413 35191539 refGene_gn=HDHD5;refGene_tx=XM_003988298.4
B4 35198480 35198684 refGene_gn=HDHD5;refGene_tx=XM_003988298.4
B4 35199979 35200092 refGene_gn=HDHD5;refGene_tx=XM_003988298.4
B4 35201903 35201997 refGene_gn=HDHD5;refGene_tx=XM_003988298.4
B4 35203322 35203356 refGene_gn=HDHD5;refGene_tx=XM_003988298.4
B4 35205565 35205740 refGene_gn=HDHD5;refGene_tx=XM_003988298.4
B4 35209449 35209638 refGene_gn=HDHD5;refGene_tx=XM_003988298.4
B4 35209826 35210160 refGene_gn=HDHD5;refGene_tx=XM_003988298.4
B4 35237064 35237124 refGene_gn=TMEM121B;refGene_tx=XM_019834246.2
B4 35237125 35238844 refGene_gn=TMEM121B;refGene_tx=XM_019834246.2
B4 35248035 35249534
refGene_gn=IL17RA;refGene_tx=XM_023256595.1;refGene_gn=IL17RA;refGene_tx=XM_023256596.1;refGene_gn=IL17RA;refGene_tx=XM_023256597.1;refGene_gn=IL17RA;refGene_tx=XM_023256598.1
B4 35250133 35250172
refGene_gn=IL17RA;refGene_tx=XM_023256595.1;refGene_gn=IL17RA;refGene_tx=XM_023256596.1;refGene_gn=IL17RA;refGene_tx=XM_023256597.1;refGene_gn=IL17RA;refGene_tx=XM_023256598.1
B4 35252131 35252233
refGene_gn=IL17RA;refGene_tx=XM_023256595.1;refGene_gn=IL17RA;refGene_tx=XM_023256596.1
B4 35252501 35252513
refGene_gn=IL17RA;refGene_tx=XM_023256595.1;refGene_gn=IL17RA;refGene_tx=XM_023256597.1
B4 35253329 35253414
refGene_gn=IL17RA;refGene_tx=XM_023256595.1;refGene_gn=IL17RA;refGene_tx=XM_023256596.1;refGene_gn=IL17RA;refGene_tx=XM_023256597.1;refGene_gn=IL17RA;refGene_tx=XM_023256598.1
B4 35253776 35253860
refGene_gn=IL17RA;refGene_tx=XM_023256595.1;refGene_gn=IL17RA;refGene_tx=XM_023256596.1;refGene_gn=IL17RA;refGene_tx=XM_023256597.1;refGene_gn=IL17RA;refGene_tx=XM_023256598.1
B4 35254551 35254715
refGene_gn=IL17RA;refGene_tx=XM_023256595.1;refGene_gn=IL17RA;refGene_tx=XM_023256596.1;refGene_gn=IL17RA;refGene_tx=XM_023256597.1;refGene_gn=IL17RA;refGene_tx=XM_023256598.1
B4 35254817 35254865
refGene_gn=IL17RA;refGene_tx=XM_023256595.1;refGene_gn=IL17RA;refGene_tx=XM_023256596.1;refGene_gn=IL17RA;refGene_tx=XM_023256597.1;refGene_gn=IL17RA;refGene_tx=XM_023256598.1
B4 35256447 35256574
refGene_gn=IL17RA;refGene_tx=XM_023256595.1;refGene_gn=IL17RA;refGene_tx=XM_023256596.1;refGene_gn=IL17RA;refGene_tx=XM_023256597.1;refGene_gn=IL17RA;refGene_tx=XM_023256598.1
B4 35257944 35258057
refGene_gn=IL17RA;refGene_tx=XM_023256595.1;refGene_gn=IL17RA;refGene_tx=XM_023256596.1;refGene_gn=IL17RA;refGene_tx=XM_023256597.1;refGene_

gn=IL17RA;refGene_tx=XM_023256598.1
B4 35259031 35259178
refGene_gn=IL17RA;refGene_tx=XM_023256595.1;refGene_gn=IL17RA;refGene_tx=XM_023256596.1;refGene_gn=IL17RA;refGene_tx=XM_023256597.1;refGene_gn=IL17RA;refGene_tx=XM_023256598.1
B4 35259904 35259929
refGene_gn=IL17RA;refGene_tx=XM_023256595.1;refGene_gn=IL17RA;refGene_tx=XM_023256596.1;refGene_gn=IL17RA;refGene_tx=XM_023256597.1;refGene_gn=IL17RA;refGene_tx=XM_023256598.1
B4 35269644 35269788
refGene_gn=IL17RA;refGene_tx=XM_023256595.1;refGene_gn=IL17RA;refGene_tx=XM_023256596.1;refGene_gn=IL17RA;refGene_tx=XM_023256597.1;refGene_gn=IL17RA;refGene_tx=XM_023256598.1
B4 35300723 35301464
refGene_gn=LOC102899890;refGene_tx=XM_006933384.3;refGene_gn=LOC102899890;refGene_tx=XM_006933386.3
B4 35321646 35321949 refGene_gn=CACNA1C;refGene_tx=XM_023256599.1
B4 35323874 35324210 refGene_gn=CACNA1C;refGene_tx=XM_023256599.1
B4 35326014 35326118 refGene_gn=CACNA1C;refGene_tx=XM_023256599.1
B4 35326444 35326551 refGene_gn=CACNA1C;refGene_tx=XM_023256599.1
B4 35330058 35330187 refGene_gn=CACNA1C;refGene_tx=XM_023256599.1
B4 35332538 35332565 refGene_gn=CACNA1C;refGene_tx=XM_023256599.1
B4 35333165 35333518 refGene_gn=CACNA1C;refGene_tx=XM_023256599.1
B4 35335068 35335203 refGene_gn=CACNA1C;refGene_tx=XM_023256599.1
B4 35335628 35335756 refGene_gn=CACNA1C;refGene_tx=XM_023256599.1
B4 35338802 35338904 refGene_gn=CACNA1C;refGene_tx=XM_023256599.1
B4 35349039 35349142 refGene_gn=CACNA1C;refGene_tx=XM_023256599.1
B4 35350956 35351053 refGene_gn=CACNA1C;refGene_tx=XM_023256599.1
B4 35351706 35351834 refGene_gn=CACNA1C;refGene_tx=XM_023256599.1
B4 35352431 35352597 refGene_gn=CACNA1C;refGene_tx=XM_023256599.1
B4 35360663 35360755 refGene_gn=CACNA1C;refGene_tx=XM_023256599.1
B4 35362317 35362383 refGene_gn=CACNA1C;refGene_tx=XM_023256599.1
B4 35364871 35365000 refGene_gn=CACNA1C;refGene_tx=XM_023256599.1
B4 35368295 35368328 refGene_gn=CACNA1C;refGene_tx=XM_023256599.1
B4 35382363 35382447 refGene_gn=CACNA1C;refGene_tx=XM_023256599.1
B4 35383000 35383084 refGene_gn=CACNA1C;refGene_tx=XM_023256599.1
B4 35399774 35399885 refGene_gn=CACNA1C;refGene_tx=XM_023256599.1
B4 35400504 35400663 refGene_gn=CACNA1C;refGene_tx=XM_023256599.1
B4 35402234 35402436 refGene_gn=CACNA1C;refGene_tx=XM_023256599.1
B4 35403803 35403950 refGene_gn=CACNA1C;refGene_tx=XM_023256599.1
B4 35404274 35404327 refGene_gn=CACNA1C;refGene_tx=XM_023256599.1
B4 35405179 35405287 refGene_gn=CACNA1C;refGene_tx=XM_023256599.1
B4 35406070 35406158 refGene_gn=CACNA1C;refGene_tx=XM_023256599.1
B4 35408320 35408411 refGene_gn=CACNA1C;refGene_tx=XM_023256599.1
B4 35408413 35408426 refGene_gn=CACNA1C;refGene_tx=XM_023256599.1
B4 35412812 35412872 refGene_gn=CACNA1C;refGene_tx=XM_023256599.1
B4 35413013 35413073 refGene_gn=CACNA1C;refGene_tx=XM_023256599.1
B4 35414266 35414396 refGene_gn=CACNA1C;refGene_tx=XM_023256599.1
B4 35418587 35418720 refGene_gn=CACNA1C;refGene_tx=XM_023256599.1
B4 35426128 35426198 refGene_gn=CACNA1C;refGene_tx=XM_023256599.1

B4	35426544	35426665	refGene_gn=CACNA1C;refGene_tx=XM_023256599.1
B4	35427467	35427582	refGene_gn=CACNA1C;refGene_tx=XM_023256599.1
B4	35429272	35429393	refGene_gn=CACNA1C;refGene_tx=XM_023256599.1
B4	35430146	35430354	refGene_gn=CACNA1C;refGene_tx=XM_023256599.1
B4	35444677	35444903	refGene_gn=CACNA1C;refGene_tx=XM_023256599.1
B4	35445858	35446019	refGene_gn=CACNA1C;refGene_tx=XM_023256599.1
B4	35460813	35460840	refGene_gn=CACNA1C;refGene_tx=XM_023256599.1
B4	35468296	35468387	refGene_gn=CACNA1C;refGene_tx=XM_023256599.1
B4	35500668	35500841	refGene_gn=CACNA1C;refGene_tx=XM_023256599.1
B4	35508172	35508276	refGene_gn=CACNA1C;refGene_tx=XM_023256599.1
B4	35519359	35519556	refGene_gn=CACNA1C;refGene_tx=XM_023256599.1
B4	35526220	35526379	refGene_gn=CACNA1C;refGene_tx=XM_023256599.1
B4	35563535	35563675	refGene_gn=CACNA1C;refGene_tx=XM_023256599.1
B4	35570734	35570874	refGene_gn=CACNA1C;refGene_tx=XM_023256599.1
B4	35916378	35916484	refGene_gn=CACNA1C;refGene_tx=XM_023256599.1
B4	35921100	35921422	refGene_gn=CACNA1C;refGene_tx=XM_023256599.1
B4	36033664	36033814	refGene_gn=DCP1B;refGene_tx=XM_003988262.4
B4	36040371	36040412	refGene_gn=DCP1B;refGene_tx=XM_003988262.4
B4	36044738	36044866	refGene_gn=DCP1B;refGene_tx=XM_003988262.4
B4	36071325	36071464	refGene_gn=CACNA1C;refGene_tx=XM_023256599.1
B4	36074413	36074480	refGene_gn=DCP1B;refGene_tx=XM_003988262.4
B4	36076637	36076773	refGene_gn=DCP1B;refGene_tx=XM_003988262.4
B4	36084674	36084803	refGene_gn=DCP1B;refGene_tx=XM_003988262.4
B4	36086302	36087166	refGene_gn=DCP1B;refGene_tx=XM_003988262.4
B4	36089647	36089896	refGene_gn=DCP1B;refGene_tx=XM_003988262.4
B4	36092283	36092364	refGene_gn=DCP1B;refGene_tx=XM_003988262.4
B4	36121069	36121245
refGene_gn=CACNA2D4;refGene_tx=XM_003988301.2;refGene_gn=CACNA2D4;refGene_tx=XM_019834293.1;refGene_gn=CACNA2D4;refGene_tx=XM_023256600.1;refGene_gn=CACNA2D4;refGene_tx=XM_023256601.1;refGene_gn=CACNA2D4;refGene_tx=XM_023256602.1;refGene_gn=CACNA2D4;refGene_tx=XM_023256603.1;refGene_gn=CACNA2D4;refGene_tx=XM_023256604.1
B4	36125041	36125123
refGene_gn=CACNA2D4;refGene_tx=XM_003988301.2;refGene_gn=CACNA2D4;refGene_tx=XM_019834293.1;refGene_gn=CACNA2D4;refGene_tx=XM_023256600.1;refGene_gn=CACNA2D4;refGene_tx=XM_023256601.1;refGene_gn=CACNA2D4;refGene_tx=XM_023256602.1;refGene_gn=CACNA2D4;refGene_tx=XM_023256603.1;refGene_gn=CACNA2D4;refGene_tx=XM_023256604.1
B4	36126895	36127012
refGene_gn=CACNA2D4;refGene_tx=XM_003988301.2;refGene_gn=CACNA2D4;refGene_tx=XM_019834293.1;refGene_gn=CACNA2D4;refGene_tx=XM_023256600.1;refGene_gn=CACNA2D4;refGene_tx=XM_023256601.1;refGene_gn=CACNA2D4;refGene_tx=XM_023256602.1;refGene_gn=CACNA2D4;refGene_tx=XM_023256603.1;refGene_gn=CACNA2D4;refGene_tx=XM_023256604.1
B4	36132268	36132328
refGene_gn=CACNA2D4;refGene_tx=XM_003988301.2;refGene_gn=CACNA2D4;refGene_tx=XM_019834293.1;refGene_gn=CACNA2D4;refGene_tx=XM_023256600.1;refGene_gn=CACNA2D4;refGene_tx=XM_023256601.1;refGene_gn=CACNA2D4;refGene_tx=XM_023256602.1;refGene_gn=CACNA2D4;refGene_tx=XM_023256603.1;refGene_gn=CACNA2D4;refGene_tx=XM_023256604.1

B4 36134481 36134644
refGene_gn=CACNA2D4;refGene_tx=XM_003988301.2;refGene_gn=CACNA2D4;refGene_tx=XM_019834293.1;refGene_gn=CACNA2D4;refGene_tx=XM_023256600.1;refGene_gn=CACNA2D4;refGene_tx=XM_023256601.1;refGene_gn=CACNA2D4;refGene_tx=XM_023256602.1;refGene_gn=CACNA2D4;refGene_tx=XM_023256603.1;refGene_gn=CACNA2D4;refGene_tx=XM_023256604.1
B4 36135005 36135137
refGene_gn=CACNA2D4;refGene_tx=XM_003988301.2;refGene_gn=CACNA2D4;refGene_tx=XM_019834293.1;refGene_gn=CACNA2D4;refGene_tx=XM_023256600.1;refGene_gn=CACNA2D4;refGene_tx=XM_023256601.1;refGene_gn=CACNA2D4;refGene_tx=XM_023256602.1;refGene_gn=CACNA2D4;refGene_tx=XM_023256603.1;refGene_gn=CACNA2D4;refGene_tx=XM_023256604.1
B4 36144955 36145016
refGene_gn=CACNA2D4;refGene_tx=XM_003988301.2;refGene_gn=CACNA2D4;refGene_tx=XM_019834293.1;refGene_gn=CACNA2D4;refGene_tx=XM_023256600.1;refGene_gn=CACNA2D4;refGene_tx=XM_023256601.1;refGene_gn=CACNA2D4;refGene_tx=XM_023256602.1;refGene_gn=CACNA2D4;refGene_tx=XM_023256603.1;refGene_gn=CACNA2D4;refGene_tx=XM_023256604.1
B4 36145675 36145826
refGene_gn=CACNA2D4;refGene_tx=XM_003988301.2;refGene_gn=CACNA2D4;refGene_tx=XM_019834293.1;refGene_gn=CACNA2D4;refGene_tx=XM_023256600.1;refGene_gn=CACNA2D4;refGene_tx=XM_023256601.1;refGene_gn=CACNA2D4;refGene_tx=XM_023256602.1;refGene_gn=CACNA2D4;refGene_tx=XM_023256603.1;refGene_gn=CACNA2D4;refGene_tx=XM_023256604.1
B4 36147575 36147650
refGene_gn=CACNA2D4;refGene_tx=XM_003988301.2;refGene_gn=CACNA2D4;refGene_tx=XM_019834293.1;refGene_gn=CACNA2D4;refGene_tx=XM_023256600.1;refGene_gn=CACNA2D4;refGene_tx=XM_023256601.1;refGene_gn=CACNA2D4;refGene_tx=XM_023256602.1;refGene_gn=CACNA2D4;refGene_tx=XM_023256603.1;refGene_gn=CACNA2D4;refGene_tx=XM_023256604.1
B4 36148560 36148650
refGene_gn=CACNA2D4;refGene_tx=XM_003988301.2;refGene_gn=CACNA2D4;refGene_tx=XM_019834293.1;refGene_gn=CACNA2D4;refGene_tx=XM_023256600.1;refGene_gn=CACNA2D4;refGene_tx=XM_023256601.1;refGene_gn=CACNA2D4;refGene_tx=XM_023256602.1;refGene_gn=CACNA2D4;refGene_tx=XM_023256603.1;refGene_gn=CACNA2D4;refGene_tx=XM_023256604.1
B4 36148759 36148873
refGene_gn=CACNA2D4;refGene_tx=XM_003988301.2;refGene_gn=CACNA2D4;refGene_tx=XM_019834293.1;refGene_gn=CACNA2D4;refGene_tx=XM_023256600.1;refGene_gn=CACNA2D4;refGene_tx=XM_023256601.1;refGene_gn=CACNA2D4;refGene_tx=XM_023256602.1;refGene_gn=CACNA2D4;refGene_tx=XM_023256603.1;refGene_gn=CACNA2D4;refGene_tx=XM_023256604.1
B4 36149349 36149428
refGene_gn=CACNA2D4;refGene_tx=XM_003988301.2;refGene_gn=CACNA2D4;refGene_tx=XM_019834293.1;refGene_gn=CACNA2D4;refGene_tx=XM_023256600.1;refGene_gn=CACNA2D4;refGene_tx=XM_023256601.1;refGene_gn=CACNA2D4;refGene_tx=XM_023256602.1;refGene_gn=CACNA2D4;refGene_tx=XM_023256603.1;refGene_gn=CACNA2D4;refGene_tx=XM_023256604.1
B4 36150909 36151043
refGene_gn=CACNA2D4;refGene_tx=XM_003988301.2;refGene_gn=CACNA2D4;refG

ene_tx=XM_019834293.1;refGene_gn=CACNA2D4;refGene_tx=XM_023256600.1;refGene_gn=CACNA2D4;refGene_tx=XM_023256601.1;refGene_gn=CACNA2D4;refGene_tx=XM_023256602.1;refGene_gn=CACNA2D4;refGene_tx=XM_023256603.1;refGene_gn=CACNA2D4;refGene_tx=XM_023256604.1
B4 36152787 36152865
refGene_gn=CACNA2D4;refGene_tx=XM_003988301.2;refGene_gn=CACNA2D4;refGene_tx=XM_019834293.1;refGene_gn=CACNA2D4;refGene_tx=XM_023256600.1;refGene_gn=CACNA2D4;refGene_tx=XM_023256601.1;refGene_gn=CACNA2D4;refGene_tx=XM_023256602.1;refGene_gn=CACNA2D4;refGene_tx=XM_023256603.1;refGene_gn=CACNA2D4;refGene_tx=XM_023256604.1
B4 36153653 36153734
refGene_gn=CACNA2D4;refGene_tx=XM_003988301.2;refGene_gn=CACNA2D4;refGene_tx=XM_019834293.1;refGene_gn=CACNA2D4;refGene_tx=XM_023256600.1;refGene_gn=CACNA2D4;refGene_tx=XM_023256601.1;refGene_gn=CACNA2D4;refGene_tx=XM_023256602.1;refGene_gn=CACNA2D4;refGene_tx=XM_023256604.1
B4 36154309 36154384
refGene_gn=CACNA2D4;refGene_tx=XM_003988301.2;refGene_gn=CACNA2D4;refGene_tx=XM_019834293.1;refGene_gn=CACNA2D4;refGene_tx=XM_023256600.1;refGene_gn=CACNA2D4;refGene_tx=XM_023256601.1;refGene_gn=CACNA2D4;refGene_tx=XM_023256602.1;refGene_gn=CACNA2D4;refGene_tx=XM_023256603.1;refGene_gn=CACNA2D4;refGene_tx=XM_023256604.1
B4 36157489 36157576
refGene_gn=CACNA2D4;refGene_tx=XM_003988301.2;refGene_gn=CACNA2D4;refGene_tx=XM_019834293.1;refGene_gn=CACNA2D4;refGene_tx=XM_023256600.1;refGene_gn=CACNA2D4;refGene_tx=XM_023256601.1;refGene_gn=CACNA2D4;refGene_tx=XM_023256602.1;refGene_gn=CACNA2D4;refGene_tx=XM_023256603.1;refGene_gn=CACNA2D4;refGene_tx=XM_023256604.1
B4 36158118 36158190
refGene_gn=CACNA2D4;refGene_tx=XM_003988301.2;refGene_gn=CACNA2D4;refGene_tx=XM_019834293.1;refGene_gn=CACNA2D4;refGene_tx=XM_023256600.1;refGene_gn=CACNA2D4;refGene_tx=XM_023256601.1;refGene_gn=CACNA2D4;refGene_tx=XM_023256603.1;refGene_gn=CACNA2D4;refGene_tx=XM_023256604.1
B4 36169469 36169531
refGene_gn=CACNA2D4;refGene_tx=XM_003988301.2;refGene_gn=CACNA2D4;refGene_tx=XM_019834293.1;refGene_gn=CACNA2D4;refGene_tx=XM_023256600.1;refGene_gn=CACNA2D4;refGene_tx=XM_023256601.1;refGene_gn=CACNA2D4;refGene_tx=XM_023256602.1;refGene_gn=CACNA2D4;refGene_tx=XM_023256603.1;refGene_gn=CACNA2D4;refGene_tx=XM_023256604.1
B4 36170804 36170872
refGene_gn=CACNA2D4;refGene_tx=XM_003988301.2;refGene_gn=CACNA2D4;refGene_tx=XM_019834293.1;refGene_gn=CACNA2D4;refGene_tx=XM_023256600.1;refGene_gn=CACNA2D4;refGene_tx=XM_023256601.1;refGene_gn=CACNA2D4;refGene_tx=XM_023256602.1;refGene_gn=CACNA2D4;refGene_tx=XM_023256603.1;refGene_gn=CACNA2D4;refGene_tx=XM_023256604.1
B4 36173120 36173166
refGene_gn=CACNA2D4;refGene_tx=XM_003988301.2;refGene_gn=CACNA2D4;refGene_tx=XM_019834293.1;refGene_gn=CACNA2D4;refGene_tx=XM_023256600.1;refGene_gn=CACNA2D4;refGene_tx=XM_023256601.1;refGene_gn=CACNA2D4;refGene_tx=XM_023256602.1;refGene_gn=CACNA2D4;refGene_tx=XM_023256603.1;refGene_gn=CACNA2D4;refGene_tx=XM_023256604.1

B4 36173259 36173357
refGene_gn=CACNA2D4;refGene_tx=XM_003988301.2;refGene_gn=CACNA2D4;refGene_tx=XM_019834293.1;refGene_gn=CACNA2D4;refGene_tx=XM_023256600.1;refGene_gn=CACNA2D4;refGene_tx=XM_023256601.1;refGene_gn=CACNA2D4;refGene_tx=XM_023256602.1;refGene_gn=CACNA2D4;refGene_tx=XM_023256603.1;refGene_gn=CACNA2D4;refGene_tx=XM_023256604.1
B4 36175656 36175750
refGene_gn=CACNA2D4;refGene_tx=XM_003988301.2;refGene_gn=CACNA2D4;refGene_tx=XM_019834293.1;refGene_gn=CACNA2D4;refGene_tx=XM_023256600.1;refGene_gn=CACNA2D4;refGene_tx=XM_023256601.1;refGene_gn=CACNA2D4;refGene_tx=XM_023256602.1;refGene_gn=CACNA2D4;refGene_tx=XM_023256603.1;refGene_gn=CACNA2D4;refGene_tx=XM_023256604.1
B4 36183884 36183980
refGene_gn=CACNA2D4;refGene_tx=XM_003988301.2;refGene_gn=CACNA2D4;refGene_tx=XM_019834293.1;refGene_gn=CACNA2D4;refGene_tx=XM_023256600.1;refGene_gn=CACNA2D4;refGene_tx=XM_023256601.1;refGene_gn=CACNA2D4;refGene_tx=XM_023256602.1;refGene_gn=CACNA2D4;refGene_tx=XM_023256603.1
B4 36186025 36186153
refGene_gn=CACNA2D4;refGene_tx=XM_003988301.2;refGene_gn=CACNA2D4;refGene_tx=XM_019834293.1;refGene_gn=CACNA2D4;refGene_tx=XM_023256600.1;refGene_gn=CACNA2D4;refGene_tx=XM_023256601.1;refGene_gn=CACNA2D4;refGene_tx=XM_023256602.1;refGene_gn=CACNA2D4;refGene_tx=XM_023256603.1;refGene_gn=CACNA2D4;refGene_tx=XM_023256604.1
B4 36189798 36189879
refGene_gn=CACNA2D4;refGene_tx=XM_003988301.2;refGene_gn=CACNA2D4;refGene_tx=XM_019834293.1;refGene_gn=CACNA2D4;refGene_tx=XM_023256600.1;refGene_gn=CACNA2D4;refGene_tx=XM_023256601.1;refGene_gn=CACNA2D4;refGene_tx=XM_023256602.1;refGene_gn=CACNA2D4;refGene_tx=XM_023256603.1;refGene_gn=CACNA2D4;refGene_tx=XM_023256604.1
B4 36194180 36194203 refGene_gn=LRTM2;refGene_tx=XM_019834298.2
B4 36195237 36195692
refGene_gn=LRTM2;refGene_tx=XM_003988263.5;refGene_gn=LRTM2;refGene_tx=XM_019834295.2;refGene_gn=LRTM2;refGene_tx=XM_019834297.2;refGene_gn=LRTM2;refGene_tx=XM_019834298.2
B4 36198651 36199242
refGene_gn=LRTM2;refGene_tx=XM_003988263.5;refGene_gn=LRTM2;refGene_tx=XM_019834295.2;refGene_gn=LRTM2;refGene_tx=XM_019834297.2;refGene_gn=LRTM2;refGene_tx=XM_019834298.2
B4 36201864 36201931
refGene_gn=LRTM2;refGene_tx=XM_003988263.5;refGene_gn=LRTM2;refGene_tx=XM_019834295.2;refGene_gn=LRTM2;refGene_tx=XM_019834297.2;refGene_gn=LRTM2;refGene_tx=XM_019834298.2
B4 36216698 36216760
refGene_gn=CACNA2D4;refGene_tx=XM_003988301.2;refGene_gn=CACNA2D4;refGene_tx=XM_019834293.1;refGene_gn=CACNA2D4;refGene_tx=XM_023256600.1;refGene_gn=CACNA2D4;refGene_tx=XM_023256601.1;refGene_gn=CACNA2D4;refGene_tx=XM_023256602.1;refGene_gn=CACNA2D4;refGene_tx=XM_023256603.1;refGene_gn=CACNA2D4;refGene_tx=XM_023256604.1
B4 36217740 36217785
refGene_gn=CACNA2D4;refGene_tx=XM_003988301.2;refGene_gn=CACNA2D4;refG

ene_tx=XM_019834293.1;refGene_gn=CACNA2D4;refGene_tx=XM_023256600.1;refGene_gn=CACNA2D4;refGene_tx=XM_023256601.1;refGene_gn=CACNA2D4;refGene_tx=XM_023256602.1;refGene_gn=CACNA2D4;refGene_tx=XM_023256603.1;refGene_gn=CACNA2D4;refGene_tx=XM_023256604.1
B4 36217949 36218012
refGene_gn=CACNA2D4;refGene_tx=XM_003988301.2;refGene_gn=CACNA2D4;refGene_tx=XM_019834293.1;refGene_gn=CACNA2D4;refGene_tx=XM_023256600.1;refGene_gn=CACNA2D4;refGene_tx=XM_023256602.1;refGene_gn=CACNA2D4;refGene_tx=XM_023256603.1;refGene_gn=CACNA2D4;refGene_tx=XM_023256604.1
B4 36225274 36225345
refGene_gn=CACNA2D4;refGene_tx=XM_003988301.2;refGene_gn=CACNA2D4;refGene_tx=XM_019834293.1;refGene_gn=CACNA2D4;refGene_tx=XM_023256600.1;refGene_gn=CACNA2D4;refGene_tx=XM_023256601.1;refGene_gn=CACNA2D4;refGene_tx=XM_023256602.1;refGene_gn=CACNA2D4;refGene_tx=XM_023256603.1;refGene_gn=CACNA2D4;refGene_tx=XM_023256604.1
B4 36225732 36225808
refGene_gn=CACNA2D4;refGene_tx=XM_003988301.2;refGene_gn=CACNA2D4;refGene_tx=XM_019834293.1;refGene_gn=CACNA2D4;refGene_tx=XM_023256600.1;refGene_gn=CACNA2D4;refGene_tx=XM_023256601.1;refGene_gn=CACNA2D4;refGene_tx=XM_023256602.1;refGene_gn=CACNA2D4;refGene_tx=XM_023256603.1;refGene_gn=CACNA2D4;refGene_tx=XM_023256604.1
B4 36226332 36226385
refGene_gn=CACNA2D4;refGene_tx=XM_003988301.2;refGene_gn=CACNA2D4;refGene_tx=XM_019834293.1;refGene_gn=CACNA2D4;refGene_tx=XM_023256600.1;refGene_gn=CACNA2D4;refGene_tx=XM_023256601.1;refGene_gn=CACNA2D4;refGene_tx=XM_023256602.1;refGene_gn=CACNA2D4;refGene_tx=XM_023256603.1;refGene_gn=CACNA2D4;refGene_tx=XM_023256604.1
B4 36226738 36226794
refGene_gn=CACNA2D4;refGene_tx=XM_003988301.2;refGene_gn=CACNA2D4;refGene_tx=XM_019834293.1;refGene_gn=CACNA2D4;refGene_tx=XM_023256600.1;refGene_gn=CACNA2D4;refGene_tx=XM_023256601.1;refGene_gn=CACNA2D4;refGene_tx=XM_023256602.1;refGene_gn=CACNA2D4;refGene_tx=XM_023256603.1;refGene_gn=CACNA2D4;refGene_tx=XM_023256604.1
B4 36227110 36227131
refGene_gn=CACNA2D4;refGene_tx=XM_003988301.2;refGene_gn=CACNA2D4;refGene_tx=XM_019834293.1;refGene_gn=CACNA2D4;refGene_tx=XM_023256601.1;refGene_gn=CACNA2D4;refGene_tx=XM_023256602.1;refGene_gn=CACNA2D4;refGene_tx=XM_023256603.1;refGene_gn=CACNA2D4;refGene_tx=XM_023256604.1
B4 36229648 36229766
refGene_gn=CACNA2D4;refGene_tx=XM_003988301.2;refGene_gn=CACNA2D4;refGene_tx=XM_023256600.1;refGene_gn=CACNA2D4;refGene_tx=XM_023256601.1;refGene_gn=CACNA2D4;refGene_tx=XM_023256602.1;refGene_gn=CACNA2D4;refGene_tx=XM_023256603.1;refGene_gn=CACNA2D4;refGene_tx=XM_023256604.1
B4 36231401 36231514
refGene_gn=CACNA2D4;refGene_tx=XM_003988301.2;refGene_gn=CACNA2D4;refGene_tx=XM_019834293.1;refGene_gn=CACNA2D4;refGene_tx=XM_023256600.1;refGene_gn=CACNA2D4;refGene_tx=XM_023256601.1;refGene_gn=CACNA2D4;refGene_tx=XM_023256602.1;refGene_gn=CACNA2D4;refGene_tx=XM_023256603.1;refGene_gn=CACNA2D4;refGene_tx=XM_023256604.1
B4 36231835 36231918

refGene_gn=CACNA2D4;refGene_tx=XM_003988301.2;refGene_gn=CACNA2D4;refGene_tx=XM_023256600.1;refGene_gn=CACNA2D4;refGene_tx=XM_023256601.1;refGene_gn=CACNA2D4;refGene_tx=XM_023256602.1;refGene_gn=CACNA2D4;refGene_tx=XM_023256603.1;refGene_gn=CACNA2D4;refGene_tx=XM_023256604.1
B4 36232705 36232810
refGene_gn=CACNA2D4;refGene_tx=XM_003988301.2;refGene_gn=CACNA2D4;refGene_tx=XM_023256600.1;refGene_gn=CACNA2D4;refGene_tx=XM_023256601.1;refGene_gn=CACNA2D4;refGene_tx=XM_023256602.1;refGene_gn=CACNA2D4;refGene_tx=XM_023256603.1;refGene_gn=CACNA2D4;refGene_tx=XM_023256604.1
B4 36239878 36240007
refGene_gn=ADIPOR2;refGene_tx=XM_003988264.5;refGene_gn=ADIPOR2;refGene_tx=XM_006933391.4;refGene_gn=ADIPOR2;refGene_tx=XM_011283752.3;refGene_gn=ADIPOR2;refGene_tx=XM_019834302.1
B4 36241788 36241982
refGene_gn=ADIPOR2;refGene_tx=XM_003988264.5;refGene_gn=ADIPOR2;refGene_tx=XM_006933391.4;refGene_gn=ADIPOR2;refGene_tx=XM_011283752.3;refGene_gn=ADIPOR2;refGene_tx=XM_019834302.1
B4 36244277 36244465
refGene_gn=ADIPOR2;refGene_tx=XM_003988264.5;refGene_gn=ADIPOR2;refGene_tx=XM_006933391.4;refGene_gn=ADIPOR2;refGene_tx=XM_011283752.3;refGene_gn=ADIPOR2;refGene_tx=XM_019834302.1
B4 36244723 36244910
refGene_gn=ADIPOR2;refGene_tx=XM_003988264.5;refGene_gn=ADIPOR2;refGene_tx=XM_006933391.4;refGene_gn=ADIPOR2;refGene_tx=XM_011283752.3;refGene_gn=ADIPOR2;refGene_tx=XM_019834302.1
B4 36248460 36248632
refGene_gn=ADIPOR2;refGene_tx=XM_003988264.5;refGene_gn=ADIPOR2;refGene_tx=XM_006933391.4;refGene_gn=ADIPOR2;refGene_tx=XM_011283752.3;refGene_gn=ADIPOR2;refGene_tx=XM_019834302.1
B4 36253299 36253419
refGene_gn=ADIPOR2;refGene_tx=XM_003988264.5;refGene_gn=ADIPOR2;refGene_tx=XM_006933391.4;refGene_gn=ADIPOR2;refGene_tx=XM_011283752.3;refGene_gn=ADIPOR2;refGene_tx=XM_019834302.1
B4 36275301 36275472
refGene_gn=ADIPOR2;refGene_tx=XM_003988264.5;refGene_gn=ADIPOR2;refGene_tx=XM_006933391.4;refGene_gn=ADIPOR2;refGene_tx=XM_011283752.3;refGene_gn=ADIPOR2;refGene_tx=XM_019834302.1
B4 36373227 36373686 refGene_gn=WNT5B;refGene_tx=XM_023256605.1
B4 36378861 36379154 refGene_gn=WNT5B;refGene_tx=XM_023256605.1
B4 36384308 36384556 refGene_gn=WNT5B;refGene_tx=XM_023256605.1
B4 36385784 36385942 refGene_gn=WNT5B;refGene_tx=XM_023256605.1
B4 36422404 36423607 refGene_gn=FBXL14;refGene_tx=XM_019834300.2
B4 36476377 36476383 refGene_gn=WNT5B;refGene_tx=XM_023256605.1
B4 36511281 36511419
refGene_gn=ERC1;refGene_tx=XM_023256606.1;refGene_gn=ERC1;refGene_tx=XM_023256607.1;refGene_gn=ERC1;refGene_tx=XM_023256608.1;refGene_gn=ERC1;refGene_tx=XM_023256609.1;refGene_gn=ERC1;refGene_tx=XM_023256610.1;refGene_gn=ERC1;refGene_tx=XM_023256611.1;refGene_gn=ERC1;refGene_tx=XM_023256612.1;refGene_gn=ERC1;refGene_tx=XM_023256613.1
B4 36551208 36551397

refGene_gn=ERC1;refGene_tx=XM_023256606.1;refGene_gn=ERC1;refGene_tx=XM_023256607.1;refGene_gn=ERC1;refGene_tx=XM_023256608.1;refGene_gn=ERC1;refGene_tx=XM_023256609.1;refGene_gn=ERC1;refGene_tx=XM_023256610.1;refGene_gn=ERC1;refGene_tx=XM_023256611.1;refGene_gn=ERC1;refGene_tx=XM_023256612.1;refGene_gn=ERC1;refGene_tx=XM_023256613.1
B4 36588739 36588766
refGene_gn=ERC1;refGene_tx=XM_023256614.1;refGene_gn=ERC1;refGene_tx=XM_023256615.1;refGene_gn=ERC1;refGene_tx=XM_023256616.1;refGene_gn=ERC1;refGene_tx=XM_023256617.1
B4 36590254 36590353
refGene_gn=ERC1;refGene_tx=XM_023256606.1;refGene_gn=ERC1;refGene_tx=XM_023256607.1;refGene_gn=ERC1;refGene_tx=XM_023256608.1;refGene_gn=ERC1;refGene_tx=XM_023256609.1;refGene_gn=ERC1;refGene_tx=XM_023256610.1;refGene_gn=ERC1;refGene_tx=XM_023256611.1;refGene_gn=ERC1;refGene_tx=XM_023256612.1;refGene_gn=ERC1;refGene_tx=XM_023256613.1;refGene_gn=ERC1;refGene_tx=XM_023256614.1;refGene_gn=ERC1;refGene_tx=XM_023256615.1;refGene_gn=ERC1;refGene_tx=XM_023256616.1;refGene_gn=ERC1;refGene_tx=XM_023256617.1
B4 36631418 36631563
refGene_gn=ERC1;refGene_tx=XM_023256606.1;refGene_gn=ERC1;refGene_tx=XM_023256607.1;refGene_gn=ERC1;refGene_tx=XM_023256608.1;refGene_gn=ERC1;refGene_tx=XM_023256609.1;refGene_gn=ERC1;refGene_tx=XM_023256610.1;refGene_gn=ERC1;refGene_tx=XM_023256611.1;refGene_gn=ERC1;refGene_tx=XM_023256612.1;refGene_gn=ERC1;refGene_tx=XM_023256613.1;refGene_gn=ERC1;refGene_tx=XM_023256614.1;refGene_gn=ERC1;refGene_tx=XM_023256615.1;refGene_gn=ERC1;refGene_tx=XM_023256616.1;refGene_gn=ERC1;refGene_tx=XM_023256617.1
B4 36735177 36735338
refGene_gn=ERC1;refGene_tx=XM_023256606.1;refGene_gn=ERC1;refGene_tx=XM_023256607.1;refGene_gn=ERC1;refGene_tx=XM_023256608.1;refGene_gn=ERC1;refGene_tx=XM_023256609.1;refGene_gn=ERC1;refGene_tx=XM_023256610.1;refGene_gn=ERC1;refGene_tx=XM_023256611.1;refGene_gn=ERC1;refGene_tx=XM_023256612.1;refGene_gn=ERC1;refGene_tx=XM_023256613.1;refGene_gn=ERC1;refGene_tx=XM_023256614.1;refGene_gn=ERC1;refGene_tx=XM_023256615.1;refGene_gn=ERC1;refGene_tx=XM_023256616.1;refGene_gn=ERC1;refGene_tx=XM_023256617.1
B4 36773932 36774064
refGene_gn=ERC1;refGene_tx=XM_023256606.1;refGene_gn=ERC1;refGene_tx=XM_023256607.1;refGene_gn=ERC1;refGene_tx=XM_023256608.1;refGene_gn=ERC1;refGene_tx=XM_023256609.1;refGene_gn=ERC1;refGene_tx=XM_023256610.1;refGene_gn=ERC1;refGene_tx=XM_023256611.1;refGene_gn=ERC1;refGene_tx=XM_023256612.1;refGene_gn=ERC1;refGene_tx=XM_023256614.1;refGene_gn=ERC1;refGene_tx=XM_023256615.1
B4 36793242 36793378
refGene_gn=ERC1;refGene_tx=XM_023256606.1;refGene_gn=ERC1;refGene_tx=XM_023256607.1;refGene_gn=ERC1;refGene_tx=XM_023256608.1;refGene_gn=ERC1;refGene_tx=XM_023256609.1;refGene_gn=ERC1;refGene_tx=XM_023256610.1;refGene_gn=ERC1;refGene_tx=XM_023256611.1;refGene_gn=ERC1;refGene_tx=XM_023256612.1;refGene_gn=ERC1;refGene_tx=XM_023256613.1;refGene_gn=ERC1;refGene_tx=XM_023256614.1;refGene_gn=ERC1;refGene_tx=XM_023256615.1;

refGene_gn=ERC1;refGene_tx=XM_023256616.1;refGene_gn=ERC1;refGene_tx=XM_023256617.1
B4 36810907 36810919
refGene_gn=ERC1;refGene_tx=XM_023256606.1;refGene_gn=ERC1;refGene_tx=XM_023256607.1;refGene_gn=ERC1;refGene_tx=XM_023256608.1;refGene_gn=ERC1;refGene_tx=XM_023256611.1;refGene_gn=ERC1;refGene_tx=XM_023256613.1;refGene_gn=ERC1;refGene_tx=XM_023256614.1;refGene_gn=ERC1;refGene_tx=XM_023256615.1;refGene_gn=ERC1;refGene_tx=XM_023256616.1;refGene_gn=ERC1;refGene_tx=XM_023256617.1
B4 36830929 36831123
refGene_gn=ERC1;refGene_tx=XM_023256606.1;refGene_gn=ERC1;refGene_tx=XM_023256607.1;refGene_gn=ERC1;refGene_tx=XM_023256608.1;refGene_gn=ERC1;refGene_tx=XM_023256609.1;refGene_gn=ERC1;refGene_tx=XM_023256610.1;refGene_gn=ERC1;refGene_tx=XM_023256611.1;refGene_gn=ERC1;refGene_tx=XM_023256612.1;refGene_gn=ERC1;refGene_tx=XM_023256613.1;refGene_gn=ERC1;refGene_tx=XM_023256614.1;refGene_gn=ERC1;refGene_tx=XM_023256615.1;refGene_gn=ERC1;refGene_tx=XM_023256616.1;refGene_gn=ERC1;refGene_tx=XM_023256617.1
B4 36840793 36840934
refGene_gn=ERC1;refGene_tx=XM_023256606.1;refGene_gn=ERC1;refGene_tx=XM_023256607.1;refGene_gn=ERC1;refGene_tx=XM_023256608.1;refGene_gn=ERC1;refGene_tx=XM_023256609.1;refGene_gn=ERC1;refGene_tx=XM_023256610.1;refGene_gn=ERC1;refGene_tx=XM_023256611.1;refGene_gn=ERC1;refGene_tx=XM_023256612.1;refGene_gn=ERC1;refGene_tx=XM_023256613.1;refGene_gn=ERC1;refGene_tx=XM_023256614.1;refGene_gn=ERC1;refGene_tx=XM_023256615.1;refGene_gn=ERC1;refGene_tx=XM_023256616.1;refGene_gn=ERC1;refGene_tx=XM_023256617.1
B4 36843382 36843523
refGene_gn=ERC1;refGene_tx=XM_023256606.1;refGene_gn=ERC1;refGene_tx=XM_023256607.1;refGene_gn=ERC1;refGene_tx=XM_023256608.1;refGene_gn=ERC1;refGene_tx=XM_023256609.1;refGene_gn=ERC1;refGene_tx=XM_023256610.1;refGene_gn=ERC1;refGene_tx=XM_023256611.1;refGene_gn=ERC1;refGene_tx=XM_023256612.1;refGene_gn=ERC1;refGene_tx=XM_023256613.1;refGene_gn=ERC1;refGene_tx=XM_023256614.1;refGene_gn=ERC1;refGene_tx=XM_023256615.1;refGene_gn=ERC1;refGene_tx=XM_023256616.1;refGene_gn=ERC1;refGene_tx=XM_023256617.1
B4 36844882 36845020
refGene_gn=ERC1;refGene_tx=XM_023256606.1;refGene_gn=ERC1;refGene_tx=XM_023256607.1;refGene_gn=ERC1;refGene_tx=XM_023256608.1;refGene_gn=ERC1;refGene_tx=XM_023256609.1;refGene_gn=ERC1;refGene_tx=XM_023256610.1;refGene_gn=ERC1;refGene_tx=XM_023256611.1;refGene_gn=ERC1;refGene_tx=XM_023256612.1;refGene_gn=ERC1;refGene_tx=XM_023256613.1;refGene_gn=ERC1;refGene_tx=XM_023256614.1;refGene_gn=ERC1;refGene_tx=XM_023256615.1;refGene_gn=ERC1;refGene_tx=XM_023256616.1;refGene_gn=ERC1;refGene_tx=XM_023256617.1
B4 36874339 36874507
refGene_gn=ERC1;refGene_tx=XM_023256606.1;refGene_gn=ERC1;refGene_tx=XM_023256607.1;refGene_gn=ERC1;refGene_tx=XM_023256608.1;refGene_gn=ERC1;refGene_tx=XM_023256609.1;refGene_gn=ERC1;refGene_tx=XM_023256610.1;refGene_gn=ERC1;refGene_tx=XM_023256611.1;refGene_gn=ERC1;refGene_tx=X

M_023256612.1;refGene_gn=ERC1;refGene_tx=XM_023256613.1;refGene_gn=ERC1;refGene_tx=XM_023256614.1;refGene_gn=ERC1;refGene_tx=XM_023256615.1;refGene_gn=ERC1;refGene_tx=XM_023256616.1;refGene_gn=ERC1;refGene_tx=XM_023256617.1
B4 36896198 36896366
refGene_gn=ERC1;refGene_tx=XM_023256606.1;refGene_gn=ERC1;refGene_tx=XM_023256607.1;refGene_gn=ERC1;refGene_tx=XM_023256608.1;refGene_gn=ERC1;refGene_tx=XM_023256609.1;refGene_gn=ERC1;refGene_tx=XM_023256610.1;refGene_gn=ERC1;refGene_tx=XM_023256611.1;refGene_gn=ERC1;refGene_tx=XM_023256612.1;refGene_gn=ERC1;refGene_tx=XM_023256613.1;refGene_gn=ERC1;refGene_tx=XM_023256614.1;refGene_gn=ERC1;refGene_tx=XM_023256615.1;refGene_gn=ERC1;refGene_tx=XM_023256616.1;refGene_gn=ERC1;refGene_tx=XM_023256617.1
B4 36899918 36900011
refGene_gn=ERC1;refGene_tx=XM_023256606.1;refGene_gn=ERC1;refGene_tx=XM_023256607.1;refGene_gn=ERC1;refGene_tx=XM_023256608.1;refGene_gn=ERC1;refGene_tx=XM_023256609.1;refGene_gn=ERC1;refGene_tx=XM_023256610.1;refGene_gn=ERC1;refGene_tx=XM_023256613.1;refGene_gn=ERC1;refGene_tx=XM_023256614.1;refGene_gn=ERC1;refGene_tx=XM_023256616.1
B4 36901272 36901428
refGene_gn=ERC1;refGene_tx=XM_023256606.1;refGene_gn=ERC1;refGene_tx=XM_023256607.1;refGene_gn=ERC1;refGene_tx=XM_023256608.1;refGene_gn=ERC1;refGene_tx=XM_023256609.1;refGene_gn=ERC1;refGene_tx=XM_023256610.1;refGene_gn=ERC1;refGene_tx=XM_023256611.1;refGene_gn=ERC1;refGene_tx=XM_023256612.1;refGene_gn=ERC1;refGene_tx=XM_023256613.1;refGene_gn=ERC1;refGene_tx=XM_023256614.1;refGene_gn=ERC1;refGene_tx=XM_023256615.1;refGene_gn=ERC1;refGene_tx=XM_023256616.1;refGene_gn=ERC1;refGene_tx=XM_023256617.1
B4 36908172 36908247
refGene_gn=ERC1;refGene_tx=XM_023256606.1;refGene_gn=ERC1;refGene_tx=XM_023256607.1;refGene_gn=ERC1;refGene_tx=XM_023256608.1;refGene_gn=ERC1;refGene_tx=XM_023256609.1;refGene_gn=ERC1;refGene_tx=XM_023256610.1;refGene_gn=ERC1;refGene_tx=XM_023256611.1;refGene_gn=ERC1;refGene_tx=XM_023256612.1;refGene_gn=ERC1;refGene_tx=XM_023256613.1;refGene_gn=ERC1;refGene_tx=XM_023256614.1;refGene_gn=ERC1;refGene_tx=XM_023256615.1;refGene_gn=ERC1;refGene_tx=XM_023256616.1;refGene_gn=ERC1;refGene_tx=XM_023256617.1
B4 36925901 36926318
refGene_gn=ERC1;refGene_tx=XM_023256606.1;refGene_gn=ERC1;refGene_tx=XM_023256607.1;refGene_gn=ERC1;refGene_tx=XM_023256608.1;refGene_gn=ERC1;refGene_tx=XM_023256609.1;refGene_gn=ERC1;refGene_tx=XM_023256610.1;refGene_gn=ERC1;refGene_tx=XM_023256611.1;refGene_gn=ERC1;refGene_tx=XM_023256612.1;refGene_gn=ERC1;refGene_tx=XM_023256613.1;refGene_gn=ERC1;refGene_tx=XM_023256614.1;refGene_gn=ERC1;refGene_tx=XM_023256615.1;refGene_gn=ERC1;refGene_tx=XM_023256616.1;refGene_gn=ERC1;refGene_tx=XM_023256617.1
B4 36980496 36981165
refGene_gn=ERC1;refGene_tx=XM_023256606.1;refGene_gn=ERC1;refGene_tx=XM_023256607.1;refGene_gn=ERC1;refGene_tx=XM_023256608.1;refGene_gn=ERC1;refGene_tx=XM_023256609.1;refGene_gn=ERC1;refGene_tx=XM_023256610.1;

refGene_gn=ERC1;refGene_tx=XM_023256611.1;refGene_gn=ERC1;refGene_tx=XM_023256612.1;refGene_gn=ERC1;refGene_tx=XM_023256613.1;refGene_gn=ERC1;refGene_tx=XM_023256614.1;refGene_gn=ERC1;refGene_tx=XM_023256615.1;refGene_gn=ERC1;refGene_tx=XM_023256616.1;refGene_gn=ERC1;refGene_tx=XM_023256617.1
B4 37091477 37091561
refGene_gn=RAD52;refGene_tx=XM_003988268.3;refGene_gn=RAD52;refGene_tx=XM_006933396.2;refGene_gn=RAD52;refGene_tx=XM_006933397.2;refGene_gn=RAD52;refGene_tx=XM_006933398.2;refGene_gn=RAD52;refGene_tx=XM_019834330.2
B4 37092654 37092756
refGene_gn=RAD52;refGene_tx=XM_003988268.3;refGene_gn=RAD52;refGene_tx=XM_006933396.2;refGene_gn=RAD52;refGene_tx=XM_006933397.2;refGene_gn=RAD52;refGene_tx=XM_006933398.2;refGene_gn=RAD52;refGene_tx=XM_019834330.2
B4 37093336 37093430
refGene_gn=RAD52;refGene_tx=XM_003988268.3;refGene_gn=RAD52;refGene_tx=XM_006933396.2;refGene_gn=RAD52;refGene_tx=XM_006933397.2;refGene_gn=RAD52;refGene_tx=XM_006933398.2
B4 37093600 37093668
refGene_gn=RAD52;refGene_tx=XM_003988268.3;refGene_gn=RAD52;refGene_tx=XM_006933396.2;refGene_gn=RAD52;refGene_tx=XM_006933397.2;refGene_gn=RAD52;refGene_tx=XM_006933398.2
B4 37096699 37096818
refGene_gn=RAD52;refGene_tx=XM_003988268.3;refGene_gn=RAD52;refGene_tx=XM_006933396.2;refGene_gn=RAD52;refGene_tx=XM_006933397.2;refGene_gn=RAD52;refGene_tx=XM_019834330.2
B4 37097680 37097756
refGene_gn=RAD52;refGene_tx=XM_003988268.3;refGene_gn=RAD52;refGene_tx=XM_006933396.2;refGene_gn=RAD52;refGene_tx=XM_006933397.2;refGene_gn=RAD52;refGene_tx=XM_019834330.2
B4 37105015 37105212
refGene_gn=RAD52;refGene_tx=XM_003988268.3;refGene_gn=RAD52;refGene_tx=XM_006933396.2;refGene_gn=RAD52;refGene_tx=XM_006933397.2;refGene_gn=RAD52;refGene_tx=XM_006933398.2;refGene_gn=RAD52;refGene_tx=XM_019834330.2
B4 37105355 37105495
refGene_gn=RAD52;refGene_tx=XM_003988268.3;refGene_gn=RAD52;refGene_tx=XM_006933396.2;refGene_gn=RAD52;refGene_tx=XM_006933397.2;refGene_gn=RAD52;refGene_tx=XM_006933398.2;refGene_gn=RAD52;refGene_tx=XM_019834330.2
B4 37106135 37106216
refGene_gn=RAD52;refGene_tx=XM_003988268.3;refGene_gn=RAD52;refGene_tx=XM_006933396.2;refGene_gn=RAD52;refGene_tx=XM_006933397.2;refGene_gn=RAD52;refGene_tx=XM_006933398.2;refGene_gn=RAD52;refGene_tx=XM_019834330.2
B4 37106817 37107033
refGene_gn=RAD52;refGene_tx=XM_003988268.3;refGene_gn=RAD52;refGene_tx=XM_006933396.2;refGene_gn=RAD52;refGene_tx=XM_006933397.2;refGene_gn=RAD52;refGene_tx=XM_006933398.2;refGene_gn=RAD52;refGene_tx=XM_0198343

30.2
B4 37107640 37107702
refGene_gn=RAD52;refGene_tx=XM_003988268.3;refGene_gn=RAD52;refGene_tx=XM_006933396.2;refGene_gn=RAD52;refGene_tx=XM_006933397.2;refGene_gn=RAD52;refGene_tx=XM_006933398.2;refGene_gn=RAD52;refGene_tx=XM_019834330.2
B4 37111579 37111897
refGene_gn=WNK1;refGene_tx=XM_023256618.1;refGene_gn=WNK1;refGene_tx=XM_023256619.1;refGene_gn=WNK1;refGene_tx=XM_023256620.1;refGene_gn=WNK1;refGene_tx=XM_023256621.1;refGene_gn=WNK1;refGene_tx=XM_023256622.1;refGene_gn=WNK1;refGene_tx=XM_023256623.1;refGene_gn=WNK1;refGene_tx=XM_023256624.1;refGene_gn=WNK1;refGene_tx=XM_023256625.1;refGene_gn=WNK1;refGene_tx=XM_023256626.1;refGene_gn=WNK1;refGene_tx=XM_023256627.1;refGene_gn=WNK1;refGene_tx=XM_023256628.1;refGene_gn=WNK1;refGene_tx=XM_023256629.1;refGene_gn=WNK1;refGene_tx=XM_023256630.1;refGene_gn=WNK1;refGene_tx=XM_023256631.1;refGene_gn=WNK1;refGene_tx=XM_023256632.1;refGene_gn=WNK1;refGene_tx=XM_023256633.1;refGene_gn=WNK1;refGene_tx=XM_023256634.1;refGene_gn=WNK1;refGene_tx=XM_023256635.1;refGene_gn=WNK1;refGene_tx=XM_023256636.1;refGene_gn=WNK1;refGene_tx=XM_023256637.1;refGene_gn=WNK1;refGene_tx=XM_023256638.1;refGene_gn=WNK1;refGene_tx=XM_023256639.1;refGene_gn=WNK1;refGene_tx=XM_023256640.1
B4 37112403 37112591
refGene_gn=WNK1;refGene_tx=XM_023256618.1;refGene_gn=WNK1;refGene_tx=XM_023256619.1;refGene_gn=WNK1;refGene_tx=XM_023256620.1;refGene_gn=WNK1;refGene_tx=XM_023256621.1;refGene_gn=WNK1;refGene_tx=XM_023256622.1;refGene_gn=WNK1;refGene_tx=XM_023256623.1;refGene_gn=WNK1;refGene_tx=XM_023256624.1;refGene_gn=WNK1;refGene_tx=XM_023256625.1;refGene_gn=WNK1;refGene_tx=XM_023256626.1;refGene_gn=WNK1;refGene_tx=XM_023256627.1;refGene_gn=WNK1;refGene_tx=XM_023256628.1;refGene_gn=WNK1;refGene_tx=XM_023256629.1;refGene_gn=WNK1;refGene_tx=XM_023256630.1;refGene_gn=WNK1;refGene_tx=XM_023256631.1;refGene_gn=WNK1;refGene_tx=XM_023256632.1;refGene_gn=WNK1;refGene_tx=XM_023256633.1;refGene_gn=WNK1;refGene_tx=XM_023256634.1;refGene_gn=WNK1;refGene_tx=XM_023256635.1;refGene_gn=WNK1;refGene_tx=XM_023256636.1;refGene_gn=WNK1;refGene_tx=XM_023256637.1;refGene_gn=WNK1;refGene_tx=XM_023256638.1;refGene_gn=WNK1;refGene_tx=XM_023256639.1;refGene_gn=WNK1;refGene_tx=XM_023256640.1
B4 37113543 37113570
refGene_gn=WNK1;refGene_tx=XM_023256618.1;refGene_gn=WNK1;refGene_tx=XM_023256619.1;refGene_gn=WNK1;refGene_tx=XM_023256620.1;refGene_gn=WNK1;refGene_tx=XM_023256621.1;refGene_gn=WNK1;refGene_tx=XM_023256624.1;refGene_gn=WNK1;refGene_tx=XM_023256625.1;refGene_gn=WNK1;refGene_tx=XM_023256628.1;refGene_gn=WNK1;refGene_tx=XM_023256630.1;refGene_gn=WNK1;refGene_tx=XM_023256631.1;refGene_gn=WNK1;refGene_tx=XM_023256632.1;refGene_gn=WNK1;refGene_tx=XM_023256634.1;refGene_gn=WNK1;refGene_tx=XM_023256635.1
B4 37115502 37115544
refGene_gn=WNK1;refGene_tx=XM_023256618.1;refGene_gn=WNK1;refGene_tx=XM_023256619.1;refGene_gn=WNK1;refGene_tx=XM_023256620.1;refGene_gn=WNK1;refGene_tx=XM_023256621.1;refGene_gn=WNK1;refGene_tx=XM_023256622.1;refGene_gn=WNK1;refGene_tx=XM_023256623.1;refGene_gn=WNK1;refGene_tx=X

M_023256624.1;refGene_gn=WNK1;refGene_tx=XM_023256625.1;refGene_gn=WNK1;refGene_tx=XM_023256626.1;refGene_gn=WNK1;refGene_tx=XM_023256628.1;refGene_gn=WNK1;refGene_tx=XM_023256629.1;refGene_gn=WNK1;refGene_tx=XM_023256630.1;refGene_gn=WNK1;refGene_tx=XM_023256631.1;refGene_gn=WNK1;refGene_tx=XM_023256632.1;refGene_gn=WNK1;refGene_tx=XM_023256634.1;refGene_gn=WNK1;refGene_tx=XM_023256635.1
B4 37117782 37117875
refGene_gn=WNK1;refGene_tx=XM_023256618.1;refGene_gn=WNK1;refGene_tx=XM_023256619.1;refGene_gn=WNK1;refGene_tx=XM_023256620.1;refGene_gn=WNK1;refGene_tx=XM_023256621.1;refGene_gn=WNK1;refGene_tx=XM_023256622.1;refGene_gn=WNK1;refGene_tx=XM_023256623.1;refGene_gn=WNK1;refGene_tx=XM_023256624.1;refGene_gn=WNK1;refGene_tx=XM_023256628.1;refGene_gn=WNK1;refGene_tx=XM_023256629.1;refGene_gn=WNK1;refGene_tx=XM_023256630.1;refGene_gn=WNK1;refGene_tx=XM_023256631.1;refGene_gn=WNK1;refGene_tx=XM_023256632.1;refGene_gn=WNK1;refGene_tx=XM_023256634.1;refGene_gn=WNK1;refGene_tx=XM_023256635.1
B4 37118799 37118994
refGene_gn=WNK1;refGene_tx=XM_023256618.1;refGene_gn=WNK1;refGene_tx=XM_023256619.1;refGene_gn=WNK1;refGene_tx=XM_023256620.1;refGene_gn=WNK1;refGene_tx=XM_023256621.1;refGene_gn=WNK1;refGene_tx=XM_023256622.1;refGene_gn=WNK1;refGene_tx=XM_023256623.1;refGene_gn=WNK1;refGene_tx=XM_023256624.1;refGene_gn=WNK1;refGene_tx=XM_023256625.1;refGene_gn=WNK1;refGene_tx=XM_023256626.1;refGene_gn=WNK1;refGene_tx=XM_023256627.1;refGene_gn=WNK1;refGene_tx=XM_023256628.1;refGene_gn=WNK1;refGene_tx=XM_023256629.1;refGene_gn=WNK1;refGene_tx=XM_023256630.1;refGene_gn=WNK1;refGene_tx=XM_023256631.1;refGene_gn=WNK1;refGene_tx=XM_023256632.1;refGene_gn=WNK1;refGene_tx=XM_023256633.1;refGene_gn=WNK1;refGene_tx=XM_023256634.1;refGene_gn=WNK1;refGene_tx=XM_023256635.1;refGene_gn=WNK1;refGene_tx=XM_023256636.1;refGene_gn=WNK1;refGene_tx=XM_023256637.1;refGene_gn=WNK1;refGene_tx=XM_023256638.1;refGene_gn=WNK1;refGene_tx=XM_023256639.1;refGene_gn=WNK1;refGene_tx=XM_023256640.1
B4 37120931 37121134
refGene_gn=WNK1;refGene_tx=XM_023256618.1;refGene_gn=WNK1;refGene_tx=XM_023256619.1;refGene_gn=WNK1;refGene_tx=XM_023256620.1;refGene_gn=WNK1;refGene_tx=XM_023256621.1;refGene_gn=WNK1;refGene_tx=XM_023256622.1;refGene_gn=WNK1;refGene_tx=XM_023256623.1;refGene_gn=WNK1;refGene_tx=XM_023256624.1;refGene_gn=WNK1;refGene_tx=XM_023256625.1;refGene_gn=WNK1;refGene_tx=XM_023256626.1;refGene_gn=WNK1;refGene_tx=XM_023256627.1;refGene_gn=WNK1;refGene_tx=XM_023256628.1;refGene_gn=WNK1;refGene_tx=XM_023256629.1;refGene_gn=WNK1;refGene_tx=XM_023256630.1;refGene_gn=WNK1;refGene_tx=XM_023256631.1;refGene_gn=WNK1;refGene_tx=XM_023256632.1;refGene_gn=WNK1;refGene_tx=XM_023256633.1;refGene_gn=WNK1;refGene_tx=XM_023256634.1;refGene_gn=WNK1;refGene_tx=XM_023256635.1;refGene_gn=WNK1;refGene_tx=XM_023256636.1;refGene_gn=WNK1;refGene_tx=XM_023256637.1;refGene_gn=WNK1;refGene_tx=XM_023256638.1;refGene_gn=WNK1;refGene_tx=XM_023256639.1;refGene_gn=WNK1;refGene_tx=XM_023256640.1
B4 37121794 37122450
refGene_gn=WNK1;refGene_tx=XM_023256618.1;refGene_gn=WNK1;refGene_tx=XM_023256619.1;refGene_gn=WNK1;refGene_tx=XM_023256620.1;refGene_gn=WNK1;refGene_tx=XM_023256621.1;refGene_gn=WNK1;refGene_tx=XM_023256622.1;

refGene_gn=WNK1;refGene_tx=XM_023256623.1;refGene_gn=WNK1;refGene_tx=X
M_023256624.1;refGene_gn=WNK1;refGene_tx=XM_023256625.1;refGene_gn=WNK
1;refGene_tx=XM_023256626.1;refGene_gn=WNK1;refGene_tx=XM_023256627.1;
refGene_gn=WNK1;refGene_tx=XM_023256628.1;refGene_gn=WNK1;refGene_tx=X
M_023256629.1;refGene_gn=WNK1;refGene_tx=XM_023256630.1;refGene_gn=WNK
1;refGene_tx=XM_023256631.1;refGene_gn=WNK1;refGene_tx=XM_023256632.1;
refGene_gn=WNK1;refGene_tx=XM_023256633.1;refGene_gn=WNK1;refGene_tx=X
M_023256634.1;refGene_gn=WNK1;refGene_tx=XM_023256635.1;refGene_gn=WNK
1;refGene_tx=XM_023256636.1;refGene_gn=WNK1;refGene_tx=XM_023256637.1;
refGene_gn=WNK1;refGene_tx=XM_023256638.1;refGene_gn=WNK1;refGene_tx=X
M_023256639.1;refGene_gn=WNK1;refGene_tx=XM_023256640.1
B4 37124029 37124103
refGene_gn=WNK1;refGene_tx=XM_023256618.1;refGene_gn=WNK1;refGene_tx=X
M_023256619.1;refGene_gn=WNK1;refGene_tx=XM_023256620.1;refGene_gn=WNK
1;refGene_tx=XM_023256621.1;refGene_gn=WNK1;refGene_tx=XM_023256622.1;
refGene_gn=WNK1;refGene_tx=XM_023256623.1;refGene_gn=WNK1;refGene_tx=X
M_023256624.1;refGene_gn=WNK1;refGene_tx=XM_023256625.1;refGene_gn=WNK
1;refGene_tx=XM_023256626.1;refGene_gn=WNK1;refGene_tx=XM_023256627.1;
refGene_gn=WNK1;refGene_tx=XM_023256628.1;refGene_gn=WNK1;refGene_tx=X
M_023256629.1;refGene_gn=WNK1;refGene_tx=XM_023256630.1;refGene_gn=WNK
1;refGene_tx=XM_023256631.1;refGene_gn=WNK1;refGene_tx=XM_023256632.1;
refGene_gn=WNK1;refGene_tx=XM_023256633.1;refGene_gn=WNK1;refGene_tx=X
M_023256634.1;refGene_gn=WNK1;refGene_tx=XM_023256635.1;refGene_gn=WNK
1;refGene_tx=XM_023256636.1;refGene_gn=WNK1;refGene_tx=XM_023256637.1;
refGene_gn=WNK1;refGene_tx=XM_023256638.1;refGene_gn=WNK1;refGene_tx=X
M_023256639.1;refGene_gn=WNK1;refGene_tx=XM_023256640.1
B4 37126940 37127001
refGene_gn=WNK1;refGene_tx=XM_023256618.1;refGene_gn=WNK1;refGene_tx=X
M_023256619.1;refGene_gn=WNK1;refGene_tx=XM_023256620.1;refGene_gn=WNK
1;refGene_tx=XM_023256621.1;refGene_gn=WNK1;refGene_tx=XM_023256622.1;
refGene_gn=WNK1;refGene_tx=XM_023256623.1;refGene_gn=WNK1;refGene_tx=X
M_023256624.1;refGene_gn=WNK1;refGene_tx=XM_023256625.1;refGene_gn=WNK
1;refGene_tx=XM_023256626.1;refGene_gn=WNK1;refGene_tx=XM_023256627.1;
refGene_gn=WNK1;refGene_tx=XM_023256628.1;refGene_gn=WNK1;refGene_tx=X
M_023256629.1;refGene_gn=WNK1;refGene_tx=XM_023256630.1;refGene_gn=WNK
1;refGene_tx=XM_023256631.1;refGene_gn=WNK1;refGene_tx=XM_023256632.1;
refGene_gn=WNK1;refGene_tx=XM_023256633.1;refGene_gn=WNK1;refGene_tx=X
M_023256634.1;refGene_gn=WNK1;refGene_tx=XM_023256635.1;refGene_gn=WNK
1;refGene_tx=XM_023256636.1;refGene_gn=WNK1;refGene_tx=XM_023256637.1;
refGene_gn=WNK1;refGene_tx=XM_023256638.1;refGene_gn=WNK1;refGene_tx=X
M_023256639.1;refGene_gn=WNK1;refGene_tx=XM_023256640.1
B4 37127654 37127738
refGene_gn=WNK1;refGene_tx=XM_023256618.1;refGene_gn=WNK1;refGene_tx=X
M_023256619.1;refGene_gn=WNK1;refGene_tx=XM_023256620.1;refGene_gn=WNK
1;refGene_tx=XM_023256621.1;refGene_gn=WNK1;refGene_tx=XM_023256622.1;
refGene_gn=WNK1;refGene_tx=XM_023256623.1;refGene_gn=WNK1;refGene_tx=X
M_023256624.1;refGene_gn=WNK1;refGene_tx=XM_023256625.1;refGene_gn=WNK
1;refGene_tx=XM_023256626.1;refGene_gn=WNK1;refGene_tx=XM_023256627.1;
refGene_gn=WNK1;refGene_tx=XM_023256628.1;refGene_gn=WNK1;refGene_tx=X
M_023256629.1;refGene_gn=WNK1;refGene_tx=XM_023256630.1;refGene_gn=WNK

1;refGene_tx=XM_023256631.1;refGene_gn=WNK1;refGene_tx=XM_023256632.1;
refGene_gn=WNK1;refGene_tx=XM_023256633.1;refGene_gn=WNK1;refGene_tx=X
M_023256634.1;refGene_gn=WNK1;refGene_tx=XM_023256635.1;refGene_gn=WNK
1;refGene_tx=XM_023256636.1;refGene_gn=WNK1;refGene_tx=XM_023256637.1;
refGene_gn=WNK1;refGene_tx=XM_023256638.1;refGene_gn=WNK1;refGene_tx=X
M_023256639.1;refGene_gn=WNK1;refGene_tx=XM_023256640.1
B4 37129751 37129835
refGene_gn=WNK1;refGene_tx=XM_023256618.1;refGene_gn=WNK1;refGene_tx=X
M_023256619.1;refGene_gn=WNK1;refGene_tx=XM_023256620.1;refGene_gn=WNK
1;refGene_tx=XM_023256621.1;refGene_gn=WNK1;refGene_tx=XM_023256622.1;
refGene_gn=WNK1;refGene_tx=XM_023256623.1;refGene_gn=WNK1;refGene_tx=X
M_023256625.1;refGene_gn=WNK1;refGene_tx=XM_023256626.1;refGene_gn=WNK
1;refGene_tx=XM_023256627.1;refGene_gn=WNK1;refGene_tx=XM_023256628.1;
refGene_gn=WNK1;refGene_tx=XM_023256629.1;refGene_gn=WNK1;refGene_tx=X
M_023256630.1;refGene_gn=WNK1;refGene_tx=XM_023256631.1;refGene_gn=WNK
1;refGene_tx=XM_023256632.1;refGene_gn=WNK1;refGene_tx=XM_023256633.1;
refGene_gn=WNK1;refGene_tx=XM_023256634.1;refGene_gn=WNK1;refGene_tx=X
M_023256635.1;refGene_gn=WNK1;refGene_tx=XM_023256636.1;refGene_gn=WNK
1;refGene_tx=XM_023256637.1;refGene_gn=WNK1;refGene_tx=XM_023256638.1;
refGene_gn=WNK1;refGene_tx=XM_023256639.1;refGene_gn=WNK1;refGene_tx=X
M_023256640.1
B4 37130962 37132410
refGene_gn=WNK1;refGene_tx=XM_023256618.1;refGene_gn=WNK1;refGene_tx=X
M_023256619.1;refGene_gn=WNK1;refGene_tx=XM_023256620.1;refGene_gn=WNK
1;refGene_tx=XM_023256621.1;refGene_gn=WNK1;refGene_tx=XM_023256622.1;
refGene_gn=WNK1;refGene_tx=XM_023256623.1;refGene_gn=WNK1;refGene_tx=X
M_023256624.1;refGene_gn=WNK1;refGene_tx=XM_023256625.1;refGene_gn=WNK
1;refGene_tx=XM_023256626.1;refGene_gn=WNK1;refGene_tx=XM_023256627.1;
refGene_gn=WNK1;refGene_tx=XM_023256628.1;refGene_gn=WNK1;refGene_tx=X
M_023256629.1;refGene_gn=WNK1;refGene_tx=XM_023256630.1;refGene_gn=WNK
1;refGene_tx=XM_023256631.1;refGene_gn=WNK1;refGene_tx=XM_023256632.1;
refGene_gn=WNK1;refGene_tx=XM_023256633.1;refGene_gn=WNK1;refGene_tx=X
M_023256634.1;refGene_gn=WNK1;refGene_tx=XM_023256635.1;refGene_gn=WNK
1;refGene_tx=XM_023256636.1;refGene_gn=WNK1;refGene_tx=XM_023256637.1;
refGene_gn=WNK1;refGene_tx=XM_023256638.1;refGene_gn=WNK1;refGene_tx=X
M_023256639.1;refGene_gn=WNK1;refGene_tx=XM_023256640.1
B4 37132974 37133097
refGene_gn=WNK1;refGene_tx=XM_023256618.1;refGene_gn=WNK1;refGene_tx=X
M_023256619.1;refGene_gn=WNK1;refGene_tx=XM_023256620.1;refGene_gn=WNK
1;refGene_tx=XM_023256621.1;refGene_gn=WNK1;refGene_tx=XM_023256622.1;
refGene_gn=WNK1;refGene_tx=XM_023256623.1;refGene_gn=WNK1;refGene_tx=X
M_023256624.1;refGene_gn=WNK1;refGene_tx=XM_023256625.1;refGene_gn=WNK
1;refGene_tx=XM_023256626.1;refGene_gn=WNK1;refGene_tx=XM_023256627.1;
refGene_gn=WNK1;refGene_tx=XM_023256628.1;refGene_gn=WNK1;refGene_tx=X
M_023256629.1;refGene_gn=WNK1;refGene_tx=XM_023256630.1;refGene_gn=WNK
1;refGene_tx=XM_023256631.1;refGene_gn=WNK1;refGene_tx=XM_023256632.1;
refGene_gn=WNK1;refGene_tx=XM_023256633.1;refGene_gn=WNK1;refGene_tx=X
M_023256634.1;refGene_gn=WNK1;refGene_tx=XM_023256635.1;refGene_gn=WNK
1;refGene_tx=XM_023256636.1;refGene_gn=WNK1;refGene_tx=XM_023256637.1;
refGene_gn=WNK1;refGene_tx=XM_023256638.1;refGene_gn=WNK1;refGene_tx=X

M_023256639.1;refGene_gn=WNK1;refGene_tx=XM_023256640.1
B4 37133223 37133281
refGene_gn=WNK1;refGene_tx=XM_023256618.1;refGene_gn=WNK1;refGene_tx=XM_023256619.1;refGene_gn=WNK1;refGene_tx=XM_023256620.1;refGene_gn=WNK1;refGene_tx=XM_023256621.1;refGene_gn=WNK1;refGene_tx=XM_023256622.1;refGene_gn=WNK1;refGene_tx=XM_023256623.1;refGene_gn=WNK1;refGene_tx=XM_023256624.1;refGene_gn=WNK1;refGene_tx=XM_023256625.1;refGene_gn=WNK1;refGene_tx=XM_023256626.1;refGene_gn=WNK1;refGene_tx=XM_023256627.1;refGene_gn=WNK1;refGene_tx=XM_023256628.1;refGene_gn=WNK1;refGene_tx=XM_023256629.1;refGene_gn=WNK1;refGene_tx=XM_023256630.1;refGene_gn=WNK1;refGene_tx=XM_023256631.1;refGene_gn=WNK1;refGene_tx=XM_023256632.1;refGene_gn=WNK1;refGene_tx=XM_023256633.1;refGene_gn=WNK1;refGene_tx=XM_023256634.1;refGene_gn=WNK1;refGene_tx=XM_023256635.1;refGene_gn=WNK1;refGene_tx=XM_023256636.1;refGene_gn=WNK1;refGene_tx=XM_023256637.1;refGene_gn=WNK1;refGene_tx=XM_023256638.1;refGene_gn=WNK1;refGene_tx=XM_023256639.1;refGene_gn=WNK1;refGene_tx=XM_023256640.1
B4 37133513 37133687
refGene_gn=WNK1;refGene_tx=XM_023256618.1;refGene_gn=WNK1;refGene_tx=XM_023256619.1;refGene_gn=WNK1;refGene_tx=XM_023256620.1;refGene_gn=WNK1;refGene_tx=XM_023256621.1;refGene_gn=WNK1;refGene_tx=XM_023256622.1;refGene_gn=WNK1;refGene_tx=XM_023256623.1;refGene_gn=WNK1;refGene_tx=XM_023256624.1;refGene_gn=WNK1;refGene_tx=XM_023256625.1;refGene_gn=WNK1;refGene_tx=XM_023256626.1;refGene_gn=WNK1;refGene_tx=XM_023256627.1;refGene_gn=WNK1;refGene_tx=XM_023256628.1;refGene_gn=WNK1;refGene_tx=XM_023256629.1;refGene_gn=WNK1;refGene_tx=XM_023256630.1;refGene_gn=WNK1;refGene_tx=XM_023256631.1;refGene_gn=WNK1;refGene_tx=XM_023256632.1;refGene_gn=WNK1;refGene_tx=XM_023256633.1;refGene_gn=WNK1;refGene_tx=XM_023256634.1;refGene_gn=WNK1;refGene_tx=XM_023256635.1;refGene_gn=WNK1;refGene_tx=XM_023256636.1;refGene_gn=WNK1;refGene_tx=XM_023256637.1;refGene_gn=WNK1;refGene_tx=XM_023256638.1;refGene_gn=WNK1;refGene_tx=XM_023256639.1;refGene_gn=WNK1;refGene_tx=XM_023256640.1
B4 37134008 37134125
refGene_gn=WNK1;refGene_tx=XM_023256618.1;refGene_gn=WNK1;refGene_tx=XM_023256619.1;refGene_gn=WNK1;refGene_tx=XM_023256620.1;refGene_gn=WNK1;refGene_tx=XM_023256621.1;refGene_gn=WNK1;refGene_tx=XM_023256622.1;refGene_gn=WNK1;refGene_tx=XM_023256623.1;refGene_gn=WNK1;refGene_tx=XM_023256624.1;refGene_gn=WNK1;refGene_tx=XM_023256625.1;refGene_gn=WNK1;refGene_tx=XM_023256626.1;refGene_gn=WNK1;refGene_tx=XM_023256627.1;refGene_gn=WNK1;refGene_tx=XM_023256628.1;refGene_gn=WNK1;refGene_tx=XM_023256629.1;refGene_gn=WNK1;refGene_tx=XM_023256630.1;refGene_gn=WNK1;refGene_tx=XM_023256631.1;refGene_gn=WNK1;refGene_tx=XM_023256632.1;refGene_gn=WNK1;refGene_tx=XM_023256633.1;refGene_gn=WNK1;refGene_tx=XM_023256634.1;refGene_gn=WNK1;refGene_tx=XM_023256635.1;refGene_gn=WNK1;refGene_tx=XM_023256636.1;refGene_gn=WNK1;refGene_tx=XM_023256637.1;refGene_gn=WNK1;refGene_tx=XM_023256638.1;refGene_gn=WNK1;refGene_tx=XM_023256639.1;refGene_gn=WNK1;refGene_tx=XM_023256640.1
B4 37134550 37134713
refGene_gn=WNK1;refGene_tx=XM_023256618.1;refGene_gn=WNK1;refGene_tx=XM_023256619.1;refGene_gn=WNK1;refGene_tx=XM_023256620.1;refGene_gn=WNK1;refGene_tx=XM_023256621.1;refGene_gn=WNK1;refGene_tx=XM_023256622.1;

refGene_gn=WNK1;refGene_tx=XM_023256623.1;refGene_gn=WNK1;refGene_tx=XM_023256624.1;refGene_gn=WNK1;refGene_tx=XM_023256625.1;refGene_gn=WNK1;refGene_tx=XM_023256626.1;refGene_gn=WNK1;refGene_tx=XM_023256627.1;refGene_gn=WNK1;refGene_tx=XM_023256628.1;refGene_gn=WNK1;refGene_tx=XM_023256629.1;refGene_gn=WNK1;refGene_tx=XM_023256630.1;refGene_gn=WNK1;refGene_tx=XM_023256631.1;refGene_gn=WNK1;refGene_tx=XM_023256632.1;refGene_gn=WNK1;refGene_tx=XM_023256633.1;refGene_gn=WNK1;refGene_tx=XM_023256634.1;refGene_gn=WNK1;refGene_tx=XM_023256635.1;refGene_gn=WNK1;refGene_tx=XM_023256636.1;refGene_gn=WNK1;refGene_tx=XM_023256637.1;refGene_gn=WNK1;refGene_tx=XM_023256638.1;refGene_gn=WNK1;refGene_tx=XM_023256639.1;refGene_gn=WNK1;refGene_tx=XM_023256640.1
B4 37134806 37134895
refGene_gn=WNK1;refGene_tx=XM_023256618.1;refGene_gn=WNK1;refGene_tx=XM_023256619.1;refGene_gn=WNK1;refGene_tx=XM_023256620.1;refGene_gn=WNK1;refGene_tx=XM_023256621.1;refGene_gn=WNK1;refGene_tx=XM_023256622.1;refGene_gn=WNK1;refGene_tx=XM_023256623.1;refGene_gn=WNK1;refGene_tx=XM_023256624.1;refGene_gn=WNK1;refGene_tx=XM_023256625.1;refGene_gn=WNK1;refGene_tx=XM_023256626.1;refGene_gn=WNK1;refGene_tx=XM_023256627.1;refGene_gn=WNK1;refGene_tx=XM_023256628.1;refGene_gn=WNK1;refGene_tx=XM_023256629.1;refGene_gn=WNK1;refGene_tx=XM_023256630.1;refGene_gn=WNK1;refGene_tx=XM_023256631.1;refGene_gn=WNK1;refGene_tx=XM_023256632.1;refGene_gn=WNK1;refGene_tx=XM_023256633.1;refGene_gn=WNK1;refGene_tx=XM_023256634.1;refGene_gn=WNK1;refGene_tx=XM_023256635.1;refGene_gn=WNK1;refGene_tx=XM_023256636.1;refGene_gn=WNK1;refGene_tx=XM_023256637.1;refGene_gn=WNK1;refGene_tx=XM_023256638.1;refGene_gn=WNK1;refGene_tx=XM_023256639.1;refGene_gn=WNK1;refGene_tx=XM_023256640.1
B4 37135599 37135881
refGene_gn=WNK1;refGene_tx=XM_023256618.1;refGene_gn=WNK1;refGene_tx=XM_023256619.1;refGene_gn=WNK1;refGene_tx=XM_023256620.1;refGene_gn=WNK1;refGene_tx=XM_023256621.1;refGene_gn=WNK1;refGene_tx=XM_023256622.1;refGene_gn=WNK1;refGene_tx=XM_023256623.1;refGene_gn=WNK1;refGene_tx=XM_023256624.1;refGene_gn=WNK1;refGene_tx=XM_023256625.1;refGene_gn=WNK1;refGene_tx=XM_023256626.1;refGene_gn=WNK1;refGene_tx=XM_023256627.1;refGene_gn=WNK1;refGene_tx=XM_023256628.1;refGene_gn=WNK1;refGene_tx=XM_023256629.1;refGene_gn=WNK1;refGene_tx=XM_023256630.1;refGene_gn=WNK1;refGene_tx=XM_023256631.1;refGene_gn=WNK1;refGene_tx=XM_023256633.1;refGene_gn=WNK1;refGene_tx=XM_023256634.1;refGene_gn=WNK1;refGene_tx=XM_023256635.1;refGene_gn=WNK1;refGene_tx=XM_023256636.1;refGene_gn=WNK1;refGene_tx=XM_023256639.1;refGene_gn=WNK1;refGene_tx=XM_023256640.1
B4 37136652 37137111
refGene_gn=WNK1;refGene_tx=XM_023256618.1;refGene_gn=WNK1;refGene_tx=XM_023256619.1;refGene_gn=WNK1;refGene_tx=XM_023256620.1;refGene_gn=WNK1;refGene_tx=XM_023256621.1;refGene_gn=WNK1;refGene_tx=XM_023256622.1;refGene_gn=WNK1;refGene_tx=XM_023256623.1;refGene_gn=WNK1;refGene_tx=XM_023256624.1;refGene_gn=WNK1;refGene_tx=XM_023256625.1;refGene_gn=WNK1;refGene_tx=XM_023256626.1;refGene_gn=WNK1;refGene_tx=XM_023256627.1;refGene_gn=WNK1;refGene_tx=XM_023256628.1;refGene_gn=WNK1;refGene_tx=XM_023256629.1;refGene_gn=WNK1;refGene_tx=XM_023256634.1;refGene_gn=WNK1;refGene_tx=XM_023256635.1;refGene_gn=WNK1;refGene_tx=XM_023256636.1;refGene_gn=WNK1;refGene_tx=XM_023256640.1

B4 37138365 37138515
refGene_gn=WNK1;refGene_tx=XM_023256618.1;refGene_gn=WNK1;refGene_tx=X
M_023256619.1;refGene_gn=WNK1;refGene_tx=XM_023256620.1;refGene_gn=WNK
1;refGene_tx=XM_023256621.1;refGene_gn=WNK1;refGene_tx=XM_023256622.1;
refGene_gn=WNK1;refGene_tx=XM_023256623.1;refGene_gn=WNK1;refGene_tx=X
M_023256624.1;refGene_gn=WNK1;refGene_tx=XM_023256625.1;refGene_gn=WNK
1;refGene_tx=XM_023256626.1;refGene_gn=WNK1;refGene_tx=XM_023256627.1;
refGene_gn=WNK1;refGene_tx=XM_023256628.1;refGene_gn=WNK1;refGene_tx=X
M_023256629.1;refGene_gn=WNK1;refGene_tx=XM_023256630.1;refGene_gn=WNK
1;refGene_tx=XM_023256631.1;refGene_gn=WNK1;refGene_tx=XM_023256632.1;
refGene_gn=WNK1;refGene_tx=XM_023256633.1;refGene_gn=WNK1;refGene_tx=X
M_023256634.1;refGene_gn=WNK1;refGene_tx=XM_023256635.1;refGene_gn=WNK
1;refGene_tx=XM_023256636.1;refGene_gn=WNK1;refGene_tx=XM_023256637.1;
refGene_gn=WNK1;refGene_tx=XM_023256638.1;refGene_gn=WNK1;refGene_tx=X
M_023256639.1;refGene_gn=WNK1;refGene_tx=XM_023256640.1
B4 37144964 37145048
refGene_gn=WNK1;refGene_tx=XM_023256618.1;refGene_gn=WNK1;refGene_tx=X
M_023256619.1;refGene_gn=WNK1;refGene_tx=XM_023256620.1;refGene_gn=WNK
1;refGene_tx=XM_023256621.1;refGene_gn=WNK1;refGene_tx=XM_023256622.1;
refGene_gn=WNK1;refGene_tx=XM_023256623.1;refGene_gn=WNK1;refGene_tx=X
M_023256624.1;refGene_gn=WNK1;refGene_tx=XM_023256625.1;refGene_gn=WNK
1;refGene_tx=XM_023256626.1;refGene_gn=WNK1;refGene_tx=XM_023256627.1;
refGene_gn=WNK1;refGene_tx=XM_023256628.1;refGene_gn=WNK1;refGene_tx=X
M_023256629.1;refGene_gn=WNK1;refGene_tx=XM_023256630.1;refGene_gn=WNK
1;refGene_tx=XM_023256631.1;refGene_gn=WNK1;refGene_tx=XM_023256632.1;
refGene_gn=WNK1;refGene_tx=XM_023256633.1;refGene_gn=WNK1;refGene_tx=X
M_023256634.1;refGene_gn=WNK1;refGene_tx=XM_023256635.1;refGene_gn=WNK
1;refGene_tx=XM_023256636.1;refGene_gn=WNK1;refGene_tx=XM_023256637.1;
refGene_gn=WNK1;refGene_tx=XM_023256638.1;refGene_gn=WNK1;refGene_tx=X
M_023256639.1;refGene_gn=WNK1;refGene_tx=XM_023256640.1
B4 37147209 37148439
refGene_gn=WNK1;refGene_tx=XM_023256618.1;refGene_gn=WNK1;refGene_tx=X
M_023256619.1;refGene_gn=WNK1;refGene_tx=XM_023256620.1;refGene_gn=WNK
1;refGene_tx=XM_023256621.1;refGene_gn=WNK1;refGene_tx=XM_023256622.1;
refGene_gn=WNK1;refGene_tx=XM_023256623.1;refGene_gn=WNK1;refGene_tx=X
M_023256624.1;refGene_gn=WNK1;refGene_tx=XM_023256625.1;refGene_gn=WNK
1;refGene_tx=XM_023256626.1;refGene_gn=WNK1;refGene_tx=XM_023256627.1;
refGene_gn=WNK1;refGene_tx=XM_023256628.1;refGene_gn=WNK1;refGene_tx=X
M_023256629.1;refGene_gn=WNK1;refGene_tx=XM_023256630.1;refGene_gn=WNK
1;refGene_tx=XM_023256631.1;refGene_gn=WNK1;refGene_tx=XM_023256632.1;
refGene_gn=WNK1;refGene_tx=XM_023256633.1;refGene_gn=WNK1;refGene_tx=X
M_023256639.1;refGene_gn=WNK1;refGene_tx=XM_023256640.1
B4 37150351 37150609
refGene_gn=WNK1;refGene_tx=XM_023256618.1;refGene_gn=WNK1;refGene_tx=X
M_023256619.1;refGene_gn=WNK1;refGene_tx=XM_023256620.1;refGene_gn=WNK
1;refGene_tx=XM_023256621.1;refGene_gn=WNK1;refGene_tx=XM_023256622.1;
refGene_gn=WNK1;refGene_tx=XM_023256623.1;refGene_gn=WNK1;refGene_tx=X
M_023256624.1;refGene_gn=WNK1;refGene_tx=XM_023256625.1;refGene_gn=WNK
1;refGene_tx=XM_023256626.1;refGene_gn=WNK1;refGene_tx=XM_023256627.1;
refGene_gn=WNK1;refGene_tx=XM_023256630.1;refGene_gn=WNK1;refGene_tx=X

M_023256632.1;refGene_gn=WNK1;refGene_tx=XM_023256640.1
B4 37153261 37153449
refGene_gn=WNK1;refGene_tx=XM_023256618.1;refGene_gn=WNK1;refGene_tx=XM_023256619.1;refGene_gn=WNK1;refGene_tx=XM_023256620.1;refGene_gn=WNK1;refGene_tx=XM_023256621.1;refGene_gn=WNK1;refGene_tx=XM_023256622.1;refGene_gn=WNK1;refGene_tx=XM_023256623.1;refGene_gn=WNK1;refGene_tx=XM_023256624.1;refGene_gn=WNK1;refGene_tx=XM_023256625.1;refGene_gn=WNK1;refGene_tx=XM_023256626.1;refGene_gn=WNK1;refGene_tx=XM_023256627.1;refGene_gn=WNK1;refGene_tx=XM_023256628.1;refGene_gn=WNK1;refGene_tx=XM_023256629.1;refGene_gn=WNK1;refGene_tx=XM_023256630.1;refGene_gn=WNK1;refGene_tx=XM_023256631.1;refGene_gn=WNK1;refGene_tx=XM_023256632.1;refGene_gn=WNK1;refGene_tx=XM_023256633.1;refGene_gn=WNK1;refGene_tx=XM_023256634.1;refGene_gn=WNK1;refGene_tx=XM_023256635.1;refGene_gn=WNK1;refGene_tx=XM_023256636.1;refGene_gn=WNK1;refGene_tx=XM_023256637.1;refGene_gn=WNK1;refGene_tx=XM_023256638.1;refGene_gn=WNK1;refGene_tx=XM_023256639.1;refGene_gn=WNK1;refGene_tx=XM_023256640.1
B4 37154795 37155129
refGene_gn=WNK1;refGene_tx=XM_023256618.1;refGene_gn=WNK1;refGene_tx=XM_023256619.1;refGene_gn=WNK1;refGene_tx=XM_023256620.1;refGene_gn=WNK1;refGene_tx=XM_023256621.1;refGene_gn=WNK1;refGene_tx=XM_023256622.1;refGene_gn=WNK1;refGene_tx=XM_023256623.1;refGene_gn=WNK1;refGene_tx=XM_023256624.1;refGene_gn=WNK1;refGene_tx=XM_023256625.1;refGene_gn=WNK1;refGene_tx=XM_023256626.1;refGene_gn=WNK1;refGene_tx=XM_023256627.1;refGene_gn=WNK1;refGene_tx=XM_023256628.1;refGene_gn=WNK1;refGene_tx=XM_023256629.1;refGene_gn=WNK1;refGene_tx=XM_023256630.1;refGene_gn=WNK1;refGene_tx=XM_023256631.1;refGene_gn=WNK1;refGene_tx=XM_023256632.1;refGene_gn=WNK1;refGene_tx=XM_023256633.1;refGene_gn=WNK1;refGene_tx=XM_023256634.1;refGene_gn=WNK1;refGene_tx=XM_023256635.1;refGene_gn=WNK1;refGene_tx=XM_023256636.1;refGene_gn=WNK1;refGene_tx=XM_023256637.1;refGene_gn=WNK1;refGene_tx=XM_023256638.1;refGene_gn=WNK1;refGene_tx=XM_023256639.1;refGene_gn=WNK1;refGene_tx=XM_023256640.1
B4 37156477 37156697
refGene_gn=WNK1;refGene_tx=XM_023256618.1;refGene_gn=WNK1;refGene_tx=XM_023256619.1;refGene_gn=WNK1;refGene_tx=XM_023256620.1;refGene_gn=WNK1;refGene_tx=XM_023256621.1;refGene_gn=WNK1;refGene_tx=XM_023256622.1;refGene_gn=WNK1;refGene_tx=XM_023256623.1;refGene_gn=WNK1;refGene_tx=XM_023256624.1;refGene_gn=WNK1;refGene_tx=XM_023256625.1;refGene_gn=WNK1;refGene_tx=XM_023256626.1;refGene_gn=WNK1;refGene_tx=XM_023256627.1;refGene_gn=WNK1;refGene_tx=XM_023256628.1;refGene_gn=WNK1;refGene_tx=XM_023256629.1;refGene_gn=WNK1;refGene_tx=XM_023256630.1;refGene_gn=WNK1;refGene_tx=XM_023256631.1;refGene_gn=WNK1;refGene_tx=XM_023256632.1;refGene_gn=WNK1;refGene_tx=XM_023256633.1;refGene_gn=WNK1;refGene_tx=XM_023256634.1;refGene_gn=WNK1;refGene_tx=XM_023256635.1;refGene_gn=WNK1;refGene_tx=XM_023256636.1;refGene_gn=WNK1;refGene_tx=XM_023256637.1;refGene_gn=WNK1;refGene_tx=XM_023256638.1;refGene_gn=WNK1;refGene_tx=XM_023256639.1;refGene_gn=WNK1;refGene_tx=XM_023256640.1
B4 37158749 37158838
refGene_gn=WNK1;refGene_tx=XM_023256618.1;refGene_gn=WNK1;refGene_tx=XM_023256619.1;refGene_gn=WNK1;refGene_tx=XM_023256620.1;refGene_gn=WNK1;refGene_tx=XM_023256621.1;refGene_gn=WNK1;refGene_tx=XM_023256622.1;

refGene_gn=WNK1;refGene_tx=XM_023256623.1;refGene_gn=WNK1;refGene_tx=XM_023256624.1;refGene_gn=WNK1;refGene_tx=XM_023256625.1;refGene_gn=WNK1;refGene_tx=XM_023256626.1;refGene_gn=WNK1;refGene_tx=XM_023256627.1;refGene_gn=WNK1;refGene_tx=XM_023256628.1;refGene_gn=WNK1;refGene_tx=XM_023256629.1;refGene_gn=WNK1;refGene_tx=XM_023256630.1;refGene_gn=WNK1;refGene_tx=XM_023256631.1;refGene_gn=WNK1;refGene_tx=XM_023256632.1;refGene_gn=WNK1;refGene_tx=XM_023256633.1;refGene_gn=WNK1;refGene_tx=XM_023256634.1;refGene_gn=WNK1;refGene_tx=XM_023256635.1;refGene_gn=WNK1;refGene_tx=XM_023256636.1;refGene_gn=WNK1;refGene_tx=XM_023256637.1;refGene_gn=WNK1;refGene_tx=XM_023256638.1;refGene_gn=WNK1;refGene_tx=XM_023256639.1;refGene_gn=WNK1;refGene_tx=XM_023256640.1
B4 37182953 37183111
refGene_gn=WNK1;refGene_tx=XM_023256618.1;refGene_gn=WNK1;refGene_tx=XM_023256619.1;refGene_gn=WNK1;refGene_tx=XM_023256620.1;refGene_gn=WNK1;refGene_tx=XM_023256621.1;refGene_gn=WNK1;refGene_tx=XM_023256622.1;refGene_gn=WNK1;refGene_tx=XM_023256623.1;refGene_gn=WNK1;refGene_tx=XM_023256624.1;refGene_gn=WNK1;refGene_tx=XM_023256625.1;refGene_gn=WNK1;refGene_tx=XM_023256626.1;refGene_gn=WNK1;refGene_tx=XM_023256627.1;refGene_gn=WNK1;refGene_tx=XM_023256628.1;refGene_gn=WNK1;refGene_tx=XM_023256629.1;refGene_gn=WNK1;refGene_tx=XM_023256630.1;refGene_gn=WNK1;refGene_tx=XM_023256631.1;refGene_gn=WNK1;refGene_tx=XM_023256632.1;refGene_gn=WNK1;refGene_tx=XM_023256633.1;refGene_gn=WNK1;refGene_tx=XM_023256634.1;refGene_gn=WNK1;refGene_tx=XM_023256635.1;refGene_gn=WNK1;refGene_tx=XM_023256636.1;refGene_gn=WNK1;refGene_tx=XM_023256637.1;refGene_gn=WNK1;refGene_tx=XM_023256638.1;refGene_gn=WNK1;refGene_tx=XM_023256639.1;refGene_gn=WNK1;refGene_tx=XM_023256640.1
B4 37185733 37185954
refGene_gn=WNK1;refGene_tx=XM_023256618.1;refGene_gn=WNK1;refGene_tx=XM_023256619.1;refGene_gn=WNK1;refGene_tx=XM_023256620.1;refGene_gn=WNK1;refGene_tx=XM_023256621.1;refGene_gn=WNK1;refGene_tx=XM_023256622.1;refGene_gn=WNK1;refGene_tx=XM_023256623.1;refGene_gn=WNK1;refGene_tx=XM_023256624.1;refGene_gn=WNK1;refGene_tx=XM_023256625.1;refGene_gn=WNK1;refGene_tx=XM_023256626.1;refGene_gn=WNK1;refGene_tx=XM_023256627.1;refGene_gn=WNK1;refGene_tx=XM_023256628.1;refGene_gn=WNK1;refGene_tx=XM_023256629.1;refGene_gn=WNK1;refGene_tx=XM_023256630.1;refGene_gn=WNK1;refGene_tx=XM_023256631.1;refGene_gn=WNK1;refGene_tx=XM_023256632.1;refGene_gn=WNK1;refGene_tx=XM_023256633.1;refGene_gn=WNK1;refGene_tx=XM_023256634.1;refGene_gn=WNK1;refGene_tx=XM_023256635.1;refGene_gn=WNK1;refGene_tx=XM_023256636.1;refGene_gn=WNK1;refGene_tx=XM_023256637.1;refGene_gn=WNK1;refGene_tx=XM_023256638.1;refGene_gn=WNK1;refGene_tx=XM_023256639.1;refGene_gn=WNK1;refGene_tx=XM_023256640.1
B4 37200465 37200638
refGene_gn=WNK1;refGene_tx=XM_023256618.1;refGene_gn=WNK1;refGene_tx=XM_023256619.1;refGene_gn=WNK1;refGene_tx=XM_023256620.1;refGene_gn=WNK1;refGene_tx=XM_023256621.1;refGene_gn=WNK1;refGene_tx=XM_023256622.1;refGene_gn=WNK1;refGene_tx=XM_023256623.1;refGene_gn=WNK1;refGene_tx=XM_023256624.1;refGene_gn=WNK1;refGene_tx=XM_023256625.1;refGene_gn=WNK1;refGene_tx=XM_023256626.1;refGene_gn=WNK1;refGene_tx=XM_023256627.1;refGene_gn=WNK1;refGene_tx=XM_023256628.1;refGene_gn=WNK1;refGene_tx=XM_023256629.1;refGene_gn=WNK1;refGene_tx=XM_023256630.1;refGene_gn=WNK

1;refGene_tx=XM_023256631.1;refGene_gn=WNK1;refGene_tx=XM_023256632.1;refGene_gn=WNK1;refGene_tx=XM_023256633.1;refGene_gn=WNK1;refGene_tx=XM_023256634.1;refGene_gn=WNK1;refGene_tx=XM_023256635.1;refGene_gn=WNK1;refGene_tx=XM_023256636.1;refGene_gn=WNK1;refGene_tx=XM_023256637.1;refGene_gn=WNK1;refGene_tx=XM_023256638.1;refGene_gn=WNK1;refGene_tx=XM_023256639.1;refGene_gn=WNK1;refGene_tx=XM_023256640.1
B4 37268046 37268808 refGene_gn=WNK1;refGene_tx=XM_023256618.1;refGene_gn=WNK1;refGene_tx=XM_023256619.1;refGene_gn=WNK1;refGene_tx=XM_023256620.1;refGene_gn=WNK1;refGene_tx=XM_023256621.1;refGene_gn=WNK1;refGene_tx=XM_023256622.1;refGene_gn=WNK1;refGene_tx=XM_023256623.1;refGene_gn=WNK1;refGene_tx=XM_023256624.1;refGene_gn=WNK1;refGene_tx=XM_023256625.1;refGene_gn=WNK1;refGene_tx=XM_023256626.1;refGene_gn=WNK1;refGene_tx=XM_023256627.1;refGene_gn=WNK1;refGene_tx=XM_023256628.1;refGene_gn=WNK1;refGene_tx=XM_023256629.1;refGene_gn=WNK1;refGene_tx=XM_023256630.1;refGene_gn=WNK1;refGene_tx=XM_023256631.1;refGene_gn=WNK1;refGene_tx=XM_023256632.1;refGene_gn=WNK1;refGene_tx=XM_023256633.1;refGene_gn=WNK1;refGene_tx=XM_023256634.1;refGene_gn=WNK1;refGene_tx=XM_023256635.1;refGene_gn=WNK1;refGene_tx=XM_023256636.1;refGene_gn=WNK1;refGene_tx=XM_023256637.1;refGene_gn=WNK1;refGene_tx=XM_023256638.1;refGene_gn=WNK1;refGene_tx=XM_023256639.1;refGene_gn=WNK1;refGene_tx=XM_023256640.1
B4 37338232 37338406 refGene_gn=NINJ2;refGene_tx=XM_019834331.2
B4 37346469 37346502 refGene_gn=NINJ2;refGene_tx=XM_019834332.2
B4 37382288 37382523 refGene_gn=B4GALNT3;refGene_tx=XM_023256641.1
B4 37417292 37417524 refGene_gn=NINJ2;refGene_tx=XM_011283747.3;refGene_gn=NINJ2;refGene_tx=XM_019834331.2;refGene_gn=NINJ2;refGene_tx=XM_019834332.2
B4 37418053 37418220 refGene_gn=NINJ2;refGene_tx=XM_011283747.3;refGene_gn=NINJ2;refGene_tx=XM_019834331.2;refGene_gn=NINJ2;refGene_tx=XM_019834332.2
B4 37422080 37422189 refGene_gn=B4GALNT3;refGene_tx=XM_011283761.3
B4 37424482 37424609 refGene_gn=B4GALNT3;refGene_tx=XM_011283761.3;refGene_gn=B4GALNT3;refGene_tx=XM_023256641.1
B4 37425212 37425366 refGene_gn=B4GALNT3;refGene_tx=XM_011283761.3;refGene_gn=B4GALNT3;refGene_tx=XM_023256641.1
B4 37425778 37425851 refGene_gn=B4GALNT3;refGene_tx=XM_011283761.3;refGene_gn=B4GALNT3;refGene_tx=XM_023256641.1
B4 37426110 37426264 refGene_gn=B4GALNT3;refGene_tx=XM_011283761.3;refGene_gn=B4GALNT3;refGene_tx=XM_023256641.1
B4 37426924 37427244 refGene_gn=B4GALNT3;refGene_tx=XM_011283761.3;refGene_gn=B4GALNT3;refGene_tx=XM_023256641.1
B4 37429165 37429961 refGene_gn=B4GALNT3;refGene_tx=XM_011283761.3;refGene_gn=B4GALNT3;refGene_tx=XM_023256641.1
B4 37430631 37430693

refGene_gn=B4GALNT3;refGene_tx=XM_011283761.3;refGene_gn=B4GALNT3;refGene_tx=XM_023256641.1
B4 37430980 37431081
refGene_gn=B4GALNT3;refGene_tx=XM_011283761.3;refGene_gn=B4GALNT3;refGene_tx=XM_023256641.1
B4 37432029 37432139
refGene_gn=B4GALNT3;refGene_tx=XM_011283761.3;refGene_gn=B4GALNT3;refGene_tx=XM_023256641.1
B4 37432946 37433090
refGene_gn=B4GALNT3;refGene_tx=XM_011283761.3;refGene_gn=B4GALNT3;refGene_tx=XM_023256641.1
B4 37434982 37435049
refGene_gn=B4GALNT3;refGene_tx=XM_011283761.3;refGene_gn=B4GALNT3;refGene_tx=XM_023256641.1
B4 37435175 37435254
refGene_gn=B4GALNT3;refGene_tx=XM_011283761.3;refGene_gn=B4GALNT3;refGene_tx=XM_023256641.1
B4 37436879 37436947
refGene_gn=B4GALNT3;refGene_tx=XM_011283761.3;refGene_gn=B4GALNT3;refGene_tx=XM_023256641.1
B4 37437491 37437592
refGene_gn=B4GALNT3;refGene_tx=XM_011283761.3;refGene_gn=B4GALNT3;refGene_tx=XM_023256641.1
B4 37437972 37438063
refGene_gn=B4GALNT3;refGene_tx=XM_011283761.3;refGene_gn=B4GALNT3;refGene_tx=XM_023256641.1
B4 37438568 37438664
refGene_gn=B4GALNT3;refGene_tx=XM_011283761.3;refGene_gn=B4GALNT3;refGene_tx=XM_023256641.1
B4 37446283 37446361
refGene_gn=B4GALNT3;refGene_tx=XM_011283761.3;refGene_gn=B4GALNT3;refGene_tx=XM_023256641.1
B4 37447358 37447462
refGene_gn=B4GALNT3;refGene_tx=XM_011283761.3;refGene_gn=B4GALNT3;refGene_tx=XM_023256641.1
B4 37520743 37520912
refGene_gn=B4GALNT3;refGene_tx=XM_011283761.3;refGene_gn=B4GALNT3;refGene_tx=XM_023256641.1
B4 37532690 37532837
refGene_gn=CCDC77;refGene_tx=XM_003988269.5;refGene_gn=CCDC77;refGene_tx=XM_006933400.4;refGene_gn=CCDC77;refGene_tx=XM_019834333.2;refGene_gn=CCDC77;refGene_tx=XM_019834336.2;refGene_gn=CCDC77;refGene_tx=XM_019834337.2
B4 37533593 37533746
refGene_gn=CCDC77;refGene_tx=XM_003988269.5;refGene_gn=CCDC77;refGene_tx=XM_006933400.4;refGene_gn=CCDC77;refGene_tx=XM_019834333.2;refGene_gn=CCDC77;refGene_tx=XM_019834336.2;refGene_gn=CCDC77;refGene_tx=XM_019834337.2
B4 37533847 37533973
refGene_gn=CCDC77;refGene_tx=XM_003988269.5;refGene_gn=CCDC77;refGene_

tx=XM_006933400.4;refGene_gn=CCDC77;refGene_tx=XM_019834333.2;refGene_gn=CCDC77;refGene_tx=XM_019834336.2;refGene_gn=CCDC77;refGene_tx=XM_019834337.2
B4 37536245 37536465 refGene_gn=CCDC77;refGene_tx=XM_003988269.5;refGene_gn=CCDC77;refGene_tx=XM_006933400.4;refGene_gn=CCDC77;refGene_tx=XM_019834333.2;refGene_gn=CCDC77;refGene_tx=XM_019834336.2;refGene_gn=CCDC77;refGene_tx=XM_019834337.2
B4 37539109 37539258 refGene_gn=CCDC77;refGene_tx=XM_003988269.5;refGene_gn=CCDC77;refGene_tx=XM_006933400.4;refGene_gn=CCDC77;refGene_tx=XM_019834333.2;refGene_gn=CCDC77;refGene_tx=XM_019834336.2;refGene_gn=CCDC77;refGene_tx=XM_019834337.2
B4 37540437 37540529 refGene_gn=CCDC77;refGene_tx=XM_003988269.5;refGene_gn=CCDC77;refGene_tx=XM_006933400.4;refGene_gn=CCDC77;refGene_tx=XM_019834333.2;refGene_gn=CCDC77;refGene_tx=XM_019834336.2;refGene_gn=CCDC77;refGene_tx=XM_019834337.2
B4 37547831 37547907 refGene_gn=CCDC77;refGene_tx=XM_003988269.5;refGene_gn=CCDC77;refGene_tx=XM_006933400.4;refGene_gn=CCDC77;refGene_tx=XM_019834333.2;refGene_gn=CCDC77;refGene_tx=XM_019834336.2;refGene_gn=CCDC77;refGene_tx=XM_019834337.2
B4 37549222 37549319 refGene_gn=CCDC77;refGene_tx=XM_003988269.5;refGene_gn=CCDC77;refGene_tx=XM_006933400.4;refGene_gn=CCDC77;refGene_tx=XM_019834333.2;refGene_gn=CCDC77;refGene_tx=XM_019834336.2;refGene_gn=CCDC77;refGene_tx=XM_019834337.2
B4 37551106 37551249 refGene_gn=CCDC77;refGene_tx=XM_003988269.5;refGene_gn=CCDC77;refGene_tx=XM_006933400.4;refGene_gn=CCDC77;refGene_tx=XM_019834333.2;refGene_gn=CCDC77;refGene_tx=XM_019834336.2;refGene_gn=CCDC77;refGene_tx=XM_019834337.2
B4 37554069 37554313 refGene_gn=CCDC77;refGene_tx=XM_003988269.5;refGene_gn=CCDC77;refGene_tx=XM_006933400.4;refGene_gn=CCDC77;refGene_tx=XM_019834333.2;refGene_gn=CCDC77;refGene_tx=XM_019834336.2;refGene_gn=CCDC77;refGene_tx=XM_019834337.2
B4 37556124 37556162 refGene_gn=CCDC77;refGene_tx=XM_003988269.5;refGene_gn=CCDC77;refGene_tx=XM_006933400.4;refGene_gn=CCDC77;refGene_tx=XM_019834333.2;refGene_gn=CCDC77;refGene_tx=XM_019834336.2;refGene_gn=CCDC77;refGene_tx=XM_019834337.2
B4 37576605 37576770 refGene_gn=KDM5A;refGene_tx=XM_006933402.4
B4 37579265 37579343 refGene_gn=KDM5A;refGene_tx=XM_006933402.4
B4 37580432 37580555 refGene_gn=KDM5A;refGene_tx=XM_006933402.4
B4 37591046 37591217 refGene_gn=KDM5A;refGene_tx=XM_006933402.4
B4 37593407 37593542 refGene_gn=KDM5A;refGene_tx=XM_006933402.4
B4 37602026 37602132 refGene_gn=KDM5A;refGene_tx=XM_006933402.4
B4 37603253 37603345 refGene_gn=KDM5A;refGene_tx=XM_006933402.4

B4 37603931 37604090 refGene_gn=KDM5A;refGene_tx=XM_006933402.4
B4 37606273 37606393 refGene_gn=KDM5A;refGene_tx=XM_006933402.4
B4 37607086 37607245 refGene_gn=KDM5A;refGene_tx=XM_006933402.4
B4 37617602 37617784 refGene_gn=KDM5A;refGene_tx=XM_006933402.4
B4 37619678 37619841 refGene_gn=KDM5A;refGene_tx=XM_006933402.4
B4 37621395 37621515 refGene_gn=KDM5A;refGene_tx=XM_006933402.4
B4 37626674 37626869 refGene_gn=KDM5A;refGene_tx=XM_006933402.4
B4 37634030 37634212 refGene_gn=KDM5A;refGene_tx=XM_006933402.4
B4 37634575 37634700 refGene_gn=KDM5A;refGene_tx=XM_006933402.4
B4 37635585 37635736 refGene_gn=KDM5A;refGene_tx=XM_006933402.4
B4 37637069 37637184 refGene_gn=KDM5A;refGene_tx=XM_006933402.4
B4 37639809 37640165 refGene_gn=KDM5A;refGene_tx=XM_006933402.4
B4 37644317 37644456 refGene_gn=KDM5A;refGene_tx=XM_006933402.4
B4 37647071 37647251 refGene_gn=KDM5A;refGene_tx=XM_006933402.4
B4 37647844 37648006 refGene_gn=KDM5A;refGene_tx=XM_006933402.4
B4 37652215 37652767 refGene_gn=KDM5A;refGene_tx=XM_006933402.4
B4 37653582 37653726 refGene_gn=KDM5A;refGene_tx=XM_006933402.4
B4 37660773 37660933 refGene_gn=KDM5A;refGene_tx=XM_006933402.4
B4 37661816 37662037 refGene_gn=KDM5A;refGene_tx=XM_006933402.4
B4 37664562 37664973 refGene_gn=KDM5A;refGene_tx=XM_006933402.4
B4 37671756 37671963 refGene_gn=KDM5A;refGene_tx=XM_006933402.4
B4 37702639 37702841
refGene_gn=SLC6A13;refGene_tx=XM_003988271.4;refGene_gn=SLC6A13;refGene_tx=XM_006933404.3
B4 37704899 37704975
refGene_gn=SLC6A13;refGene_tx=XM_019834338.2;refGene_gn=SLC6A13;refGene_tx=XM_019834339.2;refGene_gn=SLC6A13;refGene_tx=XM_019834340.2;refGene_gn=SLC6A13;refGene_tx=XM_019834341.2
B4 37711755 37711890
refGene_gn=SLC6A13;refGene_tx=XM_003988271.4;refGene_gn=SLC6A13;refGene_tx=XM_019834338.2;refGene_gn=SLC6A13;refGene_tx=XM_019834339.2;refGene_gn=SLC6A13;refGene_tx=XM_019834340.2;refGene_gn=SLC6A13;refGene_tx=XM_019834341.2
B4 37714494 37714635
refGene_gn=SLC6A13;refGene_tx=XM_003988271.4;refGene_gn=SLC6A13;refGene_tx=XM_006933404.3;refGene_gn=SLC6A13;refGene_tx=XM_019834338.2;refGene_gn=SLC6A13;refGene_tx=XM_019834339.2;refGene_gn=SLC6A13;refGene_tx=XM_019834340.2;refGene_gn=SLC6A13;refGene_tx=XM_019834341.2;refGene_gn=SLC6A13;refGene_tx=XM_023256642.1
B4 37718975 37719060
refGene_gn=SLC6A13;refGene_tx=XM_003988271.4;refGene_gn=SLC6A13;refGene_tx=XM_006933404.3;refGene_gn=SLC6A13;refGene_tx=XM_019834338.2;refGene_gn=SLC6A13;refGene_tx=XM_019834339.2;refGene_gn=SLC6A13;refGene_tx=XM_019834340.2;refGene_gn=SLC6A13;refGene_tx=XM_019834341.2;refGene_gn=SLC6A13;refGene_tx=XM_023256642.1
B4 37719730 37719863
refGene_gn=SLC6A13;refGene_tx=XM_003988271.4;refGene_gn=SLC6A13;refGene_tx=XM_006933404.3;refGene_gn=SLC6A13;refGene_tx=XM_019834338.2;refGene_gn=SLC6A13;refGene_tx=XM_019834339.2;refGene_gn=SLC6A13;refGene_tx=XM_019834340.2;refGene_gn=SLC6A13;refGene_tx=XM_019834341.2;refGene_gn

=SLC6A13;refGene_tx=XM_023256642.1
B4	37720765 37720900
refGene_gn=SLC6A13;refGene_tx=XM_003988271.4;refGene_gn=SLC6A13;refGene_tx=XM_006933404.3;refGene_gn=SLC6A13;refGene_tx=XM_019834338.2;refGene_gn=SLC6A13;refGene_tx=XM_019834339.2;refGene_gn=SLC6A13;refGene_tx=XM_019834340.2;refGene_gn=SLC6A13;refGene_tx=XM_019834341.2;refGene_gn=SLC6A13;refGene_tx=XM_023256642.1
B4	37727453 37727557
refGene_gn=SLC6A13;refGene_tx=XM_003988271.4;refGene_gn=SLC6A13;refGene_tx=XM_006933404.3;refGene_gn=SLC6A13;refGene_tx=XM_019834338.2;refGene_gn=SLC6A13;refGene_tx=XM_019834339.2;refGene_gn=SLC6A13;refGene_tx=XM_019834340.2;refGene_gn=SLC6A13;refGene_tx=XM_019834341.2;refGene_gn=SLC6A13;refGene_tx=XM_023256642.1
B4	37728476 37728601
refGene_gn=SLC6A13;refGene_tx=XM_003988271.4;refGene_gn=SLC6A13;refGene_tx=XM_006933404.3;refGene_gn=SLC6A13;refGene_tx=XM_019834338.2;refGene_gn=SLC6A13;refGene_tx=XM_019834339.2;refGene_gn=SLC6A13;refGene_tx=XM_019834340.2;refGene_gn=SLC6A13;refGene_tx=XM_019834341.2;refGene_gn=SLC6A13;refGene_tx=XM_023256642.1
B4	37730050 37730163
refGene_gn=SLC6A13;refGene_tx=XM_003988271.4;refGene_gn=SLC6A13;refGene_tx=XM_006933404.3;refGene_gn=SLC6A13;refGene_tx=XM_019834338.2;refGene_gn=SLC6A13;refGene_tx=XM_019834339.2;refGene_gn=SLC6A13;refGene_tx=XM_019834340.2;refGene_gn=SLC6A13;refGene_tx=XM_019834341.2;refGene_gn=SLC6A13;refGene_tx=XM_023256642.1
B4	37730485 37730623
refGene_gn=SLC6A13;refGene_tx=XM_003988271.4;refGene_gn=SLC6A13;refGene_tx=XM_006933404.3;refGene_gn=SLC6A13;refGene_tx=XM_019834338.2;refGene_gn=SLC6A13;refGene_tx=XM_019834339.2;refGene_gn=SLC6A13;refGene_tx=XM_019834340.2;refGene_gn=SLC6A13;refGene_tx=XM_019834341.2;refGene_gn=SLC6A13;refGene_tx=XM_023256642.1
B4	37731396 37731499
refGene_gn=SLC6A13;refGene_tx=XM_003988271.4;refGene_gn=SLC6A13;refGene_tx=XM_006933404.3;refGene_gn=SLC6A13;refGene_tx=XM_019834338.2;refGene_gn=SLC6A13;refGene_tx=XM_019834339.2;refGene_gn=SLC6A13;refGene_tx=XM_019834340.2;refGene_gn=SLC6A13;refGene_tx=XM_019834341.2;refGene_gn=SLC6A13;refGene_tx=XM_023256642.1
B4	37731951 37732052
refGene_gn=SLC6A13;refGene_tx=XM_003988271.4;refGene_gn=SLC6A13;refGene_tx=XM_006933404.3;refGene_gn=SLC6A13;refGene_tx=XM_019834338.2;refGene_gn=SLC6A13;refGene_tx=XM_019834339.2;refGene_gn=SLC6A13;refGene_tx=XM_019834340.2;refGene_gn=SLC6A13;refGene_tx=XM_019834341.2;refGene_gn=SLC6A13;refGene_tx=XM_023256642.1
B4	37733037 37733208
refGene_gn=SLC6A13;refGene_tx=XM_003988271.4;refGene_gn=SLC6A13;refGene_tx=XM_006933404.3;refGene_gn=SLC6A13;refGene_tx=XM_019834338.2;refGene_gn=SLC6A13;refGene_tx=XM_019834339.2;refGene_gn=SLC6A13;refGene_tx=XM_019834340.2;refGene_gn=SLC6A13;refGene_tx=XM_019834341.2;refGene_gn=SLC6A13;refGene_tx=XM_023256642.1
B4	37733503 37733626

refGene_gn=SLC6A13;refGene_tx=XM_003988271.4;refGene_gn=SLC6A13;refGen
e_tx=XM_006933404.3;refGene_gn=SLC6A13;refGene_tx=XM_019834338.2;refGe
ne_gn=SLC6A13;refGene_tx=XM_019834339.2;refGene_gn=SLC6A13;refGene_tx=
XM_019834340.2;refGene_gn=SLC6A13;refGene_tx=XM_019834341.2;refGene_gn
=SLC6A13;refGene_tx=XM_023256642.1
B4 37743934 37744148
refGene_gn=LOC101088338;refGene_tx=XM_019834344.1;refGene_gn=LOC101088
338;refGene_tx=XM_019834345.1;refGene_gn=LOC101088338;refGene_tx=XM_01
9834346.1;refGene_gn=LOC101088338;refGene_tx=XM_019834347.2;refGene_gn
=LOC101088338;refGene_tx=XM_019834348.1;refGene_gn=LOC101088338;refGen
e_tx=XM_019834349.1;refGene_gn=LOC101088338;refGene_tx=XM_019834351.1
B4 37750088 37750223
refGene_gn=LOC101088338;refGene_tx=XM_019834344.1;refGene_gn=LOC101088
338;refGene_tx=XM_019834345.1;refGene_gn=LOC101088338;refGene_tx=XM_01
9834346.1;refGene_gn=LOC101088338;refGene_tx=XM_019834347.2;refGene_gn
=LOC101088338;refGene_tx=XM_019834348.1;refGene_gn=LOC101088338;refGen
e_tx=XM_019834349.1;refGene_gn=LOC101088338;refGene_tx=XM_019834351.1
B4 37753218 37753359
refGene_gn=LOC101088338;refGene_tx=XM_019834344.1;refGene_gn=LOC101088
338;refGene_tx=XM_019834345.1;refGene_gn=LOC101088338;refGene_tx=XM_01
9834346.1;refGene_gn=LOC101088338;refGene_tx=XM_019834347.2;refGene_gn
=LOC101088338;refGene_tx=XM_019834348.1;refGene_gn=LOC101088338;refGen
e_tx=XM_019834349.1;refGene_gn=LOC101088338;refGene_tx=XM_019834351.1
B4 37754179 37754267
refGene_gn=LOC101088338;refGene_tx=XM_019834344.1;refGene_gn=LOC101088
338;refGene_tx=XM_019834345.1;refGene_gn=LOC101088338;refGene_tx=XM_01
9834346.1;refGene_gn=LOC101088338;refGene_tx=XM_019834347.2;refGene_gn
=LOC101088338;refGene_tx=XM_019834348.1;refGene_gn=LOC101088338;refGen
e_tx=XM_019834349.1;refGene_gn=LOC101088338;refGene_tx=XM_019834351.1
B4 37755241 37755374
refGene_gn=LOC101088338;refGene_tx=XM_019834344.1;refGene_gn=LOC101088
338;refGene_tx=XM_019834345.1;refGene_gn=LOC101088338;refGene_tx=XM_01
9834346.1;refGene_gn=LOC101088338;refGene_tx=XM_019834347.2;refGene_gn
=LOC101088338;refGene_tx=XM_019834348.1;refGene_gn=LOC101088338;refGen
e_tx=XM_019834349.1;refGene_gn=LOC101088338;refGene_tx=XM_019834351.1
B4 37756824 37756959
refGene_gn=LOC101088338;refGene_tx=XM_019834344.1;refGene_gn=LOC101088
338;refGene_tx=XM_019834345.1;refGene_gn=LOC101088338;refGene_tx=XM_01
9834346.1;refGene_gn=LOC101088338;refGene_tx=XM_019834347.2;refGene_gn
=LOC101088338;refGene_tx=XM_019834348.1;refGene_gn=LOC101088338;refGen
e_tx=XM_019834349.1;refGene_gn=LOC101088338;refGene_tx=XM_019834351.1
B4 37758758 37758862
refGene_gn=LOC101088338;refGene_tx=XM_019834344.1;refGene_gn=LOC101088
338;refGene_tx=XM_019834345.1;refGene_gn=LOC101088338;refGene_tx=XM_01
9834346.1;refGene_gn=LOC101088338;refGene_tx=XM_019834347.2;refGene_gn
=LOC101088338;refGene_tx=XM_019834348.1;refGene_gn=LOC101088338;refGen
e_tx=XM_019834349.1;refGene_gn=LOC101088338;refGene_tx=XM_019834351.1
B4 37759177 37759302
refGene_gn=LOC101088338;refGene_tx=XM_019834344.1;refGene_gn=LOC101088
338;refGene_tx=XM_019834345.1;refGene_gn=LOC101088338;refGene_tx=XM_01

9834346.1;refGene_gn=LOC101088338;refGene_tx=XM_019834347.2;refGene_gn=LOC101088338;refGene_tx=XM_019834348.1;refGene_gn=LOC101088338;refGene_tx=XM_019834349.1;refGene_gn=LOC101088338;refGene_tx=XM_019834351.1
B4 37759712 37759825
refGene_gn=LOC101088338;refGene_tx=XM_019834344.1;refGene_gn=LOC101088338;refGene_tx=XM_019834345.1;refGene_gn=LOC101088338;refGene_tx=XM_019834346.1;refGene_gn=LOC101088338;refGene_tx=XM_019834347.2;refGene_gn=LOC101088338;refGene_tx=XM_019834348.1;refGene_gn=LOC101088338;refGene_tx=XM_019834349.1;refGene_gn=LOC101088338;refGene_tx=XM_019834351.1
B4 37760281 37760419
refGene_gn=LOC101088338;refGene_tx=XM_019834344.1;refGene_gn=LOC101088338;refGene_tx=XM_019834345.1;refGene_gn=LOC101088338;refGene_tx=XM_019834346.1;refGene_gn=LOC101088338;refGene_tx=XM_019834347.2;refGene_gn=LOC101088338;refGene_tx=XM_019834348.1;refGene_gn=LOC101088338;refGene_tx=XM_019834349.1;refGene_gn=LOC101088338;refGene_tx=XM_019834351.1
B4 37761132 37761235
refGene_gn=LOC101088338;refGene_tx=XM_019834344.1;refGene_gn=LOC101088338;refGene_tx=XM_019834345.1;refGene_gn=LOC101088338;refGene_tx=XM_019834346.1;refGene_gn=LOC101088338;refGene_tx=XM_019834347.2;refGene_gn=LOC101088338;refGene_tx=XM_019834348.1;refGene_gn=LOC101088338;refGene_tx=XM_019834349.1;refGene_gn=LOC101088338;refGene_tx=XM_019834351.1
B4 37763089 37763190
refGene_gn=LOC101088338;refGene_tx=XM_019834344.1;refGene_gn=LOC101088338;refGene_tx=XM_019834345.1;refGene_gn=LOC101088338;refGene_tx=XM_019834346.1;refGene_gn=LOC101088338;refGene_tx=XM_019834347.2;refGene_gn=LOC101088338;refGene_tx=XM_019834348.1;refGene_gn=LOC101088338;refGene_tx=XM_019834349.1;refGene_gn=LOC101088338;refGene_tx=XM_019834351.1
B4 37763897 37764068
refGene_gn=LOC101088338;refGene_tx=XM_019834344.1;refGene_gn=LOC101088338;refGene_tx=XM_019834345.1;refGene_gn=LOC101088338;refGene_tx=XM_019834346.1;refGene_gn=LOC101088338;refGene_tx=XM_019834347.2;refGene_gn=LOC101088338;refGene_tx=XM_019834348.1;refGene_gn=LOC101088338;refGene_tx=XM_019834349.1;refGene_gn=LOC101088338;refGene_tx=XM_019834351.1
B4 37765222 37765366
refGene_gn=LOC101088338;refGene_tx=XM_019834344.1;refGene_gn=LOC101088338;refGene_tx=XM_019834345.1;refGene_gn=LOC101088338;refGene_tx=XM_019834346.1;refGene_gn=LOC101088338;refGene_tx=XM_019834347.2;refGene_gn=LOC101088338;refGene_tx=XM_019834348.1;refGene_gn=LOC101088338;refGene_tx=XM_019834349.1;refGene_gn=LOC101088338;refGene_tx=XM_019834351.1
B4 37780218 37780653
refGene_gn=IQSEC3;refGene_tx=XM_023256643.1;refGene_gn=IQSEC3;refGene_tx=XM_023256644.1;refGene_gn=IQSEC3;refGene_tx=XM_023256645.1;refGene_gn=IQSEC3;refGene_tx=XM_023256648.1
B4 37783768 37783843
refGene_gn=IQSEC3;refGene_tx=XM_023256646.1;refGene_gn=IQSEC3;refGene_tx=XM_023256647.1;refGene_gn=IQSEC3;refGene_tx=XM_023256649.1;refGene_gn=IQSEC3;refGene_tx=XM_023256650.1
B4 37783931 37783981
refGene_gn=IQSEC3;refGene_tx=XM_023256643.1;refGene_gn=IQSEC3;refGene_tx=XM_023256644.1;refGene_gn=IQSEC3;refGene_tx=XM_023256645.1;refGene_

gn=IQSEC3;refGene_tx=XM_023256646.1;refGene_gn=IQSEC3;refGene_tx=XM_02
3256647.1
B4 37786284 37786377
refGene_gn=IQSEC3;refGene_tx=XM_023256643.1;refGene_gn=IQSEC3;refGene_
tx=XM_023256644.1;refGene_gn=IQSEC3;refGene_tx=XM_023256645.1;refGene_
gn=IQSEC3;refGene_tx=XM_023256646.1;refGene_gn=IQSEC3;refGene_tx=XM_02
3256647.1;refGene_gn=IQSEC3;refGene_tx=XM_023256648.1;refGene_gn=IQSEC
3;refGene_tx=XM_023256649.1;refGene_gn=IQSEC3;refGene_tx=XM_023256650.
1
B4 37789206 37789368
refGene_gn=IQSEC3;refGene_tx=XM_023256643.1;refGene_gn=IQSEC3;refGene_
tx=XM_023256644.1;refGene_gn=IQSEC3;refGene_tx=XM_023256645.1;refGene_
gn=IQSEC3;refGene_tx=XM_023256646.1;refGene_gn=IQSEC3;refGene_tx=XM_02
3256647.1;refGene_gn=IQSEC3;refGene_tx=XM_023256648.1;refGene_gn=IQSEC
3;refGene_tx=XM_023256649.1;refGene_gn=IQSEC3;refGene_tx=XM_023256650.
1
B4 37789544 37789644
refGene_gn=IQSEC3;refGene_tx=XM_023256643.1;refGene_gn=IQSEC3;refGene_
tx=XM_023256644.1;refGene_gn=IQSEC3;refGene_tx=XM_023256645.1;refGene_
gn=IQSEC3;refGene_tx=XM_023256646.1;refGene_gn=IQSEC3;refGene_tx=XM_02
3256647.1;refGene_gn=IQSEC3;refGene_tx=XM_023256648.1;refGene_gn=IQSEC
3;refGene_tx=XM_023256649.1;refGene_gn=IQSEC3;refGene_tx=XM_023256650.
1
B4 37791613 37791739
refGene_gn=IQSEC3;refGene_tx=XM_023256643.1;refGene_gn=IQSEC3;refGene_
tx=XM_023256644.1;refGene_gn=IQSEC3;refGene_tx=XM_023256645.1;refGene_
gn=IQSEC3;refGene_tx=XM_023256646.1;refGene_gn=IQSEC3;refGene_tx=XM_02
3256647.1;refGene_gn=IQSEC3;refGene_tx=XM_023256648.1;refGene_gn=IQSEC
3;refGene_tx=XM_023256649.1;refGene_gn=IQSEC3;refGene_tx=XM_023256650.
1
B4 37792853 37792993
refGene_gn=IQSEC3;refGene_tx=XM_023256643.1;refGene_gn=IQSEC3;refGene_
tx=XM_023256644.1;refGene_gn=IQSEC3;refGene_tx=XM_023256645.1;refGene_
gn=IQSEC3;refGene_tx=XM_023256646.1;refGene_gn=IQSEC3;refGene_tx=XM_02
3256647.1;refGene_gn=IQSEC3;refGene_tx=XM_023256648.1;refGene_gn=IQSEC
3;refGene_tx=XM_023256649.1;refGene_gn=IQSEC3;refGene_tx=XM_023256650.
1
B4 37798335 37798502
refGene_gn=IQSEC3;refGene_tx=XM_023256643.1;refGene_gn=IQSEC3;refGene_
tx=XM_023256644.1;refGene_gn=IQSEC3;refGene_tx=XM_023256645.1;refGene_
gn=IQSEC3;refGene_tx=XM_023256646.1;refGene_gn=IQSEC3;refGene_tx=XM_02
3256647.1;refGene_gn=IQSEC3;refGene_tx=XM_023256648.1;refGene_gn=IQSEC
3;refGene_tx=XM_023256649.1;refGene_gn=IQSEC3;refGene_tx=XM_023256650.
1
B4 37798957 37799080
refGene_gn=IQSEC3;refGene_tx=XM_023256643.1;refGene_gn=IQSEC3;refGene_
tx=XM_023256644.1;refGene_gn=IQSEC3;refGene_tx=XM_023256645.1;refGene_
gn=IQSEC3;refGene_tx=XM_023256646.1;refGene_gn=IQSEC3;refGene_tx=XM_02
3256647.1;refGene_gn=IQSEC3;refGene_tx=XM_023256648.1;refGene_gn=IQSEC
3;refGene_tx=XM_023256649.1;refGene_gn=IQSEC3;refGene_tx=XM_023256650.

1
B4	37814521	37814683
refGene_gn=IQSEC3;refGene_tx=XM_023256643.1;refGene_gn=IQSEC3;refGene_tx=XM_023256644.1;refGene_gn=IQSEC3;refGene_tx=XM_023256645.1;refGene_gn=IQSEC3;refGene_tx=XM_023256646.1;refGene_gn=IQSEC3;refGene_tx=XM_023256647.1;refGene_gn=IQSEC3;refGene_tx=XM_023256648.1;refGene_gn=IQSEC3;refGene_tx=XM_023256649.1;refGene_gn=IQSEC3;refGene_tx=XM_023256650.1
B4	37816688	37817818
refGene_gn=IQSEC3;refGene_tx=XM_023256643.1;refGene_gn=IQSEC3;refGene_tx=XM_023256644.1;refGene_gn=IQSEC3;refGene_tx=XM_023256645.1;refGene_gn=IQSEC3;refGene_tx=XM_023256646.1;refGene_gn=IQSEC3;refGene_tx=XM_023256647.1;refGene_gn=IQSEC3;refGene_tx=XM_023256648.1;refGene_gn=IQSEC3;refGene_tx=XM_023256649.1;refGene_gn=IQSEC3;refGene_tx=XM_023256650.1
B4	37829224	37829504
refGene_gn=IQSEC3;refGene_tx=XM_023256643.1;refGene_gn=IQSEC3;refGene_tx=XM_023256644.1;refGene_gn=IQSEC3;refGene_tx=XM_023256645.1;refGene_gn=IQSEC3;refGene_tx=XM_023256646.1;refGene_gn=IQSEC3;refGene_tx=XM_023256647.1;refGene_gn=IQSEC3;refGene_tx=XM_023256648.1;refGene_gn=IQSEC3;refGene_tx=XM_023256649.1;refGene_gn=IQSEC3;refGene_tx=XM_023256650.1
B4	37851538	37851622
refGene_gn=IQSEC3;refGene_tx=XM_023256643.1;refGene_gn=IQSEC3;refGene_tx=XM_023256644.1;refGene_gn=IQSEC3;refGene_tx=XM_023256645.1;refGene_gn=IQSEC3;refGene_tx=XM_023256646.1;refGene_gn=IQSEC3;refGene_tx=XM_023256647.1;refGene_gn=IQSEC3;refGene_tx=XM_023256648.1;refGene_gn=IQSEC3;refGene_tx=XM_023256649.1;refGene_gn=IQSEC3;refGene_tx=XM_023256650.1
B4	37881805	37881910
refGene_gn=IQSEC3;refGene_tx=XM_023256643.1;refGene_gn=IQSEC3;refGene_tx=XM_023256645.1;refGene_gn=IQSEC3;refGene_tx=XM_023256646.1;refGene_gn=IQSEC3;refGene_tx=XM_023256647.1;refGene_gn=IQSEC3;refGene_tx=XM_023256648.1;refGene_gn=IQSEC3;refGene_tx=XM_023256649.1;refGene_gn=IQSEC3;refGene_tx=XM_023256650.1
B4	37886471	37887028
refGene_gn=IQSEC3;refGene_tx=XM_023256643.1;refGene_gn=IQSEC3;refGene_tx=XM_023256644.1;refGene_gn=IQSEC3;refGene_tx=XM_023256645.1;refGene_gn=IQSEC3;refGene_tx=XM_023256646.1;refGene_gn=IQSEC3;refGene_tx=XM_023256647.1;refGene_gn=IQSEC3;refGene_tx=XM_023256648.1;refGene_gn=IQSEC3;refGene_tx=XM_023256649.1;refGene_gn=IQSEC3;refGene_tx=XM_023256650.1
B4	37924487	37924636
refGene_gn=WASHC1;refGene_tx=XM_003988273.5;refGene_gn=WASHC1;refGene_tx=XM_023256651.1
B4	37934965	37935077
refGene_gn=WASHC1;refGene_tx=XM_003988273.5;refGene_gn=WASHC1;refGene_tx=XM_023256651.1
B4	37935286	37935433
refGene_gn=WASHC1;refGene_tx=XM_003988273.5;refGene_gn=WASHC1;refGene_

tx=XM_023256651.1
B4	37935596	37935733
refGene_gn=WASHC1;refGene_tx=XM_003988273.5;refGene_gn=WASHC1;refGene_tx=XM_023256651.1
B4	37935991	37936127
refGene_gn=WASHC1;refGene_tx=XM_003988273.5;refGene_gn=WASHC1;refGene_tx=XM_023256651.1
B4	37936320	37936522
refGene_gn=WASHC1;refGene_tx=XM_003988273.5;refGene_gn=WASHC1;refGene_tx=XM_023256651.1
B4	37936611	37936776
refGene_gn=WASHC1;refGene_tx=XM_003988273.5;refGene_gn=WASHC1;refGene_tx=XM_023256651.1
B4	37937488	37937647
refGene_gn=WASHC1;refGene_tx=XM_003988273.5;refGene_gn=WASHC1;refGene_tx=XM_023256651.1
B4	37938416	37938485
refGene_gn=WASHC1;refGene_tx=XM_003988273.5;refGene_gn=WASHC1;refGene_tx=XM_023256651.1
B4	37938643	37938783
refGene_gn=WASHC1;refGene_tx=XM_003988273.5;refGene_gn=WASHC1;refGene_tx=XM_023256651.1
B4	37940273	37940303
refGene_gn=DDX11;refGene_tx=XM_011283741.3;refGene_gn=DDX11;refGene_tx=XM_019834354.2
B4	37940385	37940540
refGene_gn=DDX11;refGene_tx=XM_011283741.3;refGene_gn=DDX11;refGene_tx=XM_019834354.2
B4	37940700	37940779
refGene_gn=DDX11;refGene_tx=XM_011283741.3;refGene_gn=DDX11;refGene_tx=XM_019834354.2
B4	37941105	37941190
refGene_gn=DDX11;refGene_tx=XM_011283741.3;refGene_gn=DDX11;refGene_tx=XM_019834354.2
B4	37941609	37941710
refGene_gn=DDX11;refGene_tx=XM_011283741.3;refGene_gn=DDX11;refGene_tx=XM_019834354.2
B4	37941824	37941893
refGene_gn=DDX11;refGene_tx=XM_011283741.3;refGene_gn=DDX11;refGene_tx=XM_019834354.2
B4	37942181	37942331
refGene_gn=DDX11;refGene_tx=XM_011283741.3;refGene_gn=DDX11;refGene_tx=XM_019834354.2
B4	37942996	37943100
refGene_gn=DDX11;refGene_tx=XM_011283741.3;refGene_gn=DDX11;refGene_tx=XM_019834354.2
B4	37943406	37943479
refGene_gn=DDX11;refGene_tx=XM_011283741.3;refGene_gn=DDX11;refGene_tx=XM_019834354.2
B4	37950979	37951092

refGene_gn=DDX11;refGene_tx=XM_011283741.3;refGene_gn=DDX11;refGene_tx=XM_019834354.2
B4 37951590 37951722
refGene_gn=DDX11;refGene_tx=XM_011283741.3;refGene_gn=DDX11;refGene_tx=XM_019834354.2
B4 37951841 37951953
refGene_gn=DDX11;refGene_tx=XM_011283741.3;refGene_gn=DDX11;refGene_tx=XM_019834354.2
B4 37952219 37952258
refGene_gn=DDX11;refGene_tx=XM_011283741.3;refGene_gn=DDX11;refGene_tx=XM_019834354.2
B4 37953789 37953857
refGene_gn=DDX11;refGene_tx=XM_011283741.3;refGene_gn=DDX11;refGene_tx=XM_019834354.2
B4 37953970 37954015
refGene_gn=DDX11;refGene_tx=XM_011283741.3;refGene_gn=DDX11;refGene_tx=XM_019834354.2
B4 37954923 37955003
refGene_gn=DDX11;refGene_tx=XM_011283741.3;refGene_gn=DDX11;refGene_tx=XM_019834354.2
B4 37955341 37955388
refGene_gn=DDX11;refGene_tx=XM_011283741.3;refGene_gn=DDX11;refGene_tx=XM_019834354.2
B4 37956040 37956193 refGene_gn=DDX11;refGene_tx=XM_011283741.3
B4 37957782 37957988
refGene_gn=DDX11;refGene_tx=XM_011283741.3;refGene_gn=DDX11;refGene_tx=XM_019834354.2
B4 37958397 37958485
refGene_gn=DDX11;refGene_tx=XM_011283741.3;refGene_gn=DDX11;refGene_tx=XM_019834354.2
B4 37958704 37958812
refGene_gn=DDX11;refGene_tx=XM_011283741.3;refGene_gn=DDX11;refGene_tx=XM_019834354.2
B4 37959733 37959785
refGene_gn=DDX11;refGene_tx=XM_011283741.3;refGene_gn=DDX11;refGene_tx=XM_019834354.2
B4 37966289 37966447
refGene_gn=DDX11;refGene_tx=XM_011283741.3;refGene_gn=DDX11;refGene_tx=XM_019834354.2
B4 37966766 37966853
refGene_gn=DDX11;refGene_tx=XM_011283741.3;refGene_gn=DDX11;refGene_tx=XM_019834354.2
B4 37967551 37967800
refGene_gn=DDX11;refGene_tx=XM_011283741.3;refGene_gn=DDX11;refGene_tx=XM_019834354.2
B4 37974707 37974851
refGene_gn=DDX11;refGene_tx=XM_011283741.3;refGene_gn=DDX11;refGene_tx=XM_019834354.2
B4 37989954 37990283
refGene_gn=LOC101098061;refGene_tx=XM_023256652.1;refGene_gn=LOC101098

061;refGene_tx=XM_023256653.1;refGene_gn=LOC101098061;refGene_tx=XM_023256654.1;refGene_gn=LOC101098061;refGene_tx=XM_023256655.1;refGene_gn=LOC101098061;refGene_tx=XM_023256656.1;refGene_gn=LOC101098061;refGene_tx=XM_023256657.1;refGene_gn=LOC101098061;refGene_tx=XM_023256659.1;refGene_gn=LOC101098061;refGene_tx=XM_023256660.1
B4 37996145 37996211
refGene_gn=LOC101098061;refGene_tx=XM_023256652.1;refGene_gn=LOC101098061;refGene_tx=XM_023256653.1;refGene_gn=LOC101098061;refGene_tx=XM_023256654.1;refGene_gn=LOC101098061;refGene_tx=XM_023256655.1;refGene_gn=LOC101098061;refGene_tx=XM_023256656.1;refGene_gn=LOC101098061;refGene_tx=XM_023256657.1;refGene_gn=LOC101098061;refGene_tx=XM_023256659.1;refGene_gn=LOC101098061;refGene_tx=XM_023256660.1
B4 37998018 37998124
refGene_gn=LOC101098061;refGene_tx=XM_023256652.1;refGene_gn=LOC101098061;refGene_tx=XM_023256653.1;refGene_gn=LOC101098061;refGene_tx=XM_023256654.1;refGene_gn=LOC101098061;refGene_tx=XM_023256655.1;refGene_gn=LOC101098061;refGene_tx=XM_023256656.1;refGene_gn=LOC101098061;refGene_tx=XM_023256657.1;refGene_gn=LOC101098061;refGene_tx=XM_023256659.1;refGene_gn=LOC101098061;refGene_tx=XM_023256660.1
B4 37999874 37999997
refGene_gn=LOC101098061;refGene_tx=XM_023256652.1;refGene_gn=LOC101098061;refGene_tx=XM_023256653.1;refGene_gn=LOC101098061;refGene_tx=XM_023256654.1;refGene_gn=LOC101098061;refGene_tx=XM_023256655.1;refGene_gn=LOC101098061;refGene_tx=XM_023256656.1;refGene_gn=LOC101098061;refGene_tx=XM_023256657.1;refGene_gn=LOC101098061;refGene_tx=XM_023256659.1
B4 38000382 38000541
refGene_gn=LOC101098061;refGene_tx=XM_023256652.1;refGene_gn=LOC101098061;refGene_tx=XM_023256653.1;refGene_gn=LOC101098061;refGene_tx=XM_023256654.1;refGene_gn=LOC101098061;refGene_tx=XM_023256655.1;refGene_gn=LOC101098061;refGene_tx=XM_023256656.1;refGene_gn=LOC101098061;refGene_tx=XM_023256657.1;refGene_gn=LOC101098061;refGene_tx=XM_023256659.1
B4 38004742 38004916
refGene_gn=LOC101098061;refGene_tx=XM_023256652.1;refGene_gn=LOC101098061;refGene_tx=XM_023256653.1;refGene_gn=LOC101098061;refGene_tx=XM_023256654.1;refGene_gn=LOC101098061;refGene_tx=XM_023256655.1;refGene_gn=LOC101098061;refGene_tx=XM_023256657.1;refGene_gn=LOC101098061;refGene_tx=XM_023256659.1
B4 38004937 38004967
refGene_gn=LOC101098061;refGene_tx=XM_023256661.1
B4 38005281 38005444
refGene_gn=LOC101098061;refGene_tx=XM_023256652.1;refGene_gn=LOC101098061;refGene_tx=XM_023256653.1;refGene_gn=LOC101098061;refGene_tx=XM_023256654.1;refGene_gn=LOC101098061;refGene_tx=XM_023256655.1;refGene_gn=LOC101098061;refGene_tx=XM_023256656.1;refGene_gn=LOC101098061;refGene_tx=XM_023256657.1;refGene_gn=LOC101098061;refGene_tx=XM_023256659.1;refGene_gn=LOC101098061;refGene_tx=XM_023256660.1;refGene_gn=LOC101098061;refGene_tx=XM_023256661.1;refGene_gn=LOC101098061;refGene_tx=XM_023256662.1;refGene_gn=LOC101098061;refGene_tx=XM_023256663.1;refGene_gn=LOC101098061;refGene_tx=XM_023256664.1;refGene_gn=LOC101098061;refGene_tx=XM_023256665.1

B4 38006069 38006187
refGene_gn=LOC101098061;refGene_tx=XM_023256652.1;refGene_gn=LOC101098061;refGene_tx=XM_023256653.1;refGene_gn=LOC101098061;refGene_tx=XM_023256654.1;refGene_gn=LOC101098061;refGene_tx=XM_023256655.1;refGene_gn=LOC101098061;refGene_tx=XM_023256656.1;refGene_gn=LOC101098061;refGene_tx=XM_023256657.1;refGene_gn=LOC101098061;refGene_tx=XM_023256659.1;refGene_gn=LOC101098061;refGene_tx=XM_023256660.1;refGene_gn=LOC101098061;refGene_tx=XM_023256661.1;refGene_gn=LOC101098061;refGene_tx=XM_023256662.1;refGene_gn=LOC101098061;refGene_tx=XM_023256663.1;refGene_gn=LOC101098061;refGene_tx=XM_023256664.1;refGene_gn=LOC101098061;refGene_tx=XM_023256665.1
B4 38006379 38006517
refGene_gn=LOC101098061;refGene_tx=XM_023256652.1;refGene_gn=LOC101098061;refGene_tx=XM_023256653.1;refGene_gn=LOC101098061;refGene_tx=XM_023256654.1;refGene_gn=LOC101098061;refGene_tx=XM_023256655.1;refGene_gn=LOC101098061;refGene_tx=XM_023256656.1;refGene_gn=LOC101098061;refGene_tx=XM_023256657.1;refGene_gn=LOC101098061;refGene_tx=XM_023256659.1;refGene_gn=LOC101098061;refGene_tx=XM_023256660.1;refGene_gn=LOC101098061;refGene_tx=XM_023256661.1;refGene_gn=LOC101098061;refGene_tx=XM_023256662.1;refGene_gn=LOC101098061;refGene_tx=XM_023256663.1;refGene_gn=LOC101098061;refGene_tx=XM_023256664.1;refGene_gn=LOC101098061;refGene_tx=XM_023256665.1
B4 38007394 38007588
refGene_gn=LOC101098061;refGene_tx=XM_023256652.1;refGene_gn=LOC101098061;refGene_tx=XM_023256653.1;refGene_gn=LOC101098061;refGene_tx=XM_023256654.1;refGene_gn=LOC101098061;refGene_tx=XM_023256655.1;refGene_gn=LOC101098061;refGene_tx=XM_023256656.1;refGene_gn=LOC101098061;refGene_tx=XM_023256657.1;refGene_gn=LOC101098061;refGene_tx=XM_023256659.1;refGene_gn=LOC101098061;refGene_tx=XM_023256660.1;refGene_gn=LOC101098061;refGene_tx=XM_023256661.1;refGene_gn=LOC101098061;refGene_tx=XM_023256662.1;refGene_gn=LOC101098061;refGene_tx=XM_023256663.1;refGene_gn=LOC101098061;refGene_tx=XM_023256664.1;refGene_gn=LOC101098061;refGene_tx=XM_023256665.1
B4 38008197 38008333
refGene_gn=LOC101098061;refGene_tx=XM_023256652.1;refGene_gn=LOC101098061;refGene_tx=XM_023256653.1;refGene_gn=LOC101098061;refGene_tx=XM_023256654.1;refGene_gn=LOC101098061;refGene_tx=XM_023256655.1;refGene_gn=LOC101098061;refGene_tx=XM_023256656.1;refGene_gn=LOC101098061;refGene_tx=XM_023256657.1;refGene_gn=LOC101098061;refGene_tx=XM_023256659.1;refGene_gn=LOC101098061;refGene_tx=XM_023256660.1;refGene_gn=LOC101098061;refGene_tx=XM_023256661.1;refGene_gn=LOC101098061;refGene_tx=XM_023256662.1;refGene_gn=LOC101098061;refGene_tx=XM_023256663.1;refGene_gn=LOC101098061;refGene_tx=XM_023256664.1;refGene_gn=LOC101098061;refGene_tx=XM_023256665.1
B4 38008577 38008732
refGene_gn=LOC101098061;refGene_tx=XM_023256652.1;refGene_gn=LOC101098061;refGene_tx=XM_023256653.1;refGene_gn=LOC101098061;refGene_tx=XM_023256654.1;refGene_gn=LOC101098061;refGene_tx=XM_023256655.1;refGene_gn=LOC101098061;refGene_tx=XM_023256656.1;refGene_gn=LOC101098061;refGene_tx=XM_023256657.1;refGene_gn=LOC101098061;refGene_tx=XM_023256659.1;

refGene_gn=LOC101098061;refGene_tx=XM_023256660.1;refGene_gn=LOC101098061;refGene_tx=XM_023256661.1;refGene_gn=LOC101098061;refGene_tx=XM_023256662.1;refGene_gn=LOC101098061;refGene_tx=XM_023256663.1;refGene_gn=LOC101098061;refGene_tx=XM_023256664.1;refGene_gn=LOC101098061;refGene_tx=XM_023256665.1
B4 38009835 38009985
refGene_gn=LOC101098061;refGene_tx=XM_023256652.1;refGene_gn=LOC101098061;refGene_tx=XM_023256653.1;refGene_gn=LOC101098061;refGene_tx=XM_023256654.1;refGene_gn=LOC101098061;refGene_tx=XM_023256655.1;refGene_gn=LOC101098061;refGene_tx=XM_023256656.1;refGene_gn=LOC101098061;refGene_tx=XM_023256657.1;refGene_gn=LOC101098061;refGene_tx=XM_023256659.1;refGene_gn=LOC101098061;refGene_tx=XM_023256660.1;refGene_gn=LOC101098061;refGene_tx=XM_023256661.1;refGene_gn=LOC101098061;refGene_tx=XM_023256662.1;refGene_gn=LOC101098061;refGene_tx=XM_023256663.1;refGene_gn=LOC101098061;refGene_tx=XM_023256664.1;refGene_gn=LOC101098061;refGene_tx=XM_023256665.1
B4 38010540 38010638
refGene_gn=LOC101098061;refGene_tx=XM_023256652.1;refGene_gn=LOC101098061;refGene_tx=XM_023256653.1;refGene_gn=LOC101098061;refGene_tx=XM_023256654.1;refGene_gn=LOC101098061;refGene_tx=XM_023256655.1;refGene_gn=LOC101098061;refGene_tx=XM_023256656.1;refGene_gn=LOC101098061;refGene_tx=XM_023256657.1;refGene_gn=LOC101098061;refGene_tx=XM_023256659.1;refGene_gn=LOC101098061;refGene_tx=XM_023256660.1;refGene_gn=LOC101098061;refGene_tx=XM_023256661.1;refGene_gn=LOC101098061;refGene_tx=XM_023256662.1;refGene_gn=LOC101098061;refGene_tx=XM_023256663.1;refGene_gn=LOC101098061;refGene_tx=XM_023256664.1;refGene_gn=LOC101098061;refGene_tx=XM_023256665.1
B4 38011092 38011228
refGene_gn=LOC101098061;refGene_tx=XM_023256652.1;refGene_gn=LOC101098061;refGene_tx=XM_023256653.1;refGene_gn=LOC101098061;refGene_tx=XM_023256654.1;refGene_gn=LOC101098061;refGene_tx=XM_023256655.1;refGene_gn=LOC101098061;refGene_tx=XM_023256656.1;refGene_gn=LOC101098061;refGene_tx=XM_023256657.1;refGene_gn=LOC101098061;refGene_tx=XM_023256659.1;refGene_gn=LOC101098061;refGene_tx=XM_023256660.1;refGene_gn=LOC101098061;refGene_tx=XM_023256661.1;refGene_gn=LOC101098061;refGene_tx=XM_023256662.1;refGene_gn=LOC101098061;refGene_tx=XM_023256663.1;refGene_gn=LOC101098061;refGene_tx=XM_023256664.1;refGene_gn=LOC101098061;refGene_tx=XM_023256665.1
B4 38011891 38011996
refGene_gn=LOC101098061;refGene_tx=XM_023256652.1;refGene_gn=LOC101098061;refGene_tx=XM_023256653.1;refGene_gn=LOC101098061;refGene_tx=XM_023256654.1;refGene_gn=LOC101098061;refGene_tx=XM_023256655.1;refGene_gn=LOC101098061;refGene_tx=XM_023256656.1;refGene_gn=LOC101098061;refGene_tx=XM_023256657.1;refGene_gn=LOC101098061;refGene_tx=XM_023256659.1;refGene_gn=LOC101098061;refGene_tx=XM_023256660.1;refGene_gn=LOC101098061;refGene_tx=XM_023256661.1;refGene_gn=LOC101098061;refGene_tx=XM_023256662.1;refGene_gn=LOC101098061;refGene_tx=XM_023256663.1;refGene_gn=LOC101098061;refGene_tx=XM_023256664.1;refGene_gn=LOC101098061;refGene_tx=XM_023256665.1
B4 38034650 38034740

refGene_gn=LOC101098061;refGene_tx=XM_023256652.1;refGene_gn=LOC101098061;refGene_tx=XM_023256653.1;refGene_gn=LOC101098061;refGene_tx=XM_023256654.1;refGene_gn=LOC101098061;refGene_tx=XM_023256655.1;refGene_gn=LOC101098061;refGene_tx=XM_023256656.1;refGene_gn=LOC101098061;refGene_tx=XM_023256657.1;refGene_gn=LOC101098061;refGene_tx=XM_023256659.1;refGene_gn=LOC101098061;refGene_tx=XM_023256660.1;refGene_gn=LOC101098061;refGene_tx=XM_023256661.1;refGene_gn=LOC101098061;refGene_tx=XM_023256662.1;refGene_gn=LOC101098061;refGene_tx=XM_023256663.1;refGene_gn=LOC101098061;refGene_tx=XM_023256664.1;refGene_gn=LOC101098061;refGene_tx=XM_023256665.1
B4 38035796 38035932
refGene_gn=LOC101098061;refGene_tx=XM_023256652.1;refGene_gn=LOC101098061;refGene_tx=XM_023256653.1;refGene_gn=LOC101098061;refGene_tx=XM_023256654.1;refGene_gn=LOC101098061;refGene_tx=XM_023256655.1;refGene_gn=LOC101098061;refGene_tx=XM_023256656.1;refGene_gn=LOC101098061;refGene_tx=XM_023256657.1;refGene_gn=LOC101098061;refGene_tx=XM_023256659.1;refGene_gn=LOC101098061;refGene_tx=XM_023256660.1;refGene_gn=LOC101098061;refGene_tx=XM_023256661.1;refGene_gn=LOC101098061;refGene_tx=XM_023256662.1;refGene_gn=LOC101098061;refGene_tx=XM_023256663.1;refGene_gn=LOC101098061;refGene_tx=XM_023256664.1;refGene_gn=LOC101098061;refGene_tx=XM_023256665.1
B4 38036533 38036613
refGene_gn=LOC101098061;refGene_tx=XM_023256652.1;refGene_gn=LOC101098061;refGene_tx=XM_023256653.1;refGene_gn=LOC101098061;refGene_tx=XM_023256654.1;refGene_gn=LOC101098061;refGene_tx=XM_023256655.1;refGene_gn=LOC101098061;refGene_tx=XM_023256656.1;refGene_gn=LOC101098061;refGene_tx=XM_023256657.1;refGene_gn=LOC101098061;refGene_tx=XM_023256659.1;refGene_gn=LOC101098061;refGene_tx=XM_023256660.1;refGene_gn=LOC101098061;refGene_tx=XM_023256661.1;refGene_gn=LOC101098061;refGene_tx=XM_023256662.1;refGene_gn=LOC101098061;refGene_tx=XM_023256663.1;refGene_gn=LOC101098061;refGene_tx=XM_023256664.1;refGene_gn=LOC101098061;refGene_tx=XM_023256665.1
B4 38036979 38037102
refGene_gn=LOC101098061;refGene_tx=XM_023256652.1;refGene_gn=LOC101098061;refGene_tx=XM_023256653.1;refGene_gn=LOC101098061;refGene_tx=XM_023256654.1;refGene_gn=LOC101098061;refGene_tx=XM_023256655.1;refGene_gn=LOC101098061;refGene_tx=XM_023256656.1;refGene_gn=LOC101098061;refGene_tx=XM_023256657.1;refGene_gn=LOC101098061;refGene_tx=XM_023256660.1;refGene_gn=LOC101098061;refGene_tx=XM_023256661.1;refGene_gn=LOC101098061;refGene_tx=XM_023256662.1;refGene_gn=LOC101098061;refGene_tx=XM_023256663.1;refGene_gn=LOC101098061;refGene_tx=XM_023256664.1;refGene_gn=LOC101098061;refGene_tx=XM_023256665.1
B4 38037560 38037713
refGene_gn=LOC101098061;refGene_tx=XM_023256652.1;refGene_gn=LOC101098061;refGene_tx=XM_023256653.1;refGene_gn=LOC101098061;refGene_tx=XM_023256654.1;refGene_gn=LOC101098061;refGene_tx=XM_023256655.1;refGene_gn=LOC101098061;refGene_tx=XM_023256656.1;refGene_gn=LOC101098061;refGene_tx=XM_023256657.1;refGene_gn=LOC101098061;refGene_tx=XM_023256660.1;refGene_gn=LOC101098061;refGene_tx=XM_023256661.1;refGene_gn=LOC101098061;refGene_tx=XM_023256662.1;refGene_gn=LOC101098061;refGene_tx=XM_02

3256663.1;refGene_gn=LOC101098061;refGene_tx=XM_023256664.1;refGene_gn=LOC101098061;refGene_tx=XM_023256665.1
B4 38038071 38038194
refGene_gn=LOC101098061;refGene_tx=XM_023256652.1;refGene_gn=LOC101098061;refGene_tx=XM_023256653.1;refGene_gn=LOC101098061;refGene_tx=XM_023256654.1;refGene_gn=LOC101098061;refGene_tx=XM_023256655.1;refGene_gn=LOC101098061;refGene_tx=XM_023256656.1;refGene_gn=LOC101098061;refGene_tx=XM_023256657.1;refGene_gn=LOC101098061;refGene_tx=XM_023256660.1;refGene_gn=LOC101098061;refGene_tx=XM_023256661.1;refGene_gn=LOC101098061;refGene_tx=XM_023256662.1;refGene_gn=LOC101098061;refGene_tx=XM_023256663.1;refGene_gn=LOC101098061;refGene_tx=XM_023256664.1;refGene_gn=LOC101098061;refGene_tx=XM_023256665.1
B4 38048113 38048230
refGene_gn=LOC101098061;refGene_tx=XM_023256652.1;refGene_gn=LOC101098061;refGene_tx=XM_023256653.1;refGene_gn=LOC101098061;refGene_tx=XM_023256654.1;refGene_gn=LOC101098061;refGene_tx=XM_023256655.1;refGene_gn=LOC101098061;refGene_tx=XM_023256656.1;refGene_gn=LOC101098061;refGene_tx=XM_023256657.1;refGene_gn=LOC101098061;refGene_tx=XM_023256659.1;refGene_gn=LOC101098061;refGene_tx=XM_023256660.1;refGene_gn=LOC101098061;refGene_tx=XM_023256661.1;refGene_gn=LOC101098061;refGene_tx=XM_023256662.1;refGene_gn=LOC101098061;refGene_tx=XM_023256663.1;refGene_gn=LOC101098061;refGene_tx=XM_023256664.1;refGene_gn=LOC101098061;refGene_tx=XM_023256665.1
B4 38053458 38053566
refGene_gn=LOC101098061;refGene_tx=XM_023256652.1;refGene_gn=LOC101098061;refGene_tx=XM_023256653.1;refGene_gn=LOC101098061;refGene_tx=XM_023256654.1;refGene_gn=LOC101098061;refGene_tx=XM_023256655.1;refGene_gn=LOC101098061;refGene_tx=XM_023256656.1;refGene_gn=LOC101098061;refGene_tx=XM_023256659.1;refGene_gn=LOC101098061;refGene_tx=XM_023256660.1;refGene_gn=LOC101098061;refGene_tx=XM_023256661.1;refGene_gn=LOC101098061;refGene_tx=XM_023256662.1;refGene_gn=LOC101098061;refGene_tx=XM_023256663.1;refGene_gn=LOC101098061;refGene_tx=XM_023256664.1;refGene_gn=LOC101098061;refGene_tx=XM_023256665.1
B4 38054165 38054218
refGene_gn=LOC101098061;refGene_tx=XM_023256652.1;refGene_gn=LOC101098061;refGene_tx=XM_023256653.1;refGene_gn=LOC101098061;refGene_tx=XM_023256654.1;refGene_gn=LOC101098061;refGene_tx=XM_023256655.1;refGene_gn=LOC101098061;refGene_tx=XM_023256656.1;refGene_gn=LOC101098061;refGene_tx=XM_023256659.1;refGene_gn=LOC101098061;refGene_tx=XM_023256660.1;refGene_gn=LOC101098061;refGene_tx=XM_023256661.1;refGene_gn=LOC101098061;refGene_tx=XM_023256662.1;refGene_gn=LOC101098061;refGene_tx=XM_023256663.1;refGene_gn=LOC101098061;refGene_tx=XM_023256664.1;refGene_gn=LOC101098061;refGene_tx=XM_023256665.1
B4 38054733 38054893
refGene_gn=LOC101098061;refGene_tx=XM_023256652.1;refGene_gn=LOC101098061;refGene_tx=XM_023256653.1;refGene_gn=LOC101098061;refGene_tx=XM_023256654.1;refGene_gn=LOC101098061;refGene_tx=XM_023256655.1;refGene_gn=LOC101098061;refGene_tx=XM_023256656.1;refGene_gn=LOC101098061;refGene_tx=XM_023256659.1;refGene_gn=LOC101098061;refGene_tx=XM_023256660.1;refGene_gn=LOC101098061;refGene_tx=XM_023256661.1;refGene_gn=LOC101098

061;refGene_tx=XM_023256662.1;refGene_gn=LOC101098061;refGene_tx=XM_02
3256663.1;refGene_gn=LOC101098061;refGene_tx=XM_023256664.1;refGene_gn
=LOC101098061;refGene_tx=XM_023256665.1
B4 38055623 38055667
refGene_gn=LOC101098061;refGene_tx=XM_023256652.1;refGene_gn=LOC101098
061;refGene_tx=XM_023256653.1;refGene_gn=LOC101098061;refGene_tx=XM_02
3256654.1;refGene_gn=LOC101098061;refGene_tx=XM_023256655.1;refGene_gn
=LOC101098061;refGene_tx=XM_023256656.1;refGene_gn=LOC101098061;refGen
e_tx=XM_023256657.1;refGene_gn=LOC101098061;refGene_tx=XM_023256659.1;
refGene_gn=LOC101098061;refGene_tx=XM_023256660.1;refGene_gn=LOC101098
061;refGene_tx=XM_023256661.1;refGene_gn=LOC101098061;refGene_tx=XM_02
3256662.1;refGene_gn=LOC101098061;refGene_tx=XM_023256663.1;refGene_gn
=LOC101098061;refGene_tx=XM_023256664.1;refGene_gn=LOC101098061;refGen
e_tx=XM_023256665.1
B4 38057230 38057375
refGene_gn=LOC101098061;refGene_tx=XM_023256652.1;refGene_gn=LOC101098
061;refGene_tx=XM_023256653.1;refGene_gn=LOC101098061;refGene_tx=XM_02
3256654.1;refGene_gn=LOC101098061;refGene_tx=XM_023256655.1;refGene_gn
=LOC101098061;refGene_tx=XM_023256656.1;refGene_gn=LOC101098061;refGen
e_tx=XM_023256657.1;refGene_gn=LOC101098061;refGene_tx=XM_023256659.1;
refGene_gn=LOC101098061;refGene_tx=XM_023256660.1;refGene_gn=LOC101098
061;refGene_tx=XM_023256661.1;refGene_gn=LOC101098061;refGene_tx=XM_02
3256662.1;refGene_gn=LOC101098061;refGene_tx=XM_023256663.1;refGene_gn
=LOC101098061;refGene_tx=XM_023256664.1;refGene_gn=LOC101098061;refGen
e_tx=XM_023256665.1
B4 38058421 38058466
refGene_gn=LOC101098061;refGene_tx=XM_023256652.1;refGene_gn=LOC101098
061;refGene_tx=XM_023256653.1;refGene_gn=LOC101098061;refGene_tx=XM_02
3256654.1;refGene_gn=LOC101098061;refGene_tx=XM_023256655.1;refGene_gn
=LOC101098061;refGene_tx=XM_023256656.1;refGene_gn=LOC101098061;refGen
e_tx=XM_023256657.1;refGene_gn=LOC101098061;refGene_tx=XM_023256659.1;
refGene_gn=LOC101098061;refGene_tx=XM_023256660.1;refGene_gn=LOC101098
061;refGene_tx=XM_023256661.1;refGene_gn=LOC101098061;refGene_tx=XM_02
3256662.1;refGene_gn=LOC101098061;refGene_tx=XM_023256663.1;refGene_gn
=LOC101098061;refGene_tx=XM_023256664.1;refGene_gn=LOC101098061;refGen
e_tx=XM_023256665.1
B4 38058677 38058794
refGene_gn=LOC101098061;refGene_tx=XM_023256652.1;refGene_gn=LOC101098
061;refGene_tx=XM_023256653.1;refGene_gn=LOC101098061;refGene_tx=XM_02
3256654.1;refGene_gn=LOC101098061;refGene_tx=XM_023256655.1;refGene_gn
=LOC101098061;refGene_tx=XM_023256656.1;refGene_gn=LOC101098061;refGen
e_tx=XM_023256657.1;refGene_gn=LOC101098061;refGene_tx=XM_023256659.1;
refGene_gn=LOC101098061;refGene_tx=XM_023256660.1;refGene_gn=LOC101098
061;refGene_tx=XM_023256661.1;refGene_gn=LOC101098061;refGene_tx=XM_02
3256662.1;refGene_gn=LOC101098061;refGene_tx=XM_023256663.1;refGene_gn
=LOC101098061;refGene_tx=XM_023256664.1;refGene_gn=LOC101098061;refGen
e_tx=XM_023256665.1
B4 38059935 38060009
refGene_gn=LOC101098061;refGene_tx=XM_023256652.1;refGene_gn=LOC101098
061;refGene_tx=XM_023256653.1;refGene_gn=LOC101098061;refGene_tx=XM_02

3256654.1;refGene_gn=LOC101098061;refGene_tx=XM_023256655.1;refGene_gn=LOC101098061;refGene_tx=XM_023256656.1;refGene_gn=LOC101098061;refGene_tx=XM_023256657.1;refGene_gn=LOC101098061;refGene_tx=XM_023256659.1;refGene_gn=LOC101098061;refGene_tx=XM_023256660.1;refGene_gn=LOC101098061;refGene_tx=XM_023256661.1;refGene_gn=LOC101098061;refGene_tx=XM_023256662.1;refGene_gn=LOC101098061;refGene_tx=XM_023256663.1;refGene_gn=LOC101098061;refGene_tx=XM_023256664.1;refGene_gn=LOC101098061;refGene_tx=XM_023256665.1
B4 38060998 38061109
refGene_gn=LOC101098061;refGene_tx=XM_023256652.1;refGene_gn=LOC101098061;refGene_tx=XM_023256653.1;refGene_gn=LOC101098061;refGene_tx=XM_023256654.1;refGene_gn=LOC101098061;refGene_tx=XM_023256655.1;refGene_gn=LOC101098061;refGene_tx=XM_023256656.1;refGene_gn=LOC101098061;refGene_tx=XM_023256657.1;refGene_gn=LOC101098061;refGene_tx=XM_023256659.1;refGene_gn=LOC101098061;refGene_tx=XM_023256660.1;refGene_gn=LOC101098061;refGene_tx=XM_023256661.1;refGene_gn=LOC101098061;refGene_tx=XM_023256662.1;refGene_gn=LOC101098061;refGene_tx=XM_023256663.1;refGene_gn=LOC101098061;refGene_tx=XM_023256664.1;refGene_gn=LOC101098061;refGene_tx=XM_023256665.1
B4 38061894 38061973
refGene_gn=LOC101098061;refGene_tx=XM_023256652.1;refGene_gn=LOC101098061;refGene_tx=XM_023256653.1;refGene_gn=LOC101098061;refGene_tx=XM_023256654.1;refGene_gn=LOC101098061;refGene_tx=XM_023256655.1;refGene_gn=LOC101098061;refGene_tx=XM_023256656.1;refGene_gn=LOC101098061;refGene_tx=XM_023256657.1;refGene_gn=LOC101098061;refGene_tx=XM_023256659.1;refGene_gn=LOC101098061;refGene_tx=XM_023256660.1;refGene_gn=LOC101098061;refGene_tx=XM_023256661.1;refGene_gn=LOC101098061;refGene_tx=XM_023256662.1;refGene_gn=LOC101098061;refGene_tx=XM_023256663.1;refGene_gn=LOC101098061;refGene_tx=XM_023256664.1;refGene_gn=LOC101098061;refGene_tx=XM_023256665.1
B4 38062546 38062645
refGene_gn=LOC101098061;refGene_tx=XM_023256652.1;refGene_gn=LOC101098061;refGene_tx=XM_023256653.1;refGene_gn=LOC101098061;refGene_tx=XM_023256654.1;refGene_gn=LOC101098061;refGene_tx=XM_023256655.1;refGene_gn=LOC101098061;refGene_tx=XM_023256656.1;refGene_gn=LOC101098061;refGene_tx=XM_023256657.1;refGene_gn=LOC101098061;refGene_tx=XM_023256659.1;refGene_gn=LOC101098061;refGene_tx=XM_023256660.1;refGene_gn=LOC101098061;refGene_tx=XM_023256661.1;refGene_gn=LOC101098061;refGene_tx=XM_023256662.1;refGene_gn=LOC101098061;refGene_tx=XM_023256663.1;refGene_gn=LOC101098061;refGene_tx=XM_023256664.1;refGene_gn=LOC101098061;refGene_tx=XM_023256665.1
B4 38064660 38064801
refGene_gn=LOC101098061;refGene_tx=XM_023256652.1;refGene_gn=LOC101098061;refGene_tx=XM_023256653.1;refGene_gn=LOC101098061;refGene_tx=XM_023256654.1;refGene_gn=LOC101098061;refGene_tx=XM_023256655.1;refGene_gn=LOC101098061;refGene_tx=XM_023256656.1;refGene_gn=LOC101098061;refGene_tx=XM_023256657.1;refGene_gn=LOC101098061;refGene_tx=XM_023256659.1;refGene_gn=LOC101098061;refGene_tx=XM_023256660.1;refGene_gn=LOC101098061;refGene_tx=XM_023256661.1;refGene_gn=LOC101098061;refGene_tx=XM_023256662.1;refGene_gn=LOC101098061;refGene_tx=XM_023256663.1;refGene_gn

=LOC101098061;refGene_tx=XM_023256664.1;refGene_gn=LOC101098061;refGene_tx=XM_023256665.1
B4 38065170 38065325
refGene_gn=LOC101098061;refGene_tx=XM_023256652.1;refGene_gn=LOC101098061;refGene_tx=XM_023256653.1;refGene_gn=LOC101098061;refGene_tx=XM_023256654.1;refGene_gn=LOC101098061;refGene_tx=XM_023256655.1;refGene_gn=LOC101098061;refGene_tx=XM_023256656.1;refGene_gn=LOC101098061;refGene_tx=XM_023256657.1;refGene_gn=LOC101098061;refGene_tx=XM_023256659.1;refGene_gn=LOC101098061;refGene_tx=XM_023256660.1;refGene_gn=LOC101098061;refGene_tx=XM_023256661.1;refGene_gn=LOC101098061;refGene_tx=XM_023256662.1;refGene_gn=LOC101098061;refGene_tx=XM_023256663.1;refGene_gn=LOC101098061;refGene_tx=XM_023256664.1;refGene_gn=LOC101098061;refGene_tx=XM_023256665.1
B4 38065754 38065847
refGene_gn=LOC101098061;refGene_tx=XM_023256652.1;refGene_gn=LOC101098061;refGene_tx=XM_023256653.1;refGene_gn=LOC101098061;refGene_tx=XM_023256654.1;refGene_gn=LOC101098061;refGene_tx=XM_023256655.1;refGene_gn=LOC101098061;refGene_tx=XM_023256656.1;refGene_gn=LOC101098061;refGene_tx=XM_023256657.1;refGene_gn=LOC101098061;refGene_tx=XM_023256659.1;refGene_gn=LOC101098061;refGene_tx=XM_023256660.1;refGene_gn=LOC101098061;refGene_tx=XM_023256661.1;refGene_gn=LOC101098061;refGene_tx=XM_023256662.1;refGene_gn=LOC101098061;refGene_tx=XM_023256663.1;refGene_gn=LOC101098061;refGene_tx=XM_023256664.1;refGene_gn=LOC101098061;refGene_tx=XM_023256665.1
B4 38066919 38067082
refGene_gn=LOC101098061;refGene_tx=XM_023256652.1;refGene_gn=LOC101098061;refGene_tx=XM_023256653.1;refGene_gn=LOC101098061;refGene_tx=XM_023256654.1;refGene_gn=LOC101098061;refGene_tx=XM_023256655.1;refGene_gn=LOC101098061;refGene_tx=XM_023256656.1;refGene_gn=LOC101098061;refGene_tx=XM_023256657.1;refGene_gn=LOC101098061;refGene_tx=XM_023256659.1;refGene_gn=LOC101098061;refGene_tx=XM_023256660.1;refGene_gn=LOC101098061;refGene_tx=XM_023256661.1;refGene_gn=LOC101098061;refGene_tx=XM_023256662.1;refGene_gn=LOC101098061;refGene_tx=XM_023256663.1;refGene_gn=LOC101098061;refGene_tx=XM_023256664.1;refGene_gn=LOC101098061;refGene_tx=XM_023256665.1
B4 38068005 38068128
refGene_gn=LOC101098061;refGene_tx=XM_023256652.1;refGene_gn=LOC101098061;refGene_tx=XM_023256653.1;refGene_gn=LOC101098061;refGene_tx=XM_023256654.1;refGene_gn=LOC101098061;refGene_tx=XM_023256655.1;refGene_gn=LOC101098061;refGene_tx=XM_023256656.1;refGene_gn=LOC101098061;refGene_tx=XM_023256657.1;refGene_gn=LOC101098061;refGene_tx=XM_023256659.1;refGene_gn=LOC101098061;refGene_tx=XM_023256660.1;refGene_gn=LOC101098061;refGene_tx=XM_023256661.1;refGene_gn=LOC101098061;refGene_tx=XM_023256662.1;refGene_gn=LOC101098061;refGene_tx=XM_023256663.1;refGene_gn=LOC101098061;refGene_tx=XM_023256664.1;refGene_gn=LOC101098061;refGene_tx=XM_023256665.1
B4 38068456 38068561
refGene_gn=LOC101098061;refGene_tx=XM_023256652.1;refGene_gn=LOC101098061;refGene_tx=XM_023256653.1;refGene_gn=LOC101098061;refGene_tx=XM_023256654.1;refGene_gn=LOC101098061;refGene_tx=XM_023256656.1;refGene_gn

=LOC101098061;refGene_tx=XM_023256657.1;refGene_gn=LOC101098061;refGene_tx=XM_023256659.1;refGene_gn=LOC101098061;refGene_tx=XM_023256660.1;refGene_gn=LOC101098061;refGene_tx=XM_023256661.1;refGene_gn=LOC101098061;refGene_tx=XM_023256662.1;refGene_gn=LOC101098061;refGene_tx=XM_023256663.1;refGene_gn=LOC101098061;refGene_tx=XM_023256664.1;refGene_gn=LOC101098061;refGene_tx=XM_023256665.1
B4 38109462 38109567 refGene_gn=FKBP4;refGene_tx=XM_011283738.3
B4 38111569 38111714 refGene_gn=FKBP4;refGene_tx=XM_011283738.3
B4 38112192 38112335 refGene_gn=FKBP4;refGene_tx=XM_011283738.3
B4 38113146 38113267 refGene_gn=FKBP4;refGene_tx=XM_011283738.3
B4 38113777 38113934 refGene_gn=FKBP4;refGene_tx=XM_011283738.3
B4 38114582 38114673 refGene_gn=FKBP4;refGene_tx=XM_011283738.3
B4 38114765 38114849 refGene_gn=FKBP4;refGene_tx=XM_011283738.3
B4 38115112 38115298 refGene_gn=FKBP4;refGene_tx=XM_011283738.3
B4 38115852 38116092 refGene_gn=FKBP4;refGene_tx=XM_011283738.3
B4 38117695 38117803 refGene_gn=FKBP4;refGene_tx=XM_011283738.3
B4 38124647 38124743
refGene_gn=ITFG2;refGene_tx=XM_019834359.2;refGene_gn=ITFG2;refGene_tx=XM_019834360.2
B4 38124751 38124790 refGene_gn=ITFG2;refGene_tx=XM_019834361.2
B4 38129617 38129713
refGene_gn=ITFG2;refGene_tx=XM_019834359.2;refGene_gn=ITFG2;refGene_tx=XM_019834360.2;refGene_gn=ITFG2;refGene_tx=XM_019834361.2
B4 38130269 38130311
refGene_gn=ITFG2;refGene_tx=XM_019834359.2;refGene_gn=ITFG2;refGene_tx=XM_019834360.2;refGene_gn=ITFG2;refGene_tx=XM_019834361.2
B4 38130467 38130669
refGene_gn=ITFG2;refGene_tx=XM_019834359.2;refGene_gn=ITFG2;refGene_tx=XM_019834360.2;refGene_gn=ITFG2;refGene_tx=XM_019834361.2
B4 38131379 38131519
refGene_gn=ITFG2;refGene_tx=XM_019834359.2;refGene_gn=ITFG2;refGene_tx=XM_019834360.2;refGene_gn=ITFG2;refGene_tx=XM_019834361.2
B4 38131990 38132139
refGene_gn=ITFG2;refGene_tx=XM_019834359.2;refGene_gn=ITFG2;refGene_tx=XM_019834360.2;refGene_gn=ITFG2;refGene_tx=XM_019834361.2
B4 38132530 38132628
refGene_gn=ITFG2;refGene_tx=XM_019834359.2;refGene_gn=ITFG2;refGene_tx=XM_019834360.2;refGene_gn=ITFG2;refGene_tx=XM_019834361.2
B4 38132806 38132860
refGene_gn=ITFG2;refGene_tx=XM_019834359.2;refGene_gn=ITFG2;refGene_tx=XM_019834360.2;refGene_gn=ITFG2;refGene_tx=XM_019834361.2
B4 38132952 38133044
refGene_gn=ITFG2;refGene_tx=XM_019834359.2;refGene_gn=ITFG2;refGene_tx=XM_019834360.2;refGene_gn=ITFG2;refGene_tx=XM_019834361.2
B4 38133779 38133897
refGene_gn=ITFG2;refGene_tx=XM_019834359.2;refGene_gn=ITFG2;refGene_tx=XM_019834360.2;refGene_gn=ITFG2;refGene_tx=XM_019834361.2
B4 38135500 38135674
refGene_gn=ITFG2;refGene_tx=XM_019834359.2;refGene_gn=ITFG2;refGene_tx=XM_019834360.2;refGene_gn=ITFG2;refGene_tx=XM_019834361.2

B4 38135764 38135868
refGene_gn=ITFG2;refGene_tx=XM_019834359.2;refGene_gn=ITFG2;refGene_tx=XM_019834360.2;refGene_gn=ITFG2;refGene_tx=XM_019834361.2
B4 38138141 38138244 refGene_gn=NRIP2;refGene_tx=XM_011283736.3
B4 38138337 38138512
refGene_gn=NRIP2;refGene_tx=XM_004001512.5;refGene_gn=NRIP2;refGene_tx=XM_011283736.3
B4 38138849 38138902
refGene_gn=NRIP2;refGene_tx=XM_004001512.5;refGene_gn=NRIP2;refGene_tx=XM_011283736.3
B4 38139093 38139212
refGene_gn=NRIP2;refGene_tx=XM_004001512.5;refGene_gn=NRIP2;refGene_tx=XM_011283736.3
B4 38139565 38139648
refGene_gn=NRIP2;refGene_tx=XM_004001512.5;refGene_gn=NRIP2;refGene_tx=XM_011283736.3
B4 38140748 38140901
refGene_gn=NRIP2;refGene_tx=XM_004001512.5;refGene_gn=NRIP2;refGene_tx=XM_011283736.3
B4 38144786 38145038
refGene_gn=NRIP2;refGene_tx=XM_004001512.5;refGene_gn=NRIP2;refGene_tx=XM_011283736.3
B4 38157384 38157679 refGene_gn=TEX52;refGene_tx=XM_011283735.3
B4 38161117 38161668 refGene_gn=TEX52;refGene_tx=XM_011283735.3
B4 38162728 38162863 refGene_gn=TEX52;refGene_tx=XM_011283735.3
B4 38164435 38165461
refGene_gn=FOXM1;refGene_tx=XM_011283734.3;refGene_gn=FOXM1;refGene_tx=XM_019834356.2;refGene_gn=FOXM1;refGene_tx=XM_019834357.2;refGene_gn=FOXM1;refGene_tx=XM_019834358.2
B4 38168177 38168353
refGene_gn=FOXM1;refGene_tx=XM_011283734.3;refGene_gn=FOXM1;refGene_tx=XM_019834356.2;refGene_gn=FOXM1;refGene_tx=XM_019834357.2;refGene_gn=FOXM1;refGene_tx=XM_019834358.2
B4 38168500 38168573
refGene_gn=FOXM1;refGene_tx=XM_011283734.3;refGene_gn=FOXM1;refGene_tx=XM_019834356.2;refGene_gn=FOXM1;refGene_tx=XM_019834357.2;refGene_gn=FOXM1;refGene_tx=XM_019834358.2
B4 38169214 38169259
refGene_gn=FOXM1;refGene_tx=XM_011283734.3;refGene_gn=FOXM1;refGene_tx=XM_019834356.2;refGene_gn=FOXM1;refGene_tx=XM_019834358.2
B4 38170159 38170288
refGene_gn=FOXM1;refGene_tx=XM_011283734.3;refGene_gn=FOXM1;refGene_tx=XM_019834356.2;refGene_gn=FOXM1;refGene_tx=XM_019834357.2;refGene_gn=FOXM1;refGene_tx=XM_019834358.2
B4 38171180 38171372
refGene_gn=FOXM1;refGene_tx=XM_011283734.3;refGene_gn=FOXM1;refGene_tx=XM_019834356.2;refGene_gn=FOXM1;refGene_tx=XM_019834357.2
B4 38172472 38172618
refGene_gn=FOXM1;refGene_tx=XM_011283734.3;refGene_gn=FOXM1;refGene_tx=XM_019834356.2;refGene_gn=FOXM1;refGene_tx=XM_019834357.2;refGene_gn=

FOXM1;refGene_tx=XM_019834358.2
B4 38173213 38173715
refGene_gn=FOXM1;refGene_tx=XM_011283734.3;refGene_gn=FOXM1;refGene_tx=XM_019834356.2;refGene_gn=FOXM1;refGene_tx=XM_019834357.2;refGene_gn=FOXM1;refGene_tx=XM_019834358.2
B4 38180736 38180904
refGene_gn=RHNO1;refGene_tx=XM_003988277.4;refGene_gn=RHNO1;refGene_tx=XM_023256673.1;refGene_gn=RHNO1;refGene_tx=XM_023256674.1
B4 38182180 38182777
refGene_gn=RHNO1;refGene_tx=XM_003988277.4;refGene_gn=RHNO1;refGene_tx=XM_023256673.1;refGene_gn=RHNO1;refGene_tx=XM_023256674.1
B4 38184718 38184759
refGene_gn=TULP3;refGene_tx=XM_023256666.1;refGene_gn=TULP3;refGene_tx=XM_023256668.1;refGene_gn=TULP3;refGene_tx=XM_023256669.1;refGene_gn=TULP3;refGene_tx=XM_023256671.1;refGene_gn=TULP3;refGene_tx=XM_023256672.1
B4 38219582 38219634
refGene_gn=TULP3;refGene_tx=XM_023256666.1;refGene_gn=TULP3;refGene_tx=XM_023256667.1;refGene_gn=TULP3;refGene_tx=XM_023256668.1;refGene_gn=TULP3;refGene_tx=XM_023256669.1;refGene_gn=TULP3;refGene_tx=XM_023256671.1;refGene_gn=TULP3;refGene_tx=XM_023256672.1
B4 38229130 38229290
refGene_gn=TULP3;refGene_tx=XM_023256666.1;refGene_gn=TULP3;refGene_tx=XM_023256667.1;refGene_gn=TULP3;refGene_tx=XM_023256668.1;refGene_gn=TULP3;refGene_tx=XM_023256669.1;refGene_gn=TULP3;refGene_tx=XM_023256671.1;refGene_gn=TULP3;refGene_tx=XM_023256672.1
B4 38229952 38230129
refGene_gn=TULP3;refGene_tx=XM_023256666.1;refGene_gn=TULP3;refGene_tx=XM_023256667.1;refGene_gn=TULP3;refGene_tx=XM_023256668.1;refGene_gn=TULP3;refGene_tx=XM_023256669.1;refGene_gn=TULP3;refGene_tx=XM_023256671.1;refGene_gn=TULP3;refGene_tx=XM_023256672.1
B4 38242261 38242389
refGene_gn=TULP3;refGene_tx=XM_023256666.1;refGene_gn=TULP3;refGene_tx=XM_023256667.1;refGene_gn=TULP3;refGene_tx=XM_023256668.1;refGene_gn=TULP3;refGene_tx=XM_023256669.1;refGene_gn=TULP3;refGene_tx=XM_023256671.1;refGene_gn=TULP3;refGene_tx=XM_023256672.1
B4 38242993 38243191
refGene_gn=TULP3;refGene_tx=XM_023256666.1;refGene_gn=TULP3;refGene_tx=XM_023256667.1;refGene_gn=TULP3;refGene_tx=XM_023256668.1;refGene_gn=TULP3;refGene_tx=XM_023256669.1;refGene_gn=TULP3;refGene_tx=XM_023256671.1
B4 38245784 38245897
refGene_gn=TULP3;refGene_tx=XM_023256666.1;refGene_gn=TULP3;refGene_tx=XM_023256667.1;refGene_gn=TULP3;refGene_tx=XM_023256668.1;refGene_gn=TULP3;refGene_tx=XM_023256669.1;refGene_gn=TULP3;refGene_tx=XM_023256671.1;refGene_gn=TULP3;refGene_tx=XM_023256672.1
B4 38246452 38246567
refGene_gn=TULP3;refGene_tx=XM_023256666.1;refGene_gn=TULP3;refGene_tx=XM_023256667.1;refGene_gn=TULP3;refGene_tx=XM_023256668.1;refGene_gn=TULP3;refGene_tx=XM_023256669.1;refGene_gn=TULP3;refGene_tx=XM_0232566

71.1;refGene_gn=TULP3;refGene_tx=XM_023256672.1
B4	38249345	38249444
refGene_gn=TULP3;refGene_tx=XM_023256666.1;refGene_gn=TULP3;refGene_tx=XM_023256667.1;refGene_gn=TULP3;refGene_tx=XM_023256668.1;refGene_gn=TULP3;refGene_tx=XM_023256669.1;refGene_gn=TULP3;refGene_tx=XM_023256671.1;refGene_gn=TULP3;refGene_tx=XM_023256672.1
B4	38249845	38250017
refGene_gn=TULP3;refGene_tx=XM_023256666.1;refGene_gn=TULP3;refGene_tx=XM_023256667.1;refGene_gn=TULP3;refGene_tx=XM_023256668.1;refGene_gn=TULP3;refGene_tx=XM_023256669.1;refGene_gn=TULP3;refGene_tx=XM_023256671.1;refGene_gn=TULP3;refGene_tx=XM_023256672.1
B4	38250936	38251070
refGene_gn=TULP3;refGene_tx=XM_023256666.1;refGene_gn=TULP3;refGene_tx=XM_023256667.1;refGene_gn=TULP3;refGene_tx=XM_023256668.1;refGene_gn=TULP3;refGene_tx=XM_023256669.1;refGene_gn=TULP3;refGene_tx=XM_023256671.1;refGene_gn=TULP3;refGene_tx=XM_023256672.1
B4	38264235	38264377
refGene_gn=TEAD4;refGene_tx=XM_006933413.4;refGene_gn=TEAD4;refGene_tx=XM_023256676.1;refGene_gn=TEAD4;refGene_tx=XM_023256678.1;refGene_gn=TEAD4;refGene_tx=XM_023256679.1
B4	38285977	38286032
refGene_gn=TEAD4;refGene_tx=XM_006933414.4;refGene_gn=TEAD4;refGene_tx=XM_023256677.1
B4	38290680	38290935
refGene_gn=TEAD4;refGene_tx=XM_006933413.4;refGene_gn=TEAD4;refGene_tx=XM_006933414.4;refGene_gn=TEAD4;refGene_tx=XM_023256676.1;refGene_gn=TEAD4;refGene_tx=XM_023256677.1;refGene_gn=TEAD4;refGene_tx=XM_023256678.1;refGene_gn=TEAD4;refGene_tx=XM_023256679.1
B4	38304541	38304606
refGene_gn=TEAD4;refGene_tx=XM_006933413.4;refGene_gn=TEAD4;refGene_tx=XM_006933414.4;refGene_gn=TEAD4;refGene_tx=XM_023256676.1;refGene_gn=TEAD4;refGene_tx=XM_023256677.1;refGene_gn=TEAD4;refGene_tx=XM_023256678.1;refGene_gn=TEAD4;refGene_tx=XM_023256679.1
B4	38305077	38305140
refGene_gn=TEAD4;refGene_tx=XM_006933413.4;refGene_gn=TEAD4;refGene_tx=XM_006933414.4;refGene_gn=TEAD4;refGene_tx=XM_023256676.1;refGene_gn=TEAD4;refGene_tx=XM_023256677.1;refGene_gn=TEAD4;refGene_tx=XM_023256678.1;refGene_gn=TEAD4;refGene_tx=XM_023256679.1
B4	38308990	38309119
refGene_gn=TEAD4;refGene_tx=XM_006933413.4;refGene_gn=TEAD4;refGene_tx=XM_006933414.4;refGene_gn=TEAD4;refGene_tx=XM_023256676.1;refGene_gn=TEAD4;refGene_tx=XM_023256677.1
B4	38310209	38310253
refGene_gn=TEAD4;refGene_tx=XM_006933413.4;refGene_gn=TEAD4;refGene_tx=XM_006933414.4;refGene_gn=TEAD4;refGene_tx=XM_023256676.1;refGene_gn=TEAD4;refGene_tx=XM_023256677.1;refGene_gn=TEAD4;refGene_tx=XM_023256678.1;refGene_gn=TEAD4;refGene_tx=XM_023256679.1
B4	38310771	38310827
refGene_gn=TEAD4;refGene_tx=XM_006933413.4;refGene_gn=TEAD4;refGene_tx=XM_006933414.4;refGene_gn=TEAD4;refGene_tx=XM_023256676.1;refGene_gn=

TEAD4;refGene_tx=XM_023256677.1;refGene_gn=TEAD4;refGene_tx=XM_0232566
78.1;refGene_gn=TEAD4;refGene_tx=XM_023256679.1
B4 38313182 38313370
refGene_gn=TEAD4;refGene_tx=XM_006933413.4;refGene_gn=TEAD4;refGene_tx
=XM_006933414.4;refGene_gn=TEAD4;refGene_tx=XM_023256676.1;refGene_gn=
TEAD4;refGene_tx=XM_023256677.1;refGene_gn=TEAD4;refGene_tx=XM_0232566
78.1;refGene_gn=TEAD4;refGene_tx=XM_023256679.1
B4 38314467 38314641
refGene_gn=TEAD4;refGene_tx=XM_006933413.4;refGene_gn=TEAD4;refGene_tx
=XM_006933414.4;refGene_gn=TEAD4;refGene_tx=XM_023256676.1;refGene_gn=
TEAD4;refGene_tx=XM_023256677.1;refGene_gn=TEAD4;refGene_tx=XM_0232566
78.1;refGene_gn=TEAD4;refGene_tx=XM_023256679.1
B4 38326527 38326668
refGene_gn=TEAD4;refGene_tx=XM_006933413.4;refGene_gn=TEAD4;refGene_tx
=XM_006933414.4;refGene_gn=TEAD4;refGene_tx=XM_023256676.1;refGene_gn=
TEAD4;refGene_tx=XM_023256677.1;refGene_gn=TEAD4;refGene_tx=XM_0232566
78.1;refGene_gn=TEAD4;refGene_tx=XM_023256679.1
B4 38327448 38327601
refGene_gn=TEAD4;refGene_tx=XM_006933413.4;refGene_gn=TEAD4;refGene_tx
=XM_006933414.4;refGene_gn=TEAD4;refGene_tx=XM_023256676.1;refGene_gn=
TEAD4;refGene_tx=XM_023256677.1;refGene_gn=TEAD4;refGene_tx=XM_0232566
78.1;refGene_gn=TEAD4;refGene_tx=XM_023256679.1
B4 38327722 38327836
refGene_gn=TEAD4;refGene_tx=XM_006933413.4;refGene_gn=TEAD4;refGene_tx
=XM_006933414.4;refGene_gn=TEAD4;refGene_tx=XM_023256676.1;refGene_gn=
TEAD4;refGene_tx=XM_023256677.1;refGene_gn=TEAD4;refGene_tx=XM_0232566
78.1;refGene_gn=TEAD4;refGene_tx=XM_023256679.1
B4 38471560 38471896
refGene_gn=TSPAN9;refGene_tx=XM_023256680.1;refGene_gn=TSPAN9;refGene_
tx=XM_023256681.1;refGene_gn=TSPAN9;refGene_tx=XM_023256682.1
B4 38493770 38493872
refGene_gn=TSPAN9;refGene_tx=XM_023256680.1;refGene_gn=TSPAN9;refGene_
tx=XM_023256682.1
B4 38550206 38550398
refGene_gn=TSPAN9;refGene_tx=XM_023256680.1;refGene_gn=TSPAN9;refGene_
tx=XM_023256681.1;refGene_gn=TSPAN9;refGene_tx=XM_023256682.1
B4 38550772 38550847
refGene_gn=TSPAN9;refGene_tx=XM_023256680.1;refGene_gn=TSPAN9;refGene_
tx=XM_023256681.1;refGene_gn=TSPAN9;refGene_tx=XM_023256682.1
B4 38551172 38551274
refGene_gn=TSPAN9;refGene_tx=XM_023256680.1;refGene_gn=TSPAN9;refGene_
tx=XM_023256681.1;refGene_gn=TSPAN9;refGene_tx=XM_023256682.1
B4 38551906 38552038
refGene_gn=TSPAN9;refGene_tx=XM_023256680.1;refGene_gn=TSPAN9;refGene_
tx=XM_023256681.1;refGene_gn=TSPAN9;refGene_tx=XM_023256682.1
B4 38552468 38552552
refGene_gn=TSPAN9;refGene_tx=XM_023256680.1;refGene_gn=TSPAN9;refGene_
tx=XM_023256681.1;refGene_gn=TSPAN9;refGene_tx=XM_023256682.1
B4 38554227 38554299
refGene_gn=TSPAN9;refGene_tx=XM_023256680.1;refGene_gn=TSPAN9;refGene_

tx=XM_023256681.1;refGene_gn=TSPAN9;refGene_tx=XM_023256682.1
B4	38568111 38568171
refGene_gn=TSPAN11;refGene_tx=XM_023256684.1;refGene_gn=TSPAN11;refGene_tx=XM_023256685.1
B4	38574551 38574629 refGene_gn=TSPAN11;refGene_tx=XM_023256683.1
B4	38574782 38574869
refGene_gn=TSPAN11;refGene_tx=XM_023256683.1;refGene_gn=TSPAN11;refGene_tx=XM_023256684.1
B4	38575234 38575393
refGene_gn=TSPAN11;refGene_tx=XM_023256683.1;refGene_gn=TSPAN11;refGene_tx=XM_023256684.1;refGene_gn=TSPAN11;refGene_tx=XM_023256685.1
B4	38578234 38578339
refGene_gn=TSPAN11;refGene_tx=XM_023256683.1;refGene_gn=TSPAN11;refGene_tx=XM_023256684.1;refGene_gn=TSPAN11;refGene_tx=XM_023256685.1
B4	38579231 38579306
refGene_gn=TSPAN11;refGene_tx=XM_023256683.1;refGene_gn=TSPAN11;refGene_tx=XM_023256684.1;refGene_gn=TSPAN11;refGene_tx=XM_023256685.1
B4	38595274 38595466
refGene_gn=TSPAN11;refGene_tx=XM_023256683.1;refGene_gn=TSPAN11;refGene_tx=XM_023256684.1;refGene_gn=TSPAN11;refGene_tx=XM_023256685.1
B4	38608290 38608374
refGene_gn=TSPAN11;refGene_tx=XM_023256683.1;refGene_gn=TSPAN11;refGene_tx=XM_023256684.1;refGene_gn=TSPAN11;refGene_tx=XM_023256685.1
B4	38779667 38779730 refGene_gn=PRMT8;refGene_tx=XM_023256686.1
B4	38788993 38789179 refGene_gn=PRMT8;refGene_tx=XM_023256686.1
B4	38802711 38802867 refGene_gn=PRMT8;refGene_tx=XM_023256686.1
B4	38806028 38806092 refGene_gn=PRMT8;refGene_tx=XM_023256686.1
B4	38820882 38820971 refGene_gn=PRMT8;refGene_tx=XM_023256686.1
B4	38820972 38821026 refGene_gn=PRMT8;refGene_tx=XM_023256686.1
B4	38821649 38821737 refGene_gn=PRMT8;refGene_tx=XM_023256686.1
B4	38828975 38829091 refGene_gn=PRMT8;refGene_tx=XM_023256686.1
B4	38834745 38834896 refGene_gn=PRMT8;refGene_tx=XM_023256686.1
B4	38843699 38843821 refGene_gn=PRMT8;refGene_tx=XM_023256686.1
B4	38844597 38844681 refGene_gn=PRMT8;refGene_tx=XM_023256686.1
B4	38870097 38870182
refGene_gn=CRACR2A;refGene_tx=XM_011283726.3;refGene_gn=CRACR2A;refGene_tx=XM_019834378.2
B4	38871817 38871894
refGene_gn=CRACR2A;refGene_tx=XM_011283726.3;refGene_gn=CRACR2A;refGene_tx=XM_019834378.2
B4	38874583 38874685
refGene_gn=CRACR2A;refGene_tx=XM_011283726.3;refGene_gn=CRACR2A;refGene_tx=XM_019834378.2
B4	38882992 38883107
refGene_gn=CRACR2A;refGene_tx=XM_011283726.3;refGene_gn=CRACR2A;refGene_tx=XM_019834378.2
B4	38883231 38883313
refGene_gn=CRACR2A;refGene_tx=XM_011283726.3;refGene_gn=CRACR2A;refGene_tx=XM_019834378.2
B4	38889318 38889451

refGene_gn=CRACR2A;refGene_tx=XM_011283726.3;refGene_gn=CRACR2A;refGene_tx=XM_019834378.2
B4 38893508 38893827
refGene_gn=CRACR2A;refGene_tx=XM_011283726.3;refGene_gn=CRACR2A;refGene_tx=XM_019834378.2
B4 38897618 38897725
refGene_gn=CRACR2A;refGene_tx=XM_011283726.3;refGene_gn=CRACR2A;refGene_tx=XM_019834378.2
B4 38899657 38899703
refGene_gn=CRACR2A;refGene_tx=XM_011283726.3;refGene_gn=CRACR2A;refGene_tx=XM_019834378.2
B4 38903675 38903747
refGene_gn=CRACR2A;refGene_tx=XM_011283726.3;refGene_gn=CRACR2A;refGene_tx=XM_019834378.2
B4 38908867 38909055
refGene_gn=CRACR2A;refGene_tx=XM_011283726.3;refGene_gn=CRACR2A;refGene_tx=XM_019834378.2
B4 38911255 38911351
refGene_gn=CRACR2A;refGene_tx=XM_011283726.3;refGene_gn=CRACR2A;refGene_tx=XM_019834378.2
B4 38914914 38915005
refGene_gn=CRACR2A;refGene_tx=XM_011283726.3;refGene_gn=CRACR2A;refGene_tx=XM_019834378.2
B4 38929212 38929359
refGene_gn=CRACR2A;refGene_tx=XM_011283726.3;refGene_gn=CRACR2A;refGene_tx=XM_019834378.2
B4 38932883 38933067
refGene_gn=CRACR2A;refGene_tx=XM_011283726.3;refGene_gn=CRACR2A;refGene_tx=XM_019834378.2
B4 38934199 38934311
refGene_gn=CRACR2A;refGene_tx=XM_011283726.3;refGene_gn=CRACR2A;refGene_tx=XM_019834378.2
B4 38943288 38943504
refGene_gn=CRACR2A;refGene_tx=XM_011283726.3;refGene_gn=CRACR2A;refGene_tx=XM_019834378.2
B4 39064890 39065207
refGene_gn=PARP11;refGene_tx=XM_006933421.3;refGene_gn=PARP11;refGene_tx=XM_023256687.1;refGene_gn=PARP11;refGene_tx=XM_023256688.1;refGene_gn=PARP11;refGene_tx=XM_023256689.1;refGene_gn=PARP11;refGene_tx=XM_023256690.1
B4 39066731 39066883
refGene_gn=PARP11;refGene_tx=XM_006933421.3;refGene_gn=PARP11;refGene_tx=XM_023256687.1;refGene_gn=PARP11;refGene_tx=XM_023256688.1;refGene_gn=PARP11;refGene_tx=XM_023256689.1;refGene_gn=PARP11;refGene_tx=XM_023256690.1
B4 39076029 39076160
refGene_gn=PARP11;refGene_tx=XM_006933421.3;refGene_gn=PARP11;refGene_tx=XM_023256687.1;refGene_gn=PARP11;refGene_tx=XM_023256688.1;refGene_gn=PARP11;refGene_tx=XM_023256689.1;refGene_gn=PARP11;refGene_tx=XM_023256690.1

B4 39076238 39076311
refGene_gn=PARP11;refGene_tx=XM_006933421.3;refGene_gn=PARP11;refGene_tx=XM_023256687.1;refGene_gn=PARP11;refGene_tx=XM_023256688.1;refGene_gn=PARP11;refGene_tx=XM_023256689.1;refGene_gn=PARP11;refGene_tx=XM_023256690.1
B4 39079401 39079477
refGene_gn=PARP11;refGene_tx=XM_006933421.3;refGene_gn=PARP11;refGene_tx=XM_023256687.1;refGene_gn=PARP11;refGene_tx=XM_023256688.1;refGene_gn=PARP11;refGene_tx=XM_023256689.1;refGene_gn=PARP11;refGene_tx=XM_023256690.1
B4 39082665 39082786
refGene_gn=PARP11;refGene_tx=XM_006933421.3;refGene_gn=PARP11;refGene_tx=XM_023256687.1;refGene_gn=PARP11;refGene_tx=XM_023256688.1;refGene_gn=PARP11;refGene_tx=XM_023256689.1
B4 39083856 39083985
refGene_gn=PARP11;refGene_tx=XM_006933421.3;refGene_gn=PARP11;refGene_tx=XM_023256687.1;refGene_gn=PARP11;refGene_tx=XM_023256688.1;refGene_gn=PARP11;refGene_tx=XM_023256689.1
B4 39100990 39101014 refGene_gn=PARP11;refGene_tx=XM_006933421.3
B4 39109515 39109533 refGene_gn=PARP11;refGene_tx=XM_023256687.1
B4 39524508 39524703 refGene_gn=CCND2;refGene_tx=XM_003988283.5
B4 39526507 39526723 refGene_gn=CCND2;refGene_tx=XM_003988283.5
B4 39529390 39529550 refGene_gn=CCND2;refGene_tx=XM_003988283.5
B4 39538787 39538936 refGene_gn=CCND2;refGene_tx=XM_003988283.5
B4 39549007 39549157 refGene_gn=CCND2;refGene_tx=XM_003988283.5
B4 39568391 39568423 refGene_gn=TIGAR;refGene_tx=XM_003988284.5
B4 39575843 39575881 refGene_gn=TIGAR;refGene_tx=XM_003988284.5
B4 39580017 39580139 refGene_gn=TIGAR;refGene_tx=XM_003988284.5
B4 39587490 39587568 refGene_gn=TIGAR;refGene_tx=XM_003988284.5
B4 39590690 39590801 refGene_gn=TIGAR;refGene_tx=XM_003988284.5
B4 39591653 39592085 refGene_gn=TIGAR;refGene_tx=XM_003988284.5
B4 39607795 39608233 refGene_gn=FGF23;refGene_tx=XM_011283757.3
B4 39609438 39609542 refGene_gn=FGF23;refGene_tx=XM_011283757.3
B4 39615476 39615672 refGene_gn=FGF23;refGene_tx=XM_011283757.3
B4 39667610 39667787 refGene_gn=FGF6;refGene_tx=XM_003988285.4
B4 39677307 39677411 refGene_gn=FGF6;refGene_tx=XM_003988285.4
B4 39678372 39678718 refGene_gn=FGF6;refGene_tx=XM_003988285.4
B4 39720209 39720302
refGene_gn=CB4H12orf4;refGene_tx=XM_003988286.4;refGene_gn=CB4H12orf4;refGene_tx=XM_019834385.2;refGene_gn=CB4H12orf4;refGene_tx=XM_023256691.1;refGene_gn=CB4H12orf4;refGene_tx=XM_023256692.1;refGene_gn=CB4H12orf4;refGene_tx=XM_023256693.1;refGene_gn=CB4H12orf4;refGene_tx=XM_023256694.1
B4 39720893 39720961
refGene_gn=CB4H12orf4;refGene_tx=XM_003988286.4;refGene_gn=CB4H12orf4;refGene_tx=XM_019834385.2;refGene_gn=CB4H12orf4;refGene_tx=XM_023256691.1;refGene_gn=CB4H12orf4;refGene_tx=XM_023256692.1;refGene_gn=CB4H12orf4;refGene_tx=XM_023256693.1;refGene_gn=CB4H12orf4;refGene_tx=XM_023256694.1
B4 39721547 39721605

refGene_gn=CB4H12orf4;refGene_tx=XM_003988286.4;refGene_gn=CB4H12orf4;
refGene_tx=XM_019834385.2;refGene_gn=CB4H12orf4;refGene_tx=XM_02325669
1.1;refGene_gn=CB4H12orf4;refGene_tx=XM_023256692.1;refGene_gn=CB4H12o
rf4;refGene_tx=XM_023256693.1;refGene_gn=CB4H12orf4;refGene_tx=XM_0232
56694.1
B4 39729092 39729295
refGene_gn=CB4H12orf4;refGene_tx=XM_003988286.4;refGene_gn=CB4H12orf4;
refGene_tx=XM_019834385.2;refGene_gn=CB4H12orf4;refGene_tx=XM_02325669
1.1;refGene_gn=CB4H12orf4;refGene_tx=XM_023256692.1;refGene_gn=CB4H12o
rf4;refGene_tx=XM_023256693.1;refGene_gn=CB4H12orf4;refGene_tx=XM_0232
56694.1
B4 39732879 39732954
refGene_gn=CB4H12orf4;refGene_tx=XM_003988286.4;refGene_gn=CB4H12orf4;
refGene_tx=XM_019834385.2;refGene_gn=CB4H12orf4;refGene_tx=XM_02325669
1.1;refGene_gn=CB4H12orf4;refGene_tx=XM_023256692.1;refGene_gn=CB4H12o
rf4;refGene_tx=XM_023256693.1;refGene_gn=CB4H12orf4;refGene_tx=XM_0232
56694.1
B4 39737417 39737546
refGene_gn=CB4H12orf4;refGene_tx=XM_003988286.4;refGene_gn=CB4H12orf4;
refGene_tx=XM_019834385.2;refGene_gn=CB4H12orf4;refGene_tx=XM_02325669
1.1;refGene_gn=CB4H12orf4;refGene_tx=XM_023256692.1;refGene_gn=CB4H12o
rf4;refGene_tx=XM_023256693.1;refGene_gn=CB4H12orf4;refGene_tx=XM_0232
56694.1
B4 39738378 39738576
refGene_gn=CB4H12orf4;refGene_tx=XM_003988286.4;refGene_gn=CB4H12orf4;
refGene_tx=XM_019834385.2;refGene_gn=CB4H12orf4;refGene_tx=XM_02325669
1.1;refGene_gn=CB4H12orf4;refGene_tx=XM_023256692.1;refGene_gn=CB4H12o
rf4;refGene_tx=XM_023256693.1;refGene_gn=CB4H12orf4;refGene_tx=XM_0232
56694.1
B4 39739235 39739413
refGene_gn=CB4H12orf4;refGene_tx=XM_003988286.4;refGene_gn=CB4H12orf4;
refGene_tx=XM_019834385.2;refGene_gn=CB4H12orf4;refGene_tx=XM_02325669
1.1;refGene_gn=CB4H12orf4;refGene_tx=XM_023256692.1;refGene_gn=CB4H12o
rf4;refGene_tx=XM_023256693.1
B4 39740480 39740684
refGene_gn=CB4H12orf4;refGene_tx=XM_023256692.1;refGene_gn=CB4H12orf4;
refGene_tx=XM_023256693.1
B4 39746718 39746840
refGene_gn=CB4H12orf4;refGene_tx=XM_003988286.4;refGene_gn=CB4H12orf4;
refGene_tx=XM_019834385.2;refGene_gn=CB4H12orf4;refGene_tx=XM_02325669
1.1
B4 39746941 39747041
refGene_gn=CB4H12orf4;refGene_tx=XM_003988286.4;refGene_gn=CB4H12orf4;
refGene_tx=XM_019834385.2;refGene_gn=CB4H12orf4;refGene_tx=XM_02325669
1.1
B4 39751655 39751837
refGene_gn=CB4H12orf4;refGene_tx=XM_003988286.4;refGene_gn=CB4H12orf4;
refGene_tx=XM_019834385.2;refGene_gn=CB4H12orf4;refGene_tx=XM_02325669
1.1
B4 39764518 39764664

refGene_gn=CB4H12orf4;refGene_tx=XM_003988286.4;refGene_gn=CB4H12orf4;
refGene_tx=XM_019834385.2;refGene_gn=CB4H12orf4;refGene_tx=XM_02325669
1.1
B4	39766338	39766516
refGene_gn=CB4H12orf4;refGene_tx=XM_003988286.4;refGene_gn=CB4H12orf4;
refGene_tx=XM_019834385.2;refGene_gn=CB4H12orf4;refGene_tx=XM_02325669
1.1
B4	39768914	39769030
refGene_gn=RAD51AP1;refGene_tx=XM_023256701.1;refGene_gn=RAD51AP1;refG
ene_tx=XM_023256702.1;refGene_gn=RAD51AP1;refGene_tx=XM_023256703.1
B4	39771813	39771863
refGene_gn=RAD51AP1;refGene_tx=XM_023256701.1;refGene_gn=RAD51AP1;refG
ene_tx=XM_023256702.1;refGene_gn=RAD51AP1;refGene_tx=XM_023256703.1
B4	39774087	39774229
refGene_gn=RAD51AP1;refGene_tx=XM_023256701.1;refGene_gn=RAD51AP1;refG
ene_tx=XM_023256702.1;refGene_gn=RAD51AP1;refGene_tx=XM_023256703.1
B4	39776819	39776929
refGene_gn=RAD51AP1;refGene_tx=XM_023256701.1;refGene_gn=RAD51AP1;refG
ene_tx=XM_023256702.1;refGene_gn=RAD51AP1;refGene_tx=XM_023256703.1
B4	39778300	39778387
refGene_gn=RAD51AP1;refGene_tx=XM_023256701.1;refGene_gn=RAD51AP1;refG
ene_tx=XM_023256702.1;refGene_gn=RAD51AP1;refGene_tx=XM_023256703.1
B4	39778912	39779062
refGene_gn=RAD51AP1;refGene_tx=XM_023256701.1;refGene_gn=RAD51AP1;refG
ene_tx=XM_023256702.1;refGene_gn=RAD51AP1;refGene_tx=XM_023256703.1
B4	39784086	39784251
refGene_gn=RAD51AP1;refGene_tx=XM_023256701.1;refGene_gn=RAD51AP1;refG
ene_tx=XM_023256702.1;refGene_gn=RAD51AP1;refGene_tx=XM_023256703.1
B4	39787460	39787610
refGene_gn=RAD51AP1;refGene_tx=XM_023256701.1;refGene_gn=RAD51AP1;refG
ene_tx=XM_023256702.1;refGene_gn=RAD51AP1;refGene_tx=XM_023256703.1
B4	39788071	39788085	refGene_gn=RAD51AP1;refGene_tx=XM_023256703.1
B4	39789968	39790102
refGene_gn=RAD51AP1;refGene_tx=XM_023256701.1;refGene_gn=RAD51AP1;refG
ene_tx=XM_023256702.1
B4	39799759	39799846
refGene_gn=DYRK4;refGene_tx=XM_006933425.4;refGene_gn=DYRK4;refGene_tx
=XM_019834386.2;refGene_gn=DYRK4;refGene_tx=XM_023256697.1;refGene_gn=
DYRK4;refGene_tx=XM_023256698.1;refGene_gn=DYRK4;refGene_tx=XM_0232566
99.1;refGene_gn=DYRK4;refGene_tx=XM_023256700.1
B4	39817062	39817143
refGene_gn=DYRK4;refGene_tx=XM_006933425.4;refGene_gn=DYRK4;refGene_tx
=XM_019834386.2;refGene_gn=DYRK4;refGene_tx=XM_023256697.1;refGene_gn=
DYRK4;refGene_tx=XM_023256698.1;refGene_gn=DYRK4;refGene_tx=XM_0232566
99.1
B4	39817870	39817987
refGene_gn=DYRK4;refGene_tx=XM_006933425.4;refGene_gn=DYRK4;refGene_tx
=XM_019834386.2;refGene_gn=DYRK4;refGene_tx=XM_023256697.1;refGene_gn=
DYRK4;refGene_tx=XM_023256698.1
B4	39818720	39818859

refGene_gn=DYRK4;refGene_tx=XM_006933425.4;refGene_gn=DYRK4;refGene_tx=XM_019834386.2;refGene_gn=DYRK4;refGene_tx=XM_023256697.1;refGene_gn=DYRK4;refGene_tx=XM_023256698.1;refGene_gn=DYRK4;refGene_tx=XM_023256699.1;refGene_gn=DYRK4;refGene_tx=XM_023256700.1
B4 39819914 39820075
refGene_gn=DYRK4;refGene_tx=XM_006933425.4;refGene_gn=DYRK4;refGene_tx=XM_019834386.2;refGene_gn=DYRK4;refGene_tx=XM_023256697.1;refGene_gn=DYRK4;refGene_tx=XM_023256698.1;refGene_gn=DYRK4;refGene_tx=XM_023256699.1;refGene_gn=DYRK4;refGene_tx=XM_023256700.1
B4 39823502 39823639
refGene_gn=DYRK4;refGene_tx=XM_006933425.4;refGene_gn=DYRK4;refGene_tx=XM_019834386.2;refGene_gn=DYRK4;refGene_tx=XM_023256697.1;refGene_gn=DYRK4;refGene_tx=XM_023256698.1;refGene_gn=DYRK4;refGene_tx=XM_023256699.1;refGene_gn=DYRK4;refGene_tx=XM_023256700.1
B4 39823974 39824115
refGene_gn=DYRK4;refGene_tx=XM_006933425.4;refGene_gn=DYRK4;refGene_tx=XM_019834386.2;refGene_gn=DYRK4;refGene_tx=XM_023256697.1;refGene_gn=DYRK4;refGene_tx=XM_023256698.1;refGene_gn=DYRK4;refGene_tx=XM_023256699.1;refGene_gn=DYRK4;refGene_tx=XM_023256700.1
B4 39825976 39826115
refGene_gn=DYRK4;refGene_tx=XM_006933425.4;refGene_gn=DYRK4;refGene_tx=XM_019834386.2;refGene_gn=DYRK4;refGene_tx=XM_023256697.1;refGene_gn=DYRK4;refGene_tx=XM_023256698.1;refGene_gn=DYRK4;refGene_tx=XM_023256699.1;refGene_gn=DYRK4;refGene_tx=XM_023256700.1
B4 39828748 39828830
refGene_gn=DYRK4;refGene_tx=XM_006933425.4;refGene_gn=DYRK4;refGene_tx=XM_019834386.2;refGene_gn=DYRK4;refGene_tx=XM_023256697.1;refGene_gn=DYRK4;refGene_tx=XM_023256698.1;refGene_gn=DYRK4;refGene_tx=XM_023256699.1;refGene_gn=DYRK4;refGene_tx=XM_023256700.1
B4 39831559 39831732
refGene_gn=DYRK4;refGene_tx=XM_006933425.4;refGene_gn=DYRK4;refGene_tx=XM_019834386.2;refGene_gn=DYRK4;refGene_tx=XM_023256697.1;refGene_gn=DYRK4;refGene_tx=XM_023256698.1;refGene_gn=DYRK4;refGene_tx=XM_023256699.1;refGene_gn=DYRK4;refGene_tx=XM_023256700.1
B4 39835058 39835119
refGene_gn=DYRK4;refGene_tx=XM_006933425.4;refGene_gn=DYRK4;refGene_tx=XM_019834386.2;refGene_gn=DYRK4;refGene_tx=XM_023256697.1;refGene_gn=DYRK4;refGene_tx=XM_023256698.1;refGene_gn=DYRK4;refGene_tx=XM_023256699.1;refGene_gn=DYRK4;refGene_tx=XM_023256700.1
B4 39838924 39839054
refGene_gn=DYRK4;refGene_tx=XM_006933425.4;refGene_gn=DYRK4;refGene_tx=XM_019834386.2;refGene_gn=DYRK4;refGene_tx=XM_023256697.1;refGene_gn=DYRK4;refGene_tx=XM_023256698.1;refGene_gn=DYRK4;refGene_tx=XM_023256699.1;refGene_gn=DYRK4;refGene_tx=XM_023256700.1
B4 39840702 39840859 refGene_gn=DYRK4;refGene_tx=XM_019834386.2
B4 39844568 39844744
refGene_gn=DYRK4;refGene_tx=XM_006933425.4;refGene_gn=DYRK4;refGene_tx=XM_019834386.2;refGene_gn=DYRK4;refGene_tx=XM_023256697.1;refGene_gn=DYRK4;refGene_tx=XM_023256698.1;refGene_gn=DYRK4;refGene_tx=XM_023256699.1;refGene_gn=DYRK4;refGene_tx=XM_023256700.1

B4 39845425 39845670
refGene_gn=DYRK4;refGene_tx=XM_006933425.4;refGene_gn=DYRK4;refGene_tx=XM_023256697.1;refGene_gn=DYRK4;refGene_tx=XM_023256698.1;refGene_gn=DYRK4;refGene_tx=XM_023256699.1;refGene_gn=DYRK4;refGene_tx=XM_023256700.1
B4 39847521 39847677
refGene_gn=AKAP3;refGene_tx=XM_003988289.5;refGene_gn=AKAP3;refGene_tx=XM_006933427.4;refGene_gn=AKAP3;refGene_tx=XM_019834387.2;refGene_gn=AKAP3;refGene_tx=XM_019834388.2;refGene_gn=AKAP3;refGene_tx=XM_023256695.1;refGene_gn=AKAP3;refGene_tx=XM_023256696.1
B4 39867607 39869956
refGene_gn=AKAP3;refGene_tx=XM_003988289.5;refGene_gn=AKAP3;refGene_tx=XM_006933427.4;refGene_gn=AKAP3;refGene_tx=XM_019834387.2;refGene_gn=AKAP3;refGene_tx=XM_019834388.2;refGene_gn=AKAP3;refGene_tx=XM_023256695.1;refGene_gn=AKAP3;refGene_tx=XM_023256696.1
B4 39885696 39885792
refGene_gn=AKAP3;refGene_tx=XM_003988289.5;refGene_gn=AKAP3;refGene_tx=XM_006933427.4;refGene_gn=AKAP3;refGene_tx=XM_019834387.2;refGene_gn=AKAP3;refGene_tx=XM_019834388.2;refGene_gn=AKAP3;refGene_tx=XM_023256695.1;refGene_gn=AKAP3;refGene_tx=XM_023256696.1
B4 39894476 39894525 refGene_gn=NDUFA9;refGene_tx=XM_003988290.4
B4 39901199 39901370 refGene_gn=NDUFA9;refGene_tx=XM_003988290.4
B4 39901730 39901828 refGene_gn=NDUFA9;refGene_tx=XM_003988290.4
B4 39906292 39906384 refGene_gn=NDUFA9;refGene_tx=XM_003988290.4
B4 39908805 39908947 refGene_gn=NDUFA9;refGene_tx=XM_003988290.4
B4 39913500 39913603 refGene_gn=NDUFA9;refGene_tx=XM_003988290.4
B4 39919667 39919735 refGene_gn=NDUFA9;refGene_tx=XM_003988290.4
B4 39920849 39920926 refGene_gn=NDUFA9;refGene_tx=XM_003988290.4
B4 39925291 39925387 refGene_gn=NDUFA9;refGene_tx=XM_003988290.4
B4 39928006 39928073 refGene_gn=NDUFA9;refGene_tx=XM_003988290.4
B4 39930890 39931061 refGene_gn=NDUFA9;refGene_tx=XM_003988290.4
B4 39968592 39968803 refGene_gn=GALNT8;refGene_tx=XM_011283725.2
B4 39975823 39976001 refGene_gn=GALNT8;refGene_tx=XM_011283725.2
B4 39976002 39976110 refGene_gn=GALNT8;refGene_tx=XM_011283725.2
B4 39992641 39992808 refGene_gn=GALNT8;refGene_tx=XM_011283725.2
B4 39998463 39998485 refGene_gn=GALNT8;refGene_tx=XM_011283725.2
B4 39998729 39998892 refGene_gn=GALNT8;refGene_tx=XM_011283725.2
B4 39999386 39999388 refGene_gn=GALNT8;refGene_tx=XM_011283725.2
B4 39999873 40000071 refGene_gn=GALNT8;refGene_tx=XM_011283725.2
B4 40000824 40000939 refGene_gn=GALNT8;refGene_tx=XM_011283725.2
B4 40006243 40006299 refGene_gn=GALNT8;refGene_tx=XM_011283725.2
B4 40006300 40006427 refGene_gn=GALNT8;refGene_tx=XM_011283725.2
B4 40007386 40007524 refGene_gn=GALNT8;refGene_tx=XM_011283725.2
B4 40008075 40008171 refGene_gn=GALNT8;refGene_tx=XM_011283725.2
B4 40009199 40009367 refGene_gn=GALNT8;refGene_tx=XM_011283725.2
B4 40017243 40017369 refGene_gn=GALNT8;refGene_tx=XM_011283725.2
B4 40048662 40050249
refGene_gn=KCNA6;refGene_tx=XM_003988291.5;refGene_gn=KCNA6;refGene_tx=XM_006933429.4;refGene_gn=KCNA6;refGene_tx=XM_019834391.2;refGene_gn=KCNA6;refGene_tx=XM_023256704.1

B4	40156756 40158244 refGene_gn=KCNA1;refGene_tx=XM_019834392.2
B4	40255992 40256110
refGene_gn=LOC102900369;refGene_tx=XM_019834393.2;refGene_gn=LOC102900369;refGene_tx=XM_019834395.2;refGene_gn=LOC102900369;refGene_tx=XM_019834396.2
B4	40256291 40256476
refGene_gn=LOC102900369;refGene_tx=XM_019834393.2;refGene_gn=LOC102900369;refGene_tx=XM_019834395.2;refGene_gn=LOC102900369;refGene_tx=XM_019834396.2
B4	40257396 40257488
refGene_gn=LOC102900369;refGene_tx=XM_019834393.2;refGene_gn=LOC102900369;refGene_tx=XM_019834395.2;refGene_gn=LOC102900369;refGene_tx=XM_019834396.2
B4	40260163 40260301
refGene_gn=LOC102900369;refGene_tx=XM_019834393.2;refGene_gn=LOC102900369;refGene_tx=XM_019834395.2;refGene_gn=LOC102900369;refGene_tx=XM_019834396.2
B4	40261419 40261523
refGene_gn=LOC102900369;refGene_tx=XM_019834393.2;refGene_gn=LOC102900369;refGene_tx=XM_019834395.2;refGene_gn=LOC102900369;refGene_tx=XM_019834396.2
B4	40261705 40261764
refGene_gn=LOC102900369;refGene_tx=XM_019834396.2
B4	40262748 40262936
refGene_gn=LOC102900369;refGene_tx=XM_019834393.2;refGene_gn=LOC102900369;refGene_tx=XM_019834395.2
B4	40282823 40284635 refGene_gn=KCNA5;refGene_tx=XM_006933430.4
B4	40711633 40712405 refGene_gn=NTF3;refGene_tx=NM_001009367.1
B4	40775291 40775561
refGene_gn=ANO2;refGene_tx=XM_019834397.2;refGene_gn=ANO2;refGene_tx=XM_019834398.2;refGene_gn=ANO2;refGene_tx=XM_019834399.2
B4	40777459 40777565
refGene_gn=ANO2;refGene_tx=XM_019834397.2;refGene_gn=ANO2;refGene_tx=XM_019834398.2;refGene_gn=ANO2;refGene_tx=XM_019834399.2
B4	40785318 40785500
refGene_gn=ANO2;refGene_tx=XM_019834397.2;refGene_gn=ANO2;refGene_tx=XM_019834398.2;refGene_gn=ANO2;refGene_tx=XM_019834399.2
B4	40787383 40787436
refGene_gn=ANO2;refGene_tx=XM_019834397.2;refGene_gn=ANO2;refGene_tx=XM_019834398.2;refGene_gn=ANO2;refGene_tx=XM_019834399.2
B4	40787797 40787950
refGene_gn=ANO2;refGene_tx=XM_019834397.2;refGene_gn=ANO2;refGene_tx=XM_019834398.2;refGene_gn=ANO2;refGene_tx=XM_019834399.2
B4	40806392 40806538
refGene_gn=ANO2;refGene_tx=XM_019834397.2;refGene_gn=ANO2;refGene_tx=XM_019834398.2;refGene_gn=ANO2;refGene_tx=XM_019834399.2
B4	40815971 40816072
refGene_gn=ANO2;refGene_tx=XM_019834397.2;refGene_gn=ANO2;refGene_tx=XM_019834398.2;refGene_gn=ANO2;refGene_tx=XM_019834399.2
B4	40816236 40816294

refGene_gn=ANO2;refGene_tx=XM_019834397.2;refGene_gn=ANO2;refGene_tx=XM_019834398.2;refGene_gn=ANO2;refGene_tx=XM_019834399.2
B4 40818477 40818589
refGene_gn=ANO2;refGene_tx=XM_019834397.2;refGene_gn=ANO2;refGene_tx=XM_019834398.2;refGene_gn=ANO2;refGene_tx=XM_019834399.2
B4 40836285 40836481
refGene_gn=ANO2;refGene_tx=XM_019834397.2;refGene_gn=ANO2;refGene_tx=XM_019834398.2;refGene_gn=ANO2;refGene_tx=XM_019834399.2
B4 40846895 40846970
refGene_gn=ANO2;refGene_tx=XM_019834397.2;refGene_gn=ANO2;refGene_tx=XM_019834398.2;refGene_gn=ANO2;refGene_tx=XM_019834399.2
B4 40919303 40919414
refGene_gn=ANO2;refGene_tx=XM_019834397.2;refGene_gn=ANO2;refGene_tx=XM_019834398.2;refGene_gn=ANO2;refGene_tx=XM_019834399.2
B4 40919635 40919647
refGene_gn=ANO2;refGene_tx=XM_019834397.2;refGene_gn=ANO2;refGene_tx=XM_019834398.2
B4 40925913 40925996
refGene_gn=ANO2;refGene_tx=XM_019834397.2;refGene_gn=ANO2;refGene_tx=XM_019834398.2;refGene_gn=ANO2;refGene_tx=XM_019834399.2
B4 40930855 40931016
refGene_gn=ANO2;refGene_tx=XM_019834397.2;refGene_gn=ANO2;refGene_tx=XM_019834398.2;refGene_gn=ANO2;refGene_tx=XM_019834399.2
B4 40937354 40937489
refGene_gn=ANO2;refGene_tx=XM_019834397.2;refGene_gn=ANO2;refGene_tx=XM_019834398.2;refGene_gn=ANO2;refGene_tx=XM_019834399.2
B4 40983339 40983404
refGene_gn=ANO2;refGene_tx=XM_019834397.2;refGene_gn=ANO2;refGene_tx=XM_019834398.2;refGene_gn=ANO2;refGene_tx=XM_019834399.2
B4 40990967 40991009
refGene_gn=ANO2;refGene_tx=XM_019834397.2;refGene_gn=ANO2;refGene_tx=XM_019834398.2;refGene_gn=ANO2;refGene_tx=XM_019834399.2
B4 40992164 40992220
refGene_gn=ANO2;refGene_tx=XM_019834397.2;refGene_gn=ANO2;refGene_tx=XM_019834398.2;refGene_gn=ANO2;refGene_tx=XM_019834399.2
B4 41005888 41005940
refGene_gn=ANO2;refGene_tx=XM_019834397.2;refGene_gn=ANO2;refGene_tx=XM_019834398.2;refGene_gn=ANO2;refGene_tx=XM_019834399.2
B4 41008391 41008446
refGene_gn=ANO2;refGene_tx=XM_019834397.2;refGene_gn=ANO2;refGene_tx=XM_019834398.2;refGene_gn=ANO2;refGene_tx=XM_019834399.2
B4 41011610 41011762
refGene_gn=ANO2;refGene_tx=XM_019834397.2;refGene_gn=ANO2;refGene_tx=XM_019834398.2;refGene_gn=ANO2;refGene_tx=XM_019834399.2
B4 41031041 41031044
refGene_gn=ANO2;refGene_tx=XM_019834397.2;refGene_gn=ANO2;refGene_tx=XM_019834399.2
B4 41031551 41031650
refGene_gn=ANO2;refGene_tx=XM_019834397.2;refGene_gn=ANO2;refGene_tx=XM_019834398.2;refGene_gn=ANO2;refGene_tx=XM_019834399.2

B4 41092327 41092654
refGene_gn=ANO2;refGene_tx=XM_019834397.2;refGene_gn=ANO2;refGene_tx=XM_019834398.2;refGene_gn=ANO2;refGene_tx=XM_019834399.2
B4 41093952 41094143
refGene_gn=ANO2;refGene_tx=XM_019834397.2;refGene_gn=ANO2;refGene_tx=XM_019834398.2;refGene_gn=ANO2;refGene_tx=XM_019834399.2
B4 41116101 41116123
refGene_gn=ANO2;refGene_tx=XM_019834397.2;refGene_gn=ANO2;refGene_tx=XM_019834398.2;refGene_gn=ANO2;refGene_tx=XM_019834399.2
B4 41119991 41120180 refGene_gn=VWF;refGene_tx=NM_001246279.1
B4 41120611 41120709 refGene_gn=VWF;refGene_tx=NM_001246279.1
B4 41122846 41122886 refGene_gn=VWF;refGene_tx=NM_001246279.1
B4 41123401 41123530 refGene_gn=VWF;refGene_tx=NM_001246279.1
B4 41124206 41124305 refGene_gn=VWF;refGene_tx=NM_001246279.1
B4 41130840 41130957 refGene_gn=VWF;refGene_tx=NM_001246279.1
B4 41131464 41131505 refGene_gn=VWF;refGene_tx=NM_001246279.1
B4 41132216 41132397 refGene_gn=VWF;refGene_tx=NM_001246279.1
B4 41133304 41133415 refGene_gn=VWF;refGene_tx=NM_001246279.1
B4 41137920 41138070 refGene_gn=VWF;refGene_tx=NM_001246279.1
B4 41140671 41140877 refGene_gn=VWF;refGene_tx=NM_001246279.1
B4 41142164 41142269 refGene_gn=VWF;refGene_tx=NM_001246279.1
B4 41143467 41143542 refGene_gn=VWF;refGene_tx=NM_001246279.1
B4 41143961 41144064 refGene_gn=VWF;refGene_tx=NM_001246279.1
B4 41150405 41150605 refGene_gn=VWF;refGene_tx=NM_001246279.1
B4 41152976 41153318 refGene_gn=VWF;refGene_tx=NM_001246279.1
B4 41153515 41153708 refGene_gn=VWF;refGene_tx=NM_001246279.1
B4 41154892 41155113 refGene_gn=VWF;refGene_tx=NM_001246279.1
B4 41159329 41159507 refGene_gn=VWF;refGene_tx=NM_001246279.1
B4 41159809 41159853 refGene_gn=VWF;refGene_tx=NM_001246279.1
B4 41160782 41160947 refGene_gn=VWF;refGene_tx=NM_001246279.1
B4 41162213 41162357 refGene_gn=VWF;refGene_tx=NM_001246279.1
B4 41162629 41162770 refGene_gn=VWF;refGene_tx=NM_001246279.1
B4 41162865 41162982 refGene_gn=VWF;refGene_tx=NM_001246279.1
B4 41165105 41166484 refGene_gn=VWF;refGene_tx=NM_001246279.1
B4 41169006 41169142 refGene_gn=VWF;refGene_tx=NM_001246279.1
B4 41169819 41169978 refGene_gn=VWF;refGene_tx=NM_001246279.1
B4 41170755 41170912 refGene_gn=VWF;refGene_tx=NM_001246279.1
B4 41172123 41172237 refGene_gn=VWF;refGene_tx=NM_001246279.1
B4 41172441 41172582 refGene_gn=VWF;refGene_tx=NM_001246279.1
B4 41175224 41175371 refGene_gn=VWF;refGene_tx=NM_001246279.1
B4 41177556 41177691 refGene_gn=VWF;refGene_tx=NM_001246279.1
B4 41180033 41180172 refGene_gn=VWF;refGene_tx=NM_001246279.1
B4 41181465 41181569 refGene_gn=VWF;refGene_tx=NM_001246279.1
B4 41189225 41189386 refGene_gn=VWF;refGene_tx=NM_001246279.1
B4 41193244 41193339 refGene_gn=VWF;refGene_tx=NM_001246279.1
B4 41199150 41199388 refGene_gn=VWF;refGene_tx=NM_001246279.1
B4 41203089 41203305 refGene_gn=VWF;refGene_tx=NM_001246279.1
B4 41203939 41204135 refGene_gn=VWF;refGene_tx=NM_001246279.1
B4 41207630 41207731 refGene_gn=VWF;refGene_tx=NM_001246279.1
B4 41208842 41208981 refGene_gn=VWF;refGene_tx=NM_001246279.1

B4 41209416 41209553 refGene_gn=VWF;refGene_tx=NM_001246279.1
B4 41218224 41218271 refGene_gn=VWF;refGene_tx=NM_001246279.1
B4 41219048 41219160 refGene_gn=VWF;refGene_tx=NM_001246279.1
B4 41220042 41220165 refGene_gn=VWF;refGene_tx=NM_001246279.1
B4 41221838 41222055 refGene_gn=VWF;refGene_tx=NM_001246279.1
B4 41235476 41235601 refGene_gn=VWF;refGene_tx=NM_001246279.1
B4 41243526 41243735 refGene_gn=VWF;refGene_tx=NM_001246279.1
B4 41244030 41244133 refGene_gn=VWF;refGene_tx=NM_001246279.1
B4 41253791 41253956 refGene_gn=VWF;refGene_tx=NM_001246279.1
B4 41255662 41255717 refGene_gn=VWF;refGene_tx=NM_001246279.1
B4 41321949 41322015 refGene_gn=CD9;refGene_tx=NM_001009305.1
B4 41342847 41342950 refGene_gn=CD9;refGene_tx=NM_001009305.1
B4 41350145 41350243 refGene_gn=CD9;refGene_tx=NM_001009305.1
B4 41350967 41351042 refGene_gn=CD9;refGene_tx=NM_001009305.1
B4 41352590 41352689 refGene_gn=CD9;refGene_tx=NM_001009305.1
B4 41352842 41352932 refGene_gn=CD9;refGene_tx=NM_001009305.1
B4 41353592 41353676 refGene_gn=CD9;refGene_tx=NM_001009305.1
B4 41355014 41355080 refGene_gn=CD9;refGene_tx=NM_001009305.1
B4 41425604 41425742 refGene_gn=PLEKHG6;refGene_tx=XM_006933439.4
B4 41427045 41427201 refGene_gn=PLEKHG6;refGene_tx=XM_006933439.4
B4 41427773 41427938 refGene_gn=PLEKHG6;refGene_tx=XM_006933439.4
B4 41428290 41428386 refGene_gn=PLEKHG6;refGene_tx=XM_006933439.4
B4 41428676 41428727 refGene_gn=PLEKHG6;refGene_tx=XM_006933439.4
B4 41429085 41429235 refGene_gn=PLEKHG6;refGene_tx=XM_006933439.4
B4 41430402 41430513 refGene_gn=PLEKHG6;refGene_tx=XM_006933439.4
B4 41430667 41430817 refGene_gn=PLEKHG6;refGene_tx=XM_006933439.4
B4 41431028 41431166 refGene_gn=PLEKHG6;refGene_tx=XM_006933439.4
B4 41431464 41431584 refGene_gn=PLEKHG6;refGene_tx=XM_006933439.4
B4 41431944 41432077 refGene_gn=PLEKHG6;refGene_tx=XM_006933439.4
B4 41432197 41432313 refGene_gn=PLEKHG6;refGene_tx=XM_006933439.4
B4 41438873 41439019 refGene_gn=PLEKHG6;refGene_tx=XM_006933439.4
B4 41439668 41440364 refGene_gn=PLEKHG6;refGene_tx=XM_006933439.4
B4 41440561 41440571 refGene_gn=PLEKHG6;refGene_tx=XM_006933439.4
B4 41441581 41441898 refGene_gn=TNFRSF1A;refGene_tx=NM_001009361.1
B4 41442009 41442262 refGene_gn=TNFRSF1A;refGene_tx=NM_001009361.1
B4 41442433 41442462 refGene_gn=TNFRSF1A;refGene_tx=NM_001009361.1
B4 41442768 41442879 refGene_gn=TNFRSF1A;refGene_tx=NM_001009361.1
B4 41443005 41443079 refGene_gn=TNFRSF1A;refGene_tx=NM_001009361.1
B4 41444589 41444668 refGene_gn=TNFRSF1A;refGene_tx=NM_001009361.1
B4 41444903 41445053 refGene_gn=TNFRSF1A;refGene_tx=NM_001009361.1
B4 41445258 41445387 refGene_gn=TNFRSF1A;refGene_tx=NM_001009361.1
B4 41445636 41445790 refGene_gn=TNFRSF1A;refGene_tx=NM_001009361.1
B4 41453184 41453223 refGene_gn=TNFRSF1A;refGene_tx=NM_001009361.1
B4 41458626 41458995
refGene_gn=SCNN1A;refGene_tx=XM_003988358.5;refGene_gn=SCNN1A;refGene_tx=XM_019834403.2;refGene_gn=SCNN1A;refGene_tx=XM_019834404.2
B4 41459356 41459432
refGene_gn=SCNN1A;refGene_tx=XM_003988358.5;refGene_gn=SCNN1A;refGene_tx=XM_019834403.2;refGene_gn=SCNN1A;refGene_tx=XM_019834404.2
B4 41459570 41459626

refGene_gn=SCNN1A;refGene_tx=XM_003988358.5;refGene_gn=SCNN1A;refGene_tx=XM_019834403.2;refGene_gn=SCNN1A;refGene_tx=XM_019834404.2
B4 41459818 41459876
refGene_gn=SCNN1A;refGene_tx=XM_003988358.5;refGene_gn=SCNN1A;refGene_tx=XM_019834403.2;refGene_gn=SCNN1A;refGene_tx=XM_019834404.2
B4 41459976 41460055
refGene_gn=SCNN1A;refGene_tx=XM_003988358.5;refGene_gn=SCNN1A;refGene_tx=XM_019834403.2;refGene_gn=SCNN1A;refGene_tx=XM_019834404.2
B4 41462924 41463042
refGene_gn=SCNN1A;refGene_tx=XM_003988358.5;refGene_gn=SCNN1A;refGene_tx=XM_019834403.2;refGene_gn=SCNN1A;refGene_tx=XM_019834404.2
B4 41463281 41463380
refGene_gn=SCNN1A;refGene_tx=XM_003988358.5;refGene_gn=SCNN1A;refGene_tx=XM_019834403.2;refGene_gn=SCNN1A;refGene_tx=XM_019834404.2
B4 41463795 41463959
refGene_gn=SCNN1A;refGene_tx=XM_003988358.5;refGene_gn=SCNN1A;refGene_tx=XM_019834403.2;refGene_gn=SCNN1A;refGene_tx=XM_019834404.2
B4 41464345 41464449
refGene_gn=SCNN1A;refGene_tx=XM_003988358.5;refGene_gn=SCNN1A;refGene_tx=XM_019834403.2;refGene_gn=SCNN1A;refGene_tx=XM_019834404.2
B4 41468245 41468436
refGene_gn=SCNN1A;refGene_tx=XM_003988358.5;refGene_gn=SCNN1A;refGene_tx=XM_019834403.2;refGene_gn=SCNN1A;refGene_tx=XM_019834404.2
B4 41469253 41469521
refGene_gn=SCNN1A;refGene_tx=XM_003988358.5;refGene_gn=SCNN1A;refGene_tx=XM_019834403.2;refGene_gn=SCNN1A;refGene_tx=XM_019834404.2
B4 41480969 41481325
refGene_gn=SCNN1A;refGene_tx=XM_003988358.5;refGene_gn=SCNN1A;refGene_tx=XM_019834403.2;refGene_gn=SCNN1A;refGene_tx=XM_019834404.2
B4 41489058 41489175 refGene_gn=LTBR;refGene_tx=XM_003988317.5
B4 41489293 41489390 refGene_gn=LTBR;refGene_tx=XM_003988317.5
B4 41489650 41489776 refGene_gn=LTBR;refGene_tx=XM_003988317.5
B4 41489859 41490012 refGene_gn=LTBR;refGene_tx=XM_003988317.5
B4 41490379 41490476 refGene_gn=LTBR;refGene_tx=XM_003988317.5
B4 41490598 41490690 refGene_gn=LTBR;refGene_tx=XM_003988317.5
B4 41493651 41493759 refGene_gn=LTBR;refGene_tx=XM_003988317.5
B4 41494074 41494100 refGene_gn=LTBR;refGene_tx=XM_003988317.5
B4 41494672 41494901 refGene_gn=LTBR;refGene_tx=XM_003988317.5
B4 41495060 41495278 refGene_gn=LTBR;refGene_tx=XM_003988317.5
B4 41495279 41495302 refGene_gn=LTBR;refGene_tx=XM_003988317.5
B4 41495303 41495415 refGene_gn=LTBR;refGene_tx=XM_003988317.5
B4 41559649 41559785
refGene_gn=CD27;refGene_tx=XM_011283723.3;refGene_gn=CD27;refGene_tx=XM_019834412.2
B4 41559931 41560063 refGene_gn=CD27;refGene_tx=XM_011283723.3
B4 41564436 41564616
refGene_gn=CD27;refGene_tx=XM_011283723.3;refGene_gn=CD27;refGene_tx=XM_019834412.2
B4 41564817 41564907
refGene_gn=CD27;refGene_tx=XM_011283723.3;refGene_gn=CD27;refGene_tx=X

M_019834412.2
B4 41565172 41565292
refGene_gn=CD27;refGene_tx=XM_011283723.3;refGene_gn=CD27;refGene_tx=XM_019834412.2
B4 41565539 41565664
refGene_gn=CD27;refGene_tx=XM_011283723.3;refGene_gn=CD27;refGene_tx=XM_019834412.2
B4 41566199 41566245
refGene_gn=TAPBPL;refGene_tx=XM_006933446.4;refGene_gn=TAPBPL;refGene_tx=XM_023256705.1
B4 41566428 41566677
refGene_gn=TAPBPL;refGene_tx=XM_023256705.1;refGene_gn=TAPBPL;refGene_tx=XM_023256706.1
B4 41567461 41567680
refGene_gn=TAPBPL;refGene_tx=XM_006933446.4;refGene_gn=TAPBPL;refGene_tx=XM_023256705.1;refGene_gn=TAPBPL;refGene_tx=XM_023256706.1
B4 41567826 41568081
refGene_gn=TAPBPL;refGene_tx=XM_006933446.4;refGene_gn=TAPBPL;refGene_tx=XM_023256705.1;refGene_gn=TAPBPL;refGene_tx=XM_023256706.1
B4 41570629 41570968
refGene_gn=TAPBPL;refGene_tx=XM_006933446.4;refGene_gn=TAPBPL;refGene_tx=XM_023256705.1;refGene_gn=TAPBPL;refGene_tx=XM_023256706.1
B4 41571541 41571844
refGene_gn=TAPBPL;refGene_tx=XM_006933446.4;refGene_gn=TAPBPL;refGene_tx=XM_023256705.1;refGene_gn=TAPBPL;refGene_tx=XM_023256706.1
B4 41573024 41573108
refGene_gn=TAPBPL;refGene_tx=XM_006933446.4;refGene_gn=TAPBPL;refGene_tx=XM_023256705.1;refGene_gn=TAPBPL;refGene_tx=XM_023256706.1
B4 41573292 41573336
refGene_gn=TAPBPL;refGene_tx=XM_006933446.4;refGene_gn=TAPBPL;refGene_tx=XM_023256705.1;refGene_gn=TAPBPL;refGene_tx=XM_023256706.1
B4 41576589 41576606 refGene_gn=VAMP1;refGene_tx=XM_006933444.4
B4 41577016 41577068 refGene_gn=VAMP1;refGene_tx=XM_006933444.4
B4 41577699 41577858 refGene_gn=VAMP1;refGene_tx=XM_006933444.4
B4 41578058 41578185 refGene_gn=VAMP1;refGene_tx=XM_006933444.4
B4 41580910 41580912 refGene_gn=VAMP1;refGene_tx=XM_006933444.4
B4 41594782 41594979 refGene_gn=MRPL51;refGene_tx=XM_023256717.1
B4 41595428 41595539 refGene_gn=MRPL51;refGene_tx=XM_023256717.1
B4 41595622 41595655 refGene_gn=MRPL51;refGene_tx=XM_023256717.1
B4 41595657 41595700 refGene_gn=MRPL51;refGene_tx=XM_023256717.1
B4 41597795 41597922
refGene_gn=NCAPD2;refGene_tx=XM_019834405.2;refGene_gn=NCAPD2;refGene_tx=XM_019834406.2;refGene_gn=NCAPD2;refGene_tx=XM_019834407.2;refGene_gn=NCAPD2;refGene_tx=XM_019834408.2
B4 41606781 41606857
refGene_gn=NCAPD2;refGene_tx=XM_019834405.2;refGene_gn=NCAPD2;refGene_tx=XM_019834406.2;refGene_gn=NCAPD2;refGene_tx=XM_019834407.2;refGene_gn=NCAPD2;refGene_tx=XM_019834408.2
B4 41607122 41607181
refGene_gn=NCAPD2;refGene_tx=XM_019834405.2;refGene_gn=NCAPD2;refGene_

tx=XM_019834406.2;refGene_gn=NCAPD2;refGene_tx=XM_019834407.2;refGene_gn=NCAPD2;refGene_tx=XM_019834408.2
B4 41607663 41607842
refGene_gn=NCAPD2;refGene_tx=XM_019834405.2;refGene_gn=NCAPD2;refGene_tx=XM_019834406.2;refGene_gn=NCAPD2;refGene_tx=XM_019834407.2;refGene_gn=NCAPD2;refGene_tx=XM_019834408.2
B4 41608152 41608295
refGene_gn=NCAPD2;refGene_tx=XM_019834405.2;refGene_gn=NCAPD2;refGene_tx=XM_019834406.2;refGene_gn=NCAPD2;refGene_tx=XM_019834407.2;refGene_gn=NCAPD2;refGene_tx=XM_019834408.2
B4 41610425 41610553
refGene_gn=NCAPD2;refGene_tx=XM_019834405.2;refGene_gn=NCAPD2;refGene_tx=XM_019834406.2;refGene_gn=NCAPD2;refGene_tx=XM_019834407.2;refGene_gn=NCAPD2;refGene_tx=XM_019834408.2
B4 41610625 41610749
refGene_gn=NCAPD2;refGene_tx=XM_019834405.2;refGene_gn=NCAPD2;refGene_tx=XM_019834406.2;refGene_gn=NCAPD2;refGene_tx=XM_019834407.2;refGene_gn=NCAPD2;refGene_tx=XM_019834408.2
B4 41610922 41611070
refGene_gn=NCAPD2;refGene_tx=XM_019834405.2;refGene_gn=NCAPD2;refGene_tx=XM_019834406.2;refGene_gn=NCAPD2;refGene_tx=XM_019834407.2;refGene_gn=NCAPD2;refGene_tx=XM_019834408.2
B4 41612005 41612203
refGene_gn=NCAPD2;refGene_tx=XM_019834405.2;refGene_gn=NCAPD2;refGene_tx=XM_019834406.2;refGene_gn=NCAPD2;refGene_tx=XM_019834407.2;refGene_gn=NCAPD2;refGene_tx=XM_019834408.2
B4 41612538 41612673
refGene_gn=NCAPD2;refGene_tx=XM_019834405.2;refGene_gn=NCAPD2;refGene_tx=XM_019834406.2;refGene_gn=NCAPD2;refGene_tx=XM_019834407.2;refGene_gn=NCAPD2;refGene_tx=XM_019834408.2
B4 41612770 41612858
refGene_gn=NCAPD2;refGene_tx=XM_019834405.2;refGene_gn=NCAPD2;refGene_tx=XM_019834406.2;refGene_gn=NCAPD2;refGene_tx=XM_019834407.2;refGene_gn=NCAPD2;refGene_tx=XM_019834408.2
B4 41612950 41613131
refGene_gn=NCAPD2;refGene_tx=XM_019834405.2;refGene_gn=NCAPD2;refGene_tx=XM_019834406.2;refGene_gn=NCAPD2;refGene_tx=XM_019834407.2;refGene_gn=NCAPD2;refGene_tx=XM_019834408.2
B4 41617192 41617317
refGene_gn=NCAPD2;refGene_tx=XM_019834405.2;refGene_gn=NCAPD2;refGene_tx=XM_019834406.2;refGene_gn=NCAPD2;refGene_tx=XM_019834407.2;refGene_gn=NCAPD2;refGene_tx=XM_019834408.2
B4 41617687 41617924
refGene_gn=NCAPD2;refGene_tx=XM_019834405.2;refGene_gn=NCAPD2;refGene_tx=XM_019834406.2;refGene_gn=NCAPD2;refGene_tx=XM_019834407.2;refGene_gn=NCAPD2;refGene_tx=XM_019834408.2
B4 41618233 41618408
refGene_gn=NCAPD2;refGene_tx=XM_019834405.2;refGene_gn=NCAPD2;refGene_tx=XM_019834406.2;refGene_gn=NCAPD2;refGene_tx=XM_019834407.2;refGene_gn=NCAPD2;refGene_tx=XM_019834408.2

B4	41618657 41618742
refGene_gn=NCAPD2;refGene_tx=XM_019834405.2;refGene_gn=NCAPD2;refGene_tx=XM_019834406.2;refGene_gn=NCAPD2;refGene_tx=XM_019834407.2;refGene_gn=NCAPD2;refGene_tx=XM_019834408.2
B4	41620818 41620952
refGene_gn=NCAPD2;refGene_tx=XM_019834405.2;refGene_gn=NCAPD2;refGene_tx=XM_019834406.2;refGene_gn=NCAPD2;refGene_tx=XM_019834407.2;refGene_gn=NCAPD2;refGene_tx=XM_019834408.2
B4	41621304 41621437
refGene_gn=NCAPD2;refGene_tx=XM_019834405.2;refGene_gn=NCAPD2;refGene_tx=XM_019834406.2;refGene_gn=NCAPD2;refGene_tx=XM_019834407.2;refGene_gn=NCAPD2;refGene_tx=XM_019834408.2
B4	41621526 41621611
refGene_gn=NCAPD2;refGene_tx=XM_019834405.2;refGene_gn=NCAPD2;refGene_tx=XM_019834406.2;refGene_gn=NCAPD2;refGene_tx=XM_019834407.2;refGene_gn=NCAPD2;refGene_tx=XM_019834408.2
B4	41621684 41621852
refGene_gn=NCAPD2;refGene_tx=XM_019834405.2;refGene_gn=NCAPD2;refGene_tx=XM_019834406.2;refGene_gn=NCAPD2;refGene_tx=XM_019834407.2;refGene_gn=NCAPD2;refGene_tx=XM_019834408.2
B4	41622176 41622346
refGene_gn=NCAPD2;refGene_tx=XM_019834405.2;refGene_gn=NCAPD2;refGene_tx=XM_019834406.2;refGene_gn=NCAPD2;refGene_tx=XM_019834407.2;refGene_gn=NCAPD2;refGene_tx=XM_019834408.2
B4	41623358 41623470
refGene_gn=NCAPD2;refGene_tx=XM_019834405.2;refGene_gn=NCAPD2;refGene_tx=XM_019834406.2;refGene_gn=NCAPD2;refGene_tx=XM_019834407.2;refGene_gn=NCAPD2;refGene_tx=XM_019834408.2
B4	41623545 41623669
refGene_gn=NCAPD2;refGene_tx=XM_019834405.2;refGene_gn=NCAPD2;refGene_tx=XM_019834406.2;refGene_gn=NCAPD2;refGene_tx=XM_019834407.2;refGene_gn=NCAPD2;refGene_tx=XM_019834408.2
B4	41623749 41623905
refGene_gn=NCAPD2;refGene_tx=XM_019834405.2;refGene_gn=NCAPD2;refGene_tx=XM_019834406.2;refGene_gn=NCAPD2;refGene_tx=XM_019834407.2;refGene_gn=NCAPD2;refGene_tx=XM_019834408.2
B4	41624381 41624559
refGene_gn=NCAPD2;refGene_tx=XM_019834405.2;refGene_gn=NCAPD2;refGene_tx=XM_019834406.2;refGene_gn=NCAPD2;refGene_tx=XM_019834407.2;refGene_gn=NCAPD2;refGene_tx=XM_019834408.2
B4	41624650 41624745
refGene_gn=NCAPD2;refGene_tx=XM_019834405.2;refGene_gn=NCAPD2;refGene_tx=XM_019834406.2;refGene_gn=NCAPD2;refGene_tx=XM_019834407.2;refGene_gn=NCAPD2;refGene_tx=XM_019834408.2
B4	41625182 41625263
refGene_gn=NCAPD2;refGene_tx=XM_019834405.2;refGene_gn=NCAPD2;refGene_tx=XM_019834406.2;refGene_gn=NCAPD2;refGene_tx=XM_019834407.2;refGene_gn=NCAPD2;refGene_tx=XM_019834408.2
B4	41625373 41625557
refGene_gn=NCAPD2;refGene_tx=XM_019834405.2;refGene_gn=NCAPD2;refGene_

tx=XM_019834406.2;refGene_gn=NCAPD2;refGene_tx=XM_019834407.2;refGene_gn=NCAPD2;refGene_tx=XM_019834408.2
B4	41626512	41626639
refGene_gn=NCAPD2;refGene_tx=XM_019834405.2;refGene_gn=NCAPD2;refGene_tx=XM_019834406.2;refGene_gn=NCAPD2;refGene_tx=XM_019834407.2;refGene_gn=NCAPD2;refGene_tx=XM_019834408.2
B4	41626726	41626888
refGene_gn=NCAPD2;refGene_tx=XM_019834405.2;refGene_gn=NCAPD2;refGene_tx=XM_019834406.2;refGene_gn=NCAPD2;refGene_tx=XM_019834407.2;refGene_gn=NCAPD2;refGene_tx=XM_019834408.2
B4	41627140	41627226
refGene_gn=NCAPD2;refGene_tx=XM_019834405.2;refGene_gn=NCAPD2;refGene_tx=XM_019834406.2;refGene_gn=NCAPD2;refGene_tx=XM_019834407.2;refGene_gn=NCAPD2;refGene_tx=XM_019834408.2
B4	41630334	41630357
refGene_gn=GAPDH;refGene_tx=NM_001009307.1;refGene_gn=GAPDH;refGene_tx=XM_006933438.3
B4	41632154	41632254
refGene_gn=GAPDH;refGene_tx=NM_001009307.1;refGene_gn=GAPDH;refGene_tx=XM_006933438.3
B4	41632337	41632444
refGene_gn=GAPDH;refGene_tx=NM_001009307.1;refGene_gn=GAPDH;refGene_tx=XM_006933438.3
B4	41632566	41632657
refGene_gn=GAPDH;refGene_tx=NM_001009307.1;refGene_gn=GAPDH;refGene_tx=XM_006933438.3
B4	41632766	41632882
refGene_gn=GAPDH;refGene_tx=NM_001009307.1;refGene_gn=GAPDH;refGene_tx=XM_006933438.3
B4	41632984	41633066
refGene_gn=GAPDH;refGene_tx=NM_001009307.1;refGene_gn=GAPDH;refGene_tx=XM_006933438.3
B4	41633292	41633444
refGene_gn=GAPDH;refGene_tx=NM_001009307.1;refGene_gn=GAPDH;refGene_tx=XM_006933438.3
B4	41633528	41633607
refGene_gn=GAPDH;refGene_tx=NM_001009307.1;refGene_gn=GAPDH;refGene_tx=XM_006933438.3
B4	41633710	41633892
refGene_gn=GAPDH;refGene_tx=NM_001009307.1;refGene_gn=GAPDH;refGene_tx=XM_006933438.3
B4	41633995	41634065
refGene_gn=GAPDH;refGene_tx=NM_001009307.1;refGene_gn=GAPDH;refGene_tx=XM_006933438.3
B4	41636317	41636426
refGene_gn=IFFO1;refGene_tx=XM_019834411.2;refGene_gn=IFFO1;refGene_tx=XM_023256707.1;refGene_gn=IFFO1;refGene_tx=XM_023256708.1;refGene_gn=IFFO1;refGene_tx=XM_023256709.1;refGene_gn=IFFO1;refGene_tx=XM_023256710.1;refGene_gn=IFFO1;refGene_tx=XM_023256711.1;refGene_gn=IFFO1;refGene_tx=XM_023256712.1;refGene_gn=IFFO1;refGene_tx=XM_023256713.1;refGen

e_gn=IFFO1;refGene_tx=XM_023256714.1;refGene_gn=IFFO1;refGene_tx=XM_023256715.1;refGene_gn=IFFO1;refGene_tx=XM_023256716.1
B4 41636994 41637125
refGene_gn=IFFO1;refGene_tx=XM_019834411.2;refGene_gn=IFFO1;refGene_tx=XM_023256707.1;refGene_gn=IFFO1;refGene_tx=XM_023256708.1;refGene_gn=IFFO1;refGene_tx=XM_023256709.1;refGene_gn=IFFO1;refGene_tx=XM_023256710.1;refGene_gn=IFFO1;refGene_tx=XM_023256711.1;refGene_gn=IFFO1;refGene_tx=XM_023256712.1;refGene_gn=IFFO1;refGene_tx=XM_023256713.1;refGene_gn=IFFO1;refGene_tx=XM_023256714.1;refGene_gn=IFFO1;refGene_tx=XM_023256715.1;refGene_gn=IFFO1;refGene_tx=XM_023256716.1
B4 41640822 41640918
refGene_gn=IFFO1;refGene_tx=XM_019834411.2;refGene_gn=IFFO1;refGene_tx=XM_023256707.1;refGene_gn=IFFO1;refGene_tx=XM_023256708.1;refGene_gn=IFFO1;refGene_tx=XM_023256709.1;refGene_gn=IFFO1;refGene_tx=XM_023256710.1;refGene_gn=IFFO1;refGene_tx=XM_023256711.1;refGene_gn=IFFO1;refGene_tx=XM_023256712.1;refGene_gn=IFFO1;refGene_tx=XM_023256713.1;refGene_gn=IFFO1;refGene_tx=XM_023256714.1;refGene_gn=IFFO1;refGene_tx=XM_023256715.1;refGene_gn=IFFO1;refGene_tx=XM_023256716.1
B4 41641182 41641303
refGene_gn=IFFO1;refGene_tx=XM_019834411.2;refGene_gn=IFFO1;refGene_tx=XM_023256707.1;refGene_gn=IFFO1;refGene_tx=XM_023256708.1;refGene_gn=IFFO1;refGene_tx=XM_023256709.1;refGene_gn=IFFO1;refGene_tx=XM_023256710.1;refGene_gn=IFFO1;refGene_tx=XM_023256711.1;refGene_gn=IFFO1;refGene_tx=XM_023256712.1;refGene_gn=IFFO1;refGene_tx=XM_023256713.1;refGene_gn=IFFO1;refGene_tx=XM_023256714.1;refGene_gn=IFFO1;refGene_tx=XM_023256715.1
B4 41641426 41641608
refGene_gn=IFFO1;refGene_tx=XM_019834411.2;refGene_gn=IFFO1;refGene_tx=XM_023256707.1;refGene_gn=IFFO1;refGene_tx=XM_023256708.1;refGene_gn=IFFO1;refGene_tx=XM_023256709.1;refGene_gn=IFFO1;refGene_tx=XM_023256710.1;refGene_gn=IFFO1;refGene_tx=XM_023256711.1;refGene_gn=IFFO1;refGene_tx=XM_023256712.1;refGene_gn=IFFO1;refGene_tx=XM_023256713.1;refGene_gn=IFFO1;refGene_tx=XM_023256714.1;refGene_gn=IFFO1;refGene_tx=XM_023256715.1
B4 41642235 41642244
refGene_gn=IFFO1;refGene_tx=XM_019834411.2;refGene_gn=IFFO1;refGene_tx=XM_023256707.1;refGene_gn=IFFO1;refGene_tx=XM_023256708.1;refGene_gn=IFFO1;refGene_tx=XM_023256709.1;refGene_gn=IFFO1;refGene_tx=XM_023256710.1;refGene_gn=IFFO1;refGene_tx=XM_023256711.1;refGene_gn=IFFO1;refGene_tx=XM_023256712.1;refGene_gn=IFFO1;refGene_tx=XM_023256713.1
B4 41642585 41642726
refGene_gn=IFFO1;refGene_tx=XM_019834411.2;refGene_gn=IFFO1;refGene_tx=XM_023256707.1;refGene_gn=IFFO1;refGene_tx=XM_023256708.1;refGene_gn=IFFO1;refGene_tx=XM_023256709.1;refGene_gn=IFFO1;refGene_tx=XM_023256710.1;refGene_gn=IFFO1;refGene_tx=XM_023256711.1;refGene_gn=IFFO1;refGene_tx=XM_023256712.1;refGene_gn=IFFO1;refGene_tx=XM_023256713.1;refGene_gn=IFFO1;refGene_tx=XM_023256714.1;refGene_gn=IFFO1;refGene_tx=XM_023256715.1;refGene_gn=IFFO1;refGene_tx=XM_023256716.1
B4 41643416 41643512
refGene_gn=IFFO1;refGene_tx=XM_019834411.2;refGene_gn=IFFO1;refGene_tx

=XM_023256707.1;refGene_gn=IFFO1;refGene_tx=XM_023256708.1;refGene_gn=IFFO1;refGene_tx=XM_023256709.1;refGene_gn=IFFO1;refGene_tx=XM_023256710.1;refGene_gn=IFFO1;refGene_tx=XM_023256711.1;refGene_gn=IFFO1;refGene_tx=XM_023256712.1;refGene_gn=IFFO1;refGene_tx=XM_023256713.1;refGene_gn=IFFO1;refGene_tx=XM_023256714.1;refGene_gn=IFFO1;refGene_tx=XM_023256715.1;refGene_gn=IFFO1;refGene_tx=XM_023256716.1
B4 41643659 41643720
refGene_gn=IFFO1;refGene_tx=XM_019834411.2;refGene_gn=IFFO1;refGene_tx=XM_023256707.1;refGene_gn=IFFO1;refGene_tx=XM_023256708.1;refGene_gn=IFFO1;refGene_tx=XM_023256709.1;refGene_gn=IFFO1;refGene_tx=XM_023256710.1;refGene_gn=IFFO1;refGene_tx=XM_023256711.1;refGene_gn=IFFO1;refGene_tx=XM_023256712.1;refGene_gn=IFFO1;refGene_tx=XM_023256713.1;refGene_gn=IFFO1;refGene_tx=XM_023256714.1;refGene_gn=IFFO1;refGene_tx=XM_023256715.1;refGene_gn=IFFO1;refGene_tx=XM_023256716.1
B4 41647691 41648464
refGene_gn=IFFO1;refGene_tx=XM_019834411.2;refGene_gn=IFFO1;refGene_tx=XM_023256707.1;refGene_gn=IFFO1;refGene_tx=XM_023256708.1;refGene_gn=IFFO1;refGene_tx=XM_023256709.1;refGene_gn=IFFO1;refGene_tx=XM_023256710.1;refGene_gn=IFFO1;refGene_tx=XM_023256711.1;refGene_gn=IFFO1;refGene_tx=XM_023256712.1;refGene_gn=IFFO1;refGene_tx=XM_023256713.1;refGene_gn=IFFO1;refGene_tx=XM_023256714.1;refGene_gn=IFFO1;refGene_tx=XM_023256715.1;refGene_gn=IFFO1;refGene_tx=XM_023256716.1
B4 41649120 41649779 refGene_gn=NOP2;refGene_tx=XM_006933447.4
B4 41651087 41651316 refGene_gn=NOP2;refGene_tx=XM_006933447.4
B4 41651437 41651560 refGene_gn=NOP2;refGene_tx=XM_006933447.4
B4 41651738 41651828 refGene_gn=NOP2;refGene_tx=XM_006933447.4
B4 41651974 41651992 refGene_gn=NOP2;refGene_tx=XM_006933447.4
B4 41651993 41652115 refGene_gn=NOP2;refGene_tx=XM_006933447.4
B4 41652693 41652834 refGene_gn=NOP2;refGene_tx=XM_006933447.4
B4 41652910 41652998 refGene_gn=NOP2;refGene_tx=XM_006933447.4
B4 41653844 41653934 refGene_gn=NOP2;refGene_tx=XM_006933447.4
B4 41654089 41654289 refGene_gn=NOP2;refGene_tx=XM_006933447.4
B4 41654388 41654555 refGene_gn=NOP2;refGene_tx=XM_006933447.4
B4 41654670 41654726 refGene_gn=NOP2;refGene_tx=XM_006933447.4
B4 41656945 41657196 refGene_gn=NOP2;refGene_tx=XM_006933447.4
B4 41657372 41657461 refGene_gn=NOP2;refGene_tx=XM_006933447.4
B4 41657633 41657679 refGene_gn=NOP2;refGene_tx=XM_006933447.4
B4 41659353 41659456 refGene_gn=NOP2;refGene_tx=XM_006933447.4
B4 41661352 41661370
refGene_gn=CHD4;refGene_tx=XM_023256718.1;refGene_gn=CHD4;refGene_tx=XM_023256719.1;refGene_gn=CHD4;refGene_tx=XM_023256720.1;refGene_gn=CHD4;refGene_tx=XM_023256721.1;refGene_gn=CHD4;refGene_tx=XM_023256722.1;refGene_gn=CHD4;refGene_tx=XM_023256723.1;refGene_gn=CHD4;refGene_tx=XM_023256724.1
B4 41661534 41661698
refGene_gn=CHD4;refGene_tx=XM_023256718.1;refGene_gn=CHD4;refGene_tx=XM_023256719.1;refGene_gn=CHD4;refGene_tx=XM_023256720.1;refGene_gn=CHD4;refGene_tx=XM_023256721.1;refGene_gn=CHD4;refGene_tx=XM_023256722.1;refGene_gn=CHD4;refGene_tx=XM_023256723.1;refGene_gn=CHD4;refGene_tx=XM_023256724.1

B4 41662416 41662615
refGene_gn=CHD4;refGene_tx=XM_023256718.1;refGene_gn=CHD4;refGene_tx=XM_023256719.1;refGene_gn=CHD4;refGene_tx=XM_023256720.1;refGene_gn=CHD4;refGene_tx=XM_023256721.1;refGene_gn=CHD4;refGene_tx=XM_023256722.1;refGene_gn=CHD4;refGene_tx=XM_023256723.1;refGene_gn=CHD4;refGene_tx=XM_023256724.1
B4 41666532 41666665
refGene_gn=CHD4;refGene_tx=XM_023256718.1;refGene_gn=CHD4;refGene_tx=XM_023256719.1;refGene_gn=CHD4;refGene_tx=XM_023256720.1;refGene_gn=CHD4;refGene_tx=XM_023256721.1;refGene_gn=CHD4;refGene_tx=XM_023256722.1;refGene_gn=CHD4;refGene_tx=XM_023256723.1;refGene_gn=CHD4;refGene_tx=XM_023256724.1
B4 41666771 41666880
refGene_gn=CHD4;refGene_tx=XM_023256718.1;refGene_gn=CHD4;refGene_tx=XM_023256719.1;refGene_gn=CHD4;refGene_tx=XM_023256720.1;refGene_gn=CHD4;refGene_tx=XM_023256721.1;refGene_gn=CHD4;refGene_tx=XM_023256722.1;refGene_gn=CHD4;refGene_tx=XM_023256723.1;refGene_gn=CHD4;refGene_tx=XM_023256724.1
B4 41667109 41667247
refGene_gn=CHD4;refGene_tx=XM_023256718.1;refGene_gn=CHD4;refGene_tx=XM_023256719.1;refGene_gn=CHD4;refGene_tx=XM_023256720.1;refGene_gn=CHD4;refGene_tx=XM_023256721.1;refGene_gn=CHD4;refGene_tx=XM_023256722.1;refGene_gn=CHD4;refGene_tx=XM_023256723.1;refGene_gn=CHD4;refGene_tx=XM_023256724.1
B4 41667555 41667627
refGene_gn=CHD4;refGene_tx=XM_023256718.1;refGene_gn=CHD4;refGene_tx=XM_023256719.1;refGene_gn=CHD4;refGene_tx=XM_023256720.1;refGene_gn=CHD4;refGene_tx=XM_023256721.1;refGene_gn=CHD4;refGene_tx=XM_023256722.1;refGene_gn=CHD4;refGene_tx=XM_023256723.1;refGene_gn=CHD4;refGene_tx=XM_023256724.1
B4 41668894 41669024
refGene_gn=CHD4;refGene_tx=XM_023256718.1;refGene_gn=CHD4;refGene_tx=XM_023256719.1;refGene_gn=CHD4;refGene_tx=XM_023256720.1;refGene_gn=CHD4;refGene_tx=XM_023256721.1;refGene_gn=CHD4;refGene_tx=XM_023256722.1;refGene_gn=CHD4;refGene_tx=XM_023256723.1;refGene_gn=CHD4;refGene_tx=XM_023256724.1
B4 41669128 41669226
refGene_gn=CHD4;refGene_tx=XM_023256718.1;refGene_gn=CHD4;refGene_tx=XM_023256719.1;refGene_gn=CHD4;refGene_tx=XM_023256720.1;refGene_gn=CHD4;refGene_tx=XM_023256721.1;refGene_gn=CHD4;refGene_tx=XM_023256722.1;refGene_gn=CHD4;refGene_tx=XM_023256723.1;refGene_gn=CHD4;refGene_tx=XM_023256724.1
B4 41669481 41669647
refGene_gn=CHD4;refGene_tx=XM_023256718.1;refGene_gn=CHD4;refGene_tx=XM_023256719.1;refGene_gn=CHD4;refGene_tx=XM_023256720.1;refGene_gn=CHD4;refGene_tx=XM_023256721.1;refGene_gn=CHD4;refGene_tx=XM_023256722.1;refGene_gn=CHD4;refGene_tx=XM_023256723.1;refGene_gn=CHD4;refGene_tx=XM_023256724.1
B4 41669761 41669906
refGene_gn=CHD4;refGene_tx=XM_023256718.1;refGene_gn=CHD4;refGene_tx=X

M_023256719.1;refGene_gn=CHD4;refGene_tx=XM_023256720.1;refGene_gn=CHD
4;refGene_tx=XM_023256721.1;refGene_gn=CHD4;refGene_tx=XM_023256722.1;
refGene_gn=CHD4;refGene_tx=XM_023256723.1;refGene_gn=CHD4;refGene_tx=X
M_023256724.1
B4 41670247 41670381
refGene_gn=CHD4;refGene_tx=XM_023256718.1;refGene_gn=CHD4;refGene_tx=X
M_023256719.1;refGene_gn=CHD4;refGene_tx=XM_023256720.1;refGene_gn=CHD
4;refGene_tx=XM_023256721.1;refGene_gn=CHD4;refGene_tx=XM_023256722.1;
refGene_gn=CHD4;refGene_tx=XM_023256723.1;refGene_gn=CHD4;refGene_tx=X
M_023256724.1
B4 41670487 41670576
refGene_gn=CHD4;refGene_tx=XM_023256718.1;refGene_gn=CHD4;refGene_tx=X
M_023256719.1;refGene_gn=CHD4;refGene_tx=XM_023256720.1;refGene_gn=CHD
4;refGene_tx=XM_023256721.1;refGene_gn=CHD4;refGene_tx=XM_023256722.1;
refGene_gn=CHD4;refGene_tx=XM_023256723.1;refGene_gn=CHD4;refGene_tx=X
M_023256724.1
B4 41670665 41671017
refGene_gn=CHD4;refGene_tx=XM_023256718.1;refGene_gn=CHD4;refGene_tx=X
M_023256719.1;refGene_gn=CHD4;refGene_tx=XM_023256720.1;refGene_gn=CHD
4;refGene_tx=XM_023256721.1;refGene_gn=CHD4;refGene_tx=XM_023256722.1;
refGene_gn=CHD4;refGene_tx=XM_023256723.1;refGene_gn=CHD4;refGene_tx=X
M_023256724.1
B4 41674147 41674323
refGene_gn=CHD4;refGene_tx=XM_023256718.1;refGene_gn=CHD4;refGene_tx=X
M_023256719.1;refGene_gn=CHD4;refGene_tx=XM_023256720.1;refGene_gn=CHD
4;refGene_tx=XM_023256721.1;refGene_gn=CHD4;refGene_tx=XM_023256722.1;
refGene_gn=CHD4;refGene_tx=XM_023256723.1;refGene_gn=CHD4;refGene_tx=X
M_023256724.1
B4 41674486 41674724
refGene_gn=CHD4;refGene_tx=XM_023256718.1;refGene_gn=CHD4;refGene_tx=X
M_023256719.1;refGene_gn=CHD4;refGene_tx=XM_023256720.1;refGene_gn=CHD
4;refGene_tx=XM_023256721.1;refGene_gn=CHD4;refGene_tx=XM_023256722.1;
refGene_gn=CHD4;refGene_tx=XM_023256723.1;refGene_gn=CHD4;refGene_tx=X
M_023256724.1
B4 41675009 41675134
refGene_gn=CHD4;refGene_tx=XM_023256718.1;refGene_gn=CHD4;refGene_tx=X
M_023256719.1;refGene_gn=CHD4;refGene_tx=XM_023256720.1;refGene_gn=CHD
4;refGene_tx=XM_023256721.1;refGene_gn=CHD4;refGene_tx=XM_023256722.1;
refGene_gn=CHD4;refGene_tx=XM_023256723.1;refGene_gn=CHD4;refGene_tx=X
M_023256724.1
B4 41677407 41677525
refGene_gn=CHD4;refGene_tx=XM_023256718.1;refGene_gn=CHD4;refGene_tx=X
M_023256719.1;refGene_gn=CHD4;refGene_tx=XM_023256720.1;refGene_gn=CHD
4;refGene_tx=XM_023256721.1;refGene_gn=CHD4;refGene_tx=XM_023256722.1;
refGene_gn=CHD4;refGene_tx=XM_023256723.1;refGene_gn=CHD4;refGene_tx=X
M_023256724.1
B4 41677640 41677772
refGene_gn=CHD4;refGene_tx=XM_023256718.1;refGene_gn=CHD4;refGene_tx=X
M_023256719.1;refGene_gn=CHD4;refGene_tx=XM_023256720.1;refGene_gn=CHD
4;refGene_tx=XM_023256721.1;refGene_gn=CHD4;refGene_tx=XM_023256722.1;

refGene_gn=CHD4;refGene_tx=XM_023256723.1;refGene_gn=CHD4;refGene_tx=XM_023256724.1
B4 41677857 41677999
refGene_gn=CHD4;refGene_tx=XM_023256718.1;refGene_gn=CHD4;refGene_tx=XM_023256719.1;refGene_gn=CHD4;refGene_tx=XM_023256720.1;refGene_gn=CHD4;refGene_tx=XM_023256721.1;refGene_gn=CHD4;refGene_tx=XM_023256722.1;refGene_gn=CHD4;refGene_tx=XM_023256723.1;refGene_gn=CHD4;refGene_tx=XM_023256724.1
B4 41678357 41678531
refGene_gn=CHD4;refGene_tx=XM_023256718.1;refGene_gn=CHD4;refGene_tx=XM_023256719.1;refGene_gn=CHD4;refGene_tx=XM_023256720.1;refGene_gn=CHD4;refGene_tx=XM_023256721.1;refGene_gn=CHD4;refGene_tx=XM_023256722.1;refGene_gn=CHD4;refGene_tx=XM_023256723.1;refGene_gn=CHD4;refGene_tx=XM_023256724.1
B4 41678625 41678747
refGene_gn=CHD4;refGene_tx=XM_023256718.1;refGene_gn=CHD4;refGene_tx=XM_023256719.1;refGene_gn=CHD4;refGene_tx=XM_023256720.1;refGene_gn=CHD4;refGene_tx=XM_023256721.1;refGene_gn=CHD4;refGene_tx=XM_023256722.1;refGene_gn=CHD4;refGene_tx=XM_023256723.1;refGene_gn=CHD4;refGene_tx=XM_023256724.1
B4 41679023 41679161
refGene_gn=CHD4;refGene_tx=XM_023256718.1;refGene_gn=CHD4;refGene_tx=XM_023256719.1;refGene_gn=CHD4;refGene_tx=XM_023256720.1;refGene_gn=CHD4;refGene_tx=XM_023256721.1;refGene_gn=CHD4;refGene_tx=XM_023256722.1;refGene_gn=CHD4;refGene_tx=XM_023256723.1;refGene_gn=CHD4;refGene_tx=XM_023256724.1
B4 41679326 41679527
refGene_gn=CHD4;refGene_tx=XM_023256718.1;refGene_gn=CHD4;refGene_tx=XM_023256719.1;refGene_gn=CHD4;refGene_tx=XM_023256720.1;refGene_gn=CHD4;refGene_tx=XM_023256721.1;refGene_gn=CHD4;refGene_tx=XM_023256722.1;refGene_gn=CHD4;refGene_tx=XM_023256723.1;refGene_gn=CHD4;refGene_tx=XM_023256724.1
B4 41680230 41680422
refGene_gn=CHD4;refGene_tx=XM_023256718.1;refGene_gn=CHD4;refGene_tx=XM_023256719.1;refGene_gn=CHD4;refGene_tx=XM_023256720.1;refGene_gn=CHD4;refGene_tx=XM_023256721.1;refGene_gn=CHD4;refGene_tx=XM_023256722.1;refGene_gn=CHD4;refGene_tx=XM_023256723.1;refGene_gn=CHD4;refGene_tx=XM_023256724.1
B4 41681290 41681387
refGene_gn=CHD4;refGene_tx=XM_023256718.1;refGene_gn=CHD4;refGene_tx=XM_023256719.1;refGene_gn=CHD4;refGene_tx=XM_023256720.1;refGene_gn=CHD4;refGene_tx=XM_023256721.1;refGene_gn=CHD4;refGene_tx=XM_023256722.1;refGene_gn=CHD4;refGene_tx=XM_023256723.1;refGene_gn=CHD4;refGene_tx=XM_023256724.1
B4 41681467 41681599
refGene_gn=CHD4;refGene_tx=XM_023256718.1;refGene_gn=CHD4;refGene_tx=XM_023256719.1;refGene_gn=CHD4;refGene_tx=XM_023256720.1;refGene_gn=CHD4;refGene_tx=XM_023256721.1;refGene_gn=CHD4;refGene_tx=XM_023256722.1;refGene_gn=CHD4;refGene_tx=XM_023256723.1;refGene_gn=CHD4;refGene_tx=XM_023256724.1

B4 41681973 41682179
refGene_gn=CHD4;refGene_tx=XM_023256718.1;refGene_gn=CHD4;refGene_tx=XM_023256719.1;refGene_gn=CHD4;refGene_tx=XM_023256720.1;refGene_gn=CHD4;refGene_tx=XM_023256721.1;refGene_gn=CHD4;refGene_tx=XM_023256722.1;refGene_gn=CHD4;refGene_tx=XM_023256723.1;refGene_gn=CHD4;refGene_tx=XM_023256724.1
B4 41682301 41682505
refGene_gn=CHD4;refGene_tx=XM_023256718.1;refGene_gn=CHD4;refGene_tx=XM_023256719.1;refGene_gn=CHD4;refGene_tx=XM_023256720.1;refGene_gn=CHD4;refGene_tx=XM_023256721.1;refGene_gn=CHD4;refGene_tx=XM_023256722.1;refGene_gn=CHD4;refGene_tx=XM_023256723.1;refGene_gn=CHD4;refGene_tx=XM_023256724.1
B4 41683022 41683262
refGene_gn=CHD4;refGene_tx=XM_023256718.1;refGene_gn=CHD4;refGene_tx=XM_023256719.1;refGene_gn=CHD4;refGene_tx=XM_023256720.1;refGene_gn=CHD4;refGene_tx=XM_023256721.1;refGene_gn=CHD4;refGene_tx=XM_023256722.1;refGene_gn=CHD4;refGene_tx=XM_023256723.1;refGene_gn=CHD4;refGene_tx=XM_023256724.1
B4 41683451 41683630
refGene_gn=CHD4;refGene_tx=XM_023256718.1;refGene_gn=CHD4;refGene_tx=XM_023256719.1;refGene_gn=CHD4;refGene_tx=XM_023256720.1;refGene_gn=CHD4;refGene_tx=XM_023256721.1;refGene_gn=CHD4;refGene_tx=XM_023256722.1;refGene_gn=CHD4;refGene_tx=XM_023256723.1;refGene_gn=CHD4;refGene_tx=XM_023256724.1
B4 41683782 41683918
refGene_gn=CHD4;refGene_tx=XM_023256718.1;refGene_gn=CHD4;refGene_tx=XM_023256719.1;refGene_gn=CHD4;refGene_tx=XM_023256720.1;refGene_gn=CHD4;refGene_tx=XM_023256721.1;refGene_gn=CHD4;refGene_tx=XM_023256722.1;refGene_gn=CHD4;refGene_tx=XM_023256723.1;refGene_gn=CHD4;refGene_tx=XM_023256724.1
B4 41684164 41684292
refGene_gn=CHD4;refGene_tx=XM_023256718.1;refGene_gn=CHD4;refGene_tx=XM_023256719.1;refGene_gn=CHD4;refGene_tx=XM_023256720.1;refGene_gn=CHD4;refGene_tx=XM_023256721.1;refGene_gn=CHD4;refGene_tx=XM_023256722.1;refGene_gn=CHD4;refGene_tx=XM_023256723.1;refGene_gn=CHD4;refGene_tx=XM_023256724.1
B4 41684541 41684783
refGene_gn=CHD4;refGene_tx=XM_023256718.1;refGene_gn=CHD4;refGene_tx=XM_023256719.1;refGene_gn=CHD4;refGene_tx=XM_023256720.1;refGene_gn=CHD4;refGene_tx=XM_023256721.1;refGene_gn=CHD4;refGene_tx=XM_023256722.1;refGene_gn=CHD4;refGene_tx=XM_023256723.1;refGene_gn=CHD4;refGene_tx=XM_023256724.1
B4 41684908 41685027
refGene_gn=CHD4;refGene_tx=XM_023256718.1;refGene_gn=CHD4;refGene_tx=XM_023256719.1;refGene_gn=CHD4;refGene_tx=XM_023256720.1;refGene_gn=CHD4;refGene_tx=XM_023256721.1;refGene_gn=CHD4;refGene_tx=XM_023256722.1;refGene_gn=CHD4;refGene_tx=XM_023256723.1;refGene_gn=CHD4;refGene_tx=XM_023256724.1
B4 41685241 41685457
refGene_gn=CHD4;refGene_tx=XM_023256718.1;refGene_gn=CHD4;refGene_tx=X

M_023256719.1;refGene_gn=CHD4;refGene_tx=XM_023256720.1;refGene_gn=CHD4;refGene_tx=XM_023256721.1;refGene_gn=CHD4;refGene_tx=XM_023256722.1;refGene_gn=CHD4;refGene_tx=XM_023256723.1;refGene_gn=CHD4;refGene_tx=XM_023256724.1
B4 41685679 41685801
refGene_gn=CHD4;refGene_tx=XM_023256718.1;refGene_gn=CHD4;refGene_tx=XM_023256719.1;refGene_gn=CHD4;refGene_tx=XM_023256720.1;refGene_gn=CHD4;refGene_tx=XM_023256721.1;refGene_gn=CHD4;refGene_tx=XM_023256722.1;refGene_gn=CHD4;refGene_tx=XM_023256723.1;refGene_gn=CHD4;refGene_tx=XM_023256724.1
B4 41689445 41689545
refGene_gn=CHD4;refGene_tx=XM_023256718.1;refGene_gn=CHD4;refGene_tx=XM_023256719.1;refGene_gn=CHD4;refGene_tx=XM_023256720.1;refGene_gn=CHD4;refGene_tx=XM_023256721.1;refGene_gn=CHD4;refGene_tx=XM_023256722.1;refGene_gn=CHD4;refGene_tx=XM_023256723.1;refGene_gn=CHD4;refGene_tx=XM_023256724.1
B4 41701280 41702402
refGene_gn=LPAR5;refGene_tx=XM_011283722.3;refGene_gn=LPAR5;refGene_tx=XM_019834421.2;refGene_gn=LPAR5;refGene_tx=XM_019834422.2;refGene_gn=LPAR5;refGene_tx=XM_019834425.2;refGene_gn=LPAR5;refGene_tx=XM_019834426.2;refGene_gn=LPAR5;refGene_tx=XM_019834427.2;refGene_gn=LPAR5;refGene_tx=XM_019834429.2
B4 41712745 41712868
refGene_gn=ACRBP;refGene_tx=XM_003988322.4;refGene_gn=ACRBP;refGene_tx=XM_019834430.2
B4 41713445 41713529
refGene_gn=ACRBP;refGene_tx=XM_003988322.4;refGene_gn=ACRBP;refGene_tx=XM_019834430.2
B4 41714576 41714744
refGene_gn=ACRBP;refGene_tx=XM_003988322.4;refGene_gn=ACRBP;refGene_tx=XM_019834430.2
B4 41714882 41715062
refGene_gn=ACRBP;refGene_tx=XM_003988322.4;refGene_gn=ACRBP;refGene_tx=XM_019834430.2
B4 41717034 41717167
refGene_gn=ACRBP;refGene_tx=XM_003988322.4;refGene_gn=ACRBP;refGene_tx=XM_019834430.2
B4 41717649 41718118
refGene_gn=ACRBP;refGene_tx=XM_003988322.4;refGene_gn=ACRBP;refGene_tx=XM_019834430.2
B4 41718762 41718880
refGene_gn=ACRBP;refGene_tx=XM_003988322.4;refGene_gn=ACRBP;refGene_tx=XM_019834430.2
B4 41719371 41719466
refGene_gn=ACRBP;refGene_tx=XM_003988322.4;refGene_gn=ACRBP;refGene_tx=XM_019834430.2
B4 41719641 41719860 refGene_gn=ACRBP;refGene_tx=XM_003988322.4
B4 41720162 41720205
refGene_gn=ACRBP;refGene_tx=XM_003988322.4;refGene_gn=ACRBP;refGene_tx=XM_019834430.2

B4 41721773 41721813
refGene_gn=ING4;refGene_tx=XM_003988323.5;refGene_gn=ING4;refGene_tx=XM_006933453.4;refGene_gn=ING4;refGene_tx=XM_006933454.4;refGene_gn=ING4;refGene_tx=XM_006933455.3;refGene_gn=ING4;refGene_tx=XM_019834431.2
B4 41721924 41721986
refGene_gn=ING4;refGene_tx=XM_003988323.5;refGene_gn=ING4;refGene_tx=XM_006933453.4;refGene_gn=ING4;refGene_tx=XM_006933454.4;refGene_gn=ING4;refGene_tx=XM_006933455.3;refGene_gn=ING4;refGene_tx=XM_019834431.2;refGene_gn=ING4;refGene_tx=XM_019834433.2
B4 41722250 41722398
refGene_gn=ING4;refGene_tx=XM_003988323.5;refGene_gn=ING4;refGene_tx=XM_006933453.4;refGene_gn=ING4;refGene_tx=XM_006933454.4;refGene_gn=ING4;refGene_tx=XM_006933455.3;refGene_gn=ING4;refGene_tx=XM_019834431.2
B4 41722642 41722751
refGene_gn=ING4;refGene_tx=XM_003988323.5;refGene_gn=ING4;refGene_tx=XM_006933453.4;refGene_gn=ING4;refGene_tx=XM_006933454.4;refGene_gn=ING4;refGene_tx=XM_006933455.3;refGene_gn=ING4;refGene_tx=XM_019834431.2;refGene_gn=ING4;refGene_tx=XM_019834433.2
B4 41722914 41723029
refGene_gn=ING4;refGene_tx=XM_003988323.5;refGene_gn=ING4;refGene_tx=XM_006933453.4;refGene_gn=ING4;refGene_tx=XM_006933454.4;refGene_gn=ING4;refGene_tx=XM_006933455.3;refGene_gn=ING4;refGene_tx=XM_019834431.2;refGene_gn=ING4;refGene_tx=XM_019834433.2
B4 41723208 41723375
refGene_gn=ING4;refGene_tx=XM_003988323.5;refGene_gn=ING4;refGene_tx=XM_006933453.4;refGene_gn=ING4;refGene_tx=XM_006933454.4;refGene_gn=ING4;refGene_tx=XM_006933455.3;refGene_gn=ING4;refGene_tx=XM_019834431.2;refGene_gn=ING4;refGene_tx=XM_019834433.2
B4 41725101 41725173
refGene_gn=ING4;refGene_tx=XM_003988323.5;refGene_gn=ING4;refGene_tx=XM_006933453.4;refGene_gn=ING4;refGene_tx=XM_006933454.4;refGene_gn=ING4;refGene_tx=XM_019834431.2;refGene_gn=ING4;refGene_tx=XM_019834433.2
B4 41727943 41727980
refGene_gn=ING4;refGene_tx=XM_003988323.5;refGene_gn=ING4;refGene_tx=XM_006933453.4;refGene_gn=ING4;refGene_tx=XM_006933454.4;refGene_gn=ING4;refGene_tx=XM_019834431.2;refGene_gn=ING4;refGene_tx=XM_019834433.2
B4 41731449 41731836
refGene_gn=ZNF384;refGene_tx=XM_011283720.3;refGene_gn=ZNF384;refGene_tx=XM_019834434.2;refGene_gn=ZNF384;refGene_tx=XM_019834435.2;refGene_gn=ZNF384;refGene_tx=XM_019834436.2;refGene_gn=ZNF384;refGene_tx=XM_019834438.2;refGene_gn=ZNF384;refGene_tx=XM_019834439.2;refGene_gn=ZNF384;refGene_tx=XM_023256725.1;refGene_gn=ZNF384;refGene_tx=XM_023256726.1
B4 41732514 41732673
refGene_gn=ZNF384;refGene_tx=XM_011283720.3;refGene_gn=ZNF384;refGene_tx=XM_019834434.2;refGene_gn=ZNF384;refGene_tx=XM_019834435.2;refGene_gn=ZNF384;refGene_tx=XM_019834436.2;refGene_gn=ZNF384;refGene_tx=XM_019834438.2;refGene_gn=ZNF384;refGene_tx=XM_019834439.2;refGene_gn=ZNF384;refGene_tx=XM_023256725.1;refGene_gn=ZNF384;refGene_tx=XM_023256726.1

B4	41733634	41733713
refGene_gn=ZNF384;refGene_tx=XM_011283720.3;refGene_gn=ZNF384;refGene_tx=XM_019834434.2;refGene_gn=ZNF384;refGene_tx=XM_019834435.2;refGene_gn=ZNF384;refGene_tx=XM_019834436.2;refGene_gn=ZNF384;refGene_tx=XM_019834438.2;refGene_gn=ZNF384;refGene_tx=XM_019834439.2;refGene_gn=ZNF384;refGene_tx=XM_023256725.1;refGene_gn=ZNF384;refGene_tx=XM_023256726.1
B4	41735220	41735403
refGene_gn=ZNF384;refGene_tx=XM_011283720.3;refGene_gn=ZNF384;refGene_tx=XM_019834434.2;refGene_gn=ZNF384;refGene_tx=XM_019834435.2;refGene_gn=ZNF384;refGene_tx=XM_019834436.2;refGene_gn=ZNF384;refGene_tx=XM_023256725.1;refGene_gn=ZNF384;refGene_tx=XM_023256726.1
B4	41736108	41736333
refGene_gn=ZNF384;refGene_tx=XM_011283720.3;refGene_gn=ZNF384;refGene_tx=XM_019834434.2;refGene_gn=ZNF384;refGene_tx=XM_019834435.2;refGene_gn=ZNF384;refGene_tx=XM_019834436.2;refGene_gn=ZNF384;refGene_tx=XM_019834438.2;refGene_gn=ZNF384;refGene_tx=XM_019834439.2;refGene_gn=ZNF384;refGene_tx=XM_023256725.1;refGene_gn=ZNF384;refGene_tx=XM_023256726.1
B4	41740070	41740163
refGene_gn=ZNF384;refGene_tx=XM_011283720.3;refGene_gn=ZNF384;refGene_tx=XM_019834434.2;refGene_gn=ZNF384;refGene_tx=XM_019834435.2;refGene_gn=ZNF384;refGene_tx=XM_019834438.2;refGene_gn=ZNF384;refGene_tx=XM_023256725.1;refGene_gn=ZNF384;refGene_tx=XM_023256726.1
B4	41741009	41741343
refGene_gn=ZNF384;refGene_tx=XM_011283720.3;refGene_gn=ZNF384;refGene_tx=XM_019834434.2;refGene_gn=ZNF384;refGene_tx=XM_019834435.2;refGene_gn=ZNF384;refGene_tx=XM_019834436.2;refGene_gn=ZNF384;refGene_tx=XM_019834438.2;refGene_gn=ZNF384;refGene_tx=XM_019834439.2;refGene_gn=ZNF384;refGene_tx=XM_023256725.1;refGene_gn=ZNF384;refGene_tx=XM_023256726.1
B4	41741547	41741595
refGene_gn=ZNF384;refGene_tx=XM_011283720.3;refGene_gn=ZNF384;refGene_tx=XM_019834434.2;refGene_gn=ZNF384;refGene_tx=XM_019834436.2;refGene_gn=ZNF384;refGene_tx=XM_019834438.2;refGene_gn=ZNF384;refGene_tx=XM_019834439.2;refGene_gn=ZNF384;refGene_tx=XM_023256725.1;refGene_gn=ZNF384;refGene_tx=XM_023256726.1
B4	41741830	41742068
refGene_gn=ZNF384;refGene_tx=XM_011283720.3;refGene_gn=ZNF384;refGene_tx=XM_019834434.2;refGene_gn=ZNF384;refGene_tx=XM_019834435.2;refGene_gn=ZNF384;refGene_tx=XM_019834436.2;refGene_gn=ZNF384;refGene_tx=XM_019834438.2;refGene_gn=ZNF384;refGene_tx=XM_019834439.2;refGene_gn=ZNF384;refGene_tx=XM_023256725.1;refGene_gn=ZNF384;refGene_tx=XM_023256726.1
B4	41742352	41742418
refGene_gn=ZNF384;refGene_tx=XM_011283720.3;refGene_gn=ZNF384;refGene_tx=XM_019834434.2;refGene_gn=ZNF384;refGene_tx=XM_019834435.2;refGene_gn=ZNF384;refGene_tx=XM_019834436.2;refGene_gn=ZNF384;refGene_tx=XM_019834438.2;refGene_gn=ZNF384;refGene_tx=XM_019834439.2;refGene_gn=ZNF384;refGene_tx=XM_023256725.1;refGene_gn=ZNF384;refGene_tx=XM_023256726.

1
B4 41754954 41754976 refGene_gn=PIANP;refGene_tx=XM_004001495.5;refGene_gn=PIANP;refGene_tx=XM_019834440.2;refGene_gn=PIANP;refGene_tx=XM_023256727.1
B4 41755287 41755509 refGene_gn=PIANP;refGene_tx=XM_004001495.5;refGene_gn=PIANP;refGene_tx=XM_019834440.2;refGene_gn=PIANP;refGene_tx=XM_023256727.1
B4 41756322 41756404 refGene_gn=PIANP;refGene_tx=XM_004001495.5;refGene_gn=PIANP;refGene_tx=XM_019834440.2;refGene_gn=PIANP;refGene_tx=XM_023256727.1
B4 41757166 41757678 refGene_gn=PIANP;refGene_tx=XM_004001495.5;refGene_gn=PIANP;refGene_tx=XM_019834440.2;refGene_gn=PIANP;refGene_tx=XM_023256727.1
B4 41757941 41757958 refGene_gn=PIANP;refGene_tx=XM_004001495.5;refGene_gn=PIANP;refGene_tx=XM_019834440.2;refGene_gn=PIANP;refGene_tx=XM_023256727.1
B4 41778197 41778359 refGene_gn=COPS7A;refGene_tx=XM_003988328.5;refGene_gn=COPS7A;refGene_tx=XM_003988329.5;refGene_gn=COPS7A;refGene_tx=XM_006933458.4
B4 41780082 41780158 refGene_gn=COPS7A;refGene_tx=XM_003988328.5;refGene_gn=COPS7A;refGene_tx=XM_003988329.5;refGene_gn=COPS7A;refGene_tx=XM_006933458.4
B4 41780618 41780707 refGene_gn=COPS7A;refGene_tx=XM_003988328.5;refGene_gn=COPS7A;refGene_tx=XM_003988329.5;refGene_gn=COPS7A;refGene_tx=XM_006933458.4
B4 41781665 41781868 refGene_gn=COPS7A;refGene_tx=XM_003988328.5;refGene_gn=COPS7A;refGene_tx=XM_003988329.5;refGene_gn=COPS7A;refGene_tx=XM_006933458.4
B4 41782478 41782584 refGene_gn=COPS7A;refGene_tx=XM_003988328.5;refGene_gn=COPS7A;refGene_tx=XM_003988329.5;refGene_gn=COPS7A;refGene_tx=XM_006933458.4
B4 41782748 41782900 refGene_gn=COPS7A;refGene_tx=XM_003988328.5;refGene_gn=COPS7A;refGene_tx=XM_003988329.5;refGene_gn=COPS7A;refGene_tx=XM_006933458.4
B4 41783069 41783109 refGene_gn=COPS7A;refGene_tx=XM_003988328.5;refGene_gn=COPS7A;refGene_tx=XM_003988329.5;refGene_gn=COPS7A;refGene_tx=XM_006933458.4
B4 41797574 41797692 refGene_gn=LOC101092863;refGene_tx=XM_019834441.2
B4 41797817 41797993 refGene_gn=LOC101092863;refGene_tx=XM_019834441.2
B4 41798072 41798145 refGene_gn=LOC101092863;refGene_tx=XM_019834441.2
B4 41798725 41798927 refGene_gn=LOC101092863;refGene_tx=XM_019834441.2
B4 41801776 41801867 refGene_gn=LOC101092863;refGene_tx=XM_019834441.2
B4 41802795 41802983 refGene_gn=MLF2;refGene_tx=XM_023256728.1
B4 41803803 41803963 refGene_gn=MLF2;refGene_tx=XM_023256728.1
B4 41804130 41804259 refGene_gn=MLF2;refGene_tx=XM_023256728.1

| | | | |
|---|---|---|---|
| B4 | 41804662 | 41804716 | refGene_gn=MLF2;refGene_tx=XM_023256728.1 |
| B4 | 41805537 | 41805573 | refGene_gn=MLF2;refGene_tx=XM_023256728.1 |
| B4 | 41805814 | 41805941 | refGene_gn=MLF2;refGene_tx=XM_023256728.1 |
| B4 | 41806203 | 41806253 | refGene_gn=MLF2;refGene_tx=XM_023256728.1 |
| B4 | 41816580 | 41816631 | refGene_gn=PTMS;refGene_tx=XM_023257737.1 |
| B4 | 41817765 | 41818449 | refGene_gn=PTMS;refGene_tx=XM_023257737.1 |
| B4 | 41821416 | 41821488 | refGene_gn=PTMS;refGene_tx=XM_023257737.1 |
| B4 | 41821683 | 41821762 | refGene_gn=PTMS;refGene_tx=XM_023257737.1 |
| B4 | 41821904 | 41821966 | refGene_gn=PTMS;refGene_tx=XM_023257737.1 |
| B4 | 41822136 | 41822187 | refGene_gn=PTMS;refGene_tx=XM_023257737.1 |
| B4 | 41824527 | 41824585 | refGene_gn=LAG3;refGene_tx=XM_023256729.1 |
| B4 | 41824884 | 41825032 | refGene_gn=LAG3;refGene_tx=XM_023256729.1 |
| B4 | 41825403 | 41825690 | refGene_gn=LAG3;refGene_tx=XM_023256729.1 |
| B4 | 41826417 | 41826675 | refGene_gn=LAG3;refGene_tx=XM_023256729.1 |
| B4 | 41827088 | 41827364 | refGene_gn=LAG3;refGene_tx=XM_023256729.1 |
| B4 | 41829245 | 41829488 | refGene_gn=LAG3;refGene_tx=XM_023256729.1 |
| B4 | 41829762 | 41829893 | refGene_gn=LAG3;refGene_tx=XM_023256729.1 |
| B4 | 41830227 | 41830368 | refGene_gn=LAG3;refGene_tx=XM_023256729.1 |
| B4 | 41845990 | 41846039 | refGene_gn=CD4;refGene_tx=NM_001009250.1 |
| B4 | 41846155 | 41846326 | refGene_gn=CD4;refGene_tx=NM_001009250.1 |
| B4 | 41879189 | 41879348 | refGene_gn=CD4;refGene_tx=NM_001009250.1 |
| B4 | 41879753 | 41880038 | refGene_gn=CD4;refGene_tx=NM_001009250.1 |
| B4 | 41881402 | 41881738 | refGene_gn=CD4;refGene_tx=NM_001009250.1 |
| B4 | 41882465 | 41882666 | refGene_gn=CD4;refGene_tx=NM_001009250.1 |
| B4 | 41883230 | 41883355 | refGene_gn=CD4;refGene_tx=NM_001009250.1 |
| B4 | 41883688 | 41883756 | refGene_gn=CD4;refGene_tx=NM_001009250.1 |
| B4 | 41884101 | 41884132 | refGene_gn=CD4;refGene_tx=NM_001009250.1 |
| B4 | 41888754 | 41889621 | refGene_gn=GPR162;refGene_tx=XM_003988331.3 |
| B4 | 41890321 | 41890511 | refGene_gn=GPR162;refGene_tx=XM_003988331.3 |
| B4 | 41891142 | 41891300 | refGene_gn=GPR162;refGene_tx=XM_003988331.3 |
| B4 | 41891803 | 41892355 | refGene_gn=GPR162;refGene_tx=XM_003988331.3 |
| B4 | 41893479 | 41893971 | refGene_gn=P3H3;refGene_tx=XM_011283712.3 |
| B4 | 41894895 | 41895048 | refGene_gn=P3H3;refGene_tx=XM_011283712.3 |
| B4 | 41895369 | 41895571 | refGene_gn=P3H3;refGene_tx=XM_011283712.3 |
| B4 | 41895645 | 41895777 | refGene_gn=P3H3;refGene_tx=XM_011283712.3 |
| B4 | 41896286 | 41896423 | refGene_gn=P3H3;refGene_tx=XM_011283712.3 |
| B4 | 41896775 | 41896865 | refGene_gn=P3H3;refGene_tx=XM_011283712.3 |
| B4 | 41899011 | 41899064 | refGene_gn=P3H3;refGene_tx=XM_011283712.3 |
| B4 | 41899177 | 41899245 | refGene_gn=P3H3;refGene_tx=XM_011283712.3 |
| B4 | 41899361 | 41899486 | refGene_gn=P3H3;refGene_tx=XM_011283712.3 |
| B4 | 41902344 | 41902446 | refGene_gn=P3H3;refGene_tx=XM_011283712.3 |
| B4 | 41902791 | 41902942 | refGene_gn=P3H3;refGene_tx=XM_011283712.3 |
| B4 | 41903160 | 41903278 | refGene_gn=P3H3;refGene_tx=XM_011283712.3 |
| B4 | 41903361 | 41903437 | refGene_gn=P3H3;refGene_tx=XM_011283712.3 |
| B4 | 41904227 | 41904368 | refGene_gn=P3H3;refGene_tx=XM_011283712.3 |
| B4 | 41904508 | 41904679 | refGene_gn=P3H3;refGene_tx=XM_011283712.3 |
| B4 | 41906534 | 41906591 | refGene_gn=GNB3;refGene_tx=XM_003988332.5 |
| B4 | 41906883 | 41906922 | refGene_gn=GNB3;refGene_tx=XM_003988332.5 |
| B4 | 41908145 | 41908252 | refGene_gn=GNB3;refGene_tx=XM_003988332.5 |
| B4 | 41908335 | 41908399 | refGene_gn=GNB3;refGene_tx=XM_003988332.5 |

B4 41908522 41908685 refGene_gn=GNB3;refGene_tx=XM_003988332.5
B4 41908793 41908860 refGene_gn=GNB3;refGene_tx=XM_003988332.5
B4 41908952 41909154 refGene_gn=GNB3;refGene_tx=XM_003988332.5
B4 41910226 41910281 refGene_gn=GNB3;refGene_tx=XM_003988332.5
B4 41910283 41910396 refGene_gn=GNB3;refGene_tx=XM_003988332.5
B4 41910398 41910444 refGene_gn=GNB3;refGene_tx=XM_003988332.5
B4 41911316 41911423 refGene_gn=GNB3;refGene_tx=XM_003988332.5
B4 41913363 41913522
refGene_gn=CDCA3;refGene_tx=XM_003988333.5;refGene_gn=CDCA3;refGene_tx=XM_006933463.3;refGene_gn=CDCA3;refGene_tx=XM_011283713.3;refGene_gn=CDCA3;refGene_tx=XM_023256731.1;refGene_gn=CDCA3;refGene_tx=XM_023256732.1
B4 41913658 41913762
refGene_gn=CDCA3;refGene_tx=XM_003988333.5;refGene_gn=CDCA3;refGene_tx=XM_006933463.3;refGene_gn=CDCA3;refGene_tx=XM_023256731.1;refGene_gn=CDCA3;refGene_tx=XM_023256732.1
B4 41913884 41914178
refGene_gn=CDCA3;refGene_tx=XM_003988333.5;refGene_gn=CDCA3;refGene_tx=XM_006933463.3;refGene_gn=CDCA3;refGene_tx=XM_011283713.3;refGene_gn=CDCA3;refGene_tx=XM_023256731.1;refGene_gn=CDCA3;refGene_tx=XM_023256732.1
B4 41914451 41914581
refGene_gn=CDCA3;refGene_tx=XM_003988333.5;refGene_gn=CDCA3;refGene_tx=XM_006933463.3;refGene_gn=CDCA3;refGene_tx=XM_011283713.3;refGene_gn=CDCA3;refGene_tx=XM_023256731.1;refGene_gn=CDCA3;refGene_tx=XM_023256732.1
B4 41914834 41914954
refGene_gn=CDCA3;refGene_tx=XM_003988333.5;refGene_gn=CDCA3;refGene_tx=XM_006933463.3;refGene_gn=CDCA3;refGene_tx=XM_011283713.3;refGene_gn=CDCA3;refGene_tx=XM_023256731.1;refGene_gn=CDCA3;refGene_tx=XM_023256732.1
B4 41916069 41916279
refGene_gn=USP5;refGene_tx=XM_003988334.5;refGene_gn=USP5;refGene_tx=XM_003988335.5;refGene_gn=USP5;refGene_tx=XM_019834444.2;refGene_gn=USP5;refGene_tx=XM_019834445.2
B4 41919043 41919169
refGene_gn=USP5;refGene_tx=XM_003988334.5;refGene_gn=USP5;refGene_tx=XM_003988335.5;refGene_gn=USP5;refGene_tx=XM_019834444.2;refGene_gn=USP5;refGene_tx=XM_019834445.2
B4 41919432 41919499
refGene_gn=USP5;refGene_tx=XM_003988334.5;refGene_gn=USP5;refGene_tx=XM_003988335.5;refGene_gn=USP5;refGene_tx=XM_019834444.2;refGene_gn=USP5;refGene_tx=XM_019834445.2
B4 41919693 41919827
refGene_gn=USP5;refGene_tx=XM_003988334.5;refGene_gn=USP5;refGene_tx=XM_003988335.5;refGene_gn=USP5;refGene_tx=XM_019834444.2;refGene_gn=USP5;refGene_tx=XM_019834445.2
B4 41919982 41920128
refGene_gn=USP5;refGene_tx=XM_003988334.5;refGene_gn=USP5;refGene_tx=XM_003988335.5;refGene_gn=USP5;refGene_tx=XM_019834444.2;refGene_gn=USP

5;refGene_tx=XM_019834445.2
B4 41920368 41920553
refGene_gn=USP5;refGene_tx=XM_003988334.5;refGene_gn=USP5;refGene_tx=XM_003988335.5;refGene_gn=USP5;refGene_tx=XM_019834444.2;refGene_gn=USP5;refGene_tx=XM_019834445.2
B4 41921023 41921118
refGene_gn=USP5;refGene_tx=XM_003988334.5;refGene_gn=USP5;refGene_tx=XM_003988335.5;refGene_gn=USP5;refGene_tx=XM_019834444.2;refGene_gn=USP5;refGene_tx=XM_019834445.2
B4 41921781 41921975
refGene_gn=USP5;refGene_tx=XM_003988334.5;refGene_gn=USP5;refGene_tx=XM_003988335.5;refGene_gn=USP5;refGene_tx=XM_019834444.2;refGene_gn=USP5;refGene_tx=XM_019834445.2
B4 41922701 41922773
refGene_gn=USP5;refGene_tx=XM_003988334.5;refGene_gn=USP5;refGene_tx=XM_003988335.5;refGene_gn=USP5;refGene_tx=XM_019834444.2;refGene_gn=USP5;refGene_tx=XM_019834445.2
B4 41923045 41923133
refGene_gn=USP5;refGene_tx=XM_003988334.5;refGene_gn=USP5;refGene_tx=XM_003988335.5;refGene_gn=USP5;refGene_tx=XM_019834444.2;refGene_gn=USP5;refGene_tx=XM_019834445.2
B4 41923262 41923388
refGene_gn=USP5;refGene_tx=XM_003988334.5;refGene_gn=USP5;refGene_tx=XM_003988335.5;refGene_gn=USP5;refGene_tx=XM_019834444.2;refGene_gn=USP5;refGene_tx=XM_019834445.2
B4 41923890 41924044
refGene_gn=USP5;refGene_tx=XM_003988334.5;refGene_gn=USP5;refGene_tx=XM_003988335.5;refGene_gn=USP5;refGene_tx=XM_019834444.2;refGene_gn=USP5;refGene_tx=XM_019834445.2
B4 41924369 41924544
refGene_gn=USP5;refGene_tx=XM_003988334.5;refGene_gn=USP5;refGene_tx=XM_003988335.5;refGene_gn=USP5;refGene_tx=XM_019834444.2;refGene_gn=USP5;refGene_tx=XM_019834445.2
B4 41925188 41925277
refGene_gn=USP5;refGene_tx=XM_003988334.5;refGene_gn=USP5;refGene_tx=XM_003988335.5;refGene_gn=USP5;refGene_tx=XM_019834444.2;refGene_gn=USP5;refGene_tx=XM_019834445.2
B4 41925901 41926093
refGene_gn=USP5;refGene_tx=XM_003988334.5;refGene_gn=USP5;refGene_tx=XM_003988335.5;refGene_gn=USP5;refGene_tx=XM_019834444.2;refGene_gn=USP5;refGene_tx=XM_019834445.2
B4 41926622 41926766
refGene_gn=USP5;refGene_tx=XM_003988334.5;refGene_gn=USP5;refGene_tx=XM_003988335.5;refGene_gn=USP5;refGene_tx=XM_019834444.2;refGene_gn=USP5;refGene_tx=XM_019834445.2
B4 41926843 41926989
refGene_gn=USP5;refGene_tx=XM_003988334.5;refGene_gn=USP5;refGene_tx=XM_003988335.5;refGene_gn=USP5;refGene_tx=XM_019834444.2;refGene_gn=USP5;refGene_tx=XM_019834445.2
B4 41927185 41927363

refGene_gn=USP5;refGene_tx=XM_003988334.5;refGene_gn=USP5;refGene_tx=XM_003988335.5;refGene_gn=USP5;refGene_tx=XM_019834444.2;refGene_gn=USP5;refGene_tx=XM_019834445.2
B4 41927660 41927745
refGene_gn=USP5;refGene_tx=XM_003988334.5;refGene_gn=USP5;refGene_tx=XM_003988335.5;refGene_gn=USP5;refGene_tx=XM_019834444.2;refGene_gn=USP5;refGene_tx=XM_019834445.2
B4 41928604 41928698
refGene_gn=USP5;refGene_tx=XM_003988334.5;refGene_gn=USP5;refGene_tx=XM_003988335.5;refGene_gn=USP5;refGene_tx=XM_019834444.2;refGene_gn=USP5;refGene_tx=XM_019834445.2
B4 41930025 41930251 refGene_gn=TPI1;refGene_tx=XM_006933465.3
B4 41931456 41931580 refGene_gn=TPI1;refGene_tx=XM_006933465.3
B4 41931695 41931780 refGene_gn=TPI1;refGene_tx=XM_006933465.3
B4 41931854 41931987 refGene_gn=TPI1;refGene_tx=XM_006933465.3
B4 41932290 41932376 refGene_gn=TPI1;refGene_tx=XM_006933465.3
B4 41932832 41932920 refGene_gn=TPI1;refGene_tx=XM_006933465.3
B4 41933010 41933129 refGene_gn=TPI1;refGene_tx=XM_006933465.3
B4 41934003 41934131 refGene_gn=SPSB2;refGene_tx=XM_003988338.5
B4 41935028 41935281 refGene_gn=SPSB2;refGene_tx=XM_003988338.5
B4 41935282 41935693 refGene_gn=SPSB2;refGene_tx=XM_003988338.5
B4 41957701 41957827
refGene_gn=LRRC23;refGene_tx=XM_003988339.5;refGene_gn=LRRC23;refGene_tx=XM_023256734.1
B4 41957912 41958022
refGene_gn=LRRC23;refGene_tx=XM_003988339.5;refGene_gn=LRRC23;refGene_tx=XM_023256734.1
B4 41958465 41958719
refGene_gn=LRRC23;refGene_tx=XM_003988339.5;refGene_gn=LRRC23;refGene_tx=XM_023256734.1
B4 41959382 41959513
refGene_gn=LRRC23;refGene_tx=XM_003988339.5;refGene_gn=LRRC23;refGene_tx=XM_023256734.1
B4 41960146 41960283
refGene_gn=LRRC23;refGene_tx=XM_003988339.5;refGene_gn=LRRC23;refGene_tx=XM_023256734.1
B4 41961917 41962191
refGene_gn=LRRC23;refGene_tx=XM_003988339.5;refGene_gn=LRRC23;refGene_tx=XM_023256734.1
B4 41964788 41964873
refGene_gn=ENO2;refGene_tx=XM_011283710.3;refGene_gn=ENO2;refGene_tx=XM_019834450.2
B4 41965356 41965452
refGene_gn=ENO2;refGene_tx=XM_011283710.3;refGene_gn=ENO2;refGene_tx=XM_019834450.2
B4 41965596 41965655
refGene_gn=ENO2;refGene_tx=XM_011283710.3;refGene_gn=ENO2;refGene_tx=XM_019834450.2
B4 41965945 41966015
refGene_gn=ENO2;refGene_tx=XM_011283710.3;refGene_gn=ENO2;refGene_tx=X

M_019834450.2;refGene_gn=ENO2;refGene_tx=XM_023256733.1
B4 41966503 41966637
refGene_gn=ENO2;refGene_tx=XM_011283710.3;refGene_gn=ENO2;refGene_tx=XM_019834450.2;refGene_gn=ENO2;refGene_tx=XM_023256733.1
B4 41966870 41967093
refGene_gn=ENO2;refGene_tx=XM_011283710.3;refGene_gn=ENO2;refGene_tx=XM_019834450.2;refGene_gn=ENO2;refGene_tx=XM_023256733.1
B4 41968137 41968335
refGene_gn=ENO2;refGene_tx=XM_011283710.3;refGene_gn=ENO2;refGene_tx=XM_019834450.2;refGene_gn=ENO2;refGene_tx=XM_023256733.1
B4 41969190 41969392
refGene_gn=ENO2;refGene_tx=XM_011283710.3;refGene_gn=ENO2;refGene_tx=XM_019834450.2;refGene_gn=ENO2;refGene_tx=XM_023256733.1
B4 41969922 41970031
refGene_gn=ENO2;refGene_tx=XM_011283710.3;refGene_gn=ENO2;refGene_tx=XM_019834450.2;refGene_gn=ENO2;refGene_tx=XM_023256733.1
B4 41970191 41970250
refGene_gn=ENO2;refGene_tx=XM_011283710.3;refGene_gn=ENO2;refGene_tx=XM_019834450.2;refGene_gn=ENO2;refGene_tx=XM_023256733.1
B4 41970538 41970608
refGene_gn=ENO2;refGene_tx=XM_011283710.3;refGene_gn=ENO2;refGene_tx=XM_019834450.2;refGene_gn=ENO2;refGene_tx=XM_023256733.1
B4 41980466 41980493
refGene_gn=ATN1;refGene_tx=XM_023256735.1;refGene_gn=ATN1;refGene_tx=XM_023256736.1;refGene_gn=ATN1;refGene_tx=XM_023256737.1
B4 41980696 41980834
refGene_gn=ATN1;refGene_tx=XM_023256735.1;refGene_gn=ATN1;refGene_tx=XM_023256736.1;refGene_gn=ATN1;refGene_tx=XM_023256737.1
B4 41980983 41981097
refGene_gn=ATN1;refGene_tx=XM_023256735.1;refGene_gn=ATN1;refGene_tx=XM_023256736.1
B4 41981808 41983832
refGene_gn=ATN1;refGene_tx=XM_023256735.1;refGene_gn=ATN1;refGene_tx=XM_023256736.1;refGene_gn=ATN1;refGene_tx=XM_023256737.1
B4 41984079 41984302
refGene_gn=ATN1;refGene_tx=XM_023256735.1;refGene_gn=ATN1;refGene_tx=XM_023256736.1;refGene_gn=ATN1;refGene_tx=XM_023256737.1
B4 41984713 41985410
refGene_gn=ATN1;refGene_tx=XM_023256735.1;refGene_gn=ATN1;refGene_tx=XM_023256736.1;refGene_gn=ATN1;refGene_tx=XM_023256737.1
B4 41986275 41986419
refGene_gn=ATN1;refGene_tx=XM_023256735.1;refGene_gn=ATN1;refGene_tx=XM_023256736.1;refGene_gn=ATN1;refGene_tx=XM_023256737.1
B4 41986749 41986930
refGene_gn=ATN1;refGene_tx=XM_023256735.1;refGene_gn=ATN1;refGene_tx=XM_023256736.1;refGene_gn=ATN1;refGene_tx=XM_023256737.1
B4 41987128 41987162
refGene_gn=ATN1;refGene_tx=XM_023256735.1;refGene_gn=ATN1;refGene_tx=XM_023256736.1;refGene_gn=ATN1;refGene_tx=XM_023256737.1
B4 41988915 41988967

refGene_gn=CB4H12orf57;refGene_tx=XM_003988345.5
B4 41989251 41989428
refGene_gn=CB4H12orf57;refGene_tx=XM_003988345.5
B4 41990206 41990358
refGene_gn=CB4H12orf57;refGene_tx=XM_003988345.5
B4 41991125 41991139 refGene_gn=PTPN6;refGene_tx=XM_019834452.2
B4 41995624 41995632 refGene_gn=PTPN6;refGene_tx=XM_003988343.5
B4 41995718 41995841
refGene_gn=PTPN6;refGene_tx=XM_003988343.5;refGene_gn=PTPN6;refGene_tx
=XM_019834452.2
B4 41996053 41996248
refGene_gn=PTPN6;refGene_tx=XM_003988343.5;refGene_gn=PTPN6;refGene_tx
=XM_019834452.2
B4 41998792 41998982
refGene_gn=PTPN6;refGene_tx=XM_003988343.5;refGene_gn=PTPN6;refGene_tx
=XM_019834452.2
B4 41999141 41999258
refGene_gn=PTPN6;refGene_tx=XM_003988343.5;refGene_gn=PTPN6;refGene_tx
=XM_019834452.2
B4 41999344 41999458
refGene_gn=PTPN6;refGene_tx=XM_003988343.5;refGene_gn=PTPN6;refGene_tx
=XM_019834452.2
B4 41999581 41999678
refGene_gn=PTPN6;refGene_tx=XM_003988343.5;refGene_gn=PTPN6;refGene_tx
=XM_019834452.2
B4 42000068 42000148
refGene_gn=PTPN6;refGene_tx=XM_003988343.5;refGene_gn=PTPN6;refGene_tx
=XM_019834452.2
B4 42000313 42000463
refGene_gn=PTPN6;refGene_tx=XM_003988343.5;refGene_gn=PTPN6;refGene_tx
=XM_019834452.2
B4 42001826 42001958
refGene_gn=PTPN6;refGene_tx=XM_003988343.5;refGene_gn=PTPN6;refGene_tx
=XM_019834452.2
B4 42002083 42002238
refGene_gn=PTPN6;refGene_tx=XM_003988343.5;refGene_gn=PTPN6;refGene_tx
=XM_019834452.2
B4 42004271 42004339
refGene_gn=PTPN6;refGene_tx=XM_003988343.5;refGene_gn=PTPN6;refGene_tx
=XM_019834452.2
B4 42004440 42004592
refGene_gn=PTPN6;refGene_tx=XM_003988343.5;refGene_gn=PTPN6;refGene_tx
=XM_019834452.2
B4 42004690 42004782
refGene_gn=PTPN6;refGene_tx=XM_003988343.5;refGene_gn=PTPN6;refGene_tx
=XM_019834452.2
B4 42005175 42005290
refGene_gn=PTPN6;refGene_tx=XM_003988343.5;refGene_gn=PTPN6;refGene_tx
=XM_019834452.2
B4 42010418 42010446 refGene_gn=PHB2;refGene_tx=XM_003988346.5

B4 42010634 42010640 refGene_gn=PHB2;refGene_tx=XM_003988346.5
B4 42011097 42011174 refGene_gn=PHB2;refGene_tx=XM_003988346.5
B4 42011525 42011603 refGene_gn=PHB2;refGene_tx=XM_003988346.5
B4 42012023 42012127 refGene_gn=PHB2;refGene_tx=XM_003988346.5
B4 42012225 42012355 refGene_gn=PHB2;refGene_tx=XM_003988346.5
B4 42012783 42012968 refGene_gn=PHB2;refGene_tx=XM_003988346.5
B4 42014122 42014202 refGene_gn=PHB2;refGene_tx=XM_003988346.5
B4 42014513 42014598 refGene_gn=PHB2;refGene_tx=XM_003988346.5
B4 42014767 42014894 refGene_gn=PHB2;refGene_tx=XM_003988346.5
B4 42015273 42015441 refGene_gn=EMG1;refGene_tx=XM_003988347.5
B4 42019135 42019237 refGene_gn=EMG1;refGene_tx=XM_003988347.5
B4 42019346 42019488 refGene_gn=EMG1;refGene_tx=XM_003988347.5
B4 42019854 42019913 refGene_gn=EMG1;refGene_tx=XM_003988347.5
B4 42020000 42020150 refGene_gn=EMG1;refGene_tx=XM_003988347.5
B4 42020813 42020927 refGene_gn=EMG1;refGene_tx=XM_003988347.5
B4 42022045 42022162
refGene_gn=LPCAT3;refGene_tx=XM_003988348.5;refGene_gn=LPCAT3;refGene_tx=XM_023256738.1
B4 42022253 42022412
refGene_gn=LPCAT3;refGene_tx=XM_003988348.5;refGene_gn=LPCAT3;refGene_tx=XM_023256738.1
B4 42022487 42022635
refGene_gn=LPCAT3;refGene_tx=XM_003988348.5;refGene_gn=LPCAT3;refGene_tx=XM_023256738.1
B4 42023694 42023861
refGene_gn=LPCAT3;refGene_tx=XM_003988348.5;refGene_gn=LPCAT3;refGene_tx=XM_023256738.1
B4 42023960 42024047
refGene_gn=LPCAT3;refGene_tx=XM_003988348.5;refGene_gn=LPCAT3;refGene_tx=XM_023256738.1
B4 42024788 42024897
refGene_gn=LPCAT3;refGene_tx=XM_003988348.5;refGene_gn=LPCAT3;refGene_tx=XM_023256738.1
B4 42025752 42025931
refGene_gn=LPCAT3;refGene_tx=XM_003988348.5;refGene_gn=LPCAT3;refGene_tx=XM_023256738.1
B4 42026365 42026403
refGene_gn=LPCAT3;refGene_tx=XM_003988348.5;refGene_gn=LPCAT3;refGene_tx=XM_023256738.1
B4 42026568 42026662
refGene_gn=LPCAT3;refGene_tx=XM_003988348.5;refGene_gn=LPCAT3;refGene_tx=XM_023256738.1
B4 42027458 42027565
refGene_gn=LPCAT3;refGene_tx=XM_003988348.5;refGene_gn=LPCAT3;refGene_tx=XM_023256738.1
B4 42028099 42028207
refGene_gn=LPCAT3;refGene_tx=XM_003988348.5;refGene_gn=LPCAT3;refGene_tx=XM_023256738.1
B4 42061583 42061734 refGene_gn=LPCAT3;refGene_tx=XM_003988348.5
B4 42112866 42112889

refGene_gn=C1S;refGene_tx=XM_006933467.4;refGene_gn=C1S;refGene_tx=XM_006933468.4
B4 42113139 42113347
refGene_gn=C1S;refGene_tx=XM_006933467.4;refGene_gn=C1S;refGene_tx=XM_006933468.4
B4 42113478 42113656
refGene_gn=C1S;refGene_tx=XM_006933467.4;refGene_gn=C1S;refGene_tx=XM_006933468.4
B4 42114771 42114897
refGene_gn=C1S;refGene_tx=XM_006933467.4;refGene_gn=C1S;refGene_tx=XM_006933468.4
B4 42115341 42115541
refGene_gn=C1S;refGene_tx=XM_006933467.4;refGene_gn=C1S;refGene_tx=XM_006933468.4
B4 42115753 42115907
refGene_gn=C1S;refGene_tx=XM_006933467.4;refGene_gn=C1S;refGene_tx=XM_006933468.4
B4 42116346 42116462
refGene_gn=C1S;refGene_tx=XM_006933467.4;refGene_gn=C1S;refGene_tx=XM_006933468.4
B4 42116875 42116954
refGene_gn=C1S;refGene_tx=XM_006933467.4;refGene_gn=C1S;refGene_tx=XM_006933468.4
B4 42117442 42117571
refGene_gn=C1S;refGene_tx=XM_006933467.4;refGene_gn=C1S;refGene_tx=XM_006933468.4
B4 42118599 42118674
refGene_gn=C1S;refGene_tx=XM_006933467.4;refGene_gn=C1S;refGene_tx=XM_006933468.4
B4 42120003 42120803
refGene_gn=C1S;refGene_tx=XM_006933467.4;refGene_gn=C1S;refGene_tx=XM_006933468.4
B4 42141718 42142497 refGene_gn=C1R;refGene_tx=XM_023256739.1
B4 42142974 42143049 refGene_gn=C1R;refGene_tx=XM_023256739.1
B4 42144855 42145007 refGene_gn=C1R;refGene_tx=XM_023256739.1
B4 42145370 42145428 refGene_gn=C1R;refGene_tx=XM_023256739.1
B4 42145430 42145455 refGene_gn=C1R;refGene_tx=XM_023256739.1
B4 42147412 42147534 refGene_gn=C1R;refGene_tx=XM_023256739.1
B4 42147641 42147789 refGene_gn=C1R;refGene_tx=XM_023256739.1
B4 42148075 42148167 refGene_gn=C1R;refGene_tx=XM_023256739.1
B4 42148169 42148271 refGene_gn=C1R;refGene_tx=XM_023256739.1
B4 42148371 42148518 refGene_gn=C1R;refGene_tx=XM_023256739.1
B4 42148842 42149006 refGene_gn=C1R;refGene_tx=XM_023256739.1
B4 42149007 42149033 refGene_gn=C1R;refGene_tx=XM_023256739.1
B4 42150588 42150817 refGene_gn=C1R;refGene_tx=XM_023256739.1
B4 42150911 42150937 refGene_gn=C1R;refGene_tx=XM_023256739.1
B4 42156295 42157001 refGene_gn=C1RL;refGene_tx=XM_023257739.1
B4 42157002 42157081 refGene_gn=C1RL;refGene_tx=XM_023257739.1
B4 42160491 42160566 refGene_gn=C1RL;refGene_tx=XM_023257739.1
B4 42160703 42160817 refGene_gn=C1RL;refGene_tx=XM_023257739.1

B4 42162745 42162935 refGene_gn=C1RL;refGene_tx=XM_023257739.1
B4 42168879 42169107 refGene_gn=C1RL;refGene_tx=XM_023257739.1
B4 42170337 42170505 refGene_gn=C1RL;refGene_tx=XM_023257739.1
B4 42170783 42170822 refGene_gn=C1RL;refGene_tx=XM_023257739.1
B4 42196904 42196958 refGene_gn=RBP5;refGene_tx=XM_003988352.4
B4 42197190 42197295
refGene_gn=RBP5;refGene_tx=XM_019834455.2;refGene_gn=RBP5;refGene_tx=XM_019834456.2
B4 42197407 42197509
refGene_gn=RBP5;refGene_tx=XM_003988352.4;refGene_gn=RBP5;refGene_tx=XM_019834455.2;refGene_gn=RBP5;refGene_tx=XM_019834456.2
B4 42199805 42199984
refGene_gn=RBP5;refGene_tx=XM_003988352.4;refGene_gn=RBP5;refGene_tx=XM_019834455.2;refGene_gn=RBP5;refGene_tx=XM_019834456.2
B4 42200270 42200343
refGene_gn=RBP5;refGene_tx=XM_003988352.4;refGene_gn=RBP5;refGene_tx=XM_019834455.2
B4 42202442 42202506 refGene_gn=CLSTN3;refGene_tx=XM_023256740.1
B4 42204921 42205044 refGene_gn=CLSTN3;refGene_tx=XM_023256740.1
B4 42205479 42205675 refGene_gn=CLSTN3;refGene_tx=XM_023256740.1
B4 42207075 42207284 refGene_gn=CLSTN3;refGene_tx=XM_023256740.1
B4 42207556 42207706 refGene_gn=CLSTN3;refGene_tx=XM_023256740.1
B4 42207950 42208136 refGene_gn=CLSTN3;refGene_tx=XM_023256740.1
B4 42208583 42208865 refGene_gn=CLSTN3;refGene_tx=XM_023256740.1
B4 42209634 42209747 refGene_gn=CLSTN3;refGene_tx=XM_023256740.1
B4 42212707 42212870 refGene_gn=CLSTN3;refGene_tx=XM_023256740.1
B4 42213516 42213573 refGene_gn=CLSTN3;refGene_tx=XM_023256740.1
B4 42214304 42214462 refGene_gn=CLSTN3;refGene_tx=XM_023256740.1
B4 42214594 42214634 refGene_gn=CLSTN3;refGene_tx=XM_023256740.1
B4 42214636 42214742 refGene_gn=CLSTN3;refGene_tx=XM_023256740.1
B4 42219819 42220046 refGene_gn=CLSTN3;refGene_tx=XM_023256740.1
B4 42220311 42220482 refGene_gn=CLSTN3;refGene_tx=XM_023256740.1
B4 42221584 42221730 refGene_gn=CLSTN3;refGene_tx=XM_023256740.1
B4 42221970 42222106 refGene_gn=CLSTN3;refGene_tx=XM_023256740.1
B4 42228305 42228508 refGene_gn=CLSTN3;refGene_tx=XM_023256740.1
B4 42228743 42228884 refGene_gn=CLSTN3;refGene_tx=XM_023256740.1
B4 42246386 42246533
refGene_gn=PEX5;refGene_tx=XM_003988355.4;refGene_gn=PEX5;refGene_tx=XM_023256742.1;refGene_gn=PEX5;refGene_tx=XM_023256743.1;refGene_gn=PEX5;refGene_tx=XM_023256744.1;refGene_gn=PEX5;refGene_tx=XM_023256745.1;refGene_gn=PEX5;refGene_tx=XM_023256746.1;refGene_gn=PEX5;refGene_tx=XM_023256747.1
B4 42246874 42246910
refGene_gn=PEX5;refGene_tx=XM_003988355.4;refGene_gn=PEX5;refGene_tx=XM_023256742.1;refGene_gn=PEX5;refGene_tx=XM_023256743.1;refGene_gn=PEX5;refGene_tx=XM_023256745.1;refGene_gn=PEX5;refGene_tx=XM_023256746.1;refGene_gn=PEX5;refGene_tx=XM_023256747.1
B4 42247172 42247305
refGene_gn=PEX5;refGene_tx=XM_003988355.4;refGene_gn=PEX5;refGene_tx=XM_023256742.1;refGene_gn=PEX5;refGene_tx=XM_023256743.1;refGene_gn=PEX

5;refGene_tx=XM_023256744.1;refGene_gn=PEX5;refGene_tx=XM_023256745.1;
refGene_gn=PEX5;refGene_tx=XM_023256746.1;refGene_gn=PEX5;refGene_tx=X
M_023256747.1
B4 42247519 42247651
refGene_gn=PEX5;refGene_tx=XM_003988355.4;refGene_gn=PEX5;refGene_tx=X
M_023256742.1;refGene_gn=PEX5;refGene_tx=XM_023256743.1;refGene_gn=PEX
5;refGene_tx=XM_023256744.1;refGene_gn=PEX5;refGene_tx=XM_023256745.1;
refGene_gn=PEX5;refGene_tx=XM_023256746.1;refGene_gn=PEX5;refGene_tx=X
M_023256747.1
B4 42253607 42253713
refGene_gn=PEX5;refGene_tx=XM_003988355.4;refGene_gn=PEX5;refGene_tx=X
M_023256742.1;refGene_gn=PEX5;refGene_tx=XM_023256743.1;refGene_gn=PEX
5;refGene_tx=XM_023256744.1;refGene_gn=PEX5;refGene_tx=XM_023256745.1;
refGene_gn=PEX5;refGene_tx=XM_023256746.1;refGene_gn=PEX5;refGene_tx=X
M_023256747.1
B4 42254843 42254934
refGene_gn=PEX5;refGene_tx=XM_003988355.4;refGene_gn=PEX5;refGene_tx=X
M_023256742.1;refGene_gn=PEX5;refGene_tx=XM_023256743.1;refGene_gn=PEX
5;refGene_tx=XM_023256744.1;refGene_gn=PEX5;refGene_tx=XM_023256745.1;
refGene_gn=PEX5;refGene_tx=XM_023256746.1;refGene_gn=PEX5;refGene_tx=X
M_023256747.1
B4 42255332 42255443
refGene_gn=PEX5;refGene_tx=XM_003988355.4;refGene_gn=PEX5;refGene_tx=X
M_023256742.1;refGene_gn=PEX5;refGene_tx=XM_023256743.1;refGene_gn=PEX
5;refGene_tx=XM_023256744.1
B4 42255689 42255782
refGene_gn=PEX5;refGene_tx=XM_003988355.4;refGene_gn=PEX5;refGene_tx=X
M_023256742.1;refGene_gn=PEX5;refGene_tx=XM_023256743.1;refGene_gn=PEX
5;refGene_tx=XM_023256744.1;refGene_gn=PEX5;refGene_tx=XM_023256745.1;
refGene_gn=PEX5;refGene_tx=XM_023256746.1
B4 42256123 42256243
refGene_gn=PEX5;refGene_tx=XM_003988355.4;refGene_gn=PEX5;refGene_tx=X
M_023256742.1;refGene_gn=PEX5;refGene_tx=XM_023256743.1;refGene_gn=PEX
5;refGene_tx=XM_023256744.1;refGene_gn=PEX5;refGene_tx=XM_023256745.1;
refGene_gn=PEX5;refGene_tx=XM_023256746.1;refGene_gn=PEX5;refGene_tx=X
M_023256747.1
B4 42261102 42261246
refGene_gn=PEX5;refGene_tx=XM_003988355.4;refGene_gn=PEX5;refGene_tx=X
M_023256742.1;refGene_gn=PEX5;refGene_tx=XM_023256743.1;refGene_gn=PEX
5;refGene_tx=XM_023256744.1;refGene_gn=PEX5;refGene_tx=XM_023256745.1;
refGene_gn=PEX5;refGene_tx=XM_023256746.1;refGene_gn=PEX5;refGene_tx=X
M_023256747.1
B4 42261445 42261516
refGene_gn=PEX5;refGene_tx=XM_003988355.4;refGene_gn=PEX5;refGene_tx=X
M_023256742.1;refGene_gn=PEX5;refGene_tx=XM_023256743.1;refGene_gn=PEX
5;refGene_tx=XM_023256744.1;refGene_gn=PEX5;refGene_tx=XM_023256745.1;
refGene_gn=PEX5;refGene_tx=XM_023256746.1;refGene_gn=PEX5;refGene_tx=X
M_023256747.1
B4 42261909 42262122
refGene_gn=PEX5;refGene_tx=XM_003988355.4;refGene_gn=PEX5;refGene_tx=X

M_023256742.1;refGene_gn=PEX5;refGene_tx=XM_023256743.1;refGene_gn=PEX5;refGene_tx=XM_023256744.1;refGene_gn=PEX5;refGene_tx=XM_023256745.1;refGene_gn=PEX5;refGene_tx=XM_023256746.1;refGene_gn=PEX5;refGene_tx=XM_023256747.1
B4 42262696 42262862
refGene_gn=PEX5;refGene_tx=XM_003988355.4;refGene_gn=PEX5;refGene_tx=XM_023256742.1;refGene_gn=PEX5;refGene_tx=XM_023256743.1;refGene_gn=PEX5;refGene_tx=XM_023256744.1;refGene_gn=PEX5;refGene_tx=XM_023256745.1;refGene_gn=PEX5;refGene_tx=XM_023256746.1;refGene_gn=PEX5;refGene_tx=XM_023256747.1
B4 42263190 42263348
refGene_gn=PEX5;refGene_tx=XM_003988355.4;refGene_gn=PEX5;refGene_tx=XM_023256742.1;refGene_gn=PEX5;refGene_tx=XM_023256743.1;refGene_gn=PEX5;refGene_tx=XM_023256744.1;refGene_gn=PEX5;refGene_tx=XM_023256745.1;refGene_gn=PEX5;refGene_tx=XM_023256746.1;refGene_gn=PEX5;refGene_tx=XM_023256747.1
B4 42263524 42263726
refGene_gn=PEX5;refGene_tx=XM_003988355.4;refGene_gn=PEX5;refGene_tx=XM_023256742.1;refGene_gn=PEX5;refGene_tx=XM_023256743.1;refGene_gn=PEX5;refGene_tx=XM_023256744.1;refGene_gn=PEX5;refGene_tx=XM_023256745.1;refGene_gn=PEX5;refGene_tx=XM_023256746.1;refGene_gn=PEX5;refGene_tx=XM_023256747.1
B4 42313382 42313689
refGene_gn=LOC101090625;refGene_tx=XM_023257789.1
B4 42313755 42313900
refGene_gn=LOC101090625;refGene_tx=XM_023257789.1
B4 42314138 42314177
refGene_gn=LOC101090625;refGene_tx=XM_023257789.1
B4 42314179 42314246
refGene_gn=LOC101090625;refGene_tx=XM_023257789.1
B4 42344448 42344509
refGene_gn=LOC101094355;refGene_tx=XM_023257740.1
B4 42350248 42350341
refGene_gn=LOC101094355;refGene_tx=XM_023257740.1
B4 42350417 42350732
refGene_gn=LOC101094355;refGene_tx=XM_023257740.1
B4 42352121 42352436
refGene_gn=LOC101094355;refGene_tx=XM_023257740.1
B4 42357273 42357315
refGene_gn=LOC101094355;refGene_tx=XM_023257740.1
B4 42357397 42357712
refGene_gn=LOC101094355;refGene_tx=XM_023257740.1
B4 42357941 42357974
refGene_gn=LOC101094355;refGene_tx=XM_023257740.1
B4 42358545 42358683
refGene_gn=LOC101094355;refGene_tx=XM_023257740.1
B4 42359075 42359168
refGene_gn=LOC101094355;refGene_tx=XM_023257740.1
B4 42360053 42360188
refGene_gn=LOC101094355;refGene_tx=XM_023257740.1

B4	42361268	42361387
refGene_gn=LOC101094355;refGene_tx=XM_023257740.1
B4	42476624	42476631	refGene_gn=CD163;refGene_tx=XM_011283764.3
B4	42479828	42479890	refGene_gn=CD163;refGene_tx=XM_011283764.3
B4	42479892	42479966	refGene_gn=CD163;refGene_tx=XM_011283764.3
B4	42480831	42480927	refGene_gn=CD163;refGene_tx=XM_011283764.3
B4	42482164	42482272	refGene_gn=CD163;refGene_tx=XM_011283764.3
B4	42482473	42482509	refGene_gn=CD163;refGene_tx=XM_011283764.3
B4	42482868	42483177	refGene_gn=CD163;refGene_tx=XM_011283764.3
B4	42483880	42484198	refGene_gn=CD163;refGene_tx=XM_011283764.3
B4	42485253	42485568	refGene_gn=CD163;refGene_tx=XM_011283764.3
B4	42485650	42485743	refGene_gn=CD163;refGene_tx=XM_011283764.3
B4	42486258	42486573	refGene_gn=CD163;refGene_tx=XM_011283764.3
B4	42486671	42486986	refGene_gn=CD163;refGene_tx=XM_011283764.3
B4	42497165	42497436	refGene_gn=CD163;refGene_tx=XM_011283764.3
B4	42497437	42497487	refGene_gn=CD163;refGene_tx=XM_011283764.3
B4	42498191	42498512	refGene_gn=CD163;refGene_tx=XM_011283764.3
B4	42500318	42500639	refGene_gn=CD163;refGene_tx=XM_011283764.3
B4	42503615	42503783	refGene_gn=CD163;refGene_tx=XM_011283764.3
B4	42503785	42503935	refGene_gn=CD163;refGene_tx=XM_011283764.3
B4	42504931	42505018	refGene_gn=CD163;refGene_tx=XM_011283764.3
B4	42506144	42506175	refGene_gn=CD163;refGene_tx=XM_011283764.3
B4	42559624	42560344
refGene_gn=LOC102899560;refGene_tx=XM_023257790.1
B4	42625258	42625325	refGene_gn=CLEC4E;refGene_tx=XM_003988370.4
B4	42626397	42626490	refGene_gn=CLEC4E;refGene_tx=XM_003988370.4
B4	42627097	42627187	refGene_gn=CLEC4E;refGene_tx=XM_003988370.4
B4	42628602	42628754	refGene_gn=CLEC4E;refGene_tx=XM_003988370.4
B4	42629670	42629786	refGene_gn=CLEC4E;refGene_tx=XM_003988370.4
B4	42630795	42630964	refGene_gn=CLEC4E;refGene_tx=XM_003988370.4
B4	42637427	42637519	refGene_gn=CLEC4D;refGene_tx=XM_023256748.1
B4	42637521	42637577	refGene_gn=CLEC4D;refGene_tx=XM_023256748.1
B4	42637858	42637974	refGene_gn=CLEC4D;refGene_tx=XM_023256748.1
B4	42639080	42639232	refGene_gn=CLEC4D;refGene_tx=XM_023256748.1
B4	42640391	42640496	refGene_gn=CLEC4D;refGene_tx=XM_023256748.1
B4	42643181	42643271	refGene_gn=CLEC4D;refGene_tx=XM_023256748.1
B4	42644957	42644982	refGene_gn=CLEC4D;refGene_tx=XM_023256748.1
B4	42716317	42716462
refGene_gn=CLEC6A;refGene_tx=XM_019834458.2;refGene_gn=CLEC6A;refGene_tx=XM_023256749.1
B4	42718083	42718199
refGene_gn=CLEC6A;refGene_tx=XM_019834458.2;refGene_gn=CLEC6A;refGene_tx=XM_023256749.1
B4	42728532	42728684
refGene_gn=CLEC6A;refGene_tx=XM_019834458.2;refGene_gn=CLEC6A;refGene_tx=XM_023256749.1
B4	42733835	42733937
refGene_gn=CLEC6A;refGene_tx=XM_019834458.2;refGene_gn=CLEC6A;refGene_tx=XM_023256749.1
B4	42735396	42735489	refGene_gn=CLEC6A;refGene_tx=XM_019834458.2

B4	42736217	42736248
refGene_gn=CLEC6A;refGene_tx=XM_019834458.2;refGene_gn=CLEC6A;refGene_tx=XM_023256749.1
B4	42743836	42743984
refGene_gn=CLEC4A;refGene_tx=XM_003988372.5;refGene_gn=CLEC4A;refGene_tx=XM_011283789.3;refGene_gn=CLEC4A;refGene_tx=XM_011283790.3;refGene_gn=CLEC4A;refGene_tx=XM_019834461.2
B4	42745008	42745124
refGene_gn=CLEC4A;refGene_tx=XM_003988372.5;refGene_gn=CLEC4A;refGene_tx=XM_011283789.3;refGene_gn=CLEC4A;refGene_tx=XM_011283790.3;refGene_gn=CLEC4A;refGene_tx=XM_019834461.2
B4	42746215	42746367
refGene_gn=CLEC4A;refGene_tx=XM_003988372.5;refGene_gn=CLEC4A;refGene_tx=XM_011283789.3;refGene_gn=CLEC4A;refGene_tx=XM_011283790.3;refGene_gn=CLEC4A;refGene_tx=XM_019834461.2
B4	42759714	42759816
refGene_gn=CLEC4A;refGene_tx=XM_003988372.5;refGene_gn=CLEC4A;refGene_tx=XM_011283789.3;refGene_gn=CLEC4A;refGene_tx=XM_011283790.3;refGene_gn=CLEC4A;refGene_tx=XM_019834461.2
B4	42761047	42761164
refGene_gn=CLEC4A;refGene_tx=XM_003988372.5;refGene_gn=CLEC4A;refGene_tx=XM_011283789.3;refGene_gn=CLEC4A;refGene_tx=XM_011283790.3;refGene_gn=CLEC4A;refGene_tx=XM_019834461.2
B4	42765781	42765863
refGene_gn=CLEC4A;refGene_tx=XM_003988372.5;refGene_gn=CLEC4A;refGene_tx=XM_011283789.3;refGene_gn=CLEC4A;refGene_tx=XM_011283790.3;refGene_gn=CLEC4A;refGene_tx=XM_019834461.2
B4	42797787	42797836	refGene_gn=NECAP1;refGene_tx=XM_003988373.3
B4	42798131	42798234	refGene_gn=NECAP1;refGene_tx=XM_003988373.3
B4	42800865	42801049	refGene_gn=NECAP1;refGene_tx=XM_003988373.3
B4	42801137	42801246	refGene_gn=NECAP1;refGene_tx=XM_003988373.3
B4	42802100	42802182	refGene_gn=NECAP1;refGene_tx=XM_003988373.3
B4	42803099	42803204	refGene_gn=NECAP1;refGene_tx=XM_003988373.3
B4	42803616	42803717	refGene_gn=NECAP1;refGene_tx=XM_003988373.3
B4	42809444	42809539	refGene_gn=NECAP1;refGene_tx=XM_003988373.3
B4	42838658	42840086	refGene_gn=C3AR1;refGene_tx=XM_003988384.5
B4	42846303	42846392	refGene_gn=FOXJ2;refGene_tx=XM_011283788.3
B4	42847549	42847648	refGene_gn=FOXJ2;refGene_tx=XM_011283788.3
B4	42848275	42848485	refGene_gn=FOXJ2;refGene_tx=XM_011283788.3
B4	42849303	42849405	refGene_gn=FOXJ2;refGene_tx=XM_011283788.3
B4	42849784	42850189	refGene_gn=FOXJ2;refGene_tx=XM_011283788.3
B4	42853448	42853644	refGene_gn=FOXJ2;refGene_tx=XM_011283788.3
B4	42854324	42854465	refGene_gn=FOXJ2;refGene_tx=XM_011283788.3
B4	42854648	42854717	refGene_gn=FOXJ2;refGene_tx=XM_011283788.3
B4	42856760	42856835	refGene_gn=FOXJ2;refGene_tx=XM_011283788.3
B4	42858596	42858929	refGene_gn=FOXJ2;refGene_tx=XM_011283788.3
B4	42880133	42880178	refGene_gn=SLC2A3;refGene_tx=XM_019834462.2
B4	42882112	42882133
refGene_gn=SLC2A3;refGene_tx=XM_006933484.4;refGene_gn=SLC2A3;refGene_tx=XM_006933485.4;refGene_gn=SLC2A3;refGene_tx=XM_019834463.2;refGene_

gn=SLC2A3;refGene_tx=XM_019834464.2
B4 42954928 42954943 refGene_gn=SLC2A3;refGene_tx=XM_003988375.5
B4 42956510 42956603
refGene_gn=SLC2A3;refGene_tx=XM_003988375.5;refGene_gn=SLC2A3;refGene_tx=XM_006933484.4;refGene_gn=SLC2A3;refGene_tx=XM_006933485.4;refGene_gn=SLC2A3;refGene_tx=XM_019834462.2;refGene_gn=SLC2A3;refGene_tx=XM_019834463.2;refGene_gn=SLC2A3;refGene_tx=XM_019834464.2
B4 42957124 42957285
refGene_gn=SLC2A3;refGene_tx=XM_003988375.5;refGene_gn=SLC2A3;refGene_tx=XM_006933484.4;refGene_gn=SLC2A3;refGene_tx=XM_006933485.4;refGene_gn=SLC2A3;refGene_tx=XM_019834462.2;refGene_gn=SLC2A3;refGene_tx=XM_019834463.2;refGene_gn=SLC2A3;refGene_tx=XM_019834464.2
B4 42959191 42959432
refGene_gn=SLC2A3;refGene_tx=XM_003988375.5;refGene_gn=SLC2A3;refGene_tx=XM_006933484.4;refGene_gn=SLC2A3;refGene_tx=XM_006933485.4;refGene_gn=SLC2A3;refGene_tx=XM_019834462.2;refGene_gn=SLC2A3;refGene_tx=XM_019834463.2;refGene_gn=SLC2A3;refGene_tx=XM_019834464.2
B4 42959703 42959866
refGene_gn=SLC2A3;refGene_tx=XM_003988375.5;refGene_gn=SLC2A3;refGene_tx=XM_006933484.4;refGene_gn=SLC2A3;refGene_tx=XM_006933485.4;refGene_gn=SLC2A3;refGene_tx=XM_019834462.2;refGene_gn=SLC2A3;refGene_tx=XM_019834463.2;refGene_gn=SLC2A3;refGene_tx=XM_019834464.2
B4 42960571 42960759
refGene_gn=SLC2A3;refGene_tx=XM_003988375.5;refGene_gn=SLC2A3;refGene_tx=XM_006933484.4;refGene_gn=SLC2A3;refGene_tx=XM_006933485.4;refGene_gn=SLC2A3;refGene_tx=XM_019834462.2;refGene_gn=SLC2A3;refGene_tx=XM_019834463.2;refGene_gn=SLC2A3;refGene_tx=XM_019834464.2
B4 42965351 42965456
refGene_gn=SLC2A3;refGene_tx=XM_003988375.5;refGene_gn=SLC2A3;refGene_tx=XM_006933484.4;refGene_gn=SLC2A3;refGene_tx=XM_006933485.4;refGene_gn=SLC2A3;refGene_tx=XM_019834462.2;refGene_gn=SLC2A3;refGene_tx=XM_019834463.2;refGene_gn=SLC2A3;refGene_tx=XM_019834464.2
B4 42965819 42965921
refGene_gn=SLC2A3;refGene_tx=XM_003988375.5;refGene_gn=SLC2A3;refGene_tx=XM_006933484.4;refGene_gn=SLC2A3;refGene_tx=XM_006933485.4;refGene_gn=SLC2A3;refGene_tx=XM_019834462.2;refGene_gn=SLC2A3;refGene_tx=XM_019834463.2;refGene_gn=SLC2A3;refGene_tx=XM_019834464.2
B4 42966392 42966596
refGene_gn=SLC2A3;refGene_tx=XM_003988375.5;refGene_gn=SLC2A3;refGene_tx=XM_006933484.4;refGene_gn=SLC2A3;refGene_tx=XM_006933485.4;refGene_gn=SLC2A3;refGene_tx=XM_019834462.2;refGene_gn=SLC2A3;refGene_tx=XM_019834463.2;refGene_gn=SLC2A3;refGene_tx=XM_019834464.2
B4 42968037 42968256
refGene_gn=SLC2A3;refGene_tx=XM_003988375.5;refGene_gn=SLC2A3;refGene_tx=XM_006933484.4;refGene_gn=SLC2A3;refGene_tx=XM_006933485.4;refGene_gn=SLC2A3;refGene_tx=XM_019834462.2;refGene_gn=SLC2A3;refGene_tx=XM_019834463.2;refGene_gn=SLC2A3;refGene_tx=XM_019834464.2
B4 42996380 42996812 refGene_gn=NANOG;refGene_tx=NM_001173442.1
B4 42997300 42997387 refGene_gn=NANOG;refGene_tx=NM_001173442.1
B4 42998827 42999093 refGene_gn=NANOG;refGene_tx=NM_001173442.1

B4	43001253 43001404 refGene_gn=NANOG;refGene_tx=NM_001173442.1
B4	43039500 43039507 refGene_gn=NANOGNB;refGene_tx=XM_019835817.1
B4	43040553 43040633 refGene_gn=NANOGNB;refGene_tx=XM_019835817.1
B4	43040743 43041060 refGene_gn=NANOGNB;refGene_tx=XM_019835817.1
B4	43045647 43045708 refGene_gn=NANOGNB;refGene_tx=XM_019835817.1
B4	43123502 43123610 refGene_gn=DPPA3;refGene_tx=XM_023256750.1
B4	43124739 43125002 refGene_gn=DPPA3;refGene_tx=XM_023256750.1
B4	43127459 43127544 refGene_gn=DPPA3;refGene_tx=XM_023256750.1
B4	43155637 43155905 refGene_gn=GDF3;refGene_tx=XM_003988376.3
B4	43162171 43163001 refGene_gn=GDF3;refGene_tx=XM_003988376.3
B4	43166469 43166514 refGene_gn=APOBEC1;refGene_tx=XM_023257742.1
B4	43184346 43184794 refGene_gn=APOBEC1;refGene_tx=XM_023257742.1
B4	43185624 43185743 refGene_gn=APOBEC1;refGene_tx=XM_023257742.1
B4	43186609 43186759 refGene_gn=APOBEC1;refGene_tx=XM_023257742.1
B4	43209304 43209358 refGene_gn=AICDA;refGene_tx=XM_023256751.1
B4	43209778 43209894 refGene_gn=AICDA;refGene_tx=XM_023256751.1
B4	43210164 43210435 refGene_gn=AICDA;refGene_tx=XM_023256751.1
B4	43212507 43212655 refGene_gn=AICDA;refGene_tx=XM_023256751.1
B4	43219340 43219348 refGene_gn=AICDA;refGene_tx=XM_023256751.1
B4	43243276 43243389
refGene_gn=MFAP5;refGene_tx=XM_003988377.4;refGene_gn=MFAP5;refGene_tx
=XM_006933487.3;refGene_gn=MFAP5;refGene_tx=XM_006933488.3;refGene_gn=
MFAP5;refGene_tx=XM_006933490.3;refGene_gn=MFAP5;refGene_tx=XM_0112837
87.2
B4	43245037 43245179
refGene_gn=MFAP5;refGene_tx=XM_003988377.4;refGene_gn=MFAP5;refGene_tx
=XM_006933487.3;refGene_gn=MFAP5;refGene_tx=XM_006933488.3;refGene_gn=
MFAP5;refGene_tx=XM_006933489.3;refGene_gn=MFAP5;refGene_tx=XM_0069334
90.3;refGene_gn=MFAP5;refGene_tx=XM_011283787.2
B4	43245706 43245794
refGene_gn=MFAP5;refGene_tx=XM_003988377.4;refGene_gn=MFAP5;refGene_tx
=XM_006933487.3;refGene_gn=MFAP5;refGene_tx=XM_006933488.3;refGene_gn=
MFAP5;refGene_tx=XM_006933489.3;refGene_gn=MFAP5;refGene_tx=XM_0069334
90.3;refGene_gn=MFAP5;refGene_tx=XM_011283787.2
B4	43246514 43246544
refGene_gn=MFAP5;refGene_tx=XM_003988377.4;refGene_gn=MFAP5;refGene_tx
=XM_006933487.3;refGene_gn=MFAP5;refGene_tx=XM_006933488.3;refGene_gn=
MFAP5;refGene_tx=XM_006933489.3;refGene_gn=MFAP5;refGene_tx=XM_0069334
90.3;refGene_gn=MFAP5;refGene_tx=XM_011283787.2
B4	43250315 43250351
refGene_gn=MFAP5;refGene_tx=XM_003988377.4;refGene_gn=MFAP5;refGene_tx
=XM_006933487.3;refGene_gn=MFAP5;refGene_tx=XM_006933489.3;refGene_gn=
MFAP5;refGene_tx=XM_011283787.2
B4	43250771 43250804
refGene_gn=MFAP5;refGene_tx=XM_003988377.4;refGene_gn=MFAP5;refGene_tx
=XM_006933488.3;refGene_gn=MFAP5;refGene_tx=XM_006933489.3;refGene_gn=
MFAP5;refGene_tx=XM_011283787.2
B4	43251053 43251098
refGene_gn=MFAP5;refGene_tx=XM_003988377.4;refGene_gn=MFAP5;refGene_tx
=XM_006933487.3;refGene_gn=MFAP5;refGene_tx=XM_006933488.3;refGene_gn=

MFAP5;refGene_tx=XM_006933489.3;refGene_gn=MFAP5;refGene_tx=XM_0069334
90.3;refGene_gn=MFAP5;refGene_tx=XM_011283787.2
B4 43255034 43255070
refGene_gn=MFAP5;refGene_tx=XM_003988377.4;refGene_gn=MFAP5;refGene_tx
=XM_006933487.3;refGene_gn=MFAP5;refGene_tx=XM_006933488.3;refGene_gn=
MFAP5;refGene_tx=XM_006933489.3;refGene_gn=MFAP5;refGene_tx=XM_0069334
90.3;refGene_gn=MFAP5;refGene_tx=XM_011283787.2
B4 43256369 43256427
refGene_gn=MFAP5;refGene_tx=XM_003988377.4;refGene_gn=MFAP5;refGene_tx
=XM_006933487.3;refGene_gn=MFAP5;refGene_tx=XM_006933488.3;refGene_gn=
MFAP5;refGene_tx=XM_006933489.3;refGene_gn=MFAP5;refGene_tx=XM_0069334
90.3;refGene_gn=MFAP5;refGene_tx=XM_011283787.2
B4 43307257 43307432
refGene_gn=RIMKLB;refGene_tx=XM_019834465.2;refGene_gn=RIMKLB;refGene_
tx=XM_019834467.2;refGene_gn=RIMKLB;refGene_tx=XM_023256752.1;refGene_
gn=RIMKLB;refGene_tx=XM_023256753.1;refGene_gn=RIMKLB;refGene_tx=XM_02
3256754.1;refGene_gn=RIMKLB;refGene_tx=XM_023256755.1;refGene_gn=RIMKL
B;refGene_tx=XM_023256756.1
B4 43342955 43343186
refGene_gn=RIMKLB;refGene_tx=XM_019834465.2;refGene_gn=RIMKLB;refGene_
tx=XM_019834467.2;refGene_gn=RIMKLB;refGene_tx=XM_023256752.1;refGene_
gn=RIMKLB;refGene_tx=XM_023256753.1;refGene_gn=RIMKLB;refGene_tx=XM_02
3256754.1;refGene_gn=RIMKLB;refGene_tx=XM_023256755.1;refGene_gn=RIMKL
B;refGene_tx=XM_023256756.1
B4 43345064 43345151
refGene_gn=RIMKLB;refGene_tx=XM_019834465.2;refGene_gn=RIMKLB;refGene_
tx=XM_019834467.2;refGene_gn=RIMKLB;refGene_tx=XM_023256752.1;refGene_
gn=RIMKLB;refGene_tx=XM_023256753.1;refGene_gn=RIMKLB;refGene_tx=XM_02
3256754.1;refGene_gn=RIMKLB;refGene_tx=XM_023256755.1;refGene_gn=RIMKL
B;refGene_tx=XM_023256756.1
B4 43346223 43346427
refGene_gn=RIMKLB;refGene_tx=XM_019834465.2;refGene_gn=RIMKLB;refGene_
tx=XM_019834467.2;refGene_gn=RIMKLB;refGene_tx=XM_023256752.1;refGene_
gn=RIMKLB;refGene_tx=XM_023256753.1;refGene_gn=RIMKLB;refGene_tx=XM_02
3256754.1;refGene_gn=RIMKLB;refGene_tx=XM_023256755.1;refGene_gn=RIMKL
B;refGene_tx=XM_023256756.1
B4 43350454 43350918
refGene_gn=RIMKLB;refGene_tx=XM_019834465.2;refGene_gn=RIMKLB;refGene_
tx=XM_019834467.2;refGene_gn=RIMKLB;refGene_tx=XM_023256752.1;refGene_
gn=RIMKLB;refGene_tx=XM_023256753.1;refGene_gn=RIMKLB;refGene_tx=XM_02
3256754.1;refGene_gn=RIMKLB;refGene_tx=XM_023256755.1;refGene_gn=RIMKL
B;refGene_tx=XM_023256756.1
B4 43401506 43401571
refGene_gn=A2ML1;refGene_tx=XM_019834470.1;refGene_gn=A2ML1;refGene_tx
=XM_023256757.1
B4 43402314 43402498
refGene_gn=A2ML1;refGene_tx=XM_019834470.1;refGene_gn=A2ML1;refGene_tx
=XM_023256757.1
B4 43402843 43403006
refGene_gn=A2ML1;refGene_tx=XM_019834470.1;refGene_gn=A2ML1;refGene_tx

=XM_023256757.1
B4	43403600	43403653
refGene_gn=A2ML1;refGene_tx=XM_019834470.1;refGene_gn=A2ML1;refGene_tx
=XM_023256757.1
B4	43411345	43411366
refGene_gn=A2ML1;refGene_tx=XM_019834470.1;refGene_gn=A2ML1;refGene_tx
=XM_023256757.1
B4	43412210	43412370
refGene_gn=A2ML1;refGene_tx=XM_019834470.1;refGene_gn=A2ML1;refGene_tx
=XM_023256757.1
B4	43412638	43412723
refGene_gn=A2ML1;refGene_tx=XM_019834470.1;refGene_gn=A2ML1;refGene_tx
=XM_023256757.1
B4	43413654	43413781
refGene_gn=A2ML1;refGene_tx=XM_019834470.1;refGene_gn=A2ML1;refGene_tx
=XM_023256757.1
B4	43414223	43414338
refGene_gn=A2ML1;refGene_tx=XM_019834470.1;refGene_gn=A2ML1;refGene_tx
=XM_023256757.1
B4	43414663	43414773
refGene_gn=A2ML1;refGene_tx=XM_019834470.1;refGene_gn=A2ML1;refGene_tx
=XM_023256757.1
B4	43416302	43416470
refGene_gn=A2ML1;refGene_tx=XM_019834470.1;refGene_gn=A2ML1;refGene_tx
=XM_023256757.1
B4	43419243	43419471
refGene_gn=A2ML1;refGene_tx=XM_019834470.1;refGene_gn=A2ML1;refGene_tx
=XM_023256757.1
B4	43423601	43423662
refGene_gn=A2ML1;refGene_tx=XM_019834470.1;refGene_gn=A2ML1;refGene_tx
=XM_023256757.1
B4	43423893	43424039
refGene_gn=A2ML1;refGene_tx=XM_019834470.1;refGene_gn=A2ML1;refGene_tx
=XM_023256757.1
B4	43424779	43424929
refGene_gn=A2ML1;refGene_tx=XM_019834470.1;refGene_gn=A2ML1;refGene_tx
=XM_023256757.1
B4	43425578	43425737
refGene_gn=A2ML1;refGene_tx=XM_019834470.1;refGene_gn=A2ML1;refGene_tx
=XM_023256757.1
B4	43426899	43426993
refGene_gn=A2ML1;refGene_tx=XM_019834470.1;refGene_gn=A2ML1;refGene_tx
=XM_023256757.1
B4	43427575	43427696
refGene_gn=A2ML1;refGene_tx=XM_019834470.1;refGene_gn=A2ML1;refGene_tx
=XM_023256757.1
B4	43428806	43429035
refGene_gn=A2ML1;refGene_tx=XM_019834470.1;refGene_gn=A2ML1;refGene_tx
=XM_023256757.1
B4	43429159	43429286

refGene_gn=A2ML1;refGene_tx=XM_019834470.1;refGene_gn=A2ML1;refGene_tx=XM_023256757.1
B4 43430483 43430605 refGene_gn=A2ML1;refGene_tx=XM_019834470.1;refGene_gn=A2ML1;refGene_tx=XM_023256757.1
B4 43431729 43431781 refGene_gn=A2ML1;refGene_tx=XM_019834470.1;refGene_gn=A2ML1;refGene_tx=XM_023256757.1
B4 43432147 43432231 refGene_gn=A2ML1;refGene_tx=XM_019834470.1;refGene_gn=A2ML1;refGene_tx=XM_023256757.1
B4 43434135 43434312 refGene_gn=A2ML1;refGene_tx=XM_019834470.1;refGene_gn=A2ML1;refGene_tx=XM_023256757.1
B4 43434477 43434559 refGene_gn=A2ML1;refGene_tx=XM_019834470.1;refGene_gn=A2ML1;refGene_tx=XM_023256757.1
B4 43435161 43435318 refGene_gn=A2ML1;refGene_tx=XM_019834470.1;refGene_gn=A2ML1;refGene_tx=XM_023256757.1
B4 43437055 43437130 refGene_gn=A2ML1;refGene_tx=XM_019834470.1;refGene_gn=A2ML1;refGene_tx=XM_023256757.1
B4 43437301 43437464 refGene_gn=A2ML1;refGene_tx=XM_019834470.1;refGene_gn=A2ML1;refGene_tx=XM_023256757.1
B4 43439543 43439758 refGene_gn=A2ML1;refGene_tx=XM_019834470.1;refGene_gn=A2ML1;refGene_tx=XM_023256757.1
B4 43443536 43443752 refGene_gn=A2ML1;refGene_tx=XM_019834470.1;refGene_gn=A2ML1;refGene_tx=XM_023256757.1
B4 43444171 43444299 refGene_gn=A2ML1;refGene_tx=XM_019834470.1;refGene_gn=A2ML1;refGene_tx=XM_023256757.1
B4 43444458 43444549 refGene_gn=A2ML1;refGene_tx=XM_019834470.1;refGene_gn=A2ML1;refGene_tx=XM_023256757.1
B4 43444901 43444970 refGene_gn=A2ML1;refGene_tx=XM_019834470.1;refGene_gn=A2ML1;refGene_tx=XM_023256757.1
B4 43446311 43446414 refGene_gn=A2ML1;refGene_tx=XM_019834470.1;refGene_gn=A2ML1;refGene_tx=XM_023256757.1
B4 43446854 43446895 refGene_gn=A2ML1;refGene_tx=XM_019834470.1;refGene_gn=A2ML1;refGene_tx=XM_023256757.1
B4 43481600 43481714 refGene_gn=PHC1;refGene_tx=XM_003988379.4
B4 43483391 43483502 refGene_gn=PHC1;refGene_tx=XM_003988379.4
B4 43484292 43484370 refGene_gn=PHC1;refGene_tx=XM_003988379.4

B4	43484916 43485066 refGene_gn=PHC1;refGene_tx=XM_003988379.4
B4	43485666 43485822 refGene_gn=PHC1;refGene_tx=XM_003988379.4
B4	43492667 43493163 refGene_gn=PHC1;refGene_tx=XM_003988379.4
B4	43494992 43495621 refGene_gn=PHC1;refGene_tx=XM_003988379.4
B4	43495622 43495658 refGene_gn=PHC1;refGene_tx=XM_003988379.4
B4	43495659 43495797 refGene_gn=PHC1;refGene_tx=XM_003988379.4
B4	43496819 43496967 refGene_gn=PHC1;refGene_tx=XM_003988379.4
B4	43497215 43497427 refGene_gn=PHC1;refGene_tx=XM_003988379.4
B4	43498005 43498120 refGene_gn=PHC1;refGene_tx=XM_003988379.4
B4	43499502 43499611 refGene_gn=PHC1;refGene_tx=XM_003988379.4
B4	43499843 43499994 refGene_gn=PHC1;refGene_tx=XM_003988379.4
B4	43500450 43500682 refGene_gn=PHC1;refGene_tx=XM_003988379.4
B4	43501809 43501964 refGene_gn=PHC1;refGene_tx=XM_003988379.4
B4	43504377 43504500
refGene_gn=M6PR;refGene_tx=XM_003988381.5;refGene_gn=M6PR;refGene_tx=XM_006933495.4;refGene_gn=M6PR;refGene_tx=XM_006933496.2;refGene_gn=M6PR;refGene_tx=XM_019834472.1
B4	43504981 43505108
refGene_gn=M6PR;refGene_tx=XM_003988381.5;refGene_gn=M6PR;refGene_tx=XM_006933495.4;refGene_gn=M6PR;refGene_tx=XM_006933496.2;refGene_gn=M6PR;refGene_tx=XM_019834472.1
B4	43505654 43505785
refGene_gn=M6PR;refGene_tx=XM_003988381.5;refGene_gn=M6PR;refGene_tx=XM_006933495.4;refGene_gn=M6PR;refGene_tx=XM_006933496.2;refGene_gn=M6PR;refGene_tx=XM_019834472.1
B4	43506065 43506175
refGene_gn=M6PR;refGene_tx=XM_003988381.5;refGene_gn=M6PR;refGene_tx=XM_006933495.4;refGene_gn=M6PR;refGene_tx=XM_006933496.2;refGene_gn=M6PR;refGene_tx=XM_019834472.1
B4	43508007 43508174
refGene_gn=M6PR;refGene_tx=XM_003988381.5;refGene_gn=M6PR;refGene_tx=XM_006933495.4;refGene_gn=M6PR;refGene_tx=XM_006933496.2;refGene_gn=M6PR;refGene_tx=XM_019834472.1
B4	43508960 43509139
refGene_gn=M6PR;refGene_tx=XM_003988381.5;refGene_gn=M6PR;refGene_tx=XM_006933495.4;refGene_gn=M6PR;refGene_tx=XM_006933496.2;refGene_gn=M6PR;refGene_tx=XM_019834472.1
B4	43532260 43532342 refGene_gn=KLRG1;refGene_tx=XM_023256758.1
B4	43536165 43536270 refGene_gn=KLRG1;refGene_tx=XM_023256758.1
B4	43538705 43538866 refGene_gn=KLRG1;refGene_tx=XM_023256758.1
B4	43549393 43549494 refGene_gn=KLRG1;refGene_tx=XM_023256758.1
B4	43550329 43550441 refGene_gn=KLRG1;refGene_tx=XM_023256758.1
B4	43585249 43585266 refGene_gn=A2M;refGene_tx=XM_006933497.3
B4	43585602 43585644 refGene_gn=A2M;refGene_tx=XM_006933497.3
B4	43586342 43586445 refGene_gn=A2M;refGene_tx=XM_006933497.3
B4	43587946 43588015 refGene_gn=A2M;refGene_tx=XM_006933497.3
B4	43588664 43588755 refGene_gn=A2M;refGene_tx=XM_006933497.3
B4	43589839 43589967 refGene_gn=A2M;refGene_tx=XM_006933497.3
B4	43590391 43590610 refGene_gn=A2M;refGene_tx=XM_006933497.3
B4	43591938 43592162 refGene_gn=A2M;refGene_tx=XM_006933497.3

B4 43593714 43593895 refGene_gn=A2M;refGene_tx=XM_006933497.3
B4 43594580 43594655 refGene_gn=A2M;refGene_tx=XM_006933497.3
B4 43594930 43595087 refGene_gn=A2M;refGene_tx=XM_006933497.3
B4 43596569 43596657 refGene_gn=A2M;refGene_tx=XM_006933497.3
B4 43597411 43597588 refGene_gn=A2M;refGene_tx=XM_006933497.3
B4 43597872 43597956 refGene_gn=A2M;refGene_tx=XM_006933497.3
B4 43602649 43602701 refGene_gn=A2M;refGene_tx=XM_006933497.3
B4 43603031 43603153 refGene_gn=A2M;refGene_tx=XM_006933497.3
B4 43603733 43603854 refGene_gn=A2M;refGene_tx=XM_006933497.3
B4 43604574 43604803 refGene_gn=A2M;refGene_tx=XM_006933497.3
B4 43605478 43605599 refGene_gn=A2M;refGene_tx=XM_006933497.3
B4 43607933 43608042 refGene_gn=A2M;refGene_tx=XM_006933497.3
B4 43608666 43608834 refGene_gn=A2M;refGene_tx=XM_006933497.3
B4 43611010 43611160 refGene_gn=A2M;refGene_tx=XM_006933497.3
B4 43611738 43611881 refGene_gn=A2M;refGene_tx=XM_006933497.3
B4 43613115 43613179 refGene_gn=A2M;refGene_tx=XM_006933497.3
B4 43613428 43613656 refGene_gn=A2M;refGene_tx=XM_006933497.3
B4 43618480 43618642 refGene_gn=A2M;refGene_tx=XM_006933497.3
B4 43620484 43620594 refGene_gn=A2M;refGene_tx=XM_006933497.3
B4 43621043 43621158 refGene_gn=A2M;refGene_tx=XM_006933497.3
B4 43621912 43622033 refGene_gn=A2M;refGene_tx=XM_006933497.3
B4 43623611 43623696 refGene_gn=A2M;refGene_tx=XM_006933497.3
B4 43624133 43624302 refGene_gn=A2M;refGene_tx=XM_006933497.3
B4 43624578 43624599 refGene_gn=A2M;refGene_tx=XM_006933497.3
B4 43626519 43626572 refGene_gn=A2M;refGene_tx=XM_006933497.3
B4 43626740 43626900 refGene_gn=A2M;refGene_tx=XM_006933497.3
B4 43627994 43628178 refGene_gn=A2M;refGene_tx=XM_006933497.3
B4 43630459 43630545 refGene_gn=A2M;refGene_tx=XM_006933497.3
B4 43663229 43663252 refGene_gn=PZP;refGene_tx=XM_023256759.1
B4 43663808 43663850 refGene_gn=PZP;refGene_tx=XM_023256759.1
B4 43664327 43664430 refGene_gn=PZP;refGene_tx=XM_023256759.1
B4 43665184 43665253
refGene_gn=PZP;refGene_tx=XM_023256759.1;refGene_gn=PZP;refGene_tx=XM_
023256760.1
B4 43666853 43666944 refGene_gn=PZP;refGene_tx=XM_023256759.1
B4 43668204 43668332
refGene_gn=PZP;refGene_tx=XM_023256759.1;refGene_gn=PZP;refGene_tx=XM_
023256760.1
B4 43668496 43668715
refGene_gn=PZP;refGene_tx=XM_023256759.1;refGene_gn=PZP;refGene_tx=XM_
023256760.1
B4 43670302 43670526
refGene_gn=PZP;refGene_tx=XM_023256759.1;refGene_gn=PZP;refGene_tx=XM_
023256760.1
B4 43671429 43671610
refGene_gn=PZP;refGene_tx=XM_023256759.1;refGene_gn=PZP;refGene_tx=XM_
023256760.1
B4 43672024 43672099
refGene_gn=PZP;refGene_tx=XM_023256759.1;refGene_gn=PZP;refGene_tx=XM_
023256760.1

B4 43672662 43672819
refGene_gn=PZP;refGene_tx=XM_023256759.1;refGene_gn=PZP;refGene_tx=XM_023256760.1
B4 43674178 43674266
refGene_gn=PZP;refGene_tx=XM_023256759.1;refGene_gn=PZP;refGene_tx=XM_023256760.1
B4 43674543 43674720
refGene_gn=PZP;refGene_tx=XM_023256759.1;refGene_gn=PZP;refGene_tx=XM_023256760.1
B4 43674996 43675080
refGene_gn=PZP;refGene_tx=XM_023256759.1;refGene_gn=PZP;refGene_tx=XM_023256760.1
B4 43676280 43676332
refGene_gn=PZP;refGene_tx=XM_023256759.1;refGene_gn=PZP;refGene_tx=XM_023256760.1
B4 43677013 43677135
refGene_gn=PZP;refGene_tx=XM_023256759.1;refGene_gn=PZP;refGene_tx=XM_023256760.1
B4 43677559 43677686
refGene_gn=PZP;refGene_tx=XM_023256759.1;refGene_gn=PZP;refGene_tx=XM_023256760.1
B4 43678576 43678805
refGene_gn=PZP;refGene_tx=XM_023256759.1;refGene_gn=PZP;refGene_tx=XM_023256760.1
B4 43679451 43679587
refGene_gn=PZP;refGene_tx=XM_023256759.1;refGene_gn=PZP;refGene_tx=XM_023256760.1
B4 43684853 43684956
refGene_gn=PZP;refGene_tx=XM_023256759.1;refGene_gn=PZP;refGene_tx=XM_023256760.1
B4 43685634 43685799
refGene_gn=PZP;refGene_tx=XM_023256759.1;refGene_gn=PZP;refGene_tx=XM_023256760.1
B4 43689950 43690100
refGene_gn=PZP;refGene_tx=XM_023256759.1;refGene_gn=PZP;refGene_tx=XM_023256760.1
B4 43690690 43690833
refGene_gn=PZP;refGene_tx=XM_023256759.1;refGene_gn=PZP;refGene_tx=XM_023256760.1
B4 43694293 43694357
refGene_gn=PZP;refGene_tx=XM_023256759.1;refGene_gn=PZP;refGene_tx=XM_023256760.1
B4 43694609 43694837
refGene_gn=PZP;refGene_tx=XM_023256759.1;refGene_gn=PZP;refGene_tx=XM_023256760.1
B4 43696444 43696606
refGene_gn=PZP;refGene_tx=XM_023256759.1;refGene_gn=PZP;refGene_tx=XM_023256760.1
B4 43697470 43697580
refGene_gn=PZP;refGene_tx=XM_023256759.1;refGene_gn=PZP;refGene_tx=XM_

023256760.1
B4 43697730 43697845
refGene_gn=PZP;refGene_tx=XM_023256759.1;refGene_gn=PZP;refGene_tx=XM_
023256760.1
B4 43698164 43698276
refGene_gn=PZP;refGene_tx=XM_023256759.1;refGene_gn=PZP;refGene_tx=XM_
023256760.1
B4 43704746 43704831
refGene_gn=PZP;refGene_tx=XM_023256759.1;refGene_gn=PZP;refGene_tx=XM_
023256760.1
B4 43705446 43705615
refGene_gn=PZP;refGene_tx=XM_023256759.1;refGene_gn=PZP;refGene_tx=XM_
023256760.1
B4 43705873 43705894
refGene_gn=PZP;refGene_tx=XM_023256759.1;refGene_gn=PZP;refGene_tx=XM_
023256760.1
B4 43706916 43706969
refGene_gn=PZP;refGene_tx=XM_023256759.1;refGene_gn=PZP;refGene_tx=XM_
023256760.1
B4 43707143 43707303
refGene_gn=PZP;refGene_tx=XM_023256759.1;refGene_gn=PZP;refGene_tx=XM_
023256760.1
B4 43708690 43708874
refGene_gn=PZP;refGene_tx=XM_023256759.1;refGene_gn=PZP;refGene_tx=XM_
023256760.1
B4 43713870 43713956
refGene_gn=PZP;refGene_tx=XM_023256759.1;refGene_gn=PZP;refGene_tx=XM_
023256760.1
B4 43728963 43728986
refGene_gn=LOC101080976;refGene_tx=XM_019834473.2
B4 43729886 43729928
refGene_gn=LOC101080976;refGene_tx=XM_019834473.2
B4 43730308 43730411
refGene_gn=LOC101080976;refGene_tx=XM_019834473.2
B4 43730743 43730812
refGene_gn=LOC101080976;refGene_tx=XM_019834473.2
B4 43731497 43731588
refGene_gn=LOC101080976;refGene_tx=XM_019834473.2
B4 43731962 43732093
refGene_gn=LOC101080976;refGene_tx=XM_019834473.2
B4 43732246 43732465
refGene_gn=LOC101080976;refGene_tx=XM_019834473.2
B4 43733299 43733523
refGene_gn=LOC101080976;refGene_tx=XM_019834473.2
B4 43733936 43734120
refGene_gn=LOC101080976;refGene_tx=XM_019834473.2
B4 43734408 43734483
refGene_gn=LOC101080976;refGene_tx=XM_019834473.2
B4 43734868 43735025
refGene_gn=LOC101080976;refGene_tx=XM_019834473.2

B4 43735675 43735763
refGene_gn=LOC101080976;refGene_tx=XM_019834473.2
B4 43736053 43736230
refGene_gn=LOC101080976;refGene_tx=XM_019834473.2
B4 43736752 43736836
refGene_gn=LOC101080976;refGene_tx=XM_019834473.2
B4 43739500 43739552
refGene_gn=LOC101080976;refGene_tx=XM_019834473.2
B4 43739774 43739896
refGene_gn=LOC101080976;refGene_tx=XM_019834473.2
B4 43740106 43740233
refGene_gn=LOC101080976;refGene_tx=XM_019834473.2
B4 43740538 43740767
refGene_gn=LOC101080976;refGene_tx=XM_019834473.2
B4 43741149 43741309
refGene_gn=LOC101080976;refGene_tx=XM_019834473.2
B4 43744532 43744623
refGene_gn=LOC101080976;refGene_tx=XM_019834473.2
B4 43746784 43746949
refGene_gn=LOC101080976;refGene_tx=XM_019834473.2
B4 43750238 43750388
refGene_gn=LOC101080976;refGene_tx=XM_019834473.2
B4 43754082 43754225
refGene_gn=LOC101080976;refGene_tx=XM_019834473.2
B4 43758595 43758659
refGene_gn=LOC101080976;refGene_tx=XM_019834473.2
B4 43760390 43760618
refGene_gn=LOC101080976;refGene_tx=XM_019834473.2
B4 43761433 43761595
refGene_gn=LOC101080976;refGene_tx=XM_019834473.2
B4 43764204 43764314
refGene_gn=LOC101080976;refGene_tx=XM_019834473.2
B4 43766131 43766246
refGene_gn=LOC101080976;refGene_tx=XM_019834473.2
B4 43768206 43768330
refGene_gn=LOC101080976;refGene_tx=XM_019834473.2
B4 43770679 43770764
refGene_gn=LOC101080976;refGene_tx=XM_019834473.2
B4 43772026 43772195
refGene_gn=LOC101080976;refGene_tx=XM_019834473.2
B4 43772668 43772689
refGene_gn=LOC101080976;refGene_tx=XM_019834473.2
B4 43773977 43774030
refGene_gn=LOC101080976;refGene_tx=XM_019834473.2
B4 43774188 43774342
refGene_gn=LOC101080976;refGene_tx=XM_019834473.2
B4 43775051 43775235
refGene_gn=LOC101080976;refGene_tx=XM_019834473.2
B4 43777573 43777656
refGene_gn=LOC101080976;refGene_tx=XM_019834473.2

B4	43806741	43807803
refGene_gn=LOC101081229;refGene_tx=XM_019834474.2
B4	45851867	45851920
refGene_gn=LOC101081475;refGene_tx=XM_019834475.1;refGene_gn=LOC101081475;refGene_tx=XM_019834476.1
B4	45852625	45852728
refGene_gn=LOC101081475;refGene_tx=XM_019834475.1;refGene_gn=LOC101081475;refGene_tx=XM_019834476.1
B4	45854230	45854299
refGene_gn=LOC101081475;refGene_tx=XM_019834475.1;refGene_gn=LOC101081475;refGene_tx=XM_019834476.1
B4	45855121	45855212
refGene_gn=LOC101081475;refGene_tx=XM_019834475.1;refGene_gn=LOC101081475;refGene_tx=XM_019834476.1
B4	45856547	45856672
refGene_gn=LOC101081475;refGene_tx=XM_019834475.1;refGene_gn=LOC101081475;refGene_tx=XM_019834476.1
B4	45857342	45857552
refGene_gn=LOC101081475;refGene_tx=XM_019834475.1;refGene_gn=LOC101081475;refGene_tx=XM_019834476.1
B4	45858737	45858949
refGene_gn=LOC101081475;refGene_tx=XM_019834475.1;refGene_gn=LOC101081475;refGene_tx=XM_019834476.1
B4	45859968	45860134
refGene_gn=LOC101081475;refGene_tx=XM_019834475.1;refGene_gn=LOC101081475;refGene_tx=XM_019834476.1
B4	45860862	45860937
refGene_gn=LOC101081475;refGene_tx=XM_019834475.1;refGene_gn=LOC101081475;refGene_tx=XM_019834476.1
B4	45863880	45864037
refGene_gn=LOC101081475;refGene_tx=XM_019834475.1;refGene_gn=LOC101081475;refGene_tx=XM_019834476.1
B4	45864674	45864756
refGene_gn=LOC101081475;refGene_tx=XM_019834475.1;refGene_gn=LOC101081475;refGene_tx=XM_019834476.1
B4	45865553	45865730
refGene_gn=LOC101081475;refGene_tx=XM_019834475.1;refGene_gn=LOC101081475;refGene_tx=XM_019834476.1
B4	45868367	45868451
refGene_gn=LOC101081475;refGene_tx=XM_019834475.1;refGene_gn=LOC101081475;refGene_tx=XM_019834476.1
B4	45869212	45869264
refGene_gn=LOC101081475;refGene_tx=XM_019834475.1;refGene_gn=LOC101081475;refGene_tx=XM_019834476.1
B4	45870437	45870556
refGene_gn=LOC101081475;refGene_tx=XM_019834475.1;refGene_gn=LOC101081475;refGene_tx=XM_019834476.1
B4	45872862	45872992
refGene_gn=LOC101081475;refGene_tx=XM_019834475.1;refGene_gn=LOC101081475;refGene_tx=XM_019834476.1

B4 45873242 45873471
refGene_gn=LOC101081475;refGene_tx=XM_019834475.1;refGene_gn=LOC101081475;refGene_tx=XM_019834476.1
B4 45874135 45874241
refGene_gn=LOC101081475;refGene_tx=XM_019834475.1;refGene_gn=LOC101081475;refGene_tx=XM_019834476.1
B4 45876927 45877072
refGene_gn=LOC101081475;refGene_tx=XM_019834475.1;refGene_gn=LOC101081475;refGene_tx=XM_019834476.1
B4 45878063 45878234
refGene_gn=LOC101081475;refGene_tx=XM_019834475.1;refGene_gn=LOC101081475;refGene_tx=XM_019834476.1
B4 45887182 45887332
refGene_gn=LOC101081475;refGene_tx=XM_019834475.1;refGene_gn=LOC101081475;refGene_tx=XM_019834476.1
B4 45887715 45887861
refGene_gn=LOC101081475;refGene_tx=XM_019834475.1;refGene_gn=LOC101081475;refGene_tx=XM_019834476.1
B4 45888300 45888355
refGene_gn=LOC101081475;refGene_tx=XM_019834475.1;refGene_gn=LOC101081475;refGene_tx=XM_019834476.1
B4 45889154 45889382
refGene_gn=LOC101081475;refGene_tx=XM_019834475.1;refGene_gn=LOC101081475;refGene_tx=XM_019834476.1
B4 45890294 45890453
refGene_gn=LOC101081475;refGene_tx=XM_019834475.1;refGene_gn=LOC101081475;refGene_tx=XM_019834476.1
B4 45890791 45890901
refGene_gn=LOC101081475;refGene_tx=XM_019834475.1;refGene_gn=LOC101081475;refGene_tx=XM_019834476.1
B4 45892191 45892303
refGene_gn=LOC101081475;refGene_tx=XM_019834475.1;refGene_gn=LOC101081475;refGene_tx=XM_019834476.1
B4 45897144 45897262
refGene_gn=LOC101081475;refGene_tx=XM_019834475.1;refGene_gn=LOC101081475;refGene_tx=XM_019834476.1
B4 45901554 45901639
refGene_gn=LOC101081475;refGene_tx=XM_019834475.1;refGene_gn=LOC101081475;refGene_tx=XM_019834476.1
B4 45902131 45902300
refGene_gn=LOC101081475;refGene_tx=XM_019834475.1;refGene_gn=LOC101081475;refGene_tx=XM_019834476.1
B4 45904903 45904924
refGene_gn=LOC101081475;refGene_tx=XM_019834475.1;refGene_gn=LOC101081475;refGene_tx=XM_019834476.1
B4 45905360 45905413
refGene_gn=LOC101081475;refGene_tx=XM_019834475.1;refGene_gn=LOC101081475;refGene_tx=XM_019834476.1
B4 45906940 45907097
refGene_gn=LOC101081475;refGene_tx=XM_019834475.1;refGene_gn=LOC101081

475;refGene_tx=XM_019834476.1
B4	45907849	45908033
refGene_gn=LOC101081475;refGene_tx=XM_019834475.1;refGene_gn=LOC101081475;refGene_tx=XM_019834476.1
B4	45908350	45908415
refGene_gn=LOC101081475;refGene_tx=XM_019834475.1;refGene_gn=LOC101081475;refGene_tx=XM_019834476.1
B4	45951428	45951576	refGene_gn=KLRB1;refGene_tx=XM_006933504.2
B4	45955188	45955304	refGene_gn=KLRB1;refGene_tx=XM_006933504.2
B4	45955670	45955825	refGene_gn=KLRB1;refGene_tx=XM_006933504.2
B4	45957267	45957342	refGene_gn=KLRB1;refGene_tx=XM_006933504.2
B4	45958004	45958103	refGene_gn=KLRB1;refGene_tx=XM_006933504.2
B4	45964341	45964426	refGene_gn=KLRB1;refGene_tx=XM_006933504.2
B4	46019731	46019792
refGene_gn=CLEC2D;refGene_tx=XM_003988394.5;refGene_gn=CLEC2D;refGene_tx=XM_006933502.3
B4	46022649	46022760	refGene_gn=CLEC2D;refGene_tx=XM_006933502.3
B4	46029365	46029550
refGene_gn=CLEC2D;refGene_tx=XM_003988394.5;refGene_gn=CLEC2D;refGene_tx=XM_006933502.3
B4	46033561	46033665
refGene_gn=CLEC2D;refGene_tx=XM_003988394.5;refGene_gn=CLEC2D;refGene_tx=XM_006933502.3
B4	46035398	46035537
refGene_gn=CLEC2D;refGene_tx=XM_003988394.5;refGene_gn=CLEC2D;refGene_tx=XM_006933502.3
B4	46045316	46045441
refGene_gn=LOC109501446;refGene_tx=XM_019834478.1
B4	46059918	46060010
refGene_gn=LOC109501446;refGene_tx=XM_019834478.1
B4	46060924	46061023
refGene_gn=LOC109501446;refGene_tx=XM_019834478.1
B4	46062424	46062514
refGene_gn=LOC109501446;refGene_tx=XM_019834478.1
B4	46064748	46064903
refGene_gn=LOC109501446;refGene_tx=XM_019834478.1
B4	46065561	46065674
refGene_gn=LOC109501446;refGene_tx=XM_019834478.1
B4	46068223	46068356
refGene_gn=LOC109501446;refGene_tx=XM_019834478.1
B4	46111144	46111259
refGene_gn=CLECL1;refGene_tx=XM_006933505.4;refGene_gn=CLECL1;refGene_tx=XM_019834481.2
B4	46112060	46112173
refGene_gn=CLECL1;refGene_tx=XM_006933505.4;refGene_gn=CLECL1;refGene_tx=XM_019834481.2
B4	46117713	46117883
refGene_gn=CLECL1;refGene_tx=XM_006933505.4;refGene_gn=CLECL1;refGene_tx=XM_019834481.2
B4	46129368	46129467

refGene_gn=CLECL1;refGene_tx=XM_006933505.4;refGene_gn=CLECL1;refGene_tx=XM_019834481.2
B4 46131498 46131585 refGene_gn=CLECL1;refGene_tx=XM_006933505.4
B4 46136310 46136392
refGene_gn=CLECL1;refGene_tx=XM_006933505.4;refGene_gn=CLECL1;refGene_tx=XM_019834481.2
B4 46164160 46164269 refGene_gn=CD69;refGene_tx=XM_003988395.5
B4 46165291 46165395 refGene_gn=CD69;refGene_tx=XM_003988395.5
B4 46165739 46165939 refGene_gn=CD69;refGene_tx=XM_003988395.5
B4 46167222 46167345 refGene_gn=CD69;refGene_tx=XM_003988395.5
B4 46171727 46171791 refGene_gn=CD69;refGene_tx=XM_003988395.5
B4 46217439 46217524 refGene_gn=KLRF1;refGene_tx=XM_003988396.4
B4 46224278 46224377 refGene_gn=KLRF1;refGene_tx=XM_003988396.4
B4 46225274 46225430 refGene_gn=KLRF1;refGene_tx=XM_003988396.4
B4 46227601 46227741 refGene_gn=KLRF1;refGene_tx=XM_003988396.4
B4 46228358 46228471 refGene_gn=KLRF1;refGene_tx=XM_003988396.4
B4 46229226 46229335 refGene_gn=KLRF1;refGene_tx=XM_003988396.4
B4 46237889 46237998
refGene_gn=CLEC2B;refGene_tx=XM_023256762.1;refGene_gn=CLEC2B;refGene_tx=XM_023256763.1
B4 46239480 46239584
refGene_gn=CLEC2B;refGene_tx=XM_023256762.1;refGene_gn=CLEC2B;refGene_tx=XM_023256763.1
B4 46244474 46244638
refGene_gn=CLEC2B;refGene_tx=XM_023256762.1;refGene_gn=CLEC2B;refGene_tx=XM_023256763.1
B4 46247575 46247654
refGene_gn=CLEC2B;refGene_tx=XM_023256762.1;refGene_gn=CLEC2B;refGene_tx=XM_023256763.1
B4 46280595 46280659 refGene_gn=KLRF2;refGene_tx=XM_023256764.1
B4 46283896 46283995 refGene_gn=KLRF2;refGene_tx=XM_023256764.1
B4 46288018 46288066 refGene_gn=KLRF2;refGene_tx=XM_023256764.1
B4 46292448 46292597 refGene_gn=KLRF2;refGene_tx=XM_023256764.1
B4 46292862 46292978 refGene_gn=KLRF2;refGene_tx=XM_023256764.1
B4 46295762 46295880 refGene_gn=KLRF2;refGene_tx=XM_023256764.1
B4 46303217 46303326
refGene_gn=CLEC2A;refGene_tx=XM_019834485.2;refGene_gn=CLEC2A;refGene_tx=XM_019834487.2;refGene_gn=CLEC2A;refGene_tx=XM_019834493.1
B4 46305332 46305436
refGene_gn=CLEC2A;refGene_tx=XM_019834485.2;refGene_gn=CLEC2A;refGene_tx=XM_019834487.2;refGene_gn=CLEC2A;refGene_tx=XM_019834488.1;refGene_gn=CLEC2A;refGene_tx=XM_019834493.1
B4 46307569 46307739
refGene_gn=CLEC2A;refGene_tx=XM_019834485.2;refGene_gn=CLEC2A;refGene_tx=XM_019834487.2;refGene_gn=CLEC2A;refGene_tx=XM_019834488.1;refGene_gn=CLEC2A;refGene_tx=XM_019834493.1
B4 46310599 46310674
refGene_gn=CLEC2A;refGene_tx=XM_019834485.2;refGene_gn=CLEC2A;refGene_tx=XM_019834487.2;refGene_gn=CLEC2A;refGene_tx=XM_019834488.1;refGene_gn=CLEC2A;refGene_tx=XM_019834493.1

B4 46317300 46317386
refGene_gn=CLEC2A;refGene_tx=XM_019834485.2;refGene_gn=CLEC2A;refGene_tx=XM_019834488.1
B4 46319946 46320001
refGene_gn=CLEC2A;refGene_tx=XM_019834485.2;refGene_gn=CLEC2A;refGene_tx=XM_019834487.2;refGene_gn=CLEC2A;refGene_tx=XM_019834488.1;refGene_gn=CLEC2A;refGene_tx=XM_019834493.1
B4 46322883 46322998
refGene_gn=LOC102902233;refGene_tx=XM_006933507.3;refGene_gn=LOC102902233;refGene_tx=XM_006933508.3;refGene_gn=LOC102902233;refGene_tx=XM_019834494.2
B4 46325170 46325277
refGene_gn=LOC102902233;refGene_tx=XM_006933507.3;refGene_gn=LOC102902233;refGene_tx=XM_006933508.3;refGene_gn=LOC102902233;refGene_tx=XM_019834494.2
B4 46326175 46326363
refGene_gn=LOC102902233;refGene_tx=XM_006933507.3;refGene_gn=LOC102902233;refGene_tx=XM_006933508.3;refGene_gn=LOC102902233;refGene_tx=XM_019834494.2
B4 46332140 46332230
refGene_gn=LOC102902233;refGene_tx=XM_006933507.3;refGene_gn=LOC102902233;refGene_tx=XM_019834494.2
B4 46332758 46332834
refGene_gn=LOC102902233;refGene_tx=XM_006933507.3;refGene_gn=LOC102902233;refGene_tx=XM_006933508.3
B4 46344531 46344622 refGene_gn=CLEC12A;refGene_tx=XM_003988405.3
B4 46346925 46346944 refGene_gn=CLEC12A;refGene_tx=XM_023256766.1
B4 46352854 46352953
refGene_gn=CLEC12A;refGene_tx=XM_003988405.3;refGene_gn=CLEC12A;refGene_tx=XM_023256766.1
B4 46353249 46353438
refGene_gn=CLEC12A;refGene_tx=XM_003988405.3;refGene_gn=CLEC12A;refGene_tx=XM_023256766.1
B4 46354617 46354769
refGene_gn=CLEC12A;refGene_tx=XM_003988405.3;refGene_gn=CLEC12A;refGene_tx=XM_023256766.1
B4 46356147 46356254
refGene_gn=CLEC12A;refGene_tx=XM_003988405.3;refGene_gn=CLEC12A;refGene_tx=XM_023256766.1
B4 46359028 46359206
refGene_gn=CLEC12A;refGene_tx=XM_003988405.3;refGene_gn=CLEC12A;refGene_tx=XM_023256766.1
B4 46410591 46410742
refGene_gn=CLEC12B;refGene_tx=XM_003988397.4;refGene_gn=CLEC12B;refGene_tx=XM_019834496.2
B4 46415536 46415652
refGene_gn=CLEC12B;refGene_tx=XM_003988397.4;refGene_gn=CLEC12B;refGene_tx=XM_019834496.2
B4 46415828 46415983
refGene_gn=CLEC12B;refGene_tx=XM_003988397.4;refGene_gn=CLEC12B;refGen

e_tx=XM_019834496.2
B4	46416741 46416960
refGene_gn=CLEC12B;refGene_tx=XM_003988397.4;refGene_gn=CLEC12B;refGene_tx=XM_019834496.2
B4	46418988 46419087
refGene_gn=CLEC12B;refGene_tx=XM_003988397.4;refGene_gn=CLEC12B;refGene_tx=XM_019834496.2
B4	46421183 46421274 refGene_gn=CLEC12B;refGene_tx=XM_003988397.4
B4	46433103 46433290 refGene_gn=CLEC1B;refGene_tx=XM_003988398.4
B4	46433893 46433992 refGene_gn=CLEC1B;refGene_tx=XM_003988398.4
B4	46435500 46435620 refGene_gn=CLEC1B;refGene_tx=XM_003988398.4
B4	46435782 46435934 refGene_gn=CLEC1B;refGene_tx=XM_003988398.4
B4	46439115 46439222 refGene_gn=CLEC1B;refGene_tx=XM_003988398.4
B4	46443501 46443646 refGene_gn=CLEC1B;refGene_tx=XM_003988398.4
B4	46447669 46447760 refGene_gn=CLEC9A;refGene_tx=XM_019835818.1
B4	46449715 46449796 refGene_gn=CLEC9A;refGene_tx=XM_019835818.1
B4	46456981 46457128 refGene_gn=CLEC9A;refGene_tx=XM_019835818.1
B4	46459063 46459215 refGene_gn=CLEC9A;refGene_tx=XM_019835818.1
B4	46460623 46460745 refGene_gn=CLEC9A;refGene_tx=XM_019835818.1
B4	46461406 46461539 refGene_gn=CLEC9A;refGene_tx=XM_019835818.1
B4	46468493 46468644
refGene_gn=CLEC1A;refGene_tx=XM_011283779.2;refGene_gn=CLEC1A;refGene_tx=XM_019834497.2;refGene_gn=CLEC1A;refGene_tx=XM_019834498.2;refGene_gn=CLEC1A;refGene_tx=XM_019834499.1;refGene_gn=CLEC1A;refGene_tx=XM_023256768.1;refGene_gn=CLEC1A;refGene_tx=XM_023256769.1
B4	46469703 46469822
refGene_gn=CLEC1A;refGene_tx=XM_011283779.2;refGene_gn=CLEC1A;refGene_tx=XM_019834497.2;refGene_gn=CLEC1A;refGene_tx=XM_019834498.2;refGene_gn=CLEC1A;refGene_tx=XM_019834499.1;refGene_gn=CLEC1A;refGene_tx=XM_023256768.1;refGene_gn=CLEC1A;refGene_tx=XM_023256769.1
B4	46471748 46471900
refGene_gn=CLEC1A;refGene_tx=XM_011283779.2;refGene_gn=CLEC1A;refGene_tx=XM_019834497.2;refGene_gn=CLEC1A;refGene_tx=XM_019834498.2;refGene_gn=CLEC1A;refGene_tx=XM_019834499.1;refGene_gn=CLEC1A;refGene_tx=XM_023256768.1;refGene_gn=CLEC1A;refGene_tx=XM_023256769.1
B4	46476466 46476643
refGene_gn=CLEC1A;refGene_tx=XM_011283779.2;refGene_gn=CLEC1A;refGene_tx=XM_019834497.2;refGene_gn=CLEC1A;refGene_tx=XM_019834498.2;refGene_gn=CLEC1A;refGene_tx=XM_019834499.1;refGene_gn=CLEC1A;refGene_tx=XM_023256768.1;refGene_gn=CLEC1A;refGene_tx=XM_023256769.1
B4	46477326 46477365
refGene_gn=CLEC1A;refGene_tx=XM_019834499.1;refGene_gn=CLEC1A;refGene_tx=XM_023256768.1;refGene_gn=CLEC1A;refGene_tx=XM_023256769.1
B4	46477979 46478078
refGene_gn=CLEC1A;refGene_tx=XM_011283779.2;refGene_gn=CLEC1A;refGene_tx=XM_019834497.2;refGene_gn=CLEC1A;refGene_tx=XM_019834498.2
B4	46490316 46490431
refGene_gn=CLEC1A;refGene_tx=XM_011283779.2;refGene_gn=CLEC1A;refGene_tx=XM_019834497.2;refGene_gn=CLEC1A;refGene_tx=XM_019834498.2
B4	46502669 46502805

refGene_gn=CLEC7A;refGene_tx=XM_003988400.5;refGene_gn=CLEC7A;refGene_tx=XM_006933509.3;refGene_gn=CLEC7A;refGene_tx=XM_006933510.3;refGene_gn=CLEC7A;refGene_tx=XM_023256771.1
B4 46508243 46508362
refGene_gn=CLEC7A;refGene_tx=XM_006933509.3;refGene_gn=CLEC7A;refGene_tx=XM_006933510.3
B4 46510379 46510531
refGene_gn=CLEC7A;refGene_tx=XM_003988400.5;refGene_gn=CLEC7A;refGene_tx=XM_006933509.3;refGene_gn=CLEC7A;refGene_tx=XM_006933510.3;refGene_gn=CLEC7A;refGene_tx=XM_023256771.1
B4 46512103 46512241
refGene_gn=CLEC7A;refGene_tx=XM_003988400.5;refGene_gn=CLEC7A;refGene_tx=XM_006933509.3
B4 46513359 46513458
refGene_gn=CLEC7A;refGene_tx=XM_003988400.5;refGene_gn=CLEC7A;refGene_tx=XM_006933509.3;refGene_gn=CLEC7A;refGene_tx=XM_006933510.3;refGene_gn=CLEC7A;refGene_tx=XM_023256771.1
B4 46515295 46515398
refGene_gn=CLEC7A;refGene_tx=XM_003988400.5;refGene_gn=CLEC7A;refGene_tx=XM_006933509.3;refGene_gn=CLEC7A;refGene_tx=XM_006933510.3;refGene_gn=CLEC7A;refGene_tx=XM_023256771.1
B4 46542955 46543100
refGene_gn=OLR1;refGene_tx=XM_006933511.4;refGene_gn=OLR1;refGene_tx=XM_019834502.2;refGene_gn=OLR1;refGene_tx=XM_019834503.2
B4 46543591 46543707
refGene_gn=OLR1;refGene_tx=XM_006933511.4;refGene_gn=OLR1;refGene_tx=XM_019834502.2;refGene_gn=OLR1;refGene_tx=XM_019834503.2
B4 46544005 46544145
refGene_gn=OLR1;refGene_tx=XM_006933511.4;refGene_gn=OLR1;refGene_tx=XM_019834502.2;refGene_gn=OLR1;refGene_tx=XM_019834503.2
B4 46547919 46548165
refGene_gn=OLR1;refGene_tx=XM_006933511.4;refGene_gn=OLR1;refGene_tx=XM_019834502.2;refGene_gn=OLR1;refGene_tx=XM_019834503.2
B4 46550103 46550205
refGene_gn=OLR1;refGene_tx=XM_006933511.4;refGene_gn=OLR1;refGene_tx=XM_019834502.2
B4 46552737 46552813
refGene_gn=OLR1;refGene_tx=XM_006933511.4;refGene_gn=OLR1;refGene_tx=XM_019834502.2;refGene_gn=OLR1;refGene_tx=XM_019834503.2
B4 46563638 46563692
refGene_gn=TMEM52B;refGene_tx=XM_003988409.2;refGene_gn=TMEM52B;refGene_tx=XM_019834504.1;refGene_gn=TMEM52B;refGene_tx=XM_019834505.1;refGene_gn=TMEM52B;refGene_tx=XM_019834506.1
B4 46565756 46565800
refGene_gn=TMEM52B;refGene_tx=XM_003988409.2;refGene_gn=TMEM52B;refGene_tx=XM_019834504.1;refGene_gn=TMEM52B;refGene_tx=XM_019834505.1;refGene_gn=TMEM52B;refGene_tx=XM_019834506.1
B4 46568590 46568629
refGene_gn=TMEM52B;refGene_tx=XM_003988409.2;refGene_gn=TMEM52B;refGene_tx=XM_019834504.1;refGene_gn=TMEM52B;refGene_tx=XM_019834505.1;refGe

ne_gn=TMEM52B;refGene_tx=XM_019834506.1
B4 46569167 46569334
refGene_gn=TMEM52B;refGene_tx=XM_003988409.2;refGene_gn=TMEM52B;refGene_tx=XM_019834504.1;refGene_gn=TMEM52B;refGene_tx=XM_019834505.1;refGene_gn=TMEM52B;refGene_tx=XM_019834506.1
B4 46570546 46570788
refGene_gn=TMEM52B;refGene_tx=XM_003988409.2;refGene_gn=TMEM52B;refGene_tx=XM_019834504.1;refGene_gn=TMEM52B;refGene_tx=XM_019834505.1;refGene_gn=TMEM52B;refGene_tx=XM_019834506.1
B4 46588927 46589017
refGene_gn=GABARAPL1;refGene_tx=XM_003988401.5
B4 46594187 46594266
refGene_gn=GABARAPL1;refGene_tx=XM_003988401.5
B4 46595290 46595409
refGene_gn=GABARAPL1;refGene_tx=XM_003988401.5
B4 46596282 46596348
refGene_gn=GABARAPL1;refGene_tx=XM_003988401.5
B4 46682464 46682471 refGene_gn=KLRD1;refGene_tx=XM_006933512.3
B4 46683763 46683856 refGene_gn=KLRD1;refGene_tx=XM_006933512.3
B4 46684282 46684345 refGene_gn=KLRD1;refGene_tx=XM_006933512.3
B4 46685866 46686018 refGene_gn=KLRD1;refGene_tx=XM_006933512.3
B4 46686846 46686950 refGene_gn=KLRD1;refGene_tx=XM_006933512.3
B4 46687752 46687894 refGene_gn=KLRD1;refGene_tx=XM_006933512.3
B4 46703439 46703557
refGene_gn=LOC101086984;refGene_tx=XM_003988411.3
B4 46707795 46707899
refGene_gn=LOC101086984;refGene_tx=XM_003988411.3
B4 46708181 46708333
refGene_gn=LOC101086984;refGene_tx=XM_003988411.3
B4 46708706 46708742
refGene_gn=LOC101086984;refGene_tx=XM_003988411.3
B4 46710002 46710095
refGene_gn=LOC101086984;refGene_tx=XM_003988411.3
B4 46711534 46711639
refGene_gn=LOC101086984;refGene_tx=XM_003988411.3
B4 46713844 46713884
refGene_gn=LOC101086984;refGene_tx=XM_003988411.3
B4 46784771 46785029
refGene_gn=LOC101087492;refGene_tx=XM_023257744.1
B4 46785491 46785577
refGene_gn=LOC101087492;refGene_tx=XM_023257744.1
B4 46786041 46786171
refGene_gn=LOC101087492;refGene_tx=XM_023257744.1
B4 46835026 46835134
refGene_gn=LOC111561447;refGene_tx=XM_023257745.1
B4 46839995 46840147
refGene_gn=LOC111561447;refGene_tx=XM_023257745.1
B4 46840724 46840796
refGene_gn=LOC111561447;refGene_tx=XM_023257745.1
B4 46841272 46841346

refGene_gn=LOC111561447;refGene_tx=XM_023257745.1
B4 46841699 46841815
refGene_gn=LOC111561447;refGene_tx=XM_023257745.1
B4 46854024 46854233
refGene_gn=LOC101100397;refGene_tx=XM_023257746.1
B4 46854883 46854945
refGene_gn=LOC101100397;refGene_tx=XM_023257746.1
B4 46862951 46863141
refGene_gn=LOC101100397;refGene_tx=XM_023257746.1
B4 46863614 46863668
refGene_gn=LOC101100397;refGene_tx=XM_023257746.1
B4 46864122 46864200
refGene_gn=LOC101100397;refGene_tx=XM_023257746.1
B4 46864531 46864718
refGene_gn=LOC101100397;refGene_tx=XM_023257746.1
B4 46923736 46923866
refGene_gn=LOC101100897;refGene_tx=XM_019834512.1
B4 46924351 46924455
refGene_gn=LOC101100897;refGene_tx=XM_019834512.1
B4 46947558 46947707
refGene_gn=LOC101100897;refGene_tx=XM_019834512.1
B4 46948239 46948293
refGene_gn=LOC101100897;refGene_tx=XM_019834512.1
B4 46948758 46948836
refGene_gn=LOC101100897;refGene_tx=XM_019834512.1
B4 46949241 46949428
refGene_gn=LOC101100897;refGene_tx=XM_019834512.1
B4 47118014 47118141
refGene_gn=LOC102899579;refGene_tx=XM_023256772.1;refGene_gn=LOC102899579;refGene_tx=XM_023256773.1
B4 47120841 47120945
refGene_gn=LOC102899579;refGene_tx=XM_023256772.1;refGene_gn=LOC102899579;refGene_tx=XM_023256773.1
B4 47125234 47125380
refGene_gn=LOC102899579;refGene_tx=XM_023256772.1;refGene_gn=LOC102899579;refGene_tx=XM_023256773.1
B4 47127724 47127838
refGene_gn=LOC102899579;refGene_tx=XM_023256772.1;refGene_gn=LOC102899579;refGene_tx=XM_023256773.1
B4 47130642 47130726
refGene_gn=LOC102899579;refGene_tx=XM_023256772.1;refGene_gn=LOC102899579;refGene_tx=XM_023256773.1
B4 47131004 47131038
refGene_gn=LOC102899579;refGene_tx=XM_023256772.1;refGene_gn=LOC102899579;refGene_tx=XM_023256773.1
B4 47186251 47186411
refGene_gn=KLRAP1;refGene_tx=NM_001009834.1;refGene_gn=KLRAP1;refGene_tx=XM_006933516.4;refGene_gn=KLRAP1;refGene_tx=XM_019833856.2
B4 47189038 47189145
refGene_gn=KLRAP1;refGene_tx=NM_001009834.1;refGene_gn=KLRAP1;refGene_

tx=XM_006933516.4;refGene_gn=KLRAP1;refGene_tx=XM_019833856.2
B4 47191220 47191363
refGene_gn=KLRAP1;refGene_tx=NM_001009834.1;refGene_gn=KLRAP1;refGene_tx=XM_006933516.4;refGene_gn=KLRAP1;refGene_tx=XM_019833856.2
B4 47193262 47193469
refGene_gn=KLRAP1;refGene_tx=NM_001009834.1;refGene_gn=KLRAP1;refGene_tx=XM_006933516.4;refGene_gn=KLRAP1;refGene_tx=XM_019833856.2
B4 47194628 47194718
refGene_gn=KLRAP1;refGene_tx=NM_001009834.1;refGene_gn=KLRAP1;refGene_tx=XM_006933516.4;refGene_gn=KLRAP1;refGene_tx=XM_019833856.2
B4 47196278 47196393
refGene_gn=KLRAP1;refGene_tx=NM_001009834.1;refGene_gn=KLRAP1;refGene_tx=XM_006933516.4;refGene_gn=KLRAP1;refGene_tx=XM_019833856.2
B4 47216626 47216726
refGene_gn=MAGOHB;refGene_tx=XM_019834516.2;refGene_gn=MAGOHB;refGene_tx=XM_023256774.1;refGene_gn=MAGOHB;refGene_tx=XM_023256775.1
B4 47218426 47218509
refGene_gn=MAGOHB;refGene_tx=XM_019834516.2;refGene_gn=MAGOHB;refGene_tx=XM_023256774.1;refGene_gn=MAGOHB;refGene_tx=XM_023256775.1
B4 47219738 47219780 refGene_gn=MAGOHB;refGene_tx=XM_023256776.1
B4 47220471 47220582
refGene_gn=MAGOHB;refGene_tx=XM_019834516.2;refGene_gn=MAGOHB;refGene_tx=XM_023256774.1;refGene_gn=MAGOHB;refGene_tx=XM_023256775.1;refGene_gn=MAGOHB;refGene_tx=XM_023256776.1
B4 47221561 47221664
refGene_gn=MAGOHB;refGene_tx=XM_019834516.2;refGene_gn=MAGOHB;refGene_tx=XM_023256774.1;refGene_gn=MAGOHB;refGene_tx=XM_023256775.1;refGene_gn=MAGOHB;refGene_tx=XM_023256776.1
B4 47225369 47225530
refGene_gn=MAGOHB;refGene_tx=XM_023256774.1;refGene_gn=MAGOHB;refGene_tx=XM_023256775.1;refGene_gn=MAGOHB;refGene_tx=XM_023256776.1
B4 47235194 47235399
refGene_gn=STYK1;refGene_tx=XM_003988414.5;refGene_gn=STYK1;refGene_tx=XM_019834514.2
B4 47237909 47238006
refGene_gn=STYK1;refGene_tx=XM_003988414.5;refGene_gn=STYK1;refGene_tx=XM_019834514.2
B4 47238612 47238653
refGene_gn=STYK1;refGene_tx=XM_003988414.5;refGene_gn=STYK1;refGene_tx=XM_019834514.2
B4 47240443 47240652
refGene_gn=STYK1;refGene_tx=XM_003988414.5;refGene_gn=STYK1;refGene_tx=XM_019834514.2
B4 47243005 47243089
refGene_gn=STYK1;refGene_tx=XM_003988414.5;refGene_gn=STYK1;refGene_tx=XM_019834514.2
B4 47245662 47245844
refGene_gn=STYK1;refGene_tx=XM_003988414.5;refGene_gn=STYK1;refGene_tx=XM_019834514.2
B4 47246658 47246925

refGene_gn=STYK1;refGene_tx=XM_003988414.5;refGene_gn=STYK1;refGene_tx=XM_019834514.2
B4 47249330 47249465
refGene_gn=STYK1;refGene_tx=XM_003988414.5;refGene_gn=STYK1;refGene_tx=XM_019834514.2
B4 47250159 47250205
refGene_gn=STYK1;refGene_tx=XM_003988414.5;refGene_gn=STYK1;refGene_tx=XM_019834514.2
B4 47296096 47296162
refGene_gn=YBX3;refGene_tx=XM_023256777.1;refGene_gn=YBX3;refGene_tx=XM_023256778.1
B4 47296865 47297037
refGene_gn=YBX3;refGene_tx=XM_023256777.1;refGene_gn=YBX3;refGene_tx=XM_023256778.1
B4 47298865 47298963
refGene_gn=YBX3;refGene_tx=XM_023256777.1;refGene_gn=YBX3;refGene_tx=XM_023256778.1
B4 47305009 47305213 refGene_gn=YBX3;refGene_tx=XM_023256777.1
B4 47310135 47310258
refGene_gn=YBX3;refGene_tx=XM_023256777.1;refGene_gn=YBX3;refGene_tx=XM_023256778.1
B4 47311796 47311886
refGene_gn=YBX3;refGene_tx=XM_023256777.1;refGene_gn=YBX3;refGene_tx=XM_023256778.1
B4 47314368 47314402
refGene_gn=YBX3;refGene_tx=XM_023256777.1;refGene_gn=YBX3;refGene_tx=XM_023256778.1
B4 47315449 47315513
refGene_gn=YBX3;refGene_tx=XM_023256777.1;refGene_gn=YBX3;refGene_tx=XM_023256778.1
B4 47318894 47319212
refGene_gn=LOC109501454;refGene_tx=XM_023256779.1;refGene_gn=LOC109501454;refGene_tx=XM_023256780.1
B4 47319460 47319725
refGene_gn=YBX3;refGene_tx=XM_023256777.1;refGene_gn=YBX3;refGene_tx=XM_023256778.1
B4 47320153 47320240
refGene_gn=LOC109501454;refGene_tx=XM_023256779.1
B4 47376632 47376659
refGene_gn=LOC109501454;refGene_tx=XM_023256780.1
B4 47396378 47397314
refGene_gn=LOC101089022;refGene_tx=XM_003988415.3
B4 47401462 47402041 refGene_gn=TAS2R8;refGene_tx=XM_011283793.3
B4 47402043 47402172 refGene_gn=TAS2R8;refGene_tx=XM_011283793.3
B4 47402173 47402206 refGene_gn=TAS2R8;refGene_tx=XM_011283793.3
B4 47402207 47402417 refGene_gn=TAS2R8;refGene_tx=XM_011283793.3
B4 47404517 47405531 refGene_gn=TAS2R9;refGene_tx=XM_003988465.4
B4 47419700 47419772
refGene_gn=LOC101089276;refGene_tx=XM_003988416.4
B4 47419773 47419905

refGene_gn=LOC101089276;refGene_tx=XM_003988416.4
B4	47419906	47420674
refGene_gn=LOC101089276;refGene_tx=XM_003988416.4
B4	47441269	47442199
refGene_gn=LOC101080977;refGene_tx=XM_003988466.1
B4	47476954	47477881
refGene_gn=LOC101089527;refGene_tx=XM_023257791.1
B4	47497729	47497799
refGene_gn=LOC101090025;refGene_tx=XM_011283776.2
B4	47498028	47499005
refGene_gn=LOC101090025;refGene_tx=XM_011283776.2
B4	47535749	47535986	refGene_gn=SMIM10L1;refGene_tx=XM_006933518.4
B4	47545388	47546327
refGene_gn=LOC101090527;refGene_tx=XM_003988420.2
B4	47553082	47554051
refGene_gn=LOC101090778;refGene_tx=XM_003988421.3
B4	47578431	47578472
refGene_gn=LOC102900024;refGene_tx=XM_019834520.2;refGene_gn=LOC102900024;refGene_tx=XM_019834521.2;refGene_gn=LOC102900024;refGene_tx=XM_019834522.2;refGene_gn=LOC102900024;refGene_tx=XM_019834523.2;refGene_gn=LOC102900024;refGene_tx=XM_019834524.2
B4	47578929	47578977
refGene_gn=LOC102900024;refGene_tx=XM_019834520.2;refGene_gn=LOC102900024;refGene_tx=XM_019834521.2;refGene_gn=LOC102900024;refGene_tx=XM_019834522.2;refGene_gn=LOC102900024;refGene_tx=XM_019834523.2;refGene_gn=LOC102900024;refGene_tx=XM_019834524.2
B4	47580546	47580609
refGene_gn=LOC102900024;refGene_tx=XM_019834520.2;refGene_gn=LOC102900024;refGene_tx=XM_019834521.2;refGene_gn=LOC102900024;refGene_tx=XM_019834522.2;refGene_gn=LOC102900024;refGene_tx=XM_019834523.2;refGene_gn=LOC102900024;refGene_tx=XM_019834524.2
B4	47580758	47581175
refGene_gn=LOC102900024;refGene_tx=XM_019834520.2;refGene_gn=LOC102900024;refGene_tx=XM_019834521.2;refGene_gn=LOC102900024;refGene_tx=XM_019834522.2;refGene_gn=LOC102900024;refGene_tx=XM_019834523.2;refGene_gn=LOC102900024;refGene_tx=XM_019834524.2
B4	47583017	47583087
refGene_gn=LOC102900024;refGene_tx=XM_019834520.2;refGene_gn=LOC102900024;refGene_tx=XM_019834521.2;refGene_gn=LOC102900024;refGene_tx=XM_019834522.2;refGene_gn=LOC102900024;refGene_tx=XM_019834523.2;refGene_gn=LOC102900024;refGene_tx=XM_019834524.2
B4	47719734	47719767
refGene_gn=ETV6;refGene_tx=XM_023256781.1;refGene_gn=ETV6;refGene_tx=XM_023256782.1
B4	47818390	47818520
refGene_gn=ETV6;refGene_tx=XM_023256781.1;refGene_gn=ETV6;refGene_tx=XM_023256782.1
B4	47919498	47919663
refGene_gn=ETV6;refGene_tx=XM_023256781.1;refGene_gn=ETV6;refGene_tx=XM_023256782.1

B4 47934268 47934403
refGene_gn=ETV6;refGene_tx=XM_023256781.1;refGene_gn=ETV6;refGene_tx=XM_023256782.1
B4 47949663 47950209
refGene_gn=ETV6;refGene_tx=XM_023256781.1;refGene_gn=ETV6;refGene_tx=XM_023256782.1
B4 47963643 47963924
refGene_gn=ETV6;refGene_tx=XM_023256781.1;refGene_gn=ETV6;refGene_tx=XM_023256782.1
B4 48114252 48114284 refGene_gn=BCL2L14;refGene_tx=XM_023256783.1
B4 48115956 48115988 refGene_gn=BCL2L14;refGene_tx=XM_019834525.2
B4 48135650 48136087
refGene_gn=BCL2L14;refGene_tx=XM_003988422.4;refGene_gn=BCL2L14;refGene_tx=XM_019834525.2;refGene_gn=BCL2L14;refGene_tx=XM_019834526.1;refGene_gn=BCL2L14;refGene_tx=XM_023256783.1
B4 48140576 48140750
refGene_gn=BCL2L14;refGene_tx=XM_003988422.4;refGene_gn=BCL2L14;refGene_tx=XM_019834525.2;refGene_gn=BCL2L14;refGene_tx=XM_019834526.1;refGene_gn=BCL2L14;refGene_tx=XM_023256783.1
B4 48142780 48142851
refGene_gn=BCL2L14;refGene_tx=XM_003988422.4;refGene_gn=BCL2L14;refGene_tx=XM_019834525.2;refGene_gn=BCL2L14;refGene_tx=XM_019834526.1;refGene_gn=BCL2L14;refGene_tx=XM_023256783.1
B4 48144960 48145227
refGene_gn=BCL2L14;refGene_tx=XM_003988422.4;refGene_gn=BCL2L14;refGene_tx=XM_019834525.2;refGene_gn=BCL2L14;refGene_tx=XM_019834526.1;refGene_gn=BCL2L14;refGene_tx=XM_023256783.1
B4 48149962 48150001
refGene_gn=BCL2L14;refGene_tx=XM_003988422.4;refGene_gn=BCL2L14;refGene_tx=XM_019834525.2;refGene_gn=BCL2L14;refGene_tx=XM_019834526.1;refGene_gn=BCL2L14;refGene_tx=XM_023256783.1
B4 48165360 48165655 refGene_gn=LRP6;refGene_tx=XM_023256784.1
B4 48167726 48167824 refGene_gn=LRP6;refGene_tx=XM_023256784.1
B4 48168496 48168633 refGene_gn=LRP6;refGene_tx=XM_023256784.1
B4 48169893 48170124 refGene_gn=LRP6;refGene_tx=XM_023256784.1
B4 48174372 48174483 refGene_gn=LRP6;refGene_tx=XM_023256784.1
B4 48174957 48175194 refGene_gn=LRP6;refGene_tx=XM_023256784.1
B4 48180488 48180614 refGene_gn=LRP6;refGene_tx=XM_023256784.1
B4 48183366 48183576 refGene_gn=LRP6;refGene_tx=XM_023256784.1
B4 48192675 48192866 refGene_gn=LRP6;refGene_tx=XM_023256784.1
B4 48193801 48194013 refGene_gn=LRP6;refGene_tx=XM_023256784.1
B4 48195792 48195995 refGene_gn=LRP6;refGene_tx=XM_023256784.1
B4 48204685 48205012 refGene_gn=LRP6;refGene_tx=XM_023256784.1
B4 48205633 48205818 refGene_gn=LRP6;refGene_tx=XM_023256784.1
B4 48207778 48208005 refGene_gn=LRP6;refGene_tx=XM_023256784.1
B4 48209520 48209810 refGene_gn=LRP6;refGene_tx=XM_023256784.1
B4 48210570 48210787 refGene_gn=LRP6;refGene_tx=XM_023256784.1
B4 48229272 48229444 refGene_gn=LRP6;refGene_tx=XM_023256784.1
B4 48230582 48230979 refGene_gn=LRP6;refGene_tx=XM_023256784.1
B4 48233995 48234127 refGene_gn=LRP6;refGene_tx=XM_023256784.1

B4 48235926 48236123 refGene_gn=LRP6;refGene_tx=XM_023256784.1
B4 48267478 48267676 refGene_gn=LRP6;refGene_tx=XM_023256784.1
B4 48316386 48316780 refGene_gn=LRP6;refGene_tx=XM_023256784.1
B4 48339862 48339917 refGene_gn=LRP6;refGene_tx=XM_023256784.1
B4 48340017 48340767
refGene_gn=LOC109501459;refGene_tx=XM_023256785.1
B4 48382636 48383572 refGene_gn=MANSC1;refGene_tx=XM_003988423.5
B4 48383573 48383599 refGene_gn=MANSC1;refGene_tx=XM_003988423.5
B4 48390928 48391069 refGene_gn=MANSC1;refGene_tx=XM_003988423.5
B4 48395108 48395331 refGene_gn=MANSC1;refGene_tx=XM_003988423.5
B4 48409741 48409742 refGene_gn=BORCS5;refGene_tx=XM_011283772.2
B4 48410077 48410135 refGene_gn=BORCS5;refGene_tx=XM_003988424.3
B4 48412503 48412647
refGene_gn=BORCS5;refGene_tx=XM_003988424.3;refGene_gn=BORCS5;refGene_tx=XM_011283772.2;refGene_gn=BORCS5;refGene_tx=XM_019834528.2
B4 48482069 48482227
refGene_gn=BORCS5;refGene_tx=XM_003988424.3;refGene_gn=BORCS5;refGene_tx=XM_011283772.2;refGene_gn=BORCS5;refGene_tx=XM_019834528.2
B4 48507102 48507333
refGene_gn=BORCS5;refGene_tx=XM_003988424.3;refGene_gn=BORCS5;refGene_tx=XM_011283772.2;refGene_gn=BORCS5;refGene_tx=XM_019834528.2
B4 48516626 48517803
refGene_gn=DUSP16;refGene_tx=XM_011283771.3;refGene_gn=DUSP16;refGene_tx=XM_019834529.2;refGene_gn=DUSP16;refGene_tx=XM_019834530.2;refGene_gn=DUSP16;refGene_tx=XM_023256786.1;refGene_gn=DUSP16;refGene_tx=XM_023256788.1
B4 48519614 48519738
refGene_gn=DUSP16;refGene_tx=XM_011283771.3;refGene_gn=DUSP16;refGene_tx=XM_019834529.2;refGene_gn=DUSP16;refGene_tx=XM_019834530.2;refGene_gn=DUSP16;refGene_tx=XM_023256786.1;refGene_gn=DUSP16;refGene_tx=XM_023256788.1
B4 48527660 48527820
refGene_gn=DUSP16;refGene_tx=XM_011283771.3;refGene_gn=DUSP16;refGene_tx=XM_019834529.2;refGene_gn=DUSP16;refGene_tx=XM_019834530.2;refGene_gn=DUSP16;refGene_tx=XM_023256786.1;refGene_gn=DUSP16;refGene_tx=XM_023256788.1
B4 48539948 48540112
refGene_gn=DUSP16;refGene_tx=XM_011283771.3;refGene_gn=DUSP16;refGene_tx=XM_019834529.2;refGene_gn=DUSP16;refGene_tx=XM_019834530.2;refGene_gn=DUSP16;refGene_tx=XM_023256786.1;refGene_gn=DUSP16;refGene_tx=XM_023256788.1
B4 48556953 48557092
refGene_gn=DUSP16;refGene_tx=XM_011283771.3;refGene_gn=DUSP16;refGene_tx=XM_019834529.2;refGene_gn=DUSP16;refGene_tx=XM_019834530.2;refGene_gn=DUSP16;refGene_tx=XM_023256786.1;refGene_gn=DUSP16;refGene_tx=XM_023256788.1
B4 48558035 48558263
refGene_gn=DUSP16;refGene_tx=XM_011283771.3;refGene_gn=DUSP16;refGene_tx=XM_019834529.2;refGene_gn=DUSP16;refGene_tx=XM_019834530.2;refGene_gn=DUSP16;refGene_tx=XM_023256786.1;refGene_gn=DUSP16;refGene_tx=XM_02

3256788.1
B4	48641603	48641618	refGene_gn=CREBL2;refGene_tx=XM_003988426.5
B4	48660772	48660970	refGene_gn=CREBL2;refGene_tx=XM_003988426.5
B4	48662378	48662523	refGene_gn=CREBL2;refGene_tx=XM_003988426.5
B4	48664878	48664883	refGene_gn=CREBL2;refGene_tx=XM_003988426.5
B4	48678979	48680227
refGene_gn=GPR19;refGene_tx=XM_006933523.4;refGene_gn=GPR19;refGene_tx=XM_019834532.2;refGene_gn=GPR19;refGene_tx=XM_019834534.1
B4	48736047	48736522	refGene_gn=CDKN1B;refGene_tx=NM_001009870.1
B4	48737091	48737213	refGene_gn=CDKN1B;refGene_tx=NM_001009870.1
B4	48744456	48744559	refGene_gn=APOLD1;refGene_tx=XM_019834544.1
B4	48751647	48751669	refGene_gn=APOLD1;refGene_tx=XM_006933525.3
B4	48768718	48768855
refGene_gn=APOLD1;refGene_tx=XM_006933525.3;refGene_gn=APOLD1;refGene_tx=XM_019834544.1
B4	48772420	48772891
refGene_gn=APOLD1;refGene_tx=XM_006933525.3;refGene_gn=APOLD1;refGene_tx=XM_019834544.1
B4	48797175	48797178
refGene_gn=APOLD1;refGene_tx=XM_003988429.4;refGene_gn=APOLD1;refGene_tx=XM_019834535.2;refGene_gn=APOLD1;refGene_tx=XM_019834536.2;refGene_gn=APOLD1;refGene_tx=XM_019834538.1;refGene_gn=APOLD1;refGene_tx=XM_019834539.2;refGene_gn=APOLD1;refGene_tx=XM_019834540.2;refGene_gn=APOLD1;refGene_tx=XM_023256790.1
B4	48798423	48799167
refGene_gn=APOLD1;refGene_tx=XM_003988429.4;refGene_gn=APOLD1;refGene_tx=XM_019834535.2;refGene_gn=APOLD1;refGene_tx=XM_019834536.2;refGene_gn=APOLD1;refGene_tx=XM_019834538.1;refGene_gn=APOLD1;refGene_tx=XM_019834539.2;refGene_gn=APOLD1;refGene_tx=XM_019834540.2;refGene_gn=APOLD1;refGene_tx=XM_019834543.2;refGene_gn=APOLD1;refGene_tx=XM_023256790.1
B4	48822366	48822427	refGene_gn=DDX47;refGene_tx=XM_003988430.5
B4	48822429	48822452	refGene_gn=DDX47;refGene_tx=XM_003988430.5
B4	48823337	48823431	refGene_gn=DDX47;refGene_tx=XM_003988430.5
B4	48828990	48829179	refGene_gn=DDX47;refGene_tx=XM_003988430.5
B4	48829427	48829499	refGene_gn=DDX47;refGene_tx=XM_003988430.5
B4	48829730	48829849	refGene_gn=DDX47;refGene_tx=XM_003988430.5
B4	48830426	48830498	refGene_gn=DDX47;refGene_tx=XM_003988430.5
B4	48830594	48830711	refGene_gn=DDX47;refGene_tx=XM_003988430.5
B4	48831460	48831607	refGene_gn=DDX47;refGene_tx=XM_003988430.5
B4	48832154	48832292	refGene_gn=DDX47;refGene_tx=XM_003988430.5
B4	48834248	48834319	refGene_gn=DDX47;refGene_tx=XM_003988430.5
B4	48835330	48835460	refGene_gn=DDX47;refGene_tx=XM_003988430.5
B4	48837007	48837139	refGene_gn=DDX47;refGene_tx=XM_003988430.5
B4	48908259	48909181
refGene_gn=GPRC5A;refGene_tx=XM_003988432.4;refGene_gn=GPRC5A;refGene_tx=XM_019834547.1
B4	48911492	48911551
refGene_gn=GPRC5A;refGene_tx=XM_003988432.4;refGene_gn=GPRC5A;refGene_tx=XM_019834547.1

B4 48911844 48911937
refGene_gn=GPRC5A;refGene_tx=XM_003988432.4;refGene_gn=GPRC5A;refGene_tx=XM_019834547.1
B4 48936107 48936188
refGene_gn=GPRC5D;refGene_tx=XM_003988470.4;refGene_gn=GPRC5D;refGene_tx=XM_011283806.2;refGene_gn=GPRC5D;refGene_tx=XM_019834548.1
B4 48937285 48937353
refGene_gn=GPRC5D;refGene_tx=XM_003988470.4;refGene_gn=GPRC5D;refGene_tx=XM_019834548.1
B4 48942878 48943860
refGene_gn=GPRC5D;refGene_tx=XM_003988470.4;refGene_gn=GPRC5D;refGene_tx=XM_011283806.2;refGene_gn=GPRC5D;refGene_tx=XM_019834548.1
B4 48964156 48964328
refGene_gn=HEBP1;refGene_tx=XM_003988433.4;refGene_gn=HEBP1;refGene_tx=XM_019834549.2;refGene_gn=HEBP1;refGene_tx=XM_019834550.2
B4 48977383 48977491
refGene_gn=HEBP1;refGene_tx=XM_019834549.2;refGene_gn=HEBP1;refGene_tx=XM_019834550.2
B4 48978124 48978305
refGene_gn=HEBP1;refGene_tx=XM_003988433.4;refGene_gn=HEBP1;refGene_tx=XM_019834549.2;refGene_gn=HEBP1;refGene_tx=XM_019834550.2
B4 48980070 48980209
refGene_gn=HEBP1;refGene_tx=XM_003988433.4;refGene_gn=HEBP1;refGene_tx=XM_019834549.2;refGene_gn=HEBP1;refGene_tx=XM_019834550.2
B4 48988891 48988969
refGene_gn=HEBP1;refGene_tx=XM_003988433.4;refGene_gn=HEBP1;refGene_tx=XM_019834549.2
B4 49018546 49018583 refGene_gn=FAM234B;refGene_tx=XM_023256791.1
B4 49029644 49030043 refGene_gn=FAM234B;refGene_tx=XM_023256791.1
B4 49033417 49033516 refGene_gn=FAM234B;refGene_tx=XM_023256791.1
B4 49036872 49037061 refGene_gn=FAM234B;refGene_tx=XM_023256791.1
B4 49038100 49038231 refGene_gn=FAM234B;refGene_tx=XM_023256791.1
B4 49041858 49042006 refGene_gn=FAM234B;refGene_tx=XM_023256791.1
B4 49042366 49042508 refGene_gn=FAM234B;refGene_tx=XM_023256791.1
B4 49043510 49043654 refGene_gn=FAM234B;refGene_tx=XM_023256791.1
B4 49043817 49043899 refGene_gn=FAM234B;refGene_tx=XM_023256791.1
B4 49046883 49047039 refGene_gn=FAM234B;refGene_tx=XM_023256791.1
B4 49049379 49049497 refGene_gn=FAM234B;refGene_tx=XM_023256791.1
B4 49051052 49051273 refGene_gn=FAM234B;refGene_tx=XM_023256791.1
B4 49051854 49051860 refGene_gn=FAM234B;refGene_tx=XM_023256791.1
B4 49055163 49055506
refGene_gn=GSG1;refGene_tx=XM_006933531.4;refGene_gn=GSG1;refGene_tx=XM_006933532.4;refGene_gn=GSG1;refGene_tx=XM_019834552.2
B4 49057458 49057570
refGene_gn=GSG1;refGene_tx=XM_006933531.4;refGene_gn=GSG1;refGene_tx=XM_006933532.4;refGene_gn=GSG1;refGene_tx=XM_019834552.2
B4 49058241 49058394
refGene_gn=GSG1;refGene_tx=XM_006933531.4;refGene_gn=GSG1;refGene_tx=XM_006933532.4;refGene_gn=GSG1;refGene_tx=XM_019834552.2
B4 49059111 49059282

refGene_gn=GSG1;refGene_tx=XM_006933531.4;refGene_gn=GSG1;refGene_tx=XM_006933532.4
B4 49060929 49061251
refGene_gn=GSG1;refGene_tx=XM_006933531.4;refGene_gn=GSG1;refGene_tx=XM_006933532.4;refGene_gn=GSG1;refGene_tx=XM_019834552.2
B4 49070836 49070884
refGene_gn=GSG1;refGene_tx=XM_006933531.4;refGene_gn=GSG1;refGene_tx=XM_006933532.4;refGene_gn=GSG1;refGene_tx=XM_019834552.2
B4 49173213 49173291 refGene_gn=EMP1;refGene_tx=XM_003988435.4
B4 49174877 49174974 refGene_gn=EMP1;refGene_tx=XM_003988435.4
B4 49175088 49175229 refGene_gn=EMP1;refGene_tx=XM_003988435.4
B4 49176166 49176333 refGene_gn=EMP1;refGene_tx=XM_003988435.4
B4 49484503 49486366
refGene_gn=GRIN2B;refGene_tx=XM_019834554.2;refGene_gn=GRIN2B;refGene_tx=XM_019834555.1
B4 49488723 49488962
refGene_gn=GRIN2B;refGene_tx=XM_019834554.2;refGene_gn=GRIN2B;refGene_tx=XM_019834555.1
B4 49491930 49492118
refGene_gn=GRIN2B;refGene_tx=XM_019834554.2;refGene_gn=GRIN2B;refGene_tx=XM_019834555.1
B4 49493980 49494141
refGene_gn=GRIN2B;refGene_tx=XM_019834554.2;refGene_gn=GRIN2B;refGene_tx=XM_019834555.1
B4 49528378 49528608
refGene_gn=GRIN2B;refGene_tx=XM_019834554.2;refGene_gn=GRIN2B;refGene_tx=XM_019834555.1
B4 49531142 49531268
refGene_gn=GRIN2B;refGene_tx=XM_019834554.2;refGene_gn=GRIN2B;refGene_tx=XM_019834555.1
B4 49534643 49534797
refGene_gn=GRIN2B;refGene_tx=XM_019834554.2;refGene_gn=GRIN2B;refGene_tx=XM_019834555.1
B4 49535015 49535187
refGene_gn=GRIN2B;refGene_tx=XM_019834554.2;refGene_gn=GRIN2B;refGene_tx=XM_019834555.1
B4 49535966 49536169
refGene_gn=GRIN2B;refGene_tx=XM_019834554.2;refGene_gn=GRIN2B;refGene_tx=XM_019834555.1
B4 49593821 49593936
refGene_gn=GRIN2B;refGene_tx=XM_019834554.2;refGene_gn=GRIN2B;refGene_tx=XM_019834555.1
B4 49665542 49666141
refGene_gn=GRIN2B;refGene_tx=XM_019834554.2;refGene_gn=GRIN2B;refGene_tx=XM_019834555.1
B4 49772728 49773139
refGene_gn=GRIN2B;refGene_tx=XM_019834554.2;refGene_gn=GRIN2B;refGene_tx=XM_019834555.1
B4 50185928 50185945
refGene_gn=ATF7IP;refGene_tx=XM_019834556.2;refGene_gn=ATF7IP;refGene_

tx=XM_019834557.2
B4 50222051 50223592
refGene_gn=ATF7IP;refGene_tx=XM_011283810.3;refGene_gn=ATF7IP;refGene_tx=XM_019834556.2;refGene_gn=ATF7IP;refGene_tx=XM_019834557.2;refGene_gn=ATF7IP;refGene_tx=XM_019834558.2
B4 50229215 50229299
refGene_gn=ATF7IP;refGene_tx=XM_011283810.3;refGene_gn=ATF7IP;refGene_tx=XM_019834556.2;refGene_gn=ATF7IP;refGene_tx=XM_019834557.2;refGene_gn=ATF7IP;refGene_tx=XM_019834558.2
B4 50230797 50230943
refGene_gn=ATF7IP;refGene_tx=XM_011283810.3;refGene_gn=ATF7IP;refGene_tx=XM_019834556.2;refGene_gn=ATF7IP;refGene_tx=XM_019834557.2;refGene_gn=ATF7IP;refGene_tx=XM_019834558.2
B4 50232268 50232406
refGene_gn=ATF7IP;refGene_tx=XM_011283810.3;refGene_gn=ATF7IP;refGene_tx=XM_019834556.2;refGene_gn=ATF7IP;refGene_tx=XM_019834557.2;refGene_gn=ATF7IP;refGene_tx=XM_019834558.2
B4 50249721 50249787
refGene_gn=ATF7IP;refGene_tx=XM_011283810.3;refGene_gn=ATF7IP;refGene_tx=XM_019834556.2;refGene_gn=ATF7IP;refGene_tx=XM_019834557.2;refGene_gn=ATF7IP;refGene_tx=XM_019834558.2
B4 50261401 50261478
refGene_gn=ATF7IP;refGene_tx=XM_011283810.3;refGene_gn=ATF7IP;refGene_tx=XM_019834556.2;refGene_gn=ATF7IP;refGene_tx=XM_019834557.2;refGene_gn=ATF7IP;refGene_tx=XM_019834558.2
B4 50262088 50262177
refGene_gn=ATF7IP;refGene_tx=XM_011283810.3;refGene_gn=ATF7IP;refGene_tx=XM_019834556.2;refGene_gn=ATF7IP;refGene_tx=XM_019834557.2;refGene_gn=ATF7IP;refGene_tx=XM_019834558.2
B4 50265897 50266536
refGene_gn=ATF7IP;refGene_tx=XM_011283810.3;refGene_gn=ATF7IP;refGene_tx=XM_019834556.2;refGene_gn=ATF7IP;refGene_tx=XM_019834557.2;refGene_gn=ATF7IP;refGene_tx=XM_019834558.2
B4 50271118 50271183
refGene_gn=ATF7IP;refGene_tx=XM_011283810.3;refGene_gn=ATF7IP;refGene_tx=XM_019834556.2;refGene_gn=ATF7IP;refGene_tx=XM_019834557.2;refGene_gn=ATF7IP;refGene_tx=XM_019834558.2
B4 50273092 50273171
refGene_gn=ATF7IP;refGene_tx=XM_011283810.3;refGene_gn=ATF7IP;refGene_tx=XM_019834556.2;refGene_gn=ATF7IP;refGene_tx=XM_019834557.2;refGene_gn=ATF7IP;refGene_tx=XM_019834558.2
B4 50281173 50281329
refGene_gn=ATF7IP;refGene_tx=XM_011283810.3;refGene_gn=ATF7IP;refGene_tx=XM_019834556.2;refGene_gn=ATF7IP;refGene_tx=XM_019834557.2;refGene_gn=ATF7IP;refGene_tx=XM_019834558.2
B4 50282977 50283160
refGene_gn=ATF7IP;refGene_tx=XM_011283810.3;refGene_gn=ATF7IP;refGene_tx=XM_019834556.2;refGene_gn=ATF7IP;refGene_tx=XM_019834557.2;refGene_gn=ATF7IP;refGene_tx=XM_019834558.2
B4 50288695 50288808

refGene_gn=ATF7IP;refGene_tx=XM_011283810.3;refGene_gn=ATF7IP;refGene_tx=XM_019834556.2;refGene_gn=ATF7IP;refGene_tx=XM_019834557.2;refGene_gn=ATF7IP;refGene_tx=XM_019834558.2
B4 50292898 50293318
refGene_gn=ATF7IP;refGene_tx=XM_011283810.3;refGene_gn=ATF7IP;refGene_tx=XM_019834556.2;refGene_gn=ATF7IP;refGene_tx=XM_019834557.2;refGene_gn=ATF7IP;refGene_tx=XM_019834558.2
B4 50298374 50298557 refGene_gn=PLBD1;refGene_tx=XM_003988438.4
B4 50300071 50300178 refGene_gn=PLBD1;refGene_tx=XM_003988438.4
B4 50300795 50300981 refGene_gn=PLBD1;refGene_tx=XM_003988438.4
B4 50305766 50305907 refGene_gn=PLBD1;refGene_tx=XM_003988438.4
B4 50306013 50306214 refGene_gn=PLBD1;refGene_tx=XM_003988438.4
B4 50320813 50320958 refGene_gn=PLBD1;refGene_tx=XM_003988438.4
B4 50321401 50321542 refGene_gn=PLBD1;refGene_tx=XM_003988438.4
B4 50327364 50327503 refGene_gn=PLBD1;refGene_tx=XM_003988438.4
B4 50328636 50328720 refGene_gn=PLBD1;refGene_tx=XM_003988438.4
B4 50339838 50340058 refGene_gn=PLBD1;refGene_tx=XM_003988438.4
B4 50352166 50352275 refGene_gn=PLBD1;refGene_tx=XM_003988438.4
B4 50409670 50409845
refGene_gn=GUCY2C;refGene_tx=XM_003988439.4;refGene_gn=GUCY2C;refGene_tx=XM_019834559.2;refGene_gn=GUCY2C;refGene_tx=XM_019834560.2;refGene_gn=GUCY2C;refGene_tx=XM_019834562.2;refGene_gn=GUCY2C;refGene_tx=XM_019834564.2;refGene_gn=GUCY2C;refGene_tx=XM_019834565.2;refGene_gn=GUCY2C;refGene_tx=XM_023256792.1
B4 50410944 50411021
refGene_gn=GUCY2C;refGene_tx=XM_003988439.4;refGene_gn=GUCY2C;refGene_tx=XM_019834559.2;refGene_gn=GUCY2C;refGene_tx=XM_019834560.2;refGene_gn=GUCY2C;refGene_tx=XM_019834562.2;refGene_gn=GUCY2C;refGene_tx=XM_019834564.2;refGene_gn=GUCY2C;refGene_tx=XM_019834565.2;refGene_gn=GUCY2C;refGene_tx=XM_023256792.1
B4 50412460 50412555
refGene_gn=GUCY2C;refGene_tx=XM_003988439.4;refGene_gn=GUCY2C;refGene_tx=XM_019834559.2;refGene_gn=GUCY2C;refGene_tx=XM_019834560.2;refGene_gn=GUCY2C;refGene_tx=XM_019834562.2;refGene_gn=GUCY2C;refGene_tx=XM_019834564.2;refGene_gn=GUCY2C;refGene_tx=XM_019834565.2;refGene_gn=GUCY2C;refGene_tx=XM_023256792.1
B4 50414204 50414303
refGene_gn=GUCY2C;refGene_tx=XM_003988439.4;refGene_gn=GUCY2C;refGene_tx=XM_019834559.2;refGene_gn=GUCY2C;refGene_tx=XM_019834560.2;refGene_gn=GUCY2C;refGene_tx=XM_019834562.2;refGene_gn=GUCY2C;refGene_tx=XM_019834564.2;refGene_gn=GUCY2C;refGene_tx=XM_019834565.2;refGene_gn=GUCY2C;refGene_tx=XM_023256792.1
B4 50416754 50416929
refGene_gn=GUCY2C;refGene_tx=XM_003988439.4;refGene_gn=GUCY2C;refGene_tx=XM_019834559.2;refGene_gn=GUCY2C;refGene_tx=XM_019834560.2;refGene_gn=GUCY2C;refGene_tx=XM_019834562.2;refGene_gn=GUCY2C;refGene_tx=XM_019834564.2;refGene_gn=GUCY2C;refGene_tx=XM_019834565.2;refGene_gn=GUCY2C;refGene_tx=XM_023256792.1
B4 50418434 50418627
refGene_gn=GUCY2C;refGene_tx=XM_003988439.4;refGene_gn=GUCY2C;refGene_

tx=XM_019834559.2;refGene_gn=GUCY2C;refGene_tx=XM_019834560.2;refGene_gn=GUCY2C;refGene_tx=XM_019834562.2;refGene_gn=GUCY2C;refGene_tx=XM_019834564.2;refGene_gn=GUCY2C;refGene_tx=XM_019834565.2;refGene_gn=GUCY2C;refGene_tx=XM_023256792.1
B4 50422783 50422942
refGene_gn=GUCY2C;refGene_tx=XM_003988439.4;refGene_gn=GUCY2C;refGene_tx=XM_019834559.2;refGene_gn=GUCY2C;refGene_tx=XM_019834560.2;refGene_gn=GUCY2C;refGene_tx=XM_019834562.2;refGene_gn=GUCY2C;refGene_tx=XM_019834564.2;refGene_gn=GUCY2C;refGene_tx=XM_019834565.2;refGene_gn=GUCY2C;refGene_tx=XM_023256792.1
B4 50425713 50425805
refGene_gn=GUCY2C;refGene_tx=XM_003988439.4;refGene_gn=GUCY2C;refGene_tx=XM_019834559.2;refGene_gn=GUCY2C;refGene_tx=XM_019834560.2;refGene_gn=GUCY2C;refGene_tx=XM_019834562.2;refGene_gn=GUCY2C;refGene_tx=XM_019834564.2;refGene_gn=GUCY2C;refGene_tx=XM_019834565.2;refGene_gn=GUCY2C;refGene_tx=XM_023256792.1
B4 50427478 50427567
refGene_gn=GUCY2C;refGene_tx=XM_003988439.4;refGene_gn=GUCY2C;refGene_tx=XM_019834559.2;refGene_gn=GUCY2C;refGene_tx=XM_019834560.2;refGene_gn=GUCY2C;refGene_tx=XM_019834562.2;refGene_gn=GUCY2C;refGene_tx=XM_019834564.2;refGene_gn=GUCY2C;refGene_tx=XM_019834565.2;refGene_gn=GUCY2C;refGene_tx=XM_023256792.1
B4 50428381 50428519
refGene_gn=GUCY2C;refGene_tx=XM_003988439.4;refGene_gn=GUCY2C;refGene_tx=XM_019834559.2;refGene_gn=GUCY2C;refGene_tx=XM_019834560.2;refGene_gn=GUCY2C;refGene_tx=XM_019834562.2;refGene_gn=GUCY2C;refGene_tx=XM_019834564.2;refGene_gn=GUCY2C;refGene_tx=XM_019834565.2;refGene_gn=GUCY2C;refGene_tx=XM_023256792.1
B4 50430552 50430685
refGene_gn=GUCY2C;refGene_tx=XM_003988439.4;refGene_gn=GUCY2C;refGene_tx=XM_019834559.2;refGene_gn=GUCY2C;refGene_tx=XM_019834560.2;refGene_gn=GUCY2C;refGene_tx=XM_019834562.2;refGene_gn=GUCY2C;refGene_tx=XM_019834564.2;refGene_gn=GUCY2C;refGene_tx=XM_019834565.2;refGene_gn=GUCY2C;refGene_tx=XM_023256792.1
B4 50431758 50431845
refGene_gn=GUCY2C;refGene_tx=XM_003988439.4;refGene_gn=GUCY2C;refGene_tx=XM_019834559.2;refGene_gn=GUCY2C;refGene_tx=XM_019834560.2;refGene_gn=GUCY2C;refGene_tx=XM_019834562.2;refGene_gn=GUCY2C;refGene_tx=XM_019834564.2;refGene_gn=GUCY2C;refGene_tx=XM_019834565.2;refGene_gn=GUCY2C;refGene_tx=XM_023256792.1
B4 50436661 50436766
refGene_gn=GUCY2C;refGene_tx=XM_003988439.4;refGene_gn=GUCY2C;refGene_tx=XM_019834559.2;refGene_gn=GUCY2C;refGene_tx=XM_019834560.2;refGene_gn=GUCY2C;refGene_tx=XM_019834562.2;refGene_gn=GUCY2C;refGene_tx=XM_019834564.2;refGene_gn=GUCY2C;refGene_tx=XM_019834565.2;refGene_gn=GUCY2C;refGene_tx=XM_023256792.1
B4 50437232 50437304
refGene_gn=GUCY2C;refGene_tx=XM_003988439.4;refGene_gn=GUCY2C;refGene_tx=XM_019834559.2;refGene_gn=GUCY2C;refGene_tx=XM_019834560.2;refGene_gn=GUCY2C;refGene_tx=XM_019834562.2;refGene_gn=GUCY2C;refGene_tx=XM_01

9834564.2;refGene_gn=GUCY2C;refGene_tx=XM_019834565.2;refGene_gn=GUCY2C;refGene_tx=XM_023256792.1
B4 50437876 50437939
refGene_gn=GUCY2C;refGene_tx=XM_003988439.4;refGene_gn=GUCY2C;refGene_tx=XM_019834559.2;refGene_gn=GUCY2C;refGene_tx=XM_019834560.2;refGene_gn=GUCY2C;refGene_tx=XM_019834562.2;refGene_gn=GUCY2C;refGene_tx=XM_019834564.2;refGene_gn=GUCY2C;refGene_tx=XM_019834565.2;refGene_gn=GUCY2C;refGene_tx=XM_023256792.1
B4 50440814 50440920
refGene_gn=GUCY2C;refGene_tx=XM_003988439.4;refGene_gn=GUCY2C;refGene_tx=XM_019834559.2;refGene_gn=GUCY2C;refGene_tx=XM_019834560.2;refGene_gn=GUCY2C;refGene_tx=XM_019834562.2;refGene_gn=GUCY2C;refGene_tx=XM_019834564.2;refGene_gn=GUCY2C;refGene_tx=XM_019834565.2;refGene_gn=GUCY2C;refGene_tx=XM_023256792.1
B4 50444033 50444115
refGene_gn=GUCY2C;refGene_tx=XM_003988439.4;refGene_gn=GUCY2C;refGene_tx=XM_019834559.2;refGene_gn=GUCY2C;refGene_tx=XM_019834560.2;refGene_gn=GUCY2C;refGene_tx=XM_019834562.2;refGene_gn=GUCY2C;refGene_tx=XM_019834564.2;refGene_gn=GUCY2C;refGene_tx=XM_019834565.2;refGene_gn=GUCY2C;refGene_tx=XM_023256792.1
B4 50449883 50449995
refGene_gn=GUCY2C;refGene_tx=XM_003988439.4;refGene_gn=GUCY2C;refGene_tx=XM_019834559.2;refGene_gn=GUCY2C;refGene_tx=XM_019834560.2;refGene_gn=GUCY2C;refGene_tx=XM_019834562.2;refGene_gn=GUCY2C;refGene_tx=XM_019834564.2;refGene_gn=GUCY2C;refGene_tx=XM_019834565.2;refGene_gn=GUCY2C;refGene_tx=XM_023256792.1
B4 50452648 50452734
refGene_gn=GUCY2C;refGene_tx=XM_003988439.4;refGene_gn=GUCY2C;refGene_tx=XM_019834559.2;refGene_gn=GUCY2C;refGene_tx=XM_019834560.2;refGene_gn=GUCY2C;refGene_tx=XM_019834562.2;refGene_gn=GUCY2C;refGene_tx=XM_019834564.2;refGene_gn=GUCY2C;refGene_tx=XM_019834565.2;refGene_gn=GUCY2C;refGene_tx=XM_023256792.1
B4 50459544 50459680
refGene_gn=GUCY2C;refGene_tx=XM_003988439.4;refGene_gn=GUCY2C;refGene_tx=XM_019834559.2;refGene_gn=GUCY2C;refGene_tx=XM_019834560.2;refGene_gn=GUCY2C;refGene_tx=XM_019834562.2;refGene_gn=GUCY2C;refGene_tx=XM_019834564.2;refGene_gn=GUCY2C;refGene_tx=XM_019834565.2;refGene_gn=GUCY2C;refGene_tx=XM_023256792.1
B4 50461887 50462005
refGene_gn=GUCY2C;refGene_tx=XM_003988439.4;refGene_gn=GUCY2C;refGene_tx=XM_019834559.2;refGene_gn=GUCY2C;refGene_tx=XM_019834560.2;refGene_gn=GUCY2C;refGene_tx=XM_019834562.2;refGene_gn=GUCY2C;refGene_tx=XM_019834564.2;refGene_gn=GUCY2C;refGene_tx=XM_019834565.2;refGene_gn=GUCY2C;refGene_tx=XM_023256792.1
B4 50464090 50464187
refGene_gn=GUCY2C;refGene_tx=XM_003988439.4;refGene_gn=GUCY2C;refGene_tx=XM_019834559.2;refGene_gn=GUCY2C;refGene_tx=XM_019834560.2;refGene_gn=GUCY2C;refGene_tx=XM_019834562.2;refGene_gn=GUCY2C;refGene_tx=XM_019834564.2;refGene_gn=GUCY2C;refGene_tx=XM_019834565.2;refGene_gn=GUCY2C;refGene_tx=XM_023256792.1

B4 50465873 50465995
refGene_gn=GUCY2C;refGene_tx=XM_003988439.4;refGene_gn=GUCY2C;refGene_tx=XM_019834559.2;refGene_gn=GUCY2C;refGene_tx=XM_019834560.2;refGene_gn=GUCY2C;refGene_tx=XM_019834562.2;refGene_gn=GUCY2C;refGene_tx=XM_019834564.2;refGene_gn=GUCY2C;refGene_tx=XM_019834565.2;refGene_gn=GUCY2C;refGene_tx=XM_023256792.1
B4 50467087 50467303
refGene_gn=GUCY2C;refGene_tx=XM_003988439.4;refGene_gn=GUCY2C;refGene_tx=XM_019834559.2;refGene_gn=GUCY2C;refGene_tx=XM_019834560.2;refGene_gn=GUCY2C;refGene_tx=XM_019834562.2;refGene_gn=GUCY2C;refGene_tx=XM_019834564.2;refGene_gn=GUCY2C;refGene_tx=XM_019834565.2;refGene_gn=GUCY2C;refGene_tx=XM_023256792.1
B4 50470107 50470172
refGene_gn=GUCY2C;refGene_tx=XM_003988439.4;refGene_gn=GUCY2C;refGene_tx=XM_019834559.2;refGene_gn=GUCY2C;refGene_tx=XM_019834560.2;refGene_gn=GUCY2C;refGene_tx=XM_019834562.2;refGene_gn=GUCY2C;refGene_tx=XM_019834564.2;refGene_gn=GUCY2C;refGene_tx=XM_019834565.2;refGene_gn=GUCY2C;refGene_tx=XM_023256792.1
B4 50471528 50471641
refGene_gn=GUCY2C;refGene_tx=XM_003988439.4;refGene_gn=GUCY2C;refGene_tx=XM_019834559.2;refGene_gn=GUCY2C;refGene_tx=XM_019834560.2;refGene_gn=GUCY2C;refGene_tx=XM_019834562.2;refGene_gn=GUCY2C;refGene_tx=XM_019834564.2;refGene_gn=GUCY2C;refGene_tx=XM_019834565.2;refGene_gn=GUCY2C;refGene_tx=XM_023256792.1
B4 50478121 50478338
refGene_gn=GUCY2C;refGene_tx=XM_003988439.4;refGene_gn=GUCY2C;refGene_tx=XM_019834559.2;refGene_gn=GUCY2C;refGene_tx=XM_019834560.2;refGene_gn=GUCY2C;refGene_tx=XM_019834562.2;refGene_gn=GUCY2C;refGene_tx=XM_019834564.2;refGene_gn=GUCY2C;refGene_tx=XM_023256792.1
B4 50480341 50480550
refGene_gn=GUCY2C;refGene_tx=XM_019834559.2;refGene_gn=GUCY2C;refGene_tx=XM_019834560.2;refGene_gn=GUCY2C;refGene_tx=XM_019834562.2;refGene_gn=GUCY2C;refGene_tx=XM_023256792.1
B4 50491759 50491839
refGene_gn=GUCY2C;refGene_tx=XM_019834559.2;refGene_gn=GUCY2C;refGene_tx=XM_019834560.2;refGene_gn=GUCY2C;refGene_tx=XM_019834562.2;refGene_gn=GUCY2C;refGene_tx=XM_019834564.2;refGene_gn=GUCY2C;refGene_tx=XM_019834565.2;refGene_gn=GUCY2C;refGene_tx=XM_023256792.1
B4 50550050 50550362
refGene_gn=LOC101095347;refGene_tx=XM_006933534.3
B4 50553556 50553946
refGene_gn=LOC101095750;refGene_tx=XM_003988442.5
B4 50571227 50571661 refGene_gn=WBP11;refGene_tx=XM_011283816.3
B4 50573061 50573244 refGene_gn=WBP11;refGene_tx=XM_011283816.3
B4 50574212 50574506 refGene_gn=WBP11;refGene_tx=XM_011283816.3
B4 50574928 50575030 refGene_gn=WBP11;refGene_tx=XM_011283816.3
B4 50577266 50577458 refGene_gn=WBP11;refGene_tx=XM_011283816.3
B4 50578117 50578317 refGene_gn=WBP11;refGene_tx=XM_011283816.3
B4 50578528 50578662 refGene_gn=WBP11;refGene_tx=XM_011283816.3
B4 50579943 50580140 refGene_gn=WBP11;refGene_tx=XM_011283816.3

B4	50583067	50583161	refGene_gn=WBP11;refGene_tx=XM_011283816.3
B4	50583995	50584027	refGene_gn=WBP11;refGene_tx=XM_011283816.3
B4	50584513	50584577	refGene_gn=WBP11;refGene_tx=XM_011283816.3
B4	50589035	50589713
refGene_gn=SMCO3;refGene_tx=XM_006933536.4;refGene_gn=SMCO3;refGene_tx
=XM_006933537.4;refGene_gn=SMCO3;refGene_tx=XM_019834573.2
B4	50602031	50602766
refGene_gn=CB4H12orf60;refGene_tx=XM_003988472.3;refGene_gn=CB4H12orf6
0;refGene_tx=XM_019834568.1;refGene_gn=CB4H12orf60;refGene_tx=XM_01983
4569.1;refGene_gn=CB4H12orf60;refGene_tx=XM_019834570.1;refGene_gn=CB4
H12orf60;refGene_tx=XM_019834571.1;refGene_gn=CB4H12orf60;refGene_tx=X
M_019834572.1
B4	50608971	50609063	refGene_gn=ART4;refGene_tx=XM_003988444.5
B4	50616633	50617342	refGene_gn=ART4;refGene_tx=XM_003988444.5
B4	50619119	50619305	refGene_gn=ART4;refGene_tx=XM_003988444.5
B4	50660845	50660987	refGene_gn=MGP;refGene_tx=XM_003988445.4
B4	50661400	50661476	refGene_gn=MGP;refGene_tx=XM_003988445.4
B4	50662329	50662362	refGene_gn=MGP;refGene_tx=XM_003988445.4
B4	50663825	50663886	refGene_gn=MGP;refGene_tx=XM_003988445.4
B4	50698477	50698534
refGene_gn=ERP27;refGene_tx=XM_019834566.2;refGene_gn=ERP27;refGene_tx
=XM_019834567.1
B4	50701039	50701087	refGene_gn=ERP27;refGene_tx=XM_003988446.5
B4	50701492	50701690
refGene_gn=ERP27;refGene_tx=XM_003988446.5;refGene_gn=ERP27;refGene_tx
=XM_019834566.2;refGene_gn=ERP27;refGene_tx=XM_019834567.1
B4	50703828	50703954
refGene_gn=ERP27;refGene_tx=XM_003988446.5;refGene_gn=ERP27;refGene_tx
=XM_019834566.2;refGene_gn=ERP27;refGene_tx=XM_019834567.1
B4	50707956	50708073
refGene_gn=ERP27;refGene_tx=XM_003988446.5;refGene_gn=ERP27;refGene_tx
=XM_019834566.2;refGene_gn=ERP27;refGene_tx=XM_019834567.1
B4	50720948	50721086
refGene_gn=ERP27;refGene_tx=XM_003988446.5;refGene_gn=ERP27;refGene_tx
=XM_019834566.2;refGene_gn=ERP27;refGene_tx=XM_019834567.1
B4	50722967	50723068
refGene_gn=ERP27;refGene_tx=XM_003988446.5;refGene_gn=ERP27;refGene_tx
=XM_019834566.2;refGene_gn=ERP27;refGene_tx=XM_019834567.1
B4	50723444	50723538
refGene_gn=ERP27;refGene_tx=XM_003988446.5;refGene_gn=ERP27;refGene_tx
=XM_019834566.2
B4	50725302	50725502	refGene_gn=ARHGDIB;refGene_tx=XM_003988447.4
B4	50726360	50726424	refGene_gn=ARHGDIB;refGene_tx=XM_003988447.4
B4	50729539	50729616	refGene_gn=ARHGDIB;refGene_tx=XM_003988447.4
B4	50732412	50732496	refGene_gn=ARHGDIB;refGene_tx=XM_003988447.4
B4	50733149	50733327	refGene_gn=ARHGDIB;refGene_tx=XM_003988447.4
B4	50762050	50762184
refGene_gn=PDE6H;refGene_tx=XM_006933535.3;refGene_gn=PDE6H;refGene_tx
=XM_019834575.2;refGene_gn=PDE6H;refGene_tx=XM_019834576.1;refGene_gn=
PDE6H;refGene_tx=XM_019834577.2;refGene_gn=PDE6H;refGene_tx=XM_0198345

78.1;refGene_gn=PDE6H;refGene_tx=XM_023256794.1;refGene_gn=PDE6H;refGene_tx=XM_023256795.1;refGene_gn=PDE6H;refGene_tx=XM_023256796.1
B4 50763210 50763251
refGene_gn=PDE6H;refGene_tx=XM_006933535.3;refGene_gn=PDE6H;refGene_tx=XM_019834575.2;refGene_gn=PDE6H;refGene_tx=XM_019834576.1;refGene_gn=PDE6H;refGene_tx=XM_019834577.2;refGene_gn=PDE6H;refGene_tx=XM_019834578.1;refGene_gn=PDE6H;refGene_tx=XM_023256794.1;refGene_gn=PDE6H;refGene_tx=XM_023256795.1;refGene_gn=PDE6H;refGene_tx=XM_023256796.1
B4 50765479 50765556
refGene_gn=PDE6H;refGene_tx=XM_006933535.3;refGene_gn=PDE6H;refGene_tx=XM_019834575.2;refGene_gn=PDE6H;refGene_tx=XM_019834576.1;refGene_gn=PDE6H;refGene_tx=XM_019834577.2;refGene_gn=PDE6H;refGene_tx=XM_019834578.1;refGene_gn=PDE6H;refGene_tx=XM_023256794.1;refGene_gn=PDE6H;refGene_tx=XM_023256795.1;refGene_gn=PDE6H;refGene_tx=XM_023256796.1
B4 50873720 50874128 refGene_gn=RERG;refGene_tx=XM_003988449.5
B4 50875698 50875772 refGene_gn=RERG;refGene_tx=XM_003988449.5
B4 50886385 50886442 refGene_gn=RERG;refGene_tx=XM_003988449.5
B4 50985032 50985093 refGene_gn=RERG;refGene_tx=XM_003988449.5
B4 51078006 51078081
refGene_gn=PTPRO;refGene_tx=XM_023256797.1;refGene_gn=PTPRO;refGene_tx=XM_023256798.1
B4 51211804 51212078
refGene_gn=PTPRO;refGene_tx=XM_023256797.1;refGene_gn=PTPRO;refGene_tx=XM_023256798.1
B4 51226497 51226656
refGene_gn=PTPRO;refGene_tx=XM_023256797.1;refGene_gn=PTPRO;refGene_tx=XM_023256798.1
B4 51228384 51228537
refGene_gn=PTPRO;refGene_tx=XM_023256797.1;refGene_gn=PTPRO;refGene_tx=XM_023256798.1
B4 51230253 51230697
refGene_gn=PTPRO;refGene_tx=XM_023256797.1;refGene_gn=PTPRO;refGene_tx=XM_023256798.1
B4 51232531 51232693
refGene_gn=PTPRO;refGene_tx=XM_023256797.1;refGene_gn=PTPRO;refGene_tx=XM_023256798.1
B4 51236266 51236463
refGene_gn=PTPRO;refGene_tx=XM_023256797.1;refGene_gn=PTPRO;refGene_tx=XM_023256798.1
B4 51243487 51243608
refGene_gn=PTPRO;refGene_tx=XM_023256797.1;refGene_gn=PTPRO;refGene_tx=XM_023256798.1
B4 51243957 51244151
refGene_gn=PTPRO;refGene_tx=XM_023256797.1;refGene_gn=PTPRO;refGene_tx=XM_023256798.1
B4 51245328 51245440
refGene_gn=PTPRO;refGene_tx=XM_023256797.1;refGene_gn=PTPRO;refGene_tx=XM_023256798.1
B4 51249284 51249436
refGene_gn=PTPRO;refGene_tx=XM_023256797.1;refGene_gn=PTPRO;refGene_tx

=XM_023256798.1
B4 51250612 51250733
refGene_gn=PTPRO;refGene_tx=XM_023256797.1;refGene_gn=PTPRO;refGene_tx=XM_023256798.1
B4 51267713 51267853
refGene_gn=PTPRO;refGene_tx=XM_023256797.1;refGene_gn=PTPRO;refGene_tx=XM_023256798.1
B4 51270454 51270587
refGene_gn=PTPRO;refGene_tx=XM_023256797.1;refGene_gn=PTPRO;refGene_tx=XM_023256798.1
B4 51272709 51272830
refGene_gn=PTPRO;refGene_tx=XM_023256797.1;refGene_gn=PTPRO;refGene_tx=XM_023256798.1
B4 51278517 51278586
refGene_gn=PTPRO;refGene_tx=XM_023256797.1;refGene_gn=PTPRO;refGene_tx=XM_023256798.1
B4 51280928 51281012 refGene_gn=PTPRO;refGene_tx=XM_023256797.1
B4 51287976 51288012
refGene_gn=PTPRO;refGene_tx=XM_023256797.1;refGene_gn=PTPRO;refGene_tx=XM_023256798.1
B4 51291254 51291336
refGene_gn=PTPRO;refGene_tx=XM_023256797.1;refGene_gn=PTPRO;refGene_tx=XM_023256798.1
B4 51301705 51301796
refGene_gn=PTPRO;refGene_tx=XM_023256797.1;refGene_gn=PTPRO;refGene_tx=XM_023256798.1
B4 51302859 51302936
refGene_gn=PTPRO;refGene_tx=XM_023256797.1;refGene_gn=PTPRO;refGene_tx=XM_023256798.1
B4 51303775 51303910
refGene_gn=PTPRO;refGene_tx=XM_023256797.1;refGene_gn=PTPRO;refGene_tx=XM_023256798.1
B4 51305185 51305308
refGene_gn=PTPRO;refGene_tx=XM_023256797.1;refGene_gn=PTPRO;refGene_tx=XM_023256798.1
B4 51310132 51310287
refGene_gn=PTPRO;refGene_tx=XM_023256797.1;refGene_gn=PTPRO;refGene_tx=XM_023256798.1
B4 51312394 51312530
refGene_gn=PTPRO;refGene_tx=XM_023256797.1;refGene_gn=PTPRO;refGene_tx=XM_023256798.1
B4 51317500 51317605
refGene_gn=PTPRO;refGene_tx=XM_023256797.1;refGene_gn=PTPRO;refGene_tx=XM_023256798.1
B4 51334145 51334259
refGene_gn=EPS8;refGene_tx=XM_011283819.3;refGene_gn=EPS8;refGene_tx=XM_023256799.1;refGene_gn=EPS8;refGene_tx=XM_023256800.1;refGene_gn=EPS8;refGene_tx=XM_023256801.1
B4 51336038 51336168
refGene_gn=EPS8;refGene_tx=XM_011283819.3;refGene_gn=EPS8;refGene_tx=X

M_023256799.1;refGene_gn=EPS8;refGene_tx=XM_023256800.1;refGene_gn=EPS8;refGene_tx=XM_023256801.1
B4 51337309 51337490 refGene_gn=EPS8;refGene_tx=XM_011283819.3;refGene_gn=EPS8;refGene_tx=XM_023256799.1;refGene_gn=EPS8;refGene_tx=XM_023256800.1;refGene_gn=EPS8;refGene_tx=XM_023256801.1
B4 51344518 51344741 refGene_gn=EPS8;refGene_tx=XM_011283819.3;refGene_gn=EPS8;refGene_tx=XM_023256799.1;refGene_gn=EPS8;refGene_tx=XM_023256800.1;refGene_gn=EPS8;refGene_tx=XM_023256801.1
B4 51354627 51354771 refGene_gn=EPS8;refGene_tx=XM_011283819.3;refGene_gn=EPS8;refGene_tx=XM_023256799.1;refGene_gn=EPS8;refGene_tx=XM_023256800.1;refGene_gn=EPS8;refGene_tx=XM_023256801.1
B4 51355659 51355768 refGene_gn=EPS8;refGene_tx=XM_011283819.3;refGene_gn=EPS8;refGene_tx=XM_023256799.1;refGene_gn=EPS8;refGene_tx=XM_023256800.1;refGene_gn=EPS8;refGene_tx=XM_023256801.1
B4 51361024 51361155 refGene_gn=EPS8;refGene_tx=XM_011283819.3;refGene_gn=EPS8;refGene_tx=XM_023256799.1;refGene_gn=EPS8;refGene_tx=XM_023256800.1;refGene_gn=EPS8;refGene_tx=XM_023256801.1
B4 51366174 51366358 refGene_gn=EPS8;refGene_tx=XM_011283819.3;refGene_gn=EPS8;refGene_tx=XM_023256799.1;refGene_gn=EPS8;refGene_tx=XM_023256800.1;refGene_gn=EPS8;refGene_tx=XM_023256801.1
B4 51369096 51369245 refGene_gn=EPS8;refGene_tx=XM_011283819.3;refGene_gn=EPS8;refGene_tx=XM_023256799.1;refGene_gn=EPS8;refGene_tx=XM_023256800.1;refGene_gn=EPS8;refGene_tx=XM_023256801.1
B4 51374812 51374887 refGene_gn=EPS8;refGene_tx=XM_011283819.3;refGene_gn=EPS8;refGene_tx=XM_023256799.1;refGene_gn=EPS8;refGene_tx=XM_023256800.1;refGene_gn=EPS8;refGene_tx=XM_023256801.1
B4 51375263 51375352 refGene_gn=EPS8;refGene_tx=XM_011283819.3;refGene_gn=EPS8;refGene_tx=XM_023256799.1;refGene_gn=EPS8;refGene_tx=XM_023256800.1;refGene_gn=EPS8;refGene_tx=XM_023256801.1
B4 51377527 51377654 refGene_gn=EPS8;refGene_tx=XM_011283819.3;refGene_gn=EPS8;refGene_tx=XM_023256799.1;refGene_gn=EPS8;refGene_tx=XM_023256800.1;refGene_gn=EPS8;refGene_tx=XM_023256801.1
B4 51378907 51378981 refGene_gn=EPS8;refGene_tx=XM_011283819.3;refGene_gn=EPS8;refGene_tx=XM_023256799.1;refGene_gn=EPS8;refGene_tx=XM_023256800.1;refGene_gn=EPS8;refGene_tx=XM_023256801.1
B4 51382510 51382647 refGene_gn=EPS8;refGene_tx=XM_011283819.3;refGene_gn=EPS8;refGene_tx=XM_023256799.1;refGene_gn=EPS8;refGene_tx=XM_023256800.1;refGene_gn=EPS8;refGene_tx=XM_023256801.1

B4 51383247 51383330
refGene_gn=EPS8;refGene_tx=XM_011283819.3;refGene_gn=EPS8;refGene_tx=XM_023256799.1;refGene_gn=EPS8;refGene_tx=XM_023256800.1;refGene_gn=EPS8;refGene_tx=XM_023256801.1
B4 51389557 51389707
refGene_gn=EPS8;refGene_tx=XM_011283819.3;refGene_gn=EPS8;refGene_tx=XM_023256799.1;refGene_gn=EPS8;refGene_tx=XM_023256800.1;refGene_gn=EPS8;refGene_tx=XM_023256801.1
B4 51391482 51391644
refGene_gn=EPS8;refGene_tx=XM_011283819.3;refGene_gn=EPS8;refGene_tx=XM_023256799.1;refGene_gn=EPS8;refGene_tx=XM_023256800.1
B4 51393013 51393081
refGene_gn=EPS8;refGene_tx=XM_011283819.3;refGene_gn=EPS8;refGene_tx=XM_023256799.1;refGene_gn=EPS8;refGene_tx=XM_023256800.1
B4 51403532 51403609
refGene_gn=EPS8;refGene_tx=XM_011283819.3;refGene_gn=EPS8;refGene_tx=XM_023256799.1;refGene_gn=EPS8;refGene_tx=XM_023256800.1
B4 51405497 51405556
refGene_gn=EPS8;refGene_tx=XM_011283819.3;refGene_gn=EPS8;refGene_tx=XM_023256799.1;refGene_gn=EPS8;refGene_tx=XM_023256800.1
B4 51523922 51523984
refGene_gn=LOC111561448;refGene_tx=XM_023257748.1
B4 51528676 51529167
refGene_gn=LOC111561448;refGene_tx=XM_023257748.1
B4 51529281 51529328
refGene_gn=LOC111561448;refGene_tx=XM_023257748.1
B4 51604610 51604722
refGene_gn=STRAP;refGene_tx=XM_003988454.5;refGene_gn=STRAP;refGene_tx=XM_019834580.2
B4 51605431 51605567
refGene_gn=STRAP;refGene_tx=XM_003988454.5;refGene_gn=STRAP;refGene_tx=XM_019834580.2
B4 51611890 51611972
refGene_gn=STRAP;refGene_tx=XM_003988454.5;refGene_gn=STRAP;refGene_tx=XM_019834580.2
B4 51614233 51614306
refGene_gn=STRAP;refGene_tx=XM_003988454.5;refGene_gn=STRAP;refGene_tx=XM_019834580.2
B4 51617546 51617643
refGene_gn=STRAP;refGene_tx=XM_003988454.5;refGene_gn=STRAP;refGene_tx=XM_019834580.2
B4 51618238 51618376
refGene_gn=STRAP;refGene_tx=XM_003988454.5;refGene_gn=STRAP;refGene_tx=XM_019834580.2
B4 51620807 51620944
refGene_gn=STRAP;refGene_tx=XM_003988454.5;refGene_gn=STRAP;refGene_tx=XM_019834580.2
B4 51623502 51623652
refGene_gn=STRAP;refGene_tx=XM_003988454.5;refGene_gn=STRAP;refGene_tx=XM_019834580.2

B4 51624327 51624393
refGene_gn=STRAP;refGene_tx=XM_003988454.5;refGene_gn=STRAP;refGene_tx=XM_019834580.2
B4 51625105 51625167
refGene_gn=STRAP;refGene_tx=XM_003988454.5;refGene_gn=STRAP;refGene_tx=XM_019834580.2
B4 51635375 51635406 refGene_gn=DERA;refGene_tx=XM_023256802.1
B4 51685890 51685988 refGene_gn=DERA;refGene_tx=XM_023256802.1
B4 51687144 51687292 refGene_gn=DERA;refGene_tx=XM_023256802.1
B4 51690237 51690333 refGene_gn=DERA;refGene_tx=XM_023256802.1
B4 51692331 51692466 refGene_gn=DERA;refGene_tx=XM_023256802.1
B4 51708792 51708921 refGene_gn=DERA;refGene_tx=XM_023256802.1
B4 51751288 51751401 refGene_gn=DERA;refGene_tx=XM_023256802.1
B4 51754941 51755091 refGene_gn=DERA;refGene_tx=XM_023256802.1
B4 51755420 51755477 refGene_gn=DERA;refGene_tx=XM_023256802.1
B4 51908422 51908570 refGene_gn=SLC15A5;refGene_tx=XM_003988456.5
B4 51913637 51913746 refGene_gn=SLC15A5;refGene_tx=XM_003988456.5
B4 51938648 51938777 refGene_gn=SLC15A5;refGene_tx=XM_003988456.5
B4 51948936 51949125 refGene_gn=SLC15A5;refGene_tx=XM_003988456.5
B4 51964109 51964296 refGene_gn=SLC15A5;refGene_tx=XM_003988456.5
B4 51968525 51968746 refGene_gn=SLC15A5;refGene_tx=XM_003988456.5
B4 51978962 51979132 refGene_gn=SLC15A5;refGene_tx=XM_003988456.5
B4 51994644 51994867 refGene_gn=SLC15A5;refGene_tx=XM_003988456.5
B4 51999016 51999377 refGene_gn=SLC15A5;refGene_tx=XM_003988456.5
B4 52070808 52070934
refGene_gn=MGST1;refGene_tx=XM_003988457.5;refGene_gn=MGST1;refGene_tx=XM_006933539.3;refGene_gn=MGST1;refGene_tx=XM_006933540.3;refGene_gn=MGST1;refGene_tx=XM_006933541.3;refGene_gn=MGST1;refGene_tx=XM_006933542.4;refGene_gn=MGST1;refGene_tx=XM_006933543.4;refGene_gn=MGST1;refGene_tx=XM_019834584.2
B4 52073391 52073486
refGene_gn=MGST1;refGene_tx=XM_003988457.5;refGene_gn=MGST1;refGene_tx=XM_006933539.3;refGene_gn=MGST1;refGene_tx=XM_006933540.3;refGene_gn=MGST1;refGene_tx=XM_006933541.3;refGene_gn=MGST1;refGene_tx=XM_006933542.4;refGene_gn=MGST1;refGene_tx=XM_006933543.4;refGene_gn=MGST1;refGene_tx=XM_019834584.2
B4 52077746 52077945 refGene_gn=MGST1;refGene_tx=XM_019834584.2
B4 52084175 52084422
refGene_gn=MGST1;refGene_tx=XM_003988457.5;refGene_gn=MGST1;refGene_tx=XM_006933539.3;refGene_gn=MGST1;refGene_tx=XM_006933540.3;refGene_gn=MGST1;refGene_tx=XM_006933541.3;refGene_gn=MGST1;refGene_tx=XM_006933542.4;refGene_gn=MGST1;refGene_tx=XM_006933543.4
B4 52258146 52258252
refGene_gn=LMO3;refGene_tx=XM_006933544.3;refGene_gn=LMO3;refGene_tx=XM_006933546.4;refGene_gn=LMO3;refGene_tx=XM_019834586.2;refGene_gn=LMO3;refGene_tx=XM_019834587.2;refGene_gn=LMO3;refGene_tx=XM_019834588.2;refGene_gn=LMO3;refGene_tx=XM_019834590.2;refGene_gn=LMO3;refGene_tx=XM_023256803.1;refGene_gn=LMO3;refGene_tx=XM_023256804.1
B4 52264126 52264181 refGene_gn=LMO3;refGene_tx=XM_019834589.2
B4 52267950 52268076

refGene_gn=LMO3;refGene_tx=XM_006933544.3;refGene_gn=LMO3;refGene_tx=XM_006933546.4;refGene_gn=LMO3;refGene_tx=XM_019834586.2;refGene_gn=LMO3;refGene_tx=XM_019834587.2;refGene_gn=LMO3;refGene_tx=XM_019834588.2;refGene_gn=LMO3;refGene_tx=XM_019834589.2;refGene_gn=LMO3;refGene_tx=XM_019834590.2;refGene_gn=LMO3;refGene_tx=XM_023256803.1;refGene_gn=LMO3;refGene_tx=XM_023256804.1
B4 52305284 52305498
refGene_gn=LMO3;refGene_tx=XM_006933544.3;refGene_gn=LMO3;refGene_tx=XM_006933546.4;refGene_gn=LMO3;refGene_tx=XM_019834586.2;refGene_gn=LMO3;refGene_tx=XM_019834587.2;refGene_gn=LMO3;refGene_tx=XM_019834588.2;refGene_gn=LMO3;refGene_tx=XM_019834589.2;refGene_gn=LMO3;refGene_tx=XM_019834590.2;refGene_gn=LMO3;refGene_tx=XM_023256803.1;refGene_gn=LMO3;refGene_tx=XM_023256804.1
B4 52309541 52309566
refGene_gn=LMO3;refGene_tx=XM_019834586.2;refGene_gn=LMO3;refGene_tx=XM_019834589.2
B4 53524975 53525276
refGene_gn=RERGL;refGene_tx=XM_003988473.5;refGene_gn=RERGL;refGene_tx=XM_011283826.3;refGene_gn=RERGL;refGene_tx=XM_019834606.2;refGene_gn=RERGL;refGene_tx=XM_023256806.1
B4 53527777 53527926
refGene_gn=RERGL;refGene_tx=XM_003988473.5;refGene_gn=RERGL;refGene_tx=XM_011283826.3;refGene_gn=RERGL;refGene_tx=XM_019834606.2;refGene_gn=RERGL;refGene_tx=XM_023256806.1
B4 53528184 53528258
refGene_gn=RERGL;refGene_tx=XM_003988473.5;refGene_gn=RERGL;refGene_tx=XM_011283826.3;refGene_gn=RERGL;refGene_tx=XM_019834606.2;refGene_gn=RERGL;refGene_tx=XM_023256806.1
B4 53531987 53532044
refGene_gn=RERGL;refGene_tx=XM_003988473.5;refGene_gn=RERGL;refGene_tx=XM_011283826.3;refGene_gn=RERGL;refGene_tx=XM_023256806.1
B4 53533090 53533142
refGene_gn=RERGL;refGene_tx=XM_003988473.5;refGene_gn=RERGL;refGene_tx=XM_011283826.3;refGene_gn=RERGL;refGene_tx=XM_019834606.2
B4 53535341 53535354 refGene_gn=RERGL;refGene_tx=XM_023256806.1
B4 53693040 53693718
refGene_gn=PIK3C2G;refGene_tx=XM_006933551.3;refGene_gn=PIK3C2G;refGene_tx=XM_011283827.2;refGene_gn=PIK3C2G;refGene_tx=XM_019834592.2;refGene_gn=PIK3C2G;refGene_tx=XM_019834593.1;refGene_gn=PIK3C2G;refGene_tx=XM_019834594.2
B4 53699172 53699255
refGene_gn=PIK3C2G;refGene_tx=XM_006933551.3;refGene_gn=PIK3C2G;refGene_tx=XM_011283827.2;refGene_gn=PIK3C2G;refGene_tx=XM_019834592.2;refGene_gn=PIK3C2G;refGene_tx=XM_019834593.1;refGene_gn=PIK3C2G;refGene_tx=XM_019834594.2
B4 53702695 53702853
refGene_gn=PIK3C2G;refGene_tx=XM_006933551.3;refGene_gn=PIK3C2G;refGene_tx=XM_011283827.2;refGene_gn=PIK3C2G;refGene_tx=XM_019834592.2;refGene_gn=PIK3C2G;refGene_tx=XM_019834593.1;refGene_gn=PIK3C2G;refGene_tx=XM_019834594.2

B4 53708233 53708348
refGene_gn=PIK3C2G;refGene_tx=XM_006933551.3;refGene_gn=PIK3C2G;refGene_tx=XM_011283827.2;refGene_gn=PIK3C2G;refGene_tx=XM_019834592.2;refGene_gn=PIK3C2G;refGene_tx=XM_019834593.1;refGene_gn=PIK3C2G;refGene_tx=XM_019834594.2
B4 53727005 53727108
refGene_gn=PIK3C2G;refGene_tx=XM_006933551.3;refGene_gn=PIK3C2G;refGene_tx=XM_011283827.2;refGene_gn=PIK3C2G;refGene_tx=XM_019834592.2;refGene_gn=PIK3C2G;refGene_tx=XM_019834593.1;refGene_gn=PIK3C2G;refGene_tx=XM_019834594.2
B4 53732732 53732803
refGene_gn=PIK3C2G;refGene_tx=XM_006933551.3;refGene_gn=PIK3C2G;refGene_tx=XM_011283827.2;refGene_gn=PIK3C2G;refGene_tx=XM_019834592.2;refGene_gn=PIK3C2G;refGene_tx=XM_019834593.1;refGene_gn=PIK3C2G;refGene_tx=XM_019834594.2
B4 53741534 53741598
refGene_gn=PIK3C2G;refGene_tx=XM_006933551.3;refGene_gn=PIK3C2G;refGene_tx=XM_011283827.2;refGene_gn=PIK3C2G;refGene_tx=XM_019834592.2;refGene_gn=PIK3C2G;refGene_tx=XM_019834593.1;refGene_gn=PIK3C2G;refGene_tx=XM_019834594.2
B4 53753954 53754077
refGene_gn=PIK3C2G;refGene_tx=XM_006933551.3;refGene_gn=PIK3C2G;refGene_tx=XM_011283827.2;refGene_gn=PIK3C2G;refGene_tx=XM_019834592.2;refGene_gn=PIK3C2G;refGene_tx=XM_019834594.2
B4 53758971 53759005
refGene_gn=PIK3C2G;refGene_tx=XM_006933551.3;refGene_gn=PIK3C2G;refGene_tx=XM_011283827.2;refGene_gn=PIK3C2G;refGene_tx=XM_019834592.2;refGene_gn=PIK3C2G;refGene_tx=XM_019834593.1;refGene_gn=PIK3C2G;refGene_tx=XM_019834594.2
B4 53764109 53764305
refGene_gn=PIK3C2G;refGene_tx=XM_006933551.3;refGene_gn=PIK3C2G;refGene_tx=XM_011283827.2;refGene_gn=PIK3C2G;refGene_tx=XM_019834592.2;refGene_gn=PIK3C2G;refGene_tx=XM_019834593.1;refGene_gn=PIK3C2G;refGene_tx=XM_019834594.2
B4 53766797 53766920
refGene_gn=PIK3C2G;refGene_tx=XM_006933551.3;refGene_gn=PIK3C2G;refGene_tx=XM_019834592.2;refGene_gn=PIK3C2G;refGene_tx=XM_019834593.1;refGene_gn=PIK3C2G;refGene_tx=XM_019834594.2
B4 53773536 53773668
refGene_gn=PIK3C2G;refGene_tx=XM_006933551.3;refGene_gn=PIK3C2G;refGene_tx=XM_011283827.2;refGene_gn=PIK3C2G;refGene_tx=XM_019834592.2;refGene_gn=PIK3C2G;refGene_tx=XM_019834593.1;refGene_gn=PIK3C2G;refGene_tx=XM_019834594.2
B4 53787394 53787509
refGene_gn=PIK3C2G;refGene_tx=XM_006933551.3;refGene_gn=PIK3C2G;refGene_tx=XM_011283827.2;refGene_gn=PIK3C2G;refGene_tx=XM_019834592.2;refGene_gn=PIK3C2G;refGene_tx=XM_019834593.1;refGene_gn=PIK3C2G;refGene_tx=XM_019834594.2
B4 53794775 53794906
refGene_gn=PIK3C2G;refGene_tx=XM_006933551.3;refGene_gn=PIK3C2G;refGen

e_tx=XM_011283827.2;refGene_gn=PIK3C2G;refGene_tx=XM_019834592.2;refGene_gn=PIK3C2G;refGene_tx=XM_019834593.1;refGene_gn=PIK3C2G;refGene_tx=XM_019834594.2
B4 53802822 53803011
refGene_gn=PIK3C2G;refGene_tx=XM_006933551.3;refGene_gn=PIK3C2G;refGene_tx=XM_011283827.2;refGene_gn=PIK3C2G;refGene_tx=XM_019834592.2;refGene_gn=PIK3C2G;refGene_tx=XM_019834593.1;refGene_gn=PIK3C2G;refGene_tx=XM_019834594.2
B4 53820533 53820627
refGene_gn=PIK3C2G;refGene_tx=XM_006933551.3;refGene_gn=PIK3C2G;refGene_tx=XM_011283827.2;refGene_gn=PIK3C2G;refGene_tx=XM_019834592.2;refGene_gn=PIK3C2G;refGene_tx=XM_019834593.1;refGene_gn=PIK3C2G;refGene_tx=XM_019834594.2
B4 53824636 53824731
refGene_gn=PIK3C2G;refGene_tx=XM_006933551.3;refGene_gn=PIK3C2G;refGene_tx=XM_011283827.2;refGene_gn=PIK3C2G;refGene_tx=XM_019834592.2;refGene_gn=PIK3C2G;refGene_tx=XM_019834593.1;refGene_gn=PIK3C2G;refGene_tx=XM_019834594.2
B4 53887342 53887523
refGene_gn=PIK3C2G;refGene_tx=XM_006933551.3;refGene_gn=PIK3C2G;refGene_tx=XM_011283827.2;refGene_gn=PIK3C2G;refGene_tx=XM_019834592.2;refGene_gn=PIK3C2G;refGene_tx=XM_019834593.1;refGene_gn=PIK3C2G;refGene_tx=XM_019834594.2
B4 53889304 53889412
refGene_gn=PIK3C2G;refGene_tx=XM_006933551.3;refGene_gn=PIK3C2G;refGene_tx=XM_011283827.2;refGene_gn=PIK3C2G;refGene_tx=XM_019834592.2;refGene_gn=PIK3C2G;refGene_tx=XM_019834593.1;refGene_gn=PIK3C2G;refGene_tx=XM_019834594.2
B4 53895144 53895237
refGene_gn=PIK3C2G;refGene_tx=XM_006933551.3;refGene_gn=PIK3C2G;refGene_tx=XM_011283827.2;refGene_gn=PIK3C2G;refGene_tx=XM_019834592.2;refGene_gn=PIK3C2G;refGene_tx=XM_019834593.1;refGene_gn=PIK3C2G;refGene_tx=XM_019834594.2
B4 53896696 53896826
refGene_gn=PIK3C2G;refGene_tx=XM_006933551.3;refGene_gn=PIK3C2G;refGene_tx=XM_011283827.2;refGene_gn=PIK3C2G;refGene_tx=XM_019834592.2;refGene_gn=PIK3C2G;refGene_tx=XM_019834593.1;refGene_gn=PIK3C2G;refGene_tx=XM_019834594.2
B4 53901774 53901911
refGene_gn=PIK3C2G;refGene_tx=XM_006933551.3;refGene_gn=PIK3C2G;refGene_tx=XM_011283827.2;refGene_gn=PIK3C2G;refGene_tx=XM_019834592.2;refGene_gn=PIK3C2G;refGene_tx=XM_019834593.1;refGene_gn=PIK3C2G;refGene_tx=XM_019834594.2
B4 53903743 53903913
refGene_gn=PIK3C2G;refGene_tx=XM_006933551.3;refGene_gn=PIK3C2G;refGene_tx=XM_011283827.2;refGene_gn=PIK3C2G;refGene_tx=XM_019834592.2;refGene_gn=PIK3C2G;refGene_tx=XM_019834593.1;refGene_gn=PIK3C2G;refGene_tx=XM_019834594.2
B4 53923403 53923560
refGene_gn=PIK3C2G;refGene_tx=XM_006933551.3;refGene_gn=PIK3C2G;refGen

e_tx=XM_011283827.2;refGene_gn=PIK3C2G;refGene_tx=XM_019834592.2;refGene_gn=PIK3C2G;refGene_tx=XM_019834593.1;refGene_gn=PIK3C2G;refGene_tx=XM_019834594.2
B4 53930557 53930667
refGene_gn=PIK3C2G;refGene_tx=XM_006933551.3;refGene_gn=PIK3C2G;refGene_tx=XM_011283827.2;refGene_gn=PIK3C2G;refGene_tx=XM_019834592.2;refGene_gn=PIK3C2G;refGene_tx=XM_019834593.1;refGene_gn=PIK3C2G;refGene_tx=XM_019834594.2
B4 53949877 53950067
refGene_gn=PIK3C2G;refGene_tx=XM_006933551.3;refGene_gn=PIK3C2G;refGene_tx=XM_011283827.2;refGene_gn=PIK3C2G;refGene_tx=XM_019834592.2;refGene_gn=PIK3C2G;refGene_tx=XM_019834593.1;refGene_gn=PIK3C2G;refGene_tx=XM_019834594.2
B4 53950558 53950680
refGene_gn=PIK3C2G;refGene_tx=XM_006933551.3;refGene_gn=PIK3C2G;refGene_tx=XM_011283827.2;refGene_gn=PIK3C2G;refGene_tx=XM_019834592.2;refGene_gn=PIK3C2G;refGene_tx=XM_019834593.1;refGene_gn=PIK3C2G;refGene_tx=XM_019834594.2
B4 53954491 53954600
refGene_gn=PIK3C2G;refGene_tx=XM_006933551.3;refGene_gn=PIK3C2G;refGene_tx=XM_011283827.2;refGene_gn=PIK3C2G;refGene_tx=XM_019834592.2;refGene_gn=PIK3C2G;refGene_tx=XM_019834593.1;refGene_gn=PIK3C2G;refGene_tx=XM_019834594.2
B4 53964253 53964268 refGene_gn=PIK3C2G;refGene_tx=XM_019834594.2
B4 53975526 53975602
refGene_gn=PIK3C2G;refGene_tx=XM_006933551.3;refGene_gn=PIK3C2G;refGene_tx=XM_011283827.2;refGene_gn=PIK3C2G;refGene_tx=XM_019834592.2;refGene_gn=PIK3C2G;refGene_tx=XM_019834593.1
B4 53987995 53988090
refGene_gn=PIK3C2G;refGene_tx=XM_006933551.3;refGene_gn=PIK3C2G;refGene_tx=XM_011283827.2;refGene_gn=PIK3C2G;refGene_tx=XM_019834592.2;refGene_gn=PIK3C2G;refGene_tx=XM_019834593.1
B4 54005370 54005496
refGene_gn=PIK3C2G;refGene_tx=XM_006933551.3;refGene_gn=PIK3C2G;refGene_tx=XM_011283827.2;refGene_gn=PIK3C2G;refGene_tx=XM_019834592.2;refGene_gn=PIK3C2G;refGene_tx=XM_019834593.1
B4 54014353 54014506
refGene_gn=PIK3C2G;refGene_tx=XM_006933551.3;refGene_gn=PIK3C2G;refGene_tx=XM_011283827.2;refGene_gn=PIK3C2G;refGene_tx=XM_019834592.2;refGene_gn=PIK3C2G;refGene_tx=XM_019834593.1
B4 54047417 54047503
refGene_gn=PLCZ1;refGene_tx=XM_006933552.4;refGene_gn=PLCZ1;refGene_tx=XM_019834608.2;refGene_gn=PLCZ1;refGene_tx=XM_019834609.2;refGene_gn=PLCZ1;refGene_tx=XM_019834612.2;refGene_gn=PLCZ1;refGene_tx=XM_019834613.2;refGene_gn=PLCZ1;refGene_tx=XM_019834614.2;refGene_gn=PLCZ1;refGene_tx=XM_023256807.1;refGene_gn=PLCZ1;refGene_tx=XM_023256808.1;refGene_gn=PLCZ1;refGene_tx=XM_023256809.1
B4 54048293 54048443
refGene_gn=PLCZ1;refGene_tx=XM_006933552.4;refGene_gn=PLCZ1;refGene_tx=XM_019834608.2;refGene_gn=PLCZ1;refGene_tx=XM_019834609.2;refGene_gn=

PLCZ1;refGene_tx=XM_019834612.2;refGene_gn=PLCZ1;refGene_tx=XM_0198346
13.2;refGene_gn=PLCZ1;refGene_tx=XM_019834614.2;refGene_gn=PLCZ1;refGe
ne_tx=XM_023256807.1;refGene_gn=PLCZ1;refGene_tx=XM_023256808.1;refGen
e_gn=PLCZ1;refGene_tx=XM_023256809.1
B4 54053028 54053161
refGene_gn=PLCZ1;refGene_tx=XM_006933552.4;refGene_gn=PLCZ1;refGene_tx
=XM_019834608.2;refGene_gn=PLCZ1;refGene_tx=XM_019834609.2;refGene_gn=
PLCZ1;refGene_tx=XM_019834612.2;refGene_gn=PLCZ1;refGene_tx=XM_0198346
13.2;refGene_gn=PLCZ1;refGene_tx=XM_019834614.2;refGene_gn=PLCZ1;refGe
ne_tx=XM_023256807.1;refGene_gn=PLCZ1;refGene_tx=XM_023256808.1;refGen
e_gn=PLCZ1;refGene_tx=XM_023256809.1
B4 54059363 54059536
refGene_gn=PLCZ1;refGene_tx=XM_006933552.4;refGene_gn=PLCZ1;refGene_tx
=XM_019834608.2;refGene_gn=PLCZ1;refGene_tx=XM_019834609.2;refGene_gn=
PLCZ1;refGene_tx=XM_019834612.2;refGene_gn=PLCZ1;refGene_tx=XM_0198346
13.2;refGene_gn=PLCZ1;refGene_tx=XM_019834614.2;refGene_gn=PLCZ1;refGe
ne_tx=XM_023256807.1;refGene_gn=PLCZ1;refGene_tx=XM_023256808.1;refGen
e_gn=PLCZ1;refGene_tx=XM_023256809.1
B4 54061084 54061201
refGene_gn=PLCZ1;refGene_tx=XM_006933552.4;refGene_gn=PLCZ1;refGene_tx
=XM_019834608.2;refGene_gn=PLCZ1;refGene_tx=XM_019834609.2;refGene_gn=
PLCZ1;refGene_tx=XM_019834612.2;refGene_gn=PLCZ1;refGene_tx=XM_0198346
13.2;refGene_gn=PLCZ1;refGene_tx=XM_019834614.2;refGene_gn=PLCZ1;refGe
ne_tx=XM_023256808.1;refGene_gn=PLCZ1;refGene_tx=XM_023256809.1
B4 54065120 54065268
refGene_gn=PLCZ1;refGene_tx=XM_006933552.4;refGene_gn=PLCZ1;refGene_tx
=XM_019834608.2;refGene_gn=PLCZ1;refGene_tx=XM_019834609.2;refGene_gn=
PLCZ1;refGene_tx=XM_019834612.2;refGene_gn=PLCZ1;refGene_tx=XM_0198346
13.2;refGene_gn=PLCZ1;refGene_tx=XM_019834614.2;refGene_gn=PLCZ1;refGe
ne_tx=XM_023256807.1;refGene_gn=PLCZ1;refGene_tx=XM_023256808.1;refGen
e_gn=PLCZ1;refGene_tx=XM_023256809.1
B4 54066516 54066738
refGene_gn=PLCZ1;refGene_tx=XM_006933552.4;refGene_gn=PLCZ1;refGene_tx
=XM_019834608.2;refGene_gn=PLCZ1;refGene_tx=XM_019834609.2;refGene_gn=
PLCZ1;refGene_tx=XM_019834612.2;refGene_gn=PLCZ1;refGene_tx=XM_0198346
13.2;refGene_gn=PLCZ1;refGene_tx=XM_019834614.2;refGene_gn=PLCZ1;refGe
ne_tx=XM_023256807.1;refGene_gn=PLCZ1;refGene_tx=XM_023256808.1;refGen
e_gn=PLCZ1;refGene_tx=XM_023256809.1
B4 54069276 54069426
refGene_gn=PLCZ1;refGene_tx=XM_006933552.4;refGene_gn=PLCZ1;refGene_tx
=XM_019834608.2;refGene_gn=PLCZ1;refGene_tx=XM_019834612.2;refGene_gn=
PLCZ1;refGene_tx=XM_019834613.2;refGene_gn=PLCZ1;refGene_tx=XM_0198346
14.2;refGene_gn=PLCZ1;refGene_tx=XM_023256807.1;refGene_gn=PLCZ1;refGe
ne_tx=XM_023256808.1;refGene_gn=PLCZ1;refGene_tx=XM_023256809.1
B4 54074146 54074291
refGene_gn=PLCZ1;refGene_tx=XM_006933552.4;refGene_gn=PLCZ1;refGene_tx
=XM_019834608.2;refGene_gn=PLCZ1;refGene_tx=XM_019834609.2;refGene_gn=
PLCZ1;refGene_tx=XM_019834612.2;refGene_gn=PLCZ1;refGene_tx=XM_0198346
13.2;refGene_gn=PLCZ1;refGene_tx=XM_019834614.2;refGene_gn=PLCZ1;refGe
ne_tx=XM_023256807.1;refGene_gn=PLCZ1;refGene_tx=XM_023256808.1;refGen

e_gn=PLCZ1;refGene_tx=XM_023256809.1
B4 54078684 54078886
refGene_gn=PLCZ1;refGene_tx=XM_006933552.4;refGene_gn=PLCZ1;refGene_tx=XM_019834608.2;refGene_gn=PLCZ1;refGene_tx=XM_019834609.2;refGene_gn=PLCZ1;refGene_tx=XM_019834612.2;refGene_gn=PLCZ1;refGene_tx=XM_019834613.2;refGene_gn=PLCZ1;refGene_tx=XM_019834614.2;refGene_gn=PLCZ1;refGene_tx=XM_023256807.1;refGene_gn=PLCZ1;refGene_tx=XM_023256808.1;refGene_gn=PLCZ1;refGene_tx=XM_023256809.1
B4 54079919 54080151
refGene_gn=PLCZ1;refGene_tx=XM_006933552.4;refGene_gn=PLCZ1;refGene_tx=XM_019834608.2;refGene_gn=PLCZ1;refGene_tx=XM_019834609.2;refGene_gn=PLCZ1;refGene_tx=XM_019834612.2;refGene_gn=PLCZ1;refGene_tx=XM_019834613.2;refGene_gn=PLCZ1;refGene_tx=XM_019834614.2;refGene_gn=PLCZ1;refGene_tx=XM_023256807.1;refGene_gn=PLCZ1;refGene_tx=XM_023256808.1;refGene_gn=PLCZ1;refGene_tx=XM_023256809.1
B4 54091640 54091764
refGene_gn=PLCZ1;refGene_tx=XM_006933552.4;refGene_gn=PLCZ1;refGene_tx=XM_019834608.2;refGene_gn=PLCZ1;refGene_tx=XM_019834609.2;refGene_gn=PLCZ1;refGene_tx=XM_023256807.1
B4 54092792 54092803
refGene_gn=PLCZ1;refGene_tx=XM_006933552.4;refGene_gn=PLCZ1;refGene_tx=XM_019834608.2;refGene_gn=PLCZ1;refGene_tx=XM_019834609.2;refGene_gn=PLCZ1;refGene_tx=XM_023256807.1
B4 54093717 54094617 refGene_gn=CAPZA3;refGene_tx=XM_019834618.2
B4 54437192 54437284
refGene_gn=PLEKHA5;refGene_tx=XM_023256810.1;refGene_gn=PLEKHA5;refGene_tx=XM_023256811.1;refGene_gn=PLEKHA5;refGene_tx=XM_023256812.1;refGene_gn=PLEKHA5;refGene_tx=XM_023256813.1;refGene_gn=PLEKHA5;refGene_tx=XM_023256814.1;refGene_gn=PLEKHA5;refGene_tx=XM_023256815.1;refGene_gn=PLEKHA5;refGene_tx=XM_023256816.1;refGene_gn=PLEKHA5;refGene_tx=XM_023256817.1;refGene_gn=PLEKHA5;refGene_tx=XM_023256818.1;refGene_gn=PLEKHA5;refGene_tx=XM_023256819.1;refGene_gn=PLEKHA5;refGene_tx=XM_023256821.1;refGene_gn=PLEKHA5;refGene_tx=XM_023256822.1;refGene_gn=PLEKHA5;refGene_tx=XM_023256823.1
B4 54437435 54437515
refGene_gn=PLEKHA5;refGene_tx=XM_023256810.1;refGene_gn=PLEKHA5;refGene_tx=XM_023256811.1;refGene_gn=PLEKHA5;refGene_tx=XM_023256812.1;refGene_gn=PLEKHA5;refGene_tx=XM_023256813.1;refGene_gn=PLEKHA5;refGene_tx=XM_023256814.1;refGene_gn=PLEKHA5;refGene_tx=XM_023256815.1;refGene_gn=PLEKHA5;refGene_tx=XM_023256816.1;refGene_gn=PLEKHA5;refGene_tx=XM_023256817.1;refGene_gn=PLEKHA5;refGene_tx=XM_023256818.1;refGene_gn=PLEKHA5;refGene_tx=XM_023256819.1;refGene_gn=PLEKHA5;refGene_tx=XM_023256821.1;refGene_gn=PLEKHA5;refGene_tx=XM_023256822.1;refGene_gn=PLEKHA5;refGene_tx=XM_023256823.1
B4 54439866 54439924
refGene_gn=PLEKHA5;refGene_tx=XM_023256810.1;refGene_gn=PLEKHA5;refGene_tx=XM_023256811.1;refGene_gn=PLEKHA5;refGene_tx=XM_023256812.1;refGene_gn=PLEKHA5;refGene_tx=XM_023256813.1;refGene_gn=PLEKHA5;refGene_tx=XM_023256814.1;refGene_gn=PLEKHA5;refGene_tx=XM_023256815.1;refGene_gn=PLEKHA5;refGene_tx=XM_023256816.1;refGene_gn=PLEKHA5;refGene_tx=XM_02

3256817.1;refGene_gn=PLEKHA5;refGene_tx=XM_023256818.1;refGene_gn=PLEKHA5;refGene_tx=XM_023256819.1;refGene_gn=PLEKHA5;refGene_tx=XM_023256821.1;refGene_gn=PLEKHA5;refGene_tx=XM_023256822.1;refGene_gn=PLEKHA5;refGene_tx=XM_023256823.1
B4	54447459 54447466 refGene_gn=PLEKHA5;refGene_tx=XM_023256822.1
B4	54578620 54578704
refGene_gn=PLEKHA5;refGene_tx=XM_023256810.1;refGene_gn=PLEKHA5;refGene_tx=XM_023256811.1;refGene_gn=PLEKHA5;refGene_tx=XM_023256812.1;refGene_gn=PLEKHA5;refGene_tx=XM_023256813.1;refGene_gn=PLEKHA5;refGene_tx=XM_023256814.1;refGene_gn=PLEKHA5;refGene_tx=XM_023256815.1;refGene_gn=PLEKHA5;refGene_tx=XM_023256816.1;refGene_gn=PLEKHA5;refGene_tx=XM_023256817.1;refGene_gn=PLEKHA5;refGene_tx=XM_023256818.1;refGene_gn=PLEKHA5;refGene_tx=XM_023256819.1;refGene_gn=PLEKHA5;refGene_tx=XM_023256821.1;refGene_gn=PLEKHA5;refGene_tx=XM_023256823.1
B4	54579632 54579753
refGene_gn=PLEKHA5;refGene_tx=XM_019834625.2;refGene_gn=PLEKHA5;refGene_tx=XM_023256810.1;refGene_gn=PLEKHA5;refGene_tx=XM_023256811.1;refGene_gn=PLEKHA5;refGene_tx=XM_023256812.1;refGene_gn=PLEKHA5;refGene_tx=XM_023256813.1;refGene_gn=PLEKHA5;refGene_tx=XM_023256814.1;refGene_gn=PLEKHA5;refGene_tx=XM_023256815.1;refGene_gn=PLEKHA5;refGene_tx=XM_023256816.1;refGene_gn=PLEKHA5;refGene_tx=XM_023256817.1;refGene_gn=PLEKHA5;refGene_tx=XM_023256818.1;refGene_gn=PLEKHA5;refGene_tx=XM_023256819.1;refGene_gn=PLEKHA5;refGene_tx=XM_023256821.1;refGene_gn=PLEKHA5;refGene_tx=XM_023256823.1
B4	54582707 54582812
refGene_gn=PLEKHA5;refGene_tx=XM_019834625.2;refGene_gn=PLEKHA5;refGene_tx=XM_023256810.1;refGene_gn=PLEKHA5;refGene_tx=XM_023256811.1;refGene_gn=PLEKHA5;refGene_tx=XM_023256812.1;refGene_gn=PLEKHA5;refGene_tx=XM_023256813.1;refGene_gn=PLEKHA5;refGene_tx=XM_023256814.1;refGene_gn=PLEKHA5;refGene_tx=XM_023256815.1;refGene_gn=PLEKHA5;refGene_tx=XM_023256816.1;refGene_gn=PLEKHA5;refGene_tx=XM_023256817.1;refGene_gn=PLEKHA5;refGene_tx=XM_023256818.1;refGene_gn=PLEKHA5;refGene_tx=XM_023256819.1;refGene_gn=PLEKHA5;refGene_tx=XM_023256821.1;refGene_gn=PLEKHA5;refGene_tx=XM_023256823.1
B4	54585820 54585893
refGene_gn=PLEKHA5;refGene_tx=XM_019834625.2;refGene_gn=PLEKHA5;refGene_tx=XM_023256810.1;refGene_gn=PLEKHA5;refGene_tx=XM_023256811.1;refGene_gn=PLEKHA5;refGene_tx=XM_023256812.1;refGene_gn=PLEKHA5;refGene_tx=XM_023256813.1;refGene_gn=PLEKHA5;refGene_tx=XM_023256814.1;refGene_gn=PLEKHA5;refGene_tx=XM_023256815.1;refGene_gn=PLEKHA5;refGene_tx=XM_023256816.1;refGene_gn=PLEKHA5;refGene_tx=XM_023256817.1;refGene_gn=PLEKHA5;refGene_tx=XM_023256818.1;refGene_gn=PLEKHA5;refGene_tx=XM_023256819.1;refGene_gn=PLEKHA5;refGene_tx=XM_023256821.1;refGene_gn=PLEKHA5;refGene_tx=XM_023256823.1
B4	54592278 54592379
refGene_gn=PLEKHA5;refGene_tx=XM_019834625.2;refGene_gn=PLEKHA5;refGene_tx=XM_023256810.1;refGene_gn=PLEKHA5;refGene_tx=XM_023256811.1;refGene_gn=PLEKHA5;refGene_tx=XM_023256812.1;refGene_gn=PLEKHA5;refGene_tx=XM_023256813.1;refGene_gn=PLEKHA5;refGene_tx=XM_023256814.1;refGene_gn=PLEKHA5;refGene_tx=XM_023256815.1;refGene_gn=PLEKHA5;refGene_tx=XM_02

3256816.1;refGene_gn=PLEKHA5;refGene_tx=XM_023256817.1;refGene_gn=PLEKHA5;refGene_tx=XM_023256818.1;refGene_gn=PLEKHA5;refGene_tx=XM_023256819.1;refGene_gn=PLEKHA5;refGene_tx=XM_023256821.1;refGene_gn=PLEKHA5;refGene_tx=XM_023256823.1
B4 54596380 54596496
refGene_gn=PLEKHA5;refGene_tx=XM_019834625.2;refGene_gn=PLEKHA5;refGene_tx=XM_023256810.1;refGene_gn=PLEKHA5;refGene_tx=XM_023256811.1;refGene_gn=PLEKHA5;refGene_tx=XM_023256812.1;refGene_gn=PLEKHA5;refGene_tx=XM_023256813.1;refGene_gn=PLEKHA5;refGene_tx=XM_023256814.1;refGene_gn=PLEKHA5;refGene_tx=XM_023256815.1;refGene_gn=PLEKHA5;refGene_tx=XM_023256816.1;refGene_gn=PLEKHA5;refGene_tx=XM_023256817.1;refGene_gn=PLEKHA5;refGene_tx=XM_023256818.1;refGene_gn=PLEKHA5;refGene_tx=XM_023256819.1;refGene_gn=PLEKHA5;refGene_tx=XM_023256820.1;refGene_gn=PLEKHA5;refGene_tx=XM_023256821.1;refGene_gn=PLEKHA5;refGene_tx=XM_023256823.1
B4 54596789 54596807
refGene_gn=PLEKHA5;refGene_tx=XM_019834625.2;refGene_gn=PLEKHA5;refGene_tx=XM_023256810.1;refGene_gn=PLEKHA5;refGene_tx=XM_023256812.1;refGene_gn=PLEKHA5;refGene_tx=XM_023256813.1;refGene_gn=PLEKHA5;refGene_tx=XM_023256814.1;refGene_gn=PLEKHA5;refGene_tx=XM_023256816.1;refGene_gn=PLEKHA5;refGene_tx=XM_023256817.1;refGene_gn=PLEKHA5;refGene_tx=XM_023256819.1;refGene_gn=PLEKHA5;refGene_tx=XM_023256820.1;refGene_gn=PLEKHA5;refGene_tx=XM_023256821.1;refGene_gn=PLEKHA5;refGene_tx=XM_023256823.1
B4 54600238 54600706
refGene_gn=PLEKHA5;refGene_tx=XM_019834625.2;refGene_gn=PLEKHA5;refGene_tx=XM_023256810.1;refGene_gn=PLEKHA5;refGene_tx=XM_023256811.1;refGene_gn=PLEKHA5;refGene_tx=XM_023256812.1;refGene_gn=PLEKHA5;refGene_tx=XM_023256813.1;refGene_gn=PLEKHA5;refGene_tx=XM_023256814.1;refGene_gn=PLEKHA5;refGene_tx=XM_023256815.1;refGene_gn=PLEKHA5;refGene_tx=XM_023256816.1;refGene_gn=PLEKHA5;refGene_tx=XM_023256817.1;refGene_gn=PLEKHA5;refGene_tx=XM_023256818.1;refGene_gn=PLEKHA5;refGene_tx=XM_023256819.1;refGene_gn=PLEKHA5;refGene_tx=XM_023256820.1;refGene_gn=PLEKHA5;refGene_tx=XM_023256821.1;refGene_gn=PLEKHA5;refGene_tx=XM_023256823.1
B4 54606544 54607010
refGene_gn=PLEKHA5;refGene_tx=XM_019834625.2;refGene_gn=PLEKHA5;refGene_tx=XM_023256810.1;refGene_gn=PLEKHA5;refGene_tx=XM_023256811.1;refGene_gn=PLEKHA5;refGene_tx=XM_023256812.1;refGene_gn=PLEKHA5;refGene_tx=XM_023256813.1;refGene_gn=PLEKHA5;refGene_tx=XM_023256814.1;refGene_gn=PLEKHA5;refGene_tx=XM_023256815.1;refGene_gn=PLEKHA5;refGene_tx=XM_023256816.1;refGene_gn=PLEKHA5;refGene_tx=XM_023256817.1;refGene_gn=PLEKHA5;refGene_tx=XM_023256818.1;refGene_gn=PLEKHA5;refGene_tx=XM_023256819.1;refGene_gn=PLEKHA5;refGene_tx=XM_023256820.1;refGene_gn=PLEKHA5;refGene_tx=XM_023256821.1;refGene_gn=PLEKHA5;refGene_tx=XM_023256823.1
B4 54610283 54610367
refGene_gn=PLEKHA5;refGene_tx=XM_019834625.2;refGene_gn=PLEKHA5;refGene_tx=XM_023256810.1;refGene_gn=PLEKHA5;refGene_tx=XM_023256811.1;refGene_gn=PLEKHA5;refGene_tx=XM_023256812.1;refGene_gn=PLEKHA5;refGene_tx=XM_023256813.1;refGene_gn=PLEKHA5;refGene_tx=XM_023256814.1;refGene_gn=PLEKHA5;refGene_tx=XM_023256815.1;refGene_gn=PLEKHA5;refGene_tx=XM_023256816.1;refGene_gn=PLEKHA5;refGene_tx=XM_023256817.1;refGene_gn=PLEK

HA5;refGene_tx=XM_023256818.1;refGene_gn=PLEKHA5;refGene_tx=XM_0232568
19.1;refGene_gn=PLEKHA5;refGene_tx=XM_023256820.1;refGene_gn=PLEKHA5;r
efGene_tx=XM_023256821.1;refGene_gn=PLEKHA5;refGene_tx=XM_023256823.1
B4 54613383 54613503
refGene_gn=PLEKHA5;refGene_tx=XM_019834625.2;refGene_gn=PLEKHA5;refGen
e_tx=XM_023256810.1;refGene_gn=PLEKHA5;refGene_tx=XM_023256811.1;refGe
ne_gn=PLEKHA5;refGene_tx=XM_023256812.1;refGene_gn=PLEKHA5;refGene_tx=
XM_023256813.1;refGene_gn=PLEKHA5;refGene_tx=XM_023256815.1;refGene_gn
=PLEKHA5;refGene_tx=XM_023256819.1;refGene_gn=PLEKHA5;refGene_tx=XM_02
3256820.1;refGene_gn=PLEKHA5;refGene_tx=XM_023256821.1;refGene_gn=PLEK
HA5;refGene_tx=XM_023256823.1
B4 54614310 54614364
refGene_gn=PLEKHA5;refGene_tx=XM_019834625.2;refGene_gn=PLEKHA5;refGen
e_tx=XM_023256810.1;refGene_gn=PLEKHA5;refGene_tx=XM_023256811.1;refGe
ne_gn=PLEKHA5;refGene_tx=XM_023256812.1;refGene_gn=PLEKHA5;refGene_tx=
XM_023256813.1;refGene_gn=PLEKHA5;refGene_tx=XM_023256814.1;refGene_gn
=PLEKHA5;refGene_tx=XM_023256815.1;refGene_gn=PLEKHA5;refGene_tx=XM_02
3256816.1;refGene_gn=PLEKHA5;refGene_tx=XM_023256819.1;refGene_gn=PLEK
HA5;refGene_tx=XM_023256820.1;refGene_gn=PLEKHA5;refGene_tx=XM_0232568
21.1;refGene_gn=PLEKHA5;refGene_tx=XM_023256823.1
B4 54630946 54630985 refGene_gn=PLEKHA5;refGene_tx=XM_023256821.1
B4 54630995 54631076
refGene_gn=PLEKHA5;refGene_tx=XM_019834625.2;refGene_gn=PLEKHA5;refGen
e_tx=XM_023256810.1;refGene_gn=PLEKHA5;refGene_tx=XM_023256811.1;refGe
ne_gn=PLEKHA5;refGene_tx=XM_023256812.1;refGene_gn=PLEKHA5;refGene_tx=
XM_023256814.1;refGene_gn=PLEKHA5;refGene_tx=XM_023256819.1;refGene_gn
=PLEKHA5;refGene_tx=XM_023256820.1;refGene_gn=PLEKHA5;refGene_tx=XM_02
3256823.1
B4 54636403 54636439
refGene_gn=PLEKHA5;refGene_tx=XM_019834625.2;refGene_gn=PLEKHA5;refGen
e_tx=XM_023256810.1;refGene_gn=PLEKHA5;refGene_tx=XM_023256811.1;refGe
ne_gn=PLEKHA5;refGene_tx=XM_023256814.1;refGene_gn=PLEKHA5;refGene_tx=
XM_023256820.1;refGene_gn=PLEKHA5;refGene_tx=XM_023256823.1
B4 54636947 54637010
refGene_gn=PLEKHA5;refGene_tx=XM_019834625.2;refGene_gn=PLEKHA5;refGen
e_tx=XM_023256810.1;refGene_gn=PLEKHA5;refGene_tx=XM_023256811.1;refGe
ne_gn=PLEKHA5;refGene_tx=XM_023256812.1;refGene_gn=PLEKHA5;refGene_tx=
XM_023256813.1;refGene_gn=PLEKHA5;refGene_tx=XM_023256814.1;refGene_gn
=PLEKHA5;refGene_tx=XM_023256815.1;refGene_gn=PLEKHA5;refGene_tx=XM_02
3256816.1;refGene_gn=PLEKHA5;refGene_tx=XM_023256817.1;refGene_gn=PLEK
HA5;refGene_tx=XM_023256818.1;refGene_gn=PLEKHA5;refGene_tx=XM_0232568
19.1;refGene_gn=PLEKHA5;refGene_tx=XM_023256820.1;refGene_gn=PLEKHA5;r
efGene_tx=XM_023256823.1
B4 54638595 54638676
refGene_gn=PLEKHA5;refGene_tx=XM_019834625.2;refGene_gn=PLEKHA5;refGen
e_tx=XM_023256810.1;refGene_gn=PLEKHA5;refGene_tx=XM_023256811.1;refGe
ne_gn=PLEKHA5;refGene_tx=XM_023256812.1;refGene_gn=PLEKHA5;refGene_tx=
XM_023256813.1;refGene_gn=PLEKHA5;refGene_tx=XM_023256814.1;refGene_gn
=PLEKHA5;refGene_tx=XM_023256815.1;refGene_gn=PLEKHA5;refGene_tx=XM_02
3256816.1;refGene_gn=PLEKHA5;refGene_tx=XM_023256817.1;refGene_gn=PLEK

HA5;refGene_tx=XM_023256818.1;refGene_gn=PLEKHA5;refGene_tx=XM_0232568
19.1;refGene_gn=PLEKHA5;refGene_tx=XM_023256820.1;refGene_gn=PLEKHA5;r
efGene_tx=XM_023256823.1
B4 54638807 54638957
refGene_gn=PLEKHA5;refGene_tx=XM_019834625.2;refGene_gn=PLEKHA5;refGen
e_tx=XM_023256810.1;refGene_gn=PLEKHA5;refGene_tx=XM_023256811.1;refGe
ne_gn=PLEKHA5;refGene_tx=XM_023256812.1;refGene_gn=PLEKHA5;refGene_tx=
XM_023256813.1;refGene_gn=PLEKHA5;refGene_tx=XM_023256814.1;refGene_gn
=PLEKHA5;refGene_tx=XM_023256815.1;refGene_gn=PLEKHA5;refGene_tx=XM_02
3256816.1;refGene_gn=PLEKHA5;refGene_tx=XM_023256817.1;refGene_gn=PLEK
HA5;refGene_tx=XM_023256818.1;refGene_gn=PLEKHA5;refGene_tx=XM_0232568
19.1;refGene_gn=PLEKHA5;refGene_tx=XM_023256820.1;refGene_gn=PLEKHA5;r
efGene_tx=XM_023256823.1
B4 54648389 54648491
refGene_gn=PLEKHA5;refGene_tx=XM_019834625.2;refGene_gn=PLEKHA5;refGen
e_tx=XM_023256810.1;refGene_gn=PLEKHA5;refGene_tx=XM_023256811.1;refGe
ne_gn=PLEKHA5;refGene_tx=XM_023256812.1;refGene_gn=PLEKHA5;refGene_tx=
XM_023256813.1;refGene_gn=PLEKHA5;refGene_tx=XM_023256814.1;refGene_gn
=PLEKHA5;refGene_tx=XM_023256815.1;refGene_gn=PLEKHA5;refGene_tx=XM_02
3256816.1;refGene_gn=PLEKHA5;refGene_tx=XM_023256817.1;refGene_gn=PLEK
HA5;refGene_tx=XM_023256818.1;refGene_gn=PLEKHA5;refGene_tx=XM_0232568
19.1;refGene_gn=PLEKHA5;refGene_tx=XM_023256820.1;refGene_gn=PLEKHA5;r
efGene_tx=XM_023256823.1
B4 54651658 54651770
refGene_gn=PLEKHA5;refGene_tx=XM_019834625.2;refGene_gn=PLEKHA5;refGen
e_tx=XM_023256810.1;refGene_gn=PLEKHA5;refGene_tx=XM_023256811.1;refGe
ne_gn=PLEKHA5;refGene_tx=XM_023256812.1;refGene_gn=PLEKHA5;refGene_tx=
XM_023256813.1;refGene_gn=PLEKHA5;refGene_tx=XM_023256814.1;refGene_gn
=PLEKHA5;refGene_tx=XM_023256815.1;refGene_gn=PLEKHA5;refGene_tx=XM_02
3256816.1;refGene_gn=PLEKHA5;refGene_tx=XM_023256817.1;refGene_gn=PLEK
HA5;refGene_tx=XM_023256818.1;refGene_gn=PLEKHA5;refGene_tx=XM_0232568
19.1;refGene_gn=PLEKHA5;refGene_tx=XM_023256820.1;refGene_gn=PLEKHA5;r
efGene_tx=XM_023256823.1
B4 54653519 54653566
refGene_gn=PLEKHA5;refGene_tx=XM_019834625.2;refGene_gn=PLEKHA5;refGen
e_tx=XM_023256810.1;refGene_gn=PLEKHA5;refGene_tx=XM_023256811.1;refGe
ne_gn=PLEKHA5;refGene_tx=XM_023256812.1;refGene_gn=PLEKHA5;refGene_tx=
XM_023256813.1;refGene_gn=PLEKHA5;refGene_tx=XM_023256814.1;refGene_gn
=PLEKHA5;refGene_tx=XM_023256815.1;refGene_gn=PLEKHA5;refGene_tx=XM_02
3256816.1;refGene_gn=PLEKHA5;refGene_tx=XM_023256817.1;refGene_gn=PLEK
HA5;refGene_tx=XM_023256818.1;refGene_gn=PLEKHA5;refGene_tx=XM_0232568
19.1;refGene_gn=PLEKHA5;refGene_tx=XM_023256820.1;refGene_gn=PLEKHA5;r
efGene_tx=XM_023256823.1
B4 54654367 54654655
refGene_gn=PLEKHA5;refGene_tx=XM_023256819.1;refGene_gn=PLEKHA5;refGen
e_tx=XM_023256823.1
B4 54655391 54655509
refGene_gn=PLEKHA5;refGene_tx=XM_019834625.2;refGene_gn=PLEKHA5;refGen
e_tx=XM_023256810.1;refGene_gn=PLEKHA5;refGene_tx=XM_023256811.1;refGe
ne_gn=PLEKHA5;refGene_tx=XM_023256812.1;refGene_gn=PLEKHA5;refGene_tx=

XM_023256813.1;refGene_gn=PLEKHA5;refGene_tx=XM_023256814.1;refGene_gn=PLEKHA5;refGene_tx=XM_023256815.1;refGene_gn=PLEKHA5;refGene_tx=XM_023256816.1;refGene_gn=PLEKHA5;refGene_tx=XM_023256817.1;refGene_gn=PLEKHA5;refGene_tx=XM_023256818.1;refGene_gn=PLEKHA5;refGene_tx=XM_023256820.1;refGene_gn=PLEKHA5;refGene_tx=XM_023256823.1
B4 54663951 54664070
refGene_gn=PLEKHA5;refGene_tx=XM_019834625.2;refGene_gn=PLEKHA5;refGene_tx=XM_023256810.1;refGene_gn=PLEKHA5;refGene_tx=XM_023256811.1;refGene_gn=PLEKHA5;refGene_tx=XM_023256812.1;refGene_gn=PLEKHA5;refGene_tx=XM_023256813.1;refGene_gn=PLEKHA5;refGene_tx=XM_023256814.1;refGene_gn=PLEKHA5;refGene_tx=XM_023256815.1;refGene_gn=PLEKHA5;refGene_tx=XM_023256816.1;refGene_gn=PLEKHA5;refGene_tx=XM_023256817.1;refGene_gn=PLEKHA5;refGene_tx=XM_023256818.1;refGene_gn=PLEKHA5;refGene_tx=XM_023256820.1;refGene_gn=PLEKHA5;refGene_tx=XM_023256823.1
B4 54667822 54668032
refGene_gn=PLEKHA5;refGene_tx=XM_019834625.2;refGene_gn=PLEKHA5;refGene_tx=XM_023256810.1;refGene_gn=PLEKHA5;refGene_tx=XM_023256811.1;refGene_gn=PLEKHA5;refGene_tx=XM_023256812.1;refGene_gn=PLEKHA5;refGene_tx=XM_023256813.1;refGene_gn=PLEKHA5;refGene_tx=XM_023256814.1;refGene_gn=PLEKHA5;refGene_tx=XM_023256815.1;refGene_gn=PLEKHA5;refGene_tx=XM_023256816.1;refGene_gn=PLEKHA5;refGene_tx=XM_023256817.1;refGene_gn=PLEKHA5;refGene_tx=XM_023256818.1;refGene_gn=PLEKHA5;refGene_tx=XM_023256820.1;refGene_gn=PLEKHA5;refGene_tx=XM_023256823.1
B4 54668941 54669082
refGene_gn=PLEKHA5;refGene_tx=XM_019834625.2;refGene_gn=PLEKHA5;refGene_tx=XM_023256810.1;refGene_gn=PLEKHA5;refGene_tx=XM_023256811.1;refGene_gn=PLEKHA5;refGene_tx=XM_023256812.1;refGene_gn=PLEKHA5;refGene_tx=XM_023256813.1;refGene_gn=PLEKHA5;refGene_tx=XM_023256814.1;refGene_gn=PLEKHA5;refGene_tx=XM_023256815.1;refGene_gn=PLEKHA5;refGene_tx=XM_023256816.1;refGene_gn=PLEKHA5;refGene_tx=XM_023256817.1;refGene_gn=PLEKHA5;refGene_tx=XM_023256818.1;refGene_gn=PLEKHA5;refGene_tx=XM_023256820.1;refGene_gn=PLEKHA5;refGene_tx=XM_023256823.1
B4 54669966 54670091
refGene_gn=PLEKHA5;refGene_tx=XM_019834625.2;refGene_gn=PLEKHA5;refGene_tx=XM_023256810.1;refGene_gn=PLEKHA5;refGene_tx=XM_023256811.1;refGene_gn=PLEKHA5;refGene_tx=XM_023256812.1;refGene_gn=PLEKHA5;refGene_tx=XM_023256813.1;refGene_gn=PLEKHA5;refGene_tx=XM_023256814.1;refGene_gn=PLEKHA5;refGene_tx=XM_023256815.1;refGene_gn=PLEKHA5;refGene_tx=XM_023256816.1;refGene_gn=PLEKHA5;refGene_tx=XM_023256817.1;refGene_gn=PLEKHA5;refGene_tx=XM_023256818.1;refGene_gn=PLEKHA5;refGene_tx=XM_023256820.1;refGene_gn=PLEKHA5;refGene_tx=XM_023256823.1
B4 54672389 54672535
refGene_gn=PLEKHA5;refGene_tx=XM_019834625.2;refGene_gn=PLEKHA5;refGene_tx=XM_023256810.1;refGene_gn=PLEKHA5;refGene_tx=XM_023256811.1;refGene_gn=PLEKHA5;refGene_tx=XM_023256812.1;refGene_gn=PLEKHA5;refGene_tx=XM_023256813.1;refGene_gn=PLEKHA5;refGene_tx=XM_023256814.1;refGene_gn=PLEKHA5;refGene_tx=XM_023256815.1;refGene_gn=PLEKHA5;refGene_tx=XM_023256816.1;refGene_gn=PLEKHA5;refGene_tx=XM_023256817.1;refGene_gn=PLEKHA5;refGene_tx=XM_023256818.1;refGene_gn=PLEKHA5;refGene_tx=XM_023256820.1;refGene_gn=PLEKHA5;refGene_tx=XM_023256823.1

B4 54674200 54674295
refGene_gn=PLEKHA5;refGene_tx=XM_019834625.2;refGene_gn=PLEKHA5;refGen
e_tx=XM_023256810.1;refGene_gn=PLEKHA5;refGene_tx=XM_023256811.1;refGe
ne_gn=PLEKHA5;refGene_tx=XM_023256812.1;refGene_gn=PLEKHA5;refGene_tx=
XM_023256813.1;refGene_gn=PLEKHA5;refGene_tx=XM_023256814.1;refGene_gn
=PLEKHA5;refGene_tx=XM_023256815.1;refGene_gn=PLEKHA5;refGene_tx=XM_02
3256816.1;refGene_gn=PLEKHA5;refGene_tx=XM_023256817.1;refGene_gn=PLEK
HA5;refGene_tx=XM_023256818.1;refGene_gn=PLEKHA5;refGene_tx=XM_0232568
20.1;refGene_gn=PLEKHA5;refGene_tx=XM_023256823.1
B4 54729054 54729722
refGene_gn=AEBP2;refGene_tx=XM_023256824.1;refGene_gn=AEBP2;refGene_tx
=XM_023256825.1;refGene_gn=AEBP2;refGene_tx=XM_023256826.1
B4 54751539 54751747
refGene_gn=AEBP2;refGene_tx=XM_023256824.1;refGene_gn=AEBP2;refGene_tx
=XM_023256825.1;refGene_gn=AEBP2;refGene_tx=XM_023256826.1
B4 54759504 54759612
refGene_gn=AEBP2;refGene_tx=XM_023256824.1;refGene_gn=AEBP2;refGene_tx
=XM_023256825.1;refGene_gn=AEBP2;refGene_tx=XM_023256826.1
B4 54781586 54781773
refGene_gn=AEBP2;refGene_tx=XM_023256824.1;refGene_gn=AEBP2;refGene_tx
=XM_023256825.1;refGene_gn=AEBP2;refGene_tx=XM_023256826.1
B4 54785592 54785717
refGene_gn=AEBP2;refGene_tx=XM_023256824.1;refGene_gn=AEBP2;refGene_tx
=XM_023256825.1;refGene_gn=AEBP2;refGene_tx=XM_023256826.1
B4 54791951 54792019
refGene_gn=AEBP2;refGene_tx=XM_023256824.1;refGene_gn=AEBP2;refGene_tx
=XM_023256825.1;refGene_gn=AEBP2;refGene_tx=XM_023256826.1
B4 54794578 54794692
refGene_gn=AEBP2;refGene_tx=XM_023256824.1;refGene_gn=AEBP2;refGene_tx
=XM_023256825.1;refGene_gn=AEBP2;refGene_tx=XM_023256826.1
B4 54795186 54795202 refGene_gn=AEBP2;refGene_tx=XM_023256826.1
B4 54797303 54797330
refGene_gn=AEBP2;refGene_tx=XM_023256824.1;refGene_gn=AEBP2;refGene_tx
=XM_023256825.1
B4 54797867 54797913 refGene_gn=AEBP2;refGene_tx=XM_023256824.1
B4 54822849 54822889 refGene_gn=AEBP2;refGene_tx=XM_023256825.1
B4 55636463 55637423 refGene_gn=PDE3A;refGene_tx=XM_019834634.2
B4 55822248 55822299 refGene_gn=PDE3A;refGene_tx=XM_019834634.2
B4 55884182 55884440 refGene_gn=PDE3A;refGene_tx=XM_019834634.2
B4 55885883 55886038 refGene_gn=PDE3A;refGene_tx=XM_019834634.2
B4 55892333 55892458 refGene_gn=PDE3A;refGene_tx=XM_019834634.2
B4 55908264 55908484 refGene_gn=PDE3A;refGene_tx=XM_019834634.2
B4 55911849 55911935 refGene_gn=PDE3A;refGene_tx=XM_019834634.2
B4 55914982 55915137 refGene_gn=PDE3A;refGene_tx=XM_019834634.2
B4 55916506 55916644 refGene_gn=PDE3A;refGene_tx=XM_019834634.2
B4 55923475 55923587 refGene_gn=PDE3A;refGene_tx=XM_019834634.2
B4 55930881 55930995 refGene_gn=PDE3A;refGene_tx=XM_019834634.2
B4 55931131 55931331 refGene_gn=PDE3A;refGene_tx=XM_019834634.2
B4 55932963 55933135 refGene_gn=PDE3A;refGene_tx=XM_019834634.2
B4 55933137 55933166 refGene_gn=PDE3A;refGene_tx=XM_019834634.2

B4 55934637 55934793 refGene_gn=PDE3A;refGene_tx=XM_019834634.2
B4 55936955 55937184 refGene_gn=PDE3A;refGene_tx=XM_019834634.2
B4 55949021 55949263 refGene_gn=PDE3A;refGene_tx=XM_019834634.2
B4 55969637 55969766
refGene_gn=SLCO1C1;refGene_tx=XM_019834637.2;refGene_gn=SLCO1C1;refGen
e_tx=XM_019834639.2;refGene_gn=SLCO1C1;refGene_tx=XM_019834640.2;refGe
ne_gn=SLCO1C1;refGene_tx=XM_023256827.1
B4 55971200 55971342
refGene_gn=SLCO1C1;refGene_tx=XM_019834637.2;refGene_gn=SLCO1C1;refGen
e_tx=XM_019834639.2;refGene_gn=SLCO1C1;refGene_tx=XM_019834640.2;refGe
ne_gn=SLCO1C1;refGene_tx=XM_023256827.1
B4 55982458 55982591
refGene_gn=SLCO1C1;refGene_tx=XM_019834637.2;refGene_gn=SLCO1C1;refGen
e_tx=XM_019834639.2;refGene_gn=SLCO1C1;refGene_tx=XM_019834640.2;refGe
ne_gn=SLCO1C1;refGene_tx=XM_023256827.1
B4 55987048 55987077 refGene_gn=SLCO1C1;refGene_tx=XM_019834641.1
B4 55994151 55994276
refGene_gn=SLCO1C1;refGene_tx=XM_019834637.2;refGene_gn=SLCO1C1;refGen
e_tx=XM_019834639.2;refGene_gn=SLCO1C1;refGene_tx=XM_019834640.2;refGe
ne_gn=SLCO1C1;refGene_tx=XM_019834641.1;refGene_gn=SLCO1C1;refGene_tx=
XM_023256827.1
B4 55997672 55997819
refGene_gn=SLCO1C1;refGene_tx=XM_019834637.2;refGene_gn=SLCO1C1;refGen
e_tx=XM_019834639.2;refGene_gn=SLCO1C1;refGene_tx=XM_019834641.1;refGe
ne_gn=SLCO1C1;refGene_tx=XM_023256827.1
B4 55999721 55999820
refGene_gn=SLCO1C1;refGene_tx=XM_019834637.2;refGene_gn=SLCO1C1;refGen
e_tx=XM_019834641.1;refGene_gn=SLCO1C1;refGene_tx=XM_023256827.1
B4 56001805 56002051
refGene_gn=SLCO1C1;refGene_tx=XM_019834637.2;refGene_gn=SLCO1C1;refGen
e_tx=XM_019834639.2;refGene_gn=SLCO1C1;refGene_tx=XM_019834640.2;refGe
ne_gn=SLCO1C1;refGene_tx=XM_019834641.1;refGene_gn=SLCO1C1;refGene_tx=
XM_023256827.1
B4 56003329 56003494
refGene_gn=SLCO1C1;refGene_tx=XM_019834637.2;refGene_gn=SLCO1C1;refGen
e_tx=XM_019834639.2;refGene_gn=SLCO1C1;refGene_tx=XM_019834640.2;refGe
ne_gn=SLCO1C1;refGene_tx=XM_019834641.1;refGene_gn=SLCO1C1;refGene_tx=
XM_023256827.1
B4 56011205 56011401
refGene_gn=SLCO1C1;refGene_tx=XM_019834637.2;refGene_gn=SLCO1C1;refGen
e_tx=XM_019834639.2;refGene_gn=SLCO1C1;refGene_tx=XM_019834640.2;refGe
ne_gn=SLCO1C1;refGene_tx=XM_019834641.1;refGene_gn=SLCO1C1;refGene_tx=
XM_023256827.1
B4 56015427 56015593
refGene_gn=SLCO1C1;refGene_tx=XM_019834637.2;refGene_gn=SLCO1C1;refGen
e_tx=XM_019834639.2;refGene_gn=SLCO1C1;refGene_tx=XM_019834640.2;refGe
ne_gn=SLCO1C1;refGene_tx=XM_019834641.1;refGene_gn=SLCO1C1;refGene_tx=
XM_023256827.1
B4 56018689 56018874
refGene_gn=SLCO1C1;refGene_tx=XM_019834637.2;refGene_gn=SLCO1C1;refGen

e_tx=XM_019834639.2;refGene_gn=SLCO1C1;refGene_tx=XM_019834640.2;refGene_gn=SLCO1C1;refGene_tx=XM_019834641.1;refGene_gn=SLCO1C1;refGene_tx=XM_023256827.1
B4 56020656 56020721
refGene_gn=SLCO1C1;refGene_tx=XM_019834637.2;refGene_gn=SLCO1C1;refGene_tx=XM_019834639.2;refGene_gn=SLCO1C1;refGene_tx=XM_019834640.2;refGene_gn=SLCO1C1;refGene_tx=XM_019834641.1;refGene_gn=SLCO1C1;refGene_tx=XM_023256827.1
B4 56024425 56024543
refGene_gn=SLCO1C1;refGene_tx=XM_019834637.2;refGene_gn=SLCO1C1;refGene_tx=XM_019834639.2;refGene_gn=SLCO1C1;refGene_tx=XM_019834640.2;refGene_gn=SLCO1C1;refGene_tx=XM_019834641.1;refGene_gn=SLCO1C1;refGene_tx=XM_023256827.1
B4 56026057 56026280
refGene_gn=SLCO1C1;refGene_tx=XM_019834637.2;refGene_gn=SLCO1C1;refGene_tx=XM_019834639.2;refGene_gn=SLCO1C1;refGene_tx=XM_019834640.2;refGene_gn=SLCO1C1;refGene_tx=XM_019834641.1;refGene_gn=SLCO1C1;refGene_tx=XM_023256827.1
B4 56053558 56053645
refGene_gn=SLCO1B3;refGene_tx=XM_019834642.1;refGene_gn=SLCO1B3;refGene_tx=XM_019834643.2
B4 56080998 56081140
refGene_gn=SLCO1B3;refGene_tx=XM_019834642.1;refGene_gn=SLCO1B3;refGene_tx=XM_019834643.2
B4 56084658 56084791
refGene_gn=SLCO1B3;refGene_tx=XM_019834642.1;refGene_gn=SLCO1B3;refGene_tx=XM_019834643.2
B4 56087524 56087646
refGene_gn=SLCO1B3;refGene_tx=XM_019834642.1;refGene_gn=SLCO1B3;refGene_tx=XM_019834643.2
B4 56089001 56089148
refGene_gn=SLCO1B3;refGene_tx=XM_019834642.1;refGene_gn=SLCO1B3;refGene_tx=XM_019834643.2
B4 56089309 56089408
refGene_gn=SLCO1B3;refGene_tx=XM_019834642.1;refGene_gn=SLCO1B3;refGene_tx=XM_019834643.2
B4 56089909 56090152
refGene_gn=SLCO1B3;refGene_tx=XM_019834642.1;refGene_gn=SLCO1B3;refGene_tx=XM_019834643.2
B4 56091189 56091354
refGene_gn=SLCO1B3;refGene_tx=XM_019834642.1;refGene_gn=SLCO1B3;refGene_tx=XM_019834643.2
B4 56092968 56093164
refGene_gn=SLCO1B3;refGene_tx=XM_019834642.1;refGene_gn=SLCO1B3;refGene_tx=XM_019834643.2
B4 56096838 56097004
refGene_gn=SLCO1B3;refGene_tx=XM_019834642.1;refGene_gn=SLCO1B3;refGene_tx=XM_019834643.2
B4 56099099 56099284
refGene_gn=SLCO1B3;refGene_tx=XM_019834642.1;refGene_gn=SLCO1B3;refGen

e_tx=XM_019834643.2
B4 56104996 56105061
refGene_gn=SLCO1B3;refGene_tx=XM_019834642.1;refGene_gn=SLCO1B3;refGene_tx=XM_019834643.2
B4 56108152 56108270
refGene_gn=SLCO1B3;refGene_tx=XM_019834642.1;refGene_gn=SLCO1B3;refGene_tx=XM_019834643.2
B4 56118063 56118271
refGene_gn=SLCO1B3;refGene_tx=XM_019834642.1;refGene_gn=SLCO1B3;refGene_tx=XM_019834643.2
B4 56129025 56129245
refGene_gn=SLCO1A2;refGene_tx=XM_003988480.5;refGene_gn=SLCO1A2;refGene_tx=XM_019834644.2;refGene_gn=SLCO1A2;refGene_tx=XM_019834645.2;refGene_gn=SLCO1A2;refGene_tx=XM_019834646.2;refGene_gn=SLCO1A2;refGene_tx=XM_019834647.2;refGene_gn=SLCO1A2;refGene_tx=XM_019834648.2;refGene_gn=SLCO1A2;refGene_tx=XM_019834649.2;refGene_gn=SLCO1A2;refGene_tx=XM_019834650.2;refGene_gn=SLCO1A2;refGene_tx=XM_019834651.2;refGene_gn=SLCO1A2;refGene_tx=XM_019834652.2
B4 56142097 56142215
refGene_gn=SLCO1A2;refGene_tx=XM_003988480.5;refGene_gn=SLCO1A2;refGene_tx=XM_019834644.2;refGene_gn=SLCO1A2;refGene_tx=XM_019834645.2;refGene_gn=SLCO1A2;refGene_tx=XM_019834646.2;refGene_gn=SLCO1A2;refGene_tx=XM_019834647.2;refGene_gn=SLCO1A2;refGene_tx=XM_019834648.2;refGene_gn=SLCO1A2;refGene_tx=XM_019834649.2;refGene_gn=SLCO1A2;refGene_tx=XM_019834650.2;refGene_gn=SLCO1A2;refGene_tx=XM_019834651.2;refGene_gn=SLCO1A2;refGene_tx=XM_019834652.2
B4 56142779 56142844
refGene_gn=SLCO1A2;refGene_tx=XM_003988480.5;refGene_gn=SLCO1A2;refGene_tx=XM_019834644.2;refGene_gn=SLCO1A2;refGene_tx=XM_019834645.2;refGene_gn=SLCO1A2;refGene_tx=XM_019834646.2;refGene_gn=SLCO1A2;refGene_tx=XM_019834647.2;refGene_gn=SLCO1A2;refGene_tx=XM_019834648.2;refGene_gn=SLCO1A2;refGene_tx=XM_019834649.2;refGene_gn=SLCO1A2;refGene_tx=XM_019834650.2;refGene_gn=SLCO1A2;refGene_tx=XM_019834651.2;refGene_gn=SLCO1A2;refGene_tx=XM_019834652.2
B4 56149209 56149382
refGene_gn=SLCO1A2;refGene_tx=XM_003988480.5;refGene_gn=SLCO1A2;refGene_tx=XM_019834644.2;refGene_gn=SLCO1A2;refGene_tx=XM_019834645.2;refGene_gn=SLCO1A2;refGene_tx=XM_019834646.2;refGene_gn=SLCO1A2;refGene_tx=XM_019834647.2;refGene_gn=SLCO1A2;refGene_tx=XM_019834648.2;refGene_gn=SLCO1A2;refGene_tx=XM_019834650.2
B4 56150325 56150491
refGene_gn=SLCO1A2;refGene_tx=XM_003988480.5;refGene_gn=SLCO1A2;refGene_tx=XM_019834644.2;refGene_gn=SLCO1A2;refGene_tx=XM_019834645.2;refGene_gn=SLCO1A2;refGene_tx=XM_019834646.2;refGene_gn=SLCO1A2;refGene_tx=XM_019834647.2;refGene_gn=SLCO1A2;refGene_tx=XM_019834648.2;refGene_gn=SLCO1A2;refGene_tx=XM_019834650.2
B4 56156577 56156773
refGene_gn=SLCO1A2;refGene_tx=XM_003988480.5;refGene_gn=SLCO1A2;refGene_tx=XM_019834644.2;refGene_gn=SLCO1A2;refGene_tx=XM_019834645.2;refGene_gn=SLCO1A2;refGene_tx=XM_019834646.2;refGene_gn=SLCO1A2;refGene_tx=

XM_019834647.2;refGene_gn=SLCO1A2;refGene_tx=XM_019834648.2;refGene_gn=SLCO1A2;refGene_tx=XM_019834649.2;refGene_gn=SLCO1A2;refGene_tx=XM_019834650.2;refGene_gn=SLCO1A2;refGene_tx=XM_019834651.2;refGene_gn=SLCO1A2;refGene_tx=XM_019834652.2
B4 56157967 56158132
refGene_gn=SLCO1A2;refGene_tx=XM_003988480.5;refGene_gn=SLCO1A2;refGene_tx=XM_019834644.2;refGene_gn=SLCO1A2;refGene_tx=XM_019834645.2;refGene_gn=SLCO1A2;refGene_tx=XM_019834646.2;refGene_gn=SLCO1A2;refGene_tx=XM_019834647.2;refGene_gn=SLCO1A2;refGene_tx=XM_019834649.2;refGene_gn=SLCO1A2;refGene_tx=XM_019834650.2;refGene_gn=SLCO1A2;refGene_tx=XM_019834651.2
B4 56160690 56160912
refGene_gn=SLCO1A2;refGene_tx=XM_003988480.5;refGene_gn=SLCO1A2;refGene_tx=XM_019834644.2;refGene_gn=SLCO1A2;refGene_tx=XM_019834645.2;refGene_gn=SLCO1A2;refGene_tx=XM_019834646.2;refGene_gn=SLCO1A2;refGene_tx=XM_019834647.2;refGene_gn=SLCO1A2;refGene_tx=XM_019834648.2;refGene_gn=SLCO1A2;refGene_tx=XM_019834649.2;refGene_gn=SLCO1A2;refGene_tx=XM_019834650.2;refGene_gn=SLCO1A2;refGene_tx=XM_019834651.2;refGene_gn=SLCO1A2;refGene_tx=XM_019834652.2
B4 56162259 56162358
refGene_gn=SLCO1A2;refGene_tx=XM_003988480.5;refGene_gn=SLCO1A2;refGene_tx=XM_019834644.2;refGene_gn=SLCO1A2;refGene_tx=XM_019834645.2;refGene_gn=SLCO1A2;refGene_tx=XM_019834646.2;refGene_gn=SLCO1A2;refGene_tx=XM_019834647.2;refGene_gn=SLCO1A2;refGene_tx=XM_019834648.2;refGene_gn=SLCO1A2;refGene_tx=XM_019834649.2;refGene_gn=SLCO1A2;refGene_tx=XM_019834650.2;refGene_gn=SLCO1A2;refGene_tx=XM_019834651.2;refGene_gn=SLCO1A2;refGene_tx=XM_019834652.2
B4 56165382 56165529
refGene_gn=SLCO1A2;refGene_tx=XM_003988480.5;refGene_gn=SLCO1A2;refGene_tx=XM_019834644.2;refGene_gn=SLCO1A2;refGene_tx=XM_019834645.2;refGene_gn=SLCO1A2;refGene_tx=XM_019834646.2;refGene_gn=SLCO1A2;refGene_tx=XM_019834647.2;refGene_gn=SLCO1A2;refGene_tx=XM_019834648.2;refGene_gn=SLCO1A2;refGene_tx=XM_019834649.2;refGene_gn=SLCO1A2;refGene_tx=XM_019834650.2;refGene_gn=SLCO1A2;refGene_tx=XM_019834651.2;refGene_gn=SLCO1A2;refGene_tx=XM_019834652.2
B4 56167576 56167683
refGene_gn=SLCO1A2;refGene_tx=XM_003988480.5;refGene_gn=SLCO1A2;refGene_tx=XM_019834644.2;refGene_gn=SLCO1A2;refGene_tx=XM_019834645.2;refGene_gn=SLCO1A2;refGene_tx=XM_019834646.2;refGene_gn=SLCO1A2;refGene_tx=XM_019834647.2;refGene_gn=SLCO1A2;refGene_tx=XM_019834648.2;refGene_gn=SLCO1A2;refGene_tx=XM_019834649.2;refGene_gn=SLCO1A2;refGene_tx=XM_019834650.2;refGene_gn=SLCO1A2;refGene_tx=XM_019834651.2;refGene_gn=SLCO1A2;refGene_tx=XM_019834652.2
B4 56171210 56171343
refGene_gn=SLCO1A2;refGene_tx=XM_003988480.5;refGene_gn=SLCO1A2;refGene_tx=XM_019834644.2;refGene_gn=SLCO1A2;refGene_tx=XM_019834645.2;refGene_gn=SLCO1A2;refGene_tx=XM_019834646.2;refGene_gn=SLCO1A2;refGene_tx=XM_019834647.2;refGene_gn=SLCO1A2;refGene_tx=XM_019834648.2;refGene_gn=SLCO1A2;refGene_tx=XM_019834649.2;refGene_gn=SLCO1A2;refGene_tx=XM_019834651.2;refGene_gn=SLCO1A2;refGene_tx=XM_019834652.2

B4 56175968 56176110
refGene_gn=SLCO1A2;refGene_tx=XM_003988480.5;refGene_gn=SLCO1A2;refGene_tx=XM_019834644.2;refGene_gn=SLCO1A2;refGene_tx=XM_019834645.2;refGene_gn=SLCO1A2;refGene_tx=XM_019834646.2;refGene_gn=SLCO1A2;refGene_tx=XM_019834647.2;refGene_gn=SLCO1A2;refGene_tx=XM_019834648.2;refGene_gn=SLCO1A2;refGene_tx=XM_019834649.2;refGene_gn=SLCO1A2;refGene_tx=XM_019834651.2;refGene_gn=SLCO1A2;refGene_tx=XM_019834652.2
B4 56180999 56181059
refGene_gn=SLCO1A2;refGene_tx=XM_003988480.5;refGene_gn=SLCO1A2;refGene_tx=XM_019834644.2;refGene_gn=SLCO1A2;refGene_tx=XM_019834645.2;refGene_gn=SLCO1A2;refGene_tx=XM_019834647.2;refGene_gn=SLCO1A2;refGene_tx=XM_019834648.2;refGene_gn=SLCO1A2;refGene_tx=XM_019834649.2;refGene_gn=SLCO1A2;refGene_tx=XM_019834651.2;refGene_gn=SLCO1A2;refGene_tx=XM_019834652.2
B4 56232111 56232191
refGene_gn=IAPP;refGene_tx=NM_001043338.1;refGene_gn=IAPP;refGene_tx=XM_019833865.1
B4 56237835 56238025
refGene_gn=IAPP;refGene_tx=NM_001043338.1;refGene_gn=IAPP;refGene_tx=XM_019833865.1
B4 56269654 56269738 refGene_gn=PYROXD1;refGene_tx=XM_006933557.3
B4 56274091 56274172 refGene_gn=PYROXD1;refGene_tx=XM_006933557.3
B4 56279450 56279570
refGene_gn=PYROXD1;refGene_tx=XM_006933557.3;refGene_gn=PYROXD1;refGene_tx=XM_023256828.1
B4 56282572 56282701
refGene_gn=PYROXD1;refGene_tx=XM_006933557.3;refGene_gn=PYROXD1;refGene_tx=XM_023256828.1
B4 56284753 56284827
refGene_gn=PYROXD1;refGene_tx=XM_006933557.3;refGene_gn=PYROXD1;refGene_tx=XM_023256828.1
B4 56287678 56287839
refGene_gn=PYROXD1;refGene_tx=XM_006933557.3;refGene_gn=PYROXD1;refGene_tx=XM_023256828.1
B4 56290023 56290124
refGene_gn=PYROXD1;refGene_tx=XM_006933557.3;refGene_gn=PYROXD1;refGene_tx=XM_023256828.1
B4 56291847 56291983
refGene_gn=PYROXD1;refGene_tx=XM_006933557.3;refGene_gn=PYROXD1;refGene_tx=XM_023256828.1
B4 56293026 56293139
refGene_gn=PYROXD1;refGene_tx=XM_006933557.3;refGene_gn=PYROXD1;refGene_tx=XM_023256828.1
B4 56293997 56294120
refGene_gn=PYROXD1;refGene_tx=XM_006933557.3;refGene_gn=PYROXD1;refGene_tx=XM_023256828.1
B4 56296274 56296412
refGene_gn=PYROXD1;refGene_tx=XM_006933557.3;refGene_gn=PYROXD1;refGene_tx=XM_023256828.1
B4 56297008 56297257

refGene_gn=PYROXD1;refGene_tx=XM_006933557.3;refGene_gn=PYROXD1;refGene_tx=XM_023256828.1
B4 57200718 57200877 refGene_gn=RECQL;refGene_tx=XM_006933559.3
B4 57201477 57201607 refGene_gn=RECQL;refGene_tx=XM_006933559.3
B4 57201908 57202128 refGene_gn=RECQL;refGene_tx=XM_006933559.3
B4 57210529 57210621 refGene_gn=RECQL;refGene_tx=XM_006933559.3
B4 57212539 57212678 refGene_gn=RECQL;refGene_tx=XM_006933559.3
B4 57213203 57213321 refGene_gn=RECQL;refGene_tx=XM_006933559.3
B4 57213410 57213559 refGene_gn=RECQL;refGene_tx=XM_006933559.3
B4 57214760 57214842 refGene_gn=RECQL;refGene_tx=XM_006933559.3
B4 57215578 57215745 refGene_gn=RECQL;refGene_tx=XM_006933559.3
B4 57226648 57226847 refGene_gn=RECQL;refGene_tx=XM_006933559.3
B4 57230678 57230785 refGene_gn=RECQL;refGene_tx=XM_006933559.3
B4 57233416 57233596 refGene_gn=RECQL;refGene_tx=XM_006933559.3
B4 57234994 57235192 refGene_gn=RECQL;refGene_tx=XM_006933559.3
B4 57250505 57250524 refGene_gn=RECQL;refGene_tx=XM_006933559.3
B4 57257718 57257743 refGene_gn=GOLT1B;refGene_tx=XM_003988485.5
B4 57263722 57263814 refGene_gn=GOLT1B;refGene_tx=XM_003988485.5
B4 57265437 57265616 refGene_gn=GOLT1B;refGene_tx=XM_003988485.5
B4 57266733 57266815 refGene_gn=GOLT1B;refGene_tx=XM_003988485.5
B4 57269027 57269066 refGene_gn=GOLT1B;refGene_tx=XM_003988485.5
B4 57278188 57278194 refGene_gn=SPX;refGene_tx=XM_023257750.1
B4 57278597 57278678 refGene_gn=SPX;refGene_tx=XM_023257750.1
B4 57279181 57279233 refGene_gn=SPX;refGene_tx=XM_023257750.1
B4 57279532 57279595 refGene_gn=SPX;refGene_tx=XM_023257750.1
B4 57280597 57280681 refGene_gn=SPX;refGene_tx=XM_023257750.1
B4 57286580 57286627 refGene_gn=SPX;refGene_tx=XM_023257750.1
B4 57292594 57292813
refGene_gn=GYS2;refGene_tx=XM_003988486.4;refGene_gn=GYS2;refGene_tx=XM_019834666.1
B4 57294987 57295068
refGene_gn=GYS2;refGene_tx=XM_003988486.4;refGene_gn=GYS2;refGene_tx=XM_019834666.1
B4 57296544 57296708
refGene_gn=GYS2;refGene_tx=XM_003988486.4;refGene_gn=GYS2;refGene_tx=XM_019834666.1
B4 57297875 57297971
refGene_gn=GYS2;refGene_tx=XM_003988486.4;refGene_gn=GYS2;refGene_tx=XM_019834666.1
B4 57301279 57301406
refGene_gn=GYS2;refGene_tx=XM_003988486.4;refGene_gn=GYS2;refGene_tx=XM_019834666.1
B4 57312385 57312499
refGene_gn=GYS2;refGene_tx=XM_003988486.4;refGene_gn=GYS2;refGene_tx=XM_019834666.1
B4 57313478 57313557
refGene_gn=GYS2;refGene_tx=XM_003988486.4;refGene_gn=GYS2;refGene_tx=XM_019834666.1
B4 57314492 57314552
refGene_gn=GYS2;refGene_tx=XM_003988486.4;refGene_gn=GYS2;refGene_tx=X

M_019834666.1
B4	57315438	57315545
refGene_gn=GYS2;refGene_tx=XM_003988486.4;refGene_gn=GYS2;refGene_tx=XM_019834666.1
B4	57318493	57318614
refGene_gn=GYS2;refGene_tx=XM_003988486.4;refGene_gn=GYS2;refGene_tx=XM_019834666.1
B4	57319018	57319136
refGene_gn=GYS2;refGene_tx=XM_003988486.4;refGene_gn=GYS2;refGene_tx=XM_019834666.1
B4	57322574	57322719
refGene_gn=GYS2;refGene_tx=XM_003988486.4;refGene_gn=GYS2;refGene_tx=XM_019834666.1
B4	57328715	57328898
refGene_gn=GYS2;refGene_tx=XM_003988486.4;refGene_gn=GYS2;refGene_tx=XM_019834666.1
B4	57331033	57331225
refGene_gn=GYS2;refGene_tx=XM_003988486.4;refGene_gn=GYS2;refGene_tx=XM_019834666.1
B4	57335126	57335308
refGene_gn=GYS2;refGene_tx=XM_003988486.4;refGene_gn=GYS2;refGene_tx=XM_019834666.1
B4	57343764	57343885
refGene_gn=GYS2;refGene_tx=XM_003988486.4;refGene_gn=GYS2;refGene_tx=XM_019834666.1
B4	57378060	57378228	refGene_gn=LDHB;refGene_tx=XM_003988487.3
B4	57380061	57380185	refGene_gn=LDHB;refGene_tx=XM_003988487.3
B4	57381624	57381742	refGene_gn=LDHB;refGene_tx=XM_003988487.3
B4	57385431	57385605	refGene_gn=LDHB;refGene_tx=XM_003988487.3
B4	57387259	57387433	refGene_gn=LDHB;refGene_tx=XM_003988487.3
B4	57388653	57388771	refGene_gn=LDHB;refGene_tx=XM_003988487.3
B4	57396132	57396261	refGene_gn=LDHB;refGene_tx=XM_003988487.3
B4	57515472	57516373
refGene_gn=KCNJ8;refGene_tx=XM_003988489.5;refGene_gn=KCNJ8;refGene_tx=XM_006933561.3;refGene_gn=KCNJ8;refGene_tx=XM_019834667.2
B4	57520786	57521160
refGene_gn=KCNJ8;refGene_tx=XM_003988489.5;refGene_gn=KCNJ8;refGene_tx=XM_006933561.3;refGene_gn=KCNJ8;refGene_tx=XM_019834667.2
B4	57538828	57538966
refGene_gn=ABCC9;refGene_tx=XM_011283836.3;refGene_gn=ABCC9;refGene_tx=XM_019834668.2;refGene_gn=ABCC9;refGene_tx=XM_019834669.2;refGene_gn=ABCC9;refGene_tx=XM_019834670.2;refGene_gn=ABCC9;refGene_tx=XM_019834671.2;refGene_gn=ABCC9;refGene_tx=XM_019834672.2
B4	57543137	57543275
refGene_gn=ABCC9;refGene_tx=XM_011283834.3;refGene_gn=ABCC9;refGene_tx=XM_011283837.3
B4	57543949	57544012
refGene_gn=ABCC9;refGene_tx=XM_011283834.3;refGene_gn=ABCC9;refGene_tx=XM_011283836.3;refGene_gn=ABCC9;refGene_tx=XM_011283837.3;refGene_gn=ABCC9;refGene_tx=XM_019834668.2;refGene_gn=ABCC9;refGene_tx=XM_0198346

69.2;refGene_gn=ABCC9;refGene_tx=XM_019834670.2;refGene_gn=ABCC9;refGene_tx=XM_019834671.2;refGene_gn=ABCC9;refGene_tx=XM_019834672.2
B4 57545989 57546123
refGene_gn=ABCC9;refGene_tx=XM_011283834.3;refGene_gn=ABCC9;refGene_tx=XM_011283836.3;refGene_gn=ABCC9;refGene_tx=XM_011283837.3;refGene_gn=ABCC9;refGene_tx=XM_019834668.2;refGene_gn=ABCC9;refGene_tx=XM_019834669.2;refGene_gn=ABCC9;refGene_tx=XM_019834670.2;refGene_gn=ABCC9;refGene_tx=XM_019834671.2;refGene_gn=ABCC9;refGene_tx=XM_019834672.2
B4 57548697 57548801
refGene_gn=ABCC9;refGene_tx=XM_011283834.3;refGene_gn=ABCC9;refGene_tx=XM_011283836.3;refGene_gn=ABCC9;refGene_tx=XM_011283837.3;refGene_gn=ABCC9;refGene_tx=XM_019834668.2;refGene_gn=ABCC9;refGene_tx=XM_019834669.2;refGene_gn=ABCC9;refGene_tx=XM_019834670.2;refGene_gn=ABCC9;refGene_tx=XM_019834671.2;refGene_gn=ABCC9;refGene_tx=XM_019834672.2
B4 57550724 57550833
refGene_gn=ABCC9;refGene_tx=XM_011283834.3;refGene_gn=ABCC9;refGene_tx=XM_011283836.3;refGene_gn=ABCC9;refGene_tx=XM_011283837.3;refGene_gn=ABCC9;refGene_tx=XM_019834668.2;refGene_gn=ABCC9;refGene_tx=XM_019834669.2;refGene_gn=ABCC9;refGene_tx=XM_019834670.2;refGene_gn=ABCC9;refGene_tx=XM_019834671.2;refGene_gn=ABCC9;refGene_tx=XM_019834672.2
B4 57552722 57552801
refGene_gn=ABCC9;refGene_tx=XM_011283834.3;refGene_gn=ABCC9;refGene_tx=XM_011283836.3;refGene_gn=ABCC9;refGene_tx=XM_011283837.3;refGene_gn=ABCC9;refGene_tx=XM_019834668.2;refGene_gn=ABCC9;refGene_tx=XM_019834669.2;refGene_gn=ABCC9;refGene_tx=XM_019834670.2;refGene_gn=ABCC9;refGene_tx=XM_019834671.2;refGene_gn=ABCC9;refGene_tx=XM_019834672.2
B4 57553516 57553647
refGene_gn=ABCC9;refGene_tx=XM_011283834.3;refGene_gn=ABCC9;refGene_tx=XM_011283836.3;refGene_gn=ABCC9;refGene_tx=XM_011283837.3;refGene_gn=ABCC9;refGene_tx=XM_019834668.2;refGene_gn=ABCC9;refGene_tx=XM_019834669.2;refGene_gn=ABCC9;refGene_tx=XM_019834670.2;refGene_gn=ABCC9;refGene_tx=XM_019834671.2;refGene_gn=ABCC9;refGene_tx=XM_019834672.2
B4 57555182 57555303
refGene_gn=ABCC9;refGene_tx=XM_011283834.3;refGene_gn=ABCC9;refGene_tx=XM_011283836.3;refGene_gn=ABCC9;refGene_tx=XM_011283837.3;refGene_gn=ABCC9;refGene_tx=XM_019834668.2;refGene_gn=ABCC9;refGene_tx=XM_019834669.2;refGene_gn=ABCC9;refGene_tx=XM_019834670.2;refGene_gn=ABCC9;refGene_tx=XM_019834671.2;refGene_gn=ABCC9;refGene_tx=XM_019834672.2
B4 57555883 57555985
refGene_gn=ABCC9;refGene_tx=XM_011283834.3;refGene_gn=ABCC9;refGene_tx=XM_011283836.3;refGene_gn=ABCC9;refGene_tx=XM_011283837.3;refGene_gn=ABCC9;refGene_tx=XM_019834668.2;refGene_gn=ABCC9;refGene_tx=XM_019834669.2;refGene_gn=ABCC9;refGene_tx=XM_019834670.2;refGene_gn=ABCC9;refGene_tx=XM_019834671.2;refGene_gn=ABCC9;refGene_tx=XM_019834672.2
B4 57565058 57565161
refGene_gn=ABCC9;refGene_tx=XM_011283834.3;refGene_gn=ABCC9;refGene_tx=XM_011283836.3;refGene_gn=ABCC9;refGene_tx=XM_011283837.3;refGene_gn=ABCC9;refGene_tx=XM_019834668.2;refGene_gn=ABCC9;refGene_tx=XM_019834669.2;refGene_gn=ABCC9;refGene_tx=XM_019834670.2;refGene_gn=ABCC9;refGene_tx=XM_019834671.2;refGene_gn=ABCC9;refGene_tx=XM_019834672.2

B4 57569765 57569858
refGene_gn=ABCC9;refGene_tx=XM_011283834.3;refGene_gn=ABCC9;refGene_tx=XM_011283836.3;refGene_gn=ABCC9;refGene_tx=XM_011283837.3;refGene_gn=ABCC9;refGene_tx=XM_019834668.2;refGene_gn=ABCC9;refGene_tx=XM_019834669.2;refGene_gn=ABCC9;refGene_tx=XM_019834670.2;refGene_gn=ABCC9;refGene_tx=XM_019834671.2;refGene_gn=ABCC9;refGene_tx=XM_019834672.2
B4 57573687 57573845
refGene_gn=ABCC9;refGene_tx=XM_011283834.3;refGene_gn=ABCC9;refGene_tx=XM_011283836.3;refGene_gn=ABCC9;refGene_tx=XM_011283837.3;refGene_gn=ABCC9;refGene_tx=XM_019834668.2;refGene_gn=ABCC9;refGene_tx=XM_019834669.2;refGene_gn=ABCC9;refGene_tx=XM_019834670.2;refGene_gn=ABCC9;refGene_tx=XM_019834671.2;refGene_gn=ABCC9;refGene_tx=XM_019834672.2
B4 57575863 57575933
refGene_gn=ABCC9;refGene_tx=XM_011283834.3;refGene_gn=ABCC9;refGene_tx=XM_011283836.3;refGene_gn=ABCC9;refGene_tx=XM_011283837.3;refGene_gn=ABCC9;refGene_tx=XM_019834668.2;refGene_gn=ABCC9;refGene_tx=XM_019834669.2;refGene_gn=ABCC9;refGene_tx=XM_019834670.2;refGene_gn=ABCC9;refGene_tx=XM_019834671.2;refGene_gn=ABCC9;refGene_tx=XM_019834672.2
B4 57576166 57576315
refGene_gn=ABCC9;refGene_tx=XM_011283834.3;refGene_gn=ABCC9;refGene_tx=XM_011283836.3;refGene_gn=ABCC9;refGene_tx=XM_011283837.3;refGene_gn=ABCC9;refGene_tx=XM_019834668.2;refGene_gn=ABCC9;refGene_tx=XM_019834669.2;refGene_gn=ABCC9;refGene_tx=XM_019834670.2;refGene_gn=ABCC9;refGene_tx=XM_019834671.2;refGene_gn=ABCC9;refGene_tx=XM_019834672.2
B4 57576993 57577223
refGene_gn=ABCC9;refGene_tx=XM_011283834.3;refGene_gn=ABCC9;refGene_tx=XM_011283836.3;refGene_gn=ABCC9;refGene_tx=XM_011283837.3;refGene_gn=ABCC9;refGene_tx=XM_019834668.2;refGene_gn=ABCC9;refGene_tx=XM_019834669.2;refGene_gn=ABCC9;refGene_tx=XM_019834670.2;refGene_gn=ABCC9;refGene_tx=XM_019834671.2;refGene_gn=ABCC9;refGene_tx=XM_019834672.2
B4 57579310 57579407
refGene_gn=ABCC9;refGene_tx=XM_011283834.3;refGene_gn=ABCC9;refGene_tx=XM_011283836.3;refGene_gn=ABCC9;refGene_tx=XM_011283837.3;refGene_gn=ABCC9;refGene_tx=XM_019834668.2;refGene_gn=ABCC9;refGene_tx=XM_019834669.2;refGene_gn=ABCC9;refGene_tx=XM_019834670.2;refGene_gn=ABCC9;refGene_tx=XM_019834671.2;refGene_gn=ABCC9;refGene_tx=XM_019834672.2
B4 57583256 57583382
refGene_gn=ABCC9;refGene_tx=XM_011283834.3;refGene_gn=ABCC9;refGene_tx=XM_011283836.3;refGene_gn=ABCC9;refGene_tx=XM_011283837.3;refGene_gn=ABCC9;refGene_tx=XM_019834668.2;refGene_gn=ABCC9;refGene_tx=XM_019834669.2;refGene_gn=ABCC9;refGene_tx=XM_019834670.2;refGene_gn=ABCC9;refGene_tx=XM_019834671.2;refGene_gn=ABCC9;refGene_tx=XM_019834672.2
B4 57583713 57583851
refGene_gn=ABCC9;refGene_tx=XM_011283834.3;refGene_gn=ABCC9;refGene_tx=XM_011283836.3;refGene_gn=ABCC9;refGene_tx=XM_011283837.3;refGene_gn=ABCC9;refGene_tx=XM_019834668.2;refGene_gn=ABCC9;refGene_tx=XM_019834669.2;refGene_gn=ABCC9;refGene_tx=XM_019834670.2;refGene_gn=ABCC9;refGene_tx=XM_019834671.2;refGene_gn=ABCC9;refGene_tx=XM_019834672.2
B4 57594484 57594565
refGene_gn=ABCC9;refGene_tx=XM_011283834.3;refGene_gn=ABCC9;refGene_tx

=XM_011283836.3;refGene_gn=ABCC9;refGene_tx=XM_011283837.3;refGene_gn=
ABCC9;refGene_tx=XM_019834668.2;refGene_gn=ABCC9;refGene_tx=XM_0198346
69.2;refGene_gn=ABCC9;refGene_tx=XM_019834670.2;refGene_gn=ABCC9;refGe
ne_tx=XM_019834671.2;refGene_gn=ABCC9;refGene_tx=XM_019834672.2
B4 57595771 57595856
refGene_gn=ABCC9;refGene_tx=XM_011283834.3;refGene_gn=ABCC9;refGene_tx
=XM_011283836.3;refGene_gn=ABCC9;refGene_tx=XM_011283837.3;refGene_gn=
ABCC9;refGene_tx=XM_019834668.2;refGene_gn=ABCC9;refGene_tx=XM_0198346
69.2;refGene_gn=ABCC9;refGene_tx=XM_019834670.2;refGene_gn=ABCC9;refGe
ne_tx=XM_019834671.2;refGene_gn=ABCC9;refGene_tx=XM_019834672.2
B4 57597529 57597631
refGene_gn=ABCC9;refGene_tx=XM_011283834.3;refGene_gn=ABCC9;refGene_tx
=XM_011283836.3;refGene_gn=ABCC9;refGene_tx=XM_011283837.3;refGene_gn=
ABCC9;refGene_tx=XM_019834668.2;refGene_gn=ABCC9;refGene_tx=XM_0198346
69.2;refGene_gn=ABCC9;refGene_tx=XM_019834670.2;refGene_gn=ABCC9;refGe
ne_tx=XM_019834671.2;refGene_gn=ABCC9;refGene_tx=XM_019834672.2
B4 57599013 57599052
refGene_gn=ABCC9;refGene_tx=XM_011283834.3;refGene_gn=ABCC9;refGene_tx
=XM_011283836.3;refGene_gn=ABCC9;refGene_tx=XM_019834668.2;refGene_gn=
ABCC9;refGene_tx=XM_019834669.2;refGene_gn=ABCC9;refGene_tx=XM_0198346
70.2;refGene_gn=ABCC9;refGene_tx=XM_019834671.2
B4 57607018 57607124
refGene_gn=ABCC9;refGene_tx=XM_011283834.3;refGene_gn=ABCC9;refGene_tx
=XM_011283836.3;refGene_gn=ABCC9;refGene_tx=XM_011283837.3;refGene_gn=
ABCC9;refGene_tx=XM_019834668.2;refGene_gn=ABCC9;refGene_tx=XM_0198346
69.2;refGene_gn=ABCC9;refGene_tx=XM_019834670.2;refGene_gn=ABCC9;refGe
ne_tx=XM_019834671.2;refGene_gn=ABCC9;refGene_tx=XM_019834672.2
B4 57608016 57608089
refGene_gn=ABCC9;refGene_tx=XM_011283834.3;refGene_gn=ABCC9;refGene_tx
=XM_011283836.3;refGene_gn=ABCC9;refGene_tx=XM_011283837.3;refGene_gn=
ABCC9;refGene_tx=XM_019834668.2;refGene_gn=ABCC9;refGene_tx=XM_0198346
69.2;refGene_gn=ABCC9;refGene_tx=XM_019834670.2;refGene_gn=ABCC9;refGe
ne_tx=XM_019834671.2;refGene_gn=ABCC9;refGene_tx=XM_019834672.2
B4 57614100 57614208
refGene_gn=ABCC9;refGene_tx=XM_011283834.3;refGene_gn=ABCC9;refGene_tx
=XM_011283836.3;refGene_gn=ABCC9;refGene_tx=XM_011283837.3;refGene_gn=
ABCC9;refGene_tx=XM_019834668.2;refGene_gn=ABCC9;refGene_tx=XM_0198346
69.2;refGene_gn=ABCC9;refGene_tx=XM_019834670.2;refGene_gn=ABCC9;refGe
ne_tx=XM_019834671.2;refGene_gn=ABCC9;refGene_tx=XM_019834672.2
B4 57618888 57618997
refGene_gn=ABCC9;refGene_tx=XM_011283834.3;refGene_gn=ABCC9;refGene_tx
=XM_011283836.3;refGene_gn=ABCC9;refGene_tx=XM_011283837.3;refGene_gn=
ABCC9;refGene_tx=XM_019834668.2;refGene_gn=ABCC9;refGene_tx=XM_0198346
69.2;refGene_gn=ABCC9;refGene_tx=XM_019834670.2;refGene_gn=ABCC9;refGe
ne_tx=XM_019834671.2;refGene_gn=ABCC9;refGene_tx=XM_019834672.2
B4 57627898 57628041
refGene_gn=ABCC9;refGene_tx=XM_011283834.3;refGene_gn=ABCC9;refGene_tx
=XM_011283836.3;refGene_gn=ABCC9;refGene_tx=XM_011283837.3;refGene_gn=
ABCC9;refGene_tx=XM_019834668.2;refGene_gn=ABCC9;refGene_tx=XM_0198346
69.2;refGene_gn=ABCC9;refGene_tx=XM_019834670.2;refGene_gn=ABCC9;refGe

ne_tx=XM_019834671.2;refGene_gn=ABCC9;refGene_tx=XM_019834672.2
B4 57628943 57628984
refGene_gn=ABCC9;refGene_tx=XM_011283834.3;refGene_gn=ABCC9;refGene_tx=XM_011283836.3;refGene_gn=ABCC9;refGene_tx=XM_011283837.3;refGene_gn=ABCC9;refGene_tx=XM_019834668.2;refGene_gn=ABCC9;refGene_tx=XM_019834669.2;refGene_gn=ABCC9;refGene_tx=XM_019834670.2;refGene_gn=ABCC9;refGene_tx=XM_019834671.2;refGene_gn=ABCC9;refGene_tx=XM_019834672.2
B4 57633366 57633529
refGene_gn=ABCC9;refGene_tx=XM_011283834.3;refGene_gn=ABCC9;refGene_tx=XM_011283836.3;refGene_gn=ABCC9;refGene_tx=XM_011283837.3;refGene_gn=ABCC9;refGene_tx=XM_019834668.2;refGene_gn=ABCC9;refGene_tx=XM_019834669.2;refGene_gn=ABCC9;refGene_tx=XM_019834670.2;refGene_gn=ABCC9;refGene_tx=XM_019834671.2;refGene_gn=ABCC9;refGene_tx=XM_019834672.2
B4 57635575 57635710
refGene_gn=ABCC9;refGene_tx=XM_011283834.3;refGene_gn=ABCC9;refGene_tx=XM_011283836.3;refGene_gn=ABCC9;refGene_tx=XM_011283837.3;refGene_gn=ABCC9;refGene_tx=XM_019834668.2;refGene_gn=ABCC9;refGene_tx=XM_019834669.2;refGene_gn=ABCC9;refGene_tx=XM_019834670.2;refGene_gn=ABCC9;refGene_tx=XM_019834671.2;refGene_gn=ABCC9;refGene_tx=XM_019834672.2
B4 57638742 57638898
refGene_gn=ABCC9;refGene_tx=XM_011283834.3;refGene_gn=ABCC9;refGene_tx=XM_011283836.3;refGene_gn=ABCC9;refGene_tx=XM_011283837.3;refGene_gn=ABCC9;refGene_tx=XM_019834668.2;refGene_gn=ABCC9;refGene_tx=XM_019834669.2;refGene_gn=ABCC9;refGene_tx=XM_019834670.2;refGene_gn=ABCC9;refGene_tx=XM_019834671.2;refGene_gn=ABCC9;refGene_tx=XM_019834672.2
B4 57639370 57639523
refGene_gn=ABCC9;refGene_tx=XM_011283834.3;refGene_gn=ABCC9;refGene_tx=XM_011283836.3;refGene_gn=ABCC9;refGene_tx=XM_011283837.3;refGene_gn=ABCC9;refGene_tx=XM_019834668.2;refGene_gn=ABCC9;refGene_tx=XM_019834669.2;refGene_gn=ABCC9;refGene_tx=XM_019834670.2;refGene_gn=ABCC9;refGene_tx=XM_019834671.2;refGene_gn=ABCC9;refGene_tx=XM_019834672.2
B4 57641371 57641566
refGene_gn=ABCC9;refGene_tx=XM_011283834.3;refGene_gn=ABCC9;refGene_tx=XM_011283836.3;refGene_gn=ABCC9;refGene_tx=XM_011283837.3;refGene_gn=ABCC9;refGene_tx=XM_019834668.2;refGene_gn=ABCC9;refGene_tx=XM_019834669.2;refGene_gn=ABCC9;refGene_tx=XM_019834670.2;refGene_gn=ABCC9;refGene_tx=XM_019834671.2;refGene_gn=ABCC9;refGene_tx=XM_019834672.2
B4 57643447 57643690
refGene_gn=ABCC9;refGene_tx=XM_011283834.3;refGene_gn=ABCC9;refGene_tx=XM_011283836.3;refGene_gn=ABCC9;refGene_tx=XM_011283837.3;refGene_gn=ABCC9;refGene_tx=XM_019834668.2;refGene_gn=ABCC9;refGene_tx=XM_019834669.2;refGene_gn=ABCC9;refGene_tx=XM_019834670.2;refGene_gn=ABCC9;refGene_tx=XM_019834671.2;refGene_gn=ABCC9;refGene_tx=XM_019834672.2
B4 57644728 57644895
refGene_gn=ABCC9;refGene_tx=XM_011283834.3;refGene_gn=ABCC9;refGene_tx=XM_011283836.3;refGene_gn=ABCC9;refGene_tx=XM_011283837.3;refGene_gn=ABCC9;refGene_tx=XM_019834668.2;refGene_gn=ABCC9;refGene_tx=XM_019834669.2;refGene_gn=ABCC9;refGene_tx=XM_019834670.2;refGene_gn=ABCC9;refGene_tx=XM_019834671.2;refGene_gn=ABCC9;refGene_tx=XM_019834672.2
B4 57653554 57653676

refGene_gn=ABCC9;refGene_tx=XM_011283834.3;refGene_gn=ABCC9;refGene_tx=XM_011283836.3;refGene_gn=ABCC9;refGene_tx=XM_011283837.3;refGene_gn=ABCC9;refGene_tx=XM_019834668.2;refGene_gn=ABCC9;refGene_tx=XM_019834669.2;refGene_gn=ABCC9;refGene_tx=XM_019834670.2;refGene_gn=ABCC9;refGene_tx=XM_019834671.2;refGene_gn=ABCC9;refGene_tx=XM_019834672.2
B4 57659425 57659567
refGene_gn=ABCC9;refGene_tx=XM_011283834.3;refGene_gn=ABCC9;refGene_tx=XM_011283836.3;refGene_gn=ABCC9;refGene_tx=XM_011283837.3;refGene_gn=ABCC9;refGene_tx=XM_019834668.2;refGene_gn=ABCC9;refGene_tx=XM_019834669.2;refGene_gn=ABCC9;refGene_tx=XM_019834670.2;refGene_gn=ABCC9;refGene_tx=XM_019834671.2;refGene_gn=ABCC9;refGene_tx=XM_019834672.2
B4 57661939 57662081
refGene_gn=ABCC9;refGene_tx=XM_011283834.3;refGene_gn=ABCC9;refGene_tx=XM_011283836.3;refGene_gn=ABCC9;refGene_tx=XM_011283837.3;refGene_gn=ABCC9;refGene_tx=XM_019834668.2;refGene_gn=ABCC9;refGene_tx=XM_019834669.2;refGene_gn=ABCC9;refGene_tx=XM_019834670.2;refGene_gn=ABCC9;refGene_tx=XM_019834671.2;refGene_gn=ABCC9;refGene_tx=XM_019834672.2
B4 57738883 57739143 refGene_gn=CMAS;refGene_tx=XM_011283838.3
B4 57746488 57746631 refGene_gn=CMAS;refGene_tx=XM_011283838.3
B4 57746777 57746933 refGene_gn=CMAS;refGene_tx=XM_011283838.3
B4 57752178 57752312 refGene_gn=CMAS;refGene_tx=XM_011283838.3
B4 57754027 57754122 refGene_gn=CMAS;refGene_tx=XM_011283838.3
B4 57754489 57754661 refGene_gn=CMAS;refGene_tx=XM_011283838.3
B4 57755456 57755610 refGene_gn=CMAS;refGene_tx=XM_011283838.3
B4 57758879 57759070 refGene_gn=CMAS;refGene_tx=XM_011283838.3
B4 57774121 57774240
refGene_gn=LOC102901968;refGene_tx=XM_006933565.3
B4 57777182 57777369
refGene_gn=LOC102901968;refGene_tx=XM_006933565.3
B4 57777519 57777614
refGene_gn=LOC102901968;refGene_tx=XM_006933565.3
B4 57783394 57783521
refGene_gn=LOC102901968;refGene_tx=XM_006933565.3
B4 57785720 57785905
refGene_gn=LOC102901968;refGene_tx=XM_006933565.3
B4 57788231 57788376
refGene_gn=LOC102901968;refGene_tx=XM_006933565.3
B4 57823752 57824239
refGene_gn=ST8SIA1;refGene_tx=XM_003988490.5;refGene_gn=ST8SIA1;refGene_tx=XM_019834673.2
B4 57869821 57869914
refGene_gn=ST8SIA1;refGene_tx=XM_003988490.5;refGene_gn=ST8SIA1;refGene_tx=XM_019834673.2
B4 57881045 57881155 refGene_gn=ST8SIA1;refGene_tx=XM_003988490.5
B4 57916129 57916274 refGene_gn=ST8SIA1;refGene_tx=XM_003988490.5
B4 57966304 57966540
refGene_gn=ST8SIA1;refGene_tx=XM_003988490.5;refGene_gn=ST8SIA1;refGene_tx=XM_019834673.2
B4 58055143 58055275
refGene_gn=C2CD5;refGene_tx=XM_011283840.3;refGene_gn=C2CD5;refGene_tx

=XM_011283841.3;refGene_gn=C2CD5;refGene_tx=XM_011283842.3;refGene_gn=C2CD5;refGene_tx=XM_019834674.2;refGene_gn=C2CD5;refGene_tx=XM_019834675.2;refGene_gn=C2CD5;refGene_tx=XM_019834676.2;refGene_gn=C2CD5;refGene_tx=XM_019834677.2;refGene_gn=C2CD5;refGene_tx=XM_019834678.2;refGene_gn=C2CD5;refGene_tx=XM_019834679.1;refGene_gn=C2CD5;refGene_tx=XM_019834680.2;refGene_gn=C2CD5;refGene_tx=XM_019834681.2;refGene_gn=C2CD5;refGene_tx=XM_019834682.2;refGene_gn=C2CD5;refGene_tx=XM_019834683.2;refGene_gn=C2CD5;refGene_tx=XM_019834684.2;refGene_gn=C2CD5;refGene_tx=XM_019834685.2;refGene_gn=C2CD5;refGene_tx=XM_019834686.2;refGene_gn=C2CD5;refGene_tx=XM_019834687.2
B4 58056010 58056157
refGene_gn=C2CD5;refGene_tx=XM_011283840.3;refGene_gn=C2CD5;refGene_tx=XM_011283841.3;refGene_gn=C2CD5;refGene_tx=XM_011283842.3;refGene_gn=C2CD5;refGene_tx=XM_019834674.2;refGene_gn=C2CD5;refGene_tx=XM_019834675.2;refGene_gn=C2CD5;refGene_tx=XM_019834676.2;refGene_gn=C2CD5;refGene_tx=XM_019834677.2;refGene_gn=C2CD5;refGene_tx=XM_019834678.2;refGene_gn=C2CD5;refGene_tx=XM_019834679.1;refGene_gn=C2CD5;refGene_tx=XM_019834680.2;refGene_gn=C2CD5;refGene_tx=XM_019834681.2;refGene_gn=C2CD5;refGene_tx=XM_019834682.2;refGene_gn=C2CD5;refGene_tx=XM_019834683.2;refGene_gn=C2CD5;refGene_tx=XM_019834684.2;refGene_gn=C2CD5;refGene_tx=XM_019834685.2;refGene_gn=C2CD5;refGene_tx=XM_019834686.2;refGene_gn=C2CD5;refGene_tx=XM_019834687.2
B4 58059493 58059684
refGene_gn=C2CD5;refGene_tx=XM_011283840.3;refGene_gn=C2CD5;refGene_tx=XM_011283841.3;refGene_gn=C2CD5;refGene_tx=XM_011283842.3;refGene_gn=C2CD5;refGene_tx=XM_019834674.2;refGene_gn=C2CD5;refGene_tx=XM_019834675.2;refGene_gn=C2CD5;refGene_tx=XM_019834676.2;refGene_gn=C2CD5;refGene_tx=XM_019834677.2;refGene_gn=C2CD5;refGene_tx=XM_019834678.2;refGene_gn=C2CD5;refGene_tx=XM_019834679.1;refGene_gn=C2CD5;refGene_tx=XM_019834680.2;refGene_gn=C2CD5;refGene_tx=XM_019834681.2;refGene_gn=C2CD5;refGene_tx=XM_019834682.2;refGene_gn=C2CD5;refGene_tx=XM_019834683.2;refGene_gn=C2CD5;refGene_tx=XM_019834684.2;refGene_gn=C2CD5;refGene_tx=XM_019834685.2;refGene_gn=C2CD5;refGene_tx=XM_019834686.2;refGene_gn=C2CD5;refGene_tx=XM_019834687.2
B4 58060986 58061088
refGene_gn=C2CD5;refGene_tx=XM_011283840.3;refGene_gn=C2CD5;refGene_tx=XM_019834674.2;refGene_gn=C2CD5;refGene_tx=XM_019834675.2;refGene_gn=C2CD5;refGene_tx=XM_019834676.2;refGene_gn=C2CD5;refGene_tx=XM_019834677.2;refGene_gn=C2CD5;refGene_tx=XM_019834678.2;refGene_gn=C2CD5;refGene_tx=XM_019834681.2;refGene_gn=C2CD5;refGene_tx=XM_019834682.2;refGene_gn=C2CD5;refGene_tx=XM_019834683.2;refGene_gn=C2CD5;refGene_tx=XM_019834684.2;refGene_gn=C2CD5;refGene_tx=XM_019834685.2;refGene_gn=C2CD5;refGene_tx=XM_019834686.2;refGene_gn=C2CD5;refGene_tx=XM_019834687.2
B4 58061963 58062014
refGene_gn=C2CD5;refGene_tx=XM_011283840.3;refGene_gn=C2CD5;refGene_tx=XM_011283841.3;refGene_gn=C2CD5;refGene_tx=XM_019834674.2;refGene_gn=C2CD5;refGene_tx=XM_019834675.2;refGene_gn=C2CD5;refGene_tx=XM_019834676.2;refGene_gn=C2CD5;refGene_tx=XM_019834678.2;refGene_gn=C2CD5;refGene_tx=XM_019834679.1;refGene_gn=C2CD5;refGene_tx=XM_019834681.2;refGene_gn=C2CD5;refGene_tx=XM_019834682.2;refGene_gn=C2CD5;refGene_tx=XM_01

9834683.2;refGene_gn=C2CD5;refGene_tx=XM_019834684.2;refGene_gn=C2CD5;
refGene_tx=XM_019834685.2;refGene_gn=C2CD5;refGene_tx=XM_019834686.2;r
efGene_gn=C2CD5;refGene_tx=XM_019834687.2
B4 58070709 58070796
refGene_gn=C2CD5;refGene_tx=XM_011283840.3;refGene_gn=C2CD5;refGene_tx
=XM_011283841.3;refGene_gn=C2CD5;refGene_tx=XM_011283842.3;refGene_gn=
C2CD5;refGene_tx=XM_019834674.2;refGene_gn=C2CD5;refGene_tx=XM_0198346
75.2;refGene_gn=C2CD5;refGene_tx=XM_019834676.2;refGene_gn=C2CD5;refGe
ne_tx=XM_019834677.2;refGene_gn=C2CD5;refGene_tx=XM_019834678.2;refGen
e_gn=C2CD5;refGene_tx=XM_019834679.1;refGene_gn=C2CD5;refGene_tx=XM_01
9834680.2;refGene_gn=C2CD5;refGene_tx=XM_019834681.2;refGene_gn=C2CD5;
refGene_tx=XM_019834682.2;refGene_gn=C2CD5;refGene_tx=XM_019834683.2;r
efGene_gn=C2CD5;refGene_tx=XM_019834684.2;refGene_gn=C2CD5;refGene_tx=
XM_019834685.2;refGene_gn=C2CD5;refGene_tx=XM_019834686.2;refGene_gn=C
2CD5;refGene_tx=XM_019834687.2
B4 58071760 58071848
refGene_gn=C2CD5;refGene_tx=XM_011283840.3;refGene_gn=C2CD5;refGene_tx
=XM_011283841.3;refGene_gn=C2CD5;refGene_tx=XM_011283842.3;refGene_gn=
C2CD5;refGene_tx=XM_019834674.2;refGene_gn=C2CD5;refGene_tx=XM_0198346
75.2;refGene_gn=C2CD5;refGene_tx=XM_019834676.2;refGene_gn=C2CD5;refGe
ne_tx=XM_019834677.2;refGene_gn=C2CD5;refGene_tx=XM_019834678.2;refGen
e_gn=C2CD5;refGene_tx=XM_019834679.1;refGene_gn=C2CD5;refGene_tx=XM_01
9834680.2;refGene_gn=C2CD5;refGene_tx=XM_019834681.2;refGene_gn=C2CD5;
refGene_tx=XM_019834682.2;refGene_gn=C2CD5;refGene_tx=XM_019834683.2;r
efGene_gn=C2CD5;refGene_tx=XM_019834684.2;refGene_gn=C2CD5;refGene_tx=
XM_019834685.2;refGene_gn=C2CD5;refGene_tx=XM_019834686.2;refGene_gn=C
2CD5;refGene_tx=XM_019834687.2
B4 58072543 58072633
refGene_gn=C2CD5;refGene_tx=XM_011283840.3;refGene_gn=C2CD5;refGene_tx
=XM_011283841.3;refGene_gn=C2CD5;refGene_tx=XM_011283842.3;refGene_gn=
C2CD5;refGene_tx=XM_019834674.2;refGene_gn=C2CD5;refGene_tx=XM_0198346
75.2;refGene_gn=C2CD5;refGene_tx=XM_019834676.2;refGene_gn=C2CD5;refGe
ne_tx=XM_019834677.2;refGene_gn=C2CD5;refGene_tx=XM_019834678.2;refGen
e_gn=C2CD5;refGene_tx=XM_019834679.1;refGene_gn=C2CD5;refGene_tx=XM_01
9834680.2;refGene_gn=C2CD5;refGene_tx=XM_019834681.2;refGene_gn=C2CD5;
refGene_tx=XM_019834682.2;refGene_gn=C2CD5;refGene_tx=XM_019834683.2;r
efGene_gn=C2CD5;refGene_tx=XM_019834684.2;refGene_gn=C2CD5;refGene_tx=
XM_019834685.2;refGene_gn=C2CD5;refGene_tx=XM_019834686.2;refGene_gn=C
2CD5;refGene_tx=XM_019834687.2
B4 58073144 58073243
refGene_gn=C2CD5;refGene_tx=XM_011283840.3;refGene_gn=C2CD5;refGene_tx
=XM_011283841.3;refGene_gn=C2CD5;refGene_tx=XM_011283842.3;refGene_gn=
C2CD5;refGene_tx=XM_019834674.2;refGene_gn=C2CD5;refGene_tx=XM_0198346
75.2;refGene_gn=C2CD5;refGene_tx=XM_019834676.2;refGene_gn=C2CD5;refGe
ne_tx=XM_019834677.2;refGene_gn=C2CD5;refGene_tx=XM_019834678.2;refGen
e_gn=C2CD5;refGene_tx=XM_019834679.1;refGene_gn=C2CD5;refGene_tx=XM_01
9834680.2;refGene_gn=C2CD5;refGene_tx=XM_019834681.2;refGene_gn=C2CD5;
refGene_tx=XM_019834682.2;refGene_gn=C2CD5;refGene_tx=XM_019834683.2;r
efGene_gn=C2CD5;refGene_tx=XM_019834684.2;refGene_gn=C2CD5;refGene_tx=
XM_019834685.2;refGene_gn=C2CD5;refGene_tx=XM_019834686.2;refGene_gn=C

2CD5;refGene_tx=XM_019834687.2
B4 58073499 58073561
refGene_gn=C2CD5;refGene_tx=XM_011283840.3;refGene_gn=C2CD5;refGene_tx=XM_011283841.3;refGene_gn=C2CD5;refGene_tx=XM_011283842.3;refGene_gn=C2CD5;refGene_tx=XM_019834674.2;refGene_gn=C2CD5;refGene_tx=XM_019834675.2;refGene_gn=C2CD5;refGene_tx=XM_019834676.2;refGene_gn=C2CD5;refGene_tx=XM_019834677.2;refGene_gn=C2CD5;refGene_tx=XM_019834678.2;refGene_gn=C2CD5;refGene_tx=XM_019834679.1;refGene_gn=C2CD5;refGene_tx=XM_019834680.2;refGene_gn=C2CD5;refGene_tx=XM_019834681.2;refGene_gn=C2CD5;refGene_tx=XM_019834682.2;refGene_gn=C2CD5;refGene_tx=XM_019834683.2;refGene_gn=C2CD5;refGene_tx=XM_019834684.2;refGene_gn=C2CD5;refGene_tx=XM_019834685.2;refGene_gn=C2CD5;refGene_tx=XM_019834686.2;refGene_gn=C2CD5;refGene_tx=XM_019834687.2
B4 58074380 58074444
refGene_gn=C2CD5;refGene_tx=XM_011283840.3;refGene_gn=C2CD5;refGene_tx=XM_011283841.3;refGene_gn=C2CD5;refGene_tx=XM_011283842.3;refGene_gn=C2CD5;refGene_tx=XM_019834674.2;refGene_gn=C2CD5;refGene_tx=XM_019834675.2;refGene_gn=C2CD5;refGene_tx=XM_019834676.2;refGene_gn=C2CD5;refGene_tx=XM_019834677.2;refGene_gn=C2CD5;refGene_tx=XM_019834678.2;refGene_gn=C2CD5;refGene_tx=XM_019834679.1;refGene_gn=C2CD5;refGene_tx=XM_019834680.2;refGene_gn=C2CD5;refGene_tx=XM_019834681.2;refGene_gn=C2CD5;refGene_tx=XM_019834682.2;refGene_gn=C2CD5;refGene_tx=XM_019834683.2;refGene_gn=C2CD5;refGene_tx=XM_019834684.2;refGene_gn=C2CD5;refGene_tx=XM_019834685.2;refGene_gn=C2CD5;refGene_tx=XM_019834686.2;refGene_gn=C2CD5;refGene_tx=XM_019834687.2
B4 58076774 58076915
refGene_gn=C2CD5;refGene_tx=XM_011283840.3;refGene_gn=C2CD5;refGene_tx=XM_011283841.3;refGene_gn=C2CD5;refGene_tx=XM_011283842.3;refGene_gn=C2CD5;refGene_tx=XM_019834674.2;refGene_gn=C2CD5;refGene_tx=XM_019834675.2;refGene_gn=C2CD5;refGene_tx=XM_019834676.2;refGene_gn=C2CD5;refGene_tx=XM_019834677.2;refGene_gn=C2CD5;refGene_tx=XM_019834678.2;refGene_gn=C2CD5;refGene_tx=XM_019834679.1;refGene_gn=C2CD5;refGene_tx=XM_019834680.2;refGene_gn=C2CD5;refGene_tx=XM_019834681.2;refGene_gn=C2CD5;refGene_tx=XM_019834682.2;refGene_gn=C2CD5;refGene_tx=XM_019834683.2;refGene_gn=C2CD5;refGene_tx=XM_019834684.2;refGene_gn=C2CD5;refGene_tx=XM_019834685.2;refGene_gn=C2CD5;refGene_tx=XM_019834686.2;refGene_gn=C2CD5;refGene_tx=XM_019834687.2
B4 58080531 58080696
refGene_gn=C2CD5;refGene_tx=XM_011283840.3;refGene_gn=C2CD5;refGene_tx=XM_011283841.3;refGene_gn=C2CD5;refGene_tx=XM_011283842.3;refGene_gn=C2CD5;refGene_tx=XM_019834674.2;refGene_gn=C2CD5;refGene_tx=XM_019834675.2;refGene_gn=C2CD5;refGene_tx=XM_019834676.2;refGene_gn=C2CD5;refGene_tx=XM_019834677.2;refGene_gn=C2CD5;refGene_tx=XM_019834678.2;refGene_gn=C2CD5;refGene_tx=XM_019834679.1;refGene_gn=C2CD5;refGene_tx=XM_019834680.2;refGene_gn=C2CD5;refGene_tx=XM_019834681.2;refGene_gn=C2CD5;refGene_tx=XM_019834682.2;refGene_gn=C2CD5;refGene_tx=XM_019834683.2;refGene_gn=C2CD5;refGene_tx=XM_019834684.2;refGene_gn=C2CD5;refGene_tx=XM_019834685.2;refGene_gn=C2CD5;refGene_tx=XM_019834686.2;refGene_gn=C2CD5;refGene_tx=XM_019834687.2
B4 58083648 58083835

refGene_gn=C2CD5;refGene_tx=XM_011283840.3;refGene_gn=C2CD5;refGene_tx=XM_011283841.3;refGene_gn=C2CD5;refGene_tx=XM_011283842.3;refGene_gn=C2CD5;refGene_tx=XM_019834674.2;refGene_gn=C2CD5;refGene_tx=XM_019834675.2;refGene_gn=C2CD5;refGene_tx=XM_019834676.2;refGene_gn=C2CD5;refGene_tx=XM_019834677.2;refGene_gn=C2CD5;refGene_tx=XM_019834678.2;refGene_gn=C2CD5;refGene_tx=XM_019834679.1;refGene_gn=C2CD5;refGene_tx=XM_019834680.2;refGene_gn=C2CD5;refGene_tx=XM_019834681.2;refGene_gn=C2CD5;refGene_tx=XM_019834682.2;refGene_gn=C2CD5;refGene_tx=XM_019834683.2;refGene_gn=C2CD5;refGene_tx=XM_019834684.2;refGene_gn=C2CD5;refGene_tx=XM_019834685.2;refGene_gn=C2CD5;refGene_tx=XM_019834686.2;refGene_gn=C2CD5;refGene_tx=XM_019834687.2
B4 58088626 58088818
refGene_gn=C2CD5;refGene_tx=XM_011283840.3;refGene_gn=C2CD5;refGene_tx=XM_011283841.3;refGene_gn=C2CD5;refGene_tx=XM_011283842.3;refGene_gn=C2CD5;refGene_tx=XM_019834674.2;refGene_gn=C2CD5;refGene_tx=XM_019834675.2;refGene_gn=C2CD5;refGene_tx=XM_019834676.2;refGene_gn=C2CD5;refGene_tx=XM_019834677.2;refGene_gn=C2CD5;refGene_tx=XM_019834678.2;refGene_gn=C2CD5;refGene_tx=XM_019834679.1;refGene_gn=C2CD5;refGene_tx=XM_019834680.2;refGene_gn=C2CD5;refGene_tx=XM_019834681.2;refGene_gn=C2CD5;refGene_tx=XM_019834682.2;refGene_gn=C2CD5;refGene_tx=XM_019834683.2;refGene_gn=C2CD5;refGene_tx=XM_019834684.2;refGene_gn=C2CD5;refGene_tx=XM_019834685.2;refGene_gn=C2CD5;refGene_tx=XM_019834686.2;refGene_gn=C2CD5;refGene_tx=XM_019834687.2
B4 58091627 58091723
refGene_gn=C2CD5;refGene_tx=XM_011283840.3;refGene_gn=C2CD5;refGene_tx=XM_011283841.3;refGene_gn=C2CD5;refGene_tx=XM_011283842.3;refGene_gn=C2CD5;refGene_tx=XM_019834674.2;refGene_gn=C2CD5;refGene_tx=XM_019834675.2;refGene_gn=C2CD5;refGene_tx=XM_019834676.2;refGene_gn=C2CD5;refGene_tx=XM_019834677.2;refGene_gn=C2CD5;refGene_tx=XM_019834678.2;refGene_gn=C2CD5;refGene_tx=XM_019834679.1;refGene_gn=C2CD5;refGene_tx=XM_019834680.2;refGene_gn=C2CD5;refGene_tx=XM_019834681.2;refGene_gn=C2CD5;refGene_tx=XM_019834682.2;refGene_gn=C2CD5;refGene_tx=XM_019834683.2;refGene_gn=C2CD5;refGene_tx=XM_019834684.2;refGene_gn=C2CD5;refGene_tx=XM_019834685.2;refGene_gn=C2CD5;refGene_tx=XM_019834686.2;refGene_gn=C2CD5;refGene_tx=XM_019834687.2
B4 58094784 58094899
refGene_gn=C2CD5;refGene_tx=XM_011283840.3;refGene_gn=C2CD5;refGene_tx=XM_011283841.3;refGene_gn=C2CD5;refGene_tx=XM_011283842.3;refGene_gn=C2CD5;refGene_tx=XM_019834674.2;refGene_gn=C2CD5;refGene_tx=XM_019834675.2;refGene_gn=C2CD5;refGene_tx=XM_019834676.2;refGene_gn=C2CD5;refGene_tx=XM_019834677.2;refGene_gn=C2CD5;refGene_tx=XM_019834678.2;refGene_gn=C2CD5;refGene_tx=XM_019834679.1;refGene_gn=C2CD5;refGene_tx=XM_019834680.2;refGene_gn=C2CD5;refGene_tx=XM_019834681.2;refGene_gn=C2CD5;refGene_tx=XM_019834682.2;refGene_gn=C2CD5;refGene_tx=XM_019834683.2;refGene_gn=C2CD5;refGene_tx=XM_019834684.2;refGene_gn=C2CD5;refGene_tx=XM_019834685.2;refGene_gn=C2CD5;refGene_tx=XM_019834686.2;refGene_gn=C2CD5;refGene_tx=XM_019834687.2
B4 58104906 58104972
refGene_gn=C2CD5;refGene_tx=XM_019834674.2;refGene_gn=C2CD5;refGene_tx=XM_019834675.2;refGene_gn=C2CD5;refGene_tx=XM_019834676.2;refGene_gn=

C2CD5;refGene_tx=XM_019834677.2;refGene_gn=C2CD5;refGene_tx=XM_019834679.1;refGene_gn=C2CD5;refGene_tx=XM_019834680.2;refGene_gn=C2CD5;refGene_tx=XM_019834681.2;refGene_gn=C2CD5;refGene_tx=XM_019834682.2;refGene_gn=C2CD5;refGene_tx=XM_019834683.2;refGene_gn=C2CD5;refGene_tx=XM_019834684.2;refGene_gn=C2CD5;refGene_tx=XM_019834685.2
B4 58107723 58107832
refGene_gn=C2CD5;refGene_tx=XM_011283840.3;refGene_gn=C2CD5;refGene_tx=XM_011283841.3;refGene_gn=C2CD5;refGene_tx=XM_011283842.3;refGene_gn=C2CD5;refGene_tx=XM_019834674.2;refGene_gn=C2CD5;refGene_tx=XM_019834675.2;refGene_gn=C2CD5;refGene_tx=XM_019834676.2;refGene_gn=C2CD5;refGene_tx=XM_019834677.2;refGene_gn=C2CD5;refGene_tx=XM_019834678.2;refGene_gn=C2CD5;refGene_tx=XM_019834679.1;refGene_gn=C2CD5;refGene_tx=XM_019834680.2;refGene_gn=C2CD5;refGene_tx=XM_019834681.2;refGene_gn=C2CD5;refGene_tx=XM_019834682.2;refGene_gn=C2CD5;refGene_tx=XM_019834683.2;refGene_gn=C2CD5;refGene_tx=XM_019834684.2;refGene_gn=C2CD5;refGene_tx=XM_019834685.2;refGene_gn=C2CD5;refGene_tx=XM_019834686.2;refGene_gn=C2CD5;refGene_tx=XM_019834687.2
B4 58113384 58113417
refGene_gn=C2CD5;refGene_tx=XM_019834674.2;refGene_gn=C2CD5;refGene_tx=XM_019834675.2;refGene_gn=C2CD5;refGene_tx=XM_019834677.2;refGene_gn=C2CD5;refGene_tx=XM_019834678.2;refGene_gn=C2CD5;refGene_tx=XM_019834679.1;refGene_gn=C2CD5;refGene_tx=XM_019834680.2;refGene_gn=C2CD5;refGene_tx=XM_019834681.2;refGene_gn=C2CD5;refGene_tx=XM_019834682.2;refGene_gn=C2CD5;refGene_tx=XM_019834686.2
B4 58114029 58114115
refGene_gn=C2CD5;refGene_tx=XM_011283840.3;refGene_gn=C2CD5;refGene_tx=XM_011283841.3;refGene_gn=C2CD5;refGene_tx=XM_011283842.3;refGene_gn=C2CD5;refGene_tx=XM_019834674.2;refGene_gn=C2CD5;refGene_tx=XM_019834675.2;refGene_gn=C2CD5;refGene_tx=XM_019834676.2;refGene_gn=C2CD5;refGene_tx=XM_019834677.2;refGene_gn=C2CD5;refGene_tx=XM_019834678.2;refGene_gn=C2CD5;refGene_tx=XM_019834679.1;refGene_gn=C2CD5;refGene_tx=XM_019834680.2;refGene_gn=C2CD5;refGene_tx=XM_019834681.2;refGene_gn=C2CD5;refGene_tx=XM_019834682.2;refGene_gn=C2CD5;refGene_tx=XM_019834683.2;refGene_gn=C2CD5;refGene_tx=XM_019834684.2;refGene_gn=C2CD5;refGene_tx=XM_019834685.2;refGene_gn=C2CD5;refGene_tx=XM_019834686.2;refGene_gn=C2CD5;refGene_tx=XM_019834687.2
B4 58116049 58116241
refGene_gn=C2CD5;refGene_tx=XM_019834674.2;refGene_gn=C2CD5;refGene_tx=XM_019834675.2;refGene_gn=C2CD5;refGene_tx=XM_019834676.2;refGene_gn=C2CD5;refGene_tx=XM_019834677.2;refGene_gn=C2CD5;refGene_tx=XM_019834678.2;refGene_gn=C2CD5;refGene_tx=XM_019834679.1;refGene_gn=C2CD5;refGene_tx=XM_019834680.2
B4 58119369 58119521
refGene_gn=C2CD5;refGene_tx=XM_011283840.3;refGene_gn=C2CD5;refGene_tx=XM_011283841.3;refGene_gn=C2CD5;refGene_tx=XM_011283842.3;refGene_gn=C2CD5;refGene_tx=XM_019834674.2;refGene_gn=C2CD5;refGene_tx=XM_019834675.2;refGene_gn=C2CD5;refGene_tx=XM_019834676.2;refGene_gn=C2CD5;refGene_tx=XM_019834677.2;refGene_gn=C2CD5;refGene_tx=XM_019834678.2;refGene_gn=C2CD5;refGene_tx=XM_019834679.1;refGene_gn=C2CD5;refGene_tx=XM_019834680.2;refGene_gn=C2CD5;refGene_tx=XM_019834681.2;refGene_gn=C2CD5;

refGene_tx=XM_019834683.2
B4	58120516	58120641
refGene_gn=C2CD5;refGene_tx=XM_019834682.2;refGene_gn=C2CD5;refGene_tx=XM_019834684.2;refGene_gn=C2CD5;refGene_tx=XM_019834685.2;refGene_gn=C2CD5;refGene_tx=XM_019834686.2;refGene_gn=C2CD5;refGene_tx=XM_019834687.2
B4	58126212	58126411
refGene_gn=C2CD5;refGene_tx=XM_011283840.3;refGene_gn=C2CD5;refGene_tx=XM_011283841.3;refGene_gn=C2CD5;refGene_tx=XM_011283842.3;refGene_gn=C2CD5;refGene_tx=XM_019834674.2;refGene_gn=C2CD5;refGene_tx=XM_019834675.2;refGene_gn=C2CD5;refGene_tx=XM_019834676.2;refGene_gn=C2CD5;refGene_tx=XM_019834677.2;refGene_gn=C2CD5;refGene_tx=XM_019834678.2;refGene_gn=C2CD5;refGene_tx=XM_019834679.1;refGene_gn=C2CD5;refGene_tx=XM_019834680.2;refGene_gn=C2CD5;refGene_tx=XM_019834681.2;refGene_gn=C2CD5;refGene_tx=XM_019834682.2;refGene_gn=C2CD5;refGene_tx=XM_019834683.2;refGene_gn=C2CD5;refGene_tx=XM_019834684.2;refGene_gn=C2CD5;refGene_tx=XM_019834685.2;refGene_gn=C2CD5;refGene_tx=XM_019834686.2;refGene_gn=C2CD5;refGene_tx=XM_019834687.2
B4	58127458	58127614
refGene_gn=C2CD5;refGene_tx=XM_011283840.3;refGene_gn=C2CD5;refGene_tx=XM_011283841.3;refGene_gn=C2CD5;refGene_tx=XM_011283842.3;refGene_gn=C2CD5;refGene_tx=XM_019834674.2;refGene_gn=C2CD5;refGene_tx=XM_019834675.2;refGene_gn=C2CD5;refGene_tx=XM_019834676.2;refGene_gn=C2CD5;refGene_tx=XM_019834677.2;refGene_gn=C2CD5;refGene_tx=XM_019834678.2;refGene_gn=C2CD5;refGene_tx=XM_019834679.1;refGene_gn=C2CD5;refGene_tx=XM_019834680.2;refGene_gn=C2CD5;refGene_tx=XM_019834681.2;refGene_gn=C2CD5;refGene_tx=XM_019834682.2;refGene_gn=C2CD5;refGene_tx=XM_019834683.2;refGene_gn=C2CD5;refGene_tx=XM_019834684.2;refGene_gn=C2CD5;refGene_tx=XM_019834685.2;refGene_gn=C2CD5;refGene_tx=XM_019834686.2;refGene_gn=C2CD5;refGene_tx=XM_019834687.2
B4	58129970	58130066
refGene_gn=C2CD5;refGene_tx=XM_011283840.3;refGene_gn=C2CD5;refGene_tx=XM_011283841.3;refGene_gn=C2CD5;refGene_tx=XM_011283842.3;refGene_gn=C2CD5;refGene_tx=XM_019834674.2;refGene_gn=C2CD5;refGene_tx=XM_019834675.2;refGene_gn=C2CD5;refGene_tx=XM_019834676.2;refGene_gn=C2CD5;refGene_tx=XM_019834677.2;refGene_gn=C2CD5;refGene_tx=XM_019834678.2;refGene_gn=C2CD5;refGene_tx=XM_019834679.1;refGene_gn=C2CD5;refGene_tx=XM_019834680.2;refGene_gn=C2CD5;refGene_tx=XM_019834681.2;refGene_gn=C2CD5;refGene_tx=XM_019834682.2;refGene_gn=C2CD5;refGene_tx=XM_019834683.2;refGene_gn=C2CD5;refGene_tx=XM_019834684.2;refGene_gn=C2CD5;refGene_tx=XM_019834685.2;refGene_gn=C2CD5;refGene_tx=XM_019834686.2;refGene_gn=C2CD5;refGene_tx=XM_019834687.2
B4	58132559	58132731
refGene_gn=C2CD5;refGene_tx=XM_011283840.3;refGene_gn=C2CD5;refGene_tx=XM_011283841.3;refGene_gn=C2CD5;refGene_tx=XM_011283842.3;refGene_gn=C2CD5;refGene_tx=XM_019834674.2;refGene_gn=C2CD5;refGene_tx=XM_019834675.2;refGene_gn=C2CD5;refGene_tx=XM_019834676.2;refGene_gn=C2CD5;refGene_tx=XM_019834677.2;refGene_gn=C2CD5;refGene_tx=XM_019834678.2;refGene_gn=C2CD5;refGene_tx=XM_019834679.1;refGene_gn=C2CD5;refGene_tx=XM_019834680.2;refGene_gn=C2CD5;refGene_tx=XM_019834681.2;refGene_gn=C2CD5;

refGene_tx=XM_019834682.2;refGene_gn=C2CD5;refGene_tx=XM_019834683.2;refGene_gn=C2CD5;refGene_tx=XM_019834684.2;refGene_gn=C2CD5;refGene_tx=XM_019834685.2;refGene_gn=C2CD5;refGene_tx=XM_019834686.2;refGene_gn=C2CD5;refGene_tx=XM_019834687.2
B4 58140411 58140498
refGene_gn=C2CD5;refGene_tx=XM_011283840.3;refGene_gn=C2CD5;refGene_tx=XM_011283841.3;refGene_gn=C2CD5;refGene_tx=XM_011283842.3;refGene_gn=C2CD5;refGene_tx=XM_019834674.2;refGene_gn=C2CD5;refGene_tx=XM_019834675.2;refGene_gn=C2CD5;refGene_tx=XM_019834676.2;refGene_gn=C2CD5;refGene_tx=XM_019834677.2;refGene_gn=C2CD5;refGene_tx=XM_019834678.2;refGene_gn=C2CD5;refGene_tx=XM_019834679.1;refGene_gn=C2CD5;refGene_tx=XM_019834680.2;refGene_gn=C2CD5;refGene_tx=XM_019834681.2;refGene_gn=C2CD5;refGene_tx=XM_019834682.2;refGene_gn=C2CD5;refGene_tx=XM_019834683.2;refGene_gn=C2CD5;refGene_tx=XM_019834684.2;refGene_gn=C2CD5;refGene_tx=XM_019834685.2;refGene_gn=C2CD5;refGene_tx=XM_019834686.2;refGene_gn=C2CD5;refGene_tx=XM_019834687.2
B4 58155061 58155151
refGene_gn=C2CD5;refGene_tx=XM_011283840.3;refGene_gn=C2CD5;refGene_tx=XM_011283841.3;refGene_gn=C2CD5;refGene_tx=XM_011283842.3;refGene_gn=C2CD5;refGene_tx=XM_019834674.2;refGene_gn=C2CD5;refGene_tx=XM_019834675.2;refGene_gn=C2CD5;refGene_tx=XM_019834676.2;refGene_gn=C2CD5;refGene_tx=XM_019834677.2;refGene_gn=C2CD5;refGene_tx=XM_019834678.2;refGene_gn=C2CD5;refGene_tx=XM_019834679.1;refGene_gn=C2CD5;refGene_tx=XM_019834680.2;refGene_gn=C2CD5;refGene_tx=XM_019834681.2;refGene_gn=C2CD5;refGene_tx=XM_019834682.2;refGene_gn=C2CD5;refGene_tx=XM_019834683.2;refGene_gn=C2CD5;refGene_tx=XM_019834684.2;refGene_gn=C2CD5;refGene_tx=XM_019834685.2;refGene_gn=C2CD5;refGene_tx=XM_019834686.2;refGene_gn=C2CD5;refGene_tx=XM_019834687.2
B4 58234549 58234972 refGene_gn=ETNK1;refGene_tx=XM_023256829.1
B4 58254623 58254883 refGene_gn=ETNK1;refGene_tx=XM_023256829.1
B4 58266728 58266869 refGene_gn=ETNK1;refGene_tx=XM_023256829.1
B4 58269894 58270037 refGene_gn=ETNK1;refGene_tx=XM_023256829.1
B4 58279199 58279283 refGene_gn=ETNK1;refGene_tx=XM_023256829.1
B4 58282499 58282660 refGene_gn=ETNK1;refGene_tx=XM_023256829.1
B4 58297157 58297231 refGene_gn=ETNK1;refGene_tx=XM_023256829.1
B4 58297570 58297643 refGene_gn=ETNK1;refGene_tx=XM_023256829.1
B4 59029108 59029412
refGene_gn=SOX5;refGene_tx=XM_023256830.1;refGene_gn=SOX5;refGene_tx=XM_023256831.1;refGene_gn=SOX5;refGene_tx=XM_023256832.1;refGene_gn=SOX5;refGene_tx=XM_023256833.1;refGene_gn=SOX5;refGene_tx=XM_023256834.1;refGene_gn=SOX5;refGene_tx=XM_023256835.1;refGene_gn=SOX5;refGene_tx=XM_023256836.1;refGene_gn=SOX5;refGene_tx=XM_023256837.1;refGene_gn=SOX5;refGene_tx=XM_023256838.1;refGene_gn=SOX5;refGene_tx=XM_023256839.1;refGene_gn=SOX5;refGene_tx=XM_023256841.1;refGene_gn=SOX5;refGene_tx=XM_023256842.1;refGene_gn=SOX5;refGene_tx=XM_023256843.1;refGene_gn=SOX5;refGene_tx=XM_023256844.1;refGene_gn=SOX5;refGene_tx=XM_023256845.1
B4 59031004 59031221
refGene_gn=SOX5;refGene_tx=XM_023256830.1;refGene_gn=SOX5;refGene_tx=XM_023256831.1;refGene_gn=SOX5;refGene_tx=XM_023256832.1;refGene_gn=SOX5;refGene_tx=XM_023256833.1;refGene_gn=SOX5;refGene_tx=XM_023256834.1;

refGene_gn=SOX5;refGene_tx=XM_023256835.1;refGene_gn=SOX5;refGene_tx=XM_023256836.1;refGene_gn=SOX5;refGene_tx=XM_023256837.1;refGene_gn=SOX5;refGene_tx=XM_023256838.1;refGene_gn=SOX5;refGene_tx=XM_023256839.1;refGene_gn=SOX5;refGene_tx=XM_023256841.1;refGene_gn=SOX5;refGene_tx=XM_023256842.1;refGene_gn=SOX5;refGene_tx=XM_023256843.1;refGene_gn=SOX5;refGene_tx=XM_023256844.1;refGene_gn=SOX5;refGene_tx=XM_023256845.1
B4 59037338 59037512
refGene_gn=SOX5;refGene_tx=XM_023256830.1;refGene_gn=SOX5;refGene_tx=XM_023256831.1;refGene_gn=SOX5;refGene_tx=XM_023256832.1;refGene_gn=SOX5;refGene_tx=XM_023256833.1;refGene_gn=SOX5;refGene_tx=XM_023256834.1;refGene_gn=SOX5;refGene_tx=XM_023256835.1;refGene_gn=SOX5;refGene_tx=XM_023256836.1;refGene_gn=SOX5;refGene_tx=XM_023256837.1;refGene_gn=SOX5;refGene_tx=XM_023256838.1;refGene_gn=SOX5;refGene_tx=XM_023256839.1;refGene_gn=SOX5;refGene_tx=XM_023256841.1;refGene_gn=SOX5;refGene_tx=XM_023256842.1;refGene_gn=SOX5;refGene_tx=XM_023256843.1;refGene_gn=SOX5;refGene_tx=XM_023256844.1;refGene_gn=SOX5;refGene_tx=XM_023256845.1
B4 59040810 59040919
refGene_gn=SOX5;refGene_tx=XM_023256830.1;refGene_gn=SOX5;refGene_tx=XM_023256831.1;refGene_gn=SOX5;refGene_tx=XM_023256832.1;refGene_gn=SOX5;refGene_tx=XM_023256833.1;refGene_gn=SOX5;refGene_tx=XM_023256834.1;refGene_gn=SOX5;refGene_tx=XM_023256835.1;refGene_gn=SOX5;refGene_tx=XM_023256836.1;refGene_gn=SOX5;refGene_tx=XM_023256837.1;refGene_gn=SOX5;refGene_tx=XM_023256838.1;refGene_gn=SOX5;refGene_tx=XM_023256839.1;refGene_gn=SOX5;refGene_tx=XM_023256841.1;refGene_gn=SOX5;refGene_tx=XM_023256842.1;refGene_gn=SOX5;refGene_tx=XM_023256843.1;refGene_gn=SOX5;refGene_tx=XM_023256844.1;refGene_gn=SOX5;refGene_tx=XM_023256845.1
B4 59058534 59058680
refGene_gn=SOX5;refGene_tx=XM_023256830.1;refGene_gn=SOX5;refGene_tx=XM_023256831.1;refGene_gn=SOX5;refGene_tx=XM_023256832.1;refGene_gn=SOX5;refGene_tx=XM_023256833.1;refGene_gn=SOX5;refGene_tx=XM_023256834.1;refGene_gn=SOX5;refGene_tx=XM_023256835.1;refGene_gn=SOX5;refGene_tx=XM_023256836.1;refGene_gn=SOX5;refGene_tx=XM_023256837.1;refGene_gn=SOX5;refGene_tx=XM_023256838.1;refGene_gn=SOX5;refGene_tx=XM_023256839.1;refGene_gn=SOX5;refGene_tx=XM_023256841.1;refGene_gn=SOX5;refGene_tx=XM_023256842.1;refGene_gn=SOX5;refGene_tx=XM_023256843.1;refGene_gn=SOX5;refGene_tx=XM_023256844.1;refGene_gn=SOX5;refGene_tx=XM_023256845.1
B4 59070181 59070359
refGene_gn=SOX5;refGene_tx=XM_023256830.1;refGene_gn=SOX5;refGene_tx=XM_023256831.1;refGene_gn=SOX5;refGene_tx=XM_023256832.1;refGene_gn=SOX5;refGene_tx=XM_023256833.1;refGene_gn=SOX5;refGene_tx=XM_023256834.1;refGene_gn=SOX5;refGene_tx=XM_023256835.1;refGene_gn=SOX5;refGene_tx=XM_023256836.1;refGene_gn=SOX5;refGene_tx=XM_023256837.1;refGene_gn=SOX5;refGene_tx=XM_023256838.1;refGene_gn=SOX5;refGene_tx=XM_023256839.1;refGene_gn=SOX5;refGene_tx=XM_023256841.1;refGene_gn=SOX5;refGene_tx=XM_023256842.1;refGene_gn=SOX5;refGene_tx=XM_023256843.1;refGene_gn=SOX5;refGene_tx=XM_023256844.1;refGene_gn=SOX5;refGene_tx=XM_023256845.1
B4 59099028 59099175
refGene_gn=SOX5;refGene_tx=XM_023256830.1;refGene_gn=SOX5;refGene_tx=XM_023256831.1;refGene_gn=SOX5;refGene_tx=XM_023256832.1;refGene_gn=SOX5;refGene_tx=XM_023256833.1;refGene_gn=SOX5;refGene_tx=XM_023256834.1;

refGene_gn=SOX5;refGene_tx=XM_023256835.1;refGene_gn=SOX5;refGene_tx=XM_023256836.1;refGene_gn=SOX5;refGene_tx=XM_023256837.1;refGene_gn=SOX5;refGene_tx=XM_023256838.1;refGene_gn=SOX5;refGene_tx=XM_023256839.1;refGene_gn=SOX5;refGene_tx=XM_023256842.1;refGene_gn=SOX5;refGene_tx=XM_023256843.1;refGene_gn=SOX5;refGene_tx=XM_023256844.1;refGene_gn=SOX5;refGene_tx=XM_023256845.1
B4 59133266 59133352
refGene_gn=SOX5;refGene_tx=XM_023256830.1;refGene_gn=SOX5;refGene_tx=XM_023256831.1;refGene_gn=SOX5;refGene_tx=XM_023256832.1;refGene_gn=SOX5;refGene_tx=XM_023256833.1;refGene_gn=SOX5;refGene_tx=XM_023256834.1;refGene_gn=SOX5;refGene_tx=XM_023256835.1;refGene_gn=SOX5;refGene_tx=XM_023256836.1;refGene_gn=SOX5;refGene_tx=XM_023256837.1;refGene_gn=SOX5;refGene_tx=XM_023256838.1;refGene_gn=SOX5;refGene_tx=XM_023256839.1;refGene_gn=SOX5;refGene_tx=XM_023256841.1;refGene_gn=SOX5;refGene_tx=XM_023256842.1;refGene_gn=SOX5;refGene_tx=XM_023256843.1;refGene_gn=SOX5;refGene_tx=XM_023256844.1;refGene_gn=SOX5;refGene_tx=XM_023256845.1
B4 59160202 59160326
refGene_gn=SOX5;refGene_tx=XM_023256830.1;refGene_gn=SOX5;refGene_tx=XM_023256831.1;refGene_gn=SOX5;refGene_tx=XM_023256832.1;refGene_gn=SOX5;refGene_tx=XM_023256833.1;refGene_gn=SOX5;refGene_tx=XM_023256834.1;refGene_gn=SOX5;refGene_tx=XM_023256835.1;refGene_gn=SOX5;refGene_tx=XM_023256836.1;refGene_gn=SOX5;refGene_tx=XM_023256837.1;refGene_gn=SOX5;refGene_tx=XM_023256838.1;refGene_gn=SOX5;refGene_tx=XM_023256839.1;refGene_gn=SOX5;refGene_tx=XM_023256841.1;refGene_gn=SOX5;refGene_tx=XM_023256842.1;refGene_gn=SOX5;refGene_tx=XM_023256843.1;refGene_gn=SOX5;refGene_tx=XM_023256844.1;refGene_gn=SOX5;refGene_tx=XM_023256845.1
B4 59231315 59231384
refGene_gn=SOX5;refGene_tx=XM_023256830.1;refGene_gn=SOX5;refGene_tx=XM_023256831.1;refGene_gn=SOX5;refGene_tx=XM_023256832.1;refGene_gn=SOX5;refGene_tx=XM_023256833.1;refGene_gn=SOX5;refGene_tx=XM_023256834.1;refGene_gn=SOX5;refGene_tx=XM_023256835.1;refGene_gn=SOX5;refGene_tx=XM_023256836.1;refGene_gn=SOX5;refGene_tx=XM_023256837.1;refGene_gn=SOX5;refGene_tx=XM_023256838.1;refGene_gn=SOX5;refGene_tx=XM_023256839.1;refGene_gn=SOX5;refGene_tx=XM_023256841.1;refGene_gn=SOX5;refGene_tx=XM_023256842.1;refGene_gn=SOX5;refGene_tx=XM_023256843.1
B4 59236668 59236841
refGene_gn=SOX5;refGene_tx=XM_023256830.1;refGene_gn=SOX5;refGene_tx=XM_023256831.1;refGene_gn=SOX5;refGene_tx=XM_023256832.1;refGene_gn=SOX5;refGene_tx=XM_023256833.1;refGene_gn=SOX5;refGene_tx=XM_023256834.1;refGene_gn=SOX5;refGene_tx=XM_023256835.1;refGene_gn=SOX5;refGene_tx=XM_023256836.1;refGene_gn=SOX5;refGene_tx=XM_023256837.1;refGene_gn=SOX5;refGene_tx=XM_023256838.1;refGene_gn=SOX5;refGene_tx=XM_023256839.1;refGene_gn=SOX5;refGene_tx=XM_023256841.1;refGene_gn=SOX5;refGene_tx=XM_023256842.1;refGene_gn=SOX5;refGene_tx=XM_023256843.1
B4 59251940 59252027
refGene_gn=SOX5;refGene_tx=XM_023256830.1;refGene_gn=SOX5;refGene_tx=XM_023256831.1;refGene_gn=SOX5;refGene_tx=XM_023256832.1;refGene_gn=SOX5;refGene_tx=XM_023256833.1;refGene_gn=SOX5;refGene_tx=XM_023256834.1;refGene_gn=SOX5;refGene_tx=XM_023256835.1;refGene_gn=SOX5;refGene_tx=XM_023256836.1;refGene_gn=SOX5;refGene_tx=XM_023256837.1;refGene_gn=SOX

5;refGene_tx=XM_023256838.1;refGene_gn=SOX5;refGene_tx=XM_023256839.1;
refGene_gn=SOX5;refGene_tx=XM_023256841.1;refGene_gn=SOX5;refGene_tx=X
M_023256842.1;refGene_gn=SOX5;refGene_tx=XM_023256843.1
B4 59338391 59338602
refGene_gn=SOX5;refGene_tx=XM_023256830.1;refGene_gn=SOX5;refGene_tx=X
M_023256831.1;refGene_gn=SOX5;refGene_tx=XM_023256832.1;refGene_gn=SOX
5;refGene_tx=XM_023256833.1;refGene_gn=SOX5;refGene_tx=XM_023256834.1;
refGene_gn=SOX5;refGene_tx=XM_023256835.1;refGene_gn=SOX5;refGene_tx=X
M_023256836.1;refGene_gn=SOX5;refGene_tx=XM_023256837.1;refGene_gn=SOX
5;refGene_tx=XM_023256838.1;refGene_gn=SOX5;refGene_tx=XM_023256839.1;
refGene_gn=SOX5;refGene_tx=XM_023256841.1
B4 59384431 59384663
refGene_gn=SOX5;refGene_tx=XM_023256830.1;refGene_gn=SOX5;refGene_tx=X
M_023256831.1;refGene_gn=SOX5;refGene_tx=XM_023256832.1;refGene_gn=SOX
5;refGene_tx=XM_023256833.1;refGene_gn=SOX5;refGene_tx=XM_023256834.1;
refGene_gn=SOX5;refGene_tx=XM_023256835.1;refGene_gn=SOX5;refGene_tx=X
M_023256836.1;refGene_gn=SOX5;refGene_tx=XM_023256837.1;refGene_gn=SOX
5;refGene_tx=XM_023256838.1;refGene_gn=SOX5;refGene_tx=XM_023256839.1;
refGene_gn=SOX5;refGene_tx=XM_023256841.1
B4 59439460 59439498
refGene_gn=SOX5;refGene_tx=XM_023256830.1;refGene_gn=SOX5;refGene_tx=X
M_023256831.1;refGene_gn=SOX5;refGene_tx=XM_023256838.1;refGene_gn=SOX
5;refGene_tx=XM_023256839.1;refGene_gn=SOX5;refGene_tx=XM_023256841.1
B4 59586805 59586807
refGene_gn=SOX5;refGene_tx=XM_023256832.1;refGene_gn=SOX5;refGene_tx=X
M_023256833.1;refGene_gn=SOX5;refGene_tx=XM_023256834.1;refGene_gn=SOX
5;refGene_tx=XM_023256835.1;refGene_gn=SOX5;refGene_tx=XM_023256836.1;
refGene_gn=SOX5;refGene_tx=XM_023256837.1
B4 60268696 60268738
refGene_gn=BCAT1;refGene_tx=XM_006933567.3;refGene_gn=BCAT1;refGene_tx
=XM_006933568.4;refGene_gn=BCAT1;refGene_tx=XM_019834701.2
B4 60279198 60279273
refGene_gn=BCAT1;refGene_tx=XM_006933567.3;refGene_gn=BCAT1;refGene_tx
=XM_006933568.4;refGene_gn=BCAT1;refGene_tx=XM_019834701.2
B4 60281395 60281536
refGene_gn=BCAT1;refGene_tx=XM_006933567.3;refGene_gn=BCAT1;refGene_tx
=XM_006933568.4;refGene_gn=BCAT1;refGene_tx=XM_019834701.2
B4 60284192 60284278
refGene_gn=BCAT1;refGene_tx=XM_006933567.3;refGene_gn=BCAT1;refGene_tx
=XM_006933568.4;refGene_gn=BCAT1;refGene_tx=XM_019834701.2
B4 60289282 60289425
refGene_gn=BCAT1;refGene_tx=XM_006933567.3;refGene_gn=BCAT1;refGene_tx
=XM_006933568.4;refGene_gn=BCAT1;refGene_tx=XM_019834701.2
B4 60294939 60295103
refGene_gn=BCAT1;refGene_tx=XM_006933567.3;refGene_gn=BCAT1;refGene_tx
=XM_006933568.4;refGene_gn=BCAT1;refGene_tx=XM_019834701.2
B4 60314803 60314923
refGene_gn=BCAT1;refGene_tx=XM_006933567.3;refGene_gn=BCAT1;refGene_tx
=XM_006933568.4;refGene_gn=BCAT1;refGene_tx=XM_019834701.2
B4 60317282 60317393

refGene_gn=BCAT1;refGene_tx=XM_006933567.3;refGene_gn=BCAT1;refGene_tx
=XM_006933568.4;refGene_gn=BCAT1;refGene_tx=XM_019834701.2
B4	60324004	60324205
refGene_gn=BCAT1;refGene_tx=XM_006933567.3;refGene_gn=BCAT1;refGene_tx
=XM_006933568.4;refGene_gn=BCAT1;refGene_tx=XM_019834701.2
B4	60334375	60334447
refGene_gn=BCAT1;refGene_tx=XM_006933567.3;refGene_gn=BCAT1;refGene_tx
=XM_006933568.4;refGene_gn=BCAT1;refGene_tx=XM_019834701.2
B4	60334521	60334524	refGene_gn=BCAT1;refGene_tx=XM_006933568.4
B4	60335537	60335579	refGene_gn=BCAT1;refGene_tx=XM_006933567.3
B4	60370203	60370209	refGene_gn=BCAT1;refGene_tx=XM_019834701.2
B4	60415468	60416039
refGene_gn=LOC109501496;refGene_tx=XM_023256855.1
B4	60422343	60422454
refGene_gn=LOC109501496;refGene_tx=XM_023256855.1
B4	60450787	60450968
refGene_gn=LOC109501496;refGene_tx=XM_023256855.1
B4	60453495	60453575
refGene_gn=LOC109501496;refGene_tx=XM_023256855.1
B4	60453665	60453744
refGene_gn=LOC109501496;refGene_tx=XM_023256855.1
B4	60455113	60455272
refGene_gn=LOC109501496;refGene_tx=XM_023256855.1
B4	60459053	60459171
refGene_gn=LOC109501496;refGene_tx=XM_023256855.1
B4	60461242	60461293
refGene_gn=LOC109501496;refGene_tx=XM_023256855.1
B4	60464506	60464584
refGene_gn=LOC109501496;refGene_tx=XM_023256855.1
B4	60466813	60466870
refGene_gn=LOC109501496;refGene_tx=XM_023256855.1
B4	60468575	60468635
refGene_gn=LOC109501496;refGene_tx=XM_023256855.1
B4	60468734	60468799
refGene_gn=LOC109501496;refGene_tx=XM_023256855.1
B4	60469508	60469608
refGene_gn=LOC109501496;refGene_tx=XM_023256855.1
B4	60471293	60471425
refGene_gn=LOC109501496;refGene_tx=XM_023256855.1
B4	60472163	60472265
refGene_gn=LOC109501496;refGene_tx=XM_023256855.1
B4	60473439	60473603
refGene_gn=LOC109501496;refGene_tx=XM_023256855.1
B4	60494129	60494274
refGene_gn=LOC109501496;refGene_tx=XM_023256855.1
B4	60494275	60494314
refGene_gn=LOC109501496;refGene_tx=XM_023256855.1
B4	60494662	60494852	refGene_gn=LRMP;refGene_tx=XM_023257751.1
B4	60501752	60501814	refGene_gn=LRMP;refGene_tx=XM_023257751.1
B4	60502960	60503041	refGene_gn=LRMP;refGene_tx=XM_023257751.1

B4 60503616 60503713 refGene_gn=LRMP;refGene_tx=XM_023257751.1
B4 60506119 60506267 refGene_gn=LRMP;refGene_tx=XM_023257751.1
B4 60507000 60507082 refGene_gn=LRMP;refGene_tx=XM_023257751.1
B4 60518057 60518104 refGene_gn=LRMP;refGene_tx=XM_023257751.1
B4 60518206 60518271 refGene_gn=LRMP;refGene_tx=XM_023257751.1
B4 60518428 60518536 refGene_gn=LRMP;refGene_tx=XM_023257751.1
B4 60521549 60521620 refGene_gn=LRMP;refGene_tx=XM_023257751.1
B4 60524473 60524531 refGene_gn=LRMP;refGene_tx=XM_023257751.1
B4 60526312 60526376 refGene_gn=LRMP;refGene_tx=XM_023257751.1
B4 60526462 60526491 refGene_gn=LRMP;refGene_tx=XM_023257751.1
B4 60526749 60526889 refGene_gn=LRMP;refGene_tx=XM_023257751.1
B4 60533522 60533657 refGene_gn=LRMP;refGene_tx=XM_023257751.1
B4 60540078 60540226 refGene_gn=LRMP;refGene_tx=XM_023257751.1
B4 60541073 60541117 refGene_gn=LRMP;refGene_tx=XM_023257751.1
B4 60542611 60542674 refGene_gn=LRMP;refGene_tx=XM_023257751.1
B4 60542789 60542836 refGene_gn=LRMP;refGene_tx=XM_023257751.1
B4 60543094 60543196 refGene_gn=LRMP;refGene_tx=XM_023257751.1
B4 60545018 60545126 refGene_gn=LRMP;refGene_tx=XM_023257751.1
B4 60545971 60546215 refGene_gn=LRMP;refGene_tx=XM_023257751.1
B4 60546679 60546958
refGene_gn=CASC1;refGene_tx=XM_023256846.1;refGene_gn=CASC1;refGene_tx=XM_023256847.1;refGene_gn=CASC1;refGene_tx=XM_023256848.1;refGene_gn=CASC1;refGene_tx=XM_023256849.1;refGene_gn=CASC1;refGene_tx=XM_023256850.1;refGene_gn=CASC1;refGene_tx=XM_023256851.1;refGene_gn=CASC1;refGene_tx=XM_023256852.1
B4 60547693 60547807
refGene_gn=CASC1;refGene_tx=XM_023256846.1;refGene_gn=CASC1;refGene_tx=XM_023256847.1;refGene_gn=CASC1;refGene_tx=XM_023256848.1;refGene_gn=CASC1;refGene_tx=XM_023256849.1;refGene_gn=CASC1;refGene_tx=XM_023256850.1;refGene_gn=CASC1;refGene_tx=XM_023256851.1;refGene_gn=CASC1;refGene_tx=XM_023256852.1
B4 60549187 60549355
refGene_gn=CASC1;refGene_tx=XM_023256846.1;refGene_gn=CASC1;refGene_tx=XM_023256847.1;refGene_gn=CASC1;refGene_tx=XM_023256848.1;refGene_gn=CASC1;refGene_tx=XM_023256849.1;refGene_gn=CASC1;refGene_tx=XM_023256850.1;refGene_gn=CASC1;refGene_tx=XM_023256851.1;refGene_gn=CASC1;refGene_tx=XM_023256852.1
B4 60550288 60550503
refGene_gn=CASC1;refGene_tx=XM_023256846.1;refGene_gn=CASC1;refGene_tx=XM_023256847.1;refGene_gn=CASC1;refGene_tx=XM_023256848.1;refGene_gn=CASC1;refGene_tx=XM_023256849.1;refGene_gn=CASC1;refGene_tx=XM_023256850.1;refGene_gn=CASC1;refGene_tx=XM_023256851.1;refGene_gn=CASC1;refGene_tx=XM_023256852.1
B4 60553171 60553328
refGene_gn=CASC1;refGene_tx=XM_023256846.1;refGene_gn=CASC1;refGene_tx=XM_023256847.1;refGene_gn=CASC1;refGene_tx=XM_023256848.1;refGene_gn=CASC1;refGene_tx=XM_023256849.1;refGene_gn=CASC1;refGene_tx=XM_023256850.1;refGene_gn=CASC1;refGene_tx=XM_023256851.1;refGene_gn=CASC1;refGene_tx=XM_023256852.1
B4 60556199 60556360

refGene_gn=CASC1;refGene_tx=XM_023256846.1;refGene_gn=CASC1;refGene_tx
=XM_023256847.1;refGene_gn=CASC1;refGene_tx=XM_023256848.1;refGene_gn=
CASC1;refGene_tx=XM_023256849.1;refGene_gn=CASC1;refGene_tx=XM_0232568
50.1;refGene_gn=CASC1;refGene_tx=XM_023256851.1
B4 60558965 60559041
refGene_gn=CASC1;refGene_tx=XM_023256846.1;refGene_gn=CASC1;refGene_tx
=XM_023256847.1;refGene_gn=CASC1;refGene_tx=XM_023256848.1;refGene_gn=
CASC1;refGene_tx=XM_023256849.1;refGene_gn=CASC1;refGene_tx=XM_0232568
50.1;refGene_gn=CASC1;refGene_tx=XM_023256851.1
B4 60577310 60577620
refGene_gn=CASC1;refGene_tx=XM_023256846.1;refGene_gn=CASC1;refGene_tx
=XM_023256847.1;refGene_gn=CASC1;refGene_tx=XM_023256848.1;refGene_gn=
CASC1;refGene_tx=XM_023256849.1;refGene_gn=CASC1;refGene_tx=XM_0232568
50.1;refGene_gn=CASC1;refGene_tx=XM_023256851.1;refGene_gn=CASC1;refGe
ne_tx=XM_023256852.1
B4 60579636 60579740
refGene_gn=CASC1;refGene_tx=XM_023256846.1;refGene_gn=CASC1;refGene_tx
=XM_023256847.1;refGene_gn=CASC1;refGene_tx=XM_023256848.1;refGene_gn=
CASC1;refGene_tx=XM_023256849.1;refGene_gn=CASC1;refGene_tx=XM_0232568
50.1;refGene_gn=CASC1;refGene_tx=XM_023256851.1;refGene_gn=CASC1;refGe
ne_tx=XM_023256852.1
B4 60582313 60582460
refGene_gn=CASC1;refGene_tx=XM_023256846.1;refGene_gn=CASC1;refGene_tx
=XM_023256847.1;refGene_gn=CASC1;refGene_tx=XM_023256848.1;refGene_gn=
CASC1;refGene_tx=XM_023256849.1;refGene_gn=CASC1;refGene_tx=XM_0232568
50.1;refGene_gn=CASC1;refGene_tx=XM_023256851.1;refGene_gn=CASC1;refGe
ne_tx=XM_023256852.1
B4 60587390 60587528
refGene_gn=CASC1;refGene_tx=XM_023256846.1;refGene_gn=CASC1;refGene_tx
=XM_023256847.1;refGene_gn=CASC1;refGene_tx=XM_023256848.1;refGene_gn=
CASC1;refGene_tx=XM_023256849.1;refGene_gn=CASC1;refGene_tx=XM_0232568
50.1;refGene_gn=CASC1;refGene_tx=XM_023256852.1
B4 60588381 60588483
refGene_gn=CASC1;refGene_tx=XM_023256846.1;refGene_gn=CASC1;refGene_tx
=XM_023256847.1;refGene_gn=CASC1;refGene_tx=XM_023256848.1;refGene_gn=
CASC1;refGene_tx=XM_023256849.1;refGene_gn=CASC1;refGene_tx=XM_0232568
50.1;refGene_gn=CASC1;refGene_tx=XM_023256851.1;refGene_gn=CASC1;refGe
ne_tx=XM_023256852.1
B4 60590479 60590571
refGene_gn=CASC1;refGene_tx=XM_023256846.1;refGene_gn=CASC1;refGene_tx
=XM_023256847.1;refGene_gn=CASC1;refGene_tx=XM_023256848.1;refGene_gn=
CASC1;refGene_tx=XM_023256849.1;refGene_gn=CASC1;refGene_tx=XM_0232568
50.1;refGene_gn=CASC1;refGene_tx=XM_023256851.1;refGene_gn=CASC1;refGe
ne_tx=XM_023256852.1
B4 60592806 60592888
refGene_gn=CASC1;refGene_tx=XM_023256846.1;refGene_gn=CASC1;refGene_tx
=XM_023256847.1;refGene_gn=CASC1;refGene_tx=XM_023256851.1;refGene_gn=
CASC1;refGene_tx=XM_023256852.1
B4 60619296 60619314
refGene_gn=CASC1;refGene_tx=XM_023256846.1;refGene_gn=CASC1;refGene_tx

=XM_023256851.1;refGene_gn=CASC1;refGene_tx=XM_023256852.1
B4 60623002 60623005
refGene_gn=CASC1;refGene_tx=XM_023256846.1;refGene_gn=CASC1;refGene_tx=XM_023256847.1;refGene_gn=CASC1;refGene_tx=XM_023256851.1;refGene_gn=CASC1;refGene_tx=XM_023256852.1
B4 60628437 60628488
refGene_gn=ETFRF1;refGene_tx=XM_003988501.5;refGene_gn=ETFRF1;refGene_tx=XM_011283847.2
B4 60628569 60628791
refGene_gn=ETFRF1;refGene_tx=XM_003988501.5;refGene_gn=ETFRF1;refGene_tx=XM_011283847.2
B4 60634154 60634271
refGene_gn=KRAS;refGene_tx=XM_023256853.1;refGene_gn=KRAS;refGene_tx=XM_023256854.1
B4 60646361 60646521
refGene_gn=KRAS;refGene_tx=XM_023256853.1;refGene_gn=KRAS;refGene_tx=XM_023256854.1
B4 60648028 60648207
refGene_gn=KRAS;refGene_tx=XM_023256853.1;refGene_gn=KRAS;refGene_tx=XM_023256854.1
B4 60666117 60666228
refGene_gn=KRAS;refGene_tx=XM_023256853.1;refGene_gn=KRAS;refGene_tx=XM_023256854.1
B4 60806664 60806697 refGene_gn=LMNTD1;refGene_tx=XM_023256865.1
B4 60879888 60879942
refGene_gn=LMNTD1;refGene_tx=XM_023256860.1;refGene_gn=LMNTD1;refGene_tx=XM_023256862.1;refGene_gn=LMNTD1;refGene_tx=XM_023256863.1
B4 60889488 60889578
refGene_gn=LMNTD1;refGene_tx=XM_019834724.2;refGene_gn=LMNTD1;refGene_tx=XM_019834725.2;refGene_gn=LMNTD1;refGene_tx=XM_019834726.1;refGene_gn=LMNTD1;refGene_tx=XM_023256856.1;refGene_gn=LMNTD1;refGene_tx=XM_023256857.1;refGene_gn=LMNTD1;refGene_tx=XM_023256858.1;refGene_gn=LMNTD1;refGene_tx=XM_023256859.1;refGene_gn=LMNTD1;refGene_tx=XM_023256861.1;refGene_gn=LMNTD1;refGene_tx=XM_023256864.1;refGene_gn=LMNTD1;refGene_tx=XM_023256866.1;refGene_gn=LMNTD1;refGene_tx=XM_023256867.1;refGene_gn=LMNTD1;refGene_tx=XM_023256868.1
B4 60898248 60898420
refGene_gn=LMNTD1;refGene_tx=XM_019834724.2;refGene_gn=LMNTD1;refGene_tx=XM_019834725.2;refGene_gn=LMNTD1;refGene_tx=XM_019834726.1;refGene_gn=LMNTD1;refGene_tx=XM_023256856.1;refGene_gn=LMNTD1;refGene_tx=XM_023256857.1;refGene_gn=LMNTD1;refGene_tx=XM_023256858.1;refGene_gn=LMNTD1;refGene_tx=XM_023256859.1;refGene_gn=LMNTD1;refGene_tx=XM_023256860.1;refGene_gn=LMNTD1;refGene_tx=XM_023256861.1;refGene_gn=LMNTD1;refGene_tx=XM_023256862.1;refGene_gn=LMNTD1;refGene_tx=XM_023256863.1;refGene_gn=LMNTD1;refGene_tx=XM_023256864.1;refGene_gn=LMNTD1;refGene_tx=XM_023256865.1;refGene_gn=LMNTD1;refGene_tx=XM_023256866.1;refGene_gn=LMNTD1;refGene_tx=XM_023256867.1;refGene_gn=LMNTD1;refGene_tx=XM_023256868.1
B4 60898987 60899202
refGene_gn=LMNTD1;refGene_tx=XM_019834724.2;refGene_gn=LMNTD1;refGene_

tx=XM_019834725.2;refGene_gn=LMNTD1;refGene_tx=XM_019834726.1;refGene_gn=LMNTD1;refGene_tx=XM_023256856.1;refGene_gn=LMNTD1;refGene_tx=XM_023256857.1;refGene_gn=LMNTD1;refGene_tx=XM_023256858.1;refGene_gn=LMNTD1;refGene_tx=XM_023256859.1;refGene_gn=LMNTD1;refGene_tx=XM_023256860.1;refGene_gn=LMNTD1;refGene_tx=XM_023256861.1;refGene_gn=LMNTD1;refGene_tx=XM_023256862.1;refGene_gn=LMNTD1;refGene_tx=XM_023256863.1;refGene_gn=LMNTD1;refGene_tx=XM_023256864.1;refGene_gn=LMNTD1;refGene_tx=XM_023256865.1;refGene_gn=LMNTD1;refGene_tx=XM_023256866.1;refGene_gn=LMNTD1;refGene_tx=XM_023256867.1;refGene_gn=LMNTD1;refGene_tx=XM_023256868.1
B4 60900870 60900990
refGene_gn=LMNTD1;refGene_tx=XM_019834724.2;refGene_gn=LMNTD1;refGene_tx=XM_019834725.2;refGene_gn=LMNTD1;refGene_tx=XM_019834726.1;refGene_gn=LMNTD1;refGene_tx=XM_023256856.1;refGene_gn=LMNTD1;refGene_tx=XM_023256857.1;refGene_gn=LMNTD1;refGene_tx=XM_023256858.1;refGene_gn=LMNTD1;refGene_tx=XM_023256859.1;refGene_gn=LMNTD1;refGene_tx=XM_023256860.1;refGene_gn=LMNTD1;refGene_tx=XM_023256861.1;refGene_gn=LMNTD1;refGene_tx=XM_023256862.1;refGene_gn=LMNTD1;refGene_tx=XM_023256863.1;refGene_gn=LMNTD1;refGene_tx=XM_023256864.1;refGene_gn=LMNTD1;refGene_tx=XM_023256865.1;refGene_gn=LMNTD1;refGene_tx=XM_023256866.1;refGene_gn=LMNTD1;refGene_tx=XM_023256867.1;refGene_gn=LMNTD1;refGene_tx=XM_023256868.1
B4 60901535 60901722
refGene_gn=LMNTD1;refGene_tx=XM_019834724.2;refGene_gn=LMNTD1;refGene_tx=XM_019834725.2;refGene_gn=LMNTD1;refGene_tx=XM_019834726.1;refGene_gn=LMNTD1;refGene_tx=XM_023256856.1;refGene_gn=LMNTD1;refGene_tx=XM_023256857.1;refGene_gn=LMNTD1;refGene_tx=XM_023256858.1;refGene_gn=LMNTD1;refGene_tx=XM_023256859.1;refGene_gn=LMNTD1;refGene_tx=XM_023256860.1;refGene_gn=LMNTD1;refGene_tx=XM_023256861.1;refGene_gn=LMNTD1;refGene_tx=XM_023256862.1;refGene_gn=LMNTD1;refGene_tx=XM_023256863.1;refGene_gn=LMNTD1;refGene_tx=XM_023256864.1;refGene_gn=LMNTD1;refGene_tx=XM_023256865.1;refGene_gn=LMNTD1;refGene_tx=XM_023256866.1;refGene_gn=LMNTD1;refGene_tx=XM_023256867.1;refGene_gn=LMNTD1;refGene_tx=XM_023256868.1
B4 60911628 60911827
refGene_gn=LMNTD1;refGene_tx=XM_019834724.2;refGene_gn=LMNTD1;refGene_tx=XM_019834725.2;refGene_gn=LMNTD1;refGene_tx=XM_019834726.1;refGene_gn=LMNTD1;refGene_tx=XM_023256856.1;refGene_gn=LMNTD1;refGene_tx=XM_023256857.1;refGene_gn=LMNTD1;refGene_tx=XM_023256858.1;refGene_gn=LMNTD1;refGene_tx=XM_023256859.1;refGene_gn=LMNTD1;refGene_tx=XM_023256860.1;refGene_gn=LMNTD1;refGene_tx=XM_023256861.1;refGene_gn=LMNTD1;refGene_tx=XM_023256862.1;refGene_gn=LMNTD1;refGene_tx=XM_023256863.1;refGene_gn=LMNTD1;refGene_tx=XM_023256864.1;refGene_gn=LMNTD1;refGene_tx=XM_023256865.1;refGene_gn=LMNTD1;refGene_tx=XM_023256866.1;refGene_gn=LMNTD1;refGene_tx=XM_023256867.1;refGene_gn=LMNTD1;refGene_tx=XM_023256868.1
B4 60914486 60914716
refGene_gn=LMNTD1;refGene_tx=XM_019834724.2;refGene_gn=LMNTD1;refGene_tx=XM_019834725.2;refGene_gn=LMNTD1;refGene_tx=XM_019834726.1;refGene_gn=LMNTD1;refGene_tx=XM_023256856.1;refGene_gn=LMNTD1;refGene_tx=XM_02

3256857.1;refGene_gn=LMNTD1;refGene_tx=XM_023256858.1;refGene_gn=LMNTD1;refGene_tx=XM_023256859.1;refGene_gn=LMNTD1;refGene_tx=XM_023256860.1;refGene_gn=LMNTD1;refGene_tx=XM_023256861.1;refGene_gn=LMNTD1;refGene_tx=XM_023256862.1;refGene_gn=LMNTD1;refGene_tx=XM_023256863.1;refGene_gn=LMNTD1;refGene_tx=XM_023256864.1;refGene_gn=LMNTD1;refGene_tx=XM_023256865.1;refGene_gn=LMNTD1;refGene_tx=XM_023256866.1;refGene_gn=LMNTD1;refGene_tx=XM_023256867.1;refGene_gn=LMNTD1;refGene_tx=XM_023256868.1

B4 60917708 60917827 refGene_gn=LMNTD1;refGene_tx=XM_019834724.2;refGene_gn=LMNTD1;refGene_tx=XM_019834725.2;refGene_gn=LMNTD1;refGene_tx=XM_019834726.1;refGene_gn=LMNTD1;refGene_tx=XM_023256856.1;refGene_gn=LMNTD1;refGene_tx=XM_023256857.1;refGene_gn=LMNTD1;refGene_tx=XM_023256858.1;refGene_gn=LMNTD1;refGene_tx=XM_023256859.1;refGene_gn=LMNTD1;refGene_tx=XM_023256860.1;refGene_gn=LMNTD1;refGene_tx=XM_023256861.1;refGene_gn=LMNTD1;refGene_tx=XM_023256862.1;refGene_gn=LMNTD1;refGene_tx=XM_023256863.1;refGene_gn=LMNTD1;refGene_tx=XM_023256864.1;refGene_gn=LMNTD1;refGene_tx=XM_023256865.1;refGene_gn=LMNTD1;refGene_tx=XM_023256866.1;refGene_gn=LMNTD1;refGene_tx=XM_023256867.1

B4 60919656 60919737 refGene_gn=LMNTD1;refGene_tx=XM_023256864.1

B4 61032104 61032188 refGene_gn=LMNTD1;refGene_tx=XM_023256861.1

B4 61034868 61034970 refGene_gn=LMNTD1;refGene_tx=XM_023256856.1;refGene_gn=LMNTD1;refGene_tx=XM_023256857.1;refGene_gn=LMNTD1;refGene_tx=XM_023256858.1;refGene_gn=LMNTD1;refGene_tx=XM_023256859.1;refGene_gn=LMNTD1;refGene_tx=XM_023256860.1;refGene_gn=LMNTD1;refGene_tx=XM_023256862.1;refGene_gn=LMNTD1;refGene_tx=XM_023256863.1;refGene_gn=LMNTD1;refGene_tx=XM_023256865.1;refGene_gn=LMNTD1;refGene_tx=XM_023256866.1;refGene_gn=LMNTD1;refGene_tx=XM_023256867.1

B4 61043633 61043651 refGene_gn=LMNTD1;refGene_tx=XM_023256859.1

B4 61198266 61198284 refGene_gn=LMNTD1;refGene_tx=XM_023256856.1;refGene_gn=LMNTD1;refGene_tx=XM_023256857.1;refGene_gn=LMNTD1;refGene_tx=XM_023256858.1;refGene_gn=LMNTD1;refGene_tx=XM_023256860.1;refGene_gn=LMNTD1;refGene_tx=XM_023256862.1;refGene_gn=LMNTD1;refGene_tx=XM_023256863.1;refGene_gn=LMNTD1;refGene_tx=XM_023256865.1;refGene_gn=LMNTD1;refGene_tx=XM_023256866.1

B4 61426859 61426962 refGene_gn=RASSF8;refGene_tx=XM_003988504.5;refGene_gn=RASSF8;refGene_tx=XM_023256869.1

B4 61437346 61438236 refGene_gn=RASSF8;refGene_tx=XM_003988504.5;refGene_gn=RASSF8;refGene_tx=XM_023256869.1

B4 61440330 61440475 refGene_gn=RASSF8;refGene_tx=XM_003988504.5;refGene_gn=RASSF8;refGene_tx=XM_023256869.1

B4 61441082 61441204 refGene_gn=RASSF8;refGene_tx=XM_003988504.5;refGene_gn=RASSF8;refGene_tx=XM_023256869.1

B4 61486150 61487259 refGene_gn=BHLHE41;refGene_tx=XM_023256870.1

B4 61487694 61487806 refGene_gn=BHLHE41;refGene_tx=XM_023256870.1
B4 61488135 61488243 refGene_gn=BHLHE41;refGene_tx=XM_023256870.1
B4 61488564 61488628 refGene_gn=BHLHE41;refGene_tx=XM_023256870.1
B4 61488772 61489164 refGene_gn=BHLHE41;refGene_tx=XM_023256870.1
B4 61563099 61563375
refGene_gn=SSPN;refGene_tx=XM_003988505.4;refGene_gn=SSPN;refGene_tx=XM_019834728.1
B4 61588516 61588567 refGene_gn=SSPN;refGene_tx=XM_006933572.4
B4 61589074 61589161
refGene_gn=SSPN;refGene_tx=XM_003988505.4;refGene_gn=SSPN;refGene_tx=XM_006933572.4
B4 61595841 61596207
refGene_gn=SSPN;refGene_tx=XM_003988505.4;refGene_gn=SSPN;refGene_tx=XM_006933572.4;refGene_gn=SSPN;refGene_tx=XM_019834728.1
B4 61678559 61678646
refGene_gn=ITPR2;refGene_tx=XM_019834731.2;refGene_gn=ITPR2;refGene_tx=XM_019834732.2;refGene_gn=ITPR2;refGene_tx=XM_019834733.2;refGene_gn=ITPR2;refGene_tx=XM_019834734.1;refGene_gn=ITPR2;refGene_tx=XM_023256871.1
B4 61679376 61679538
refGene_gn=ITPR2;refGene_tx=XM_019834731.2;refGene_gn=ITPR2;refGene_tx=XM_019834732.2;refGene_gn=ITPR2;refGene_tx=XM_019834733.2;refGene_gn=ITPR2;refGene_tx=XM_019834734.1;refGene_gn=ITPR2;refGene_tx=XM_023256871.1
B4 61724606 61724767
refGene_gn=ITPR2;refGene_tx=XM_019834731.2;refGene_gn=ITPR2;refGene_tx=XM_019834732.2;refGene_gn=ITPR2;refGene_tx=XM_019834733.2;refGene_gn=ITPR2;refGene_tx=XM_019834734.1;refGene_gn=ITPR2;refGene_tx=XM_023256871.1
B4 61737565 61737731
refGene_gn=ITPR2;refGene_tx=XM_019834731.2;refGene_gn=ITPR2;refGene_tx=XM_019834732.2;refGene_gn=ITPR2;refGene_tx=XM_019834733.2;refGene_gn=ITPR2;refGene_tx=XM_019834734.1;refGene_gn=ITPR2;refGene_tx=XM_023256871.1
B4 61739055 61739186
refGene_gn=ITPR2;refGene_tx=XM_019834731.2;refGene_gn=ITPR2;refGene_tx=XM_019834732.2;refGene_gn=ITPR2;refGene_tx=XM_019834733.2;refGene_gn=ITPR2;refGene_tx=XM_019834734.1;refGene_gn=ITPR2;refGene_tx=XM_023256871.1
B4 61749720 61749813
refGene_gn=ITPR2;refGene_tx=XM_019834731.2;refGene_gn=ITPR2;refGene_tx=XM_019834732.2;refGene_gn=ITPR2;refGene_tx=XM_019834733.2;refGene_gn=ITPR2;refGene_tx=XM_019834734.1;refGene_gn=ITPR2;refGene_tx=XM_023256871.1
B4 61754486 61754682
refGene_gn=ITPR2;refGene_tx=XM_019834731.2;refGene_gn=ITPR2;refGene_tx=XM_019834732.2;refGene_gn=ITPR2;refGene_tx=XM_019834733.2;refGene_gn=ITPR2;refGene_tx=XM_019834734.1;refGene_gn=ITPR2;refGene_tx=XM_023256871.1
B4 61758098 61758263

refGene_gn=ITPR2;refGene_tx=XM_019834731.2;refGene_gn=ITPR2;refGene_tx=XM_019834732.2;refGene_gn=ITPR2;refGene_tx=XM_019834733.2;refGene_gn=ITPR2;refGene_tx=XM_019834734.1;refGene_gn=ITPR2;refGene_tx=XM_023256871.1
B4 61766261 61766437
refGene_gn=ITPR2;refGene_tx=XM_019834731.2;refGene_gn=ITPR2;refGene_tx=XM_019834732.2;refGene_gn=ITPR2;refGene_tx=XM_019834733.2;refGene_gn=ITPR2;refGene_tx=XM_019834734.1;refGene_gn=ITPR2;refGene_tx=XM_023256871.1
B4 61778874 61779000
refGene_gn=ITPR2;refGene_tx=XM_019834731.2;refGene_gn=ITPR2;refGene_tx=XM_019834732.2;refGene_gn=ITPR2;refGene_tx=XM_019834733.2;refGene_gn=ITPR2;refGene_tx=XM_019834734.1;refGene_gn=ITPR2;refGene_tx=XM_023256871.1
B4 61782527 61782720
refGene_gn=ITPR2;refGene_tx=XM_019834731.2;refGene_gn=ITPR2;refGene_tx=XM_019834732.2;refGene_gn=ITPR2;refGene_tx=XM_019834733.2;refGene_gn=ITPR2;refGene_tx=XM_019834734.1;refGene_gn=ITPR2;refGene_tx=XM_023256871.1
B4 61790332 61790440
refGene_gn=ITPR2;refGene_tx=XM_019834731.2;refGene_gn=ITPR2;refGene_tx=XM_019834732.2;refGene_gn=ITPR2;refGene_tx=XM_019834733.2;refGene_gn=ITPR2;refGene_tx=XM_019834734.1;refGene_gn=ITPR2;refGene_tx=XM_023256871.1
B4 61824087 61824210
refGene_gn=ITPR2;refGene_tx=XM_019834731.2;refGene_gn=ITPR2;refGene_tx=XM_019834732.2;refGene_gn=ITPR2;refGene_tx=XM_019834733.2;refGene_gn=ITPR2;refGene_tx=XM_019834734.1;refGene_gn=ITPR2;refGene_tx=XM_023256871.1
B4 61826511 61826607
refGene_gn=ITPR2;refGene_tx=XM_019834731.2;refGene_gn=ITPR2;refGene_tx=XM_019834732.2;refGene_gn=ITPR2;refGene_tx=XM_019834733.2;refGene_gn=ITPR2;refGene_tx=XM_019834734.1;refGene_gn=ITPR2;refGene_tx=XM_023256871.1
B4 61831027 61831138
refGene_gn=ITPR2;refGene_tx=XM_019834731.2;refGene_gn=ITPR2;refGene_tx=XM_019834732.2;refGene_gn=ITPR2;refGene_tx=XM_019834733.2;refGene_gn=ITPR2;refGene_tx=XM_019834734.1;refGene_gn=ITPR2;refGene_tx=XM_023256871.1
B4 61834088 61834289
refGene_gn=ITPR2;refGene_tx=XM_019834731.2;refGene_gn=ITPR2;refGene_tx=XM_019834732.2;refGene_gn=ITPR2;refGene_tx=XM_019834733.2;refGene_gn=ITPR2;refGene_tx=XM_019834734.1;refGene_gn=ITPR2;refGene_tx=XM_023256871.1
B4 61836881 61837138
refGene_gn=ITPR2;refGene_tx=XM_019834731.2;refGene_gn=ITPR2;refGene_tx=XM_019834732.2;refGene_gn=ITPR2;refGene_tx=XM_019834733.2;refGene_gn=ITPR2;refGene_tx=XM_019834734.1;refGene_gn=ITPR2;refGene_tx=XM_023256871.1
B4 61837820 61838004

refGene_gn=ITPR2;refGene_tx=XM_019834731.2;refGene_gn=ITPR2;refGene_tx=XM_019834732.2;refGene_gn=ITPR2;refGene_tx=XM_019834733.2;refGene_gn=ITPR2;refGene_tx=XM_019834734.1;refGene_gn=ITPR2;refGene_tx=XM_023256871.1
B4 61843848 61844036
refGene_gn=ITPR2;refGene_tx=XM_019834731.2;refGene_gn=ITPR2;refGene_tx=XM_019834732.2;refGene_gn=ITPR2;refGene_tx=XM_019834733.2;refGene_gn=ITPR2;refGene_tx=XM_019834734.1;refGene_gn=ITPR2;refGene_tx=XM_023256871.1
B4 61844765 61844874
refGene_gn=ITPR2;refGene_tx=XM_019834731.2;refGene_gn=ITPR2;refGene_tx=XM_019834732.2;refGene_gn=ITPR2;refGene_tx=XM_019834733.2;refGene_gn=ITPR2;refGene_tx=XM_019834734.1;refGene_gn=ITPR2;refGene_tx=XM_023256871.1
B4 61898336 61898445
refGene_gn=ITPR2;refGene_tx=XM_019834731.2;refGene_gn=ITPR2;refGene_tx=XM_019834732.2;refGene_gn=ITPR2;refGene_tx=XM_019834733.2;refGene_gn=ITPR2;refGene_tx=XM_019834734.1;refGene_gn=ITPR2;refGene_tx=XM_023256871.1
B4 61903703 61903846
refGene_gn=ITPR2;refGene_tx=XM_019834731.2;refGene_gn=ITPR2;refGene_tx=XM_019834732.2;refGene_gn=ITPR2;refGene_tx=XM_019834733.2;refGene_gn=ITPR2;refGene_tx=XM_019834734.1;refGene_gn=ITPR2;refGene_tx=XM_023256871.1
B4 61909546 61909737
refGene_gn=ITPR2;refGene_tx=XM_019834731.2;refGene_gn=ITPR2;refGene_tx=XM_019834732.2;refGene_gn=ITPR2;refGene_tx=XM_019834733.2;refGene_gn=ITPR2;refGene_tx=XM_019834734.1;refGene_gn=ITPR2;refGene_tx=XM_023256871.1
B4 61926544 61926665
refGene_gn=ITPR2;refGene_tx=XM_019834731.2;refGene_gn=ITPR2;refGene_tx=XM_019834732.2;refGene_gn=ITPR2;refGene_tx=XM_019834733.2;refGene_gn=ITPR2;refGene_tx=XM_019834734.1;refGene_gn=ITPR2;refGene_tx=XM_023256871.1
B4 61928115 61928244
refGene_gn=ITPR2;refGene_tx=XM_019834731.2;refGene_gn=ITPR2;refGene_tx=XM_019834732.2;refGene_gn=ITPR2;refGene_tx=XM_019834733.2;refGene_gn=ITPR2;refGene_tx=XM_019834734.1;refGene_gn=ITPR2;refGene_tx=XM_023256871.1
B4 61941031 61941157
refGene_gn=ITPR2;refGene_tx=XM_019834731.2;refGene_gn=ITPR2;refGene_tx=XM_019834732.2;refGene_gn=ITPR2;refGene_tx=XM_019834733.2;refGene_gn=ITPR2;refGene_tx=XM_019834734.1;refGene_gn=ITPR2;refGene_tx=XM_023256871.1
B4 61944789 61945041
refGene_gn=ITPR2;refGene_tx=XM_019834731.2;refGene_gn=ITPR2;refGene_tx=XM_019834732.2;refGene_gn=ITPR2;refGene_tx=XM_019834733.2;refGene_gn=ITPR2;refGene_tx=XM_019834734.1;refGene_gn=ITPR2;refGene_tx=XM_023256871.1
B4 61947128 61947329

refGene_gn=ITPR2;refGene_tx=XM_019834731.2;refGene_gn=ITPR2;refGene_tx=XM_019834732.2;refGene_gn=ITPR2;refGene_tx=XM_019834733.2;refGene_gn=ITPR2;refGene_tx=XM_019834734.1;refGene_gn=ITPR2;refGene_tx=XM_023256871.1
B4 61947959 61948082
refGene_gn=ITPR2;refGene_tx=XM_019834731.2;refGene_gn=ITPR2;refGene_tx=XM_019834732.2;refGene_gn=ITPR2;refGene_tx=XM_019834733.2;refGene_gn=ITPR2;refGene_tx=XM_019834734.1;refGene_gn=ITPR2;refGene_tx=XM_023256871.1
B4 61951094 61951220
refGene_gn=ITPR2;refGene_tx=XM_019834731.2;refGene_gn=ITPR2;refGene_tx=XM_019834732.2;refGene_gn=ITPR2;refGene_tx=XM_019834733.2;refGene_gn=ITPR2;refGene_tx=XM_019834734.1;refGene_gn=ITPR2;refGene_tx=XM_023256871.1
B4 61951337 61951427
refGene_gn=ITPR2;refGene_tx=XM_019834731.2;refGene_gn=ITPR2;refGene_tx=XM_019834732.2;refGene_gn=ITPR2;refGene_tx=XM_019834733.2;refGene_gn=ITPR2;refGene_tx=XM_019834734.1;refGene_gn=ITPR2;refGene_tx=XM_023256871.1
B4 61971990 61972164
refGene_gn=ITPR2;refGene_tx=XM_019834731.2;refGene_gn=ITPR2;refGene_tx=XM_019834732.2;refGene_gn=ITPR2;refGene_tx=XM_019834733.2;refGene_gn=ITPR2;refGene_tx=XM_019834734.1;refGene_gn=ITPR2;refGene_tx=XM_023256871.1
B4 61973164 61973330
refGene_gn=ITPR2;refGene_tx=XM_019834731.2;refGene_gn=ITPR2;refGene_tx=XM_019834732.2;refGene_gn=ITPR2;refGene_tx=XM_019834733.2;refGene_gn=ITPR2;refGene_tx=XM_019834734.1;refGene_gn=ITPR2;refGene_tx=XM_023256871.1
B4 61974828 61974886
refGene_gn=ITPR2;refGene_tx=XM_019834731.2;refGene_gn=ITPR2;refGene_tx=XM_019834732.2;refGene_gn=ITPR2;refGene_tx=XM_019834733.2;refGene_gn=ITPR2;refGene_tx=XM_019834734.1;refGene_gn=ITPR2;refGene_tx=XM_023256871.1
B4 61977143 61977273
refGene_gn=ITPR2;refGene_tx=XM_019834731.2;refGene_gn=ITPR2;refGene_tx=XM_019834732.2;refGene_gn=ITPR2;refGene_tx=XM_019834733.2;refGene_gn=ITPR2;refGene_tx=XM_019834734.1;refGene_gn=ITPR2;refGene_tx=XM_023256871.1
B4 61981098 61981292
refGene_gn=ITPR2;refGene_tx=XM_019834731.2;refGene_gn=ITPR2;refGene_tx=XM_019834732.2;refGene_gn=ITPR2;refGene_tx=XM_019834733.2;refGene_gn=ITPR2;refGene_tx=XM_019834734.1;refGene_gn=ITPR2;refGene_tx=XM_023256871.1
B4 62002392 62002543
refGene_gn=ITPR2;refGene_tx=XM_019834731.2;refGene_gn=ITPR2;refGene_tx=XM_019834732.2;refGene_gn=ITPR2;refGene_tx=XM_019834733.2;refGene_gn=ITPR2;refGene_tx=XM_019834734.1;refGene_gn=ITPR2;refGene_tx=XM_023256871.1
B4 62003998 62004143

refGene_gn=ITPR2;refGene_tx=XM_019834731.2;refGene_gn=ITPR2;refGene_tx=XM_019834732.2;refGene_gn=ITPR2;refGene_tx=XM_019834733.2;refGene_gn=ITPR2;refGene_tx=XM_019834734.1;refGene_gn=ITPR2;refGene_tx=XM_023256871.1
B4 62005412 62005664
refGene_gn=ITPR2;refGene_tx=XM_019834731.2;refGene_gn=ITPR2;refGene_tx=XM_019834732.2;refGene_gn=ITPR2;refGene_tx=XM_019834733.2;refGene_gn=ITPR2;refGene_tx=XM_019834734.1;refGene_gn=ITPR2;refGene_tx=XM_023256871.1
B4 62006787 62006973
refGene_gn=ITPR2;refGene_tx=XM_019834731.2;refGene_gn=ITPR2;refGene_tx=XM_019834732.2;refGene_gn=ITPR2;refGene_tx=XM_019834733.2;refGene_gn=ITPR2;refGene_tx=XM_019834734.1;refGene_gn=ITPR2;refGene_tx=XM_023256871.1
B4 62007338 62007458
refGene_gn=ITPR2;refGene_tx=XM_019834731.2;refGene_gn=ITPR2;refGene_tx=XM_019834732.2;refGene_gn=ITPR2;refGene_tx=XM_019834733.2;refGene_gn=ITPR2;refGene_tx=XM_019834734.1;refGene_gn=ITPR2;refGene_tx=XM_023256871.1
B4 62008441 62008614
refGene_gn=ITPR2;refGene_tx=XM_019834731.2;refGene_gn=ITPR2;refGene_tx=XM_019834732.2;refGene_gn=ITPR2;refGene_tx=XM_019834733.2;refGene_gn=ITPR2;refGene_tx=XM_019834734.1;refGene_gn=ITPR2;refGene_tx=XM_023256871.1
B4 62014415 62014577
refGene_gn=ITPR2;refGene_tx=XM_019834731.2;refGene_gn=ITPR2;refGene_tx=XM_019834732.2;refGene_gn=ITPR2;refGene_tx=XM_019834733.2;refGene_gn=ITPR2;refGene_tx=XM_019834734.1;refGene_gn=ITPR2;refGene_tx=XM_023256871.1
B4 62016845 62016987
refGene_gn=ITPR2;refGene_tx=XM_019834731.2;refGene_gn=ITPR2;refGene_tx=XM_019834732.2;refGene_gn=ITPR2;refGene_tx=XM_019834733.2;refGene_gn=ITPR2;refGene_tx=XM_019834734.1;refGene_gn=ITPR2;refGene_tx=XM_023256871.1
B4 62025027 62025188
refGene_gn=ITPR2;refGene_tx=XM_019834731.2;refGene_gn=ITPR2;refGene_tx=XM_019834732.2;refGene_gn=ITPR2;refGene_tx=XM_019834733.2;refGene_gn=ITPR2;refGene_tx=XM_019834734.1;refGene_gn=ITPR2;refGene_tx=XM_023256871.1
B4 62026140 62026240
refGene_gn=ITPR2;refGene_tx=XM_019834731.2;refGene_gn=ITPR2;refGene_tx=XM_019834732.2;refGene_gn=ITPR2;refGene_tx=XM_019834733.2;refGene_gn=ITPR2;refGene_tx=XM_019834734.1;refGene_gn=ITPR2;refGene_tx=XM_023256871.1
B4 62029285 62029437
refGene_gn=ITPR2;refGene_tx=XM_019834731.2;refGene_gn=ITPR2;refGene_tx=XM_019834732.2;refGene_gn=ITPR2;refGene_tx=XM_019834733.2;refGene_gn=ITPR2;refGene_tx=XM_019834734.1;refGene_gn=ITPR2;refGene_tx=XM_023256871.1
B4 62039685 62039730

refGene_gn=ITPR2;refGene_tx=XM_019834731.2;refGene_gn=ITPR2;refGene_tx=XM_019834732.2;refGene_gn=ITPR2;refGene_tx=XM_019834733.2;refGene_gn=ITPR2;refGene_tx=XM_019834734.1;refGene_gn=ITPR2;refGene_tx=XM_023256871.1
B4 62054758 62054854
refGene_gn=ITPR2;refGene_tx=XM_019834731.2;refGene_gn=ITPR2;refGene_tx=XM_019834732.2;refGene_gn=ITPR2;refGene_tx=XM_019834733.2;refGene_gn=ITPR2;refGene_tx=XM_019834734.1;refGene_gn=ITPR2;refGene_tx=XM_023256871.1
B4 62058099 62058246
refGene_gn=ITPR2;refGene_tx=XM_019834731.2;refGene_gn=ITPR2;refGene_tx=XM_019834732.2;refGene_gn=ITPR2;refGene_tx=XM_019834733.2;refGene_gn=ITPR2;refGene_tx=XM_019834734.1;refGene_gn=ITPR2;refGene_tx=XM_023256871.1
B4 62058545 62058629
refGene_gn=ITPR2;refGene_tx=XM_019834731.2;refGene_gn=ITPR2;refGene_tx=XM_019834732.2;refGene_gn=ITPR2;refGene_tx=XM_019834733.2;refGene_gn=ITPR2;refGene_tx=XM_019834734.1;refGene_gn=ITPR2;refGene_tx=XM_023256871.1
B4 62058911 62059010
refGene_gn=ITPR2;refGene_tx=XM_019834731.2;refGene_gn=ITPR2;refGene_tx=XM_019834732.2;refGene_gn=ITPR2;refGene_tx=XM_019834733.2;refGene_gn=ITPR2;refGene_tx=XM_019834734.1;refGene_gn=ITPR2;refGene_tx=XM_023256871.1
B4 62064928 62065087
refGene_gn=ITPR2;refGene_tx=XM_019834731.2;refGene_gn=ITPR2;refGene_tx=XM_019834732.2;refGene_gn=ITPR2;refGene_tx=XM_019834733.2;refGene_gn=ITPR2;refGene_tx=XM_023256871.1
B4 62067071 62067158
refGene_gn=ITPR2;refGene_tx=XM_019834731.2;refGene_gn=ITPR2;refGene_tx=XM_019834732.2;refGene_gn=ITPR2;refGene_tx=XM_019834733.2;refGene_gn=ITPR2;refGene_tx=XM_023256871.1
B4 62067949 62068065
refGene_gn=ITPR2;refGene_tx=XM_019834731.2;refGene_gn=ITPR2;refGene_tx=XM_019834732.2;refGene_gn=ITPR2;refGene_tx=XM_019834733.2;refGene_gn=ITPR2;refGene_tx=XM_023256871.1
B4 62126150 62126221
refGene_gn=ITPR2;refGene_tx=XM_019834731.2;refGene_gn=ITPR2;refGene_tx=XM_019834732.2
B4 62163549 62163641
refGene_gn=ITPR2;refGene_tx=XM_019834731.2;refGene_gn=ITPR2;refGene_tx=XM_019834732.2
B4 62239442 62239482
refGene_gn=INTS13;refGene_tx=XM_011283852.3;refGene_gn=INTS13;refGene_tx=XM_019834735.2
B4 62240251 62240387
refGene_gn=INTS13;refGene_tx=XM_011283852.3;refGene_gn=INTS13;refGene_tx=XM_019834735.2
B4 62245303 62245443
refGene_gn=INTS13;refGene_tx=XM_011283852.3;refGene_gn=INTS13;refGene_

tx=XM_019834735.2
B4	62247229 62247460
refGene_gn=INTS13;refGene_tx=XM_011283852.3;refGene_gn=INTS13;refGene_tx=XM_019834735.2
B4	62247825 62247980
refGene_gn=INTS13;refGene_tx=XM_011283852.3;refGene_gn=INTS13;refGene_tx=XM_019834735.2
B4	62248221 62248392
refGene_gn=INTS13;refGene_tx=XM_011283852.3;refGene_gn=INTS13;refGene_tx=XM_019834735.2
B4	62249492 62249671
refGene_gn=INTS13;refGene_tx=XM_011283852.3;refGene_gn=INTS13;refGene_tx=XM_019834735.2
B4	62250742 62250850
refGene_gn=INTS13;refGene_tx=XM_011283852.3;refGene_gn=INTS13;refGene_tx=XM_019834735.2
B4	62251277 62251367
refGene_gn=INTS13;refGene_tx=XM_011283852.3;refGene_gn=INTS13;refGene_tx=XM_019834735.2
B4	62253827 62253912
refGene_gn=INTS13;refGene_tx=XM_011283852.3;refGene_gn=INTS13;refGene_tx=XM_019834735.2
B4	62255310 62255439
refGene_gn=INTS13;refGene_tx=XM_011283852.3;refGene_gn=INTS13;refGene_tx=XM_019834735.2
B4	62256885 62256976
refGene_gn=INTS13;refGene_tx=XM_011283852.3;refGene_gn=INTS13;refGene_tx=XM_019834735.2
B4	62257753 62257834
refGene_gn=INTS13;refGene_tx=XM_011283852.3;refGene_gn=INTS13;refGene_tx=XM_019834735.2
B4	62258200 62258403
refGene_gn=INTS13;refGene_tx=XM_011283852.3;refGene_gn=INTS13;refGene_tx=XM_019834735.2
B4	62262008 62262083
refGene_gn=INTS13;refGene_tx=XM_011283852.3;refGene_gn=INTS13;refGene_tx=XM_019834735.2
B4	62263993 62264218
refGene_gn=INTS13;refGene_tx=XM_011283852.3;refGene_gn=INTS13;refGene_tx=XM_019834735.2
B4	62288285 62288420
refGene_gn=FGFR1OP2;refGene_tx=XM_003988513.5;refGene_gn=FGFR1OP2;refGene_tx=XM_003988514.5;refGene_gn=FGFR1OP2;refGene_tx=XM_006933577.4
B4	62291171 62291289
refGene_gn=FGFR1OP2;refGene_tx=XM_003988513.5;refGene_gn=FGFR1OP2;refGene_tx=XM_003988514.5;refGene_gn=FGFR1OP2;refGene_tx=XM_006933577.4
B4	62292619 62292762
refGene_gn=FGFR1OP2;refGene_tx=XM_003988513.5;refGene_gn=FGFR1OP2;refGene_tx=XM_003988514.5;refGene_gn=FGFR1OP2;refGene_tx=XM_006933577.4
B4	62295125 62295239

refGene_gn=FGFR1OP2;refGene_tx=XM_003988514.5;refGene_gn=FGFR1OP2;refGene_tx=XM_006933577.4
B4 62297726 62297840
refGene_gn=FGFR1OP2;refGene_tx=XM_003988513.5;refGene_gn=FGFR1OP2;refGene_tx=XM_003988514.5;refGene_gn=FGFR1OP2;refGene_tx=XM_006933577.4
B4 62299442 62299580
refGene_gn=FGFR1OP2;refGene_tx=XM_003988513.5;refGene_gn=FGFR1OP2;refGene_tx=XM_003988514.5;refGene_gn=FGFR1OP2;refGene_tx=XM_006933577.4
B4 62310121 62310384 refGene_gn=TM7SF3;refGene_tx=XM_003988516.5
B4 62311662 62311825 refGene_gn=TM7SF3;refGene_tx=XM_003988516.5
B4 62312409 62312507 refGene_gn=TM7SF3;refGene_tx=XM_003988516.5
B4 62315743 62315896 refGene_gn=TM7SF3;refGene_tx=XM_003988516.5
B4 62316523 62316604 refGene_gn=TM7SF3;refGene_tx=XM_003988516.5
B4 62318396 62318483 refGene_gn=TM7SF3;refGene_tx=XM_003988516.5
B4 62324478 62324656 refGene_gn=TM7SF3;refGene_tx=XM_003988516.5
B4 62328879 62329051 refGene_gn=TM7SF3;refGene_tx=XM_003988516.5
B4 62329812 62329933 refGene_gn=TM7SF3;refGene_tx=XM_003988516.5
B4 62332390 62332541 refGene_gn=TM7SF3;refGene_tx=XM_003988516.5
B4 62335796 62335951 refGene_gn=TM7SF3;refGene_tx=XM_003988516.5
B4 62345760 62345851 refGene_gn=TM7SF3;refGene_tx=XM_003988516.5
B4 62353128 62353170
refGene_gn=MED21;refGene_tx=XM_003988517.3;refGene_gn=MED21;refGene_tx=XM_011283854.2;refGene_gn=MED21;refGene_tx=XM_019834739.1
B4 62357686 62357801
refGene_gn=MED21;refGene_tx=XM_003988517.3;refGene_gn=MED21;refGene_tx=XM_011283854.2;refGene_gn=MED21;refGene_tx=XM_019834739.1
B4 62358334 62358451
refGene_gn=MED21;refGene_tx=XM_003988517.3;refGene_gn=MED21;refGene_tx=XM_011283854.2;refGene_gn=MED21;refGene_tx=XM_019834739.1
B4 62359573 62359774
refGene_gn=MED21;refGene_tx=XM_003988517.3;refGene_gn=MED21;refGene_tx=XM_019834739.1
B4 62473982 62474615
refGene_gn=LOC111561451;refGene_tx=XM_023257752.1
B4 62479951 62479952
refGene_gn=STK38L;refGene_tx=XM_023256872.1;refGene_gn=STK38L;refGene_tx=XM_023256874.1
B4 62526677 62526822
refGene_gn=STK38L;refGene_tx=XM_023256872.1;refGene_gn=STK38L;refGene_tx=XM_023256873.1;refGene_gn=STK38L;refGene_tx=XM_023256874.1
B4 62530332 62530384
refGene_gn=STK38L;refGene_tx=XM_023256872.1;refGene_gn=STK38L;refGene_tx=XM_023256873.1;refGene_gn=STK38L;refGene_tx=XM_023256874.1
B4 62537093 62537216
refGene_gn=STK38L;refGene_tx=XM_023256872.1;refGene_gn=STK38L;refGene_tx=XM_023256873.1;refGene_gn=STK38L;refGene_tx=XM_023256874.1
B4 62537654 62537738
refGene_gn=STK38L;refGene_tx=XM_023256872.1;refGene_gn=STK38L;refGene_tx=XM_023256873.1;refGene_gn=STK38L;refGene_tx=XM_023256874.1
B4 62545266 62545390

refGene_gn=STK38L;refGene_tx=XM_023256872.1;refGene_gn=STK38L;refGene_tx=XM_023256873.1;refGene_gn=STK38L;refGene_tx=XM_023256874.1
B4 62547207 62547362
refGene_gn=STK38L;refGene_tx=XM_023256872.1;refGene_gn=STK38L;refGene_tx=XM_023256873.1;refGene_gn=STK38L;refGene_tx=XM_023256874.1
B4 62547728 62547831
refGene_gn=STK38L;refGene_tx=XM_023256872.1;refGene_gn=STK38L;refGene_tx=XM_023256873.1;refGene_gn=STK38L;refGene_tx=XM_023256874.1
B4 62548009 62548071
refGene_gn=STK38L;refGene_tx=XM_023256872.1;refGene_gn=STK38L;refGene_tx=XM_023256873.1;refGene_gn=STK38L;refGene_tx=XM_023256874.1
B4 62550394 62550512
refGene_gn=STK38L;refGene_tx=XM_023256872.1;refGene_gn=STK38L;refGene_tx=XM_023256873.1;refGene_gn=STK38L;refGene_tx=XM_023256874.1
B4 62550937 62551065
refGene_gn=STK38L;refGene_tx=XM_023256872.1;refGene_gn=STK38L;refGene_tx=XM_023256873.1;refGene_gn=STK38L;refGene_tx=XM_023256874.1
B4 62552250 62552345
refGene_gn=STK38L;refGene_tx=XM_023256872.1;refGene_gn=STK38L;refGene_tx=XM_023256873.1;refGene_gn=STK38L;refGene_tx=XM_023256874.1
B4 62555641 62555733 refGene_gn=STK38L;refGene_tx=XM_023256874.1
B4 62555837 62555965 refGene_gn=STK38L;refGene_tx=XM_023256874.1
B4 62566066 62566143
refGene_gn=ARNTL2;refGene_tx=XM_023256875.1;refGene_gn=ARNTL2;refGene_tx=XM_023256876.1;refGene_gn=ARNTL2;refGene_tx=XM_023256877.1;refGene_gn=ARNTL2;refGene_tx=XM_023256878.1;refGene_gn=ARNTL2;refGene_tx=XM_023256879.1;refGene_gn=ARNTL2;refGene_tx=XM_023256882.1;refGene_gn=ARNTL2;refGene_tx=XM_023256883.1;refGene_gn=ARNTL2;refGene_tx=XM_023256886.1
B4 62616473 62616624
refGene_gn=ARNTL2;refGene_tx=XM_023256875.1;refGene_gn=ARNTL2;refGene_tx=XM_023256876.1;refGene_gn=ARNTL2;refGene_tx=XM_023256877.1;refGene_gn=ARNTL2;refGene_tx=XM_023256878.1;refGene_gn=ARNTL2;refGene_tx=XM_023256879.1;refGene_gn=ARNTL2;refGene_tx=XM_023256880.1;refGene_gn=ARNTL2;refGene_tx=XM_023256881.1;refGene_gn=ARNTL2;refGene_tx=XM_023256882.1;refGene_gn=ARNTL2;refGene_tx=XM_023256883.1;refGene_gn=ARNTL2;refGene_tx=XM_023256886.1
B4 62632947 62632989
refGene_gn=ARNTL2;refGene_tx=XM_023256875.1;refGene_gn=ARNTL2;refGene_tx=XM_023256876.1;refGene_gn=ARNTL2;refGene_tx=XM_023256877.1;refGene_gn=ARNTL2;refGene_tx=XM_023256880.1;refGene_gn=ARNTL2;refGene_tx=XM_023256881.1;refGene_gn=ARNTL2;refGene_tx=XM_023256882.1;refGene_gn=ARNTL2;refGene_tx=XM_023256883.1;refGene_gn=ARNTL2;refGene_tx=XM_023256884.1;refGene_gn=ARNTL2;refGene_tx=XM_023256885.1
B4 62636615 62636773
refGene_gn=ARNTL2;refGene_tx=XM_023256875.1;refGene_gn=ARNTL2;refGene_tx=XM_023256876.1;refGene_gn=ARNTL2;refGene_tx=XM_023256877.1;refGene_gn=ARNTL2;refGene_tx=XM_023256878.1;refGene_gn=ARNTL2;refGene_tx=XM_023256879.1;refGene_gn=ARNTL2;refGene_tx=XM_023256880.1;refGene_gn=ARNTL2;refGene_tx=XM_023256881.1;refGene_gn=ARNTL2;refGene_tx=XM_023256882.

1;refGene_gn=ARNTL2;refGene_tx=XM_023256883.1;refGene_gn=ARNTL2;refGen
e_tx=XM_023256884.1;refGene_gn=ARNTL2;refGene_tx=XM_023256885.1;refGen
e_gn=ARNTL2;refGene_tx=XM_023256886.1
B4 62643004 62643084
refGene_gn=ARNTL2;refGene_tx=XM_023256875.1;refGene_gn=ARNTL2;refGene_
tx=XM_023256876.1;refGene_gn=ARNTL2;refGene_tx=XM_023256877.1;refGene_
gn=ARNTL2;refGene_tx=XM_023256878.1;refGene_gn=ARNTL2;refGene_tx=XM_02
3256879.1;refGene_gn=ARNTL2;refGene_tx=XM_023256880.1;refGene_gn=ARNTL
2;refGene_tx=XM_023256881.1;refGene_gn=ARNTL2;refGene_tx=XM_023256882.
1;refGene_gn=ARNTL2;refGene_tx=XM_023256883.1;refGene_gn=ARNTL2;refGen
e_tx=XM_023256884.1;refGene_gn=ARNTL2;refGene_tx=XM_023256885.1;refGen
e_gn=ARNTL2;refGene_tx=XM_023256886.1
B4 62646086 62646179
refGene_gn=ARNTL2;refGene_tx=XM_023256875.1;refGene_gn=ARNTL2;refGene_
tx=XM_023256876.1;refGene_gn=ARNTL2;refGene_tx=XM_023256877.1;refGene_
gn=ARNTL2;refGene_tx=XM_023256878.1;refGene_gn=ARNTL2;refGene_tx=XM_02
3256879.1;refGene_gn=ARNTL2;refGene_tx=XM_023256880.1;refGene_gn=ARNTL
2;refGene_tx=XM_023256881.1;refGene_gn=ARNTL2;refGene_tx=XM_023256882.
1;refGene_gn=ARNTL2;refGene_tx=XM_023256883.1;refGene_gn=ARNTL2;refGen
e_tx=XM_023256884.1;refGene_gn=ARNTL2;refGene_tx=XM_023256885.1;refGen
e_gn=ARNTL2;refGene_tx=XM_023256886.1
B4 62648370 62648485
refGene_gn=ARNTL2;refGene_tx=XM_023256875.1;refGene_gn=ARNTL2;refGene_
tx=XM_023256876.1;refGene_gn=ARNTL2;refGene_tx=XM_023256877.1;refGene_
gn=ARNTL2;refGene_tx=XM_023256878.1;refGene_gn=ARNTL2;refGene_tx=XM_02
3256879.1;refGene_gn=ARNTL2;refGene_tx=XM_023256880.1;refGene_gn=ARNTL
2;refGene_tx=XM_023256881.1;refGene_gn=ARNTL2;refGene_tx=XM_023256882.
1;refGene_gn=ARNTL2;refGene_tx=XM_023256883.1;refGene_gn=ARNTL2;refGen
e_tx=XM_023256884.1;refGene_gn=ARNTL2;refGene_tx=XM_023256885.1;refGen
e_gn=ARNTL2;refGene_tx=XM_023256886.1
B4 62649281 62649425
refGene_gn=ARNTL2;refGene_tx=XM_023256875.1;refGene_gn=ARNTL2;refGene_
tx=XM_023256876.1;refGene_gn=ARNTL2;refGene_tx=XM_023256877.1;refGene_
gn=ARNTL2;refGene_tx=XM_023256878.1;refGene_gn=ARNTL2;refGene_tx=XM_02
3256879.1;refGene_gn=ARNTL2;refGene_tx=XM_023256880.1;refGene_gn=ARNTL
2;refGene_tx=XM_023256881.1;refGene_gn=ARNTL2;refGene_tx=XM_023256883.
1;refGene_gn=ARNTL2;refGene_tx=XM_023256884.1;refGene_gn=ARNTL2;refGen
e_tx=XM_023256885.1;refGene_gn=ARNTL2;refGene_tx=XM_023256886.1
B4 62677466 62677707
refGene_gn=ARNTL2;refGene_tx=XM_023256875.1;refGene_gn=ARNTL2;refGene_
tx=XM_023256876.1;refGene_gn=ARNTL2;refGene_tx=XM_023256877.1;refGene_
gn=ARNTL2;refGene_tx=XM_023256878.1;refGene_gn=ARNTL2;refGene_tx=XM_02
3256879.1;refGene_gn=ARNTL2;refGene_tx=XM_023256880.1;refGene_gn=ARNTL
2;refGene_tx=XM_023256881.1;refGene_gn=ARNTL2;refGene_tx=XM_023256882.
1;refGene_gn=ARNTL2;refGene_tx=XM_023256883.1;refGene_gn=ARNTL2;refGen
e_tx=XM_023256884.1;refGene_gn=ARNTL2;refGene_tx=XM_023256885.1;refGen
e_gn=ARNTL2;refGene_tx=XM_023256886.1
B4 62678103 62678210
refGene_gn=ARNTL2;refGene_tx=XM_023256875.1;refGene_gn=ARNTL2;refGene_
tx=XM_023256876.1;refGene_gn=ARNTL2;refGene_tx=XM_023256877.1;refGene_

gn=ARNTL2;refGene_tx=XM_023256878.1;refGene_gn=ARNTL2;refGene_tx=XM_023256879.1;refGene_gn=ARNTL2;refGene_tx=XM_023256880.1;refGene_gn=ARNTL2;refGene_tx=XM_023256881.1;refGene_gn=ARNTL2;refGene_tx=XM_023256882.1;refGene_gn=ARNTL2;refGene_tx=XM_023256883.1;refGene_gn=ARNTL2;refGene_tx=XM_023256884.1;refGene_gn=ARNTL2;refGene_tx=XM_023256885.1;refGene_gn=ARNTL2;refGene_tx=XM_023256886.1
B4 62678360 62678511
refGene_gn=ARNTL2;refGene_tx=XM_023256875.1;refGene_gn=ARNTL2;refGene_tx=XM_023256876.1;refGene_gn=ARNTL2;refGene_tx=XM_023256877.1;refGene_gn=ARNTL2;refGene_tx=XM_023256878.1;refGene_gn=ARNTL2;refGene_tx=XM_023256879.1;refGene_gn=ARNTL2;refGene_tx=XM_023256880.1;refGene_gn=ARNTL2;refGene_tx=XM_023256881.1;refGene_gn=ARNTL2;refGene_tx=XM_023256882.1;refGene_gn=ARNTL2;refGene_tx=XM_023256883.1;refGene_gn=ARNTL2;refGene_tx=XM_023256884.1;refGene_gn=ARNTL2;refGene_tx=XM_023256885.1;refGene_gn=ARNTL2;refGene_tx=XM_023256886.1
B4 62679439 62679498
refGene_gn=ARNTL2;refGene_tx=XM_023256875.1;refGene_gn=ARNTL2;refGene_tx=XM_023256876.1;refGene_gn=ARNTL2;refGene_tx=XM_023256877.1;refGene_gn=ARNTL2;refGene_tx=XM_023256878.1;refGene_gn=ARNTL2;refGene_tx=XM_023256879.1;refGene_gn=ARNTL2;refGene_tx=XM_023256880.1;refGene_gn=ARNTL2;refGene_tx=XM_023256881.1;refGene_gn=ARNTL2;refGene_tx=XM_023256882.1;refGene_gn=ARNTL2;refGene_tx=XM_023256883.1;refGene_gn=ARNTL2;refGene_tx=XM_023256884.1;refGene_gn=ARNTL2;refGene_tx=XM_023256885.1;refGene_gn=ARNTL2;refGene_tx=XM_023256886.1
B4 62680247 62680356
refGene_gn=ARNTL2;refGene_tx=XM_023256875.1;refGene_gn=ARNTL2;refGene_tx=XM_023256876.1;refGene_gn=ARNTL2;refGene_tx=XM_023256877.1;refGene_gn=ARNTL2;refGene_tx=XM_023256878.1;refGene_gn=ARNTL2;refGene_tx=XM_023256879.1;refGene_gn=ARNTL2;refGene_tx=XM_023256880.1;refGene_gn=ARNTL2;refGene_tx=XM_023256881.1;refGene_gn=ARNTL2;refGene_tx=XM_023256882.1;refGene_gn=ARNTL2;refGene_tx=XM_023256883.1;refGene_gn=ARNTL2;refGene_tx=XM_023256884.1;refGene_gn=ARNTL2;refGene_tx=XM_023256885.1;refGene_gn=ARNTL2;refGene_tx=XM_023256886.1
B4 62690844 62690932
refGene_gn=ARNTL2;refGene_tx=XM_023256875.1;refGene_gn=ARNTL2;refGene_tx=XM_023256876.1;refGene_gn=ARNTL2;refGene_tx=XM_023256877.1;refGene_gn=ARNTL2;refGene_tx=XM_023256878.1;refGene_gn=ARNTL2;refGene_tx=XM_023256879.1;refGene_gn=ARNTL2;refGene_tx=XM_023256880.1;refGene_gn=ARNTL2;refGene_tx=XM_023256881.1;refGene_gn=ARNTL2;refGene_tx=XM_023256882.1;refGene_gn=ARNTL2;refGene_tx=XM_023256883.1;refGene_gn=ARNTL2;refGene_tx=XM_023256884.1;refGene_gn=ARNTL2;refGene_tx=XM_023256885.1;refGene_gn=ARNTL2;refGene_tx=XM_023256886.1
B4 62692712 62692821
refGene_gn=ARNTL2;refGene_tx=XM_023256875.1;refGene_gn=ARNTL2;refGene_tx=XM_023256876.1;refGene_gn=ARNTL2;refGene_tx=XM_023256877.1;refGene_gn=ARNTL2;refGene_tx=XM_023256878.1;refGene_gn=ARNTL2;refGene_tx=XM_023256879.1;refGene_gn=ARNTL2;refGene_tx=XM_023256880.1;refGene_gn=ARNTL2;refGene_tx=XM_023256881.1;refGene_gn=ARNTL2;refGene_tx=XM_023256882.1;refGene_gn=ARNTL2;refGene_tx=XM_023256884.1;refGene_gn=ARNTL2;refGene_tx=XM_023256885.1

B4 62694166 62694303
refGene_gn=ARNTL2;refGene_tx=XM_023256875.1;refGene_gn=ARNTL2;refGene_tx=XM_023256876.1;refGene_gn=ARNTL2;refGene_tx=XM_023256877.1;refGene_gn=ARNTL2;refGene_tx=XM_023256878.1;refGene_gn=ARNTL2;refGene_tx=XM_023256879.1;refGene_gn=ARNTL2;refGene_tx=XM_023256880.1;refGene_gn=ARNTL2;refGene_tx=XM_023256881.1;refGene_gn=ARNTL2;refGene_tx=XM_023256882.1;refGene_gn=ARNTL2;refGene_tx=XM_023256883.1;refGene_gn=ARNTL2;refGene_tx=XM_023256884.1;refGene_gn=ARNTL2;refGene_tx=XM_023256885.1;refGene_gn=ARNTL2;refGene_tx=XM_023256886.1
B4 62756394 62756522
refGene_gn=SMCO2;refGene_tx=XM_006933580.4;refGene_gn=SMCO2;refGene_tx=XM_006933581.4
B4 62758897 62759015
refGene_gn=SMCO2;refGene_tx=XM_006933580.4;refGene_gn=SMCO2;refGene_tx=XM_006933581.4;refGene_gn=SMCO2;refGene_tx=XM_006933582.4;refGene_gn=SMCO2;refGene_tx=XM_019834741.2
B4 62769138 62769275
refGene_gn=SMCO2;refGene_tx=XM_006933580.4;refGene_gn=SMCO2;refGene_tx=XM_006933581.4;refGene_gn=SMCO2;refGene_tx=XM_006933582.4;refGene_gn=SMCO2;refGene_tx=XM_019834741.2
B4 62770499 62770649
refGene_gn=SMCO2;refGene_tx=XM_006933580.4;refGene_gn=SMCO2;refGene_tx=XM_006933581.4;refGene_gn=SMCO2;refGene_tx=XM_006933582.4;refGene_gn=SMCO2;refGene_tx=XM_019834741.2
B4 62774114 62774205
refGene_gn=SMCO2;refGene_tx=XM_006933580.4;refGene_gn=SMCO2;refGene_tx=XM_006933581.4;refGene_gn=SMCO2;refGene_tx=XM_006933582.4;refGene_gn=SMCO2;refGene_tx=XM_019834741.2
B4 62778569 62778626
refGene_gn=SMCO2;refGene_tx=XM_006933580.4;refGene_gn=SMCO2;refGene_tx=XM_006933581.4;refGene_gn=SMCO2;refGene_tx=XM_006933582.4;refGene_gn=SMCO2;refGene_tx=XM_019834741.2
B4 62780277 62780334
refGene_gn=SMCO2;refGene_tx=XM_006933580.4;refGene_gn=SMCO2;refGene_tx=XM_006933581.4;refGene_gn=SMCO2;refGene_tx=XM_006933582.4;refGene_gn=SMCO2;refGene_tx=XM_019834741.2
B4 62781387 62781563
refGene_gn=SMCO2;refGene_tx=XM_006933580.4;refGene_gn=SMCO2;refGene_tx=XM_006933581.4;refGene_gn=SMCO2;refGene_tx=XM_006933582.4;refGene_gn=SMCO2;refGene_tx=XM_019834741.2
B4 62785957 62786156
refGene_gn=SMCO2;refGene_tx=XM_006933580.4;refGene_gn=SMCO2;refGene_tx=XM_006933581.4;refGene_gn=SMCO2;refGene_tx=XM_006933582.4;refGene_gn=SMCO2;refGene_tx=XM_019834741.2
B4 62885986 62886325
refGene_gn=CB4H12orf71;refGene_tx=XM_006933583.3;refGene_gn=CB4H12orf71;refGene_tx=XM_019834746.2
B4 62887008 62887647
refGene_gn=CB4H12orf71;refGene_tx=XM_006933583.3;refGene_gn=CB4H12orf71;refGene_tx=XM_019834746.2

B4 62923881 62923945
refGene_gn=PPFIBP1;refGene_tx=XM_003988520.4;refGene_gn=PPFIBP1;refGene_tx=XM_011283855.2;refGene_gn=PPFIBP1;refGene_tx=XM_011283856.2;refGene_gn=PPFIBP1;refGene_tx=XM_011283857.3;refGene_gn=PPFIBP1;refGene_tx=XM_011283858.2;refGene_gn=PPFIBP1;refGene_tx=XM_011283859.2;refGene_gn=PPFIBP1;refGene_tx=XM_011283860.2;refGene_gn=PPFIBP1;refGene_tx=XM_011283861.2;refGene_gn=PPFIBP1;refGene_tx=XM_011283862.2
B4 62927316 62927522
refGene_gn=PPFIBP1;refGene_tx=XM_003988520.4;refGene_gn=PPFIBP1;refGene_tx=XM_011283855.2;refGene_gn=PPFIBP1;refGene_tx=XM_011283856.2;refGene_gn=PPFIBP1;refGene_tx=XM_011283857.3;refGene_gn=PPFIBP1;refGene_tx=XM_011283858.2;refGene_gn=PPFIBP1;refGene_tx=XM_011283859.2;refGene_gn=PPFIBP1;refGene_tx=XM_011283860.2;refGene_gn=PPFIBP1;refGene_tx=XM_011283861.2;refGene_gn=PPFIBP1;refGene_tx=XM_011283862.2
B4 62936843 62936930
refGene_gn=PPFIBP1;refGene_tx=XM_003988520.4;refGene_gn=PPFIBP1;refGene_tx=XM_011283855.2;refGene_gn=PPFIBP1;refGene_tx=XM_011283856.2;refGene_gn=PPFIBP1;refGene_tx=XM_011283857.3;refGene_gn=PPFIBP1;refGene_tx=XM_011283858.2;refGene_gn=PPFIBP1;refGene_tx=XM_011283859.2;refGene_gn=PPFIBP1;refGene_tx=XM_011283860.2;refGene_gn=PPFIBP1;refGene_tx=XM_011283861.2;refGene_gn=PPFIBP1;refGene_tx=XM_011283862.2
B4 62938131 62938245
refGene_gn=PPFIBP1;refGene_tx=XM_003988520.4;refGene_gn=PPFIBP1;refGene_tx=XM_011283855.2;refGene_gn=PPFIBP1;refGene_tx=XM_011283856.2;refGene_gn=PPFIBP1;refGene_tx=XM_011283857.3;refGene_gn=PPFIBP1;refGene_tx=XM_011283858.2;refGene_gn=PPFIBP1;refGene_tx=XM_011283859.2;refGene_gn=PPFIBP1;refGene_tx=XM_011283860.2;refGene_gn=PPFIBP1;refGene_tx=XM_011283861.2;refGene_gn=PPFIBP1;refGene_tx=XM_011283862.2
B4 62939172 62939304
refGene_gn=PPFIBP1;refGene_tx=XM_003988520.4;refGene_gn=PPFIBP1;refGene_tx=XM_011283855.2;refGene_gn=PPFIBP1;refGene_tx=XM_011283856.2;refGene_gn=PPFIBP1;refGene_tx=XM_011283857.3;refGene_gn=PPFIBP1;refGene_tx=XM_011283858.2;refGene_gn=PPFIBP1;refGene_tx=XM_011283859.2;refGene_gn=PPFIBP1;refGene_tx=XM_011283860.2;refGene_gn=PPFIBP1;refGene_tx=XM_011283861.2;refGene_gn=PPFIBP1;refGene_tx=XM_011283862.2
B4 62943800 62943893
refGene_gn=PPFIBP1;refGene_tx=XM_003988520.4;refGene_gn=PPFIBP1;refGene_tx=XM_011283855.2;refGene_gn=PPFIBP1;refGene_tx=XM_011283856.2;refGene_gn=PPFIBP1;refGene_tx=XM_011283857.3;refGene_gn=PPFIBP1;refGene_tx=XM_011283858.2;refGene_gn=PPFIBP1;refGene_tx=XM_011283859.2;refGene_gn=PPFIBP1;refGene_tx=XM_011283860.2;refGene_gn=PPFIBP1;refGene_tx=XM_011283861.2;refGene_gn=PPFIBP1;refGene_tx=XM_011283862.2
B4 62944217 62944310
refGene_gn=PPFIBP1;refGene_tx=XM_011283855.2;refGene_gn=PPFIBP1;refGene_tx=XM_011283856.2;refGene_gn=PPFIBP1;refGene_tx=XM_011283857.3;refGene_gn=PPFIBP1;refGene_tx=XM_011283858.2;refGene_gn=PPFIBP1;refGene_tx=XM_011283859.2;refGene_gn=PPFIBP1;refGene_tx=XM_011283860.2;refGene_gn=PPFIBP1;refGene_tx=XM_011283861.2
B4 62945667 62945782
refGene_gn=PPFIBP1;refGene_tx=XM_003988520.4;refGene_gn=PPFIBP1;refGen

e_tx=XM_011283855.2;refGene_gn=PPFIBP1;refGene_tx=XM_011283856.2;refGene_gn=PPFIBP1;refGene_tx=XM_011283857.3;refGene_gn=PPFIBP1;refGene_tx=XM_011283858.2;refGene_gn=PPFIBP1;refGene_tx=XM_011283859.2;refGene_gn=PPFIBP1;refGene_tx=XM_011283860.2;refGene_gn=PPFIBP1;refGene_tx=XM_011283861.2;refGene_gn=PPFIBP1;refGene_tx=XM_011283862.2
B4 62947748 62947781
refGene_gn=PPFIBP1;refGene_tx=XM_003988520.4;refGene_gn=PPFIBP1;refGene_tx=XM_011283855.2;refGene_gn=PPFIBP1;refGene_tx=XM_011283856.2;refGene_gn=PPFIBP1;refGene_tx=XM_011283857.3;refGene_gn=PPFIBP1;refGene_tx=XM_011283858.2;refGene_gn=PPFIBP1;refGene_tx=XM_011283860.2
B4 62949598 62949660
refGene_gn=PPFIBP1;refGene_tx=XM_003988520.4;refGene_gn=PPFIBP1;refGene_tx=XM_011283855.2;refGene_gn=PPFIBP1;refGene_tx=XM_011283856.2;refGene_gn=PPFIBP1;refGene_tx=XM_011283857.3;refGene_gn=PPFIBP1;refGene_tx=XM_011283858.2;refGene_gn=PPFIBP1;refGene_tx=XM_011283859.2;refGene_gn=PPFIBP1;refGene_tx=XM_011283860.2;refGene_gn=PPFIBP1;refGene_tx=XM_011283861.2;refGene_gn=PPFIBP1;refGene_tx=XM_011283862.2
B4 62952731 62952816
refGene_gn=PPFIBP1;refGene_tx=XM_003988520.4;refGene_gn=PPFIBP1;refGene_tx=XM_011283855.2;refGene_gn=PPFIBP1;refGene_tx=XM_011283856.2;refGene_gn=PPFIBP1;refGene_tx=XM_011283857.3;refGene_gn=PPFIBP1;refGene_tx=XM_011283858.2;refGene_gn=PPFIBP1;refGene_tx=XM_011283859.2;refGene_gn=PPFIBP1;refGene_tx=XM_011283860.2;refGene_gn=PPFIBP1;refGene_tx=XM_011283861.2;refGene_gn=PPFIBP1;refGene_tx=XM_011283862.2
B4 62955398 62955553
refGene_gn=PPFIBP1;refGene_tx=XM_003988520.4;refGene_gn=PPFIBP1;refGene_tx=XM_011283855.2;refGene_gn=PPFIBP1;refGene_tx=XM_011283856.2;refGene_gn=PPFIBP1;refGene_tx=XM_011283857.3;refGene_gn=PPFIBP1;refGene_tx=XM_011283858.2;refGene_gn=PPFIBP1;refGene_tx=XM_011283859.2;refGene_gn=PPFIBP1;refGene_tx=XM_011283860.2;refGene_gn=PPFIBP1;refGene_tx=XM_011283861.2;refGene_gn=PPFIBP1;refGene_tx=XM_011283862.2
B4 62961026 62961139
refGene_gn=PPFIBP1;refGene_tx=XM_003988520.4;refGene_gn=PPFIBP1;refGene_tx=XM_011283855.2;refGene_gn=PPFIBP1;refGene_tx=XM_011283856.2;refGene_gn=PPFIBP1;refGene_tx=XM_011283857.3;refGene_gn=PPFIBP1;refGene_tx=XM_011283858.2;refGene_gn=PPFIBP1;refGene_tx=XM_011283859.2;refGene_gn=PPFIBP1;refGene_tx=XM_011283860.2;refGene_gn=PPFIBP1;refGene_tx=XM_011283861.2;refGene_gn=PPFIBP1;refGene_tx=XM_011283862.2
B4 62961747 62961804
refGene_gn=PPFIBP1;refGene_tx=XM_003988520.4;refGene_gn=PPFIBP1;refGene_tx=XM_011283855.2;refGene_gn=PPFIBP1;refGene_tx=XM_011283856.2;refGene_gn=PPFIBP1;refGene_tx=XM_011283857.3;refGene_gn=PPFIBP1;refGene_tx=XM_011283858.2;refGene_gn=PPFIBP1;refGene_tx=XM_011283859.2;refGene_gn=PPFIBP1;refGene_tx=XM_011283860.2;refGene_gn=PPFIBP1;refGene_tx=XM_011283861.2;refGene_gn=PPFIBP1;refGene_tx=XM_011283862.2
B4 62964355 62964416
refGene_gn=PPFIBP1;refGene_tx=XM_003988520.4;refGene_gn=PPFIBP1;refGene_tx=XM_011283855.2;refGene_gn=PPFIBP1;refGene_tx=XM_011283856.2;refGene_gn=PPFIBP1;refGene_tx=XM_011283857.3;refGene_gn=PPFIBP1;refGene_tx=XM_011283858.2;refGene_gn=PPFIBP1;refGene_tx=XM_011283859.2;refGene_gn

=PPFIBP1;refGene_tx=XM_011283860.2;refGene_gn=PPFIBP1;refGene_tx=XM_011283861.2;refGene_gn=PPFIBP1;refGene_tx=XM_011283862.2
B4 62964798 62964828
refGene_gn=PPFIBP1;refGene_tx=XM_003988520.4;refGene_gn=PPFIBP1;refGene_tx=XM_011283855.2;refGene_gn=PPFIBP1;refGene_tx=XM_011283856.2;refGene_gn=PPFIBP1;refGene_tx=XM_011283857.3;refGene_gn=PPFIBP1;refGene_tx=XM_011283858.2;refGene_gn=PPFIBP1;refGene_tx=XM_011283859.2;refGene_gn=PPFIBP1;refGene_tx=XM_011283860.2;refGene_gn=PPFIBP1;refGene_tx=XM_011283861.2;refGene_gn=PPFIBP1;refGene_tx=XM_011283862.2
B4 62968033 62968205
refGene_gn=PPFIBP1;refGene_tx=XM_003988520.4;refGene_gn=PPFIBP1;refGene_tx=XM_011283855.2;refGene_gn=PPFIBP1;refGene_tx=XM_011283856.2;refGene_gn=PPFIBP1;refGene_tx=XM_011283857.3;refGene_gn=PPFIBP1;refGene_tx=XM_011283858.2;refGene_gn=PPFIBP1;refGene_tx=XM_011283859.2;refGene_gn=PPFIBP1;refGene_tx=XM_011283860.2;refGene_gn=PPFIBP1;refGene_tx=XM_011283861.2;refGene_gn=PPFIBP1;refGene_tx=XM_011283862.2
B4 62968690 62968723
refGene_gn=PPFIBP1;refGene_tx=XM_003988520.4;refGene_gn=PPFIBP1;refGene_tx=XM_011283855.2;refGene_gn=PPFIBP1;refGene_tx=XM_011283856.2;refGene_gn=PPFIBP1;refGene_tx=XM_011283857.3;refGene_gn=PPFIBP1;refGene_tx=XM_011283858.2;refGene_gn=PPFIBP1;refGene_tx=XM_011283859.2
B4 62970844 62970995
refGene_gn=PPFIBP1;refGene_tx=XM_003988520.4;refGene_gn=PPFIBP1;refGene_tx=XM_011283855.2;refGene_gn=PPFIBP1;refGene_tx=XM_011283856.2;refGene_gn=PPFIBP1;refGene_tx=XM_011283857.3;refGene_gn=PPFIBP1;refGene_tx=XM_011283858.2;refGene_gn=PPFIBP1;refGene_tx=XM_011283859.2;refGene_gn=PPFIBP1;refGene_tx=XM_011283860.2;refGene_gn=PPFIBP1;refGene_tx=XM_011283861.2;refGene_gn=PPFIBP1;refGene_tx=XM_011283862.2
B4 62971275 62971404
refGene_gn=PPFIBP1;refGene_tx=XM_003988520.4;refGene_gn=PPFIBP1;refGene_tx=XM_011283855.2;refGene_gn=PPFIBP1;refGene_tx=XM_011283856.2;refGene_gn=PPFIBP1;refGene_tx=XM_011283857.3;refGene_gn=PPFIBP1;refGene_tx=XM_011283858.2;refGene_gn=PPFIBP1;refGene_tx=XM_011283859.2;refGene_gn=PPFIBP1;refGene_tx=XM_011283860.2;refGene_gn=PPFIBP1;refGene_tx=XM_011283861.2;refGene_gn=PPFIBP1;refGene_tx=XM_011283862.2
B4 62972902 62973053
refGene_gn=PPFIBP1;refGene_tx=XM_003988520.4;refGene_gn=PPFIBP1;refGene_tx=XM_011283855.2;refGene_gn=PPFIBP1;refGene_tx=XM_011283856.2;refGene_gn=PPFIBP1;refGene_tx=XM_011283857.3;refGene_gn=PPFIBP1;refGene_tx=XM_011283858.2;refGene_gn=PPFIBP1;refGene_tx=XM_011283859.2;refGene_gn=PPFIBP1;refGene_tx=XM_011283860.2;refGene_gn=PPFIBP1;refGene_tx=XM_011283861.2;refGene_gn=PPFIBP1;refGene_tx=XM_011283862.2
B4 62973743 62973855
refGene_gn=PPFIBP1;refGene_tx=XM_003988520.4;refGene_gn=PPFIBP1;refGene_tx=XM_011283855.2;refGene_gn=PPFIBP1;refGene_tx=XM_011283856.2;refGene_gn=PPFIBP1;refGene_tx=XM_011283857.3;refGene_gn=PPFIBP1;refGene_tx=XM_011283858.2;refGene_gn=PPFIBP1;refGene_tx=XM_011283859.2;refGene_gn=PPFIBP1;refGene_tx=XM_011283860.2;refGene_gn=PPFIBP1;refGene_tx=XM_011283861.2;refGene_gn=PPFIBP1;refGene_tx=XM_011283862.2
B4 62973992 62974081

refGene_gn=PPFIBP1;refGene_tx=XM_003988520.4;refGene_gn=PPFIBP1;refGene_tx=XM_011283855.2;refGene_gn=PPFIBP1;refGene_tx=XM_011283856.2;refGene_gn=PPFIBP1;refGene_tx=XM_011283857.3;refGene_gn=PPFIBP1;refGene_tx=XM_011283858.2;refGene_gn=PPFIBP1;refGene_tx=XM_011283859.2;refGene_gn=PPFIBP1;refGene_tx=XM_011283860.2;refGene_gn=PPFIBP1;refGene_tx=XM_011283861.2;refGene_gn=PPFIBP1;refGene_tx=XM_011283862.2
B4 62977145 62977268
refGene_gn=PPFIBP1;refGene_tx=XM_003988520.4;refGene_gn=PPFIBP1;refGene_tx=XM_011283855.2;refGene_gn=PPFIBP1;refGene_tx=XM_011283856.2;refGene_gn=PPFIBP1;refGene_tx=XM_011283857.3;refGene_gn=PPFIBP1;refGene_tx=XM_011283858.2;refGene_gn=PPFIBP1;refGene_tx=XM_011283859.2;refGene_gn=PPFIBP1;refGene_tx=XM_011283860.2;refGene_gn=PPFIBP1;refGene_tx=XM_011283861.2;refGene_gn=PPFIBP1;refGene_tx=XM_011283862.2
B4 62978292 62978418
refGene_gn=PPFIBP1;refGene_tx=XM_003988520.4;refGene_gn=PPFIBP1;refGene_tx=XM_011283855.2;refGene_gn=PPFIBP1;refGene_tx=XM_011283856.2;refGene_gn=PPFIBP1;refGene_tx=XM_011283857.3;refGene_gn=PPFIBP1;refGene_tx=XM_011283858.2;refGene_gn=PPFIBP1;refGene_tx=XM_011283859.2;refGene_gn=PPFIBP1;refGene_tx=XM_011283860.2;refGene_gn=PPFIBP1;refGene_tx=XM_011283861.2;refGene_gn=PPFIBP1;refGene_tx=XM_011283862.2
B4 62979019 62979208
refGene_gn=PPFIBP1;refGene_tx=XM_003988520.4;refGene_gn=PPFIBP1;refGene_tx=XM_011283855.2;refGene_gn=PPFIBP1;refGene_tx=XM_011283856.2;refGene_gn=PPFIBP1;refGene_tx=XM_011283857.3;refGene_gn=PPFIBP1;refGene_tx=XM_011283858.2;refGene_gn=PPFIBP1;refGene_tx=XM_011283859.2;refGene_gn=PPFIBP1;refGene_tx=XM_011283860.2;refGene_gn=PPFIBP1;refGene_tx=XM_011283861.2;refGene_gn=PPFIBP1;refGene_tx=XM_011283862.2
B4 62981226 62981406
refGene_gn=PPFIBP1;refGene_tx=XM_003988520.4;refGene_gn=PPFIBP1;refGene_tx=XM_011283855.2;refGene_gn=PPFIBP1;refGene_tx=XM_011283856.2;refGene_gn=PPFIBP1;refGene_tx=XM_011283857.3;refGene_gn=PPFIBP1;refGene_tx=XM_011283858.2;refGene_gn=PPFIBP1;refGene_tx=XM_011283859.2;refGene_gn=PPFIBP1;refGene_tx=XM_011283860.2;refGene_gn=PPFIBP1;refGene_tx=XM_011283861.2;refGene_gn=PPFIBP1;refGene_tx=XM_011283862.2
B4 62982595 62982661
refGene_gn=PPFIBP1;refGene_tx=XM_003988520.4;refGene_gn=PPFIBP1;refGene_tx=XM_011283855.2;refGene_gn=PPFIBP1;refGene_tx=XM_011283856.2;refGene_gn=PPFIBP1;refGene_tx=XM_011283857.3;refGene_gn=PPFIBP1;refGene_tx=XM_011283858.2;refGene_gn=PPFIBP1;refGene_tx=XM_011283859.2;refGene_gn=PPFIBP1;refGene_tx=XM_011283860.2;refGene_gn=PPFIBP1;refGene_tx=XM_011283861.2;refGene_gn=PPFIBP1;refGene_tx=XM_011283862.2
B4 62983043 62983130
refGene_gn=PPFIBP1;refGene_tx=XM_003988520.4;refGene_gn=PPFIBP1;refGene_tx=XM_011283855.2;refGene_gn=PPFIBP1;refGene_tx=XM_011283856.2;refGene_gn=PPFIBP1;refGene_tx=XM_011283857.3;refGene_gn=PPFIBP1;refGene_tx=XM_011283858.2;refGene_gn=PPFIBP1;refGene_tx=XM_011283859.2;refGene_gn=PPFIBP1;refGene_tx=XM_011283860.2;refGene_gn=PPFIBP1;refGene_tx=XM_011283861.2;refGene_gn=PPFIBP1;refGene_tx=XM_011283862.2
B4 62985578 62986289 refGene_gn=REP15;refGene_tx=XM_003988521.5
B4 62991263 62991375

refGene_gn=MRPS35;refGene_tx=XM_006933585.4;refGene_gn=MRPS35;refGene_tx=XM_019834743.2
B4 62994943 62994990
refGene_gn=MRPS35;refGene_tx=XM_006933585.4;refGene_gn=MRPS35;refGene_tx=XM_019834743.2
B4 62996174 62996177 refGene_gn=MRPS35;refGene_tx=XM_023256887.1
B4 62996856 62997024
refGene_gn=MRPS35;refGene_tx=XM_006933585.4;refGene_gn=MRPS35;refGene_tx=XM_006933586.4;refGene_gn=MRPS35;refGene_tx=XM_019834743.2;refGene_gn=MRPS35;refGene_tx=XM_023256887.1;refGene_gn=MRPS35;refGene_tx=XM_023256888.1
B4 62999740 62999801
refGene_gn=MRPS35;refGene_tx=XM_006933585.4;refGene_gn=MRPS35;refGene_tx=XM_006933586.4;refGene_gn=MRPS35;refGene_tx=XM_019834743.2;refGene_gn=MRPS35;refGene_tx=XM_023256887.1;refGene_gn=MRPS35;refGene_tx=XM_023256888.1
B4 63007374 63007514
refGene_gn=MRPS35;refGene_tx=XM_006933585.4;refGene_gn=MRPS35;refGene_tx=XM_006933586.4;refGene_gn=MRPS35;refGene_tx=XM_019834743.2;refGene_gn=MRPS35;refGene_tx=XM_023256887.1;refGene_gn=MRPS35;refGene_tx=XM_023256888.1
B4 63016235 63016345
refGene_gn=MRPS35;refGene_tx=XM_006933585.4;refGene_gn=MRPS35;refGene_tx=XM_006933586.4;refGene_gn=MRPS35;refGene_tx=XM_019834743.2;refGene_gn=MRPS35;refGene_tx=XM_023256887.1;refGene_gn=MRPS35;refGene_tx=XM_023256888.1
B4 63018060 63018130
refGene_gn=MRPS35;refGene_tx=XM_006933585.4;refGene_gn=MRPS35;refGene_tx=XM_006933586.4;refGene_gn=MRPS35;refGene_tx=XM_019834743.2;refGene_gn=MRPS35;refGene_tx=XM_023256887.1;refGene_gn=MRPS35;refGene_tx=XM_023256888.1
B4 63026126 63026206 refGene_gn=MRPS35;refGene_tx=XM_019834743.2
B4 63037423 63037693
refGene_gn=MRPS35;refGene_tx=XM_006933585.4;refGene_gn=MRPS35;refGene_tx=XM_006933586.4;refGene_gn=MRPS35;refGene_tx=XM_019834743.2;refGene_gn=MRPS35;refGene_tx=XM_023256887.1;refGene_gn=MRPS35;refGene_tx=XM_023256888.1
B4 63042793 63043434 refGene_gn=MANSC4;refGene_tx=XM_019834742.2
B4 63046052 63046187 refGene_gn=MANSC4;refGene_tx=XM_019834742.2
B4 63048701 63048930 refGene_gn=MANSC4;refGene_tx=XM_019834742.2
B4 63055458 63056330 refGene_gn=KLHL42;refGene_tx=XM_011283990.2
B4 63066409 63066603 refGene_gn=KLHL42;refGene_tx=XM_011283990.2
B4 63071092 63071544 refGene_gn=KLHL42;refGene_tx=XM_011283990.2
B4 63235724 63235734
refGene_gn=PTHLH;refGene_tx=XM_006933587.3;refGene_gn=PTHLH;refGene_tx=XM_006933588.3;refGene_gn=PTHLH;refGene_tx=XM_006933589.3;refGene_gn=PTHLH;refGene_tx=XM_019834748.1
B4 63240072 63240495
refGene_gn=PTHLH;refGene_tx=XM_006933587.3;refGene_gn=PTHLH;refGene_tx=XM_006933588.3;refGene_gn=PTHLH;refGene_tx=XM_006933589.3;refGene_gn=

PTHLH;refGene_tx=XM_019834748.1
B4 63246241 63246342
refGene_gn=PTHLH;refGene_tx=XM_006933587.3;refGene_gn=PTHLH;refGene_tx=XM_006933588.3;refGene_gn=PTHLH;refGene_tx=XM_006933589.3;refGene_gn=PTHLH;refGene_tx=XM_019834748.1
B4 63457460 63457467 refGene_gn=CCDC91;refGene_tx=XM_019834752.2
B4 63505423 63505487 refGene_gn=CCDC91;refGene_tx=XM_019834749.2
B4 63536181 63536224
refGene_gn=CCDC91;refGene_tx=XM_006933592.4;refGene_gn=CCDC91;refGene_tx=XM_006933593.2;refGene_gn=CCDC91;refGene_tx=XM_006933595.4;refGene_gn=CCDC91;refGene_tx=XM_006933596.3;refGene_gn=CCDC91;refGene_tx=XM_011283863.3;refGene_gn=CCDC91;refGene_tx=XM_019834749.2;refGene_gn=CCDC91;refGene_tx=XM_019834752.2;refGene_gn=CCDC91;refGene_tx=XM_019834753.2;refGene_gn=CCDC91;refGene_tx=XM_019834754.2;refGene_gn=CCDC91;refGene_tx=XM_019834757.2;refGene_gn=CCDC91;refGene_tx=XM_019834758.2;refGene_gn=CCDC91;refGene_tx=XM_019834759.2;refGene_gn=CCDC91;refGene_tx=XM_019834764.1;refGene_gn=CCDC91;refGene_tx=XM_023256889.1
B4 63538372 63538464
refGene_gn=CCDC91;refGene_tx=XM_006933592.4;refGene_gn=CCDC91;refGene_tx=XM_006933593.2;refGene_gn=CCDC91;refGene_tx=XM_006933595.4;refGene_gn=CCDC91;refGene_tx=XM_006933596.3;refGene_gn=CCDC91;refGene_tx=XM_011283863.3;refGene_gn=CCDC91;refGene_tx=XM_019834749.2;refGene_gn=CCDC91;refGene_tx=XM_019834752.2;refGene_gn=CCDC91;refGene_tx=XM_019834753.2;refGene_gn=CCDC91;refGene_tx=XM_019834754.2;refGene_gn=CCDC91;refGene_tx=XM_019834757.2;refGene_gn=CCDC91;refGene_tx=XM_019834758.2;refGene_gn=CCDC91;refGene_tx=XM_019834759.2;refGene_gn=CCDC91;refGene_tx=XM_019834764.1;refGene_gn=CCDC91;refGene_tx=XM_023256889.1
B4 63581959 63581978
refGene_gn=CCDC91;refGene_tx=XM_019834761.2;refGene_gn=CCDC91;refGene_tx=XM_019834762.1;refGene_gn=CCDC91;refGene_tx=XM_019834763.2
B4 63582831 63582992
refGene_gn=CCDC91;refGene_tx=XM_006933592.4;refGene_gn=CCDC91;refGene_tx=XM_006933593.2;refGene_gn=CCDC91;refGene_tx=XM_006933595.4;refGene_gn=CCDC91;refGene_tx=XM_006933596.3;refGene_gn=CCDC91;refGene_tx=XM_011283863.3;refGene_gn=CCDC91;refGene_tx=XM_019834749.2;refGene_gn=CCDC91;refGene_tx=XM_019834752.2;refGene_gn=CCDC91;refGene_tx=XM_019834753.2;refGene_gn=CCDC91;refGene_tx=XM_019834754.2;refGene_gn=CCDC91;refGene_tx=XM_019834757.2;refGene_gn=CCDC91;refGene_tx=XM_019834758.2;refGene_gn=CCDC91;refGene_tx=XM_019834759.2;refGene_gn=CCDC91;refGene_tx=XM_019834761.2;refGene_gn=CCDC91;refGene_tx=XM_019834762.1;refGene_gn=CCDC91;refGene_tx=XM_019834763.2;refGene_gn=CCDC91;refGene_tx=XM_019834764.1;refGene_gn=CCDC91;refGene_tx=XM_023256889.1
B4 63583921 63584125
refGene_gn=CCDC91;refGene_tx=XM_006933592.4;refGene_gn=CCDC91;refGene_tx=XM_006933593.2;refGene_gn=CCDC91;refGene_tx=XM_006933595.4;refGene_gn=CCDC91;refGene_tx=XM_006933596.3;refGene_gn=CCDC91;refGene_tx=XM_011283863.3;refGene_gn=CCDC91;refGene_tx=XM_019834749.2;refGene_gn=CCDC91;refGene_tx=XM_019834752.2;refGene_gn=CCDC91;refGene_tx=XM_019834753.2;refGene_gn=CCDC91;refGene_tx=XM_019834754.2;refGene_gn=CCDC91;refGene_tx=XM_019834757.2;refGene_gn=CCDC91;refGene_tx=XM_019834758.2;refGen

e_gn=CCDC91;refGene_tx=XM_019834759.2;refGene_gn=CCDC91;refGene_tx=XM_019834761.2;refGene_gn=CCDC91;refGene_tx=XM_019834762.1;refGene_gn=CCDC91;refGene_tx=XM_019834763.2;refGene_gn=CCDC91;refGene_tx=XM_019834764.1;refGene_gn=CCDC91;refGene_tx=XM_023256889.1
B4 63585054 63585159
refGene_gn=CCDC91;refGene_tx=XM_006933592.4;refGene_gn=CCDC91;refGene_tx=XM_006933593.2;refGene_gn=CCDC91;refGene_tx=XM_006933595.4;refGene_gn=CCDC91;refGene_tx=XM_006933596.3;refGene_gn=CCDC91;refGene_tx=XM_011283863.3;refGene_gn=CCDC91;refGene_tx=XM_019834749.2;refGene_gn=CCDC91;refGene_tx=XM_019834752.2;refGene_gn=CCDC91;refGene_tx=XM_019834753.2;refGene_gn=CCDC91;refGene_tx=XM_019834754.2;refGene_gn=CCDC91;refGene_tx=XM_019834757.2;refGene_gn=CCDC91;refGene_tx=XM_019834758.2;refGene_gn=CCDC91;refGene_tx=XM_019834759.2;refGene_gn=CCDC91;refGene_tx=XM_019834761.2;refGene_gn=CCDC91;refGene_tx=XM_019834762.1;refGene_gn=CCDC91;refGene_tx=XM_019834763.2;refGene_gn=CCDC91;refGene_tx=XM_019834764.1;refGene_gn=CCDC91;refGene_tx=XM_023256889.1
B4 63639205 63639283
refGene_gn=CCDC91;refGene_tx=XM_006933592.4;refGene_gn=CCDC91;refGene_tx=XM_006933593.2;refGene_gn=CCDC91;refGene_tx=XM_006933595.4;refGene_gn=CCDC91;refGene_tx=XM_006933596.3;refGene_gn=CCDC91;refGene_tx=XM_011283863.3;refGene_gn=CCDC91;refGene_tx=XM_019834749.2;refGene_gn=CCDC91;refGene_tx=XM_019834752.2;refGene_gn=CCDC91;refGene_tx=XM_019834753.2;refGene_gn=CCDC91;refGene_tx=XM_019834754.2;refGene_gn=CCDC91;refGene_tx=XM_019834757.2;refGene_gn=CCDC91;refGene_tx=XM_019834758.2;refGene_gn=CCDC91;refGene_tx=XM_019834759.2;refGene_gn=CCDC91;refGene_tx=XM_019834761.2;refGene_gn=CCDC91;refGene_tx=XM_019834762.1;refGene_gn=CCDC91;refGene_tx=XM_019834763.2;refGene_gn=CCDC91;refGene_tx=XM_019834764.1;refGene_gn=CCDC91;refGene_tx=XM_023256889.1
B4 63666021 63666129
refGene_gn=CCDC91;refGene_tx=XM_006933592.4;refGene_gn=CCDC91;refGene_tx=XM_006933593.2;refGene_gn=CCDC91;refGene_tx=XM_006933595.4;refGene_gn=CCDC91;refGene_tx=XM_006933596.3;refGene_gn=CCDC91;refGene_tx=XM_011283863.3;refGene_gn=CCDC91;refGene_tx=XM_019834749.2;refGene_gn=CCDC91;refGene_tx=XM_019834752.2;refGene_gn=CCDC91;refGene_tx=XM_019834753.2;refGene_gn=CCDC91;refGene_tx=XM_019834754.2;refGene_gn=CCDC91;refGene_tx=XM_019834757.2;refGene_gn=CCDC91;refGene_tx=XM_019834758.2;refGene_gn=CCDC91;refGene_tx=XM_019834759.2;refGene_gn=CCDC91;refGene_tx=XM_019834761.2;refGene_gn=CCDC91;refGene_tx=XM_019834762.1;refGene_gn=CCDC91;refGene_tx=XM_019834763.2
B4 63719945 63720038
refGene_gn=CCDC91;refGene_tx=XM_006933592.4;refGene_gn=CCDC91;refGene_tx=XM_006933593.2;refGene_gn=CCDC91;refGene_tx=XM_006933595.4;refGene_gn=CCDC91;refGene_tx=XM_006933596.3;refGene_gn=CCDC91;refGene_tx=XM_011283863.3;refGene_gn=CCDC91;refGene_tx=XM_019834749.2;refGene_gn=CCDC91;refGene_tx=XM_019834752.2;refGene_gn=CCDC91;refGene_tx=XM_019834753.2;refGene_gn=CCDC91;refGene_tx=XM_019834754.2;refGene_gn=CCDC91;refGene_tx=XM_019834757.2;refGene_gn=CCDC91;refGene_tx=XM_019834758.2;refGene_gn=CCDC91;refGene_tx=XM_019834759.2;refGene_gn=CCDC91;refGene_tx=XM_019834761.2;refGene_gn=CCDC91;refGene_tx=XM_019834762.1;refGene_gn=CCDC91;refGene_tx=XM_019834763.2;refGene_gn=CCDC91;refGene_tx=XM_01983476

4.1;refGene_gn=CCDC91;refGene_tx=XM_023256889.1
B4 63720146 63720215
refGene_gn=CCDC91;refGene_tx=XM_006933592.4;refGene_gn=CCDC91;refGene_tx=XM_006933593.2;refGene_gn=CCDC91;refGene_tx=XM_006933595.4;refGene_gn=CCDC91;refGene_tx=XM_006933596.3;refGene_gn=CCDC91;refGene_tx=XM_011283863.3;refGene_gn=CCDC91;refGene_tx=XM_019834749.2;refGene_gn=CCDC91;refGene_tx=XM_019834752.2;refGene_gn=CCDC91;refGene_tx=XM_019834753.2;refGene_gn=CCDC91;refGene_tx=XM_019834754.2;refGene_gn=CCDC91;refGene_tx=XM_019834757.2;refGene_gn=CCDC91;refGene_tx=XM_019834758.2;refGene_gn=CCDC91;refGene_tx=XM_019834759.2;refGene_gn=CCDC91;refGene_tx=XM_019834761.2;refGene_gn=CCDC91;refGene_tx=XM_019834762.1;refGene_gn=CCDC91;refGene_tx=XM_019834763.2;refGene_gn=CCDC91;refGene_tx=XM_019834764.1;refGene_gn=CCDC91;refGene_tx=XM_023256889.1
B4 63723298 63723475
refGene_gn=CCDC91;refGene_tx=XM_006933592.4;refGene_gn=CCDC91;refGene_tx=XM_006933593.2;refGene_gn=CCDC91;refGene_tx=XM_006933595.4;refGene_gn=CCDC91;refGene_tx=XM_006933596.3;refGene_gn=CCDC91;refGene_tx=XM_011283863.3;refGene_gn=CCDC91;refGene_tx=XM_019834749.2;refGene_gn=CCDC91;refGene_tx=XM_019834752.2;refGene_gn=CCDC91;refGene_tx=XM_019834753.2;refGene_gn=CCDC91;refGene_tx=XM_019834754.2;refGene_gn=CCDC91;refGene_tx=XM_019834757.2;refGene_gn=CCDC91;refGene_tx=XM_019834758.2;refGene_gn=CCDC91;refGene_tx=XM_019834759.2;refGene_gn=CCDC91;refGene_tx=XM_019834761.2;refGene_gn=CCDC91;refGene_tx=XM_019834762.1;refGene_gn=CCDC91;refGene_tx=XM_019834763.2;refGene_gn=CCDC91;refGene_tx=XM_019834764.1;refGene_gn=CCDC91;refGene_tx=XM_023256889.1
B4 63749755 63749869
refGene_gn=CCDC91;refGene_tx=XM_006933592.4;refGene_gn=CCDC91;refGene_tx=XM_006933593.2;refGene_gn=CCDC91;refGene_tx=XM_006933595.4;refGene_gn=CCDC91;refGene_tx=XM_006933596.3;refGene_gn=CCDC91;refGene_tx=XM_011283863.3;refGene_gn=CCDC91;refGene_tx=XM_019834749.2;refGene_gn=CCDC91;refGene_tx=XM_019834752.2;refGene_gn=CCDC91;refGene_tx=XM_019834753.2;refGene_gn=CCDC91;refGene_tx=XM_019834754.2;refGene_gn=CCDC91;refGene_tx=XM_019834757.2;refGene_gn=CCDC91;refGene_tx=XM_019834758.2;refGene_gn=CCDC91;refGene_tx=XM_019834759.2;refGene_gn=CCDC91;refGene_tx=XM_019834761.2;refGene_gn=CCDC91;refGene_tx=XM_019834762.1;refGene_gn=CCDC91;refGene_tx=XM_019834763.2;refGene_gn=CCDC91;refGene_tx=XM_019834764.1;refGene_gn=CCDC91;refGene_tx=XM_023256889.1
B4 63805516 63805627
refGene_gn=CCDC91;refGene_tx=XM_006933592.4;refGene_gn=CCDC91;refGene_tx=XM_006933593.2;refGene_gn=CCDC91;refGene_tx=XM_006933595.4;refGene_gn=CCDC91;refGene_tx=XM_006933596.3;refGene_gn=CCDC91;refGene_tx=XM_011283863.3;refGene_gn=CCDC91;refGene_tx=XM_019834749.2;refGene_gn=CCDC91;refGene_tx=XM_019834752.2;refGene_gn=CCDC91;refGene_tx=XM_019834753.2;refGene_gn=CCDC91;refGene_tx=XM_019834754.2;refGene_gn=CCDC91;refGene_tx=XM_019834757.2;refGene_gn=CCDC91;refGene_tx=XM_019834758.2;refGene_gn=CCDC91;refGene_tx=XM_019834759.2;refGene_gn=CCDC91;refGene_tx=XM_019834761.2;refGene_gn=CCDC91;refGene_tx=XM_019834762.1;refGene_gn=CCDC91;refGene_tx=XM_019834763.2;refGene_gn=CCDC91;refGene_tx=XM_019834764.1;refGene_gn=CCDC91;refGene_tx=XM_023256889.1
B4 64201909 64201985

refGene_gn=LOC101085811;refGene_tx=XM_011283864.3
B4 64201987 64203700
refGene_gn=LOC101085811;refGene_tx=XM_011283864.3
B4 64203702 64203998
refGene_gn=LOC101085811;refGene_tx=XM_011283864.3
B4 64418074 64418263
refGene_gn=FAR2;refGene_tx=XM_003988531.5;refGene_gn=FAR2;refGene_tx=XM_023256890.1
B4 64434168 64434344
refGene_gn=FAR2;refGene_tx=XM_003988531.5;refGene_gn=FAR2;refGene_tx=XM_023256890.1
B4 64437231 64437411
refGene_gn=FAR2;refGene_tx=XM_003988531.5;refGene_gn=FAR2;refGene_tx=XM_023256890.1
B4 64447506 64447684
refGene_gn=FAR2;refGene_tx=XM_003988531.5;refGene_gn=FAR2;refGene_tx=XM_023256890.1
B4 64448995 64449040
refGene_gn=FAR2;refGene_tx=XM_003988531.5;refGene_gn=FAR2;refGene_tx=XM_023256890.1
B4 64451239 64451358
refGene_gn=FAR2;refGene_tx=XM_003988531.5;refGene_gn=FAR2;refGene_tx=XM_023256890.1
B4 64452235 64452303
refGene_gn=FAR2;refGene_tx=XM_003988531.5;refGene_gn=FAR2;refGene_tx=XM_023256890.1
B4 64454625 64454797
refGene_gn=FAR2;refGene_tx=XM_003988531.5;refGene_gn=FAR2;refGene_tx=XM_023256890.1
B4 64458714 64458844
refGene_gn=FAR2;refGene_tx=XM_003988531.5;refGene_gn=FAR2;refGene_tx=XM_023256890.1
B4 64462166 64462294
refGene_gn=FAR2;refGene_tx=XM_003988531.5;refGene_gn=FAR2;refGene_tx=XM_023256890.1
B4 64463144 64463307
refGene_gn=FAR2;refGene_tx=XM_003988531.5;refGene_gn=FAR2;refGene_tx=XM_023256890.1
B4 64470280 64470343
refGene_gn=ERGIC2;refGene_tx=XM_003988532.4;refGene_gn=ERGIC2;refGene_tx=XM_006933600.3
B4 64470509 64470592
refGene_gn=ERGIC2;refGene_tx=XM_003988532.4;refGene_gn=ERGIC2;refGene_tx=XM_006933600.3
B4 64471685 64471848
refGene_gn=ERGIC2;refGene_tx=XM_003988532.4;refGene_gn=ERGIC2;refGene_tx=XM_006933600.3
B4 64475017 64475115
refGene_gn=ERGIC2;refGene_tx=XM_003988532.4;refGene_gn=ERGIC2;refGene_tx=XM_006933600.3

B4 64478650 64478749
refGene_gn=ERGIC2;refGene_tx=XM_003988532.4;refGene_gn=ERGIC2;refGene_tx=XM_006933600.3
B4 64479626 64479682
refGene_gn=ERGIC2;refGene_tx=XM_003988532.4;refGene_gn=ERGIC2;refGene_tx=XM_006933600.3
B4 64485334 64485430
refGene_gn=ERGIC2;refGene_tx=XM_003988532.4;refGene_gn=ERGIC2;refGene_tx=XM_006933600.3
B4 64486332 64486434
refGene_gn=ERGIC2;refGene_tx=XM_003988532.4;refGene_gn=ERGIC2;refGene_tx=XM_006933600.3
B4 64490098 64490139
refGene_gn=ERGIC2;refGene_tx=XM_003988532.4;refGene_gn=ERGIC2;refGene_tx=XM_006933600.3
B4 64495984 64496055
refGene_gn=ERGIC2;refGene_tx=XM_003988532.4;refGene_gn=ERGIC2;refGene_tx=XM_006933600.3
B4 64497565 64497612
refGene_gn=ERGIC2;refGene_tx=XM_003988532.4;refGene_gn=ERGIC2;refGene_tx=XM_006933600.3
B4 64499927 64500036
refGene_gn=ERGIC2;refGene_tx=XM_003988532.4;refGene_gn=ERGIC2;refGene_tx=XM_006933600.3
B4 64501573 64501679
refGene_gn=ERGIC2;refGene_tx=XM_003988532.4;refGene_gn=ERGIC2;refGene_tx=XM_006933600.3
B4 64523625 64523790
refGene_gn=LOC111561341;refGene_tx=XM_023256891.1
B4 64524446 64524580
refGene_gn=LOC111561341;refGene_tx=XM_023256891.1
B4 64525725 64525824
refGene_gn=LOC111561341;refGene_tx=XM_023256891.1
B4 64531480 64531641
refGene_gn=LOC111561341;refGene_tx=XM_023256891.1
B4 64532934 64533202
refGene_gn=LOC111561341;refGene_tx=XM_023256891.1
B4 64534749 64534923
refGene_gn=LOC111561341;refGene_tx=XM_023256891.1
B4 64541454 64541602
refGene_gn=LOC111561341;refGene_tx=XM_023256891.1
B4 64544348 64544647 refGene_gn=OVCH1;refGene_tx=XM_023257753.1
B4 64550129 64550269 refGene_gn=OVCH1;refGene_tx=XM_023257753.1
B4 64551068 64551204 refGene_gn=OVCH1;refGene_tx=XM_023257753.1
B4 64552445 64552571 refGene_gn=OVCH1;refGene_tx=XM_023257753.1
B4 64557979 64558204 refGene_gn=OVCH1;refGene_tx=XM_023257753.1
B4 64561810 64561912 refGene_gn=OVCH1;refGene_tx=XM_023257753.1
B4 64562239 64562396 refGene_gn=OVCH1;refGene_tx=XM_023257753.1
B4 64569672 64569824 refGene_gn=OVCH1;refGene_tx=XM_023257753.1
B4 64573394 64573590 refGene_gn=OVCH1;refGene_tx=XM_023257753.1

B4	64574264	64574337	refGene_gn=OVCH1;refGene_tx=XM_023257753.1
B4	64580472	64580653	refGene_gn=OVCH1;refGene_tx=XM_023257753.1
B4	64581583	64581658	refGene_gn=OVCH1;refGene_tx=XM_023257753.1
B4	64583718	64583847	refGene_gn=OVCH1;refGene_tx=XM_023257753.1
B4	64585349	64585443	refGene_gn=OVCH1;refGene_tx=XM_023257753.1
B4	64586409	64586540	refGene_gn=OVCH1;refGene_tx=XM_023257753.1
B4	64586643	64586784	refGene_gn=OVCH1;refGene_tx=XM_023257753.1
B4	64587969	64588103	refGene_gn=OVCH1;refGene_tx=XM_023257753.1
B4	64594480	64594568	refGene_gn=OVCH1;refGene_tx=XM_023257753.1
B4	64596523	64596710	refGene_gn=OVCH1;refGene_tx=XM_023257753.1
B4	64598437	64598589	refGene_gn=OVCH1;refGene_tx=XM_023257753.1
B4	64599442	64599538	refGene_gn=OVCH1;refGene_tx=XM_023257753.1
B4	64603629	64603799	refGene_gn=OVCH1;refGene_tx=XM_023257753.1
B4	64604563	64604661	refGene_gn=OVCH1;refGene_tx=XM_023257753.1
B4	64604935	64605074	refGene_gn=OVCH1;refGene_tx=XM_023257753.1
B4	64605654	64606110	refGene_gn=OVCH1;refGene_tx=XM_023257753.1
B4	64607669	64607695	refGene_gn=OVCH1;refGene_tx=XM_023257753.1
B4	64614815	64614956	refGene_gn=TMTC1;refGene_tx=XM_023256892.1
B4	64619608	64619686	refGene_gn=TMTC1;refGene_tx=XM_023256892.1
B4	64622562	64622685	refGene_gn=TMTC1;refGene_tx=XM_023256892.1
B4	64624592	64624730	refGene_gn=TMTC1;refGene_tx=XM_023256892.1
B4	64625635	64625780	refGene_gn=TMTC1;refGene_tx=XM_023256892.1
B4	64626590	64626726	refGene_gn=TMTC1;refGene_tx=XM_023256892.1
B4	64628729	64628832	refGene_gn=TMTC1;refGene_tx=XM_023256892.1
B4	64647116	64647225	refGene_gn=TMTC1;refGene_tx=XM_023256892.1
B4	64667918	64668062	refGene_gn=TMTC1;refGene_tx=XM_023256892.1
B4	64686097	64686211	refGene_gn=TMTC1;refGene_tx=XM_023256892.1
B4	64693974	64694142	refGene_gn=TMTC1;refGene_tx=XM_023256892.1
B4	64710915	64711037	refGene_gn=TMTC1;refGene_tx=XM_023256892.1
B4	64739108	64739298	refGene_gn=TMTC1;refGene_tx=XM_023256892.1
B4	64762617	64762721	refGene_gn=TMTC1;refGene_tx=XM_023256892.1
B4	64765318	64765400	refGene_gn=TMTC1;refGene_tx=XM_023256892.1
B4	64851953	64852160	refGene_gn=TMTC1;refGene_tx=XM_023256892.1
B4	64856181	64856358	refGene_gn=TMTC1;refGene_tx=XM_023256892.1
B4	64859346	64859420	refGene_gn=TMTC1;refGene_tx=XM_023256892.1
B4	64868881	64869059	refGene_gn=TMTC1;refGene_tx=XM_023256892.1
B4	64889187	64889495	refGene_gn=TMTC1;refGene_tx=XM_023256892.1
B4	65501822	65501920
refGene_gn=IPO8;refGene_tx=XM_003988535.5;refGene_gn=IPO8;refGene_tx=XM_019834767.2;refGene_gn=IPO8;refGene_tx=XM_019834768.2;refGene_gn=IPO8;refGene_tx=XM_023256893.1
B4	65513392	65513509
refGene_gn=IPO8;refGene_tx=XM_003988535.5;refGene_gn=IPO8;refGene_tx=XM_019834767.2;refGene_gn=IPO8;refGene_tx=XM_019834768.2;refGene_gn=IPO8;refGene_tx=XM_023256893.1
B4	65514065	65514269
refGene_gn=IPO8;refGene_tx=XM_003988535.5;refGene_gn=IPO8;refGene_tx=XM_019834767.2;refGene_gn=IPO8;refGene_tx=XM_019834768.2;refGene_gn=IPO8;refGene_tx=XM_023256893.1
B4	65517107	65517313

refGene_gn=IPO8;refGene_tx=XM_003988535.5;refGene_gn=IPO8;refGene_tx=XM_019834767.2;refGene_gn=IPO8;refGene_tx=XM_019834768.2;refGene_gn=IPO8;refGene_tx=XM_023256893.1
B4 65521377 65521598
refGene_gn=IPO8;refGene_tx=XM_003988535.5;refGene_gn=IPO8;refGene_tx=XM_019834767.2;refGene_gn=IPO8;refGene_tx=XM_019834768.2;refGene_gn=IPO8;refGene_tx=XM_023256893.1
B4 65539017 65539113
refGene_gn=IPO8;refGene_tx=XM_003988535.5;refGene_gn=IPO8;refGene_tx=XM_019834767.2;refGene_gn=IPO8;refGene_tx=XM_019834768.2;refGene_gn=IPO8;refGene_tx=XM_023256893.1
B4 65542096 65542194
refGene_gn=IPO8;refGene_tx=XM_003988535.5;refGene_gn=IPO8;refGene_tx=XM_019834767.2;refGene_gn=IPO8;refGene_tx=XM_019834768.2;refGene_gn=IPO8;refGene_tx=XM_023256893.1
B4 65542870 65542996
refGene_gn=IPO8;refGene_tx=XM_003988535.5;refGene_gn=IPO8;refGene_tx=XM_019834767.2;refGene_gn=IPO8;refGene_tx=XM_019834768.2;refGene_gn=IPO8;refGene_tx=XM_023256893.1
B4 65546698 65546765
refGene_gn=IPO8;refGene_tx=XM_003988535.5;refGene_gn=IPO8;refGene_tx=XM_019834767.2;refGene_gn=IPO8;refGene_tx=XM_019834768.2;refGene_gn=IPO8;refGene_tx=XM_023256893.1
B4 65550207 65550333
refGene_gn=IPO8;refGene_tx=XM_003988535.5;refGene_gn=IPO8;refGene_tx=XM_019834767.2;refGene_gn=IPO8;refGene_tx=XM_019834768.2;refGene_gn=IPO8;refGene_tx=XM_023256893.1
B4 65551167 65551328
refGene_gn=IPO8;refGene_tx=XM_003988535.5;refGene_gn=IPO8;refGene_tx=XM_019834767.2;refGene_gn=IPO8;refGene_tx=XM_019834768.2;refGene_gn=IPO8;refGene_tx=XM_023256893.1
B4 65552505 65552671
refGene_gn=IPO8;refGene_tx=XM_003988535.5;refGene_gn=IPO8;refGene_tx=XM_019834767.2;refGene_gn=IPO8;refGene_tx=XM_019834768.2;refGene_gn=IPO8;refGene_tx=XM_023256893.1
B4 65554353 65554443
refGene_gn=IPO8;refGene_tx=XM_003988535.5;refGene_gn=IPO8;refGene_tx=XM_019834767.2;refGene_gn=IPO8;refGene_tx=XM_019834768.2;refGene_gn=IPO8;refGene_tx=XM_023256893.1
B4 65554874 65554991
refGene_gn=IPO8;refGene_tx=XM_003988535.5;refGene_gn=IPO8;refGene_tx=XM_019834767.2;refGene_gn=IPO8;refGene_tx=XM_019834768.2;refGene_gn=IPO8;refGene_tx=XM_023256893.1
B4 65555278 65555355
refGene_gn=IPO8;refGene_tx=XM_003988535.5;refGene_gn=IPO8;refGene_tx=XM_019834767.2;refGene_gn=IPO8;refGene_tx=XM_019834768.2;refGene_gn=IPO8;refGene_tx=XM_023256893.1
B4 65558362 65558462
refGene_gn=IPO8;refGene_tx=XM_003988535.5;refGene_gn=IPO8;refGene_tx=XM_019834767.2;refGene_gn=IPO8;refGene_tx=XM_019834768.2;refGene_gn=IPO

8;refGene_tx=XM_023256893.1
B4 65560277 65560412
refGene_gn=IPO8;refGene_tx=XM_003988535.5;refGene_gn=IPO8;refGene_tx=XM_019834767.2;refGene_gn=IPO8;refGene_tx=XM_019834768.2;refGene_gn=IPO8;refGene_tx=XM_023256893.1
B4 65562704 65562789
refGene_gn=IPO8;refGene_tx=XM_003988535.5;refGene_gn=IPO8;refGene_tx=XM_019834767.2;refGene_gn=IPO8;refGene_tx=XM_019834768.2;refGene_gn=IPO8;refGene_tx=XM_023256893.1
B4 65563645 65563740
refGene_gn=IPO8;refGene_tx=XM_003988535.5;refGene_gn=IPO8;refGene_tx=XM_019834767.2;refGene_gn=IPO8;refGene_tx=XM_019834768.2;refGene_gn=IPO8;refGene_tx=XM_023256893.1
B4 65565325 65565415
refGene_gn=IPO8;refGene_tx=XM_003988535.5;refGene_gn=IPO8;refGene_tx=XM_019834767.2;refGene_gn=IPO8;refGene_tx=XM_019834768.2;refGene_gn=IPO8;refGene_tx=XM_023256893.1
B4 65569132 65569289
refGene_gn=IPO8;refGene_tx=XM_003988535.5;refGene_gn=IPO8;refGene_tx=XM_019834767.2;refGene_gn=IPO8;refGene_tx=XM_019834768.2;refGene_gn=IPO8;refGene_tx=XM_023256893.1
B4 65570287 65570446
refGene_gn=IPO8;refGene_tx=XM_003988535.5;refGene_gn=IPO8;refGene_tx=XM_019834768.2;refGene_gn=IPO8;refGene_tx=XM_023256893.1
B4 65573499 65573656
refGene_gn=IPO8;refGene_tx=XM_003988535.5;refGene_gn=IPO8;refGene_tx=XM_019834767.2;refGene_gn=IPO8;refGene_tx=XM_023256893.1
B4 65580068 65580150
refGene_gn=IPO8;refGene_tx=XM_003988535.5;refGene_gn=IPO8;refGene_tx=XM_019834767.2
B4 65586397 65586481
refGene_gn=IPO8;refGene_tx=XM_003988535.5;refGene_gn=IPO8;refGene_tx=XM_019834767.2
B4 65598318 65598887
refGene_gn=CAPRIN2;refGene_tx=XM_023256894.1;refGene_gn=CAPRIN2;refGene_tx=XM_023256895.1;refGene_gn=CAPRIN2;refGene_tx=XM_023256896.1;refGene_gn=CAPRIN2;refGene_tx=XM_023256897.1;refGene_gn=CAPRIN2;refGene_tx=XM_023256898.1;refGene_gn=CAPRIN2;refGene_tx=XM_023256899.1;refGene_gn=CAPRIN2;refGene_tx=XM_023256900.1;refGene_gn=CAPRIN2;refGene_tx=XM_023256901.1
B4 65600054 65600118
refGene_gn=CAPRIN2;refGene_tx=XM_023256894.1;refGene_gn=CAPRIN2;refGene_tx=XM_023256895.1;refGene_gn=CAPRIN2;refGene_tx=XM_023256896.1;refGene_gn=CAPRIN2;refGene_tx=XM_023256897.1;refGene_gn=CAPRIN2;refGene_tx=XM_023256898.1;refGene_gn=CAPRIN2;refGene_tx=XM_023256899.1;refGene_gn=CAPRIN2;refGene_tx=XM_023256900.1;refGene_gn=CAPRIN2;refGene_tx=XM_023256901.1
B4 65602647 65602748
refGene_gn=CAPRIN2;refGene_tx=XM_023256894.1;refGene_gn=CAPRIN2;refGene_tx=XM_023256895.1;refGene_gn=CAPRIN2;refGene_tx=XM_023256896.1;refGe

ne_gn=CAPRIN2;refGene_tx=XM_023256897.1;refGene_gn=CAPRIN2;refGene_tx=XM_023256898.1;refGene_gn=CAPRIN2;refGene_tx=XM_023256899.1;refGene_gn=CAPRIN2;refGene_tx=XM_023256900.1;refGene_gn=CAPRIN2;refGene_tx=XM_023256901.1
B4 65603511 65603694
refGene_gn=CAPRIN2;refGene_tx=XM_023256894.1;refGene_gn=CAPRIN2;refGene_tx=XM_023256895.1;refGene_gn=CAPRIN2;refGene_tx=XM_023256896.1;refGene_gn=CAPRIN2;refGene_tx=XM_023256897.1;refGene_gn=CAPRIN2;refGene_tx=XM_023256898.1;refGene_gn=CAPRIN2;refGene_tx=XM_023256900.1;refGene_gn=CAPRIN2;refGene_tx=XM_023256901.1
B4 65605124 65605293
refGene_gn=CAPRIN2;refGene_tx=XM_023256894.1;refGene_gn=CAPRIN2;refGene_tx=XM_023256895.1;refGene_gn=CAPRIN2;refGene_tx=XM_023256896.1;refGene_gn=CAPRIN2;refGene_tx=XM_023256897.1;refGene_gn=CAPRIN2;refGene_tx=XM_023256898.1;refGene_gn=CAPRIN2;refGene_tx=XM_023256899.1;refGene_gn=CAPRIN2;refGene_tx=XM_023256900.1;refGene_gn=CAPRIN2;refGene_tx=XM_023256901.1
B4 65608054 65608201
refGene_gn=CAPRIN2;refGene_tx=XM_023256894.1;refGene_gn=CAPRIN2;refGene_tx=XM_023256895.1;refGene_gn=CAPRIN2;refGene_tx=XM_023256896.1;refGene_gn=CAPRIN2;refGene_tx=XM_023256897.1;refGene_gn=CAPRIN2;refGene_tx=XM_023256898.1;refGene_gn=CAPRIN2;refGene_tx=XM_023256899.1;refGene_gn=CAPRIN2;refGene_tx=XM_023256901.1
B4 65610306 65610411
refGene_gn=CAPRIN2;refGene_tx=XM_023256894.1;refGene_gn=CAPRIN2;refGene_tx=XM_023256895.1;refGene_gn=CAPRIN2;refGene_tx=XM_023256896.1;refGene_gn=CAPRIN2;refGene_tx=XM_023256897.1;refGene_gn=CAPRIN2;refGene_tx=XM_023256899.1;refGene_gn=CAPRIN2;refGene_tx=XM_023256901.1
B4 65613415 65613471
refGene_gn=CAPRIN2;refGene_tx=XM_023256894.1;refGene_gn=CAPRIN2;refGene_tx=XM_023256895.1;refGene_gn=CAPRIN2;refGene_tx=XM_023256896.1;refGene_gn=CAPRIN2;refGene_tx=XM_023256897.1;refGene_gn=CAPRIN2;refGene_tx=XM_023256898.1;refGene_gn=CAPRIN2;refGene_tx=XM_023256899.1;refGene_gn=CAPRIN2;refGene_tx=XM_023256900.1;refGene_gn=CAPRIN2;refGene_tx=XM_023256901.1
B4 65614488 65614570
refGene_gn=CAPRIN2;refGene_tx=XM_023256894.1;refGene_gn=CAPRIN2;refGene_tx=XM_023256895.1;refGene_gn=CAPRIN2;refGene_tx=XM_023256896.1;refGene_gn=CAPRIN2;refGene_tx=XM_023256897.1;refGene_gn=CAPRIN2;refGene_tx=XM_023256898.1;refGene_gn=CAPRIN2;refGene_tx=XM_023256899.1;refGene_gn=CAPRIN2;refGene_tx=XM_023256900.1;refGene_gn=CAPRIN2;refGene_tx=XM_023256901.1
B4 65616389 65616512
refGene_gn=CAPRIN2;refGene_tx=XM_023256894.1;refGene_gn=CAPRIN2;refGene_tx=XM_023256895.1;refGene_gn=CAPRIN2;refGene_tx=XM_023256896.1;refGene_gn=CAPRIN2;refGene_tx=XM_023256897.1;refGene_gn=CAPRIN2;refGene_tx=XM_023256898.1;refGene_gn=CAPRIN2;refGene_tx=XM_023256899.1;refGene_gn=CAPRIN2;refGene_tx=XM_023256900.1;refGene_gn=CAPRIN2;refGene_tx=XM_023256901.1
B4 65618999 65619674

refGene_gn=CAPRIN2;refGene_tx=XM_023256894.1;refGene_gn=CAPRIN2;refGene_tx=XM_023256895.1;refGene_gn=CAPRIN2;refGene_tx=XM_023256896.1;refGene_gn=CAPRIN2;refGene_tx=XM_023256897.1;refGene_gn=CAPRIN2;refGene_tx=XM_023256898.1;refGene_gn=CAPRIN2;refGene_tx=XM_023256899.1;refGene_gn=CAPRIN2;refGene_tx=XM_023256900.1
B4 65620582 65620626
refGene_gn=CAPRIN2;refGene_tx=XM_023256894.1;refGene_gn=CAPRIN2;refGene_tx=XM_023256895.1;refGene_gn=CAPRIN2;refGene_tx=XM_023256896.1;refGene_gn=CAPRIN2;refGene_tx=XM_023256897.1;refGene_gn=CAPRIN2;refGene_tx=XM_023256898.1;refGene_gn=CAPRIN2;refGene_tx=XM_023256899.1;refGene_gn=CAPRIN2;refGene_tx=XM_023256900.1;refGene_gn=CAPRIN2;refGene_tx=XM_023256901.1
B4 65621652 65621820
refGene_gn=CAPRIN2;refGene_tx=XM_023256894.1;refGene_gn=CAPRIN2;refGene_tx=XM_023256895.1;refGene_gn=CAPRIN2;refGene_tx=XM_023256896.1;refGene_gn=CAPRIN2;refGene_tx=XM_023256897.1;refGene_gn=CAPRIN2;refGene_tx=XM_023256898.1;refGene_gn=CAPRIN2;refGene_tx=XM_023256899.1;refGene_gn=CAPRIN2;refGene_tx=XM_023256900.1;refGene_gn=CAPRIN2;refGene_tx=XM_023256901.1
B4 65623996 65624079
refGene_gn=CAPRIN2;refGene_tx=XM_023256894.1;refGene_gn=CAPRIN2;refGene_tx=XM_023256895.1;refGene_gn=CAPRIN2;refGene_tx=XM_023256896.1;refGene_gn=CAPRIN2;refGene_tx=XM_023256897.1;refGene_gn=CAPRIN2;refGene_tx=XM_023256898.1;refGene_gn=CAPRIN2;refGene_tx=XM_023256899.1;refGene_gn=CAPRIN2;refGene_tx=XM_023256900.1;refGene_gn=CAPRIN2;refGene_tx=XM_023256901.1
B4 65625579 65625818
refGene_gn=CAPRIN2;refGene_tx=XM_023256894.1;refGene_gn=CAPRIN2;refGene_tx=XM_023256895.1;refGene_gn=CAPRIN2;refGene_tx=XM_023256896.1;refGene_gn=CAPRIN2;refGene_tx=XM_023256897.1;refGene_gn=CAPRIN2;refGene_tx=XM_023256898.1;refGene_gn=CAPRIN2;refGene_tx=XM_023256899.1;refGene_gn=CAPRIN2;refGene_tx=XM_023256900.1;refGene_gn=CAPRIN2;refGene_tx=XM_023256901.1
B4 65630484 65630571
refGene_gn=CAPRIN2;refGene_tx=XM_023256894.1;refGene_gn=CAPRIN2;refGene_tx=XM_023256895.1;refGene_gn=CAPRIN2;refGene_tx=XM_023256896.1;refGene_gn=CAPRIN2;refGene_tx=XM_023256897.1;refGene_gn=CAPRIN2;refGene_tx=XM_023256898.1;refGene_gn=CAPRIN2;refGene_tx=XM_023256899.1;refGene_gn=CAPRIN2;refGene_tx=XM_023256900.1;refGene_gn=CAPRIN2;refGene_tx=XM_023256901.1
B4 65639792 65639855
refGene_gn=CAPRIN2;refGene_tx=XM_023256894.1;refGene_gn=CAPRIN2;refGene_tx=XM_023256895.1;refGene_gn=CAPRIN2;refGene_tx=XM_023256896.1;refGene_gn=CAPRIN2;refGene_tx=XM_023256897.1;refGene_gn=CAPRIN2;refGene_tx=XM_023256898.1;refGene_gn=CAPRIN2;refGene_tx=XM_023256899.1;refGene_gn=CAPRIN2;refGene_tx=XM_023256900.1;refGene_gn=CAPRIN2;refGene_tx=XM_023256901.1
B4 65641844 65642096
refGene_gn=CAPRIN2;refGene_tx=XM_023256894.1;refGene_gn=CAPRIN2;refGene_tx=XM_023256895.1;refGene_gn=CAPRIN2;refGene_tx=XM_023256896.1;refGe

ne_gn=CAPRIN2;refGene_tx=XM_023256897.1;refGene_gn=CAPRIN2;refGene_tx=XM_023256898.1;refGene_gn=CAPRIN2;refGene_tx=XM_023256899.1;refGene_gn=CAPRIN2;refGene_tx=XM_023256900.1;refGene_gn=CAPRIN2;refGene_tx=XM_023256901.1
B4 65713430 65713590
refGene_gn=SINHCAF;refGene_tx=XM_003988537.5;refGene_gn=SINHCAF;refGene_tx=XM_006933602.4;refGene_gn=SINHCAF;refGene_tx=XM_019834775.2
B4 65716679 65716830
refGene_gn=SINHCAF;refGene_tx=XM_003988537.5;refGene_gn=SINHCAF;refGene_tx=XM_006933602.4;refGene_gn=SINHCAF;refGene_tx=XM_019834775.2
B4 65723305 65723432
refGene_gn=SINHCAF;refGene_tx=XM_003988537.5;refGene_gn=SINHCAF;refGene_tx=XM_006933602.4;refGene_gn=SINHCAF;refGene_tx=XM_019834775.2
B4 65725347 65725447
refGene_gn=SINHCAF;refGene_tx=XM_003988537.5;refGene_gn=SINHCAF;refGene_tx=XM_006933602.4;refGene_gn=SINHCAF;refGene_tx=XM_019834775.2
B4 65726724 65726852
refGene_gn=SINHCAF;refGene_tx=XM_003988537.5;refGene_gn=SINHCAF;refGene_tx=XM_006933602.4;refGene_gn=SINHCAF;refGene_tx=XM_019834775.2
B4 65795383 65795567
refGene_gn=DENND5B;refGene_tx=XM_003988538.5;refGene_gn=DENND5B;refGene_tx=XM_003988539.5;refGene_gn=DENND5B;refGene_tx=XM_019834776.2;refGene_gn=DENND5B;refGene_tx=XM_019834778.2;refGene_gn=DENND5B;refGene_tx=XM_019834779.2;refGene_gn=DENND5B;refGene_tx=XM_019834780.2
B4 65797251 65797426
refGene_gn=DENND5B;refGene_tx=XM_003988538.5;refGene_gn=DENND5B;refGene_tx=XM_003988539.5;refGene_gn=DENND5B;refGene_tx=XM_019834776.2;refGene_gn=DENND5B;refGene_tx=XM_019834778.2;refGene_gn=DENND5B;refGene_tx=XM_019834779.2;refGene_gn=DENND5B;refGene_tx=XM_019834780.2
B4 65799420 65799547
refGene_gn=DENND5B;refGene_tx=XM_003988538.5;refGene_gn=DENND5B;refGene_tx=XM_003988539.5;refGene_gn=DENND5B;refGene_tx=XM_019834776.2;refGene_gn=DENND5B;refGene_tx=XM_019834778.2;refGene_gn=DENND5B;refGene_tx=XM_019834779.2;refGene_gn=DENND5B;refGene_tx=XM_019834780.2
B4 65800140 65800223
refGene_gn=DENND5B;refGene_tx=XM_003988538.5;refGene_gn=DENND5B;refGene_tx=XM_003988539.5;refGene_gn=DENND5B;refGene_tx=XM_019834776.2;refGene_gn=DENND5B;refGene_tx=XM_019834778.2;refGene_gn=DENND5B;refGene_tx=XM_019834779.2;refGene_gn=DENND5B;refGene_tx=XM_019834780.2
B4 65805614 65805802
refGene_gn=DENND5B;refGene_tx=XM_003988538.5;refGene_gn=DENND5B;refGene_tx=XM_003988539.5;refGene_gn=DENND5B;refGene_tx=XM_019834776.2;refGene_gn=DENND5B;refGene_tx=XM_019834778.2;refGene_gn=DENND5B;refGene_tx=XM_019834779.2;refGene_gn=DENND5B;refGene_tx=XM_019834780.2
B4 65806832 65806951
refGene_gn=DENND5B;refGene_tx=XM_003988538.5;refGene_gn=DENND5B;refGene_tx=XM_003988539.5;refGene_gn=DENND5B;refGene_tx=XM_019834776.2;refGene_gn=DENND5B;refGene_tx=XM_019834778.2;refGene_gn=DENND5B;refGene_tx=XM_019834779.2;refGene_gn=DENND5B;refGene_tx=XM_019834780.2
B4 65809006 65809152

refGene_gn=DENND5B;refGene_tx=XM_003988538.5;refGene_gn=DENND5B;refGene_tx=XM_003988539.5;refGene_gn=DENND5B;refGene_tx=XM_019834776.2;refGene_gn=DENND5B;refGene_tx=XM_019834778.2;refGene_gn=DENND5B;refGene_tx=XM_019834779.2;refGene_gn=DENND5B;refGene_tx=XM_019834780.2
B4 65813108 65813230
refGene_gn=DENND5B;refGene_tx=XM_003988538.5;refGene_gn=DENND5B;refGene_tx=XM_003988539.5;refGene_gn=DENND5B;refGene_tx=XM_019834776.2;refGene_gn=DENND5B;refGene_tx=XM_019834778.2;refGene_gn=DENND5B;refGene_tx=XM_019834779.2;refGene_gn=DENND5B;refGene_tx=XM_019834780.2
B4 65816710 65816839
refGene_gn=DENND5B;refGene_tx=XM_003988538.5;refGene_gn=DENND5B;refGene_tx=XM_003988539.5;refGene_gn=DENND5B;refGene_tx=XM_019834776.2;refGene_gn=DENND5B;refGene_tx=XM_019834778.2;refGene_gn=DENND5B;refGene_tx=XM_019834779.2;refGene_gn=DENND5B;refGene_tx=XM_019834780.2
B4 65829657 65829739
refGene_gn=DENND5B;refGene_tx=XM_003988538.5;refGene_gn=DENND5B;refGene_tx=XM_003988539.5;refGene_gn=DENND5B;refGene_tx=XM_019834776.2;refGene_gn=DENND5B;refGene_tx=XM_019834778.2;refGene_gn=DENND5B;refGene_tx=XM_019834779.2;refGene_gn=DENND5B;refGene_tx=XM_019834780.2
B4 65841265 65841344
refGene_gn=DENND5B;refGene_tx=XM_003988538.5;refGene_gn=DENND5B;refGene_tx=XM_003988539.5;refGene_gn=DENND5B;refGene_tx=XM_019834776.2;refGene_gn=DENND5B;refGene_tx=XM_019834778.2;refGene_gn=DENND5B;refGene_tx=XM_019834779.2;refGene_gn=DENND5B;refGene_tx=XM_019834780.2
B4 65842211 65842364
refGene_gn=DENND5B;refGene_tx=XM_003988538.5;refGene_gn=DENND5B;refGene_tx=XM_003988539.5;refGene_gn=DENND5B;refGene_tx=XM_019834776.2;refGene_gn=DENND5B;refGene_tx=XM_019834778.2;refGene_gn=DENND5B;refGene_tx=XM_019834779.2;refGene_gn=DENND5B;refGene_tx=XM_019834780.2
B4 65844062 65844194
refGene_gn=DENND5B;refGene_tx=XM_003988538.5;refGene_gn=DENND5B;refGene_tx=XM_003988539.5;refGene_gn=DENND5B;refGene_tx=XM_019834776.2;refGene_gn=DENND5B;refGene_tx=XM_019834778.2;refGene_gn=DENND5B;refGene_tx=XM_019834779.2;refGene_gn=DENND5B;refGene_tx=XM_019834780.2
B4 65847337 65847448
refGene_gn=DENND5B;refGene_tx=XM_019834776.2;refGene_gn=DENND5B;refGene_tx=XM_019834778.2;refGene_gn=DENND5B;refGene_tx=XM_019834779.2
B4 65848270 65848364
refGene_gn=DENND5B;refGene_tx=XM_003988538.5;refGene_gn=DENND5B;refGene_tx=XM_003988539.5;refGene_gn=DENND5B;refGene_tx=XM_019834776.2;refGene_gn=DENND5B;refGene_tx=XM_019834778.2;refGene_gn=DENND5B;refGene_tx=XM_019834779.2;refGene_gn=DENND5B;refGene_tx=XM_019834780.2
B4 65857619 65857770
refGene_gn=DENND5B;refGene_tx=XM_003988538.5;refGene_gn=DENND5B;refGene_tx=XM_003988539.5;refGene_gn=DENND5B;refGene_tx=XM_019834776.2;refGene_gn=DENND5B;refGene_tx=XM_019834778.2;refGene_gn=DENND5B;refGene_tx=XM_019834779.2;refGene_gn=DENND5B;refGene_tx=XM_019834780.2
B4 65859763 65859847
refGene_gn=DENND5B;refGene_tx=XM_019834776.2;refGene_gn=DENND5B;refGene_tx=XM_019834778.2;refGene_gn=DENND5B;refGene_tx=XM_019834780.2

B4	65862463	65862695
refGene_gn=DENND5B;refGene_tx=XM_003988538.5;refGene_gn=DENND5B;refGene_tx=XM_003988539.5;refGene_gn=DENND5B;refGene_tx=XM_019834776.2;refGene_gn=DENND5B;refGene_tx=XM_019834778.2;refGene_gn=DENND5B;refGene_tx=XM_019834779.2;refGene_gn=DENND5B;refGene_tx=XM_019834780.2
B4	65865048	65865585
refGene_gn=DENND5B;refGene_tx=XM_003988538.5;refGene_gn=DENND5B;refGene_tx=XM_003988539.5;refGene_gn=DENND5B;refGene_tx=XM_019834776.2;refGene_gn=DENND5B;refGene_tx=XM_019834778.2;refGene_gn=DENND5B;refGene_tx=XM_019834779.2;refGene_gn=DENND5B;refGene_tx=XM_019834780.2
B4	65871791	65871979
refGene_gn=DENND5B;refGene_tx=XM_003988538.5;refGene_gn=DENND5B;refGene_tx=XM_003988539.5;refGene_gn=DENND5B;refGene_tx=XM_019834776.2;refGene_gn=DENND5B;refGene_tx=XM_019834778.2;refGene_gn=DENND5B;refGene_tx=XM_019834779.2;refGene_gn=DENND5B;refGene_tx=XM_019834780.2
B4	65894690	65895357
refGene_gn=DENND5B;refGene_tx=XM_003988538.5;refGene_gn=DENND5B;refGene_tx=XM_003988539.5;refGene_gn=DENND5B;refGene_tx=XM_019834776.2;refGene_gn=DENND5B;refGene_tx=XM_019834778.2;refGene_gn=DENND5B;refGene_tx=XM_019834779.2;refGene_gn=DENND5B;refGene_tx=XM_019834780.2
B4	65906045	65906155
refGene_gn=DENND5B;refGene_tx=XM_003988538.5;refGene_gn=DENND5B;refGene_tx=XM_003988539.5;refGene_gn=DENND5B;refGene_tx=XM_019834776.2;refGene_gn=DENND5B;refGene_tx=XM_019834778.2;refGene_gn=DENND5B;refGene_tx=XM_019834779.2;refGene_gn=DENND5B;refGene_tx=XM_019834780.2
B4	65909275	65909341
refGene_gn=DENND5B;refGene_tx=XM_003988538.5;refGene_gn=DENND5B;refGene_tx=XM_019834776.2;refGene_gn=DENND5B;refGene_tx=XM_019834779.2;refGene_gn=DENND5B;refGene_tx=XM_019834780.2
B4	65982356	65982477
refGene_gn=DENND5B;refGene_tx=XM_003988538.5;refGene_gn=DENND5B;refGene_tx=XM_003988539.5;refGene_gn=DENND5B;refGene_tx=XM_019834776.2;refGene_gn=DENND5B;refGene_tx=XM_019834778.2;refGene_gn=DENND5B;refGene_tx=XM_019834779.2;refGene_gn=DENND5B;refGene_tx=XM_019834780.2
B4	66038171	66038497	refGene_gn=ETFBKMT;refGene_tx=XM_019834783.2
B4	66040804	66040910	refGene_gn=ETFBKMT;refGene_tx=XM_019834783.2
B4	66040911	66040936	refGene_gn=ETFBKMT;refGene_tx=XM_019834783.2
B4	66041726	66042070	refGene_gn=ETFBKMT;refGene_tx=XM_019834783.2
B4	66047211	66047285
refGene_gn=AMN1;refGene_tx=XM_003988553.5;refGene_gn=AMN1;refGene_tx=XM_019834785.2;refGene_gn=AMN1;refGene_tx=XM_023256903.1
B4	66062279	66062391
refGene_gn=AMN1;refGene_tx=XM_003988553.5;refGene_gn=AMN1;refGene_tx=XM_019834785.2;refGene_gn=AMN1;refGene_tx=XM_023256903.1
B4	66069914	66069971
refGene_gn=AMN1;refGene_tx=XM_003988553.5;refGene_gn=AMN1;refGene_tx=XM_019834785.2;refGene_gn=AMN1;refGene_tx=XM_023256903.1
B4	66070295	66070513
refGene_gn=AMN1;refGene_tx=XM_003988553.5;refGene_gn=AMN1;refGene_tx=XM_019834785.2;refGene_gn=AMN1;refGene_tx=XM_023256903.1

B4 66072366 66072511
refGene_gn=AMN1;refGene_tx=XM_003988553.5;refGene_gn=AMN1;refGene_tx=XM_019834785.2;refGene_gn=AMN1;refGene_tx=XM_023256903.1
B4 66076018 66076151
refGene_gn=AMN1;refGene_tx=XM_003988553.5;refGene_gn=AMN1;refGene_tx=XM_019834785.2;refGene_gn=AMN1;refGene_tx=XM_023256903.1
B4 66085168 66085203 refGene_gn=AMN1;refGene_tx=XM_019834785.2
B4 66096647 66096685 refGene_gn=AMN1;refGene_tx=XM_003988553.5
B4 66319890 66324901
refGene_gn=KIAA1551;refGene_tx=XM_023256904.1;refGene_gn=KIAA1551;refGene_tx=XM_023256905.1;refGene_gn=KIAA1551;refGene_tx=XM_023256906.1
B4 66326281 66326365
refGene_gn=KIAA1551;refGene_tx=XM_023256904.1;refGene_gn=KIAA1551;refGene_tx=XM_023256905.1;refGene_gn=KIAA1551;refGene_tx=XM_023256906.1
B4 66328828 66328995
refGene_gn=KIAA1551;refGene_tx=XM_023256904.1;refGene_gn=KIAA1551;refGene_tx=XM_023256905.1;refGene_gn=KIAA1551;refGene_tx=XM_023256906.1
B4 66452263 66452551
refGene_gn=LOC101093947;refGene_tx=XM_019835914.2
B4 66452608 66452779
refGene_gn=LOC101093947;refGene_tx=XM_019835914.2
B4 66471501 66471714
refGene_gn=BICD1;refGene_tx=XM_023256907.1;refGene_gn=BICD1;refGene_tx=XM_023256908.1;refGene_gn=BICD1;refGene_tx=XM_023256909.1;refGene_gn=BICD1;refGene_tx=XM_023256910.1;refGene_gn=BICD1;refGene_tx=XM_023256911.1;refGene_gn=BICD1;refGene_tx=XM_023256912.1;refGene_gn=BICD1;refGene_tx=XM_023256913.1
B4 66589521 66589734
refGene_gn=BICD1;refGene_tx=XM_023256907.1;refGene_gn=BICD1;refGene_tx=XM_023256908.1;refGene_gn=BICD1;refGene_tx=XM_023256909.1;refGene_gn=BICD1;refGene_tx=XM_023256910.1;refGene_gn=BICD1;refGene_tx=XM_023256911.1;refGene_gn=BICD1;refGene_tx=XM_023256912.1;refGene_gn=BICD1;refGene_tx=XM_023256913.1
B4 66665639 66665792
refGene_gn=BICD1;refGene_tx=XM_023256907.1;refGene_gn=BICD1;refGene_tx=XM_023256908.1;refGene_gn=BICD1;refGene_tx=XM_023256909.1;refGene_gn=BICD1;refGene_tx=XM_023256910.1;refGene_gn=BICD1;refGene_tx=XM_023256911.1;refGene_gn=BICD1;refGene_tx=XM_023256912.1;refGene_gn=BICD1;refGene_tx=XM_023256913.1
B4 66673768 66674194
refGene_gn=BICD1;refGene_tx=XM_023256907.1;refGene_gn=BICD1;refGene_tx=XM_023256908.1;refGene_gn=BICD1;refGene_tx=XM_023256909.1;refGene_gn=BICD1;refGene_tx=XM_023256910.1;refGene_gn=BICD1;refGene_tx=XM_023256911.1;refGene_gn=BICD1;refGene_tx=XM_023256912.1;refGene_gn=BICD1;refGene_tx=XM_023256913.1
B4 66686737 66687832
refGene_gn=BICD1;refGene_tx=XM_023256907.1;refGene_gn=BICD1;refGene_tx=XM_023256908.1;refGene_gn=BICD1;refGene_tx=XM_023256909.1;refGene_gn=BICD1;refGene_tx=XM_023256910.1;refGene_gn=BICD1;refGene_tx=XM_023256911.1;refGene_gn=BICD1;refGene_tx=XM_023256912.1;refGene_gn=BICD1;refGe

ne_tx=XM_023256913.1
B4 66692470 66692622
refGene_gn=BICD1;refGene_tx=XM_023256907.1;refGene_gn=BICD1;refGene_tx=XM_023256908.1;refGene_gn=BICD1;refGene_tx=XM_023256909.1;refGene_gn=BICD1;refGene_tx=XM_023256910.1;refGene_gn=BICD1;refGene_tx=XM_023256911.1;refGene_gn=BICD1;refGene_tx=XM_023256912.1;refGene_gn=BICD1;refGene_tx=XM_023256913.1
B4 66695212 66695530
refGene_gn=BICD1;refGene_tx=XM_023256907.1;refGene_gn=BICD1;refGene_tx=XM_023256908.1;refGene_gn=BICD1;refGene_tx=XM_023256909.1;refGene_gn=BICD1;refGene_tx=XM_023256910.1;refGene_gn=BICD1;refGene_tx=XM_023256911.1;refGene_gn=BICD1;refGene_tx=XM_023256912.1;refGene_gn=BICD1;refGene_tx=XM_023256913.1
B4 66696499 66696693
refGene_gn=BICD1;refGene_tx=XM_023256907.1;refGene_gn=BICD1;refGene_tx=XM_023256908.1;refGene_gn=BICD1;refGene_tx=XM_023256912.1
B4 66709792 66709801
refGene_gn=BICD1;refGene_tx=XM_023256907.1;refGene_gn=BICD1;refGene_tx=XM_023256910.1
B4 66720698 66720784
refGene_gn=BICD1;refGene_tx=XM_023256907.1;refGene_gn=BICD1;refGene_tx=XM_023256908.1;refGene_gn=BICD1;refGene_tx=XM_023256909.1;refGene_gn=BICD1;refGene_tx=XM_023256910.1;refGene_gn=BICD1;refGene_tx=XM_023256911.1
B4 66726318 66726406
refGene_gn=BICD1;refGene_tx=XM_023256907.1;refGene_gn=BICD1;refGene_tx=XM_023256908.1;refGene_gn=BICD1;refGene_tx=XM_023256909.1;refGene_gn=BICD1;refGene_tx=XM_023256913.1
B4 66746589 66746755
refGene_gn=FGD4;refGene_tx=XM_019834793.2;refGene_gn=FGD4;refGene_tx=XM_023256914.1
B4 66819188 66819198 refGene_gn=FGD4;refGene_tx=XM_019834798.2
B4 66842023 66842129 refGene_gn=FGD4;refGene_tx=XM_019834795.2
B4 66863174 66863265 refGene_gn=FGD4;refGene_tx=XM_011283867.3
B4 66867988 66868070 refGene_gn=FGD4;refGene_tx=XM_019834797.2
B4 66891001 66891154
refGene_gn=FGD4;refGene_tx=XM_011283867.3;refGene_gn=FGD4;refGene_tx=XM_019834793.2;refGene_gn=FGD4;refGene_tx=XM_019834795.2;refGene_gn=FGD4;refGene_tx=XM_019834797.2;refGene_gn=FGD4;refGene_tx=XM_019834798.2;refGene_gn=FGD4;refGene_tx=XM_023256914.1
B4 66900005 66900189
refGene_gn=FGD4;refGene_tx=XM_011283867.3;refGene_gn=FGD4;refGene_tx=XM_019834793.2;refGene_gn=FGD4;refGene_tx=XM_019834795.2;refGene_gn=FGD4;refGene_tx=XM_019834797.2;refGene_gn=FGD4;refGene_tx=XM_019834798.2;refGene_gn=FGD4;refGene_tx=XM_019834799.2;refGene_gn=FGD4;refGene_tx=XM_019834800.2;refGene_gn=FGD4;refGene_tx=XM_023256914.1
B4 66905293 66905804
refGene_gn=FGD4;refGene_tx=XM_011283867.3;refGene_gn=FGD4;refGene_tx=XM_019834793.2;refGene_gn=FGD4;refGene_tx=XM_019834795.2;refGene_gn=FGD4;refGene_tx=XM_019834797.2;refGene_gn=FGD4;refGene_tx=XM_019834798.2;

refGene_gn=FGD4;refGene_tx=XM_019834799.2;refGene_gn=FGD4;refGene_tx=XM_019834800.2;refGene_gn=FGD4;refGene_tx=XM_023256914.1
B4 66908235 66908265 refGene_gn=FGD4;refGene_tx=XM_019834802.2
B4 66922475 66922565
refGene_gn=FGD4;refGene_tx=XM_011283867.3;refGene_gn=FGD4;refGene_tx=XM_019834793.2;refGene_gn=FGD4;refGene_tx=XM_019834795.2;refGene_gn=FGD4;refGene_tx=XM_019834797.2;refGene_gn=FGD4;refGene_tx=XM_019834798.2;refGene_gn=FGD4;refGene_tx=XM_019834799.2;refGene_gn=FGD4;refGene_tx=XM_019834800.2;refGene_gn=FGD4;refGene_tx=XM_019834802.2;refGene_gn=FGD4;refGene_tx=XM_023256914.1
B4 66924545 66924691
refGene_gn=FGD4;refGene_tx=XM_011283867.3;refGene_gn=FGD4;refGene_tx=XM_019834793.2;refGene_gn=FGD4;refGene_tx=XM_019834795.2;refGene_gn=FGD4;refGene_tx=XM_019834797.2;refGene_gn=FGD4;refGene_tx=XM_019834798.2;refGene_gn=FGD4;refGene_tx=XM_019834799.2;refGene_gn=FGD4;refGene_tx=XM_019834800.2;refGene_gn=FGD4;refGene_tx=XM_019834802.2;refGene_gn=FGD4;refGene_tx=XM_019834803.2;refGene_gn=FGD4;refGene_tx=XM_023256914.1
B4 66925326 66925483
refGene_gn=FGD4;refGene_tx=XM_011283867.3;refGene_gn=FGD4;refGene_tx=XM_019834793.2;refGene_gn=FGD4;refGene_tx=XM_019834795.2;refGene_gn=FGD4;refGene_tx=XM_019834797.2;refGene_gn=FGD4;refGene_tx=XM_019834798.2;refGene_gn=FGD4;refGene_tx=XM_019834799.2;refGene_gn=FGD4;refGene_tx=XM_019834800.2;refGene_gn=FGD4;refGene_tx=XM_019834802.2;refGene_gn=FGD4;refGene_tx=XM_019834803.2;refGene_gn=FGD4;refGene_tx=XM_023256914.1
B4 66931126 66931265
refGene_gn=FGD4;refGene_tx=XM_011283867.3;refGene_gn=FGD4;refGene_tx=XM_019834793.2;refGene_gn=FGD4;refGene_tx=XM_019834795.2;refGene_gn=FGD4;refGene_tx=XM_019834797.2;refGene_gn=FGD4;refGene_tx=XM_019834798.2;refGene_gn=FGD4;refGene_tx=XM_019834799.2;refGene_gn=FGD4;refGene_tx=XM_019834800.2;refGene_gn=FGD4;refGene_tx=XM_019834802.2;refGene_gn=FGD4;refGene_tx=XM_019834803.2;refGene_gn=FGD4;refGene_tx=XM_023256914.1
B4 66933548 66933607
refGene_gn=FGD4;refGene_tx=XM_011283867.3;refGene_gn=FGD4;refGene_tx=XM_019834793.2;refGene_gn=FGD4;refGene_tx=XM_019834795.2;refGene_gn=FGD4;refGene_tx=XM_019834797.2;refGene_gn=FGD4;refGene_tx=XM_019834798.2;refGene_gn=FGD4;refGene_tx=XM_019834799.2;refGene_gn=FGD4;refGene_tx=XM_019834800.2;refGene_gn=FGD4;refGene_tx=XM_019834802.2;refGene_gn=FGD4;refGene_tx=XM_019834803.2;refGene_gn=FGD4;refGene_tx=XM_023256914.1
B4 66933941 66934088
refGene_gn=FGD4;refGene_tx=XM_011283867.3;refGene_gn=FGD4;refGene_tx=XM_019834793.2;refGene_gn=FGD4;refGene_tx=XM_019834795.2;refGene_gn=FGD4;refGene_tx=XM_019834797.2;refGene_gn=FGD4;refGene_tx=XM_019834798.2;refGene_gn=FGD4;refGene_tx=XM_019834799.2;refGene_gn=FGD4;refGene_tx=XM_019834800.2;refGene_gn=FGD4;refGene_tx=XM_019834802.2;refGene_gn=FGD4;refGene_tx=XM_019834803.2;refGene_gn=FGD4;refGene_tx=XM_023256914.1
B4 66940310 66940483
refGene_gn=FGD4;refGene_tx=XM_011283867.3;refGene_gn=FGD4;refGene_tx=XM_019834793.2;refGene_gn=FGD4;refGene_tx=XM_019834795.2;refGene_gn=FGD4;refGene_tx=XM_019834797.2;refGene_gn=FGD4;refGene_tx=XM_019834798.2;refGene_gn=FGD4;refGene_tx=XM_019834799.2;refGene_gn=FGD4;refGene_tx=X

M_019834800.2;refGene_gn=FGD4;refGene_tx=XM_019834802.2;refGene_gn=FGD4;refGene_tx=XM_019834803.2;refGene_gn=FGD4;refGene_tx=XM_023256914.1
B4 66944465 66944496
refGene_gn=FGD4;refGene_tx=XM_011283867.3;refGene_gn=FGD4;refGene_tx=XM_019834793.2;refGene_gn=FGD4;refGene_tx=XM_019834795.2;refGene_gn=FGD4;refGene_tx=XM_019834797.2;refGene_gn=FGD4;refGene_tx=XM_019834798.2;refGene_gn=FGD4;refGene_tx=XM_019834799.2;refGene_gn=FGD4;refGene_tx=XM_019834800.2;refGene_gn=FGD4;refGene_tx=XM_019834802.2;refGene_gn=FGD4;refGene_tx=XM_019834803.2;refGene_gn=FGD4;refGene_tx=XM_023256914.1
B4 66944717 66944810
refGene_gn=FGD4;refGene_tx=XM_011283867.3;refGene_gn=FGD4;refGene_tx=XM_019834793.2;refGene_gn=FGD4;refGene_tx=XM_019834795.2;refGene_gn=FGD4;refGene_tx=XM_019834797.2;refGene_gn=FGD4;refGene_tx=XM_019834798.2;refGene_gn=FGD4;refGene_tx=XM_019834799.2;refGene_gn=FGD4;refGene_tx=XM_019834800.2;refGene_gn=FGD4;refGene_tx=XM_019834802.2;refGene_gn=FGD4;refGene_tx=XM_019834803.2;refGene_gn=FGD4;refGene_tx=XM_023256914.1
B4 66945080 66945206
refGene_gn=FGD4;refGene_tx=XM_011283867.3;refGene_gn=FGD4;refGene_tx=XM_019834793.2;refGene_gn=FGD4;refGene_tx=XM_019834795.2;refGene_gn=FGD4;refGene_tx=XM_019834797.2;refGene_gn=FGD4;refGene_tx=XM_019834798.2;refGene_gn=FGD4;refGene_tx=XM_019834799.2;refGene_gn=FGD4;refGene_tx=XM_019834800.2;refGene_gn=FGD4;refGene_tx=XM_019834802.2;refGene_gn=FGD4;refGene_tx=XM_019834803.2;refGene_gn=FGD4;refGene_tx=XM_023256914.1
B4 66950234 66950375
refGene_gn=FGD4;refGene_tx=XM_011283867.3;refGene_gn=FGD4;refGene_tx=XM_019834793.2;refGene_gn=FGD4;refGene_tx=XM_019834795.2;refGene_gn=FGD4;refGene_tx=XM_019834797.2;refGene_gn=FGD4;refGene_tx=XM_019834798.2;refGene_gn=FGD4;refGene_tx=XM_019834799.2;refGene_gn=FGD4;refGene_tx=XM_019834800.2;refGene_gn=FGD4;refGene_tx=XM_019834802.2;refGene_gn=FGD4;refGene_tx=XM_019834803.2;refGene_gn=FGD4;refGene_tx=XM_023256914.1
B4 66957697 66957838
refGene_gn=FGD4;refGene_tx=XM_011283867.3;refGene_gn=FGD4;refGene_tx=XM_019834793.2;refGene_gn=FGD4;refGene_tx=XM_019834795.2;refGene_gn=FGD4;refGene_tx=XM_019834797.2;refGene_gn=FGD4;refGene_tx=XM_019834798.2;refGene_gn=FGD4;refGene_tx=XM_019834799.2;refGene_gn=FGD4;refGene_tx=XM_019834800.2;refGene_gn=FGD4;refGene_tx=XM_019834802.2;refGene_gn=FGD4;refGene_tx=XM_019834803.2;refGene_gn=FGD4;refGene_tx=XM_023256914.1
B4 66959019 66959277
refGene_gn=FGD4;refGene_tx=XM_011283867.3;refGene_gn=FGD4;refGene_tx=XM_019834793.2;refGene_gn=FGD4;refGene_tx=XM_019834795.2;refGene_gn=FGD4;refGene_tx=XM_019834797.2;refGene_gn=FGD4;refGene_tx=XM_019834798.2;refGene_gn=FGD4;refGene_tx=XM_019834799.2;refGene_gn=FGD4;refGene_tx=XM_019834800.2;refGene_gn=FGD4;refGene_tx=XM_019834802.2;refGene_gn=FGD4;refGene_tx=XM_019834803.2;refGene_gn=FGD4;refGene_tx=XM_023256914.1
B4 66990276 66990287 refGene_gn=FGD4;refGene_tx=XM_023256914.1
B4 67007351 67007568
refGene_gn=DNM1L;refGene_tx=XM_023256915.1;refGene_gn=DNM1L;refGene_tx=XM_023256916.1;refGene_gn=DNM1L;refGene_tx=XM_023256917.1;refGene_gn=DNM1L;refGene_tx=XM_023256918.1;refGene_gn=DNM1L;refGene_tx=XM_023256919.1;refGene_gn=DNM1L;refGene_tx=XM_023256920.1;refGene_gn=DNM1L;refGe

ne_tx=XM_023256921.1;refGene_gn=DNM1L;refGene_tx=XM_023256922.1;refGene_gn=DNM1L;refGene_tx=XM_023256923.1
B4 67025524 67025672
refGene_gn=DNM1L;refGene_tx=XM_023256915.1;refGene_gn=DNM1L;refGene_tx=XM_023256916.1;refGene_gn=DNM1L;refGene_tx=XM_023256917.1;refGene_gn=DNM1L;refGene_tx=XM_023256918.1;refGene_gn=DNM1L;refGene_tx=XM_023256919.1;refGene_gn=DNM1L;refGene_tx=XM_023256920.1;refGene_gn=DNM1L;refGene_tx=XM_023256921.1;refGene_gn=DNM1L;refGene_tx=XM_023256922.1
B4 67029015 67029054
refGene_gn=DNM1L;refGene_tx=XM_023256915.1;refGene_gn=DNM1L;refGene_tx=XM_023256916.1;refGene_gn=DNM1L;refGene_tx=XM_023256919.1;refGene_gn=DNM1L;refGene_tx=XM_023256920.1
B4 67030528 67030575
refGene_gn=DNM1L;refGene_tx=XM_023256915.1;refGene_gn=DNM1L;refGene_tx=XM_023256916.1;refGene_gn=DNM1L;refGene_tx=XM_023256917.1;refGene_gn=DNM1L;refGene_tx=XM_023256918.1;refGene_gn=DNM1L;refGene_tx=XM_023256919.1;refGene_gn=DNM1L;refGene_tx=XM_023256920.1;refGene_gn=DNM1L;refGene_tx=XM_023256921.1;refGene_gn=DNM1L;refGene_tx=XM_023256922.1;refGene_gn=DNM1L;refGene_tx=XM_023256923.1
B4 67030988 67031060
refGene_gn=DNM1L;refGene_tx=XM_023256915.1;refGene_gn=DNM1L;refGene_tx=XM_023256916.1;refGene_gn=DNM1L;refGene_tx=XM_023256917.1;refGene_gn=DNM1L;refGene_tx=XM_023256918.1;refGene_gn=DNM1L;refGene_tx=XM_023256919.1;refGene_gn=DNM1L;refGene_tx=XM_023256920.1;refGene_gn=DNM1L;refGene_tx=XM_023256921.1;refGene_gn=DNM1L;refGene_tx=XM_023256922.1;refGene_gn=DNM1L;refGene_tx=XM_023256923.1
B4 67033627 67033714
refGene_gn=DNM1L;refGene_tx=XM_023256915.1;refGene_gn=DNM1L;refGene_tx=XM_023256916.1;refGene_gn=DNM1L;refGene_tx=XM_023256917.1;refGene_gn=DNM1L;refGene_tx=XM_023256918.1;refGene_gn=DNM1L;refGene_tx=XM_023256919.1;refGene_gn=DNM1L;refGene_tx=XM_023256920.1;refGene_gn=DNM1L;refGene_tx=XM_023256921.1;refGene_gn=DNM1L;refGene_tx=XM_023256922.1;refGene_gn=DNM1L;refGene_tx=XM_023256923.1
B4 67035189 67035352
refGene_gn=DNM1L;refGene_tx=XM_023256915.1;refGene_gn=DNM1L;refGene_tx=XM_023256916.1;refGene_gn=DNM1L;refGene_tx=XM_023256917.1;refGene_gn=DNM1L;refGene_tx=XM_023256918.1;refGene_gn=DNM1L;refGene_tx=XM_023256919.1;refGene_gn=DNM1L;refGene_tx=XM_023256920.1;refGene_gn=DNM1L;refGene_tx=XM_023256921.1;refGene_gn=DNM1L;refGene_tx=XM_023256922.1;refGene_gn=DNM1L;refGene_tx=XM_023256923.1
B4 67038733 67038854
refGene_gn=DNM1L;refGene_tx=XM_023256915.1;refGene_gn=DNM1L;refGene_tx=XM_023256916.1;refGene_gn=DNM1L;refGene_tx=XM_023256917.1;refGene_gn=DNM1L;refGene_tx=XM_023256918.1;refGene_gn=DNM1L;refGene_tx=XM_023256919.1;refGene_gn=DNM1L;refGene_tx=XM_023256920.1;refGene_gn=DNM1L;refGene_tx=XM_023256921.1;refGene_gn=DNM1L;refGene_tx=XM_023256922.1;refGene_gn=DNM1L;refGene_tx=XM_023256923.1
B4 67040585 67040717
refGene_gn=DNM1L;refGene_tx=XM_023256915.1;refGene_gn=DNM1L;refGene_tx=XM_023256916.1;refGene_gn=DNM1L;refGene_tx=XM_023256917.1;refGene_gn=

DNM1L;refGene_tx=XM_023256918.1;refGene_gn=DNM1L;refGene_tx=XM_023256919.1;refGene_gn=DNM1L;refGene_tx=XM_023256920.1;refGene_gn=DNM1L;refGene_tx=XM_023256921.1;refGene_gn=DNM1L;refGene_tx=XM_023256922.1;refGene_gn=DNM1L;refGene_tx=XM_023256923.1
B4 67041618 67041825
refGene_gn=DNM1L;refGene_tx=XM_023256915.1;refGene_gn=DNM1L;refGene_tx=XM_023256916.1;refGene_gn=DNM1L;refGene_tx=XM_023256917.1;refGene_gn=DNM1L;refGene_tx=XM_023256918.1;refGene_gn=DNM1L;refGene_tx=XM_023256919.1;refGene_gn=DNM1L;refGene_tx=XM_023256920.1;refGene_gn=DNM1L;refGene_tx=XM_023256921.1;refGene_gn=DNM1L;refGene_tx=XM_023256922.1;refGene_gn=DNM1L;refGene_tx=XM_023256923.1
B4 67044931 67045052
refGene_gn=DNM1L;refGene_tx=XM_023256915.1;refGene_gn=DNM1L;refGene_tx=XM_023256916.1;refGene_gn=DNM1L;refGene_tx=XM_023256917.1;refGene_gn=DNM1L;refGene_tx=XM_023256918.1;refGene_gn=DNM1L;refGene_tx=XM_023256919.1;refGene_gn=DNM1L;refGene_tx=XM_023256920.1;refGene_gn=DNM1L;refGene_tx=XM_023256921.1;refGene_gn=DNM1L;refGene_tx=XM_023256922.1;refGene_gn=DNM1L;refGene_tx=XM_023256923.1
B4 67045285 67045441
refGene_gn=DNM1L;refGene_tx=XM_023256915.1;refGene_gn=DNM1L;refGene_tx=XM_023256916.1;refGene_gn=DNM1L;refGene_tx=XM_023256917.1;refGene_gn=DNM1L;refGene_tx=XM_023256918.1;refGene_gn=DNM1L;refGene_tx=XM_023256919.1;refGene_gn=DNM1L;refGene_tx=XM_023256920.1;refGene_gn=DNM1L;refGene_tx=XM_023256921.1;refGene_gn=DNM1L;refGene_tx=XM_023256922.1;refGene_gn=DNM1L;refGene_tx=XM_023256923.1
B4 67045682 67045772
refGene_gn=DNM1L;refGene_tx=XM_023256915.1;refGene_gn=DNM1L;refGene_tx=XM_023256916.1;refGene_gn=DNM1L;refGene_tx=XM_023256917.1;refGene_gn=DNM1L;refGene_tx=XM_023256918.1;refGene_gn=DNM1L;refGene_tx=XM_023256919.1;refGene_gn=DNM1L;refGene_tx=XM_023256920.1;refGene_gn=DNM1L;refGene_tx=XM_023256921.1;refGene_gn=DNM1L;refGene_tx=XM_023256922.1;refGene_gn=DNM1L;refGene_tx=XM_023256923.1
B4 67047360 67047453
refGene_gn=DNM1L;refGene_tx=XM_023256915.1;refGene_gn=DNM1L;refGene_tx=XM_023256916.1;refGene_gn=DNM1L;refGene_tx=XM_023256917.1;refGene_gn=DNM1L;refGene_tx=XM_023256918.1;refGene_gn=DNM1L;refGene_tx=XM_023256919.1;refGene_gn=DNM1L;refGene_tx=XM_023256920.1;refGene_gn=DNM1L;refGene_tx=XM_023256921.1;refGene_gn=DNM1L;refGene_tx=XM_023256922.1;refGene_gn=DNM1L;refGene_tx=XM_023256923.1
B4 67050048 67050105
refGene_gn=DNM1L;refGene_tx=XM_023256915.1;refGene_gn=DNM1L;refGene_tx=XM_023256916.1;refGene_gn=DNM1L;refGene_tx=XM_023256917.1;refGene_gn=DNM1L;refGene_tx=XM_023256918.1;refGene_gn=DNM1L;refGene_tx=XM_023256919.1;refGene_gn=DNM1L;refGene_tx=XM_023256920.1;refGene_gn=DNM1L;refGene_tx=XM_023256921.1;refGene_gn=DNM1L;refGene_tx=XM_023256922.1;refGene_gn=DNM1L;refGene_tx=XM_023256923.1
B4 67051008 67051086
refGene_gn=DNM1L;refGene_tx=XM_023256915.1;refGene_gn=DNM1L;refGene_tx=XM_023256916.1;refGene_gn=DNM1L;refGene_tx=XM_023256917.1;refGene_gn=DNM1L;refGene_tx=XM_023256918.1;refGene_gn=DNM1L;refGene_tx=XM_0232569

23.1
B4	67051399 67051432
refGene_gn=DNM1L;refGene_tx=XM_023256915.1;refGene_gn=DNM1L;refGene_tx=XM_023256917.1;refGene_gn=DNM1L;refGene_tx=XM_023256919.1;refGene_gn=DNM1L;refGene_tx=XM_023256921.1;refGene_gn=DNM1L;refGene_tx=XM_023256923.1
B4	67053212 67053389
refGene_gn=DNM1L;refGene_tx=XM_023256915.1;refGene_gn=DNM1L;refGene_tx=XM_023256916.1;refGene_gn=DNM1L;refGene_tx=XM_023256917.1;refGene_gn=DNM1L;refGene_tx=XM_023256918.1;refGene_gn=DNM1L;refGene_tx=XM_023256919.1;refGene_gn=DNM1L;refGene_tx=XM_023256920.1;refGene_gn=DNM1L;refGene_tx=XM_023256921.1;refGene_gn=DNM1L;refGene_tx=XM_023256922.1;refGene_gn=DNM1L;refGene_tx=XM_023256923.1
B4	67053728 67053838
refGene_gn=DNM1L;refGene_tx=XM_023256915.1;refGene_gn=DNM1L;refGene_tx=XM_023256916.1;refGene_gn=DNM1L;refGene_tx=XM_023256917.1;refGene_gn=DNM1L;refGene_tx=XM_023256918.1;refGene_gn=DNM1L;refGene_tx=XM_023256919.1;refGene_gn=DNM1L;refGene_tx=XM_023256920.1;refGene_gn=DNM1L;refGene_tx=XM_023256921.1;refGene_gn=DNM1L;refGene_tx=XM_023256922.1;refGene_gn=DNM1L;refGene_tx=XM_023256923.1
B4	67053972 67054132
refGene_gn=DNM1L;refGene_tx=XM_023256915.1;refGene_gn=DNM1L;refGene_tx=XM_023256916.1;refGene_gn=DNM1L;refGene_tx=XM_023256917.1;refGene_gn=DNM1L;refGene_tx=XM_023256918.1;refGene_gn=DNM1L;refGene_tx=XM_023256919.1;refGene_gn=DNM1L;refGene_tx=XM_023256920.1;refGene_gn=DNM1L;refGene_tx=XM_023256921.1;refGene_gn=DNM1L;refGene_tx=XM_023256922.1;refGene_gn=DNM1L;refGene_tx=XM_023256923.1
B4	67054380 67054437
refGene_gn=DNM1L;refGene_tx=XM_023256915.1;refGene_gn=DNM1L;refGene_tx=XM_023256916.1;refGene_gn=DNM1L;refGene_tx=XM_023256917.1;refGene_gn=DNM1L;refGene_tx=XM_023256918.1;refGene_gn=DNM1L;refGene_tx=XM_023256919.1;refGene_gn=DNM1L;refGene_tx=XM_023256920.1;refGene_gn=DNM1L;refGene_tx=XM_023256921.1;refGene_gn=DNM1L;refGene_tx=XM_023256922.1;refGene_gn=DNM1L;refGene_tx=XM_023256923.1
B4	67057482 67057642 refGene_gn=YARS2;refGene_tx=XM_003988547.5
B4	67059161 67059332 refGene_gn=YARS2;refGene_tx=XM_003988547.5
B4	67059631 67059787 refGene_gn=YARS2;refGene_tx=XM_003988547.5
B4	67075474 67075642 refGene_gn=YARS2;refGene_tx=XM_003988547.5
B4	67076439 67077218 refGene_gn=YARS2;refGene_tx=XM_003988547.5
B4	67117712 67117810 refGene_gn=PKP2;refGene_tx=XM_023256924.1
B4	67118299 67118352 refGene_gn=PKP2;refGene_tx=XM_023256924.1
B4	67121651 67121841 refGene_gn=PKP2;refGene_tx=XM_023256924.1
B4	67128871 67129025 refGene_gn=PKP2;refGene_tx=XM_023256924.1
B4	67135855 67136029 refGene_gn=PKP2;refGene_tx=XM_023256924.1
B4	67136935 67137100 refGene_gn=PKP2;refGene_tx=XM_023256924.1
B4	67137962 67138080 refGene_gn=PKP2;refGene_tx=XM_023256924.1
B4	67156008 67156186 refGene_gn=PKP2;refGene_tx=XM_023256924.1
B4	67165905 67166113 refGene_gn=PKP2;refGene_tx=XM_023256924.1
B4	67182781 67182917 refGene_gn=PKP2;refGene_tx=XM_023256924.1
B4	67195215 67195916 refGene_gn=PKP2;refGene_tx=XM_023256924.1

B4 67196637 67196750 refGene_gn=PKP2;refGene_tx=XM_023256924.1
B4 67212274 67212497 refGene_gn=PKP2;refGene_tx=XM_023256924.1
B4 67567832 67567983
refGene_gn=SYT10;refGene_tx=XM_023256925.1;refGene_gn=SYT10;refGene_tx=XM_023256926.1
B4 67570988 67571118 refGene_gn=SYT10;refGene_tx=XM_023256925.1
B4 67573634 67573806
refGene_gn=SYT10;refGene_tx=XM_023256925.1;refGene_gn=SYT10;refGene_tx=XM_023256926.1
B4 67577147 67577268
refGene_gn=SYT10;refGene_tx=XM_023256925.1;refGene_gn=SYT10;refGene_tx=XM_023256926.1
B4 67598703 67599271
refGene_gn=SYT10;refGene_tx=XM_023256925.1;refGene_gn=SYT10;refGene_tx=XM_023256926.1
B4 67612783 67613141
refGene_gn=SYT10;refGene_tx=XM_023256925.1;refGene_gn=SYT10;refGene_tx=XM_023256926.1
B4 67631720 67631871
refGene_gn=SYT10;refGene_tx=XM_023256925.1;refGene_gn=SYT10;refGene_tx=XM_023256926.1
B4 68146872 68147055 refGene_gn=ALG10;refGene_tx=XM_003988550.5
B4 68148313 68148511 refGene_gn=ALG10;refGene_tx=XM_003988550.5
B4 68150190 68151246 refGene_gn=ALG10;refGene_tx=XM_003988550.5
B4 68449360 68449549 refGene_gn=CPNE8;refGene_tx=XM_003988551.5
B4 68463962 68464036 refGene_gn=CPNE8;refGene_tx=XM_003988551.5
B4 68468845 68468903 refGene_gn=CPNE8;refGene_tx=XM_003988551.5
B4 68470653 68470756 refGene_gn=CPNE8;refGene_tx=XM_003988551.5
B4 68479223 68479227 refGene_gn=CPNE8;refGene_tx=XM_019834813.1
B4 68487383 68487511
refGene_gn=CPNE8;refGene_tx=XM_003988551.5;refGene_gn=CPNE8;refGene_tx=XM_019834813.1
B4 68494291 68494473
refGene_gn=CPNE8;refGene_tx=XM_003988551.5;refGene_gn=CPNE8;refGene_tx=XM_019834813.1
B4 68500984 68501031
refGene_gn=CPNE8;refGene_tx=XM_003988551.5;refGene_gn=CPNE8;refGene_tx=XM_019834813.1
B4 68525027 68525089
refGene_gn=CPNE8;refGene_tx=XM_003988551.5;refGene_gn=CPNE8;refGene_tx=XM_019834813.1
B4 68526439 68526493
refGene_gn=CPNE8;refGene_tx=XM_003988551.5;refGene_gn=CPNE8;refGene_tx=XM_019834813.1
B4 68529597 68529673
refGene_gn=CPNE8;refGene_tx=XM_003988551.5;refGene_gn=CPNE8;refGene_tx=XM_019834813.1
B4 68555325 68555367
refGene_gn=CPNE8;refGene_tx=XM_003988551.5;refGene_gn=CPNE8;refGene_tx=XM_019834813.1

B4 68556864 68556969
refGene_gn=CPNE8;refGene_tx=XM_003988551.5;refGene_gn=CPNE8;refGene_tx=XM_019834813.1
B4 68562208 68562312
refGene_gn=CPNE8;refGene_tx=XM_003988551.5;refGene_gn=CPNE8;refGene_tx=XM_019834813.1
B4 68569377 68569441
refGene_gn=CPNE8;refGene_tx=XM_003988551.5;refGene_gn=CPNE8;refGene_tx=XM_019834813.1
B4 68595083 68595160
refGene_gn=CPNE8;refGene_tx=XM_003988551.5;refGene_gn=CPNE8;refGene_tx=XM_019834813.1
B4 68610110 68610150
refGene_gn=CPNE8;refGene_tx=XM_003988551.5;refGene_gn=CPNE8;refGene_tx=XM_019834813.1
B4 68621800 68621904
refGene_gn=CPNE8;refGene_tx=XM_003988551.5;refGene_gn=CPNE8;refGene_tx=XM_019834813.1
B4 68646290 68646337
refGene_gn=CPNE8;refGene_tx=XM_003988551.5;refGene_gn=CPNE8;refGene_tx=XM_019834813.1
B4 68648331 68648372
refGene_gn=CPNE8;refGene_tx=XM_003988551.5;refGene_gn=CPNE8;refGene_tx=XM_019834813.1
B4 68679260 68679397
refGene_gn=CPNE8;refGene_tx=XM_003988551.5;refGene_gn=CPNE8;refGene_tx=XM_019834813.1
B4 69014756 69014850
refGene_gn=KIF21A;refGene_tx=XM_023256927.1;refGene_gn=KIF21A;refGene_tx=XM_023256928.1;refGene_gn=KIF21A;refGene_tx=XM_023256929.1;refGene_gn=KIF21A;refGene_tx=XM_023256930.1;refGene_gn=KIF21A;refGene_tx=XM_023256932.1;refGene_gn=KIF21A;refGene_tx=XM_023256933.1;refGene_gn=KIF21A;refGene_tx=XM_023256934.1;refGene_gn=KIF21A;refGene_tx=XM_023256935.1;refGene_gn=KIF21A;refGene_tx=XM_023256936.1;refGene_gn=KIF21A;refGene_tx=XM_023256937.1;refGene_gn=KIF21A;refGene_tx=XM_023256938.1;refGene_gn=KIF21A;refGene_tx=XM_023256939.1;refGene_gn=KIF21A;refGene_tx=XM_023256940.1
B4 69019894 69020094
refGene_gn=KIF21A;refGene_tx=XM_023256927.1;refGene_gn=KIF21A;refGene_tx=XM_023256928.1;refGene_gn=KIF21A;refGene_tx=XM_023256929.1;refGene_gn=KIF21A;refGene_tx=XM_023256930.1;refGene_gn=KIF21A;refGene_tx=XM_023256932.1;refGene_gn=KIF21A;refGene_tx=XM_023256933.1;refGene_gn=KIF21A;refGene_tx=XM_023256934.1;refGene_gn=KIF21A;refGene_tx=XM_023256935.1;refGene_gn=KIF21A;refGene_tx=XM_023256936.1;refGene_gn=KIF21A;refGene_tx=XM_023256937.1;refGene_gn=KIF21A;refGene_tx=XM_023256938.1;refGene_gn=KIF21A;refGene_tx=XM_023256939.1;refGene_gn=KIF21A;refGene_tx=XM_023256940.1
B4 69021374 69021545
refGene_gn=KIF21A;refGene_tx=XM_023256927.1;refGene_gn=KIF21A;refGene_tx=XM_023256928.1;refGene_gn=KIF21A;refGene_tx=XM_023256929.1;refGene_

gn=KIF21A;refGene_tx=XM_023256930.1;refGene_gn=KIF21A;refGene_tx=XM_02
3256932.1;refGene_gn=KIF21A;refGene_tx=XM_023256933.1;refGene_gn=KIF21
A;refGene_tx=XM_023256934.1;refGene_gn=KIF21A;refGene_tx=XM_023256935.
1;refGene_gn=KIF21A;refGene_tx=XM_023256936.1;refGene_gn=KIF21A;refGen
e_tx=XM_023256937.1;refGene_gn=KIF21A;refGene_tx=XM_023256938.1;refGen
e_gn=KIF21A;refGene_tx=XM_023256939.1;refGene_gn=KIF21A;refGene_tx=XM_
023256940.1
B4 69022961 69023079
refGene_gn=KIF21A;refGene_tx=XM_023256927.1;refGene_gn=KIF21A;refGene_
tx=XM_023256928.1;refGene_gn=KIF21A;refGene_tx=XM_023256929.1;refGene_
gn=KIF21A;refGene_tx=XM_023256930.1;refGene_gn=KIF21A;refGene_tx=XM_02
3256932.1;refGene_gn=KIF21A;refGene_tx=XM_023256933.1;refGene_gn=KIF21
A;refGene_tx=XM_023256934.1;refGene_gn=KIF21A;refGene_tx=XM_023256935.
1;refGene_gn=KIF21A;refGene_tx=XM_023256936.1;refGene_gn=KIF21A;refGen
e_tx=XM_023256937.1;refGene_gn=KIF21A;refGene_tx=XM_023256938.1;refGen
e_gn=KIF21A;refGene_tx=XM_023256939.1;refGene_gn=KIF21A;refGene_tx=XM_
023256940.1
B4 69025363 69025528
refGene_gn=KIF21A;refGene_tx=XM_023256927.1;refGene_gn=KIF21A;refGene_
tx=XM_023256928.1;refGene_gn=KIF21A;refGene_tx=XM_023256929.1;refGene_
gn=KIF21A;refGene_tx=XM_023256930.1;refGene_gn=KIF21A;refGene_tx=XM_02
3256932.1;refGene_gn=KIF21A;refGene_tx=XM_023256933.1;refGene_gn=KIF21
A;refGene_tx=XM_023256934.1;refGene_gn=KIF21A;refGene_tx=XM_023256935.
1;refGene_gn=KIF21A;refGene_tx=XM_023256936.1;refGene_gn=KIF21A;refGen
e_tx=XM_023256937.1;refGene_gn=KIF21A;refGene_tx=XM_023256938.1;refGen
e_gn=KIF21A;refGene_tx=XM_023256939.1;refGene_gn=KIF21A;refGene_tx=XM_
023256940.1
B4 69027489 69027670
refGene_gn=KIF21A;refGene_tx=XM_023256927.1;refGene_gn=KIF21A;refGene_
tx=XM_023256928.1;refGene_gn=KIF21A;refGene_tx=XM_023256929.1;refGene_
gn=KIF21A;refGene_tx=XM_023256930.1;refGene_gn=KIF21A;refGene_tx=XM_02
3256932.1;refGene_gn=KIF21A;refGene_tx=XM_023256933.1;refGene_gn=KIF21
A;refGene_tx=XM_023256934.1;refGene_gn=KIF21A;refGene_tx=XM_023256935.
1;refGene_gn=KIF21A;refGene_tx=XM_023256936.1;refGene_gn=KIF21A;refGen
e_tx=XM_023256937.1;refGene_gn=KIF21A;refGene_tx=XM_023256938.1;refGen
e_gn=KIF21A;refGene_tx=XM_023256939.1;refGene_gn=KIF21A;refGene_tx=XM_
023256940.1
B4 69029610 69029747
refGene_gn=KIF21A;refGene_tx=XM_023256927.1;refGene_gn=KIF21A;refGene_
tx=XM_023256928.1;refGene_gn=KIF21A;refGene_tx=XM_023256929.1;refGene_
gn=KIF21A;refGene_tx=XM_023256930.1;refGene_gn=KIF21A;refGene_tx=XM_02
3256932.1;refGene_gn=KIF21A;refGene_tx=XM_023256933.1;refGene_gn=KIF21
A;refGene_tx=XM_023256934.1;refGene_gn=KIF21A;refGene_tx=XM_023256935.
1;refGene_gn=KIF21A;refGene_tx=XM_023256936.1;refGene_gn=KIF21A;refGen
e_tx=XM_023256937.1;refGene_gn=KIF21A;refGene_tx=XM_023256938.1;refGen
e_gn=KIF21A;refGene_tx=XM_023256939.1;refGene_gn=KIF21A;refGene_tx=XM_
023256940.1
B4 69033149 69033164
refGene_gn=KIF21A;refGene_tx=XM_023256927.1;refGene_gn=KIF21A;refGene_
tx=XM_023256928.1;refGene_gn=KIF21A;refGene_tx=XM_023256929.1;refGene_

gn=KIF21A;refGene_tx=XM_023256930.1;refGene_gn=KIF21A;refGene_tx=XM_023256932.1;refGene_gn=KIF21A;refGene_tx=XM_023256934.1;refGene_gn=KIF21A;refGene_tx=XM_023256935.1;refGene_gn=KIF21A;refGene_tx=XM_023256937.1;refGene_gn=KIF21A;refGene_tx=XM_023256938.1
B4 69033853 69033892
refGene_gn=KIF21A;refGene_tx=XM_023256927.1;refGene_gn=KIF21A;refGene_tx=XM_023256928.1;refGene_gn=KIF21A;refGene_tx=XM_023256929.1;refGene_gn=KIF21A;refGene_tx=XM_023256930.1;refGene_gn=KIF21A;refGene_tx=XM_023256932.1;refGene_gn=KIF21A;refGene_tx=XM_023256934.1;refGene_gn=KIF21A;refGene_tx=XM_023256935.1;refGene_gn=KIF21A;refGene_tx=XM_023256937.1;refGene_gn=KIF21A;refGene_tx=XM_023256938.1
B4 69036228 69036357
refGene_gn=KIF21A;refGene_tx=XM_023256927.1;refGene_gn=KIF21A;refGene_tx=XM_023256928.1;refGene_gn=KIF21A;refGene_tx=XM_023256929.1;refGene_gn=KIF21A;refGene_tx=XM_023256930.1;refGene_gn=KIF21A;refGene_tx=XM_023256932.1;refGene_gn=KIF21A;refGene_tx=XM_023256933.1;refGene_gn=KIF21A;refGene_tx=XM_023256934.1;refGene_gn=KIF21A;refGene_tx=XM_023256935.1;refGene_gn=KIF21A;refGene_tx=XM_023256936.1;refGene_gn=KIF21A;refGene_tx=XM_023256940.1
B4 69037746 69037851
refGene_gn=KIF21A;refGene_tx=XM_023256927.1;refGene_gn=KIF21A;refGene_tx=XM_023256928.1;refGene_gn=KIF21A;refGene_tx=XM_023256929.1;refGene_gn=KIF21A;refGene_tx=XM_023256930.1;refGene_gn=KIF21A;refGene_tx=XM_023256932.1;refGene_gn=KIF21A;refGene_tx=XM_023256933.1;refGene_gn=KIF21A;refGene_tx=XM_023256934.1;refGene_gn=KIF21A;refGene_tx=XM_023256935.1;refGene_gn=KIF21A;refGene_tx=XM_023256936.1;refGene_gn=KIF21A;refGene_tx=XM_023256937.1;refGene_gn=KIF21A;refGene_tx=XM_023256938.1;refGene_gn=KIF21A;refGene_tx=XM_023256939.1;refGene_gn=KIF21A;refGene_tx=XM_023256940.1
B4 69040407 69040622
refGene_gn=KIF21A;refGene_tx=XM_023256927.1;refGene_gn=KIF21A;refGene_tx=XM_023256928.1;refGene_gn=KIF21A;refGene_tx=XM_023256929.1;refGene_gn=KIF21A;refGene_tx=XM_023256930.1;refGene_gn=KIF21A;refGene_tx=XM_023256932.1;refGene_gn=KIF21A;refGene_tx=XM_023256933.1;refGene_gn=KIF21A;refGene_tx=XM_023256934.1;refGene_gn=KIF21A;refGene_tx=XM_023256935.1;refGene_gn=KIF21A;refGene_tx=XM_023256936.1;refGene_gn=KIF21A;refGene_tx=XM_023256937.1;refGene_gn=KIF21A;refGene_tx=XM_023256938.1;refGene_gn=KIF21A;refGene_tx=XM_023256939.1;refGene_gn=KIF21A;refGene_tx=XM_023256940.1
B4 69043310 69043365
refGene_gn=KIF21A;refGene_tx=XM_023256927.1;refGene_gn=KIF21A;refGene_tx=XM_023256928.1;refGene_gn=KIF21A;refGene_tx=XM_023256929.1;refGene_gn=KIF21A;refGene_tx=XM_023256930.1;refGene_gn=KIF21A;refGene_tx=XM_023256932.1;refGene_gn=KIF21A;refGene_tx=XM_023256933.1;refGene_gn=KIF21A;refGene_tx=XM_023256934.1;refGene_gn=KIF21A;refGene_tx=XM_023256935.1;refGene_gn=KIF21A;refGene_tx=XM_023256936.1;refGene_gn=KIF21A;refGene_tx=XM_023256937.1;refGene_gn=KIF21A;refGene_tx=XM_023256938.1;refGene_gn=KIF21A;refGene_tx=XM_023256939.1;refGene_gn=KIF21A;refGene_tx=XM_023256940.1
B4 69043719 69043780

refGene_gn=KIF21A;refGene_tx=XM_023256927.1;refGene_gn=KIF21A;refGene_
tx=XM_023256928.1;refGene_gn=KIF21A;refGene_tx=XM_023256929.1;refGene_
gn=KIF21A;refGene_tx=XM_023256930.1;refGene_gn=KIF21A;refGene_tx=XM_02
3256932.1;refGene_gn=KIF21A;refGene_tx=XM_023256933.1;refGene_gn=KIF21
A;refGene_tx=XM_023256934.1;refGene_gn=KIF21A;refGene_tx=XM_023256935.
1;refGene_gn=KIF21A;refGene_tx=XM_023256936.1;refGene_gn=KIF21A;refGen
e_tx=XM_023256937.1;refGene_gn=KIF21A;refGene_tx=XM_023256938.1;refGen
e_gn=KIF21A;refGene_tx=XM_023256939.1;refGene_gn=KIF21A;refGene_tx=XM_
023256940.1
B4 69048490 69048511
refGene_gn=KIF21A;refGene_tx=XM_023256927.1;refGene_gn=KIF21A;refGene_
tx=XM_023256928.1;refGene_gn=KIF21A;refGene_tx=XM_023256930.1;refGene_
gn=KIF21A;refGene_tx=XM_023256932.1;refGene_gn=KIF21A;refGene_tx=XM_02
3256933.1;refGene_gn=KIF21A;refGene_tx=XM_023256937.1;refGene_gn=KIF21
A;refGene_tx=XM_023256938.1;refGene_gn=KIF21A;refGene_tx=XM_023256939.
1;refGene_gn=KIF21A;refGene_tx=XM_023256940.1
B4 69048986 69049152
refGene_gn=KIF21A;refGene_tx=XM_023256927.1;refGene_gn=KIF21A;refGene_
tx=XM_023256928.1;refGene_gn=KIF21A;refGene_tx=XM_023256929.1;refGene_
gn=KIF21A;refGene_tx=XM_023256930.1;refGene_gn=KIF21A;refGene_tx=XM_02
3256932.1;refGene_gn=KIF21A;refGene_tx=XM_023256933.1;refGene_gn=KIF21
A;refGene_tx=XM_023256934.1;refGene_gn=KIF21A;refGene_tx=XM_023256935.
1;refGene_gn=KIF21A;refGene_tx=XM_023256936.1;refGene_gn=KIF21A;refGen
e_tx=XM_023256937.1;refGene_gn=KIF21A;refGene_tx=XM_023256938.1;refGen
e_gn=KIF21A;refGene_tx=XM_023256939.1;refGene_gn=KIF21A;refGene_tx=XM_
023256940.1
B4 69049950 69050052
refGene_gn=KIF21A;refGene_tx=XM_023256927.1;refGene_gn=KIF21A;refGene_
tx=XM_023256928.1;refGene_gn=KIF21A;refGene_tx=XM_023256929.1;refGene_
gn=KIF21A;refGene_tx=XM_023256930.1;refGene_gn=KIF21A;refGene_tx=XM_02
3256932.1;refGene_gn=KIF21A;refGene_tx=XM_023256933.1;refGene_gn=KIF21
A;refGene_tx=XM_023256934.1;refGene_gn=KIF21A;refGene_tx=XM_023256935.
1;refGene_gn=KIF21A;refGene_tx=XM_023256936.1;refGene_gn=KIF21A;refGen
e_tx=XM_023256937.1;refGene_gn=KIF21A;refGene_tx=XM_023256938.1;refGen
e_gn=KIF21A;refGene_tx=XM_023256939.1;refGene_gn=KIF21A;refGene_tx=XM_
023256940.1
B4 69050722 69050917
refGene_gn=KIF21A;refGene_tx=XM_023256927.1;refGene_gn=KIF21A;refGene_
tx=XM_023256928.1;refGene_gn=KIF21A;refGene_tx=XM_023256929.1;refGene_
gn=KIF21A;refGene_tx=XM_023256930.1;refGene_gn=KIF21A;refGene_tx=XM_02
3256932.1;refGene_gn=KIF21A;refGene_tx=XM_023256933.1;refGene_gn=KIF21
A;refGene_tx=XM_023256934.1;refGene_gn=KIF21A;refGene_tx=XM_023256935.
1;refGene_gn=KIF21A;refGene_tx=XM_023256936.1;refGene_gn=KIF21A;refGen
e_tx=XM_023256937.1;refGene_gn=KIF21A;refGene_tx=XM_023256938.1;refGen
e_gn=KIF21A;refGene_tx=XM_023256939.1;refGene_gn=KIF21A;refGene_tx=XM_
023256940.1
B4 69051448 69051602
refGene_gn=KIF21A;refGene_tx=XM_023256927.1;refGene_gn=KIF21A;refGene_
tx=XM_023256928.1;refGene_gn=KIF21A;refGene_tx=XM_023256929.1;refGene_
gn=KIF21A;refGene_tx=XM_023256930.1;refGene_gn=KIF21A;refGene_tx=XM_02

3256932.1;refGene_gn=KIF21A;refGene_tx=XM_023256933.1;refGene_gn=KIF21A;refGene_tx=XM_023256934.1;refGene_gn=KIF21A;refGene_tx=XM_023256935.1;refGene_gn=KIF21A;refGene_tx=XM_023256936.1;refGene_gn=KIF21A;refGene_tx=XM_023256937.1;refGene_gn=KIF21A;refGene_tx=XM_023256938.1;refGene_gn=KIF21A;refGene_tx=XM_023256939.1;refGene_gn=KIF21A;refGene_tx=XM_023256940.1
B4 69051756 69051971
refGene_gn=KIF21A;refGene_tx=XM_023256927.1;refGene_gn=KIF21A;refGene_tx=XM_023256928.1;refGene_gn=KIF21A;refGene_tx=XM_023256929.1;refGene_gn=KIF21A;refGene_tx=XM_023256930.1;refGene_gn=KIF21A;refGene_tx=XM_023256932.1;refGene_gn=KIF21A;refGene_tx=XM_023256933.1;refGene_gn=KIF21A;refGene_tx=XM_023256934.1;refGene_gn=KIF21A;refGene_tx=XM_023256935.1;refGene_gn=KIF21A;refGene_tx=XM_023256936.1;refGene_gn=KIF21A;refGene_tx=XM_023256937.1;refGene_gn=KIF21A;refGene_tx=XM_023256938.1;refGene_gn=KIF21A;refGene_tx=XM_023256939.1;refGene_gn=KIF21A;refGene_tx=XM_023256940.1
B4 69052065 69052134
refGene_gn=KIF21A;refGene_tx=XM_023256927.1;refGene_gn=KIF21A;refGene_tx=XM_023256928.1;refGene_gn=KIF21A;refGene_tx=XM_023256929.1;refGene_gn=KIF21A;refGene_tx=XM_023256930.1;refGene_gn=KIF21A;refGene_tx=XM_023256932.1;refGene_gn=KIF21A;refGene_tx=XM_023256933.1;refGene_gn=KIF21A;refGene_tx=XM_023256934.1;refGene_gn=KIF21A;refGene_tx=XM_023256935.1;refGene_gn=KIF21A;refGene_tx=XM_023256936.1;refGene_gn=KIF21A;refGene_tx=XM_023256937.1;refGene_gn=KIF21A;refGene_tx=XM_023256938.1;refGene_gn=KIF21A;refGene_tx=XM_023256939.1;refGene_gn=KIF21A;refGene_tx=XM_023256940.1
B4 69055653 69055761
refGene_gn=KIF21A;refGene_tx=XM_023256927.1;refGene_gn=KIF21A;refGene_tx=XM_023256928.1;refGene_gn=KIF21A;refGene_tx=XM_023256929.1;refGene_gn=KIF21A;refGene_tx=XM_023256930.1;refGene_gn=KIF21A;refGene_tx=XM_023256932.1;refGene_gn=KIF21A;refGene_tx=XM_023256933.1;refGene_gn=KIF21A;refGene_tx=XM_023256934.1;refGene_gn=KIF21A;refGene_tx=XM_023256935.1;refGene_gn=KIF21A;refGene_tx=XM_023256936.1;refGene_gn=KIF21A;refGene_tx=XM_023256937.1;refGene_gn=KIF21A;refGene_tx=XM_023256938.1;refGene_gn=KIF21A;refGene_tx=XM_023256939.1;refGene_gn=KIF21A;refGene_tx=XM_023256940.1
B4 69056817 69057017
refGene_gn=KIF21A;refGene_tx=XM_023256927.1;refGene_gn=KIF21A;refGene_tx=XM_023256928.1;refGene_gn=KIF21A;refGene_tx=XM_023256929.1;refGene_gn=KIF21A;refGene_tx=XM_023256930.1;refGene_gn=KIF21A;refGene_tx=XM_023256932.1;refGene_gn=KIF21A;refGene_tx=XM_023256933.1;refGene_gn=KIF21A;refGene_tx=XM_023256934.1;refGene_gn=KIF21A;refGene_tx=XM_023256935.1;refGene_gn=KIF21A;refGene_tx=XM_023256936.1;refGene_gn=KIF21A;refGene_tx=XM_023256937.1;refGene_gn=KIF21A;refGene_tx=XM_023256938.1;refGene_gn=KIF21A;refGene_tx=XM_023256939.1;refGene_gn=KIF21A;refGene_tx=XM_023256940.1
B4 69057440 69057629
refGene_gn=KIF21A;refGene_tx=XM_023256927.1;refGene_gn=KIF21A;refGene_tx=XM_023256928.1;refGene_gn=KIF21A;refGene_tx=XM_023256929.1;refGene_gn=KIF21A;refGene_tx=XM_023256930.1;refGene_gn=KIF21A;refGene_tx=XM_02

3256932.1;refGene_gn=KIF21A;refGene_tx=XM_023256933.1;refGene_gn=KIF21A;refGene_tx=XM_023256934.1;refGene_gn=KIF21A;refGene_tx=XM_023256935.1;refGene_gn=KIF21A;refGene_tx=XM_023256936.1;refGene_gn=KIF21A;refGene_tx=XM_023256937.1;refGene_gn=KIF21A;refGene_tx=XM_023256938.1;refGene_gn=KIF21A;refGene_tx=XM_023256939.1;refGene_gn=KIF21A;refGene_tx=XM_023256940.1
B4 69058023 69058141
refGene_gn=KIF21A;refGene_tx=XM_023256927.1;refGene_gn=KIF21A;refGene_tx=XM_023256928.1;refGene_gn=KIF21A;refGene_tx=XM_023256929.1;refGene_gn=KIF21A;refGene_tx=XM_023256930.1;refGene_gn=KIF21A;refGene_tx=XM_023256932.1;refGene_gn=KIF21A;refGene_tx=XM_023256933.1;refGene_gn=KIF21A;refGene_tx=XM_023256934.1;refGene_gn=KIF21A;refGene_tx=XM_023256935.1;refGene_gn=KIF21A;refGene_tx=XM_023256936.1;refGene_gn=KIF21A;refGene_tx=XM_023256937.1;refGene_gn=KIF21A;refGene_tx=XM_023256938.1;refGene_gn=KIF21A;refGene_tx=XM_023256939.1;refGene_gn=KIF21A;refGene_tx=XM_023256940.1
B4 69058551 69058642
refGene_gn=KIF21A;refGene_tx=XM_023256927.1;refGene_gn=KIF21A;refGene_tx=XM_023256928.1;refGene_gn=KIF21A;refGene_tx=XM_023256929.1;refGene_gn=KIF21A;refGene_tx=XM_023256930.1;refGene_gn=KIF21A;refGene_tx=XM_023256932.1;refGene_gn=KIF21A;refGene_tx=XM_023256933.1;refGene_gn=KIF21A;refGene_tx=XM_023256934.1;refGene_gn=KIF21A;refGene_tx=XM_023256935.1;refGene_gn=KIF21A;refGene_tx=XM_023256936.1;refGene_gn=KIF21A;refGene_tx=XM_023256937.1;refGene_gn=KIF21A;refGene_tx=XM_023256938.1;refGene_gn=KIF21A;refGene_tx=XM_023256939.1;refGene_gn=KIF21A;refGene_tx=XM_023256940.1
B4 69062715 69062754
refGene_gn=KIF21A;refGene_tx=XM_023256927.1;refGene_gn=KIF21A;refGene_tx=XM_023256928.1;refGene_gn=KIF21A;refGene_tx=XM_023256929.1;refGene_gn=KIF21A;refGene_tx=XM_023256933.1;refGene_gn=KIF21A;refGene_tx=XM_023256937.1;refGene_gn=KIF21A;refGene_tx=XM_023256938.1;refGene_gn=KIF21A;refGene_tx=XM_023256939.1
B4 69068316 69068520
refGene_gn=KIF21A;refGene_tx=XM_023256927.1;refGene_gn=KIF21A;refGene_tx=XM_023256928.1;refGene_gn=KIF21A;refGene_tx=XM_023256929.1;refGene_gn=KIF21A;refGene_tx=XM_023256930.1;refGene_gn=KIF21A;refGene_tx=XM_023256932.1;refGene_gn=KIF21A;refGene_tx=XM_023256933.1;refGene_gn=KIF21A;refGene_tx=XM_023256934.1;refGene_gn=KIF21A;refGene_tx=XM_023256935.1;refGene_gn=KIF21A;refGene_tx=XM_023256936.1;refGene_gn=KIF21A;refGene_tx=XM_023256937.1;refGene_gn=KIF21A;refGene_tx=XM_023256938.1;refGene_gn=KIF21A;refGene_tx=XM_023256939.1;refGene_gn=KIF21A;refGene_tx=XM_023256940.1
B4 69073546 69073610
refGene_gn=KIF21A;refGene_tx=XM_023256927.1;refGene_gn=KIF21A;refGene_tx=XM_023256928.1;refGene_gn=KIF21A;refGene_tx=XM_023256929.1;refGene_gn=KIF21A;refGene_tx=XM_023256930.1;refGene_gn=KIF21A;refGene_tx=XM_023256932.1;refGene_gn=KIF21A;refGene_tx=XM_023256933.1;refGene_gn=KIF21A;refGene_tx=XM_023256934.1;refGene_gn=KIF21A;refGene_tx=XM_023256935.1;refGene_gn=KIF21A;refGene_tx=XM_023256936.1;refGene_gn=KIF21A;refGene_tx=XM_023256937.1;refGene_gn=KIF21A;refGene_tx=XM_023256938.1;refGen

e_gn=KIF21A;refGene_tx=XM_023256939.1;refGene_gn=KIF21A;refGene_tx=XM_023256940.1
B4 69073982 69074172
refGene_gn=KIF21A;refGene_tx=XM_023256927.1;refGene_gn=KIF21A;refGene_tx=XM_023256928.1;refGene_gn=KIF21A;refGene_tx=XM_023256929.1;refGene_gn=KIF21A;refGene_tx=XM_023256930.1;refGene_gn=KIF21A;refGene_tx=XM_023256932.1;refGene_gn=KIF21A;refGene_tx=XM_023256933.1;refGene_gn=KIF21A;refGene_tx=XM_023256934.1;refGene_gn=KIF21A;refGene_tx=XM_023256935.1;refGene_gn=KIF21A;refGene_tx=XM_023256936.1;refGene_gn=KIF21A;refGene_tx=XM_023256937.1;refGene_gn=KIF21A;refGene_tx=XM_023256938.1;refGene_gn=KIF21A;refGene_tx=XM_023256939.1;refGene_gn=KIF21A;refGene_tx=XM_023256940.1
B4 69074568 69074764
refGene_gn=KIF21A;refGene_tx=XM_023256927.1;refGene_gn=KIF21A;refGene_tx=XM_023256928.1;refGene_gn=KIF21A;refGene_tx=XM_023256929.1;refGene_gn=KIF21A;refGene_tx=XM_023256930.1;refGene_gn=KIF21A;refGene_tx=XM_023256932.1;refGene_gn=KIF21A;refGene_tx=XM_023256933.1;refGene_gn=KIF21A;refGene_tx=XM_023256934.1;refGene_gn=KIF21A;refGene_tx=XM_023256935.1;refGene_gn=KIF21A;refGene_tx=XM_023256936.1;refGene_gn=KIF21A;refGene_tx=XM_023256937.1;refGene_gn=KIF21A;refGene_tx=XM_023256938.1;refGene_gn=KIF21A;refGene_tx=XM_023256939.1;refGene_gn=KIF21A;refGene_tx=XM_023256940.1
B4 69079017 69079133
refGene_gn=KIF21A;refGene_tx=XM_023256927.1;refGene_gn=KIF21A;refGene_tx=XM_023256928.1;refGene_gn=KIF21A;refGene_tx=XM_023256929.1;refGene_gn=KIF21A;refGene_tx=XM_023256930.1;refGene_gn=KIF21A;refGene_tx=XM_023256932.1;refGene_gn=KIF21A;refGene_tx=XM_023256933.1;refGene_gn=KIF21A;refGene_tx=XM_023256934.1;refGene_gn=KIF21A;refGene_tx=XM_023256935.1;refGene_gn=KIF21A;refGene_tx=XM_023256936.1;refGene_gn=KIF21A;refGene_tx=XM_023256937.1;refGene_gn=KIF21A;refGene_tx=XM_023256938.1;refGene_gn=KIF21A;refGene_tx=XM_023256939.1;refGene_gn=KIF21A;refGene_tx=XM_023256940.1
B4 69081484 69081652
refGene_gn=KIF21A;refGene_tx=XM_023256927.1;refGene_gn=KIF21A;refGene_tx=XM_023256928.1;refGene_gn=KIF21A;refGene_tx=XM_023256929.1;refGene_gn=KIF21A;refGene_tx=XM_023256930.1;refGene_gn=KIF21A;refGene_tx=XM_023256932.1;refGene_gn=KIF21A;refGene_tx=XM_023256933.1;refGene_gn=KIF21A;refGene_tx=XM_023256934.1;refGene_gn=KIF21A;refGene_tx=XM_023256935.1;refGene_gn=KIF21A;refGene_tx=XM_023256936.1;refGene_gn=KIF21A;refGene_tx=XM_023256937.1;refGene_gn=KIF21A;refGene_tx=XM_023256938.1;refGene_gn=KIF21A;refGene_tx=XM_023256939.1;refGene_gn=KIF21A;refGene_tx=XM_023256940.1
B4 69082708 69082843
refGene_gn=KIF21A;refGene_tx=XM_023256927.1;refGene_gn=KIF21A;refGene_tx=XM_023256928.1;refGene_gn=KIF21A;refGene_tx=XM_023256929.1;refGene_gn=KIF21A;refGene_tx=XM_023256930.1;refGene_gn=KIF21A;refGene_tx=XM_023256932.1;refGene_gn=KIF21A;refGene_tx=XM_023256933.1;refGene_gn=KIF21A;refGene_tx=XM_023256934.1;refGene_gn=KIF21A;refGene_tx=XM_023256935.1;refGene_gn=KIF21A;refGene_tx=XM_023256936.1;refGene_gn=KIF21A;refGene_tx=XM_023256937.1;refGene_gn=KIF21A;refGene_tx=XM_023256938.1;refGen

e_gn=KIF21A;refGene_tx=XM_023256939.1;refGene_gn=KIF21A;refGene_tx=XM_023256940.1
B4 69083538 69083688 refGene_gn=KIF21A;refGene_tx=XM_023256927.1;refGene_gn=KIF21A;refGene_tx=XM_023256928.1;refGene_gn=KIF21A;refGene_tx=XM_023256929.1;refGene_gn=KIF21A;refGene_tx=XM_023256930.1;refGene_gn=KIF21A;refGene_tx=XM_023256932.1;refGene_gn=KIF21A;refGene_tx=XM_023256933.1;refGene_gn=KIF21A;refGene_tx=XM_023256934.1;refGene_gn=KIF21A;refGene_tx=XM_023256935.1;refGene_gn=KIF21A;refGene_tx=XM_023256936.1;refGene_gn=KIF21A;refGene_tx=XM_023256937.1;refGene_gn=KIF21A;refGene_tx=XM_023256938.1;refGene_gn=KIF21A;refGene_tx=XM_023256939.1;refGene_gn=KIF21A;refGene_tx=XM_023256940.1
B4 69085072 69085255 refGene_gn=KIF21A;refGene_tx=XM_023256927.1;refGene_gn=KIF21A;refGene_tx=XM_023256928.1;refGene_gn=KIF21A;refGene_tx=XM_023256929.1;refGene_gn=KIF21A;refGene_tx=XM_023256930.1;refGene_gn=KIF21A;refGene_tx=XM_023256932.1;refGene_gn=KIF21A;refGene_tx=XM_023256933.1;refGene_gn=KIF21A;refGene_tx=XM_023256934.1;refGene_gn=KIF21A;refGene_tx=XM_023256935.1;refGene_gn=KIF21A;refGene_tx=XM_023256936.1;refGene_gn=KIF21A;refGene_tx=XM_023256937.1;refGene_gn=KIF21A;refGene_tx=XM_023256938.1;refGene_gn=KIF21A;refGene_tx=XM_023256939.1;refGene_gn=KIF21A;refGene_tx=XM_023256940.1
B4 69085389 69085612 refGene_gn=KIF21A;refGene_tx=XM_023256927.1;refGene_gn=KIF21A;refGene_tx=XM_023256928.1;refGene_gn=KIF21A;refGene_tx=XM_023256929.1;refGene_gn=KIF21A;refGene_tx=XM_023256930.1;refGene_gn=KIF21A;refGene_tx=XM_023256932.1;refGene_gn=KIF21A;refGene_tx=XM_023256933.1;refGene_gn=KIF21A;refGene_tx=XM_023256934.1;refGene_gn=KIF21A;refGene_tx=XM_023256935.1;refGene_gn=KIF21A;refGene_tx=XM_023256936.1;refGene_gn=KIF21A;refGene_tx=XM_023256937.1;refGene_gn=KIF21A;refGene_tx=XM_023256938.1;refGene_gn=KIF21A;refGene_tx=XM_023256939.1;refGene_gn=KIF21A;refGene_tx=XM_023256940.1
B4 69165577 69165621 refGene_gn=KIF21A;refGene_tx=XM_023256927.1;refGene_gn=KIF21A;refGene_tx=XM_023256928.1;refGene_gn=KIF21A;refGene_tx=XM_023256929.1;refGene_gn=KIF21A;refGene_tx=XM_023256930.1;refGene_gn=KIF21A;refGene_tx=XM_023256932.1;refGene_gn=KIF21A;refGene_tx=XM_023256933.1;refGene_gn=KIF21A;refGene_tx=XM_023256934.1;refGene_gn=KIF21A;refGene_tx=XM_023256935.1;refGene_gn=KIF21A;refGene_tx=XM_023256936.1;refGene_gn=KIF21A;refGene_tx=XM_023256937.1;refGene_gn=KIF21A;refGene_tx=XM_023256938.1;refGene_gn=KIF21A;refGene_tx=XM_023256939.1;refGene_gn=KIF21A;refGene_tx=XM_023256940.1
B4 69261728 69261948 refGene_gn=ABCD2;refGene_tx=XM_011283869.3
B4 69267506 69267594 refGene_gn=ABCD2;refGene_tx=XM_011283869.3
B4 69267596 69267631 refGene_gn=ABCD2;refGene_tx=XM_011283869.3
B4 69272004 69272089 refGene_gn=ABCD2;refGene_tx=XM_011283869.3
B4 69279730 69279876 refGene_gn=ABCD2;refGene_tx=XM_011283869.3
B4 69303549 69303695 refGene_gn=ABCD2;refGene_tx=XM_011283869.3
B4 69307983 69308078 refGene_gn=ABCD2;refGene_tx=XM_011283869.3
B4 69308718 69308887 refGene_gn=ABCD2;refGene_tx=XM_011283869.3

B4 69312168 69312290 refGene_gn=ABCD2;refGene_tx=XM_011283869.3
B4 69323151 69323332 refGene_gn=ABCD2;refGene_tx=XM_011283869.3
B4 69324905 69325844 refGene_gn=ABCD2;refGene_tx=XM_011283869.3
B4 69369509 69369615
refGene_gn=CB4H12orf40;refGene_tx=XM_023256942.1
B4 69374102 69374235
refGene_gn=CB4H12orf40;refGene_tx=XM_023256942.1
B4 69374638 69374672
refGene_gn=CB4H12orf40;refGene_tx=XM_023256942.1
B4 69376024 69376213
refGene_gn=CB4H12orf40;refGene_tx=XM_023256942.1;refGene_gn=CB4H12orf4
0;refGene_tx=XM_023256943.1
B4 69378816 69378932
refGene_gn=CB4H12orf40;refGene_tx=XM_023256942.1;refGene_gn=CB4H12orf4
0;refGene_tx=XM_023256943.1
B4 69391912 69392430
refGene_gn=CB4H12orf40;refGene_tx=XM_023256942.1;refGene_gn=CB4H12orf4
0;refGene_tx=XM_023256943.1
B4 69392735 69392805
refGene_gn=CB4H12orf40;refGene_tx=XM_023256942.1;refGene_gn=CB4H12orf4
0;refGene_tx=XM_023256943.1
B4 69394121 69394182
refGene_gn=CB4H12orf40;refGene_tx=XM_023256942.1;refGene_gn=CB4H12orf4
0;refGene_tx=XM_023256943.1
B4 69408378 69408439
refGene_gn=CB4H12orf40;refGene_tx=XM_023256942.1;refGene_gn=CB4H12orf4
0;refGene_tx=XM_023256943.1
B4 69419224 69419307
refGene_gn=CB4H12orf40;refGene_tx=XM_023256942.1;refGene_gn=CB4H12orf4
0;refGene_tx=XM_023256943.1
B4 69423378 69423741
refGene_gn=CB4H12orf40;refGene_tx=XM_023256942.1;refGene_gn=CB4H12orf4
0;refGene_tx=XM_023256943.1
B4 69463986 69464213 refGene_gn=SLC2A13;refGene_tx=XM_023256944.1
B4 69476281 69476434 refGene_gn=SLC2A13;refGene_tx=XM_023256944.1
B4 69476547 69476669 refGene_gn=SLC2A13;refGene_tx=XM_023256944.1
B4 69550079 69550205 refGene_gn=SLC2A13;refGene_tx=XM_023256944.1
B4 69595667 69595788 refGene_gn=SLC2A13;refGene_tx=XM_023256944.1
B4 69603202 69603366 refGene_gn=SLC2A13;refGene_tx=XM_023256944.1
B4 69683569 69683678 refGene_gn=SLC2A13;refGene_tx=XM_023256944.1
B4 69761653 69761862 refGene_gn=SLC2A13;refGene_tx=XM_023256944.1
B4 69789032 69789192 refGene_gn=SLC2A13;refGene_tx=XM_023256944.1
B4 69844714 69845273 refGene_gn=SLC2A13;refGene_tx=XM_023256944.1
B4 69965118 69965272 refGene_gn=LRRK2;refGene_tx=XM_023256946.1
B4 69965527 69965613 refGene_gn=LRRK2;refGene_tx=XM_023256946.1
B4 69970454 69970564 refGene_gn=LRRK2;refGene_tx=XM_023256946.1
B4 69972814 69972903 refGene_gn=LRRK2;refGene_tx=XM_023256946.1
B4 69975901 69976036 refGene_gn=LRRK2;refGene_tx=XM_023256946.1
B4 69978577 69978712 refGene_gn=LRRK2;refGene_tx=XM_023256946.1
B4 69982084 69982216 refGene_gn=LRRK2;refGene_tx=XM_023256946.1

| | | | |
|---|---|---|---|
| B4 | 69983962 | 69984082 | refGene_gn=LRRK2;refGene_tx=XM_023256946.1 |
| B4 | 69984781 | 69984921 | refGene_gn=LRRK2;refGene_tx=XM_023256946.1 |
| B4 | 69985014 | 69985094 | refGene_gn=LRRK2;refGene_tx=XM_023256946.1 |
| B4 | 69986658 | 69986765 | refGene_gn=LRRK2;refGene_tx=XM_023256946.1 |
| B4 | 69990995 | 69991125 | refGene_gn=LRRK2;refGene_tx=XM_023256946.1 |
| B4 | 69993305 | 69993430 | refGene_gn=LRRK2;refGene_tx=XM_023256946.1 |
| B4 | 69999685 | 69999798 | refGene_gn=LRRK2;refGene_tx=XM_023256946.1 |
| B4 | 70004947 | 70005092 | refGene_gn=LRRK2;refGene_tx=XM_023256946.1 |
| B4 | 70005657 | 70005797 | refGene_gn=LRRK2;refGene_tx=XM_023256946.1 |
| B4 | 70010230 | 70010359 | refGene_gn=LRRK2;refGene_tx=XM_023256946.1 |
| B4 | 70010434 | 70010605 | refGene_gn=LRRK2;refGene_tx=XM_023256946.1 |
| B4 | 70016211 | 70016470 | refGene_gn=LRRK2;refGene_tx=XM_023256946.1 |
| B4 | 70019922 | 70019951 | refGene_gn=LRRK2;refGene_tx=XM_023256946.1 |
| B4 | 70019953 | 70020113 | refGene_gn=LRRK2;refGene_tx=XM_023256946.1 |
| B4 | 70027233 | 70027352 | refGene_gn=LRRK2;refGene_tx=XM_023256946.1 |
| B4 | 70027829 | 70027899 | refGene_gn=LRRK2;refGene_tx=XM_023256946.1 |
| B4 | 70028411 | 70028629 | refGene_gn=LRRK2;refGene_tx=XM_023256946.1 |
| B4 | 70031134 | 70031385 | refGene_gn=LRRK2;refGene_tx=XM_023256946.1 |
| B4 | 70031676 | 70031825 | refGene_gn=LRRK2;refGene_tx=XM_023256946.1 |
| B4 | 70034770 | 70034864 | refGene_gn=LRRK2;refGene_tx=XM_023256946.1 |
| B4 | 70037428 | 70037615 | refGene_gn=LRRK2;refGene_tx=XM_023256946.1 |
| B4 | 70038934 | 70039116 | refGene_gn=LRRK2;refGene_tx=XM_023256946.1 |
| B4 | 70041922 | 70042152 | refGene_gn=LRRK2;refGene_tx=XM_023256946.1 |
| B4 | 70042525 | 70042653 | refGene_gn=LRRK2;refGene_tx=XM_023256946.1 |
| B4 | 70044337 | 70044556 | refGene_gn=LRRK2;refGene_tx=XM_023256946.1 |
| B4 | 70047933 | 70048135 | refGene_gn=LRRK2;refGene_tx=XM_023256946.1 |
| B4 | 70049172 | 70049261 | refGene_gn=LRRK2;refGene_tx=XM_023256946.1 |
| B4 | 70054278 | 70054466 | refGene_gn=LRRK2;refGene_tx=XM_023256946.1 |
| B4 | 70055313 | 70055468 | refGene_gn=LRRK2;refGene_tx=XM_023256946.1 |
| B4 | 70056575 | 70056722 | refGene_gn=LRRK2;refGene_tx=XM_023256946.1 |
| B4 | 70056864 | 70057056 | refGene_gn=LRRK2;refGene_tx=XM_023256946.1 |
| B4 | 70058518 | 70058665 | refGene_gn=LRRK2;refGene_tx=XM_023256946.1 |
| B4 | 70062314 | 70062415 | refGene_gn=LRRK2;refGene_tx=XM_023256946.1 |
| B4 | 70065197 | 70065388 | refGene_gn=LRRK2;refGene_tx=XM_023256946.1 |
| B4 | 70071088 | 70071249 | refGene_gn=LRRK2;refGene_tx=XM_023256946.1 |
| B4 | 70081878 | 70082049 | refGene_gn=LRRK2;refGene_tx=XM_023256946.1 |
| B4 | 70083284 | 70083385 | refGene_gn=LRRK2;refGene_tx=XM_023256946.1 |
| B4 | 70088940 | 70089135 | refGene_gn=LRRK2;refGene_tx=XM_023256946.1 |
| B4 | 70090186 | 70090380 | refGene_gn=LRRK2;refGene_tx=XM_023256946.1 |
| B4 | 70091902 | 70091975 | refGene_gn=LRRK2;refGene_tx=XM_023256946.1 |
| B4 | 70092760 | 70092945 | refGene_gn=LRRK2;refGene_tx=XM_023256946.1 |
| B4 | 70097693 | 70097846 | refGene_gn=LRRK2;refGene_tx=XM_023256946.1 |
| B4 | 70099513 | 70099722 | refGene_gn=LRRK2;refGene_tx=XM_023256946.1 |
| B4 | 70101802 | 70101874 | refGene_gn=LRRK2;refGene_tx=XM_023256946.1 |
| B4 | 70102464 | 70102586 | refGene_gn=LRRK2;refGene_tx=XM_023256946.1 |
| B4 | 70111470 | 70111817 | refGene_gn=MUC19;refGene_tx=XM_023257754.1 |
| B4 | 70117780 | 70117914 | refGene_gn=MUC19;refGene_tx=XM_023257754.1 |
| B4 | 70138535 | 70138556 | refGene_gn=MUC19;refGene_tx=XM_023257754.1 |
| B4 | 70138952 | 70139117 | refGene_gn=MUC19;refGene_tx=XM_023257754.1 |
| B4 | 70141440 | 70141623 | refGene_gn=MUC19;refGene_tx=XM_023257754.1 |

| | | | |
|---|---|---|---|
| B4 | 70141757 | 70141925 | refGene_gn=MUC19;refGene_tx=XM_023257754.1 |
| B4 | 70142378 | 70142528 | refGene_gn=MUC19;refGene_tx=XM_023257754.1 |
| B4 | 70142618 | 70142741 | refGene_gn=MUC19;refGene_tx=XM_023257754.1 |
| B4 | 70143616 | 70143799 | refGene_gn=MUC19;refGene_tx=XM_023257754.1 |
| B4 | 70143900 | 70143948 | refGene_gn=MUC19;refGene_tx=XM_023257754.1 |
| B4 | 70145800 | 70145851 | refGene_gn=MUC19;refGene_tx=XM_023257754.1 |
| B4 | 70146901 | 70146961 | refGene_gn=MUC19;refGene_tx=XM_023257754.1 |
| B4 | 70147589 | 70147818 | refGene_gn=MUC19;refGene_tx=XM_023257754.1 |
| B4 | 70148803 | 70148921 | refGene_gn=MUC19;refGene_tx=XM_023257754.1 |
| B4 | 70149371 | 70149432 | refGene_gn=MUC19;refGene_tx=XM_023257754.1 |
| B4 | 70149657 | 70149755 | refGene_gn=MUC19;refGene_tx=XM_023257754.1 |
| B4 | 70153389 | 70153600 | refGene_gn=MUC19;refGene_tx=XM_023257754.1 |
| B4 | 70154317 | 70154440 | refGene_gn=MUC19;refGene_tx=XM_023257754.1 |
| B4 | 70155111 | 70155235 | refGene_gn=MUC19;refGene_tx=XM_023257754.1 |
| B4 | 70156815 | 70156951 | refGene_gn=MUC19;refGene_tx=XM_023257754.1 |
| B4 | 70158194 | 70158287 | refGene_gn=MUC19;refGene_tx=XM_023257754.1 |
| B4 | 70159281 | 70159366 | refGene_gn=MUC19;refGene_tx=XM_023257754.1 |
| B4 | 70159532 | 70159670 | refGene_gn=MUC19;refGene_tx=XM_023257754.1 |
| B4 | 70160630 | 70160780 | refGene_gn=MUC19;refGene_tx=XM_023257754.1 |
| B4 | 70163685 | 70163780 | refGene_gn=MUC19;refGene_tx=XM_023257754.1 |
| B4 | 70164401 | 70164528 | refGene_gn=MUC19;refGene_tx=XM_023257754.1 |
| B4 | 70166298 | 70166545 | refGene_gn=MUC19;refGene_tx=XM_023257754.1 |
| B4 | 70168254 | 70168307 | refGene_gn=MUC19;refGene_tx=XM_023257754.1 |
| B4 | 70168404 | 70168505 | refGene_gn=MUC19;refGene_tx=XM_023257754.1 |
| B4 | 70168745 | 70168897 | refGene_gn=MUC19;refGene_tx=XM_023257754.1 |
| B4 | 70169615 | 70169754 | refGene_gn=MUC19;refGene_tx=XM_023257754.1 |
| B4 | 70170791 | 70170896 | refGene_gn=MUC19;refGene_tx=XM_023257754.1 |
| B4 | 70170990 | 70171185 | refGene_gn=MUC19;refGene_tx=XM_023257754.1 |
| B4 | 70172174 | 70172414 | refGene_gn=MUC19;refGene_tx=XM_023257754.1 |
| B4 | 70172484 | 70172641 | refGene_gn=MUC19;refGene_tx=XM_023257754.1 |
| B4 | 70173592 | 70173718 | refGene_gn=MUC19;refGene_tx=XM_023257754.1 |
| B4 | 70175303 | 70175445 | refGene_gn=MUC19;refGene_tx=XM_023257754.1 |
| B4 | 70176074 | 70176106 | refGene_gn=MUC19;refGene_tx=XM_023257754.1 |
| B4 | 70176656 | 70176686 | refGene_gn=MUC19;refGene_tx=XM_023257754.1 |
| B4 | 70177945 | 70177987 | refGene_gn=MUC19;refGene_tx=XM_023257754.1 |
| B4 | 70178532 | 70178565 | refGene_gn=MUC19;refGene_tx=XM_023257754.1 |
| B4 | 70178969 | 70179014 | refGene_gn=MUC19;refGene_tx=XM_023257754.1 |
| B4 | 70189209 | 70189248 | refGene_gn=MUC19;refGene_tx=XM_023257754.1 |
| B4 | 70190009 | 70190105 | refGene_gn=MUC19;refGene_tx=XM_023257754.1 |
| B4 | 70192093 | 70192189 | refGene_gn=MUC19;refGene_tx=XM_023257754.1 |
| B4 | 70192631 | 70192730 | refGene_gn=MUC19;refGene_tx=XM_023257754.1 |
| B4 | 70193970 | 70194036 | refGene_gn=MUC19;refGene_tx=XM_023257754.1 |
| B4 | 70202047 | 70215472 | refGene_gn=MUC19;refGene_tx=XM_023257754.1 |
| B4 | 70215557 | 70220681 | refGene_gn=MUC19;refGene_tx=XM_023257754.1 |
| B4 | 70221777 | 70221861 | refGene_gn=MUC19;refGene_tx=XM_023257754.1 |
| B4 | 70223677 | 70223713 | refGene_gn=MUC19;refGene_tx=XM_023257754.1 |
| B4 | 70225549 | 70225603 | refGene_gn=MUC19;refGene_tx=XM_023257754.1 |
| B4 | 70225748 | 70225802 | refGene_gn=MUC19;refGene_tx=XM_023257754.1 |
| B4 | 70226693 | 70226747 | refGene_gn=MUC19;refGene_tx=XM_023257754.1 |
| B4 | 70227439 | 70227493 | refGene_gn=MUC19;refGene_tx=XM_023257754.1 |

B4 70227878 70227932 refGene_gn=MUC19;refGene_tx=XM_023257754.1
B4 70228069 70228123 refGene_gn=MUC19;refGene_tx=XM_023257754.1
B4 70228550 70228595 refGene_gn=MUC19;refGene_tx=XM_023257754.1
B4 70228856 70228910 refGene_gn=MUC19;refGene_tx=XM_023257754.1
B4 70229032 70229083 refGene_gn=MUC19;refGene_tx=XM_023257754.1
B4 70230879 70230924 refGene_gn=MUC19;refGene_tx=XM_023257754.1
B4 70231358 70231403 refGene_gn=MUC19;refGene_tx=XM_023257754.1
B4 70232109 70232163 refGene_gn=MUC19;refGene_tx=XM_023257754.1
B4 70233209 70233263 refGene_gn=MUC19;refGene_tx=XM_023257754.1
B4 70234738 70234792 refGene_gn=MUC19;refGene_tx=XM_023257754.1
B4 70235203 70235239 refGene_gn=MUC19;refGene_tx=XM_023257754.1
B4 70235484 70235538 refGene_gn=MUC19;refGene_tx=XM_023257754.1
B4 70236301 70236349 refGene_gn=MUC19;refGene_tx=XM_023257754.1
B4 70236597 70236648 refGene_gn=MUC19;refGene_tx=XM_023257754.1
B4 70237228 70237282 refGene_gn=MUC19;refGene_tx=XM_023257754.1
B4 70237371 70237425 refGene_gn=MUC19;refGene_tx=XM_023257754.1
B4 70237670 70237727 refGene_gn=MUC19;refGene_tx=XM_023257754.1
B4 70238412 70238466 refGene_gn=MUC19;refGene_tx=XM_023257754.1
B4 70240507 70240561 refGene_gn=MUC19;refGene_tx=XM_023257754.1
B4 70241001 70241055 refGene_gn=MUC19;refGene_tx=XM_023257754.1
B4 70241687 70241741 refGene_gn=MUC19;refGene_tx=XM_023257754.1
B4 70242354 70242408 refGene_gn=MUC19;refGene_tx=XM_023257754.1
B4 70243411 70243465 refGene_gn=MUC19;refGene_tx=XM_023257754.1
B4 70243590 70243641 refGene_gn=MUC19;refGene_tx=XM_023257754.1
B4 70243979 70244033 refGene_gn=MUC19;refGene_tx=XM_023257754.1
B4 70244206 70244254 refGene_gn=MUC19;refGene_tx=XM_023257754.1
B4 70246474 70246504 refGene_gn=MUC19;refGene_tx=XM_023257754.1
B4 70248606 70248638 refGene_gn=MUC19;refGene_tx=XM_023257754.1
B4 70248724 70248887 refGene_gn=MUC19;refGene_tx=XM_023257754.1
B4 70249232 70249270 refGene_gn=MUC19;refGene_tx=XM_023257754.1
B4 70251080 70251195 refGene_gn=MUC19;refGene_tx=XM_023257754.1
B4 70252184 70252220 refGene_gn=MUC19;refGene_tx=XM_023257754.1
B4 70252867 70252976 refGene_gn=MUC19;refGene_tx=XM_023257754.1
B4 70253642 70253826 refGene_gn=MUC19;refGene_tx=XM_023257754.1
B4 70549607 70549668
refGene_gn=CNTN1;refGene_tx=XM_003988561.4;refGene_gn=CNTN1;refGene_tx
=XM_011283870.2;refGene_gn=CNTN1;refGene_tx=XM_011283871.3;refGene_gn=
CNTN1;refGene_tx=XM_011283872.2
B4 70551068 70551101
refGene_gn=CNTN1;refGene_tx=XM_003988561.4;refGene_gn=CNTN1;refGene_tx
=XM_011283870.2;refGene_gn=CNTN1;refGene_tx=XM_011283871.3;refGene_gn=
CNTN1;refGene_tx=XM_011283872.2
B4 70561034 70561068 refGene_gn=CNTN1;refGene_tx=XM_019834827.2
B4 70561536 70561669
refGene_gn=CNTN1;refGene_tx=XM_003988561.4;refGene_gn=CNTN1;refGene_tx
=XM_011283870.2;refGene_gn=CNTN1;refGene_tx=XM_011283871.3;refGene_gn=
CNTN1;refGene_tx=XM_011283872.2;refGene_gn=CNTN1;refGene_tx=XM_0198348
27.2
B4 70565275 70565448
refGene_gn=CNTN1;refGene_tx=XM_003988561.4;refGene_gn=CNTN1;refGene_tx

=XM_011283870.2;refGene_gn=CNTN1;refGene_tx=XM_011283871.3;refGene_gn=CNTN1;refGene_tx=XM_011283872.2;refGene_gn=CNTN1;refGene_tx=XM_019834827.2
B4	70567878	70567974
refGene_gn=CNTN1;refGene_tx=XM_003988561.4;refGene_gn=CNTN1;refGene_tx=XM_011283870.2;refGene_gn=CNTN1;refGene_tx=XM_011283871.3;refGene_gn=CNTN1;refGene_tx=XM_011283872.2;refGene_gn=CNTN1;refGene_tx=XM_019834827.2
B4	70570094	70570301
refGene_gn=CNTN1;refGene_tx=XM_003988561.4;refGene_gn=CNTN1;refGene_tx=XM_011283870.2;refGene_gn=CNTN1;refGene_tx=XM_011283871.3;refGene_gn=CNTN1;refGene_tx=XM_011283872.2;refGene_gn=CNTN1;refGene_tx=XM_019834827.2
B4	70573851	70573951
refGene_gn=CNTN1;refGene_tx=XM_003988561.4;refGene_gn=CNTN1;refGene_tx=XM_011283870.2;refGene_gn=CNTN1;refGene_tx=XM_011283871.3;refGene_gn=CNTN1;refGene_tx=XM_011283872.2;refGene_gn=CNTN1;refGene_tx=XM_019834827.2
B4	70574107	70574289
refGene_gn=CNTN1;refGene_tx=XM_003988561.4;refGene_gn=CNTN1;refGene_tx=XM_011283870.2;refGene_gn=CNTN1;refGene_tx=XM_011283871.3;refGene_gn=CNTN1;refGene_tx=XM_011283872.2;refGene_gn=CNTN1;refGene_tx=XM_019834827.2
B4	70579962	70580087
refGene_gn=CNTN1;refGene_tx=XM_003988561.4;refGene_gn=CNTN1;refGene_tx=XM_011283870.2;refGene_gn=CNTN1;refGene_tx=XM_011283871.3;refGene_gn=CNTN1;refGene_tx=XM_011283872.2;refGene_gn=CNTN1;refGene_tx=XM_019834827.2
B4	70580779	70580897
refGene_gn=CNTN1;refGene_tx=XM_003988561.4;refGene_gn=CNTN1;refGene_tx=XM_011283870.2;refGene_gn=CNTN1;refGene_tx=XM_011283871.3;refGene_gn=CNTN1;refGene_tx=XM_011283872.2;refGene_gn=CNTN1;refGene_tx=XM_019834827.2
B4	70587005	70587156
refGene_gn=CNTN1;refGene_tx=XM_003988561.4;refGene_gn=CNTN1;refGene_tx=XM_011283870.2;refGene_gn=CNTN1;refGene_tx=XM_011283871.3;refGene_gn=CNTN1;refGene_tx=XM_011283872.2;refGene_gn=CNTN1;refGene_tx=XM_019834827.2
B4	70592799	70592927
refGene_gn=CNTN1;refGene_tx=XM_003988561.4;refGene_gn=CNTN1;refGene_tx=XM_011283870.2;refGene_gn=CNTN1;refGene_tx=XM_011283871.3;refGene_gn=CNTN1;refGene_tx=XM_011283872.2;refGene_gn=CNTN1;refGene_tx=XM_019834827.2
B4	70593470	70593646
refGene_gn=CNTN1;refGene_tx=XM_003988561.4;refGene_gn=CNTN1;refGene_tx=XM_011283870.2;refGene_gn=CNTN1;refGene_tx=XM_011283871.3;refGene_gn=CNTN1;refGene_tx=XM_011283872.2;refGene_gn=CNTN1;refGene_tx=XM_019834827.2
B4	70612083	70612204
refGene_gn=CNTN1;refGene_tx=XM_003988561.4;refGene_gn=CNTN1;refGene_tx

=XM_011283870.2;refGene_gn=CNTN1;refGene_tx=XM_011283871.3;refGene_gn=CNTN1;refGene_tx=XM_011283872.2;refGene_gn=CNTN1;refGene_tx=XM_019834827.2
B4 70629445 70629525 refGene_gn=CNTN1;refGene_tx=XM_011283872.2
B4 70638320 70638479
refGene_gn=CNTN1;refGene_tx=XM_003988561.4;refGene_gn=CNTN1;refGene_tx=XM_011283870.2;refGene_gn=CNTN1;refGene_tx=XM_011283871.3;refGene_gn=CNTN1;refGene_tx=XM_019834827.2
B4 70650843 70650993
refGene_gn=CNTN1;refGene_tx=XM_003988561.4;refGene_gn=CNTN1;refGene_tx=XM_011283870.2;refGene_gn=CNTN1;refGene_tx=XM_011283871.3;refGene_gn=CNTN1;refGene_tx=XM_019834827.2
B4 70661432 70661503
refGene_gn=CNTN1;refGene_tx=XM_003988561.4;refGene_gn=CNTN1;refGene_tx=XM_011283870.2;refGene_gn=CNTN1;refGene_tx=XM_011283871.3;refGene_gn=CNTN1;refGene_tx=XM_019834827.2
B4 70666678 70666913
refGene_gn=CNTN1;refGene_tx=XM_003988561.4;refGene_gn=CNTN1;refGene_tx=XM_011283870.2;refGene_gn=CNTN1;refGene_tx=XM_011283871.3;refGene_gn=CNTN1;refGene_tx=XM_019834827.2
B4 70669431 70669535
refGene_gn=CNTN1;refGene_tx=XM_003988561.4;refGene_gn=CNTN1;refGene_tx=XM_011283870.2;refGene_gn=CNTN1;refGene_tx=XM_011283871.3;refGene_gn=CNTN1;refGene_tx=XM_019834827.2
B4 70677933 70678120
refGene_gn=CNTN1;refGene_tx=XM_003988561.4;refGene_gn=CNTN1;refGene_tx=XM_011283870.2;refGene_gn=CNTN1;refGene_tx=XM_011283871.3;refGene_gn=CNTN1;refGene_tx=XM_019834827.2
B4 70680570 70680683
refGene_gn=CNTN1;refGene_tx=XM_003988561.4;refGene_gn=CNTN1;refGene_tx=XM_011283870.2;refGene_gn=CNTN1;refGene_tx=XM_011283871.3;refGene_gn=CNTN1;refGene_tx=XM_019834827.2
B4 70681492 70681649
refGene_gn=CNTN1;refGene_tx=XM_003988561.4;refGene_gn=CNTN1;refGene_tx=XM_011283870.2;refGene_gn=CNTN1;refGene_tx=XM_011283871.3;refGene_gn=CNTN1;refGene_tx=XM_019834827.2
B4 70723953 70724030
refGene_gn=CNTN1;refGene_tx=XM_003988561.4;refGene_gn=CNTN1;refGene_tx=XM_011283870.2;refGene_gn=CNTN1;refGene_tx=XM_011283871.3;refGene_gn=CNTN1;refGene_tx=XM_019834827.2
B4 70815996 70816647 refGene_gn=PDZRN4;refGene_tx=XM_023256948.1
B4 70819264 70819351 refGene_gn=PDZRN4;refGene_tx=XM_023256948.1
B4 70821194 70821302 refGene_gn=PDZRN4;refGene_tx=XM_023256948.1
B4 71042529 71042598 refGene_gn=PDZRN4;refGene_tx=XM_019834828.2
B4 71112307 71112564
refGene_gn=PDZRN4;refGene_tx=XM_019834828.2;refGene_gn=PDZRN4;refGene_tx=XM_023256948.1
B4 71115716 71115819
refGene_gn=PDZRN4;refGene_tx=XM_019834828.2;refGene_gn=PDZRN4;refGene_tx=XM_023256948.1

B4 71161455 71161554
refGene_gn=PDZRN4;refGene_tx=XM_019834828.2;refGene_gn=PDZRN4;refGene_
tx=XM_023256948.1
B4 71164079 71164142
refGene_gn=PDZRN4;refGene_tx=XM_019834828.2;refGene_gn=PDZRN4;refGene_
tx=XM_023256948.1
B4 71171159 71171261
refGene_gn=PDZRN4;refGene_tx=XM_019834828.2;refGene_gn=PDZRN4;refGene_
tx=XM_023256948.1
B4 71175611 71175728
refGene_gn=PDZRN4;refGene_tx=XM_019834828.2;refGene_gn=PDZRN4;refGene_
tx=XM_023256948.1
B4 71180477 71182004
refGene_gn=PDZRN4;refGene_tx=XM_019834828.2;refGene_gn=PDZRN4;refGene_
tx=XM_023256948.1
B4 71661657 71661663 refGene_gn=GXYLT1;refGene_tx=XM_023256951.1
B4 71661903 71662065
refGene_gn=GXYLT1;refGene_tx=XM_023256949.1;refGene_gn=GXYLT1;refGene_
tx=XM_023256950.1
B4 71674444 71674617
refGene_gn=GXYLT1;refGene_tx=XM_023256949.1;refGene_gn=GXYLT1;refGene_
tx=XM_023256950.1;refGene_gn=GXYLT1;refGene_tx=XM_023256951.1
B4 71674931 71675055
refGene_gn=GXYLT1;refGene_tx=XM_023256949.1;refGene_gn=GXYLT1;refGene_
tx=XM_023256950.1;refGene_gn=GXYLT1;refGene_tx=XM_023256951.1
B4 71681813 71682065
refGene_gn=GXYLT1;refGene_tx=XM_023256949.1;refGene_gn=GXYLT1;refGene_
tx=XM_023256950.1;refGene_gn=GXYLT1;refGene_tx=XM_023256951.1
B4 71685853 71685979
refGene_gn=GXYLT1;refGene_tx=XM_023256949.1;refGene_gn=GXYLT1;refGene_
tx=XM_023256950.1;refGene_gn=GXYLT1;refGene_tx=XM_023256951.1
B4 71687391 71687563
refGene_gn=GXYLT1;refGene_tx=XM_023256949.1;refGene_gn=GXYLT1;refGene_
tx=XM_023256950.1;refGene_gn=GXYLT1;refGene_tx=XM_023256951.1
B4 71699899 71699992
refGene_gn=GXYLT1;refGene_tx=XM_023256949.1;refGene_gn=GXYLT1;refGene_
tx=XM_023256951.1
B4 71709349 71709570
refGene_gn=GXYLT1;refGene_tx=XM_023256949.1;refGene_gn=GXYLT1;refGene_
tx=XM_023256950.1;refGene_gn=GXYLT1;refGene_tx=XM_023256951.1
B4 71719756 71719994
refGene_gn=YAF2;refGene_tx=XM_023256952.1;refGene_gn=YAF2;refGene_tx=X
M_023256953.1;refGene_gn=YAF2;refGene_tx=XM_023256954.1
B4 71720897 71721050
refGene_gn=YAF2;refGene_tx=XM_023256952.1;refGene_gn=YAF2;refGene_tx=X
M_023256953.1;refGene_gn=YAF2;refGene_tx=XM_023256954.1
B4 71769549 71769682
refGene_gn=YAF2;refGene_tx=XM_023256952.1;refGene_gn=YAF2;refGene_tx=X
M_023256953.1
B4 71792187 71792389 refGene_gn=YAF2;refGene_tx=XM_023256957.1

B4 71794026 71794152
refGene_gn=YAF2;refGene_tx=XM_023256952.1;refGene_gn=YAF2;refGene_tx=XM_023256953.1;refGene_gn=YAF2;refGene_tx=XM_023256954.1;refGene_gn=YAF2;refGene_tx=XM_023256955.1;refGene_gn=YAF2;refGene_tx=XM_023256956.1;refGene_gn=YAF2;refGene_tx=XM_023256957.1
B4 71794587 71795032
refGene_gn=YAF2;refGene_tx=XM_023256952.1;refGene_gn=YAF2;refGene_tx=XM_023256954.1;refGene_gn=YAF2;refGene_tx=XM_023256955.1;refGene_gn=YAF2;refGene_tx=XM_023256956.1;refGene_gn=YAF2;refGene_tx=XM_023256957.1
B4 71872191 71872323
refGene_gn=ZCRB1;refGene_tx=XM_011283873.3;refGene_gn=ZCRB1;refGene_tx=XM_011283874.3
B4 71873328 71873404
refGene_gn=ZCRB1;refGene_tx=XM_011283873.3;refGene_gn=ZCRB1;refGene_tx=XM_011283874.3
B4 71873502 71873615
refGene_gn=ZCRB1;refGene_tx=XM_011283873.3;refGene_gn=ZCRB1;refGene_tx=XM_011283874.3
B4 71876565 71876673
refGene_gn=ZCRB1;refGene_tx=XM_011283873.3;refGene_gn=ZCRB1;refGene_tx=XM_011283874.3
B4 71877017 71877129
refGene_gn=ZCRB1;refGene_tx=XM_011283873.3;refGene_gn=ZCRB1;refGene_tx=XM_011283874.3
B4 71885909 71885938 refGene_gn=ZCRB1;refGene_tx=XM_011283873.3
B4 71888049 71888133 refGene_gn=ZCRB1;refGene_tx=XM_011283873.3
B4 71899315 71899316
refGene_gn=PPHLN1;refGene_tx=XM_006933619.3;refGene_gn=PPHLN1;refGene_tx=XM_006933622.3;refGene_gn=PPHLN1;refGene_tx=XM_011283875.2;refGene_gn=PPHLN1;refGene_tx=XM_011283876.3;refGene_gn=PPHLN1;refGene_tx=XM_011283879.2;refGene_gn=PPHLN1;refGene_tx=XM_023256959.1
B4 71902122 71902214
refGene_gn=PPHLN1;refGene_tx=XM_006933619.3;refGene_gn=PPHLN1;refGene_tx=XM_006933622.3;refGene_gn=PPHLN1;refGene_tx=XM_011283875.2;refGene_gn=PPHLN1;refGene_tx=XM_011283876.3;refGene_gn=PPHLN1;refGene_tx=XM_011283878.2;refGene_gn=PPHLN1;refGene_tx=XM_011283879.2;refGene_gn=PPHLN1;refGene_tx=XM_019834831.2;refGene_gn=PPHLN1;refGene_tx=XM_023256959.1
B4 71914710 71914875
refGene_gn=PPHLN1;refGene_tx=XM_006933619.3;refGene_gn=PPHLN1;refGene_tx=XM_011283875.2;refGene_gn=PPHLN1;refGene_tx=XM_011283876.3;refGene_gn=PPHLN1;refGene_tx=XM_011283878.2;refGene_gn=PPHLN1;refGene_tx=XM_023256959.1
B4 71917190 71917252
refGene_gn=PPHLN1;refGene_tx=XM_006933619.3;refGene_gn=PPHLN1;refGene_tx=XM_006933622.3;refGene_gn=PPHLN1;refGene_tx=XM_011283875.2;refGene_gn=PPHLN1;refGene_tx=XM_011283876.3;refGene_gn=PPHLN1;refGene_tx=XM_011283878.2;refGene_gn=PPHLN1;refGene_tx=XM_011283879.2;refGene_gn=PPHLN1;refGene_tx=XM_019834831.2;refGene_gn=PPHLN1;refGene_tx=XM_023256959.1

B4 71945974 71946186
refGene_gn=PPHLN1;refGene_tx=XM_006933619.3;refGene_gn=PPHLN1;refGene_tx=XM_006933622.3;refGene_gn=PPHLN1;refGene_tx=XM_011283875.2;refGene_gn=PPHLN1;refGene_tx=XM_011283876.3;refGene_gn=PPHLN1;refGene_tx=XM_011283878.2;refGene_gn=PPHLN1;refGene_tx=XM_011283879.2;refGene_gn=PPHLN1;refGene_tx=XM_019834831.2;refGene_gn=PPHLN1;refGene_tx=XM_023256959.1
B4 71956052 71956109
refGene_gn=PPHLN1;refGene_tx=XM_011283875.2;refGene_gn=PPHLN1;refGene_tx=XM_011283876.3;refGene_gn=PPHLN1;refGene_tx=XM_011283878.2;refGene_gn=PPHLN1;refGene_tx=XM_011283879.2;refGene_gn=PPHLN1;refGene_tx=XM_019834831.2;refGene_gn=PPHLN1;refGene_tx=XM_023256959.1
B4 71958807 71958887
refGene_gn=PPHLN1;refGene_tx=XM_006933619.3;refGene_gn=PPHLN1;refGene_tx=XM_006933622.3;refGene_gn=PPHLN1;refGene_tx=XM_011283875.2;refGene_gn=PPHLN1;refGene_tx=XM_011283876.3;refGene_gn=PPHLN1;refGene_tx=XM_011283878.2;refGene_gn=PPHLN1;refGene_tx=XM_011283879.2;refGene_gn=PPHLN1;refGene_tx=XM_019834831.2;refGene_gn=PPHLN1;refGene_tx=XM_023256959.1
B4 71961495 71961615
refGene_gn=PPHLN1;refGene_tx=XM_006933619.3;refGene_gn=PPHLN1;refGene_tx=XM_006933622.3;refGene_gn=PPHLN1;refGene_tx=XM_011283875.2;refGene_gn=PPHLN1;refGene_tx=XM_011283876.3;refGene_gn=PPHLN1;refGene_tx=XM_011283878.2;refGene_gn=PPHLN1;refGene_tx=XM_011283879.2;refGene_gn=PPHLN1;refGene_tx=XM_019834831.2;refGene_gn=PPHLN1;refGene_tx=XM_023256959.1
B4 71984433 71984574
refGene_gn=PPHLN1;refGene_tx=XM_006933619.3;refGene_gn=PPHLN1;refGene_tx=XM_006933622.3;refGene_gn=PPHLN1;refGene_tx=XM_011283875.2;refGene_gn=PPHLN1;refGene_tx=XM_011283876.3;refGene_gn=PPHLN1;refGene_tx=XM_011283878.2;refGene_gn=PPHLN1;refGene_tx=XM_011283879.2;refGene_gn=PPHLN1;refGene_tx=XM_019834831.2;refGene_gn=PPHLN1;refGene_tx=XM_023256959.1
B4 72030191 72030386
refGene_gn=PPHLN1;refGene_tx=XM_006933619.3;refGene_gn=PPHLN1;refGene_tx=XM_006933622.3;refGene_gn=PPHLN1;refGene_tx=XM_011283875.2;refGene_gn=PPHLN1;refGene_tx=XM_011283876.3;refGene_gn=PPHLN1;refGene_tx=XM_011283878.2;refGene_gn=PPHLN1;refGene_tx=XM_011283879.2;refGene_gn=PPHLN1;refGene_tx=XM_019834831.2;refGene_gn=PPHLN1;refGene_tx=XM_023256959.1
B4 72041750 72042613
refGene_gn=PRICKLE1;refGene_tx=XM_003988570.5;refGene_gn=PRICKLE1;refGene_tx=XM_006933623.3;refGene_gn=PRICKLE1;refGene_tx=XM_011283880.3;refGene_gn=PRICKLE1;refGene_tx=XM_023256958.1
B4 72044466 72045327
refGene_gn=PRICKLE1;refGene_tx=XM_003988570.5;refGene_gn=PRICKLE1;refGene_tx=XM_006933623.3;refGene_gn=PRICKLE1;refGene_tx=XM_011283880.3;refGene_gn=PRICKLE1;refGene_tx=XM_023256958.1
B4 72046239 72046426
refGene_gn=PRICKLE1;refGene_tx=XM_003988570.5;refGene_gn=PRICKLE1;refG

ene_tx=XM_006933623.3;refGene_gn=PRICKLE1;refGene_tx=XM_011283880.3;refGene_gn=PRICKLE1;refGene_tx=XM_023256958.1
B4 72047711 72047915
refGene_gn=PRICKLE1;refGene_tx=XM_003988570.5;refGene_gn=PRICKLE1;refGene_tx=XM_006933623.3;refGene_gn=PRICKLE1;refGene_tx=XM_011283880.3;refGene_gn=PRICKLE1;refGene_tx=XM_023256958.1
B4 72048489 72048627
refGene_gn=PRICKLE1;refGene_tx=XM_003988570.5;refGene_gn=PRICKLE1;refGene_tx=XM_006933623.3;refGene_gn=PRICKLE1;refGene_tx=XM_011283880.3;refGene_gn=PRICKLE1;refGene_tx=XM_023256958.1
B4 72049279 72049393
refGene_gn=PRICKLE1;refGene_tx=XM_003988570.5;refGene_gn=PRICKLE1;refGene_tx=XM_006933623.3;refGene_gn=PRICKLE1;refGene_tx=XM_011283880.3;refGene_gn=PRICKLE1;refGene_tx=XM_023256958.1
B4 72050973 72051105
refGene_gn=PRICKLE1;refGene_tx=XM_003988570.5;refGene_gn=PRICKLE1;refGene_tx=XM_006933623.3;refGene_gn=PRICKLE1;refGene_tx=XM_011283880.3;refGene_gn=PRICKLE1;refGene_tx=XM_023256958.1
B4 72847043 72847133
refGene_gn=ADAMTS20;refGene_tx=XM_023256960.1;refGene_gn=ADAMTS20;refGene_tx=XM_023256961.1;refGene_gn=ADAMTS20;refGene_tx=XM_023256963.1
B4 72849533 72849638
refGene_gn=ADAMTS20;refGene_tx=XM_023256960.1;refGene_gn=ADAMTS20;refGene_tx=XM_023256961.1;refGene_gn=ADAMTS20;refGene_tx=XM_023256963.1
B4 72867310 72867402
refGene_gn=ADAMTS20;refGene_tx=XM_023256960.1;refGene_gn=ADAMTS20;refGene_tx=XM_023256961.1;refGene_gn=ADAMTS20;refGene_tx=XM_023256963.1
B4 72874756 72874890
refGene_gn=ADAMTS20;refGene_tx=XM_023256960.1;refGene_gn=ADAMTS20;refGene_tx=XM_023256961.1;refGene_gn=ADAMTS20;refGene_tx=XM_023256963.1
B4 72875631 72875726
refGene_gn=ADAMTS20;refGene_tx=XM_023256960.1;refGene_gn=ADAMTS20;refGene_tx=XM_023256961.1;refGene_gn=ADAMTS20;refGene_tx=XM_023256963.1
B4 72875811 72875906
refGene_gn=ADAMTS20;refGene_tx=XM_023256960.1;refGene_gn=ADAMTS20;refGene_tx=XM_023256961.1;refGene_gn=ADAMTS20;refGene_tx=XM_023256963.1
B4 72876557 72876687
refGene_gn=ADAMTS20;refGene_tx=XM_023256960.1;refGene_gn=ADAMTS20;refGene_tx=XM_023256961.1;refGene_gn=ADAMTS20;refGene_tx=XM_023256963.1
B4 72877394 72877595
refGene_gn=ADAMTS20;refGene_tx=XM_023256960.1;refGene_gn=ADAMTS20;refGene_tx=XM_023256961.1;refGene_gn=ADAMTS20;refGene_tx=XM_023256963.1
B4 72881767 72881938
refGene_gn=ADAMTS20;refGene_tx=XM_023256960.1;refGene_gn=ADAMTS20;refGene_tx=XM_023256961.1;refGene_gn=ADAMTS20;refGene_tx=XM_023256963.1
B4 72882021 72882195
refGene_gn=ADAMTS20;refGene_tx=XM_023256960.1;refGene_gn=ADAMTS20;refGene_tx=XM_023256961.1;refGene_gn=ADAMTS20;refGene_tx=XM_023256963.1
B4 72889481 72889649
refGene_gn=ADAMTS20;refGene_tx=XM_023256960.1;refGene_gn=ADAMTS20;refG

ene_tx=XM_023256961.1;refGene_gn=ADAMTS20;refGene_tx=XM_023256963.1
B4	72916640	72916817
refGene_gn=ADAMTS20;refGene_tx=XM_023256960.1;refGene_gn=ADAMTS20;refGene_tx=XM_023256961.1
B4	72917988	72918150
refGene_gn=ADAMTS20;refGene_tx=XM_023256960.1;refGene_gn=ADAMTS20;refGene_tx=XM_023256961.1;refGene_gn=ADAMTS20;refGene_tx=XM_023256963.1
B4	72918855	72919146
refGene_gn=ADAMTS20;refGene_tx=XM_023256960.1;refGene_gn=ADAMTS20;refGene_tx=XM_023256961.1;refGene_gn=ADAMTS20;refGene_tx=XM_023256963.1
B4	72919230	72919395
refGene_gn=ADAMTS20;refGene_tx=XM_023256960.1;refGene_gn=ADAMTS20;refGene_tx=XM_023256961.1;refGene_gn=ADAMTS20;refGene_tx=XM_023256963.1
B4	72920330	72920438
refGene_gn=ADAMTS20;refGene_tx=XM_023256960.1;refGene_gn=ADAMTS20;refGene_tx=XM_023256961.1;refGene_gn=ADAMTS20;refGene_tx=XM_023256963.1
B4	72921012	72921132
refGene_gn=ADAMTS20;refGene_tx=XM_023256960.1;refGene_gn=ADAMTS20;refGene_tx=XM_023256961.1;refGene_gn=ADAMTS20;refGene_tx=XM_023256963.1
B4	72922282	72922447
refGene_gn=ADAMTS20;refGene_tx=XM_023256960.1;refGene_gn=ADAMTS20;refGene_tx=XM_023256961.1;refGene_gn=ADAMTS20;refGene_tx=XM_023256963.1
B4	72922905	72923070
refGene_gn=ADAMTS20;refGene_tx=XM_023256960.1;refGene_gn=ADAMTS20;refGene_tx=XM_023256961.1;refGene_gn=ADAMTS20;refGene_tx=XM_023256963.1
B4	72923227	72923438
refGene_gn=ADAMTS20;refGene_tx=XM_023256960.1;refGene_gn=ADAMTS20;refGene_tx=XM_023256961.1;refGene_gn=ADAMTS20;refGene_tx=XM_023256963.1
B4	72925149	72925276
refGene_gn=ADAMTS20;refGene_tx=XM_023256960.1;refGene_gn=ADAMTS20;refGene_tx=XM_023256961.1;refGene_gn=ADAMTS20;refGene_tx=XM_023256963.1
B4	72931057	72931187
refGene_gn=ADAMTS20;refGene_tx=XM_023256960.1;refGene_gn=ADAMTS20;refGene_tx=XM_023256961.1;refGene_gn=ADAMTS20;refGene_tx=XM_023256963.1
B4	72931335	72931502
refGene_gn=ADAMTS20;refGene_tx=XM_023256960.1;refGene_gn=ADAMTS20;refGene_tx=XM_023256961.1;refGene_gn=ADAMTS20;refGene_tx=XM_023256963.1
B4	72934661	72934757
refGene_gn=ADAMTS20;refGene_tx=XM_023256960.1;refGene_gn=ADAMTS20;refGene_tx=XM_023256961.1;refGene_gn=ADAMTS20;refGene_tx=XM_023256963.1
B4	72936994	72937112
refGene_gn=ADAMTS20;refGene_tx=XM_023256960.1;refGene_gn=ADAMTS20;refGene_tx=XM_023256961.1;refGene_gn=ADAMTS20;refGene_tx=XM_023256963.1
B4	72939334	72939471
refGene_gn=ADAMTS20;refGene_tx=XM_023256960.1;refGene_gn=ADAMTS20;refGene_tx=XM_023256961.1;refGene_gn=ADAMTS20;refGene_tx=XM_023256963.1
B4	72939569	72939751
refGene_gn=ADAMTS20;refGene_tx=XM_023256960.1;refGene_gn=ADAMTS20;refGene_tx=XM_023256961.1;refGene_gn=ADAMTS20;refGene_tx=XM_023256963.1
B4	72941041	72941187

refGene_gn=ADAMTS20;refGene_tx=XM_023256960.1;refGene_gn=ADAMTS20;refGene_tx=XM_023256961.1;refGene_gn=ADAMTS20;refGene_tx=XM_023256963.1
B4 72947457 72947562
refGene_gn=ADAMTS20;refGene_tx=XM_023256960.1;refGene_gn=ADAMTS20;refGene_tx=XM_023256961.1;refGene_gn=ADAMTS20;refGene_tx=XM_023256963.1
B4 72949689 72949831
refGene_gn=ADAMTS20;refGene_tx=XM_023256960.1;refGene_gn=ADAMTS20;refGene_tx=XM_023256961.1;refGene_gn=ADAMTS20;refGene_tx=XM_023256963.1
B4 72952248 72952395
refGene_gn=ADAMTS20;refGene_tx=XM_023256960.1;refGene_gn=ADAMTS20;refGene_tx=XM_023256961.1;refGene_gn=ADAMTS20;refGene_tx=XM_023256963.1
B4 72954177 72954283
refGene_gn=ADAMTS20;refGene_tx=XM_023256960.1;refGene_gn=ADAMTS20;refGene_tx=XM_023256961.1;refGene_gn=ADAMTS20;refGene_tx=XM_023256963.1
B4 72959491 72959532
refGene_gn=ADAMTS20;refGene_tx=XM_023256960.1;refGene_gn=ADAMTS20;refGene_tx=XM_023256961.1;refGene_gn=ADAMTS20;refGene_tx=XM_023256963.1
B4 72960703 72960828
refGene_gn=ADAMTS20;refGene_tx=XM_023256960.1;refGene_gn=ADAMTS20;refGene_tx=XM_023256961.1;refGene_gn=ADAMTS20;refGene_tx=XM_023256963.1
B4 72961297 72961381
refGene_gn=ADAMTS20;refGene_tx=XM_023256960.1;refGene_gn=ADAMTS20;refGene_tx=XM_023256961.1;refGene_gn=ADAMTS20;refGene_tx=XM_023256963.1
B4 72974539 72974799
refGene_gn=ADAMTS20;refGene_tx=XM_023256960.1;refGene_gn=ADAMTS20;refGene_tx=XM_023256961.1;refGene_gn=ADAMTS20;refGene_tx=XM_023256963.1
B4 73005126 73005289
refGene_gn=ADAMTS20;refGene_tx=XM_023256960.1;refGene_gn=ADAMTS20;refGene_tx=XM_023256961.1;refGene_gn=ADAMTS20;refGene_tx=XM_023256963.1
B4 73023189 73023551
refGene_gn=ADAMTS20;refGene_tx=XM_023256960.1;refGene_gn=ADAMTS20;refGene_tx=XM_023256961.1;refGene_gn=ADAMTS20;refGene_tx=XM_023256963.1
B4 73024066 73024157
refGene_gn=ADAMTS20;refGene_tx=XM_023256960.1;refGene_gn=ADAMTS20;refGene_tx=XM_023256961.1;refGene_gn=ADAMTS20;refGene_tx=XM_023256963.1
B4 73161059 73161386 refGene_gn=PUS7L;refGene_tx=XM_023256964.1
B4 73162424 73162478 refGene_gn=PUS7L;refGene_tx=XM_023256964.1
B4 73166139 73166420 refGene_gn=PUS7L;refGene_tx=XM_023256964.1
B4 73167560 73167642 refGene_gn=PUS7L;refGene_tx=XM_023256964.1
B4 73169997 73170096 refGene_gn=PUS7L;refGene_tx=XM_023256964.1
B4 73172755 73172948 refGene_gn=PUS7L;refGene_tx=XM_023256964.1
B4 73175486 73175614 refGene_gn=PUS7L;refGene_tx=XM_023256964.1
B4 73175615 73175647 refGene_gn=PUS7L;refGene_tx=XM_023256964.1
B4 73188622 73189526 refGene_gn=PUS7L;refGene_tx=XM_023256964.1
B4 73199162 73199323 refGene_gn=IRAK4;refGene_tx=XM_006933627.4
B4 73202572 73202718 refGene_gn=IRAK4;refGene_tx=XM_006933627.4
B4 73203539 73203722
refGene_gn=IRAK4;refGene_tx=XM_006933627.4;refGene_gn=IRAK4;refGene_tx=XM_023256965.1
B4 73204208 73204369

refGene_gn=IRAK4;refGene_tx=XM_006933627.4;refGene_gn=IRAK4;refGene_tx=XM_023256965.1
B4 73205287 73205352
refGene_gn=IRAK4;refGene_tx=XM_006933627.4;refGene_gn=IRAK4;refGene_tx=XM_023256965.1
B4 73208068 73208183
refGene_gn=IRAK4;refGene_tx=XM_006933627.4;refGene_gn=IRAK4;refGene_tx=XM_023256965.1
B4 73208577 73208687
refGene_gn=IRAK4;refGene_tx=XM_006933627.4;refGene_gn=IRAK4;refGene_tx=XM_023256965.1
B4 73212371 73212555
refGene_gn=IRAK4;refGene_tx=XM_006933627.4;refGene_gn=IRAK4;refGene_tx=XM_023256965.1
B4 73213410 73213473
refGene_gn=IRAK4;refGene_tx=XM_006933627.4;refGene_gn=IRAK4;refGene_tx=XM_023256965.1
B4 73217079 73217238
refGene_gn=IRAK4;refGene_tx=XM_006933627.4;refGene_gn=IRAK4;refGene_tx=XM_023256965.1
B4 73217366 73217402
refGene_gn=IRAK4;refGene_tx=XM_006933627.4;refGene_gn=IRAK4;refGene_tx=XM_023256965.1
B4 73243951 73244122
refGene_gn=TWF1;refGene_tx=XM_023256966.1;refGene_gn=TWF1;refGene_tx=XM_023256967.1
B4 73245853 73245975
refGene_gn=TWF1;refGene_tx=XM_023256966.1;refGene_gn=TWF1;refGene_tx=XM_023256967.1
B4 73246128 73246279
refGene_gn=TWF1;refGene_tx=XM_023256966.1;refGene_gn=TWF1;refGene_tx=XM_023256967.1
B4 73246735 73246861
refGene_gn=TWF1;refGene_tx=XM_023256966.1;refGene_gn=TWF1;refGene_tx=XM_023256967.1
B4 73247641 73247662 refGene_gn=TWF1;refGene_tx=XM_023256967.1
B4 73248515 73248620
refGene_gn=TWF1;refGene_tx=XM_023256966.1;refGene_gn=TWF1;refGene_tx=XM_023256967.1
B4 73249716 73249812
refGene_gn=TWF1;refGene_tx=XM_023256966.1;refGene_gn=TWF1;refGene_tx=XM_023256967.1
B4 73251034 73251213
refGene_gn=TWF1;refGene_tx=XM_023256966.1;refGene_gn=TWF1;refGene_tx=XM_023256967.1
B4 73252829 73252907
refGene_gn=TWF1;refGene_tx=XM_023256966.1;refGene_gn=TWF1;refGene_tx=XM_023256967.1
B4 73254529 73254554
refGene_gn=TWF1;refGene_tx=XM_023256966.1;refGene_gn=TWF1;refGene_tx=X

M_023256967.1
B4	73300479 73300756
refGene_gn=TMEM117;refGene_tx=XM_023256968.1;refGene_gn=TMEM117;refGene_tx=XM_023256969.1;refGene_gn=TMEM117;refGene_tx=XM_023256971.1;refGene_gn=TMEM117;refGene_tx=XM_023256972.1
B4	73382453 73382586
refGene_gn=TMEM117;refGene_tx=XM_023256968.1;refGene_gn=TMEM117;refGene_tx=XM_023256969.1;refGene_gn=TMEM117;refGene_tx=XM_023256971.1;refGene_gn=TMEM117;refGene_tx=XM_023256972.1
B4	73507373 73507408 refGene_gn=TMEM117;refGene_tx=XM_023256970.1
B4	73544922 73545022
refGene_gn=TMEM117;refGene_tx=XM_023256968.1;refGene_gn=TMEM117;refGene_tx=XM_023256969.1;refGene_gn=TMEM117;refGene_tx=XM_023256970.1;refGene_gn=TMEM117;refGene_tx=XM_023256971.1;refGene_gn=TMEM117;refGene_tx=XM_023256972.1
B4	73623819 73623917
refGene_gn=TMEM117;refGene_tx=XM_023256968.1;refGene_gn=TMEM117;refGene_tx=XM_023256969.1;refGene_gn=TMEM117;refGene_tx=XM_023256970.1;refGene_gn=TMEM117;refGene_tx=XM_023256971.1;refGene_gn=TMEM117;refGene_tx=XM_023256972.1
B4	73709883 73710043
refGene_gn=TMEM117;refGene_tx=XM_023256968.1;refGene_gn=TMEM117;refGene_tx=XM_023256969.1;refGene_gn=TMEM117;refGene_tx=XM_023256970.1;refGene_gn=TMEM117;refGene_tx=XM_023256971.1;refGene_gn=TMEM117;refGene_tx=XM_023256972.1
B4	73729301 73729376 refGene_gn=TMEM117;refGene_tx=XM_023256972.1
B4	73776533 73776663
refGene_gn=TMEM117;refGene_tx=XM_023256968.1;refGene_gn=TMEM117;refGene_tx=XM_023256969.1;refGene_gn=TMEM117;refGene_tx=XM_023256970.1;refGene_gn=TMEM117;refGene_tx=XM_023256971.1
B4	73788025 73788672
refGene_gn=TMEM117;refGene_tx=XM_023256968.1;refGene_gn=TMEM117;refGene_tx=XM_023256969.1;refGene_gn=TMEM117;refGene_tx=XM_023256970.1
B4	73802128 73802139 refGene_gn=TMEM117;refGene_tx=XM_023256971.1
B4	73900552 73900603
refGene_gn=NELL2;refGene_tx=XM_011283881.3;refGene_gn=NELL2;refGene_tx=XM_019834835.1;refGene_gn=NELL2;refGene_tx=XM_019834836.1;refGene_gn=NELL2;refGene_tx=XM_023256973.1
B4	73910299 73910524
refGene_gn=NELL2;refGene_tx=XM_011283881.3;refGene_gn=NELL2;refGene_tx=XM_019834835.1;refGene_gn=NELL2;refGene_tx=XM_019834836.1;refGene_gn=NELL2;refGene_tx=XM_023256973.1
B4	73912009 73912186
refGene_gn=NELL2;refGene_tx=XM_011283881.3;refGene_gn=NELL2;refGene_tx=XM_019834835.1;refGene_gn=NELL2;refGene_tx=XM_019834836.1;refGene_gn=NELL2;refGene_tx=XM_023256973.1
B4	73913219 73913413
refGene_gn=NELL2;refGene_tx=XM_011283881.3;refGene_gn=NELL2;refGene_tx=XM_019834835.1;refGene_gn=NELL2;refGene_tx=XM_019834836.1;refGene_gn=NELL2;refGene_tx=XM_023256973.1

B4 73923032 73923173
refGene_gn=NELL2;refGene_tx=XM_011283881.3;refGene_gn=NELL2;refGene_tx=XM_019834835.1;refGene_gn=NELL2;refGene_tx=XM_019834836.1;refGene_gn=NELL2;refGene_tx=XM_023256973.1
B4 73992932 73993028
refGene_gn=NELL2;refGene_tx=XM_011283881.3;refGene_gn=NELL2;refGene_tx=XM_019834835.1;refGene_gn=NELL2;refGene_tx=XM_019834836.1;refGene_gn=NELL2;refGene_tx=XM_023256973.1
B4 73996025 73996148
refGene_gn=NELL2;refGene_tx=XM_011283881.3;refGene_gn=NELL2;refGene_tx=XM_019834835.1;refGene_gn=NELL2;refGene_tx=XM_019834836.1;refGene_gn=NELL2;refGene_tx=XM_023256973.1
B4 74046140 74046266
refGene_gn=NELL2;refGene_tx=XM_011283881.3;refGene_gn=NELL2;refGene_tx=XM_019834835.1;refGene_gn=NELL2;refGene_tx=XM_019834836.1;refGene_gn=NELL2;refGene_tx=XM_023256973.1
B4 74080110 74080239
refGene_gn=NELL2;refGene_tx=XM_011283881.3;refGene_gn=NELL2;refGene_tx=XM_019834835.1;refGene_gn=NELL2;refGene_tx=XM_019834836.1;refGene_gn=NELL2;refGene_tx=XM_023256973.1
B4 74089303 74089406
refGene_gn=NELL2;refGene_tx=XM_011283881.3;refGene_gn=NELL2;refGene_tx=XM_019834835.1;refGene_gn=NELL2;refGene_tx=XM_019834836.1;refGene_gn=NELL2;refGene_tx=XM_023256973.1
B4 74093527 74093619
refGene_gn=NELL2;refGene_tx=XM_011283881.3;refGene_gn=NELL2;refGene_tx=XM_019834835.1;refGene_gn=NELL2;refGene_tx=XM_019834836.1;refGene_gn=NELL2;refGene_tx=XM_023256973.1
B4 74139859 74139962
refGene_gn=NELL2;refGene_tx=XM_011283881.3;refGene_gn=NELL2;refGene_tx=XM_019834835.1;refGene_gn=NELL2;refGene_tx=XM_019834836.1;refGene_gn=NELL2;refGene_tx=XM_023256973.1
B4 74141439 74141568
refGene_gn=NELL2;refGene_tx=XM_011283881.3;refGene_gn=NELL2;refGene_tx=XM_019834835.1;refGene_gn=NELL2;refGene_tx=XM_019834836.1;refGene_gn=NELL2;refGene_tx=XM_023256973.1
B4 74142443 74142526
refGene_gn=NELL2;refGene_tx=XM_011283881.3;refGene_gn=NELL2;refGene_tx=XM_019834835.1;refGene_gn=NELL2;refGene_tx=XM_019834836.1;refGene_gn=NELL2;refGene_tx=XM_023256973.1
B4 74142696 74142769
refGene_gn=NELL2;refGene_tx=XM_011283881.3;refGene_gn=NELL2;refGene_tx=XM_019834835.1;refGene_gn=NELL2;refGene_tx=XM_019834836.1;refGene_gn=NELL2;refGene_tx=XM_023256973.1
B4 74145570 74145667
refGene_gn=NELL2;refGene_tx=XM_011283881.3;refGene_gn=NELL2;refGene_tx=XM_019834835.1;refGene_gn=NELL2;refGene_tx=XM_019834836.1;refGene_gn=NELL2;refGene_tx=XM_023256973.1
B4 74145753 74145927
refGene_gn=NELL2;refGene_tx=XM_011283881.3;refGene_gn=NELL2;refGene_tx

=XM_019834835.1;refGene_gn=NELL2;refGene_tx=XM_019834836.1;refGene_gn=NELL2;refGene_tx=XM_023256973.1
B4 74179039 74179190 refGene_gn=NELL2;refGene_tx=XM_011283881.3;refGene_gn=NELL2;refGene_tx=XM_019834835.1;refGene_gn=NELL2;refGene_tx=XM_019834836.1;refGene_gn=NELL2;refGene_tx=XM_023256973.1
B4 74228606 74228735 refGene_gn=NELL2;refGene_tx=XM_011283881.3;refGene_gn=NELL2;refGene_tx=XM_019834835.1;refGene_gn=NELL2;refGene_tx=XM_019834836.1;refGene_gn=NELL2;refGene_tx=XM_023256973.1
B4 74228878 74228924 refGene_gn=NELL2;refGene_tx=XM_019834836.1
B4 74229176 74229231 refGene_gn=NELL2;refGene_tx=XM_011283881.3;refGene_gn=NELL2;refGene_tx=XM_019834835.1;refGene_gn=NELL2;refGene_tx=XM_023256973.1
B4 74346207 74346537 refGene_gn=DBX2;refGene_tx=XM_023256974.1
B4 74352174 74352362 refGene_gn=DBX2;refGene_tx=XM_023256974.1
B4 74364562 74364658 refGene_gn=DBX2;refGene_tx=XM_023256974.1
B4 74379102 74379523 refGene_gn=DBX2;refGene_tx=XM_023256974.1
B4 74530906 74530979 refGene_gn=ANO6;refGene_tx=XM_023256976.1;refGene_gn=ANO6;refGene_tx=XM_023256977.1
B4 74577691 74577754 refGene_gn=ANO6;refGene_tx=XM_023256976.1
B4 74608466 74608546 refGene_gn=ANO6;refGene_tx=XM_023256976.1;refGene_gn=ANO6;refGene_tx=XM_023256977.1
B4 74643051 74643180 refGene_gn=ANO6;refGene_tx=XM_023256976.1;refGene_gn=ANO6;refGene_tx=XM_023256977.1
B4 74657836 74657902 refGene_gn=ANO6;refGene_tx=XM_023256976.1;refGene_gn=ANO6;refGene_tx=XM_023256977.1
B4 74658901 74659189 refGene_gn=ANO6;refGene_tx=XM_023256976.1;refGene_gn=ANO6;refGene_tx=XM_023256977.1
B4 74659391 74659505 refGene_gn=ANO6;refGene_tx=XM_023256976.1;refGene_gn=ANO6;refGene_tx=XM_023256977.1
B4 74661507 74661623 refGene_gn=ANO6;refGene_tx=XM_023256976.1;refGene_gn=ANO6;refGene_tx=XM_023256977.1
B4 74668235 74668370 refGene_gn=ANO6;refGene_tx=XM_023256976.1;refGene_gn=ANO6;refGene_tx=XM_023256977.1
B4 74677082 74677188 refGene_gn=ANO6;refGene_tx=XM_023256976.1;refGene_gn=ANO6;refGene_tx=XM_023256977.1
B4 74681879 74681940 refGene_gn=ANO6;refGene_tx=XM_023256976.1;refGene_gn=ANO6;refGene_tx=XM_023256977.1
B4 74688729 74688872

refGene_gn=ANO6;refGene_tx=XM_023256976.1;refGene_gn=ANO6;refGene_tx=XM_023256977.1
B4	74691180	74691258
refGene_gn=ANO6;refGene_tx=XM_023256976.1;refGene_gn=ANO6;refGene_tx=XM_023256977.1
B4	74696243	74696469
refGene_gn=ANO6;refGene_tx=XM_023256976.1;refGene_gn=ANO6;refGene_tx=XM_023256977.1
B4	74697512	74697682
refGene_gn=ANO6;refGene_tx=XM_023256976.1;refGene_gn=ANO6;refGene_tx=XM_023256977.1
B4	74697889	74697987
refGene_gn=ANO6;refGene_tx=XM_023256976.1;refGene_gn=ANO6;refGene_tx=XM_023256977.1
B4	74705240	74705371
refGene_gn=ANO6;refGene_tx=XM_023256976.1;refGene_gn=ANO6;refGene_tx=XM_023256977.1
B4	74712906	74713112
refGene_gn=ANO6;refGene_tx=XM_023256976.1;refGene_gn=ANO6;refGene_tx=XM_023256977.1
B4	74717539	74717742
refGene_gn=ANO6;refGene_tx=XM_023256976.1;refGene_gn=ANO6;refGene_tx=XM_023256977.1
B4	74719466	74719572
refGene_gn=ANO6;refGene_tx=XM_023256976.1;refGene_gn=ANO6;refGene_tx=XM_023256977.1
B4	74724829	74725036
refGene_gn=ANO6;refGene_tx=XM_023256976.1;refGene_gn=ANO6;refGene_tx=XM_023256977.1
B4	75057671	75057763
refGene_gn=ARID2;refGene_tx=XM_023256978.1;refGene_gn=ARID2;refGene_tx=XM_023256979.1
B4	75057878	75057972
refGene_gn=ARID2;refGene_tx=XM_023256978.1;refGene_gn=ARID2;refGene_tx=XM_023256979.1
B4	75059041	75059139
refGene_gn=ARID2;refGene_tx=XM_023256978.1;refGene_gn=ARID2;refGene_tx=XM_023256979.1
B4	75147843	75147977
refGene_gn=ARID2;refGene_tx=XM_023256978.1;refGene_gn=ARID2;refGene_tx=XM_023256979.1
B4	75152064	75152283
refGene_gn=ARID2;refGene_tx=XM_023256978.1;refGene_gn=ARID2;refGene_tx=XM_023256979.1
B4	75155741	75155809
refGene_gn=ARID2;refGene_tx=XM_023256978.1;refGene_gn=ARID2;refGene_tx=XM_023256979.1
B4	75164882	75164949
refGene_gn=ARID2;refGene_tx=XM_023256978.1;refGene_gn=ARID2;refGene_tx=XM_023256979.1

B4	75165041	75165292
refGene_gn=ARID2;refGene_tx=XM_023256978.1;refGene_gn=ARID2;refGene_tx=XM_023256979.1
B4	75165618	75165715
refGene_gn=ARID2;refGene_tx=XM_023256978.1;refGene_gn=ARID2;refGene_tx=XM_023256979.1
B4	75165811	75166021
refGene_gn=ARID2;refGene_tx=XM_023256978.1;refGene_gn=ARID2;refGene_tx=XM_023256979.1
B4	75166876	75167044
refGene_gn=ARID2;refGene_tx=XM_023256978.1;refGene_gn=ARID2;refGene_tx=XM_023256979.1
B4	75173112	75173194
refGene_gn=ARID2;refGene_tx=XM_023256978.1;refGene_gn=ARID2;refGene_tx=XM_023256979.1
B4	75175120	75175255
refGene_gn=ARID2;refGene_tx=XM_023256978.1;refGene_gn=ARID2;refGene_tx=XM_023256979.1
B4	75175895	75176092
refGene_gn=ARID2;refGene_tx=XM_023256978.1;refGene_gn=ARID2;refGene_tx=XM_023256979.1
B4	75176362	75179226
refGene_gn=ARID2;refGene_tx=XM_023256978.1;refGene_gn=ARID2;refGene_tx=XM_023256979.1
B4	75181948	75182062	refGene_gn=ARID2;refGene_tx=XM_023256978.1
B4	75188618	75188767
refGene_gn=ARID2;refGene_tx=XM_023256978.1;refGene_gn=ARID2;refGene_tx=XM_023256979.1
B4	75206855	75206994
refGene_gn=ARID2;refGene_tx=XM_023256978.1;refGene_gn=ARID2;refGene_tx=XM_023256979.1
B4	75207087	75207173
refGene_gn=ARID2;refGene_tx=XM_023256978.1;refGene_gn=ARID2;refGene_tx=XM_023256979.1
B4	75208907	75209031
refGene_gn=ARID2;refGene_tx=XM_023256978.1;refGene_gn=ARID2;refGene_tx=XM_023256979.1
B4	75209118	75209210
refGene_gn=ARID2;refGene_tx=XM_023256978.1;refGene_gn=ARID2;refGene_tx=XM_023256979.1
B4	75223727	75223872
refGene_gn=ARID2;refGene_tx=XM_023256978.1;refGene_gn=ARID2;refGene_tx=XM_023256979.1
B4	75238311	75238458
refGene_gn=SCAF11;refGene_tx=XM_023256980.1;refGene_gn=SCAF11;refGene_tx=XM_023256981.1;refGene_gn=SCAF11;refGene_tx=XM_023256982.1;refGene_gn=SCAF11;refGene_tx=XM_023256983.1
B4	75238722	75238842
refGene_gn=SCAF11;refGene_tx=XM_023256980.1;refGene_gn=SCAF11;refGene_tx=XM_023256981.1;refGene_gn=SCAF11;refGene_tx=XM_023256982.1;refGene_

gn=SCAF11;refGene_tx=XM_023256983.1
B4 75239191 75239410
refGene_gn=SCAF11;refGene_tx=XM_023256980.1;refGene_gn=SCAF11;refGene_tx=XM_023256981.1;refGene_gn=SCAF11;refGene_tx=XM_023256982.1;refGene_gn=SCAF11;refGene_tx=XM_023256983.1
B4 75240379 75240726
refGene_gn=SCAF11;refGene_tx=XM_023256980.1;refGene_gn=SCAF11;refGene_tx=XM_023256981.1;refGene_gn=SCAF11;refGene_tx=XM_023256982.1;refGene_gn=SCAF11;refGene_tx=XM_023256983.1
B4 75245816 75248534
refGene_gn=SCAF11;refGene_tx=XM_023256980.1;refGene_gn=SCAF11;refGene_tx=XM_023256981.1;refGene_gn=SCAF11;refGene_tx=XM_023256982.1;refGene_gn=SCAF11;refGene_tx=XM_023256983.1
B4 75249210 75249317
refGene_gn=SCAF11;refGene_tx=XM_023256980.1;refGene_gn=SCAF11;refGene_tx=XM_023256981.1;refGene_gn=SCAF11;refGene_tx=XM_023256982.1;refGene_gn=SCAF11;refGene_tx=XM_023256983.1
B4 75251075 75251177
refGene_gn=SCAF11;refGene_tx=XM_023256980.1;refGene_gn=SCAF11;refGene_tx=XM_023256981.1;refGene_gn=SCAF11;refGene_tx=XM_023256982.1;refGene_gn=SCAF11;refGene_tx=XM_023256983.1
B4 75252159 75252269
refGene_gn=SCAF11;refGene_tx=XM_023256980.1;refGene_gn=SCAF11;refGene_tx=XM_023256981.1;refGene_gn=SCAF11;refGene_tx=XM_023256982.1;refGene_gn=SCAF11;refGene_tx=XM_023256983.1
B4 75252648 75252707
refGene_gn=SCAF11;refGene_tx=XM_023256980.1;refGene_gn=SCAF11;refGene_tx=XM_023256981.1;refGene_gn=SCAF11;refGene_tx=XM_023256982.1;refGene_gn=SCAF11;refGene_tx=XM_023256983.1
B4 75259264 75259326
refGene_gn=SCAF11;refGene_tx=XM_023256980.1;refGene_gn=SCAF11;refGene_tx=XM_023256981.1;refGene_gn=SCAF11;refGene_tx=XM_023256982.1;refGene_gn=SCAF11;refGene_tx=XM_023256983.1
B4 75261658 75261759
refGene_gn=SCAF11;refGene_tx=XM_023256980.1;refGene_gn=SCAF11;refGene_tx=XM_023256981.1;refGene_gn=SCAF11;refGene_tx=XM_023256982.1;refGene_gn=SCAF11;refGene_tx=XM_023256983.1
B4 75265336 75265414
refGene_gn=SCAF11;refGene_tx=XM_023256980.1;refGene_gn=SCAF11;refGene_tx=XM_023256981.1;refGene_gn=SCAF11;refGene_tx=XM_023256982.1;refGene_gn=SCAF11;refGene_tx=XM_023256983.1
B4 75275804 75275962
refGene_gn=SCAF11;refGene_tx=XM_023256980.1;refGene_gn=SCAF11;refGene_tx=XM_023256981.1;refGene_gn=SCAF11;refGene_tx=XM_023256982.1;refGene_gn=SCAF11;refGene_tx=XM_023256983.1
B4 75279353 75279414
refGene_gn=SCAF11;refGene_tx=XM_023256980.1;refGene_gn=SCAF11;refGene_tx=XM_023256981.1;refGene_gn=SCAF11;refGene_tx=XM_023256982.1;refGene_gn=SCAF11;refGene_tx=XM_023256983.1
B4 75475791 75475893

refGene_gn=SLC38A1;refGene_tx=XM_006933636.4;refGene_gn=SLC38A1;refGene_tx=XM_011283882.3
B4 75477636 75477734
refGene_gn=SLC38A1;refGene_tx=XM_006933636.4;refGene_gn=SLC38A1;refGene_tx=XM_011283882.3
B4 75477834 75477976
refGene_gn=SLC38A1;refGene_tx=XM_006933636.4;refGene_gn=SLC38A1;refGene_tx=XM_011283882.3
B4 75479170 75479289
refGene_gn=SLC38A1;refGene_tx=XM_006933636.4;refGene_gn=SLC38A1;refGene_tx=XM_011283882.3
B4 75481367 75481468
refGene_gn=SLC38A1;refGene_tx=XM_006933636.4;refGene_gn=SLC38A1;refGene_tx=XM_011283882.3
B4 75483079 75483159
refGene_gn=SLC38A1;refGene_tx=XM_006933636.4;refGene_gn=SLC38A1;refGene_tx=XM_011283882.3
B4 75484526 75484643
refGene_gn=SLC38A1;refGene_tx=XM_006933636.4;refGene_gn=SLC38A1;refGene_tx=XM_011283882.3
B4 75484765 75484824
refGene_gn=SLC38A1;refGene_tx=XM_006933636.4;refGene_gn=SLC38A1;refGene_tx=XM_011283882.3
B4 75486973 75487056
refGene_gn=SLC38A1;refGene_tx=XM_006933636.4;refGene_gn=SLC38A1;refGene_tx=XM_011283882.3
B4 75488073 75488155
refGene_gn=SLC38A1;refGene_tx=XM_006933636.4;refGene_gn=SLC38A1;refGene_tx=XM_011283882.3
B4 75488461 75488554
refGene_gn=SLC38A1;refGene_tx=XM_006933636.4;refGene_gn=SLC38A1;refGene_tx=XM_011283882.3
B4 75489961 75490035
refGene_gn=SLC38A1;refGene_tx=XM_006933636.4;refGene_gn=SLC38A1;refGene_tx=XM_011283882.3
B4 75511130 75511246
refGene_gn=SLC38A1;refGene_tx=XM_006933636.4;refGene_gn=SLC38A1;refGene_tx=XM_011283882.3
B4 75511534 75511610
refGene_gn=SLC38A1;refGene_tx=XM_006933636.4;refGene_gn=SLC38A1;refGene_tx=XM_011283882.3
B4 75523400 75523522
refGene_gn=SLC38A1;refGene_tx=XM_006933636.4;refGene_gn=SLC38A1;refGene_tx=XM_011283882.3
B4 75593249 75593348
refGene_gn=SLC38A2;refGene_tx=XM_003988587.5;refGene_gn=SLC38A2;refGene_tx=XM_003988588.3;refGene_gn=SLC38A2;refGene_tx=XM_006933638.4
B4 75594404 75594502
refGene_gn=SLC38A2;refGene_tx=XM_003988587.5;refGene_gn=SLC38A2;refGene_tx=XM_003988588.3;refGene_gn=SLC38A2;refGene_tx=XM_006933638.4

B4	75594623 75594768
refGene_gn=SLC38A2;refGene_tx=XM_003988587.5;refGene_gn=SLC38A2;refGene_tx=XM_003988588.3;refGene_gn=SLC38A2;refGene_tx=XM_006933638.4
B4	75595108 75595233
refGene_gn=SLC38A2;refGene_tx=XM_003988587.5;refGene_gn=SLC38A2;refGene_tx=XM_003988588.3;refGene_gn=SLC38A2;refGene_tx=XM_006933638.4
B4	75595816 75595917
refGene_gn=SLC38A2;refGene_tx=XM_003988587.5;refGene_gn=SLC38A2;refGene_tx=XM_003988588.3;refGene_gn=SLC38A2;refGene_tx=XM_006933638.4
B4	75596031 75596111
refGene_gn=SLC38A2;refGene_tx=XM_003988587.5;refGene_gn=SLC38A2;refGene_tx=XM_003988588.3;refGene_gn=SLC38A2;refGene_tx=XM_006933638.4
B4	75596518 75596686
refGene_gn=SLC38A2;refGene_tx=XM_003988587.5;refGene_gn=SLC38A2;refGene_tx=XM_003988588.3;refGene_gn=SLC38A2;refGene_tx=XM_006933638.4
B4	75596798 75596857
refGene_gn=SLC38A2;refGene_tx=XM_003988587.5;refGene_gn=SLC38A2;refGene_tx=XM_003988588.3;refGene_gn=SLC38A2;refGene_tx=XM_006933638.4
B4	75597253 75597336
refGene_gn=SLC38A2;refGene_tx=XM_003988587.5;refGene_gn=SLC38A2;refGene_tx=XM_003988588.3;refGene_gn=SLC38A2;refGene_tx=XM_006933638.4
B4	75598738 75598820
refGene_gn=SLC38A2;refGene_tx=XM_003988587.5;refGene_gn=SLC38A2;refGene_tx=XM_003988588.3;refGene_gn=SLC38A2;refGene_tx=XM_006933638.4
B4	75598949 75599042
refGene_gn=SLC38A2;refGene_tx=XM_003988587.5;refGene_gn=SLC38A2;refGene_tx=XM_003988588.3;refGene_gn=SLC38A2;refGene_tx=XM_006933638.4
B4	75599128 75599202
refGene_gn=SLC38A2;refGene_tx=XM_003988587.5;refGene_gn=SLC38A2;refGene_tx=XM_003988588.3;refGene_gn=SLC38A2;refGene_tx=XM_006933638.4
B4	75602085 75602201
refGene_gn=SLC38A2;refGene_tx=XM_003988587.5;refGene_gn=SLC38A2;refGene_tx=XM_006933638.4
B4	75602348 75602430
refGene_gn=SLC38A2;refGene_tx=XM_003988587.5;refGene_gn=SLC38A2;refGene_tx=XM_003988588.3;refGene_gn=SLC38A2;refGene_tx=XM_006933638.4
B4	75602739 75602855
refGene_gn=SLC38A2;refGene_tx=XM_003988587.5;refGene_gn=SLC38A2;refGene_tx=XM_006933638.4
B4	75916646 75916748
refGene_gn=SLC38A4;refGene_tx=XM_006933639.2;refGene_gn=SLC38A4;refGene_tx=XM_019834842.1
B4	75918674 75918772
refGene_gn=SLC38A4;refGene_tx=XM_006933639.2;refGene_gn=SLC38A4;refGene_tx=XM_019834842.1
B4	75920129 75920274
refGene_gn=SLC38A4;refGene_tx=XM_006933639.2;refGene_gn=SLC38A4;refGene_tx=XM_019834842.1
B4	75925978 75926103
refGene_gn=SLC38A4;refGene_tx=XM_006933639.2;refGene_gn=SLC38A4;refGen

e_tx=XM_019834842.1
B4	75927853 75927954
refGene_gn=SLC38A4;refGene_tx=XM_006933639.2;refGene_gn=SLC38A4;refGen
e_tx=XM_019834842.1
B4	75929300 75929380
refGene_gn=SLC38A4;refGene_tx=XM_006933639.2;refGene_gn=SLC38A4;refGen
e_tx=XM_019834842.1
B4	75929492 75929768
refGene_gn=SLC38A4;refGene_tx=XM_006933639.2;refGene_gn=SLC38A4;refGen
e_tx=XM_019834842.1
B4	75930620 75930679
refGene_gn=SLC38A4;refGene_tx=XM_006933639.2;refGene_gn=SLC38A4;refGen
e_tx=XM_019834842.1
B4	75930779 75930862
refGene_gn=SLC38A4;refGene_tx=XM_006933639.2;refGene_gn=SLC38A4;refGen
e_tx=XM_019834842.1
B4	75930952 75931034
refGene_gn=SLC38A4;refGene_tx=XM_006933639.2;refGene_gn=SLC38A4;refGen
e_tx=XM_019834842.1
B4	75935755 75935848
refGene_gn=SLC38A4;refGene_tx=XM_006933639.2;refGene_gn=SLC38A4;refGen
e_tx=XM_019834842.1
B4	75936471 75936545
refGene_gn=SLC38A4;refGene_tx=XM_006933639.2;refGene_gn=SLC38A4;refGen
e_tx=XM_019834842.1
B4	75939355 75939471
refGene_gn=SLC38A4;refGene_tx=XM_006933639.2;refGene_gn=SLC38A4;refGen
e_tx=XM_019834842.1
B4	75939973 75940064
refGene_gn=SLC38A4;refGene_tx=XM_006933639.2;refGene_gn=SLC38A4;refGen
e_tx=XM_019834842.1
B4	75943955 75944074
refGene_gn=SLC38A4;refGene_tx=XM_006933639.2;refGene_gn=SLC38A4;refGen
e_tx=XM_019834842.1
B4	76163552 76165118 refGene_gn=AMIGO2;refGene_tx=XM_003988590.4
B4	76301054 76302353
refGene_gn=PCED1B;refGene_tx=XM_019834846.2;refGene_gn=PCED1B;refGene_
tx=XM_023256986.1
B4	76565136 76565222 refGene_gn=RPAP3;refGene_tx=XM_011283884.3
B4	76568031 76568132 refGene_gn=RPAP3;refGene_tx=XM_011283884.3
B4	76568852 76568950 refGene_gn=RPAP3;refGene_tx=XM_011283884.3
B4	76569779 76569966 refGene_gn=RPAP3;refGene_tx=XM_011283884.3
B4	76571230 76571451 refGene_gn=RPAP3;refGene_tx=XM_011283884.3
B4	76578160 76578262 refGene_gn=RPAP3;refGene_tx=XM_011283884.3
B4	76578379 76578450 refGene_gn=RPAP3;refGene_tx=XM_011283884.3
B4	76580150 76580272 refGene_gn=RPAP3;refGene_tx=XM_011283884.3
B4	76584192 76584230 refGene_gn=RPAP3;refGene_tx=XM_011283884.3
B4	76584231 76584321 refGene_gn=RPAP3;refGene_tx=XM_011283884.3
B4	76585246 76585372 refGene_gn=RPAP3;refGene_tx=XM_011283884.3
B4	76586510 76586581 refGene_gn=RPAP3;refGene_tx=XM_011283884.3

B4 76587298 76587420 refGene_gn=RPAP3;refGene_tx=XM_011283884.3
B4 76596712 76596840 refGene_gn=RPAP3;refGene_tx=XM_011283884.3
B4 76597669 76597792 refGene_gn=RPAP3;refGene_tx=XM_011283884.3
B4 76601311 76601452 refGene_gn=RPAP3;refGene_tx=XM_011283884.3
B4 76602676 76602829 refGene_gn=RPAP3;refGene_tx=XM_011283884.3
B4 76615683 76615804
refGene_gn=ENDOU;refGene_tx=XM_003988593.4;refGene_gn=ENDOU;refGene_tx=XM_019834847.2
B4 76616984 76617127
refGene_gn=ENDOU;refGene_tx=XM_003988593.4;refGene_gn=ENDOU;refGene_tx=XM_019834847.2
B4 76617868 76617975
refGene_gn=ENDOU;refGene_tx=XM_003988593.4;refGene_gn=ENDOU;refGene_tx=XM_019834847.2
B4 76618581 76618695
refGene_gn=ENDOU;refGene_tx=XM_003988593.4;refGene_gn=ENDOU;refGene_tx=XM_019834847.2
B4 76621670 76621870
refGene_gn=ENDOU;refGene_tx=XM_003988593.4;refGene_gn=ENDOU;refGene_tx=XM_019834847.2
B4 76622279 76622451
refGene_gn=ENDOU;refGene_tx=XM_003988593.4;refGene_gn=ENDOU;refGene_tx=XM_019834847.2
B4 76622924 76623059
refGene_gn=ENDOU;refGene_tx=XM_003988593.4;refGene_gn=ENDOU;refGene_tx=XM_019834847.2
B4 76623634 76623703
refGene_gn=ENDOU;refGene_tx=XM_003988593.4;refGene_gn=ENDOU;refGene_tx=XM_019834847.2
B4 76625986 76626109
refGene_gn=ENDOU;refGene_tx=XM_003988593.4;refGene_gn=ENDOU;refGene_tx=XM_019834847.2
B4 76631034 76631089 refGene_gn=ENDOU;refGene_tx=XM_003988593.4
B4 76643142 76643261
refGene_gn=RAPGEF3;refGene_tx=XM_003988595.5;refGene_gn=RAPGEF3;refGene_tx=XM_023256988.1
B4 76644083 76644155
refGene_gn=RAPGEF3;refGene_tx=XM_003988595.5;refGene_gn=RAPGEF3;refGene_tx=XM_023256988.1
B4 76644263 76644318
refGene_gn=RAPGEF3;refGene_tx=XM_003988595.5;refGene_gn=RAPGEF3;refGene_tx=XM_023256988.1
B4 76644954 76645019
refGene_gn=RAPGEF3;refGene_tx=XM_003988595.5;refGene_gn=RAPGEF3;refGene_tx=XM_023256988.1
B4 76645377 76645465
refGene_gn=RAPGEF3;refGene_tx=XM_003988595.5;refGene_gn=RAPGEF3;refGene_tx=XM_023256988.1
B4 76646393 76646444
refGene_gn=RAPGEF3;refGene_tx=XM_003988595.5;refGene_gn=RAPGEF3;refGen

e_tx=XM_023256988.1
B4	76646558 76646649
refGene_gn=RAPGEF3;refGene_tx=XM_003988595.5;refGene_gn=RAPGEF3;refGene_tx=XM_023256988.1
B4	76646863 76647045
refGene_gn=RAPGEF3;refGene_tx=XM_003988595.5;refGene_gn=RAPGEF3;refGene_tx=XM_023256988.1
B4	76647135 76647261
refGene_gn=RAPGEF3;refGene_tx=XM_003988595.5;refGene_gn=RAPGEF3;refGene_tx=XM_023256988.1
B4	76647732 76647830
refGene_gn=RAPGEF3;refGene_tx=XM_003988595.5;refGene_gn=RAPGEF3;refGene_tx=XM_023256988.1
B4	76649749 76649896
refGene_gn=RAPGEF3;refGene_tx=XM_003988595.5;refGene_gn=RAPGEF3;refGene_tx=XM_023256988.1
B4	76650220 76650302
refGene_gn=RAPGEF3;refGene_tx=XM_003988595.5;refGene_gn=RAPGEF3;refGene_tx=XM_023256988.1
B4	76652747 76652787
refGene_gn=RAPGEF3;refGene_tx=XM_003988595.5;refGene_gn=RAPGEF3;refGene_tx=XM_023256988.1
B4	76653607 76653690
refGene_gn=RAPGEF3;refGene_tx=XM_003988595.5;refGene_gn=RAPGEF3;refGene_tx=XM_023256988.1
B4	76653774 76653925
refGene_gn=RAPGEF3;refGene_tx=XM_003988595.5;refGene_gn=RAPGEF3;refGene_tx=XM_023256988.1
B4	76654131 76654210
refGene_gn=RAPGEF3;refGene_tx=XM_003988595.5;refGene_gn=RAPGEF3;refGene_tx=XM_023256988.1
B4	76654514 76654603
refGene_gn=RAPGEF3;refGene_tx=XM_003988595.5;refGene_gn=RAPGEF3;refGene_tx=XM_023256988.1
B4	76654909 76655022
refGene_gn=RAPGEF3;refGene_tx=XM_003988595.5;refGene_gn=RAPGEF3;refGene_tx=XM_023256988.1
B4	76655547 76655694
refGene_gn=RAPGEF3;refGene_tx=XM_003988595.5;refGene_gn=RAPGEF3;refGene_tx=XM_023256988.1
B4	76655884 76655961 refGene_gn=RAPGEF3;refGene_tx=XM_003988595.5
B4	76656069 76656130 refGene_gn=RAPGEF3;refGene_tx=XM_003988595.5
B4	76656437 76656522
refGene_gn=RAPGEF3;refGene_tx=XM_003988595.5;refGene_gn=RAPGEF3;refGene_tx=XM_023256988.1
B4	76657077 76657246
refGene_gn=RAPGEF3;refGene_tx=XM_003988595.5;refGene_gn=RAPGEF3;refGene_tx=XM_023256988.1
B4	76657390 76657512
refGene_gn=RAPGEF3;refGene_tx=XM_003988595.5;refGene_gn=RAPGEF3;refGen

e_tx=XM_023256988.1
B4	76657867	76657974
refGene_gn=RAPGEF3;refGene_tx=XM_003988595.5;refGene_gn=RAPGEF3;refGen
e_tx=XM_023256988.1
B4	76658058	76658112
refGene_gn=RAPGEF3;refGene_tx=XM_003988595.5;refGene_gn=RAPGEF3;refGen
e_tx=XM_023256988.1
B4	76663761	76663974
refGene_gn=RAPGEF3;refGene_tx=XM_003988595.5;refGene_gn=RAPGEF3;refGen
e_tx=XM_023256988.1
B4	76664455	76664461
refGene_gn=RAPGEF3;refGene_tx=XM_003988595.5;refGene_gn=RAPGEF3;refGen
e_tx=XM_023256988.1
B4	76678845	76678981	refGene_gn=SLC48A1;refGene_tx=XM_023256992.1
B4	76684372	76684540	refGene_gn=SLC48A1;refGene_tx=XM_023256992.1
B4	76685166	76685303	refGene_gn=SLC48A1;refGene_tx=XM_023256992.1
B4	76688606	76688652
refGene_gn=HDAC7;refGene_tx=XM_006933642.4;refGene_gn=HDAC7;refGene_tx
=XM_011283885.2;refGene_gn=HDAC7;refGene_tx=XM_011283886.2;refGene_gn=
HDAC7;refGene_tx=XM_011283887.2;refGene_gn=HDAC7;refGene_tx=XM_0198348
49.1;refGene_gn=HDAC7;refGene_tx=XM_023256989.1;refGene_gn=HDAC7;refGe
ne_tx=XM_023256990.1;refGene_gn=HDAC7;refGene_tx=XM_023256991.1
B4	76688824	76688963
refGene_gn=HDAC7;refGene_tx=XM_006933642.4;refGene_gn=HDAC7;refGene_tx
=XM_011283885.2;refGene_gn=HDAC7;refGene_tx=XM_011283886.2;refGene_gn=
HDAC7;refGene_tx=XM_011283887.2;refGene_gn=HDAC7;refGene_tx=XM_0198348
49.1;refGene_gn=HDAC7;refGene_tx=XM_023256989.1;refGene_gn=HDAC7;refGe
ne_tx=XM_023256990.1;refGene_gn=HDAC7;refGene_tx=XM_023256991.1
B4	76690236	76690321
refGene_gn=HDAC7;refGene_tx=XM_006933642.4;refGene_gn=HDAC7;refGene_tx
=XM_011283885.2;refGene_gn=HDAC7;refGene_tx=XM_011283886.2;refGene_gn=
HDAC7;refGene_tx=XM_011283887.2;refGene_gn=HDAC7;refGene_tx=XM_0198348
49.1;refGene_gn=HDAC7;refGene_tx=XM_023256989.1;refGene_gn=HDAC7;refGe
ne_tx=XM_023256990.1;refGene_gn=HDAC7;refGene_tx=XM_023256991.1
B4	76690630	76690764
refGene_gn=HDAC7;refGene_tx=XM_006933642.4;refGene_gn=HDAC7;refGene_tx
=XM_011283885.2;refGene_gn=HDAC7;refGene_tx=XM_011283886.2;refGene_gn=
HDAC7;refGene_tx=XM_011283887.2;refGene_gn=HDAC7;refGene_tx=XM_0198348
49.1;refGene_gn=HDAC7;refGene_tx=XM_023256989.1;refGene_gn=HDAC7;refGe
ne_tx=XM_023256990.1;refGene_gn=HDAC7;refGene_tx=XM_023256991.1
B4	76691246	76691365
refGene_gn=HDAC7;refGene_tx=XM_006933642.4;refGene_gn=HDAC7;refGene_tx
=XM_011283885.2;refGene_gn=HDAC7;refGene_tx=XM_011283886.2;refGene_gn=
HDAC7;refGene_tx=XM_011283887.2;refGene_gn=HDAC7;refGene_tx=XM_0198348
49.1;refGene_gn=HDAC7;refGene_tx=XM_023256989.1;refGene_gn=HDAC7;refGe
ne_tx=XM_023256990.1;refGene_gn=HDAC7;refGene_tx=XM_023256991.1
B4	76692538	76692636
refGene_gn=HDAC7;refGene_tx=XM_006933642.4;refGene_gn=HDAC7;refGene_tx
=XM_011283885.2;refGene_gn=HDAC7;refGene_tx=XM_011283886.2;refGene_gn=
HDAC7;refGene_tx=XM_011283887.2;refGene_gn=HDAC7;refGene_tx=XM_0198348

49.1;refGene_gn=HDAC7;refGene_tx=XM_023256989.1;refGene_gn=HDAC7;refGene_tx=XM_023256990.1;refGene_gn=HDAC7;refGene_tx=XM_023256991.1
B4 76692865 76692985
refGene_gn=HDAC7;refGene_tx=XM_006933642.4;refGene_gn=HDAC7;refGene_tx=XM_011283885.2;refGene_gn=HDAC7;refGene_tx=XM_011283886.2;refGene_gn=HDAC7;refGene_tx=XM_011283887.2;refGene_gn=HDAC7;refGene_tx=XM_019834849.1;refGene_gn=HDAC7;refGene_tx=XM_023256989.1;refGene_gn=HDAC7;refGene_tx=XM_023256990.1;refGene_gn=HDAC7;refGene_tx=XM_023256991.1
B4 76694063 76694151
refGene_gn=HDAC7;refGene_tx=XM_006933642.4;refGene_gn=HDAC7;refGene_tx=XM_011283885.2;refGene_gn=HDAC7;refGene_tx=XM_011283886.2;refGene_gn=HDAC7;refGene_tx=XM_011283887.2;refGene_gn=HDAC7;refGene_tx=XM_019834849.1;refGene_gn=HDAC7;refGene_tx=XM_023256989.1;refGene_gn=HDAC7;refGene_tx=XM_023256990.1;refGene_gn=HDAC7;refGene_tx=XM_023256991.1
B4 76694326 76694382
refGene_gn=HDAC7;refGene_tx=XM_006933642.4;refGene_gn=HDAC7;refGene_tx=XM_011283885.2;refGene_gn=HDAC7;refGene_tx=XM_011283886.2;refGene_gn=HDAC7;refGene_tx=XM_011283887.2;refGene_gn=HDAC7;refGene_tx=XM_019834849.1;refGene_gn=HDAC7;refGene_tx=XM_023256989.1;refGene_gn=HDAC7;refGene_tx=XM_023256990.1;refGene_gn=HDAC7;refGene_tx=XM_023256991.1
B4 76695398 76695506
refGene_gn=HDAC7;refGene_tx=XM_006933642.4;refGene_gn=HDAC7;refGene_tx=XM_011283885.2;refGene_gn=HDAC7;refGene_tx=XM_011283886.2;refGene_gn=HDAC7;refGene_tx=XM_011283887.2;refGene_gn=HDAC7;refGene_tx=XM_019834849.1;refGene_gn=HDAC7;refGene_tx=XM_023256989.1;refGene_gn=HDAC7;refGene_tx=XM_023256990.1;refGene_gn=HDAC7;refGene_tx=XM_023256991.1
B4 76696731 76696781
refGene_gn=HDAC7;refGene_tx=XM_006933642.4;refGene_gn=HDAC7;refGene_tx=XM_011283885.2;refGene_gn=HDAC7;refGene_tx=XM_011283886.2;refGene_gn=HDAC7;refGene_tx=XM_011283887.2;refGene_gn=HDAC7;refGene_tx=XM_019834849.1;refGene_gn=HDAC7;refGene_tx=XM_023256989.1;refGene_gn=HDAC7;refGene_tx=XM_023256990.1;refGene_gn=HDAC7;refGene_tx=XM_023256991.1
B4 76697041 76697162
refGene_gn=HDAC7;refGene_tx=XM_006933642.4;refGene_gn=HDAC7;refGene_tx=XM_011283885.2;refGene_gn=HDAC7;refGene_tx=XM_011283886.2;refGene_gn=HDAC7;refGene_tx=XM_011283887.2;refGene_gn=HDAC7;refGene_tx=XM_019834849.1;refGene_gn=HDAC7;refGene_tx=XM_023256989.1;refGene_gn=HDAC7;refGene_tx=XM_023256990.1;refGene_gn=HDAC7;refGene_tx=XM_023256991.1
B4 76697370 76697504
refGene_gn=HDAC7;refGene_tx=XM_006933642.4;refGene_gn=HDAC7;refGene_tx=XM_011283885.2;refGene_gn=HDAC7;refGene_tx=XM_011283886.2;refGene_gn=HDAC7;refGene_tx=XM_011283887.2;refGene_gn=HDAC7;refGene_tx=XM_019834849.1;refGene_gn=HDAC7;refGene_tx=XM_023256989.1;refGene_gn=HDAC7;refGene_tx=XM_023256990.1;refGene_gn=HDAC7;refGene_tx=XM_023256991.1
B4 76698838 76699058
refGene_gn=HDAC7;refGene_tx=XM_006933642.4;refGene_gn=HDAC7;refGene_tx=XM_011283885.2;refGene_gn=HDAC7;refGene_tx=XM_011283886.2;refGene_gn=HDAC7;refGene_tx=XM_011283887.2;refGene_gn=HDAC7;refGene_tx=XM_019834849.1;refGene_gn=HDAC7;refGene_tx=XM_023256989.1;refGene_gn=HDAC7;refGene_tx=XM_023256990.1;refGene_gn=HDAC7;refGene_tx=XM_023256991.1

B4 76700215 76700389
refGene_gn=HDAC7;refGene_tx=XM_006933642.4;refGene_gn=HDAC7;refGene_tx=XM_011283885.2;refGene_gn=HDAC7;refGene_tx=XM_011283886.2;refGene_gn=HDAC7;refGene_tx=XM_011283887.2;refGene_gn=HDAC7;refGene_tx=XM_019834849.1;refGene_gn=HDAC7;refGene_tx=XM_023256989.1;refGene_gn=HDAC7;refGene_tx=XM_023256990.1;refGene_gn=HDAC7;refGene_tx=XM_023256991.1
B4 76700635 76700832
refGene_gn=HDAC7;refGene_tx=XM_006933642.4;refGene_gn=HDAC7;refGene_tx=XM_011283885.2;refGene_gn=HDAC7;refGene_tx=XM_011283886.2;refGene_gn=HDAC7;refGene_tx=XM_011283887.2;refGene_gn=HDAC7;refGene_tx=XM_019834849.1;refGene_gn=HDAC7;refGene_tx=XM_023256989.1;refGene_gn=HDAC7;refGene_tx=XM_023256990.1;refGene_gn=HDAC7;refGene_tx=XM_023256991.1
B4 76701041 76701222
refGene_gn=HDAC7;refGene_tx=XM_006933642.4;refGene_gn=HDAC7;refGene_tx=XM_011283885.2;refGene_gn=HDAC7;refGene_tx=XM_011283886.2;refGene_gn=HDAC7;refGene_tx=XM_011283887.2;refGene_gn=HDAC7;refGene_tx=XM_019834849.1;refGene_gn=HDAC7;refGene_tx=XM_023256989.1;refGene_gn=HDAC7;refGene_tx=XM_023256990.1;refGene_gn=HDAC7;refGene_tx=XM_023256991.1
B4 76701363 76701474
refGene_gn=HDAC7;refGene_tx=XM_006933642.4;refGene_gn=HDAC7;refGene_tx=XM_011283885.2;refGene_gn=HDAC7;refGene_tx=XM_011283886.2;refGene_gn=HDAC7;refGene_tx=XM_019834849.1;refGene_gn=HDAC7;refGene_tx=XM_023256989.1;refGene_gn=HDAC7;refGene_tx=XM_023256990.1;refGene_gn=HDAC7;refGene_tx=XM_023256991.1
B4 76701676 76701768
refGene_gn=HDAC7;refGene_tx=XM_006933642.4;refGene_gn=HDAC7;refGene_tx=XM_011283885.2;refGene_gn=HDAC7;refGene_tx=XM_011283886.2;refGene_gn=HDAC7;refGene_tx=XM_011283887.2;refGene_gn=HDAC7;refGene_tx=XM_019834849.1;refGene_gn=HDAC7;refGene_tx=XM_023256989.1;refGene_gn=HDAC7;refGene_tx=XM_023256990.1;refGene_gn=HDAC7;refGene_tx=XM_023256991.1
B4 76702102 76702228
refGene_gn=HDAC7;refGene_tx=XM_006933642.4;refGene_gn=HDAC7;refGene_tx=XM_011283885.2;refGene_gn=HDAC7;refGene_tx=XM_011283886.2;refGene_gn=HDAC7;refGene_tx=XM_011283887.2;refGene_gn=HDAC7;refGene_tx=XM_019834849.1;refGene_gn=HDAC7;refGene_tx=XM_023256989.1;refGene_gn=HDAC7;refGene_tx=XM_023256990.1;refGene_gn=HDAC7;refGene_tx=XM_023256991.1
B4 76702442 76702558
refGene_gn=HDAC7;refGene_tx=XM_006933642.4;refGene_gn=HDAC7;refGene_tx=XM_011283885.2;refGene_gn=HDAC7;refGene_tx=XM_011283886.2;refGene_gn=HDAC7;refGene_tx=XM_011283887.2;refGene_gn=HDAC7;refGene_tx=XM_019834849.1;refGene_gn=HDAC7;refGene_tx=XM_023256989.1;refGene_gn=HDAC7;refGene_tx=XM_023256990.1;refGene_gn=HDAC7;refGene_tx=XM_023256991.1
B4 76703187 76703299
refGene_gn=HDAC7;refGene_tx=XM_006933642.4;refGene_gn=HDAC7;refGene_tx=XM_011283885.2;refGene_gn=HDAC7;refGene_tx=XM_011283886.2;refGene_gn=HDAC7;refGene_tx=XM_011283887.2;refGene_gn=HDAC7;refGene_tx=XM_019834849.1;refGene_gn=HDAC7;refGene_tx=XM_023256989.1;refGene_gn=HDAC7;refGene_tx=XM_023256990.1;refGene_gn=HDAC7;refGene_tx=XM_023256991.1
B4 76703512 76703603
refGene_gn=HDAC7;refGene_tx=XM_006933642.4;refGene_gn=HDAC7;refGene_tx

=XM_011283885.2;refGene_gn=HDAC7;refGene_tx=XM_011283886.2;refGene_gn=HDAC7;refGene_tx=XM_011283887.2;refGene_gn=HDAC7;refGene_tx=XM_019834849.1;refGene_gn=HDAC7;refGene_tx=XM_023256989.1;refGene_gn=HDAC7;refGene_tx=XM_023256990.1;refGene_gn=HDAC7;refGene_tx=XM_023256991.1
B4 76703740 76703928
refGene_gn=HDAC7;refGene_tx=XM_006933642.4;refGene_gn=HDAC7;refGene_tx=XM_011283885.2;refGene_gn=HDAC7;refGene_tx=XM_011283886.2;refGene_gn=HDAC7;refGene_tx=XM_011283887.2;refGene_gn=HDAC7;refGene_tx=XM_019834849.1;refGene_gn=HDAC7;refGene_tx=XM_023256989.1;refGene_gn=HDAC7;refGene_tx=XM_023256990.1;refGene_gn=HDAC7;refGene_tx=XM_023256991.1
B4 76707411 76707462
refGene_gn=HDAC7;refGene_tx=XM_006933642.4;refGene_gn=HDAC7;refGene_tx=XM_011283885.2;refGene_gn=HDAC7;refGene_tx=XM_011283886.2;refGene_gn=HDAC7;refGene_tx=XM_011283887.2;refGene_gn=HDAC7;refGene_tx=XM_019834849.1;refGene_gn=HDAC7;refGene_tx=XM_023256989.1
B4 76707702 76707718 refGene_gn=HDAC7;refGene_tx=XM_011283886.2
B4 76713461 76713513 refGene_gn=HDAC7;refGene_tx=XM_006933642.4
B4 76723405 76723424
refGene_gn=HDAC7;refGene_tx=XM_023256989.1;refGene_gn=HDAC7;refGene_tx=XM_023256990.1
B4 76723866 76723936
refGene_gn=HDAC7;refGene_tx=XM_011283885.2;refGene_gn=HDAC7;refGene_tx=XM_011283887.2;refGene_gn=HDAC7;refGene_tx=XM_019834849.1
B4 76747762 76748022
refGene_gn=VDR;refGene_tx=XM_003988596.5;refGene_gn=VDR;refGene_tx=XM_011283890.2;refGene_gn=VDR;refGene_tx=XM_019834851.2;refGene_gn=VDR;refGene_tx=XM_019834852.2
B4 76749333 76749450
refGene_gn=VDR;refGene_tx=XM_003988596.5;refGene_gn=VDR;refGene_tx=XM_011283890.2;refGene_gn=VDR;refGene_tx=XM_019834851.2;refGene_gn=VDR;refGene_tx=XM_019834852.2
B4 76749666 76749818
refGene_gn=VDR;refGene_tx=XM_003988596.5;refGene_gn=VDR;refGene_tx=XM_011283890.2;refGene_gn=VDR;refGene_tx=XM_019834851.2;refGene_gn=VDR;refGene_tx=XM_019834852.2
B4 76756018 76756190
refGene_gn=VDR;refGene_tx=XM_003988596.5;refGene_gn=VDR;refGene_tx=XM_011283890.2;refGene_gn=VDR;refGene_tx=XM_019834851.2;refGene_gn=VDR;refGene_tx=XM_019834852.2
B4 76758904 76759025
refGene_gn=VDR;refGene_tx=XM_003988596.5;refGene_gn=VDR;refGene_tx=XM_011283890.2;refGene_gn=VDR;refGene_tx=XM_019834851.2;refGene_gn=VDR;refGene_tx=XM_019834852.2
B4 76759295 76759480
refGene_gn=VDR;refGene_tx=XM_003988596.5;refGene_gn=VDR;refGene_tx=XM_011283890.2;refGene_gn=VDR;refGene_tx=XM_019834851.2;refGene_gn=VDR;refGene_tx=XM_019834852.2
B4 76767347 76767478
refGene_gn=VDR;refGene_tx=XM_003988596.5;refGene_gn=VDR;refGene_tx=XM_011283890.2;refGene_gn=VDR;refGene_tx=XM_019834851.2;refGene_gn=VDR;re

fGene_tx=XM_019834852.2
B4	76777538	76777686
refGene_gn=VDR;refGene_tx=XM_003988596.5;refGene_gn=VDR;refGene_tx=XM_011283890.2;refGene_gn=VDR;refGene_tx=XM_019834851.2;refGene_gn=VDR;refGene_tx=XM_019834852.2
B4	76801955	76801995	refGene_gn=VDR;refGene_tx=XM_019834851.2
B4	76848101	76848285
refGene_gn=TMEM106C;refGene_tx=XM_003988597.4;refGene_gn=TMEM106C;refGene_tx=XM_003988598.3;refGene_gn=TMEM106C;refGene_tx=XM_006933646.2;refGene_gn=TMEM106C;refGene_tx=XM_006933647.4;refGene_gn=TMEM106C;refGene_tx=XM_006933648.3;refGene_gn=TMEM106C;refGene_tx=XM_006933649.4;refGene_gn=TMEM106C;refGene_tx=XM_019834853.2
B4	76849142	76849206
refGene_gn=TMEM106C;refGene_tx=XM_003988597.4;refGene_gn=TMEM106C;refGene_tx=XM_003988598.3;refGene_gn=TMEM106C;refGene_tx=XM_006933646.2;refGene_gn=TMEM106C;refGene_tx=XM_006933647.4;refGene_gn=TMEM106C;refGene_tx=XM_006933648.3;refGene_gn=TMEM106C;refGene_tx=XM_006933649.4;refGene_gn=TMEM106C;refGene_tx=XM_019834853.2
B4	76849710	76849870
refGene_gn=TMEM106C;refGene_tx=XM_003988597.4;refGene_gn=TMEM106C;refGene_tx=XM_003988598.3;refGene_gn=TMEM106C;refGene_tx=XM_006933646.2;refGene_gn=TMEM106C;refGene_tx=XM_006933647.4;refGene_gn=TMEM106C;refGene_tx=XM_006933648.3;refGene_gn=TMEM106C;refGene_tx=XM_006933649.4;refGene_gn=TMEM106C;refGene_tx=XM_019834853.2
B4	76849961	76850102
refGene_gn=TMEM106C;refGene_tx=XM_003988597.4;refGene_gn=TMEM106C;refGene_tx=XM_003988598.3;refGene_gn=TMEM106C;refGene_tx=XM_006933646.2;refGene_gn=TMEM106C;refGene_tx=XM_006933647.4;refGene_gn=TMEM106C;refGene_tx=XM_006933648.3;refGene_gn=TMEM106C;refGene_tx=XM_006933649.4;refGene_gn=TMEM106C;refGene_tx=XM_019834853.2
B4	76850555	76850605
refGene_gn=TMEM106C;refGene_tx=XM_003988597.4;refGene_gn=TMEM106C;refGene_tx=XM_003988598.3;refGene_gn=TMEM106C;refGene_tx=XM_006933646.2;refGene_gn=TMEM106C;refGene_tx=XM_006933647.4;refGene_gn=TMEM106C;refGene_tx=XM_006933648.3;refGene_gn=TMEM106C;refGene_tx=XM_006933649.4;refGene_gn=TMEM106C;refGene_tx=XM_019834853.2
B4	76851057	76851111
refGene_gn=TMEM106C;refGene_tx=XM_003988597.4;refGene_gn=TMEM106C;refGene_tx=XM_003988598.3;refGene_gn=TMEM106C;refGene_tx=XM_006933646.2;refGene_gn=TMEM106C;refGene_tx=XM_006933647.4;refGene_gn=TMEM106C;refGene_tx=XM_006933648.3;refGene_gn=TMEM106C;refGene_tx=XM_006933649.4;refGene_gn=TMEM106C;refGene_tx=XM_019834853.2
B4	76852039	76852136
refGene_gn=TMEM106C;refGene_tx=XM_003988597.4;refGene_gn=TMEM106C;refGene_tx=XM_003988598.3;refGene_gn=TMEM106C;refGene_tx=XM_006933646.2;refGene_gn=TMEM106C;refGene_tx=XM_006933647.4;refGene_gn=TMEM106C;refGene_tx=XM_006933648.3;refGene_gn=TMEM106C;refGene_tx=XM_006933649.4;refGene_gn=TMEM106C;refGene_tx=XM_019834853.2
B4	76868304	76868451	refGene_gn=COL2A1;refGene_tx=XM_023256993.1
B4	76868910	76869153	refGene_gn=COL2A1;refGene_tx=XM_023256993.1

| | | | |
|---|---|---|---|
| B4 | 76869349 | 76869537 | refGene_gn=COL2A1;refGene_tx=XM_023256993.1 |
| B4 | 76870015 | 76870304 | refGene_gn=COL2A1;refGene_tx=XM_023256993.1 |
| B4 | 76870658 | 76870766 | refGene_gn=COL2A1;refGene_tx=XM_023256993.1 |
| B4 | 76871170 | 76871224 | refGene_gn=COL2A1;refGene_tx=XM_023256993.1 |
| B4 | 76871452 | 76871560 | refGene_gn=COL2A1;refGene_tx=XM_023256993.1 |
| B4 | 76871707 | 76871761 | refGene_gn=COL2A1;refGene_tx=XM_023256993.1 |
| B4 | 76871949 | 76872057 | refGene_gn=COL2A1;refGene_tx=XM_023256993.1 |
| B4 | 76872230 | 76872284 | refGene_gn=COL2A1;refGene_tx=XM_023256993.1 |
| B4 | 76872621 | 76872729 | refGene_gn=COL2A1;refGene_tx=XM_023256993.1 |
| B4 | 76872969 | 76873077 | refGene_gn=COL2A1;refGene_tx=XM_023256993.1 |
| B4 | 76873267 | 76873429 | refGene_gn=COL2A1;refGene_tx=XM_023256993.1 |
| B4 | 76874165 | 76874219 | refGene_gn=COL2A1;refGene_tx=XM_023256993.1 |
| B4 | 76874713 | 76874767 | refGene_gn=COL2A1;refGene_tx=XM_023256993.1 |
| B4 | 76875359 | 76875467 | refGene_gn=COL2A1;refGene_tx=XM_023256993.1 |
| B4 | 76875764 | 76875818 | refGene_gn=COL2A1;refGene_tx=XM_023256993.1 |
| B4 | 76876166 | 76876220 | refGene_gn=COL2A1;refGene_tx=XM_023256993.1 |
| B4 | 76876598 | 76876652 | refGene_gn=COL2A1;refGene_tx=XM_023256993.1 |
| B4 | 76876891 | 76876945 | refGene_gn=COL2A1;refGene_tx=XM_023256993.1 |
| B4 | 76877272 | 76877380 | refGene_gn=COL2A1;refGene_tx=XM_023256993.1 |
| B4 | 76877575 | 76877674 | refGene_gn=COL2A1;refGene_tx=XM_023256993.1 |
| B4 | 76877858 | 76877903 | refGene_gn=COL2A1;refGene_tx=XM_023256993.1 |
| B4 | 76878102 | 76878156 | refGene_gn=COL2A1;refGene_tx=XM_023256993.1 |
| B4 | 76878369 | 76878423 | refGene_gn=COL2A1;refGene_tx=XM_023256993.1 |
| B4 | 76878740 | 76878797 | refGene_gn=COL2A1;refGene_tx=XM_023256993.1 |
| B4 | 76879173 | 76879227 | refGene_gn=COL2A1;refGene_tx=XM_023256993.1 |
| B4 | 76879615 | 76879714 | refGene_gn=COL2A1;refGene_tx=XM_023256993.1 |
| B4 | 76880150 | 76880204 | refGene_gn=COL2A1;refGene_tx=XM_023256993.1 |
| B4 | 76880360 | 76880459 | refGene_gn=COL2A1;refGene_tx=XM_023256993.1 |
| B4 | 76880540 | 76880594 | refGene_gn=COL2A1;refGene_tx=XM_023256993.1 |
| B4 | 76880990 | 76881098 | refGene_gn=COL2A1;refGene_tx=XM_023256993.1 |
| B4 | 76881448 | 76881502 | refGene_gn=COL2A1;refGene_tx=XM_023256993.1 |
| B4 | 76881694 | 76881793 | refGene_gn=COL2A1;refGene_tx=XM_023256993.1 |
| B4 | 76881886 | 76881931 | refGene_gn=COL2A1;refGene_tx=XM_023256993.1 |
| B4 | 76882240 | 76882339 | refGene_gn=COL2A1;refGene_tx=XM_023256993.1 |
| B4 | 76883815 | 76883869 | refGene_gn=COL2A1;refGene_tx=XM_023256993.1 |
| B4 | 76884302 | 76884347 | refGene_gn=COL2A1;refGene_tx=XM_023256993.1 |
| B4 | 76887005 | 76887059 | refGene_gn=COL2A1;refGene_tx=XM_023256993.1 |
| B4 | 76887604 | 76887649 | refGene_gn=COL2A1;refGene_tx=XM_023256993.1 |
| B4 | 76887938 | 76887992 | refGene_gn=COL2A1;refGene_tx=XM_023256993.1 |
| B4 | 76888110 | 76888164 | refGene_gn=COL2A1;refGene_tx=XM_023256993.1 |
| B4 | 76888581 | 76888635 | refGene_gn=COL2A1;refGene_tx=XM_023256993.1 |
| B4 | 76889400 | 76889454 | refGene_gn=COL2A1;refGene_tx=XM_023256993.1 |
| B4 | 76889885 | 76889939 | refGene_gn=COL2A1;refGene_tx=XM_023256993.1 |
| B4 | 76890033 | 76890078 | refGene_gn=COL2A1;refGene_tx=XM_023256993.1 |
| B4 | 76890723 | 76890801 | refGene_gn=COL2A1;refGene_tx=XM_023256993.1 |
| B4 | 76892016 | 76892118 | refGene_gn=COL2A1;refGene_tx=XM_023256993.1 |
| B4 | 76892285 | 76892339 | refGene_gn=COL2A1;refGene_tx=XM_023256993.1 |
| B4 | 76892437 | 76892470 | refGene_gn=COL2A1;refGene_tx=XM_023256993.1 |
| B4 | 76892576 | 76892609 | refGene_gn=COL2A1;refGene_tx=XM_023256993.1 |
| B4 | 76893050 | 76893067 | refGene_gn=COL2A1;refGene_tx=XM_023256993.1 |

B4	76894494 76894701 refGene_gn=COL2A1;refGene_tx=XM_023256993.1
B4	76898804 76898889 refGene_gn=COL2A1;refGene_tx=XM_023256993.1
B4	76938998 76939007 refGene_gn=SENP1;refGene_tx=XM_023256997.1
B4	76941569 76941632
refGene_gn=SENP1;refGene_tx=XM_019834858.2;refGene_gn=SENP1;refGene_tx
=XM_023256995.1;refGene_gn=SENP1;refGene_tx=XM_023256996.1;refGene_gn=
SENP1;refGene_tx=XM_023256998.1;refGene_gn=SENP1;refGene_tx=XM_0232569
99.1;refGene_gn=SENP1;refGene_tx=XM_023257001.1
B4	76942581 76942677
refGene_gn=SENP1;refGene_tx=XM_019834858.2;refGene_gn=SENP1;refGene_tx
=XM_023256995.1;refGene_gn=SENP1;refGene_tx=XM_023256996.1;refGene_gn=
SENP1;refGene_tx=XM_023256997.1;refGene_gn=SENP1;refGene_tx=XM_0232569
98.1;refGene_gn=SENP1;refGene_tx=XM_023256999.1;refGene_gn=SENP1;refGe
ne_tx=XM_023257001.1
B4	76943195 76943280
refGene_gn=SENP1;refGene_tx=XM_019834858.2;refGene_gn=SENP1;refGene_tx
=XM_023256995.1;refGene_gn=SENP1;refGene_tx=XM_023256996.1;refGene_gn=
SENP1;refGene_tx=XM_023256997.1;refGene_gn=SENP1;refGene_tx=XM_0232569
98.1;refGene_gn=SENP1;refGene_tx=XM_023256999.1;refGene_gn=SENP1;refGe
ne_tx=XM_023257001.1
B4	76944221 76944301
refGene_gn=SENP1;refGene_tx=XM_019834858.2;refGene_gn=SENP1;refGene_tx
=XM_023256995.1;refGene_gn=SENP1;refGene_tx=XM_023256996.1;refGene_gn=
SENP1;refGene_tx=XM_023256997.1;refGene_gn=SENP1;refGene_tx=XM_0232569
98.1;refGene_gn=SENP1;refGene_tx=XM_023256999.1;refGene_gn=SENP1;refGe
ne_tx=XM_023257001.1
B4	76945129 76945333
refGene_gn=SENP1;refGene_tx=XM_019834858.2;refGene_gn=SENP1;refGene_tx
=XM_023256995.1;refGene_gn=SENP1;refGene_tx=XM_023256996.1;refGene_gn=
SENP1;refGene_tx=XM_023256997.1;refGene_gn=SENP1;refGene_tx=XM_0232569
98.1;refGene_gn=SENP1;refGene_tx=XM_023256999.1;refGene_gn=SENP1;refGe
ne_tx=XM_023257001.1
B4	76958150 76958282
refGene_gn=SENP1;refGene_tx=XM_019834858.2;refGene_gn=SENP1;refGene_tx
=XM_023256995.1;refGene_gn=SENP1;refGene_tx=XM_023256996.1;refGene_gn=
SENP1;refGene_tx=XM_023256997.1;refGene_gn=SENP1;refGene_tx=XM_0232569
98.1;refGene_gn=SENP1;refGene_tx=XM_023256999.1;refGene_gn=SENP1;refGe
ne_tx=XM_023257001.1
B4	76959360 76959516
refGene_gn=SENP1;refGene_tx=XM_019834858.2;refGene_gn=SENP1;refGene_tx
=XM_023256995.1;refGene_gn=SENP1;refGene_tx=XM_023256996.1;refGene_gn=
SENP1;refGene_tx=XM_023256997.1;refGene_gn=SENP1;refGene_tx=XM_0232569
98.1;refGene_gn=SENP1;refGene_tx=XM_023256999.1;refGene_gn=SENP1;refGe
ne_tx=XM_023257001.1
B4	76959751 76959836
refGene_gn=SENP1;refGene_tx=XM_019834858.2;refGene_gn=SENP1;refGene_tx
=XM_023256995.1;refGene_gn=SENP1;refGene_tx=XM_023256996.1;refGene_gn=
SENP1;refGene_tx=XM_023256997.1;refGene_gn=SENP1;refGene_tx=XM_0232569
98.1;refGene_gn=SENP1;refGene_tx=XM_023256999.1;refGene_gn=SENP1;refGe
ne_tx=XM_023257001.1

B4 76960699 76960738
refGene_gn=SENP1;refGene_tx=XM_019834858.2;refGene_gn=SENP1;refGene_tx=XM_023256995.1;refGene_gn=SENP1;refGene_tx=XM_023256996.1;refGene_gn=SENP1;refGene_tx=XM_023256997.1;refGene_gn=SENP1;refGene_tx=XM_023256998.1;refGene_gn=SENP1;refGene_tx=XM_023256999.1;refGene_gn=SENP1;refGene_tx=XM_023257001.1
B4 76964950 76965005
refGene_gn=SENP1;refGene_tx=XM_019834858.2;refGene_gn=SENP1;refGene_tx=XM_023256995.1;refGene_gn=SENP1;refGene_tx=XM_023256996.1;refGene_gn=SENP1;refGene_tx=XM_023256997.1;refGene_gn=SENP1;refGene_tx=XM_023256998.1;refGene_gn=SENP1;refGene_tx=XM_023256999.1;refGene_gn=SENP1;refGene_tx=XM_023257001.1
B4 76966509 76966796
refGene_gn=SENP1;refGene_tx=XM_019834858.2;refGene_gn=SENP1;refGene_tx=XM_023256995.1;refGene_gn=SENP1;refGene_tx=XM_023256996.1;refGene_gn=SENP1;refGene_tx=XM_023256997.1;refGene_gn=SENP1;refGene_tx=XM_023256998.1;refGene_gn=SENP1;refGene_tx=XM_023256999.1;refGene_gn=SENP1;refGene_tx=XM_023257001.1
B4 76966878 76966982
refGene_gn=SENP1;refGene_tx=XM_019834858.2;refGene_gn=SENP1;refGene_tx=XM_023256995.1;refGene_gn=SENP1;refGene_tx=XM_023256996.1;refGene_gn=SENP1;refGene_tx=XM_023256997.1;refGene_gn=SENP1;refGene_tx=XM_023256998.1;refGene_gn=SENP1;refGene_tx=XM_023256999.1;refGene_gn=SENP1;refGene_tx=XM_023257001.1
B4 76974171 76974343
refGene_gn=SENP1;refGene_tx=XM_019834858.2;refGene_gn=SENP1;refGene_tx=XM_023256995.1;refGene_gn=SENP1;refGene_tx=XM_023256996.1;refGene_gn=SENP1;refGene_tx=XM_023256997.1;refGene_gn=SENP1;refGene_tx=XM_023256998.1;refGene_gn=SENP1;refGene_tx=XM_023256999.1;refGene_gn=SENP1;refGene_tx=XM_023257001.1
B4 76978453 76978613
refGene_gn=SENP1;refGene_tx=XM_019834858.2;refGene_gn=SENP1;refGene_tx=XM_023256995.1;refGene_gn=SENP1;refGene_tx=XM_023256996.1;refGene_gn=SENP1;refGene_tx=XM_023256997.1;refGene_gn=SENP1;refGene_tx=XM_023256998.1;refGene_gn=SENP1;refGene_tx=XM_023256999.1;refGene_gn=SENP1;refGene_tx=XM_023257001.1
B4 76983138 76983180
refGene_gn=SENP1;refGene_tx=XM_023256995.1;refGene_gn=SENP1;refGene_tx=XM_023256996.1;refGene_gn=SENP1;refGene_tx=XM_023256997.1;refGene_gn=SENP1;refGene_tx=XM_023256998.1;refGene_gn=SENP1;refGene_tx=XM_023256999.1;refGene_gn=SENP1;refGene_tx=XM_023257001.1
B4 76984034 76984119
refGene_gn=SENP1;refGene_tx=XM_019834858.2;refGene_gn=SENP1;refGene_tx=XM_023256995.1;refGene_gn=SENP1;refGene_tx=XM_023256996.1;refGene_gn=SENP1;refGene_tx=XM_023256997.1;refGene_gn=SENP1;refGene_tx=XM_023256998.1;refGene_gn=SENP1;refGene_tx=XM_023256999.1;refGene_gn=SENP1;refGene_tx=XM_023257001.1
B4 76985827 76985958
refGene_gn=SENP1;refGene_tx=XM_019834858.2;refGene_gn=SENP1;refGene_tx=XM_023256995.1;refGene_gn=SENP1;refGene_tx=XM_023256996.1;refGene_gn=

SENP1;refGene_tx=XM_023256997.1;refGene_gn=SENP1;refGene_tx=XM_0232569
98.1;refGene_gn=SENP1;refGene_tx=XM_023256999.1;refGene_gn=SENP1;refGe
ne_tx=XM_023257001.1
B4	76987418	76987466
refGene_gn=SENP1;refGene_tx=XM_019834858.2;refGene_gn=SENP1;refGene_tx
=XM_023256995.1;refGene_gn=SENP1;refGene_tx=XM_023256997.1;refGene_gn=
SENP1;refGene_tx=XM_023256998.1;refGene_gn=SENP1;refGene_tx=XM_0232569
99.1
B4	76990820	76990872	refGene_gn=SENP1;refGene_tx=XM_023256998.1
B4	76991455	76991669
refGene_gn=SENP1;refGene_tx=XM_019834858.2;refGene_gn=SENP1;refGene_tx
=XM_023256995.1;refGene_gn=SENP1;refGene_tx=XM_023256996.1;refGene_gn=
SENP1;refGene_tx=XM_023256997.1
B4	76992786	76992868
refGene_gn=PFKM;refGene_tx=XM_019834856.1;refGene_gn=PFKM;refGene_tx=X
M_019834857.1
B4	76993519	76993642
refGene_gn=PFKM;refGene_tx=XM_019834856.1;refGene_gn=PFKM;refGene_tx=X
M_019834857.1
B4	77015753	77015846
refGene_gn=PFKM;refGene_tx=XM_006933653.2;refGene_gn=PFKM;refGene_tx=X
M_019834856.1;refGene_gn=PFKM;refGene_tx=XM_019834857.1
B4	77022022	77022096
refGene_gn=PFKM;refGene_tx=XM_006933653.2;refGene_gn=PFKM;refGene_tx=X
M_019834856.1;refGene_gn=PFKM;refGene_tx=XM_019834857.1
B4	77022937	77023015
refGene_gn=PFKM;refGene_tx=XM_006933653.2;refGene_gn=PFKM;refGene_tx=X
M_019834856.1;refGene_gn=PFKM;refGene_tx=XM_019834857.1
B4	77024885	77025075
refGene_gn=PFKM;refGene_tx=XM_006933653.2;refGene_gn=PFKM;refGene_tx=X
M_019834856.1;refGene_gn=PFKM;refGene_tx=XM_019834857.1
B4	77025286	77025452
refGene_gn=PFKM;refGene_tx=XM_006933653.2;refGene_gn=PFKM;refGene_tx=X
M_019834856.1;refGene_gn=PFKM;refGene_tx=XM_019834857.1
B4	77026187	77026232
refGene_gn=PFKM;refGene_tx=XM_006933653.2;refGene_gn=PFKM;refGene_tx=X
M_019834856.1;refGene_gn=PFKM;refGene_tx=XM_019834857.1
B4	77026655	77026764
refGene_gn=PFKM;refGene_tx=XM_006933653.2;refGene_gn=PFKM;refGene_tx=X
M_019834856.1;refGene_gn=PFKM;refGene_tx=XM_019834857.1
B4	77026876	77026972
refGene_gn=PFKM;refGene_tx=XM_006933653.2;refGene_gn=PFKM;refGene_tx=X
M_019834856.1;refGene_gn=PFKM;refGene_tx=XM_019834857.1
B4	77027205	77027298
refGene_gn=PFKM;refGene_tx=XM_006933653.2;refGene_gn=PFKM;refGene_tx=X
M_019834856.1;refGene_gn=PFKM;refGene_tx=XM_019834857.1
B4	77027916	77028042
refGene_gn=PFKM;refGene_tx=XM_006933653.2;refGene_gn=PFKM;refGene_tx=X
M_019834856.1;refGene_gn=PFKM;refGene_tx=XM_019834857.1
B4	77029196	77029261

refGene_gn=PFKM;refGene_tx=XM_006933653.2;refGene_gn=PFKM;refGene_tx=XM_019834856.1;refGene_gn=PFKM;refGene_tx=XM_019834857.1
B4 77029752 77029822
refGene_gn=PFKM;refGene_tx=XM_006933653.2;refGene_gn=PFKM;refGene_tx=XM_019834856.1;refGene_gn=PFKM;refGene_tx=XM_019834857.1
B4 77030574 77030724
refGene_gn=PFKM;refGene_tx=XM_006933653.2;refGene_gn=PFKM;refGene_tx=XM_019834856.1;refGene_gn=PFKM;refGene_tx=XM_019834857.1
B4 77031141 77031212
refGene_gn=PFKM;refGene_tx=XM_006933653.2;refGene_gn=PFKM;refGene_tx=XM_019834856.1;refGene_gn=PFKM;refGene_tx=XM_019834857.1
B4 77031566 77031654
refGene_gn=PFKM;refGene_tx=XM_006933653.2;refGene_gn=PFKM;refGene_tx=XM_019834856.1;refGene_gn=PFKM;refGene_tx=XM_019834857.1
B4 77031728 77031881
refGene_gn=PFKM;refGene_tx=XM_006933653.2;refGene_gn=PFKM;refGene_tx=XM_019834856.1;refGene_gn=PFKM;refGene_tx=XM_019834857.1
B4 77032367 77032532
refGene_gn=PFKM;refGene_tx=XM_006933653.2;refGene_gn=PFKM;refGene_tx=XM_019834856.1;refGene_gn=PFKM;refGene_tx=XM_019834857.1
B4 77033606 77033668
refGene_gn=PFKM;refGene_tx=XM_006933653.2;refGene_gn=PFKM;refGene_tx=XM_019834856.1;refGene_gn=PFKM;refGene_tx=XM_019834857.1
B4 77033897 77034009
refGene_gn=PFKM;refGene_tx=XM_006933653.2;refGene_gn=PFKM;refGene_tx=XM_019834856.1;refGene_gn=PFKM;refGene_tx=XM_019834857.1
B4 77035314 77035414
refGene_gn=PFKM;refGene_tx=XM_006933653.2;refGene_gn=PFKM;refGene_tx=XM_019834856.1;refGene_gn=PFKM;refGene_tx=XM_019834857.1
B4 77035495 77035601
refGene_gn=PFKM;refGene_tx=XM_006933653.2;refGene_gn=PFKM;refGene_tx=XM_019834856.1;refGene_gn=PFKM;refGene_tx=XM_019834857.1
B4 77035825 77035970
refGene_gn=PFKM;refGene_tx=XM_006933653.2;refGene_gn=PFKM;refGene_tx=XM_019834856.1;refGene_gn=PFKM;refGene_tx=XM_019834857.1
B4 77038751 77039384
refGene_gn=ASB8;refGene_tx=XM_003988612.5;refGene_gn=ASB8;refGene_tx=XM_006933652.4;refGene_gn=ASB8;refGene_tx=XM_019834860.2;refGene_gn=ASB8;refGene_tx=XM_019834861.2;refGene_gn=ASB8;refGene_tx=XM_019834862.2
B4 77041224 77041329
refGene_gn=ASB8;refGene_tx=XM_003988612.5;refGene_gn=ASB8;refGene_tx=XM_006933652.4;refGene_gn=ASB8;refGene_tx=XM_019834860.2;refGene_gn=ASB8;refGene_tx=XM_019834862.2
B4 77042781 77042910
refGene_gn=ASB8;refGene_tx=XM_003988612.5;refGene_gn=ASB8;refGene_tx=XM_006933652.4;refGene_gn=ASB8;refGene_tx=XM_019834860.2;refGene_gn=ASB8;refGene_tx=XM_019834861.2
B4 77073571 77074159 refGene_gn=CCDC184;refGene_tx=XM_019834863.2
B4 77091251 77092208
refGene_gn=LOC101097405;refGene_tx=XM_023257003.1

B4 77114365 77115322
refGene_gn=LOC101097653;refGene_tx=XM_011284010.3
B4 77215179 77215206
refGene_gn=LOC101097901;refGene_tx=XM_019835841.1
B4 77215999 77216147
refGene_gn=LOC101097901;refGene_tx=XM_019835841.1
B4 77216149 77216922
refGene_gn=LOC101097901;refGene_tx=XM_019835841.1
B4 77227670 77228425 refGene_gn=ZNF641;refGene_tx=XM_023257002.1
B4 77229253 77229367 refGene_gn=ZNF641;refGene_tx=XM_023257002.1
B4 77230253 77230383 refGene_gn=ZNF641;refGene_tx=XM_023257002.1
B4 77231518 77231610 refGene_gn=ZNF641;refGene_tx=XM_023257002.1
B4 77232234 77232418 refGene_gn=ZNF641;refGene_tx=XM_023257002.1
B4 77284824 77285736
refGene_gn=LOC101098390;refGene_tx=XM_003988649.3
B4 77308790 77309702
refGene_gn=LOC101098641;refGene_tx=XM_003988650.1
B4 77362423 77363458
refGene_gn=LOC101098877;refGene_tx=XM_011284012.2
B4 77386162 77387098
refGene_gn=LOC101099132;refGene_tx=XM_023257755.1
B4 77421473 77422391
refGene_gn=LOC101099384;refGene_tx=XM_003988652.2
B4 77503277 77504213
refGene_gn=LOC101100140;refGene_tx=XM_003988653.1
B4 77508038 77508977
refGene_gn=LOC101100398;refGene_tx=XM_011284015.1
B4 77542029 77542965
refGene_gn=LOC101100898;refGene_tx=XM_003988654.1
B4 77556389 77557316
refGene_gn=LOC101101145;refGene_tx=XM_003988655.1
B4 77569545 77570508
refGene_gn=LOC101101393;refGene_tx=XM_003988656.1
B4 77585606 77586554
refGene_gn=LOC101101645;refGene_tx=XM_003988657.1
B4 77597533 77598487
refGene_gn=LOC101080462;refGene_tx=XM_003988658.3
B4 77600387 77602514
refGene_gn=LOC109501563;refGene_tx=XM_023257756.1
B4 77603069 77603605
refGene_gn=LOC109501563;refGene_tx=XM_023257756.1
B4 77603740 77605824
refGene_gn=LOC109501563;refGene_tx=XM_023257756.1
B4 77614576 77615548
refGene_gn=LOC101080978;refGene_tx=XM_023257005.1
B4 77630957 77631893
refGene_gn=LOC101081231;refGene_tx=XM_023257796.1
B4 77654797 77654858 refGene_gn=LALBA;refGene_tx=XM_003988615.4
B4 77655379 77655455 refGene_gn=LALBA;refGene_tx=XM_003988615.4
B4 77655925 77656084 refGene_gn=LALBA;refGene_tx=XM_003988615.4

B4 77656663 77656796 refGene_gn=LALBA;refGene_tx=XM_003988615.4
B4 77678654 77679707
refGene_gn=LOC101081477;refGene_tx=XM_003988661.2
B4 77685120 77685259
refGene_gn=LOC102900869;refGene_tx=XM_011283893.3
B4 77686500 77686662
refGene_gn=LOC102900869;refGene_tx=XM_011283893.3
B4 77687210 77687289
refGene_gn=LOC102900869;refGene_tx=XM_011283893.3
B4 77687634 77687701
refGene_gn=LOC102900869;refGene_tx=XM_011283893.3
B4 77710403 77710535
refGene_gn=KANSL2;refGene_tx=XM_006933655.4;refGene_gn=KANSL2;refGene_tx=XM_006933656.4;refGene_gn=KANSL2;refGene_tx=XM_011283894.3
B4 77711156 77711330
refGene_gn=KANSL2;refGene_tx=XM_006933655.4;refGene_gn=KANSL2;refGene_tx=XM_006933656.4;refGene_gn=KANSL2;refGene_tx=XM_011283894.3;refGene_gn=KANSL2;refGene_tx=XM_011283895.3
B4 77717476 77717730
refGene_gn=KANSL2;refGene_tx=XM_006933655.4;refGene_gn=KANSL2;refGene_tx=XM_006933656.4;refGene_gn=KANSL2;refGene_tx=XM_011283894.3;refGene_gn=KANSL2;refGene_tx=XM_011283895.3
B4 77721287 77721384
refGene_gn=KANSL2;refGene_tx=XM_006933655.4;refGene_gn=KANSL2;refGene_tx=XM_006933656.4;refGene_gn=KANSL2;refGene_tx=XM_011283894.3;refGene_gn=KANSL2;refGene_tx=XM_011283895.3
B4 77722479 77722646
refGene_gn=KANSL2;refGene_tx=XM_006933655.4;refGene_gn=KANSL2;refGene_tx=XM_006933656.4;refGene_gn=KANSL2;refGene_tx=XM_011283894.3;refGene_gn=KANSL2;refGene_tx=XM_011283895.3
B4 77722871 77723035
refGene_gn=KANSL2;refGene_tx=XM_006933655.4;refGene_gn=KANSL2;refGene_tx=XM_006933656.4;refGene_gn=KANSL2;refGene_tx=XM_011283894.3;refGene_gn=KANSL2;refGene_tx=XM_011283895.3
B4 77728250 77728365
refGene_gn=KANSL2;refGene_tx=XM_006933655.4;refGene_gn=KANSL2;refGene_tx=XM_006933656.4;refGene_gn=KANSL2;refGene_tx=XM_011283894.3;refGene_gn=KANSL2;refGene_tx=XM_011283895.3
B4 77728907 77729086
refGene_gn=KANSL2;refGene_tx=XM_006933655.4;refGene_gn=KANSL2;refGene_tx=XM_006933656.4;refGene_gn=KANSL2;refGene_tx=XM_011283894.3;refGene_gn=KANSL2;refGene_tx=XM_011283895.3
B4 77729877 77730128
refGene_gn=KANSL2;refGene_tx=XM_006933655.4;refGene_gn=KANSL2;refGene_tx=XM_006933656.4;refGene_gn=KANSL2;refGene_tx=XM_011283894.3;refGene_gn=KANSL2;refGene_tx=XM_011283895.3
B4 77740627 77741266 refGene_gn=CCNT1;refGene_tx=XM_003988618.5
B4 77750389 77750460 refGene_gn=CCNT1;refGene_tx=XM_003988618.5
B4 77750618 77750782 refGene_gn=CCNT1;refGene_tx=XM_003988618.5
B4 77751937 77751983 refGene_gn=CCNT1;refGene_tx=XM_003988618.5

B4 77753891 77753954 refGene_gn=CCNT1;refGene_tx=XM_003988618.5
B4 77756394 77756455 refGene_gn=CCNT1;refGene_tx=XM_003988618.5
B4 77759603 77759732 refGene_gn=CCNT1;refGene_tx=XM_003988618.5
B4 77766498 77766580 refGene_gn=CCNT1;refGene_tx=XM_003988618.5
B4 77768350 77768511 refGene_gn=CCNT1;refGene_tx=XM_003988618.5
B4 77776097 77776277
refGene_gn=LOC111561359;refGene_tx=XM_023257008.1;refGene_gn=LOC111561
359;refGene_tx=XM_023257009.1
B4 77785187 77785245
refGene_gn=LOC111561359;refGene_tx=XM_023257008.1
B4 77800831 77800954
refGene_gn=LOC111561359;refGene_tx=XM_023257008.1;refGene_gn=LOC111561
359;refGene_tx=XM_023257009.1
B4 77802607 77802752
refGene_gn=LOC111561359;refGene_tx=XM_023257008.1;refGene_gn=LOC111561
359;refGene_tx=XM_023257009.1
B4 77803193 77803242
refGene_gn=LOC111561359;refGene_tx=XM_023257008.1;refGene_gn=LOC111561
359;refGene_tx=XM_023257009.1
B4 77806120 77806246
refGene_gn=ADCY6;refGene_tx=XM_006933657.4;refGene_gn=ADCY6;refGene_tx
=XM_011283897.3;refGene_gn=ADCY6;refGene_tx=XM_019834866.2;refGene_gn=
ADCY6;refGene_tx=XM_019834867.2;refGene_gn=ADCY6;refGene_tx=XM_0232570
06.1;refGene_gn=ADCY6;refGene_tx=XM_023257007.1
B4 77806446 77806571
refGene_gn=ADCY6;refGene_tx=XM_006933657.4;refGene_gn=ADCY6;refGene_tx
=XM_011283897.3;refGene_gn=ADCY6;refGene_tx=XM_019834866.2;refGene_gn=
ADCY6;refGene_tx=XM_019834867.2;refGene_gn=ADCY6;refGene_tx=XM_0232570
06.1;refGene_gn=ADCY6;refGene_tx=XM_023257007.1
B4 77807396 77807601
refGene_gn=ADCY6;refGene_tx=XM_006933657.4;refGene_gn=ADCY6;refGene_tx
=XM_011283897.3;refGene_gn=ADCY6;refGene_tx=XM_019834866.2;refGene_gn=
ADCY6;refGene_tx=XM_019834867.2;refGene_gn=ADCY6;refGene_tx=XM_0232570
06.1;refGene_gn=ADCY6;refGene_tx=XM_023257007.1
B4 77808417 77808681
refGene_gn=ADCY6;refGene_tx=XM_006933657.4;refGene_gn=ADCY6;refGene_tx
=XM_011283897.3;refGene_gn=ADCY6;refGene_tx=XM_019834866.2;refGene_gn=
ADCY6;refGene_tx=XM_019834867.2;refGene_gn=ADCY6;refGene_tx=XM_0232570
06.1;refGene_gn=ADCY6;refGene_tx=XM_023257007.1
B4 77808998 77809128
refGene_gn=ADCY6;refGene_tx=XM_006933657.4;refGene_gn=ADCY6;refGene_tx
=XM_011283897.3;refGene_gn=ADCY6;refGene_tx=XM_019834866.2;refGene_gn=
ADCY6;refGene_tx=XM_019834867.2;refGene_gn=ADCY6;refGene_tx=XM_0232570
06.1;refGene_gn=ADCY6;refGene_tx=XM_023257007.1
B4 77809211 77809247
refGene_gn=ADCY6;refGene_tx=XM_006933657.4;refGene_gn=ADCY6;refGene_tx
=XM_011283897.3;refGene_gn=ADCY6;refGene_tx=XM_019834866.2;refGene_gn=
ADCY6;refGene_tx=XM_019834867.2;refGene_gn=ADCY6;refGene_tx=XM_0232570
06.1;refGene_gn=ADCY6;refGene_tx=XM_023257007.1
B4 77810133 77810312

refGene_gn=ADCY6;refGene_tx=XM_006933657.4;refGene_gn=ADCY6;refGene_tx=XM_011283897.3;refGene_gn=ADCY6;refGene_tx=XM_019834866.2;refGene_gn=ADCY6;refGene_tx=XM_019834867.2;refGene_gn=ADCY6;refGene_tx=XM_023257006.1;refGene_gn=ADCY6;refGene_tx=XM_023257007.1
B4 77810617 77810776
refGene_gn=ADCY6;refGene_tx=XM_006933657.4;refGene_gn=ADCY6;refGene_tx=XM_011283897.3;refGene_gn=ADCY6;refGene_tx=XM_019834866.2;refGene_gn=ADCY6;refGene_tx=XM_019834867.2;refGene_gn=ADCY6;refGene_tx=XM_023257006.1;refGene_gn=ADCY6;refGene_tx=XM_023257007.1
B4 77811057 77811174
refGene_gn=ADCY6;refGene_tx=XM_006933657.4;refGene_gn=ADCY6;refGene_tx=XM_011283897.3;refGene_gn=ADCY6;refGene_tx=XM_019834866.2;refGene_gn=ADCY6;refGene_tx=XM_019834867.2;refGene_gn=ADCY6;refGene_tx=XM_023257006.1;refGene_gn=ADCY6;refGene_tx=XM_023257007.1
B4 77811350 77811438
refGene_gn=ADCY6;refGene_tx=XM_006933657.4;refGene_gn=ADCY6;refGene_tx=XM_011283897.3;refGene_gn=ADCY6;refGene_tx=XM_019834866.2;refGene_gn=ADCY6;refGene_tx=XM_019834867.2;refGene_gn=ADCY6;refGene_tx=XM_023257006.1;refGene_gn=ADCY6;refGene_tx=XM_023257007.1
B4 77811615 77811713
refGene_gn=ADCY6;refGene_tx=XM_006933657.4;refGene_gn=ADCY6;refGene_tx=XM_011283897.3;refGene_gn=ADCY6;refGene_tx=XM_019834866.2;refGene_gn=ADCY6;refGene_tx=XM_019834867.2;refGene_gn=ADCY6;refGene_tx=XM_023257006.1;refGene_gn=ADCY6;refGene_tx=XM_023257007.1
B4 77811954 77812102
refGene_gn=ADCY6;refGene_tx=XM_006933657.4;refGene_gn=ADCY6;refGene_tx=XM_011283897.3;refGene_gn=ADCY6;refGene_tx=XM_019834866.2;refGene_gn=ADCY6;refGene_tx=XM_019834867.2;refGene_gn=ADCY6;refGene_tx=XM_023257006.1;refGene_gn=ADCY6;refGene_tx=XM_023257007.1
B4 77812335 77812361
refGene_gn=ADCY6;refGene_tx=XM_006933657.4;refGene_gn=ADCY6;refGene_tx=XM_011283897.3;refGene_gn=ADCY6;refGene_tx=XM_019834866.2;refGene_gn=ADCY6;refGene_tx=XM_019834867.2;refGene_gn=ADCY6;refGene_tx=XM_023257006.1;refGene_gn=ADCY6;refGene_tx=XM_023257007.1
B4 77812632 77812761
refGene_gn=ADCY6;refGene_tx=XM_006933657.4;refGene_gn=ADCY6;refGene_tx=XM_011283897.3;refGene_gn=ADCY6;refGene_tx=XM_019834866.2;refGene_gn=ADCY6;refGene_tx=XM_019834867.2;refGene_gn=ADCY6;refGene_tx=XM_023257006.1;refGene_gn=ADCY6;refGene_tx=XM_023257007.1
B4 77812877 77813019
refGene_gn=ADCY6;refGene_tx=XM_006933657.4;refGene_gn=ADCY6;refGene_tx=XM_011283897.3;refGene_gn=ADCY6;refGene_tx=XM_019834866.2;refGene_gn=ADCY6;refGene_tx=XM_019834867.2;refGene_gn=ADCY6;refGene_tx=XM_023257006.1;refGene_gn=ADCY6;refGene_tx=XM_023257007.1
B4 77813096 77813255
refGene_gn=ADCY6;refGene_tx=XM_006933657.4;refGene_gn=ADCY6;refGene_tx=XM_019834866.2;refGene_gn=ADCY6;refGene_tx=XM_019834867.2;refGene_gn=ADCY6;refGene_tx=XM_023257006.1;refGene_gn=ADCY6;refGene_tx=XM_023257007.1
B4 77813747 77813875

refGene_gn=ADCY6;refGene_tx=XM_006933657.4;refGene_gn=ADCY6;refGene_tx=XM_011283897.3;refGene_gn=ADCY6;refGene_tx=XM_019834866.2;refGene_gn=ADCY6;refGene_tx=XM_019834867.2;refGene_gn=ADCY6;refGene_tx=XM_023257006.1;refGene_gn=ADCY6;refGene_tx=XM_023257007.1
B4 77814028 77814140
refGene_gn=ADCY6;refGene_tx=XM_006933657.4;refGene_gn=ADCY6;refGene_tx=XM_011283897.3;refGene_gn=ADCY6;refGene_tx=XM_019834866.2;refGene_gn=ADCY6;refGene_tx=XM_019834867.2;refGene_gn=ADCY6;refGene_tx=XM_023257006.1;refGene_gn=ADCY6;refGene_tx=XM_023257007.1
B4 77814236 77814358
refGene_gn=ADCY6;refGene_tx=XM_006933657.4;refGene_gn=ADCY6;refGene_tx=XM_011283897.3;refGene_gn=ADCY6;refGene_tx=XM_019834866.2;refGene_gn=ADCY6;refGene_tx=XM_019834867.2;refGene_gn=ADCY6;refGene_tx=XM_023257006.1;refGene_gn=ADCY6;refGene_tx=XM_023257007.1
B4 77814742 77814892
refGene_gn=ADCY6;refGene_tx=XM_006933657.4;refGene_gn=ADCY6;refGene_tx=XM_011283897.3;refGene_gn=ADCY6;refGene_tx=XM_019834866.2;refGene_gn=ADCY6;refGene_tx=XM_019834867.2;refGene_gn=ADCY6;refGene_tx=XM_023257006.1;refGene_gn=ADCY6;refGene_tx=XM_023257007.1
B4 77818492 77819348
refGene_gn=ADCY6;refGene_tx=XM_006933657.4;refGene_gn=ADCY6;refGene_tx=XM_011283897.3;refGene_gn=ADCY6;refGene_tx=XM_019834866.2;refGene_gn=ADCY6;refGene_tx=XM_019834867.2;refGene_gn=ADCY6;refGene_tx=XM_023257006.1;refGene_gn=ADCY6;refGene_tx=XM_023257007.1
B4 77819463 77819756
refGene_gn=ADCY6;refGene_tx=XM_006933657.4;refGene_gn=ADCY6;refGene_tx=XM_011283897.3;refGene_gn=ADCY6;refGene_tx=XM_019834866.2
B4 77844155 77844197 refGene_gn=CACNB3;refGene_tx=XM_003988623.3
B4 77847574 77847619 refGene_gn=CACNB3;refGene_tx=XM_003988622.4
B4 77851264 77851387
refGene_gn=CACNB3;refGene_tx=XM_003988622.4;refGene_gn=CACNB3;refGene_tx=XM_003988623.3
B4 77851603 77851726
refGene_gn=CACNB3;refGene_tx=XM_003988622.4;refGene_gn=CACNB3;refGene_tx=XM_003988623.3
B4 77852173 77852289
refGene_gn=CACNB3;refGene_tx=XM_003988622.4;refGene_gn=CACNB3;refGene_tx=XM_003988623.3
B4 77852516 77852581
refGene_gn=CACNB3;refGene_tx=XM_003988622.4;refGene_gn=CACNB3;refGene_tx=XM_003988623.3
B4 77852800 77852820
refGene_gn=CACNB3;refGene_tx=XM_003988622.4;refGene_gn=CACNB3;refGene_tx=XM_003988623.3
B4 77853018 77853099
refGene_gn=CACNB3;refGene_tx=XM_003988622.4;refGene_gn=CACNB3;refGene_tx=XM_003988623.3
B4 77853300 77853359
refGene_gn=CACNB3;refGene_tx=XM_003988622.4;refGene_gn=CACNB3;refGene_tx=XM_003988623.3

B4	77853529	77853639
refGene_gn=CACNB3;refGene_tx=XM_003988622.4;refGene_gn=CACNB3;refGene_tx=XM_003988623.3
B4	77853942	77854094
refGene_gn=CACNB3;refGene_tx=XM_003988622.4;refGene_gn=CACNB3;refGene_tx=XM_003988623.3
B4	77854296	77854392
refGene_gn=CACNB3;refGene_tx=XM_003988622.4;refGene_gn=CACNB3;refGene_tx=XM_003988623.3
B4	77854521	77854671
refGene_gn=CACNB3;refGene_tx=XM_003988622.4;refGene_gn=CACNB3;refGene_tx=XM_003988623.3
B4	77855129	77855444
refGene_gn=CACNB3;refGene_tx=XM_003988622.4;refGene_gn=CACNB3;refGene_tx=XM_003988623.3
B4	77858033	77858257
refGene_gn=DDX23;refGene_tx=XM_003988626.4;refGene_gn=DDX23;refGene_tx=XM_019834872.2
B4	77858645	77858820
refGene_gn=DDX23;refGene_tx=XM_003988626.4;refGene_gn=DDX23;refGene_tx=XM_019834872.2
B4	77859578	77859687
refGene_gn=DDX23;refGene_tx=XM_003988626.4;refGene_gn=DDX23;refGene_tx=XM_019834872.2
B4	77859902	77860054
refGene_gn=DDX23;refGene_tx=XM_003988626.4;refGene_gn=DDX23;refGene_tx=XM_019834872.2
B4	77860466	77860709
refGene_gn=DDX23;refGene_tx=XM_003988626.4;refGene_gn=DDX23;refGene_tx=XM_019834872.2
B4	77861449	77861627
refGene_gn=DDX23;refGene_tx=XM_003988626.4;refGene_gn=DDX23;refGene_tx=XM_019834872.2
B4	77861989	77862135
refGene_gn=DDX23;refGene_tx=XM_003988626.4;refGene_gn=DDX23;refGene_tx=XM_019834872.2
B4	77862432	77862658
refGene_gn=DDX23;refGene_tx=XM_003988626.4;refGene_gn=DDX23;refGene_tx=XM_019834872.2
B4	77862759	77862903
refGene_gn=DDX23;refGene_tx=XM_003988626.4;refGene_gn=DDX23;refGene_tx=XM_019834872.2
B4	77863129	77863242
refGene_gn=DDX23;refGene_tx=XM_003988626.4;refGene_gn=DDX23;refGene_tx=XM_019834872.2
B4	77863379	77863513
refGene_gn=DDX23;refGene_tx=XM_003988626.4;refGene_gn=DDX23;refGene_tx=XM_019834872.2
B4	77863788	77863927
refGene_gn=DDX23;refGene_tx=XM_003988626.4;refGene_gn=DDX23;refGene_tx

=XM_019834872.2
B4 77864736 77864802
refGene_gn=DDX23;refGene_tx=XM_003988626.4;refGene_gn=DDX23;refGene_tx=XM_019834872.2
B4 77864916 77865010
refGene_gn=DDX23;refGene_tx=XM_003988626.4;refGene_gn=DDX23;refGene_tx=XM_019834872.2
B4 77865908 77866019
refGene_gn=DDX23;refGene_tx=XM_003988626.4;refGene_gn=DDX23;refGene_tx=XM_019834872.2
B4 77866646 77866855
refGene_gn=DDX23;refGene_tx=XM_003988626.4;refGene_gn=DDX23;refGene_tx=XM_019834872.2
B4 77875036 77875282 refGene_gn=RND1;refGene_tx=XM_003988627.4
B4 77876799 77876934 refGene_gn=RND1;refGene_tx=XM_003988627.4
B4 77877798 77877908 refGene_gn=RND1;refGene_tx=XM_003988627.4
B4 77880521 77880609 refGene_gn=RND1;refGene_tx=XM_003988627.4
B4 77881388 77881508 refGene_gn=RND1;refGene_tx=XM_003988627.4
B4 77917194 77917326 refGene_gn=CCDC65;refGene_tx=XM_003988628.4
B4 77918160 77918328 refGene_gn=CCDC65;refGene_tx=XM_003988628.4
B4 77924708 77924878 refGene_gn=CCDC65;refGene_tx=XM_003988628.4
B4 77925549 77925688 refGene_gn=CCDC65;refGene_tx=XM_003988628.4
B4 77926631 77926829 refGene_gn=CCDC65;refGene_tx=XM_003988628.4
B4 77927024 77927243 refGene_gn=CCDC65;refGene_tx=XM_003988628.4
B4 77928152 77928305 refGene_gn=CCDC65;refGene_tx=XM_003988628.4
B4 77928420 77928738 refGene_gn=CCDC65;refGene_tx=XM_003988628.4
B4 77929224 77929442 refGene_gn=FKBP11;refGene_tx=XM_003988629.4
B4 77930757 77930828 refGene_gn=FKBP11;refGene_tx=XM_003988629.4
B4 77931187 77931221 refGene_gn=FKBP11;refGene_tx=XM_003988629.4
B4 77931485 77931573 refGene_gn=FKBP11;refGene_tx=XM_003988629.4
B4 77931976 77932042 refGene_gn=FKBP11;refGene_tx=XM_003988629.4
B4 77932225 77932351 refGene_gn=FKBP11;refGene_tx=XM_003988629.4
B4 77942569 77942731
refGene_gn=ARF3;refGene_tx=XM_003988630.5;refGene_gn=ARF3;refGene_tx=XM_006933660.4;refGene_gn=ARF3;refGene_tx=XM_006933661.3
B4 77943286 77943411
refGene_gn=ARF3;refGene_tx=XM_003988630.5;refGene_gn=ARF3;refGene_tx=XM_006933660.4;refGene_gn=ARF3;refGene_tx=XM_006933661.3
B4 77943652 77943763
refGene_gn=ARF3;refGene_tx=XM_003988630.5;refGene_gn=ARF3;refGene_tx=XM_006933660.4;refGene_gn=ARF3;refGene_tx=XM_006933661.3
B4 77944565 77944713
refGene_gn=ARF3;refGene_tx=XM_003988630.5;refGene_gn=ARF3;refGene_tx=XM_006933660.4;refGene_gn=ARF3;refGene_tx=XM_006933661.3
B4 77963966 77964425
refGene_gn=WNT10B;refGene_tx=XM_023257010.1;refGene_gn=WNT10B;refGene_tx=XM_023257011.1
B4 77965080 77965454
refGene_gn=WNT10B;refGene_tx=XM_023257010.1;refGene_gn=WNT10B;refGene_tx=XM_023257011.1

B4 77968135 77968398
refGene_gn=WNT10B;refGene_tx=XM_023257010.1;refGene_gn=WNT10B;refGene_tx=XM_023257011.1
B4 77968528 77968602
refGene_gn=WNT10B;refGene_tx=XM_023257010.1;refGene_gn=WNT10B;refGene_tx=XM_023257011.1
B4 77977937 77978041 refGene_gn=WNT1;refGene_tx=XM_023257012.1
B4 77978766 77979020 refGene_gn=WNT1;refGene_tx=XM_023257012.1
B4 77979682 77979948 refGene_gn=WNT1;refGene_tx=XM_023257012.1
B4 77980422 77980911 refGene_gn=WNT1;refGene_tx=XM_023257012.1
B4 77993330 77995245 refGene_gn=DDN;refGene_tx=XM_003988635.4
B4 77995656 77995865 refGene_gn=DDN;refGene_tx=XM_003988635.4
B4 77999417 77999522
refGene_gn=PRKAG1;refGene_tx=XM_003988636.5;refGene_gn=PRKAG1;refGene_tx=XM_006933663.4;refGene_gn=PRKAG1;refGene_tx=XM_019834873.2
B4 77999661 77999808
refGene_gn=PRKAG1;refGene_tx=XM_003988636.5;refGene_gn=PRKAG1;refGene_tx=XM_006933663.4;refGene_gn=PRKAG1;refGene_tx=XM_019834873.2
B4 78000075 78000113
refGene_gn=PRKAG1;refGene_tx=XM_003988636.5;refGene_gn=PRKAG1;refGene_tx=XM_006933663.4;refGene_gn=PRKAG1;refGene_tx=XM_019834873.2
B4 78000267 78000433
refGene_gn=PRKAG1;refGene_tx=XM_003988636.5;refGene_gn=PRKAG1;refGene_tx=XM_006933663.4;refGene_gn=PRKAG1;refGene_tx=XM_019834873.2
B4 78000538 78000665
refGene_gn=PRKAG1;refGene_tx=XM_003988636.5;refGene_gn=PRKAG1;refGene_tx=XM_006933663.4;refGene_gn=PRKAG1;refGene_tx=XM_019834873.2
B4 78000981 78001036
refGene_gn=PRKAG1;refGene_tx=XM_003988636.5;refGene_gn=PRKAG1;refGene_tx=XM_006933663.4;refGene_gn=PRKAG1;refGene_tx=XM_019834873.2
B4 78001138 78001184
refGene_gn=PRKAG1;refGene_tx=XM_003988636.5;refGene_gn=PRKAG1;refGene_tx=XM_006933663.4;refGene_gn=PRKAG1;refGene_tx=XM_019834873.2
B4 78001324 78001383
refGene_gn=PRKAG1;refGene_tx=XM_003988636.5;refGene_gn=PRKAG1;refGene_tx=XM_006933663.4;refGene_gn=PRKAG1;refGene_tx=XM_019834873.2
B4 78001481 78001563
refGene_gn=PRKAG1;refGene_tx=XM_003988636.5;refGene_gn=PRKAG1;refGene_tx=XM_006933663.4;refGene_gn=PRKAG1;refGene_tx=XM_019834873.2
B4 78001763 78001916
refGene_gn=PRKAG1;refGene_tx=XM_003988636.5;refGene_gn=PRKAG1;refGene_tx=XM_006933663.4
B4 78007564 78007613
refGene_gn=PRKAG1;refGene_tx=XM_003988636.5;refGene_gn=PRKAG1;refGene_tx=XM_006933663.4;refGene_gn=PRKAG1;refGene_tx=XM_019834873.2
B4 78011566 78011664
refGene_gn=PRKAG1;refGene_tx=XM_003988636.5;refGene_gn=PRKAG1;refGene_tx=XM_006933663.4;refGene_gn=PRKAG1;refGene_tx=XM_019834873.2
B4 78014647 78014740 refGene_gn=KMT2D;refGene_tx=XM_023257013.1
B4 78014898 78015007 refGene_gn=KMT2D;refGene_tx=XM_023257013.1

| | | | |
|---|---|---|---|
| B4 | 78015131 | 78015205 | refGene_gn=KMT2D;refGene_tx=XM_023257013.1 |
| B4 | 78015444 | 78015730 | refGene_gn=KMT2D;refGene_tx=XM_023257013.1 |
| B4 | 78017409 | 78017540 | refGene_gn=KMT2D;refGene_tx=XM_023257013.1 |
| B4 | 78017632 | 78017769 | refGene_gn=KMT2D;refGene_tx=XM_023257013.1 |
| B4 | 78018620 | 78019761 | refGene_gn=KMT2D;refGene_tx=XM_023257013.1 |
| B4 | 78019911 | 78020039 | refGene_gn=KMT2D;refGene_tx=XM_023257013.1 |
| B4 | 78020114 | 78020247 | refGene_gn=KMT2D;refGene_tx=XM_023257013.1 |
| B4 | 78020916 | 78021047 | refGene_gn=KMT2D;refGene_tx=XM_023257013.1 |
| B4 | 78021144 | 78021320 | refGene_gn=KMT2D;refGene_tx=XM_023257013.1 |
| B4 | 78021496 | 78021572 | refGene_gn=KMT2D;refGene_tx=XM_023257013.1 |
| B4 | 78022205 | 78022365 | refGene_gn=KMT2D;refGene_tx=XM_023257013.1 |
| B4 | 78022521 | 78022689 | refGene_gn=KMT2D;refGene_tx=XM_023257013.1 |
| B4 | 78022812 | 78022953 | refGene_gn=KMT2D;refGene_tx=XM_023257013.1 |
| B4 | 78023087 | 78024641 | refGene_gn=KMT2D;refGene_tx=XM_023257013.1 |
| B4 | 78024706 | 78025285 | refGene_gn=KMT2D;refGene_tx=XM_023257013.1 |
| B4 | 78025576 | 78026239 | refGene_gn=KMT2D;refGene_tx=XM_023257013.1 |
| B4 | 78026339 | 78026572 | refGene_gn=KMT2D;refGene_tx=XM_023257013.1 |
| B4 | 78026677 | 78026744 | refGene_gn=KMT2D;refGene_tx=XM_023257013.1 |
| B4 | 78026849 | 78026934 | refGene_gn=KMT2D;refGene_tx=XM_023257013.1 |
| B4 | 78027082 | 78027206 | refGene_gn=KMT2D;refGene_tx=XM_023257013.1 |
| B4 | 78028851 | 78030752 | refGene_gn=KMT2D;refGene_tx=XM_023257013.1 |
| B4 | 78030967 | 78031104 | refGene_gn=KMT2D;refGene_tx=XM_023257013.1 |
| B4 | 78031184 | 78031367 | refGene_gn=KMT2D;refGene_tx=XM_023257013.1 |
| B4 | 78031471 | 78033283 | refGene_gn=KMT2D;refGene_tx=XM_023257013.1 |
| B4 | 78033406 | 78033457 | refGene_gn=KMT2D;refGene_tx=XM_023257013.1 |
| B4 | 78033663 | 78033737 | refGene_gn=KMT2D;refGene_tx=XM_023257013.1 |
| B4 | 78033835 | 78034077 | refGene_gn=KMT2D;refGene_tx=XM_023257013.1 |
| B4 | 78034315 | 78034400 | refGene_gn=KMT2D;refGene_tx=XM_023257013.1 |
| B4 | 78034505 | 78034643 | refGene_gn=KMT2D;refGene_tx=XM_023257013.1 |
| B4 | 78034848 | 78034959 | refGene_gn=KMT2D;refGene_tx=XM_023257013.1 |
| B4 | 78035136 | 78035202 | refGene_gn=KMT2D;refGene_tx=XM_023257013.1 |
| B4 | 78035403 | 78035551 | refGene_gn=KMT2D;refGene_tx=XM_023257013.1 |
| B4 | 78035634 | 78035765 | refGene_gn=KMT2D;refGene_tx=XM_023257013.1 |
| B4 | 78035994 | 78036099 | refGene_gn=KMT2D;refGene_tx=XM_023257013.1 |
| B4 | 78036186 | 78036306 | refGene_gn=KMT2D;refGene_tx=XM_023257013.1 |
| B4 | 78036538 | 78036760 | refGene_gn=KMT2D;refGene_tx=XM_023257013.1 |
| B4 | 78037394 | 78037442 | refGene_gn=KMT2D;refGene_tx=XM_023257013.1 |
| B4 | 78037545 | 78037655 | refGene_gn=KMT2D;refGene_tx=XM_023257013.1 |
| B4 | 78037747 | 78037912 | refGene_gn=KMT2D;refGene_tx=XM_023257013.1 |
| B4 | 78038090 | 78038272 | refGene_gn=KMT2D;refGene_tx=XM_023257013.1 |
| B4 | 78039043 | 78039148 | refGene_gn=KMT2D;refGene_tx=XM_023257013.1 |
| B4 | 78039785 | 78039896 | refGene_gn=KMT2D;refGene_tx=XM_023257013.1 |
| B4 | 78040245 | 78040359 | refGene_gn=KMT2D;refGene_tx=XM_023257013.1 |
| B4 | 78040839 | 78041948 | refGene_gn=KMT2D;refGene_tx=XM_023257013.1 |
| B4 | 78042037 | 78043684 | refGene_gn=KMT2D;refGene_tx=XM_023257013.1 |
| B4 | 78043822 | 78043968 | refGene_gn=KMT2D;refGene_tx=XM_023257013.1 |
| B4 | 78044177 | 78044335 | refGene_gn=KMT2D;refGene_tx=XM_023257013.1 |
| B4 | 78044474 | 78044589 | refGene_gn=KMT2D;refGene_tx=XM_023257013.1 |
| B4 | 78044688 | 78044854 | refGene_gn=KMT2D;refGene_tx=XM_023257013.1 |
| B4 | 78045155 | 78045318 | refGene_gn=KMT2D;refGene_tx=XM_023257013.1 |

B4 78045489 78045599 refGene_gn=KMT2D;refGene_tx=XM_023257013.1
B4 78045724 78045948 refGene_gn=KMT2D;refGene_tx=XM_023257013.1
B4 78046127 78046254 refGene_gn=KMT2D;refGene_tx=XM_023257013.1
B4 78046752 78046801 refGene_gn=KMT2D;refGene_tx=XM_023257013.1
B4 78055973 78056063 refGene_gn=RHEBL1;refGene_tx=XM_003988640.5
B4 78056223 78056305 refGene_gn=RHEBL1;refGene_tx=XM_003988640.5
B4 78056867 78056915 refGene_gn=RHEBL1;refGene_tx=XM_003988640.5
B4 78057107 78057164 refGene_gn=RHEBL1;refGene_tx=XM_003988640.5
B4 78057253 78057336 refGene_gn=RHEBL1;refGene_tx=XM_003988640.5
B4 78057611 78057679 refGene_gn=RHEBL1;refGene_tx=XM_003988640.5
B4 78058028 78058100 refGene_gn=RHEBL1;refGene_tx=XM_003988640.5
B4 78058737 78058789 refGene_gn=RHEBL1;refGene_tx=XM_003988640.5
B4 78074093 78074719 refGene_gn=DHH;refGene_tx=XM_003988641.5
B4 78074998 78075260 refGene_gn=DHH;refGene_tx=XM_003988641.5
B4 78077935 78077989 refGene_gn=DHH;refGene_tx=XM_003988641.5
B4 78077990 78078239 refGene_gn=DHH;refGene_tx=XM_003988641.5
B4 78080782 78080850 refGene_gn=LMBR1L;refGene_tx=XM_003988642.4
B4 78081052 78081214 refGene_gn=LMBR1L;refGene_tx=XM_003988642.4
B4 78082834 78082901 refGene_gn=LMBR1L;refGene_tx=XM_003988642.4
B4 78083007 78083098 refGene_gn=LMBR1L;refGene_tx=XM_003988642.4
B4 78083320 78083394 refGene_gn=LMBR1L;refGene_tx=XM_003988642.4
B4 78083543 78083621 refGene_gn=LMBR1L;refGene_tx=XM_003988642.4
B4 78084165 78084242 refGene_gn=LMBR1L;refGene_tx=XM_003988642.4
B4 78084336 78084420 refGene_gn=LMBR1L;refGene_tx=XM_003988642.4
B4 78084511 78084584 refGene_gn=LMBR1L;refGene_tx=XM_003988642.4
B4 78084908 78084973 refGene_gn=LMBR1L;refGene_tx=XM_003988642.4
B4 78085113 78085182 refGene_gn=LMBR1L;refGene_tx=XM_003988642.4
B4 78085733 78085860 refGene_gn=LMBR1L;refGene_tx=XM_003988642.4
B4 78086425 78086529 refGene_gn=LMBR1L;refGene_tx=XM_003988642.4
B4 78086717 78086857 refGene_gn=LMBR1L;refGene_tx=XM_003988642.4
B4 78087883 78087917 refGene_gn=LMBR1L;refGene_tx=XM_003988642.4
B4 78088951 78089036 refGene_gn=LMBR1L;refGene_tx=XM_003988642.4
B4 78092203 78092275 refGene_gn=LMBR1L;refGene_tx=XM_003988642.4
B4 78103552 78104533
refGene_gn=LOC101096402;refGene_tx=XM_003988644.5
B4 78104803 78104952
refGene_gn=LOC101096402;refGene_tx=XM_003988644.5
B4 78105095 78105318
refGene_gn=LOC101096402;refGene_tx=XM_003988644.5
B4 78106680 78106683
refGene_gn=LOC101096402;refGene_tx=XM_003988644.5
B4 78154515 78154961
refGene_gn=LOC101096902;refGene_tx=XM_023257014.1
B4 78155231 78155380
refGene_gn=LOC101096902;refGene_tx=XM_023257014.1
B4 78155523 78155746
refGene_gn=LOC101096902;refGene_tx=XM_023257014.1
B4 78157829 78157832
refGene_gn=LOC101096902;refGene_tx=XM_023257014.1
B4 78236268 78236271

refGene_gn=LOC101081992;refGene_tx=XM_011283901.3
B4 78238896 78239119
refGene_gn=LOC101081992;refGene_tx=XM_011283901.3
B4 78239262 78239411
refGene_gn=LOC101081992;refGene_tx=XM_011283901.3
B4 78242590 78243565
refGene_gn=LOC101081992;refGene_tx=XM_011283901.3
B4 78259371 78259913
refGene_gn=PRPH;refGene_tx=XM_011283902.3;refGene_gn=PRPH;refGene_tx=XM_011283903.3
B4 78260271 78260332
refGene_gn=PRPH;refGene_tx=XM_011283902.3;refGene_gn=PRPH;refGene_tx=XM_011283903.3
B4 78260513 78260609
refGene_gn=PRPH;refGene_tx=XM_011283902.3;refGene_gn=PRPH;refGene_tx=XM_011283903.3
B4 78261012 78261180
refGene_gn=PRPH;refGene_tx=XM_011283902.3;refGene_gn=PRPH;refGene_tx=XM_011283903.3
B4 78261280 78261406
refGene_gn=PRPH;refGene_tx=XM_011283902.3;refGene_gn=PRPH;refGene_tx=XM_011283903.3
B4 78261503 78261724
refGene_gn=PRPH;refGene_tx=XM_011283902.3;refGene_gn=PRPH;refGene_tx=XM_011283903.3
B4 78261837 78261887
refGene_gn=PRPH;refGene_tx=XM_011283902.3;refGene_gn=PRPH;refGene_tx=XM_011283903.3
B4 78262131 78262211
refGene_gn=PRPH;refGene_tx=XM_011283902.3;refGene_gn=PRPH;refGene_tx=XM_011283903.3
B4 78262460 78262529
refGene_gn=PRPH;refGene_tx=XM_011283902.3;refGene_gn=PRPH;refGene_tx=XM_011283903.3
B4 78281603 78281747 refGene_gn=TROAP;refGene_tx=XM_003988664.3
B4 78281851 78282035 refGene_gn=TROAP;refGene_tx=XM_003988664.3
B4 78282549 78282707 refGene_gn=TROAP;refGene_tx=XM_003988664.3
B4 78282796 78282934 refGene_gn=TROAP;refGene_tx=XM_003988664.3
B4 78283123 78283206 refGene_gn=TROAP;refGene_tx=XM_003988664.3
B4 78283703 78283756 refGene_gn=TROAP;refGene_tx=XM_003988664.3
B4 78284242 78284355 refGene_gn=TROAP;refGene_tx=XM_003988664.3
B4 78285238 78285355 refGene_gn=TROAP;refGene_tx=XM_003988664.3
B4 78285462 78285546 refGene_gn=TROAP;refGene_tx=XM_003988664.3
B4 78285655 78285715 refGene_gn=TROAP;refGene_tx=XM_003988664.3
B4 78285970 78286105 refGene_gn=TROAP;refGene_tx=XM_003988664.3
B4 78286257 78286891 refGene_gn=TROAP;refGene_tx=XM_003988664.3
B4 78287153 78287347 refGene_gn=TROAP;refGene_tx=XM_003988664.3
B4 78287525 78287573 refGene_gn=TROAP;refGene_tx=XM_003988664.3
B4 78289404 78289584 refGene_gn=C1QL4;refGene_tx=XM_023257016.1
B4 78291711 78292248 refGene_gn=C1QL4;refGene_tx=XM_023257016.1

B4	78297988 78298201
refGene_gn=LOC111561454;refGene_tx=XM_023257757.1
B4	78298900 78299017
refGene_gn=LOC111561454;refGene_tx=XM_023257757.1
B4	78300599 78300770
refGene_gn=LOC111561454;refGene_tx=XM_023257757.1
B4	78300877 78300952
refGene_gn=LOC111561454;refGene_tx=XM_023257757.1
B4	78304588 78305428 refGene_gn=DNAJC22;refGene_tx=XM_019834879.2
B4	78307141 78307327 refGene_gn=DNAJC22;refGene_tx=XM_019834879.2
B4	78392974 78392999
refGene_gn=SPATS2;refGene_tx=XM_003988666.5;refGene_gn=SPATS2;refGene_tx=XM_019834880.2
B4	78427667 78427747
refGene_gn=SPATS2;refGene_tx=XM_003988666.5;refGene_gn=SPATS2;refGene_tx=XM_006933672.4;refGene_gn=SPATS2;refGene_tx=XM_019834880.2
B4	78432052 78432161
refGene_gn=SPATS2;refGene_tx=XM_003988666.5;refGene_gn=SPATS2;refGene_tx=XM_006933672.4;refGene_gn=SPATS2;refGene_tx=XM_019834880.2
B4	78433302 78433352
refGene_gn=SPATS2;refGene_tx=XM_003988666.5;refGene_gn=SPATS2;refGene_tx=XM_006933672.4;refGene_gn=SPATS2;refGene_tx=XM_019834880.2
B4	78442755 78443017
refGene_gn=SPATS2;refGene_tx=XM_003988666.5;refGene_gn=SPATS2;refGene_tx=XM_006933672.4;refGene_gn=SPATS2;refGene_tx=XM_019834880.2
B4	78444932 78445112
refGene_gn=SPATS2;refGene_tx=XM_003988666.5;refGene_gn=SPATS2;refGene_tx=XM_006933672.4;refGene_gn=SPATS2;refGene_tx=XM_019834880.2
B4	78447485 78447621
refGene_gn=SPATS2;refGene_tx=XM_003988666.5;refGene_gn=SPATS2;refGene_tx=XM_006933672.4;refGene_gn=SPATS2;refGene_tx=XM_019834880.2
B4	78460419 78460478
refGene_gn=SPATS2;refGene_tx=XM_003988666.5;refGene_gn=SPATS2;refGene_tx=XM_006933672.4;refGene_gn=SPATS2;refGene_tx=XM_019834880.2
B4	78464022 78464132
refGene_gn=SPATS2;refGene_tx=XM_003988666.5;refGene_gn=SPATS2;refGene_tx=XM_006933672.4;refGene_gn=SPATS2;refGene_tx=XM_019834880.2
B4	78467072 78467175
refGene_gn=SPATS2;refGene_tx=XM_003988666.5;refGene_gn=SPATS2;refGene_tx=XM_006933672.4;refGene_gn=SPATS2;refGene_tx=XM_019834880.2
B4	78468454 78468669
refGene_gn=SPATS2;refGene_tx=XM_003988666.5;refGene_gn=SPATS2;refGene_tx=XM_006933672.4;refGene_gn=SPATS2;refGene_tx=XM_019834880.2
B4	78469732 78470044
refGene_gn=SPATS2;refGene_tx=XM_003988666.5;refGene_gn=SPATS2;refGene_tx=XM_006933672.4;refGene_gn=SPATS2;refGene_tx=XM_019834880.2
B4	78480283 78480359
refGene_gn=KCNH3;refGene_tx=XM_003988667.5;refGene_gn=KCNH3;refGene_tx=XM_019834881.2
B4	78481804 78482038

refGene_gn=KCNH3;refGene_tx=XM_003988667.5;refGene_gn=KCNH3;refGene_tx=XM_019834881.2
B4 78482521 78482656 refGene_gn=KCNH3;refGene_tx=XM_003988667.5
B4 78483428 78483562 refGene_gn=KCNH3;refGene_tx=XM_003988667.5;refGene_gn=KCNH3;refGene_tx=XM_019834881.2
B4 78484001 78484245 refGene_gn=KCNH3;refGene_tx=XM_003988667.5;refGene_gn=KCNH3;refGene_tx=XM_019834881.2
B4 78484634 78484792 refGene_gn=KCNH3;refGene_tx=XM_003988667.5;refGene_gn=KCNH3;refGene_tx=XM_019834881.2
B4 78484877 78485085 refGene_gn=KCNH3;refGene_tx=XM_003988667.5;refGene_gn=KCNH3;refGene_tx=XM_019834881.2
B4 78488990 78489272 refGene_gn=KCNH3;refGene_tx=XM_003988667.5;refGene_gn=KCNH3;refGene_tx=XM_019834881.2
B4 78489565 78489765 refGene_gn=KCNH3;refGene_tx=XM_003988667.5;refGene_gn=KCNH3;refGene_tx=XM_019834881.2
B4 78490214 78490464 refGene_gn=KCNH3;refGene_tx=XM_003988667.5;refGene_gn=KCNH3;refGene_tx=XM_019834881.2
B4 78493717 78493935 refGene_gn=KCNH3;refGene_tx=XM_003988667.5;refGene_gn=KCNH3;refGene_tx=XM_019834881.2
B4 78494696 78495028 refGene_gn=KCNH3;refGene_tx=XM_003988667.5;refGene_gn=KCNH3;refGene_tx=XM_019834881.2
B4 78495416 78495523 refGene_gn=KCNH3;refGene_tx=XM_003988667.5;refGene_gn=KCNH3;refGene_tx=XM_019834881.2
B4 78496155 78496232 refGene_gn=KCNH3;refGene_tx=XM_003988667.5;refGene_gn=KCNH3;refGene_tx=XM_019834881.2
B4 78496353 78496950 refGene_gn=KCNH3;refGene_tx=XM_003988667.5;refGene_gn=KCNH3;refGene_tx=XM_019834881.2
B4 78497653 78497740 refGene_gn=MCRS1;refGene_tx=XM_003988668.4;refGene_gn=MCRS1;refGene_tx=XM_003988669.5;refGene_gn=MCRS1;refGene_tx=XM_006933673.2;refGene_gn=MCRS1;refGene_tx=XM_019834884.2
B4 78497857 78497985 refGene_gn=MCRS1;refGene_tx=XM_003988668.4;refGene_gn=MCRS1;refGene_tx=XM_003988669.5;refGene_gn=MCRS1;refGene_tx=XM_006933673.2;refGene_gn=MCRS1;refGene_tx=XM_019834884.2
B4 78498220 78498308 refGene_gn=MCRS1;refGene_tx=XM_003988668.4;refGene_gn=MCRS1;refGene_tx=XM_003988669.5;refGene_gn=MCRS1;refGene_tx=XM_006933673.2;refGene_gn=

MCRS1;refGene_tx=XM_019834884.2
B4 78498465 78498548
refGene_gn=MCRS1;refGene_tx=XM_003988668.4;refGene_gn=MCRS1;refGene_tx=XM_003988669.5;refGene_gn=MCRS1;refGene_tx=XM_006933673.2;refGene_gn=MCRS1;refGene_tx=XM_019834884.2
B4 78498732 78498825
refGene_gn=MCRS1;refGene_tx=XM_003988668.4;refGene_gn=MCRS1;refGene_tx=XM_003988669.5;refGene_gn=MCRS1;refGene_tx=XM_006933673.2;refGene_gn=MCRS1;refGene_tx=XM_019834884.2
B4 78498941 78498970
refGene_gn=MCRS1;refGene_tx=XM_003988668.4;refGene_gn=MCRS1;refGene_tx=XM_003988669.5;refGene_gn=MCRS1;refGene_tx=XM_006933673.2;refGene_gn=MCRS1;refGene_tx=XM_019834884.2
B4 78499333 78499409
refGene_gn=MCRS1;refGene_tx=XM_003988668.4;refGene_gn=MCRS1;refGene_tx=XM_003988669.5;refGene_gn=MCRS1;refGene_tx=XM_006933673.2;refGene_gn=MCRS1;refGene_tx=XM_019834884.2
B4 78501606 78501745
refGene_gn=MCRS1;refGene_tx=XM_003988668.4;refGene_gn=MCRS1;refGene_tx=XM_003988669.5;refGene_gn=MCRS1;refGene_tx=XM_006933673.2;refGene_gn=MCRS1;refGene_tx=XM_019834884.2
B4 78502008 78502117
refGene_gn=MCRS1;refGene_tx=XM_003988668.4;refGene_gn=MCRS1;refGene_tx=XM_003988669.5;refGene_gn=MCRS1;refGene_tx=XM_006933673.2;refGene_gn=MCRS1;refGene_tx=XM_019834884.2
B4 78502956 78503066
refGene_gn=MCRS1;refGene_tx=XM_003988668.4;refGene_gn=MCRS1;refGene_tx=XM_003988669.5;refGene_gn=MCRS1;refGene_tx=XM_006933673.2;refGene_gn=MCRS1;refGene_tx=XM_019834884.2
B4 78503216 78503375
refGene_gn=MCRS1;refGene_tx=XM_003988668.4;refGene_gn=MCRS1;refGene_tx=XM_003988669.5;refGene_gn=MCRS1;refGene_tx=XM_006933673.2;refGene_gn=MCRS1;refGene_tx=XM_019834884.2
B4 78503989 78504128
refGene_gn=MCRS1;refGene_tx=XM_003988668.4;refGene_gn=MCRS1;refGene_tx=XM_003988669.5;refGene_gn=MCRS1;refGene_tx=XM_006933673.2;refGene_gn=MCRS1;refGene_tx=XM_019834884.2
B4 78504495 78504634
refGene_gn=MCRS1;refGene_tx=XM_003988668.4;refGene_gn=MCRS1;refGene_tx=XM_003988669.5;refGene_gn=MCRS1;refGene_tx=XM_006933673.2;refGene_gn=MCRS1;refGene_tx=XM_019834884.2
B4 78504822 78504871 refGene_gn=MCRS1;refGene_tx=XM_003988669.5
B4 78505154 78505164 refGene_gn=MCRS1;refGene_tx=XM_003988668.4
B4 78518640 78518662 refGene_gn=FAM186B;refGene_tx=XM_006933676.3
B4 78524116 78524285
refGene_gn=FAM186B;refGene_tx=XM_006933675.3;refGene_gn=FAM186B;refGene_tx=XM_006933677.2
B4 78526995 78527185
refGene_gn=FAM186B;refGene_tx=XM_006933675.3;refGene_gn=FAM186B;refGene_tx=XM_006933676.3;refGene_gn=FAM186B;refGene_tx=XM_006933677.2;refGe

ne_gn=FAM186B;refGene_tx=XM_019834886.2
B4 78527887 78528066
refGene_gn=FAM186B;refGene_tx=XM_006933675.3;refGene_gn=FAM186B;refGene_tx=XM_006933676.3;refGene_gn=FAM186B;refGene_tx=XM_006933677.2;refGene_gn=FAM186B;refGene_tx=XM_019834886.2
B4 78529713 78529906
refGene_gn=FAM186B;refGene_tx=XM_006933675.3;refGene_gn=FAM186B;refGene_tx=XM_006933676.3;refGene_gn=FAM186B;refGene_tx=XM_006933677.2;refGene_gn=FAM186B;refGene_tx=XM_019834886.2
B4 78530421 78532096
refGene_gn=FAM186B;refGene_tx=XM_006933675.3;refGene_gn=FAM186B;refGene_tx=XM_006933676.3;refGene_gn=FAM186B;refGene_tx=XM_006933677.2;refGene_gn=FAM186B;refGene_tx=XM_019834886.2
B4 78534507 78534690
refGene_gn=FAM186B;refGene_tx=XM_006933675.3;refGene_gn=FAM186B;refGene_tx=XM_006933676.3;refGene_gn=FAM186B;refGene_tx=XM_006933677.2;refGene_gn=FAM186B;refGene_tx=XM_019834886.2
B4 78538941 78539167
refGene_gn=FAM186B;refGene_tx=XM_006933675.3;refGene_gn=FAM186B;refGene_tx=XM_006933676.3;refGene_gn=FAM186B;refGene_tx=XM_006933677.2;refGene_gn=FAM186B;refGene_tx=XM_019834886.2
B4 78540000 78540096
refGene_gn=FAM186B;refGene_tx=XM_006933675.3;refGene_gn=FAM186B;refGene_tx=XM_006933676.3;refGene_gn=FAM186B;refGene_tx=XM_019834886.2
B4 78548434 78548437
refGene_gn=PRPF40B;refGene_tx=XM_023257018.1;refGene_gn=PRPF40B;refGene_tx=XM_023257019.1;refGene_gn=PRPF40B;refGene_tx=XM_023257020.1;refGene_gn=PRPF40B;refGene_tx=XM_023257021.1;refGene_gn=PRPF40B;refGene_tx=XM_023257022.1;refGene_gn=PRPF40B;refGene_tx=XM_023257023.1
B4 78554938 78555019
refGene_gn=PRPF40B;refGene_tx=XM_011283909.3;refGene_gn=PRPF40B;refGene_tx=XM_023257018.1;refGene_gn=PRPF40B;refGene_tx=XM_023257019.1;refGene_gn=PRPF40B;refGene_tx=XM_023257020.1;refGene_gn=PRPF40B;refGene_tx=XM_023257022.1;refGene_gn=PRPF40B;refGene_tx=XM_023257023.1
B4 78555791 78555935
refGene_gn=PRPF40B;refGene_tx=XM_011283909.3;refGene_gn=PRPF40B;refGene_tx=XM_023257018.1;refGene_gn=PRPF40B;refGene_tx=XM_023257019.1;refGene_gn=PRPF40B;refGene_tx=XM_023257020.1;refGene_gn=PRPF40B;refGene_tx=XM_023257021.1;refGene_gn=PRPF40B;refGene_tx=XM_023257022.1;refGene_gn=PRPF40B;refGene_tx=XM_023257023.1
B4 78556246 78556312
refGene_gn=PRPF40B;refGene_tx=XM_011283909.3;refGene_gn=PRPF40B;refGene_tx=XM_023257018.1;refGene_gn=PRPF40B;refGene_tx=XM_023257019.1;refGene_gn=PRPF40B;refGene_tx=XM_023257020.1;refGene_gn=PRPF40B;refGene_tx=XM_023257021.1;refGene_gn=PRPF40B;refGene_tx=XM_023257022.1;refGene_gn=PRPF40B;refGene_tx=XM_023257023.1
B4 78556991 78557019
refGene_gn=PRPF40B;refGene_tx=XM_011283909.3;refGene_gn=PRPF40B;refGene_tx=XM_023257018.1;refGene_gn=PRPF40B;refGene_tx=XM_023257019.1;refGene_gn=PRPF40B;refGene_tx=XM_023257020.1;refGene_gn=PRPF40B;refGene_tx=

XM_023257021.1;refGene_gn=PRPF40B;refGene_tx=XM_023257022.1;refGene_gn=PRPF40B;refGene_tx=XM_023257023.1
B4	78557247	78557273
refGene_gn=PRPF40B;refGene_tx=XM_011283909.3;refGene_gn=PRPF40B;refGene_tx=XM_023257018.1;refGene_gn=PRPF40B;refGene_tx=XM_023257019.1;refGene_gn=PRPF40B;refGene_tx=XM_023257020.1;refGene_gn=PRPF40B;refGene_tx=XM_023257021.1;refGene_gn=PRPF40B;refGene_tx=XM_023257022.1;refGene_gn=PRPF40B;refGene_tx=XM_023257023.1
B4	78557412	78557523
refGene_gn=PRPF40B;refGene_tx=XM_011283909.3;refGene_gn=PRPF40B;refGene_tx=XM_023257018.1;refGene_gn=PRPF40B;refGene_tx=XM_023257019.1;refGene_gn=PRPF40B;refGene_tx=XM_023257020.1;refGene_gn=PRPF40B;refGene_tx=XM_023257021.1
B4	78557811	78557932
refGene_gn=PRPF40B;refGene_tx=XM_011283909.3;refGene_gn=PRPF40B;refGene_tx=XM_023257018.1;refGene_gn=PRPF40B;refGene_tx=XM_023257019.1;refGene_gn=PRPF40B;refGene_tx=XM_023257020.1;refGene_gn=PRPF40B;refGene_tx=XM_023257021.1;refGene_gn=PRPF40B;refGene_tx=XM_023257022.1;refGene_gn=PRPF40B;refGene_tx=XM_023257023.1
B4	78558014	78558039
refGene_gn=PRPF40B;refGene_tx=XM_011283909.3;refGene_gn=PRPF40B;refGene_tx=XM_023257018.1;refGene_gn=PRPF40B;refGene_tx=XM_023257019.1;refGene_gn=PRPF40B;refGene_tx=XM_023257020.1;refGene_gn=PRPF40B;refGene_tx=XM_023257021.1;refGene_gn=PRPF40B;refGene_tx=XM_023257022.1;refGene_gn=PRPF40B;refGene_tx=XM_023257023.1
B4	78558257	78558440
refGene_gn=PRPF40B;refGene_tx=XM_011283909.3;refGene_gn=PRPF40B;refGene_tx=XM_023257018.1;refGene_gn=PRPF40B;refGene_tx=XM_023257019.1;refGene_gn=PRPF40B;refGene_tx=XM_023257020.1;refGene_gn=PRPF40B;refGene_tx=XM_023257021.1;refGene_gn=PRPF40B;refGene_tx=XM_023257022.1;refGene_gn=PRPF40B;refGene_tx=XM_023257023.1
B4	78558675	78558805
refGene_gn=PRPF40B;refGene_tx=XM_011283909.3;refGene_gn=PRPF40B;refGene_tx=XM_023257018.1;refGene_gn=PRPF40B;refGene_tx=XM_023257019.1;refGene_gn=PRPF40B;refGene_tx=XM_023257020.1;refGene_gn=PRPF40B;refGene_tx=XM_023257021.1;refGene_gn=PRPF40B;refGene_tx=XM_023257022.1;refGene_gn=PRPF40B;refGene_tx=XM_023257023.1
B4	78558887	78558952
refGene_gn=PRPF40B;refGene_tx=XM_011283909.3;refGene_gn=PRPF40B;refGene_tx=XM_023257018.1;refGene_gn=PRPF40B;refGene_tx=XM_023257019.1;refGene_gn=PRPF40B;refGene_tx=XM_023257020.1;refGene_gn=PRPF40B;refGene_tx=XM_023257021.1;refGene_gn=PRPF40B;refGene_tx=XM_023257022.1;refGene_gn=PRPF40B;refGene_tx=XM_023257023.1
B4	78559467	78559632
refGene_gn=PRPF40B;refGene_tx=XM_011283909.3;refGene_gn=PRPF40B;refGene_tx=XM_023257018.1;refGene_gn=PRPF40B;refGene_tx=XM_023257019.1;refGene_gn=PRPF40B;refGene_tx=XM_023257020.1;refGene_gn=PRPF40B;refGene_tx=XM_023257021.1;refGene_gn=PRPF40B;refGene_tx=XM_023257022.1;refGene_gn=PRPF40B;refGene_tx=XM_023257023.1
B4	78559733	78559842

refGene_gn=PRPF40B;refGene_tx=XM_011283909.3;refGene_gn=PRPF40B;refGene_tx=XM_023257018.1;refGene_gn=PRPF40B;refGene_tx=XM_023257019.1;refGene_gn=PRPF40B;refGene_tx=XM_023257020.1;refGene_gn=PRPF40B;refGene_tx=XM_023257021.1;refGene_gn=PRPF40B;refGene_tx=XM_023257022.1;refGene_gn=PRPF40B;refGene_tx=XM_023257023.1
B4 78560219 78560370
refGene_gn=PRPF40B;refGene_tx=XM_011283909.3;refGene_gn=PRPF40B;refGene_tx=XM_023257018.1;refGene_gn=PRPF40B;refGene_tx=XM_023257019.1;refGene_gn=PRPF40B;refGene_tx=XM_023257020.1;refGene_gn=PRPF40B;refGene_tx=XM_023257021.1;refGene_gn=PRPF40B;refGene_tx=XM_023257022.1;refGene_gn=PRPF40B;refGene_tx=XM_023257023.1
B4 78561006 78561140
refGene_gn=PRPF40B;refGene_tx=XM_011283909.3;refGene_gn=PRPF40B;refGene_tx=XM_023257018.1;refGene_gn=PRPF40B;refGene_tx=XM_023257019.1;refGene_gn=PRPF40B;refGene_tx=XM_023257020.1;refGene_gn=PRPF40B;refGene_tx=XM_023257021.1;refGene_gn=PRPF40B;refGene_tx=XM_023257022.1;refGene_gn=PRPF40B;refGene_tx=XM_023257023.1
B4 78561807 78561922
refGene_gn=PRPF40B;refGene_tx=XM_011283909.3;refGene_gn=PRPF40B;refGene_tx=XM_023257018.1;refGene_gn=PRPF40B;refGene_tx=XM_023257019.1;refGene_gn=PRPF40B;refGene_tx=XM_023257020.1;refGene_gn=PRPF40B;refGene_tx=XM_023257021.1;refGene_gn=PRPF40B;refGene_tx=XM_023257022.1;refGene_gn=PRPF40B;refGene_tx=XM_023257023.1
B4 78562071 78562163
refGene_gn=PRPF40B;refGene_tx=XM_011283909.3;refGene_gn=PRPF40B;refGene_tx=XM_023257018.1;refGene_gn=PRPF40B;refGene_tx=XM_023257019.1;refGene_gn=PRPF40B;refGene_tx=XM_023257020.1;refGene_gn=PRPF40B;refGene_tx=XM_023257021.1;refGene_gn=PRPF40B;refGene_tx=XM_023257022.1;refGene_gn=PRPF40B;refGene_tx=XM_023257023.1
B4 78566581 78566698
refGene_gn=PRPF40B;refGene_tx=XM_011283909.3;refGene_gn=PRPF40B;refGene_tx=XM_023257018.1;refGene_gn=PRPF40B;refGene_tx=XM_023257019.1;refGene_gn=PRPF40B;refGene_tx=XM_023257020.1;refGene_gn=PRPF40B;refGene_tx=XM_023257021.1;refGene_gn=PRPF40B;refGene_tx=XM_023257022.1;refGene_gn=PRPF40B;refGene_tx=XM_023257023.1
B4 78566926 78567064
refGene_gn=PRPF40B;refGene_tx=XM_011283909.3;refGene_gn=PRPF40B;refGene_tx=XM_023257018.1;refGene_gn=PRPF40B;refGene_tx=XM_023257019.1;refGene_gn=PRPF40B;refGene_tx=XM_023257020.1;refGene_gn=PRPF40B;refGene_tx=XM_023257021.1;refGene_gn=PRPF40B;refGene_tx=XM_023257022.1;refGene_gn=PRPF40B;refGene_tx=XM_023257023.1
B4 78567272 78567368
refGene_gn=PRPF40B;refGene_tx=XM_011283909.3;refGene_gn=PRPF40B;refGene_tx=XM_023257018.1;refGene_gn=PRPF40B;refGene_tx=XM_023257019.1;refGene_gn=PRPF40B;refGene_tx=XM_023257020.1;refGene_gn=PRPF40B;refGene_tx=XM_023257021.1;refGene_gn=PRPF40B;refGene_tx=XM_023257022.1;refGene_gn=PRPF40B;refGene_tx=XM_023257023.1
B4 78567626 78567713
refGene_gn=PRPF40B;refGene_tx=XM_011283909.3;refGene_gn=PRPF40B;refGene_tx=XM_023257018.1;refGene_gn=PRPF40B;refGene_tx=XM_023257019.1;refGe

ne_gn=PRPF40B;refGene_tx=XM_023257020.1;refGene_gn=PRPF40B;refGene_tx=XM_023257021.1;refGene_gn=PRPF40B;refGene_tx=XM_023257022.1;refGene_gn=PRPF40B;refGene_tx=XM_023257023.1
B4 78567930 78568108
refGene_gn=PRPF40B;refGene_tx=XM_011283909.3;refGene_gn=PRPF40B;refGene_tx=XM_023257018.1;refGene_gn=PRPF40B;refGene_tx=XM_023257019.1;refGene_gn=PRPF40B;refGene_tx=XM_023257020.1;refGene_gn=PRPF40B;refGene_tx=XM_023257021.1;refGene_gn=PRPF40B;refGene_tx=XM_023257022.1;refGene_gn=PRPF40B;refGene_tx=XM_023257023.1
B4 78568370 78568432
refGene_gn=PRPF40B;refGene_tx=XM_011283909.3;refGene_gn=PRPF40B;refGene_tx=XM_023257018.1;refGene_gn=PRPF40B;refGene_tx=XM_023257019.1;refGene_gn=PRPF40B;refGene_tx=XM_023257020.1;refGene_gn=PRPF40B;refGene_tx=XM_023257021.1;refGene_gn=PRPF40B;refGene_tx=XM_023257022.1;refGene_gn=PRPF40B;refGene_tx=XM_023257023.1
B4 78568545 78568689
refGene_gn=PRPF40B;refGene_tx=XM_011283909.3;refGene_gn=PRPF40B;refGene_tx=XM_023257018.1;refGene_gn=PRPF40B;refGene_tx=XM_023257019.1;refGene_gn=PRPF40B;refGene_tx=XM_023257020.1;refGene_gn=PRPF40B;refGene_tx=XM_023257021.1;refGene_gn=PRPF40B;refGene_tx=XM_023257022.1;refGene_gn=PRPF40B;refGene_tx=XM_023257023.1
B4 78568789 78568882
refGene_gn=PRPF40B;refGene_tx=XM_011283909.3;refGene_gn=PRPF40B;refGene_tx=XM_023257018.1;refGene_gn=PRPF40B;refGene_tx=XM_023257019.1;refGene_gn=PRPF40B;refGene_tx=XM_023257020.1;refGene_gn=PRPF40B;refGene_tx=XM_023257021.1;refGene_gn=PRPF40B;refGene_tx=XM_023257022.1;refGene_gn=PRPF40B;refGene_tx=XM_023257023.1
B4 78570507 78570596
refGene_gn=FMNL3;refGene_tx=XM_011283912.2;refGene_gn=FMNL3;refGene_tx=XM_023257017.1
B4 78571326 78571418
refGene_gn=FMNL3;refGene_tx=XM_011283910.3;refGene_gn=FMNL3;refGene_tx=XM_011283911.3;refGene_gn=FMNL3;refGene_tx=XM_011283913.3;refGene_gn=FMNL3;refGene_tx=XM_019834885.2
B4 78571549 78571673
refGene_gn=FMNL3;refGene_tx=XM_011283910.3;refGene_gn=FMNL3;refGene_tx=XM_011283911.3;refGene_gn=FMNL3;refGene_tx=XM_011283912.2;refGene_gn=FMNL3;refGene_tx=XM_011283913.3;refGene_gn=FMNL3;refGene_tx=XM_019834885.2;refGene_gn=FMNL3;refGene_tx=XM_023257017.1
B4 78571959 78572052
refGene_gn=FMNL3;refGene_tx=XM_011283910.3;refGene_gn=FMNL3;refGene_tx=XM_011283911.3;refGene_gn=FMNL3;refGene_tx=XM_011283912.2;refGene_gn=FMNL3;refGene_tx=XM_011283913.3;refGene_gn=FMNL3;refGene_tx=XM_019834885.2;refGene_gn=FMNL3;refGene_tx=XM_023257017.1
B4 78572371 78572473
refGene_gn=FMNL3;refGene_tx=XM_011283910.3;refGene_gn=FMNL3;refGene_tx=XM_011283911.3;refGene_gn=FMNL3;refGene_tx=XM_011283912.2;refGene_gn=FMNL3;refGene_tx=XM_011283913.3;refGene_gn=FMNL3;refGene_tx=XM_019834885.2;refGene_gn=FMNL3;refGene_tx=XM_023257017.1
B4 78572824 78572985

refGene_gn=FMNL3;refGene_tx=XM_011283910.3;refGene_gn=FMNL3;refGene_tx=XM_011283911.3;refGene_gn=FMNL3;refGene_tx=XM_011283912.2;refGene_gn=FMNL3;refGene_tx=XM_011283913.3;refGene_gn=FMNL3;refGene_tx=XM_019834885.2;refGene_gn=FMNL3;refGene_tx=XM_023257017.1
B4 78573626 78573756
refGene_gn=FMNL3;refGene_tx=XM_011283910.3;refGene_gn=FMNL3;refGene_tx=XM_011283911.3;refGene_gn=FMNL3;refGene_tx=XM_011283912.2;refGene_gn=FMNL3;refGene_tx=XM_011283913.3;refGene_gn=FMNL3;refGene_tx=XM_019834885.2;refGene_gn=FMNL3;refGene_tx=XM_023257017.1
B4 78573845 78573926
refGene_gn=FMNL3;refGene_tx=XM_011283910.3;refGene_gn=FMNL3;refGene_tx=XM_011283911.3;refGene_gn=FMNL3;refGene_tx=XM_011283912.2;refGene_gn=FMNL3;refGene_tx=XM_011283913.3;refGene_gn=FMNL3;refGene_tx=XM_019834885.2;refGene_gn=FMNL3;refGene_tx=XM_023257017.1
B4 78574030 78574099
refGene_gn=FMNL3;refGene_tx=XM_011283910.3;refGene_gn=FMNL3;refGene_tx=XM_011283911.3;refGene_gn=FMNL3;refGene_tx=XM_011283912.2;refGene_gn=FMNL3;refGene_tx=XM_011283913.3;refGene_gn=FMNL3;refGene_tx=XM_019834885.2;refGene_gn=FMNL3;refGene_tx=XM_023257017.1
B4 78574260 78574495
refGene_gn=FMNL3;refGene_tx=XM_011283910.3;refGene_gn=FMNL3;refGene_tx=XM_011283911.3;refGene_gn=FMNL3;refGene_tx=XM_011283912.2;refGene_gn=FMNL3;refGene_tx=XM_011283913.3;refGene_gn=FMNL3;refGene_tx=XM_019834885.2;refGene_gn=FMNL3;refGene_tx=XM_023257017.1
B4 78575242 78575445
refGene_gn=FMNL3;refGene_tx=XM_011283910.3;refGene_gn=FMNL3;refGene_tx=XM_011283911.3;refGene_gn=FMNL3;refGene_tx=XM_011283912.2;refGene_gn=FMNL3;refGene_tx=XM_011283913.3;refGene_gn=FMNL3;refGene_tx=XM_019834885.2;refGene_gn=FMNL3;refGene_tx=XM_023257017.1
B4 78575644 78575769
refGene_gn=FMNL3;refGene_tx=XM_011283910.3;refGene_gn=FMNL3;refGene_tx=XM_011283911.3;refGene_gn=FMNL3;refGene_tx=XM_011283912.2;refGene_gn=FMNL3;refGene_tx=XM_011283913.3;refGene_gn=FMNL3;refGene_tx=XM_019834885.2;refGene_gn=FMNL3;refGene_tx=XM_023257017.1
B4 78575858 78575927
refGene_gn=FMNL3;refGene_tx=XM_011283910.3;refGene_gn=FMNL3;refGene_tx=XM_011283911.3;refGene_gn=FMNL3;refGene_tx=XM_011283912.2;refGene_gn=FMNL3;refGene_tx=XM_011283913.3;refGene_gn=FMNL3;refGene_tx=XM_019834885.2;refGene_gn=FMNL3;refGene_tx=XM_023257017.1
B4 78576390 78576670
refGene_gn=FMNL3;refGene_tx=XM_011283910.3;refGene_gn=FMNL3;refGene_tx=XM_011283911.3;refGene_gn=FMNL3;refGene_tx=XM_011283912.2;refGene_gn=FMNL3;refGene_tx=XM_011283913.3;refGene_gn=FMNL3;refGene_tx=XM_019834885.2;refGene_gn=FMNL3;refGene_tx=XM_023257017.1
B4 78577958 78578060
refGene_gn=FMNL3;refGene_tx=XM_011283910.3;refGene_gn=FMNL3;refGene_tx=XM_011283911.3;refGene_gn=FMNL3;refGene_tx=XM_011283912.2;refGene_gn=FMNL3;refGene_tx=XM_011283913.3;refGene_gn=FMNL3;refGene_tx=XM_019834885.2;refGene_gn=FMNL3;refGene_tx=XM_023257017.1
B4 78578458 78578608

refGene_gn=FMNL3;refGene_tx=XM_011283910.3;refGene_gn=FMNL3;refGene_tx
=XM_011283911.3;refGene_gn=FMNL3;refGene_tx=XM_011283912.2;refGene_gn=
FMNL3;refGene_tx=XM_011283913.3;refGene_gn=FMNL3;refGene_tx=XM_0198348
85.2;refGene_gn=FMNL3;refGene_tx=XM_023257017.1
B4 78578924 78579035
refGene_gn=FMNL3;refGene_tx=XM_011283910.3;refGene_gn=FMNL3;refGene_tx
=XM_011283911.3;refGene_gn=FMNL3;refGene_tx=XM_011283912.2;refGene_gn=
FMNL3;refGene_tx=XM_011283913.3;refGene_gn=FMNL3;refGene_tx=XM_0198348
85.2;refGene_gn=FMNL3;refGene_tx=XM_023257017.1
B4 78579815 78579890
refGene_gn=FMNL3;refGene_tx=XM_011283910.3;refGene_gn=FMNL3;refGene_tx
=XM_011283911.3;refGene_gn=FMNL3;refGene_tx=XM_011283912.2;refGene_gn=
FMNL3;refGene_tx=XM_011283913.3;refGene_gn=FMNL3;refGene_tx=XM_0198348
85.2;refGene_gn=FMNL3;refGene_tx=XM_023257017.1
B4 78581268 78581362
refGene_gn=FMNL3;refGene_tx=XM_011283910.3;refGene_gn=FMNL3;refGene_tx
=XM_011283911.3;refGene_gn=FMNL3;refGene_tx=XM_011283912.2;refGene_gn=
FMNL3;refGene_tx=XM_011283913.3;refGene_gn=FMNL3;refGene_tx=XM_0198348
85.2;refGene_gn=FMNL3;refGene_tx=XM_023257017.1
B4 78581712 78581789
refGene_gn=FMNL3;refGene_tx=XM_011283910.3;refGene_gn=FMNL3;refGene_tx
=XM_011283911.3;refGene_gn=FMNL3;refGene_tx=XM_011283912.2;refGene_gn=
FMNL3;refGene_tx=XM_011283913.3;refGene_gn=FMNL3;refGene_tx=XM_0198348
85.2;refGene_gn=FMNL3;refGene_tx=XM_023257017.1
B4 78582008 78582117
refGene_gn=FMNL3;refGene_tx=XM_011283910.3;refGene_gn=FMNL3;refGene_tx
=XM_011283911.3;refGene_gn=FMNL3;refGene_tx=XM_011283912.2;refGene_gn=
FMNL3;refGene_tx=XM_011283913.3;refGene_gn=FMNL3;refGene_tx=XM_0198348
85.2;refGene_gn=FMNL3;refGene_tx=XM_023257017.1
B4 78583336 78583489
refGene_gn=FMNL3;refGene_tx=XM_011283910.3;refGene_gn=FMNL3;refGene_tx
=XM_011283911.3;refGene_gn=FMNL3;refGene_tx=XM_011283912.2;refGene_gn=
FMNL3;refGene_tx=XM_019834885.2
B4 78584068 78584079 refGene_gn=FMNL3;refGene_tx=XM_019834885.2
B4 78586253 78586337
refGene_gn=FMNL3;refGene_tx=XM_011283910.3;refGene_gn=FMNL3;refGene_tx
=XM_011283911.3;refGene_gn=FMNL3;refGene_tx=XM_011283912.2;refGene_gn=
FMNL3;refGene_tx=XM_011283913.3;refGene_gn=FMNL3;refGene_tx=XM_0232570
17.1
B4 78589528 78589605
refGene_gn=FMNL3;refGene_tx=XM_011283910.3;refGene_gn=FMNL3;refGene_tx
=XM_011283911.3;refGene_gn=FMNL3;refGene_tx=XM_011283912.2;refGene_gn=
FMNL3;refGene_tx=XM_011283913.3;refGene_gn=FMNL3;refGene_tx=XM_0232570
17.1
B4 78589885 78589966
refGene_gn=FMNL3;refGene_tx=XM_011283910.3;refGene_gn=FMNL3;refGene_tx
=XM_011283911.3;refGene_gn=FMNL3;refGene_tx=XM_011283912.2;refGene_gn=
FMNL3;refGene_tx=XM_011283913.3;refGene_gn=FMNL3;refGene_tx=XM_0232570
17.1
B4 78591956 78592040

refGene_gn=FMNL3;refGene_tx=XM_011283910.3;refGene_gn=FMNL3;refGene_tx=XM_011283911.3;refGene_gn=FMNL3;refGene_tx=XM_011283912.2;refGene_gn=FMNL3;refGene_tx=XM_011283913.3;refGene_gn=FMNL3;refGene_tx=XM_023257017.1
B4 78623283 78623409
refGene_gn=FMNL3;refGene_tx=XM_011283910.3;refGene_gn=FMNL3;refGene_tx=XM_011283911.3;refGene_gn=FMNL3;refGene_tx=XM_011283912.2;refGene_gn=FMNL3;refGene_tx=XM_011283913.3;refGene_gn=FMNL3;refGene_tx=XM_023257017.1
B4 78649027 78649083
refGene_gn=TMBIM6;refGene_tx=XM_003988673.4;refGene_gn=TMBIM6;refGene_tx=XM_006933679.3
B4 78650165 78650274
refGene_gn=TMBIM6;refGene_tx=XM_003988673.4;refGene_gn=TMBIM6;refGene_tx=XM_006933679.3
B4 78653215 78653336
refGene_gn=TMBIM6;refGene_tx=XM_003988673.4;refGene_gn=TMBIM6;refGene_tx=XM_006933679.3
B4 78654724 78654773
refGene_gn=TMBIM6;refGene_tx=XM_003988673.4;refGene_gn=TMBIM6;refGene_tx=XM_006933679.3
B4 78654879 78654977
refGene_gn=TMBIM6;refGene_tx=XM_003988673.4;refGene_gn=TMBIM6;refGene_tx=XM_006933679.3
B4 78655182 78655262
refGene_gn=TMBIM6;refGene_tx=XM_003988673.4;refGene_gn=TMBIM6;refGene_tx=XM_006933679.3
B4 78655572 78655673
refGene_gn=TMBIM6;refGene_tx=XM_003988673.4;refGene_gn=TMBIM6;refGene_tx=XM_006933679.3
B4 78656511 78656587
refGene_gn=TMBIM6;refGene_tx=XM_003988673.4;refGene_gn=TMBIM6;refGene_tx=XM_006933679.3
B4 78657798 78657822
refGene_gn=TMBIM6;refGene_tx=XM_003988673.4;refGene_gn=TMBIM6;refGene_tx=XM_006933679.3
B4 78679284 78679497
refGene_gn=NCKAP5L;refGene_tx=XM_011283915.3;refGene_gn=NCKAP5L;refGene_tx=XM_011283916.3;refGene_gn=NCKAP5L;refGene_tx=XM_011283917.2;refGene_gn=NCKAP5L;refGene_tx=XM_011283918.3;refGene_gn=NCKAP5L;refGene_tx=XM_011283919.2;refGene_gn=NCKAP5L;refGene_tx=XM_011283920.2;refGene_gn=NCKAP5L;refGene_tx=XM_019834896.2
B4 78679868 78680011
refGene_gn=NCKAP5L;refGene_tx=XM_011283915.3;refGene_gn=NCKAP5L;refGene_tx=XM_011283916.3;refGene_gn=NCKAP5L;refGene_tx=XM_011283917.2;refGene_gn=NCKAP5L;refGene_tx=XM_011283918.3;refGene_gn=NCKAP5L;refGene_tx=XM_011283919.2;refGene_gn=NCKAP5L;refGene_tx=XM_011283920.2;refGene_gn=NCKAP5L;refGene_tx=XM_019834896.2
B4 78680089 78680398
refGene_gn=NCKAP5L;refGene_tx=XM_011283915.3;refGene_gn=NCKAP5L;refGen

e_tx=XM_011283916.3;refGene_gn=NCKAP5L;refGene_tx=XM_011283917.2;refGene_gn=NCKAP5L;refGene_tx=XM_011283918.3;refGene_gn=NCKAP5L;refGene_tx=XM_011283919.2;refGene_gn=NCKAP5L;refGene_tx=XM_011283920.2;refGene_gn=NCKAP5L;refGene_tx=XM_019834896.2
B4 78680750 78680832
refGene_gn=NCKAP5L;refGene_tx=XM_011283915.3;refGene_gn=NCKAP5L;refGene_tx=XM_011283916.3;refGene_gn=NCKAP5L;refGene_tx=XM_011283917.2;refGene_gn=NCKAP5L;refGene_tx=XM_011283918.3;refGene_gn=NCKAP5L;refGene_tx=XM_011283919.2;refGene_gn=NCKAP5L;refGene_tx=XM_011283920.2;refGene_gn=NCKAP5L;refGene_tx=XM_019834896.2
B4 78681042 78681205
refGene_gn=NCKAP5L;refGene_tx=XM_011283915.3;refGene_gn=NCKAP5L;refGene_tx=XM_011283916.3;refGene_gn=NCKAP5L;refGene_tx=XM_011283917.2;refGene_gn=NCKAP5L;refGene_tx=XM_011283918.3;refGene_gn=NCKAP5L;refGene_tx=XM_011283919.2;refGene_gn=NCKAP5L;refGene_tx=XM_011283920.2;refGene_gn=NCKAP5L;refGene_tx=XM_019834896.2
B4 78682110 78684731
refGene_gn=NCKAP5L;refGene_tx=XM_011283915.3;refGene_gn=NCKAP5L;refGene_tx=XM_011283916.3;refGene_gn=NCKAP5L;refGene_tx=XM_011283917.2;refGene_gn=NCKAP5L;refGene_tx=XM_011283918.3;refGene_gn=NCKAP5L;refGene_tx=XM_011283919.2;refGene_gn=NCKAP5L;refGene_tx=XM_011283920.2;refGene_gn=NCKAP5L;refGene_tx=XM_019834896.2
B4 78686045 78686159
refGene_gn=NCKAP5L;refGene_tx=XM_011283915.3;refGene_gn=NCKAP5L;refGene_tx=XM_011283916.3;refGene_gn=NCKAP5L;refGene_tx=XM_011283917.2;refGene_gn=NCKAP5L;refGene_tx=XM_011283918.3;refGene_gn=NCKAP5L;refGene_tx=XM_011283919.2;refGene_gn=NCKAP5L;refGene_tx=XM_011283920.2;refGene_gn=NCKAP5L;refGene_tx=XM_019834896.2
B4 78688815 78688935
refGene_gn=NCKAP5L;refGene_tx=XM_011283915.3;refGene_gn=NCKAP5L;refGene_tx=XM_011283916.3;refGene_gn=NCKAP5L;refGene_tx=XM_011283917.2;refGene_gn=NCKAP5L;refGene_tx=XM_011283918.3;refGene_gn=NCKAP5L;refGene_tx=XM_011283919.2;refGene_gn=NCKAP5L;refGene_tx=XM_011283920.2;refGene_gn=NCKAP5L;refGene_tx=XM_019834896.2
B4 78689663 78689702
refGene_gn=NCKAP5L;refGene_tx=XM_011283915.3;refGene_gn=NCKAP5L;refGene_tx=XM_011283916.3;refGene_gn=NCKAP5L;refGene_tx=XM_011283917.2;refGene_gn=NCKAP5L;refGene_tx=XM_011283918.3;refGene_gn=NCKAP5L;refGene_tx=XM_011283919.2;refGene_gn=NCKAP5L;refGene_tx=XM_011283920.2;refGene_gn=NCKAP5L;refGene_tx=XM_019834896.2
B4 78689826 78689895
refGene_gn=NCKAP5L;refGene_tx=XM_011283915.3;refGene_gn=NCKAP5L;refGene_tx=XM_011283916.3;refGene_gn=NCKAP5L;refGene_tx=XM_011283917.2;refGene_gn=NCKAP5L;refGene_tx=XM_011283918.3;refGene_gn=NCKAP5L;refGene_tx=XM_011283919.2;refGene_gn=NCKAP5L;refGene_tx=XM_011283920.2;refGene_gn=NCKAP5L;refGene_tx=XM_019834896.2
B4 78691039 78691369
refGene_gn=NCKAP5L;refGene_tx=XM_011283915.3;refGene_gn=NCKAP5L;refGene_tx=XM_011283916.3;refGene_gn=NCKAP5L;refGene_tx=XM_011283917.2;refGene_gn=NCKAP5L;refGene_tx=XM_011283918.3;refGene_gn=NCKAP5L;refGene_tx=

XM_011283919.2;refGene_gn=NCKAP5L;refGene_tx=XM_011283920.2;refGene_gn=NCKAP5L;refGene_tx=XM_019834896.2
B4 78692273 78692333 refGene_gn=NCKAP5L;refGene_tx=XM_011283915.3
B4 78694929 78695008 refGene_gn=NCKAP5L;refGene_tx=XM_011283915.3
B4 78728770 78729415 refGene_gn=BCDIN3D;refGene_tx=XM_003988675.4
B4 78731969 78732203 refGene_gn=BCDIN3D;refGene_tx=XM_003988675.4
B4 78752898 78753048
refGene_gn=FAIM2;refGene_tx=XM_006933680.4;refGene_gn=FAIM2;refGene_tx=XM_006933681.4
B4 78766383 78766437
refGene_gn=FAIM2;refGene_tx=XM_006933680.4;refGene_gn=FAIM2;refGene_tx=XM_006933681.4
B4 78768103 78768199
refGene_gn=FAIM2;refGene_tx=XM_006933680.4;refGene_gn=FAIM2;refGene_tx=XM_006933681.4
B4 78768477 78768565
refGene_gn=FAIM2;refGene_tx=XM_006933680.4;refGene_gn=FAIM2;refGene_tx=XM_006933681.4
B4 78769045 78769083
refGene_gn=FAIM2;refGene_tx=XM_006933680.4;refGene_gn=FAIM2;refGene_tx=XM_006933681.4
B4 78769598 78769638
refGene_gn=FAIM2;refGene_tx=XM_006933680.4;refGene_gn=FAIM2;refGene_tx=XM_006933681.4
B4 78769970 78770021
refGene_gn=FAIM2;refGene_tx=XM_006933680.4;refGene_gn=FAIM2;refGene_tx=XM_006933681.4
B4 78775767 78775821
refGene_gn=FAIM2;refGene_tx=XM_006933680.4;refGene_gn=FAIM2;refGene_tx=XM_006933681.4
B4 78776247 78776312
refGene_gn=FAIM2;refGene_tx=XM_006933680.4;refGene_gn=FAIM2;refGene_tx=XM_006933681.4
B4 78776761 78776862
refGene_gn=FAIM2;refGene_tx=XM_006933680.4;refGene_gn=FAIM2;refGene_tx=XM_006933681.4
B4 78778549 78778634 refGene_gn=FAIM2;refGene_tx=XM_006933681.4
B4 78779553 78779749 refGene_gn=FAIM2;refGene_tx=XM_006933680.4
B4 78783315 78783330 refGene_gn=FAIM2;refGene_tx=XM_006933680.4
B4 78833391 78833751
refGene_gn=AQP2;refGene_tx=XM_023257024.1;refGene_gn=AQP2;refGene_tx=XM_023257026.1
B4 78836535 78836700
refGene_gn=AQP2;refGene_tx=XM_023257024.1;refGene_gn=AQP2;refGene_tx=XM_023257026.1
B4 78837110 78837191
refGene_gn=AQP2;refGene_tx=XM_023257024.1;refGene_gn=AQP2;refGene_tx=XM_023257026.1
B4 78837898 78838269
refGene_gn=AQP2;refGene_tx=XM_023257024.1;refGene_gn=AQP2;refGene_tx=X

M_023257026.1
B4 78845546 78845909 refGene_gn=AQP5;refGene_tx=XM_003988711.5
B4 78846977 78847142 refGene_gn=AQP5;refGene_tx=XM_003988711.5
B4 78847532 78847616 refGene_gn=AQP5;refGene_tx=XM_003988711.5
B4 78848376 78848562 refGene_gn=AQP5;refGene_tx=XM_003988711.5
B4 78855635 78855998 refGene_gn=AQP6;refGene_tx=XM_023257027.1
B4 78856807 78856966 refGene_gn=AQP6;refGene_tx=XM_023257027.1
B4 78857221 78857302 refGene_gn=AQP6;refGene_tx=XM_023257027.1
B4 78857959 78858172 refGene_gn=AQP6;refGene_tx=XM_023257027.1
B4 78869537 78869613
refGene_gn=RACGAP1;refGene_tx=XM_006933682.4;refGene_gn=RACGAP1;refGene_tx=XM_006933683.4;refGene_gn=RACGAP1;refGene_tx=XM_019834897.2;refGene_gn=RACGAP1;refGene_tx=XM_023257028.1;refGene_gn=RACGAP1;refGene_tx=XM_023257029.1;refGene_gn=RACGAP1;refGene_tx=XM_023257030.1
B4 78869934 78870043
refGene_gn=RACGAP1;refGene_tx=XM_006933682.4;refGene_gn=RACGAP1;refGene_tx=XM_006933683.4;refGene_gn=RACGAP1;refGene_tx=XM_019834897.2;refGene_gn=RACGAP1;refGene_tx=XM_023257028.1;refGene_gn=RACGAP1;refGene_tx=XM_023257029.1;refGene_gn=RACGAP1;refGene_tx=XM_023257030.1
B4 78871291 78871427
refGene_gn=RACGAP1;refGene_tx=XM_006933682.4;refGene_gn=RACGAP1;refGene_tx=XM_006933683.4;refGene_gn=RACGAP1;refGene_tx=XM_019834897.2;refGene_gn=RACGAP1;refGene_tx=XM_023257028.1;refGene_gn=RACGAP1;refGene_tx=XM_023257029.1;refGene_gn=RACGAP1;refGene_tx=XM_023257030.1
B4 78871643 78871776
refGene_gn=RACGAP1;refGene_tx=XM_006933682.4;refGene_gn=RACGAP1;refGene_tx=XM_006933683.4;refGene_gn=RACGAP1;refGene_tx=XM_019834897.2;refGene_gn=RACGAP1;refGene_tx=XM_023257028.1;refGene_gn=RACGAP1;refGene_tx=XM_023257029.1;refGene_gn=RACGAP1;refGene_tx=XM_023257030.1
B4 78871952 78872058
refGene_gn=RACGAP1;refGene_tx=XM_006933682.4;refGene_gn=RACGAP1;refGene_tx=XM_006933683.4;refGene_gn=RACGAP1;refGene_tx=XM_019834897.2;refGene_gn=RACGAP1;refGene_tx=XM_023257028.1;refGene_gn=RACGAP1;refGene_tx=XM_023257029.1;refGene_gn=RACGAP1;refGene_tx=XM_023257030.1
B4 78873368 78873567
refGene_gn=RACGAP1;refGene_tx=XM_006933682.4;refGene_gn=RACGAP1;refGene_tx=XM_006933683.4;refGene_gn=RACGAP1;refGene_tx=XM_019834897.2;refGene_gn=RACGAP1;refGene_tx=XM_023257028.1;refGene_gn=RACGAP1;refGene_tx=XM_023257029.1;refGene_gn=RACGAP1;refGene_tx=XM_023257030.1
B4 78873654 78873750
refGene_gn=RACGAP1;refGene_tx=XM_006933682.4;refGene_gn=RACGAP1;refGene_tx=XM_006933683.4;refGene_gn=RACGAP1;refGene_tx=XM_019834897.2;refGene_gn=RACGAP1;refGene_tx=XM_023257028.1;refGene_gn=RACGAP1;refGene_tx=XM_023257029.1;refGene_gn=RACGAP1;refGene_tx=XM_023257030.1
B4 78874998 78875163
refGene_gn=RACGAP1;refGene_tx=XM_006933682.4;refGene_gn=RACGAP1;refGene_tx=XM_006933683.4;refGene_gn=RACGAP1;refGene_tx=XM_019834897.2;refGene_gn=RACGAP1;refGene_tx=XM_023257028.1;refGene_gn=RACGAP1;refGene_tx=XM_023257029.1;refGene_gn=RACGAP1;refGene_tx=XM_023257030.1
B4 78876080 78876211

refGene_gn=RACGAP1;refGene_tx=XM_006933682.4;refGene_gn=RACGAP1;refGene_tx=XM_006933683.4;refGene_gn=RACGAP1;refGene_tx=XM_019834897.2;refGene_gn=RACGAP1;refGene_tx=XM_023257028.1;refGene_gn=RACGAP1;refGene_tx=XM_023257029.1;refGene_gn=RACGAP1;refGene_tx=XM_023257030.1
B4	78876528	78876646
refGene_gn=RACGAP1;refGene_tx=XM_006933682.4;refGene_gn=RACGAP1;refGene_tx=XM_006933683.4;refGene_gn=RACGAP1;refGene_tx=XM_019834897.2;refGene_gn=RACGAP1;refGene_tx=XM_023257028.1;refGene_gn=RACGAP1;refGene_tx=XM_023257029.1;refGene_gn=RACGAP1;refGene_tx=XM_023257030.1
B4	78877515	78877596
refGene_gn=RACGAP1;refGene_tx=XM_006933682.4;refGene_gn=RACGAP1;refGene_tx=XM_006933683.4;refGene_gn=RACGAP1;refGene_tx=XM_019834897.2;refGene_gn=RACGAP1;refGene_tx=XM_023257028.1;refGene_gn=RACGAP1;refGene_tx=XM_023257029.1;refGene_gn=RACGAP1;refGene_tx=XM_023257030.1
B4	78878270	78878324
refGene_gn=RACGAP1;refGene_tx=XM_006933682.4;refGene_gn=RACGAP1;refGene_tx=XM_006933683.4;refGene_gn=RACGAP1;refGene_tx=XM_019834897.2;refGene_gn=RACGAP1;refGene_tx=XM_023257028.1;refGene_gn=RACGAP1;refGene_tx=XM_023257029.1;refGene_gn=RACGAP1;refGene_tx=XM_023257030.1
B4	78880196	78880266
refGene_gn=RACGAP1;refGene_tx=XM_006933682.4;refGene_gn=RACGAP1;refGene_tx=XM_006933683.4;refGene_gn=RACGAP1;refGene_tx=XM_019834897.2;refGene_gn=RACGAP1;refGene_tx=XM_023257028.1;refGene_gn=RACGAP1;refGene_tx=XM_023257029.1
B4	78881219	78881356
refGene_gn=RACGAP1;refGene_tx=XM_006933682.4;refGene_gn=RACGAP1;refGene_tx=XM_006933683.4;refGene_gn=RACGAP1;refGene_tx=XM_019834897.2;refGene_gn=RACGAP1;refGene_tx=XM_023257028.1;refGene_gn=RACGAP1;refGene_tx=XM_023257029.1
B4	78882367	78882570
refGene_gn=RACGAP1;refGene_tx=XM_006933682.4;refGene_gn=RACGAP1;refGene_tx=XM_006933683.4;refGene_gn=RACGAP1;refGene_tx=XM_019834897.2;refGene_gn=RACGAP1;refGene_tx=XM_023257028.1;refGene_gn=RACGAP1;refGene_tx=XM_023257029.1;refGene_gn=RACGAP1;refGene_tx=XM_023257030.1
B4	78891282	78891367
refGene_gn=RACGAP1;refGene_tx=XM_006933682.4;refGene_gn=RACGAP1;refGene_tx=XM_006933683.4;refGene_gn=RACGAP1;refGene_tx=XM_019834897.2;refGene_gn=RACGAP1;refGene_tx=XM_023257028.1;refGene_gn=RACGAP1;refGene_tx=XM_023257029.1;refGene_gn=RACGAP1;refGene_tx=XM_023257030.1
B4	78922967	78923329
refGene_gn=ASIC1;refGene_tx=XM_023257031.1;refGene_gn=ASIC1;refGene_tx=XM_023257032.1
B4	78923955	78924151
refGene_gn=ASIC1;refGene_tx=XM_023257031.1;refGene_gn=ASIC1;refGene_tx=XM_023257032.1
B4	78937802	78938453
refGene_gn=ASIC1;refGene_tx=XM_003988679.5;refGene_gn=ASIC1;refGene_tx=XM_019834899.2
B4	78941154	78941305
refGene_gn=ASIC1;refGene_tx=XM_003988679.5;refGene_gn=ASIC1;refGene_tx

=XM_019834899.2;refGene_gn=ASIC1;refGene_tx=XM_023257031.1;refGene_gn=ASIC1;refGene_tx=XM_023257032.1
B4	78941920	78942048	refGene_gn=ASIC1;refGene_tx=XM_003988679.5;refGene_gn=ASIC1;refGene_tx=XM_019834899.2;refGene_gn=ASIC1;refGene_tx=XM_023257031.1;refGene_gn=ASIC1;refGene_tx=XM_023257032.1
B4	78942327	78942481	refGene_gn=ASIC1;refGene_tx=XM_003988679.5;refGene_gn=ASIC1;refGene_tx=XM_019834899.2;refGene_gn=ASIC1;refGene_tx=XM_023257031.1;refGene_gn=ASIC1;refGene_tx=XM_023257032.1
B4	78942861	78942918	refGene_gn=ASIC1;refGene_tx=XM_003988679.5;refGene_gn=ASIC1;refGene_tx=XM_019834899.2;refGene_gn=ASIC1;refGene_tx=XM_023257031.1;refGene_gn=ASIC1;refGene_tx=XM_023257032.1
B4	78943388	78943542	refGene_gn=ASIC1;refGene_tx=XM_003988679.5;refGene_gn=ASIC1;refGene_tx=XM_019834899.2;refGene_gn=ASIC1;refGene_tx=XM_023257031.1;refGene_gn=ASIC1;refGene_tx=XM_023257032.1
B4	78943981	78944073	refGene_gn=ASIC1;refGene_tx=XM_003988679.5;refGene_gn=ASIC1;refGene_tx=XM_019834899.2;refGene_gn=ASIC1;refGene_tx=XM_023257031.1;refGene_gn=ASIC1;refGene_tx=XM_023257032.1
B4	78944573	78944653	refGene_gn=ASIC1;refGene_tx=XM_003988679.5;refGene_gn=ASIC1;refGene_tx=XM_019834899.2;refGene_gn=ASIC1;refGene_tx=XM_023257031.1;refGene_gn=ASIC1;refGene_tx=XM_023257032.1
B4	78944741	78944846	refGene_gn=ASIC1;refGene_tx=XM_003988679.5;refGene_gn=ASIC1;refGene_tx=XM_019834899.2;refGene_gn=ASIC1;refGene_tx=XM_023257031.1;refGene_gn=ASIC1;refGene_tx=XM_023257032.1
B4	78945006	78945111	refGene_gn=ASIC1;refGene_tx=XM_003988679.5;refGene_gn=ASIC1;refGene_tx=XM_019834899.2;refGene_gn=ASIC1;refGene_tx=XM_023257031.1;refGene_gn=ASIC1;refGene_tx=XM_023257032.1
B4	78948068	78948245	refGene_gn=SMARCD1;refGene_tx=XM_003988680.5;refGene_gn=SMARCD1;refGene_tx=XM_019834901.2
B4	78948941	78949129	refGene_gn=SMARCD1;refGene_tx=XM_003988680.5;refGene_gn=SMARCD1;refGene_tx=XM_019834901.2
B4	78949435	78949478	refGene_gn=SMARCD1;refGene_tx=XM_003988680.5;refGene_gn=SMARCD1;refGene_tx=XM_019834901.2
B4	78949572	78949695	refGene_gn=SMARCD1;refGene_tx=XM_003988680.5;refGene_gn=SMARCD1;refGene_tx=XM_019834901.2
B4	78950162	78950285	refGene_gn=SMARCD1;refGene_tx=XM_003988680.5
B4	78951338	78951455	refGene_gn=SMARCD1;refGene_tx=XM_003988680.5;refGene_gn=SMARCD1;refGene_tx=XM_019834901.2

B4	78952729	78952831
refGene_gn=SMARCD1;refGene_tx=XM_003988680.5;refGene_gn=SMARCD1;refGene_tx=XM_019834901.2
B4	78953125	78953287
refGene_gn=SMARCD1;refGene_tx=XM_003988680.5;refGene_gn=SMARCD1;refGene_tx=XM_019834901.2
B4	78953608	78953706
refGene_gn=SMARCD1;refGene_tx=XM_003988680.5;refGene_gn=SMARCD1;refGene_tx=XM_019834901.2
B4	78955669	78955805
refGene_gn=SMARCD1;refGene_tx=XM_003988680.5;refGene_gn=SMARCD1;refGene_tx=XM_019834901.2
B4	78957360	78957483
refGene_gn=SMARCD1;refGene_tx=XM_003988680.5;refGene_gn=SMARCD1;refGene_tx=XM_019834901.2
B4	78959086	78959188
refGene_gn=SMARCD1;refGene_tx=XM_003988680.5;refGene_gn=SMARCD1;refGene_tx=XM_019834901.2
B4	78959310	78959364
refGene_gn=SMARCD1;refGene_tx=XM_003988680.5;refGene_gn=SMARCD1;refGene_tx=XM_019834901.2
B4	78964179	78964220	refGene_gn=GPD1;refGene_tx=XM_003988682.4
B4	78964726	78964904	refGene_gn=GPD1;refGene_tx=XM_003988682.4
B4	78965718	78965859	refGene_gn=GPD1;refGene_tx=XM_003988682.4
B4	78966436	78966575	refGene_gn=GPD1;refGene_tx=XM_003988682.4
B4	78966819	78966932	refGene_gn=GPD1;refGene_tx=XM_003988682.4
B4	78967590	78967824	refGene_gn=GPD1;refGene_tx=XM_003988682.4
B4	78967968	78968075	refGene_gn=GPD1;refGene_tx=XM_003988682.4
B4	78969311	78969408	refGene_gn=GPD1;refGene_tx=XM_003988682.4
B4	78975983	78976157
refGene_gn=LOC101086986;refGene_tx=XM_003988684.5;refGene_gn=LOC101086986;refGene_tx=XM_006933687.4;refGene_gn=LOC101086986;refGene_tx=XM_006933688.4
B4	78993304	78993454
refGene_gn=CERS5;refGene_tx=XM_003988685.5;refGene_gn=CERS5;refGene_tx=XM_019834902.2;refGene_gn=CERS5;refGene_tx=XM_023257039.1;refGene_gn=CERS5;refGene_tx=XM_023257040.1
B4	78995555	78995664	refGene_gn=CERS5;refGene_tx=XM_019834902.2
B4	78996678	78996835
refGene_gn=CERS5;refGene_tx=XM_003988685.5;refGene_gn=CERS5;refGene_tx=XM_019834902.2;refGene_gn=CERS5;refGene_tx=XM_023257039.1
B4	78997604	78997711
refGene_gn=CERS5;refGene_tx=XM_003988685.5;refGene_gn=CERS5;refGene_tx=XM_019834902.2;refGene_gn=CERS5;refGene_tx=XM_023257039.1;refGene_gn=CERS5;refGene_tx=XM_023257040.1
B4	78997828	78997957
refGene_gn=CERS5;refGene_tx=XM_003988685.5;refGene_gn=CERS5;refGene_tx=XM_019834902.2;refGene_gn=CERS5;refGene_tx=XM_023257039.1;refGene_gn=CERS5;refGene_tx=XM_023257040.1
B4	78999807	78999900

refGene_gn=CERS5;refGene_tx=XM_003988685.5;refGene_gn=CERS5;refGene_tx=XM_019834902.2;refGene_gn=CERS5;refGene_tx=XM_023257039.1;refGene_gn=CERS5;refGene_tx=XM_023257040.1
B4 79000584 79000635
refGene_gn=CERS5;refGene_tx=XM_003988685.5;refGene_gn=CERS5;refGene_tx=XM_019834902.2;refGene_gn=CERS5;refGene_tx=XM_023257039.1;refGene_gn=CERS5;refGene_tx=XM_023257040.1
B4 79002433 79002491
refGene_gn=CERS5;refGene_tx=XM_003988685.5;refGene_gn=CERS5;refGene_tx=XM_019834902.2;refGene_gn=CERS5;refGene_tx=XM_023257039.1;refGene_gn=CERS5;refGene_tx=XM_023257040.1
B4 79003329 79003460
refGene_gn=CERS5;refGene_tx=XM_003988685.5;refGene_gn=CERS5;refGene_tx=XM_019834902.2;refGene_gn=CERS5;refGene_tx=XM_023257039.1;refGene_gn=CERS5;refGene_tx=XM_023257040.1
B4 79004296 79004402
refGene_gn=CERS5;refGene_tx=XM_003988685.5;refGene_gn=CERS5;refGene_tx=XM_019834902.2;refGene_gn=CERS5;refGene_tx=XM_023257039.1;refGene_gn=CERS5;refGene_tx=XM_023257040.1
B4 79006571 79006582 refGene_gn=CERS5;refGene_tx=XM_023257039.1
B4 79023541 79023738
refGene_gn=CERS5;refGene_tx=XM_003988685.5;refGene_gn=CERS5;refGene_tx=XM_019834902.2;refGene_gn=CERS5;refGene_tx=XM_023257040.1
B4 79029081 79030090
refGene_gn=LIMA1;refGene_tx=XM_019834904.2;refGene_gn=LIMA1;refGene_tx=XM_019834905.2;refGene_gn=LIMA1;refGene_tx=XM_019834906.2;refGene_gn=LIMA1;refGene_tx=XM_023257033.1;refGene_gn=LIMA1;refGene_tx=XM_023257034.1;refGene_gn=LIMA1;refGene_tx=XM_023257035.1;refGene_gn=LIMA1;refGene_tx=XM_023257036.1;refGene_gn=LIMA1;refGene_tx=XM_023257037.1;refGene_gn=LIMA1;refGene_tx=XM_023257038.1
B4 79032204 79032338
refGene_gn=LIMA1;refGene_tx=XM_019834904.2;refGene_gn=LIMA1;refGene_tx=XM_019834905.2;refGene_gn=LIMA1;refGene_tx=XM_019834906.2;refGene_gn=LIMA1;refGene_tx=XM_023257033.1;refGene_gn=LIMA1;refGene_tx=XM_023257034.1;refGene_gn=LIMA1;refGene_tx=XM_023257035.1;refGene_gn=LIMA1;refGene_tx=XM_023257036.1;refGene_gn=LIMA1;refGene_tx=XM_023257037.1;refGene_gn=LIMA1;refGene_tx=XM_023257038.1
B4 79036624 79036738
refGene_gn=LIMA1;refGene_tx=XM_023257033.1;refGene_gn=LIMA1;refGene_tx=XM_023257034.1;refGene_gn=LIMA1;refGene_tx=XM_023257035.1;refGene_gn=LIMA1;refGene_tx=XM_023257036.1;refGene_gn=LIMA1;refGene_tx=XM_023257038.1
B4 79039329 79039442
refGene_gn=LIMA1;refGene_tx=XM_019834904.2;refGene_gn=LIMA1;refGene_tx=XM_019834905.2;refGene_gn=LIMA1;refGene_tx=XM_019834906.2;refGene_gn=LIMA1;refGene_tx=XM_023257033.1;refGene_gn=LIMA1;refGene_tx=XM_023257034.1;refGene_gn=LIMA1;refGene_tx=XM_023257035.1;refGene_gn=LIMA1;refGene_tx=XM_023257036.1;refGene_gn=LIMA1;refGene_tx=XM_023257037.1;refGene_gn=LIMA1;refGene_tx=XM_023257038.1
B4 79040945 79041003

refGene_gn=LIMA1;refGene_tx=XM_019834904.2;refGene_gn=LIMA1;refGene_tx
=XM_019834905.2;refGene_gn=LIMA1;refGene_tx=XM_019834906.2;refGene_gn=
LIMA1;refGene_tx=XM_023257033.1;refGene_gn=LIMA1;refGene_tx=XM_0232570
34.1;refGene_gn=LIMA1;refGene_tx=XM_023257035.1;refGene_gn=LIMA1;refGe
ne_tx=XM_023257036.1;refGene_gn=LIMA1;refGene_tx=XM_023257037.1;refGen
e_gn=LIMA1;refGene_tx=XM_023257038.1
B4 79045580 79045688
refGene_gn=LIMA1;refGene_tx=XM_019834904.2;refGene_gn=LIMA1;refGene_tx
=XM_019834905.2;refGene_gn=LIMA1;refGene_tx=XM_019834906.2;refGene_gn=
LIMA1;refGene_tx=XM_023257033.1;refGene_gn=LIMA1;refGene_tx=XM_0232570
34.1;refGene_gn=LIMA1;refGene_tx=XM_023257035.1;refGene_gn=LIMA1;refGe
ne_tx=XM_023257036.1;refGene_gn=LIMA1;refGene_tx=XM_023257037.1;refGen
e_gn=LIMA1;refGene_tx=XM_023257038.1
B4 79048707 79048856
refGene_gn=LIMA1;refGene_tx=XM_019834904.2;refGene_gn=LIMA1;refGene_tx
=XM_019834905.2;refGene_gn=LIMA1;refGene_tx=XM_019834906.2;refGene_gn=
LIMA1;refGene_tx=XM_023257033.1;refGene_gn=LIMA1;refGene_tx=XM_0232570
34.1;refGene_gn=LIMA1;refGene_tx=XM_023257035.1;refGene_gn=LIMA1;refGe
ne_tx=XM_023257036.1;refGene_gn=LIMA1;refGene_tx=XM_023257037.1;refGen
e_gn=LIMA1;refGene_tx=XM_023257038.1
B4 79050299 79050387
refGene_gn=LIMA1;refGene_tx=XM_019834904.2;refGene_gn=LIMA1;refGene_tx
=XM_019834905.2;refGene_gn=LIMA1;refGene_tx=XM_019834906.2;refGene_gn=
LIMA1;refGene_tx=XM_023257033.1;refGene_gn=LIMA1;refGene_tx=XM_0232570
34.1;refGene_gn=LIMA1;refGene_tx=XM_023257035.1;refGene_gn=LIMA1;refGe
ne_tx=XM_023257036.1;refGene_gn=LIMA1;refGene_tx=XM_023257037.1;refGen
e_gn=LIMA1;refGene_tx=XM_023257038.1
B4 79063376 79063841
refGene_gn=LIMA1;refGene_tx=XM_019834904.2;refGene_gn=LIMA1;refGene_tx
=XM_019834905.2;refGene_gn=LIMA1;refGene_tx=XM_019834906.2;refGene_gn=
LIMA1;refGene_tx=XM_023257033.1;refGene_gn=LIMA1;refGene_tx=XM_0232570
34.1;refGene_gn=LIMA1;refGene_tx=XM_023257035.1;refGene_gn=LIMA1;refGe
ne_tx=XM_023257036.1;refGene_gn=LIMA1;refGene_tx=XM_023257037.1;refGen
e_gn=LIMA1;refGene_tx=XM_023257038.1
B4 79073370 79073416
refGene_gn=LIMA1;refGene_tx=XM_019834904.2;refGene_gn=LIMA1;refGene_tx
=XM_019834905.2;refGene_gn=LIMA1;refGene_tx=XM_019834906.2;refGene_gn=
LIMA1;refGene_tx=XM_023257033.1;refGene_gn=LIMA1;refGene_tx=XM_0232570
34.1;refGene_gn=LIMA1;refGene_tx=XM_023257035.1;refGene_gn=LIMA1;refGe
ne_tx=XM_023257036.1;refGene_gn=LIMA1;refGene_tx=XM_023257037.1
B4 79087621 79087763
refGene_gn=LIMA1;refGene_tx=XM_019834904.2;refGene_gn=LIMA1;refGene_tx
=XM_019834905.2;refGene_gn=LIMA1;refGene_tx=XM_019834906.2;refGene_gn=
LIMA1;refGene_tx=XM_023257033.1;refGene_gn=LIMA1;refGene_tx=XM_0232570
34.1;refGene_gn=LIMA1;refGene_tx=XM_023257035.1;refGene_gn=LIMA1;refGe
ne_tx=XM_023257036.1;refGene_gn=LIMA1;refGene_tx=XM_023257037.1;refGen
e_gn=LIMA1;refGene_tx=XM_023257038.1
B4 79088751 79088788
refGene_gn=LIMA1;refGene_tx=XM_019834904.2;refGene_gn=LIMA1;refGene_tx
=XM_019834905.2;refGene_gn=LIMA1;refGene_tx=XM_019834906.2;refGene_gn=

LIMA1;refGene_tx=XM_023257033.1;refGene_gn=LIMA1;refGene_tx=XM_0232570
34.1;refGene_gn=LIMA1;refGene_tx=XM_023257035.1;refGene_gn=LIMA1;refGe
ne_tx=XM_023257037.1
B4 79154830 79154852 refGene_gn=FAM186A;refGene_tx=XM_023257041.1
B4 79161283 79161466 refGene_gn=FAM186A;refGene_tx=XM_023257041.1
B4 79163064 79163216 refGene_gn=FAM186A;refGene_tx=XM_023257041.1
B4 79165952 79166145 refGene_gn=FAM186A;refGene_tx=XM_023257041.1
B4 79173699 79176290 refGene_gn=FAM186A;refGene_tx=XM_023257041.1
B4 79176361 79176499 refGene_gn=FAM186A;refGene_tx=XM_023257041.1
B4 79176859 79177501 refGene_gn=FAM186A;refGene_tx=XM_023257041.1
B4 79177616 79177802 refGene_gn=FAM186A;refGene_tx=XM_023257041.1
B4 79178587 79181313 refGene_gn=FAM186A;refGene_tx=XM_023257041.1
B4 79194717 79194888 refGene_gn=FAM186A;refGene_tx=XM_023257041.1
B4 79195834 79196051 refGene_gn=FAM186A;refGene_tx=XM_023257041.1
B4 79237673 79237880 refGene_gn=FAM186A;refGene_tx=XM_023257041.1
B4 79240082 79240094
refGene_gn=LARP4;refGene_tx=XM_011283925.3;refGene_gn=LARP4;refGene_tx
=XM_019834914.2;refGene_gn=LARP4;refGene_tx=XM_019834917.2;refGene_gn=
LARP4;refGene_tx=XM_023257042.1
B4 79241403 79241421
refGene_gn=LARP4;refGene_tx=XM_011283923.3;refGene_gn=LARP4;refGene_tx
=XM_011283924.3;refGene_gn=LARP4;refGene_tx=XM_011283926.3;refGene_gn=
LARP4;refGene_tx=XM_011283927.3;refGene_gn=LARP4;refGene_tx=XM_0112839
28.3;refGene_gn=LARP4;refGene_tx=XM_011283929.3;refGene_gn=LARP4;refGe
ne_tx=XM_023257043.1
B4 79259166 79259314
refGene_gn=LARP4;refGene_tx=XM_011283923.3;refGene_gn=LARP4;refGene_tx
=XM_011283924.3;refGene_gn=LARP4;refGene_tx=XM_011283925.3;refGene_gn=
LARP4;refGene_tx=XM_011283926.3;refGene_gn=LARP4;refGene_tx=XM_0112839
27.3;refGene_gn=LARP4;refGene_tx=XM_011283928.3;refGene_gn=LARP4;refGe
ne_tx=XM_011283929.3;refGene_gn=LARP4;refGene_tx=XM_019834914.2;refGen
e_gn=LARP4;refGene_tx=XM_019834917.2;refGene_gn=LARP4;refGene_tx=XM_02
3257042.1;refGene_gn=LARP4;refGene_tx=XM_023257043.1
B4 79259942 79260116
refGene_gn=LARP4;refGene_tx=XM_011283923.3;refGene_gn=LARP4;refGene_tx
=XM_011283924.3;refGene_gn=LARP4;refGene_tx=XM_011283925.3;refGene_gn=
LARP4;refGene_tx=XM_011283926.3;refGene_gn=LARP4;refGene_tx=XM_0112839
27.3;refGene_gn=LARP4;refGene_tx=XM_011283928.3;refGene_gn=LARP4;refGe
ne_tx=XM_011283929.3;refGene_gn=LARP4;refGene_tx=XM_019834914.2;refGen
e_gn=LARP4;refGene_tx=XM_019834917.2;refGene_gn=LARP4;refGene_tx=XM_02
3257042.1;refGene_gn=LARP4;refGene_tx=XM_023257043.1
B4 79262657 79262733
refGene_gn=LARP4;refGene_tx=XM_011283923.3;refGene_gn=LARP4;refGene_tx
=XM_011283924.3;refGene_gn=LARP4;refGene_tx=XM_011283925.3;refGene_gn=
LARP4;refGene_tx=XM_011283926.3;refGene_gn=LARP4;refGene_tx=XM_0112839
27.3;refGene_gn=LARP4;refGene_tx=XM_011283928.3;refGene_gn=LARP4;refGe
ne_tx=XM_011283929.3;refGene_gn=LARP4;refGene_tx=XM_019834914.2;refGen
e_gn=LARP4;refGene_tx=XM_019834917.2;refGene_gn=LARP4;refGene_tx=XM_02
3257042.1;refGene_gn=LARP4;refGene_tx=XM_023257043.1
B4 79266499 79266636

refGene_gn=LARP4;refGene_tx=XM_011283923.3;refGene_gn=LARP4;refGene_tx=XM_011283924.3;refGene_gn=LARP4;refGene_tx=XM_011283925.3;refGene_gn=LARP4;refGene_tx=XM_011283926.3;refGene_gn=LARP4;refGene_tx=XM_011283927.3;refGene_gn=LARP4;refGene_tx=XM_011283928.3;refGene_gn=LARP4;refGene_tx=XM_011283929.3;refGene_gn=LARP4;refGene_tx=XM_019834914.2;refGene_gn=LARP4;refGene_tx=XM_019834917.2;refGene_gn=LARP4;refGene_tx=XM_023257042.1;refGene_gn=LARP4;refGene_tx=XM_023257043.1
B4 79269410 79269514
refGene_gn=LARP4;refGene_tx=XM_011283923.3;refGene_gn=LARP4;refGene_tx=XM_011283924.3;refGene_gn=LARP4;refGene_tx=XM_011283925.3;refGene_gn=LARP4;refGene_tx=XM_011283926.3;refGene_gn=LARP4;refGene_tx=XM_011283927.3;refGene_gn=LARP4;refGene_tx=XM_011283928.3;refGene_gn=LARP4;refGene_tx=XM_011283929.3;refGene_gn=LARP4;refGene_tx=XM_019834914.2;refGene_gn=LARP4;refGene_tx=XM_019834917.2;refGene_gn=LARP4;refGene_tx=XM_023257042.1;refGene_gn=LARP4;refGene_tx=XM_023257043.1
B4 79272948 79273059
refGene_gn=LARP4;refGene_tx=XM_011283923.3;refGene_gn=LARP4;refGene_tx=XM_011283924.3;refGene_gn=LARP4;refGene_tx=XM_011283925.3;refGene_gn=LARP4;refGene_tx=XM_011283926.3;refGene_gn=LARP4;refGene_tx=XM_011283927.3;refGene_gn=LARP4;refGene_tx=XM_011283928.3;refGene_gn=LARP4;refGene_tx=XM_011283929.3;refGene_gn=LARP4;refGene_tx=XM_019834914.2;refGene_gn=LARP4;refGene_tx=XM_019834917.2;refGene_gn=LARP4;refGene_tx=XM_023257042.1;refGene_gn=LARP4;refGene_tx=XM_023257043.1
B4 79273927 79273981
refGene_gn=LARP4;refGene_tx=XM_011283923.3;refGene_gn=LARP4;refGene_tx=XM_011283924.3;refGene_gn=LARP4;refGene_tx=XM_011283925.3;refGene_gn=LARP4;refGene_tx=XM_011283926.3;refGene_gn=LARP4;refGene_tx=XM_011283927.3;refGene_gn=LARP4;refGene_tx=XM_011283928.3;refGene_gn=LARP4;refGene_tx=XM_011283929.3;refGene_gn=LARP4;refGene_tx=XM_019834914.2;refGene_gn=LARP4;refGene_tx=XM_019834917.2;refGene_gn=LARP4;refGene_tx=XM_023257042.1;refGene_gn=LARP4;refGene_tx=XM_023257043.1
B4 79286150 79286363
refGene_gn=LARP4;refGene_tx=XM_011283923.3;refGene_gn=LARP4;refGene_tx=XM_011283924.3;refGene_gn=LARP4;refGene_tx=XM_011283925.3;refGene_gn=LARP4;refGene_tx=XM_011283926.3;refGene_gn=LARP4;refGene_tx=XM_011283927.3;refGene_gn=LARP4;refGene_tx=XM_011283928.3;refGene_gn=LARP4;refGene_tx=XM_011283929.3;refGene_gn=LARP4;refGene_tx=XM_019834914.2;refGene_gn=LARP4;refGene_tx=XM_019834917.2;refGene_gn=LARP4;refGene_tx=XM_023257042.1;refGene_gn=LARP4;refGene_tx=XM_023257043.1
B4 79287217 79287321
refGene_gn=LARP4;refGene_tx=XM_011283923.3;refGene_gn=LARP4;refGene_tx=XM_011283924.3;refGene_gn=LARP4;refGene_tx=XM_011283925.3;refGene_gn=LARP4;refGene_tx=XM_011283926.3;refGene_gn=LARP4;refGene_tx=XM_011283927.3;refGene_gn=LARP4;refGene_tx=XM_011283928.3;refGene_gn=LARP4;refGene_tx=XM_011283929.3;refGene_gn=LARP4;refGene_tx=XM_019834914.2;refGene_gn=LARP4;refGene_tx=XM_019834917.2;refGene_gn=LARP4;refGene_tx=XM_023257042.1;refGene_gn=LARP4;refGene_tx=XM_023257043.1
B4 79291754 79291971
refGene_gn=LARP4;refGene_tx=XM_011283923.3;refGene_gn=LARP4;refGene_tx=XM_011283924.3;refGene_gn=LARP4;refGene_tx=XM_011283925.3;refGene_gn=

LARP4;refGene_tx=XM_011283926.3;refGene_gn=LARP4;refGene_tx=XM_0112839
27.3;refGene_gn=LARP4;refGene_tx=XM_011283928.3;refGene_gn=LARP4;refGe
ne_tx=XM_011283929.3;refGene_gn=LARP4;refGene_tx=XM_019834914.2;refGen
e_gn=LARP4;refGene_tx=XM_019834917.2;refGene_gn=LARP4;refGene_tx=XM_02
3257042.1;refGene_gn=LARP4;refGene_tx=XM_023257043.1
B4 79292676 79292725
refGene_gn=LARP4;refGene_tx=XM_011283923.3;refGene_gn=LARP4;refGene_tx
=XM_011283924.3;refGene_gn=LARP4;refGene_tx=XM_011283925.3;refGene_gn=
LARP4;refGene_tx=XM_011283926.3;refGene_gn=LARP4;refGene_tx=XM_0112839
27.3;refGene_gn=LARP4;refGene_tx=XM_011283928.3;refGene_gn=LARP4;refGe
ne_tx=XM_011283929.3;refGene_gn=LARP4;refGene_tx=XM_019834914.2;refGen
e_gn=LARP4;refGene_tx=XM_019834917.2;refGene_gn=LARP4;refGene_tx=XM_02
3257042.1
B4 79295643 79295805
refGene_gn=LARP4;refGene_tx=XM_011283923.3;refGene_gn=LARP4;refGene_tx
=XM_011283924.3;refGene_gn=LARP4;refGene_tx=XM_011283925.3;refGene_gn=
LARP4;refGene_tx=XM_011283926.3;refGene_gn=LARP4;refGene_tx=XM_0112839
27.3;refGene_gn=LARP4;refGene_tx=XM_019834914.2;refGene_gn=LARP4;refGe
ne_tx=XM_019834917.2;refGene_gn=LARP4;refGene_tx=XM_023257042.1
B4 79300621 79300746
refGene_gn=LARP4;refGene_tx=XM_011283923.3;refGene_gn=LARP4;refGene_tx
=XM_011283924.3;refGene_gn=LARP4;refGene_tx=XM_011283925.3;refGene_gn=
LARP4;refGene_tx=XM_011283926.3;refGene_gn=LARP4;refGene_tx=XM_0112839
27.3;refGene_gn=LARP4;refGene_tx=XM_011283928.3;refGene_gn=LARP4;refGe
ne_tx=XM_011283929.3;refGene_gn=LARP4;refGene_tx=XM_019834914.2;refGen
e_gn=LARP4;refGene_tx=XM_019834917.2;refGene_gn=LARP4;refGene_tx=XM_02
3257042.1
B4 79300860 79301029
refGene_gn=LARP4;refGene_tx=XM_011283923.3;refGene_gn=LARP4;refGene_tx
=XM_011283924.3;refGene_gn=LARP4;refGene_tx=XM_011283925.3;refGene_gn=
LARP4;refGene_tx=XM_011283926.3;refGene_gn=LARP4;refGene_tx=XM_0112839
27.3;refGene_gn=LARP4;refGene_tx=XM_011283928.3;refGene_gn=LARP4;refGe
ne_tx=XM_011283929.3;refGene_gn=LARP4;refGene_tx=XM_019834914.2;refGen
e_gn=LARP4;refGene_tx=XM_019834917.2;refGene_gn=LARP4;refGene_tx=XM_02
3257042.1
B4 79302612 79302951
refGene_gn=LARP4;refGene_tx=XM_011283923.3;refGene_gn=LARP4;refGene_tx
=XM_011283924.3;refGene_gn=LARP4;refGene_tx=XM_011283925.3;refGene_gn=
LARP4;refGene_tx=XM_011283926.3;refGene_gn=LARP4;refGene_tx=XM_0112839
27.3;refGene_gn=LARP4;refGene_tx=XM_011283928.3;refGene_gn=LARP4;refGe
ne_tx=XM_011283929.3;refGene_gn=LARP4;refGene_tx=XM_019834914.2;refGen
e_gn=LARP4;refGene_tx=XM_019834917.2;refGene_gn=LARP4;refGene_tx=XM_02
3257042.1
B4 79330133 79330233
refGene_gn=DIP2B;refGene_tx=XM_023257044.1;refGene_gn=DIP2B;refGene_tx
=XM_023257045.1;refGene_gn=DIP2B;refGene_tx=XM_023257046.1;refGene_gn=
DIP2B;refGene_tx=XM_023257047.1
B4 79439129 79439201
refGene_gn=DIP2B;refGene_tx=XM_023257044.1;refGene_gn=DIP2B;refGene_tx
=XM_023257045.1;refGene_gn=DIP2B;refGene_tx=XM_023257046.1;refGene_gn=

DIP2B;refGene_tx=XM_023257047.1
B4 79459033 79459162
refGene_gn=DIP2B;refGene_tx=XM_023257044.1;refGene_gn=DIP2B;refGene_tx=XM_023257045.1;refGene_gn=DIP2B;refGene_tx=XM_023257046.1;refGene_gn=DIP2B;refGene_tx=XM_023257047.1
B4 79477768 79477894
refGene_gn=DIP2B;refGene_tx=XM_023257044.1;refGene_gn=DIP2B;refGene_tx=XM_023257045.1;refGene_gn=DIP2B;refGene_tx=XM_023257046.1;refGene_gn=DIP2B;refGene_tx=XM_023257047.1
B4 79488479 79488692
refGene_gn=DIP2B;refGene_tx=XM_023257044.1;refGene_gn=DIP2B;refGene_tx=XM_023257045.1;refGene_gn=DIP2B;refGene_tx=XM_023257046.1
B4 79491591 79491747
refGene_gn=DIP2B;refGene_tx=XM_023257044.1;refGene_gn=DIP2B;refGene_tx=XM_023257045.1;refGene_gn=DIP2B;refGene_tx=XM_023257046.1;refGene_gn=DIP2B;refGene_tx=XM_023257047.1
B4 79492218 79492341
refGene_gn=DIP2B;refGene_tx=XM_023257044.1;refGene_gn=DIP2B;refGene_tx=XM_023257045.1;refGene_gn=DIP2B;refGene_tx=XM_023257046.1;refGene_gn=DIP2B;refGene_tx=XM_023257047.1
B4 79495645 79495843
refGene_gn=DIP2B;refGene_tx=XM_023257044.1;refGene_gn=DIP2B;refGene_tx=XM_023257045.1;refGene_gn=DIP2B;refGene_tx=XM_023257046.1;refGene_gn=DIP2B;refGene_tx=XM_023257047.1
B4 79498649 79498741
refGene_gn=DIP2B;refGene_tx=XM_023257044.1;refGene_gn=DIP2B;refGene_tx=XM_023257045.1;refGene_gn=DIP2B;refGene_tx=XM_023257046.1;refGene_gn=DIP2B;refGene_tx=XM_023257047.1
B4 79500484 79500595
refGene_gn=DIP2B;refGene_tx=XM_023257044.1;refGene_gn=DIP2B;refGene_tx=XM_023257045.1;refGene_gn=DIP2B;refGene_tx=XM_023257046.1;refGene_gn=DIP2B;refGene_tx=XM_023257047.1
B4 79502402 79502526
refGene_gn=DIP2B;refGene_tx=XM_023257044.1;refGene_gn=DIP2B;refGene_tx=XM_023257045.1;refGene_gn=DIP2B;refGene_tx=XM_023257046.1;refGene_gn=DIP2B;refGene_tx=XM_023257047.1
B4 79503143 79503253
refGene_gn=DIP2B;refGene_tx=XM_023257044.1;refGene_gn=DIP2B;refGene_tx=XM_023257045.1;refGene_gn=DIP2B;refGene_tx=XM_023257046.1;refGene_gn=DIP2B;refGene_tx=XM_023257047.1
B4 79506351 79506454
refGene_gn=DIP2B;refGene_tx=XM_023257044.1;refGene_gn=DIP2B;refGene_tx=XM_023257045.1;refGene_gn=DIP2B;refGene_tx=XM_023257046.1;refGene_gn=DIP2B;refGene_tx=XM_023257047.1
B4 79507599 79507664
refGene_gn=DIP2B;refGene_tx=XM_023257044.1;refGene_gn=DIP2B;refGene_tx=XM_023257045.1;refGene_gn=DIP2B;refGene_tx=XM_023257046.1;refGene_gn=DIP2B;refGene_tx=XM_023257047.1
B4 79509704 79509798
refGene_gn=DIP2B;refGene_tx=XM_023257044.1;refGene_gn=DIP2B;refGene_tx

=XM_023257045.1;refGene_gn=DIP2B;refGene_tx=XM_023257046.1;refGene_gn=DIP2B;refGene_tx=XM_023257047.1
B4	79510286 79510406
refGene_gn=DIP2B;refGene_tx=XM_023257044.1;refGene_gn=DIP2B;refGene_tx=XM_023257045.1;refGene_gn=DIP2B;refGene_tx=XM_023257046.1;refGene_gn=DIP2B;refGene_tx=XM_023257047.1
B4	79512009 79512124
refGene_gn=DIP2B;refGene_tx=XM_023257044.1;refGene_gn=DIP2B;refGene_tx=XM_023257045.1;refGene_gn=DIP2B;refGene_tx=XM_023257046.1;refGene_gn=DIP2B;refGene_tx=XM_023257047.1
B4	79512957 79513097
refGene_gn=DIP2B;refGene_tx=XM_023257044.1;refGene_gn=DIP2B;refGene_tx=XM_023257045.1;refGene_gn=DIP2B;refGene_tx=XM_023257046.1;refGene_gn=DIP2B;refGene_tx=XM_023257047.1
B4	79513756 79513893
refGene_gn=DIP2B;refGene_tx=XM_023257044.1;refGene_gn=DIP2B;refGene_tx=XM_023257045.1;refGene_gn=DIP2B;refGene_tx=XM_023257046.1;refGene_gn=DIP2B;refGene_tx=XM_023257047.1
B4	79516932 79517013
refGene_gn=DIP2B;refGene_tx=XM_023257044.1;refGene_gn=DIP2B;refGene_tx=XM_023257045.1;refGene_gn=DIP2B;refGene_tx=XM_023257046.1;refGene_gn=DIP2B;refGene_tx=XM_023257047.1
B4	79520438 79520566
refGene_gn=DIP2B;refGene_tx=XM_023257044.1;refGene_gn=DIP2B;refGene_tx=XM_023257045.1;refGene_gn=DIP2B;refGene_tx=XM_023257046.1;refGene_gn=DIP2B;refGene_tx=XM_023257047.1
B4	79522360 79522475
refGene_gn=DIP2B;refGene_tx=XM_023257044.1;refGene_gn=DIP2B;refGene_tx=XM_023257045.1;refGene_gn=DIP2B;refGene_tx=XM_023257046.1;refGene_gn=DIP2B;refGene_tx=XM_023257047.1
B4	79526016 79526218
refGene_gn=DIP2B;refGene_tx=XM_023257044.1;refGene_gn=DIP2B;refGene_tx=XM_023257045.1;refGene_gn=DIP2B;refGene_tx=XM_023257046.1;refGene_gn=DIP2B;refGene_tx=XM_023257047.1
B4	79531134 79531244
refGene_gn=DIP2B;refGene_tx=XM_023257044.1;refGene_gn=DIP2B;refGene_tx=XM_023257045.1;refGene_gn=DIP2B;refGene_tx=XM_023257046.1;refGene_gn=DIP2B;refGene_tx=XM_023257047.1
B4	79531366 79531447
refGene_gn=DIP2B;refGene_tx=XM_023257044.1;refGene_gn=DIP2B;refGene_tx=XM_023257045.1;refGene_gn=DIP2B;refGene_tx=XM_023257046.1;refGene_gn=DIP2B;refGene_tx=XM_023257047.1
B4	79534792 79534916
refGene_gn=DIP2B;refGene_tx=XM_023257044.1;refGene_gn=DIP2B;refGene_tx=XM_023257045.1;refGene_gn=DIP2B;refGene_tx=XM_023257046.1;refGene_gn=DIP2B;refGene_tx=XM_023257047.1
B4	79536368 79536490
refGene_gn=DIP2B;refGene_tx=XM_023257044.1;refGene_gn=DIP2B;refGene_tx=XM_023257045.1;refGene_gn=DIP2B;refGene_tx=XM_023257046.1;refGene_gn=DIP2B;refGene_tx=XM_023257047.1

B4 79537975 79538087
refGene_gn=DIP2B;refGene_tx=XM_023257044.1;refGene_gn=DIP2B;refGene_tx=XM_023257045.1;refGene_gn=DIP2B;refGene_tx=XM_023257046.1;refGene_gn=DIP2B;refGene_tx=XM_023257047.1
B4 79540277 79540387
refGene_gn=DIP2B;refGene_tx=XM_023257044.1;refGene_gn=DIP2B;refGene_tx=XM_023257045.1;refGene_gn=DIP2B;refGene_tx=XM_023257046.1;refGene_gn=DIP2B;refGene_tx=XM_023257047.1
B4 79540875 79541006
refGene_gn=DIP2B;refGene_tx=XM_023257044.1;refGene_gn=DIP2B;refGene_tx=XM_023257045.1;refGene_gn=DIP2B;refGene_tx=XM_023257046.1;refGene_gn=DIP2B;refGene_tx=XM_023257047.1
B4 79542625 79542794
refGene_gn=DIP2B;refGene_tx=XM_023257044.1;refGene_gn=DIP2B;refGene_tx=XM_023257045.1;refGene_gn=DIP2B;refGene_tx=XM_023257046.1;refGene_gn=DIP2B;refGene_tx=XM_023257047.1
B4 79543825 79543996
refGene_gn=DIP2B;refGene_tx=XM_023257044.1;refGene_gn=DIP2B;refGene_tx=XM_023257045.1;refGene_gn=DIP2B;refGene_tx=XM_023257046.1;refGene_gn=DIP2B;refGene_tx=XM_023257047.1
B4 79545011 79545073
refGene_gn=DIP2B;refGene_tx=XM_023257044.1;refGene_gn=DIP2B;refGene_tx=XM_023257045.1;refGene_gn=DIP2B;refGene_tx=XM_023257046.1;refGene_gn=DIP2B;refGene_tx=XM_023257047.1
B4 79546269 79546327
refGene_gn=DIP2B;refGene_tx=XM_023257044.1;refGene_gn=DIP2B;refGene_tx=XM_023257045.1;refGene_gn=DIP2B;refGene_tx=XM_023257046.1;refGene_gn=DIP2B;refGene_tx=XM_023257047.1
B4 79548434 79548509
refGene_gn=DIP2B;refGene_tx=XM_023257044.1;refGene_gn=DIP2B;refGene_tx=XM_023257045.1;refGene_gn=DIP2B;refGene_tx=XM_023257046.1;refGene_gn=DIP2B;refGene_tx=XM_023257047.1
B4 79549990 79550168
refGene_gn=DIP2B;refGene_tx=XM_023257044.1;refGene_gn=DIP2B;refGene_tx=XM_023257045.1;refGene_gn=DIP2B;refGene_tx=XM_023257046.1;refGene_gn=DIP2B;refGene_tx=XM_023257047.1
B4 79551995 79552119
refGene_gn=DIP2B;refGene_tx=XM_023257044.1;refGene_gn=DIP2B;refGene_tx=XM_023257045.1;refGene_gn=DIP2B;refGene_tx=XM_023257046.1;refGene_gn=DIP2B;refGene_tx=XM_023257047.1
B4 79554008 79554261
refGene_gn=DIP2B;refGene_tx=XM_023257044.1;refGene_gn=DIP2B;refGene_tx=XM_023257045.1;refGene_gn=DIP2B;refGene_tx=XM_023257046.1;refGene_gn=DIP2B;refGene_tx=XM_023257047.1
B4 79571396 79571573 refGene_gn=ATF1;refGene_tx=XM_019834924.2
B4 79578056 79578155
refGene_gn=ATF1;refGene_tx=XM_019834924.2;refGene_gn=ATF1;refGene_tx=XM_019834925.1
B4 79615553 79615673 refGene_gn=ATF1;refGene_tx=XM_019834924.2
B4 79618155 79618256

refGene_gn=ATF1;refGene_tx=XM_019834924.2;refGene_gn=ATF1;refGene_tx=XM_019834925.1
B4 79634627 79634758
refGene_gn=ATF1;refGene_tx=XM_019834924.2;refGene_gn=ATF1;refGene_tx=XM_019834925.1
B4 79638860 79639043
refGene_gn=ATF1;refGene_tx=XM_019834924.2;refGene_gn=ATF1;refGene_tx=XM_019834925.1
B4 79639128 79639288
refGene_gn=ATF1;refGene_tx=XM_019834924.2;refGene_gn=ATF1;refGene_tx=XM_019834925.1
B4 79646308 79646453
refGene_gn=ATF1;refGene_tx=XM_019834924.2;refGene_gn=ATF1;refGene_tx=XM_019834925.1
B4 79665112 79665149 refGene_gn=TMPRSS12;refGene_tx=XM_019834926.1
B4 79665537 79665769 refGene_gn=TMPRSS12;refGene_tx=XM_011283930.2
B4 79666443 79666639
refGene_gn=TMPRSS12;refGene_tx=XM_011283930.2;refGene_gn=TMPRSS12;refGene_tx=XM_019834926.1
B4 79674283 79674552
refGene_gn=TMPRSS12;refGene_tx=XM_011283930.2;refGene_gn=TMPRSS12;refGene_tx=XM_019834926.1
B4 79683685 79683828
refGene_gn=TMPRSS12;refGene_tx=XM_011283930.2;refGene_gn=TMPRSS12;refGene_tx=XM_019834926.1
B4 79687160 79687412
refGene_gn=TMPRSS12;refGene_tx=XM_011283930.2;refGene_gn=TMPRSS12;refGene_tx=XM_019834926.1
B4 79698932 79699430 refGene_gn=METTL7A;refGene_tx=XM_003988690.5
B4 79706555 79706792 refGene_gn=METTL7A;refGene_tx=XM_003988690.5
B4 79724054 79724226 refGene_gn=HIGD1C;refGene_tx=XM_023257069.1
B4 79728756 79728891 refGene_gn=HIGD1C;refGene_tx=XM_023257069.1
B4 79738758 79738823 refGene_gn=HIGD1C;refGene_tx=XM_023257069.1
B4 79744221 79744299
refGene_gn=SLC11A2;refGene_tx=XM_011283933.3;refGene_gn=SLC11A2;refGene_tx=XM_023257049.1;refGene_gn=SLC11A2;refGene_tx=XM_023257050.1;refGene_gn=SLC11A2;refGene_tx=XM_023257051.1;refGene_gn=SLC11A2;refGene_tx=XM_023257052.1;refGene_gn=SLC11A2;refGene_tx=XM_023257053.1;refGene_gn=SLC11A2;refGene_tx=XM_023257054.1;refGene_gn=SLC11A2;refGene_tx=XM_023257055.1;refGene_gn=SLC11A2;refGene_tx=XM_023257056.1;refGene_gn=SLC11A2;refGene_tx=XM_023257058.1
B4 79749768 79749822
refGene_gn=SLC11A2;refGene_tx=XM_011283933.3;refGene_gn=SLC11A2;refGene_tx=XM_023257049.1;refGene_gn=SLC11A2;refGene_tx=XM_023257050.1;refGene_gn=SLC11A2;refGene_tx=XM_023257051.1;refGene_gn=SLC11A2;refGene_tx=XM_023257052.1;refGene_gn=SLC11A2;refGene_tx=XM_023257053.1;refGene_gn=SLC11A2;refGene_tx=XM_023257054.1;refGene_gn=SLC11A2;refGene_tx=XM_023257055.1;refGene_gn=SLC11A2;refGene_tx=XM_023257056.1;refGene_gn=SLC11A2;refGene_tx=XM_023257057.1;refGene_gn=SLC11A2;refGene_tx=XM_023257058.1

B4 79751618 79751772
refGene_gn=SLC11A2;refGene_tx=XM_011283933.3;refGene_gn=SLC11A2;refGene_tx=XM_023257049.1;refGene_gn=SLC11A2;refGene_tx=XM_023257050.1;refGene_gn=SLC11A2;refGene_tx=XM_023257051.1;refGene_gn=SLC11A2;refGene_tx=XM_023257052.1;refGene_gn=SLC11A2;refGene_tx=XM_023257053.1;refGene_gn=SLC11A2;refGene_tx=XM_023257054.1;refGene_gn=SLC11A2;refGene_tx=XM_023257055.1;refGene_gn=SLC11A2;refGene_tx=XM_023257056.1;refGene_gn=SLC11A2;refGene_tx=XM_023257058.1
B4 79752391 79752465
refGene_gn=SLC11A2;refGene_tx=XM_011283933.3;refGene_gn=SLC11A2;refGene_tx=XM_023257049.1;refGene_gn=SLC11A2;refGene_tx=XM_023257050.1;refGene_gn=SLC11A2;refGene_tx=XM_023257051.1;refGene_gn=SLC11A2;refGene_tx=XM_023257052.1;refGene_gn=SLC11A2;refGene_tx=XM_023257053.1;refGene_gn=SLC11A2;refGene_tx=XM_023257054.1;refGene_gn=SLC11A2;refGene_tx=XM_023257055.1;refGene_gn=SLC11A2;refGene_tx=XM_023257056.1;refGene_gn=SLC11A2;refGene_tx=XM_023257057.1;refGene_gn=SLC11A2;refGene_tx=XM_023257058.1
B4 79753733 79753883
refGene_gn=SLC11A2;refGene_tx=XM_011283933.3;refGene_gn=SLC11A2;refGene_tx=XM_023257049.1;refGene_gn=SLC11A2;refGene_tx=XM_023257050.1;refGene_gn=SLC11A2;refGene_tx=XM_023257051.1;refGene_gn=SLC11A2;refGene_tx=XM_023257052.1;refGene_gn=SLC11A2;refGene_tx=XM_023257053.1;refGene_gn=SLC11A2;refGene_tx=XM_023257054.1;refGene_gn=SLC11A2;refGene_tx=XM_023257055.1;refGene_gn=SLC11A2;refGene_tx=XM_023257056.1;refGene_gn=SLC11A2;refGene_tx=XM_023257057.1;refGene_gn=SLC11A2;refGene_tx=XM_023257058.1
B4 79754025 79754145
refGene_gn=SLC11A2;refGene_tx=XM_011283933.3;refGene_gn=SLC11A2;refGene_tx=XM_023257049.1;refGene_gn=SLC11A2;refGene_tx=XM_023257050.1;refGene_gn=SLC11A2;refGene_tx=XM_023257051.1;refGene_gn=SLC11A2;refGene_tx=XM_023257052.1;refGene_gn=SLC11A2;refGene_tx=XM_023257053.1;refGene_gn=SLC11A2;refGene_tx=XM_023257054.1;refGene_gn=SLC11A2;refGene_tx=XM_023257055.1;refGene_gn=SLC11A2;refGene_tx=XM_023257056.1;refGene_gn=SLC11A2;refGene_tx=XM_023257057.1;refGene_gn=SLC11A2;refGene_tx=XM_023257058.1
B4 79755098 79755185
refGene_gn=SLC11A2;refGene_tx=XM_011283933.3;refGene_gn=SLC11A2;refGene_tx=XM_023257049.1;refGene_gn=SLC11A2;refGene_tx=XM_023257050.1;refGene_gn=SLC11A2;refGene_tx=XM_023257051.1;refGene_gn=SLC11A2;refGene_tx=XM_023257052.1;refGene_gn=SLC11A2;refGene_tx=XM_023257053.1;refGene_gn=SLC11A2;refGene_tx=XM_023257054.1;refGene_gn=SLC11A2;refGene_tx=XM_023257055.1;refGene_gn=SLC11A2;refGene_tx=XM_023257056.1;refGene_gn=SLC11A2;refGene_tx=XM_023257057.1;refGene_gn=SLC11A2;refGene_tx=XM_023257058.1
B4 79756003 79756162
refGene_gn=SLC11A2;refGene_tx=XM_011283933.3;refGene_gn=SLC11A2;refGene_tx=XM_023257049.1;refGene_gn=SLC11A2;refGene_tx=XM_023257050.1;refGene_gn=SLC11A2;refGene_tx=XM_023257051.1;refGene_gn=SLC11A2;refGene_tx=XM_023257052.1;refGene_gn=SLC11A2;refGene_tx=XM_023257053.1;refGene_gn=SLC11A2;refGene_tx=XM_023257054.1;refGene_gn=SLC11A2;refGene_tx=XM_02

3257055.1;refGene_gn=SLC11A2;refGene_tx=XM_023257056.1;refGene_gn=SLC11A2;refGene_tx=XM_023257057.1;refGene_gn=SLC11A2;refGene_tx=XM_023257058.1
B4 79756812 79756968
refGene_gn=SLC11A2;refGene_tx=XM_011283933.3;refGene_gn=SLC11A2;refGene_tx=XM_023257049.1;refGene_gn=SLC11A2;refGene_tx=XM_023257050.1;refGene_gn=SLC11A2;refGene_tx=XM_023257051.1;refGene_gn=SLC11A2;refGene_tx=XM_023257052.1;refGene_gn=SLC11A2;refGene_tx=XM_023257053.1;refGene_gn=SLC11A2;refGene_tx=XM_023257054.1;refGene_gn=SLC11A2;refGene_tx=XM_023257055.1;refGene_gn=SLC11A2;refGene_tx=XM_023257056.1;refGene_gn=SLC11A2;refGene_tx=XM_023257057.1;refGene_gn=SLC11A2;refGene_tx=XM_023257058.1
B4 79758283 79758351
refGene_gn=SLC11A2;refGene_tx=XM_011283933.3;refGene_gn=SLC11A2;refGene_tx=XM_023257049.1;refGene_gn=SLC11A2;refGene_tx=XM_023257050.1;refGene_gn=SLC11A2;refGene_tx=XM_023257051.1;refGene_gn=SLC11A2;refGene_tx=XM_023257052.1;refGene_gn=SLC11A2;refGene_tx=XM_023257053.1;refGene_gn=SLC11A2;refGene_tx=XM_023257054.1;refGene_gn=SLC11A2;refGene_tx=XM_023257055.1;refGene_gn=SLC11A2;refGene_tx=XM_023257056.1;refGene_gn=SLC11A2;refGene_tx=XM_023257057.1;refGene_gn=SLC11A2;refGene_tx=XM_023257058.1
B4 79758454 79758525
refGene_gn=SLC11A2;refGene_tx=XM_011283933.3;refGene_gn=SLC11A2;refGene_tx=XM_023257049.1;refGene_gn=SLC11A2;refGene_tx=XM_023257050.1;refGene_gn=SLC11A2;refGene_tx=XM_023257051.1;refGene_gn=SLC11A2;refGene_tx=XM_023257052.1;refGene_gn=SLC11A2;refGene_tx=XM_023257053.1;refGene_gn=SLC11A2;refGene_tx=XM_023257054.1;refGene_gn=SLC11A2;refGene_tx=XM_023257055.1;refGene_gn=SLC11A2;refGene_tx=XM_023257056.1;refGene_gn=SLC11A2;refGene_tx=XM_023257057.1;refGene_gn=SLC11A2;refGene_tx=XM_023257058.1
B4 79759056 79759163
refGene_gn=SLC11A2;refGene_tx=XM_011283933.3;refGene_gn=SLC11A2;refGene_tx=XM_023257049.1;refGene_gn=SLC11A2;refGene_tx=XM_023257050.1;refGene_gn=SLC11A2;refGene_tx=XM_023257051.1;refGene_gn=SLC11A2;refGene_tx=XM_023257052.1;refGene_gn=SLC11A2;refGene_tx=XM_023257053.1;refGene_gn=SLC11A2;refGene_tx=XM_023257054.1;refGene_gn=SLC11A2;refGene_tx=XM_023257055.1;refGene_gn=SLC11A2;refGene_tx=XM_023257056.1;refGene_gn=SLC11A2;refGene_tx=XM_023257057.1;refGene_gn=SLC11A2;refGene_tx=XM_023257058.1
B4 79760817 79760937
refGene_gn=SLC11A2;refGene_tx=XM_011283933.3;refGene_gn=SLC11A2;refGene_tx=XM_023257049.1;refGene_gn=SLC11A2;refGene_tx=XM_023257050.1;refGene_gn=SLC11A2;refGene_tx=XM_023257051.1;refGene_gn=SLC11A2;refGene_tx=XM_023257052.1;refGene_gn=SLC11A2;refGene_tx=XM_023257053.1;refGene_gn=SLC11A2;refGene_tx=XM_023257054.1;refGene_gn=SLC11A2;refGene_tx=XM_023257055.1;refGene_gn=SLC11A2;refGene_tx=XM_023257056.1;refGene_gn=SLC11A2;refGene_tx=XM_023257057.1;refGene_gn=SLC11A2;refGene_tx=XM_023257058.1
B4 79761815 79761941
refGene_gn=SLC11A2;refGene_tx=XM_011283933.3;refGene_gn=SLC11A2;refGen

e_tx=XM_023257049.1;refGene_gn=SLC11A2;refGene_tx=XM_023257050.1;refGene_gn=SLC11A2;refGene_tx=XM_023257051.1;refGene_gn=SLC11A2;refGene_tx=XM_023257052.1;refGene_gn=SLC11A2;refGene_tx=XM_023257053.1;refGene_gn=SLC11A2;refGene_tx=XM_023257054.1;refGene_gn=SLC11A2;refGene_tx=XM_023257055.1;refGene_gn=SLC11A2;refGene_tx=XM_023257056.1;refGene_gn=SLC11A2;refGene_tx=XM_023257057.1;refGene_gn=SLC11A2;refGene_tx=XM_023257058.1
B4 79762481 79762630
refGene_gn=SLC11A2;refGene_tx=XM_011283933.3;refGene_gn=SLC11A2;refGene_tx=XM_023257049.1;refGene_gn=SLC11A2;refGene_tx=XM_023257050.1;refGene_gn=SLC11A2;refGene_tx=XM_023257051.1;refGene_gn=SLC11A2;refGene_tx=XM_023257052.1;refGene_gn=SLC11A2;refGene_tx=XM_023257053.1;refGene_gn=SLC11A2;refGene_tx=XM_023257054.1;refGene_gn=SLC11A2;refGene_tx=XM_023257055.1;refGene_gn=SLC11A2;refGene_tx=XM_023257056.1;refGene_gn=SLC11A2;refGene_tx=XM_023257057.1;refGene_gn=SLC11A2;refGene_tx=XM_023257058.1
B4 79763336 79763358 refGene_gn=SLC11A2;refGene_tx=XM_023257058.1
B4 79764550 79764622
refGene_gn=SLC11A2;refGene_tx=XM_011283933.3;refGene_gn=SLC11A2;refGene_tx=XM_023257049.1;refGene_gn=SLC11A2;refGene_tx=XM_023257050.1;refGene_gn=SLC11A2;refGene_tx=XM_023257051.1;refGene_gn=SLC11A2;refGene_tx=XM_023257052.1;refGene_gn=SLC11A2;refGene_tx=XM_023257053.1;refGene_gn=SLC11A2;refGene_tx=XM_023257054.1;refGene_gn=SLC11A2;refGene_tx=XM_023257055.1;refGene_gn=SLC11A2;refGene_tx=XM_023257056.1;refGene_gn=SLC11A2;refGene_tx=XM_023257057.1
B4 79776796 79777022
refGene_gn=SLC11A2;refGene_tx=XM_023257049.1;refGene_gn=SLC11A2;refGene_tx=XM_023257050.1;refGene_gn=SLC11A2;refGene_tx=XM_023257051.1;refGene_gn=SLC11A2;refGene_tx=XM_023257052.1;refGene_gn=SLC11A2;refGene_tx=XM_023257053.1;refGene_gn=SLC11A2;refGene_tx=XM_023257054.1;refGene_gn=SLC11A2;refGene_tx=XM_023257055.1;refGene_gn=SLC11A2;refGene_tx=XM_023257057.1
B4 79794934 79795062
refGene_gn=LETMD1;refGene_tx=XM_003988697.4;refGene_gn=LETMD1;refGene_tx=XM_006933693.3;refGene_gn=LETMD1;refGene_tx=XM_019834923.1;refGene_gn=LETMD1;refGene_tx=XM_023257060.1;refGene_gn=LETMD1;refGene_tx=XM_023257061.1;refGene_gn=LETMD1;refGene_tx=XM_023257064.1;refGene_gn=LETMD1;refGene_tx=XM_023257066.1;refGene_gn=LETMD1;refGene_tx=XM_023257067.1;refGene_gn=LETMD1;refGene_tx=XM_023257068.1
B4 79795590 79795763
refGene_gn=LETMD1;refGene_tx=XM_003988697.4;refGene_gn=LETMD1;refGene_tx=XM_006933693.3;refGene_gn=LETMD1;refGene_tx=XM_019834923.1;refGene_gn=LETMD1;refGene_tx=XM_023257060.1;refGene_gn=LETMD1;refGene_tx=XM_023257061.1;refGene_gn=LETMD1;refGene_tx=XM_023257062.1;refGene_gn=LETMD1;refGene_tx=XM_023257064.1;refGene_gn=LETMD1;refGene_tx=XM_023257066.1;refGene_gn=LETMD1;refGene_tx=XM_023257067.1;refGene_gn=LETMD1;refGene_tx=XM_023257068.1
B4 79797598 79797714
refGene_gn=LETMD1;refGene_tx=XM_003988697.4;refGene_gn=LETMD1;refGene_tx=XM_006933693.3;refGene_gn=LETMD1;refGene_tx=XM_019834923.1;refGene_

gn=LETMD1;refGene_tx=XM_023257060.1;refGene_gn=LETMD1;refGene_tx=XM_02
3257062.1;refGene_gn=LETMD1;refGene_tx=XM_023257063.1;refGene_gn=LETMD
1;refGene_tx=XM_023257064.1;refGene_gn=LETMD1;refGene_tx=XM_023257066.
1;refGene_gn=LETMD1;refGene_tx=XM_023257068.1
B4 79798377 79798460
refGene_gn=LETMD1;refGene_tx=XM_003988697.4;refGene_gn=LETMD1;refGene_
tx=XM_006933693.3;refGene_gn=LETMD1;refGene_tx=XM_023257060.1;refGene_
gn=LETMD1;refGene_tx=XM_023257061.1;refGene_gn=LETMD1;refGene_tx=XM_02
3257062.1;refGene_gn=LETMD1;refGene_tx=XM_023257063.1;refGene_gn=LETMD
1;refGene_tx=XM_023257065.1;refGene_gn=LETMD1;refGene_tx=XM_023257066.
1;refGene_gn=LETMD1;refGene_tx=XM_023257067.1
B4 79800011 79800198
refGene_gn=LETMD1;refGene_tx=XM_003988697.4;refGene_gn=LETMD1;refGene_
tx=XM_006933693.3;refGene_gn=LETMD1;refGene_tx=XM_023257060.1;refGene_
gn=LETMD1;refGene_tx=XM_023257061.1;refGene_gn=LETMD1;refGene_tx=XM_02
3257062.1;refGene_gn=LETMD1;refGene_tx=XM_023257063.1;refGene_gn=LETMD
1;refGene_tx=XM_023257065.1;refGene_gn=LETMD1;refGene_tx=XM_023257066.
1
B4 79800321 79800423
refGene_gn=LETMD1;refGene_tx=XM_003988697.4;refGene_gn=LETMD1;refGene_
tx=XM_006933693.3;refGene_gn=LETMD1;refGene_tx=XM_023257061.1;refGene_
gn=LETMD1;refGene_tx=XM_023257062.1;refGene_gn=LETMD1;refGene_tx=XM_02
3257063.1;refGene_gn=LETMD1;refGene_tx=XM_023257065.1;refGene_gn=LETMD
1;refGene_tx=XM_023257066.1;refGene_gn=LETMD1;refGene_tx=XM_023257067.
1
B4 79800521 79800674
refGene_gn=LETMD1;refGene_tx=XM_003988697.4;refGene_gn=LETMD1;refGene_
tx=XM_006933693.3;refGene_gn=LETMD1;refGene_tx=XM_019834923.1;refGene_
gn=LETMD1;refGene_tx=XM_023257060.1;refGene_gn=LETMD1;refGene_tx=XM_02
3257061.1;refGene_gn=LETMD1;refGene_tx=XM_023257062.1;refGene_gn=LETMD
1;refGene_tx=XM_023257063.1;refGene_gn=LETMD1;refGene_tx=XM_023257064.
1;refGene_gn=LETMD1;refGene_tx=XM_023257065.1;refGene_gn=LETMD1;refGen
e_tx=XM_023257066.1;refGene_gn=LETMD1;refGene_tx=XM_023257067.1;refGen
e_gn=LETMD1;refGene_tx=XM_023257068.1
B4 79801100 79801197
refGene_gn=LETMD1;refGene_tx=XM_003988697.4;refGene_gn=LETMD1;refGene_
tx=XM_006933693.3;refGene_gn=LETMD1;refGene_tx=XM_019834923.1;refGene_
gn=LETMD1;refGene_tx=XM_023257060.1;refGene_gn=LETMD1;refGene_tx=XM_02
3257061.1;refGene_gn=LETMD1;refGene_tx=XM_023257062.1;refGene_gn=LETMD
1;refGene_tx=XM_023257063.1;refGene_gn=LETMD1;refGene_tx=XM_023257064.
1;refGene_gn=LETMD1;refGene_tx=XM_023257065.1;refGene_gn=LETMD1;refGen
e_tx=XM_023257067.1
B4 79802736 79802811
refGene_gn=LETMD1;refGene_tx=XM_003988697.4;refGene_gn=LETMD1;refGene_
tx=XM_006933693.3;refGene_gn=LETMD1;refGene_tx=XM_019834923.1;refGene_
gn=LETMD1;refGene_tx=XM_023257060.1;refGene_gn=LETMD1;refGene_tx=XM_02
3257061.1;refGene_gn=LETMD1;refGene_tx=XM_023257062.1;refGene_gn=LETMD
1;refGene_tx=XM_023257063.1;refGene_gn=LETMD1;refGene_tx=XM_023257064.
1;refGene_gn=LETMD1;refGene_tx=XM_023257065.1;refGene_gn=LETMD1;refGen
e_tx=XM_023257066.1;refGene_gn=LETMD1;refGene_tx=XM_023257067.1;refGen

e_gn=LETMD1;refGene_tx=XM_023257068.1
B4 79806798 79807722
refGene_gn=CSRNP2;refGene_tx=XM_011283934.3;refGene_gn=CSRNP2;refGene_tx=XM_011283935.2;refGene_gn=CSRNP2;refGene_tx=XM_023257059.1
B4 79809502 79809799
refGene_gn=CSRNP2;refGene_tx=XM_011283934.3;refGene_gn=CSRNP2;refGene_tx=XM_011283935.2;refGene_gn=CSRNP2;refGene_tx=XM_023257059.1
B4 79813306 79813566
refGene_gn=CSRNP2;refGene_tx=XM_011283934.3;refGene_gn=CSRNP2;refGene_tx=XM_011283935.2;refGene_gn=CSRNP2;refGene_tx=XM_023257059.1
B4 79815313 79815464
refGene_gn=CSRNP2;refGene_tx=XM_011283934.3;refGene_gn=CSRNP2;refGene_tx=XM_011283935.2;refGene_gn=CSRNP2;refGene_tx=XM_023257059.1
B4 79828958 79828996
refGene_gn=TFCP2;refGene_tx=XM_003988700.5;refGene_gn=TFCP2;refGene_tx=XM_006933695.4;refGene_gn=TFCP2;refGene_tx=XM_006933696.4;refGene_gn=TFCP2;refGene_tx=XM_006933697.4
B4 79829417 79829469
refGene_gn=TFCP2;refGene_tx=XM_003988700.5;refGene_gn=TFCP2;refGene_tx=XM_006933695.4;refGene_gn=TFCP2;refGene_tx=XM_006933696.4;refGene_gn=TFCP2;refGene_tx=XM_006933697.4
B4 79831226 79831369
refGene_gn=TFCP2;refGene_tx=XM_003988700.5;refGene_gn=TFCP2;refGene_tx=XM_006933695.4;refGene_gn=TFCP2;refGene_tx=XM_006933696.4;refGene_gn=TFCP2;refGene_tx=XM_006933697.4
B4 79831798 79831917
refGene_gn=TFCP2;refGene_tx=XM_003988700.5;refGene_gn=TFCP2;refGene_tx=XM_006933695.4;refGene_gn=TFCP2;refGene_tx=XM_006933696.4;refGene_gn=TFCP2;refGene_tx=XM_006933697.4
B4 79833440 79833531
refGene_gn=TFCP2;refGene_tx=XM_003988700.5;refGene_gn=TFCP2;refGene_tx=XM_006933695.4;refGene_gn=TFCP2;refGene_tx=XM_006933696.4;refGene_gn=TFCP2;refGene_tx=XM_006933697.4
B4 79834403 79834497
refGene_gn=TFCP2;refGene_tx=XM_003988700.5;refGene_gn=TFCP2;refGene_tx=XM_006933695.4;refGene_gn=TFCP2;refGene_tx=XM_006933696.4;refGene_gn=TFCP2;refGene_tx=XM_006933697.4
B4 79834905 79834972
refGene_gn=TFCP2;refGene_tx=XM_003988700.5;refGene_gn=TFCP2;refGene_tx=XM_006933695.4;refGene_gn=TFCP2;refGene_tx=XM_006933696.4;refGene_gn=TFCP2;refGene_tx=XM_006933697.4
B4 79837087 79837176
refGene_gn=TFCP2;refGene_tx=XM_003988700.5;refGene_gn=TFCP2;refGene_tx=XM_006933695.4;refGene_gn=TFCP2;refGene_tx=XM_006933696.4;refGene_gn=TFCP2;refGene_tx=XM_006933697.4
B4 79837528 79837639
refGene_gn=TFCP2;refGene_tx=XM_003988700.5;refGene_gn=TFCP2;refGene_tx=XM_006933695.4;refGene_gn=TFCP2;refGene_tx=XM_006933696.4;refGene_gn=TFCP2;refGene_tx=XM_006933697.4
B4 79838479 79838632

refGene_gn=TFCP2;refGene_tx=XM_003988700.5;refGene_gn=TFCP2;refGene_tx=XM_006933695.4;refGene_gn=TFCP2;refGene_tx=XM_006933696.4;refGene_gn=TFCP2;refGene_tx=XM_006933697.4
B4 79839830 79839937
refGene_gn=TFCP2;refGene_tx=XM_003988700.5;refGene_gn=TFCP2;refGene_tx=XM_006933695.4;refGene_gn=TFCP2;refGene_tx=XM_006933696.4;refGene_gn=TFCP2;refGene_tx=XM_006933697.4
B4 79843499 79843605
refGene_gn=TFCP2;refGene_tx=XM_003988700.5;refGene_gn=TFCP2;refGene_tx=XM_006933695.4;refGene_gn=TFCP2;refGene_tx=XM_006933696.4;refGene_gn=TFCP2;refGene_tx=XM_006933697.4
B4 79845869 79845946
refGene_gn=TFCP2;refGene_tx=XM_003988700.5;refGene_gn=TFCP2;refGene_tx=XM_006933695.4;refGene_gn=TFCP2;refGene_tx=XM_006933696.4;refGene_gn=TFCP2;refGene_tx=XM_006933697.4
B4 79846516 79846668
refGene_gn=TFCP2;refGene_tx=XM_003988700.5;refGene_gn=TFCP2;refGene_tx=XM_006933695.4;refGene_gn=TFCP2;refGene_tx=XM_006933696.4;refGene_gn=TFCP2;refGene_tx=XM_006933697.4
B4 79872129 79872251
refGene_gn=TFCP2;refGene_tx=XM_003988700.5;refGene_gn=TFCP2;refGene_tx=XM_006933695.4;refGene_gn=TFCP2;refGene_tx=XM_006933696.4;refGene_gn=TFCP2;refGene_tx=XM_006933697.4
B4 79884420 79884766
refGene_gn=POU6F1;refGene_tx=XM_019834932.1;refGene_gn=POU6F1;refGene_tx=XM_023257070.1;refGene_gn=POU6F1;refGene_tx=XM_023257071.1;refGene_gn=POU6F1;refGene_tx=XM_023257072.1;refGene_gn=POU6F1;refGene_tx=XM_023257073.1;refGene_gn=POU6F1;refGene_tx=XM_023257074.1;refGene_gn=POU6F1;refGene_tx=XM_023257075.1
B4 79885774 79885943
refGene_gn=POU6F1;refGene_tx=XM_019834932.1;refGene_gn=POU6F1;refGene_tx=XM_023257070.1;refGene_gn=POU6F1;refGene_tx=XM_023257071.1;refGene_gn=POU6F1;refGene_tx=XM_023257072.1;refGene_gn=POU6F1;refGene_tx=XM_023257073.1;refGene_gn=POU6F1;refGene_tx=XM_023257074.1;refGene_gn=POU6F1;refGene_tx=XM_023257075.1
B4 79886495 79886637
refGene_gn=POU6F1;refGene_tx=XM_019834932.1;refGene_gn=POU6F1;refGene_tx=XM_023257070.1;refGene_gn=POU6F1;refGene_tx=XM_023257071.1;refGene_gn=POU6F1;refGene_tx=XM_023257072.1;refGene_gn=POU6F1;refGene_tx=XM_023257073.1;refGene_gn=POU6F1;refGene_tx=XM_023257074.1;refGene_gn=POU6F1;refGene_tx=XM_023257075.1
B4 79889740 79889944
refGene_gn=POU6F1;refGene_tx=XM_019834932.1;refGene_gn=POU6F1;refGene_tx=XM_023257070.1;refGene_gn=POU6F1;refGene_tx=XM_023257071.1;refGene_gn=POU6F1;refGene_tx=XM_023257072.1;refGene_gn=POU6F1;refGene_tx=XM_023257073.1;refGene_gn=POU6F1;refGene_tx=XM_023257074.1;refGene_gn=POU6F1;refGene_tx=XM_023257075.1
B4 79890555 79890684
refGene_gn=POU6F1;refGene_tx=XM_019834932.1;refGene_gn=POU6F1;refGene_tx=XM_023257070.1;refGene_gn=POU6F1;refGene_tx=XM_023257071.1;refGene_

gn=POU6F1;refGene_tx=XM_023257072.1;refGene_gn=POU6F1;refGene_tx=XM_023257073.1;refGene_gn=POU6F1;refGene_tx=XM_023257074.1;refGene_gn=POU6F1;refGene_tx=XM_023257075.1
B4 79891354 79891608
refGene_gn=POU6F1;refGene_tx=XM_019834932.1;refGene_gn=POU6F1;refGene_tx=XM_023257070.1;refGene_gn=POU6F1;refGene_tx=XM_023257071.1;refGene_gn=POU6F1;refGene_tx=XM_023257072.1;refGene_gn=POU6F1;refGene_tx=XM_023257073.1;refGene_gn=POU6F1;refGene_tx=XM_023257074.1;refGene_gn=POU6F1;refGene_tx=XM_023257075.1
B4 79892157 79892383
refGene_gn=POU6F1;refGene_tx=XM_019834932.1;refGene_gn=POU6F1;refGene_tx=XM_023257070.1;refGene_gn=POU6F1;refGene_tx=XM_023257071.1;refGene_gn=POU6F1;refGene_tx=XM_023257072.1;refGene_gn=POU6F1;refGene_tx=XM_023257073.1;refGene_gn=POU6F1;refGene_tx=XM_023257074.1;refGene_gn=POU6F1;refGene_tx=XM_023257075.1
B4 79893351 79893473
refGene_gn=POU6F1;refGene_tx=XM_019834932.1;refGene_gn=POU6F1;refGene_tx=XM_023257070.1;refGene_gn=POU6F1;refGene_tx=XM_023257071.1;refGene_gn=POU6F1;refGene_tx=XM_023257072.1;refGene_gn=POU6F1;refGene_tx=XM_023257073.1;refGene_gn=POU6F1;refGene_tx=XM_023257074.1;refGene_gn=POU6F1;refGene_tx=XM_023257075.1
B4 79897791 79897987
refGene_gn=POU6F1;refGene_tx=XM_019834932.1;refGene_gn=POU6F1;refGene_tx=XM_023257070.1;refGene_gn=POU6F1;refGene_tx=XM_023257071.1;refGene_gn=POU6F1;refGene_tx=XM_023257072.1;refGene_gn=POU6F1;refGene_tx=XM_023257073.1;refGene_gn=POU6F1;refGene_tx=XM_023257074.1;refGene_gn=POU6F1;refGene_tx=XM_023257075.1
B4 79900178 79900226
refGene_gn=POU6F1;refGene_tx=XM_019834932.1;refGene_gn=POU6F1;refGene_tx=XM_023257070.1;refGene_gn=POU6F1;refGene_tx=XM_023257071.1;refGene_gn=POU6F1;refGene_tx=XM_023257072.1;refGene_gn=POU6F1;refGene_tx=XM_023257073.1;refGene_gn=POU6F1;refGene_tx=XM_023257074.1;refGene_gn=POU6F1;refGene_tx=XM_023257075.1
B4 79927900 79927913
refGene_gn=DAZAP2;refGene_tx=XM_003988703.5;refGene_gn=DAZAP2;refGene_tx=XM_011283942.2
B4 79928948 79929067
refGene_gn=DAZAP2;refGene_tx=XM_003988703.5;refGene_gn=DAZAP2;refGene_tx=XM_011283942.2
B4 79929459 79929705
refGene_gn=DAZAP2;refGene_tx=XM_003988703.5;refGene_gn=DAZAP2;refGene_tx=XM_011283942.2
B4 79930382 79930705
refGene_gn=DAZAP2;refGene_tx=XM_003988703.5;refGene_gn=DAZAP2;refGene_tx=XM_011283942.2
B4 79933840 79934019 refGene_gn=SMAGP;refGene_tx=XM_023257076.1
B4 79934741 79934822 refGene_gn=SMAGP;refGene_tx=XM_023257076.1
B4 79948861 79948920 refGene_gn=SMAGP;refGene_tx=XM_023257076.1
B4 79949833 79950135 refGene_gn=SMAGP;refGene_tx=XM_023257076.1
B4 79954958 79954982

refGene_gn=BIN2;refGene_tx=XM_011283945.3;refGene_gn=BIN2;refGene_tx=XM_019834937.2;refGene_gn=BIN2;refGene_tx=XM_019834939.2;refGene_gn=BIN2;refGene_tx=XM_019834941.2;refGene_gn=BIN2;refGene_tx=XM_023257077.1
B4 79955773 79955845
refGene_gn=BIN2;refGene_tx=XM_011283945.3;refGene_gn=BIN2;refGene_tx=XM_019834937.2;refGene_gn=BIN2;refGene_tx=XM_019834939.2;refGene_gn=BIN2;refGene_tx=XM_019834941.2;refGene_gn=BIN2;refGene_tx=XM_023257077.1
B4 79957452 79957533
refGene_gn=BIN2;refGene_tx=XM_011283945.3;refGene_gn=BIN2;refGene_tx=XM_019834937.2;refGene_gn=BIN2;refGene_tx=XM_019834939.2;refGene_gn=BIN2;refGene_tx=XM_019834941.2;refGene_gn=BIN2;refGene_tx=XM_023257077.1
B4 79959975 79960696
refGene_gn=BIN2;refGene_tx=XM_011283945.3;refGene_gn=BIN2;refGene_tx=XM_019834937.2;refGene_gn=BIN2;refGene_tx=XM_019834939.2;refGene_gn=BIN2;refGene_tx=XM_019834941.2;refGene_gn=BIN2;refGene_tx=XM_023257077.1
B4 79964625 79964708
refGene_gn=BIN2;refGene_tx=XM_011283945.3;refGene_gn=BIN2;refGene_tx=XM_019834937.2;refGene_gn=BIN2;refGene_tx=XM_019834939.2;refGene_gn=BIN2;refGene_tx=XM_019834941.2;refGene_gn=BIN2;refGene_tx=XM_023257077.1
B4 79966161 79966237
refGene_gn=BIN2;refGene_tx=XM_011283945.3;refGene_gn=BIN2;refGene_tx=XM_019834937.2;refGene_gn=BIN2;refGene_tx=XM_019834939.2;refGene_gn=BIN2;refGene_tx=XM_019834941.2;refGene_gn=BIN2;refGene_tx=XM_023257077.1
B4 79967834 79967920
refGene_gn=BIN2;refGene_tx=XM_011283945.3;refGene_gn=BIN2;refGene_tx=XM_019834937.2;refGene_gn=BIN2;refGene_tx=XM_019834939.2;refGene_gn=BIN2;refGene_tx=XM_019834941.2;refGene_gn=BIN2;refGene_tx=XM_023257077.1
B4 79968223 79968331
refGene_gn=BIN2;refGene_tx=XM_011283945.3;refGene_gn=BIN2;refGene_tx=XM_019834937.2;refGene_gn=BIN2;refGene_tx=XM_019834939.2;refGene_gn=BIN2;refGene_tx=XM_019834941.2;refGene_gn=BIN2;refGene_tx=XM_023257077.1
B4 79970232 79970328
refGene_gn=BIN2;refGene_tx=XM_011283945.3;refGene_gn=BIN2;refGene_tx=XM_019834937.2;refGene_gn=BIN2;refGene_tx=XM_019834939.2;refGene_gn=BIN2;refGene_tx=XM_023257077.1
B4 79970908 79971003
refGene_gn=BIN2;refGene_tx=XM_011283945.3;refGene_gn=BIN2;refGene_tx=XM_019834937.2;refGene_gn=BIN2;refGene_tx=XM_019834939.2;refGene_gn=BIN2;refGene_tx=XM_023257077.1
B4 79971272 79971327
refGene_gn=BIN2;refGene_tx=XM_019834937.2;refGene_gn=BIN2;refGene_tx=XM_019834939.2
B4 79983075 79983159 refGene_gn=BIN2;refGene_tx=XM_019834939.2
B4 79983603 79983684
refGene_gn=BIN2;refGene_tx=XM_019834937.2;refGene_gn=BIN2;refGene_tx=XM_019834941.2
B4 79992461 79992542
refGene_gn=BIN2;refGene_tx=XM_019834937.2;refGene_gn=BIN2;refGene_tx=XM_019834941.2
B4 79995739 79995757 refGene_gn=CELA1;refGene_tx=NM_001009389.2

B4 79996551 79996701 refGene_gn=CELA1;refGene_tx=NM_001009389.2
B4 80007339 80007485 refGene_gn=CELA1;refGene_tx=NM_001009389.2
B4 80008756 80008893 refGene_gn=CELA1;refGene_tx=NM_001009389.2
B4 80009679 80009805 refGene_gn=CELA1;refGene_tx=NM_001009389.2
B4 80010944 80011045 refGene_gn=CELA1;refGene_tx=NM_001009389.2
B4 80012204 80012287 refGene_gn=CELA1;refGene_tx=NM_001009389.2
B4 80012966 80013006 refGene_gn=CELA1;refGene_tx=NM_001009389.2
B4 80020887 80021001
refGene_gn=GALNT6;refGene_tx=XM_006933704.3;refGene_gn=GALNT6;refGene_tx=XM_019834943.2;refGene_gn=GALNT6;refGene_tx=XM_019834945.2;refGene_gn=GALNT6;refGene_tx=XM_023257078.1;refGene_gn=GALNT6;refGene_tx=XM_023257079.1
B4 80022272 80022425
refGene_gn=GALNT6;refGene_tx=XM_006933704.3;refGene_gn=GALNT6;refGene_tx=XM_019834943.2;refGene_gn=GALNT6;refGene_tx=XM_019834945.2;refGene_gn=GALNT6;refGene_tx=XM_023257078.1;refGene_gn=GALNT6;refGene_tx=XM_023257079.1
B4 80023448 80023550
refGene_gn=GALNT6;refGene_tx=XM_006933704.3;refGene_gn=GALNT6;refGene_tx=XM_019834943.2;refGene_gn=GALNT6;refGene_tx=XM_019834945.2;refGene_gn=GALNT6;refGene_tx=XM_023257078.1;refGene_gn=GALNT6;refGene_tx=XM_023257079.1
B4 80024366 80024498
refGene_gn=GALNT6;refGene_tx=XM_006933704.3;refGene_gn=GALNT6;refGene_tx=XM_019834943.2;refGene_gn=GALNT6;refGene_tx=XM_019834945.2;refGene_gn=GALNT6;refGene_tx=XM_023257078.1;refGene_gn=GALNT6;refGene_tx=XM_023257079.1
B4 80025004 80025205
refGene_gn=GALNT6;refGene_tx=XM_006933704.3;refGene_gn=GALNT6;refGene_tx=XM_019834943.2;refGene_gn=GALNT6;refGene_tx=XM_019834945.2;refGene_gn=GALNT6;refGene_tx=XM_023257078.1;refGene_gn=GALNT6;refGene_tx=XM_023257079.1
B4 80026510 80026628
refGene_gn=GALNT6;refGene_tx=XM_006933704.3;refGene_gn=GALNT6;refGene_tx=XM_019834943.2;refGene_gn=GALNT6;refGene_tx=XM_019834945.2;refGene_gn=GALNT6;refGene_tx=XM_023257078.1;refGene_gn=GALNT6;refGene_tx=XM_023257079.1
B4 80028675 80028910
refGene_gn=GALNT6;refGene_tx=XM_006933704.3;refGene_gn=GALNT6;refGene_tx=XM_019834943.2;refGene_gn=GALNT6;refGene_tx=XM_019834945.2;refGene_gn=GALNT6;refGene_tx=XM_023257078.1;refGene_gn=GALNT6;refGene_tx=XM_023257079.1
B4 80029959 80030109
refGene_gn=GALNT6;refGene_tx=XM_006933704.3;refGene_gn=GALNT6;refGene_tx=XM_019834943.2;refGene_gn=GALNT6;refGene_tx=XM_019834945.2;refGene_gn=GALNT6;refGene_tx=XM_023257078.1;refGene_gn=GALNT6;refGene_tx=XM_023257079.1
B4 80041344 80041517
refGene_gn=GALNT6;refGene_tx=XM_006933704.3;refGene_gn=GALNT6;refGene_tx=XM_019834943.2;refGene_gn=GALNT6;refGene_tx=XM_019834945.2;refGene_

gn=GALNT6;refGene_tx=XM_023257078.1;refGene_gn=GALNT6;refGene_tx=XM_02
3257079.1
B4 80043092 80043583
refGene_gn=GALNT6;refGene_tx=XM_006933704.3;refGene_gn=GALNT6;refGene_
tx=XM_019834943.2;refGene_gn=GALNT6;refGene_tx=XM_019834945.2;refGene_
gn=GALNT6;refGene_tx=XM_023257078.1;refGene_gn=GALNT6;refGene_tx=XM_02
3257079.1
B4 80084383 80084428 refGene_gn=SLC4A8;refGene_tx=XM_019834951.2
B4 80098592 80098643
refGene_gn=SLC4A8;refGene_tx=XM_006933705.4;refGene_gn=SLC4A8;refGene_
tx=XM_019834949.2;refGene_gn=SLC4A8;refGene_tx=XM_019834952.2;refGene_
gn=SLC4A8;refGene_tx=XM_019834953.2;refGene_gn=SLC4A8;refGene_tx=XM_01
9834955.2;refGene_gn=SLC4A8;refGene_tx=XM_019834956.2;refGene_gn=SLC4A
8;refGene_tx=XM_019834957.2
B4 80098769 80098851
refGene_gn=SLC4A8;refGene_tx=XM_006933705.4;refGene_gn=SLC4A8;refGene_
tx=XM_019834949.2;refGene_gn=SLC4A8;refGene_tx=XM_019834951.2;refGene_
gn=SLC4A8;refGene_tx=XM_019834952.2;refGene_gn=SLC4A8;refGene_tx=XM_01
9834953.2;refGene_gn=SLC4A8;refGene_tx=XM_019834954.2;refGene_gn=SLC4A
8;refGene_tx=XM_019834955.2;refGene_gn=SLC4A8;refGene_tx=XM_019834956.
2;refGene_gn=SLC4A8;refGene_tx=XM_019834957.2
B4 80109333 80109520
refGene_gn=SLC4A8;refGene_tx=XM_006933705.4;refGene_gn=SLC4A8;refGene_
tx=XM_019834949.2;refGene_gn=SLC4A8;refGene_tx=XM_019834951.2;refGene_
gn=SLC4A8;refGene_tx=XM_019834952.2;refGene_gn=SLC4A8;refGene_tx=XM_01
9834953.2;refGene_gn=SLC4A8;refGene_tx=XM_019834954.2;refGene_gn=SLC4A
8;refGene_tx=XM_019834955.2;refGene_gn=SLC4A8;refGene_tx=XM_019834956.
2;refGene_gn=SLC4A8;refGene_tx=XM_019834957.2
B4 80110020 80110156
refGene_gn=SLC4A8;refGene_tx=XM_006933705.4;refGene_gn=SLC4A8;refGene_
tx=XM_019834949.2;refGene_gn=SLC4A8;refGene_tx=XM_019834951.2;refGene_
gn=SLC4A8;refGene_tx=XM_019834952.2;refGene_gn=SLC4A8;refGene_tx=XM_01
9834953.2;refGene_gn=SLC4A8;refGene_tx=XM_019834954.2;refGene_gn=SLC4A
8;refGene_tx=XM_019834955.2;refGene_gn=SLC4A8;refGene_tx=XM_019834956.
2;refGene_gn=SLC4A8;refGene_tx=XM_019834957.2
B4 80111889 80112050
refGene_gn=SLC4A8;refGene_tx=XM_006933705.4;refGene_gn=SLC4A8;refGene_
tx=XM_019834949.2;refGene_gn=SLC4A8;refGene_tx=XM_019834951.2;refGene_
gn=SLC4A8;refGene_tx=XM_019834952.2;refGene_gn=SLC4A8;refGene_tx=XM_01
9834953.2;refGene_gn=SLC4A8;refGene_tx=XM_019834954.2;refGene_gn=SLC4A
8;refGene_tx=XM_019834955.2;refGene_gn=SLC4A8;refGene_tx=XM_019834956.
2;refGene_gn=SLC4A8;refGene_tx=XM_019834957.2
B4 80115550 80115739
refGene_gn=SLC4A8;refGene_tx=XM_006933705.4;refGene_gn=SLC4A8;refGene_
tx=XM_019834949.2;refGene_gn=SLC4A8;refGene_tx=XM_019834951.2;refGene_
gn=SLC4A8;refGene_tx=XM_019834952.2;refGene_gn=SLC4A8;refGene_tx=XM_01
9834953.2;refGene_gn=SLC4A8;refGene_tx=XM_019834954.2;refGene_gn=SLC4A
8;refGene_tx=XM_019834955.2;refGene_gn=SLC4A8;refGene_tx=XM_019834956.
2;refGene_gn=SLC4A8;refGene_tx=XM_019834957.2
B4 80116615 80116707

refGene_gn=SLC4A8;refGene_tx=XM_006933705.4;refGene_gn=SLC4A8;refGene_tx=XM_019834949.2;refGene_gn=SLC4A8;refGene_tx=XM_019834951.2;refGene_gn=SLC4A8;refGene_tx=XM_019834952.2;refGene_gn=SLC4A8;refGene_tx=XM_019834953.2;refGene_gn=SLC4A8;refGene_tx=XM_019834954.2;refGene_gn=SLC4A8;refGene_tx=XM_019834955.2;refGene_gn=SLC4A8;refGene_tx=XM_019834956.2;refGene_gn=SLC4A8;refGene_tx=XM_019834957.2
B4 80117668 80117826
refGene_gn=SLC4A8;refGene_tx=XM_006933705.4;refGene_gn=SLC4A8;refGene_tx=XM_019834949.2;refGene_gn=SLC4A8;refGene_tx=XM_019834951.2;refGene_gn=SLC4A8;refGene_tx=XM_019834952.2;refGene_gn=SLC4A8;refGene_tx=XM_019834953.2;refGene_gn=SLC4A8;refGene_tx=XM_019834954.2;refGene_gn=SLC4A8;refGene_tx=XM_019834955.2;refGene_gn=SLC4A8;refGene_tx=XM_019834956.2;refGene_gn=SLC4A8;refGene_tx=XM_019834957.2
B4 80118758 80118846
refGene_gn=SLC4A8;refGene_tx=XM_006933705.4;refGene_gn=SLC4A8;refGene_tx=XM_019834949.2;refGene_gn=SLC4A8;refGene_tx=XM_019834951.2;refGene_gn=SLC4A8;refGene_tx=XM_019834952.2;refGene_gn=SLC4A8;refGene_tx=XM_019834953.2;refGene_gn=SLC4A8;refGene_tx=XM_019834954.2;refGene_gn=SLC4A8;refGene_tx=XM_019834955.2;refGene_gn=SLC4A8;refGene_tx=XM_019834956.2;refGene_gn=SLC4A8;refGene_tx=XM_019834957.2
B4 80120242 80120389
refGene_gn=SLC4A8;refGene_tx=XM_006933705.4;refGene_gn=SLC4A8;refGene_tx=XM_019834949.2;refGene_gn=SLC4A8;refGene_tx=XM_019834951.2;refGene_gn=SLC4A8;refGene_tx=XM_019834952.2;refGene_gn=SLC4A8;refGene_tx=XM_019834953.2;refGene_gn=SLC4A8;refGene_tx=XM_019834954.2;refGene_gn=SLC4A8;refGene_tx=XM_019834955.2;refGene_gn=SLC4A8;refGene_tx=XM_019834956.2;refGene_gn=SLC4A8;refGene_tx=XM_019834957.2
B4 80121227 80121328
refGene_gn=SLC4A8;refGene_tx=XM_006933705.4;refGene_gn=SLC4A8;refGene_tx=XM_019834949.2;refGene_gn=SLC4A8;refGene_tx=XM_019834951.2;refGene_gn=SLC4A8;refGene_tx=XM_019834952.2;refGene_gn=SLC4A8;refGene_tx=XM_019834953.2;refGene_gn=SLC4A8;refGene_tx=XM_019834954.2;refGene_gn=SLC4A8;refGene_tx=XM_019834955.2;refGene_gn=SLC4A8;refGene_tx=XM_019834956.2;refGene_gn=SLC4A8;refGene_tx=XM_019834957.2
B4 80127162 80127337
refGene_gn=SLC4A8;refGene_tx=XM_006933705.4;refGene_gn=SLC4A8;refGene_tx=XM_019834949.2;refGene_gn=SLC4A8;refGene_tx=XM_019834951.2;refGene_gn=SLC4A8;refGene_tx=XM_019834952.2;refGene_gn=SLC4A8;refGene_tx=XM_019834953.2;refGene_gn=SLC4A8;refGene_tx=XM_019834954.2;refGene_gn=SLC4A8;refGene_tx=XM_019834955.2;refGene_gn=SLC4A8;refGene_tx=XM_019834956.2
B4 80127936 80128070
refGene_gn=SLC4A8;refGene_tx=XM_006933705.4;refGene_gn=SLC4A8;refGene_tx=XM_019834949.2;refGene_gn=SLC4A8;refGene_tx=XM_019834951.2;refGene_gn=SLC4A8;refGene_tx=XM_019834952.2;refGene_gn=SLC4A8;refGene_tx=XM_019834953.2;refGene_gn=SLC4A8;refGene_tx=XM_019834954.2;refGene_gn=SLC4A8;refGene_tx=XM_019834955.2;refGene_gn=SLC4A8;refGene_tx=XM_019834956.2
B4 80129096 80129342
refGene_gn=SLC4A8;refGene_tx=XM_006933705.4;refGene_gn=SLC4A8;refGene_

tx=XM_019834949.2;refGene_gn=SLC4A8;refGene_tx=XM_019834951.2;refGene_gn=SLC4A8;refGene_tx=XM_019834952.2;refGene_gn=SLC4A8;refGene_tx=XM_019834953.2;refGene_gn=SLC4A8;refGene_tx=XM_019834954.2;refGene_gn=SLC4A8;refGene_tx=XM_019834955.2;refGene_gn=SLC4A8;refGene_tx=XM_019834956.2;refGene_gn=SLC4A8;refGene_tx=XM_019834957.2
B4 80132998 80133104
refGene_gn=SLC4A8;refGene_tx=XM_006933705.4;refGene_gn=SLC4A8;refGene_tx=XM_019834949.2;refGene_gn=SLC4A8;refGene_tx=XM_019834951.2;refGene_gn=SLC4A8;refGene_tx=XM_019834952.2;refGene_gn=SLC4A8;refGene_tx=XM_019834953.2;refGene_gn=SLC4A8;refGene_tx=XM_019834954.2;refGene_gn=SLC4A8;refGene_tx=XM_019834955.2;refGene_gn=SLC4A8;refGene_tx=XM_019834956.2;refGene_gn=SLC4A8;refGene_tx=XM_019834957.2
B4 80133709 80133871
refGene_gn=SLC4A8;refGene_tx=XM_006933705.4;refGene_gn=SLC4A8;refGene_tx=XM_019834949.2;refGene_gn=SLC4A8;refGene_tx=XM_019834951.2;refGene_gn=SLC4A8;refGene_tx=XM_019834952.2;refGene_gn=SLC4A8;refGene_tx=XM_019834953.2;refGene_gn=SLC4A8;refGene_tx=XM_019834954.2;refGene_gn=SLC4A8;refGene_tx=XM_019834955.2;refGene_gn=SLC4A8;refGene_tx=XM_019834956.2;refGene_gn=SLC4A8;refGene_tx=XM_019834957.2
B4 80136982 80137096
refGene_gn=SLC4A8;refGene_tx=XM_006933705.4;refGene_gn=SLC4A8;refGene_tx=XM_019834949.2;refGene_gn=SLC4A8;refGene_tx=XM_019834951.2;refGene_gn=SLC4A8;refGene_tx=XM_019834952.2;refGene_gn=SLC4A8;refGene_tx=XM_019834953.2;refGene_gn=SLC4A8;refGene_tx=XM_019834954.2;refGene_gn=SLC4A8;refGene_tx=XM_019834955.2;refGene_gn=SLC4A8;refGene_tx=XM_019834956.2;refGene_gn=SLC4A8;refGene_tx=XM_019834957.2
B4 80139464 80139626
refGene_gn=SLC4A8;refGene_tx=XM_006933705.4;refGene_gn=SLC4A8;refGene_tx=XM_019834949.2;refGene_gn=SLC4A8;refGene_tx=XM_019834951.2;refGene_gn=SLC4A8;refGene_tx=XM_019834952.2;refGene_gn=SLC4A8;refGene_tx=XM_019834953.2;refGene_gn=SLC4A8;refGene_tx=XM_019834954.2;refGene_gn=SLC4A8;refGene_tx=XM_019834955.2;refGene_gn=SLC4A8;refGene_tx=XM_019834956.2;refGene_gn=SLC4A8;refGene_tx=XM_019834957.2
B4 80140518 80140770
refGene_gn=SLC4A8;refGene_tx=XM_006933705.4;refGene_gn=SLC4A8;refGene_tx=XM_019834949.2;refGene_gn=SLC4A8;refGene_tx=XM_019834951.2;refGene_gn=SLC4A8;refGene_tx=XM_019834952.2;refGene_gn=SLC4A8;refGene_tx=XM_019834953.2;refGene_gn=SLC4A8;refGene_tx=XM_019834954.2;refGene_gn=SLC4A8;refGene_tx=XM_019834955.2;refGene_gn=SLC4A8;refGene_tx=XM_019834956.2;refGene_gn=SLC4A8;refGene_tx=XM_019834957.2
B4 80144370 80144439
refGene_gn=SLC4A8;refGene_tx=XM_006933705.4;refGene_gn=SLC4A8;refGene_tx=XM_019834949.2;refGene_gn=SLC4A8;refGene_tx=XM_019834951.2;refGene_gn=SLC4A8;refGene_tx=XM_019834952.2;refGene_gn=SLC4A8;refGene_tx=XM_019834953.2;refGene_gn=SLC4A8;refGene_tx=XM_019834954.2;refGene_gn=SLC4A8;refGene_tx=XM_019834955.2;refGene_gn=SLC4A8;refGene_tx=XM_019834956.2;refGene_gn=SLC4A8;refGene_tx=XM_019834957.2
B4 80145726 80145900
refGene_gn=SLC4A8;refGene_tx=XM_006933705.4;refGene_gn=SLC4A8;refGene_tx=XM_019834949.2;refGene_gn=SLC4A8;refGene_tx=XM_019834951.2;refGene_

gn=SLC4A8;refGene_tx=XM_019834952.2;refGene_gn=SLC4A8;refGene_tx=XM_01
9834953.2;refGene_gn=SLC4A8;refGene_tx=XM_019834954.2;refGene_gn=SLC4A
8;refGene_tx=XM_019834955.2;refGene_gn=SLC4A8;refGene_tx=XM_019834956.
2;refGene_gn=SLC4A8;refGene_tx=XM_019834957.2
B4 80148969 80149107
refGene_gn=SLC4A8;refGene_tx=XM_006933705.4;refGene_gn=SLC4A8;refGene_
tx=XM_019834949.2;refGene_gn=SLC4A8;refGene_tx=XM_019834951.2;refGene_
gn=SLC4A8;refGene_tx=XM_019834952.2;refGene_gn=SLC4A8;refGene_tx=XM_01
9834953.2;refGene_gn=SLC4A8;refGene_tx=XM_019834954.2;refGene_gn=SLC4A
8;refGene_tx=XM_019834957.2
B4 80155120 80155212
refGene_gn=SLC4A8;refGene_tx=XM_006933705.4;refGene_gn=SLC4A8;refGene_
tx=XM_019834949.2;refGene_gn=SLC4A8;refGene_tx=XM_019834951.2;refGene_
gn=SLC4A8;refGene_tx=XM_019834952.2;refGene_gn=SLC4A8;refGene_tx=XM_01
9834953.2;refGene_gn=SLC4A8;refGene_tx=XM_019834954.2;refGene_gn=SLC4A
8;refGene_tx=XM_019834955.2;refGene_gn=SLC4A8;refGene_tx=XM_019834956.
2;refGene_gn=SLC4A8;refGene_tx=XM_019834957.2
B4 80157340 80157448
refGene_gn=SLC4A8;refGene_tx=XM_006933705.4;refGene_gn=SLC4A8;refGene_
tx=XM_019834949.2;refGene_gn=SLC4A8;refGene_tx=XM_019834951.2;refGene_
gn=SLC4A8;refGene_tx=XM_019834952.2;refGene_gn=SLC4A8;refGene_tx=XM_01
9834953.2;refGene_gn=SLC4A8;refGene_tx=XM_019834954.2;refGene_gn=SLC4A
8;refGene_tx=XM_019834955.2;refGene_gn=SLC4A8;refGene_tx=XM_019834956.
2;refGene_gn=SLC4A8;refGene_tx=XM_019834957.2
B4 80158615 80158628
refGene_gn=SLC4A8;refGene_tx=XM_006933705.4;refGene_gn=SLC4A8;refGene_
tx=XM_019834949.2;refGene_gn=SLC4A8;refGene_tx=XM_019834951.2;refGene_
gn=SLC4A8;refGene_tx=XM_019834952.2;refGene_gn=SLC4A8;refGene_tx=XM_01
9834953.2;refGene_gn=SLC4A8;refGene_tx=XM_019834954.2;refGene_gn=SLC4A
8;refGene_tx=XM_019834955.2;refGene_gn=SLC4A8;refGene_tx=XM_019834956.
2;refGene_gn=SLC4A8;refGene_tx=XM_019834957.2
B4 80274609 80274885
refGene_gn=SCN8A;refGene_tx=XM_023257080.1;refGene_gn=SCN8A;refGene_tx
=XM_023257081.1;refGene_gn=SCN8A;refGene_tx=XM_023257082.1;refGene_gn=
SCN8A;refGene_tx=XM_023257083.1;refGene_gn=SCN8A;refGene_tx=XM_0232570
84.1;refGene_gn=SCN8A;refGene_tx=XM_023257085.1;refGene_gn=SCN8A;refGe
ne_tx=XM_023257086.1;refGene_gn=SCN8A;refGene_tx=XM_023257087.1
B4 80300823 80300942
refGene_gn=SCN8A;refGene_tx=XM_023257080.1;refGene_gn=SCN8A;refGene_tx
=XM_023257081.1;refGene_gn=SCN8A;refGene_tx=XM_023257082.1;refGene_gn=
SCN8A;refGene_tx=XM_023257083.1;refGene_gn=SCN8A;refGene_tx=XM_0232570
84.1;refGene_gn=SCN8A;refGene_tx=XM_023257085.1;refGene_gn=SCN8A;refGe
ne_tx=XM_023257086.1;refGene_gn=SCN8A;refGene_tx=XM_023257087.1
B4 80302840 80302930
refGene_gn=SCN8A;refGene_tx=XM_023257080.1;refGene_gn=SCN8A;refGene_tx
=XM_023257081.1;refGene_gn=SCN8A;refGene_tx=XM_023257082.1;refGene_gn=
SCN8A;refGene_tx=XM_023257083.1;refGene_gn=SCN8A;refGene_tx=XM_0232570
84.1;refGene_gn=SCN8A;refGene_tx=XM_023257085.1;refGene_gn=SCN8A;refGe
ne_tx=XM_023257086.1;refGene_gn=SCN8A;refGene_tx=XM_023257087.1
B4 80303604 80303733

refGene_gn=SCN8A;refGene_tx=XM_023257080.1;refGene_gn=SCN8A;refGene_tx=XM_023257081.1;refGene_gn=SCN8A;refGene_tx=XM_023257082.1;refGene_gn=SCN8A;refGene_tx=XM_023257083.1;refGene_gn=SCN8A;refGene_tx=XM_023257084.1;refGene_gn=SCN8A;refGene_tx=XM_023257085.1;refGene_gn=SCN8A;refGene_tx=XM_023257086.1;refGene_gn=SCN8A;refGene_tx=XM_023257087.1
B4 80305336 80305428
refGene_gn=SCN8A;refGene_tx=XM_023257082.1;refGene_gn=SCN8A;refGene_tx=XM_023257084.1;refGene_gn=SCN8A;refGene_tx=XM_023257086.1
B4 80305585 80305677
refGene_gn=SCN8A;refGene_tx=XM_023257080.1;refGene_gn=SCN8A;refGene_tx=XM_023257081.1;refGene_gn=SCN8A;refGene_tx=XM_023257083.1;refGene_gn=SCN8A;refGene_tx=XM_023257085.1;refGene_gn=SCN8A;refGene_tx=XM_023257087.1
B4 80316220 80316442
refGene_gn=SCN8A;refGene_tx=XM_023257080.1;refGene_gn=SCN8A;refGene_tx=XM_023257081.1;refGene_gn=SCN8A;refGene_tx=XM_023257082.1;refGene_gn=SCN8A;refGene_tx=XM_023257083.1;refGene_gn=SCN8A;refGene_tx=XM_023257084.1;refGene_gn=SCN8A;refGene_tx=XM_023257085.1;refGene_gn=SCN8A;refGene_tx=XM_023257086.1;refGene_gn=SCN8A;refGene_tx=XM_023257087.1
B4 80317782 80317846
refGene_gn=SCN8A;refGene_tx=XM_023257080.1;refGene_gn=SCN8A;refGene_tx=XM_023257081.1;refGene_gn=SCN8A;refGene_tx=XM_023257082.1;refGene_gn=SCN8A;refGene_tx=XM_023257083.1;refGene_gn=SCN8A;refGene_tx=XM_023257084.1;refGene_gn=SCN8A;refGene_tx=XM_023257085.1;refGene_gn=SCN8A;refGene_tx=XM_023257086.1;refGene_gn=SCN8A;refGene_tx=XM_023257087.1
B4 80319992 80320134
refGene_gn=SCN8A;refGene_tx=XM_023257080.1;refGene_gn=SCN8A;refGene_tx=XM_023257081.1;refGene_gn=SCN8A;refGene_tx=XM_023257082.1;refGene_gn=SCN8A;refGene_tx=XM_023257083.1;refGene_gn=SCN8A;refGene_tx=XM_023257084.1;refGene_gn=SCN8A;refGene_tx=XM_023257085.1;refGene_gn=SCN8A;refGene_tx=XM_023257086.1;refGene_gn=SCN8A;refGene_tx=XM_023257087.1
B4 80322089 80322296
refGene_gn=SCN8A;refGene_tx=XM_023257080.1;refGene_gn=SCN8A;refGene_tx=XM_023257081.1;refGene_gn=SCN8A;refGene_tx=XM_023257082.1;refGene_gn=SCN8A;refGene_tx=XM_023257083.1;refGene_gn=SCN8A;refGene_tx=XM_023257084.1;refGene_gn=SCN8A;refGene_tx=XM_023257085.1;refGene_gn=SCN8A;refGene_tx=XM_023257086.1;refGene_gn=SCN8A;refGene_tx=XM_023257087.1
B4 80323179 80323473
refGene_gn=SCN8A;refGene_tx=XM_023257080.1;refGene_gn=SCN8A;refGene_tx=XM_023257081.1;refGene_gn=SCN8A;refGene_tx=XM_023257082.1;refGene_gn=SCN8A;refGene_tx=XM_023257083.1;refGene_gn=SCN8A;refGene_tx=XM_023257084.1;refGene_gn=SCN8A;refGene_tx=XM_023257085.1;refGene_gn=SCN8A;refGene_tx=XM_023257086.1;refGene_gn=SCN8A;refGene_tx=XM_023257087.1
B4 80334841 80335237
refGene_gn=SCN8A;refGene_tx=XM_023257080.1;refGene_gn=SCN8A;refGene_tx=XM_023257081.1;refGene_gn=SCN8A;refGene_tx=XM_023257082.1;refGene_gn=SCN8A;refGene_tx=XM_023257083.1;refGene_gn=SCN8A;refGene_tx=XM_023257084.1;refGene_gn=SCN8A;refGene_tx=XM_023257085.1;refGene_gn=SCN8A;refGene_tx=XM_023257086.1;refGene_gn=SCN8A;refGene_tx=XM_023257087.1
B4 80348768 80348901

refGene_gn=SCN8A;refGene_tx=XM_023257080.1;refGene_gn=SCN8A;refGene_tx=XM_023257081.1;refGene_gn=SCN8A;refGene_tx=XM_023257082.1;refGene_gn=SCN8A;refGene_tx=XM_023257083.1;refGene_gn=SCN8A;refGene_tx=XM_023257084.1;refGene_gn=SCN8A;refGene_tx=XM_023257085.1;refGene_gn=SCN8A;refGene_tx=XM_023257086.1;refGene_gn=SCN8A;refGene_tx=XM_023257087.1
B4 80354360 80354599
refGene_gn=SCN8A;refGene_tx=XM_023257080.1;refGene_gn=SCN8A;refGene_tx=XM_023257081.1;refGene_gn=SCN8A;refGene_tx=XM_023257082.1;refGene_gn=SCN8A;refGene_tx=XM_023257083.1;refGene_gn=SCN8A;refGene_tx=XM_023257084.1;refGene_gn=SCN8A;refGene_tx=XM_023257085.1;refGene_gn=SCN8A;refGene_tx=XM_023257086.1;refGene_gn=SCN8A;refGene_tx=XM_023257087.1
B4 80357983 80358157
refGene_gn=SCN8A;refGene_tx=XM_023257080.1;refGene_gn=SCN8A;refGene_tx=XM_023257081.1;refGene_gn=SCN8A;refGene_tx=XM_023257082.1;refGene_gn=SCN8A;refGene_tx=XM_023257083.1;refGene_gn=SCN8A;refGene_tx=XM_023257084.1;refGene_gn=SCN8A;refGene_tx=XM_023257085.1;refGene_gn=SCN8A;refGene_tx=XM_023257086.1;refGene_gn=SCN8A;refGene_tx=XM_023257087.1
B4 80360330 80360687
refGene_gn=SCN8A;refGene_tx=XM_023257080.1;refGene_gn=SCN8A;refGene_tx=XM_023257081.1;refGene_gn=SCN8A;refGene_tx=XM_023257082.1;refGene_gn=SCN8A;refGene_tx=XM_023257083.1;refGene_gn=SCN8A;refGene_tx=XM_023257084.1;refGene_gn=SCN8A;refGene_tx=XM_023257085.1;refGene_gn=SCN8A;refGene_tx=XM_023257086.1;refGene_gn=SCN8A;refGene_tx=XM_023257087.1
B4 80363992 80364463
refGene_gn=SCN8A;refGene_tx=XM_023257080.1;refGene_gn=SCN8A;refGene_tx=XM_023257081.1;refGene_gn=SCN8A;refGene_tx=XM_023257082.1;refGene_gn=SCN8A;refGene_tx=XM_023257083.1;refGene_gn=SCN8A;refGene_tx=XM_023257084.1;refGene_gn=SCN8A;refGene_tx=XM_023257085.1;refGene_gn=SCN8A;refGene_tx=XM_023257086.1;refGene_gn=SCN8A;refGene_tx=XM_023257087.1
B4 80364995 80365113
refGene_gn=SCN8A;refGene_tx=XM_023257080.1;refGene_gn=SCN8A;refGene_tx=XM_023257081.1;refGene_gn=SCN8A;refGene_tx=XM_023257082.1;refGene_gn=SCN8A;refGene_tx=XM_023257083.1;refGene_gn=SCN8A;refGene_tx=XM_023257084.1;refGene_gn=SCN8A;refGene_tx=XM_023257085.1;refGene_gn=SCN8A;refGene_tx=XM_023257086.1;refGene_gn=SCN8A;refGene_tx=XM_023257087.1
B4 80365634 80365789
refGene_gn=SCN8A;refGene_tx=XM_023257080.1;refGene_gn=SCN8A;refGene_tx=XM_023257081.1;refGene_gn=SCN8A;refGene_tx=XM_023257082.1;refGene_gn=SCN8A;refGene_tx=XM_023257083.1;refGene_gn=SCN8A;refGene_tx=XM_023257084.1;refGene_gn=SCN8A;refGene_tx=XM_023257085.1;refGene_gn=SCN8A;refGene_tx=XM_023257086.1;refGene_gn=SCN8A;refGene_tx=XM_023257087.1
B4 80368043 80368217
refGene_gn=SCN8A;refGene_tx=XM_023257080.1;refGene_gn=SCN8A;refGene_tx=XM_023257081.1;refGene_gn=SCN8A;refGene_tx=XM_023257082.1;refGene_gn=SCN8A;refGene_tx=XM_023257083.1;refGene_gn=SCN8A;refGene_tx=XM_023257084.1;refGene_gn=SCN8A;refGene_tx=XM_023257085.1;refGene_gn=SCN8A;refGene_tx=XM_023257086.1;refGene_gn=SCN8A;refGene_tx=XM_023257087.1
B4 80374043 80374166
refGene_gn=SCN8A;refGene_tx=XM_023257080.1;refGene_gn=SCN8A;refGene_tx=XM_023257081.1;refGene_gn=SCN8A;refGene_tx=XM_023257082.1;refGene_gn=

SCN8A;refGene_tx=XM_023257083.1;refGene_gn=SCN8A;refGene_tx=XM_0232570
84.1;refGene_gn=SCN8A;refGene_tx=XM_023257087.1
B4 80379949 80380234
refGene_gn=SCN8A;refGene_tx=XM_023257080.1;refGene_gn=SCN8A;refGene_tx
=XM_023257081.1;refGene_gn=SCN8A;refGene_tx=XM_023257082.1;refGene_gn=
SCN8A;refGene_tx=XM_023257083.1;refGene_gn=SCN8A;refGene_tx=XM_0232570
84.1;refGene_gn=SCN8A;refGene_tx=XM_023257085.1;refGene_gn=SCN8A;refGe
ne_tx=XM_023257086.1;refGene_gn=SCN8A;refGene_tx=XM_023257087.1
B4 80382109 80382163
refGene_gn=SCN8A;refGene_tx=XM_023257080.1;refGene_gn=SCN8A;refGene_tx
=XM_023257081.1;refGene_gn=SCN8A;refGene_tx=XM_023257082.1;refGene_gn=
SCN8A;refGene_tx=XM_023257083.1;refGene_gn=SCN8A;refGene_tx=XM_0232570
84.1;refGene_gn=SCN8A;refGene_tx=XM_023257085.1;refGene_gn=SCN8A;refGe
ne_tx=XM_023257086.1;refGene_gn=SCN8A;refGene_tx=XM_023257087.1
B4 80382648 80382786
refGene_gn=SCN8A;refGene_tx=XM_023257080.1;refGene_gn=SCN8A;refGene_tx
=XM_023257081.1;refGene_gn=SCN8A;refGene_tx=XM_023257082.1;refGene_gn=
SCN8A;refGene_tx=XM_023257083.1;refGene_gn=SCN8A;refGene_tx=XM_0232570
84.1;refGene_gn=SCN8A;refGene_tx=XM_023257085.1;refGene_gn=SCN8A;refGe
ne_tx=XM_023257086.1;refGene_gn=SCN8A;refGene_tx=XM_023257087.1
B4 80383763 80383868
refGene_gn=SCN8A;refGene_tx=XM_023257080.1;refGene_gn=SCN8A;refGene_tx
=XM_023257081.1;refGene_gn=SCN8A;refGene_tx=XM_023257082.1;refGene_gn=
SCN8A;refGene_tx=XM_023257083.1;refGene_gn=SCN8A;refGene_tx=XM_0232570
84.1;refGene_gn=SCN8A;refGene_tx=XM_023257085.1;refGene_gn=SCN8A;refGe
ne_tx=XM_023257086.1;refGene_gn=SCN8A;refGene_tx=XM_023257087.1
B4 80386281 80386552
refGene_gn=SCN8A;refGene_tx=XM_023257080.1;refGene_gn=SCN8A;refGene_tx
=XM_023257081.1;refGene_gn=SCN8A;refGene_tx=XM_023257082.1;refGene_gn=
SCN8A;refGene_tx=XM_023257083.1;refGene_gn=SCN8A;refGene_tx=XM_0232570
84.1;refGene_gn=SCN8A;refGene_tx=XM_023257085.1;refGene_gn=SCN8A;refGe
ne_tx=XM_023257086.1;refGene_gn=SCN8A;refGene_tx=XM_023257087.1
B4 80390209 80391357
refGene_gn=SCN8A;refGene_tx=XM_023257080.1;refGene_gn=SCN8A;refGene_tx
=XM_023257081.1;refGene_gn=SCN8A;refGene_tx=XM_023257082.1;refGene_gn=
SCN8A;refGene_tx=XM_023257083.1;refGene_gn=SCN8A;refGene_tx=XM_0232570
84.1;refGene_gn=SCN8A;refGene_tx=XM_023257085.1;refGene_gn=SCN8A;refGe
ne_tx=XM_023257086.1;refGene_gn=SCN8A;refGene_tx=XM_023257087.1
B4 80403651 80405631 refGene_gn=FIGNL2;refGene_tx=XM_023257088.1
B4 80469914 80470079
refGene_gn=ANKRD33;refGene_tx=XM_011283946.3;refGene_gn=ANKRD33;refGen
e_tx=XM_019834966.2
B4 80470294 80470424
refGene_gn=ANKRD33;refGene_tx=XM_011283946.3;refGene_gn=ANKRD33;refGen
e_tx=XM_019834966.2;refGene_gn=ANKRD33;refGene_tx=XM_019834967.1;refGe
ne_gn=ANKRD33;refGene_tx=XM_019834969.1
B4 80470608 80470749
refGene_gn=ANKRD33;refGene_tx=XM_011283946.3;refGene_gn=ANKRD33;refGen
e_tx=XM_019834966.2;refGene_gn=ANKRD33;refGene_tx=XM_019834967.1;refGe
ne_gn=ANKRD33;refGene_tx=XM_019834969.1

B4 80472418 80473149
refGene_gn=ANKRD33;refGene_tx=XM_011283946.3;refGene_gn=ANKRD33;refGen
e_tx=XM_019834966.2;refGene_gn=ANKRD33;refGene_tx=XM_019834967.1;refGe
ne_gn=ANKRD33;refGene_tx=XM_019834969.1
B4 80490840 80490904
refGene_gn=ACVRL1;refGene_tx=XM_011283947.2;refGene_gn=ACVRL1;refGene_
tx=XM_023257089.1
B4 80491460 80491712
refGene_gn=ACVRL1;refGene_tx=XM_011283947.2;refGene_gn=ACVRL1;refGene_
tx=XM_023257089.1
B4 80491906 80492121
refGene_gn=ACVRL1;refGene_tx=XM_011283947.2;refGene_gn=ACVRL1;refGene_
tx=XM_023257089.1
B4 80492390 80492490
refGene_gn=ACVRL1;refGene_tx=XM_011283947.2;refGene_gn=ACVRL1;refGene_
tx=XM_023257089.1
B4 80492847 80492994
refGene_gn=ACVRL1;refGene_tx=XM_011283947.2;refGene_gn=ACVRL1;refGene_
tx=XM_023257089.1
B4 80493257 80493533
refGene_gn=ACVRL1;refGene_tx=XM_011283947.2;refGene_gn=ACVRL1;refGene_
tx=XM_023257089.1
B4 80493669 80494118 refGene_gn=ACVRL1;refGene_tx=XM_023257089.1
B4 80494136 80494334 refGene_gn=ACVRL1;refGene_tx=XM_011283947.2
B4 80497217 80497348 refGene_gn=ACVRL1;refGene_tx=XM_011283947.2
B4 80498552 80498687 refGene_gn=ACVRL1;refGene_tx=XM_011283947.2
B4 80531405 80531496 refGene_gn=ACVR1B;refGene_tx=XM_023257090.1
B4 80547743 80547983
refGene_gn=ACVR1B;refGene_tx=XM_023257090.1;refGene_gn=ACVR1B;refGene_
tx=XM_023257091.1
B4 80548780 80549029
refGene_gn=ACVR1B;refGene_tx=XM_023257090.1;refGene_gn=ACVR1B;refGene_
tx=XM_023257091.1
B4 80552205 80552436
refGene_gn=ACVR1B;refGene_tx=XM_023257090.1;refGene_gn=ACVR1B;refGene_
tx=XM_023257091.1
B4 80555328 80555496
refGene_gn=ACVR1B;refGene_tx=XM_023257090.1;refGene_gn=ACVR1B;refGene_
tx=XM_023257091.1
B4 80556549 80556706
refGene_gn=ACVR1B;refGene_tx=XM_023257090.1;refGene_gn=ACVR1B;refGene_
tx=XM_023257091.1
B4 80557938 80558063
refGene_gn=ACVR1B;refGene_tx=XM_023257090.1;refGene_gn=ACVR1B;refGene_
tx=XM_023257091.1
B4 80561293 80561424
refGene_gn=ACVR1B;refGene_tx=XM_023257090.1;refGene_gn=ACVR1B;refGene_
tx=XM_023257091.1
B4 80562941 80563067
refGene_gn=ACVR1B;refGene_tx=XM_023257090.1;refGene_gn=ACVR1B;refGene_

tx=XM_023257091.1
B4 80574921 80575173 refGene_gn=GRASP;refGene_tx=XM_023257092.1
B4 80577323 80577373 refGene_gn=GRASP;refGene_tx=XM_023257092.1
B4 80578992 80579047 refGene_gn=GRASP;refGene_tx=XM_023257092.1
B4 80579151 80579254
refGene_gn=GRASP;refGene_tx=XM_023257092.1;refGene_gn=GRASP;refGene_tx
=XM_023257093.1
B4 80580403 80580497
refGene_gn=GRASP;refGene_tx=XM_023257092.1;refGene_gn=GRASP;refGene_tx
=XM_023257093.1
B4 80580600 80580667
refGene_gn=GRASP;refGene_tx=XM_023257092.1;refGene_gn=GRASP;refGene_tx
=XM_023257093.1
B4 80580882 80580949
refGene_gn=GRASP;refGene_tx=XM_023257092.1;refGene_gn=GRASP;refGene_tx
=XM_023257093.1
B4 80581388 80581888
refGene_gn=GRASP;refGene_tx=XM_023257092.1;refGene_gn=GRASP;refGene_tx
=XM_023257093.1
B4 80623061 80623937
refGene_gn=NR4A1;refGene_tx=XM_003988726.5;refGene_gn=NR4A1;refGene_tx
=XM_006933714.2;refGene_gn=NR4A1;refGene_tx=XM_006933716.2;refGene_gn=
NR4A1;refGene_tx=XM_019834972.2
B4 80624778 80624908
refGene_gn=NR4A1;refGene_tx=XM_003988726.5;refGene_gn=NR4A1;refGene_tx
=XM_006933714.2;refGene_gn=NR4A1;refGene_tx=XM_006933716.2;refGene_gn=
NR4A1;refGene_tx=XM_019834972.2
B4 80625194 80625346
refGene_gn=NR4A1;refGene_tx=XM_003988726.5;refGene_gn=NR4A1;refGene_tx
=XM_006933714.2;refGene_gn=NR4A1;refGene_tx=XM_006933716.2;refGene_gn=
NR4A1;refGene_tx=XM_019834972.2
B4 80625686 80625889
refGene_gn=NR4A1;refGene_tx=XM_003988726.5;refGene_gn=NR4A1;refGene_tx
=XM_006933714.2;refGene_gn=NR4A1;refGene_tx=XM_006933716.2;refGene_gn=
NR4A1;refGene_tx=XM_019834972.2
B4 80625975 80626154
refGene_gn=NR4A1;refGene_tx=XM_003988726.5;refGene_gn=NR4A1;refGene_tx
=XM_006933714.2;refGene_gn=NR4A1;refGene_tx=XM_006933716.2;refGene_gn=
NR4A1;refGene_tx=XM_019834972.2
B4 80627216 80627473
refGene_gn=NR4A1;refGene_tx=XM_003988726.5;refGene_gn=NR4A1;refGene_tx
=XM_006933714.2;refGene_gn=NR4A1;refGene_tx=XM_006933716.2;refGene_gn=
NR4A1;refGene_tx=XM_019834972.2
B4 80640329 80640581
refGene_gn=ATG101;refGene_tx=XM_003988728.3;refGene_gn=ATG101;refGene_
tx=XM_023257094.1;refGene_gn=ATG101;refGene_tx=XM_023257095.1
B4 80642277 80642682
refGene_gn=ATG101;refGene_tx=XM_003988728.3;refGene_gn=ATG101;refGene_
tx=XM_023257094.1;refGene_gn=ATG101;refGene_tx=XM_023257095.1
B4 80693331 80693491

refGene_gn=KRT80;refGene_tx=XM_003988729.5;refGene_gn=KRT80;refGene_tx=XM_019834973.2
B4 80693587 80693643
refGene_gn=KRT80;refGene_tx=XM_003988729.5;refGene_gn=KRT80;refGene_tx=XM_019834973.2
B4 80694111 80694332
refGene_gn=KRT80;refGene_tx=XM_003988729.5;refGene_gn=KRT80;refGene_tx=XM_019834973.2
B4 80694976 80695102
refGene_gn=KRT80;refGene_tx=XM_003988729.5;refGene_gn=KRT80;refGene_tx=XM_019834973.2
B4 80695521 80695686
refGene_gn=KRT80;refGene_tx=XM_003988729.5;refGene_gn=KRT80;refGene_tx=XM_019834973.2
B4 80702203 80702299
refGene_gn=KRT80;refGene_tx=XM_003988729.5;refGene_gn=KRT80;refGene_tx=XM_019834973.2
B4 80702561 80702622
refGene_gn=KRT80;refGene_tx=XM_003988729.5;refGene_gn=KRT80;refGene_tx=XM_019834973.2
B4 80707212 80707421
refGene_gn=KRT80;refGene_tx=XM_003988729.5;refGene_gn=KRT80;refGene_tx=XM_019834973.2
B4 80713267 80713702
refGene_gn=KRT80;refGene_tx=XM_003988729.5;refGene_gn=KRT80;refGene_tx=XM_019834973.2
B4 80725087 80725149
refGene_gn=CB4H12orf80;refGene_tx=XM_023257758.1
B4 80725490 80725607
refGene_gn=CB4H12orf80;refGene_tx=XM_023257758.1
B4 80726381 80726514
refGene_gn=CB4H12orf80;refGene_tx=XM_023257758.1
B4 80727013 80727294
refGene_gn=CB4H12orf80;refGene_tx=XM_023257758.1
B4 80733110 80733156
refGene_gn=CB4H12orf80;refGene_tx=XM_023257758.1
B4 80751098 80751428 refGene_gn=KRT7;refGene_tx=XM_019834974.2
B4 80752899 80753111 refGene_gn=KRT7;refGene_tx=XM_019834974.2
B4 80755229 80755290 refGene_gn=KRT7;refGene_tx=XM_019834974.2
B4 80756490 80756586 refGene_gn=KRT7;refGene_tx=XM_019834974.2
B4 80758311 80758476 refGene_gn=KRT7;refGene_tx=XM_019834974.2
B4 80759449 80759575 refGene_gn=KRT7;refGene_tx=XM_019834974.2
B4 80761432 80761653 refGene_gn=KRT7;refGene_tx=XM_019834974.2
B4 80764245 80764280 refGene_gn=KRT7;refGene_tx=XM_019834974.2
B4 80764668 80764835 refGene_gn=KRT7;refGene_tx=XM_019834974.2
B4 80766794 80766970
refGene_gn=LOC101096492;refGene_tx=XM_023257096.1
B4 80767480 80767512
refGene_gn=LOC101096492;refGene_tx=XM_023257096.1
B4 80767662 80767883

refGene_gn=LOC101096492;refGene_tx=XM_023257096.1
B4	80768128	80768254
refGene_gn=LOC101096492;refGene_tx=XM_023257096.1
B4	80768518	80768683
refGene_gn=LOC101096492;refGene_tx=XM_023257096.1
B4	80768917	80769013
refGene_gn=LOC101096492;refGene_tx=XM_023257096.1
B4	80769979	80770040
refGene_gn=LOC101096492;refGene_tx=XM_023257096.1
B4	80770675	80770884
refGene_gn=LOC101096492;refGene_tx=XM_023257096.1
B4	80773304	80773865
refGene_gn=LOC101096492;refGene_tx=XM_023257096.1
B4	80786132	80786135
refGene_gn=LOC101096742;refGene_tx=XM_019834975.2;refGene_gn=LOC101096742;refGene_tx=XM_019834976.2;refGene_gn=LOC101096742;refGene_tx=XM_019834977.2;refGene_gn=LOC101096742;refGene_tx=XM_019834978.2;refGene_gn=LOC101096742;refGene_tx=XM_023257097.1
B4	80786570	80786666
refGene_gn=LOC101096742;refGene_tx=XM_019834975.2;refGene_gn=LOC101096742;refGene_tx=XM_019834976.2;refGene_gn=LOC101096742;refGene_tx=XM_019834977.2;refGene_gn=LOC101096742;refGene_tx=XM_019834978.2
B4	80787286	80787412
refGene_gn=LOC101096742;refGene_tx=XM_019834975.2;refGene_gn=LOC101096742;refGene_tx=XM_019834976.2;refGene_gn=LOC101096742;refGene_tx=XM_019834977.2;refGene_gn=LOC101096742;refGene_tx=XM_019834978.2;refGene_gn=LOC101096742;refGene_tx=XM_023257097.1
B4	80789228	80789449
refGene_gn=LOC101096742;refGene_tx=XM_019834975.2;refGene_gn=LOC101096742;refGene_tx=XM_019834976.2;refGene_gn=LOC101096742;refGene_tx=XM_019834977.2;refGene_gn=LOC101096742;refGene_tx=XM_019834978.2;refGene_gn=LOC101096742;refGene_tx=XM_023257097.1
B4	80790129	80790232
refGene_gn=LOC101096742;refGene_tx=XM_019834977.2
B4	80790399	80790431
refGene_gn=LOC101096742;refGene_tx=XM_019834975.2;refGene_gn=LOC101096742;refGene_tx=XM_019834976.2;refGene_gn=LOC101096742;refGene_tx=XM_019834978.2;refGene_gn=LOC101096742;refGene_tx=XM_023257097.1
B4	80790948	80791112
refGene_gn=LOC101096742;refGene_tx=XM_019834975.2;refGene_gn=LOC101096742;refGene_tx=XM_019834976.2;refGene_gn=LOC101096742;refGene_tx=XM_023257097.1
B4	80791295	80791300
refGene_gn=LOC101096742;refGene_tx=XM_019834978.2
B4	80796767	80797012
refGene_gn=LOC101098967;refGene_tx=XM_003988731.4
B4	80797308	80797340
refGene_gn=LOC101098967;refGene_tx=XM_003988731.4
B4	80797605	80797826
refGene_gn=LOC101098967;refGene_tx=XM_003988731.4

B4 80798079 80798205
refGene_gn=LOC101098967;refGene_tx=XM_003988731.4
B4 80798470 80798635
refGene_gn=LOC101098967;refGene_tx=XM_003988731.4
B4 80798849 80798945
refGene_gn=LOC101098967;refGene_tx=XM_003988731.4
B4 80799698 80799759
refGene_gn=LOC101098967;refGene_tx=XM_003988731.4
B4 80800548 80800757
refGene_gn=LOC101098967;refGene_tx=XM_003988731.4
B4 80801500 80801869
refGene_gn=LOC101098967;refGene_tx=XM_003988731.4
B4 80811130 80811499
refGene_gn=LOC101096995;refGene_tx=XM_011283950.3
B4 80812242 80812362
refGene_gn=LOC101096995;refGene_tx=XM_011283950.3
B4 80812364 80812468
refGene_gn=LOC101096995;refGene_tx=XM_011283950.3
B4 80813257 80813318
refGene_gn=LOC101096995;refGene_tx=XM_011283950.3
B4 80814072 80814168
refGene_gn=LOC101096995;refGene_tx=XM_011283950.3
B4 80814383 80814548
refGene_gn=LOC101096995;refGene_tx=XM_011283950.3
B4 80814808 80814934
refGene_gn=LOC101096995;refGene_tx=XM_011283950.3
B4 80815187 80815408
refGene_gn=LOC101096995;refGene_tx=XM_011283950.3
B4 80817012 80817044
refGene_gn=LOC101096995;refGene_tx=XM_011283950.3
B4 80817175 80817336
refGene_gn=LOC101096995;refGene_tx=XM_011283950.3
B4 80825002 80825190
refGene_gn=LOC101099222;refGene_tx=XM_003988732.4
B4 80825670 80825702
refGene_gn=LOC101099222;refGene_tx=XM_003988732.4
B4 80826385 80826606
refGene_gn=LOC101099222;refGene_tx=XM_003988732.4
B4 80826851 80826977
refGene_gn=LOC101099222;refGene_tx=XM_003988732.4
B4 80827242 80827407
refGene_gn=LOC101099222;refGene_tx=XM_003988732.4
B4 80827992 80828088
refGene_gn=LOC101099222;refGene_tx=XM_003988732.4
B4 80828218 80828279
refGene_gn=LOC101099222;refGene_tx=XM_003988732.4
B4 80829340 80829549
refGene_gn=LOC101099222;refGene_tx=XM_003988732.4
B4 80831074 80831458
refGene_gn=LOC101099222;refGene_tx=XM_003988732.4

B4 80845004 80845427
refGene_gn=LOC101099477;refGene_tx=XM_003988733.4
B4 80848119 80848328
refGene_gn=LOC101099477;refGene_tx=XM_003988733.4
B4 80849811 80849872
refGene_gn=LOC101099477;refGene_tx=XM_003988733.4
B4 80850462 80850558
refGene_gn=LOC101099477;refGene_tx=XM_003988733.4
B4 80851301 80851466
refGene_gn=LOC101099477;refGene_tx=XM_003988733.4
B4 80851701 80851827
refGene_gn=LOC101099477;refGene_tx=XM_003988733.4
B4 80852728 80852949
refGene_gn=LOC101099477;refGene_tx=XM_003988733.4
B4 80855415 80855447
refGene_gn=LOC101099477;refGene_tx=XM_003988733.4
B4 80856156 80856308
refGene_gn=LOC101099477;refGene_tx=XM_003988733.4
B4 80862523 80862717
refGene_gn=LOC101099731;refGene_tx=XM_003988734.4
B4 80863119 80863151
refGene_gn=LOC101099731;refGene_tx=XM_003988734.4
B4 80863872 80864093
refGene_gn=LOC101099731;refGene_tx=XM_003988734.4
B4 80864516 80864642
refGene_gn=LOC101099731;refGene_tx=XM_003988734.4
B4 80864903 80865068
refGene_gn=LOC101099731;refGene_tx=XM_003988734.4
B4 80865783 80865879
refGene_gn=LOC101099731;refGene_tx=XM_003988734.4
B4 80866008 80866069
refGene_gn=LOC101099731;refGene_tx=XM_003988734.4
B4 80866648 80866857
refGene_gn=LOC101099731;refGene_tx=XM_003988734.4
B4 80869093 80869513
refGene_gn=LOC101099731;refGene_tx=XM_003988734.4
B4 80882364 80882406 refGene_gn=KRT84;refGene_tx=XM_003988735.3
B4 80882407 80882557 refGene_gn=KRT84;refGene_tx=XM_003988735.3
B4 80882559 80882711 refGene_gn=KRT84;refGene_tx=XM_003988735.3
B4 80884359 80884391 refGene_gn=KRT84;refGene_tx=XM_003988735.3
B4 80884845 80885066 refGene_gn=KRT84;refGene_tx=XM_003988735.3
B4 80885171 80885297 refGene_gn=KRT84;refGene_tx=XM_003988735.3
B4 80885467 80885632 refGene_gn=KRT84;refGene_tx=XM_003988735.3
B4 80886557 80886653 refGene_gn=KRT84;refGene_tx=XM_003988735.3
B4 80887137 80887198 refGene_gn=KRT84;refGene_tx=XM_003988735.3
B4 80887801 80888010 refGene_gn=KRT84;refGene_tx=XM_003988735.3
B4 80889189 80889741 refGene_gn=KRT84;refGene_tx=XM_003988735.3
B4 80899069 80899311 refGene_gn=KRT82;refGene_tx=XM_003988811.3
B4 80899858 80899890 refGene_gn=KRT82;refGene_tx=XM_003988811.3
B4 80900113 80900334 refGene_gn=KRT82;refGene_tx=XM_003988811.3

B4 80901861 80901987 refGene_gn=KRT82;refGene_tx=XM_003988811.3
B4 80903750 80903915 refGene_gn=KRT82;refGene_tx=XM_003988811.3
B4 80904272 80904368 refGene_gn=KRT82;refGene_tx=XM_003988811.3
B4 80905049 80905110 refGene_gn=KRT82;refGene_tx=XM_003988811.3
B4 80907406 80907615 refGene_gn=KRT82;refGene_tx=XM_003988811.3
B4 80909604 80910009 refGene_gn=KRT82;refGene_tx=XM_003988811.3
B4 80913930 80914133
refGene_gn=LOC101097497;refGene_tx=XM_006933719.2
B4 80915413 80915448
refGene_gn=LOC101097497;refGene_tx=XM_006933719.2
B4 80916010 80916231
refGene_gn=LOC101097497;refGene_tx=XM_006933719.2
B4 80917134 80917260
refGene_gn=LOC101097497;refGene_tx=XM_006933719.2
B4 80918023 80918188
refGene_gn=LOC101097497;refGene_tx=XM_006933719.2
B4 80920241 80920337
refGene_gn=LOC101097497;refGene_tx=XM_006933719.2
B4 80921427 80921488
refGene_gn=LOC101097497;refGene_tx=XM_006933719.2
B4 80924209 80924418
refGene_gn=LOC101097497;refGene_tx=XM_006933719.2
B4 80926212 80926692
refGene_gn=LOC101097497;refGene_tx=XM_006933719.2
B4 80928816 80929055
refGene_gn=LOC101100226;refGene_tx=XM_003988736.4
B4 80931238 80931273
refGene_gn=LOC101100226;refGene_tx=XM_003988736.4
B4 80932621 80932842
refGene_gn=LOC101100226;refGene_tx=XM_003988736.4
B4 80932984 80933110
refGene_gn=LOC101100226;refGene_tx=XM_003988736.4
B4 80934999 80935164
refGene_gn=LOC101100226;refGene_tx=XM_003988736.4
B4 80936010 80936106
refGene_gn=LOC101100226;refGene_tx=XM_003988736.4
B4 80936461 80936522
refGene_gn=LOC101100226;refGene_tx=XM_003988736.4
B4 80937645 80937860
refGene_gn=LOC101100226;refGene_tx=XM_003988736.4
B4 80938375 80938873
refGene_gn=LOC101100226;refGene_tx=XM_003988736.4
B4 80957059 80957313
refGene_gn=LOC101100486;refGene_tx=XM_023257098.1
B4 80957645 80957900
refGene_gn=LOC105260741;refGene_tx=XM_023257099.1
B4 80958746 80958872
refGene_gn=LOC105260741;refGene_tx=XM_023257099.1
B4 80959361 80959526
refGene_gn=LOC105260741;refGene_tx=XM_023257099.1

B4 80959648 80959807
refGene_gn=LOC105260741;refGene_tx=XM_023257099.1
B4 80961047 80961587
refGene_gn=LOC105260741;refGene_tx=XM_023257099.1
B4 80977081 80977335
refGene_gn=LOC101100486;refGene_tx=XM_011279741.3
B4 80977446 80977481
refGene_gn=LOC101100486;refGene_tx=XM_011279741.3;refGene_gn=LOC101100486;refGene_tx=XM_023257098.1
B4 80977701 80977922
refGene_gn=LOC101100486;refGene_tx=XM_011279741.3;refGene_gn=LOC101100486;refGene_tx=XM_023257098.1
B4 80978779 80978905
refGene_gn=LOC101100486;refGene_tx=XM_011279741.3;refGene_gn=LOC101100486;refGene_tx=XM_023257098.1
B4 80979401 80979566
refGene_gn=LOC101100486;refGene_tx=XM_011279741.3;refGene_gn=LOC101100486;refGene_tx=XM_023257098.1
B4 80979687 80979783
refGene_gn=LOC101100486;refGene_tx=XM_011279741.3;refGene_gn=LOC101100486;refGene_tx=XM_023257098.1
B4 80979940 80980001
refGene_gn=LOC101100486;refGene_tx=XM_011279741.3;refGene_gn=LOC101100486;refGene_tx=XM_023257098.1
B4 80980348 80980563
refGene_gn=LOC101100486;refGene_tx=XM_011279741.3;refGene_gn=LOC101100486;refGene_tx=XM_023257098.1
B4 80981751 80982291
refGene_gn=LOC101100486;refGene_tx=XM_011279741.3;refGene_gn=LOC101100486;refGene_tx=XM_023257098.1
B4 80993388 80993642
refGene_gn=LOC105259896;refGene_tx=XM_023257100.1
B4 80993753 80993788
refGene_gn=LOC105259896;refGene_tx=XM_023257100.1
B4 80994008 80994229
refGene_gn=LOC105259896;refGene_tx=XM_023257100.1
B4 80995089 80995215
refGene_gn=LOC105259896;refGene_tx=XM_023257100.1
B4 80995711 80995876
refGene_gn=LOC105259896;refGene_tx=XM_023257100.1
B4 80995997 80996093
refGene_gn=LOC105259896;refGene_tx=XM_023257100.1
B4 80996250 80996311
refGene_gn=LOC105259896;refGene_tx=XM_023257100.1
B4 80996658 80996873
refGene_gn=LOC105259896;refGene_tx=XM_023257100.1
B4 80998064 80998604
refGene_gn=LOC105259896;refGene_tx=XM_023257100.1
B4 81013486 81013809 refGene_gn=KRT5;refGene_tx=XM_004001288.5
B4 81014272 81014307 refGene_gn=KRT5;refGene_tx=XM_004001288.5

B4 81015044 81015265 refGene_gn=KRT5;refGene_tx=XM_004001288.5
B4 81015510 81015636 refGene_gn=KRT5;refGene_tx=XM_004001288.5
B4 81016025 81016190 refGene_gn=KRT5;refGene_tx=XM_004001288.5
B4 81016773 81016869 refGene_gn=KRT5;refGene_tx=XM_004001288.5
B4 81016979 81017040 refGene_gn=KRT5;refGene_tx=XM_004001288.5
B4 81017831 81018046 refGene_gn=KRT5;refGene_tx=XM_004001288.5
B4 81018626 81019178 refGene_gn=KRT5;refGene_tx=XM_004001288.5
B4 81044518 81044730 refGene_gn=KRT71;refGene_tx=NM_001195239.1
B4 81045615 81045650 refGene_gn=KRT71;refGene_tx=NM_001195239.1
B4 81046226 81046447 refGene_gn=KRT71;refGene_tx=NM_001195239.1
B4 81047819 81047945 refGene_gn=KRT71;refGene_tx=NM_001195239.1
B4 81048146 81048311 refGene_gn=KRT71;refGene_tx=NM_001195239.1
B4 81048680 81048776 refGene_gn=KRT71;refGene_tx=NM_001195239.1
B4 81049300 81049361 refGene_gn=KRT71;refGene_tx=NM_001195239.1
B4 81050048 81050263 refGene_gn=KRT71;refGene_tx=NM_001195239.1
B4 81052967 81053411 refGene_gn=KRT71;refGene_tx=NM_001195239.1
B4 81064677 81064868 refGene_gn=KRT74;refGene_tx=NM_001278857.1
B4 81065808 81065843 refGene_gn=KRT74;refGene_tx=NM_001278857.1
B4 81066299 81066520 refGene_gn=KRT74;refGene_tx=NM_001278857.1
B4 81067983 81068109 refGene_gn=KRT74;refGene_tx=NM_001278857.1
B4 81069303 81069468 refGene_gn=KRT74;refGene_tx=NM_001278857.1
B4 81069943 81070039 refGene_gn=KRT74;refGene_tx=NM_001278857.1
B4 81070531 81070592 refGene_gn=KRT74;refGene_tx=NM_001278857.1
B4 81071039 81071254 refGene_gn=KRT74;refGene_tx=NM_001278857.1
B4 81071862 81072333 refGene_gn=KRT74;refGene_tx=NM_001278857.1
B4 81083164 81083358 refGene_gn=KRT72;refGene_tx=XM_011283953.3
B4 81084108 81084143 refGene_gn=KRT72;refGene_tx=XM_011283953.3
B4 81084774 81084995 refGene_gn=KRT72;refGene_tx=XM_011283953.3
B4 81087364 81087490 refGene_gn=KRT72;refGene_tx=XM_011283953.3
B4 81088069 81088234 refGene_gn=KRT72;refGene_tx=XM_011283953.3
B4 81088920 81089016 refGene_gn=KRT72;refGene_tx=XM_011283953.3
B4 81089369 81089430 refGene_gn=KRT72;refGene_tx=XM_011283953.3
B4 81092310 81092525 refGene_gn=KRT72;refGene_tx=XM_011283953.3
B4 81094214 81094673 refGene_gn=KRT72;refGene_tx=XM_011283953.3
B4 81101786 81102043
refGene_gn=KRT73;refGene_tx=NM_001290235.1;refGene_gn=KRT73;refGene_tx
=XM_019833854.1
B4 81102818 81102853
refGene_gn=KRT73;refGene_tx=NM_001290235.1;refGene_gn=KRT73;refGene_tx
=XM_019833854.1
B4 81104081 81104302
refGene_gn=KRT73;refGene_tx=NM_001290235.1;refGene_gn=KRT73;refGene_tx
=XM_019833854.1
B4 81104696 81104822
refGene_gn=KRT73;refGene_tx=NM_001290235.1;refGene_gn=KRT73;refGene_tx
=XM_019833854.1
B4 81107312 81107477
refGene_gn=KRT73;refGene_tx=NM_001290235.1;refGene_gn=KRT73;refGene_tx
=XM_019833854.1
B4 81108193 81108289

refGene_gn=KRT73;refGene_tx=NM_001290235.1;refGene_gn=KRT73;refGene_tx=XM_019833854.1
B4 81109156 81109217 refGene_gn=KRT73;refGene_tx=NM_001290235.1;refGene_gn=KRT73;refGene_tx=XM_019833854.1
B4 81110013 81110228 refGene_gn=KRT73;refGene_tx=NM_001290235.1;refGene_gn=KRT73;refGene_tx=XM_019833854.1
B4 81111722 81112169 refGene_gn=KRT73;refGene_tx=NM_001290235.1;refGene_gn=KRT73;refGene_tx=XM_019833854.1
B4 81132923 81133390 refGene_gn=KRT2;refGene_tx=XM_003988739.3
B4 81133484 81133519 refGene_gn=KRT2;refGene_tx=XM_003988739.3
B4 81134730 81134951 refGene_gn=KRT2;refGene_tx=XM_003988739.3
B4 81135761 81135887 refGene_gn=KRT2;refGene_tx=XM_003988739.3
B4 81136189 81136354 refGene_gn=KRT2;refGene_tx=XM_003988739.3
B4 81137025 81137121 refGene_gn=KRT2;refGene_tx=XM_003988739.3
B4 81137360 81137421 refGene_gn=KRT2;refGene_tx=XM_003988739.3
B4 81137767 81137982 refGene_gn=KRT2;refGene_tx=XM_003988739.3
B4 81138974 81139589 refGene_gn=KRT2;refGene_tx=XM_003988739.3
B4 81161181 81161546 refGene_gn=KRT1;refGene_tx=XM_003988740.4
B4 81161637 81161672 refGene_gn=KRT1;refGene_tx=XM_003988740.4
B4 81162187 81162408 refGene_gn=KRT1;refGene_tx=XM_003988740.4
B4 81162980 81163106 refGene_gn=KRT1;refGene_tx=XM_003988740.4
B4 81163238 81163403 refGene_gn=KRT1;refGene_tx=XM_003988740.4
B4 81163566 81163662 refGene_gn=KRT1;refGene_tx=XM_003988740.4
B4 81164070 81164131 refGene_gn=KRT1;refGene_tx=XM_003988740.4
B4 81164444 81164659 refGene_gn=KRT1;refGene_tx=XM_003988740.4
B4 81165593 81166217 refGene_gn=KRT1;refGene_tx=XM_003988740.4
B4 81175759 81176019 refGene_gn=LOC101080303;refGene_tx=XM_003988741.4
B4 81176502 81176537 refGene_gn=LOC101080303;refGene_tx=XM_003988741.4
B4 81176957 81177178 refGene_gn=LOC101080303;refGene_tx=XM_003988741.4
B4 81177293 81177419 refGene_gn=LOC101080303;refGene_tx=XM_003988741.4
B4 81179135 81179300 refGene_gn=LOC101080303;refGene_tx=XM_003988741.4
B4 81180255 81180351 refGene_gn=LOC101080303;refGene_tx=XM_003988741.4
B4 81180879 81180940 refGene_gn=LOC101080303;refGene_tx=XM_003988741.4
B4 81182035 81182250 refGene_gn=LOC101080303;refGene_tx=XM_003988741.4
B4 81186133 81186679 refGene_gn=LOC101080303;refGene_tx=XM_003988741.4
B4 81195543 81195776 refGene_gn=LOC101098240;refGene_tx=XM_023257759.1
B4 81196073 81196108

refGene_gn=LOC101098240;refGene_tx=XM_023257759.1
B4 81196747 81196968
refGene_gn=LOC101098240;refGene_tx=XM_023257759.1
B4 81197578 81197704
refGene_gn=LOC101098240;refGene_tx=XM_023257759.1
B4 81198809 81198963
refGene_gn=LOC101098240;refGene_tx=XM_023257759.1
B4 81199183 81199296
refGene_gn=LOC101098240;refGene_tx=XM_023257759.1
B4 81199936 81199997
refGene_gn=LOC101098240;refGene_tx=XM_023257759.1
B4 81200969 81201190
refGene_gn=LOC101098240;refGene_tx=XM_023257759.1
B4 81222508 81222663
refGene_gn=LOC101098240;refGene_tx=XM_023257759.1
B4 81226431 81226444
refGene_gn=LOC101098240;refGene_tx=XM_023257759.1
B4 81237054 81237395
refGene_gn=LOC101098725;refGene_tx=XM_023257797.1
B4 81237816 81237851
refGene_gn=LOC101098725;refGene_tx=XM_023257797.1
B4 81239494 81239715
refGene_gn=LOC101098725;refGene_tx=XM_023257797.1
B4 81240574 81240700
refGene_gn=LOC101098725;refGene_tx=XM_023257797.1
B4 81240809 81240956
refGene_gn=LOC101098725;refGene_tx=XM_023257797.1
B4 81241574 81241670
refGene_gn=LOC101098725;refGene_tx=XM_023257797.1
B4 81242360 81242421
refGene_gn=LOC101098725;refGene_tx=XM_023257797.1
B4 81243828 81244049
refGene_gn=LOC101098725;refGene_tx=XM_023257797.1
B4 81245606 81246290
refGene_gn=LOC101098725;refGene_tx=XM_023257797.1
B4 81257940 81258317 refGene_gn=KRT3;refGene_tx=XM_023257101.1
B4 81258805 81258840 refGene_gn=KRT3;refGene_tx=XM_023257101.1
B4 81259220 81259441 refGene_gn=KRT3;refGene_tx=XM_023257101.1
B4 81259772 81259898 refGene_gn=KRT3;refGene_tx=XM_023257101.1
B4 81260321 81260486 refGene_gn=KRT3;refGene_tx=XM_023257101.1
B4 81260767 81260863 refGene_gn=KRT3;refGene_tx=XM_023257101.1
B4 81261212 81261273 refGene_gn=KRT3;refGene_tx=XM_023257101.1
B4 81261807 81262028 refGene_gn=KRT3;refGene_tx=XM_023257101.1
B4 81263065 81263725 refGene_gn=KRT3;refGene_tx=XM_023257101.1
B4 81275742 81275930 refGene_gn=KRT4;refGene_tx=XM_003988742.4
B4 81276029 81276064 refGene_gn=KRT4;refGene_tx=XM_003988742.4
B4 81276316 81276537 refGene_gn=KRT4;refGene_tx=XM_003988742.4
B4 81276956 81277082 refGene_gn=KRT4;refGene_tx=XM_003988742.4
B4 81277341 81277506 refGene_gn=KRT4;refGene_tx=XM_003988742.4
B4 81278023 81278119 refGene_gn=KRT4;refGene_tx=XM_003988742.4

B4 81278505 81278566 refGene_gn=KRT4;refGene_tx=XM_003988742.4
B4 81279512 81279727 refGene_gn=KRT4;refGene_tx=XM_003988742.4
B4 81281556 81282018 refGene_gn=KRT4;refGene_tx=XM_003988742.4
B4 81294084 81294287 refGene_gn=KRT79;refGene_tx=XM_003988743.4
B4 81294563 81294598 refGene_gn=KRT79;refGene_tx=XM_003988743.4
B4 81294926 81295041 refGene_gn=KRT79;refGene_tx=XM_019834983.2
B4 81295149 81295370
refGene_gn=KRT79;refGene_tx=XM_003988743.4;refGene_gn=KRT79;refGene_tx
=XM_019834983.2
B4 81295999 81296125
refGene_gn=KRT79;refGene_tx=XM_003988743.4;refGene_gn=KRT79;refGene_tx
=XM_019834983.2
B4 81296287 81296452
refGene_gn=KRT79;refGene_tx=XM_003988743.4;refGene_gn=KRT79;refGene_tx
=XM_019834983.2
B4 81300572 81300668
refGene_gn=KRT79;refGene_tx=XM_003988743.4;refGene_gn=KRT79;refGene_tx
=XM_019834983.2
B4 81300792 81300853
refGene_gn=KRT79;refGene_tx=XM_003988743.4;refGene_gn=KRT79;refGene_tx
=XM_019834983.2
B4 81302158 81302379
refGene_gn=KRT79;refGene_tx=XM_003988743.4;refGene_gn=KRT79;refGene_tx
=XM_019834983.2
B4 81305097 81305574
refGene_gn=KRT79;refGene_tx=XM_003988743.4;refGene_gn=KRT79;refGene_tx
=XM_019834983.2
B4 81311563 81311835 refGene_gn=KRT78;refGene_tx=XM_003988744.3
B4 81311916 81311951 refGene_gn=KRT78;refGene_tx=XM_003988744.3
B4 81312227 81312448 refGene_gn=KRT78;refGene_tx=XM_003988744.3
B4 81315450 81315576 refGene_gn=KRT78;refGene_tx=XM_003988744.3
B4 81315930 81316095 refGene_gn=KRT78;refGene_tx=XM_003988744.3
B4 81317194 81317290 refGene_gn=KRT78;refGene_tx=XM_003988744.3
B4 81317767 81317828 refGene_gn=KRT78;refGene_tx=XM_003988744.3
B4 81318872 81319087 refGene_gn=KRT78;refGene_tx=XM_003988744.3
B4 81319747 81320113 refGene_gn=KRT78;refGene_tx=XM_003988744.3
B4 81356700 81356891 refGene_gn=KRT8;refGene_tx=XM_003988745.4
B4 81357196 81357255 refGene_gn=KRT8;refGene_tx=XM_003988745.4
B4 81357400 81357621 refGene_gn=KRT8;refGene_tx=XM_003988745.4
B4 81358316 81358442 refGene_gn=KRT8;refGene_tx=XM_003988745.4
B4 81358714 81358879 refGene_gn=KRT8;refGene_tx=XM_003988745.4
B4 81359500 81359596 refGene_gn=KRT8;refGene_tx=XM_003988745.4
B4 81360045 81360106 refGene_gn=KRT8;refGene_tx=XM_003988745.4
B4 81360721 81360930 refGene_gn=KRT8;refGene_tx=XM_003988745.4
B4 81363619 81363961 refGene_gn=KRT8;refGene_tx=XM_003988745.4
B4 81398359 81398776
refGene_gn=KRT18;refGene_tx=XM_003988746.3;refGene_gn=KRT18;refGene_tx
=XM_019834984.2
B4 81399540 81399623 refGene_gn=KRT18;refGene_tx=XM_003988746.3
B4 81399979 81400136 refGene_gn=KRT18;refGene_tx=XM_003988746.3

B4 81400689 81400854
refGene_gn=KRT18;refGene_tx=XM_003988746.3;refGene_gn=KRT18;refGene_tx=XM_019834984.2
B4 81400960 81401086
refGene_gn=KRT18;refGene_tx=XM_003988746.3;refGene_gn=KRT18;refGene_tx=XM_019834984.2
B4 81401407 81401631
refGene_gn=KRT18;refGene_tx=XM_003988746.3;refGene_gn=KRT18;refGene_tx=XM_019834984.2
B4 81402024 81402145
refGene_gn=KRT18;refGene_tx=XM_003988746.3;refGene_gn=KRT18;refGene_tx=XM_019834984.2
B4 81446429 81446442
refGene_gn=EIF4B;refGene_tx=XM_003988747.3;refGene_gn=EIF4B;refGene_tx=XM_006933722.2
B4 81454821 81454959
refGene_gn=EIF4B;refGene_tx=XM_003988747.3;refGene_gn=EIF4B;refGene_tx=XM_006933722.2
B4 81455989 81456198
refGene_gn=EIF4B;refGene_tx=XM_003988747.3;refGene_gn=EIF4B;refGene_tx=XM_006933722.2
B4 81459165 81459282
refGene_gn=EIF4B;refGene_tx=XM_003988747.3;refGene_gn=EIF4B;refGene_tx=XM_006933722.2
B4 81461420 81461475
refGene_gn=EIF4B;refGene_tx=XM_003988747.3;refGene_gn=EIF4B;refGene_tx=XM_006933722.2
B4 81462124 81462259
refGene_gn=EIF4B;refGene_tx=XM_003988747.3;refGene_gn=EIF4B;refGene_tx=XM_006933722.2
B4 81465701 81465839
refGene_gn=EIF4B;refGene_tx=XM_003988747.3;refGene_gn=EIF4B;refGene_tx=XM_006933722.2
B4 81465940 81466114
refGene_gn=EIF4B;refGene_tx=XM_003988747.3;refGene_gn=EIF4B;refGene_tx=XM_006933722.2
B4 81469100 81469329
refGene_gn=EIF4B;refGene_tx=XM_003988747.3;refGene_gn=EIF4B;refGene_tx=XM_006933722.2
B4 81470513 81470626
refGene_gn=EIF4B;refGene_tx=XM_003988747.3;refGene_gn=EIF4B;refGene_tx=XM_006933722.2
B4 81471844 81472058
refGene_gn=EIF4B;refGene_tx=XM_003988747.3;refGene_gn=EIF4B;refGene_tx=XM_006933722.2
B4 81472678 81472734
refGene_gn=EIF4B;refGene_tx=XM_003988747.3;refGene_gn=EIF4B;refGene_tx=XM_006933722.2
B4 81473299 81473405
refGene_gn=EIF4B;refGene_tx=XM_003988747.3;refGene_gn=EIF4B;refGene_tx

=XM_006933722.2
B4	81473689 81473762
refGene_gn=EIF4B;refGene_tx=XM_003988747.3;refGene_gn=EIF4B;refGene_tx
=XM_006933722.2
B4	81474156 81474237
refGene_gn=EIF4B;refGene_tx=XM_003988747.3;refGene_gn=EIF4B;refGene_tx
=XM_006933722.2
B4	81478108 81478183 refGene_gn=TNS2;refGene_tx=XM_006933723.4
B4	81480552 81480627 refGene_gn=TNS2;refGene_tx=XM_006933725.3
B4	81483307 81483382 refGene_gn=TNS2;refGene_tx=XM_003988749.5
B4	81485007 81485116
refGene_gn=TNS2;refGene_tx=XM_003988749.5;refGene_gn=TNS2;refGene_tx=X
M_006933723.4;refGene_gn=TNS2;refGene_tx=XM_006933725.3
B4	81485613 81485651
refGene_gn=TNS2;refGene_tx=XM_003988749.5;refGene_gn=TNS2;refGene_tx=X
M_006933723.4;refGene_gn=TNS2;refGene_tx=XM_006933725.3
B4	81486535 81486574
refGene_gn=TNS2;refGene_tx=XM_003988749.5;refGene_gn=TNS2;refGene_tx=X
M_006933723.4;refGene_gn=TNS2;refGene_tx=XM_006933725.3
B4	81486889 81486928
refGene_gn=TNS2;refGene_tx=XM_003988749.5;refGene_gn=TNS2;refGene_tx=X
M_006933723.4;refGene_gn=TNS2;refGene_tx=XM_006933725.3
B4	81487081 81487131
refGene_gn=TNS2;refGene_tx=XM_003988749.5;refGene_gn=TNS2;refGene_tx=X
M_006933723.4;refGene_gn=TNS2;refGene_tx=XM_006933725.3
B4	81487336 81487508
refGene_gn=TNS2;refGene_tx=XM_003988749.5;refGene_gn=TNS2;refGene_tx=X
M_006933723.4;refGene_gn=TNS2;refGene_tx=XM_006933725.3
B4	81487791 81487842
refGene_gn=TNS2;refGene_tx=XM_003988749.5;refGene_gn=TNS2;refGene_tx=X
M_006933723.4;refGene_gn=TNS2;refGene_tx=XM_006933725.3
B4	81488153 81488276
refGene_gn=TNS2;refGene_tx=XM_003988749.5;refGene_gn=TNS2;refGene_tx=X
M_006933723.4;refGene_gn=TNS2;refGene_tx=XM_006933725.3
B4	81488363 81488428
refGene_gn=TNS2;refGene_tx=XM_003988749.5;refGene_gn=TNS2;refGene_tx=X
M_006933723.4;refGene_gn=TNS2;refGene_tx=XM_006933725.3
B4	81489284 81489368
refGene_gn=TNS2;refGene_tx=XM_003988749.5;refGene_gn=TNS2;refGene_tx=X
M_006933723.4;refGene_gn=TNS2;refGene_tx=XM_006933725.3;refGene_gn=TNS
2;refGene_tx=XM_006933726.3
B4	81489831 81489944
refGene_gn=TNS2;refGene_tx=XM_003988749.5;refGene_gn=TNS2;refGene_tx=X
M_006933723.4;refGene_gn=TNS2;refGene_tx=XM_006933725.3;refGene_gn=TNS
2;refGene_tx=XM_006933726.3
B4	81490037 81490098
refGene_gn=TNS2;refGene_tx=XM_003988749.5;refGene_gn=TNS2;refGene_tx=X
M_006933723.4;refGene_gn=TNS2;refGene_tx=XM_006933725.3;refGene_gn=TNS
2;refGene_tx=XM_006933726.3
B4	81490366 81490442

refGene_gn=TNS2;refGene_tx=XM_003988749.5;refGene_gn=TNS2;refGene_tx=XM_006933723.4;refGene_gn=TNS2;refGene_tx=XM_006933725.3;refGene_gn=TNS2;refGene_tx=XM_006933726.3
B4 81490695 81490825
refGene_gn=TNS2;refGene_tx=XM_003988749.5;refGene_gn=TNS2;refGene_tx=XM_006933723.4;refGene_gn=TNS2;refGene_tx=XM_006933725.3;refGene_gn=TNS2;refGene_tx=XM_006933726.3
B4 81490905 81490971
refGene_gn=TNS2;refGene_tx=XM_003988749.5;refGene_gn=TNS2;refGene_tx=XM_006933723.4;refGene_gn=TNS2;refGene_tx=XM_006933725.3;refGene_gn=TNS2;refGene_tx=XM_006933726.3
B4 81491045 81491159
refGene_gn=TNS2;refGene_tx=XM_003988749.5;refGene_gn=TNS2;refGene_tx=XM_006933723.4;refGene_gn=TNS2;refGene_tx=XM_006933725.3;refGene_gn=TNS2;refGene_tx=XM_006933726.3
B4 81491345 81492557
refGene_gn=TNS2;refGene_tx=XM_003988749.5;refGene_gn=TNS2;refGene_tx=XM_006933723.4;refGene_gn=TNS2;refGene_tx=XM_006933725.3;refGene_gn=TNS2;refGene_tx=XM_006933726.3
B4 81492697 81492848
refGene_gn=TNS2;refGene_tx=XM_003988749.5;refGene_gn=TNS2;refGene_tx=XM_006933723.4;refGene_gn=TNS2;refGene_tx=XM_006933725.3;refGene_gn=TNS2;refGene_tx=XM_006933726.3
B4 81492964 81493554
refGene_gn=TNS2;refGene_tx=XM_003988749.5;refGene_gn=TNS2;refGene_tx=XM_006933723.4;refGene_gn=TNS2;refGene_tx=XM_006933725.3;refGene_gn=TNS2;refGene_tx=XM_006933726.3
B4 81493669 81493759
refGene_gn=TNS2;refGene_tx=XM_003988749.5;refGene_gn=TNS2;refGene_tx=XM_006933723.4;refGene_gn=TNS2;refGene_tx=XM_006933725.3;refGene_gn=TNS2;refGene_tx=XM_006933726.3
B4 81494102 81494228
refGene_gn=TNS2;refGene_tx=XM_003988749.5;refGene_gn=TNS2;refGene_tx=XM_006933723.4;refGene_gn=TNS2;refGene_tx=XM_006933725.3;refGene_gn=TNS2;refGene_tx=XM_006933726.3
B4 81494450 81494543
refGene_gn=TNS2;refGene_tx=XM_003988749.5;refGene_gn=TNS2;refGene_tx=XM_006933723.4;refGene_gn=TNS2;refGene_tx=XM_006933725.3;refGene_gn=TNS2;refGene_tx=XM_006933726.3
B4 81494645 81494723
refGene_gn=TNS2;refGene_tx=XM_003988749.5;refGene_gn=TNS2;refGene_tx=XM_006933723.4;refGene_gn=TNS2;refGene_tx=XM_006933725.3;refGene_gn=TNS2;refGene_tx=XM_006933726.3
B4 81494891 81494969
refGene_gn=TNS2;refGene_tx=XM_003988749.5;refGene_gn=TNS2;refGene_tx=XM_006933723.4;refGene_gn=TNS2;refGene_tx=XM_006933725.3;refGene_gn=TNS2;refGene_tx=XM_006933726.3
B4 81495371 81495540
refGene_gn=TNS2;refGene_tx=XM_003988749.5;refGene_gn=TNS2;refGene_tx=XM_006933723.4;refGene_gn=TNS2;refGene_tx=XM_006933725.3;refGene_gn=TNS

2;refGene_tx=XM_006933726.3
B4 81495621 81495690
refGene_gn=TNS2;refGene_tx=XM_003988749.5;refGene_gn=TNS2;refGene_tx=XM_006933723.4;refGene_gn=TNS2;refGene_tx=XM_006933725.3;refGene_gn=TNS2;refGene_tx=XM_006933726.3
B4 81495837 81495867
refGene_gn=TNS2;refGene_tx=XM_003988749.5;refGene_gn=TNS2;refGene_tx=XM_006933723.4;refGene_gn=TNS2;refGene_tx=XM_006933725.3;refGene_gn=TNS2;refGene_tx=XM_006933726.3
B4 81495998 81496137
refGene_gn=TNS2;refGene_tx=XM_003988749.5;refGene_gn=TNS2;refGene_tx=XM_006933723.4;refGene_gn=TNS2;refGene_tx=XM_006933725.3;refGene_gn=TNS2;refGene_tx=XM_006933726.3
B4 81498062 81498197
refGene_gn=SPRYD3;refGene_tx=XM_003988750.5;refGene_gn=SPRYD3;refGene_tx=XM_011283954.3
B4 81498533 81498706
refGene_gn=SPRYD3;refGene_tx=XM_003988750.5;refGene_gn=SPRYD3;refGene_tx=XM_011283954.3
B4 81498790 81498910
refGene_gn=SPRYD3;refGene_tx=XM_003988750.5;refGene_gn=SPRYD3;refGene_tx=XM_011283954.3
B4 81499589 81499647
refGene_gn=SPRYD3;refGene_tx=XM_003988750.5;refGene_gn=SPRYD3;refGene_tx=XM_011283954.3
B4 81500103 81500253
refGene_gn=SPRYD3;refGene_tx=XM_003988750.5;refGene_gn=SPRYD3;refGene_tx=XM_011283954.3
B4 81506399 81506585
refGene_gn=SPRYD3;refGene_tx=XM_003988750.5;refGene_gn=SPRYD3;refGene_tx=XM_011283954.3
B4 81507422 81507558
refGene_gn=SPRYD3;refGene_tx=XM_003988750.5;refGene_gn=SPRYD3;refGene_tx=XM_011283954.3
B4 81507877 81508002
refGene_gn=SPRYD3;refGene_tx=XM_003988750.5;refGene_gn=SPRYD3;refGene_tx=XM_011283954.3
B4 81508506 81508582
refGene_gn=SPRYD3;refGene_tx=XM_003988750.5;refGene_gn=SPRYD3;refGene_tx=XM_011283954.3
B4 81509568 81509715
refGene_gn=SPRYD3;refGene_tx=XM_003988750.5;refGene_gn=SPRYD3;refGene_tx=XM_011283954.3
B4 81509836 81509838 refGene_gn=SPRYD3;refGene_tx=XM_011283954.3
B4 81511420 81511443 refGene_gn=SPRYD3;refGene_tx=XM_003988750.5
B4 81523957 81524285 refGene_gn=IGFBP6;refGene_tx=XM_023257102.1
B4 81526278 81526418 refGene_gn=IGFBP6;refGene_tx=XM_023257102.1
B4 81526603 81526723 refGene_gn=IGFBP6;refGene_tx=XM_023257102.1
B4 81527518 81527641 refGene_gn=IGFBP6;refGene_tx=XM_023257102.1
B4 81529161 81529237

refGene_gn=SOAT2;refGene_tx=XM_003988817.5;refGene_gn=SOAT2;refGene_tx
=XM_006933728.3
B4 81529713 81529769
refGene_gn=SOAT2;refGene_tx=XM_003988817.5;refGene_gn=SOAT2;refGene_tx
=XM_006933728.3
B4 81530293 81530430
refGene_gn=SOAT2;refGene_tx=XM_003988817.5;refGene_gn=SOAT2;refGene_tx
=XM_006933728.3
B4 81530784 81530844
refGene_gn=SOAT2;refGene_tx=XM_003988817.5;refGene_gn=SOAT2;refGene_tx
=XM_006933728.3
B4 81531131 81531239
refGene_gn=SOAT2;refGene_tx=XM_003988817.5;refGene_gn=SOAT2;refGene_tx
=XM_006933728.3
B4 81532703 81532980
refGene_gn=SOAT2;refGene_tx=XM_003988817.5;refGene_gn=SOAT2;refGene_tx
=XM_006933728.3
B4 81533411 81533481
refGene_gn=SOAT2;refGene_tx=XM_003988817.5;refGene_gn=SOAT2;refGene_tx
=XM_006933728.3
B4 81534249 81534334
refGene_gn=SOAT2;refGene_tx=XM_003988817.5;refGene_gn=SOAT2;refGene_tx
=XM_006933728.3
B4 81534839 81534885
refGene_gn=SOAT2;refGene_tx=XM_003988817.5;refGene_gn=SOAT2;refGene_tx
=XM_006933728.3
B4 81535092 81535222
refGene_gn=SOAT2;refGene_tx=XM_003988817.5;refGene_gn=SOAT2;refGene_tx
=XM_006933728.3
B4 81536186 81536284
refGene_gn=SOAT2;refGene_tx=XM_003988817.5;refGene_gn=SOAT2;refGene_tx
=XM_006933728.3
B4 81536712 81536811
refGene_gn=SOAT2;refGene_tx=XM_003988817.5;refGene_gn=SOAT2;refGene_tx
=XM_006933728.3
B4 81537316 81537452
refGene_gn=SOAT2;refGene_tx=XM_003988817.5;refGene_gn=SOAT2;refGene_tx
=XM_006933728.3
B4 81537946 81538092
refGene_gn=SOAT2;refGene_tx=XM_003988817.5;refGene_gn=SOAT2;refGene_tx
=XM_006933728.3
B4 81538293 81538344 refGene_gn=SOAT2;refGene_tx=XM_003988817.5
B4 81554446 81554893
refGene_gn=LOC101099478;refGene_tx=XM_019834986.2
B4 81556267 81556441 refGene_gn=CSAD;refGene_tx=XM_019834985.2
B4 81557333 81557423 refGene_gn=CSAD;refGene_tx=XM_019834985.2
B4 81557571 81557623 refGene_gn=CSAD;refGene_tx=XM_019834985.2
B4 81557828 81558028 refGene_gn=CSAD;refGene_tx=XM_019834985.2
B4 81558472 81558554 refGene_gn=CSAD;refGene_tx=XM_019834985.2
B4 81558815 81558880 refGene_gn=CSAD;refGene_tx=XM_019834985.2

B4 81558956 81559073 refGene_gn=CSAD;refGene_tx=XM_019834985.2
B4 81567307 81567362 refGene_gn=CSAD;refGene_tx=XM_019834985.2
B4 81567665 81567745 refGene_gn=CSAD;refGene_tx=XM_019834985.2
B4 81568226 81568342 refGene_gn=CSAD;refGene_tx=XM_019834985.2
B4 81568771 81568878 refGene_gn=CSAD;refGene_tx=XM_019834985.2
B4 81569236 81569327 refGene_gn=CSAD;refGene_tx=XM_019834985.2
B4 81569409 81569536 refGene_gn=CSAD;refGene_tx=XM_019834985.2
B4 81569797 81569934 refGene_gn=CSAD;refGene_tx=XM_019834985.2
B4 81570180 81570310 refGene_gn=CSAD;refGene_tx=XM_019834985.2
B4 81587398 81587407 refGene_gn=ZNF740;refGene_tx=XM_003988752.5
B4 81587598 81587607 refGene_gn=ZNF740;refGene_tx=XM_006933729.3
B4 81590025 81590175
refGene_gn=ZNF740;refGene_tx=XM_003988752.5;refGene_gn=ZNF740;refGene_tx=XM_006933729.3
B4 81590523 81590613
refGene_gn=ZNF740;refGene_tx=XM_003988752.5;refGene_gn=ZNF740;refGene_tx=XM_006933729.3
B4 81591061 81591185
refGene_gn=ZNF740;refGene_tx=XM_003988752.5;refGene_gn=ZNF740;refGene_tx=XM_006933729.3
B4 81591495 81591614
refGene_gn=ZNF740;refGene_tx=XM_003988752.5;refGene_gn=ZNF740;refGene_tx=XM_006933729.3
B4 81592582 81592672
refGene_gn=ZNF740;refGene_tx=XM_003988752.5;refGene_gn=ZNF740;refGene_tx=XM_006933729.3
B4 81596578 81596659
refGene_gn=ITGB7;refGene_tx=XM_006933733.4;refGene_gn=ITGB7;refGene_tx=XM_019834987.2;refGene_gn=ITGB7;refGene_tx=XM_023257103.1;refGene_gn=ITGB7;refGene_tx=XM_023257104.1;refGene_gn=ITGB7;refGene_tx=XM_023257105.1;refGene_gn=ITGB7;refGene_tx=XM_023257106.1
B4 81596927 81597085
refGene_gn=ITGB7;refGene_tx=XM_006933733.4;refGene_gn=ITGB7;refGene_tx=XM_019834987.2;refGene_gn=ITGB7;refGene_tx=XM_023257103.1;refGene_gn=ITGB7;refGene_tx=XM_023257104.1;refGene_gn=ITGB7;refGene_tx=XM_023257105.1;refGene_gn=ITGB7;refGene_tx=XM_023257106.1
B4 81597413 81597622
refGene_gn=ITGB7;refGene_tx=XM_006933733.4;refGene_gn=ITGB7;refGene_tx=XM_019834987.2;refGene_gn=ITGB7;refGene_tx=XM_023257103.1;refGene_gn=ITGB7;refGene_tx=XM_023257104.1;refGene_gn=ITGB7;refGene_tx=XM_023257105.1
B4 81597762 81597982
refGene_gn=ITGB7;refGene_tx=XM_006933733.4;refGene_gn=ITGB7;refGene_tx=XM_019834987.2;refGene_gn=ITGB7;refGene_tx=XM_023257103.1;refGene_gn=ITGB7;refGene_tx=XM_023257104.1;refGene_gn=ITGB7;refGene_tx=XM_023257105.1
B4 81598222 81598446
refGene_gn=ITGB7;refGene_tx=XM_006933733.4;refGene_gn=ITGB7;refGene_tx=XM_019834987.2;refGene_gn=ITGB7;refGene_tx=XM_023257103.1;refGene_gn=ITGB7;refGene_tx=XM_023257104.1;refGene_gn=ITGB7;refGene_tx=XM_0232571

05.1;refGene_gn=ITGB7;refGene_tx=XM_023257106.1
B4 81598808 81599002
refGene_gn=ITGB7;refGene_tx=XM_006933733.4;refGene_gn=ITGB7;refGene_tx=XM_019834987.2;refGene_gn=ITGB7;refGene_tx=XM_023257103.1;refGene_gn=ITGB7;refGene_tx=XM_023257104.1;refGene_gn=ITGB7;refGene_tx=XM_023257105.1;refGene_gn=ITGB7;refGene_tx=XM_023257106.1
B4 81599307 81599454
refGene_gn=ITGB7;refGene_tx=XM_006933733.4;refGene_gn=ITGB7;refGene_tx=XM_019834987.2;refGene_gn=ITGB7;refGene_tx=XM_023257103.1;refGene_gn=ITGB7;refGene_tx=XM_023257104.1;refGene_gn=ITGB7;refGene_tx=XM_023257105.1;refGene_gn=ITGB7;refGene_tx=XM_023257106.1
B4 81600452 81600542
refGene_gn=ITGB7;refGene_tx=XM_006933733.4;refGene_gn=ITGB7;refGene_tx=XM_019834987.2;refGene_gn=ITGB7;refGene_tx=XM_023257103.1;refGene_gn=ITGB7;refGene_tx=XM_023257104.1;refGene_gn=ITGB7;refGene_tx=XM_023257105.1;refGene_gn=ITGB7;refGene_tx=XM_023257106.1
B4 81600707 81600803
refGene_gn=ITGB7;refGene_tx=XM_006933733.4;refGene_gn=ITGB7;refGene_tx=XM_019834987.2;refGene_gn=ITGB7;refGene_tx=XM_023257103.1;refGene_gn=ITGB7;refGene_tx=XM_023257104.1;refGene_gn=ITGB7;refGene_tx=XM_023257105.1;refGene_gn=ITGB7;refGene_tx=XM_023257106.1
B4 81601143 81601302
refGene_gn=ITGB7;refGene_tx=XM_006933733.4;refGene_gn=ITGB7;refGene_tx=XM_019834987.2;refGene_gn=ITGB7;refGene_tx=XM_023257103.1;refGene_gn=ITGB7;refGene_tx=XM_023257104.1;refGene_gn=ITGB7;refGene_tx=XM_023257105.1;refGene_gn=ITGB7;refGene_tx=XM_023257106.1
B4 81601692 81601934
refGene_gn=ITGB7;refGene_tx=XM_006933733.4;refGene_gn=ITGB7;refGene_tx=XM_019834987.2;refGene_gn=ITGB7;refGene_tx=XM_023257103.1;refGene_gn=ITGB7;refGene_tx=XM_023257104.1;refGene_gn=ITGB7;refGene_tx=XM_023257105.1;refGene_gn=ITGB7;refGene_tx=XM_023257106.1
B4 81602654 81602825
refGene_gn=ITGB7;refGene_tx=XM_006933733.4;refGene_gn=ITGB7;refGene_tx=XM_019834987.2;refGene_gn=ITGB7;refGene_tx=XM_023257103.1;refGene_gn=ITGB7;refGene_tx=XM_023257104.1;refGene_gn=ITGB7;refGene_tx=XM_023257105.1;refGene_gn=ITGB7;refGene_tx=XM_023257106.1
B4 81602917 81603119
refGene_gn=ITGB7;refGene_tx=XM_006933733.4;refGene_gn=ITGB7;refGene_tx=XM_019834987.2;refGene_gn=ITGB7;refGene_tx=XM_023257103.1;refGene_gn=ITGB7;refGene_tx=XM_023257104.1;refGene_gn=ITGB7;refGene_tx=XM_023257105.1;refGene_gn=ITGB7;refGene_tx=XM_023257106.1
B4 81604818 81605019
refGene_gn=ITGB7;refGene_tx=XM_006933733.4;refGene_gn=ITGB7;refGene_tx=XM_019834987.2;refGene_gn=ITGB7;refGene_tx=XM_023257103.1;refGene_gn=ITGB7;refGene_tx=XM_023257104.1;refGene_gn=ITGB7;refGene_tx=XM_023257105.1;refGene_gn=ITGB7;refGene_tx=XM_023257106.1
B4 81614187 81614387
refGene_gn=RARG;refGene_tx=XM_011283963.3;refGene_gn=RARG;refGene_tx=XM_023257107.1;refGene_gn=RARG;refGene_tx=XM_023257108.1;refGene_gn=RARG;refGene_tx=XM_023257109.1

B4 81615184 81615343
refGene_gn=RARG;refGene_tx=XM_011283963.3;refGene_gn=RARG;refGene_tx=XM_023257107.1;refGene_gn=RARG;refGene_tx=XM_023257108.1;refGene_gn=RARG;refGene_tx=XM_023257109.1
B4 81615786 81615991
refGene_gn=RARG;refGene_tx=XM_011283963.3;refGene_gn=RARG;refGene_tx=XM_023257107.1;refGene_gn=RARG;refGene_tx=XM_023257108.1;refGene_gn=RARG;refGene_tx=XM_023257109.1
B4 81616392 81616569
refGene_gn=RARG;refGene_tx=XM_011283963.3;refGene_gn=RARG;refGene_tx=XM_023257107.1;refGene_gn=RARG;refGene_tx=XM_023257108.1;refGene_gn=RARG;refGene_tx=XM_023257109.1
B4 81616757 81616918
refGene_gn=RARG;refGene_tx=XM_011283963.3;refGene_gn=RARG;refGene_tx=XM_023257107.1;refGene_gn=RARG;refGene_tx=XM_023257108.1;refGene_gn=RARG;refGene_tx=XM_023257109.1
B4 81617544 81617686
refGene_gn=RARG;refGene_tx=XM_011283963.3;refGene_gn=RARG;refGene_tx=XM_023257107.1;refGene_gn=RARG;refGene_tx=XM_023257108.1;refGene_gn=RARG;refGene_tx=XM_023257109.1
B4 81617907 81618056
refGene_gn=RARG;refGene_tx=XM_011283963.3;refGene_gn=RARG;refGene_tx=XM_023257107.1;refGene_gn=RARG;refGene_tx=XM_023257108.1;refGene_gn=RARG;refGene_tx=XM_023257109.1
B4 81622463 81622614 refGene_gn=RARG;refGene_tx=XM_011283963.3
B4 81629083 81629270
refGene_gn=RARG;refGene_tx=XM_023257107.1;refGene_gn=RARG;refGene_tx=XM_023257108.1
B4 81650935 81652288 refGene_gn=MFSD5;refGene_tx=XM_003988756.4
B4 81661451 81661532
refGene_gn=ESPL1;refGene_tx=XM_003988757.4;refGene_gn=ESPL1;refGene_tx=XM_019834990.2
B4 81661697 81662756
refGene_gn=ESPL1;refGene_tx=XM_003988757.4;refGene_gn=ESPL1;refGene_tx=XM_019834990.2
B4 81663047 81663152
refGene_gn=ESPL1;refGene_tx=XM_003988757.4;refGene_gn=ESPL1;refGene_tx=XM_019834990.2
B4 81663312 81663433
refGene_gn=ESPL1;refGene_tx=XM_003988757.4;refGene_gn=ESPL1;refGene_tx=XM_019834990.2
B4 81666645 81666782
refGene_gn=ESPL1;refGene_tx=XM_003988757.4;refGene_gn=ESPL1;refGene_tx=XM_019834990.2
B4 81667781 81667975
refGene_gn=ESPL1;refGene_tx=XM_003988757.4;refGene_gn=ESPL1;refGene_tx=XM_019834990.2
B4 81668288 81668528
refGene_gn=ESPL1;refGene_tx=XM_003988757.4;refGene_gn=ESPL1;refGene_tx=XM_019834990.2

B4	81668752	81668897
refGene_gn=ESPL1;refGene_tx=XM_003988757.4;refGene_gn=ESPL1;refGene_tx=XM_019834990.2
B4	81669008	81669147
refGene_gn=ESPL1;refGene_tx=XM_003988757.4;refGene_gn=ESPL1;refGene_tx=XM_019834990.2
B4	81669343	81669483
refGene_gn=ESPL1;refGene_tx=XM_003988757.4;refGene_gn=ESPL1;refGene_tx=XM_019834990.2
B4	81671170	81671305
refGene_gn=ESPL1;refGene_tx=XM_003988757.4;refGene_gn=ESPL1;refGene_tx=XM_019834990.2
B4	81672362	81672482
refGene_gn=ESPL1;refGene_tx=XM_003988757.4;refGene_gn=ESPL1;refGene_tx=XM_019834990.2
B4	81672937	81673109
refGene_gn=ESPL1;refGene_tx=XM_003988757.4;refGene_gn=ESPL1;refGene_tx=XM_019834990.2
B4	81673502	81673631
refGene_gn=ESPL1;refGene_tx=XM_003988757.4;refGene_gn=ESPL1;refGene_tx=XM_019834990.2
B4	81673760	81673917
refGene_gn=ESPL1;refGene_tx=XM_003988757.4;refGene_gn=ESPL1;refGene_tx=XM_019834990.2
B4	81674432	81674542
refGene_gn=ESPL1;refGene_tx=XM_003988757.4;refGene_gn=ESPL1;refGene_tx=XM_019834990.2
B4	81675617	81676606
refGene_gn=ESPL1;refGene_tx=XM_003988757.4;refGene_gn=ESPL1;refGene_tx=XM_019834990.2
B4	81676997	81677346
refGene_gn=ESPL1;refGene_tx=XM_003988757.4;refGene_gn=ESPL1;refGene_tx=XM_019834990.2
B4	81677546	81677708
refGene_gn=ESPL1;refGene_tx=XM_003988757.4;refGene_gn=ESPL1;refGene_tx=XM_019834990.2
B4	81678147	81678361
refGene_gn=ESPL1;refGene_tx=XM_003988757.4;refGene_gn=ESPL1;refGene_tx=XM_019834990.2
B4	81678455	81678646
refGene_gn=ESPL1;refGene_tx=XM_003988757.4;refGene_gn=ESPL1;refGene_tx=XM_019834990.2
B4	81679185	81679313
refGene_gn=ESPL1;refGene_tx=XM_003988757.4;refGene_gn=ESPL1;refGene_tx=XM_019834990.2
B4	81679449	81679572
refGene_gn=ESPL1;refGene_tx=XM_003988757.4;refGene_gn=ESPL1;refGene_tx=XM_019834990.2
B4	81680042	81680198
refGene_gn=ESPL1;refGene_tx=XM_003988757.4;refGene_gn=ESPL1;refGene_tx

=XM_019834990.2
B4 81680308 81680479
refGene_gn=ESPL1;refGene_tx=XM_003988757.4;refGene_gn=ESPL1;refGene_tx
=XM_019834990.2
B4 81680605 81680710
refGene_gn=ESPL1;refGene_tx=XM_003988757.4;refGene_gn=ESPL1;refGene_tx
=XM_019834990.2
B4 81680843 81680993
refGene_gn=ESPL1;refGene_tx=XM_003988757.4;refGene_gn=ESPL1;refGene_tx
=XM_019834990.2
B4 81681152 81681236 refGene_gn=ESPL1;refGene_tx=XM_003988757.4
B4 81681392 81681557 refGene_gn=ESPL1;refGene_tx=XM_003988757.4
B4 81681679 81681881 refGene_gn=ESPL1;refGene_tx=XM_003988757.4
B4 81683315 81683387 refGene_gn=PFDN5;refGene_tx=XM_003988758.3
B4 81683589 81683692 refGene_gn=PFDN5;refGene_tx=XM_003988758.3
B4 81683985 81684017 refGene_gn=PFDN5;refGene_tx=XM_003988758.3
B4 81685154 81685229 refGene_gn=PFDN5;refGene_tx=XM_003988758.3
B4 81685345 81685451 refGene_gn=PFDN5;refGene_tx=XM_003988758.3
B4 81685815 81685892 refGene_gn=PFDN5;refGene_tx=XM_003988758.3
B4 81686310 81686529
refGene_gn=CB4H12orf10;refGene_tx=XM_011283964.3;refGene_gn=CB4H12orf1
0;refGene_tx=XM_019834992.1
B4 81686627 81686740
refGene_gn=CB4H12orf10;refGene_tx=XM_011283964.3;refGene_gn=CB4H12orf1
0;refGene_tx=XM_019834992.1
B4 81689488 81689648
refGene_gn=CB4H12orf10;refGene_tx=XM_011283964.3;refGene_gn=CB4H12orf1
0;refGene_tx=XM_019834992.1
B4 81691374 81691527
refGene_gn=CB4H12orf10;refGene_tx=XM_011283964.3;refGene_gn=CB4H12orf1
0;refGene_tx=XM_019834992.1
B4 81691665 81691788
refGene_gn=CB4H12orf10;refGene_tx=XM_011283964.3;refGene_gn=CB4H12orf1
0;refGene_tx=XM_019834992.1
B4 81691902 81692084
refGene_gn=CB4H12orf10;refGene_tx=XM_011283964.3;refGene_gn=CB4H12orf1
0;refGene_tx=XM_019834992.1
B4 81692239 81692435
refGene_gn=CB4H12orf10;refGene_tx=XM_011283964.3
B4 81692768 81692993
refGene_gn=AAAS;refGene_tx=XM_003988761.5;refGene_gn=AAAS;refGene_tx=X
M_019834991.1
B4 81693111 81693196
refGene_gn=AAAS;refGene_tx=XM_003988761.5;refGene_gn=AAAS;refGene_tx=X
M_019834991.1
B4 81693308 81693390
refGene_gn=AAAS;refGene_tx=XM_003988761.5;refGene_gn=AAAS;refGene_tx=X
M_019834991.1
B4 81693531 81693599
refGene_gn=AAAS;refGene_tx=XM_003988761.5;refGene_gn=AAAS;refGene_tx=X

M_019834991.1
B4 81693681 81693775
refGene_gn=AAAS;refGene_tx=XM_003988761.5;refGene_gn=AAAS;refGene_tx=XM_019834991.1
B4 81693978 81694069
refGene_gn=AAAS;refGene_tx=XM_003988761.5;refGene_gn=AAAS;refGene_tx=XM_019834991.1
B4 81694188 81694249
refGene_gn=AAAS;refGene_tx=XM_003988761.5;refGene_gn=AAAS;refGene_tx=XM_019834991.1
B4 81694439 81694564
refGene_gn=AAAS;refGene_tx=XM_003988761.5;refGene_gn=AAAS;refGene_tx=XM_019834991.1
B4 81694907 81695028
refGene_gn=AAAS;refGene_tx=XM_003988761.5;refGene_gn=AAAS;refGene_tx=XM_019834991.1
B4 81700385 81700470
refGene_gn=CB4H12orf10;refGene_tx=XM_019834992.1
B4 81700496 81700640
refGene_gn=AAAS;refGene_tx=XM_003988761.5;refGene_gn=AAAS;refGene_tx=XM_019834991.1
B4 81700953 81701052
refGene_gn=AAAS;refGene_tx=XM_003988761.5;refGene_gn=AAAS;refGene_tx=XM_019834991.1
B4 81701310 81701357
refGene_gn=AAAS;refGene_tx=XM_003988761.5;refGene_gn=AAAS;refGene_tx=XM_019834991.1
B4 81701545 81701637
refGene_gn=AAAS;refGene_tx=XM_003988761.5;refGene_gn=AAAS;refGene_tx=XM_019834991.1
B4 81701930 81701986
refGene_gn=AAAS;refGene_tx=XM_003988761.5;refGene_gn=AAAS;refGene_tx=XM_019834991.1
B4 81704670 81704798 refGene_gn=AAAS;refGene_tx=XM_003988761.5
B4 81705113 81705128
refGene_gn=CB4H12orf10;refGene_tx=XM_019834992.1
B4 81705450 81705573 refGene_gn=AAAS;refGene_tx=XM_003988761.5
B4 81711040 81712315 refGene_gn=SP7;refGene_tx=XM_023257110.1
B4 81717148 81717169 refGene_gn=SP7;refGene_tx=XM_023257110.1
B4 81752184 81752191 refGene_gn=SP1;refGene_tx=XM_023257111.1
B4 81753229 81753384 refGene_gn=SP1;refGene_tx=XM_023257111.1
B4 81753701 81755217 refGene_gn=SP1;refGene_tx=XM_023257111.1
B4 81783467 81783636 refGene_gn=SP1;refGene_tx=XM_023257111.1
B4 81786453 81786653 refGene_gn=SP1;refGene_tx=XM_023257111.1
B4 81787051 81787365 refGene_gn=SP1;refGene_tx=XM_023257111.1
B4 81798174 81798247
refGene_gn=AMHR2;refGene_tx=XM_023257112.1;refGene_gn=AMHR2;refGene_tx=XM_023257113.1
B4 81798382 81798731
refGene_gn=AMHR2;refGene_tx=XM_006933741.2;refGene_gn=AMHR2;refGene_tx

=XM_023257112.1
B4 81799016 81799208
refGene_gn=AMHR2;refGene_tx=XM_006933741.2;refGene_gn=AMHR2;refGene_tx=XM_023257112.1;refGene_gn=AMHR2;refGene_tx=XM_023257113.1
B4 81799474 81799552
refGene_gn=AMHR2;refGene_tx=XM_006933741.2;refGene_gn=AMHR2;refGene_tx=XM_023257112.1;refGene_gn=AMHR2;refGene_tx=XM_023257113.1
B4 81799738 81799872
refGene_gn=AMHR2;refGene_tx=XM_006933741.2;refGene_gn=AMHR2;refGene_tx=XM_023257112.1;refGene_gn=AMHR2;refGene_tx=XM_023257113.1
B4 81799986 81800217
refGene_gn=AMHR2;refGene_tx=XM_006933741.2;refGene_gn=AMHR2;refGene_tx=XM_023257112.1;refGene_gn=AMHR2;refGene_tx=XM_023257113.1
B4 81801810 81801925
refGene_gn=AMHR2;refGene_tx=XM_006933741.2;refGene_gn=AMHR2;refGene_tx=XM_023257112.1;refGene_gn=AMHR2;refGene_tx=XM_023257113.1
B4 81802069 81802242
refGene_gn=AMHR2;refGene_tx=XM_006933741.2;refGene_gn=AMHR2;refGene_tx=XM_023257112.1;refGene_gn=AMHR2;refGene_tx=XM_023257113.1
B4 81802445 81802593
refGene_gn=AMHR2;refGene_tx=XM_006933741.2;refGene_gn=AMHR2;refGene_tx=XM_023257112.1;refGene_gn=AMHR2;refGene_tx=XM_023257113.1
B4 81802764 81802901
refGene_gn=AMHR2;refGene_tx=XM_006933741.2;refGene_gn=AMHR2;refGene_tx=XM_023257112.1;refGene_gn=AMHR2;refGene_tx=XM_023257113.1
B4 81803786 81803996
refGene_gn=AMHR2;refGene_tx=XM_006933741.2;refGene_gn=AMHR2;refGene_tx=XM_023257112.1;refGene_gn=AMHR2;refGene_tx=XM_023257113.1
B4 81817647 81817669 refGene_gn=PRR13;refGene_tx=XM_023257760.1
B4 81818030 81818395 refGene_gn=PRR13;refGene_tx=XM_023257760.1
B4 81819625 81819670 refGene_gn=PRR13;refGene_tx=XM_023257760.1
B4 81831226 81831295
refGene_gn=PCBP2;refGene_tx=XM_003988768.3;refGene_gn=PCBP2;refGene_tx=XM_003988769.4;refGene_gn=PCBP2;refGene_tx=XM_006933744.3;refGene_gn=PCBP2;refGene_tx=XM_006933745.3;refGene_gn=PCBP2;refGene_tx=XM_006933746.2;refGene_gn=PCBP2;refGene_tx=XM_006933747.2;refGene_gn=PCBP2;refGene_tx=XM_006933748.2;refGene_gn=PCBP2;refGene_tx=XM_006933750.2;refGene_gn=PCBP2;refGene_tx=XM_011283967.2;refGene_gn=PCBP2;refGene_tx=XM_011283968.3;refGene_gn=PCBP2;refGene_tx=XM_019835000.1;refGene_gn=PCBP2;refGene_tx=XM_019835001.2;refGene_gn=PCBP2;refGene_tx=XM_019835002.1;refGene_gn=PCBP2;refGene_tx=XM_019835003.1;refGene_gn=PCBP2;refGene_tx=XM_019835004.1;refGene_gn=PCBP2;refGene_tx=XM_019835005.1;refGene_gn=PCBP2;refGene_tx=XM_019835006.2;refGene_gn=PCBP2;refGene_tx=XM_019835007.2;refGene_gn=PCBP2;refGene_tx=XM_019835008.2;refGene_gn=PCBP2;refGene_tx=XM_019835009.2;refGene_gn=PCBP2;refGene_tx=XM_019835010.2;refGene_gn=PCBP2;refGene_tx=XM_019835011.2;refGene_gn=PCBP2;refGene_tx=XM_019835012.2
B4 81831853 81831877
refGene_gn=PCBP2;refGene_tx=XM_003988768.3;refGene_gn=PCBP2;refGene_tx=XM_003988769.4;refGene_gn=PCBP2;refGene_tx=XM_006933744.3;refGene_gn=

PCBP2;refGene_tx=XM_006933745.3;refGene_gn=PCBP2;refGene_tx=XM_0069337
46.2;refGene_gn=PCBP2;refGene_tx=XM_006933747.2;refGene_gn=PCBP2;refGe
ne_tx=XM_006933748.2;refGene_gn=PCBP2;refGene_tx=XM_006933750.2;refGen
e_gn=PCBP2;refGene_tx=XM_011283967.2;refGene_gn=PCBP2;refGene_tx=XM_01
1283968.3;refGene_gn=PCBP2;refGene_tx=XM_019835000.1;refGene_gn=PCBP2;
refGene_tx=XM_019835001.2;refGene_gn=PCBP2;refGene_tx=XM_019835002.1;r
efGene_gn=PCBP2;refGene_tx=XM_019835003.1;refGene_gn=PCBP2;refGene_tx=
XM_019835004.1;refGene_gn=PCBP2;refGene_tx=XM_019835005.1;refGene_gn=P
CBP2;refGene_tx=XM_019835006.2;refGene_gn=PCBP2;refGene_tx=XM_01983500
7.2;refGene_gn=PCBP2;refGene_tx=XM_019835008.2;refGene_gn=PCBP2;refGen
e_tx=XM_019835009.2;refGene_gn=PCBP2;refGene_tx=XM_019835010.2;refGene
_gn=PCBP2;refGene_tx=XM_019835011.2;refGene_gn=PCBP2;refGene_tx=XM_019
835012.2
B4 81831968 81832001
refGene_gn=PCBP2;refGene_tx=XM_003988768.3;refGene_gn=PCBP2;refGene_tx
=XM_003988769.4;refGene_gn=PCBP2;refGene_tx=XM_006933744.3;refGene_gn=
PCBP2;refGene_tx=XM_006933745.3;refGene_gn=PCBP2;refGene_tx=XM_0069337
46.2;refGene_gn=PCBP2;refGene_tx=XM_006933747.2;refGene_gn=PCBP2;refGe
ne_tx=XM_006933748.2;refGene_gn=PCBP2;refGene_tx=XM_006933750.2;refGen
e_gn=PCBP2;refGene_tx=XM_011283967.2;refGene_gn=PCBP2;refGene_tx=XM_01
1283968.3;refGene_gn=PCBP2;refGene_tx=XM_019835000.1;refGene_gn=PCBP2;
refGene_tx=XM_019835001.2;refGene_gn=PCBP2;refGene_tx=XM_019835002.1;r
efGene_gn=PCBP2;refGene_tx=XM_019835003.1;refGene_gn=PCBP2;refGene_tx=
XM_019835004.1;refGene_gn=PCBP2;refGene_tx=XM_019835005.1;refGene_gn=P
CBP2;refGene_tx=XM_019835006.2;refGene_gn=PCBP2;refGene_tx=XM_01983500
7.2;refGene_gn=PCBP2;refGene_tx=XM_019835008.2;refGene_gn=PCBP2;refGen
e_tx=XM_019835009.2;refGene_gn=PCBP2;refGene_tx=XM_019835010.2;refGene
_gn=PCBP2;refGene_tx=XM_019835011.2;refGene_gn=PCBP2;refGene_tx=XM_019
835012.2
B4 81832374 81832491
refGene_gn=PCBP2;refGene_tx=XM_003988768.3;refGene_gn=PCBP2;refGene_tx
=XM_003988769.4;refGene_gn=PCBP2;refGene_tx=XM_006933744.3;refGene_gn=
PCBP2;refGene_tx=XM_006933745.3;refGene_gn=PCBP2;refGene_tx=XM_0069337
46.2;refGene_gn=PCBP2;refGene_tx=XM_006933747.2;refGene_gn=PCBP2;refGe
ne_tx=XM_006933748.2;refGene_gn=PCBP2;refGene_tx=XM_006933750.2;refGen
e_gn=PCBP2;refGene_tx=XM_011283967.2;refGene_gn=PCBP2;refGene_tx=XM_01
1283968.3;refGene_gn=PCBP2;refGene_tx=XM_019835000.1;refGene_gn=PCBP2;
refGene_tx=XM_019835001.2;refGene_gn=PCBP2;refGene_tx=XM_019835002.1;r
efGene_gn=PCBP2;refGene_tx=XM_019835003.1;refGene_gn=PCBP2;refGene_tx=
XM_019835004.1;refGene_gn=PCBP2;refGene_tx=XM_019835005.1;refGene_gn=P
CBP2;refGene_tx=XM_019835006.2;refGene_gn=PCBP2;refGene_tx=XM_01983500
7.2;refGene_gn=PCBP2;refGene_tx=XM_019835008.2;refGene_gn=PCBP2;refGen
e_tx=XM_019835009.2;refGene_gn=PCBP2;refGene_tx=XM_019835010.2;refGene
_gn=PCBP2;refGene_tx=XM_019835011.2;refGene_gn=PCBP2;refGene_tx=XM_019
835012.2
B4 81833996 81834128
refGene_gn=PCBP2;refGene_tx=XM_003988768.3;refGene_gn=PCBP2;refGene_tx
=XM_003988769.4;refGene_gn=PCBP2;refGene_tx=XM_006933744.3;refGene_gn=
PCBP2;refGene_tx=XM_006933745.3;refGene_gn=PCBP2;refGene_tx=XM_0069337
46.2;refGene_gn=PCBP2;refGene_tx=XM_006933747.2;refGene_gn=PCBP2;refGe

ne_tx=XM_006933748.2;refGene_gn=PCBP2;refGene_tx=XM_006933750.2;refGen
e_gn=PCBP2;refGene_tx=XM_011283967.2;refGene_gn=PCBP2;refGene_tx=XM_01
1283968.3;refGene_gn=PCBP2;refGene_tx=XM_019835000.1;refGene_gn=PCBP2;
refGene_tx=XM_019835001.2;refGene_gn=PCBP2;refGene_tx=XM_019835002.1;r
efGene_gn=PCBP2;refGene_tx=XM_019835003.1;refGene_gn=PCBP2;refGene_tx=
XM_019835004.1;refGene_gn=PCBP2;refGene_tx=XM_019835005.1;refGene_gn=P
CBP2;refGene_tx=XM_019835006.2;refGene_gn=PCBP2;refGene_tx=XM_01983500
7.2;refGene_gn=PCBP2;refGene_tx=XM_019835008.2;refGene_gn=PCBP2;refGen
e_tx=XM_019835009.2;refGene_gn=PCBP2;refGene_tx=XM_019835010.2;refGene
_gn=PCBP2;refGene_tx=XM_019835011.2;refGene_gn=PCBP2;refGene_tx=XM_019
835012.2
B4 81835136 81835265
refGene_gn=PCBP2;refGene_tx=XM_003988768.3;refGene_gn=PCBP2;refGene_tx
=XM_003988769.4;refGene_gn=PCBP2;refGene_tx=XM_006933744.3;refGene_gn=
PCBP2;refGene_tx=XM_006933745.3;refGene_gn=PCBP2;refGene_tx=XM_0069337
46.2;refGene_gn=PCBP2;refGene_tx=XM_006933747.2;refGene_gn=PCBP2;refGe
ne_tx=XM_006933748.2;refGene_gn=PCBP2;refGene_tx=XM_006933750.2;refGen
e_gn=PCBP2;refGene_tx=XM_011283967.2;refGene_gn=PCBP2;refGene_tx=XM_01
1283968.3;refGene_gn=PCBP2;refGene_tx=XM_019835000.1;refGene_gn=PCBP2;
refGene_tx=XM_019835001.2;refGene_gn=PCBP2;refGene_tx=XM_019835002.1;r
efGene_gn=PCBP2;refGene_tx=XM_019835003.1;refGene_gn=PCBP2;refGene_tx=
XM_019835004.1;refGene_gn=PCBP2;refGene_tx=XM_019835005.1;refGene_gn=P
CBP2;refGene_tx=XM_019835006.2;refGene_gn=PCBP2;refGene_tx=XM_01983500
7.2;refGene_gn=PCBP2;refGene_tx=XM_019835008.2;refGene_gn=PCBP2;refGen
e_tx=XM_019835009.2;refGene_gn=PCBP2;refGene_tx=XM_019835010.2;refGene
_gn=PCBP2;refGene_tx=XM_019835011.2;refGene_gn=PCBP2;refGene_tx=XM_019
835012.2
B4 81836279 81836366
refGene_gn=PCBP2;refGene_tx=XM_003988768.3;refGene_gn=PCBP2;refGene_tx
=XM_003988769.4;refGene_gn=PCBP2;refGene_tx=XM_006933744.3;refGene_gn=
PCBP2;refGene_tx=XM_006933745.3;refGene_gn=PCBP2;refGene_tx=XM_0069337
46.2;refGene_gn=PCBP2;refGene_tx=XM_006933747.2;refGene_gn=PCBP2;refGe
ne_tx=XM_006933748.2;refGene_gn=PCBP2;refGene_tx=XM_006933750.2;refGen
e_gn=PCBP2;refGene_tx=XM_011283967.2;refGene_gn=PCBP2;refGene_tx=XM_01
1283968.3;refGene_gn=PCBP2;refGene_tx=XM_019835000.1;refGene_gn=PCBP2;
refGene_tx=XM_019835001.2;refGene_gn=PCBP2;refGene_tx=XM_019835002.1;r
efGene_gn=PCBP2;refGene_tx=XM_019835003.1;refGene_gn=PCBP2;refGene_tx=
XM_019835004.1;refGene_gn=PCBP2;refGene_tx=XM_019835005.1;refGene_gn=P
CBP2;refGene_tx=XM_019835006.2;refGene_gn=PCBP2;refGene_tx=XM_01983500
7.2;refGene_gn=PCBP2;refGene_tx=XM_019835008.2;refGene_gn=PCBP2;refGen
e_tx=XM_019835009.2;refGene_gn=PCBP2;refGene_tx=XM_019835010.2;refGene
_gn=PCBP2;refGene_tx=XM_019835011.2;refGene_gn=PCBP2;refGene_tx=XM_019
835012.2
B4 81838213 81838306
refGene_gn=PCBP2;refGene_tx=XM_003988769.4;refGene_gn=PCBP2;refGene_tx
=XM_006933744.3;refGene_gn=PCBP2;refGene_tx=XM_006933745.3;refGene_gn=
PCBP2;refGene_tx=XM_006933746.2;refGene_gn=PCBP2;refGene_tx=XM_0112839
67.2;refGene_gn=PCBP2;refGene_tx=XM_011283968.3;refGene_gn=PCBP2;refGe
ne_tx=XM_019835000.1;refGene_gn=PCBP2;refGene_tx=XM_019835001.2;refGen
e_gn=PCBP2;refGene_tx=XM_019835002.1;refGene_gn=PCBP2;refGene_tx=XM_01

9835003.1;refGene_gn=PCBP2;refGene_tx=XM_019835004.1;refGene_gn=PCBP2;
refGene_tx=XM_019835005.1;refGene_gn=PCBP2;refGene_tx=XM_019835012.2
B4 81839428 81839470
refGene_gn=PCBP2;refGene_tx=XM_003988768.3;refGene_gn=PCBP2;refGene_tx
=XM_003988769.4;refGene_gn=PCBP2;refGene_tx=XM_006933744.3;refGene_gn=
PCBP2;refGene_tx=XM_006933745.3;refGene_gn=PCBP2;refGene_tx=XM_0069337
46.2;refGene_gn=PCBP2;refGene_tx=XM_006933747.2;refGene_gn=PCBP2;refGe
ne_tx=XM_006933748.2;refGene_gn=PCBP2;refGene_tx=XM_006933750.2;refGen
e_gn=PCBP2;refGene_tx=XM_011283967.2;refGene_gn=PCBP2;refGene_tx=XM_01
1283968.3;refGene_gn=PCBP2;refGene_tx=XM_019835000.1;refGene_gn=PCBP2;
refGene_tx=XM_019835001.2;refGene_gn=PCBP2;refGene_tx=XM_019835002.1;r
efGene_gn=PCBP2;refGene_tx=XM_019835003.1;refGene_gn=PCBP2;refGene_tx=
XM_019835004.1;refGene_gn=PCBP2;refGene_tx=XM_019835005.1;refGene_gn=P
CBP2;refGene_tx=XM_019835006.2;refGene_gn=PCBP2;refGene_tx=XM_01983500
7.2;refGene_gn=PCBP2;refGene_tx=XM_019835012.2
B4 81840723 81840796
refGene_gn=PCBP2;refGene_tx=XM_003988768.3;refGene_gn=PCBP2;refGene_tx
=XM_003988769.4;refGene_gn=PCBP2;refGene_tx=XM_006933744.3;refGene_gn=
PCBP2;refGene_tx=XM_006933745.3;refGene_gn=PCBP2;refGene_tx=XM_0069337
46.2;refGene_gn=PCBP2;refGene_tx=XM_006933747.2;refGene_gn=PCBP2;refGe
ne_tx=XM_006933748.2;refGene_gn=PCBP2;refGene_tx=XM_006933750.2;refGen
e_gn=PCBP2;refGene_tx=XM_011283967.2;refGene_gn=PCBP2;refGene_tx=XM_01
1283968.3;refGene_gn=PCBP2;refGene_tx=XM_019835000.1;refGene_gn=PCBP2;
refGene_tx=XM_019835001.2;refGene_gn=PCBP2;refGene_tx=XM_019835002.1;r
efGene_gn=PCBP2;refGene_tx=XM_019835003.1;refGene_gn=PCBP2;refGene_tx=
XM_019835004.1;refGene_gn=PCBP2;refGene_tx=XM_019835005.1;refGene_gn=P
CBP2;refGene_tx=XM_019835006.2;refGene_gn=PCBP2;refGene_tx=XM_01983500
7.2;refGene_gn=PCBP2;refGene_tx=XM_019835008.2;refGene_gn=PCBP2;refGen
e_tx=XM_019835009.2;refGene_gn=PCBP2;refGene_tx=XM_019835010.2;refGene
_gn=PCBP2;refGene_tx=XM_019835011.2;refGene_gn=PCBP2;refGene_tx=XM_019
835012.2
B4 81841293 81841332
refGene_gn=PCBP2;refGene_tx=XM_003988768.3;refGene_gn=PCBP2;refGene_tx
=XM_003988769.4;refGene_gn=PCBP2;refGene_tx=XM_006933744.3;refGene_gn=
PCBP2;refGene_tx=XM_006933747.2;refGene_gn=PCBP2;refGene_tx=XM_0112839
67.2;refGene_gn=PCBP2;refGene_tx=XM_011283968.3;refGene_gn=PCBP2;refGe
ne_tx=XM_019835000.1;refGene_gn=PCBP2;refGene_tx=XM_019835001.2;refGen
e_gn=PCBP2;refGene_tx=XM_019835002.1;refGene_gn=PCBP2;refGene_tx=XM_01
9835003.1;refGene_gn=PCBP2;refGene_tx=XM_019835006.2;refGene_gn=PCBP2;
refGene_tx=XM_019835007.2;refGene_gn=PCBP2;refGene_tx=XM_019835008.2;r
efGene_gn=PCBP2;refGene_tx=XM_019835009.2;refGene_gn=PCBP2;refGene_tx=
XM_019835012.2
B4 81842275 81842331
refGene_gn=PCBP2;refGene_tx=XM_003988768.3;refGene_gn=PCBP2;refGene_tx
=XM_003988769.4;refGene_gn=PCBP2;refGene_tx=XM_006933744.3;refGene_gn=
PCBP2;refGene_tx=XM_006933745.3;refGene_gn=PCBP2;refGene_tx=XM_0069337
46.2;refGene_gn=PCBP2;refGene_tx=XM_006933747.2;refGene_gn=PCBP2;refGe
ne_tx=XM_006933748.2;refGene_gn=PCBP2;refGene_tx=XM_006933750.2;refGen
e_gn=PCBP2;refGene_tx=XM_011283967.2;refGene_gn=PCBP2;refGene_tx=XM_01
1283968.3;refGene_gn=PCBP2;refGene_tx=XM_019835000.1;refGene_gn=PCBP2;

refGene_tx=XM_019835001.2;refGene_gn=PCBP2;refGene_tx=XM_019835002.1;refGene_gn=PCBP2;refGene_tx=XM_019835003.1;refGene_gn=PCBP2;refGene_tx=XM_019835004.1;refGene_gn=PCBP2;refGene_tx=XM_019835005.1;refGene_gn=PCBP2;refGene_tx=XM_019835006.2;refGene_gn=PCBP2;refGene_tx=XM_019835007.2;refGene_gn=PCBP2;refGene_tx=XM_019835008.2;refGene_gn=PCBP2;refGene_tx=XM_019835009.2;refGene_gn=PCBP2;refGene_tx=XM_019835010.2;refGene_gn=PCBP2;refGene_tx=XM_019835011.2;refGene_gn=PCBP2;refGene_tx=XM_019835012.2
B4 81843241 81843411 refGene_gn=PCBP2;refGene_tx=XM_003988768.3;refGene_gn=PCBP2;refGene_tx=XM_003988769.4;refGene_gn=PCBP2;refGene_tx=XM_006933744.3;refGene_gn=PCBP2;refGene_tx=XM_006933745.3;refGene_gn=PCBP2;refGene_tx=XM_006933746.2;refGene_gn=PCBP2;refGene_tx=XM_006933747.2;refGene_gn=PCBP2;refGene_tx=XM_006933748.2;refGene_gn=PCBP2;refGene_tx=XM_006933750.2;refGene_gn=PCBP2;refGene_tx=XM_011283967.2;refGene_gn=PCBP2;refGene_tx=XM_011283968.3;refGene_gn=PCBP2;refGene_tx=XM_019835000.1;refGene_gn=PCBP2;refGene_tx=XM_019835001.2;refGene_gn=PCBP2;refGene_tx=XM_019835002.1;refGene_gn=PCBP2;refGene_tx=XM_019835003.1;refGene_gn=PCBP2;refGene_tx=XM_019835004.1;refGene_gn=PCBP2;refGene_tx=XM_019835005.1;refGene_gn=PCBP2;refGene_tx=XM_019835006.2;refGene_gn=PCBP2;refGene_tx=XM_019835007.2;refGene_gn=PCBP2;refGene_tx=XM_019835008.2;refGene_gn=PCBP2;refGene_tx=XM_019835009.2;refGene_gn=PCBP2;refGene_tx=XM_019835010.2;refGene_gn=PCBP2;refGene_tx=XM_019835011.2;refGene_gn=PCBP2;refGene_tx=XM_019835012.2
B4 81846051 81846403 refGene_gn=PCBP2;refGene_tx=XM_019835000.1;refGene_gn=PCBP2;refGene_tx=XM_019835001.2;refGene_gn=PCBP2;refGene_tx=XM_019835002.1;refGene_gn=PCBP2;refGene_tx=XM_019835003.1;refGene_gn=PCBP2;refGene_tx=XM_019835004.1;refGene_gn=PCBP2;refGene_tx=XM_019835005.1;refGene_gn=PCBP2;refGene_tx=XM_019835006.2;refGene_gn=PCBP2;refGene_tx=XM_019835007.2;refGene_gn=PCBP2;refGene_tx=XM_019835008.2;refGene_gn=PCBP2;refGene_tx=XM_019835009.2;refGene_gn=PCBP2;refGene_tx=XM_019835010.2;refGene_gn=PCBP2;refGene_tx=XM_019835011.2
B4 81847895 81847908 refGene_gn=PCBP2;refGene_tx=XM_019835012.2
B4 81849694 81849731 refGene_gn=PCBP2;refGene_tx=XM_003988768.3;refGene_gn=PCBP2;refGene_tx=XM_003988769.4;refGene_gn=PCBP2;refGene_tx=XM_006933744.3;refGene_gn=PCBP2;refGene_tx=XM_006933745.3;refGene_gn=PCBP2;refGene_tx=XM_006933746.2;refGene_gn=PCBP2;refGene_tx=XM_006933747.2;refGene_gn=PCBP2;refGene_tx=XM_006933748.2;refGene_gn=PCBP2;refGene_tx=XM_006933750.2;refGene_gn=PCBP2;refGene_tx=XM_011283967.2;refGene_gn=PCBP2;refGene_tx=XM_011283968.3
B4 81851505 81851604 refGene_gn=MAP3K12;refGene_tx=XM_003988770.5
B4 81852028 81852302 refGene_gn=MAP3K12;refGene_tx=XM_003988770.5
B4 81852393 81852464 refGene_gn=MAP3K12;refGene_tx=XM_003988770.5
B4 81852650 81853272 refGene_gn=MAP3K12;refGene_tx=XM_003988770.5
B4 81853466 81853604 refGene_gn=MAP3K12;refGene_tx=XM_003988770.5
B4 81853728 81853845 refGene_gn=MAP3K12;refGene_tx=XM_003988770.5
B4 81853979 81854089 refGene_gn=MAP3K12;refGene_tx=XM_003988770.5
B4 81854314 81854423 refGene_gn=MAP3K12;refGene_tx=XM_003988770.5

B4 81855094 81855253 refGene_gn=MAP3K12;refGene_tx=XM_003988770.5
B4 81855404 81855563 refGene_gn=MAP3K12;refGene_tx=XM_003988770.5
B4 81855836 81856028 refGene_gn=MAP3K12;refGene_tx=XM_003988770.5
B4 81856207 81856391 refGene_gn=MAP3K12;refGene_tx=XM_003988770.5
B4 81856713 81857158 refGene_gn=MAP3K12;refGene_tx=XM_003988770.5
B4 81867808 81867861
refGene_gn=TARBP2;refGene_tx=XM_003988771.5;refGene_gn=TARBP2;refGene_tx=XM_019835014.2
B4 81868415 81868585
refGene_gn=TARBP2;refGene_tx=XM_003988771.5;refGene_gn=TARBP2;refGene_tx=XM_003988773.5;refGene_gn=TARBP2;refGene_tx=XM_019835014.2
B4 81869397 81869500
refGene_gn=TARBP2;refGene_tx=XM_003988771.5;refGene_gn=TARBP2;refGene_tx=XM_003988773.5
B4 81870081 81870177
refGene_gn=TARBP2;refGene_tx=XM_003988771.5;refGene_gn=TARBP2;refGene_tx=XM_003988773.5;refGene_gn=TARBP2;refGene_tx=XM_019835014.2
B4 81870768 81870841
refGene_gn=TARBP2;refGene_tx=XM_003988771.5;refGene_gn=TARBP2;refGene_tx=XM_003988773.5;refGene_gn=TARBP2;refGene_tx=XM_019835014.2
B4 81871053 81871171
refGene_gn=TARBP2;refGene_tx=XM_003988771.5;refGene_gn=TARBP2;refGene_tx=XM_003988773.5;refGene_gn=TARBP2;refGene_tx=XM_019835014.2
B4 81871480 81871608
refGene_gn=TARBP2;refGene_tx=XM_003988771.5;refGene_gn=TARBP2;refGene_tx=XM_003988773.5;refGene_gn=TARBP2;refGene_tx=XM_019835014.2
B4 81872027 81872229
refGene_gn=TARBP2;refGene_tx=XM_003988771.5;refGene_gn=TARBP2;refGene_tx=XM_003988773.5;refGene_gn=TARBP2;refGene_tx=XM_019835014.2
B4 81872388 81872546
refGene_gn=TARBP2;refGene_tx=XM_003988771.5;refGene_gn=TARBP2;refGene_tx=XM_003988773.5;refGene_gn=TARBP2;refGene_tx=XM_019835014.2
B4 81873204 81873322 refGene_gn=NPFF;refGene_tx=XM_019835015.2
B4 81873435 81873557 refGene_gn=NPFF;refGene_tx=XM_019835015.2
B4 81873757 81874084 refGene_gn=NPFF;refGene_tx=XM_019835015.2
B4 81878929 81879105 refGene_gn=ATF7;refGene_tx=XM_006933754.3
B4 81884187 81884405 refGene_gn=ATF7;refGene_tx=XM_006933753.4
B4 81888001 81888110
refGene_gn=ATF7;refGene_tx=XM_006933753.4;refGene_gn=ATF7;refGene_tx=XM_006933754.3
B4 81889401 81889599
refGene_gn=ATF7;refGene_tx=XM_006933753.4;refGene_gn=ATF7;refGene_tx=XM_006933754.3
B4 81892310 81892463
refGene_gn=ATF7;refGene_tx=XM_006933753.4;refGene_gn=ATF7;refGene_tx=XM_006933754.3
B4 81893075 81893189
refGene_gn=ATF7;refGene_tx=XM_006933753.4;refGene_gn=ATF7;refGene_tx=XM_006933754.3
B4 81893720 81893820

refGene_gn=ATF7;refGene_tx=XM_006933753.4;refGene_gn=ATF7;refGene_tx=XM_006933754.3
B4 81894852 81895010
refGene_gn=ATF7;refGene_tx=XM_006933753.4;refGene_gn=ATF7;refGene_tx=XM_006933754.3
B4 81897305 81897443
refGene_gn=ATF7;refGene_tx=XM_006933753.4;refGene_gn=ATF7;refGene_tx=XM_006933754.3
B4 81900531 81900650
refGene_gn=ATF7;refGene_tx=XM_006933753.4;refGene_gn=ATF7;refGene_tx=XM_006933754.3
B4 81904937 81905034
refGene_gn=ATF7;refGene_tx=XM_006933753.4;refGene_gn=ATF7;refGene_tx=XM_006933754.3
B4 81945881 81945929
refGene_gn=ATF7;refGene_tx=XM_006933753.4;refGene_gn=ATF7;refGene_tx=XM_006933754.3
B4 82008848 82008963
refGene_gn=ATP5MC2;refGene_tx=XM_006933755.3;refGene_gn=ATP5MC2;refGene_tx=XM_019835019.1
B4 82010783 82010977
refGene_gn=ATP5MC2;refGene_tx=XM_006933755.3;refGene_gn=ATP5MC2;refGene_tx=XM_019835019.1
B4 82011533 82011611
refGene_gn=ATP5MC2;refGene_tx=XM_006933755.3;refGene_gn=ATP5MC2;refGene_tx=XM_019835019.1
B4 82012842 82012912
refGene_gn=ATP5MC2;refGene_tx=XM_006933755.3;refGene_gn=ATP5MC2;refGene_tx=XM_019835019.1
B4 82015469 82015602 refGene_gn=ATP5MC2;refGene_tx=XM_019835019.1
B4 82016272 82016372
refGene_gn=ATP5MC2;refGene_tx=XM_006933755.3;refGene_gn=ATP5MC2;refGene_tx=XM_019835019.1
B4 82016988 82017045 refGene_gn=ATP5MC2;refGene_tx=XM_019835019.1
B4 82018054 82018160 refGene_gn=ATP5MC2;refGene_tx=XM_006933755.3
B4 82048042 82048168 refGene_gn=CALCOCO1;refGene_tx=XM_003988778.5
B4 82048169 82048221 refGene_gn=CALCOCO1;refGene_tx=XM_003988778.5
B4 82049205 82049312 refGene_gn=CALCOCO1;refGene_tx=XM_003988778.5
B4 82049719 82049919 refGene_gn=CALCOCO1;refGene_tx=XM_003988778.5
B4 82050144 82050253 refGene_gn=CALCOCO1;refGene_tx=XM_003988778.5
B4 82050658 82050754 refGene_gn=CALCOCO1;refGene_tx=XM_003988778.5
B4 82051249 82051375 refGene_gn=CALCOCO1;refGene_tx=XM_003988778.5
B4 82051842 82052097 refGene_gn=CALCOCO1;refGene_tx=XM_003988778.5
B4 82052290 82052446 refGene_gn=CALCOCO1;refGene_tx=XM_003988778.5
B4 82055137 82055228 refGene_gn=CALCOCO1;refGene_tx=XM_003988778.5
B4 82056892 82057041 refGene_gn=CALCOCO1;refGene_tx=XM_003988778.5
B4 82057448 82057607 refGene_gn=CALCOCO1;refGene_tx=XM_003988778.5
B4 82060316 82060507 refGene_gn=CALCOCO1;refGene_tx=XM_003988778.5
B4 82061515 82061618 refGene_gn=CALCOCO1;refGene_tx=XM_003988778.5
B4 82061954 82062110 refGene_gn=CALCOCO1;refGene_tx=XM_003988778.5

B4	82272354	82273087	refGene_gn=HOXC13;refGene_tx=XM_023257114.1
B4	82278611	82278868	refGene_gn=HOXC13;refGene_tx=XM_023257114.1
B4	82288243	82288850	refGene_gn=HOXC12;refGene_tx=XM_003988780.2
B4	82289641	82289880	refGene_gn=HOXC12;refGene_tx=XM_003988780.2
B4	82306678	82307360	refGene_gn=HOXC11;refGene_tx=XM_003988781.4
B4	82308629	82308862	refGene_gn=HOXC11;refGene_tx=XM_003988781.4
B4	82318914	82319665	refGene_gn=HOXC10;refGene_tx=XM_003988782.4
B4	82322854	82323132	refGene_gn=HOXC10;refGene_tx=XM_003988782.4
B4	82328049	82328840
refGene_gn=LOC109501596;refGene_tx=XM_019835022.2;refGene_gn=LOC109501596;refGene_tx=XM_019835023.2;refGene_gn=LOC109501596;refGene_tx=XM_019835024.2
B4	82329670	82329878
refGene_gn=LOC109501596;refGene_tx=XM_019835022.2;refGene_gn=LOC109501596;refGene_tx=XM_019835023.2;refGene_gn=LOC109501596;refGene_tx=XM_019835024.2
B4	82334051	82334589	refGene_gn=HOXC9;refGene_tx=XM_003988783.4
B4	82336341	82336586	refGene_gn=HOXC9;refGene_tx=XM_003988783.4
B4	82343396	82343832	refGene_gn=HOXC8;refGene_tx=XM_003988784.3
B4	82345204	82345497	refGene_gn=HOXC8;refGene_tx=XM_003988784.3
B4	82362502	82362902
refGene_gn=HOXC6;refGene_tx=XM_003988785.5;refGene_gn=HOXC6;refGene_tx=XM_019835029.1;refGene_gn=HOXC6;refGene_tx=XM_023257116.1
B4	82363609	82363941
refGene_gn=HOXC6;refGene_tx=XM_003988785.5;refGene_gn=HOXC6;refGene_tx=XM_019835029.1;refGene_gn=HOXC6;refGene_tx=XM_023257116.1
B4	82367178	82367632	refGene_gn=HOXC5;refGene_tx=XM_003988787.5
B4	82368348	82368563	refGene_gn=HOXC5;refGene_tx=XM_003988787.5
B4	82387710	82388149
refGene_gn=HOXC4;refGene_tx=XM_011283972.3;refGene_gn=HOXC4;refGene_tx=XM_019835026.2;refGene_gn=HOXC4;refGene_tx=XM_019835027.2;refGene_gn=HOXC4;refGene_tx=XM_023257115.1
B4	82388644	82389000
refGene_gn=HOXC4;refGene_tx=XM_011283972.3;refGene_gn=HOXC4;refGene_tx=XM_019835026.2;refGene_gn=HOXC4;refGene_tx=XM_019835027.2;refGene_gn=HOXC4;refGene_tx=XM_023257115.1
B4	82507911	82508439
refGene_gn=SMUG1;refGene_tx=XM_003988788.4;refGene_gn=SMUG1;refGene_tx=XM_006933759.3;refGene_gn=SMUG1;refGene_tx=XM_006933760.3;refGene_gn=SMUG1;refGene_tx=XM_011283973.3;refGene_gn=SMUG1;refGene_tx=XM_019835031.2
B4	82509483	82509774
refGene_gn=SMUG1;refGene_tx=XM_003988788.4;refGene_gn=SMUG1;refGene_tx=XM_006933759.3;refGene_gn=SMUG1;refGene_tx=XM_006933760.3;refGene_gn=SMUG1;refGene_tx=XM_011283973.3;refGene_gn=SMUG1;refGene_tx=XM_019835031.2
B4	82555246	82555397
refGene_gn=CBX5;refGene_tx=XM_003988789.5;refGene_gn=CBX5;refGene_tx=XM_003988790.4;refGene_gn=CBX5;refGene_tx=XM_006933761.4;refGene_gn=CBX5;refGene_tx=XM_006933762.4;refGene_gn=CBX5;refGene_tx=XM_019835033.2

B4 82557064 82557165
refGene_gn=CBX5;refGene_tx=XM_003988789.5;refGene_gn=CBX5;refGene_tx=XM_003988790.4;refGene_gn=CBX5;refGene_tx=XM_006933761.4;refGene_gn=CBX5;refGene_tx=XM_006933762.4;refGene_gn=CBX5;refGene_tx=XM_019835033.2
B4 82560186 82560373
refGene_gn=CBX5;refGene_tx=XM_003988789.5;refGene_gn=CBX5;refGene_tx=XM_003988790.4;refGene_gn=CBX5;refGene_tx=XM_006933761.4;refGene_gn=CBX5;refGene_tx=XM_006933762.4;refGene_gn=CBX5;refGene_tx=XM_019835033.2
B4 82565668 82565805
refGene_gn=CBX5;refGene_tx=XM_003988789.5;refGene_gn=CBX5;refGene_tx=XM_003988790.4;refGene_gn=CBX5;refGene_tx=XM_006933761.4;refGene_gn=CBX5;refGene_tx=XM_006933762.4;refGene_gn=CBX5;refGene_tx=XM_019835033.2
B4 82592094 82592109
refGene_gn=HNRNPA1;refGene_tx=XM_003988792.4;refGene_gn=HNRNPA1;refGene_tx=XM_003988793.4;refGene_gn=HNRNPA1;refGene_tx=XM_019835032.1
B4 82592676 82592793
refGene_gn=HNRNPA1;refGene_tx=XM_003988792.4;refGene_gn=HNRNPA1;refGene_tx=XM_003988793.4;refGene_gn=HNRNPA1;refGene_tx=XM_019835032.1
B4 82593044 82593191
refGene_gn=HNRNPA1;refGene_tx=XM_003988792.4;refGene_gn=HNRNPA1;refGene_tx=XM_003988793.4;refGene_gn=HNRNPA1;refGene_tx=XM_019835032.1
B4 82593330 82593541
refGene_gn=HNRNPA1;refGene_tx=XM_003988792.4;refGene_gn=HNRNPA1;refGene_tx=XM_003988793.4;refGene_gn=HNRNPA1;refGene_tx=XM_019835032.1
B4 82593634 82593727
refGene_gn=HNRNPA1;refGene_tx=XM_003988792.4;refGene_gn=HNRNPA1;refGene_tx=XM_003988793.4;refGene_gn=HNRNPA1;refGene_tx=XM_019835032.1
B4 82593807 82593900
refGene_gn=HNRNPA1;refGene_tx=XM_003988792.4;refGene_gn=HNRNPA1;refGene_tx=XM_003988793.4;refGene_gn=HNRNPA1;refGene_tx=XM_019835032.1
B4 82594190 82594265
refGene_gn=HNRNPA1;refGene_tx=XM_003988792.4;refGene_gn=HNRNPA1;refGene_tx=XM_003988793.4;refGene_gn=HNRNPA1;refGene_tx=XM_019835032.1
B4 82594461 82594620
refGene_gn=HNRNPA1;refGene_tx=XM_003988793.4;refGene_gn=HNRNPA1;refGene_tx=XM_019835032.1
B4 82595181 82595337
refGene_gn=HNRNPA1;refGene_tx=XM_003988792.4;refGene_gn=HNRNPA1;refGene_tx=XM_003988793.4;refGene_gn=HNRNPA1;refGene_tx=XM_019835032.1
B4 82595568 82595624
refGene_gn=HNRNPA1;refGene_tx=XM_003988792.4;refGene_gn=HNRNPA1;refGene_tx=XM_003988793.4;refGene_gn=HNRNPA1;refGene_tx=XM_019835032.1
B4 82601836 82602844
refGene_gn=NFE2;refGene_tx=XM_011283975.3;refGene_gn=NFE2;refGene_tx=XM_011283976.3;refGene_gn=NFE2;refGene_tx=XM_019835035.1
B4 82604034 82604148
refGene_gn=NFE2;refGene_tx=XM_011283975.3;refGene_gn=NFE2;refGene_tx=XM_011283976.3;refGene_gn=NFE2;refGene_tx=XM_019835035.1
B4 82630767 82630785 refGene_gn=COPZ1;refGene_tx=XM_003988794.4
B4 82641983 82642052 refGene_gn=COPZ1;refGene_tx=XM_003988794.4

| | | | |
|---|---|---|---|
| B4 | 82643406 | 82643488 | refGene_gn=COPZ1;refGene_tx=XM_003988794.4 |
| B4 | 82644103 | 82644195 | refGene_gn=COPZ1;refGene_tx=XM_003988794.4 |
| B4 | 82645768 | 82645824 | refGene_gn=COPZ1;refGene_tx=XM_003988794.4 |
| B4 | 82647158 | 82647236 | refGene_gn=COPZ1;refGene_tx=XM_003988794.4 |
| B4 | 82647396 | 82647448 | refGene_gn=COPZ1;refGene_tx=XM_003988794.4 |
| B4 | 82648627 | 82648666 | refGene_gn=COPZ1;refGene_tx=XM_003988794.4 |
| B4 | 82649533 | 82649581 | refGene_gn=COPZ1;refGene_tx=XM_003988794.4 |
| B4 | 82660749 | 82661934 | refGene_gn=GPR84;refGene_tx=XM_006933782.3 |
| B4 | 82669821 | 82670052 | refGene_gn=ZNF385A;refGene_tx=XM_003988798.5 |
| B4 | 82670199 | 82670295 | refGene_gn=ZNF385A;refGene_tx=XM_003988798.5 |
| B4 | 82670516 | 82670686 | refGene_gn=ZNF385A;refGene_tx=XM_003988798.5 |
| B4 | 82671049 | 82671182 | refGene_gn=ZNF385A;refGene_tx=XM_003988798.5 |
| B4 | 82671183 | 82671293 | refGene_gn=ZNF385A;refGene_tx=XM_003988798.5 |
| B4 | 82673170 | 82673333 | refGene_gn=ZNF385A;refGene_tx=XM_003988798.5 |
| B4 | 82674953 | 82675064 | refGene_gn=ZNF385A;refGene_tx=XM_003988798.5 |
| B4 | 82682564 | 82682701 | refGene_gn=ZNF385A;refGene_tx=XM_003988798.5 |
| B4 | 82689352 | 82689362 | refGene_gn=ZNF385A;refGene_tx=XM_003988798.5 |
| B4 | 82694148 | 82694232 | refGene_gn=ITGA5;refGene_tx=XM_003988800.4 |
| B4 | 82695339 | 82695462 | refGene_gn=ITGA5;refGene_tx=XM_003988800.4 |
| B4 | 82696057 | 82696159 | refGene_gn=ITGA5;refGene_tx=XM_003988800.4 |
| B4 | 82697010 | 82697124 | refGene_gn=ITGA5;refGene_tx=XM_003988800.4 |
| B4 | 82697203 | 82697290 | refGene_gn=ITGA5;refGene_tx=XM_003988800.4 |
| B4 | 82698449 | 82698602 | refGene_gn=ITGA5;refGene_tx=XM_003988800.4 |
| B4 | 82699034 | 82699140 | refGene_gn=ITGA5;refGene_tx=XM_003988800.4 |
| B4 | 82699248 | 82699329 | refGene_gn=ITGA5;refGene_tx=XM_003988800.4 |
| B4 | 82699446 | 82699526 | refGene_gn=ITGA5;refGene_tx=XM_003988800.4 |
| B4 | 82699672 | 82699765 | refGene_gn=ITGA5;refGene_tx=XM_003988800.4 |
| B4 | 82699850 | 82700001 | refGene_gn=ITGA5;refGene_tx=XM_003988800.4 |
| B4 | 82700382 | 82700450 | refGene_gn=ITGA5;refGene_tx=XM_003988800.4 |
| B4 | 82700583 | 82700721 | refGene_gn=ITGA5;refGene_tx=XM_003988800.4 |
| B4 | 82701053 | 82701208 | refGene_gn=ITGA5;refGene_tx=XM_003988800.4 |
| B4 | 82701348 | 82701404 | refGene_gn=ITGA5;refGene_tx=XM_003988800.4 |
| B4 | 82701595 | 82701697 | refGene_gn=ITGA5;refGene_tx=XM_003988800.4 |
| B4 | 82701863 | 82701909 | refGene_gn=ITGA5;refGene_tx=XM_003988800.4 |
| B4 | 82702136 | 82702328 | refGene_gn=ITGA5;refGene_tx=XM_003988800.4 |
| B4 | 82702646 | 82702855 | refGene_gn=ITGA5;refGene_tx=XM_003988800.4 |
| B4 | 82703162 | 82703215 | refGene_gn=ITGA5;refGene_tx=XM_003988800.4 |
| B4 | 82703394 | 82703451 | refGene_gn=ITGA5;refGene_tx=XM_003988800.4 |
| B4 | 82705546 | 82705590 | refGene_gn=ITGA5;refGene_tx=XM_003988800.4 |
| B4 | 82705727 | 82705772 | refGene_gn=ITGA5;refGene_tx=XM_003988800.4 |
| B4 | 82706018 | 82706144 | refGene_gn=ITGA5;refGene_tx=XM_003988800.4 |
| B4 | 82706408 | 82706454 | refGene_gn=ITGA5;refGene_tx=XM_003988800.4 |
| B4 | 82706543 | 82706605 | refGene_gn=ITGA5;refGene_tx=XM_003988800.4 |
| B4 | 82706782 | 82706903 | refGene_gn=ITGA5;refGene_tx=XM_003988800.4 |
| B4 | 82707038 | 82707151 | refGene_gn=ITGA5;refGene_tx=XM_003988800.4 |
| B4 | 82708923 | 82709054 | refGene_gn=ITGA5;refGene_tx=XM_003988800.4 |
| B4 | 82715440 | 82715817 | refGene_gn=ITGA5;refGene_tx=XM_003988800.4 |
| B4 | 82753885 | 82753902 | refGene_gn=GTSF1;refGene_tx=XM_003988801.4;refGene_gn=GTSF1;refGene_tx=XM_019835039.2 |

B4 82754813 82754908
refGene_gn=GTSF1;refGene_tx=XM_003988801.4;refGene_gn=GTSF1;refGene_tx=XM_019835039.2
B4 82756268 82756332
refGene_gn=GTSF1;refGene_tx=XM_003988801.4;refGene_gn=GTSF1;refGene_tx=XM_019835039.2
B4 82756780 82756864
refGene_gn=GTSF1;refGene_tx=XM_003988801.4;refGene_gn=GTSF1;refGene_tx=XM_019835039.2
B4 82757302 82757429
refGene_gn=GTSF1;refGene_tx=XM_003988801.4;refGene_gn=GTSF1;refGene_tx=XM_019835039.2
B4 82759219 82759320
refGene_gn=GTSF1;refGene_tx=XM_003988801.4;refGene_gn=GTSF1;refGene_tx=XM_019835039.2
B4 82764756 82764801
refGene_gn=GTSF1;refGene_tx=XM_003988801.4;refGene_gn=GTSF1;refGene_tx=XM_019835039.2
B4 82765379 82765461 refGene_gn=GTSF1;refGene_tx=XM_019835039.2
B4 82774712 82774726
refGene_gn=LOC102899071;refGene_tx=XM_011283979.3
B4 82775485 82775556
refGene_gn=LOC102899071;refGene_tx=XM_011283979.3
B4 82777022 82777089
refGene_gn=LOC102899071;refGene_tx=XM_011283979.3
B4 82778263 82778365
refGene_gn=LOC102899071;refGene_tx=XM_011283979.3
B4 82778533 82778654
refGene_gn=LOC102899071;refGene_tx=XM_011283979.3
B4 82779328 82779421
refGene_gn=LOC102899071;refGene_tx=XM_011283979.3
B4 82784913 82785015
refGene_gn=NCKAP1L;refGene_tx=XM_003988802.5;refGene_gn=NCKAP1L;refGene_tx=XM_023257118.1
B4 82786123 82786234
refGene_gn=NCKAP1L;refGene_tx=XM_003988802.5;refGene_gn=NCKAP1L;refGene_tx=XM_023257118.1
B4 82786965 82787058
refGene_gn=NCKAP1L;refGene_tx=XM_003988802.5;refGene_gn=NCKAP1L;refGene_tx=XM_023257118.1
B4 82792838 82792895
refGene_gn=NCKAP1L;refGene_tx=XM_003988802.5;refGene_gn=NCKAP1L;refGene_tx=XM_023257118.1
B4 82793388 82793531
refGene_gn=NCKAP1L;refGene_tx=XM_003988802.5;refGene_gn=NCKAP1L;refGene_tx=XM_023257118.1
B4 82794693 82794784
refGene_gn=NCKAP1L;refGene_tx=XM_003988802.5;refGene_gn=NCKAP1L;refGene_tx=XM_023257118.1
B4 82794874 82795012

refGene_gn=NCKAP1L;refGene_tx=XM_003988802.5;refGene_gn=NCKAP1L;refGene_tx=XM_023257118.1
B4 82795824 82795873
refGene_gn=NCKAP1L;refGene_tx=XM_003988802.5;refGene_gn=NCKAP1L;refGene_tx=XM_023257118.1
B4 82795971 82796128
refGene_gn=NCKAP1L;refGene_tx=XM_003988802.5;refGene_gn=NCKAP1L;refGene_tx=XM_023257118.1
B4 82798848 82798905
refGene_gn=NCKAP1L;refGene_tx=XM_003988802.5;refGene_gn=NCKAP1L;refGene_tx=XM_023257118.1
B4 82799162 82799259
refGene_gn=NCKAP1L;refGene_tx=XM_003988802.5;refGene_gn=NCKAP1L;refGene_tx=XM_023257118.1
B4 82799814 82799924
refGene_gn=NCKAP1L;refGene_tx=XM_003988802.5;refGene_gn=NCKAP1L;refGene_tx=XM_023257118.1
B4 82800094 82800227
refGene_gn=NCKAP1L;refGene_tx=XM_003988802.5;refGene_gn=NCKAP1L;refGene_tx=XM_023257118.1
B4 82800694 82800776
refGene_gn=NCKAP1L;refGene_tx=XM_003988802.5;refGene_gn=NCKAP1L;refGene_tx=XM_023257118.1
B4 82800957 82801016
refGene_gn=NCKAP1L;refGene_tx=XM_003988802.5;refGene_gn=NCKAP1L;refGene_tx=XM_023257118.1
B4 82801231 82801377
refGene_gn=NCKAP1L;refGene_tx=XM_003988802.5;refGene_gn=NCKAP1L;refGene_tx=XM_023257118.1
B4 82802755 82802888
refGene_gn=NCKAP1L;refGene_tx=XM_003988802.5;refGene_gn=NCKAP1L;refGene_tx=XM_023257118.1
B4 82803157 82803277
refGene_gn=NCKAP1L;refGene_tx=XM_003988802.5;refGene_gn=NCKAP1L;refGene_tx=XM_023257118.1
B4 82805360 82805506
refGene_gn=NCKAP1L;refGene_tx=XM_003988802.5;refGene_gn=NCKAP1L;refGene_tx=XM_023257118.1
B4 82805789 82805921
refGene_gn=NCKAP1L;refGene_tx=XM_003988802.5;refGene_gn=NCKAP1L;refGene_tx=XM_023257118.1
B4 82807363 82807582 refGene_gn=NCKAP1L;refGene_tx=XM_003988802.5
B4 82809487 82809618
refGene_gn=NCKAP1L;refGene_tx=XM_003988802.5;refGene_gn=NCKAP1L;refGene_tx=XM_023257118.1
B4 82812318 82812416
refGene_gn=NCKAP1L;refGene_tx=XM_003988802.5;refGene_gn=NCKAP1L;refGene_tx=XM_023257118.1
B4 82812551 82812645
refGene_gn=NCKAP1L;refGene_tx=XM_003988802.5;refGene_gn=NCKAP1L;refGen

e_tx=XM_023257118.1
B4 82812801 82812884
refGene_gn=NCKAP1L;refGene_tx=XM_003988802.5;refGene_gn=NCKAP1L;refGene_tx=XM_023257118.1
B4 82813183 82813264
refGene_gn=NCKAP1L;refGene_tx=XM_003988802.5;refGene_gn=NCKAP1L;refGene_tx=XM_023257118.1
B4 82816220 82816314
refGene_gn=NCKAP1L;refGene_tx=XM_003988802.5;refGene_gn=NCKAP1L;refGene_tx=XM_023257118.1
B4 82817469 82817586
refGene_gn=NCKAP1L;refGene_tx=XM_003988802.5;refGene_gn=NCKAP1L;refGene_tx=XM_023257118.1
B4 82817961 82818071
refGene_gn=NCKAP1L;refGene_tx=XM_003988802.5;refGene_gn=NCKAP1L;refGene_tx=XM_023257118.1
B4 82819550 82819640
refGene_gn=NCKAP1L;refGene_tx=XM_003988802.5;refGene_gn=NCKAP1L;refGene_tx=XM_023257118.1
B4 82823798 82823930
refGene_gn=NCKAP1L;refGene_tx=XM_003988802.5;refGene_gn=NCKAP1L;refGene_tx=XM_023257118.1
B4 82830283 82830393 refGene_gn=PDE1B;refGene_tx=XM_011283982.3
B4 82842283 82842336 refGene_gn=PDE1B;refGene_tx=XM_019835041.2
B4 82846759 82846873
refGene_gn=PDE1B;refGene_tx=XM_011283982.3;refGene_gn=PDE1B;refGene_tx=XM_019835041.2
B4 82848225 82848408
refGene_gn=PDE1B;refGene_tx=XM_011283982.3;refGene_gn=PDE1B;refGene_tx=XM_019835041.2
B4 82848604 82848671
refGene_gn=PDE1B;refGene_tx=XM_011283982.3;refGene_gn=PDE1B;refGene_tx=XM_019835041.2
B4 82849018 82849135
refGene_gn=PDE1B;refGene_tx=XM_011283982.3;refGene_gn=PDE1B;refGene_tx=XM_019835041.2
B4 82851307 82851448
refGene_gn=PDE1B;refGene_tx=XM_011283982.3;refGene_gn=PDE1B;refGene_tx=XM_019835041.2
B4 82851857 82851958
refGene_gn=PDE1B;refGene_tx=XM_011283982.3;refGene_gn=PDE1B;refGene_tx=XM_019835041.2
B4 82852066 82852192
refGene_gn=PDE1B;refGene_tx=XM_011283982.3;refGene_gn=PDE1B;refGene_tx=XM_019835041.2
B4 82852316 82852418
refGene_gn=PDE1B;refGene_tx=XM_011283982.3;refGene_gn=PDE1B;refGene_tx=XM_019835041.2
B4 82853507 82853628
refGene_gn=PDE1B;refGene_tx=XM_011283982.3;refGene_gn=PDE1B;refGene_tx

=XM_019835041.2
B4 82853929 82854011
refGene_gn=PDE1B;refGene_tx=XM_011283982.3;refGene_gn=PDE1B;refGene_tx=XM_019835041.2
B4 82854341 82854450
refGene_gn=PDE1B;refGene_tx=XM_011283982.3;refGene_gn=PDE1B;refGene_tx=XM_019835041.2
B4 82854927 82855058
refGene_gn=PDE1B;refGene_tx=XM_011283982.3;refGene_gn=PDE1B;refGene_tx=XM_019835041.2
B4 82855566 82855670
refGene_gn=PDE1B;refGene_tx=XM_011283982.3;refGene_gn=PDE1B;refGene_tx=XM_019835041.2
B4 82858669 82858675 refGene_gn=PPP1R1A;refGene_tx=XM_023257119.1
B4 82859209 82859316 refGene_gn=PPP1R1A;refGene_tx=XM_023257119.1
B4 82860229 82860385 refGene_gn=PPP1R1A;refGene_tx=XM_023257119.1
B4 82860963 82861027 refGene_gn=PPP1R1A;refGene_tx=XM_023257119.1
B4 82861352 82861390 refGene_gn=PPP1R1A;refGene_tx=XM_023257119.1
B4 82862394 82862455 refGene_gn=PPP1R1A;refGene_tx=XM_023257119.1
B4 82866247 82866331 refGene_gn=PPP1R1A;refGene_tx=XM_023257119.1
B4 82905737 82905799
refGene_gn=LOC102901478;refGene_tx=XM_006933768.3;refGene_gn=LOC102901478;refGene_tx=XM_006933769.3
B4 82906246 82906333
refGene_gn=LOC102901478;refGene_tx=XM_006933768.3;refGene_gn=LOC102901478;refGene_tx=XM_006933769.3
B4 82906710 82906815
refGene_gn=LOC102901478;refGene_tx=XM_006933768.3;refGene_gn=LOC102901478;refGene_tx=XM_006933769.3
B4 82907956 82908004
refGene_gn=LOC102901478;refGene_tx=XM_006933768.3
B4 82909252 82909310
refGene_gn=LOC102901478;refGene_tx=XM_006933768.3;refGene_gn=LOC102901478;refGene_tx=XM_006933769.3
B4 83024500 83024642
refGene_gn=LOC102901584;refGene_tx=XM_019835043.1;refGene_gn=LOC102901584;refGene_tx=XM_019835045.1;refGene_gn=LOC102901584;refGene_tx=XM_019835046.1
B4 83025165 83025198
refGene_gn=LOC102901584;refGene_tx=XM_019835046.1
B4 83027112 83027202
refGene_gn=LOC102901584;refGene_tx=XM_019835046.1
B4 83030145 83030183
refGene_gn=LOC102901584;refGene_tx=XM_019835046.1
B4 83051530 83051672
refGene_gn=LOC102901584;refGene_tx=XM_019835042.1;refGene_gn=LOC102901584;refGene_tx=XM_019835044.1
B4 83052172 83052208
refGene_gn=LOC102901584;refGene_tx=XM_019835042.1;refGene_gn=LOC102901584;refGene_tx=XM_019835043.1;refGene_gn=LOC102901584;refGene_tx=XM_01

9835044.1;refGene_gn=LOC102901584;refGene_tx=XM_019835045.1
B4 83054124 83054255
refGene_gn=LOC102901584;refGene_tx=XM_019835042.1;refGene_gn=LOC102901584;refGene_tx=XM_019835043.1;refGene_gn=LOC102901584;refGene_tx=XM_019835044.1;refGene_gn=LOC102901584;refGene_tx=XM_019835045.1
B4 83055981 83056011
refGene_gn=LOC102901584;refGene_tx=XM_019835042.1;refGene_gn=LOC102901584;refGene_tx=XM_019835043.1;refGene_gn=LOC102901584;refGene_tx=XM_019835044.1;refGene_gn=LOC102901584;refGene_tx=XM_019835045.1
B4 83100195 83101145 refGene_gn=TESPA1;refGene_tx=XM_019835047.2
B4 83103166 83103375 refGene_gn=TESPA1;refGene_tx=XM_019835047.2
B4 83104895 83104994 refGene_gn=TESPA1;refGene_tx=XM_019835047.2
B4 83105268 83105305 refGene_gn=TESPA1;refGene_tx=XM_019835047.2
B4 83106070 83106124 refGene_gn=TESPA1;refGene_tx=XM_019835047.2
B4 83106750 83106800 refGene_gn=TESPA1;refGene_tx=XM_019835047.2
B4 83112459 83112502 refGene_gn=TESPA1;refGene_tx=XM_019835047.2
B4 83113448 83113611 refGene_gn=TESPA1;refGene_tx=XM_019835047.2
B4 83159883 83160879 refGene_gn=NEUROD4;refGene_tx=XM_019835048.1
B4 83243127 83244069
refGene_gn=LOC101081393;refGene_tx=XM_003988828.1
B4 83280376 83281318
refGene_gn=LOC101081906;refGene_tx=XM_003988830.4
B4 83314139 83315081
refGene_gn=LOC101082164;refGene_tx=XM_003988831.1
B4 83377961 83378614
refGene_gn=LOC101090598;refGene_tx=XM_023257798.1
B4 83378615 83378901
refGene_gn=LOC101090598;refGene_tx=XM_023257798.1
B4 83384028 83384871
refGene_gn=LOC111561480;refGene_tx=XM_023257799.1
B4 83409731 83410670
refGene_gn=LOC101081645;refGene_tx=XM_003988829.1
B4 83450798 83451740
refGene_gn=LOC101083947;refGene_tx=XM_006933788.3
B4 83463143 83464106
refGene_gn=LOC101084207;refGene_tx=XM_003988836.4
B4 83472385 83473324
refGene_gn=LOC101084462;refGene_tx=XM_003988837.1
B4 83484755 83485703
refGene_gn=LOC101084720;refGene_tx=XM_003988838.2
B4 83520312 83521254
refGene_gn=LOC101100488;refGene_tx=XM_003988897.1
B4 83532138 83533089
refGene_gn=LOC111561481;refGene_tx=XM_023257801.1
B4 83548411 83549350
refGene_gn=LOC101100737;refGene_tx=XM_011284052.1
B4 83684465 83684740
refGene_gn=LOC101100980;refGene_tx=XM_023257124.1
B4 83684742 83685173
refGene_gn=LOC101100980;refGene_tx=XM_023257124.1

B4 83685174 83685401
refGene_gn=LOC101100980;refGene_tx=XM_023257124.1
B4 83702614 83703553
refGene_gn=LOC101085730;refGene_tx=XM_011284064.2
B4 83721415 83722354
refGene_gn=LOC101085987;refGene_tx=XM_003988841.2
B4 83741014 83741599
refGene_gn=LOC101101234;refGene_tx=XM_023257761.1
B4 83755544 83756483
refGene_gn=LOC101101485;refGene_tx=XM_011284062.2
B4 83789170 83790109
refGene_gn=LOC105260754;refGene_tx=XM_023257123.1
B4 83796399 83797335
refGene_gn=LOC111561371;refGene_tx=XM_023257125.1
B4 83813842 83814790
refGene_gn=LOC111561369;refGene_tx=XM_023257121.1
B4 83820182 83821121
refGene_gn=LOC105260753;refGene_tx=XM_023257122.1
B4 83853229 83854165
refGene_gn=LOC101081060;refGene_tx=XM_006933820.2
B4 83862885 83863824
refGene_gn=LOC105260758;refGene_tx=XM_023257802.1
B4 83870392 83871331
refGene_gn=LOC105260757;refGene_tx=XM_011284059.2
B4 83902918 83903854
refGene_gn=LOC105260751;refGene_tx=XM_011284049.1
B4 83912574 83913513
refGene_gn=LOC105260756;refGene_tx=XM_011284058.2
B4 83920080 83921019
refGene_gn=LOC101081313;refGene_tx=XM_011284057.2
B4 83935123 83936062
refGene_gn=LOC101086734;refGene_tx=XM_011284048.1
B4 83950959 83951895
refGene_gn=LOC101086987;refGene_tx=XM_003988844.1
B4 83979388 83980336
refGene_gn=LOC101081560;refGene_tx=XM_003988900.1
B4 84004388 84005327
refGene_gn=LOC101081818;refGene_tx=XM_011284056.1
B4 84063029 84063977
refGene_gn=LOC101082325;refGene_tx=XM_011284055.1
B4 84075876 84076848
refGene_gn=LOC101082580;refGene_tx=XM_006933821.1
B4 84097933 84098872
refGene_gn=LOC101087755;refGene_tx=XM_011284047.1
B4 84119335 84120274
refGene_gn=LOC101082834;refGene_tx=XM_011284054.1
B4 84155529 84156468
refGene_gn=LOC101088007;refGene_tx=XM_003988846.1
B4 84187014 84187941
refGene_gn=LOC101088501;refGene_tx=XM_011284046.1

B4 84195307 84195369
refGene_gn=LOC101083349;refGene_tx=XM_003988902.2
B4 84196291 84197201
refGene_gn=LOC101083349;refGene_tx=XM_003988902.2
B4 84226462 84227392
refGene_gn=LOC101088763;refGene_tx=XM_003988848.1
B4 84259277 84260222
refGene_gn=LOC101083860;refGene_tx=XM_003988904.3
B4 84285523 84286450
refGene_gn=LOC101089023;refGene_tx=XM_003988849.1
B4 84312113 84313049
refGene_gn=LOC101084119;refGene_tx=XM_006933822.1
B4 84329658 84330597
refGene_gn=LOC101089279;refGene_tx=XM_003988850.2
B4 84343000 84344041
refGene_gn=LOC101084373;refGene_tx=XM_003988906.1
B4 84379178 84379676 refGene_gn=METTL7B;refGene_tx=XM_003988851.4
B4 84380769 84381006 refGene_gn=METTL7B;refGene_tx=XM_003988851.4
B4 84382125 84382356
refGene_gn=ITGA7;refGene_tx=XM_003988852.3;refGene_gn=ITGA7;refGene_tx=XM_003988853.3;refGene_gn=ITGA7;refGene_tx=XM_006933789.2;refGene_gn=ITGA7;refGene_tx=XM_006933790.2;refGene_gn=ITGA7;refGene_tx=XM_011284045.2
B4 84383297 84383410 refGene_gn=ITGA7;refGene_tx=XM_006933789.2
B4 84384386 84384512
refGene_gn=ITGA7;refGene_tx=XM_003988852.3;refGene_gn=ITGA7;refGene_tx=XM_003988853.3;refGene_gn=ITGA7;refGene_tx=XM_006933789.2;refGene_gn=ITGA7;refGene_tx=XM_006933790.2;refGene_gn=ITGA7;refGene_tx=XM_011284045.2
B4 84384600 84384699
refGene_gn=ITGA7;refGene_tx=XM_003988852.3;refGene_gn=ITGA7;refGene_tx=XM_003988853.3;refGene_gn=ITGA7;refGene_tx=XM_006933789.2;refGene_gn=ITGA7;refGene_tx=XM_006933790.2;refGene_gn=ITGA7;refGene_tx=XM_011284045.2
B4 84384923 84385037
refGene_gn=ITGA7;refGene_tx=XM_003988852.3;refGene_gn=ITGA7;refGene_tx=XM_003988853.3;refGene_gn=ITGA7;refGene_tx=XM_006933789.2;refGene_gn=ITGA7;refGene_tx=XM_006933790.2;refGene_gn=ITGA7;refGene_tx=XM_011284045.2
B4 84387334 84387466
refGene_gn=ITGA7;refGene_tx=XM_003988852.3;refGene_gn=ITGA7;refGene_tx=XM_003988853.3;refGene_gn=ITGA7;refGene_tx=XM_006933789.2;refGene_gn=ITGA7;refGene_tx=XM_006933790.2;refGene_gn=ITGA7;refGene_tx=XM_011284045.2
B4 84387644 84387821
refGene_gn=ITGA7;refGene_tx=XM_003988852.3;refGene_gn=ITGA7;refGene_tx=XM_003988853.3;refGene_gn=ITGA7;refGene_tx=XM_006933789.2;refGene_gn=ITGA7;refGene_tx=XM_006933790.2;refGene_gn=ITGA7;refGene_tx=XM_011284045.2
B4 84388356 84388459

refGene_gn=ITGA7;refGene_tx=XM_003988852.3;refGene_gn=ITGA7;refGene_tx=XM_003988853.3;refGene_gn=ITGA7;refGene_tx=XM_006933789.2;refGene_gn=ITGA7;refGene_tx=XM_006933790.2;refGene_gn=ITGA7;refGene_tx=XM_011284045.2
B4 84388602 84388677
refGene_gn=ITGA7;refGene_tx=XM_003988852.3;refGene_gn=ITGA7;refGene_tx=XM_003988853.3;refGene_gn=ITGA7;refGene_tx=XM_006933789.2;refGene_gn=ITGA7;refGene_tx=XM_006933790.2;refGene_gn=ITGA7;refGene_tx=XM_011284045.2
B4 84388819 84388899
refGene_gn=ITGA7;refGene_tx=XM_003988852.3;refGene_gn=ITGA7;refGene_tx=XM_003988853.3;refGene_gn=ITGA7;refGene_tx=XM_006933789.2;refGene_gn=ITGA7;refGene_tx=XM_006933790.2;refGene_gn=ITGA7;refGene_tx=XM_011284045.2
B4 84389005 84389086
refGene_gn=ITGA7;refGene_tx=XM_003988852.3;refGene_gn=ITGA7;refGene_tx=XM_003988853.3;refGene_gn=ITGA7;refGene_tx=XM_006933789.2;refGene_gn=ITGA7;refGene_tx=XM_006933790.2;refGene_gn=ITGA7;refGene_tx=XM_011284045.2
B4 84389168 84389361
refGene_gn=ITGA7;refGene_tx=XM_003988852.3;refGene_gn=ITGA7;refGene_tx=XM_003988853.3;refGene_gn=ITGA7;refGene_tx=XM_006933789.2;refGene_gn=ITGA7;refGene_tx=XM_006933790.2;refGene_gn=ITGA7;refGene_tx=XM_011284045.2
B4 84389924 84390040
refGene_gn=ITGA7;refGene_tx=XM_003988852.3;refGene_gn=ITGA7;refGene_tx=XM_003988853.3;refGene_gn=ITGA7;refGene_tx=XM_006933789.2;refGene_gn=ITGA7;refGene_tx=XM_006933790.2;refGene_gn=ITGA7;refGene_tx=XM_011284045.2
B4 84390717 84390867
refGene_gn=ITGA7;refGene_tx=XM_003988852.3;refGene_gn=ITGA7;refGene_tx=XM_003988853.3;refGene_gn=ITGA7;refGene_tx=XM_006933789.2;refGene_gn=ITGA7;refGene_tx=XM_006933790.2;refGene_gn=ITGA7;refGene_tx=XM_011284045.2
B4 84391291 84391461
refGene_gn=ITGA7;refGene_tx=XM_003988852.3;refGene_gn=ITGA7;refGene_tx=XM_003988853.3;refGene_gn=ITGA7;refGene_tx=XM_006933789.2;refGene_gn=ITGA7;refGene_tx=XM_006933790.2;refGene_gn=ITGA7;refGene_tx=XM_011284045.2
B4 84391606 84391668
refGene_gn=ITGA7;refGene_tx=XM_003988852.3;refGene_gn=ITGA7;refGene_tx=XM_003988853.3;refGene_gn=ITGA7;refGene_tx=XM_006933789.2;refGene_gn=ITGA7;refGene_tx=XM_006933790.2;refGene_gn=ITGA7;refGene_tx=XM_011284045.2
B4 84391824 84391920
refGene_gn=ITGA7;refGene_tx=XM_003988852.3;refGene_gn=ITGA7;refGene_tx=XM_003988853.3;refGene_gn=ITGA7;refGene_tx=XM_006933789.2;refGene_gn=ITGA7;refGene_tx=XM_006933790.2;refGene_gn=ITGA7;refGene_tx=XM_011284045.2
B4 84392029 84392157

refGene_gn=ITGA7;refGene_tx=XM_003988852.3;refGene_gn=ITGA7;refGene_tx=XM_003988853.3;refGene_gn=ITGA7;refGene_tx=XM_006933789.2;refGene_gn=ITGA7;refGene_tx=XM_006933790.2;refGene_gn=ITGA7;refGene_tx=XM_011284045.2
B4 84392285 84392374
refGene_gn=ITGA7;refGene_tx=XM_003988852.3;refGene_gn=ITGA7;refGene_tx=XM_003988853.3;refGene_gn=ITGA7;refGene_tx=XM_006933789.2;refGene_gn=ITGA7;refGene_tx=XM_006933790.2;refGene_gn=ITGA7;refGene_tx=XM_011284045.2
B4 84392737 84392931
refGene_gn=ITGA7;refGene_tx=XM_003988852.3;refGene_gn=ITGA7;refGene_tx=XM_003988853.3;refGene_gn=ITGA7;refGene_tx=XM_006933789.2;refGene_gn=ITGA7;refGene_tx=XM_006933790.2;refGene_gn=ITGA7;refGene_tx=XM_011284045.2
B4 84393059 84393267
refGene_gn=ITGA7;refGene_tx=XM_003988852.3;refGene_gn=ITGA7;refGene_tx=XM_003988853.3;refGene_gn=ITGA7;refGene_tx=XM_006933789.2;refGene_gn=ITGA7;refGene_tx=XM_006933790.2;refGene_gn=ITGA7;refGene_tx=XM_011284045.2
B4 84394115 84394235
refGene_gn=ITGA7;refGene_tx=XM_003988853.3;refGene_gn=ITGA7;refGene_tx=XM_006933789.2;refGene_gn=ITGA7;refGene_tx=XM_006933790.2;refGene_gn=ITGA7;refGene_tx=XM_011284045.2
B4 84394527 84394659
refGene_gn=ITGA7;refGene_tx=XM_003988852.3;refGene_gn=ITGA7;refGene_tx=XM_003988853.3;refGene_gn=ITGA7;refGene_tx=XM_006933789.2;refGene_gn=ITGA7;refGene_tx=XM_011284045.2
B4 84395139 84395395
refGene_gn=ITGA7;refGene_tx=XM_003988852.3;refGene_gn=ITGA7;refGene_tx=XM_003988853.3;refGene_gn=ITGA7;refGene_tx=XM_006933789.2;refGene_gn=ITGA7;refGene_tx=XM_006933790.2;refGene_gn=ITGA7;refGene_tx=XM_011284045.2
B4 84396588 84396668
refGene_gn=ITGA7;refGene_tx=XM_003988852.3;refGene_gn=ITGA7;refGene_tx=XM_003988853.3;refGene_gn=ITGA7;refGene_tx=XM_006933789.2;refGene_gn=ITGA7;refGene_tx=XM_006933790.2;refGene_gn=ITGA7;refGene_tx=XM_011284045.2
B4 84396769 84396897
refGene_gn=ITGA7;refGene_tx=XM_003988852.3;refGene_gn=ITGA7;refGene_tx=XM_003988853.3;refGene_gn=ITGA7;refGene_tx=XM_006933789.2;refGene_gn=ITGA7;refGene_tx=XM_006933790.2;refGene_gn=ITGA7;refGene_tx=XM_011284045.2
B4 84401174 84401380
refGene_gn=ITGA7;refGene_tx=XM_003988852.3;refGene_gn=ITGA7;refGene_tx=XM_003988853.3;refGene_gn=ITGA7;refGene_tx=XM_006933789.2;refGene_gn=ITGA7;refGene_tx=XM_006933790.2;refGene_gn=ITGA7;refGene_tx=XM_011284045.2
B4 84411561 84411622 refGene_gn=BLOC1S1;refGene_tx=XM_003988854.5
B4 84412414 84412487 refGene_gn=BLOC1S1;refGene_tx=XM_003988854.5
B4 84414569 84414702 refGene_gn=BLOC1S1;refGene_tx=XM_003988854.5

B4 84414977 84415088 refGene_gn=BLOC1S1;refGene_tx=XM_003988854.5
B4 84416567 84416877
refGene_gn=RDH5;refGene_tx=XM_006933791.3;refGene_gn=RDH5;refGene_tx=XM_019835050.1
B4 84417032 84417291
refGene_gn=RDH5;refGene_tx=XM_006933791.3;refGene_gn=RDH5;refGene_tx=XM_019835050.1
B4 84419201 84419365
refGene_gn=RDH5;refGene_tx=XM_006933791.3;refGene_gn=RDH5;refGene_tx=XM_019835050.1
B4 84419652 84419876
refGene_gn=RDH5;refGene_tx=XM_006933791.3;refGene_gn=RDH5;refGene_tx=XM_019835050.1
B4 84421157 84421223
refGene_gn=CD63;refGene_tx=NM_001009855.1;refGene_gn=CD63;refGene_tx=XM_023256248.1
B4 84421400 84421484
refGene_gn=CD63;refGene_tx=NM_001009855.1;refGene_gn=CD63;refGene_tx=XM_023256248.1
B4 84421725 84421866
refGene_gn=CD63;refGene_tx=NM_001009855.1;refGene_gn=CD63;refGene_tx=XM_023256248.1
B4 84421978 84422074
refGene_gn=CD63;refGene_tx=NM_001009855.1;refGene_gn=CD63;refGene_tx=XM_023256248.1
B4 84422173 84422248
refGene_gn=CD63;refGene_tx=NM_001009855.1;refGene_gn=CD63;refGene_tx=XM_023256248.1
B4 84422423 84422612
refGene_gn=CD63;refGene_tx=NM_001009855.1;refGene_gn=CD63;refGene_tx=XM_023256248.1
B4 84423487 84423553
refGene_gn=CD63;refGene_tx=NM_001009855.1;refGene_gn=CD63;refGene_tx=XM_023256248.1
B4 84435408 84435844 refGene_gn=GDF11;refGene_tx=XM_003988856.3
B4 84440770 84441168 refGene_gn=GDF11;refGene_tx=XM_003988856.3
B4 84441693 84442074 refGene_gn=GDF11;refGene_tx=XM_003988856.3
B4 84449592 84449634 refGene_gn=SARNP;refGene_tx=XM_003988857.4
B4 84453905 84453995 refGene_gn=SARNP;refGene_tx=XM_003988857.4
B4 84477084 84477153 refGene_gn=SARNP;refGene_tx=XM_003988857.4
B4 84478296 84478322 refGene_gn=SARNP;refGene_tx=XM_003988857.4
B4 84480707 84480736 refGene_gn=SARNP;refGene_tx=XM_003988857.4
B4 84481179 84481253 refGene_gn=SARNP;refGene_tx=XM_003988857.4
B4 84482309 84482361 refGene_gn=SARNP;refGene_tx=XM_003988857.4
B4 84486596 84486664 refGene_gn=SARNP;refGene_tx=XM_003988857.4
B4 84486876 84486923 refGene_gn=SARNP;refGene_tx=XM_003988857.4
B4 84488586 84488686 refGene_gn=SARNP;refGene_tx=XM_003988857.4
B4 84499891 84499927 refGene_gn=SARNP;refGene_tx=XM_003988857.4
B4 84500256 84500342 refGene_gn=ORMDL2;refGene_tx=XM_003988859.4
B4 84501268 84501443

refGene_gn=ORMDL2;refGene_tx=XM_003988859.4;refGene_gn=ORMDL2;refGene_tx=XM_019835055.2;refGene_gn=ORMDL2;refGene_tx=XM_019835056.2
B4 84501608 84501760
refGene_gn=ORMDL2;refGene_tx=XM_003988859.4;refGene_gn=ORMDL2;refGene_tx=XM_019835055.2;refGene_gn=ORMDL2;refGene_tx=XM_019835056.2
B4 84502335 84502471
refGene_gn=ORMDL2;refGene_tx=XM_003988859.4;refGene_gn=ORMDL2;refGene_tx=XM_019835055.2;refGene_gn=ORMDL2;refGene_tx=XM_019835056.2
B4 84503812 84504023
refGene_gn=DNAJC14;refGene_tx=XM_003988861.5;refGene_gn=DNAJC14;refGene_tx=XM_006933792.3;refGene_gn=DNAJC14;refGene_tx=XM_019835052.2
B4 84504223 84504326
refGene_gn=DNAJC14;refGene_tx=XM_003988861.5;refGene_gn=DNAJC14;refGene_tx=XM_006933792.3;refGene_gn=DNAJC14;refGene_tx=XM_019835052.2
B4 84504408 84504569
refGene_gn=DNAJC14;refGene_tx=XM_003988861.5;refGene_gn=DNAJC14;refGene_tx=XM_006933792.3;refGene_gn=DNAJC14;refGene_tx=XM_019835052.2
B4 84505179 84505299
refGene_gn=DNAJC14;refGene_tx=XM_003988861.5;refGene_gn=DNAJC14;refGene_tx=XM_006933792.3;refGene_gn=DNAJC14;refGene_tx=XM_019835052.2
B4 84505527 84505634
refGene_gn=DNAJC14;refGene_tx=XM_003988861.5;refGene_gn=DNAJC14;refGene_tx=XM_006933792.3;refGene_gn=DNAJC14;refGene_tx=XM_019835052.2
B4 84508162 84509572
refGene_gn=DNAJC14;refGene_tx=XM_003988861.5;refGene_gn=DNAJC14;refGene_tx=XM_006933792.3;refGene_gn=DNAJC14;refGene_tx=XM_019835052.2
B4 84512749 84512924
refGene_gn=LOC101084636;refGene_tx=XM_011284042.3
B4 84513523 84514105
refGene_gn=LOC101084636;refGene_tx=XM_011284042.3;refGene_gn=LOC101084636;refGene_tx=XM_019835053.2;refGene_gn=LOC101084636;refGene_tx=XM_019835054.1
B4 84514522 84514680
refGene_gn=LOC101084636;refGene_tx=XM_011284042.3;refGene_gn=LOC101084636;refGene_tx=XM_019835053.2;refGene_gn=LOC101084636;refGene_tx=XM_019835054.1
B4 84514801 84514939
refGene_gn=LOC101084636;refGene_tx=XM_011284042.3;refGene_gn=LOC101084636;refGene_tx=XM_019835053.2;refGene_gn=LOC101084636;refGene_tx=XM_019835054.1
B4 84516611 84517067 refGene_gn=MMP19;refGene_tx=XM_003988862.3
B4 84517246 84517374 refGene_gn=MMP19;refGene_tx=XM_003988862.3
B4 84517536 84517701 refGene_gn=MMP19;refGene_tx=XM_003988862.3
B4 84518298 84518424 refGene_gn=MMP19;refGene_tx=XM_003988862.3
B4 84518857 84519103 refGene_gn=MMP19;refGene_tx=XM_003988862.3
B4 84519588 84519804 refGene_gn=MMP19;refGene_tx=XM_003988862.3
B4 84520040 84520171 refGene_gn=MMP19;refGene_tx=XM_003988862.3
B4 84521078 84521164 refGene_gn=MMP19;refGene_tx=XM_003988862.3
B4 84521473 84521563 refGene_gn=MMP19;refGene_tx=XM_003988862.3
B4 84552114 84552598 refGene_gn=PYM1;refGene_tx=XM_003988863.5

B4 84552989 84553083 refGene_gn=PYM1;refGene_tx=XM_003988863.5
B4 84569750 84569784 refGene_gn=PYM1;refGene_tx=XM_003988863.5
B4 84578088 84578142 refGene_gn=DGKA;refGene_tx=XM_019835058.2
B4 84578645 84578787
refGene_gn=DGKA;refGene_tx=XM_011284040.3;refGene_gn=DGKA;refGene_tx=XM_019835057.2;refGene_gn=DGKA;refGene_tx=XM_019835058.2;refGene_gn=DGKA;refGene_tx=XM_023257126.1;refGene_gn=DGKA;refGene_tx=XM_023257127.1
B4 84579202 84579276
refGene_gn=DGKA;refGene_tx=XM_011284040.3;refGene_gn=DGKA;refGene_tx=XM_019835057.2;refGene_gn=DGKA;refGene_tx=XM_019835058.2;refGene_gn=DGKA;refGene_tx=XM_023257126.1;refGene_gn=DGKA;refGene_tx=XM_023257127.1
B4 84579607 84579740
refGene_gn=DGKA;refGene_tx=XM_011284040.3;refGene_gn=DGKA;refGene_tx=XM_019835057.2;refGene_gn=DGKA;refGene_tx=XM_019835058.2;refGene_gn=DGKA;refGene_tx=XM_023257126.1;refGene_gn=DGKA;refGene_tx=XM_023257127.1
B4 84579987 84580062
refGene_gn=DGKA;refGene_tx=XM_011284040.3;refGene_gn=DGKA;refGene_tx=XM_019835057.2;refGene_gn=DGKA;refGene_tx=XM_019835058.2;refGene_gn=DGKA;refGene_tx=XM_023257126.1;refGene_gn=DGKA;refGene_tx=XM_023257127.1
B4 84580525 84580575
refGene_gn=DGKA;refGene_tx=XM_011284040.3;refGene_gn=DGKA;refGene_tx=XM_019835057.2;refGene_gn=DGKA;refGene_tx=XM_019835058.2;refGene_gn=DGKA;refGene_tx=XM_023257126.1;refGene_gn=DGKA;refGene_tx=XM_023257127.1
B4 84580938 84581013
refGene_gn=DGKA;refGene_tx=XM_011284040.3;refGene_gn=DGKA;refGene_tx=XM_019835057.2;refGene_gn=DGKA;refGene_tx=XM_019835058.2;refGene_gn=DGKA;refGene_tx=XM_023257126.1;refGene_gn=DGKA;refGene_tx=XM_023257127.1
B4 84581211 84581331
refGene_gn=DGKA;refGene_tx=XM_011284040.3;refGene_gn=DGKA;refGene_tx=XM_019835057.2;refGene_gn=DGKA;refGene_tx=XM_019835058.2;refGene_gn=DGKA;refGene_tx=XM_023257126.1;refGene_gn=DGKA;refGene_tx=XM_023257127.1
B4 84581443 84581558
refGene_gn=DGKA;refGene_tx=XM_011284040.3;refGene_gn=DGKA;refGene_tx=XM_019835057.2;refGene_gn=DGKA;refGene_tx=XM_019835058.2;refGene_gn=DGKA;refGene_tx=XM_023257126.1;refGene_gn=DGKA;refGene_tx=XM_023257127.1
B4 84582288 84582377
refGene_gn=DGKA;refGene_tx=XM_011284040.3;refGene_gn=DGKA;refGene_tx=XM_019835057.2;refGene_gn=DGKA;refGene_tx=XM_019835058.2;refGene_gn=DGKA;refGene_tx=XM_023257126.1;refGene_gn=DGKA;refGene_tx=XM_023257127.1
B4 84582514 84582634
refGene_gn=DGKA;refGene_tx=XM_011284040.3;refGene_gn=DGKA;refGene_tx=XM_019835057.2;refGene_gn=DGKA;refGene_tx=XM_019835058.2;refGene_gn=DGKA;refGene_tx=XM_023257126.1;refGene_gn=DGKA;refGene_tx=XM_023257127.1
B4 84582835 84582934
refGene_gn=DGKA;refGene_tx=XM_011284040.3;refGene_gn=DGKA;refGene_tx=XM_019835057.2;refGene_gn=DGKA;refGene_tx=XM_019835058.2;refGene_gn=DGKA;refGene_tx=XM_023257126.1;refGene_gn=DGKA;refGene_tx=XM_023257127.1
B4 84583132 84583216
refGene_gn=DGKA;refGene_tx=XM_011284040.3;refGene_gn=DGKA;refGene_tx=XM_019835057.2;refGene_gn=DGKA;refGene_tx=XM_019835058.2;refGene_gn=DGK

A;refGene_tx=XM_023257126.1;refGene_gn=DGKA;refGene_tx=XM_023257127.1
B4 84583433 84583507
refGene_gn=DGKA;refGene_tx=XM_011284040.3;refGene_gn=DGKA;refGene_tx=XM_019835057.2;refGene_gn=DGKA;refGene_tx=XM_019835058.2;refGene_gn=DGKA;refGene_tx=XM_023257126.1;refGene_gn=DGKA;refGene_tx=XM_023257127.1
B4 84583667 84583742
refGene_gn=DGKA;refGene_tx=XM_011284040.3;refGene_gn=DGKA;refGene_tx=XM_019835057.2;refGene_gn=DGKA;refGene_tx=XM_019835058.2;refGene_gn=DGKA;refGene_tx=XM_023257126.1;refGene_gn=DGKA;refGene_tx=XM_023257127.1
B4 84584359 84584445
refGene_gn=DGKA;refGene_tx=XM_011284040.3;refGene_gn=DGKA;refGene_tx=XM_019835057.2;refGene_gn=DGKA;refGene_tx=XM_019835058.2;refGene_gn=DGKA;refGene_tx=XM_023257126.1;refGene_gn=DGKA;refGene_tx=XM_023257127.1
B4 84584536 84584626
refGene_gn=DGKA;refGene_tx=XM_011284040.3;refGene_gn=DGKA;refGene_tx=XM_019835057.2;refGene_gn=DGKA;refGene_tx=XM_019835058.2;refGene_gn=DGKA;refGene_tx=XM_023257126.1;refGene_gn=DGKA;refGene_tx=XM_023257127.1
B4 84591751 84591912
refGene_gn=DGKA;refGene_tx=XM_011284040.3;refGene_gn=DGKA;refGene_tx=XM_019835057.2;refGene_gn=DGKA;refGene_tx=XM_019835058.2;refGene_gn=DGKA;refGene_tx=XM_023257126.1;refGene_gn=DGKA;refGene_tx=XM_023257127.1
B4 84592169 84592234
refGene_gn=DGKA;refGene_tx=XM_011284040.3;refGene_gn=DGKA;refGene_tx=XM_019835057.2;refGene_gn=DGKA;refGene_tx=XM_019835058.2;refGene_gn=DGKA;refGene_tx=XM_023257126.1;refGene_gn=DGKA;refGene_tx=XM_023257127.1
B4 84592946 84593037
refGene_gn=DGKA;refGene_tx=XM_011284040.3;refGene_gn=DGKA;refGene_tx=XM_019835057.2;refGene_gn=DGKA;refGene_tx=XM_019835058.2;refGene_gn=DGKA;refGene_tx=XM_023257126.1;refGene_gn=DGKA;refGene_tx=XM_023257127.1
B4 84593326 84593525
refGene_gn=DGKA;refGene_tx=XM_011284040.3;refGene_gn=DGKA;refGene_tx=XM_019835057.2;refGene_gn=DGKA;refGene_tx=XM_019835058.2;refGene_gn=DGKA;refGene_tx=XM_023257126.1;refGene_gn=DGKA;refGene_tx=XM_023257127.1
B4 84593635 84593756
refGene_gn=DGKA;refGene_tx=XM_011284040.3;refGene_gn=DGKA;refGene_tx=XM_019835057.2;refGene_gn=DGKA;refGene_tx=XM_019835058.2;refGene_gn=DGKA;refGene_tx=XM_023257126.1;refGene_gn=DGKA;refGene_tx=XM_023257127.1
B4 84593950 84594011
refGene_gn=DGKA;refGene_tx=XM_011284040.3;refGene_gn=DGKA;refGene_tx=XM_019835057.2;refGene_gn=DGKA;refGene_tx=XM_019835058.2;refGene_gn=DGKA;refGene_tx=XM_023257126.1;refGene_gn=DGKA;refGene_tx=XM_023257127.1
B4 84594284 84594368
refGene_gn=DGKA;refGene_tx=XM_011284040.3;refGene_gn=DGKA;refGene_tx=XM_019835057.2;refGene_gn=DGKA;refGene_tx=XM_019835058.2;refGene_gn=DGKA;refGene_tx=XM_023257126.1;refGene_gn=DGKA;refGene_tx=XM_023257127.1
B4 84594857 84595002
refGene_gn=PMEL;refGene_tx=XM_003988865.4;refGene_gn=PMEL;refGene_tx=XM_019835059.2
B4 84595621 84595709 refGene_gn=PMEL;refGene_tx=XM_003988865.4
B4 84595811 84596017

refGene_gn=PMEL;refGene_tx=XM_003988865.4;refGene_gn=PMEL;refGene_tx=XM_019835059.2
B4 84596127 84596212
refGene_gn=PMEL;refGene_tx=XM_003988865.4;refGene_gn=PMEL;refGene_tx=XM_019835059.2
B4 84596460 84596577
refGene_gn=PMEL;refGene_tx=XM_003988865.4;refGene_gn=PMEL;refGene_tx=XM_019835059.2
B4 84597326 84598046
refGene_gn=PMEL;refGene_tx=XM_003988865.4;refGene_gn=PMEL;refGene_tx=XM_019835059.2
B4 84598295 84598457
refGene_gn=PMEL;refGene_tx=XM_003988865.4;refGene_gn=PMEL;refGene_tx=XM_019835059.2
B4 84598784 84598919
refGene_gn=PMEL;refGene_tx=XM_003988865.4;refGene_gn=PMEL;refGene_tx=XM_019835059.2
B4 84599632 84599779
refGene_gn=PMEL;refGene_tx=XM_003988865.4;refGene_gn=PMEL;refGene_tx=XM_019835059.2
B4 84599925 84600036
refGene_gn=PMEL;refGene_tx=XM_003988865.4;refGene_gn=PMEL;refGene_tx=XM_019835059.2
B4 84602271 84602347
refGene_gn=PMEL;refGene_tx=XM_003988865.4;refGene_gn=PMEL;refGene_tx=XM_019835059.2
B4 84603350 84603466
refGene_gn=CDK2;refGene_tx=XM_003988868.5;refGene_gn=CDK2;refGene_tx=XM_003988869.5;refGene_gn=CDK2;refGene_tx=XM_019835062.2
B4 84604176 84604254
refGene_gn=CDK2;refGene_tx=XM_003988868.5;refGene_gn=CDK2;refGene_tx=XM_003988869.5;refGene_gn=CDK2;refGene_tx=XM_019835062.2
B4 84604358 84604479
refGene_gn=CDK2;refGene_tx=XM_003988868.5;refGene_gn=CDK2;refGene_tx=XM_003988869.5;refGene_gn=CDK2;refGene_tx=XM_019835062.2
B4 84605142 84605313
refGene_gn=CDK2;refGene_tx=XM_003988868.5;refGene_gn=CDK2;refGene_tx=XM_003988869.5;refGene_gn=CDK2;refGene_tx=XM_019835062.2
B4 84605792 84605894
refGene_gn=CDK2;refGene_tx=XM_003988868.5;refGene_gn=CDK2;refGene_tx=XM_019835062.2
B4 84606563 84606707 refGene_gn=CDK2;refGene_tx=XM_019835062.2
B4 84607010 84607214
refGene_gn=CDK2;refGene_tx=XM_003988868.5;refGene_gn=CDK2;refGene_tx=XM_003988869.5;refGene_gn=CDK2;refGene_tx=XM_019835062.2
B4 84607495 84607600
refGene_gn=CDK2;refGene_tx=XM_003988868.5;refGene_gn=CDK2;refGene_tx=XM_003988869.5;refGene_gn=CDK2;refGene_tx=XM_019835062.2
B4 84620315 84620478
refGene_gn=RAB5B;refGene_tx=XM_003988870.5;refGene_gn=RAB5B;refGene_tx

=XM_006933797.4
B4	84621548	84621700
refGene_gn=RAB5B;refGene_tx=XM_003988870.5;refGene_gn=RAB5B;refGene_tx
=XM_006933797.4
B4	84622091	84622214
refGene_gn=RAB5B;refGene_tx=XM_003988870.5;refGene_gn=RAB5B;refGene_tx
=XM_006933797.4
B4	84622761	84622855
refGene_gn=RAB5B;refGene_tx=XM_003988870.5;refGene_gn=RAB5B;refGene_tx
=XM_006933797.4
B4	84623191	84623307
refGene_gn=RAB5B;refGene_tx=XM_003988870.5;refGene_gn=RAB5B;refGene_tx
=XM_006933797.4
B4	84627855	84627945	refGene_gn=SUOX;refGene_tx=XM_019835061.2
B4	84629129	84629465
refGene_gn=SUOX;refGene_tx=XM_003988872.5;refGene_gn=SUOX;refGene_tx=X
M_006933798.4;refGene_gn=SUOX;refGene_tx=XM_019835060.2;refGene_gn=SUO
X;refGene_tx=XM_019835061.2;refGene_gn=SUOX;refGene_tx=XM_023257134.1
B4	84629735	84629913
refGene_gn=SUOX;refGene_tx=XM_003988872.5;refGene_gn=SUOX;refGene_tx=X
M_006933798.4;refGene_gn=SUOX;refGene_tx=XM_023257134.1
B4	84630151	84631561
refGene_gn=SUOX;refGene_tx=XM_003988872.5;refGene_gn=SUOX;refGene_tx=X
M_006933798.4;refGene_gn=SUOX;refGene_tx=XM_019835060.2;refGene_gn=SUO
X;refGene_tx=XM_019835061.2;refGene_gn=SUOX;refGene_tx=XM_023257134.1
B4	84641500	84641540
refGene_gn=IKZF4;refGene_tx=XM_006933802.4;refGene_gn=IKZF4;refGene_tx
=XM_006933803.4;refGene_gn=IKZF4;refGene_tx=XM_023257128.1;refGene_gn=
IKZF4;refGene_tx=XM_023257129.1
B4	84641666	84641682	refGene_gn=IKZF4;refGene_tx=XM_023257131.1
B4	84645056	84645143
refGene_gn=IKZF4;refGene_tx=XM_006933799.4;refGene_gn=IKZF4;refGene_tx
=XM_023257132.1
B4	84646902	84646996
refGene_gn=IKZF4;refGene_tx=XM_006933799.4;refGene_gn=IKZF4;refGene_tx
=XM_023257128.1;refGene_gn=IKZF4;refGene_tx=XM_023257129.1;refGene_gn=
IKZF4;refGene_tx=XM_023257130.1;refGene_gn=IKZF4;refGene_tx=XM_0232571
32.1
B4	84649054	84649159
refGene_gn=IKZF4;refGene_tx=XM_006933799.4;refGene_gn=IKZF4;refGene_tx
=XM_006933802.4;refGene_gn=IKZF4;refGene_tx=XM_006933803.4;refGene_gn=
IKZF4;refGene_tx=XM_023257128.1;refGene_gn=IKZF4;refGene_tx=XM_0232571
29.1;refGene_gn=IKZF4;refGene_tx=XM_023257130.1;refGene_gn=IKZF4;refGe
ne_tx=XM_023257131.1;refGene_gn=IKZF4;refGene_tx=XM_023257132.1
B4	84650171	84650432
refGene_gn=IKZF4;refGene_tx=XM_006933799.4;refGene_gn=IKZF4;refGene_tx
=XM_006933802.4;refGene_gn=IKZF4;refGene_tx=XM_006933803.4;refGene_gn=
IKZF4;refGene_tx=XM_006933804.4;refGene_gn=IKZF4;refGene_tx=XM_0198350
69.2;refGene_gn=IKZF4;refGene_tx=XM_023257128.1;refGene_gn=IKZF4;refGe
ne_tx=XM_023257129.1;refGene_gn=IKZF4;refGene_tx=XM_023257130.1;refGen

e_gn=IKZF4;refGene_tx=XM_023257131.1;refGene_gn=IKZF4;refGene_tx=XM_023257133.1
B4 84651102 84651270
refGene_gn=IKZF4;refGene_tx=XM_006933799.4;refGene_gn=IKZF4;refGene_tx=XM_006933802.4;refGene_gn=IKZF4;refGene_tx=XM_006933803.4;refGene_gn=IKZF4;refGene_tx=XM_006933804.4;refGene_gn=IKZF4;refGene_tx=XM_019835069.2;refGene_gn=IKZF4;refGene_tx=XM_023257128.1;refGene_gn=IKZF4;refGene_tx=XM_023257129.1;refGene_gn=IKZF4;refGene_tx=XM_023257130.1;refGene_gn=IKZF4;refGene_tx=XM_023257131.1;refGene_gn=IKZF4;refGene_tx=XM_023257132.1;refGene_gn=IKZF4;refGene_tx=XM_023257133.1
B4 84654307 84654457
refGene_gn=IKZF4;refGene_tx=XM_006933799.4;refGene_gn=IKZF4;refGene_tx=XM_006933802.4;refGene_gn=IKZF4;refGene_tx=XM_006933803.4;refGene_gn=IKZF4;refGene_tx=XM_006933804.4;refGene_gn=IKZF4;refGene_tx=XM_019835069.2;refGene_gn=IKZF4;refGene_tx=XM_023257128.1;refGene_gn=IKZF4;refGene_tx=XM_023257129.1;refGene_gn=IKZF4;refGene_tx=XM_023257130.1;refGene_gn=IKZF4;refGene_tx=XM_023257131.1;refGene_gn=IKZF4;refGene_tx=XM_023257132.1;refGene_gn=IKZF4;refGene_tx=XM_023257133.1
B4 84654927 84655059
refGene_gn=IKZF4;refGene_tx=XM_006933799.4;refGene_gn=IKZF4;refGene_tx=XM_006933802.4;refGene_gn=IKZF4;refGene_tx=XM_006933803.4;refGene_gn=IKZF4;refGene_tx=XM_006933804.4;refGene_gn=IKZF4;refGene_tx=XM_019835069.2;refGene_gn=IKZF4;refGene_tx=XM_023257128.1;refGene_gn=IKZF4;refGene_tx=XM_023257129.1;refGene_gn=IKZF4;refGene_tx=XM_023257130.1;refGene_gn=IKZF4;refGene_tx=XM_023257131.1;refGene_gn=IKZF4;refGene_tx=XM_023257132.1;refGene_gn=IKZF4;refGene_tx=XM_023257133.1
B4 84655676 84656437
refGene_gn=IKZF4;refGene_tx=XM_006933799.4;refGene_gn=IKZF4;refGene_tx=XM_006933802.4;refGene_gn=IKZF4;refGene_tx=XM_006933803.4;refGene_gn=IKZF4;refGene_tx=XM_006933804.4;refGene_gn=IKZF4;refGene_tx=XM_019835069.2;refGene_gn=IKZF4;refGene_tx=XM_023257128.1;refGene_gn=IKZF4;refGene_tx=XM_023257129.1;refGene_gn=IKZF4;refGene_tx=XM_023257130.1;refGene_gn=IKZF4;refGene_tx=XM_023257131.1;refGene_gn=IKZF4;refGene_tx=XM_023257132.1;refGene_gn=IKZF4;refGene_tx=XM_023257133.1
B4 84662148 84662151 refGene_gn=RPS26;refGene_tx=XM_003988875.4
B4 84662415 84662593 refGene_gn=RPS26;refGene_tx=XM_003988875.4
B4 84663359 84663490 refGene_gn=RPS26;refGene_tx=XM_003988875.4
B4 84663815 84663851 refGene_gn=RPS26;refGene_tx=XM_003988875.4
B4 84715308 84715390
refGene_gn=ERBB3;refGene_tx=NM_001319274.1;refGene_gn=ERBB3;refGene_tx=XM_019833853.1
B4 84718876 84719028
refGene_gn=ERBB3;refGene_tx=NM_001319274.1;refGene_gn=ERBB3;refGene_tx=XM_019833853.1
B4 84719496 84719683
refGene_gn=ERBB3;refGene_tx=NM_001319274.1;refGene_gn=ERBB3;refGene_tx=XM_019833853.1
B4 84720900 84721026
refGene_gn=ERBB3;refGene_tx=NM_001319274.1;refGene_gn=ERBB3;refGene_tx=XM_019833853.1

B4	84721663	84721729
refGene_gn=ERBB3;refGene_tx=NM_001319274.1;refGene_gn=ERBB3;refGene_tx=XM_019833853.1
B4	84722136	84722255
refGene_gn=ERBB3;refGene_tx=NM_001319274.1;refGene_gn=ERBB3;refGene_tx=XM_019833853.1
B4	84722359	84722501
refGene_gn=ERBB3;refGene_tx=NM_001319274.1;refGene_gn=ERBB3;refGene_tx=XM_019833853.1
B4	84722855	84722969
refGene_gn=ERBB3;refGene_tx=NM_001319274.1;refGene_gn=ERBB3;refGene_tx=XM_019833853.1
B4	84723061	84723182
refGene_gn=ERBB3;refGene_tx=NM_001319274.1;refGene_gn=ERBB3;refGene_tx=XM_019833853.1
B4	84724161	84724235
refGene_gn=ERBB3;refGene_tx=NM_001319274.1;refGene_gn=ERBB3;refGene_tx=XM_019833853.1
B4	84724395	84724486
refGene_gn=ERBB3;refGene_tx=NM_001319274.1;refGene_gn=ERBB3;refGene_tx=XM_019833853.1
B4	84724738	84724944
refGene_gn=ERBB3;refGene_tx=NM_001319274.1;refGene_gn=ERBB3;refGene_tx=XM_019833853.1
B4	84725148	84725281
refGene_gn=ERBB3;refGene_tx=NM_001319274.1;refGene_gn=ERBB3;refGene_tx=XM_019833853.1
B4	84725490	84725581
refGene_gn=ERBB3;refGene_tx=NM_001319274.1;refGene_gn=ERBB3;refGene_tx=XM_019833853.1
B4	84725799	84725954
refGene_gn=ERBB3;refGene_tx=NM_001319274.1;refGene_gn=ERBB3;refGene_tx=XM_019833853.1
B4	84726353	84726407
refGene_gn=ERBB3;refGene_tx=NM_001319274.1;refGene_gn=ERBB3;refGene_tx=XM_019833853.1
B4	84727009	84727151
refGene_gn=ERBB3;refGene_tx=NM_001319274.1;refGene_gn=ERBB3;refGene_tx=XM_019833853.1
B4	84727751	84727871
refGene_gn=ERBB3;refGene_tx=NM_001319274.1;refGene_gn=ERBB3;refGene_tx=XM_019833853.1
B4	84727998	84728097
refGene_gn=ERBB3;refGene_tx=NM_001319274.1;refGene_gn=ERBB3;refGene_tx=XM_019833853.1
B4	84728290	84728476
refGene_gn=ERBB3;refGene_tx=NM_001319274.1;refGene_gn=ERBB3;refGene_tx=XM_019833853.1
B4	84728626	84728782	refGene_gn=ERBB3;refGene_tx=NM_001319274.1
B4	84729021	84729097

refGene_gn=ERBB3;refGene_tx=NM_001319274.1;refGene_gn=ERBB3;refGene_tx=XM_019833853.1
B4	84729281	84729428
refGene_gn=ERBB3;refGene_tx=NM_001319274.1;refGene_gn=ERBB3;refGene_tx=XM_019833853.1
B4	84729748	84729846
refGene_gn=ERBB3;refGene_tx=NM_001319274.1;refGene_gn=ERBB3;refGene_tx=XM_019833853.1
B4	84729943	84730132
refGene_gn=ERBB3;refGene_tx=NM_001319274.1;refGene_gn=ERBB3;refGene_tx=XM_019833853.1
B4	84730294	84730366
refGene_gn=ERBB3;refGene_tx=NM_001319274.1;refGene_gn=ERBB3;refGene_tx=XM_019833853.1
B4	84730810	84731105
refGene_gn=ERBB3;refGene_tx=NM_001319274.1;refGene_gn=ERBB3;refGene_tx=XM_019833853.1
B4	84731504	84732031
refGene_gn=ERBB3;refGene_tx=NM_001319274.1;refGene_gn=ERBB3;refGene_tx=XM_019833853.1
B4	84734311	84734399	refGene_gn=PA2G4;refGene_tx=XM_003988876.5
B4	84736274	84736403	refGene_gn=PA2G4;refGene_tx=XM_003988876.5
B4	84736684	84736790	refGene_gn=PA2G4;refGene_tx=XM_003988876.5
B4	84736881	84736951	refGene_gn=PA2G4;refGene_tx=XM_003988876.5
B4	84737151	84737244	refGene_gn=PA2G4;refGene_tx=XM_003988876.5
B4	84739777	84739841	refGene_gn=PA2G4;refGene_tx=XM_003988876.5
B4	84740062	84740141	refGene_gn=PA2G4;refGene_tx=XM_003988876.5
B4	84740293	84740372	refGene_gn=PA2G4;refGene_tx=XM_003988876.5
B4	84740467	84740601	refGene_gn=PA2G4;refGene_tx=XM_003988876.5
B4	84740735	84740830	refGene_gn=PA2G4;refGene_tx=XM_003988876.5
B4	84740948	84741076	refGene_gn=PA2G4;refGene_tx=XM_003988876.5
B4	84741236	84741290	refGene_gn=PA2G4;refGene_tx=XM_003988876.5
B4	84741502	84741568	refGene_gn=PA2G4;refGene_tx=XM_003988876.5
B4	84744985	84744997	refGene_gn=RPL41;refGene_tx=NM_001048157.1
B4	84745399	84745422	refGene_gn=RPL41;refGene_tx=NM_001048157.1
B4	84745693	84745736	refGene_gn=RPL41;refGene_tx=NM_001048157.1
B4	84748917	84750225	refGene_gn=ZC3H10;refGene_tx=XM_006933806.4
B4	84755631	84756021	refGene_gn=ESYT1;refGene_tx=XM_003988877.4
B4	84757664	84757706	refGene_gn=ESYT1;refGene_tx=XM_003988877.4
B4	84757884	84758019	refGene_gn=ESYT1;refGene_tx=XM_003988877.4
B4	84758136	84758210	refGene_gn=ESYT1;refGene_tx=XM_003988877.4
B4	84758331	84758404	refGene_gn=ESYT1;refGene_tx=XM_003988877.4
B4	84758563	84758653	refGene_gn=ESYT1;refGene_tx=XM_003988877.4
B4	84758829	84758885	refGene_gn=ESYT1;refGene_tx=XM_003988877.4
B4	84759317	84759441	refGene_gn=ESYT1;refGene_tx=XM_003988877.4
B4	84759529	84759706	refGene_gn=ESYT1;refGene_tx=XM_003988877.4
B4	84759833	84759916	refGene_gn=ESYT1;refGene_tx=XM_003988877.4
B4	84760286	84760335	refGene_gn=ESYT1;refGene_tx=XM_003988877.4
B4	84760465	84760552	refGene_gn=ESYT1;refGene_tx=XM_003988877.4
B4	84760654	84760747	refGene_gn=ESYT1;refGene_tx=XM_003988877.4

B4	84760976 84761048 refGene_gn=ESYT1;refGene_tx=XM_003988877.4
B4	84761259 84761346 refGene_gn=ESYT1;refGene_tx=XM_003988877.4
B4	84762781 84762931 refGene_gn=ESYT1;refGene_tx=XM_003988877.4
B4	84763246 84763402 refGene_gn=ESYT1;refGene_tx=XM_003988877.4
B4	84763526 84763703 refGene_gn=ESYT1;refGene_tx=XM_003988877.4
B4	84763875 84763958 refGene_gn=ESYT1;refGene_tx=XM_003988877.4
B4	84764084 84764133 refGene_gn=ESYT1;refGene_tx=XM_003988877.4
B4	84764242 84764332 refGene_gn=ESYT1;refGene_tx=XM_003988877.4
B4	84764446 84764542 refGene_gn=ESYT1;refGene_tx=XM_003988877.4
B4	84764824 84764896 refGene_gn=ESYT1;refGene_tx=XM_003988877.4
B4	84764988 84765075 refGene_gn=ESYT1;refGene_tx=XM_003988877.4
B4	84767496 84767637 refGene_gn=ESYT1;refGene_tx=XM_003988877.4
B4	84767749 84767904 refGene_gn=ESYT1;refGene_tx=XM_003988877.4
B4	84768010 84768109 refGene_gn=ESYT1;refGene_tx=XM_003988877.4
B4	84768187 84768319 refGene_gn=ESYT1;refGene_tx=XM_003988877.4
B4	84768393 84768499 refGene_gn=ESYT1;refGene_tx=XM_003988877.4
B4	84768731 84768781 refGene_gn=ESYT1;refGene_tx=XM_003988877.4
B4	84768972 84769012 refGene_gn=ESYT1;refGene_tx=XM_003988877.4
B4	84789596 84789779
refGene_gn=MYL6B;refGene_tx=XM_006933807.4;refGene_gn=MYL6B;refGene_tx
=XM_019835075.2;refGene_gn=MYL6B;refGene_tx=XM_023257142.1
B4	84790519 84790547
refGene_gn=MYL6B;refGene_tx=XM_006933807.4;refGene_gn=MYL6B;refGene_tx
=XM_019835075.2;refGene_gn=MYL6B;refGene_tx=XM_023257142.1
B4	84790783 84790927
refGene_gn=MYL6B;refGene_tx=XM_006933807.4;refGene_gn=MYL6B;refGene_tx
=XM_019835075.2;refGene_gn=MYL6B;refGene_tx=XM_023257142.1
B4	84791157 84791331
refGene_gn=MYL6B;refGene_tx=XM_006933807.4;refGene_gn=MYL6B;refGene_tx
=XM_019835075.2;refGene_gn=MYL6B;refGene_tx=XM_023257142.1
B4	84791828 84791906
refGene_gn=MYL6B;refGene_tx=XM_006933807.4;refGene_gn=MYL6B;refGene_tx
=XM_019835075.2;refGene_gn=MYL6B;refGene_tx=XM_023257142.1
B4	84792037 84792066
refGene_gn=MYL6B;refGene_tx=XM_006933807.4;refGene_gn=MYL6B;refGene_tx
=XM_019835075.2;refGene_gn=MYL6B;refGene_tx=XM_023257142.1
B4	84792680 84792683
refGene_gn=MYL6;refGene_tx=XM_003988881.3;refGene_gn=MYL6;refGene_tx=X
M_023257143.1
B4	84792935 84792963
refGene_gn=MYL6;refGene_tx=XM_003988881.3;refGene_gn=MYL6;refGene_tx=X
M_023257143.1
B4	84793828 84793972
refGene_gn=MYL6;refGene_tx=XM_003988881.3;refGene_gn=MYL6;refGene_tx=X
M_023257143.1
B4	84794200 84794374
refGene_gn=MYL6;refGene_tx=XM_003988881.3;refGene_gn=MYL6;refGene_tx=X
M_023257143.1
B4	84794462 84794540
refGene_gn=MYL6;refGene_tx=XM_003988881.3;refGene_gn=MYL6;refGene_tx=X

M_023257143.1
B4	84794834 84794863 refGene_gn=MYL6;refGene_tx=XM_003988881.3
B4	84795581 84795610 refGene_gn=MYL6;refGene_tx=XM_023257143.1
B4	84797871 84797948
refGene_gn=SMARCC2;refGene_tx=XM_023257135.1;refGene_gn=SMARCC2;refGene_tx=XM_023257136.1;refGene_gn=SMARCC2;refGene_tx=XM_023257137.1;refGene_gn=SMARCC2;refGene_tx=XM_023257138.1;refGene_gn=SMARCC2;refGene_tx=XM_023257139.1
B4	84798499 84798565
refGene_gn=SMARCC2;refGene_tx=XM_023257135.1;refGene_gn=SMARCC2;refGene_tx=XM_023257136.1;refGene_gn=SMARCC2;refGene_tx=XM_023257138.1
B4	84798845 84798929
refGene_gn=SMARCC2;refGene_tx=XM_023257135.1;refGene_gn=SMARCC2;refGene_tx=XM_023257136.1;refGene_gn=SMARCC2;refGene_tx=XM_023257137.1;refGene_gn=SMARCC2;refGene_tx=XM_023257138.1;refGene_gn=SMARCC2;refGene_tx=XM_023257139.1
B4	84799223 84799605
refGene_gn=SMARCC2;refGene_tx=XM_023257135.1;refGene_gn=SMARCC2;refGene_tx=XM_023257136.1;refGene_gn=SMARCC2;refGene_tx=XM_023257137.1;refGene_gn=SMARCC2;refGene_tx=XM_023257138.1;refGene_gn=SMARCC2;refGene_tx=XM_023257139.1
B4	84801461 84801596
refGene_gn=SMARCC2;refGene_tx=XM_023257135.1;refGene_gn=SMARCC2;refGene_tx=XM_023257136.1;refGene_gn=SMARCC2;refGene_tx=XM_023257137.1;refGene_gn=SMARCC2;refGene_tx=XM_023257138.1;refGene_gn=SMARCC2;refGene_tx=XM_023257139.1
B4	84802009 84802176
refGene_gn=SMARCC2;refGene_tx=XM_023257135.1;refGene_gn=SMARCC2;refGene_tx=XM_023257136.1;refGene_gn=SMARCC2;refGene_tx=XM_023257137.1;refGene_gn=SMARCC2;refGene_tx=XM_023257138.1;refGene_gn=SMARCC2;refGene_tx=XM_023257139.1
B4	84802258 84802394
refGene_gn=SMARCC2;refGene_tx=XM_023257135.1;refGene_gn=SMARCC2;refGene_tx=XM_023257136.1;refGene_gn=SMARCC2;refGene_tx=XM_023257137.1;refGene_gn=SMARCC2;refGene_tx=XM_023257138.1;refGene_gn=SMARCC2;refGene_tx=XM_023257139.1
B4	84802597 84802662
refGene_gn=SMARCC2;refGene_tx=XM_023257135.1;refGene_gn=SMARCC2;refGene_tx=XM_023257136.1;refGene_gn=SMARCC2;refGene_tx=XM_023257137.1;refGene_gn=SMARCC2;refGene_tx=XM_023257138.1;refGene_gn=SMARCC2;refGene_tx=XM_023257139.1
B4	84802924 84803086
refGene_gn=SMARCC2;refGene_tx=XM_023257135.1;refGene_gn=SMARCC2;refGene_tx=XM_023257136.1;refGene_gn=SMARCC2;refGene_tx=XM_023257137.1;refGene_gn=SMARCC2;refGene_tx=XM_023257138.1;refGene_gn=SMARCC2;refGene_tx=XM_023257139.1
B4	84803320 84803579
refGene_gn=SMARCC2;refGene_tx=XM_023257135.1;refGene_gn=SMARCC2;refGene_tx=XM_023257136.1;refGene_gn=SMARCC2;refGene_tx=XM_023257137.1;refGene_gn=SMARCC2;refGene_tx=XM_023257138.1;refGene_gn=SMARCC2;refGene_tx=

XM_023257139.1
B4 84803920 84803983
refGene_gn=SMARCC2;refGene_tx=XM_023257135.1;refGene_gn=SMARCC2;refGene_tx=XM_023257136.1;refGene_gn=SMARCC2;refGene_tx=XM_023257137.1;refGene_gn=SMARCC2;refGene_tx=XM_023257138.1;refGene_gn=SMARCC2;refGene_tx=XM_023257139.1
B4 84804075 84804195
refGene_gn=SMARCC2;refGene_tx=XM_023257135.1;refGene_gn=SMARCC2;refGene_tx=XM_023257136.1;refGene_gn=SMARCC2;refGene_tx=XM_023257137.1;refGene_gn=SMARCC2;refGene_tx=XM_023257138.1;refGene_gn=SMARCC2;refGene_tx=XM_023257139.1
B4 84804428 84804521
refGene_gn=SMARCC2;refGene_tx=XM_023257135.1;refGene_gn=SMARCC2;refGene_tx=XM_023257136.1;refGene_gn=SMARCC2;refGene_tx=XM_023257137.1
B4 84805206 84805360
refGene_gn=SMARCC2;refGene_tx=XM_023257135.1;refGene_gn=SMARCC2;refGene_tx=XM_023257136.1;refGene_gn=SMARCC2;refGene_tx=XM_023257137.1;refGene_gn=SMARCC2;refGene_tx=XM_023257138.1;refGene_gn=SMARCC2;refGene_tx=XM_023257139.1
B4 84805835 84805949
refGene_gn=SMARCC2;refGene_tx=XM_023257135.1;refGene_gn=SMARCC2;refGene_tx=XM_023257136.1;refGene_gn=SMARCC2;refGene_tx=XM_023257137.1;refGene_gn=SMARCC2;refGene_tx=XM_023257138.1;refGene_gn=SMARCC2;refGene_tx=XM_023257139.1
B4 84808075 84808147
refGene_gn=SMARCC2;refGene_tx=XM_023257135.1;refGene_gn=SMARCC2;refGene_tx=XM_023257136.1;refGene_gn=SMARCC2;refGene_tx=XM_023257137.1;refGene_gn=SMARCC2;refGene_tx=XM_023257138.1;refGene_gn=SMARCC2;refGene_tx=XM_023257139.1
B4 84808415 84808546
refGene_gn=SMARCC2;refGene_tx=XM_023257135.1;refGene_gn=SMARCC2;refGene_tx=XM_023257136.1;refGene_gn=SMARCC2;refGene_tx=XM_023257137.1;refGene_gn=SMARCC2;refGene_tx=XM_023257138.1;refGene_gn=SMARCC2;refGene_tx=XM_023257139.1
B4 84808831 84808869
refGene_gn=SMARCC2;refGene_tx=XM_023257135.1;refGene_gn=SMARCC2;refGene_tx=XM_023257136.1;refGene_gn=SMARCC2;refGene_tx=XM_023257137.1;refGene_gn=SMARCC2;refGene_tx=XM_023257138.1;refGene_gn=SMARCC2;refGene_tx=XM_023257139.1
B4 84808990 84809050
refGene_gn=SMARCC2;refGene_tx=XM_023257135.1;refGene_gn=SMARCC2;refGene_tx=XM_023257136.1;refGene_gn=SMARCC2;refGene_tx=XM_023257137.1;refGene_gn=SMARCC2;refGene_tx=XM_023257138.1;refGene_gn=SMARCC2;refGene_tx=XM_023257139.1
B4 84810095 84810220
refGene_gn=SMARCC2;refGene_tx=XM_023257135.1;refGene_gn=SMARCC2;refGene_tx=XM_023257136.1;refGene_gn=SMARCC2;refGene_tx=XM_023257137.1;refGene_gn=SMARCC2;refGene_tx=XM_023257138.1;refGene_gn=SMARCC2;refGene_tx=XM_023257139.1
B4 84810599 84810715

refGene_gn=SMARCC2;refGene_tx=XM_023257135.1;refGene_gn=SMARCC2;refGen
e_tx=XM_023257136.1;refGene_gn=SMARCC2;refGene_tx=XM_023257137.1;refGe
ne_gn=SMARCC2;refGene_tx=XM_023257138.1;refGene_gn=SMARCC2;refGene_tx=
XM_023257139.1
B4 84810814 84810946
refGene_gn=SMARCC2;refGene_tx=XM_023257135.1;refGene_gn=SMARCC2;refGen
e_tx=XM_023257136.1;refGene_gn=SMARCC2;refGene_tx=XM_023257137.1;refGe
ne_gn=SMARCC2;refGene_tx=XM_023257138.1;refGene_gn=SMARCC2;refGene_tx=
XM_023257139.1
B4 84811484 84811560
refGene_gn=SMARCC2;refGene_tx=XM_023257135.1;refGene_gn=SMARCC2;refGen
e_tx=XM_023257136.1;refGene_gn=SMARCC2;refGene_tx=XM_023257137.1;refGe
ne_gn=SMARCC2;refGene_tx=XM_023257138.1;refGene_gn=SMARCC2;refGene_tx=
XM_023257139.1
B4 84812248 84812318
refGene_gn=SMARCC2;refGene_tx=XM_023257135.1;refGene_gn=SMARCC2;refGen
e_tx=XM_023257136.1;refGene_gn=SMARCC2;refGene_tx=XM_023257137.1;refGe
ne_gn=SMARCC2;refGene_tx=XM_023257138.1;refGene_gn=SMARCC2;refGene_tx=
XM_023257139.1
B4 84812548 84812618
refGene_gn=SMARCC2;refGene_tx=XM_023257135.1;refGene_gn=SMARCC2;refGen
e_tx=XM_023257136.1;refGene_gn=SMARCC2;refGene_tx=XM_023257137.1;refGe
ne_gn=SMARCC2;refGene_tx=XM_023257138.1;refGene_gn=SMARCC2;refGene_tx=
XM_023257139.1
B4 84813219 84813312
refGene_gn=SMARCC2;refGene_tx=XM_023257135.1;refGene_gn=SMARCC2;refGen
e_tx=XM_023257136.1;refGene_gn=SMARCC2;refGene_tx=XM_023257137.1;refGe
ne_gn=SMARCC2;refGene_tx=XM_023257138.1;refGene_gn=SMARCC2;refGene_tx=
XM_023257139.1
B4 84813413 84813495
refGene_gn=SMARCC2;refGene_tx=XM_023257135.1;refGene_gn=SMARCC2;refGen
e_tx=XM_023257136.1;refGene_gn=SMARCC2;refGene_tx=XM_023257137.1;refGe
ne_gn=SMARCC2;refGene_tx=XM_023257138.1;refGene_gn=SMARCC2;refGene_tx=
XM_023257139.1
B4 84813955 84814041
refGene_gn=SMARCC2;refGene_tx=XM_023257135.1;refGene_gn=SMARCC2;refGen
e_tx=XM_023257136.1;refGene_gn=SMARCC2;refGene_tx=XM_023257137.1;refGe
ne_gn=SMARCC2;refGene_tx=XM_023257138.1;refGene_gn=SMARCC2;refGene_tx=
XM_023257139.1
B4 84815170 84815290
refGene_gn=SMARCC2;refGene_tx=XM_023257135.1;refGene_gn=SMARCC2;refGen
e_tx=XM_023257136.1;refGene_gn=SMARCC2;refGene_tx=XM_023257137.1;refGe
ne_gn=SMARCC2;refGene_tx=XM_023257138.1;refGene_gn=SMARCC2;refGene_tx=
XM_023257139.1
B4 84817383 84817494
refGene_gn=SMARCC2;refGene_tx=XM_023257135.1;refGene_gn=SMARCC2;refGen
e_tx=XM_023257136.1;refGene_gn=SMARCC2;refGene_tx=XM_023257137.1;refGe
ne_gn=SMARCC2;refGene_tx=XM_023257138.1;refGene_gn=SMARCC2;refGene_tx=
XM_023257139.1
B4 84832197 84832549

refGene_gn=RNF41;refGene_tx=XM_003988883.5;refGene_gn=RNF41;refGene_tx=XM_006933808.4;refGene_gn=RNF41;refGene_tx=XM_006933809.4;refGene_gn=RNF41;refGene_tx=XM_019835071.2;refGene_gn=RNF41;refGene_tx=XM_019835072.2
B4 84833386 84833490
refGene_gn=RNF41;refGene_tx=XM_003988883.5;refGene_gn=RNF41;refGene_tx=XM_006933808.4;refGene_gn=RNF41;refGene_tx=XM_006933809.4;refGene_gn=RNF41;refGene_tx=XM_019835071.2;refGene_gn=RNF41;refGene_tx=XM_019835072.2
B4 84833962 84834098
refGene_gn=RNF41;refGene_tx=XM_003988883.5;refGene_gn=RNF41;refGene_tx=XM_006933808.4;refGene_gn=RNF41;refGene_tx=XM_006933809.4;refGene_gn=RNF41;refGene_tx=XM_019835071.2;refGene_gn=RNF41;refGene_tx=XM_019835072.2
B4 84834794 84835066
refGene_gn=RNF41;refGene_tx=XM_003988883.5;refGene_gn=RNF41;refGene_tx=XM_006933808.4;refGene_gn=RNF41;refGene_tx=XM_006933809.4;refGene_gn=RNF41;refGene_tx=XM_019835071.2;refGene_gn=RNF41;refGene_tx=XM_019835072.2
B4 84838511 84838601
refGene_gn=RNF41;refGene_tx=XM_003988883.5;refGene_gn=RNF41;refGene_tx=XM_006933808.4;refGene_gn=RNF41;refGene_tx=XM_006933809.4;refGene_gn=RNF41;refGene_tx=XM_019835071.2;refGene_gn=RNF41;refGene_tx=XM_019835072.2
B4 84863659 84863738
refGene_gn=NABP2;refGene_tx=XM_003988885.5;refGene_gn=NABP2;refGene_tx=XM_006933812.4;refGene_gn=NABP2;refGene_tx=XM_006933813.4;refGene_gn=NABP2;refGene_tx=XM_023257141.1
B4 84864086 84864225
refGene_gn=NABP2;refGene_tx=XM_003988885.5;refGene_gn=NABP2;refGene_tx=XM_006933812.4;refGene_gn=NABP2;refGene_tx=XM_006933813.4;refGene_gn=NABP2;refGene_tx=XM_023257141.1
B4 84864339 84864411
refGene_gn=NABP2;refGene_tx=XM_003988885.5;refGene_gn=NABP2;refGene_tx=XM_006933812.4;refGene_gn=NABP2;refGene_tx=XM_006933813.4;refGene_gn=NABP2;refGene_tx=XM_023257141.1
B4 84864543 84864625
refGene_gn=NABP2;refGene_tx=XM_003988885.5;refGene_gn=NABP2;refGene_tx=XM_006933812.4;refGene_gn=NABP2;refGene_tx=XM_006933813.4;refGene_gn=NABP2;refGene_tx=XM_023257141.1
B4 84864719 84864783
refGene_gn=NABP2;refGene_tx=XM_003988885.5;refGene_gn=NABP2;refGene_tx=XM_006933812.4;refGene_gn=NABP2;refGene_tx=XM_006933813.4;refGene_gn=NABP2;refGene_tx=XM_023257141.1
B4 84866726 84866926
refGene_gn=NABP2;refGene_tx=XM_003988885.5;refGene_gn=NABP2;refGene_tx=XM_006933812.4;refGene_gn=NABP2;refGene_tx=XM_006933813.4;refGene_gn=NABP2;refGene_tx=XM_023257141.1
B4 84868578 84868862 refGene_gn=SLC39A5;refGene_tx=XM_019835070.2
B4 84869374 84869558 refGene_gn=SLC39A5;refGene_tx=XM_019835070.2

B4 84870390 84870553 refGene_gn=SLC39A5;refGene_tx=XM_019835070.2
B4 84870730 84870900 refGene_gn=SLC39A5;refGene_tx=XM_019835070.2
B4 84871143 84871284 refGene_gn=SLC39A5;refGene_tx=XM_019835070.2
B4 84871667 84871764 refGene_gn=SLC39A5;refGene_tx=XM_019835070.2
B4 84871849 84872023 refGene_gn=SLC39A5;refGene_tx=XM_019835070.2
B4 84872138 84872219 refGene_gn=SLC39A5;refGene_tx=XM_019835070.2
B4 84872362 84872553 refGene_gn=SLC39A5;refGene_tx=XM_019835070.2
B4 84872809 84872953 refGene_gn=SLC39A5;refGene_tx=XM_019835070.2
B4 84878447 84878698 refGene_gn=ANKRD52;refGene_tx=XM_023257140.1
B4 84879135 84879227 refGene_gn=ANKRD52;refGene_tx=XM_023257140.1
B4 84879409 84879492 refGene_gn=ANKRD52;refGene_tx=XM_023257140.1
B4 84879690 84879773 refGene_gn=ANKRD52;refGene_tx=XM_023257140.1
B4 84879992 84880138 refGene_gn=ANKRD52;refGene_tx=XM_023257140.1
B4 84880246 84880330 refGene_gn=ANKRD52;refGene_tx=XM_023257140.1
B4 84880438 84880526 refGene_gn=ANKRD52;refGene_tx=XM_023257140.1
B4 84880747 84880967 refGene_gn=ANKRD52;refGene_tx=XM_023257140.1
B4 84881574 84881692 refGene_gn=ANKRD52;refGene_tx=XM_023257140.1
B4 84881768 84881856 refGene_gn=ANKRD52;refGene_tx=XM_023257140.1
B4 84882117 84882319 refGene_gn=ANKRD52;refGene_tx=XM_023257140.1
B4 84882596 84882668 refGene_gn=ANKRD52;refGene_tx=XM_023257140.1
B4 84882873 84882985 refGene_gn=ANKRD52;refGene_tx=XM_023257140.1
B4 84884196 84884277 refGene_gn=ANKRD52;refGene_tx=XM_023257140.1
B4 84884353 84884494 refGene_gn=ANKRD52;refGene_tx=XM_023257140.1
B4 84884668 84884737 refGene_gn=ANKRD52;refGene_tx=XM_023257140.1
B4 84884943 84885061 refGene_gn=ANKRD52;refGene_tx=XM_023257140.1
B4 84885167 84885250 refGene_gn=ANKRD52;refGene_tx=XM_023257140.1
B4 84885431 84885546 refGene_gn=ANKRD52;refGene_tx=XM_023257140.1
B4 84885860 84885939 refGene_gn=ANKRD52;refGene_tx=XM_023257140.1
B4 84886183 84886396 refGene_gn=ANKRD52;refGene_tx=XM_023257140.1
B4 84886697 84886840 refGene_gn=ANKRD52;refGene_tx=XM_023257140.1
B4 84886982 84887070 refGene_gn=ANKRD52;refGene_tx=XM_023257140.1
B4 84887872 84888073 refGene_gn=ANKRD52;refGene_tx=XM_023257140.1
B4 84889091 84889162 refGene_gn=ANKRD52;refGene_tx=XM_023257140.1
B4 84889385 84889464 refGene_gn=ANKRD52;refGene_tx=XM_023257140.1
B4 84889947 84890031 refGene_gn=ANKRD52;refGene_tx=XM_023257140.1
B4 84890355 84890382 refGene_gn=ANKRD52;refGene_tx=XM_023257140.1
B4 84896299 84896433
refGene_gn=COQ10A;refGene_tx=XM_003988887.5;refGene_gn=COQ10A;refGene_tx=XM_006933814.4;refGene_gn=COQ10A;refGene_tx=XM_019835083.2;refGene_gn=COQ10A;refGene_tx=XM_019835084.2
B4 84897002 84897149
refGene_gn=COQ10A;refGene_tx=XM_003988887.5;refGene_gn=COQ10A;refGene_tx=XM_006933814.4;refGene_gn=COQ10A;refGene_tx=XM_019835083.2;refGene_gn=COQ10A;refGene_tx=XM_019835084.2
B4 84898023 84898216
refGene_gn=COQ10A;refGene_tx=XM_003988887.5;refGene_gn=COQ10A;refGene_tx=XM_006933814.4;refGene_gn=COQ10A;refGene_tx=XM_019835083.2;refGene_gn=COQ10A;refGene_tx=XM_019835084.2
B4 84898452 84898554
refGene_gn=COQ10A;refGene_tx=XM_003988887.5;refGene_gn=COQ10A;refGene_

tx=XM_006933814.4;refGene_gn=COQ10A;refGene_tx=XM_019835083.2;refGene_gn=COQ10A;refGene_tx=XM_019835084.2
B4 84898839 84899007 refGene_gn=COQ10A;refGene_tx=XM_003988887.5
B4 84901814 84901985 refGene_gn=CS;refGene_tx=XM_003988888.4;refGene_gn=CS;refGene_tx=XM_019835081.2
B4 84902516 84902726 refGene_gn=CS;refGene_tx=XM_003988888.4;refGene_gn=CS;refGene_tx=XM_019835081.2
B4 84902937 84903039 refGene_gn=CS;refGene_tx=XM_003988888.4;refGene_gn=CS;refGene_tx=XM_019835081.2
B4 84903163 84903293 refGene_gn=CS;refGene_tx=XM_003988888.4;refGene_gn=CS;refGene_tx=XM_019835081.2
B4 84903488 84903620 refGene_gn=COQ10A;refGene_tx=XM_006933814.4;refGene_gn=COQ10A;refGene_tx=XM_019835083.2
B4 84903842 84904042 refGene_gn=CS;refGene_tx=XM_003988888.4;refGene_gn=CS;refGene_tx=XM_019835081.2
B4 84911328 84911397 refGene_gn=COQ10A;refGene_tx=XM_019835083.2;refGene_gn=COQ10A;refGene_tx=XM_019835084.2
B4 84911715 84911904 refGene_gn=CS;refGene_tx=XM_003988888.4;refGene_gn=CS;refGene_tx=XM_019835081.2
B4 84912161 84912293 refGene_gn=CS;refGene_tx=XM_003988888.4;refGene_gn=CS;refGene_tx=XM_019835081.2
B4 84912727 84912793 refGene_gn=CS;refGene_tx=XM_003988888.4;refGene_gn=CS;refGene_tx=XM_019835081.2
B4 84914338 84914446 refGene_gn=CS;refGene_tx=XM_003988888.4;refGene_gn=CS;refGene_tx=XM_019835081.2
B4 84914999 84915050 refGene_gn=CS;refGene_tx=XM_003988888.4;refGene_gn=CS;refGene_tx=XM_019835081.2
B4 84928201 84928243 refGene_gn=CS;refGene_tx=XM_003988888.4
B4 84932452 84932488 refGene_gn=CS;refGene_tx=XM_019835081.2
B4 84933840 84933884 refGene_gn=CNPY2;refGene_tx=XM_003988889.5
B4 84934250 84934347 refGene_gn=CNPY2;refGene_tx=XM_003988889.5
B4 84934505 84934709 refGene_gn=CNPY2;refGene_tx=XM_003988889.5
B4 84935542 84935658 refGene_gn=CNPY2;refGene_tx=XM_003988889.5
B4 84935810 84935898 refGene_gn=CNPY2;refGene_tx=XM_003988889.5
B4 84938086 84938121 refGene_gn=PAN2;refGene_tx=XM_003988891.5;refGene_gn=PAN2;refGene_tx=XM_006933817.3;refGene_gn=PAN2;refGene_tx=XM_019835076.2;refGene_gn=PAN2;refGene_tx=XM_019835077.2;refGene_gn=PAN2;refGene_tx=XM_019835078.2;

refGene_gn=PAN2;refGene_tx=XM_019835079.1
B4 84938462 84938660
refGene_gn=PAN2;refGene_tx=XM_003988891.5;refGene_gn=PAN2;refGene_tx=XM_006933817.3;refGene_gn=PAN2;refGene_tx=XM_019835076.2;refGene_gn=PAN2;refGene_tx=XM_019835077.2;refGene_gn=PAN2;refGene_tx=XM_019835078.2;refGene_gn=PAN2;refGene_tx=XM_019835079.1
B4 84938936 84939030
refGene_gn=PAN2;refGene_tx=XM_003988891.5;refGene_gn=PAN2;refGene_tx=XM_006933817.3;refGene_gn=PAN2;refGene_tx=XM_019835076.2;refGene_gn=PAN2;refGene_tx=XM_019835077.2;refGene_gn=PAN2;refGene_tx=XM_019835078.2;refGene_gn=PAN2;refGene_tx=XM_019835079.1
B4 84939157 84939337
refGene_gn=PAN2;refGene_tx=XM_003988891.5;refGene_gn=PAN2;refGene_tx=XM_006933817.3;refGene_gn=PAN2;refGene_tx=XM_019835076.2;refGene_gn=PAN2;refGene_tx=XM_019835077.2;refGene_gn=PAN2;refGene_tx=XM_019835079.1
B4 84939478 84939622
refGene_gn=PAN2;refGene_tx=XM_003988891.5;refGene_gn=PAN2;refGene_tx=XM_006933817.3;refGene_gn=PAN2;refGene_tx=XM_019835076.2;refGene_gn=PAN2;refGene_tx=XM_019835077.2;refGene_gn=PAN2;refGene_tx=XM_019835078.2;refGene_gn=PAN2;refGene_tx=XM_019835079.1
B4 84939721 84939879
refGene_gn=PAN2;refGene_tx=XM_003988891.5;refGene_gn=PAN2;refGene_tx=XM_006933817.3;refGene_gn=PAN2;refGene_tx=XM_019835076.2;refGene_gn=PAN2;refGene_tx=XM_019835077.2;refGene_gn=PAN2;refGene_tx=XM_019835078.2;refGene_gn=PAN2;refGene_tx=XM_019835079.1
B4 84940473 84940564
refGene_gn=PAN2;refGene_tx=XM_003988891.5;refGene_gn=PAN2;refGene_tx=XM_006933817.3;refGene_gn=PAN2;refGene_tx=XM_019835076.2;refGene_gn=PAN2;refGene_tx=XM_019835077.2;refGene_gn=PAN2;refGene_tx=XM_019835078.2;refGene_gn=PAN2;refGene_tx=XM_019835079.1
B4 84940789 84940849
refGene_gn=PAN2;refGene_tx=XM_003988891.5;refGene_gn=PAN2;refGene_tx=XM_006933817.3;refGene_gn=PAN2;refGene_tx=XM_019835076.2;refGene_gn=PAN2;refGene_tx=XM_019835077.2;refGene_gn=PAN2;refGene_tx=XM_019835078.2;refGene_gn=PAN2;refGene_tx=XM_019835079.1
B4 84940980 84941124
refGene_gn=PAN2;refGene_tx=XM_003988891.5;refGene_gn=PAN2;refGene_tx=XM_006933817.3;refGene_gn=PAN2;refGene_tx=XM_019835076.2;refGene_gn=PAN2;refGene_tx=XM_019835077.2;refGene_gn=PAN2;refGene_tx=XM_019835078.2;refGene_gn=PAN2;refGene_tx=XM_019835079.1
B4 84941422 84941570
refGene_gn=PAN2;refGene_tx=XM_003988891.5;refGene_gn=PAN2;refGene_tx=XM_006933817.3;refGene_gn=PAN2;refGene_tx=XM_019835076.2;refGene_gn=PAN2;refGene_tx=XM_019835077.2;refGene_gn=PAN2;refGene_tx=XM_019835078.2;refGene_gn=PAN2;refGene_tx=XM_019835079.1
B4 84941674 84941748
refGene_gn=PAN2;refGene_tx=XM_003988891.5;refGene_gn=PAN2;refGene_tx=XM_006933817.3;refGene_gn=PAN2;refGene_tx=XM_019835076.2;refGene_gn=PAN2;refGene_tx=XM_019835077.2;refGene_gn=PAN2;refGene_tx=XM_019835078.2;refGene_gn=PAN2;refGene_tx=XM_019835079.1

B4 84941865 84941964
refGene_gn=PAN2;refGene_tx=XM_003988891.5;refGene_gn=PAN2;refGene_tx=XM_006933817.3;refGene_gn=PAN2;refGene_tx=XM_019835076.2;refGene_gn=PAN2;refGene_tx=XM_019835077.2;refGene_gn=PAN2;refGene_tx=XM_019835078.2;refGene_gn=PAN2;refGene_tx=XM_019835079.1
B4 84942153 84942260
refGene_gn=PAN2;refGene_tx=XM_003988891.5;refGene_gn=PAN2;refGene_tx=XM_006933817.3;refGene_gn=PAN2;refGene_tx=XM_019835076.2;refGene_gn=PAN2;refGene_tx=XM_019835077.2;refGene_gn=PAN2;refGene_tx=XM_019835078.2;refGene_gn=PAN2;refGene_tx=XM_019835079.1
B4 84942384 84942533
refGene_gn=PAN2;refGene_tx=XM_003988891.5;refGene_gn=PAN2;refGene_tx=XM_006933817.3;refGene_gn=PAN2;refGene_tx=XM_019835076.2;refGene_gn=PAN2;refGene_tx=XM_019835077.2;refGene_gn=PAN2;refGene_tx=XM_019835078.2;refGene_gn=PAN2;refGene_tx=XM_019835079.1
B4 84942641 84942841
refGene_gn=PAN2;refGene_tx=XM_003988891.5;refGene_gn=PAN2;refGene_tx=XM_006933817.3;refGene_gn=PAN2;refGene_tx=XM_019835076.2;refGene_gn=PAN2;refGene_tx=XM_019835077.2;refGene_gn=PAN2;refGene_tx=XM_019835078.2;refGene_gn=PAN2;refGene_tx=XM_019835079.1
B4 84942968 84943097
refGene_gn=PAN2;refGene_tx=XM_003988891.5;refGene_gn=PAN2;refGene_tx=XM_006933817.3;refGene_gn=PAN2;refGene_tx=XM_019835076.2;refGene_gn=PAN2;refGene_tx=XM_019835077.2;refGene_gn=PAN2;refGene_tx=XM_019835078.2;refGene_gn=PAN2;refGene_tx=XM_019835079.1
B4 84943369 84943489
refGene_gn=PAN2;refGene_tx=XM_003988891.5;refGene_gn=PAN2;refGene_tx=XM_006933817.3;refGene_gn=PAN2;refGene_tx=XM_019835076.2;refGene_gn=PAN2;refGene_tx=XM_019835077.2;refGene_gn=PAN2;refGene_tx=XM_019835078.2;refGene_gn=PAN2;refGene_tx=XM_019835079.1
B4 84943705 84943825
refGene_gn=PAN2;refGene_tx=XM_003988891.5;refGene_gn=PAN2;refGene_tx=XM_006933817.3;refGene_gn=PAN2;refGene_tx=XM_019835076.2;refGene_gn=PAN2;refGene_tx=XM_019835077.2;refGene_gn=PAN2;refGene_tx=XM_019835078.2;refGene_gn=PAN2;refGene_tx=XM_019835079.1
B4 84944668 84944765
refGene_gn=PAN2;refGene_tx=XM_003988891.5;refGene_gn=PAN2;refGene_tx=XM_006933817.3;refGene_gn=PAN2;refGene_tx=XM_019835076.2;refGene_gn=PAN2;refGene_tx=XM_019835077.2;refGene_gn=PAN2;refGene_tx=XM_019835078.2;refGene_gn=PAN2;refGene_tx=XM_019835079.1
B4 84944960 84945303
refGene_gn=PAN2;refGene_tx=XM_003988891.5;refGene_gn=PAN2;refGene_tx=XM_006933817.3;refGene_gn=PAN2;refGene_tx=XM_019835076.2;refGene_gn=PAN2;refGene_tx=XM_019835077.2;refGene_gn=PAN2;refGene_tx=XM_019835078.2;refGene_gn=PAN2;refGene_tx=XM_019835079.1
B4 84945646 84945914
refGene_gn=PAN2;refGene_tx=XM_003988891.5;refGene_gn=PAN2;refGene_tx=XM_006933817.3;refGene_gn=PAN2;refGene_tx=XM_019835076.2;refGene_gn=PAN2;refGene_tx=XM_019835077.2;refGene_gn=PAN2;refGene_tx=XM_019835078.2;refGene_gn=PAN2;refGene_tx=XM_019835079.1

B4 84946297 84946375
refGene_gn=PAN2;refGene_tx=XM_003988891.5;refGene_gn=PAN2;refGene_tx=XM_006933817.3;refGene_gn=PAN2;refGene_tx=XM_019835076.2;refGene_gn=PAN2;refGene_tx=XM_019835077.2;refGene_gn=PAN2;refGene_tx=XM_019835078.2;refGene_gn=PAN2;refGene_tx=XM_019835079.1
B4 84946537 84946658
refGene_gn=PAN2;refGene_tx=XM_003988891.5;refGene_gn=PAN2;refGene_tx=XM_006933817.3;refGene_gn=PAN2;refGene_tx=XM_019835076.2;refGene_gn=PAN2;refGene_tx=XM_019835077.2;refGene_gn=PAN2;refGene_tx=XM_019835078.2;refGene_gn=PAN2;refGene_tx=XM_019835079.1
B4 84946771 84946941
refGene_gn=PAN2;refGene_tx=XM_003988891.5;refGene_gn=PAN2;refGene_tx=XM_006933817.3;refGene_gn=PAN2;refGene_tx=XM_019835076.2;refGene_gn=PAN2;refGene_tx=XM_019835077.2;refGene_gn=PAN2;refGene_tx=XM_019835078.2;refGene_gn=PAN2;refGene_tx=XM_019835079.1
B4 84949635 84949917
refGene_gn=PAN2;refGene_tx=XM_003988891.5;refGene_gn=PAN2;refGene_tx=XM_006933817.3;refGene_gn=PAN2;refGene_tx=XM_019835076.2;refGene_gn=PAN2;refGene_tx=XM_019835077.2;refGene_gn=PAN2;refGene_tx=XM_019835078.2
B4 84954355 84954535 refGene_gn=IL23A;refGene_tx=XM_003988892.3
B4 84954763 84954865 refGene_gn=IL23A;refGene_tx=XM_003988892.3
B4 84955370 84955517 refGene_gn=IL23A;refGene_tx=XM_003988892.3
B4 84955621 84955783 refGene_gn=IL23A;refGene_tx=XM_003988892.3
B4 84958117 84958260
refGene_gn=STAT2;refGene_tx=XM_003988893.5;refGene_gn=STAT2;refGene_tx=XM_019835080.1
B4 84958539 84958892
refGene_gn=STAT2;refGene_tx=XM_003988893.5;refGene_gn=STAT2;refGene_tx=XM_019835080.1
B4 84959699 84959757
refGene_gn=STAT2;refGene_tx=XM_003988893.5;refGene_gn=STAT2;refGene_tx=XM_019835080.1
B4 84959993 84960176
refGene_gn=STAT2;refGene_tx=XM_003988893.5;refGene_gn=STAT2;refGene_tx=XM_019835080.1
B4 84960353 84960490
refGene_gn=STAT2;refGene_tx=XM_003988893.5;refGene_gn=STAT2;refGene_tx=XM_019835080.1
B4 84961068 84961163
refGene_gn=STAT2;refGene_tx=XM_003988893.5;refGene_gn=STAT2;refGene_tx=XM_019835080.1
B4 84961298 84961351
refGene_gn=STAT2;refGene_tx=XM_003988893.5;refGene_gn=STAT2;refGene_tx=XM_019835080.1
B4 84961472 84961608
refGene_gn=STAT2;refGene_tx=XM_003988893.5;refGene_gn=STAT2;refGene_tx=XM_019835080.1
B4 84961711 84961810
refGene_gn=STAT2;refGene_tx=XM_003988893.5;refGene_gn=STAT2;refGene_tx=XM_019835080.1

B4 84961967 84962051
refGene_gn=STAT2;refGene_tx=XM_003988893.5;refGene_gn=STAT2;refGene_tx=XM_019835080.1
B4 84962134 84962182
refGene_gn=STAT2;refGene_tx=XM_003988893.5;refGene_gn=STAT2;refGene_tx=XM_019835080.1
B4 84962328 84962422
refGene_gn=STAT2;refGene_tx=XM_003988893.5;refGene_gn=STAT2;refGene_tx=XM_019835080.1
B4 84962631 84962652
refGene_gn=STAT2;refGene_tx=XM_003988893.5;refGene_gn=STAT2;refGene_tx=XM_019835080.1
B4 84963042 84963102
refGene_gn=STAT2;refGene_tx=XM_003988893.5;refGene_gn=STAT2;refGene_tx=XM_019835080.1
B4 84963315 84963408
refGene_gn=STAT2;refGene_tx=XM_003988893.5;refGene_gn=STAT2;refGene_tx=XM_019835080.1
B4 84963514 84963673
refGene_gn=STAT2;refGene_tx=XM_003988893.5;refGene_gn=STAT2;refGene_tx=XM_019835080.1
B4 84970166 84970315
refGene_gn=STAT2;refGene_tx=XM_003988893.5;refGene_gn=STAT2;refGene_tx=XM_019835080.1
B4 84970476 84970562
refGene_gn=STAT2;refGene_tx=XM_003988893.5;refGene_gn=STAT2;refGene_tx=XM_019835080.1
B4 84971002 84971078
refGene_gn=STAT2;refGene_tx=XM_003988893.5;refGene_gn=STAT2;refGene_tx=XM_019835080.1
B4 84971169 84971259
refGene_gn=STAT2;refGene_tx=XM_003988893.5;refGene_gn=STAT2;refGene_tx=XM_019835080.1
B4 84971460 84971556
refGene_gn=STAT2;refGene_tx=XM_003988893.5;refGene_gn=STAT2;refGene_tx=XM_019835080.1
B4 84971900 84972054 refGene_gn=STAT2;refGene_tx=XM_003988893.5
B4 84972449 84972580 refGene_gn=STAT2;refGene_tx=XM_003988893.5
B4 84977047 84977914
refGene_gn=APOF;refGene_tx=XM_006933826.2;refGene_gn=APOF;refGene_tx=XM_019835086.1
B4 84978152 84978155 refGene_gn=APOF;refGene_tx=XM_019835086.1
B4 84991751 84991825
refGene_gn=TIMELESS;refGene_tx=XM_006933819.3;refGene_gn=TIMELESS;refGene_tx=XM_019835087.2;refGene_gn=TIMELESS;refGene_tx=XM_019835088.2
B4 84991966 84992071
refGene_gn=TIMELESS;refGene_tx=XM_006933819.3;refGene_gn=TIMELESS;refGene_tx=XM_019835087.2;refGene_gn=TIMELESS;refGene_tx=XM_019835088.2
B4 84992195 84992421
refGene_gn=TIMELESS;refGene_tx=XM_006933819.3;refGene_gn=TIMELESS;refG

ene_tx=XM_019835087.2;refGene_gn=TIMELESS;refGene_tx=XM_019835088.2
B4 84993804 84993923
refGene_gn=TIMELESS;refGene_tx=XM_006933819.3;refGene_gn=TIMELESS;refGene_tx=XM_019835087.2;refGene_gn=TIMELESS;refGene_tx=XM_019835088.2
B4 84994045 84994114
refGene_gn=TIMELESS;refGene_tx=XM_006933819.3;refGene_gn=TIMELESS;refGene_tx=XM_019835087.2;refGene_gn=TIMELESS;refGene_tx=XM_019835088.2
B4 84994196 84994374
refGene_gn=TIMELESS;refGene_tx=XM_006933819.3;refGene_gn=TIMELESS;refGene_tx=XM_019835087.2;refGene_gn=TIMELESS;refGene_tx=XM_019835088.2
B4 84994574 84994717
refGene_gn=TIMELESS;refGene_tx=XM_006933819.3;refGene_gn=TIMELESS;refGene_tx=XM_019835087.2;refGene_gn=TIMELESS;refGene_tx=XM_019835088.2
B4 84994945 84995028
refGene_gn=TIMELESS;refGene_tx=XM_006933819.3;refGene_gn=TIMELESS;refGene_tx=XM_019835087.2;refGene_gn=TIMELESS;refGene_tx=XM_019835088.2
B4 84995124 84995239
refGene_gn=TIMELESS;refGene_tx=XM_006933819.3;refGene_gn=TIMELESS;refGene_tx=XM_019835087.2;refGene_gn=TIMELESS;refGene_tx=XM_019835088.2
B4 84995327 84995413
refGene_gn=TIMELESS;refGene_tx=XM_006933819.3;refGene_gn=TIMELESS;refGene_tx=XM_019835087.2;refGene_gn=TIMELESS;refGene_tx=XM_019835088.2
B4 84996569 84996715
refGene_gn=TIMELESS;refGene_tx=XM_006933819.3;refGene_gn=TIMELESS;refGene_tx=XM_019835087.2;refGene_gn=TIMELESS;refGene_tx=XM_019835088.2
B4 84996985 84997187
refGene_gn=TIMELESS;refGene_tx=XM_006933819.3;refGene_gn=TIMELESS;refGene_tx=XM_019835087.2;refGene_gn=TIMELESS;refGene_tx=XM_019835088.2
B4 84997295 84997422
refGene_gn=TIMELESS;refGene_tx=XM_006933819.3;refGene_gn=TIMELESS;refGene_tx=XM_019835087.2;refGene_gn=TIMELESS;refGene_tx=XM_019835088.2
B4 84997765 84997863
refGene_gn=TIMELESS;refGene_tx=XM_006933819.3;refGene_gn=TIMELESS;refGene_tx=XM_019835087.2;refGene_gn=TIMELESS;refGene_tx=XM_019835088.2
B4 84999219 84999371
refGene_gn=TIMELESS;refGene_tx=XM_006933819.3;refGene_gn=TIMELESS;refGene_tx=XM_019835087.2;refGene_gn=TIMELESS;refGene_tx=XM_019835088.2
B4 84999469 84999604
refGene_gn=TIMELESS;refGene_tx=XM_006933819.3;refGene_gn=TIMELESS;refGene_tx=XM_019835087.2;refGene_gn=TIMELESS;refGene_tx=XM_019835088.2
B4 85000691 85000861
refGene_gn=TIMELESS;refGene_tx=XM_006933819.3;refGene_gn=TIMELESS;refGene_tx=XM_019835087.2;refGene_gn=TIMELESS;refGene_tx=XM_019835088.2
B4 85001006 85001110
refGene_gn=TIMELESS;refGene_tx=XM_006933819.3;refGene_gn=TIMELESS;refGene_tx=XM_019835087.2;refGene_gn=TIMELESS;refGene_tx=XM_019835088.2
B4 85001353 85001571
refGene_gn=TIMELESS;refGene_tx=XM_006933819.3;refGene_gn=TIMELESS;refGene_tx=XM_019835087.2;refGene_gn=TIMELESS;refGene_tx=XM_019835088.2
B4 85001947 85002124

refGene_gn=TIMELESS;refGene_tx=XM_006933819.3;refGene_gn=TIMELESS;refGene_tx=XM_019835087.2;refGene_gn=TIMELESS;refGene_tx=XM_019835088.2
B4 85003095 85003183
refGene_gn=TIMELESS;refGene_tx=XM_006933819.3;refGene_gn=TIMELESS;refGene_tx=XM_019835087.2;refGene_gn=TIMELESS;refGene_tx=XM_019835088.2
B4 85003399 85003533
refGene_gn=TIMELESS;refGene_tx=XM_006933819.3;refGene_gn=TIMELESS;refGene_tx=XM_019835087.2;refGene_gn=TIMELESS;refGene_tx=XM_019835088.2
B4 85003775 85003931
refGene_gn=TIMELESS;refGene_tx=XM_006933819.3;refGene_gn=TIMELESS;refGene_tx=XM_019835087.2;refGene_gn=TIMELESS;refGene_tx=XM_019835088.2
B4 85004372 85004474
refGene_gn=TIMELESS;refGene_tx=XM_006933819.3;refGene_gn=TIMELESS;refGene_tx=XM_019835087.2;refGene_gn=TIMELESS;refGene_tx=XM_019835088.2
B4 85004996 85005059
refGene_gn=TIMELESS;refGene_tx=XM_006933819.3;refGene_gn=TIMELESS;refGene_tx=XM_019835087.2;refGene_gn=TIMELESS;refGene_tx=XM_019835088.2
B4 85005154 85005269
refGene_gn=TIMELESS;refGene_tx=XM_006933819.3;refGene_gn=TIMELESS;refGene_tx=XM_019835087.2;refGene_gn=TIMELESS;refGene_tx=XM_019835088.2
B4 85009512 85009666
refGene_gn=TIMELESS;refGene_tx=XM_006933819.3;refGene_gn=TIMELESS;refGene_tx=XM_019835087.2;refGene_gn=TIMELESS;refGene_tx=XM_019835088.2
B4 85009813 85009898
refGene_gn=TIMELESS;refGene_tx=XM_006933819.3;refGene_gn=TIMELESS;refGene_tx=XM_019835087.2;refGene_gn=TIMELESS;refGene_tx=XM_019835088.2
B4 85026393 85026579 refGene_gn=MIP;refGene_tx=XM_003988894.4
B4 85028396 85028477 refGene_gn=MIP;refGene_tx=XM_003988894.4
B4 85028907 85029072 refGene_gn=MIP;refGene_tx=XM_003988894.4
B4 85029582 85029942 refGene_gn=MIP;refGene_tx=XM_003988894.4
B4 85041311 85041396 refGene_gn=SPRYD4;refGene_tx=XM_003988895.5
B4 85041711 85042250 refGene_gn=SPRYD4;refGene_tx=XM_003988895.5
B4 85043796 85043953
refGene_gn=GLS2;refGene_tx=XM_003988896.5;refGene_gn=GLS2;refGene_tx=XM_011284104.3;refGene_gn=GLS2;refGene_tx=XM_019835093.2;refGene_gn=GLS2;refGene_tx=XM_019835094.1
B4 85044086 85044150
refGene_gn=GLS2;refGene_tx=XM_003988896.5;refGene_gn=GLS2;refGene_tx=XM_011284104.3;refGene_gn=GLS2;refGene_tx=XM_019835093.2;refGene_gn=GLS2;refGene_tx=XM_019835094.1
B4 85044561 85044638
refGene_gn=GLS2;refGene_tx=XM_003988896.5;refGene_gn=GLS2;refGene_tx=XM_011284104.3;refGene_gn=GLS2;refGene_tx=XM_019835093.2;refGene_gn=GLS2;refGene_tx=XM_019835094.1
B4 85045340 85045402
refGene_gn=GLS2;refGene_tx=XM_003988896.5;refGene_gn=GLS2;refGene_tx=XM_011284104.3;refGene_gn=GLS2;refGene_tx=XM_019835093.2;refGene_gn=GLS2;refGene_tx=XM_019835094.1
B4 85045571 85045664
refGene_gn=GLS2;refGene_tx=XM_003988896.5;refGene_gn=GLS2;refGene_tx=X

M_011284104.3;refGene_gn=GLS2;refGene_tx=XM_019835093.2;refGene_gn=GLS2;refGene_tx=XM_019835094.1
B4 85045777 85045909 refGene_gn=GLS2;refGene_tx=XM_003988896.5;refGene_gn=GLS2;refGene_tx=XM_011284104.3;refGene_gn=GLS2;refGene_tx=XM_019835093.2;refGene_gn=GLS2;refGene_tx=XM_019835094.1
B4 85046584 85046761 refGene_gn=GLS2;refGene_tx=XM_003988896.5;refGene_gn=GLS2;refGene_tx=XM_011284104.3;refGene_gn=GLS2;refGene_tx=XM_019835094.1
B4 85046982 85047033 refGene_gn=GLS2;refGene_tx=XM_003988896.5;refGene_gn=GLS2;refGene_tx=XM_011284104.3;refGene_gn=GLS2;refGene_tx=XM_019835093.2;refGene_gn=GLS2;refGene_tx=XM_019835094.1
B4 85047185 85047252 refGene_gn=GLS2;refGene_tx=XM_003988896.5;refGene_gn=GLS2;refGene_tx=XM_011284104.3;refGene_gn=GLS2;refGene_tx=XM_019835093.2;refGene_gn=GLS2;refGene_tx=XM_019835094.1
B4 85047974 85048033 refGene_gn=GLS2;refGene_tx=XM_003988896.5;refGene_gn=GLS2;refGene_tx=XM_011284104.3;refGene_gn=GLS2;refGene_tx=XM_019835093.2;refGene_gn=GLS2;refGene_tx=XM_019835094.1
B4 85048316 85048349 refGene_gn=GLS2;refGene_tx=XM_003988896.5;refGene_gn=GLS2;refGene_tx=XM_011284104.3;refGene_gn=GLS2;refGene_tx=XM_019835093.2;refGene_gn=GLS2;refGene_tx=XM_019835094.1
B4 85049647 85049706 refGene_gn=GLS2;refGene_tx=XM_003988896.5;refGene_gn=GLS2;refGene_tx=XM_011284104.3;refGene_gn=GLS2;refGene_tx=XM_019835093.2;refGene_gn=GLS2;refGene_tx=XM_019835094.1
B4 85049930 85050094 refGene_gn=GLS2;refGene_tx=XM_003988896.5;refGene_gn=GLS2;refGene_tx=XM_011284104.3;refGene_gn=GLS2;refGene_tx=XM_019835093.2;refGene_gn=GLS2;refGene_tx=XM_019835094.1
B4 85050180 85050260 refGene_gn=GLS2;refGene_tx=XM_003988896.5;refGene_gn=GLS2;refGene_tx=XM_011284104.3;refGene_gn=GLS2;refGene_tx=XM_019835093.2;refGene_gn=GLS2;refGene_tx=XM_019835094.1
B4 85050765 85050895 refGene_gn=GLS2;refGene_tx=XM_003988896.5;refGene_gn=GLS2;refGene_tx=XM_011284104.3;refGene_gn=GLS2;refGene_tx=XM_019835093.2;refGene_gn=GLS2;refGene_tx=XM_019835094.1
B4 85051511 85051633 refGene_gn=GLS2;refGene_tx=XM_003988896.5;refGene_gn=GLS2;refGene_tx=XM_011284104.3;refGene_gn=GLS2;refGene_tx=XM_019835093.2;refGene_gn=GLS2;refGene_tx=XM_019835094.1
B4 85052041 85052141 refGene_gn=GLS2;refGene_tx=XM_003988896.5;refGene_gn=GLS2;refGene_tx=XM_011284104.3;refGene_gn=GLS2;refGene_tx=XM_019835093.2;refGene_gn=GLS2;refGene_tx=XM_019835094.1
B4 85057105 85057218 refGene_gn=GLS2;refGene_tx=XM_011284104.3

B4 85057348 85057530
refGene_gn=GLS2;refGene_tx=XM_003988896.5;refGene_gn=GLS2;refGene_tx=XM_019835093.2
B4 85081690 85081756 refGene_gn=RBMS2;refGene_tx=XM_011284101.3
B4 85121396 85121563
refGene_gn=RBMS2;refGene_tx=XM_011284101.3;refGene_gn=RBMS2;refGene_tx=XM_011284102.3;refGene_gn=RBMS2;refGene_tx=XM_019835096.2
B4 85129708 85129767
refGene_gn=RBMS2;refGene_tx=XM_011284101.3;refGene_gn=RBMS2;refGene_tx=XM_011284102.3;refGene_gn=RBMS2;refGene_tx=XM_019835096.2
B4 85130610 85130702
refGene_gn=RBMS2;refGene_tx=XM_011284101.3;refGene_gn=RBMS2;refGene_tx=XM_011284102.3;refGene_gn=RBMS2;refGene_tx=XM_019835096.2
B4 85132155 85132313
refGene_gn=RBMS2;refGene_tx=XM_011284101.3;refGene_gn=RBMS2;refGene_tx=XM_011284102.3;refGene_gn=RBMS2;refGene_tx=XM_019835096.2
B4 85138653 85138733
refGene_gn=RBMS2;refGene_tx=XM_011284101.3;refGene_gn=RBMS2;refGene_tx=XM_011284102.3;refGene_gn=RBMS2;refGene_tx=XM_019835096.2
B4 85138868 85138978
refGene_gn=RBMS2;refGene_tx=XM_011284101.3;refGene_gn=RBMS2;refGene_tx=XM_011284102.3;refGene_gn=RBMS2;refGene_tx=XM_019835096.2
B4 85139313 85139360
refGene_gn=RBMS2;refGene_tx=XM_011284101.3;refGene_gn=RBMS2;refGene_tx=XM_011284102.3;refGene_gn=RBMS2;refGene_tx=XM_019835096.2
B4 85139529 85139623
refGene_gn=RBMS2;refGene_tx=XM_011284101.3;refGene_gn=RBMS2;refGene_tx=XM_011284102.3;refGene_gn=RBMS2;refGene_tx=XM_019835096.2
B4 85146382 85146460
refGene_gn=RBMS2;refGene_tx=XM_011284101.3;refGene_gn=RBMS2;refGene_tx=XM_011284102.3;refGene_gn=RBMS2;refGene_tx=XM_019835096.2
B4 85146748 85146859
refGene_gn=RBMS2;refGene_tx=XM_011284101.3;refGene_gn=RBMS2;refGene_tx=XM_011284102.3;refGene_gn=RBMS2;refGene_tx=XM_019835096.2
B4 85147446 85147527
refGene_gn=RBMS2;refGene_tx=XM_011284101.3;refGene_gn=RBMS2;refGene_tx=XM_011284102.3;refGene_gn=RBMS2;refGene_tx=XM_019835096.2
B4 85147929 85148010
refGene_gn=RBMS2;refGene_tx=XM_011284101.3;refGene_gn=RBMS2;refGene_tx=XM_011284102.3;refGene_gn=RBMS2;refGene_tx=XM_019835096.2
B4 85156837 85157003
refGene_gn=BAZ2A;refGene_tx=XM_003988914.5;refGene_gn=BAZ2A;refGene_tx=XM_006933827.4;refGene_gn=BAZ2A;refGene_tx=XM_019835089.2;refGene_gn=BAZ2A;refGene_tx=XM_019835090.2;refGene_gn=BAZ2A;refGene_tx=XM_019835091.2;refGene_gn=BAZ2A;refGene_tx=XM_019835092.2;refGene_gn=BAZ2A;refGene_tx=XM_023257145.1
B4 85157105 85157249
refGene_gn=BAZ2A;refGene_tx=XM_003988914.5;refGene_gn=BAZ2A;refGene_tx=XM_006933827.4;refGene_gn=BAZ2A;refGene_tx=XM_019835089.2;refGene_gn=BAZ2A;refGene_tx=XM_019835090.2;refGene_gn=BAZ2A;refGene_tx=XM_0198350

91.2;refGene_gn=BAZ2A;refGene_tx=XM_019835092.2;refGene_gn=BAZ2A;refGene_tx=XM_023257145.1
B4	85157369	85157596
refGene_gn=BAZ2A;refGene_tx=XM_003988914.5;refGene_gn=BAZ2A;refGene_tx=XM_006933827.4;refGene_gn=BAZ2A;refGene_tx=XM_019835089.2;refGene_gn=BAZ2A;refGene_tx=XM_019835090.2;refGene_gn=BAZ2A;refGene_tx=XM_019835091.2;refGene_gn=BAZ2A;refGene_tx=XM_019835092.2;refGene_gn=BAZ2A;refGene_tx=XM_023257145.1
B4	85157974	85158121
refGene_gn=BAZ2A;refGene_tx=XM_003988914.5;refGene_gn=BAZ2A;refGene_tx=XM_006933827.4;refGene_gn=BAZ2A;refGene_tx=XM_019835089.2;refGene_gn=BAZ2A;refGene_tx=XM_019835090.2;refGene_gn=BAZ2A;refGene_tx=XM_019835091.2;refGene_gn=BAZ2A;refGene_tx=XM_019835092.2;refGene_gn=BAZ2A;refGene_tx=XM_023257145.1
B4	85158212	85158345
refGene_gn=BAZ2A;refGene_tx=XM_003988914.5;refGene_gn=BAZ2A;refGene_tx=XM_006933827.4;refGene_gn=BAZ2A;refGene_tx=XM_019835089.2;refGene_gn=BAZ2A;refGene_tx=XM_019835090.2;refGene_gn=BAZ2A;refGene_tx=XM_019835091.2;refGene_gn=BAZ2A;refGene_tx=XM_019835092.2;refGene_gn=BAZ2A;refGene_tx=XM_023257145.1
B4	85158442	85158732
refGene_gn=BAZ2A;refGene_tx=XM_003988914.5;refGene_gn=BAZ2A;refGene_tx=XM_006933827.4;refGene_gn=BAZ2A;refGene_tx=XM_019835089.2;refGene_gn=BAZ2A;refGene_tx=XM_019835090.2;refGene_gn=BAZ2A;refGene_tx=XM_019835091.2;refGene_gn=BAZ2A;refGene_tx=XM_019835092.2;refGene_gn=BAZ2A;refGene_tx=XM_023257145.1
B4	85158897	85159049
refGene_gn=BAZ2A;refGene_tx=XM_003988914.5;refGene_gn=BAZ2A;refGene_tx=XM_006933827.4;refGene_gn=BAZ2A;refGene_tx=XM_019835089.2;refGene_gn=BAZ2A;refGene_tx=XM_019835090.2;refGene_gn=BAZ2A;refGene_tx=XM_019835091.2;refGene_gn=BAZ2A;refGene_tx=XM_019835092.2;refGene_gn=BAZ2A;refGene_tx=XM_023257145.1
B4	85159156	85159312
refGene_gn=BAZ2A;refGene_tx=XM_003988914.5;refGene_gn=BAZ2A;refGene_tx=XM_006933827.4;refGene_gn=BAZ2A;refGene_tx=XM_019835089.2;refGene_gn=BAZ2A;refGene_tx=XM_019835090.2;refGene_gn=BAZ2A;refGene_tx=XM_019835091.2;refGene_gn=BAZ2A;refGene_tx=XM_019835092.2;refGene_gn=BAZ2A;refGene_tx=XM_023257145.1
B4	85159400	85159623
refGene_gn=BAZ2A;refGene_tx=XM_003988914.5;refGene_gn=BAZ2A;refGene_tx=XM_006933827.4;refGene_gn=BAZ2A;refGene_tx=XM_019835089.2;refGene_gn=BAZ2A;refGene_tx=XM_019835090.2;refGene_gn=BAZ2A;refGene_tx=XM_019835091.2;refGene_gn=BAZ2A;refGene_tx=XM_019835092.2;refGene_gn=BAZ2A;refGene_tx=XM_023257145.1
B4	85159779	85160438
refGene_gn=BAZ2A;refGene_tx=XM_003988914.5;refGene_gn=BAZ2A;refGene_tx=XM_006933827.4;refGene_gn=BAZ2A;refGene_tx=XM_019835089.2;refGene_gn=BAZ2A;refGene_tx=XM_019835090.2;refGene_gn=BAZ2A;refGene_tx=XM_019835091.2;refGene_gn=BAZ2A;refGene_tx=XM_019835092.2;refGene_gn=BAZ2A;refGene_tx=XM_023257145.1

B4 85160976 85161121
refGene_gn=BAZ2A;refGene_tx=XM_003988914.5;refGene_gn=BAZ2A;refGene_tx=XM_006933827.4;refGene_gn=BAZ2A;refGene_tx=XM_019835089.2;refGene_gn=BAZ2A;refGene_tx=XM_019835090.2;refGene_gn=BAZ2A;refGene_tx=XM_019835091.2;refGene_gn=BAZ2A;refGene_tx=XM_019835092.2;refGene_gn=BAZ2A;refGene_tx=XM_023257145.1
B4 85161313 85161373
refGene_gn=BAZ2A;refGene_tx=XM_003988914.5;refGene_gn=BAZ2A;refGene_tx=XM_006933827.4;refGene_gn=BAZ2A;refGene_tx=XM_019835089.2;refGene_gn=BAZ2A;refGene_tx=XM_019835090.2;refGene_gn=BAZ2A;refGene_tx=XM_019835091.2;refGene_gn=BAZ2A;refGene_tx=XM_019835092.2;refGene_gn=BAZ2A;refGene_tx=XM_023257145.1
B4 85161481 85161662
refGene_gn=BAZ2A;refGene_tx=XM_003988914.5;refGene_gn=BAZ2A;refGene_tx=XM_006933827.4;refGene_gn=BAZ2A;refGene_tx=XM_019835089.2;refGene_gn=BAZ2A;refGene_tx=XM_019835090.2;refGene_gn=BAZ2A;refGene_tx=XM_019835091.2;refGene_gn=BAZ2A;refGene_tx=XM_019835092.2;refGene_gn=BAZ2A;refGene_tx=XM_023257145.1
B4 85161877 85161952
refGene_gn=BAZ2A;refGene_tx=XM_003988914.5;refGene_gn=BAZ2A;refGene_tx=XM_006933827.4;refGene_gn=BAZ2A;refGene_tx=XM_019835089.2;refGene_gn=BAZ2A;refGene_tx=XM_019835090.2;refGene_gn=BAZ2A;refGene_tx=XM_019835091.2;refGene_gn=BAZ2A;refGene_tx=XM_019835092.2;refGene_gn=BAZ2A;refGene_tx=XM_023257145.1
B4 85162736 85162951
refGene_gn=BAZ2A;refGene_tx=XM_003988914.5;refGene_gn=BAZ2A;refGene_tx=XM_006933827.4;refGene_gn=BAZ2A;refGene_tx=XM_019835089.2;refGene_gn=BAZ2A;refGene_tx=XM_019835090.2;refGene_gn=BAZ2A;refGene_tx=XM_019835091.2;refGene_gn=BAZ2A;refGene_tx=XM_019835092.2;refGene_gn=BAZ2A;refGene_tx=XM_023257145.1
B4 85163326 85163581
refGene_gn=BAZ2A;refGene_tx=XM_003988914.5;refGene_gn=BAZ2A;refGene_tx=XM_006933827.4;refGene_gn=BAZ2A;refGene_tx=XM_019835089.2;refGene_gn=BAZ2A;refGene_tx=XM_019835090.2;refGene_gn=BAZ2A;refGene_tx=XM_019835091.2;refGene_gn=BAZ2A;refGene_tx=XM_019835092.2;refGene_gn=BAZ2A;refGene_tx=XM_023257145.1
B4 85163728 85163962
refGene_gn=BAZ2A;refGene_tx=XM_003988914.5;refGene_gn=BAZ2A;refGene_tx=XM_006933827.4;refGene_gn=BAZ2A;refGene_tx=XM_019835089.2;refGene_gn=BAZ2A;refGene_tx=XM_019835090.2;refGene_gn=BAZ2A;refGene_tx=XM_019835091.2;refGene_gn=BAZ2A;refGene_tx=XM_019835092.2;refGene_gn=BAZ2A;refGene_tx=XM_023257145.1
B4 85164146 85164212
refGene_gn=BAZ2A;refGene_tx=XM_003988914.5;refGene_gn=BAZ2A;refGene_tx=XM_006933827.4;refGene_gn=BAZ2A;refGene_tx=XM_019835089.2;refGene_gn=BAZ2A;refGene_tx=XM_019835090.2;refGene_gn=BAZ2A;refGene_tx=XM_019835091.2;refGene_gn=BAZ2A;refGene_tx=XM_019835092.2;refGene_gn=BAZ2A;refGene_tx=XM_023257145.1
B4 85164541 85164642
refGene_gn=BAZ2A;refGene_tx=XM_003988914.5;refGene_gn=BAZ2A;refGene_tx

=XM_006933827.4;refGene_gn=BAZ2A;refGene_tx=XM_019835089.2;refGene_gn=
BAZ2A;refGene_tx=XM_019835090.2;refGene_gn=BAZ2A;refGene_tx=XM_0198350
91.2;refGene_gn=BAZ2A;refGene_tx=XM_019835092.2;refGene_gn=BAZ2A;refGe
ne_tx=XM_023257145.1
B4 85166149 85166360
refGene_gn=BAZ2A;refGene_tx=XM_003988914.5;refGene_gn=BAZ2A;refGene_tx
=XM_006933827.4;refGene_gn=BAZ2A;refGene_tx=XM_019835089.2;refGene_gn=
BAZ2A;refGene_tx=XM_019835090.2;refGene_gn=BAZ2A;refGene_tx=XM_0198350
91.2;refGene_gn=BAZ2A;refGene_tx=XM_019835092.2;refGene_gn=BAZ2A;refGe
ne_tx=XM_023257145.1
B4 85166519 85166621
refGene_gn=BAZ2A;refGene_tx=XM_003988914.5;refGene_gn=BAZ2A;refGene_tx
=XM_006933827.4;refGene_gn=BAZ2A;refGene_tx=XM_019835089.2;refGene_gn=
BAZ2A;refGene_tx=XM_019835090.2;refGene_gn=BAZ2A;refGene_tx=XM_0198350
91.2;refGene_gn=BAZ2A;refGene_tx=XM_019835092.2;refGene_gn=BAZ2A;refGe
ne_tx=XM_023257145.1
B4 85166787 85166896
refGene_gn=BAZ2A;refGene_tx=XM_003988914.5;refGene_gn=BAZ2A;refGene_tx
=XM_006933827.4;refGene_gn=BAZ2A;refGene_tx=XM_019835089.2;refGene_gn=
BAZ2A;refGene_tx=XM_019835090.2;refGene_gn=BAZ2A;refGene_tx=XM_0198350
91.2;refGene_gn=BAZ2A;refGene_tx=XM_019835092.2;refGene_gn=BAZ2A;refGe
ne_tx=XM_023257145.1
B4 85168187 85168248
refGene_gn=BAZ2A;refGene_tx=XM_003988914.5;refGene_gn=BAZ2A;refGene_tx
=XM_006933827.4;refGene_gn=BAZ2A;refGene_tx=XM_019835089.2;refGene_gn=
BAZ2A;refGene_tx=XM_019835090.2;refGene_gn=BAZ2A;refGene_tx=XM_0198350
91.2;refGene_gn=BAZ2A;refGene_tx=XM_019835092.2;refGene_gn=BAZ2A;refGe
ne_tx=XM_023257145.1
B4 85168385 85168886
refGene_gn=BAZ2A;refGene_tx=XM_003988914.5;refGene_gn=BAZ2A;refGene_tx
=XM_006933827.4;refGene_gn=BAZ2A;refGene_tx=XM_019835089.2;refGene_gn=
BAZ2A;refGene_tx=XM_019835090.2;refGene_gn=BAZ2A;refGene_tx=XM_0198350
91.2;refGene_gn=BAZ2A;refGene_tx=XM_019835092.2;refGene_gn=BAZ2A;refGe
ne_tx=XM_023257145.1
B4 85169312 85169531
refGene_gn=BAZ2A;refGene_tx=XM_003988914.5;refGene_gn=BAZ2A;refGene_tx
=XM_006933827.4;refGene_gn=BAZ2A;refGene_tx=XM_019835089.2;refGene_gn=
BAZ2A;refGene_tx=XM_019835090.2;refGene_gn=BAZ2A;refGene_tx=XM_0198350
91.2;refGene_gn=BAZ2A;refGene_tx=XM_019835092.2;refGene_gn=BAZ2A;refGe
ne_tx=XM_023257145.1
B4 85170129 85170315
refGene_gn=BAZ2A;refGene_tx=XM_003988914.5;refGene_gn=BAZ2A;refGene_tx
=XM_006933827.4;refGene_gn=BAZ2A;refGene_tx=XM_019835089.2;refGene_gn=
BAZ2A;refGene_tx=XM_019835090.2;refGene_gn=BAZ2A;refGene_tx=XM_0198350
91.2;refGene_gn=BAZ2A;refGene_tx=XM_019835092.2;refGene_gn=BAZ2A;refGe
ne_tx=XM_023257145.1
B4 85171193 85171787
refGene_gn=BAZ2A;refGene_tx=XM_003988914.5;refGene_gn=BAZ2A;refGene_tx
=XM_006933827.4;refGene_gn=BAZ2A;refGene_tx=XM_019835089.2;refGene_gn=
BAZ2A;refGene_tx=XM_019835090.2;refGene_gn=BAZ2A;refGene_tx=XM_0198350

91.2;refGene_gn=BAZ2A;refGene_tx=XM_019835092.2;refGene_gn=BAZ2A;refGene_tx=XM_023257145.1
B4 85173365 85173501 refGene_gn=BAZ2A;refGene_tx=XM_003988914.5;refGene_gn=BAZ2A;refGene_tx=XM_006933827.4;refGene_gn=BAZ2A;refGene_tx=XM_019835089.2;refGene_gn=BAZ2A;refGene_tx=XM_019835090.2;refGene_gn=BAZ2A;refGene_tx=XM_019835091.2;refGene_gn=BAZ2A;refGene_tx=XM_019835092.2;refGene_gn=BAZ2A;refGene_tx=XM_023257145.1
B4 85190706 85190804 refGene_gn=ATP5F1B;refGene_tx=XM_003988915.4
B4 85191381 85191583 refGene_gn=ATP5F1B;refGene_tx=XM_003988915.4
B4 85192003 85192216 refGene_gn=ATP5F1B;refGene_tx=XM_003988915.4
B4 85193128 85193251 refGene_gn=ATP5F1B;refGene_tx=XM_003988915.4
B4 85193382 85193541 refGene_gn=ATP5F1B;refGene_tx=XM_003988915.4
B4 85194279 85194464 refGene_gn=ATP5F1B;refGene_tx=XM_003988915.4
B4 85194713 85194835 refGene_gn=ATP5F1B;refGene_tx=XM_003988915.4
B4 85195955 85196130 refGene_gn=ATP5F1B;refGene_tx=XM_003988915.4
B4 85196360 85196543 refGene_gn=ATP5F1B;refGene_tx=XM_003988915.4
B4 85197017 85197144 refGene_gn=ATP5F1B;refGene_tx=XM_003988915.4
B4 85207406 85207426 refGene_gn=PTGES3;refGene_tx=XM_003988916.5
B4 85207737 85207762 refGene_gn=PTGES3;refGene_tx=XM_003988916.5
B4 85208604 85208667 refGene_gn=PTGES3;refGene_tx=XM_003988916.5
B4 85210578 85210668 refGene_gn=PTGES3;refGene_tx=XM_003988916.5
B4 85211225 85211324 refGene_gn=PTGES3;refGene_tx=XM_003988916.5
B4 85211822 85211892 refGene_gn=PTGES3;refGene_tx=XM_003988916.5
B4 85212038 85212152 refGene_gn=PTGES3;refGene_tx=XM_003988916.5
B4 85227189 85227191 refGene_gn=PTGES3;refGene_tx=XM_003988916.5
B4 85236987 85238442 refGene_gn=LOC109496917;refGene_tx=XM_023257146.1
B4 85238630 85240585 refGene_gn=LOC111561459;refGene_tx=XM_023257762.1
B4 85240587 85241987 refGene_gn=LOC111561459;refGene_tx=XM_023257762.1
B4 85241988 85243760 refGene_gn=LOC111561459;refGene_tx=XM_023257762.1
B4 85267941 85267956 refGene_gn=NACA;refGene_tx=XM_003988918.4;refGene_gn=NACA;refGene_tx=XM_023257147.1
B4 85268157 85268280 refGene_gn=NACA;refGene_tx=XM_003988918.4;refGene_gn=NACA;refGene_tx=XM_023257147.1
B4 85268934 85269063 refGene_gn=NACA;refGene_tx=XM_003988918.4;refGene_gn=NACA;refGene_tx=XM_023257147.1
B4 85269493 85269640 refGene_gn=NACA;refGene_tx=XM_003988918.4;refGene_gn=NACA;refGene_tx=XM_023257147.1
B4 85269930 85270008 refGene_gn=NACA;refGene_tx=XM_003988918.4;refGene_gn=NACA;refGene_tx=XM_023257147.1
B4 85270196 85270282

refGene_gn=NACA;refGene_tx=XM_003988918.4;refGene_gn=NACA;refGene_tx=XM_023257147.1
B4 85271472 85276524 refGene_gn=NACA;refGene_tx=XM_023257147.1
B4 85279207 85279277
refGene_gn=NACA;refGene_tx=XM_003988918.4;refGene_gn=NACA;refGene_tx=XM_023257147.1
B4 85283746 85283766
refGene_gn=PRIM1;refGene_tx=XM_023257148.1;refGene_gn=PRIM1;refGene_tx=XM_023257149.1
B4 85285299 85285398
refGene_gn=PRIM1;refGene_tx=XM_023257148.1;refGene_gn=PRIM1;refGene_tx=XM_023257149.1
B4 85293019 85293108
refGene_gn=PRIM1;refGene_tx=XM_023257148.1;refGene_gn=PRIM1;refGene_tx=XM_023257149.1
B4 85293903 85293973
refGene_gn=PRIM1;refGene_tx=XM_023257148.1;refGene_gn=PRIM1;refGene_tx=XM_023257149.1
B4 85295469 85295611
refGene_gn=PRIM1;refGene_tx=XM_023257148.1;refGene_gn=PRIM1;refGene_tx=XM_023257149.1
B4 85295783 85295872
refGene_gn=PRIM1;refGene_tx=XM_023257148.1;refGene_gn=PRIM1;refGene_tx=XM_023257149.1
B4 85296589 85296699
refGene_gn=PRIM1;refGene_tx=XM_023257148.1;refGene_gn=PRIM1;refGene_tx=XM_023257149.1
B4 85298022 85298081
refGene_gn=PRIM1;refGene_tx=XM_023257148.1;refGene_gn=PRIM1;refGene_tx=XM_023257149.1
B4 85298454 85298591
refGene_gn=PRIM1;refGene_tx=XM_023257148.1;refGene_gn=PRIM1;refGene_tx=XM_023257149.1
B4 85299685 85299759
refGene_gn=PRIM1;refGene_tx=XM_023257148.1;refGene_gn=PRIM1;refGene_tx=XM_023257149.1
B4 85299830 85299937
refGene_gn=PRIM1;refGene_tx=XM_023257148.1;refGene_gn=PRIM1;refGene_tx=XM_023257149.1
B4 85308941 85309099
refGene_gn=PRIM1;refGene_tx=XM_023257148.1;refGene_gn=PRIM1;refGene_tx=XM_023257149.1
B4 85309730 85309854 refGene_gn=PRIM1;refGene_tx=XM_023257148.1
B4 85345684 85345997
refGene_gn=HSD17B6;refGene_tx=XM_003988920.2;refGene_gn=HSD17B6;refGene_tx=XM_006933829.2;refGene_gn=HSD17B6;refGene_tx=XM_006933830.3;refGene_gn=HSD17B6;refGene_tx=XM_019835097.1;refGene_gn=HSD17B6;refGene_tx=XM_019835098.1
B4 85350743 85351002
refGene_gn=HSD17B6;refGene_tx=XM_003988920.2;refGene_gn=HSD17B6;refGen

e_tx=XM_006933829.2;refGene_gn=HSD17B6;refGene_tx=XM_006933830.3;refGene_gn=HSD17B6;refGene_tx=XM_019835097.1;refGene_gn=HSD17B6;refGene_tx=XM_019835098.1
B4 85355620 85355784
refGene_gn=HSD17B6;refGene_tx=XM_003988920.2;refGene_gn=HSD17B6;refGene_tx=XM_006933829.2;refGene_gn=HSD17B6;refGene_tx=XM_006933830.3;refGene_gn=HSD17B6;refGene_tx=XM_019835097.1;refGene_gn=HSD17B6;refGene_tx=XM_019835098.1
B4 85358782 85359000
refGene_gn=HSD17B6;refGene_tx=XM_003988920.2;refGene_gn=HSD17B6;refGene_tx=XM_006933829.2;refGene_gn=HSD17B6;refGene_tx=XM_006933830.3;refGene_gn=HSD17B6;refGene_tx=XM_019835097.1;refGene_gn=HSD17B6;refGene_tx=XM_019835098.1
B4 85471071 85471289 refGene_gn=SDR9C7;refGene_tx=XM_003988921.3
B4 85472813 85472977 refGene_gn=SDR9C7;refGene_tx=XM_003988921.3
B4 85473647 85473906 refGene_gn=SDR9C7;refGene_tx=XM_003988921.3
B4 85477141 85477442 refGene_gn=SDR9C7;refGene_tx=XM_003988921.3
B4 85481425 85481643
refGene_gn=RDH16;refGene_tx=XM_003988922.5;refGene_gn=RDH16;refGene_tx=XM_006933833.4;refGene_gn=RDH16;refGene_tx=XM_019835102.1;refGene_gn=RDH16;refGene_tx=XM_019835103.1
B4 85482190 85482354
refGene_gn=RDH16;refGene_tx=XM_003988922.5;refGene_gn=RDH16;refGene_tx=XM_006933833.4;refGene_gn=RDH16;refGene_tx=XM_019835102.1;refGene_gn=RDH16;refGene_tx=XM_019835103.1
B4 85483902 85484161
refGene_gn=RDH16;refGene_tx=XM_003988922.5;refGene_gn=RDH16;refGene_tx=XM_006933833.4;refGene_gn=RDH16;refGene_tx=XM_019835102.1;refGene_gn=RDH16;refGene_tx=XM_019835103.1
B4 85485320 85485655
refGene_gn=RDH16;refGene_tx=XM_003988922.5;refGene_gn=RDH16;refGene_tx=XM_019835102.1;refGene_gn=RDH16;refGene_tx=XM_019835103.1
B4 85488162 85488221 refGene_gn=RDH16;refGene_tx=XM_003988922.5
B4 85489099 85489412 refGene_gn=RDH16;refGene_tx=XM_006933833.4
B4 85497354 85498500 refGene_gn=GPR182;refGene_tx=XM_003988979.5
B4 85505722 85507858
refGene_gn=ZBTB39;refGene_tx=XM_003988923.4;refGene_gn=ZBTB39;refGene_tx=XM_023257150.1
B4 85516208 85516282
refGene_gn=TAC3;refGene_tx=XM_003988924.5;refGene_gn=TAC3;refGene_tx=XM_019835104.2
B4 85516678 85516732
refGene_gn=TAC3;refGene_tx=XM_003988924.5;refGene_gn=TAC3;refGene_tx=XM_019835104.2
B4 85517224 85517254
refGene_gn=TAC3;refGene_tx=XM_003988924.5;refGene_gn=TAC3;refGene_tx=XM_019835104.2
B4 85517447 85517556
refGene_gn=TAC3;refGene_tx=XM_003988924.5;refGene_gn=TAC3;refGene_tx=XM_019835104.2

B4	85519951	85520065
refGene_gn=TAC3;refGene_tx=XM_003988924.5;refGene_gn=TAC3;refGene_tx=XM_019835104.2
B4	85526252	85526376
refGene_gn=MYO1A;refGene_tx=XM_003988925.4;refGene_gn=MYO1A;refGene_tx=XM_019835108.1
B4	85526671	85526799
refGene_gn=MYO1A;refGene_tx=XM_003988925.4;refGene_gn=MYO1A;refGene_tx=XM_019835108.1
B4	85526976	85527129
refGene_gn=MYO1A;refGene_tx=XM_003988925.4;refGene_gn=MYO1A;refGene_tx=XM_019835108.1;refGene_gn=MYO1A;refGene_tx=XM_023257151.1
B4	85527245	85527378
refGene_gn=MYO1A;refGene_tx=XM_003988925.4;refGene_gn=MYO1A;refGene_tx=XM_019835108.1
B4	85527747	85527854
refGene_gn=MYO1A;refGene_tx=XM_003988925.4;refGene_gn=MYO1A;refGene_tx=XM_019835108.1;refGene_gn=MYO1A;refGene_tx=XM_023257151.1
B4	85528554	85528689
refGene_gn=MYO1A;refGene_tx=XM_003988925.4;refGene_gn=MYO1A;refGene_tx=XM_019835108.1;refGene_gn=MYO1A;refGene_tx=XM_023257151.1
B4	85531983	85532058
refGene_gn=MYO1A;refGene_tx=XM_003988925.4;refGene_gn=MYO1A;refGene_tx=XM_019835108.1;refGene_gn=MYO1A;refGene_tx=XM_023257151.1
B4	85532464	85532533
refGene_gn=MYO1A;refGene_tx=XM_003988925.4;refGene_gn=MYO1A;refGene_tx=XM_019835108.1;refGene_gn=MYO1A;refGene_tx=XM_023257151.1
B4	85532845	85532995
refGene_gn=MYO1A;refGene_tx=XM_003988925.4;refGene_gn=MYO1A;refGene_tx=XM_019835108.1;refGene_gn=MYO1A;refGene_tx=XM_023257151.1
B4	85533485	85533579
refGene_gn=MYO1A;refGene_tx=XM_003988925.4;refGene_gn=MYO1A;refGene_tx=XM_019835108.1;refGene_gn=MYO1A;refGene_tx=XM_023257151.1
B4	85533839	85534040
refGene_gn=MYO1A;refGene_tx=XM_003988925.4;refGene_gn=MYO1A;refGene_tx=XM_019835108.1;refGene_gn=MYO1A;refGene_tx=XM_023257151.1
B4	85534379	85534606
refGene_gn=MYO1A;refGene_tx=XM_003988925.4;refGene_gn=MYO1A;refGene_tx=XM_019835108.1;refGene_gn=MYO1A;refGene_tx=XM_023257151.1
B4	85534773	85534974
refGene_gn=MYO1A;refGene_tx=XM_003988925.4;refGene_gn=MYO1A;refGene_tx=XM_019835108.1;refGene_gn=MYO1A;refGene_tx=XM_023257151.1
B4	85535167	85535230
refGene_gn=MYO1A;refGene_tx=XM_003988925.4;refGene_gn=MYO1A;refGene_tx=XM_019835108.1;refGene_gn=MYO1A;refGene_tx=XM_023257151.1
B4	85537184	85537289
refGene_gn=MYO1A;refGene_tx=XM_003988925.4;refGene_gn=MYO1A;refGene_tx=XM_019835108.1;refGene_gn=MYO1A;refGene_tx=XM_023257151.1
B4	85537447	85537513
refGene_gn=MYO1A;refGene_tx=XM_003988925.4;refGene_gn=MYO1A;refGene_tx

=XM_019835108.1;refGene_gn=MYO1A;refGene_tx=XM_023257151.1
B4 85539295 85539382
refGene_gn=MYO1A;refGene_tx=XM_003988925.4;refGene_gn=MYO1A;refGene_tx=XM_019835108.1;refGene_gn=MYO1A;refGene_tx=XM_023257151.1
B4 85539460 85539579
refGene_gn=MYO1A;refGene_tx=XM_003988925.4;refGene_gn=MYO1A;refGene_tx=XM_019835108.1;refGene_gn=MYO1A;refGene_tx=XM_023257151.1
B4 85540183 85540331
refGene_gn=MYO1A;refGene_tx=XM_003988925.4;refGene_gn=MYO1A;refGene_tx=XM_019835108.1;refGene_gn=MYO1A;refGene_tx=XM_023257151.1
B4 85540445 85540549
refGene_gn=MYO1A;refGene_tx=XM_003988925.4;refGene_gn=MYO1A;refGene_tx=XM_019835108.1;refGene_gn=MYO1A;refGene_tx=XM_023257151.1
B4 85542849 85542948
refGene_gn=MYO1A;refGene_tx=XM_003988925.4;refGene_gn=MYO1A;refGene_tx=XM_019835108.1;refGene_gn=MYO1A;refGene_tx=XM_023257151.1
B4 85543153 85543217
refGene_gn=MYO1A;refGene_tx=XM_003988925.4;refGene_gn=MYO1A;refGene_tx=XM_019835108.1;refGene_gn=MYO1A;refGene_tx=XM_023257151.1
B4 85543353 85543400
refGene_gn=MYO1A;refGene_tx=XM_003988925.4;refGene_gn=MYO1A;refGene_tx=XM_019835108.1;refGene_gn=MYO1A;refGene_tx=XM_023257151.1
B4 85543598 85543703
refGene_gn=MYO1A;refGene_tx=XM_003988925.4;refGene_gn=MYO1A;refGene_tx=XM_019835108.1;refGene_gn=MYO1A;refGene_tx=XM_023257151.1
B4 85543895 85543990
refGene_gn=MYO1A;refGene_tx=XM_003988925.4;refGene_gn=MYO1A;refGene_tx=XM_023257151.1
B4 85544252 85544368
refGene_gn=MYO1A;refGene_tx=XM_003988925.4;refGene_gn=MYO1A;refGene_tx=XM_023257151.1
B4 85544497 85544611
refGene_gn=MYO1A;refGene_tx=XM_003988925.4;refGene_gn=MYO1A;refGene_tx=XM_023257151.1
B4 85554657 85554838 refGene_gn=NEMP1;refGene_tx=XM_023257763.1
B4 85555588 85555762 refGene_gn=NEMP1;refGene_tx=XM_023257763.1
B4 85556554 85556780 refGene_gn=NEMP1;refGene_tx=XM_023257763.1
B4 85557917 85558032 refGene_gn=NEMP1;refGene_tx=XM_023257763.1
B4 85558487 85558581 refGene_gn=NEMP1;refGene_tx=XM_023257763.1
B4 85559913 85559986 refGene_gn=NEMP1;refGene_tx=XM_023257763.1
B4 85561700 85561920 refGene_gn=NEMP1;refGene_tx=XM_023257763.1
B4 85563176 85563301 refGene_gn=NEMP1;refGene_tx=XM_023257763.1
B4 85572291 85572547 refGene_gn=NEMP1;refGene_tx=XM_023257763.1
B4 85581712 85582242 refGene_gn=NEMP1;refGene_tx=XM_023257763.1
B4 85582336 85582523
refGene_gn=NAB2;refGene_tx=XM_023257152.1;refGene_gn=NAB2;refGene_tx=XM_023257153.1;refGene_gn=NEMP1;refGene_tx=XM_023257763.1
B4 85583967 85584841
refGene_gn=NAB2;refGene_tx=XM_023257152.1;refGene_gn=NAB2;refGene_tx=XM_023257153.1

B4 85585255 85585389
refGene_gn=NAB2;refGene_tx=XM_023257152.1;refGene_gn=NAB2;refGene_tx=XM_023257153.1
B4 85585716 85585768
refGene_gn=NAB2;refGene_tx=XM_023257152.1;refGene_gn=NAB2;refGene_tx=XM_023257153.1
B4 85585862 85585995
refGene_gn=NAB2;refGene_tx=XM_023257152.1;refGene_gn=NAB2;refGene_tx=XM_023257153.1
B4 85586200 85586392 refGene_gn=NAB2;refGene_tx=XM_023257152.1
B4 85587384 85587494
refGene_gn=NAB2;refGene_tx=XM_023257152.1;refGene_gn=NAB2;refGene_tx=XM_023257153.1
B4 85589365 85589555
refGene_gn=STAT6;refGene_tx=XM_006933834.4;refGene_gn=STAT6;refGene_tx=XM_019835110.2
B4 85589645 85589774
refGene_gn=STAT6;refGene_tx=XM_006933834.4;refGene_gn=STAT6;refGene_tx=XM_019835110.2
B4 85589865 85589931
refGene_gn=STAT6;refGene_tx=XM_006933834.4;refGene_gn=STAT6;refGene_tx=XM_019835110.2
B4 85591218 85591302
refGene_gn=STAT6;refGene_tx=XM_006933834.4;refGene_gn=STAT6;refGene_tx=XM_019835110.2
B4 85591496 85591607
refGene_gn=STAT6;refGene_tx=XM_006933834.4;refGene_gn=STAT6;refGene_tx=XM_019835110.2
B4 85591712 85591776
refGene_gn=STAT6;refGene_tx=XM_006933834.4;refGene_gn=STAT6;refGene_tx=XM_019835110.2
B4 85591993 85592140
refGene_gn=STAT6;refGene_tx=XM_006933834.4;refGene_gn=STAT6;refGene_tx=XM_019835110.2
B4 85592468 85592605
refGene_gn=STAT6;refGene_tx=XM_006933834.4;refGene_gn=STAT6;refGene_tx=XM_019835110.2
B4 85592700 85592795
refGene_gn=STAT6;refGene_tx=XM_006933834.4;refGene_gn=STAT6;refGene_tx=XM_019835110.2
B4 85594027 85594234
refGene_gn=STAT6;refGene_tx=XM_006933834.4;refGene_gn=STAT6;refGene_tx=XM_019835110.2
B4 85594606 85594699
refGene_gn=STAT6;refGene_tx=XM_006933834.4;refGene_gn=STAT6;refGene_tx=XM_019835110.2
B4 85595281 85595404
refGene_gn=STAT6;refGene_tx=XM_006933834.4;refGene_gn=STAT6;refGene_tx=XM_019835110.2
B4 85595541 85595629

refGene_gn=STAT6;refGene_tx=XM_006933834.4;refGene_gn=STAT6;refGene_tx=XM_019835110.2
B4 85595940 85596129
refGene_gn=STAT6;refGene_tx=XM_006933834.4;refGene_gn=STAT6;refGene_tx=XM_019835110.2
B4 85596255 85596387
refGene_gn=STAT6;refGene_tx=XM_006933834.4;refGene_gn=STAT6;refGene_tx=XM_019835110.2
B4 85596945 85597094
refGene_gn=STAT6;refGene_tx=XM_006933834.4;refGene_gn=STAT6;refGene_tx=XM_019835110.2
B4 85597301 85597354
refGene_gn=STAT6;refGene_tx=XM_006933834.4;refGene_gn=STAT6;refGene_tx=XM_019835110.2
B4 85597472 85597611
refGene_gn=STAT6;refGene_tx=XM_006933834.4;refGene_gn=STAT6;refGene_tx=XM_019835110.2
B4 85598048 85598132
refGene_gn=STAT6;refGene_tx=XM_006933834.4;refGene_gn=STAT6;refGene_tx=XM_019835110.2
B4 85598462 85598601
refGene_gn=STAT6;refGene_tx=XM_006933834.4;refGene_gn=STAT6;refGene_tx=XM_019835110.2
B4 85599004 85599120
refGene_gn=STAT6;refGene_tx=XM_006933834.4;refGene_gn=STAT6;refGene_tx=XM_019835110.2
B4 85616441 85616508
refGene_gn=LRP1;refGene_tx=XM_003988931.4;refGene_gn=LRP1;refGene_tx=XM_019835109.1
B4 85625078 85625201
refGene_gn=LRP1;refGene_tx=XM_003988931.4;refGene_gn=LRP1;refGene_tx=XM_019835109.1
B4 85628158 85628296
refGene_gn=LRP1;refGene_tx=XM_003988931.4;refGene_gn=LRP1;refGene_tx=XM_019835109.1
B4 85630429 85630549
refGene_gn=LRP1;refGene_tx=XM_003988931.4;refGene_gn=LRP1;refGene_tx=XM_019835109.1
B4 85631999 85632128
refGene_gn=LRP1;refGene_tx=XM_003988931.4;refGene_gn=LRP1;refGene_tx=XM_019835109.1
B4 85632248 85632512
refGene_gn=LRP1;refGene_tx=XM_003988931.4;refGene_gn=LRP1;refGene_tx=XM_019835109.1
B4 85641413 85641576
refGene_gn=LRP1;refGene_tx=XM_003988931.4;refGene_gn=LRP1;refGene_tx=XM_019835109.1
B4 85641655 85641878
refGene_gn=LRP1;refGene_tx=XM_003988931.4;refGene_gn=LRP1;refGene_tx=XM_019835109.1

B4 85642883 85643073
refGene_gn=LRP1;refGene_tx=XM_003988931.4;refGene_gn=LRP1;refGene_tx=XM_019835109.1
B4 85643710 85643854
refGene_gn=LRP1;refGene_tx=XM_003988931.4;refGene_gn=LRP1;refGene_tx=XM_019835109.1
B4 85644976 85645213
refGene_gn=LRP1;refGene_tx=XM_003988931.4;refGene_gn=LRP1;refGene_tx=XM_019835109.1
B4 85646410 85646591
refGene_gn=LRP1;refGene_tx=XM_003988931.4;refGene_gn=LRP1;refGene_tx=XM_019835109.1
B4 85647464 85647687
refGene_gn=LRP1;refGene_tx=XM_003988931.4;refGene_gn=LRP1;refGene_tx=XM_019835109.1
B4 85648822 85649024
refGene_gn=LRP1;refGene_tx=XM_003988931.4;refGene_gn=LRP1;refGene_tx=XM_019835109.1
B4 85649378 85649504
refGene_gn=LRP1;refGene_tx=XM_003988931.4;refGene_gn=LRP1;refGene_tx=XM_019835109.1
B4 85651993 85652134
refGene_gn=LRP1;refGene_tx=XM_003988931.4;refGene_gn=LRP1;refGene_tx=XM_019835109.1
B4 85652230 85652356
refGene_gn=LRP1;refGene_tx=XM_003988931.4;refGene_gn=LRP1;refGene_tx=XM_019835109.1
B4 85652825 85652942
refGene_gn=LRP1;refGene_tx=XM_003988931.4;refGene_gn=LRP1;refGene_tx=XM_019835109.1
B4 85653404 85653485
refGene_gn=LRP1;refGene_tx=XM_003988931.4;refGene_gn=LRP1;refGene_tx=XM_019835109.1
B4 85654131 85654182 refGene_gn=LRP1;refGene_tx=XM_019835109.1
B4 85655098 85655266
refGene_gn=LRP1;refGene_tx=XM_003988931.4;refGene_gn=LRP1;refGene_tx=XM_019835109.1
B4 85660464 85660647
refGene_gn=LRP1;refGene_tx=XM_003988931.4;refGene_gn=LRP1;refGene_tx=XM_019835109.1
B4 85661078 85661279
refGene_gn=LRP1;refGene_tx=XM_003988931.4;refGene_gn=LRP1;refGene_tx=XM_019835109.1
B4 85662396 85662642
refGene_gn=LRP1;refGene_tx=XM_003988931.4;refGene_gn=LRP1;refGene_tx=XM_019835109.1
B4 85662856 85663054
refGene_gn=LRP1;refGene_tx=XM_003988931.4;refGene_gn=LRP1;refGene_tx=XM_019835109.1
B4 85664319 85664524

refGene_gn=LRP1;refGene_tx=XM_003988931.4;refGene_gn=LRP1;refGene_tx=XM_019835109.1
B4 85664707 85664872
refGene_gn=LRP1;refGene_tx=XM_003988931.4;refGene_gn=LRP1;refGene_tx=XM_019835109.1
B4 85665556 85665801
refGene_gn=LRP1;refGene_tx=XM_003988931.4;refGene_gn=LRP1;refGene_tx=XM_019835109.1
B4 85666026 85666158
refGene_gn=LRP1;refGene_tx=XM_003988931.4;refGene_gn=LRP1;refGene_tx=XM_019835109.1
B4 85666467 85666695
refGene_gn=LRP1;refGene_tx=XM_003988931.4;refGene_gn=LRP1;refGene_tx=XM_019835109.1
B4 85666947 85667122
refGene_gn=LRP1;refGene_tx=XM_003988931.4;refGene_gn=LRP1;refGene_tx=XM_019835109.1
B4 85667210 85667305
refGene_gn=LRP1;refGene_tx=XM_003988931.4;refGene_gn=LRP1;refGene_tx=XM_019835109.1
B4 85667504 85667654
refGene_gn=LRP1;refGene_tx=XM_003988931.4;refGene_gn=LRP1;refGene_tx=XM_019835109.1
B4 85667757 85667898
refGene_gn=LRP1;refGene_tx=XM_003988931.4;refGene_gn=LRP1;refGene_tx=XM_019835109.1
B4 85668265 85668400
refGene_gn=LRP1;refGene_tx=XM_003988931.4;refGene_gn=LRP1;refGene_tx=XM_019835109.1
B4 85669999 85670131
refGene_gn=LRP1;refGene_tx=XM_003988931.4;refGene_gn=LRP1;refGene_tx=XM_019835109.1
B4 85670392 85670527
refGene_gn=LRP1;refGene_tx=XM_003988931.4;refGene_gn=LRP1;refGene_tx=XM_019835109.1
B4 85670728 85670858
refGene_gn=LRP1;refGene_tx=XM_003988931.4;refGene_gn=LRP1;refGene_tx=XM_019835109.1
B4 85670999 85671126
refGene_gn=LRP1;refGene_tx=XM_003988931.4;refGene_gn=LRP1;refGene_tx=XM_019835109.1
B4 85671547 85671699
refGene_gn=LRP1;refGene_tx=XM_003988931.4;refGene_gn=LRP1;refGene_tx=XM_019835109.1
B4 85671797 85671922
refGene_gn=LRP1;refGene_tx=XM_003988931.4;refGene_gn=LRP1;refGene_tx=XM_019835109.1
B4 85672207 85672585
refGene_gn=LRP1;refGene_tx=XM_003988931.4;refGene_gn=LRP1;refGene_tx=XM_019835109.1

B4 85673927 85674117
refGene_gn=LRP1;refGene_tx=XM_003988931.4;refGene_gn=LRP1;refGene_tx=XM_019835109.1
B4 85677358 85677563
refGene_gn=LRP1;refGene_tx=XM_003988931.4;refGene_gn=LRP1;refGene_tx=XM_019835109.1
B4 85677954 85678147
refGene_gn=LRP1;refGene_tx=XM_003988931.4;refGene_gn=LRP1;refGene_tx=XM_019835109.1
B4 85678897 85679023
refGene_gn=LRP1;refGene_tx=XM_003988931.4;refGene_gn=LRP1;refGene_tx=XM_019835109.1
B4 85679235 85679364
refGene_gn=LRP1;refGene_tx=XM_003988931.4;refGene_gn=LRP1;refGene_tx=XM_019835109.1
B4 85679641 85679761
refGene_gn=LRP1;refGene_tx=XM_003988931.4;refGene_gn=LRP1;refGene_tx=XM_019835109.1
B4 85679973 85680087
refGene_gn=LRP1;refGene_tx=XM_003988931.4;refGene_gn=LRP1;refGene_tx=XM_019835109.1
B4 85680392 85680542
refGene_gn=LRP1;refGene_tx=XM_003988931.4;refGene_gn=LRP1;refGene_tx=XM_019835109.1
B4 85680626 85680749
refGene_gn=LRP1;refGene_tx=XM_003988931.4;refGene_gn=LRP1;refGene_tx=XM_019835109.1
B4 85681051 85681168
refGene_gn=LRP1;refGene_tx=XM_003988931.4;refGene_gn=LRP1;refGene_tx=XM_019835109.1
B4 85681347 85681476
refGene_gn=LRP1;refGene_tx=XM_003988931.4;refGene_gn=LRP1;refGene_tx=XM_019835109.1
B4 85681749 85681872
refGene_gn=LRP1;refGene_tx=XM_003988931.4;refGene_gn=LRP1;refGene_tx=XM_019835109.1
B4 85681966 85682107
refGene_gn=LRP1;refGene_tx=XM_003988931.4;refGene_gn=LRP1;refGene_tx=XM_019835109.1
B4 85682187 85682378
refGene_gn=LRP1;refGene_tx=XM_003988931.4;refGene_gn=LRP1;refGene_tx=XM_019835109.1
B4 85683144 85683328
refGene_gn=LRP1;refGene_tx=XM_003988931.4;refGene_gn=LRP1;refGene_tx=XM_019835109.1
B4 85683432 85683518
refGene_gn=LRP1;refGene_tx=XM_003988931.4;refGene_gn=LRP1;refGene_tx=XM_019835109.1
B4 85683692 85683812
refGene_gn=LRP1;refGene_tx=XM_003988931.4;refGene_gn=LRP1;refGene_tx=X

M_019835109.1
B4 85684337 85684525
refGene_gn=LRP1;refGene_tx=XM_003988931.4;refGene_gn=LRP1;refGene_tx=X
M_019835109.1
B4 85684661 85684867
refGene_gn=LRP1;refGene_tx=XM_003988931.4;refGene_gn=LRP1;refGene_tx=X
M_019835109.1
B4 85685609 85685798
refGene_gn=LRP1;refGene_tx=XM_003988931.4;refGene_gn=LRP1;refGene_tx=X
M_019835109.1
B4 85686355 85686504
refGene_gn=LRP1;refGene_tx=XM_003988931.4;refGene_gn=LRP1;refGene_tx=X
M_019835109.1
B4 85686929 85687023
refGene_gn=LRP1;refGene_tx=XM_003988931.4;refGene_gn=LRP1;refGene_tx=X
M_019835109.1
B4 85687143 85687260
refGene_gn=LRP1;refGene_tx=XM_003988931.4;refGene_gn=LRP1;refGene_tx=X
M_019835109.1
B4 85687509 85687629
refGene_gn=LRP1;refGene_tx=XM_003988931.4;refGene_gn=LRP1;refGene_tx=X
M_019835109.1
B4 85687963 85688086
refGene_gn=LRP1;refGene_tx=XM_003988931.4;refGene_gn=LRP1;refGene_tx=X
M_019835109.1
B4 85688240 85688366
refGene_gn=LRP1;refGene_tx=XM_003988931.4;refGene_gn=LRP1;refGene_tx=X
M_019835109.1
B4 85688773 85688890
refGene_gn=LRP1;refGene_tx=XM_003988931.4;refGene_gn=LRP1;refGene_tx=X
M_019835109.1
B4 85689565 85689672
refGene_gn=LRP1;refGene_tx=XM_003988931.4;refGene_gn=LRP1;refGene_tx=X
M_019835109.1
B4 85689776 85689909
refGene_gn=LRP1;refGene_tx=XM_003988931.4;refGene_gn=LRP1;refGene_tx=X
M_019835109.1
B4 85690762 85690882
refGene_gn=LRP1;refGene_tx=XM_003988931.4;refGene_gn=LRP1;refGene_tx=X
M_019835109.1
B4 85690994 85691117
refGene_gn=LRP1;refGene_tx=XM_003988931.4;refGene_gn=LRP1;refGene_tx=X
M_019835109.1
B4 85691463 85691604
refGene_gn=LRP1;refGene_tx=XM_003988931.4;refGene_gn=LRP1;refGene_tx=X
M_019835109.1
B4 85691714 85691849
refGene_gn=LRP1;refGene_tx=XM_003988931.4;refGene_gn=LRP1;refGene_tx=X
M_019835109.1
B4 85691921 85692041

refGene_gn=LRP1;refGene_tx=XM_003988931.4;refGene_gn=LRP1;refGene_tx=XM_019835109.1
B4 85692961 85693230
refGene_gn=LRP1;refGene_tx=XM_003988931.4;refGene_gn=LRP1;refGene_tx=XM_019835109.1
B4 85693780 85693959
refGene_gn=LRP1;refGene_tx=XM_003988931.4;refGene_gn=LRP1;refGene_tx=XM_019835109.1
B4 85694466 85694573
refGene_gn=LRP1;refGene_tx=XM_003988931.4;refGene_gn=LRP1;refGene_tx=XM_019835109.1
B4 85694820 85694937
refGene_gn=LRP1;refGene_tx=XM_003988931.4;refGene_gn=LRP1;refGene_tx=XM_019835109.1
B4 85695397 85695574
refGene_gn=LRP1;refGene_tx=XM_003988931.4;refGene_gn=LRP1;refGene_tx=XM_019835109.1
B4 85695720 85695861
refGene_gn=LRP1;refGene_tx=XM_003988931.4;refGene_gn=LRP1;refGene_tx=XM_019835109.1
B4 85695963 85696137
refGene_gn=LRP1;refGene_tx=XM_003988931.4;refGene_gn=LRP1;refGene_tx=XM_019835109.1
B4 85696407 85696569
refGene_gn=LRP1;refGene_tx=XM_003988931.4;refGene_gn=LRP1;refGene_tx=XM_019835109.1
B4 85696985 85697160
refGene_gn=LRP1;refGene_tx=XM_003988931.4;refGene_gn=LRP1;refGene_tx=XM_019835109.1
B4 85697343 85697445
refGene_gn=LRP1;refGene_tx=XM_003988931.4;refGene_gn=LRP1;refGene_tx=XM_019835109.1
B4 85697596 85697661
refGene_gn=LRP1;refGene_tx=XM_003988931.4;refGene_gn=LRP1;refGene_tx=XM_019835109.1
B4 85697770 85697861
refGene_gn=LRP1;refGene_tx=XM_003988931.4;refGene_gn=LRP1;refGene_tx=XM_019835109.1
B4 85697959 85698104
refGene_gn=LRP1;refGene_tx=XM_003988931.4;refGene_gn=LRP1;refGene_tx=XM_019835109.1
B4 85698272 85698413
refGene_gn=LRP1;refGene_tx=XM_003988931.4;refGene_gn=LRP1;refGene_tx=XM_019835109.1
B4 85702953 85703010 refGene_gn=NXPH4;refGene_tx=XM_003988932.4
B4 85710678 85711533 refGene_gn=NXPH4;refGene_tx=XM_003988932.4
B4 85714918 85714951 refGene_gn=SHMT2;refGene_tx=XM_003988933.4
B4 85715354 85715369 refGene_gn=SHMT2;refGene_tx=XM_019835111.2
B4 85715933 85716131
refGene_gn=SHMT2;refGene_tx=XM_003988933.4;refGene_gn=SHMT2;refGene_tx

=XM_019835111.2
B4 85717158 85717238
refGene_gn=SHMT2;refGene_tx=XM_003988933.4;refGene_gn=SHMT2;refGene_tx
=XM_019835111.2
B4 85717403 85717604
refGene_gn=SHMT2;refGene_tx=XM_003988933.4;refGene_gn=SHMT2;refGene_tx
=XM_019835111.2
B4 85717915 85717997
refGene_gn=SHMT2;refGene_tx=XM_003988933.4;refGene_gn=SHMT2;refGene_tx
=XM_019835111.2
B4 85718133 85718256
refGene_gn=SHMT2;refGene_tx=XM_003988933.4;refGene_gn=SHMT2;refGene_tx
=XM_019835111.2
B4 85718382 85718522
refGene_gn=SHMT2;refGene_tx=XM_003988933.4;refGene_gn=SHMT2;refGene_tx
=XM_019835111.2
B4 85718867 85719033
refGene_gn=SHMT2;refGene_tx=XM_003988933.4;refGene_gn=SHMT2;refGene_tx
=XM_019835111.2
B4 85719224 85719324
refGene_gn=SHMT2;refGene_tx=XM_003988933.4;refGene_gn=SHMT2;refGene_tx
=XM_019835111.2
B4 85719411 85719567
refGene_gn=SHMT2;refGene_tx=XM_003988933.4;refGene_gn=SHMT2;refGene_tx
=XM_019835111.2
B4 85719647 85719755
refGene_gn=SHMT2;refGene_tx=XM_003988933.4;refGene_gn=SHMT2;refGene_tx
=XM_019835111.2
B4 85719873 85720001
refGene_gn=SHMT2;refGene_tx=XM_003988933.4;refGene_gn=SHMT2;refGene_tx
=XM_019835111.2
B4 85721199 85721256 refGene_gn=NDUFA4L2;refGene_tx=XM_003988935.5
B4 85721374 85721435 refGene_gn=NDUFA4L2;refGene_tx=XM_003988935.5
B4 85722259 85722348 refGene_gn=NDUFA4L2;refGene_tx=XM_003988935.5
B4 85722711 85722768 refGene_gn=NDUFA4L2;refGene_tx=XM_003988935.5
B4 85729620 85729719 refGene_gn=STAC3;refGene_tx=XM_011284099.3
B4 85729895 85730033 refGene_gn=STAC3;refGene_tx=XM_011284099.3
B4 85730122 85730174 refGene_gn=STAC3;refGene_tx=XM_011284099.3
B4 85730379 85730465 refGene_gn=STAC3;refGene_tx=XM_011284099.3
B4 85730788 85730838 refGene_gn=STAC3;refGene_tx=XM_011284099.3
B4 85731218 85731285 refGene_gn=STAC3;refGene_tx=XM_011284099.3
B4 85732846 85732944 refGene_gn=STAC3;refGene_tx=XM_011284099.3
B4 85733839 85733912 refGene_gn=STAC3;refGene_tx=XM_011284099.3
B4 85734123 85734221 refGene_gn=STAC3;refGene_tx=XM_011284099.3
B4 85734460 85734728 refGene_gn=STAC3;refGene_tx=XM_011284099.3
B4 85734996 85735062 refGene_gn=STAC3;refGene_tx=XM_011284099.3
B4 85738677 85739018
refGene_gn=R3HDM2;refGene_tx=XM_006933843.3;refGene_gn=R3HDM2;refGene_
tx=XM_011284097.3;refGene_gn=R3HDM2;refGene_tx=XM_011284098.3;refGene_
gn=R3HDM2;refGene_tx=XM_019835112.2;refGene_gn=R3HDM2;refGene_tx=XM_01

9835113.2;refGene_gn=R3HDM2;refGene_tx=XM_019835114.2;refGene_gn=R3HDM
2;refGene_tx=XM_019835115.2;refGene_gn=R3HDM2;refGene_tx=XM_019835117.
2;refGene_gn=R3HDM2;refGene_tx=XM_019835118.2;refGene_gn=R3HDM2;refGen
e_tx=XM_019835119.2;refGene_gn=R3HDM2;refGene_tx=XM_019835120.2;refGen
e_gn=R3HDM2;refGene_tx=XM_019835121.2;refGene_gn=R3HDM2;refGene_tx=XM_
019835122.2;refGene_gn=R3HDM2;refGene_tx=XM_019835123.2;refGene_gn=R3H
DM2;refGene_tx=XM_019835124.2;refGene_gn=R3HDM2;refGene_tx=XM_01983512
5.2;refGene_gn=R3HDM2;refGene_tx=XM_019835126.2;refGene_gn=R3HDM2;refG
ene_tx=XM_023257154.1;refGene_gn=R3HDM2;refGene_tx=XM_023257155.1;refG
ene_gn=R3HDM2;refGene_tx=XM_023257156.1;refGene_gn=R3HDM2;refGene_tx=X
M_023257157.1;refGene_gn=R3HDM2;refGene_tx=XM_023257158.1;refGene_gn=R
3HDM2;refGene_tx=XM_023257159.1;refGene_gn=R3HDM2;refGene_tx=XM_023257
160.1
B4 85740086 85740171
refGene_gn=R3HDM2;refGene_tx=XM_006933843.3;refGene_gn=R3HDM2;refGene_
tx=XM_011284097.3;refGene_gn=R3HDM2;refGene_tx=XM_011284098.3;refGene_
gn=R3HDM2;refGene_tx=XM_019835112.2;refGene_gn=R3HDM2;refGene_tx=XM_01
9835113.2;refGene_gn=R3HDM2;refGene_tx=XM_019835114.2;refGene_gn=R3HDM
2;refGene_tx=XM_019835115.2;refGene_gn=R3HDM2;refGene_tx=XM_019835117.
2;refGene_gn=R3HDM2;refGene_tx=XM_019835118.2;refGene_gn=R3HDM2;refGen
e_tx=XM_019835119.2;refGene_gn=R3HDM2;refGene_tx=XM_019835120.2;refGen
e_gn=R3HDM2;refGene_tx=XM_019835121.2;refGene_gn=R3HDM2;refGene_tx=XM_
019835122.2;refGene_gn=R3HDM2;refGene_tx=XM_019835123.2;refGene_gn=R3H
DM2;refGene_tx=XM_019835124.2;refGene_gn=R3HDM2;refGene_tx=XM_01983512
5.2;refGene_gn=R3HDM2;refGene_tx=XM_019835126.2;refGene_gn=R3HDM2;refG
ene_tx=XM_023257154.1;refGene_gn=R3HDM2;refGene_tx=XM_023257155.1;refG
ene_gn=R3HDM2;refGene_tx=XM_023257156.1;refGene_gn=R3HDM2;refGene_tx=X
M_023257157.1;refGene_gn=R3HDM2;refGene_tx=XM_023257158.1;refGene_gn=R
3HDM2;refGene_tx=XM_023257159.1;refGene_gn=R3HDM2;refGene_tx=XM_023257
160.1
B4 85740544 85740642
refGene_gn=R3HDM2;refGene_tx=XM_006933843.3;refGene_gn=R3HDM2;refGene_
tx=XM_011284097.3;refGene_gn=R3HDM2;refGene_tx=XM_011284098.3;refGene_
gn=R3HDM2;refGene_tx=XM_019835112.2;refGene_gn=R3HDM2;refGene_tx=XM_01
9835113.2;refGene_gn=R3HDM2;refGene_tx=XM_019835114.2;refGene_gn=R3HDM
2;refGene_tx=XM_019835115.2;refGene_gn=R3HDM2;refGene_tx=XM_019835117.
2;refGene_gn=R3HDM2;refGene_tx=XM_019835118.2;refGene_gn=R3HDM2;refGen
e_tx=XM_019835119.2;refGene_gn=R3HDM2;refGene_tx=XM_019835120.2;refGen
e_gn=R3HDM2;refGene_tx=XM_019835121.2;refGene_gn=R3HDM2;refGene_tx=XM_
019835122.2;refGene_gn=R3HDM2;refGene_tx=XM_019835123.2;refGene_gn=R3H
DM2;refGene_tx=XM_019835124.2;refGene_gn=R3HDM2;refGene_tx=XM_01983512
5.2;refGene_gn=R3HDM2;refGene_tx=XM_019835126.2;refGene_gn=R3HDM2;refG
ene_tx=XM_023257154.1;refGene_gn=R3HDM2;refGene_tx=XM_023257155.1;refG
ene_gn=R3HDM2;refGene_tx=XM_023257156.1;refGene_gn=R3HDM2;refGene_tx=X
M_023257157.1;refGene_gn=R3HDM2;refGene_tx=XM_023257158.1;refGene_gn=R
3HDM2;refGene_tx=XM_023257159.1;refGene_gn=R3HDM2;refGene_tx=XM_023257
160.1
B4 85742048 85742196
refGene_gn=R3HDM2;refGene_tx=XM_006933843.3;refGene_gn=R3HDM2;refGene_
tx=XM_011284097.3;refGene_gn=R3HDM2;refGene_tx=XM_011284098.3;refGene_

gn=R3HDM2;refGene_tx=XM_019835112.2;refGene_gn=R3HDM2;refGene_tx=XM_01
9835113.2;refGene_gn=R3HDM2;refGene_tx=XM_019835114.2;refGene_gn=R3HDM
2;refGene_tx=XM_019835115.2;refGene_gn=R3HDM2;refGene_tx=XM_019835117.
2;refGene_gn=R3HDM2;refGene_tx=XM_019835118.2;refGene_gn=R3HDM2;refGen
e_tx=XM_019835119.2;refGene_gn=R3HDM2;refGene_tx=XM_019835120.2;refGen
e_gn=R3HDM2;refGene_tx=XM_019835121.2;refGene_gn=R3HDM2;refGene_tx=XM_
019835122.2;refGene_gn=R3HDM2;refGene_tx=XM_019835123.2;refGene_gn=R3H
DM2;refGene_tx=XM_019835124.2;refGene_gn=R3HDM2;refGene_tx=XM_01983512
5.2;refGene_gn=R3HDM2;refGene_tx=XM_019835126.2;refGene_gn=R3HDM2;refG
ene_tx=XM_023257154.1;refGene_gn=R3HDM2;refGene_tx=XM_023257155.1;refG
ene_gn=R3HDM2;refGene_tx=XM_023257156.1;refGene_gn=R3HDM2;refGene_tx=X
M_023257157.1;refGene_gn=R3HDM2;refGene_tx=XM_023257158.1;refGene_gn=R
3HDM2;refGene_tx=XM_023257159.1;refGene_gn=R3HDM2;refGene_tx=XM_023257
160.1
B4 85742830 85743000
refGene_gn=R3HDM2;refGene_tx=XM_006933843.3;refGene_gn=R3HDM2;refGene_
tx=XM_011284097.3;refGene_gn=R3HDM2;refGene_tx=XM_011284098.3;refGene_
gn=R3HDM2;refGene_tx=XM_019835112.2;refGene_gn=R3HDM2;refGene_tx=XM_01
9835113.2;refGene_gn=R3HDM2;refGene_tx=XM_019835114.2;refGene_gn=R3HDM
2;refGene_tx=XM_019835115.2;refGene_gn=R3HDM2;refGene_tx=XM_019835117.
2;refGene_gn=R3HDM2;refGene_tx=XM_019835118.2;refGene_gn=R3HDM2;refGen
e_tx=XM_019835119.2;refGene_gn=R3HDM2;refGene_tx=XM_019835120.2;refGen
e_gn=R3HDM2;refGene_tx=XM_019835121.2;refGene_gn=R3HDM2;refGene_tx=XM_
019835122.2;refGene_gn=R3HDM2;refGene_tx=XM_019835123.2;refGene_gn=R3H
DM2;refGene_tx=XM_019835124.2;refGene_gn=R3HDM2;refGene_tx=XM_01983512
5.2;refGene_gn=R3HDM2;refGene_tx=XM_019835126.2;refGene_gn=R3HDM2;refG
ene_tx=XM_023257154.1;refGene_gn=R3HDM2;refGene_tx=XM_023257155.1;refG
ene_gn=R3HDM2;refGene_tx=XM_023257156.1;refGene_gn=R3HDM2;refGene_tx=X
M_023257157.1;refGene_gn=R3HDM2;refGene_tx=XM_023257158.1;refGene_gn=R
3HDM2;refGene_tx=XM_023257159.1;refGene_gn=R3HDM2;refGene_tx=XM_023257
160.1
B4 85748181 85748282
refGene_gn=R3HDM2;refGene_tx=XM_006933843.3;refGene_gn=R3HDM2;refGene_
tx=XM_011284097.3;refGene_gn=R3HDM2;refGene_tx=XM_011284098.3;refGene_
gn=R3HDM2;refGene_tx=XM_019835112.2;refGene_gn=R3HDM2;refGene_tx=XM_01
9835113.2;refGene_gn=R3HDM2;refGene_tx=XM_019835114.2;refGene_gn=R3HDM
2;refGene_tx=XM_019835115.2;refGene_gn=R3HDM2;refGene_tx=XM_019835117.
2;refGene_gn=R3HDM2;refGene_tx=XM_019835118.2;refGene_gn=R3HDM2;refGen
e_tx=XM_019835119.2;refGene_gn=R3HDM2;refGene_tx=XM_019835120.2;refGen
e_gn=R3HDM2;refGene_tx=XM_019835121.2;refGene_gn=R3HDM2;refGene_tx=XM_
019835122.2;refGene_gn=R3HDM2;refGene_tx=XM_019835123.2;refGene_gn=R3H
DM2;refGene_tx=XM_019835124.2;refGene_gn=R3HDM2;refGene_tx=XM_01983512
5.2;refGene_gn=R3HDM2;refGene_tx=XM_019835126.2;refGene_gn=R3HDM2;refG
ene_tx=XM_023257154.1;refGene_gn=R3HDM2;refGene_tx=XM_023257155.1;refG
ene_gn=R3HDM2;refGene_tx=XM_023257156.1;refGene_gn=R3HDM2;refGene_tx=X
M_023257157.1;refGene_gn=R3HDM2;refGene_tx=XM_023257158.1;refGene_gn=R
3HDM2;refGene_tx=XM_023257159.1;refGene_gn=R3HDM2;refGene_tx=XM_023257
160.1
B4 85749613 85749768
refGene_gn=R3HDM2;refGene_tx=XM_006933843.3;refGene_gn=R3HDM2;refGene_

tx=XM_011284097.3;refGene_gn=R3HDM2;refGene_tx=XM_011284098.3;refGene_
gn=R3HDM2;refGene_tx=XM_019835112.2;refGene_gn=R3HDM2;refGene_tx=XM_01
9835113.2;refGene_gn=R3HDM2;refGene_tx=XM_019835114.2;refGene_gn=R3HDM
2;refGene_tx=XM_019835115.2;refGene_gn=R3HDM2;refGene_tx=XM_019835117.
2;refGene_gn=R3HDM2;refGene_tx=XM_019835118.2;refGene_gn=R3HDM2;refGen
e_tx=XM_019835119.2;refGene_gn=R3HDM2;refGene_tx=XM_019835120.2;refGen
e_gn=R3HDM2;refGene_tx=XM_019835121.2;refGene_gn=R3HDM2;refGene_tx=XM_
019835122.2;refGene_gn=R3HDM2;refGene_tx=XM_019835123.2;refGene_gn=R3H
DM2;refGene_tx=XM_019835124.2;refGene_gn=R3HDM2;refGene_tx=XM_01983512
5.2;refGene_gn=R3HDM2;refGene_tx=XM_019835126.2;refGene_gn=R3HDM2;refG
ene_tx=XM_023257154.1;refGene_gn=R3HDM2;refGene_tx=XM_023257155.1;refG
ene_gn=R3HDM2;refGene_tx=XM_023257156.1;refGene_gn=R3HDM2;refGene_tx=X
M_023257157.1;refGene_gn=R3HDM2;refGene_tx=XM_023257158.1;refGene_gn=R
3HDM2;refGene_tx=XM_023257159.1;refGene_gn=R3HDM2;refGene_tx=XM_023257
160.1
B4 85750217 85750378
refGene_gn=R3HDM2;refGene_tx=XM_006933843.3;refGene_gn=R3HDM2;refGene_
tx=XM_011284097.3;refGene_gn=R3HDM2;refGene_tx=XM_011284098.3;refGene_
gn=R3HDM2;refGene_tx=XM_019835112.2;refGene_gn=R3HDM2;refGene_tx=XM_01
9835113.2;refGene_gn=R3HDM2;refGene_tx=XM_019835114.2;refGene_gn=R3HDM
2;refGene_tx=XM_019835115.2;refGene_gn=R3HDM2;refGene_tx=XM_019835117.
2;refGene_gn=R3HDM2;refGene_tx=XM_019835118.2;refGene_gn=R3HDM2;refGen
e_tx=XM_019835119.2;refGene_gn=R3HDM2;refGene_tx=XM_019835120.2;refGen
e_gn=R3HDM2;refGene_tx=XM_019835121.2;refGene_gn=R3HDM2;refGene_tx=XM_
019835122.2;refGene_gn=R3HDM2;refGene_tx=XM_019835123.2;refGene_gn=R3H
DM2;refGene_tx=XM_019835124.2;refGene_gn=R3HDM2;refGene_tx=XM_01983512
5.2;refGene_gn=R3HDM2;refGene_tx=XM_019835126.2;refGene_gn=R3HDM2;refG
ene_tx=XM_023257154.1;refGene_gn=R3HDM2;refGene_tx=XM_023257155.1;refG
ene_gn=R3HDM2;refGene_tx=XM_023257156.1;refGene_gn=R3HDM2;refGene_tx=X
M_023257157.1;refGene_gn=R3HDM2;refGene_tx=XM_023257158.1;refGene_gn=R
3HDM2;refGene_tx=XM_023257159.1;refGene_gn=R3HDM2;refGene_tx=XM_023257
160.1
B4 85750638 85750765
refGene_gn=R3HDM2;refGene_tx=XM_006933843.3;refGene_gn=R3HDM2;refGene_
tx=XM_011284097.3;refGene_gn=R3HDM2;refGene_tx=XM_011284098.3;refGene_
gn=R3HDM2;refGene_tx=XM_019835112.2;refGene_gn=R3HDM2;refGene_tx=XM_01
9835113.2;refGene_gn=R3HDM2;refGene_tx=XM_019835114.2;refGene_gn=R3HDM
2;refGene_tx=XM_019835115.2;refGene_gn=R3HDM2;refGene_tx=XM_019835117.
2;refGene_gn=R3HDM2;refGene_tx=XM_019835118.2;refGene_gn=R3HDM2;refGen
e_tx=XM_019835119.2;refGene_gn=R3HDM2;refGene_tx=XM_019835120.2;refGen
e_gn=R3HDM2;refGene_tx=XM_019835121.2;refGene_gn=R3HDM2;refGene_tx=XM_
019835122.2;refGene_gn=R3HDM2;refGene_tx=XM_019835123.2;refGene_gn=R3H
DM2;refGene_tx=XM_019835124.2;refGene_gn=R3HDM2;refGene_tx=XM_01983512
5.2;refGene_gn=R3HDM2;refGene_tx=XM_019835126.2;refGene_gn=R3HDM2;refG
ene_tx=XM_023257154.1;refGene_gn=R3HDM2;refGene_tx=XM_023257155.1;refG
ene_gn=R3HDM2;refGene_tx=XM_023257156.1;refGene_gn=R3HDM2;refGene_tx=X
M_023257157.1;refGene_gn=R3HDM2;refGene_tx=XM_023257158.1;refGene_gn=R
3HDM2;refGene_tx=XM_023257159.1;refGene_gn=R3HDM2;refGene_tx=XM_023257
160.1
B4 85751099 85751342

refGene_gn=R3HDM2;refGene_tx=XM_006933843.3;refGene_gn=R3HDM2;refGene_tx=XM_011284097.3;refGene_gn=R3HDM2;refGene_tx=XM_011284098.3;refGene_gn=R3HDM2;refGene_tx=XM_019835112.2;refGene_gn=R3HDM2;refGene_tx=XM_019835113.2;refGene_gn=R3HDM2;refGene_tx=XM_019835114.2;refGene_gn=R3HDM2;refGene_tx=XM_019835115.2;refGene_gn=R3HDM2;refGene_tx=XM_019835117.2;refGene_gn=R3HDM2;refGene_tx=XM_019835118.2;refGene_gn=R3HDM2;refGene_tx=XM_019835119.2;refGene_gn=R3HDM2;refGene_tx=XM_019835120.2;refGene_gn=R3HDM2;refGene_tx=XM_019835121.2;refGene_gn=R3HDM2;refGene_tx=XM_019835122.2;refGene_gn=R3HDM2;refGene_tx=XM_019835123.2;refGene_gn=R3HDM2;refGene_tx=XM_019835124.2;refGene_gn=R3HDM2;refGene_tx=XM_019835125.2;refGene_gn=R3HDM2;refGene_tx=XM_019835126.2;refGene_gn=R3HDM2;refGene_tx=XM_023257154.1;refGene_gn=R3HDM2;refGene_tx=XM_023257155.1;refGene_gn=R3HDM2;refGene_tx=XM_023257156.1;refGene_gn=R3HDM2;refGene_tx=XM_023257157.1;refGene_gn=R3HDM2;refGene_tx=XM_023257158.1;refGene_gn=R3HDM2;refGene_tx=XM_023257159.1;refGene_gn=R3HDM2;refGene_tx=XM_023257160.1
B4 85753682 85753784
refGene_gn=R3HDM2;refGene_tx=XM_006933843.3;refGene_gn=R3HDM2;refGene_tx=XM_011284097.3;refGene_gn=R3HDM2;refGene_tx=XM_011284098.3;refGene_gn=R3HDM2;refGene_tx=XM_019835112.2;refGene_gn=R3HDM2;refGene_tx=XM_019835113.2;refGene_gn=R3HDM2;refGene_tx=XM_019835114.2;refGene_gn=R3HDM2;refGene_tx=XM_019835115.2;refGene_gn=R3HDM2;refGene_tx=XM_019835117.2;refGene_gn=R3HDM2;refGene_tx=XM_019835118.2;refGene_gn=R3HDM2;refGene_tx=XM_019835120.2;refGene_gn=R3HDM2;refGene_tx=XM_019835121.2;refGene_gn=R3HDM2;refGene_tx=XM_019835122.2;refGene_gn=R3HDM2;refGene_tx=XM_019835123.2;refGene_gn=R3HDM2;refGene_tx=XM_023257154.1;refGene_gn=R3HDM2;refGene_tx=XM_023257156.1;refGene_gn=R3HDM2;refGene_tx=XM_023257158.1
B4 85758124 85758300
refGene_gn=R3HDM2;refGene_tx=XM_006933843.3;refGene_gn=R3HDM2;refGene_tx=XM_011284097.3;refGene_gn=R3HDM2;refGene_tx=XM_011284098.3;refGene_gn=R3HDM2;refGene_tx=XM_019835112.2;refGene_gn=R3HDM2;refGene_tx=XM_019835113.2;refGene_gn=R3HDM2;refGene_tx=XM_019835114.2;refGene_gn=R3HDM2;refGene_tx=XM_019835115.2;refGene_gn=R3HDM2;refGene_tx=XM_019835117.2;refGene_gn=R3HDM2;refGene_tx=XM_019835118.2;refGene_gn=R3HDM2;refGene_tx=XM_019835119.2;refGene_gn=R3HDM2;refGene_tx=XM_019835120.2;refGene_gn=R3HDM2;refGene_tx=XM_019835121.2;refGene_gn=R3HDM2;refGene_tx=XM_019835122.2;refGene_gn=R3HDM2;refGene_tx=XM_019835123.2;refGene_gn=R3HDM2;refGene_tx=XM_019835124.2;refGene_gn=R3HDM2;refGene_tx=XM_019835125.2;refGene_gn=R3HDM2;refGene_tx=XM_019835126.2;refGene_gn=R3HDM2;refGene_tx=XM_023257154.1;refGene_gn=R3HDM2;refGene_tx=XM_023257155.1;refGene_gn=R3HDM2;refGene_tx=XM_023257156.1;refGene_gn=R3HDM2;refGene_tx=XM_023257157.1;refGene_gn=R3HDM2;refGene_tx=XM_023257158.1;refGene_gn=R3HDM2;refGene_tx=XM_023257159.1;refGene_gn=R3HDM2;refGene_tx=XM_023257160.1
B4 85760652 85760885
refGene_gn=R3HDM2;refGene_tx=XM_006933843.3;refGene_gn=R3HDM2;refGene_tx=XM_011284097.3;refGene_gn=R3HDM2;refGene_tx=XM_011284098.3;refGene_gn=R3HDM2;refGene_tx=XM_019835112.2;refGene_gn=R3HDM2;refGene_tx=XM_019835113.2;refGene_gn=R3HDM2;refGene_tx=XM_019835114.2;refGene_gn=R3HDM

2;refGene_tx=XM_019835115.2;refGene_gn=R3HDM2;refGene_tx=XM_019835117.2;refGene_gn=R3HDM2;refGene_tx=XM_019835118.2;refGene_gn=R3HDM2;refGene_tx=XM_019835119.2;refGene_gn=R3HDM2;refGene_tx=XM_019835120.2;refGene_gn=R3HDM2;refGene_tx=XM_019835121.2;refGene_gn=R3HDM2;refGene_tx=XM_019835122.2;refGene_gn=R3HDM2;refGene_tx=XM_019835123.2;refGene_gn=R3HDM2;refGene_tx=XM_019835124.2;refGene_gn=R3HDM2;refGene_tx=XM_019835125.2;refGene_gn=R3HDM2;refGene_tx=XM_019835126.2;refGene_gn=R3HDM2;refGene_tx=XM_023257154.1;refGene_gn=R3HDM2;refGene_tx=XM_023257155.1;refGene_gn=R3HDM2;refGene_tx=XM_023257156.1;refGene_gn=R3HDM2;refGene_tx=XM_023257157.1;refGene_gn=R3HDM2;refGene_tx=XM_023257158.1;refGene_gn=R3HDM2;refGene_tx=XM_023257159.1;refGene_gn=R3HDM2;refGene_tx=XM_023257160.1

B4 85765124 85765178
refGene_gn=R3HDM2;refGene_tx=XM_006933843.3;refGene_gn=R3HDM2;refGene_tx=XM_011284097.3;refGene_gn=R3HDM2;refGene_tx=XM_011284098.3;refGene_gn=R3HDM2;refGene_tx=XM_019835112.2;refGene_gn=R3HDM2;refGene_tx=XM_019835113.2;refGene_gn=R3HDM2;refGene_tx=XM_019835114.2;refGene_gn=R3HDM2;refGene_tx=XM_019835117.2;refGene_gn=R3HDM2;refGene_tx=XM_019835118.2;refGene_gn=R3HDM2;refGene_tx=XM_019835119.2;refGene_gn=R3HDM2;refGene_tx=XM_019835120.2;refGene_gn=R3HDM2;refGene_tx=XM_019835121.2;refGene_gn=R3HDM2;refGene_tx=XM_019835122.2;refGene_gn=R3HDM2;refGene_tx=XM_019835125.2;refGene_gn=R3HDM2;refGene_tx=XM_023257155.1;refGene_gn=R3HDM2;refGene_tx=XM_023257156.1;refGene_gn=R3HDM2;refGene_tx=XM_023257159.1

B4 85765276 85765308
refGene_gn=R3HDM2;refGene_tx=XM_006933843.3;refGene_gn=R3HDM2;refGene_tx=XM_011284097.3;refGene_gn=R3HDM2;refGene_tx=XM_011284098.3;refGene_gn=R3HDM2;refGene_tx=XM_019835112.2;refGene_gn=R3HDM2;refGene_tx=XM_019835113.2;refGene_gn=R3HDM2;refGene_tx=XM_019835114.2;refGene_gn=R3HDM2;refGene_tx=XM_019835115.2;refGene_gn=R3HDM2;refGene_tx=XM_019835117.2;refGene_gn=R3HDM2;refGene_tx=XM_019835118.2;refGene_gn=R3HDM2;refGene_tx=XM_019835119.2;refGene_gn=R3HDM2;refGene_tx=XM_019835120.2;refGene_gn=R3HDM2;refGene_tx=XM_019835121.2;refGene_gn=R3HDM2;refGene_tx=XM_019835122.2;refGene_gn=R3HDM2;refGene_tx=XM_019835123.2;refGene_gn=R3HDM2;refGene_tx=XM_019835124.2;refGene_gn=R3HDM2;refGene_tx=XM_019835125.2;refGene_gn=R3HDM2;refGene_tx=XM_019835126.2;refGene_gn=R3HDM2;refGene_tx=XM_023257154.1;refGene_gn=R3HDM2;refGene_tx=XM_023257155.1;refGene_gn=R3HDM2;refGene_tx=XM_023257156.1;refGene_gn=R3HDM2;refGene_tx=XM_023257157.1;refGene_gn=R3HDM2;refGene_tx=XM_023257158.1;refGene_gn=R3HDM2;refGene_tx=XM_023257159.1;refGene_gn=R3HDM2;refGene_tx=XM_023257160.1

B4 85768202 85768298
refGene_gn=R3HDM2;refGene_tx=XM_006933843.3;refGene_gn=R3HDM2;refGene_tx=XM_011284097.3;refGene_gn=R3HDM2;refGene_tx=XM_011284098.3;refGene_gn=R3HDM2;refGene_tx=XM_019835112.2;refGene_gn=R3HDM2;refGene_tx=XM_019835113.2;refGene_gn=R3HDM2;refGene_tx=XM_019835114.2;refGene_gn=R3HDM2;refGene_tx=XM_019835115.2;refGene_gn=R3HDM2;refGene_tx=XM_019835117.2;refGene_gn=R3HDM2;refGene_tx=XM_019835119.2;refGene_gn=R3HDM2;refGene_tx=XM_019835121.2;refGene_gn=R3HDM2;refGene_tx=XM_019835122.2;refGene_gn=R3HDM2;refGene_tx=XM_019835124.2;refGene_gn=R3HDM2;refGene_tx=XM_

023257154.1;refGene_gn=R3HDM2;refGene_tx=XM_023257155.1;refGene_gn=R3HDM2;refGene_tx=XM_023257156.1;refGene_gn=R3HDM2;refGene_tx=XM_023257157.1;refGene_gn=R3HDM2;refGene_tx=XM_023257159.1
B4 85771053 85771162
refGene_gn=R3HDM2;refGene_tx=XM_006933843.3;refGene_gn=R3HDM2;refGene_tx=XM_011284097.3;refGene_gn=R3HDM2;refGene_tx=XM_011284098.3;refGene_gn=R3HDM2;refGene_tx=XM_019835112.2;refGene_gn=R3HDM2;refGene_tx=XM_019835113.2;refGene_gn=R3HDM2;refGene_tx=XM_019835114.2;refGene_gn=R3HDM2;refGene_tx=XM_019835115.2;refGene_gn=R3HDM2;refGene_tx=XM_019835117.2;refGene_gn=R3HDM2;refGene_tx=XM_019835118.2;refGene_gn=R3HDM2;refGene_tx=XM_019835119.2;refGene_gn=R3HDM2;refGene_tx=XM_019835120.2;refGene_gn=R3HDM2;refGene_tx=XM_019835121.2;refGene_gn=R3HDM2;refGene_tx=XM_019835122.2;refGene_gn=R3HDM2;refGene_tx=XM_019835123.2;refGene_gn=R3HDM2;refGene_tx=XM_019835124.2;refGene_gn=R3HDM2;refGene_tx=XM_019835125.2;refGene_gn=R3HDM2;refGene_tx=XM_019835126.2;refGene_gn=R3HDM2;refGene_tx=XM_023257154.1;refGene_gn=R3HDM2;refGene_tx=XM_023257155.1;refGene_gn=R3HDM2;refGene_tx=XM_023257156.1;refGene_gn=R3HDM2;refGene_tx=XM_023257157.1;refGene_gn=R3HDM2;refGene_tx=XM_023257158.1;refGene_gn=R3HDM2;refGene_tx=XM_023257159.1;refGene_gn=R3HDM2;refGene_tx=XM_023257160.1
B4 85771569 85771710
refGene_gn=R3HDM2;refGene_tx=XM_006933843.3;refGene_gn=R3HDM2;refGene_tx=XM_011284097.3;refGene_gn=R3HDM2;refGene_tx=XM_011284098.3;refGene_gn=R3HDM2;refGene_tx=XM_019835112.2;refGene_gn=R3HDM2;refGene_tx=XM_019835113.2;refGene_gn=R3HDM2;refGene_tx=XM_019835114.2;refGene_gn=R3HDM2;refGene_tx=XM_019835115.2;refGene_gn=R3HDM2;refGene_tx=XM_019835117.2;refGene_gn=R3HDM2;refGene_tx=XM_019835118.2;refGene_gn=R3HDM2;refGene_tx=XM_019835119.2;refGene_gn=R3HDM2;refGene_tx=XM_019835120.2;refGene_gn=R3HDM2;refGene_tx=XM_019835121.2;refGene_gn=R3HDM2;refGene_tx=XM_019835122.2;refGene_gn=R3HDM2;refGene_tx=XM_019835123.2;refGene_gn=R3HDM2;refGene_tx=XM_019835124.2;refGene_gn=R3HDM2;refGene_tx=XM_019835125.2;refGene_gn=R3HDM2;refGene_tx=XM_019835126.2;refGene_gn=R3HDM2;refGene_tx=XM_023257154.1;refGene_gn=R3HDM2;refGene_tx=XM_023257155.1;refGene_gn=R3HDM2;refGene_tx=XM_023257156.1;refGene_gn=R3HDM2;refGene_tx=XM_023257157.1;refGene_gn=R3HDM2;refGene_tx=XM_023257158.1;refGene_gn=R3HDM2;refGene_tx=XM_023257159.1;refGene_gn=R3HDM2;refGene_tx=XM_023257160.1
B4 85772273 85772333
refGene_gn=R3HDM2;refGene_tx=XM_006933843.3;refGene_gn=R3HDM2;refGene_tx=XM_011284097.3;refGene_gn=R3HDM2;refGene_tx=XM_011284098.3;refGene_gn=R3HDM2;refGene_tx=XM_019835112.2;refGene_gn=R3HDM2;refGene_tx=XM_019835113.2;refGene_gn=R3HDM2;refGene_tx=XM_019835114.2;refGene_gn=R3HDM2;refGene_tx=XM_019835115.2;refGene_gn=R3HDM2;refGene_tx=XM_019835118.2;refGene_gn=R3HDM2;refGene_tx=XM_019835119.2;refGene_gn=R3HDM2;refGene_tx=XM_019835120.2;refGene_gn=R3HDM2;refGene_tx=XM_019835121.2;refGene_gn=R3HDM2;refGene_tx=XM_019835122.2;refGene_gn=R3HDM2;refGene_tx=XM_019835123.2;refGene_gn=R3HDM2;refGene_tx=XM_019835124.2;refGene_gn=R3HDM2;refGene_tx=XM_019835125.2;refGene_gn=R3HDM2;refGene_tx=XM_019835126.2;refGene_gn=R3HDM2;refGene_tx=XM_023257154.1;refGene_gn=R3HDM2;refGene_tx=XM_023257155.1;refGene_gn=R3HDM2;refGene_tx=XM_023257156.1;refG

ene_gn=R3HDM2;refGene_tx=XM_023257157.1;refGene_gn=R3HDM2;refGene_tx=XM_023257158.1;refGene_gn=R3HDM2;refGene_tx=XM_023257160.1
B4 85772977 85773056
refGene_gn=R3HDM2;refGene_tx=XM_006933843.3;refGene_gn=R3HDM2;refGene_tx=XM_011284097.3;refGene_gn=R3HDM2;refGene_tx=XM_011284098.3;refGene_gn=R3HDM2;refGene_tx=XM_019835112.2;refGene_gn=R3HDM2;refGene_tx=XM_019835113.2;refGene_gn=R3HDM2;refGene_tx=XM_019835114.2;refGene_gn=R3HDM2;refGene_tx=XM_019835115.2;refGene_gn=R3HDM2;refGene_tx=XM_019835117.2;refGene_gn=R3HDM2;refGene_tx=XM_019835118.2;refGene_gn=R3HDM2;refGene_tx=XM_019835119.2;refGene_gn=R3HDM2;refGene_tx=XM_019835120.2;refGene_gn=R3HDM2;refGene_tx=XM_019835121.2;refGene_gn=R3HDM2;refGene_tx=XM_019835122.2;refGene_gn=R3HDM2;refGene_tx=XM_019835123.2;refGene_gn=R3HDM2;refGene_tx=XM_019835124.2;refGene_gn=R3HDM2;refGene_tx=XM_019835125.2;refGene_gn=R3HDM2;refGene_tx=XM_019835126.2;refGene_gn=R3HDM2;refGene_tx=XM_023257154.1;refGene_gn=R3HDM2;refGene_tx=XM_023257155.1;refGene_gn=R3HDM2;refGene_tx=XM_023257156.1;refGene_gn=R3HDM2;refGene_tx=XM_023257157.1;refGene_gn=R3HDM2;refGene_tx=XM_023257158.1;refGene_gn=R3HDM2;refGene_tx=XM_023257159.1;refGene_gn=R3HDM2;refGene_tx=XM_023257160.1
B4 85773904 85774031
refGene_gn=R3HDM2;refGene_tx=XM_006933843.3;refGene_gn=R3HDM2;refGene_tx=XM_011284097.3;refGene_gn=R3HDM2;refGene_tx=XM_011284098.3;refGene_gn=R3HDM2;refGene_tx=XM_019835112.2;refGene_gn=R3HDM2;refGene_tx=XM_019835113.2;refGene_gn=R3HDM2;refGene_tx=XM_019835114.2;refGene_gn=R3HDM2;refGene_tx=XM_019835115.2;refGene_gn=R3HDM2;refGene_tx=XM_019835117.2;refGene_gn=R3HDM2;refGene_tx=XM_019835118.2;refGene_gn=R3HDM2;refGene_tx=XM_019835119.2;refGene_gn=R3HDM2;refGene_tx=XM_019835120.2;refGene_gn=R3HDM2;refGene_tx=XM_019835121.2;refGene_gn=R3HDM2;refGene_tx=XM_019835122.2;refGene_gn=R3HDM2;refGene_tx=XM_019835123.2;refGene_gn=R3HDM2;refGene_tx=XM_019835124.2;refGene_gn=R3HDM2;refGene_tx=XM_019835125.2;refGene_gn=R3HDM2;refGene_tx=XM_019835126.2;refGene_gn=R3HDM2;refGene_tx=XM_023257154.1;refGene_gn=R3HDM2;refGene_tx=XM_023257155.1;refGene_gn=R3HDM2;refGene_tx=XM_023257156.1;refGene_gn=R3HDM2;refGene_tx=XM_023257157.1;refGene_gn=R3HDM2;refGene_tx=XM_023257158.1;refGene_gn=R3HDM2;refGene_tx=XM_023257159.1;refGene_gn=R3HDM2;refGene_tx=XM_023257160.1
B4 85774554 85774641
refGene_gn=R3HDM2;refGene_tx=XM_006933843.3;refGene_gn=R3HDM2;refGene_tx=XM_011284097.3;refGene_gn=R3HDM2;refGene_tx=XM_011284098.3;refGene_gn=R3HDM2;refGene_tx=XM_019835112.2;refGene_gn=R3HDM2;refGene_tx=XM_019835113.2;refGene_gn=R3HDM2;refGene_tx=XM_019835114.2;refGene_gn=R3HDM2;refGene_tx=XM_019835115.2;refGene_gn=R3HDM2;refGene_tx=XM_019835117.2;refGene_gn=R3HDM2;refGene_tx=XM_019835118.2;refGene_gn=R3HDM2;refGene_tx=XM_019835119.2;refGene_gn=R3HDM2;refGene_tx=XM_019835120.2;refGene_gn=R3HDM2;refGene_tx=XM_019835121.2;refGene_gn=R3HDM2;refGene_tx=XM_019835122.2;refGene_gn=R3HDM2;refGene_tx=XM_019835123.2;refGene_gn=R3HDM2;refGene_tx=XM_019835124.2;refGene_gn=R3HDM2;refGene_tx=XM_019835125.2;refGene_gn=R3HDM2;refGene_tx=XM_019835126.2;refGene_gn=R3HDM2;refGene_tx=XM_023257154.1;refGene_gn=R3HDM2;refGene_tx=XM_023257155.1;refGene_gn=R3HDM2;refGene_tx=XM_023257156.1;refGene_gn=R3HDM2;refGene_tx=X

M_023257157.1;refGene_gn=R3HDM2;refGene_tx=XM_023257158.1;refGene_gn=R3HDM2;refGene_tx=XM_023257159.1;refGene_gn=R3HDM2;refGene_tx=XM_023257160.1
B4 85777227 85777269
refGene_gn=R3HDM2;refGene_tx=XM_006933843.3;refGene_gn=R3HDM2;refGene_tx=XM_011284097.3;refGene_gn=R3HDM2;refGene_tx=XM_011284098.3;refGene_gn=R3HDM2;refGene_tx=XM_019835112.2;refGene_gn=R3HDM2;refGene_tx=XM_019835113.2;refGene_gn=R3HDM2;refGene_tx=XM_019835114.2;refGene_gn=R3HDM2;refGene_tx=XM_019835115.2;refGene_gn=R3HDM2;refGene_tx=XM_019835117.2;refGene_gn=R3HDM2;refGene_tx=XM_019835118.2;refGene_gn=R3HDM2;refGene_tx=XM_019835119.2;refGene_gn=R3HDM2;refGene_tx=XM_019835120.2;refGene_gn=R3HDM2;refGene_tx=XM_019835121.2;refGene_gn=R3HDM2;refGene_tx=XM_019835122.2;refGene_gn=R3HDM2;refGene_tx=XM_019835123.2;refGene_gn=R3HDM2;refGene_tx=XM_019835124.2;refGene_gn=R3HDM2;refGene_tx=XM_019835125.2;refGene_gn=R3HDM2;refGene_tx=XM_019835126.2;refGene_gn=R3HDM2;refGene_tx=XM_023257154.1;refGene_gn=R3HDM2;refGene_tx=XM_023257155.1;refGene_gn=R3HDM2;refGene_tx=XM_023257156.1;refGene_gn=R3HDM2;refGene_tx=XM_023257157.1;refGene_gn=R3HDM2;refGene_tx=XM_023257158.1;refGene_gn=R3HDM2;refGene_tx=XM_023257159.1;refGene_gn=R3HDM2;refGene_tx=XM_023257160.1
B4 85779080 85779104
refGene_gn=R3HDM2;refGene_tx=XM_006933843.3;refGene_gn=R3HDM2;refGene_tx=XM_011284097.3;refGene_gn=R3HDM2;refGene_tx=XM_011284098.3;refGene_gn=R3HDM2;refGene_tx=XM_019835112.2;refGene_gn=R3HDM2;refGene_tx=XM_019835113.2;refGene_gn=R3HDM2;refGene_tx=XM_019835115.2;refGene_gn=R3HDM2;refGene_tx=XM_019835117.2;refGene_gn=R3HDM2;refGene_tx=XM_019835118.2;refGene_gn=R3HDM2;refGene_tx=XM_019835119.2;refGene_gn=R3HDM2;refGene_tx=XM_019835121.2;refGene_gn=R3HDM2;refGene_tx=XM_019835122.2;refGene_gn=R3HDM2;refGene_tx=XM_019835123.2;refGene_gn=R3HDM2;refGene_tx=XM_019835126.2;refGene_gn=R3HDM2;refGene_tx=XM_023257156.1;refGene_gn=R3HDM2;refGene_tx=XM_023257157.1
B4 85781632 85781797
refGene_gn=R3HDM2;refGene_tx=XM_006933843.3;refGene_gn=R3HDM2;refGene_tx=XM_011284097.3;refGene_gn=R3HDM2;refGene_tx=XM_011284098.3;refGene_gn=R3HDM2;refGene_tx=XM_019835112.2;refGene_gn=R3HDM2;refGene_tx=XM_019835113.2;refGene_gn=R3HDM2;refGene_tx=XM_019835114.2;refGene_gn=R3HDM2;refGene_tx=XM_019835115.2;refGene_gn=R3HDM2;refGene_tx=XM_019835117.2;refGene_gn=R3HDM2;refGene_tx=XM_019835118.2;refGene_gn=R3HDM2;refGene_tx=XM_019835119.2;refGene_gn=R3HDM2;refGene_tx=XM_019835120.2;refGene_gn=R3HDM2;refGene_tx=XM_019835121.2;refGene_gn=R3HDM2;refGene_tx=XM_019835122.2;refGene_gn=R3HDM2;refGene_tx=XM_019835123.2;refGene_gn=R3HDM2;refGene_tx=XM_019835124.2;refGene_gn=R3HDM2;refGene_tx=XM_019835125.2;refGene_gn=R3HDM2;refGene_tx=XM_019835126.2;refGene_gn=R3HDM2;refGene_tx=XM_023257154.1;refGene_gn=R3HDM2;refGene_tx=XM_023257155.1;refGene_gn=R3HDM2;refGene_tx=XM_023257156.1;refGene_gn=R3HDM2;refGene_tx=XM_023257157.1;refGene_gn=R3HDM2;refGene_tx=XM_023257158.1;refGene_gn=R3HDM2;refGene_tx=XM_023257159.1;refGene_gn=R3HDM2;refGene_tx=XM_023257160.1
B4 85899270 85899583 refGene_gn=INHBC;refGene_tx=XM_023257161.1
B4 85906579 85907328 refGene_gn=INHBC;refGene_tx=XM_023257161.1

B4 85911870 85912168 refGene_gn=INHBE;refGene_tx=XM_003988938.2
B4 85912429 85913181 refGene_gn=INHBE;refGene_tx=XM_003988938.2
B4 85919214 85919311
refGene_gn=GLI1;refGene_tx=XM_003988939.4;refGene_gn=GLI1;refGene_tx=XM_006933846.3;refGene_gn=GLI1;refGene_tx=XM_006933847.3;refGene_gn=GLI1;refGene_tx=XM_019835128.2;refGene_gn=GLI1;refGene_tx=XM_019835129.2;refGene_gn=GLI1;refGene_tx=XM_019835130.2;refGene_gn=GLI1;refGene_tx=XM_019835131.2
B4 85919542 85919635
refGene_gn=GLI1;refGene_tx=XM_003988939.4;refGene_gn=GLI1;refGene_tx=XM_006933846.3;refGene_gn=GLI1;refGene_tx=XM_006933847.3;refGene_gn=GLI1;refGene_tx=XM_019835128.2;refGene_gn=GLI1;refGene_tx=XM_019835129.2;refGene_gn=GLI1;refGene_tx=XM_019835130.2;refGene_gn=GLI1;refGene_tx=XM_019835131.2
B4 85919910 85920106
refGene_gn=GLI1;refGene_tx=XM_003988939.4;refGene_gn=GLI1;refGene_tx=XM_006933846.3;refGene_gn=GLI1;refGene_tx=XM_006933847.3;refGene_gn=GLI1;refGene_tx=XM_019835128.2;refGene_gn=GLI1;refGene_tx=XM_019835129.2;refGene_gn=GLI1;refGene_tx=XM_019835130.2;refGene_gn=GLI1;refGene_tx=XM_019835131.2
B4 85920314 85920459
refGene_gn=GLI1;refGene_tx=XM_003988939.4;refGene_gn=GLI1;refGene_tx=XM_006933846.3;refGene_gn=GLI1;refGene_tx=XM_006933847.3;refGene_gn=GLI1;refGene_tx=XM_019835128.2;refGene_gn=GLI1;refGene_tx=XM_019835129.2;refGene_gn=GLI1;refGene_tx=XM_019835130.2;refGene_gn=GLI1;refGene_tx=XM_019835131.2;refGene_gn=GLI1;refGene_tx=XM_019835133.1
B4 85920630 85920894
refGene_gn=GLI1;refGene_tx=XM_003988939.4;refGene_gn=GLI1;refGene_tx=XM_006933846.3;refGene_gn=GLI1;refGene_tx=XM_006933847.3;refGene_gn=GLI1;refGene_tx=XM_019835128.2;refGene_gn=GLI1;refGene_tx=XM_019835129.2;refGene_gn=GLI1;refGene_tx=XM_019835130.2;refGene_gn=GLI1;refGene_tx=XM_019835131.2;refGene_gn=GLI1;refGene_tx=XM_019835132.1;refGene_gn=GLI1;refGene_tx=XM_019835133.1
B4 85920980 85921118
refGene_gn=GLI1;refGene_tx=XM_003988939.4;refGene_gn=GLI1;refGene_tx=XM_006933846.3;refGene_gn=GLI1;refGene_tx=XM_006933847.3;refGene_gn=GLI1;refGene_tx=XM_019835128.2;refGene_gn=GLI1;refGene_tx=XM_019835129.2;refGene_gn=GLI1;refGene_tx=XM_019835130.2;refGene_gn=GLI1;refGene_tx=XM_019835131.2;refGene_gn=GLI1;refGene_tx=XM_019835132.1;refGene_gn=GLI1;refGene_tx=XM_019835133.1
B4 85921445 85921595
refGene_gn=GLI1;refGene_tx=XM_003988939.4;refGene_gn=GLI1;refGene_tx=XM_006933846.3;refGene_gn=GLI1;refGene_tx=XM_006933847.3;refGene_gn=GLI1;refGene_tx=XM_019835128.2;refGene_gn=GLI1;refGene_tx=XM_019835129.2;refGene_gn=GLI1;refGene_tx=XM_019835130.2;refGene_gn=GLI1;refGene_tx=XM_019835131.2;refGene_gn=GLI1;refGene_tx=XM_019835132.1;refGene_gn=GLI1;refGene_tx=XM_019835133.1
B4 85922695 85922860
refGene_gn=GLI1;refGene_tx=XM_003988939.4;refGene_gn=GLI1;refGene_tx=XM_006933846.3;refGene_gn=GLI1;refGene_tx=XM_006933847.3;refGene_gn=GLI

1;refGene_tx=XM_019835128.2;refGene_gn=GLI1;refGene_tx=XM_019835129.2;
refGene_gn=GLI1;refGene_tx=XM_019835130.2;refGene_gn=GLI1;refGene_tx=X
M_019835131.2;refGene_gn=GLI1;refGene_tx=XM_019835132.1;refGene_gn=GLI
1;refGene_tx=XM_019835133.1
B4 85923351 85923582
refGene_gn=GLI1;refGene_tx=XM_003988939.4;refGene_gn=GLI1;refGene_tx=X
M_006933846.3;refGene_gn=GLI1;refGene_tx=XM_006933847.3;refGene_gn=GLI
1;refGene_tx=XM_019835128.2;refGene_gn=GLI1;refGene_tx=XM_019835129.2;
refGene_gn=GLI1;refGene_tx=XM_019835130.2;refGene_gn=GLI1;refGene_tx=X
M_019835131.2;refGene_gn=GLI1;refGene_tx=XM_019835132.1;refGene_gn=GLI
1;refGene_tx=XM_019835133.1
B4 85924052 85924320
refGene_gn=GLI1;refGene_tx=XM_003988939.4;refGene_gn=GLI1;refGene_tx=X
M_006933846.3;refGene_gn=GLI1;refGene_tx=XM_006933847.3;refGene_gn=GLI
1;refGene_tx=XM_019835128.2;refGene_gn=GLI1;refGene_tx=XM_019835129.2;
refGene_gn=GLI1;refGene_tx=XM_019835130.2;refGene_gn=GLI1;refGene_tx=X
M_019835131.2;refGene_gn=GLI1;refGene_tx=XM_019835132.1;refGene_gn=GLI
1;refGene_tx=XM_019835133.1
B4 85924731 85926476
refGene_gn=GLI1;refGene_tx=XM_003988939.4;refGene_gn=GLI1;refGene_tx=X
M_006933846.3;refGene_gn=GLI1;refGene_tx=XM_006933847.3;refGene_gn=GLI
1;refGene_tx=XM_019835128.2;refGene_gn=GLI1;refGene_tx=XM_019835129.2;
refGene_gn=GLI1;refGene_tx=XM_019835130.2;refGene_gn=GLI1;refGene_tx=X
M_019835131.2;refGene_gn=GLI1;refGene_tx=XM_019835132.1;refGene_gn=GLI
1;refGene_tx=XM_019835133.1
B4 85926944 85927116
refGene_gn=ARHGAP9;refGene_tx=XM_003988942.5;refGene_gn=ARHGAP9;refGen
e_tx=XM_011284095.3;refGene_gn=ARHGAP9;refGene_tx=XM_019835134.2;refGe
ne_gn=ARHGAP9;refGene_tx=XM_019835135.2;refGene_gn=ARHGAP9;refGene_tx=
XM_019835140.1;refGene_gn=ARHGAP9;refGene_tx=XM_023257165.1;refGene_gn
=ARHGAP9;refGene_tx=XM_023257166.1;refGene_gn=ARHGAP9;refGene_tx=XM_02
3257167.1
B4 85927722 85927828
refGene_gn=ARHGAP9;refGene_tx=XM_003988942.5;refGene_gn=ARHGAP9;refGen
e_tx=XM_011284095.3;refGene_gn=ARHGAP9;refGene_tx=XM_019835134.2;refGe
ne_gn=ARHGAP9;refGene_tx=XM_019835135.2;refGene_gn=ARHGAP9;refGene_tx=
XM_019835140.1;refGene_gn=ARHGAP9;refGene_tx=XM_023257165.1;refGene_gn
=ARHGAP9;refGene_tx=XM_023257166.1;refGene_gn=ARHGAP9;refGene_tx=XM_02
3257167.1
B4 85928491 85928626
refGene_gn=ARHGAP9;refGene_tx=XM_003988942.5;refGene_gn=ARHGAP9;refGen
e_tx=XM_011284095.3;refGene_gn=ARHGAP9;refGene_tx=XM_019835134.2;refGe
ne_gn=ARHGAP9;refGene_tx=XM_019835135.2;refGene_gn=ARHGAP9;refGene_tx=
XM_019835140.1;refGene_gn=ARHGAP9;refGene_tx=XM_023257165.1;refGene_gn
=ARHGAP9;refGene_tx=XM_023257166.1;refGene_gn=ARHGAP9;refGene_tx=XM_02
3257167.1
B4 85928881 85928935
refGene_gn=ARHGAP9;refGene_tx=XM_003988942.5;refGene_gn=ARHGAP9;refGen
e_tx=XM_011284095.3;refGene_gn=ARHGAP9;refGene_tx=XM_019835134.2;refGe
ne_gn=ARHGAP9;refGene_tx=XM_019835135.2;refGene_gn=ARHGAP9;refGene_tx=

XM_019835140.1;refGene_gn=ARHGAP9;refGene_tx=XM_023257165.1;refGene_gn=ARHGAP9;refGene_tx=XM_023257166.1;refGene_gn=ARHGAP9;refGene_tx=XM_023257167.1
B4 85929081 85929159
refGene_gn=ARHGAP9;refGene_tx=XM_003988942.5;refGene_gn=ARHGAP9;refGene_tx=XM_011284095.3;refGene_gn=ARHGAP9;refGene_tx=XM_019835134.2;refGene_gn=ARHGAP9;refGene_tx=XM_019835135.2;refGene_gn=ARHGAP9;refGene_tx=XM_019835140.1;refGene_gn=ARHGAP9;refGene_tx=XM_023257165.1;refGene_gn=ARHGAP9;refGene_tx=XM_023257166.1;refGene_gn=ARHGAP9;refGene_tx=XM_023257167.1
B4 85929339 85929438
refGene_gn=ARHGAP9;refGene_tx=XM_003988942.5;refGene_gn=ARHGAP9;refGene_tx=XM_011284095.3;refGene_gn=ARHGAP9;refGene_tx=XM_019835134.2;refGene_gn=ARHGAP9;refGene_tx=XM_019835135.2;refGene_gn=ARHGAP9;refGene_tx=XM_019835140.1;refGene_gn=ARHGAP9;refGene_tx=XM_023257165.1;refGene_gn=ARHGAP9;refGene_tx=XM_023257166.1
B4 85929775 85929883
refGene_gn=ARHGAP9;refGene_tx=XM_003988942.5;refGene_gn=ARHGAP9;refGene_tx=XM_011284095.3;refGene_gn=ARHGAP9;refGene_tx=XM_019835134.2;refGene_gn=ARHGAP9;refGene_tx=XM_019835135.2;refGene_gn=ARHGAP9;refGene_tx=XM_019835140.1;refGene_gn=ARHGAP9;refGene_tx=XM_023257165.1;refGene_gn=ARHGAP9;refGene_tx=XM_023257166.1;refGene_gn=ARHGAP9;refGene_tx=XM_023257167.1
B4 85929987 85930129
refGene_gn=ARHGAP9;refGene_tx=XM_003988942.5;refGene_gn=ARHGAP9;refGene_tx=XM_011284095.3;refGene_gn=ARHGAP9;refGene_tx=XM_019835134.2;refGene_gn=ARHGAP9;refGene_tx=XM_019835135.2;refGene_gn=ARHGAP9;refGene_tx=XM_019835140.1;refGene_gn=ARHGAP9;refGene_tx=XM_023257165.1;refGene_gn=ARHGAP9;refGene_tx=XM_023257166.1;refGene_gn=ARHGAP9;refGene_tx=XM_023257167.1
B4 85930384 85930483
refGene_gn=ARHGAP9;refGene_tx=XM_003988942.5;refGene_gn=ARHGAP9;refGene_tx=XM_011284095.3;refGene_gn=ARHGAP9;refGene_tx=XM_019835134.2;refGene_gn=ARHGAP9;refGene_tx=XM_019835135.2;refGene_gn=ARHGAP9;refGene_tx=XM_019835140.1;refGene_gn=ARHGAP9;refGene_tx=XM_023257165.1;refGene_gn=ARHGAP9;refGene_tx=XM_023257166.1;refGene_gn=ARHGAP9;refGene_tx=XM_023257167.1
B4 85930625 85930721
refGene_gn=ARHGAP9;refGene_tx=XM_003988942.5;refGene_gn=ARHGAP9;refGene_tx=XM_011284095.3;refGene_gn=ARHGAP9;refGene_tx=XM_019835134.2;refGene_gn=ARHGAP9;refGene_tx=XM_019835135.2;refGene_gn=ARHGAP9;refGene_tx=XM_019835140.1;refGene_gn=ARHGAP9;refGene_tx=XM_023257165.1;refGene_gn=ARHGAP9;refGene_tx=XM_023257166.1;refGene_gn=ARHGAP9;refGene_tx=XM_023257167.1
B4 85930915 85931006
refGene_gn=ARHGAP9;refGene_tx=XM_003988942.5;refGene_gn=ARHGAP9;refGene_tx=XM_011284095.3;refGene_gn=ARHGAP9;refGene_tx=XM_019835134.2;refGene_gn=ARHGAP9;refGene_tx=XM_019835135.2;refGene_gn=ARHGAP9;refGene_tx=XM_019835140.1;refGene_gn=ARHGAP9;refGene_tx=XM_023257165.1;refGene_gn=ARHGAP9;refGene_tx=XM_023257166.1;refGene_gn=ARHGAP9;refGene_tx=XM_02

3257167.1
B4	85931144 85931206
refGene_gn=ARHGAP9;refGene_tx=XM_003988942.5;refGene_gn=ARHGAP9;refGene_tx=XM_011284095.3;refGene_gn=ARHGAP9;refGene_tx=XM_019835134.2;refGene_gn=ARHGAP9;refGene_tx=XM_019835135.2;refGene_gn=ARHGAP9;refGene_tx=XM_019835140.1;refGene_gn=ARHGAP9;refGene_tx=XM_023257165.1;refGene_gn=ARHGAP9;refGene_tx=XM_023257166.1;refGene_gn=ARHGAP9;refGene_tx=XM_023257167.1
B4	85931448 85931553
refGene_gn=ARHGAP9;refGene_tx=XM_003988942.5;refGene_gn=ARHGAP9;refGene_tx=XM_011284095.3;refGene_gn=ARHGAP9;refGene_tx=XM_019835134.2;refGene_gn=ARHGAP9;refGene_tx=XM_019835135.2;refGene_gn=ARHGAP9;refGene_tx=XM_019835140.1;refGene_gn=ARHGAP9;refGene_tx=XM_023257165.1;refGene_gn=ARHGAP9;refGene_tx=XM_023257166.1;refGene_gn=ARHGAP9;refGene_tx=XM_023257167.1
B4	85931745 85931859
refGene_gn=ARHGAP9;refGene_tx=XM_003988942.5;refGene_gn=ARHGAP9;refGene_tx=XM_011284095.3;refGene_gn=ARHGAP9;refGene_tx=XM_019835134.2;refGene_gn=ARHGAP9;refGene_tx=XM_019835135.2;refGene_gn=ARHGAP9;refGene_tx=XM_019835140.1;refGene_gn=ARHGAP9;refGene_tx=XM_023257165.1;refGene_gn=ARHGAP9;refGene_tx=XM_023257166.1;refGene_gn=ARHGAP9;refGene_tx=XM_023257167.1
B4	85932036 85932261
refGene_gn=ARHGAP9;refGene_tx=XM_003988942.5;refGene_gn=ARHGAP9;refGene_tx=XM_011284095.3;refGene_gn=ARHGAP9;refGene_tx=XM_019835134.2;refGene_gn=ARHGAP9;refGene_tx=XM_019835135.2;refGene_gn=ARHGAP9;refGene_tx=XM_019835140.1;refGene_gn=ARHGAP9;refGene_tx=XM_023257165.1;refGene_gn=ARHGAP9;refGene_tx=XM_023257166.1;refGene_gn=ARHGAP9;refGene_tx=XM_023257167.1
B4	85933224 85933448
refGene_gn=ARHGAP9;refGene_tx=XM_003988942.5;refGene_gn=ARHGAP9;refGene_tx=XM_019835134.2;refGene_gn=ARHGAP9;refGene_tx=XM_019835135.2;refGene_gn=ARHGAP9;refGene_tx=XM_023257165.1;refGene_gn=ARHGAP9;refGene_tx=XM_023257166.1;refGene_gn=ARHGAP9;refGene_tx=XM_023257167.1
B4	85933746 85934062
refGene_gn=ARHGAP9;refGene_tx=XM_003988942.5;refGene_gn=ARHGAP9;refGene_tx=XM_019835134.2;refGene_gn=ARHGAP9;refGene_tx=XM_019835135.2;refGene_gn=ARHGAP9;refGene_tx=XM_023257165.1;refGene_gn=ARHGAP9;refGene_tx=XM_023257166.1;refGene_gn=ARHGAP9;refGene_tx=XM_023257167.1
B4	85938421 85938524 refGene_gn=MARS;refGene_tx=XM_023257168.1
B4	85939356 85939453 refGene_gn=MARS;refGene_tx=XM_023257168.1
B4	85939612 85939691 refGene_gn=MARS;refGene_tx=XM_023257168.1
B4	85939769 85939904 refGene_gn=MARS;refGene_tx=XM_023257168.1
B4	85940248 85940324 refGene_gn=MARS;refGene_tx=XM_023257168.1
B4	85940575 85940748 refGene_gn=MARS;refGene_tx=XM_023257168.1
B4	85940926 85941033 refGene_gn=MARS;refGene_tx=XM_023257168.1
B4	85942946 85943063 refGene_gn=MARS;refGene_tx=XM_023257168.1
B4	85943202 85943406 refGene_gn=MARS;refGene_tx=XM_023257168.1
B4	85944184 85944386 refGene_gn=MARS;refGene_tx=XM_023257168.1
B4	85946615 85946690 refGene_gn=MARS;refGene_tx=XM_023257168.1

B4 85947876 85948047 refGene_gn=MARS;refGene_tx=XM_023257168.1
B4 85948216 85948312 refGene_gn=MARS;refGene_tx=XM_023257168.1
B4 85948450 85948568 refGene_gn=MARS;refGene_tx=XM_023257168.1
B4 85948932 85949146 refGene_gn=MARS;refGene_tx=XM_023257168.1
B4 85949935 85950067 refGene_gn=MARS;refGene_tx=XM_023257168.1
B4 85950175 85950280 refGene_gn=MARS;refGene_tx=XM_023257168.1
B4 85950352 85950539 refGene_gn=MARS;refGene_tx=XM_023257168.1
B4 85951139 85951211 refGene_gn=MARS;refGene_tx=XM_023257168.1
B4 85951462 85951555 refGene_gn=MARS;refGene_tx=XM_023257168.1
B4 85951654 85951801 refGene_gn=MARS;refGene_tx=XM_023257168.1
B4 85952035 85952404 refGene_gn=DDIT3;refGene_tx=XM_006933848.4
B4 85952500 85952635 refGene_gn=DDIT3;refGene_tx=XM_006933848.4
B4 85959350 85959463
refGene_gn=MBD6;refGene_tx=XM_003988948.5;refGene_gn=MBD6;refGene_tx=XM_023257162.1;refGene_gn=MBD6;refGene_tx=XM_023257163.1;refGene_gn=MBD6;refGene_tx=XM_023257164.1
B4 85959759 85959862
refGene_gn=MBD6;refGene_tx=XM_003988948.5;refGene_gn=MBD6;refGene_tx=XM_023257162.1;refGene_gn=MBD6;refGene_tx=XM_023257163.1;refGene_gn=MBD6;refGene_tx=XM_023257164.1
B4 85959992 85960155
refGene_gn=MBD6;refGene_tx=XM_003988948.5;refGene_gn=MBD6;refGene_tx=XM_023257162.1;refGene_gn=MBD6;refGene_tx=XM_023257163.1;refGene_gn=MBD6;refGene_tx=XM_023257164.1
B4 85960384 85961437
refGene_gn=MBD6;refGene_tx=XM_003988948.5;refGene_gn=MBD6;refGene_tx=XM_023257162.1;refGene_gn=MBD6;refGene_tx=XM_023257163.1;refGene_gn=MBD6;refGene_tx=XM_023257164.1
B4 85961616 85962278
refGene_gn=MBD6;refGene_tx=XM_003988948.5;refGene_gn=MBD6;refGene_tx=XM_023257162.1;refGene_gn=MBD6;refGene_tx=XM_023257163.1;refGene_gn=MBD6;refGene_tx=XM_023257164.1
B4 85962553 85962707
refGene_gn=MBD6;refGene_tx=XM_003988948.5;refGene_gn=MBD6;refGene_tx=XM_023257162.1;refGene_gn=MBD6;refGene_tx=XM_023257163.1;refGene_gn=MBD6;refGene_tx=XM_023257164.1
B4 85962884 85963054
refGene_gn=MBD6;refGene_tx=XM_003988948.5;refGene_gn=MBD6;refGene_tx=XM_023257162.1;refGene_gn=MBD6;refGene_tx=XM_023257163.1;refGene_gn=MBD6;refGene_tx=XM_023257164.1
B4 85963177 85963591
refGene_gn=MBD6;refGene_tx=XM_003988948.5;refGene_gn=MBD6;refGene_tx=XM_023257162.1;refGene_gn=MBD6;refGene_tx=XM_023257163.1;refGene_gn=MBD6;refGene_tx=XM_023257164.1
B4 85963682 85963750
refGene_gn=MBD6;refGene_tx=XM_003988948.5;refGene_gn=MBD6;refGene_tx=XM_023257162.1;refGene_gn=MBD6;refGene_tx=XM_023257163.1;refGene_gn=MBD6;refGene_tx=XM_023257164.1
B4 85963965 85964029
refGene_gn=MBD6;refGene_tx=XM_003988948.5;refGene_gn=MBD6;refGene_tx=X

M_023257162.1;refGene_gn=MBD6;refGene_tx=XM_023257163.1;refGene_gn=MBD6;refGene_tx=XM_023257164.1
B4 85964182 85964257
refGene_gn=MBD6;refGene_tx=XM_003988948.5;refGene_gn=MBD6;refGene_tx=XM_023257162.1;refGene_gn=MBD6;refGene_tx=XM_023257163.1;refGene_gn=MBD6;refGene_tx=XM_023257164.1
B4 85965673 85965760
refGene_gn=DCTN2;refGene_tx=XM_003988945.4;refGene_gn=DCTN2;refGene_tx=XM_003988947.4;refGene_gn=DCTN2;refGene_tx=XM_019835141.1
B4 85966624 85966716
refGene_gn=DCTN2;refGene_tx=XM_003988945.4;refGene_gn=DCTN2;refGene_tx=XM_003988947.4;refGene_gn=DCTN2;refGene_tx=XM_019835141.1
B4 85966829 85966932
refGene_gn=DCTN2;refGene_tx=XM_003988945.4;refGene_gn=DCTN2;refGene_tx=XM_003988947.4;refGene_gn=DCTN2;refGene_tx=XM_019835141.1
B4 85967172 85967244
refGene_gn=DCTN2;refGene_tx=XM_003988945.4;refGene_gn=DCTN2;refGene_tx=XM_003988947.4;refGene_gn=DCTN2;refGene_tx=XM_019835141.1
B4 85967332 85967410
refGene_gn=DCTN2;refGene_tx=XM_003988945.4;refGene_gn=DCTN2;refGene_tx=XM_003988947.4;refGene_gn=DCTN2;refGene_tx=XM_019835141.1
B4 85967587 85967626
refGene_gn=DCTN2;refGene_tx=XM_003988945.4;refGene_gn=DCTN2;refGene_tx=XM_003988947.4;refGene_gn=DCTN2;refGene_tx=XM_019835141.1
B4 85967840 85967906
refGene_gn=DCTN2;refGene_tx=XM_003988945.4;refGene_gn=DCTN2;refGene_tx=XM_003988947.4;refGene_gn=DCTN2;refGene_tx=XM_019835141.1
B4 85968234 85968385
refGene_gn=DCTN2;refGene_tx=XM_003988945.4;refGene_gn=DCTN2;refGene_tx=XM_003988947.4;refGene_gn=DCTN2;refGene_tx=XM_019835141.1
B4 85968577 85968738
refGene_gn=DCTN2;refGene_tx=XM_003988945.4;refGene_gn=DCTN2;refGene_tx=XM_003988947.4;refGene_gn=DCTN2;refGene_tx=XM_019835141.1
B4 85969226 85969325
refGene_gn=DCTN2;refGene_tx=XM_003988945.4;refGene_gn=DCTN2;refGene_tx=XM_003988947.4;refGene_gn=DCTN2;refGene_tx=XM_019835141.1
B4 85969884 85969946
refGene_gn=DCTN2;refGene_tx=XM_003988945.4;refGene_gn=DCTN2;refGene_tx=XM_003988947.4;refGene_gn=DCTN2;refGene_tx=XM_019835141.1
B4 85970126 85970223
refGene_gn=DCTN2;refGene_tx=XM_003988945.4;refGene_gn=DCTN2;refGene_tx=XM_003988947.4;refGene_gn=DCTN2;refGene_tx=XM_019835141.1
B4 85972553 85972559
refGene_gn=DCTN2;refGene_tx=XM_003988947.4;refGene_gn=DCTN2;refGene_tx=XM_019835141.1
B4 85975059 85975068 refGene_gn=DCTN2;refGene_tx=XM_019835141.1
B4 85978202 85978271
refGene_gn=DCTN2;refGene_tx=XM_003988945.4;refGene_gn=DCTN2;refGene_tx=XM_003988947.4;refGene_gn=DCTN2;refGene_tx=XM_019835141.1
B4 85979209 85979245

refGene_gn=DCTN2;refGene_tx=XM_003988945.4;refGene_gn=DCTN2;refGene_tx=XM_003988947.4;refGene_gn=DCTN2;refGene_tx=XM_019835141.1
B4 85982688 85982817
refGene_gn=KIF5A;refGene_tx=XM_006933850.3;refGene_gn=KIF5A;refGene_tx=XM_011284094.3
B4 85995005 85995093
refGene_gn=KIF5A;refGene_tx=XM_006933850.3;refGene_gn=KIF5A;refGene_tx=XM_011284094.3
B4 85995186 85995260
refGene_gn=KIF5A;refGene_tx=XM_006933850.3;refGene_gn=KIF5A;refGene_tx=XM_011284094.3
B4 85995666 85995771
refGene_gn=KIF5A;refGene_tx=XM_006933850.3;refGene_gn=KIF5A;refGene_tx=XM_011284094.3
B4 85996016 85996065
refGene_gn=KIF5A;refGene_tx=XM_006933850.3;refGene_gn=KIF5A;refGene_tx=XM_011284094.3
B4 85996727 85996783
refGene_gn=KIF5A;refGene_tx=XM_006933850.3;refGene_gn=KIF5A;refGene_tx=XM_011284094.3
B4 85997028 85997116
refGene_gn=KIF5A;refGene_tx=XM_006933850.3;refGene_gn=KIF5A;refGene_tx=XM_011284094.3
B4 85997396 85997521
refGene_gn=KIF5A;refGene_tx=XM_006933850.3;refGene_gn=KIF5A;refGene_tx=XM_011284094.3
B4 85997982 85998087
refGene_gn=KIF5A;refGene_tx=XM_006933850.3;refGene_gn=KIF5A;refGene_tx=XM_011284094.3
B4 85998271 85998420
refGene_gn=KIF5A;refGene_tx=XM_006933850.3;refGene_gn=KIF5A;refGene_tx=XM_011284094.3
B4 85998525 85998674
refGene_gn=KIF5A;refGene_tx=XM_006933850.3;refGene_gn=KIF5A;refGene_tx=XM_011284094.3
B4 85998999 85999175
refGene_gn=KIF5A;refGene_tx=XM_006933850.3;refGene_gn=KIF5A;refGene_tx=XM_011284094.3
B4 86000564 86000633
refGene_gn=KIF5A;refGene_tx=XM_006933850.3;refGene_gn=KIF5A;refGene_tx=XM_011284094.3
B4 86000988 86001195
refGene_gn=KIF5A;refGene_tx=XM_006933850.3;refGene_gn=KIF5A;refGene_tx=XM_011284094.3
B4 86001487 86001634
refGene_gn=KIF5A;refGene_tx=XM_006933850.3;refGene_gn=KIF5A;refGene_tx=XM_011284094.3
B4 86002425 86002614
refGene_gn=KIF5A;refGene_tx=XM_006933850.3;refGene_gn=KIF5A;refGene_tx=XM_011284094.3

B4	86002979	86003097
refGene_gn=KIF5A;refGene_tx=XM_006933850.3;refGene_gn=KIF5A;refGene_tx=XM_011284094.3
B4	86003473	86003538
refGene_gn=KIF5A;refGene_tx=XM_006933850.3;refGene_gn=KIF5A;refGene_tx=XM_011284094.3
B4	86003648	86003758
refGene_gn=KIF5A;refGene_tx=XM_006933850.3;refGene_gn=KIF5A;refGene_tx=XM_011284094.3
B4	86004135	86004237
refGene_gn=KIF5A;refGene_tx=XM_006933850.3;refGene_gn=KIF5A;refGene_tx=XM_011284094.3
B4	86004998	86005059
refGene_gn=KIF5A;refGene_tx=XM_006933850.3;refGene_gn=KIF5A;refGene_tx=XM_011284094.3
B4	86005296	86005368
refGene_gn=KIF5A;refGene_tx=XM_006933850.3;refGene_gn=KIF5A;refGene_tx=XM_011284094.3
B4	86005451	86005556
refGene_gn=KIF5A;refGene_tx=XM_006933850.3;refGene_gn=KIF5A;refGene_tx=XM_011284094.3
B4	86006098	86006119	refGene_gn=KIF5A;refGene_tx=XM_011284094.3
B4	86008319	86008536
refGene_gn=KIF5A;refGene_tx=XM_006933850.3;refGene_gn=KIF5A;refGene_tx=XM_011284094.3
B4	86008800	86008954
refGene_gn=KIF5A;refGene_tx=XM_006933850.3;refGene_gn=KIF5A;refGene_tx=XM_011284094.3
B4	86009253	86009336
refGene_gn=KIF5A;refGene_tx=XM_006933850.3;refGene_gn=KIF5A;refGene_tx=XM_011284094.3
B4	86009736	86009764
refGene_gn=KIF5A;refGene_tx=XM_006933850.3;refGene_gn=KIF5A;refGene_tx=XM_011284094.3
B4	86010230	86010309
refGene_gn=KIF5A;refGene_tx=XM_006933850.3;refGene_gn=KIF5A;refGene_tx=XM_011284094.3
B4	86016898	86017072	refGene_gn=PIP4K2C;refGene_tx=XM_003988950.5
B4	86019624	86019722	refGene_gn=PIP4K2C;refGene_tx=XM_003988950.5
B4	86020424	86020521	refGene_gn=PIP4K2C;refGene_tx=XM_003988950.5
B4	86020889	86021033	refGene_gn=PIP4K2C;refGene_tx=XM_003988950.5
B4	86022850	86022997	refGene_gn=PIP4K2C;refGene_tx=XM_003988950.5
B4	86023188	86023227	refGene_gn=PIP4K2C;refGene_tx=XM_003988950.5
B4	86023822	86023936	refGene_gn=PIP4K2C;refGene_tx=XM_003988950.5
B4	86024307	86024575	refGene_gn=PIP4K2C;refGene_tx=XM_003988950.5
B4	86024774	86024878	refGene_gn=PIP4K2C;refGene_tx=XM_003988950.5
B4	86025058	86025139	refGene_gn=PIP4K2C;refGene_tx=XM_003988950.5
B4	86030064	86030093
refGene_gn=DTX3;refGene_tx=XM_011284087.3;refGene_gn=DTX3;refGene_tx=XM_011284089.3;refGene_gn=DTX3;refGene_tx=XM_011284090.2;refGene_gn=DTX

3;refGene_tx=XM_011284091.3;refGene_gn=DTX3;refGene_tx=XM_011284092.3;
refGene_gn=DTX3;refGene_tx=XM_019835143.1
B4 86030447 86031202
refGene_gn=DTX3;refGene_tx=XM_011284087.3;refGene_gn=DTX3;refGene_tx=X
M_011284089.3;refGene_gn=DTX3;refGene_tx=XM_011284090.2;refGene_gn=DTX
3;refGene_tx=XM_011284091.3;refGene_gn=DTX3;refGene_tx=XM_011284092.3;
refGene_gn=DTX3;refGene_tx=XM_019835143.1
B4 86032160 86032538
refGene_gn=DTX3;refGene_tx=XM_011284087.3;refGene_gn=DTX3;refGene_tx=X
M_011284089.3;refGene_gn=DTX3;refGene_tx=XM_011284090.2;refGene_gn=DTX
3;refGene_tx=XM_011284091.3;refGene_gn=DTX3;refGene_tx=XM_011284092.3;
refGene_gn=DTX3;refGene_tx=XM_019835143.1
B4 86032726 86032802
refGene_gn=DTX3;refGene_tx=XM_011284092.3;refGene_gn=DTX3;refGene_tx=X
M_019835143.1
B4 86033818 86033917 refGene_gn=ARHGEF25;refGene_tx=XM_003988953.5
B4 86034216 86034331 refGene_gn=ARHGEF25;refGene_tx=XM_003988953.5
B4 86035563 86035660 refGene_gn=ARHGEF25;refGene_tx=XM_003988954.5
B4 86036253 86036401 refGene_gn=ARHGEF25;refGene_tx=XM_006933851.4
B4 86036601 86036816
refGene_gn=ARHGEF25;refGene_tx=XM_003988953.5;refGene_gn=ARHGEF25;refG
ene_tx=XM_003988954.5;refGene_gn=ARHGEF25;refGene_tx=XM_006933851.4
B4 86036952 86037048
refGene_gn=ARHGEF25;refGene_tx=XM_003988953.5;refGene_gn=ARHGEF25;refG
ene_tx=XM_003988954.5;refGene_gn=ARHGEF25;refGene_tx=XM_006933851.4
B4 86037122 86037199
refGene_gn=ARHGEF25;refGene_tx=XM_003988953.5;refGene_gn=ARHGEF25;refG
ene_tx=XM_003988954.5;refGene_gn=ARHGEF25;refGene_tx=XM_006933851.4
B4 86037398 86037465
refGene_gn=ARHGEF25;refGene_tx=XM_003988953.5;refGene_gn=ARHGEF25;refG
ene_tx=XM_003988954.5;refGene_gn=ARHGEF25;refGene_tx=XM_006933851.4
B4 86037661 86037765
refGene_gn=ARHGEF25;refGene_tx=XM_003988953.5;refGene_gn=ARHGEF25;refG
ene_tx=XM_003988954.5;refGene_gn=ARHGEF25;refGene_tx=XM_006933851.4
B4 86037952 86038022
refGene_gn=ARHGEF25;refGene_tx=XM_003988953.5;refGene_gn=ARHGEF25;refG
ene_tx=XM_003988954.5;refGene_gn=ARHGEF25;refGene_tx=XM_006933851.4
B4 86038140 86038230
refGene_gn=ARHGEF25;refGene_tx=XM_003988953.5;refGene_gn=ARHGEF25;refG
ene_tx=XM_003988954.5;refGene_gn=ARHGEF25;refGene_tx=XM_006933851.4
B4 86038314 86038407
refGene_gn=ARHGEF25;refGene_tx=XM_003988953.5;refGene_gn=ARHGEF25;refG
ene_tx=XM_003988954.5;refGene_gn=ARHGEF25;refGene_tx=XM_006933851.4
B4 86038594 86038645
refGene_gn=ARHGEF25;refGene_tx=XM_003988953.5;refGene_gn=ARHGEF25;refG
ene_tx=XM_003988954.5;refGene_gn=ARHGEF25;refGene_tx=XM_006933851.4
B4 86038844 86038922
refGene_gn=ARHGEF25;refGene_tx=XM_003988953.5;refGene_gn=ARHGEF25;refG
ene_tx=XM_003988954.5;refGene_gn=ARHGEF25;refGene_tx=XM_006933851.4
B4 86039122 86039323

refGene_gn=ARHGEF25;refGene_tx=XM_003988953.5;refGene_gn=ARHGEF25;refGene_tx=XM_003988954.5;refGene_gn=ARHGEF25;refGene_tx=XM_006933851.4
B4 86039467 86039648
refGene_gn=ARHGEF25;refGene_tx=XM_003988953.5;refGene_gn=ARHGEF25;refGene_tx=XM_003988954.5;refGene_gn=ARHGEF25;refGene_tx=XM_006933851.4
B4 86039888 86040100
refGene_gn=ARHGEF25;refGene_tx=XM_003988953.5;refGene_gn=ARHGEF25;refGene_tx=XM_003988954.5;refGene_gn=ARHGEF25;refGene_tx=XM_006933851.4
B4 86040399 86040510
refGene_gn=ARHGEF25;refGene_tx=XM_003988953.5;refGene_gn=ARHGEF25;refGene_tx=XM_003988954.5;refGene_gn=ARHGEF25;refGene_tx=XM_006933851.4
B4 86042571 86042914
refGene_gn=SLC26A10;refGene_tx=XM_006933854.3;refGene_gn=SLC26A10;refGene_tx=XM_019835144.2;refGene_gn=SLC26A10;refGene_tx=XM_019835145.2;refGene_gn=SLC26A10;refGene_tx=XM_019835146.2;refGene_gn=SLC26A10;refGene_tx=XM_023257169.1
B4 86043371 86043482
refGene_gn=SLC26A10;refGene_tx=XM_006933854.3;refGene_gn=SLC26A10;refGene_tx=XM_019835144.2;refGene_gn=SLC26A10;refGene_tx=XM_019835145.2;refGene_gn=SLC26A10;refGene_tx=XM_019835146.2;refGene_gn=SLC26A10;refGene_tx=XM_023257169.1
B4 86044048 86044206
refGene_gn=SLC26A10;refGene_tx=XM_006933854.3;refGene_gn=SLC26A10;refGene_tx=XM_019835144.2;refGene_gn=SLC26A10;refGene_tx=XM_019835145.2;refGene_gn=SLC26A10;refGene_tx=XM_019835146.2;refGene_gn=SLC26A10;refGene_tx=XM_023257169.1
B4 86044382 86044547
refGene_gn=SLC26A10;refGene_tx=XM_006933854.3;refGene_gn=SLC26A10;refGene_tx=XM_019835144.2;refGene_gn=SLC26A10;refGene_tx=XM_019835146.2;refGene_gn=SLC26A10;refGene_tx=XM_023257169.1
B4 86044804 86044957
refGene_gn=SLC26A10;refGene_tx=XM_006933854.3;refGene_gn=SLC26A10;refGene_tx=XM_019835144.2;refGene_gn=SLC26A10;refGene_tx=XM_019835145.2;refGene_gn=SLC26A10;refGene_tx=XM_019835146.2;refGene_gn=SLC26A10;refGene_tx=XM_023257169.1
B4 86045707 86045790
refGene_gn=SLC26A10;refGene_tx=XM_006933854.3;refGene_gn=SLC26A10;refGene_tx=XM_019835144.2;refGene_gn=SLC26A10;refGene_tx=XM_019835145.2;refGene_gn=SLC26A10;refGene_tx=XM_019835146.2;refGene_gn=SLC26A10;refGene_tx=XM_023257169.1
B4 86045917 86046065
refGene_gn=SLC26A10;refGene_tx=XM_006933854.3;refGene_gn=SLC26A10;refGene_tx=XM_019835144.2;refGene_gn=SLC26A10;refGene_tx=XM_019835145.2;refGene_gn=SLC26A10;refGene_tx=XM_019835146.2;refGene_gn=SLC26A10;refGene_tx=XM_023257169.1
B4 86046204 86046318
refGene_gn=SLC26A10;refGene_tx=XM_006933854.3;refGene_gn=SLC26A10;refGene_tx=XM_019835144.2;refGene_gn=SLC26A10;refGene_tx=XM_019835145.2;refGene_gn=SLC26A10;refGene_tx=XM_019835146.2;refGene_gn=SLC26A10;refGene_tx=XM_023257169.1

B4 86047022 86047289
refGene_gn=SLC26A10;refGene_tx=XM_006933854.3;refGene_gn=SLC26A10;refGene_tx=XM_019835144.2;refGene_gn=SLC26A10;refGene_tx=XM_019835145.2;refGene_gn=SLC26A10;refGene_tx=XM_019835146.2;refGene_gn=SLC26A10;refGene_tx=XM_023257169.1
B4 86047710 86047817
refGene_gn=SLC26A10;refGene_tx=XM_006933854.3;refGene_gn=SLC26A10;refGene_tx=XM_019835144.2;refGene_gn=SLC26A10;refGene_tx=XM_019835145.2
B4 86047927 86047997
refGene_gn=SLC26A10;refGene_tx=XM_006933854.3;refGene_gn=SLC26A10;refGene_tx=XM_019835144.2;refGene_gn=SLC26A10;refGene_tx=XM_019835145.2;refGene_gn=SLC26A10;refGene_tx=XM_019835146.2;refGene_gn=SLC26A10;refGene_tx=XM_023257169.1
B4 86048070 86048187
refGene_gn=SLC26A10;refGene_tx=XM_006933854.3;refGene_gn=SLC26A10;refGene_tx=XM_019835144.2;refGene_gn=SLC26A10;refGene_tx=XM_019835145.2
B4 86048323 86048357
refGene_gn=SLC26A10;refGene_tx=XM_006933854.3;refGene_gn=SLC26A10;refGene_tx=XM_019835144.2;refGene_gn=SLC26A10;refGene_tx=XM_019835145.2
B4 86048436 86048519
refGene_gn=SLC26A10;refGene_tx=XM_006933854.3;refGene_gn=SLC26A10;refGene_tx=XM_019835144.2;refGene_gn=SLC26A10;refGene_tx=XM_019835145.2
B4 86048609 86048664
refGene_gn=SLC26A10;refGene_tx=XM_006933854.3;refGene_gn=SLC26A10;refGene_tx=XM_019835144.2;refGene_gn=SLC26A10;refGene_tx=XM_019835145.2
B4 86048811 86048918
refGene_gn=SLC26A10;refGene_tx=XM_006933854.3;refGene_gn=SLC26A10;refGene_tx=XM_019835144.2;refGene_gn=SLC26A10;refGene_tx=XM_019835145.2
B4 86049229 86049268
refGene_gn=SLC26A10;refGene_tx=XM_006933854.3;refGene_gn=SLC26A10;refGene_tx=XM_019835144.2;refGene_gn=SLC26A10;refGene_tx=XM_019835145.2
B4 86049952 86050170
refGene_gn=B4GALNT1;refGene_tx=XM_019835147.1;refGene_gn=B4GALNT1;refGene_tx=XM_019835148.2;refGene_gn=B4GALNT1;refGene_tx=XM_023257170.1;refGene_gn=B4GALNT1;refGene_tx=XM_023257171.1
B4 86050758 86050999
refGene_gn=B4GALNT1;refGene_tx=XM_019835147.1;refGene_gn=B4GALNT1;refGene_tx=XM_019835148.2;refGene_gn=B4GALNT1;refGene_tx=XM_023257170.1;refGene_gn=B4GALNT1;refGene_tx=XM_023257171.1
B4 86051258 86051399
refGene_gn=B4GALNT1;refGene_tx=XM_019835147.1;refGene_gn=B4GALNT1;refGene_tx=XM_019835148.2;refGene_gn=B4GALNT1;refGene_tx=XM_023257170.1;refGene_gn=B4GALNT1;refGene_tx=XM_023257171.1
B4 86051867 86052058
refGene_gn=B4GALNT1;refGene_tx=XM_019835147.1;refGene_gn=B4GALNT1;refGene_tx=XM_019835148.2;refGene_gn=B4GALNT1;refGene_tx=XM_023257170.1;refGene_gn=B4GALNT1;refGene_tx=XM_023257171.1
B4 86052184 86052292
refGene_gn=B4GALNT1;refGene_tx=XM_019835147.1;refGene_gn=B4GALNT1;refGene_tx=XM_019835148.2;refGene_gn=B4GALNT1;refGene_tx=XM_023257170.1;re

fGene_gn=B4GALNT1;refGene_tx=XM_023257171.1
B4 86053305 86053486
refGene_gn=B4GALNT1;refGene_tx=XM_019835147.1;refGene_gn=B4GALNT1;refGene_tx=XM_019835148.2;refGene_gn=B4GALNT1;refGene_tx=XM_023257170.1;refGene_gn=B4GALNT1;refGene_tx=XM_023257171.1
B4 86054295 86054336
refGene_gn=B4GALNT1;refGene_tx=XM_019835147.1;refGene_gn=B4GALNT1;refGene_tx=XM_019835148.2;refGene_gn=B4GALNT1;refGene_tx=XM_023257170.1;refGene_gn=B4GALNT1;refGene_tx=XM_023257171.1
B4 86054807 86054914
refGene_gn=B4GALNT1;refGene_tx=XM_019835147.1;refGene_gn=B4GALNT1;refGene_tx=XM_019835148.2;refGene_gn=B4GALNT1;refGene_tx=XM_023257170.1;refGene_gn=B4GALNT1;refGene_tx=XM_023257171.1
B4 86055035 86055200
refGene_gn=B4GALNT1;refGene_tx=XM_019835147.1;refGene_gn=B4GALNT1;refGene_tx=XM_019835148.2;refGene_gn=B4GALNT1;refGene_tx=XM_023257170.1;refGene_gn=B4GALNT1;refGene_tx=XM_023257171.1
B4 86055749 86055968
refGene_gn=B4GALNT1;refGene_tx=XM_019835147.1;refGene_gn=B4GALNT1;refGene_tx=XM_019835148.2;refGene_gn=B4GALNT1;refGene_tx=XM_023257170.1;refGene_gn=B4GALNT1;refGene_tx=XM_023257171.1
B4 86056569 86056795 refGene_gn=B4GALNT1;refGene_tx=XM_023257170.1
B4 86106691 86106853
refGene_gn=OS9;refGene_tx=XM_003988955.4;refGene_gn=OS9;refGene_tx=XM_003988956.4;refGene_gn=OS9;refGene_tx=XM_006933858.3;refGene_gn=OS9;refGene_tx=XM_006933859.3
B4 86107266 86107443
refGene_gn=OS9;refGene_tx=XM_003988955.4;refGene_gn=OS9;refGene_tx=XM_003988956.4;refGene_gn=OS9;refGene_tx=XM_006933858.3;refGene_gn=OS9;refGene_tx=XM_006933859.3
B4 86108311 86108375
refGene_gn=OS9;refGene_tx=XM_003988955.4;refGene_gn=OS9;refGene_tx=XM_003988956.4;refGene_gn=OS9;refGene_tx=XM_006933858.3;refGene_gn=OS9;refGene_tx=XM_006933859.3
B4 86108505 86108582
refGene_gn=OS9;refGene_tx=XM_003988955.4;refGene_gn=OS9;refGene_tx=XM_003988956.4;refGene_gn=OS9;refGene_tx=XM_006933858.3;refGene_gn=OS9;refGene_tx=XM_006933859.3
B4 86108768 86108867
refGene_gn=OS9;refGene_tx=XM_003988955.4;refGene_gn=OS9;refGene_tx=XM_003988956.4;refGene_gn=OS9;refGene_tx=XM_006933858.3;refGene_gn=OS9;refGene_tx=XM_006933859.3
B4 86134126 86134337
refGene_gn=OS9;refGene_tx=XM_003988955.4;refGene_gn=OS9;refGene_tx=XM_003988956.4;refGene_gn=OS9;refGene_tx=XM_006933858.3;refGene_gn=OS9;refGene_tx=XM_006933859.3
B4 86134449 86134554
refGene_gn=OS9;refGene_tx=XM_003988955.4;refGene_gn=OS9;refGene_tx=XM_003988956.4;refGene_gn=OS9;refGene_tx=XM_006933858.3;refGene_gn=OS9;refGene_tx=XM_006933859.3

B4 86134768 86134869
refGene_gn=OS9;refGene_tx=XM_003988955.4;refGene_gn=OS9;refGene_tx=XM_003988956.4;refGene_gn=OS9;refGene_tx=XM_006933858.3;refGene_gn=OS9;refGene_tx=XM_006933859.3
B4 86135047 86135099
refGene_gn=OS9;refGene_tx=XM_003988955.4;refGene_gn=OS9;refGene_tx=XM_003988956.4;refGene_gn=OS9;refGene_tx=XM_006933858.3;refGene_gn=OS9;refGene_tx=XM_006933859.3
B4 86135787 86135876
refGene_gn=OS9;refGene_tx=XM_003988955.4;refGene_gn=OS9;refGene_tx=XM_003988956.4;refGene_gn=OS9;refGene_tx=XM_006933858.3;refGene_gn=OS9;refGene_tx=XM_006933859.3
B4 86136102 86136381
refGene_gn=OS9;refGene_tx=XM_003988955.4;refGene_gn=OS9;refGene_tx=XM_003988956.4;refGene_gn=OS9;refGene_tx=XM_006933858.3;refGene_gn=OS9;refGene_tx=XM_006933859.3
B4 86136614 86136657 refGene_gn=B4GALNT1;refGene_tx=XM_023257170.1
B4 86136662 86136852
refGene_gn=OS9;refGene_tx=XM_003988955.4;refGene_gn=OS9;refGene_tx=XM_003988956.4;refGene_gn=OS9;refGene_tx=XM_006933858.3;refGene_gn=OS9;refGene_tx=XM_006933859.3
B4 86137792 86137957
refGene_gn=OS9;refGene_tx=XM_003988956.4;refGene_gn=OS9;refGene_tx=XM_006933858.3
B4 86138123 86138236
refGene_gn=OS9;refGene_tx=XM_003988955.4;refGene_gn=OS9;refGene_tx=XM_003988956.4;refGene_gn=OS9;refGene_tx=XM_006933858.3;refGene_gn=OS9;refGene_tx=XM_006933859.3
B4 86138549 86138675
refGene_gn=OS9;refGene_tx=XM_003988955.4;refGene_gn=OS9;refGene_tx=XM_003988956.4;refGene_gn=OS9;refGene_tx=XM_006933858.3;refGene_gn=OS9;refGene_tx=XM_006933859.3
B4 86144268 86144511
refGene_gn=AGAP2;refGene_tx=XM_006933860.4;refGene_gn=AGAP2;refGene_tx=XM_011284083.3;refGene_gn=AGAP2;refGene_tx=XM_011284084.3;refGene_gn=AGAP2;refGene_tx=XM_019835149.2;refGene_gn=AGAP2;refGene_tx=XM_019835150.2
B4 86144675 86144931
refGene_gn=AGAP2;refGene_tx=XM_006933860.4;refGene_gn=AGAP2;refGene_tx=XM_011284083.3;refGene_gn=AGAP2;refGene_tx=XM_011284084.3;refGene_gn=AGAP2;refGene_tx=XM_019835149.2;refGene_gn=AGAP2;refGene_tx=XM_019835150.2
B4 86145052 86145275
refGene_gn=AGAP2;refGene_tx=XM_006933860.4;refGene_gn=AGAP2;refGene_tx=XM_011284083.3;refGene_gn=AGAP2;refGene_tx=XM_011284084.3;refGene_gn=AGAP2;refGene_tx=XM_019835149.2;refGene_gn=AGAP2;refGene_tx=XM_019835150.2
B4 86145381 86145472
refGene_gn=AGAP2;refGene_tx=XM_006933860.4;refGene_gn=AGAP2;refGene_tx=XM_011284083.3;refGene_gn=AGAP2;refGene_tx=XM_011284084.3;refGene_gn=

AGAP2;refGene_tx=XM_019835149.2;refGene_gn=AGAP2;refGene_tx=XM_0198351
50.2
B4 86145576 86145725
refGene_gn=AGAP2;refGene_tx=XM_006933860.4;refGene_gn=AGAP2;refGene_tx
=XM_011284083.3;refGene_gn=AGAP2;refGene_tx=XM_011284084.3;refGene_gn=
AGAP2;refGene_tx=XM_019835149.2;refGene_gn=AGAP2;refGene_tx=XM_0198351
50.2
B4 86145957 86146017
refGene_gn=AGAP2;refGene_tx=XM_011284083.3;refGene_gn=AGAP2;refGene_tx
=XM_019835149.2;refGene_gn=AGAP2;refGene_tx=XM_019835150.2
B4 86146920 86147049
refGene_gn=AGAP2;refGene_tx=XM_006933860.4;refGene_gn=AGAP2;refGene_tx
=XM_011284083.3;refGene_gn=AGAP2;refGene_tx=XM_011284084.3;refGene_gn=
AGAP2;refGene_tx=XM_019835149.2;refGene_gn=AGAP2;refGene_tx=XM_0198351
50.2
B4 86147771 86147891
refGene_gn=AGAP2;refGene_tx=XM_006933860.4;refGene_gn=AGAP2;refGene_tx
=XM_011284083.3;refGene_gn=AGAP2;refGene_tx=XM_011284084.3;refGene_gn=
AGAP2;refGene_tx=XM_019835149.2;refGene_gn=AGAP2;refGene_tx=XM_0198351
50.2
B4 86148070 86148233
refGene_gn=AGAP2;refGene_tx=XM_006933860.4;refGene_gn=AGAP2;refGene_tx
=XM_011284083.3;refGene_gn=AGAP2;refGene_tx=XM_011284084.3;refGene_gn=
AGAP2;refGene_tx=XM_019835149.2;refGene_gn=AGAP2;refGene_tx=XM_0198351
50.2
B4 86148677 86148782
refGene_gn=AGAP2;refGene_tx=XM_006933860.4;refGene_gn=AGAP2;refGene_tx
=XM_011284083.3;refGene_gn=AGAP2;refGene_tx=XM_011284084.3;refGene_gn=
AGAP2;refGene_tx=XM_019835149.2;refGene_gn=AGAP2;refGene_tx=XM_0198351
50.2
B4 86148869 86148956
refGene_gn=AGAP2;refGene_tx=XM_006933860.4;refGene_gn=AGAP2;refGene_tx
=XM_011284083.3;refGene_gn=AGAP2;refGene_tx=XM_011284084.3;refGene_gn=
AGAP2;refGene_tx=XM_019835149.2;refGene_gn=AGAP2;refGene_tx=XM_0198351
50.2
B4 86149117 86149276
refGene_gn=AGAP2;refGene_tx=XM_006933860.4;refGene_gn=AGAP2;refGene_tx
=XM_011284083.3;refGene_gn=AGAP2;refGene_tx=XM_011284084.3;refGene_gn=
AGAP2;refGene_tx=XM_019835149.2;refGene_gn=AGAP2;refGene_tx=XM_0198351
50.2
B4 86149676 86149786
refGene_gn=AGAP2;refGene_tx=XM_006933860.4;refGene_gn=AGAP2;refGene_tx
=XM_011284083.3;refGene_gn=AGAP2;refGene_tx=XM_011284084.3;refGene_gn=
AGAP2;refGene_tx=XM_019835149.2;refGene_gn=AGAP2;refGene_tx=XM_0198351
50.2
B4 86150111 86150246
refGene_gn=AGAP2;refGene_tx=XM_006933860.4;refGene_gn=AGAP2;refGene_tx
=XM_011284083.3;refGene_gn=AGAP2;refGene_tx=XM_011284084.3;refGene_gn=
AGAP2;refGene_tx=XM_019835149.2;refGene_gn=AGAP2;refGene_tx=XM_0198351
50.2

B4 86151271 86151419
refGene_gn=AGAP2;refGene_tx=XM_006933860.4;refGene_gn=AGAP2;refGene_tx=XM_011284083.3;refGene_gn=AGAP2;refGene_tx=XM_011284084.3;refGene_gn=AGAP2;refGene_tx=XM_019835149.2;refGene_gn=AGAP2;refGene_tx=XM_019835150.2
B4 86151566 86151652
refGene_gn=AGAP2;refGene_tx=XM_006933860.4;refGene_gn=AGAP2;refGene_tx=XM_011284083.3;refGene_gn=AGAP2;refGene_tx=XM_011284084.3;refGene_gn=AGAP2;refGene_tx=XM_019835149.2;refGene_gn=AGAP2;refGene_tx=XM_019835150.2
B4 86151836 86151924
refGene_gn=AGAP2;refGene_tx=XM_006933860.4;refGene_gn=AGAP2;refGene_tx=XM_011284083.3;refGene_gn=AGAP2;refGene_tx=XM_011284084.3;refGene_gn=AGAP2;refGene_tx=XM_019835149.2;refGene_gn=AGAP2;refGene_tx=XM_019835150.2
B4 86152520 86152579
refGene_gn=AGAP2;refGene_tx=XM_006933860.4;refGene_gn=AGAP2;refGene_tx=XM_011284083.3;refGene_gn=AGAP2;refGene_tx=XM_011284084.3;refGene_gn=AGAP2;refGene_tx=XM_019835149.2;refGene_gn=AGAP2;refGene_tx=XM_019835150.2
B4 86154259 86155436
refGene_gn=AGAP2;refGene_tx=XM_006933860.4;refGene_gn=AGAP2;refGene_tx=XM_011284083.3;refGene_gn=AGAP2;refGene_tx=XM_019835149.2
B4 86159145 86159305
refGene_gn=AGAP2;refGene_tx=XM_011284084.3;refGene_gn=AGAP2;refGene_tx=XM_019835150.2
B4 86161962 86162025
refGene_gn=TSPAN31;refGene_tx=XM_003988964.5;refGene_gn=TSPAN31;refGene_tx=XM_019835151.2
B4 86162523 86162691
refGene_gn=TSPAN31;refGene_tx=XM_003988964.5;refGene_gn=TSPAN31;refGene_tx=XM_019835151.2
B4 86162965 86163046
refGene_gn=TSPAN31;refGene_tx=XM_003988964.5;refGene_gn=TSPAN31;refGene_tx=XM_019835151.2;refGene_gn=TSPAN31;refGene_tx=XM_019835152.1
B4 86163350 86163482
refGene_gn=TSPAN31;refGene_tx=XM_003988964.5;refGene_gn=TSPAN31;refGene_tx=XM_019835151.2;refGene_gn=TSPAN31;refGene_tx=XM_019835152.1
B4 86163806 86163920
refGene_gn=TSPAN31;refGene_tx=XM_003988964.5;refGene_gn=TSPAN31;refGene_tx=XM_019835152.1
B4 86164032 86164107
refGene_gn=TSPAN31;refGene_tx=XM_003988964.5;refGene_gn=TSPAN31;refGene_tx=XM_019835151.2;refGene_gn=TSPAN31;refGene_tx=XM_019835152.1
B4 86165083 86165176
refGene_gn=CDK4;refGene_tx=XM_003988965.5;refGene_gn=CDK4;refGene_tx=XM_011284081.3
B4 86165448 86165584
refGene_gn=CDK4;refGene_tx=XM_003988965.5;refGene_gn=CDK4;refGene_tx=XM_011284081.3

B4 86165720 86165771
refGene_gn=CDK4;refGene_tx=XM_003988965.5;refGene_gn=CDK4;refGene_tx=XM_011284081.3
B4 86166026 86166136
refGene_gn=CDK4;refGene_tx=XM_003988965.5;refGene_gn=CDK4;refGene_tx=XM_011284081.3
B4 86166258 86166438
refGene_gn=CDK4;refGene_tx=XM_003988965.5;refGene_gn=CDK4;refGene_tx=XM_011284081.3
B4 86166555 86166691
refGene_gn=CDK4;refGene_tx=XM_003988965.5;refGene_gn=CDK4;refGene_tx=XM_011284081.3
B4 86166849 86167067
refGene_gn=CDK4;refGene_tx=XM_003988965.5;refGene_gn=CDK4;refGene_tx=XM_011284081.3
B4 86170897 86171254 refGene_gn=MARCH9;refGene_tx=XM_003988987.4
B4 86172281 86172437 refGene_gn=MARCH9;refGene_tx=XM_003988987.4
B4 86173390 86173583 refGene_gn=MARCH9;refGene_tx=XM_003988987.4
B4 86173845 86174180 refGene_gn=MARCH9;refGene_tx=XM_003988987.4
B4 86178389 86178503
refGene_gn=CYP27B1;refGene_tx=XM_003988966.3;refGene_gn=CYP27B1;refGene_tx=XM_019835154.1;refGene_gn=CYP27B1;refGene_tx=XM_019835155.1
B4 86178819 86179017
refGene_gn=CYP27B1;refGene_tx=XM_003988966.3;refGene_gn=CYP27B1;refGene_tx=XM_019835154.1;refGene_gn=CYP27B1;refGene_tx=XM_019835155.1
B4 86179299 86179378
refGene_gn=CYP27B1;refGene_tx=XM_003988966.3;refGene_gn=CYP27B1;refGene_tx=XM_019835154.1
B4 86179584 86179757
refGene_gn=CYP27B1;refGene_tx=XM_003988966.3;refGene_gn=CYP27B1;refGene_tx=XM_019835154.1;refGene_gn=CYP27B1;refGene_tx=XM_019835155.1
B4 86179963 86180136
refGene_gn=CYP27B1;refGene_tx=XM_003988966.3;refGene_gn=CYP27B1;refGene_tx=XM_019835154.1;refGene_gn=CYP27B1;refGene_tx=XM_019835155.1
B4 86180221 86180422
refGene_gn=CYP27B1;refGene_tx=XM_003988966.3;refGene_gn=CYP27B1;refGene_tx=XM_019835154.1;refGene_gn=CYP27B1;refGene_tx=XM_019835155.1
B4 86180502 86180705
refGene_gn=CYP27B1;refGene_tx=XM_003988966.3;refGene_gn=CYP27B1;refGene_tx=XM_019835154.1;refGene_gn=CYP27B1;refGene_tx=XM_019835155.1
B4 86181216 86181650
refGene_gn=CYP27B1;refGene_tx=XM_003988966.3;refGene_gn=CYP27B1;refGene_tx=XM_019835154.1;refGene_gn=CYP27B1;refGene_tx=XM_019835155.1
B4 86182065 86182260 refGene_gn=CYP27B1;refGene_tx=XM_003988966.3
B4 86183754 86183910
refGene_gn=METTL1;refGene_tx=XM_003988967.3;refGene_gn=METTL1;refGene_tx=XM_023257174.1
B4 86184073 86184175
refGene_gn=METTL1;refGene_tx=XM_003988967.3;refGene_gn=METTL1;refGene_tx=XM_023257174.1

B4 86184334 86184448
refGene_gn=METTL1;refGene_tx=XM_003988967.3;refGene_gn=METTL1;refGene_tx=XM_023257174.1
B4 86184538 86184723
refGene_gn=METTL1;refGene_tx=XM_003988967.3;refGene_gn=METTL1;refGene_tx=XM_023257174.1
B4 86185698 86185862
refGene_gn=METTL1;refGene_tx=XM_003988967.3;refGene_gn=METTL1;refGene_tx=XM_023257174.1
B4 86186583 86186696
refGene_gn=METTL1;refGene_tx=XM_003988967.3;refGene_gn=METTL1;refGene_tx=XM_023257174.1
B4 86187316 86187493
refGene_gn=EEF1AKMT3;refGene_tx=XM_023257175.1;refGene_gn=EEF1AKMT3;refGene_tx=XM_023257176.1
B4 86187616 86187728
refGene_gn=EEF1AKMT3;refGene_tx=XM_023257175.1;refGene_gn=EEF1AKMT3;refGene_tx=XM_023257176.1
B4 86194385 86194777
refGene_gn=EEF1AKMT3;refGene_tx=XM_023257175.1;refGene_gn=EEF1AKMT3;refGene_tx=XM_023257176.1
B4 86196557 86196614
refGene_gn=TSFM;refGene_tx=XM_003988969.4;refGene_gn=TSFM;refGene_tx=XM_019835156.1;refGene_gn=TSFM;refGene_tx=XM_023257173.1
B4 86196889 86197063
refGene_gn=TSFM;refGene_tx=XM_003988969.4;refGene_gn=TSFM;refGene_tx=XM_019835156.1;refGene_gn=TSFM;refGene_tx=XM_023257173.1
B4 86197667 86197796
refGene_gn=TSFM;refGene_tx=XM_003988969.4;refGene_gn=TSFM;refGene_tx=XM_019835156.1;refGene_gn=TSFM;refGene_tx=XM_023257173.1
B4 86201108 86201231
refGene_gn=TSFM;refGene_tx=XM_003988969.4;refGene_gn=TSFM;refGene_tx=XM_023257173.1
B4 86205694 86205782
refGene_gn=TSFM;refGene_tx=XM_003988969.4;refGene_gn=TSFM;refGene_tx=XM_019835156.1;refGene_gn=TSFM;refGene_tx=XM_023257173.1
B4 86208115 86208522
refGene_gn=TSFM;refGene_tx=XM_003988969.4;refGene_gn=TSFM;refGene_tx=XM_019835156.1
B4 86210034 86210148
refGene_gn=AVIL;refGene_tx=XM_011284079.3;refGene_gn=AVIL;refGene_tx=XM_011284080.3
B4 86211776 86211893
refGene_gn=AVIL;refGene_tx=XM_011284079.3;refGene_gn=AVIL;refGene_tx=XM_011284080.3
B4 86214083 86214152
refGene_gn=AVIL;refGene_tx=XM_011284079.3;refGene_gn=AVIL;refGene_tx=XM_011284080.3
B4 86214161 86214214 refGene_gn=TSFM;refGene_tx=XM_023257173.1
B4 86214805 86214994

refGene_gn=AVIL;refGene_tx=XM_011284079.3;refGene_gn=AVIL;refGene_tx=XM_011284080.3
B4 86216386 86216531 refGene_gn=AVIL;refGene_tx=XM_011284079.3;refGene_gn=AVIL;refGene_tx=XM_011284080.3
B4 86216669 86216815 refGene_gn=AVIL;refGene_tx=XM_011284079.3;refGene_gn=AVIL;refGene_tx=XM_011284080.3
B4 86218235 86218415 refGene_gn=AVIL;refGene_tx=XM_011284079.3;refGene_gn=AVIL;refGene_tx=XM_011284080.3
B4 86219439 86219598 refGene_gn=AVIL;refGene_tx=XM_011284079.3;refGene_gn=AVIL;refGene_tx=XM_011284080.3
B4 86219704 86219842 refGene_gn=AVIL;refGene_tx=XM_011284079.3;refGene_gn=AVIL;refGene_tx=XM_011284080.3
B4 86220334 86220435 refGene_gn=AVIL;refGene_tx=XM_011284079.3;refGene_gn=AVIL;refGene_tx=XM_011284080.3
B4 86220538 86220692 refGene_gn=AVIL;refGene_tx=XM_011284079.3;refGene_gn=AVIL;refGene_tx=XM_011284080.3
B4 86221383 86221482 refGene_gn=AVIL;refGene_tx=XM_011284079.3;refGene_gn=AVIL;refGene_tx=XM_011284080.3
B4 86221588 86221667 refGene_gn=AVIL;refGene_tx=XM_011284079.3;refGene_gn=AVIL;refGene_tx=XM_011284080.3
B4 86222131 86222334 refGene_gn=AVIL;refGene_tx=XM_011284079.3;refGene_gn=AVIL;refGene_tx=XM_011284080.3
B4 86222598 86222709 refGene_gn=AVIL;refGene_tx=XM_011284079.3;refGene_gn=AVIL;refGene_tx=XM_011284080.3
B4 86222784 86222893 refGene_gn=AVIL;refGene_tx=XM_011284079.3;refGene_gn=AVIL;refGene_tx=XM_011284080.3
B4 86224358 86224555 refGene_gn=AVIL;refGene_tx=XM_011284079.3;refGene_gn=AVIL;refGene_tx=XM_011284080.3
B4 86225229 86225304 refGene_gn=AVIL;refGene_tx=XM_011284079.3;refGene_gn=AVIL;refGene_tx=XM_011284080.3
B4 86226636 86226702 refGene_gn=AVIL;refGene_tx=XM_011284079.3;refGene_gn=AVIL;refGene_tx=XM_011284080.3
B4 86233765 86233891 refGene_gn=CTDSP2;refGene_tx=XM_003988971.5
B4 86234093 86234279 refGene_gn=CTDSP2;refGene_tx=XM_003988971.5
B4 86234435 86234528 refGene_gn=CTDSP2;refGene_tx=XM_003988971.5

B4 86236525 86236582 refGene_gn=CTDSP2;refGene_tx=XM_003988971.5
B4 86237185 86237287 refGene_gn=CTDSP2;refGene_tx=XM_003988971.5
B4 86237703 86237742 refGene_gn=CTDSP2;refGene_tx=XM_003988971.5
B4 86239589 86239738 refGene_gn=CTDSP2;refGene_tx=XM_003988971.5
B4 86253574 86253638 refGene_gn=CTDSP2;refGene_tx=XM_003988971.5
B4 86264835 86264931
refGene_gn=LOC109501628;refGene_tx=XM_023257764.1
B4 86270583 86271178
refGene_gn=LOC109501628;refGene_tx=XM_023257764.1
B4 86295369 86295439
refGene_gn=LOC109501628;refGene_tx=XM_023257764.1
B4 86295636 86295718
refGene_gn=LOC109501628;refGene_tx=XM_023257764.1
B4 86296369 86296378
refGene_gn=LOC109501628;refGene_tx=XM_023257764.1
B4 86341571 86341758 refGene_gn=ATP23;refGene_tx=XM_006933863.3
B4 86345190 86345236 refGene_gn=ATP23;refGene_tx=XM_006933863.3
B4 86345918 86346000 refGene_gn=ATP23;refGene_tx=XM_006933863.3
B4 86349866 86350004
refGene_gn=ATP23;refGene_tx=XM_006933863.3;refGene_gn=ATP23;refGene_tx
=XM_019835157.1;refGene_gn=ATP23;refGene_tx=XM_019835159.1;refGene_gn=
ATP23;refGene_tx=XM_019835160.1
B4 86351836 86351920
refGene_gn=ATP23;refGene_tx=XM_006933863.3;refGene_gn=ATP23;refGene_tx
=XM_019835157.1;refGene_gn=ATP23;refGene_tx=XM_019835159.1;refGene_gn=
ATP23;refGene_tx=XM_019835160.1
B4 86354550 86354754
refGene_gn=ATP23;refGene_tx=XM_006933863.3;refGene_gn=ATP23;refGene_tx
=XM_019835157.1;refGene_gn=ATP23;refGene_tx=XM_019835159.1;refGene_gn=
ATP23;refGene_tx=XM_019835160.1
B4 87117089 87117334
refGene_gn=LRIG3;refGene_tx=XM_023257177.1;refGene_gn=LRIG3;refGene_tx
=XM_023257178.1;refGene_gn=LRIG3;refGene_tx=XM_023257179.1
B4 87118571 87118847
refGene_gn=LRIG3;refGene_tx=XM_023257177.1;refGene_gn=LRIG3;refGene_tx
=XM_023257179.1
B4 87118940 87119084
refGene_gn=LRIG3;refGene_tx=XM_023257177.1;refGene_gn=LRIG3;refGene_tx
=XM_023257178.1;refGene_gn=LRIG3;refGene_tx=XM_023257179.1
B4 87120918 87121077
refGene_gn=LRIG3;refGene_tx=XM_023257177.1;refGene_gn=LRIG3;refGene_tx
=XM_023257178.1;refGene_gn=LRIG3;refGene_tx=XM_023257179.1
B4 87121839 87122292
refGene_gn=LRIG3;refGene_tx=XM_023257177.1;refGene_gn=LRIG3;refGene_tx
=XM_023257178.1;refGene_gn=LRIG3;refGene_tx=XM_023257179.1
B4 87123267 87123549
refGene_gn=LRIG3;refGene_tx=XM_023257177.1;refGene_gn=LRIG3;refGene_tx
=XM_023257178.1;refGene_gn=LRIG3;refGene_tx=XM_023257179.1
B4 87125073 87125394
refGene_gn=LRIG3;refGene_tx=XM_023257177.1;refGene_gn=LRIG3;refGene_tx

=XM_023257178.1;refGene_gn=LRIG3;refGene_tx=XM_023257179.1
B4 87127703 87127867
refGene_gn=LRIG3;refGene_tx=XM_023257177.1;refGene_gn=LRIG3;refGene_tx=XM_023257178.1;refGene_gn=LRIG3;refGene_tx=XM_023257179.1
B4 87128365 87128437
refGene_gn=LRIG3;refGene_tx=XM_023257177.1;refGene_gn=LRIG3;refGene_tx=XM_023257178.1;refGene_gn=LRIG3;refGene_tx=XM_023257179.1
B4 87130825 87130897
refGene_gn=LRIG3;refGene_tx=XM_023257177.1;refGene_gn=LRIG3;refGene_tx=XM_023257178.1;refGene_gn=LRIG3;refGene_tx=XM_023257179.1
B4 87131799 87131880
refGene_gn=LRIG3;refGene_tx=XM_023257177.1;refGene_gn=LRIG3;refGene_tx=XM_023257178.1;refGene_gn=LRIG3;refGene_tx=XM_023257179.1
B4 87132805 87132949
refGene_gn=LRIG3;refGene_tx=XM_023257177.1;refGene_gn=LRIG3;refGene_tx=XM_023257178.1;refGene_gn=LRIG3;refGene_tx=XM_023257179.1
B4 87133342 87133486
refGene_gn=LRIG3;refGene_tx=XM_023257177.1;refGene_gn=LRIG3;refGene_tx=XM_023257178.1;refGene_gn=LRIG3;refGene_tx=XM_023257179.1
B4 87133787 87133931
refGene_gn=LRIG3;refGene_tx=XM_023257177.1;refGene_gn=LRIG3;refGene_tx=XM_023257178.1;refGene_gn=LRIG3;refGene_tx=XM_023257179.1
B4 87134943 87135087
refGene_gn=LRIG3;refGene_tx=XM_023257177.1;refGene_gn=LRIG3;refGene_tx=XM_023257178.1;refGene_gn=LRIG3;refGene_tx=XM_023257179.1
B4 87135877 87136009
refGene_gn=LRIG3;refGene_tx=XM_023257177.1;refGene_gn=LRIG3;refGene_tx=XM_023257178.1;refGene_gn=LRIG3;refGene_tx=XM_023257179.1
B4 87143377 87143418 refGene_gn=LRIG3;refGene_tx=XM_023257179.1
B4 87160516 87160591
refGene_gn=LRIG3;refGene_tx=XM_023257177.1;refGene_gn=LRIG3;refGene_tx=XM_023257178.1
B4 87160804 87160876
refGene_gn=LRIG3;refGene_tx=XM_023257177.1;refGene_gn=LRIG3;refGene_tx=XM_023257178.1
B4 87166469 87166714
refGene_gn=LRIG3;refGene_tx=XM_023257177.1;refGene_gn=LRIG3;refGene_tx=XM_023257178.1
B4 87870226 87870440
refGene_gn=SLC16A7;refGene_tx=XM_011284078.2;refGene_gn=SLC16A7;refGene_tx=XM_019835162.2;refGene_gn=SLC16A7;refGene_tx=XM_023257180.1;refGene_gn=SLC16A7;refGene_tx=XM_023257181.1;refGene_gn=SLC16A7;refGene_tx=XM_023257182.1
B4 87955765 87955909
refGene_gn=SLC16A7;refGene_tx=XM_011284078.2;refGene_gn=SLC16A7;refGene_tx=XM_019835162.2;refGene_gn=SLC16A7;refGene_tx=XM_023257180.1;refGene_gn=SLC16A7;refGene_tx=XM_023257181.1;refGene_gn=SLC16A7;refGene_tx=XM_023257182.1
B4 87958191 87959010
refGene_gn=SLC16A7;refGene_tx=XM_011284078.2;refGene_gn=SLC16A7;refGen

e_tx=XM_019835162.2;refGene_gn=SLC16A7;refGene_tx=XM_023257180.1;refGene_gn=SLC16A7;refGene_tx=XM_023257181.1;refGene_gn=SLC16A7;refGene_tx=XM_023257182.1
B4	87962992	87963264	refGene_gn=SLC16A7;refGene_tx=XM_011284078.2;refGene_gn=SLC16A7;refGene_tx=XM_019835162.2;refGene_gn=SLC16A7;refGene_tx=XM_023257180.1;refGene_gn=SLC16A7;refGene_tx=XM_023257181.1;refGene_gn=SLC16A7;refGene_tx=XM_023257182.1
B4	89594186	89594198	refGene_gn=FAM19A2;refGene_tx=XM_011284073.2;refGene_gn=FAM19A2;refGene_tx=XM_019835164.1;refGene_gn=FAM19A2;refGene_tx=XM_023257183.1
B4	89640805	89640930	refGene_gn=FAM19A2;refGene_tx=XM_011284073.2;refGene_gn=FAM19A2;refGene_tx=XM_019835164.1;refGene_gn=FAM19A2;refGene_tx=XM_023257183.1
B4	89642171	89642324	refGene_gn=FAM19A2;refGene_tx=XM_011284073.2;refGene_gn=FAM19A2;refGene_tx=XM_019835164.1;refGene_gn=FAM19A2;refGene_tx=XM_023257183.1
B4	89762033	89762140	refGene_gn=FAM19A2;refGene_tx=XM_011284073.2;refGene_gn=FAM19A2;refGene_tx=XM_019835164.1;refGene_gn=FAM19A2;refGene_tx=XM_023257183.1
B4	90080359	90080391	refGene_gn=FAM19A2;refGene_tx=XM_019835164.1
B4	90138537	90138626	refGene_gn=USP15;refGene_tx=XM_006933864.4
B4	90160744	90160872	refGene_gn=USP15;refGene_tx=XM_006933864.4
B4	90168514	90168645	refGene_gn=USP15;refGene_tx=XM_006933864.4
B4	90178232	90178359	refGene_gn=USP15;refGene_tx=XM_006933864.4
B4	90184616	90184762	refGene_gn=USP15;refGene_tx=XM_006933864.4
B4	90188702	90188764	refGene_gn=USP15;refGene_tx=XM_006933864.4
B4	90207171	90207258	refGene_gn=USP15;refGene_tx=XM_006933864.4
B4	90213941	90214063	refGene_gn=USP15;refGene_tx=XM_006933864.4
B4	90214065	90214085	refGene_gn=USP15;refGene_tx=XM_006933864.4
B4	90235777	90235951	refGene_gn=USP15;refGene_tx=XM_006933864.4
B4	90237390	90237549	refGene_gn=USP15;refGene_tx=XM_006933864.4
B4	90237628	90237853	refGene_gn=USP15;refGene_tx=XM_006933864.4
B4	90242816	90242900	refGene_gn=USP15;refGene_tx=XM_006933864.4
B4	90242990	90243011	refGene_gn=USP15;refGene_tx=XM_006933864.4
B4	90243012	90243039	refGene_gn=USP15;refGene_tx=XM_006933864.4
B4	90243040	90243087	refGene_gn=USP15;refGene_tx=XM_006933864.4
B4	90243185	90243377	refGene_gn=USP15;refGene_tx=XM_006933864.4
B4	90244284	90244400	refGene_gn=USP15;refGene_tx=XM_006933864.4
B4	90244573	90244846	refGene_gn=USP15;refGene_tx=XM_006933864.4
B4	90245228	90245299	refGene_gn=USP15;refGene_tx=XM_006933864.4
B4	90245615	90245675	refGene_gn=USP15;refGene_tx=XM_006933864.4
B4	90245676	90245732	refGene_gn=USP15;refGene_tx=XM_006933864.4
B4	90246350	90246500	refGene_gn=USP15;refGene_tx=XM_006933864.4
B4	90248088	90248192	refGene_gn=USP15;refGene_tx=XM_006933864.4
B4	90252204	90252293	refGene_gn=USP15;refGene_tx=XM_006933864.4
B4	90255336	90255519	refGene_gn=USP15;refGene_tx=XM_006933864.4
B4	90302037	90302148	refGene_gn=MON2;refGene_tx=XM_011284068.3;refGene_gn=MON2;refGene_tx=XM_011284069.3;refGene_gn=MON2;refGene_tx=XM_019835165.2;refGene_gn=MON

2;refGene_tx=XM_019835166.2;refGene_gn=MON2;refGene_tx=XM_019835167.2
B4	90316852 90316916
refGene_gn=MON2;refGene_tx=XM_011284068.3;refGene_gn=MON2;refGene_tx=XM_011284069.3;refGene_gn=MON2;refGene_tx=XM_019835165.2;refGene_gn=MON2;refGene_tx=XM_019835166.2;refGene_gn=MON2;refGene_tx=XM_019835167.2
B4	90319530 90319537 refGene_gn=MON2;refGene_tx=XM_019835168.2
B4	90325560 90325688
refGene_gn=MON2;refGene_tx=XM_011284068.3;refGene_gn=MON2;refGene_tx=XM_011284069.3;refGene_gn=MON2;refGene_tx=XM_019835165.2;refGene_gn=MON2;refGene_tx=XM_019835166.2;refGene_gn=MON2;refGene_tx=XM_019835167.2;refGene_gn=MON2;refGene_tx=XM_019835168.2;refGene_gn=MON2;refGene_tx=XM_019835170.2;refGene_gn=MON2;refGene_tx=XM_023257184.1
B4	90326886 90327018
refGene_gn=MON2;refGene_tx=XM_011284068.3;refGene_gn=MON2;refGene_tx=XM_011284069.3;refGene_gn=MON2;refGene_tx=XM_019835165.2;refGene_gn=MON2;refGene_tx=XM_019835166.2;refGene_gn=MON2;refGene_tx=XM_019835167.2;refGene_gn=MON2;refGene_tx=XM_019835168.2;refGene_gn=MON2;refGene_tx=XM_019835170.2;refGene_gn=MON2;refGene_tx=XM_023257184.1
B4	90328880 90329010
refGene_gn=MON2;refGene_tx=XM_011284068.3;refGene_gn=MON2;refGene_tx=XM_011284069.3;refGene_gn=MON2;refGene_tx=XM_019835165.2;refGene_gn=MON2;refGene_tx=XM_019835166.2;refGene_gn=MON2;refGene_tx=XM_019835167.2;refGene_gn=MON2;refGene_tx=XM_019835168.2;refGene_gn=MON2;refGene_tx=XM_019835170.2;refGene_gn=MON2;refGene_tx=XM_023257184.1
B4	90334902 90335000
refGene_gn=MON2;refGene_tx=XM_011284068.3;refGene_gn=MON2;refGene_tx=XM_011284069.3;refGene_gn=MON2;refGene_tx=XM_019835165.2;refGene_gn=MON2;refGene_tx=XM_019835166.2;refGene_gn=MON2;refGene_tx=XM_019835167.2;refGene_gn=MON2;refGene_tx=XM_019835168.2;refGene_gn=MON2;refGene_tx=XM_019835170.2;refGene_gn=MON2;refGene_tx=XM_023257184.1
B4	90335715 90335841
refGene_gn=MON2;refGene_tx=XM_011284068.3;refGene_gn=MON2;refGene_tx=XM_011284069.3;refGene_gn=MON2;refGene_tx=XM_019835165.2;refGene_gn=MON2;refGene_tx=XM_019835166.2;refGene_gn=MON2;refGene_tx=XM_019835167.2;refGene_gn=MON2;refGene_tx=XM_019835168.2;refGene_gn=MON2;refGene_tx=XM_019835170.2;refGene_gn=MON2;refGene_tx=XM_023257184.1
B4	90338018 90338213
refGene_gn=MON2;refGene_tx=XM_011284068.3;refGene_gn=MON2;refGene_tx=XM_011284069.3;refGene_gn=MON2;refGene_tx=XM_019835165.2;refGene_gn=MON2;refGene_tx=XM_019835166.2;refGene_gn=MON2;refGene_tx=XM_019835167.2;refGene_gn=MON2;refGene_tx=XM_019835168.2;refGene_gn=MON2;refGene_tx=XM_019835170.2;refGene_gn=MON2;refGene_tx=XM_023257184.1
B4	90348253 90348378
refGene_gn=MON2;refGene_tx=XM_011284068.3;refGene_gn=MON2;refGene_tx=XM_011284069.3;refGene_gn=MON2;refGene_tx=XM_019835165.2;refGene_gn=MON2;refGene_tx=XM_019835166.2;refGene_gn=MON2;refGene_tx=XM_019835167.2;refGene_gn=MON2;refGene_tx=XM_019835168.2;refGene_gn=MON2;refGene_tx=XM_019835170.2;refGene_gn=MON2;refGene_tx=XM_023257184.1
B4	90348796 90348933
refGene_gn=MON2;refGene_tx=XM_011284068.3;refGene_gn=MON2;refGene_tx=X

M_011284069.3;refGene_gn=MON2;refGene_tx=XM_019835165.2;refGene_gn=MON2;refGene_tx=XM_019835166.2;refGene_gn=MON2;refGene_tx=XM_019835167.2;refGene_gn=MON2;refGene_tx=XM_019835168.2;refGene_gn=MON2;refGene_tx=XM_019835170.2;refGene_gn=MON2;refGene_tx=XM_023257184.1
B4 90351233 90351387
refGene_gn=MON2;refGene_tx=XM_011284068.3;refGene_gn=MON2;refGene_tx=XM_011284069.3;refGene_gn=MON2;refGene_tx=XM_019835165.2;refGene_gn=MON2;refGene_tx=XM_019835166.2;refGene_gn=MON2;refGene_tx=XM_019835167.2;refGene_gn=MON2;refGene_tx=XM_019835168.2;refGene_gn=MON2;refGene_tx=XM_019835170.2;refGene_gn=MON2;refGene_tx=XM_023257184.1
B4 90357694 90357927
refGene_gn=MON2;refGene_tx=XM_011284068.3;refGene_gn=MON2;refGene_tx=XM_011284069.3;refGene_gn=MON2;refGene_tx=XM_019835165.2;refGene_gn=MON2;refGene_tx=XM_019835166.2;refGene_gn=MON2;refGene_tx=XM_019835167.2;refGene_gn=MON2;refGene_tx=XM_019835168.2;refGene_gn=MON2;refGene_tx=XM_019835170.2;refGene_gn=MON2;refGene_tx=XM_023257184.1
B4 90360286 90360368
refGene_gn=MON2;refGene_tx=XM_011284068.3;refGene_gn=MON2;refGene_tx=XM_011284069.3;refGene_gn=MON2;refGene_tx=XM_019835165.2;refGene_gn=MON2;refGene_tx=XM_019835166.2;refGene_gn=MON2;refGene_tx=XM_019835167.2;refGene_gn=MON2;refGene_tx=XM_019835168.2;refGene_gn=MON2;refGene_tx=XM_019835170.2;refGene_gn=MON2;refGene_tx=XM_023257184.1
B4 90360946 90361131
refGene_gn=MON2;refGene_tx=XM_011284068.3;refGene_gn=MON2;refGene_tx=XM_011284069.3;refGene_gn=MON2;refGene_tx=XM_019835165.2;refGene_gn=MON2;refGene_tx=XM_019835166.2;refGene_gn=MON2;refGene_tx=XM_019835167.2;refGene_gn=MON2;refGene_tx=XM_019835168.2;refGene_gn=MON2;refGene_tx=XM_019835170.2;refGene_gn=MON2;refGene_tx=XM_023257184.1
B4 90362297 90362410
refGene_gn=MON2;refGene_tx=XM_011284068.3;refGene_gn=MON2;refGene_tx=XM_011284069.3;refGene_gn=MON2;refGene_tx=XM_019835165.2;refGene_gn=MON2;refGene_tx=XM_019835166.2;refGene_gn=MON2;refGene_tx=XM_019835167.2;refGene_gn=MON2;refGene_tx=XM_019835168.2;refGene_gn=MON2;refGene_tx=XM_019835170.2;refGene_gn=MON2;refGene_tx=XM_023257184.1
B4 90369825 90369930
refGene_gn=MON2;refGene_tx=XM_011284068.3;refGene_gn=MON2;refGene_tx=XM_011284069.3;refGene_gn=MON2;refGene_tx=XM_019835165.2;refGene_gn=MON2;refGene_tx=XM_019835166.2;refGene_gn=MON2;refGene_tx=XM_019835167.2;refGene_gn=MON2;refGene_tx=XM_019835168.2;refGene_gn=MON2;refGene_tx=XM_019835170.2;refGene_gn=MON2;refGene_tx=XM_023257184.1
B4 90370286 90370367
refGene_gn=MON2;refGene_tx=XM_011284068.3;refGene_gn=MON2;refGene_tx=XM_011284069.3;refGene_gn=MON2;refGene_tx=XM_019835165.2;refGene_gn=MON2;refGene_tx=XM_019835166.2;refGene_gn=MON2;refGene_tx=XM_019835167.2;refGene_gn=MON2;refGene_tx=XM_019835168.2;refGene_gn=MON2;refGene_tx=XM_019835170.2;refGene_gn=MON2;refGene_tx=XM_023257184.1
B4 90370445 90370519
refGene_gn=MON2;refGene_tx=XM_011284068.3;refGene_gn=MON2;refGene_tx=XM_011284069.3;refGene_gn=MON2;refGene_tx=XM_019835165.2;refGene_gn=MON2;refGene_tx=XM_019835166.2;refGene_gn=MON2;refGene_tx=XM_019835167.2;

refGene_gn=MON2;refGene_tx=XM_019835168.2;refGene_gn=MON2;refGene_tx=XM_019835170.2;refGene_gn=MON2;refGene_tx=XM_023257184.1
B4 90370787 90370878
refGene_gn=MON2;refGene_tx=XM_011284068.3;refGene_gn=MON2;refGene_tx=XM_011284069.3;refGene_gn=MON2;refGene_tx=XM_019835165.2;refGene_gn=MON2;refGene_tx=XM_019835166.2;refGene_gn=MON2;refGene_tx=XM_019835167.2;refGene_gn=MON2;refGene_tx=XM_019835168.2;refGene_gn=MON2;refGene_tx=XM_019835170.2;refGene_gn=MON2;refGene_tx=XM_023257184.1
B4 90375586 90375688
refGene_gn=MON2;refGene_tx=XM_011284068.3;refGene_gn=MON2;refGene_tx=XM_011284069.3;refGene_gn=MON2;refGene_tx=XM_019835165.2;refGene_gn=MON2;refGene_tx=XM_019835166.2;refGene_gn=MON2;refGene_tx=XM_019835167.2;refGene_gn=MON2;refGene_tx=XM_019835168.2;refGene_gn=MON2;refGene_tx=XM_019835170.2;refGene_gn=MON2;refGene_tx=XM_023257184.1
B4 90376856 90376859
refGene_gn=MON2;refGene_tx=XM_019835165.2;refGene_gn=MON2;refGene_tx=XM_019835166.2;refGene_gn=MON2;refGene_tx=XM_019835167.2;refGene_gn=MON2;refGene_tx=XM_019835168.2;refGene_gn=MON2;refGene_tx=XM_019835170.2;refGene_gn=MON2;refGene_tx=XM_023257184.1
B4 90377047 90377158
refGene_gn=MON2;refGene_tx=XM_011284068.3;refGene_gn=MON2;refGene_tx=XM_011284069.3;refGene_gn=MON2;refGene_tx=XM_019835165.2;refGene_gn=MON2;refGene_tx=XM_019835166.2;refGene_gn=MON2;refGene_tx=XM_019835167.2;refGene_gn=MON2;refGene_tx=XM_019835168.2;refGene_gn=MON2;refGene_tx=XM_019835170.2;refGene_gn=MON2;refGene_tx=XM_023257184.1
B4 90378706 90378882
refGene_gn=MON2;refGene_tx=XM_011284068.3;refGene_gn=MON2;refGene_tx=XM_011284069.3;refGene_gn=MON2;refGene_tx=XM_019835165.2;refGene_gn=MON2;refGene_tx=XM_019835166.2;refGene_gn=MON2;refGene_tx=XM_019835167.2;refGene_gn=MON2;refGene_tx=XM_019835168.2;refGene_gn=MON2;refGene_tx=XM_019835170.2;refGene_gn=MON2;refGene_tx=XM_023257184.1
B4 90381618 90381781
refGene_gn=MON2;refGene_tx=XM_011284068.3;refGene_gn=MON2;refGene_tx=XM_011284069.3;refGene_gn=MON2;refGene_tx=XM_019835165.2;refGene_gn=MON2;refGene_tx=XM_019835166.2;refGene_gn=MON2;refGene_tx=XM_019835167.2;refGene_gn=MON2;refGene_tx=XM_019835168.2;refGene_gn=MON2;refGene_tx=XM_019835170.2;refGene_gn=MON2;refGene_tx=XM_023257184.1
B4 90383408 90383702
refGene_gn=MON2;refGene_tx=XM_011284068.3;refGene_gn=MON2;refGene_tx=XM_011284069.3;refGene_gn=MON2;refGene_tx=XM_019835165.2;refGene_gn=MON2;refGene_tx=XM_019835166.2;refGene_gn=MON2;refGene_tx=XM_019835167.2;refGene_gn=MON2;refGene_tx=XM_019835168.2;refGene_gn=MON2;refGene_tx=XM_019835170.2;refGene_gn=MON2;refGene_tx=XM_023257184.1
B4 90385855 90386054
refGene_gn=MON2;refGene_tx=XM_011284068.3;refGene_gn=MON2;refGene_tx=XM_011284069.3;refGene_gn=MON2;refGene_tx=XM_019835165.2;refGene_gn=MON2;refGene_tx=XM_019835166.2;refGene_gn=MON2;refGene_tx=XM_019835167.2;refGene_gn=MON2;refGene_tx=XM_019835168.2;refGene_gn=MON2;refGene_tx=XM_019835170.2;refGene_gn=MON2;refGene_tx=XM_023257184.1
B4 90390328 90390957

refGene_gn=MON2;refGene_tx=XM_011284068.3;refGene_gn=MON2;refGene_tx=XM_011284069.3;refGene_gn=MON2;refGene_tx=XM_019835165.2;refGene_gn=MON2;refGene_tx=XM_019835166.2;refGene_gn=MON2;refGene_tx=XM_019835167.2;refGene_gn=MON2;refGene_tx=XM_019835168.2;refGene_gn=MON2;refGene_tx=XM_019835170.2;refGene_gn=MON2;refGene_tx=XM_023257184.1
B4 90395396 90395540
refGene_gn=MON2;refGene_tx=XM_011284068.3;refGene_gn=MON2;refGene_tx=XM_011284069.3;refGene_gn=MON2;refGene_tx=XM_019835165.2;refGene_gn=MON2;refGene_tx=XM_019835166.2;refGene_gn=MON2;refGene_tx=XM_019835167.2;refGene_gn=MON2;refGene_tx=XM_019835168.2;refGene_gn=MON2;refGene_tx=XM_019835170.2;refGene_gn=MON2;refGene_tx=XM_023257184.1
B4 90396132 90396150
refGene_gn=MON2;refGene_tx=XM_011284068.3;refGene_gn=MON2;refGene_tx=XM_019835165.2;refGene_gn=MON2;refGene_tx=XM_019835167.2;refGene_gn=MON2;refGene_tx=XM_019835168.2;refGene_gn=MON2;refGene_tx=XM_019835170.2;refGene_gn=MON2;refGene_tx=XM_023257184.1
B4 90396433 90396562
refGene_gn=MON2;refGene_tx=XM_011284068.3;refGene_gn=MON2;refGene_tx=XM_011284069.3;refGene_gn=MON2;refGene_tx=XM_019835165.2;refGene_gn=MON2;refGene_tx=XM_019835166.2;refGene_gn=MON2;refGene_tx=XM_019835168.2;refGene_gn=MON2;refGene_tx=XM_019835170.2;refGene_gn=MON2;refGene_tx=XM_023257184.1
B4 90400609 90400800
refGene_gn=MON2;refGene_tx=XM_011284068.3;refGene_gn=MON2;refGene_tx=XM_011284069.3;refGene_gn=MON2;refGene_tx=XM_019835165.2;refGene_gn=MON2;refGene_tx=XM_019835166.2;refGene_gn=MON2;refGene_tx=XM_019835167.2;refGene_gn=MON2;refGene_tx=XM_019835168.2;refGene_gn=MON2;refGene_tx=XM_019835170.2;refGene_gn=MON2;refGene_tx=XM_023257184.1
B4 90407029 90407090
refGene_gn=MON2;refGene_tx=XM_011284068.3;refGene_gn=MON2;refGene_tx=XM_011284069.3;refGene_gn=MON2;refGene_tx=XM_019835165.2;refGene_gn=MON2;refGene_tx=XM_019835166.2;refGene_gn=MON2;refGene_tx=XM_019835167.2;refGene_gn=MON2;refGene_tx=XM_019835168.2;refGene_gn=MON2;refGene_tx=XM_019835170.2;refGene_gn=MON2;refGene_tx=XM_023257184.1
B4 90408176 90408300
refGene_gn=MON2;refGene_tx=XM_011284068.3;refGene_gn=MON2;refGene_tx=XM_011284069.3;refGene_gn=MON2;refGene_tx=XM_019835165.2;refGene_gn=MON2;refGene_tx=XM_019835166.2;refGene_gn=MON2;refGene_tx=XM_019835167.2;refGene_gn=MON2;refGene_tx=XM_019835168.2;refGene_gn=MON2;refGene_tx=XM_019835170.2;refGene_gn=MON2;refGene_tx=XM_023257184.1
B4 90410587 90410795
refGene_gn=MON2;refGene_tx=XM_011284068.3;refGene_gn=MON2;refGene_tx=XM_011284069.3;refGene_gn=MON2;refGene_tx=XM_019835165.2;refGene_gn=MON2;refGene_tx=XM_019835166.2;refGene_gn=MON2;refGene_tx=XM_019835167.2;refGene_gn=MON2;refGene_tx=XM_019835168.2;refGene_gn=MON2;refGene_tx=XM_019835170.2;refGene_gn=MON2;refGene_tx=XM_023257184.1
B4 90413458 90413541
refGene_gn=MON2;refGene_tx=XM_011284068.3;refGene_gn=MON2;refGene_tx=XM_011284069.3;refGene_gn=MON2;refGene_tx=XM_019835165.2;refGene_gn=MON2;refGene_tx=XM_019835166.2;refGene_gn=MON2;refGene_tx=XM_019835167.2;

refGene_gn=MON2;refGene_tx=XM_019835168.2;refGene_gn=MON2;refGene_tx=XM_019835170.2;refGene_gn=MON2;refGene_tx=XM_023257184.1
B4 90419785 90419949
refGene_gn=MON2;refGene_tx=XM_011284068.3;refGene_gn=MON2;refGene_tx=XM_011284069.3;refGene_gn=MON2;refGene_tx=XM_019835165.2;refGene_gn=MON2;refGene_tx=XM_019835166.2;refGene_gn=MON2;refGene_tx=XM_019835167.2;refGene_gn=MON2;refGene_tx=XM_019835168.2;refGene_gn=MON2;refGene_tx=XM_019835170.2;refGene_gn=MON2;refGene_tx=XM_023257184.1
B4 90484366 90484514 refGene_gn=PPM1H;refGene_tx=XM_003988974.5
B4 90501653 90501805 refGene_gn=PPM1H;refGene_tx=XM_003988974.5
B4 90515724 90515745 refGene_gn=PPM1H;refGene_tx=XM_023257185.1
B4 90521827 90521935
refGene_gn=PPM1H;refGene_tx=XM_003988974.5;refGene_gn=PPM1H;refGene_tx=XM_023257185.1
B4 90525511 90525575
refGene_gn=PPM1H;refGene_tx=XM_003988974.5;refGene_gn=PPM1H;refGene_tx=XM_023257185.1
B4 90549932 90550051
refGene_gn=PPM1H;refGene_tx=XM_003988974.5;refGene_gn=PPM1H;refGene_tx=XM_023257185.1
B4 90566649 90566734
refGene_gn=PPM1H;refGene_tx=XM_003988974.5;refGene_gn=PPM1H;refGene_tx=XM_023257185.1
B4 90604535 90604648
refGene_gn=PPM1H;refGene_tx=XM_003988974.5;refGene_gn=PPM1H;refGene_tx=XM_023257185.1
B4 90613797 90614142
refGene_gn=PPM1H;refGene_tx=XM_003988974.5;refGene_gn=PPM1H;refGene_tx=XM_023257185.1
B4 90637138 90637304
refGene_gn=PPM1H;refGene_tx=XM_003988974.5;refGene_gn=PPM1H;refGene_tx=XM_023257185.1
B4 90740814 90741059
refGene_gn=PPM1H;refGene_tx=XM_003988974.5;refGene_gn=PPM1H;refGene_tx=XM_023257185.1
B4 90835237 90835393
refGene_gn=LOC111561460;refGene_tx=XM_023257766.1
B4 90838138 90838192
refGene_gn=LOC111561461;refGene_tx=XM_023257767.1
B4 90951554 90951841 refGene_gn=AVPR1A;refGene_tx=XM_003988975.5
B4 90954042 90955021 refGene_gn=AVPR1A;refGene_tx=XM_003988975.5
B4 91370026 91370195 refGene_gn=TMEM5;refGene_tx=XM_003988976.5
B4 91370989 91371145 refGene_gn=TMEM5;refGene_tx=XM_003988976.5
B4 91375492 91375595 refGene_gn=TMEM5;refGene_tx=XM_003988976.5
B4 91390190 91390505
refGene_gn=TMEM5;refGene_tx=XM_003988976.5;refGene_gn=TMEM5;refGene_tx=XM_019835171.2
B4 91392979 91393150
refGene_gn=TMEM5;refGene_tx=XM_003988976.5;refGene_gn=TMEM5;refGene_tx=XM_019835171.2

B4 91396104 91396522
refGene_gn=TMEM5;refGene_tx=XM_003988976.5;refGene_gn=TMEM5;refGene_tx
=XM_019835171.2
B4 91425719 91425786
refGene_gn=SRGAP1;refGene_tx=XM_011284066.3;refGene_gn=SRGAP1;refGene_
tx=XM_011284067.3
B4 91564664 91564860
refGene_gn=SRGAP1;refGene_tx=XM_011284066.3;refGene_gn=SRGAP1;refGene_
tx=XM_011284067.3
B4 91569247 91569410
refGene_gn=SRGAP1;refGene_tx=XM_011284066.3;refGene_gn=SRGAP1;refGene_
tx=XM_011284067.3
B4 91598220 91598283
refGene_gn=SRGAP1;refGene_tx=XM_011284066.3;refGene_gn=SRGAP1;refGene_
tx=XM_011284067.3
B4 91622242 91622425
refGene_gn=SRGAP1;refGene_tx=XM_011284066.3;refGene_gn=SRGAP1;refGene_
tx=XM_011284067.3
B4 91622856 91622985
refGene_gn=SRGAP1;refGene_tx=XM_011284066.3;refGene_gn=SRGAP1;refGene_
tx=XM_011284067.3
B4 91639782 91640004
refGene_gn=SRGAP1;refGene_tx=XM_011284066.3;refGene_gn=SRGAP1;refGene_
tx=XM_011284067.3
B4 91642219 91642321
refGene_gn=SRGAP1;refGene_tx=XM_011284066.3;refGene_gn=SRGAP1;refGene_
tx=XM_011284067.3
B4 91653757 91653955
refGene_gn=SRGAP1;refGene_tx=XM_011284066.3;refGene_gn=SRGAP1;refGene_
tx=XM_011284067.3
B4 91654696 91654781
refGene_gn=SRGAP1;refGene_tx=XM_011284066.3;refGene_gn=SRGAP1;refGene_
tx=XM_011284067.3
B4 91662062 91662090
refGene_gn=SRGAP1;refGene_tx=XM_011284066.3;refGene_gn=SRGAP1;refGene_
tx=XM_011284067.3
B4 91667857 91667960 refGene_gn=SRGAP1;refGene_tx=XM_011284066.3
B4 91668531 91668565 refGene_gn=SRGAP1;refGene_tx=XM_011284067.3
B4 91671687 91671748
refGene_gn=SRGAP1;refGene_tx=XM_011284066.3;refGene_gn=SRGAP1;refGene_
tx=XM_011284067.3
B4 91672275 91672353
refGene_gn=SRGAP1;refGene_tx=XM_011284066.3;refGene_gn=SRGAP1;refGene_
tx=XM_011284067.3
B4 91674360 91674495
refGene_gn=SRGAP1;refGene_tx=XM_011284066.3;refGene_gn=SRGAP1;refGene_
tx=XM_011284067.3
B4 91686642 91686748
refGene_gn=SRGAP1;refGene_tx=XM_011284066.3;refGene_gn=SRGAP1;refGene_
tx=XM_011284067.3

B4 91690661 91690886
refGene_gn=SRGAP1;refGene_tx=XM_011284066.3;refGene_gn=SRGAP1;refGene_tx=XM_011284067.3
B4 91692846 91692926
refGene_gn=SRGAP1;refGene_tx=XM_011284066.3;refGene_gn=SRGAP1;refGene_tx=XM_011284067.3
B4 91703201 91703382
refGene_gn=SRGAP1;refGene_tx=XM_011284066.3;refGene_gn=SRGAP1;refGene_tx=XM_011284067.3
B4 91704833 91704968
refGene_gn=SRGAP1;refGene_tx=XM_011284066.3;refGene_gn=SRGAP1;refGene_tx=XM_011284067.3
B4 91705105 91705445
refGene_gn=SRGAP1;refGene_tx=XM_011284066.3;refGene_gn=SRGAP1;refGene_tx=XM_011284067.3
B4 91709347 91709725
refGene_gn=SRGAP1;refGene_tx=XM_011284066.3;refGene_gn=SRGAP1;refGene_tx=XM_011284067.3
B4 91751438 91752241
refGene_gn=CB4H12orf66;refGene_tx=XM_003988995.4
B4 91752243 91752254
refGene_gn=CB4H12orf66;refGene_tx=XM_003988995.4
B4 91763747 91764033
refGene_gn=CB4H12orf66;refGene_tx=XM_003988995.4
B4 91769867 91770102
refGene_gn=CB4H12orf66;refGene_tx=XM_003988995.4
B4 91774081 91774103
refGene_gn=CB4H12orf56;refGene_tx=XM_023257190.1
B4 91801424 91801569
refGene_gn=CB4H12orf56;refGene_tx=XM_023257187.1
B4 91802743 91802870
refGene_gn=CB4H12orf56;refGene_tx=XM_023257188.1
B4 91803109 91803293
refGene_gn=CB4H12orf56;refGene_tx=XM_023257186.1;refGene_gn=CB4H12orf56;refGene_tx=XM_023257189.1;refGene_gn=CB4H12orf56;refGene_tx=XM_023257191.1
B4 91805125 91805304
refGene_gn=CB4H12orf56;refGene_tx=XM_023257186.1;refGene_gn=CB4H12orf56;refGene_tx=XM_023257187.1;refGene_gn=CB4H12orf56;refGene_tx=XM_023257188.1;refGene_gn=CB4H12orf56;refGene_tx=XM_023257189.1;refGene_gn=CB4H12orf56;refGene_tx=XM_023257190.1;refGene_gn=CB4H12orf56;refGene_tx=XM_023257191.1
B4 91808694 91808769
refGene_gn=CB4H12orf56;refGene_tx=XM_023257186.1;refGene_gn=CB4H12orf56;refGene_tx=XM_023257187.1;refGene_gn=CB4H12orf56;refGene_tx=XM_023257188.1;refGene_gn=CB4H12orf56;refGene_tx=XM_023257189.1;refGene_gn=CB4H12orf56;refGene_tx=XM_023257190.1;refGene_gn=CB4H12orf56;refGene_tx=XM_023257191.1
B4 91809065 91809140
refGene_gn=CB4H12orf56;refGene_tx=XM_023257186.1;refGene_gn=CB4H12orf5

6;refGene_tx=XM_023257187.1;refGene_gn=CB4H12orf56;refGene_tx=XM_02325
7188.1;refGene_gn=CB4H12orf56;refGene_tx=XM_023257189.1;refGene_gn=CB4
H12orf56;refGene_tx=XM_023257190.1;refGene_gn=CB4H12orf56;refGene_tx=X
M_023257191.1
B4 91812091 91812215
refGene_gn=CB4H12orf56;refGene_tx=XM_023257186.1;refGene_gn=CB4H12orf5
6;refGene_tx=XM_023257187.1;refGene_gn=CB4H12orf56;refGene_tx=XM_02325
7188.1;refGene_gn=CB4H12orf56;refGene_tx=XM_023257189.1;refGene_gn=CB4
H12orf56;refGene_tx=XM_023257190.1;refGene_gn=CB4H12orf56;refGene_tx=X
M_023257191.1
B4 91820024 91820114
refGene_gn=CB4H12orf56;refGene_tx=XM_023257186.1;refGene_gn=CB4H12orf5
6;refGene_tx=XM_023257187.1;refGene_gn=CB4H12orf56;refGene_tx=XM_02325
7188.1;refGene_gn=CB4H12orf56;refGene_tx=XM_023257189.1;refGene_gn=CB4
H12orf56;refGene_tx=XM_023257190.1;refGene_gn=CB4H12orf56;refGene_tx=X
M_023257191.1
B4 91820944 91821051
refGene_gn=CB4H12orf56;refGene_tx=XM_023257186.1;refGene_gn=CB4H12orf5
6;refGene_tx=XM_023257187.1;refGene_gn=CB4H12orf56;refGene_tx=XM_02325
7188.1;refGene_gn=CB4H12orf56;refGene_tx=XM_023257189.1;refGene_gn=CB4
H12orf56;refGene_tx=XM_023257190.1;refGene_gn=CB4H12orf56;refGene_tx=X
M_023257191.1
B4 91827383 91827528
refGene_gn=CB4H12orf56;refGene_tx=XM_023257186.1;refGene_gn=CB4H12orf5
6;refGene_tx=XM_023257187.1;refGene_gn=CB4H12orf56;refGene_tx=XM_02325
7188.1;refGene_gn=CB4H12orf56;refGene_tx=XM_023257189.1;refGene_gn=CB4
H12orf56;refGene_tx=XM_023257190.1;refGene_gn=CB4H12orf56;refGene_tx=X
M_023257191.1
B4 91833279 91833350
refGene_gn=CB4H12orf56;refGene_tx=XM_023257186.1;refGene_gn=CB4H12orf5
6;refGene_tx=XM_023257187.1;refGene_gn=CB4H12orf56;refGene_tx=XM_02325
7188.1;refGene_gn=CB4H12orf56;refGene_tx=XM_023257189.1;refGene_gn=CB4
H12orf56;refGene_tx=XM_023257190.1;refGene_gn=CB4H12orf56;refGene_tx=X
M_023257191.1
B4 91838527 91838936
refGene_gn=CB4H12orf56;refGene_tx=XM_023257186.1;refGene_gn=CB4H12orf5
6;refGene_tx=XM_023257187.1;refGene_gn=CB4H12orf56;refGene_tx=XM_02325
7188.1;refGene_gn=CB4H12orf56;refGene_tx=XM_023257189.1;refGene_gn=CB4
H12orf56;refGene_tx=XM_023257190.1;refGene_gn=CB4H12orf56;refGene_tx=X
M_023257191.1
B4 91852039 91852112
refGene_gn=CB4H12orf56;refGene_tx=XM_023257186.1;refGene_gn=CB4H12orf5
6;refGene_tx=XM_023257187.1;refGene_gn=CB4H12orf56;refGene_tx=XM_02325
7188.1;refGene_gn=CB4H12orf56;refGene_tx=XM_023257189.1;refGene_gn=CB4
H12orf56;refGene_tx=XM_023257190.1;refGene_gn=CB4H12orf56;refGene_tx=X
M_023257191.1
B4 91861050 91861231
refGene_gn=CB4H12orf56;refGene_tx=XM_023257186.1;refGene_gn=CB4H12orf5
6;refGene_tx=XM_023257187.1;refGene_gn=CB4H12orf56;refGene_tx=XM_02325
7188.1;refGene_gn=CB4H12orf56;refGene_tx=XM_023257189.1;refGene_gn=CB4

H12orf56;refGene_tx=XM_023257190.1;refGene_gn=CB4H12orf56;refGene_tx=XM_023257191.1
B4	91887578	91888049
refGene_gn=CB4H12orf56;refGene_tx=XM_023257186.1;refGene_gn=CB4H12orf56;refGene_tx=XM_023257187.1;refGene_gn=CB4H12orf56;refGene_tx=XM_023257188.1;refGene_gn=CB4H12orf56;refGene_tx=XM_023257189.1;refGene_gn=CB4H12orf56;refGene_tx=XM_023257190.1;refGene_gn=CB4H12orf56;refGene_tx=XM_023257191.1
B4	91898774	91898834
refGene_gn=XPOT;refGene_tx=XM_011284111.3;refGene_gn=XPOT;refGene_tx=XM_011284112.3
B4	91902596	91902679
refGene_gn=XPOT;refGene_tx=XM_011284111.3;refGene_gn=XPOT;refGene_tx=XM_011284112.3
B4	91905843	91905900
refGene_gn=XPOT;refGene_tx=XM_011284111.3;refGene_gn=XPOT;refGene_tx=XM_011284112.3
B4	91906582	91906652
refGene_gn=XPOT;refGene_tx=XM_011284111.3;refGene_gn=XPOT;refGene_tx=XM_011284112.3
B4	91907382	91907601
refGene_gn=XPOT;refGene_tx=XM_011284111.3;refGene_gn=XPOT;refGene_tx=XM_011284112.3
B4	91910068	91910253
refGene_gn=XPOT;refGene_tx=XM_011284111.3;refGene_gn=XPOT;refGene_tx=XM_011284112.3
B4	91910353	91910522
refGene_gn=XPOT;refGene_tx=XM_011284111.3;refGene_gn=XPOT;refGene_tx=XM_011284112.3
B4	91912039	91912276
refGene_gn=XPOT;refGene_tx=XM_011284111.3;refGene_gn=XPOT;refGene_tx=XM_011284112.3
B4	91914403	91914442
refGene_gn=XPOT;refGene_tx=XM_011284111.3;refGene_gn=XPOT;refGene_tx=XM_011284112.3
B4	91914568	91914631
refGene_gn=XPOT;refGene_tx=XM_011284111.3;refGene_gn=XPOT;refGene_tx=XM_011284112.3
B4	91915559	91915684
refGene_gn=XPOT;refGene_tx=XM_011284111.3;refGene_gn=XPOT;refGene_tx=XM_011284112.3
B4	91916414	91916559
refGene_gn=XPOT;refGene_tx=XM_011284111.3;refGene_gn=XPOT;refGene_tx=XM_011284112.3
B4	91916709	91916829
refGene_gn=XPOT;refGene_tx=XM_011284111.3;refGene_gn=XPOT;refGene_tx=XM_011284112.3
B4	91917190	91917285
refGene_gn=XPOT;refGene_tx=XM_011284111.3;refGene_gn=XPOT;refGene_tx=XM_011284112.3

B4 91918072 91918142
refGene_gn=XPOT;refGene_tx=XM_011284111.3;refGene_gn=XPOT;refGene_tx=XM_011284112.3
B4 91920729 91920968
refGene_gn=XPOT;refGene_tx=XM_011284111.3;refGene_gn=XPOT;refGene_tx=XM_011284112.3
B4 91922414 91922700
refGene_gn=XPOT;refGene_tx=XM_011284111.3;refGene_gn=XPOT;refGene_tx=XM_011284112.3
B4 91923594 91923784
refGene_gn=XPOT;refGene_tx=XM_011284111.3;refGene_gn=XPOT;refGene_tx=XM_011284112.3
B4 91924585 91924702
refGene_gn=XPOT;refGene_tx=XM_011284111.3;refGene_gn=XPOT;refGene_tx=XM_011284112.3
B4 91924874 91924990
refGene_gn=XPOT;refGene_tx=XM_011284111.3;refGene_gn=XPOT;refGene_tx=XM_011284112.3
B4 91925668 91925716
refGene_gn=XPOT;refGene_tx=XM_011284111.3;refGene_gn=XPOT;refGene_tx=XM_011284112.3
B4 91926256 91926328
refGene_gn=XPOT;refGene_tx=XM_011284111.3;refGene_gn=XPOT;refGene_tx=XM_011284112.3
B4 91929486 91929543
refGene_gn=XPOT;refGene_tx=XM_011284111.3;refGene_gn=XPOT;refGene_tx=XM_011284112.3
B4 91932146 91932173
refGene_gn=XPOT;refGene_tx=XM_011284111.3;refGene_gn=XPOT;refGene_tx=XM_011284112.3
B4 91938595 91938682
refGene_gn=TBK1;refGene_tx=XM_023257192.1;refGene_gn=TBK1;refGene_tx=XM_023257193.1;refGene_gn=TBK1;refGene_tx=XM_023257194.1
B4 91944133 91944274
refGene_gn=TBK1;refGene_tx=XM_023257192.1;refGene_gn=TBK1;refGene_tx=XM_023257193.1;refGene_gn=TBK1;refGene_tx=XM_023257194.1
B4 91947377 91947507
refGene_gn=TBK1;refGene_tx=XM_023257192.1;refGene_gn=TBK1;refGene_tx=XM_023257193.1;refGene_gn=TBK1;refGene_tx=XM_023257194.1
B4 91950694 91950876
refGene_gn=TBK1;refGene_tx=XM_023257192.1;refGene_gn=TBK1;refGene_tx=XM_023257193.1;refGene_gn=TBK1;refGene_tx=XM_023257194.1
B4 91958185 91958346
refGene_gn=TBK1;refGene_tx=XM_023257192.1;refGene_gn=TBK1;refGene_tx=XM_023257193.1;refGene_gn=TBK1;refGene_tx=XM_023257194.1
B4 91961023 91961134
refGene_gn=TBK1;refGene_tx=XM_023257192.1;refGene_gn=TBK1;refGene_tx=XM_023257193.1;refGene_gn=TBK1;refGene_tx=XM_023257194.1
B4 91966168 91966348
refGene_gn=TBK1;refGene_tx=XM_023257192.1;refGene_gn=TBK1;refGene_tx=X

M_023257193.1;refGene_gn=TBK1;refGene_tx=XM_023257194.1
B4 91968562 91968759
refGene_gn=TBK1;refGene_tx=XM_023257192.1;refGene_gn=TBK1;refGene_tx=XM_023257193.1;refGene_gn=TBK1;refGene_tx=XM_023257194.1
B4 91969516 91969575
refGene_gn=TBK1;refGene_tx=XM_023257192.1;refGene_gn=TBK1;refGene_tx=XM_023257193.1;refGene_gn=TBK1;refGene_tx=XM_023257194.1
B4 91970431 91970523
refGene_gn=TBK1;refGene_tx=XM_023257192.1;refGene_gn=TBK1;refGene_tx=XM_023257193.1;refGene_gn=TBK1;refGene_tx=XM_023257194.1
B4 91974748 91974850
refGene_gn=TBK1;refGene_tx=XM_023257192.1;refGene_gn=TBK1;refGene_tx=XM_023257193.1;refGene_gn=TBK1;refGene_tx=XM_023257194.1
B4 91975632 91975711
refGene_gn=TBK1;refGene_tx=XM_023257192.1;refGene_gn=TBK1;refGene_tx=XM_023257193.1;refGene_gn=TBK1;refGene_tx=XM_023257194.1
B4 91976305 91976427
refGene_gn=TBK1;refGene_tx=XM_023257192.1;refGene_gn=TBK1;refGene_tx=XM_023257193.1;refGene_gn=TBK1;refGene_tx=XM_023257194.1
B4 91976526 91976603
refGene_gn=TBK1;refGene_tx=XM_023257192.1;refGene_gn=TBK1;refGene_tx=XM_023257193.1;refGene_gn=TBK1;refGene_tx=XM_023257194.1
B4 91977456 91977496
refGene_gn=TBK1;refGene_tx=XM_023257192.1;refGene_gn=TBK1;refGene_tx=XM_023257193.1;refGene_gn=TBK1;refGene_tx=XM_023257194.1
B4 91978006 91978108
refGene_gn=TBK1;refGene_tx=XM_023257192.1;refGene_gn=TBK1;refGene_tx=XM_023257193.1;refGene_gn=TBK1;refGene_tx=XM_023257194.1
B4 91978213 91978310
refGene_gn=TBK1;refGene_tx=XM_023257192.1;refGene_gn=TBK1;refGene_tx=XM_023257193.1;refGene_gn=TBK1;refGene_tx=XM_023257194.1
B4 91979187 91979294
refGene_gn=TBK1;refGene_tx=XM_023257192.1;refGene_gn=TBK1;refGene_tx=XM_023257193.1;refGene_gn=TBK1;refGene_tx=XM_023257194.1
B4 91979510 91979582
refGene_gn=TBK1;refGene_tx=XM_023257192.1;refGene_gn=TBK1;refGene_tx=XM_023257193.1;refGene_gn=TBK1;refGene_tx=XM_023257194.1
B4 91985646 91986711
refGene_gn=LOC101087675;refGene_tx=XM_003988997.3
B4 91987683 91987735
refGene_gn=TBK1;refGene_tx=XM_023257192.1;refGene_gn=TBK1;refGene_tx=XM_023257193.1
B4 91990580 91990590 refGene_gn=TBK1;refGene_tx=XM_023257194.1
B4 92073978 92074089 refGene_gn=RASSF3;refGene_tx=XM_011284114.3
B4 92135060 92135168 refGene_gn=RASSF3;refGene_tx=XM_011284114.3
B4 92137252 92137490 refGene_gn=RASSF3;refGene_tx=XM_011284114.3
B4 92139488 92139598 refGene_gn=RASSF3;refGene_tx=XM_011284114.3
B4 92143687 92143837 refGene_gn=RASSF3;refGene_tx=XM_011284114.3
B4 92160808 92160887 refGene_gn=GNS;refGene_tx=XM_023257195.1
B4 92163904 92164065 refGene_gn=GNS;refGene_tx=XM_023257195.1

| | | | |
|---|---|---|---|
| B4 | 92165854 | 92165965 | refGene_gn=GNS;refGene_tx=XM_023257195.1 |
| B4 | 92166878 | 92166986 | refGene_gn=GNS;refGene_tx=XM_023257195.1 |
| B4 | 92188258 | 92188360 | refGene_gn=GNS;refGene_tx=XM_023257195.1 |
| B4 | 92195261 | 92195365 | refGene_gn=GNS;refGene_tx=XM_023257195.1 |
| B4 | 92197244 | 92197363 | refGene_gn=GNS;refGene_tx=XM_023257195.1 |
| B4 | 92198614 | 92198697 | refGene_gn=GNS;refGene_tx=XM_023257195.1 |
| B4 | 92200544 | 92200712 | refGene_gn=GNS;refGene_tx=XM_023257195.1 |
| B4 | 92202361 | 92202460 | refGene_gn=GNS;refGene_tx=XM_023257195.1 |
| B4 | 92204751 | 92204817 | refGene_gn=GNS;refGene_tx=XM_023257195.1 |
| B4 | 92208676 | 92208883 | refGene_gn=GNS;refGene_tx=XM_023257195.1 |
| B4 | 92211653 | 92211713 | refGene_gn=GNS;refGene_tx=XM_023257195.1 |
| B4 | 92217146 | 92217341 | refGene_gn=GNS;refGene_tx=XM_023257195.1 |
| B4 | 92230286 | 92230740 | refGene_gn=TBC1D30;refGene_tx=XM_023257196.1 |
| B4 | 92235372 | 92235537 | refGene_gn=TBC1D30;refGene_tx=XM_023257196.1 |
| B4 | 92278696 | 92278758 | refGene_gn=TBC1D30;refGene_tx=XM_023257196.1 |
| B4 | 92279285 | 92279351 | refGene_gn=TBC1D30;refGene_tx=XM_023257196.1 |
| B4 | 92281630 | 92281642 | refGene_gn=TBC1D30;refGene_tx=XM_023257196.1 |
| B4 | 92281643 | 92281757 | refGene_gn=TBC1D30;refGene_tx=XM_023257196.1 |
| B4 | 92284438 | 92284624 | refGene_gn=TBC1D30;refGene_tx=XM_023257196.1 |
| B4 | 92290033 | 92290202 | refGene_gn=TBC1D30;refGene_tx=XM_023257196.1 |
| B4 | 92292114 | 92292283 | refGene_gn=TBC1D30;refGene_tx=XM_023257196.1 |
| B4 | 92296364 | 92296470 | refGene_gn=TBC1D30;refGene_tx=XM_023257196.1 |
| B4 | 92305212 | 92305325 | refGene_gn=TBC1D30;refGene_tx=XM_023257196.1 |
| B4 | 92307080 | 92307220 | refGene_gn=TBC1D30;refGene_tx=XM_023257196.1 |
| B4 | 92309588 | 92309795 | refGene_gn=TBC1D30;refGene_tx=XM_023257196.1 |
| B4 | 92312964 | 92313737 | refGene_gn=TBC1D30;refGene_tx=XM_023257196.1 |
| B4 | 92456275 | 92456397 | refGene_gn=WIF1;refGene_tx=XM_003989002.5 |
| B4 | 92459011 | 92459107 | refGene_gn=WIF1;refGene_tx=XM_003989002.5 |
| B4 | 92460135 | 92460231 | refGene_gn=WIF1;refGene_tx=XM_003989002.5 |
| B4 | 92466142 | 92466238 | refGene_gn=WIF1;refGene_tx=XM_003989002.5 |
| B4 | 92470186 | 92470282 | refGene_gn=WIF1;refGene_tx=XM_003989002.5 |
| B4 | 92471834 | 92471930 | refGene_gn=WIF1;refGene_tx=XM_003989002.5 |
| B4 | 92473238 | 92473379 | refGene_gn=WIF1;refGene_tx=XM_003989002.5 |
| B4 | 92482635 | 92482744 | refGene_gn=WIF1;refGene_tx=XM_003989002.5 |
| B4 | 92523680 | 92523820 | refGene_gn=WIF1;refGene_tx=XM_003989002.5 |
| B4 | 92524304 | 92524452 | refGene_gn=WIF1;refGene_tx=XM_003989002.5 |
| B4 | 92577111 | 92578645 | refGene_gn=LEMD3;refGene_tx=XM_023257197.1 |
| B4 | 92605903 | 92605941 | refGene_gn=LEMD3;refGene_tx=XM_023257197.1 |
| B4 | 92609856 | 92609923 | refGene_gn=LEMD3;refGene_tx=XM_023257197.1 |
| B4 | 92612102 | 92612170 | refGene_gn=LEMD3;refGene_tx=XM_023257197.1 |
| B4 | 92628235 | 92628315 | refGene_gn=LEMD3;refGene_tx=XM_023257197.1 |
| B4 | 92628400 | 92628546 | refGene_gn=LEMD3;refGene_tx=XM_023257197.1 |
| B4 | 92629664 | 92629766 | refGene_gn=LEMD3;refGene_tx=XM_023257197.1 |
| B4 | 92629866 | 92629969 | refGene_gn=LEMD3;refGene_tx=XM_023257197.1 |
| B4 | 92630663 | 92630842 | refGene_gn=LEMD3;refGene_tx=XM_023257197.1 |
| B4 | 92636569 | 92636651 | refGene_gn=LEMD3;refGene_tx=XM_023257197.1 |
| B4 | 92638193 | 92638299 | refGene_gn=LEMD3;refGene_tx=XM_023257197.1 |
| B4 | 92638416 | 92638495 | refGene_gn=LEMD3;refGene_tx=XM_023257197.1 |
| B4 | 92638720 | 92638884 | refGene_gn=LEMD3;refGene_tx=XM_023257197.1 |
| B4 | 92668885 | 92669297 | refGene_gn=MSRB3;refGene_tx=XM_023257198.1 |

B4	92701437	92701513
refGene_gn=MSRB3;refGene_tx=XM_023257199.1;refGene_gn=MSRB3;refGene_tx=XM_023257200.1;refGene_gn=MSRB3;refGene_tx=XM_023257201.1
B4	92719267	92719376
refGene_gn=MSRB3;refGene_tx=XM_023257198.1;refGene_gn=MSRB3;refGene_tx=XM_023257199.1;refGene_gn=MSRB3;refGene_tx=XM_023257200.1;refGene_gn=MSRB3;refGene_tx=XM_023257201.1
B4	92721007	92721085
refGene_gn=MSRB3;refGene_tx=XM_023257198.1;refGene_gn=MSRB3;refGene_tx=XM_023257199.1;refGene_gn=MSRB3;refGene_tx=XM_023257200.1;refGene_gn=MSRB3;refGene_tx=XM_023257201.1
B4	92753446	92753475
refGene_gn=MSRB3;refGene_tx=XM_023257198.1;refGene_gn=MSRB3;refGene_tx=XM_023257199.1;refGene_gn=MSRB3;refGene_tx=XM_023257200.1;refGene_gn=MSRB3;refGene_tx=XM_023257201.1
B4	92832160	92832258
refGene_gn=MSRB3;refGene_tx=XM_023257198.1;refGene_gn=MSRB3;refGene_tx=XM_023257199.1;refGene_gn=MSRB3;refGene_tx=XM_023257200.1;refGene_gn=MSRB3;refGene_tx=XM_023257201.1
B4	92840382	92840550
refGene_gn=MSRB3;refGene_tx=XM_023257198.1;refGene_gn=MSRB3;refGene_tx=XM_023257199.1;refGene_gn=MSRB3;refGene_tx=XM_023257200.1;refGene_gn=MSRB3;refGene_tx=XM_023257201.1
B4	93165668	93165779
refGene_gn=HMGA2;refGene_tx=XM_023257202.1;refGene_gn=HMGA2;refGene_tx=XM_023257203.1
B4	93168431	93168518
refGene_gn=HMGA2;refGene_tx=XM_023257202.1;refGene_gn=HMGA2;refGene_tx=XM_023257203.1
B4	93178700	93178751	refGene_gn=HMGA2;refGene_tx=XM_023257202.1
B4	93292777	93292810
refGene_gn=HMGA2;refGene_tx=XM_023257202.1;refGene_gn=HMGA2;refGene_tx=XM_023257203.1
B4	93304404	93304452
refGene_gn=HMGA2;refGene_tx=XM_023257202.1;refGene_gn=HMGA2;refGene_tx=XM_023257203.1
B4	93460622	93460801
refGene_gn=LLPH;refGene_tx=XM_003989004.4;refGene_gn=LLPH;refGene_tx=XM_019835179.2
B4	93466122	93466327	refGene_gn=LLPH;refGene_tx=XM_003989004.4
B4	93468443	93468705	refGene_gn=LLPH;refGene_tx=XM_019835179.2
B4	93474107	93474314	refGene_gn=TMBIM4;refGene_tx=XM_003989005.5
B4	93474872	93474918	refGene_gn=TMBIM4;refGene_tx=XM_003989005.5
B4	93481115	93481233	refGene_gn=TMBIM4;refGene_tx=XM_003989005.5
B4	93484122	93484156	refGene_gn=TMBIM4;refGene_tx=XM_003989005.5
B4	93488303	93488409	refGene_gn=TMBIM4;refGene_tx=XM_003989005.5
B4	93489797	93489864	refGene_gn=TMBIM4;refGene_tx=XM_003989005.5
B4	93489866	93489905	refGene_gn=TMBIM4;refGene_tx=XM_003989005.5
B4	93493407	93493504	refGene_gn=TMBIM4;refGene_tx=XM_003989005.5
B4	93506656	93506808	refGene_gn=IRAK3;refGene_tx=XM_023257205.1

B4 93523187 93523366
refGene_gn=IRAK3;refGene_tx=XM_023257204.1;refGene_gn=IRAK3;refGene_tx=XM_023257205.1
B4 93528498 93528557
refGene_gn=IRAK3;refGene_tx=XM_023257204.1;refGene_gn=IRAK3;refGene_tx=XM_023257205.1
B4 93529181 93529236
refGene_gn=IRAK3;refGene_tx=XM_023257204.1;refGene_gn=IRAK3;refGene_tx=XM_023257205.1
B4 93530523 93530675
refGene_gn=IRAK3;refGene_tx=XM_023257204.1;refGene_gn=IRAK3;refGene_tx=XM_023257205.1
B4 93534495 93534560
refGene_gn=IRAK3;refGene_tx=XM_023257204.1;refGene_gn=IRAK3;refGene_tx=XM_023257205.1
B4 93540860 93540975
refGene_gn=IRAK3;refGene_tx=XM_023257204.1;refGene_gn=IRAK3;refGene_tx=XM_023257205.1
B4 93541788 93541907
refGene_gn=IRAK3;refGene_tx=XM_023257204.1;refGene_gn=IRAK3;refGene_tx=XM_023257205.1
B4 93555502 93555701
refGene_gn=IRAK3;refGene_tx=XM_023257204.1;refGene_gn=IRAK3;refGene_tx=XM_023257205.1
B4 93555963 93556026
refGene_gn=IRAK3;refGene_tx=XM_023257204.1;refGene_gn=IRAK3;refGene_tx=XM_023257205.1
B4 93556120 93556285
refGene_gn=IRAK3;refGene_tx=XM_023257204.1;refGene_gn=IRAK3;refGene_tx=XM_023257205.1
B4 93558074 93558590
refGene_gn=IRAK3;refGene_tx=XM_023257204.1;refGene_gn=IRAK3;refGene_tx=XM_023257205.1
B4 93598530 93598714 refGene_gn=HELB;refGene_tx=XM_003989007.5
B4 93600180 93600591
refGene_gn=HELB;refGene_tx=XM_003989007.5;refGene_gn=HELB;refGene_tx=XM_019835183.2
B4 93601505 93601675
refGene_gn=HELB;refGene_tx=XM_003989007.5;refGene_gn=HELB;refGene_tx=XM_019835183.2
B4 93603958 93604837
refGene_gn=HELB;refGene_tx=XM_003989007.5;refGene_gn=HELB;refGene_tx=XM_019835183.2
B4 93608774 93608952
refGene_gn=HELB;refGene_tx=XM_003989007.5;refGene_gn=HELB;refGene_tx=XM_019835183.2
B4 93610371 93610513
refGene_gn=HELB;refGene_tx=XM_003989007.5;refGene_gn=HELB;refGene_tx=XM_019835183.2
B4 93613004 93613159

refGene_gn=HELB;refGene_tx=XM_003989007.5;refGene_gn=HELB;refGene_tx=XM_019835183.2
B4	93616605	93616687
refGene_gn=HELB;refGene_tx=XM_003989007.5;refGene_gn=HELB;refGene_tx=XM_019835183.2
B4	93617180	93617240
refGene_gn=HELB;refGene_tx=XM_003989007.5;refGene_gn=HELB;refGene_tx=XM_019835183.2
B4	93619517	93619746
refGene_gn=HELB;refGene_tx=XM_003989007.5;refGene_gn=HELB;refGene_tx=XM_019835183.2
B4	93620539	93620683
refGene_gn=HELB;refGene_tx=XM_003989007.5;refGene_gn=HELB;refGene_tx=XM_019835183.2
B4	93626195	93626681
refGene_gn=HELB;refGene_tx=XM_003989007.5;refGene_gn=HELB;refGene_tx=XM_019835183.2
B4	93629887	93629992
refGene_gn=HELB;refGene_tx=XM_003989007.5;refGene_gn=HELB;refGene_tx=XM_019835183.2
B4	93635092	93635320
refGene_gn=GRIP1;refGene_tx=XM_019835185.2;refGene_gn=GRIP1;refGene_tx=XM_019835187.2;refGene_gn=GRIP1;refGene_tx=XM_019835188.2;refGene_gn=GRIP1;refGene_tx=XM_023257206.1;refGene_gn=GRIP1;refGene_tx=XM_023257207.1;refGene_gn=GRIP1;refGene_tx=XM_023257208.1;refGene_gn=GRIP1;refGene_tx=XM_023257209.1;refGene_gn=GRIP1;refGene_tx=XM_023257210.1;refGene_gn=GRIP1;refGene_tx=XM_023257211.1;refGene_gn=GRIP1;refGene_tx=XM_023257212.1;refGene_gn=GRIP1;refGene_tx=XM_023257213.1;refGene_gn=GRIP1;refGene_tx=XM_023257214.1
B4	93637281	93637428
refGene_gn=GRIP1;refGene_tx=XM_019835185.2;refGene_gn=GRIP1;refGene_tx=XM_019835187.2;refGene_gn=GRIP1;refGene_tx=XM_019835188.2;refGene_gn=GRIP1;refGene_tx=XM_023257206.1;refGene_gn=GRIP1;refGene_tx=XM_023257207.1;refGene_gn=GRIP1;refGene_tx=XM_023257208.1;refGene_gn=GRIP1;refGene_tx=XM_023257209.1;refGene_gn=GRIP1;refGene_tx=XM_023257210.1;refGene_gn=GRIP1;refGene_tx=XM_023257211.1;refGene_gn=GRIP1;refGene_tx=XM_023257212.1;refGene_gn=GRIP1;refGene_tx=XM_023257213.1;refGene_gn=GRIP1;refGene_tx=XM_023257214.1
B4	93654698	93654932
refGene_gn=GRIP1;refGene_tx=XM_019835185.2;refGene_gn=GRIP1;refGene_tx=XM_019835187.2;refGene_gn=GRIP1;refGene_tx=XM_019835188.2;refGene_gn=GRIP1;refGene_tx=XM_023257206.1;refGene_gn=GRIP1;refGene_tx=XM_023257207.1;refGene_gn=GRIP1;refGene_tx=XM_023257208.1;refGene_gn=GRIP1;refGene_tx=XM_023257209.1;refGene_gn=GRIP1;refGene_tx=XM_023257210.1;refGene_gn=GRIP1;refGene_tx=XM_023257211.1;refGene_gn=GRIP1;refGene_tx=XM_023257212.1;refGene_gn=GRIP1;refGene_tx=XM_023257214.1
B4	93659747	93659792
refGene_gn=GRIP1;refGene_tx=XM_019835185.2;refGene_gn=GRIP1;refGene_tx=XM_019835187.2;refGene_gn=GRIP1;refGene_tx=XM_019835188.2;refGene_gn=GRIP1;refGene_tx=XM_023257206.1;refGene_gn=GRIP1;refGene_tx=XM_0232572

08.1;refGene_gn=GRIP1;refGene_tx=XM_023257209.1;refGene_gn=GRIP1;refGene_tx=XM_023257210.1;refGene_gn=GRIP1;refGene_tx=XM_023257211.1;refGene_gn=GRIP1;refGene_tx=XM_023257214.1
B4 93659911 93660023
refGene_gn=GRIP1;refGene_tx=XM_019835185.2;refGene_gn=GRIP1;refGene_tx=XM_019835187.2;refGene_gn=GRIP1;refGene_tx=XM_019835188.2;refGene_gn=GRIP1;refGene_tx=XM_023257206.1;refGene_gn=GRIP1;refGene_tx=XM_023257207.1;refGene_gn=GRIP1;refGene_tx=XM_023257208.1;refGene_gn=GRIP1;refGene_tx=XM_023257209.1;refGene_gn=GRIP1;refGene_tx=XM_023257210.1;refGene_gn=GRIP1;refGene_tx=XM_023257211.1;refGene_gn=GRIP1;refGene_tx=XM_023257212.1;refGene_gn=GRIP1;refGene_tx=XM_023257213.1;refGene_gn=GRIP1;refGene_tx=XM_023257214.1
B4 93660998 93661155
refGene_gn=GRIP1;refGene_tx=XM_019835185.2;refGene_gn=GRIP1;refGene_tx=XM_019835187.2;refGene_gn=GRIP1;refGene_tx=XM_019835188.2;refGene_gn=GRIP1;refGene_tx=XM_023257206.1;refGene_gn=GRIP1;refGene_tx=XM_023257207.1;refGene_gn=GRIP1;refGene_tx=XM_023257208.1;refGene_gn=GRIP1;refGene_tx=XM_023257209.1;refGene_gn=GRIP1;refGene_tx=XM_023257210.1;refGene_gn=GRIP1;refGene_tx=XM_023257211.1;refGene_gn=GRIP1;refGene_tx=XM_023257212.1;refGene_gn=GRIP1;refGene_tx=XM_023257213.1;refGene_gn=GRIP1;refGene_tx=XM_023257214.1
B4 93664415 93664522 refGene_gn=GRIP1;refGene_tx=XM_023257215.1
B4 93675218 93675413
refGene_gn=GRIP1;refGene_tx=XM_019835185.2;refGene_gn=GRIP1;refGene_tx=XM_019835187.2;refGene_gn=GRIP1;refGene_tx=XM_019835188.2;refGene_gn=GRIP1;refGene_tx=XM_023257206.1;refGene_gn=GRIP1;refGene_tx=XM_023257207.1;refGene_gn=GRIP1;refGene_tx=XM_023257208.1;refGene_gn=GRIP1;refGene_tx=XM_023257209.1;refGene_gn=GRIP1;refGene_tx=XM_023257210.1;refGene_gn=GRIP1;refGene_tx=XM_023257211.1;refGene_gn=GRIP1;refGene_tx=XM_023257212.1;refGene_gn=GRIP1;refGene_tx=XM_023257213.1;refGene_gn=GRIP1;refGene_tx=XM_023257215.1
B4 93675567 93675707
refGene_gn=GRIP1;refGene_tx=XM_019835185.2;refGene_gn=GRIP1;refGene_tx=XM_019835187.2;refGene_gn=GRIP1;refGene_tx=XM_019835188.2;refGene_gn=GRIP1;refGene_tx=XM_023257206.1;refGene_gn=GRIP1;refGene_tx=XM_023257207.1;refGene_gn=GRIP1;refGene_tx=XM_023257208.1;refGene_gn=GRIP1;refGene_tx=XM_023257209.1;refGene_gn=GRIP1;refGene_tx=XM_023257210.1;refGene_gn=GRIP1;refGene_tx=XM_023257211.1;refGene_gn=GRIP1;refGene_tx=XM_023257212.1;refGene_gn=GRIP1;refGene_tx=XM_023257213.1;refGene_gn=GRIP1;refGene_tx=XM_023257215.1
B4 93677853 93677998
refGene_gn=GRIP1;refGene_tx=XM_019835185.2;refGene_gn=GRIP1;refGene_tx=XM_019835187.2;refGene_gn=GRIP1;refGene_tx=XM_019835188.2;refGene_gn=GRIP1;refGene_tx=XM_023257206.1;refGene_gn=GRIP1;refGene_tx=XM_023257207.1;refGene_gn=GRIP1;refGene_tx=XM_023257208.1;refGene_gn=GRIP1;refGene_tx=XM_023257209.1;refGene_gn=GRIP1;refGene_tx=XM_023257210.1;refGene_gn=GRIP1;refGene_tx=XM_023257211.1;refGene_gn=GRIP1;refGene_tx=XM_023257212.1;refGene_gn=GRIP1;refGene_tx=XM_023257213.1;refGene_gn=GRIP1;refGene_tx=XM_023257215.1
B4 93691121 93691267

refGene_gn=GRIP1;refGene_tx=XM_019835185.2;refGene_gn=GRIP1;refGene_tx=XM_019835187.2;refGene_gn=GRIP1;refGene_tx=XM_019835188.2;refGene_gn=GRIP1;refGene_tx=XM_023257206.1;refGene_gn=GRIP1;refGene_tx=XM_023257207.1;refGene_gn=GRIP1;refGene_tx=XM_023257208.1;refGene_gn=GRIP1;refGene_tx=XM_023257209.1;refGene_gn=GRIP1;refGene_tx=XM_023257210.1;refGene_gn=GRIP1;refGene_tx=XM_023257211.1;refGene_gn=GRIP1;refGene_tx=XM_023257212.1;refGene_gn=GRIP1;refGene_tx=XM_023257213.1;refGene_gn=GRIP1;refGene_tx=XM_023257214.1;refGene_gn=GRIP1;refGene_tx=XM_023257215.1
B4 93693019 93693089
refGene_gn=GRIP1;refGene_tx=XM_019835185.2;refGene_gn=GRIP1;refGene_tx=XM_019835187.2;refGene_gn=GRIP1;refGene_tx=XM_019835188.2;refGene_gn=GRIP1;refGene_tx=XM_023257206.1;refGene_gn=GRIP1;refGene_tx=XM_023257207.1;refGene_gn=GRIP1;refGene_tx=XM_023257208.1;refGene_gn=GRIP1;refGene_tx=XM_023257209.1;refGene_gn=GRIP1;refGene_tx=XM_023257210.1;refGene_gn=GRIP1;refGene_tx=XM_023257211.1;refGene_gn=GRIP1;refGene_tx=XM_023257212.1;refGene_gn=GRIP1;refGene_tx=XM_023257213.1;refGene_gn=GRIP1;refGene_tx=XM_023257214.1;refGene_gn=GRIP1;refGene_tx=XM_023257215.1
B4 93705910 93705991
refGene_gn=GRIP1;refGene_tx=XM_019835185.2;refGene_gn=GRIP1;refGene_tx=XM_019835187.2;refGene_gn=GRIP1;refGene_tx=XM_019835188.2;refGene_gn=GRIP1;refGene_tx=XM_023257206.1;refGene_gn=GRIP1;refGene_tx=XM_023257207.1;refGene_gn=GRIP1;refGene_tx=XM_023257208.1;refGene_gn=GRIP1;refGene_tx=XM_023257209.1;refGene_gn=GRIP1;refGene_tx=XM_023257210.1;refGene_gn=GRIP1;refGene_tx=XM_023257211.1;refGene_gn=GRIP1;refGene_tx=XM_023257212.1;refGene_gn=GRIP1;refGene_tx=XM_023257213.1;refGene_gn=GRIP1;refGene_tx=XM_023257214.1;refGene_gn=GRIP1;refGene_tx=XM_023257215.1
B4 93718804 93718950
refGene_gn=GRIP1;refGene_tx=XM_019835185.2;refGene_gn=GRIP1;refGene_tx=XM_019835187.2;refGene_gn=GRIP1;refGene_tx=XM_019835188.2;refGene_gn=GRIP1;refGene_tx=XM_023257206.1;refGene_gn=GRIP1;refGene_tx=XM_023257207.1;refGene_gn=GRIP1;refGene_tx=XM_023257208.1;refGene_gn=GRIP1;refGene_tx=XM_023257209.1;refGene_gn=GRIP1;refGene_tx=XM_023257210.1;refGene_gn=GRIP1;refGene_tx=XM_023257211.1;refGene_gn=GRIP1;refGene_tx=XM_023257212.1;refGene_gn=GRIP1;refGene_tx=XM_023257213.1;refGene_gn=GRIP1;refGene_tx=XM_023257214.1;refGene_gn=GRIP1;refGene_tx=XM_023257215.1
B4 93719532 93719719
refGene_gn=GRIP1;refGene_tx=XM_019835185.2;refGene_gn=GRIP1;refGene_tx=XM_019835187.2;refGene_gn=GRIP1;refGene_tx=XM_019835188.2;refGene_gn=GRIP1;refGene_tx=XM_023257206.1;refGene_gn=GRIP1;refGene_tx=XM_023257207.1;refGene_gn=GRIP1;refGene_tx=XM_023257208.1;refGene_gn=GRIP1;refGene_tx=XM_023257209.1;refGene_gn=GRIP1;refGene_tx=XM_023257210.1;refGene_gn=GRIP1;refGene_tx=XM_023257211.1;refGene_gn=GRIP1;refGene_tx=XM_023257212.1;refGene_gn=GRIP1;refGene_tx=XM_023257213.1;refGene_gn=GRIP1;refGene_tx=XM_023257214.1;refGene_gn=GRIP1;refGene_tx=XM_023257215.1
B4 93730542 93730698
refGene_gn=GRIP1;refGene_tx=XM_019835185.2;refGene_gn=GRIP1;refGene_tx=XM_019835187.2;refGene_gn=GRIP1;refGene_tx=XM_019835188.2;refGene_gn=GRIP1;refGene_tx=XM_023257206.1;refGene_gn=GRIP1;refGene_tx=XM_023257207.1;refGene_gn=GRIP1;refGene_tx=XM_023257208.1;refGene_gn=GRIP1;refGene_tx=XM_023257209.1;refGene_gn=GRIP1;refGene_tx=XM_023257210.1;refGen

e_gn=GRIP1;refGene_tx=XM_023257211.1;refGene_gn=GRIP1;refGene_tx=XM_02
3257212.1;refGene_gn=GRIP1;refGene_tx=XM_023257213.1;refGene_gn=GRIP1;
refGene_tx=XM_023257214.1;refGene_gn=GRIP1;refGene_tx=XM_023257215.1
B4 93731374 93731530
refGene_gn=GRIP1;refGene_tx=XM_019835185.2;refGene_gn=GRIP1;refGene_tx
=XM_019835187.2;refGene_gn=GRIP1;refGene_tx=XM_019835188.2;refGene_gn=
GRIP1;refGene_tx=XM_023257206.1;refGene_gn=GRIP1;refGene_tx=XM_0232572
07.1;refGene_gn=GRIP1;refGene_tx=XM_023257208.1;refGene_gn=GRIP1;refGe
ne_tx=XM_023257209.1;refGene_gn=GRIP1;refGene_tx=XM_023257213.1;refGen
e_gn=GRIP1;refGene_tx=XM_023257214.1;refGene_gn=GRIP1;refGene_tx=XM_02
3257215.1
B4 93737006 93737176
refGene_gn=GRIP1;refGene_tx=XM_019835185.2;refGene_gn=GRIP1;refGene_tx
=XM_019835187.2;refGene_gn=GRIP1;refGene_tx=XM_019835188.2;refGene_gn=
GRIP1;refGene_tx=XM_023257206.1;refGene_gn=GRIP1;refGene_tx=XM_0232572
07.1;refGene_gn=GRIP1;refGene_tx=XM_023257208.1;refGene_gn=GRIP1;refGe
ne_tx=XM_023257209.1;refGene_gn=GRIP1;refGene_tx=XM_023257210.1;refGen
e_gn=GRIP1;refGene_tx=XM_023257211.1;refGene_gn=GRIP1;refGene_tx=XM_02
3257212.1;refGene_gn=GRIP1;refGene_tx=XM_023257213.1;refGene_gn=GRIP1;
refGene_tx=XM_023257214.1;refGene_gn=GRIP1;refGene_tx=XM_023257215.1
B4 93739059 93739207
refGene_gn=GRIP1;refGene_tx=XM_019835185.2;refGene_gn=GRIP1;refGene_tx
=XM_019835187.2;refGene_gn=GRIP1;refGene_tx=XM_019835188.2;refGene_gn=
GRIP1;refGene_tx=XM_023257206.1;refGene_gn=GRIP1;refGene_tx=XM_0232572
07.1;refGene_gn=GRIP1;refGene_tx=XM_023257208.1;refGene_gn=GRIP1;refGe
ne_tx=XM_023257209.1;refGene_gn=GRIP1;refGene_tx=XM_023257210.1;refGen
e_gn=GRIP1;refGene_tx=XM_023257211.1;refGene_gn=GRIP1;refGene_tx=XM_02
3257212.1;refGene_gn=GRIP1;refGene_tx=XM_023257213.1;refGene_gn=GRIP1;
refGene_tx=XM_023257214.1;refGene_gn=GRIP1;refGene_tx=XM_023257215.1
B4 93773079 93773225
refGene_gn=GRIP1;refGene_tx=XM_019835185.2;refGene_gn=GRIP1;refGene_tx
=XM_019835187.2;refGene_gn=GRIP1;refGene_tx=XM_019835188.2;refGene_gn=
GRIP1;refGene_tx=XM_023257206.1;refGene_gn=GRIP1;refGene_tx=XM_0232572
07.1;refGene_gn=GRIP1;refGene_tx=XM_023257208.1;refGene_gn=GRIP1;refGe
ne_tx=XM_023257209.1;refGene_gn=GRIP1;refGene_tx=XM_023257210.1;refGen
e_gn=GRIP1;refGene_tx=XM_023257211.1;refGene_gn=GRIP1;refGene_tx=XM_02
3257212.1;refGene_gn=GRIP1;refGene_tx=XM_023257213.1;refGene_gn=GRIP1;
refGene_tx=XM_023257214.1;refGene_gn=GRIP1;refGene_tx=XM_023257215.1
B4 93775389 93775465
refGene_gn=GRIP1;refGene_tx=XM_019835185.2;refGene_gn=GRIP1;refGene_tx
=XM_019835187.2;refGene_gn=GRIP1;refGene_tx=XM_019835188.2;refGene_gn=
GRIP1;refGene_tx=XM_023257206.1;refGene_gn=GRIP1;refGene_tx=XM_0232572
07.1;refGene_gn=GRIP1;refGene_tx=XM_023257208.1;refGene_gn=GRIP1;refGe
ne_tx=XM_023257209.1;refGene_gn=GRIP1;refGene_tx=XM_023257210.1;refGen
e_gn=GRIP1;refGene_tx=XM_023257211.1;refGene_gn=GRIP1;refGene_tx=XM_02
3257212.1;refGene_gn=GRIP1;refGene_tx=XM_023257213.1;refGene_gn=GRIP1;
refGene_tx=XM_023257214.1;refGene_gn=GRIP1;refGene_tx=XM_023257215.1
B4 93779944 93780028
refGene_gn=GRIP1;refGene_tx=XM_019835185.2;refGene_gn=GRIP1;refGene_tx
=XM_019835187.2;refGene_gn=GRIP1;refGene_tx=XM_019835188.2;refGene_gn=

GRIP1;refGene_tx=XM_023257206.1;refGene_gn=GRIP1;refGene_tx=XM_0232572
07.1;refGene_gn=GRIP1;refGene_tx=XM_023257208.1;refGene_gn=GRIP1;refGe
ne_tx=XM_023257209.1;refGene_gn=GRIP1;refGene_tx=XM_023257210.1;refGen
e_gn=GRIP1;refGene_tx=XM_023257211.1;refGene_gn=GRIP1;refGene_tx=XM_02
3257212.1;refGene_gn=GRIP1;refGene_tx=XM_023257213.1;refGene_gn=GRIP1;
refGene_tx=XM_023257214.1;refGene_gn=GRIP1;refGene_tx=XM_023257215.1
B4 93788234 93788380
refGene_gn=GRIP1;refGene_tx=XM_019835185.2;refGene_gn=GRIP1;refGene_tx
=XM_019835187.2;refGene_gn=GRIP1;refGene_tx=XM_019835188.2;refGene_gn=
GRIP1;refGene_tx=XM_023257206.1;refGene_gn=GRIP1;refGene_tx=XM_0232572
07.1;refGene_gn=GRIP1;refGene_tx=XM_023257208.1;refGene_gn=GRIP1;refGe
ne_tx=XM_023257209.1;refGene_gn=GRIP1;refGene_tx=XM_023257210.1;refGen
e_gn=GRIP1;refGene_tx=XM_023257211.1;refGene_gn=GRIP1;refGene_tx=XM_02
3257212.1;refGene_gn=GRIP1;refGene_tx=XM_023257213.1;refGene_gn=GRIP1;
refGene_tx=XM_023257214.1;refGene_gn=GRIP1;refGene_tx=XM_023257215.1
B4 93791729 93791865
refGene_gn=GRIP1;refGene_tx=XM_019835185.2;refGene_gn=GRIP1;refGene_tx
=XM_019835187.2;refGene_gn=GRIP1;refGene_tx=XM_019835188.2;refGene_gn=
GRIP1;refGene_tx=XM_023257206.1;refGene_gn=GRIP1;refGene_tx=XM_0232572
07.1;refGene_gn=GRIP1;refGene_tx=XM_023257208.1;refGene_gn=GRIP1;refGe
ne_tx=XM_023257209.1;refGene_gn=GRIP1;refGene_tx=XM_023257210.1;refGen
e_gn=GRIP1;refGene_tx=XM_023257211.1;refGene_gn=GRIP1;refGene_tx=XM_02
3257212.1;refGene_gn=GRIP1;refGene_tx=XM_023257213.1;refGene_gn=GRIP1;
refGene_tx=XM_023257214.1;refGene_gn=GRIP1;refGene_tx=XM_023257215.1
B4 93825593 93825674
refGene_gn=GRIP1;refGene_tx=XM_019835185.2;refGene_gn=GRIP1;refGene_tx
=XM_019835187.2;refGene_gn=GRIP1;refGene_tx=XM_023257206.1;refGene_gn=
GRIP1;refGene_tx=XM_023257207.1;refGene_gn=GRIP1;refGene_tx=XM_0232572
08.1;refGene_gn=GRIP1;refGene_tx=XM_023257210.1;refGene_gn=GRIP1;refGe
ne_tx=XM_023257211.1;refGene_gn=GRIP1;refGene_tx=XM_023257212.1;refGen
e_gn=GRIP1;refGene_tx=XM_023257213.1;refGene_gn=GRIP1;refGene_tx=XM_02
3257214.1;refGene_gn=GRIP1;refGene_tx=XM_023257215.1
B4 93913936 93913991 refGene_gn=GRIP1;refGene_tx=XM_019835187.2
B4 94044576 94044709
refGene_gn=GRIP1;refGene_tx=XM_019835185.2;refGene_gn=GRIP1;refGene_tx
=XM_019835188.2;refGene_gn=GRIP1;refGene_tx=XM_023257206.1;refGene_gn=
GRIP1;refGene_tx=XM_023257207.1;refGene_gn=GRIP1;refGene_tx=XM_0232572
10.1;refGene_gn=GRIP1;refGene_tx=XM_023257211.1;refGene_gn=GRIP1;refGe
ne_tx=XM_023257212.1;refGene_gn=GRIP1;refGene_tx=XM_023257213.1;refGen
e_gn=GRIP1;refGene_tx=XM_023257214.1;refGene_gn=GRIP1;refGene_tx=XM_02
3257215.1
B4 94326086 94326144
refGene_gn=GRIP1;refGene_tx=XM_023257208.1;refGene_gn=GRIP1;refGene_tx
=XM_023257209.1
B4 94525394 94525462 refGene_gn=CAND1;refGene_tx=XM_003989017.5
B4 94536731 94536875 refGene_gn=CAND1;refGene_tx=XM_003989017.5
B4 94546630 94546785 refGene_gn=CAND1;refGene_tx=XM_003989017.5
B4 94548566 94548690
refGene_gn=CAND1;refGene_tx=XM_003989017.5;refGene_gn=CAND1;refGene_tx
=XM_019835191.1

B4 94550294 94550551
refGene_gn=CAND1;refGene_tx=XM_003989017.5;refGene_gn=CAND1;refGene_tx
=XM_019835191.1
B4 94550630 94550736
refGene_gn=CAND1;refGene_tx=XM_003989017.5;refGene_gn=CAND1;refGene_tx
=XM_019835191.1
B4 94552601 94552747
refGene_gn=CAND1;refGene_tx=XM_003989017.5;refGene_gn=CAND1;refGene_tx
=XM_019835191.1
B4 94555869 94556162
refGene_gn=CAND1;refGene_tx=XM_003989017.5;refGene_gn=CAND1;refGene_tx
=XM_019835191.1
B4 94557911 94558053
refGene_gn=CAND1;refGene_tx=XM_003989017.5;refGene_gn=CAND1;refGene_tx
=XM_019835191.1
B4 94558386 94559880
refGene_gn=CAND1;refGene_tx=XM_003989017.5;refGene_gn=CAND1;refGene_tx
=XM_019835191.1
B4 94560578 94560674
refGene_gn=CAND1;refGene_tx=XM_003989017.5;refGene_gn=CAND1;refGene_tx
=XM_019835191.1
B4 94563575 94563745
refGene_gn=CAND1;refGene_tx=XM_003989017.5;refGene_gn=CAND1;refGene_tx
=XM_019835191.1
B4 94563826 94563991
refGene_gn=CAND1;refGene_tx=XM_003989017.5;refGene_gn=CAND1;refGene_tx
=XM_019835191.1
B4 94565559 94565667
refGene_gn=CAND1;refGene_tx=XM_003989017.5;refGene_gn=CAND1;refGene_tx
=XM_019835191.1
B4 94566633 94566858
refGene_gn=CAND1;refGene_tx=XM_003989017.5;refGene_gn=CAND1;refGene_tx
=XM_019835191.1
B4 94891480 94891529 refGene_gn=DYRK2;refGene_tx=XM_023257216.1
B4 94892145 94892294 refGene_gn=DYRK2;refGene_tx=XM_023257216.1
B4 94899283 94900891
refGene_gn=DYRK2;refGene_tx=XM_006933882.4;refGene_gn=DYRK2;refGene_tx
=XM_023257216.1
B4 95329502 95329637 refGene_gn=IFNG;refGene_tx=NM_001009873.1
B4 95332368 95332554 refGene_gn=IFNG;refGene_tx=NM_001009873.1
B4 95332649 95332718 refGene_gn=IFNG;refGene_tx=NM_001009873.1
B4 95333917 95334031 refGene_gn=IFNG;refGene_tx=NM_001009873.1
B4 95382466 95382553 refGene_gn=IL26;refGene_tx=XM_006933883.2
B4 95382643 95382709 refGene_gn=IL26;refGene_tx=XM_006933883.2
B4 95398833 95398968 refGene_gn=IL26;refGene_tx=XM_006933883.2
B4 95399146 95399203 refGene_gn=IL26;refGene_tx=XM_006933883.2
B4 95399288 95399459 refGene_gn=IL26;refGene_tx=XM_006933883.2
B4 95406752 95406850 refGene_gn=IL22;refGene_tx=XM_003989020.2
B4 95416669 95416739 refGene_gn=IL22;refGene_tx=XM_003989020.2
B4 95419426 95419492 refGene_gn=IL22;refGene_tx=XM_003989020.2

B4 95420395 95420539 refGene_gn=IL22;refGene_tx=XM_003989020.2
B4 95420663 95420729 refGene_gn=IL22;refGene_tx=XM_003989020.2
B4 95421160 95421346 refGene_gn=IL22;refGene_tx=XM_003989020.2
B4 95458393 95458506
refGene_gn=MDM1;refGene_tx=XM_003989021.5;refGene_gn=MDM1;refGene_tx=XM_006933884.4
B4 95459783 95459843
refGene_gn=MDM1;refGene_tx=XM_003989021.5;refGene_gn=MDM1;refGene_tx=XM_006933884.4
B4 95462202 95462455
refGene_gn=MDM1;refGene_tx=XM_003989021.5;refGene_gn=MDM1;refGene_tx=XM_006933884.4
B4 95469509 95469619
refGene_gn=MDM1;refGene_tx=XM_003989021.5;refGene_gn=MDM1;refGene_tx=XM_006933884.4
B4 95469708 95469821
refGene_gn=MDM1;refGene_tx=XM_003989021.5;refGene_gn=MDM1;refGene_tx=XM_006933884.4
B4 95470577 95470877
refGene_gn=MDM1;refGene_tx=XM_003989021.5;refGene_gn=MDM1;refGene_tx=XM_006933884.4
B4 95472379 95472555
refGene_gn=MDM1;refGene_tx=XM_003989021.5;refGene_gn=MDM1;refGene_tx=XM_006933884.4
B4 95472905 95472935 refGene_gn=MDM1;refGene_tx=XM_006933884.4
B4 95478331 95478431
refGene_gn=MDM1;refGene_tx=XM_003989021.5;refGene_gn=MDM1;refGene_tx=XM_006933884.4
B4 95478512 95478616
refGene_gn=MDM1;refGene_tx=XM_003989021.5;refGene_gn=MDM1;refGene_tx=XM_006933884.4
B4 95481070 95481238
refGene_gn=MDM1;refGene_tx=XM_003989021.5;refGene_gn=MDM1;refGene_tx=XM_006933884.4
B4 95485724 95485859
refGene_gn=MDM1;refGene_tx=XM_003989021.5;refGene_gn=MDM1;refGene_tx=XM_006933884.4
B4 95487186 95487554
refGene_gn=MDM1;refGene_tx=XM_003989021.5;refGene_gn=MDM1;refGene_tx=XM_006933884.4
B4 95491100 95491215
refGene_gn=MDM1;refGene_tx=XM_003989021.5;refGene_gn=MDM1;refGene_tx=XM_006933884.4
B4 95492245 95492263
refGene_gn=MDM1;refGene_tx=XM_003989021.5;refGene_gn=MDM1;refGene_tx=XM_006933884.4
B4 95746964 95747021 refGene_gn=RAP1B;refGene_tx=XM_023257217.1
B4 95748077 95748146 refGene_gn=RAP1B;refGene_tx=XM_023257217.1
B4 95749626 95749683 refGene_gn=RAP1B;refGene_tx=XM_023257217.1
B4 95750724 95750865 refGene_gn=RAP1B;refGene_tx=XM_023257217.1

B4 95752502 95752646 refGene_gn=RAP1B;refGene_tx=XM_023257217.1
B4 95753567 95753600 refGene_gn=RAP1B;refGene_tx=XM_023257217.1
B4 95753601 95753655 refGene_gn=RAP1B;refGene_tx=XM_023257217.1
B4 95780484 95780492 refGene_gn=NUP107;refGene_tx=XM_003989029.5
B4 95782509 95782601
refGene_gn=NUP107;refGene_tx=XM_003989029.5;refGene_gn=NUP107;refGene_tx=XM_011284115.3
B4 95783033 95783124
refGene_gn=NUP107;refGene_tx=XM_003989029.5;refGene_gn=NUP107;refGene_tx=XM_011284115.3;refGene_gn=NUP107;refGene_tx=XM_019835195.2
B4 95784140 95784280
refGene_gn=NUP107;refGene_tx=XM_003989029.5;refGene_gn=NUP107;refGene_tx=XM_011284115.3;refGene_gn=NUP107;refGene_tx=XM_019835195.2
B4 95786915 95787060
refGene_gn=NUP107;refGene_tx=XM_003989029.5;refGene_gn=NUP107;refGene_tx=XM_011284115.3;refGene_gn=NUP107;refGene_tx=XM_019835194.2;refGene_gn=NUP107;refGene_tx=XM_019835195.2
B4 95789923 95790027
refGene_gn=NUP107;refGene_tx=XM_003989029.5;refGene_gn=NUP107;refGene_tx=XM_011284115.3;refGene_gn=NUP107;refGene_tx=XM_019835194.2;refGene_gn=NUP107;refGene_tx=XM_019835195.2
B4 95792451 95792579
refGene_gn=NUP107;refGene_tx=XM_003989029.5;refGene_gn=NUP107;refGene_tx=XM_011284115.3;refGene_gn=NUP107;refGene_tx=XM_019835194.2;refGene_gn=NUP107;refGene_tx=XM_019835195.2
B4 95793568 95793617
refGene_gn=NUP107;refGene_tx=XM_003989029.5;refGene_gn=NUP107;refGene_tx=XM_011284115.3;refGene_gn=NUP107;refGene_tx=XM_019835194.2;refGene_gn=NUP107;refGene_tx=XM_019835195.2
B4 95795815 95795887
refGene_gn=NUP107;refGene_tx=XM_003989029.5;refGene_gn=NUP107;refGene_tx=XM_011284115.3;refGene_gn=NUP107;refGene_tx=XM_019835194.2;refGene_gn=NUP107;refGene_tx=XM_019835195.2
B4 95796293 95796382
refGene_gn=NUP107;refGene_tx=XM_003989029.5;refGene_gn=NUP107;refGene_tx=XM_011284115.3;refGene_gn=NUP107;refGene_tx=XM_019835194.2;refGene_gn=NUP107;refGene_tx=XM_019835195.2
B4 95798428 95798507
refGene_gn=NUP107;refGene_tx=XM_003989029.5;refGene_gn=NUP107;refGene_tx=XM_011284115.3;refGene_gn=NUP107;refGene_tx=XM_019835194.2;refGene_gn=NUP107;refGene_tx=XM_019835195.2
B4 95800273 95800387
refGene_gn=NUP107;refGene_tx=XM_003989029.5;refGene_gn=NUP107;refGene_tx=XM_011284115.3;refGene_gn=NUP107;refGene_tx=XM_019835194.2;refGene_gn=NUP107;refGene_tx=XM_019835195.2
B4 95804907 95804998
refGene_gn=NUP107;refGene_tx=XM_003989029.5;refGene_gn=NUP107;refGene_tx=XM_011284115.3;refGene_gn=NUP107;refGene_tx=XM_019835194.2;refGene_gn=NUP107;refGene_tx=XM_019835195.2
B4 95805156 95805233

refGene_gn=NUP107;refGene_tx=XM_003989029.5;refGene_gn=NUP107;refGene_tx=XM_011284115.3;refGene_gn=NUP107;refGene_tx=XM_019835194.2;refGene_gn=NUP107;refGene_tx=XM_019835195.2
B4 95807432 95807492
refGene_gn=NUP107;refGene_tx=XM_003989029.5;refGene_gn=NUP107;refGene_tx=XM_011284115.3;refGene_gn=NUP107;refGene_tx=XM_019835194.2;refGene_gn=NUP107;refGene_tx=XM_019835195.2
B4 95808348 95808494
refGene_gn=NUP107;refGene_tx=XM_003989029.5;refGene_gn=NUP107;refGene_tx=XM_011284115.3;refGene_gn=NUP107;refGene_tx=XM_019835194.2;refGene_gn=NUP107;refGene_tx=XM_019835195.2
B4 95808617 95808666
refGene_gn=NUP107;refGene_tx=XM_003989029.5;refGene_gn=NUP107;refGene_tx=XM_011284115.3;refGene_gn=NUP107;refGene_tx=XM_019835194.2;refGene_gn=NUP107;refGene_tx=XM_019835195.2
B4 95811924 95811994
refGene_gn=NUP107;refGene_tx=XM_003989029.5;refGene_gn=NUP107;refGene_tx=XM_011284115.3;refGene_gn=NUP107;refGene_tx=XM_019835194.2;refGene_gn=NUP107;refGene_tx=XM_019835195.2
B4 95812310 95812429
refGene_gn=NUP107;refGene_tx=XM_003989029.5;refGene_gn=NUP107;refGene_tx=XM_011284115.3;refGene_gn=NUP107;refGene_tx=XM_019835194.2;refGene_gn=NUP107;refGene_tx=XM_019835195.2
B4 95813192 95813231
refGene_gn=NUP107;refGene_tx=XM_003989029.5;refGene_gn=NUP107;refGene_tx=XM_011284115.3;refGene_gn=NUP107;refGene_tx=XM_019835194.2;refGene_gn=NUP107;refGene_tx=XM_019835195.2
B4 95813535 95813686
refGene_gn=NUP107;refGene_tx=XM_003989029.5;refGene_gn=NUP107;refGene_tx=XM_011284115.3;refGene_gn=NUP107;refGene_tx=XM_019835194.2;refGene_gn=NUP107;refGene_tx=XM_019835195.2
B4 95813999 95814112
refGene_gn=NUP107;refGene_tx=XM_003989029.5;refGene_gn=NUP107;refGene_tx=XM_011284115.3;refGene_gn=NUP107;refGene_tx=XM_019835194.2;refGene_gn=NUP107;refGene_tx=XM_019835195.2
B4 95815410 95815513
refGene_gn=NUP107;refGene_tx=XM_003989029.5;refGene_gn=NUP107;refGene_tx=XM_011284115.3;refGene_gn=NUP107;refGene_tx=XM_019835194.2;refGene_gn=NUP107;refGene_tx=XM_019835195.2
B4 95815970 95816131
refGene_gn=NUP107;refGene_tx=XM_003989029.5;refGene_gn=NUP107;refGene_tx=XM_011284115.3;refGene_gn=NUP107;refGene_tx=XM_019835194.2;refGene_gn=NUP107;refGene_tx=XM_019835195.2
B4 95816920 95817046
refGene_gn=NUP107;refGene_tx=XM_003989029.5;refGene_gn=NUP107;refGene_tx=XM_011284115.3;refGene_gn=NUP107;refGene_tx=XM_019835194.2;refGene_gn=NUP107;refGene_tx=XM_019835195.2
B4 95819930 95820044
refGene_gn=NUP107;refGene_tx=XM_003989029.5;refGene_gn=NUP107;refGene_tx=XM_011284115.3;refGene_gn=NUP107;refGene_tx=XM_019835194.2;refGene_

gn=NUP107;refGene_tx=XM_019835195.2
B4 95823491 95823659
refGene_gn=NUP107;refGene_tx=XM_003989029.5;refGene_gn=NUP107;refGene_tx=XM_011284115.3;refGene_gn=NUP107;refGene_tx=XM_019835194.2;refGene_gn=NUP107;refGene_tx=XM_019835195.2
B4 95824009 95824117
refGene_gn=NUP107;refGene_tx=XM_003989029.5;refGene_gn=NUP107;refGene_tx=XM_011284115.3;refGene_gn=NUP107;refGene_tx=XM_019835194.2;refGene_gn=NUP107;refGene_tx=XM_019835195.2
B4 95830997 95831399 refGene_gn=SLC35E3;refGene_tx=XM_003989030.3
B4 95832262 95832373 refGene_gn=SLC35E3;refGene_tx=XM_003989030.3
B4 95836229 95836388 refGene_gn=SLC35E3;refGene_tx=XM_003989030.3
B4 95839852 95839935 refGene_gn=SLC35E3;refGene_tx=XM_003989030.3
B4 95845523 95845710 refGene_gn=SLC35E3;refGene_tx=XM_003989030.3
B4 95878994 95879075
refGene_gn=MDM2;refGene_tx=NM_001009346.1;refGene_gn=MDM2;refGene_tx=XM_006933879.3;refGene_gn=MDM2;refGene_tx=XM_006933880.3;refGene_gn=MDM2;refGene_tx=XM_006933881.4;refGene_gn=MDM2;refGene_tx=XM_019833860.2;refGene_gn=MDM2;refGene_tx=XM_019833861.2
B4 95882190 95882265
refGene_gn=MDM2;refGene_tx=NM_001009346.1;refGene_gn=MDM2;refGene_tx=XM_006933879.3;refGene_gn=MDM2;refGene_tx=XM_006933880.3;refGene_gn=MDM2;refGene_tx=XM_006933881.4;refGene_gn=MDM2;refGene_tx=XM_019833860.2;refGene_gn=MDM2;refGene_tx=XM_019833861.2
B4 95883844 95883978
refGene_gn=MDM2;refGene_tx=NM_001009346.1;refGene_gn=MDM2;refGene_tx=XM_006933879.3;refGene_gn=MDM2;refGene_tx=XM_006933880.3;refGene_gn=MDM2;refGene_tx=XM_006933881.4;refGene_gn=MDM2;refGene_tx=XM_011284110.2;refGene_gn=MDM2;refGene_tx=XM_019833860.2;refGene_gn=MDM2;refGene_tx=XM_019833861.2
B4 95887311 95887361
refGene_gn=MDM2;refGene_tx=NM_001009346.1;refGene_gn=MDM2;refGene_tx=XM_006933879.3;refGene_gn=MDM2;refGene_tx=XM_006933880.3;refGene_gn=MDM2;refGene_tx=XM_006933881.4;refGene_gn=MDM2;refGene_tx=XM_011284110.2;refGene_gn=MDM2;refGene_tx=XM_019833860.2;refGene_gn=MDM2;refGene_tx=XM_019833861.2
B4 95890732 95890800
refGene_gn=MDM2;refGene_tx=NM_001009346.1;refGene_gn=MDM2;refGene_tx=XM_006933879.3;refGene_gn=MDM2;refGene_tx=XM_006933880.3;refGene_gn=MDM2;refGene_tx=XM_006933881.4;refGene_gn=MDM2;refGene_tx=XM_011284110.2;refGene_gn=MDM2;refGene_tx=XM_019833861.2
B4 95890929 95891026
refGene_gn=MDM2;refGene_tx=NM_001009346.1;refGene_gn=MDM2;refGene_tx=XM_006933879.3;refGene_gn=MDM2;refGene_tx=XM_006933880.3;refGene_gn=MDM2;refGene_tx=XM_006933881.4;refGene_gn=MDM2;refGene_tx=XM_011284110.2;refGene_gn=MDM2;refGene_tx=XM_019833861.2
B4 95895603 95895764
refGene_gn=MDM2;refGene_tx=NM_001009346.1;refGene_gn=MDM2;refGene_tx=XM_006933879.3;refGene_gn=MDM2;refGene_tx=XM_006933880.3;refGene_gn=MDM2;refGene_tx=XM_006933881.4;refGene_gn=MDM2;refGene_tx=XM_011284110.2;

refGene_gn=MDM2;refGene_tx=XM_019833860.2;refGene_gn=MDM2;refGene_tx=XM_019833861.2
B4 95900141 95900297
refGene_gn=MDM2;refGene_tx=NM_001009346.1;refGene_gn=MDM2;refGene_tx=XM_006933879.3;refGene_gn=MDM2;refGene_tx=XM_006933880.3;refGene_gn=MDM2;refGene_tx=XM_006933881.4;refGene_gn=MDM2;refGene_tx=XM_011284110.2;refGene_gn=MDM2;refGene_tx=XM_019833860.2;refGene_gn=MDM2;refGene_tx=XM_019833861.2
B4 95901253 95901331
refGene_gn=MDM2;refGene_tx=NM_001009346.1;refGene_gn=MDM2;refGene_tx=XM_006933879.3;refGene_gn=MDM2;refGene_tx=XM_006933880.3;refGene_gn=MDM2;refGene_tx=XM_006933881.4;refGene_gn=MDM2;refGene_tx=XM_011284110.2;refGene_gn=MDM2;refGene_tx=XM_019833860.2;refGene_gn=MDM2;refGene_tx=XM_019833861.2
B4 95903930 95904506
refGene_gn=MDM2;refGene_tx=NM_001009346.1;refGene_gn=MDM2;refGene_tx=XM_006933879.3;refGene_gn=MDM2;refGene_tx=XM_006933880.3;refGene_gn=MDM2;refGene_tx=XM_006933881.4;refGene_gn=MDM2;refGene_tx=XM_011284110.2;refGene_gn=MDM2;refGene_tx=XM_019833860.2;refGene_gn=MDM2;refGene_tx=XM_019833861.2
B4 95913885 95914128
refGene_gn=CPM;refGene_tx=XM_003989031.5;refGene_gn=CPM;refGene_tx=XM_023257218.1;refGene_gn=CPM;refGene_tx=XM_023257219.1
B4 95916093 95916242
refGene_gn=CPM;refGene_tx=XM_003989031.5;refGene_gn=CPM;refGene_tx=XM_023257218.1;refGene_gn=CPM;refGene_tx=XM_023257219.1
B4 95925957 95926110
refGene_gn=CPM;refGene_tx=XM_003989031.5;refGene_gn=CPM;refGene_tx=XM_023257218.1;refGene_gn=CPM;refGene_tx=XM_023257219.1
B4 95929508 95929679
refGene_gn=CPM;refGene_tx=XM_003989031.5;refGene_gn=CPM;refGene_tx=XM_023257218.1;refGene_gn=CPM;refGene_tx=XM_023257219.1
B4 95930382 95930567
refGene_gn=CPM;refGene_tx=XM_003989031.5;refGene_gn=CPM;refGene_tx=XM_023257218.1;refGene_gn=CPM;refGene_tx=XM_023257219.1
B4 95932404 95932577
refGene_gn=CPM;refGene_tx=XM_003989031.5;refGene_gn=CPM;refGene_tx=XM_023257218.1;refGene_gn=CPM;refGene_tx=XM_023257219.1
B4 95945945 95946043
refGene_gn=CPM;refGene_tx=XM_003989031.5;refGene_gn=CPM;refGene_tx=XM_023257218.1;refGene_gn=CPM;refGene_tx=XM_023257219.1
B4 95996503 95996663
refGene_gn=CPM;refGene_tx=XM_003989031.5;refGene_gn=CPM;refGene_tx=XM_023257218.1;refGene_gn=CPM;refGene_tx=XM_023257219.1
B4 96232735 96232795
refGene_gn=CPSF6;refGene_tx=XM_023257220.1;refGene_gn=CPSF6;refGene_tx=XM_023257221.1;refGene_gn=CPSF6;refGene_tx=XM_023257222.1;refGene_gn=CPSF6;refGene_tx=XM_023257223.1
B4 96240802 96241012
refGene_gn=CPSF6;refGene_tx=XM_023257220.1;refGene_gn=CPSF6;refGene_tx

=XM_023257221.1;refGene_gn=CPSF6;refGene_tx=XM_023257222.1;refGene_gn=CPSF6;refGene_tx=XM_023257223.1
B4 96243109 96243213 refGene_gn=CPSF6;refGene_tx=XM_023257220.1;refGene_gn=CPSF6;refGene_tx=XM_023257221.1;refGene_gn=CPSF6;refGene_tx=XM_023257222.1;refGene_gn=CPSF6;refGene_tx=XM_023257223.1
B4 96246323 96246469 refGene_gn=CPSF6;refGene_tx=XM_023257220.1;refGene_gn=CPSF6;refGene_tx=XM_023257221.1;refGene_gn=CPSF6;refGene_tx=XM_023257222.1;refGene_gn=CPSF6;refGene_tx=XM_023257223.1
B4 96247612 96247786 refGene_gn=CPSF6;refGene_tx=XM_023257220.1;refGene_gn=CPSF6;refGene_tx=XM_023257221.1;refGene_gn=CPSF6;refGene_tx=XM_023257222.1;refGene_gn=CPSF6;refGene_tx=XM_023257223.1
B4 96247916 96248027 refGene_gn=CPSF6;refGene_tx=XM_023257220.1;refGene_gn=CPSF6;refGene_tx=XM_023257221.1
B4 96248472 96248977 refGene_gn=CPSF6;refGene_tx=XM_023257220.1;refGene_gn=CPSF6;refGene_tx=XM_023257221.1;refGene_gn=CPSF6;refGene_tx=XM_023257222.1;refGene_gn=CPSF6;refGene_tx=XM_023257223.1
B4 96251195 96251311 refGene_gn=CPSF6;refGene_tx=XM_023257220.1;refGene_gn=CPSF6;refGene_tx=XM_023257221.1;refGene_gn=CPSF6;refGene_tx=XM_023257222.1;refGene_gn=CPSF6;refGene_tx=XM_023257223.1
B4 96251806 96251960 refGene_gn=CPSF6;refGene_tx=XM_023257220.1;refGene_gn=CPSF6;refGene_tx=XM_023257221.1;refGene_gn=CPSF6;refGene_tx=XM_023257222.1;refGene_gn=CPSF6;refGene_tx=XM_023257223.1
B4 96254707 96254897 refGene_gn=CPSF6;refGene_tx=XM_023257220.1;refGene_gn=CPSF6;refGene_tx=XM_023257221.1;refGene_gn=CPSF6;refGene_tx=XM_023257222.1;refGene_gn=CPSF6;refGene_tx=XM_023257223.1
B4 96330718 96330854 refGene_gn=LYZ;refGene_tx=XM_023257226.1
B4 96332830 96332995 refGene_gn=LYZ;refGene_tx=XM_023257226.1
B4 96334631 96334710 refGene_gn=LYZ;refGene_tx=XM_023257226.1
B4 96335799 96335866 refGene_gn=LYZ;refGene_tx=XM_023257226.1
B4 96346759 96346810 refGene_gn=YEATS4;refGene_tx=XM_003989033.5;refGene_gn=YEATS4;refGene_tx=XM_023257224.1;refGene_gn=YEATS4;refGene_tx=XM_023257225.1
B4 96349507 96349627 refGene_gn=YEATS4;refGene_tx=XM_003989033.5;refGene_gn=YEATS4;refGene_tx=XM_023257224.1;refGene_gn=YEATS4;refGene_tx=XM_023257225.1
B4 96350709 96350776 refGene_gn=YEATS4;refGene_tx=XM_003989033.5;refGene_gn=YEATS4;refGene_tx=XM_023257224.1;refGene_gn=YEATS4;refGene_tx=XM_023257225.1
B4 96350883 96350978 refGene_gn=YEATS4;refGene_tx=XM_003989033.5;refGene_gn=YEATS4;refGene_tx=XM_023257224.1;refGene_gn=YEATS4;refGene_tx=XM_023257225.1
B4 96355892 96355985

refGene_gn=YEATS4;refGene_tx=XM_003989033.5;refGene_gn=YEATS4;refGene_tx=XM_023257224.1;refGene_gn=YEATS4;refGene_tx=XM_023257225.1
B4 96356072 96356160
refGene_gn=YEATS4;refGene_tx=XM_003989033.5;refGene_gn=YEATS4;refGene_tx=XM_023257224.1;refGene_gn=YEATS4;refGene_tx=XM_023257225.1
B4 96367505 96367675 refGene_gn=YEATS4;refGene_tx=XM_003989033.5
B4 96387140 96387208 refGene_gn=YEATS4;refGene_tx=XM_023257225.1
B4 96406390 96406439 refGene_gn=YEATS4;refGene_tx=XM_023257224.1
B4 96411898 96411989 refGene_gn=YEATS4;refGene_tx=XM_023257224.1
B4 96533516 96533582
refGene_gn=FRS2;refGene_tx=XM_023257227.1;refGene_gn=FRS2;refGene_tx=XM_023257228.1
B4 96534831 96535018
refGene_gn=FRS2;refGene_tx=XM_023257227.1;refGene_gn=FRS2;refGene_tx=XM_023257228.1
B4 96536081 96536240
refGene_gn=FRS2;refGene_tx=XM_023257227.1;refGene_gn=FRS2;refGene_tx=XM_023257228.1
B4 96536627 96536791
refGene_gn=FRS2;refGene_tx=XM_023257227.1;refGene_gn=FRS2;refGene_tx=XM_023257228.1
B4 96538188 96539139
refGene_gn=FRS2;refGene_tx=XM_023257227.1;refGene_gn=FRS2;refGene_tx=XM_023257228.1
B4 96547522 96547525 refGene_gn=CCT2;refGene_tx=XM_003989035.4
B4 96548279 96548354 refGene_gn=CCT2;refGene_tx=XM_003989035.4
B4 96548459 96548525 refGene_gn=CCT2;refGene_tx=XM_003989035.4
B4 96549183 96549295 refGene_gn=CCT2;refGene_tx=XM_003989035.4
B4 96549617 96549694 refGene_gn=CCT2;refGene_tx=XM_003989035.4
B4 96549998 96550111 refGene_gn=CCT2;refGene_tx=XM_003989035.4
B4 96551094 96551297 refGene_gn=CCT2;refGene_tx=XM_003989035.4
B4 96552788 96552889 refGene_gn=CCT2;refGene_tx=XM_003989035.4
B4 96553330 96553458 refGene_gn=CCT2;refGene_tx=XM_003989035.4
B4 96556946 96557050 refGene_gn=CCT2;refGene_tx=XM_003989035.4
B4 96559649 96559769 refGene_gn=CCT2;refGene_tx=XM_003989035.4
B4 96560158 96560287 refGene_gn=CCT2;refGene_tx=XM_003989035.4
B4 96560483 96560587 refGene_gn=CCT2;refGene_tx=XM_003989035.4
B4 96560866 96560966 refGene_gn=CCT2;refGene_tx=XM_003989035.4
B4 96562634 96562776 refGene_gn=CCT2;refGene_tx=XM_003989035.4
B4 96565429 96565460 refGene_gn=CCT2;refGene_tx=XM_003989035.4
B4 96571819 96572659 refGene_gn=LRRC10;refGene_tx=XM_003989036.3
B4 96601597 96602540
refGene_gn=BEST3;refGene_tx=XM_019835201.2;refGene_gn=BEST3;refGene_tx=XM_019835207.2;refGene_gn=BEST3;refGene_tx=XM_023257229.1
B4 96615662 96615814
refGene_gn=BEST3;refGene_tx=XM_019835201.2;refGene_gn=BEST3;refGene_tx=XM_019835207.2;refGene_gn=BEST3;refGene_tx=XM_023257229.1
B4 96618423 96618504
refGene_gn=BEST3;refGene_tx=XM_019835201.2;refGene_gn=BEST3;refGene_tx=XM_019835207.2;refGene_gn=BEST3;refGene_tx=XM_023257229.1

B4 96621618 96621771
refGene_gn=BEST3;refGene_tx=XM_019835201.2;refGene_gn=BEST3;refGene_tx
=XM_019835207.2;refGene_gn=BEST3;refGene_tx=XM_023257229.1
B4 96622136 96622214
refGene_gn=BEST3;refGene_tx=XM_019835207.2;refGene_gn=BEST3;refGene_tx
=XM_023257229.1
B4 96623432 96623587
refGene_gn=BEST3;refGene_tx=XM_019835201.2;refGene_gn=BEST3;refGene_tx
=XM_019835207.2;refGene_gn=BEST3;refGene_tx=XM_023257229.1
B4 96635118 96635352
refGene_gn=BEST3;refGene_tx=XM_019835201.2;refGene_gn=BEST3;refGene_tx
=XM_019835207.2;refGene_gn=BEST3;refGene_tx=XM_023257229.1
B4 96635817 96635912
refGene_gn=BEST3;refGene_tx=XM_019835201.2;refGene_gn=BEST3;refGene_tx
=XM_023257229.1
B4 96639471 96639623
refGene_gn=BEST3;refGene_tx=XM_019835201.2;refGene_gn=BEST3;refGene_tx
=XM_023257229.1
B4 96678879 96679435
refGene_gn=LOC111561462;refGene_tx=XM_023257769.1
B4 96679565 96679640
refGene_gn=LOC111561462;refGene_tx=XM_023257769.1
B4 96679841 96679852
refGene_gn=RAB3IP;refGene_tx=XM_023257230.1;refGene_gn=RAB3IP;refGene_
tx=XM_023257231.1;refGene_gn=RAB3IP;refGene_tx=XM_023257234.1;refGene_
gn=RAB3IP;refGene_tx=XM_023257235.1;refGene_gn=RAB3IP;refGene_tx=XM_02
3257237.1
B4 96695183 96695459
refGene_gn=RAB3IP;refGene_tx=XM_023257230.1;refGene_gn=RAB3IP;refGene_
tx=XM_023257231.1;refGene_gn=RAB3IP;refGene_tx=XM_023257232.1;refGene_
gn=RAB3IP;refGene_tx=XM_023257233.1;refGene_gn=RAB3IP;refGene_tx=XM_02
3257234.1;refGene_gn=RAB3IP;refGene_tx=XM_023257235.1;refGene_gn=RAB3I
P;refGene_tx=XM_023257236.1;refGene_gn=RAB3IP;refGene_tx=XM_023257237.
1
B4 96696176 96696435
refGene_gn=RAB3IP;refGene_tx=XM_023257230.1;refGene_gn=RAB3IP;refGene_
tx=XM_023257231.1;refGene_gn=RAB3IP;refGene_tx=XM_023257232.1;refGene_
gn=RAB3IP;refGene_tx=XM_023257233.1;refGene_gn=RAB3IP;refGene_tx=XM_02
3257234.1;refGene_gn=RAB3IP;refGene_tx=XM_023257235.1;refGene_gn=RAB3I
P;refGene_tx=XM_023257236.1;refGene_gn=RAB3IP;refGene_tx=XM_023257237.
1
B4 96707808 96707904
refGene_gn=RAB3IP;refGene_tx=XM_023257230.1;refGene_gn=RAB3IP;refGene_
tx=XM_023257231.1;refGene_gn=RAB3IP;refGene_tx=XM_023257232.1;refGene_
gn=RAB3IP;refGene_tx=XM_023257233.1;refGene_gn=RAB3IP;refGene_tx=XM_02
3257234.1;refGene_gn=RAB3IP;refGene_tx=XM_023257236.1;refGene_gn=RAB3I
P;refGene_tx=XM_023257237.1
B4 96713950 96714028
refGene_gn=RAB3IP;refGene_tx=XM_023257230.1;refGene_gn=RAB3IP;refGene_
tx=XM_023257231.1;refGene_gn=RAB3IP;refGene_tx=XM_023257232.1;refGene_

gn=RAB3IP;refGene_tx=XM_023257233.1;refGene_gn=RAB3IP;refGene_tx=XM_02
3257234.1;refGene_gn=RAB3IP;refGene_tx=XM_023257235.1;refGene_gn=RAB3I
P;refGene_tx=XM_023257236.1;refGene_gn=RAB3IP;refGene_tx=XM_023257237.
1
B4 96714658 96714862
refGene_gn=RAB3IP;refGene_tx=XM_023257230.1;refGene_gn=RAB3IP;refGene_
tx=XM_023257231.1;refGene_gn=RAB3IP;refGene_tx=XM_023257232.1;refGene_
gn=RAB3IP;refGene_tx=XM_023257233.1;refGene_gn=RAB3IP;refGene_tx=XM_02
3257234.1;refGene_gn=RAB3IP;refGene_tx=XM_023257235.1;refGene_gn=RAB3I
P;refGene_tx=XM_023257236.1;refGene_gn=RAB3IP;refGene_tx=XM_023257237.
1
B4 96715015 96715072
refGene_gn=RAB3IP;refGene_tx=XM_023257230.1;refGene_gn=RAB3IP;refGene_
tx=XM_023257231.1;refGene_gn=RAB3IP;refGene_tx=XM_023257232.1;refGene_
gn=RAB3IP;refGene_tx=XM_023257233.1;refGene_gn=RAB3IP;refGene_tx=XM_02
3257235.1;refGene_gn=RAB3IP;refGene_tx=XM_023257236.1
B4 96717621 96717750
refGene_gn=RAB3IP;refGene_tx=XM_023257230.1;refGene_gn=RAB3IP;refGene_
tx=XM_023257231.1;refGene_gn=RAB3IP;refGene_tx=XM_023257232.1;refGene_
gn=RAB3IP;refGene_tx=XM_023257233.1;refGene_gn=RAB3IP;refGene_tx=XM_02
3257234.1;refGene_gn=RAB3IP;refGene_tx=XM_023257235.1;refGene_gn=RAB3I
P;refGene_tx=XM_023257236.1
B4 96719009 96719122
refGene_gn=RAB3IP;refGene_tx=XM_023257230.1;refGene_gn=RAB3IP;refGene_
tx=XM_023257231.1;refGene_gn=RAB3IP;refGene_tx=XM_023257232.1;refGene_
gn=RAB3IP;refGene_tx=XM_023257233.1;refGene_gn=RAB3IP;refGene_tx=XM_02
3257234.1;refGene_gn=RAB3IP;refGene_tx=XM_023257235.1;refGene_gn=RAB3I
P;refGene_tx=XM_023257236.1;refGene_gn=RAB3IP;refGene_tx=XM_023257237.
1
B4 96722922 96723022
refGene_gn=RAB3IP;refGene_tx=XM_023257230.1;refGene_gn=RAB3IP;refGene_
tx=XM_023257231.1;refGene_gn=RAB3IP;refGene_tx=XM_023257232.1;refGene_
gn=RAB3IP;refGene_tx=XM_023257233.1;refGene_gn=RAB3IP;refGene_tx=XM_02
3257234.1;refGene_gn=RAB3IP;refGene_tx=XM_023257235.1;refGene_gn=RAB3I
P;refGene_tx=XM_023257236.1;refGene_gn=RAB3IP;refGene_tx=XM_023257237.
1
B4 96723105 96723175
refGene_gn=RAB3IP;refGene_tx=XM_023257230.1;refGene_gn=RAB3IP;refGene_
tx=XM_023257231.1;refGene_gn=RAB3IP;refGene_tx=XM_023257232.1;refGene_
gn=RAB3IP;refGene_tx=XM_023257233.1;refGene_gn=RAB3IP;refGene_tx=XM_02
3257234.1;refGene_gn=RAB3IP;refGene_tx=XM_023257235.1;refGene_gn=RAB3I
P;refGene_tx=XM_023257236.1;refGene_gn=RAB3IP;refGene_tx=XM_023257237.
1
B4 96725416 96725499
refGene_gn=RAB3IP;refGene_tx=XM_023257230.1;refGene_gn=RAB3IP;refGene_
tx=XM_023257231.1;refGene_gn=RAB3IP;refGene_tx=XM_023257232.1;refGene_
gn=RAB3IP;refGene_tx=XM_023257233.1;refGene_gn=RAB3IP;refGene_tx=XM_02
3257234.1;refGene_gn=RAB3IP;refGene_tx=XM_023257235.1;refGene_gn=RAB3I
P;refGene_tx=XM_023257236.1;refGene_gn=RAB3IP;refGene_tx=XM_023257237.
1

B4 96737772 96737818 refGene_gn=MYRFL;refGene_tx=XM_023257770.1
B4 96785346 96785770 refGene_gn=MYRFL;refGene_tx=XM_023257770.1
B4 96786647 96786739 refGene_gn=MYRFL;refGene_tx=XM_023257770.1
B4 96792392 96792543 refGene_gn=MYRFL;refGene_tx=XM_023257770.1
B4 96796019 96796215 refGene_gn=MYRFL;refGene_tx=XM_023257770.1
B4 96798495 96798572 refGene_gn=MYRFL;refGene_tx=XM_023257770.1
B4 96800065 96800176 refGene_gn=MYRFL;refGene_tx=XM_023257770.1
B4 96801736 96801827 refGene_gn=MYRFL;refGene_tx=XM_023257770.1
B4 96811248 96811449 refGene_gn=MYRFL;refGene_tx=XM_023257770.1
B4 96820427 96820536 refGene_gn=MYRFL;refGene_tx=XM_023257770.1
B4 96821273 96821383 refGene_gn=MYRFL;refGene_tx=XM_023257770.1
B4 96839697 96839861 refGene_gn=MYRFL;refGene_tx=XM_023257770.1
B4 96840477 96840541 refGene_gn=MYRFL;refGene_tx=XM_023257770.1
B4 96845460 96845546 refGene_gn=MYRFL;refGene_tx=XM_023257770.1
B4 96848264 96848339 refGene_gn=MYRFL;refGene_tx=XM_023257770.1
B4 96848422 96848475 refGene_gn=MYRFL;refGene_tx=XM_023257770.1
B4 96848596 96848776 refGene_gn=MYRFL;refGene_tx=XM_023257770.1
B4 96856366 96856429 refGene_gn=MYRFL;refGene_tx=XM_023257770.1
B4 96857078 96857166 refGene_gn=MYRFL;refGene_tx=XM_023257770.1
B4 96859133 96859208 refGene_gn=MYRFL;refGene_tx=XM_023257770.1
B4 96861997 96862118 refGene_gn=MYRFL;refGene_tx=XM_023257770.1
B4 96862450 96862525 refGene_gn=MYRFL;refGene_tx=XM_023257770.1
B4 96862684 96862771 refGene_gn=MYRFL;refGene_tx=XM_023257770.1
B4 97071651 97071656
refGene_gn=CNOT2;refGene_tx=XM_006933889.4;refGene_gn=CNOT2;refGene_tx=XM_023257238.1
B4 97124238 97124311
refGene_gn=CNOT2;refGene_tx=XM_003989040.5;refGene_gn=CNOT2;refGene_tx=XM_006933889.4;refGene_gn=CNOT2;refGene_tx=XM_023257238.1
B4 97156939 97157062
refGene_gn=CNOT2;refGene_tx=XM_003989040.5;refGene_gn=CNOT2;refGene_tx=XM_006933889.4;refGene_gn=CNOT2;refGene_tx=XM_023257238.1
B4 97167851 97167918
refGene_gn=CNOT2;refGene_tx=XM_003989040.5;refGene_gn=CNOT2;refGene_tx=XM_006933889.4;refGene_gn=CNOT2;refGene_tx=XM_023257238.1
B4 97178255 97178403
refGene_gn=CNOT2;refGene_tx=XM_003989040.5;refGene_gn=CNOT2;refGene_tx=XM_006933889.4;refGene_gn=CNOT2;refGene_tx=XM_019835217.1;refGene_gn=CNOT2;refGene_tx=XM_023257238.1
B4 97179136 97179319
refGene_gn=CNOT2;refGene_tx=XM_003989040.5;refGene_gn=CNOT2;refGene_tx=XM_006933889.4;refGene_gn=CNOT2;refGene_tx=XM_019835217.1
B4 97181634 97181714
refGene_gn=CNOT2;refGene_tx=XM_003989040.5;refGene_gn=CNOT2;refGene_tx=XM_006933889.4;refGene_gn=CNOT2;refGene_tx=XM_019835217.1;refGene_gn=CNOT2;refGene_tx=XM_023257238.1
B4 97184583 97184709
refGene_gn=CNOT2;refGene_tx=XM_003989040.5;refGene_gn=CNOT2;refGene_tx=XM_006933889.4;refGene_gn=CNOT2;refGene_tx=XM_019835217.1;refGene_gn=CNOT2;refGene_tx=XM_023257238.1

B4 97186517 97186642
refGene_gn=CNOT2;refGene_tx=XM_003989040.5;refGene_gn=CNOT2;refGene_tx=XM_006933889.4;refGene_gn=CNOT2;refGene_tx=XM_019835217.1;refGene_gn=CNOT2;refGene_tx=XM_023257238.1
B4 97187594 97187715
refGene_gn=CNOT2;refGene_tx=XM_003989040.5;refGene_gn=CNOT2;refGene_tx=XM_006933889.4;refGene_gn=CNOT2;refGene_tx=XM_019835217.1;refGene_gn=CNOT2;refGene_tx=XM_023257238.1
B4 97187806 97187963
refGene_gn=CNOT2;refGene_tx=XM_003989040.5;refGene_gn=CNOT2;refGene_tx=XM_006933889.4;refGene_gn=CNOT2;refGene_tx=XM_019835217.1;refGene_gn=CNOT2;refGene_tx=XM_023257238.1
B4 97190490 97190552
refGene_gn=CNOT2;refGene_tx=XM_003989040.5;refGene_gn=CNOT2;refGene_tx=XM_006933889.4;refGene_gn=CNOT2;refGene_tx=XM_019835217.1;refGene_gn=CNOT2;refGene_tx=XM_023257238.1
B4 97190641 97190691
refGene_gn=CNOT2;refGene_tx=XM_003989040.5;refGene_gn=CNOT2;refGene_tx=XM_006933889.4;refGene_gn=CNOT2;refGene_tx=XM_019835217.1;refGene_gn=CNOT2;refGene_tx=XM_023257238.1
B4 97192594 97192695
refGene_gn=CNOT2;refGene_tx=XM_003989040.5;refGene_gn=CNOT2;refGene_tx=XM_006933889.4;refGene_gn=CNOT2;refGene_tx=XM_019835217.1;refGene_gn=CNOT2;refGene_tx=XM_023257238.1
B4 97194549 97194694
refGene_gn=CNOT2;refGene_tx=XM_003989040.5;refGene_gn=CNOT2;refGene_tx=XM_006933889.4;refGene_gn=CNOT2;refGene_tx=XM_019835217.1;refGene_gn=CNOT2;refGene_tx=XM_023257238.1
B4 97201113 97201200
refGene_gn=CNOT2;refGene_tx=XM_003989040.5;refGene_gn=CNOT2;refGene_tx=XM_006933889.4;refGene_gn=CNOT2;refGene_tx=XM_019835217.1;refGene_gn=CNOT2;refGene_tx=XM_023257238.1
B4 97215514 97215850 refGene_gn=KCNMB4;refGene_tx=XM_006933893.3
B4 97244180 97244308 refGene_gn=KCNMB4;refGene_tx=XM_006933893.3
B4 97278529 97278698 refGene_gn=KCNMB4;refGene_tx=XM_006933893.3
B4 97307237 97307378
refGene_gn=PTPRB;refGene_tx=XM_003989041.4;refGene_gn=PTPRB;refGene_tx=XM_006933890.3;refGene_gn=PTPRB;refGene_tx=XM_006933891.4;refGene_gn=PTPRB;refGene_tx=XM_019835219.1;refGene_gn=PTPRB;refGene_tx=XM_019835220.2;refGene_gn=PTPRB;refGene_tx=XM_019835221.2
B4 97309248 97309369
refGene_gn=PTPRB;refGene_tx=XM_003989041.4;refGene_gn=PTPRB;refGene_tx=XM_006933891.4;refGene_gn=PTPRB;refGene_tx=XM_019835219.1;refGene_gn=PTPRB;refGene_tx=XM_019835220.2;refGene_gn=PTPRB;refGene_tx=XM_019835221.2
B4 97315552 97315688
refGene_gn=PTPRB;refGene_tx=XM_003989041.4;refGene_gn=PTPRB;refGene_tx=XM_006933890.3;refGene_gn=PTPRB;refGene_tx=XM_006933891.4;refGene_gn=PTPRB;refGene_tx=XM_019835219.1;refGene_gn=PTPRB;refGene_tx=XM_019835220.2;refGene_gn=PTPRB;refGene_tx=XM_019835221.2

B4 97317421 97317585
refGene_gn=PTPRB;refGene_tx=XM_003989041.4;refGene_gn=PTPRB;refGene_tx=XM_006933890.3;refGene_gn=PTPRB;refGene_tx=XM_006933891.4;refGene_gn=PTPRB;refGene_tx=XM_019835219.1;refGene_gn=PTPRB;refGene_tx=XM_019835220.2;refGene_gn=PTPRB;refGene_tx=XM_019835221.2
B4 97317812 97317935
refGene_gn=PTPRB;refGene_tx=XM_003989041.4;refGene_gn=PTPRB;refGene_tx=XM_006933890.3;refGene_gn=PTPRB;refGene_tx=XM_006933891.4;refGene_gn=PTPRB;refGene_tx=XM_019835219.1;refGene_gn=PTPRB;refGene_tx=XM_019835220.2;refGene_gn=PTPRB;refGene_tx=XM_019835221.2
B4 97319202 97319337
refGene_gn=PTPRB;refGene_tx=XM_003989041.4;refGene_gn=PTPRB;refGene_tx=XM_006933890.3;refGene_gn=PTPRB;refGene_tx=XM_006933891.4;refGene_gn=PTPRB;refGene_tx=XM_019835219.1;refGene_gn=PTPRB;refGene_tx=XM_019835220.2;refGene_gn=PTPRB;refGene_tx=XM_019835221.2
B4 97321009 97321086
refGene_gn=PTPRB;refGene_tx=XM_003989041.4;refGene_gn=PTPRB;refGene_tx=XM_006933890.3;refGene_gn=PTPRB;refGene_tx=XM_006933891.4;refGene_gn=PTPRB;refGene_tx=XM_019835219.1;refGene_gn=PTPRB;refGene_tx=XM_019835220.2;refGene_gn=PTPRB;refGene_tx=XM_019835221.2
B4 97321771 97321862
refGene_gn=PTPRB;refGene_tx=XM_003989041.4;refGene_gn=PTPRB;refGene_tx=XM_006933890.3;refGene_gn=PTPRB;refGene_tx=XM_006933891.4;refGene_gn=PTPRB;refGene_tx=XM_019835219.1;refGene_gn=PTPRB;refGene_tx=XM_019835220.2;refGene_gn=PTPRB;refGene_tx=XM_019835221.2
B4 97322460 97322542
refGene_gn=PTPRB;refGene_tx=XM_003989041.4;refGene_gn=PTPRB;refGene_tx=XM_006933890.3;refGene_gn=PTPRB;refGene_tx=XM_006933891.4;refGene_gn=PTPRB;refGene_tx=XM_019835219.1;refGene_gn=PTPRB;refGene_tx=XM_019835220.2;refGene_gn=PTPRB;refGene_tx=XM_019835221.2
B4 97322663 97322681
refGene_gn=PTPRB;refGene_tx=XM_003989041.4;refGene_gn=PTPRB;refGene_tx=XM_006933890.3;refGene_gn=PTPRB;refGene_tx=XM_006933891.4;refGene_gn=PTPRB;refGene_tx=XM_019835219.1;refGene_gn=PTPRB;refGene_tx=XM_019835220.2;refGene_gn=PTPRB;refGene_tx=XM_019835221.2
B4 97322783 97322867
refGene_gn=PTPRB;refGene_tx=XM_003989041.4;refGene_gn=PTPRB;refGene_tx=XM_006933890.3;refGene_gn=PTPRB;refGene_tx=XM_006933891.4;refGene_gn=PTPRB;refGene_tx=XM_019835219.1;refGene_gn=PTPRB;refGene_tx=XM_019835220.2;refGene_gn=PTPRB;refGene_tx=XM_019835221.2
B4 97323661 97323761
refGene_gn=PTPRB;refGene_tx=XM_003989041.4;refGene_gn=PTPRB;refGene_tx=XM_006933890.3;refGene_gn=PTPRB;refGene_tx=XM_006933891.4;refGene_gn=PTPRB;refGene_tx=XM_019835219.1;refGene_gn=PTPRB;refGene_tx=XM_019835220.2;refGene_gn=PTPRB;refGene_tx=XM_019835221.2
B4 97327681 97327788
refGene_gn=PTPRB;refGene_tx=XM_003989041.4;refGene_gn=PTPRB;refGene_tx=XM_006933890.3;refGene_gn=PTPRB;refGene_tx=XM_006933891.4;refGene_gn=PTPRB;refGene_tx=XM_019835219.1;refGene_gn=PTPRB;refGene_tx=XM_019835220.2;refGene_gn=PTPRB;refGene_tx=XM_019835221.2

B4 97333725 97333969
refGene_gn=PTPRB;refGene_tx=XM_003989041.4;refGene_gn=PTPRB;refGene_tx=XM_006933890.3;refGene_gn=PTPRB;refGene_tx=XM_006933891.4;refGene_gn=PTPRB;refGene_tx=XM_019835219.1;refGene_gn=PTPRB;refGene_tx=XM_019835220.2;refGene_gn=PTPRB;refGene_tx=XM_019835221.2
B4 97336337 97336487
refGene_gn=PTPRB;refGene_tx=XM_003989041.4;refGene_gn=PTPRB;refGene_tx=XM_006933890.3;refGene_gn=PTPRB;refGene_tx=XM_006933891.4;refGene_gn=PTPRB;refGene_tx=XM_019835219.1;refGene_gn=PTPRB;refGene_tx=XM_019835220.2;refGene_gn=PTPRB;refGene_tx=XM_019835221.2
B4 97337014 97337293
refGene_gn=PTPRB;refGene_tx=XM_003989041.4;refGene_gn=PTPRB;refGene_tx=XM_006933890.3;refGene_gn=PTPRB;refGene_tx=XM_006933891.4;refGene_gn=PTPRB;refGene_tx=XM_019835219.1;refGene_gn=PTPRB;refGene_tx=XM_019835220.2;refGene_gn=PTPRB;refGene_tx=XM_019835221.2
B4 97341769 97342051
refGene_gn=PTPRB;refGene_tx=XM_003989041.4;refGene_gn=PTPRB;refGene_tx=XM_006933890.3;refGene_gn=PTPRB;refGene_tx=XM_006933891.4;refGene_gn=PTPRB;refGene_tx=XM_019835219.1;refGene_gn=PTPRB;refGene_tx=XM_019835220.2;refGene_gn=PTPRB;refGene_tx=XM_019835221.2
B4 97342800 97343064
refGene_gn=PTPRB;refGene_tx=XM_003989041.4;refGene_gn=PTPRB;refGene_tx=XM_006933890.3;refGene_gn=PTPRB;refGene_tx=XM_006933891.4;refGene_gn=PTPRB;refGene_tx=XM_019835219.1;refGene_gn=PTPRB;refGene_tx=XM_019835220.2;refGene_gn=PTPRB;refGene_tx=XM_019835221.2
B4 97344684 97344948
refGene_gn=PTPRB;refGene_tx=XM_003989041.4;refGene_gn=PTPRB;refGene_tx=XM_006933890.3;refGene_gn=PTPRB;refGene_tx=XM_006933891.4;refGene_gn=PTPRB;refGene_tx=XM_019835219.1;refGene_gn=PTPRB;refGene_tx=XM_019835220.2;refGene_gn=PTPRB;refGene_tx=XM_019835221.2
B4 97348893 97349163
refGene_gn=PTPRB;refGene_tx=XM_003989041.4;refGene_gn=PTPRB;refGene_tx=XM_006933890.3;refGene_gn=PTPRB;refGene_tx=XM_006933891.4;refGene_gn=PTPRB;refGene_tx=XM_019835219.1;refGene_gn=PTPRB;refGene_tx=XM_019835220.2;refGene_gn=PTPRB;refGene_tx=XM_019835221.2
B4 97351360 97351624
refGene_gn=PTPRB;refGene_tx=XM_003989041.4;refGene_gn=PTPRB;refGene_tx=XM_006933890.3;refGene_gn=PTPRB;refGene_tx=XM_006933891.4;refGene_gn=PTPRB;refGene_tx=XM_019835219.1;refGene_gn=PTPRB;refGene_tx=XM_019835220.2;refGene_gn=PTPRB;refGene_tx=XM_019835221.2
B4 97352386 97352650
refGene_gn=PTPRB;refGene_tx=XM_003989041.4;refGene_gn=PTPRB;refGene_tx=XM_006933890.3;refGene_gn=PTPRB;refGene_tx=XM_006933891.4;refGene_gn=PTPRB;refGene_tx=XM_019835219.1;refGene_gn=PTPRB;refGene_tx=XM_019835220.2;refGene_gn=PTPRB;refGene_tx=XM_019835221.2
B4 97353195 97353459
refGene_gn=PTPRB;refGene_tx=XM_003989041.4;refGene_gn=PTPRB;refGene_tx=XM_006933890.3;refGene_gn=PTPRB;refGene_tx=XM_006933891.4;refGene_gn=PTPRB;refGene_tx=XM_019835219.1;refGene_gn=PTPRB;refGene_tx=XM_019835220.2;refGene_gn=PTPRB;refGene_tx=XM_019835221.2

B4	97359518 97359782
refGene_gn=PTPRB;refGene_tx=XM_003989041.4;refGene_gn=PTPRB;refGene_tx=XM_006933890.3;refGene_gn=PTPRB;refGene_tx=XM_006933891.4;refGene_gn=PTPRB;refGene_tx=XM_019835219.1;refGene_gn=PTPRB;refGene_tx=XM_019835220.2;refGene_gn=PTPRB;refGene_tx=XM_019835221.2
B4	97363830 97364097
refGene_gn=PTPRB;refGene_tx=XM_003989041.4;refGene_gn=PTPRB;refGene_tx=XM_006933890.3;refGene_gn=PTPRB;refGene_tx=XM_006933891.4;refGene_gn=PTPRB;refGene_tx=XM_019835219.1;refGene_gn=PTPRB;refGene_tx=XM_019835220.2;refGene_gn=PTPRB;refGene_tx=XM_019835221.2
B4	97368598 97368859
refGene_gn=PTPRB;refGene_tx=XM_003989041.4;refGene_gn=PTPRB;refGene_tx=XM_006933890.3;refGene_gn=PTPRB;refGene_tx=XM_006933891.4;refGene_gn=PTPRB;refGene_tx=XM_019835219.1;refGene_gn=PTPRB;refGene_tx=XM_019835220.2;refGene_gn=PTPRB;refGene_tx=XM_019835221.2
B4	97370916 97371186
refGene_gn=PTPRB;refGene_tx=XM_003989041.4;refGene_gn=PTPRB;refGene_tx=XM_006933890.3;refGene_gn=PTPRB;refGene_tx=XM_006933891.4;refGene_gn=PTPRB;refGene_tx=XM_019835219.1;refGene_gn=PTPRB;refGene_tx=XM_019835220.2;refGene_gn=PTPRB;refGene_tx=XM_019835221.2
B4	97372959 97373223
refGene_gn=PTPRB;refGene_tx=XM_003989041.4;refGene_gn=PTPRB;refGene_tx=XM_006933890.3;refGene_gn=PTPRB;refGene_tx=XM_006933891.4;refGene_gn=PTPRB;refGene_tx=XM_019835219.1;refGene_gn=PTPRB;refGene_tx=XM_019835220.2;refGene_gn=PTPRB;refGene_tx=XM_019835221.2
B4	97375373 97375631
refGene_gn=PTPRB;refGene_tx=XM_003989041.4;refGene_gn=PTPRB;refGene_tx=XM_006933890.3;refGene_gn=PTPRB;refGene_tx=XM_006933891.4;refGene_gn=PTPRB;refGene_tx=XM_019835220.2;refGene_gn=PTPRB;refGene_tx=XM_019835221.2
B4	97376471 97376747
refGene_gn=PTPRB;refGene_tx=XM_003989041.4;refGene_gn=PTPRB;refGene_tx=XM_006933890.3;refGene_gn=PTPRB;refGene_tx=XM_006933891.4;refGene_gn=PTPRB;refGene_tx=XM_019835219.1;refGene_gn=PTPRB;refGene_tx=XM_019835220.2;refGene_gn=PTPRB;refGene_tx=XM_019835221.2
B4	97389580 97389854
refGene_gn=PTPRB;refGene_tx=XM_003989041.4;refGene_gn=PTPRB;refGene_tx=XM_006933890.3;refGene_gn=PTPRB;refGene_tx=XM_006933891.4;refGene_gn=PTPRB;refGene_tx=XM_019835219.1;refGene_gn=PTPRB;refGene_tx=XM_019835220.2;refGene_gn=PTPRB;refGene_tx=XM_019835221.2
B4	97390260 97390314 refGene_gn=PTPRB;refGene_tx=XM_006933891.4
B4	97408466 97408723
refGene_gn=PTPRB;refGene_tx=XM_003989041.4;refGene_gn=PTPRB;refGene_tx=XM_006933890.3;refGene_gn=PTPRB;refGene_tx=XM_019835219.1;refGene_gn=PTPRB;refGene_tx=XM_019835220.2;refGene_gn=PTPRB;refGene_tx=XM_019835221.2
B4	97418168 97418564
refGene_gn=PTPRB;refGene_tx=XM_003989041.4;refGene_gn=PTPRB;refGene_tx=XM_006933890.3;refGene_gn=PTPRB;refGene_tx=XM_019835219.1;refGene_gn=PTPRB;refGene_tx=XM_019835220.2;refGene_gn=PTPRB;refGene_tx=XM_0198352

21.2
B4 97419838 97419893
refGene_gn=PTPRB;refGene_tx=XM_003989041.4;refGene_gn=PTPRB;refGene_tx=XM_006933890.3;refGene_gn=PTPRB;refGene_tx=XM_019835219.1
B4 97421686 97421780
refGene_gn=PTPRR;refGene_tx=XM_011284123.3;refGene_gn=PTPRR;refGene_tx=XM_019835222.2
B4 97439216 97439330
refGene_gn=PTPRR;refGene_tx=XM_011284123.3;refGene_gn=PTPRR;refGene_tx=XM_019835222.2
B4 97443919 97444077
refGene_gn=PTPRR;refGene_tx=XM_011284123.3;refGene_gn=PTPRR;refGene_tx=XM_019835222.2
B4 97446128 97446239
refGene_gn=PTPRR;refGene_tx=XM_011284123.3;refGene_gn=PTPRR;refGene_tx=XM_019835222.2
B4 97455219 97455357
refGene_gn=PTPRR;refGene_tx=XM_011284123.3;refGene_gn=PTPRR;refGene_tx=XM_019835222.2
B4 97455790 97455870
refGene_gn=PTPRR;refGene_tx=XM_011284123.3;refGene_gn=PTPRR;refGene_tx=XM_019835222.2
B4 97464415 97464500
refGene_gn=PTPRR;refGene_tx=XM_011284123.3;refGene_gn=PTPRR;refGene_tx=XM_019835222.2
B4 97467098 97467285
refGene_gn=PTPRR;refGene_tx=XM_011284123.3;refGene_gn=PTPRR;refGene_tx=XM_019835222.2
B4 97515122 97515385
refGene_gn=PTPRR;refGene_tx=XM_011284123.3;refGene_gn=PTPRR;refGene_tx=XM_019835222.2
B4 97527890 97528001
refGene_gn=PTPRR;refGene_tx=XM_011284123.3;refGene_gn=PTPRR;refGene_tx=XM_019835222.2
B4 97534783 97534939
refGene_gn=PTPRR;refGene_tx=XM_011284123.3;refGene_gn=PTPRR;refGene_tx=XM_019835222.2
B4 97537988 97538102
refGene_gn=PTPRR;refGene_tx=XM_011284123.3;refGene_gn=PTPRR;refGene_tx=XM_019835222.2
B4 97644244 97644543 refGene_gn=PTPRR;refGene_tx=XM_019835222.2
B4 97664345 97664403 refGene_gn=PTPRR;refGene_tx=XM_019835222.2
B4 97845335 97845389
refGene_gn=TSPAN8;refGene_tx=XM_003989042.5;refGene_gn=TSPAN8;refGene_tx=XM_023257239.1
B4 97849789 97849873
refGene_gn=TSPAN8;refGene_tx=XM_003989042.5;refGene_gn=TSPAN8;refGene_tx=XM_023257239.1
B4 97852651 97852786
refGene_gn=TSPAN8;refGene_tx=XM_003989042.5;refGene_gn=TSPAN8;refGene_

tx=XM_023257239.1
B4 97857079 97857187
refGene_gn=TSPAN8;refGene_tx=XM_003989042.5;refGene_gn=TSPAN8;refGene_tx=XM_023257239.1
B4 97857284 97857359
refGene_gn=TSPAN8;refGene_tx=XM_003989042.5;refGene_gn=TSPAN8;refGene_tx=XM_023257239.1
B4 97858823 97858961
refGene_gn=TSPAN8;refGene_tx=XM_003989042.5;refGene_gn=TSPAN8;refGene_tx=XM_023257239.1
B4 97863988 97864051
refGene_gn=TSPAN8;refGene_tx=XM_003989042.5;refGene_gn=TSPAN8;refGene_tx=XM_023257239.1
B4 97877205 97877265
refGene_gn=TSPAN8;refGene_tx=XM_003989042.5;refGene_gn=TSPAN8;refGene_tx=XM_023257239.1
B4 98085832 98086044
refGene_gn=LGR5;refGene_tx=XM_023257240.1;refGene_gn=LGR5;refGene_tx=XM_023257241.1;refGene_gn=LGR5;refGene_tx=XM_023257242.1
B4 98142674 98142746
refGene_gn=LGR5;refGene_tx=XM_023257240.1;refGene_gn=LGR5;refGene_tx=XM_023257241.1;refGene_gn=LGR5;refGene_tx=XM_023257242.1
B4 98161528 98161600
refGene_gn=LGR5;refGene_tx=XM_023257240.1;refGene_gn=LGR5;refGene_tx=XM_023257242.1
B4 98172801 98172873
refGene_gn=LGR5;refGene_tx=XM_023257240.1;refGene_gn=LGR5;refGene_tx=XM_023257241.1;refGene_gn=LGR5;refGene_tx=XM_023257242.1
B4 98190845 98191061
refGene_gn=LGR5;refGene_tx=XM_023257240.1;refGene_gn=LGR5;refGene_tx=XM_023257241.1;refGene_gn=LGR5;refGene_tx=XM_023257242.1
B4 98192976 98193048
refGene_gn=LGR5;refGene_tx=XM_023257240.1;refGene_gn=LGR5;refGene_tx=XM_023257241.1;refGene_gn=LGR5;refGene_tx=XM_023257242.1
B4 98195929 98195998
refGene_gn=LGR5;refGene_tx=XM_023257240.1;refGene_gn=LGR5;refGene_tx=XM_023257241.1;refGene_gn=LGR5;refGene_tx=XM_023257242.1
B4 98197608 98197680
refGene_gn=LGR5;refGene_tx=XM_023257240.1;refGene_gn=LGR5;refGene_tx=XM_023257241.1
B4 98201060 98201132
refGene_gn=LGR5;refGene_tx=XM_023257240.1;refGene_gn=LGR5;refGene_tx=XM_023257241.1;refGene_gn=LGR5;refGene_tx=XM_023257242.1
B4 98201281 98201350
refGene_gn=LGR5;refGene_tx=XM_023257240.1;refGene_gn=LGR5;refGene_tx=XM_023257241.1;refGene_gn=LGR5;refGene_tx=XM_023257242.1
B4 98201491 98201563
refGene_gn=LGR5;refGene_tx=XM_023257240.1;refGene_gn=LGR5;refGene_tx=XM_023257241.1;refGene_gn=LGR5;refGene_tx=XM_023257242.1
B4 98204917 98204983

refGene_gn=LGR5;refGene_tx=XM_023257240.1;refGene_gn=LGR5;refGene_tx=XM_023257241.1;refGene_gn=LGR5;refGene_tx=XM_023257242.1
B4 98206368 98206440
refGene_gn=LGR5;refGene_tx=XM_023257240.1;refGene_gn=LGR5;refGene_tx=XM_023257241.1;refGene_gn=LGR5;refGene_tx=XM_023257242.1
B4 98211306 98211378
refGene_gn=LGR5;refGene_tx=XM_023257240.1;refGene_gn=LGR5;refGene_tx=XM_023257241.1;refGene_gn=LGR5;refGene_tx=XM_023257242.1
B4 98212180 98212306
refGene_gn=LGR5;refGene_tx=XM_023257240.1;refGene_gn=LGR5;refGene_tx=XM_023257241.1;refGene_gn=LGR5;refGene_tx=XM_023257242.1
B4 98213389 98213535
refGene_gn=LGR5;refGene_tx=XM_023257240.1;refGene_gn=LGR5;refGene_tx=XM_023257241.1;refGene_gn=LGR5;refGene_tx=XM_023257242.1
B4 98215290 98215374
refGene_gn=LGR5;refGene_tx=XM_023257240.1;refGene_gn=LGR5;refGene_tx=XM_023257241.1;refGene_gn=LGR5;refGene_tx=XM_023257242.1
B4 98216440 98217528
refGene_gn=LGR5;refGene_tx=XM_023257240.1;refGene_gn=LGR5;refGene_tx=XM_023257241.1;refGene_gn=LGR5;refGene_tx=XM_023257242.1
B4 98241080 98241218
refGene_gn=ZFC3H1;refGene_tx=XM_011284124.3;refGene_gn=ZFC3H1;refGene_tx=XM_019835224.2
B4 98241347 98241410
refGene_gn=ZFC3H1;refGene_tx=XM_011284124.3;refGene_gn=ZFC3H1;refGene_tx=XM_019835224.2
B4 98241650 98241690
refGene_gn=ZFC3H1;refGene_tx=XM_011284124.3;refGene_gn=ZFC3H1;refGene_tx=XM_019835224.2
B4 98242331 98242433
refGene_gn=ZFC3H1;refGene_tx=XM_011284124.3;refGene_gn=ZFC3H1;refGene_tx=XM_019835224.2
B4 98244048 98244149
refGene_gn=ZFC3H1;refGene_tx=XM_011284124.3;refGene_gn=ZFC3H1;refGene_tx=XM_019835224.2
B4 98245227 98245393
refGene_gn=ZFC3H1;refGene_tx=XM_011284124.3;refGene_gn=ZFC3H1;refGene_tx=XM_019835224.2
B4 98245527 98245632
refGene_gn=ZFC3H1;refGene_tx=XM_011284124.3;refGene_gn=ZFC3H1;refGene_tx=XM_019835224.2
B4 98245914 98246025
refGene_gn=ZFC3H1;refGene_tx=XM_011284124.3;refGene_gn=ZFC3H1;refGene_tx=XM_019835224.2
B4 98249696 98249791
refGene_gn=ZFC3H1;refGene_tx=XM_011284124.3;refGene_gn=ZFC3H1;refGene_tx=XM_019835224.2
B4 98250554 98250753
refGene_gn=ZFC3H1;refGene_tx=XM_011284124.3;refGene_gn=ZFC3H1;refGene_tx=XM_019835224.2

B4 98250840 98250946
refGene_gn=ZFC3H1;refGene_tx=XM_011284124.3;refGene_gn=ZFC3H1;refGene_tx=XM_019835224.2
B4 98252860 98253098
refGene_gn=ZFC3H1;refGene_tx=XM_011284124.3;refGene_gn=ZFC3H1;refGene_tx=XM_019835224.2
B4 98253742 98253931 refGene_gn=ZFC3H1;refGene_tx=XM_011284124.3
B4 98255403 98255590
refGene_gn=ZFC3H1;refGene_tx=XM_011284124.3;refGene_gn=ZFC3H1;refGene_tx=XM_019835224.2
B4 98257174 98257358
refGene_gn=ZFC3H1;refGene_tx=XM_011284124.3;refGene_gn=ZFC3H1;refGene_tx=XM_019835224.2
B4 98258517 98258637
refGene_gn=ZFC3H1;refGene_tx=XM_011284124.3;refGene_gn=ZFC3H1;refGene_tx=XM_019835224.2
B4 98258918 98259020
refGene_gn=ZFC3H1;refGene_tx=XM_011284124.3;refGene_gn=ZFC3H1;refGene_tx=XM_019835224.2
B4 98259987 98260109
refGene_gn=ZFC3H1;refGene_tx=XM_011284124.3;refGene_gn=ZFC3H1;refGene_tx=XM_019835224.2
B4 98260213 98260345
refGene_gn=ZFC3H1;refGene_tx=XM_011284124.3;refGene_gn=ZFC3H1;refGene_tx=XM_019835224.2
B4 98261393 98261503
refGene_gn=ZFC3H1;refGene_tx=XM_011284124.3;refGene_gn=ZFC3H1;refGene_tx=XM_019835224.2
B4 98261588 98262131
refGene_gn=ZFC3H1;refGene_tx=XM_011284124.3;refGene_gn=ZFC3H1;refGene_tx=XM_019835224.2
B4 98262488 98262620
refGene_gn=ZFC3H1;refGene_tx=XM_011284124.3;refGene_gn=ZFC3H1;refGene_tx=XM_019835224.2
B4 98262870 98263045
refGene_gn=ZFC3H1;refGene_tx=XM_011284124.3;refGene_gn=ZFC3H1;refGene_tx=XM_019835224.2
B4 98265752 98265902
refGene_gn=ZFC3H1;refGene_tx=XM_011284124.3;refGene_gn=ZFC3H1;refGene_tx=XM_019835224.2
B4 98266479 98266601
refGene_gn=ZFC3H1;refGene_tx=XM_011284124.3;refGene_gn=ZFC3H1;refGene_tx=XM_019835224.2
B4 98267436 98267574
refGene_gn=ZFC3H1;refGene_tx=XM_011284124.3;refGene_gn=ZFC3H1;refGene_tx=XM_019835224.2
B4 98268086 98268251
refGene_gn=ZFC3H1;refGene_tx=XM_011284124.3;refGene_gn=ZFC3H1;refGene_tx=XM_019835224.2
B4 98268421 98268631

refGene_gn=ZFC3H1;refGene_tx=XM_011284124.3;refGene_gn=ZFC3H1;refGene_tx=XM_019835224.2
B4 98269949 98270047
refGene_gn=ZFC3H1;refGene_tx=XM_011284124.3;refGene_gn=ZFC3H1;refGene_tx=XM_019835224.2
B4 98272752 98272876
refGene_gn=ZFC3H1;refGene_tx=XM_011284124.3;refGene_gn=ZFC3H1;refGene_tx=XM_019835224.2
B4 98274080 98274304
refGene_gn=ZFC3H1;refGene_tx=XM_011284124.3;refGene_gn=ZFC3H1;refGene_tx=XM_019835224.2
B4 98275009 98275208
refGene_gn=ZFC3H1;refGene_tx=XM_011284124.3;refGene_gn=ZFC3H1;refGene_tx=XM_019835224.2
B4 98279415 98279480
refGene_gn=ZFC3H1;refGene_tx=XM_011284124.3;refGene_gn=ZFC3H1;refGene_tx=XM_019835224.2
B4 98288131 98288548
refGene_gn=ZFC3H1;refGene_tx=XM_011284124.3;refGene_gn=ZFC3H1;refGene_tx=XM_019835224.2
B4 98293180 98293778
refGene_gn=ZFC3H1;refGene_tx=XM_011284124.3;refGene_gn=ZFC3H1;refGene_tx=XM_019835224.2
B4 98294654 98294725 refGene_gn=THAP2;refGene_tx=XM_011284125.3
B4 98300968 98301164 refGene_gn=THAP2;refGene_tx=XM_011284125.3
B4 98302840 98303230 refGene_gn=THAP2;refGene_tx=XM_011284125.3
B4 98311744 98311874 refGene_gn=TMEM19;refGene_tx=XM_023257243.1
B4 98313589 98313703 refGene_gn=TMEM19;refGene_tx=XM_023257243.1
B4 98328042 98328180 refGene_gn=TMEM19;refGene_tx=XM_023257243.1
B4 98328555 98328810 refGene_gn=TMEM19;refGene_tx=XM_023257243.1
B4 98329632 98329842 refGene_gn=TMEM19;refGene_tx=XM_023257243.1
B4 98331731 98331895 refGene_gn=TMEM19;refGene_tx=XM_023257243.1
B4 98363790 98363940 refGene_gn=RAB21;refGene_tx=XM_003989049.4
B4 98375749 98375809 refGene_gn=RAB21;refGene_tx=XM_003989049.4
B4 98376591 98376699 refGene_gn=RAB21;refGene_tx=XM_003989049.4
B4 98380139 98380203 refGene_gn=RAB21;refGene_tx=XM_003989049.4
B4 98386876 98386931 refGene_gn=RAB21;refGene_tx=XM_003989049.4
B4 98387395 98387484 refGene_gn=RAB21;refGene_tx=XM_003989049.4
B4 98402328 98402471 refGene_gn=RAB21;refGene_tx=XM_003989049.4
B4 98434688 98434718
refGene_gn=TBC1D15;refGene_tx=XM_003989050.5;refGene_gn=TBC1D15;refGene_tx=XM_019835226.1
B4 98457606 98457705
refGene_gn=TBC1D15;refGene_tx=XM_003989050.5;refGene_gn=TBC1D15;refGene_tx=XM_019835226.1
B4 98458242 98458317
refGene_gn=TBC1D15;refGene_tx=XM_003989050.5;refGene_gn=TBC1D15;refGene_tx=XM_019835226.1
B4 98463611 98463750
refGene_gn=TBC1D15;refGene_tx=XM_003989050.5;refGene_gn=TBC1D15;refGen

e_tx=XM_019835226.1
B4	98468582	98468793
refGene_gn=TBC1D15;refGene_tx=XM_003989050.5;refGene_gn=TBC1D15;refGene_tx=XM_019835226.1
B4	98476413	98476516
refGene_gn=TBC1D15;refGene_tx=XM_003989050.5;refGene_gn=TBC1D15;refGene_tx=XM_019835226.1
B4	98478179	98478230	refGene_gn=TBC1D15;refGene_tx=XM_019835226.1
B4	98478530	98478728
refGene_gn=TBC1D15;refGene_tx=XM_003989050.5;refGene_gn=TBC1D15;refGene_tx=XM_019835226.1
B4	98479796	98479925
refGene_gn=TBC1D15;refGene_tx=XM_003989050.5;refGene_gn=TBC1D15;refGene_tx=XM_019835226.1
B4	98480468	98480572
refGene_gn=TBC1D15;refGene_tx=XM_003989050.5;refGene_gn=TBC1D15;refGene_tx=XM_019835226.1
B4	98481587	98481682
refGene_gn=TBC1D15;refGene_tx=XM_003989050.5;refGene_gn=TBC1D15;refGene_tx=XM_019835226.1
B4	98489549	98489666
refGene_gn=TBC1D15;refGene_tx=XM_003989050.5;refGene_gn=TBC1D15;refGene_tx=XM_019835226.1
B4	98495243	98495344
refGene_gn=TBC1D15;refGene_tx=XM_003989050.5;refGene_gn=TBC1D15;refGene_tx=XM_019835226.1
B4	98499383	98499483
refGene_gn=TBC1D15;refGene_tx=XM_003989050.5;refGene_gn=TBC1D15;refGene_tx=XM_019835226.1
B4	98499866	98499964
refGene_gn=TBC1D15;refGene_tx=XM_003989050.5;refGene_gn=TBC1D15;refGene_tx=XM_019835226.1
B4	98503994	98504111
refGene_gn=TBC1D15;refGene_tx=XM_003989050.5;refGene_gn=TBC1D15;refGene_tx=XM_019835226.1
B4	98504627	98504714
refGene_gn=TBC1D15;refGene_tx=XM_003989050.5;refGene_gn=TBC1D15;refGene_tx=XM_019835226.1
B4	98505713	98505935
refGene_gn=TBC1D15;refGene_tx=XM_003989050.5;refGene_gn=TBC1D15;refGene_tx=XM_019835226.1
B4	98524707	98524812	refGene_gn=TPH2;refGene_tx=XM_003989052.3
B4	98527372	98527525	refGene_gn=TPH2;refGene_tx=XM_003989052.3
B4	98530286	98530470	refGene_gn=TPH2;refGene_tx=XM_003989052.3
B4	98530580	98530681	refGene_gn=TPH2;refGene_tx=XM_003989052.3
B4	98535667	98535735	refGene_gn=TPH2;refGene_tx=XM_003989052.3
B4	98557616	98557813	refGene_gn=TPH2;refGene_tx=XM_003989052.3
B4	98562534	98562670	refGene_gn=TPH2;refGene_tx=XM_003989052.3
B4	98588122	98588249	refGene_gn=TPH2;refGene_tx=XM_003989052.3
B4	98618007	98618103	refGene_gn=TPH2;refGene_tx=XM_003989052.3

B4 98627410 98627544 refGene_gn=TPH2;refGene_tx=XM_003989052.3
B4 98627674 98627849 refGene_gn=TPH2;refGene_tx=XM_003989052.3
B4 98867456 98868373 refGene_gn=TRHDE;refGene_tx=XM_011284128.3
B4 98882053 98882327 refGene_gn=TRHDE;refGene_tx=XM_011284128.3
B4 98981129 98981256 refGene_gn=TRHDE;refGene_tx=XM_011284128.3
B4 99072050 99072205 refGene_gn=TRHDE;refGene_tx=XM_011284128.3
B4 99074955 99075069 refGene_gn=TRHDE;refGene_tx=XM_011284128.3
B4 99096058 99096196 refGene_gn=TRHDE;refGene_tx=XM_011284128.3
B4 99139610 99139676 refGene_gn=TRHDE;refGene_tx=XM_011284128.3
B4 99159859 99159925 refGene_gn=TRHDE;refGene_tx=XM_011284128.3
B4 99160568 99160756 refGene_gn=TRHDE;refGene_tx=XM_011284128.3
B4 99165901 99165990 refGene_gn=TRHDE;refGene_tx=XM_011284128.3
B4 99172851 99172982 refGene_gn=TRHDE;refGene_tx=XM_011284128.3
B4 99173090 99173146 refGene_gn=TRHDE;refGene_tx=XM_011284128.3
B4 99219295 99219443 refGene_gn=TRHDE;refGene_tx=XM_011284128.3
B4 99221349 99221447 refGene_gn=TRHDE;refGene_tx=XM_011284128.3
B4 99221917 99222025 refGene_gn=TRHDE;refGene_tx=XM_011284128.3
B4 99243140 99243308 refGene_gn=TRHDE;refGene_tx=XM_011284128.3
B4 99243819 99243960 refGene_gn=TRHDE;refGene_tx=XM_011284128.3
B4 99246538 99246620 refGene_gn=TRHDE;refGene_tx=XM_011284128.3
B4 99252132 99252276 refGene_gn=TRHDE;refGene_tx=XM_011284128.3
B4 100666947 100667241
refGene_gn=ATXN7L3B;refGene_tx=XM_006933896.3
B4 100993229 100993339
refGene_gn=KCNC2;refGene_tx=XM_023257256.1
B4 100994441 100994503
refGene_gn=KCNC2;refGene_tx=XM_023257250.1;refGene_gn=KCNC2;refGene_tx=XM_023257251.1;refGene_gn=KCNC2;refGene_tx=XM_023257257.1;refGene_gn=KCNC2;refGene_tx=XM_023257258.1
B4 100995211 100995348
refGene_gn=KCNC2;refGene_tx=XM_023257244.1;refGene_gn=KCNC2;refGene_tx=XM_023257245.1;refGene_gn=KCNC2;refGene_tx=XM_023257246.1;refGene_gn=KCNC2;refGene_tx=XM_023257247.1;refGene_gn=KCNC2;refGene_tx=XM_023257248.1;refGene_gn=KCNC2;refGene_tx=XM_023257249.1;refGene_gn=KCNC2;refGene_tx=XM_023257255.1
B4 101000499 101000664
refGene_gn=KCNC2;refGene_tx=XM_023257244.1;refGene_gn=KCNC2;refGene_tx=XM_023257245.1;refGene_gn=KCNC2;refGene_tx=XM_023257246.1;refGene_gn=KCNC2;refGene_tx=XM_023257247.1;refGene_gn=KCNC2;refGene_tx=XM_023257248.1;refGene_gn=KCNC2;refGene_tx=XM_023257249.1;refGene_gn=KCNC2;refGene_tx=XM_023257250.1;refGene_gn=KCNC2;refGene_tx=XM_023257251.1;refGene_gn=KCNC2;refGene_tx=XM_023257252.1;refGene_gn=KCNC2;refGene_tx=XM_023257253.1;refGene_gn=KCNC2;refGene_tx=XM_023257254.1
B4 101002226 101002330
refGene_gn=KCNC2;refGene_tx=XM_023257252.1;refGene_gn=KCNC2;refGene_tx=XM_023257253.1;refGene_gn=KCNC2;refGene_tx=XM_023257254.1
B4 101002736 101003664
refGene_gn=KCNC2;refGene_tx=XM_023257244.1;refGene_gn=KCNC2;refGene_tx=XM_023257245.1;refGene_gn=KCNC2;refGene_tx=XM_023257246.1;refGene_gn=KCNC2;refGene_tx=XM_023257247.1;refGene_gn=KCNC2;refGene_tx=XM_0232572

48.1;refGene_gn=KCNC2;refGene_tx=XM_023257249.1;refGene_gn=KCNC2;refGene_tx=XM_023257250.1;refGene_gn=KCNC2;refGene_tx=XM_023257251.1;refGene_gn=KCNC2;refGene_tx=XM_023257252.1;refGene_gn=KCNC2;refGene_tx=XM_023257253.1;refGene_gn=KCNC2;refGene_tx=XM_023257254.1;refGene_gn=KCNC2;refGene_tx=XM_023257255.1;refGene_gn=KCNC2;refGene_tx=XM_023257256.1;refGene_gn=KCNC2;refGene_tx=XM_023257257.1;refGene_gn=KCNC2;refGene_tx=XM_023257258.1
B4 101187770 101188433
refGene_gn=KCNC2;refGene_tx=XM_023257244.1;refGene_gn=KCNC2;refGene_tx=XM_023257245.1;refGene_gn=KCNC2;refGene_tx=XM_023257246.1;refGene_gn=KCNC2;refGene_tx=XM_023257247.1;refGene_gn=KCNC2;refGene_tx=XM_023257248.1;refGene_gn=KCNC2;refGene_tx=XM_023257249.1;refGene_gn=KCNC2;refGene_tx=XM_023257250.1;refGene_gn=KCNC2;refGene_tx=XM_023257251.1;refGene_gn=KCNC2;refGene_tx=XM_023257252.1;refGene_gn=KCNC2;refGene_tx=XM_023257253.1;refGene_gn=KCNC2;refGene_tx=XM_023257254.1;refGene_gn=KCNC2;refGene_tx=XM_023257255.1;refGene_gn=KCNC2;refGene_tx=XM_023257256.1;refGene_gn=KCNC2;refGene_tx=XM_023257257.1;refGene_gn=KCNC2;refGene_tx=XM_023257258.1
B4 101235769 101235945
refGene_gn=CAPS2;refGene_tx=XM_019835231.2;refGene_gn=CAPS2;refGene_tx=XM_019835232.2;refGene_gn=CAPS2;refGene_tx=XM_019835234.2;refGene_gn=CAPS2;refGene_tx=XM_019835236.2;refGene_gn=CAPS2;refGene_tx=XM_019835237.2;refGene_gn=CAPS2;refGene_tx=XM_023257259.1
B4 101240506 101240603
refGene_gn=CAPS2;refGene_tx=XM_019835231.2;refGene_gn=CAPS2;refGene_tx=XM_019835232.2;refGene_gn=CAPS2;refGene_tx=XM_019835234.2;refGene_gn=CAPS2;refGene_tx=XM_019835236.2;refGene_gn=CAPS2;refGene_tx=XM_019835237.2;refGene_gn=CAPS2;refGene_tx=XM_023257259.1
B4 101243533 101243653
refGene_gn=CAPS2;refGene_tx=XM_019835231.2;refGene_gn=CAPS2;refGene_tx=XM_019835232.2;refGene_gn=CAPS2;refGene_tx=XM_019835234.2;refGene_gn=CAPS2;refGene_tx=XM_019835236.2;refGene_gn=CAPS2;refGene_tx=XM_019835237.2;refGene_gn=CAPS2;refGene_tx=XM_023257259.1
B4 101246058 101246213
refGene_gn=CAPS2;refGene_tx=XM_019835231.2;refGene_gn=CAPS2;refGene_tx=XM_019835232.2;refGene_gn=CAPS2;refGene_tx=XM_019835234.2;refGene_gn=CAPS2;refGene_tx=XM_019835236.2;refGene_gn=CAPS2;refGene_tx=XM_019835237.2;refGene_gn=CAPS2;refGene_tx=XM_023257259.1
B4 101251133 101251207
refGene_gn=CAPS2;refGene_tx=XM_019835231.2;refGene_gn=CAPS2;refGene_tx=XM_019835232.2;refGene_gn=CAPS2;refGene_tx=XM_019835234.2;refGene_gn=CAPS2;refGene_tx=XM_019835236.2;refGene_gn=CAPS2;refGene_tx=XM_019835237.2;refGene_gn=CAPS2;refGene_tx=XM_023257259.1
B4 101252884 101253003
refGene_gn=CAPS2;refGene_tx=XM_019835231.2;refGene_gn=CAPS2;refGene_tx=XM_019835232.2;refGene_gn=CAPS2;refGene_tx=XM_019835234.2;refGene_gn=CAPS2;refGene_tx=XM_019835236.2;refGene_gn=CAPS2;refGene_tx=XM_019835237.2;refGene_gn=CAPS2;refGene_tx=XM_023257259.1
B4 101257936 101258030
refGene_gn=CAPS2;refGene_tx=XM_019835231.2;refGene_gn=CAPS2;refGene_tx

=XM_019835232.2;refGene_gn=CAPS2;refGene_tx=XM_019835234.2;refGene_gn=CAPS2;refGene_tx=XM_019835236.2;refGene_gn=CAPS2;refGene_tx=XM_019835237.2;refGene_gn=CAPS2;refGene_tx=XM_023257259.1
B4 101258118 101258214
refGene_gn=CAPS2;refGene_tx=XM_019835231.2;refGene_gn=CAPS2;refGene_tx=XM_019835232.2;refGene_gn=CAPS2;refGene_tx=XM_019835234.2;refGene_gn=CAPS2;refGene_tx=XM_019835236.2;refGene_gn=CAPS2;refGene_tx=XM_019835237.2;refGene_gn=CAPS2;refGene_tx=XM_023257259.1
B4 101259395 101259470
refGene_gn=CAPS2;refGene_tx=XM_019835231.2;refGene_gn=CAPS2;refGene_tx=XM_019835232.2;refGene_gn=CAPS2;refGene_tx=XM_019835234.2;refGene_gn=CAPS2;refGene_tx=XM_019835236.2;refGene_gn=CAPS2;refGene_tx=XM_019835237.2;refGene_gn=CAPS2;refGene_tx=XM_023257259.1
B4 101270568 101270688
refGene_gn=CAPS2;refGene_tx=XM_019835231.2;refGene_gn=CAPS2;refGene_tx=XM_019835232.2;refGene_gn=CAPS2;refGene_tx=XM_019835234.2;refGene_gn=CAPS2;refGene_tx=XM_019835236.2;refGene_gn=CAPS2;refGene_tx=XM_019835237.2;refGene_gn=CAPS2;refGene_tx=XM_023257259.1
B4 101274455 101274523
refGene_gn=CAPS2;refGene_tx=XM_019835231.2;refGene_gn=CAPS2;refGene_tx=XM_019835232.2;refGene_gn=CAPS2;refGene_tx=XM_019835234.2;refGene_gn=CAPS2;refGene_tx=XM_019835236.2;refGene_gn=CAPS2;refGene_tx=XM_019835237.2;refGene_gn=CAPS2;refGene_tx=XM_023257259.1
B4 101275266 101275281
refGene_gn=CAPS2;refGene_tx=XM_019835237.2
B4 101277032 101277155
refGene_gn=CAPS2;refGene_tx=XM_019835231.2;refGene_gn=CAPS2;refGene_tx=XM_019835232.2;refGene_gn=CAPS2;refGene_tx=XM_019835234.2;refGene_gn=CAPS2;refGene_tx=XM_019835236.2;refGene_gn=CAPS2;refGene_tx=XM_023257259.1
B4 101281547 101281723
refGene_gn=CAPS2;refGene_tx=XM_019835231.2;refGene_gn=CAPS2;refGene_tx=XM_019835232.2;refGene_gn=CAPS2;refGene_tx=XM_019835234.2;refGene_gn=CAPS2;refGene_tx=XM_019835236.2;refGene_gn=CAPS2;refGene_tx=XM_023257259.1
B4 101282160 101282204
refGene_gn=CAPS2;refGene_tx=XM_019835231.2;refGene_gn=CAPS2;refGene_tx=XM_019835232.2
B4 101283853 101283903
refGene_gn=CAPS2;refGene_tx=XM_019835231.2;refGene_gn=CAPS2;refGene_tx=XM_019835232.2
B4 101289266 101289344
refGene_gn=CAPS2;refGene_tx=XM_019835231.2;refGene_gn=CAPS2;refGene_tx=XM_019835232.2;refGene_gn=CAPS2;refGene_tx=XM_019835234.2
B4 101300460 101300736
refGene_gn=GLIPR1L1;refGene_tx=XM_023257260.1
B4 101318261 101318507
refGene_gn=GLIPR1L1;refGene_tx=XM_023257260.1
B4 101322207 101322308
refGene_gn=GLIPR1L1;refGene_tx=XM_023257260.1

B4	101331502	101331591
refGene_gn=GLIPR1L1;refGene_tx=XM_023257260.1
B4	101336428	101336455
refGene_gn=GLIPR1L1;refGene_tx=XM_023257260.1
B4	101336543	101336635
refGene_gn=GLIPR1L1;refGene_tx=XM_023257260.1
B4	101351973	101352207
refGene_gn=GLIPR1L2;refGene_tx=XM_019835241.2
B4	101354591	101354837
refGene_gn=GLIPR1L2;refGene_tx=XM_019835241.2
B4	101357177	101357281
refGene_gn=GLIPR1L2;refGene_tx=XM_019835241.2
B4	101373222	101373308
refGene_gn=GLIPR1L2;refGene_tx=XM_019835241.2
B4	101381200	101381227
refGene_gn=GLIPR1L2;refGene_tx=XM_019835241.2
B4	101381310	101381684
refGene_gn=GLIPR1L2;refGene_tx=XM_019835241.2
B4	101423233	101423425
refGene_gn=GLIPR1;refGene_tx=XM_003989068.4;refGene_gn=GLIPR1;refGene_tx=XM_019835243.2;refGene_gn=GLIPR1;refGene_tx=XM_023257262.1
B4	101424614	101424860
refGene_gn=GLIPR1;refGene_tx=XM_003989068.4;refGene_gn=GLIPR1;refGene_tx=XM_019835243.2;refGene_gn=GLIPR1;refGene_tx=XM_023257262.1
B4	101433801	101433911
refGene_gn=GLIPR1;refGene_tx=XM_003989068.4;refGene_gn=GLIPR1;refGene_tx=XM_019835243.2;refGene_gn=GLIPR1;refGene_tx=XM_023257262.1
B4	101439506	101439589
refGene_gn=GLIPR1;refGene_tx=XM_003989068.4;refGene_gn=GLIPR1;refGene_tx=XM_019835243.2;refGene_gn=GLIPR1;refGene_tx=XM_023257262.1
B4	101442333	101442360
refGene_gn=GLIPR1;refGene_tx=XM_003989068.4;refGene_gn=GLIPR1;refGene_tx=XM_019835243.2;refGene_gn=GLIPR1;refGene_tx=XM_023257262.1
B4	101442463	101442609
refGene_gn=GLIPR1;refGene_tx=XM_003989068.4;refGene_gn=GLIPR1;refGene_tx=XM_019835243.2;refGene_gn=GLIPR1;refGene_tx=XM_023257262.1
B4	101443615	101443758
refGene_gn=KRR1;refGene_tx=XM_019835242.2;refGene_gn=KRR1;refGene_tx=XM_023257261.1
B4	101445479	101445573
refGene_gn=KRR1;refGene_tx=XM_019835242.2;refGene_gn=KRR1;refGene_tx=XM_023257261.1
B4	101445683	101445761
refGene_gn=KRR1;refGene_tx=XM_019835242.2;refGene_gn=KRR1;refGene_tx=XM_023257261.1
B4	101446953	101447124
refGene_gn=KRR1;refGene_tx=XM_019835242.2;refGene_gn=KRR1;refGene_tx=XM_023257261.1
B4	101449015	101449072
refGene_gn=KRR1;refGene_tx=XM_019835242.2;refGene_gn=KRR1;refGene_tx=X

M_023257261.1
B4	101450200	101450284
refGene_gn=KRR1;refGene_tx=XM_019835242.2;refGene_gn=KRR1;refGene_tx=X
M_023257261.1
B4	101450376	101450502
refGene_gn=KRR1;refGene_tx=XM_019835242.2;refGene_gn=KRR1;refGene_tx=X
M_023257261.1
B4	101450681	101450816
refGene_gn=KRR1;refGene_tx=XM_019835242.2;refGene_gn=KRR1;refGene_tx=X
M_023257261.1
B4	101452670	101452843
refGene_gn=KRR1;refGene_tx=XM_019835242.2;refGene_gn=KRR1;refGene_tx=X
M_023257261.1
B4	101455881	101455966
refGene_gn=KRR1;refGene_tx=XM_023257261.1
B4	101557287	101557312
refGene_gn=KRR1;refGene_tx=XM_019835242.2
B4	101922105	101923275
refGene_gn=PHLDA1;refGene_tx=XM_004001594.4
B4	101939228	101939264
refGene_gn=NAP1L1;refGene_tx=XM_003989056.4;refGene_gn=NAP1L1;refGene_
tx=XM_006933898.3;refGene_gn=NAP1L1;refGene_tx=XM_006933899.3;refGene_
gn=NAP1L1;refGene_tx=XM_006933900.3
B4	101940331	101940382
refGene_gn=NAP1L1;refGene_tx=XM_003989056.4;refGene_gn=NAP1L1;refGene_
tx=XM_006933898.3;refGene_gn=NAP1L1;refGene_tx=XM_006933899.3;refGene_
gn=NAP1L1;refGene_tx=XM_006933900.3
B4	101940597	101940660
refGene_gn=NAP1L1;refGene_tx=XM_011284132.2
B4	101940892	101940922
refGene_gn=NAP1L1;refGene_tx=XM_003989056.4;refGene_gn=NAP1L1;refGene_
tx=XM_006933898.3;refGene_gn=NAP1L1;refGene_tx=XM_006933899.3;refGene_
gn=NAP1L1;refGene_tx=XM_006933900.3;refGene_gn=NAP1L1;refGene_tx=XM_01
1284132.2
B4	101941627	101941750
refGene_gn=NAP1L1;refGene_tx=XM_003989056.4;refGene_gn=NAP1L1;refGene_
tx=XM_006933898.3;refGene_gn=NAP1L1;refGene_tx=XM_006933899.3;refGene_
gn=NAP1L1;refGene_tx=XM_006933900.3;refGene_gn=NAP1L1;refGene_tx=XM_01
1284132.2
B4	101943599	101943619
refGene_gn=NAP1L1;refGene_tx=XM_003989056.4;refGene_gn=NAP1L1;refGene_
tx=XM_006933898.3;refGene_gn=NAP1L1;refGene_tx=XM_006933899.3;refGene_
gn=NAP1L1;refGene_tx=XM_006933900.3;refGene_gn=NAP1L1;refGene_tx=XM_01
1284132.2
B4	101943713	101943859
refGene_gn=NAP1L1;refGene_tx=XM_003989056.4;refGene_gn=NAP1L1;refGene_
tx=XM_006933898.3;refGene_gn=NAP1L1;refGene_tx=XM_006933899.3;refGene_
gn=NAP1L1;refGene_tx=XM_006933900.3;refGene_gn=NAP1L1;refGene_tx=XM_01
1284132.2
B4	101944302	101944442

refGene_gn=NAP1L1;refGene_tx=XM_003989056.4;refGene_gn=NAP1L1;refGene_tx=XM_006933898.3;refGene_gn=NAP1L1;refGene_tx=XM_006933899.3;refGene_gn=NAP1L1;refGene_tx=XM_006933900.3;refGene_gn=NAP1L1;refGene_tx=XM_011284132.2
B4 101945586 101945658
refGene_gn=NAP1L1;refGene_tx=XM_003989056.4;refGene_gn=NAP1L1;refGene_tx=XM_006933898.3;refGene_gn=NAP1L1;refGene_tx=XM_006933899.3;refGene_gn=NAP1L1;refGene_tx=XM_006933900.3;refGene_gn=NAP1L1;refGene_tx=XM_011284132.2
B4 101946627 101946756
refGene_gn=NAP1L1;refGene_tx=XM_003989056.4;refGene_gn=NAP1L1;refGene_tx=XM_006933898.3;refGene_gn=NAP1L1;refGene_tx=XM_006933899.3;refGene_gn=NAP1L1;refGene_tx=XM_006933900.3;refGene_gn=NAP1L1;refGene_tx=XM_011284132.2
B4 101947678 101947759
refGene_gn=NAP1L1;refGene_tx=XM_003989056.4;refGene_gn=NAP1L1;refGene_tx=XM_006933898.3;refGene_gn=NAP1L1;refGene_tx=XM_006933899.3;refGene_gn=NAP1L1;refGene_tx=XM_006933900.3;refGene_gn=NAP1L1;refGene_tx=XM_011284132.2
B4 101948012 101948154
refGene_gn=NAP1L1;refGene_tx=XM_003989056.4;refGene_gn=NAP1L1;refGene_tx=XM_006933898.3;refGene_gn=NAP1L1;refGene_tx=XM_006933899.3;refGene_gn=NAP1L1;refGene_tx=XM_006933900.3;refGene_gn=NAP1L1;refGene_tx=XM_011284132.2
B4 101953753 101953856
refGene_gn=NAP1L1;refGene_tx=XM_003989056.4;refGene_gn=NAP1L1;refGene_tx=XM_006933898.3;refGene_gn=NAP1L1;refGene_tx=XM_006933899.3;refGene_gn=NAP1L1;refGene_tx=XM_011284132.2
B4 101955338 101955424
refGene_gn=NAP1L1;refGene_tx=XM_003989056.4;refGene_gn=NAP1L1;refGene_tx=XM_006933898.3;refGene_gn=NAP1L1;refGene_tx=XM_006933899.3;refGene_gn=NAP1L1;refGene_tx=XM_011284132.2
B4 101959928 101959945
refGene_gn=NAP1L1;refGene_tx=XM_003989056.4;refGene_gn=NAP1L1;refGene_tx=XM_006933898.3;refGene_gn=NAP1L1;refGene_tx=XM_006933899.3;refGene_gn=NAP1L1;refGene_tx=XM_006933900.3;refGene_gn=NAP1L1;refGene_tx=XM_011284132.2
B4 101969578 101970303
refGene_gn=LOC111561464;refGene_tx=XM_023257771.1
B4 102164826 102166807
refGene_gn=BBS10;refGene_tx=XM_003989057.5
B4 102167235 102167542
refGene_gn=BBS10;refGene_tx=XM_003989057.5
B4 102167930 102167991
refGene_gn=BBS10;refGene_tx=XM_003989057.5
B4 102173712 102173845
refGene_gn=OSBPL8;refGene_tx=XM_019835250.2;refGene_gn=OSBPL8;refGene_tx=XM_023257263.1;refGene_gn=OSBPL8;refGene_tx=XM_023257264.1;refGene_gn=OSBPL8;refGene_tx=XM_023257265.1;refGene_gn=OSBPL8;refGene_tx=XM_023257266.1;refGene_gn=OSBPL8;refGene_tx=XM_023257267.1;refGene_gn=OSBPL

8;refGene_tx=XM_023257268.1;refGene_gn=OSBPL8;refGene_tx=XM_023257269.
1;refGene_gn=OSBPL8;refGene_tx=XM_023257270.1;refGene_gn=OSBPL8;refGen
e_tx=XM_023257271.1;refGene_gn=OSBPL8;refGene_tx=XM_023257272.1;refGen
e_gn=OSBPL8;refGene_tx=XM_023257273.1
B4 102175879 102175982
refGene_gn=OSBPL8;refGene_tx=XM_019835250.2;refGene_gn=OSBPL8;refGene_
tx=XM_023257263.1;refGene_gn=OSBPL8;refGene_tx=XM_023257264.1;refGene_
gn=OSBPL8;refGene_tx=XM_023257265.1;refGene_gn=OSBPL8;refGene_tx=XM_02
3257266.1;refGene_gn=OSBPL8;refGene_tx=XM_023257267.1;refGene_gn=OSBPL
8;refGene_tx=XM_023257268.1;refGene_gn=OSBPL8;refGene_tx=XM_023257269.
1;refGene_gn=OSBPL8;refGene_tx=XM_023257270.1;refGene_gn=OSBPL8;refGen
e_tx=XM_023257271.1;refGene_gn=OSBPL8;refGene_tx=XM_023257272.1;refGen
e_gn=OSBPL8;refGene_tx=XM_023257273.1
B4 102177813 102177919
refGene_gn=OSBPL8;refGene_tx=XM_019835250.2;refGene_gn=OSBPL8;refGene_
tx=XM_023257263.1;refGene_gn=OSBPL8;refGene_tx=XM_023257264.1;refGene_
gn=OSBPL8;refGene_tx=XM_023257265.1;refGene_gn=OSBPL8;refGene_tx=XM_02
3257266.1;refGene_gn=OSBPL8;refGene_tx=XM_023257267.1;refGene_gn=OSBPL
8;refGene_tx=XM_023257268.1;refGene_gn=OSBPL8;refGene_tx=XM_023257269.
1;refGene_gn=OSBPL8;refGene_tx=XM_023257270.1;refGene_gn=OSBPL8;refGen
e_tx=XM_023257271.1;refGene_gn=OSBPL8;refGene_tx=XM_023257272.1;refGen
e_gn=OSBPL8;refGene_tx=XM_023257273.1
B4 102180968 102181056
refGene_gn=OSBPL8;refGene_tx=XM_019835250.2;refGene_gn=OSBPL8;refGene_
tx=XM_023257263.1;refGene_gn=OSBPL8;refGene_tx=XM_023257264.1;refGene_
gn=OSBPL8;refGene_tx=XM_023257265.1;refGene_gn=OSBPL8;refGene_tx=XM_02
3257266.1;refGene_gn=OSBPL8;refGene_tx=XM_023257267.1;refGene_gn=OSBPL
8;refGene_tx=XM_023257268.1;refGene_gn=OSBPL8;refGene_tx=XM_023257269.
1;refGene_gn=OSBPL8;refGene_tx=XM_023257270.1;refGene_gn=OSBPL8;refGen
e_tx=XM_023257271.1;refGene_gn=OSBPL8;refGene_tx=XM_023257272.1;refGen
e_gn=OSBPL8;refGene_tx=XM_023257273.1
B4 102181390 102181576
refGene_gn=OSBPL8;refGene_tx=XM_019835250.2;refGene_gn=OSBPL8;refGene_
tx=XM_023257263.1;refGene_gn=OSBPL8;refGene_tx=XM_023257264.1;refGene_
gn=OSBPL8;refGene_tx=XM_023257265.1;refGene_gn=OSBPL8;refGene_tx=XM_02
3257266.1;refGene_gn=OSBPL8;refGene_tx=XM_023257267.1;refGene_gn=OSBPL
8;refGene_tx=XM_023257268.1;refGene_gn=OSBPL8;refGene_tx=XM_023257269.
1;refGene_gn=OSBPL8;refGene_tx=XM_023257270.1;refGene_gn=OSBPL8;refGen
e_tx=XM_023257271.1;refGene_gn=OSBPL8;refGene_tx=XM_023257272.1;refGen
e_gn=OSBPL8;refGene_tx=XM_023257273.1
B4 102182940 102183077
refGene_gn=OSBPL8;refGene_tx=XM_019835250.2;refGene_gn=OSBPL8;refGene_
tx=XM_023257263.1;refGene_gn=OSBPL8;refGene_tx=XM_023257264.1;refGene_
gn=OSBPL8;refGene_tx=XM_023257265.1;refGene_gn=OSBPL8;refGene_tx=XM_02
3257266.1;refGene_gn=OSBPL8;refGene_tx=XM_023257267.1;refGene_gn=OSBPL
8;refGene_tx=XM_023257268.1;refGene_gn=OSBPL8;refGene_tx=XM_023257269.
1;refGene_gn=OSBPL8;refGene_tx=XM_023257270.1;refGene_gn=OSBPL8;refGen
e_tx=XM_023257271.1;refGene_gn=OSBPL8;refGene_tx=XM_023257272.1;refGen
e_gn=OSBPL8;refGene_tx=XM_023257273.1
B4 102184588 102184678

refGene_gn=OSBPL8;refGene_tx=XM_019835250.2;refGene_gn=OSBPL8;refGene_tx=XM_023257263.1;refGene_gn=OSBPL8;refGene_tx=XM_023257264.1;refGene_gn=OSBPL8;refGene_tx=XM_023257265.1;refGene_gn=OSBPL8;refGene_tx=XM_023257266.1;refGene_gn=OSBPL8;refGene_tx=XM_023257267.1;refGene_gn=OSBPL8;refGene_tx=XM_023257268.1;refGene_gn=OSBPL8;refGene_tx=XM_023257269.1;refGene_gn=OSBPL8;refGene_tx=XM_023257270.1;refGene_gn=OSBPL8;refGene_tx=XM_023257271.1;refGene_gn=OSBPL8;refGene_tx=XM_023257272.1;refGene_gn=OSBPL8;refGene_tx=XM_023257273.1
B4 102186541 102186639
refGene_gn=OSBPL8;refGene_tx=XM_019835250.2;refGene_gn=OSBPL8;refGene_tx=XM_023257263.1;refGene_gn=OSBPL8;refGene_tx=XM_023257264.1;refGene_gn=OSBPL8;refGene_tx=XM_023257265.1;refGene_gn=OSBPL8;refGene_tx=XM_023257266.1;refGene_gn=OSBPL8;refGene_tx=XM_023257267.1;refGene_gn=OSBPL8;refGene_tx=XM_023257268.1;refGene_gn=OSBPL8;refGene_tx=XM_023257269.1;refGene_gn=OSBPL8;refGene_tx=XM_023257270.1;refGene_gn=OSBPL8;refGene_tx=XM_023257271.1;refGene_gn=OSBPL8;refGene_tx=XM_023257272.1;refGene_gn=OSBPL8;refGene_tx=XM_023257273.1
B4 102188314 102188413
refGene_gn=OSBPL8;refGene_tx=XM_019835250.2;refGene_gn=OSBPL8;refGene_tx=XM_023257263.1;refGene_gn=OSBPL8;refGene_tx=XM_023257264.1;refGene_gn=OSBPL8;refGene_tx=XM_023257265.1;refGene_gn=OSBPL8;refGene_tx=XM_023257266.1;refGene_gn=OSBPL8;refGene_tx=XM_023257267.1;refGene_gn=OSBPL8;refGene_tx=XM_023257268.1;refGene_gn=OSBPL8;refGene_tx=XM_023257269.1;refGene_gn=OSBPL8;refGene_tx=XM_023257270.1;refGene_gn=OSBPL8;refGene_tx=XM_023257271.1;refGene_gn=OSBPL8;refGene_tx=XM_023257272.1;refGene_gn=OSBPL8;refGene_tx=XM_023257273.1
B4 102193627 102193724
refGene_gn=OSBPL8;refGene_tx=XM_019835250.2;refGene_gn=OSBPL8;refGene_tx=XM_023257263.1;refGene_gn=OSBPL8;refGene_tx=XM_023257264.1;refGene_gn=OSBPL8;refGene_tx=XM_023257265.1;refGene_gn=OSBPL8;refGene_tx=XM_023257266.1;refGene_gn=OSBPL8;refGene_tx=XM_023257267.1;refGene_gn=OSBPL8;refGene_tx=XM_023257268.1;refGene_gn=OSBPL8;refGene_tx=XM_023257269.1;refGene_gn=OSBPL8;refGene_tx=XM_023257270.1;refGene_gn=OSBPL8;refGene_tx=XM_023257271.1;refGene_gn=OSBPL8;refGene_tx=XM_023257272.1;refGene_gn=OSBPL8;refGene_tx=XM_023257273.1
B4 102195640 102195739
refGene_gn=OSBPL8;refGene_tx=XM_019835250.2;refGene_gn=OSBPL8;refGene_tx=XM_023257263.1;refGene_gn=OSBPL8;refGene_tx=XM_023257264.1;refGene_gn=OSBPL8;refGene_tx=XM_023257265.1;refGene_gn=OSBPL8;refGene_tx=XM_023257266.1;refGene_gn=OSBPL8;refGene_tx=XM_023257267.1;refGene_gn=OSBPL8;refGene_tx=XM_023257268.1;refGene_gn=OSBPL8;refGene_tx=XM_023257269.1;refGene_gn=OSBPL8;refGene_tx=XM_023257270.1;refGene_gn=OSBPL8;refGene_tx=XM_023257271.1;refGene_gn=OSBPL8;refGene_tx=XM_023257272.1;refGene_gn=OSBPL8;refGene_tx=XM_023257273.1
B4 102196008 102196090
refGene_gn=OSBPL8;refGene_tx=XM_019835250.2;refGene_gn=OSBPL8;refGene_tx=XM_023257263.1;refGene_gn=OSBPL8;refGene_tx=XM_023257264.1;refGene_gn=OSBPL8;refGene_tx=XM_023257265.1;refGene_gn=OSBPL8;refGene_tx=XM_023257266.1;refGene_gn=OSBPL8;refGene_tx=XM_023257267.1;refGene_gn=OSBPL8;refGene_tx=XM_023257268.1;refGene_gn=OSBPL8;refGene_tx=XM_023257269.

1;refGene_gn=OSBPL8;refGene_tx=XM_023257270.1;refGene_gn=OSBPL8;refGene_tx=XM_023257271.1;refGene_gn=OSBPL8;refGene_tx=XM_023257272.1;refGene_gn=OSBPL8;refGene_tx=XM_023257273.1
B4 102199265 102199450
refGene_gn=OSBPL8;refGene_tx=XM_019835250.2;refGene_gn=OSBPL8;refGene_tx=XM_023257263.1;refGene_gn=OSBPL8;refGene_tx=XM_023257264.1;refGene_gn=OSBPL8;refGene_tx=XM_023257265.1;refGene_gn=OSBPL8;refGene_tx=XM_023257266.1;refGene_gn=OSBPL8;refGene_tx=XM_023257267.1;refGene_gn=OSBPL8;refGene_tx=XM_023257268.1;refGene_gn=OSBPL8;refGene_tx=XM_023257269.1;refGene_gn=OSBPL8;refGene_tx=XM_023257270.1;refGene_gn=OSBPL8;refGene_tx=XM_023257271.1;refGene_gn=OSBPL8;refGene_tx=XM_023257272.1;refGene_gn=OSBPL8;refGene_tx=XM_023257273.1
B4 102200063 102200301
refGene_gn=OSBPL8;refGene_tx=XM_019835250.2;refGene_gn=OSBPL8;refGene_tx=XM_023257263.1;refGene_gn=OSBPL8;refGene_tx=XM_023257264.1;refGene_gn=OSBPL8;refGene_tx=XM_023257265.1;refGene_gn=OSBPL8;refGene_tx=XM_023257266.1;refGene_gn=OSBPL8;refGene_tx=XM_023257267.1;refGene_gn=OSBPL8;refGene_tx=XM_023257268.1;refGene_gn=OSBPL8;refGene_tx=XM_023257269.1;refGene_gn=OSBPL8;refGene_tx=XM_023257270.1;refGene_gn=OSBPL8;refGene_tx=XM_023257271.1;refGene_gn=OSBPL8;refGene_tx=XM_023257272.1;refGene_gn=OSBPL8;refGene_tx=XM_023257273.1
B4 102203484 102203656
refGene_gn=OSBPL8;refGene_tx=XM_019835250.2;refGene_gn=OSBPL8;refGene_tx=XM_023257263.1;refGene_gn=OSBPL8;refGene_tx=XM_023257264.1;refGene_gn=OSBPL8;refGene_tx=XM_023257265.1;refGene_gn=OSBPL8;refGene_tx=XM_023257266.1;refGene_gn=OSBPL8;refGene_tx=XM_023257267.1;refGene_gn=OSBPL8;refGene_tx=XM_023257268.1;refGene_gn=OSBPL8;refGene_tx=XM_023257269.1;refGene_gn=OSBPL8;refGene_tx=XM_023257270.1;refGene_gn=OSBPL8;refGene_tx=XM_023257271.1;refGene_gn=OSBPL8;refGene_tx=XM_023257272.1;refGene_gn=OSBPL8;refGene_tx=XM_023257273.1
B4 102205193 102205278
refGene_gn=OSBPL8;refGene_tx=XM_019835250.2;refGene_gn=OSBPL8;refGene_tx=XM_023257263.1;refGene_gn=OSBPL8;refGene_tx=XM_023257264.1;refGene_gn=OSBPL8;refGene_tx=XM_023257265.1;refGene_gn=OSBPL8;refGene_tx=XM_023257266.1;refGene_gn=OSBPL8;refGene_tx=XM_023257267.1;refGene_gn=OSBPL8;refGene_tx=XM_023257268.1;refGene_gn=OSBPL8;refGene_tx=XM_023257269.1;refGene_gn=OSBPL8;refGene_tx=XM_023257270.1;refGene_gn=OSBPL8;refGene_tx=XM_023257271.1;refGene_gn=OSBPL8;refGene_tx=XM_023257272.1;refGene_gn=OSBPL8;refGene_tx=XM_023257273.1
B4 102208052 102208256
refGene_gn=OSBPL8;refGene_tx=XM_019835250.2;refGene_gn=OSBPL8;refGene_tx=XM_023257263.1;refGene_gn=OSBPL8;refGene_tx=XM_023257264.1;refGene_gn=OSBPL8;refGene_tx=XM_023257265.1;refGene_gn=OSBPL8;refGene_tx=XM_023257266.1;refGene_gn=OSBPL8;refGene_tx=XM_023257267.1;refGene_gn=OSBPL8;refGene_tx=XM_023257268.1;refGene_gn=OSBPL8;refGene_tx=XM_023257269.1;refGene_gn=OSBPL8;refGene_tx=XM_023257270.1;refGene_gn=OSBPL8;refGene_tx=XM_023257271.1;refGene_gn=OSBPL8;refGene_tx=XM_023257272.1;refGene_gn=OSBPL8;refGene_tx=XM_023257273.1
B4 102210254 102210356
refGene_gn=OSBPL8;refGene_tx=XM_019835250.2;refGene_gn=OSBPL8;refGene_

tx=XM_023257263.1;refGene_gn=OSBPL8;refGene_tx=XM_023257264.1;refGene_gn=OSBPL8;refGene_tx=XM_023257265.1;refGene_gn=OSBPL8;refGene_tx=XM_023257266.1;refGene_gn=OSBPL8;refGene_tx=XM_023257267.1;refGene_gn=OSBPL8;refGene_tx=XM_023257268.1;refGene_gn=OSBPL8;refGene_tx=XM_023257269.1;refGene_gn=OSBPL8;refGene_tx=XM_023257270.1;refGene_gn=OSBPL8;refGene_tx=XM_023257271.1;refGene_gn=OSBPL8;refGene_tx=XM_023257272.1;refGene_gn=OSBPL8;refGene_tx=XM_023257273.1
B4 102212116 102212194
refGene_gn=OSBPL8;refGene_tx=XM_019835250.2;refGene_gn=OSBPL8;refGene_tx=XM_023257263.1;refGene_gn=OSBPL8;refGene_tx=XM_023257264.1;refGene_gn=OSBPL8;refGene_tx=XM_023257265.1;refGene_gn=OSBPL8;refGene_tx=XM_023257266.1;refGene_gn=OSBPL8;refGene_tx=XM_023257267.1;refGene_gn=OSBPL8;refGene_tx=XM_023257268.1;refGene_gn=OSBPL8;refGene_tx=XM_023257269.1;refGene_gn=OSBPL8;refGene_tx=XM_023257270.1;refGene_gn=OSBPL8;refGene_tx=XM_023257271.1;refGene_gn=OSBPL8;refGene_tx=XM_023257272.1;refGene_gn=OSBPL8;refGene_tx=XM_023257273.1
B4 102218000 102218071
refGene_gn=OSBPL8;refGene_tx=XM_019835250.2;refGene_gn=OSBPL8;refGene_tx=XM_023257263.1;refGene_gn=OSBPL8;refGene_tx=XM_023257264.1;refGene_gn=OSBPL8;refGene_tx=XM_023257265.1;refGene_gn=OSBPL8;refGene_tx=XM_023257266.1;refGene_gn=OSBPL8;refGene_tx=XM_023257267.1;refGene_gn=OSBPL8;refGene_tx=XM_023257268.1;refGene_gn=OSBPL8;refGene_tx=XM_023257269.1;refGene_gn=OSBPL8;refGene_tx=XM_023257270.1;refGene_gn=OSBPL8;refGene_tx=XM_023257271.1;refGene_gn=OSBPL8;refGene_tx=XM_023257272.1;refGene_gn=OSBPL8;refGene_tx=XM_023257273.1
B4 102226598 102226743
refGene_gn=OSBPL8;refGene_tx=XM_019835250.2
B4 102244723 102244861
refGene_gn=OSBPL8;refGene_tx=XM_023257263.1;refGene_gn=OSBPL8;refGene_tx=XM_023257264.1;refGene_gn=OSBPL8;refGene_tx=XM_023257265.1;refGene_gn=OSBPL8;refGene_tx=XM_023257266.1;refGene_gn=OSBPL8;refGene_tx=XM_023257267.1;refGene_gn=OSBPL8;refGene_tx=XM_023257268.1;refGene_gn=OSBPL8;refGene_tx=XM_023257269.1;refGene_gn=OSBPL8;refGene_tx=XM_023257270.1;refGene_gn=OSBPL8;refGene_tx=XM_023257271.1;refGene_gn=OSBPL8;refGene_tx=XM_023257272.1;refGene_gn=OSBPL8;refGene_tx=XM_023257273.1
B4 102254985 102255022
refGene_gn=OSBPL8;refGene_tx=XM_023257263.1;refGene_gn=OSBPL8;refGene_tx=XM_023257264.1;refGene_gn=OSBPL8;refGene_tx=XM_023257265.1;refGene_gn=OSBPL8;refGene_tx=XM_023257266.1;refGene_gn=OSBPL8;refGene_tx=XM_023257267.1;refGene_gn=OSBPL8;refGene_tx=XM_023257268.1
B4 102261665 102261704
refGene_gn=OSBPL8;refGene_tx=XM_023257263.1;refGene_gn=OSBPL8;refGene_tx=XM_023257264.1;refGene_gn=OSBPL8;refGene_tx=XM_023257265.1;refGene_gn=OSBPL8;refGene_tx=XM_023257266.1
B4 102263868 102263892
refGene_gn=OSBPL8;refGene_tx=XM_023257265.1;refGene_gn=OSBPL8;refGene_tx=XM_023257266.1
B4 102275577 102275619
refGene_gn=OSBPL8;refGene_tx=XM_023257263.1;refGene_gn=OSBPL8;refGene_tx=XM_023257264.1;refGene_gn=OSBPL8;refGene_tx=XM_023257267.1;refGene_

gn=OSBPL8;refGene_tx=XM_023257268.1
B4 102491315 102491408
refGene_gn=ZDHHC17;refGene_tx=XM_003989060.5;refGene_gn=ZDHHC17;refGene_tx=XM_006933902.3;refGene_gn=ZDHHC17;refGene_tx=XM_019835254.1;refGene_gn=ZDHHC17;refGene_tx=XM_019835255.1
B4 102513281 102513377
refGene_gn=ZDHHC17;refGene_tx=XM_006933902.3;refGene_gn=ZDHHC17;refGene_tx=XM_019835254.1;refGene_gn=ZDHHC17;refGene_tx=XM_019835255.1
B4 102516422 102516526
refGene_gn=ZDHHC17;refGene_tx=XM_003989060.5;refGene_gn=ZDHHC17;refGene_tx=XM_006933902.3;refGene_gn=ZDHHC17;refGene_tx=XM_019835254.1;refGene_gn=ZDHHC17;refGene_tx=XM_019835255.1
B4 102524242 102524365
refGene_gn=ZDHHC17;refGene_tx=XM_003989060.5;refGene_gn=ZDHHC17;refGene_tx=XM_006933902.3;refGene_gn=ZDHHC17;refGene_tx=XM_019835254.1;refGene_gn=ZDHHC17;refGene_tx=XM_019835255.1
B4 102532321 102532399
refGene_gn=ZDHHC17;refGene_tx=XM_003989060.5;refGene_gn=ZDHHC17;refGene_tx=XM_006933902.3;refGene_gn=ZDHHC17;refGene_tx=XM_019835254.1;refGene_gn=ZDHHC17;refGene_tx=XM_019835255.1
B4 102532984 102533129
refGene_gn=ZDHHC17;refGene_tx=XM_003989060.5;refGene_gn=ZDHHC17;refGene_tx=XM_006933902.3;refGene_gn=ZDHHC17;refGene_tx=XM_019835254.1;refGene_gn=ZDHHC17;refGene_tx=XM_019835255.1
B4 102538105 102538170
refGene_gn=ZDHHC17;refGene_tx=XM_003989060.5;refGene_gn=ZDHHC17;refGene_tx=XM_006933902.3;refGene_gn=ZDHHC17;refGene_tx=XM_019835254.1;refGene_gn=ZDHHC17;refGene_tx=XM_019835255.1
B4 102538810 102538973
refGene_gn=ZDHHC17;refGene_tx=XM_003989060.5;refGene_gn=ZDHHC17;refGene_tx=XM_006933902.3;refGene_gn=ZDHHC17;refGene_tx=XM_019835254.1;refGene_gn=ZDHHC17;refGene_tx=XM_019835255.1
B4 102546766 102546892
refGene_gn=ZDHHC17;refGene_tx=XM_003989060.5;refGene_gn=ZDHHC17;refGene_tx=XM_006933902.3;refGene_gn=ZDHHC17;refGene_tx=XM_019835254.1;refGene_gn=ZDHHC17;refGene_tx=XM_019835255.1
B4 102558824 102558967
refGene_gn=ZDHHC17;refGene_tx=XM_003989060.5;refGene_gn=ZDHHC17;refGene_tx=XM_006933902.3;refGene_gn=ZDHHC17;refGene_tx=XM_019835254.1;refGene_gn=ZDHHC17;refGene_tx=XM_019835255.1
B4 102560307 102560408
refGene_gn=ZDHHC17;refGene_tx=XM_003989060.5;refGene_gn=ZDHHC17;refGene_tx=XM_006933902.3;refGene_gn=ZDHHC17;refGene_tx=XM_019835254.1;refGene_gn=ZDHHC17;refGene_tx=XM_019835255.1
B4 102568469 102568594
refGene_gn=ZDHHC17;refGene_tx=XM_003989060.5;refGene_gn=ZDHHC17;refGene_tx=XM_006933902.3;refGene_gn=ZDHHC17;refGene_tx=XM_019835254.1;refGene_gn=ZDHHC17;refGene_tx=XM_019835255.1
B4 102569667 102569730
refGene_gn=ZDHHC17;refGene_tx=XM_003989060.5;refGene_gn=ZDHHC17;refGen

e_tx=XM_006933902.3;refGene_gn=ZDHHC17;refGene_tx=XM_019835254.1;refGene_gn=ZDHHC17;refGene_tx=XM_019835255.1
B4 102572644 102572738
refGene_gn=ZDHHC17;refGene_tx=XM_003989060.5;refGene_gn=ZDHHC17;refGene_tx=XM_006933902.3;refGene_gn=ZDHHC17;refGene_tx=XM_019835254.1;refGene_gn=ZDHHC17;refGene_tx=XM_019835255.1
B4 102573616 102573700
refGene_gn=ZDHHC17;refGene_tx=XM_003989060.5;refGene_gn=ZDHHC17;refGene_tx=XM_006933902.3;refGene_gn=ZDHHC17;refGene_tx=XM_019835254.1;refGene_gn=ZDHHC17;refGene_tx=XM_019835255.1
B4 102574945 102575103
refGene_gn=ZDHHC17;refGene_tx=XM_003989060.5;refGene_gn=ZDHHC17;refGene_tx=XM_006933902.3;refGene_gn=ZDHHC17;refGene_tx=XM_019835254.1;refGene_gn=ZDHHC17;refGene_tx=XM_019835255.1
B4 102575800 102575895
refGene_gn=ZDHHC17;refGene_tx=XM_003989060.5;refGene_gn=ZDHHC17;refGene_tx=XM_006933902.3;refGene_gn=ZDHHC17;refGene_tx=XM_019835254.1;refGene_gn=ZDHHC17;refGene_tx=XM_019835255.1
B4 102578352 102578428
refGene_gn=ZDHHC17;refGene_tx=XM_019835254.1
B4 102578703 102578842
refGene_gn=ZDHHC17;refGene_tx=XM_003989060.5;refGene_gn=ZDHHC17;refGene_tx=XM_006933902.3
B4 102587258 102587335
refGene_gn=CSRP2;refGene_tx=XM_003989061.3
B4 102587883 102587977
refGene_gn=CSRP2;refGene_tx=XM_003989061.3
B4 102588664 102588794
refGene_gn=CSRP2;refGene_tx=XM_003989061.3
B4 102592002 102592171
refGene_gn=CSRP2;refGene_tx=XM_003989061.3
B4 102595014 102595018
refGene_gn=ZDHHC17;refGene_tx=XM_019835255.1
B4 102595083 102595195
refGene_gn=CSRP2;refGene_tx=XM_003989061.3
B4 102720500 102720671
refGene_gn=E2F7;refGene_tx=XM_006933903.3
B4 102721949 102722371
refGene_gn=E2F7;refGene_tx=XM_006933903.3
B4 102723862 102724118
refGene_gn=E2F7;refGene_tx=XM_006933903.3
B4 102725484 102725986
refGene_gn=E2F7;refGene_tx=XM_006933903.3
B4 102728503 102728576
refGene_gn=E2F7;refGene_tx=XM_006933903.3
B4 102729059 102729245
refGene_gn=E2F7;refGene_tx=XM_006933903.3
B4 102738406 102738541
refGene_gn=E2F7;refGene_tx=XM_006933903.3
B4 102740025 102740184

refGene_gn=E2F7;refGene_tx=XM_006933903.3
B4 102741574 102741865
refGene_gn=E2F7;refGene_tx=XM_006933903.3
B4 102745659 102745828
refGene_gn=E2F7;refGene_tx=XM_006933903.3
B4 102749813 102750089
refGene_gn=E2F7;refGene_tx=XM_006933903.3
B4 102757022 102757115
refGene_gn=E2F7;refGene_tx=XM_006933903.3
B4 103223171 103223243
refGene_gn=NAV3;refGene_tx=XM_011284133.3
B4 103461846 103462089
refGene_gn=NAV3;refGene_tx=XM_019835257.2;refGene_gn=NAV3;refGene_tx=XM_019835258.2;refGene_gn=NAV3;refGene_tx=XM_019835259.2;refGene_gn=NAV3;refGene_tx=XM_019835261.2;refGene_gn=NAV3;refGene_tx=XM_019835262.2;refGene_gn=NAV3;refGene_tx=XM_019835263.2
B4 103565542 103565660
refGene_gn=NAV3;refGene_tx=XM_011284133.3;refGene_gn=NAV3;refGene_tx=XM_019835257.2;refGene_gn=NAV3;refGene_tx=XM_019835258.2;refGene_gn=NAV3;refGene_tx=XM_019835259.2;refGene_gn=NAV3;refGene_tx=XM_019835261.2;refGene_gn=NAV3;refGene_tx=XM_019835262.2;refGene_gn=NAV3;refGene_tx=XM_019835263.2
B4 103566300 103566353
refGene_gn=NAV3;refGene_tx=XM_011284133.3;refGene_gn=NAV3;refGene_tx=XM_019835257.2;refGene_gn=NAV3;refGene_tx=XM_019835258.2;refGene_gn=NAV3;refGene_tx=XM_019835259.2;refGene_gn=NAV3;refGene_tx=XM_019835261.2;refGene_gn=NAV3;refGene_tx=XM_019835262.2;refGene_gn=NAV3;refGene_tx=XM_019835263.2
B4 103584925 103584998
refGene_gn=NAV3;refGene_tx=XM_011284133.3;refGene_gn=NAV3;refGene_tx=XM_019835257.2;refGene_gn=NAV3;refGene_tx=XM_019835258.2;refGene_gn=NAV3;refGene_tx=XM_019835259.2;refGene_gn=NAV3;refGene_tx=XM_019835261.2;refGene_gn=NAV3;refGene_tx=XM_019835262.2;refGene_gn=NAV3;refGene_tx=XM_019835263.2
B4 103587489 103587673
refGene_gn=NAV3;refGene_tx=XM_011284133.3;refGene_gn=NAV3;refGene_tx=XM_019835257.2;refGene_gn=NAV3;refGene_tx=XM_019835258.2;refGene_gn=NAV3;refGene_tx=XM_019835259.2;refGene_gn=NAV3;refGene_tx=XM_019835261.2;refGene_gn=NAV3;refGene_tx=XM_019835262.2;refGene_gn=NAV3;refGene_tx=XM_019835263.2
B4 103610650 103610719
refGene_gn=NAV3;refGene_tx=XM_011284133.3;refGene_gn=NAV3;refGene_tx=XM_019835257.2;refGene_gn=NAV3;refGene_tx=XM_019835258.2;refGene_gn=NAV3;refGene_tx=XM_019835259.2;refGene_gn=NAV3;refGene_tx=XM_019835261.2;refGene_gn=NAV3;refGene_tx=XM_019835262.2;refGene_gn=NAV3;refGene_tx=XM_019835263.2
B4 103613916 103614056
refGene_gn=NAV3;refGene_tx=XM_011284133.3;refGene_gn=NAV3;refGene_tx=XM_019835257.2;refGene_gn=NAV3;refGene_tx=XM_019835258.2;refGene_gn=NAV3;refGene_tx=XM_019835259.2;refGene_gn=NAV3;refGene_tx=XM_019835261.2;

refGene_gn=NAV3;refGene_tx=XM_019835262.2;refGene_gn=NAV3;refGene_tx=X
M_019835263.2
B4 103624008 103625035
refGene_gn=NAV3;refGene_tx=XM_011284133.3;refGene_gn=NAV3;refGene_tx=X
M_019835257.2;refGene_gn=NAV3;refGene_tx=XM_019835258.2;refGene_gn=NAV
3;refGene_tx=XM_019835259.2;refGene_gn=NAV3;refGene_tx=XM_019835261.2;
refGene_gn=NAV3;refGene_tx=XM_019835262.2;refGene_gn=NAV3;refGene_tx=X
M_019835263.2
B4 103640997 103641113
refGene_gn=NAV3;refGene_tx=XM_011284133.3;refGene_gn=NAV3;refGene_tx=X
M_019835257.2;refGene_gn=NAV3;refGene_tx=XM_019835258.2;refGene_gn=NAV
3;refGene_tx=XM_019835259.2;refGene_gn=NAV3;refGene_tx=XM_019835261.2;
refGene_gn=NAV3;refGene_tx=XM_019835262.2;refGene_gn=NAV3;refGene_tx=X
M_019835263.2
B4 103657955 103658064
refGene_gn=NAV3;refGene_tx=XM_011284133.3;refGene_gn=NAV3;refGene_tx=X
M_019835257.2;refGene_gn=NAV3;refGene_tx=XM_019835258.2;refGene_gn=NAV
3;refGene_tx=XM_019835259.2;refGene_gn=NAV3;refGene_tx=XM_019835261.2;
refGene_gn=NAV3;refGene_tx=XM_019835262.2;refGene_gn=NAV3;refGene_tx=X
M_019835263.2
B4 103658771 103659155
refGene_gn=NAV3;refGene_tx=XM_011284133.3;refGene_gn=NAV3;refGene_tx=X
M_019835257.2;refGene_gn=NAV3;refGene_tx=XM_019835258.2;refGene_gn=NAV
3;refGene_tx=XM_019835259.2;refGene_gn=NAV3;refGene_tx=XM_019835261.2;
refGene_gn=NAV3;refGene_tx=XM_019835262.2;refGene_gn=NAV3;refGene_tx=X
M_019835263.2
B4 103671134 103671254
refGene_gn=NAV3;refGene_tx=XM_011284133.3;refGene_gn=NAV3;refGene_tx=X
M_019835257.2;refGene_gn=NAV3;refGene_tx=XM_019835258.2;refGene_gn=NAV
3;refGene_tx=XM_019835259.2;refGene_gn=NAV3;refGene_tx=XM_019835261.2;
refGene_gn=NAV3;refGene_tx=XM_019835262.2;refGene_gn=NAV3;refGene_tx=X
M_019835263.2
B4 103726472 103726605
refGene_gn=NAV3;refGene_tx=XM_011284133.3;refGene_gn=NAV3;refGene_tx=X
M_019835257.2;refGene_gn=NAV3;refGene_tx=XM_019835258.2;refGene_gn=NAV
3;refGene_tx=XM_019835259.2;refGene_gn=NAV3;refGene_tx=XM_019835261.2;
refGene_gn=NAV3;refGene_tx=XM_019835262.2;refGene_gn=NAV3;refGene_tx=X
M_019835263.2
B4 103727538 103727809
refGene_gn=NAV3;refGene_tx=XM_011284133.3;refGene_gn=NAV3;refGene_tx=X
M_019835257.2;refGene_gn=NAV3;refGene_tx=XM_019835258.2;refGene_gn=NAV
3;refGene_tx=XM_019835259.2;refGene_gn=NAV3;refGene_tx=XM_019835261.2;
refGene_gn=NAV3;refGene_tx=XM_019835262.2;refGene_gn=NAV3;refGene_tx=X
M_019835263.2
B4 103728986 103729695
refGene_gn=NAV3;refGene_tx=XM_011284133.3;refGene_gn=NAV3;refGene_tx=X
M_019835257.2;refGene_gn=NAV3;refGene_tx=XM_019835258.2;refGene_gn=NAV
3;refGene_tx=XM_019835259.2;refGene_gn=NAV3;refGene_tx=XM_019835261.2;
refGene_gn=NAV3;refGene_tx=XM_019835262.2;refGene_gn=NAV3;refGene_tx=X
M_019835263.2

B4 103731732 103732221
refGene_gn=NAV3;refGene_tx=XM_011284133.3;refGene_gn=NAV3;refGene_tx=XM_019835257.2;refGene_gn=NAV3;refGene_tx=XM_019835258.2;refGene_gn=NAV3;refGene_tx=XM_019835259.2;refGene_gn=NAV3;refGene_tx=XM_019835263.2
B4 103737532 103737574
refGene_gn=NAV3;refGene_tx=XM_011284133.3;refGene_gn=NAV3;refGene_tx=XM_019835257.2;refGene_gn=NAV3;refGene_tx=XM_019835258.2;refGene_gn=NAV3;refGene_tx=XM_019835261.2;refGene_gn=NAV3;refGene_tx=XM_019835263.2
B4 103738974 103739135
refGene_gn=NAV3;refGene_tx=XM_011284133.3;refGene_gn=NAV3;refGene_tx=XM_019835257.2;refGene_gn=NAV3;refGene_tx=XM_019835258.2;refGene_gn=NAV3;refGene_tx=XM_019835259.2;refGene_gn=NAV3;refGene_tx=XM_019835261.2;refGene_gn=NAV3;refGene_tx=XM_019835262.2;refGene_gn=NAV3;refGene_tx=XM_019835263.2
B4 103744421 103744610
refGene_gn=NAV3;refGene_tx=XM_011284133.3;refGene_gn=NAV3;refGene_tx=XM_019835257.2;refGene_gn=NAV3;refGene_tx=XM_019835258.2;refGene_gn=NAV3;refGene_tx=XM_019835259.2;refGene_gn=NAV3;refGene_tx=XM_019835261.2;refGene_gn=NAV3;refGene_tx=XM_019835262.2;refGene_gn=NAV3;refGene_tx=XM_019835263.2
B4 103747977 103748030
refGene_gn=NAV3;refGene_tx=XM_011284133.3;refGene_gn=NAV3;refGene_tx=XM_019835257.2;refGene_gn=NAV3;refGene_tx=XM_019835258.2;refGene_gn=NAV3;refGene_tx=XM_019835259.2;refGene_gn=NAV3;refGene_tx=XM_019835261.2;refGene_gn=NAV3;refGene_tx=XM_019835262.2;refGene_gn=NAV3;refGene_tx=XM_019835263.2
B4 103753734 103753758
refGene_gn=NAV3;refGene_tx=XM_011284133.3;refGene_gn=NAV3;refGene_tx=XM_019835257.2;refGene_gn=NAV3;refGene_tx=XM_019835258.2;refGene_gn=NAV3;refGene_tx=XM_019835259.2;refGene_gn=NAV3;refGene_tx=XM_019835261.2;refGene_gn=NAV3;refGene_tx=XM_019835262.2;refGene_gn=NAV3;refGene_tx=XM_019835263.2
B4 103756319 103756397
refGene_gn=NAV3;refGene_tx=XM_011284133.3;refGene_gn=NAV3;refGene_tx=XM_019835257.2;refGene_gn=NAV3;refGene_tx=XM_019835258.2;refGene_gn=NAV3;refGene_tx=XM_019835259.2;refGene_gn=NAV3;refGene_tx=XM_019835261.2;refGene_gn=NAV3;refGene_tx=XM_019835262.2;refGene_gn=NAV3;refGene_tx=XM_019835263.2
B4 103763983 103764067
refGene_gn=NAV3;refGene_tx=XM_011284133.3;refGene_gn=NAV3;refGene_tx=XM_019835257.2;refGene_gn=NAV3;refGene_tx=XM_019835258.2;refGene_gn=NAV3;refGene_tx=XM_019835259.2;refGene_gn=NAV3;refGene_tx=XM_019835261.2;refGene_gn=NAV3;refGene_tx=XM_019835262.2;refGene_gn=NAV3;refGene_tx=XM_019835263.2
B4 103771648 103771760
refGene_gn=NAV3;refGene_tx=XM_011284133.3;refGene_gn=NAV3;refGene_tx=XM_019835257.2;refGene_gn=NAV3;refGene_tx=XM_019835258.2;refGene_gn=NAV3;refGene_tx=XM_019835259.2;refGene_gn=NAV3;refGene_tx=XM_019835261.2;refGene_gn=NAV3;refGene_tx=XM_019835262.2;refGene_gn=NAV3;refGene_tx=XM_019835263.2

B4	103779097	103779219
refGene_gn=NAV3;refGene_tx=XM_011284133.3;refGene_gn=NAV3;refGene_tx=XM_019835257.2;refGene_gn=NAV3;refGene_tx=XM_019835258.2;refGene_gn=NAV3;refGene_tx=XM_019835259.2;refGene_gn=NAV3;refGene_tx=XM_019835261.2;refGene_gn=NAV3;refGene_tx=XM_019835262.2;refGene_gn=NAV3;refGene_tx=XM_019835263.2
B4	103780231	103780252
refGene_gn=NAV3;refGene_tx=XM_011284133.3;refGene_gn=NAV3;refGene_tx=XM_019835257.2;refGene_gn=NAV3;refGene_tx=XM_019835259.2;refGene_gn=NAV3;refGene_tx=XM_019835261.2;refGene_gn=NAV3;refGene_tx=XM_019835262.2;refGene_gn=NAV3;refGene_tx=XM_019835263.2
B4	103780937	103781110
refGene_gn=NAV3;refGene_tx=XM_011284133.3;refGene_gn=NAV3;refGene_tx=XM_019835257.2;refGene_gn=NAV3;refGene_tx=XM_019835258.2;refGene_gn=NAV3;refGene_tx=XM_019835259.2;refGene_gn=NAV3;refGene_tx=XM_019835261.2;refGene_gn=NAV3;refGene_tx=XM_019835262.2;refGene_gn=NAV3;refGene_tx=XM_019835263.2
B4	103781404	103781470
refGene_gn=NAV3;refGene_tx=XM_019835263.2
B4	103782732	103782886
refGene_gn=NAV3;refGene_tx=XM_011284133.3;refGene_gn=NAV3;refGene_tx=XM_019835257.2;refGene_gn=NAV3;refGene_tx=XM_019835258.2;refGene_gn=NAV3;refGene_tx=XM_019835259.2;refGene_gn=NAV3;refGene_tx=XM_019835261.2;refGene_gn=NAV3;refGene_tx=XM_019835262.2;refGene_gn=NAV3;refGene_tx=XM_019835263.2
B4	103784093	103784268
refGene_gn=NAV3;refGene_tx=XM_011284133.3;refGene_gn=NAV3;refGene_tx=XM_019835257.2;refGene_gn=NAV3;refGene_tx=XM_019835258.2;refGene_gn=NAV3;refGene_tx=XM_019835259.2;refGene_gn=NAV3;refGene_tx=XM_019835261.2;refGene_gn=NAV3;refGene_tx=XM_019835262.2;refGene_gn=NAV3;refGene_tx=XM_019835263.2
B4	103788947	103789045
refGene_gn=NAV3;refGene_tx=XM_011284133.3;refGene_gn=NAV3;refGene_tx=XM_019835257.2;refGene_gn=NAV3;refGene_tx=XM_019835258.2;refGene_gn=NAV3;refGene_tx=XM_019835259.2;refGene_gn=NAV3;refGene_tx=XM_019835261.2;refGene_gn=NAV3;refGene_tx=XM_019835262.2;refGene_gn=NAV3;refGene_tx=XM_019835263.2
B4	103791800	103791896
refGene_gn=NAV3;refGene_tx=XM_011284133.3;refGene_gn=NAV3;refGene_tx=XM_019835257.2;refGene_gn=NAV3;refGene_tx=XM_019835258.2;refGene_gn=NAV3;refGene_tx=XM_019835259.2;refGene_gn=NAV3;refGene_tx=XM_019835261.2;refGene_gn=NAV3;refGene_tx=XM_019835262.2;refGene_gn=NAV3;refGene_tx=XM_019835263.2
B4	103792163	103792332
refGene_gn=NAV3;refGene_tx=XM_011284133.3;refGene_gn=NAV3;refGene_tx=XM_019835257.2;refGene_gn=NAV3;refGene_tx=XM_019835258.2;refGene_gn=NAV3;refGene_tx=XM_019835259.2;refGene_gn=NAV3;refGene_tx=XM_019835261.2;refGene_gn=NAV3;refGene_tx=XM_019835262.2;refGene_gn=NAV3;refGene_tx=XM_019835263.2
B4	103793625	103793861

refGene_gn=NAV3;refGene_tx=XM_011284133.3;refGene_gn=NAV3;refGene_tx=X
M_019835257.2;refGene_gn=NAV3;refGene_tx=XM_019835258.2;refGene_gn=NAV
3;refGene_tx=XM_019835259.2;refGene_gn=NAV3;refGene_tx=XM_019835261.2;
refGene_gn=NAV3;refGene_tx=XM_019835262.2;refGene_gn=NAV3;refGene_tx=X
M_019835263.2
B4 103804753 103804908
refGene_gn=NAV3;refGene_tx=XM_011284133.3;refGene_gn=NAV3;refGene_tx=X
M_019835257.2;refGene_gn=NAV3;refGene_tx=XM_019835258.2;refGene_gn=NAV
3;refGene_tx=XM_019835259.2;refGene_gn=NAV3;refGene_tx=XM_019835261.2;
refGene_gn=NAV3;refGene_tx=XM_019835262.2;refGene_gn=NAV3;refGene_tx=X
M_019835263.2
B4 103805943 103806015
refGene_gn=NAV3;refGene_tx=XM_011284133.3;refGene_gn=NAV3;refGene_tx=X
M_019835257.2;refGene_gn=NAV3;refGene_tx=XM_019835258.2;refGene_gn=NAV
3;refGene_tx=XM_019835259.2;refGene_gn=NAV3;refGene_tx=XM_019835261.2;
refGene_gn=NAV3;refGene_tx=XM_019835262.2;refGene_gn=NAV3;refGene_tx=X
M_019835263.2
B4 103806629 103806826
refGene_gn=NAV3;refGene_tx=XM_011284133.3;refGene_gn=NAV3;refGene_tx=X
M_019835257.2;refGene_gn=NAV3;refGene_tx=XM_019835258.2;refGene_gn=NAV
3;refGene_tx=XM_019835259.2;refGene_gn=NAV3;refGene_tx=XM_019835261.2;
refGene_gn=NAV3;refGene_tx=XM_019835262.2;refGene_gn=NAV3;refGene_tx=X
M_019835263.2
B4 103808043 103808162
refGene_gn=NAV3;refGene_tx=XM_011284133.3;refGene_gn=NAV3;refGene_tx=X
M_019835257.2;refGene_gn=NAV3;refGene_tx=XM_019835258.2;refGene_gn=NAV
3;refGene_tx=XM_019835259.2;refGene_gn=NAV3;refGene_tx=XM_019835261.2;
refGene_gn=NAV3;refGene_tx=XM_019835262.2;refGene_gn=NAV3;refGene_tx=X
M_019835263.2
B4 103812388 103812592
refGene_gn=NAV3;refGene_tx=XM_011284133.3;refGene_gn=NAV3;refGene_tx=X
M_019835257.2;refGene_gn=NAV3;refGene_tx=XM_019835258.2;refGene_gn=NAV
3;refGene_tx=XM_019835259.2;refGene_gn=NAV3;refGene_tx=XM_019835261.2;
refGene_gn=NAV3;refGene_tx=XM_019835262.2;refGene_gn=NAV3;refGene_tx=X
M_019835263.2
B4 103817051 103817171
refGene_gn=NAV3;refGene_tx=XM_011284133.3;refGene_gn=NAV3;refGene_tx=X
M_019835257.2;refGene_gn=NAV3;refGene_tx=XM_019835258.2;refGene_gn=NAV
3;refGene_tx=XM_019835259.2;refGene_gn=NAV3;refGene_tx=XM_019835261.2;
refGene_gn=NAV3;refGene_tx=XM_019835262.2;refGene_gn=NAV3;refGene_tx=X
M_019835263.2
B4 104766913 104767079
refGene_gn=SYT1;refGene_tx=XM_011284134.2;refGene_gn=SYT1;refGene_tx=X
M_011284135.2;refGene_gn=SYT1;refGene_tx=XM_011284136.2;refGene_gn=SYT
1;refGene_tx=XM_019835264.1
B4 104819751 104819936
refGene_gn=SYT1;refGene_tx=XM_011284134.2;refGene_gn=SYT1;refGene_tx=X
M_011284135.2;refGene_gn=SYT1;refGene_tx=XM_011284136.2;refGene_gn=SYT
1;refGene_tx=XM_019835264.1
B4 104825914 104826037

refGene_gn=SYT1;refGene_tx=XM_011284134.2;refGene_gn=SYT1;refGene_tx=XM_011284135.2;refGene_gn=SYT1;refGene_tx=XM_011284136.2;refGene_gn=SYT1;refGene_tx=XM_019835264.1
B4 104830239 104830407
refGene_gn=SYT1;refGene_tx=XM_011284134.2;refGene_gn=SYT1;refGene_tx=XM_011284135.2;refGene_gn=SYT1;refGene_tx=XM_011284136.2;refGene_gn=SYT1;refGene_tx=XM_019835264.1
B4 104833593 104833761
refGene_gn=SYT1;refGene_tx=XM_011284134.2;refGene_gn=SYT1;refGene_tx=XM_011284135.2;refGene_gn=SYT1;refGene_tx=XM_011284136.2;refGene_gn=SYT1;refGene_tx=XM_019835264.1
B4 104878287 104878405
refGene_gn=SYT1;refGene_tx=XM_011284134.2;refGene_gn=SYT1;refGene_tx=XM_011284135.2;refGene_gn=SYT1;refGene_tx=XM_011284136.2;refGene_gn=SYT1;refGene_tx=XM_019835264.1
B4 104971622 104971756
refGene_gn=SYT1;refGene_tx=XM_011284134.2;refGene_gn=SYT1;refGene_tx=XM_011284135.2;refGene_gn=SYT1;refGene_tx=XM_011284136.2;refGene_gn=SYT1;refGene_tx=XM_019835264.1
B4 104976345 104976552
refGene_gn=SYT1;refGene_tx=XM_011284134.2;refGene_gn=SYT1;refGene_tx=XM_011284135.2;refGene_gn=SYT1;refGene_tx=XM_011284136.2;refGene_gn=SYT1;refGene_tx=XM_019835264.1
B4 105038225 105038241
refGene_gn=PAWR;refGene_tx=XM_023257274.1
B4 105052435 105052522
refGene_gn=PAWR;refGene_tx=XM_023257274.1;refGene_gn=PAWR;refGene_tx=XM_023257275.1
B4 105054637 105054742
refGene_gn=PAWR;refGene_tx=XM_023257274.1;refGene_gn=PAWR;refGene_tx=XM_023257275.1
B4 105057043 105057191
refGene_gn=PAWR;refGene_tx=XM_023257274.1;refGene_gn=PAWR;refGene_tx=XM_023257275.1
B4 105078819 105078854
refGene_gn=PAWR;refGene_tx=XM_023257274.1;refGene_gn=PAWR;refGene_tx=XM_023257275.1
B4 105093263 105093395
refGene_gn=PAWR;refGene_tx=XM_023257274.1;refGene_gn=PAWR;refGene_tx=XM_023257275.1
B4 105177272 105177797
refGene_gn=PAWR;refGene_tx=XM_023257274.1;refGene_gn=PAWR;refGene_tx=XM_023257275.1
B4 105243699 105243786
refGene_gn=PPP1R12A;refGene_tx=XM_006933907.4;refGene_gn=PPP1R12A;refGene_tx=XM_006933908.3
B4 105245290 105245425
refGene_gn=PPP1R12A;refGene_tx=XM_019835266.2;refGene_gn=PPP1R12A;refGene_tx=XM_019835267.2;refGene_gn=PPP1R12A;refGene_tx=XM_019835270.2;refGene_gn=PPP1R12A;refGene_tx=XM_019835271.2

B4	105246798	105246849
refGene_gn=PPP1R12A;refGene_tx=XM_006933907.4;refGene_gn=PPP1R12A;refGene_tx=XM_006933908.3;refGene_gn=PPP1R12A;refGene_tx=XM_019835266.2;refGene_gn=PPP1R12A;refGene_tx=XM_019835267.2;refGene_gn=PPP1R12A;refGene_tx=XM_019835268.2;refGene_gn=PPP1R12A;refGene_tx=XM_019835269.2;refGene_gn=PPP1R12A;refGene_tx=XM_019835270.2;refGene_gn=PPP1R12A;refGene_tx=XM_019835271.2
B4	105247578	105247609
refGene_gn=PPP1R12A;refGene_tx=XM_019835268.2;refGene_gn=PPP1R12A;refGene_tx=XM_019835269.2
B4	105249297	105249345
refGene_gn=PPP1R12A;refGene_tx=XM_006933907.4;refGene_gn=PPP1R12A;refGene_tx=XM_006933908.3;refGene_gn=PPP1R12A;refGene_tx=XM_019835266.2;refGene_gn=PPP1R12A;refGene_tx=XM_019835267.2;refGene_gn=PPP1R12A;refGene_tx=XM_019835268.2;refGene_gn=PPP1R12A;refGene_tx=XM_019835269.2;refGene_gn=PPP1R12A;refGene_tx=XM_019835270.2;refGene_gn=PPP1R12A;refGene_tx=XM_019835271.2
B4	105254625	105254730
refGene_gn=PPP1R12A;refGene_tx=XM_006933907.4;refGene_gn=PPP1R12A;refGene_tx=XM_006933908.3;refGene_gn=PPP1R12A;refGene_tx=XM_019835266.2;refGene_gn=PPP1R12A;refGene_tx=XM_019835267.2;refGene_gn=PPP1R12A;refGene_tx=XM_019835268.2;refGene_gn=PPP1R12A;refGene_tx=XM_019835269.2;refGene_gn=PPP1R12A;refGene_tx=XM_019835270.2;refGene_gn=PPP1R12A;refGene_tx=XM_019835271.2
B4	105256655	105256791
refGene_gn=PPP1R12A;refGene_tx=XM_006933907.4;refGene_gn=PPP1R12A;refGene_tx=XM_006933908.3;refGene_gn=PPP1R12A;refGene_tx=XM_019835266.2;refGene_gn=PPP1R12A;refGene_tx=XM_019835267.2;refGene_gn=PPP1R12A;refGene_tx=XM_019835268.2;refGene_gn=PPP1R12A;refGene_tx=XM_019835269.2;refGene_gn=PPP1R12A;refGene_tx=XM_019835270.2;refGene_gn=PPP1R12A;refGene_tx=XM_019835271.2
B4	105257840	105257857
refGene_gn=PPP1R12A;refGene_tx=XM_006933907.4;refGene_gn=PPP1R12A;refGene_tx=XM_006933908.3;refGene_gn=PPP1R12A;refGene_tx=XM_019835266.2;refGene_gn=PPP1R12A;refGene_tx=XM_019835267.2;refGene_gn=PPP1R12A;refGene_tx=XM_019835268.2;refGene_gn=PPP1R12A;refGene_tx=XM_019835269.2;refGene_gn=PPP1R12A;refGene_tx=XM_019835270.2;refGene_gn=PPP1R12A;refGene_tx=XM_019835271.2
B4	105264081	105264147
refGene_gn=PPP1R12A;refGene_tx=XM_006933907.4;refGene_gn=PPP1R12A;refGene_tx=XM_006933908.3;refGene_gn=PPP1R12A;refGene_tx=XM_019835266.2;refGene_gn=PPP1R12A;refGene_tx=XM_019835267.2;refGene_gn=PPP1R12A;refGene_tx=XM_019835268.2;refGene_gn=PPP1R12A;refGene_tx=XM_019835269.2;refGene_gn=PPP1R12A;refGene_tx=XM_019835270.2;refGene_gn=PPP1R12A;refGene_tx=XM_019835271.2
B4	105276812	105276934
refGene_gn=PPP1R12A;refGene_tx=XM_006933907.4;refGene_gn=PPP1R12A;refGene_tx=XM_006933908.3;refGene_gn=PPP1R12A;refGene_tx=XM_019835266.2;refGene_gn=PPP1R12A;refGene_tx=XM_019835267.2;refGene_gn=PPP1R12A;refGene_tx=XM_019835268.2;refGene_gn=PPP1R12A;refGene_tx=XM_019835269.2;refG

ene_gn=PPP1R12A;refGene_tx=XM_019835270.2;refGene_gn=PPP1R12A;refGene_tx=XM_019835271.2
B4 105279173 105279342
refGene_gn=PPP1R12A;refGene_tx=XM_006933907.4;refGene_gn=PPP1R12A;refGene_tx=XM_006933908.3;refGene_gn=PPP1R12A;refGene_tx=XM_019835266.2;refGene_gn=PPP1R12A;refGene_tx=XM_019835267.2;refGene_gn=PPP1R12A;refGene_tx=XM_019835268.2;refGene_gn=PPP1R12A;refGene_tx=XM_019835269.2;refGene_gn=PPP1R12A;refGene_tx=XM_019835270.2;refGene_gn=PPP1R12A;refGene_tx=XM_019835271.2
B4 105279576 105279777
refGene_gn=PPP1R12A;refGene_tx=XM_006933907.4;refGene_gn=PPP1R12A;refGene_tx=XM_006933908.3;refGene_gn=PPP1R12A;refGene_tx=XM_019835266.2;refGene_gn=PPP1R12A;refGene_tx=XM_019835267.2;refGene_gn=PPP1R12A;refGene_tx=XM_019835268.2;refGene_gn=PPP1R12A;refGene_tx=XM_019835269.2;refGene_gn=PPP1R12A;refGene_tx=XM_019835270.2;refGene_gn=PPP1R12A;refGene_tx=XM_019835271.2
B4 105281071 105281162
refGene_gn=PPP1R12A;refGene_tx=XM_006933907.4;refGene_gn=PPP1R12A;refGene_tx=XM_006933908.3;refGene_gn=PPP1R12A;refGene_tx=XM_019835266.2;refGene_gn=PPP1R12A;refGene_tx=XM_019835267.2;refGene_gn=PPP1R12A;refGene_tx=XM_019835268.2;refGene_gn=PPP1R12A;refGene_tx=XM_019835269.2;refGene_gn=PPP1R12A;refGene_tx=XM_019835270.2;refGene_gn=PPP1R12A;refGene_tx=XM_019835271.2
B4 105288652 105288829
refGene_gn=PPP1R12A;refGene_tx=XM_006933907.4;refGene_gn=PPP1R12A;refGene_tx=XM_006933908.3;refGene_gn=PPP1R12A;refGene_tx=XM_019835266.2;refGene_gn=PPP1R12A;refGene_tx=XM_019835267.2;refGene_gn=PPP1R12A;refGene_tx=XM_019835268.2;refGene_gn=PPP1R12A;refGene_tx=XM_019835269.2;refGene_gn=PPP1R12A;refGene_tx=XM_019835270.2
B4 105289223 105289391
refGene_gn=PPP1R12A;refGene_tx=XM_006933907.4;refGene_gn=PPP1R12A;refGene_tx=XM_019835266.2;refGene_gn=PPP1R12A;refGene_tx=XM_019835267.2;refGene_gn=PPP1R12A;refGene_tx=XM_019835268.2;refGene_gn=PPP1R12A;refGene_tx=XM_019835269.2;refGene_gn=PPP1R12A;refGene_tx=XM_019835271.2
B4 105290275 105290380
refGene_gn=PPP1R12A;refGene_tx=XM_006933907.4;refGene_gn=PPP1R12A;refGene_tx=XM_006933908.3;refGene_gn=PPP1R12A;refGene_tx=XM_019835266.2;refGene_gn=PPP1R12A;refGene_tx=XM_019835267.2;refGene_gn=PPP1R12A;refGene_tx=XM_019835268.2;refGene_gn=PPP1R12A;refGene_tx=XM_019835269.2;refGene_gn=PPP1R12A;refGene_tx=XM_019835270.2;refGene_gn=PPP1R12A;refGene_tx=XM_019835271.2
B4 105291191 105291286
refGene_gn=PPP1R12A;refGene_tx=XM_006933907.4;refGene_gn=PPP1R12A;refGene_tx=XM_006933908.3;refGene_gn=PPP1R12A;refGene_tx=XM_019835266.2;refGene_gn=PPP1R12A;refGene_tx=XM_019835267.2;refGene_gn=PPP1R12A;refGene_tx=XM_019835268.2;refGene_gn=PPP1R12A;refGene_tx=XM_019835269.2;refGene_gn=PPP1R12A;refGene_tx=XM_019835270.2;refGene_gn=PPP1R12A;refGene_tx=XM_019835271.2
B4 105293230 105293446
refGene_gn=PPP1R12A;refGene_tx=XM_006933907.4;refGene_gn=PPP1R12A;refG

ene_tx=XM_006933908.3;refGene_gn=PPP1R12A;refGene_tx=XM_019835266.2;re
fGene_gn=PPP1R12A;refGene_tx=XM_019835267.2;refGene_gn=PPP1R12A;refGen
e_tx=XM_019835268.2;refGene_gn=PPP1R12A;refGene_tx=XM_019835269.2;refG
ene_gn=PPP1R12A;refGene_tx=XM_019835270.2;refGene_gn=PPP1R12A;refGene_
tx=XM_019835271.2
B4 105302002 105302124
refGene_gn=PPP1R12A;refGene_tx=XM_006933907.4;refGene_gn=PPP1R12A;refG
ene_tx=XM_006933908.3;refGene_gn=PPP1R12A;refGene_tx=XM_019835266.2;re
fGene_gn=PPP1R12A;refGene_tx=XM_019835267.2;refGene_gn=PPP1R12A;refGen
e_tx=XM_019835268.2;refGene_gn=PPP1R12A;refGene_tx=XM_019835269.2;refG
ene_gn=PPP1R12A;refGene_tx=XM_019835270.2;refGene_gn=PPP1R12A;refGene_
tx=XM_019835271.2
B4 105309416 105309574
refGene_gn=PPP1R12A;refGene_tx=XM_006933907.4;refGene_gn=PPP1R12A;refG
ene_tx=XM_006933908.3;refGene_gn=PPP1R12A;refGene_tx=XM_019835266.2;re
fGene_gn=PPP1R12A;refGene_tx=XM_019835267.2;refGene_gn=PPP1R12A;refGen
e_tx=XM_019835268.2;refGene_gn=PPP1R12A;refGene_tx=XM_019835269.2;refG
ene_gn=PPP1R12A;refGene_tx=XM_019835270.2;refGene_gn=PPP1R12A;refGene_
tx=XM_019835271.2
B4 105309727 105309816
refGene_gn=PPP1R12A;refGene_tx=XM_006933907.4;refGene_gn=PPP1R12A;refG
ene_tx=XM_006933908.3;refGene_gn=PPP1R12A;refGene_tx=XM_019835266.2;re
fGene_gn=PPP1R12A;refGene_tx=XM_019835267.2;refGene_gn=PPP1R12A;refGen
e_tx=XM_019835268.2;refGene_gn=PPP1R12A;refGene_tx=XM_019835269.2;refG
ene_gn=PPP1R12A;refGene_tx=XM_019835270.2;refGene_gn=PPP1R12A;refGene_
tx=XM_019835271.2
B4 105310487 105310562
refGene_gn=PPP1R12A;refGene_tx=XM_006933907.4;refGene_gn=PPP1R12A;refG
ene_tx=XM_006933908.3;refGene_gn=PPP1R12A;refGene_tx=XM_019835266.2;re
fGene_gn=PPP1R12A;refGene_tx=XM_019835267.2;refGene_gn=PPP1R12A;refGen
e_tx=XM_019835268.2;refGene_gn=PPP1R12A;refGene_tx=XM_019835269.2;refG
ene_gn=PPP1R12A;refGene_tx=XM_019835270.2;refGene_gn=PPP1R12A;refGene_
tx=XM_019835271.2
B4 105317236 105317381
refGene_gn=PPP1R12A;refGene_tx=XM_006933907.4;refGene_gn=PPP1R12A;refG
ene_tx=XM_006933908.3;refGene_gn=PPP1R12A;refGene_tx=XM_019835266.2;re
fGene_gn=PPP1R12A;refGene_tx=XM_019835267.2;refGene_gn=PPP1R12A;refGen
e_tx=XM_019835268.2;refGene_gn=PPP1R12A;refGene_tx=XM_019835269.2;refG
ene_gn=PPP1R12A;refGene_tx=XM_019835270.2;refGene_gn=PPP1R12A;refGene_
tx=XM_019835271.2
B4 105321926 105322086
refGene_gn=PPP1R12A;refGene_tx=XM_006933907.4;refGene_gn=PPP1R12A;refG
ene_tx=XM_006933908.3;refGene_gn=PPP1R12A;refGene_tx=XM_019835266.2;re
fGene_gn=PPP1R12A;refGene_tx=XM_019835267.2;refGene_gn=PPP1R12A;refGen
e_tx=XM_019835268.2;refGene_gn=PPP1R12A;refGene_tx=XM_019835269.2;refG
ene_gn=PPP1R12A;refGene_tx=XM_019835270.2;refGene_gn=PPP1R12A;refGene_
tx=XM_019835271.2
B4 105330886 105331005
refGene_gn=PPP1R12A;refGene_tx=XM_006933907.4;refGene_gn=PPP1R12A;refG
ene_tx=XM_006933908.3;refGene_gn=PPP1R12A;refGene_tx=XM_019835266.2;re

fGene_gn=PPP1R12A;refGene_tx=XM_019835267.2;refGene_gn=PPP1R12A;refGene_tx=XM_019835268.2;refGene_gn=PPP1R12A;refGene_tx=XM_019835269.2;refGene_gn=PPP1R12A;refGene_tx=XM_019835270.2;refGene_gn=PPP1R12A;refGene_tx=XM_019835271.2
B4 105353035 105353166
refGene_gn=PPP1R12A;refGene_tx=XM_006933907.4;refGene_gn=PPP1R12A;refGene_tx=XM_006933908.3;refGene_gn=PPP1R12A;refGene_tx=XM_019835266.2;refGene_gn=PPP1R12A;refGene_tx=XM_019835267.2;refGene_gn=PPP1R12A;refGene_tx=XM_019835268.2;refGene_gn=PPP1R12A;refGene_tx=XM_019835269.2;refGene_gn=PPP1R12A;refGene_tx=XM_019835270.2;refGene_gn=PPP1R12A;refGene_tx=XM_019835271.2
B4 105405892 105406129
refGene_gn=PPP1R12A;refGene_tx=XM_006933907.4;refGene_gn=PPP1R12A;refGene_tx=XM_006933908.3;refGene_gn=PPP1R12A;refGene_tx=XM_019835266.2;refGene_gn=PPP1R12A;refGene_tx=XM_019835267.2;refGene_gn=PPP1R12A;refGene_tx=XM_019835268.2;refGene_gn=PPP1R12A;refGene_tx=XM_019835269.2;refGene_gn=PPP1R12A;refGene_tx=XM_019835270.2;refGene_gn=PPP1R12A;refGene_tx=XM_019835271.2
B4 105631573 105631649
refGene_gn=OTOGL;refGene_tx=XM_019835272.2
B4 105632750 105632790
refGene_gn=OTOGL;refGene_tx=XM_019835272.2
B4 105633747 105633796
refGene_gn=OTOGL;refGene_tx=XM_019835272.2
B4 105638500 105638567
refGene_gn=OTOGL;refGene_tx=XM_019835272.2
B4 105640207 105640306
refGene_gn=OTOGL;refGene_tx=XM_019835272.2
B4 105641854 105642009
refGene_gn=OTOGL;refGene_tx=XM_019835272.2
B4 105646006 105646128
refGene_gn=OTOGL;refGene_tx=XM_019835272.2
B4 105650431 105650637
refGene_gn=OTOGL;refGene_tx=XM_019835272.2
B4 105656424 105656552
refGene_gn=OTOGL;refGene_tx=XM_019835272.2
B4 105657161 105657268
refGene_gn=OTOGL;refGene_tx=XM_019835272.2
B4 105661775 105661882
refGene_gn=OTOGL;refGene_tx=XM_019835272.2
B4 105662098 105662224
refGene_gn=OTOGL;refGene_tx=XM_019835272.2
B4 105663811 105663920
refGene_gn=OTOGL;refGene_tx=XM_019835272.2
B4 105665104 105665151
refGene_gn=OTOGL;refGene_tx=XM_019835272.2
B4 105665620 105665766
refGene_gn=OTOGL;refGene_tx=XM_019835272.2
B4 105666938 105667062
refGene_gn=OTOGL;refGene_tx=XM_019835272.2

B4 105669191 105669369
refGene_gn=OTOGL;refGene_tx=XM_019835272.2
B4 105673667 105673792
refGene_gn=OTOGL;refGene_tx=XM_019835272.2
B4 105677108 105677318
refGene_gn=OTOGL;refGene_tx=XM_019835272.2
B4 105679799 105679965
refGene_gn=OTOGL;refGene_tx=XM_019835272.2
B4 105680779 105680854
refGene_gn=OTOGL;refGene_tx=XM_019835272.2
B4 105683433 105683486
refGene_gn=OTOGL;refGene_tx=XM_019835272.2
B4 105685065 105685228
refGene_gn=OTOGL;refGene_tx=XM_019835272.2
B4 105689215 105689323
refGene_gn=OTOGL;refGene_tx=XM_019835272.2
B4 105690077 105690216
refGene_gn=OTOGL;refGene_tx=XM_019835272.2
B4 105697754 105697889
refGene_gn=OTOGL;refGene_tx=XM_019835272.2
B4 105702876 105703026
refGene_gn=OTOGL;refGene_tx=XM_019835272.2
B4 105705239 105705359
refGene_gn=OTOGL;refGene_tx=XM_019835272.2
B4 105708098 105708215
refGene_gn=OTOGL;refGene_tx=XM_019835272.2
B4 105709327 105709484
refGene_gn=OTOGL;refGene_tx=XM_019835272.2
B4 105710157 105710184
refGene_gn=OTOGL;refGene_tx=XM_019835272.2
B4 105722007 105722175
refGene_gn=OTOGL;refGene_tx=XM_019835272.2
B4 105724353 105724632
refGene_gn=OTOGL;refGene_tx=XM_019835272.2
B4 105726558 105726676
refGene_gn=OTOGL;refGene_tx=XM_019835272.2
B4 105731561 105731641
refGene_gn=OTOGL;refGene_tx=XM_019835272.2
B4 105731949 105731991
refGene_gn=OTOGL;refGene_tx=XM_019835272.2
B4 105734808 105734882
refGene_gn=OTOGL;refGene_tx=XM_019835272.2
B4 105738121 105738299
refGene_gn=OTOGL;refGene_tx=XM_019835272.2
B4 105738817 105738960
refGene_gn=OTOGL;refGene_tx=XM_019835272.2
B4 105739203 105739227
refGene_gn=OTOGL;refGene_tx=XM_019835272.2
B4 105739322 105739415
refGene_gn=OTOGL;refGene_tx=XM_019835272.2

B4 105741613 105741803
refGene_gn=OTOGL;refGene_tx=XM_019835272.2
B4 105744671 105744886
refGene_gn=OTOGL;refGene_tx=XM_019835272.2
B4 105748200 105748342
refGene_gn=OTOGL;refGene_tx=XM_019835272.2
B4 105750042 105750228
refGene_gn=OTOGL;refGene_tx=XM_019835272.2
B4 105752310 105752523
refGene_gn=OTOGL;refGene_tx=XM_019835272.2
B4 105752960 105753065
refGene_gn=OTOGL;refGene_tx=XM_019835272.2
B4 105754131 105754239
refGene_gn=OTOGL;refGene_tx=XM_019835272.2
B4 105756031 105756133
refGene_gn=OTOGL;refGene_tx=XM_019835272.2
B4 105756636 105756741
refGene_gn=OTOGL;refGene_tx=XM_019835272.2
B4 105756826 105756867
refGene_gn=OTOGL;refGene_tx=XM_019835272.2
B4 105761501 105761565
refGene_gn=OTOGL;refGene_tx=XM_019835272.2
B4 105762620 105762799
refGene_gn=OTOGL;refGene_tx=XM_019835272.2
B4 105763233 105763338
refGene_gn=OTOGL;refGene_tx=XM_019835272.2
B4 105766553 105766673
refGene_gn=OTOGL;refGene_tx=XM_019835272.2
B4 105768064 105768110
refGene_gn=OTOGL;refGene_tx=XM_019835272.2
B4 105770878 105770958
refGene_gn=OTOGL;refGene_tx=XM_019835272.2
B4 105771603 105771804
refGene_gn=OTOGL;refGene_tx=XM_019835272.2
B4 105791474 105791664
refGene_gn=LOC109501922;refGene_tx=XM_023257276.1
B4 105792140 105792398
refGene_gn=LOC109501922;refGene_tx=XM_023257276.1
B4 105793704 105793919
refGene_gn=LOC109501922;refGene_tx=XM_023257276.1
B4 105796747 105796937
refGene_gn=LOC109501922;refGene_tx=XM_023257276.1
B4 105797044 105797350
refGene_gn=LOC109501922;refGene_tx=XM_023257276.1
B4 105799222 105799498
refGene_gn=LOC109501922;refGene_tx=XM_023257276.1
B4 105804734 105805004
refGene_gn=LOC109501922;refGene_tx=XM_023257276.1
B4 105809868 105810129
refGene_gn=LOC109501922;refGene_tx=XM_023257276.1

B4 105811769 105811900
refGene_gn=LOC109501922;refGene_tx=XM_023257276.1
B4 105812080 105812210
refGene_gn=LOC109501922;refGene_tx=XM_023257276.1
B4 105813836 105813987
refGene_gn=LOC109501922;refGene_tx=XM_023257276.1
B4 105814441 105814613
refGene_gn=LOC109501922;refGene_tx=XM_023257276.1
B4 105842353 105842410
refGene_gn=LOC101091541;refGene_tx=XM_023257277.1
B4 105842756 105842868
refGene_gn=LOC101091541;refGene_tx=XM_023257277.1
B4 105843548 105843775
refGene_gn=LOC101091541;refGene_tx=XM_023257277.1
B4 105846791 105846861
refGene_gn=LOC101091541;refGene_tx=XM_023257277.1
B4 105849854 105850054
refGene_gn=LOC101091541;refGene_tx=XM_023257277.1
B4 105851206 105851456
refGene_gn=LOC101091541;refGene_tx=XM_023257277.1
B4 105852999 105853128
refGene_gn=LOC101091541;refGene_tx=XM_023257277.1
B4 105855003 105855150
refGene_gn=LOC101091541;refGene_tx=XM_023257277.1
B4 105861398 105861571
refGene_gn=LOC101091541;refGene_tx=XM_023257277.1
B4 105871080 105871258
refGene_gn=LOC101091541;refGene_tx=XM_023257277.1
B4 105872549 105872711
refGene_gn=LOC101091541;refGene_tx=XM_023257277.1
B4 105872817 105872997
refGene_gn=LOC101091541;refGene_tx=XM_023257277.1
B4 105873804 105873912
refGene_gn=LOC101091541;refGene_tx=XM_023257277.1
B4 105874051 105874333
refGene_gn=LOC101091541;refGene_tx=XM_023257277.1
B4 105885272 105885455
refGene_gn=LOC101091541;refGene_tx=XM_023257277.1
B4 105885819 105885921
refGene_gn=LOC101091541;refGene_tx=XM_023257277.1
B4 105889632 105889753
refGene_gn=LOC101091541;refGene_tx=XM_023257277.1
B4 105908872 105909031
refGene_gn=LOC101091541;refGene_tx=XM_023257277.1
B4 105909754 105909902
refGene_gn=LOC101091541;refGene_tx=XM_023257277.1
B4 105914515 105914684
refGene_gn=LOC101091541;refGene_tx=XM_023257277.1
B4 105916088 105916379
refGene_gn=LOC101091541;refGene_tx=XM_023257277.1

B4 105916640 105916916
refGene_gn=LOC101091541;refGene_tx=XM_023257277.1
B4 105917291 105917443
refGene_gn=LOC101091541;refGene_tx=XM_023257277.1
B4 105920699 105920841
refGene_gn=LOC101091541;refGene_tx=XM_023257277.1
B4 105924001 105924271
refGene_gn=LOC101091541;refGene_tx=XM_023257277.1
B4 105952054 105952378
refGene_gn=LOC101091541;refGene_tx=XM_023257277.1
B4 105971985 105972107
refGene_gn=LOC101091541;refGene_tx=XM_023257277.1
B4 105977712 105977899
refGene_gn=LOC101091541;refGene_tx=XM_023257277.1
B4 105981152 105981397
refGene_gn=LOC101091541;refGene_tx=XM_023257277.1
B4 105983845 105983912
refGene_gn=LOC101091541;refGene_tx=XM_023257277.1
B4 105987384 105987543
refGene_gn=LOC101091541;refGene_tx=XM_023257277.1
B4 105988100 105988323
refGene_gn=LOC101091541;refGene_tx=XM_023257277.1
B4 105989961 105990035
refGene_gn=LOC101091541;refGene_tx=XM_023257277.1
B4 105999937 106000037
refGene_gn=LOC101091541;refGene_tx=XM_023257277.1
B4 106002564 106002693
refGene_gn=LOC101091541;refGene_tx=XM_023257277.1
B4 106015956 106015983
refGene_gn=LOC101091541;refGene_tx=XM_023257277.1
B4 106016641 106016723
refGene_gn=LOC101091541;refGene_tx=XM_023257277.1
B4 106020078 106020169
refGene_gn=LOC101091541;refGene_tx=XM_023257277.1
B4 106025355 106025432
refGene_gn=LOC101091541;refGene_tx=XM_023257277.1
B4 106031876 106032011
refGene_gn=LOC101091541;refGene_tx=XM_023257277.1
B4 106032209 106032335
refGene_gn=LOC101091541;refGene_tx=XM_023257277.1
B4 106033454 106033603
refGene_gn=LOC101091541;refGene_tx=XM_023257277.1
B4 106042800 106042936
refGene_gn=LOC101091541;refGene_tx=XM_023257277.1
B4 106047894 106048018
refGene_gn=LOC101091541;refGene_tx=XM_023257277.1
B4 106048248 106048286
refGene_gn=LOC101091541;refGene_tx=XM_023257277.1
B4 106072998 106073517
refGene_gn=MYF6;refGene_tx=XM_003989075.2

B4 106073808 106073899
refGene_gn=MYF6;refGene_tx=XM_003989075.2
B4 106074140 106074259
refGene_gn=MYF6;refGene_tx=XM_003989075.2
B4 106082370 106082871
refGene_gn=MYF5;refGene_tx=XM_003989076.3
B4 106083678 106083754
refGene_gn=MYF5;refGene_tx=XM_003989076.3
B4 106084191 106084382
refGene_gn=MYF5;refGene_tx=XM_003989076.3
B4 106146791 106147010
refGene_gn=LIN7A;refGene_tx=XM_003989077.5;refGene_gn=LIN7A;refGene_tx
=XM_023257278.1
B4 106174028 106174238
refGene_gn=LIN7A;refGene_tx=XM_003989077.5;refGene_gn=LIN7A;refGene_tx
=XM_023257278.1
B4 106176570 106176642
refGene_gn=LIN7A;refGene_tx=XM_003989077.5;refGene_gn=LIN7A;refGene_tx
=XM_023257278.1
B4 106219631 106219750
refGene_gn=LIN7A;refGene_tx=XM_003989077.5;refGene_gn=LIN7A;refGene_tx
=XM_023257278.1
B4 106261564 106261646
refGene_gn=LIN7A;refGene_tx=XM_003989077.5;refGene_gn=LIN7A;refGene_tx
=XM_023257278.1
B4 106377780 106378382
refGene_gn=ACSS3;refGene_tx=XM_011284138.3
B4 106412966 106413111
refGene_gn=ACSS3;refGene_tx=XM_011284138.3
B4 106426813 106427002
refGene_gn=ACSS3;refGene_tx=XM_011284138.3
B4 106430973 106431108
refGene_gn=ACSS3;refGene_tx=XM_011284138.3
B4 106435180 106435321
refGene_gn=ACSS3;refGene_tx=XM_011284138.3
B4 106444943 106445024
refGene_gn=ACSS3;refGene_tx=XM_011284138.3
B4 106445105 106445201
refGene_gn=ACSS3;refGene_tx=XM_011284138.3
B4 106470163 106470315
refGene_gn=ACSS3;refGene_tx=XM_011284138.3
B4 106484898 106485002
refGene_gn=ACSS3;refGene_tx=XM_011284138.3
B4 106501857 106501953
refGene_gn=ACSS3;refGene_tx=XM_011284138.3
B4 106504752 106504816
refGene_gn=ACSS3;refGene_tx=XM_011284138.3
B4 106515094 106515176
refGene_gn=ACSS3;refGene_tx=XM_011284138.3
B4 106517339 106517462

refGene_gn=ACSS3;refGene_tx=XM_011284138.3
B4	106530933	106531033
refGene_gn=ACSS3;refGene_tx=XM_011284138.3
B4	106531123	106531299
refGene_gn=ACSS3;refGene_tx=XM_011284138.3
B4	106532663	106532732
refGene_gn=ACSS3;refGene_tx=XM_011284138.3
B4	106536936	106536965
refGene_gn=PPFIA2;refGene_tx=XM_019835279.2
B4	106539246	106539305
refGene_gn=PPFIA2;refGene_tx=XM_006933911.4;refGene_gn=PPFIA2;refGene_tx=XM_019835275.2;refGene_gn=PPFIA2;refGene_tx=XM_019835276.2;refGene_gn=PPFIA2;refGene_tx=XM_019835277.2;refGene_gn=PPFIA2;refGene_tx=XM_019835278.2;refGene_gn=PPFIA2;refGene_tx=XM_019835280.2;refGene_gn=PPFIA2;refGene_tx=XM_019835281.2;refGene_gn=PPFIA2;refGene_tx=XM_019835282.2;refGene_gn=PPFIA2;refGene_tx=XM_019835283.2;refGene_gn=PPFIA2;refGene_tx=XM_019835285.2;refGene_gn=PPFIA2;refGene_tx=XM_019835286.2;refGene_gn=PPFIA2;refGene_tx=XM_019835287.2;refGene_gn=PPFIA2;refGene_tx=XM_019835288.2;refGene_gn=PPFIA2;refGene_tx=XM_019835289.2;refGene_gn=PPFIA2;refGene_tx=XM_019835290.2;refGene_gn=PPFIA2;refGene_tx=XM_019835291.2;refGene_gn=PPFIA2;refGene_tx=XM_019835292.2;refGene_gn=PPFIA2;refGene_tx=XM_019835293.2
B4	106540519	106540679
refGene_gn=PPFIA2;refGene_tx=XM_006933911.4;refGene_gn=PPFIA2;refGene_tx=XM_019835275.2;refGene_gn=PPFIA2;refGene_tx=XM_019835276.2;refGene_gn=PPFIA2;refGene_tx=XM_019835277.2;refGene_gn=PPFIA2;refGene_tx=XM_019835278.2;refGene_gn=PPFIA2;refGene_tx=XM_019835279.2;refGene_gn=PPFIA2;refGene_tx=XM_019835280.2;refGene_gn=PPFIA2;refGene_tx=XM_019835281.2;refGene_gn=PPFIA2;refGene_tx=XM_019835282.2;refGene_gn=PPFIA2;refGene_tx=XM_019835283.2;refGene_gn=PPFIA2;refGene_tx=XM_019835285.2;refGene_gn=PPFIA2;refGene_tx=XM_019835286.2;refGene_gn=PPFIA2;refGene_tx=XM_019835287.2;refGene_gn=PPFIA2;refGene_tx=XM_019835288.2;refGene_gn=PPFIA2;refGene_tx=XM_019835289.2;refGene_gn=PPFIA2;refGene_tx=XM_019835290.2;refGene_gn=PPFIA2;refGene_tx=XM_019835291.2;refGene_gn=PPFIA2;refGene_tx=XM_019835292.2;refGene_gn=PPFIA2;refGene_tx=XM_019835293.2
B4	106544259	106544328
refGene_gn=PPFIA2;refGene_tx=XM_006933911.4;refGene_gn=PPFIA2;refGene_tx=XM_019835275.2;refGene_gn=PPFIA2;refGene_tx=XM_019835276.2;refGene_gn=PPFIA2;refGene_tx=XM_019835277.2;refGene_gn=PPFIA2;refGene_tx=XM_019835278.2;refGene_gn=PPFIA2;refGene_tx=XM_019835279.2;refGene_gn=PPFIA2;refGene_tx=XM_019835280.2;refGene_gn=PPFIA2;refGene_tx=XM_019835281.2;refGene_gn=PPFIA2;refGene_tx=XM_019835282.2;refGene_gn=PPFIA2;refGene_tx=XM_019835283.2;refGene_gn=PPFIA2;refGene_tx=XM_019835285.2;refGene_gn=PPFIA2;refGene_tx=XM_019835286.2;refGene_gn=PPFIA2;refGene_tx=XM_019835287.2;refGene_gn=PPFIA2;refGene_tx=XM_019835288.2;refGene_gn=PPFIA2;refGene_tx=XM_019835289.2;refGene_gn=PPFIA2;refGene_tx=XM_019835290.2;refGene_gn=PPFIA2;refGene_tx=XM_019835291.2;refGene_gn=PPFIA2;refGene_tx=XM_019835292.2;refGene_gn=PPFIA2;refGene_tx=XM_019835293.2
B4	106545217	106545393
refGene_gn=PPFIA2;refGene_tx=XM_006933911.4;refGene_gn=PPFIA2;refGene_

tx=XM_019835275.2;refGene_gn=PPFIA2;refGene_tx=XM_019835276.2;refGene_
gn=PPFIA2;refGene_tx=XM_019835277.2;refGene_gn=PPFIA2;refGene_tx=XM_01
9835278.2;refGene_gn=PPFIA2;refGene_tx=XM_019835279.2;refGene_gn=PPFIA
2;refGene_tx=XM_019835280.2;refGene_gn=PPFIA2;refGene_tx=XM_019835281.
2;refGene_gn=PPFIA2;refGene_tx=XM_019835282.2;refGene_gn=PPFIA2;refGen
e_tx=XM_019835283.2;refGene_gn=PPFIA2;refGene_tx=XM_019835285.2;refGen
e_gn=PPFIA2;refGene_tx=XM_019835286.2;refGene_gn=PPFIA2;refGene_tx=XM_
019835287.2;refGene_gn=PPFIA2;refGene_tx=XM_019835288.2;refGene_gn=PPF
IA2;refGene_tx=XM_019835289.2;refGene_gn=PPFIA2;refGene_tx=XM_01983529
0.2;refGene_gn=PPFIA2;refGene_tx=XM_019835291.2;refGene_gn=PPFIA2;refG
ene_tx=XM_019835292.2;refGene_gn=PPFIA2;refGene_tx=XM_019835293.2
B4 106551807 106551905
refGene_gn=PPFIA2;refGene_tx=XM_006933911.4;refGene_gn=PPFIA2;refGene_
tx=XM_019835275.2;refGene_gn=PPFIA2;refGene_tx=XM_019835276.2;refGene_
gn=PPFIA2;refGene_tx=XM_019835277.2;refGene_gn=PPFIA2;refGene_tx=XM_01
9835278.2;refGene_gn=PPFIA2;refGene_tx=XM_019835279.2;refGene_gn=PPFIA
2;refGene_tx=XM_019835280.2;refGene_gn=PPFIA2;refGene_tx=XM_019835281.
2;refGene_gn=PPFIA2;refGene_tx=XM_019835282.2;refGene_gn=PPFIA2;refGen
e_tx=XM_019835283.2;refGene_gn=PPFIA2;refGene_tx=XM_019835285.2;refGen
e_gn=PPFIA2;refGene_tx=XM_019835286.2;refGene_gn=PPFIA2;refGene_tx=XM_
019835287.2;refGene_gn=PPFIA2;refGene_tx=XM_019835288.2;refGene_gn=PPF
IA2;refGene_tx=XM_019835289.2;refGene_gn=PPFIA2;refGene_tx=XM_01983529
0.2;refGene_gn=PPFIA2;refGene_tx=XM_019835291.2;refGene_gn=PPFIA2;refG
ene_tx=XM_019835292.2;refGene_gn=PPFIA2;refGene_tx=XM_019835293.2
B4 106556349 106556543
refGene_gn=PPFIA2;refGene_tx=XM_006933911.4;refGene_gn=PPFIA2;refGene_
tx=XM_019835275.2;refGene_gn=PPFIA2;refGene_tx=XM_019835276.2;refGene_
gn=PPFIA2;refGene_tx=XM_019835277.2;refGene_gn=PPFIA2;refGene_tx=XM_01
9835278.2;refGene_gn=PPFIA2;refGene_tx=XM_019835279.2;refGene_gn=PPFIA
2;refGene_tx=XM_019835280.2;refGene_gn=PPFIA2;refGene_tx=XM_019835281.
2;refGene_gn=PPFIA2;refGene_tx=XM_019835282.2;refGene_gn=PPFIA2;refGen
e_tx=XM_019835283.2;refGene_gn=PPFIA2;refGene_tx=XM_019835285.2;refGen
e_gn=PPFIA2;refGene_tx=XM_019835286.2;refGene_gn=PPFIA2;refGene_tx=XM_
019835287.2;refGene_gn=PPFIA2;refGene_tx=XM_019835288.2;refGene_gn=PPF
IA2;refGene_tx=XM_019835289.2;refGene_gn=PPFIA2;refGene_tx=XM_01983529
0.2;refGene_gn=PPFIA2;refGene_tx=XM_019835291.2;refGene_gn=PPFIA2;refG
ene_tx=XM_019835292.2;refGene_gn=PPFIA2;refGene_tx=XM_019835293.2
B4 106558135 106558165
refGene_gn=PPFIA2;refGene_tx=XM_006933911.4;refGene_gn=PPFIA2;refGene_
tx=XM_019835275.2;refGene_gn=PPFIA2;refGene_tx=XM_019835276.2;refGene_
gn=PPFIA2;refGene_tx=XM_019835277.2;refGene_gn=PPFIA2;refGene_tx=XM_01
9835278.2;refGene_gn=PPFIA2;refGene_tx=XM_019835279.2;refGene_gn=PPFIA
2;refGene_tx=XM_019835281.2;refGene_gn=PPFIA2;refGene_tx=XM_019835282.
2;refGene_gn=PPFIA2;refGene_tx=XM_019835283.2;refGene_gn=PPFIA2;refGen
e_tx=XM_019835285.2;refGene_gn=PPFIA2;refGene_tx=XM_019835286.2;refGen
e_gn=PPFIA2;refGene_tx=XM_019835287.2;refGene_gn=PPFIA2;refGene_tx=XM_
019835288.2;refGene_gn=PPFIA2;refGene_tx=XM_019835289.2;refGene_gn=PPF
IA2;refGene_tx=XM_019835290.2;refGene_gn=PPFIA2;refGene_tx=XM_01983529
1.2;refGene_gn=PPFIA2;refGene_tx=XM_019835292.2;refGene_gn=PPFIA2;refG
ene_tx=XM_019835293.2

B4 106559719 106559782
refGene_gn=PPFIA2;refGene_tx=XM_006933911.4;refGene_gn=PPFIA2;refGene_tx=XM_019835275.2;refGene_gn=PPFIA2;refGene_tx=XM_019835276.2;refGene_gn=PPFIA2;refGene_tx=XM_019835277.2;refGene_gn=PPFIA2;refGene_tx=XM_019835278.2;refGene_gn=PPFIA2;refGene_tx=XM_019835279.2;refGene_gn=PPFIA2;refGene_tx=XM_019835280.2;refGene_gn=PPFIA2;refGene_tx=XM_019835281.2;refGene_gn=PPFIA2;refGene_tx=XM_019835282.2;refGene_gn=PPFIA2;refGene_tx=XM_019835283.2;refGene_gn=PPFIA2;refGene_tx=XM_019835285.2;refGene_gn=PPFIA2;refGene_tx=XM_019835286.2;refGene_gn=PPFIA2;refGene_tx=XM_019835287.2;refGene_gn=PPFIA2;refGene_tx=XM_019835288.2;refGene_gn=PPFIA2;refGene_tx=XM_019835289.2;refGene_gn=PPFIA2;refGene_tx=XM_019835290.2;refGene_gn=PPFIA2;refGene_tx=XM_019835291.2;refGene_gn=PPFIA2;refGene_tx=XM_019835292.2;refGene_gn=PPFIA2;refGene_tx=XM_019835293.2
B4 106559890 106559947
refGene_gn=PPFIA2;refGene_tx=XM_023257280.1
B4 106567653 106567854
refGene_gn=PPFIA2;refGene_tx=XM_006933911.4;refGene_gn=PPFIA2;refGene_tx=XM_019835275.2;refGene_gn=PPFIA2;refGene_tx=XM_019835276.2;refGene_gn=PPFIA2;refGene_tx=XM_019835277.2;refGene_gn=PPFIA2;refGene_tx=XM_019835278.2;refGene_gn=PPFIA2;refGene_tx=XM_019835279.2;refGene_gn=PPFIA2;refGene_tx=XM_019835280.2;refGene_gn=PPFIA2;refGene_tx=XM_019835281.2;refGene_gn=PPFIA2;refGene_tx=XM_019835282.2;refGene_gn=PPFIA2;refGene_tx=XM_019835283.2;refGene_gn=PPFIA2;refGene_tx=XM_019835285.2;refGene_gn=PPFIA2;refGene_tx=XM_019835286.2;refGene_gn=PPFIA2;refGene_tx=XM_019835287.2;refGene_gn=PPFIA2;refGene_tx=XM_019835288.2;refGene_gn=PPFIA2;refGene_tx=XM_019835289.2;refGene_gn=PPFIA2;refGene_tx=XM_019835290.2;refGene_gn=PPFIA2;refGene_tx=XM_019835291.2;refGene_gn=PPFIA2;refGene_tx=XM_019835292.2;refGene_gn=PPFIA2;refGene_tx=XM_019835293.2;refGene_gn=PPFIA2;refGene_tx=XM_023257280.1
B4 106571781 106571863
refGene_gn=PPFIA2;refGene_tx=XM_006933911.4;refGene_gn=PPFIA2;refGene_tx=XM_019835275.2;refGene_gn=PPFIA2;refGene_tx=XM_019835276.2;refGene_gn=PPFIA2;refGene_tx=XM_019835277.2;refGene_gn=PPFIA2;refGene_tx=XM_019835278.2;refGene_gn=PPFIA2;refGene_tx=XM_019835279.2;refGene_gn=PPFIA2;refGene_tx=XM_019835280.2;refGene_gn=PPFIA2;refGene_tx=XM_019835281.2;refGene_gn=PPFIA2;refGene_tx=XM_019835282.2;refGene_gn=PPFIA2;refGene_tx=XM_019835283.2;refGene_gn=PPFIA2;refGene_tx=XM_019835285.2;refGene_gn=PPFIA2;refGene_tx=XM_019835286.2;refGene_gn=PPFIA2;refGene_tx=XM_019835287.2;refGene_gn=PPFIA2;refGene_tx=XM_019835288.2;refGene_gn=PPFIA2;refGene_tx=XM_019835289.2;refGene_gn=PPFIA2;refGene_tx=XM_019835290.2;refGene_gn=PPFIA2;refGene_tx=XM_019835291.2;refGene_gn=PPFIA2;refGene_tx=XM_019835292.2;refGene_gn=PPFIA2;refGene_tx=XM_019835293.2;refGene_gn=PPFIA2;refGene_tx=XM_023257280.1
B4 106583099 106583108
refGene_gn=PPFIA2;refGene_tx=XM_019835275.2;refGene_gn=PPFIA2;refGene_tx=XM_019835277.2;refGene_gn=PPFIA2;refGene_tx=XM_019835279.2;refGene_gn=PPFIA2;refGene_tx=XM_019835280.2;refGene_gn=PPFIA2;refGene_tx=XM_019835281.2;refGene_gn=PPFIA2;refGene_tx=XM_019835282.2;refGene_gn=PPFIA2;refGene_tx=XM_019835283.2;refGene_gn=PPFIA2;refGene_tx=XM_019835286.2;refGene_gn=PPFIA2;refGene_tx=XM_019835287.2;refGene_gn=PPFIA2;refGen

e_tx=XM_019835288.2;refGene_gn=PPFIA2;refGene_tx=XM_023257280.1
B4 106596848 106596942
refGene_gn=PPFIA2;refGene_tx=XM_006933911.4;refGene_gn=PPFIA2;refGene_tx=XM_019835275.2;refGene_gn=PPFIA2;refGene_tx=XM_019835276.2;refGene_gn=PPFIA2;refGene_tx=XM_019835277.2;refGene_gn=PPFIA2;refGene_tx=XM_019835278.2;refGene_gn=PPFIA2;refGene_tx=XM_019835279.2;refGene_gn=PPFIA2;refGene_tx=XM_019835280.2;refGene_gn=PPFIA2;refGene_tx=XM_019835281.2;refGene_gn=PPFIA2;refGene_tx=XM_019835282.2;refGene_gn=PPFIA2;refGene_tx=XM_019835283.2;refGene_gn=PPFIA2;refGene_tx=XM_019835285.2;refGene_gn=PPFIA2;refGene_tx=XM_019835286.2;refGene_gn=PPFIA2;refGene_tx=XM_019835287.2;refGene_gn=PPFIA2;refGene_tx=XM_019835288.2;refGene_gn=PPFIA2;refGene_tx=XM_019835289.2;refGene_gn=PPFIA2;refGene_tx=XM_019835290.2;refGene_gn=PPFIA2;refGene_tx=XM_019835291.2;refGene_gn=PPFIA2;refGene_tx=XM_019835292.2;refGene_gn=PPFIA2;refGene_tx=XM_019835293.2;refGene_gn=PPFIA2;refGene_tx=XM_023257280.1
B4 106606063 106606218
refGene_gn=PPFIA2;refGene_tx=XM_006933911.4;refGene_gn=PPFIA2;refGene_tx=XM_019835275.2;refGene_gn=PPFIA2;refGene_tx=XM_019835276.2;refGene_gn=PPFIA2;refGene_tx=XM_019835277.2;refGene_gn=PPFIA2;refGene_tx=XM_019835278.2;refGene_gn=PPFIA2;refGene_tx=XM_019835279.2;refGene_gn=PPFIA2;refGene_tx=XM_019835280.2;refGene_gn=PPFIA2;refGene_tx=XM_019835281.2;refGene_gn=PPFIA2;refGene_tx=XM_019835282.2;refGene_gn=PPFIA2;refGene_tx=XM_019835283.2;refGene_gn=PPFIA2;refGene_tx=XM_019835285.2;refGene_gn=PPFIA2;refGene_tx=XM_019835286.2;refGene_gn=PPFIA2;refGene_tx=XM_019835287.2;refGene_gn=PPFIA2;refGene_tx=XM_019835288.2;refGene_gn=PPFIA2;refGene_tx=XM_019835289.2;refGene_gn=PPFIA2;refGene_tx=XM_019835290.2;refGene_gn=PPFIA2;refGene_tx=XM_019835291.2;refGene_gn=PPFIA2;refGene_tx=XM_019835292.2;refGene_gn=PPFIA2;refGene_tx=XM_019835293.2;refGene_gn=PPFIA2;refGene_tx=XM_023257280.1
B4 106609363 106609494
refGene_gn=PPFIA2;refGene_tx=XM_006933911.4;refGene_gn=PPFIA2;refGene_tx=XM_019835275.2;refGene_gn=PPFIA2;refGene_tx=XM_019835276.2;refGene_gn=PPFIA2;refGene_tx=XM_019835277.2;refGene_gn=PPFIA2;refGene_tx=XM_019835278.2;refGene_gn=PPFIA2;refGene_tx=XM_019835279.2;refGene_gn=PPFIA2;refGene_tx=XM_019835280.2;refGene_gn=PPFIA2;refGene_tx=XM_019835281.2;refGene_gn=PPFIA2;refGene_tx=XM_019835282.2;refGene_gn=PPFIA2;refGene_tx=XM_019835283.2;refGene_gn=PPFIA2;refGene_tx=XM_019835285.2;refGene_gn=PPFIA2;refGene_tx=XM_019835286.2;refGene_gn=PPFIA2;refGene_tx=XM_019835287.2;refGene_gn=PPFIA2;refGene_tx=XM_019835288.2;refGene_gn=PPFIA2;refGene_tx=XM_019835289.2;refGene_gn=PPFIA2;refGene_tx=XM_019835290.2;refGene_gn=PPFIA2;refGene_tx=XM_019835291.2;refGene_gn=PPFIA2;refGene_tx=XM_019835292.2;refGene_gn=PPFIA2;refGene_tx=XM_019835293.2;refGene_gn=PPFIA2;refGene_tx=XM_023257280.1
B4 106613352 106613382
refGene_gn=PPFIA2;refGene_tx=XM_006933911.4;refGene_gn=PPFIA2;refGene_tx=XM_019835275.2;refGene_gn=PPFIA2;refGene_tx=XM_019835276.2;refGene_gn=PPFIA2;refGene_tx=XM_019835277.2;refGene_gn=PPFIA2;refGene_tx=XM_019835278.2;refGene_gn=PPFIA2;refGene_tx=XM_019835279.2;refGene_gn=PPFIA2;refGene_tx=XM_019835280.2;refGene_gn=PPFIA2;refGene_tx=XM_019835281.2;refGene_gn=PPFIA2;refGene_tx=XM_019835282.2;refGene_gn=PPFIA2;refGen

e_tx=XM_019835283.2;refGene_gn=PPFIA2;refGene_tx=XM_019835285.2;refGene_gn=PPFIA2;refGene_tx=XM_019835286.2;refGene_gn=PPFIA2;refGene_tx=XM_019835287.2;refGene_gn=PPFIA2;refGene_tx=XM_019835288.2;refGene_gn=PPFIA2;refGene_tx=XM_019835289.2;refGene_gn=PPFIA2;refGene_tx=XM_019835290.2;refGene_gn=PPFIA2;refGene_tx=XM_019835291.2;refGene_gn=PPFIA2;refGene_tx=XM_019835292.2;refGene_gn=PPFIA2;refGene_tx=XM_019835293.2;refGene_gn=PPFIA2;refGene_tx=XM_023257280.1
B4 106616686 106616924
refGene_gn=PPFIA2;refGene_tx=XM_006933911.4;refGene_gn=PPFIA2;refGene_tx=XM_019835275.2;refGene_gn=PPFIA2;refGene_tx=XM_019835276.2;refGene_gn=PPFIA2;refGene_tx=XM_019835277.2;refGene_gn=PPFIA2;refGene_tx=XM_019835278.2;refGene_gn=PPFIA2;refGene_tx=XM_019835279.2;refGene_gn=PPFIA2;refGene_tx=XM_019835280.2;refGene_gn=PPFIA2;refGene_tx=XM_019835281.2;refGene_gn=PPFIA2;refGene_tx=XM_019835282.2;refGene_gn=PPFIA2;refGene_tx=XM_019835283.2;refGene_gn=PPFIA2;refGene_tx=XM_019835285.2;refGene_gn=PPFIA2;refGene_tx=XM_019835286.2;refGene_gn=PPFIA2;refGene_tx=XM_019835287.2;refGene_gn=PPFIA2;refGene_tx=XM_019835288.2;refGene_gn=PPFIA2;refGene_tx=XM_019835289.2;refGene_gn=PPFIA2;refGene_tx=XM_019835290.2;refGene_gn=PPFIA2;refGene_tx=XM_019835291.2;refGene_gn=PPFIA2;refGene_tx=XM_019835292.2;refGene_gn=PPFIA2;refGene_tx=XM_019835293.2;refGene_gn=PPFIA2;refGene_tx=XM_023257280.1
B4 106629411 106629632
refGene_gn=PPFIA2;refGene_tx=XM_006933911.4;refGene_gn=PPFIA2;refGene_tx=XM_019835275.2;refGene_gn=PPFIA2;refGene_tx=XM_019835276.2;refGene_gn=PPFIA2;refGene_tx=XM_019835277.2;refGene_gn=PPFIA2;refGene_tx=XM_019835278.2;refGene_gn=PPFIA2;refGene_tx=XM_019835279.2;refGene_gn=PPFIA2;refGene_tx=XM_019835280.2;refGene_gn=PPFIA2;refGene_tx=XM_019835281.2;refGene_gn=PPFIA2;refGene_tx=XM_019835282.2;refGene_gn=PPFIA2;refGene_tx=XM_019835283.2;refGene_gn=PPFIA2;refGene_tx=XM_019835285.2;refGene_gn=PPFIA2;refGene_tx=XM_019835286.2;refGene_gn=PPFIA2;refGene_tx=XM_019835287.2;refGene_gn=PPFIA2;refGene_tx=XM_019835288.2;refGene_gn=PPFIA2;refGene_tx=XM_019835289.2;refGene_gn=PPFIA2;refGene_tx=XM_019835290.2;refGene_gn=PPFIA2;refGene_tx=XM_019835291.2;refGene_gn=PPFIA2;refGene_tx=XM_019835292.2;refGene_gn=PPFIA2;refGene_tx=XM_019835293.2;refGene_gn=PPFIA2;refGene_tx=XM_023257280.1
B4 106631466 106631602
refGene_gn=PPFIA2;refGene_tx=XM_006933911.4;refGene_gn=PPFIA2;refGene_tx=XM_019835275.2;refGene_gn=PPFIA2;refGene_tx=XM_019835276.2;refGene_gn=PPFIA2;refGene_tx=XM_019835277.2;refGene_gn=PPFIA2;refGene_tx=XM_019835278.2;refGene_gn=PPFIA2;refGene_tx=XM_019835279.2;refGene_gn=PPFIA2;refGene_tx=XM_019835280.2;refGene_gn=PPFIA2;refGene_tx=XM_019835281.2;refGene_gn=PPFIA2;refGene_tx=XM_019835282.2;refGene_gn=PPFIA2;refGene_tx=XM_019835283.2;refGene_gn=PPFIA2;refGene_tx=XM_019835285.2;refGene_gn=PPFIA2;refGene_tx=XM_019835286.2;refGene_gn=PPFIA2;refGene_tx=XM_019835287.2;refGene_gn=PPFIA2;refGene_tx=XM_019835288.2;refGene_gn=PPFIA2;refGene_tx=XM_019835289.2;refGene_gn=PPFIA2;refGene_tx=XM_019835290.2;refGene_gn=PPFIA2;refGene_tx=XM_019835291.2;refGene_gn=PPFIA2;refGene_tx=XM_019835292.2;refGene_gn=PPFIA2;refGene_tx=XM_019835293.2;refGene_gn=PPFIA2;refGene_tx=XM_023257280.1
B4 106635819 106635911

refGene_gn=PPFIA2;refGene_tx=XM_006933911.4;refGene_gn=PPFIA2;refGene_tx=XM_019835275.2;refGene_gn=PPFIA2;refGene_tx=XM_019835276.2;refGene_gn=PPFIA2;refGene_tx=XM_019835277.2;refGene_gn=PPFIA2;refGene_tx=XM_019835278.2;refGene_gn=PPFIA2;refGene_tx=XM_019835279.2;refGene_gn=PPFIA2;refGene_tx=XM_019835280.2;refGene_gn=PPFIA2;refGene_tx=XM_019835281.2;refGene_gn=PPFIA2;refGene_tx=XM_019835282.2;refGene_gn=PPFIA2;refGene_tx=XM_019835283.2;refGene_gn=PPFIA2;refGene_tx=XM_019835285.2;refGene_gn=PPFIA2;refGene_tx=XM_019835286.2;refGene_gn=PPFIA2;refGene_tx=XM_019835287.2;refGene_gn=PPFIA2;refGene_tx=XM_019835288.2;refGene_gn=PPFIA2;refGene_tx=XM_019835289.2;refGene_gn=PPFIA2;refGene_tx=XM_019835290.2;refGene_gn=PPFIA2;refGene_tx=XM_019835291.2;refGene_gn=PPFIA2;refGene_tx=XM_019835292.2;refGene_gn=PPFIA2;refGene_tx=XM_019835293.2;refGene_gn=PPFIA2;refGene_tx=XM_023257280.1
B4 106640688 106640751
refGene_gn=PPFIA2;refGene_tx=XM_006933911.4;refGene_gn=PPFIA2;refGene_tx=XM_019835275.2;refGene_gn=PPFIA2;refGene_tx=XM_019835276.2;refGene_gn=PPFIA2;refGene_tx=XM_019835277.2;refGene_gn=PPFIA2;refGene_tx=XM_019835278.2;refGene_gn=PPFIA2;refGene_tx=XM_019835279.2;refGene_gn=PPFIA2;refGene_tx=XM_019835280.2;refGene_gn=PPFIA2;refGene_tx=XM_019835281.2;refGene_gn=PPFIA2;refGene_tx=XM_019835282.2;refGene_gn=PPFIA2;refGene_tx=XM_019835283.2;refGene_gn=PPFIA2;refGene_tx=XM_019835285.2;refGene_gn=PPFIA2;refGene_tx=XM_019835286.2;refGene_gn=PPFIA2;refGene_tx=XM_019835287.2;refGene_gn=PPFIA2;refGene_tx=XM_019835288.2;refGene_gn=PPFIA2;refGene_tx=XM_019835289.2;refGene_gn=PPFIA2;refGene_tx=XM_019835290.2;refGene_gn=PPFIA2;refGene_tx=XM_019835291.2;refGene_gn=PPFIA2;refGene_tx=XM_019835292.2;refGene_gn=PPFIA2;refGene_tx=XM_019835293.2;refGene_gn=PPFIA2;refGene_tx=XM_023257280.1
B4 106642285 106642417
refGene_gn=PPFIA2;refGene_tx=XM_006933911.4;refGene_gn=PPFIA2;refGene_tx=XM_019835275.2;refGene_gn=PPFIA2;refGene_tx=XM_019835276.2;refGene_gn=PPFIA2;refGene_tx=XM_019835277.2;refGene_gn=PPFIA2;refGene_tx=XM_019835278.2;refGene_gn=PPFIA2;refGene_tx=XM_019835279.2;refGene_gn=PPFIA2;refGene_tx=XM_019835280.2;refGene_gn=PPFIA2;refGene_tx=XM_019835281.2;refGene_gn=PPFIA2;refGene_tx=XM_019835282.2;refGene_gn=PPFIA2;refGene_tx=XM_019835283.2;refGene_gn=PPFIA2;refGene_tx=XM_019835285.2;refGene_gn=PPFIA2;refGene_tx=XM_019835286.2;refGene_gn=PPFIA2;refGene_tx=XM_019835287.2;refGene_gn=PPFIA2;refGene_tx=XM_019835288.2;refGene_gn=PPFIA2;refGene_tx=XM_019835289.2;refGene_gn=PPFIA2;refGene_tx=XM_019835290.2;refGene_gn=PPFIA2;refGene_tx=XM_019835291.2;refGene_gn=PPFIA2;refGene_tx=XM_019835292.2;refGene_gn=PPFIA2;refGene_tx=XM_019835293.2;refGene_gn=PPFIA2;refGene_tx=XM_023257280.1
B4 106642678 106642762
refGene_gn=PPFIA2;refGene_tx=XM_006933911.4;refGene_gn=PPFIA2;refGene_tx=XM_019835275.2;refGene_gn=PPFIA2;refGene_tx=XM_019835276.2;refGene_gn=PPFIA2;refGene_tx=XM_019835277.2;refGene_gn=PPFIA2;refGene_tx=XM_019835278.2;refGene_gn=PPFIA2;refGene_tx=XM_019835279.2;refGene_gn=PPFIA2;refGene_tx=XM_019835280.2;refGene_gn=PPFIA2;refGene_tx=XM_019835281.2;refGene_gn=PPFIA2;refGene_tx=XM_019835282.2;refGene_gn=PPFIA2;refGene_tx=XM_019835283.2;refGene_gn=PPFIA2;refGene_tx=XM_019835285.2;refGene_gn=PPFIA2;refGene_tx=XM_019835286.2;refGene_gn=PPFIA2;refGene_tx=XM_

019835287.2;refGene_gn=PPFIA2;refGene_tx=XM_019835288.2;refGene_gn=PPFIA2;refGene_tx=XM_019835289.2;refGene_gn=PPFIA2;refGene_tx=XM_019835290.2;refGene_gn=PPFIA2;refGene_tx=XM_019835291.2;refGene_gn=PPFIA2;refGene_tx=XM_019835292.2;refGene_gn=PPFIA2;refGene_tx=XM_019835293.2;refGene_gn=PPFIA2;refGene_tx=XM_023257280.1
B4 106643189 106643276
refGene_gn=PPFIA2;refGene_tx=XM_019835275.2;refGene_gn=PPFIA2;refGene_tx=XM_019835276.2;refGene_gn=PPFIA2;refGene_tx=XM_019835277.2;refGene_gn=PPFIA2;refGene_tx=XM_019835278.2;refGene_gn=PPFIA2;refGene_tx=XM_019835279.2;refGene_gn=PPFIA2;refGene_tx=XM_019835280.2;refGene_gn=PPFIA2;refGene_tx=XM_019835281.2;refGene_gn=PPFIA2;refGene_tx=XM_019835282.2;refGene_gn=PPFIA2;refGene_tx=XM_019835287.2;refGene_gn=PPFIA2;refGene_tx=XM_023257280.1
B4 106648149 106648284
refGene_gn=PPFIA2;refGene_tx=XM_006933911.4;refGene_gn=PPFIA2;refGene_tx=XM_019835275.2;refGene_gn=PPFIA2;refGene_tx=XM_019835276.2;refGene_gn=PPFIA2;refGene_tx=XM_019835277.2;refGene_gn=PPFIA2;refGene_tx=XM_019835278.2;refGene_gn=PPFIA2;refGene_tx=XM_019835279.2;refGene_gn=PPFIA2;refGene_tx=XM_019835280.2;refGene_gn=PPFIA2;refGene_tx=XM_019835281.2;refGene_gn=PPFIA2;refGene_tx=XM_019835282.2;refGene_gn=PPFIA2;refGene_tx=XM_019835283.2;refGene_gn=PPFIA2;refGene_tx=XM_019835285.2;refGene_gn=PPFIA2;refGene_tx=XM_019835286.2;refGene_gn=PPFIA2;refGene_tx=XM_019835287.2;refGene_gn=PPFIA2;refGene_tx=XM_019835288.2;refGene_gn=PPFIA2;refGene_tx=XM_019835289.2;refGene_gn=PPFIA2;refGene_tx=XM_019835290.2;refGene_gn=PPFIA2;refGene_tx=XM_019835291.2;refGene_gn=PPFIA2;refGene_tx=XM_019835292.2;refGene_gn=PPFIA2;refGene_tx=XM_019835293.2;refGene_gn=PPFIA2;refGene_tx=XM_023257280.1
B4 106649299 106649446
refGene_gn=PPFIA2;refGene_tx=XM_006933911.4;refGene_gn=PPFIA2;refGene_tx=XM_019835275.2;refGene_gn=PPFIA2;refGene_tx=XM_019835276.2;refGene_gn=PPFIA2;refGene_tx=XM_019835277.2;refGene_gn=PPFIA2;refGene_tx=XM_019835278.2;refGene_gn=PPFIA2;refGene_tx=XM_019835279.2;refGene_gn=PPFIA2;refGene_tx=XM_019835280.2;refGene_gn=PPFIA2;refGene_tx=XM_019835281.2;refGene_gn=PPFIA2;refGene_tx=XM_019835282.2;refGene_gn=PPFIA2;refGene_tx=XM_019835283.2;refGene_gn=PPFIA2;refGene_tx=XM_019835285.2;refGene_gn=PPFIA2;refGene_tx=XM_019835286.2;refGene_gn=PPFIA2;refGene_tx=XM_019835287.2;refGene_gn=PPFIA2;refGene_tx=XM_019835288.2;refGene_gn=PPFIA2;refGene_tx=XM_019835289.2;refGene_gn=PPFIA2;refGene_tx=XM_019835290.2;refGene_gn=PPFIA2;refGene_tx=XM_019835291.2;refGene_gn=PPFIA2;refGene_tx=XM_019835292.2;refGene_gn=PPFIA2;refGene_tx=XM_019835293.2;refGene_gn=PPFIA2;refGene_tx=XM_023257280.1
B4 106658373 106658595
refGene_gn=PPFIA2;refGene_tx=XM_006933911.4;refGene_gn=PPFIA2;refGene_tx=XM_019835275.2;refGene_gn=PPFIA2;refGene_tx=XM_019835276.2;refGene_gn=PPFIA2;refGene_tx=XM_019835277.2;refGene_gn=PPFIA2;refGene_tx=XM_019835278.2;refGene_gn=PPFIA2;refGene_tx=XM_019835279.2;refGene_gn=PPFIA2;refGene_tx=XM_019835280.2;refGene_gn=PPFIA2;refGene_tx=XM_019835281.2;refGene_gn=PPFIA2;refGene_tx=XM_019835282.2;refGene_gn=PPFIA2;refGene_tx=XM_019835283.2;refGene_gn=PPFIA2;refGene_tx=XM_019835285.2;refGene_gn=PPFIA2;refGene_tx=XM_019835286.2;refGene_gn=PPFIA2;refGene_tx=XM_

019835287.2;refGene_gn=PPFIA2;refGene_tx=XM_019835288.2;refGene_gn=PPFIA2;refGene_tx=XM_019835289.2;refGene_gn=PPFIA2;refGene_tx=XM_019835290.2;refGene_gn=PPFIA2;refGene_tx=XM_019835291.2;refGene_gn=PPFIA2;refGene_tx=XM_019835292.2;refGene_gn=PPFIA2;refGene_tx=XM_019835293.2;refGene_gn=PPFIA2;refGene_tx=XM_023257280.1
B4 106684887 106685004
refGene_gn=PPFIA2;refGene_tx=XM_006933911.4;refGene_gn=PPFIA2;refGene_tx=XM_019835275.2;refGene_gn=PPFIA2;refGene_tx=XM_019835276.2;refGene_gn=PPFIA2;refGene_tx=XM_019835277.2;refGene_gn=PPFIA2;refGene_tx=XM_019835278.2;refGene_gn=PPFIA2;refGene_tx=XM_019835279.2;refGene_gn=PPFIA2;refGene_tx=XM_019835280.2;refGene_gn=PPFIA2;refGene_tx=XM_019835281.2;refGene_gn=PPFIA2;refGene_tx=XM_019835282.2;refGene_gn=PPFIA2;refGene_tx=XM_019835283.2;refGene_gn=PPFIA2;refGene_tx=XM_019835285.2;refGene_gn=PPFIA2;refGene_tx=XM_019835286.2;refGene_gn=PPFIA2;refGene_tx=XM_019835287.2;refGene_gn=PPFIA2;refGene_tx=XM_019835288.2;refGene_gn=PPFIA2;refGene_tx=XM_019835289.2;refGene_gn=PPFIA2;refGene_tx=XM_019835290.2;refGene_gn=PPFIA2;refGene_tx=XM_019835291.2;refGene_gn=PPFIA2;refGene_tx=XM_019835292.2;refGene_gn=PPFIA2;refGene_tx=XM_019835293.2;refGene_gn=PPFIA2;refGene_tx=XM_023257280.1
B4 106717244 106717319
refGene_gn=PPFIA2;refGene_tx=XM_006933911.4;refGene_gn=PPFIA2;refGene_tx=XM_019835275.2;refGene_gn=PPFIA2;refGene_tx=XM_019835276.2;refGene_gn=PPFIA2;refGene_tx=XM_019835277.2;refGene_gn=PPFIA2;refGene_tx=XM_019835278.2;refGene_gn=PPFIA2;refGene_tx=XM_019835279.2;refGene_gn=PPFIA2;refGene_tx=XM_019835280.2;refGene_gn=PPFIA2;refGene_tx=XM_019835281.2;refGene_gn=PPFIA2;refGene_tx=XM_019835283.2;refGene_gn=PPFIA2;refGene_tx=XM_019835285.2;refGene_gn=PPFIA2;refGene_tx=XM_019835286.2;refGene_gn=PPFIA2;refGene_tx=XM_019835287.2;refGene_gn=PPFIA2;refGene_tx=XM_019835288.2;refGene_gn=PPFIA2;refGene_tx=XM_019835289.2;refGene_gn=PPFIA2;refGene_tx=XM_019835291.2;refGene_gn=PPFIA2;refGene_tx=XM_019835292.2;refGene_gn=PPFIA2;refGene_tx=XM_019835293.2;refGene_gn=PPFIA2;refGene_tx=XM_023257280.1
B4 106723351 106723516
refGene_gn=PPFIA2;refGene_tx=XM_006933911.4;refGene_gn=PPFIA2;refGene_tx=XM_019835275.2;refGene_gn=PPFIA2;refGene_tx=XM_019835276.2;refGene_gn=PPFIA2;refGene_tx=XM_019835277.2;refGene_gn=PPFIA2;refGene_tx=XM_019835278.2;refGene_gn=PPFIA2;refGene_tx=XM_019835279.2;refGene_gn=PPFIA2;refGene_tx=XM_019835280.2;refGene_gn=PPFIA2;refGene_tx=XM_019835281.2;refGene_gn=PPFIA2;refGene_tx=XM_019835282.2;refGene_gn=PPFIA2;refGene_tx=XM_019835283.2;refGene_gn=PPFIA2;refGene_tx=XM_019835285.2;refGene_gn=PPFIA2;refGene_tx=XM_019835286.2;refGene_gn=PPFIA2;refGene_tx=XM_019835287.2;refGene_gn=PPFIA2;refGene_tx=XM_019835288.2;refGene_gn=PPFIA2;refGene_tx=XM_019835289.2;refGene_gn=PPFIA2;refGene_tx=XM_019835290.2;refGene_gn=PPFIA2;refGene_tx=XM_019835291.2;refGene_gn=PPFIA2;refGene_tx=XM_019835292.2;refGene_gn=PPFIA2;refGene_tx=XM_019835293.2;refGene_gn=PPFIA2;refGene_tx=XM_023257280.1
B4 106733974 106734076
refGene_gn=PPFIA2;refGene_tx=XM_006933911.4;refGene_gn=PPFIA2;refGene_tx=XM_019835275.2;refGene_gn=PPFIA2;refGene_tx=XM_019835276.2;refGene_gn=PPFIA2;refGene_tx=XM_019835277.2;refGene_gn=PPFIA2;refGene_tx=XM_01

9835278.2;refGene_gn=PPFIA2;refGene_tx=XM_019835279.2;refGene_gn=PPFIA2;refGene_tx=XM_019835280.2;refGene_gn=PPFIA2;refGene_tx=XM_019835281.2;refGene_gn=PPFIA2;refGene_tx=XM_019835282.2;refGene_gn=PPFIA2;refGene_tx=XM_019835283.2;refGene_gn=PPFIA2;refGene_tx=XM_019835285.2;refGene_gn=PPFIA2;refGene_tx=XM_019835286.2;refGene_gn=PPFIA2;refGene_tx=XM_019835287.2;refGene_gn=PPFIA2;refGene_tx=XM_019835288.2;refGene_gn=PPFIA2;refGene_tx=XM_019835289.2;refGene_gn=PPFIA2;refGene_tx=XM_019835290.2;refGene_gn=PPFIA2;refGene_tx=XM_019835291.2;refGene_gn=PPFIA2;refGene_tx=XM_019835292.2;refGene_gn=PPFIA2;refGene_tx=XM_019835293.2;refGene_gn=PPFIA2;refGene_tx=XM_023257280.1
B4 106856665 106856746
refGene_gn=PPFIA2;refGene_tx=XM_019835287.2;refGene_gn=PPFIA2;refGene_tx=XM_019835288.2;refGene_gn=PPFIA2;refGene_tx=XM_019835289.2;refGene_gn=PPFIA2;refGene_tx=XM_019835290.2;refGene_gn=PPFIA2;refGene_tx=XM_019835291.2
B4 106929560 106929614
refGene_gn=PPFIA2;refGene_tx=XM_006933911.4;refGene_gn=PPFIA2;refGene_tx=XM_019835275.2;refGene_gn=PPFIA2;refGene_tx=XM_019835276.2;refGene_gn=PPFIA2;refGene_tx=XM_019835277.2;refGene_gn=PPFIA2;refGene_tx=XM_019835278.2;refGene_gn=PPFIA2;refGene_tx=XM_019835279.2;refGene_gn=PPFIA2;refGene_tx=XM_019835280.2;refGene_gn=PPFIA2;refGene_tx=XM_019835282.2;refGene_gn=PPFIA2;refGene_tx=XM_019835283.2;refGene_gn=PPFIA2;refGene_tx=XM_019835285.2;refGene_gn=PPFIA2;refGene_tx=XM_023257280.1
B4 107008314 107008563
refGene_gn=PPFIA2;refGene_tx=XM_006933911.4;refGene_gn=PPFIA2;refGene_tx=XM_019835275.2;refGene_gn=PPFIA2;refGene_tx=XM_019835276.2;refGene_gn=PPFIA2;refGene_tx=XM_019835277.2;refGene_gn=PPFIA2;refGene_tx=XM_019835278.2;refGene_gn=PPFIA2;refGene_tx=XM_019835279.2;refGene_gn=PPFIA2;refGene_tx=XM_019835280.2;refGene_gn=PPFIA2;refGene_tx=XM_019835281.2;refGene_gn=PPFIA2;refGene_tx=XM_019835282.2;refGene_gn=PPFIA2;refGene_tx=XM_019835283.2;refGene_gn=PPFIA2;refGene_tx=XM_019835285.2;refGene_gn=PPFIA2;refGene_tx=XM_019835286.2;refGene_gn=PPFIA2;refGene_tx=XM_019835292.2;refGene_gn=PPFIA2;refGene_tx=XM_019835293.2;refGene_gn=PPFIA2;refGene_tx=XM_023257280.1
B4 107471750 107471912
refGene_gn=CCDC59;refGene_tx=XM_003989086.5
B4 107474032 107474132
refGene_gn=CCDC59;refGene_tx=XM_003989086.5
B4 107476614 107476927
refGene_gn=CCDC59;refGene_tx=XM_003989086.5
B4 107478159 107478370
refGene_gn=CCDC59;refGene_tx=XM_003989086.5
B4 107478501 107478760
refGene_gn=METTL25;refGene_tx=XM_003989085.3
B4 107503268 107503433
refGene_gn=METTL25;refGene_tx=XM_003989085.3
B4 107503910 107504017
refGene_gn=METTL25;refGene_tx=XM_003989085.3
B4 107518505 107519102
refGene_gn=METTL25;refGene_tx=XM_003989085.3

B4 107521477 107521625
refGene_gn=METTL25;refGene_tx=XM_003989085.3
B4 107558154 107558249
refGene_gn=METTL25;refGene_tx=XM_003989085.3
B4 107562096 107562126
refGene_gn=METTL25;refGene_tx=XM_003989085.3
B4 107565849 107565923
refGene_gn=METTL25;refGene_tx=XM_003989085.3
B4 107579576 107579670
refGene_gn=METTL25;refGene_tx=XM_003989085.3
B4 107604376 107604451
refGene_gn=METTL25;refGene_tx=XM_003989085.3
B4 107605014 107605086
refGene_gn=METTL25;refGene_tx=XM_003989085.3
B4 107607003 107607096
refGene_gn=METTL25;refGene_tx=XM_003989085.3
B4 107774371 107774454
refGene_gn=TMTC2;refGene_tx=XM_019835295.2;refGene_gn=TMTC2;refGene_tx=XM_023257281.1
B4 107927662 107928233
refGene_gn=TMTC2;refGene_tx=XM_019835295.2;refGene_gn=TMTC2;refGene_tx=XM_023257281.1
B4 107964700 107965529
refGene_gn=TMTC2;refGene_tx=XM_019835295.2;refGene_gn=TMTC2;refGene_tx=XM_023257281.1
B4 107989386 107989501
refGene_gn=TMTC2;refGene_tx=XM_019835295.2;refGene_gn=TMTC2;refGene_tx=XM_023257281.1
B4 108025604 108025690
refGene_gn=TMTC2;refGene_tx=XM_019835295.2;refGene_gn=TMTC2;refGene_tx=XM_023257281.1
B4 108026425 108026610
refGene_gn=TMTC2;refGene_tx=XM_019835295.2;refGene_gn=TMTC2;refGene_tx=XM_023257281.1
B4 108027689 108027768
refGene_gn=TMTC2;refGene_tx=XM_019835295.2;refGene_gn=TMTC2;refGene_tx=XM_023257281.1
B4 108046967 108047089
refGene_gn=TMTC2;refGene_tx=XM_019835295.2;refGene_gn=TMTC2;refGene_tx=XM_023257281.1
B4 108086118 108086200
refGene_gn=TMTC2;refGene_tx=XM_019835295.2;refGene_gn=TMTC2;refGene_tx=XM_023257281.1
B4 108104617 108104732
refGene_gn=TMTC2;refGene_tx=XM_019835295.2;refGene_gn=TMTC2;refGene_tx=XM_023257281.1
B4 108117765 108117829
refGene_gn=TMTC2;refGene_tx=XM_019835295.2;refGene_gn=TMTC2;refGene_tx=XM_023257281.1
B4 108171343 108171523

refGene_gn=TMTC2;refGene_tx=XM_023257281.1
B4	108177821	108177851
refGene_gn=TMTC2;refGene_tx=XM_019835295.2
B4	109720210	109720582
refGene_gn=SLC6A15;refGene_tx=XM_011284141.3;refGene_gn=SLC6A15;refGene_tx=XM_019835296.2;refGene_gn=SLC6A15;refGene_tx=XM_023257282.1
B4	109721909	109722072
refGene_gn=SLC6A15;refGene_tx=XM_011284141.3;refGene_gn=SLC6A15;refGene_tx=XM_019835296.2;refGene_gn=SLC6A15;refGene_tx=XM_023257282.1
B4	109725424	109725584
refGene_gn=SLC6A15;refGene_tx=XM_011284141.3;refGene_gn=SLC6A15;refGene_tx=XM_019835296.2;refGene_gn=SLC6A15;refGene_tx=XM_023257282.1
B4	109727989	109728182
refGene_gn=SLC6A15;refGene_tx=XM_011284141.3;refGene_gn=SLC6A15;refGene_tx=XM_019835296.2;refGene_gn=SLC6A15;refGene_tx=XM_023257282.1
B4	109729218	109729411
refGene_gn=SLC6A15;refGene_tx=XM_011284141.3;refGene_gn=SLC6A15;refGene_tx=XM_019835296.2;refGene_gn=SLC6A15;refGene_tx=XM_023257282.1
B4	109729514	109729756
refGene_gn=SLC6A15;refGene_tx=XM_011284141.3;refGene_gn=SLC6A15;refGene_tx=XM_019835296.2;refGene_gn=SLC6A15;refGene_tx=XM_023257282.1
B4	109732806	109732917
refGene_gn=SLC6A15;refGene_tx=XM_011284141.3;refGene_gn=SLC6A15;refGene_tx=XM_019835296.2;refGene_gn=SLC6A15;refGene_tx=XM_023257282.1
B4	109734287	109734469
refGene_gn=SLC6A15;refGene_tx=XM_011284141.3;refGene_gn=SLC6A15;refGene_tx=XM_019835296.2;refGene_gn=SLC6A15;refGene_tx=XM_023257282.1
B4	109734978	109735105
refGene_gn=SLC6A15;refGene_tx=XM_011284141.3;refGene_gn=SLC6A15;refGene_tx=XM_019835296.2;refGene_gn=SLC6A15;refGene_tx=XM_023257282.1
B4	109735454	109735612
refGene_gn=SLC6A15;refGene_tx=XM_011284141.3;refGene_gn=SLC6A15;refGene_tx=XM_019835296.2;refGene_gn=SLC6A15;refGene_tx=XM_023257282.1
B4	109739595	109739884
refGene_gn=SLC6A15;refGene_tx=XM_011284141.3;refGene_gn=SLC6A15;refGene_tx=XM_019835296.2;refGene_gn=SLC6A15;refGene_tx=XM_023257282.1
B4	109828744	109828813
refGene_gn=TSPAN19;refGene_tx=XM_023257804.1
B4	109830379	109830463
refGene_gn=TSPAN19;refGene_tx=XM_023257804.1
B4	109831376	109831520
refGene_gn=TSPAN19;refGene_tx=XM_023257804.1
B4	109833680	109833791
refGene_gn=TSPAN19;refGene_tx=XM_023257804.1
B4	109837770	109837845
refGene_gn=TSPAN19;refGene_tx=XM_023257804.1
B4	109843921	109844046
refGene_gn=TSPAN19;refGene_tx=XM_023257804.1
B4	109846801	109846874
refGene_gn=TSPAN19;refGene_tx=XM_023257804.1

B4 109846960 109847026
refGene_gn=TSPAN19;refGene_tx=XM_023257804.1
B4 109859417 109859561
refGene_gn=LRRIQ1;refGene_tx=XM_011284143.3;refGene_gn=LRRIQ1;refGene_tx=XM_019835297.2;refGene_gn=LRRIQ1;refGene_tx=XM_019835298.2;refGene_gn=LRRIQ1;refGene_tx=XM_019835299.2;refGene_gn=LRRIQ1;refGene_tx=XM_019835300.2;refGene_gn=LRRIQ1;refGene_tx=XM_023257283.1
B4 109861176 109861288
refGene_gn=LRRIQ1;refGene_tx=XM_011284143.3;refGene_gn=LRRIQ1;refGene_tx=XM_019835297.2;refGene_gn=LRRIQ1;refGene_tx=XM_019835298.2;refGene_gn=LRRIQ1;refGene_tx=XM_019835299.2;refGene_gn=LRRIQ1;refGene_tx=XM_019835300.2;refGene_gn=LRRIQ1;refGene_tx=XM_023257283.1
B4 109865356 109865448
refGene_gn=LRRIQ1;refGene_tx=XM_011284143.3;refGene_gn=LRRIQ1;refGene_tx=XM_019835297.2;refGene_gn=LRRIQ1;refGene_tx=XM_019835298.2;refGene_gn=LRRIQ1;refGene_tx=XM_019835299.2;refGene_gn=LRRIQ1;refGene_tx=XM_019835300.2;refGene_gn=LRRIQ1;refGene_tx=XM_023257283.1
B4 109866814 109866932
refGene_gn=LRRIQ1;refGene_tx=XM_011284143.3;refGene_gn=LRRIQ1;refGene_tx=XM_019835297.2;refGene_gn=LRRIQ1;refGene_tx=XM_019835298.2;refGene_gn=LRRIQ1;refGene_tx=XM_019835299.2;refGene_gn=LRRIQ1;refGene_tx=XM_019835300.2;refGene_gn=LRRIQ1;refGene_tx=XM_023257283.1
B4 109867966 109868190
refGene_gn=LRRIQ1;refGene_tx=XM_011284143.3;refGene_gn=LRRIQ1;refGene_tx=XM_019835297.2;refGene_gn=LRRIQ1;refGene_tx=XM_019835298.2;refGene_gn=LRRIQ1;refGene_tx=XM_019835299.2;refGene_gn=LRRIQ1;refGene_tx=XM_019835300.2;refGene_gn=LRRIQ1;refGene_tx=XM_023257283.1
B4 109872640 109872715
refGene_gn=LRRIQ1;refGene_tx=XM_011284143.3;refGene_gn=LRRIQ1;refGene_tx=XM_019835297.2;refGene_gn=LRRIQ1;refGene_tx=XM_019835298.2;refGene_gn=LRRIQ1;refGene_tx=XM_019835299.2;refGene_gn=LRRIQ1;refGene_tx=XM_019835300.2;refGene_gn=LRRIQ1;refGene_tx=XM_023257283.1
B4 109876190 109877822
refGene_gn=LRRIQ1;refGene_tx=XM_011284143.3;refGene_gn=LRRIQ1;refGene_tx=XM_019835297.2;refGene_gn=LRRIQ1;refGene_tx=XM_019835298.2;refGene_gn=LRRIQ1;refGene_tx=XM_019835299.2;refGene_gn=LRRIQ1;refGene_tx=XM_019835300.2;refGene_gn=LRRIQ1;refGene_tx=XM_023257283.1
B4 109884537 109884690
refGene_gn=LRRIQ1;refGene_tx=XM_011284143.3;refGene_gn=LRRIQ1;refGene_tx=XM_019835297.2;refGene_gn=LRRIQ1;refGene_tx=XM_019835298.2;refGene_gn=LRRIQ1;refGene_tx=XM_019835299.2;refGene_gn=LRRIQ1;refGene_tx=XM_019835300.2;refGene_gn=LRRIQ1;refGene_tx=XM_023257283.1
B4 109886910 109887061
refGene_gn=LRRIQ1;refGene_tx=XM_011284143.3;refGene_gn=LRRIQ1;refGene_tx=XM_019835297.2;refGene_gn=LRRIQ1;refGene_tx=XM_019835298.2;refGene_gn=LRRIQ1;refGene_tx=XM_019835299.2;refGene_gn=LRRIQ1;refGene_tx=XM_019835300.2;refGene_gn=LRRIQ1;refGene_tx=XM_023257283.1
B4 109905870 109906064
refGene_gn=LRRIQ1;refGene_tx=XM_011284143.3;refGene_gn=LRRIQ1;refGene_tx=XM_019835297.2;refGene_gn=LRRIQ1;refGene_tx=XM_019835298.2;refGene_

gn=LRRIQ1;refGene_tx=XM_019835299.2;refGene_gn=LRRIQ1;refGene_tx=XM_019835300.2;refGene_gn=LRRIQ1;refGene_tx=XM_023257283.1
B4 109906586 109906714
refGene_gn=LRRIQ1;refGene_tx=XM_011284143.3;refGene_gn=LRRIQ1;refGene_tx=XM_019835297.2;refGene_gn=LRRIQ1;refGene_tx=XM_019835298.2;refGene_gn=LRRIQ1;refGene_tx=XM_019835299.2;refGene_gn=LRRIQ1;refGene_tx=XM_019835300.2;refGene_gn=LRRIQ1;refGene_tx=XM_023257283.1
B4 109911553 109911627
refGene_gn=LRRIQ1;refGene_tx=XM_011284143.3;refGene_gn=LRRIQ1;refGene_tx=XM_019835297.2;refGene_gn=LRRIQ1;refGene_tx=XM_019835298.2;refGene_gn=LRRIQ1;refGene_tx=XM_019835299.2;refGene_gn=LRRIQ1;refGene_tx=XM_019835300.2;refGene_gn=LRRIQ1;refGene_tx=XM_023257283.1
B4 109920378 109920472
refGene_gn=LRRIQ1;refGene_tx=XM_011284143.3;refGene_gn=LRRIQ1;refGene_tx=XM_019835297.2;refGene_gn=LRRIQ1;refGene_tx=XM_019835298.2;refGene_gn=LRRIQ1;refGene_tx=XM_019835299.2;refGene_gn=LRRIQ1;refGene_tx=XM_019835300.2;refGene_gn=LRRIQ1;refGene_tx=XM_023257283.1
B4 109936729 109936909
refGene_gn=LRRIQ1;refGene_tx=XM_011284143.3;refGene_gn=LRRIQ1;refGene_tx=XM_019835297.2;refGene_gn=LRRIQ1;refGene_tx=XM_019835298.2;refGene_gn=LRRIQ1;refGene_tx=XM_019835299.2;refGene_gn=LRRIQ1;refGene_tx=XM_019835300.2;refGene_gn=LRRIQ1;refGene_tx=XM_023257283.1
B4 109939676 109940123
refGene_gn=LRRIQ1;refGene_tx=XM_011284143.3;refGene_gn=LRRIQ1;refGene_tx=XM_019835297.2;refGene_gn=LRRIQ1;refGene_tx=XM_019835298.2;refGene_gn=LRRIQ1;refGene_tx=XM_019835299.2;refGene_gn=LRRIQ1;refGene_tx=XM_019835300.2;refGene_gn=LRRIQ1;refGene_tx=XM_023257283.1
B4 109944059 109944270
refGene_gn=LRRIQ1;refGene_tx=XM_011284143.3;refGene_gn=LRRIQ1;refGene_tx=XM_019835297.2;refGene_gn=LRRIQ1;refGene_tx=XM_019835298.2;refGene_gn=LRRIQ1;refGene_tx=XM_019835299.2;refGene_gn=LRRIQ1;refGene_tx=XM_019835300.2;refGene_gn=LRRIQ1;refGene_tx=XM_023257283.1
B4 109950587 109950707
refGene_gn=LRRIQ1;refGene_tx=XM_011284143.3;refGene_gn=LRRIQ1;refGene_tx=XM_019835297.2;refGene_gn=LRRIQ1;refGene_tx=XM_019835298.2;refGene_gn=LRRIQ1;refGene_tx=XM_019835299.2;refGene_gn=LRRIQ1;refGene_tx=XM_019835300.2;refGene_gn=LRRIQ1;refGene_tx=XM_023257283.1
B4 109967292 109967382
refGene_gn=LRRIQ1;refGene_tx=XM_011284143.3;refGene_gn=LRRIQ1;refGene_tx=XM_019835297.2;refGene_gn=LRRIQ1;refGene_tx=XM_019835298.2;refGene_gn=LRRIQ1;refGene_tx=XM_019835299.2;refGene_gn=LRRIQ1;refGene_tx=XM_019835300.2;refGene_gn=LRRIQ1;refGene_tx=XM_023257283.1
B4 109968041 109968163
refGene_gn=LRRIQ1;refGene_tx=XM_011284143.3;refGene_gn=LRRIQ1;refGene_tx=XM_019835297.2;refGene_gn=LRRIQ1;refGene_tx=XM_019835298.2;refGene_gn=LRRIQ1;refGene_tx=XM_019835299.2;refGene_gn=LRRIQ1;refGene_tx=XM_019835300.2;refGene_gn=LRRIQ1;refGene_tx=XM_023257283.1
B4 109968677 109968773
refGene_gn=LRRIQ1;refGene_tx=XM_011284143.3;refGene_gn=LRRIQ1;refGene_tx=XM_019835297.2;refGene_gn=LRRIQ1;refGene_tx=XM_019835298.2;refGene_

gn=LRRIQ1;refGene_tx=XM_019835299.2;refGene_gn=LRRIQ1;refGene_tx=XM_01
9835300.2;refGene_gn=LRRIQ1;refGene_tx=XM_023257283.1
B4 109969017 109969100
refGene_gn=LRRIQ1;refGene_tx=XM_011284143.3;refGene_gn=LRRIQ1;refGene_
tx=XM_019835297.2;refGene_gn=LRRIQ1;refGene_tx=XM_019835298.2;refGene_
gn=LRRIQ1;refGene_tx=XM_019835299.2;refGene_gn=LRRIQ1;refGene_tx=XM_01
9835300.2;refGene_gn=LRRIQ1;refGene_tx=XM_023257283.1
B4 109972081 109972108
refGene_gn=LRRIQ1;refGene_tx=XM_019835297.2;refGene_gn=LRRIQ1;refGene_
tx=XM_019835298.2;refGene_gn=LRRIQ1;refGene_tx=XM_019835300.2;refGene_
gn=LRRIQ1;refGene_tx=XM_023257283.1
B4 109977206 109977308
refGene_gn=LRRIQ1;refGene_tx=XM_011284143.3;refGene_gn=LRRIQ1;refGene_
tx=XM_019835297.2;refGene_gn=LRRIQ1;refGene_tx=XM_019835298.2;refGene_
gn=LRRIQ1;refGene_tx=XM_019835299.2;refGene_gn=LRRIQ1;refGene_tx=XM_01
9835300.2;refGene_gn=LRRIQ1;refGene_tx=XM_023257283.1
B4 110029140 110029273
refGene_gn=LRRIQ1;refGene_tx=XM_011284143.3;refGene_gn=LRRIQ1;refGene_
tx=XM_019835297.2;refGene_gn=LRRIQ1;refGene_tx=XM_019835298.2;refGene_
gn=LRRIQ1;refGene_tx=XM_019835299.2;refGene_gn=LRRIQ1;refGene_tx=XM_01
9835300.2;refGene_gn=LRRIQ1;refGene_tx=XM_023257283.1
B4 110031843 110031904
refGene_gn=LRRIQ1;refGene_tx=XM_011284143.3;refGene_gn=LRRIQ1;refGene_
tx=XM_019835297.2;refGene_gn=LRRIQ1;refGene_tx=XM_019835298.2;refGene_
gn=LRRIQ1;refGene_tx=XM_019835299.2;refGene_gn=LRRIQ1;refGene_tx=XM_01
9835300.2;refGene_gn=LRRIQ1;refGene_tx=XM_023257283.1
B4 110042736 110042889
refGene_gn=LRRIQ1;refGene_tx=XM_011284143.3;refGene_gn=LRRIQ1;refGene_
tx=XM_019835297.2;refGene_gn=LRRIQ1;refGene_tx=XM_019835298.2;refGene_
gn=LRRIQ1;refGene_tx=XM_019835299.2;refGene_gn=LRRIQ1;refGene_tx=XM_01
9835300.2;refGene_gn=LRRIQ1;refGene_tx=XM_023257283.1
B4 110065093 110065236
refGene_gn=LRRIQ1;refGene_tx=XM_019835297.2;refGene_gn=LRRIQ1;refGene_
tx=XM_019835298.2;refGene_gn=LRRIQ1;refGene_tx=XM_019835299.2
B4 110070144 110070179
refGene_gn=LRRIQ1;refGene_tx=XM_023257283.1
B4 110084910 110085136
refGene_gn=ALX1;refGene_tx=XM_003989090.4
B4 110087974 110088279
refGene_gn=ALX1;refGene_tx=XM_003989090.4
B4 110091285 110091414
refGene_gn=ALX1;refGene_tx=XM_003989090.4
B4 110104756 110105077
refGene_gn=ALX1;refGene_tx=XM_003989090.4
B4 110557377 110558638
refGene_gn=RASSF9;refGene_tx=XM_019835305.2
B4 110585691 110585738
refGene_gn=RASSF9;refGene_tx=XM_019835305.2
B4 110613753 110613835
refGene_gn=NTS;refGene_tx=XM_003989092.5

B4	110616089	110616151
refGene_gn=NTS;refGene_tx=XM_003989092.5
B4	110622571	110622796
refGene_gn=NTS;refGene_tx=XM_003989092.5
B4	110625426	110625579
refGene_gn=NTS;refGene_tx=XM_003989092.5
B4	110712399	110713541
refGene_gn=MGAT4C;refGene_tx=XM_006933915.4;refGene_gn=MGAT4C;refGene_tx=XM_019835308.2;refGene_gn=MGAT4C;refGene_tx=XM_019835311.2;refGene_gn=MGAT4C;refGene_tx=XM_023257285.1;refGene_gn=MGAT4C;refGene_tx=XM_023257286.1;refGene_gn=MGAT4C;refGene_tx=XM_023257287.1
B4	110717010	110717158
refGene_gn=MGAT4C;refGene_tx=XM_006933915.4;refGene_gn=MGAT4C;refGene_tx=XM_019835308.2;refGene_gn=MGAT4C;refGene_tx=XM_019835311.2;refGene_gn=MGAT4C;refGene_tx=XM_023257285.1;refGene_gn=MGAT4C;refGene_tx=XM_023257286.1;refGene_gn=MGAT4C;refGene_tx=XM_023257287.1
B4	110722378	110722525
refGene_gn=MGAT4C;refGene_tx=XM_006933915.4;refGene_gn=MGAT4C;refGene_tx=XM_019835308.2;refGene_gn=MGAT4C;refGene_tx=XM_019835311.2;refGene_gn=MGAT4C;refGene_tx=XM_023257285.1;refGene_gn=MGAT4C;refGene_tx=XM_023257286.1;refGene_gn=MGAT4C;refGene_tx=XM_023257287.1
B4	112460978	112461004
refGene_gn=CB4H12orf50;refGene_tx=XM_003989095.4;refGene_gn=CB4H12orf50;refGene_tx=XM_019835314.2;refGene_gn=CB4H12orf50;refGene_tx=XM_019835315.2;refGene_gn=CB4H12orf50;refGene_tx=XM_019835316.2;refGene_gn=CB4H12orf50;refGene_tx=XM_019835317.2;refGene_gn=CB4H12orf50;refGene_tx=XM_019835318.2;refGene_gn=CB4H12orf50;refGene_tx=XM_023257288.1
B4	112464090	112464183
refGene_gn=CB4H12orf50;refGene_tx=XM_003989095.4;refGene_gn=CB4H12orf50;refGene_tx=XM_019835314.2;refGene_gn=CB4H12orf50;refGene_tx=XM_019835315.2;refGene_gn=CB4H12orf50;refGene_tx=XM_019835316.2;refGene_gn=CB4H12orf50;refGene_tx=XM_019835317.2;refGene_gn=CB4H12orf50;refGene_tx=XM_019835318.2;refGene_gn=CB4H12orf50;refGene_tx=XM_023257288.1
B4	112466046	112466250
refGene_gn=CB4H12orf50;refGene_tx=XM_003989095.4;refGene_gn=CB4H12orf50;refGene_tx=XM_019835314.2;refGene_gn=CB4H12orf50;refGene_tx=XM_019835315.2;refGene_gn=CB4H12orf50;refGene_tx=XM_019835316.2;refGene_gn=CB4H12orf50;refGene_tx=XM_019835317.2;refGene_gn=CB4H12orf50;refGene_tx=XM_019835318.2;refGene_gn=CB4H12orf50;refGene_tx=XM_023257288.1
B4	112466504	112466609
refGene_gn=CB4H12orf50;refGene_tx=XM_003989095.4;refGene_gn=CB4H12orf50;refGene_tx=XM_019835314.2;refGene_gn=CB4H12orf50;refGene_tx=XM_019835315.2;refGene_gn=CB4H12orf50;refGene_tx=XM_019835316.2;refGene_gn=CB4H12orf50;refGene_tx=XM_019835317.2;refGene_gn=CB4H12orf50;refGene_tx=XM_019835318.2;refGene_gn=CB4H12orf50;refGene_tx=XM_023257288.1
B4	112468122	112468239
refGene_gn=CB4H12orf50;refGene_tx=XM_003989095.4;refGene_gn=CB4H12orf50;refGene_tx=XM_019835314.2;refGene_gn=CB4H12orf50;refGene_tx=XM_019835315.2;refGene_gn=CB4H12orf50;refGene_tx=XM_019835317.2;refGene_gn=CB4H12orf50;refGene_tx=XM_019835318.2;refGene_gn=CB4H12orf50;refGene_tx=X

M_023257288.1
B4	112469596	112469704
refGene_gn=CB4H12orf50;refGene_tx=XM_003989095.4;refGene_gn=CB4H12orf50;refGene_tx=XM_019835314.2;refGene_gn=CB4H12orf50;refGene_tx=XM_019835315.2;refGene_gn=CB4H12orf50;refGene_tx=XM_019835316.2;refGene_gn=CB4H12orf50;refGene_tx=XM_019835317.2;refGene_gn=CB4H12orf50;refGene_tx=XM_019835318.2;refGene_gn=CB4H12orf50;refGene_tx=XM_023257288.1
B4	112471120	112471231
refGene_gn=CB4H12orf50;refGene_tx=XM_003989095.4;refGene_gn=CB4H12orf50;refGene_tx=XM_019835314.2;refGene_gn=CB4H12orf50;refGene_tx=XM_019835315.2;refGene_gn=CB4H12orf50;refGene_tx=XM_019835316.2;refGene_gn=CB4H12orf50;refGene_tx=XM_019835317.2;refGene_gn=CB4H12orf50;refGene_tx=XM_019835318.2;refGene_gn=CB4H12orf50;refGene_tx=XM_023257288.1
B4	112473149	112473260
refGene_gn=CB4H12orf50;refGene_tx=XM_003989095.4;refGene_gn=CB4H12orf50;refGene_tx=XM_019835314.2;refGene_gn=CB4H12orf50;refGene_tx=XM_019835315.2;refGene_gn=CB4H12orf50;refGene_tx=XM_019835316.2;refGene_gn=CB4H12orf50;refGene_tx=XM_019835317.2;refGene_gn=CB4H12orf50;refGene_tx=XM_019835318.2;refGene_gn=CB4H12orf50;refGene_tx=XM_023257288.1
B4	112473341	112473419
refGene_gn=CB4H12orf50;refGene_tx=XM_003989095.4;refGene_gn=CB4H12orf50;refGene_tx=XM_019835314.2;refGene_gn=CB4H12orf50;refGene_tx=XM_019835315.2;refGene_gn=CB4H12orf50;refGene_tx=XM_019835316.2;refGene_gn=CB4H12orf50;refGene_tx=XM_019835317.2;refGene_gn=CB4H12orf50;refGene_tx=XM_019835318.2;refGene_gn=CB4H12orf50;refGene_tx=XM_023257288.1
B4	112473869	112474028
refGene_gn=CB4H12orf50;refGene_tx=XM_003989095.4;refGene_gn=CB4H12orf50;refGene_tx=XM_019835314.2;refGene_gn=CB4H12orf50;refGene_tx=XM_019835315.2;refGene_gn=CB4H12orf50;refGene_tx=XM_019835316.2;refGene_gn=CB4H12orf50;refGene_tx=XM_019835317.2;refGene_gn=CB4H12orf50;refGene_tx=XM_019835318.2;refGene_gn=CB4H12orf50;refGene_tx=XM_023257288.1
B4	112497085	112497206
refGene_gn=CB4H12orf50;refGene_tx=XM_003989095.4;refGene_gn=CB4H12orf50;refGene_tx=XM_019835314.2;refGene_gn=CB4H12orf50;refGene_tx=XM_019835315.2;refGene_gn=CB4H12orf50;refGene_tx=XM_019835316.2;refGene_gn=CB4H12orf50;refGene_tx=XM_019835317.2;refGene_gn=CB4H12orf50;refGene_tx=XM_019835318.2;refGene_gn=CB4H12orf50;refGene_tx=XM_023257288.1
B4	112497546	112497666
refGene_gn=CB4H12orf50;refGene_tx=XM_003989095.4;refGene_gn=CB4H12orf50;refGene_tx=XM_019835314.2;refGene_gn=CB4H12orf50;refGene_tx=XM_019835315.2;refGene_gn=CB4H12orf50;refGene_tx=XM_019835316.2;refGene_gn=CB4H12orf50;refGene_tx=XM_019835317.2;refGene_gn=CB4H12orf50;refGene_tx=XM_019835318.2;refGene_gn=CB4H12orf50;refGene_tx=XM_023257288.1
B4	112499665	112499719
refGene_gn=CB4H12orf50;refGene_tx=XM_019835314.2;refGene_gn=CB4H12orf50;refGene_tx=XM_019835315.2;refGene_gn=CB4H12orf50;refGene_tx=XM_019835316.2
B4	112503451	112503610
refGene_gn=CB4H12orf29;refGene_tx=XM_003989096.5;refGene_gn=CB4H12orf29;refGene_tx=XM_019835319.2;refGene_gn=CB4H12orf29;refGene_tx=XM_01983

5321.2
B4	112506191	112506309
refGene_gn=CB4H12orf29;refGene_tx=XM_003989096.5;refGene_gn=CB4H12orf29;refGene_tx=XM_019835319.2;refGene_gn=CB4H12orf29;refGene_tx=XM_019835321.2
B4	112509280	112509376
refGene_gn=CB4H12orf29;refGene_tx=XM_003989096.5;refGene_gn=CB4H12orf29;refGene_tx=XM_019835319.2;refGene_gn=CB4H12orf29;refGene_tx=XM_019835321.2
B4	112510123	112510240
refGene_gn=CB4H12orf29;refGene_tx=XM_003989096.5;refGene_gn=CB4H12orf29;refGene_tx=XM_019835319.2
B4	112511670	112511868
refGene_gn=CB4H12orf29;refGene_tx=XM_003989096.5;refGene_gn=CB4H12orf29;refGene_tx=XM_019835319.2;refGene_gn=CB4H12orf29;refGene_tx=XM_019835321.2
B4	112513191	112513361
refGene_gn=CB4H12orf29;refGene_tx=XM_003989096.5;refGene_gn=CB4H12orf29;refGene_tx=XM_019835319.2;refGene_gn=CB4H12orf29;refGene_tx=XM_019835321.2
B4	112514615	112514804
refGene_gn=CB4H12orf29;refGene_tx=XM_003989096.5;refGene_gn=CB4H12orf29;refGene_tx=XM_019835319.2;refGene_gn=CB4H12orf29;refGene_tx=XM_019835321.2
B4	112515598	112515829
refGene_gn=CEP290;refGene_tx=NM_001101658.1;refGene_gn=CEP290;refGene_tx=XM_023256243.1;refGene_gn=CEP290;refGene_tx=XM_023256244.1;refGene_gn=CEP290;refGene_tx=XM_023256245.1
B4	112516840	112516920
refGene_gn=CEP290;refGene_tx=NM_001101658.1;refGene_gn=CEP290;refGene_tx=XM_023256243.1;refGene_gn=CEP290;refGene_tx=XM_023256244.1;refGene_gn=CEP290;refGene_tx=XM_023256245.1
B4	112520731	112520826
refGene_gn=CEP290;refGene_tx=NM_001101658.1;refGene_gn=CEP290;refGene_tx=XM_023256243.1;refGene_gn=CEP290;refGene_tx=XM_023256244.1;refGene_gn=CEP290;refGene_tx=XM_023256245.1
B4	112521353	112521427
refGene_gn=CEP290;refGene_tx=NM_001101658.1;refGene_gn=CEP290;refGene_tx=XM_023256243.1;refGene_gn=CEP290;refGene_tx=XM_023256244.1;refGene_gn=CEP290;refGene_tx=XM_023256245.1
B4	112522826	112522968
refGene_gn=CEP290;refGene_tx=NM_001101658.1;refGene_gn=CEP290;refGene_tx=XM_023256243.1;refGene_gn=CEP290;refGene_tx=XM_023256244.1;refGene_gn=CEP290;refGene_tx=XM_023256245.1
B4	112525577	112525750
refGene_gn=CEP290;refGene_tx=NM_001101658.1;refGene_gn=CEP290;refGene_tx=XM_023256243.1;refGene_gn=CEP290;refGene_tx=XM_023256244.1;refGene_gn=CEP290;refGene_tx=XM_023256245.1
B4	112527212	112527335
refGene_gn=CEP290;refGene_tx=NM_001101658.1;refGene_gn=CEP290;refGene_

tx=XM_023256243.1;refGene_gn=CEP290;refGene_tx=XM_023256244.1;refGene_gn=CEP290;refGene_tx=XM_023256245.1
B4	112528333	112528498
refGene_gn=CEP290;refGene_tx=NM_001101658.1;refGene_gn=CEP290;refGene_tx=XM_023256243.1;refGene_gn=CEP290;refGene_tx=XM_023256244.1;refGene_gn=CEP290;refGene_tx=XM_023256245.1
B4	112529995	112530019
refGene_gn=CEP290;refGene_tx=XM_023256246.1
B4	112530161	112530248
refGene_gn=CEP290;refGene_tx=NM_001101658.1;refGene_gn=CEP290;refGene_tx=XM_023256243.1;refGene_gn=CEP290;refGene_tx=XM_023256244.1;refGene_gn=CEP290;refGene_tx=XM_023256245.1;refGene_gn=CEP290;refGene_tx=XM_023256246.1
B4	112531623	112531758
refGene_gn=CEP290;refGene_tx=NM_001101658.1;refGene_gn=CEP290;refGene_tx=XM_023256243.1;refGene_gn=CEP290;refGene_tx=XM_023256244.1;refGene_gn=CEP290;refGene_tx=XM_023256245.1;refGene_gn=CEP290;refGene_tx=XM_023256246.1
B4	112534911	112535535
refGene_gn=CEP290;refGene_tx=XM_023256243.1;refGene_gn=CEP290;refGene_tx=XM_023256244.1;refGene_gn=CEP290;refGene_tx=XM_023256245.1;refGene_gn=CEP290;refGene_tx=XM_023256246.1
B4	112535983	112536107
refGene_gn=CEP290;refGene_tx=NM_001101658.1;refGene_gn=CEP290;refGene_tx=XM_023256243.1;refGene_gn=CEP290;refGene_tx=XM_023256244.1;refGene_gn=CEP290;refGene_tx=XM_023256245.1;refGene_gn=CEP290;refGene_tx=XM_023256246.1
B4	112538787	112538943
refGene_gn=CEP290;refGene_tx=NM_001101658.1;refGene_gn=CEP290;refGene_tx=XM_023256243.1;refGene_gn=CEP290;refGene_tx=XM_023256244.1;refGene_gn=CEP290;refGene_tx=XM_023256245.1;refGene_gn=CEP290;refGene_tx=XM_023256246.1
B4	112539304	112539450
refGene_gn=CEP290;refGene_tx=NM_001101658.1;refGene_gn=CEP290;refGene_tx=XM_023256243.1;refGene_gn=CEP290;refGene_tx=XM_023256244.1;refGene_gn=CEP290;refGene_tx=XM_023256245.1;refGene_gn=CEP290;refGene_tx=XM_023256246.1
B4	112543054	112543177
refGene_gn=CEP290;refGene_tx=NM_001101658.1;refGene_gn=CEP290;refGene_tx=XM_023256243.1;refGene_gn=CEP290;refGene_tx=XM_023256244.1;refGene_gn=CEP290;refGene_tx=XM_023256245.1;refGene_gn=CEP290;refGene_tx=XM_023256246.1
B4	112543503	112543725
refGene_gn=CEP290;refGene_tx=NM_001101658.1;refGene_gn=CEP290;refGene_tx=XM_023256243.1;refGene_gn=CEP290;refGene_tx=XM_023256244.1;refGene_gn=CEP290;refGene_tx=XM_023256245.1;refGene_gn=CEP290;refGene_tx=XM_023256246.1
B4	112545688	112545826
refGene_gn=CEP290;refGene_tx=NM_001101658.1;refGene_gn=CEP290;refGene_tx=XM_023256243.1;refGene_gn=CEP290;refGene_tx=XM_023256244.1;refGene_

gn=CEP290;refGene_tx=XM_023256245.1;refGene_gn=CEP290;refGene_tx=XM_023256246.1
B4	112546958	112547172
refGene_gn=CEP290;refGene_tx=NM_001101658.1;refGene_gn=CEP290;refGene_tx=XM_023256243.1;refGene_gn=CEP290;refGene_tx=XM_023256244.1;refGene_gn=CEP290;refGene_tx=XM_023256245.1;refGene_gn=CEP290;refGene_tx=XM_023256246.1
B4	112549335	112549535
refGene_gn=CEP290;refGene_tx=NM_001101658.1;refGene_gn=CEP290;refGene_tx=XM_023256243.1;refGene_gn=CEP290;refGene_tx=XM_023256244.1;refGene_gn=CEP290;refGene_tx=XM_023256245.1;refGene_gn=CEP290;refGene_tx=XM_023256246.1
B4	112550578	112550686
refGene_gn=CEP290;refGene_tx=NM_001101658.1;refGene_gn=CEP290;refGene_tx=XM_023256243.1;refGene_gn=CEP290;refGene_tx=XM_023256244.1;refGene_gn=CEP290;refGene_tx=XM_023256245.1;refGene_gn=CEP290;refGene_tx=XM_023256246.1
B4	112551326	112551593
refGene_gn=CEP290;refGene_tx=NM_001101658.1;refGene_gn=CEP290;refGene_tx=XM_023256243.1;refGene_gn=CEP290;refGene_tx=XM_023256244.1;refGene_gn=CEP290;refGene_tx=XM_023256245.1;refGene_gn=CEP290;refGene_tx=XM_023256246.1
B4	112552777	112552912
refGene_gn=CEP290;refGene_tx=NM_001101658.1;refGene_gn=CEP290;refGene_tx=XM_023256243.1;refGene_gn=CEP290;refGene_tx=XM_023256244.1;refGene_gn=CEP290;refGene_tx=XM_023256245.1;refGene_gn=CEP290;refGene_tx=XM_023256246.1
B4	112553122	112553230
refGene_gn=CEP290;refGene_tx=NM_001101658.1;refGene_gn=CEP290;refGene_tx=XM_023256243.1;refGene_gn=CEP290;refGene_tx=XM_023256244.1;refGene_gn=CEP290;refGene_tx=XM_023256245.1;refGene_gn=CEP290;refGene_tx=XM_023256246.1
B4	112554545	112554710
refGene_gn=CEP290;refGene_tx=NM_001101658.1;refGene_gn=CEP290;refGene_tx=XM_023256243.1;refGene_gn=CEP290;refGene_tx=XM_023256244.1;refGene_gn=CEP290;refGene_tx=XM_023256245.1;refGene_gn=CEP290;refGene_tx=XM_023256246.1
B4	112555483	112555939
refGene_gn=CEP290;refGene_tx=NM_001101658.1;refGene_gn=CEP290;refGene_tx=XM_023256243.1;refGene_gn=CEP290;refGene_tx=XM_023256244.1;refGene_gn=CEP290;refGene_tx=XM_023256245.1;refGene_gn=CEP290;refGene_tx=XM_023256246.1
B4	112556570	112556682
refGene_gn=CEP290;refGene_tx=NM_001101658.1;refGene_gn=CEP290;refGene_tx=XM_023256243.1;refGene_gn=CEP290;refGene_tx=XM_023256244.1;refGene_gn=CEP290;refGene_tx=XM_023256245.1;refGene_gn=CEP290;refGene_tx=XM_023256246.1
B4	112559647	112559799
refGene_gn=CEP290;refGene_tx=NM_001101658.1;refGene_gn=CEP290;refGene_tx=XM_023256243.1;refGene_gn=CEP290;refGene_tx=XM_023256244.1;refGene_

gn=CEP290;refGene_tx=XM_023256245.1;refGene_gn=CEP290;refGene_tx=XM_02
3256246.1
B4 112560705 112560911
refGene_gn=CEP290;refGene_tx=NM_001101658.1;refGene_gn=CEP290;refGene_
tx=XM_023256243.1;refGene_gn=CEP290;refGene_tx=XM_023256244.1;refGene_
gn=CEP290;refGene_tx=XM_023256245.1;refGene_gn=CEP290;refGene_tx=XM_02
3256246.1
B4 112562994 112563106
refGene_gn=CEP290;refGene_tx=NM_001101658.1;refGene_gn=CEP290;refGene_
tx=XM_023256243.1;refGene_gn=CEP290;refGene_tx=XM_023256244.1;refGene_
gn=CEP290;refGene_tx=XM_023256245.1;refGene_gn=CEP290;refGene_tx=XM_02
3256246.1
B4 112568095 112568269
refGene_gn=CEP290;refGene_tx=NM_001101658.1;refGene_gn=CEP290;refGene_
tx=XM_023256243.1;refGene_gn=CEP290;refGene_tx=XM_023256244.1;refGene_
gn=CEP290;refGene_tx=XM_023256245.1;refGene_gn=CEP290;refGene_tx=XM_02
3256246.1
B4 112572492 112572723
refGene_gn=CEP290;refGene_tx=NM_001101658.1;refGene_gn=CEP290;refGene_
tx=XM_023256243.1;refGene_gn=CEP290;refGene_tx=XM_023256244.1;refGene_
gn=CEP290;refGene_tx=XM_023256245.1;refGene_gn=CEP290;refGene_tx=XM_02
3256246.1
B4 112572813 112572916
refGene_gn=CEP290;refGene_tx=NM_001101658.1;refGene_gn=CEP290;refGene_
tx=XM_023256243.1;refGene_gn=CEP290;refGene_tx=XM_023256244.1;refGene_
gn=CEP290;refGene_tx=XM_023256245.1;refGene_gn=CEP290;refGene_tx=XM_02
3256246.1
B4 112574165 112574281
refGene_gn=CEP290;refGene_tx=NM_001101658.1;refGene_gn=CEP290;refGene_
tx=XM_023256243.1;refGene_gn=CEP290;refGene_tx=XM_023256244.1;refGene_
gn=CEP290;refGene_tx=XM_023256245.1;refGene_gn=CEP290;refGene_tx=XM_02
3256246.1
B4 112576301 112576451
refGene_gn=CEP290;refGene_tx=NM_001101658.1;refGene_gn=CEP290;refGene_
tx=XM_023256243.1;refGene_gn=CEP290;refGene_tx=XM_023256244.1;refGene_
gn=CEP290;refGene_tx=XM_023256245.1;refGene_gn=CEP290;refGene_tx=XM_02
3256246.1
B4 112576819 112576984
refGene_gn=CEP290;refGene_tx=NM_001101658.1;refGene_gn=CEP290;refGene_
tx=XM_023256243.1;refGene_gn=CEP290;refGene_tx=XM_023256244.1;refGene_
gn=CEP290;refGene_tx=XM_023256245.1;refGene_gn=CEP290;refGene_tx=XM_02
3256246.1
B4 112581054 112581194
refGene_gn=CEP290;refGene_tx=NM_001101658.1;refGene_gn=CEP290;refGene_
tx=XM_023256243.1;refGene_gn=CEP290;refGene_tx=XM_023256244.1;refGene_
gn=CEP290;refGene_tx=XM_023256245.1;refGene_gn=CEP290;refGene_tx=XM_02
3256246.1
B4 112581749 112581834
refGene_gn=CEP290;refGene_tx=NM_001101658.1;refGene_gn=CEP290;refGene_
tx=XM_023256243.1;refGene_gn=CEP290;refGene_tx=XM_023256244.1;refGene_

gn=CEP290;refGene_tx=XM_023256245.1;refGene_gn=CEP290;refGene_tx=XM_023256246.1
B4 112583619 112583732
refGene_gn=CEP290;refGene_tx=NM_001101658.1;refGene_gn=CEP290;refGene_tx=XM_023256243.1;refGene_gn=CEP290;refGene_tx=XM_023256244.1;refGene_gn=CEP290;refGene_tx=XM_023256245.1;refGene_gn=CEP290;refGene_tx=XM_023256246.1
B4 112584972 112585060
refGene_gn=CEP290;refGene_tx=NM_001101658.1;refGene_gn=CEP290;refGene_tx=XM_023256243.1;refGene_gn=CEP290;refGene_tx=XM_023256244.1;refGene_gn=CEP290;refGene_tx=XM_023256245.1;refGene_gn=CEP290;refGene_tx=XM_023256246.1
B4 112585131 112585232
refGene_gn=CEP290;refGene_tx=NM_001101658.1;refGene_gn=CEP290;refGene_tx=XM_023256243.1;refGene_gn=CEP290;refGene_tx=XM_023256244.1;refGene_gn=CEP290;refGene_tx=XM_023256245.1;refGene_gn=CEP290;refGene_tx=XM_023256246.1
B4 112586685 112586848
refGene_gn=CEP290;refGene_tx=NM_001101658.1;refGene_gn=CEP290;refGene_tx=XM_023256243.1;refGene_gn=CEP290;refGene_tx=XM_023256244.1;refGene_gn=CEP290;refGene_tx=XM_023256245.1;refGene_gn=CEP290;refGene_tx=XM_023256246.1
B4 112587528 112587701
refGene_gn=CEP290;refGene_tx=NM_001101658.1;refGene_gn=CEP290;refGene_tx=XM_023256243.1;refGene_gn=CEP290;refGene_tx=XM_023256244.1;refGene_gn=CEP290;refGene_tx=XM_023256245.1;refGene_gn=CEP290;refGene_tx=XM_023256246.1
B4 112591793 112591917
refGene_gn=CEP290;refGene_tx=NM_001101658.1;refGene_gn=CEP290;refGene_tx=XM_023256243.1;refGene_gn=CEP290;refGene_tx=XM_023256244.1;refGene_gn=CEP290;refGene_tx=XM_023256245.1;refGene_gn=CEP290;refGene_tx=XM_023256246.1
B4 112592980 112593103
refGene_gn=CEP290;refGene_tx=NM_001101658.1;refGene_gn=CEP290;refGene_tx=XM_023256243.1;refGene_gn=CEP290;refGene_tx=XM_023256244.1;refGene_gn=CEP290;refGene_tx=XM_023256245.1;refGene_gn=CEP290;refGene_tx=XM_023256246.1
B4 112596092 112596182
refGene_gn=CEP290;refGene_tx=NM_001101658.1;refGene_gn=CEP290;refGene_tx=XM_023256243.1;refGene_gn=CEP290;refGene_tx=XM_023256244.1;refGene_gn=CEP290;refGene_tx=XM_023256245.1;refGene_gn=CEP290;refGene_tx=XM_023256246.1
B4 112596897 112597080
refGene_gn=CEP290;refGene_tx=NM_001101658.1;refGene_gn=CEP290;refGene_tx=XM_023256243.1;refGene_gn=CEP290;refGene_tx=XM_023256244.1;refGene_gn=CEP290;refGene_tx=XM_023256245.1;refGene_gn=CEP290;refGene_tx=XM_023256246.1
B4 112597497 112597650
refGene_gn=CEP290;refGene_tx=NM_001101658.1;refGene_gn=CEP290;refGene_tx=XM_023256243.1;refGene_gn=CEP290;refGene_tx=XM_023256244.1;refGene_

gn=CEP290;refGene_tx=XM_023256245.1;refGene_gn=CEP290;refGene_tx=XM_02
3256246.1
B4 112597779 112597800
refGene_gn=CEP290;refGene_tx=NM_001101658.1;refGene_gn=CEP290;refGene_
tx=XM_023256243.1;refGene_gn=CEP290;refGene_tx=XM_023256245.1;refGene_
gn=CEP290;refGene_tx=XM_023256246.1
B4 112598390 112598444
refGene_gn=CEP290;refGene_tx=NM_001101658.1;refGene_gn=CEP290;refGene_
tx=XM_023256243.1;refGene_gn=CEP290;refGene_tx=XM_023256244.1;refGene_
gn=CEP290;refGene_tx=XM_023256245.1;refGene_gn=CEP290;refGene_tx=XM_02
3256246.1
B4 112602721 112602865
refGene_gn=CEP290;refGene_tx=NM_001101658.1;refGene_gn=CEP290;refGene_
tx=XM_023256243.1;refGene_gn=CEP290;refGene_tx=XM_023256244.1;refGene_
gn=CEP290;refGene_tx=XM_023256245.1;refGene_gn=CEP290;refGene_tx=XM_02
3256246.1
B4 112605308 112605355
refGene_gn=CEP290;refGene_tx=NM_001101658.1;refGene_gn=CEP290;refGene_
tx=XM_023256243.1;refGene_gn=CEP290;refGene_tx=XM_023256244.1;refGene_
gn=CEP290;refGene_tx=XM_023256245.1;refGene_gn=CEP290;refGene_tx=XM_02
3256246.1
B4 112605620 112605690
refGene_gn=CEP290;refGene_tx=NM_001101658.1;refGene_gn=CEP290;refGene_
tx=XM_023256243.1;refGene_gn=CEP290;refGene_tx=XM_023256244.1;refGene_
gn=CEP290;refGene_tx=XM_023256245.1;refGene_gn=CEP290;refGene_tx=XM_02
3256246.1
B4 112606655 112606733
refGene_gn=CEP290;refGene_tx=NM_001101658.1;refGene_gn=CEP290;refGene_
tx=XM_023256243.1;refGene_gn=CEP290;refGene_tx=XM_023256244.1;refGene_
gn=CEP290;refGene_tx=XM_023256245.1;refGene_gn=CEP290;refGene_tx=XM_02
3256246.1
B4 112606889 112606991
refGene_gn=CEP290;refGene_tx=NM_001101658.1;refGene_gn=CEP290;refGene_
tx=XM_023256243.1;refGene_gn=CEP290;refGene_tx=XM_023256244.1;refGene_
gn=CEP290;refGene_tx=XM_023256245.1;refGene_gn=CEP290;refGene_tx=XM_02
3256246.1
B4 112617828 112618017
refGene_gn=TMTC3;refGene_tx=XM_006933916.4
B4 112629289 112629508
refGene_gn=TMTC3;refGene_tx=XM_006933916.4
B4 112631414 112631514
refGene_gn=TMTC3;refGene_tx=XM_006933916.4
B4 112635206 112635322
refGene_gn=TMTC3;refGene_tx=XM_006933916.4
B4 112635768 112635941
refGene_gn=TMTC3;refGene_tx=XM_006933916.4
B4 112641674 112641927
refGene_gn=TMTC3;refGene_tx=XM_006933916.4
B4 112646935 112647084
refGene_gn=TMTC3;refGene_tx=XM_006933916.4

B4 112648862 112648983
refGene_gn=TMTC3;refGene_tx=XM_006933916.4
B4 112651450 112651562
refGene_gn=TMTC3;refGene_tx=XM_006933916.4
B4 112655962 112656066
refGene_gn=TMTC3;refGene_tx=XM_006933916.4
B4 112658051 112658221
refGene_gn=TMTC3;refGene_tx=XM_006933916.4
B4 112660140 112660367
refGene_gn=TMTC3;refGene_tx=XM_006933916.4
B4 112662114 112662929
refGene_gn=TMTC3;refGene_tx=XM_006933916.4
B4 112965574 112965614
refGene_gn=KITLG;refGene_tx=NM_001009343.1
B4 112966544 112966612
refGene_gn=KITLG;refGene_tx=NM_001009343.1
B4 112967564 112967674
refGene_gn=KITLG;refGene_tx=NM_001009343.1
B4 112976284 112976368
refGene_gn=KITLG;refGene_tx=NM_001009343.1
B4 112977100 112977248
refGene_gn=KITLG;refGene_tx=NM_001009343.1
B4 112977249 112977261
refGene_gn=KITLG;refGene_tx=NM_001009343.1
B4 112979402 112979573
refGene_gn=KITLG;refGene_tx=NM_001009343.1
B4 112993581 112993644
refGene_gn=KITLG;refGene_tx=NM_001009343.1
B4 113004556 113004670
refGene_gn=KITLG;refGene_tx=NM_001009343.1
B4 113038349 113038364
refGene_gn=KITLG;refGene_tx=NM_001009343.1
B4 113704928 113705236
refGene_gn=DUSP6;refGene_tx=XM_003989097.5
B4 113706343 113706781
refGene_gn=DUSP6;refGene_tx=XM_003989097.5
B4 113707441 113707841
refGene_gn=DUSP6;refGene_tx=XM_003989097.5
B4 113771040 113771145
refGene_gn=POC1B;refGene_tx=XM_023257289.1;refGene_gn=POC1B;refGene_tx=XM_023257290.1;refGene_gn=POC1B;refGene_tx=XM_023257291.1
B4 113775081 113775300
refGene_gn=POC1B;refGene_tx=XM_023257289.1;refGene_gn=POC1B;refGene_tx=XM_023257290.1;refGene_gn=POC1B;refGene_tx=XM_023257291.1
B4 113812641 113812722
refGene_gn=POC1B;refGene_tx=XM_023257289.1;refGene_gn=POC1B;refGene_tx=XM_023257290.1
B4 113820173 113820326
refGene_gn=POC1B;refGene_tx=XM_023257289.1;refGene_gn=POC1B;refGene_tx=XM_023257290.1;refGene_gn=POC1B;refGene_tx=XM_023257291.1

B4	113821090	113821159
refGene_gn=POC1B;refGene_tx=XM_023257289.1;refGene_gn=POC1B;refGene_tx=XM_023257290.1;refGene_gn=POC1B;refGene_tx=XM_023257291.1
B4	113823783	113823917
refGene_gn=POC1B;refGene_tx=XM_023257289.1;refGene_gn=POC1B;refGene_tx=XM_023257290.1;refGene_gn=POC1B;refGene_tx=XM_023257291.1
B4	113825058	113825174
refGene_gn=POC1B;refGene_tx=XM_023257289.1;refGene_gn=POC1B;refGene_tx=XM_023257290.1;refGene_gn=POC1B;refGene_tx=XM_023257291.1
B4	113825308	113825416
refGene_gn=POC1B;refGene_tx=XM_023257289.1;refGene_gn=POC1B;refGene_tx=XM_023257290.1;refGene_gn=POC1B;refGene_tx=XM_023257291.1
B4	113843307	113843487
refGene_gn=POC1B;refGene_tx=XM_023257289.1;refGene_gn=POC1B;refGene_tx=XM_023257290.1;refGene_gn=POC1B;refGene_tx=XM_023257291.1
B4	113848139	113848311
refGene_gn=POC1B;refGene_tx=XM_023257289.1;refGene_gn=POC1B;refGene_tx=XM_023257290.1;refGene_gn=POC1B;refGene_tx=XM_023257291.1
B4	113852254	113852300
refGene_gn=POC1B;refGene_tx=XM_023257290.1
B4	113866910	113868659
refGene_gn=GALNT4;refGene_tx=XM_006933918.4
B4	113869204	113869289
refGene_gn=POC1B;refGene_tx=XM_023257289.1;refGene_gn=POC1B;refGene_tx=XM_023257291.1
B4	113869949	113869964
refGene_gn=POC1B;refGene_tx=XM_023257289.1;refGene_gn=POC1B;refGene_tx=XM_023257291.1
B4	113935189	113935501
refGene_gn=ATP2B1;refGene_tx=XM_003989101.4;refGene_gn=ATP2B1;refGene_tx=XM_006933920.3;refGene_gn=ATP2B1;refGene_tx=XM_019835323.1;refGene_gn=ATP2B1;refGene_tx=XM_019835324.1;refGene_gn=ATP2B1;refGene_tx=XM_023257292.1
B4	113940404	113940556
refGene_gn=ATP2B1;refGene_tx=XM_006933920.3;refGene_gn=ATP2B1;refGene_tx=XM_019835323.1;refGene_gn=ATP2B1;refGene_tx=XM_019835324.1;refGene_gn=ATP2B1;refGene_tx=XM_023257292.1
B4	113940941	113941124
refGene_gn=ATP2B1;refGene_tx=XM_003989101.4;refGene_gn=ATP2B1;refGene_tx=XM_006933920.3;refGene_gn=ATP2B1;refGene_tx=XM_019835323.1;refGene_gn=ATP2B1;refGene_tx=XM_019835324.1;refGene_gn=ATP2B1;refGene_tx=XM_023257292.1
B4	113943042	113943150
refGene_gn=ATP2B1;refGene_tx=XM_003989101.4;refGene_gn=ATP2B1;refGene_tx=XM_006933920.3;refGene_gn=ATP2B1;refGene_tx=XM_019835323.1;refGene_gn=ATP2B1;refGene_tx=XM_023257292.1
B4	113947713	113947925
refGene_gn=ATP2B1;refGene_tx=XM_003989101.4;refGene_gn=ATP2B1;refGene_tx=XM_006933920.3;refGene_gn=ATP2B1;refGene_tx=XM_019835323.1;refGene_gn=ATP2B1;refGene_tx=XM_023257292.1

B4	113948366	113948580
refGene_gn=ATP2B1;refGene_tx=XM_003989101.4;refGene_gn=ATP2B1;refGene_tx=XM_006933920.3;refGene_gn=ATP2B1;refGene_tx=XM_019835323.1;refGene_gn=ATP2B1;refGene_tx=XM_023257292.1
B4	113948817	113949009
refGene_gn=ATP2B1;refGene_tx=XM_003989101.4;refGene_gn=ATP2B1;refGene_tx=XM_006933920.3;refGene_gn=ATP2B1;refGene_tx=XM_019835323.1;refGene_gn=ATP2B1;refGene_tx=XM_019835324.1;refGene_gn=ATP2B1;refGene_tx=XM_023257292.1
B4	113955734	113955841
refGene_gn=ATP2B1;refGene_tx=XM_003989101.4;refGene_gn=ATP2B1;refGene_tx=XM_006933920.3;refGene_gn=ATP2B1;refGene_tx=XM_019835323.1;refGene_gn=ATP2B1;refGene_tx=XM_019835324.1;refGene_gn=ATP2B1;refGene_tx=XM_023257292.1
B4	113956447	113956535
refGene_gn=ATP2B1;refGene_tx=XM_003989101.4;refGene_gn=ATP2B1;refGene_tx=XM_006933920.3;refGene_gn=ATP2B1;refGene_tx=XM_019835323.1;refGene_gn=ATP2B1;refGene_tx=XM_019835324.1;refGene_gn=ATP2B1;refGene_tx=XM_023257292.1
B4	113957205	113957385
refGene_gn=ATP2B1;refGene_tx=XM_003989101.4;refGene_gn=ATP2B1;refGene_tx=XM_006933920.3;refGene_gn=ATP2B1;refGene_tx=XM_019835323.1;refGene_gn=ATP2B1;refGene_tx=XM_019835324.1;refGene_gn=ATP2B1;refGene_tx=XM_023257292.1
B4	113962020	113962258
refGene_gn=ATP2B1;refGene_tx=XM_003989101.4;refGene_gn=ATP2B1;refGene_tx=XM_006933920.3;refGene_gn=ATP2B1;refGene_tx=XM_019835323.1;refGene_gn=ATP2B1;refGene_tx=XM_019835324.1;refGene_gn=ATP2B1;refGene_tx=XM_023257292.1
B4	113964815	113965057
refGene_gn=ATP2B1;refGene_tx=XM_003989101.4;refGene_gn=ATP2B1;refGene_tx=XM_006933920.3;refGene_gn=ATP2B1;refGene_tx=XM_019835323.1;refGene_gn=ATP2B1;refGene_tx=XM_019835324.1;refGene_gn=ATP2B1;refGene_tx=XM_023257292.1
B4	113966400	113966643
refGene_gn=ATP2B1;refGene_tx=XM_003989101.4;refGene_gn=ATP2B1;refGene_tx=XM_006933920.3;refGene_gn=ATP2B1;refGene_tx=XM_019835323.1;refGene_gn=ATP2B1;refGene_tx=XM_019835324.1;refGene_gn=ATP2B1;refGene_tx=XM_023257292.1
B4	113969108	113969323
refGene_gn=ATP2B1;refGene_tx=XM_003989101.4;refGene_gn=ATP2B1;refGene_tx=XM_006933920.3;refGene_gn=ATP2B1;refGene_tx=XM_019835323.1;refGene_gn=ATP2B1;refGene_tx=XM_019835324.1;refGene_gn=ATP2B1;refGene_tx=XM_023257292.1
B4	113971090	113971252
refGene_gn=ATP2B1;refGene_tx=XM_003989101.4;refGene_gn=ATP2B1;refGene_tx=XM_006933920.3;refGene_gn=ATP2B1;refGene_tx=XM_019835323.1;refGene_gn=ATP2B1;refGene_tx=XM_019835324.1;refGene_gn=ATP2B1;refGene_tx=XM_023257292.1
B4	113972044	113972083

refGene_gn=ATP2B1;refGene_tx=XM_003989101.4;refGene_gn=ATP2B1;refGene_tx=XM_006933920.3;refGene_gn=ATP2B1;refGene_tx=XM_019835323.1;refGene_gn=ATP2B1;refGene_tx=XM_019835324.1;refGene_gn=ATP2B1;refGene_tx=XM_023257292.1
B4 113974602 113974743
refGene_gn=ATP2B1;refGene_tx=XM_003989101.4;refGene_gn=ATP2B1;refGene_tx=XM_006933920.3;refGene_gn=ATP2B1;refGene_tx=XM_019835323.1;refGene_gn=ATP2B1;refGene_tx=XM_019835324.1;refGene_gn=ATP2B1;refGene_tx=XM_023257292.1
B4 113978594 113978720
refGene_gn=ATP2B1;refGene_tx=XM_003989101.4;refGene_gn=ATP2B1;refGene_tx=XM_006933920.3;refGene_gn=ATP2B1;refGene_tx=XM_019835323.1;refGene_gn=ATP2B1;refGene_tx=XM_019835324.1;refGene_gn=ATP2B1;refGene_tx=XM_023257292.1
B4 113978810 113979065
refGene_gn=ATP2B1;refGene_tx=XM_003989101.4;refGene_gn=ATP2B1;refGene_tx=XM_006933920.3;refGene_gn=ATP2B1;refGene_tx=XM_019835323.1;refGene_gn=ATP2B1;refGene_tx=XM_019835324.1;refGene_gn=ATP2B1;refGene_tx=XM_023257292.1
B4 113985302 113985500
refGene_gn=ATP2B1;refGene_tx=XM_003989101.4;refGene_gn=ATP2B1;refGene_tx=XM_006933920.3;refGene_gn=ATP2B1;refGene_tx=XM_019835323.1;refGene_gn=ATP2B1;refGene_tx=XM_019835324.1;refGene_gn=ATP2B1;refGene_tx=XM_023257292.1
B4 114003720 114003928
refGene_gn=ATP2B1;refGene_tx=XM_003989101.4;refGene_gn=ATP2B1;refGene_tx=XM_006933920.3;refGene_gn=ATP2B1;refGene_tx=XM_019835323.1;refGene_gn=ATP2B1;refGene_tx=XM_019835324.1;refGene_gn=ATP2B1;refGene_tx=XM_023257292.1
B4 115068580 115069825
refGene_gn=CCER1;refGene_tx=XM_023257293.1
B4 115077644 115077815
refGene_gn=EPYC;refGene_tx=XM_003989107.4
B4 115082987 115083083
refGene_gn=EPYC;refGene_tx=XM_003989107.4
B4 115084436 115084639
refGene_gn=EPYC;refGene_tx=XM_003989107.4
B4 115085734 115085893
refGene_gn=EPYC;refGene_tx=XM_003989107.4
B4 115090903 115091072
refGene_gn=EPYC;refGene_tx=XM_003989107.4
B4 115111665 115111830
refGene_gn=EPYC;refGene_tx=XM_003989107.4
B4 115161430 115161597
refGene_gn=KERA;refGene_tx=XM_003989108.3;refGene_gn=KERA;refGene_tx=XM_006933924.3
B4 115165545 115166431
refGene_gn=KERA;refGene_tx=XM_003989108.3;refGene_gn=KERA;refGene_tx=XM_006933924.3
B4 115206291 115206446

refGene_gn=LUM;refGene_tx=XM_003989109.5
B4 115210079 115210944
refGene_gn=LUM;refGene_tx=XM_003989109.5
B4 115250345 115250540
refGene_gn=DCN;refGene_tx=XM_003989110.5;refGene_gn=DCN;refGene_tx=XM_003989111.5
B4 115255902 115256041
refGene_gn=DCN;refGene_tx=XM_003989110.5;refGene_gn=DCN;refGene_tx=XM_003989111.5
B4 115259493 115259587
refGene_gn=DCN;refGene_tx=XM_003989110.5;refGene_gn=DCN;refGene_tx=XM_003989111.5
B4 115264627 115264741
refGene_gn=DCN;refGene_tx=XM_003989110.5;refGene_gn=DCN;refGene_tx=XM_003989111.5
B4 115265742 115265956
refGene_gn=DCN;refGene_tx=XM_003989110.5;refGene_gn=DCN;refGene_tx=XM_003989111.5
B4 115272473 115272586
refGene_gn=DCN;refGene_tx=XM_003989110.5;refGene_gn=DCN;refGene_tx=XM_003989111.5
B4 115287475 115287689
refGene_gn=DCN;refGene_tx=XM_003989110.5;refGene_gn=DCN;refGene_tx=XM_003989111.5
B4 116115978 116116346
refGene_gn=BTG1;refGene_tx=XM_023257294.1
B4 116117379 116117527
refGene_gn=BTG1;refGene_tx=XM_023257294.1
B4 116563906 116564092
refGene_gn=CB4H12orf74;refGene_tx=XM_019835868.1
B4 116564094 116564355
refGene_gn=CB4H12orf74;refGene_tx=XM_019835868.1
B4 116565466 116565530
refGene_gn=PLEKHG7;refGene_tx=XM_023257773.1
B4 116565744 116565812
refGene_gn=PLEKHG7;refGene_tx=XM_023257773.1
B4 116593453 116593549
refGene_gn=PLEKHG7;refGene_tx=XM_023257773.1
B4 116593878 116593980
refGene_gn=PLEKHG7;refGene_tx=XM_023257773.1
B4 116594321 116594337
refGene_gn=PLEKHG7;refGene_tx=XM_023257773.1
B4 116598001 116598103
refGene_gn=PLEKHG7;refGene_tx=XM_023257773.1
B4 116604000 116604083
refGene_gn=PLEKHG7;refGene_tx=XM_023257773.1
B4 116604085 116604171
refGene_gn=PLEKHG7;refGene_tx=XM_023257773.1
B4 116604386 116604460
refGene_gn=PLEKHG7;refGene_tx=XM_023257773.1

B4 116605704 116605805
refGene_gn=PLEKHG7;refGene_tx=XM_023257773.1
B4 116606155 116606264
refGene_gn=PLEKHG7;refGene_tx=XM_023257773.1
B4 116611458 116611537
refGene_gn=PLEKHG7;refGene_tx=XM_023257773.1
B4 116618567 116618722
refGene_gn=PLEKHG7;refGene_tx=XM_023257773.1
B4 116623640 116623699
refGene_gn=PLEKHG7;refGene_tx=XM_023257773.1
B4 116623701 116623740
refGene_gn=PLEKHG7;refGene_tx=XM_023257773.1
B4 116624714 116624822
refGene_gn=PLEKHG7;refGene_tx=XM_023257773.1
B4 116631902 116632025
refGene_gn=EEA1;refGene_tx=XM_023257295.1;refGene_gn=EEA1;refGene_tx=XM_023257296.1;refGene_gn=EEA1;refGene_tx=XM_023257297.1;refGene_gn=EEA1;refGene_tx=XM_023257298.1;refGene_gn=EEA1;refGene_tx=XM_023257299.1
B4 116632732 116632831
refGene_gn=EEA1;refGene_tx=XM_023257295.1;refGene_gn=EEA1;refGene_tx=XM_023257296.1;refGene_gn=EEA1;refGene_tx=XM_023257297.1;refGene_gn=EEA1;refGene_tx=XM_023257298.1;refGene_gn=EEA1;refGene_tx=XM_023257299.1
B4 116633897 116634018
refGene_gn=EEA1;refGene_tx=XM_023257295.1;refGene_gn=EEA1;refGene_tx=XM_023257296.1;refGene_gn=EEA1;refGene_tx=XM_023257297.1;refGene_gn=EEA1;refGene_tx=XM_023257298.1;refGene_gn=EEA1;refGene_tx=XM_023257299.1
B4 116634290 116634529
refGene_gn=EEA1;refGene_tx=XM_023257295.1;refGene_gn=EEA1;refGene_tx=XM_023257296.1;refGene_gn=EEA1;refGene_tx=XM_023257297.1;refGene_gn=EEA1;refGene_tx=XM_023257298.1;refGene_gn=EEA1;refGene_tx=XM_023257299.1
B4 116635474 116635660
refGene_gn=EEA1;refGene_tx=XM_023257295.1;refGene_gn=EEA1;refGene_tx=XM_023257296.1;refGene_gn=EEA1;refGene_tx=XM_023257297.1;refGene_gn=EEA1;refGene_tx=XM_023257298.1;refGene_gn=EEA1;refGene_tx=XM_023257299.1
B4 116636830 116636962
refGene_gn=EEA1;refGene_tx=XM_023257295.1;refGene_gn=EEA1;refGene_tx=XM_023257296.1;refGene_gn=EEA1;refGene_tx=XM_023257297.1;refGene_gn=EEA1;refGene_tx=XM_023257298.1;refGene_gn=EEA1;refGene_tx=XM_023257299.1
B4 116638450 116638636
refGene_gn=EEA1;refGene_tx=XM_023257295.1;refGene_gn=EEA1;refGene_tx=XM_023257296.1;refGene_gn=EEA1;refGene_tx=XM_023257297.1;refGene_gn=EEA1;refGene_tx=XM_023257298.1;refGene_gn=EEA1;refGene_tx=XM_023257299.1
B4 116644424 116644607
refGene_gn=EEA1;refGene_tx=XM_023257295.1;refGene_gn=EEA1;refGene_tx=XM_023257296.1;refGene_gn=EEA1;refGene_tx=XM_023257297.1;refGene_gn=EEA1;refGene_tx=XM_023257298.1;refGene_gn=EEA1;refGene_tx=XM_023257299.1
B4 116650500 116650695
refGene_gn=EEA1;refGene_tx=XM_023257295.1;refGene_gn=EEA1;refGene_tx=XM_023257296.1;refGene_gn=EEA1;refGene_tx=XM_023257297.1;refGene_gn=EEA1;refGene_tx=XM_023257298.1;refGene_gn=EEA1;refGene_tx=XM_023257299.1

B4 116652456 116652558
refGene_gn=EEA1;refGene_tx=XM_023257295.1;refGene_gn=EEA1;refGene_tx=XM_023257296.1;refGene_gn=EEA1;refGene_tx=XM_023257297.1;refGene_gn=EEA1;refGene_tx=XM_023257298.1;refGene_gn=EEA1;refGene_tx=XM_023257299.1
B4 116653634 116653965
refGene_gn=EEA1;refGene_tx=XM_023257295.1;refGene_gn=EEA1;refGene_tx=XM_023257296.1;refGene_gn=EEA1;refGene_tx=XM_023257297.1;refGene_gn=EEA1;refGene_tx=XM_023257298.1;refGene_gn=EEA1;refGene_tx=XM_023257299.1
B4 116661352 116661492
refGene_gn=EEA1;refGene_tx=XM_023257295.1;refGene_gn=EEA1;refGene_tx=XM_023257296.1;refGene_gn=EEA1;refGene_tx=XM_023257297.1;refGene_gn=EEA1;refGene_tx=XM_023257298.1;refGene_gn=EEA1;refGene_tx=XM_023257299.1
B4 116663470 116663626
refGene_gn=EEA1;refGene_tx=XM_023257295.1;refGene_gn=EEA1;refGene_tx=XM_023257296.1;refGene_gn=EEA1;refGene_tx=XM_023257297.1;refGene_gn=EEA1;refGene_tx=XM_023257298.1;refGene_gn=EEA1;refGene_tx=XM_023257299.1
B4 116666486 116666600
refGene_gn=EEA1;refGene_tx=XM_023257295.1;refGene_gn=EEA1;refGene_tx=XM_023257296.1;refGene_gn=EEA1;refGene_tx=XM_023257297.1;refGene_gn=EEA1;refGene_tx=XM_023257298.1;refGene_gn=EEA1;refGene_tx=XM_023257299.1
B4 116668765 116668966
refGene_gn=EEA1;refGene_tx=XM_023257295.1;refGene_gn=EEA1;refGene_tx=XM_023257296.1;refGene_gn=EEA1;refGene_tx=XM_023257297.1;refGene_gn=EEA1;refGene_tx=XM_023257298.1;refGene_gn=EEA1;refGene_tx=XM_023257299.1
B4 116671888 116672092
refGene_gn=EEA1;refGene_tx=XM_023257295.1;refGene_gn=EEA1;refGene_tx=XM_023257296.1;refGene_gn=EEA1;refGene_tx=XM_023257297.1;refGene_gn=EEA1;refGene_tx=XM_023257298.1;refGene_gn=EEA1;refGene_tx=XM_023257299.1
B4 116680512 116680632
refGene_gn=EEA1;refGene_tx=XM_023257295.1;refGene_gn=EEA1;refGene_tx=XM_023257296.1;refGene_gn=EEA1;refGene_tx=XM_023257297.1;refGene_gn=EEA1;refGene_tx=XM_023257298.1;refGene_gn=EEA1;refGene_tx=XM_023257299.1
B4 116681897 116682047
refGene_gn=EEA1;refGene_tx=XM_023257295.1;refGene_gn=EEA1;refGene_tx=XM_023257296.1;refGene_gn=EEA1;refGene_tx=XM_023257297.1;refGene_gn=EEA1;refGene_tx=XM_023257298.1;refGene_gn=EEA1;refGene_tx=XM_023257299.1
B4 116688041 116688380
refGene_gn=EEA1;refGene_tx=XM_023257295.1;refGene_gn=EEA1;refGene_tx=XM_023257296.1;refGene_gn=EEA1;refGene_tx=XM_023257297.1;refGene_gn=EEA1;refGene_tx=XM_023257298.1;refGene_gn=EEA1;refGene_tx=XM_023257299.1
B4 116697573 116697690
refGene_gn=EEA1;refGene_tx=XM_023257295.1;refGene_gn=EEA1;refGene_tx=XM_023257296.1;refGene_gn=EEA1;refGene_tx=XM_023257297.1;refGene_gn=EEA1;refGene_tx=XM_023257298.1;refGene_gn=EEA1;refGene_tx=XM_023257299.1
B4 116701195 116701210
refGene_gn=EEA1;refGene_tx=XM_023257295.1;refGene_gn=EEA1;refGene_tx=XM_023257297.1;refGene_gn=EEA1;refGene_tx=XM_023257299.1
B4 116704768 116704924
refGene_gn=EEA1;refGene_tx=XM_023257295.1;refGene_gn=EEA1;refGene_tx=XM_023257296.1;refGene_gn=EEA1;refGene_tx=XM_023257297.1;refGene_gn=EEA

1;refGene_tx=XM_023257298.1;refGene_gn=EEA1;refGene_tx=XM_023257299.1
B4 116706067 116706189
refGene_gn=EEA1;refGene_tx=XM_023257295.1;refGene_gn=EEA1;refGene_tx=XM_023257296.1;refGene_gn=EEA1;refGene_tx=XM_023257297.1;refGene_gn=EEA1;refGene_tx=XM_023257298.1;refGene_gn=EEA1;refGene_tx=XM_023257299.1
B4 116706898 116707012
refGene_gn=EEA1;refGene_tx=XM_023257295.1;refGene_gn=EEA1;refGene_tx=XM_023257296.1;refGene_gn=EEA1;refGene_tx=XM_023257297.1;refGene_gn=EEA1;refGene_tx=XM_023257298.1;refGene_gn=EEA1;refGene_tx=XM_023257299.1
B4 116708419 116708459
refGene_gn=EEA1;refGene_tx=XM_023257295.1;refGene_gn=EEA1;refGene_tx=XM_023257296.1;refGene_gn=EEA1;refGene_tx=XM_023257297.1;refGene_gn=EEA1;refGene_tx=XM_023257298.1;refGene_gn=EEA1;refGene_tx=XM_023257299.1
B4 116710583 116710649
refGene_gn=EEA1;refGene_tx=XM_023257295.1;refGene_gn=EEA1;refGene_tx=XM_023257296.1;refGene_gn=EEA1;refGene_tx=XM_023257297.1;refGene_gn=EEA1;refGene_tx=XM_023257298.1;refGene_gn=EEA1;refGene_tx=XM_023257299.1
B4 116710742 116710797
refGene_gn=EEA1;refGene_tx=XM_023257295.1;refGene_gn=EEA1;refGene_tx=XM_023257296.1;refGene_gn=EEA1;refGene_tx=XM_023257297.1;refGene_gn=EEA1;refGene_tx=XM_023257298.1;refGene_gn=EEA1;refGene_tx=XM_023257299.1
B4 116711385 116711510
refGene_gn=EEA1;refGene_tx=XM_023257295.1;refGene_gn=EEA1;refGene_tx=XM_023257296.1;refGene_gn=EEA1;refGene_tx=XM_023257297.1;refGene_gn=EEA1;refGene_tx=XM_023257298.1;refGene_gn=EEA1;refGene_tx=XM_023257299.1
B4 116716040 116716166
refGene_gn=EEA1;refGene_tx=XM_023257295.1;refGene_gn=EEA1;refGene_tx=XM_023257296.1
B4 116723227 116723320
refGene_gn=EEA1;refGene_tx=XM_023257295.1;refGene_gn=EEA1;refGene_tx=XM_023257296.1;refGene_gn=EEA1;refGene_tx=XM_023257297.1;refGene_gn=EEA1;refGene_tx=XM_023257298.1
B4 116744533 116744557
refGene_gn=EEA1;refGene_tx=XM_023257295.1;refGene_gn=EEA1;refGene_tx=XM_023257296.1;refGene_gn=EEA1;refGene_tx=XM_023257297.1;refGene_gn=EEA1;refGene_tx=XM_023257298.1
B4 117128518 117128617
refGene_gn=NUDT4;refGene_tx=XM_006933953.4;refGene_gn=NUDT4;refGene_tx=XM_011284150.3
B4 117138870 117138981
refGene_gn=NUDT4;refGene_tx=XM_006933953.4;refGene_gn=NUDT4;refGene_tx=XM_011284150.3
B4 117140216 117140261
refGene_gn=NUDT4;refGene_tx=XM_006933953.4;refGene_gn=NUDT4;refGene_tx=XM_011284150.3
B4 117142630 117142718
refGene_gn=NUDT4;refGene_tx=XM_006933953.4;refGene_gn=NUDT4;refGene_tx=XM_011284150.3
B4 117143009 117143212
refGene_gn=NUDT4;refGene_tx=XM_006933953.4;refGene_gn=NUDT4;refGene_tx

=XM_011284150.3
B4	117153450	117153491
refGene_gn=UBE2N;refGene_tx=XM_023257300.1
B4	117154174	117154315
refGene_gn=UBE2N;refGene_tx=XM_023257300.1
B4	117154508	117154755
refGene_gn=UBE2N;refGene_tx=XM_023257300.1
B4	117189298	117189328
refGene_gn=UBE2N;refGene_tx=XM_023257300.1
B4	117209874	117209944
refGene_gn=MRPL42;refGene_tx=XM_011284151.3;refGene_gn=MRPL42;refGene_tx=XM_019835333.2;refGene_gn=MRPL42;refGene_tx=XM_023257301.1
B4	117214857	117214921
refGene_gn=MRPL42;refGene_tx=XM_011284151.3;refGene_gn=MRPL42;refGene_tx=XM_019835333.2;refGene_gn=MRPL42;refGene_tx=XM_023257301.1
B4	117216582	117216667
refGene_gn=MRPL42;refGene_tx=XM_011284151.3;refGene_gn=MRPL42;refGene_tx=XM_019835333.2;refGene_gn=MRPL42;refGene_tx=XM_023257301.1
B4	117222858	117223022
refGene_gn=MRPL42;refGene_tx=XM_011284151.3;refGene_gn=MRPL42;refGene_tx=XM_019835333.2;refGene_gn=MRPL42;refGene_tx=XM_023257301.1
B4	117226017	117226141
refGene_gn=MRPL42;refGene_tx=XM_019835333.2;refGene_gn=MRPL42;refGene_tx=XM_023257301.1
B4	117239895	117239941
refGene_gn=MRPL42;refGene_tx=XM_011284151.3
B4	117304302	117304441
refGene_gn=SOCS2;refGene_tx=XM_011284154.3;refGene_gn=SOCS2;refGene_tx=XM_019835335.2
B4	117306151	117306609
refGene_gn=SOCS2;refGene_tx=XM_011284154.3;refGene_gn=SOCS2;refGene_tx=XM_019835335.2
B4	117396917	117397163
refGene_gn=CRADD;refGene_tx=XM_006933926.4
B4	117398281	117398389
refGene_gn=CRADD;refGene_tx=XM_003989115.5
B4	117398487	117398791
refGene_gn=CRADD;refGene_tx=XM_003989115.5;refGene_gn=CRADD;refGene_tx=XM_006933926.4
B4	117571913	117572215
refGene_gn=CRADD;refGene_tx=XM_003989115.5;refGene_gn=CRADD;refGene_tx=XM_006933926.4
B4	117803842	117804907
refGene_gn=PLXNC1;refGene_tx=XM_023257304.1;refGene_gn=PLXNC1;refGene_tx=XM_023257305.1
B4	117827916	117828057
refGene_gn=PLXNC1;refGene_tx=XM_023257304.1;refGene_gn=PLXNC1;refGene_tx=XM_023257305.1
B4	117840019	117840154
refGene_gn=PLXNC1;refGene_tx=XM_023257304.1;refGene_gn=PLXNC1;refGene_

tx=XM_023257305.1
B4 117843568 117843669
refGene_gn=PLXNC1;refGene_tx=XM_023257304.1;refGene_gn=PLXNC1;refGene_tx=XM_023257305.1
B4 117864526 117864641
refGene_gn=PLXNC1;refGene_tx=XM_023257304.1;refGene_gn=PLXNC1;refGene_tx=XM_023257305.1
B4 117873093 117873247
refGene_gn=PLXNC1;refGene_tx=XM_023257304.1;refGene_gn=PLXNC1;refGene_tx=XM_023257305.1
B4 117876746 117876834
refGene_gn=PLXNC1;refGene_tx=XM_023257304.1;refGene_gn=PLXNC1;refGene_tx=XM_023257305.1
B4 117878983 117879086
refGene_gn=PLXNC1;refGene_tx=XM_023257304.1;refGene_gn=PLXNC1;refGene_tx=XM_023257305.1
B4 117879227 117879329
refGene_gn=PLXNC1;refGene_tx=XM_023257304.1;refGene_gn=PLXNC1;refGene_tx=XM_023257305.1
B4 117890315 117890458
refGene_gn=PLXNC1;refGene_tx=XM_023257304.1;refGene_gn=PLXNC1;refGene_tx=XM_023257305.1
B4 117893156 117893336
refGene_gn=PLXNC1;refGene_tx=XM_023257304.1;refGene_gn=PLXNC1;refGene_tx=XM_023257305.1
B4 117896170 117896258
refGene_gn=PLXNC1;refGene_tx=XM_023257304.1;refGene_gn=PLXNC1;refGene_tx=XM_023257305.1
B4 117898574 117898778
refGene_gn=PLXNC1;refGene_tx=XM_023257304.1;refGene_gn=PLXNC1;refGene_tx=XM_023257305.1
B4 117898901 117899087
refGene_gn=PLXNC1;refGene_tx=XM_023257304.1;refGene_gn=PLXNC1;refGene_tx=XM_023257305.1
B4 117900901 117901004
refGene_gn=PLXNC1;refGene_tx=XM_023257304.1;refGene_gn=PLXNC1;refGene_tx=XM_023257305.1
B4 117904292 117904394
refGene_gn=PLXNC1;refGene_tx=XM_023257304.1;refGene_gn=PLXNC1;refGene_tx=XM_023257305.1
B4 117904703 117904807
refGene_gn=PLXNC1;refGene_tx=XM_023257304.1;refGene_gn=PLXNC1;refGene_tx=XM_023257305.1
B4 117908989 117909028
refGene_gn=PLXNC1;refGene_tx=XM_023257304.1;refGene_gn=PLXNC1;refGene_tx=XM_023257305.1
B4 117909267 117909392
refGene_gn=PLXNC1;refGene_tx=XM_023257304.1;refGene_gn=PLXNC1;refGene_tx=XM_023257305.1
B4 117910677 117910876

refGene_gn=PLXNC1;refGene_tx=XM_023257304.1;refGene_gn=PLXNC1;refGene_tx=XM_023257305.1
B4 117915819 117915966
refGene_gn=PLXNC1;refGene_tx=XM_023257304.1;refGene_gn=PLXNC1;refGene_tx=XM_023257305.1
B4 117929218 117929396
refGene_gn=PLXNC1;refGene_tx=XM_023257304.1;refGene_gn=PLXNC1;refGene_tx=XM_023257305.1
B4 117931898 117932002
refGene_gn=PLXNC1;refGene_tx=XM_023257304.1;refGene_gn=PLXNC1;refGene_tx=XM_023257305.1
B4 117941118 117941173
refGene_gn=PLXNC1;refGene_tx=XM_023257304.1;refGene_gn=PLXNC1;refGene_tx=XM_023257305.1
B4 117943623 117943655
refGene_gn=PLXNC1;refGene_tx=XM_023257304.1;refGene_gn=PLXNC1;refGene_tx=XM_023257305.1
B4 117943752 117943860
refGene_gn=PLXNC1;refGene_tx=XM_023257304.1;refGene_gn=PLXNC1;refGene_tx=XM_023257305.1
B4 117945001 117945165
refGene_gn=PLXNC1;refGene_tx=XM_023257304.1;refGene_gn=PLXNC1;refGene_tx=XM_023257305.1
B4 117947085 117947233
refGene_gn=PLXNC1;refGene_tx=XM_023257304.1;refGene_gn=PLXNC1;refGene_tx=XM_023257305.1
B4 117950072 117950213
refGene_gn=PLXNC1;refGene_tx=XM_023257304.1;refGene_gn=PLXNC1;refGene_tx=XM_023257305.1
B4 117950305 117950380
refGene_gn=PLXNC1;refGene_tx=XM_023257304.1;refGene_gn=PLXNC1;refGene_tx=XM_023257305.1
B4 117951560 117951665
refGene_gn=PLXNC1;refGene_tx=XM_023257304.1;refGene_gn=PLXNC1;refGene_tx=XM_023257305.1
B4 117955329 117955434
refGene_gn=CEP83;refGene_tx=XM_011284155.3;refGene_gn=CEP83;refGene_tx=XM_011284158.3;refGene_gn=CEP83;refGene_tx=XM_019835338.2;refGene_gn=CEP83;refGene_tx=XM_019835339.2;refGene_gn=CEP83;refGene_tx=XM_023257306.1;refGene_gn=CEP83;refGene_tx=XM_023257307.1
B4 117956383 117956573
refGene_gn=CEP83;refGene_tx=XM_011284155.3;refGene_gn=CEP83;refGene_tx=XM_011284158.3;refGene_gn=CEP83;refGene_tx=XM_019835338.2;refGene_gn=CEP83;refGene_tx=XM_019835339.2;refGene_gn=CEP83;refGene_tx=XM_023257306.1;refGene_gn=CEP83;refGene_tx=XM_023257307.1
B4 117959485 117959589
refGene_gn=CEP83;refGene_tx=XM_011284155.3;refGene_gn=CEP83;refGene_tx=XM_011284158.3;refGene_gn=CEP83;refGene_tx=XM_019835338.2;refGene_gn=CEP83;refGene_tx=XM_019835339.2;refGene_gn=CEP83;refGene_tx=XM_023257306.1;refGene_gn=CEP83;refGene_tx=XM_023257307.1

B4 117974229 117974359
refGene_gn=CEP83;refGene_tx=XM_011284155.3;refGene_gn=CEP83;refGene_tx=XM_011284158.3;refGene_gn=CEP83;refGene_tx=XM_019835338.2;refGene_gn=CEP83;refGene_tx=XM_019835339.2;refGene_gn=CEP83;refGene_tx=XM_023257306.1;refGene_gn=CEP83;refGene_tx=XM_023257307.1
B4 117976231 117976389
refGene_gn=CEP83;refGene_tx=XM_011284155.3;refGene_gn=CEP83;refGene_tx=XM_011284158.3;refGene_gn=CEP83;refGene_tx=XM_019835338.2;refGene_gn=CEP83;refGene_tx=XM_019835339.2;refGene_gn=CEP83;refGene_tx=XM_023257306.1;refGene_gn=CEP83;refGene_tx=XM_023257307.1
B4 117979184 117979260
refGene_gn=CEP83;refGene_tx=XM_011284155.3;refGene_gn=CEP83;refGene_tx=XM_011284158.3;refGene_gn=CEP83;refGene_tx=XM_019835338.2;refGene_gn=CEP83;refGene_tx=XM_019835339.2;refGene_gn=CEP83;refGene_tx=XM_023257306.1;refGene_gn=CEP83;refGene_tx=XM_023257307.1
B4 117992258 117992408
refGene_gn=CEP83;refGene_tx=XM_011284155.3;refGene_gn=CEP83;refGene_tx=XM_011284158.3;refGene_gn=CEP83;refGene_tx=XM_019835338.2;refGene_gn=CEP83;refGene_tx=XM_019835339.2;refGene_gn=CEP83;refGene_tx=XM_023257306.1;refGene_gn=CEP83;refGene_tx=XM_023257307.1
B4 117992521 117992666
refGene_gn=CEP83;refGene_tx=XM_011284155.3;refGene_gn=CEP83;refGene_tx=XM_011284158.3;refGene_gn=CEP83;refGene_tx=XM_019835338.2;refGene_gn=CEP83;refGene_tx=XM_019835339.2;refGene_gn=CEP83;refGene_tx=XM_023257306.1;refGene_gn=CEP83;refGene_tx=XM_023257307.1
B4 117993927 117994042
refGene_gn=CEP83;refGene_tx=XM_011284155.3;refGene_gn=CEP83;refGene_tx=XM_011284158.3;refGene_gn=CEP83;refGene_tx=XM_019835338.2;refGene_gn=CEP83;refGene_tx=XM_019835339.2;refGene_gn=CEP83;refGene_tx=XM_023257306.1;refGene_gn=CEP83;refGene_tx=XM_023257307.1
B4 118000293 118000425
refGene_gn=CEP83;refGene_tx=XM_011284155.3;refGene_gn=CEP83;refGene_tx=XM_019835338.2;refGene_gn=CEP83;refGene_tx=XM_019835339.2;refGene_gn=CEP83;refGene_tx=XM_023257306.1;refGene_gn=CEP83;refGene_tx=XM_023257307.1
B4 118003836 118004088
refGene_gn=CEP83;refGene_tx=XM_011284155.3;refGene_gn=CEP83;refGene_tx=XM_011284158.3;refGene_gn=CEP83;refGene_tx=XM_019835338.2;refGene_gn=CEP83;refGene_tx=XM_019835339.2;refGene_gn=CEP83;refGene_tx=XM_023257306.1;refGene_gn=CEP83;refGene_tx=XM_023257307.1
B4 118029026 118029158
refGene_gn=CEP83;refGene_tx=XM_011284155.3;refGene_gn=CEP83;refGene_tx=XM_011284158.3;refGene_gn=CEP83;refGene_tx=XM_019835338.2;refGene_gn=CEP83;refGene_tx=XM_019835339.2;refGene_gn=CEP83;refGene_tx=XM_023257306.1;refGene_gn=CEP83;refGene_tx=XM_023257307.1
B4 118031264 118031357
refGene_gn=CEP83;refGene_tx=XM_011284155.3;refGene_gn=CEP83;refGene_tx=XM_011284158.3;refGene_gn=CEP83;refGene_tx=XM_019835338.2;refGene_gn=CEP83;refGene_tx=XM_019835339.2;refGene_gn=CEP83;refGene_tx=XM_023257306.1;refGene_gn=CEP83;refGene_tx=XM_023257307.1

B4	118039912	118040063
refGene_gn=CEP83;refGene_tx=XM_011284155.3;refGene_gn=CEP83;refGene_tx=XM_011284158.3;refGene_gn=CEP83;refGene_tx=XM_019835338.2;refGene_gn=CEP83;refGene_tx=XM_019835339.2;refGene_gn=CEP83;refGene_tx=XM_023257306.1;refGene_gn=CEP83;refGene_tx=XM_023257307.1
B4	118040699	118040848
refGene_gn=CEP83;refGene_tx=XM_011284155.3;refGene_gn=CEP83;refGene_tx=XM_011284158.3;refGene_gn=CEP83;refGene_tx=XM_019835338.2;refGene_gn=CEP83;refGene_tx=XM_019835339.2;refGene_gn=CEP83;refGene_tx=XM_023257306.1;refGene_gn=CEP83;refGene_tx=XM_023257307.1
B4	118126561	118126816
refGene_gn=TMCC3;refGene_tx=XM_023257310.1
B4	118157996	118158161
refGene_gn=TMCC3;refGene_tx=XM_023257314.1
B4	118167040	118167343
refGene_gn=TMCC3;refGene_tx=XM_006933927.4;refGene_gn=TMCC3;refGene_tx=XM_019835343.2;refGene_gn=TMCC3;refGene_tx=XM_019835344.2;refGene_gn=TMCC3;refGene_tx=XM_019835348.2;refGene_gn=TMCC3;refGene_tx=XM_023257309.1;refGene_gn=TMCC3;refGene_tx=XM_023257311.1;refGene_gn=TMCC3;refGene_tx=XM_023257313.1
B4	118172641	118172777
refGene_gn=TMCC3;refGene_tx=XM_006933927.4;refGene_gn=TMCC3;refGene_tx=XM_019835343.2;refGene_gn=TMCC3;refGene_tx=XM_019835344.2;refGene_gn=TMCC3;refGene_tx=XM_019835348.2;refGene_gn=TMCC3;refGene_tx=XM_023257309.1;refGene_gn=TMCC3;refGene_tx=XM_023257310.1;refGene_gn=TMCC3;refGene_tx=XM_023257311.1;refGene_gn=TMCC3;refGene_tx=XM_023257313.1;refGene_gn=TMCC3;refGene_tx=XM_023257314.1
B4	118176140	118177057
refGene_gn=TMCC3;refGene_tx=XM_006933927.4;refGene_gn=TMCC3;refGene_tx=XM_019835343.2;refGene_gn=TMCC3;refGene_tx=XM_019835344.2;refGene_gn=TMCC3;refGene_tx=XM_019835348.2;refGene_gn=TMCC3;refGene_tx=XM_023257309.1;refGene_gn=TMCC3;refGene_tx=XM_023257310.1;refGene_gn=TMCC3;refGene_tx=XM_023257311.1;refGene_gn=TMCC3;refGene_tx=XM_023257313.1;refGene_gn=TMCC3;refGene_tx=XM_023257314.1
B4	118243074	118243152
refGene_gn=TMCC3;refGene_tx=XM_023257309.1;refGene_gn=TMCC3;refGene_tx=XM_023257310.1;refGene_gn=TMCC3;refGene_tx=XM_023257314.1
B4	118455262	118455271
refGene_gn=TMCC3;refGene_tx=XM_023257311.1
B4	118549389	118549570
refGene_gn=NDUFA12;refGene_tx=XM_003989116.5
B4	118574610	118574698
refGene_gn=NDUFA12;refGene_tx=XM_003989116.5
B4	118577986	118578069
refGene_gn=NDUFA12;refGene_tx=XM_003989116.5
B4	118578966	118579052
refGene_gn=NDUFA12;refGene_tx=XM_003989116.5
B4	118594157	118594332
refGene_gn=NR2C1;refGene_tx=XM_003989118.5;refGene_gn=NR2C1;refGene_tx=XM_006933928.4;refGene_gn=NR2C1;refGene_tx=XM_006933930.4;refGene_gn=

NR2C1;refGene_tx=XM_006933931.4;refGene_gn=NR2C1;refGene_tx=XM_0198353
54.2;refGene_gn=NR2C1;refGene_tx=XM_019835355.2;refGene_gn=NR2C1;refGe
ne_tx=XM_019835356.2;refGene_gn=NR2C1;refGene_tx=XM_019835357.2;refGen
e_gn=NR2C1;refGene_tx=XM_019835358.2;refGene_gn=NR2C1;refGene_tx=XM_01
9835359.2;refGene_gn=NR2C1;refGene_tx=XM_019835360.2
B4 118597248 118597354
refGene_gn=NR2C1;refGene_tx=XM_003989118.5;refGene_gn=NR2C1;refGene_tx
=XM_006933928.4;refGene_gn=NR2C1;refGene_tx=XM_006933930.4;refGene_gn=
NR2C1;refGene_tx=XM_006933931.4;refGene_gn=NR2C1;refGene_tx=XM_0198353
54.2;refGene_gn=NR2C1;refGene_tx=XM_019835355.2;refGene_gn=NR2C1;refGe
ne_tx=XM_019835356.2;refGene_gn=NR2C1;refGene_tx=XM_019835357.2;refGen
e_gn=NR2C1;refGene_tx=XM_019835358.2;refGene_gn=NR2C1;refGene_tx=XM_01
9835359.2;refGene_gn=NR2C1;refGene_tx=XM_019835360.2
B4 118598680 118598818
refGene_gn=NR2C1;refGene_tx=XM_003989118.5;refGene_gn=NR2C1;refGene_tx
=XM_006933928.4;refGene_gn=NR2C1;refGene_tx=XM_006933930.4;refGene_gn=
NR2C1;refGene_tx=XM_006933931.4;refGene_gn=NR2C1;refGene_tx=XM_0198353
54.2;refGene_gn=NR2C1;refGene_tx=XM_019835355.2;refGene_gn=NR2C1;refGe
ne_tx=XM_019835356.2;refGene_gn=NR2C1;refGene_tx=XM_019835357.2;refGen
e_gn=NR2C1;refGene_tx=XM_019835358.2;refGene_gn=NR2C1;refGene_tx=XM_01
9835359.2;refGene_gn=NR2C1;refGene_tx=XM_019835360.2
B4 118600652 118600792
refGene_gn=NR2C1;refGene_tx=XM_003989118.5;refGene_gn=NR2C1;refGene_tx
=XM_006933928.4;refGene_gn=NR2C1;refGene_tx=XM_006933930.4;refGene_gn=
NR2C1;refGene_tx=XM_006933931.4;refGene_gn=NR2C1;refGene_tx=XM_0198353
54.2;refGene_gn=NR2C1;refGene_tx=XM_019835355.2;refGene_gn=NR2C1;refGe
ne_tx=XM_019835356.2;refGene_gn=NR2C1;refGene_tx=XM_019835357.2;refGen
e_gn=NR2C1;refGene_tx=XM_019835358.2;refGene_gn=NR2C1;refGene_tx=XM_01
9835359.2;refGene_gn=NR2C1;refGene_tx=XM_019835360.2
B4 118607244 118607366
refGene_gn=NR2C1;refGene_tx=XM_003989118.5;refGene_gn=NR2C1;refGene_tx
=XM_006933928.4;refGene_gn=NR2C1;refGene_tx=XM_006933930.4;refGene_gn=
NR2C1;refGene_tx=XM_006933931.4;refGene_gn=NR2C1;refGene_tx=XM_0198353
54.2;refGene_gn=NR2C1;refGene_tx=XM_019835355.2;refGene_gn=NR2C1;refGe
ne_tx=XM_019835356.2;refGene_gn=NR2C1;refGene_tx=XM_019835357.2;refGen
e_gn=NR2C1;refGene_tx=XM_019835358.2;refGene_gn=NR2C1;refGene_tx=XM_01
9835359.2;refGene_gn=NR2C1;refGene_tx=XM_019835360.2
B4 118613776 118613942
refGene_gn=NR2C1;refGene_tx=XM_003989118.5;refGene_gn=NR2C1;refGene_tx
=XM_006933928.4;refGene_gn=NR2C1;refGene_tx=XM_006933930.4;refGene_gn=
NR2C1;refGene_tx=XM_006933931.4;refGene_gn=NR2C1;refGene_tx=XM_0198353
54.2;refGene_gn=NR2C1;refGene_tx=XM_019835355.2;refGene_gn=NR2C1;refGe
ne_tx=XM_019835356.2;refGene_gn=NR2C1;refGene_tx=XM_019835357.2;refGen
e_gn=NR2C1;refGene_tx=XM_019835358.2;refGene_gn=NR2C1;refGene_tx=XM_01
9835359.2;refGene_gn=NR2C1;refGene_tx=XM_019835360.2
B4 118615658 118615843
refGene_gn=NR2C1;refGene_tx=XM_003989118.5;refGene_gn=NR2C1;refGene_tx
=XM_006933928.4;refGene_gn=NR2C1;refGene_tx=XM_006933930.4;refGene_gn=
NR2C1;refGene_tx=XM_006933931.4;refGene_gn=NR2C1;refGene_tx=XM_0198353
54.2;refGene_gn=NR2C1;refGene_tx=XM_019835355.2;refGene_gn=NR2C1;refGe

ne_tx=XM_019835356.2;refGene_gn=NR2C1;refGene_tx=XM_019835357.2;refGen
e_gn=NR2C1;refGene_tx=XM_019835358.2;refGene_gn=NR2C1;refGene_tx=XM_01
9835359.2;refGene_gn=NR2C1;refGene_tx=XM_019835360.2
B4	118619032	118619123
refGene_gn=NR2C1;refGene_tx=XM_003989118.5;refGene_gn=NR2C1;refGene_tx
=XM_006933928.4;refGene_gn=NR2C1;refGene_tx=XM_006933930.4;refGene_gn=
NR2C1;refGene_tx=XM_006933931.4;refGene_gn=NR2C1;refGene_tx=XM_0198353
54.2;refGene_gn=NR2C1;refGene_tx=XM_019835355.2;refGene_gn=NR2C1;refGe
ne_tx=XM_019835356.2;refGene_gn=NR2C1;refGene_tx=XM_019835357.2;refGen
e_gn=NR2C1;refGene_tx=XM_019835358.2;refGene_gn=NR2C1;refGene_tx=XM_01
9835359.2;refGene_gn=NR2C1;refGene_tx=XM_019835360.2
B4	118619260	118619408
refGene_gn=NR2C1;refGene_tx=XM_003989118.5;refGene_gn=NR2C1;refGene_tx
=XM_006933928.4;refGene_gn=NR2C1;refGene_tx=XM_006933930.4;refGene_gn=
NR2C1;refGene_tx=XM_006933931.4;refGene_gn=NR2C1;refGene_tx=XM_0198353
54.2;refGene_gn=NR2C1;refGene_tx=XM_019835355.2;refGene_gn=NR2C1;refGe
ne_tx=XM_019835356.2;refGene_gn=NR2C1;refGene_tx=XM_019835357.2;refGen
e_gn=NR2C1;refGene_tx=XM_019835358.2;refGene_gn=NR2C1;refGene_tx=XM_01
9835359.2;refGene_gn=NR2C1;refGene_tx=XM_019835360.2
B4	118620424	118620604
refGene_gn=NR2C1;refGene_tx=XM_003989118.5;refGene_gn=NR2C1;refGene_tx
=XM_006933928.4;refGene_gn=NR2C1;refGene_tx=XM_006933930.4;refGene_gn=
NR2C1;refGene_tx=XM_006933931.4;refGene_gn=NR2C1;refGene_tx=XM_0198353
54.2;refGene_gn=NR2C1;refGene_tx=XM_019835355.2;refGene_gn=NR2C1;refGe
ne_tx=XM_019835356.2;refGene_gn=NR2C1;refGene_tx=XM_019835357.2;refGen
e_gn=NR2C1;refGene_tx=XM_019835358.2;refGene_gn=NR2C1;refGene_tx=XM_01
9835359.2;refGene_gn=NR2C1;refGene_tx=XM_019835360.2
B4	118622465	118622547
refGene_gn=NR2C1;refGene_tx=XM_003989118.5;refGene_gn=NR2C1;refGene_tx
=XM_006933928.4;refGene_gn=NR2C1;refGene_tx=XM_006933930.4;refGene_gn=
NR2C1;refGene_tx=XM_006933931.4;refGene_gn=NR2C1;refGene_tx=XM_0198353
54.2;refGene_gn=NR2C1;refGene_tx=XM_019835355.2;refGene_gn=NR2C1;refGe
ne_tx=XM_019835356.2;refGene_gn=NR2C1;refGene_tx=XM_019835357.2;refGen
e_gn=NR2C1;refGene_tx=XM_019835358.2;refGene_gn=NR2C1;refGene_tx=XM_01
9835359.2;refGene_gn=NR2C1;refGene_tx=XM_019835360.2
B4	118625544	118625775
refGene_gn=NR2C1;refGene_tx=XM_003989118.5;refGene_gn=NR2C1;refGene_tx
=XM_006933928.4;refGene_gn=NR2C1;refGene_tx=XM_006933930.4;refGene_gn=
NR2C1;refGene_tx=XM_019835354.2;refGene_gn=NR2C1;refGene_tx=XM_0198353
55.2;refGene_gn=NR2C1;refGene_tx=XM_019835356.2;refGene_gn=NR2C1;refGe
ne_tx=XM_019835357.2
B4	118628424	118628478
refGene_gn=NR2C1;refGene_tx=XM_003989118.5;refGene_gn=NR2C1;refGene_tx
=XM_006933928.4;refGene_gn=NR2C1;refGene_tx=XM_006933930.4;refGene_gn=
NR2C1;refGene_tx=XM_006933931.4;refGene_gn=NR2C1;refGene_tx=XM_0198353
54.2;refGene_gn=NR2C1;refGene_tx=XM_019835355.2;refGene_gn=NR2C1;refGe
ne_tx=XM_019835356.2;refGene_gn=NR2C1;refGene_tx=XM_019835357.2;refGen
e_gn=NR2C1;refGene_tx=XM_019835358.2;refGene_gn=NR2C1;refGene_tx=XM_01
9835359.2;refGene_gn=NR2C1;refGene_tx=XM_019835360.2
B4	118646690	118646727

refGene_gn=FGD6;refGene_tx=XM_003989119.5;refGene_gn=FGD6;refGene_tx=XM_019835362.2;refGene_gn=FGD6;refGene_tx=XM_019835363.2;refGene_gn=FGD6;refGene_tx=XM_019835364.2
B4 118649759 118649908
refGene_gn=FGD6;refGene_tx=XM_003989119.5;refGene_gn=FGD6;refGene_tx=XM_019835362.2;refGene_gn=FGD6;refGene_tx=XM_019835363.2;refGene_gn=FGD6;refGene_tx=XM_019835364.2
B4 118650681 118650810
refGene_gn=FGD6;refGene_tx=XM_003989119.5;refGene_gn=FGD6;refGene_tx=XM_019835362.2;refGene_gn=FGD6;refGene_tx=XM_019835363.2;refGene_gn=FGD6;refGene_tx=XM_019835364.2
B4 118656249 118656377
refGene_gn=FGD6;refGene_tx=XM_003989119.5;refGene_gn=FGD6;refGene_tx=XM_019835362.2;refGene_gn=FGD6;refGene_tx=XM_019835363.2;refGene_gn=FGD6;refGene_tx=XM_019835364.2
B4 118658268 118658371
refGene_gn=FGD6;refGene_tx=XM_003989119.5;refGene_gn=FGD6;refGene_tx=XM_019835362.2;refGene_gn=FGD6;refGene_tx=XM_019835363.2;refGene_gn=FGD6;refGene_tx=XM_019835364.2
B4 118659209 118659356
refGene_gn=FGD6;refGene_tx=XM_003989119.5;refGene_gn=FGD6;refGene_tx=XM_019835362.2;refGene_gn=FGD6;refGene_tx=XM_019835363.2;refGene_gn=FGD6;refGene_tx=XM_019835364.2
B4 118659833 118659936
refGene_gn=FGD6;refGene_tx=XM_003989119.5;refGene_gn=FGD6;refGene_tx=XM_019835362.2;refGene_gn=FGD6;refGene_tx=XM_019835363.2;refGene_gn=FGD6;refGene_tx=XM_019835364.2
B4 118667463 118667543
refGene_gn=FGD6;refGene_tx=XM_003989119.5;refGene_gn=FGD6;refGene_tx=XM_019835362.2;refGene_gn=FGD6;refGene_tx=XM_019835363.2;refGene_gn=FGD6;refGene_tx=XM_019835364.2
B4 118668287 118668371
refGene_gn=FGD6;refGene_tx=XM_003989119.5;refGene_gn=FGD6;refGene_tx=XM_019835362.2;refGene_gn=FGD6;refGene_tx=XM_019835363.2;refGene_gn=FGD6;refGene_tx=XM_019835364.2
B4 118669791 118669860
refGene_gn=FGD6;refGene_tx=XM_003989119.5;refGene_gn=FGD6;refGene_tx=XM_019835362.2;refGene_gn=FGD6;refGene_tx=XM_019835363.2;refGene_gn=FGD6;refGene_tx=XM_019835364.2
B4 118670557 118670629
refGene_gn=FGD6;refGene_tx=XM_003989119.5;refGene_gn=FGD6;refGene_tx=XM_019835362.2;refGene_gn=FGD6;refGene_tx=XM_019835363.2;refGene_gn=FGD6;refGene_tx=XM_019835364.2
B4 118670712 118670771
refGene_gn=FGD6;refGene_tx=XM_003989119.5;refGene_gn=FGD6;refGene_tx=XM_019835362.2;refGene_gn=FGD6;refGene_tx=XM_019835363.2;refGene_gn=FGD6;refGene_tx=XM_019835364.2
B4 118674041 118674092
refGene_gn=FGD6;refGene_tx=XM_003989119.5;refGene_gn=FGD6;refGene_tx=XM_019835362.2;refGene_gn=FGD6;refGene_tx=XM_019835363.2;refGene_gn=FGD

6;refGene_tx=XM_019835364.2
B4 118692028 118692116
refGene_gn=FGD6;refGene_tx=XM_003989119.5;refGene_gn=FGD6;refGene_tx=XM_019835362.2;refGene_gn=FGD6;refGene_tx=XM_019835363.2;refGene_gn=FGD6;refGene_tx=XM_019835364.2
B4 118694083 118694240
refGene_gn=FGD6;refGene_tx=XM_003989119.5;refGene_gn=FGD6;refGene_tx=XM_019835362.2;refGene_gn=FGD6;refGene_tx=XM_019835363.2;refGene_gn=FGD6;refGene_tx=XM_019835364.2
B4 118696230 118696382
refGene_gn=FGD6;refGene_tx=XM_003989119.5;refGene_gn=FGD6;refGene_tx=XM_019835362.2;refGene_gn=FGD6;refGene_tx=XM_019835363.2;refGene_gn=FGD6;refGene_tx=XM_019835364.2
B4 118709143 118709174
refGene_gn=FGD6;refGene_tx=XM_003989119.5;refGene_gn=FGD6;refGene_tx=XM_019835362.2;refGene_gn=FGD6;refGene_tx=XM_019835363.2
B4 118709275 118709355
refGene_gn=FGD6;refGene_tx=XM_003989119.5;refGene_gn=FGD6;refGene_tx=XM_019835362.2;refGene_gn=FGD6;refGene_tx=XM_019835363.2
B4 118725068 118725216
refGene_gn=FGD6;refGene_tx=XM_003989119.5;refGene_gn=FGD6;refGene_tx=XM_019835362.2;refGene_gn=FGD6;refGene_tx=XM_019835363.2;refGene_gn=FGD6;refGene_tx=XM_019835364.2
B4 118753375 118755776
refGene_gn=FGD6;refGene_tx=XM_003989119.5;refGene_gn=FGD6;refGene_tx=XM_019835362.2;refGene_gn=FGD6;refGene_tx=XM_019835363.2;refGene_gn=FGD6;refGene_tx=XM_019835364.2
B4 118761183 118761199
refGene_gn=FGD6;refGene_tx=XM_003989119.5;refGene_gn=FGD6;refGene_tx=XM_019835362.2;refGene_gn=FGD6;refGene_tx=XM_019835363.2;refGene_gn=FGD6;refGene_tx=XM_019835364.2
B4 118761839 118761875
refGene_gn=VEZT;refGene_tx=XM_003989120.5;refGene_gn=VEZT;refGene_tx=XM_023257318.1;refGene_gn=VEZT;refGene_tx=XM_023257319.1;refGene_gn=VEZT;refGene_tx=XM_023257321.1;refGene_gn=VEZT;refGene_tx=XM_023257322.1;refGene_gn=VEZT;refGene_tx=XM_023257323.1;refGene_gn=VEZT;refGene_tx=XM_023257324.1;refGene_gn=VEZT;refGene_tx=XM_023257325.1;refGene_gn=VEZT;refGene_tx=XM_023257326.1
B4 118770746 118770755
refGene_gn=VEZT;refGene_tx=XM_023257320.1
B4 118785056 118785191
refGene_gn=VEZT;refGene_tx=XM_003989120.5;refGene_gn=VEZT;refGene_tx=XM_023257318.1;refGene_gn=VEZT;refGene_tx=XM_023257319.1;refGene_gn=VEZT;refGene_tx=XM_023257320.1;refGene_gn=VEZT;refGene_tx=XM_023257321.1;refGene_gn=VEZT;refGene_tx=XM_023257322.1;refGene_gn=VEZT;refGene_tx=XM_023257323.1;refGene_gn=VEZT;refGene_tx=XM_023257324.1;refGene_gn=VEZT;refGene_tx=XM_023257325.1;refGene_gn=VEZT;refGene_tx=XM_023257326.1;refGene_gn=VEZT;refGene_tx=XM_023257327.1;refGene_gn=VEZT;refGene_tx=XM_023257328.1
B4 118788062 118788065

refGene_gn=VEZT;refGene_tx=XM_023257327.1;refGene_gn=VEZT;refGene_tx=XM_023257328.1
B4 118789988 118790075
refGene_gn=VEZT;refGene_tx=XM_003989120.5;refGene_gn=VEZT;refGene_tx=XM_023257318.1;refGene_gn=VEZT;refGene_tx=XM_023257319.1;refGene_gn=VEZT;refGene_tx=XM_023257320.1;refGene_gn=VEZT;refGene_tx=XM_023257321.1;refGene_gn=VEZT;refGene_tx=XM_023257322.1;refGene_gn=VEZT;refGene_tx=XM_023257323.1;refGene_gn=VEZT;refGene_tx=XM_023257324.1;refGene_gn=VEZT;refGene_tx=XM_023257325.1;refGene_gn=VEZT;refGene_tx=XM_023257326.1;refGene_gn=VEZT;refGene_tx=XM_023257327.1;refGene_gn=VEZT;refGene_tx=XM_023257328.1
B4 118791410 118791449
refGene_gn=VEZT;refGene_tx=XM_023257318.1;refGene_gn=VEZT;refGene_tx=XM_023257319.1;refGene_gn=VEZT;refGene_tx=XM_023257320.1;refGene_gn=VEZT;refGene_tx=XM_023257322.1;refGene_gn=VEZT;refGene_tx=XM_023257323.1;refGene_gn=VEZT;refGene_tx=XM_023257324.1
B4 118797368 118797544
refGene_gn=VEZT;refGene_tx=XM_003989120.5;refGene_gn=VEZT;refGene_tx=XM_023257318.1;refGene_gn=VEZT;refGene_tx=XM_023257319.1;refGene_gn=VEZT;refGene_tx=XM_023257320.1;refGene_gn=VEZT;refGene_tx=XM_023257321.1;refGene_gn=VEZT;refGene_tx=XM_023257322.1;refGene_gn=VEZT;refGene_tx=XM_023257323.1;refGene_gn=VEZT;refGene_tx=XM_023257324.1;refGene_gn=VEZT;refGene_tx=XM_023257325.1;refGene_gn=VEZT;refGene_tx=XM_023257326.1;refGene_gn=VEZT;refGene_tx=XM_023257327.1;refGene_gn=VEZT;refGene_tx=XM_023257328.1
B4 118799957 118800233
refGene_gn=VEZT;refGene_tx=XM_003989120.5;refGene_gn=VEZT;refGene_tx=XM_023257318.1;refGene_gn=VEZT;refGene_tx=XM_023257319.1;refGene_gn=VEZT;refGene_tx=XM_023257320.1;refGene_gn=VEZT;refGene_tx=XM_023257321.1;refGene_gn=VEZT;refGene_tx=XM_023257322.1;refGene_gn=VEZT;refGene_tx=XM_023257323.1;refGene_gn=VEZT;refGene_tx=XM_023257324.1;refGene_gn=VEZT;refGene_tx=XM_023257325.1;refGene_gn=VEZT;refGene_tx=XM_023257326.1;refGene_gn=VEZT;refGene_tx=XM_023257327.1;refGene_gn=VEZT;refGene_tx=XM_023257328.1
B4 118802969 118803119
refGene_gn=VEZT;refGene_tx=XM_003989120.5;refGene_gn=VEZT;refGene_tx=XM_023257318.1;refGene_gn=VEZT;refGene_tx=XM_023257319.1;refGene_gn=VEZT;refGene_tx=XM_023257320.1;refGene_gn=VEZT;refGene_tx=XM_023257321.1;refGene_gn=VEZT;refGene_tx=XM_023257322.1;refGene_gn=VEZT;refGene_tx=XM_023257323.1;refGene_gn=VEZT;refGene_tx=XM_023257324.1;refGene_gn=VEZT;refGene_tx=XM_023257325.1;refGene_gn=VEZT;refGene_tx=XM_023257326.1;refGene_gn=VEZT;refGene_tx=XM_023257327.1;refGene_gn=VEZT;refGene_tx=XM_023257328.1
B4 118807612 118807760
refGene_gn=VEZT;refGene_tx=XM_003989120.5;refGene_gn=VEZT;refGene_tx=XM_023257318.1;refGene_gn=VEZT;refGene_tx=XM_023257319.1;refGene_gn=VEZT;refGene_tx=XM_023257320.1;refGene_gn=VEZT;refGene_tx=XM_023257321.1;refGene_gn=VEZT;refGene_tx=XM_023257322.1;refGene_gn=VEZT;refGene_tx=XM_023257323.1;refGene_gn=VEZT;refGene_tx=XM_023257324.1;refGene_gn=VEZT;refGene_tx=XM_023257325.1;refGene_gn=VEZT;refGene_tx=XM_023257326.1;

refGene_gn=VEZT;refGene_tx=XM_023257327.1;refGene_gn=VEZT;refGene_tx=X
M_023257328.1
B4 118809592 118809924
refGene_gn=VEZT;refGene_tx=XM_003989120.5;refGene_gn=VEZT;refGene_tx=X
M_023257318.1;refGene_gn=VEZT;refGene_tx=XM_023257319.1;refGene_gn=VEZ
T;refGene_tx=XM_023257320.1;refGene_gn=VEZT;refGene_tx=XM_023257321.1;
refGene_gn=VEZT;refGene_tx=XM_023257322.1;refGene_gn=VEZT;refGene_tx=X
M_023257323.1;refGene_gn=VEZT;refGene_tx=XM_023257324.1;refGene_gn=VEZ
T;refGene_tx=XM_023257325.1;refGene_gn=VEZT;refGene_tx=XM_023257326.1;
refGene_gn=VEZT;refGene_tx=XM_023257327.1;refGene_gn=VEZT;refGene_tx=X
M_023257328.1
B4 118812338 118812532
refGene_gn=VEZT;refGene_tx=XM_003989120.5;refGene_gn=VEZT;refGene_tx=X
M_023257318.1;refGene_gn=VEZT;refGene_tx=XM_023257319.1;refGene_gn=VEZ
T;refGene_tx=XM_023257320.1;refGene_gn=VEZT;refGene_tx=XM_023257321.1;
refGene_gn=VEZT;refGene_tx=XM_023257322.1;refGene_gn=VEZT;refGene_tx=X
M_023257323.1;refGene_gn=VEZT;refGene_tx=XM_023257324.1;refGene_gn=VEZ
T;refGene_tx=XM_023257325.1;refGene_gn=VEZT;refGene_tx=XM_023257326.1;
refGene_gn=VEZT;refGene_tx=XM_023257327.1;refGene_gn=VEZT;refGene_tx=X
M_023257328.1
B4 118817348 118817449
refGene_gn=VEZT;refGene_tx=XM_003989120.5;refGene_gn=VEZT;refGene_tx=X
M_023257318.1;refGene_gn=VEZT;refGene_tx=XM_023257319.1;refGene_gn=VEZ
T;refGene_tx=XM_023257320.1;refGene_gn=VEZT;refGene_tx=XM_023257321.1;
refGene_gn=VEZT;refGene_tx=XM_023257322.1;refGene_gn=VEZT;refGene_tx=X
M_023257323.1;refGene_gn=VEZT;refGene_tx=XM_023257324.1;refGene_gn=VEZ
T;refGene_tx=XM_023257325.1;refGene_gn=VEZT;refGene_tx=XM_023257326.1;
refGene_gn=VEZT;refGene_tx=XM_023257327.1;refGene_gn=VEZT;refGene_tx=X
M_023257328.1
B4 118819348 118819556
refGene_gn=VEZT;refGene_tx=XM_003989120.5;refGene_gn=VEZT;refGene_tx=X
M_023257318.1;refGene_gn=VEZT;refGene_tx=XM_023257319.1;refGene_gn=VEZ
T;refGene_tx=XM_023257320.1;refGene_gn=VEZT;refGene_tx=XM_023257321.1;
refGene_gn=VEZT;refGene_tx=XM_023257322.1;refGene_gn=VEZT;refGene_tx=X
M_023257323.1;refGene_gn=VEZT;refGene_tx=XM_023257324.1;refGene_gn=VEZ
T;refGene_tx=XM_023257325.1;refGene_gn=VEZT;refGene_tx=XM_023257326.1;
refGene_gn=VEZT;refGene_tx=XM_023257327.1;refGene_gn=VEZT;refGene_tx=X
M_023257328.1
B4 118822288 118822311
refGene_gn=VEZT;refGene_tx=XM_023257323.1;refGene_gn=VEZT;refGene_tx=X
M_023257324.1;refGene_gn=VEZT;refGene_tx=XM_023257325.1;refGene_gn=VEZ
T;refGene_tx=XM_023257326.1;refGene_gn=VEZT;refGene_tx=XM_023257328.1
B4 118823362 118823877
refGene_gn=VEZT;refGene_tx=XM_003989120.5;refGene_gn=VEZT;refGene_tx=X
M_023257318.1;refGene_gn=VEZT;refGene_tx=XM_023257319.1;refGene_gn=VEZ
T;refGene_tx=XM_023257320.1;refGene_gn=VEZT;refGene_tx=XM_023257321.1;
refGene_gn=VEZT;refGene_tx=XM_023257327.1
B4 118834459 118834606
refGene_gn=LOC102901233;refGene_tx=XM_023257317.1
B4 118839920 118839949

refGene_gn=VEZT;refGene_tx=XM_023257322.1
B4 118863107 118863224
refGene_gn=LOC102901233;refGene_tx=XM_023257317.1
B4 118868749 118869047
refGene_gn=LOC102901233;refGene_tx=XM_023257317.1
B4 118879542 118879768
refGene_gn=LOC102901233;refGene_tx=XM_023257317.1
B4 118881628 118881788
refGene_gn=LOC102901233;refGene_tx=XM_023257317.1
B4 118895098 118895200
refGene_gn=LOC102901233;refGene_tx=XM_023257317.1
B4 118896304 118896439
refGene_gn=LOC102901233;refGene_tx=XM_023257317.1
B4 118898903 118899024
refGene_gn=LOC102901233;refGene_tx=XM_023257317.1
B4 118915527 118915677
refGene_gn=LOC102901233;refGene_tx=XM_023257317.1
B4 118917021 118917249
refGene_gn=LOC102901233;refGene_tx=XM_023257317.1
B4 118927682 118927782
refGene_gn=LOC102901233;refGene_tx=XM_023257317.1
B4 118929663 118929762
refGene_gn=LOC102901233;refGene_tx=XM_023257317.1
B4 118937021 118937482
refGene_gn=LOC102901233;refGene_tx=XM_023257317.1
B4 118947229 118947315
refGene_gn=LOC102901233;refGene_tx=XM_023257317.1
B4 118947426 118947486
refGene_gn=LOC102901233;refGene_tx=XM_023257317.1
B4 118961867 118962018
refGene_gn=METAP2;refGene_tx=XM_003989121.5;refGene_gn=METAP2;refGene_tx=XM_006933934.4
B4 118963806 118963914
refGene_gn=METAP2;refGene_tx=XM_003989121.5;refGene_gn=METAP2;refGene_tx=XM_006933934.4
B4 118972066 118972132
refGene_gn=METAP2;refGene_tx=XM_003989121.5;refGene_gn=METAP2;refGene_tx=XM_006933934.4
B4 118975960 118976063
refGene_gn=METAP2;refGene_tx=XM_003989121.5;refGene_gn=METAP2;refGene_tx=XM_006933934.4
B4 118983906 118984068
refGene_gn=METAP2;refGene_tx=XM_003989121.5;refGene_gn=METAP2;refGene_tx=XM_006933934.4
B4 118984806 118984988
refGene_gn=METAP2;refGene_tx=XM_003989121.5;refGene_gn=METAP2;refGene_tx=XM_006933934.4
B4 118985843 118985938
refGene_gn=METAP2;refGene_tx=XM_003989121.5;refGene_gn=METAP2;refGene_tx=XM_006933934.4

B4 118986197 118986294
refGene_gn=METAP2;refGene_tx=XM_003989121.5;refGene_gn=METAP2;refGene_tx=XM_006933934.4
B4 118988497 118988601
refGene_gn=METAP2;refGene_tx=XM_003989121.5;refGene_gn=METAP2;refGene_tx=XM_006933934.4
B4 118989109 118989225
refGene_gn=METAP2;refGene_tx=XM_003989121.5;refGene_gn=METAP2;refGene_tx=XM_006933934.4
B4 118989928 118990181
refGene_gn=METAP2;refGene_tx=XM_003989121.5;refGene_gn=METAP2;refGene_tx=XM_006933934.4
B4 118994439 118994639
refGene_gn=USP44;refGene_tx=XM_023257329.1;refGene_gn=USP44;refGene_tx=XM_023257330.1;refGene_gn=USP44;refGene_tx=XM_023257331.1;refGene_gn=USP44;refGene_tx=XM_023257332.1;refGene_gn=USP44;refGene_tx=XM_023257333.1;refGene_gn=USP44;refGene_tx=XM_023257334.1;refGene_gn=USP44;refGene_tx=XM_023257335.1;refGene_gn=USP44;refGene_tx=XM_023257336.1
B4 118995812 118996018
refGene_gn=USP44;refGene_tx=XM_023257329.1;refGene_gn=USP44;refGene_tx=XM_023257330.1;refGene_gn=USP44;refGene_tx=XM_023257331.1;refGene_gn=USP44;refGene_tx=XM_023257332.1;refGene_gn=USP44;refGene_tx=XM_023257333.1;refGene_gn=USP44;refGene_tx=XM_023257334.1;refGene_gn=USP44;refGene_tx=XM_023257335.1;refGene_gn=USP44;refGene_tx=XM_023257336.1
B4 119001056 119001165
refGene_gn=USP44;refGene_tx=XM_023257329.1;refGene_gn=USP44;refGene_tx=XM_023257330.1;refGene_gn=USP44;refGene_tx=XM_023257331.1;refGene_gn=USP44;refGene_tx=XM_023257332.1;refGene_gn=USP44;refGene_tx=XM_023257333.1;refGene_gn=USP44;refGene_tx=XM_023257334.1;refGene_gn=USP44;refGene_tx=XM_023257335.1;refGene_gn=USP44;refGene_tx=XM_023257336.1
B4 119003531 119003727
refGene_gn=USP44;refGene_tx=XM_023257329.1;refGene_gn=USP44;refGene_tx=XM_023257330.1;refGene_gn=USP44;refGene_tx=XM_023257331.1;refGene_gn=USP44;refGene_tx=XM_023257332.1;refGene_gn=USP44;refGene_tx=XM_023257333.1;refGene_gn=USP44;refGene_tx=XM_023257334.1;refGene_gn=USP44;refGene_tx=XM_023257335.1;refGene_gn=USP44;refGene_tx=XM_023257336.1
B4 119007637 119009062
refGene_gn=USP44;refGene_tx=XM_023257329.1;refGene_gn=USP44;refGene_tx=XM_023257330.1;refGene_gn=USP44;refGene_tx=XM_023257331.1;refGene_gn=USP44;refGene_tx=XM_023257332.1;refGene_gn=USP44;refGene_tx=XM_023257333.1;refGene_gn=USP44;refGene_tx=XM_023257334.1;refGene_gn=USP44;refGene_tx=XM_023257335.1;refGene_gn=USP44;refGene_tx=XM_023257336.1
B4 119120009 119120146
refGene_gn=NTN4;refGene_tx=XM_011284159.3;refGene_gn=NTN4;refGene_tx=XM_023257337.1
B4 119126506 119126677
refGene_gn=NTN4;refGene_tx=XM_011284159.3;refGene_gn=NTN4;refGene_tx=XM_023257337.1
B4 119129772 119129841
refGene_gn=NTN4;refGene_tx=XM_011284159.3;refGene_gn=NTN4;refGene_tx=X

M_023257337.1
B4	119137200	119137316
refGene_gn=NTN4;refGene_tx=XM_011284159.3;refGene_gn=NTN4;refGene_tx=X
M_023257337.1
B4	119137653	119137867
refGene_gn=NTN4;refGene_tx=XM_011284159.3;refGene_gn=NTN4;refGene_tx=X
M_023257337.1
B4	119158383	119158572
refGene_gn=NTN4;refGene_tx=XM_011284159.3;refGene_gn=NTN4;refGene_tx=X
M_023257337.1
B4	119160588	119160715
refGene_gn=NTN4;refGene_tx=XM_011284159.3;refGene_gn=NTN4;refGene_tx=X
M_023257337.1
B4	119195671	119195950
refGene_gn=NTN4;refGene_tx=XM_011284159.3;refGene_gn=NTN4;refGene_tx=X
M_023257337.1
B4	119239913	119240443
refGene_gn=NTN4;refGene_tx=XM_011284159.3;refGene_gn=NTN4;refGene_tx=X
M_023257337.1
B4	119243266	119243321
refGene_gn=NTN4;refGene_tx=XM_011284159.3;refGene_gn=NTN4;refGene_tx=X
M_023257337.1
B4	119285394	119285397
refGene_gn=SNRPF;refGene_tx=XM_006933937.4
B4	119287271	119287397
refGene_gn=SNRPF;refGene_tx=XM_006933937.4
B4	119290797	119290862
refGene_gn=SNRPF;refGene_tx=XM_006933937.4
B4	119291419	119291486
refGene_gn=SNRPF;refGene_tx=XM_006933937.4
B4	119292399	119292513
refGene_gn=CCDC38;refGene_tx=XM_011284160.3
B4	119296223	119296317
refGene_gn=CCDC38;refGene_tx=XM_011284160.3
B4	119298765	119298971
refGene_gn=CCDC38;refGene_tx=XM_011284160.3
B4	119303655	119303791
refGene_gn=CCDC38;refGene_tx=XM_011284160.3
B4	119305068	119305220
refGene_gn=CCDC38;refGene_tx=XM_011284160.3
B4	119306268	119306338
refGene_gn=CCDC38;refGene_tx=XM_011284160.3
B4	119320952	119321010
refGene_gn=CCDC38;refGene_tx=XM_011284160.3
B4	119321688	119321769
refGene_gn=CCDC38;refGene_tx=XM_011284160.3
B4	119323569	119323730
refGene_gn=CCDC38;refGene_tx=XM_011284160.3
B4	119326128	119326209
refGene_gn=CCDC38;refGene_tx=XM_011284160.3

B4	119326319	119326483
refGene_gn=CCDC38;refGene_tx=XM_011284160.3
B4	119329174	119329239
refGene_gn=CCDC38;refGene_tx=XM_011284160.3
B4	119334664	119334830
refGene_gn=CCDC38;refGene_tx=XM_011284160.3
B4	119335990	119336091
refGene_gn=CCDC38;refGene_tx=XM_011284160.3
B4	119339782	119339822
refGene_gn=CCDC38;refGene_tx=XM_011284160.3
B4	119349124	119349261
refGene_gn=AMDHD1;refGene_tx=XM_003989123.5
B4	119350484	119350591
refGene_gn=AMDHD1;refGene_tx=XM_003989123.5
B4	119352237	119352302
refGene_gn=AMDHD1;refGene_tx=XM_003989123.5
B4	119354996	119355274
refGene_gn=AMDHD1;refGene_tx=XM_003989123.5
B4	119357520	119357746
refGene_gn=AMDHD1;refGene_tx=XM_003989123.5
B4	119360394	119360519
refGene_gn=AMDHD1;refGene_tx=XM_003989123.5
B4	119364363	119364457
refGene_gn=AMDHD1;refGene_tx=XM_003989123.5
B4	119365036	119365197
refGene_gn=AMDHD1;refGene_tx=XM_003989123.5
B4	119366059	119366147
refGene_gn=AMDHD1;refGene_tx=XM_003989123.5
B4	119373336	119373477
refGene_gn=HAL;refGene_tx=XM_006933958.3
B4	119376655	119376725
refGene_gn=HAL;refGene_tx=XM_006933958.3
B4	119376824	119376933
refGene_gn=HAL;refGene_tx=XM_006933958.3
B4	119382546	119382681
refGene_gn=HAL;refGene_tx=XM_006933958.3
B4	119385185	119385351
refGene_gn=HAL;refGene_tx=XM_006933958.3
B4	119385434	119385500
refGene_gn=HAL;refGene_tx=XM_006933958.3
B4	119388339	119388420
refGene_gn=HAL;refGene_tx=XM_006933958.3
B4	119389495	119389554
refGene_gn=HAL;refGene_tx=XM_006933958.3
B4	119389655	119389751
refGene_gn=HAL;refGene_tx=XM_006933958.3
B4	119390505	119390653
refGene_gn=HAL;refGene_tx=XM_006933958.3
B4	119391294	119391342
refGene_gn=HAL;refGene_tx=XM_006933958.3

B4 119392683 119392823
refGene_gn=HAL;refGene_tx=XM_006933958.3
B4 119397028 119397154
refGene_gn=HAL;refGene_tx=XM_006933958.3
B4 119397748 119397786
refGene_gn=HAL;refGene_tx=XM_006933958.3
B4 119398094 119398161
refGene_gn=HAL;refGene_tx=XM_006933958.3
B4 119398253 119398326
refGene_gn=HAL;refGene_tx=XM_006933958.3
B4 119398419 119398494
refGene_gn=HAL;refGene_tx=XM_006933958.3
B4 119398828 119398856
refGene_gn=HAL;refGene_tx=XM_006933958.3
B4 119398963 119399024
refGene_gn=HAL;refGene_tx=XM_006933958.3
B4 119399622 119399869
refGene_gn=HAL;refGene_tx=XM_006933958.3
B4 119406594 119406712
refGene_gn=LTA4H;refGene_tx=XM_011284161.3
B4 119407653 119407758
refGene_gn=LTA4H;refGene_tx=XM_011284161.3;refGene_gn=LTA4H;refGene_tx
=XM_019835367.2
B4 119408435 119408518
refGene_gn=LTA4H;refGene_tx=XM_011284161.3
B4 119410462 119410558
refGene_gn=LTA4H;refGene_tx=XM_011284161.3;refGene_gn=LTA4H;refGene_tx
=XM_019835367.2
B4 119417280 119417335
refGene_gn=LTA4H;refGene_tx=XM_011284161.3;refGene_gn=LTA4H;refGene_tx
=XM_019835367.2
B4 119419433 119419504
refGene_gn=LTA4H;refGene_tx=XM_011284161.3;refGene_gn=LTA4H;refGene_tx
=XM_019835367.2
B4 119419858 119419962
refGene_gn=LTA4H;refGene_tx=XM_011284161.3;refGene_gn=LTA4H;refGene_tx
=XM_019835367.2
B4 119421114 119421259
refGene_gn=LTA4H;refGene_tx=XM_011284161.3;refGene_gn=LTA4H;refGene_tx
=XM_019835367.2
B4 119421808 119421920
refGene_gn=LTA4H;refGene_tx=XM_011284161.3;refGene_gn=LTA4H;refGene_tx
=XM_019835367.2
B4 119422604 119422675
refGene_gn=LTA4H;refGene_tx=XM_011284161.3;refGene_gn=LTA4H;refGene_tx
=XM_019835367.2
B4 119423356 119423380
refGene_gn=LTA4H;refGene_tx=XM_011284161.3;refGene_gn=LTA4H;refGene_tx
=XM_019835367.2
B4 119424165 119424306

refGene_gn=LTA4H;refGene_tx=XM_011284161.3;refGene_gn=LTA4H;refGene_tx=XM_019835367.2
B4 119424818 119424891
refGene_gn=LTA4H;refGene_tx=XM_011284161.3;refGene_gn=LTA4H;refGene_tx=XM_019835367.2
B4 119426769 119426822
refGene_gn=LTA4H;refGene_tx=XM_011284161.3;refGene_gn=LTA4H;refGene_tx=XM_019835367.2
B4 119428093 119428198
refGene_gn=LTA4H;refGene_tx=XM_011284161.3;refGene_gn=LTA4H;refGene_tx=XM_019835367.2
B4 119429815 119429884
refGene_gn=LTA4H;refGene_tx=XM_011284161.3;refGene_gn=LTA4H;refGene_tx=XM_019835367.2
B4 119432583 119432704
refGene_gn=LTA4H;refGene_tx=XM_011284161.3;refGene_gn=LTA4H;refGene_tx=XM_019835367.2
B4 119434210 119434341
refGene_gn=LTA4H;refGene_tx=XM_011284161.3;refGene_gn=LTA4H;refGene_tx=XM_019835367.2
B4 119440771 119440930
refGene_gn=LTA4H;refGene_tx=XM_011284161.3;refGene_gn=LTA4H;refGene_tx=XM_019835367.2
B4 119593822 119594029
refGene_gn=ELK3;refGene_tx=XM_023257338.1;refGene_gn=ELK3;refGene_tx=XM_023257339.1
B4 119614788 119615568
refGene_gn=ELK3;refGene_tx=XM_023257338.1;refGene_gn=ELK3;refGene_tx=XM_023257339.1
B4 119625579 119625702
refGene_gn=ELK3;refGene_tx=XM_023257338.1
B4 119631573 119631672
refGene_gn=ELK3;refGene_tx=XM_023257338.1;refGene_gn=ELK3;refGene_tx=XM_023257339.1
B4 119641865 119641903
refGene_gn=CDK17;refGene_tx=XM_011284162.3;refGene_gn=CDK17;refGene_tx=XM_023257340.1;refGene_gn=CDK17;refGene_tx=XM_023257341.1;refGene_gn=CDK17;refGene_tx=XM_023257342.1
B4 119642249 119642257
refGene_gn=CDK17;refGene_tx=XM_023257343.1
B4 119642429 119642507
refGene_gn=CDK17;refGene_tx=XM_011284162.3;refGene_gn=CDK17;refGene_tx=XM_023257340.1;refGene_gn=CDK17;refGene_tx=XM_023257341.1;refGene_gn=CDK17;refGene_tx=XM_023257342.1;refGene_gn=CDK17;refGene_tx=XM_023257343.1
B4 119643813 119643904
refGene_gn=CDK17;refGene_tx=XM_011284162.3;refGene_gn=CDK17;refGene_tx=XM_023257340.1;refGene_gn=CDK17;refGene_tx=XM_023257341.1;refGene_gn=CDK17;refGene_tx=XM_023257342.1;refGene_gn=CDK17;refGene_tx=XM_023257343.1

B4	119644892	119644935
refGene_gn=CDK17;refGene_tx=XM_011284162.3;refGene_gn=CDK17;refGene_tx=XM_023257340.1;refGene_gn=CDK17;refGene_tx=XM_023257341.1;refGene_gn=CDK17;refGene_tx=XM_023257342.1;refGene_gn=CDK17;refGene_tx=XM_023257343.1
B4	119647812	119647918
refGene_gn=CDK17;refGene_tx=XM_011284162.3;refGene_gn=CDK17;refGene_tx=XM_023257340.1;refGene_gn=CDK17;refGene_tx=XM_023257341.1;refGene_gn=CDK17;refGene_tx=XM_023257342.1;refGene_gn=CDK17;refGene_tx=XM_023257343.1
B4	119648560	119648658
refGene_gn=CDK17;refGene_tx=XM_011284162.3;refGene_gn=CDK17;refGene_tx=XM_023257340.1;refGene_gn=CDK17;refGene_tx=XM_023257341.1;refGene_gn=CDK17;refGene_tx=XM_023257342.1;refGene_gn=CDK17;refGene_tx=XM_023257343.1
B4	119649259	119649380
refGene_gn=CDK17;refGene_tx=XM_011284162.3;refGene_gn=CDK17;refGene_tx=XM_023257340.1;refGene_gn=CDK17;refGene_tx=XM_023257341.1;refGene_gn=CDK17;refGene_tx=XM_023257342.1;refGene_gn=CDK17;refGene_tx=XM_023257343.1
B4	119654021	119654145
refGene_gn=CDK17;refGene_tx=XM_011284162.3;refGene_gn=CDK17;refGene_tx=XM_023257340.1;refGene_gn=CDK17;refGene_tx=XM_023257341.1;refGene_gn=CDK17;refGene_tx=XM_023257342.1;refGene_gn=CDK17;refGene_tx=XM_023257343.1
B4	119655131	119655194
refGene_gn=CDK17;refGene_tx=XM_011284162.3;refGene_gn=CDK17;refGene_tx=XM_023257340.1;refGene_gn=CDK17;refGene_tx=XM_023257341.1;refGene_gn=CDK17;refGene_tx=XM_023257342.1;refGene_gn=CDK17;refGene_tx=XM_023257343.1
B4	119655432	119655527
refGene_gn=CDK17;refGene_tx=XM_011284162.3;refGene_gn=CDK17;refGene_tx=XM_023257340.1;refGene_gn=CDK17;refGene_tx=XM_023257341.1;refGene_gn=CDK17;refGene_tx=XM_023257342.1;refGene_gn=CDK17;refGene_tx=XM_023257343.1
B4	119656500	119656615
refGene_gn=CDK17;refGene_tx=XM_011284162.3;refGene_gn=CDK17;refGene_tx=XM_023257340.1;refGene_gn=CDK17;refGene_tx=XM_023257341.1;refGene_gn=CDK17;refGene_tx=XM_023257342.1;refGene_gn=CDK17;refGene_tx=XM_023257343.1
B4	119657656	119657713
refGene_gn=CDK17;refGene_tx=XM_011284162.3;refGene_gn=CDK17;refGene_tx=XM_023257340.1;refGene_gn=CDK17;refGene_tx=XM_023257341.1;refGene_gn=CDK17;refGene_tx=XM_023257342.1;refGene_gn=CDK17;refGene_tx=XM_023257343.1
B4	119666231	119666357
refGene_gn=CDK17;refGene_tx=XM_011284162.3;refGene_gn=CDK17;refGene_tx=XM_023257340.1;refGene_gn=CDK17;refGene_tx=XM_023257341.1;refGene_gn=CDK17;refGene_tx=XM_023257342.1;refGene_gn=CDK17;refGene_tx=XM_023257343.1

B4 119669027 119669161
refGene_gn=CDK17;refGene_tx=XM_011284162.3;refGene_gn=CDK17;refGene_tx=XM_023257340.1;refGene_gn=CDK17;refGene_tx=XM_023257341.1;refGene_gn=CDK17;refGene_tx=XM_023257342.1;refGene_gn=CDK17;refGene_tx=XM_023257343.1
B4 119673322 119673487
refGene_gn=CDK17;refGene_tx=XM_011284162.3;refGene_gn=CDK17;refGene_tx=XM_023257340.1;refGene_gn=CDK17;refGene_tx=XM_023257341.1;refGene_gn=CDK17;refGene_tx=XM_023257342.1;refGene_gn=CDK17;refGene_tx=XM_023257343.1
B4 119678167 119678285
refGene_gn=CDK17;refGene_tx=XM_011284162.3;refGene_gn=CDK17;refGene_tx=XM_023257340.1;refGene_gn=CDK17;refGene_tx=XM_023257341.1;refGene_gn=CDK17;refGene_tx=XM_023257342.1;refGene_gn=CDK17;refGene_tx=XM_023257343.1
B4 119820864 119821193
refGene_gn=CFAP54;refGene_tx=XM_023257344.1
B4 119829039 119829145
refGene_gn=CFAP54;refGene_tx=XM_023257344.1
B4 119832041 119832185
refGene_gn=CFAP54;refGene_tx=XM_023257344.1
B4 119836634 119836806
refGene_gn=CFAP54;refGene_tx=XM_023257344.1
B4 119840781 119840840
refGene_gn=CFAP54;refGene_tx=XM_023257344.1
B4 119848010 119848154
refGene_gn=CFAP54;refGene_tx=XM_023257344.1
B4 119849920 119850034
refGene_gn=CFAP54;refGene_tx=XM_023257344.1
B4 119850157 119850259
refGene_gn=CFAP54;refGene_tx=XM_023257344.1
B4 119854864 119855063
refGene_gn=CFAP54;refGene_tx=XM_023257344.1
B4 119858041 119858204
refGene_gn=CFAP54;refGene_tx=XM_023257344.1
B4 119858287 119858472
refGene_gn=CFAP54;refGene_tx=XM_023257344.1
B4 119859381 119859464
refGene_gn=CFAP54;refGene_tx=XM_023257344.1
B4 119865157 119865292
refGene_gn=CFAP54;refGene_tx=XM_023257344.1
B4 119866291 119866442
refGene_gn=CFAP54;refGene_tx=XM_023257344.1
B4 119870580 119870657
refGene_gn=CFAP54;refGene_tx=XM_023257344.1
B4 119876781 119876910
refGene_gn=CFAP54;refGene_tx=XM_023257344.1
B4 119877275 119877402
refGene_gn=CFAP54;refGene_tx=XM_023257344.1
B4 119889918 119890005

refGene_gn=CFAP54;refGene_tx=XM_023257344.1
B4 119890097 119890219
refGene_gn=CFAP54;refGene_tx=XM_023257344.1
B4 119905165 119905342
refGene_gn=CFAP54;refGene_tx=XM_023257344.1
B4 119908657 119908750
refGene_gn=CFAP54;refGene_tx=XM_023257344.1
B4 119909006 119909192
refGene_gn=CFAP54;refGene_tx=XM_023257344.1
B4 119919628 119919765
refGene_gn=CFAP54;refGene_tx=XM_023257344.1
B4 119923296 119923444
refGene_gn=CFAP54;refGene_tx=XM_023257344.1
B4 119925827 119925983
refGene_gn=CFAP54;refGene_tx=XM_023257344.1
B4 119930849 119930972
refGene_gn=CFAP54;refGene_tx=XM_023257344.1
B4 119939715 119939844
refGene_gn=CFAP54;refGene_tx=XM_023257344.1
B4 119941515 119941630
refGene_gn=CFAP54;refGene_tx=XM_023257344.1
B4 119943407 119943497
refGene_gn=CFAP54;refGene_tx=XM_023257344.1
B4 119944027 119944154
refGene_gn=CFAP54;refGene_tx=XM_023257344.1
B4 119949330 119949445
refGene_gn=CFAP54;refGene_tx=XM_023257344.1
B4 119950449 119950550
refGene_gn=CFAP54;refGene_tx=XM_023257344.1
B4 119959442 119959673
refGene_gn=CFAP54;refGene_tx=XM_023257344.1
B4 119961462 119961520
refGene_gn=CFAP54;refGene_tx=XM_023257344.1
B4 119961521 119961606
refGene_gn=CFAP54;refGene_tx=XM_023257344.1
B4 119963635 119963817
refGene_gn=CFAP54;refGene_tx=XM_023257344.1
B4 119965359 119965587
refGene_gn=CFAP54;refGene_tx=XM_023257344.1
B4 119975913 119976137
refGene_gn=CFAP54;refGene_tx=XM_023257344.1
B4 119976237 119976373
refGene_gn=CFAP54;refGene_tx=XM_023257344.1
B4 119979863 119979972
refGene_gn=CFAP54;refGene_tx=XM_023257344.1
B4 119988113 119988266
refGene_gn=CFAP54;refGene_tx=XM_023257344.1
B4 119990029 119990224
refGene_gn=CFAP54;refGene_tx=XM_023257344.1
B4 119992141 119992229

refGene_gn=CFAP54;refGene_tx=XM_023257344.1
B4	119992531	119992741
refGene_gn=CFAP54;refGene_tx=XM_023257344.1
B4	119994947	119995014
refGene_gn=CFAP54;refGene_tx=XM_023257344.1
B4	119996768	119996951
refGene_gn=CFAP54;refGene_tx=XM_023257344.1
B4	119999355	119999442
refGene_gn=CFAP54;refGene_tx=XM_023257344.1
B4	120004148	120004271
refGene_gn=CFAP54;refGene_tx=XM_023257344.1
B4	120008251	120008305
refGene_gn=CFAP54;refGene_tx=XM_023257344.1
B4	120013577	120013785
refGene_gn=CFAP54;refGene_tx=XM_023257344.1
B4	120015901	120015981
refGene_gn=CFAP54;refGene_tx=XM_023257344.1
B4	120017611	120017781
refGene_gn=CFAP54;refGene_tx=XM_023257344.1
B4	120025330	120025436
refGene_gn=CFAP54;refGene_tx=XM_023257344.1
B4	120027571	120027719
refGene_gn=CFAP54;refGene_tx=XM_023257344.1
B4	120028519	120028677
refGene_gn=CFAP54;refGene_tx=XM_023257344.1
B4	120028843	120029023
refGene_gn=CFAP54;refGene_tx=XM_023257344.1
B4	120029123	120029250
refGene_gn=CFAP54;refGene_tx=XM_023257344.1
B4	120033131	120033287
refGene_gn=CFAP54;refGene_tx=XM_023257344.1
B4	120034596	120034702
refGene_gn=CFAP54;refGene_tx=XM_023257344.1
B4	120035625	120035719
refGene_gn=CFAP54;refGene_tx=XM_023257344.1
B4	120038846	120038945
refGene_gn=CFAP54;refGene_tx=XM_023257344.1
B4	120039481	120039623
refGene_gn=CFAP54;refGene_tx=XM_023257344.1
B4	120055362	120055536
refGene_gn=CFAP54;refGene_tx=XM_023257344.1
B4	120057073	120057294
refGene_gn=CFAP54;refGene_tx=XM_023257344.1
B4	120061010	120061178
refGene_gn=CFAP54;refGene_tx=XM_023257344.1
B4	120074993	120075103
refGene_gn=CFAP54;refGene_tx=XM_023257344.1
B4	120078977	120079119
refGene_gn=CFAP54;refGene_tx=XM_023257344.1
B4	120089727	120089802

refGene_gn=CFAP54;refGene_tx=XM_023257344.1
B4	120123237	120123357
refGene_gn=CFAP54;refGene_tx=XM_023257344.1
B4	120166105	120166241
refGene_gn=NEDD1;refGene_tx=XM_003989128.5;refGene_gn=NEDD1;refGene_tx=XM_006933941.4;refGene_gn=NEDD1;refGene_tx=XM_006933942.4
B4	120168982	120169077
refGene_gn=NEDD1;refGene_tx=XM_003989128.5;refGene_gn=NEDD1;refGene_tx=XM_006933941.4;refGene_gn=NEDD1;refGene_tx=XM_006933942.4
B4	120173164	120173281
refGene_gn=NEDD1;refGene_tx=XM_003989128.5;refGene_gn=NEDD1;refGene_tx=XM_006933941.4;refGene_gn=NEDD1;refGene_tx=XM_006933942.4;refGene_gn=NEDD1;refGene_tx=XM_019835369.2
B4	120176326	120176467
refGene_gn=NEDD1;refGene_tx=XM_003989128.5;refGene_gn=NEDD1;refGene_tx=XM_006933941.4;refGene_gn=NEDD1;refGene_tx=XM_006933942.4;refGene_gn=NEDD1;refGene_tx=XM_019835369.2
B4	120189743	120189973
refGene_gn=NEDD1;refGene_tx=XM_003989128.5;refGene_gn=NEDD1;refGene_tx=XM_006933941.4;refGene_gn=NEDD1;refGene_tx=XM_006933942.4;refGene_gn=NEDD1;refGene_tx=XM_019835369.2
B4	120191377	120191579
refGene_gn=NEDD1;refGene_tx=XM_003989128.5;refGene_gn=NEDD1;refGene_tx=XM_006933941.4;refGene_gn=NEDD1;refGene_tx=XM_006933942.4;refGene_gn=NEDD1;refGene_tx=XM_019835369.2
B4	120191960	120192156
refGene_gn=NEDD1;refGene_tx=XM_003989128.5;refGene_gn=NEDD1;refGene_tx=XM_006933941.4;refGene_gn=NEDD1;refGene_tx=XM_006933942.4;refGene_gn=NEDD1;refGene_tx=XM_019835369.2
B4	120196732	120196861
refGene_gn=NEDD1;refGene_tx=XM_003989128.5;refGene_gn=NEDD1;refGene_tx=XM_006933941.4;refGene_gn=NEDD1;refGene_tx=XM_006933942.4;refGene_gn=NEDD1;refGene_tx=XM_019835369.2
B4	120199581	120199626
refGene_gn=NEDD1;refGene_tx=XM_003989128.5;refGene_gn=NEDD1;refGene_tx=XM_006933941.4;refGene_gn=NEDD1;refGene_tx=XM_006933942.4;refGene_gn=NEDD1;refGene_tx=XM_019835369.2
B4	120200519	120200722
refGene_gn=NEDD1;refGene_tx=XM_003989128.5;refGene_gn=NEDD1;refGene_tx=XM_006933941.4;refGene_gn=NEDD1;refGene_tx=XM_006933942.4;refGene_gn=NEDD1;refGene_tx=XM_019835369.2
B4	120201643	120201797
refGene_gn=NEDD1;refGene_tx=XM_003989128.5;refGene_gn=NEDD1;refGene_tx=XM_006933941.4;refGene_gn=NEDD1;refGene_tx=XM_006933942.4;refGene_gn=NEDD1;refGene_tx=XM_019835369.2
B4	120202938	120203095
refGene_gn=NEDD1;refGene_tx=XM_003989128.5;refGene_gn=NEDD1;refGene_tx=XM_006933941.4;refGene_gn=NEDD1;refGene_tx=XM_006933942.4;refGene_gn=NEDD1;refGene_tx=XM_019835369.2
B4	120209970	120210037

refGene_gn=NEDD1;refGene_tx=XM_003989128.5;refGene_gn=NEDD1;refGene_tx=XM_006933941.4;refGene_gn=NEDD1;refGene_tx=XM_006933942.4;refGene_gn=NEDD1;refGene_tx=XM_019835369.2
B4 120210504 120210609
refGene_gn=NEDD1;refGene_tx=XM_003989128.5;refGene_gn=NEDD1;refGene_tx=XM_006933941.4;refGene_gn=NEDD1;refGene_tx=XM_006933942.4;refGene_gn=NEDD1;refGene_tx=XM_019835369.2
B4 121565705 121565984
refGene_gn=TMPO;refGene_tx=XM_023257345.1;refGene_gn=TMPO;refGene_tx=XM_023257346.1;refGene_gn=TMPO;refGene_tx=XM_023257347.1;refGene_gn=TMPO;refGene_tx=XM_023257348.1;refGene_gn=TMPO;refGene_tx=XM_023257349.1
B4 121574778 121574905
refGene_gn=TMPO;refGene_tx=XM_023257345.1;refGene_gn=TMPO;refGene_tx=XM_023257346.1;refGene_gn=TMPO;refGene_tx=XM_023257347.1;refGene_gn=TMPO;refGene_tx=XM_023257348.1;refGene_gn=TMPO;refGene_tx=XM_023257349.1
B4 121578480 121578639
refGene_gn=TMPO;refGene_tx=XM_023257345.1;refGene_gn=TMPO;refGene_tx=XM_023257346.1;refGene_gn=TMPO;refGene_tx=XM_023257347.1;refGene_gn=TMPO;refGene_tx=XM_023257348.1;refGene_gn=TMPO;refGene_tx=XM_023257349.1
B4 121579478 121580995
refGene_gn=TMPO;refGene_tx=XM_023257345.1
B4 121584119 121584217
refGene_gn=TMPO;refGene_tx=XM_023257346.1;refGene_gn=TMPO;refGene_tx=XM_023257347.1;refGene_gn=TMPO;refGene_tx=XM_023257348.1;refGene_gn=TMPO;refGene_tx=XM_023257349.1
B4 121591232 121591352
refGene_gn=TMPO;refGene_tx=XM_023257346.1
B4 121591451 121591547
refGene_gn=TMPO;refGene_tx=XM_023257346.1;refGene_gn=TMPO;refGene_tx=XM_023257347.1
B4 121591961 121592072
refGene_gn=TMPO;refGene_tx=XM_023257346.1;refGene_gn=TMPO;refGene_tx=XM_023257347.1;refGene_gn=TMPO;refGene_tx=XM_023257348.1
B4 121593114 121593203
refGene_gn=TMPO;refGene_tx=XM_023257346.1;refGene_gn=TMPO;refGene_tx=XM_023257347.1;refGene_gn=TMPO;refGene_tx=XM_023257348.1;refGene_gn=TMPO;refGene_tx=XM_023257349.1
B4 121594021 121594307
refGene_gn=TMPO;refGene_tx=XM_023257346.1;refGene_gn=TMPO;refGene_tx=XM_023257347.1;refGene_gn=TMPO;refGene_tx=XM_023257348.1;refGene_gn=TMPO;refGene_tx=XM_023257349.1
B4 121633837 121633994
refGene_gn=SLC25A3;refGene_tx=XM_006933943.3;refGene_gn=SLC25A3;refGene_tx=XM_006933944.3
B4 121635241 121635366
refGene_gn=SLC25A3;refGene_tx=XM_006933943.3
B4 121635535 121635657
refGene_gn=SLC25A3;refGene_tx=XM_006933944.3
B4 121637467 121637647
refGene_gn=SLC25A3;refGene_tx=XM_006933943.3;refGene_gn=SLC25A3;refGen

e_tx=XM_006933944.3
B4 121638110 121638292
refGene_gn=SLC25A3;refGene_tx=XM_006933943.3;refGene_gn=SLC25A3;refGene_tx=XM_006933944.3
B4 121638919 121639092
refGene_gn=SLC25A3;refGene_tx=XM_006933943.3;refGene_gn=SLC25A3;refGene_tx=XM_006933944.3
B4 121639754 121639865
refGene_gn=SLC25A3;refGene_tx=XM_006933943.3;refGene_gn=SLC25A3;refGene_tx=XM_006933944.3
B4 121639948 121640109
refGene_gn=SLC25A3;refGene_tx=XM_006933943.3;refGene_gn=SLC25A3;refGene_tx=XM_006933944.3
B4 121647855 121648611
refGene_gn=IKBIP;refGene_tx=XM_023257351.1
B4 121654807 121655644
refGene_gn=IKBIP;refGene_tx=XM_023257350.1;refGene_gn=IKBIP;refGene_tx=XM_023257352.1
B4 121662440 121662558
refGene_gn=IKBIP;refGene_tx=XM_023257350.1;refGene_gn=IKBIP;refGene_tx=XM_023257351.1;refGene_gn=IKBIP;refGene_tx=XM_023257352.1
B4 121669977 121670375
refGene_gn=IKBIP;refGene_tx=XM_023257350.1;refGene_gn=IKBIP;refGene_tx=XM_023257351.1;refGene_gn=IKBIP;refGene_tx=XM_023257352.1
B4 121674438 121674576
refGene_gn=APAF1;refGene_tx=XM_019835370.2;refGene_gn=APAF1;refGene_tx=XM_023257353.1;refGene_gn=APAF1;refGene_tx=XM_023257354.1;refGene_gn=APAF1;refGene_tx=XM_023257355.1
B4 121674768 121674961
refGene_gn=APAF1;refGene_tx=XM_019835370.2;refGene_gn=APAF1;refGene_tx=XM_023257353.1;refGene_gn=APAF1;refGene_tx=XM_023257354.1;refGene_gn=APAF1;refGene_tx=XM_023257355.1
B4 121675619 121675817
refGene_gn=APAF1;refGene_tx=XM_019835370.2;refGene_gn=APAF1;refGene_tx=XM_023257353.1;refGene_gn=APAF1;refGene_tx=XM_023257354.1;refGene_gn=APAF1;refGene_tx=XM_023257355.1
B4 121689478 121689662
refGene_gn=APAF1;refGene_tx=XM_019835370.2;refGene_gn=APAF1;refGene_tx=XM_023257353.1;refGene_gn=APAF1;refGene_tx=XM_023257354.1;refGene_gn=APAF1;refGene_tx=XM_023257355.1
B4 121692544 121692657
refGene_gn=APAF1;refGene_tx=XM_019835370.2;refGene_gn=APAF1;refGene_tx=XM_023257353.1;refGene_gn=APAF1;refGene_tx=XM_023257354.1;refGene_gn=APAF1;refGene_tx=XM_023257355.1
B4 121692755 121692887
refGene_gn=APAF1;refGene_tx=XM_019835370.2;refGene_gn=APAF1;refGene_tx=XM_023257353.1;refGene_gn=APAF1;refGene_tx=XM_023257354.1;refGene_gn=APAF1;refGene_tx=XM_023257355.1
B4 121695350 121695589
refGene_gn=APAF1;refGene_tx=XM_019835370.2;refGene_gn=APAF1;refGene_tx

=XM_023257353.1;refGene_gn=APAF1;refGene_tx=XM_023257354.1;refGene_gn=APAF1;refGene_tx=XM_023257355.1
B4 121695996 121696164
refGene_gn=APAF1;refGene_tx=XM_019835370.2;refGene_gn=APAF1;refGene_tx=XM_023257353.1;refGene_gn=APAF1;refGene_tx=XM_023257354.1;refGene_gn=APAF1;refGene_tx=XM_023257355.1
B4 121696876 121697008
refGene_gn=APAF1;refGene_tx=XM_019835370.2;refGene_gn=APAF1;refGene_tx=XM_023257353.1;refGene_gn=APAF1;refGene_tx=XM_023257354.1;refGene_gn=APAF1;refGene_tx=XM_023257355.1
B4 121700371 121700485
refGene_gn=APAF1;refGene_tx=XM_019835370.2;refGene_gn=APAF1;refGene_tx=XM_023257353.1;refGene_gn=APAF1;refGene_tx=XM_023257354.1;refGene_gn=APAF1;refGene_tx=XM_023257355.1
B4 121701328 121701513
refGene_gn=APAF1;refGene_tx=XM_019835370.2;refGene_gn=APAF1;refGene_tx=XM_023257353.1;refGene_gn=APAF1;refGene_tx=XM_023257354.1;refGene_gn=APAF1;refGene_tx=XM_023257355.1
B4 121706071 121706198
refGene_gn=APAF1;refGene_tx=XM_019835370.2;refGene_gn=APAF1;refGene_tx=XM_023257353.1;refGene_gn=APAF1;refGene_tx=XM_023257354.1
B4 121707077 121707203
refGene_gn=APAF1;refGene_tx=XM_019835370.2;refGene_gn=APAF1;refGene_tx=XM_023257353.1;refGene_gn=APAF1;refGene_tx=XM_023257354.1;refGene_gn=APAF1;refGene_tx=XM_023257355.1
B4 121715371 121715503
refGene_gn=APAF1;refGene_tx=XM_019835370.2;refGene_gn=APAF1;refGene_tx=XM_023257353.1;refGene_gn=APAF1;refGene_tx=XM_023257354.1
B4 121719723 121719849
refGene_gn=APAF1;refGene_tx=XM_019835370.2;refGene_gn=APAF1;refGene_tx=XM_023257353.1;refGene_gn=APAF1;refGene_tx=XM_023257354.1
B4 121726788 121726950
refGene_gn=APAF1;refGene_tx=XM_019835370.2;refGene_gn=APAF1;refGene_tx=XM_023257353.1;refGene_gn=APAF1;refGene_tx=XM_023257354.1
B4 121730919 121731048
refGene_gn=APAF1;refGene_tx=XM_019835370.2;refGene_gn=APAF1;refGene_tx=XM_023257353.1;refGene_gn=APAF1;refGene_tx=XM_023257354.1
B4 121733313 121733439
refGene_gn=APAF1;refGene_tx=XM_019835370.2;refGene_gn=APAF1;refGene_tx=XM_023257353.1;refGene_gn=APAF1;refGene_tx=XM_023257354.1
B4 121735967 121736087
refGene_gn=APAF1;refGene_tx=XM_019835370.2;refGene_gn=APAF1;refGene_tx=XM_023257353.1;refGene_gn=APAF1;refGene_tx=XM_023257354.1
B4 121738915 121739032
refGene_gn=APAF1;refGene_tx=XM_019835370.2;refGene_gn=APAF1;refGene_tx=XM_023257353.1;refGene_gn=APAF1;refGene_tx=XM_023257354.1
B4 121741294 121741420
refGene_gn=APAF1;refGene_tx=XM_019835370.2;refGene_gn=APAF1;refGene_tx=XM_023257353.1;refGene_gn=APAF1;refGene_tx=XM_023257354.1
B4 121742372 121742411

refGene_gn=APAF1;refGene_tx=XM_023257354.1
B4 121750077 121750197
refGene_gn=APAF1;refGene_tx=XM_019835370.2;refGene_gn=APAF1;refGene_tx=XM_023257353.1
B4 121750729 121750855
refGene_gn=APAF1;refGene_tx=XM_019835370.2;refGene_gn=APAF1;refGene_tx=XM_023257353.1
B4 121752461 121752587
refGene_gn=APAF1;refGene_tx=XM_019835370.2;refGene_gn=APAF1;refGene_tx=XM_023257353.1
B4 121754786 121754933
refGene_gn=APAF1;refGene_tx=XM_019835370.2;refGene_gn=APAF1;refGene_tx=XM_023257353.1
B4 121760067 121760214
refGene_gn=APAF1;refGene_tx=XM_019835370.2;refGene_gn=APAF1;refGene_tx=XM_023257353.1
B4 121775259 121775386
refGene_gn=ANKS1B;refGene_tx=XM_011284164.3;refGene_gn=ANKS1B;refGene_tx=XM_011284166.3;refGene_gn=ANKS1B;refGene_tx=XM_011284168.3;refGene_gn=ANKS1B;refGene_tx=XM_019835371.2;refGene_gn=ANKS1B;refGene_tx=XM_019835372.2;refGene_gn=ANKS1B;refGene_tx=XM_019835373.2;refGene_gn=ANKS1B;refGene_tx=XM_019835374.2;refGene_gn=ANKS1B;refGene_tx=XM_019835377.2;refGene_gn=ANKS1B;refGene_tx=XM_019835378.2;refGene_gn=ANKS1B;refGene_tx=XM_023257356.1;refGene_gn=ANKS1B;refGene_tx=XM_023257357.1;refGene_gn=ANKS1B;refGene_tx=XM_023257358.1;refGene_gn=ANKS1B;refGene_tx=XM_023257359.1;refGene_gn=ANKS1B;refGene_tx=XM_023257360.1;refGene_gn=ANKS1B;refGene_tx=XM_023257361.1;refGene_gn=ANKS1B;refGene_tx=XM_023257362.1
B4 121786386 121786554
refGene_gn=ANKS1B;refGene_tx=XM_011284164.3;refGene_gn=ANKS1B;refGene_tx=XM_011284166.3;refGene_gn=ANKS1B;refGene_tx=XM_011284168.3;refGene_gn=ANKS1B;refGene_tx=XM_019835371.2;refGene_gn=ANKS1B;refGene_tx=XM_019835372.2;refGene_gn=ANKS1B;refGene_tx=XM_019835373.2;refGene_gn=ANKS1B;refGene_tx=XM_019835374.2;refGene_gn=ANKS1B;refGene_tx=XM_019835376.1;refGene_gn=ANKS1B;refGene_tx=XM_019835377.2;refGene_gn=ANKS1B;refGene_tx=XM_019835378.2;refGene_gn=ANKS1B;refGene_tx=XM_023257356.1;refGene_gn=ANKS1B;refGene_tx=XM_023257357.1;refGene_gn=ANKS1B;refGene_tx=XM_023257358.1;refGene_gn=ANKS1B;refGene_tx=XM_023257359.1;refGene_gn=ANKS1B;refGene_tx=XM_023257360.1;refGene_gn=ANKS1B;refGene_tx=XM_023257361.1;refGene_gn=ANKS1B;refGene_tx=XM_023257362.1
B4 121802812 121802950
refGene_gn=ANKS1B;refGene_tx=XM_011284164.3;refGene_gn=ANKS1B;refGene_tx=XM_011284166.3;refGene_gn=ANKS1B;refGene_tx=XM_011284168.3;refGene_gn=ANKS1B;refGene_tx=XM_019835371.2;refGene_gn=ANKS1B;refGene_tx=XM_019835372.2;refGene_gn=ANKS1B;refGene_tx=XM_019835373.2;refGene_gn=ANKS1B;refGene_tx=XM_019835374.2;refGene_gn=ANKS1B;refGene_tx=XM_019835376.1;refGene_gn=ANKS1B;refGene_tx=XM_019835377.2;refGene_gn=ANKS1B;refGene_tx=XM_019835378.2;refGene_gn=ANKS1B;refGene_tx=XM_023257356.1;refGene_gn=ANKS1B;refGene_tx=XM_023257357.1;refGene_gn=ANKS1B;refGene_tx=XM_023257358.1;refGene_gn=ANKS1B;refGene_tx=XM_023257359.1;refGene_gn=ANK

S1B;refGene_tx=XM_023257360.1;refGene_gn=ANKS1B;refGene_tx=XM_02325736
1.1;refGene_gn=ANKS1B;refGene_tx=XM_023257362.1
B4 121812071 121812158
refGene_gn=ANKS1B;refGene_tx=XM_011284164.3;refGene_gn=ANKS1B;refGene_
tx=XM_011284166.3;refGene_gn=ANKS1B;refGene_tx=XM_011284168.3;refGene_
gn=ANKS1B;refGene_tx=XM_019835371.2;refGene_gn=ANKS1B;refGene_tx=XM_01
9835372.2;refGene_gn=ANKS1B;refGene_tx=XM_019835373.2;refGene_gn=ANKS1
B;refGene_tx=XM_019835374.2;refGene_gn=ANKS1B;refGene_tx=XM_019835376.
1;refGene_gn=ANKS1B;refGene_tx=XM_019835377.2;refGene_gn=ANKS1B;refGen
e_tx=XM_019835378.2;refGene_gn=ANKS1B;refGene_tx=XM_023257356.1;refGen
e_gn=ANKS1B;refGene_tx=XM_023257357.1;refGene_gn=ANKS1B;refGene_tx=XM_
023257358.1;refGene_gn=ANKS1B;refGene_tx=XM_023257359.1;refGene_gn=ANK
S1B;refGene_tx=XM_023257360.1;refGene_gn=ANKS1B;refGene_tx=XM_02325736
1.1;refGene_gn=ANKS1B;refGene_tx=XM_023257362.1
B4 121813049 121813061
refGene_gn=ANKS1B;refGene_tx=XM_011284164.3;refGene_gn=ANKS1B;refGene_
tx=XM_011284166.3;refGene_gn=ANKS1B;refGene_tx=XM_011284168.3;refGene_
gn=ANKS1B;refGene_tx=XM_019835371.2;refGene_gn=ANKS1B;refGene_tx=XM_01
9835372.2;refGene_gn=ANKS1B;refGene_tx=XM_019835374.2;refGene_gn=ANKS1
B;refGene_tx=XM_019835376.1;refGene_gn=ANKS1B;refGene_tx=XM_019835377.
2;refGene_gn=ANKS1B;refGene_tx=XM_019835378.2;refGene_gn=ANKS1B;refGen
e_tx=XM_023257356.1;refGene_gn=ANKS1B;refGene_tx=XM_023257357.1;refGen
e_gn=ANKS1B;refGene_tx=XM_023257358.1;refGene_gn=ANKS1B;refGene_tx=XM_
023257359.1;refGene_gn=ANKS1B;refGene_tx=XM_023257360.1;refGene_gn=ANK
S1B;refGene_tx=XM_023257361.1;refGene_gn=ANKS1B;refGene_tx=XM_02325736
2.1
B4 121830197 121830269
refGene_gn=ANKS1B;refGene_tx=XM_011284164.3;refGene_gn=ANKS1B;refGene_
tx=XM_011284166.3;refGene_gn=ANKS1B;refGene_tx=XM_011284168.3;refGene_
gn=ANKS1B;refGene_tx=XM_019835371.2;refGene_gn=ANKS1B;refGene_tx=XM_01
9835372.2;refGene_gn=ANKS1B;refGene_tx=XM_019835373.2;refGene_gn=ANKS1
B;refGene_tx=XM_019835374.2;refGene_gn=ANKS1B;refGene_tx=XM_019835376.
1;refGene_gn=ANKS1B;refGene_tx=XM_019835377.2;refGene_gn=ANKS1B;refGen
e_tx=XM_019835378.2;refGene_gn=ANKS1B;refGene_tx=XM_023257356.1;refGen
e_gn=ANKS1B;refGene_tx=XM_023257357.1;refGene_gn=ANKS1B;refGene_tx=XM_
023257358.1;refGene_gn=ANKS1B;refGene_tx=XM_023257359.1;refGene_gn=ANK
S1B;refGene_tx=XM_023257360.1;refGene_gn=ANKS1B;refGene_tx=XM_02325736
1.1;refGene_gn=ANKS1B;refGene_tx=XM_023257362.1
B4 121832399 121832528
refGene_gn=ANKS1B;refGene_tx=XM_011284164.3;refGene_gn=ANKS1B;refGene_
tx=XM_011284166.3;refGene_gn=ANKS1B;refGene_tx=XM_011284168.3;refGene_
gn=ANKS1B;refGene_tx=XM_019835371.2;refGene_gn=ANKS1B;refGene_tx=XM_01
9835372.2;refGene_gn=ANKS1B;refGene_tx=XM_019835373.2;refGene_gn=ANKS1
B;refGene_tx=XM_019835374.2;refGene_gn=ANKS1B;refGene_tx=XM_019835376.
1;refGene_gn=ANKS1B;refGene_tx=XM_019835377.2;refGene_gn=ANKS1B;refGen
e_tx=XM_019835378.2;refGene_gn=ANKS1B;refGene_tx=XM_023257356.1;refGen
e_gn=ANKS1B;refGene_tx=XM_023257357.1;refGene_gn=ANKS1B;refGene_tx=XM_
023257358.1;refGene_gn=ANKS1B;refGene_tx=XM_023257359.1;refGene_gn=ANK
S1B;refGene_tx=XM_023257360.1
B4 121838760 121838835

refGene_gn=ANKS1B;refGene_tx=XM_011284164.3;refGene_gn=ANKS1B;refGene_tx=XM_019835371.2;refGene_gn=ANKS1B;refGene_tx=XM_019835372.2;refGene_gn=ANKS1B;refGene_tx=XM_019835373.2;refGene_gn=ANKS1B;refGene_tx=XM_019835374.2;refGene_gn=ANKS1B;refGene_tx=XM_019835376.1;refGene_gn=ANKS1B;refGene_tx=XM_023257356.1;refGene_gn=ANKS1B;refGene_tx=XM_023257358.1;refGene_gn=ANKS1B;refGene_tx=XM_023257359.1;refGene_gn=ANKS1B;refGene_tx=XM_023257360.1
B4 121857848 121858028
refGene_gn=ANKS1B;refGene_tx=XM_011284164.3;refGene_gn=ANKS1B;refGene_tx=XM_011284168.3;refGene_gn=ANKS1B;refGene_tx=XM_019835371.2;refGene_gn=ANKS1B;refGene_tx=XM_019835372.2;refGene_gn=ANKS1B;refGene_tx=XM_019835373.2;refGene_gn=ANKS1B;refGene_tx=XM_019835374.2;refGene_gn=ANKS1B;refGene_tx=XM_019835376.1;refGene_gn=ANKS1B;refGene_tx=XM_019835378.2;refGene_gn=ANKS1B;refGene_tx=XM_023257356.1;refGene_gn=ANKS1B;refGene_tx=XM_023257357.1;refGene_gn=ANKS1B;refGene_tx=XM_023257358.1;refGene_gn=ANKS1B;refGene_tx=XM_023257359.1;refGene_gn=ANKS1B;refGene_tx=XM_023257360.1
B4 121860110 121860218
refGene_gn=ANKS1B;refGene_tx=XM_011284164.3;refGene_gn=ANKS1B;refGene_tx=XM_011284166.3;refGene_gn=ANKS1B;refGene_tx=XM_011284168.3;refGene_gn=ANKS1B;refGene_tx=XM_019835371.2;refGene_gn=ANKS1B;refGene_tx=XM_019835372.2;refGene_gn=ANKS1B;refGene_tx=XM_019835373.2;refGene_gn=ANKS1B;refGene_tx=XM_019835374.2;refGene_gn=ANKS1B;refGene_tx=XM_019835376.1;refGene_gn=ANKS1B;refGene_tx=XM_019835377.2;refGene_gn=ANKS1B;refGene_tx=XM_019835378.2;refGene_gn=ANKS1B;refGene_tx=XM_023257356.1;refGene_gn=ANKS1B;refGene_tx=XM_023257357.1
B4 122061901 122062054
refGene_gn=ANKS1B;refGene_tx=XM_011284164.3;refGene_gn=ANKS1B;refGene_tx=XM_011284166.3;refGene_gn=ANKS1B;refGene_tx=XM_011284168.3;refGene_gn=ANKS1B;refGene_tx=XM_019835371.2;refGene_gn=ANKS1B;refGene_tx=XM_019835372.2;refGene_gn=ANKS1B;refGene_tx=XM_019835373.2;refGene_gn=ANKS1B;refGene_tx=XM_019835374.2;refGene_gn=ANKS1B;refGene_tx=XM_019835376.1;refGene_gn=ANKS1B;refGene_tx=XM_019835377.2;refGene_gn=ANKS1B;refGene_tx=XM_019835378.2;refGene_gn=ANKS1B;refGene_tx=XM_023257356.1;refGene_gn=ANKS1B;refGene_tx=XM_023257357.1
B4 122088644 122088743
refGene_gn=ANKS1B;refGene_tx=XM_011284164.3;refGene_gn=ANKS1B;refGene_tx=XM_011284166.3;refGene_gn=ANKS1B;refGene_tx=XM_011284168.3;refGene_gn=ANKS1B;refGene_tx=XM_019835371.2;refGene_gn=ANKS1B;refGene_tx=XM_019835372.2;refGene_gn=ANKS1B;refGene_tx=XM_019835373.2;refGene_gn=ANKS1B;refGene_tx=XM_019835374.2;refGene_gn=ANKS1B;refGene_tx=XM_019835376.1;refGene_gn=ANKS1B;refGene_tx=XM_019835377.2;refGene_gn=ANKS1B;refGene_tx=XM_019835378.2;refGene_gn=ANKS1B;refGene_tx=XM_023257356.1;refGene_gn=ANKS1B;refGene_tx=XM_023257357.1
B4 122153682 122153886
refGene_gn=ANKS1B;refGene_tx=XM_011284164.3;refGene_gn=ANKS1B;refGene_tx=XM_011284166.3;refGene_gn=ANKS1B;refGene_tx=XM_011284168.3;refGene_gn=ANKS1B;refGene_tx=XM_019835371.2;refGene_gn=ANKS1B;refGene_tx=XM_019835372.2;refGene_gn=ANKS1B;refGene_tx=XM_019835373.2;refGene_gn=ANKS1B;refGene_tx=XM_019835374.2;refGene_gn=ANKS1B;refGene_tx=XM_019835376.

1;refGene_gn=ANKS1B;refGene_tx=XM_019835377.2;refGene_gn=ANKS1B;refGene_tx=XM_019835378.2;refGene_gn=ANKS1B;refGene_tx=XM_023257356.1;refGene_gn=ANKS1B;refGene_tx=XM_023257357.1
B4 122226084 122226157
refGene_gn=ANKS1B;refGene_tx=XM_011284164.3;refGene_gn=ANKS1B;refGene_tx=XM_011284166.3;refGene_gn=ANKS1B;refGene_tx=XM_011284168.3;refGene_gn=ANKS1B;refGene_tx=XM_019835371.2;refGene_gn=ANKS1B;refGene_tx=XM_019835372.2;refGene_gn=ANKS1B;refGene_tx=XM_019835373.2;refGene_gn=ANKS1B;refGene_tx=XM_019835374.2;refGene_gn=ANKS1B;refGene_tx=XM_019835376.1;refGene_gn=ANKS1B;refGene_tx=XM_019835377.2;refGene_gn=ANKS1B;refGene_tx=XM_023257356.1;refGene_gn=ANKS1B;refGene_tx=XM_023257357.1
B4 122228089 122228679
refGene_gn=ANKS1B;refGene_tx=XM_011284164.3;refGene_gn=ANKS1B;refGene_tx=XM_011284166.3;refGene_gn=ANKS1B;refGene_tx=XM_011284168.3;refGene_gn=ANKS1B;refGene_tx=XM_019835371.2;refGene_gn=ANKS1B;refGene_tx=XM_019835372.2;refGene_gn=ANKS1B;refGene_tx=XM_019835373.2;refGene_gn=ANKS1B;refGene_tx=XM_019835374.2;refGene_gn=ANKS1B;refGene_tx=XM_019835376.1;refGene_gn=ANKS1B;refGene_tx=XM_019835377.2;refGene_gn=ANKS1B;refGene_tx=XM_023257356.1;refGene_gn=ANKS1B;refGene_tx=XM_023257357.1
B4 122376268 122376449
refGene_gn=ANKS1B;refGene_tx=XM_011284164.3;refGene_gn=ANKS1B;refGene_tx=XM_011284166.3;refGene_gn=ANKS1B;refGene_tx=XM_011284168.3;refGene_gn=ANKS1B;refGene_tx=XM_019835371.2;refGene_gn=ANKS1B;refGene_tx=XM_019835372.2;refGene_gn=ANKS1B;refGene_tx=XM_019835373.2;refGene_gn=ANKS1B;refGene_tx=XM_019835374.2;refGene_gn=ANKS1B;refGene_tx=XM_019835376.1;refGene_gn=ANKS1B;refGene_tx=XM_019835377.2;refGene_gn=ANKS1B;refGene_tx=XM_023257356.1;refGene_gn=ANKS1B;refGene_tx=XM_023257357.1
B4 122420929 122421066
refGene_gn=ANKS1B;refGene_tx=XM_011284164.3;refGene_gn=ANKS1B;refGene_tx=XM_011284166.3;refGene_gn=ANKS1B;refGene_tx=XM_011284168.3;refGene_gn=ANKS1B;refGene_tx=XM_019835371.2;refGene_gn=ANKS1B;refGene_tx=XM_019835372.2;refGene_gn=ANKS1B;refGene_tx=XM_019835373.2;refGene_gn=ANKS1B;refGene_tx=XM_019835374.2;refGene_gn=ANKS1B;refGene_tx=XM_019835376.1;refGene_gn=ANKS1B;refGene_tx=XM_019835377.2;refGene_gn=ANKS1B;refGene_tx=XM_023257356.1;refGene_gn=ANKS1B;refGene_tx=XM_023257357.1
B4 122449598 122449764
refGene_gn=ANKS1B;refGene_tx=XM_011284164.3;refGene_gn=ANKS1B;refGene_tx=XM_011284166.3;refGene_gn=ANKS1B;refGene_tx=XM_011284168.3;refGene_gn=ANKS1B;refGene_tx=XM_019835371.2;refGene_gn=ANKS1B;refGene_tx=XM_019835372.2;refGene_gn=ANKS1B;refGene_tx=XM_019835373.2;refGene_gn=ANKS1B;refGene_tx=XM_019835374.2;refGene_gn=ANKS1B;refGene_tx=XM_019835376.1;refGene_gn=ANKS1B;refGene_tx=XM_019835377.2;refGene_gn=ANKS1B;refGene_tx=XM_023257356.1;refGene_gn=ANKS1B;refGene_tx=XM_023257357.1
B4 122551465 122552080
refGene_gn=FAM71C;refGene_tx=XM_019835379.1
B4 122552793 122552907
refGene_gn=FAM71C;refGene_tx=XM_019835379.1
B4 122556802 122556946
refGene_gn=ANKS1B;refGene_tx=XM_011284164.3;refGene_gn=ANKS1B;refGene_tx=XM_011284166.3;refGene_gn=ANKS1B;refGene_tx=XM_011284168.3;refGene_

gn=ANKS1B;refGene_tx=XM_019835371.2;refGene_gn=ANKS1B;refGene_tx=XM_01
9835372.2;refGene_gn=ANKS1B;refGene_tx=XM_019835373.2;refGene_gn=ANKS1
B;refGene_tx=XM_019835374.2;refGene_gn=ANKS1B;refGene_tx=XM_019835376.
1;refGene_gn=ANKS1B;refGene_tx=XM_019835377.2;refGene_gn=ANKS1B;refGen
e_tx=XM_023257356.1;refGene_gn=ANKS1B;refGene_tx=XM_023257357.1
B4 122652383 122652550
refGene_gn=ANKS1B;refGene_tx=XM_011284164.3;refGene_gn=ANKS1B;refGene_
tx=XM_011284166.3;refGene_gn=ANKS1B;refGene_tx=XM_011284168.3;refGene_
gn=ANKS1B;refGene_tx=XM_019835371.2;refGene_gn=ANKS1B;refGene_tx=XM_01
9835372.2;refGene_gn=ANKS1B;refGene_tx=XM_019835373.2;refGene_gn=ANKS1
B;refGene_tx=XM_019835374.2;refGene_gn=ANKS1B;refGene_tx=XM_019835376.
1;refGene_gn=ANKS1B;refGene_tx=XM_019835377.2;refGene_gn=ANKS1B;refGen
e_tx=XM_023257356.1;refGene_gn=ANKS1B;refGene_tx=XM_023257357.1
B4 122653491 122653605
refGene_gn=ANKS1B;refGene_tx=XM_011284164.3;refGene_gn=ANKS1B;refGene_
tx=XM_011284166.3;refGene_gn=ANKS1B;refGene_tx=XM_011284168.3;refGene_
gn=ANKS1B;refGene_tx=XM_019835371.2;refGene_gn=ANKS1B;refGene_tx=XM_01
9835372.2;refGene_gn=ANKS1B;refGene_tx=XM_019835373.2;refGene_gn=ANKS1
B;refGene_tx=XM_019835374.2;refGene_gn=ANKS1B;refGene_tx=XM_019835376.
1;refGene_gn=ANKS1B;refGene_tx=XM_019835377.2;refGene_gn=ANKS1B;refGen
e_tx=XM_023257356.1;refGene_gn=ANKS1B;refGene_tx=XM_023257357.1
B4 122658106 122658208
refGene_gn=ANKS1B;refGene_tx=XM_011284164.3;refGene_gn=ANKS1B;refGene_
tx=XM_011284166.3;refGene_gn=ANKS1B;refGene_tx=XM_011284168.3;refGene_
gn=ANKS1B;refGene_tx=XM_019835371.2;refGene_gn=ANKS1B;refGene_tx=XM_01
9835372.2;refGene_gn=ANKS1B;refGene_tx=XM_019835373.2;refGene_gn=ANKS1
B;refGene_tx=XM_019835374.2;refGene_gn=ANKS1B;refGene_tx=XM_019835376.
1;refGene_gn=ANKS1B;refGene_tx=XM_019835377.2;refGene_gn=ANKS1B;refGen
e_tx=XM_023257356.1;refGene_gn=ANKS1B;refGene_tx=XM_023257357.1
B4 122660375 122660451
refGene_gn=ANKS1B;refGene_tx=XM_011284164.3;refGene_gn=ANKS1B;refGene_
tx=XM_011284166.3;refGene_gn=ANKS1B;refGene_tx=XM_011284168.3;refGene_
gn=ANKS1B;refGene_tx=XM_019835371.2;refGene_gn=ANKS1B;refGene_tx=XM_01
9835372.2;refGene_gn=ANKS1B;refGene_tx=XM_019835373.2;refGene_gn=ANKS1
B;refGene_tx=XM_019835374.2;refGene_gn=ANKS1B;refGene_tx=XM_019835376.
1;refGene_gn=ANKS1B;refGene_tx=XM_019835377.2;refGene_gn=ANKS1B;refGen
e_tx=XM_023257356.1;refGene_gn=ANKS1B;refGene_tx=XM_023257357.1
B4 122674572 122674869
refGene_gn=ANKS1B;refGene_tx=XM_011284164.3;refGene_gn=ANKS1B;refGene_
tx=XM_011284166.3;refGene_gn=ANKS1B;refGene_tx=XM_011284168.3;refGene_
gn=ANKS1B;refGene_tx=XM_019835371.2;refGene_gn=ANKS1B;refGene_tx=XM_01
9835372.2;refGene_gn=ANKS1B;refGene_tx=XM_019835373.2;refGene_gn=ANKS1
B;refGene_tx=XM_019835374.2;refGene_gn=ANKS1B;refGene_tx=XM_019835376.
1;refGene_gn=ANKS1B;refGene_tx=XM_019835377.2;refGene_gn=ANKS1B;refGen
e_tx=XM_023257356.1;refGene_gn=ANKS1B;refGene_tx=XM_023257357.1
B4 122681850 122682007
refGene_gn=ANKS1B;refGene_tx=XM_011284164.3;refGene_gn=ANKS1B;refGene_
tx=XM_011284166.3;refGene_gn=ANKS1B;refGene_tx=XM_011284168.3;refGene_
gn=ANKS1B;refGene_tx=XM_019835371.2;refGene_gn=ANKS1B;refGene_tx=XM_01
9835372.2;refGene_gn=ANKS1B;refGene_tx=XM_019835373.2;refGene_gn=ANKS1

B;refGene_tx=XM_019835374.2;refGene_gn=ANKS1B;refGene_tx=XM_019835376.1;refGene_gn=ANKS1B;refGene_tx=XM_019835377.2;refGene_gn=ANKS1B;refGene_tx=XM_023257356.1;refGene_gn=ANKS1B;refGene_tx=XM_023257357.1
B4 122691198 122691279
refGene_gn=ANKS1B;refGene_tx=XM_011284164.3;refGene_gn=ANKS1B;refGene_tx=XM_011284166.3;refGene_gn=ANKS1B;refGene_tx=XM_011284168.3;refGene_gn=ANKS1B;refGene_tx=XM_019835371.2;refGene_gn=ANKS1B;refGene_tx=XM_019835372.2;refGene_gn=ANKS1B;refGene_tx=XM_019835373.2;refGene_gn=ANKS1B;refGene_tx=XM_019835374.2;refGene_gn=ANKS1B;refGene_tx=XM_019835376.1;refGene_gn=ANKS1B;refGene_tx=XM_019835377.2;refGene_gn=ANKS1B;refGene_tx=XM_023257356.1;refGene_gn=ANKS1B;refGene_tx=XM_023257357.1
B4 122852792 122852926
refGene_gn=ANKS1B;refGene_tx=XM_011284164.3;refGene_gn=ANKS1B;refGene_tx=XM_011284166.3;refGene_gn=ANKS1B;refGene_tx=XM_011284168.3;refGene_gn=ANKS1B;refGene_tx=XM_019835371.2;refGene_gn=ANKS1B;refGene_tx=XM_019835372.2;refGene_gn=ANKS1B;refGene_tx=XM_019835373.2;refGene_gn=ANKS1B;refGene_tx=XM_019835374.2;refGene_gn=ANKS1B;refGene_tx=XM_019835376.1;refGene_gn=ANKS1B;refGene_tx=XM_019835377.2;refGene_gn=ANKS1B;refGene_tx=XM_023257356.1;refGene_gn=ANKS1B;refGene_tx=XM_023257357.1
B4 122902389 122902509
refGene_gn=UHRF1BP1L;refGene_tx=XM_011284169.3;refGene_gn=UHRF1BP1L;refGene_tx=XM_011284170.2
B4 122904319 122904513
refGene_gn=UHRF1BP1L;refGene_tx=XM_011284169.3;refGene_gn=UHRF1BP1L;refGene_tx=XM_011284170.2
B4 122920377 122920486
refGene_gn=UHRF1BP1L;refGene_tx=XM_011284169.3;refGene_gn=UHRF1BP1L;refGene_tx=XM_011284170.2
B4 122920708 122920962
refGene_gn=UHRF1BP1L;refGene_tx=XM_011284169.3;refGene_gn=UHRF1BP1L;refGene_tx=XM_011284170.2
B4 122924718 122924869
refGene_gn=UHRF1BP1L;refGene_tx=XM_011284169.3;refGene_gn=UHRF1BP1L;refGene_tx=XM_011284170.2
B4 122925527 122925667
refGene_gn=UHRF1BP1L;refGene_tx=XM_011284169.3;refGene_gn=UHRF1BP1L;refGene_tx=XM_011284170.2
B4 122926371 122926547
refGene_gn=UHRF1BP1L;refGene_tx=XM_011284169.3;refGene_gn=UHRF1BP1L;refGene_tx=XM_011284170.2
B4 122926720 122928219
refGene_gn=UHRF1BP1L;refGene_tx=XM_011284169.3;refGene_gn=UHRF1BP1L;refGene_tx=XM_011284170.2
B4 122929829 122930023
refGene_gn=UHRF1BP1L;refGene_tx=XM_011284169.3;refGene_gn=UHRF1BP1L;refGene_tx=XM_011284170.2
B4 122933977 122934119
refGene_gn=UHRF1BP1L;refGene_tx=XM_011284169.3;refGene_gn=UHRF1BP1L;refGene_tx=XM_011284170.2
B4 122937606 122937725

refGene_gn=UHRF1BP1L;refGene_tx=XM_011284169.3;refGene_gn=UHRF1BP1L;refGene_tx=XM_011284170.2
B4	122938244	122938427
refGene_gn=UHRF1BP1L;refGene_tx=XM_011284169.3;refGene_gn=UHRF1BP1L;refGene_tx=XM_011284170.2
B4	122952951	122953035
refGene_gn=UHRF1BP1L;refGene_tx=XM_011284169.3;refGene_gn=UHRF1BP1L;refGene_tx=XM_011284170.2
B4	122955045	122955226
refGene_gn=UHRF1BP1L;refGene_tx=XM_011284169.3;refGene_gn=UHRF1BP1L;refGene_tx=XM_011284170.2
B4	122959520	122959691
refGene_gn=UHRF1BP1L;refGene_tx=XM_011284169.3;refGene_gn=UHRF1BP1L;refGene_tx=XM_011284170.2
B4	122961086	122961227
refGene_gn=UHRF1BP1L;refGene_tx=XM_011284169.3;refGene_gn=UHRF1BP1L;refGene_tx=XM_011284170.2
B4	122961799	122961989
refGene_gn=UHRF1BP1L;refGene_tx=XM_011284169.3;refGene_gn=UHRF1BP1L;refGene_tx=XM_011284170.2
B4	122968276	122968368
refGene_gn=UHRF1BP1L;refGene_tx=XM_011284169.3;refGene_gn=UHRF1BP1L;refGene_tx=XM_011284170.2
B4	122969474	122969513
refGene_gn=UHRF1BP1L;refGene_tx=XM_011284169.3;refGene_gn=UHRF1BP1L;refGene_tx=XM_011284170.2
B4	122971822	122971985
refGene_gn=UHRF1BP1L;refGene_tx=XM_011284169.3;refGene_gn=UHRF1BP1L;refGene_tx=XM_011284170.2
B4	123015768	123015812
refGene_gn=UHRF1BP1L;refGene_tx=XM_011284169.3
B4	123024520	123024588
refGene_gn=ACTR6;refGene_tx=XM_003989157.4
B4	123026144	123026262
refGene_gn=ACTR6;refGene_tx=XM_003989157.4
B4	123026746	123026815
refGene_gn=ACTR6;refGene_tx=XM_003989157.4
B4	123028441	123028565
refGene_gn=ACTR6;refGene_tx=XM_003989157.4
B4	123030247	123030383
refGene_gn=ACTR6;refGene_tx=XM_003989157.4
B4	123030474	123030531
refGene_gn=ACTR6;refGene_tx=XM_003989157.4
B4	123038407	123038506
refGene_gn=ACTR6;refGene_tx=XM_003989157.4
B4	123038600	123038679
refGene_gn=ACTR6;refGene_tx=XM_003989157.4
B4	123044796	123044968
refGene_gn=ACTR6;refGene_tx=XM_003989157.4
B4	123046859	123046998

refGene_gn=ACTR6;refGene_tx=XM_003989157.4
B4	123051310	123051440
refGene_gn=ACTR6;refGene_tx=XM_003989157.4
B4	123086105	123086282
refGene_gn=SCYL2;refGene_tx=XM_011284172.3;refGene_gn=SCYL2;refGene_tx=XM_019835381.2;refGene_gn=SCYL2;refGene_tx=XM_019835382.2;refGene_gn=SCYL2;refGene_tx=XM_019835383.2
B4	123095251	123095409
refGene_gn=SCYL2;refGene_tx=XM_011284172.3;refGene_gn=SCYL2;refGene_tx=XM_019835381.2;refGene_gn=SCYL2;refGene_tx=XM_019835382.2;refGene_gn=SCYL2;refGene_tx=XM_019835383.2
B4	123099508	123099653
refGene_gn=SCYL2;refGene_tx=XM_011284172.3;refGene_gn=SCYL2;refGene_tx=XM_019835381.2;refGene_gn=SCYL2;refGene_tx=XM_019835382.2;refGene_gn=SCYL2;refGene_tx=XM_019835383.2
B4	123105909	123106059
refGene_gn=SCYL2;refGene_tx=XM_011284172.3;refGene_gn=SCYL2;refGene_tx=XM_019835381.2;refGene_gn=SCYL2;refGene_tx=XM_019835382.2;refGene_gn=SCYL2;refGene_tx=XM_019835383.2
B4	123107372	123107594
refGene_gn=SCYL2;refGene_tx=XM_011284172.3;refGene_gn=SCYL2;refGene_tx=XM_019835381.2;refGene_gn=SCYL2;refGene_tx=XM_019835382.2;refGene_gn=SCYL2;refGene_tx=XM_019835383.2
B4	123108917	123109034
refGene_gn=SCYL2;refGene_tx=XM_011284172.3;refGene_gn=SCYL2;refGene_tx=XM_019835381.2;refGene_gn=SCYL2;refGene_tx=XM_019835382.2;refGene_gn=SCYL2;refGene_tx=XM_019835383.2
B4	123109608	123109734
refGene_gn=SCYL2;refGene_tx=XM_011284172.3;refGene_gn=SCYL2;refGene_tx=XM_019835381.2;refGene_gn=SCYL2;refGene_tx=XM_019835382.2;refGene_gn=SCYL2;refGene_tx=XM_019835383.2
B4	123110549	123110726
refGene_gn=SCYL2;refGene_tx=XM_011284172.3;refGene_gn=SCYL2;refGene_tx=XM_019835381.2;refGene_gn=SCYL2;refGene_tx=XM_019835382.2;refGene_gn=SCYL2;refGene_tx=XM_019835383.2
B4	123112578	123112590
refGene_gn=SCYL2;refGene_tx=XM_019835381.2;refGene_gn=SCYL2;refGene_tx=XM_019835382.2;refGene_gn=SCYL2;refGene_tx=XM_019835383.2
B4	123112701	123112824
refGene_gn=SCYL2;refGene_tx=XM_011284172.3;refGene_gn=SCYL2;refGene_tx=XM_019835381.2;refGene_gn=SCYL2;refGene_tx=XM_019835382.2;refGene_gn=SCYL2;refGene_tx=XM_019835383.2
B4	123116806	123116920
refGene_gn=SCYL2;refGene_tx=XM_011284172.3;refGene_gn=SCYL2;refGene_tx=XM_019835381.2;refGene_gn=SCYL2;refGene_tx=XM_019835382.2;refGene_gn=SCYL2;refGene_tx=XM_019835383.2
B4	123120829	123120962
refGene_gn=SCYL2;refGene_tx=XM_011284172.3;refGene_gn=SCYL2;refGene_tx=XM_019835381.2;refGene_gn=SCYL2;refGene_tx=XM_019835382.2;refGene_gn=SCYL2;refGene_tx=XM_019835383.2

B4 123122774 123122893
refGene_gn=SCYL2;refGene_tx=XM_011284172.3;refGene_gn=SCYL2;refGene_tx=XM_019835381.2;refGene_gn=SCYL2;refGene_tx=XM_019835382.2;refGene_gn=SCYL2;refGene_tx=XM_019835383.2
B4 123132848 123132949
refGene_gn=SCYL2;refGene_tx=XM_011284172.3;refGene_gn=SCYL2;refGene_tx=XM_019835381.2;refGene_gn=SCYL2;refGene_tx=XM_019835382.2;refGene_gn=SCYL2;refGene_tx=XM_019835383.2
B4 123134527 123134594
refGene_gn=SCYL2;refGene_tx=XM_011284172.3;refGene_gn=SCYL2;refGene_tx=XM_019835381.2;refGene_gn=SCYL2;refGene_tx=XM_019835382.2;refGene_gn=SCYL2;refGene_tx=XM_019835383.2
B4 123134729 123134825
refGene_gn=SCYL2;refGene_tx=XM_011284172.3;refGene_gn=SCYL2;refGene_tx=XM_019835381.2;refGene_gn=SCYL2;refGene_tx=XM_019835382.2;refGene_gn=SCYL2;refGene_tx=XM_019835383.2
B4 123135851 123135971
refGene_gn=SCYL2;refGene_tx=XM_011284172.3;refGene_gn=SCYL2;refGene_tx=XM_019835381.2;refGene_gn=SCYL2;refGene_tx=XM_019835382.2
B4 123137275 123137920
refGene_gn=SCYL2;refGene_tx=XM_011284172.3;refGene_gn=SCYL2;refGene_tx=XM_019835381.2;refGene_gn=SCYL2;refGene_tx=XM_019835382.2;refGene_gn=SCYL2;refGene_tx=XM_019835383.2
B4 123152909 123153010
refGene_gn=SLC17A8;refGene_tx=XM_003989158.4
B4 123168270 123168523
refGene_gn=SLC17A8;refGene_tx=XM_003989158.4
B4 123175861 123175980
refGene_gn=SLC17A8;refGene_tx=XM_003989158.4
B4 123177830 123177945
refGene_gn=SLC17A8;refGene_tx=XM_003989158.4
B4 123180511 123180599
refGene_gn=SLC17A8;refGene_tx=XM_003989158.4
B4 123184528 123184615
refGene_gn=SLC17A8;refGene_tx=XM_003989158.4
B4 123185205 123185345
refGene_gn=SLC17A8;refGene_tx=XM_003989158.4
B4 123185473 123185623
refGene_gn=SLC17A8;refGene_tx=XM_003989158.4
B4 123186847 123186980
refGene_gn=SLC17A8;refGene_tx=XM_003989158.4
B4 123193433 123193544
refGene_gn=SLC17A8;refGene_tx=XM_003989158.4
B4 123198128 123198256
refGene_gn=SLC17A8;refGene_tx=XM_003989158.4
B4 123200274 123200619
refGene_gn=SLC17A8;refGene_tx=XM_003989158.4
B4 123264183 123264262
refGene_gn=NR1H4;refGene_tx=XM_011284174.3;refGene_gn=NR1H4;refGene_tx=XM_019835387.2

B4 123273175 123273284
refGene_gn=NR1H4;refGene_tx=XM_019835385.2;refGene_gn=NR1H4;refGene_tx
=XM_019835386.1;refGene_gn=NR1H4;refGene_tx=XM_019835388.2
B4 123280497 123280863
refGene_gn=NR1H4;refGene_tx=XM_011284174.3;refGene_gn=NR1H4;refGene_tx
=XM_019835385.2;refGene_gn=NR1H4;refGene_tx=XM_019835386.1;refGene_gn=
NR1H4;refGene_tx=XM_019835387.2
B4 123295833 123295986
refGene_gn=NR1H4;refGene_tx=XM_011284174.3;refGene_gn=NR1H4;refGene_tx
=XM_019835385.2;refGene_gn=NR1H4;refGene_tx=XM_019835386.1;refGene_gn=
NR1H4;refGene_tx=XM_019835387.2;refGene_gn=NR1H4;refGene_tx=XM_0198353
88.2
B4 123297866 123298000
refGene_gn=NR1H4;refGene_tx=XM_011284174.3;refGene_gn=NR1H4;refGene_tx
=XM_019835385.2;refGene_gn=NR1H4;refGene_tx=XM_019835386.1;refGene_gn=
NR1H4;refGene_tx=XM_019835387.2;refGene_gn=NR1H4;refGene_tx=XM_0198353
88.2
B4 123300049 123300148
refGene_gn=NR1H4;refGene_tx=XM_011284174.3;refGene_gn=NR1H4;refGene_tx
=XM_019835385.2;refGene_gn=NR1H4;refGene_tx=XM_019835386.1;refGene_gn=
NR1H4;refGene_tx=XM_019835387.2;refGene_gn=NR1H4;refGene_tx=XM_0198353
88.2
B4 123300459 123300559
refGene_gn=NR1H4;refGene_tx=XM_011284174.3;refGene_gn=NR1H4;refGene_tx
=XM_019835385.2;refGene_gn=NR1H4;refGene_tx=XM_019835386.1;refGene_gn=
NR1H4;refGene_tx=XM_019835387.2;refGene_gn=NR1H4;refGene_tx=XM_0198353
88.2
B4 123304238 123304385
refGene_gn=NR1H4;refGene_tx=XM_011284174.3;refGene_gn=NR1H4;refGene_tx
=XM_019835385.2;refGene_gn=NR1H4;refGene_tx=XM_019835386.1;refGene_gn=
NR1H4;refGene_tx=XM_019835387.2;refGene_gn=NR1H4;refGene_tx=XM_0198353
88.2
B4 123319862 123319976
refGene_gn=NR1H4;refGene_tx=XM_011284174.3;refGene_gn=NR1H4;refGene_tx
=XM_019835385.2;refGene_gn=NR1H4;refGene_tx=XM_019835386.1;refGene_gn=
NR1H4;refGene_tx=XM_019835387.2;refGene_gn=NR1H4;refGene_tx=XM_0198353
88.2
B4 123321416 123321655
refGene_gn=NR1H4;refGene_tx=XM_011284174.3;refGene_gn=NR1H4;refGene_tx
=XM_019835385.2;refGene_gn=NR1H4;refGene_tx=XM_019835386.1;refGene_gn=
NR1H4;refGene_tx=XM_019835387.2;refGene_gn=NR1H4;refGene_tx=XM_0198353
88.2
B4 123344429 123344453
refGene_gn=GAS2L3;refGene_tx=XM_003989163.3;refGene_gn=GAS2L3;refGene_
tx=XM_019835389.2;refGene_gn=GAS2L3;refGene_tx=XM_019835392.1
B4 123350343 123350512
refGene_gn=GAS2L3;refGene_tx=XM_003989163.3;refGene_gn=GAS2L3;refGene_
tx=XM_019835389.2;refGene_gn=GAS2L3;refGene_tx=XM_019835392.1
B4 123351878 123351994
refGene_gn=GAS2L3;refGene_tx=XM_003989163.3;refGene_gn=GAS2L3;refGene_

tx=XM_019835389.2;refGene_gn=GAS2L3;refGene_tx=XM_019835392.1
B4 123353901 123354043
refGene_gn=GAS2L3;refGene_tx=XM_003989163.3;refGene_gn=GAS2L3;refGene_tx=XM_019835389.2;refGene_gn=GAS2L3;refGene_tx=XM_019835392.1
B4 123358848 123358912
refGene_gn=GAS2L3;refGene_tx=XM_003989163.3;refGene_gn=GAS2L3;refGene_tx=XM_019835389.2;refGene_gn=GAS2L3;refGene_tx=XM_019835392.1
B4 123359849 123359988
refGene_gn=GAS2L3;refGene_tx=XM_003989163.3;refGene_gn=GAS2L3;refGene_tx=XM_019835389.2;refGene_gn=GAS2L3;refGene_tx=XM_019835392.1
B4 123364383 123364491
refGene_gn=GAS2L3;refGene_tx=XM_003989163.3;refGene_gn=GAS2L3;refGene_tx=XM_019835389.2;refGene_gn=GAS2L3;refGene_tx=XM_019835392.1
B4 123366481 123367795
refGene_gn=GAS2L3;refGene_tx=XM_003989163.3;refGene_gn=GAS2L3;refGene_tx=XM_019835389.2;refGene_gn=GAS2L3;refGene_tx=XM_019835392.1
B4 123445475 123445596
refGene_gn=ANO4;refGene_tx=XM_023257364.1;refGene_gn=ANO4;refGene_tx=XM_023257365.1;refGene_gn=ANO4;refGene_tx=XM_023257366.1
B4 123458252 123458336
refGene_gn=ANO4;refGene_tx=XM_023257364.1;refGene_gn=ANO4;refGene_tx=XM_023257365.1;refGene_gn=ANO4;refGene_tx=XM_023257366.1
B4 123466030 123466282
refGene_gn=ANO4;refGene_tx=XM_023257364.1;refGene_gn=ANO4;refGene_tx=XM_023257365.1;refGene_gn=ANO4;refGene_tx=XM_023257366.1
B4 123630733 123630925
refGene_gn=ANO4;refGene_tx=XM_019835393.2;refGene_gn=ANO4;refGene_tx=XM_023257364.1;refGene_gn=ANO4;refGene_tx=XM_023257365.1;refGene_gn=ANO4;refGene_tx=XM_023257366.1
B4 123649068 123649173
refGene_gn=ANO4;refGene_tx=XM_019835393.2;refGene_gn=ANO4;refGene_tx=XM_023257364.1;refGene_gn=ANO4;refGene_tx=XM_023257365.1
B4 123661370 123661507
refGene_gn=ANO4;refGene_tx=XM_019835393.2;refGene_gn=ANO4;refGene_tx=XM_023257364.1;refGene_gn=ANO4;refGene_tx=XM_023257365.1;refGene_gn=ANO4;refGene_tx=XM_023257366.1
B4 123664548 123664707
refGene_gn=ANO4;refGene_tx=XM_019835393.2;refGene_gn=ANO4;refGene_tx=XM_023257364.1;refGene_gn=ANO4;refGene_tx=XM_023257365.1;refGene_gn=ANO4;refGene_tx=XM_023257366.1
B4 123709462 123709563
refGene_gn=ANO4;refGene_tx=XM_019835393.2;refGene_gn=ANO4;refGene_tx=XM_023257364.1;refGene_gn=ANO4;refGene_tx=XM_023257365.1;refGene_gn=ANO4;refGene_tx=XM_023257366.1
B4 123712958 123713047
refGene_gn=ANO4;refGene_tx=XM_019835393.2;refGene_gn=ANO4;refGene_tx=XM_019835401.1;refGene_gn=ANO4;refGene_tx=XM_019835402.1;refGene_gn=ANO4;refGene_tx=XM_023257364.1;refGene_gn=ANO4;refGene_tx=XM_023257366.1
B4 123722780 123722912
refGene_gn=ANO4;refGene_tx=XM_019835393.2;refGene_gn=ANO4;refGene_tx=X

M_019835401.1;refGene_gn=ANO4;refGene_tx=XM_019835402.1;refGene_gn=ANO4;refGene_tx=XM_023257364.1;refGene_gn=ANO4;refGene_tx=XM_023257365.1;refGene_gn=ANO4;refGene_tx=XM_023257366.1
B4 123752226 123752333
refGene_gn=ANO4;refGene_tx=XM_019835393.2;refGene_gn=ANO4;refGene_tx=XM_019835401.1;refGene_gn=ANO4;refGene_tx=XM_019835402.1;refGene_gn=ANO4;refGene_tx=XM_023257364.1;refGene_gn=ANO4;refGene_tx=XM_023257365.1;refGene_gn=ANO4;refGene_tx=XM_023257366.1
B4 123764109 123764165
refGene_gn=ANO4;refGene_tx=XM_019835393.2;refGene_gn=ANO4;refGene_tx=XM_019835401.1;refGene_gn=ANO4;refGene_tx=XM_019835402.1;refGene_gn=ANO4;refGene_tx=XM_023257364.1;refGene_gn=ANO4;refGene_tx=XM_023257365.1;refGene_gn=ANO4;refGene_tx=XM_023257366.1
B4 123766458 123766580
refGene_gn=ANO4;refGene_tx=XM_019835393.2;refGene_gn=ANO4;refGene_tx=XM_019835401.1;refGene_gn=ANO4;refGene_tx=XM_019835402.1;refGene_gn=ANO4;refGene_tx=XM_023257364.1;refGene_gn=ANO4;refGene_tx=XM_023257365.1;refGene_gn=ANO4;refGene_tx=XM_023257366.1
B4 123769918 123770053
refGene_gn=ANO4;refGene_tx=XM_019835393.2;refGene_gn=ANO4;refGene_tx=XM_019835401.1;refGene_gn=ANO4;refGene_tx=XM_019835402.1;refGene_gn=ANO4;refGene_tx=XM_023257364.1;refGene_gn=ANO4;refGene_tx=XM_023257365.1;refGene_gn=ANO4;refGene_tx=XM_023257366.1
B4 123771114 123771211
refGene_gn=ANO4;refGene_tx=XM_019835393.2;refGene_gn=ANO4;refGene_tx=XM_019835401.1;refGene_gn=ANO4;refGene_tx=XM_019835402.1;refGene_gn=ANO4;refGene_tx=XM_023257364.1;refGene_gn=ANO4;refGene_tx=XM_023257365.1;refGene_gn=ANO4;refGene_tx=XM_023257366.1
B4 123776418 123776479
refGene_gn=ANO4;refGene_tx=XM_019835393.2;refGene_gn=ANO4;refGene_tx=XM_019835401.1;refGene_gn=ANO4;refGene_tx=XM_019835402.1;refGene_gn=ANO4;refGene_tx=XM_023257364.1;refGene_gn=ANO4;refGene_tx=XM_023257365.1;refGene_gn=ANO4;refGene_tx=XM_023257366.1
B4 123803236 123803319
refGene_gn=ANO4;refGene_tx=XM_019835393.2;refGene_gn=ANO4;refGene_tx=XM_019835401.1;refGene_gn=ANO4;refGene_tx=XM_019835402.1;refGene_gn=ANO4;refGene_tx=XM_023257364.1;refGene_gn=ANO4;refGene_tx=XM_023257365.1;refGene_gn=ANO4;refGene_tx=XM_023257366.1
B4 123806149 123806290
refGene_gn=ANO4;refGene_tx=XM_019835393.2;refGene_gn=ANO4;refGene_tx=XM_019835401.1;refGene_gn=ANO4;refGene_tx=XM_019835402.1;refGene_gn=ANO4;refGene_tx=XM_023257364.1;refGene_gn=ANO4;refGene_tx=XM_023257365.1;refGene_gn=ANO4;refGene_tx=XM_023257366.1
B4 123809079 123809244
refGene_gn=ANO4;refGene_tx=XM_019835393.2;refGene_gn=ANO4;refGene_tx=XM_019835401.1;refGene_gn=ANO4;refGene_tx=XM_019835402.1;refGene_gn=ANO4;refGene_tx=XM_023257364.1;refGene_gn=ANO4;refGene_tx=XM_023257365.1;refGene_gn=ANO4;refGene_tx=XM_023257366.1
B4 123815282 123815319
refGene_gn=ANO4;refGene_tx=XM_019835393.2;refGene_gn=ANO4;refGene_tx=X

M_019835401.1;refGene_gn=ANO4;refGene_tx=XM_019835402.1;refGene_gn=ANO4;refGene_tx=XM_023257364.1;refGene_gn=ANO4;refGene_tx=XM_023257365.1;refGene_gn=ANO4;refGene_tx=XM_023257366.1
B4	123817762	123817874	refGene_gn=ANO4;refGene_tx=XM_019835393.2;refGene_gn=ANO4;refGene_tx=XM_019835401.1;refGene_gn=ANO4;refGene_tx=XM_019835402.1;refGene_gn=ANO4;refGene_tx=XM_023257364.1;refGene_gn=ANO4;refGene_tx=XM_023257365.1;refGene_gn=ANO4;refGene_tx=XM_023257366.1
B4	123818849	123818907	refGene_gn=ANO4;refGene_tx=XM_019835393.2;refGene_gn=ANO4;refGene_tx=XM_019835401.1;refGene_gn=ANO4;refGene_tx=XM_019835402.1;refGene_gn=ANO4;refGene_tx=XM_023257364.1;refGene_gn=ANO4;refGene_tx=XM_023257365.1;refGene_gn=ANO4;refGene_tx=XM_023257366.1
B4	123819045	123819143	refGene_gn=ANO4;refGene_tx=XM_019835393.2;refGene_gn=ANO4;refGene_tx=XM_019835401.1;refGene_gn=ANO4;refGene_tx=XM_019835402.1;refGene_gn=ANO4;refGene_tx=XM_023257364.1;refGene_gn=ANO4;refGene_tx=XM_023257365.1;refGene_gn=ANO4;refGene_tx=XM_023257366.1
B4	123821437	123821580	refGene_gn=ANO4;refGene_tx=XM_019835393.2;refGene_gn=ANO4;refGene_tx=XM_019835401.1;refGene_gn=ANO4;refGene_tx=XM_019835402.1;refGene_gn=ANO4;refGene_tx=XM_023257364.1;refGene_gn=ANO4;refGene_tx=XM_023257365.1;refGene_gn=ANO4;refGene_tx=XM_023257366.1
B4	123837625	123837778	refGene_gn=ANO4;refGene_tx=XM_019835393.2;refGene_gn=ANO4;refGene_tx=XM_019835401.1;refGene_gn=ANO4;refGene_tx=XM_019835402.1;refGene_gn=ANO4;refGene_tx=XM_023257364.1;refGene_gn=ANO4;refGene_tx=XM_023257365.1;refGene_gn=ANO4;refGene_tx=XM_023257366.1
B4	123838759	123838907	refGene_gn=ANO4;refGene_tx=XM_019835393.2;refGene_gn=ANO4;refGene_tx=XM_019835401.1;refGene_gn=ANO4;refGene_tx=XM_019835402.1;refGene_gn=ANO4;refGene_tx=XM_023257364.1;refGene_gn=ANO4;refGene_tx=XM_023257365.1;refGene_gn=ANO4;refGene_tx=XM_023257366.1
B4	123844891	123845011	refGene_gn=ANO4;refGene_tx=XM_019835393.2;refGene_gn=ANO4;refGene_tx=XM_019835401.1;refGene_gn=ANO4;refGene_tx=XM_019835402.1;refGene_gn=ANO4;refGene_tx=XM_023257364.1;refGene_gn=ANO4;refGene_tx=XM_023257365.1;refGene_gn=ANO4;refGene_tx=XM_023257366.1
B4	123857110	123857216	refGene_gn=ANO4;refGene_tx=XM_019835393.2;refGene_gn=ANO4;refGene_tx=XM_019835401.1;refGene_gn=ANO4;refGene_tx=XM_019835402.1;refGene_gn=ANO4;refGene_tx=XM_023257364.1;refGene_gn=ANO4;refGene_tx=XM_023257365.1;refGene_gn=ANO4;refGene_tx=XM_023257366.1
B4	123858759	123858951	refGene_gn=ANO4;refGene_tx=XM_019835393.2;refGene_gn=ANO4;refGene_tx=XM_019835401.1;refGene_gn=ANO4;refGene_tx=XM_019835402.1;refGene_gn=ANO4;refGene_tx=XM_023257364.1;refGene_gn=ANO4;refGene_tx=XM_023257365.1;refGene_gn=ANO4;refGene_tx=XM_023257366.1
B4	123887964	123888090	refGene_gn=SLC5A8;refGene_tx=XM_023257368.1

B4 123888348 123888428
refGene_gn=SLC5A8;refGene_tx=XM_023257368.1
B4 123893037 123893141
refGene_gn=SLC5A8;refGene_tx=XM_023257368.1
B4 123897439 123897645
refGene_gn=SLC5A8;refGene_tx=XM_023257368.1
B4 123900961 123901048
refGene_gn=SLC5A8;refGene_tx=XM_023257368.1
B4 123912527 123912595
refGene_gn=SLC5A8;refGene_tx=XM_023257368.1
B4 123915308 123915421
refGene_gn=SLC5A8;refGene_tx=XM_023257368.1
B4 123922448 123922537
refGene_gn=SLC5A8;refGene_tx=XM_023257368.1
B4 123926399 123926529
refGene_gn=SLC5A8;refGene_tx=XM_023257368.1
B4 123929911 123930052
refGene_gn=SLC5A8;refGene_tx=XM_023257368.1
B4 123932720 123932875
refGene_gn=SLC5A8;refGene_tx=XM_023257368.1
B4 123933895 123933963
refGene_gn=SLC5A8;refGene_tx=XM_023257368.1
B4 123938538 123938590
refGene_gn=SLC5A8;refGene_tx=XM_023257368.1
B4 123940734 123940800
refGene_gn=SLC5A8;refGene_tx=XM_023257368.1
B4 123945878 123946229
refGene_gn=SLC5A8;refGene_tx=XM_023257368.1
B4 124009613 124009658
refGene_gn=UTP20;refGene_tx=XM_006933978.3;refGene_gn=UTP20;refGene_tx
=XM_019835403.1
B4 124010608 124010689
refGene_gn=UTP20;refGene_tx=XM_006933978.3;refGene_gn=UTP20;refGene_tx
=XM_019835403.1
B4 124014252 124014319
refGene_gn=UTP20;refGene_tx=XM_006933978.3;refGene_gn=UTP20;refGene_tx
=XM_019835403.1
B4 124014456 124014589
refGene_gn=UTP20;refGene_tx=XM_006933978.3;refGene_gn=UTP20;refGene_tx
=XM_019835403.1
B4 124015041 124015230
refGene_gn=UTP20;refGene_tx=XM_006933978.3;refGene_gn=UTP20;refGene_tx
=XM_019835403.1
B4 124018758 124018840
refGene_gn=UTP20;refGene_tx=XM_006933978.3;refGene_gn=UTP20;refGene_tx
=XM_019835403.1
B4 124019525 124019663
refGene_gn=UTP20;refGene_tx=XM_006933978.3;refGene_gn=UTP20;refGene_tx
=XM_019835403.1
B4 124020134 124020290

refGene_gn=UTP20;refGene_tx=XM_006933978.3;refGene_gn=UTP20;refGene_tx=XM_019835403.1
B4 124020888 124021035
refGene_gn=UTP20;refGene_tx=XM_006933978.3;refGene_gn=UTP20;refGene_tx=XM_019835403.1
B4 124021118 124021253
refGene_gn=UTP20;refGene_tx=XM_006933978.3;refGene_gn=UTP20;refGene_tx=XM_019835403.1
B4 124022236 124022314
refGene_gn=UTP20;refGene_tx=XM_006933978.3;refGene_gn=UTP20;refGene_tx=XM_019835403.1
B4 124024143 124024322
refGene_gn=UTP20;refGene_tx=XM_006933978.3;refGene_gn=UTP20;refGene_tx=XM_019835403.1
B4 124028685 124028841
refGene_gn=UTP20;refGene_tx=XM_006933978.3;refGene_gn=UTP20;refGene_tx=XM_019835403.1
B4 124028983 124029072
refGene_gn=UTP20;refGene_tx=XM_006933978.3;refGene_gn=UTP20;refGene_tx=XM_019835403.1
B4 124030529 124030635
refGene_gn=UTP20;refGene_tx=XM_006933978.3;refGene_gn=UTP20;refGene_tx=XM_019835403.1
B4 124034961 124035112
refGene_gn=UTP20;refGene_tx=XM_006933978.3;refGene_gn=UTP20;refGene_tx=XM_019835403.1
B4 124035769 124035832
refGene_gn=UTP20;refGene_tx=XM_006933978.3;refGene_gn=UTP20;refGene_tx=XM_019835403.1
B4 124036616 124036775
refGene_gn=UTP20;refGene_tx=XM_006933978.3;refGene_gn=UTP20;refGene_tx=XM_019835403.1
B4 124037837 124037914
refGene_gn=UTP20;refGene_tx=XM_006933978.3;refGene_gn=UTP20;refGene_tx=XM_019835403.1
B4 124048055 124048135
refGene_gn=UTP20;refGene_tx=XM_006933978.3;refGene_gn=UTP20;refGene_tx=XM_019835403.1
B4 124048377 124048618
refGene_gn=UTP20;refGene_tx=XM_006933978.3;refGene_gn=UTP20;refGene_tx=XM_019835403.1
B4 124053215 124053401
refGene_gn=UTP20;refGene_tx=XM_006933978.3;refGene_gn=UTP20;refGene_tx=XM_019835403.1
B4 124057448 124057539
refGene_gn=UTP20;refGene_tx=XM_006933978.3;refGene_gn=UTP20;refGene_tx=XM_019835403.1
B4 124058151 124058237
refGene_gn=UTP20;refGene_tx=XM_006933978.3;refGene_gn=UTP20;refGene_tx=XM_019835403.1

B4 124058798 124058924
refGene_gn=UTP20;refGene_tx=XM_006933978.3;refGene_gn=UTP20;refGene_tx
=XM_019835403.1
B4 124062500 124062667
refGene_gn=UTP20;refGene_tx=XM_006933978.3;refGene_gn=UTP20;refGene_tx
=XM_019835403.1
B4 124064809 124065018
refGene_gn=UTP20;refGene_tx=XM_006933978.3;refGene_gn=UTP20;refGene_tx
=XM_019835403.1
B4 124071191 124071335
refGene_gn=UTP20;refGene_tx=XM_006933978.3;refGene_gn=UTP20;refGene_tx
=XM_019835403.1
B4 124072043 124072123
refGene_gn=UTP20;refGene_tx=XM_006933978.3;refGene_gn=UTP20;refGene_tx
=XM_019835403.1
B4 124076874 124077092
refGene_gn=UTP20;refGene_tx=XM_006933978.3;refGene_gn=UTP20;refGene_tx
=XM_019835403.1
B4 124077635 124077780
refGene_gn=UTP20;refGene_tx=XM_006933978.3;refGene_gn=UTP20;refGene_tx
=XM_019835403.1
B4 124078997 124079085
refGene_gn=UTP20;refGene_tx=XM_006933978.3;refGene_gn=UTP20;refGene_tx
=XM_019835403.1
B4 124080188 124080332
refGene_gn=UTP20;refGene_tx=XM_006933978.3;refGene_gn=UTP20;refGene_tx
=XM_019835403.1
B4 124080531 124080582
refGene_gn=UTP20;refGene_tx=XM_006933978.3;refGene_gn=UTP20;refGene_tx
=XM_019835403.1
B4 124080694 124080847
refGene_gn=UTP20;refGene_tx=XM_006933978.3;refGene_gn=UTP20;refGene_tx
=XM_019835403.1
B4 124082940 124083096
refGene_gn=UTP20;refGene_tx=XM_006933978.3;refGene_gn=UTP20;refGene_tx
=XM_019835403.1
B4 124083437 124083578
refGene_gn=UTP20;refGene_tx=XM_006933978.3;refGene_gn=UTP20;refGene_tx
=XM_019835403.1
B4 124084244 124084382
refGene_gn=UTP20;refGene_tx=XM_006933978.3;refGene_gn=UTP20;refGene_tx
=XM_019835403.1
B4 124088792 124088932
refGene_gn=UTP20;refGene_tx=XM_006933978.3;refGene_gn=UTP20;refGene_tx
=XM_019835403.1
B4 124089182 124089265
refGene_gn=UTP20;refGene_tx=XM_006933978.3;refGene_gn=UTP20;refGene_tx
=XM_019835403.1
B4 124091387 124091704
refGene_gn=UTP20;refGene_tx=XM_006933978.3;refGene_gn=UTP20;refGene_tx

=XM_019835403.1
B4	124093244	124093384
refGene_gn=UTP20;refGene_tx=XM_006933978.3;refGene_gn=UTP20;refGene_tx=XM_019835403.1
B4	124093830	124093987
refGene_gn=UTP20;refGene_tx=XM_006933978.3;refGene_gn=UTP20;refGene_tx=XM_019835403.1
B4	124098892	124098991
refGene_gn=UTP20;refGene_tx=XM_006933978.3;refGene_gn=UTP20;refGene_tx=XM_019835403.1
B4	124099832	124100000
refGene_gn=UTP20;refGene_tx=XM_006933978.3;refGene_gn=UTP20;refGene_tx=XM_019835403.1
B4	124101373	124101540
refGene_gn=UTP20;refGene_tx=XM_006933978.3;refGene_gn=UTP20;refGene_tx=XM_019835403.1
B4	124102584	124102726
refGene_gn=UTP20;refGene_tx=XM_006933978.3;refGene_gn=UTP20;refGene_tx=XM_019835403.1
B4	124103820	124103937
refGene_gn=UTP20;refGene_tx=XM_006933978.3;refGene_gn=UTP20;refGene_tx=XM_019835403.1
B4	124105407	124105578
refGene_gn=UTP20;refGene_tx=XM_006933978.3;refGene_gn=UTP20;refGene_tx=XM_019835403.1
B4	124105811	124105943
refGene_gn=UTP20;refGene_tx=XM_006933978.3;refGene_gn=UTP20;refGene_tx=XM_019835403.1
B4	124106405	124106516
refGene_gn=UTP20;refGene_tx=XM_006933978.3;refGene_gn=UTP20;refGene_tx=XM_019835403.1
B4	124108109	124108189
refGene_gn=UTP20;refGene_tx=XM_006933978.3;refGene_gn=UTP20;refGene_tx=XM_019835403.1
B4	124108615	124108685
refGene_gn=UTP20;refGene_tx=XM_006933978.3;refGene_gn=UTP20;refGene_tx=XM_019835403.1
B4	124108813	124108996
refGene_gn=UTP20;refGene_tx=XM_006933978.3;refGene_gn=UTP20;refGene_tx=XM_019835403.1
B4	124110220	124110352
refGene_gn=UTP20;refGene_tx=XM_006933978.3;refGene_gn=UTP20;refGene_tx=XM_019835403.1
B4	124110801	124110934
refGene_gn=UTP20;refGene_tx=XM_006933978.3;refGene_gn=UTP20;refGene_tx=XM_019835403.1
B4	124112502	124112690
refGene_gn=UTP20;refGene_tx=XM_006933978.3;refGene_gn=UTP20;refGene_tx=XM_019835403.1
B4	124113413	124113485

refGene_gn=UTP20;refGene_tx=XM_006933978.3;refGene_gn=UTP20;refGene_tx=XM_019835403.1
B4 124115436 124115736
refGene_gn=UTP20;refGene_tx=XM_006933978.3;refGene_gn=UTP20;refGene_tx=XM_019835403.1
B4 124115948 124116075
refGene_gn=UTP20;refGene_tx=XM_006933978.3;refGene_gn=UTP20;refGene_tx=XM_019835403.1
B4 124117354 124117500
refGene_gn=UTP20;refGene_tx=XM_006933978.3;refGene_gn=UTP20;refGene_tx=XM_019835403.1
B4 124117779 124117935
refGene_gn=UTP20;refGene_tx=XM_006933978.3;refGene_gn=UTP20;refGene_tx=XM_019835403.1
B4 124122365 124122396
refGene_gn=ARL1;refGene_tx=XM_003989168.5
B4 124123154 124123333
refGene_gn=ARL1;refGene_tx=XM_003989168.5
B4 124127024 124127136
refGene_gn=ARL1;refGene_tx=XM_003989168.5
B4 124128535 124128617
refGene_gn=ARL1;refGene_tx=XM_003989168.5
B4 124131047 124131185
refGene_gn=ARL1;refGene_tx=XM_003989168.5
B4 124133976 124133980
refGene_gn=ARL1;refGene_tx=XM_003989168.5
B4 124177297 124177300
refGene_gn=SPIC;refGene_tx=XM_019835404.1;refGene_gn=SPIC;refGene_tx=XM_019835405.1;refGene_gn=SPIC;refGene_tx=XM_019835406.1;refGene_gn=SPIC;refGene_tx=XM_019835407.1
B4 124178085 124178182
refGene_gn=SPIC;refGene_tx=XM_019835404.1;refGene_gn=SPIC;refGene_tx=XM_019835405.1;refGene_gn=SPIC;refGene_tx=XM_019835406.1;refGene_gn=SPIC;refGene_tx=XM_019835407.1
B4 124179097 124179234
refGene_gn=SPIC;refGene_tx=XM_019835404.1;refGene_gn=SPIC;refGene_tx=XM_019835405.1;refGene_gn=SPIC;refGene_tx=XM_019835406.1;refGene_gn=SPIC;refGene_tx=XM_019835407.1
B4 124180674 124180783
refGene_gn=SPIC;refGene_tx=XM_019835404.1;refGene_gn=SPIC;refGene_tx=XM_019835405.1;refGene_gn=SPIC;refGene_tx=XM_019835406.1;refGene_gn=SPIC;refGene_tx=XM_019835407.1
B4 124184101 124184529
refGene_gn=SPIC;refGene_tx=XM_019835404.1;refGene_gn=SPIC;refGene_tx=XM_019835405.1;refGene_gn=SPIC;refGene_tx=XM_019835406.1;refGene_gn=SPIC;refGene_tx=XM_019835407.1
B4 124242815 124242840
refGene_gn=MYBPC1;refGene_tx=XM_023257370.1
B4 124259988 124260039
refGene_gn=MYBPC1;refGene_tx=XM_023257370.1

B4 124262404 124262446
refGene_gn=MYBPC1;refGene_tx=XM_023257370.1
B4 124271036 124271075
refGene_gn=MYBPC1;refGene_tx=XM_023257370.1
B4 124271575 124271587
refGene_gn=MYBPC1;refGene_tx=XM_023257370.1
B4 124271589 124271610
refGene_gn=MYBPC1;refGene_tx=XM_023257370.1
B4 124273243 124273354
refGene_gn=MYBPC1;refGene_tx=XM_023257370.1
B4 124275108 124275257
refGene_gn=MYBPC1;refGene_tx=XM_023257370.1
B4 124275578 124275696
refGene_gn=MYBPC1;refGene_tx=XM_023257370.1
B4 124277170 124277222
refGene_gn=MYBPC1;refGene_tx=XM_023257370.1
B4 124279258 124279315
refGene_gn=MYBPC1;refGene_tx=XM_023257370.1
B4 124285022 124285189
refGene_gn=MYBPC1;refGene_tx=XM_023257370.1
B4 124287416 124287549
refGene_gn=MYBPC1;refGene_tx=XM_023257370.1
B4 124289384 124289509
refGene_gn=MYBPC1;refGene_tx=XM_023257370.1
B4 124290417 124290523
refGene_gn=MYBPC1;refGene_tx=XM_023257370.1
B4 124291704 124291871
refGene_gn=MYBPC1;refGene_tx=XM_023257370.1
B4 124293307 124293470
refGene_gn=MYBPC1;refGene_tx=XM_023257370.1
B4 124295150 124295257
refGene_gn=MYBPC1;refGene_tx=XM_023257370.1
B4 124295568 124295702
refGene_gn=MYBPC1;refGene_tx=XM_023257370.1
B4 124300614 124300774
refGene_gn=MYBPC1;refGene_tx=XM_023257370.1
B4 124301824 124301929
refGene_gn=MYBPC1;refGene_tx=XM_023257370.1
B4 124302708 124302897
refGene_gn=MYBPC1;refGene_tx=XM_023257370.1
B4 124303761 124303896
refGene_gn=MYBPC1;refGene_tx=XM_023257370.1
B4 124307191 124307245
refGene_gn=MYBPC1;refGene_tx=XM_023257370.1
B4 124308224 124308392
refGene_gn=MYBPC1;refGene_tx=XM_023257370.1
B4 124312769 124312858
refGene_gn=MYBPC1;refGene_tx=XM_023257370.1
B4 124315783 124315979
refGene_gn=MYBPC1;refGene_tx=XM_023257370.1

B4 124317976 124318116
refGene_gn=MYBPC1;refGene_tx=XM_023257370.1
B4 124321053 124321213
refGene_gn=MYBPC1;refGene_tx=XM_023257370.1
B4 124322589 124322726
refGene_gn=MYBPC1;refGene_tx=XM_023257370.1
B4 124324575 124324762
refGene_gn=MYBPC1;refGene_tx=XM_023257370.1
B4 124329136 124329237
refGene_gn=MYBPC1;refGene_tx=XM_023257370.1
B4 124344277 124344550
refGene_gn=CHPT1;refGene_tx=XM_019835408.2
B4 124360201 124360240
refGene_gn=CHPT1;refGene_tx=XM_019835408.2
B4 124360241 124360350
refGene_gn=CHPT1;refGene_tx=XM_019835408.2
B4 124360662 124360804
refGene_gn=CHPT1;refGene_tx=XM_019835408.2
B4 124367978 124368063
refGene_gn=CHPT1;refGene_tx=XM_019835408.2
B4 124370678 124370810
refGene_gn=CHPT1;refGene_tx=XM_019835408.2
B4 124372836 124372857
refGene_gn=SYCP3;refGene_tx=XM_011284177.3;refGene_gn=SYCP3;refGene_tx=XM_019835409.2;refGene_gn=SYCP3;refGene_tx=XM_019835410.2
B4 124374010 124374169
refGene_gn=CHPT1;refGene_tx=XM_019835408.2
B4 124374622 124374748
refGene_gn=CHPT1;refGene_tx=XM_019835408.2
B4 124377341 124377452
refGene_gn=CHPT1;refGene_tx=XM_019835408.2
B4 124380171 124380251
refGene_gn=CHPT1;refGene_tx=XM_019835408.2;refGene_gn=SYCP3;refGene_tx=XM_023257371.1
B4 124380382 124380487
refGene_gn=SYCP3;refGene_tx=XM_011284177.3;refGene_gn=SYCP3;refGene_tx=XM_019835409.2;refGene_gn=SYCP3;refGene_tx=XM_019835410.2;refGene_gn=SYCP3;refGene_tx=XM_023257371.1
B4 124383262 124383361
refGene_gn=SYCP3;refGene_tx=XM_011284177.3;refGene_gn=SYCP3;refGene_tx=XM_019835409.2;refGene_gn=SYCP3;refGene_tx=XM_023257371.1
B4 124386028 124386128
refGene_gn=SYCP3;refGene_tx=XM_011284177.3;refGene_gn=SYCP3;refGene_tx=XM_019835409.2;refGene_gn=SYCP3;refGene_tx=XM_019835410.2;refGene_gn=SYCP3;refGene_tx=XM_023257371.1
B4 124386972 124387090
refGene_gn=SYCP3;refGene_tx=XM_011284177.3;refGene_gn=SYCP3;refGene_tx=XM_019835409.2;refGene_gn=SYCP3;refGene_tx=XM_019835410.2;refGene_gn=SYCP3;refGene_tx=XM_023257371.1
B4 124387366 124387401

refGene_gn=SYCP3;refGene_tx=XM_011284177.3;refGene_gn=SYCP3;refGene_tx=XM_019835409.2;refGene_gn=SYCP3;refGene_tx=XM_019835410.2;refGene_gn=SYCP3;refGene_tx=XM_023257371.1
B4 124387540 124387607
refGene_gn=SYCP3;refGene_tx=XM_011284177.3;refGene_gn=SYCP3;refGene_tx=XM_019835409.2;refGene_gn=SYCP3;refGene_tx=XM_019835410.2;refGene_gn=SYCP3;refGene_tx=XM_023257371.1
B4 124388115 124388248
refGene_gn=SYCP3;refGene_tx=XM_011284177.3;refGene_gn=SYCP3;refGene_tx=XM_019835409.2;refGene_gn=SYCP3;refGene_tx=XM_019835410.2;refGene_gn=SYCP3;refGene_tx=XM_023257371.1
B4 124415551 124415629
refGene_gn=GNPTAB;refGene_tx=XM_003989173.5
B4 124417828 124417919
refGene_gn=GNPTAB;refGene_tx=XM_003989173.5
B4 124420907 124421075
refGene_gn=GNPTAB;refGene_tx=XM_003989173.5
B4 124425090 124425189
refGene_gn=GNPTAB;refGene_tx=XM_003989173.5
B4 124425435 124425521
refGene_gn=GNPTAB;refGene_tx=XM_003989173.5
B4 124426647 124426761
refGene_gn=GNPTAB;refGene_tx=XM_003989173.5
B4 124427911 124428131
refGene_gn=GNPTAB;refGene_tx=XM_003989173.5
B4 124428379 124428579
refGene_gn=GNPTAB;refGene_tx=XM_003989173.5
B4 124431079 124431491
refGene_gn=GNPTAB;refGene_tx=XM_003989173.5
B4 124431492 124431830
refGene_gn=GNPTAB;refGene_tx=XM_003989173.5
B4 124431831 124432181
refGene_gn=GNPTAB;refGene_tx=XM_003989173.5
B4 124433110 124433314
refGene_gn=GNPTAB;refGene_tx=XM_003989173.5
B4 124434616 124434740
refGene_gn=GNPTAB;refGene_tx=XM_003989173.5
B4 124436721 124436892
refGene_gn=GNPTAB;refGene_tx=XM_003989173.5
B4 124437126 124437306
refGene_gn=GNPTAB;refGene_tx=XM_003989173.5
B4 124438023 124438185
refGene_gn=GNPTAB;refGene_tx=XM_003989173.5
B4 124446716 124446851
refGene_gn=GNPTAB;refGene_tx=XM_003989173.5
B4 124447119 124447184
refGene_gn=GNPTAB;refGene_tx=XM_003989173.5
B4 124452669 124452875
refGene_gn=GNPTAB;refGene_tx=XM_003989173.5
B4 124454928 124454970

refGene_gn=GNPTAB;refGene_tx=XM_003989173.5
B4 124456283 124456403
refGene_gn=GNPTAB;refGene_tx=XM_003989173.5
B4 124463663 124463749
refGene_gn=GNPTAB;refGene_tx=XM_003989173.5
B4 124487693 124487810
refGene_gn=GNPTAB;refGene_tx=XM_003989173.5
B4 124525209 124525340
refGene_gn=DRAM1;refGene_tx=XM_023257372.1;refGene_gn=DRAM1;refGene_tx=XM_023257373.1
B4 124544756 124544824
refGene_gn=DRAM1;refGene_tx=XM_023257372.1;refGene_gn=DRAM1;refGene_tx=XM_023257373.1
B4 124548730 124548873
refGene_gn=DRAM1;refGene_tx=XM_023257372.1;refGene_gn=DRAM1;refGene_tx=XM_023257373.1
B4 124554672 124555086
refGene_gn=DRAM1;refGene_tx=XM_023257372.1;refGene_gn=DRAM1;refGene_tx=XM_023257373.1
B4 124559048 124559107
refGene_gn=DRAM1;refGene_tx=XM_023257373.1
B4 124563117 124563210
refGene_gn=DRAM1;refGene_tx=XM_023257373.1
B4 124563792 124563837
refGene_gn=DRAM1;refGene_tx=XM_023257373.1
B4 124645427 124645464
refGene_gn=WASHC3;refGene_tx=XM_019835412.2
B4 124647258 124647343
refGene_gn=WASHC3;refGene_tx=XM_003989174.5;refGene_gn=WASHC3;refGene_tx=XM_006933972.4
B4 124651386 124651426
refGene_gn=WASHC3;refGene_tx=XM_019835411.2
B4 124661822 124661887
refGene_gn=WASHC3;refGene_tx=XM_003989174.5;refGene_gn=WASHC3;refGene_tx=XM_006933972.4;refGene_gn=WASHC3;refGene_tx=XM_019835411.2;refGene_gn=WASHC3;refGene_tx=XM_019835412.2
B4 124680735 124680846
refGene_gn=WASHC3;refGene_tx=XM_003989174.5;refGene_gn=WASHC3;refGene_tx=XM_006933972.4;refGene_gn=WASHC3;refGene_tx=XM_019835411.2;refGene_gn=WASHC3;refGene_tx=XM_019835412.2
B4 124684588 124684696
refGene_gn=WASHC3;refGene_tx=XM_003989174.5;refGene_gn=WASHC3;refGene_tx=XM_006933972.4;refGene_gn=WASHC3;refGene_tx=XM_019835411.2;refGene_gn=WASHC3;refGene_tx=XM_019835412.2
B4 124686532 124686598
refGene_gn=WASHC3;refGene_tx=XM_003989174.5;refGene_gn=WASHC3;refGene_tx=XM_006933972.4;refGene_gn=WASHC3;refGene_tx=XM_019835411.2;refGene_gn=WASHC3;refGene_tx=XM_019835412.2
B4 124693303 124693402
refGene_gn=WASHC3;refGene_tx=XM_003989174.5;refGene_gn=WASHC3;refGene_

tx=XM_006933972.4;refGene_gn=WASHC3;refGene_tx=XM_019835411.2;refGene_gn=WASHC3;refGene_tx=XM_019835412.2
B4 124693977 124694028
refGene_gn=WASHC3;refGene_tx=XM_003989174.5;refGene_gn=WASHC3;refGene_tx=XM_006933972.4;refGene_gn=WASHC3;refGene_tx=XM_019835411.2;refGene_gn=WASHC3;refGene_tx=XM_019835412.2
B4 124707180 124707294
refGene_gn=NUP37;refGene_tx=XM_003989175.5;refGene_gn=NUP37;refGene_tx=XM_006933973.2
B4 124707805 124707899
refGene_gn=NUP37;refGene_tx=XM_003989175.5;refGene_gn=NUP37;refGene_tx=XM_006933973.2
B4 124709745 124709796
refGene_gn=NUP37;refGene_tx=XM_003989175.5;refGene_gn=NUP37;refGene_tx=XM_006933973.2
B4 124710284 124710466
refGene_gn=NUP37;refGene_tx=XM_003989175.5;refGene_gn=NUP37;refGene_tx=XM_006933973.2
B4 124716442 124716533
refGene_gn=NUP37;refGene_tx=XM_003989175.5;refGene_gn=NUP37;refGene_tx=XM_006933973.2
B4 124731964 124732059
refGene_gn=NUP37;refGene_tx=XM_003989175.5;refGene_gn=NUP37;refGene_tx=XM_006933973.2
B4 124733744 124733817
refGene_gn=NUP37;refGene_tx=XM_003989175.5;refGene_gn=NUP37;refGene_tx=XM_006933973.2
B4 124737584 124737709
refGene_gn=NUP37;refGene_tx=XM_003989175.5;refGene_gn=NUP37;refGene_tx=XM_006933973.2
B4 124746369 124746525
refGene_gn=NUP37;refGene_tx=XM_003989175.5;refGene_gn=NUP37;refGene_tx=XM_006933973.2
B4 124747986 124748064
refGene_gn=PARPBP;refGene_tx=XM_011284179.3
B4 124752171 124752327
refGene_gn=PARPBP;refGene_tx=XM_011284179.3
B4 124765879 124766113
refGene_gn=PARPBP;refGene_tx=XM_011284179.3
B4 124772180 124772288
refGene_gn=PARPBP;refGene_tx=XM_011284179.3
B4 124781445 124781616
refGene_gn=PARPBP;refGene_tx=XM_011284179.3
B4 124782698 124782853
refGene_gn=PARPBP;refGene_tx=XM_011284179.3
B4 124793005 124793189
refGene_gn=PARPBP;refGene_tx=XM_011284179.3
B4 124796465 124796644
refGene_gn=PARPBP;refGene_tx=XM_011284179.3
B4 124798584 124798666

refGene_gn=PARPBP;refGene_tx=XM_011284179.3
B4 124812054 124812190
refGene_gn=PARPBP;refGene_tx=XM_011284179.3
B4 124812672 124813013
refGene_gn=PARPBP;refGene_tx=XM_011284179.3
B4 124813601 124813651
refGene_gn=PMCH;refGene_tx=XM_003989176.3;refGene_gn=PMCH;refGene_tx=XM_019835414.2
B4 124813935 124814134
refGene_gn=PMCH;refGene_tx=XM_003989176.3;refGene_gn=PMCH;refGene_tx=XM_019835414.2
B4 124814467 124814716
refGene_gn=PMCH;refGene_tx=XM_003989176.3;refGene_gn=PMCH;refGene_tx=XM_019835414.2
B4 124823881 124823909
refGene_gn=PMCH;refGene_tx=XM_019835414.2
B4 124984253 124984313
refGene_gn=IGF1;refGene_tx=XM_003989177.5;refGene_gn=IGF1;refGene_tx=XM_003989178.5;refGene_gn=IGF1;refGene_tx=XM_006933975.4
B4 124998064 124998226
refGene_gn=IGF1;refGene_tx=XM_006933975.4;refGene_gn=IGF1;refGene_tx=XM_023257374.1
B4 124999731 124999913
refGene_gn=IGF1;refGene_tx=XM_003989177.5;refGene_gn=IGF1;refGene_tx=XM_003989178.5;refGene_gn=IGF1;refGene_tx=XM_006933975.4;refGene_gn=IGF1;refGene_tx=XM_023257374.1
B4 125049970 125050127
refGene_gn=IGF1;refGene_tx=XM_003989177.5;refGene_gn=IGF1;refGene_tx=XM_003989178.5;refGene_gn=IGF1;refGene_tx=XM_006933975.4;refGene_gn=IGF1;refGene_tx=XM_023257374.1
B4 125052814 125052829
refGene_gn=IGF1;refGene_tx=XM_003989178.5;refGene_gn=IGF1;refGene_tx=XM_023257374.1
B4 125054742 125054805
refGene_gn=IGF1;refGene_tx=XM_003989177.5;refGene_gn=IGF1;refGene_tx=XM_006933975.4
B4 125352393 125352437
refGene_gn=PAH;refGene_tx=XM_019835417.2;refGene_gn=PAH;refGene_tx=XM_019835418.2;refGene_gn=PAH;refGene_tx=XM_019835419.2
B4 125354452 125354568
refGene_gn=PAH;refGene_tx=XM_019835417.2;refGene_gn=PAH;refGene_tx=XM_019835418.2;refGene_gn=PAH;refGene_tx=XM_019835419.2
B4 125358956 125359090
refGene_gn=PAH;refGene_tx=XM_019835417.2;refGene_gn=PAH;refGene_tx=XM_019835418.2;refGene_gn=PAH;refGene_tx=XM_019835419.2
B4 125359748 125359844
refGene_gn=PAH;refGene_tx=XM_019835417.2;refGene_gn=PAH;refGene_tx=XM_019835418.2;refGene_gn=PAH;refGene_tx=XM_019835419.2
B4 125361170 125361227
refGene_gn=PAH;refGene_tx=XM_019835417.2;refGene_gn=PAH;refGene_tx=XM_

019835418.2;refGene_gn=PAH;refGene_tx=XM_019835419.2
B4	125363423	125363493
refGene_gn=PAH;refGene_tx=XM_019835417.2;refGene_gn=PAH;refGene_tx=XM_
019835418.2;refGene_gn=PAH;refGene_tx=XM_019835419.2
B4	125364792	125364928
refGene_gn=PAH;refGene_tx=XM_019835417.2;refGene_gn=PAH;refGene_tx=XM_
019835418.2;refGene_gn=PAH;refGene_tx=XM_019835419.2
B4	125367072	125367269
refGene_gn=PAH;refGene_tx=XM_019835417.2;refGene_gn=PAH;refGene_tx=XM_
019835418.2;refGene_gn=PAH;refGene_tx=XM_019835419.2
B4	125374088	125374156
refGene_gn=PAH;refGene_tx=XM_019835417.2;refGene_gn=PAH;refGene_tx=XM_
019835418.2;refGene_gn=PAH;refGene_tx=XM_019835419.2
B4	125385641	125385730
refGene_gn=PAH;refGene_tx=XM_019835417.2;refGene_gn=PAH;refGene_tx=XM_
019835418.2;refGene_gn=PAH;refGene_tx=XM_019835419.2
B4	125400171	125400355
refGene_gn=PAH;refGene_tx=XM_019835417.2;refGene_gn=PAH;refGene_tx=XM_
019835418.2;refGene_gn=PAH;refGene_tx=XM_019835419.2
B4	125416942	125417050
refGene_gn=PAH;refGene_tx=XM_019835417.2;refGene_gn=PAH;refGene_tx=XM_
019835418.2;refGene_gn=PAH;refGene_tx=XM_019835419.2
B4	125423307	125423364
refGene_gn=PAH;refGene_tx=XM_019835417.2;refGene_gn=PAH;refGene_tx=XM_
019835418.2;refGene_gn=PAH;refGene_tx=XM_019835419.2
B4	125454243	125454984
refGene_gn=ASCL1;refGene_tx=XM_006933981.3;refGene_gn=ASCL1;refGene_tx
=XM_019835421.2
B4	125728246	125728321
refGene_gn=CB4H12orf42;refGene_tx=XM_023257775.1
B4	125728323	125728473
refGene_gn=CB4H12orf42;refGene_tx=XM_023257775.1
B4	125728474	125728540
refGene_gn=CB4H12orf42;refGene_tx=XM_023257775.1
B4	125728541	125728723
refGene_gn=CB4H12orf42;refGene_tx=XM_023257775.1
B4	125731732	125731793
refGene_gn=CB4H12orf42;refGene_tx=XM_023257775.1
B4	125732503	125732895
refGene_gn=CB4H12orf42;refGene_tx=XM_023257775.1
B4	125745922	125745986
refGene_gn=CB4H12orf42;refGene_tx=XM_023257775.1
B4	125963560	125963638
refGene_gn=STAB2;refGene_tx=XM_011284182.3;refGene_gn=STAB2;refGene_tx
=XM_023257375.1;refGene_gn=STAB2;refGene_tx=XM_023257376.1
B4	125968368	125968502
refGene_gn=STAB2;refGene_tx=XM_011284182.3;refGene_gn=STAB2;refGene_tx
=XM_023257375.1;refGene_gn=STAB2;refGene_tx=XM_023257376.1
B4	125970905	125971021
refGene_gn=STAB2;refGene_tx=XM_011284182.3;refGene_gn=STAB2;refGene_tx

=XM_023257375.1;refGene_gn=STAB2;refGene_tx=XM_023257376.1
B4	125994514	125994600
refGene_gn=STAB2;refGene_tx=XM_011284182.3;refGene_gn=STAB2;refGene_tx=XM_023257375.1;refGene_gn=STAB2;refGene_tx=XM_023257376.1
B4	125995959	125996029
refGene_gn=STAB2;refGene_tx=XM_011284182.3;refGene_gn=STAB2;refGene_tx=XM_023257375.1;refGene_gn=STAB2;refGene_tx=XM_023257376.1
B4	126004051	126004147
refGene_gn=STAB2;refGene_tx=XM_011284182.3;refGene_gn=STAB2;refGene_tx=XM_023257375.1;refGene_gn=STAB2;refGene_tx=XM_023257376.1
B4	126009533	126009659
refGene_gn=STAB2;refGene_tx=XM_011284182.3;refGene_gn=STAB2;refGene_tx=XM_023257375.1;refGene_gn=STAB2;refGene_tx=XM_023257376.1
B4	126010546	126010743
refGene_gn=STAB2;refGene_tx=XM_011284182.3;refGene_gn=STAB2;refGene_tx=XM_023257375.1;refGene_gn=STAB2;refGene_tx=XM_023257376.1
B4	126012357	126012491
refGene_gn=STAB2;refGene_tx=XM_011284182.3;refGene_gn=STAB2;refGene_tx=XM_023257375.1;refGene_gn=STAB2;refGene_tx=XM_023257376.1
B4	126019283	126019417
refGene_gn=STAB2;refGene_tx=XM_011284182.3;refGene_gn=STAB2;refGene_tx=XM_023257375.1;refGene_gn=STAB2;refGene_tx=XM_023257376.1
B4	126021065	126021148
refGene_gn=STAB2;refGene_tx=XM_011284182.3;refGene_gn=STAB2;refGene_tx=XM_023257375.1;refGene_gn=STAB2;refGene_tx=XM_023257376.1
B4	126022362	126022512
refGene_gn=STAB2;refGene_tx=XM_011284182.3;refGene_gn=STAB2;refGene_tx=XM_023257375.1;refGene_gn=STAB2;refGene_tx=XM_023257376.1
B4	126023888	126024032
refGene_gn=STAB2;refGene_tx=XM_011284182.3;refGene_gn=STAB2;refGene_tx=XM_023257375.1;refGene_gn=STAB2;refGene_tx=XM_023257376.1
B4	126024575	126024632
refGene_gn=STAB2;refGene_tx=XM_011284182.3;refGene_gn=STAB2;refGene_tx=XM_023257375.1;refGene_gn=STAB2;refGene_tx=XM_023257376.1
B4	126024785	126024911
refGene_gn=STAB2;refGene_tx=XM_011284182.3;refGene_gn=STAB2;refGene_tx=XM_023257375.1;refGene_gn=STAB2;refGene_tx=XM_023257376.1
B4	126029566	126029620
refGene_gn=STAB2;refGene_tx=XM_011284182.3;refGene_gn=STAB2;refGene_tx=XM_023257375.1;refGene_gn=STAB2;refGene_tx=XM_023257376.1
B4	126029920	126030001
refGene_gn=STAB2;refGene_tx=XM_011284182.3;refGene_gn=STAB2;refGene_tx=XM_023257375.1;refGene_gn=STAB2;refGene_tx=XM_023257376.1
B4	126031882	126032035
refGene_gn=STAB2;refGene_tx=XM_011284182.3;refGene_gn=STAB2;refGene_tx=XM_023257375.1
B4	126034749	126034812
refGene_gn=STAB2;refGene_tx=XM_011284182.3;refGene_gn=STAB2;refGene_tx=XM_023257375.1;refGene_gn=STAB2;refGene_tx=XM_023257376.1
B4	126037674	126037761

refGene_gn=STAB2;refGene_tx=XM_011284182.3;refGene_gn=STAB2;refGene_tx=XM_023257375.1;refGene_gn=STAB2;refGene_tx=XM_023257376.1
B4 126039124 126039211
refGene_gn=STAB2;refGene_tx=XM_011284182.3;refGene_gn=STAB2;refGene_tx=XM_023257375.1;refGene_gn=STAB2;refGene_tx=XM_023257376.1
B4 126040545 126040657
refGene_gn=STAB2;refGene_tx=XM_011284182.3;refGene_gn=STAB2;refGene_tx=XM_023257375.1;refGene_gn=STAB2;refGene_tx=XM_023257376.1
B4 126042569 126042750
refGene_gn=STAB2;refGene_tx=XM_011284182.3;refGene_gn=STAB2;refGene_tx=XM_023257375.1;refGene_gn=STAB2;refGene_tx=XM_023257376.1
B4 126044792 126044886
refGene_gn=STAB2;refGene_tx=XM_011284182.3;refGene_gn=STAB2;refGene_tx=XM_023257375.1;refGene_gn=STAB2;refGene_tx=XM_023257376.1
B4 126046123 126046282
refGene_gn=STAB2;refGene_tx=XM_011284182.3;refGene_gn=STAB2;refGene_tx=XM_023257375.1;refGene_gn=STAB2;refGene_tx=XM_023257376.1
B4 126049174 126049270
refGene_gn=STAB2;refGene_tx=XM_011284182.3;refGene_gn=STAB2;refGene_tx=XM_023257375.1;refGene_gn=STAB2;refGene_tx=XM_023257376.1
B4 126051254 126051350
refGene_gn=STAB2;refGene_tx=XM_011284182.3;refGene_gn=STAB2;refGene_tx=XM_023257375.1;refGene_gn=STAB2;refGene_tx=XM_023257376.1
B4 126053668 126053716
refGene_gn=STAB2;refGene_tx=XM_011284182.3;refGene_gn=STAB2;refGene_tx=XM_023257375.1;refGene_gn=STAB2;refGene_tx=XM_023257376.1
B4 126055152 126055289
refGene_gn=STAB2;refGene_tx=XM_011284182.3;refGene_gn=STAB2;refGene_tx=XM_023257375.1;refGene_gn=STAB2;refGene_tx=XM_023257376.1
B4 126055698 126055813
refGene_gn=STAB2;refGene_tx=XM_011284182.3;refGene_gn=STAB2;refGene_tx=XM_023257375.1;refGene_gn=STAB2;refGene_tx=XM_023257376.1
B4 126058191 126058269
refGene_gn=STAB2;refGene_tx=XM_011284182.3;refGene_gn=STAB2;refGene_tx=XM_023257376.1
B4 126060853 126060952
refGene_gn=STAB2;refGene_tx=XM_011284182.3;refGene_gn=STAB2;refGene_tx=XM_023257375.1;refGene_gn=STAB2;refGene_tx=XM_023257376.1
B4 126061040 126061148
refGene_gn=STAB2;refGene_tx=XM_011284182.3;refGene_gn=STAB2;refGene_tx=XM_023257375.1;refGene_gn=STAB2;refGene_tx=XM_023257376.1
B4 126064110 126064242
refGene_gn=STAB2;refGene_tx=XM_011284182.3;refGene_gn=STAB2;refGene_tx=XM_023257375.1;refGene_gn=STAB2;refGene_tx=XM_023257376.1
B4 126068040 126068169
refGene_gn=STAB2;refGene_tx=XM_011284182.3;refGene_gn=STAB2;refGene_tx=XM_023257375.1;refGene_gn=STAB2;refGene_tx=XM_023257376.1
B4 126069428 126069485
refGene_gn=STAB2;refGene_tx=XM_011284182.3;refGene_gn=STAB2;refGene_tx=XM_023257375.1;refGene_gn=STAB2;refGene_tx=XM_023257376.1

B4	126071175	126071271
refGene_gn=STAB2;refGene_tx=XM_011284182.3;refGene_gn=STAB2;refGene_tx=XM_023257375.1;refGene_gn=STAB2;refGene_tx=XM_023257376.1
B4	126073027	126073223
refGene_gn=STAB2;refGene_tx=XM_011284182.3;refGene_gn=STAB2;refGene_tx=XM_023257375.1;refGene_gn=STAB2;refGene_tx=XM_023257376.1
B4	126074644	126074740
refGene_gn=STAB2;refGene_tx=XM_011284182.3;refGene_gn=STAB2;refGene_tx=XM_023257375.1;refGene_gn=STAB2;refGene_tx=XM_023257376.1
B4	126078211	126078257
refGene_gn=STAB2;refGene_tx=XM_011284182.3;refGene_gn=STAB2;refGene_tx=XM_023257375.1;refGene_gn=STAB2;refGene_tx=XM_023257376.1
B4	126079089	126079166
refGene_gn=STAB2;refGene_tx=XM_011284182.3;refGene_gn=STAB2;refGene_tx=XM_023257375.1;refGene_gn=STAB2;refGene_tx=XM_023257376.1
B4	126081126	126081252
refGene_gn=STAB2;refGene_tx=XM_011284182.3;refGene_gn=STAB2;refGene_tx=XM_023257375.1;refGene_gn=STAB2;refGene_tx=XM_023257376.1
B4	126083209	126083283
refGene_gn=STAB2;refGene_tx=XM_011284182.3;refGene_gn=STAB2;refGene_tx=XM_023257375.1;refGene_gn=STAB2;refGene_tx=XM_023257376.1
B4	126085288	126085360
refGene_gn=STAB2;refGene_tx=XM_011284182.3;refGene_gn=STAB2;refGene_tx=XM_023257375.1;refGene_gn=STAB2;refGene_tx=XM_023257376.1
B4	126088696	126088816
refGene_gn=STAB2;refGene_tx=XM_011284182.3;refGene_gn=STAB2;refGene_tx=XM_023257375.1;refGene_gn=STAB2;refGene_tx=XM_023257376.1
B4	126089734	126089782
refGene_gn=STAB2;refGene_tx=XM_011284182.3;refGene_gn=STAB2;refGene_tx=XM_023257375.1;refGene_gn=STAB2;refGene_tx=XM_023257376.1
B4	126090940	126091024
refGene_gn=STAB2;refGene_tx=XM_011284182.3;refGene_gn=STAB2;refGene_tx=XM_023257375.1;refGene_gn=STAB2;refGene_tx=XM_023257376.1
B4	126092306	126092453
refGene_gn=STAB2;refGene_tx=XM_011284182.3;refGene_gn=STAB2;refGene_tx=XM_023257375.1;refGene_gn=STAB2;refGene_tx=XM_023257376.1
B4	126093243	126093384
refGene_gn=STAB2;refGene_tx=XM_011284182.3;refGene_gn=STAB2;refGene_tx=XM_023257375.1;refGene_gn=STAB2;refGene_tx=XM_023257376.1
B4	126094384	126094444
refGene_gn=STAB2;refGene_tx=XM_011284182.3;refGene_gn=STAB2;refGene_tx=XM_023257375.1;refGene_gn=STAB2;refGene_tx=XM_023257376.1
B4	126096646	126096823
refGene_gn=STAB2;refGene_tx=XM_011284182.3;refGene_gn=STAB2;refGene_tx=XM_023257375.1;refGene_gn=STAB2;refGene_tx=XM_023257376.1
B4	126098166	126098256
refGene_gn=STAB2;refGene_tx=XM_011284182.3;refGene_gn=STAB2;refGene_tx=XM_023257375.1;refGene_gn=STAB2;refGene_tx=XM_023257376.1
B4	126100240	126100387
refGene_gn=STAB2;refGene_tx=XM_011284182.3;refGene_gn=STAB2;refGene_tx

=XM_023257375.1;refGene_gn=STAB2;refGene_tx=XM_023257376.1
B4	126102000	126102057
refGene_gn=STAB2;refGene_tx=XM_011284182.3;refGene_gn=STAB2;refGene_tx=XM_023257375.1;refGene_gn=STAB2;refGene_tx=XM_023257376.1
B4	126103230	126103357
refGene_gn=STAB2;refGene_tx=XM_011284182.3;refGene_gn=STAB2;refGene_tx=XM_023257375.1;refGene_gn=STAB2;refGene_tx=XM_023257376.1
B4	126104912	126105062
refGene_gn=STAB2;refGene_tx=XM_011284182.3;refGene_gn=STAB2;refGene_tx=XM_023257375.1;refGene_gn=STAB2;refGene_tx=XM_023257376.1
B4	126106801	126106906
refGene_gn=STAB2;refGene_tx=XM_011284182.3;refGene_gn=STAB2;refGene_tx=XM_023257375.1;refGene_gn=STAB2;refGene_tx=XM_023257376.1
B4	126108855	126108969
refGene_gn=NT5DC3;refGene_tx=XM_023257377.1;refGene_gn=STAB2;refGene_tx=XM_011284182.3;refGene_gn=STAB2;refGene_tx=XM_023257375.1;refGene_gn=STAB2;refGene_tx=XM_023257376.1
B4	126110883	126111077
refGene_gn=STAB2;refGene_tx=XM_011284182.3;refGene_gn=STAB2;refGene_tx=XM_023257375.1;refGene_gn=STAB2;refGene_tx=XM_023257376.1
B4	126112530	126112672
refGene_gn=STAB2;refGene_tx=XM_011284182.3;refGene_gn=STAB2;refGene_tx=XM_023257375.1;refGene_gn=STAB2;refGene_tx=XM_023257376.1
B4	126117421	126117555
refGene_gn=STAB2;refGene_tx=XM_011284182.3;refGene_gn=STAB2;refGene_tx=XM_023257375.1;refGene_gn=STAB2;refGene_tx=XM_023257376.1
B4	126118415	126118581
refGene_gn=STAB2;refGene_tx=XM_011284182.3;refGene_gn=STAB2;refGene_tx=XM_023257375.1;refGene_gn=STAB2;refGene_tx=XM_023257376.1
B4	126118725	126118832
refGene_gn=STAB2;refGene_tx=XM_011284182.3;refGene_gn=STAB2;refGene_tx=XM_023257375.1;refGene_gn=STAB2;refGene_tx=XM_023257376.1
B4	126120890	126121010
refGene_gn=STAB2;refGene_tx=XM_011284182.3;refGene_gn=STAB2;refGene_tx=XM_023257375.1;refGene_gn=STAB2;refGene_tx=XM_023257376.1
B4	126121865	126122006
refGene_gn=STAB2;refGene_tx=XM_011284182.3;refGene_gn=STAB2;refGene_tx=XM_023257375.1;refGene_gn=STAB2;refGene_tx=XM_023257376.1
B4	126123263	126123371
refGene_gn=STAB2;refGene_tx=XM_011284182.3;refGene_gn=STAB2;refGene_tx=XM_023257375.1;refGene_gn=STAB2;refGene_tx=XM_023257376.1
B4	126124520	126124649
refGene_gn=STAB2;refGene_tx=XM_011284182.3;refGene_gn=STAB2;refGene_tx=XM_023257375.1;refGene_gn=STAB2;refGene_tx=XM_023257376.1
B4	126125727	126125844
refGene_gn=STAB2;refGene_tx=XM_011284182.3;refGene_gn=STAB2;refGene_tx=XM_023257375.1;refGene_gn=STAB2;refGene_tx=XM_023257376.1
B4	126129881	126129932
refGene_gn=STAB2;refGene_tx=XM_011284182.3;refGene_gn=STAB2;refGene_tx=XM_023257375.1;refGene_gn=STAB2;refGene_tx=XM_023257376.1

B4 126136769 126137022
refGene_gn=NT5DC3;refGene_tx=XM_011284289.3
B4 126141803 126141868
refGene_gn=NT5DC3;refGene_tx=XM_011284289.3;refGene_gn=NT5DC3;refGene_tx=XM_023257377.1
B4 126147471 126147612
refGene_gn=NT5DC3;refGene_tx=XM_011284289.3;refGene_gn=NT5DC3;refGene_tx=XM_023257377.1
B4 126152587 126152674
refGene_gn=NT5DC3;refGene_tx=XM_011284289.3;refGene_gn=NT5DC3;refGene_tx=XM_023257377.1
B4 126153773 126153855
refGene_gn=NT5DC3;refGene_tx=XM_011284289.3;refGene_gn=NT5DC3;refGene_tx=XM_023257377.1
B4 126156799 126156901
refGene_gn=NT5DC3;refGene_tx=XM_011284289.3;refGene_gn=NT5DC3;refGene_tx=XM_023257377.1
B4 126157045 126157148
refGene_gn=NT5DC3;refGene_tx=XM_011284289.3;refGene_gn=NT5DC3;refGene_tx=XM_023257377.1
B4 126157560 126157621
refGene_gn=NT5DC3;refGene_tx=XM_011284289.3;refGene_gn=NT5DC3;refGene_tx=XM_023257377.1
B4 126160005 126160143
refGene_gn=NT5DC3;refGene_tx=XM_011284289.3;refGene_gn=NT5DC3;refGene_tx=XM_023257377.1
B4 126161020 126161111
refGene_gn=NT5DC3;refGene_tx=XM_011284289.3;refGene_gn=NT5DC3;refGene_tx=XM_023257377.1
B4 126166290 126166346
refGene_gn=NT5DC3;refGene_tx=XM_011284289.3;refGene_gn=NT5DC3;refGene_tx=XM_023257377.1
B4 126166794 126166869
refGene_gn=NT5DC3;refGene_tx=XM_011284289.3;refGene_gn=NT5DC3;refGene_tx=XM_023257377.1
B4 126176213 126176398
refGene_gn=NT5DC3;refGene_tx=XM_011284289.3;refGene_gn=NT5DC3;refGene_tx=XM_023257377.1
B4 126197637 126197836
refGene_gn=NT5DC3;refGene_tx=XM_011284289.3;refGene_gn=NT5DC3;refGene_tx=XM_023257377.1
B4 126200022 126200701
refGene_gn=LOC101083779;refGene_tx=XM_023257776.1
B4 126200792 126200922
refGene_gn=LOC101083779;refGene_tx=XM_023257776.1
B4 126209740 126209874
refGene_gn=LOC101083779;refGene_tx=XM_023257776.1
B4 126214663 126214816
refGene_gn=LOC101083779;refGene_tx=XM_023257776.1
B4 126215799 126215947

refGene_gn=LOC101083779;refGene_tx=XM_023257776.1
B4 126216783 126216823
refGene_gn=LOC101083779;refGene_tx=XM_023257776.1
B4 126221195 126221335
refGene_gn=LOC101083779;refGene_tx=XM_023257776.1
B4 126230672 126230851
refGene_gn=LOC101083779;refGene_tx=XM_023257776.1
B4 126231695 126231727
refGene_gn=LOC101083779;refGene_tx=XM_023257776.1
B4 126231728 126231895
refGene_gn=LOC101083779;refGene_tx=XM_023257776.1
B4 126238228 126238252
refGene_gn=LOC101083779;refGene_tx=XM_023257776.1
B4 126238254 126238467
refGene_gn=LOC101083779;refGene_tx=XM_023257776.1
B4 126240918 126241357
refGene_gn=LOC101083779;refGene_tx=XM_023257776.1
B4 126244167 126244328
refGene_gn=LOC101083779;refGene_tx=XM_023257776.1
B4 126253366 126253581
refGene_gn=LOC101083779;refGene_tx=XM_023257776.1
B4 126257217 126257335
refGene_gn=LOC101083779;refGene_tx=XM_023257776.1
B4 126257337 126257394
refGene_gn=LOC101083779;refGene_tx=XM_023257776.1
B4 126261256 126261973
refGene_gn=LOC101083779;refGene_tx=XM_023257776.1
B4 126261975 126262057
refGene_gn=LOC101083779;refGene_tx=XM_023257776.1
B4 126278324 126278373
refGene_gn=HSP90B1;refGene_tx=XM_019835423.2
B4 126279391 126279494
refGene_gn=HSP90B1;refGene_tx=XM_019835423.2
B4 126280302 126280444
refGene_gn=HSP90B1;refGene_tx=XM_019835423.2
B4 126280945 126281062
refGene_gn=HSP90B1;refGene_tx=XM_019835423.2
B4 126282089 126282421
refGene_gn=HSP90B1;refGene_tx=XM_019835423.2
B4 126287105 126287217
refGene_gn=HSP90B1;refGene_tx=XM_019835423.2
B4 126287427 126287547
refGene_gn=HSP90B1;refGene_tx=XM_019835423.2
B4 126288106 126288223
refGene_gn=HSP90B1;refGene_tx=XM_019835423.2
B4 126290447 126290585
refGene_gn=HSP90B1;refGene_tx=XM_019835423.2
B4 126290668 126290746
refGene_gn=HSP90B1;refGene_tx=XM_019835423.2
B4 126291068 126291134

refGene_gn=HSP90B1;refGene_tx=XM_019835423.2
B4 126291735 126292005
refGene_gn=HSP90B1;refGene_tx=XM_019835423.2
B4 126292254 126292500
refGene_gn=HSP90B1;refGene_tx=XM_019835423.2
B4 126293613 126293750
refGene_gn=HSP90B1;refGene_tx=XM_019835423.2
B4 126295106 126295185
refGene_gn=HSP90B1;refGene_tx=XM_019835423.2
B4 126295273 126295429
refGene_gn=HSP90B1;refGene_tx=XM_019835423.2
B4 126295695 126295818
refGene_gn=HSP90B1;refGene_tx=XM_019835423.2
B4 126296373 126296403
refGene_gn=HSP90B1;refGene_tx=XM_019835423.2
B4 126313887 126313910
refGene_gn=TDG;refGene_tx=XM_006933983.3
B4 126322085 126322228
refGene_gn=TDG;refGene_tx=XM_006933983.3
B4 126324635 126324877
refGene_gn=TDG;refGene_tx=XM_006933983.3
B4 126326005 126326075
refGene_gn=TDG;refGene_tx=XM_006933983.3
B4 126327626 126327762
refGene_gn=TDG;refGene_tx=XM_006933983.3
B4 126327956 126328039
refGene_gn=TDG;refGene_tx=XM_006933983.3
B4 126328118 126328213
refGene_gn=TDG;refGene_tx=XM_006933983.3
B4 126331292 126331464
refGene_gn=TDG;refGene_tx=XM_006933983.3
B4 126331895 126332021
refGene_gn=TDG;refGene_tx=XM_006933983.3
B4 126332620 126332751
refGene_gn=TDG;refGene_tx=XM_006933983.3
B4 126335095 126335268
refGene_gn=GLT8D2;refGene_tx=XM_003989187.5;refGene_gn=GLT8D2;refGene_tx=XM_006933984.4;refGene_gn=GLT8D2;refGene_tx=XM_019835424.2
B4 126338854 126338967
refGene_gn=GLT8D2;refGene_tx=XM_003989187.5;refGene_gn=GLT8D2;refGene_tx=XM_006933984.4;refGene_gn=GLT8D2;refGene_tx=XM_019835424.2
B4 126340015 126340182
refGene_gn=GLT8D2;refGene_tx=XM_003989187.5;refGene_gn=GLT8D2;refGene_tx=XM_006933984.4;refGene_gn=GLT8D2;refGene_tx=XM_019835424.2
B4 126342302 126342415
refGene_gn=GLT8D2;refGene_tx=XM_003989187.5;refGene_gn=GLT8D2;refGene_tx=XM_006933984.4;refGene_gn=GLT8D2;refGene_tx=XM_019835424.2
B4 126343046 126343131
refGene_gn=GLT8D2;refGene_tx=XM_003989187.5;refGene_gn=GLT8D2;refGene_tx=XM_006933984.4;refGene_gn=GLT8D2;refGene_tx=XM_019835424.2

B4 126344345 126344463
refGene_gn=GLT8D2;refGene_tx=XM_003989187.5;refGene_gn=GLT8D2;refGene_tx=XM_006933984.4;refGene_gn=GLT8D2;refGene_tx=XM_019835424.2
B4 126350105 126350277
refGene_gn=GLT8D2;refGene_tx=XM_003989187.5;refGene_gn=GLT8D2;refGene_tx=XM_006933984.4;refGene_gn=GLT8D2;refGene_tx=XM_019835424.2
B4 126357875 126357968
refGene_gn=GLT8D2;refGene_tx=XM_003989187.5;refGene_gn=GLT8D2;refGene_tx=XM_006933984.4;refGene_gn=GLT8D2;refGene_tx=XM_019835424.2
B4 126361271 126361290
refGene_gn=GLT8D2;refGene_tx=XM_003989187.5;refGene_gn=GLT8D2;refGene_tx=XM_006933984.4;refGene_gn=GLT8D2;refGene_tx=XM_019835424.2
B4 126394535 126394698
refGene_gn=HCFC2;refGene_tx=XM_023257378.1;refGene_gn=HCFC2;refGene_tx=XM_023257379.1;refGene_gn=HCFC2;refGene_tx=XM_023257380.1;refGene_gn=HCFC2;refGene_tx=XM_023257381.1;refGene_gn=HCFC2;refGene_tx=XM_023257382.1;refGene_gn=HCFC2;refGene_tx=XM_023257383.1
B4 126396175 126396324
refGene_gn=HCFC2;refGene_tx=XM_023257378.1;refGene_gn=HCFC2;refGene_tx=XM_023257379.1;refGene_gn=HCFC2;refGene_tx=XM_023257380.1;refGene_gn=HCFC2;refGene_tx=XM_023257381.1;refGene_gn=HCFC2;refGene_tx=XM_023257382.1;refGene_gn=HCFC2;refGene_tx=XM_023257383.1
B4 126397768 126397929
refGene_gn=HCFC2;refGene_tx=XM_023257378.1;refGene_gn=HCFC2;refGene_tx=XM_023257379.1;refGene_gn=HCFC2;refGene_tx=XM_023257380.1;refGene_gn=HCFC2;refGene_tx=XM_023257381.1;refGene_gn=HCFC2;refGene_tx=XM_023257382.1;refGene_gn=HCFC2;refGene_tx=XM_023257383.1
B4 126408614 126408823
refGene_gn=HCFC2;refGene_tx=XM_023257378.1;refGene_gn=HCFC2;refGene_tx=XM_023257379.1;refGene_gn=HCFC2;refGene_tx=XM_023257380.1;refGene_gn=HCFC2;refGene_tx=XM_023257381.1;refGene_gn=HCFC2;refGene_tx=XM_023257382.1;refGene_gn=HCFC2;refGene_tx=XM_023257383.1
B4 126409622 126409707
refGene_gn=HCFC2;refGene_tx=XM_023257378.1;refGene_gn=HCFC2;refGene_tx=XM_023257379.1;refGene_gn=HCFC2;refGene_tx=XM_023257380.1;refGene_gn=HCFC2;refGene_tx=XM_023257381.1;refGene_gn=HCFC2;refGene_tx=XM_023257382.1;refGene_gn=HCFC2;refGene_tx=XM_023257383.1
B4 126411340 126411447
refGene_gn=HCFC2;refGene_tx=XM_023257378.1;refGene_gn=HCFC2;refGene_tx=XM_023257379.1;refGene_gn=HCFC2;refGene_tx=XM_023257380.1;refGene_gn=HCFC2;refGene_tx=XM_023257381.1;refGene_gn=HCFC2;refGene_tx=XM_023257382.1;refGene_gn=HCFC2;refGene_tx=XM_023257383.1
B4 126411556 126411745
refGene_gn=HCFC2;refGene_tx=XM_023257378.1;refGene_gn=HCFC2;refGene_tx=XM_023257379.1;refGene_gn=HCFC2;refGene_tx=XM_023257380.1;refGene_gn=HCFC2;refGene_tx=XM_023257381.1;refGene_gn=HCFC2;refGene_tx=XM_023257382.1;refGene_gn=HCFC2;refGene_tx=XM_023257383.1
B4 126421082 126421250
refGene_gn=HCFC2;refGene_tx=XM_023257378.1;refGene_gn=HCFC2;refGene_tx=XM_023257379.1;refGene_gn=HCFC2;refGene_tx=XM_023257380.1;refGene_gn=

HCFC2;refGene_tx=XM_023257381.1;refGene_gn=HCFC2;refGene_tx=XM_0232573
82.1;refGene_gn=HCFC2;refGene_tx=XM_023257383.1
B4 126421794 126421847
refGene_gn=HCFC2;refGene_tx=XM_023257378.1;refGene_gn=HCFC2;refGene_tx
=XM_023257379.1;refGene_gn=HCFC2;refGene_tx=XM_023257380.1;refGene_gn=
HCFC2;refGene_tx=XM_023257381.1;refGene_gn=HCFC2;refGene_tx=XM_0232573
82.1;refGene_gn=HCFC2;refGene_tx=XM_023257383.1
B4 126423314 126423492
refGene_gn=HCFC2;refGene_tx=XM_023257378.1;refGene_gn=HCFC2;refGene_tx
=XM_023257379.1;refGene_gn=HCFC2;refGene_tx=XM_023257380.1;refGene_gn=
HCFC2;refGene_tx=XM_023257381.1;refGene_gn=HCFC2;refGene_tx=XM_0232573
82.1;refGene_gn=HCFC2;refGene_tx=XM_023257383.1
B4 126427567 126427771
refGene_gn=HCFC2;refGene_tx=XM_023257378.1;refGene_gn=HCFC2;refGene_tx
=XM_023257379.1;refGene_gn=HCFC2;refGene_tx=XM_023257380.1;refGene_gn=
HCFC2;refGene_tx=XM_023257381.1;refGene_gn=HCFC2;refGene_tx=XM_0232573
82.1;refGene_gn=HCFC2;refGene_tx=XM_023257383.1
B4 126428338 126428412
refGene_gn=HCFC2;refGene_tx=XM_023257378.1;refGene_gn=HCFC2;refGene_tx
=XM_023257379.1;refGene_gn=HCFC2;refGene_tx=XM_023257380.1;refGene_gn=
HCFC2;refGene_tx=XM_023257381.1;refGene_gn=HCFC2;refGene_tx=XM_0232573
82.1;refGene_gn=HCFC2;refGene_tx=XM_023257383.1
B4 126429885 126429901
refGene_gn=HCFC2;refGene_tx=XM_023257380.1
B4 126430362 126430500
refGene_gn=HCFC2;refGene_tx=XM_023257378.1;refGene_gn=HCFC2;refGene_tx
=XM_023257379.1;refGene_gn=HCFC2;refGene_tx=XM_023257380.1;refGene_gn=
HCFC2;refGene_tx=XM_023257381.1;refGene_gn=HCFC2;refGene_tx=XM_0232573
82.1;refGene_gn=HCFC2;refGene_tx=XM_023257383.1
B4 126432299 126432485
refGene_gn=HCFC2;refGene_tx=XM_023257378.1;refGene_gn=HCFC2;refGene_tx
=XM_023257379.1;refGene_gn=HCFC2;refGene_tx=XM_023257380.1;refGene_gn=
HCFC2;refGene_tx=XM_023257381.1;refGene_gn=HCFC2;refGene_tx=XM_0232573
82.1;refGene_gn=HCFC2;refGene_tx=XM_023257383.1
B4 126432883 126432922
refGene_gn=HCFC2;refGene_tx=XM_023257378.1;refGene_gn=HCFC2;refGene_tx
=XM_023257380.1;refGene_gn=HCFC2;refGene_tx=XM_023257381.1;refGene_gn=
HCFC2;refGene_tx=XM_023257382.1
B4 126433189 126433504
refGene_gn=HCFC2;refGene_tx=XM_023257378.1;refGene_gn=HCFC2;refGene_tx
=XM_023257379.1;refGene_gn=HCFC2;refGene_tx=XM_023257380.1;refGene_gn=
HCFC2;refGene_tx=XM_023257381.1;refGene_gn=HCFC2;refGene_tx=XM_0232573
82.1;refGene_gn=HCFC2;refGene_tx=XM_023257383.1
B4 126434446 126434487
refGene_gn=HCFC2;refGene_tx=XM_023257382.1
B4 126434685 126434870
refGene_gn=HCFC2;refGene_tx=XM_023257378.1;refGene_gn=HCFC2;refGene_tx
=XM_023257379.1;refGene_gn=HCFC2;refGene_tx=XM_023257380.1
B4 126446307 126446399
refGene_gn=HCFC2;refGene_tx=XM_023257381.1

B4 126449891 126449924
refGene_gn=NFYB;refGene_tx=XM_003989188.4;refGene_gn=NFYB;refGene_tx=XM_011284183.3;refGene_gn=NFYB;refGene_tx=XM_023257384.1;refGene_gn=NFYB;refGene_tx=XM_023257385.1
B4 126451046 126451126
refGene_gn=NFYB;refGene_tx=XM_003989188.4;refGene_gn=NFYB;refGene_tx=XM_011284183.3;refGene_gn=NFYB;refGene_tx=XM_023257384.1;refGene_gn=NFYB;refGene_tx=XM_023257385.1
B4 126452494 126452576
refGene_gn=NFYB;refGene_tx=XM_003989188.4;refGene_gn=NFYB;refGene_tx=XM_011284183.3;refGene_gn=NFYB;refGene_tx=XM_023257384.1;refGene_gn=NFYB;refGene_tx=XM_023257385.1
B4 126453830 126454028
refGene_gn=NFYB;refGene_tx=XM_003989188.4;refGene_gn=NFYB;refGene_tx=XM_011284183.3;refGene_gn=NFYB;refGene_tx=XM_023257384.1;refGene_gn=NFYB;refGene_tx=XM_023257385.1
B4 126457321 126457452
refGene_gn=NFYB;refGene_tx=XM_003989188.4;refGene_gn=NFYB;refGene_tx=XM_011284183.3;refGene_gn=NFYB;refGene_tx=XM_023257384.1;refGene_gn=NFYB;refGene_tx=XM_023257385.1
B4 126458839 126458933
refGene_gn=NFYB;refGene_tx=XM_003989188.4;refGene_gn=NFYB;refGene_tx=XM_011284183.3;refGene_gn=NFYB;refGene_tx=XM_023257384.1;refGene_gn=NFYB;refGene_tx=XM_023257385.1
B4 126538502 126538542
refGene_gn=TXNRD1;refGene_tx=XM_006933988.4;refGene_gn=TXNRD1;refGene_tx=XM_023257386.1
B4 126559145 126559255
refGene_gn=TXNRD1;refGene_tx=XM_006933988.4;refGene_gn=TXNRD1;refGene_tx=XM_023257386.1
B4 126570608 126571766
refGene_gn=EID3;refGene_tx=XM_003989195.5
B4 126578668 126578974
refGene_gn=TXNRD1;refGene_tx=XM_019835431.2
B4 126580719 126580842
refGene_gn=TXNRD1;refGene_tx=XM_006933988.4;refGene_gn=TXNRD1;refGene_tx=XM_006933989.4;refGene_gn=TXNRD1;refGene_tx=XM_019835431.2;refGene_gn=TXNRD1;refGene_tx=XM_023257386.1
B4 126582959 126583032
refGene_gn=TXNRD1;refGene_tx=XM_006933988.4;refGene_gn=TXNRD1;refGene_tx=XM_006933989.4;refGene_gn=TXNRD1;refGene_tx=XM_019835431.2;refGene_gn=TXNRD1;refGene_tx=XM_023257386.1
B4 126585513 126585633
refGene_gn=TXNRD1;refGene_tx=XM_006933988.4;refGene_gn=TXNRD1;refGene_tx=XM_006933989.4;refGene_gn=TXNRD1;refGene_tx=XM_019835431.2;refGene_gn=TXNRD1;refGene_tx=XM_023257386.1
B4 126590935 126591078
refGene_gn=TXNRD1;refGene_tx=XM_006933988.4;refGene_gn=TXNRD1;refGene_tx=XM_006933989.4;refGene_gn=TXNRD1;refGene_tx=XM_019835431.2;refGene_gn=TXNRD1;refGene_tx=XM_023257386.1

B4 126591492 126591608
refGene_gn=TXNRD1;refGene_tx=XM_006933988.4;refGene_gn=TXNRD1;refGene_tx=XM_006933989.4;refGene_gn=TXNRD1;refGene_tx=XM_019835431.2;refGene_gn=TXNRD1;refGene_tx=XM_023257386.1
B4 126592603 126592829
refGene_gn=TXNRD1;refGene_tx=XM_006933988.4;refGene_gn=TXNRD1;refGene_tx=XM_006933989.4;refGene_gn=TXNRD1;refGene_tx=XM_019835431.2;refGene_gn=TXNRD1;refGene_tx=XM_023257386.1
B4 126597612 126597705
refGene_gn=TXNRD1;refGene_tx=XM_006933988.4;refGene_gn=TXNRD1;refGene_tx=XM_006933989.4;refGene_gn=TXNRD1;refGene_tx=XM_019835431.2;refGene_gn=TXNRD1;refGene_tx=XM_023257386.1
B4 126598733 126598810
refGene_gn=TXNRD1;refGene_tx=XM_006933988.4;refGene_gn=TXNRD1;refGene_tx=XM_006933989.4;refGene_gn=TXNRD1;refGene_tx=XM_019835431.2;refGene_gn=TXNRD1;refGene_tx=XM_023257386.1
B4 126600123 126600280
refGene_gn=TXNRD1;refGene_tx=XM_006933988.4;refGene_gn=TXNRD1;refGene_tx=XM_006933989.4;refGene_gn=TXNRD1;refGene_tx=XM_019835431.2;refGene_gn=TXNRD1;refGene_tx=XM_023257386.1
B4 126606827 126606935
refGene_gn=TXNRD1;refGene_tx=XM_006933988.4;refGene_gn=TXNRD1;refGene_tx=XM_006933989.4;refGene_gn=TXNRD1;refGene_tx=XM_019835431.2;refGene_gn=TXNRD1;refGene_tx=XM_023257386.1
B4 126609793 126609889
refGene_gn=TXNRD1;refGene_tx=XM_006933988.4;refGene_gn=TXNRD1;refGene_tx=XM_006933989.4;refGene_gn=TXNRD1;refGene_tx=XM_019835431.2;refGene_gn=TXNRD1;refGene_tx=XM_023257386.1
B4 126613889 126614024
refGene_gn=TXNRD1;refGene_tx=XM_006933988.4;refGene_gn=TXNRD1;refGene_tx=XM_006933989.4;refGene_gn=TXNRD1;refGene_tx=XM_019835431.2;refGene_gn=TXNRD1;refGene_tx=XM_023257386.1
B4 126626410 126626473
refGene_gn=TXNRD1;refGene_tx=XM_006933988.4;refGene_gn=TXNRD1;refGene_tx=XM_006933989.4;refGene_gn=TXNRD1;refGene_tx=XM_019835431.2
B4 126658822 126658885
refGene_gn=TXNRD1;refGene_tx=XM_023257386.1
B4 126717688 126717806
refGene_gn=CHST11;refGene_tx=XM_011284184.3;refGene_gn=CHST11;refGene_tx=XM_011284185.3
B4 126846008 126846094
refGene_gn=CHST11;refGene_tx=XM_011284184.3;refGene_gn=CHST11;refGene_tx=XM_011284185.3;refGene_gn=CHST11;refGene_tx=XM_019835432.2
B4 126984341 126985196
refGene_gn=CHST11;refGene_tx=XM_011284184.3;refGene_gn=CHST11;refGene_tx=XM_011284185.3;refGene_gn=CHST11;refGene_tx=XM_019835432.2
B4 127027674 127027860
refGene_gn=SLC41A2;refGene_tx=XM_003989197.5;refGene_gn=SLC41A2;refGene_tx=XM_019835434.2;refGene_gn=SLC41A2;refGene_tx=XM_019835435.2;refGene_gn=SLC41A2;refGene_tx=XM_019835436.2;refGene_gn=SLC41A2;refGene_tx=

XM_023257387.1;refGene_gn=SLC41A2;refGene_tx=XM_023257388.1
B4 127056999 127057398
refGene_gn=LOC101086240;refGene_tx=XM_011284186.3
B4 127065848 127065997
refGene_gn=SLC41A2;refGene_tx=XM_003989197.5;refGene_gn=SLC41A2;refGene_tx=XM_019835434.2;refGene_gn=SLC41A2;refGene_tx=XM_019835435.2;refGene_gn=SLC41A2;refGene_tx=XM_019835436.2;refGene_gn=SLC41A2;refGene_tx=XM_023257387.1;refGene_gn=SLC41A2;refGene_tx=XM_023257388.1
B4 127067558 127067690
refGene_gn=SLC41A2;refGene_tx=XM_003989197.5;refGene_gn=SLC41A2;refGene_tx=XM_019835434.2;refGene_gn=SLC41A2;refGene_tx=XM_019835435.2;refGene_gn=SLC41A2;refGene_tx=XM_019835436.2;refGene_gn=SLC41A2;refGene_tx=XM_023257387.1;refGene_gn=SLC41A2;refGene_tx=XM_023257388.1
B4 127071364 127071444
refGene_gn=SLC41A2;refGene_tx=XM_003989197.5;refGene_gn=SLC41A2;refGene_tx=XM_019835434.2;refGene_gn=SLC41A2;refGene_tx=XM_019835435.2;refGene_gn=SLC41A2;refGene_tx=XM_019835436.2;refGene_gn=SLC41A2;refGene_tx=XM_023257387.1;refGene_gn=SLC41A2;refGene_tx=XM_023257388.1
B4 127076695 127076843
refGene_gn=SLC41A2;refGene_tx=XM_003989197.5;refGene_gn=SLC41A2;refGene_tx=XM_019835434.2;refGene_gn=SLC41A2;refGene_tx=XM_019835435.2;refGene_gn=SLC41A2;refGene_tx=XM_019835436.2;refGene_gn=SLC41A2;refGene_tx=XM_023257387.1;refGene_gn=SLC41A2;refGene_tx=XM_023257388.1
B4 127086710 127086857
refGene_gn=SLC41A2;refGene_tx=XM_003989197.5;refGene_gn=SLC41A2;refGene_tx=XM_019835434.2;refGene_gn=SLC41A2;refGene_tx=XM_019835435.2;refGene_gn=SLC41A2;refGene_tx=XM_019835436.2;refGene_gn=SLC41A2;refGene_tx=XM_023257387.1;refGene_gn=SLC41A2;refGene_tx=XM_023257388.1
B4 127091128 127091273
refGene_gn=SLC41A2;refGene_tx=XM_003989197.5;refGene_gn=SLC41A2;refGene_tx=XM_019835434.2;refGene_gn=SLC41A2;refGene_tx=XM_019835435.2;refGene_gn=SLC41A2;refGene_tx=XM_019835436.2;refGene_gn=SLC41A2;refGene_tx=XM_023257387.1;refGene_gn=SLC41A2;refGene_tx=XM_023257388.1
B4 127093415 127093487
refGene_gn=SLC41A2;refGene_tx=XM_003989197.5;refGene_gn=SLC41A2;refGene_tx=XM_019835434.2;refGene_gn=SLC41A2;refGene_tx=XM_019835435.2;refGene_gn=SLC41A2;refGene_tx=XM_019835436.2;refGene_gn=SLC41A2;refGene_tx=XM_023257387.1;refGene_gn=SLC41A2;refGene_tx=XM_023257388.1
B4 127105394 127105502
refGene_gn=SLC41A2;refGene_tx=XM_003989197.5;refGene_gn=SLC41A2;refGene_tx=XM_019835434.2;refGene_gn=SLC41A2;refGene_tx=XM_019835435.2;refGene_gn=SLC41A2;refGene_tx=XM_019835436.2;refGene_gn=SLC41A2;refGene_tx=XM_023257387.1;refGene_gn=SLC41A2;refGene_tx=XM_023257388.1
B4 127124745 127125300
refGene_gn=SLC41A2;refGene_tx=XM_003989197.5;refGene_gn=SLC41A2;refGene_tx=XM_019835434.2;refGene_gn=SLC41A2;refGene_tx=XM_019835435.2;refGene_gn=SLC41A2;refGene_tx=XM_019835436.2;refGene_gn=SLC41A2;refGene_tx=XM_023257387.1;refGene_gn=SLC41A2;refGene_tx=XM_023257388.1
B4 127171625 127171737
refGene_gn=CB4H12orf45;refGene_tx=XM_003989198.4

B4	127173563	127173650
refGene_gn=CB4H12orf45;refGene_tx=XM_003989198.4
B4	127176097	127176234
refGene_gn=CB4H12orf45;refGene_tx=XM_003989198.4
B4	127178195	127178414
refGene_gn=CB4H12orf45;refGene_tx=XM_003989198.4
B4	127187669	127187725
refGene_gn=ALDH1L2;refGene_tx=XM_023257389.1;refGene_gn=ALDH1L2;refGene_tx=XM_023257390.1
B4	127189591	127189791
refGene_gn=ALDH1L2;refGene_tx=XM_023257389.1;refGene_gn=ALDH1L2;refGene_tx=XM_023257390.1
B4	127192480	127192586
refGene_gn=ALDH1L2;refGene_tx=XM_023257389.1;refGene_gn=ALDH1L2;refGene_tx=XM_023257390.1
B4	127194334	127194500
refGene_gn=ALDH1L2;refGene_tx=XM_023257389.1;refGene_gn=ALDH1L2;refGene_tx=XM_023257390.1
B4	127196976	127197075
refGene_gn=ALDH1L2;refGene_tx=XM_023257389.1;refGene_gn=ALDH1L2;refGene_tx=XM_023257390.1
B4	127197684	127197784
refGene_gn=ALDH1L2;refGene_tx=XM_023257389.1;refGene_gn=ALDH1L2;refGene_tx=XM_023257390.1
B4	127199386	127199480
refGene_gn=ALDH1L2;refGene_tx=XM_023257389.1;refGene_gn=ALDH1L2;refGene_tx=XM_023257390.1
B4	127199958	127200046
refGene_gn=ALDH1L2;refGene_tx=XM_023257389.1;refGene_gn=ALDH1L2;refGene_tx=XM_023257390.1
B4	127208392	127208498
refGene_gn=ALDH1L2;refGene_tx=XM_023257389.1;refGene_gn=ALDH1L2;refGene_tx=XM_023257390.1
B4	127208584	127208655
refGene_gn=ALDH1L2;refGene_tx=XM_023257389.1;refGene_gn=ALDH1L2;refGene_tx=XM_023257390.1
B4	127211273	127211424
refGene_gn=ALDH1L2;refGene_tx=XM_023257389.1;refGene_gn=ALDH1L2;refGene_tx=XM_023257390.1
B4	127213127	127213255
refGene_gn=ALDH1L2;refGene_tx=XM_023257389.1;refGene_gn=ALDH1L2;refGene_tx=XM_023257390.1
B4	127213852	127213972
refGene_gn=ALDH1L2;refGene_tx=XM_023257389.1;refGene_gn=ALDH1L2;refGene_tx=XM_023257390.1
B4	127225978	127226126
refGene_gn=ALDH1L2;refGene_tx=XM_023257389.1;refGene_gn=ALDH1L2;refGene_tx=XM_023257390.1
B4	127227422	127227514
refGene_gn=ALDH1L2;refGene_tx=XM_023257389.1;refGene_gn=ALDH1L2;refGen

e_tx=XM_023257390.1
B4 127228079 127228205
refGene_gn=ALDH1L2;refGene_tx=XM_023257389.1;refGene_gn=ALDH1L2;refGen
e_tx=XM_023257390.1
B4 127229366 127229501
refGene_gn=ALDH1L2;refGene_tx=XM_023257389.1;refGene_gn=ALDH1L2;refGen
e_tx=XM_023257390.1
B4 127231342 127231432
refGene_gn=ALDH1L2;refGene_tx=XM_023257389.1;refGene_gn=ALDH1L2;refGen
e_tx=XM_023257390.1
B4 127231642 127231744
refGene_gn=ALDH1L2;refGene_tx=XM_023257389.1;refGene_gn=ALDH1L2;refGen
e_tx=XM_023257390.1
B4 127233460 127233626
refGene_gn=ALDH1L2;refGene_tx=XM_023257389.1;refGene_gn=ALDH1L2;refGen
e_tx=XM_023257390.1
B4 127236457 127236692
refGene_gn=ALDH1L2;refGene_tx=XM_023257389.1;refGene_gn=ALDH1L2;refGen
e_tx=XM_023257390.1
B4 127241659 127241804
refGene_gn=ALDH1L2;refGene_tx=XM_023257389.1;refGene_gn=ALDH1L2;refGen
e_tx=XM_023257390.1
B4 127253778 127253826
refGene_gn=ALDH1L2;refGene_tx=XM_023257389.1
B4 127266462 127266523
refGene_gn=WASHC4;refGene_tx=XM_023257391.1;refGene_gn=WASHC4;refGene_
tx=XM_023257392.1;refGene_gn=WASHC4;refGene_tx=XM_023257393.1
B4 127269393 127269533
refGene_gn=WASHC4;refGene_tx=XM_023257391.1;refGene_gn=WASHC4;refGene_
tx=XM_023257392.1;refGene_gn=WASHC4;refGene_tx=XM_023257393.1
B4 127270869 127270923
refGene_gn=WASHC4;refGene_tx=XM_023257391.1;refGene_gn=WASHC4;refGene_
tx=XM_023257392.1;refGene_gn=WASHC4;refGene_tx=XM_023257393.1
B4 127271017 127271083
refGene_gn=WASHC4;refGene_tx=XM_023257391.1;refGene_gn=WASHC4;refGene_
tx=XM_023257392.1;refGene_gn=WASHC4;refGene_tx=XM_023257393.1
B4 127271980 127272026
refGene_gn=WASHC4;refGene_tx=XM_023257391.1;refGene_gn=WASHC4;refGene_
tx=XM_023257392.1;refGene_gn=WASHC4;refGene_tx=XM_023257393.1
B4 127272492 127272560
refGene_gn=WASHC4;refGene_tx=XM_023257391.1;refGene_gn=WASHC4;refGene_
tx=XM_023257392.1;refGene_gn=WASHC4;refGene_tx=XM_023257393.1
B4 127275671 127275754
refGene_gn=WASHC4;refGene_tx=XM_023257391.1;refGene_gn=WASHC4;refGene_
tx=XM_023257392.1;refGene_gn=WASHC4;refGene_tx=XM_023257393.1
B4 127277312 127277355
refGene_gn=WASHC4;refGene_tx=XM_023257391.1;refGene_gn=WASHC4;refGene_
tx=XM_023257392.1;refGene_gn=WASHC4;refGene_tx=XM_023257393.1
B4 127277816 127277920
refGene_gn=WASHC4;refGene_tx=XM_023257391.1;refGene_gn=WASHC4;refGene_

tx=XM_023257392.1;refGene_gn=WASHC4;refGene_tx=XM_023257393.1
B4	127279382	127279503
refGene_gn=WASHC4;refGene_tx=XM_023257391.1;refGene_gn=WASHC4;refGene_tx=XM_023257392.1;refGene_gn=WASHC4;refGene_tx=XM_023257393.1
B4	127281006	127281130
refGene_gn=WASHC4;refGene_tx=XM_023257391.1;refGene_gn=WASHC4;refGene_tx=XM_023257392.1;refGene_gn=WASHC4;refGene_tx=XM_023257393.1
B4	127281229	127281357
refGene_gn=WASHC4;refGene_tx=XM_023257391.1;refGene_gn=WASHC4;refGene_tx=XM_023257392.1;refGene_gn=WASHC4;refGene_tx=XM_023257393.1
B4	127282354	127282515
refGene_gn=WASHC4;refGene_tx=XM_023257391.1;refGene_gn=WASHC4;refGene_tx=XM_023257392.1;refGene_gn=WASHC4;refGene_tx=XM_023257393.1
B4	127288243	127288370
refGene_gn=WASHC4;refGene_tx=XM_023257391.1;refGene_gn=WASHC4;refGene_tx=XM_023257392.1;refGene_gn=WASHC4;refGene_tx=XM_023257393.1
B4	127293091	127293217
refGene_gn=WASHC4;refGene_tx=XM_023257391.1;refGene_gn=WASHC4;refGene_tx=XM_023257392.1;refGene_gn=WASHC4;refGene_tx=XM_023257393.1
B4	127296891	127296999
refGene_gn=WASHC4;refGene_tx=XM_023257391.1;refGene_gn=WASHC4;refGene_tx=XM_023257393.1
B4	127298092	127298239
refGene_gn=WASHC4;refGene_tx=XM_023257391.1;refGene_gn=WASHC4;refGene_tx=XM_023257392.1;refGene_gn=WASHC4;refGene_tx=XM_023257393.1
B4	127298364	127298444
refGene_gn=WASHC4;refGene_tx=XM_023257391.1;refGene_gn=WASHC4;refGene_tx=XM_023257392.1;refGene_gn=WASHC4;refGene_tx=XM_023257393.1
B4	127299575	127299681
refGene_gn=WASHC4;refGene_tx=XM_023257391.1;refGene_gn=WASHC4;refGene_tx=XM_023257392.1;refGene_gn=WASHC4;refGene_tx=XM_023257393.1
B4	127300270	127300387
refGene_gn=WASHC4;refGene_tx=XM_023257391.1;refGene_gn=WASHC4;refGene_tx=XM_023257392.1;refGene_gn=WASHC4;refGene_tx=XM_023257393.1
B4	127302002	127302171
refGene_gn=WASHC4;refGene_tx=XM_023257391.1;refGene_gn=WASHC4;refGene_tx=XM_023257392.1;refGene_gn=WASHC4;refGene_tx=XM_023257393.1
B4	127302452	127302607
refGene_gn=WASHC4;refGene_tx=XM_023257391.1;refGene_gn=WASHC4;refGene_tx=XM_023257392.1;refGene_gn=WASHC4;refGene_tx=XM_023257393.1
B4	127304109	127304184
refGene_gn=WASHC4;refGene_tx=XM_023257391.1;refGene_gn=WASHC4;refGene_tx=XM_023257392.1;refGene_gn=WASHC4;refGene_tx=XM_023257393.1
B4	127304452	127304557
refGene_gn=WASHC4;refGene_tx=XM_023257391.1;refGene_gn=WASHC4;refGene_tx=XM_023257392.1;refGene_gn=WASHC4;refGene_tx=XM_023257393.1
B4	127307453	127307588
refGene_gn=WASHC4;refGene_tx=XM_023257391.1;refGene_gn=WASHC4;refGene_tx=XM_023257392.1;refGene_gn=WASHC4;refGene_tx=XM_023257393.1
B4	127309246	127309355

refGene_gn=WASHC4;refGene_tx=XM_023257391.1;refGene_gn=WASHC4;refGene_tx=XM_023257392.1;refGene_gn=WASHC4;refGene_tx=XM_023257393.1
B4 127313838 127313905
refGene_gn=WASHC4;refGene_tx=XM_023257391.1;refGene_gn=WASHC4;refGene_tx=XM_023257392.1;refGene_gn=WASHC4;refGene_tx=XM_023257393.1
B4 127314348 127314435
refGene_gn=WASHC4;refGene_tx=XM_023257391.1;refGene_gn=WASHC4;refGene_tx=XM_023257392.1;refGene_gn=WASHC4;refGene_tx=XM_023257393.1
B4 127318211 127318359
refGene_gn=WASHC4;refGene_tx=XM_023257391.1;refGene_gn=WASHC4;refGene_tx=XM_023257392.1;refGene_gn=WASHC4;refGene_tx=XM_023257393.1
B4 127320738 127320835
refGene_gn=WASHC4;refGene_tx=XM_023257391.1;refGene_gn=WASHC4;refGene_tx=XM_023257392.1;refGene_gn=WASHC4;refGene_tx=XM_023257393.1
B4 127321349 127321546
refGene_gn=WASHC4;refGene_tx=XM_023257391.1;refGene_gn=WASHC4;refGene_tx=XM_023257392.1;refGene_gn=WASHC4;refGene_tx=XM_023257393.1
B4 127321806 127321903
refGene_gn=WASHC4;refGene_tx=XM_023257391.1;refGene_gn=WASHC4;refGene_tx=XM_023257392.1;refGene_gn=WASHC4;refGene_tx=XM_023257393.1
B4 127323815 127323883
refGene_gn=WASHC4;refGene_tx=XM_023257391.1;refGene_gn=WASHC4;refGene_tx=XM_023257392.1
B4 127331942 127331969
refGene_gn=APPL2;refGene_tx=XM_023257396.1
B4 127332048 127332183
refGene_gn=APPL2;refGene_tx=XM_023257394.1;refGene_gn=APPL2;refGene_tx=XM_023257395.1
B4 127333411 127333459
refGene_gn=APPL2;refGene_tx=XM_023257394.1;refGene_gn=APPL2;refGene_tx=XM_023257395.1;refGene_gn=APPL2;refGene_tx=XM_023257396.1
B4 127334196 127334337
refGene_gn=APPL2;refGene_tx=XM_023257394.1;refGene_gn=APPL2;refGene_tx=XM_023257395.1;refGene_gn=APPL2;refGene_tx=XM_023257396.1
B4 127334522 127334559
refGene_gn=APPL2;refGene_tx=XM_023257394.1;refGene_gn=APPL2;refGene_tx=XM_023257395.1;refGene_gn=APPL2;refGene_tx=XM_023257396.1
B4 127338535 127338710
refGene_gn=APPL2;refGene_tx=XM_023257394.1;refGene_gn=APPL2;refGene_tx=XM_023257395.1;refGene_gn=APPL2;refGene_tx=XM_023257396.1
B4 127340264 127340317
refGene_gn=APPL2;refGene_tx=XM_023257394.1;refGene_gn=APPL2;refGene_tx=XM_023257395.1;refGene_gn=APPL2;refGene_tx=XM_023257396.1
B4 127340455 127340614
refGene_gn=APPL2;refGene_tx=XM_023257394.1;refGene_gn=APPL2;refGene_tx=XM_023257395.1;refGene_gn=APPL2;refGene_tx=XM_023257396.1
B4 127345155 127345244
refGene_gn=APPL2;refGene_tx=XM_023257394.1;refGene_gn=APPL2;refGene_tx=XM_023257395.1;refGene_gn=APPL2;refGene_tx=XM_023257396.1
B4 127345340 127345397

refGene_gn=APPL2;refGene_tx=XM_023257394.1;refGene_gn=APPL2;refGene_tx=XM_023257395.1;refGene_gn=APPL2;refGene_tx=XM_023257396.1
B4 127345479 127345522
refGene_gn=APPL2;refGene_tx=XM_023257394.1;refGene_gn=APPL2;refGene_tx=XM_023257395.1;refGene_gn=APPL2;refGene_tx=XM_023257396.1
B4 127346778 127346967
refGene_gn=APPL2;refGene_tx=XM_023257394.1;refGene_gn=APPL2;refGene_tx=XM_023257395.1;refGene_gn=APPL2;refGene_tx=XM_023257396.1
B4 127348347 127348506
refGene_gn=APPL2;refGene_tx=XM_023257394.1;refGene_gn=APPL2;refGene_tx=XM_023257395.1;refGene_gn=APPL2;refGene_tx=XM_023257396.1
B4 127352156 127352239
refGene_gn=APPL2;refGene_tx=XM_023257394.1;refGene_gn=APPL2;refGene_tx=XM_023257395.1;refGene_gn=APPL2;refGene_tx=XM_023257396.1
B4 127356645 127356792
refGene_gn=APPL2;refGene_tx=XM_023257394.1;refGene_gn=APPL2;refGene_tx=XM_023257395.1;refGene_gn=APPL2;refGene_tx=XM_023257396.1
B4 127357944 127358003
refGene_gn=APPL2;refGene_tx=XM_023257394.1;refGene_gn=APPL2;refGene_tx=XM_023257395.1;refGene_gn=APPL2;refGene_tx=XM_023257396.1
B4 127358365 127358407
refGene_gn=APPL2;refGene_tx=XM_023257394.1;refGene_gn=APPL2;refGene_tx=XM_023257395.1;refGene_gn=APPL2;refGene_tx=XM_023257396.1
B4 127361413 127361501
refGene_gn=APPL2;refGene_tx=XM_023257394.1;refGene_gn=APPL2;refGene_tx=XM_023257395.1;refGene_gn=APPL2;refGene_tx=XM_023257396.1
B4 127365751 127365823
refGene_gn=APPL2;refGene_tx=XM_023257394.1;refGene_gn=APPL2;refGene_tx=XM_023257395.1;refGene_gn=APPL2;refGene_tx=XM_023257396.1
B4 127366374 127366434
refGene_gn=APPL2;refGene_tx=XM_023257394.1;refGene_gn=APPL2;refGene_tx=XM_023257395.1;refGene_gn=APPL2;refGene_tx=XM_023257396.1
B4 127377703 127377802
refGene_gn=APPL2;refGene_tx=XM_023257394.1;refGene_gn=APPL2;refGene_tx=XM_023257396.1
B4 127384726 127384780
refGene_gn=APPL2;refGene_tx=XM_023257394.1;refGene_gn=APPL2;refGene_tx=XM_023257395.1;refGene_gn=APPL2;refGene_tx=XM_023257396.1
B4 127453438 127454061
refGene_gn=CB4H12orf75;refGene_tx=XM_023257397.1
B4 127485683 127485719
refGene_gn=CB4H12orf75;refGene_tx=XM_023257397.1
B4 127486774 127486854
refGene_gn=CB4H12orf75;refGene_tx=XM_023257397.1
B4 127487666 127487671
refGene_gn=CB4H12orf75;refGene_tx=XM_023257397.1
B4 128080872 128082026
refGene_gn=NUAK1;refGene_tx=XM_023257398.1
B4 128085777 128085910
refGene_gn=NUAK1;refGene_tx=XM_023257398.1

B4 128087716 128087836
refGene_gn=NUAK1;refGene_tx=XM_023257398.1
B4 128097105 128097171
refGene_gn=NUAK1;refGene_tx=XM_023257398.1
B4 128100229 128100381
refGene_gn=NUAK1;refGene_tx=XM_023257398.1
B4 128117716 128117837
refGene_gn=NUAK1;refGene_tx=XM_023257398.1
B4 128149587 128149845
refGene_gn=NUAK1;refGene_tx=XM_023257398.1
B4 128226078 128227401
refGene_gn=CKAP4;refGene_tx=XM_023257399.1
B4 128234092 128235260
refGene_gn=CKAP4;refGene_tx=XM_023257399.1;refGene_gn=LOC109501766;refGene_tx=XM_023257400.1
B4 128280997 128281154
refGene_gn=TCP11L2;refGene_tx=XM_006933990.4;refGene_gn=TCP11L2;refGene_tx=XM_019835444.2
B4 128283787 128283923
refGene_gn=TCP11L2;refGene_tx=XM_006933990.4;refGene_gn=TCP11L2;refGene_tx=XM_019835444.2
B4 128287552 128287673
refGene_gn=TCP11L2;refGene_tx=XM_006933990.4;refGene_gn=TCP11L2;refGene_tx=XM_019835444.2
B4 128288966 128289187
refGene_gn=TCP11L2;refGene_tx=XM_006933990.4;refGene_gn=TCP11L2;refGene_tx=XM_019835444.2
B4 128291341 128291478
refGene_gn=TCP11L2;refGene_tx=XM_006933990.4;refGene_gn=TCP11L2;refGene_tx=XM_019835444.2
B4 128302279 128302464
refGene_gn=TCP11L2;refGene_tx=XM_006933990.4;refGene_gn=TCP11L2;refGene_tx=XM_019835444.2
B4 128302659 128302841
refGene_gn=TCP11L2;refGene_tx=XM_006933990.4
B4 128305199 128305372
refGene_gn=TCP11L2;refGene_tx=XM_006933990.4;refGene_gn=TCP11L2;refGene_tx=XM_019835444.2
B4 128311089 128311331
refGene_gn=TCP11L2;refGene_tx=XM_006933990.4
B4 128320846 128320918
refGene_gn=POLR3B;refGene_tx=XM_003989200.5;refGene_gn=POLR3B;refGene_tx=XM_023257401.1
B4 128323707 128323740
refGene_gn=POLR3B;refGene_tx=XM_003989200.5;refGene_gn=POLR3B;refGene_tx=XM_023257401.1
B4 128326311 128326368
refGene_gn=POLR3B;refGene_tx=XM_003989200.5;refGene_gn=POLR3B;refGene_tx=XM_023257401.1
B4 128326559 128326624

refGene_gn=POLR3B;refGene_tx=XM_003989200.5;refGene_gn=POLR3B;refGene_tx=XM_023257401.1
B4 128329375 128329451
refGene_gn=POLR3B;refGene_tx=XM_003989200.5;refGene_gn=POLR3B;refGene_tx=XM_023257401.1
B4 128332547 128332648
refGene_gn=POLR3B;refGene_tx=XM_003989200.5;refGene_gn=POLR3B;refGene_tx=XM_023257401.1
B4 128334952 128335044
refGene_gn=POLR3B;refGene_tx=XM_003989200.5;refGene_gn=POLR3B;refGene_tx=XM_023257401.1
B4 128336184 128336302
refGene_gn=POLR3B;refGene_tx=XM_003989200.5;refGene_gn=POLR3B;refGene_tx=XM_023257401.1
B4 128338381 128338490
refGene_gn=POLR3B;refGene_tx=XM_003989200.5;refGene_gn=POLR3B;refGene_tx=XM_023257401.1
B4 128346510 128346633
refGene_gn=POLR3B;refGene_tx=XM_003989200.5;refGene_gn=POLR3B;refGene_tx=XM_023257401.1
B4 128351401 128351521
refGene_gn=POLR3B;refGene_tx=XM_003989200.5;refGene_gn=POLR3B;refGene_tx=XM_023257401.1
B4 128354064 128354199
refGene_gn=POLR3B;refGene_tx=XM_003989200.5;refGene_gn=POLR3B;refGene_tx=XM_023257401.1
B4 128363027 128363189
refGene_gn=POLR3B;refGene_tx=XM_003989200.5;refGene_gn=POLR3B;refGene_tx=XM_023257401.1
B4 128365878 128366079
refGene_gn=POLR3B;refGene_tx=XM_003989200.5;refGene_gn=POLR3B;refGene_tx=XM_023257401.1
B4 128367871 128368034
refGene_gn=POLR3B;refGene_tx=XM_003989200.5;refGene_gn=POLR3B;refGene_tx=XM_023257401.1
B4 128369187 128369341
refGene_gn=POLR3B;refGene_tx=XM_003989200.5;refGene_gn=POLR3B;refGene_tx=XM_023257401.1
B4 128372897 128372972
refGene_gn=POLR3B;refGene_tx=XM_003989200.5;refGene_gn=POLR3B;refGene_tx=XM_023257401.1
B4 128373538 128373637
refGene_gn=POLR3B;refGene_tx=XM_003989200.5;refGene_gn=POLR3B;refGene_tx=XM_023257401.1
B4 128378266 128378394
refGene_gn=POLR3B;refGene_tx=XM_003989200.5;refGene_gn=POLR3B;refGene_tx=XM_023257401.1
B4 128386339 128386549
refGene_gn=POLR3B;refGene_tx=XM_003989200.5;refGene_gn=POLR3B;refGene_tx=XM_023257401.1

B4       128388864        128389023
refGene_gn=POLR3B;refGene_tx=XM_003989200.5;refGene_gn=POLR3B;refGene_tx=XM_023257401.1
B4       128390723        128390841
refGene_gn=POLR3B;refGene_tx=XM_003989200.5;refGene_gn=POLR3B;refGene_tx=XM_023257401.1
B4       128392722        128392865
refGene_gn=POLR3B;refGene_tx=XM_003989200.5;refGene_gn=POLR3B;refGene_tx=XM_023257401.1
B4       128393107        128393145
refGene_gn=POLR3B;refGene_tx=XM_023257401.1
B4       128410733        128410837
refGene_gn=POLR3B;refGene_tx=XM_003989200.5
B4       128411420        128411587
refGene_gn=POLR3B;refGene_tx=XM_003989200.5
B4       128415124        128415238
refGene_gn=POLR3B;refGene_tx=XM_003989200.5
B4       128418141        128418315
refGene_gn=POLR3B;refGene_tx=XM_003989200.5
B4       128423645        128423775
refGene_gn=POLR3B;refGene_tx=XM_003989200.5
B4       128469132        128469175
refGene_gn=RFX4;refGene_tx=XM_019835449.2
B4       128495613        128495700
refGene_gn=RFX4;refGene_tx=XM_019835449.2
B4       128517191        128517252
refGene_gn=RFX4;refGene_tx=XM_019835449.2
B4       128530708        128530832
refGene_gn=RFX4;refGene_tx=XM_019835449.2
B4       128549779        128549841
refGene_gn=RFX4;refGene_tx=XM_019835449.2
B4       128552351        128552446
refGene_gn=RFX4;refGene_tx=XM_006933991.4
B4       128554334        128554548
refGene_gn=RFX4;refGene_tx=XM_006933991.4;refGene_gn=RFX4;refGene_tx=XM_019835449.2
B4       128555879        128555957
refGene_gn=RFX4;refGene_tx=XM_006933991.4;refGene_gn=RFX4;refGene_tx=XM_019835449.2
B4       128562238        128562402
refGene_gn=RFX4;refGene_tx=XM_006933991.4;refGene_gn=RFX4;refGene_tx=XM_019835449.2
B4       128585077        128585178
refGene_gn=RFX4;refGene_tx=XM_006933991.4;refGene_gn=RFX4;refGene_tx=XM_019835449.2
B4       128587328        128587387
refGene_gn=RFX4;refGene_tx=XM_006933991.4;refGene_gn=RFX4;refGene_tx=XM_019835449.2
B4       128591256        128591401
refGene_gn=RFX4;refGene_tx=XM_006933991.4;refGene_gn=RFX4;refGene_tx=X

M_019835449.2
B4	128595598	128595693
refGene_gn=RFX4;refGene_tx=XM_006933991.4;refGene_gn=RFX4;refGene_tx=XM_019835449.2
B4	128596121	128596239
refGene_gn=RFX4;refGene_tx=XM_006933991.4;refGene_gn=RFX4;refGene_tx=XM_019835449.2
B4	128608512	128608632
refGene_gn=RFX4;refGene_tx=XM_006933991.4;refGene_gn=RFX4;refGene_tx=XM_019835449.2
B4	128608973	128609135
refGene_gn=RFX4;refGene_tx=XM_006933991.4;refGene_gn=RFX4;refGene_tx=XM_019835449.2
B4	128621328	128621491
refGene_gn=RFX4;refGene_tx=XM_006933991.4;refGene_gn=RFX4;refGene_tx=XM_019835449.2
B4	128623435	128623574
refGene_gn=RFX4;refGene_tx=XM_006933991.4;refGene_gn=RFX4;refGene_tx=XM_019835449.2
B4	128632068	128632341
refGene_gn=RFX4;refGene_tx=XM_006933991.4;refGene_gn=RFX4;refGene_tx=XM_019835449.2
B4	128644845	128644929
refGene_gn=RIC8B;refGene_tx=XM_003989204.4;refGene_gn=RIC8B;refGene_tx=XM_003989206.4;refGene_gn=RIC8B;refGene_tx=XM_019835451.2;refGene_gn=RIC8B;refGene_tx=XM_023257402.1
B4	128653746	128653794
refGene_gn=RIC8B;refGene_tx=XM_003989204.4;refGene_gn=RIC8B;refGene_tx=XM_003989206.4;refGene_gn=RIC8B;refGene_tx=XM_019835451.2;refGene_gn=RIC8B;refGene_tx=XM_023257402.1
B4	128685159	128685768
refGene_gn=RIC8B;refGene_tx=XM_003989204.4;refGene_gn=RIC8B;refGene_tx=XM_003989206.4;refGene_gn=RIC8B;refGene_tx=XM_019835451.2;refGene_gn=RIC8B;refGene_tx=XM_023257402.1
B4	128695808	128695903
refGene_gn=RIC8B;refGene_tx=XM_003989204.4;refGene_gn=RIC8B;refGene_tx=XM_003989206.4;refGene_gn=RIC8B;refGene_tx=XM_019835451.2;refGene_gn=RIC8B;refGene_tx=XM_023257402.1
B4	128707042	128707271
refGene_gn=RIC8B;refGene_tx=XM_003989204.4;refGene_gn=RIC8B;refGene_tx=XM_003989206.4;refGene_gn=RIC8B;refGene_tx=XM_019835451.2;refGene_gn=RIC8B;refGene_tx=XM_023257402.1
B4	128708021	128708117
refGene_gn=RIC8B;refGene_tx=XM_003989204.4;refGene_gn=RIC8B;refGene_tx=XM_003989206.4;refGene_gn=RIC8B;refGene_tx=XM_019835451.2;refGene_gn=RIC8B;refGene_tx=XM_023257402.1
B4	128713364	128713509
refGene_gn=RIC8B;refGene_tx=XM_003989204.4;refGene_gn=RIC8B;refGene_tx=XM_003989206.4;refGene_gn=RIC8B;refGene_tx=XM_019835451.2;refGene_gn=RIC8B;refGene_tx=XM_023257402.1

B4 128724264 128724409
refGene_gn=RIC8B;refGene_tx=XM_003989204.4;refGene_gn=RIC8B;refGene_tx=XM_003989206.4;refGene_gn=RIC8B;refGene_tx=XM_019835451.2;refGene_gn=RIC8B;refGene_tx=XM_023257402.1
B4 128731475 128731595
refGene_gn=RIC8B;refGene_tx=XM_003989206.4;refGene_gn=RIC8B;refGene_tx=XM_023257402.1
B4 128734793 128734833
refGene_gn=RIC8B;refGene_tx=XM_019835451.2;refGene_gn=RIC8B;refGene_tx=XM_023257402.1
B4 128745863 128745975
refGene_gn=RIC8B;refGene_tx=XM_003989204.4;refGene_gn=RIC8B;refGene_tx=XM_003989206.4
B4 128761430 128761534
refGene_gn=LOC111561417;refGene_tx=XM_023257403.1
B4 128831866 128831927
refGene_gn=TMEM263;refGene_tx=XM_003989207.5
B4 128835907 128836194
refGene_gn=TMEM263;refGene_tx=XM_003989207.5
B4 128842554 128843712
refGene_gn=MTERF2;refGene_tx=XM_006933994.4;refGene_gn=MTERF2;refGene_tx=XM_011284188.3;refGene_gn=MTERF2;refGene_tx=XM_011284190.3;refGene_gn=MTERF2;refGene_tx=XM_011284191.2;refGene_gn=MTERF2;refGene_tx=XM_011284192.2;refGene_gn=MTERF2;refGene_tx=XM_011284193.3;refGene_gn=MTERF2;refGene_tx=XM_011284194.3
B4 128856064 128856168
refGene_gn=CRY1;refGene_tx=XM_003989209.5;refGene_gn=CRY1;refGene_tx=XM_019835452.1
B4 128856244 128856316
refGene_gn=CRY1;refGene_tx=XM_003989209.5;refGene_gn=CRY1;refGene_tx=XM_019835452.1
B4 128861235 128861331
refGene_gn=CRY1;refGene_tx=XM_003989209.5;refGene_gn=CRY1;refGene_tx=XM_019835452.1
B4 128861437 128861640
refGene_gn=CRY1;refGene_tx=XM_003989209.5;refGene_gn=CRY1;refGene_tx=XM_019835452.1
B4 128861873 128862025
refGene_gn=CRY1;refGene_tx=XM_003989209.5;refGene_gn=CRY1;refGene_tx=XM_019835452.1
B4 128864017 128864329
refGene_gn=CRY1;refGene_tx=XM_003989209.5;refGene_gn=CRY1;refGene_tx=XM_019835452.1
B4 128864410 128864551
refGene_gn=CRY1;refGene_tx=XM_003989209.5;refGene_gn=CRY1;refGene_tx=XM_019835452.1
B4 128865333 128865422
refGene_gn=CRY1;refGene_tx=XM_003989209.5;refGene_gn=CRY1;refGene_tx=XM_019835452.1
B4 128865827 128866012

refGene_gn=CRY1;refGene_tx=XM_003989209.5;refGene_gn=CRY1;refGene_tx=XM_019835452.1
B4 128869266 128869409
refGene_gn=CRY1;refGene_tx=XM_003989209.5;refGene_gn=CRY1;refGene_tx=XM_019835452.1
B4 128886075 128886184
refGene_gn=CRY1;refGene_tx=XM_003989209.5;refGene_gn=CRY1;refGene_tx=XM_019835452.1
B4 128909448 128909549
refGene_gn=CRY1;refGene_tx=XM_019835452.1
B4 128947089 128947247
refGene_gn=CRY1;refGene_tx=XM_003989209.5
B4 129138067 129139205
refGene_gn=BTBD11;refGene_tx=XM_023257404.1
B4 129321348 129321498
refGene_gn=BTBD11;refGene_tx=XM_023257404.1
B4 129337904 129338109
refGene_gn=BTBD11;refGene_tx=XM_023257404.1
B4 129375808 129375961
refGene_gn=BTBD11;refGene_tx=XM_023257404.1
B4 129404396 129404562
refGene_gn=BTBD11;refGene_tx=XM_023257404.1
B4 129406963 129407053
refGene_gn=BTBD11;refGene_tx=XM_023257404.1
B4 129408998 129409100
refGene_gn=BTBD11;refGene_tx=XM_023257404.1
B4 129411052 129411146
refGene_gn=BTBD11;refGene_tx=XM_023257404.1
B4 129411262 129411398
refGene_gn=BTBD11;refGene_tx=XM_023257404.1
B4 129412118 129412269
refGene_gn=BTBD11;refGene_tx=XM_023257404.1
B4 129412270 129412340
refGene_gn=BTBD11;refGene_tx=XM_023257404.1
B4 129413886 129414065
refGene_gn=BTBD11;refGene_tx=XM_023257404.1
B4 129428857 129428963
refGene_gn=BTBD11;refGene_tx=XM_023257404.1
B4 129433776 129433879
refGene_gn=BTBD11;refGene_tx=XM_023257404.1
B4 129436182 129436264
refGene_gn=BTBD11;refGene_tx=XM_023257404.1
B4 129442994 129443063
refGene_gn=BTBD11;refGene_tx=XM_023257404.1
B4 129444838 129444952
refGene_gn=BTBD11;refGene_tx=XM_023257404.1
B4 129453012 129453222
refGene_gn=BTBD11;refGene_tx=XM_023257404.1
B4 129476185 129476257
refGene_gn=PWP1;refGene_tx=XM_003989210.4

B4	129478748	129478807
refGene_gn=PWP1;refGene_tx=XM_003989210.4
B4	129479175	129479360
refGene_gn=PWP1;refGene_tx=XM_003989210.4
B4	129481172	129481258
refGene_gn=PWP1;refGene_tx=XM_003989210.4
B4	129481358	129481455
refGene_gn=PWP1;refGene_tx=XM_003989210.4
B4	129486916	129487027
refGene_gn=PWP1;refGene_tx=XM_003989210.4
B4	129487874	129488005
refGene_gn=PWP1;refGene_tx=XM_003989210.4
B4	129488623	129488685
refGene_gn=PWP1;refGene_tx=XM_003989210.4
B4	129493921	129494018
refGene_gn=PWP1;refGene_tx=XM_003989210.4
B4	129494672	129494734
refGene_gn=PWP1;refGene_tx=XM_003989210.4
B4	129496146	129496258
refGene_gn=PWP1;refGene_tx=XM_003989210.4
B4	129497821	129497912
refGene_gn=PWP1;refGene_tx=XM_003989210.4
B4	129498002	129498124
refGene_gn=PWP1;refGene_tx=XM_003989210.4
B4	129499121	129499227
refGene_gn=PWP1;refGene_tx=XM_003989210.4
B4	129500018	129500128
refGene_gn=PWP1;refGene_tx=XM_003989210.4
B4	129520757	129521093
refGene_gn=PRDM4;refGene_tx=XM_023257406.1;refGene_gn=PRDM4;refGene_tx=XM_023257407.1;refGene_gn=PRDM4;refGene_tx=XM_023257408.1;refGene_gn=PRDM4;refGene_tx=XM_023257409.1
B4	129526565	129526682
refGene_gn=PRDM4;refGene_tx=XM_023257406.1;refGene_gn=PRDM4;refGene_tx=XM_023257407.1
B4	129526957	129527126
refGene_gn=PRDM4;refGene_tx=XM_023257406.1;refGene_gn=PRDM4;refGene_tx=XM_023257407.1;refGene_gn=PRDM4;refGene_tx=XM_023257408.1;refGene_gn=PRDM4;refGene_tx=XM_023257409.1
B4	129528505	129528820
refGene_gn=PRDM4;refGene_tx=XM_023257406.1;refGene_gn=PRDM4;refGene_tx=XM_023257407.1;refGene_gn=PRDM4;refGene_tx=XM_023257408.1;refGene_gn=PRDM4;refGene_tx=XM_023257409.1
B4	129529681	129529809
refGene_gn=PRDM4;refGene_tx=XM_023257406.1;refGene_gn=PRDM4;refGene_tx=XM_023257407.1;refGene_gn=PRDM4;refGene_tx=XM_023257408.1;refGene_gn=PRDM4;refGene_tx=XM_023257409.1
B4	129530419	129530505
refGene_gn=PRDM4;refGene_tx=XM_023257406.1;refGene_gn=PRDM4;refGene_tx=XM_023257407.1;refGene_gn=PRDM4;refGene_tx=XM_023257408.1;refGene_gn=

PRDM4;refGene_tx=XM_023257409.1
B4 129531502 129531621
refGene_gn=PRDM4;refGene_tx=XM_023257406.1;refGene_gn=PRDM4;refGene_tx=XM_023257407.1;refGene_gn=PRDM4;refGene_tx=XM_023257408.1;refGene_gn=PRDM4;refGene_tx=XM_023257409.1
B4 129532573 129532723
refGene_gn=PRDM4;refGene_tx=XM_023257406.1;refGene_gn=PRDM4;refGene_tx=XM_023257407.1;refGene_gn=PRDM4;refGene_tx=XM_023257408.1;refGene_gn=PRDM4;refGene_tx=XM_023257409.1
B4 129536404 129537199
refGene_gn=PRDM4;refGene_tx=XM_023257406.1;refGene_gn=PRDM4;refGene_tx=XM_023257407.1;refGene_gn=PRDM4;refGene_tx=XM_023257408.1;refGene_gn=PRDM4;refGene_tx=XM_023257409.1
B4 129539273 129539459
refGene_gn=PRDM4;refGene_tx=XM_023257406.1;refGene_gn=PRDM4;refGene_tx=XM_023257407.1;refGene_gn=PRDM4;refGene_tx=XM_023257408.1;refGene_gn=PRDM4;refGene_tx=XM_023257409.1
B4 129542199 129542333
refGene_gn=PRDM4;refGene_tx=XM_023257406.1;refGene_gn=PRDM4;refGene_tx=XM_023257407.1;refGene_gn=PRDM4;refGene_tx=XM_023257408.1;refGene_gn=PRDM4;refGene_tx=XM_023257409.1
B4 129544458 129544469
refGene_gn=PRDM4;refGene_tx=XM_023257406.1;refGene_gn=PRDM4;refGene_tx=XM_023257407.1;refGene_gn=PRDM4;refGene_tx=XM_023257408.1;refGene_gn=PRDM4;refGene_tx=XM_023257409.1
B4 129562116 129562638
refGene_gn=ASCL4;refGene_tx=XM_023257410.1
B4 129585640 129585748
refGene_gn=RTCB;refGene_tx=XM_003989212.5;refGene_gn=RTCB;refGene_tx=XM_023257411.1
B4 129590376 129590496
refGene_gn=RTCB;refGene_tx=XM_003989212.5;refGene_gn=RTCB;refGene_tx=XM_023257411.1
B4 129592094 129592205
refGene_gn=RTCB;refGene_tx=XM_003989212.5;refGene_gn=RTCB;refGene_tx=XM_023257411.1
B4 129593154 129593343
refGene_gn=RTCB;refGene_tx=XM_003989212.5;refGene_gn=RTCB;refGene_tx=XM_023257411.1
B4 129593897 129594073
refGene_gn=RTCB;refGene_tx=XM_003989212.5;refGene_gn=RTCB;refGene_tx=XM_023257411.1
B4 129595197 129595357
refGene_gn=RTCB;refGene_tx=XM_003989212.5;refGene_gn=RTCB;refGene_tx=XM_023257411.1
B4 129596356 129596513
refGene_gn=RTCB;refGene_tx=XM_003989212.5;refGene_gn=RTCB;refGene_tx=XM_023257411.1
B4 129597558 129597715
refGene_gn=RTCB;refGene_tx=XM_003989212.5;refGene_gn=RTCB;refGene_tx=X

M_023257411.1
B4	129600733	129600833
refGene_gn=RTCB;refGene_tx=XM_003989212.5;refGene_gn=RTCB;refGene_tx=XM_023257411.1
B4	129602144	129602212
refGene_gn=RTCB;refGene_tx=XM_003989212.5;refGene_gn=RTCB;refGene_tx=XM_023257411.1
B4	129603385	129603464
refGene_gn=RTCB;refGene_tx=XM_003989212.5;refGene_gn=RTCB;refGene_tx=XM_023257411.1
B4	129606479	129606572
refGene_gn=RTCB;refGene_tx=XM_003989212.5
B4	129608826	129608949
refGene_gn=BPIFC;refGene_tx=XM_019835460.2;refGene_gn=BPIFC;refGene_tx=XM_019835461.2;refGene_gn=BPIFC;refGene_tx=XM_023257416.1
B4	129609654	129609731
refGene_gn=BPIFC;refGene_tx=XM_019835460.2;refGene_gn=BPIFC;refGene_tx=XM_019835461.2;refGene_gn=BPIFC;refGene_tx=XM_023257416.1
B4	129610726	129610790
refGene_gn=BPIFC;refGene_tx=XM_019835460.2;refGene_gn=BPIFC;refGene_tx=XM_019835461.2;refGene_gn=BPIFC;refGene_tx=XM_023257416.1
B4	129613287	129613330
refGene_gn=BPIFC;refGene_tx=XM_019835460.2;refGene_gn=BPIFC;refGene_tx=XM_019835461.2;refGene_gn=BPIFC;refGene_tx=XM_023257416.1
B4	129622303	129622371
refGene_gn=BPIFC;refGene_tx=XM_019835460.2;refGene_gn=BPIFC;refGene_tx=XM_019835461.2;refGene_gn=BPIFC;refGene_tx=XM_023257416.1
B4	129623049	129623220
refGene_gn=BPIFC;refGene_tx=XM_019835460.2;refGene_gn=BPIFC;refGene_tx=XM_019835461.2;refGene_gn=BPIFC;refGene_tx=XM_023257416.1
B4	129623961	129624015
refGene_gn=BPIFC;refGene_tx=XM_019835460.2;refGene_gn=BPIFC;refGene_tx=XM_019835461.2;refGene_gn=BPIFC;refGene_tx=XM_023257416.1
B4	129625199	129625376
refGene_gn=BPIFC;refGene_tx=XM_019835460.2;refGene_gn=BPIFC;refGene_tx=XM_019835461.2;refGene_gn=BPIFC;refGene_tx=XM_023257416.1
B4	129626373	129626465
refGene_gn=BPIFC;refGene_tx=XM_019835460.2;refGene_gn=BPIFC;refGene_tx=XM_019835461.2;refGene_gn=BPIFC;refGene_tx=XM_023257416.1
B4	129630218	129630279
refGene_gn=BPIFC;refGene_tx=XM_019835460.2;refGene_gn=BPIFC;refGene_tx=XM_019835461.2;refGene_gn=BPIFC;refGene_tx=XM_023257416.1
B4	129632112	129632176
refGene_gn=BPIFC;refGene_tx=XM_019835460.2;refGene_gn=BPIFC;refGene_tx=XM_019835461.2;refGene_gn=BPIFC;refGene_tx=XM_023257416.1
B4	129632299	129632455
refGene_gn=BPIFC;refGene_tx=XM_019835460.2;refGene_gn=BPIFC;refGene_tx=XM_019835461.2;refGene_gn=BPIFC;refGene_tx=XM_023257416.1
B4	129633605	129633734
refGene_gn=BPIFC;refGene_tx=XM_019835460.2;refGene_gn=BPIFC;refGene_tx

=XM_019835461.2;refGene_gn=BPIFC;refGene_tx=XM_023257416.1
B4	129637451	129637572
refGene_gn=BPIFC;refGene_tx=XM_019835460.2;refGene_gn=BPIFC;refGene_tx
=XM_019835461.2;refGene_gn=BPIFC;refGene_tx=XM_023257416.1
B4	129641525	129641649
refGene_gn=BPIFC;refGene_tx=XM_019835460.2;refGene_gn=BPIFC;refGene_tx
=XM_019835461.2;refGene_gn=BPIFC;refGene_tx=XM_023257416.1
B4	129663163	129663285
refGene_gn=FBXO7;refGene_tx=XM_023257414.1
B4	129663438	129663475
refGene_gn=FBXO7;refGene_tx=XM_023257415.1
B4	129666901	129667196
refGene_gn=FBXO7;refGene_tx=XM_023257414.1;refGene_gn=FBXO7;refGene_tx
=XM_023257415.1
B4	129672135	129672363
refGene_gn=FBXO7;refGene_tx=XM_023257414.1;refGene_gn=FBXO7;refGene_tx
=XM_023257415.1
B4	129673672	129673814
refGene_gn=FBXO7;refGene_tx=XM_023257414.1;refGene_gn=FBXO7;refGene_tx
=XM_023257415.1
B4	129675932	129676016
refGene_gn=FBXO7;refGene_tx=XM_023257414.1;refGene_gn=FBXO7;refGene_tx
=XM_023257415.1
B4	129679076	129679172
refGene_gn=FBXO7;refGene_tx=XM_023257414.1;refGene_gn=FBXO7;refGene_tx
=XM_023257415.1
B4	129680964	129681141
refGene_gn=FBXO7;refGene_tx=XM_023257414.1;refGene_gn=FBXO7;refGene_tx
=XM_023257415.1
B4	129683249	129683287
refGene_gn=FBXO7;refGene_tx=XM_023257414.1;refGene_gn=FBXO7;refGene_tx
=XM_023257415.1
B4	129684266	129684656
refGene_gn=FBXO7;refGene_tx=XM_023257414.1;refGene_gn=FBXO7;refGene_tx
=XM_023257415.1
B4	129691078	129691211
refGene_gn=SYN3;refGene_tx=XM_019835455.1;refGene_gn=SYN3;refGene_tx=X
M_023257413.1
B4	129694196	129694488
refGene_gn=SYN3;refGene_tx=XM_019835455.1;refGene_gn=SYN3;refGene_tx=X
M_023257413.1
B4	129700332	129700420
refGene_gn=SYN3;refGene_tx=XM_019835455.1;refGene_gn=SYN3;refGene_tx=X
M_023257413.1
B4	129701259	129701394
refGene_gn=SYN3;refGene_tx=XM_019835455.1;refGene_gn=SYN3;refGene_tx=X
M_023257413.1
B4	129705753	129705856
refGene_gn=SYN3;refGene_tx=XM_019835455.1;refGene_gn=SYN3;refGene_tx=X
M_023257413.1

B4 129710197 129710272
refGene_gn=SYN3;refGene_tx=XM_019835455.1;refGene_gn=SYN3;refGene_tx=XM_023257413.1
B4 129714986 129715129
refGene_gn=SYN3;refGene_tx=XM_019835455.1;refGene_gn=SYN3;refGene_tx=XM_023257413.1
B4 129761275 129761338
refGene_gn=SYN3;refGene_tx=XM_019835455.1;refGene_gn=SYN3;refGene_tx=XM_023257413.1
B4 129932263 129932384
refGene_gn=TIMP3;refGene_tx=XM_003989216.5
B4 129979177 129979260
refGene_gn=TIMP3;refGene_tx=XM_003989216.5
B4 129987104 129987216
refGene_gn=TIMP3;refGene_tx=XM_003989216.5
B4 129987919 129988041
refGene_gn=TIMP3;refGene_tx=XM_003989216.5
B4 129989119 129989317
refGene_gn=TIMP3;refGene_tx=XM_003989216.5
B4 129995094 129995184
refGene_gn=SYN3;refGene_tx=XM_019835455.1;refGene_gn=SYN3;refGene_tx=XM_023257413.1
B4 129998673 129998830
refGene_gn=SYN3;refGene_tx=XM_019835455.1;refGene_gn=SYN3;refGene_tx=XM_023257413.1
B4 130060275 130060367
refGene_gn=SYN3;refGene_tx=XM_019835455.1;refGene_gn=SYN3;refGene_tx=XM_023257413.1
B4 130093723 130093781
refGene_gn=SYN3;refGene_tx=XM_019835455.1;refGene_gn=SYN3;refGene_tx=XM_023257413.1
B4 130112601 130112912
refGene_gn=SYN3;refGene_tx=XM_019835455.1;refGene_gn=SYN3;refGene_tx=XM_023257413.1
B4 130286021 130286219
refGene_gn=LARGE1;refGene_tx=XM_006933997.3;refGene_gn=LARGE1;refGene_tx=XM_023257418.1
B4 130289045 130289241
refGene_gn=LARGE1;refGene_tx=XM_006933997.3;refGene_gn=LARGE1;refGene_tx=XM_023257418.1
B4 130295022 130295169
refGene_gn=LARGE1;refGene_tx=XM_006933997.3;refGene_gn=LARGE1;refGene_tx=XM_023257418.1
B4 130315918 130316197
refGene_gn=LARGE1;refGene_tx=XM_006933997.3;refGene_gn=LARGE1;refGene_tx=XM_023257418.1
B4 130326629 130326793
refGene_gn=LARGE1;refGene_tx=XM_006933997.3;refGene_gn=LARGE1;refGene_tx=XM_023257418.1
B4 130341437 130341593

refGene_gn=LARGE1;refGene_tx=XM_006933997.3
B4 130365246 130365372
refGene_gn=LARGE1;refGene_tx=XM_006933997.3;refGene_gn=LARGE1;refGene_tx=XM_023257418.1
B4 130367076 130367189
refGene_gn=LARGE1;refGene_tx=XM_006933997.3;refGene_gn=LARGE1;refGene_tx=XM_023257418.1
B4 130403208 130403313
refGene_gn=LARGE1;refGene_tx=XM_006933997.3;refGene_gn=LARGE1;refGene_tx=XM_023257418.1
B4 130514597 130514769
refGene_gn=LARGE1;refGene_tx=XM_006933997.3;refGene_gn=LARGE1;refGene_tx=XM_023257418.1
B4 130549365 130549489
refGene_gn=LARGE1;refGene_tx=XM_006933997.3;refGene_gn=LARGE1;refGene_tx=XM_023257418.1
B4 130566921 130567004
refGene_gn=LARGE1;refGene_tx=XM_006933997.3;refGene_gn=LARGE1;refGene_tx=XM_023257418.1
B4 130583836 130584138
refGene_gn=LARGE1;refGene_tx=XM_006933997.3;refGene_gn=LARGE1;refGene_tx=XM_023257418.1
B4 130685626 130685732
refGene_gn=LARGE1;refGene_tx=XM_006933997.3;refGene_gn=LARGE1;refGene_tx=XM_023257418.1
B4 131719749 131719780
refGene_gn=ISX;refGene_tx=XM_011284195.3
B4 131720033 131720228
refGene_gn=ISX;refGene_tx=XM_011284195.3
B4 131736406 131736558
refGene_gn=ISX;refGene_tx=XM_011284195.3
B4 131738467 131738581
refGene_gn=ISX;refGene_tx=XM_011284195.3
B4 131739719 131739959
refGene_gn=ISX;refGene_tx=XM_011284195.3
B4 131881784 131881944
refGene_gn=HMGXB4;refGene_tx=XM_019835464.2
B4 131886457 131886488
refGene_gn=HMGXB4;refGene_tx=XM_003989219.5
B4 131887136 131887285
refGene_gn=HMGXB4;refGene_tx=XM_003989219.5;refGene_gn=HMGXB4;refGene_tx=XM_019835464.2
B4 131887499 131887578
refGene_gn=HMGXB4;refGene_tx=XM_003989219.5;refGene_gn=HMGXB4;refGene_tx=XM_019835464.2
B4 131888348 131889304
refGene_gn=HMGXB4;refGene_tx=XM_003989219.5;refGene_gn=HMGXB4;refGene_tx=XM_019835464.2;refGene_gn=HMGXB4;refGene_tx=XM_019835465.2
B4 131900962 131901044
refGene_gn=HMGXB4;refGene_tx=XM_003989219.5;refGene_gn=HMGXB4;refGene_

tx=XM_019835464.2;refGene_gn=HMGXB4;refGene_tx=XM_019835465.2
B4	131902045	131902110
refGene_gn=HMGXB4;refGene_tx=XM_003989219.5;refGene_gn=HMGXB4;refGene_tx=XM_019835464.2;refGene_gn=HMGXB4;refGene_tx=XM_019835465.2
B4	131903599	131903705
refGene_gn=HMGXB4;refGene_tx=XM_003989219.5;refGene_gn=HMGXB4;refGene_tx=XM_019835464.2;refGene_gn=HMGXB4;refGene_tx=XM_019835465.2
B4	131904154	131904324
refGene_gn=HMGXB4;refGene_tx=XM_003989219.5;refGene_gn=HMGXB4;refGene_tx=XM_019835464.2;refGene_gn=HMGXB4;refGene_tx=XM_019835465.2
B4	131909127	131909250
refGene_gn=HMGXB4;refGene_tx=XM_003989219.5;refGene_gn=HMGXB4;refGene_tx=XM_019835464.2;refGene_gn=HMGXB4;refGene_tx=XM_019835465.2
B4	131909727	131909772
refGene_gn=HMGXB4;refGene_tx=XM_003989219.5;refGene_gn=HMGXB4;refGene_tx=XM_019835464.2;refGene_gn=HMGXB4;refGene_tx=XM_019835465.2
B4	131915392	131915444
refGene_gn=TOM1;refGene_tx=XM_019835467.2
B4	131922319	131922370
refGene_gn=TOM1;refGene_tx=XM_023257419.1;refGene_gn=TOM1;refGene_tx=XM_023257420.1
B4	131923000	131923148
refGene_gn=TOM1;refGene_tx=XM_019835466.2;refGene_gn=TOM1;refGene_tx=XM_023257419.1;refGene_gn=TOM1;refGene_tx=XM_023257420.1;refGene_gn=TOM1;refGene_tx=XM_023257421.1
B4	131935047	131935132
refGene_gn=TOM1;refGene_tx=XM_019835466.2;refGene_gn=TOM1;refGene_tx=XM_019835467.2;refGene_gn=TOM1;refGene_tx=XM_023257419.1;refGene_gn=TOM1;refGene_tx=XM_023257420.1;refGene_gn=TOM1;refGene_tx=XM_023257421.1;refGene_gn=TOM1;refGene_tx=XM_023257422.1;refGene_gn=TOM1;refGene_tx=XM_023257423.1
B4	131938231	131938310
refGene_gn=TOM1;refGene_tx=XM_019835466.2;refGene_gn=TOM1;refGene_tx=XM_019835467.2;refGene_gn=TOM1;refGene_tx=XM_023257419.1;refGene_gn=TOM1;refGene_tx=XM_023257420.1;refGene_gn=TOM1;refGene_tx=XM_023257421.1;refGene_gn=TOM1;refGene_tx=XM_023257422.1;refGene_gn=TOM1;refGene_tx=XM_023257423.1
B4	131939177	131939327
refGene_gn=TOM1;refGene_tx=XM_019835466.2;refGene_gn=TOM1;refGene_tx=XM_019835467.2;refGene_gn=TOM1;refGene_tx=XM_023257419.1;refGene_gn=TOM1;refGene_tx=XM_023257420.1;refGene_gn=TOM1;refGene_tx=XM_023257421.1;refGene_gn=TOM1;refGene_tx=XM_023257422.1;refGene_gn=TOM1;refGene_tx=XM_023257423.1
B4	131939649	131939784
refGene_gn=TOM1;refGene_tx=XM_019835466.2;refGene_gn=TOM1;refGene_tx=XM_019835467.2;refGene_gn=TOM1;refGene_tx=XM_023257419.1;refGene_gn=TOM1;refGene_tx=XM_023257420.1;refGene_gn=TOM1;refGene_tx=XM_023257421.1;refGene_gn=TOM1;refGene_tx=XM_023257422.1;refGene_gn=TOM1;refGene_tx=XM_023257423.1
B4	131939904	131940051

refGene_gn=TOM1;refGene_tx=XM_019835466.2;refGene_gn=TOM1;refGene_tx=XM_019835467.2;refGene_gn=TOM1;refGene_tx=XM_023257419.1;refGene_gn=TOM1;refGene_tx=XM_023257420.1;refGene_gn=TOM1;refGene_tx=XM_023257421.1;refGene_gn=TOM1;refGene_tx=XM_023257422.1;refGene_gn=TOM1;refGene_tx=XM_023257423.1
B4 131942859 131942976
refGene_gn=TOM1;refGene_tx=XM_019835466.2;refGene_gn=TOM1;refGene_tx=XM_019835467.2;refGene_gn=TOM1;refGene_tx=XM_023257419.1;refGene_gn=TOM1;refGene_tx=XM_023257420.1;refGene_gn=TOM1;refGene_tx=XM_023257421.1;refGene_gn=TOM1;refGene_tx=XM_023257422.1;refGene_gn=TOM1;refGene_tx=XM_023257423.1
B4 131944956 131945090
refGene_gn=TOM1;refGene_tx=XM_019835466.2;refGene_gn=TOM1;refGene_tx=XM_019835467.2;refGene_gn=TOM1;refGene_tx=XM_023257419.1;refGene_gn=TOM1;refGene_tx=XM_023257420.1;refGene_gn=TOM1;refGene_tx=XM_023257421.1;refGene_gn=TOM1;refGene_tx=XM_023257422.1;refGene_gn=TOM1;refGene_tx=XM_023257423.1
B4 131946658 131946692
refGene_gn=TOM1;refGene_tx=XM_019835466.2;refGene_gn=TOM1;refGene_tx=XM_019835467.2;refGene_gn=TOM1;refGene_tx=XM_023257419.1;refGene_gn=TOM1;refGene_tx=XM_023257420.1;refGene_gn=TOM1;refGene_tx=XM_023257421.1;refGene_gn=TOM1;refGene_tx=XM_023257422.1;refGene_gn=TOM1;refGene_tx=XM_023257423.1
B4 131947001 131947095
refGene_gn=TOM1;refGene_tx=XM_019835466.2;refGene_gn=TOM1;refGene_tx=XM_019835467.2;refGene_gn=TOM1;refGene_tx=XM_023257419.1;refGene_gn=TOM1;refGene_tx=XM_023257420.1;refGene_gn=TOM1;refGene_tx=XM_023257421.1;refGene_gn=TOM1;refGene_tx=XM_023257422.1;refGene_gn=TOM1;refGene_tx=XM_023257423.1
B4 131947966 131948087
refGene_gn=TOM1;refGene_tx=XM_019835466.2;refGene_gn=TOM1;refGene_tx=XM_019835467.2;refGene_gn=TOM1;refGene_tx=XM_023257419.1;refGene_gn=TOM1;refGene_tx=XM_023257420.1;refGene_gn=TOM1;refGene_tx=XM_023257421.1;refGene_gn=TOM1;refGene_tx=XM_023257422.1;refGene_gn=TOM1;refGene_tx=XM_023257423.1
B4 131952275 131952351
refGene_gn=TOM1;refGene_tx=XM_019835466.2;refGene_gn=TOM1;refGene_tx=XM_019835467.2;refGene_gn=TOM1;refGene_tx=XM_023257419.1;refGene_gn=TOM1;refGene_tx=XM_023257420.1;refGene_gn=TOM1;refGene_tx=XM_023257421.1;refGene_gn=TOM1;refGene_tx=XM_023257422.1;refGene_gn=TOM1;refGene_tx=XM_023257423.1
B4 131954986 131955040
refGene_gn=TOM1;refGene_tx=XM_019835467.2;refGene_gn=TOM1;refGene_tx=XM_023257419.1;refGene_gn=TOM1;refGene_tx=XM_023257420.1;refGene_gn=TOM1;refGene_tx=XM_023257421.1;refGene_gn=TOM1;refGene_tx=XM_023257422.1;refGene_gn=TOM1;refGene_tx=XM_023257423.1
B4 131956148 131956208
refGene_gn=TOM1;refGene_tx=XM_019835466.2;refGene_gn=TOM1;refGene_tx=XM_019835467.2;refGene_gn=TOM1;refGene_tx=XM_023257419.1;refGene_gn=TOM1;refGene_tx=XM_023257420.1;refGene_gn=TOM1;refGene_tx=XM_023257421.1;

refGene_gn=TOM1;refGene_tx=XM_023257422.1;refGene_gn=TOM1;refGene_tx=XM_023257423.1
B4 131957168 131957208
refGene_gn=TOM1;refGene_tx=XM_019835466.2;refGene_gn=TOM1;refGene_tx=XM_019835467.2;refGene_gn=TOM1;refGene_tx=XM_023257419.1;refGene_gn=TOM1;refGene_tx=XM_023257420.1;refGene_gn=TOM1;refGene_tx=XM_023257421.1;refGene_gn=TOM1;refGene_tx=XM_023257422.1;refGene_gn=TOM1;refGene_tx=XM_023257423.1
B4 131957289 131957444
refGene_gn=TOM1;refGene_tx=XM_019835466.2;refGene_gn=TOM1;refGene_tx=XM_019835467.2;refGene_gn=TOM1;refGene_tx=XM_023257419.1;refGene_gn=TOM1;refGene_tx=XM_023257420.1;refGene_gn=TOM1;refGene_tx=XM_023257421.1;refGene_gn=TOM1;refGene_tx=XM_023257422.1;refGene_gn=TOM1;refGene_tx=XM_023257423.1
B4 131988920 131988943
refGene_gn=HMOX1;refGene_tx=XM_003989222.5
B4 131990368 131990489
refGene_gn=HMOX1;refGene_tx=XM_003989222.5
B4 131992665 131993157
refGene_gn=HMOX1;refGene_tx=XM_003989222.5
B4 131994518 131994618
refGene_gn=HMOX1;refGene_tx=XM_003989222.5
B4 131995621 131995752
refGene_gn=HMOX1;refGene_tx=XM_003989222.5
B4 131998902 131999069
refGene_gn=MCM5;refGene_tx=XM_003989223.5;refGene_gn=MCM5;refGene_tx=XM_006934004.4;refGene_gn=MCM5;refGene_tx=XM_006934005.4
B4 132000960 132001087
refGene_gn=MCM5;refGene_tx=XM_003989223.5;refGene_gn=MCM5;refGene_tx=XM_006934004.4;refGene_gn=MCM5;refGene_tx=XM_006934005.4
B4 132001175 132001304
refGene_gn=MCM5;refGene_tx=XM_003989223.5;refGene_gn=MCM5;refGene_tx=XM_006934004.4;refGene_gn=MCM5;refGene_tx=XM_006934005.4
B4 132004165 132004338
refGene_gn=MCM5;refGene_tx=XM_003989223.5;refGene_gn=MCM5;refGene_tx=XM_006934004.4;refGene_gn=MCM5;refGene_tx=XM_006934005.4
B4 132006214 132006370
refGene_gn=MCM5;refGene_tx=XM_003989223.5;refGene_gn=MCM5;refGene_tx=XM_006934004.4;refGene_gn=MCM5;refGene_tx=XM_006934005.4
B4 132007766 132007933
refGene_gn=MCM5;refGene_tx=XM_003989223.5;refGene_gn=MCM5;refGene_tx=XM_006934004.4;refGene_gn=MCM5;refGene_tx=XM_006934005.4
B4 132008277 132008449
refGene_gn=MCM5;refGene_tx=XM_003989223.5;refGene_gn=MCM5;refGene_tx=XM_006934004.4;refGene_gn=MCM5;refGene_tx=XM_006934005.4
B4 132008898 132009010
refGene_gn=MCM5;refGene_tx=XM_003989223.5;refGene_gn=MCM5;refGene_tx=XM_006934004.4;refGene_gn=MCM5;refGene_tx=XM_006934005.4
B4 132010599 132010743
refGene_gn=MCM5;refGene_tx=XM_003989223.5;refGene_gn=MCM5;refGene_tx=X

M_006934004.4;refGene_gn=MCM5;refGene_tx=XM_006934005.4
B4	132011045	132011111
refGene_gn=MCM5;refGene_tx=XM_003989223.5;refGene_gn=MCM5;refGene_tx=XM_006934004.4;refGene_gn=MCM5;refGene_tx=XM_006934005.4
B4	132011301	132011478
refGene_gn=MCM5;refGene_tx=XM_003989223.5;refGene_gn=MCM5;refGene_tx=XM_006934004.4;refGene_gn=MCM5;refGene_tx=XM_006934005.4
B4	132012133	132012246
refGene_gn=MCM5;refGene_tx=XM_003989223.5;refGene_gn=MCM5;refGene_tx=XM_006934004.4;refGene_gn=MCM5;refGene_tx=XM_006934005.4
B4	132013432	132013561
refGene_gn=MCM5;refGene_tx=XM_003989223.5;refGene_gn=MCM5;refGene_tx=XM_006934004.4;refGene_gn=MCM5;refGene_tx=XM_006934005.4
B4	132014782	132014925
refGene_gn=MCM5;refGene_tx=XM_003989223.5;refGene_gn=MCM5;refGene_tx=XM_006934004.4;refGene_gn=MCM5;refGene_tx=XM_006934005.4
B4	132016748	132016876
refGene_gn=MCM5;refGene_tx=XM_003989223.5;refGene_gn=MCM5;refGene_tx=XM_006934004.4;refGene_gn=MCM5;refGene_tx=XM_006934005.4
B4	132018026	132018128
refGene_gn=MCM5;refGene_tx=XM_003989223.5;refGene_gn=MCM5;refGene_tx=XM_006934004.4;refGene_gn=MCM5;refGene_tx=XM_006934005.4
B4	132114555	132114826
refGene_gn=RASD2;refGene_tx=XM_023257425.1
B4	132118534	132119064
refGene_gn=RASD2;refGene_tx=XM_023257425.1
B4	132166046	132166193
refGene_gn=MB;refGene_tx=XM_003989225.4
B4	132169717	132169940
refGene_gn=MB;refGene_tx=XM_003989225.4
B4	132174342	132174437
refGene_gn=MB;refGene_tx=XM_003989225.4
B4	132206546	132206647
refGene_gn=APOL6;refGene_tx=XM_006934006.4;refGene_gn=APOL6;refGene_tx=XM_006934007.4;refGene_gn=APOL6;refGene_tx=XM_019835470.2;refGene_gn=APOL6;refGene_tx=XM_019835473.2
B4	132208453	132208625
refGene_gn=APOL6;refGene_tx=XM_006934006.4;refGene_gn=APOL6;refGene_tx=XM_006934007.4;refGene_gn=APOL6;refGene_tx=XM_006934008.4;refGene_gn=APOL6;refGene_tx=XM_019835470.2
B4	132209517	132209531
refGene_gn=APOL6;refGene_tx=XM_006934009.4
B4	132210416	132211251
refGene_gn=APOL6;refGene_tx=XM_006934006.4;refGene_gn=APOL6;refGene_tx=XM_006934007.4;refGene_gn=APOL6;refGene_tx=XM_006934008.4;refGene_gn=APOL6;refGene_tx=XM_006934009.4;refGene_gn=APOL6;refGene_tx=XM_019835470.2;refGene_gn=APOL6;refGene_tx=XM_019835473.2
B4	132212090	132212243
refGene_gn=APOL6;refGene_tx=XM_006934006.4;refGene_gn=APOL6;refGene_tx=XM_006934007.4;refGene_gn=APOL6;refGene_tx=XM_006934008.4;refGene_gn=

APOL6;refGene_tx=XM_006934009.4;refGene_gn=APOL6;refGene_tx=XM_0198354
70.2;refGene_gn=APOL6;refGene_tx=XM_019835473.2
B4 132246489 132246557
refGene_gn=APOL5;refGene_tx=XM_023257777.1
B4 132251692 132252274
refGene_gn=APOL5;refGene_tx=XM_023257777.1
B4 132252275 132252630
refGene_gn=APOL5;refGene_tx=XM_023257777.1
B4 132272421 132272476
refGene_gn=RBFOX2;refGene_tx=XM_019835484.2;refGene_gn=RBFOX2;refGene_
tx=XM_019835485.2;refGene_gn=RBFOX2;refGene_tx=XM_019835486.2;refGene_
gn=RBFOX2;refGene_tx=XM_019835487.2;refGene_gn=RBFOX2;refGene_tx=XM_01
9835488.2;refGene_gn=RBFOX2;refGene_tx=XM_019835489.2;refGene_gn=RBFOX
2;refGene_tx=XM_023257426.1;refGene_gn=RBFOX2;refGene_tx=XM_023257427.
1;refGene_gn=RBFOX2;refGene_tx=XM_023257428.1;refGene_gn=RBFOX2;refGen
e_tx=XM_023257429.1;refGene_gn=RBFOX2;refGene_tx=XM_023257430.1;refGen
e_gn=RBFOX2;refGene_tx=XM_023257431.1;refGene_gn=RBFOX2;refGene_tx=XM_
023257432.1;refGene_gn=RBFOX2;refGene_tx=XM_023257433.1;refGene_gn=RBF
OX2;refGene_tx=XM_023257434.1;refGene_gn=RBFOX2;refGene_tx=XM_02325743
5.1;refGene_gn=RBFOX2;refGene_tx=XM_023257436.1;refGene_gn=RBFOX2;refG
ene_tx=XM_023257437.1;refGene_gn=RBFOX2;refGene_tx=XM_023257438.1;refG
ene_gn=RBFOX2;refGene_tx=XM_023257439.1;refGene_gn=RBFOX2;refGene_tx=X
M_023257440.1;refGene_gn=RBFOX2;refGene_tx=XM_023257442.1;refGene_gn=R
BFOX2;refGene_tx=XM_023257443.1;refGene_gn=RBFOX2;refGene_tx=XM_023257
444.1;refGene_gn=RBFOX2;refGene_tx=XM_023257445.1;refGene_gn=RBFOX2;re
fGene_tx=XM_023257446.1
B4 132274141 132274214
refGene_gn=RBFOX2;refGene_tx=XM_019835484.2;refGene_gn=RBFOX2;refGene_
tx=XM_019835485.2;refGene_gn=RBFOX2;refGene_tx=XM_019835486.2;refGene_
gn=RBFOX2;refGene_tx=XM_019835487.2;refGene_gn=RBFOX2;refGene_tx=XM_01
9835488.2;refGene_gn=RBFOX2;refGene_tx=XM_019835489.2;refGene_gn=RBFOX
2;refGene_tx=XM_023257426.1;refGene_gn=RBFOX2;refGene_tx=XM_023257427.
1;refGene_gn=RBFOX2;refGene_tx=XM_023257428.1;refGene_gn=RBFOX2;refGen
e_tx=XM_023257429.1;refGene_gn=RBFOX2;refGene_tx=XM_023257430.1;refGen
e_gn=RBFOX2;refGene_tx=XM_023257431.1;refGene_gn=RBFOX2;refGene_tx=XM_
023257432.1;refGene_gn=RBFOX2;refGene_tx=XM_023257433.1;refGene_gn=RBF
OX2;refGene_tx=XM_023257434.1;refGene_gn=RBFOX2;refGene_tx=XM_02325743
5.1;refGene_gn=RBFOX2;refGene_tx=XM_023257436.1;refGene_gn=RBFOX2;refG
ene_tx=XM_023257437.1;refGene_gn=RBFOX2;refGene_tx=XM_023257438.1;refG
ene_gn=RBFOX2;refGene_tx=XM_023257439.1;refGene_gn=RBFOX2;refGene_tx=X
M_023257440.1;refGene_gn=RBFOX2;refGene_tx=XM_023257442.1;refGene_gn=R
BFOX2;refGene_tx=XM_023257443.1;refGene_gn=RBFOX2;refGene_tx=XM_023257
444.1;refGene_gn=RBFOX2;refGene_tx=XM_023257445.1;refGene_gn=RBFOX2;re
fGene_tx=XM_023257446.1
B4 132274638 132274727
refGene_gn=RBFOX2;refGene_tx=XM_019835484.2;refGene_gn=RBFOX2;refGene_
tx=XM_019835485.2;refGene_gn=RBFOX2;refGene_tx=XM_019835486.2;refGene_
gn=RBFOX2;refGene_tx=XM_019835487.2;refGene_gn=RBFOX2;refGene_tx=XM_01
9835488.2;refGene_gn=RBFOX2;refGene_tx=XM_019835489.2;refGene_gn=RBFOX
2;refGene_tx=XM_023257426.1;refGene_gn=RBFOX2;refGene_tx=XM_023257427.

1;refGene_gn=RBFOX2;refGene_tx=XM_023257428.1;refGene_gn=RBFOX2;refGene_tx=XM_023257429.1;refGene_gn=RBFOX2;refGene_tx=XM_023257430.1;refGene_gn=RBFOX2;refGene_tx=XM_023257431.1;refGene_gn=RBFOX2;refGene_tx=XM_023257432.1;refGene_gn=RBFOX2;refGene_tx=XM_023257433.1;refGene_gn=RBFOX2;refGene_tx=XM_023257434.1;refGene_gn=RBFOX2;refGene_tx=XM_023257435.1;refGene_gn=RBFOX2;refGene_tx=XM_023257436.1;refGene_gn=RBFOX2;refGene_tx=XM_023257437.1;refGene_gn=RBFOX2;refGene_tx=XM_023257438.1;refGene_gn=RBFOX2;refGene_tx=XM_023257439.1;refGene_gn=RBFOX2;refGene_tx=XM_023257440.1;refGene_gn=RBFOX2;refGene_tx=XM_023257442.1;refGene_gn=RBFOX2;refGene_tx=XM_023257443.1;refGene_gn=RBFOX2;refGene_tx=XM_023257444.1;refGene_gn=RBFOX2;refGene_tx=XM_023257445.1;refGene_gn=RBFOX2;refGene_tx=XM_023257446.1
B4 132278300 132278332
refGene_gn=RBFOX2;refGene_tx=XM_019835484.2;refGene_gn=RBFOX2;refGene_tx=XM_019835485.2;refGene_gn=RBFOX2;refGene_tx=XM_019835486.2;refGene_gn=RBFOX2;refGene_tx=XM_019835487.2;refGene_gn=RBFOX2;refGene_tx=XM_019835488.2;refGene_gn=RBFOX2;refGene_tx=XM_019835489.2;refGene_gn=RBFOX2;refGene_tx=XM_023257426.1;refGene_gn=RBFOX2;refGene_tx=XM_023257427.1;refGene_gn=RBFOX2;refGene_tx=XM_023257428.1;refGene_gn=RBFOX2;refGene_tx=XM_023257429.1;refGene_gn=RBFOX2;refGene_tx=XM_023257433.1;refGene_gn=RBFOX2;refGene_tx=XM_023257437.1;refGene_gn=RBFOX2;refGene_tx=XM_023257438.1;refGene_gn=RBFOX2;refGene_tx=XM_023257439.1;refGene_gn=RBFOX2;refGene_tx=XM_023257440.1;refGene_gn=RBFOX2;refGene_tx=XM_023257442.1;refGene_gn=RBFOX2;refGene_tx=XM_023257443.1;refGene_gn=RBFOX2;refGene_tx=XM_023257444.1
B4 132280473 132280516
refGene_gn=RBFOX2;refGene_tx=XM_023257430.1;refGene_gn=RBFOX2;refGene_tx=XM_023257432.1
B4 132283504 132283544
refGene_gn=RBFOX2;refGene_tx=XM_019835484.2;refGene_gn=RBFOX2;refGene_tx=XM_019835485.2;refGene_gn=RBFOX2;refGene_tx=XM_019835486.2;refGene_gn=RBFOX2;refGene_tx=XM_019835487.2;refGene_gn=RBFOX2;refGene_tx=XM_019835488.2;refGene_gn=RBFOX2;refGene_tx=XM_019835489.2;refGene_gn=RBFOX2;refGene_tx=XM_023257426.1;refGene_gn=RBFOX2;refGene_tx=XM_023257427.1;refGene_gn=RBFOX2;refGene_tx=XM_023257428.1;refGene_gn=RBFOX2;refGene_tx=XM_023257429.1;refGene_gn=RBFOX2;refGene_tx=XM_023257431.1;refGene_gn=RBFOX2;refGene_tx=XM_023257433.1;refGene_gn=RBFOX2;refGene_tx=XM_023257434.1;refGene_gn=RBFOX2;refGene_tx=XM_023257435.1;refGene_gn=RBFOX2;refGene_tx=XM_023257437.1;refGene_gn=RBFOX2;refGene_tx=XM_023257438.1;refGene_gn=RBFOX2;refGene_tx=XM_023257439.1;refGene_gn=RBFOX2;refGene_tx=XM_023257440.1;refGene_gn=RBFOX2;refGene_tx=XM_023257442.1;refGene_gn=RBFOX2;refGene_tx=XM_023257443.1;refGene_gn=RBFOX2;refGene_tx=XM_023257444.1;refGene_gn=RBFOX2;refGene_tx=XM_023257445.1;refGene_gn=RBFOX2;refGene_tx=XM_023257446.1
B4 132287356 132287501
refGene_gn=RBFOX2;refGene_tx=XM_019835484.2;refGene_gn=RBFOX2;refGene_tx=XM_019835485.2;refGene_gn=RBFOX2;refGene_tx=XM_019835486.2;refGene_gn=RBFOX2;refGene_tx=XM_019835487.2;refGene_gn=RBFOX2;refGene_tx=XM_019835488.2;refGene_gn=RBFOX2;refGene_tx=XM_019835489.2;refGene_gn=RBFOX2;refGene_tx=XM_023257426.1;refGene_gn=RBFOX2;refGene_tx=XM_023257427.

1;refGene_gn=RBFOX2;refGene_tx=XM_023257428.1;refGene_gn=RBFOX2;refGene_tx=XM_023257429.1;refGene_gn=RBFOX2;refGene_tx=XM_023257430.1;refGene_gn=RBFOX2;refGene_tx=XM_023257431.1;refGene_gn=RBFOX2;refGene_tx=XM_023257432.1;refGene_gn=RBFOX2;refGene_tx=XM_023257433.1;refGene_gn=RBFOX2;refGene_tx=XM_023257434.1;refGene_gn=RBFOX2;refGene_tx=XM_023257435.1;refGene_gn=RBFOX2;refGene_tx=XM_023257436.1;refGene_gn=RBFOX2;refGene_tx=XM_023257437.1;refGene_gn=RBFOX2;refGene_tx=XM_023257438.1;refGene_gn=RBFOX2;refGene_tx=XM_023257439.1;refGene_gn=RBFOX2;refGene_tx=XM_023257440.1;refGene_gn=RBFOX2;refGene_tx=XM_023257442.1;refGene_gn=RBFOX2;refGene_tx=XM_023257443.1;refGene_gn=RBFOX2;refGene_tx=XM_023257444.1;refGene_gn=RBFOX2;refGene_tx=XM_023257445.1;refGene_gn=RBFOX2;refGene_tx=XM_023257446.1
B4 132288597 132288690
refGene_gn=RBFOX2;refGene_tx=XM_019835484.2;refGene_gn=RBFOX2;refGene_tx=XM_019835485.2;refGene_gn=RBFOX2;refGene_tx=XM_019835486.2;refGene_gn=RBFOX2;refGene_tx=XM_019835487.2;refGene_gn=RBFOX2;refGene_tx=XM_019835488.2;refGene_gn=RBFOX2;refGene_tx=XM_019835489.2;refGene_gn=RBFOX2;refGene_tx=XM_023257426.1;refGene_gn=RBFOX2;refGene_tx=XM_023257427.1;refGene_gn=RBFOX2;refGene_tx=XM_023257428.1;refGene_gn=RBFOX2;refGene_tx=XM_023257429.1;refGene_gn=RBFOX2;refGene_tx=XM_023257430.1;refGene_gn=RBFOX2;refGene_tx=XM_023257431.1;refGene_gn=RBFOX2;refGene_tx=XM_023257432.1;refGene_gn=RBFOX2;refGene_tx=XM_023257433.1;refGene_gn=RBFOX2;refGene_tx=XM_023257434.1;refGene_gn=RBFOX2;refGene_tx=XM_023257435.1;refGene_gn=RBFOX2;refGene_tx=XM_023257436.1;refGene_gn=RBFOX2;refGene_tx=XM_023257437.1;refGene_gn=RBFOX2;refGene_tx=XM_023257438.1;refGene_gn=RBFOX2;refGene_tx=XM_023257439.1;refGene_gn=RBFOX2;refGene_tx=XM_023257440.1;refGene_gn=RBFOX2;refGene_tx=XM_023257442.1;refGene_gn=RBFOX2;refGene_tx=XM_023257443.1;refGene_gn=RBFOX2;refGene_tx=XM_023257444.1;refGene_gn=RBFOX2;refGene_tx=XM_023257445.1;refGene_gn=RBFOX2;refGene_tx=XM_023257446.1
B4 132288812 132288866
refGene_gn=RBFOX2;refGene_tx=XM_019835484.2;refGene_gn=RBFOX2;refGene_tx=XM_019835485.2;refGene_gn=RBFOX2;refGene_tx=XM_019835486.2;refGene_gn=RBFOX2;refGene_tx=XM_019835487.2;refGene_gn=RBFOX2;refGene_tx=XM_019835488.2;refGene_gn=RBFOX2;refGene_tx=XM_019835489.2;refGene_gn=RBFOX2;refGene_tx=XM_023257426.1;refGene_gn=RBFOX2;refGene_tx=XM_023257427.1;refGene_gn=RBFOX2;refGene_tx=XM_023257428.1;refGene_gn=RBFOX2;refGene_tx=XM_023257429.1;refGene_gn=RBFOX2;refGene_tx=XM_023257430.1;refGene_gn=RBFOX2;refGene_tx=XM_023257431.1;refGene_gn=RBFOX2;refGene_tx=XM_023257432.1;refGene_gn=RBFOX2;refGene_tx=XM_023257433.1;refGene_gn=RBFOX2;refGene_tx=XM_023257434.1;refGene_gn=RBFOX2;refGene_tx=XM_023257435.1;refGene_gn=RBFOX2;refGene_tx=XM_023257436.1;refGene_gn=RBFOX2;refGene_tx=XM_023257437.1;refGene_gn=RBFOX2;refGene_tx=XM_023257438.1;refGene_gn=RBFOX2;refGene_tx=XM_023257439.1;refGene_gn=RBFOX2;refGene_tx=XM_023257440.1;refGene_gn=RBFOX2;refGene_tx=XM_023257442.1;refGene_gn=RBFOX2;refGene_tx=XM_023257443.1;refGene_gn=RBFOX2;refGene_tx=XM_023257444.1;refGene_gn=RBFOX2;refGene_tx=XM_023257445.1;refGene_gn=RBFOX2;refGene_tx=XM_023257446.1
B4 132292224 132292285
refGene_gn=RBFOX2;refGene_tx=XM_019835484.2;refGene_gn=RBFOX2;refGene_

tx=XM_019835485.2;refGene_gn=RBFOX2;refGene_tx=XM_019835486.2;refGene_gn=RBFOX2;refGene_tx=XM_019835487.2;refGene_gn=RBFOX2;refGene_tx=XM_019835488.2;refGene_gn=RBFOX2;refGene_tx=XM_019835489.2;refGene_gn=RBFOX2;refGene_tx=XM_023257426.1;refGene_gn=RBFOX2;refGene_tx=XM_023257427.1;refGene_gn=RBFOX2;refGene_tx=XM_023257428.1;refGene_gn=RBFOX2;refGene_tx=XM_023257429.1;refGene_gn=RBFOX2;refGene_tx=XM_023257430.1;refGene_gn=RBFOX2;refGene_tx=XM_023257431.1;refGene_gn=RBFOX2;refGene_tx=XM_023257432.1;refGene_gn=RBFOX2;refGene_tx=XM_023257433.1;refGene_gn=RBFOX2;refGene_tx=XM_023257434.1;refGene_gn=RBFOX2;refGene_tx=XM_023257435.1;refGene_gn=RBFOX2;refGene_tx=XM_023257436.1;refGene_gn=RBFOX2;refGene_tx=XM_023257437.1;refGene_gn=RBFOX2;refGene_tx=XM_023257438.1;refGene_gn=RBFOX2;refGene_tx=XM_023257439.1;refGene_gn=RBFOX2;refGene_tx=XM_023257440.1;refGene_gn=RBFOX2;refGene_tx=XM_023257442.1;refGene_gn=RBFOX2;refGene_tx=XM_023257443.1;refGene_gn=RBFOX2;refGene_tx=XM_023257444.1;refGene_gn=RBFOX2;refGene_tx=XM_023257445.1;refGene_gn=RBFOX2;refGene_tx=XM_023257446.1
B4 132295112 132295205
refGene_gn=RBFOX2;refGene_tx=XM_019835484.2;refGene_gn=RBFOX2;refGene_tx=XM_019835485.2;refGene_gn=RBFOX2;refGene_tx=XM_019835486.2;refGene_gn=RBFOX2;refGene_tx=XM_019835487.2;refGene_gn=RBFOX2;refGene_tx=XM_019835488.2;refGene_gn=RBFOX2;refGene_tx=XM_019835489.2;refGene_gn=RBFOX2;refGene_tx=XM_023257426.1;refGene_gn=RBFOX2;refGene_tx=XM_023257427.1;refGene_gn=RBFOX2;refGene_tx=XM_023257428.1;refGene_gn=RBFOX2;refGene_tx=XM_023257429.1;refGene_gn=RBFOX2;refGene_tx=XM_023257430.1;refGene_gn=RBFOX2;refGene_tx=XM_023257431.1;refGene_gn=RBFOX2;refGene_tx=XM_023257432.1;refGene_gn=RBFOX2;refGene_tx=XM_023257433.1;refGene_gn=RBFOX2;refGene_tx=XM_023257434.1;refGene_gn=RBFOX2;refGene_tx=XM_023257435.1;refGene_gn=RBFOX2;refGene_tx=XM_023257436.1;refGene_gn=RBFOX2;refGene_tx=XM_023257439.1;refGene_gn=RBFOX2;refGene_tx=XM_023257440.1;refGene_gn=RBFOX2;refGene_tx=XM_023257442.1;refGene_gn=RBFOX2;refGene_tx=XM_023257443.1;refGene_gn=RBFOX2;refGene_tx=XM_023257444.1;refGene_gn=RBFOX2;refGene_tx=XM_023257445.1;refGene_gn=RBFOX2;refGene_tx=XM_023257446.1
B4 132304836 132304890
refGene_gn=RBFOX2;refGene_tx=XM_019835484.2;refGene_gn=RBFOX2;refGene_tx=XM_019835485.2;refGene_gn=RBFOX2;refGene_tx=XM_019835486.2;refGene_gn=RBFOX2;refGene_tx=XM_019835487.2;refGene_gn=RBFOX2;refGene_tx=XM_019835488.2;refGene_gn=RBFOX2;refGene_tx=XM_019835489.2;refGene_gn=RBFOX2;refGene_tx=XM_023257426.1;refGene_gn=RBFOX2;refGene_tx=XM_023257427.1;refGene_gn=RBFOX2;refGene_tx=XM_023257428.1;refGene_gn=RBFOX2;refGene_tx=XM_023257429.1;refGene_gn=RBFOX2;refGene_tx=XM_023257430.1;refGene_gn=RBFOX2;refGene_tx=XM_023257431.1;refGene_gn=RBFOX2;refGene_tx=XM_023257432.1;refGene_gn=RBFOX2;refGene_tx=XM_023257433.1;refGene_gn=RBFOX2;refGene_tx=XM_023257434.1;refGene_gn=RBFOX2;refGene_tx=XM_023257435.1;refGene_gn=RBFOX2;refGene_tx=XM_023257436.1;refGene_gn=RBFOX2;refGene_tx=XM_023257437.1;refGene_gn=RBFOX2;refGene_tx=XM_023257438.1;refGene_gn=RBFOX2;refGene_tx=XM_023257439.1;refGene_gn=RBFOX2;refGene_tx=XM_023257440.1;refGene_gn=RBFOX2;refGene_tx=XM_023257442.1;refGene_gn=RBFOX2;refGene_tx=XM_023257443.1;refGene_gn=RBFOX2;refGene_tx=XM_023257444.1;refGene_gn=RBFOX2;refGene_tx=XM_023257445.1;refGene_gn=RBFOX2;re

fGene_tx=XM_023257446.1
B4 132307899 132308046
refGene_gn=RBFOX2;refGene_tx=XM_019835484.2;refGene_gn=RBFOX2;refGene_tx=XM_019835485.2;refGene_gn=RBFOX2;refGene_tx=XM_019835486.2;refGene_gn=RBFOX2;refGene_tx=XM_019835487.2;refGene_gn=RBFOX2;refGene_tx=XM_019835488.2;refGene_gn=RBFOX2;refGene_tx=XM_019835489.2;refGene_gn=RBFOX2;refGene_tx=XM_023257426.1;refGene_gn=RBFOX2;refGene_tx=XM_023257427.1;refGene_gn=RBFOX2;refGene_tx=XM_023257428.1;refGene_gn=RBFOX2;refGene_tx=XM_023257429.1;refGene_gn=RBFOX2;refGene_tx=XM_023257430.1;refGene_gn=RBFOX2;refGene_tx=XM_023257431.1;refGene_gn=RBFOX2;refGene_tx=XM_023257432.1;refGene_gn=RBFOX2;refGene_tx=XM_023257433.1;refGene_gn=RBFOX2;refGene_tx=XM_023257434.1;refGene_gn=RBFOX2;refGene_tx=XM_023257435.1;refGene_gn=RBFOX2;refGene_tx=XM_023257436.1;refGene_gn=RBFOX2;refGene_tx=XM_023257437.1;refGene_gn=RBFOX2;refGene_tx=XM_023257438.1;refGene_gn=RBFOX2;refGene_tx=XM_023257439.1;refGene_gn=RBFOX2;refGene_tx=XM_023257440.1;refGene_gn=RBFOX2;refGene_tx=XM_023257442.1;refGene_gn=RBFOX2;refGene_tx=XM_023257443.1;refGene_gn=RBFOX2;refGene_tx=XM_023257444.1;refGene_gn=RBFOX2;refGene_tx=XM_023257445.1;refGene_gn=RBFOX2;refGene_tx=XM_023257446.1
B4 132332433 132332658
refGene_gn=RBFOX2;refGene_tx=XM_019835484.2;refGene_gn=RBFOX2;refGene_tx=XM_019835485.2;refGene_gn=RBFOX2;refGene_tx=XM_019835486.2;refGene_gn=RBFOX2;refGene_tx=XM_019835487.2;refGene_gn=RBFOX2;refGene_tx=XM_019835488.2;refGene_gn=RBFOX2;refGene_tx=XM_019835489.2;refGene_gn=RBFOX2;refGene_tx=XM_023257426.1;refGene_gn=RBFOX2;refGene_tx=XM_023257427.1;refGene_gn=RBFOX2;refGene_tx=XM_023257428.1;refGene_gn=RBFOX2;refGene_tx=XM_023257429.1;refGene_gn=RBFOX2;refGene_tx=XM_023257430.1;refGene_gn=RBFOX2;refGene_tx=XM_023257431.1;refGene_gn=RBFOX2;refGene_tx=XM_023257432.1;refGene_gn=RBFOX2;refGene_tx=XM_023257433.1;refGene_gn=RBFOX2;refGene_tx=XM_023257434.1;refGene_gn=RBFOX2;refGene_tx=XM_023257435.1;refGene_gn=RBFOX2;refGene_tx=XM_023257436.1;refGene_gn=RBFOX2;refGene_tx=XM_023257437.1;refGene_gn=RBFOX2;refGene_tx=XM_023257438.1;refGene_gn=RBFOX2;refGene_tx=XM_023257439.1;refGene_gn=RBFOX2;refGene_tx=XM_023257440.1;refGene_gn=RBFOX2;refGene_tx=XM_023257442.1;refGene_gn=RBFOX2;refGene_tx=XM_023257443.1;refGene_gn=RBFOX2;refGene_tx=XM_023257444.1;refGene_gn=RBFOX2;refGene_tx=XM_023257445.1;refGene_gn=RBFOX2;refGene_tx=XM_023257446.1
B4 132346575 132346582
refGene_gn=RBFOX2;refGene_tx=XM_019835488.2
B4 132362551 132362605
refGene_gn=RBFOX2;refGene_tx=XM_019835485.2
B4 132370809 132370836
refGene_gn=RBFOX2;refGene_tx=XM_019835487.2
B4 132382002 132382047
refGene_gn=RBFOX2;refGene_tx=XM_019835486.2
B4 132425499 132425514
refGene_gn=RBFOX2;refGene_tx=XM_019835489.2
B4 132443778 132443979
refGene_gn=LOC111561439;refGene_tx=XM_023257680.1
B4 132444440 132444661

refGene_gn=LOC111561439;refGene_tx=XM_023257680.1
B4	132449521	132449632
refGene_gn=LOC111561439;refGene_tx=XM_023257680.1
B4	132451466	132451692
refGene_gn=LOC111561439;refGene_tx=XM_023257680.1
B4	132451693	132452035
refGene_gn=LOC111561439;refGene_tx=XM_023257680.1
B4	132465928	132465979
refGene_gn=RBFOX2;refGene_tx=XM_019835484.2;refGene_gn=RBFOX2;refGene_tx=XM_023257426.1;refGene_gn=RBFOX2;refGene_tx=XM_023257427.1;refGene_gn=RBFOX2;refGene_tx=XM_023257428.1;refGene_gn=RBFOX2;refGene_tx=XM_023257429.1;refGene_gn=RBFOX2;refGene_tx=XM_023257430.1;refGene_gn=RBFOX2;refGene_tx=XM_023257431.1;refGene_gn=RBFOX2;refGene_tx=XM_023257432.1;refGene_gn=RBFOX2;refGene_tx=XM_023257434.1;refGene_gn=RBFOX2;refGene_tx=XM_023257435.1;refGene_gn=RBFOX2;refGene_tx=XM_023257436.1;refGene_gn=RBFOX2;refGene_tx=XM_023257437.1;refGene_gn=RBFOX2;refGene_tx=XM_023257438.1;refGene_gn=RBFOX2;refGene_tx=XM_023257445.1
B4	132489840	132489882
refGene_gn=RBFOX2;refGene_tx=XM_019835484.2
B4	132556638	132556824
refGene_gn=RBFOX2;refGene_tx=XM_023257426.1;refGene_gn=RBFOX2;refGene_tx=XM_023257427.1;refGene_gn=RBFOX2;refGene_tx=XM_023257428.1;refGene_gn=RBFOX2;refGene_tx=XM_023257429.1;refGene_gn=RBFOX2;refGene_tx=XM_023257430.1;refGene_gn=RBFOX2;refGene_tx=XM_023257431.1;refGene_gn=RBFOX2;refGene_tx=XM_023257432.1;refGene_gn=RBFOX2;refGene_tx=XM_023257433.1;refGene_gn=RBFOX2;refGene_tx=XM_023257434.1;refGene_gn=RBFOX2;refGene_tx=XM_023257435.1;refGene_gn=RBFOX2;refGene_tx=XM_023257436.1;refGene_gn=RBFOX2;refGene_tx=XM_023257437.1;refGene_gn=RBFOX2;refGene_tx=XM_023257438.1;refGene_gn=RBFOX2;refGene_tx=XM_023257445.1;refGene_gn=RBFOX2;refGene_tx=XM_023257446.1
B4	132626989	132627848
refGene_gn=LOC111556223;refGene_tx=XM_023257447.1
B4	132631199	132631312
refGene_gn=LOC111556223;refGene_tx=XM_023257447.1
B4	132650448	132650566
refGene_gn=MYH9;refGene_tx=XM_023257448.1;refGene_gn=MYH9;refGene_tx=XM_023257450.1
B4	132651850	132652023
refGene_gn=MYH9;refGene_tx=XM_023257448.1;refGene_gn=MYH9;refGene_tx=XM_023257450.1
B4	132652126	132652235
refGene_gn=MYH9;refGene_tx=XM_023257448.1;refGene_gn=MYH9;refGene_tx=XM_023257450.1
B4	132652700	132652909
refGene_gn=MYH9;refGene_tx=XM_023257448.1;refGene_gn=MYH9;refGene_tx=XM_023257450.1
B4	132653175	132653299
refGene_gn=MYH9;refGene_tx=XM_023257448.1;refGene_gn=MYH9;refGene_tx=XM_023257450.1
B4	132653389	132653478

refGene_gn=MYH9;refGene_tx=XM_023257448.1;refGene_gn=MYH9;refGene_tx=XM_023257450.1
B4 132654438 132654567
refGene_gn=MYH9;refGene_tx=XM_023257448.1;refGene_gn=MYH9;refGene_tx=XM_023257450.1
B4 132656257 132656419
refGene_gn=MYH9;refGene_tx=XM_023257448.1;refGene_gn=MYH9;refGene_tx=XM_023257450.1
B4 132656703 132656916
refGene_gn=MYH9;refGene_tx=XM_023257448.1;refGene_gn=MYH9;refGene_tx=XM_023257450.1
B4 132657035 132657248
refGene_gn=MYH9;refGene_tx=XM_023257448.1;refGene_gn=MYH9;refGene_tx=XM_023257450.1
B4 132658524 132658773
refGene_gn=MYH9;refGene_tx=XM_023257448.1;refGene_gn=MYH9;refGene_tx=XM_023257450.1
B4 132659817 132659970
refGene_gn=MYH9;refGene_tx=XM_023257448.1;refGene_gn=MYH9;refGene_tx=XM_023257450.1
B4 132660221 132660326
refGene_gn=MYH9;refGene_tx=XM_023257448.1;refGene_gn=MYH9;refGene_tx=XM_023257450.1
B4 132660460 132660667
refGene_gn=MYH9;refGene_tx=XM_023257448.1;refGene_gn=MYH9;refGene_tx=XM_023257450.1
B4 132661926 132662071
refGene_gn=MYH9;refGene_tx=XM_023257448.1;refGene_gn=MYH9;refGene_tx=XM_023257450.1
B4 132662445 132662658
refGene_gn=MYH9;refGene_tx=XM_023257448.1;refGene_gn=MYH9;refGene_tx=XM_023257450.1
B4 132663319 132663491
refGene_gn=MYH9;refGene_tx=XM_023257448.1;refGene_gn=MYH9;refGene_tx=XM_023257450.1
B4 132665420 132665544
refGene_gn=MYH9;refGene_tx=XM_023257448.1;refGene_gn=MYH9;refGene_tx=XM_023257450.1
B4 132666587 132666725
refGene_gn=MYH9;refGene_tx=XM_023257448.1;refGene_gn=MYH9;refGene_tx=XM_023257450.1
B4 132667039 132667246
refGene_gn=MYH9;refGene_tx=XM_023257448.1;refGene_gn=MYH9;refGene_tx=XM_023257450.1
B4 132667662 132667794
refGene_gn=MYH9;refGene_tx=XM_023257448.1;refGene_gn=MYH9;refGene_tx=XM_023257450.1
B4 132669150 132669259
refGene_gn=MYH9;refGene_tx=XM_023257448.1;refGene_gn=MYH9;refGene_tx=XM_023257450.1

B4	132670112	132670273
refGene_gn=MYH9;refGene_tx=XM_023257448.1;refGene_gn=MYH9;refGene_tx=XM_023257450.1
B4	132671963	132672033
refGene_gn=MYH9;refGene_tx=XM_023257448.1;refGene_gn=MYH9;refGene_tx=XM_023257450.1
B4	132672775	132672897
refGene_gn=MYH9;refGene_tx=XM_023257448.1;refGene_gn=MYH9;refGene_tx=XM_023257450.1
B4	132673214	132673408
refGene_gn=MYH9;refGene_tx=XM_023257448.1;refGene_gn=MYH9;refGene_tx=XM_023257450.1
B4	132675555	132675670
refGene_gn=MYH9;refGene_tx=XM_023257448.1;refGene_gn=MYH9;refGene_tx=XM_023257450.1
B4	132677522	132677696
refGene_gn=MYH9;refGene_tx=XM_023257448.1;refGene_gn=MYH9;refGene_tx=XM_023257450.1
B4	132679336	132679510
refGene_gn=MYH9;refGene_tx=XM_023257448.1;refGene_gn=MYH9;refGene_tx=XM_023257450.1
B4	132680430	132680583
refGene_gn=MYH9;refGene_tx=XM_023257448.1;refGene_gn=MYH9;refGene_tx=XM_023257450.1
B4	132682665	132682784
refGene_gn=MYH9;refGene_tx=XM_023257448.1;refGene_gn=MYH9;refGene_tx=XM_023257450.1
B4	132683622	132683718
refGene_gn=MYH9;refGene_tx=XM_023257448.1;refGene_gn=MYH9;refGene_tx=XM_023257450.1
B4	132684508	132684652
refGene_gn=MYH9;refGene_tx=XM_023257448.1;refGene_gn=MYH9;refGene_tx=XM_023257450.1
B4	132684997	132685096
refGene_gn=MYH9;refGene_tx=XM_023257448.1;refGene_gn=MYH9;refGene_tx=XM_023257450.1
B4	132685702	132685766
refGene_gn=MYH9;refGene_tx=XM_023257448.1;refGene_gn=MYH9;refGene_tx=XM_023257450.1
B4	132686289	132686382
refGene_gn=MYH9;refGene_tx=XM_023257448.1;refGene_gn=MYH9;refGene_tx=XM_023257450.1
B4	132690391	132690485
refGene_gn=MYH9;refGene_tx=XM_023257448.1;refGene_gn=MYH9;refGene_tx=XM_023257450.1
B4	132691266	132691294
refGene_gn=MYH9;refGene_tx=XM_023257448.1;refGene_gn=MYH9;refGene_tx=XM_023257450.1
B4	132704467	132704624
refGene_gn=MYH9;refGene_tx=XM_023257448.1;refGene_gn=MYH9;refGene_tx=X

M_023257450.1
B4	132709905	132710238
refGene_gn=MYH9;refGene_tx=XM_023257448.1;refGene_gn=MYH9;refGene_tx=XM_023257450.1
B4	132808905	132809019
refGene_gn=TXN2;refGene_tx=XM_003989228.5;refGene_gn=TXN2;refGene_tx=XM_019835506.2
B4	132816648	132816772
refGene_gn=TXN2;refGene_tx=XM_003989228.5;refGene_gn=TXN2;refGene_tx=XM_019835506.2
B4	132818615	132818869
refGene_gn=TXN2;refGene_tx=XM_003989228.5;refGene_gn=TXN2;refGene_tx=XM_019835506.2
B4	132834988	132835125
refGene_gn=FOXRED2;refGene_tx=XM_023257453.1;refGene_gn=FOXRED2;refGene_tx=XM_023257454.1
B4	132837628	132837876
refGene_gn=FOXRED2;refGene_tx=XM_023257451.1;refGene_gn=FOXRED2;refGene_tx=XM_023257452.1;refGene_gn=FOXRED2;refGene_tx=XM_023257455.1
B4	132841043	132841149
refGene_gn=FOXRED2;refGene_tx=XM_023257452.1
B4	132841569	132841740
refGene_gn=FOXRED2;refGene_tx=XM_023257451.1;refGene_gn=FOXRED2;refGene_tx=XM_023257452.1;refGene_gn=FOXRED2;refGene_tx=XM_023257453.1;refGene_gn=FOXRED2;refGene_tx=XM_023257454.1;refGene_gn=FOXRED2;refGene_tx=XM_023257455.1
B4	132843592	132843834
refGene_gn=FOXRED2;refGene_tx=XM_023257451.1;refGene_gn=FOXRED2;refGene_tx=XM_023257452.1;refGene_gn=FOXRED2;refGene_tx=XM_023257453.1
B4	132845102	132845268
refGene_gn=FOXRED2;refGene_tx=XM_023257451.1;refGene_gn=FOXRED2;refGene_tx=XM_023257452.1;refGene_gn=FOXRED2;refGene_tx=XM_023257453.1;refGene_gn=FOXRED2;refGene_tx=XM_023257454.1;refGene_gn=FOXRED2;refGene_tx=XM_023257455.1
B4	132848199	132848366
refGene_gn=FOXRED2;refGene_tx=XM_023257451.1;refGene_gn=FOXRED2;refGene_tx=XM_023257452.1;refGene_gn=FOXRED2;refGene_tx=XM_023257453.1;refGene_gn=FOXRED2;refGene_tx=XM_023257454.1;refGene_gn=FOXRED2;refGene_tx=XM_023257455.1
B4	132849302	132849566
refGene_gn=FOXRED2;refGene_tx=XM_023257451.1;refGene_gn=FOXRED2;refGene_tx=XM_023257452.1;refGene_gn=FOXRED2;refGene_tx=XM_023257453.1;refGene_gn=FOXRED2;refGene_tx=XM_023257454.1;refGene_gn=FOXRED2;refGene_tx=XM_023257455.1
B4	132849696	132849948
refGene_gn=FOXRED2;refGene_tx=XM_023257451.1;refGene_gn=FOXRED2;refGene_tx=XM_023257452.1;refGene_gn=FOXRED2;refGene_tx=XM_023257453.1;refGene_gn=FOXRED2;refGene_tx=XM_023257454.1;refGene_gn=FOXRED2;refGene_tx=XM_023257455.1
B4	132851140	132851688

refGene_gn=FOXRED2;refGene_tx=XM_023257451.1;refGene_gn=FOXRED2;refGene_tx=XM_023257452.1;refGene_gn=FOXRED2;refGene_tx=XM_023257453.1;refGene_gn=FOXRED2;refGene_tx=XM_023257454.1;refGene_gn=FOXRED2;refGene_tx=XM_023257455.1
B4 132855193 132855207
refGene_gn=EIF3D;refGene_tx=XM_003989229.5;refGene_gn=EIF3D;refGene_tx=XM_019835507.2
B4 132855573 132855857
refGene_gn=EIF3D;refGene_tx=XM_003989229.5;refGene_gn=EIF3D;refGene_tx=XM_019835507.2
B4 132856043 132856186
refGene_gn=EIF3D;refGene_tx=XM_003989229.5;refGene_gn=EIF3D;refGene_tx=XM_019835507.2
B4 132858754 132858884
refGene_gn=EIF3D;refGene_tx=XM_003989229.5;refGene_gn=EIF3D;refGene_tx=XM_019835507.2
B4 132858965 132859051
refGene_gn=EIF3D;refGene_tx=XM_003989229.5;refGene_gn=EIF3D;refGene_tx=XM_019835507.2
B4 132859750 132859881
refGene_gn=EIF3D;refGene_tx=XM_003989229.5;refGene_gn=EIF3D;refGene_tx=XM_019835507.2
B4 132860657 132860805
refGene_gn=EIF3D;refGene_tx=XM_003989229.5;refGene_gn=EIF3D;refGene_tx=XM_019835507.2
B4 132861840 132861973
refGene_gn=EIF3D;refGene_tx=XM_003989229.5;refGene_gn=EIF3D;refGene_tx=XM_019835507.2
B4 132862586 132862699
refGene_gn=EIF3D;refGene_tx=XM_003989229.5;refGene_gn=EIF3D;refGene_tx=XM_019835507.2
B4 132863836 132863909
refGene_gn=EIF3D;refGene_tx=XM_003989229.5;refGene_gn=EIF3D;refGene_tx=XM_019835507.2
B4 132864452 132864538
refGene_gn=EIF3D;refGene_tx=XM_003989229.5;refGene_gn=EIF3D;refGene_tx=XM_019835507.2
B4 132865049 132865186
refGene_gn=EIF3D;refGene_tx=XM_003989229.5;refGene_gn=EIF3D;refGene_tx=XM_019835507.2
B4 132866320 132866366
refGene_gn=EIF3D;refGene_tx=XM_003989229.5;refGene_gn=EIF3D;refGene_tx=XM_019835507.2
B4 132866655 132866778
refGene_gn=EIF3D;refGene_tx=XM_003989229.5;refGene_gn=EIF3D;refGene_tx=XM_019835507.2
B4 132899075 132899611
refGene_gn=CACNG2;refGene_tx=XM_011284199.3
B4 132900707 132900848
refGene_gn=CACNG2;refGene_tx=XM_011284199.3

B4 132912735 132912819
refGene_gn=CACNG2;refGene_tx=XM_011284199.3
B4 133005810 133006021
refGene_gn=CACNG2;refGene_tx=XM_011284199.3
B4 133049817 133049916
refGene_gn=IFT27;refGene_tx=XM_003989232.5
B4 133055504 133055614
refGene_gn=IFT27;refGene_tx=XM_003989232.5
B4 133056526 133056644
refGene_gn=IFT27;refGene_tx=XM_003989232.5
B4 133058550 133058610
refGene_gn=IFT27;refGene_tx=XM_003989232.5
B4 133059749 133059809
refGene_gn=IFT27;refGene_tx=XM_003989232.5
B4 133060395 133060475
refGene_gn=IFT27;refGene_tx=XM_003989232.5
B4 133066530 133066564
refGene_gn=IFT27;refGene_tx=XM_003989232.5
B4 133085927 133085956
refGene_gn=PVALB;refGene_tx=XM_003989233.4;refGene_gn=PVALB;refGene_tx=XM_019835510.2;refGene_gn=PVALB;refGene_tx=XM_019835511.2
B4 133097135 133097245
refGene_gn=PVALB;refGene_tx=XM_003989233.4;refGene_gn=PVALB;refGene_tx=XM_019835510.2;refGene_gn=PVALB;refGene_tx=XM_019835511.2
B4 133098427 133098560
refGene_gn=PVALB;refGene_tx=XM_003989233.4;refGene_gn=PVALB;refGene_tx=XM_019835510.2;refGene_gn=PVALB;refGene_tx=XM_019835511.2
B4 133100194 133100255
refGene_gn=PVALB;refGene_tx=XM_003989233.4;refGene_gn=PVALB;refGene_tx=XM_019835510.2;refGene_gn=PVALB;refGene_tx=XM_019835511.2
B4 133147131 133147163
refGene_gn=NCF4;refGene_tx=XM_003989234.5
B4 133152130 133152215
refGene_gn=NCF4;refGene_tx=XM_003989234.5
B4 133152868 133153022
refGene_gn=NCF4;refGene_tx=XM_003989234.5
B4 133154659 133154730
refGene_gn=NCF4;refGene_tx=XM_003989234.5
B4 133158081 133158209
refGene_gn=NCF4;refGene_tx=XM_003989234.5
B4 133159347 133159405
refGene_gn=NCF4;refGene_tx=XM_003989234.5
B4 133159973 133160072
refGene_gn=NCF4;refGene_tx=XM_003989234.5
B4 133165717 133165848
refGene_gn=NCF4;refGene_tx=XM_003989234.5
B4 133166075 133166141
refGene_gn=NCF4;refGene_tx=XM_003989234.5
B4 133167748 133167818
refGene_gn=NCF4;refGene_tx=XM_003989234.5

B4 133167819 133167945
refGene_gn=NCF4;refGene_tx=XM_003989234.5
B4 133206300 133206376
refGene_gn=CSF2RB;refGene_tx=XM_011284200.3
B4 133207541 133207665
refGene_gn=CSF2RB;refGene_tx=XM_011284200.3
B4 133209980 133210174
refGene_gn=CSF2RB;refGene_tx=XM_011284200.3
B4 133213397 133213549
refGene_gn=CSF2RB;refGene_tx=XM_011284200.3
B4 133213631 133213800
refGene_gn=CSF2RB;refGene_tx=XM_011284200.3
B4 133214384 133214520
refGene_gn=CSF2RB;refGene_tx=XM_011284200.3
B4 133214681 133214839
refGene_gn=CSF2RB;refGene_tx=XM_011284200.3
B4 133215989 133216129
refGene_gn=CSF2RB;refGene_tx=XM_011284200.3
B4 133217121 133217284
refGene_gn=CSF2RB;refGene_tx=XM_011284200.3
B4 133218369 133218460
refGene_gn=CSF2RB;refGene_tx=XM_011284200.3
B4 133218669 133218727
refGene_gn=CSF2RB;refGene_tx=XM_011284200.3
B4 133219611 133219715
refGene_gn=CSF2RB;refGene_tx=XM_011284200.3
B4 133220313 133221418
refGene_gn=CSF2RB;refGene_tx=XM_011284200.3
B4 133256166 133256264
refGene_gn=TEX33;refGene_tx=XM_003989235.4
B4 133256465 133256576
refGene_gn=TEX33;refGene_tx=XM_003989235.4
B4 133260216 133260361
refGene_gn=TEX33;refGene_tx=XM_003989235.4
B4 133262216 133262659
refGene_gn=TEX33;refGene_tx=XM_003989235.4
B4 133265676 133265698
refGene_gn=TEX33;refGene_tx=XM_003989235.4
B4 133272186 133272485
refGene_gn=TST;refGene_tx=XM_011284201.3;refGene_gn=TST;refGene_tx=XM_
019835514.2
B4 133282874 133283760
refGene_gn=TST;refGene_tx=XM_011284201.3;refGene_gn=TST;refGene_tx=XM_
019835514.2
B4 133284159 133284255
refGene_gn=TST;refGene_tx=XM_011284201.3
B4 133284596 133284632
refGene_gn=MPST;refGene_tx=XM_023257458.1
B4 133288019 133288650
refGene_gn=MPST;refGene_tx=XM_023257457.1;refGene_gn=MPST;refGene_tx=X

M_023257458.1;refGene_gn=MPST;refGene_tx=XM_023257459.1
B4 133292214 133292513
refGene_gn=MPST;refGene_tx=XM_023257457.1;refGene_gn=MPST;refGene_tx=XM_023257458.1;refGene_gn=MPST;refGene_tx=XM_023257459.1
B4 133304202 133304412
refGene_gn=KCTD17;refGene_tx=XM_023257465.1;refGene_gn=KCTD17;refGene_tx=XM_023257466.1;refGene_gn=KCTD17;refGene_tx=XM_023257467.1;refGene_gn=KCTD17;refGene_tx=XM_023257468.1;refGene_gn=KCTD17;refGene_tx=XM_023257469.1;refGene_gn=KCTD17;refGene_tx=XM_023257470.1;refGene_gn=KCTD17;refGene_tx=XM_023257471.1
B4 133305558 133305667
refGene_gn=KCTD17;refGene_tx=XM_023257465.1;refGene_gn=KCTD17;refGene_tx=XM_023257466.1;refGene_gn=KCTD17;refGene_tx=XM_023257467.1;refGene_gn=KCTD17;refGene_tx=XM_023257468.1;refGene_gn=KCTD17;refGene_tx=XM_023257469.1;refGene_gn=KCTD17;refGene_tx=XM_023257470.1;refGene_gn=KCTD17;refGene_tx=XM_023257471.1
B4 133307857 133307949
refGene_gn=KCTD17;refGene_tx=XM_023257465.1;refGene_gn=KCTD17;refGene_tx=XM_023257466.1;refGene_gn=KCTD17;refGene_tx=XM_023257467.1;refGene_gn=KCTD17;refGene_tx=XM_023257468.1;refGene_gn=KCTD17;refGene_tx=XM_023257469.1;refGene_gn=KCTD17;refGene_tx=XM_023257470.1;refGene_gn=KCTD17;refGene_tx=XM_023257471.1
B4 133308769 133308865
refGene_gn=KCTD17;refGene_tx=XM_023257465.1;refGene_gn=KCTD17;refGene_tx=XM_023257466.1;refGene_gn=KCTD17;refGene_tx=XM_023257467.1;refGene_gn=KCTD17;refGene_tx=XM_023257468.1;refGene_gn=KCTD17;refGene_tx=XM_023257469.1;refGene_gn=KCTD17;refGene_tx=XM_023257470.1;refGene_gn=KCTD17;refGene_tx=XM_023257471.1
B4 133310657 133310783
refGene_gn=KCTD17;refGene_tx=XM_023257465.1;refGene_gn=KCTD17;refGene_tx=XM_023257466.1;refGene_gn=KCTD17;refGene_tx=XM_023257467.1;refGene_gn=KCTD17;refGene_tx=XM_023257468.1;refGene_gn=KCTD17;refGene_tx=XM_023257469.1;refGene_gn=KCTD17;refGene_tx=XM_023257470.1;refGene_gn=KCTD17;refGene_tx=XM_023257471.1
B4 133312110 133312189
refGene_gn=KCTD17;refGene_tx=XM_023257465.1;refGene_gn=KCTD17;refGene_tx=XM_023257466.1;refGene_gn=KCTD17;refGene_tx=XM_023257467.1;refGene_gn=KCTD17;refGene_tx=XM_023257468.1;refGene_gn=KCTD17;refGene_tx=XM_023257469.1
B4 133312378 133312743
refGene_gn=KCTD17;refGene_tx=XM_023257465.1;refGene_gn=KCTD17;refGene_tx=XM_023257467.1
B4 133312811 133313053
refGene_gn=KCTD17;refGene_tx=XM_023257468.1;refGene_gn=KCTD17;refGene_tx=XM_023257469.1;refGene_gn=KCTD17;refGene_tx=XM_023257471.1
B4 133313833 133314210
refGene_gn=KCTD17;refGene_tx=XM_023257466.1;refGene_gn=KCTD17;refGene_tx=XM_023257469.1;refGene_gn=KCTD17;refGene_tx=XM_023257470.1
B4 133317781 133317940
refGene_gn=TMPRSS6;refGene_tx=XM_023257460.1;refGene_gn=TMPRSS6;refGen

e_tx=XM_023257461.1;refGene_gn=TMPRSS6;refGene_tx=XM_023257462.1;refGene_gn=TMPRSS6;refGene_tx=XM_023257463.1;refGene_gn=TMPRSS6;refGene_tx=XM_023257464.1
B4 133318492 133318629
refGene_gn=TMPRSS6;refGene_tx=XM_023257460.1;refGene_gn=TMPRSS6;refGene_tx=XM_023257461.1;refGene_gn=TMPRSS6;refGene_tx=XM_023257462.1;refGene_gn=TMPRSS6;refGene_tx=XM_023257463.1;refGene_gn=TMPRSS6;refGene_tx=XM_023257464.1
B4 133322006 133322278
refGene_gn=TMPRSS6;refGene_tx=XM_023257460.1;refGene_gn=TMPRSS6;refGene_tx=XM_023257461.1;refGene_gn=TMPRSS6;refGene_tx=XM_023257462.1;refGene_gn=TMPRSS6;refGene_tx=XM_023257463.1;refGene_gn=TMPRSS6;refGene_tx=XM_023257464.1
B4 133323660 133323829
refGene_gn=TMPRSS6;refGene_tx=XM_023257460.1;refGene_gn=TMPRSS6;refGene_tx=XM_023257461.1;refGene_gn=TMPRSS6;refGene_tx=XM_023257462.1;refGene_gn=TMPRSS6;refGene_tx=XM_023257463.1;refGene_gn=TMPRSS6;refGene_tx=XM_023257464.1
B4 133324134 133324251
refGene_gn=TMPRSS6;refGene_tx=XM_023257460.1;refGene_gn=TMPRSS6;refGene_tx=XM_023257461.1;refGene_gn=TMPRSS6;refGene_tx=XM_023257462.1;refGene_gn=TMPRSS6;refGene_tx=XM_023257463.1;refGene_gn=TMPRSS6;refGene_tx=XM_023257464.1
B4 133331079 133331193
refGene_gn=TMPRSS6;refGene_tx=XM_023257460.1;refGene_gn=TMPRSS6;refGene_tx=XM_023257461.1;refGene_gn=TMPRSS6;refGene_tx=XM_023257462.1;refGene_gn=TMPRSS6;refGene_tx=XM_023257463.1;refGene_gn=TMPRSS6;refGene_tx=XM_023257464.1
B4 133332079 133332178
refGene_gn=TMPRSS6;refGene_tx=XM_023257460.1;refGene_gn=TMPRSS6;refGene_tx=XM_023257461.1;refGene_gn=TMPRSS6;refGene_tx=XM_023257462.1;refGene_gn=TMPRSS6;refGene_tx=XM_023257463.1;refGene_gn=TMPRSS6;refGene_tx=XM_023257464.1
B4 133332504 133332650
refGene_gn=TMPRSS6;refGene_tx=XM_023257460.1;refGene_gn=TMPRSS6;refGene_tx=XM_023257461.1;refGene_gn=TMPRSS6;refGene_tx=XM_023257462.1;refGene_gn=TMPRSS6;refGene_tx=XM_023257463.1;refGene_gn=TMPRSS6;refGene_tx=XM_023257464.1
B4 133339486 133339596
refGene_gn=TMPRSS6;refGene_tx=XM_023257460.1;refGene_gn=TMPRSS6;refGene_tx=XM_023257461.1;refGene_gn=TMPRSS6;refGene_tx=XM_023257462.1;refGene_gn=TMPRSS6;refGene_tx=XM_023257463.1;refGene_gn=TMPRSS6;refGene_tx=XM_023257464.1
B4 133339929 133340042
refGene_gn=TMPRSS6;refGene_tx=XM_023257460.1;refGene_gn=TMPRSS6;refGene_tx=XM_023257461.1;refGene_gn=TMPRSS6;refGene_tx=XM_023257462.1;refGene_gn=TMPRSS6;refGene_tx=XM_023257463.1;refGene_gn=TMPRSS6;refGene_tx=XM_023257464.1
B4 133341446 133341583
refGene_gn=TMPRSS6;refGene_tx=XM_023257460.1;refGene_gn=TMPRSS6;refGen

e_tx=XM_023257461.1;refGene_gn=TMPRSS6;refGene_tx=XM_023257462.1;refGene_gn=TMPRSS6;refGene_tx=XM_023257463.1;refGene_gn=TMPRSS6;refGene_tx=XM_023257464.1
B4 133343430 133343635
refGene_gn=TMPRSS6;refGene_tx=XM_023257460.1;refGene_gn=TMPRSS6;refGene_tx=XM_023257461.1;refGene_gn=TMPRSS6;refGene_tx=XM_023257462.1;refGene_gn=TMPRSS6;refGene_tx=XM_023257463.1;refGene_gn=TMPRSS6;refGene_tx=XM_023257464.1
B4 133347813 133347855
refGene_gn=TMPRSS6;refGene_tx=XM_023257460.1;refGene_gn=TMPRSS6;refGene_tx=XM_023257461.1;refGene_gn=TMPRSS6;refGene_tx=XM_023257462.1;refGene_gn=TMPRSS6;refGene_tx=XM_023257464.1
B4 133348138 133348323
refGene_gn=TMPRSS6;refGene_tx=XM_023257460.1;refGene_gn=TMPRSS6;refGene_tx=XM_023257461.1;refGene_gn=TMPRSS6;refGene_tx=XM_023257462.1;refGene_gn=TMPRSS6;refGene_tx=XM_023257463.1;refGene_gn=TMPRSS6;refGene_tx=XM_023257464.1
B4 133348806 133348874
refGene_gn=TMPRSS6;refGene_tx=XM_023257460.1;refGene_gn=TMPRSS6;refGene_tx=XM_023257461.1;refGene_gn=TMPRSS6;refGene_tx=XM_023257462.1;refGene_gn=TMPRSS6;refGene_tx=XM_023257463.1;refGene_gn=TMPRSS6;refGene_tx=XM_023257464.1
B4 133350031 133350165
refGene_gn=TMPRSS6;refGene_tx=XM_023257460.1;refGene_gn=TMPRSS6;refGene_tx=XM_023257461.1;refGene_gn=TMPRSS6;refGene_tx=XM_023257462.1;refGene_gn=TMPRSS6;refGene_tx=XM_023257463.1;refGene_gn=TMPRSS6;refGene_tx=XM_023257464.1
B4 133353581 133353777
refGene_gn=TMPRSS6;refGene_tx=XM_023257460.1;refGene_gn=TMPRSS6;refGene_tx=XM_023257461.1;refGene_gn=TMPRSS6;refGene_tx=XM_023257462.1;refGene_gn=TMPRSS6;refGene_tx=XM_023257463.1;refGene_gn=TMPRSS6;refGene_tx=XM_023257464.1
B4 133369830 133370568
refGene_gn=IL2RB;refGene_tx=XM_023257472.1;refGene_gn=IL2RB;refGene_tx=XM_023257473.1
B4 133372986 133373071
refGene_gn=IL2RB;refGene_tx=XM_023257472.1;refGene_gn=IL2RB;refGene_tx=XM_023257473.1
B4 133374339 133374460
refGene_gn=IL2RB;refGene_tx=XM_023257472.1;refGene_gn=IL2RB;refGene_tx=XM_023257473.1
B4 133374660 133374826
refGene_gn=IL2RB;refGene_tx=XM_023257472.1;refGene_gn=IL2RB;refGene_tx=XM_023257473.1
B4 133376005 133376154
refGene_gn=IL2RB;refGene_tx=XM_023257472.1;refGene_gn=IL2RB;refGene_tx=XM_023257473.1
B4 133377022 133377128
refGene_gn=IL2RB;refGene_tx=XM_023257472.1;refGene_gn=IL2RB;refGene_tx=XM_023257473.1

B4 133377889 133377968
refGene_gn=IL2RB;refGene_tx=XM_023257472.1;refGene_gn=IL2RB;refGene_tx=XM_023257473.1
B4 133378691 133378806
refGene_gn=IL2RB;refGene_tx=XM_023257472.1;refGene_gn=IL2RB;refGene_tx=XM_023257473.1
B4 133379237 133379334
refGene_gn=IL2RB;refGene_tx=XM_023257472.1;refGene_gn=IL2RB;refGene_tx=XM_023257473.1
B4 133386562 133386586
refGene_gn=LOC111561466;refGene_tx=XM_023257778.1
B4 133391261 133392302
refGene_gn=LOC111561466;refGene_tx=XM_023257778.1
B4 133392583 133393058
refGene_gn=LOC111561466;refGene_tx=XM_023257778.1
B4 133396211 133396258
refGene_gn=LOC111561466;refGene_tx=XM_023257778.1
B4 133406663 133407211
refGene_gn=C1QTNF6;refGene_tx=XM_003989237.5
B4 133409607 133409860
refGene_gn=C1QTNF6;refGene_tx=XM_003989237.5
B4 133422501 133423764
refGene_gn=SSTR3;refGene_tx=XM_019835518.2;refGene_gn=SSTR3;refGene_tx=XM_019835519.2;refGene_gn=SSTR3;refGene_tx=XM_019835523.2;refGene_gn=SSTR3;refGene_tx=XM_023257474.1
B4 133440237 133440368
refGene_gn=RAC2;refGene_tx=XM_023257475.1;refGene_gn=RAC2;refGene_tx=XM_023257476.1;refGene_gn=RAC2;refGene_tx=XM_023257477.1;refGene_gn=RAC2;refGene_tx=XM_023257478.1
B4 133443642 133443802
refGene_gn=RAC2;refGene_tx=XM_023257475.1;refGene_gn=RAC2;refGene_tx=XM_023257476.1;refGene_gn=RAC2;refGene_tx=XM_023257477.1;refGene_gn=RAC2;refGene_tx=XM_023257478.1
B4 133444142 133444205
refGene_gn=RAC2;refGene_tx=XM_023257475.1;refGene_gn=RAC2;refGene_tx=XM_023257476.1;refGene_gn=RAC2;refGene_tx=XM_023257477.1;refGene_gn=RAC2;refGene_tx=XM_023257478.1
B4 133444810 133444928
refGene_gn=RAC2;refGene_tx=XM_023257475.1;refGene_gn=RAC2;refGene_tx=XM_023257476.1;refGene_gn=RAC2;refGene_tx=XM_023257477.1;refGene_gn=RAC2;refGene_tx=XM_023257478.1
B4 133450550 133450622
refGene_gn=RAC2;refGene_tx=XM_023257475.1;refGene_gn=RAC2;refGene_tx=XM_023257476.1;refGene_gn=RAC2;refGene_tx=XM_023257477.1;refGene_gn=RAC2;refGene_tx=XM_023257478.1
B4 133451020 133451181
refGene_gn=RAC2;refGene_tx=XM_023257475.1;refGene_gn=RAC2;refGene_tx=XM_023257476.1
B4 133452480 133452515
refGene_gn=RAC2;refGene_tx=XM_023257477.1;refGene_gn=RAC2;refGene_tx=X

M_023257478.1
B4 133478526 133478545
refGene_gn=CYTH4;refGene_tx=XM_003989240.5
B4 133486040 133486123
refGene_gn=CYTH4;refGene_tx=XM_003989240.5
B4 133488458 133488523
refGene_gn=CYTH4;refGene_tx=XM_003989240.5
B4 133489639 133489706
refGene_gn=CYTH4;refGene_tx=XM_003989240.5
B4 133491244 133491363
refGene_gn=CYTH4;refGene_tx=XM_003989240.5
B4 133492602 133492683
refGene_gn=CYTH4;refGene_tx=XM_003989240.5
B4 133494116 133494229
refGene_gn=CYTH4;refGene_tx=XM_003989240.5
B4 133495849 133495998
refGene_gn=CYTH4;refGene_tx=XM_003989240.5
B4 133501786 133501898
refGene_gn=CYTH4;refGene_tx=XM_003989240.5
B4 133503261 133503338
refGene_gn=CYTH4;refGene_tx=XM_003989240.5
B4 133503750 133503822
refGene_gn=CYTH4;refGene_tx=XM_003989240.5
B4 133504378 133504533
refGene_gn=CYTH4;refGene_tx=XM_003989240.5
B4 133505726 133505799
refGene_gn=CYTH4;refGene_tx=XM_003989240.5
B4 133554816 133557303
refGene_gn=ELFN2;refGene_tx=XM_023257479.1;refGene_gn=ELFN2;refGene_tx
=XM_023257480.1;refGene_gn=ELFN2;refGene_tx=XM_023257481.1
B4 133642426 133642493
refGene_gn=MFNG;refGene_tx=XM_003989241.4
B4 133644364 133644450
refGene_gn=MFNG;refGene_tx=XM_003989241.4
B4 133646027 133646193
refGene_gn=MFNG;refGene_tx=XM_003989241.4
B4 133647912 133647998
refGene_gn=MFNG;refGene_tx=XM_003989241.4
B4 133650180 133650334
refGene_gn=MFNG;refGene_tx=XM_003989241.4
B4 133650757 133650860
refGene_gn=MFNG;refGene_tx=XM_003989241.4
B4 133651180 133651229
refGene_gn=MFNG;refGene_tx=XM_003989241.4
B4 133656373 133656628
refGene_gn=MFNG;refGene_tx=XM_003989241.4
B4 133660962 133661197
refGene_gn=CARD10;refGene_tx=XM_023257779.1
B4 133661314 133661427
refGene_gn=CARD10;refGene_tx=XM_023257779.1

B4	133661917	133662033
refGene_gn=CARD10;refGene_tx=XM_023257779.1
B4	133662128	133662287
refGene_gn=CARD10;refGene_tx=XM_023257779.1
B4	133663634	133663737
refGene_gn=CARD10;refGene_tx=XM_023257779.1
B4	133664751	133664821
refGene_gn=CARD10;refGene_tx=XM_023257779.1
B4	133665019	133665263
refGene_gn=CARD10;refGene_tx=XM_023257779.1
B4	133665698	133665810
refGene_gn=CARD10;refGene_tx=XM_023257779.1
B4	133666239	133666399
refGene_gn=CARD10;refGene_tx=XM_023257779.1
B4	133671024	133671148
refGene_gn=CARD10;refGene_tx=XM_023257779.1
B4	133671612	133671641
refGene_gn=CARD10;refGene_tx=XM_023257779.1
B4	133672219	133672335
refGene_gn=CARD10;refGene_tx=XM_023257779.1
B4	133672683	133672818
refGene_gn=CARD10;refGene_tx=XM_023257779.1
B4	133674022	133674214
refGene_gn=CARD10;refGene_tx=XM_023257779.1
B4	133675291	133675417
refGene_gn=CARD10;refGene_tx=XM_023257779.1
B4	133676289	133676445
refGene_gn=CARD10;refGene_tx=XM_023257779.1
B4	133677352	133677550
refGene_gn=CARD10;refGene_tx=XM_023257779.1
B4	133681797	133682295
refGene_gn=CARD10;refGene_tx=XM_023257779.1
B4	133683987	133684034
refGene_gn=CARD10;refGene_tx=XM_023257779.1
B4	133684901	133685136
refGene_gn=CARD10;refGene_tx=XM_023257779.1
B4	133717910	133718373
refGene_gn=CDC42EP1;refGene_tx=XM_023257482.1;refGene_gn=CDC42EP1;refGene_tx=XM_023257483.1
B4	133719602	133720243
refGene_gn=CDC42EP1;refGene_tx=XM_023257482.1;refGene_gn=CDC42EP1;refGene_tx=XM_023257483.1
B4	133721519	133721666
refGene_gn=LGALS2;refGene_tx=XM_011284202.3;refGene_gn=LGALS2;refGene_tx=XM_019835531.1
B4	133721840	133721997
refGene_gn=LGALS2;refGene_tx=XM_011284202.3;refGene_gn=LGALS2;refGene_tx=XM_019835531.1
B4	133722145	133722303
refGene_gn=LGALS2;refGene_tx=XM_011284202.3

B4 133723152 133723235
refGene_gn=LGALS2;refGene_tx=XM_019835531.1
B4 133727200 133727206
refGene_gn=LGALS2;refGene_tx=XM_019835531.1
B4 133749143 133749186
refGene_gn=GGA1;refGene_tx=XM_023257484.1;refGene_gn=GGA1;refGene_tx=XM_023257488.1
B4 133752883 133752968
refGene_gn=GGA1;refGene_tx=XM_023257484.1;refGene_gn=GGA1;refGene_tx=XM_023257488.1
B4 133754161 133754237
refGene_gn=GGA1;refGene_tx=XM_023257484.1;refGene_gn=GGA1;refGene_tx=XM_023257488.1
B4 133755657 133755756
refGene_gn=GGA1;refGene_tx=XM_023257484.1;refGene_gn=GGA1;refGene_tx=XM_023257485.1;refGene_gn=GGA1;refGene_tx=XM_023257486.1;refGene_gn=GGA1;refGene_tx=XM_023257487.1;refGene_gn=GGA1;refGene_tx=XM_023257488.1
B4 133756834 133756958
refGene_gn=GGA1;refGene_tx=XM_023257484.1;refGene_gn=GGA1;refGene_tx=XM_023257485.1;refGene_gn=GGA1;refGene_tx=XM_023257486.1;refGene_gn=GGA1;refGene_tx=XM_023257487.1;refGene_gn=GGA1;refGene_tx=XM_023257488.1
B4 133757393 133757494
refGene_gn=GGA1;refGene_tx=XM_023257484.1;refGene_gn=GGA1;refGene_tx=XM_023257485.1;refGene_gn=GGA1;refGene_tx=XM_023257486.1;refGene_gn=GGA1;refGene_tx=XM_023257487.1;refGene_gn=GGA1;refGene_tx=XM_023257488.1
B4 133758162 133758243
refGene_gn=GGA1;refGene_tx=XM_023257484.1;refGene_gn=GGA1;refGene_tx=XM_023257485.1;refGene_gn=GGA1;refGene_tx=XM_023257486.1;refGene_gn=GGA1;refGene_tx=XM_023257487.1;refGene_gn=GGA1;refGene_tx=XM_023257488.1
B4 133759117 133759258
refGene_gn=GGA1;refGene_tx=XM_023257484.1;refGene_gn=GGA1;refGene_tx=XM_023257485.1;refGene_gn=GGA1;refGene_tx=XM_023257486.1;refGene_gn=GGA1;refGene_tx=XM_023257487.1;refGene_gn=GGA1;refGene_tx=XM_023257488.1
B4 133759336 133759418
refGene_gn=GGA1;refGene_tx=XM_023257484.1;refGene_gn=GGA1;refGene_tx=XM_023257485.1;refGene_gn=GGA1;refGene_tx=XM_023257486.1;refGene_gn=GGA1;refGene_tx=XM_023257487.1;refGene_gn=GGA1;refGene_tx=XM_023257488.1
B4 133760887 133760995
refGene_gn=GGA1;refGene_tx=XM_023257484.1;refGene_gn=GGA1;refGene_tx=XM_023257485.1;refGene_gn=GGA1;refGene_tx=XM_023257486.1;refGene_gn=GGA1;refGene_tx=XM_023257487.1;refGene_gn=GGA1;refGene_tx=XM_023257488.1
B4 133761743 133761896
refGene_gn=GGA1;refGene_tx=XM_023257484.1;refGene_gn=GGA1;refGene_tx=XM_023257485.1;refGene_gn=GGA1;refGene_tx=XM_023257486.1;refGene_gn=GGA1;refGene_tx=XM_023257487.1;refGene_gn=GGA1;refGene_tx=XM_023257488.1
B4 133764707 133764772
refGene_gn=GGA1;refGene_tx=XM_023257484.1;refGene_gn=GGA1;refGene_tx=XM_023257485.1;refGene_gn=GGA1;refGene_tx=XM_023257486.1;refGene_gn=GGA1;refGene_tx=XM_023257487.1;refGene_gn=GGA1;refGene_tx=XM_023257488.1
B4 133765223 133765390

refGene_gn=GGA1;refGene_tx=XM_023257484.1;refGene_gn=GGA1;refGene_tx=XM_023257485.1;refGene_gn=GGA1;refGene_tx=XM_023257486.1;refGene_gn=GGA1;refGene_tx=XM_023257487.1;refGene_gn=GGA1;refGene_tx=XM_023257488.1
B4 133765698 133765909
refGene_gn=GGA1;refGene_tx=XM_023257484.1;refGene_gn=GGA1;refGene_tx=XM_023257485.1;refGene_gn=GGA1;refGene_tx=XM_023257486.1;refGene_gn=GGA1;refGene_tx=XM_023257487.1;refGene_gn=GGA1;refGene_tx=XM_023257488.1
B4 133766790 133766960
refGene_gn=GGA1;refGene_tx=XM_023257484.1;refGene_gn=GGA1;refGene_tx=XM_023257485.1;refGene_gn=GGA1;refGene_tx=XM_023257486.1;refGene_gn=GGA1;refGene_tx=XM_023257487.1;refGene_gn=GGA1;refGene_tx=XM_023257488.1
B4 133767180 133767291
refGene_gn=GGA1;refGene_tx=XM_023257484.1;refGene_gn=GGA1;refGene_tx=XM_023257485.1;refGene_gn=GGA1;refGene_tx=XM_023257486.1;refGene_gn=GGA1;refGene_tx=XM_023257487.1
B4 133767384 133767495
refGene_gn=GGA1;refGene_tx=XM_023257484.1;refGene_gn=GGA1;refGene_tx=XM_023257485.1;refGene_gn=GGA1;refGene_tx=XM_023257486.1;refGene_gn=GGA1;refGene_tx=XM_023257487.1
B4 133774952 133775011
refGene_gn=SH3BP1;refGene_tx=XM_023257489.1;refGene_gn=SH3BP1;refGene_tx=XM_023257490.1
B4 133776248 133776291
refGene_gn=SH3BP1;refGene_tx=XM_023257489.1;refGene_gn=SH3BP1;refGene_tx=XM_023257490.1
B4 133776530 133776635
refGene_gn=SH3BP1;refGene_tx=XM_023257489.1;refGene_gn=SH3BP1;refGene_tx=XM_023257490.1
B4 133777715 133777792
refGene_gn=SH3BP1;refGene_tx=XM_023257489.1;refGene_gn=SH3BP1;refGene_tx=XM_023257490.1
B4 133778072 133778184
refGene_gn=SH3BP1;refGene_tx=XM_023257489.1;refGene_gn=SH3BP1;refGene_tx=XM_023257490.1
B4 133778287 133778364
refGene_gn=SH3BP1;refGene_tx=XM_023257489.1;refGene_gn=SH3BP1;refGene_tx=XM_023257490.1
B4 133778867 133779012
refGene_gn=SH3BP1;refGene_tx=XM_023257489.1;refGene_gn=SH3BP1;refGene_tx=XM_023257490.1
B4 133779729 133779798
refGene_gn=SH3BP1;refGene_tx=XM_023257489.1;refGene_gn=SH3BP1;refGene_tx=XM_023257490.1
B4 133779962 133780053
refGene_gn=SH3BP1;refGene_tx=XM_023257489.1;refGene_gn=SH3BP1;refGene_tx=XM_023257490.1
B4 133780442 133780588
refGene_gn=SH3BP1;refGene_tx=XM_023257489.1;refGene_gn=SH3BP1;refGene_tx=XM_023257490.1
B4 133781279 133781391

refGene_gn=SH3BP1;refGene_tx=XM_023257489.1;refGene_gn=SH3BP1;refGene_tx=XM_023257490.1
B4 133781765 133781847
refGene_gn=SH3BP1;refGene_tx=XM_023257489.1;refGene_gn=SH3BP1;refGene_tx=XM_023257490.1
B4 133781933 133782014
refGene_gn=SH3BP1;refGene_tx=XM_023257489.1;refGene_gn=SH3BP1;refGene_tx=XM_023257490.1
B4 133782964 133783081
refGene_gn=SH3BP1;refGene_tx=XM_023257489.1;refGene_gn=SH3BP1;refGene_tx=XM_023257490.1
B4 133783777 133783875
refGene_gn=SH3BP1;refGene_tx=XM_023257489.1;refGene_gn=SH3BP1;refGene_tx=XM_023257490.1
B4 133784200 133784354
refGene_gn=SH3BP1;refGene_tx=XM_023257489.1;refGene_gn=SH3BP1;refGene_tx=XM_023257490.1
B4 133785383 133785478
refGene_gn=SH3BP1;refGene_tx=XM_023257489.1;refGene_gn=SH3BP1;refGene_tx=XM_023257490.1
B4 133786845 133787243
refGene_gn=SH3BP1;refGene_tx=XM_023257489.1
B4 133787471 133787620
refGene_gn=SH3BP1;refGene_tx=XM_023257490.1
B4 133789880 133790454
refGene_gn=PDXP;refGene_tx=XM_023257491.1
B4 133794268 133794585
refGene_gn=PDXP;refGene_tx=XM_023257491.1
B4 133804820 133804829
refGene_gn=LGALS1;refGene_tx=XM_003989245.5
B4 133806069 133806149
refGene_gn=LGALS1;refGene_tx=XM_003989245.5
B4 133807446 133807618
refGene_gn=LGALS1;refGene_tx=XM_003989245.5
B4 133807912 133808059
refGene_gn=LGALS1;refGene_tx=XM_003989245.5
B4 133812754 133812837
refGene_gn=NOL12;refGene_tx=XM_003989246.5
B4 133814186 133814292
refGene_gn=NOL12;refGene_tx=XM_003989246.5
B4 133814782 133814831
refGene_gn=NOL12;refGene_tx=XM_003989246.5
B4 133815380 133815523
refGene_gn=NOL12;refGene_tx=XM_003989246.5
B4 133816669 133816767
refGene_gn=NOL12;refGene_tx=XM_003989246.5
B4 133817188 133817354
refGene_gn=NOL12;refGene_tx=XM_003989246.5
B4 133826872 133826986
refGene_gn=TRIOBP;refGene_tx=XM_023257492.1

B4	133831065	133831205
refGene_gn=TRIOBP;refGene_tx=XM_023257492.1
B4	133832408	133832598
refGene_gn=TRIOBP;refGene_tx=XM_023257492.1
B4	133833615	133833784
refGene_gn=TRIOBP;refGene_tx=XM_023257492.1
B4	133837794	133838366
refGene_gn=TRIOBP;refGene_tx=XM_023257492.1
B4	133838920	133840538
refGene_gn=TRIOBP;refGene_tx=XM_023257492.1
B4	133845420	133845535
refGene_gn=TRIOBP;refGene_tx=XM_023257492.1
B4	133846222	133847107
refGene_gn=TRIOBP;refGene_tx=XM_023257492.1
B4	133849673	133849751
refGene_gn=TRIOBP;refGene_tx=XM_023257492.1
B4	133851943	133852081
refGene_gn=TRIOBP;refGene_tx=XM_023257492.1
B4	133861206	133861263
refGene_gn=TRIOBP;refGene_tx=XM_023257492.1
B4	133863694	133863763
refGene_gn=TRIOBP;refGene_tx=XM_023257492.1
B4	133863887	133863977
refGene_gn=TRIOBP;refGene_tx=XM_023257492.1
B4	133864292	133864402
refGene_gn=TRIOBP;refGene_tx=XM_023257492.1
B4	133865843	133866363
refGene_gn=TRIOBP;refGene_tx=XM_023257492.1
B4	133867020	133867131
refGene_gn=TRIOBP;refGene_tx=XM_023257492.1
B4	133872145	133872293
refGene_gn=TRIOBP;refGene_tx=XM_023257492.1
B4	133873998	133874101
refGene_gn=TRIOBP;refGene_tx=XM_023257492.1
B4	133874643	133874803
refGene_gn=TRIOBP;refGene_tx=XM_023257492.1
B4	133874878	133874992
refGene_gn=TRIOBP;refGene_tx=XM_023257492.1
B4	133875522	133875609
refGene_gn=TRIOBP;refGene_tx=XM_023257492.1
B4	133876374	133876407
refGene_gn=TRIOBP;refGene_tx=XM_023257492.1
B4	133876408	133876540
refGene_gn=TRIOBP;refGene_tx=XM_023257492.1
B4	133906917	133907502
refGene_gn=LOC101098644;refGene_tx=XM_006934030.4
B4	133909390	133909586
refGene_gn=GCAT;refGene_tx=XM_003989248.4;refGene_gn=GCAT;refGene_tx=X
M_019835542.2
B4	133911275	133911406

refGene_gn=GCAT;refGene_tx=XM_003989248.4;refGene_gn=GCAT;refGene_tx=XM_019835542.2
B4 133912350 133912452
refGene_gn=GCAT;refGene_tx=XM_003989248.4;refGene_gn=GCAT;refGene_tx=XM_019835542.2
B4 133912920 133913067
refGene_gn=GCAT;refGene_tx=XM_003989248.4;refGene_gn=GCAT;refGene_tx=XM_019835542.2
B4 133913533 133913688
refGene_gn=GCAT;refGene_tx=XM_003989248.4;refGene_gn=GCAT;refGene_tx=XM_019835542.2
B4 133913820 133913903
refGene_gn=GCAT;refGene_tx=XM_003989248.4;refGene_gn=GCAT;refGene_tx=XM_019835542.2
B4 133914081 133914253
refGene_gn=GCAT;refGene_tx=XM_003989248.4;refGene_gn=GCAT;refGene_tx=XM_019835542.2
B4 133914599 133914675
refGene_gn=GCAT;refGene_tx=XM_019835542.2
B4 133914695 133914817
refGene_gn=GCAT;refGene_tx=XM_003989248.4
B4 133915099 133915251
refGene_gn=GCAT;refGene_tx=XM_003989248.4
B4 133920945 133921304
refGene_gn=GALR3;refGene_tx=XM_023257781.1
B4 133921711 133922501
refGene_gn=GALR3;refGene_tx=XM_023257781.1
B4 133932990 133933065
refGene_gn=ANKRD54;refGene_tx=XM_003989249.5
B4 133933436 133933544
refGene_gn=ANKRD54;refGene_tx=XM_003989249.5
B4 133933782 133933907
refGene_gn=ANKRD54;refGene_tx=XM_003989249.5
B4 133933982 133934030
refGene_gn=ANKRD54;refGene_tx=XM_003989249.5
B4 133934478 133934550
refGene_gn=ANKRD54;refGene_tx=XM_003989249.5
B4 133938864 133938963
refGene_gn=ANKRD54;refGene_tx=XM_003989249.5
B4 133944249 133944297
refGene_gn=ANKRD54;refGene_tx=XM_003989249.5
B4 133945251 133945576
refGene_gn=ANKRD54;refGene_tx=XM_003989249.5
B4 133954148 133954181
refGene_gn=EIF3L;refGene_tx=XM_003989250.5
B4 133954742 133954791
refGene_gn=EIF3L;refGene_tx=XM_003989250.5
B4 133955699 133955910
refGene_gn=EIF3L;refGene_tx=XM_003989250.5
B4 133959808 133959888

refGene_gn=EIF3L;refGene_tx=XM_003989250.5
B4	133963949	133964011
refGene_gn=EIF3L;refGene_tx=XM_003989250.5
B4	133967827	133967897
refGene_gn=EIF3L;refGene_tx=XM_003989250.5
B4	133968108	133968182
refGene_gn=EIF3L;refGene_tx=XM_003989250.5
B4	133971215	133971387
refGene_gn=EIF3L;refGene_tx=XM_003989250.5
B4	133973478	133973633
refGene_gn=EIF3L;refGene_tx=XM_003989250.5
B4	133974389	133974560
refGene_gn=EIF3L;refGene_tx=XM_003989250.5
B4	133975335	133975833
refGene_gn=EIF3L;refGene_tx=XM_003989250.5
B4	133980694	133980775
refGene_gn=EIF3L;refGene_tx=XM_003989250.5
B4	133981937	133981976
refGene_gn=EIF3L;refGene_tx=XM_003989250.5
B4	133996222	133996368
refGene_gn=MICALL1;refGene_tx=XM_023257682.1;refGene_gn=MICALL1;refGen
e_tx=XM_023257683.1;refGene_gn=MICALL1;refGene_tx=XM_023257684.1
B4	134000966	134001015
refGene_gn=MICALL1;refGene_tx=XM_023257682.1;refGene_gn=MICALL1;refGen
e_tx=XM_023257683.1;refGene_gn=MICALL1;refGene_tx=XM_023257684.1
B4	134001502	134001644
refGene_gn=MICALL1;refGene_tx=XM_023257682.1;refGene_gn=MICALL1;refGen
e_tx=XM_023257683.1;refGene_gn=MICALL1;refGene_tx=XM_023257684.1
B4	134006102	134006191
refGene_gn=MICALL1;refGene_tx=XM_023257682.1;refGene_gn=MICALL1;refGen
e_tx=XM_023257683.1;refGene_gn=MICALL1;refGene_tx=XM_023257684.1
B4	134007534	134007677
refGene_gn=MICALL1;refGene_tx=XM_023257682.1;refGene_gn=MICALL1;refGen
e_tx=XM_023257683.1;refGene_gn=MICALL1;refGene_tx=XM_023257684.1
B4	134009009	134009455
refGene_gn=MICALL1;refGene_tx=XM_023257682.1;refGene_gn=MICALL1;refGen
e_tx=XM_023257683.1;refGene_gn=MICALL1;refGene_tx=XM_023257684.1
B4	134010078	134010136
refGene_gn=MICALL1;refGene_tx=XM_023257682.1;refGene_gn=MICALL1;refGen
e_tx=XM_023257683.1;refGene_gn=MICALL1;refGene_tx=XM_023257684.1
B4	134011032	134011421
refGene_gn=MICALL1;refGene_tx=XM_023257682.1;refGene_gn=MICALL1;refGen
e_tx=XM_023257683.1;refGene_gn=MICALL1;refGene_tx=XM_023257684.1
B4	134012854	134013270
refGene_gn=MICALL1;refGene_tx=XM_023257682.1;refGene_gn=MICALL1;refGen
e_tx=XM_023257683.1;refGene_gn=MICALL1;refGene_tx=XM_023257684.1
B4	134015117	134015252
refGene_gn=MICALL1;refGene_tx=XM_023257682.1;refGene_gn=MICALL1;refGen
e_tx=XM_023257683.1;refGene_gn=MICALL1;refGene_tx=XM_023257684.1
B4	134015874	134016001

refGene_gn=MICALL1;refGene_tx=XM_023257682.1;refGene_gn=MICALL1;refGene_tx=XM_023257683.1;refGene_gn=MICALL1;refGene_tx=XM_023257684.1
B4	134016140	134016231
refGene_gn=MICALL1;refGene_tx=XM_023257682.1;refGene_gn=MICALL1;refGene_tx=XM_023257683.1;refGene_gn=MICALL1;refGene_tx=XM_023257684.1
B4	134016390	134016464
refGene_gn=MICALL1;refGene_tx=XM_023257682.1;refGene_gn=MICALL1;refGene_tx=XM_023257683.1;refGene_gn=MICALL1;refGene_tx=XM_023257684.1
B4	134019122	134019237
refGene_gn=MICALL1;refGene_tx=XM_023257682.1;refGene_gn=MICALL1;refGene_tx=XM_023257683.1;refGene_gn=MICALL1;refGene_tx=XM_023257684.1
B4	134019440	134019487
refGene_gn=MICALL1;refGene_tx=XM_023257682.1;refGene_gn=MICALL1;refGene_tx=XM_023257683.1;refGene_gn=MICALL1;refGene_tx=XM_023257684.1
B4	134020437	134020559
refGene_gn=MICALL1;refGene_tx=XM_023257682.1;refGene_gn=MICALL1;refGene_tx=XM_023257683.1;refGene_gn=MICALL1;refGene_tx=XM_023257684.1
B4	134021637	134021643
refGene_gn=MICALL1;refGene_tx=XM_023257684.1
B4	134023952	134024024
refGene_gn=CB4H22orf23;refGene_tx=XM_003989252.4;refGene_gn=CB4H22orf23;refGene_tx=XM_019835551.2;refGene_gn=CB4H22orf23;refGene_tx=XM_019835553.2;refGene_gn=CB4H22orf23;refGene_tx=XM_019835554.2;refGene_gn=CB4H22orf23;refGene_tx=XM_023257718.1
B4	134024165	134024266
refGene_gn=CB4H22orf23;refGene_tx=XM_003989252.4;refGene_gn=CB4H22orf23;refGene_tx=XM_019835551.2;refGene_gn=CB4H22orf23;refGene_tx=XM_019835553.2;refGene_gn=CB4H22orf23;refGene_tx=XM_019835554.2;refGene_gn=CB4H22orf23;refGene_tx=XM_023257718.1
B4	134024482	134024614
refGene_gn=CB4H22orf23;refGene_tx=XM_003989252.4;refGene_gn=CB4H22orf23;refGene_tx=XM_019835551.2;refGene_gn=CB4H22orf23;refGene_tx=XM_019835553.2;refGene_gn=CB4H22orf23;refGene_tx=XM_019835554.2;refGene_gn=CB4H22orf23;refGene_tx=XM_023257718.1
B4	134025259	134025442
refGene_gn=CB4H22orf23;refGene_tx=XM_003989252.4;refGene_gn=CB4H22orf23;refGene_tx=XM_019835551.2;refGene_gn=CB4H22orf23;refGene_tx=XM_019835553.2;refGene_gn=CB4H22orf23;refGene_tx=XM_019835554.2;refGene_gn=CB4H22orf23;refGene_tx=XM_023257718.1
B4	134028989	134029052
refGene_gn=CB4H22orf23;refGene_tx=XM_003989252.4;refGene_gn=CB4H22orf23;refGene_tx=XM_019835551.2;refGene_gn=CB4H22orf23;refGene_tx=XM_019835553.2;refGene_gn=CB4H22orf23;refGene_tx=XM_019835554.2;refGene_gn=CB4H22orf23;refGene_tx=XM_023257718.1
B4	134029845	134029846
refGene_gn=CB4H22orf23;refGene_tx=XM_019835554.2
B4	134030086	134030239
refGene_gn=CB4H22orf23;refGene_tx=XM_003989252.4;refGene_gn=CB4H22orf23;refGene_tx=XM_019835551.2;refGene_gn=CB4H22orf23;refGene_tx=XM_019835553.2;refGene_gn=CB4H22orf23;refGene_tx=XM_023257718.1

B4	134030434	134030453
refGene_gn=CB4H22orf23;refGene_tx=XM_019835551.2
B4	134030912	134030932
refGene_gn=POLR2F;refGene_tx=XM_023257730.1
B4	134032528	134032598
refGene_gn=POLR2F;refGene_tx=XM_023257730.1
B4	134035199	134035330
refGene_gn=POLR2F;refGene_tx=XM_023257730.1
B4	134039317	134039389
refGene_gn=POLR2F;refGene_tx=XM_023257730.1
B4	134039880	134039971
refGene_gn=POLR2F;refGene_tx=XM_023257730.1
B4	134045032	134045736
refGene_gn=SOX10;refGene_tx=XM_023257695.1
B4	134047983	134048252
refGene_gn=SOX10;refGene_tx=XM_023257695.1
B4	134052868	134053332
refGene_gn=SOX10;refGene_tx=XM_023257695.1
B4	134115243	134115284
refGene_gn=PICK1;refGene_tx=XM_023257706.1;refGene_gn=PICK1;refGene_tx=XM_023257707.1;refGene_gn=PICK1;refGene_tx=XM_023257708.1
B4	134115980	134116092
refGene_gn=PICK1;refGene_tx=XM_023257706.1;refGene_gn=PICK1;refGene_tx=XM_023257707.1;refGene_gn=PICK1;refGene_tx=XM_023257708.1
B4	134127936	134128065
refGene_gn=PICK1;refGene_tx=XM_023257706.1;refGene_gn=PICK1;refGene_tx=XM_023257707.1;refGene_gn=PICK1;refGene_tx=XM_023257708.1
B4	134129549	134129616
refGene_gn=PICK1;refGene_tx=XM_023257706.1;refGene_gn=PICK1;refGene_tx=XM_023257707.1;refGene_gn=PICK1;refGene_tx=XM_023257708.1
B4	134130747	134130837
refGene_gn=PICK1;refGene_tx=XM_023257706.1;refGene_gn=PICK1;refGene_tx=XM_023257707.1;refGene_gn=PICK1;refGene_tx=XM_023257708.1
B4	134132730	134132784
refGene_gn=PICK1;refGene_tx=XM_023257706.1;refGene_gn=PICK1;refGene_tx=XM_023257707.1;refGene_gn=PICK1;refGene_tx=XM_023257708.1
B4	134133288	134133351
refGene_gn=PICK1;refGene_tx=XM_023257706.1;refGene_gn=PICK1;refGene_tx=XM_023257707.1;refGene_gn=PICK1;refGene_tx=XM_023257708.1
B4	134134131	134134265
refGene_gn=PICK1;refGene_tx=XM_023257706.1;refGene_gn=PICK1;refGene_tx=XM_023257707.1;refGene_gn=PICK1;refGene_tx=XM_023257708.1
B4	134134637	134134730
refGene_gn=PICK1;refGene_tx=XM_023257706.1;refGene_gn=PICK1;refGene_tx=XM_023257707.1;refGene_gn=PICK1;refGene_tx=XM_023257708.1
B4	134135567	134135618
refGene_gn=PICK1;refGene_tx=XM_023257706.1;refGene_gn=PICK1;refGene_tx=XM_023257707.1;refGene_gn=PICK1;refGene_tx=XM_023257708.1
B4	134136450	134136595
refGene_gn=PICK1;refGene_tx=XM_023257706.1;refGene_gn=PICK1;refGene_tx

=XM_023257707.1;refGene_gn=PICK1;refGene_tx=XM_023257708.1
B4 134137059 134137334
refGene_gn=PICK1;refGene_tx=XM_023257706.1;refGene_gn=PICK1;refGene_tx
=XM_023257707.1;refGene_gn=PICK1;refGene_tx=XM_023257708.1
B4 134139721 134140038
refGene_gn=SLC16A8;refGene_tx=XM_019835547.2;refGene_gn=SLC16A8;refGen
e_tx=XM_023257694.1
B4 134141989 134142817
refGene_gn=SLC16A8;refGene_tx=XM_019835547.2;refGene_gn=SLC16A8;refGen
e_tx=XM_023257694.1
B4 134143079 134143223
refGene_gn=SLC16A8;refGene_tx=XM_019835547.2;refGene_gn=SLC16A8;refGen
e_tx=XM_023257694.1
B4 134143986 134144200
refGene_gn=SLC16A8;refGene_tx=XM_019835547.2;refGene_gn=SLC16A8;refGen
e_tx=XM_023257694.1
B4 134146944 134147020
refGene_gn=BAIAP2L2;refGene_tx=XM_023257731.1
B4 134147251 134147295
refGene_gn=BAIAP2L2;refGene_tx=XM_023257731.1
B4 134147872 134148080
refGene_gn=BAIAP2L2;refGene_tx=XM_023257731.1
B4 134148421 134148544
refGene_gn=BAIAP2L2;refGene_tx=XM_023257731.1
B4 134150218 134150456
refGene_gn=BAIAP2L2;refGene_tx=XM_023257731.1
B4 134150580 134150728
refGene_gn=BAIAP2L2;refGene_tx=XM_023257731.1
B4 134151029 134151182
refGene_gn=BAIAP2L2;refGene_tx=XM_023257731.1
B4 134159723 134159870
refGene_gn=BAIAP2L2;refGene_tx=XM_023257731.1
B4 134160895 134161012
refGene_gn=BAIAP2L2;refGene_tx=XM_023257731.1
B4 134161170 134161242
refGene_gn=BAIAP2L2;refGene_tx=XM_023257731.1
B4 134170831 134170893
refGene_gn=BAIAP2L2;refGene_tx=XM_023257731.1
B4 134171348 134171435
refGene_gn=BAIAP2L2;refGene_tx=XM_023257731.1
B4 134172184 134172260
refGene_gn=BAIAP2L2;refGene_tx=XM_023257731.1
B4 134173462 134173513
refGene_gn=BAIAP2L2;refGene_tx=XM_023257731.1
B4 134175123 134175268
refGene_gn=PLA2G6;refGene_tx=XM_011284207.3;refGene_gn=PLA2G6;refGene_
tx=XM_011284210.3;refGene_gn=PLA2G6;refGene_tx=XM_019835555.2;refGene_
gn=PLA2G6;refGene_tx=XM_019835556.2
B4 134175472 134175546
refGene_gn=PLA2G6;refGene_tx=XM_011284207.3;refGene_gn=PLA2G6;refGene_

tx=XM_011284210.3;refGene_gn=PLA2G6;refGene_tx=XM_019835555.2;refGene_gn=PLA2G6;refGene_tx=XM_019835556.2
B4 134175989 134176157
refGene_gn=PLA2G6;refGene_tx=XM_011284207.3;refGene_gn=PLA2G6;refGene_tx=XM_011284210.3;refGene_gn=PLA2G6;refGene_tx=XM_019835555.2;refGene_gn=PLA2G6;refGene_tx=XM_019835556.2
B4 134177295 134177542
refGene_gn=PLA2G6;refGene_tx=XM_011284207.3;refGene_gn=PLA2G6;refGene_tx=XM_011284210.3;refGene_gn=PLA2G6;refGene_tx=XM_019835555.2;refGene_gn=PLA2G6;refGene_tx=XM_019835556.2;refGene_gn=PLA2G6;refGene_tx=XM_019835559.2;refGene_gn=PLA2G6;refGene_tx=XM_019835560.2
B4 134177840 134177977
refGene_gn=PLA2G6;refGene_tx=XM_011284207.3;refGene_gn=PLA2G6;refGene_tx=XM_011284210.3;refGene_gn=PLA2G6;refGene_tx=XM_019835555.2;refGene_gn=PLA2G6;refGene_tx=XM_019835556.2;refGene_gn=PLA2G6;refGene_tx=XM_019835559.2;refGene_gn=PLA2G6;refGene_tx=XM_019835560.2
B4 134187293 134187444
refGene_gn=PLA2G6;refGene_tx=XM_011284207.3;refGene_gn=PLA2G6;refGene_tx=XM_011284210.3;refGene_gn=PLA2G6;refGene_tx=XM_019835555.2;refGene_gn=PLA2G6;refGene_tx=XM_019835556.2;refGene_gn=PLA2G6;refGene_tx=XM_019835559.2;refGene_gn=PLA2G6;refGene_tx=XM_019835560.2
B4 134189009 134189173
refGene_gn=PLA2G6;refGene_tx=XM_011284207.3;refGene_gn=PLA2G6;refGene_tx=XM_011284210.3;refGene_gn=PLA2G6;refGene_tx=XM_019835555.2;refGene_gn=PLA2G6;refGene_tx=XM_019835556.2;refGene_gn=PLA2G6;refGene_tx=XM_019835559.2;refGene_gn=PLA2G6;refGene_tx=XM_019835560.2
B4 134191607 134191686
refGene_gn=PLA2G6;refGene_tx=XM_011284207.3;refGene_gn=PLA2G6;refGene_tx=XM_011284210.3;refGene_gn=PLA2G6;refGene_tx=XM_019835555.2;refGene_gn=PLA2G6;refGene_tx=XM_019835556.2;refGene_gn=PLA2G6;refGene_tx=XM_019835559.2;refGene_gn=PLA2G6;refGene_tx=XM_019835560.2
B4 134193575 134193740
refGene_gn=PLA2G6;refGene_tx=XM_011284207.3;refGene_gn=PLA2G6;refGene_tx=XM_019835555.2;refGene_gn=PLA2G6;refGene_tx=XM_019835556.2;refGene_gn=PLA2G6;refGene_tx=XM_019835559.2;refGene_gn=PLA2G6;refGene_tx=XM_019835560.2
B4 134194948 134195057
refGene_gn=PLA2G6;refGene_tx=XM_011284207.3;refGene_gn=PLA2G6;refGene_tx=XM_011284210.3;refGene_gn=PLA2G6;refGene_tx=XM_019835555.2;refGene_gn=PLA2G6;refGene_tx=XM_019835556.2;refGene_gn=PLA2G6;refGene_tx=XM_019835559.2;refGene_gn=PLA2G6;refGene_tx=XM_019835560.2
B4 134199296 134199479
refGene_gn=PLA2G6;refGene_tx=XM_011284207.3;refGene_gn=PLA2G6;refGene_tx=XM_011284210.3;refGene_gn=PLA2G6;refGene_tx=XM_019835555.2;refGene_gn=PLA2G6;refGene_tx=XM_019835556.2;refGene_gn=PLA2G6;refGene_tx=XM_019835559.2;refGene_gn=PLA2G6;refGene_tx=XM_019835560.2
B4 134200633 134200730
refGene_gn=PLA2G6;refGene_tx=XM_011284207.3;refGene_gn=PLA2G6;refGene_tx=XM_011284210.3;refGene_gn=PLA2G6;refGene_tx=XM_019835555.2;refGene_gn=PLA2G6;refGene_tx=XM_019835556.2;refGene_gn=PLA2G6;refGene_tx=XM_01

9835559.2;refGene_gn=PLA2G6;refGene_tx=XM_019835560.2
B4 134202951 134203139
refGene_gn=PLA2G6;refGene_tx=XM_011284207.3;refGene_gn=PLA2G6;refGene_tx=XM_011284210.3;refGene_gn=PLA2G6;refGene_tx=XM_019835555.2;refGene_gn=PLA2G6;refGene_tx=XM_019835556.2;refGene_gn=PLA2G6;refGene_tx=XM_019835559.2;refGene_gn=PLA2G6;refGene_tx=XM_019835560.2
B4 134204000 134204184
refGene_gn=PLA2G6;refGene_tx=XM_011284207.3;refGene_gn=PLA2G6;refGene_tx=XM_011284210.3;refGene_gn=PLA2G6;refGene_tx=XM_019835555.2;refGene_gn=PLA2G6;refGene_tx=XM_019835556.2;refGene_gn=PLA2G6;refGene_tx=XM_019835559.2;refGene_gn=PLA2G6;refGene_tx=XM_019835560.2
B4 134205114 134205330
refGene_gn=PLA2G6;refGene_tx=XM_011284207.3;refGene_gn=PLA2G6;refGene_tx=XM_011284210.3;refGene_gn=PLA2G6;refGene_tx=XM_019835555.2;refGene_gn=PLA2G6;refGene_tx=XM_019835556.2;refGene_gn=PLA2G6;refGene_tx=XM_019835559.2;refGene_gn=PLA2G6;refGene_tx=XM_019835560.2
B4 134221873 134222127
refGene_gn=PLA2G6;refGene_tx=XM_011284207.3;refGene_gn=PLA2G6;refGene_tx=XM_011284210.3;refGene_gn=PLA2G6;refGene_tx=XM_019835555.2;refGene_gn=PLA2G6;refGene_tx=XM_019835556.2;refGene_gn=PLA2G6;refGene_tx=XM_019835559.2;refGene_gn=PLA2G6;refGene_tx=XM_019835560.2
B4 134225936 134225987
refGene_gn=PLA2G6;refGene_tx=XM_011284207.3;refGene_gn=PLA2G6;refGene_tx=XM_011284210.3;refGene_gn=PLA2G6;refGene_tx=XM_019835559.2;refGene_gn=PLA2G6;refGene_tx=XM_019835560.2
B4 134254918 134254954
refGene_gn=MAFF;refGene_tx=XM_023257725.1;refGene_gn=MAFF;refGene_tx=XM_023257726.1;refGene_gn=MAFF;refGene_tx=XM_023257727.1;refGene_gn=MAFF;refGene_tx=XM_023257728.1
B4 134255215 134255668
refGene_gn=MAFF;refGene_tx=XM_023257725.1;refGene_gn=MAFF;refGene_tx=XM_023257726.1;refGene_gn=MAFF;refGene_tx=XM_023257727.1;refGene_gn=MAFF;refGene_tx=XM_023257728.1
B4 134260800 134261042
refGene_gn=TMEM184B;refGene_tx=XM_023257710.1;refGene_gn=TMEM184B;refGene_tx=XM_023257711.1
B4 134261894 134261915
refGene_gn=TMEM184B;refGene_tx=XM_023257710.1
B4 134263692 134263887
refGene_gn=TMEM184B;refGene_tx=XM_023257710.1;refGene_gn=TMEM184B;refGene_tx=XM_023257711.1
B4 134264331 134264501
refGene_gn=TMEM184B;refGene_tx=XM_023257710.1;refGene_gn=TMEM184B;refGene_tx=XM_023257711.1
B4 134265471 134265563
refGene_gn=TMEM184B;refGene_tx=XM_023257710.1;refGene_gn=TMEM184B;refGene_tx=XM_023257711.1
B4 134268601 134268677
refGene_gn=TMEM184B;refGene_tx=XM_023257710.1;refGene_gn=TMEM184B;refGene_tx=XM_023257711.1

B4	134269158	134269249
refGene_gn=TMEM184B;refGene_tx=XM_023257710.1;refGene_gn=TMEM184B;refGene_tx=XM_023257711.1
B4	134281191	134281357
refGene_gn=TMEM184B;refGene_tx=XM_023257710.1;refGene_gn=TMEM184B;refGene_tx=XM_023257711.1
B4	134282806	134282998
refGene_gn=TMEM184B;refGene_tx=XM_023257710.1;refGene_gn=TMEM184B;refGene_tx=XM_023257711.1
B4	134322404	134322437
refGene_gn=LOC101101321;refGene_tx=XM_023257699.1;refGene_gn=LOC101101321;refGene_tx=XM_023257700.1;refGene_gn=LOC101101321;refGene_tx=XM_023257701.1;refGene_gn=LOC101101321;refGene_tx=XM_023257702.1;refGene_gn=LOC101101321;refGene_tx=XM_023257703.1
B4	134323175	134323315
refGene_gn=LOC101101321;refGene_tx=XM_023257699.1;refGene_gn=LOC101101321;refGene_tx=XM_023257700.1;refGene_gn=LOC101101321;refGene_tx=XM_023257701.1;refGene_gn=LOC101101321;refGene_tx=XM_023257702.1;refGene_gn=LOC101101321;refGene_tx=XM_023257703.1
B4	134323414	134323607
refGene_gn=LOC101101321;refGene_tx=XM_023257699.1;refGene_gn=LOC101101321;refGene_tx=XM_023257700.1;refGene_gn=LOC101101321;refGene_tx=XM_023257701.1;refGene_gn=LOC101101321;refGene_tx=XM_023257702.1;refGene_gn=LOC101101321;refGene_tx=XM_023257703.1
B4	134324414	134324475
refGene_gn=LOC101101321;refGene_tx=XM_023257699.1;refGene_gn=LOC101101321;refGene_tx=XM_023257700.1;refGene_gn=LOC101101321;refGene_tx=XM_023257701.1
B4	134326822	134326937
refGene_gn=LOC101101321;refGene_tx=XM_023257704.1;refGene_gn=LOC101101321;refGene_tx=XM_023257705.1
B4	134327033	134327137
refGene_gn=LOC101101321;refGene_tx=XM_023257704.1
B4	134327238	134327876
refGene_gn=LOC101101321;refGene_tx=XM_023257697.1;refGene_gn=LOC101101321;refGene_tx=XM_023257698.1;refGene_gn=LOC101101321;refGene_tx=XM_023257699.1;refGene_gn=LOC101101321;refGene_tx=XM_023257700.1;refGene_gn=LOC101101321;refGene_tx=XM_023257701.1;refGene_gn=LOC101101321;refGene_tx=XM_023257702.1;refGene_gn=LOC101101321;refGene_tx=XM_023257703.1;refGene_gn=LOC101101321;refGene_tx=XM_023257704.1;refGene_gn=LOC101101321;refGene_tx=XM_023257705.1
B4	134329335	134329506
refGene_gn=LOC101101321;refGene_tx=XM_023257697.1;refGene_gn=LOC101101321;refGene_tx=XM_023257698.1;refGene_gn=LOC101101321;refGene_tx=XM_023257699.1;refGene_gn=LOC101101321;refGene_tx=XM_023257700.1;refGene_gn=LOC101101321;refGene_tx=XM_023257701.1;refGene_gn=LOC101101321;refGene_tx=XM_023257702.1;refGene_gn=LOC101101321;refGene_tx=XM_023257703.1;refGene_gn=LOC101101321;refGene_tx=XM_023257704.1;refGene_gn=LOC101101321;refGene_tx=XM_023257705.1
B4	134330468	134330697

refGene_gn=LOC101101321;refGene_tx=XM_023257697.1;refGene_gn=LOC101101321;refGene_tx=XM_023257698.1;refGene_gn=LOC101101321;refGene_tx=XM_023257699.1;refGene_gn=LOC101101321;refGene_tx=XM_023257700.1;refGene_gn=LOC101101321;refGene_tx=XM_023257701.1;refGene_gn=LOC101101321;refGene_tx=XM_023257702.1;refGene_gn=LOC101101321;refGene_tx=XM_023257703.1;refGene_gn=LOC101101321;refGene_tx=XM_023257704.1;refGene_gn=LOC101101321;refGene_tx=XM_023257705.1
B4 134332166 134332315
refGene_gn=LOC101101321;refGene_tx=XM_023257697.1;refGene_gn=LOC101101321;refGene_tx=XM_023257698.1;refGene_gn=LOC101101321;refGene_tx=XM_023257699.1;refGene_gn=LOC101101321;refGene_tx=XM_023257700.1;refGene_gn=LOC101101321;refGene_tx=XM_023257701.1;refGene_gn=LOC101101321;refGene_tx=XM_023257702.1;refGene_gn=LOC101101321;refGene_tx=XM_023257703.1;refGene_gn=LOC101101321;refGene_tx=XM_023257704.1;refGene_gn=LOC101101321;refGene_tx=XM_023257705.1
B4 134332434 134332545
refGene_gn=LOC101101321;refGene_tx=XM_023257697.1;refGene_gn=LOC101101321;refGene_tx=XM_023257698.1;refGene_gn=LOC101101321;refGene_tx=XM_023257699.1;refGene_gn=LOC101101321;refGene_tx=XM_023257700.1;refGene_gn=LOC101101321;refGene_tx=XM_023257701.1;refGene_gn=LOC101101321;refGene_tx=XM_023257702.1;refGene_gn=LOC101101321;refGene_tx=XM_023257703.1;refGene_gn=LOC101101321;refGene_tx=XM_023257704.1;refGene_gn=LOC101101321;refGene_tx=XM_023257705.1
B4 134340671 134340747
refGene_gn=LOC101101321;refGene_tx=XM_023257697.1;refGene_gn=LOC101101321;refGene_tx=XM_023257698.1;refGene_gn=LOC101101321;refGene_tx=XM_023257699.1;refGene_gn=LOC101101321;refGene_tx=XM_023257700.1;refGene_gn=LOC101101321;refGene_tx=XM_023257701.1;refGene_gn=LOC101101321;refGene_tx=XM_023257702.1;refGene_gn=LOC101101321;refGene_tx=XM_023257703.1;refGene_gn=LOC101101321;refGene_tx=XM_023257704.1;refGene_gn=LOC101101321;refGene_tx=XM_023257705.1
B4 134386532 134387906
refGene_gn=KCNJ4;refGene_tx=XM_003989260.5;refGene_gn=KCNJ4;refGene_tx=XM_019835563.2;refGene_gn=KCNJ4;refGene_tx=XM_019835564.2;refGene_gn=KCNJ4;refGene_tx=XM_019835565.2
B4 134419906 134420018
refGene_gn=KDELR3;refGene_tx=XM_003989261.5;refGene_gn=KDELR3;refGene_tx=XM_019835585.2
B4 134424347 134424448
refGene_gn=KDELR3;refGene_tx=XM_003989261.5;refGene_gn=KDELR3;refGene_tx=XM_019835585.2
B4 134428925 134429084
refGene_gn=KDELR3;refGene_tx=XM_003989261.5;refGene_gn=KDELR3;refGene_tx=XM_019835585.2
B4 134429725 134429978
refGene_gn=KDELR3;refGene_tx=XM_003989261.5;refGene_gn=KDELR3;refGene_tx=XM_019835585.2
B4 134430755 134430796
refGene_gn=KDELR3;refGene_tx=XM_003989261.5;refGene_gn=KDELR3;refGene_tx=XM_019835585.2

B4 134434241 134434747
refGene_gn=DDX17;refGene_tx=XM_019835568.2
B4 134436141 134436384
refGene_gn=DDX17;refGene_tx=XM_019835568.2
B4 134440571 134440631
refGene_gn=DDX17;refGene_tx=XM_019835568.2
B4 134442297 134442359
refGene_gn=DDX17;refGene_tx=XM_019835568.2
B4 134442482 134442593
refGene_gn=DDX17;refGene_tx=XM_019835568.2
B4 134442912 134443085
refGene_gn=DDX17;refGene_tx=XM_019835568.2
B4 134443172 134443333
refGene_gn=DDX17;refGene_tx=XM_019835568.2
B4 134443691 134443833
refGene_gn=DDX17;refGene_tx=XM_019835568.2
B4 134445580 134445646
refGene_gn=DDX17;refGene_tx=XM_019835568.2
B4 134445942 134446076
refGene_gn=DDX17;refGene_tx=XM_019835568.2
B4 134446927 134447027
refGene_gn=DDX17;refGene_tx=XM_019835568.2
B4 134447905 134448056
refGene_gn=DDX17;refGene_tx=XM_019835568.2
B4 134452431 134452724
refGene_gn=DDX17;refGene_tx=XM_019835568.2
B4 134453141 134453393
refGene_gn=DDX17;refGene_tx=XM_019835568.2
B4 134471388 134471458
refGene_gn=DMC1;refGene_tx=XM_006934043.3;refGene_gn=DMC1;refGene_tx=XM_019835584.1
B4 134472355 134472422
refGene_gn=DMC1;refGene_tx=XM_019835579.2;refGene_gn=DMC1;refGene_tx=XM_019835580.2
B4 134473631 134473725
refGene_gn=DMC1;refGene_tx=XM_023257712.1;refGene_gn=DMC1;refGene_tx=XM_023257713.1;refGene_gn=DMC1;refGene_tx=XM_023257714.1
B4 134478539 134478656
refGene_gn=DMC1;refGene_tx=XM_006934043.3;refGene_gn=DMC1;refGene_tx=XM_019835579.2;refGene_gn=DMC1;refGene_tx=XM_019835580.2;refGene_gn=DMC1;refGene_tx=XM_019835584.1;refGene_gn=DMC1;refGene_tx=XM_023257712.1;refGene_gn=DMC1;refGene_tx=XM_023257713.1;refGene_gn=DMC1;refGene_tx=XM_023257714.1
B4 134492897 134492958
refGene_gn=DMC1;refGene_tx=XM_006934043.3;refGene_gn=DMC1;refGene_tx=XM_019835579.2;refGene_gn=DMC1;refGene_tx=XM_019835580.2;refGene_gn=DMC1;refGene_tx=XM_019835584.1;refGene_gn=DMC1;refGene_tx=XM_023257712.1;refGene_gn=DMC1;refGene_tx=XM_023257713.1;refGene_gn=DMC1;refGene_tx=XM_023257714.1
B4 134493294 134493409

refGene_gn=DMC1;refGene_tx=XM_006934043.3;refGene_gn=DMC1;refGene_tx=X
M_019835579.2;refGene_gn=DMC1;refGene_tx=XM_019835580.2;refGene_gn=DMC
1;refGene_tx=XM_019835584.1;refGene_gn=DMC1;refGene_tx=XM_023257712.1;
refGene_gn=DMC1;refGene_tx=XM_023257713.1;refGene_gn=DMC1;refGene_tx=X
M_023257714.1
B4 134493538 134493612
refGene_gn=DMC1;refGene_tx=XM_006934043.3;refGene_gn=DMC1;refGene_tx=X
M_019835579.2;refGene_gn=DMC1;refGene_tx=XM_019835580.2;refGene_gn=DMC
1;refGene_tx=XM_019835584.1;refGene_gn=DMC1;refGene_tx=XM_023257712.1;
refGene_gn=DMC1;refGene_tx=XM_023257713.1;refGene_gn=DMC1;refGene_tx=X
M_023257714.1
B4 134494425 134494517
refGene_gn=DMC1;refGene_tx=XM_006934043.3;refGene_gn=DMC1;refGene_tx=X
M_019835579.2;refGene_gn=DMC1;refGene_tx=XM_019835580.2;refGene_gn=DMC
1;refGene_tx=XM_023257712.1;refGene_gn=DMC1;refGene_tx=XM_023257713.1
B4 134499290 134499363
refGene_gn=DMC1;refGene_tx=XM_006934043.3;refGene_gn=DMC1;refGene_tx=X
M_019835579.2;refGene_gn=DMC1;refGene_tx=XM_019835580.2;refGene_gn=DMC
1;refGene_tx=XM_023257712.1;refGene_gn=DMC1;refGene_tx=XM_023257713.1
B4 134500473 134500515
refGene_gn=DMC1;refGene_tx=XM_006934043.3;refGene_gn=DMC1;refGene_tx=X
M_019835579.2;refGene_gn=DMC1;refGene_tx=XM_019835580.2;refGene_gn=DMC
1;refGene_tx=XM_019835584.1;refGene_gn=DMC1;refGene_tx=XM_023257712.1;
refGene_gn=DMC1;refGene_tx=XM_023257713.1;refGene_gn=DMC1;refGene_tx=X
M_023257714.1
B4 134502681 134502734
refGene_gn=DMC1;refGene_tx=XM_006934043.3;refGene_gn=DMC1;refGene_tx=X
M_019835579.2;refGene_gn=DMC1;refGene_tx=XM_019835580.2;refGene_gn=DMC
1;refGene_tx=XM_019835584.1;refGene_gn=DMC1;refGene_tx=XM_023257712.1;
refGene_gn=DMC1;refGene_tx=XM_023257713.1;refGene_gn=DMC1;refGene_tx=X
M_023257714.1
B4 134508264 134508347
refGene_gn=DMC1;refGene_tx=XM_006934043.3;refGene_gn=DMC1;refGene_tx=X
M_019835579.2;refGene_gn=DMC1;refGene_tx=XM_019835580.2;refGene_gn=DMC
1;refGene_tx=XM_019835584.1;refGene_gn=DMC1;refGene_tx=XM_023257712.1;
refGene_gn=DMC1;refGene_tx=XM_023257713.1;refGene_gn=DMC1;refGene_tx=X
M_023257714.1
B4 134510827 134510974
refGene_gn=DMC1;refGene_tx=XM_006934043.3;refGene_gn=DMC1;refGene_tx=X
M_019835579.2;refGene_gn=DMC1;refGene_tx=XM_019835580.2;refGene_gn=DMC
1;refGene_tx=XM_019835584.1;refGene_gn=DMC1;refGene_tx=XM_023257712.1;
refGene_gn=DMC1;refGene_tx=XM_023257713.1;refGene_gn=DMC1;refGene_tx=X
M_023257714.1
B4 134511887 134511932
refGene_gn=DMC1;refGene_tx=XM_006934043.3;refGene_gn=DMC1;refGene_tx=X
M_019835579.2;refGene_gn=DMC1;refGene_tx=XM_019835580.2;refGene_gn=DMC
1;refGene_tx=XM_019835584.1;refGene_gn=DMC1;refGene_tx=XM_023257712.1;
refGene_gn=DMC1;refGene_tx=XM_023257713.1;refGene_gn=DMC1;refGene_tx=X
M_023257714.1
B4 134512535 134512586

refGene_gn=DMC1;refGene_tx=XM_006934043.3;refGene_gn=DMC1;refGene_tx=XM_019835579.2;refGene_gn=DMC1;refGene_tx=XM_019835580.2;refGene_gn=DMC1;refGene_tx=XM_019835584.1;refGene_gn=DMC1;refGene_tx=XM_023257712.1;refGene_gn=DMC1;refGene_tx=XM_023257713.1;refGene_gn=DMC1;refGene_tx=XM_023257714.1
B4 134522066 134522148
refGene_gn=FAM227A;refGene_tx=XM_023257688.1;refGene_gn=FAM227A;refGene_tx=XM_023257689.1;refGene_gn=FAM227A;refGene_tx=XM_023257690.1;refGene_gn=FAM227A;refGene_tx=XM_023257691.1
B4 134535658 134535799
refGene_gn=FAM227A;refGene_tx=XM_023257688.1;refGene_gn=FAM227A;refGene_tx=XM_023257689.1;refGene_gn=FAM227A;refGene_tx=XM_023257690.1;refGene_gn=FAM227A;refGene_tx=XM_023257691.1
B4 134542838 134542991
refGene_gn=FAM227A;refGene_tx=XM_023257688.1;refGene_gn=FAM227A;refGene_tx=XM_023257689.1;refGene_gn=FAM227A;refGene_tx=XM_023257690.1;refGene_gn=FAM227A;refGene_tx=XM_023257691.1
B4 134544895 134545053
refGene_gn=FAM227A;refGene_tx=XM_023257688.1;refGene_gn=FAM227A;refGene_tx=XM_023257689.1;refGene_gn=FAM227A;refGene_tx=XM_023257690.1;refGene_gn=FAM227A;refGene_tx=XM_023257691.1
B4 134553182 134553277
refGene_gn=FAM227A;refGene_tx=XM_023257688.1;refGene_gn=FAM227A;refGene_tx=XM_023257689.1;refGene_gn=FAM227A;refGene_tx=XM_023257690.1;refGene_gn=FAM227A;refGene_tx=XM_023257691.1
B4 134560266 134560354
refGene_gn=FAM227A;refGene_tx=XM_023257688.1;refGene_gn=FAM227A;refGene_tx=XM_023257689.1;refGene_gn=FAM227A;refGene_tx=XM_023257690.1;refGene_gn=FAM227A;refGene_tx=XM_023257691.1
B4 134568665 134568745
refGene_gn=FAM227A;refGene_tx=XM_023257688.1;refGene_gn=FAM227A;refGene_tx=XM_023257689.1;refGene_gn=FAM227A;refGene_tx=XM_023257690.1;refGene_gn=FAM227A;refGene_tx=XM_023257691.1
B4 134570601 134570709
refGene_gn=FAM227A;refGene_tx=XM_023257688.1;refGene_gn=FAM227A;refGene_tx=XM_023257689.1;refGene_gn=FAM227A;refGene_tx=XM_023257690.1;refGene_gn=FAM227A;refGene_tx=XM_023257691.1
B4 134576168 134576292
refGene_gn=FAM227A;refGene_tx=XM_023257688.1;refGene_gn=FAM227A;refGene_tx=XM_023257689.1;refGene_gn=FAM227A;refGene_tx=XM_023257690.1;refGene_gn=FAM227A;refGene_tx=XM_023257691.1
B4 134578181 134578286
refGene_gn=FAM227A;refGene_tx=XM_023257688.1;refGene_gn=FAM227A;refGene_tx=XM_023257689.1;refGene_gn=FAM227A;refGene_tx=XM_023257690.1
B4 134578758 134578860
refGene_gn=FAM227A;refGene_tx=XM_023257688.1;refGene_gn=FAM227A;refGene_tx=XM_023257689.1;refGene_gn=FAM227A;refGene_tx=XM_023257690.1;refGene_gn=FAM227A;refGene_tx=XM_023257691.1
B4 134586165 134586315
refGene_gn=FAM227A;refGene_tx=XM_023257688.1;refGene_gn=FAM227A;refGen

e_tx=XM_023257689.1;refGene_gn=FAM227A;refGene_tx=XM_023257690.1;refGene_gn=FAM227A;refGene_tx=XM_023257691.1
B4 134587142 134587219
refGene_gn=FAM227A;refGene_tx=XM_023257688.1;refGene_gn=FAM227A;refGene_tx=XM_023257689.1;refGene_gn=FAM227A;refGene_tx=XM_023257690.1;refGene_gn=FAM227A;refGene_tx=XM_023257691.1
B4 134587703 134587773
refGene_gn=FAM227A;refGene_tx=XM_023257688.1;refGene_gn=FAM227A;refGene_tx=XM_023257689.1;refGene_gn=FAM227A;refGene_tx=XM_023257690.1;refGene_gn=FAM227A;refGene_tx=XM_023257691.1
B4 134595350 134595433
refGene_gn=FAM227A;refGene_tx=XM_023257688.1;refGene_gn=FAM227A;refGene_tx=XM_023257689.1;refGene_gn=FAM227A;refGene_tx=XM_023257690.1;refGene_gn=FAM227A;refGene_tx=XM_023257691.1
B4 134598128 134598270
refGene_gn=FAM227A;refGene_tx=XM_023257688.1;refGene_gn=FAM227A;refGene_tx=XM_023257689.1;refGene_gn=FAM227A;refGene_tx=XM_023257690.1;refGene_gn=FAM227A;refGene_tx=XM_023257691.1
B4 134602044 134602177
refGene_gn=CBY1;refGene_tx=XM_019835586.2
B4 134606042 134606158
refGene_gn=CBY1;refGene_tx=XM_003989263.5;refGene_gn=CBY1;refGene_tx=XM_006934044.3;refGene_gn=CBY1;refGene_tx=XM_019835586.2
B4 134608591 134608697
refGene_gn=CBY1;refGene_tx=XM_003989263.5;refGene_gn=CBY1;refGene_tx=XM_006934044.3;refGene_gn=CBY1;refGene_tx=XM_019835586.2
B4 134608783 134608902
refGene_gn=CBY1;refGene_tx=XM_003989263.5;refGene_gn=CBY1;refGene_tx=XM_006934044.3;refGene_gn=CBY1;refGene_tx=XM_019835586.2
B4 134610397 134610478
refGene_gn=CBY1;refGene_tx=XM_003989263.5;refGene_gn=CBY1;refGene_tx=XM_006934044.3;refGene_gn=CBY1;refGene_tx=XM_019835586.2
B4 134616856 134616973
refGene_gn=TOMM22;refGene_tx=XM_003989264.5
B4 134617219 134617338
refGene_gn=TOMM22;refGene_tx=XM_003989264.5
B4 134617812 134617930
refGene_gn=TOMM22;refGene_tx=XM_003989264.5
B4 134618140 134618215
refGene_gn=TOMM22;refGene_tx=XM_003989264.5
B4 134621933 134622033
refGene_gn=JOSD1;refGene_tx=XM_003989265.5;refGene_gn=JOSD1;refGene_tx=XM_011284223.3;refGene_gn=JOSD1;refGene_tx=XM_023257723.1
B4 134622963 134623158
refGene_gn=JOSD1;refGene_tx=XM_003989265.5;refGene_gn=JOSD1;refGene_tx=XM_011284223.3;refGene_gn=JOSD1;refGene_tx=XM_023257723.1
B4 134623278 134623407
refGene_gn=JOSD1;refGene_tx=XM_003989265.5;refGene_gn=JOSD1;refGene_tx=XM_011284223.3;refGene_gn=JOSD1;refGene_tx=XM_023257723.1
B4 134630378 134630563

refGene_gn=JOSD1;refGene_tx=XM_003989265.5;refGene_gn=JOSD1;refGene_tx=XM_011284223.3;refGene_gn=JOSD1;refGene_tx=XM_023257723.1
B4 134634956 134635148
refGene_gn=GTPBP1;refGene_tx=XM_003989266.5;refGene_gn=GTPBP1;refGene_tx=XM_019835589.2
B4 134638190 134638302
refGene_gn=GTPBP1;refGene_tx=XM_003989266.5;refGene_gn=GTPBP1;refGene_tx=XM_019835589.2;refGene_gn=GTPBP1;refGene_tx=XM_019835590.2;refGene_gn=GTPBP1;refGene_tx=XM_019835591.2
B4 134644139 134644346
refGene_gn=GTPBP1;refGene_tx=XM_003989266.5;refGene_gn=GTPBP1;refGene_tx=XM_019835589.2;refGene_gn=GTPBP1;refGene_tx=XM_019835590.2;refGene_gn=GTPBP1;refGene_tx=XM_019835591.2
B4 134644980 134645329
refGene_gn=GTPBP1;refGene_tx=XM_003989266.5;refGene_gn=GTPBP1;refGene_tx=XM_019835589.2;refGene_gn=GTPBP1;refGene_tx=XM_019835590.2;refGene_gn=GTPBP1;refGene_tx=XM_019835591.2
B4 134648601 134648725
refGene_gn=GTPBP1;refGene_tx=XM_003989266.5;refGene_gn=GTPBP1;refGene_tx=XM_019835589.2;refGene_gn=GTPBP1;refGene_tx=XM_019835590.2;refGene_gn=GTPBP1;refGene_tx=XM_019835591.2
B4 134649557 134649672
refGene_gn=GTPBP1;refGene_tx=XM_003989266.5;refGene_gn=GTPBP1;refGene_tx=XM_019835589.2;refGene_gn=GTPBP1;refGene_tx=XM_019835590.2;refGene_gn=GTPBP1;refGene_tx=XM_019835591.2
B4 134651383 134651528
refGene_gn=GTPBP1;refGene_tx=XM_003989266.5;refGene_gn=GTPBP1;refGene_tx=XM_019835589.2;refGene_gn=GTPBP1;refGene_tx=XM_019835590.2;refGene_gn=GTPBP1;refGene_tx=XM_019835591.2
B4 134651659 134651842
refGene_gn=GTPBP1;refGene_tx=XM_003989266.5;refGene_gn=GTPBP1;refGene_tx=XM_019835589.2;refGene_gn=GTPBP1;refGene_tx=XM_019835590.2;refGene_gn=GTPBP1;refGene_tx=XM_019835591.2
B4 134652666 134652802
refGene_gn=GTPBP1;refGene_tx=XM_003989266.5;refGene_gn=GTPBP1;refGene_tx=XM_019835589.2;refGene_gn=GTPBP1;refGene_tx=XM_019835590.2;refGene_gn=GTPBP1;refGene_tx=XM_019835591.2
B4 134653380 134653559
refGene_gn=GTPBP1;refGene_tx=XM_003989266.5;refGene_gn=GTPBP1;refGene_tx=XM_019835589.2;refGene_gn=GTPBP1;refGene_tx=XM_019835590.2;refGene_gn=GTPBP1;refGene_tx=XM_019835591.2
B4 134654901 134655102
refGene_gn=GTPBP1;refGene_tx=XM_003989266.5;refGene_gn=GTPBP1;refGene_tx=XM_019835589.2;refGene_gn=GTPBP1;refGene_tx=XM_019835590.2;refGene_gn=GTPBP1;refGene_tx=XM_019835591.2
B4 134656011 134656104
refGene_gn=GTPBP1;refGene_tx=XM_003989266.5
B4 134661434 134661548
refGene_gn=SUN2;refGene_tx=XM_011284225.2;refGene_gn=SUN2;refGene_tx=XM_019835588.2;refGene_gn=SUN2;refGene_tx=XM_023257686.1

B4 134663044 134663137
refGene_gn=SUN2;refGene_tx=XM_011284225.2;refGene_gn=SUN2;refGene_tx=XM_019835588.2;refGene_gn=SUN2;refGene_tx=XM_023257686.1
B4 134663456 134663624
refGene_gn=SUN2;refGene_tx=XM_011284225.2;refGene_gn=SUN2;refGene_tx=XM_019835588.2;refGene_gn=SUN2;refGene_tx=XM_023257686.1
B4 134663780 134663896
refGene_gn=SUN2;refGene_tx=XM_011284225.2;refGene_gn=SUN2;refGene_tx=XM_019835588.2;refGene_gn=SUN2;refGene_tx=XM_023257686.1
B4 134664278 134664363
refGene_gn=SUN2;refGene_tx=XM_011284225.2;refGene_gn=SUN2;refGene_tx=XM_019835588.2;refGene_gn=SUN2;refGene_tx=XM_023257686.1
B4 134664691 134664913
refGene_gn=SUN2;refGene_tx=XM_011284225.2;refGene_gn=SUN2;refGene_tx=XM_019835588.2;refGene_gn=SUN2;refGene_tx=XM_023257686.1
B4 134665252 134665418
refGene_gn=SUN2;refGene_tx=XM_011284225.2;refGene_gn=SUN2;refGene_tx=XM_019835588.2;refGene_gn=SUN2;refGene_tx=XM_023257686.1
B4 134665981 134666025
refGene_gn=SUN2;refGene_tx=XM_011284225.2;refGene_gn=SUN2;refGene_tx=XM_019835588.2;refGene_gn=SUN2;refGene_tx=XM_023257686.1
B4 134666494 134666572
refGene_gn=SUN2;refGene_tx=XM_011284225.2;refGene_gn=SUN2;refGene_tx=XM_019835588.2;refGene_gn=SUN2;refGene_tx=XM_023257686.1
B4 134666981 134667239
refGene_gn=SUN2;refGene_tx=XM_011284225.2;refGene_gn=SUN2;refGene_tx=XM_019835588.2;refGene_gn=SUN2;refGene_tx=XM_023257686.1
B4 134669441 134669549
refGene_gn=GTPBP1;refGene_tx=XM_019835589.2;refGene_gn=GTPBP1;refGene_tx=XM_019835590.2;refGene_gn=GTPBP1;refGene_tx=XM_019835591.2
B4 134669633 134669749
refGene_gn=SUN2;refGene_tx=XM_011284225.2;refGene_gn=SUN2;refGene_tx=XM_019835588.2;refGene_gn=SUN2;refGene_tx=XM_023257686.1
B4 134671986 134672057
refGene_gn=SUN2;refGene_tx=XM_011284225.2;refGene_gn=SUN2;refGene_tx=XM_019835588.2;refGene_gn=SUN2;refGene_tx=XM_023257686.1
B4 134673266 134673360
refGene_gn=SUN2;refGene_tx=XM_011284225.2;refGene_gn=SUN2;refGene_tx=XM_019835588.2;refGene_gn=SUN2;refGene_tx=XM_023257686.1
B4 134673749 134673845
refGene_gn=SUN2;refGene_tx=XM_011284225.2;refGene_gn=SUN2;refGene_tx=XM_019835588.2;refGene_gn=SUN2;refGene_tx=XM_023257686.1
B4 134674426 134674564
refGene_gn=SUN2;refGene_tx=XM_011284225.2;refGene_gn=SUN2;refGene_tx=XM_019835588.2;refGene_gn=SUN2;refGene_tx=XM_023257686.1
B4 134674924 134675115
refGene_gn=SUN2;refGene_tx=XM_011284225.2;refGene_gn=SUN2;refGene_tx=XM_019835588.2;refGene_gn=SUN2;refGene_tx=XM_023257686.1
B4 134675998 134676120
refGene_gn=SUN2;refGene_tx=XM_011284225.2;refGene_gn=SUN2;refGene_tx=X

M_019835588.2;refGene_gn=SUN2;refGene_tx=XM_023257686.1
B4	134699903	134700068
refGene_gn=DNAL4;refGene_tx=XM_003989267.4
B4	134701310	134701394
refGene_gn=DNAL4;refGene_tx=XM_003989267.4
B4	134703071	134703140
refGene_gn=DNAL4;refGene_tx=XM_003989267.4
B4	134727816	134728041
refGene_gn=NPTXR;refGene_tx=XM_023257782.1
B4	134728382	134728562
refGene_gn=NPTXR;refGene_tx=XM_023257782.1
B4	134731104	134731352
refGene_gn=NPTXR;refGene_tx=XM_023257782.1
B4	134733575	134733801
refGene_gn=NPTXR;refGene_tx=XM_023257782.1
B4	134746292	134747138
refGene_gn=NPTXR;refGene_tx=XM_023257782.1
B4	134761057	134761081
refGene_gn=CBX6;refGene_tx=XM_023257709.1
B4	134765038	134766040
refGene_gn=CBX6;refGene_tx=XM_019835592.2;refGene_gn=CBX6;refGene_tx=X
M_023257709.1
B4	134770374	134770441
refGene_gn=CBX6;refGene_tx=XM_019835592.2;refGene_gn=CBX6;refGene_tx=X
M_023257709.1
B4	134770591	134770657
refGene_gn=CBX6;refGene_tx=XM_019835592.2;refGene_gn=CBX6;refGene_tx=X
M_023257709.1
B4	134770809	134770853
refGene_gn=CBX6;refGene_tx=XM_019835592.2;refGene_gn=CBX6;refGene_tx=X
M_023257709.1
B4	134771034	134771103
refGene_gn=CBX6;refGene_tx=XM_019835592.2;refGene_gn=CBX6;refGene_tx=X
M_023257709.1
B4	134878284	134878434
refGene_gn=APOBEC3Z3;refGene_tx=NM_001112710.3
B4	134879074	134879342
refGene_gn=APOBEC3Z3;refGene_tx=NM_001112710.3
B4	134879573	134879689
refGene_gn=APOBEC3Z3;refGene_tx=NM_001112710.3
B4	134880474	134880522
refGene_gn=APOBEC3Z3;refGene_tx=NM_001112710.3
B4	134902565	134902720
refGene_gn=CBX7;refGene_tx=XM_003989269.4;refGene_gn=CBX7;refGene_tx=X
M_019835594.2
B4	134903070	134903422
refGene_gn=CBX7;refGene_tx=XM_003989269.4;refGene_gn=CBX7;refGene_tx=X
M_019835594.2
B4	134906019	134906086
refGene_gn=CBX7;refGene_tx=XM_003989269.4;refGene_gn=CBX7;refGene_tx=X

M_019835594.2
B4	134910078	134910144
refGene_gn=CBX7;refGene_tx=XM_003989269.4;refGene_gn=CBX7;refGene_tx=XM_019835594.2
B4	134917236	134917280
refGene_gn=CBX7;refGene_tx=XM_003989269.4;refGene_gn=CBX7;refGene_tx=XM_019835594.2
B4	134919465	134919534
refGene_gn=CBX7;refGene_tx=XM_003989269.4;refGene_gn=CBX7;refGene_tx=XM_019835594.2
B4	134972130	134972255
refGene_gn=PDGFB;refGene_tx=XM_003989270.5;refGene_gn=PDGFB;refGene_tx=XM_006934047.3;refGene_gn=PDGFB;refGene_tx=XM_006934048.4;refGene_gn=PDGFB;refGene_tx=XM_006934049.4;refGene_gn=PDGFB;refGene_tx=XM_006934050.4
B4	134974564	134974709
refGene_gn=PDGFB;refGene_tx=XM_003989270.5;refGene_gn=PDGFB;refGene_tx=XM_006934047.3;refGene_gn=PDGFB;refGene_tx=XM_006934048.4;refGene_gn=PDGFB;refGene_tx=XM_006934049.4;refGene_gn=PDGFB;refGene_tx=XM_006934050.4
B4	134976083	134976289
refGene_gn=PDGFB;refGene_tx=XM_003989270.5;refGene_gn=PDGFB;refGene_tx=XM_006934047.3;refGene_gn=PDGFB;refGene_tx=XM_006934048.4;refGene_gn=PDGFB;refGene_tx=XM_006934049.4;refGene_gn=PDGFB;refGene_tx=XM_006934050.4
B4	134977870	134977972
refGene_gn=PDGFB;refGene_tx=XM_003989270.5;refGene_gn=PDGFB;refGene_tx=XM_006934047.3;refGene_gn=PDGFB;refGene_tx=XM_006934048.4;refGene_gn=PDGFB;refGene_tx=XM_006934049.4;refGene_gn=PDGFB;refGene_tx=XM_006934050.4
B4	134980312	134980409
refGene_gn=PDGFB;refGene_tx=XM_003989270.5;refGene_gn=PDGFB;refGene_tx=XM_006934047.3;refGene_gn=PDGFB;refGene_tx=XM_006934048.4;refGene_gn=PDGFB;refGene_tx=XM_006934049.4;refGene_gn=PDGFB;refGene_tx=XM_006934050.4
B4	134984235	134984460
refGene_gn=PDGFB;refGene_tx=XM_006934047.3;refGene_gn=PDGFB;refGene_tx=XM_006934048.4;refGene_gn=PDGFB;refGene_tx=XM_006934049.4
B4	134986306	134986761
refGene_gn=LOC109501804;refGene_tx=XM_023257715.1;refGene_gn=LOC109501804;refGene_tx=XM_023257716.1
B4	134988258	134988321
refGene_gn=PDGFB;refGene_tx=XM_003989270.5;refGene_gn=PDGFB;refGene_tx=XM_006934050.4
B4	134989591	134989712
refGene_gn=LOC109501804;refGene_tx=XM_023257715.1;refGene_gn=LOC109501804;refGene_tx=XM_023257716.1
B4	134990322	134990676
refGene_gn=LOC109501804;refGene_tx=XM_023257715.1;refGene_gn=LOC109501804;refGene_tx=XM_023257716.1

B4 134991964 134991973
refGene_gn=LOC109501804;refGene_tx=XM_023257716.1
B4 135038519 135038564
refGene_gn=RPL3;refGene_tx=XM_003989271.3
B4 135038770 135038890
refGene_gn=RPL3;refGene_tx=XM_003989271.3
B4 135039283 135039379
refGene_gn=RPL3;refGene_tx=XM_003989271.3
B4 135039849 135039951
refGene_gn=RPL3;refGene_tx=XM_003989271.3
B4 135040391 135040552
refGene_gn=RPL3;refGene_tx=XM_003989271.3
B4 135041117 135041304
refGene_gn=RPL3;refGene_tx=XM_003989271.3
B4 135042419 135042555
refGene_gn=RPL3;refGene_tx=XM_003989271.3
B4 135043115 135043284
refGene_gn=RPL3;refGene_tx=XM_003989271.3
B4 135043817 135044010
refGene_gn=RPL3;refGene_tx=XM_003989271.3
B4 135044967 135045021
refGene_gn=RPL3;refGene_tx=XM_003989271.3
B4 135069044 135069143
refGene_gn=SYNGR1;refGene_tx=XM_023257720.1;refGene_gn=SYNGR1;refGene_tx=XM_023257721.1;refGene_gn=SYNGR1;refGene_tx=XM_023257722.1
B4 135085803 135086041
refGene_gn=SYNGR1;refGene_tx=XM_023257720.1;refGene_gn=SYNGR1;refGene_tx=XM_023257721.1;refGene_gn=SYNGR1;refGene_tx=XM_023257722.1
B4 135087482 135087628
refGene_gn=SYNGR1;refGene_tx=XM_023257720.1;refGene_gn=SYNGR1;refGene_tx=XM_023257721.1;refGene_gn=SYNGR1;refGene_tx=XM_023257722.1
B4 135089155 135089248
refGene_gn=SYNGR1;refGene_tx=XM_023257722.1
B4 135091832 135092054
refGene_gn=SYNGR1;refGene_tx=XM_023257720.1;refGene_gn=SYNGR1;refGene_tx=XM_023257721.1
B4 135120114 135120147
refGene_gn=TAB1;refGene_tx=XM_003989272.5;refGene_gn=TAB1;refGene_tx=XM_006934053.4;refGene_gn=TAB1;refGene_tx=XM_019835596.2;refGene_gn=TAB1;refGene_tx=XM_019835597.2
B4 135134478 135134615
refGene_gn=TAB1;refGene_tx=XM_003989272.5;refGene_gn=TAB1;refGene_tx=XM_006934053.4;refGene_gn=TAB1;refGene_tx=XM_019835596.2;refGene_gn=TAB1;refGene_tx=XM_019835597.2
B4 135134980 135135134
refGene_gn=TAB1;refGene_tx=XM_003989272.5;refGene_gn=TAB1;refGene_tx=XM_006934053.4;refGene_gn=TAB1;refGene_tx=XM_019835596.2;refGene_gn=TAB1;refGene_tx=XM_019835597.2
B4 135141106 135141193
refGene_gn=TAB1;refGene_tx=XM_003989272.5;refGene_gn=TAB1;refGene_tx=X

M_006934053.4;refGene_gn=TAB1;refGene_tx=XM_019835596.2;refGene_gn=TAB1;refGene_tx=XM_019835597.2
B4	135141736	135141875
refGene_gn=TAB1;refGene_tx=XM_003989272.5;refGene_gn=TAB1;refGene_tx=XM_006934053.4;refGene_gn=TAB1;refGene_tx=XM_019835596.2;refGene_gn=TAB1;refGene_tx=XM_019835597.2
B4	135142580	135142794
refGene_gn=TAB1;refGene_tx=XM_003989272.5;refGene_gn=TAB1;refGene_tx=XM_006934053.4;refGene_gn=TAB1;refGene_tx=XM_019835596.2;refGene_gn=TAB1;refGene_tx=XM_019835597.2
B4	135143355	135143499
refGene_gn=TAB1;refGene_tx=XM_003989272.5;refGene_gn=TAB1;refGene_tx=XM_006934053.4;refGene_gn=TAB1;refGene_tx=XM_019835596.2;refGene_gn=TAB1;refGene_tx=XM_019835597.2
B4	135145655	135145800
refGene_gn=TAB1;refGene_tx=XM_003989272.5;refGene_gn=TAB1;refGene_tx=XM_006934053.4;refGene_gn=TAB1;refGene_tx=XM_019835596.2;refGene_gn=TAB1;refGene_tx=XM_019835597.2
B4	135147366	135147589
refGene_gn=TAB1;refGene_tx=XM_003989272.5;refGene_gn=TAB1;refGene_tx=XM_006934053.4;refGene_gn=TAB1;refGene_tx=XM_019835596.2;refGene_gn=TAB1;refGene_tx=XM_019835597.2
B4	135148640	135148803
refGene_gn=TAB1;refGene_tx=XM_003989272.5;refGene_gn=TAB1;refGene_tx=XM_006934053.4;refGene_gn=TAB1;refGene_tx=XM_019835596.2;refGene_gn=TAB1;refGene_tx=XM_019835597.2
B4	135150881	135151089
refGene_gn=TAB1;refGene_tx=XM_003989272.5;refGene_gn=TAB1;refGene_tx=XM_006934053.4;refGene_gn=TAB1;refGene_tx=XM_019835596.2;refGene_gn=TAB1;refGene_tx=XM_019835597.2
B4	135212602	135214204
refGene_gn=MGAT3;refGene_tx=XM_019835605.2;refGene_gn=MGAT3;refGene_tx=XM_019835606.2;refGene_gn=MGAT3;refGene_tx=XM_019835607.2;refGene_gn=MGAT3;refGene_tx=XM_019835608.2;refGene_gn=MGAT3;refGene_tx=XM_023257692.1
B4	135219055	135219375
refGene_gn=LOC102900764;refGene_tx=XM_019835887.2
B4	135231420	135231568
refGene_gn=LOC102900764;refGene_tx=XM_019835887.2
B4	135231977	135232297
refGene_gn=LOC102900764;refGene_tx=XM_019835887.2
B4	135232679	135232764
refGene_gn=LOC102900764;refGene_tx=XM_019835887.2
B4	135240718	135240862
refGene_gn=MIEF1;refGene_tx=XM_003989273.5;refGene_gn=MIEF1;refGene_tx=XM_006934057.4;refGene_gn=MIEF1;refGene_tx=XM_006934059.4;refGene_gn=MIEF1;refGene_tx=XM_011284228.3;refGene_gn=MIEF1;refGene_tx=XM_019835609.2;refGene_gn=MIEF1;refGene_tx=XM_019835610.2;refGene_gn=MIEF1;refGene_tx=XM_019835611.2
B4	135241800	135241978

refGene_gn=MIEF1;refGene_tx=XM_003989273.5;refGene_gn=MIEF1;refGene_tx=XM_006934057.4;refGene_gn=MIEF1;refGene_tx=XM_006934059.4;refGene_gn=MIEF1;refGene_tx=XM_011284228.3;refGene_gn=MIEF1;refGene_tx=XM_019835609.2;refGene_gn=MIEF1;refGene_tx=XM_019835610.2;refGene_gn=MIEF1;refGene_tx=XM_019835611.2
B4 135242181 135242444
refGene_gn=MIEF1;refGene_tx=XM_003989273.5;refGene_gn=MIEF1;refGene_tx=XM_006934057.4;refGene_gn=MIEF1;refGene_tx=XM_006934059.4;refGene_gn=MIEF1;refGene_tx=XM_011284228.3;refGene_gn=MIEF1;refGene_tx=XM_019835609.2;refGene_gn=MIEF1;refGene_tx=XM_019835610.2;refGene_gn=MIEF1;refGene_tx=XM_019835611.2
B4 135243052 135243859
refGene_gn=MIEF1;refGene_tx=XM_003989273.5;refGene_gn=MIEF1;refGene_tx=XM_006934057.4;refGene_gn=MIEF1;refGene_tx=XM_006934059.4;refGene_gn=MIEF1;refGene_tx=XM_011284228.3;refGene_gn=MIEF1;refGene_tx=XM_019835609.2;refGene_gn=MIEF1;refGene_tx=XM_019835610.2;refGene_gn=MIEF1;refGene_tx=XM_019835611.2
B4 135250729 135250958
refGene_gn=ATF4;refGene_tx=XM_003989275.5
B4 135251044 135251874
refGene_gn=ATF4;refGene_tx=XM_003989275.5
B4 135255577 135255579
refGene_gn=LOC102901054;refGene_tx=XM_019835613.1
B4 135256416 135257092
refGene_gn=LOC102901054;refGene_tx=XM_019835612.2;refGene_gn=LOC102901054;refGene_tx=XM_019835613.1;refGene_gn=LOC102901054;refGene_tx=XM_019835614.2;refGene_gn=LOC102901054;refGene_tx=XM_023257493.1
B4 135257419 135257670
refGene_gn=LOC102901054;refGene_tx=XM_019835612.2;refGene_gn=LOC102901054;refGene_tx=XM_019835613.1;refGene_gn=LOC102901054;refGene_tx=XM_019835614.2;refGene_gn=LOC102901054;refGene_tx=XM_023257493.1
B4 135257775 135257958
refGene_gn=LOC102901054;refGene_tx=XM_019835612.2;refGene_gn=LOC102901054;refGene_tx=XM_019835613.1;refGene_gn=LOC102901054;refGene_tx=XM_019835614.2;refGene_gn=LOC102901054;refGene_tx=XM_023257493.1
B4 135258058 135258296
refGene_gn=LOC102901054;refGene_tx=XM_019835612.2;refGene_gn=LOC102901054;refGene_tx=XM_019835613.1;refGene_gn=LOC102901054;refGene_tx=XM_019835614.2;refGene_gn=LOC102901054;refGene_tx=XM_023257493.1
B4 135259350 135259485
refGene_gn=RPS19BP1;refGene_tx=XM_011284230.2
B4 135259791 135259889
refGene_gn=RPS19BP1;refGene_tx=XM_011284230.2
B4 135261777 135261906
refGene_gn=RPS19BP1;refGene_tx=XM_011284230.2
B4 135262073 135262125
refGene_gn=RPS19BP1;refGene_tx=XM_011284230.2
B4 135296731 135296970
refGene_gn=CACNA1I;refGene_tx=XM_023257494.1;refGene_gn=CACNA1I;refGene_tx=XM_023257495.1

B4	135316669	135316781
refGene_gn=CACNA1I;refGene_tx=XM_023257494.1;refGene_gn=CACNA1I;refGene_tx=XM_023257495.1
B4	135318481	135318615
refGene_gn=CACNA1I;refGene_tx=XM_023257494.1;refGene_gn=CACNA1I;refGene_tx=XM_023257495.1
B4	135335046	135335144
refGene_gn=CACNA1I;refGene_tx=XM_023257494.1;refGene_gn=CACNA1I;refGene_tx=XM_023257495.1;refGene_gn=CACNA1I;refGene_tx=XM_023257496.1
B4	135348034	135348194
refGene_gn=CACNA1I;refGene_tx=XM_023257494.1;refGene_gn=CACNA1I;refGene_tx=XM_023257495.1;refGene_gn=CACNA1I;refGene_tx=XM_023257496.1
B4	135354418	135354734
refGene_gn=CACNA1I;refGene_tx=XM_023257494.1;refGene_gn=CACNA1I;refGene_tx=XM_023257495.1;refGene_gn=CACNA1I;refGene_tx=XM_023257496.1
B4	135356321	135356414
refGene_gn=CACNA1I;refGene_tx=XM_023257494.1;refGene_gn=CACNA1I;refGene_tx=XM_023257495.1;refGene_gn=CACNA1I;refGene_tx=XM_023257496.1
B4	135359749	135360062
refGene_gn=CACNA1I;refGene_tx=XM_023257494.1;refGene_gn=CACNA1I;refGene_tx=XM_023257495.1;refGene_gn=CACNA1I;refGene_tx=XM_023257496.1
B4	135362638	135363060
refGene_gn=CACNA1I;refGene_tx=XM_023257494.1;refGene_gn=CACNA1I;refGene_tx=XM_023257495.1;refGene_gn=CACNA1I;refGene_tx=XM_023257496.1
B4	135370778	135370930
refGene_gn=CACNA1I;refGene_tx=XM_023257494.1;refGene_gn=CACNA1I;refGene_tx=XM_023257495.1;refGene_gn=CACNA1I;refGene_tx=XM_023257496.1
B4	135371589	135371775
refGene_gn=CACNA1I;refGene_tx=XM_023257494.1;refGene_gn=CACNA1I;refGene_tx=XM_023257495.1;refGene_gn=CACNA1I;refGene_tx=XM_023257496.1
B4	135372055	135372173
refGene_gn=CACNA1I;refGene_tx=XM_023257494.1;refGene_gn=CACNA1I;refGene_tx=XM_023257495.1;refGene_gn=CACNA1I;refGene_tx=XM_023257496.1
B4	135372902	135373058
refGene_gn=CACNA1I;refGene_tx=XM_023257494.1;refGene_gn=CACNA1I;refGene_tx=XM_023257495.1;refGene_gn=CACNA1I;refGene_tx=XM_023257496.1
B4	135373528	135373622
refGene_gn=CACNA1I;refGene_tx=XM_023257494.1;refGene_gn=CACNA1I;refGene_tx=XM_023257495.1;refGene_gn=CACNA1I;refGene_tx=XM_023257496.1
B4	135374611	135374814
refGene_gn=CACNA1I;refGene_tx=XM_023257494.1;refGene_gn=CACNA1I;refGene_tx=XM_023257495.1;refGene_gn=CACNA1I;refGene_tx=XM_023257496.1
B4	135375488	135375959
refGene_gn=CACNA1I;refGene_tx=XM_023257494.1;refGene_gn=CACNA1I;refGene_tx=XM_023257495.1;refGene_gn=CACNA1I;refGene_tx=XM_023257496.1
B4	135376263	135376364
refGene_gn=CACNA1I;refGene_tx=XM_023257494.1;refGene_gn=CACNA1I;refGene_tx=XM_023257495.1;refGene_gn=CACNA1I;refGene_tx=XM_023257496.1
B4	135377099	135377223
refGene_gn=CACNA1I;refGene_tx=XM_023257494.1;refGene_gn=CACNA1I;refGen

e_tx=XM_023257495.1;refGene_gn=CACNA1I;refGene_tx=XM_023257496.1
B4	135377506	135377575
refGene_gn=CACNA1I;refGene_tx=XM_023257494.1;refGene_gn=CACNA1I;refGene_tx=XM_023257495.1;refGene_gn=CACNA1I;refGene_tx=XM_023257496.1
B4	135378106	135378291
refGene_gn=CACNA1I;refGene_tx=XM_023257494.1;refGene_gn=CACNA1I;refGene_tx=XM_023257495.1;refGene_gn=CACNA1I;refGene_tx=XM_023257496.1
B4	135378748	135378875
refGene_gn=CACNA1I;refGene_tx=XM_023257494.1;refGene_gn=CACNA1I;refGene_tx=XM_023257495.1;refGene_gn=CACNA1I;refGene_tx=XM_023257496.1
B4	135380176	135380302
refGene_gn=CACNA1I;refGene_tx=XM_023257494.1;refGene_gn=CACNA1I;refGene_tx=XM_023257495.1;refGene_gn=CACNA1I;refGene_tx=XM_023257496.1
B4	135382686	135382776
refGene_gn=CACNA1I;refGene_tx=XM_023257494.1;refGene_gn=CACNA1I;refGene_tx=XM_023257495.1;refGene_gn=CACNA1I;refGene_tx=XM_023257496.1
B4	135383341	135383534
refGene_gn=CACNA1I;refGene_tx=XM_023257494.1;refGene_gn=CACNA1I;refGene_tx=XM_023257495.1;refGene_gn=CACNA1I;refGene_tx=XM_023257496.1
B4	135384111	135384263
refGene_gn=CACNA1I;refGene_tx=XM_023257494.1;refGene_gn=CACNA1I;refGene_tx=XM_023257495.1;refGene_gn=CACNA1I;refGene_tx=XM_023257496.1
B4	135385757	135385867
refGene_gn=CACNA1I;refGene_tx=XM_023257494.1;refGene_gn=CACNA1I;refGene_tx=XM_023257495.1;refGene_gn=CACNA1I;refGene_tx=XM_023257496.1
B4	135386517	135386651
refGene_gn=CACNA1I;refGene_tx=XM_023257494.1;refGene_gn=CACNA1I;refGene_tx=XM_023257495.1;refGene_gn=CACNA1I;refGene_tx=XM_023257496.1
B4	135387536	135387607
refGene_gn=CACNA1I;refGene_tx=XM_023257494.1;refGene_gn=CACNA1I;refGene_tx=XM_023257495.1;refGene_gn=CACNA1I;refGene_tx=XM_023257496.1
B4	135390641	135390720
refGene_gn=CACNA1I;refGene_tx=XM_023257494.1;refGene_gn=CACNA1I;refGene_tx=XM_023257495.1;refGene_gn=CACNA1I;refGene_tx=XM_023257496.1
B4	135391345	135391467
refGene_gn=CACNA1I;refGene_tx=XM_023257494.1;refGene_gn=CACNA1I;refGene_tx=XM_023257495.1;refGene_gn=CACNA1I;refGene_tx=XM_023257496.1
B4	135391973	135392312
refGene_gn=CACNA1I;refGene_tx=XM_023257494.1;refGene_gn=CACNA1I;refGene_tx=XM_023257495.1;refGene_gn=CACNA1I;refGene_tx=XM_023257496.1
B4	135392581	135392728
refGene_gn=CACNA1I;refGene_tx=XM_023257494.1;refGene_gn=CACNA1I;refGene_tx=XM_023257495.1;refGene_gn=CACNA1I;refGene_tx=XM_023257496.1
B4	135393104	135393227
refGene_gn=CACNA1I;refGene_tx=XM_023257494.1;refGene_gn=CACNA1I;refGene_tx=XM_023257495.1;refGene_gn=CACNA1I;refGene_tx=XM_023257496.1
B4	135395296	135395462
refGene_gn=CACNA1I;refGene_tx=XM_023257494.1;refGene_gn=CACNA1I;refGene_tx=XM_023257495.1;refGene_gn=CACNA1I;refGene_tx=XM_023257496.1
B4	135396984	135397184

refGene_gn=CACNA1I;refGene_tx=XM_023257494.1;refGene_gn=CACNA1I;refGene_tx=XM_023257495.1;refGene_gn=CACNA1I;refGene_tx=XM_023257496.1
B4 135398128 135398758
refGene_gn=CACNA1I;refGene_tx=XM_023257494.1;refGene_gn=CACNA1I;refGene_tx=XM_023257495.1;refGene_gn=CACNA1I;refGene_tx=XM_023257496.1
B4 135447680 135447770
refGene_gn=ENTHD1;refGene_tx=XM_006934061.4;refGene_gn=ENTHD1;refGene_tx=XM_006934062.4;refGene_gn=ENTHD1;refGene_tx=XM_006934065.4;refGene_gn=ENTHD1;refGene_tx=XM_019835616.2;refGene_gn=ENTHD1;refGene_tx=XM_019835619.2;refGene_gn=ENTHD1;refGene_tx=XM_019835620.2;refGene_gn=ENTHD1;refGene_tx=XM_019835621.2;refGene_gn=ENTHD1;refGene_tx=XM_019835622.2;refGene_gn=ENTHD1;refGene_tx=XM_023257498.1;refGene_gn=ENTHD1;refGene_tx=XM_023257499.1;refGene_gn=ENTHD1;refGene_tx=XM_023257500.1;refGene_gn=ENTHD1;refGene_tx=XM_023257502.1;refGene_gn=ENTHD1;refGene_tx=XM_023257503.1
B4 135447986 135448588
refGene_gn=ENTHD1;refGene_tx=XM_006934061.4;refGene_gn=ENTHD1;refGene_tx=XM_006934062.4;refGene_gn=ENTHD1;refGene_tx=XM_006934065.4;refGene_gn=ENTHD1;refGene_tx=XM_019835616.2;refGene_gn=ENTHD1;refGene_tx=XM_019835619.2;refGene_gn=ENTHD1;refGene_tx=XM_019835620.2;refGene_gn=ENTHD1;refGene_tx=XM_019835621.2;refGene_gn=ENTHD1;refGene_tx=XM_019835622.2;refGene_gn=ENTHD1;refGene_tx=XM_023257497.1;refGene_gn=ENTHD1;refGene_tx=XM_023257498.1;refGene_gn=ENTHD1;refGene_tx=XM_023257499.1;refGene_gn=ENTHD1;refGene_tx=XM_023257500.1;refGene_gn=ENTHD1;refGene_tx=XM_023257502.1;refGene_gn=ENTHD1;refGene_tx=XM_023257503.1
B4 135450442 135450471
refGene_gn=ENTHD1;refGene_tx=XM_023257503.1
B4 135459663 135460062
refGene_gn=ENTHD1;refGene_tx=XM_006934061.4;refGene_gn=ENTHD1;refGene_tx=XM_006934062.4;refGene_gn=ENTHD1;refGene_tx=XM_006934065.4;refGene_gn=ENTHD1;refGene_tx=XM_019835616.2;refGene_gn=ENTHD1;refGene_tx=XM_019835619.2;refGene_gn=ENTHD1;refGene_tx=XM_019835620.2;refGene_gn=ENTHD1;refGene_tx=XM_019835621.2;refGene_gn=ENTHD1;refGene_tx=XM_019835622.2;refGene_gn=ENTHD1;refGene_tx=XM_023257497.1;refGene_gn=ENTHD1;refGene_tx=XM_023257498.1;refGene_gn=ENTHD1;refGene_tx=XM_023257499.1;refGene_gn=ENTHD1;refGene_tx=XM_023257500.1;refGene_gn=ENTHD1;refGene_tx=XM_023257502.1;refGene_gn=ENTHD1;refGene_tx=XM_023257503.1
B4 135493536 135493660
refGene_gn=ENTHD1;refGene_tx=XM_006934061.4;refGene_gn=ENTHD1;refGene_tx=XM_006934062.4;refGene_gn=ENTHD1;refGene_tx=XM_006934065.4;refGene_gn=ENTHD1;refGene_tx=XM_019835616.2;refGene_gn=ENTHD1;refGene_tx=XM_019835619.2;refGene_gn=ENTHD1;refGene_tx=XM_019835620.2;refGene_gn=ENTHD1;refGene_tx=XM_019835621.2;refGene_gn=ENTHD1;refGene_tx=XM_019835622.2;refGene_gn=ENTHD1;refGene_tx=XM_023257497.1;refGene_gn=ENTHD1;refGene_tx=XM_023257498.1;refGene_gn=ENTHD1;refGene_tx=XM_023257499.1;refGene_gn=ENTHD1;refGene_tx=XM_023257500.1;refGene_gn=ENTHD1;refGene_tx=XM_023257502.1
B4 135505890 135506009
refGene_gn=ENTHD1;refGene_tx=XM_006934061.4;refGene_gn=ENTHD1;refGene_tx=XM_006934062.4;refGene_gn=ENTHD1;refGene_tx=XM_006934065.4;refGene_

gn=ENTHD1;refGene_tx=XM_019835616.2;refGene_gn=ENTHD1;refGene_tx=XM_01
9835619.2;refGene_gn=ENTHD1;refGene_tx=XM_019835620.2;refGene_gn=ENTHD
1;refGene_tx=XM_019835621.2;refGene_gn=ENTHD1;refGene_tx=XM_019835622.
2;refGene_gn=ENTHD1;refGene_tx=XM_023257497.1;refGene_gn=ENTHD1;refGen
e_tx=XM_023257498.1;refGene_gn=ENTHD1;refGene_tx=XM_023257499.1;refGen
e_gn=ENTHD1;refGene_tx=XM_023257500.1;refGene_gn=ENTHD1;refGene_tx=XM_
023257502.1
B4 135526180 135526442
refGene_gn=ENTHD1;refGene_tx=XM_006934061.4;refGene_gn=ENTHD1;refGene_
tx=XM_006934062.4;refGene_gn=ENTHD1;refGene_tx=XM_006934065.4;refGene_
gn=ENTHD1;refGene_tx=XM_019835616.2;refGene_gn=ENTHD1;refGene_tx=XM_01
9835619.2;refGene_gn=ENTHD1;refGene_tx=XM_019835620.2;refGene_gn=ENTHD
1;refGene_tx=XM_019835621.2;refGene_gn=ENTHD1;refGene_tx=XM_019835622.
2;refGene_gn=ENTHD1;refGene_tx=XM_023257497.1;refGene_gn=ENTHD1;refGen
e_tx=XM_023257498.1;refGene_gn=ENTHD1;refGene_tx=XM_023257500.1;refGen
e_gn=ENTHD1;refGene_tx=XM_023257502.1
B4 135527355 135527376
refGene_gn=ENTHD1;refGene_tx=XM_019835619.2;refGene_gn=ENTHD1;refGene_
tx=XM_019835620.2;refGene_gn=ENTHD1;refGene_tx=XM_023257500.1
B4 135545573 135545976
refGene_gn=ENTHD1;refGene_tx=XM_006934061.4;refGene_gn=ENTHD1;refGene_
tx=XM_006934062.4;refGene_gn=ENTHD1;refGene_tx=XM_019835616.2;refGene_
gn=ENTHD1;refGene_tx=XM_023257497.1;refGene_gn=ENTHD1;refGene_tx=XM_02
3257498.1;refGene_gn=ENTHD1;refGene_tx=XM_023257499.1
B4 135598753 135598831
refGene_gn=GRAP2;refGene_tx=XM_011284232.3;refGene_gn=GRAP2;refGene_tx
=XM_019835624.2;refGene_gn=GRAP2;refGene_tx=XM_023257504.1
B4 135605687 135605779
refGene_gn=GRAP2;refGene_tx=XM_011284232.3;refGene_gn=GRAP2;refGene_tx
=XM_019835624.2;refGene_gn=GRAP2;refGene_tx=XM_023257504.1
B4 135608413 135608533
refGene_gn=GRAP2;refGene_tx=XM_011284232.3;refGene_gn=GRAP2;refGene_tx
=XM_019835624.2;refGene_gn=GRAP2;refGene_tx=XM_023257504.1
B4 135611935 135612104
refGene_gn=GRAP2;refGene_tx=XM_011284232.3;refGene_gn=GRAP2;refGene_tx
=XM_019835624.2;refGene_gn=GRAP2;refGene_tx=XM_023257504.1
B4 135614404 135614590
refGene_gn=GRAP2;refGene_tx=XM_011284232.3;refGene_gn=GRAP2;refGene_tx
=XM_019835624.2;refGene_gn=GRAP2;refGene_tx=XM_023257504.1
B4 135615796 135615919
refGene_gn=GRAP2;refGene_tx=XM_011284232.3;refGene_gn=GRAP2;refGene_tx
=XM_019835624.2;refGene_gn=GRAP2;refGene_tx=XM_023257504.1
B4 135616462 135616723
refGene_gn=GRAP2;refGene_tx=XM_011284232.3;refGene_gn=GRAP2;refGene_tx
=XM_019835624.2;refGene_gn=GRAP2;refGene_tx=XM_023257504.1
B4 135637016 135637526
refGene_gn=FAM83F;refGene_tx=XM_023257505.1
B4 135658658 135658826
refGene_gn=FAM83F;refGene_tx=XM_023257505.1
B4 135659360 135659482

refGene_gn=FAM83F;refGene_tx=XM_023257505.1
B4	135662032	135662727
refGene_gn=FAM83F;refGene_tx=XM_023257505.1
B4	135669377	135669427
refGene_gn=FAM83F;refGene_tx=XM_023257505.1
B4	135752788	135752833
refGene_gn=TNRC6B;refGene_tx=XM_011284235.3;refGene_gn=TNRC6B;refGene_tx=XM_019835628.2;refGene_gn=TNRC6B;refGene_tx=XM_023257506.1
B4	135778088	135778156
refGene_gn=TNRC6B;refGene_tx=XM_011284235.3;refGene_gn=TNRC6B;refGene_tx=XM_019835628.2;refGene_gn=TNRC6B;refGene_tx=XM_023257506.1
B4	135798906	135798911
refGene_gn=TNRC6B;refGene_tx=XM_019835626.2
B4	135863669	135863757
refGene_gn=TNRC6B;refGene_tx=XM_011284235.3;refGene_gn=TNRC6B;refGene_tx=XM_019835626.2;refGene_gn=TNRC6B;refGene_tx=XM_019835628.2;refGene_gn=TNRC6B;refGene_tx=XM_023257506.1
B4	135868672	135868694
refGene_gn=TNRC6B;refGene_tx=XM_011284235.3;refGene_gn=TNRC6B;refGene_tx=XM_019835626.2;refGene_gn=TNRC6B;refGene_tx=XM_019835628.2;refGene_gn=TNRC6B;refGene_tx=XM_023257506.1
B4	135877032	135877374
refGene_gn=TNRC6B;refGene_tx=XM_011284235.3;refGene_gn=TNRC6B;refGene_tx=XM_019835626.2;refGene_gn=TNRC6B;refGene_tx=XM_019835628.2;refGene_gn=TNRC6B;refGene_tx=XM_023257506.1
B4	135879856	135882205
refGene_gn=TNRC6B;refGene_tx=XM_011284235.3;refGene_gn=TNRC6B;refGene_tx=XM_019835626.2;refGene_gn=TNRC6B;refGene_tx=XM_019835628.2;refGene_gn=TNRC6B;refGene_tx=XM_023257506.1
B4	135883779	135883938
refGene_gn=TNRC6B;refGene_tx=XM_011284235.3;refGene_gn=TNRC6B;refGene_tx=XM_019835626.2;refGene_gn=TNRC6B;refGene_tx=XM_019835628.2;refGene_gn=TNRC6B;refGene_tx=XM_023257506.1
B4	135885357	135885533
refGene_gn=TNRC6B;refGene_tx=XM_011284235.3;refGene_gn=TNRC6B;refGene_tx=XM_019835626.2;refGene_gn=TNRC6B;refGene_tx=XM_019835628.2;refGene_gn=TNRC6B;refGene_tx=XM_023257506.1
B4	135887790	135887865
refGene_gn=TNRC6B;refGene_tx=XM_011284235.3;refGene_gn=TNRC6B;refGene_tx=XM_019835626.2;refGene_gn=TNRC6B;refGene_tx=XM_019835628.2;refGene_gn=TNRC6B;refGene_tx=XM_023257506.1
B4	135888958	135889004
refGene_gn=TNRC6B;refGene_tx=XM_011284235.3;refGene_gn=TNRC6B;refGene_tx=XM_019835626.2;refGene_gn=TNRC6B;refGene_tx=XM_019835628.2;refGene_gn=TNRC6B;refGene_tx=XM_023257506.1
B4	135890459	135890608
refGene_gn=TNRC6B;refGene_tx=XM_011284235.3;refGene_gn=TNRC6B;refGene_tx=XM_019835626.2;refGene_gn=TNRC6B;refGene_tx=XM_019835628.2;refGene_gn=TNRC6B;refGene_tx=XM_023257506.1
B4	135891679	135891850

refGene_gn=TNRC6B;refGene_tx=XM_011284235.3;refGene_gn=TNRC6B;refGene_tx=XM_019835626.2;refGene_gn=TNRC6B;refGene_tx=XM_023257506.1
B4 135895818 135895944
refGene_gn=TNRC6B;refGene_tx=XM_011284235.3;refGene_gn=TNRC6B;refGene_tx=XM_019835626.2;refGene_gn=TNRC6B;refGene_tx=XM_019835628.2;refGene_gn=TNRC6B;refGene_tx=XM_023257506.1
B4 135906844 135906976
refGene_gn=TNRC6B;refGene_tx=XM_011284235.3;refGene_gn=TNRC6B;refGene_tx=XM_019835626.2;refGene_gn=TNRC6B;refGene_tx=XM_019835628.2;refGene_gn=TNRC6B;refGene_tx=XM_023257506.1
B4 135907247 135907343
refGene_gn=TNRC6B;refGene_tx=XM_011284235.3;refGene_gn=TNRC6B;refGene_tx=XM_019835626.2;refGene_gn=TNRC6B;refGene_tx=XM_019835628.2;refGene_gn=TNRC6B;refGene_tx=XM_023257506.1
B4 135907519 135907724
refGene_gn=TNRC6B;refGene_tx=XM_011284235.3;refGene_gn=TNRC6B;refGene_tx=XM_019835626.2;refGene_gn=TNRC6B;refGene_tx=XM_019835628.2;refGene_gn=TNRC6B;refGene_tx=XM_023257506.1
B4 135916535 135916673
refGene_gn=TNRC6B;refGene_tx=XM_011284235.3;refGene_gn=TNRC6B;refGene_tx=XM_019835626.2;refGene_gn=TNRC6B;refGene_tx=XM_019835628.2;refGene_gn=TNRC6B;refGene_tx=XM_023257506.1
B4 135918787 135918964
refGene_gn=TNRC6B;refGene_tx=XM_011284235.3;refGene_gn=TNRC6B;refGene_tx=XM_019835626.2;refGene_gn=TNRC6B;refGene_tx=XM_019835628.2;refGene_gn=TNRC6B;refGene_tx=XM_023257506.1
B4 135920522 135920669
refGene_gn=TNRC6B;refGene_tx=XM_011284235.3;refGene_gn=TNRC6B;refGene_tx=XM_019835626.2;refGene_gn=TNRC6B;refGene_tx=XM_019835628.2;refGene_gn=TNRC6B;refGene_tx=XM_023257506.1
B4 135920922 135921018
refGene_gn=TNRC6B;refGene_tx=XM_011284235.3;refGene_gn=TNRC6B;refGene_tx=XM_019835626.2;refGene_gn=TNRC6B;refGene_tx=XM_019835628.2;refGene_gn=TNRC6B;refGene_tx=XM_023257506.1
B4 135923510 135923735
refGene_gn=TNRC6B;refGene_tx=XM_011284235.3;refGene_gn=TNRC6B;refGene_tx=XM_019835626.2;refGene_gn=TNRC6B;refGene_tx=XM_019835628.2;refGene_gn=TNRC6B;refGene_tx=XM_023257506.1
B4 135924195 135924266
refGene_gn=TNRC6B;refGene_tx=XM_011284235.3;refGene_gn=TNRC6B;refGene_tx=XM_019835626.2;refGene_gn=TNRC6B;refGene_tx=XM_019835628.2;refGene_gn=TNRC6B;refGene_tx=XM_023257506.1
B4 135926907 135927047
refGene_gn=TNRC6B;refGene_tx=XM_011284235.3;refGene_gn=TNRC6B;refGene_tx=XM_019835626.2;refGene_gn=TNRC6B;refGene_tx=XM_019835628.2;refGene_gn=TNRC6B;refGene_tx=XM_023257506.1
B4 135928041 135928429
refGene_gn=TNRC6B;refGene_tx=XM_011284235.3;refGene_gn=TNRC6B;refGene_tx=XM_019835626.2;refGene_gn=TNRC6B;refGene_tx=XM_019835628.2;refGene_gn=TNRC6B;refGene_tx=XM_023257506.1

B4 135948829 135949000
refGene_gn=ADSL;refGene_tx=XM_003989280.5;refGene_gn=ADSL;refGene_tx=XM_023257508.1
B4 135952821 135953025
refGene_gn=ADSL;refGene_tx=XM_003989280.5;refGene_gn=ADSL;refGene_tx=XM_006934067.4;refGene_gn=ADSL;refGene_tx=XM_006934068.4;refGene_gn=ADSL;refGene_tx=XM_006934069.4;refGene_gn=ADSL;refGene_tx=XM_006934070.4;refGene_gn=ADSL;refGene_tx=XM_023257508.1
B4 135954276 135954408
refGene_gn=ADSL;refGene_tx=XM_019835629.2
B4 135954704 135954749
refGene_gn=ADSL;refGene_tx=XM_003989280.5;refGene_gn=ADSL;refGene_tx=XM_006934067.4;refGene_gn=ADSL;refGene_tx=XM_006934068.4;refGene_gn=ADSL;refGene_tx=XM_006934069.4;refGene_gn=ADSL;refGene_tx=XM_006934070.4;refGene_gn=ADSL;refGene_tx=XM_019835629.2;refGene_gn=ADSL;refGene_tx=XM_023257508.1
B4 135955489 135955569
refGene_gn=ADSL;refGene_tx=XM_003989280.5;refGene_gn=ADSL;refGene_tx=XM_006934067.4;refGene_gn=ADSL;refGene_tx=XM_006934068.4;refGene_gn=ADSL;refGene_tx=XM_006934069.4;refGene_gn=ADSL;refGene_tx=XM_006934070.4;refGene_gn=ADSL;refGene_tx=XM_019835629.2;refGene_gn=ADSL;refGene_tx=XM_023257508.1
B4 135959716 135959888
refGene_gn=ADSL;refGene_tx=XM_003989280.5;refGene_gn=ADSL;refGene_tx=XM_006934067.4;refGene_gn=ADSL;refGene_tx=XM_006934068.4;refGene_gn=ADSL;refGene_tx=XM_006934069.4;refGene_gn=ADSL;refGene_tx=XM_006934070.4;refGene_gn=ADSL;refGene_tx=XM_019835629.2;refGene_gn=ADSL;refGene_tx=XM_023257508.1
B4 135960099 135960146
refGene_gn=ADSL;refGene_tx=XM_003989280.5;refGene_gn=ADSL;refGene_tx=XM_006934067.4;refGene_gn=ADSL;refGene_tx=XM_006934068.4;refGene_gn=ADSL;refGene_tx=XM_006934069.4;refGene_gn=ADSL;refGene_tx=XM_006934070.4;refGene_gn=ADSL;refGene_tx=XM_019835629.2;refGene_gn=ADSL;refGene_tx=XM_023257508.1
B4 135960898 135960989
refGene_gn=ADSL;refGene_tx=XM_003989280.5;refGene_gn=ADSL;refGene_tx=XM_006934067.4;refGene_gn=ADSL;refGene_tx=XM_006934068.4;refGene_gn=ADSL;refGene_tx=XM_006934069.4;refGene_gn=ADSL;refGene_tx=XM_006934070.4;refGene_gn=ADSL;refGene_tx=XM_019835629.2;refGene_gn=ADSL;refGene_tx=XM_023257508.1
B4 135961387 135961457
refGene_gn=ADSL;refGene_tx=XM_003989280.5;refGene_gn=ADSL;refGene_tx=XM_006934067.4;refGene_gn=ADSL;refGene_tx=XM_006934068.4;refGene_gn=ADSL;refGene_tx=XM_006934069.4;refGene_gn=ADSL;refGene_tx=XM_006934070.4;refGene_gn=ADSL;refGene_tx=XM_019835629.2;refGene_gn=ADSL;refGene_tx=XM_023257508.1
B4 135961603 135961751
refGene_gn=ADSL;refGene_tx=XM_003989280.5;refGene_gn=ADSL;refGene_tx=XM_006934067.4;refGene_gn=ADSL;refGene_tx=XM_006934068.4;refGene_gn=ADSL;refGene_tx=XM_006934069.4;refGene_gn=ADSL;refGene_tx=XM_006934070.4;

refGene_gn=ADSL;refGene_tx=XM_019835629.2;refGene_gn=ADSL;refGene_tx=X
M_023257508.1
B4 135963692 135963783
refGene_gn=ADSL;refGene_tx=XM_003989280.5;refGene_gn=ADSL;refGene_tx=X
M_006934067.4;refGene_gn=ADSL;refGene_tx=XM_006934068.4;refGene_gn=ADS
L;refGene_tx=XM_006934069.4;refGene_gn=ADSL;refGene_tx=XM_006934070.4;
refGene_gn=ADSL;refGene_tx=XM_019835629.2;refGene_gn=ADSL;refGene_tx=X
M_023257508.1
B4 135964418 135964508
refGene_gn=ADSL;refGene_tx=XM_003989280.5;refGene_gn=ADSL;refGene_tx=X
M_006934067.4;refGene_gn=ADSL;refGene_tx=XM_006934068.4;refGene_gn=ADS
L;refGene_tx=XM_006934069.4;refGene_gn=ADSL;refGene_tx=XM_006934070.4;
refGene_gn=ADSL;refGene_tx=XM_019835629.2;refGene_gn=ADSL;refGene_tx=X
M_023257508.1
B4 135965258 135965436
refGene_gn=ADSL;refGene_tx=XM_003989280.5;refGene_gn=ADSL;refGene_tx=X
M_006934067.4;refGene_gn=ADSL;refGene_tx=XM_006934068.4;refGene_gn=ADS
L;refGene_tx=XM_006934069.4;refGene_gn=ADSL;refGene_tx=XM_006934070.4;
refGene_gn=ADSL;refGene_tx=XM_019835629.2
B4 135966962 135967049
refGene_gn=ADSL;refGene_tx=XM_003989280.5;refGene_gn=ADSL;refGene_tx=X
M_006934067.4;refGene_gn=ADSL;refGene_tx=XM_006934068.4;refGene_gn=ADS
L;refGene_tx=XM_006934069.4;refGene_gn=ADSL;refGene_tx=XM_006934070.4;
refGene_gn=ADSL;refGene_tx=XM_019835629.2;refGene_gn=ADSL;refGene_tx=X
M_023257508.1
B4 136016700 136016707
refGene_gn=SGSM3;refGene_tx=XM_023257510.1;refGene_gn=SGSM3;refGene_tx
=XM_023257511.1;refGene_gn=SGSM3;refGene_tx=XM_023257512.1;refGene_gn=
SGSM3;refGene_tx=XM_023257513.1;refGene_gn=SGSM3;refGene_tx=XM_0232575
14.1
B4 136017158 136017244
refGene_gn=SGSM3;refGene_tx=XM_023257510.1;refGene_gn=SGSM3;refGene_tx
=XM_023257511.1;refGene_gn=SGSM3;refGene_tx=XM_023257512.1;refGene_gn=
SGSM3;refGene_tx=XM_023257513.1;refGene_gn=SGSM3;refGene_tx=XM_0232575
14.1
B4 136017708 136017775
refGene_gn=SGSM3;refGene_tx=XM_023257510.1;refGene_gn=SGSM3;refGene_tx
=XM_023257511.1;refGene_gn=SGSM3;refGene_tx=XM_023257512.1;refGene_gn=
SGSM3;refGene_tx=XM_023257513.1;refGene_gn=SGSM3;refGene_tx=XM_0232575
14.1
B4 136019938 136020147
refGene_gn=SGSM3;refGene_tx=XM_023257510.1;refGene_gn=SGSM3;refGene_tx
=XM_023257511.1;refGene_gn=SGSM3;refGene_tx=XM_023257512.1;refGene_gn=
SGSM3;refGene_tx=XM_023257513.1;refGene_gn=SGSM3;refGene_tx=XM_0232575
14.1
B4 136020225 136020333
refGene_gn=SGSM3;refGene_tx=XM_023257510.1;refGene_gn=SGSM3;refGene_tx
=XM_023257511.1;refGene_gn=SGSM3;refGene_tx=XM_023257512.1;refGene_gn=
SGSM3;refGene_tx=XM_023257513.1;refGene_gn=SGSM3;refGene_tx=XM_0232575
14.1

B4	136021193	136021337
refGene_gn=SGSM3;refGene_tx=XM_023257510.1;refGene_gn=SGSM3;refGene_tx=XM_023257511.1;refGene_gn=SGSM3;refGene_tx=XM_023257512.1;refGene_gn=SGSM3;refGene_tx=XM_023257514.1
B4	136021576	136021772
refGene_gn=SGSM3;refGene_tx=XM_023257510.1;refGene_gn=SGSM3;refGene_tx=XM_023257511.1;refGene_gn=SGSM3;refGene_tx=XM_023257512.1;refGene_gn=SGSM3;refGene_tx=XM_023257513.1;refGene_gn=SGSM3;refGene_tx=XM_023257514.1
B4	136022160	136022306
refGene_gn=SGSM3;refGene_tx=XM_023257510.1;refGene_gn=SGSM3;refGene_tx=XM_023257511.1;refGene_gn=SGSM3;refGene_tx=XM_023257512.1;refGene_gn=SGSM3;refGene_tx=XM_023257513.1;refGene_gn=SGSM3;refGene_tx=XM_023257514.1
B4	136022519	136022744
refGene_gn=SGSM3;refGene_tx=XM_023257510.1;refGene_gn=SGSM3;refGene_tx=XM_023257511.1;refGene_gn=SGSM3;refGene_tx=XM_023257512.1;refGene_gn=SGSM3;refGene_tx=XM_023257513.1;refGene_gn=SGSM3;refGene_tx=XM_023257514.1
B4	136023127	136023182
refGene_gn=SGSM3;refGene_tx=XM_023257510.1;refGene_gn=SGSM3;refGene_tx=XM_023257511.1;refGene_gn=SGSM3;refGene_tx=XM_023257512.1;refGene_gn=SGSM3;refGene_tx=XM_023257513.1;refGene_gn=SGSM3;refGene_tx=XM_023257514.1
B4	136023403	136023531
refGene_gn=SGSM3;refGene_tx=XM_023257510.1;refGene_gn=SGSM3;refGene_tx=XM_023257511.1;refGene_gn=SGSM3;refGene_tx=XM_023257512.1;refGene_gn=SGSM3;refGene_tx=XM_023257513.1;refGene_gn=SGSM3;refGene_tx=XM_023257514.1
B4	136023643	136023799
refGene_gn=SGSM3;refGene_tx=XM_023257510.1;refGene_gn=SGSM3;refGene_tx=XM_023257511.1;refGene_gn=SGSM3;refGene_tx=XM_023257512.1;refGene_gn=SGSM3;refGene_tx=XM_023257513.1;refGene_gn=SGSM3;refGene_tx=XM_023257514.1
B4	136023967	136024022
refGene_gn=SGSM3;refGene_tx=XM_023257510.1;refGene_gn=SGSM3;refGene_tx=XM_023257511.1;refGene_gn=SGSM3;refGene_tx=XM_023257512.1;refGene_gn=SGSM3;refGene_tx=XM_023257513.1;refGene_gn=SGSM3;refGene_tx=XM_023257514.1
B4	136024167	136024217
refGene_gn=SGSM3;refGene_tx=XM_023257510.1;refGene_gn=SGSM3;refGene_tx=XM_023257511.1;refGene_gn=SGSM3;refGene_tx=XM_023257512.1;refGene_gn=SGSM3;refGene_tx=XM_023257513.1;refGene_gn=SGSM3;refGene_tx=XM_023257514.1
B4	136024376	136024529
refGene_gn=SGSM3;refGene_tx=XM_023257510.1;refGene_gn=SGSM3;refGene_tx=XM_023257511.1;refGene_gn=SGSM3;refGene_tx=XM_023257512.1;refGene_gn=SGSM3;refGene_tx=XM_023257513.1;refGene_gn=SGSM3;refGene_tx=XM_023257514.1
B4	136024699	136024770

refGene_gn=SGSM3;refGene_tx=XM_023257510.1;refGene_gn=SGSM3;refGene_tx=XM_023257511.1;refGene_gn=SGSM3;refGene_tx=XM_023257512.1;refGene_gn=SGSM3;refGene_tx=XM_023257513.1;refGene_gn=SGSM3;refGene_tx=XM_023257514.1
B4 136024899 136024948
refGene_gn=SGSM3;refGene_tx=XM_023257510.1;refGene_gn=SGSM3;refGene_tx=XM_023257511.1;refGene_gn=SGSM3;refGene_tx=XM_023257512.1;refGene_gn=SGSM3;refGene_tx=XM_023257513.1;refGene_gn=SGSM3;refGene_tx=XM_023257514.1
B4 136025024 136025114
refGene_gn=SGSM3;refGene_tx=XM_023257510.1;refGene_gn=SGSM3;refGene_tx=XM_023257511.1;refGene_gn=SGSM3;refGene_tx=XM_023257512.1;refGene_gn=SGSM3;refGene_tx=XM_023257513.1;refGene_gn=SGSM3;refGene_tx=XM_023257514.1
B4 136025827 136025950
refGene_gn=SGSM3;refGene_tx=XM_023257510.1;refGene_gn=SGSM3;refGene_tx=XM_023257511.1;refGene_gn=SGSM3;refGene_tx=XM_023257512.1;refGene_gn=SGSM3;refGene_tx=XM_023257513.1
B4 136026065 136026126
refGene_gn=SGSM3;refGene_tx=XM_023257510.1;refGene_gn=SGSM3;refGene_tx=XM_023257511.1;refGene_gn=SGSM3;refGene_tx=XM_023257513.1
B4 136026379 136026460
refGene_gn=SGSM3;refGene_tx=XM_023257510.1;refGene_gn=SGSM3;refGene_tx=XM_023257511.1;refGene_gn=SGSM3;refGene_tx=XM_023257512.1;refGene_gn=SGSM3;refGene_tx=XM_023257513.1
B4 136028048 136028566
refGene_gn=MKL1;refGene_tx=XM_023257509.1
B4 136031948 136032009
refGene_gn=MKL1;refGene_tx=XM_023257509.1
B4 136032329 136032485
refGene_gn=MKL1;refGene_tx=XM_023257509.1
B4 136033401 136033754
refGene_gn=MKL1;refGene_tx=XM_023257509.1
B4 136033756 136034405
refGene_gn=MKL1;refGene_tx=XM_023257509.1
B4 136035573 136035745
refGene_gn=MKL1;refGene_tx=XM_023257509.1
B4 136035922 136036176
refGene_gn=MKL1;refGene_tx=XM_023257509.1
B4 136037788 136037920
refGene_gn=MKL1;refGene_tx=XM_023257509.1
B4 136038454 136038630
refGene_gn=MKL1;refGene_tx=XM_023257509.1
B4 136042445 136042607
refGene_gn=MKL1;refGene_tx=XM_023257509.1
B4 136044355 136044431
refGene_gn=MKL1;refGene_tx=XM_023257509.1
B4 136046743 136046799
refGene_gn=MKL1;refGene_tx=XM_023257509.1
B4 136070991 136071057

refGene_gn=MKL1;refGene_tx=XM_023257509.1
B4 136089503 136089633
refGene_gn=MKL1;refGene_tx=XM_023257509.1
B4 136270481 136270563
refGene_gn=MCHR1;refGene_tx=XM_003989283.5
B4 136271869 136272849
refGene_gn=MCHR1;refGene_tx=XM_003989283.5
B4 136343150 136343298
refGene_gn=SLC25A17;refGene_tx=XM_003989284.5;refGene_gn=SLC25A17;refGene_tx=XM_019835635.2
B4 136345218 136345301
refGene_gn=SLC25A17;refGene_tx=XM_003989284.5;refGene_gn=SLC25A17;refGene_tx=XM_019835635.2
B4 136347638 136347734
refGene_gn=SLC25A17;refGene_tx=XM_003989284.5;refGene_gn=SLC25A17;refGene_tx=XM_019835635.2
B4 136347819 136347965
refGene_gn=SLC25A17;refGene_tx=XM_003989284.5;refGene_gn=SLC25A17;refGene_tx=XM_019835635.2
B4 136349595 136349712
refGene_gn=SLC25A17;refGene_tx=XM_003989284.5;refGene_gn=SLC25A17;refGene_tx=XM_019835635.2
B4 136359345 136359497
refGene_gn=SLC25A17;refGene_tx=XM_003989284.5
B4 136360909 136360976
refGene_gn=SLC25A17;refGene_tx=XM_003989284.5
B4 136363380 136363441
refGene_gn=SLC25A17;refGene_tx=XM_003989284.5;refGene_gn=SLC25A17;refGene_tx=XM_019835635.2
B4 136389066 136389120
refGene_gn=SLC25A17;refGene_tx=XM_003989284.5;refGene_gn=SLC25A17;refGene_tx=XM_019835635.2
B4 136394961 136395090
refGene_gn=ST13;refGene_tx=XM_023257521.1;refGene_gn=ST13;refGene_tx=XM_023257522.1
B4 136396354 136396491
refGene_gn=ST13;refGene_tx=XM_023257521.1;refGene_gn=ST13;refGene_tx=XM_023257522.1
B4 136398662 136398708
refGene_gn=ST13;refGene_tx=XM_023257521.1;refGene_gn=ST13;refGene_tx=XM_023257522.1
B4 136399950 136400067
refGene_gn=ST13;refGene_tx=XM_023257521.1
B4 136401728 136401831
refGene_gn=ST13;refGene_tx=XM_023257521.1;refGene_gn=ST13;refGene_tx=XM_023257522.1
B4 136405586 136405697
refGene_gn=ST13;refGene_tx=XM_023257521.1;refGene_gn=ST13;refGene_tx=XM_023257522.1
B4 136406051 136406136

refGene_gn=ST13;refGene_tx=XM_023257521.1;refGene_gn=ST13;refGene_tx=XM_023257522.1
B4	136411017	136411084
refGene_gn=ST13;refGene_tx=XM_023257521.1;refGene_gn=ST13;refGene_tx=XM_023257522.1
B4	136415112	136415183
refGene_gn=ST13;refGene_tx=XM_023257521.1;refGene_gn=ST13;refGene_tx=XM_023257522.1
B4	136417391	136417467
refGene_gn=ST13;refGene_tx=XM_023257521.1;refGene_gn=ST13;refGene_tx=XM_023257522.1
B4	136419237	136419295
refGene_gn=ST13;refGene_tx=XM_023257521.1;refGene_gn=ST13;refGene_tx=XM_023257522.1
B4	136424842	136424952
refGene_gn=ST13;refGene_tx=XM_023257521.1;refGene_gn=ST13;refGene_tx=XM_023257522.1
B4	136425593	136425657
refGene_gn=XPNPEP3;refGene_tx=XM_023257515.1;refGene_gn=XPNPEP3;refGene_tx=XM_023257516.1;refGene_gn=XPNPEP3;refGene_tx=XM_023257517.1
B4	136431740	136432381
refGene_gn=DNAJB7;refGene_tx=XM_004001541.5
B4	136432383	136432663
refGene_gn=DNAJB7;refGene_tx=XM_004001541.5
B4	136441409	136441526
refGene_gn=XPNPEP3;refGene_tx=XM_023257515.1;refGene_gn=XPNPEP3;refGene_tx=XM_023257516.1;refGene_gn=XPNPEP3;refGene_tx=XM_023257517.1;refGene_gn=XPNPEP3;refGene_tx=XM_023257518.1
B4	136453995	136454403
refGene_gn=XPNPEP3;refGene_tx=XM_023257515.1;refGene_gn=XPNPEP3;refGene_tx=XM_023257516.1;refGene_gn=XPNPEP3;refGene_tx=XM_023257517.1;refGene_gn=XPNPEP3;refGene_tx=XM_023257518.1;refGene_gn=XPNPEP3;refGene_tx=XM_023257519.1;refGene_gn=XPNPEP3;refGene_tx=XM_023257520.1
B4	136457516	136457719
refGene_gn=XPNPEP3;refGene_tx=XM_023257515.1;refGene_gn=XPNPEP3;refGene_tx=XM_023257516.1;refGene_gn=XPNPEP3;refGene_tx=XM_023257517.1;refGene_gn=XPNPEP3;refGene_tx=XM_023257518.1;refGene_gn=XPNPEP3;refGene_tx=XM_023257519.1;refGene_gn=XPNPEP3;refGene_tx=XM_023257520.1
B4	136478360	136478411
refGene_gn=XPNPEP3;refGene_tx=XM_023257515.1;refGene_gn=XPNPEP3;refGene_tx=XM_023257516.1;refGene_gn=XPNPEP3;refGene_tx=XM_023257518.1;refGene_gn=XPNPEP3;refGene_tx=XM_023257519.1;refGene_gn=XPNPEP3;refGene_tx=XM_023257520.1
B4	136479851	136479914
refGene_gn=XPNPEP3;refGene_tx=XM_023257515.1;refGene_gn=XPNPEP3;refGene_tx=XM_023257516.1;refGene_gn=XPNPEP3;refGene_tx=XM_023257517.1;refGene_gn=XPNPEP3;refGene_tx=XM_023257518.1;refGene_gn=XPNPEP3;refGene_tx=XM_023257519.1;refGene_gn=XPNPEP3;refGene_tx=XM_023257520.1
B4	136481198	136481312
refGene_gn=XPNPEP3;refGene_tx=XM_023257515.1;refGene_gn=XPNPEP3;refGen

e_tx=XM_023257516.1;refGene_gn=XPNPEP3;refGene_tx=XM_023257517.1;refGene_gn=XPNPEP3;refGene_tx=XM_023257518.1;refGene_gn=XPNPEP3;refGene_tx=XM_023257519.1;refGene_gn=XPNPEP3;refGene_tx=XM_023257520.1
B4 136487791 136487877
refGene_gn=XPNPEP3;refGene_tx=XM_023257515.1;refGene_gn=XPNPEP3;refGene_tx=XM_023257516.1;refGene_gn=XPNPEP3;refGene_tx=XM_023257517.1;refGene_gn=XPNPEP3;refGene_tx=XM_023257518.1;refGene_gn=XPNPEP3;refGene_tx=XM_023257519.1;refGene_gn=XPNPEP3;refGene_tx=XM_023257520.1
B4 136495466 136495647
refGene_gn=XPNPEP3;refGene_tx=XM_023257515.1;refGene_gn=XPNPEP3;refGene_tx=XM_023257516.1;refGene_gn=XPNPEP3;refGene_tx=XM_023257517.1;refGene_gn=XPNPEP3;refGene_tx=XM_023257518.1;refGene_gn=XPNPEP3;refGene_tx=XM_023257519.1;refGene_gn=XPNPEP3;refGene_tx=XM_023257520.1
B4 136496708 136496829
refGene_gn=XPNPEP3;refGene_tx=XM_023257515.1;refGene_gn=XPNPEP3;refGene_tx=XM_023257516.1;refGene_gn=XPNPEP3;refGene_tx=XM_023257517.1;refGene_gn=XPNPEP3;refGene_tx=XM_023257518.1;refGene_gn=XPNPEP3;refGene_tx=XM_023257519.1;refGene_gn=XPNPEP3;refGene_tx=XM_023257520.1
B4 136498879 136499046
refGene_gn=XPNPEP3;refGene_tx=XM_023257515.1;refGene_gn=XPNPEP3;refGene_tx=XM_023257516.1;refGene_gn=XPNPEP3;refGene_tx=XM_023257517.1;refGene_gn=XPNPEP3;refGene_tx=XM_023257518.1;refGene_gn=XPNPEP3;refGene_tx=XM_023257519.1;refGene_gn=XPNPEP3;refGene_tx=XM_023257520.1
B4 136528944 136529220
refGene_gn=LOC105260770;refGene_tx=XM_011284240.3
B4 136531845 136531923
refGene_gn=RBX1;refGene_tx=XM_003989287.4
B4 136533838 136533917
refGene_gn=RBX1;refGene_tx=XM_003989287.4
B4 136539025 136539096
refGene_gn=RBX1;refGene_tx=XM_003989287.4
B4 136542091 136542177
refGene_gn=RBX1;refGene_tx=XM_003989287.4
B4 136545519 136545532
refGene_gn=RBX1;refGene_tx=XM_003989287.4
B4 136643664 136643758
refGene_gn=EP300;refGene_tx=XM_003989288.5
B4 136662248 136662886
refGene_gn=EP300;refGene_tx=XM_003989288.5
B4 136673088 136673265
refGene_gn=EP300;refGene_tx=XM_003989288.5
B4 136674912 136675174
refGene_gn=EP300;refGene_tx=XM_003989288.5
B4 136675799 136675913
refGene_gn=EP300;refGene_tx=XM_003989288.5
B4 136676889 136677135
refGene_gn=EP300;refGene_tx=XM_003989288.5
B4 136682960 136683054
refGene_gn=EP300;refGene_tx=XM_003989288.5
B4 136683677 136683815

refGene_gn=EP300;refGene_tx=XM_003989288.5
B4 136685496 136685614
refGene_gn=EP300;refGene_tx=XM_003989288.5
B4 136686486 136686661
refGene_gn=EP300;refGene_tx=XM_003989288.5
B4 136692519 136692597
refGene_gn=EP300;refGene_tx=XM_003989288.5
B4 136694061 136694171
refGene_gn=EP300;refGene_tx=XM_003989288.5
B4 136696449 136696587
refGene_gn=EP300;refGene_tx=XM_003989288.5
B4 136697235 136697673
refGene_gn=EP300;refGene_tx=XM_003989288.5
B4 136699148 136699328
refGene_gn=EP300;refGene_tx=XM_003989288.5
B4 136699526 136699671
refGene_gn=EP300;refGene_tx=XM_003989288.5
B4 136701505 136701624
refGene_gn=EP300;refGene_tx=XM_003989288.5
B4 136703016 136703256
refGene_gn=EP300;refGene_tx=XM_003989288.5
B4 136704110 136704199
refGene_gn=EP300;refGene_tx=XM_003989288.5
B4 136706196 136706277
refGene_gn=EP300;refGene_tx=XM_003989288.5
B4 136708579 136708636
refGene_gn=EP300;refGene_tx=XM_003989288.5
B4 136709447 136709525
refGene_gn=EP300;refGene_tx=XM_003989288.5
B4 136711265 136711333
refGene_gn=EP300;refGene_tx=XM_003989288.5
B4 136713486 136713637
refGene_gn=EP300;refGene_tx=XM_003989288.5
B4 136713756 136713903
refGene_gn=EP300;refGene_tx=XM_003989288.5
B4 136714499 136714613
refGene_gn=EP300;refGene_tx=XM_003989288.5
B4 136715571 136715737
refGene_gn=EP300;refGene_tx=XM_003989288.5
B4 136716497 136716662
refGene_gn=EP300;refGene_tx=XM_003989288.5
B4 136717720 136717882
refGene_gn=EP300;refGene_tx=XM_003989288.5
B4 136720003 136720285
refGene_gn=EP300;refGene_tx=XM_003989288.5
B4 136720544 136722758
refGene_gn=EP300;refGene_tx=XM_003989288.5
B4 136788737 136788761
refGene_gn=L3MBTL2;refGene_tx=XM_011284241.3;refGene_gn=L3MBTL2;refGen
e_tx=XM_019835639.2

B4 136790593 136790837
refGene_gn=L3MBTL2;refGene_tx=XM_011284241.3;refGene_gn=L3MBTL2;refGene_tx=XM_019835639.2
B4 136792152 136792286
refGene_gn=L3MBTL2;refGene_tx=XM_011284241.3;refGene_gn=L3MBTL2;refGene_tx=XM_019835639.2
B4 136795277 136795401
refGene_gn=L3MBTL2;refGene_tx=XM_011284241.3;refGene_gn=L3MBTL2;refGene_tx=XM_019835639.2
B4 136796164 136796244
refGene_gn=L3MBTL2;refGene_tx=XM_011284241.3;refGene_gn=L3MBTL2;refGene_tx=XM_019835639.2
B4 136797843 136797961
refGene_gn=L3MBTL2;refGene_tx=XM_011284241.3;refGene_gn=L3MBTL2;refGene_tx=XM_019835639.2
B4 136798386 136798521
refGene_gn=L3MBTL2;refGene_tx=XM_011284241.3;refGene_gn=L3MBTL2;refGene_tx=XM_019835639.2
B4 136798792 136798881
refGene_gn=L3MBTL2;refGene_tx=XM_011284241.3;refGene_gn=L3MBTL2;refGene_tx=XM_019835639.2
B4 136801979 136802211
refGene_gn=L3MBTL2;refGene_tx=XM_011284241.3;refGene_gn=L3MBTL2;refGene_tx=XM_019835639.2
B4 136802650 136802727
refGene_gn=L3MBTL2;refGene_tx=XM_011284241.3;refGene_gn=L3MBTL2;refGene_tx=XM_019835639.2
B4 136802905 136803010
refGene_gn=L3MBTL2;refGene_tx=XM_011284241.3;refGene_gn=L3MBTL2;refGene_tx=XM_019835639.2
B4 136803656 136803804
refGene_gn=L3MBTL2;refGene_tx=XM_011284241.3;refGene_gn=L3MBTL2;refGene_tx=XM_019835639.2
B4 136804208 136804291
refGene_gn=L3MBTL2;refGene_tx=XM_011284241.3;refGene_gn=L3MBTL2;refGene_tx=XM_019835639.2
B4 136804614 136804849
refGene_gn=L3MBTL2;refGene_tx=XM_011284241.3;refGene_gn=L3MBTL2;refGene_tx=XM_019835639.2
B4 136805331 136805397
refGene_gn=L3MBTL2;refGene_tx=XM_011284241.3;refGene_gn=L3MBTL2;refGene_tx=XM_019835639.2
B4 136806360 136806477
refGene_gn=L3MBTL2;refGene_tx=XM_011284241.3;refGene_gn=L3MBTL2;refGene_tx=XM_019835639.2
B4 136806526 136806553
refGene_gn=CHADL;refGene_tx=XM_023257524.1
B4 136806862 136806975
refGene_gn=L3MBTL2;refGene_tx=XM_011284241.3
B4 136808298 136808309

refGene_gn=L3MBTL2;refGene_tx=XM_019835639.2
B4 136810938 136811370
refGene_gn=CHADL;refGene_tx=XM_023257523.1
B4 136811526 136811725
refGene_gn=CHADL;refGene_tx=XM_023257523.1;refGene_gn=CHADL;refGene_tx=XM_023257524.1
B4 136812606 136812773
refGene_gn=CHADL;refGene_tx=XM_023257523.1;refGene_gn=CHADL;refGene_tx=XM_023257524.1
B4 136813333 136815034
refGene_gn=CHADL;refGene_tx=XM_023257523.1;refGene_gn=CHADL;refGene_tx=XM_023257524.1
B4 136815577 136815746
refGene_gn=CHADL;refGene_tx=XM_023257523.1;refGene_gn=CHADL;refGene_tx=XM_023257524.1
B4 136817254 136817262
refGene_gn=CHADL;refGene_tx=XM_023257523.1;refGene_gn=CHADL;refGene_tx=XM_023257524.1
B4 136824159 136824229
refGene_gn=RANGAP1;refGene_tx=XM_003989326.4;refGene_gn=RANGAP1;refGene_tx=XM_011284242.3
B4 136826315 136826437
refGene_gn=RANGAP1;refGene_tx=XM_003989326.4;refGene_gn=RANGAP1;refGene_tx=XM_011284242.3
B4 136826694 136826783
refGene_gn=RANGAP1;refGene_tx=XM_003989326.4;refGene_gn=RANGAP1;refGene_tx=XM_011284242.3
B4 136827978 136828081
refGene_gn=RANGAP1;refGene_tx=XM_003989326.4;refGene_gn=RANGAP1;refGene_tx=XM_011284242.3
B4 136830094 136830240
refGene_gn=RANGAP1;refGene_tx=XM_003989326.4;refGene_gn=RANGAP1;refGene_tx=XM_011284242.3;refGene_gn=RANGAP1;refGene_tx=XM_023257525.1
B4 136831153 136831340
refGene_gn=RANGAP1;refGene_tx=XM_003989326.4;refGene_gn=RANGAP1;refGene_tx=XM_011284242.3;refGene_gn=RANGAP1;refGene_tx=XM_023257525.1
B4 136832328 136832413
refGene_gn=RANGAP1;refGene_tx=XM_003989326.4;refGene_gn=RANGAP1;refGene_tx=XM_011284242.3;refGene_gn=RANGAP1;refGene_tx=XM_023257525.1
B4 136832499 136832599
refGene_gn=RANGAP1;refGene_tx=XM_003989326.4;refGene_gn=RANGAP1;refGene_tx=XM_011284242.3;refGene_gn=RANGAP1;refGene_tx=XM_023257525.1
B4 136833000 136833114
refGene_gn=RANGAP1;refGene_tx=XM_003989326.4;refGene_gn=RANGAP1;refGene_tx=XM_011284242.3;refGene_gn=RANGAP1;refGene_tx=XM_023257525.1
B4 136834356 136834515
refGene_gn=RANGAP1;refGene_tx=XM_003989326.4;refGene_gn=RANGAP1;refGene_tx=XM_011284242.3;refGene_gn=RANGAP1;refGene_tx=XM_023257525.1
B4 136837253 136837388
refGene_gn=RANGAP1;refGene_tx=XM_003989326.4;refGene_gn=RANGAP1;refGen

e_tx=XM_011284242.3;refGene_gn=RANGAP1;refGene_tx=XM_023257525.1
B4 136840109 136840289
refGene_gn=RANGAP1;refGene_tx=XM_003989326.4;refGene_gn=RANGAP1;refGen
e_tx=XM_011284242.3;refGene_gn=RANGAP1;refGene_tx=XM_023257525.1
B4 136843496 136843556
refGene_gn=RANGAP1;refGene_tx=XM_003989326.4;refGene_gn=RANGAP1;refGen
e_tx=XM_011284242.3;refGene_gn=RANGAP1;refGene_tx=XM_023257525.1
B4 136846468 136846596
refGene_gn=RANGAP1;refGene_tx=XM_003989326.4;refGene_gn=RANGAP1;refGen
e_tx=XM_011284242.3;refGene_gn=RANGAP1;refGene_tx=XM_023257525.1
B4 136851692 136851842
refGene_gn=RANGAP1;refGene_tx=XM_003989326.4;refGene_gn=RANGAP1;refGen
e_tx=XM_011284242.3;refGene_gn=RANGAP1;refGene_tx=XM_023257525.1
B4 136852222 136852307
refGene_gn=RANGAP1;refGene_tx=XM_003989326.4;refGene_gn=RANGAP1;refGen
e_tx=XM_023257525.1
B4 136890386 136890439
refGene_gn=ZC3H7B;refGene_tx=XM_023257526.1
B4 136894231 136894265
refGene_gn=ZC3H7B;refGene_tx=XM_023257526.1
B4 136894399 136894597
refGene_gn=ZC3H7B;refGene_tx=XM_023257526.1
B4 136895555 136895714
refGene_gn=ZC3H7B;refGene_tx=XM_023257526.1
B4 136898511 136898592
refGene_gn=ZC3H7B;refGene_tx=XM_023257526.1
B4 136899408 136899465
refGene_gn=ZC3H7B;refGene_tx=XM_023257526.1
B4 136905847 136905890
refGene_gn=ZC3H7B;refGene_tx=XM_023257526.1
B4 136906433 136906627
refGene_gn=ZC3H7B;refGene_tx=XM_023257526.1
B4 136906975 136907297
refGene_gn=ZC3H7B;refGene_tx=XM_023257526.1
B4 136908257 136908316
refGene_gn=ZC3H7B;refGene_tx=XM_023257526.1
B4 136909141 136909241
refGene_gn=ZC3H7B;refGene_tx=XM_023257526.1
B4 136910674 136910836
refGene_gn=ZC3H7B;refGene_tx=XM_023257526.1
B4 136912881 136913087
refGene_gn=ZC3H7B;refGene_tx=XM_023257526.1
B4 136914157 136914258
refGene_gn=ZC3H7B;refGene_tx=XM_023257526.1
B4 136915217 136915399
refGene_gn=ZC3H7B;refGene_tx=XM_023257526.1
B4 136917568 136917654
refGene_gn=ZC3H7B;refGene_tx=XM_023257526.1
B4 136921179 136921313
refGene_gn=ZC3H7B;refGene_tx=XM_023257526.1

B4	136921484	136921590
refGene_gn=ZC3H7B;refGene_tx=XM_023257526.1
B4	136921732	136921841
refGene_gn=ZC3H7B;refGene_tx=XM_023257526.1
B4	136922105	136922239
refGene_gn=ZC3H7B;refGene_tx=XM_023257526.1
B4	136922357	136922521
refGene_gn=ZC3H7B;refGene_tx=XM_023257526.1
B4	136922909	136923156
refGene_gn=ZC3H7B;refGene_tx=XM_023257526.1
B4	136930149	136930252
refGene_gn=TEF;refGene_tx=XM_003989329.5;refGene_gn=TEF;refGene_tx=XM_006934109.4
B4	136939272	136939417
refGene_gn=TEF;refGene_tx=XM_006934111.4
B4	136939766	136939923
refGene_gn=TEF;refGene_tx=XM_003989328.3;refGene_gn=TEF;refGene_tx=XM_023257527.1
B4	136943386	136943719
refGene_gn=TEF;refGene_tx=XM_003989328.3;refGene_gn=TEF;refGene_tx=XM_003989329.5;refGene_gn=TEF;refGene_tx=XM_006934109.4;refGene_gn=TEF;refGene_tx=XM_006934111.4;refGene_gn=TEF;refGene_tx=XM_023257527.1
B4	136948258	136948479
refGene_gn=TEF;refGene_tx=XM_003989328.3;refGene_gn=TEF;refGene_tx=XM_003989329.5;refGene_gn=TEF;refGene_tx=XM_006934109.4;refGene_gn=TEF;refGene_tx=XM_006934111.4;refGene_gn=TEF;refGene_tx=XM_023257527.1
B4	136949692	136949908
refGene_gn=TEF;refGene_tx=XM_003989328.3;refGene_gn=TEF;refGene_tx=XM_003989329.5;refGene_gn=TEF;refGene_tx=XM_006934109.4;refGene_gn=TEF;refGene_tx=XM_006934111.4;refGene_gn=TEF;refGene_tx=XM_023257527.1
B4	136978359	136979367
refGene_gn=TOB2;refGene_tx=XM_011284244.3
B4	136996976	136997066
refGene_gn=PHF5A;refGene_tx=XM_003989331.5
B4	137004287	137004454
refGene_gn=PHF5A;refGene_tx=XM_003989331.5
B4	137004964	137004988
refGene_gn=PHF5A;refGene_tx=XM_003989331.5
B4	137005386	137005438
refGene_gn=PHF5A;refGene_tx=XM_003989331.5
B4	137005940	137005976
refGene_gn=ACO2;refGene_tx=XM_003989332.5
B4	137036415	137036552
refGene_gn=ACO2;refGene_tx=XM_003989332.5
B4	137040575	137040834
refGene_gn=ACO2;refGene_tx=XM_003989332.5
B4	137044081	137044138
refGene_gn=ACO2;refGene_tx=XM_003989332.5
B4	137044139	137044175
refGene_gn=ACO2;refGene_tx=XM_003989332.5

B4 137047653 137047812
refGene_gn=ACO2;refGene_tx=XM_003989332.5
B4 137048035 137048186
refGene_gn=ACO2;refGene_tx=XM_003989332.5
B4 137049673 137049778
refGene_gn=ACO2;refGene_tx=XM_003989332.5
B4 137050753 137050845
refGene_gn=ACO2;refGene_tx=XM_003989332.5
B4 137051858 137051964
refGene_gn=ACO2;refGene_tx=XM_003989332.5
B4 137053136 137053294
refGene_gn=ACO2;refGene_tx=XM_003989332.5
B4 137053518 137053592
refGene_gn=ACO2;refGene_tx=XM_003989332.5
B4 137054071 137054183
refGene_gn=ACO2;refGene_tx=XM_003989332.5
B4 137055261 137055384
refGene_gn=ACO2;refGene_tx=XM_003989332.5
B4 137055625 137055781
refGene_gn=ACO2;refGene_tx=XM_003989332.5
B4 137056716 137056908
refGene_gn=ACO2;refGene_tx=XM_003989332.5
B4 137057348 137057481
refGene_gn=ACO2;refGene_tx=XM_003989332.5
B4 137057958 137058080
refGene_gn=ACO2;refGene_tx=XM_003989332.5
B4 137058503 137058641
refGene_gn=ACO2;refGene_tx=XM_003989332.5
B4 137060316 137060370
refGene_gn=POLR3H;refGene_tx=XM_003989333.4
B4 137061238 137061440
refGene_gn=POLR3H;refGene_tx=XM_003989333.4
B4 137062924 137062988
refGene_gn=POLR3H;refGene_tx=XM_003989333.4
B4 137063420 137063507
refGene_gn=POLR3H;refGene_tx=XM_003989333.4
B4 137069386 137069483
refGene_gn=POLR3H;refGene_tx=XM_003989333.4
B4 137072072 137072183
refGene_gn=POLR3H;refGene_tx=XM_003989333.4
B4 137092308 137092484
refGene_gn=CSDC2;refGene_tx=XM_003989335.5;refGene_gn=CSDC2;refGene_tx
=XM_006934115.4
B4 137094133 137094256
refGene_gn=CSDC2;refGene_tx=XM_003989335.5;refGene_gn=CSDC2;refGene_tx
=XM_006934115.4
B4 137094857 137095020
refGene_gn=CSDC2;refGene_tx=XM_003989335.5;refGene_gn=CSDC2;refGene_tx
=XM_006934115.4
B4 137097516 137097639

refGene_gn=PMM1;refGene_tx=XM_003989336.5;refGene_gn=PMM1;refGene_tx=XM_011284245.2
B4	137098011	137098127
refGene_gn=PMM1;refGene_tx=XM_003989336.5;refGene_gn=PMM1;refGene_tx=XM_011284245.2
B4	137098923	137098999
refGene_gn=PMM1;refGene_tx=XM_003989336.5;refGene_gn=PMM1;refGene_tx=XM_011284245.2
B4	137102361	137102461
refGene_gn=PMM1;refGene_tx=XM_003989336.5;refGene_gn=PMM1;refGene_tx=XM_011284245.2
B4	137102701	137102793
refGene_gn=PMM1;refGene_tx=XM_003989336.5;refGene_gn=PMM1;refGene_tx=XM_011284245.2
B4	137102919	137102996
refGene_gn=PMM1;refGene_tx=XM_003989336.5;refGene_gn=PMM1;refGene_tx=XM_011284245.2
B4	137104146	137104264
refGene_gn=PMM1;refGene_tx=XM_003989336.5;refGene_gn=PMM1;refGene_tx=XM_011284245.2
B4	137107281	137107368
refGene_gn=PMM1;refGene_tx=XM_003989336.5
B4	137119320	137119414
refGene_gn=DESI1;refGene_tx=XM_003989337.5
B4	137121105	137121228
refGene_gn=DESI1;refGene_tx=XM_003989337.5
B4	137121918	137122028
refGene_gn=DESI1;refGene_tx=XM_003989337.5
B4	137124351	137124421
refGene_gn=DESI1;refGene_tx=XM_003989337.5
B4	137125527	137125549
refGene_gn=DESI1;refGene_tx=XM_003989337.5
B4	137134690	137134778
refGene_gn=DESI1;refGene_tx=XM_003989337.5
B4	137135917	137135993
refGene_gn=XRCC6;refGene_tx=XM_023257528.1;refGene_gn=XRCC6;refGene_tx=XM_023257529.1;refGene_gn=XRCC6;refGene_tx=XM_023257530.1;refGene_gn=XRCC6;refGene_tx=XM_023257532.1
B4	137140737	137140850
refGene_gn=XRCC6;refGene_tx=XM_023257528.1;refGene_gn=XRCC6;refGene_tx=XM_023257529.1;refGene_gn=XRCC6;refGene_tx=XM_023257530.1
B4	137142374	137142428
refGene_gn=XRCC6;refGene_tx=XM_023257531.1
B4	137145833	137145972
refGene_gn=XRCC6;refGene_tx=XM_023257528.1;refGene_gn=XRCC6;refGene_tx=XM_023257529.1;refGene_gn=XRCC6;refGene_tx=XM_023257530.1;refGene_gn=XRCC6;refGene_tx=XM_023257531.1;refGene_gn=XRCC6;refGene_tx=XM_023257532.1
B4	137146191	137146446
refGene_gn=XRCC6;refGene_tx=XM_023257528.1;refGene_gn=XRCC6;refGene_tx

=XM_023257529.1;refGene_gn=XRCC6;refGene_tx=XM_023257530.1;refGene_gn=XRCC6;refGene_tx=XM_023257531.1;refGene_gn=XRCC6;refGene_tx=XM_023257532.1;refGene_gn=XRCC6;refGene_tx=XM_023257533.1
B4 137147009 137147193
refGene_gn=XRCC6;refGene_tx=XM_023257528.1;refGene_gn=XRCC6;refGene_tx=XM_023257529.1;refGene_gn=XRCC6;refGene_tx=XM_023257530.1;refGene_gn=XRCC6;refGene_tx=XM_023257531.1;refGene_gn=XRCC6;refGene_tx=XM_023257532.1;refGene_gn=XRCC6;refGene_tx=XM_023257533.1
B4 137149774 137149961
refGene_gn=XRCC6;refGene_tx=XM_023257528.1;refGene_gn=XRCC6;refGene_tx=XM_023257529.1;refGene_gn=XRCC6;refGene_tx=XM_023257530.1;refGene_gn=XRCC6;refGene_tx=XM_023257531.1;refGene_gn=XRCC6;refGene_tx=XM_023257532.1;refGene_gn=XRCC6;refGene_tx=XM_023257533.1
B4 137151225 137151394
refGene_gn=XRCC6;refGene_tx=XM_023257528.1;refGene_gn=XRCC6;refGene_tx=XM_023257529.1;refGene_gn=XRCC6;refGene_tx=XM_023257530.1;refGene_gn=XRCC6;refGene_tx=XM_023257531.1;refGene_gn=XRCC6;refGene_tx=XM_023257532.1;refGene_gn=XRCC6;refGene_tx=XM_023257533.1
B4 137152107 137152269
refGene_gn=XRCC6;refGene_tx=XM_023257528.1;refGene_gn=XRCC6;refGene_tx=XM_023257529.1;refGene_gn=XRCC6;refGene_tx=XM_023257530.1;refGene_gn=XRCC6;refGene_tx=XM_023257531.1;refGene_gn=XRCC6;refGene_tx=XM_023257532.1;refGene_gn=XRCC6;refGene_tx=XM_023257533.1
B4 137154188 137154318
refGene_gn=XRCC6;refGene_tx=XM_023257528.1;refGene_gn=XRCC6;refGene_tx=XM_023257529.1;refGene_gn=XRCC6;refGene_tx=XM_023257530.1;refGene_gn=XRCC6;refGene_tx=XM_023257531.1;refGene_gn=XRCC6;refGene_tx=XM_023257532.1;refGene_gn=XRCC6;refGene_tx=XM_023257533.1
B4 137155224 137155325
refGene_gn=XRCC6;refGene_tx=XM_023257528.1;refGene_gn=XRCC6;refGene_tx=XM_023257529.1;refGene_gn=XRCC6;refGene_tx=XM_023257530.1;refGene_gn=XRCC6;refGene_tx=XM_023257531.1;refGene_gn=XRCC6;refGene_tx=XM_023257532.1;refGene_gn=XRCC6;refGene_tx=XM_023257533.1
B4 137159783 137159897
refGene_gn=XRCC6;refGene_tx=XM_023257528.1;refGene_gn=XRCC6;refGene_tx=XM_023257529.1;refGene_gn=XRCC6;refGene_tx=XM_023257530.1;refGene_gn=XRCC6;refGene_tx=XM_023257531.1;refGene_gn=XRCC6;refGene_tx=XM_023257532.1;refGene_gn=XRCC6;refGene_tx=XM_023257533.1
B4 137160998 137161195
refGene_gn=XRCC6;refGene_tx=XM_023257528.1;refGene_gn=XRCC6;refGene_tx=XM_023257529.1;refGene_gn=XRCC6;refGene_tx=XM_023257530.1;refGene_gn=XRCC6;refGene_tx=XM_023257531.1;refGene_gn=XRCC6;refGene_tx=XM_023257532.1;refGene_gn=XRCC6;refGene_tx=XM_023257533.1
B4 137165894 137166157
refGene_gn=SNU13;refGene_tx=XM_003989339.4;refGene_gn=SNU13;refGene_tx=XM_003989340.4
B4 137166948 137167069
refGene_gn=SNU13;refGene_tx=XM_003989339.4;refGene_gn=SNU13;refGene_tx=XM_003989340.4
B4 137168606 137168609

refGene_gn=SNU13;refGene_tx=XM_003989339.4
B4	137173994	137173997
refGene_gn=SNU13;refGene_tx=XM_003989340.4
B4	137178478	137178694
refGene_gn=CB4H22orf46;refGene_tx=XM_023257534.1
B4	137180941	137185498
refGene_gn=CB4H22orf46;refGene_tx=XM_023257534.1
B4	137189538	137189709
refGene_gn=MEI1;refGene_tx=XM_011284253.3;refGene_gn=MEI1;refGene_tx=XM_023257535.1;refGene_gn=MEI1;refGene_tx=XM_023257536.1;refGene_gn=MEI1;refGene_tx=XM_023257537.1
B4	137192430	137192554
refGene_gn=MEI1;refGene_tx=XM_011284253.3;refGene_gn=MEI1;refGene_tx=XM_023257535.1;refGene_gn=MEI1;refGene_tx=XM_023257536.1;refGene_gn=MEI1;refGene_tx=XM_023257537.1
B4	137193287	137193338
refGene_gn=MEI1;refGene_tx=XM_011284253.3;refGene_gn=MEI1;refGene_tx=XM_023257535.1;refGene_gn=MEI1;refGene_tx=XM_023257536.1;refGene_gn=MEI1;refGene_tx=XM_023257537.1
B4	137195187	137195261
refGene_gn=MEI1;refGene_tx=XM_011284253.3;refGene_gn=MEI1;refGene_tx=XM_023257535.1;refGene_gn=MEI1;refGene_tx=XM_023257536.1;refGene_gn=MEI1;refGene_tx=XM_023257537.1
B4	137197069	137197175
refGene_gn=MEI1;refGene_tx=XM_011284253.3;refGene_gn=MEI1;refGene_tx=XM_023257535.1;refGene_gn=MEI1;refGene_tx=XM_023257536.1;refGene_gn=MEI1;refGene_tx=XM_023257537.1
B4	137197708	137197912
refGene_gn=MEI1;refGene_tx=XM_011284253.3;refGene_gn=MEI1;refGene_tx=XM_023257535.1;refGene_gn=MEI1;refGene_tx=XM_023257536.1;refGene_gn=MEI1;refGene_tx=XM_023257537.1
B4	137199960	137200091
refGene_gn=MEI1;refGene_tx=XM_011284253.3;refGene_gn=MEI1;refGene_tx=XM_023257535.1;refGene_gn=MEI1;refGene_tx=XM_023257536.1;refGene_gn=MEI1;refGene_tx=XM_023257537.1
B4	137206172	137206287
refGene_gn=MEI1;refGene_tx=XM_011284253.3;refGene_gn=MEI1;refGene_tx=XM_023257535.1;refGene_gn=MEI1;refGene_tx=XM_023257536.1;refGene_gn=MEI1;refGene_tx=XM_023257537.1
B4	137207093	137207210
refGene_gn=MEI1;refGene_tx=XM_011284253.3;refGene_gn=MEI1;refGene_tx=XM_023257535.1;refGene_gn=MEI1;refGene_tx=XM_023257536.1;refGene_gn=MEI1;refGene_tx=XM_023257537.1
B4	137208619	137208719
refGene_gn=MEI1;refGene_tx=XM_011284253.3;refGene_gn=MEI1;refGene_tx=XM_023257535.1;refGene_gn=MEI1;refGene_tx=XM_023257536.1;refGene_gn=MEI1;refGene_tx=XM_023257537.1
B4	137208828	137208963
refGene_gn=MEI1;refGene_tx=XM_011284253.3;refGene_gn=MEI1;refGene_tx=XM_023257535.1;refGene_gn=MEI1;refGene_tx=XM_023257536.1;refGene_gn=MEI

1;refGene_tx=XM_023257537.1
B4	137214878	137214993
refGene_gn=MEI1;refGene_tx=XM_011284253.3;refGene_gn=MEI1;refGene_tx=XM_023257535.1;refGene_gn=MEI1;refGene_tx=XM_023257536.1;refGene_gn=MEI1;refGene_tx=XM_023257537.1
B4	137216002	137216094
refGene_gn=MEI1;refGene_tx=XM_011284253.3;refGene_gn=MEI1;refGene_tx=XM_023257535.1;refGene_gn=MEI1;refGene_tx=XM_023257537.1
B4	137216963	137217105
refGene_gn=MEI1;refGene_tx=XM_011284253.3;refGene_gn=MEI1;refGene_tx=XM_023257535.1;refGene_gn=MEI1;refGene_tx=XM_023257537.1
B4	137219003	137219115
refGene_gn=MEI1;refGene_tx=XM_011284253.3;refGene_gn=MEI1;refGene_tx=XM_023257535.1;refGene_gn=MEI1;refGene_tx=XM_023257536.1;refGene_gn=MEI1;refGene_tx=XM_023257537.1
B4	137222994	137223055
refGene_gn=MEI1;refGene_tx=XM_011284253.3;refGene_gn=MEI1;refGene_tx=XM_023257535.1;refGene_gn=MEI1;refGene_tx=XM_023257536.1;refGene_gn=MEI1;refGene_tx=XM_023257537.1
B4	137224010	137224108
refGene_gn=MEI1;refGene_tx=XM_011284253.3;refGene_gn=MEI1;refGene_tx=XM_023257535.1;refGene_gn=MEI1;refGene_tx=XM_023257536.1;refGene_gn=MEI1;refGene_tx=XM_023257537.1
B4	137227187	137227356
refGene_gn=MEI1;refGene_tx=XM_011284253.3;refGene_gn=MEI1;refGene_tx=XM_023257535.1;refGene_gn=MEI1;refGene_tx=XM_023257536.1;refGene_gn=MEI1;refGene_tx=XM_023257537.1
B4	137232714	137232862
refGene_gn=MEI1;refGene_tx=XM_011284253.3;refGene_gn=MEI1;refGene_tx=XM_023257535.1;refGene_gn=MEI1;refGene_tx=XM_023257536.1;refGene_gn=MEI1;refGene_tx=XM_023257537.1
B4	137241671	137241947
refGene_gn=MEI1;refGene_tx=XM_011284253.3;refGene_gn=MEI1;refGene_tx=XM_023257535.1;refGene_gn=MEI1;refGene_tx=XM_023257536.1
B4	137247431	137247597
refGene_gn=MEI1;refGene_tx=XM_011284253.3;refGene_gn=MEI1;refGene_tx=XM_023257535.1;refGene_gn=MEI1;refGene_tx=XM_023257536.1;refGene_gn=MEI1;refGene_tx=XM_023257537.1
B4	137248700	137248805
refGene_gn=MEI1;refGene_tx=XM_011284253.3;refGene_gn=MEI1;refGene_tx=XM_023257535.1;refGene_gn=MEI1;refGene_tx=XM_023257536.1;refGene_gn=MEI1;refGene_tx=XM_023257537.1
B4	137250553	137250664
refGene_gn=MEI1;refGene_tx=XM_011284253.3;refGene_gn=MEI1;refGene_tx=XM_023257535.1;refGene_gn=MEI1;refGene_tx=XM_023257536.1;refGene_gn=MEI1;refGene_tx=XM_023257537.1
B4	137250980	137251141
refGene_gn=MEI1;refGene_tx=XM_011284253.3;refGene_gn=MEI1;refGene_tx=XM_023257535.1;refGene_gn=MEI1;refGene_tx=XM_023257536.1;refGene_gn=MEI1;refGene_tx=XM_023257537.1

B4	137253661	137253743
refGene_gn=MEI1;refGene_tx=XM_011284253.3;refGene_gn=MEI1;refGene_tx=XM_023257535.1;refGene_gn=MEI1;refGene_tx=XM_023257536.1;refGene_gn=MEI1;refGene_tx=XM_023257537.1
B4	137253930	137254106
refGene_gn=MEI1;refGene_tx=XM_011284253.3;refGene_gn=MEI1;refGene_tx=XM_023257535.1;refGene_gn=MEI1;refGene_tx=XM_023257536.1;refGene_gn=MEI1;refGene_tx=XM_023257537.1
B4	137264734	137264816
refGene_gn=MEI1;refGene_tx=XM_011284253.3;refGene_gn=MEI1;refGene_tx=XM_023257535.1;refGene_gn=MEI1;refGene_tx=XM_023257536.1;refGene_gn=MEI1;refGene_tx=XM_023257537.1
B4	137265282	137265389
refGene_gn=MEI1;refGene_tx=XM_011284253.3;refGene_gn=MEI1;refGene_tx=XM_023257535.1;refGene_gn=MEI1;refGene_tx=XM_023257536.1;refGene_gn=MEI1;refGene_tx=XM_023257537.1
B4	137266691	137266823
refGene_gn=MEI1;refGene_tx=XM_011284253.3;refGene_gn=MEI1;refGene_tx=XM_023257535.1;refGene_gn=MEI1;refGene_tx=XM_023257536.1;refGene_gn=MEI1;refGene_tx=XM_023257537.1
B4	137267012	137267122
refGene_gn=MEI1;refGene_tx=XM_011284253.3;refGene_gn=MEI1;refGene_tx=XM_023257535.1;refGene_gn=MEI1;refGene_tx=XM_023257536.1;refGene_gn=MEI1;refGene_tx=XM_023257537.1
B4	137267951	137267997
refGene_gn=MEI1;refGene_tx=XM_011284253.3;refGene_gn=MEI1;refGene_tx=XM_023257535.1;refGene_gn=MEI1;refGene_tx=XM_023257536.1;refGene_gn=MEI1;refGene_tx=XM_023257537.1
B4	137271300	137271403
refGene_gn=CCDC134;refGene_tx=XM_006934121.4;refGene_gn=CCDC134;refGene_tx=XM_019835652.2
B4	137272276	137272398
refGene_gn=CCDC134;refGene_tx=XM_006934121.4;refGene_gn=CCDC134;refGene_tx=XM_019835652.2
B4	137272584	137272669
refGene_gn=CCDC134;refGene_tx=XM_006934121.4;refGene_gn=CCDC134;refGene_tx=XM_019835652.2
B4	137274933	137275115
refGene_gn=CCDC134;refGene_tx=XM_006934121.4;refGene_gn=CCDC134;refGene_tx=XM_019835652.2
B4	137275408	137275480
refGene_gn=CCDC134;refGene_tx=XM_006934121.4;refGene_gn=CCDC134;refGene_tx=XM_019835652.2
B4	137283645	137283771
refGene_gn=CCDC134;refGene_tx=XM_006934121.4;refGene_gn=CCDC134;refGene_tx=XM_019835652.2
B4	137288745	137289130
refGene_gn=SREBF2;refGene_tx=XM_011284256.3
B4	137312926	137313364
refGene_gn=SREBF2;refGene_tx=XM_003989343.5;refGene_gn=SREBF2;refGene_

tx=XM_011284256.3
B4	137314388	137314570
refGene_gn=SREBF2;refGene_tx=XM_003989343.5;refGene_gn=SREBF2;refGene_
tx=XM_011284256.3
B4	137315246	137315393
refGene_gn=SREBF2;refGene_tx=XM_003989343.5;refGene_gn=SREBF2;refGene_
tx=XM_011284256.3
B4	137316773	137316995
refGene_gn=SREBF2;refGene_tx=XM_003989343.5;refGene_gn=SREBF2;refGene_
tx=XM_011284256.3
B4	137318283	137318398
refGene_gn=SREBF2;refGene_tx=XM_003989343.5;refGene_gn=SREBF2;refGene_
tx=XM_011284256.3
B4	137318584	137318766
refGene_gn=SREBF2;refGene_tx=XM_003989343.5;refGene_gn=SREBF2;refGene_
tx=XM_011284256.3
B4	137320271	137320464
refGene_gn=SREBF2;refGene_tx=XM_003989343.5;refGene_gn=SREBF2;refGene_
tx=XM_011284256.3
B4	137321056	137321238
refGene_gn=SREBF2;refGene_tx=XM_003989343.5;refGene_gn=SREBF2;refGene_
tx=XM_011284256.3
B4	137324109	137324386
refGene_gn=SREBF2;refGene_tx=XM_003989343.5;refGene_gn=SREBF2;refGene_
tx=XM_011284256.3
B4	137326097	137326267
refGene_gn=SREBF2;refGene_tx=XM_003989343.5;refGene_gn=SREBF2;refGene_
tx=XM_011284256.3
B4	137334191	137334360
refGene_gn=SREBF2;refGene_tx=XM_003989343.5;refGene_gn=SREBF2;refGene_
tx=XM_011284256.3
B4	137335649	137335767
refGene_gn=SREBF2;refGene_tx=XM_003989343.5;refGene_gn=SREBF2;refGene_
tx=XM_011284256.3
B4	137336609	137336719
refGene_gn=SREBF2;refGene_tx=XM_003989343.5;refGene_gn=SREBF2;refGene_
tx=XM_011284256.3
B4	137337876	137338009
refGene_gn=SREBF2;refGene_tx=XM_003989343.5;refGene_gn=SREBF2;refGene_
tx=XM_011284256.3
B4	137340454	137340623
refGene_gn=SREBF2;refGene_tx=XM_003989343.5;refGene_gn=SREBF2;refGene_
tx=XM_011284256.3
B4	137342994	137343180
refGene_gn=SREBF2;refGene_tx=XM_003989343.5;refGene_gn=SREBF2;refGene_
tx=XM_011284256.3
B4	137344669	137344781
refGene_gn=SREBF2;refGene_tx=XM_003989343.5;refGene_gn=SREBF2;refGene_
tx=XM_011284256.3
B4	137345184	137345405

refGene_gn=SREBF2;refGene_tx=XM_003989343.5;refGene_gn=SREBF2;refGene_tx=XM_011284256.3
B4 137349494 137349889
refGene_gn=SHISA8;refGene_tx=XM_023257538.1;refGene_gn=SHISA8;refGene_tx=XM_023257539.1
B4 137350309 137350456
refGene_gn=SHISA8;refGene_tx=XM_023257538.1;refGene_gn=SHISA8;refGene_tx=XM_023257539.1
B4 137351323 137351529
refGene_gn=SHISA8;refGene_tx=XM_023257538.1;refGene_gn=SHISA8;refGene_tx=XM_023257539.1
B4 137354160 137354684
refGene_gn=SHISA8;refGene_tx=XM_023257538.1;refGene_gn=SHISA8;refGene_tx=XM_023257539.1
B4 137364198 137364380
refGene_gn=TNFRSF13C;refGene_tx=XM_023257540.1;refGene_gn=TNFRSF13C;refGene_tx=XM_023257541.1
B4 137364764 137365010
refGene_gn=TNFRSF13C;refGene_tx=XM_023257540.1;refGene_gn=TNFRSF13C;refGene_tx=XM_023257541.1
B4 137365361 137365506
refGene_gn=TNFRSF13C;refGene_tx=XM_023257540.1;refGene_gn=TNFRSF13C;refGene_tx=XM_023257541.1
B4 137367975 137368026
refGene_gn=CENPM;refGene_tx=XM_019835658.2
B4 137370575 137370704
refGene_gn=CENPM;refGene_tx=XM_019835657.2
B4 137374653 137374794
refGene_gn=CENPM;refGene_tx=XM_003989344.5;refGene_gn=CENPM;refGene_tx=XM_006934122.4;refGene_gn=CENPM;refGene_tx=XM_011284257.3
B4 137378890 137378982
refGene_gn=CENPM;refGene_tx=XM_003989344.5;refGene_gn=CENPM;refGene_tx=XM_006934122.4;refGene_gn=CENPM;refGene_tx=XM_011284257.3;refGene_gn=CENPM;refGene_tx=XM_019835657.2;refGene_gn=CENPM;refGene_tx=XM_019835658.2
B4 137379841 137379912
refGene_gn=CENPM;refGene_tx=XM_019835659.1;refGene_gn=CENPM;refGene_tx=XM_019835660.1
B4 137380179 137380259
refGene_gn=CENPM;refGene_tx=XM_003989344.5;refGene_gn=CENPM;refGene_tx=XM_006934122.4;refGene_gn=CENPM;refGene_tx=XM_011284257.3;refGene_gn=CENPM;refGene_tx=XM_019835657.2;refGene_gn=CENPM;refGene_tx=XM_019835658.2;refGene_gn=CENPM;refGene_tx=XM_019835659.1;refGene_gn=CENPM;refGene_tx=XM_019835660.1
B4 137382638 137382731
refGene_gn=CENPM;refGene_tx=XM_003989344.5;refGene_gn=CENPM;refGene_tx=XM_006934122.4;refGene_gn=CENPM;refGene_tx=XM_011284257.3;refGene_gn=CENPM;refGene_tx=XM_019835657.2;refGene_gn=CENPM;refGene_tx=XM_019835658.2;refGene_gn=CENPM;refGene_tx=XM_019835659.1;refGene_gn=CENPM;refGene_tx=XM_019835660.1

B4	137384028	137384108
refGene_gn=CENPM;refGene_tx=XM_003989344.5;refGene_gn=CENPM;refGene_tx=XM_006934122.4;refGene_gn=CENPM;refGene_tx=XM_011284257.3;refGene_gn=CENPM;refGene_tx=XM_019835657.2;refGene_gn=CENPM;refGene_tx=XM_019835658.2;refGene_gn=CENPM;refGene_tx=XM_019835659.1;refGene_gn=CENPM;refGene_tx=XM_019835660.1
B4	137384850	137384907
refGene_gn=CENPM;refGene_tx=XM_003989344.5;refGene_gn=CENPM;refGene_tx=XM_019835657.2;refGene_gn=CENPM;refGene_tx=XM_019835658.2;refGene_gn=CENPM;refGene_tx=XM_019835659.1;refGene_gn=CENPM;refGene_tx=XM_019835660.1
B4	137409894	137411385
refGene_gn=LOC109501929;refGene_tx=XM_019835893.2
B4	137413668	137413678
refGene_gn=LOC101083017;refGene_tx=XM_011284258.3;refGene_gn=LOC101083017;refGene_tx=XM_019835661.2
B4	137417920	137418112
refGene_gn=LOC101083017;refGene_tx=XM_011284258.3;refGene_gn=LOC101083017;refGene_tx=XM_019835661.2;refGene_gn=LOC101083017;refGene_tx=XM_023257542.1
B4	137421932	137422061
refGene_gn=LOC101083017;refGene_tx=XM_011284258.3;refGene_gn=LOC101083017;refGene_tx=XM_019835661.2;refGene_gn=LOC101083017;refGene_tx=XM_023257542.1
B4	137422482	137422564
refGene_gn=LOC101083017;refGene_tx=XM_011284258.3;refGene_gn=LOC101083017;refGene_tx=XM_019835661.2;refGene_gn=LOC101083017;refGene_tx=XM_023257542.1
B4	137423648	137423786
refGene_gn=LOC101083017;refGene_tx=XM_011284258.3;refGene_gn=LOC101083017;refGene_tx=XM_019835661.2;refGene_gn=LOC101083017;refGene_tx=XM_023257542.1
B4	137425172	137425290
refGene_gn=LOC101083017;refGene_tx=XM_011284258.3;refGene_gn=LOC101083017;refGene_tx=XM_019835661.2;refGene_gn=LOC101083017;refGene_tx=XM_023257542.1
B4	137426180	137426276
refGene_gn=LOC101083017;refGene_tx=XM_011284258.3;refGene_gn=LOC101083017;refGene_tx=XM_019835661.2;refGene_gn=LOC101083017;refGene_tx=XM_023257542.1
B4	137427878	137427978
refGene_gn=LOC101083017;refGene_tx=XM_011284258.3;refGene_gn=LOC101083017;refGene_tx=XM_019835661.2;refGene_gn=LOC101083017;refGene_tx=XM_023257542.1
B4	137429599	137429651
refGene_gn=LOC101083017;refGene_tx=XM_011284258.3;refGene_gn=LOC101083017;refGene_tx=XM_019835661.2;refGene_gn=LOC101083017;refGene_tx=XM_023257542.1
B4	137429941	137430035
refGene_gn=LOC101083017;refGene_tx=XM_011284258.3;refGene_gn=LOC101083

017;refGene_tx=XM_019835661.2;refGene_gn=LOC101083017;refGene_tx=XM_023257542.1
B4 137432190 137432256
refGene_gn=LOC101083017;refGene_tx=XM_011284258.3;refGene_gn=LOC101083017;refGene_tx=XM_019835661.2;refGene_gn=LOC101083017;refGene_tx=XM_023257542.1
B4 137437148 137437210
refGene_gn=WBP2NL;refGene_tx=XM_023257543.1;refGene_gn=WBP2NL;refGene_tx=XM_023257544.1;refGene_gn=WBP2NL;refGene_tx=XM_023257545.1;refGene_gn=WBP2NL;refGene_tx=XM_023257546.1
B4 137447771 137447880
refGene_gn=WBP2NL;refGene_tx=XM_023257543.1;refGene_gn=WBP2NL;refGene_tx=XM_023257544.1;refGene_gn=WBP2NL;refGene_tx=XM_023257545.1;refGene_gn=WBP2NL;refGene_tx=XM_023257546.1
B4 137448124 137448266
refGene_gn=WBP2NL;refGene_tx=XM_023257543.1;refGene_gn=WBP2NL;refGene_tx=XM_023257544.1;refGene_gn=WBP2NL;refGene_tx=XM_023257545.1;refGene_gn=WBP2NL;refGene_tx=XM_023257546.1
B4 137448457 137448550
refGene_gn=WBP2NL;refGene_tx=XM_023257543.1;refGene_gn=WBP2NL;refGene_tx=XM_023257544.1;refGene_gn=WBP2NL;refGene_tx=XM_023257545.1;refGene_gn=WBP2NL;refGene_tx=XM_023257546.1
B4 137453501 137453612
refGene_gn=WBP2NL;refGene_tx=XM_023257543.1;refGene_gn=WBP2NL;refGene_tx=XM_023257544.1;refGene_gn=WBP2NL;refGene_tx=XM_023257545.1;refGene_gn=WBP2NL;refGene_tx=XM_023257546.1
B4 137455930 137456370
refGene_gn=WBP2NL;refGene_tx=XM_023257543.1;refGene_gn=WBP2NL;refGene_tx=XM_023257544.1;refGene_gn=WBP2NL;refGene_tx=XM_023257545.1;refGene_gn=WBP2NL;refGene_tx=XM_023257546.1
B4 137492093 137492237
refGene_gn=NAGA;refGene_tx=XM_003989346.5;refGene_gn=NAGA;refGene_tx=XM_006934130.4
B4 137492489 137492633
refGene_gn=NAGA;refGene_tx=XM_003989346.5;refGene_gn=NAGA;refGene_tx=XM_006934130.4
B4 137493722 137493920
refGene_gn=NAGA;refGene_tx=XM_003989346.5;refGene_gn=NAGA;refGene_tx=XM_006934130.4
B4 137495244 137495406
refGene_gn=NAGA;refGene_tx=XM_003989346.5;refGene_gn=NAGA;refGene_tx=XM_006934130.4
B4 137496172 137496267
refGene_gn=NAGA;refGene_tx=XM_003989346.5;refGene_gn=NAGA;refGene_tx=XM_006934130.4
B4 137496613 137496791
refGene_gn=NAGA;refGene_tx=XM_003989346.5;refGene_gn=NAGA;refGene_tx=XM_006934130.4
B4 137497281 137497453
refGene_gn=NAGA;refGene_tx=XM_003989346.5;refGene_gn=NAGA;refGene_tx=X

M_006934130.4
B4	137497859	137497995
refGene_gn=NAGA;refGene_tx=XM_003989346.5;refGene_gn=NAGA;refGene_tx=X
M_006934130.4
B4	137498745	137498761
refGene_gn=NAGA;refGene_tx=XM_003989346.5;refGene_gn=NAGA;refGene_tx=X
M_006934130.4
B4	137503921	137504704
refGene_gn=FAM109B;refGene_tx=XM_006934133.4;refGene_gn=FAM109B;refGen
e_tx=XM_023257547.1;refGene_gn=FAM109B;refGene_tx=XM_023257548.1
B4	137506737	137506923
refGene_gn=SMDT1;refGene_tx=XM_003989348.5
B4	137508379	137508517
refGene_gn=SMDT1;refGene_tx=XM_003989348.5
B4	137511288	137511420
refGene_gn=NDUFA6;refGene_tx=XM_003989349.5
B4	137512410	137512526
refGene_gn=NDUFA6;refGene_tx=XM_003989349.5
B4	137520012	137520151
refGene_gn=NDUFA6;refGene_tx=XM_003989349.5
B4	137538511	137538624
refGene_gn=LOC105260774;refGene_tx=XM_023257549.1
B4	137540303	137540342
refGene_gn=LOC105260774;refGene_tx=XM_023257549.1
B4	137541131	137541175
refGene_gn=LOC105260774;refGene_tx=XM_023257549.1
B4	137541659	137541736
refGene_gn=LOC105260774;refGene_tx=XM_023257549.1
B4	137545318	137545384
refGene_gn=LOC105260774;refGene_tx=XM_023257549.1
B4	137545524	137545563
refGene_gn=LOC105260774;refGene_tx=XM_023257549.1
B4	137545810	137545984
refGene_gn=LOC105260774;refGene_tx=XM_023257549.1
B4	137546279	137546322
refGene_gn=LOC105260774;refGene_tx=XM_023257549.1
B4	137546653	137546775
refGene_gn=LOC105260774;refGene_tx=XM_023257549.1
B4	137548386	137548565
refGene_gn=CYP2D6;refGene_tx=NM_001195406.1;refGene_gn=CYP2D6;refGene_
tx=XM_019833852.1;refGene_gn=CYP2D6;refGene_tx=XM_023256247.1
B4	137548664	137548806
refGene_gn=CYP2D6;refGene_tx=NM_001195406.1;refGene_gn=CYP2D6;refGene_
tx=XM_019833852.1;refGene_gn=CYP2D6;refGene_tx=XM_023256247.1
B4	137549193	137549438
refGene_gn=CYP2D6;refGene_tx=NM_001195406.1;refGene_gn=CYP2D6;refGene_
tx=XM_019833852.1;refGene_gn=CYP2D6;refGene_tx=XM_023256247.1
B4	137549627	137549769
refGene_gn=CYP2D6;refGene_tx=NM_001195406.1;refGene_gn=CYP2D6;refGene_
tx=XM_019833852.1;refGene_gn=CYP2D6;refGene_tx=XM_023256247.1

B4 137549931 137550108
refGene_gn=CYP2D6;refGene_tx=NM_001195406.1;refGene_gn=CYP2D6;refGene_tx=XM_019833852.1;refGene_gn=CYP2D6;refGene_tx=XM_023256247.1
B4 137550555 137550716
refGene_gn=CYP2D6;refGene_tx=NM_001195406.1;refGene_gn=CYP2D6;refGene_tx=XM_019833852.1;refGene_gn=CYP2D6;refGene_tx=XM_023256247.1
B4 137550809 137550962
refGene_gn=CYP2D6;refGene_tx=NM_001195406.1;refGene_gn=CYP2D6;refGene_tx=XM_019833852.1;refGene_gn=CYP2D6;refGene_tx=XM_023256247.1
B4 137551544 137551716
refGene_gn=CYP2D6;refGene_tx=NM_001195406.1;refGene_gn=CYP2D6;refGene_tx=XM_019833852.1;refGene_gn=CYP2D6;refGene_tx=XM_023256247.1
B4 137552648 137552837
refGene_gn=CYP2D6;refGene_tx=NM_001195406.1;refGene_gn=CYP2D6;refGene_tx=XM_019833852.1;refGene_gn=CYP2D6;refGene_tx=XM_023256247.1
B4 137559131 137559149
refGene_gn=TCF20;refGene_tx=XM_023257551.1
B4 137564368 137564452
refGene_gn=TCF20;refGene_tx=XM_006934134.3;refGene_gn=TCF20;refGene_tx=XM_019835666.1;refGene_gn=TCF20;refGene_tx=XM_023257550.1
B4 137565590 137565640
refGene_gn=TCF20;refGene_tx=XM_006934134.3;refGene_gn=TCF20;refGene_tx=XM_019835666.1;refGene_gn=TCF20;refGene_tx=XM_023257550.1;refGene_gn=TCF20;refGene_tx=XM_023257551.1
B4 137573415 137573509
refGene_gn=TCF20;refGene_tx=XM_006934134.3;refGene_gn=TCF20;refGene_tx=XM_019835666.1;refGene_gn=TCF20;refGene_tx=XM_023257550.1;refGene_gn=TCF20;refGene_tx=XM_023257551.1
B4 137607110 137612807
refGene_gn=TCF20;refGene_tx=XM_006934134.3;refGene_gn=TCF20;refGene_tx=XM_019835666.1;refGene_gn=TCF20;refGene_tx=XM_023257550.1;refGene_gn=TCF20;refGene_tx=XM_023257551.1
B4 137669203 137669209
refGene_gn=TCF20;refGene_tx=XM_019835666.1;refGene_gn=TCF20;refGene_tx=XM_023257551.1
B4 137766458 137766484
refGene_gn=NFAM1;refGene_tx=XM_011284261.2;refGene_gn=NFAM1;refGene_tx=XM_019835668.2
B4 137768258 137768346
refGene_gn=NFAM1;refGene_tx=XM_011284261.2;refGene_gn=NFAM1;refGene_tx=XM_019835668.2
B4 137776123 137776207
refGene_gn=NFAM1;refGene_tx=XM_011284261.2;refGene_gn=NFAM1;refGene_tx=XM_019835668.2
B4 137782424 137782537
refGene_gn=NFAM1;refGene_tx=XM_011284261.2;refGene_gn=NFAM1;refGene_tx=XM_019835668.2
B4 137783740 137784070
refGene_gn=NFAM1;refGene_tx=XM_011284261.2;refGene_gn=NFAM1;refGene_tx=XM_019835668.2

B4 137786722 137786825
refGene_gn=NFAM1;refGene_tx=XM_019835668.2
B4 137797405 137797505
refGene_gn=NFAM1;refGene_tx=XM_011284261.2
B4 137892002 137892006
refGene_gn=SERHL2;refGene_tx=XM_006934140.3;refGene_gn=SERHL2;refGene_tx=XM_006934141.2
B4 137892951 137893101
refGene_gn=SERHL2;refGene_tx=XM_006934140.3;refGene_gn=SERHL2;refGene_tx=XM_006934141.2;refGene_gn=SERHL2;refGene_tx=XM_006934142.2;refGene_gn=SERHL2;refGene_tx=XM_019835671.1
B4 137893213 137893327
refGene_gn=SERHL2;refGene_tx=XM_006934140.3;refGene_gn=SERHL2;refGene_tx=XM_006934141.2;refGene_gn=SERHL2;refGene_tx=XM_006934142.2;refGene_gn=SERHL2;refGene_tx=XM_019835671.1
B4 137893711 137893753
refGene_gn=SERHL2;refGene_tx=XM_006934140.3;refGene_gn=SERHL2;refGene_tx=XM_006934141.2;refGene_gn=SERHL2;refGene_tx=XM_006934142.2;refGene_gn=SERHL2;refGene_tx=XM_019835671.1
B4 137894182 137894202
refGene_gn=SERHL2;refGene_tx=XM_006934140.3;refGene_gn=SERHL2;refGene_tx=XM_006934141.2;refGene_gn=SERHL2;refGene_tx=XM_006934142.2;refGene_gn=SERHL2;refGene_tx=XM_019835671.1
B4 137894663 137894738
refGene_gn=SERHL2;refGene_tx=XM_006934140.3;refGene_gn=SERHL2;refGene_tx=XM_006934141.2;refGene_gn=SERHL2;refGene_tx=XM_006934142.2;refGene_gn=SERHL2;refGene_tx=XM_019835671.1
B4 137896133 137896306
refGene_gn=SERHL2;refGene_tx=XM_006934140.3;refGene_gn=SERHL2;refGene_tx=XM_006934141.2;refGene_gn=SERHL2;refGene_tx=XM_006934142.2;refGene_gn=SERHL2;refGene_tx=XM_019835671.1
B4 137896967 137897047
refGene_gn=SERHL2;refGene_tx=XM_006934140.3;refGene_gn=SERHL2;refGene_tx=XM_006934141.2;refGene_gn=SERHL2;refGene_tx=XM_006934142.2;refGene_gn=SERHL2;refGene_tx=XM_019835671.1
B4 137903518 137903553
refGene_gn=SERHL2;refGene_tx=XM_006934140.3;refGene_gn=SERHL2;refGene_tx=XM_006934141.2;refGene_gn=SERHL2;refGene_tx=XM_006934142.2;refGene_gn=SERHL2;refGene_tx=XM_019835671.1
B4 137909904 137909987
refGene_gn=SERHL2;refGene_tx=XM_006934140.3;refGene_gn=SERHL2;refGene_tx=XM_006934141.2;refGene_gn=SERHL2;refGene_tx=XM_006934142.2;refGene_gn=SERHL2;refGene_tx=XM_019835671.1
B4 137911471 137911568
refGene_gn=SERHL2;refGene_tx=XM_006934140.3;refGene_gn=SERHL2;refGene_tx=XM_006934141.2;refGene_gn=SERHL2;refGene_tx=XM_006934142.2;refGene_gn=SERHL2;refGene_tx=XM_019835671.1
B4 137912385 137912499
refGene_gn=SERHL2;refGene_tx=XM_006934140.3;refGene_gn=SERHL2;refGene_tx=XM_006934141.2;refGene_gn=SERHL2;refGene_tx=XM_006934142.2;refGene_

gn=SERHL2;refGene_tx=XM_019835671.1
B4 137913225 137913311
refGene_gn=RRP7A;refGene_tx=XM_003989353.4;refGene_gn=RRP7A;refGene_tx=XM_006934138.3
B4 137914549 137914748
refGene_gn=RRP7A;refGene_tx=XM_003989353.4;refGene_gn=RRP7A;refGene_tx=XM_006934138.3
B4 137915115 137915213
refGene_gn=RRP7A;refGene_tx=XM_003989353.4;refGene_gn=RRP7A;refGene_tx=XM_006934138.3
B4 137915617 137915735
refGene_gn=RRP7A;refGene_tx=XM_003989353.4;refGene_gn=RRP7A;refGene_tx=XM_006934138.3
B4 137916402 137916528
refGene_gn=RRP7A;refGene_tx=XM_003989353.4;refGene_gn=RRP7A;refGene_tx=XM_006934138.3
B4 137918122 137918265
refGene_gn=RRP7A;refGene_tx=XM_003989353.4;refGene_gn=RRP7A;refGene_tx=XM_006934138.3
B4 137920240 137920313
refGene_gn=RRP7A;refGene_tx=XM_003989353.4;refGene_gn=RRP7A;refGene_tx=XM_006934138.3
B4 137930277 137930455
refGene_gn=POLDIP3;refGene_tx=XM_003989354.5;refGene_gn=POLDIP3;refGene_tx=XM_023257552.1
B4 137931416 137931483
refGene_gn=POLDIP3;refGene_tx=XM_003989354.5;refGene_gn=POLDIP3;refGene_tx=XM_023257552.1
B4 137933863 137933993
refGene_gn=POLDIP3;refGene_tx=XM_003989354.5;refGene_gn=POLDIP3;refGene_tx=XM_023257552.1
B4 137936553 137936631
refGene_gn=POLDIP3;refGene_tx=XM_003989354.5;refGene_gn=POLDIP3;refGene_tx=XM_023257552.1
B4 137937208 137937388
refGene_gn=POLDIP3;refGene_tx=XM_003989354.5;refGene_gn=POLDIP3;refGene_tx=XM_023257552.1
B4 137939073 137939169
refGene_gn=POLDIP3;refGene_tx=XM_003989354.5;refGene_gn=POLDIP3;refGene_tx=XM_023257552.1
B4 137940297 137940384
refGene_gn=POLDIP3;refGene_tx=XM_003989354.5
B4 137941323 137941714
refGene_gn=POLDIP3;refGene_tx=XM_003989354.5;refGene_gn=POLDIP3;refGene_tx=XM_023257552.1
B4 137949638 137949697
refGene_gn=POLDIP3;refGene_tx=XM_003989354.5;refGene_gn=POLDIP3;refGene_tx=XM_023257552.1
B4 137954796 137954969
refGene_gn=LOC101091382;refGene_tx=XM_023257553.1

B4 137962372 137962475
refGene_gn=LOC101091382;refGene_tx=XM_023257553.1
B4 137967199 137967282
refGene_gn=LOC101091382;refGene_tx=XM_023257553.1
B4 137967505 137967589
refGene_gn=LOC101091382;refGene_tx=XM_023257553.1
B4 137968083 137968213
refGene_gn=LOC101091382;refGene_tx=XM_023257553.1
B4 137970247 137970354
refGene_gn=LOC101091382;refGene_tx=XM_023257553.1
B4 137970741 137970814
refGene_gn=LOC101091382;refGene_tx=XM_023257553.1
B4 137974138 137974270
refGene_gn=LOC101091382;refGene_tx=XM_023257553.1
B4 137982181 137982202
refGene_gn=LOC101091382;refGene_tx=XM_023257553.1
B4 138027452 138028560
refGene_gn=A4GALT;refGene_tx=XM_011284316.3;refGene_gn=A4GALT;refGene_tx=XM_011284317.3;refGene_gn=A4GALT;refGene_tx=XM_011284319.3;refGene_gn=A4GALT;refGene_tx=XM_019835673.2;refGene_gn=A4GALT;refGene_tx=XM_023257554.1;refGene_gn=A4GALT;refGene_tx=XM_023257555.1;refGene_gn=A4GALT;refGene_tx=XM_023257556.1;refGene_gn=A4GALT;refGene_tx=XM_023257557.1;refGene_gn=A4GALT;refGene_tx=XM_023257558.1;refGene_gn=A4GALT;refGene_tx=XM_023257560.1
B4 138030250 138030307
refGene_gn=A4GALT;refGene_tx=XM_011284316.3
B4 138041456 138041625
refGene_gn=A4GALT;refGene_tx=XM_011284316.3
B4 138043607 138043731
refGene_gn=A4GALT;refGene_tx=XM_011284316.3
B4 138090834 138090852
refGene_gn=ARFGAP3;refGene_tx=XM_023257561.1;refGene_gn=ARFGAP3;refGene_tx=XM_023257562.1
B4 138092249 138092371
refGene_gn=ARFGAP3;refGene_tx=XM_023257561.1;refGene_gn=ARFGAP3;refGene_tx=XM_023257562.1
B4 138099183 138099274
refGene_gn=ARFGAP3;refGene_tx=XM_023257561.1;refGene_gn=ARFGAP3;refGene_tx=XM_023257562.1
B4 138100173 138100297
refGene_gn=ARFGAP3;refGene_tx=XM_023257561.1;refGene_gn=ARFGAP3;refGene_tx=XM_023257562.1
B4 138105877 138106009
refGene_gn=ARFGAP3;refGene_tx=XM_023257561.1;refGene_gn=ARFGAP3;refGene_tx=XM_023257562.1
B4 138112282 138112405
refGene_gn=ARFGAP3;refGene_tx=XM_023257561.1;refGene_gn=ARFGAP3;refGene_tx=XM_023257562.1
B4 138112836 138112965
refGene_gn=ARFGAP3;refGene_tx=XM_023257561.1;refGene_gn=ARFGAP3;refGen

e_tx=XM_023257562.1
B4	138116148	138116291
refGene_gn=ARFGAP3;refGene_tx=XM_023257561.1;refGene_gn=ARFGAP3;refGene_tx=XM_023257562.1
B4	138117258	138117305
refGene_gn=ARFGAP3;refGene_tx=XM_023257561.1;refGene_gn=ARFGAP3;refGene_tx=XM_023257562.1
B4	138119126	138119186
refGene_gn=ARFGAP3;refGene_tx=XM_023257561.1;refGene_gn=ARFGAP3;refGene_tx=XM_023257562.1
B4	138123225	138123313
refGene_gn=ARFGAP3;refGene_tx=XM_023257561.1;refGene_gn=ARFGAP3;refGene_tx=XM_023257562.1
B4	138128446	138128530
refGene_gn=ARFGAP3;refGene_tx=XM_023257561.1;refGene_gn=ARFGAP3;refGene_tx=XM_023257562.1
B4	138130362	138130494
refGene_gn=ARFGAP3;refGene_tx=XM_023257561.1;refGene_gn=ARFGAP3;refGene_tx=XM_023257562.1
B4	138131494	138131567
refGene_gn=ARFGAP3;refGene_tx=XM_023257561.1;refGene_gn=ARFGAP3;refGene_tx=XM_023257562.1
B4	138138203	138138322
refGene_gn=ARFGAP3;refGene_tx=XM_023257561.1
B4	138145305	138145374
refGene_gn=ARFGAP3;refGene_tx=XM_023257561.1
B4	138158266	138158379
refGene_gn=PACSIN2;refGene_tx=XM_019835675.2;refGene_gn=PACSIN2;refGene_tx=XM_019835676.2;refGene_gn=PACSIN2;refGene_tx=XM_019835678.2;refGene_gn=PACSIN2;refGene_tx=XM_019835679.2;refGene_gn=PACSIN2;refGene_tx=XM_023257563.1
B4	138161262	138161459
refGene_gn=PACSIN2;refGene_tx=XM_019835675.2;refGene_gn=PACSIN2;refGene_tx=XM_019835676.2;refGene_gn=PACSIN2;refGene_tx=XM_019835678.2;refGene_gn=PACSIN2;refGene_tx=XM_019835679.2;refGene_gn=PACSIN2;refGene_tx=XM_023257563.1
B4	138161870	138161993
refGene_gn=PACSIN2;refGene_tx=XM_019835675.2;refGene_gn=PACSIN2;refGene_tx=XM_019835676.2;refGene_gn=PACSIN2;refGene_tx=XM_023257563.1
B4	138164275	138164397
refGene_gn=PACSIN2;refGene_tx=XM_019835675.2;refGene_gn=PACSIN2;refGene_tx=XM_019835676.2;refGene_gn=PACSIN2;refGene_tx=XM_019835678.2;refGene_gn=PACSIN2;refGene_tx=XM_019835679.2;refGene_gn=PACSIN2;refGene_tx=XM_023257563.1
B4	138165658	138165664
refGene_gn=PACSIN2;refGene_tx=XM_019835675.2;refGene_gn=PACSIN2;refGene_tx=XM_019835678.2;refGene_gn=PACSIN2;refGene_tx=XM_023257563.1
B4	138167119	138167240
refGene_gn=PACSIN2;refGene_tx=XM_019835675.2;refGene_gn=PACSIN2;refGene_tx=XM_019835676.2;refGene_gn=PACSIN2;refGene_tx=XM_019835678.2;refGe

ne_gn=PACSIN2;refGene_tx=XM_019835679.2;refGene_gn=PACSIN2;refGene_tx=
XM_023257563.1
B4	138168786	138168962
refGene_gn=PACSIN2;refGene_tx=XM_019835675.2;refGene_gn=PACSIN2;refGen
e_tx=XM_019835676.2;refGene_gn=PACSIN2;refGene_tx=XM_019835678.2;refGe
ne_gn=PACSIN2;refGene_tx=XM_019835679.2;refGene_gn=PACSIN2;refGene_tx=
XM_023257563.1
B4	138173294	138173450
refGene_gn=PACSIN2;refGene_tx=XM_019835675.2;refGene_gn=PACSIN2;refGen
e_tx=XM_019835676.2;refGene_gn=PACSIN2;refGene_tx=XM_019835678.2;refGe
ne_gn=PACSIN2;refGene_tx=XM_019835679.2;refGene_gn=PACSIN2;refGene_tx=
XM_023257563.1
B4	138174915	138175151
refGene_gn=PACSIN2;refGene_tx=XM_019835675.2;refGene_gn=PACSIN2;refGen
e_tx=XM_019835676.2;refGene_gn=PACSIN2;refGene_tx=XM_019835678.2;refGe
ne_gn=PACSIN2;refGene_tx=XM_019835679.2;refGene_gn=PACSIN2;refGene_tx=
XM_023257563.1
B4	138177163	138177320
refGene_gn=PACSIN2;refGene_tx=XM_019835675.2;refGene_gn=PACSIN2;refGen
e_tx=XM_019835676.2;refGene_gn=PACSIN2;refGene_tx=XM_019835678.2;refGe
ne_gn=PACSIN2;refGene_tx=XM_019835679.2;refGene_gn=PACSIN2;refGene_tx=
XM_023257563.1
B4	138195994	138196054
refGene_gn=PACSIN2;refGene_tx=XM_023257563.1
B4	138316069	138316199
refGene_gn=TTLL1;refGene_tx=XM_023257564.1
B4	138320223	138320387
refGene_gn=TTLL1;refGene_tx=XM_023257564.1
B4	138322062	138322149
refGene_gn=TTLL1;refGene_tx=XM_023257564.1
B4	138324127	138324202
refGene_gn=TTLL1;refGene_tx=XM_023257564.1
B4	138324203	138324272
refGene_gn=TTLL1;refGene_tx=XM_023257564.1
B4	138329172	138329281
refGene_gn=TTLL1;refGene_tx=XM_023257564.1
B4	138329863	138329998
refGene_gn=TTLL1;refGene_tx=XM_023257564.1
B4	138330625	138330806
refGene_gn=TTLL1;refGene_tx=XM_023257564.1
B4	138331471	138331680
refGene_gn=TTLL1;refGene_tx=XM_023257564.1
B4	138337262	138337375
refGene_gn=TTLL1;refGene_tx=XM_023257564.1
B4	138346293	138346670
refGene_gn=LOC109501832;refGene_tx=XM_023257783.1
B4	138347087	138347127
refGene_gn=LOC109501832;refGene_tx=XM_023257783.1
B4	138376776	138377060
refGene_gn=BIK;refGene_tx=XM_003989379.5

B4 138385372 138385513
refGene_gn=BIK;refGene_tx=XM_003989379.5;refGene_gn=BIK;refGene_tx=XM_023257565.1;refGene_gn=BIK;refGene_tx=XM_023257566.1;refGene_gn=BIK;refGene_tx=XM_023257567.1
B4 138386912 138387020
refGene_gn=BIK;refGene_tx=XM_003989379.5;refGene_gn=BIK;refGene_tx=XM_023257565.1;refGene_gn=BIK;refGene_tx=XM_023257566.1;refGene_gn=BIK;refGene_tx=XM_023257567.1
B4 138387744 138387862
refGene_gn=BIK;refGene_tx=XM_003989379.5;refGene_gn=BIK;refGene_tx=XM_023257565.1;refGene_gn=BIK;refGene_tx=XM_023257566.1;refGene_gn=BIK;refGene_tx=XM_023257567.1
B4 138388490 138388577
refGene_gn=BIK;refGene_tx=XM_003989379.5;refGene_gn=BIK;refGene_tx=XM_023257565.1;refGene_gn=BIK;refGene_tx=XM_023257566.1;refGene_gn=BIK;refGene_tx=XM_023257567.1
B4 138393716 138394139
refGene_gn=MCAT;refGene_tx=XM_003989380.5;refGene_gn=MCAT;refGene_tx=XM_006934147.4
B4 138395398 138395440
refGene_gn=MCAT;refGene_tx=XM_019835692.2
B4 138397049 138397267
refGene_gn=MCAT;refGene_tx=XM_003989380.5;refGene_gn=MCAT;refGene_tx=XM_019835692.2
B4 138399760 138399848
refGene_gn=MCAT;refGene_tx=XM_003989380.5;refGene_gn=MCAT;refGene_tx=XM_006934147.4;refGene_gn=MCAT;refGene_tx=XM_019835692.2
B4 138401188 138401611
refGene_gn=MCAT;refGene_tx=XM_003989380.5;refGene_gn=MCAT;refGene_tx=XM_006934147.4;refGene_gn=MCAT;refGene_tx=XM_019835692.2
B4 138417144 138417326
refGene_gn=TSPO;refGene_tx=XM_023257568.1
B4 138418868 138419007
refGene_gn=TSPO;refGene_tx=XM_023257568.1
B4 138419506 138419695
refGene_gn=TSPO;refGene_tx=XM_023257568.1
B4 138424336 138424488
refGene_gn=TTLL12;refGene_tx=XM_023257569.1
B4 138424858 138424997
refGene_gn=TTLL12;refGene_tx=XM_023257569.1
B4 138425557 138425626
refGene_gn=TTLL12;refGene_tx=XM_023257569.1
B4 138427366 138427448
refGene_gn=TTLL12;refGene_tx=XM_023257569.1
B4 138427845 138427997
refGene_gn=TTLL12;refGene_tx=XM_023257569.1
B4 138428848 138428960
refGene_gn=TTLL12;refGene_tx=XM_023257569.1
B4 138429266 138429461
refGene_gn=TTLL12;refGene_tx=XM_023257569.1

B4	138429542	138429659
refGene_gn=TTLL12;refGene_tx=XM_023257569.1
B4	138430679	138430756
refGene_gn=TTLL12;refGene_tx=XM_023257569.1
B4	138432851	138432985
refGene_gn=TTLL12;refGene_tx=XM_023257569.1
B4	138433071	138433231
refGene_gn=TTLL12;refGene_tx=XM_023257569.1
B4	138434019	138434218
refGene_gn=TTLL12;refGene_tx=XM_023257569.1
B4	138436175	138436348
refGene_gn=TTLL12;refGene_tx=XM_023257569.1
B4	138439841	138440018
refGene_gn=TTLL12;refGene_tx=XM_023257569.1
B4	138453287	138453440
refGene_gn=SCUBE1;refGene_tx=XM_023257570.1;refGene_gn=SCUBE1;refGene_tx=XM_023257571.1;refGene_gn=SCUBE1;refGene_tx=XM_023257572.1
B4	138455471	138455551
refGene_gn=SCUBE1;refGene_tx=XM_023257570.1;refGene_gn=SCUBE1;refGene_tx=XM_023257571.1;refGene_gn=SCUBE1;refGene_tx=XM_023257572.1
B4	138455900	138456053
refGene_gn=SCUBE1;refGene_tx=XM_023257570.1;refGene_gn=SCUBE1;refGene_tx=XM_023257571.1;refGene_gn=SCUBE1;refGene_tx=XM_023257572.1
B4	138457454	138457652
refGene_gn=SCUBE1;refGene_tx=XM_023257570.1;refGene_gn=SCUBE1;refGene_tx=XM_023257571.1;refGene_gn=SCUBE1;refGene_tx=XM_023257572.1
B4	138459411	138459573
refGene_gn=SCUBE1;refGene_tx=XM_023257570.1;refGene_gn=SCUBE1;refGene_tx=XM_023257571.1;refGene_gn=SCUBE1;refGene_tx=XM_023257572.1
B4	138461170	138461338
refGene_gn=SCUBE1;refGene_tx=XM_023257570.1;refGene_gn=SCUBE1;refGene_tx=XM_023257571.1;refGene_gn=SCUBE1;refGene_tx=XM_023257572.1
B4	138462670	138462832
refGene_gn=SCUBE1;refGene_tx=XM_023257570.1;refGene_gn=SCUBE1;refGene_tx=XM_023257571.1;refGene_gn=SCUBE1;refGene_tx=XM_023257572.1
B4	138466565	138466769
refGene_gn=SCUBE1;refGene_tx=XM_023257570.1;refGene_gn=SCUBE1;refGene_tx=XM_023257571.1;refGene_gn=SCUBE1;refGene_tx=XM_023257572.1
B4	138467667	138467805
refGene_gn=SCUBE1;refGene_tx=XM_023257570.1;refGene_gn=SCUBE1;refGene_tx=XM_023257571.1;refGene_gn=SCUBE1;refGene_tx=XM_023257572.1
B4	138468634	138468748
refGene_gn=SCUBE1;refGene_tx=XM_023257570.1;refGene_gn=SCUBE1;refGene_tx=XM_023257571.1;refGene_gn=SCUBE1;refGene_tx=XM_023257572.1
B4	138469679	138469784
refGene_gn=SCUBE1;refGene_tx=XM_023257570.1;refGene_gn=SCUBE1;refGene_tx=XM_023257571.1;refGene_gn=SCUBE1;refGene_tx=XM_023257572.1
B4	138470112	138470232
refGene_gn=SCUBE1;refGene_tx=XM_023257570.1;refGene_gn=SCUBE1;refGene_tx=XM_023257571.1;refGene_gn=SCUBE1;refGene_tx=XM_023257572.1

B4 138478489 138478612
refGene_gn=SCUBE1;refGene_tx=XM_023257570.1;refGene_gn=SCUBE1;refGene_tx=XM_023257571.1;refGene_gn=SCUBE1;refGene_tx=XM_023257572.1
B4 138480111 138480228
refGene_gn=SCUBE1;refGene_tx=XM_023257570.1;refGene_gn=SCUBE1;refGene_tx=XM_023257571.1;refGene_gn=SCUBE1;refGene_tx=XM_023257572.1
B4 138482673 138482796
refGene_gn=SCUBE1;refGene_tx=XM_023257570.1;refGene_gn=SCUBE1;refGene_tx=XM_023257571.1;refGene_gn=SCUBE1;refGene_tx=XM_023257572.1
B4 138489180 138489297
refGene_gn=SCUBE1;refGene_tx=XM_023257570.1;refGene_gn=SCUBE1;refGene_tx=XM_023257571.1;refGene_gn=SCUBE1;refGene_tx=XM_023257572.1
B4 138502335 138502425
refGene_gn=SCUBE1;refGene_tx=XM_023257570.1;refGene_gn=SCUBE1;refGene_tx=XM_023257572.1
B4 138505094 138505211
refGene_gn=SCUBE1;refGene_tx=XM_023257570.1;refGene_gn=SCUBE1;refGene_tx=XM_023257571.1;refGene_gn=SCUBE1;refGene_tx=XM_023257572.1
B4 138509335 138509461
refGene_gn=SCUBE1;refGene_tx=XM_023257570.1;refGene_gn=SCUBE1;refGene_tx=XM_023257571.1;refGene_gn=SCUBE1;refGene_tx=XM_023257572.1
B4 138534536 138534671
refGene_gn=SCUBE1;refGene_tx=XM_023257570.1;refGene_gn=SCUBE1;refGene_tx=XM_023257571.1;refGene_gn=SCUBE1;refGene_tx=XM_023257572.1
B4 138563635 138563764
refGene_gn=SCUBE1;refGene_tx=XM_023257570.1;refGene_gn=SCUBE1;refGene_tx=XM_023257571.1;refGene_gn=SCUBE1;refGene_tx=XM_023257572.1
B4 138580971 138581103
refGene_gn=SCUBE1;refGene_tx=XM_023257570.1;refGene_gn=SCUBE1;refGene_tx=XM_023257571.1
B4 138584898 138584986
refGene_gn=SCUBE1;refGene_tx=XM_023257570.1;refGene_gn=SCUBE1;refGene_tx=XM_023257571.1
B4 138661787 138662011
refGene_gn=MPPED1;refGene_tx=XM_023257573.1
B4 138669789 138669971
refGene_gn=MPPED1;refGene_tx=XM_023257573.1
B4 138699540 138699766
refGene_gn=MPPED1;refGene_tx=XM_023257573.1
B4 138717989 138718105
refGene_gn=MPPED1;refGene_tx=XM_023257573.1
B4 138719901 138720015
refGene_gn=MPPED1;refGene_tx=XM_023257573.1
B4 138721196 138721315
refGene_gn=MPPED1;refGene_tx=XM_023257573.1
B4 138735882 138736005
refGene_gn=EFCAB6;refGene_tx=XM_023257574.1;refGene_gn=EFCAB6;refGene_tx=XM_023257575.1;refGene_gn=EFCAB6;refGene_tx=XM_023257576.1
B4 138738175 138738325
refGene_gn=EFCAB6;refGene_tx=XM_023257574.1;refGene_gn=EFCAB6;refGene_

tx=XM_023257575.1;refGene_gn=EFCAB6;refGene_tx=XM_023257576.1
B4 138742128 138742313
refGene_gn=EFCAB6;refGene_tx=XM_023257574.1;refGene_gn=EFCAB6;refGene_tx=XM_023257575.1;refGene_gn=EFCAB6;refGene_tx=XM_023257576.1
B4 138744096 138744262
refGene_gn=EFCAB6;refGene_tx=XM_023257574.1;refGene_gn=EFCAB6;refGene_tx=XM_023257575.1;refGene_gn=EFCAB6;refGene_tx=XM_023257576.1
B4 138746806 138747040
refGene_gn=EFCAB6;refGene_tx=XM_023257574.1;refGene_gn=EFCAB6;refGene_tx=XM_023257575.1;refGene_gn=EFCAB6;refGene_tx=XM_023257576.1
B4 138756266 138756494
refGene_gn=EFCAB6;refGene_tx=XM_023257574.1;refGene_gn=EFCAB6;refGene_tx=XM_023257575.1;refGene_gn=EFCAB6;refGene_tx=XM_023257576.1
B4 138777308 138777500
refGene_gn=EFCAB6;refGene_tx=XM_023257574.1;refGene_gn=EFCAB6;refGene_tx=XM_023257575.1;refGene_gn=EFCAB6;refGene_tx=XM_023257576.1
B4 138780828 138781021
refGene_gn=EFCAB6;refGene_tx=XM_023257574.1;refGene_gn=EFCAB6;refGene_tx=XM_023257575.1;refGene_gn=EFCAB6;refGene_tx=XM_023257576.1
B4 138787577 138787733
refGene_gn=EFCAB6;refGene_tx=XM_023257574.1;refGene_gn=EFCAB6;refGene_tx=XM_023257575.1;refGene_gn=EFCAB6;refGene_tx=XM_023257576.1
B4 138792952 138793147
refGene_gn=EFCAB6;refGene_tx=XM_023257574.1;refGene_gn=EFCAB6;refGene_tx=XM_023257576.1
B4 138802000 138802119
refGene_gn=EFCAB6;refGene_tx=XM_023257574.1;refGene_gn=EFCAB6;refGene_tx=XM_023257575.1;refGene_gn=EFCAB6;refGene_tx=XM_023257576.1
B4 138807806 138807903
refGene_gn=EFCAB6;refGene_tx=XM_023257574.1;refGene_gn=EFCAB6;refGene_tx=XM_023257575.1;refGene_gn=EFCAB6;refGene_tx=XM_023257576.1
B4 138818519 138818752
refGene_gn=EFCAB6;refGene_tx=XM_023257574.1;refGene_gn=EFCAB6;refGene_tx=XM_023257575.1;refGene_gn=EFCAB6;refGene_tx=XM_023257576.1
B4 138823437 138823571
refGene_gn=EFCAB6;refGene_tx=XM_023257574.1;refGene_gn=EFCAB6;refGene_tx=XM_023257575.1;refGene_gn=EFCAB6;refGene_tx=XM_023257576.1
B4 138825768 138825883
refGene_gn=EFCAB6;refGene_tx=XM_023257574.1;refGene_gn=EFCAB6;refGene_tx=XM_023257575.1;refGene_gn=EFCAB6;refGene_tx=XM_023257576.1
B4 138847642 138847811
refGene_gn=EFCAB6;refGene_tx=XM_023257574.1;refGene_gn=EFCAB6;refGene_tx=XM_023257575.1
B4 138849405 138849579
refGene_gn=EFCAB6;refGene_tx=XM_023257574.1;refGene_gn=EFCAB6;refGene_tx=XM_023257575.1
B4 138854845 138855006
refGene_gn=EFCAB6;refGene_tx=XM_023257574.1;refGene_gn=EFCAB6;refGene_tx=XM_023257575.1
B4 138858153 138858321

refGene_gn=EFCAB6;refGene_tx=XM_023257574.1;refGene_gn=EFCAB6;refGene_tx=XM_023257575.1;refGene_gn=EFCAB6;refGene_tx=XM_023257576.1
B4 138864679 138864785
refGene_gn=EFCAB6;refGene_tx=XM_023257574.1;refGene_gn=EFCAB6;refGene_tx=XM_023257575.1;refGene_gn=EFCAB6;refGene_tx=XM_023257576.1
B4 138869488 138869599
refGene_gn=EFCAB6;refGene_tx=XM_023257574.1;refGene_gn=EFCAB6;refGene_tx=XM_023257575.1;refGene_gn=EFCAB6;refGene_tx=XM_023257576.1
B4 138890197 138890346
refGene_gn=EFCAB6;refGene_tx=XM_023257574.1;refGene_gn=EFCAB6;refGene_tx=XM_023257575.1;refGene_gn=EFCAB6;refGene_tx=XM_023257576.1
B4 138898515 138898640
refGene_gn=EFCAB6;refGene_tx=XM_023257574.1;refGene_gn=EFCAB6;refGene_tx=XM_023257575.1;refGene_gn=EFCAB6;refGene_tx=XM_023257576.1
B4 138908685 138908798
refGene_gn=EFCAB6;refGene_tx=XM_023257574.1;refGene_gn=EFCAB6;refGene_tx=XM_023257575.1;refGene_gn=EFCAB6;refGene_tx=XM_023257576.1
B4 138912723 138912860
refGene_gn=EFCAB6;refGene_tx=XM_023257574.1;refGene_gn=EFCAB6;refGene_tx=XM_023257575.1;refGene_gn=EFCAB6;refGene_tx=XM_023257576.1
B4 138931049 138931116
refGene_gn=EFCAB6;refGene_tx=XM_023257574.1;refGene_gn=EFCAB6;refGene_tx=XM_023257575.1;refGene_gn=EFCAB6;refGene_tx=XM_023257576.1
B4 138936741 138936830
refGene_gn=EFCAB6;refGene_tx=XM_023257574.1;refGene_gn=EFCAB6;refGene_tx=XM_023257575.1;refGene_gn=EFCAB6;refGene_tx=XM_023257576.1
B4 138948752 138948964
refGene_gn=EFCAB6;refGene_tx=XM_023257574.1;refGene_gn=EFCAB6;refGene_tx=XM_023257575.1;refGene_gn=EFCAB6;refGene_tx=XM_023257576.1
B4 138957519 138957665
refGene_gn=EFCAB6;refGene_tx=XM_023257574.1;refGene_gn=EFCAB6;refGene_tx=XM_023257575.1;refGene_gn=EFCAB6;refGene_tx=XM_023257576.1
B4 138975048 138975119
refGene_gn=EFCAB6;refGene_tx=XM_023257574.1;refGene_gn=EFCAB6;refGene_tx=XM_023257575.1;refGene_gn=EFCAB6;refGene_tx=XM_023257576.1
B4 138997365 138997478
refGene_gn=SULT4A1;refGene_tx=XM_023257577.1
B4 139000163 139000302
refGene_gn=SULT4A1;refGene_tx=XM_023257577.1
B4 139006254 139006349
refGene_gn=SULT4A1;refGene_tx=XM_023257577.1
B4 139010656 139010783
refGene_gn=SULT4A1;refGene_tx=XM_023257577.1
B4 139011290 139011371
refGene_gn=SULT4A1;refGene_tx=XM_023257577.1
B4 139012519 139012650
refGene_gn=SULT4A1;refGene_tx=XM_023257577.1
B4 139029370 139029539
refGene_gn=SULT4A1;refGene_tx=XM_023257577.1
B4 139039924 139040009

refGene_gn=PNPLA5;refGene_tx=XM_011284348.3;refGene_gn=PNPLA5;refGene_tx=XM_019835695.2;refGene_gn=PNPLA5;refGene_tx=XM_019835697.2
B4 139040631 139040748
refGene_gn=PNPLA5;refGene_tx=XM_011284348.3;refGene_gn=PNPLA5;refGene_tx=XM_019835695.2;refGene_gn=PNPLA5;refGene_tx=XM_019835697.2
B4 139043119 139043252
refGene_gn=PNPLA5;refGene_tx=XM_011284348.3;refGene_gn=PNPLA5;refGene_tx=XM_019835695.2;refGene_gn=PNPLA5;refGene_tx=XM_019835697.2
B4 139045295 139045481
refGene_gn=PNPLA5;refGene_tx=XM_011284348.3;refGene_gn=PNPLA5;refGene_tx=XM_019835695.2;refGene_gn=PNPLA5;refGene_tx=XM_019835697.2
B4 139046781 139046842
refGene_gn=PNPLA5;refGene_tx=XM_011284348.3;refGene_gn=PNPLA5;refGene_tx=XM_019835695.2;refGene_gn=PNPLA5;refGene_tx=XM_019835697.2
B4 139048289 139048499
refGene_gn=PNPLA5;refGene_tx=XM_011284348.3;refGene_gn=PNPLA5;refGene_tx=XM_019835695.2;refGene_gn=PNPLA5;refGene_tx=XM_019835697.2
B4 139048806 139048872
refGene_gn=PNPLA5;refGene_tx=XM_011284348.3;refGene_gn=PNPLA5;refGene_tx=XM_019835695.2;refGene_gn=PNPLA5;refGene_tx=XM_019835697.2
B4 139050169 139050258
refGene_gn=PNPLA5;refGene_tx=XM_019835695.2
B4 139051130 139051363
refGene_gn=PNPLA5;refGene_tx=XM_011284348.3;refGene_gn=PNPLA5;refGene_tx=XM_019835695.2;refGene_gn=PNPLA5;refGene_tx=XM_019835697.2
B4 139051514 139051692
refGene_gn=PNPLA5;refGene_tx=XM_019835695.2;refGene_gn=PNPLA5;refGene_tx=XM_019835697.2
B4 139051747 139051940
refGene_gn=PNPLA5;refGene_tx=XM_011284348.3
B4 139078791 139078978
refGene_gn=PNPLA3;refGene_tx=XM_006934165.4
B4 139081745 139081978
refGene_gn=PNPLA3;refGene_tx=XM_006934165.4
B4 139083584 139083650
refGene_gn=PNPLA3;refGene_tx=XM_006934165.4
B4 139086180 139086390
refGene_gn=PNPLA3;refGene_tx=XM_006934165.4
B4 139087267 139087328
refGene_gn=PNPLA3;refGene_tx=XM_006934165.4
B4 139089664 139089880
refGene_gn=PNPLA3;refGene_tx=XM_006934165.4
B4 139091513 139091646
refGene_gn=PNPLA3;refGene_tx=XM_006934165.4
B4 139092618 139092724
refGene_gn=PNPLA3;refGene_tx=XM_006934165.4
B4 139093137 139093212
refGene_gn=PNPLA3;refGene_tx=XM_006934165.4
B4 139098395 139098416
refGene_gn=SAMM50;refGene_tx=XM_003989363.4

B4 139102373 139102484
refGene_gn=SAMM50;refGene_tx=XM_003989363.4
B4 139105601 139105703
refGene_gn=SAMM50;refGene_tx=XM_003989363.4
B4 139107411 139107499
refGene_gn=SAMM50;refGene_tx=XM_003989363.4
B4 139109357 139109464
refGene_gn=SAMM50;refGene_tx=XM_003989363.4
B4 139110317 139110448
refGene_gn=SAMM50;refGene_tx=XM_003989363.4
B4 139110688 139110776
refGene_gn=SAMM50;refGene_tx=XM_003989363.4
B4 139113018 139113147
refGene_gn=SAMM50;refGene_tx=XM_003989363.4
B4 139113574 139113646
refGene_gn=SAMM50;refGene_tx=XM_003989363.4
B4 139114346 139114433
refGene_gn=SAMM50;refGene_tx=XM_003989363.4
B4 139116882 139116953
refGene_gn=SAMM50;refGene_tx=XM_003989363.4
B4 139119432 139119500
refGene_gn=SAMM50;refGene_tx=XM_003989363.4
B4 139123977 139124124
refGene_gn=SAMM50;refGene_tx=XM_003989363.4
B4 139124796 139124938
refGene_gn=SAMM50;refGene_tx=XM_003989363.4
B4 139131714 139131760
refGene_gn=SAMM50;refGene_tx=XM_003989363.4
B4 139152541 139152656
refGene_gn=PARVB;refGene_tx=XM_023257578.1;refGene_gn=PARVB;refGene_tx
=XM_023257579.1
B4 139179136 139179137
refGene_gn=PARVB;refGene_tx=XM_003989364.5
B4 139202065 139202155
refGene_gn=PARVB;refGene_tx=XM_003989364.5;refGene_gn=PARVB;refGene_tx
=XM_023257578.1;refGene_gn=PARVB;refGene_tx=XM_023257579.1
B4 139204958 139205029
refGene_gn=PARVB;refGene_tx=XM_003989364.5;refGene_gn=PARVB;refGene_tx
=XM_023257578.1;refGene_gn=PARVB;refGene_tx=XM_023257579.1
B4 139218900 139219003
refGene_gn=PARVB;refGene_tx=XM_003989364.5;refGene_gn=PARVB;refGene_tx
=XM_023257578.1;refGene_gn=PARVB;refGene_tx=XM_023257579.1
B4 139225696 139225837
refGene_gn=PARVB;refGene_tx=XM_003989364.5;refGene_gn=PARVB;refGene_tx
=XM_023257578.1;refGene_gn=PARVB;refGene_tx=XM_023257579.1
B4 139227420 139227536
refGene_gn=PARVB;refGene_tx=XM_003989364.5;refGene_gn=PARVB;refGene_tx
=XM_023257578.1;refGene_gn=PARVB;refGene_tx=XM_023257579.1
B4 139230366 139230425
refGene_gn=PARVB;refGene_tx=XM_003989364.5;refGene_gn=PARVB;refGene_tx

=XM_023257578.1;refGene_gn=PARVB;refGene_tx=XM_023257579.1
B4 139233059 139233079
refGene_gn=PARVB;refGene_tx=XM_003989364.5;refGene_gn=PARVB;refGene_tx=XM_023257578.1;refGene_gn=PARVB;refGene_tx=XM_023257579.1
B4 139239020 139239082
refGene_gn=PARVB;refGene_tx=XM_003989364.5;refGene_gn=PARVB;refGene_tx=XM_023257578.1;refGene_gn=PARVB;refGene_tx=XM_023257579.1
B4 139241635 139241704
refGene_gn=PARVB;refGene_tx=XM_003989364.5;refGene_gn=PARVB;refGene_tx=XM_023257578.1;refGene_gn=PARVB;refGene_tx=XM_023257579.1
B4 139245491 139245593
refGene_gn=PARVB;refGene_tx=XM_003989364.5;refGene_gn=PARVB;refGene_tx=XM_023257578.1
B4 139250115 139250188
refGene_gn=PARVB;refGene_tx=XM_003989364.5;refGene_gn=PARVB;refGene_tx=XM_023257578.1;refGene_gn=PARVB;refGene_tx=XM_023257579.1
B4 139251672 139251749
refGene_gn=PARVB;refGene_tx=XM_003989364.5;refGene_gn=PARVB;refGene_tx=XM_023257578.1;refGene_gn=PARVB;refGene_tx=XM_023257579.1
B4 139256294 139256305
refGene_gn=PARVG;refGene_tx=XM_023257580.1
B4 139258003 139258098
refGene_gn=PARVG;refGene_tx=XM_023257580.1
B4 139260582 139260605
refGene_gn=PARVG;refGene_tx=XM_023257580.1
B4 139260606 139260648
refGene_gn=PARVG;refGene_tx=XM_023257580.1
B4 139262400 139262503
refGene_gn=PARVG;refGene_tx=XM_023257580.1
B4 139263182 139263323
refGene_gn=PARVG;refGene_tx=XM_023257580.1
B4 139264213 139264329
refGene_gn=PARVG;refGene_tx=XM_023257580.1
B4 139265549 139265605
refGene_gn=PARVG;refGene_tx=XM_023257580.1
B4 139266906 139266929
refGene_gn=PARVG;refGene_tx=XM_023257580.1
B4 139269795 139269854
refGene_gn=PARVG;refGene_tx=XM_023257580.1
B4 139270052 139270121
refGene_gn=PARVG;refGene_tx=XM_023257580.1
B4 139278181 139278254
refGene_gn=PARVG;refGene_tx=XM_023257580.1
B4 139278704 139278814
refGene_gn=PARVG;refGene_tx=XM_023257580.1
B4 139300003 139300004
refGene_gn=SHISAL1;refGene_tx=XM_023257582.1
B4 139308255 139308256
refGene_gn=SHISAL1;refGene_tx=XM_023257581.1;refGene_gn=SHISAL1;refGene_tx=XM_023257583.1;refGene_gn=SHISAL1;refGene_tx=XM_023257584.1;refGe

ne_gn=SHISAL1;refGene_tx=XM_023257585.1;refGene_gn=SHISAL1;refGene_tx=XM_023257586.1
B4 139336926 139337244
refGene_gn=SHISAL1;refGene_tx=XM_023257581.1;refGene_gn=SHISAL1;refGene_tx=XM_023257582.1;refGene_gn=SHISAL1;refGene_tx=XM_023257583.1;refGene_gn=SHISAL1;refGene_tx=XM_023257584.1;refGene_gn=SHISAL1;refGene_tx=XM_023257585.1;refGene_gn=SHISAL1;refGene_tx=XM_023257586.1
B4 139346817 139347031
refGene_gn=SHISAL1;refGene_tx=XM_023257581.1;refGene_gn=SHISAL1;refGene_tx=XM_023257582.1;refGene_gn=SHISAL1;refGene_tx=XM_023257583.1;refGene_gn=SHISAL1;refGene_tx=XM_023257584.1;refGene_gn=SHISAL1;refGene_tx=XM_023257585.1;refGene_gn=SHISAL1;refGene_tx=XM_023257586.1
B4 139350545 139350612
refGene_gn=SHISAL1;refGene_tx=XM_023257581.1;refGene_gn=SHISAL1;refGene_tx=XM_023257582.1;refGene_gn=SHISAL1;refGene_tx=XM_023257583.1;refGene_gn=SHISAL1;refGene_tx=XM_023257584.1;refGene_gn=SHISAL1;refGene_tx=XM_023257585.1;refGene_gn=SHISAL1;refGene_tx=XM_023257586.1
B4 139489854 139490574
refGene_gn=RTL6;refGene_tx=XM_023257587.1
B4 139554858 139554924
refGene_gn=LOC109501931;refGene_tx=XM_023257784.1
B4 139555189 139555274
refGene_gn=LOC109501931;refGene_tx=XM_023257784.1
B4 139555555 139555612
refGene_gn=LOC109501931;refGene_tx=XM_023257784.1
B4 139577214 139577327
refGene_gn=LOC109501931;refGene_tx=XM_023257784.1
B4 139577412 139578038
refGene_gn=LOC109501931;refGene_tx=XM_023257784.1
B4 139578179 139578594
refGene_gn=LOC109501931;refGene_tx=XM_023257784.1
B4 139625199 139625336
refGene_gn=PRR5;refGene_tx=XM_023257594.1;refGene_gn=PRR5;refGene_tx=XM_023257595.1;refGene_gn=PRR5;refGene_tx=XM_023257596.1;refGene_gn=PRR5;refGene_tx=XM_023257597.1
B4 139634228 139634309
refGene_gn=PRR5;refGene_tx=XM_023257594.1;refGene_gn=PRR5;refGene_tx=XM_023257595.1;refGene_gn=PRR5;refGene_tx=XM_023257596.1;refGene_gn=PRR5;refGene_tx=XM_023257597.1
B4 139639065 139639114
refGene_gn=PRR5;refGene_tx=XM_023257594.1;refGene_gn=PRR5;refGene_tx=XM_023257595.1;refGene_gn=PRR5;refGene_tx=XM_023257596.1;refGene_gn=PRR5;refGene_tx=XM_023257597.1
B4 139640521 139640579
refGene_gn=PRR5;refGene_tx=XM_023257594.1;refGene_gn=PRR5;refGene_tx=XM_023257595.1;refGene_gn=PRR5;refGene_tx=XM_023257596.1;refGene_gn=PRR5;refGene_tx=XM_023257597.1
B4 139644619 139644711
refGene_gn=PRR5;refGene_tx=XM_023257594.1;refGene_gn=PRR5;refGene_tx=XM_023257595.1;refGene_gn=PRR5;refGene_tx=XM_023257596.1;refGene_gn=PRR

5;refGene_tx=XM_023257597.1
B4 139645158 139645299
refGene_gn=PRR5;refGene_tx=XM_023257594.1;refGene_gn=PRR5;refGene_tx=XM_023257595.1;refGene_gn=PRR5;refGene_tx=XM_023257596.1;refGene_gn=PRR5;refGene_tx=XM_023257597.1
B4 139648468 139648604
refGene_gn=PRR5;refGene_tx=XM_023257594.1;refGene_gn=PRR5;refGene_tx=XM_023257595.1;refGene_gn=PRR5;refGene_tx=XM_023257596.1;refGene_gn=PRR5;refGene_tx=XM_023257597.1
B4 139649138 139649638
refGene_gn=PRR5;refGene_tx=XM_023257594.1;refGene_gn=PRR5;refGene_tx=XM_023257596.1;refGene_gn=PRR5;refGene_tx=XM_023257597.1
B4 139650196 139650672
refGene_gn=PRR5;refGene_tx=XM_023257595.1
B4 139687750 139687787
refGene_gn=ARHGAP8;refGene_tx=XM_023257588.1;refGene_gn=ARHGAP8;refGene_tx=XM_023257590.1;refGene_gn=ARHGAP8;refGene_tx=XM_023257591.1;refGene_gn=ARHGAP8;refGene_tx=XM_023257592.1
B4 139690280 139690424
refGene_gn=ARHGAP8;refGene_tx=XM_023257588.1;refGene_gn=ARHGAP8;refGene_tx=XM_023257590.1
B4 139698009 139698034
refGene_gn=ARHGAP8;refGene_tx=XM_023257589.1
B4 139698919 139699007
refGene_gn=ARHGAP8;refGene_tx=XM_023257588.1;refGene_gn=ARHGAP8;refGene_tx=XM_023257589.1;refGene_gn=ARHGAP8;refGene_tx=XM_023257590.1;refGene_gn=ARHGAP8;refGene_tx=XM_023257591.1;refGene_gn=ARHGAP8;refGene_tx=XM_023257592.1
B4 139703243 139703375
refGene_gn=ARHGAP8;refGene_tx=XM_023257588.1;refGene_gn=ARHGAP8;refGene_tx=XM_023257589.1;refGene_gn=ARHGAP8;refGene_tx=XM_023257591.1
B4 139707307 139707394
refGene_gn=ARHGAP8;refGene_tx=XM_023257588.1;refGene_gn=ARHGAP8;refGene_tx=XM_023257589.1;refGene_gn=ARHGAP8;refGene_tx=XM_023257590.1;refGene_gn=ARHGAP8;refGene_tx=XM_023257591.1;refGene_gn=ARHGAP8;refGene_tx=XM_023257592.1
B4 139711780 139711879
refGene_gn=ARHGAP8;refGene_tx=XM_023257588.1;refGene_gn=ARHGAP8;refGene_tx=XM_023257589.1;refGene_gn=ARHGAP8;refGene_tx=XM_023257590.1;refGene_gn=ARHGAP8;refGene_tx=XM_023257591.1;refGene_gn=ARHGAP8;refGene_tx=XM_023257592.1
B4 139713007 139713118
refGene_gn=ARHGAP8;refGene_tx=XM_023257588.1;refGene_gn=ARHGAP8;refGene_tx=XM_023257589.1;refGene_gn=ARHGAP8;refGene_tx=XM_023257590.1;refGene_gn=ARHGAP8;refGene_tx=XM_023257591.1;refGene_gn=ARHGAP8;refGene_tx=XM_023257592.1
B4 139718427 139718501
refGene_gn=ARHGAP8;refGene_tx=XM_023257588.1;refGene_gn=ARHGAP8;refGene_tx=XM_023257589.1;refGene_gn=ARHGAP8;refGene_tx=XM_023257590.1;refGene_gn=ARHGAP8;refGene_tx=XM_023257591.1;refGene_gn=ARHGAP8;refGene_tx=

XM_023257592.1
B4 139721054 139721132
refGene_gn=ARHGAP8;refGene_tx=XM_023257588.1;refGene_gn=ARHGAP8;refGene_tx=XM_023257589.1;refGene_gn=ARHGAP8;refGene_tx=XM_023257590.1;refGene_gn=ARHGAP8;refGene_tx=XM_023257591.1;refGene_gn=ARHGAP8;refGene_tx=XM_023257592.1
B4 139721799 139721928
refGene_gn=ARHGAP8;refGene_tx=XM_023257588.1;refGene_gn=ARHGAP8;refGene_tx=XM_023257589.1;refGene_gn=ARHGAP8;refGene_tx=XM_023257590.1;refGene_gn=ARHGAP8;refGene_tx=XM_023257591.1;refGene_gn=ARHGAP8;refGene_tx=XM_023257592.1
B4 139725798 139725902
refGene_gn=ARHGAP8;refGene_tx=XM_023257588.1;refGene_gn=ARHGAP8;refGene_tx=XM_023257589.1;refGene_gn=ARHGAP8;refGene_tx=XM_023257590.1;refGene_gn=ARHGAP8;refGene_tx=XM_023257591.1;refGene_gn=ARHGAP8;refGene_tx=XM_023257592.1
B4 139727663 139727996
refGene_gn=ARHGAP8;refGene_tx=XM_023257588.1;refGene_gn=ARHGAP8;refGene_tx=XM_023257589.1;refGene_gn=ARHGAP8;refGene_tx=XM_023257590.1;refGene_gn=ARHGAP8;refGene_tx=XM_023257591.1;refGene_gn=ARHGAP8;refGene_tx=XM_023257592.1
B4 139740953 139741172
refGene_gn=PHF21B;refGene_tx=XM_023257598.1;refGene_gn=PHF21B;refGene_tx=XM_023257599.1
B4 139742110 139742214
refGene_gn=PHF21B;refGene_tx=XM_023257598.1;refGene_gn=PHF21B;refGene_tx=XM_023257599.1
B4 139742530 139742606
refGene_gn=PHF21B;refGene_tx=XM_023257598.1;refGene_gn=PHF21B;refGene_tx=XM_023257599.1
B4 139743831 139743990
refGene_gn=PHF21B;refGene_tx=XM_023257598.1;refGene_gn=PHF21B;refGene_tx=XM_023257599.1
B4 139745455 139745478
refGene_gn=PHF21B;refGene_tx=XM_023257598.1;refGene_gn=PHF21B;refGene_tx=XM_023257599.1
B4 139746439 139746494
refGene_gn=PHF21B;refGene_tx=XM_023257598.1;refGene_gn=PHF21B;refGene_tx=XM_023257599.1
B4 139747418 139747495
refGene_gn=PHF21B;refGene_tx=XM_023257598.1;refGene_gn=PHF21B;refGene_tx=XM_023257599.1
B4 139748066 139748118
refGene_gn=PHF21B;refGene_tx=XM_023257598.1;refGene_gn=PHF21B;refGene_tx=XM_023257599.1
B4 139757948 139758089
refGene_gn=PHF21B;refGene_tx=XM_023257598.1;refGene_gn=PHF21B;refGene_tx=XM_023257599.1
B4 139760439 139760790
refGene_gn=PHF21B;refGene_tx=XM_023257598.1;refGene_gn=PHF21B;refGene_

tx=XM_023257599.1
B4	139762860	139762953
refGene_gn=PHF21B;refGene_tx=XM_023257598.1;refGene_gn=PHF21B;refGene_tx=XM_023257599.1
B4	139831880	139831946
refGene_gn=PHF21B;refGene_tx=XM_023257598.1;refGene_gn=PHF21B;refGene_tx=XM_023257599.1
B4	139832117	139832135
refGene_gn=PHF21B;refGene_tx=XM_023257599.1
B4	139832819	139832873
refGene_gn=PHF21B;refGene_tx=XM_023257598.1
B4	139955269	139955338
refGene_gn=NUP50;refGene_tx=XM_019835710.2
B4	139957695	139957779
refGene_gn=NUP50;refGene_tx=XM_019835710.2
B4	139960827	139961014
refGene_gn=NUP50;refGene_tx=XM_019835710.2
B4	139962034	139962132
refGene_gn=NUP50;refGene_tx=XM_019835710.2
B4	139962133	139962704
refGene_gn=NUP50;refGene_tx=XM_019835710.2
B4	139965850	139965932
refGene_gn=NUP50;refGene_tx=XM_019835710.2
B4	139967758	139967877
refGene_gn=NUP50;refGene_tx=XM_019835710.2
B4	139968882	139969085
refGene_gn=NUP50;refGene_tx=XM_019835710.2
B4	139979975	139980016
refGene_gn=KIAA0930;refGene_tx=XM_006934152.4;refGene_gn=KIAA0930;refGene_tx=XM_006934153.4;refGene_gn=KIAA0930;refGene_tx=XM_011284336.3
B4	139980531	139980690
refGene_gn=KIAA0930;refGene_tx=XM_006934152.4;refGene_gn=KIAA0930;refGene_tx=XM_006934153.4;refGene_gn=KIAA0930;refGene_tx=XM_011284336.3
B4	139981903	139982066
refGene_gn=KIAA0930;refGene_tx=XM_006934152.4;refGene_gn=KIAA0930;refGene_tx=XM_006934153.4;refGene_gn=KIAA0930;refGene_tx=XM_011284336.3
B4	139984825	139985020
refGene_gn=KIAA0930;refGene_tx=XM_006934152.4;refGene_gn=KIAA0930;refGene_tx=XM_006934153.4;refGene_gn=KIAA0930;refGene_tx=XM_011284336.3
B4	139985383	139985524
refGene_gn=KIAA0930;refGene_tx=XM_006934152.4;refGene_gn=KIAA0930;refGene_tx=XM_006934153.4;refGene_gn=KIAA0930;refGene_tx=XM_011284336.3
B4	139986767	139986869
refGene_gn=KIAA0930;refGene_tx=XM_006934152.4;refGene_gn=KIAA0930;refGene_tx=XM_006934153.4;refGene_gn=KIAA0930;refGene_tx=XM_011284336.3
B4	139987218	139987296
refGene_gn=KIAA0930;refGene_tx=XM_006934152.4;refGene_gn=KIAA0930;refGene_tx=XM_006934153.4;refGene_gn=KIAA0930;refGene_tx=XM_011284336.3
B4	139987400	139987526
refGene_gn=KIAA0930;refGene_tx=XM_006934152.4;refGene_gn=KIAA0930;refG

ene_tx=XM_006934153.4;refGene_gn=KIAA0930;refGene_tx=XM_011284336.3
B4 139991250 139991402
refGene_gn=KIAA0930;refGene_tx=XM_006934152.4;refGene_gn=KIAA0930;refGene_tx=XM_006934153.4;refGene_gn=KIAA0930;refGene_tx=XM_011284336.3
B4 140009376 140009437
refGene_gn=KIAA0930;refGene_tx=XM_006934153.4
B4 140013920 140013984
refGene_gn=KIAA0930;refGene_tx=XM_011284336.3
B4 140014939 140014988
refGene_gn=KIAA0930;refGene_tx=XM_006934152.4
B4 140046174 140046226
refGene_gn=UPK3A;refGene_tx=XM_003989368.4
B4 140047239 140047395
refGene_gn=UPK3A;refGene_tx=XM_003989368.4;refGene_gn=UPK3A;refGene_tx=XM_023257601.1
B4 140048007 140048125
refGene_gn=UPK3A;refGene_tx=XM_023257601.1
B4 140048515 140048792
refGene_gn=UPK3A;refGene_tx=XM_003989368.4;refGene_gn=UPK3A;refGene_tx=XM_023257601.1
B4 140049871 140049954
refGene_gn=UPK3A;refGene_tx=XM_003989368.4;refGene_gn=UPK3A;refGene_tx=XM_023257601.1
B4 140051803 140051936
refGene_gn=UPK3A;refGene_tx=XM_003989368.4;refGene_gn=UPK3A;refGene_tx=XM_023257601.1
B4 140053522 140053670
refGene_gn=UPK3A;refGene_tx=XM_003989368.4;refGene_gn=UPK3A;refGene_tx=XM_023257601.1
B4 140069561 140069612
refGene_gn=FAM118A;refGene_tx=XM_019835715.2;refGene_gn=FAM118A;refGene_tx=XM_019835716.2
B4 140072543 140072599
refGene_gn=FAM118A;refGene_tx=XM_006934155.4;refGene_gn=FAM118A;refGene_tx=XM_019835715.2;refGene_gn=FAM118A;refGene_tx=XM_019835716.2;refGene_gn=FAM118A;refGene_tx=XM_019835717.2;refGene_gn=FAM118A;refGene_tx=XM_023257608.1;refGene_gn=FAM118A;refGene_tx=XM_023257609.1;refGene_gn=FAM118A;refGene_tx=XM_023257611.1;refGene_gn=FAM118A;refGene_tx=XM_023257612.1
B4 140073352 140073608
refGene_gn=FAM118A;refGene_tx=XM_006934155.4;refGene_gn=FAM118A;refGene_tx=XM_019835715.2;refGene_gn=FAM118A;refGene_tx=XM_019835716.2;refGene_gn=FAM118A;refGene_tx=XM_019835717.2;refGene_gn=FAM118A;refGene_tx=XM_023257608.1;refGene_gn=FAM118A;refGene_tx=XM_023257609.1;refGene_gn=FAM118A;refGene_tx=XM_023257611.1;refGene_gn=FAM118A;refGene_tx=XM_023257612.1
B4 140076604 140076826
refGene_gn=FAM118A;refGene_tx=XM_006934155.4;refGene_gn=FAM118A;refGene_tx=XM_019835715.2;refGene_gn=FAM118A;refGene_tx=XM_019835716.2;refGene_gn=FAM118A;refGene_tx=XM_019835717.2;refGene_gn=FAM118A;refGene_tx=

XM_023257608.1;refGene_gn=FAM118A;refGene_tx=XM_023257609.1;refGene_gn=FAM118A;refGene_tx=XM_023257611.1;refGene_gn=FAM118A;refGene_tx=XM_023257612.1
B4 140078476 140078605
refGene_gn=FAM118A;refGene_tx=XM_006934155.4;refGene_gn=FAM118A;refGene_tx=XM_019835715.2;refGene_gn=FAM118A;refGene_tx=XM_019835716.2;refGene_gn=FAM118A;refGene_tx=XM_019835717.2;refGene_gn=FAM118A;refGene_tx=XM_023257608.1;refGene_gn=FAM118A;refGene_tx=XM_023257609.1;refGene_gn=FAM118A;refGene_tx=XM_023257611.1;refGene_gn=FAM118A;refGene_tx=XM_023257612.1
B4 140079886 140080172
refGene_gn=FAM118A;refGene_tx=XM_006934155.4;refGene_gn=FAM118A;refGene_tx=XM_019835715.2;refGene_gn=FAM118A;refGene_tx=XM_019835716.2;refGene_gn=FAM118A;refGene_tx=XM_019835717.2;refGene_gn=FAM118A;refGene_tx=XM_023257608.1;refGene_gn=FAM118A;refGene_tx=XM_023257609.1;refGene_gn=FAM118A;refGene_tx=XM_023257611.1;refGene_gn=FAM118A;refGene_tx=XM_023257612.1
B4 140081562 140081609
refGene_gn=FAM118A;refGene_tx=XM_019835717.2
B4 140081800 140081833
refGene_gn=FAM118A;refGene_tx=XM_006934155.4;refGene_gn=FAM118A;refGene_tx=XM_019835715.2;refGene_gn=FAM118A;refGene_tx=XM_019835716.2;refGene_gn=FAM118A;refGene_tx=XM_023257608.1;refGene_gn=FAM118A;refGene_tx=XM_023257611.1;refGene_gn=FAM118A;refGene_tx=XM_023257612.1
B4 140082629 140082713
refGene_gn=FAM118A;refGene_tx=XM_006934155.4;refGene_gn=FAM118A;refGene_tx=XM_019835715.2;refGene_gn=FAM118A;refGene_tx=XM_019835716.2;refGene_gn=FAM118A;refGene_tx=XM_023257608.1;refGene_gn=FAM118A;refGene_tx=XM_023257609.1;refGene_gn=FAM118A;refGene_tx=XM_023257611.1;refGene_gn=FAM118A;refGene_tx=XM_023257612.1
B4 140085995 140086015
refGene_gn=FAM118A;refGene_tx=XM_006934155.4;refGene_gn=FAM118A;refGene_tx=XM_019835715.2;refGene_gn=FAM118A;refGene_tx=XM_019835716.2;refGene_gn=FAM118A;refGene_tx=XM_023257608.1;refGene_gn=FAM118A;refGene_tx=XM_023257609.1;refGene_gn=FAM118A;refGene_tx=XM_023257611.1;refGene_gn=FAM118A;refGene_tx=XM_023257612.1
B4 140089085 140089187
refGene_gn=SMC1B;refGene_tx=XM_011284332.3;refGene_gn=SMC1B;refGene_tx=XM_019835711.2;refGene_gn=SMC1B;refGene_tx=XM_023257602.1;refGene_gn=SMC1B;refGene_tx=XM_023257603.1;refGene_gn=SMC1B;refGene_tx=XM_023257605.1;refGene_gn=SMC1B;refGene_tx=XM_023257606.1;refGene_gn=SMC1B;refGene_tx=XM_023257607.1
B4 140090394 140090505
refGene_gn=SMC1B;refGene_tx=XM_011284332.3;refGene_gn=SMC1B;refGene_tx=XM_019835711.2;refGene_gn=SMC1B;refGene_tx=XM_023257602.1;refGene_gn=SMC1B;refGene_tx=XM_023257603.1;refGene_gn=SMC1B;refGene_tx=XM_023257605.1;refGene_gn=SMC1B;refGene_tx=XM_023257606.1;refGene_gn=SMC1B;refGene_tx=XM_023257607.1
B4 140092617 140092687
refGene_gn=SMC1B;refGene_tx=XM_011284332.3;refGene_gn=SMC1B;refGene_tx

=XM_019835711.2;refGene_gn=SMC1B;refGene_tx=XM_023257602.1;refGene_gn=SMC1B;refGene_tx=XM_023257603.1;refGene_gn=SMC1B;refGene_tx=XM_023257605.1;refGene_gn=SMC1B;refGene_tx=XM_023257606.1;refGene_gn=SMC1B;refGene_tx=XM_023257607.1
B4 140094478 140094630
refGene_gn=SMC1B;refGene_tx=XM_011284332.3;refGene_gn=SMC1B;refGene_tx=XM_019835711.2;refGene_gn=SMC1B;refGene_tx=XM_023257602.1;refGene_gn=SMC1B;refGene_tx=XM_023257603.1;refGene_gn=SMC1B;refGene_tx=XM_023257605.1;refGene_gn=SMC1B;refGene_tx=XM_023257606.1;refGene_gn=SMC1B;refGene_tx=XM_023257607.1
B4 140095668 140095823
refGene_gn=SMC1B;refGene_tx=XM_011284332.3;refGene_gn=SMC1B;refGene_tx=XM_019835711.2;refGene_gn=SMC1B;refGene_tx=XM_023257602.1;refGene_gn=SMC1B;refGene_tx=XM_023257603.1;refGene_gn=SMC1B;refGene_tx=XM_023257605.1;refGene_gn=SMC1B;refGene_tx=XM_023257606.1;refGene_gn=SMC1B;refGene_tx=XM_023257607.1
B4 140096292 140096449
refGene_gn=SMC1B;refGene_tx=XM_011284332.3;refGene_gn=SMC1B;refGene_tx=XM_019835711.2;refGene_gn=SMC1B;refGene_tx=XM_023257602.1;refGene_gn=SMC1B;refGene_tx=XM_023257603.1;refGene_gn=SMC1B;refGene_tx=XM_023257605.1;refGene_gn=SMC1B;refGene_tx=XM_023257606.1;refGene_gn=SMC1B;refGene_tx=XM_023257607.1
B4 140098484 140098583
refGene_gn=SMC1B;refGene_tx=XM_011284332.3;refGene_gn=SMC1B;refGene_tx=XM_019835711.2;refGene_gn=SMC1B;refGene_tx=XM_023257602.1;refGene_gn=SMC1B;refGene_tx=XM_023257603.1;refGene_gn=SMC1B;refGene_tx=XM_023257605.1;refGene_gn=SMC1B;refGene_tx=XM_023257606.1;refGene_gn=SMC1B;refGene_tx=XM_023257607.1
B4 140100574 140100728
refGene_gn=SMC1B;refGene_tx=XM_011284332.3;refGene_gn=SMC1B;refGene_tx=XM_019835711.2;refGene_gn=SMC1B;refGene_tx=XM_023257602.1;refGene_gn=SMC1B;refGene_tx=XM_023257603.1;refGene_gn=SMC1B;refGene_tx=XM_023257605.1;refGene_gn=SMC1B;refGene_tx=XM_023257606.1;refGene_gn=SMC1B;refGene_tx=XM_023257607.1
B4 140102026 140102172
refGene_gn=SMC1B;refGene_tx=XM_011284332.3;refGene_gn=SMC1B;refGene_tx=XM_019835711.2;refGene_gn=SMC1B;refGene_tx=XM_023257602.1;refGene_gn=SMC1B;refGene_tx=XM_023257603.1;refGene_gn=SMC1B;refGene_tx=XM_023257605.1;refGene_gn=SMC1B;refGene_tx=XM_023257606.1;refGene_gn=SMC1B;refGene_tx=XM_023257607.1
B4 140103004 140103146
refGene_gn=SMC1B;refGene_tx=XM_011284332.3;refGene_gn=SMC1B;refGene_tx=XM_019835711.2;refGene_gn=SMC1B;refGene_tx=XM_023257602.1;refGene_gn=SMC1B;refGene_tx=XM_023257603.1;refGene_gn=SMC1B;refGene_tx=XM_023257605.1;refGene_gn=SMC1B;refGene_tx=XM_023257606.1;refGene_gn=SMC1B;refGene_tx=XM_023257607.1
B4 140113746 140113853
refGene_gn=SMC1B;refGene_tx=XM_011284332.3;refGene_gn=SMC1B;refGene_tx=XM_019835711.2;refGene_gn=SMC1B;refGene_tx=XM_023257602.1;refGene_gn=SMC1B;refGene_tx=XM_023257603.1;refGene_gn=SMC1B;refGene_tx=XM_0232576

05.1;refGene_gn=SMC1B;refGene_tx=XM_023257606.1;refGene_gn=SMC1B;refGene_tx=XM_023257607.1
B4	140115259	140115376
refGene_gn=SMC1B;refGene_tx=XM_011284332.3;refGene_gn=SMC1B;refGene_tx=XM_019835711.2;refGene_gn=SMC1B;refGene_tx=XM_023257602.1;refGene_gn=SMC1B;refGene_tx=XM_023257603.1;refGene_gn=SMC1B;refGene_tx=XM_023257605.1;refGene_gn=SMC1B;refGene_tx=XM_023257606.1;refGene_gn=SMC1B;refGene_tx=XM_023257607.1
B4	140115650	140115788
refGene_gn=SMC1B;refGene_tx=XM_011284332.3;refGene_gn=SMC1B;refGene_tx=XM_019835711.2;refGene_gn=SMC1B;refGene_tx=XM_023257602.1;refGene_gn=SMC1B;refGene_tx=XM_023257603.1;refGene_gn=SMC1B;refGene_tx=XM_023257605.1;refGene_gn=SMC1B;refGene_tx=XM_023257606.1;refGene_gn=SMC1B;refGene_tx=XM_023257607.1
B4	140123916	140124063
refGene_gn=SMC1B;refGene_tx=XM_011284332.3;refGene_gn=SMC1B;refGene_tx=XM_019835711.2;refGene_gn=SMC1B;refGene_tx=XM_023257602.1;refGene_gn=SMC1B;refGene_tx=XM_023257603.1;refGene_gn=SMC1B;refGene_tx=XM_023257605.1;refGene_gn=SMC1B;refGene_tx=XM_023257606.1;refGene_gn=SMC1B;refGene_tx=XM_023257607.1
B4	140131535	140131715
refGene_gn=SMC1B;refGene_tx=XM_011284332.3;refGene_gn=SMC1B;refGene_tx=XM_019835711.2;refGene_gn=SMC1B;refGene_tx=XM_023257602.1;refGene_gn=SMC1B;refGene_tx=XM_023257603.1;refGene_gn=SMC1B;refGene_tx=XM_023257605.1;refGene_gn=SMC1B;refGene_tx=XM_023257606.1;refGene_gn=SMC1B;refGene_tx=XM_023257607.1
B4	140132894	140133080
refGene_gn=SMC1B;refGene_tx=XM_011284332.3;refGene_gn=SMC1B;refGene_tx=XM_019835711.2;refGene_gn=SMC1B;refGene_tx=XM_023257602.1;refGene_gn=SMC1B;refGene_tx=XM_023257605.1;refGene_gn=SMC1B;refGene_tx=XM_023257607.1
B4	140135770	140135978
refGene_gn=SMC1B;refGene_tx=XM_011284332.3;refGene_gn=SMC1B;refGene_tx=XM_019835711.2;refGene_gn=SMC1B;refGene_tx=XM_023257602.1;refGene_gn=SMC1B;refGene_tx=XM_023257603.1;refGene_gn=SMC1B;refGene_tx=XM_023257607.1
B4	140136763	140136846
refGene_gn=SMC1B;refGene_tx=XM_011284332.3;refGene_gn=SMC1B;refGene_tx=XM_019835711.2;refGene_gn=SMC1B;refGene_tx=XM_023257602.1;refGene_gn=SMC1B;refGene_tx=XM_023257603.1;refGene_gn=SMC1B;refGene_tx=XM_023257607.1
B4	140141644	140141785
refGene_gn=SMC1B;refGene_tx=XM_011284332.3;refGene_gn=SMC1B;refGene_tx=XM_019835711.2;refGene_gn=SMC1B;refGene_tx=XM_023257602.1;refGene_gn=SMC1B;refGene_tx=XM_023257603.1;refGene_gn=SMC1B;refGene_tx=XM_023257605.1;refGene_gn=SMC1B;refGene_tx=XM_023257606.1;refGene_gn=SMC1B;refGene_tx=XM_023257607.1
B4	140142739	140142998
refGene_gn=SMC1B;refGene_tx=XM_011284332.3;refGene_gn=SMC1B;refGene_tx=XM_019835711.2;refGene_gn=SMC1B;refGene_tx=XM_023257602.1;refGene_gn=

SMC1B;refGene_tx=XM_023257603.1;refGene_gn=SMC1B;refGene_tx=XM_023257605.1;refGene_gn=SMC1B;refGene_tx=XM_023257606.1
B4 140144276 140144515
refGene_gn=SMC1B;refGene_tx=XM_011284332.3;refGene_gn=SMC1B;refGene_tx=XM_019835711.2;refGene_gn=SMC1B;refGene_tx=XM_023257602.1;refGene_gn=SMC1B;refGene_tx=XM_023257603.1;refGene_gn=SMC1B;refGene_tx=XM_023257605.1;refGene_gn=SMC1B;refGene_tx=XM_023257606.1
B4 140147455 140147659
refGene_gn=SMC1B;refGene_tx=XM_011284332.3;refGene_gn=SMC1B;refGene_tx=XM_019835711.2;refGene_gn=SMC1B;refGene_tx=XM_023257602.1;refGene_gn=SMC1B;refGene_tx=XM_023257603.1;refGene_gn=SMC1B;refGene_tx=XM_023257605.1;refGene_gn=SMC1B;refGene_tx=XM_023257606.1;refGene_gn=SMC1B;refGene_tx=XM_023257607.1
B4 140147746 140147859
refGene_gn=SMC1B;refGene_tx=XM_011284332.3;refGene_gn=SMC1B;refGene_tx=XM_019835711.2;refGene_gn=SMC1B;refGene_tx=XM_023257602.1;refGene_gn=SMC1B;refGene_tx=XM_023257603.1;refGene_gn=SMC1B;refGene_tx=XM_023257605.1;refGene_gn=SMC1B;refGene_tx=XM_023257606.1;refGene_gn=SMC1B;refGene_tx=XM_023257607.1
B4 140150337 140150526
refGene_gn=SMC1B;refGene_tx=XM_011284332.3;refGene_gn=SMC1B;refGene_tx=XM_019835711.2;refGene_gn=SMC1B;refGene_tx=XM_023257602.1;refGene_gn=SMC1B;refGene_tx=XM_023257603.1;refGene_gn=SMC1B;refGene_tx=XM_023257605.1;refGene_gn=SMC1B;refGene_tx=XM_023257606.1;refGene_gn=SMC1B;refGene_tx=XM_023257607.1
B4 140163210 140163319
refGene_gn=SMC1B;refGene_tx=XM_023257602.1;refGene_gn=SMC1B;refGene_tx=XM_023257603.1;refGene_gn=SMC1B;refGene_tx=XM_023257605.1;refGene_gn=SMC1B;refGene_tx=XM_023257606.1;refGene_gn=SMC1B;refGene_tx=XM_023257607.1
B4 140163646 140163760
refGene_gn=RIBC2;refGene_tx=XM_023257613.1;refGene_gn=RIBC2;refGene_tx=XM_023257614.1;refGene_gn=RIBC2;refGene_tx=XM_023257615.1;refGene_gn=RIBC2;refGene_tx=XM_023257616.1
B4 140165575 140165657
refGene_gn=RIBC2;refGene_tx=XM_023257613.1;refGene_gn=RIBC2;refGene_tx=XM_023257614.1;refGene_gn=RIBC2;refGene_tx=XM_023257615.1;refGene_gn=RIBC2;refGene_tx=XM_023257616.1
B4 140169659 140170004
refGene_gn=RIBC2;refGene_tx=XM_023257613.1;refGene_gn=RIBC2;refGene_tx=XM_023257614.1;refGene_gn=RIBC2;refGene_tx=XM_023257615.1;refGene_gn=RIBC2;refGene_tx=XM_023257616.1;refGene_gn=RIBC2;refGene_tx=XM_023257617.1;refGene_gn=RIBC2;refGene_tx=XM_023257618.1
B4 140174172 140174291
refGene_gn=RIBC2;refGene_tx=XM_023257613.1;refGene_gn=RIBC2;refGene_tx=XM_023257614.1;refGene_gn=RIBC2;refGene_tx=XM_023257615.1;refGene_gn=RIBC2;refGene_tx=XM_023257616.1;refGene_gn=RIBC2;refGene_tx=XM_023257617.1;refGene_gn=RIBC2;refGene_tx=XM_023257618.1
B4 140174510 140174786
refGene_gn=RIBC2;refGene_tx=XM_023257615.1

B4 140177304 140177532
refGene_gn=RIBC2;refGene_tx=XM_023257613.1;refGene_gn=RIBC2;refGene_tx=XM_023257614.1;refGene_gn=RIBC2;refGene_tx=XM_023257616.1;refGene_gn=RIBC2;refGene_tx=XM_023257617.1;refGene_gn=RIBC2;refGene_tx=XM_023257618.1
B4 140182163 140182330
refGene_gn=RIBC2;refGene_tx=XM_023257613.1;refGene_gn=RIBC2;refGene_tx=XM_023257614.1;refGene_gn=RIBC2;refGene_tx=XM_023257617.1;refGene_gn=RIBC2;refGene_tx=XM_023257618.1
B4 140183295 140183359
refGene_gn=RIBC2;refGene_tx=XM_023257614.1
B4 140183651 140183730
refGene_gn=RIBC2;refGene_tx=XM_023257613.1;refGene_gn=RIBC2;refGene_tx=XM_023257616.1;refGene_gn=RIBC2;refGene_tx=XM_023257617.1;refGene_gn=RIBC2;refGene_tx=XM_023257618.1
B4 140229979 140230055
refGene_gn=FBLN1;refGene_tx=XM_023257619.1;refGene_gn=FBLN1;refGene_tx=XM_023257621.1
B4 140246145 140246251
refGene_gn=FBLN1;refGene_tx=XM_023257619.1;refGene_gn=FBLN1;refGene_tx=XM_023257621.1
B4 140251908 140252044
refGene_gn=FBLN1;refGene_tx=XM_023257619.1;refGene_gn=FBLN1;refGene_tx=XM_023257621.1
B4 140254088 140254260
refGene_gn=FBLN1;refGene_tx=XM_023257619.1;refGene_gn=FBLN1;refGene_tx=XM_023257621.1
B4 140258676 140258736
refGene_gn=FBLN1;refGene_tx=XM_023257619.1;refGene_gn=FBLN1;refGene_tx=XM_023257621.1
B4 140259869 140259971
refGene_gn=FBLN1;refGene_tx=XM_023257619.1;refGene_gn=FBLN1;refGene_tx=XM_023257621.1
B4 140260787 140260925
refGene_gn=FBLN1;refGene_tx=XM_023257619.1;refGene_gn=FBLN1;refGene_tx=XM_023257621.1
B4 140262071 140262209
refGene_gn=FBLN1;refGene_tx=XM_023257619.1;refGene_gn=FBLN1;refGene_tx=XM_023257621.1
B4 140266311 140266455
refGene_gn=FBLN1;refGene_tx=XM_023257619.1;refGene_gn=FBLN1;refGene_tx=XM_023257621.1
B4 140267254 140267383
refGene_gn=FBLN1;refGene_tx=XM_023257619.1;refGene_gn=FBLN1;refGene_tx=XM_023257621.1
B4 140268036 140268162
refGene_gn=FBLN1;refGene_tx=XM_023257619.1;refGene_gn=FBLN1;refGene_tx=XM_023257621.1
B4 140271235 140271355
refGene_gn=FBLN1;refGene_tx=XM_023257619.1;refGene_gn=FBLN1;refGene_tx

=XM_023257621.1
B4 140272220 140272352
refGene_gn=FBLN1;refGene_tx=XM_023257619.1;refGene_gn=FBLN1;refGene_tx
=XM_023257621.1
B4 140274082 140274206
refGene_gn=FBLN1;refGene_tx=XM_023257619.1;refGene_gn=FBLN1;refGene_tx
=XM_023257621.1
B4 140283432 140283787
refGene_gn=FBLN1;refGene_tx=XM_023257621.1
B4 140291333 140291476
refGene_gn=FBLN1;refGene_tx=XM_023257619.1
B4 140293382 140293514
refGene_gn=FBLN1;refGene_tx=XM_023257619.1
B4 140315796 140315936
refGene_gn=FBLN1;refGene_tx=XM_023257619.1
B4 140373601 140373717
refGene_gn=ATXN10;refGene_tx=XM_023257622.1;refGene_gn=ATXN10;refGene_
tx=XM_023257623.1;refGene_gn=ATXN10;refGene_tx=XM_023257624.1
B4 140392404 140392596
refGene_gn=ATXN10;refGene_tx=XM_023257622.1;refGene_gn=ATXN10;refGene_
tx=XM_023257623.1;refGene_gn=ATXN10;refGene_tx=XM_023257624.1
B4 140399696 140399779
refGene_gn=ATXN10;refGene_tx=XM_023257622.1;refGene_gn=ATXN10;refGene_
tx=XM_023257623.1;refGene_gn=ATXN10;refGene_tx=XM_023257624.1
B4 140413839 140413936
refGene_gn=ATXN10;refGene_tx=XM_023257622.1;refGene_gn=ATXN10;refGene_
tx=XM_023257623.1;refGene_gn=ATXN10;refGene_tx=XM_023257624.1
B4 140416130 140416289
refGene_gn=ATXN10;refGene_tx=XM_023257622.1;refGene_gn=ATXN10;refGene_
tx=XM_023257623.1
B4 140429999 140430080
refGene_gn=ATXN10;refGene_tx=XM_023257622.1;refGene_gn=ATXN10;refGene_
tx=XM_023257624.1
B4 140439407 140439573
refGene_gn=ATXN10;refGene_tx=XM_023257622.1;refGene_gn=ATXN10;refGene_
tx=XM_023257623.1;refGene_gn=ATXN10;refGene_tx=XM_023257624.1
B4 140447608 140447717
refGene_gn=ATXN10;refGene_tx=XM_023257622.1;refGene_gn=ATXN10;refGene_
tx=XM_023257623.1;refGene_gn=ATXN10;refGene_tx=XM_023257624.1
B4 140449140 140449310
refGene_gn=ATXN10;refGene_tx=XM_023257622.1;refGene_gn=ATXN10;refGene_
tx=XM_023257623.1;refGene_gn=ATXN10;refGene_tx=XM_023257624.1
B4 140512119 140512183
refGene_gn=ATXN10;refGene_tx=XM_023257622.1;refGene_gn=ATXN10;refGene_
tx=XM_023257623.1;refGene_gn=ATXN10;refGene_tx=XM_023257624.1
B4 140538988 140539176
refGene_gn=ATXN10;refGene_tx=XM_023257622.1;refGene_gn=ATXN10;refGene_
tx=XM_023257623.1;refGene_gn=ATXN10;refGene_tx=XM_023257624.1
B4 140539647 140539650
refGene_gn=ATXN10;refGene_tx=XM_023257622.1;refGene_gn=ATXN10;refGene_

tx=XM_023257623.1;refGene_gn=ATXN10;refGene_tx=XM_023257624.1
B4 140610067 140610547
refGene_gn=WNT7B;refGene_tx=XM_023257625.1;refGene_gn=WNT7B;refGene_tx=XM_023257626.1;refGene_gn=WNT7B;refGene_tx=XM_023257627.1
B4 140616089 140616361
refGene_gn=WNT7B;refGene_tx=XM_023257625.1;refGene_gn=WNT7B;refGene_tx=XM_023257626.1;refGene_gn=WNT7B;refGene_tx=XM_023257627.1
B4 140632401 140632628
refGene_gn=WNT7B;refGene_tx=XM_023257625.1;refGene_gn=WNT7B;refGene_tx=XM_023257626.1;refGene_gn=WNT7B;refGene_tx=XM_023257627.1
B4 140652509 140652592
refGene_gn=WNT7B;refGene_tx=XM_023257626.1
B4 140656844 140656940
refGene_gn=WNT7B;refGene_tx=XM_023257625.1;refGene_gn=WNT7B;refGene_tx=XM_023257627.1
B4 140745624 140745846
refGene_gn=LOC111561467;refGene_tx=XM_023257785.1
B4 140746000 140746312
refGene_gn=LOC111561467;refGene_tx=XM_023257785.1
B4 140765663 140765741
refGene_gn=LOC111561467;refGene_tx=XM_023257785.1
B4 140775580 140775634
refGene_gn=LOC111561467;refGene_tx=XM_023257785.1
B4 140775935 140776337
refGene_gn=LOC111561467;refGene_tx=XM_023257785.1
B4 140849923 140850131
refGene_gn=PPARA;refGene_tx=XM_023257628.1;refGene_gn=PPARA;refGene_tx=XM_023257629.1
B4 140859530 140859691
refGene_gn=PPARA;refGene_tx=XM_023257628.1;refGene_gn=PPARA;refGene_tx=XM_023257629.1
B4 140861326 140861465
refGene_gn=PPARA;refGene_tx=XM_023257628.1;refGene_gn=PPARA;refGene_tx=XM_023257629.1
B4 140862472 140862675
refGene_gn=PPARA;refGene_tx=XM_023257628.1;refGene_gn=PPARA;refGene_tx=XM_023257629.1
B4 140870512 140870960
refGene_gn=PPARA;refGene_tx=XM_023257628.1;refGene_gn=PPARA;refGene_tx=XM_023257629.1
B4 140873851 140874099
refGene_gn=PPARA;refGene_tx=XM_023257628.1;refGene_gn=PPARA;refGene_tx=XM_023257629.1
B4 140882700 140882847
refGene_gn=CDPF1;refGene_tx=XM_006934171.4;refGene_gn=CDPF1;refGene_tx=XM_011284327.3
B4 140884829 140884941
refGene_gn=CDPF1;refGene_tx=XM_006934171.4;refGene_gn=CDPF1;refGene_tx=XM_011284327.3
B4 140885193 140885306

refGene_gn=CDPF1;refGene_tx=XM_006934171.4;refGene_gn=CDPF1;refGene_tx
=XM_011284327.3
B4	140893006	140899831
refGene_gn=PKDREJ;refGene_tx=XM_019835900.2
B4	140902652	140902685
refGene_gn=TTC38;refGene_tx=XM_023257630.1
B4	140903714	140903792
refGene_gn=TTC38;refGene_tx=XM_023257630.1
B4	140905671	140905753
refGene_gn=TTC38;refGene_tx=XM_023257630.1
B4	140907495	140907667
refGene_gn=TTC38;refGene_tx=XM_023257630.1
B4	140908740	140908914
refGene_gn=TTC38;refGene_tx=XM_023257630.1
B4	140911554	140911630
refGene_gn=TTC38;refGene_tx=XM_023257630.1
B4	140914393	140914513
refGene_gn=TTC38;refGene_tx=XM_023257630.1
B4	140916306	140916366
refGene_gn=TTC38;refGene_tx=XM_023257630.1
B4	140917708	140917747
refGene_gn=TTC38;refGene_tx=XM_023257630.1
B4	140918781	140918863
refGene_gn=TTC38;refGene_tx=XM_023257630.1
B4	140920651	140920817
refGene_gn=TTC38;refGene_tx=XM_023257630.1
B4	140922275	140922435
refGene_gn=TTC38;refGene_tx=XM_023257630.1
B4	140922629	140922703
refGene_gn=TTC38;refGene_tx=XM_023257630.1
B4	140924801	140924895
refGene_gn=TTC38;refGene_tx=XM_023257630.1
B4	140928970	140928992
refGene_gn=GTSE1;refGene_tx=XM_006934172.3;refGene_gn=GTSE1;refGene_tx
=XM_019835732.2;refGene_gn=GTSE1;refGene_tx=XM_019835733.2
B4	140935033	140935091
refGene_gn=GTSE1;refGene_tx=XM_006934172.3;refGene_gn=GTSE1;refGene_tx
=XM_019835732.2;refGene_gn=GTSE1;refGene_tx=XM_019835733.2
B4	140935194	140935798
refGene_gn=GTSE1;refGene_tx=XM_006934172.3;refGene_gn=GTSE1;refGene_tx
=XM_019835732.2;refGene_gn=GTSE1;refGene_tx=XM_019835733.2
B4	140937785	140937956
refGene_gn=GTSE1;refGene_tx=XM_006934172.3;refGene_gn=GTSE1;refGene_tx
=XM_019835733.2
B4	140938735	140938862
refGene_gn=GTSE1;refGene_tx=XM_006934172.3;refGene_gn=GTSE1;refGene_tx
=XM_019835732.2;refGene_gn=GTSE1;refGene_tx=XM_019835733.2
B4	140940425	140940806
refGene_gn=GTSE1;refGene_tx=XM_006934172.3;refGene_gn=GTSE1;refGene_tx
=XM_019835732.2

B4	140943645	140943718
refGene_gn=GTSE1;refGene_tx=XM_006934172.3;refGene_gn=GTSE1;refGene_tx
=XM_019835732.2;refGene_gn=GTSE1;refGene_tx=XM_019835733.2
B4	140944493	140944733
refGene_gn=GTSE1;refGene_tx=XM_006934172.3;refGene_gn=GTSE1;refGene_tx
=XM_019835732.2;refGene_gn=GTSE1;refGene_tx=XM_019835733.2
B4	140945659	140945864
refGene_gn=GTSE1;refGene_tx=XM_006934172.3;refGene_gn=GTSE1;refGene_tx
=XM_019835732.2;refGene_gn=GTSE1;refGene_tx=XM_019835733.2
B4	140946294	140946507
refGene_gn=GTSE1;refGene_tx=XM_006934172.3;refGene_gn=GTSE1;refGene_tx
=XM_019835732.2;refGene_gn=GTSE1;refGene_tx=XM_019835733.2
B4	140946933	140947017
refGene_gn=GTSE1;refGene_tx=XM_006934172.3;refGene_gn=GTSE1;refGene_tx
=XM_019835732.2;refGene_gn=GTSE1;refGene_tx=XM_019835733.2
B4	140951105	140951187
refGene_gn=TRMU;refGene_tx=XM_023257631.1;refGene_gn=TRMU;refGene_tx=X
M_023257632.1;refGene_gn=TRMU;refGene_tx=XM_023257633.1
B4	140954115	140954281
refGene_gn=TRMU;refGene_tx=XM_023257631.1;refGene_gn=TRMU;refGene_tx=X
M_023257632.1;refGene_gn=TRMU;refGene_tx=XM_023257633.1
B4	140956405	140956512
refGene_gn=TRMU;refGene_tx=XM_011284325.2;refGene_gn=TRMU;refGene_tx=X
M_019835734.1;refGene_gn=TRMU;refGene_tx=XM_023257631.1;refGene_gn=TRM
U;refGene_tx=XM_023257632.1;refGene_gn=TRMU;refGene_tx=XM_023257633.1
B4	140958314	140958437
refGene_gn=TRMU;refGene_tx=XM_011284325.2;refGene_gn=TRMU;refGene_tx=X
M_019835734.1;refGene_gn=TRMU;refGene_tx=XM_023257631.1;refGene_gn=TRM
U;refGene_tx=XM_023257632.1;refGene_gn=TRMU;refGene_tx=XM_023257633.1
B4	140962202	140962375
refGene_gn=TRMU;refGene_tx=XM_011284325.2;refGene_gn=TRMU;refGene_tx=X
M_019835734.1;refGene_gn=TRMU;refGene_tx=XM_023257631.1;refGene_gn=TRM
U;refGene_tx=XM_023257632.1;refGene_gn=TRMU;refGene_tx=XM_023257633.1
B4	140963905	140963959
refGene_gn=TRMU;refGene_tx=XM_011284325.2;refGene_gn=TRMU;refGene_tx=X
M_019835734.1;refGene_gn=TRMU;refGene_tx=XM_023257631.1;refGene_gn=TRM
U;refGene_tx=XM_023257632.1;refGene_gn=TRMU;refGene_tx=XM_023257633.1
B4	140964043	140964110
refGene_gn=TRMU;refGene_tx=XM_011284325.2;refGene_gn=TRMU;refGene_tx=X
M_019835734.1;refGene_gn=TRMU;refGene_tx=XM_023257631.1;refGene_gn=TRM
U;refGene_tx=XM_023257633.1
B4	140965404	140965505
refGene_gn=TRMU;refGene_tx=XM_011284325.2;refGene_gn=TRMU;refGene_tx=X
M_019835734.1;refGene_gn=TRMU;refGene_tx=XM_023257631.1
B4	140966736	140966881
refGene_gn=TRMU;refGene_tx=XM_011284325.2;refGene_gn=TRMU;refGene_tx=X
M_019835734.1;refGene_gn=TRMU;refGene_tx=XM_023257631.1;refGene_gn=TRM
U;refGene_tx=XM_023257632.1;refGene_gn=TRMU;refGene_tx=XM_023257633.1
B4	140967236	140967319
refGene_gn=TRMU;refGene_tx=XM_011284325.2;refGene_gn=TRMU;refGene_tx=X

M_019835734.1;refGene_gn=TRMU;refGene_tx=XM_023257631.1;refGene_gn=TRMU;refGene_tx=XM_023257632.1;refGene_gn=TRMU;refGene_tx=XM_023257633.1
B4 140967857 140968022
refGene_gn=TRMU;refGene_tx=XM_011284325.2;refGene_gn=TRMU;refGene_tx=XM_019835734.1;refGene_gn=TRMU;refGene_tx=XM_023257631.1;refGene_gn=TRMU;refGene_tx=XM_023257632.1;refGene_gn=TRMU;refGene_tx=XM_023257633.1
B4 140973492 140973517
refGene_gn=CELSR1;refGene_tx=XM_011284343.3
B4 140974283 140974420
refGene_gn=CELSR1;refGene_tx=XM_011284343.3
B4 140974421 140974546
refGene_gn=CELSR1;refGene_tx=XM_011284343.3
B4 140975066 140975300
refGene_gn=CELSR1;refGene_tx=XM_011284343.3
B4 140975763 140975913
refGene_gn=CELSR1;refGene_tx=XM_011284343.3
B4 140976168 140976272
refGene_gn=CELSR1;refGene_tx=XM_011284343.3
B4 140976556 140976651
refGene_gn=CELSR1;refGene_tx=XM_011284343.3
B4 140977079 140977205
refGene_gn=CELSR1;refGene_tx=XM_011284343.3
B4 140977819 140977946
refGene_gn=CELSR1;refGene_tx=XM_011284343.3
B4 140978565 140978645
refGene_gn=CELSR1;refGene_tx=XM_011284343.3
B4 140978950 140979063
refGene_gn=CELSR1;refGene_tx=XM_011284343.3
B4 140979538 140979713
refGene_gn=CELSR1;refGene_tx=XM_011284343.3
B4 140984144 140984345
refGene_gn=CELSR1;refGene_tx=XM_011284343.3
B4 140986595 140986722
refGene_gn=CELSR1;refGene_tx=XM_011284343.3
B4 140988354 140988522
refGene_gn=CELSR1;refGene_tx=XM_011284343.3
B4 140989502 140989707
refGene_gn=CELSR1;refGene_tx=XM_011284343.3
B4 140993690 140993834
refGene_gn=CELSR1;refGene_tx=XM_011284343.3
B4 140995374 140995558
refGene_gn=CELSR1;refGene_tx=XM_011284343.3
B4 140996768 140996978
refGene_gn=CELSR1;refGene_tx=XM_011284343.3
B4 140997461 140997556
refGene_gn=CELSR1;refGene_tx=XM_011284343.3
B4 140998176 140998278
refGene_gn=CELSR1;refGene_tx=XM_011284343.3
B4 140998615 140998799
refGene_gn=CELSR1;refGene_tx=XM_011284343.3

B4 141000632 141000753
refGene_gn=CELSR1;refGene_tx=XM_011284343.3
B4 141002614 141002756
refGene_gn=CELSR1;refGene_tx=XM_011284343.3
B4 141003805 141003980
refGene_gn=CELSR1;refGene_tx=XM_011284343.3
B4 141004856 141004970
refGene_gn=CELSR1;refGene_tx=XM_011284343.3
B4 141006010 141006196
refGene_gn=CELSR1;refGene_tx=XM_011284343.3
B4 141011735 141011902
refGene_gn=CELSR1;refGene_tx=XM_011284343.3
B4 141013613 141013739
refGene_gn=CELSR1;refGene_tx=XM_011284343.3
B4 141014185 141014349
refGene_gn=CELSR1;refGene_tx=XM_011284343.3
B4 141015387 141015545
refGene_gn=CELSR1;refGene_tx=XM_011284343.3
B4 141034882 141034971
refGene_gn=CELSR1;refGene_tx=XM_011284343.3
B4 141035353 141035469
refGene_gn=CELSR1;refGene_tx=XM_011284343.3
B4 141039059 141039282
refGene_gn=CELSR1;refGene_tx=XM_011284343.3
B4 141056926 141057565
refGene_gn=CELSR1;refGene_tx=XM_011284343.3
B4 141109523 141113142
refGene_gn=CELSR1;refGene_tx=XM_011284343.3
B4 141146142 141146171
refGene_gn=GRAMD4;refGene_tx=XM_023257634.1
B4 141180529 141180740
refGene_gn=GRAMD4;refGene_tx=XM_023257634.1
B4 141184449 141184570
refGene_gn=GRAMD4;refGene_tx=XM_023257634.1
B4 141200269 141200390
refGene_gn=GRAMD4;refGene_tx=XM_023257634.1
B4 141204551 141204613
refGene_gn=GRAMD4;refGene_tx=XM_023257634.1
B4 141206459 141206592
refGene_gn=GRAMD4;refGene_tx=XM_023257634.1
B4 141207037 141207063
refGene_gn=GRAMD4;refGene_tx=XM_023257634.1
B4 141207613 141207705
refGene_gn=GRAMD4;refGene_tx=XM_023257634.1
B4 141208951 141209043
refGene_gn=GRAMD4;refGene_tx=XM_023257634.1
B4 141209947 141209996
refGene_gn=GRAMD4;refGene_tx=XM_023257634.1
B4 141211023 141211095
refGene_gn=GRAMD4;refGene_tx=XM_023257634.1

B4	141211540	141211584
refGene_gn=GRAMD4;refGene_tx=XM_023257634.1
B4	141211659	141211769
refGene_gn=GRAMD4;refGene_tx=XM_023257634.1
B4	141213307	141213462
refGene_gn=GRAMD4;refGene_tx=XM_023257634.1
B4	141215130	141215275
refGene_gn=GRAMD4;refGene_tx=XM_023257634.1
B4	141215949	141215984
refGene_gn=GRAMD4;refGene_tx=XM_023257634.1
B4	141215985	141216044
refGene_gn=GRAMD4;refGene_tx=XM_023257634.1
B4	141216598	141216683
refGene_gn=GRAMD4;refGene_tx=XM_023257634.1
B4	141217493	141217562
refGene_gn=GRAMD4;refGene_tx=XM_023257634.1
B4	141218190	141218295
refGene_gn=GRAMD4;refGene_tx=XM_023257634.1
B4	141225310	141225417
refGene_gn=LOC111561468;refGene_tx=XM_023257786.1
B4	141225503	141225665
refGene_gn=LOC111561468;refGene_tx=XM_023257786.1
B4	141225710	141225866
refGene_gn=LOC111561468;refGene_tx=XM_023257786.1
B4	141226669	141226807
refGene_gn=LOC111561468;refGene_tx=XM_023257786.1
B4	141226895	141226993
refGene_gn=LOC111561468;refGene_tx=XM_023257786.1
B4	141228250	141228323
refGene_gn=CERK;refGene_tx=XM_023257635.1
B4	141230119	141230328
refGene_gn=CERK;refGene_tx=XM_023257635.1
B4	141230931	141231137
refGene_gn=CERK;refGene_tx=XM_023257635.1
B4	141231745	141231822
refGene_gn=CERK;refGene_tx=XM_023257635.1
B4	141232484	141232590
refGene_gn=CERK;refGene_tx=XM_023257635.1
B4	141234487	141234640
refGene_gn=CERK;refGene_tx=XM_023257635.1
B4	141235498	141235573
refGene_gn=CERK;refGene_tx=XM_023257635.1
B4	141241135	141241281
refGene_gn=CERK;refGene_tx=XM_023257635.1
B4	141241557	141241629
refGene_gn=CERK;refGene_tx=XM_023257635.1
B4	141243518	141243636
refGene_gn=CERK;refGene_tx=XM_023257635.1
B4	141248600	141248723
refGene_gn=CERK;refGene_tx=XM_023257635.1

B4 141249048 141249162
refGene_gn=CERK;refGene_tx=XM_023257635.1
B4 141272930 141273201
refGene_gn=CERK;refGene_tx=XM_023257635.1
B4 141301184 141301246
refGene_gn=TBC1D22A;refGene_tx=XM_011284379.3;refGene_gn=TBC1D22A;refGene_tx=XM_019835736.2;refGene_gn=TBC1D22A;refGene_tx=XM_019835737.2;refGene_gn=TBC1D22A;refGene_tx=XM_019835738.2;refGene_gn=TBC1D22A;refGene_tx=XM_019835739.2;refGene_gn=TBC1D22A;refGene_tx=XM_019835740.2;refGene_gn=TBC1D22A;refGene_tx=XM_019835741.2;refGene_gn=TBC1D22A;refGene_tx=XM_019835742.2
B4 141322731 141322788
refGene_gn=TBC1D22A;refGene_tx=XM_011284379.3;refGene_gn=TBC1D22A;refGene_tx=XM_019835736.2;refGene_gn=TBC1D22A;refGene_tx=XM_019835737.2;refGene_gn=TBC1D22A;refGene_tx=XM_019835738.2;refGene_gn=TBC1D22A;refGene_tx=XM_019835739.2;refGene_gn=TBC1D22A;refGene_tx=XM_019835740.2;refGene_gn=TBC1D22A;refGene_tx=XM_019835741.2;refGene_gn=TBC1D22A;refGene_tx=XM_019835742.2
B4 141323568 141323897
refGene_gn=TBC1D22A;refGene_tx=XM_011284379.3;refGene_gn=TBC1D22A;refGene_tx=XM_019835736.2;refGene_gn=TBC1D22A;refGene_tx=XM_019835737.2;refGene_gn=TBC1D22A;refGene_tx=XM_019835738.2;refGene_gn=TBC1D22A;refGene_tx=XM_019835739.2;refGene_gn=TBC1D22A;refGene_tx=XM_019835740.2;refGene_gn=TBC1D22A;refGene_tx=XM_019835741.2;refGene_gn=TBC1D22A;refGene_tx=XM_019835742.2
B4 141329424 141329601
refGene_gn=TBC1D22A;refGene_tx=XM_011284379.3;refGene_gn=TBC1D22A;refGene_tx=XM_019835736.2;refGene_gn=TBC1D22A;refGene_tx=XM_019835737.2;refGene_gn=TBC1D22A;refGene_tx=XM_019835738.2;refGene_gn=TBC1D22A;refGene_tx=XM_019835739.2;refGene_gn=TBC1D22A;refGene_tx=XM_019835741.2
B4 141390694 141390765
refGene_gn=TBC1D22A;refGene_tx=XM_011284379.3;refGene_gn=TBC1D22A;refGene_tx=XM_019835736.2;refGene_gn=TBC1D22A;refGene_tx=XM_019835737.2;refGene_gn=TBC1D22A;refGene_tx=XM_019835738.2;refGene_gn=TBC1D22A;refGene_tx=XM_019835739.2;refGene_gn=TBC1D22A;refGene_tx=XM_019835740.2;refGene_gn=TBC1D22A;refGene_tx=XM_019835741.2;refGene_gn=TBC1D22A;refGene_tx=XM_019835742.2
B4 141402534 141402663
refGene_gn=TBC1D22A;refGene_tx=XM_011284379.3;refGene_gn=TBC1D22A;refGene_tx=XM_019835736.2;refGene_gn=TBC1D22A;refGene_tx=XM_019835737.2;refGene_gn=TBC1D22A;refGene_tx=XM_019835738.2;refGene_gn=TBC1D22A;refGene_tx=XM_019835739.2;refGene_gn=TBC1D22A;refGene_tx=XM_019835740.2;refGene_gn=TBC1D22A;refGene_tx=XM_019835741.2;refGene_gn=TBC1D22A;refGene_tx=XM_019835742.2
B4 141405825 141405888
refGene_gn=TBC1D22A;refGene_tx=XM_011284379.3;refGene_gn=TBC1D22A;refGene_tx=XM_019835736.2;refGene_gn=TBC1D22A;refGene_tx=XM_019835737.2;refGene_gn=TBC1D22A;refGene_tx=XM_019835738.2;refGene_gn=TBC1D22A;refGene_tx=XM_019835739.2;refGene_gn=TBC1D22A;refGene_tx=XM_019835740.2;refGene_gn=TBC1D22A;refGene_tx=XM_019835741.2;refGene_gn=TBC1D22A;refGene_

tx=XM_019835742.2
B4 141425184 141425299
refGene_gn=TBC1D22A;refGene_tx=XM_011284379.3;refGene_gn=TBC1D22A;refGene_tx=XM_019835736.2;refGene_gn=TBC1D22A;refGene_tx=XM_019835737.2;refGene_gn=TBC1D22A;refGene_tx=XM_019835738.2;refGene_gn=TBC1D22A;refGene_tx=XM_019835739.2;refGene_gn=TBC1D22A;refGene_tx=XM_019835740.2;refGene_gn=TBC1D22A;refGene_tx=XM_019835741.2;refGene_gn=TBC1D22A;refGene_tx=XM_019835742.2
B4 141487052 141487162
refGene_gn=TBC1D22A;refGene_tx=XM_011284379.3;refGene_gn=TBC1D22A;refGene_tx=XM_019835736.2;refGene_gn=TBC1D22A;refGene_tx=XM_019835737.2;refGene_gn=TBC1D22A;refGene_tx=XM_019835738.2;refGene_gn=TBC1D22A;refGene_tx=XM_019835739.2;refGene_gn=TBC1D22A;refGene_tx=XM_019835740.2;refGene_gn=TBC1D22A;refGene_tx=XM_019835741.2;refGene_gn=TBC1D22A;refGene_tx=XM_019835742.2
B4 141508442 141508518
refGene_gn=TBC1D22A;refGene_tx=XM_011284379.3;refGene_gn=TBC1D22A;refGene_tx=XM_019835736.2;refGene_gn=TBC1D22A;refGene_tx=XM_019835737.2;refGene_gn=TBC1D22A;refGene_tx=XM_019835738.2;refGene_gn=TBC1D22A;refGene_tx=XM_019835739.2;refGene_gn=TBC1D22A;refGene_tx=XM_019835740.2;refGene_gn=TBC1D22A;refGene_tx=XM_019835741.2;refGene_gn=TBC1D22A;refGene_tx=XM_019835742.2
B4 141517264 141517644
refGene_gn=TBC1D22A;refGene_tx=XM_019835736.2;refGene_gn=TBC1D22A;refGene_tx=XM_019835738.2;refGene_gn=TBC1D22A;refGene_tx=XM_019835740.2
B4 141536186 141536314
refGene_gn=TBC1D22A;refGene_tx=XM_011284379.3;refGene_gn=TBC1D22A;refGene_tx=XM_019835737.2;refGene_gn=TBC1D22A;refGene_tx=XM_019835739.2;refGene_gn=TBC1D22A;refGene_tx=XM_019835741.2;refGene_gn=TBC1D22A;refGene_tx=XM_019835742.2
B4 141577543 141577573
refGene_gn=TBC1D22A;refGene_tx=XM_019835741.2
B4 141590347 141590443
refGene_gn=TBC1D22A;refGene_tx=XM_011284379.3;refGene_gn=TBC1D22A;refGene_tx=XM_019835737.2;refGene_gn=TBC1D22A;refGene_tx=XM_019835742.2
B4 141623794 141623923
refGene_gn=TBC1D22A;refGene_tx=XM_011284379.3;refGene_gn=TBC1D22A;refGene_tx=XM_019835737.2;refGene_gn=TBC1D22A;refGene_tx=XM_019835739.2;refGene_gn=TBC1D22A;refGene_tx=XM_019835742.2
B4 142601739 142602523
refGene_gn=LOC109501865;refGene_tx=XM_019835745.2
B4 142602634 142602738
refGene_gn=LOC109501865;refGene_tx=XM_019835745.2
B4 142605580 142605692
refGene_gn=FAM19A5;refGene_tx=XM_023257636.1
B4 142677243 142677334
refGene_gn=FAM19A5;refGene_tx=XM_011284349.3
B4 142722059 142722209
refGene_gn=FAM19A5;refGene_tx=XM_011284349.3;refGene_gn=FAM19A5;refGene_tx=XM_023257636.1

B4	142762077	142762205
refGene_gn=FAM19A5;refGene_tx=XM_011284349.3;refGene_gn=FAM19A5;refGen
e_tx=XM_023257636.1
B4	142795869	142795878
refGene_gn=FAM19A5;refGene_tx=XM_011284349.3;refGene_gn=FAM19A5;refGen
e_tx=XM_023257636.1
B4	143528300	143528963
refGene_gn=LOC102899362;refGene_tx=XM_023257787.1
B4	143529062	143530333
refGene_gn=LOC102899362;refGene_tx=XM_023257787.1
B4	143530792	143530985
refGene_gn=LOC102899362;refGene_tx=XM_023257787.1
B4	143578262	143578446
refGene_gn=BRD1;refGene_tx=XM_023257637.1;refGene_gn=BRD1;refGene_tx=X
M_023257638.1;refGene_gn=BRD1;refGene_tx=XM_023257639.1;refGene_gn=BRD
1;refGene_tx=XM_023257640.1;refGene_gn=BRD1;refGene_tx=XM_023257641.1;
refGene_gn=BRD1;refGene_tx=XM_023257642.1;refGene_gn=BRD1;refGene_tx=X
M_023257643.1
B4	143579623	143579778
refGene_gn=BRD1;refGene_tx=XM_023257637.1;refGene_gn=BRD1;refGene_tx=X
M_023257638.1;refGene_gn=BRD1;refGene_tx=XM_023257639.1;refGene_gn=BRD
1;refGene_tx=XM_023257640.1;refGene_gn=BRD1;refGene_tx=XM_023257641.1;
refGene_gn=BRD1;refGene_tx=XM_023257642.1;refGene_gn=BRD1;refGene_tx=X
M_023257643.1
B4	143580065	143580175
refGene_gn=BRD1;refGene_tx=XM_023257637.1;refGene_gn=BRD1;refGene_tx=X
M_023257638.1;refGene_gn=BRD1;refGene_tx=XM_023257639.1;refGene_gn=BRD
1;refGene_tx=XM_023257640.1;refGene_gn=BRD1;refGene_tx=XM_023257641.1;
refGene_gn=BRD1;refGene_tx=XM_023257642.1;refGene_gn=BRD1;refGene_tx=X
M_023257643.1
B4	143580615	143580743
refGene_gn=BRD1;refGene_tx=XM_023257637.1;refGene_gn=BRD1;refGene_tx=X
M_023257638.1;refGene_gn=BRD1;refGene_tx=XM_023257639.1;refGene_gn=BRD
1;refGene_tx=XM_023257640.1;refGene_gn=BRD1;refGene_tx=XM_023257641.1;
refGene_gn=BRD1;refGene_tx=XM_023257642.1;refGene_gn=BRD1;refGene_tx=X
M_023257643.1
B4	143581086	143581222
refGene_gn=BRD1;refGene_tx=XM_023257637.1;refGene_gn=BRD1;refGene_tx=X
M_023257638.1;refGene_gn=BRD1;refGene_tx=XM_023257639.1;refGene_gn=BRD
1;refGene_tx=XM_023257640.1;refGene_gn=BRD1;refGene_tx=XM_023257641.1;
refGene_gn=BRD1;refGene_tx=XM_023257642.1;refGene_gn=BRD1;refGene_tx=X
M_023257643.1
B4	143587534	143588032
refGene_gn=BRD1;refGene_tx=XM_023257637.1;refGene_gn=BRD1;refGene_tx=X
M_023257638.1;refGene_gn=BRD1;refGene_tx=XM_023257639.1;refGene_gn=BRD
1;refGene_tx=XM_023257640.1;refGene_gn=BRD1;refGene_tx=XM_023257641.1
B4	143593305	143593566
refGene_gn=BRD1;refGene_tx=XM_023257637.1;refGene_gn=BRD1;refGene_tx=X
M_023257638.1;refGene_gn=BRD1;refGene_tx=XM_023257639.1;refGene_gn=BRD
1;refGene_tx=XM_023257640.1;refGene_gn=BRD1;refGene_tx=XM_023257641.1;

refGene_gn=BRD1;refGene_tx=XM_023257642.1;refGene_gn=BRD1;refGene_tx=X
M_023257643.1
B4 143595317 143595630
refGene_gn=BRD1;refGene_tx=XM_023257637.1;refGene_gn=BRD1;refGene_tx=X
M_023257638.1;refGene_gn=BRD1;refGene_tx=XM_023257639.1;refGene_gn=BRD
1;refGene_tx=XM_023257640.1;refGene_gn=BRD1;refGene_tx=XM_023257641.1;
refGene_gn=BRD1;refGene_tx=XM_023257642.1;refGene_gn=BRD1;refGene_tx=X
M_023257643.1
B4 143596080 143596209
refGene_gn=BRD1;refGene_tx=XM_023257637.1;refGene_gn=BRD1;refGene_tx=X
M_023257638.1;refGene_gn=BRD1;refGene_tx=XM_023257639.1;refGene_gn=BRD
1;refGene_tx=XM_023257640.1;refGene_gn=BRD1;refGene_tx=XM_023257641.1;
refGene_gn=BRD1;refGene_tx=XM_023257642.1;refGene_gn=BRD1;refGene_tx=X
M_023257643.1
B4 143596441 143596573
refGene_gn=BRD1;refGene_tx=XM_023257637.1;refGene_gn=BRD1;refGene_tx=X
M_023257638.1;refGene_gn=BRD1;refGene_tx=XM_023257639.1;refGene_gn=BRD
1;refGene_tx=XM_023257640.1;refGene_gn=BRD1;refGene_tx=XM_023257641.1;
refGene_gn=BRD1;refGene_tx=XM_023257642.1;refGene_gn=BRD1;refGene_tx=X
M_023257643.1
B4 143602421 143602578
refGene_gn=BRD1;refGene_tx=XM_023257637.1;refGene_gn=BRD1;refGene_tx=X
M_023257638.1;refGene_gn=BRD1;refGene_tx=XM_023257639.1;refGene_gn=BRD
1;refGene_tx=XM_023257640.1;refGene_gn=BRD1;refGene_tx=XM_023257641.1;
refGene_gn=BRD1;refGene_tx=XM_023257642.1;refGene_gn=BRD1;refGene_tx=X
M_023257643.1
B4 143615298 143616665
refGene_gn=BRD1;refGene_tx=XM_023257637.1;refGene_gn=BRD1;refGene_tx=X
M_023257638.1;refGene_gn=BRD1;refGene_tx=XM_023257639.1;refGene_gn=BRD
1;refGene_tx=XM_023257640.1;refGene_gn=BRD1;refGene_tx=XM_023257641.1;
refGene_gn=BRD1;refGene_tx=XM_023257642.1;refGene_gn=BRD1;refGene_tx=X
M_023257643.1
B4 143655908 143659412
refGene_gn=ZBED4;refGene_tx=XM_011284350.3;refGene_gn=ZBED4;refGene_tx
=XM_023257644.1;refGene_gn=ZBED4;refGene_tx=XM_023257645.1
B4 143673513 143673751
refGene_gn=ALG12;refGene_tx=XM_011284351.3;refGene_gn=ALG12;refGene_tx
=XM_011284352.3;refGene_gn=ALG12;refGene_tx=XM_011284353.3;refGene_gn=
ALG12;refGene_tx=XM_019835749.2;refGene_gn=ALG12;refGene_tx=XM_0198357
50.2;refGene_gn=ALG12;refGene_tx=XM_023257646.1
B4 143673876 143673952
refGene_gn=ALG12;refGene_tx=XM_011284351.3;refGene_gn=ALG12;refGene_tx
=XM_011284352.3;refGene_gn=ALG12;refGene_tx=XM_011284353.3;refGene_gn=
ALG12;refGene_tx=XM_019835749.2;refGene_gn=ALG12;refGene_tx=XM_0198357
50.2;refGene_gn=ALG12;refGene_tx=XM_023257646.1
B4 143674049 143674219
refGene_gn=ALG12;refGene_tx=XM_011284351.3;refGene_gn=ALG12;refGene_tx
=XM_011284352.3;refGene_gn=ALG12;refGene_tx=XM_011284353.3;refGene_gn=
ALG12;refGene_tx=XM_019835749.2;refGene_gn=ALG12;refGene_tx=XM_0198357
50.2;refGene_gn=ALG12;refGene_tx=XM_023257646.1

B4	143675190	143675414
refGene_gn=ALG12;refGene_tx=XM_011284351.3;refGene_gn=ALG12;refGene_tx=XM_011284352.3;refGene_gn=ALG12;refGene_tx=XM_011284353.3;refGene_gn=ALG12;refGene_tx=XM_019835749.2;refGene_gn=ALG12;refGene_tx=XM_019835750.2;refGene_gn=ALG12;refGene_tx=XM_023257646.1
B4	143675876	143675980
refGene_gn=ALG12;refGene_tx=XM_011284351.3;refGene_gn=ALG12;refGene_tx=XM_011284352.3;refGene_gn=ALG12;refGene_tx=XM_011284353.3;refGene_gn=ALG12;refGene_tx=XM_019835749.2;refGene_gn=ALG12;refGene_tx=XM_019835750.2;refGene_gn=ALG12;refGene_tx=XM_023257646.1
B4	143676309	143676504
refGene_gn=ALG12;refGene_tx=XM_011284351.3;refGene_gn=ALG12;refGene_tx=XM_011284352.3;refGene_gn=ALG12;refGene_tx=XM_019835749.2;refGene_gn=ALG12;refGene_tx=XM_019835750.2;refGene_gn=ALG12;refGene_tx=XM_023257646.1
B4	143677151	143677325
refGene_gn=ALG12;refGene_tx=XM_011284351.3;refGene_gn=ALG12;refGene_tx=XM_011284352.3;refGene_gn=ALG12;refGene_tx=XM_011284353.3;refGene_gn=ALG12;refGene_tx=XM_019835749.2;refGene_gn=ALG12;refGene_tx=XM_019835750.2;refGene_gn=ALG12;refGene_tx=XM_023257646.1
B4	143678818	143678951
refGene_gn=ALG12;refGene_tx=XM_011284351.3;refGene_gn=ALG12;refGene_tx=XM_011284352.3;refGene_gn=ALG12;refGene_tx=XM_011284353.3;refGene_gn=ALG12;refGene_tx=XM_019835749.2;refGene_gn=ALG12;refGene_tx=XM_019835750.2;refGene_gn=ALG12;refGene_tx=XM_023257646.1
B4	143679022	143679190
refGene_gn=ALG12;refGene_tx=XM_011284351.3;refGene_gn=ALG12;refGene_tx=XM_011284352.3;refGene_gn=ALG12;refGene_tx=XM_011284353.3;refGene_gn=ALG12;refGene_tx=XM_019835749.2;refGene_gn=ALG12;refGene_tx=XM_019835750.2;refGene_gn=ALG12;refGene_tx=XM_023257646.1
B4	143683152	143683293
refGene_gn=CRELD2;refGene_tx=XM_011284354.3
B4	143683815	143683898
refGene_gn=CRELD2;refGene_tx=XM_011284354.3
B4	143684417	143684528
refGene_gn=CRELD2;refGene_tx=XM_011284354.3
B4	143685004	143685096
refGene_gn=CRELD2;refGene_tx=XM_011284354.3
B4	143686011	143686188
refGene_gn=CRELD2;refGene_tx=XM_011284354.3
B4	143687056	143687106
refGene_gn=CRELD2;refGene_tx=XM_011284354.3
B4	143687107	143687153
refGene_gn=CRELD2;refGene_tx=XM_011284354.3
B4	143688663	143688747
refGene_gn=CRELD2;refGene_tx=XM_011284354.3
B4	143690076	143690172
refGene_gn=CRELD2;refGene_tx=XM_011284354.3
B4	143690754	143690901
refGene_gn=CRELD2;refGene_tx=XM_011284354.3

B4 143691949 143691996
refGene_gn=CRELD2;refGene_tx=XM_011284354.3
B4 143722303 143722388
refGene_gn=PIM3;refGene_tx=XM_023257647.1
B4 143722502 143722612
refGene_gn=PIM3;refGene_tx=XM_023257647.1
B4 143722716 143722767
refGene_gn=PIM3;refGene_tx=XM_023257647.1
B4 143722850 143723220
refGene_gn=PIM3;refGene_tx=XM_023257647.1
B4 143723756 143723933
refGene_gn=PIM3;refGene_tx=XM_023257647.1
B4 143724017 143724205
refGene_gn=PIM3;refGene_tx=XM_023257647.1
B4 143751092 143751118
refGene_gn=LOC101100574;refGene_tx=XM_023257648.1
B4 143751381 143751570
refGene_gn=LOC101100574;refGene_tx=XM_023257648.1
B4 143752284 143752342
refGene_gn=LOC101100574;refGene_tx=XM_023257648.1
B4 143752528 143752609
refGene_gn=LOC101100574;refGene_tx=XM_023257648.1
B4 143752802 143752966
refGene_gn=LOC101100574;refGene_tx=XM_023257648.1
B4 143753170 143753228
refGene_gn=LOC101100574;refGene_tx=XM_023257648.1
B4 143753508 143753553
refGene_gn=LOC101100574;refGene_tx=XM_023257648.1
B4 143753621 143753785
refGene_gn=LOC101100574;refGene_tx=XM_023257648.1
B4 143754261 143754316
refGene_gn=LOC101100574;refGene_tx=XM_023257648.1
B4 143754396 143754468
refGene_gn=LOC101100574;refGene_tx=XM_023257648.1
B4 143754668 143754808
refGene_gn=LOC101100574;refGene_tx=XM_023257648.1
B4 143755058 143755103
refGene_gn=LOC101100574;refGene_tx=XM_023257648.1
B4 143755360 143755475
refGene_gn=LOC101100574;refGene_tx=XM_023257648.1
B4 143755705 143755815
refGene_gn=LOC101100574;refGene_tx=XM_023257648.1
B4 143755995 143756145
refGene_gn=LOC101100574;refGene_tx=XM_023257648.1;refGene_gn=LOC101100
574;refGene_tx=XM_023257652.1
B4 143768413 143768702
refGene_gn=LOC101100574;refGene_tx=XM_023257650.1
B4 143768787 143768815
refGene_gn=LOC101100574;refGene_tx=XM_023257650.1;refGene_gn=LOC101100
574;refGene_tx=XM_023257652.1

B4	143776085	143776246
refGene_gn=LOC101100574;refGene_tx=XM_023257651.1
B4	143779489	143779866
refGene_gn=LOC101100574;refGene_tx=XM_023257649.1
B4	143794732	143795213
refGene_gn=LOC101100574;refGene_tx=XM_023257648.1;refGene_gn=LOC101100574;refGene_tx=XM_023257649.1;refGene_gn=LOC101100574;refGene_tx=XM_023257650.1;refGene_gn=LOC101100574;refGene_tx=XM_023257651.1;refGene_gn=LOC101100574;refGene_tx=XM_023257652.1
B4	143795815	143796251
refGene_gn=LOC101100574;refGene_tx=XM_023257648.1;refGene_gn=LOC101100574;refGene_tx=XM_023257649.1;refGene_gn=LOC101100574;refGene_tx=XM_023257650.1;refGene_gn=LOC101100574;refGene_tx=XM_023257651.1;refGene_gn=LOC101100574;refGene_tx=XM_023257652.1
B4	143798270	143798514
refGene_gn=LOC101100574;refGene_tx=XM_023257648.1;refGene_gn=LOC101100574;refGene_tx=XM_023257649.1;refGene_gn=LOC101100574;refGene_tx=XM_023257650.1;refGene_gn=LOC101100574;refGene_tx=XM_023257651.1;refGene_gn=LOC101100574;refGene_tx=XM_023257652.1
B4	143800571	143800720
refGene_gn=LOC101100574;refGene_tx=XM_023257648.1;refGene_gn=LOC101100574;refGene_tx=XM_023257649.1;refGene_gn=LOC101100574;refGene_tx=XM_023257650.1;refGene_gn=LOC101100574;refGene_tx=XM_023257651.1;refGene_gn=LOC101100574;refGene_tx=XM_023257652.1
B4	143813188	143813375
refGene_gn=LOC101100574;refGene_tx=XM_023257648.1;refGene_gn=LOC101100574;refGene_tx=XM_023257649.1;refGene_gn=LOC101100574;refGene_tx=XM_023257650.1;refGene_gn=LOC101100574;refGene_tx=XM_023257651.1;refGene_gn=LOC101100574;refGene_tx=XM_023257652.1
B4	143819569	143819677
refGene_gn=LOC101100574;refGene_tx=XM_023257648.1;refGene_gn=LOC101100574;refGene_tx=XM_023257649.1;refGene_gn=LOC101100574;refGene_tx=XM_023257650.1;refGene_gn=LOC101100574;refGene_tx=XM_023257651.1;refGene_gn=LOC101100574;refGene_tx=XM_023257652.1
B4	143821127	143821242
refGene_gn=LOC101100574;refGene_tx=XM_023257648.1;refGene_gn=LOC101100574;refGene_tx=XM_023257649.1;refGene_gn=LOC101100574;refGene_tx=XM_023257650.1;refGene_gn=LOC101100574;refGene_tx=XM_023257651.1;refGene_gn=LOC101100574;refGene_tx=XM_023257652.1
B4	143822333	143822462
refGene_gn=LOC101100574;refGene_tx=XM_023257648.1;refGene_gn=LOC101100574;refGene_tx=XM_023257649.1;refGene_gn=LOC101100574;refGene_tx=XM_023257650.1;refGene_gn=LOC101100574;refGene_tx=XM_023257651.1;refGene_gn=LOC101100574;refGene_tx=XM_023257652.1
B4	143824227	143824301
refGene_gn=LOC101100574;refGene_tx=XM_023257648.1;refGene_gn=LOC101100574;refGene_tx=XM_023257649.1;refGene_gn=LOC101100574;refGene_tx=XM_023257650.1;refGene_gn=LOC101100574;refGene_tx=XM_023257651.1;refGene_gn=LOC101100574;refGene_tx=XM_023257652.1
B4	143824962	143825101

refGene_gn=LOC101100574;refGene_tx=XM_023257648.1;refGene_gn=LOC101100574;refGene_tx=XM_023257649.1;refGene_gn=LOC101100574;refGene_tx=XM_023257650.1;refGene_gn=LOC101100574;refGene_tx=XM_023257651.1;refGene_gn=LOC101100574;refGene_tx=XM_023257652.1
B4 143828396 143828488
refGene_gn=LOC101100574;refGene_tx=XM_023257648.1;refGene_gn=LOC101100574;refGene_tx=XM_023257649.1;refGene_gn=LOC101100574;refGene_tx=XM_023257650.1;refGene_gn=LOC101100574;refGene_tx=XM_023257651.1;refGene_gn=LOC101100574;refGene_tx=XM_023257652.1
B4 143836178 143836320
refGene_gn=LOC101100574;refGene_tx=XM_023257648.1;refGene_gn=LOC101100574;refGene_tx=XM_023257649.1;refGene_gn=LOC101100574;refGene_tx=XM_023257650.1;refGene_gn=LOC101100574;refGene_tx=XM_023257651.1;refGene_gn=LOC101100574;refGene_tx=XM_023257652.1
B4 143840061 143840232
refGene_gn=MLC1;refGene_tx=XM_006934189.3;refGene_gn=MLC1;refGene_tx=XM_019835763.2
B4 143841383 143841536
refGene_gn=MLC1;refGene_tx=XM_006934189.3;refGene_gn=MLC1;refGene_tx=XM_019835763.2
B4 143845402 143845525
refGene_gn=MLC1;refGene_tx=XM_006934189.3;refGene_gn=MLC1;refGene_tx=XM_019835763.2
B4 143846197 143846254
refGene_gn=MLC1;refGene_tx=XM_006934189.3;refGene_gn=MLC1;refGene_tx=XM_019835763.2
B4 143849545 143849662
refGene_gn=MLC1;refGene_tx=XM_006934189.3;refGene_gn=MLC1;refGene_tx=XM_019835763.2
B4 143850586 143850658
refGene_gn=MLC1;refGene_tx=XM_006934189.3;refGene_gn=MLC1;refGene_tx=XM_019835763.2
B4 143851165 143851264
refGene_gn=MLC1;refGene_tx=XM_006934189.3;refGene_gn=MLC1;refGene_tx=XM_019835763.2
B4 143857728 143857830
refGene_gn=MLC1;refGene_tx=XM_006934189.3;refGene_gn=MLC1;refGene_tx=XM_019835763.2
B4 143858212 143858266
refGene_gn=MLC1;refGene_tx=XM_006934189.3;refGene_gn=MLC1;refGene_tx=XM_019835763.2
B4 143859655 143859745
refGene_gn=MLC1;refGene_tx=XM_006934189.3;refGene_gn=MLC1;refGene_tx=XM_019835763.2
B4 143861266 143861446
refGene_gn=MLC1;refGene_tx=XM_006934189.3;refGene_gn=MLC1;refGene_tx=XM_019835763.2
B4 143865071 143865171
refGene_gn=MOV10L1;refGene_tx=XM_003989414.5;refGene_gn=MOV10L1;refGene_tx=XM_023257653.1;refGene_gn=MOV10L1;refGene_tx=XM_023257655.1

B4	143866666	143866851
refGene_gn=MOV10L1;refGene_tx=XM_003989414.5;refGene_gn=MOV10L1;refGene_tx=XM_023257653.1;refGene_gn=MOV10L1;refGene_tx=XM_023257654.1;refGene_gn=MOV10L1;refGene_tx=XM_023257655.1
B4	143877344	143877507
refGene_gn=MOV10L1;refGene_tx=XM_003989414.5;refGene_gn=MOV10L1;refGene_tx=XM_023257653.1;refGene_gn=MOV10L1;refGene_tx=XM_023257654.1;refGene_gn=MOV10L1;refGene_tx=XM_023257655.1
B4	143879133	143879164
refGene_gn=MOV10L1;refGene_tx=XM_006934191.4
B4	143884314	143884427
refGene_gn=MOV10L1;refGene_tx=XM_003989414.5;refGene_gn=MOV10L1;refGene_tx=XM_006934191.4;refGene_gn=MOV10L1;refGene_tx=XM_023257653.1;refGene_gn=MOV10L1;refGene_tx=XM_023257654.1;refGene_gn=MOV10L1;refGene_tx=XM_023257655.1
B4	143884988	143885176
refGene_gn=MOV10L1;refGene_tx=XM_003989414.5;refGene_gn=MOV10L1;refGene_tx=XM_006934191.4;refGene_gn=MOV10L1;refGene_tx=XM_023257653.1;refGene_gn=MOV10L1;refGene_tx=XM_023257654.1;refGene_gn=MOV10L1;refGene_tx=XM_023257655.1
B4	143887493	143887637
refGene_gn=MOV10L1;refGene_tx=XM_003989414.5;refGene_gn=MOV10L1;refGene_tx=XM_006934191.4;refGene_gn=MOV10L1;refGene_tx=XM_023257653.1;refGene_gn=MOV10L1;refGene_tx=XM_023257654.1;refGene_gn=MOV10L1;refGene_tx=XM_023257655.1
B4	143888129	143888371
refGene_gn=MOV10L1;refGene_tx=XM_003989414.5;refGene_gn=MOV10L1;refGene_tx=XM_006934191.4;refGene_gn=MOV10L1;refGene_tx=XM_023257653.1;refGene_gn=MOV10L1;refGene_tx=XM_023257654.1;refGene_gn=MOV10L1;refGene_tx=XM_023257655.1
B4	143888784	143888917
refGene_gn=MOV10L1;refGene_tx=XM_003989414.5;refGene_gn=MOV10L1;refGene_tx=XM_006934191.4;refGene_gn=MOV10L1;refGene_tx=XM_019835761.2;refGene_gn=MOV10L1;refGene_tx=XM_023257653.1;refGene_gn=MOV10L1;refGene_tx=XM_023257654.1;refGene_gn=MOV10L1;refGene_tx=XM_023257655.1
B4	143890243	143890438
refGene_gn=MOV10L1;refGene_tx=XM_003989414.5;refGene_gn=MOV10L1;refGene_tx=XM_006934191.4;refGene_gn=MOV10L1;refGene_tx=XM_019835761.2;refGene_gn=MOV10L1;refGene_tx=XM_023257653.1;refGene_gn=MOV10L1;refGene_tx=XM_023257654.1;refGene_gn=MOV10L1;refGene_tx=XM_023257655.1
B4	143892982	143893097
refGene_gn=MOV10L1;refGene_tx=XM_003989414.5;refGene_gn=MOV10L1;refGene_tx=XM_006934191.4;refGene_gn=MOV10L1;refGene_tx=XM_019835761.2;refGene_gn=MOV10L1;refGene_tx=XM_023257653.1;refGene_gn=MOV10L1;refGene_tx=XM_023257654.1;refGene_gn=MOV10L1;refGene_tx=XM_023257655.1
B4	143899709	143899887
refGene_gn=MOV10L1;refGene_tx=XM_003989414.5;refGene_gn=MOV10L1;refGene_tx=XM_006934191.4;refGene_gn=MOV10L1;refGene_tx=XM_019835761.2;refGene_gn=MOV10L1;refGene_tx=XM_023257653.1;refGene_gn=MOV10L1;refGene_tx=XM_023257654.1;refGene_gn=MOV10L1;refGene_tx=XM_023257655.1

B4	143900163	143900234
refGene_gn=MOV10L1;refGene_tx=XM_003989414.5;refGene_gn=MOV10L1;refGene_tx=XM_006934191.4;refGene_gn=MOV10L1;refGene_tx=XM_019835761.2;refGene_gn=MOV10L1;refGene_tx=XM_023257653.1;refGene_gn=MOV10L1;refGene_tx=XM_023257654.1;refGene_gn=MOV10L1;refGene_tx=XM_023257655.1
B4	143900351	143900363
refGene_gn=MOV10L1;refGene_tx=XM_011284356.3
B4	143903545	143903637
refGene_gn=MOV10L1;refGene_tx=XM_003989414.5;refGene_gn=MOV10L1;refGene_tx=XM_006934191.4;refGene_gn=MOV10L1;refGene_tx=XM_011284356.3;refGene_gn=MOV10L1;refGene_tx=XM_019835761.2;refGene_gn=MOV10L1;refGene_tx=XM_023257653.1;refGene_gn=MOV10L1;refGene_tx=XM_023257654.1;refGene_gn=MOV10L1;refGene_tx=XM_023257655.1
B4	143910460	143910519
refGene_gn=MOV10L1;refGene_tx=XM_003989414.5;refGene_gn=MOV10L1;refGene_tx=XM_006934191.4;refGene_gn=MOV10L1;refGene_tx=XM_011284356.3;refGene_gn=MOV10L1;refGene_tx=XM_019835761.2;refGene_gn=MOV10L1;refGene_tx=XM_023257653.1;refGene_gn=MOV10L1;refGene_tx=XM_023257654.1;refGene_gn=MOV10L1;refGene_tx=XM_023257655.1
B4	143910934	143911032
refGene_gn=MOV10L1;refGene_tx=XM_003989414.5;refGene_gn=MOV10L1;refGene_tx=XM_006934191.4;refGene_gn=MOV10L1;refGene_tx=XM_011284356.3;refGene_gn=MOV10L1;refGene_tx=XM_019835761.2;refGene_gn=MOV10L1;refGene_tx=XM_023257653.1;refGene_gn=MOV10L1;refGene_tx=XM_023257654.1;refGene_gn=MOV10L1;refGene_tx=XM_023257655.1
B4	143922722	143922831
refGene_gn=MOV10L1;refGene_tx=XM_003989414.5;refGene_gn=MOV10L1;refGene_tx=XM_006934191.4;refGene_gn=MOV10L1;refGene_tx=XM_011284356.3;refGene_gn=MOV10L1;refGene_tx=XM_019835761.2;refGene_gn=MOV10L1;refGene_tx=XM_023257653.1;refGene_gn=MOV10L1;refGene_tx=XM_023257654.1;refGene_gn=MOV10L1;refGene_tx=XM_023257655.1
B4	143923286	143923465
refGene_gn=MOV10L1;refGene_tx=XM_003989414.5;refGene_gn=MOV10L1;refGene_tx=XM_006934191.4;refGene_gn=MOV10L1;refGene_tx=XM_011284356.3;refGene_gn=MOV10L1;refGene_tx=XM_019835761.2;refGene_gn=MOV10L1;refGene_tx=XM_023257653.1;refGene_gn=MOV10L1;refGene_tx=XM_023257654.1;refGene_gn=MOV10L1;refGene_tx=XM_023257655.1
B4	143924695	143924842
refGene_gn=MOV10L1;refGene_tx=XM_003989414.5;refGene_gn=MOV10L1;refGene_tx=XM_006934191.4;refGene_gn=MOV10L1;refGene_tx=XM_011284356.3;refGene_gn=MOV10L1;refGene_tx=XM_019835761.2;refGene_gn=MOV10L1;refGene_tx=XM_019835762.2;refGene_gn=MOV10L1;refGene_tx=XM_023257653.1;refGene_gn=MOV10L1;refGene_tx=XM_023257654.1;refGene_gn=MOV10L1;refGene_tx=XM_023257655.1
B4	143925290	143925412
refGene_gn=MOV10L1;refGene_tx=XM_003989414.5;refGene_gn=MOV10L1;refGene_tx=XM_006934191.4;refGene_gn=MOV10L1;refGene_tx=XM_011284356.3;refGene_gn=MOV10L1;refGene_tx=XM_019835761.2;refGene_gn=MOV10L1;refGene_tx=XM_019835762.2;refGene_gn=MOV10L1;refGene_tx=XM_023257653.1;refGene_gn=MOV10L1;refGene_tx=XM_023257654.1;refGene_gn=MOV10L1;refGene_tx=XM_02

3257655.1
B4 143926809 143926909
refGene_gn=MOV10L1;refGene_tx=XM_003989414.5;refGene_gn=MOV10L1;refGene_tx=XM_006934191.4;refGene_gn=MOV10L1;refGene_tx=XM_011284356.3;refGene_gn=MOV10L1;refGene_tx=XM_019835761.2;refGene_gn=MOV10L1;refGene_tx=XM_019835762.2;refGene_gn=MOV10L1;refGene_tx=XM_023257653.1;refGene_gn=MOV10L1;refGene_tx=XM_023257654.1;refGene_gn=MOV10L1;refGene_tx=XM_023257655.1
B4 143927833 143927998
refGene_gn=MOV10L1;refGene_tx=XM_003989414.5;refGene_gn=MOV10L1;refGene_tx=XM_006934191.4;refGene_gn=MOV10L1;refGene_tx=XM_011284356.3;refGene_gn=MOV10L1;refGene_tx=XM_019835761.2;refGene_gn=MOV10L1;refGene_tx=XM_019835762.2;refGene_gn=MOV10L1;refGene_tx=XM_023257653.1;refGene_gn=MOV10L1;refGene_tx=XM_023257654.1;refGene_gn=MOV10L1;refGene_tx=XM_023257655.1
B4 143930731 143930919
refGene_gn=MOV10L1;refGene_tx=XM_003989414.5;refGene_gn=MOV10L1;refGene_tx=XM_006934191.4;refGene_gn=MOV10L1;refGene_tx=XM_011284356.3;refGene_gn=MOV10L1;refGene_tx=XM_019835761.2;refGene_gn=MOV10L1;refGene_tx=XM_019835762.2;refGene_gn=MOV10L1;refGene_tx=XM_023257653.1;refGene_gn=MOV10L1;refGene_tx=XM_023257654.1;refGene_gn=MOV10L1;refGene_tx=XM_023257655.1
B4 143935564 143935714
refGene_gn=MOV10L1;refGene_tx=XM_003989414.5;refGene_gn=MOV10L1;refGene_tx=XM_006934191.4;refGene_gn=MOV10L1;refGene_tx=XM_011284356.3;refGene_gn=MOV10L1;refGene_tx=XM_019835761.2;refGene_gn=MOV10L1;refGene_tx=XM_019835762.2;refGene_gn=MOV10L1;refGene_tx=XM_023257653.1;refGene_gn=MOV10L1;refGene_tx=XM_023257654.1
B4 143936546 143936654
refGene_gn=MOV10L1;refGene_tx=XM_003989414.5;refGene_gn=MOV10L1;refGene_tx=XM_006934191.4;refGene_gn=MOV10L1;refGene_tx=XM_011284356.3;refGene_gn=MOV10L1;refGene_tx=XM_019835761.2;refGene_gn=MOV10L1;refGene_tx=XM_019835762.2;refGene_gn=MOV10L1;refGene_tx=XM_023257653.1;refGene_gn=MOV10L1;refGene_tx=XM_023257654.1
B4 143937237 143937375
refGene_gn=MOV10L1;refGene_tx=XM_003989414.5;refGene_gn=MOV10L1;refGene_tx=XM_006934191.4;refGene_gn=MOV10L1;refGene_tx=XM_011284356.3;refGene_gn=MOV10L1;refGene_tx=XM_019835761.2;refGene_gn=MOV10L1;refGene_tx=XM_019835762.2;refGene_gn=MOV10L1;refGene_tx=XM_023257653.1;refGene_gn=MOV10L1;refGene_tx=XM_023257654.1
B4 143937465 143937557
refGene_gn=MOV10L1;refGene_tx=XM_003989414.5;refGene_gn=MOV10L1;refGene_tx=XM_006934191.4;refGene_gn=MOV10L1;refGene_tx=XM_011284356.3;refGene_gn=MOV10L1;refGene_tx=XM_019835761.2;refGene_gn=MOV10L1;refGene_tx=XM_019835762.2;refGene_gn=MOV10L1;refGene_tx=XM_023257653.1;refGene_gn=MOV10L1;refGene_tx=XM_023257654.1
B4 143937753 143937763
refGene_gn=MOV10L1;refGene_tx=XM_003989414.5;refGene_gn=MOV10L1;refGene_tx=XM_006934191.4;refGene_gn=MOV10L1;refGene_tx=XM_011284356.3;refGene_gn=MOV10L1;refGene_tx=XM_019835761.2;refGene_gn=MOV10L1;refGene_tx=

XM_019835762.2;refGene_gn=MOV10L1;refGene_tx=XM_023257653.1;refGene_gn=MOV10L1;refGene_tx=XM_023257654.1
B4 143946306 143946532
refGene_gn=PANX2;refGene_tx=XM_023257656.1
B4 143950507 143951965
refGene_gn=PANX2;refGene_tx=XM_023257656.1
B4 143952477 143952821
refGene_gn=PANX2;refGene_tx=XM_023257656.1
B4 143962346 143962388
refGene_gn=TRABD;refGene_tx=XM_006934195.3;refGene_gn=TRABD;refGene_tx=XM_006934196.4
B4 143963150 143963219
refGene_gn=TRABD;refGene_tx=XM_003989416.4;refGene_gn=TRABD;refGene_tx=XM_006934195.3;refGene_gn=TRABD;refGene_tx=XM_006934196.4;refGene_gn=TRABD;refGene_tx=XM_006934197.3;refGene_gn=TRABD;refGene_tx=XM_006934198.3
B4 143963626 143963705
refGene_gn=TRABD;refGene_tx=XM_003989416.4;refGene_gn=TRABD;refGene_tx=XM_006934195.3;refGene_gn=TRABD;refGene_tx=XM_006934196.4;refGene_gn=TRABD;refGene_tx=XM_006934197.3;refGene_gn=TRABD;refGene_tx=XM_006934198.3
B4 143964134 143964301
refGene_gn=TRABD;refGene_tx=XM_003989416.4;refGene_gn=TRABD;refGene_tx=XM_006934195.3;refGene_gn=TRABD;refGene_tx=XM_006934196.4;refGene_gn=TRABD;refGene_tx=XM_006934197.3;refGene_gn=TRABD;refGene_tx=XM_006934198.3;refGene_gn=TRABD;refGene_tx=XM_006934199.3
B4 143964622 143964763
refGene_gn=TRABD;refGene_tx=XM_003989416.4;refGene_gn=TRABD;refGene_tx=XM_006934195.3;refGene_gn=TRABD;refGene_tx=XM_006934196.4;refGene_gn=TRABD;refGene_tx=XM_006934197.3;refGene_gn=TRABD;refGene_tx=XM_006934198.3;refGene_gn=TRABD;refGene_tx=XM_006934199.3
B4 143965210 143965321
refGene_gn=TRABD;refGene_tx=XM_003989416.4;refGene_gn=TRABD;refGene_tx=XM_006934195.3;refGene_gn=TRABD;refGene_tx=XM_006934196.4;refGene_gn=TRABD;refGene_tx=XM_006934197.3;refGene_gn=TRABD;refGene_tx=XM_006934198.3;refGene_gn=TRABD;refGene_tx=XM_006934199.3
B4 143965413 143965553
refGene_gn=TRABD;refGene_tx=XM_003989416.4;refGene_gn=TRABD;refGene_tx=XM_006934195.3;refGene_gn=TRABD;refGene_tx=XM_006934196.4;refGene_gn=TRABD;refGene_tx=XM_006934197.3;refGene_gn=TRABD;refGene_tx=XM_006934198.3;refGene_gn=TRABD;refGene_tx=XM_006934199.3
B4 143965793 143965966
refGene_gn=TRABD;refGene_tx=XM_003989416.4;refGene_gn=TRABD;refGene_tx=XM_006934195.3;refGene_gn=TRABD;refGene_tx=XM_006934196.4;refGene_gn=TRABD;refGene_tx=XM_006934197.3;refGene_gn=TRABD;refGene_tx=XM_006934198.3;refGene_gn=TRABD;refGene_tx=XM_006934199.3
B4 143966050 143966162
refGene_gn=TRABD;refGene_tx=XM_003989416.4;refGene_gn=TRABD;refGene_tx=XM_006934195.3;refGene_gn=TRABD;refGene_tx=XM_006934196.4;refGene_gn=TRABD;refGene_tx=XM_006934197.3;refGene_gn=TRABD;refGene_tx=XM_0069341

98.3;refGene_gn=TRABD;refGene_tx=XM_006934199.3
B4 143966292 143966470
refGene_gn=TRABD;refGene_tx=XM_003989416.4;refGene_gn=TRABD;refGene_tx=XM_006934195.3;refGene_gn=TRABD;refGene_tx=XM_006934196.4;refGene_gn=TRABD;refGene_tx=XM_006934197.3;refGene_gn=TRABD;refGene_tx=XM_006934198.3;refGene_gn=TRABD;refGene_tx=XM_006934199.3
B4 143968087 143968174
refGene_gn=LOC101080391;refGene_tx=XM_003989417.4
B4 143968251 143968279
refGene_gn=LOC101080391;refGene_tx=XM_003989417.4
B4 143968806 143968898
refGene_gn=LOC101080391;refGene_tx=XM_003989417.4
B4 143969659 143969786
refGene_gn=LOC101080391;refGene_tx=XM_003989417.4
B4 143969933 143970119
refGene_gn=LOC101080391;refGene_tx=XM_003989417.4
B4 143970438 143970740
refGene_gn=LOC101080391;refGene_tx=XM_003989417.4
B4 143972515 143973096
refGene_gn=SELENOO;refGene_tx=XM_023257658.1
B4 143978115 143978319
refGene_gn=SELENOO;refGene_tx=XM_023257658.1
B4 143979649 143979830
refGene_gn=SELENOO;refGene_tx=XM_023257658.1
B4 143981084 143981215
refGene_gn=SELENOO;refGene_tx=XM_023257658.1
B4 143981586 143981867
refGene_gn=SELENOO;refGene_tx=XM_023257658.1
B4 143985108 143985259
refGene_gn=SELENOO;refGene_tx=XM_023257658.1
B4 143986168 143986354
refGene_gn=SELENOO;refGene_tx=XM_023257658.1
B4 143986526 143986683
refGene_gn=SELENOO;refGene_tx=XM_023257658.1
B4 143986785 143986950
refGene_gn=SELENOO;refGene_tx=XM_023257658.1
B4 143987372 143987464
refGene_gn=TUBGCP6;refGene_tx=XM_023257657.1
B4 143987570 143987770
refGene_gn=TUBGCP6;refGene_tx=XM_023257657.1
B4 143987930 143988144
refGene_gn=TUBGCP6;refGene_tx=XM_023257657.1
B4 143988214 143988347
refGene_gn=TUBGCP6;refGene_tx=XM_023257657.1
B4 143988429 143988624
refGene_gn=TUBGCP6;refGene_tx=XM_023257657.1
B4 143988720 143988862
refGene_gn=TUBGCP6;refGene_tx=XM_023257657.1
B4 143989018 143989187
refGene_gn=TUBGCP6;refGene_tx=XM_023257657.1

B4 143989274 143989416
refGene_gn=TUBGCP6;refGene_tx=XM_023257657.1
B4 143989619 143989672
refGene_gn=TUBGCP6;refGene_tx=XM_023257657.1
B4 143989849 143991365
refGene_gn=TUBGCP6;refGene_tx=XM_023257657.1
B4 143991573 143991642
refGene_gn=TUBGCP6;refGene_tx=XM_023257657.1
B4 143991964 143992103
refGene_gn=TUBGCP6;refGene_tx=XM_023257657.1
B4 143993030 143993146
refGene_gn=TUBGCP6;refGene_tx=XM_023257657.1
B4 143993228 143993317
refGene_gn=TUBGCP6;refGene_tx=XM_023257657.1
B4 143993413 143993495
refGene_gn=TUBGCP6;refGene_tx=XM_023257657.1
B4 143994159 143994309
refGene_gn=TUBGCP6;refGene_tx=XM_023257657.1
B4 143995192 143995332
refGene_gn=TUBGCP6;refGene_tx=XM_023257657.1
B4 143995429 143995521
refGene_gn=TUBGCP6;refGene_tx=XM_023257657.1
B4 143995660 143995734
refGene_gn=TUBGCP6;refGene_tx=XM_023257657.1
B4 143995735 143995771
refGene_gn=TUBGCP6;refGene_tx=XM_023257657.1
B4 143996122 143996180
refGene_gn=TUBGCP6;refGene_tx=XM_023257657.1
B4 143996962 143997084
refGene_gn=TUBGCP6;refGene_tx=XM_023257657.1
B4 143998082 143998256
refGene_gn=TUBGCP6;refGene_tx=XM_023257657.1
B4 144001383 144001594
refGene_gn=TUBGCP6;refGene_tx=XM_023257657.1
B4 144005439 144005603
refGene_gn=TUBGCP6;refGene_tx=XM_023257657.1
B4 144006870 144007611
refGene_gn=TUBGCP6;refGene_tx=XM_023257657.1
B4 144015935 144015952
refGene_gn=HDAC10;refGene_tx=XM_011284359.3
B4 144016113 144016273
refGene_gn=HDAC10;refGene_tx=XM_011284359.3
B4 144016369 144016510
refGene_gn=HDAC10;refGene_tx=XM_011284359.3
B4 144016511 144016553
refGene_gn=HDAC10;refGene_tx=XM_011284359.3
B4 144016694 144016773
refGene_gn=HDAC10;refGene_tx=XM_011284359.3
B4 144017096 144017153
refGene_gn=HDAC10;refGene_tx=XM_011284359.3

B4 144017294 144017386
refGene_gn=HDAC10;refGene_tx=XM_011284359.3
B4 144017747 144017832
refGene_gn=HDAC10;refGene_tx=XM_011284359.3
B4 144017929 144018194
refGene_gn=HDAC10;refGene_tx=XM_011284359.3
B4 144018273 144018341
refGene_gn=HDAC10;refGene_tx=XM_011284359.3
B4 144018413 144018520
refGene_gn=HDAC10;refGene_tx=XM_011284359.3
B4 144018660 144018750
refGene_gn=HDAC10;refGene_tx=XM_011284359.3
B4 144018827 144018887
refGene_gn=HDAC10;refGene_tx=XM_011284359.3
B4 144019074 144019140
refGene_gn=HDAC10;refGene_tx=XM_011284359.3
B4 144019330 144019457
refGene_gn=HDAC10;refGene_tx=XM_011284359.3
B4 144019548 144019617
refGene_gn=HDAC10;refGene_tx=XM_011284359.3
B4 144019773 144019878
refGene_gn=HDAC10;refGene_tx=XM_011284359.3
B4 144019977 144020075
refGene_gn=HDAC10;refGene_tx=XM_011284359.3
B4 144020306 144020403
refGene_gn=HDAC10;refGene_tx=XM_011284359.3
B4 144020632 144020767
refGene_gn=HDAC10;refGene_tx=XM_011284359.3
B4 144020837 144020896
refGene_gn=HDAC10;refGene_tx=XM_011284359.3
B4 144023384 144023464
refGene_gn=MAPK12;refGene_tx=XM_023257659.1
B4 144025563 144025737
refGene_gn=MAPK12;refGene_tx=XM_023257659.1
B4 144025820 144025899
refGene_gn=MAPK12;refGene_tx=XM_023257659.1
B4 144025980 144026060
refGene_gn=MAPK12;refGene_tx=XM_023257659.1
B4 144026180 144026252
refGene_gn=MAPK12;refGene_tx=XM_023257659.1
B4 144026482 144026597
refGene_gn=MAPK12;refGene_tx=XM_023257659.1
B4 144027174 144027222
refGene_gn=MAPK12;refGene_tx=XM_023257659.1
B4 144027501 144027531
refGene_gn=MAPK12;refGene_tx=XM_023257659.1
B4 144027662 144027774
refGene_gn=MAPK12;refGene_tx=XM_023257659.1
B4 144028666 144028725
refGene_gn=MAPK12;refGene_tx=XM_023257659.1

B4 144031196 144031326
refGene_gn=MAPK12;refGene_tx=XM_023257659.1
B4 144031415 144031540
refGene_gn=MAPK12;refGene_tx=XM_023257659.1
B4 144035017 144035097
refGene_gn=MAPK11;refGene_tx=XM_023257660.1;refGene_gn=MAPK11;refGene_tx=XM_023257661.1;refGene_gn=MAPK11;refGene_tx=XM_023257662.1
B4 144035418 144035592
refGene_gn=MAPK11;refGene_tx=XM_023257660.1;refGene_gn=MAPK11;refGene_tx=XM_023257661.1;refGene_gn=MAPK11;refGene_tx=XM_023257662.1
B4 144035680 144035759
refGene_gn=MAPK11;refGene_tx=XM_023257660.1;refGene_gn=MAPK11;refGene_tx=XM_023257661.1;refGene_gn=MAPK11;refGene_tx=XM_023257662.1
B4 144036221 144036301
refGene_gn=MAPK11;refGene_tx=XM_023257660.1;refGene_gn=MAPK11;refGene_tx=XM_023257661.1;refGene_gn=MAPK11;refGene_tx=XM_023257662.1
B4 144036496 144036568
refGene_gn=MAPK11;refGene_tx=XM_023257660.1;refGene_gn=MAPK11;refGene_tx=XM_023257661.1;refGene_gn=MAPK11;refGene_tx=XM_023257662.1
B4 144036955 144037070
refGene_gn=MAPK11;refGene_tx=XM_023257660.1;refGene_gn=MAPK11;refGene_tx=XM_023257661.1;refGene_gn=MAPK11;refGene_tx=XM_023257662.1
B4 144037160 144037208
refGene_gn=MAPK11;refGene_tx=XM_023257660.1;refGene_gn=MAPK11;refGene_tx=XM_023257661.1;refGene_gn=MAPK11;refGene_tx=XM_023257662.1
B4 144037283 144037313
refGene_gn=MAPK11;refGene_tx=XM_023257660.1;refGene_gn=MAPK11;refGene_tx=XM_023257662.1
B4 144037389 144037501
refGene_gn=MAPK11;refGene_tx=XM_023257660.1;refGene_gn=MAPK11;refGene_tx=XM_023257661.1;refGene_gn=MAPK11;refGene_tx=XM_023257662.1
B4 144037580 144037639
refGene_gn=MAPK11;refGene_tx=XM_023257660.1;refGene_gn=MAPK11;refGene_tx=XM_023257661.1;refGene_gn=MAPK11;refGene_tx=XM_023257662.1
B4 144037723 144037853
refGene_gn=MAPK11;refGene_tx=XM_023257660.1;refGene_gn=MAPK11;refGene_tx=XM_023257661.1;refGene_gn=MAPK11;refGene_tx=XM_023257662.1
B4 144040026 144040079
refGene_gn=MAPK11;refGene_tx=XM_023257662.1
B4 144040226 144040342
refGene_gn=MAPK11;refGene_tx=XM_023257660.1;refGene_gn=MAPK11;refGene_tx=XM_023257661.1
B4 144045814 144045919
refGene_gn=PLXNB2;refGene_tx=XM_019835766.1
B4 144046003 144046078
refGene_gn=PLXNB2;refGene_tx=XM_019835766.1
B4 144046439 144046515
refGene_gn=PLXNB2;refGene_tx=XM_019835766.1
B4 144046656 144046721
refGene_gn=PLXNB2;refGene_tx=XM_019835766.1

B4 144047232 144047380
refGene_gn=PLXNB2;refGene_tx=XM_019835766.1
B4 144047460 144047621
refGene_gn=PLXNB2;refGene_tx=XM_019835766.1
B4 144047792 144047947
refGene_gn=PLXNB2;refGene_tx=XM_019835766.1
B4 144048095 144048180
refGene_gn=PLXNB2;refGene_tx=XM_019835766.1
B4 144048268 144048369
refGene_gn=PLXNB2;refGene_tx=XM_019835766.1
B4 144048516 144048673
refGene_gn=PLXNB2;refGene_tx=XM_019835766.1
B4 144049103 144049250
refGene_gn=PLXNB2;refGene_tx=XM_019835766.1
B4 144049382 144049449
refGene_gn=PLXNB2;refGene_tx=XM_019835766.1
B4 144049776 144049958
refGene_gn=PLXNB2;refGene_tx=XM_019835766.1
B4 144050083 144050313
refGene_gn=PLXNB2;refGene_tx=XM_019835766.1
B4 144050497 144050598
refGene_gn=PLXNB2;refGene_tx=XM_019835766.1
B4 144050699 144050839
refGene_gn=PLXNB2;refGene_tx=XM_019835766.1
B4 144050954 144051131
refGene_gn=PLXNB2;refGene_tx=XM_019835766.1
B4 144051236 144051464
refGene_gn=PLXNB2;refGene_tx=XM_019835766.1
B4 144051574 144051704
refGene_gn=PLXNB2;refGene_tx=XM_019835766.1
B4 144052177 144052348
refGene_gn=PLXNB2;refGene_tx=XM_019835766.1
B4 144052535 144052672
refGene_gn=PLXNB2;refGene_tx=XM_019835766.1
B4 144052870 144052979
refGene_gn=PLXNB2;refGene_tx=XM_019835766.1
B4 144053122 144053271
refGene_gn=PLXNB2;refGene_tx=XM_019835766.1
B4 144053387 144053548
refGene_gn=PLXNB2;refGene_tx=XM_019835766.1
B4 144053625 144053707
refGene_gn=PLXNB2;refGene_tx=XM_019835766.1
B4 144053974 144054067
refGene_gn=PLXNB2;refGene_tx=XM_019835766.1
B4 144054560 144054662
refGene_gn=PLXNB2;refGene_tx=XM_019835766.1
B4 144054743 144054852
refGene_gn=PLXNB2;refGene_tx=XM_019835766.1
B4 144054929 144055044
refGene_gn=PLXNB2;refGene_tx=XM_019835766.1

B4	144055619	144055773
refGene_gn=PLXNB2;refGene_tx=XM_019835766.1
B4	144055964	144056091
refGene_gn=PLXNB2;refGene_tx=XM_019835766.1
B4	144056196	144056297
refGene_gn=PLXNB2;refGene_tx=XM_019835766.1
B4	144056938	144057067
refGene_gn=PLXNB2;refGene_tx=XM_019835766.1
B4	144057157	144057340
refGene_gn=PLXNB2;refGene_tx=XM_019835766.1
B4	144057643	144058648
refGene_gn=PLXNB2;refGene_tx=XM_019835766.1
B4	144058649	144058718
refGene_gn=PLXNB2;refGene_tx=XM_019835766.1
B4	144077315	144077438
refGene_gn=DENND6B;refGene_tx=XM_011284366.2;refGene_gn=DENND6B;refGene_tx=XM_011284367.3;refGene_gn=DENND6B;refGene_tx=XM_019835773.2;refGene_gn=DENND6B;refGene_tx=XM_019835774.2
B4	144077518	144077596
refGene_gn=DENND6B;refGene_tx=XM_011284366.2;refGene_gn=DENND6B;refGene_tx=XM_011284367.3;refGene_gn=DENND6B;refGene_tx=XM_019835773.2;refGene_gn=DENND6B;refGene_tx=XM_019835774.2
B4	144077695	144077798
refGene_gn=DENND6B;refGene_tx=XM_011284366.2;refGene_gn=DENND6B;refGene_tx=XM_011284367.3;refGene_gn=DENND6B;refGene_tx=XM_019835773.2;refGene_gn=DENND6B;refGene_tx=XM_019835774.2
B4	144077898	144078008
refGene_gn=DENND6B;refGene_tx=XM_011284366.2;refGene_gn=DENND6B;refGene_tx=XM_011284367.3;refGene_gn=DENND6B;refGene_tx=XM_019835773.2;refGene_gn=DENND6B;refGene_tx=XM_019835774.2
B4	144078332	144078386
refGene_gn=DENND6B;refGene_tx=XM_011284366.2;refGene_gn=DENND6B;refGene_tx=XM_011284367.3;refGene_gn=DENND6B;refGene_tx=XM_019835773.2;refGene_gn=DENND6B;refGene_tx=XM_019835774.2
B4	144078461	144078551
refGene_gn=DENND6B;refGene_tx=XM_011284366.2;refGene_gn=DENND6B;refGene_tx=XM_011284367.3;refGene_gn=DENND6B;refGene_tx=XM_019835773.2;refGene_gn=DENND6B;refGene_tx=XM_019835774.2
B4	144078639	144078704
refGene_gn=DENND6B;refGene_tx=XM_011284366.2;refGene_gn=DENND6B;refGene_tx=XM_011284367.3;refGene_gn=DENND6B;refGene_tx=XM_019835773.2;refGene_gn=DENND6B;refGene_tx=XM_019835774.2
B4	144079021	144079087
refGene_gn=DENND6B;refGene_tx=XM_011284366.2;refGene_gn=DENND6B;refGene_tx=XM_011284367.3;refGene_gn=DENND6B;refGene_tx=XM_019835773.2;refGene_gn=DENND6B;refGene_tx=XM_019835774.2
B4	144079233	144079328
refGene_gn=DENND6B;refGene_tx=XM_011284366.2;refGene_gn=DENND6B;refGene_tx=XM_011284367.3;refGene_gn=DENND6B;refGene_tx=XM_019835773.2;refGene_gn=DENND6B;refGene_tx=XM_019835774.2

B4	144079424	144079520
refGene_gn=DENND6B;refGene_tx=XM_011284366.2;refGene_gn=DENND6B;refGene_tx=XM_011284367.3;refGene_gn=DENND6B;refGene_tx=XM_019835773.2;refGene_gn=DENND6B;refGene_tx=XM_019835774.2
B4	144079622	144079745
refGene_gn=DENND6B;refGene_tx=XM_011284366.2;refGene_gn=DENND6B;refGene_tx=XM_011284367.3;refGene_gn=DENND6B;refGene_tx=XM_019835773.2;refGene_gn=DENND6B;refGene_tx=XM_019835774.2
B4	144080235	144080291
refGene_gn=DENND6B;refGene_tx=XM_011284366.2;refGene_gn=DENND6B;refGene_tx=XM_011284367.3;refGene_gn=DENND6B;refGene_tx=XM_019835773.2;refGene_gn=DENND6B;refGene_tx=XM_019835774.2
B4	144080512	144080575
refGene_gn=DENND6B;refGene_tx=XM_011284366.2;refGene_gn=DENND6B;refGene_tx=XM_011284367.3;refGene_gn=DENND6B;refGene_tx=XM_019835773.2;refGene_gn=DENND6B;refGene_tx=XM_019835774.2
B4	144080661	144080741
refGene_gn=DENND6B;refGene_tx=XM_011284366.2;refGene_gn=DENND6B;refGene_tx=XM_011284367.3;refGene_gn=DENND6B;refGene_tx=XM_019835773.2;refGene_gn=DENND6B;refGene_tx=XM_019835774.2
B4	144080860	144080966
refGene_gn=DENND6B;refGene_tx=XM_011284366.2;refGene_gn=DENND6B;refGene_tx=XM_011284367.3;refGene_gn=DENND6B;refGene_tx=XM_019835773.2;refGene_gn=DENND6B;refGene_tx=XM_019835774.2
B4	144081611	144081692
refGene_gn=DENND6B;refGene_tx=XM_011284366.2;refGene_gn=DENND6B;refGene_tx=XM_011284367.3;refGene_gn=DENND6B;refGene_tx=XM_019835773.2;refGene_gn=DENND6B;refGene_tx=XM_019835774.2
B4	144082216	144082329
refGene_gn=DENND6B;refGene_tx=XM_011284366.2;refGene_gn=DENND6B;refGene_tx=XM_011284367.3;refGene_gn=DENND6B;refGene_tx=XM_019835773.2;refGene_gn=DENND6B;refGene_tx=XM_019835774.2
B4	144082810	144082853
refGene_gn=DENND6B;refGene_tx=XM_011284366.2;refGene_gn=DENND6B;refGene_tx=XM_019835773.2;refGene_gn=DENND6B;refGene_tx=XM_019835774.2
B4	144082926	144082965
refGene_gn=DENND6B;refGene_tx=XM_011284366.2;refGene_gn=DENND6B;refGene_tx=XM_011284367.3;refGene_gn=DENND6B;refGene_tx=XM_019835773.2
B4	144087190	144087352
refGene_gn=DENND6B;refGene_tx=XM_011284366.2;refGene_gn=DENND6B;refGene_tx=XM_011284367.3;refGene_gn=DENND6B;refGene_tx=XM_019835773.2;refGene_gn=DENND6B;refGene_tx=XM_019835774.2
B4	144150037	144150264
refGene_gn=PPP6R2;refGene_tx=XM_023257665.1;refGene_gn=PPP6R2;refGene_tx=XM_023257666.1;refGene_gn=PPP6R2;refGene_tx=XM_023257667.1;refGene_gn=PPP6R2;refGene_tx=XM_023257668.1;refGene_gn=PPP6R2;refGene_tx=XM_023257669.1
B4	144157792	144157979
refGene_gn=PPP6R2;refGene_tx=XM_023257665.1;refGene_gn=PPP6R2;refGene_tx=XM_023257666.1;refGene_gn=PPP6R2;refGene_tx=XM_023257667.1;refGene_

gn=PPP6R2;refGene_tx=XM_023257668.1;refGene_gn=PPP6R2;refGene_tx=XM_023257669.1
B4 144167280 144167418
refGene_gn=PPP6R2;refGene_tx=XM_023257665.1;refGene_gn=PPP6R2;refGene_tx=XM_023257666.1;refGene_gn=PPP6R2;refGene_tx=XM_023257667.1;refGene_gn=PPP6R2;refGene_tx=XM_023257668.1;refGene_gn=PPP6R2;refGene_tx=XM_023257669.1;refGene_gn=PPP6R2;refGene_tx=XM_023257670.1
B4 144168813 144168879
refGene_gn=PPP6R2;refGene_tx=XM_023257665.1;refGene_gn=PPP6R2;refGene_tx=XM_023257666.1;refGene_gn=PPP6R2;refGene_tx=XM_023257667.1;refGene_gn=PPP6R2;refGene_tx=XM_023257668.1;refGene_gn=PPP6R2;refGene_tx=XM_023257669.1;refGene_gn=PPP6R2;refGene_tx=XM_023257670.1
B4 144170482 144170595
refGene_gn=PPP6R2;refGene_tx=XM_023257665.1;refGene_gn=PPP6R2;refGene_tx=XM_023257666.1;refGene_gn=PPP6R2;refGene_tx=XM_023257667.1;refGene_gn=PPP6R2;refGene_tx=XM_023257668.1;refGene_gn=PPP6R2;refGene_tx=XM_023257669.1;refGene_gn=PPP6R2;refGene_tx=XM_023257670.1
B4 144170982 144171096
refGene_gn=PPP6R2;refGene_tx=XM_023257665.1;refGene_gn=PPP6R2;refGene_tx=XM_023257666.1;refGene_gn=PPP6R2;refGene_tx=XM_023257667.1;refGene_gn=PPP6R2;refGene_tx=XM_023257668.1;refGene_gn=PPP6R2;refGene_tx=XM_023257669.1;refGene_gn=PPP6R2;refGene_tx=XM_023257670.1
B4 144172981 144173108
refGene_gn=PPP6R2;refGene_tx=XM_023257665.1;refGene_gn=PPP6R2;refGene_tx=XM_023257666.1;refGene_gn=PPP6R2;refGene_tx=XM_023257667.1;refGene_gn=PPP6R2;refGene_tx=XM_023257668.1;refGene_gn=PPP6R2;refGene_tx=XM_023257669.1;refGene_gn=PPP6R2;refGene_tx=XM_023257670.1
B4 144174097 144174250
refGene_gn=PPP6R2;refGene_tx=XM_023257665.1;refGene_gn=PPP6R2;refGene_tx=XM_023257666.1;refGene_gn=PPP6R2;refGene_tx=XM_023257667.1;refGene_gn=PPP6R2;refGene_tx=XM_023257668.1;refGene_gn=PPP6R2;refGene_tx=XM_023257669.1;refGene_gn=PPP6R2;refGene_tx=XM_023257670.1
B4 144176876 144177080
refGene_gn=PPP6R2;refGene_tx=XM_023257665.1;refGene_gn=PPP6R2;refGene_tx=XM_023257666.1;refGene_gn=PPP6R2;refGene_tx=XM_023257667.1;refGene_gn=PPP6R2;refGene_tx=XM_023257668.1;refGene_gn=PPP6R2;refGene_tx=XM_023257669.1;refGene_gn=PPP6R2;refGene_tx=XM_023257670.1
B4 144178225 144178290
refGene_gn=PPP6R2;refGene_tx=XM_023257665.1;refGene_gn=PPP6R2;refGene_tx=XM_023257666.1;refGene_gn=PPP6R2;refGene_tx=XM_023257667.1;refGene_gn=PPP6R2;refGene_tx=XM_023257668.1;refGene_gn=PPP6R2;refGene_tx=XM_023257669.1;refGene_gn=PPP6R2;refGene_tx=XM_023257670.1
B4 144179961 144180077
refGene_gn=PPP6R2;refGene_tx=XM_023257665.1;refGene_gn=PPP6R2;refGene_tx=XM_023257666.1;refGene_gn=PPP6R2;refGene_tx=XM_023257667.1;refGene_gn=PPP6R2;refGene_tx=XM_023257668.1;refGene_gn=PPP6R2;refGene_tx=XM_023257669.1;refGene_gn=PPP6R2;refGene_tx=XM_023257670.1
B4 144182383 144182469
refGene_gn=PPP6R2;refGene_tx=XM_023257665.1;refGene_gn=PPP6R2;refGene_tx=XM_023257666.1;refGene_gn=PPP6R2;refGene_tx=XM_023257667.1;refGene_

gn=PPP6R2;refGene_tx=XM_023257668.1;refGene_gn=PPP6R2;refGene_tx=XM_023257669.1;refGene_gn=PPP6R2;refGene_tx=XM_023257670.1
B4 144182871 144182952
refGene_gn=PPP6R2;refGene_tx=XM_023257665.1;refGene_gn=PPP6R2;refGene_tx=XM_023257666.1;refGene_gn=PPP6R2;refGene_tx=XM_023257667.1;refGene_gn=PPP6R2;refGene_tx=XM_023257670.1
B4 144183316 144183414
refGene_gn=PPP6R2;refGene_tx=XM_023257665.1;refGene_gn=PPP6R2;refGene_tx=XM_023257666.1;refGene_gn=PPP6R2;refGene_tx=XM_023257667.1;refGene_gn=PPP6R2;refGene_tx=XM_023257668.1;refGene_gn=PPP6R2;refGene_tx=XM_023257669.1;refGene_gn=PPP6R2;refGene_tx=XM_023257670.1
B4 144183843 144183901
refGene_gn=PPP6R2;refGene_tx=XM_023257665.1;refGene_gn=PPP6R2;refGene_tx=XM_023257666.1;refGene_gn=PPP6R2;refGene_tx=XM_023257667.1;refGene_gn=PPP6R2;refGene_tx=XM_023257668.1;refGene_gn=PPP6R2;refGene_tx=XM_023257669.1;refGene_gn=PPP6R2;refGene_tx=XM_023257670.1
B4 144184119 144184244
refGene_gn=PPP6R2;refGene_tx=XM_023257665.1;refGene_gn=PPP6R2;refGene_tx=XM_023257666.1;refGene_gn=PPP6R2;refGene_tx=XM_023257667.1;refGene_gn=PPP6R2;refGene_tx=XM_023257668.1;refGene_gn=PPP6R2;refGene_tx=XM_023257669.1;refGene_gn=PPP6R2;refGene_tx=XM_023257670.1
B4 144184420 144184584
refGene_gn=PPP6R2;refGene_tx=XM_023257665.1;refGene_gn=PPP6R2;refGene_tx=XM_023257666.1;refGene_gn=PPP6R2;refGene_tx=XM_023257667.1;refGene_gn=PPP6R2;refGene_tx=XM_023257668.1;refGene_gn=PPP6R2;refGene_tx=XM_023257669.1;refGene_gn=PPP6R2;refGene_tx=XM_023257670.1
B4 144184996 144185162
refGene_gn=PPP6R2;refGene_tx=XM_023257665.1;refGene_gn=PPP6R2;refGene_tx=XM_023257666.1;refGene_gn=PPP6R2;refGene_tx=XM_023257667.1;refGene_gn=PPP6R2;refGene_tx=XM_023257668.1;refGene_gn=PPP6R2;refGene_tx=XM_023257669.1;refGene_gn=PPP6R2;refGene_tx=XM_023257670.1
B4 144185247 144185336
refGene_gn=PPP6R2;refGene_tx=XM_023257665.1;refGene_gn=PPP6R2;refGene_tx=XM_023257666.1;refGene_gn=PPP6R2;refGene_tx=XM_023257667.1;refGene_gn=PPP6R2;refGene_tx=XM_023257668.1;refGene_gn=PPP6R2;refGene_tx=XM_023257669.1;refGene_gn=PPP6R2;refGene_tx=XM_023257670.1
B4 144185857 144186020
refGene_gn=PPP6R2;refGene_tx=XM_023257665.1;refGene_gn=PPP6R2;refGene_tx=XM_023257666.1;refGene_gn=PPP6R2;refGene_tx=XM_023257667.1;refGene_gn=PPP6R2;refGene_tx=XM_023257668.1;refGene_gn=PPP6R2;refGene_tx=XM_023257669.1;refGene_gn=PPP6R2;refGene_tx=XM_023257670.1
B4 144187776 144188037
refGene_gn=PPP6R2;refGene_tx=XM_023257665.1;refGene_gn=PPP6R2;refGene_tx=XM_023257666.1;refGene_gn=PPP6R2;refGene_tx=XM_023257667.1;refGene_gn=PPP6R2;refGene_tx=XM_023257668.1;refGene_gn=PPP6R2;refGene_tx=XM_023257669.1;refGene_gn=PPP6R2;refGene_tx=XM_023257670.1
B4 144188119 144188171
refGene_gn=PPP6R2;refGene_tx=XM_023257665.1;refGene_gn=PPP6R2;refGene_tx=XM_023257666.1;refGene_gn=PPP6R2;refGene_tx=XM_023257667.1;refGene_gn=PPP6R2;refGene_tx=XM_023257668.1;refGene_gn=PPP6R2;refGene_tx=XM_02

3257669.1;refGene_gn=PPP6R2;refGene_tx=XM_023257670.1
B4 144190644 144190743
refGene_gn=SBF1;refGene_tx=XM_019835767.1;refGene_gn=SBF1;refGene_tx=XM_019835768.1;refGene_gn=SBF1;refGene_tx=XM_023257663.1;refGene_gn=SBF1;refGene_tx=XM_023257664.1
B4 144190830 144190962
refGene_gn=SBF1;refGene_tx=XM_019835767.1;refGene_gn=SBF1;refGene_tx=XM_019835768.1;refGene_gn=SBF1;refGene_tx=XM_023257663.1;refGene_gn=SBF1;refGene_tx=XM_023257664.1
B4 144191068 144191156
refGene_gn=SBF1;refGene_tx=XM_019835767.1;refGene_gn=SBF1;refGene_tx=XM_019835768.1;refGene_gn=SBF1;refGene_tx=XM_023257663.1;refGene_gn=SBF1;refGene_tx=XM_023257664.1
B4 144191838 144192050
refGene_gn=SBF1;refGene_tx=XM_019835767.1;refGene_gn=SBF1;refGene_tx=XM_019835768.1;refGene_gn=SBF1;refGene_tx=XM_023257663.1;refGene_gn=SBF1;refGene_tx=XM_023257664.1
B4 144192138 144192246
refGene_gn=SBF1;refGene_tx=XM_019835767.1;refGene_gn=SBF1;refGene_tx=XM_019835768.1;refGene_gn=SBF1;refGene_tx=XM_023257663.1;refGene_gn=SBF1;refGene_tx=XM_023257664.1
B4 144196971 144197202
refGene_gn=SBF1;refGene_tx=XM_019835767.1;refGene_gn=SBF1;refGene_tx=XM_019835768.1;refGene_gn=SBF1;refGene_tx=XM_023257663.1;refGene_gn=SBF1;refGene_tx=XM_023257664.1
B4 144197272 144197403
refGene_gn=SBF1;refGene_tx=XM_019835767.1;refGene_gn=SBF1;refGene_tx=XM_019835768.1;refGene_gn=SBF1;refGene_tx=XM_023257663.1;refGene_gn=SBF1;refGene_tx=XM_023257664.1
B4 144197497 144197624
refGene_gn=SBF1;refGene_tx=XM_019835767.1;refGene_gn=SBF1;refGene_tx=XM_019835768.1;refGene_gn=SBF1;refGene_tx=XM_023257663.1;refGene_gn=SBF1;refGene_tx=XM_023257664.1
B4 144197700 144197886
refGene_gn=SBF1;refGene_tx=XM_019835767.1;refGene_gn=SBF1;refGene_tx=XM_019835768.1;refGene_gn=SBF1;refGene_tx=XM_023257663.1;refGene_gn=SBF1;refGene_tx=XM_023257664.1
B4 144197962 144198064
refGene_gn=SBF1;refGene_tx=XM_019835767.1;refGene_gn=SBF1;refGene_tx=XM_019835768.1;refGene_gn=SBF1;refGene_tx=XM_023257663.1;refGene_gn=SBF1;refGene_tx=XM_023257664.1
B4 144198336 144198516
refGene_gn=SBF1;refGene_tx=XM_019835767.1;refGene_gn=SBF1;refGene_tx=XM_019835768.1;refGene_gn=SBF1;refGene_tx=XM_023257663.1;refGene_gn=SBF1;refGene_tx=XM_023257664.1
B4 144198593 144198775
refGene_gn=SBF1;refGene_tx=XM_019835767.1;refGene_gn=SBF1;refGene_tx=XM_019835768.1;refGene_gn=SBF1;refGene_tx=XM_023257663.1;refGene_gn=SBF1;refGene_tx=XM_023257664.1
B4 144199091 144199169

refGene_gn=SBF1;refGene_tx=XM_019835767.1;refGene_gn=SBF1;refGene_tx=XM_023257663.1
B4 144202231 144202369
refGene_gn=SBF1;refGene_tx=XM_019835767.1;refGene_gn=SBF1;refGene_tx=XM_019835768.1;refGene_gn=SBF1;refGene_tx=XM_023257663.1;refGene_gn=SBF1;refGene_tx=XM_023257664.1
B4 144202538 144202735
refGene_gn=SBF1;refGene_tx=XM_019835767.1;refGene_gn=SBF1;refGene_tx=XM_019835768.1;refGene_gn=SBF1;refGene_tx=XM_023257663.1;refGene_gn=SBF1;refGene_tx=XM_023257664.1
B4 144202904 144203112
refGene_gn=SBF1;refGene_tx=XM_019835767.1;refGene_gn=SBF1;refGene_tx=XM_019835768.1;refGene_gn=SBF1;refGene_tx=XM_023257663.1;refGene_gn=SBF1;refGene_tx=XM_023257664.1
B4 144203228 144203365
refGene_gn=SBF1;refGene_tx=XM_019835767.1;refGene_gn=SBF1;refGene_tx=XM_019835768.1;refGene_gn=SBF1;refGene_tx=XM_023257663.1;refGene_gn=SBF1;refGene_tx=XM_023257664.1
B4 144203478 144203657
refGene_gn=SBF1;refGene_tx=XM_019835767.1;refGene_gn=SBF1;refGene_tx=XM_019835768.1;refGene_gn=SBF1;refGene_tx=XM_023257663.1;refGene_gn=SBF1;refGene_tx=XM_023257664.1
B4 144204129 144204257
refGene_gn=SBF1;refGene_tx=XM_019835767.1;refGene_gn=SBF1;refGene_tx=XM_019835768.1;refGene_gn=SBF1;refGene_tx=XM_023257663.1;refGene_gn=SBF1;refGene_tx=XM_023257664.1
B4 144204496 144204692
refGene_gn=SBF1;refGene_tx=XM_019835767.1;refGene_gn=SBF1;refGene_tx=XM_019835768.1;refGene_gn=SBF1;refGene_tx=XM_023257663.1;refGene_gn=SBF1;refGene_tx=XM_023257664.1
B4 144204771 144204845
refGene_gn=SBF1;refGene_tx=XM_019835767.1;refGene_gn=SBF1;refGene_tx=XM_019835768.1;refGene_gn=SBF1;refGene_tx=XM_023257663.1;refGene_gn=SBF1;refGene_tx=XM_023257664.1
B4 144204927 144205100
refGene_gn=SBF1;refGene_tx=XM_019835767.1;refGene_gn=SBF1;refGene_tx=XM_019835768.1;refGene_gn=SBF1;refGene_tx=XM_023257663.1;refGene_gn=SBF1;refGene_tx=XM_023257664.1
B4 144205195 144205461
refGene_gn=SBF1;refGene_tx=XM_019835767.1;refGene_gn=SBF1;refGene_tx=XM_019835768.1;refGene_gn=SBF1;refGene_tx=XM_023257663.1;refGene_gn=SBF1;refGene_tx=XM_023257664.1
B4 144205538 144205697
refGene_gn=SBF1;refGene_tx=XM_019835767.1;refGene_gn=SBF1;refGene_tx=XM_019835768.1;refGene_gn=SBF1;refGene_tx=XM_023257663.1;refGene_gn=SBF1;refGene_tx=XM_023257664.1
B4 144205775 144205844
refGene_gn=SBF1;refGene_tx=XM_019835767.1;refGene_gn=SBF1;refGene_tx=XM_019835768.1;refGene_gn=SBF1;refGene_tx=XM_023257663.1;refGene_gn=SBF1;refGene_tx=XM_023257664.1

B4	144206158	144206308
refGene_gn=SBF1;refGene_tx=XM_019835767.1;refGene_gn=SBF1;refGene_tx=XM_019835768.1;refGene_gn=SBF1;refGene_tx=XM_023257663.1;refGene_gn=SBF1;refGene_tx=XM_023257664.1
B4	144207708	144207821
refGene_gn=SBF1;refGene_tx=XM_019835767.1;refGene_gn=SBF1;refGene_tx=XM_019835768.1;refGene_gn=SBF1;refGene_tx=XM_023257663.1;refGene_gn=SBF1;refGene_tx=XM_023257664.1
B4	144207900	144208105
refGene_gn=SBF1;refGene_tx=XM_019835767.1;refGene_gn=SBF1;refGene_tx=XM_019835768.1;refGene_gn=SBF1;refGene_tx=XM_023257663.1;refGene_gn=SBF1;refGene_tx=XM_023257664.1
B4	144208212	144208311
refGene_gn=SBF1;refGene_tx=XM_019835767.1;refGene_gn=SBF1;refGene_tx=XM_019835768.1;refGene_gn=SBF1;refGene_tx=XM_023257663.1;refGene_gn=SBF1;refGene_tx=XM_023257664.1
B4	144208394	144208523
refGene_gn=SBF1;refGene_tx=XM_019835767.1;refGene_gn=SBF1;refGene_tx=XM_019835768.1;refGene_gn=SBF1;refGene_tx=XM_023257663.1;refGene_gn=SBF1;refGene_tx=XM_023257664.1
B4	144208600	144208714
refGene_gn=SBF1;refGene_tx=XM_019835767.1;refGene_gn=SBF1;refGene_tx=XM_019835768.1;refGene_gn=SBF1;refGene_tx=XM_023257663.1;refGene_gn=SBF1;refGene_tx=XM_023257664.1
B4	144209127	144209205
refGene_gn=SBF1;refGene_tx=XM_019835767.1;refGene_gn=SBF1;refGene_tx=XM_019835768.1;refGene_gn=SBF1;refGene_tx=XM_023257663.1;refGene_gn=SBF1;refGene_tx=XM_023257664.1
B4	144209318	144209432
refGene_gn=SBF1;refGene_tx=XM_019835767.1;refGene_gn=SBF1;refGene_tx=XM_019835768.1;refGene_gn=SBF1;refGene_tx=XM_023257663.1;refGene_gn=SBF1;refGene_tx=XM_023257664.1
B4	144209525	144209634
refGene_gn=SBF1;refGene_tx=XM_019835767.1;refGene_gn=SBF1;refGene_tx=XM_019835768.1;refGene_gn=SBF1;refGene_tx=XM_023257663.1;refGene_gn=SBF1;refGene_tx=XM_023257664.1
B4	144209710	144209843
refGene_gn=SBF1;refGene_tx=XM_019835767.1;refGene_gn=SBF1;refGene_tx=XM_019835768.1;refGene_gn=SBF1;refGene_tx=XM_023257663.1;refGene_gn=SBF1;refGene_tx=XM_023257664.1
B4	144209957	144210063
refGene_gn=SBF1;refGene_tx=XM_019835767.1;refGene_gn=SBF1;refGene_tx=XM_019835768.1;refGene_gn=SBF1;refGene_tx=XM_023257663.1;refGene_gn=SBF1;refGene_tx=XM_023257664.1
B4	144210656	144210767
refGene_gn=SBF1;refGene_tx=XM_019835767.1;refGene_gn=SBF1;refGene_tx=XM_019835768.1;refGene_gn=SBF1;refGene_tx=XM_023257663.1;refGene_gn=SBF1;refGene_tx=XM_023257664.1
B4	144210852	144211011
refGene_gn=SBF1;refGene_tx=XM_019835767.1;refGene_gn=SBF1;refGene_tx=X

M_019835768.1;refGene_gn=SBF1;refGene_tx=XM_023257663.1;refGene_gn=SBF1;refGene_tx=XM_023257664.1
B4 144211245 144211383
refGene_gn=SBF1;refGene_tx=XM_019835767.1;refGene_gn=SBF1;refGene_tx=XM_019835768.1;refGene_gn=SBF1;refGene_tx=XM_023257663.1;refGene_gn=SBF1;refGene_tx=XM_023257664.1
B4 144211724 144211810
refGene_gn=SBF1;refGene_tx=XM_019835767.1;refGene_gn=SBF1;refGene_tx=XM_019835768.1;refGene_gn=SBF1;refGene_tx=XM_023257663.1;refGene_gn=SBF1;refGene_tx=XM_023257664.1
B4 144217548 144217684
refGene_gn=SBF1;refGene_tx=XM_019835767.1;refGene_gn=SBF1;refGene_tx=XM_019835768.1
B4 144218030 144218085
refGene_gn=SBF1;refGene_tx=XM_023257663.1;refGene_gn=SBF1;refGene_tx=XM_023257664.1
B4 144223223 144223333
refGene_gn=ADM2;refGene_tx=XM_011284368.3
B4 144224012 144224331
refGene_gn=ADM2;refGene_tx=XM_011284368.3
B4 144227786 144227801
refGene_gn=MIOX;refGene_tx=XM_003989422.5
B4 144228264 144228345
refGene_gn=MIOX;refGene_tx=XM_003989422.5
B4 144228578 144228659
refGene_gn=MIOX;refGene_tx=XM_003989422.5
B4 144228833 144228996
refGene_gn=MIOX;refGene_tx=XM_003989422.5
B4 144229125 144229193
refGene_gn=MIOX;refGene_tx=XM_003989422.5
B4 144229317 144229427
refGene_gn=MIOX;refGene_tx=XM_003989422.5
B4 144229503 144229571
refGene_gn=MIOX;refGene_tx=XM_003989422.5
B4 144229665 144229715
refGene_gn=MIOX;refGene_tx=XM_003989422.5
B4 144229847 144229960
refGene_gn=MIOX;refGene_tx=XM_003989422.5
B4 144230065 144230174
refGene_gn=MIOX;refGene_tx=XM_003989422.5
B4 144242682 144242988
refGene_gn=LMF2;refGene_tx=XM_023257671.1;refGene_gn=LMF2;refGene_tx=XM_023257672.1
B4 144243079 144243176
refGene_gn=LMF2;refGene_tx=XM_023257671.1;refGene_gn=LMF2;refGene_tx=XM_023257672.1
B4 144243249 144243361
refGene_gn=LMF2;refGene_tx=XM_023257671.1;refGene_gn=LMF2;refGene_tx=XM_023257672.1
B4 144243486 144243655

refGene_gn=LMF2;refGene_tx=XM_023257671.1;refGene_gn=LMF2;refGene_tx=XM_023257672.1
B4	144243758	144243941
refGene_gn=LMF2;refGene_tx=XM_023257671.1;refGene_gn=LMF2;refGene_tx=XM_023257672.1
B4	144244010	144244107
refGene_gn=LMF2;refGene_tx=XM_023257671.1;refGene_gn=LMF2;refGene_tx=XM_023257672.1
B4	144244181	144244287
refGene_gn=LMF2;refGene_tx=XM_023257671.1;refGene_gn=LMF2;refGene_tx=XM_023257672.1
B4	144244357	144244492
refGene_gn=LMF2;refGene_tx=XM_023257671.1;refGene_gn=LMF2;refGene_tx=XM_023257672.1
B4	144244632	144244774
refGene_gn=LMF2;refGene_tx=XM_023257671.1;refGene_gn=LMF2;refGene_tx=XM_023257672.1
B4	144245222	144245401
refGene_gn=LMF2;refGene_tx=XM_023257671.1;refGene_gn=LMF2;refGene_tx=XM_023257672.1
B4	144245477	144245695
refGene_gn=LMF2;refGene_tx=XM_023257671.1;refGene_gn=LMF2;refGene_tx=XM_023257672.1
B4	144245827	144245856
refGene_gn=LMF2;refGene_tx=XM_023257671.1;refGene_gn=LMF2;refGene_tx=XM_023257672.1
B4	144246034	144246288
refGene_gn=LMF2;refGene_tx=XM_023257671.1;refGene_gn=LMF2;refGene_tx=XM_023257672.1
B4	144246777	144246871
refGene_gn=LMF2;refGene_tx=XM_023257671.1;refGene_gn=LMF2;refGene_tx=XM_023257672.1
B4	144247208	144247316
refGene_gn=NCAPH2;refGene_tx=XM_006934212.4;refGene_gn=NCAPH2;refGene_tx=XM_006934213.4;refGene_gn=NCAPH2;refGene_tx=XM_019835775.2
B4	144252340	144252442
refGene_gn=NCAPH2;refGene_tx=XM_006934212.4;refGene_gn=NCAPH2;refGene_tx=XM_006934213.4;refGene_gn=NCAPH2;refGene_tx=XM_019835775.2
B4	144255005	144255061
refGene_gn=NCAPH2;refGene_tx=XM_006934212.4;refGene_gn=NCAPH2;refGene_tx=XM_006934213.4;refGene_gn=NCAPH2;refGene_tx=XM_019835775.2
B4	144255153	144255238
refGene_gn=NCAPH2;refGene_tx=XM_006934212.4;refGene_gn=NCAPH2;refGene_tx=XM_006934213.4;refGene_gn=NCAPH2;refGene_tx=XM_019835775.2
B4	144255315	144255384
refGene_gn=NCAPH2;refGene_tx=XM_006934212.4;refGene_gn=NCAPH2;refGene_tx=XM_006934213.4;refGene_gn=NCAPH2;refGene_tx=XM_019835775.2
B4	144255509	144255589
refGene_gn=NCAPH2;refGene_tx=XM_006934212.4;refGene_gn=NCAPH2;refGene_tx=XM_006934213.4;refGene_gn=NCAPH2;refGene_tx=XM_019835775.2

B4 144255681 144255815
refGene_gn=NCAPH2;refGene_tx=XM_006934212.4;refGene_gn=NCAPH2;refGene_tx=XM_006934213.4;refGene_gn=NCAPH2;refGene_tx=XM_019835775.2
B4 144256121 144256202
refGene_gn=NCAPH2;refGene_tx=XM_006934212.4;refGene_gn=NCAPH2;refGene_tx=XM_006934213.4;refGene_gn=NCAPH2;refGene_tx=XM_019835775.2
B4 144256654 144256785
refGene_gn=NCAPH2;refGene_tx=XM_006934212.4;refGene_gn=NCAPH2;refGene_tx=XM_006934213.4;refGene_gn=NCAPH2;refGene_tx=XM_019835775.2
B4 144257423 144257486
refGene_gn=NCAPH2;refGene_tx=XM_006934212.4;refGene_gn=NCAPH2;refGene_tx=XM_019835775.2
B4 144257815 144257882
refGene_gn=NCAPH2;refGene_tx=XM_006934212.4;refGene_gn=NCAPH2;refGene_tx=XM_006934213.4;refGene_gn=NCAPH2;refGene_tx=XM_019835775.2
B4 144258066 144258174
refGene_gn=NCAPH2;refGene_tx=XM_006934212.4;refGene_gn=NCAPH2;refGene_tx=XM_006934213.4;refGene_gn=NCAPH2;refGene_tx=XM_019835775.2
B4 144258269 144258323
refGene_gn=NCAPH2;refGene_tx=XM_006934212.4;refGene_gn=NCAPH2;refGene_tx=XM_006934213.4;refGene_gn=NCAPH2;refGene_tx=XM_019835775.2
B4 144258459 144258530
refGene_gn=NCAPH2;refGene_tx=XM_006934212.4;refGene_gn=NCAPH2;refGene_tx=XM_006934213.4;refGene_gn=NCAPH2;refGene_tx=XM_019835775.2
B4 144258612 144258706
refGene_gn=NCAPH2;refGene_tx=XM_006934212.4;refGene_gn=NCAPH2;refGene_tx=XM_006934213.4;refGene_gn=NCAPH2;refGene_tx=XM_019835775.2
B4 144258796 144258865
refGene_gn=NCAPH2;refGene_tx=XM_006934212.4;refGene_gn=NCAPH2;refGene_tx=XM_006934213.4;refGene_gn=NCAPH2;refGene_tx=XM_019835775.2
B4 144258977 144259033
refGene_gn=NCAPH2;refGene_tx=XM_006934212.4;refGene_gn=NCAPH2;refGene_tx=XM_006934213.4;refGene_gn=NCAPH2;refGene_tx=XM_019835775.2
B4 144259113 144259215
refGene_gn=NCAPH2;refGene_tx=XM_006934212.4;refGene_gn=NCAPH2;refGene_tx=XM_006934213.4;refGene_gn=NCAPH2;refGene_tx=XM_019835775.2
B4 144259312 144259462
refGene_gn=NCAPH2;refGene_tx=XM_006934212.4;refGene_gn=NCAPH2;refGene_tx=XM_006934213.4;refGene_gn=NCAPH2;refGene_tx=XM_019835775.2
B4 144259527 144259668
refGene_gn=NCAPH2;refGene_tx=XM_006934212.4;refGene_gn=NCAPH2;refGene_tx=XM_006934213.4;refGene_gn=NCAPH2;refGene_tx=XM_019835775.2
B4 144259925 144260726
refGene_gn=LOC101085223;refGene_tx=XM_003989435.5
B4 144265949 144266293
refGene_gn=ODF3B;refGene_tx=XM_019835905.2;refGene_gn=ODF3B;refGene_tx=XM_023257673.1
B4 144266402 144266524
refGene_gn=ODF3B;refGene_tx=XM_019835905.2;refGene_gn=ODF3B;refGene_tx=XM_023257673.1

B4 144266670 144266775
refGene_gn=ODF3B;refGene_tx=XM_019835905.2;refGene_gn=ODF3B;refGene_tx=XM_023257673.1
B4 144267048 144267252
refGene_gn=ODF3B;refGene_tx=XM_019835905.2;refGene_gn=ODF3B;refGene_tx=XM_023257673.1
B4 144267427 144267574
refGene_gn=ODF3B;refGene_tx=XM_019835905.2;refGene_gn=ODF3B;refGene_tx=XM_023257673.1
B4 144267941 144268023
refGene_gn=ODF3B;refGene_tx=XM_019835905.2;refGene_gn=ODF3B;refGene_tx=XM_023257673.1
B4 144282111 144283522
refGene_gn=KLHDC7B;refGene_tx=XM_003989425.3
B4 144283876 144285834
refGene_gn=KLHDC7B;refGene_tx=XM_003989425.3
B4 144287272 144287430
refGene_gn=SYCE3;refGene_tx=XM_003989426.5
B4 144304969 144305078
refGene_gn=SYCE3;refGene_tx=XM_003989426.5
B4 144321787 144321871
refGene_gn=CPT1B;refGene_tx=XM_003989427.5;refGene_gn=CPT1B;refGene_tx=XM_019835777.2
B4 144322017 144322110
refGene_gn=CPT1B;refGene_tx=XM_003989427.5;refGene_gn=CPT1B;refGene_tx=XM_019835777.2
B4 144322700 144322814
refGene_gn=CPT1B;refGene_tx=XM_003989427.5;refGene_gn=CPT1B;refGene_tx=XM_019835777.2
B4 144323384 144323537
refGene_gn=CPT1B;refGene_tx=XM_003989427.5;refGene_gn=CPT1B;refGene_tx=XM_019835777.2
B4 144323636 144323771
refGene_gn=CPT1B;refGene_tx=XM_003989427.5;refGene_gn=CPT1B;refGene_tx=XM_019835777.2
B4 144323850 144324015
refGene_gn=CPT1B;refGene_tx=XM_003989427.5;refGene_gn=CPT1B;refGene_tx=XM_019835777.2
B4 144324379 144324496
refGene_gn=CPT1B;refGene_tx=XM_003989427.5;refGene_gn=CPT1B;refGene_tx=XM_019835777.2
B4 144324580 144324686
refGene_gn=CPT1B;refGene_tx=XM_003989427.5;refGene_gn=CPT1B;refGene_tx=XM_019835777.2
B4 144324772 144324958
refGene_gn=CPT1B;refGene_tx=XM_003989427.5;refGene_gn=CPT1B;refGene_tx=XM_019835777.2
B4 144325282 144325478
refGene_gn=CPT1B;refGene_tx=XM_003989427.5;refGene_gn=CPT1B;refGene_tx=XM_019835777.2

B4	144326012	144326097
refGene_gn=CPT1B;refGene_tx=XM_003989427.5;refGene_gn=CPT1B;refGene_tx=XM_019835777.2
B4	144326171	144326279
refGene_gn=CPT1B;refGene_tx=XM_003989427.5;refGene_gn=CPT1B;refGene_tx=XM_019835777.2
B4	144327066	144327144
refGene_gn=CPT1B;refGene_tx=XM_003989427.5;refGene_gn=CPT1B;refGene_tx=XM_019835777.2
B4	144327242	144327380
refGene_gn=CPT1B;refGene_tx=XM_003989427.5;refGene_gn=CPT1B;refGene_tx=XM_019835777.2
B4	144327565	144327667
refGene_gn=CPT1B;refGene_tx=XM_003989427.5;refGene_gn=CPT1B;refGene_tx=XM_019835777.2
B4	144327891	144328069
refGene_gn=CPT1B;refGene_tx=XM_003989427.5;refGene_gn=CPT1B;refGene_tx=XM_019835777.2
B4	144328366	144328506
refGene_gn=CPT1B;refGene_tx=XM_003989427.5;refGene_gn=CPT1B;refGene_tx=XM_019835777.2
B4	144328870	144329011
refGene_gn=CPT1B;refGene_tx=XM_003989427.5;refGene_gn=CPT1B;refGene_tx=XM_019835777.2
B4	144330301	144330376
refGene_gn=CHKB;refGene_tx=XM_023257674.1
B4	144330528	144330610
refGene_gn=CHKB;refGene_tx=XM_023257674.1
B4	144330894	144330998
refGene_gn=CHKB;refGene_tx=XM_023257674.1
B4	144331136	144331245
refGene_gn=CHKB;refGene_tx=XM_023257674.1
B4	144331372	144331454
refGene_gn=CHKB;refGene_tx=XM_023257674.1
B4	144331540	144331599
refGene_gn=CHKB;refGene_tx=XM_023257674.1
B4	144331768	144331857
refGene_gn=CHKB;refGene_tx=XM_023257674.1
B4	144331858	144331865
refGene_gn=CHKB;refGene_tx=XM_023257674.1
B4	144332153	144332287
refGene_gn=CHKB;refGene_tx=XM_023257674.1
B4	144332495	144332609
refGene_gn=CHKB;refGene_tx=XM_023257674.1
B4	144332838	144332947
refGene_gn=CHKB;refGene_tx=XM_023257674.1
B4	144333167	144333505
refGene_gn=CHKB;refGene_tx=XM_023257674.1
B4	144353476	144353541
refGene_gn=MAPK8IP2;refGene_tx=XM_023257675.1

B4 144354511 144354617
refGene_gn=MAPK8IP2;refGene_tx=XM_023257675.1
B4 144355911 144356190
refGene_gn=MAPK8IP2;refGene_tx=XM_023257675.1
B4 144356352 144356446
refGene_gn=MAPK8IP2;refGene_tx=XM_023257675.1
B4 144356551 144357805
refGene_gn=MAPK8IP2;refGene_tx=XM_023257675.1
B4 144358066 144358142
refGene_gn=MAPK8IP2;refGene_tx=XM_023257675.1
B4 144358270 144358443
refGene_gn=MAPK8IP2;refGene_tx=XM_023257675.1
B4 144358559 144358669
refGene_gn=MAPK8IP2;refGene_tx=XM_023257675.1
B4 144359315 144359420
refGene_gn=MAPK8IP2;refGene_tx=XM_023257675.1
B4 144359564 144359635
refGene_gn=MAPK8IP2;refGene_tx=XM_023257675.1
B4 144361573 144361672
refGene_gn=MAPK8IP2;refGene_tx=XM_023257675.1
B4 144362097 144362170
refGene_gn=MAPK8IP2;refGene_tx=XM_023257675.1
B4 144372446 144372766
refGene_gn=ARSA;refGene_tx=XM_006934215.4
B4 144372876 144372979
refGene_gn=ARSA;refGene_tx=XM_006934215.4
B4 144373186 144373314
refGene_gn=ARSA;refGene_tx=XM_006934215.4
B4 144373401 144373459
refGene_gn=ARSA;refGene_tx=XM_006934215.4
B4 144373460 144373527
refGene_gn=ARSA;refGene_tx=XM_006934215.4
B4 144373834 144374004
refGene_gn=ARSA;refGene_tx=XM_006934215.4
B4 144374078 144374297
refGene_gn=ARSA;refGene_tx=XM_006934215.4
B4 144374414 144374655
refGene_gn=ARSA;refGene_tx=XM_006934215.4
B4 144374849 144374998
refGene_gn=ARSA;refGene_tx=XM_006934215.4
B4 144399057 144399120
refGene_gn=SHANK3;refGene_tx=XM_023257676.1;refGene_gn=SHANK3;refGene_tx=XM_023257677.1;refGene_gn=SHANK3;refGene_tx=XM_023257678.1
B4 144399510 144399714
refGene_gn=SHANK3;refGene_tx=XM_023257676.1;refGene_gn=SHANK3;refGene_tx=XM_023257677.1;refGene_gn=SHANK3;refGene_tx=XM_023257678.1
B4 144400531 144400603
refGene_gn=SHANK3;refGene_tx=XM_023257676.1;refGene_gn=SHANK3;refGene_tx=XM_023257677.1;refGene_gn=SHANK3;refGene_tx=XM_023257678.1
B4 144402656 144402765

refGene_gn=SHANK3;refGene_tx=XM_023257676.1;refGene_gn=SHANK3;refGene_tx=XM_023257677.1;refGene_gn=SHANK3;refGene_tx=XM_023257678.1
B4 144402840 144402992
refGene_gn=SHANK3;refGene_tx=XM_023257676.1;refGene_gn=SHANK3;refGene_tx=XM_023257677.1;refGene_gn=SHANK3;refGene_tx=XM_023257678.1
B4 144403087 144403255
refGene_gn=SHANK3;refGene_tx=XM_023257676.1;refGene_gn=SHANK3;refGene_tx=XM_023257677.1;refGene_gn=SHANK3;refGene_tx=XM_023257678.1
B4 144403425 144403542
refGene_gn=SHANK3;refGene_tx=XM_023257676.1;refGene_gn=SHANK3;refGene_tx=XM_023257677.1;refGene_gn=SHANK3;refGene_tx=XM_023257678.1
B4 144403944 144404022
refGene_gn=SHANK3;refGene_tx=XM_023257676.1;refGene_gn=SHANK3;refGene_tx=XM_023257677.1;refGene_gn=SHANK3;refGene_tx=XM_023257678.1
B4 144404986 144405053
refGene_gn=SHANK3;refGene_tx=XM_023257676.1;refGene_gn=SHANK3;refGene_tx=XM_023257677.1;refGene_gn=SHANK3;refGene_tx=XM_023257678.1
B4 144410801 144411075
refGene_gn=SHANK3;refGene_tx=XM_023257676.1;refGene_gn=SHANK3;refGene_tx=XM_023257677.1;refGene_gn=SHANK3;refGene_tx=XM_023257678.1
B4 144414044 144414238
refGene_gn=SHANK3;refGene_tx=XM_023257676.1;refGene_gn=SHANK3;refGene_tx=XM_023257677.1;refGene_gn=SHANK3;refGene_tx=XM_023257678.1
B4 144415395 144415509
refGene_gn=SHANK3;refGene_tx=XM_023257676.1;refGene_gn=SHANK3;refGene_tx=XM_023257677.1;refGene_gn=SHANK3;refGene_tx=XM_023257678.1
B4 144420770 144420846
refGene_gn=SHANK3;refGene_tx=XM_023257676.1;refGene_gn=SHANK3;refGene_tx=XM_023257677.1;refGene_gn=SHANK3;refGene_tx=XM_023257678.1
B4 144421138 144421221
refGene_gn=SHANK3;refGene_tx=XM_023257676.1;refGene_gn=SHANK3;refGene_tx=XM_023257677.1;refGene_gn=SHANK3;refGene_tx=XM_023257678.1
B4 144421752 144421877
refGene_gn=SHANK3;refGene_tx=XM_023257676.1;refGene_gn=SHANK3;refGene_tx=XM_023257677.1;refGene_gn=SHANK3;refGene_tx=XM_023257678.1
B4 144421978 144422111
refGene_gn=SHANK3;refGene_tx=XM_023257676.1;refGene_gn=SHANK3;refGene_tx=XM_023257677.1;refGene_gn=SHANK3;refGene_tx=XM_023257678.1
B4 144423072 144423153
refGene_gn=SHANK3;refGene_tx=XM_023257676.1;refGene_gn=SHANK3;refGene_tx=XM_023257677.1;refGene_gn=SHANK3;refGene_tx=XM_023257678.1
B4 144427190 144427214
refGene_gn=SHANK3;refGene_tx=XM_023257676.1;refGene_gn=SHANK3;refGene_tx=XM_023257678.1
B4 144430810 144430941
refGene_gn=SHANK3;refGene_tx=XM_023257676.1;refGene_gn=SHANK3;refGene_tx=XM_023257677.1;refGene_gn=SHANK3;refGene_tx=XM_023257678.1
B4 144431562 144431647
refGene_gn=SHANK3;refGene_tx=XM_023257676.1;refGene_gn=SHANK3;refGene_tx=XM_023257677.1;refGene_gn=SHANK3;refGene_tx=XM_023257678.1

B4 144436257 144438514
refGene_gn=SHANK3;refGene_tx=XM_023257676.1;refGene_gn=SHANK3;refGene_tx=XM_023257677.1;refGene_gn=SHANK3;refGene_tx=XM_023257678.1
B4 144445408 144445478
refGene_gn=SHANK3;refGene_tx=XM_023257678.1
B4 144446482 144447074
refGene_gn=SHANK3;refGene_tx=XM_023257676.1;refGene_gn=SHANK3;refGene_tx=XM_023257677.1
B4 144454669 144454746
refGene_gn=ACR;refGene_tx=XM_003989430.5
B4 144455605 144455812
refGene_gn=ACR;refGene_tx=XM_003989430.5
B4 144456014 144456298
refGene_gn=ACR;refGene_tx=XM_003989430.5
B4 144459224 144459370
refGene_gn=ACR;refGene_tx=XM_003989430.5
B4 144459919 144460477
refGene_gn=ACR;refGene_tx=XM_003989430.5
B4 144489746 144489827
refGene_gn=RABL2B;refGene_tx=XM_019835786.2;refGene_gn=RABL2B;refGene_tx=XM_019835787.2
B4 144490002 144490086
refGene_gn=RABL2B;refGene_tx=XM_019835786.2;refGene_gn=RABL2B;refGene_tx=XM_019835787.2
B4 144490453 144490551
refGene_gn=RABL2B;refGene_tx=XM_019835786.2;refGene_gn=RABL2B;refGene_tx=XM_019835787.2
B4 144490883 144490995
refGene_gn=RABL2B;refGene_tx=XM_019835786.2;refGene_gn=RABL2B;refGene_tx=XM_019835787.2
B4 144493950 144494030
refGene_gn=RABL2B;refGene_tx=XM_019835786.2;refGene_gn=RABL2B;refGene_tx=XM_019835787.2
B4 144494730 144494810
refGene_gn=RABL2B;refGene_tx=XM_019835786.2;refGene_gn=RABL2B;refGene_tx=XM_019835787.2
B4 144495901 144495931
refGene_gn=RABL2B;refGene_tx=XM_019835786.2;refGene_gn=RABL2B;refGene_tx=XM_019835787.2
B4 144510105 144510212
refGene_gn=RABL2B;refGene_tx=XM_019835786.2;refGene_gn=RABL2B;refGene_tx=XM_019835787.2
C1 47800 48317
refGene_gn=SAMD11;refGene_tx=XM_011284388.3;refGene_gn=SAMD11;refGene_tx=XM_019835961.2;refGene_gn=SAMD11;refGene_tx=XM_019835962.2;refGene_gn=SAMD11;refGene_tx=XM_019835963.2;refGene_gn=SAMD11;refGene_tx=XM_019835964.2;refGene_gn=SAMD11;refGene_tx=XM_019835965.2
C1 49481 49573
refGene_gn=SAMD11;refGene_tx=XM_011284388.3;refGene_gn=SAMD11;refGene_tx=XM_019835961.2;refGene_gn=SAMD11;refGene_tx=XM_019835962.2;refGene_

gn=SAMD11;refGene_tx=XM_019835963.2;refGene_gn=SAMD11;refGene_tx=XM_01
9835964.2;refGene_gn=SAMD11;refGene_tx=XM_019835965.2;refGene_gn=SAMD1
1;refGene_tx=XM_019835966.2;refGene_gn=SAMD11;refGene_tx=XM_019835967.
2
C1 51069 51234 refGene_gn=SAMD11;refGene_tx=XM_023257879.1
C1 53520 53702
refGene_gn=SAMD11;refGene_tx=XM_011284388.3;refGene_gn=SAMD11;refGene_
tx=XM_019835961.2;refGene_gn=SAMD11;refGene_tx=XM_019835962.2;refGene_
gn=SAMD11;refGene_tx=XM_019835963.2;refGene_gn=SAMD11;refGene_tx=XM_01
9835964.2;refGene_gn=SAMD11;refGene_tx=XM_019835965.2;refGene_gn=SAMD1
1;refGene_tx=XM_019835966.2;refGene_gn=SAMD11;refGene_tx=XM_019835967.
2;refGene_gn=SAMD11;refGene_tx=XM_023257879.1
C1 55061 55112
refGene_gn=SAMD11;refGene_tx=XM_011284388.3;refGene_gn=SAMD11;refGene_
tx=XM_019835961.2;refGene_gn=SAMD11;refGene_tx=XM_019835963.2;refGene_
gn=SAMD11;refGene_tx=XM_019835964.2;refGene_gn=SAMD11;refGene_tx=XM_01
9835965.2;refGene_gn=SAMD11;refGene_tx=XM_019835966.2;refGene_gn=SAMD1
1;refGene_tx=XM_019835967.2;refGene_gn=SAMD11;refGene_tx=XM_023257879.
1
C1 59092 59217
refGene_gn=SAMD11;refGene_tx=XM_011284388.3;refGene_gn=SAMD11;refGene_
tx=XM_019835961.2;refGene_gn=SAMD11;refGene_tx=XM_019835962.2;refGene_
gn=SAMD11;refGene_tx=XM_019835963.2;refGene_gn=SAMD11;refGene_tx=XM_01
9835964.2;refGene_gn=SAMD11;refGene_tx=XM_019835965.2;refGene_gn=SAMD1
1;refGene_tx=XM_019835966.2;refGene_gn=SAMD11;refGene_tx=XM_019835967.
2;refGene_gn=SAMD11;refGene_tx=XM_023257879.1
C1 62404 62494
refGene_gn=SAMD11;refGene_tx=XM_011284388.3;refGene_gn=SAMD11;refGene_
tx=XM_019835961.2;refGene_gn=SAMD11;refGene_tx=XM_019835962.2;refGene_
gn=SAMD11;refGene_tx=XM_019835963.2;refGene_gn=SAMD11;refGene_tx=XM_01
9835964.2;refGene_gn=SAMD11;refGene_tx=XM_019835965.2;refGene_gn=SAMD1
1;refGene_tx=XM_019835966.2;refGene_gn=SAMD11;refGene_tx=XM_019835967.
2;refGene_gn=SAMD11;refGene_tx=XM_023257879.1
C1 62667 62802
refGene_gn=SAMD11;refGene_tx=XM_011284388.3;refGene_gn=SAMD11;refGene_
tx=XM_019835961.2;refGene_gn=SAMD11;refGene_tx=XM_019835962.2;refGene_
gn=SAMD11;refGene_tx=XM_019835963.2;refGene_gn=SAMD11;refGene_tx=XM_01
9835964.2;refGene_gn=SAMD11;refGene_tx=XM_019835965.2;refGene_gn=SAMD1
1;refGene_tx=XM_019835966.2;refGene_gn=SAMD11;refGene_tx=XM_019835967.
2;refGene_gn=SAMD11;refGene_tx=XM_023257879.1
C1 64763 64926
refGene_gn=SAMD11;refGene_tx=XM_011284388.3;refGene_gn=SAMD11;refGene_
tx=XM_019835961.2;refGene_gn=SAMD11;refGene_tx=XM_019835962.2;refGene_
gn=SAMD11;refGene_tx=XM_019835963.2;refGene_gn=SAMD11;refGene_tx=XM_01
9835964.2;refGene_gn=SAMD11;refGene_tx=XM_019835965.2;refGene_gn=SAMD1
1;refGene_tx=XM_019835966.2;refGene_gn=SAMD11;refGene_tx=XM_019835967.
2;refGene_gn=SAMD11;refGene_tx=XM_023257879.1
C1 66013 66129
refGene_gn=SAMD11;refGene_tx=XM_011284388.3;refGene_gn=SAMD11;refGene_
tx=XM_019835961.2;refGene_gn=SAMD11;refGene_tx=XM_019835962.2;refGene_

gn=SAMD11;refGene_tx=XM_019835963.2;refGene_gn=SAMD11;refGene_tx=XM_01
9835964.2;refGene_gn=SAMD11;refGene_tx=XM_019835965.2;refGene_gn=SAMD1
1;refGene_tx=XM_019835966.2;refGene_gn=SAMD11;refGene_tx=XM_019835967.
2;refGene_gn=SAMD11;refGene_tx=XM_023257879.1
C1 66358 66437
refGene_gn=SAMD11;refGene_tx=XM_011284388.3;refGene_gn=SAMD11;refGene_
tx=XM_019835961.2;refGene_gn=SAMD11;refGene_tx=XM_019835962.2;refGene_
gn=SAMD11;refGene_tx=XM_019835963.2;refGene_gn=SAMD11;refGene_tx=XM_01
9835964.2;refGene_gn=SAMD11;refGene_tx=XM_019835965.2;refGene_gn=SAMD1
1;refGene_tx=XM_019835966.2;refGene_gn=SAMD11;refGene_tx=XM_019835967.
2;refGene_gn=SAMD11;refGene_tx=XM_023257879.1
C1 66520 67035
refGene_gn=SAMD11;refGene_tx=XM_011284388.3;refGene_gn=SAMD11;refGene_
tx=XM_019835961.2;refGene_gn=SAMD11;refGene_tx=XM_019835962.2;refGene_
gn=SAMD11;refGene_tx=XM_019835963.2;refGene_gn=SAMD11;refGene_tx=XM_01
9835964.2;refGene_gn=SAMD11;refGene_tx=XM_019835965.2;refGene_gn=SAMD1
1;refGene_tx=XM_019835966.2;refGene_gn=SAMD11;refGene_tx=XM_019835967.
2;refGene_gn=SAMD11;refGene_tx=XM_023257879.1
C1 67278 67403
refGene_gn=SAMD11;refGene_tx=XM_011284388.3;refGene_gn=SAMD11;refGene_
tx=XM_019835961.2;refGene_gn=SAMD11;refGene_tx=XM_019835962.2;refGene_
gn=SAMD11;refGene_tx=XM_019835963.2;refGene_gn=SAMD11;refGene_tx=XM_01
9835964.2;refGene_gn=SAMD11;refGene_tx=XM_019835965.2;refGene_gn=SAMD1
1;refGene_tx=XM_019835966.2;refGene_gn=SAMD11;refGene_tx=XM_019835967.
2;refGene_gn=SAMD11;refGene_tx=XM_023257879.1
C1 67763 67874
refGene_gn=SAMD11;refGene_tx=XM_011284388.3;refGene_gn=SAMD11;refGene_
tx=XM_019835961.2;refGene_gn=SAMD11;refGene_tx=XM_019835962.2;refGene_
gn=SAMD11;refGene_tx=XM_019835963.2;refGene_gn=SAMD11;refGene_tx=XM_01
9835964.2;refGene_gn=SAMD11;refGene_tx=XM_019835965.2;refGene_gn=SAMD1
1;refGene_tx=XM_019835966.2;refGene_gn=SAMD11;refGene_tx=XM_019835967.
2;refGene_gn=SAMD11;refGene_tx=XM_023257879.1
C1 67974 68265
refGene_gn=SAMD11;refGene_tx=XM_011284388.3;refGene_gn=SAMD11;refGene_
tx=XM_019835961.2;refGene_gn=SAMD11;refGene_tx=XM_019835962.2;refGene_
gn=SAMD11;refGene_tx=XM_019835964.2;refGene_gn=SAMD11;refGene_tx=XM_01
9835965.2;refGene_gn=SAMD11;refGene_tx=XM_019835966.2;refGene_gn=SAMD1
1;refGene_tx=XM_019835967.2;refGene_gn=SAMD11;refGene_tx=XM_023257879.
1
C1 68726 69046
refGene_gn=NOC2L;refGene_tx=XM_006944179.2;refGene_gn=NOC2L;refGene_tx
=XM_011284387.2;refGene_gn=SAMD11;refGene_tx=XM_019835963.2
C1 69359 69458
refGene_gn=NOC2L;refGene_tx=XM_006944179.2;refGene_gn=NOC2L;refGene_tx
=XM_011284387.2
C1 69839 69975
refGene_gn=NOC2L;refGene_tx=XM_006944179.2;refGene_gn=NOC2L;refGene_tx
=XM_011284387.2
C1 70953 71067
refGene_gn=NOC2L;refGene_tx=XM_006944179.2;refGene_gn=NOC2L;refGene_tx

=XM_011284387.2
C1	71191	71335
refGene_gn=NOC2L;refGene_tx=XM_006944179.2;refGene_gn=NOC2L;refGene_tx
=XM_011284387.2
C1	72618	72720
refGene_gn=NOC2L;refGene_tx=XM_006944179.2;refGene_gn=NOC2L;refGene_tx
=XM_011284387.2
C1	73042	73156
refGene_gn=NOC2L;refGene_tx=XM_006944179.2;refGene_gn=NOC2L;refGene_tx
=XM_011284387.2
C1	74748	74860
refGene_gn=NOC2L;refGene_tx=XM_006944179.2;refGene_gn=NOC2L;refGene_tx
=XM_011284387.2
C1	75733	75873
refGene_gn=NOC2L;refGene_tx=XM_006944179.2;refGene_gn=NOC2L;refGene_tx
=XM_011284387.2
C1	76227	76416
refGene_gn=NOC2L;refGene_tx=XM_006944179.2;refGene_gn=NOC2L;refGene_tx
=XM_011284387.2
C1	77119	77233
refGene_gn=NOC2L;refGene_tx=XM_006944179.2;refGene_gn=NOC2L;refGene_tx
=XM_011284387.2
C1	77733	77844
refGene_gn=NOC2L;refGene_tx=XM_006944179.2;refGene_gn=NOC2L;refGene_tx
=XM_011284387.2
C1	77971	78050
refGene_gn=NOC2L;refGene_tx=XM_006944179.2;refGene_gn=NOC2L;refGene_tx
=XM_011284387.2
C1	78665	78756
refGene_gn=NOC2L;refGene_tx=XM_006944179.2;refGene_gn=NOC2L;refGene_tx
=XM_011284387.2
C1	78842	78963
refGene_gn=NOC2L;refGene_tx=XM_006944179.2;refGene_gn=NOC2L;refGene_tx
=XM_011284387.2
C1	79643	79766
refGene_gn=NOC2L;refGene_tx=XM_006944179.2;refGene_gn=NOC2L;refGene_tx
=XM_011284387.2
C1	79854	80029
refGene_gn=NOC2L;refGene_tx=XM_006944179.2;refGene_gn=NOC2L;refGene_tx
=XM_011284387.2
C1	81493	81646
refGene_gn=NOC2L;refGene_tx=XM_006944179.2;refGene_gn=NOC2L;refGene_tx
=XM_011284387.2
C1	81796	81822
refGene_gn=NOC2L;refGene_tx=XM_006944179.2;refGene_gn=NOC2L;refGene_tx
=XM_011284387.2
C1	83483	83593	refGene_gn=KLHL17;refGene_tx=XM_011284391.3
C1	84485	84745	refGene_gn=KLHL17;refGene_tx=XM_011284391.3
C1	84832	84954	refGene_gn=KLHL17;refGene_tx=XM_011284391.3
C1	85106	85328	refGene_gn=KLHL17;refGene_tx=XM_011284391.3

C1 85818 85935 refGene_gn=KLHL17;refGene_tx=XM_011284391.3
C1 86141 86355 refGene_gn=KLHL17;refGene_tx=XM_011284391.3
C1 86669 86814 refGene_gn=KLHL17;refGene_tx=XM_011284391.3
C1 86897 87065 refGene_gn=KLHL17;refGene_tx=XM_011284391.3
C1 87488 87577 refGene_gn=KLHL17;refGene_tx=XM_011284391.3
C1 87688 87762 refGene_gn=KLHL17;refGene_tx=XM_011284391.3
C1 87957 88139 refGene_gn=KLHL17;refGene_tx=XM_011284391.3
C1 88497 88726 refGene_gn=KLHL17;refGene_tx=XM_011284391.3
C1 90317 90400 refGene_gn=PLEKHN1;refGene_tx=XM_006944214.4
C1 90520 90620 refGene_gn=PLEKHN1;refGene_tx=XM_006944214.4
C1 93372 93519
refGene_gn=PLEKHN1;refGene_tx=XM_006944214.4;refGene_gn=PLEKHN1;refGen
e_tx=XM_019835968.2
C1 93606 93703
refGene_gn=PLEKHN1;refGene_tx=XM_006944214.4;refGene_gn=PLEKHN1;refGen
e_tx=XM_019835968.2
C1 93826 93899
refGene_gn=PLEKHN1;refGene_tx=XM_006944214.4;refGene_gn=PLEKHN1;refGen
e_tx=XM_019835968.2
C1 94241 94369
refGene_gn=PLEKHN1;refGene_tx=XM_006944214.4;refGene_gn=PLEKHN1;refGen
e_tx=XM_019835968.2
C1 94450 94582
refGene_gn=PLEKHN1;refGene_tx=XM_006944214.4;refGene_gn=PLEKHN1;refGen
e_tx=XM_019835968.2
C1 94772 94853
refGene_gn=PLEKHN1;refGene_tx=XM_006944214.4;refGene_gn=PLEKHN1;refGen
e_tx=XM_019835968.2
C1 95625 95701
refGene_gn=PLEKHN1;refGene_tx=XM_006944214.4;refGene_gn=PLEKHN1;refGen
e_tx=XM_019835968.2
C1 95818 95967
refGene_gn=PLEKHN1;refGene_tx=XM_006944214.4;refGene_gn=PLEKHN1;refGen
e_tx=XM_019835968.2
C1 96095 96245
refGene_gn=PLEKHN1;refGene_tx=XM_006944214.4;refGene_gn=PLEKHN1;refGen
e_tx=XM_019835968.2
C1 96466 96607
refGene_gn=PLEKHN1;refGene_tx=XM_006944214.4;refGene_gn=PLEKHN1;refGen
e_tx=XM_019835968.2
C1 96777 96918
refGene_gn=PLEKHN1;refGene_tx=XM_006944214.4;refGene_gn=PLEKHN1;refGen
e_tx=XM_019835968.2
C1 97073 97283
refGene_gn=PLEKHN1;refGene_tx=XM_006944214.4;refGene_gn=PLEKHN1;refGen
e_tx=XM_019835968.2
C1 97554 97603
refGene_gn=PLEKHN1;refGene_tx=XM_006944214.4;refGene_gn=PLEKHN1;refGen
e_tx=XM_019835968.2
C1 97693 97809

refGene_gn=PLEKHN1;refGene_tx=XM_006944214.4;refGene_gn=PLEKHN1;refGene_tx=XM_019835968.2
C1	99344	99460	refGene_gn=PERM1;refGene_tx=XM_006944180.4
C1	99698	99824	refGene_gn=PERM1;refGene_tx=XM_006944180.4
C1	100462	102464	refGene_gn=PERM1;refGene_tx=XM_006944180.4
C1	119773	120135
refGene_gn=HES4;refGene_tx=XM_023257880.1;refGene_gn=HES4;refGene_tx=XM_023257881.1
C1	120231	120319
refGene_gn=HES4;refGene_tx=XM_023257880.1;refGene_gn=HES4;refGene_tx=XM_023257881.1
C1	120425	120521
refGene_gn=HES4;refGene_tx=XM_023257880.1;refGene_gn=HES4;refGene_tx=XM_023257881.1
C1	120601	120688	refGene_gn=HES4;refGene_tx=XM_023257880.1
C1	136053	136056	refGene_gn=ISG15;refGene_tx=NM_001130843.2
C1	136548	137019	refGene_gn=ISG15;refGene_tx=NM_001130843.2
C1	139515	139722
refGene_gn=AGRN;refGene_tx=XM_023257882.1;refGene_gn=AGRN;refGene_tx=XM_023257883.1;refGene_gn=AGRN;refGene_tx=XM_023257884.1;refGene_gn=AGRN;refGene_tx=XM_023257885.1;refGene_gn=AGRN;refGene_tx=XM_023257886.1;refGene_gn=AGRN;refGene_tx=XM_023257888.1
C1	141951	142213
refGene_gn=AGRN;refGene_tx=XM_023257882.1;refGene_gn=AGRN;refGene_tx=XM_023257883.1;refGene_gn=AGRN;refGene_tx=XM_023257884.1;refGene_gn=AGRN;refGene_tx=XM_023257885.1;refGene_gn=AGRN;refGene_tx=XM_023257886.1;refGene_gn=AGRN;refGene_tx=XM_023257888.1
C1	154074	154222
refGene_gn=AGRN;refGene_tx=XM_019835969.2;refGene_gn=AGRN;refGene_tx=XM_019835970.2;refGene_gn=AGRN;refGene_tx=XM_019835971.2
C1	154858	154906
refGene_gn=AGRN;refGene_tx=XM_019835969.2;refGene_gn=AGRN;refGene_tx=XM_019835970.2;refGene_gn=AGRN;refGene_tx=XM_019835971.2;refGene_gn=AGRN;refGene_tx=XM_023257882.1;refGene_gn=AGRN;refGene_tx=XM_023257883.1;refGene_gn=AGRN;refGene_tx=XM_023257884.1;refGene_gn=AGRN;refGene_tx=XM_023257885.1;refGene_gn=AGRN;refGene_tx=XM_023257886.1;refGene_gn=AGRN;refGene_tx=XM_023257888.1
C1	159383	159599
refGene_gn=AGRN;refGene_tx=XM_019835969.2;refGene_gn=AGRN;refGene_tx=XM_019835970.2;refGene_gn=AGRN;refGene_tx=XM_019835971.2;refGene_gn=AGRN;refGene_tx=XM_023257882.1;refGene_gn=AGRN;refGene_tx=XM_023257883.1;refGene_gn=AGRN;refGene_tx=XM_023257884.1;refGene_gn=AGRN;refGene_tx=XM_023257885.1;refGene_gn=AGRN;refGene_tx=XM_023257886.1;refGene_gn=AGRN;refGene_tx=XM_023257888.1
C1	159672	159897
refGene_gn=AGRN;refGene_tx=XM_019835969.2;refGene_gn=AGRN;refGene_tx=XM_019835970.2;refGene_gn=AGRN;refGene_tx=XM_019835971.2;refGene_gn=AGRN;refGene_tx=XM_023257882.1;refGene_gn=AGRN;refGene_tx=XM_023257883.1;refGene_gn=AGRN;refGene_tx=XM_023257884.1;refGene_gn=AGRN;refGene_tx=XM_023257885.1;refGene_gn=AGRN;refGene_tx=XM_023257886.1;refGene_gn=AGR

N;refGene_tx=XM_023257888.1
C1 159983 160208
refGene_gn=AGRN;refGene_tx=XM_019835969.2;refGene_gn=AGRN;refGene_tx=XM_019835970.2;refGene_gn=AGRN;refGene_tx=XM_019835971.2;refGene_gn=AGRN;refGene_tx=XM_023257882.1;refGene_gn=AGRN;refGene_tx=XM_023257883.1;refGene_gn=AGRN;refGene_tx=XM_023257884.1;refGene_gn=AGRN;refGene_tx=XM_023257885.1;refGene_gn=AGRN;refGene_tx=XM_023257886.1;refGene_gn=AGRN;refGene_tx=XM_023257888.1
C1 160371 160578
refGene_gn=AGRN;refGene_tx=XM_019835969.2;refGene_gn=AGRN;refGene_tx=XM_019835970.2;refGene_gn=AGRN;refGene_tx=XM_019835971.2;refGene_gn=AGRN;refGene_tx=XM_023257882.1;refGene_gn=AGRN;refGene_tx=XM_023257883.1;refGene_gn=AGRN;refGene_tx=XM_023257884.1;refGene_gn=AGRN;refGene_tx=XM_023257885.1;refGene_gn=AGRN;refGene_tx=XM_023257886.1;refGene_gn=AGRN;refGene_tx=XM_023257888.1
C1 161646 161865
refGene_gn=AGRN;refGene_tx=XM_019835969.2;refGene_gn=AGRN;refGene_tx=XM_019835970.2;refGene_gn=AGRN;refGene_tx=XM_019835971.2;refGene_gn=AGRN;refGene_tx=XM_023257882.1;refGene_gn=AGRN;refGene_tx=XM_023257883.1;refGene_gn=AGRN;refGene_tx=XM_023257884.1;refGene_gn=AGRN;refGene_tx=XM_023257885.1;refGene_gn=AGRN;refGene_tx=XM_023257886.1;refGene_gn=AGRN;refGene_tx=XM_023257888.1
C1 161940 162135
refGene_gn=AGRN;refGene_tx=XM_019835969.2;refGene_gn=AGRN;refGene_tx=XM_019835970.2;refGene_gn=AGRN;refGene_tx=XM_019835971.2;refGene_gn=AGRN;refGene_tx=XM_023257882.1;refGene_gn=AGRN;refGene_tx=XM_023257883.1;refGene_gn=AGRN;refGene_tx=XM_023257884.1;refGene_gn=AGRN;refGene_tx=XM_023257885.1;refGene_gn=AGRN;refGene_tx=XM_023257886.1;refGene_gn=AGRN;refGene_tx=XM_023257888.1
C1 162227 162428
refGene_gn=AGRN;refGene_tx=XM_019835969.2;refGene_gn=AGRN;refGene_tx=XM_019835970.2;refGene_gn=AGRN;refGene_tx=XM_019835971.2;refGene_gn=AGRN;refGene_tx=XM_023257882.1;refGene_gn=AGRN;refGene_tx=XM_023257883.1;refGene_gn=AGRN;refGene_tx=XM_023257884.1;refGene_gn=AGRN;refGene_tx=XM_023257885.1;refGene_gn=AGRN;refGene_tx=XM_023257886.1;refGene_gn=AGRN;refGene_tx=XM_023257888.1
C1 162511 162660
refGene_gn=AGRN;refGene_tx=XM_019835969.2;refGene_gn=AGRN;refGene_tx=XM_019835970.2;refGene_gn=AGRN;refGene_tx=XM_019835971.2;refGene_gn=AGRN;refGene_tx=XM_023257882.1;refGene_gn=AGRN;refGene_tx=XM_023257883.1;refGene_gn=AGRN;refGene_tx=XM_023257884.1;refGene_gn=AGRN;refGene_tx=XM_023257885.1;refGene_gn=AGRN;refGene_tx=XM_023257886.1;refGene_gn=AGRN;refGene_tx=XM_023257888.1
C1 162750 162856
refGene_gn=AGRN;refGene_tx=XM_019835969.2;refGene_gn=AGRN;refGene_tx=XM_019835970.2;refGene_gn=AGRN;refGene_tx=XM_019835971.2;refGene_gn=AGRN;refGene_tx=XM_023257882.1;refGene_gn=AGRN;refGene_tx=XM_023257883.1;refGene_gn=AGRN;refGene_tx=XM_023257884.1;refGene_gn=AGRN;refGene_tx=XM_023257885.1;refGene_gn=AGRN;refGene_tx=XM_023257886.1;refGene_gn=AGRN;refGene_tx=XM_023257888.1

C1 163598 163715
refGene_gn=AGRN;refGene_tx=XM_019835969.2;refGene_gn=AGRN;refGene_tx=XM_019835970.2;refGene_gn=AGRN;refGene_tx=XM_019835971.2;refGene_gn=AGRN;refGene_tx=XM_023257882.1;refGene_gn=AGRN;refGene_tx=XM_023257883.1;refGene_gn=AGRN;refGene_tx=XM_023257884.1;refGene_gn=AGRN;refGene_tx=XM_023257885.1;refGene_gn=AGRN;refGene_tx=XM_023257886.1;refGene_gn=AGRN;refGene_tx=XM_023257888.1
C1 163797 163962
refGene_gn=AGRN;refGene_tx=XM_019835969.2;refGene_gn=AGRN;refGene_tx=XM_019835970.2;refGene_gn=AGRN;refGene_tx=XM_019835971.2;refGene_gn=AGRN;refGene_tx=XM_023257882.1;refGene_gn=AGRN;refGene_tx=XM_023257883.1;refGene_gn=AGRN;refGene_tx=XM_023257884.1;refGene_gn=AGRN;refGene_tx=XM_023257885.1;refGene_gn=AGRN;refGene_tx=XM_023257886.1;refGene_gn=AGRN;refGene_tx=XM_023257888.1
C1 164145 164289
refGene_gn=AGRN;refGene_tx=XM_019835969.2;refGene_gn=AGRN;refGene_tx=XM_019835970.2;refGene_gn=AGRN;refGene_tx=XM_019835971.2;refGene_gn=AGRN;refGene_tx=XM_023257882.1;refGene_gn=AGRN;refGene_tx=XM_023257883.1;refGene_gn=AGRN;refGene_tx=XM_023257884.1;refGene_gn=AGRN;refGene_tx=XM_023257885.1;refGene_gn=AGRN;refGene_tx=XM_023257886.1;refGene_gn=AGRN;refGene_tx=XM_023257888.1
C1 164393 164518
refGene_gn=AGRN;refGene_tx=XM_019835969.2;refGene_gn=AGRN;refGene_tx=XM_019835970.2;refGene_gn=AGRN;refGene_tx=XM_019835971.2;refGene_gn=AGRN;refGene_tx=XM_023257882.1;refGene_gn=AGRN;refGene_tx=XM_023257883.1;refGene_gn=AGRN;refGene_tx=XM_023257884.1;refGene_gn=AGRN;refGene_tx=XM_023257885.1;refGene_gn=AGRN;refGene_tx=XM_023257886.1;refGene_gn=AGRN;refGene_tx=XM_023257888.1
C1 164593 164699
refGene_gn=AGRN;refGene_tx=XM_019835969.2;refGene_gn=AGRN;refGene_tx=XM_019835970.2;refGene_gn=AGRN;refGene_tx=XM_019835971.2;refGene_gn=AGRN;refGene_tx=XM_023257882.1;refGene_gn=AGRN;refGene_tx=XM_023257883.1;refGene_gn=AGRN;refGene_tx=XM_023257884.1;refGene_gn=AGRN;refGene_tx=XM_023257885.1;refGene_gn=AGRN;refGene_tx=XM_023257886.1;refGene_gn=AGRN;refGene_tx=XM_023257888.1
C1 164779 165133
refGene_gn=AGRN;refGene_tx=XM_019835969.2;refGene_gn=AGRN;refGene_tx=XM_019835970.2;refGene_gn=AGRN;refGene_tx=XM_019835971.2;refGene_gn=AGRN;refGene_tx=XM_023257882.1;refGene_gn=AGRN;refGene_tx=XM_023257883.1;refGene_gn=AGRN;refGene_tx=XM_023257884.1;refGene_gn=AGRN;refGene_tx=XM_023257885.1;refGene_gn=AGRN;refGene_tx=XM_023257886.1;refGene_gn=AGRN;refGene_tx=XM_023257888.1
C1 165213 165351
refGene_gn=AGRN;refGene_tx=XM_019835969.2;refGene_gn=AGRN;refGene_tx=XM_019835970.2;refGene_gn=AGRN;refGene_tx=XM_019835971.2;refGene_gn=AGRN;refGene_tx=XM_023257882.1;refGene_gn=AGRN;refGene_tx=XM_023257883.1;refGene_gn=AGRN;refGene_tx=XM_023257884.1;refGene_gn=AGRN;refGene_tx=XM_023257885.1;refGene_gn=AGRN;refGene_tx=XM_023257886.1;refGene_gn=AGRN;refGene_tx=XM_023257888.1
C1 165790 165918

refGene_gn=AGRN;refGene_tx=XM_019835969.2;refGene_gn=AGRN;refGene_tx=XM_019835970.2;refGene_gn=AGRN;refGene_tx=XM_019835971.2;refGene_gn=AGRN;refGene_tx=XM_023257882.1;refGene_gn=AGRN;refGene_tx=XM_023257883.1;refGene_gn=AGRN;refGene_tx=XM_023257884.1;refGene_gn=AGRN;refGene_tx=XM_023257885.1;refGene_gn=AGRN;refGene_tx=XM_023257886.1;refGene_gn=AGRN;refGene_tx=XM_023257888.1
C1 166049 166164
refGene_gn=AGRN;refGene_tx=XM_019835969.2;refGene_gn=AGRN;refGene_tx=XM_019835970.2;refGene_gn=AGRN;refGene_tx=XM_019835971.2;refGene_gn=AGRN;refGene_tx=XM_023257882.1;refGene_gn=AGRN;refGene_tx=XM_023257883.1;refGene_gn=AGRN;refGene_tx=XM_023257884.1;refGene_gn=AGRN;refGene_tx=XM_023257885.1;refGene_gn=AGRN;refGene_tx=XM_023257886.1;refGene_gn=AGRN;refGene_tx=XM_023257888.1
C1 166249 166369
refGene_gn=AGRN;refGene_tx=XM_019835969.2;refGene_gn=AGRN;refGene_tx=XM_019835970.2;refGene_gn=AGRN;refGene_tx=XM_019835971.2;refGene_gn=AGRN;refGene_tx=XM_023257882.1;refGene_gn=AGRN;refGene_tx=XM_023257883.1;refGene_gn=AGRN;refGene_tx=XM_023257884.1;refGene_gn=AGRN;refGene_tx=XM_023257885.1;refGene_gn=AGRN;refGene_tx=XM_023257886.1;refGene_gn=AGRN;refGene_tx=XM_023257888.1
C1 166502 166859
refGene_gn=AGRN;refGene_tx=XM_019835969.2;refGene_gn=AGRN;refGene_tx=XM_019835970.2;refGene_gn=AGRN;refGene_tx=XM_019835971.2;refGene_gn=AGRN;refGene_tx=XM_023257882.1;refGene_gn=AGRN;refGene_tx=XM_023257883.1;refGene_gn=AGRN;refGene_tx=XM_023257884.1;refGene_gn=AGRN;refGene_tx=XM_023257885.1;refGene_gn=AGRN;refGene_tx=XM_023257886.1;refGene_gn=AGRN;refGene_tx=XM_023257888.1
C1 167047 167240
refGene_gn=AGRN;refGene_tx=XM_019835969.2;refGene_gn=AGRN;refGene_tx=XM_019835970.2;refGene_gn=AGRN;refGene_tx=XM_019835971.2;refGene_gn=AGRN;refGene_tx=XM_023257882.1;refGene_gn=AGRN;refGene_tx=XM_023257883.1;refGene_gn=AGRN;refGene_tx=XM_023257884.1;refGene_gn=AGRN;refGene_tx=XM_023257885.1;refGene_gn=AGRN;refGene_tx=XM_023257886.1;refGene_gn=AGRN;refGene_tx=XM_023257888.1
C1 167409 167625
refGene_gn=AGRN;refGene_tx=XM_019835969.2;refGene_gn=AGRN;refGene_tx=XM_019835970.2;refGene_gn=AGRN;refGene_tx=XM_019835971.2;refGene_gn=AGRN;refGene_tx=XM_023257882.1;refGene_gn=AGRN;refGene_tx=XM_023257883.1;refGene_gn=AGRN;refGene_tx=XM_023257884.1;refGene_gn=AGRN;refGene_tx=XM_023257885.1;refGene_gn=AGRN;refGene_tx=XM_023257886.1;refGene_gn=AGRN;refGene_tx=XM_023257888.1
C1 168048 168170
refGene_gn=AGRN;refGene_tx=XM_019835969.2;refGene_gn=AGRN;refGene_tx=XM_019835970.2;refGene_gn=AGRN;refGene_tx=XM_019835971.2;refGene_gn=AGRN;refGene_tx=XM_023257882.1;refGene_gn=AGRN;refGene_tx=XM_023257883.1;refGene_gn=AGRN;refGene_tx=XM_023257884.1;refGene_gn=AGRN;refGene_tx=XM_023257885.1;refGene_gn=AGRN;refGene_tx=XM_023257886.1;refGene_gn=AGRN;refGene_tx=XM_023257888.1
C1 168364 168499
refGene_gn=AGRN;refGene_tx=XM_019835969.2;refGene_gn=AGRN;refGene_tx=X

M_019835970.2;refGene_gn=AGRN;refGene_tx=XM_019835971.2;refGene_gn=AGR
N;refGene_tx=XM_023257882.1;refGene_gn=AGRN;refGene_tx=XM_023257883.1;
refGene_gn=AGRN;refGene_tx=XM_023257884.1;refGene_gn=AGRN;refGene_tx=X
M_023257885.1;refGene_gn=AGRN;refGene_tx=XM_023257886.1;refGene_gn=AGR
N;refGene_tx=XM_023257888.1
C1 168713 168810
refGene_gn=AGRN;refGene_tx=XM_019835969.2;refGene_gn=AGRN;refGene_tx=X
M_019835970.2;refGene_gn=AGRN;refGene_tx=XM_019835971.2;refGene_gn=AGR
N;refGene_tx=XM_023257882.1;refGene_gn=AGRN;refGene_tx=XM_023257883.1;
refGene_gn=AGRN;refGene_tx=XM_023257884.1;refGene_gn=AGRN;refGene_tx=X
M_023257885.1;refGene_gn=AGRN;refGene_tx=XM_023257886.1;refGene_gn=AGR
N;refGene_tx=XM_023257888.1
C1 168904 169069
refGene_gn=AGRN;refGene_tx=XM_019835969.2;refGene_gn=AGRN;refGene_tx=X
M_019835970.2;refGene_gn=AGRN;refGene_tx=XM_019835971.2;refGene_gn=AGR
N;refGene_tx=XM_023257882.1;refGene_gn=AGRN;refGene_tx=XM_023257883.1;
refGene_gn=AGRN;refGene_tx=XM_023257884.1;refGene_gn=AGRN;refGene_tx=X
M_023257885.1;refGene_gn=AGRN;refGene_tx=XM_023257886.1;refGene_gn=AGR
N;refGene_tx=XM_023257888.1
C1 169189 169301
refGene_gn=AGRN;refGene_tx=XM_019835969.2;refGene_gn=AGRN;refGene_tx=X
M_019835970.2;refGene_gn=AGRN;refGene_tx=XM_019835971.2;refGene_gn=AGR
N;refGene_tx=XM_023257882.1;refGene_gn=AGRN;refGene_tx=XM_023257883.1;
refGene_gn=AGRN;refGene_tx=XM_023257884.1;refGene_gn=AGRN;refGene_tx=X
M_023257885.1;refGene_gn=AGRN;refGene_tx=XM_023257886.1;refGene_gn=AGR
N;refGene_tx=XM_023257888.1
C1 169535 169547
refGene_gn=AGRN;refGene_tx=XM_019835969.2;refGene_gn=AGRN;refGene_tx=X
M_019835970.2;refGene_gn=AGRN;refGene_tx=XM_019835971.2;refGene_gn=AGR
N;refGene_tx=XM_023257882.1;refGene_gn=AGRN;refGene_tx=XM_023257884.1;
refGene_gn=AGRN;refGene_tx=XM_023257885.1
C1 169764 169881
refGene_gn=AGRN;refGene_tx=XM_019835969.2;refGene_gn=AGRN;refGene_tx=X
M_019835970.2;refGene_gn=AGRN;refGene_tx=XM_019835971.2;refGene_gn=AGR
N;refGene_tx=XM_023257882.1;refGene_gn=AGRN;refGene_tx=XM_023257883.1;
refGene_gn=AGRN;refGene_tx=XM_023257884.1;refGene_gn=AGRN;refGene_tx=X
M_023257885.1;refGene_gn=AGRN;refGene_tx=XM_023257886.1;refGene_gn=AGR
N;refGene_tx=XM_023257888.1
C1 170021 170214
refGene_gn=AGRN;refGene_tx=XM_019835969.2;refGene_gn=AGRN;refGene_tx=X
M_019835970.2;refGene_gn=AGRN;refGene_tx=XM_019835971.2;refGene_gn=AGR
N;refGene_tx=XM_023257882.1;refGene_gn=AGRN;refGene_tx=XM_023257883.1;
refGene_gn=AGRN;refGene_tx=XM_023257884.1;refGene_gn=AGRN;refGene_tx=X
M_023257885.1;refGene_gn=AGRN;refGene_tx=XM_023257886.1;refGene_gn=AGR
N;refGene_tx=XM_023257888.1
C1 170295 170383
refGene_gn=AGRN;refGene_tx=XM_019835969.2;refGene_gn=AGRN;refGene_tx=X
M_019835970.2;refGene_gn=AGRN;refGene_tx=XM_019835971.2;refGene_gn=AGR
N;refGene_tx=XM_023257882.1;refGene_gn=AGRN;refGene_tx=XM_023257883.1;
refGene_gn=AGRN;refGene_tx=XM_023257884.1;refGene_gn=AGRN;refGene_tx=X

M_023257885.1;refGene_gn=AGRN;refGene_tx=XM_023257886.1;refGene_gn=AGRN;refGene_tx=XM_023257888.1
C1 170614 170638
refGene_gn=AGRN;refGene_tx=XM_019835969.2;refGene_gn=AGRN;refGene_tx=XM_019835970.2;refGene_gn=AGRN;refGene_tx=XM_023257882.1;refGene_gn=AGRN;refGene_tx=XM_023257883.1;refGene_gn=AGRN;refGene_tx=XM_023257884.1
C1 171332 171365
refGene_gn=AGRN;refGene_tx=XM_019835969.2;refGene_gn=AGRN;refGene_tx=XM_019835970.2;refGene_gn=AGRN;refGene_tx=XM_023257882.1;refGene_gn=AGRN;refGene_tx=XM_023257883.1;refGene_gn=AGRN;refGene_tx=XM_023257884.1
C1 171611 171836
refGene_gn=AGRN;refGene_tx=XM_019835969.2;refGene_gn=AGRN;refGene_tx=XM_019835970.2;refGene_gn=AGRN;refGene_tx=XM_019835971.2;refGene_gn=AGRN;refGene_tx=XM_023257882.1;refGene_gn=AGRN;refGene_tx=XM_023257883.1;refGene_gn=AGRN;refGene_tx=XM_023257884.1;refGene_gn=AGRN;refGene_tx=XM_023257885.1;refGene_gn=AGRN;refGene_tx=XM_023257886.1;refGene_gn=AGRN;refGene_tx=XM_023257888.1
C1 172318 172422
refGene_gn=AGRN;refGene_tx=XM_019835969.2;refGene_gn=AGRN;refGene_tx=XM_019835970.2;refGene_gn=AGRN;refGene_tx=XM_019835971.2;refGene_gn=AGRN;refGene_tx=XM_023257882.1;refGene_gn=AGRN;refGene_tx=XM_023257883.1;refGene_gn=AGRN;refGene_tx=XM_023257884.1;refGene_gn=AGRN;refGene_tx=XM_023257885.1;refGene_gn=AGRN;refGene_tx=XM_023257886.1;refGene_gn=AGRN;refGene_tx=XM_023257888.1
C1 172767 172925
refGene_gn=AGRN;refGene_tx=XM_019835969.2;refGene_gn=AGRN;refGene_tx=XM_019835970.2;refGene_gn=AGRN;refGene_tx=XM_019835971.2;refGene_gn=AGRN;refGene_tx=XM_023257882.1;refGene_gn=AGRN;refGene_tx=XM_023257883.1;refGene_gn=AGRN;refGene_tx=XM_023257884.1;refGene_gn=AGRN;refGene_tx=XM_023257885.1;refGene_gn=AGRN;refGene_tx=XM_023257886.1;refGene_gn=AGRN;refGene_tx=XM_023257888.1
C1 177003 177017
refGene_gn=LOC101082222;refGene_tx=XM_023257889.1;refGene_gn=LOC101082222;refGene_tx=XM_023257890.1
C1 182686 182809
refGene_gn=LOC101082222;refGene_tx=XM_023257889.1;refGene_gn=LOC101082222;refGene_tx=XM_023257890.1
C1 183871 184069
refGene_gn=LOC101082222;refGene_tx=XM_023257889.1;refGene_gn=LOC101082222;refGene_tx=XM_023257890.1
C1 184693 184880
refGene_gn=LOC101082222;refGene_tx=XM_023257889.1;refGene_gn=LOC101082222;refGene_tx=XM_023257890.1
C1 184962 185082
refGene_gn=LOC101082222;refGene_tx=XM_023257889.1;refGene_gn=LOC101082222;refGene_tx=XM_023257890.1
C1 185291 185405
refGene_gn=LOC101082222;refGene_tx=XM_023257889.1
C1 185561 185658
refGene_gn=LOC101082222;refGene_tx=XM_023257889.1;refGene_gn=LOC101082

222;refGene_tx=XM_023257890.1
C1	185919	185999
refGene_gn=LOC101082222;refGene_tx=XM_023257889.1;refGene_gn=LOC101082
222;refGene_tx=XM_023257890.1
C1	186384	186540
refGene_gn=LOC101082222;refGene_tx=XM_023257889.1;refGene_gn=LOC101082
222;refGene_tx=XM_023257890.1
C1	186759	186891
refGene_gn=LOC101082222;refGene_tx=XM_023257889.1;refGene_gn=LOC101082
222;refGene_tx=XM_023257890.1
C1	187622	187777
refGene_gn=LOC101082222;refGene_tx=XM_023257889.1;refGene_gn=LOC101082
222;refGene_tx=XM_023257890.1
C1	187881	187949
refGene_gn=LOC101082222;refGene_tx=XM_023257889.1;refGene_gn=LOC101082
222;refGene_tx=XM_023257890.1
C1	188110	188259
refGene_gn=LOC101082222;refGene_tx=XM_023257889.1;refGene_gn=LOC101082
222;refGene_tx=XM_023257890.1
C1	188365	188457
refGene_gn=LOC101082222;refGene_tx=XM_023257889.1;refGene_gn=LOC101082
222;refGene_tx=XM_023257890.1
C1	188591	188619
refGene_gn=LOC101082222;refGene_tx=XM_023257889.1;refGene_gn=LOC101082
222;refGene_tx=XM_023257890.1
C1	190650	191418
refGene_gn=LOC101082733;refGene_tx=XM_023257892.1;refGene_gn=LOC101082
733;refGene_tx=XM_023257893.1;refGene_gn=LOC101082733;refGene_tx=XM_02
3257894.1;refGene_gn=LOC101082733;refGene_tx=XM_023257895.1
C1	197914	198012
refGene_gn=LOC101082733;refGene_tx=XM_011284393.3;refGene_gn=LOC101082
733;refGene_tx=XM_019835973.2;refGene_gn=LOC101082733;refGene_tx=XM_01
9835974.2;refGene_gn=LOC101082733;refGene_tx=XM_019835975.2;refGene_gn
=LOC101082733;refGene_tx=XM_019835976.2;refGene_gn=LOC101082733;refGen
e_tx=XM_019835978.2
C1	198651	198773
refGene_gn=LOC101082733;refGene_tx=XM_011284393.3;refGene_gn=LOC101082
733;refGene_tx=XM_019835973.2;refGene_gn=LOC101082733;refGene_tx=XM_01
9835974.2;refGene_gn=LOC101082733;refGene_tx=XM_019835975.2;refGene_gn
=LOC101082733;refGene_tx=XM_019835976.2;refGene_gn=LOC101082733;refGen
e_tx=XM_019835979.2
C1	199470	199501
refGene_gn=LOC101082733;refGene_tx=XM_011284393.3;refGene_gn=LOC101082
733;refGene_tx=XM_019835973.2;refGene_gn=LOC101082733;refGene_tx=XM_01
9835974.2;refGene_gn=LOC101082733;refGene_tx=XM_019835975.2;refGene_gn
=LOC101082733;refGene_tx=XM_019835976.2;refGene_gn=LOC101082733;refGen
e_tx=XM_019835978.2;refGene_gn=LOC101082733;refGene_tx=XM_019835979.2
C1	199597	199623
refGene_gn=LOC101082733;refGene_tx=XM_011284393.3;refGene_gn=LOC101082
733;refGene_tx=XM_019835973.2;refGene_gn=LOC101082733;refGene_tx=XM_01

9835974.2;refGene_gn=LOC101082733;refGene_tx=XM_019835975.2;refGene_gn=LOC101082733;refGene_tx=XM_019835976.2;refGene_gn=LOC101082733;refGene_tx=XM_019835978.2;refGene_gn=LOC101082733;refGene_tx=XM_019835979.2
C1 200960 201095
refGene_gn=LOC101082733;refGene_tx=XM_011284393.3;refGene_gn=LOC101082733;refGene_tx=XM_019835973.2;refGene_gn=LOC101082733;refGene_tx=XM_019835974.2;refGene_gn=LOC101082733;refGene_tx=XM_019835975.2;refGene_gn=LOC101082733;refGene_tx=XM_019835976.2;refGene_gn=LOC101082733;refGene_tx=XM_019835978.2;refGene_gn=LOC101082733;refGene_tx=XM_019835979.2
C1 202555 202621
refGene_gn=LOC101082733;refGene_tx=XM_011284393.3;refGene_gn=LOC101082733;refGene_tx=XM_019835973.2;refGene_gn=LOC101082733;refGene_tx=XM_019835974.2;refGene_gn=LOC101082733;refGene_tx=XM_019835975.2;refGene_gn=LOC101082733;refGene_tx=XM_019835976.2;refGene_gn=LOC101082733;refGene_tx=XM_019835978.2;refGene_gn=LOC101082733;refGene_tx=XM_019835979.2
C1 202932 203115
refGene_gn=LOC101082733;refGene_tx=XM_011284393.3;refGene_gn=LOC101082733;refGene_tx=XM_019835973.2;refGene_gn=LOC101082733;refGene_tx=XM_019835974.2;refGene_gn=LOC101082733;refGene_tx=XM_019835975.2;refGene_gn=LOC101082733;refGene_tx=XM_019835976.2;refGene_gn=LOC101082733;refGene_tx=XM_019835978.2;refGene_gn=LOC101082733;refGene_tx=XM_019835979.2
C1 206594 206670
refGene_gn=LOC101082733;refGene_tx=XM_011284393.3;refGene_gn=LOC101082733;refGene_tx=XM_019835973.2;refGene_gn=LOC101082733;refGene_tx=XM_019835974.2;refGene_gn=LOC101082733;refGene_tx=XM_019835975.2;refGene_gn=LOC101082733;refGene_tx=XM_019835976.2;refGene_gn=LOC101082733;refGene_tx=XM_019835978.2;refGene_gn=LOC101082733;refGene_tx=XM_019835979.2
C1 208482 208554
refGene_gn=LOC101082733;refGene_tx=XM_011284393.3;refGene_gn=LOC101082733;refGene_tx=XM_019835973.2;refGene_gn=LOC101082733;refGene_tx=XM_019835974.2;refGene_gn=LOC101082733;refGene_tx=XM_019835975.2;refGene_gn=LOC101082733;refGene_tx=XM_019835976.2;refGene_gn=LOC101082733;refGene_tx=XM_019835978.2;refGene_gn=LOC101082733;refGene_tx=XM_019835979.2
C1 284105 284118
refGene_gn=TTLL10;refGene_tx=XM_023257897.1;refGene_gn=TTLL10;refGene_tx=XM_023257898.1
C1 284535 284568
refGene_gn=TTLL10;refGene_tx=XM_023257896.1;refGene_gn=TTLL10;refGene_tx=XM_023257900.1;refGene_gn=TTLL10;refGene_tx=XM_023257901.1;refGene_gn=TTLL10;refGene_tx=XM_023257903.1;refGene_gn=TTLL10;refGene_tx=XM_023257906.1
C1 284966 285034
refGene_gn=TTLL10;refGene_tx=XM_023257896.1;refGene_gn=TTLL10;refGene_tx=XM_023257897.1;refGene_gn=TTLL10;refGene_tx=XM_023257898.1;refGene_gn=TTLL10;refGene_tx=XM_023257900.1;refGene_gn=TTLL10;refGene_tx=XM_023257901.1;refGene_gn=TTLL10;refGene_tx=XM_023257903.1;refGene_gn=TTLL10;refGene_tx=XM_023257906.1
C1 287596 287949
refGene_gn=TTLL10;refGene_tx=XM_023257896.1;refGene_gn=TTLL10;refGene_tx=XM_023257897.1;refGene_gn=TTLL10;refGene_tx=XM_023257898.1;refGene_

gn=TTLL10;refGene_tx=XM_023257900.1;refGene_gn=TTLL10;refGene_tx=XM_023257901.1;refGene_gn=TTLL10;refGene_tx=XM_023257902.1;refGene_gn=TTLL10;refGene_tx=XM_023257903.1;refGene_gn=TTLL10;refGene_tx=XM_023257906.1
C1 288617 288698
refGene_gn=TTLL10;refGene_tx=XM_023257896.1;refGene_gn=TTLL10;refGene_tx=XM_023257897.1;refGene_gn=TTLL10;refGene_tx=XM_023257898.1;refGene_gn=TTLL10;refGene_tx=XM_023257900.1;refGene_gn=TTLL10;refGene_tx=XM_023257901.1;refGene_gn=TTLL10;refGene_tx=XM_023257902.1;refGene_gn=TTLL10;refGene_tx=XM_023257903.1;refGene_gn=TTLL10;refGene_tx=XM_023257904.1;refGene_gn=TTLL10;refGene_tx=XM_023257905.1;refGene_gn=TTLL10;refGene_tx=XM_023257906.1
C1 288791 288927
refGene_gn=TTLL10;refGene_tx=XM_023257896.1;refGene_gn=TTLL10;refGene_tx=XM_023257897.1;refGene_gn=TTLL10;refGene_tx=XM_023257898.1;refGene_gn=TTLL10;refGene_tx=XM_023257900.1;refGene_gn=TTLL10;refGene_tx=XM_023257901.1;refGene_gn=TTLL10;refGene_tx=XM_023257902.1;refGene_gn=TTLL10;refGene_tx=XM_023257903.1;refGene_gn=TTLL10;refGene_tx=XM_023257904.1;refGene_gn=TTLL10;refGene_tx=XM_023257905.1;refGene_gn=TTLL10;refGene_tx=XM_023257906.1
C1 289056 289175
refGene_gn=TTLL10;refGene_tx=XM_023257896.1;refGene_gn=TTLL10;refGene_tx=XM_023257897.1;refGene_gn=TTLL10;refGene_tx=XM_023257898.1;refGene_gn=TTLL10;refGene_tx=XM_023257900.1;refGene_gn=TTLL10;refGene_tx=XM_023257901.1;refGene_gn=TTLL10;refGene_tx=XM_023257902.1;refGene_gn=TTLL10;refGene_tx=XM_023257903.1;refGene_gn=TTLL10;refGene_tx=XM_023257904.1;refGene_gn=TTLL10;refGene_tx=XM_023257905.1;refGene_gn=TTLL10;refGene_tx=XM_023257906.1
C1 289350 289480
refGene_gn=TTLL10;refGene_tx=XM_023257896.1;refGene_gn=TTLL10;refGene_tx=XM_023257897.1;refGene_gn=TTLL10;refGene_tx=XM_023257898.1;refGene_gn=TTLL10;refGene_tx=XM_023257900.1;refGene_gn=TTLL10;refGene_tx=XM_023257901.1;refGene_gn=TTLL10;refGene_tx=XM_023257902.1;refGene_gn=TTLL10;refGene_tx=XM_023257903.1;refGene_gn=TTLL10;refGene_tx=XM_023257904.1;refGene_gn=TTLL10;refGene_tx=XM_023257905.1;refGene_gn=TTLL10;refGene_tx=XM_023257906.1
C1 290243 290315
refGene_gn=TTLL10;refGene_tx=XM_023257896.1;refGene_gn=TTLL10;refGene_tx=XM_023257897.1;refGene_gn=TTLL10;refGene_tx=XM_023257898.1;refGene_gn=TTLL10;refGene_tx=XM_023257901.1;refGene_gn=TTLL10;refGene_tx=XM_023257902.1;refGene_gn=TTLL10;refGene_tx=XM_023257903.1;refGene_gn=TTLL10;refGene_tx=XM_023257904.1;refGene_gn=TTLL10;refGene_tx=XM_023257905.1;refGene_gn=TTLL10;refGene_tx=XM_023257906.1
C1 290771 290857
refGene_gn=TTLL10;refGene_tx=XM_023257896.1;refGene_gn=TTLL10;refGene_tx=XM_023257897.1;refGene_gn=TTLL10;refGene_tx=XM_023257898.1;refGene_gn=TTLL10;refGene_tx=XM_023257900.1;refGene_gn=TTLL10;refGene_tx=XM_023257901.1;refGene_gn=TTLL10;refGene_tx=XM_023257902.1;refGene_gn=TTLL10;refGene_tx=XM_023257903.1;refGene_gn=TTLL10;refGene_tx=XM_023257904.1;refGene_gn=TTLL10;refGene_tx=XM_023257905.1;refGene_gn=TTLL10;refGen

e_tx=XM_023257906.1
C1 291173 291345
refGene_gn=TTLL10;refGene_tx=XM_023257896.1;refGene_gn=TTLL10;refGene_tx=XM_023257897.1;refGene_gn=TTLL10;refGene_tx=XM_023257898.1;refGene_gn=TTLL10;refGene_tx=XM_023257900.1;refGene_gn=TTLL10;refGene_tx=XM_023257901.1;refGene_gn=TTLL10;refGene_tx=XM_023257902.1;refGene_gn=TTLL10;refGene_tx=XM_023257903.1;refGene_gn=TTLL10;refGene_tx=XM_023257904.1;refGene_gn=TTLL10;refGene_tx=XM_023257905.1;refGene_gn=TTLL10;refGene_tx=XM_023257906.1
C1 292045 292217
refGene_gn=TTLL10;refGene_tx=XM_023257896.1;refGene_gn=TTLL10;refGene_tx=XM_023257897.1;refGene_gn=TTLL10;refGene_tx=XM_023257898.1;refGene_gn=TTLL10;refGene_tx=XM_023257900.1;refGene_gn=TTLL10;refGene_tx=XM_023257901.1;refGene_gn=TTLL10;refGene_tx=XM_023257902.1;refGene_gn=TTLL10;refGene_tx=XM_023257903.1;refGene_gn=TTLL10;refGene_tx=XM_023257904.1;refGene_gn=TTLL10;refGene_tx=XM_023257905.1;refGene_gn=TTLL10;refGene_tx=XM_023257906.1
C1 292829 293010
refGene_gn=TTLL10;refGene_tx=XM_023257896.1;refGene_gn=TTLL10;refGene_tx=XM_023257897.1;refGene_gn=TTLL10;refGene_tx=XM_023257898.1;refGene_gn=TTLL10;refGene_tx=XM_023257900.1;refGene_gn=TTLL10;refGene_tx=XM_023257901.1;refGene_gn=TTLL10;refGene_tx=XM_023257902.1;refGene_gn=TTLL10;refGene_tx=XM_023257903.1;refGene_gn=TTLL10;refGene_tx=XM_023257904.1;refGene_gn=TTLL10;refGene_tx=XM_023257905.1;refGene_gn=TTLL10;refGene_tx=XM_023257906.1
C1 299091 299172 refGene_gn=TTLL10;refGene_tx=XM_023257906.1
C1 299453 299570
refGene_gn=TTLL10;refGene_tx=XM_023257896.1;refGene_gn=TTLL10;refGene_tx=XM_023257897.1;refGene_gn=TTLL10;refGene_tx=XM_023257898.1;refGene_gn=TTLL10;refGene_tx=XM_023257900.1;refGene_gn=TTLL10;refGene_tx=XM_023257901.1;refGene_gn=TTLL10;refGene_tx=XM_023257902.1;refGene_gn=TTLL10;refGene_tx=XM_023257903.1;refGene_gn=TTLL10;refGene_tx=XM_023257904.1;refGene_gn=TTLL10;refGene_tx=XM_023257905.1
C1 299895 299989
refGene_gn=TTLL10;refGene_tx=XM_023257896.1;refGene_gn=TTLL10;refGene_tx=XM_023257897.1;refGene_gn=TTLL10;refGene_tx=XM_023257898.1;refGene_gn=TTLL10;refGene_tx=XM_023257900.1;refGene_gn=TTLL10;refGene_tx=XM_023257901.1;refGene_gn=TTLL10;refGene_tx=XM_023257902.1;refGene_gn=TTLL10;refGene_tx=XM_023257904.1;refGene_gn=TTLL10;refGene_tx=XM_023257905.1
C1 300280 300657
refGene_gn=TTLL10;refGene_tx=XM_023257896.1;refGene_gn=TTLL10;refGene_tx=XM_023257897.1;refGene_gn=TTLL10;refGene_tx=XM_023257898.1;refGene_gn=TTLL10;refGene_tx=XM_023257900.1;refGene_gn=TTLL10;refGene_tx=XM_023257901.1;refGene_gn=TTLL10;refGene_tx=XM_023257902.1;refGene_gn=TTLL10;refGene_tx=XM_023257904.1;refGene_gn=TTLL10;refGene_tx=XM_023257905.1
C1 306631 306756
refGene_gn=TNFRSF18;refGene_tx=XM_006944216.3;refGene_gn=TNFRSF18;refGene_tx=XM_023260045.1

C1 306819 307026
refGene_gn=TNFRSF18;refGene_tx=XM_006944216.3;refGene_gn=TNFRSF18;refGene_tx=XM_023260045.1
C1 307194 307282
refGene_gn=TNFRSF18;refGene_tx=XM_006944216.3;refGene_gn=TNFRSF18;refGene_tx=XM_023260045.1
C1 308253 308376
refGene_gn=TNFRSF18;refGene_tx=XM_006944216.3;refGene_gn=TNFRSF18;refGene_tx=XM_023260045.1
C1 309236 309390
refGene_gn=TNFRSF18;refGene_tx=XM_006944216.3;refGene_gn=TNFRSF18;refGene_tx=XM_023260045.1
C1 314100 314171
refGene_gn=TNFRSF4;refGene_tx=NM_001009200.1;refGene_gn=TNFRSF4;refGene_tx=XM_023257814.1
C1 314256 314361
refGene_gn=TNFRSF4;refGene_tx=NM_001009200.1;refGene_gn=TNFRSF4;refGene_tx=XM_023257814.1
C1 314530 314727
refGene_gn=TNFRSF4;refGene_tx=NM_001009200.1;refGene_gn=TNFRSF4;refGene_tx=XM_023257814.1
C1 315365 315432
refGene_gn=TNFRSF4;refGene_tx=NM_001009200.1;refGene_gn=TNFRSF4;refGene_tx=XM_023257814.1
C1 315817 315919
refGene_gn=TNFRSF4;refGene_tx=NM_001009200.1;refGene_gn=TNFRSF4;refGene_tx=XM_023257814.1
C1 316367 316511
refGene_gn=TNFRSF4;refGene_tx=NM_001009200.1;refGene_gn=TNFRSF4;refGene_tx=XM_023257814.1
C1 316513 316518 refGene_gn=TNFRSF4;refGene_tx=NM_001009200.1
C1 316520 316527 refGene_gn=TNFRSF4;refGene_tx=NM_001009200.1
C1 320131 320198 refGene_gn=TNFRSF4;refGene_tx=NM_001009200.1
C1 320921 320993 refGene_gn=TNFRSF4;refGene_tx=XM_023257814.1
C1 321305 321324 refGene_gn=TNFRSF4;refGene_tx=XM_023257814.1
C1 322735 322912 refGene_gn=SDF4;refGene_tx=XM_023257907.1
C1 323980 324156 refGene_gn=SDF4;refGene_tx=XM_023257907.1
C1 324493 324652 refGene_gn=SDF4;refGene_tx=XM_023257907.1
C1 328223 328337 refGene_gn=SDF4;refGene_tx=XM_023257907.1
C1 328828 328965 refGene_gn=SDF4;refGene_tx=XM_023257907.1
C1 332535 332840 refGene_gn=SDF4;refGene_tx=XM_023257907.1
C1 336311 337301 refGene_gn=B3GALT6;refGene_tx=XM_011279777.2
C1 347130 347707
refGene_gn=C1QTNF12;refGene_tx=XM_004001145.3;refGene_gn=C1QTNF12;refGene_tx=XM_023257909.1;refGene_gn=C1QTNF12;refGene_tx=XM_023257910.1;refGene_gn=C1QTNF12;refGene_tx=XM_023257911.1
C1 347979 348058
refGene_gn=C1QTNF12;refGene_tx=XM_004001145.3;refGene_gn=C1QTNF12;refGene_tx=XM_023257909.1;refGene_gn=C1QTNF12;refGene_tx=XM_023257910.1;refGene_gn=C1QTNF12;refGene_tx=XM_023257911.1

C1	348203	348294
refGene_gn=C1QTNF12;refGene_tx=XM_004001145.3;refGene_gn=C1QTNF12;refGene_tx=XM_023257911.1
C1	348501	348685
refGene_gn=C1QTNF12;refGene_tx=XM_004001145.3;refGene_gn=C1QTNF12;refGene_tx=XM_023257909.1;refGene_gn=C1QTNF12;refGene_tx=XM_023257910.1;refGene_gn=C1QTNF12;refGene_tx=XM_023257911.1
C1	349080	349232
refGene_gn=C1QTNF12;refGene_tx=XM_004001145.3;refGene_gn=C1QTNF12;refGene_tx=XM_023257909.1;refGene_gn=C1QTNF12;refGene_tx=XM_023257910.1;refGene_gn=C1QTNF12;refGene_tx=XM_023257911.1
C1	349323	349435
refGene_gn=C1QTNF12;refGene_tx=XM_004001145.3;refGene_gn=C1QTNF12;refGene_tx=XM_023257909.1;refGene_gn=C1QTNF12;refGene_tx=XM_023257910.1;refGene_gn=C1QTNF12;refGene_tx=XM_023257911.1
C1	349516	349633
refGene_gn=C1QTNF12;refGene_tx=XM_004001145.3;refGene_gn=C1QTNF12;refGene_tx=XM_023257909.1;refGene_gn=C1QTNF12;refGene_tx=XM_023257910.1;refGene_gn=C1QTNF12;refGene_tx=XM_023257911.1
C1	351916	352081
refGene_gn=C1QTNF12;refGene_tx=XM_004001145.3;refGene_gn=C1QTNF12;refGene_tx=XM_023257909.1;refGene_gn=C1QTNF12;refGene_tx=XM_023257910.1;refGene_gn=C1QTNF12;refGene_tx=XM_023257911.1
C1	356830	357115
refGene_gn=UBE2J2;refGene_tx=XM_011284396.3;refGene_gn=UBE2J2;refGene_tx=XM_019835990.2;refGene_gn=UBE2J2;refGene_tx=XM_019835991.2;refGene_gn=UBE2J2;refGene_tx=XM_023257922.1
C1	357674	357755
refGene_gn=UBE2J2;refGene_tx=XM_011284396.3;refGene_gn=UBE2J2;refGene_tx=XM_019835990.2;refGene_gn=UBE2J2;refGene_tx=XM_019835991.2;refGene_gn=UBE2J2;refGene_tx=XM_023257922.1
C1	358102	358241
refGene_gn=UBE2J2;refGene_tx=XM_011284396.3;refGene_gn=UBE2J2;refGene_tx=XM_019835990.2;refGene_gn=UBE2J2;refGene_tx=XM_019835991.2;refGene_gn=UBE2J2;refGene_tx=XM_023257922.1
C1	358326	358429
refGene_gn=UBE2J2;refGene_tx=XM_011284396.3;refGene_gn=UBE2J2;refGene_tx=XM_019835990.2;refGene_gn=UBE2J2;refGene_tx=XM_019835991.2;refGene_gn=UBE2J2;refGene_tx=XM_023257922.1
C1	363430	363471
refGene_gn=UBE2J2;refGene_tx=XM_011284396.3;refGene_gn=UBE2J2;refGene_tx=XM_019835990.2;refGene_gn=UBE2J2;refGene_tx=XM_019835991.2;refGene_gn=UBE2J2;refGene_tx=XM_023257922.1
C1	365801	365932
refGene_gn=UBE2J2;refGene_tx=XM_011284396.3;refGene_gn=UBE2J2;refGene_tx=XM_019835990.2;refGene_gn=UBE2J2;refGene_tx=XM_019835991.2;refGene_gn=UBE2J2;refGene_tx=XM_023257922.1
C1	369673	369682	refGene_gn=UBE2J2;refGene_tx=XM_019835990.2
C1	372046	372177	refGene_gn=SCNN1D;refGene_tx=XM_023257913.1
C1	372854	373026	refGene_gn=SCNN1D;refGene_tx=XM_023257913.1

C1	375856	375919	refGene_gn=SCNN1D;refGene_tx=XM_023257913.1
C1	377767	378144	refGene_gn=SCNN1D;refGene_tx=XM_023257913.1
C1	378661	378798	refGene_gn=SCNN1D;refGene_tx=XM_023257913.1
C1	378800	378868	refGene_gn=SCNN1D;refGene_tx=XM_023257913.1
C1	378977	379168	refGene_gn=SCNN1D;refGene_tx=XM_023257913.1
C1	379340	379429	refGene_gn=SCNN1D;refGene_tx=XM_023257913.1
C1	379504	379668	refGene_gn=SCNN1D;refGene_tx=XM_023257913.1
C1	379768	379867	refGene_gn=SCNN1D;refGene_tx=XM_023257913.1
C1	380152	380270	refGene_gn=SCNN1D;refGene_tx=XM_023257913.1
C1	380690	380769	refGene_gn=SCNN1D;refGene_tx=XM_023257913.1
C1	380854	380912	refGene_gn=SCNN1D;refGene_tx=XM_023257913.1
C1	381024	381152	refGene_gn=SCNN1D;refGene_tx=XM_023257913.1
C1	381265	381341	refGene_gn=SCNN1D;refGene_tx=XM_023257913.1
C1	381460	381763	refGene_gn=SCNN1D;refGene_tx=XM_023257913.1
C1	384125	384273	refGene_gn=ACAP3;refGene_tx=XM_023257912.1
C1	384393	384504	refGene_gn=ACAP3;refGene_tx=XM_023257912.1
C1	384609	384719	refGene_gn=ACAP3;refGene_tx=XM_023257912.1
C1	384900	385121	refGene_gn=ACAP3;refGene_tx=XM_023257912.1
C1	385220	385319	refGene_gn=ACAP3;refGene_tx=XM_023257912.1
C1	386433	386541	refGene_gn=ACAP3;refGene_tx=XM_023257912.1
C1	386686	386753	refGene_gn=ACAP3;refGene_tx=XM_023257912.1
C1	386754	386890	refGene_gn=ACAP3;refGene_tx=XM_023257912.1
C1	386974	387069	refGene_gn=ACAP3;refGene_tx=XM_023257912.1
C1	387202	387272	refGene_gn=ACAP3;refGene_tx=XM_023257912.1
C1	387352	387561	refGene_gn=ACAP3;refGene_tx=XM_023257912.1
C1	387860	387972	refGene_gn=ACAP3;refGene_tx=XM_023257912.1
C1	388049	388150	refGene_gn=ACAP3;refGene_tx=XM_023257912.1
C1	388396	388448	refGene_gn=ACAP3;refGene_tx=XM_023257912.1
C1	388536	388649	refGene_gn=ACAP3;refGene_tx=XM_023257912.1
C1	389253	389265	refGene_gn=ACAP3;refGene_tx=XM_023257912.1
C1	389940	390015	refGene_gn=ACAP3;refGene_tx=XM_023257912.1
C1	390099	390195	refGene_gn=ACAP3;refGene_tx=XM_023257912.1
C1	390326	390371	refGene_gn=ACAP3;refGene_tx=XM_023257912.1
C1	390852	391036	refGene_gn=ACAP3;refGene_tx=XM_023257912.1
C1	391773	391832	refGene_gn=ACAP3;refGene_tx=XM_023257912.1
C1	392952	393006	refGene_gn=ACAP3;refGene_tx=XM_023257912.1
C1	393215	393335	refGene_gn=ACAP3;refGene_tx=XM_023257912.1
C1	394032	394090	refGene_gn=ACAP3;refGene_tx=XM_023257912.1
C1	397733	397780	refGene_gn=ACAP3;refGene_tx=XM_023257912.1
C1	397908	398698
refGene_gn=PUSL1;refGene_tx=XM_023257919.1;refGene_gn=PUSL1;refGene_tx
=XM_023257920.1;refGene_gn=PUSL1;refGene_tx=XM_023257921.1
C1	398871	398929
refGene_gn=PUSL1;refGene_tx=XM_023257919.1;refGene_gn=PUSL1;refGene_tx
=XM_023257920.1;refGene_gn=PUSL1;refGene_tx=XM_023257921.1
C1	399065	399253
refGene_gn=PUSL1;refGene_tx=XM_023257919.1;refGene_gn=PUSL1;refGene_tx
=XM_023257920.1;refGene_gn=PUSL1;refGene_tx=XM_023257921.1
C1	399421	399571
refGene_gn=PUSL1;refGene_tx=XM_023257919.1;refGene_gn=PUSL1;refGene_tx

=XM_023257920.1;refGene_gn=PUSL1;refGene_tx=XM_023257921.1
C1 399659 399830
refGene_gn=PUSL1;refGene_tx=XM_023257919.1;refGene_gn=PUSL1;refGene_tx=XM_023257920.1;refGene_gn=PUSL1;refGene_tx=XM_023257921.1
C1 400639 400691
refGene_gn=PUSL1;refGene_tx=XM_023257919.1;refGene_gn=PUSL1;refGene_tx=XM_023257920.1;refGene_gn=PUSL1;refGene_tx=XM_023257921.1
C1 400772 400935
refGene_gn=PUSL1;refGene_tx=XM_023257919.1;refGene_gn=PUSL1;refGene_tx=XM_023257920.1;refGene_gn=PUSL1;refGene_tx=XM_023257921.1
C1 401064 401300
refGene_gn=PUSL1;refGene_tx=XM_023257919.1;refGene_gn=PUSL1;refGene_tx=XM_023257920.1;refGene_gn=PUSL1;refGene_tx=XM_023257921.1
C1 401932 401998
refGene_gn=INTS11;refGene_tx=XM_023257914.1;refGene_gn=INTS11;refGene_tx=XM_023257916.1;refGene_gn=INTS11;refGene_tx=XM_023257917.1;refGene_gn=INTS11;refGene_tx=XM_023257918.1
C1 402125 402255
refGene_gn=INTS11;refGene_tx=XM_023257914.1;refGene_gn=INTS11;refGene_tx=XM_023257916.1;refGene_gn=INTS11;refGene_tx=XM_023257917.1;refGene_gn=INTS11;refGene_tx=XM_023257918.1
C1 402333 402476
refGene_gn=INTS11;refGene_tx=XM_023257914.1;refGene_gn=INTS11;refGene_tx=XM_023257916.1;refGene_gn=INTS11;refGene_tx=XM_023257917.1;refGene_gn=INTS11;refGene_tx=XM_023257918.1
C1 402548 402610
refGene_gn=INTS11;refGene_tx=XM_023257914.1;refGene_gn=INTS11;refGene_tx=XM_023257916.1;refGene_gn=INTS11;refGene_tx=XM_023257917.1;refGene_gn=INTS11;refGene_tx=XM_023257918.1
C1 402697 402805
refGene_gn=INTS11;refGene_tx=XM_023257914.1;refGene_gn=INTS11;refGene_tx=XM_023257916.1;refGene_gn=INTS11;refGene_tx=XM_023257917.1;refGene_gn=INTS11;refGene_tx=XM_023257918.1
C1 402997 403160
refGene_gn=INTS11;refGene_tx=XM_023257914.1;refGene_gn=INTS11;refGene_tx=XM_023257916.1;refGene_gn=INTS11;refGene_tx=XM_023257917.1;refGene_gn=INTS11;refGene_tx=XM_023257918.1
C1 403266 403356
refGene_gn=INTS11;refGene_tx=XM_023257914.1;refGene_gn=INTS11;refGene_tx=XM_023257917.1;refGene_gn=INTS11;refGene_tx=XM_023257918.1
C1 403756 403840
refGene_gn=INTS11;refGene_tx=XM_023257914.1;refGene_gn=INTS11;refGene_tx=XM_023257916.1;refGene_gn=INTS11;refGene_tx=XM_023257917.1;refGene_gn=INTS11;refGene_tx=XM_023257918.1
C1 404177 404367
refGene_gn=INTS11;refGene_tx=XM_023257914.1;refGene_gn=INTS11;refGene_tx=XM_023257916.1;refGene_gn=INTS11;refGene_tx=XM_023257917.1;refGene_gn=INTS11;refGene_tx=XM_023257918.1
C1 404660 404725
refGene_gn=INTS11;refGene_tx=XM_023257914.1;refGene_gn=INTS11;refGene_

tx=XM_023257916.1;refGene_gn=INTS11;refGene_tx=XM_023257917.1;refGene_gn=INTS11;refGene_tx=XM_023257918.1
C1	405243	405382	refGene_gn=INTS11;refGene_tx=XM_023257914.1;refGene_gn=INTS11;refGene_tx=XM_023257916.1;refGene_gn=INTS11;refGene_tx=XM_023257917.1;refGene_gn=INTS11;refGene_tx=XM_023257918.1
C1	405635	405670	refGene_gn=INTS11;refGene_tx=XM_023257914.1;refGene_gn=INTS11;refGene_tx=XM_023257916.1;refGene_gn=INTS11;refGene_tx=XM_023257917.1;refGene_gn=INTS11;refGene_tx=XM_023257918.1
C1	405753	405852	refGene_gn=INTS11;refGene_tx=XM_023257914.1;refGene_gn=INTS11;refGene_tx=XM_023257916.1;refGene_gn=INTS11;refGene_tx=XM_023257917.1;refGene_gn=INTS11;refGene_tx=XM_023257918.1
C1	409408	409637	refGene_gn=INTS11;refGene_tx=XM_023257914.1;refGene_gn=INTS11;refGene_tx=XM_023257916.1;refGene_gn=INTS11;refGene_tx=XM_023257917.1;refGene_gn=INTS11;refGene_tx=XM_023257918.1
C1	410884	410958	refGene_gn=INTS11;refGene_tx=XM_023257914.1;refGene_gn=INTS11;refGene_tx=XM_023257916.1;refGene_gn=INTS11;refGene_tx=XM_023257917.1
C1	411496	411594	refGene_gn=INTS11;refGene_tx=XM_023257914.1;refGene_gn=INTS11;refGene_tx=XM_023257916.1;refGene_gn=INTS11;refGene_tx=XM_023257917.1
C1	413167	413369	refGene_gn=INTS11;refGene_tx=XM_023257914.1;refGene_gn=INTS11;refGene_tx=XM_023257916.1
C1	414474	414502	refGene_gn=INTS11;refGene_tx=XM_023257917.1
C1	416947	417069	refGene_gn=CPTP;refGene_tx=XM_011284400.3;refGene_gn=CPTP;refGene_tx=XM_019835993.2;refGene_gn=CPTP;refGene_tx=XM_019835994.1;refGene_gn=CPTP;refGene_tx=XM_019835995.2;refGene_gn=CPTP;refGene_tx=XM_023257924.1;refGene_gn=CPTP;refGene_tx=XM_023257925.1
C1	417324	417847	refGene_gn=CPTP;refGene_tx=XM_011284400.3;refGene_gn=CPTP;refGene_tx=XM_019835993.2;refGene_gn=CPTP;refGene_tx=XM_019835994.1;refGene_gn=CPTP;refGene_tx=XM_019835995.2;refGene_gn=CPTP;refGene_tx=XM_023257924.1;refGene_gn=CPTP;refGene_tx=XM_023257925.1
C1	420651	420854	refGene_gn=TAS1R3;refGene_tx=NM_001114547.1
C1	420920	421259	refGene_gn=TAS1R3;refGene_tx=NM_001114547.1;refGene_gn=TAS1R3;refGene_tx=XM_023257807.1
C1	421348	422128	refGene_gn=TAS1R3;refGene_tx=NM_001114547.1;refGene_gn=TAS1R3;refGene_tx=XM_023257807.1
C1	422247	422451	refGene_gn=TAS1R3;refGene_tx=NM_001114547.1;refGene_gn=TAS1R3;refGene_tx=XM_023257807.1
C1	422606	422727	refGene_gn=TAS1R3;refGene_tx=NM_001114547.1;refGene_gn=TAS1R3;refGene_

tx=XM_023257807.1
C1 422873 423862
refGene_gn=TAS1R3;refGene_tx=NM_001114547.1;refGene_gn=TAS1R3;refGene_tx=XM_023257807.1
C1 425697 426086
refGene_gn=DVL1;refGene_tx=XM_011284401.2;refGene_gn=DVL1;refGene_tx=XM_011284402.2
C1 426641 426652 refGene_gn=DVL1;refGene_tx=XM_011284403.3
C1 427311 427518
refGene_gn=DVL1;refGene_tx=XM_011284401.2;refGene_gn=DVL1;refGene_tx=XM_011284402.2;refGene_gn=DVL1;refGene_tx=XM_011284403.3
C1 427596 427764
refGene_gn=DVL1;refGene_tx=XM_011284401.2;refGene_gn=DVL1;refGene_tx=XM_011284402.2;refGene_gn=DVL1;refGene_tx=XM_011284403.3
C1 427847 427979
refGene_gn=DVL1;refGene_tx=XM_011284401.2;refGene_gn=DVL1;refGene_tx=XM_011284402.2;refGene_gn=DVL1;refGene_tx=XM_011284403.3
C1 428676 428829
refGene_gn=DVL1;refGene_tx=XM_011284401.2;refGene_gn=DVL1;refGene_tx=XM_011284402.2;refGene_gn=DVL1;refGene_tx=XM_011284403.3
C1 428993 429061
refGene_gn=DVL1;refGene_tx=XM_011284401.2;refGene_gn=DVL1;refGene_tx=XM_011284402.2;refGene_gn=DVL1;refGene_tx=XM_011284403.3
C1 429138 429215
refGene_gn=DVL1;refGene_tx=XM_011284401.2;refGene_gn=DVL1;refGene_tx=XM_011284402.2;refGene_gn=DVL1;refGene_tx=XM_011284403.3
C1 429296 429436
refGene_gn=DVL1;refGene_tx=XM_011284401.2;refGene_gn=DVL1;refGene_tx=XM_011284402.2;refGene_gn=DVL1;refGene_tx=XM_011284403.3
C1 429648 429718
refGene_gn=DVL1;refGene_tx=XM_011284401.2;refGene_gn=DVL1;refGene_tx=XM_011284402.2;refGene_gn=DVL1;refGene_tx=XM_011284403.3
C1 429798 429892
refGene_gn=DVL1;refGene_tx=XM_011284401.2;refGene_gn=DVL1;refGene_tx=XM_011284402.2;refGene_gn=DVL1;refGene_tx=XM_011284403.3
C1 430505 430644
refGene_gn=DVL1;refGene_tx=XM_011284401.2;refGene_gn=DVL1;refGene_tx=XM_011284402.2;refGene_gn=DVL1;refGene_tx=XM_011284403.3
C1 430887 430991
refGene_gn=DVL1;refGene_tx=XM_011284401.2;refGene_gn=DVL1;refGene_tx=XM_011284402.2;refGene_gn=DVL1;refGene_tx=XM_011284403.3
C1 431228 431350
refGene_gn=DVL1;refGene_tx=XM_011284401.2;refGene_gn=DVL1;refGene_tx=XM_011284402.2;refGene_gn=DVL1;refGene_tx=XM_011284403.3
C1 431492 431562
refGene_gn=DVL1;refGene_tx=XM_011284401.2;refGene_gn=DVL1;refGene_tx=XM_011284402.2;refGene_gn=DVL1;refGene_tx=XM_011284403.3
C1 436199 436369
refGene_gn=DVL1;refGene_tx=XM_011284401.2;refGene_gn=DVL1;refGene_tx=XM_011284402.2;refGene_gn=DVL1;refGene_tx=XM_011284403.3

C1 440735 440761 refGene_gn=MXRA8;refGene_tx=XM_023257926.1
C1 440909 440990 refGene_gn=MXRA8;refGene_tx=XM_023257926.1
C1 441104 441181 refGene_gn=MXRA8;refGene_tx=XM_023257926.1
C1 441276 441316 refGene_gn=MXRA8;refGene_tx=XM_023257926.1
C1 441476 441635 refGene_gn=MXRA8;refGene_tx=XM_023257926.1
C1 441818 442289 refGene_gn=MXRA8;refGene_tx=XM_023257926.1
C1 442369 442471 refGene_gn=MXRA8;refGene_tx=XM_023257926.1
C1 442566 442872 refGene_gn=MXRA8;refGene_tx=XM_023257926.1
C1 442960 442984 refGene_gn=MXRA8;refGene_tx=XM_023257926.1
C1 444141 444190 refGene_gn=MXRA8;refGene_tx=XM_023257926.1
C1 459166 459268
refGene_gn=AURKAIP1;refGene_tx=XM_006944198.4;refGene_gn=AURKAIP1;refGene_tx=XM_006944199.4;refGene_gn=AURKAIP1;refGene_tx=XM_011284404.3
C1 459365 459811
refGene_gn=AURKAIP1;refGene_tx=XM_006944198.4;refGene_gn=AURKAIP1;refGene_tx=XM_006944199.4;refGene_gn=AURKAIP1;refGene_tx=XM_011284404.3
C1 460138 460190
refGene_gn=AURKAIP1;refGene_tx=XM_006944198.4;refGene_gn=AURKAIP1;refGene_tx=XM_006944199.4;refGene_gn=AURKAIP1;refGene_tx=XM_011284404.3
C1 463054 463409
refGene_gn=CCNL2;refGene_tx=XM_019835996.2;refGene_gn=CCNL2;refGene_tx=XM_019835997.2;refGene_gn=CCNL2;refGene_tx=XM_019835998.2;refGene_gn=CCNL2;refGene_tx=XM_019835999.1;refGene_gn=CCNL2;refGene_tx=XM_019836000.1
C1 463573 463666
refGene_gn=CCNL2;refGene_tx=XM_019835996.2;refGene_gn=CCNL2;refGene_tx=XM_019835997.2;refGene_gn=CCNL2;refGene_tx=XM_019835998.2;refGene_gn=CCNL2;refGene_tx=XM_019835999.1;refGene_gn=CCNL2;refGene_tx=XM_019836000.1
C1 463739 463851
refGene_gn=CCNL2;refGene_tx=XM_019835996.2;refGene_gn=CCNL2;refGene_tx=XM_019835997.2;refGene_gn=CCNL2;refGene_tx=XM_019835998.2;refGene_gn=CCNL2;refGene_tx=XM_019835999.1;refGene_gn=CCNL2;refGene_tx=XM_019836000.1
C1 465345 465487
refGene_gn=CCNL2;refGene_tx=XM_019835996.2;refGene_gn=CCNL2;refGene_tx=XM_019835997.2;refGene_gn=CCNL2;refGene_tx=XM_019835998.2;refGene_gn=CCNL2;refGene_tx=XM_019835999.1;refGene_gn=CCNL2;refGene_tx=XM_019836000.1
C1 465571 465676
refGene_gn=CCNL2;refGene_tx=XM_019835996.2;refGene_gn=CCNL2;refGene_tx=XM_019835997.2;refGene_gn=CCNL2;refGene_tx=XM_019835998.2;refGene_gn=CCNL2;refGene_tx=XM_019835999.1;refGene_gn=CCNL2;refGene_tx=XM_019836000.1
C1 465879 465979
refGene_gn=CCNL2;refGene_tx=XM_019835996.2;refGene_gn=CCNL2;refGene_tx=XM_019835997.2;refGene_gn=CCNL2;refGene_tx=XM_019835998.2;refGene_gn=CCNL2;refGene_tx=XM_019835999.1;refGene_gn=CCNL2;refGene_tx=XM_019836000.1
C1 466387 466539

refGene_gn=CCNL2;refGene_tx=XM_019835998.2;refGene_gn=CCNL2;refGene_tx
=XM_019835999.1;refGene_gn=CCNL2;refGene_tx=XM_019836000.1
C1	467839	467861	refGene_gn=CCNL2;refGene_tx=XM_023257928.1
C1	468451	468516
refGene_gn=CCNL2;refGene_tx=XM_019835996.2;refGene_gn=CCNL2;refGene_tx
=XM_019835997.2;refGene_gn=CCNL2;refGene_tx=XM_023257928.1
C1	469493	469614
refGene_gn=CCNL2;refGene_tx=XM_019835996.2;refGene_gn=CCNL2;refGene_tx
=XM_019835997.2;refGene_gn=CCNL2;refGene_tx=XM_023257928.1
C1	470421	470531
refGene_gn=CCNL2;refGene_tx=XM_019835996.2;refGene_gn=CCNL2;refGene_tx
=XM_019835997.2;refGene_gn=CCNL2;refGene_tx=XM_023257928.1
C1	470795	470870
refGene_gn=CCNL2;refGene_tx=XM_019835996.2;refGene_gn=CCNL2;refGene_tx
=XM_019835997.2;refGene_gn=CCNL2;refGene_tx=XM_023257928.1
C1	471215	471515
refGene_gn=CCNL2;refGene_tx=XM_019835996.2;refGene_gn=CCNL2;refGene_tx
=XM_019835997.2;refGene_gn=CCNL2;refGene_tx=XM_023257928.1
C1	474613	474787	refGene_gn=MRPL20;refGene_tx=XM_023257930.1
C1	476182	476260	refGene_gn=MRPL20;refGene_tx=XM_023257930.1
C1	479240	479351	refGene_gn=MRPL20;refGene_tx=XM_023257930.1
C1	479440	479527	refGene_gn=MRPL20;refGene_tx=XM_023257930.1
C1	484301	484742	refGene_gn=ANKRD65;refGene_tx=XM_023257931.1
C1	485271	485806	refGene_gn=ANKRD65;refGene_tx=XM_023257931.1
C1	485949	486275	refGene_gn=ANKRD65;refGene_tx=XM_023257931.1
C1	489309	489654	refGene_gn=TMEM88B;refGene_tx=XM_023257933.1
C1	490621	490882	refGene_gn=TMEM88B;refGene_tx=XM_023257933.1
C1	498333	498406
refGene_gn=VWA1;refGene_tx=XM_011284405.3;refGene_gn=VWA1;refGene_tx=X
M_023257932.1
C1	499535	500093
refGene_gn=VWA1;refGene_tx=XM_011284405.3;refGene_gn=VWA1;refGene_tx=X
M_023257932.1
C1	502053	502332
refGene_gn=VWA1;refGene_tx=XM_011284405.3;refGene_gn=VWA1;refGene_tx=X
M_023257932.1
C1	502484	502822	refGene_gn=VWA1;refGene_tx=XM_023257932.1
C1	509530	509732
refGene_gn=ATAD3A;refGene_tx=XM_004001156.5;refGene_gn=ATAD3A;refGene_
tx=XM_019836001.2
C1	513264	513341
refGene_gn=ATAD3A;refGene_tx=XM_004001156.5;refGene_gn=ATAD3A;refGene_
tx=XM_019836001.2
C1	514133	514235
refGene_gn=ATAD3A;refGene_tx=XM_004001156.5;refGene_gn=ATAD3A;refGene_
tx=XM_019836001.2
C1	514537	514597
refGene_gn=ATAD3A;refGene_tx=XM_004001156.5;refGene_gn=ATAD3A;refGene_
tx=XM_019836001.2
C1	515133	515203

refGene_gn=ATAD3A;refGene_tx=XM_004001156.5;refGene_gn=ATAD3A;refGene_tx=XM_019836001.2
C1 515538 515704
refGene_gn=ATAD3A;refGene_tx=XM_004001156.5;refGene_gn=ATAD3A;refGene_tx=XM_019836001.2
C1 515861 515931
refGene_gn=ATAD3A;refGene_tx=XM_004001156.5;refGene_gn=ATAD3A;refGene_tx=XM_019836001.2
C1 519211 519367
refGene_gn=ATAD3A;refGene_tx=XM_004001156.5;refGene_gn=ATAD3A;refGene_tx=XM_019836001.2
C1 520027 520084 refGene_gn=ATAD3A;refGene_tx=XM_004001156.5
C1 520314 520440
refGene_gn=ATAD3A;refGene_tx=XM_004001156.5;refGene_gn=ATAD3A;refGene_tx=XM_019836001.2
C1 520634 520759
refGene_gn=ATAD3A;refGene_tx=XM_004001156.5;refGene_gn=ATAD3A;refGene_tx=XM_019836001.2
C1 521731 521783
refGene_gn=ATAD3A;refGene_tx=XM_004001156.5;refGene_gn=ATAD3A;refGene_tx=XM_019836001.2
C1 522122 522193
refGene_gn=ATAD3A;refGene_tx=XM_004001156.5;refGene_gn=ATAD3A;refGene_tx=XM_019836001.2
C1 522990 523158
refGene_gn=ATAD3A;refGene_tx=XM_004001156.5;refGene_gn=ATAD3A;refGene_tx=XM_019836001.2
C1 524465 524574
refGene_gn=ATAD3A;refGene_tx=XM_004001156.5;refGene_gn=ATAD3A;refGene_tx=XM_019836001.2
C1 531667 531811
refGene_gn=ATAD3A;refGene_tx=XM_004001156.5;refGene_gn=ATAD3A;refGene_tx=XM_019836001.2
C1 533238 533387 refGene_gn=TMEM240;refGene_tx=XM_023257935.1
C1 533919 534128 refGene_gn=TMEM240;refGene_tx=XM_023257935.1
C1 538433 538540 refGene_gn=TMEM240;refGene_tx=XM_023257935.1
C1 538899 538956 refGene_gn=TMEM240;refGene_tx=XM_023257935.1
C1 540460 540562 refGene_gn=SSU72;refGene_tx=XM_023257936.1
C1 542059 542178 refGene_gn=SSU72;refGene_tx=XM_023257936.1
C1 542767 542907 refGene_gn=SSU72;refGene_tx=XM_023257936.1
C1 556758 556902 refGene_gn=SSU72;refGene_tx=XM_023257936.1
C1 566312 566392 refGene_gn=SSU72;refGene_tx=XM_023257936.1
C1 575045 575735 refGene_gn=FNDC10;refGene_tx=XM_011284427.2
C1 595807 596054
refGene_gn=MIB2;refGene_tx=XM_023257937.1;refGene_gn=MIB2;refGene_tx=XM_023257938.1;refGene_gn=MIB2;refGene_tx=XM_023257939.1;refGene_gn=MIB2;refGene_tx=XM_023257941.1
C1 596233 596405
refGene_gn=MIB2;refGene_tx=XM_023257937.1;refGene_gn=MIB2;refGene_tx=XM_023257938.1;refGene_gn=MIB2;refGene_tx=XM_023257939.1;refGene_gn=MIB

2;refGene_tx=XM_023257941.1
C1 597168 597275
refGene_gn=MIB2;refGene_tx=XM_023257937.1;refGene_gn=MIB2;refGene_tx=XM_023257938.1;refGene_gn=MIB2;refGene_tx=XM_023257939.1;refGene_gn=MIB2;refGene_tx=XM_023257941.1
C1 597360 597555
refGene_gn=MIB2;refGene_tx=XM_023257937.1;refGene_gn=MIB2;refGene_tx=XM_023257938.1;refGene_gn=MIB2;refGene_tx=XM_023257939.1;refGene_gn=MIB2;refGene_tx=XM_023257941.1
C1 597712 597855
refGene_gn=MIB2;refGene_tx=XM_023257937.1;refGene_gn=MIB2;refGene_tx=XM_023257938.1;refGene_gn=MIB2;refGene_tx=XM_023257939.1;refGene_gn=MIB2;refGene_tx=XM_023257941.1
C1 597961 598069
refGene_gn=MIB2;refGene_tx=XM_023257937.1;refGene_gn=MIB2;refGene_tx=XM_023257938.1;refGene_gn=MIB2;refGene_tx=XM_023257939.1
C1 598983 599088
refGene_gn=MIB2;refGene_tx=XM_023257937.1;refGene_gn=MIB2;refGene_tx=XM_023257938.1;refGene_gn=MIB2;refGene_tx=XM_023257939.1;refGene_gn=MIB2;refGene_tx=XM_023257941.1
C1 599164 599327
refGene_gn=MIB2;refGene_tx=XM_023257937.1;refGene_gn=MIB2;refGene_tx=XM_023257938.1;refGene_gn=MIB2;refGene_tx=XM_023257939.1;refGene_gn=MIB2;refGene_tx=XM_023257941.1
C1 599410 599544
refGene_gn=MIB2;refGene_tx=XM_023257937.1;refGene_gn=MIB2;refGene_tx=XM_023257938.1;refGene_gn=MIB2;refGene_tx=XM_023257939.1;refGene_gn=MIB2;refGene_tx=XM_023257941.1
C1 599641 599790
refGene_gn=MIB2;refGene_tx=XM_023257937.1;refGene_gn=MIB2;refGene_tx=XM_023257938.1;refGene_gn=MIB2;refGene_tx=XM_023257939.1;refGene_gn=MIB2;refGene_tx=XM_023257941.1
C1 599868 600025
refGene_gn=MIB2;refGene_tx=XM_023257937.1;refGene_gn=MIB2;refGene_tx=XM_023257938.1;refGene_gn=MIB2;refGene_tx=XM_023257939.1;refGene_gn=MIB2;refGene_tx=XM_023257941.1
C1 600133 600294
refGene_gn=MIB2;refGene_tx=XM_023257937.1;refGene_gn=MIB2;refGene_tx=XM_023257938.1;refGene_gn=MIB2;refGene_tx=XM_023257939.1;refGene_gn=MIB2;refGene_tx=XM_023257941.1
C1 600390 600517
refGene_gn=MIB2;refGene_tx=XM_023257937.1;refGene_gn=MIB2;refGene_tx=XM_023257938.1;refGene_gn=MIB2;refGene_tx=XM_023257939.1;refGene_gn=MIB2;refGene_tx=XM_023257941.1
C1 600596 600830
refGene_gn=MIB2;refGene_tx=XM_023257937.1;refGene_gn=MIB2;refGene_tx=XM_023257938.1;refGene_gn=MIB2;refGene_tx=XM_023257939.1;refGene_gn=MIB2;refGene_tx=XM_023257941.1
C1 601261 601440
refGene_gn=MIB2;refGene_tx=XM_023257937.1;refGene_gn=MIB2;refGene_tx=X

M_023257938.1;refGene_gn=MIB2;refGene_tx=XM_023257939.1;refGene_gn=MIB2;refGene_tx=XM_023257941.1
C1	601549	601737	refGene_gn=MIB2;refGene_tx=XM_023257937.1;refGene_gn=MIB2;refGene_tx=XM_023257938.1;refGene_gn=MIB2;refGene_tx=XM_023257939.1;refGene_gn=MIB2;refGene_tx=XM_023257941.1
C1	601821	601887	refGene_gn=MIB2;refGene_tx=XM_023257937.1;refGene_gn=MIB2;refGene_tx=XM_023257938.1;refGene_gn=MIB2;refGene_tx=XM_023257939.1;refGene_gn=MIB2;refGene_tx=XM_023257941.1
C1	601963	602202	refGene_gn=MIB2;refGene_tx=XM_023257937.1;refGene_gn=MIB2;refGene_tx=XM_023257938.1;refGene_gn=MIB2;refGene_tx=XM_023257939.1;refGene_gn=MIB2;refGene_tx=XM_023257941.1
C1	603962	604118	refGene_gn=MMP23B;refGene_tx=XM_023257942.1
C1	604477	604617	refGene_gn=MMP23B;refGene_tx=XM_023257942.1
C1	604694	604834	refGene_gn=MMP23B;refGene_tx=XM_023257942.1
C1	604899	605067	refGene_gn=MMP23B;refGene_tx=XM_023257942.1
C1	605447	605612	refGene_gn=MMP23B;refGene_tx=XM_023257942.1
C1	605685	605796	refGene_gn=MMP23B;refGene_tx=XM_023257942.1
C1	605889	606016	refGene_gn=MMP23B;refGene_tx=XM_023257942.1
C1	606101	606276	refGene_gn=MMP23B;refGene_tx=XM_023257942.1
C1	607251	607344	refGene_gn=LOC101080954;refGene_tx=XM_004001136.5;refGene_gn=LOC101080954;refGene_tx=XM_006944209.4;refGene_gn=LOC101080954;refGene_tx=XM_006944210.2;refGene_gn=LOC101080954;refGene_tx=XM_006944211.4
C1	607437	607627	refGene_gn=LOC101080954;refGene_tx=XM_004001136.5;refGene_gn=LOC101080954;refGene_tx=XM_006944209.4;refGene_gn=LOC101080954;refGene_tx=XM_006944210.2;refGene_gn=LOC101080954;refGene_tx=XM_006944211.4
C1	607742	607891	refGene_gn=LOC101080954;refGene_tx=XM_004001136.5;refGene_gn=LOC101080954;refGene_tx=XM_006944209.4;refGene_gn=LOC101080954;refGene_tx=XM_006944210.2;refGene_gn=LOC101080954;refGene_tx=XM_006944211.4
C1	608001	608118	refGene_gn=LOC101080954;refGene_tx=XM_004001136.5;refGene_gn=LOC101080954;refGene_tx=XM_006944209.4;refGene_gn=LOC101080954;refGene_tx=XM_006944210.2;refGene_gn=LOC101080954;refGene_tx=XM_006944211.4
C1	608191	608299	refGene_gn=LOC101080954;refGene_tx=XM_004001136.5;refGene_gn=LOC101080954;refGene_tx=XM_006944209.4;refGene_gn=LOC101080954;refGene_tx=XM_006944210.2;refGene_gn=LOC101080954;refGene_tx=XM_006944211.4
C1	608380	608502	refGene_gn=LOC101080954;refGene_tx=XM_004001136.5;refGene_gn=LOC101080954;refGene_tx=XM_006944209.4;refGene_gn=LOC101080954;refGene_tx=XM_006944210.2;refGene_gn=LOC101080954;refGene_tx=XM_006944211.4
C1	608869	608975	refGene_gn=LOC101080954;refGene_tx=XM_004001136.5;refGene_gn=LOC101080954;refGene_tx=XM_006944209.4;refGene_gn=LOC101080954;refGene_tx=XM_006944210.2;refGene_gn=LOC101080954;refGene_tx=XM_006944211.4

C1 609193 609315
refGene_gn=LOC101080954;refGene_tx=XM_004001136.5;refGene_gn=LOC101080954;refGene_tx=XM_006944209.4;refGene_gn=LOC101080954;refGene_tx=XM_006944210.2;refGene_gn=LOC101080954;refGene_tx=XM_006944211.4
C1 610229 610320
refGene_gn=LOC101080954;refGene_tx=XM_004001136.5;refGene_gn=LOC101080954;refGene_tx=XM_006944209.4;refGene_gn=LOC101080954;refGene_tx=XM_006944210.2;refGene_gn=LOC101080954;refGene_tx=XM_006944211.4
C1 610743 610919
refGene_gn=LOC101080954;refGene_tx=XM_004001136.5;refGene_gn=LOC101080954;refGene_tx=XM_006944209.4;refGene_gn=LOC101080954;refGene_tx=XM_006944210.2;refGene_gn=LOC101080954;refGene_tx=XM_006944211.4
C1 611534 611600
refGene_gn=LOC101080954;refGene_tx=XM_004001136.5;refGene_gn=LOC101080954;refGene_tx=XM_006944209.4;refGene_gn=LOC101080954;refGene_tx=XM_006944210.2;refGene_gn=LOC101080954;refGene_tx=XM_006944211.4
C1 611891 612053
refGene_gn=LOC101080954;refGene_tx=XM_004001136.5;refGene_gn=LOC101080954;refGene_tx=XM_006944209.4;refGene_gn=LOC101080954;refGene_tx=XM_006944210.2;refGene_gn=LOC101080954;refGene_tx=XM_006944211.4
C1 612135 612231
refGene_gn=LOC101080954;refGene_tx=XM_004001136.5;refGene_gn=LOC101080954;refGene_tx=XM_006944209.4;refGene_gn=LOC101080954;refGene_tx=XM_006944210.2;refGene_gn=LOC101080954;refGene_tx=XM_006944211.4
C1 612779 612902
refGene_gn=LOC101080954;refGene_tx=XM_004001136.5;refGene_gn=LOC101080954;refGene_tx=XM_006944209.4;refGene_gn=LOC101080954;refGene_tx=XM_006944210.2;refGene_gn=LOC101080954;refGene_tx=XM_006944211.4
C1 613419 613556
refGene_gn=LOC101080954;refGene_tx=XM_004001136.5;refGene_gn=LOC101080954;refGene_tx=XM_006944209.4;refGene_gn=LOC101080954;refGene_tx=XM_006944210.2;refGene_gn=LOC101080954;refGene_tx=XM_006944211.4
C1 621117 621256
refGene_gn=LOC101080954;refGene_tx=XM_004001136.5;refGene_gn=LOC101080954;refGene_tx=XM_006944209.4;refGene_gn=LOC101080954;refGene_tx=XM_006944210.2;refGene_gn=LOC101080954;refGene_tx=XM_006944211.4
C1 623249 623377
refGene_gn=LOC101080954;refGene_tx=XM_004001136.5;refGene_gn=LOC101080954;refGene_tx=XM_006944209.4;refGene_gn=LOC101080954;refGene_tx=XM_006944210.2;refGene_gn=LOC101080954;refGene_tx=XM_006944211.4
C1 624394 624510
refGene_gn=LOC101080954;refGene_tx=XM_004001136.5;refGene_gn=LOC101080954;refGene_tx=XM_006944209.4;refGene_gn=LOC101080954;refGene_tx=XM_006944210.2;refGene_gn=LOC101080954;refGene_tx=XM_006944211.4
C1 625197 625206
refGene_gn=LOC101080954;refGene_tx=XM_006944210.2;refGene_gn=LOC101080954;refGene_tx=XM_006944211.4
C1 625316 625427
refGene_gn=LOC101080954;refGene_tx=XM_004001136.5;refGene_gn=LOC101080954;refGene_tx=XM_006944209.4

C1	630612	630850
refGene_gn=SLC35E2B;refGene_tx=XM_006944212.4;refGene_gn=SLC35E2B;refGene_tx=XM_019836002.2;refGene_gn=SLC35E2B;refGene_tx=XM_023257947.1;refGene_gn=SLC35E2B;refGene_tx=XM_023257948.1;refGene_gn=SLC35E2B;refGene_tx=XM_023257949.1;refGene_gn=SLC35E2B;refGene_tx=XM_023257950.1
C1	632783	632808	refGene_gn=SLC35E2B;refGene_tx=XM_023257951.1
C1	632952	633098
refGene_gn=SLC35E2B;refGene_tx=XM_006944212.4;refGene_gn=SLC35E2B;refGene_tx=XM_019836002.2;refGene_gn=SLC35E2B;refGene_tx=XM_023257947.1;refGene_gn=SLC35E2B;refGene_tx=XM_023257948.1;refGene_gn=SLC35E2B;refGene_tx=XM_023257949.1;refGene_gn=SLC35E2B;refGene_tx=XM_023257950.1;refGene_gn=SLC35E2B;refGene_tx=XM_023257951.1
C1	635724	635797
refGene_gn=SLC35E2B;refGene_tx=XM_006944212.4;refGene_gn=SLC35E2B;refGene_tx=XM_019836002.2;refGene_gn=SLC35E2B;refGene_tx=XM_023257947.1;refGene_gn=SLC35E2B;refGene_tx=XM_023257948.1;refGene_gn=SLC35E2B;refGene_tx=XM_023257949.1;refGene_gn=SLC35E2B;refGene_tx=XM_023257950.1;refGene_gn=SLC35E2B;refGene_tx=XM_023257951.1
C1	636298	636352
refGene_gn=SLC35E2B;refGene_tx=XM_006944212.4;refGene_gn=SLC35E2B;refGene_tx=XM_019836002.2;refGene_gn=SLC35E2B;refGene_tx=XM_023257947.1;refGene_gn=SLC35E2B;refGene_tx=XM_023257948.1;refGene_gn=SLC35E2B;refGene_tx=XM_023257949.1;refGene_gn=SLC35E2B;refGene_tx=XM_023257950.1;refGene_gn=SLC35E2B;refGene_tx=XM_023257951.1
C1	638211	638332
refGene_gn=SLC35E2B;refGene_tx=XM_006944212.4;refGene_gn=SLC35E2B;refGene_tx=XM_019836002.2;refGene_gn=SLC35E2B;refGene_tx=XM_023257947.1;refGene_gn=SLC35E2B;refGene_tx=XM_023257948.1;refGene_gn=SLC35E2B;refGene_tx=XM_023257949.1;refGene_gn=SLC35E2B;refGene_tx=XM_023257950.1;refGene_gn=SLC35E2B;refGene_tx=XM_023257951.1
C1	640566	640694
refGene_gn=SLC35E2B;refGene_tx=XM_006944212.4;refGene_gn=SLC35E2B;refGene_tx=XM_019836002.2;refGene_gn=SLC35E2B;refGene_tx=XM_023257947.1;refGene_gn=SLC35E2B;refGene_tx=XM_023257948.1;refGene_gn=SLC35E2B;refGene_tx=XM_023257949.1;refGene_gn=SLC35E2B;refGene_tx=XM_023257950.1;refGene_gn=SLC35E2B;refGene_tx=XM_023257951.1
C1	641396	641532
refGene_gn=SLC35E2B;refGene_tx=XM_006944212.4;refGene_gn=SLC35E2B;refGene_tx=XM_019836002.2;refGene_gn=SLC35E2B;refGene_tx=XM_023257947.1;refGene_gn=SLC35E2B;refGene_tx=XM_023257948.1;refGene_gn=SLC35E2B;refGene_tx=XM_023257949.1;refGene_gn=SLC35E2B;refGene_tx=XM_023257950.1;refGene_gn=SLC35E2B;refGene_tx=XM_023257951.1
C1	641767	642089
refGene_gn=SLC35E2B;refGene_tx=XM_006944212.4;refGene_gn=SLC35E2B;refGene_tx=XM_019836002.2;refGene_gn=SLC35E2B;refGene_tx=XM_023257947.1;refGene_gn=SLC35E2B;refGene_tx=XM_023257948.1;refGene_gn=SLC35E2B;refGene_tx=XM_023257949.1;refGene_gn=SLC35E2B;refGene_tx=XM_023257950.1;refGene_gn=SLC35E2B;refGene_tx=XM_023257951.1
C1	662152	662180	refGene_gn=NADK;refGene_tx=XM_023257946.1
C1	662424	662629

refGene_gn=NADK;refGene_tx=XM_023257943.1;refGene_gn=NADK;refGene_tx=XM_023257944.1;refGene_gn=NADK;refGene_tx=XM_023257945.1
C1 663003 663086
refGene_gn=NADK;refGene_tx=XM_023257943.1;refGene_gn=NADK;refGene_tx=XM_023257944.1;refGene_gn=NADK;refGene_tx=XM_023257945.1;refGene_gn=NADK;refGene_tx=XM_023257946.1
C1 663791 663949
refGene_gn=NADK;refGene_tx=XM_023257943.1;refGene_gn=NADK;refGene_tx=XM_023257944.1;refGene_gn=NADK;refGene_tx=XM_023257945.1;refGene_gn=NADK;refGene_tx=XM_023257946.1
C1 664027 664127
refGene_gn=NADK;refGene_tx=XM_023257943.1;refGene_gn=NADK;refGene_tx=XM_023257944.1;refGene_gn=NADK;refGene_tx=XM_023257945.1;refGene_gn=NADK;refGene_tx=XM_023257946.1
C1 664351 664506
refGene_gn=NADK;refGene_tx=XM_023257943.1;refGene_gn=NADK;refGene_tx=XM_023257944.1;refGene_gn=NADK;refGene_tx=XM_023257945.1;refGene_gn=NADK;refGene_tx=XM_023257946.1
C1 664845 664948
refGene_gn=NADK;refGene_tx=XM_023257943.1;refGene_gn=NADK;refGene_tx=XM_023257944.1;refGene_gn=NADK;refGene_tx=XM_023257945.1;refGene_gn=NADK;refGene_tx=XM_023257946.1
C1 665748 665834
refGene_gn=NADK;refGene_tx=XM_023257943.1;refGene_gn=NADK;refGene_tx=XM_023257944.1;refGene_gn=NADK;refGene_tx=XM_023257945.1;refGene_gn=NADK;refGene_tx=XM_023257946.1
C1 665992 666101
refGene_gn=NADK;refGene_tx=XM_023257943.1;refGene_gn=NADK;refGene_tx=XM_023257944.1;refGene_gn=NADK;refGene_tx=XM_023257945.1;refGene_gn=NADK;refGene_tx=XM_023257946.1
C1 666856 666986
refGene_gn=NADK;refGene_tx=XM_023257943.1;refGene_gn=NADK;refGene_tx=XM_023257944.1;refGene_gn=NADK;refGene_tx=XM_023257945.1;refGene_gn=NADK;refGene_tx=XM_023257946.1
C1 671780 671864
refGene_gn=NADK;refGene_tx=XM_023257943.1;refGene_gn=NADK;refGene_tx=XM_023257944.1;refGene_gn=NADK;refGene_tx=XM_023257945.1;refGene_gn=NADK;refGene_tx=XM_023257946.1
C1 673294 673473
refGene_gn=NADK;refGene_tx=XM_023257943.1;refGene_gn=NADK;refGene_tx=XM_023257944.1;refGene_gn=NADK;refGene_tx=XM_023257945.1;refGene_gn=NADK;refGene_tx=XM_023257946.1
C1 698867 698974
refGene_gn=GNB1;refGene_tx=XM_019836005.2;refGene_gn=GNB1;refGene_tx=XM_023257952.1;refGene_gn=GNB1;refGene_tx=XM_023257953.1
C1 701158 701375
refGene_gn=GNB1;refGene_tx=XM_019836005.2;refGene_gn=GNB1;refGene_tx=XM_023257952.1;refGene_gn=GNB1;refGene_tx=XM_023257953.1
C1 702240 702442
refGene_gn=GNB1;refGene_tx=XM_019836005.2;refGene_gn=GNB1;refGene_tx=X

M_023257952.1;refGene_gn=GNB1;refGene_tx=XM_023257953.1
C1 703807 703874
refGene_gn=GNB1;refGene_tx=XM_019836005.2;refGene_gn=GNB1;refGene_tx=XM_023257952.1;refGene_gn=GNB1;refGene_tx=XM_023257953.1
C1 715786 715949
refGene_gn=GNB1;refGene_tx=XM_019836005.2;refGene_gn=GNB1;refGene_tx=XM_023257952.1;refGene_gn=GNB1;refGene_tx=XM_023257953.1
C1 716883 716947
refGene_gn=GNB1;refGene_tx=XM_019836005.2;refGene_gn=GNB1;refGene_tx=XM_023257952.1;refGene_gn=GNB1;refGene_tx=XM_023257953.1
C1 721810 721917
refGene_gn=GNB1;refGene_tx=XM_019836005.2;refGene_gn=GNB1;refGene_tx=XM_023257952.1;refGene_gn=GNB1;refGene_tx=XM_023257953.1
C1 723642 723681
refGene_gn=GNB1;refGene_tx=XM_019836005.2;refGene_gn=GNB1;refGene_tx=XM_023257952.1;refGene_gn=GNB1;refGene_tx=XM_023257953.1
C1 729590 729647
refGene_gn=GNB1;refGene_tx=XM_019836005.2;refGene_gn=GNB1;refGene_tx=XM_023257952.1;refGene_gn=GNB1;refGene_tx=XM_023257953.1
C1 807102 807105 refGene_gn=CALML6;refGene_tx=XM_023257961.1
C1 807783 807958
refGene_gn=CALML6;refGene_tx=XM_023257961.1;refGene_gn=CALML6;refGene_tx=XM_023257962.1
C1 808074 808219
refGene_gn=CALML6;refGene_tx=XM_023257961.1;refGene_gn=CALML6;refGene_tx=XM_023257962.1
C1 808296 808450
refGene_gn=CALML6;refGene_tx=XM_023257961.1;refGene_gn=CALML6;refGene_tx=XM_023257962.1
C1 809351 809527 refGene_gn=TMEM52;refGene_tx=XM_019836007.2
C1 809606 809785 refGene_gn=TMEM52;refGene_tx=XM_019836007.2
C1 809940 810117 refGene_gn=TMEM52;refGene_tx=XM_011284408.3
C1 810347 810407
refGene_gn=TMEM52;refGene_tx=XM_011284408.3;refGene_gn=TMEM52;refGene_tx=XM_019836007.2
C1 810500 810695
refGene_gn=TMEM52;refGene_tx=XM_011284408.3;refGene_gn=TMEM52;refGene_tx=XM_019836007.2
C1 813674 813776
refGene_gn=CFAP74;refGene_tx=XM_023257954.1;refGene_gn=CFAP74;refGene_tx=XM_023257955.1;refGene_gn=CFAP74;refGene_tx=XM_023257956.1;refGene_gn=CFAP74;refGene_tx=XM_023257957.1;refGene_gn=CFAP74;refGene_tx=XM_023257958.1;refGene_gn=CFAP74;refGene_tx=XM_023257959.1
C1 813920 814055
refGene_gn=CFAP74;refGene_tx=XM_023257954.1;refGene_gn=CFAP74;refGene_tx=XM_023257955.1;refGene_gn=CFAP74;refGene_tx=XM_023257956.1;refGene_gn=CFAP74;refGene_tx=XM_023257957.1;refGene_gn=CFAP74;refGene_tx=XM_023257958.1;refGene_gn=CFAP74;refGene_tx=XM_023257959.1
C1 814264 814395
refGene_gn=CFAP74;refGene_tx=XM_023257954.1;refGene_gn=CFAP74;refGene_

tx=XM_023257955.1;refGene_gn=CFAP74;refGene_tx=XM_023257956.1;refGene_gn=CFAP74;refGene_tx=XM_023257957.1;refGene_gn=CFAP74;refGene_tx=XM_023257958.1;refGene_gn=CFAP74;refGene_tx=XM_023257959.1;refGene_gn=CFAP74;refGene_tx=XM_023257960.1
C1 814622 814755
refGene_gn=CFAP74;refGene_tx=XM_023257954.1;refGene_gn=CFAP74;refGene_tx=XM_023257955.1;refGene_gn=CFAP74;refGene_tx=XM_023257956.1;refGene_gn=CFAP74;refGene_tx=XM_023257957.1;refGene_gn=CFAP74;refGene_tx=XM_023257958.1;refGene_gn=CFAP74;refGene_tx=XM_023257959.1
C1 814985 815140
refGene_gn=CFAP74;refGene_tx=XM_023257954.1;refGene_gn=CFAP74;refGene_tx=XM_023257955.1;refGene_gn=CFAP74;refGene_tx=XM_023257956.1;refGene_gn=CFAP74;refGene_tx=XM_023257957.1;refGene_gn=CFAP74;refGene_tx=XM_023257958.1;refGene_gn=CFAP74;refGene_tx=XM_023257959.1;refGene_gn=CFAP74;refGene_tx=XM_023257960.1
C1 815818 815951
refGene_gn=CFAP74;refGene_tx=XM_023257954.1;refGene_gn=CFAP74;refGene_tx=XM_023257955.1;refGene_gn=CFAP74;refGene_tx=XM_023257956.1;refGene_gn=CFAP74;refGene_tx=XM_023257957.1;refGene_gn=CFAP74;refGene_tx=XM_023257958.1;refGene_gn=CFAP74;refGene_tx=XM_023257959.1;refGene_gn=CFAP74;refGene_tx=XM_023257960.1
C1 816812 816968
refGene_gn=CFAP74;refGene_tx=XM_023257954.1;refGene_gn=CFAP74;refGene_tx=XM_023257955.1;refGene_gn=CFAP74;refGene_tx=XM_023257956.1;refGene_gn=CFAP74;refGene_tx=XM_023257957.1;refGene_gn=CFAP74;refGene_tx=XM_023257958.1;refGene_gn=CFAP74;refGene_tx=XM_023257959.1;refGene_gn=CFAP74;refGene_tx=XM_023257960.1
C1 817204 817324
refGene_gn=CFAP74;refGene_tx=XM_023257954.1;refGene_gn=CFAP74;refGene_tx=XM_023257955.1;refGene_gn=CFAP74;refGene_tx=XM_023257956.1;refGene_gn=CFAP74;refGene_tx=XM_023257957.1;refGene_gn=CFAP74;refGene_tx=XM_023257958.1;refGene_gn=CFAP74;refGene_tx=XM_023257959.1;refGene_gn=CFAP74;refGene_tx=XM_023257960.1
C1 817456 817512
refGene_gn=CFAP74;refGene_tx=XM_023257954.1;refGene_gn=CFAP74;refGene_tx=XM_023257955.1;refGene_gn=CFAP74;refGene_tx=XM_023257956.1;refGene_gn=CFAP74;refGene_tx=XM_023257957.1;refGene_gn=CFAP74;refGene_tx=XM_023257958.1;refGene_gn=CFAP74;refGene_tx=XM_023257959.1;refGene_gn=CFAP74;refGene_tx=XM_023257960.1
C1 817620 817874
refGene_gn=CFAP74;refGene_tx=XM_023257954.1;refGene_gn=CFAP74;refGene_tx=XM_023257955.1;refGene_gn=CFAP74;refGene_tx=XM_023257956.1;refGene_gn=CFAP74;refGene_tx=XM_023257957.1;refGene_gn=CFAP74;refGene_tx=XM_023257958.1;refGene_gn=CFAP74;refGene_tx=XM_023257959.1;refGene_gn=CFAP74;refGene_tx=XM_023257960.1
C1 817896 818031 refGene_gn=CFAP74;refGene_tx=XM_023257957.1
C1 818667 818804
refGene_gn=CFAP74;refGene_tx=XM_023257954.1;refGene_gn=CFAP74;refGene_tx=XM_023257955.1;refGene_gn=CFAP74;refGene_tx=XM_023257956.1;refGene_gn=CFAP74;refGene_tx=XM_023257957.1;refGene_gn=CFAP74;refGene_tx=XM_02

3257958.1;refGene_gn=CFAP74;refGene_tx=XM_023257959.1;refGene_gn=CFAP7
4;refGene_tx=XM_023257960.1
C1	819848	819947
refGene_gn=CFAP74;refGene_tx=XM_023257954.1;refGene_gn=CFAP74;refGene_
tx=XM_023257955.1;refGene_gn=CFAP74;refGene_tx=XM_023257956.1;refGene_
gn=CFAP74;refGene_tx=XM_023257957.1;refGene_gn=CFAP74;refGene_tx=XM_02
3257958.1;refGene_gn=CFAP74;refGene_tx=XM_023257959.1;refGene_gn=CFAP7
4;refGene_tx=XM_023257960.1
C1	820908	821185
refGene_gn=CFAP74;refGene_tx=XM_023257954.1;refGene_gn=CFAP74;refGene_
tx=XM_023257955.1;refGene_gn=CFAP74;refGene_tx=XM_023257956.1;refGene_
gn=CFAP74;refGene_tx=XM_023257957.1;refGene_gn=CFAP74;refGene_tx=XM_02
3257958.1;refGene_gn=CFAP74;refGene_tx=XM_023257959.1;refGene_gn=CFAP7
4;refGene_tx=XM_023257960.1
C1	822981	823115
refGene_gn=CFAP74;refGene_tx=XM_023257954.1;refGene_gn=CFAP74;refGene_
tx=XM_023257955.1;refGene_gn=CFAP74;refGene_tx=XM_023257956.1;refGene_
gn=CFAP74;refGene_tx=XM_023257957.1;refGene_gn=CFAP74;refGene_tx=XM_02
3257958.1;refGene_gn=CFAP74;refGene_tx=XM_023257959.1;refGene_gn=CFAP7
4;refGene_tx=XM_023257960.1
C1	823887	824061
refGene_gn=CFAP74;refGene_tx=XM_023257954.1;refGene_gn=CFAP74;refGene_
tx=XM_023257955.1;refGene_gn=CFAP74;refGene_tx=XM_023257956.1;refGene_
gn=CFAP74;refGene_tx=XM_023257957.1;refGene_gn=CFAP74;refGene_tx=XM_02
3257958.1;refGene_gn=CFAP74;refGene_tx=XM_023257959.1;refGene_gn=CFAP7
4;refGene_tx=XM_023257960.1
C1	824590	824678
refGene_gn=CFAP74;refGene_tx=XM_023257954.1;refGene_gn=CFAP74;refGene_
tx=XM_023257955.1;refGene_gn=CFAP74;refGene_tx=XM_023257956.1;refGene_
gn=CFAP74;refGene_tx=XM_023257957.1;refGene_gn=CFAP74;refGene_tx=XM_02
3257958.1;refGene_gn=CFAP74;refGene_tx=XM_023257959.1;refGene_gn=CFAP7
4;refGene_tx=XM_023257960.1
C1	824862	824991
refGene_gn=CFAP74;refGene_tx=XM_023257954.1;refGene_gn=CFAP74;refGene_
tx=XM_023257955.1;refGene_gn=CFAP74;refGene_tx=XM_023257956.1;refGene_
gn=CFAP74;refGene_tx=XM_023257957.1;refGene_gn=CFAP74;refGene_tx=XM_02
3257958.1;refGene_gn=CFAP74;refGene_tx=XM_023257959.1;refGene_gn=CFAP7
4;refGene_tx=XM_023257960.1
C1	827301	827423
refGene_gn=CFAP74;refGene_tx=XM_023257954.1;refGene_gn=CFAP74;refGene_
tx=XM_023257955.1;refGene_gn=CFAP74;refGene_tx=XM_023257956.1;refGene_
gn=CFAP74;refGene_tx=XM_023257957.1;refGene_gn=CFAP74;refGene_tx=XM_02
3257958.1;refGene_gn=CFAP74;refGene_tx=XM_023257959.1;refGene_gn=CFAP7
4;refGene_tx=XM_023257960.1
C1	832489	832612
refGene_gn=CFAP74;refGene_tx=XM_023257954.1;refGene_gn=CFAP74;refGene_
tx=XM_023257955.1;refGene_gn=CFAP74;refGene_tx=XM_023257956.1;refGene_
gn=CFAP74;refGene_tx=XM_023257957.1;refGene_gn=CFAP74;refGene_tx=XM_02
3257958.1;refGene_gn=CFAP74;refGene_tx=XM_023257959.1;refGene_gn=CFAP7
4;refGene_tx=XM_023257960.1

C1	833150	833227
refGene_gn=CFAP74;refGene_tx=XM_023257954.1;refGene_gn=CFAP74;refGene_tx=XM_023257955.1;refGene_gn=CFAP74;refGene_tx=XM_023257956.1;refGene_gn=CFAP74;refGene_tx=XM_023257957.1;refGene_gn=CFAP74;refGene_tx=XM_023257958.1;refGene_gn=CFAP74;refGene_tx=XM_023257959.1;refGene_gn=CFAP74;refGene_tx=XM_023257960.1
C1	839948	840111
refGene_gn=CFAP74;refGene_tx=XM_023257954.1;refGene_gn=CFAP74;refGene_tx=XM_023257955.1;refGene_gn=CFAP74;refGene_tx=XM_023257956.1;refGene_gn=CFAP74;refGene_tx=XM_023257957.1;refGene_gn=CFAP74;refGene_tx=XM_023257958.1;refGene_gn=CFAP74;refGene_tx=XM_023257959.1;refGene_gn=CFAP74;refGene_tx=XM_023257960.1
C1	841280	841445
refGene_gn=CFAP74;refGene_tx=XM_023257954.1;refGene_gn=CFAP74;refGene_tx=XM_023257955.1;refGene_gn=CFAP74;refGene_tx=XM_023257956.1;refGene_gn=CFAP74;refGene_tx=XM_023257957.1;refGene_gn=CFAP74;refGene_tx=XM_023257958.1;refGene_gn=CFAP74;refGene_tx=XM_023257959.1;refGene_gn=CFAP74;refGene_tx=XM_023257960.1
C1	849150	849240
refGene_gn=CFAP74;refGene_tx=XM_023257954.1;refGene_gn=CFAP74;refGene_tx=XM_023257955.1;refGene_gn=CFAP74;refGene_tx=XM_023257956.1;refGene_gn=CFAP74;refGene_tx=XM_023257957.1;refGene_gn=CFAP74;refGene_tx=XM_023257958.1;refGene_gn=CFAP74;refGene_tx=XM_023257959.1;refGene_gn=CFAP74;refGene_tx=XM_023257960.1
C1	849608	849675
refGene_gn=CFAP74;refGene_tx=XM_023257954.1;refGene_gn=CFAP74;refGene_tx=XM_023257955.1;refGene_gn=CFAP74;refGene_tx=XM_023257956.1;refGene_gn=CFAP74;refGene_tx=XM_023257957.1;refGene_gn=CFAP74;refGene_tx=XM_023257958.1;refGene_gn=CFAP74;refGene_tx=XM_023257959.1;refGene_gn=CFAP74;refGene_tx=XM_023257960.1
C1	852076	852195
refGene_gn=CFAP74;refGene_tx=XM_023257954.1;refGene_gn=CFAP74;refGene_tx=XM_023257955.1;refGene_gn=CFAP74;refGene_tx=XM_023257956.1;refGene_gn=CFAP74;refGene_tx=XM_023257957.1;refGene_gn=CFAP74;refGene_tx=XM_023257958.1;refGene_gn=CFAP74;refGene_tx=XM_023257959.1;refGene_gn=CFAP74;refGene_tx=XM_023257960.1
C1	852683	852857
refGene_gn=CFAP74;refGene_tx=XM_023257954.1;refGene_gn=CFAP74;refGene_tx=XM_023257955.1;refGene_gn=CFAP74;refGene_tx=XM_023257956.1;refGene_gn=CFAP74;refGene_tx=XM_023257957.1;refGene_gn=CFAP74;refGene_tx=XM_023257958.1;refGene_gn=CFAP74;refGene_tx=XM_023257959.1;refGene_gn=CFAP74;refGene_tx=XM_023257960.1
C1	853340	853502
refGene_gn=CFAP74;refGene_tx=XM_023257954.1;refGene_gn=CFAP74;refGene_tx=XM_023257955.1;refGene_gn=CFAP74;refGene_tx=XM_023257956.1;refGene_gn=CFAP74;refGene_tx=XM_023257957.1;refGene_gn=CFAP74;refGene_tx=XM_023257958.1;refGene_gn=CFAP74;refGene_tx=XM_023257959.1;refGene_gn=CFAP74;refGene_tx=XM_023257960.1
C1	854883	855088
refGene_gn=CFAP74;refGene_tx=XM_023257954.1;refGene_gn=CFAP74;refGene_

tx=XM_023257955.1;refGene_gn=CFAP74;refGene_tx=XM_023257956.1;refGene_
gn=CFAP74;refGene_tx=XM_023257957.1;refGene_gn=CFAP74;refGene_tx=XM_02
3257958.1;refGene_gn=CFAP74;refGene_tx=XM_023257959.1;refGene_gn=CFAP7
4;refGene_tx=XM_023257960.1
C1 856244 856402
refGene_gn=CFAP74;refGene_tx=XM_023257954.1;refGene_gn=CFAP74;refGene_
tx=XM_023257955.1;refGene_gn=CFAP74;refGene_tx=XM_023257956.1;refGene_
gn=CFAP74;refGene_tx=XM_023257957.1;refGene_gn=CFAP74;refGene_tx=XM_02
3257958.1;refGene_gn=CFAP74;refGene_tx=XM_023257959.1;refGene_gn=CFAP7
4;refGene_tx=XM_023257960.1
C1 857227 857330
refGene_gn=CFAP74;refGene_tx=XM_023257954.1;refGene_gn=CFAP74;refGene_
tx=XM_023257955.1;refGene_gn=CFAP74;refGene_tx=XM_023257956.1;refGene_
gn=CFAP74;refGene_tx=XM_023257957.1;refGene_gn=CFAP74;refGene_tx=XM_02
3257958.1;refGene_gn=CFAP74;refGene_tx=XM_023257959.1;refGene_gn=CFAP7
4;refGene_tx=XM_023257960.1
C1 857719 857830
refGene_gn=CFAP74;refGene_tx=XM_023257954.1;refGene_gn=CFAP74;refGene_
tx=XM_023257955.1;refGene_gn=CFAP74;refGene_tx=XM_023257956.1;refGene_
gn=CFAP74;refGene_tx=XM_023257957.1;refGene_gn=CFAP74;refGene_tx=XM_02
3257958.1;refGene_gn=CFAP74;refGene_tx=XM_023257959.1;refGene_gn=CFAP7
4;refGene_tx=XM_023257960.1
C1 858243 858417
refGene_gn=CFAP74;refGene_tx=XM_023257954.1;refGene_gn=CFAP74;refGene_
tx=XM_023257955.1;refGene_gn=CFAP74;refGene_tx=XM_023257956.1;refGene_
gn=CFAP74;refGene_tx=XM_023257957.1;refGene_gn=CFAP74;refGene_tx=XM_02
3257958.1;refGene_gn=CFAP74;refGene_tx=XM_023257959.1;refGene_gn=CFAP7
4;refGene_tx=XM_023257960.1
C1 863601 863706
refGene_gn=CFAP74;refGene_tx=XM_023257954.1;refGene_gn=CFAP74;refGene_
tx=XM_023257955.1;refGene_gn=CFAP74;refGene_tx=XM_023257956.1;refGene_
gn=CFAP74;refGene_tx=XM_023257957.1;refGene_gn=CFAP74;refGene_tx=XM_02
3257960.1
C1 865346 865445
refGene_gn=CFAP74;refGene_tx=XM_023257954.1;refGene_gn=CFAP74;refGene_
tx=XM_023257955.1;refGene_gn=CFAP74;refGene_tx=XM_023257956.1;refGene_
gn=CFAP74;refGene_tx=XM_023257957.1;refGene_gn=CFAP74;refGene_tx=XM_02
3257960.1
C1 867758 867896
refGene_gn=CFAP74;refGene_tx=XM_023257954.1;refGene_gn=CFAP74;refGene_
tx=XM_023257955.1;refGene_gn=CFAP74;refGene_tx=XM_023257956.1;refGene_
gn=CFAP74;refGene_tx=XM_023257957.1;refGene_gn=CFAP74;refGene_tx=XM_02
3257958.1;refGene_gn=CFAP74;refGene_tx=XM_023257960.1
C1 868015 868100
refGene_gn=CFAP74;refGene_tx=XM_023257954.1;refGene_gn=CFAP74;refGene_
tx=XM_023257955.1;refGene_gn=CFAP74;refGene_tx=XM_023257956.1;refGene_
gn=CFAP74;refGene_tx=XM_023257957.1;refGene_gn=CFAP74;refGene_tx=XM_02
3257958.1;refGene_gn=CFAP74;refGene_tx=XM_023257960.1
C1 871118 871185
refGene_gn=CFAP74;refGene_tx=XM_023257954.1;refGene_gn=CFAP74;refGene_

```
tx=XM_023257955.1;refGene_gn=CFAP74;refGene_tx=XM_023257956.1;refGene_
gn=CFAP74;refGene_tx=XM_023257957.1;refGene_gn=CFAP74;refGene_tx=XM_02
3257958.1;refGene_gn=CFAP74;refGene_tx=XM_023257960.1
C1	880441	880530	refGene_gn=CFAP74;refGene_tx=XM_023257959.1
C1	894541	894609	refGene_gn=GABRD;refGene_tx=XM_019836006.2
C1	898890	899356	
refGene_gn=GABRD;refGene_tx=XM_011284411.3;refGene_gn=GABRD;refGene_tx
=XM_019836006.2
C1	899652	899720	
refGene_gn=GABRD;refGene_tx=XM_011284411.3;refGene_gn=GABRD;refGene_tx
=XM_019836006.2
C1	899815	900036	
refGene_gn=GABRD;refGene_tx=XM_011284411.3;refGene_gn=GABRD;refGene_tx
=XM_019836006.2
C1	900779	900862	
refGene_gn=GABRD;refGene_tx=XM_011284411.3;refGene_gn=GABRD;refGene_tx
=XM_019836006.2
C1	901303	901441	
refGene_gn=GABRD;refGene_tx=XM_011284411.3;refGene_gn=GABRD;refGene_tx
=XM_019836006.2
C1	902117	902273	
refGene_gn=GABRD;refGene_tx=XM_011284411.3;refGene_gn=GABRD;refGene_tx
=XM_019836006.2
C1	902506	902718	
refGene_gn=GABRD;refGene_tx=XM_011284411.3;refGene_gn=GABRD;refGene_tx
=XM_019836006.2
C1	903029	903329	
refGene_gn=GABRD;refGene_tx=XM_011284411.3;refGene_gn=GABRD;refGene_tx
=XM_019836006.2
C1	920576	920647	refGene_gn=PRKCZ;refGene_tx=XM_023257963.1
C1	923479	923601	refGene_gn=PRKCZ;refGene_tx=XM_023257963.1
C1	924388	924478	refGene_gn=PRKCZ;refGene_tx=XM_023257963.1
C1	926335	926386	refGene_gn=PRKCZ;refGene_tx=XM_023257963.1
C1	940235	940245	refGene_gn=PRKCZ;refGene_tx=XM_019836008.2
C1	987134	987220	
refGene_gn=PRKCZ;refGene_tx=XM_019836008.2;refGene_gn=PRKCZ;refGene_tx
=XM_023257963.1
C1	991791	991923	
refGene_gn=PRKCZ;refGene_tx=XM_019836008.2;refGene_gn=PRKCZ;refGene_tx
=XM_023257963.1
C1	992886	992965	
refGene_gn=PRKCZ;refGene_tx=XM_019836008.2;refGene_gn=PRKCZ;refGene_tx
=XM_023257963.1
C1	994761	994814	
refGene_gn=PRKCZ;refGene_tx=XM_019836008.2;refGene_gn=PRKCZ;refGene_tx
=XM_023257963.1
C1	997705	997894	
refGene_gn=PRKCZ;refGene_tx=XM_019836008.2;refGene_gn=PRKCZ;refGene_tx
=XM_023257963.1
C1	1000676	1000774
```

refGene_gn=PRKCZ;refGene_tx=XM_019836008.2;refGene_gn=PRKCZ;refGene_tx=XM_023257963.1
C1 1010509 1010596
refGene_gn=PRKCZ;refGene_tx=XM_019836008.2;refGene_gn=PRKCZ;refGene_tx=XM_023257963.1
C1 1012301 1012437
refGene_gn=PRKCZ;refGene_tx=XM_019836008.2;refGene_gn=PRKCZ;refGene_tx=XM_023257963.1
C1 1012639 1012727
refGene_gn=PRKCZ;refGene_tx=XM_019836008.2;refGene_gn=PRKCZ;refGene_tx=XM_023257963.1
C1 1014580 1014700
refGene_gn=PRKCZ;refGene_tx=XM_019836008.2;refGene_gn=PRKCZ;refGene_tx=XM_023257963.1
C1 1015368 1015448
refGene_gn=PRKCZ;refGene_tx=XM_019836008.2;refGene_gn=PRKCZ;refGene_tx=XM_023257963.1
C1 1015856 1015946
refGene_gn=PRKCZ;refGene_tx=XM_019836008.2;refGene_gn=PRKCZ;refGene_tx=XM_023257963.1
C1 1019759 1019875
refGene_gn=PRKCZ;refGene_tx=XM_019836008.2;refGene_gn=PRKCZ;refGene_tx=XM_023257963.1
C1 1020180 1020268
refGene_gn=PRKCZ;refGene_tx=XM_019836008.2;refGene_gn=PRKCZ;refGene_tx=XM_023257963.1
C1 1023616 1023689 refGene_gn=FAAP20;refGene_tx=XM_023257964.1
C1 1027504 1027791 refGene_gn=FAAP20;refGene_tx=XM_023257964.1
C1 1027895 1028025 refGene_gn=FAAP20;refGene_tx=XM_023257964.1
C1 1028728 1028736 refGene_gn=FAAP20;refGene_tx=XM_023257964.1
C1 1030110 1030164
refGene_gn=LOC102901092;refGene_tx=XM_023260046.1
C1 1030776 1032305
refGene_gn=LOC102901092;refGene_tx=XM_023260046.1
C1 1032486 1032627
refGene_gn=LOC102901092;refGene_tx=XM_023260046.1
C1 1033269 1033481
refGene_gn=LOC102901092;refGene_tx=XM_023260046.1
C1 1034324 1034479
refGene_gn=LOC102901092;refGene_tx=XM_023260046.1
C1 1038795 1038928
refGene_gn=LOC102901092;refGene_tx=XM_023260046.1
C1 1041110 1041268
refGene_gn=LOC102901092;refGene_tx=XM_023260046.1
C1 1053176 1054145
refGene_gn=SKI;refGene_tx=XM_023257965.1;refGene_gn=SKI;refGene_tx=XM_023257966.1;refGene_gn=SKI;refGene_tx=XM_023257967.1;refGene_gn=SKI;refGene_tx=XM_023257968.1
C1 1116332 1116455
refGene_gn=SKI;refGene_tx=XM_023257965.1;refGene_gn=SKI;refGene_tx=XM_

023257966.1;refGene_gn=SKI;refGene_tx=XM_023257967.1;refGene_gn=SKI;refGene_tx=XM_023257968.1
C1 1116575 1116694
refGene_gn=SKI;refGene_tx=XM_023257965.1;refGene_gn=SKI;refGene_tx=XM_023257966.1;refGene_gn=SKI;refGene_tx=XM_023257967.1;refGene_gn=SKI;refGene_tx=XM_023257968.1
C1 1117247 1117516
refGene_gn=SKI;refGene_tx=XM_023257965.1;refGene_gn=SKI;refGene_tx=XM_023257966.1;refGene_gn=SKI;refGene_tx=XM_023257967.1;refGene_gn=SKI;refGene_tx=XM_023257968.1
C1 1117619 1117921
refGene_gn=SKI;refGene_tx=XM_023257965.1;refGene_gn=SKI;refGene_tx=XM_023257966.1;refGene_gn=SKI;refGene_tx=XM_023257967.1;refGene_gn=SKI;refGene_tx=XM_023257968.1
C1 1118678 1118909
refGene_gn=SKI;refGene_tx=XM_023257965.1;refGene_gn=SKI;refGene_tx=XM_023257966.1;refGene_gn=SKI;refGene_tx=XM_023257967.1;refGene_gn=SKI;refGene_tx=XM_023257968.1
C1 1119690 1119879
refGene_gn=SKI;refGene_tx=XM_023257965.1;refGene_gn=SKI;refGene_tx=XM_023257966.1;refGene_gn=SKI;refGene_tx=XM_023257967.1;refGene_gn=SKI;refGene_tx=XM_023257968.1
C1 1135265 1135414
refGene_gn=MORN1;refGene_tx=XM_006934228.4;refGene_gn=MORN1;refGene_tx=XM_023257969.1;refGene_gn=MORN1;refGene_tx=XM_023257970.1
C1 1135664 1135799
refGene_gn=MORN1;refGene_tx=XM_006934229.4;refGene_gn=MORN1;refGene_tx=XM_019836022.2
C1 1137261 1137308
refGene_gn=MORN1;refGene_tx=XM_006934228.4;refGene_gn=MORN1;refGene_tx=XM_006934229.4;refGene_gn=MORN1;refGene_tx=XM_019836022.2;refGene_gn=MORN1;refGene_tx=XM_023257969.1;refGene_gn=MORN1;refGene_tx=XM_023257970.1
C1 1147885 1147962
refGene_gn=MORN1;refGene_tx=XM_006934228.4;refGene_gn=MORN1;refGene_tx=XM_006934229.4;refGene_gn=MORN1;refGene_tx=XM_023257969.1;refGene_gn=MORN1;refGene_tx=XM_023257970.1
C1 1148070 1148210
refGene_gn=MORN1;refGene_tx=XM_006934228.4;refGene_gn=MORN1;refGene_tx=XM_006934229.4;refGene_gn=MORN1;refGene_tx=XM_023257969.1;refGene_gn=MORN1;refGene_tx=XM_023257970.1
C1 1158895 1159059
refGene_gn=MORN1;refGene_tx=XM_023257971.1;refGene_gn=MORN1;refGene_tx=XM_023257972.1
C1 1159596 1159769
refGene_gn=MORN1;refGene_tx=XM_006934228.4;refGene_gn=MORN1;refGene_tx=XM_006934229.4;refGene_gn=MORN1;refGene_tx=XM_019836022.2;refGene_gn=MORN1;refGene_tx=XM_023257969.1;refGene_gn=MORN1;refGene_tx=XM_023257970.1;refGene_gn=MORN1;refGene_tx=XM_023257971.1;refGene_gn=MORN1;refGene_tx=XM_023257972.1

C1	1160175	1160299
refGene_gn=MORN1;refGene_tx=XM_006934228.4;refGene_gn=MORN1;refGene_tx=XM_006934229.4;refGene_gn=MORN1;refGene_tx=XM_019836022.2;refGene_gn=MORN1;refGene_tx=XM_023257969.1;refGene_gn=MORN1;refGene_tx=XM_023257970.1;refGene_gn=MORN1;refGene_tx=XM_023257971.1;refGene_gn=MORN1;refGene_tx=XM_023257972.1
C1	1171967	1172078
refGene_gn=MORN1;refGene_tx=XM_006934228.4;refGene_gn=MORN1;refGene_tx=XM_006934229.4;refGene_gn=MORN1;refGene_tx=XM_019836022.2;refGene_gn=MORN1;refGene_tx=XM_023257969.1;refGene_gn=MORN1;refGene_tx=XM_023257970.1;refGene_gn=MORN1;refGene_tx=XM_023257971.1;refGene_gn=MORN1;refGene_tx=XM_023257972.1
C1	1173267	1173364
refGene_gn=MORN1;refGene_tx=XM_006934228.4;refGene_gn=MORN1;refGene_tx=XM_006934229.4;refGene_gn=MORN1;refGene_tx=XM_019836022.2;refGene_gn=MORN1;refGene_tx=XM_023257969.1;refGene_gn=MORN1;refGene_tx=XM_023257970.1;refGene_gn=MORN1;refGene_tx=XM_023257971.1;refGene_gn=MORN1;refGene_tx=XM_023257972.1
C1	1181314	1181402
refGene_gn=MORN1;refGene_tx=XM_006934228.4;refGene_gn=MORN1;refGene_tx=XM_006934229.4;refGene_gn=MORN1;refGene_tx=XM_019836022.2;refGene_gn=MORN1;refGene_tx=XM_023257969.1;refGene_gn=MORN1;refGene_tx=XM_023257970.1;refGene_gn=MORN1;refGene_tx=XM_023257971.1;refGene_gn=MORN1;refGene_tx=XM_023257972.1
C1	1181900	1181991
refGene_gn=MORN1;refGene_tx=XM_006934228.4;refGene_gn=MORN1;refGene_tx=XM_006934229.4;refGene_gn=MORN1;refGene_tx=XM_019836022.2;refGene_gn=MORN1;refGene_tx=XM_023257969.1;refGene_gn=MORN1;refGene_tx=XM_023257970.1;refGene_gn=MORN1;refGene_tx=XM_023257971.1;refGene_gn=MORN1;refGene_tx=XM_023257972.1
C1	1183162	1183273
refGene_gn=MORN1;refGene_tx=XM_006934228.4;refGene_gn=MORN1;refGene_tx=XM_006934229.4;refGene_gn=MORN1;refGene_tx=XM_019836022.2;refGene_gn=MORN1;refGene_tx=XM_023257971.1;refGene_gn=MORN1;refGene_tx=XM_023257972.1
C1	1183799	1183898
refGene_gn=MORN1;refGene_tx=XM_006934228.4;refGene_gn=MORN1;refGene_tx=XM_006934229.4;refGene_gn=MORN1;refGene_tx=XM_019836022.2;refGene_gn=MORN1;refGene_tx=XM_023257969.1;refGene_gn=MORN1;refGene_tx=XM_023257971.1;refGene_gn=MORN1;refGene_tx=XM_023257972.1
C1	1186256	1186328
refGene_gn=MORN1;refGene_tx=XM_006934228.4;refGene_gn=MORN1;refGene_tx=XM_006934229.4;refGene_gn=MORN1;refGene_tx=XM_019836022.2;refGene_gn=MORN1;refGene_tx=XM_023257969.1;refGene_gn=MORN1;refGene_tx=XM_023257970.1;refGene_gn=MORN1;refGene_tx=XM_023257971.1;refGene_gn=MORN1;refGene_tx=XM_023257972.1
C1	1186955	1187028
refGene_gn=MORN1;refGene_tx=XM_006934228.4;refGene_gn=MORN1;refGene_tx=XM_006934229.4;refGene_gn=MORN1;refGene_tx=XM_019836022.2;refGene_gn=MORN1;refGene_tx=XM_023257969.1;refGene_gn=MORN1;refGene_tx=XM_0232579

70.1;refGene_gn=MORN1;refGene_tx=XM_023257971.1;refGene_gn=MORN1;refGene_tx=XM_023257972.1
C1 1190282 1190363
refGene_gn=RER1;refGene_tx=XM_023257973.1;refGene_gn=RER1;refGene_tx=XM_023257974.1
C1 1191573 1191678 refGene_gn=RER1;refGene_tx=XM_023257973.1
C1 1193621 1193721
refGene_gn=RER1;refGene_tx=XM_023257973.1;refGene_gn=RER1;refGene_tx=XM_023257974.1
C1 1195345 1195424
refGene_gn=RER1;refGene_tx=XM_023257973.1;refGene_gn=RER1;refGene_tx=XM_023257974.1
C1 1196449 1196585
refGene_gn=RER1;refGene_tx=XM_023257973.1;refGene_gn=RER1;refGene_tx=XM_023257974.1
C1 1197217 1197307
refGene_gn=RER1;refGene_tx=XM_023257973.1;refGene_gn=RER1;refGene_tx=XM_023257974.1
C1 1199823 1199892 refGene_gn=PEX10;refGene_tx=XM_011284415.2
C1 1201152 1201288 refGene_gn=PEX10;refGene_tx=XM_011284415.2
C1 1201429 1201605 refGene_gn=PEX10;refGene_tx=XM_011284415.2
C1 1202876 1203283 refGene_gn=PEX10;refGene_tx=XM_011284415.2
C1 1203780 1203861 refGene_gn=PEX10;refGene_tx=XM_011284415.2
C1 1204528 1204640 refGene_gn=PEX10;refGene_tx=XM_011284415.2
C1 1218433 1218512 refGene_gn=PLCH2;refGene_tx=XM_023257976.1
C1 1248472 1248515 refGene_gn=PLCH2;refGene_tx=XM_023257977.1
C1 1257287 1257411 refGene_gn=PLCH2;refGene_tx=XM_023257975.1
C1 1259044 1259191
refGene_gn=PLCH2;refGene_tx=XM_023257975.1;refGene_gn=PLCH2;refGene_tx=XM_023257976.1;refGene_gn=PLCH2;refGene_tx=XM_023257977.1
C1 1261153 1261397
refGene_gn=PLCH2;refGene_tx=XM_023257975.1;refGene_gn=PLCH2;refGene_tx=XM_023257976.1;refGene_gn=PLCH2;refGene_tx=XM_023257977.1
C1 1261530 1261660
refGene_gn=PLCH2;refGene_tx=XM_023257975.1;refGene_gn=PLCH2;refGene_tx=XM_023257976.1;refGene_gn=PLCH2;refGene_tx=XM_023257977.1
C1 1265346 1265517
refGene_gn=PLCH2;refGene_tx=XM_023257975.1;refGene_gn=PLCH2;refGene_tx=XM_023257976.1;refGene_gn=PLCH2;refGene_tx=XM_023257977.1
C1 1267705 1267799
refGene_gn=PLCH2;refGene_tx=XM_023257975.1;refGene_gn=PLCH2;refGene_tx=XM_023257976.1;refGene_gn=PLCH2;refGene_tx=XM_023257977.1
C1 1267989 1268193
refGene_gn=PLCH2;refGene_tx=XM_023257975.1;refGene_gn=PLCH2;refGene_tx=XM_023257976.1;refGene_gn=PLCH2;refGene_tx=XM_023257977.1
C1 1268423 1268544
refGene_gn=PLCH2;refGene_tx=XM_023257975.1;refGene_gn=PLCH2;refGene_tx=XM_023257976.1;refGene_gn=PLCH2;refGene_tx=XM_023257977.1
C1 1270304 1270476
refGene_gn=PLCH2;refGene_tx=XM_023257975.1;refGene_gn=PLCH2;refGene_tx

=XM_023257976.1;refGene_gn=PLCH2;refGene_tx=XM_023257977.1
C1 1270869 1270977
refGene_gn=PLCH2;refGene_tx=XM_023257975.1;refGene_gn=PLCH2;refGene_tx
=XM_023257976.1;refGene_gn=PLCH2;refGene_tx=XM_023257977.1
C1 1272132 1272291
refGene_gn=PLCH2;refGene_tx=XM_023257975.1;refGene_gn=PLCH2;refGene_tx
=XM_023257976.1;refGene_gn=PLCH2;refGene_tx=XM_023257977.1
C1 1276411 1276498
refGene_gn=PLCH2;refGene_tx=XM_023257975.1;refGene_gn=PLCH2;refGene_tx
=XM_023257976.1;refGene_gn=PLCH2;refGene_tx=XM_023257977.1
C1 1276797 1276880
refGene_gn=PLCH2;refGene_tx=XM_023257975.1;refGene_gn=PLCH2;refGene_tx
=XM_023257976.1;refGene_gn=PLCH2;refGene_tx=XM_023257977.1
C1 1277246 1277344
refGene_gn=PLCH2;refGene_tx=XM_023257975.1;refGene_gn=PLCH2;refGene_tx
=XM_023257976.1;refGene_gn=PLCH2;refGene_tx=XM_023257977.1
C1 1277436 1277619
refGene_gn=PLCH2;refGene_tx=XM_023257975.1;refGene_gn=PLCH2;refGene_tx
=XM_023257976.1;refGene_gn=PLCH2;refGene_tx=XM_023257977.1
C1 1277981 1278089
refGene_gn=PLCH2;refGene_tx=XM_023257975.1;refGene_gn=PLCH2;refGene_tx
=XM_023257976.1;refGene_gn=PLCH2;refGene_tx=XM_023257977.1
C1 1278428 1278553
refGene_gn=PLCH2;refGene_tx=XM_023257975.1;refGene_gn=PLCH2;refGene_tx
=XM_023257976.1;refGene_gn=PLCH2;refGene_tx=XM_023257977.1
C1 1278632 1278717
refGene_gn=PLCH2;refGene_tx=XM_023257975.1;refGene_gn=PLCH2;refGene_tx
=XM_023257976.1;refGene_gn=PLCH2;refGene_tx=XM_023257977.1
C1 1278887 1279034
refGene_gn=PLCH2;refGene_tx=XM_023257975.1;refGene_gn=PLCH2;refGene_tx
=XM_023257976.1;refGene_gn=PLCH2;refGene_tx=XM_023257977.1
C1 1282330 1282410
refGene_gn=PLCH2;refGene_tx=XM_023257975.1;refGene_gn=PLCH2;refGene_tx
=XM_023257976.1;refGene_gn=PLCH2;refGene_tx=XM_023257977.1
C1 1284554 1284852
refGene_gn=PLCH2;refGene_tx=XM_023257975.1;refGene_gn=PLCH2;refGene_tx
=XM_023257976.1;refGene_gn=PLCH2;refGene_tx=XM_023257977.1
C1 1286394 1287617
refGene_gn=PLCH2;refGene_tx=XM_023257975.1;refGene_gn=PLCH2;refGene_tx
=XM_023257976.1;refGene_gn=PLCH2;refGene_tx=XM_023257977.1
C1 1290257 1290471
refGene_gn=PANK4;refGene_tx=XM_023257978.1;refGene_gn=PANK4;refGene_tx
=XM_023257979.1;refGene_gn=PANK4;refGene_tx=XM_023257980.1;refGene_gn=
PANK4;refGene_tx=XM_023257981.1
C1 1291031 1291100
refGene_gn=PANK4;refGene_tx=XM_023257978.1;refGene_gn=PANK4;refGene_tx
=XM_023257979.1;refGene_gn=PANK4;refGene_tx=XM_023257980.1;refGene_gn=
PANK4;refGene_tx=XM_023257981.1
C1 1291194 1291295
refGene_gn=PANK4;refGene_tx=XM_023257978.1;refGene_gn=PANK4;refGene_tx

=XM_023257979.1;refGene_gn=PANK4;refGene_tx=XM_023257980.1;refGene_gn=PANK4;refGene_tx=XM_023257981.1
C1	1291600	1291705
refGene_gn=PANK4;refGene_tx=XM_023257978.1;refGene_gn=PANK4;refGene_tx=XM_023257979.1;refGene_gn=PANK4;refGene_tx=XM_023257980.1;refGene_gn=PANK4;refGene_tx=XM_023257981.1
C1	1292071	1292121
refGene_gn=PANK4;refGene_tx=XM_023257978.1;refGene_gn=PANK4;refGene_tx=XM_023257979.1;refGene_gn=PANK4;refGene_tx=XM_023257980.1;refGene_gn=PANK4;refGene_tx=XM_023257981.1
C1	1292195	1292251
refGene_gn=PANK4;refGene_tx=XM_023257978.1;refGene_gn=PANK4;refGene_tx=XM_023257979.1;refGene_gn=PANK4;refGene_tx=XM_023257980.1;refGene_gn=PANK4;refGene_tx=XM_023257981.1
C1	1293127	1293279
refGene_gn=PANK4;refGene_tx=XM_023257978.1;refGene_gn=PANK4;refGene_tx=XM_023257979.1;refGene_gn=PANK4;refGene_tx=XM_023257980.1;refGene_gn=PANK4;refGene_tx=XM_023257981.1
C1	1293901	1293989
refGene_gn=PANK4;refGene_tx=XM_023257978.1;refGene_gn=PANK4;refGene_tx=XM_023257979.1;refGene_gn=PANK4;refGene_tx=XM_023257980.1;refGene_gn=PANK4;refGene_tx=XM_023257981.1
C1	1294241	1294354
refGene_gn=PANK4;refGene_tx=XM_023257978.1;refGene_gn=PANK4;refGene_tx=XM_023257979.1;refGene_gn=PANK4;refGene_tx=XM_023257980.1;refGene_gn=PANK4;refGene_tx=XM_023257981.1
C1	1294935	1295091
refGene_gn=PANK4;refGene_tx=XM_023257978.1;refGene_gn=PANK4;refGene_tx=XM_023257979.1;refGene_gn=PANK4;refGene_tx=XM_023257980.1;refGene_gn=PANK4;refGene_tx=XM_023257981.1
C1	1297155	1297256
refGene_gn=PANK4;refGene_tx=XM_023257978.1;refGene_gn=PANK4;refGene_tx=XM_023257979.1;refGene_gn=PANK4;refGene_tx=XM_023257980.1;refGene_gn=PANK4;refGene_tx=XM_023257981.1
C1	1297852	1297934
refGene_gn=PANK4;refGene_tx=XM_023257978.1;refGene_gn=PANK4;refGene_tx=XM_023257979.1;refGene_gn=PANK4;refGene_tx=XM_023257980.1;refGene_gn=PANK4;refGene_tx=XM_023257981.1
C1	1298452	1298634
refGene_gn=PANK4;refGene_tx=XM_023257978.1;refGene_gn=PANK4;refGene_tx=XM_023257979.1;refGene_gn=PANK4;refGene_tx=XM_023257980.1;refGene_gn=PANK4;refGene_tx=XM_023257981.1
C1	1298734	1298888
refGene_gn=PANK4;refGene_tx=XM_023257978.1;refGene_gn=PANK4;refGene_tx=XM_023257979.1;refGene_gn=PANK4;refGene_tx=XM_023257980.1;refGene_gn=PANK4;refGene_tx=XM_023257981.1
C1	1299242	1299335
refGene_gn=PANK4;refGene_tx=XM_023257978.1;refGene_gn=PANK4;refGene_tx=XM_023257979.1;refGene_gn=PANK4;refGene_tx=XM_023257980.1;refGene_gn=PANK4;refGene_tx=XM_023257981.1

C1	1299532	1299716
refGene_gn=PANK4;refGene_tx=XM_023257978.1;refGene_gn=PANK4;refGene_tx=XM_023257979.1;refGene_gn=PANK4;refGene_tx=XM_023257980.1;refGene_gn=PANK4;refGene_tx=XM_023257981.1
C1	1300181	1300396
refGene_gn=PANK4;refGene_tx=XM_023257978.1;refGene_gn=PANK4;refGene_tx=XM_023257979.1;refGene_gn=PANK4;refGene_tx=XM_023257980.1
C1	1301364	1301447
refGene_gn=PANK4;refGene_tx=XM_023257978.1;refGene_gn=PANK4;refGene_tx=XM_023257979.1;refGene_gn=PANK4;refGene_tx=XM_023257980.1
C1	1303745	1303863	refGene_gn=PANK4;refGene_tx=XM_023257979.1
C1	1305407	1305531
refGene_gn=PANK4;refGene_tx=XM_023257978.1;refGene_gn=PANK4;refGene_tx=XM_023257980.1
C1	1308246	1308527	refGene_gn=HES5;refGene_tx=XM_023257982.1
C1	1308755	1308921	refGene_gn=HES5;refGene_tx=XM_023257982.1
C1	1309006	1309060	refGene_gn=HES5;refGene_tx=XM_023257982.1
C1	1333213	1333222
refGene_gn=TNFRSF14;refGene_tx=XM_006934231.4;refGene_gn=TNFRSF14;refGene_tx=XM_006934232.4;refGene_gn=TNFRSF14;refGene_tx=XM_006934237.4
C1	1333938	1334047
refGene_gn=TNFRSF14;refGene_tx=XM_006934231.4;refGene_gn=TNFRSF14;refGene_tx=XM_006934232.4;refGene_gn=TNFRSF14;refGene_tx=XM_006934237.4
C1	1334441	1334567
refGene_gn=TNFRSF14;refGene_tx=XM_006934231.4;refGene_gn=TNFRSF14;refGene_tx=XM_006934232.4;refGene_gn=TNFRSF14;refGene_tx=XM_006934237.4
C1	1335726	1335882
refGene_gn=TNFRSF14;refGene_tx=XM_006934231.4;refGene_gn=TNFRSF14;refGene_tx=XM_006934232.4;refGene_gn=TNFRSF14;refGene_tx=XM_006934237.4
C1	1336033	1336124
refGene_gn=TNFRSF14;refGene_tx=XM_006934231.4;refGene_gn=TNFRSF14;refGene_tx=XM_006934232.4;refGene_gn=TNFRSF14;refGene_tx=XM_006934237.4
C1	1336227	1336349	refGene_gn=TNFRSF14;refGene_tx=XM_006934237.4
C1	1336912	1337091
refGene_gn=TNFRSF14;refGene_tx=XM_006934232.4;refGene_gn=TNFRSF14;refGene_tx=XM_006934237.4
C1	1337782	1337811
refGene_gn=TNFRSF14;refGene_tx=XM_006934231.4;refGene_gn=TNFRSF14;refGene_tx=XM_006934232.4
C1	1338038	1338340
refGene_gn=TNFRSF14;refGene_tx=XM_006934231.4;refGene_gn=TNFRSF14;refGene_tx=XM_006934232.4
C1	1347356	1347419
refGene_gn=FAM213B;refGene_tx=XM_023257983.1;refGene_gn=FAM213B;refGene_tx=XM_023257984.1
C1	1347607	1347812
refGene_gn=FAM213B;refGene_tx=XM_023257983.1;refGene_gn=FAM213B;refGene_tx=XM_023257984.1
C1	1348386	1348438
refGene_gn=FAM213B;refGene_tx=XM_023257983.1;refGene_gn=FAM213B;refGen

e_tx=XM_023257984.1
C1 1348934 1348998
refGene_gn=FAM213B;refGene_tx=XM_023257983.1;refGene_gn=FAM213B;refGen
e_tx=XM_023257984.1
C1 1349082 1349158
refGene_gn=FAM213B;refGene_tx=XM_023257983.1;refGene_gn=FAM213B;refGen
e_tx=XM_023257984.1
C1 1349400 1349557
refGene_gn=FAM213B;refGene_tx=XM_023257983.1;refGene_gn=FAM213B;refGen
e_tx=XM_023257984.1
C1 1350004 1350031 refGene_gn=FAM213B;refGene_tx=XM_023257983.1
C1 1351501 1351601
refGene_gn=MMEL1;refGene_tx=XM_023257985.1;refGene_gn=MMEL1;refGene_tx
=XM_023257986.1;refGene_gn=MMEL1;refGene_tx=XM_023257987.1;refGene_gn=
MMEL1;refGene_tx=XM_023257988.1;refGene_gn=MMEL1;refGene_tx=XM_0232579
89.1;refGene_gn=MMEL1;refGene_tx=XM_023257990.1
C1 1351715 1351792
refGene_gn=MMEL1;refGene_tx=XM_023257985.1;refGene_gn=MMEL1;refGene_tx
=XM_023257986.1;refGene_gn=MMEL1;refGene_tx=XM_023257987.1;refGene_gn=
MMEL1;refGene_tx=XM_023257988.1;refGene_gn=MMEL1;refGene_tx=XM_0232579
89.1;refGene_gn=MMEL1;refGene_tx=XM_023257990.1
C1 1352231 1352327
refGene_gn=MMEL1;refGene_tx=XM_023257985.1;refGene_gn=MMEL1;refGene_tx
=XM_023257986.1;refGene_gn=MMEL1;refGene_tx=XM_023257987.1;refGene_gn=
MMEL1;refGene_tx=XM_023257988.1;refGene_gn=MMEL1;refGene_tx=XM_0232579
89.1;refGene_gn=MMEL1;refGene_tx=XM_023257990.1
C1 1352938 1353004
refGene_gn=MMEL1;refGene_tx=XM_023257985.1;refGene_gn=MMEL1;refGene_tx
=XM_023257986.1;refGene_gn=MMEL1;refGene_tx=XM_023257987.1;refGene_gn=
MMEL1;refGene_tx=XM_023257988.1;refGene_gn=MMEL1;refGene_tx=XM_0232579
89.1;refGene_gn=MMEL1;refGene_tx=XM_023257990.1
C1 1353115 1353249
refGene_gn=MMEL1;refGene_tx=XM_023257985.1;refGene_gn=MMEL1;refGene_tx
=XM_023257986.1;refGene_gn=MMEL1;refGene_tx=XM_023257987.1;refGene_gn=
MMEL1;refGene_tx=XM_023257988.1;refGene_gn=MMEL1;refGene_tx=XM_0232579
89.1;refGene_gn=MMEL1;refGene_tx=XM_023257990.1
C1 1354103 1354223
refGene_gn=MMEL1;refGene_tx=XM_023257985.1;refGene_gn=MMEL1;refGene_tx
=XM_023257986.1;refGene_gn=MMEL1;refGene_tx=XM_023257987.1;refGene_gn=
MMEL1;refGene_tx=XM_023257988.1;refGene_gn=MMEL1;refGene_tx=XM_0232579
89.1;refGene_gn=MMEL1;refGene_tx=XM_023257990.1
C1 1354684 1354743
refGene_gn=MMEL1;refGene_tx=XM_023257985.1;refGene_gn=MMEL1;refGene_tx
=XM_023257986.1;refGene_gn=MMEL1;refGene_tx=XM_023257987.1;refGene_gn=
MMEL1;refGene_tx=XM_023257988.1;refGene_gn=MMEL1;refGene_tx=XM_0232579
89.1;refGene_gn=MMEL1;refGene_tx=XM_023257990.1
C1 1354990 1355094
refGene_gn=MMEL1;refGene_tx=XM_023257985.1;refGene_gn=MMEL1;refGene_tx
=XM_023257986.1;refGene_gn=MMEL1;refGene_tx=XM_023257987.1;refGene_gn=
MMEL1;refGene_tx=XM_023257988.1;refGene_gn=MMEL1;refGene_tx=XM_0232579

89.1;refGene_gn=MMEL1;refGene_tx=XM_023257990.1
C1 1355470 1355554
refGene_gn=MMEL1;refGene_tx=XM_023257985.1;refGene_gn=MMEL1;refGene_tx=XM_023257986.1;refGene_gn=MMEL1;refGene_tx=XM_023257987.1;refGene_gn=MMEL1;refGene_tx=XM_023257988.1;refGene_gn=MMEL1;refGene_tx=XM_023257989.1;refGene_gn=MMEL1;refGene_tx=XM_023257990.1
C1 1356009 1356108
refGene_gn=MMEL1;refGene_tx=XM_023257985.1;refGene_gn=MMEL1;refGene_tx=XM_023257986.1;refGene_gn=MMEL1;refGene_tx=XM_023257987.1;refGene_gn=MMEL1;refGene_tx=XM_023257988.1;refGene_gn=MMEL1;refGene_tx=XM_023257989.1;refGene_gn=MMEL1;refGene_tx=XM_023257990.1
C1 1356565 1356694
refGene_gn=MMEL1;refGene_tx=XM_023257985.1;refGene_gn=MMEL1;refGene_tx=XM_023257986.1;refGene_gn=MMEL1;refGene_tx=XM_023257987.1;refGene_gn=MMEL1;refGene_tx=XM_023257988.1;refGene_gn=MMEL1;refGene_tx=XM_023257989.1;refGene_gn=MMEL1;refGene_tx=XM_023257990.1
C1 1357929 1358023
refGene_gn=MMEL1;refGene_tx=XM_023257985.1;refGene_gn=MMEL1;refGene_tx=XM_023257986.1;refGene_gn=MMEL1;refGene_tx=XM_023257987.1;refGene_gn=MMEL1;refGene_tx=XM_023257988.1;refGene_gn=MMEL1;refGene_tx=XM_023257989.1;refGene_gn=MMEL1;refGene_tx=XM_023257990.1
C1 1358374 1358511
refGene_gn=MMEL1;refGene_tx=XM_023257985.1;refGene_gn=MMEL1;refGene_tx=XM_023257986.1;refGene_gn=MMEL1;refGene_tx=XM_023257987.1;refGene_gn=MMEL1;refGene_tx=XM_023257988.1;refGene_gn=MMEL1;refGene_tx=XM_023257989.1;refGene_gn=MMEL1;refGene_tx=XM_023257990.1
C1 1361939 1362029
refGene_gn=MMEL1;refGene_tx=XM_023257985.1;refGene_gn=MMEL1;refGene_tx=XM_023257986.1;refGene_gn=MMEL1;refGene_tx=XM_023257987.1;refGene_gn=MMEL1;refGene_tx=XM_023257988.1;refGene_gn=MMEL1;refGene_tx=XM_023257989.1;refGene_gn=MMEL1;refGene_tx=XM_023257990.1
C1 1362121 1362256
refGene_gn=MMEL1;refGene_tx=XM_023257985.1;refGene_gn=MMEL1;refGene_tx=XM_023257986.1;refGene_gn=MMEL1;refGene_tx=XM_023257987.1;refGene_gn=MMEL1;refGene_tx=XM_023257988.1;refGene_gn=MMEL1;refGene_tx=XM_023257989.1;refGene_gn=MMEL1;refGene_tx=XM_023257990.1
C1 1363508 1363574
refGene_gn=MMEL1;refGene_tx=XM_023257985.1;refGene_gn=MMEL1;refGene_tx=XM_023257986.1;refGene_gn=MMEL1;refGene_tx=XM_023257987.1;refGene_gn=MMEL1;refGene_tx=XM_023257988.1;refGene_gn=MMEL1;refGene_tx=XM_023257989.1;refGene_gn=MMEL1;refGene_tx=XM_023257990.1
C1 1364076 1364220
refGene_gn=MMEL1;refGene_tx=XM_023257985.1;refGene_gn=MMEL1;refGene_tx=XM_023257986.1;refGene_gn=MMEL1;refGene_tx=XM_023257987.1;refGene_gn=MMEL1;refGene_tx=XM_023257988.1;refGene_gn=MMEL1;refGene_tx=XM_023257989.1;refGene_gn=MMEL1;refGene_tx=XM_023257990.1
C1 1364619 1364715
refGene_gn=MMEL1;refGene_tx=XM_023257985.1;refGene_gn=MMEL1;refGene_tx=XM_023257986.1;refGene_gn=MMEL1;refGene_tx=XM_023257987.1;refGene_gn=MMEL1;refGene_tx=XM_023257988.1;refGene_gn=MMEL1;refGene_tx=XM_0232579

89.1;refGene_gn=MMEL1;refGene_tx=XM_023257990.1
C1	1365773	1365854
refGene_gn=MMEL1;refGene_tx=XM_023257985.1;refGene_gn=MMEL1;refGene_tx=XM_023257986.1;refGene_gn=MMEL1;refGene_tx=XM_023257987.1;refGene_gn=MMEL1;refGene_tx=XM_023257988.1;refGene_gn=MMEL1;refGene_tx=XM_023257989.1;refGene_gn=MMEL1;refGene_tx=XM_023257990.1
C1	1366079	1366241
refGene_gn=MMEL1;refGene_tx=XM_023257985.1;refGene_gn=MMEL1;refGene_tx=XM_023257986.1;refGene_gn=MMEL1;refGene_tx=XM_023257987.1;refGene_gn=MMEL1;refGene_tx=XM_023257988.1;refGene_gn=MMEL1;refGene_tx=XM_023257989.1;refGene_gn=MMEL1;refGene_tx=XM_023257990.1
C1	1367179	1367239
refGene_gn=MMEL1;refGene_tx=XM_023257985.1;refGene_gn=MMEL1;refGene_tx=XM_023257986.1;refGene_gn=MMEL1;refGene_tx=XM_023257987.1;refGene_gn=MMEL1;refGene_tx=XM_023257988.1;refGene_gn=MMEL1;refGene_tx=XM_023257989.1;refGene_gn=MMEL1;refGene_tx=XM_023257990.1
C1	1367840	1367853	refGene_gn=MMEL1;refGene_tx=XM_023257990.1
C1	1367878	1367956
refGene_gn=MMEL1;refGene_tx=XM_023257985.1;refGene_gn=MMEL1;refGene_tx=XM_023257986.1;refGene_gn=MMEL1;refGene_tx=XM_023257987.1;refGene_gn=MMEL1;refGene_tx=XM_023257988.1
C1	1380219	1380376
refGene_gn=MMEL1;refGene_tx=XM_023257985.1;refGene_gn=MMEL1;refGene_tx=XM_023257986.1;refGene_gn=MMEL1;refGene_tx=XM_023257987.1;refGene_gn=MMEL1;refGene_tx=XM_023257988.1;refGene_gn=MMEL1;refGene_tx=XM_023257989.1
C1	1386748	1387276	refGene_gn=TTC34;refGene_tx=XM_023260047.1
C1	1390195	1390410	refGene_gn=TTC34;refGene_tx=XM_023260047.1
C1	1391115	1391386	refGene_gn=TTC34;refGene_tx=XM_023260047.1
C1	1392391	1392558	refGene_gn=TTC34;refGene_tx=XM_023260047.1
C1	1393773	1393978	refGene_gn=TTC34;refGene_tx=XM_023260047.1
C1	1395068	1395567	refGene_gn=TTC34;refGene_tx=XM_023260047.1
C1	1396274	1397118	refGene_gn=TTC34;refGene_tx=XM_023260047.1
C1	1402365	1403146	refGene_gn=TTC34;refGene_tx=XM_023260047.1
C1	1553358	1553508
refGene_gn=LOC109502033;refGene_tx=XM_023257991.1
C1	1553640	1554270
refGene_gn=LOC109502033;refGene_tx=XM_023257991.1
C1	1567693	1568827	refGene_gn=ACTRT2;refGene_tx=XM_003989440.5
C1	1604023	1605016
refGene_gn=LOC111561665;refGene_tx=XM_023257997.1
C1	1610453	1610490
refGene_gn=PRDM16;refGene_tx=XM_023257992.1;refGene_gn=PRDM16;refGene_tx=XM_023257993.1;refGene_gn=PRDM16;refGene_tx=XM_023257994.1;refGene_gn=PRDM16;refGene_tx=XM_023257995.1;refGene_gn=PRDM16;refGene_tx=XM_023257996.1
C1	1711395	1711745
refGene_gn=PRDM16;refGene_tx=XM_023257992.1;refGene_gn=PRDM16;refGene_tx=XM_023257993.1;refGene_gn=PRDM16;refGene_tx=XM_023257994.1;refGene_gn=PRDM16;refGene_tx=XM_023257995.1;refGene_gn=PRDM16;refGene_tx=XM_02

3257996.1
C1	1763514	1763565
refGene_gn=PRDM16;refGene_tx=XM_023257992.1;refGene_gn=PRDM16;refGene_tx=XM_023257993.1;refGene_gn=PRDM16;refGene_tx=XM_023257994.1;refGene_gn=PRDM16;refGene_tx=XM_023257995.1;refGene_gn=PRDM16;refGene_tx=XM_023257996.1
C1	1875275	1875410
refGene_gn=PRDM16;refGene_tx=XM_023257992.1;refGene_gn=PRDM16;refGene_tx=XM_023257993.1;refGene_gn=PRDM16;refGene_tx=XM_023257994.1;refGene_gn=PRDM16;refGene_tx=XM_023257995.1;refGene_gn=PRDM16;refGene_tx=XM_023257996.1
C1	1884120	1884223
refGene_gn=PRDM16;refGene_tx=XM_023257992.1;refGene_gn=PRDM16;refGene_tx=XM_023257993.1;refGene_gn=PRDM16;refGene_tx=XM_023257994.1;refGene_gn=PRDM16;refGene_tx=XM_023257995.1;refGene_gn=PRDM16;refGene_tx=XM_023257996.1
C1	1890210	1890424
refGene_gn=PRDM16;refGene_tx=XM_023257992.1;refGene_gn=PRDM16;refGene_tx=XM_023257993.1;refGene_gn=PRDM16;refGene_tx=XM_023257994.1;refGene_gn=PRDM16;refGene_tx=XM_023257995.1;refGene_gn=PRDM16;refGene_tx=XM_023257996.1
C1	1892301	1892449
refGene_gn=PRDM16;refGene_tx=XM_023257992.1;refGene_gn=PRDM16;refGene_tx=XM_023257993.1;refGene_gn=PRDM16;refGene_tx=XM_023257994.1;refGene_gn=PRDM16;refGene_tx=XM_023257995.1;refGene_gn=PRDM16;refGene_tx=XM_023257996.1
C1	1892809	1892963
refGene_gn=PRDM16;refGene_tx=XM_023257992.1;refGene_gn=PRDM16;refGene_tx=XM_023257993.1;refGene_gn=PRDM16;refGene_tx=XM_023257994.1;refGene_gn=PRDM16;refGene_tx=XM_023257995.1;refGene_gn=PRDM16;refGene_tx=XM_023257996.1
C1	1899002	1900419
refGene_gn=PRDM16;refGene_tx=XM_023257992.1;refGene_gn=PRDM16;refGene_tx=XM_023257993.1;refGene_gn=PRDM16;refGene_tx=XM_023257994.1;refGene_gn=PRDM16;refGene_tx=XM_023257995.1;refGene_gn=PRDM16;refGene_tx=XM_023257996.1
C1	1903125	1903213
refGene_gn=PRDM16;refGene_tx=XM_023257992.1;refGene_gn=PRDM16;refGene_tx=XM_023257993.1;refGene_gn=PRDM16;refGene_tx=XM_023257994.1;refGene_gn=PRDM16;refGene_tx=XM_023257995.1;refGene_gn=PRDM16;refGene_tx=XM_023257996.1
C1	1905651	1905821
refGene_gn=PRDM16;refGene_tx=XM_023257992.1;refGene_gn=PRDM16;refGene_tx=XM_023257993.1;refGene_gn=PRDM16;refGene_tx=XM_023257994.1;refGene_gn=PRDM16;refGene_tx=XM_023257995.1;refGene_gn=PRDM16;refGene_tx=XM_023257996.1
C1	1906421	1906499
refGene_gn=PRDM16;refGene_tx=XM_023257992.1;refGene_gn=PRDM16;refGene_tx=XM_023257993.1;refGene_gn=PRDM16;refGene_tx=XM_023257994.1;refGene_gn=PRDM16;refGene_tx=XM_023257995.1;refGene_gn=PRDM16;refGene_tx=XM_02

3257996.1
C1      1911963  1912133
refGene_gn=PRDM16;refGene_tx=XM_023257992.1;refGene_gn=PRDM16;refGene_tx=XM_023257993.1;refGene_gn=PRDM16;refGene_tx=XM_023257994.1;refGene_gn=PRDM16;refGene_tx=XM_023257995.1;refGene_gn=PRDM16;refGene_tx=XM_023257996.1
C1      1912594  1912769
refGene_gn=PRDM16;refGene_tx=XM_023257992.1;refGene_gn=PRDM16;refGene_tx=XM_023257993.1;refGene_gn=PRDM16;refGene_tx=XM_023257994.1;refGene_gn=PRDM16;refGene_tx=XM_023257995.1;refGene_gn=PRDM16;refGene_tx=XM_023257996.1
C1      1916644  1916878
refGene_gn=PRDM16;refGene_tx=XM_023257992.1;refGene_gn=PRDM16;refGene_tx=XM_023257993.1;refGene_gn=PRDM16;refGene_tx=XM_023257994.1;refGene_gn=PRDM16;refGene_tx=XM_023257995.1;refGene_gn=PRDM16;refGene_tx=XM_023257996.1
C1      1917705  1917880
refGene_gn=PRDM16;refGene_tx=XM_023257992.1;refGene_gn=PRDM16;refGene_tx=XM_023257993.1;refGene_gn=PRDM16;refGene_tx=XM_023257994.1;refGene_gn=PRDM16;refGene_tx=XM_023257995.1
C1      1919085  1919220
refGene_gn=PRDM16;refGene_tx=XM_023257992.1;refGene_gn=PRDM16;refGene_tx=XM_023257993.1;refGene_gn=PRDM16;refGene_tx=XM_023257994.1;refGene_gn=PRDM16;refGene_tx=XM_023257995.1;refGene_gn=PRDM16;refGene_tx=XM_023257996.1
C1      1944047  1944632
refGene_gn=ARHGEF16;refGene_tx=XM_011284466.3;refGene_gn=ARHGEF16;refGene_tx=XM_023257998.1
C1      1946494  1946540
refGene_gn=ARHGEF16;refGene_tx=XM_011284466.3;refGene_gn=ARHGEF16;refGene_tx=XM_023257998.1
C1      1947664  1947834
refGene_gn=ARHGEF16;refGene_tx=XM_011284466.3;refGene_gn=ARHGEF16;refGene_tx=XM_023257998.1
C1      1949011  1949068
refGene_gn=ARHGEF16;refGene_tx=XM_011284466.3;refGene_gn=ARHGEF16;refGene_tx=XM_023257998.1
C1      1949430  1949591
refGene_gn=ARHGEF16;refGene_tx=XM_011284466.3;refGene_gn=ARHGEF16;refGene_tx=XM_023257998.1
C1      1951155  1951308
refGene_gn=ARHGEF16;refGene_tx=XM_011284466.3;refGene_gn=ARHGEF16;refGene_tx=XM_023257998.1
C1      1951430  1951560
refGene_gn=ARHGEF16;refGene_tx=XM_011284466.3;refGene_gn=ARHGEF16;refGene_tx=XM_023257998.1
C1      1954490  1954565
refGene_gn=ARHGEF16;refGene_tx=XM_011284466.3;refGene_gn=ARHGEF16;refGene_tx=XM_023257998.1
C1      1954825  1954918

refGene_gn=ARHGEF16;refGene_tx=XM_011284466.3;refGene_gn=ARHGEF16;refGene_tx=XM_023257998.1
C1	1955991	1956143
refGene_gn=ARHGEF16;refGene_tx=XM_011284466.3;refGene_gn=ARHGEF16;refGene_tx=XM_023257998.1
C1	1956515	1956707
refGene_gn=ARHGEF16;refGene_tx=XM_011284466.3;refGene_gn=ARHGEF16;refGene_tx=XM_023257998.1
C1	1957405	1957473
refGene_gn=ARHGEF16;refGene_tx=XM_011284466.3;refGene_gn=ARHGEF16;refGene_tx=XM_023257998.1
C1	1957926	1958028
refGene_gn=ARHGEF16;refGene_tx=XM_011284466.3;refGene_gn=ARHGEF16;refGene_tx=XM_023257998.1
C1	1958518	1958658
refGene_gn=ARHGEF16;refGene_tx=XM_011284466.3;refGene_gn=ARHGEF16;refGene_tx=XM_023257998.1
C1	1965832	1965897
refGene_gn=MEGF6;refGene_tx=XM_023257999.1;refGene_gn=MEGF6;refGene_tx=XM_023258000.1;refGene_gn=MEGF6;refGene_tx=XM_023258001.1;refGene_gn=MEGF6;refGene_tx=XM_023258002.1
C1	1967406	1967535
refGene_gn=MEGF6;refGene_tx=XM_023257999.1;refGene_gn=MEGF6;refGene_tx=XM_023258000.1;refGene_gn=MEGF6;refGene_tx=XM_023258001.1;refGene_gn=MEGF6;refGene_tx=XM_023258002.1
C1	1968087	1968219
refGene_gn=MEGF6;refGene_tx=XM_023257999.1;refGene_gn=MEGF6;refGene_tx=XM_023258000.1;refGene_gn=MEGF6;refGene_tx=XM_023258001.1;refGene_gn=MEGF6;refGene_tx=XM_023258002.1
C1	1968307	1968436
refGene_gn=MEGF6;refGene_tx=XM_023257999.1;refGene_gn=MEGF6;refGene_tx=XM_023258000.1;refGene_gn=MEGF6;refGene_tx=XM_023258001.1;refGene_gn=MEGF6;refGene_tx=XM_023258002.1
C1	1968749	1968878
refGene_gn=MEGF6;refGene_tx=XM_023257999.1;refGene_gn=MEGF6;refGene_tx=XM_023258000.1;refGene_gn=MEGF6;refGene_tx=XM_023258001.1;refGene_gn=MEGF6;refGene_tx=XM_023258002.1
C1	1969056	1969185
refGene_gn=MEGF6;refGene_tx=XM_023257999.1;refGene_gn=MEGF6;refGene_tx=XM_023258000.1;refGene_gn=MEGF6;refGene_tx=XM_023258001.1;refGene_gn=MEGF6;refGene_tx=XM_023258002.1
C1	1970176	1970305
refGene_gn=MEGF6;refGene_tx=XM_023257999.1;refGene_gn=MEGF6;refGene_tx=XM_023258000.1;refGene_gn=MEGF6;refGene_tx=XM_023258001.1;refGene_gn=MEGF6;refGene_tx=XM_023258002.1
C1	1970895	1971024
refGene_gn=MEGF6;refGene_tx=XM_023257999.1;refGene_gn=MEGF6;refGene_tx=XM_023258000.1;refGene_gn=MEGF6;refGene_tx=XM_023258001.1;refGene_gn=MEGF6;refGene_tx=XM_023258002.1
C1	1971127	1971259

refGene_gn=MEGF6;refGene_tx=XM_023257999.1;refGene_gn=MEGF6;refGene_tx=XM_023258000.1;refGene_gn=MEGF6;refGene_tx=XM_023258001.1;refGene_gn=MEGF6;refGene_tx=XM_023258002.1
C1 1971410 1971539
refGene_gn=MEGF6;refGene_tx=XM_023257999.1;refGene_gn=MEGF6;refGene_tx=XM_023258000.1;refGene_gn=MEGF6;refGene_tx=XM_023258001.1;refGene_gn=MEGF6;refGene_tx=XM_023258002.1
C1 1971680 1971809
refGene_gn=MEGF6;refGene_tx=XM_023257999.1;refGene_gn=MEGF6;refGene_tx=XM_023258000.1;refGene_gn=MEGF6;refGene_tx=XM_023258001.1;refGene_gn=MEGF6;refGene_tx=XM_023258002.1
C1 1972029 1972158
refGene_gn=MEGF6;refGene_tx=XM_023257999.1;refGene_gn=MEGF6;refGene_tx=XM_023258000.1;refGene_gn=MEGF6;refGene_tx=XM_023258001.1;refGene_gn=MEGF6;refGene_tx=XM_023258002.1
C1 1972423 1972552
refGene_gn=MEGF6;refGene_tx=XM_023257999.1;refGene_gn=MEGF6;refGene_tx=XM_023258000.1;refGene_gn=MEGF6;refGene_tx=XM_023258001.1;refGene_gn=MEGF6;refGene_tx=XM_023258002.1
C1 1972977 1973106
refGene_gn=MEGF6;refGene_tx=XM_023257999.1;refGene_gn=MEGF6;refGene_tx=XM_023258000.1;refGene_gn=MEGF6;refGene_tx=XM_023258001.1;refGene_gn=MEGF6;refGene_tx=XM_023258002.1
C1 1973354 1973483
refGene_gn=MEGF6;refGene_tx=XM_023257999.1;refGene_gn=MEGF6;refGene_tx=XM_023258000.1;refGene_gn=MEGF6;refGene_tx=XM_023258001.1;refGene_gn=MEGF6;refGene_tx=XM_023258002.1
C1 1973560 1973689
refGene_gn=MEGF6;refGene_tx=XM_023257999.1;refGene_gn=MEGF6;refGene_tx=XM_023258000.1;refGene_gn=MEGF6;refGene_tx=XM_023258001.1;refGene_gn=MEGF6;refGene_tx=XM_023258002.1
C1 1974208 1974340
refGene_gn=MEGF6;refGene_tx=XM_023257999.1;refGene_gn=MEGF6;refGene_tx=XM_023258000.1;refGene_gn=MEGF6;refGene_tx=XM_023258001.1;refGene_gn=MEGF6;refGene_tx=XM_023258002.1
C1 1974448 1974577
refGene_gn=MEGF6;refGene_tx=XM_023257999.1;refGene_gn=MEGF6;refGene_tx=XM_023258000.1;refGene_gn=MEGF6;refGene_tx=XM_023258001.1
C1 1974654 1974786
refGene_gn=MEGF6;refGene_tx=XM_023257999.1;refGene_gn=MEGF6;refGene_tx=XM_023258000.1;refGene_gn=MEGF6;refGene_tx=XM_023258001.1;refGene_gn=MEGF6;refGene_tx=XM_023258002.1
C1 1975361 1975487
refGene_gn=MEGF6;refGene_tx=XM_023257999.1;refGene_gn=MEGF6;refGene_tx=XM_023258000.1;refGene_gn=MEGF6;refGene_tx=XM_023258001.1;refGene_gn=MEGF6;refGene_tx=XM_023258002.1
C1 1977983 1978118
refGene_gn=MEGF6;refGene_tx=XM_023257999.1;refGene_gn=MEGF6;refGene_tx=XM_023258000.1;refGene_gn=MEGF6;refGene_tx=XM_023258001.1;refGene_gn=MEGF6;refGene_tx=XM_023258002.1

C1 1978205 1978340
refGene_gn=MEGF6;refGene_tx=XM_023257999.1;refGene_gn=MEGF6;refGene_tx=XM_023258000.1;refGene_gn=MEGF6;refGene_tx=XM_023258001.1;refGene_gn=MEGF6;refGene_tx=XM_023258002.1
C1 1979038 1979167
refGene_gn=MEGF6;refGene_tx=XM_023257999.1;refGene_gn=MEGF6;refGene_tx=XM_023258000.1;refGene_gn=MEGF6;refGene_tx=XM_023258001.1;refGene_gn=MEGF6;refGene_tx=XM_023258002.1
C1 1980523 1980652
refGene_gn=MEGF6;refGene_tx=XM_023257999.1;refGene_gn=MEGF6;refGene_tx=XM_023258000.1;refGene_gn=MEGF6;refGene_tx=XM_023258001.1;refGene_gn=MEGF6;refGene_tx=XM_023258002.1
C1 1981238 1981370
refGene_gn=MEGF6;refGene_tx=XM_023257999.1;refGene_gn=MEGF6;refGene_tx=XM_023258000.1;refGene_gn=MEGF6;refGene_tx=XM_023258001.1;refGene_gn=MEGF6;refGene_tx=XM_023258002.1
C1 1981530 1981707
refGene_gn=MEGF6;refGene_tx=XM_023257999.1;refGene_gn=MEGF6;refGene_tx=XM_023258000.1;refGene_gn=MEGF6;refGene_tx=XM_023258001.1;refGene_gn=MEGF6;refGene_tx=XM_023258002.1
C1 1982189 1982312
refGene_gn=MEGF6;refGene_tx=XM_023257999.1;refGene_gn=MEGF6;refGene_tx=XM_023258000.1;refGene_gn=MEGF6;refGene_tx=XM_023258001.1;refGene_gn=MEGF6;refGene_tx=XM_023258002.1
C1 1982633 1982753
refGene_gn=MEGF6;refGene_tx=XM_023257999.1;refGene_gn=MEGF6;refGene_tx=XM_023258000.1;refGene_gn=MEGF6;refGene_tx=XM_023258001.1;refGene_gn=MEGF6;refGene_tx=XM_023258002.1
C1 1983159 1983297
refGene_gn=MEGF6;refGene_tx=XM_023257999.1;refGene_gn=MEGF6;refGene_tx=XM_023258000.1;refGene_gn=MEGF6;refGene_tx=XM_023258001.1;refGene_gn=MEGF6;refGene_tx=XM_023258002.1
C1 1983433 1983556
refGene_gn=MEGF6;refGene_tx=XM_023257999.1;refGene_gn=MEGF6;refGene_tx=XM_023258000.1;refGene_gn=MEGF6;refGene_tx=XM_023258001.1;refGene_gn=MEGF6;refGene_tx=XM_023258002.1
C1 1984523 1984646
refGene_gn=MEGF6;refGene_tx=XM_023257999.1;refGene_gn=MEGF6;refGene_tx=XM_023258000.1;refGene_gn=MEGF6;refGene_tx=XM_023258001.1;refGene_gn=MEGF6;refGene_tx=XM_023258002.1
C1 1986707 1986833
refGene_gn=MEGF6;refGene_tx=XM_023257999.1;refGene_gn=MEGF6;refGene_tx=XM_023258000.1;refGene_gn=MEGF6;refGene_tx=XM_023258001.1;refGene_gn=MEGF6;refGene_tx=XM_023258002.1
C1 1993486 1993609
refGene_gn=MEGF6;refGene_tx=XM_023257999.1;refGene_gn=MEGF6;refGene_tx=XM_023258000.1;refGene_gn=MEGF6;refGene_tx=XM_023258001.1;refGene_gn=MEGF6;refGene_tx=XM_023258002.1
C1 2033746 2033851
refGene_gn=MEGF6;refGene_tx=XM_023257999.1;refGene_gn=MEGF6;refGene_tx

=XM_023258000.1;refGene_gn=MEGF6;refGene_tx=XM_023258001.1;refGene_gn=MEGF6;refGene_tx=XM_023258002.1
C1	2043460	2043570	refGene_gn=MEGF6;refGene_tx=XM_023257999.1;refGene_gn=MEGF6;refGene_tx=XM_023258000.1;refGene_gn=MEGF6;refGene_tx=XM_023258001.1;refGene_gn=MEGF6;refGene_tx=XM_023258002.1
C1	2047986	2048121	refGene_gn=MEGF6;refGene_tx=XM_023257999.1;refGene_gn=MEGF6;refGene_tx=XM_023258000.1;refGene_gn=MEGF6;refGene_tx=XM_023258001.1;refGene_gn=MEGF6;refGene_tx=XM_023258002.1
C1	2051484	2051594	refGene_gn=MEGF6;refGene_tx=XM_023257999.1;refGene_gn=MEGF6;refGene_tx=XM_023258000.1;refGene_gn=MEGF6;refGene_tx=XM_023258001.1;refGene_gn=MEGF6;refGene_tx=XM_023258002.1
C1	2065930	2066128	refGene_gn=TPRG1L;refGene_tx=XM_011284465.2
C1	2066230	2066322	refGene_gn=TPRG1L;refGene_tx=XM_011284465.2
C1	2066407	2066584	refGene_gn=TPRG1L;refGene_tx=XM_011284465.2
C1	2067388	2067542	refGene_gn=TPRG1L;refGene_tx=XM_011284465.2
C1	2068154	2068349	refGene_gn=TPRG1L;refGene_tx=XM_011284465.2
C1	2070237	2070380	refGene_gn=WRAP73;refGene_tx=XM_011284462.3;refGene_gn=WRAP73;refGene_tx=XM_011284463.3;refGene_gn=WRAP73;refGene_tx=XM_019836036.2
C1	2070726	2070918	refGene_gn=WRAP73;refGene_tx=XM_011284462.3;refGene_gn=WRAP73;refGene_tx=XM_011284463.3;refGene_gn=WRAP73;refGene_tx=XM_019836036.2
C1	2071343	2071469	refGene_gn=WRAP73;refGene_tx=XM_011284462.3;refGene_gn=WRAP73;refGene_tx=XM_011284463.3;refGene_gn=WRAP73;refGene_tx=XM_019836036.2
C1	2072236	2072342	refGene_gn=WRAP73;refGene_tx=XM_011284462.3;refGene_gn=WRAP73;refGene_tx=XM_011284463.3;refGene_gn=WRAP73;refGene_tx=XM_019836036.2
C1	2079933	2080011	refGene_gn=WRAP73;refGene_tx=XM_011284462.3;refGene_gn=WRAP73;refGene_tx=XM_011284463.3;refGene_gn=WRAP73;refGene_tx=XM_019836036.2
C1	2080088	2080223	refGene_gn=WRAP73;refGene_tx=XM_011284462.3;refGene_gn=WRAP73;refGene_tx=XM_011284463.3;refGene_gn=WRAP73;refGene_tx=XM_019836036.2
C1	2080523	2080610	refGene_gn=WRAP73;refGene_tx=XM_011284462.3;refGene_gn=WRAP73;refGene_tx=XM_011284463.3;refGene_gn=WRAP73;refGene_tx=XM_019836036.2
C1	2081341	2081445	refGene_gn=WRAP73;refGene_tx=XM_011284462.3;refGene_gn=WRAP73;refGene_tx=XM_011284463.3;refGene_gn=WRAP73;refGene_tx=XM_019836036.2
C1	2082686	2082759	refGene_gn=WRAP73;refGene_tx=XM_011284462.3;refGene_gn=WRAP73;refGene_tx=XM_011284463.3;refGene_gn=WRAP73;refGene_tx=XM_019836036.2
C1	2088479	2088654	refGene_gn=WRAP73;refGene_tx=XM_011284462.3;refGene_gn=WRAP73;refGene_tx=XM_011284463.3
C1	2089123	2089276

refGene_gn=WRAP73;refGene_tx=XM_011284462.3;refGene_gn=WRAP73;refGene_
tx=XM_011284463.3
C1	2092882	2092951
refGene_gn=WRAP73;refGene_tx=XM_011284462.3;refGene_gn=WRAP73;refGene_
tx=XM_019836036.2
C1	2115361	2115423
refGene_gn=TP73;refGene_tx=XM_003989443.4;refGene_gn=TP73;refGene_tx=X
M_019836037.1;refGene_gn=TP73;refGene_tx=XM_023258003.1
C1	2115751	2115872
refGene_gn=TP73;refGene_tx=XM_003989443.4;refGene_gn=TP73;refGene_tx=X
M_019836037.1;refGene_gn=TP73;refGene_tx=XM_023258003.1
C1	2136633	2136876
refGene_gn=TP73;refGene_tx=XM_003989443.4;refGene_gn=TP73;refGene_tx=X
M_019836037.1;refGene_gn=TP73;refGene_tx=XM_019836041.1;refGene_gn=TP7
3;refGene_tx=XM_023258003.1
C1	2148326	2148513
refGene_gn=TP73;refGene_tx=XM_003989443.4;refGene_gn=TP73;refGene_tx=X
M_019836037.1;refGene_gn=TP73;refGene_tx=XM_019836041.1;refGene_gn=TP7
3;refGene_tx=XM_023258003.1
C1	2148863	2148979
refGene_gn=TP73;refGene_tx=XM_003989443.4;refGene_gn=TP73;refGene_tx=X
M_019836037.1;refGene_gn=TP73;refGene_tx=XM_019836041.1;refGene_gn=TP7
3;refGene_tx=XM_023258003.1
C1	2150748	2150858
refGene_gn=TP73;refGene_tx=XM_003989443.4;refGene_gn=TP73;refGene_tx=X
M_019836037.1;refGene_gn=TP73;refGene_tx=XM_019836041.1;refGene_gn=TP7
3;refGene_tx=XM_023258003.1
C1	2151179	2151322
refGene_gn=TP73;refGene_tx=XM_003989443.4;refGene_gn=TP73;refGene_tx=X
M_019836037.1;refGene_gn=TP73;refGene_tx=XM_019836041.1;refGene_gn=TP7
3;refGene_tx=XM_023258003.1
C1	2151558	2151647
refGene_gn=TP73;refGene_tx=XM_003989443.4;refGene_gn=TP73;refGene_tx=X
M_019836037.1;refGene_gn=TP73;refGene_tx=XM_019836041.1;refGene_gn=TP7
3;refGene_tx=XM_023258003.1
C1	2151919	2152041
refGene_gn=TP73;refGene_tx=XM_003989443.4;refGene_gn=TP73;refGene_tx=X
M_019836037.1;refGene_gn=TP73;refGene_tx=XM_019836041.1;refGene_gn=TP7
3;refGene_tx=XM_023258003.1
C1	2153013	2153162
refGene_gn=TP73;refGene_tx=XM_003989443.4;refGene_gn=TP73;refGene_tx=X
M_019836037.1;refGene_gn=TP73;refGene_tx=XM_019836041.1;refGene_gn=TP7
3;refGene_tx=XM_023258003.1
C1	2153413	2153549
refGene_gn=TP73;refGene_tx=XM_003989443.4;refGene_gn=TP73;refGene_tx=X
M_019836037.1;refGene_gn=TP73;refGene_tx=XM_019836041.1;refGene_gn=TP7
3;refGene_tx=XM_023258003.1
C1	2153875	2153969
refGene_gn=TP73;refGene_tx=XM_003989443.4;refGene_gn=TP73;refGene_tx=X
M_019836037.1;refGene_gn=TP73;refGene_tx=XM_019836041.1;refGene_gn=TP7

3;refGene_tx=XM_023258003.1
C1 2155155 2155482
refGene_gn=TP73;refGene_tx=XM_003989443.4;refGene_gn=TP73;refGene_tx=XM_019836037.1;refGene_gn=TP73;refGene_tx=XM_019836041.1;refGene_gn=TP73;refGene_tx=XM_023258003.1
C1 2169259 2169592
refGene_gn=CCDC27;refGene_tx=XM_019836044.2;refGene_gn=CCDC27;refGene_tx=XM_019836045.2
C1 2169997 2170130
refGene_gn=CCDC27;refGene_tx=XM_019836044.2;refGene_gn=CCDC27;refGene_tx=XM_019836045.2
C1 2171285 2171381
refGene_gn=CCDC27;refGene_tx=XM_019836044.2;refGene_gn=CCDC27;refGene_tx=XM_019836045.2
C1 2171928 2172083
refGene_gn=CCDC27;refGene_tx=XM_019836044.2;refGene_gn=CCDC27;refGene_tx=XM_019836045.2
C1 2174903 2175065
refGene_gn=CCDC27;refGene_tx=XM_019836044.2;refGene_gn=CCDC27;refGene_tx=XM_019836045.2
C1 2175952 2176048
refGene_gn=CCDC27;refGene_tx=XM_019836044.2;refGene_gn=CCDC27;refGene_tx=XM_019836045.2
C1 2176357 2176673
refGene_gn=CCDC27;refGene_tx=XM_019836044.2;refGene_gn=CCDC27;refGene_tx=XM_019836045.2
C1 2176811 2176942
refGene_gn=CCDC27;refGene_tx=XM_019836044.2;refGene_gn=CCDC27;refGene_tx=XM_019836045.2
C1 2178830 2178908
refGene_gn=CCDC27;refGene_tx=XM_019836044.2;refGene_gn=CCDC27;refGene_tx=XM_019836045.2
C1 2179189 2179402
refGene_gn=CCDC27;refGene_tx=XM_019836044.2;refGene_gn=CCDC27;refGene_tx=XM_019836045.2
C1 2181220 2181325
refGene_gn=CCDC27;refGene_tx=XM_019836044.2;refGene_gn=CCDC27;refGene_tx=XM_019836045.2
C1 2182814 2182940
refGene_gn=CCDC27;refGene_tx=XM_019836044.2;refGene_gn=CCDC27;refGene_tx=XM_019836045.2
C1 2186049 2186159
refGene_gn=SMIM1;refGene_tx=XM_023258004.1;refGene_gn=SMIM1;refGene_tx=XM_023258005.1;refGene_gn=SMIM1;refGene_tx=XM_023258006.1
C1 2186340 2186467
refGene_gn=SMIM1;refGene_tx=XM_023258004.1;refGene_gn=SMIM1;refGene_tx=XM_023258005.1;refGene_gn=SMIM1;refGene_tx=XM_023258006.1
C1 2190722 2190971 refGene_gn=LRRC47;refGene_tx=XM_019836048.2
C1 2191126 2191216 refGene_gn=LRRC47;refGene_tx=XM_019836048.2
C1 2191525 2191628 refGene_gn=LRRC47;refGene_tx=XM_019836048.2

C1	2192171	2192287	refGene_gn=LRRC47;refGene_tx=XM_019836048.2
C1	2192819	2192936	refGene_gn=LRRC47;refGene_tx=XM_019836048.2
C1	2193789	2194248	refGene_gn=LRRC47;refGene_tx=XM_019836048.2
C1	2199579	2200206	refGene_gn=LRRC47;refGene_tx=XM_019836048.2
C1	2208821	2208937
refGene_gn=CEP104;refGene_tx=XM_011284457.3;refGene_gn=CEP104;refGene_tx=XM_011284459.3;refGene_gn=CEP104;refGene_tx=XM_011284460.3;refGene_gn=CEP104;refGene_tx=XM_019836049.2;refGene_gn=CEP104;refGene_tx=XM_019836050.2
C1	2209908	2209999
refGene_gn=CEP104;refGene_tx=XM_011284457.3;refGene_gn=CEP104;refGene_tx=XM_011284459.3;refGene_gn=CEP104;refGene_tx=XM_011284460.3;refGene_gn=CEP104;refGene_tx=XM_019836049.2;refGene_gn=CEP104;refGene_tx=XM_019836050.2
C1	2213073	2213141
refGene_gn=CEP104;refGene_tx=XM_011284457.3;refGene_gn=CEP104;refGene_tx=XM_011284459.3;refGene_gn=CEP104;refGene_tx=XM_011284460.3;refGene_gn=CEP104;refGene_tx=XM_019836049.2;refGene_gn=CEP104;refGene_tx=XM_019836050.2
C1	2213294	2213433
refGene_gn=CEP104;refGene_tx=XM_011284457.3;refGene_gn=CEP104;refGene_tx=XM_011284459.3;refGene_gn=CEP104;refGene_tx=XM_011284460.3;refGene_gn=CEP104;refGene_tx=XM_019836049.2;refGene_gn=CEP104;refGene_tx=XM_019836050.2
C1	2215433	2215542
refGene_gn=CEP104;refGene_tx=XM_011284457.3;refGene_gn=CEP104;refGene_tx=XM_011284459.3;refGene_gn=CEP104;refGene_tx=XM_011284460.3;refGene_gn=CEP104;refGene_tx=XM_019836049.2;refGene_gn=CEP104;refGene_tx=XM_019836050.2
C1	2216074	2216141
refGene_gn=CEP104;refGene_tx=XM_011284457.3;refGene_gn=CEP104;refGene_tx=XM_011284459.3;refGene_gn=CEP104;refGene_tx=XM_011284460.3;refGene_gn=CEP104;refGene_tx=XM_019836049.2;refGene_gn=CEP104;refGene_tx=XM_019836050.2
C1	2216326	2216366
refGene_gn=CEP104;refGene_tx=XM_011284457.3;refGene_gn=CEP104;refGene_tx=XM_011284459.3;refGene_gn=CEP104;refGene_tx=XM_011284460.3;refGene_gn=CEP104;refGene_tx=XM_019836049.2;refGene_gn=CEP104;refGene_tx=XM_019836050.2
C1	2218381	2218498
refGene_gn=CEP104;refGene_tx=XM_011284457.3;refGene_gn=CEP104;refGene_tx=XM_011284459.3;refGene_gn=CEP104;refGene_tx=XM_011284460.3;refGene_gn=CEP104;refGene_tx=XM_019836049.2
C1	2218860	2219067
refGene_gn=CEP104;refGene_tx=XM_011284457.3;refGene_gn=CEP104;refGene_tx=XM_011284459.3;refGene_gn=CEP104;refGene_tx=XM_011284460.3;refGene_gn=CEP104;refGene_tx=XM_019836049.2
C1	2221620	2221797
refGene_gn=CEP104;refGene_tx=XM_011284457.3;refGene_gn=CEP104;refGene_tx=XM_011284459.3;refGene_gn=CEP104;refGene_tx=XM_011284460.3;refGene_

gn=CEP104;refGene_tx=XM_019836049.2;refGene_gn=CEP104;refGene_tx=XM_019836050.2
C1	2223360	2223534
refGene_gn=CEP104;refGene_tx=XM_011284457.3;refGene_gn=CEP104;refGene_tx=XM_011284459.3;refGene_gn=CEP104;refGene_tx=XM_011284460.3;refGene_gn=CEP104;refGene_tx=XM_019836049.2;refGene_gn=CEP104;refGene_tx=XM_019836050.2
C1	2224128	2224296
refGene_gn=CEP104;refGene_tx=XM_011284457.3;refGene_gn=CEP104;refGene_tx=XM_011284459.3;refGene_gn=CEP104;refGene_tx=XM_011284460.3;refGene_gn=CEP104;refGene_tx=XM_019836049.2;refGene_gn=CEP104;refGene_tx=XM_019836050.2
C1	2224888	2225083
refGene_gn=CEP104;refGene_tx=XM_011284457.3;refGene_gn=CEP104;refGene_tx=XM_011284459.3;refGene_gn=CEP104;refGene_tx=XM_011284460.3;refGene_gn=CEP104;refGene_tx=XM_019836049.2;refGene_gn=CEP104;refGene_tx=XM_019836050.2
C1	2226812	2227034
refGene_gn=CEP104;refGene_tx=XM_011284457.3;refGene_gn=CEP104;refGene_tx=XM_011284459.3;refGene_gn=CEP104;refGene_tx=XM_011284460.3;refGene_gn=CEP104;refGene_tx=XM_019836049.2;refGene_gn=CEP104;refGene_tx=XM_019836050.2
C1	2228197	2228353
refGene_gn=CEP104;refGene_tx=XM_011284457.3;refGene_gn=CEP104;refGene_tx=XM_011284459.3;refGene_gn=CEP104;refGene_tx=XM_011284460.3;refGene_gn=CEP104;refGene_tx=XM_019836049.2;refGene_gn=CEP104;refGene_tx=XM_019836050.2
C1	2228828	2228997
refGene_gn=CEP104;refGene_tx=XM_011284457.3;refGene_gn=CEP104;refGene_tx=XM_011284459.3;refGene_gn=CEP104;refGene_tx=XM_011284460.3;refGene_gn=CEP104;refGene_tx=XM_019836049.2;refGene_gn=CEP104;refGene_tx=XM_019836050.2
C1	2231866	2231943
refGene_gn=CEP104;refGene_tx=XM_011284457.3;refGene_gn=CEP104;refGene_tx=XM_011284459.3;refGene_gn=CEP104;refGene_tx=XM_011284460.3;refGene_gn=CEP104;refGene_tx=XM_019836049.2;refGene_gn=CEP104;refGene_tx=XM_019836050.2
C1	2232250	2232313
refGene_gn=CEP104;refGene_tx=XM_011284457.3;refGene_gn=CEP104;refGene_tx=XM_011284459.3;refGene_gn=CEP104;refGene_tx=XM_011284460.3;refGene_gn=CEP104;refGene_tx=XM_019836049.2;refGene_gn=CEP104;refGene_tx=XM_019836050.2
C1	2234075	2234214
refGene_gn=CEP104;refGene_tx=XM_011284457.3;refGene_gn=CEP104;refGene_tx=XM_011284459.3;refGene_gn=CEP104;refGene_tx=XM_011284460.3;refGene_gn=CEP104;refGene_tx=XM_019836049.2;refGene_gn=CEP104;refGene_tx=XM_019836050.2
C1	2234605	2234779
refGene_gn=CEP104;refGene_tx=XM_011284457.3;refGene_gn=CEP104;refGene_tx=XM_011284459.3;refGene_gn=CEP104;refGene_tx=XM_011284460.3;refGene_

gn=CEP104;refGene_tx=XM_019836049.2;refGene_gn=CEP104;refGene_tx=XM_019836050.2
C1 2237758 2237871
refGene_gn=CEP104;refGene_tx=XM_011284457.3;refGene_gn=CEP104;refGene_tx=XM_011284459.3;refGene_gn=CEP104;refGene_tx=XM_011284460.3;refGene_gn=CEP104;refGene_tx=XM_019836049.2;refGene_gn=CEP104;refGene_tx=XM_019836050.2
C1 2242256 2242379 refGene_gn=DFFB;refGene_tx=XM_019836051.1
C1 2243336 2243529
refGene_gn=DFFB;refGene_tx=XM_019836051.1;refGene_gn=DFFB;refGene_tx=XM_019836052.1
C1 2245744 2245933
refGene_gn=DFFB;refGene_tx=XM_019836051.1;refGene_gn=DFFB;refGene_tx=XM_019836052.1
C1 2246736 2246816
refGene_gn=DFFB;refGene_tx=XM_019836051.1;refGene_gn=DFFB;refGene_tx=XM_019836052.1
C1 2248624 2248795
refGene_gn=DFFB;refGene_tx=XM_019836051.1;refGene_gn=DFFB;refGene_tx=XM_019836052.1
C1 2249558 2249659
refGene_gn=DFFB;refGene_tx=XM_019836051.1;refGene_gn=DFFB;refGene_tx=XM_019836052.1
C1 2253284 2253519
refGene_gn=DFFB;refGene_tx=XM_019836051.1;refGene_gn=DFFB;refGene_tx=XM_019836052.1
C1 2800833 2800862 refGene_gn=AJAP1;refGene_tx=XM_023258007.1
C1 2857467 2858282 refGene_gn=AJAP1;refGene_tx=XM_023258007.1
C1 2908360 2908448 refGene_gn=AJAP1;refGene_tx=XM_023258007.1
C1 2910563 2910806 refGene_gn=AJAP1;refGene_tx=XM_023258007.1
C1 2912564 2912634 refGene_gn=AJAP1;refGene_tx=XM_023258007.1
C1 3743763 3743904
refGene_gn=NPHP4;refGene_tx=XM_006934240.4;refGene_gn=NPHP4;refGene_tx=XM_019836057.2;refGene_gn=NPHP4;refGene_tx=XM_019836058.2;refGene_gn=NPHP4;refGene_tx=XM_019836060.2;refGene_gn=NPHP4;refGene_tx=XM_019836062.2;refGene_gn=NPHP4;refGene_tx=XM_023258014.1;refGene_gn=NPHP4;refGene_tx=XM_023258015.1;refGene_gn=NPHP4;refGene_tx=XM_023258016.1;refGene_gn=NPHP4;refGene_tx=XM_023258017.1
C1 3744494 3744638
refGene_gn=NPHP4;refGene_tx=XM_006934240.4;refGene_gn=NPHP4;refGene_tx=XM_019836057.2;refGene_gn=NPHP4;refGene_tx=XM_019836058.2;refGene_gn=NPHP4;refGene_tx=XM_019836060.2;refGene_gn=NPHP4;refGene_tx=XM_019836062.2;refGene_gn=NPHP4;refGene_tx=XM_023258014.1;refGene_gn=NPHP4;refGene_tx=XM_023258015.1;refGene_gn=NPHP4;refGene_tx=XM_023258016.1;refGene_gn=NPHP4;refGene_tx=XM_023258017.1
C1 3744877 3745057
refGene_gn=NPHP4;refGene_tx=XM_006934240.4;refGene_gn=NPHP4;refGene_tx=XM_019836057.2;refGene_gn=NPHP4;refGene_tx=XM_019836058.2;refGene_gn=NPHP4;refGene_tx=XM_019836060.2;refGene_gn=NPHP4;refGene_tx=XM_019836062.2;refGene_gn=NPHP4;refGene_tx=XM_023258014.1;refGene_gn=NPHP4;refGe

ne_tx=XM_023258015.1;refGene_gn=NPHP4;refGene_tx=XM_023258016.1;refGene_gn=NPHP4;refGene_tx=XM_023258017.1
C1 3745650 3745822
refGene_gn=NPHP4;refGene_tx=XM_006934240.4;refGene_gn=NPHP4;refGene_tx=XM_019836057.2;refGene_gn=NPHP4;refGene_tx=XM_019836058.2;refGene_gn=NPHP4;refGene_tx=XM_019836060.2;refGene_gn=NPHP4;refGene_tx=XM_019836062.2;refGene_gn=NPHP4;refGene_tx=XM_023258014.1;refGene_gn=NPHP4;refGene_tx=XM_023258015.1;refGene_gn=NPHP4;refGene_tx=XM_023258016.1;refGene_gn=NPHP4;refGene_tx=XM_023258017.1
C1 3746661 3746747
refGene_gn=NPHP4;refGene_tx=XM_006934240.4;refGene_gn=NPHP4;refGene_tx=XM_019836057.2;refGene_gn=NPHP4;refGene_tx=XM_019836058.2;refGene_gn=NPHP4;refGene_tx=XM_019836060.2;refGene_gn=NPHP4;refGene_tx=XM_019836062.2;refGene_gn=NPHP4;refGene_tx=XM_023258014.1;refGene_gn=NPHP4;refGene_tx=XM_023258015.1;refGene_gn=NPHP4;refGene_tx=XM_023258016.1;refGene_gn=NPHP4;refGene_tx=XM_023258017.1
C1 3747303 3747389
refGene_gn=NPHP4;refGene_tx=XM_006934240.4;refGene_gn=NPHP4;refGene_tx=XM_019836057.2;refGene_gn=NPHP4;refGene_tx=XM_019836058.2;refGene_gn=NPHP4;refGene_tx=XM_019836060.2;refGene_gn=NPHP4;refGene_tx=XM_019836062.2;refGene_gn=NPHP4;refGene_tx=XM_023258014.1;refGene_gn=NPHP4;refGene_tx=XM_023258015.1;refGene_gn=NPHP4;refGene_tx=XM_023258016.1;refGene_gn=NPHP4;refGene_tx=XM_023258017.1
C1 3748160 3748317
refGene_gn=NPHP4;refGene_tx=XM_006934240.4;refGene_gn=NPHP4;refGene_tx=XM_019836057.2;refGene_gn=NPHP4;refGene_tx=XM_019836058.2;refGene_gn=NPHP4;refGene_tx=XM_019836060.2;refGene_gn=NPHP4;refGene_tx=XM_019836062.2;refGene_gn=NPHP4;refGene_tx=XM_023258014.1;refGene_gn=NPHP4;refGene_tx=XM_023258015.1;refGene_gn=NPHP4;refGene_tx=XM_023258016.1;refGene_gn=NPHP4;refGene_tx=XM_023258017.1
C1 3751452 3751536
refGene_gn=NPHP4;refGene_tx=XM_006934240.4;refGene_gn=NPHP4;refGene_tx=XM_019836057.2;refGene_gn=NPHP4;refGene_tx=XM_019836058.2;refGene_gn=NPHP4;refGene_tx=XM_019836060.2;refGene_gn=NPHP4;refGene_tx=XM_019836062.2;refGene_gn=NPHP4;refGene_tx=XM_023258014.1;refGene_gn=NPHP4;refGene_tx=XM_023258015.1;refGene_gn=NPHP4;refGene_tx=XM_023258016.1;refGene_gn=NPHP4;refGene_tx=XM_023258017.1
C1 3752339 3752511
refGene_gn=NPHP4;refGene_tx=XM_006934240.4;refGene_gn=NPHP4;refGene_tx=XM_019836057.2;refGene_gn=NPHP4;refGene_tx=XM_019836058.2;refGene_gn=NPHP4;refGene_tx=XM_019836060.2;refGene_gn=NPHP4;refGene_tx=XM_019836062.2;refGene_gn=NPHP4;refGene_tx=XM_023258014.1;refGene_gn=NPHP4;refGene_tx=XM_023258015.1;refGene_gn=NPHP4;refGene_tx=XM_023258016.1;refGene_gn=NPHP4;refGene_tx=XM_023258017.1
C1 3752633 3752860
refGene_gn=NPHP4;refGene_tx=XM_006934240.4;refGene_gn=NPHP4;refGene_tx=XM_019836057.2;refGene_gn=NPHP4;refGene_tx=XM_019836058.2;refGene_gn=NPHP4;refGene_tx=XM_019836060.2;refGene_gn=NPHP4;refGene_tx=XM_019836062.2;refGene_gn=NPHP4;refGene_tx=XM_023258014.1;refGene_gn=NPHP4;refGene_tx=XM_023258015.1;refGene_gn=NPHP4;refGene_tx=XM_023258016.1;refGen

e_gn=NPHP4;refGene_tx=XM_023258017.1
C1 3756602 3756814
refGene_gn=NPHP4;refGene_tx=XM_006934240.4;refGene_gn=NPHP4;refGene_tx=XM_019836057.2;refGene_gn=NPHP4;refGene_tx=XM_019836058.2;refGene_gn=NPHP4;refGene_tx=XM_019836060.2;refGene_gn=NPHP4;refGene_tx=XM_019836062.2;refGene_gn=NPHP4;refGene_tx=XM_023258014.1;refGene_gn=NPHP4;refGene_tx=XM_023258015.1;refGene_gn=NPHP4;refGene_tx=XM_023258016.1;refGene_gn=NPHP4;refGene_tx=XM_023258017.1
C1 3759122 3759248
refGene_gn=NPHP4;refGene_tx=XM_006934240.4;refGene_gn=NPHP4;refGene_tx=XM_019836057.2;refGene_gn=NPHP4;refGene_tx=XM_019836058.2;refGene_gn=NPHP4;refGene_tx=XM_019836060.2;refGene_gn=NPHP4;refGene_tx=XM_019836062.2;refGene_gn=NPHP4;refGene_tx=XM_023258014.1;refGene_gn=NPHP4;refGene_tx=XM_023258015.1;refGene_gn=NPHP4;refGene_tx=XM_023258016.1;refGene_gn=NPHP4;refGene_tx=XM_023258017.1
C1 3765267 3765445
refGene_gn=NPHP4;refGene_tx=XM_006934240.4;refGene_gn=NPHP4;refGene_tx=XM_019836057.2;refGene_gn=NPHP4;refGene_tx=XM_019836058.2;refGene_gn=NPHP4;refGene_tx=XM_019836060.2;refGene_gn=NPHP4;refGene_tx=XM_019836062.2;refGene_gn=NPHP4;refGene_tx=XM_023258014.1;refGene_gn=NPHP4;refGene_tx=XM_023258015.1;refGene_gn=NPHP4;refGene_tx=XM_023258016.1;refGene_gn=NPHP4;refGene_tx=XM_023258017.1
C1 3768620 3768781
refGene_gn=NPHP4;refGene_tx=XM_006934240.4;refGene_gn=NPHP4;refGene_tx=XM_019836057.2;refGene_gn=NPHP4;refGene_tx=XM_019836058.2;refGene_gn=NPHP4;refGene_tx=XM_019836060.2;refGene_gn=NPHP4;refGene_tx=XM_019836062.2;refGene_gn=NPHP4;refGene_tx=XM_023258014.1;refGene_gn=NPHP4;refGene_tx=XM_023258015.1;refGene_gn=NPHP4;refGene_tx=XM_023258016.1;refGene_gn=NPHP4;refGene_tx=XM_023258017.1
C1 3781992 3782180
refGene_gn=NPHP4;refGene_tx=XM_006934240.4;refGene_gn=NPHP4;refGene_tx=XM_019836057.2;refGene_gn=NPHP4;refGene_tx=XM_019836058.2;refGene_gn=NPHP4;refGene_tx=XM_019836060.2;refGene_gn=NPHP4;refGene_tx=XM_019836062.2;refGene_gn=NPHP4;refGene_tx=XM_023258014.1;refGene_gn=NPHP4;refGene_tx=XM_023258015.1;refGene_gn=NPHP4;refGene_tx=XM_023258016.1;refGene_gn=NPHP4;refGene_tx=XM_023258017.1
C1 3782625 3782817
refGene_gn=NPHP4;refGene_tx=XM_006934240.4;refGene_gn=NPHP4;refGene_tx=XM_019836057.2;refGene_gn=NPHP4;refGene_tx=XM_019836058.2;refGene_gn=NPHP4;refGene_tx=XM_019836060.2;refGene_gn=NPHP4;refGene_tx=XM_019836062.2;refGene_gn=NPHP4;refGene_tx=XM_023258014.1;refGene_gn=NPHP4;refGene_tx=XM_023258015.1;refGene_gn=NPHP4;refGene_tx=XM_023258016.1;refGene_gn=NPHP4;refGene_tx=XM_023258017.1
C1 3782964 3783116
refGene_gn=NPHP4;refGene_tx=XM_006934240.4;refGene_gn=NPHP4;refGene_tx=XM_019836057.2;refGene_gn=NPHP4;refGene_tx=XM_019836058.2;refGene_gn=NPHP4;refGene_tx=XM_019836060.2;refGene_gn=NPHP4;refGene_tx=XM_019836062.2;refGene_gn=NPHP4;refGene_tx=XM_023258014.1;refGene_gn=NPHP4;refGene_tx=XM_023258015.1;refGene_gn=NPHP4;refGene_tx=XM_023258016.1;refGene_gn=NPHP4;refGene_tx=XM_023258017.1

C1 3784497 3784605
refGene_gn=NPHP4;refGene_tx=XM_006934240.4;refGene_gn=NPHP4;refGene_tx=XM_019836057.2;refGene_gn=NPHP4;refGene_tx=XM_019836058.2;refGene_gn=NPHP4;refGene_tx=XM_019836060.2;refGene_gn=NPHP4;refGene_tx=XM_019836062.2;refGene_gn=NPHP4;refGene_tx=XM_023258014.1;refGene_gn=NPHP4;refGene_tx=XM_023258015.1;refGene_gn=NPHP4;refGene_tx=XM_023258016.1;refGene_gn=NPHP4;refGene_tx=XM_023258017.1
C1 3786469 3786531
refGene_gn=NPHP4;refGene_tx=XM_006934240.4;refGene_gn=NPHP4;refGene_tx=XM_019836057.2;refGene_gn=NPHP4;refGene_tx=XM_019836058.2;refGene_gn=NPHP4;refGene_tx=XM_019836060.2;refGene_gn=NPHP4;refGene_tx=XM_019836062.2;refGene_gn=NPHP4;refGene_tx=XM_023258014.1;refGene_gn=NPHP4;refGene_tx=XM_023258015.1;refGene_gn=NPHP4;refGene_tx=XM_023258016.1;refGene_gn=NPHP4;refGene_tx=XM_023258017.1
C1 3793354 3793493
refGene_gn=NPHP4;refGene_tx=XM_006934240.4;refGene_gn=NPHP4;refGene_tx=XM_019836057.2;refGene_gn=NPHP4;refGene_tx=XM_019836058.2;refGene_gn=NPHP4;refGene_tx=XM_019836060.2;refGene_gn=NPHP4;refGene_tx=XM_019836062.2;refGene_gn=NPHP4;refGene_tx=XM_023258014.1;refGene_gn=NPHP4;refGene_tx=XM_023258015.1;refGene_gn=NPHP4;refGene_tx=XM_023258016.1;refGene_gn=NPHP4;refGene_tx=XM_023258017.1
C1 3797192 3797375
refGene_gn=NPHP4;refGene_tx=XM_006934240.4;refGene_gn=NPHP4;refGene_tx=XM_019836057.2;refGene_gn=NPHP4;refGene_tx=XM_019836058.2;refGene_gn=NPHP4;refGene_tx=XM_019836060.2;refGene_gn=NPHP4;refGene_tx=XM_019836062.2;refGene_gn=NPHP4;refGene_tx=XM_023258014.1;refGene_gn=NPHP4;refGene_tx=XM_023258015.1;refGene_gn=NPHP4;refGene_tx=XM_023258016.1;refGene_gn=NPHP4;refGene_tx=XM_023258017.1
C1 3809919 3810046
refGene_gn=NPHP4;refGene_tx=XM_006934240.4;refGene_gn=NPHP4;refGene_tx=XM_019836057.2;refGene_gn=NPHP4;refGene_tx=XM_019836058.2;refGene_gn=NPHP4;refGene_tx=XM_019836060.2;refGene_gn=NPHP4;refGene_tx=XM_019836062.2;refGene_gn=NPHP4;refGene_tx=XM_023258014.1;refGene_gn=NPHP4;refGene_tx=XM_023258015.1;refGene_gn=NPHP4;refGene_tx=XM_023258016.1;refGene_gn=NPHP4;refGene_tx=XM_023258017.1
C1 3810735 3810917
refGene_gn=NPHP4;refGene_tx=XM_006934240.4;refGene_gn=NPHP4;refGene_tx=XM_019836057.2;refGene_gn=NPHP4;refGene_tx=XM_019836058.2;refGene_gn=NPHP4;refGene_tx=XM_019836060.2;refGene_gn=NPHP4;refGene_tx=XM_019836062.2;refGene_gn=NPHP4;refGene_tx=XM_023258014.1;refGene_gn=NPHP4;refGene_tx=XM_023258015.1;refGene_gn=NPHP4;refGene_tx=XM_023258016.1;refGene_gn=NPHP4;refGene_tx=XM_023258017.1
C1 3812786 3812923
refGene_gn=NPHP4;refGene_tx=XM_006934240.4;refGene_gn=NPHP4;refGene_tx=XM_019836057.2;refGene_gn=NPHP4;refGene_tx=XM_019836058.2;refGene_gn=NPHP4;refGene_tx=XM_019836060.2;refGene_gn=NPHP4;refGene_tx=XM_019836062.2;refGene_gn=NPHP4;refGene_tx=XM_023258014.1;refGene_gn=NPHP4;refGene_tx=XM_023258015.1;refGene_gn=NPHP4;refGene_tx=XM_023258016.1;refGene_gn=NPHP4;refGene_tx=XM_023258017.1
C1 3823880 3824036

refGene_gn=NPHP4;refGene_tx=XM_006934240.4;refGene_gn=NPHP4;refGene_tx
=XM_019836057.2;refGene_gn=NPHP4;refGene_tx=XM_019836058.2;refGene_gn=
NPHP4;refGene_tx=XM_019836060.2;refGene_gn=NPHP4;refGene_tx=XM_0198360
62.2;refGene_gn=NPHP4;refGene_tx=XM_023258014.1;refGene_gn=NPHP4;refGe
ne_tx=XM_023258015.1;refGene_gn=NPHP4;refGene_tx=XM_023258016.1;refGen
e_gn=NPHP4;refGene_tx=XM_023258017.1
C1 3827233 3827298
refGene_gn=NPHP4;refGene_tx=XM_006934240.4;refGene_gn=NPHP4;refGene_tx
=XM_019836057.2;refGene_gn=NPHP4;refGene_tx=XM_019836058.2;refGene_gn=
NPHP4;refGene_tx=XM_019836060.2;refGene_gn=NPHP4;refGene_tx=XM_0232580
14.1;refGene_gn=NPHP4;refGene_tx=XM_023258015.1;refGene_gn=NPHP4;refGe
ne_tx=XM_023258016.1;refGene_gn=NPHP4;refGene_tx=XM_023258017.1
C1 3828399 3828572
refGene_gn=NPHP4;refGene_tx=XM_006934240.4;refGene_gn=NPHP4;refGene_tx
=XM_019836057.2;refGene_gn=NPHP4;refGene_tx=XM_019836058.2;refGene_gn=
NPHP4;refGene_tx=XM_019836060.2;refGene_gn=NPHP4;refGene_tx=XM_0232580
14.1;refGene_gn=NPHP4;refGene_tx=XM_023258015.1;refGene_gn=NPHP4;refGe
ne_tx=XM_023258016.1;refGene_gn=NPHP4;refGene_tx=XM_023258017.1
C1 3836461 3836614
refGene_gn=NPHP4;refGene_tx=XM_006934240.4;refGene_gn=NPHP4;refGene_tx
=XM_019836057.2;refGene_gn=NPHP4;refGene_tx=XM_019836058.2;refGene_gn=
NPHP4;refGene_tx=XM_019836060.2;refGene_gn=NPHP4;refGene_tx=XM_0232580
14.1;refGene_gn=NPHP4;refGene_tx=XM_023258015.1;refGene_gn=NPHP4;refGe
ne_tx=XM_023258016.1;refGene_gn=NPHP4;refGene_tx=XM_023258017.1
C1 3846026 3846161
refGene_gn=NPHP4;refGene_tx=XM_006934240.4;refGene_gn=NPHP4;refGene_tx
=XM_019836057.2;refGene_gn=NPHP4;refGene_tx=XM_019836058.2;refGene_gn=
NPHP4;refGene_tx=XM_019836060.2;refGene_gn=NPHP4;refGene_tx=XM_0232580
14.1;refGene_gn=NPHP4;refGene_tx=XM_023258015.1;refGene_gn=NPHP4;refGe
ne_tx=XM_023258016.1;refGene_gn=NPHP4;refGene_tx=XM_023258017.1
C1 3886525 3886602
refGene_gn=KCNAB2;refGene_tx=XM_003989449.5;refGene_gn=KCNAB2;refGene_
tx=XM_019836067.2;refGene_gn=KCNAB2;refGene_tx=XM_019836068.2;refGene_
gn=KCNAB2;refGene_tx=XM_019836069.2;refGene_gn=KCNAB2;refGene_tx=XM_01
9836070.2;refGene_gn=KCNAB2;refGene_tx=XM_019836071.2
C1 3887766 3887808
refGene_gn=KCNAB2;refGene_tx=XM_003989449.5;refGene_gn=KCNAB2;refGene_
tx=XM_019836067.2;refGene_gn=KCNAB2;refGene_tx=XM_019836068.2;refGene_
gn=KCNAB2;refGene_tx=XM_019836069.2
C1 3896329 3896547
refGene_gn=KCNAB2;refGene_tx=XM_019836063.2;refGene_gn=KCNAB2;refGene_
tx=XM_019836064.2
C1 3906884 3907096
refGene_gn=KCNAB2;refGene_tx=XM_019836065.2;refGene_gn=KCNAB2;refGene_
tx=XM_019836066.2
C1 3912315 3912359
refGene_gn=KCNAB2;refGene_tx=XM_003989449.5;refGene_gn=KCNAB2;refGene_
tx=XM_019836063.2;refGene_gn=KCNAB2;refGene_tx=XM_019836064.2;refGene_
gn=KCNAB2;refGene_tx=XM_019836065.2;refGene_gn=KCNAB2;refGene_tx=XM_01
9836066.2;refGene_gn=KCNAB2;refGene_tx=XM_019836067.2;refGene_gn=KCNAB

2;refGene_tx=XM_019836068.2;refGene_gn=KCNAB2;refGene_tx=XM_019836069.2;refGene_gn=KCNAB2;refGene_tx=XM_019836070.2;refGene_gn=KCNAB2;refGene_tx=XM_019836071.2
C1 3913160 3913198
refGene_gn=KCNAB2;refGene_tx=XM_003989449.5;refGene_gn=KCNAB2;refGene_tx=XM_019836063.2;refGene_gn=KCNAB2;refGene_tx=XM_019836064.2;refGene_gn=KCNAB2;refGene_tx=XM_019836065.2;refGene_gn=KCNAB2;refGene_tx=XM_019836066.2;refGene_gn=KCNAB2;refGene_tx=XM_019836067.2;refGene_gn=KCNAB2;refGene_tx=XM_019836068.2;refGene_gn=KCNAB2;refGene_tx=XM_019836069.2;refGene_gn=KCNAB2;refGene_tx=XM_019836070.2;refGene_gn=KCNAB2;refGene_tx=XM_019836071.2
C1 3919883 3919963
refGene_gn=KCNAB2;refGene_tx=XM_003989449.5;refGene_gn=KCNAB2;refGene_tx=XM_019836063.2;refGene_gn=KCNAB2;refGene_tx=XM_019836064.2;refGene_gn=KCNAB2;refGene_tx=XM_019836065.2;refGene_gn=KCNAB2;refGene_tx=XM_019836066.2;refGene_gn=KCNAB2;refGene_tx=XM_019836067.2;refGene_gn=KCNAB2;refGene_tx=XM_019836068.2;refGene_gn=KCNAB2;refGene_tx=XM_019836069.2;refGene_gn=KCNAB2;refGene_tx=XM_019836070.2;refGene_gn=KCNAB2;refGene_tx=XM_019836071.2
C1 3922698 3922743
refGene_gn=KCNAB2;refGene_tx=XM_003989449.5;refGene_gn=KCNAB2;refGene_tx=XM_019836063.2;refGene_gn=KCNAB2;refGene_tx=XM_019836064.2;refGene_gn=KCNAB2;refGene_tx=XM_019836065.2;refGene_gn=KCNAB2;refGene_tx=XM_019836066.2;refGene_gn=KCNAB2;refGene_tx=XM_019836067.2;refGene_gn=KCNAB2;refGene_tx=XM_019836068.2;refGene_gn=KCNAB2;refGene_tx=XM_019836069.2;refGene_gn=KCNAB2;refGene_tx=XM_019836070.2;refGene_gn=KCNAB2;refGene_tx=XM_019836071.2
C1 3924716 3924761
refGene_gn=KCNAB2;refGene_tx=XM_003989449.5;refGene_gn=KCNAB2;refGene_tx=XM_019836063.2;refGene_gn=KCNAB2;refGene_tx=XM_019836064.2;refGene_gn=KCNAB2;refGene_tx=XM_019836065.2;refGene_gn=KCNAB2;refGene_tx=XM_019836066.2;refGene_gn=KCNAB2;refGene_tx=XM_019836067.2;refGene_gn=KCNAB2;refGene_tx=XM_019836068.2;refGene_gn=KCNAB2;refGene_tx=XM_019836069.2;refGene_gn=KCNAB2;refGene_tx=XM_019836070.2;refGene_gn=KCNAB2;refGene_tx=XM_019836071.2
C1 3925466 3925510
refGene_gn=KCNAB2;refGene_tx=XM_003989449.5;refGene_gn=KCNAB2;refGene_tx=XM_019836063.2;refGene_gn=KCNAB2;refGene_tx=XM_019836064.2;refGene_gn=KCNAB2;refGene_tx=XM_019836065.2;refGene_gn=KCNAB2;refGene_tx=XM_019836066.2;refGene_gn=KCNAB2;refGene_tx=XM_019836067.2;refGene_gn=KCNAB2;refGene_tx=XM_019836068.2;refGene_gn=KCNAB2;refGene_tx=XM_019836069.2;refGene_gn=KCNAB2;refGene_tx=XM_019836070.2;refGene_gn=KCNAB2;refGene_tx=XM_019836071.2
C1 3926522 3926609
refGene_gn=KCNAB2;refGene_tx=XM_003989449.5;refGene_gn=KCNAB2;refGene_tx=XM_019836063.2;refGene_gn=KCNAB2;refGene_tx=XM_019836064.2;refGene_gn=KCNAB2;refGene_tx=XM_019836065.2;refGene_gn=KCNAB2;refGene_tx=XM_019836066.2;refGene_gn=KCNAB2;refGene_tx=XM_019836067.2;refGene_gn=KCNAB2;refGene_tx=XM_019836068.2;refGene_gn=KCNAB2;refGene_tx=XM_019836069.2;refGene_gn=KCNAB2;refGene_tx=XM_019836070.2;refGene_gn=KCNAB2;refGen

e_tx=XM_019836071.2
C1	3927341	3927386
refGene_gn=KCNAB2;refGene_tx=XM_019836063.2;refGene_gn=KCNAB2;refGene_tx=XM_019836065.2;refGene_gn=KCNAB2;refGene_tx=XM_019836067.2;refGene_gn=KCNAB2;refGene_tx=XM_019836068.2;refGene_gn=KCNAB2;refGene_tx=XM_019836069.2;refGene_gn=KCNAB2;refGene_tx=XM_019836070.2
C1	3930317	3930403
refGene_gn=KCNAB2;refGene_tx=XM_003989449.5;refGene_gn=KCNAB2;refGene_tx=XM_019836063.2;refGene_gn=KCNAB2;refGene_tx=XM_019836064.2;refGene_gn=KCNAB2;refGene_tx=XM_019836065.2;refGene_gn=KCNAB2;refGene_tx=XM_019836066.2;refGene_gn=KCNAB2;refGene_tx=XM_019836067.2;refGene_gn=KCNAB2;refGene_tx=XM_019836068.2;refGene_gn=KCNAB2;refGene_tx=XM_019836069.2;refGene_gn=KCNAB2;refGene_tx=XM_019836070.2;refGene_gn=KCNAB2;refGene_tx=XM_019836071.2
C1	3932152	3932273
refGene_gn=KCNAB2;refGene_tx=XM_003989449.5;refGene_gn=KCNAB2;refGene_tx=XM_019836063.2;refGene_gn=KCNAB2;refGene_tx=XM_019836064.2;refGene_gn=KCNAB2;refGene_tx=XM_019836065.2;refGene_gn=KCNAB2;refGene_tx=XM_019836066.2;refGene_gn=KCNAB2;refGene_tx=XM_019836067.2;refGene_gn=KCNAB2;refGene_tx=XM_019836068.2;refGene_gn=KCNAB2;refGene_tx=XM_019836069.2;refGene_gn=KCNAB2;refGene_tx=XM_019836070.2;refGene_gn=KCNAB2;refGene_tx=XM_019836071.2
C1	3932455	3932550
refGene_gn=KCNAB2;refGene_tx=XM_003989449.5;refGene_gn=KCNAB2;refGene_tx=XM_019836063.2;refGene_gn=KCNAB2;refGene_tx=XM_019836064.2;refGene_gn=KCNAB2;refGene_tx=XM_019836065.2;refGene_gn=KCNAB2;refGene_tx=XM_019836066.2;refGene_gn=KCNAB2;refGene_tx=XM_019836067.2;refGene_gn=KCNAB2;refGene_tx=XM_019836068.2;refGene_gn=KCNAB2;refGene_tx=XM_019836069.2;refGene_gn=KCNAB2;refGene_tx=XM_019836070.2;refGene_gn=KCNAB2;refGene_tx=XM_019836071.2
C1	3933508	3933629
refGene_gn=KCNAB2;refGene_tx=XM_003989449.5;refGene_gn=KCNAB2;refGene_tx=XM_019836063.2;refGene_gn=KCNAB2;refGene_tx=XM_019836064.2;refGene_gn=KCNAB2;refGene_tx=XM_019836065.2;refGene_gn=KCNAB2;refGene_tx=XM_019836066.2;refGene_gn=KCNAB2;refGene_tx=XM_019836067.2;refGene_gn=KCNAB2;refGene_tx=XM_019836068.2;refGene_gn=KCNAB2;refGene_tx=XM_019836069.2;refGene_gn=KCNAB2;refGene_tx=XM_019836070.2;refGene_gn=KCNAB2;refGene_tx=XM_019836071.2
C1	3933972	3934061
refGene_gn=KCNAB2;refGene_tx=XM_003989449.5;refGene_gn=KCNAB2;refGene_tx=XM_019836063.2;refGene_gn=KCNAB2;refGene_tx=XM_019836064.2;refGene_gn=KCNAB2;refGene_tx=XM_019836065.2;refGene_gn=KCNAB2;refGene_tx=XM_019836066.2;refGene_gn=KCNAB2;refGene_tx=XM_019836067.2;refGene_gn=KCNAB2;refGene_tx=XM_019836068.2;refGene_gn=KCNAB2;refGene_tx=XM_019836069.2;refGene_gn=KCNAB2;refGene_tx=XM_019836070.2;refGene_gn=KCNAB2;refGene_tx=XM_019836071.2
C1	3935157	3935247
refGene_gn=KCNAB2;refGene_tx=XM_003989449.5;refGene_gn=KCNAB2;refGene_tx=XM_019836063.2;refGene_gn=KCNAB2;refGene_tx=XM_019836064.2;refGene_gn=KCNAB2;refGene_tx=XM_019836065.2;refGene_gn=KCNAB2;refGene_tx=XM_01

9836066.2;refGene_gn=KCNAB2;refGene_tx=XM_019836067.2;refGene_gn=KCNAB2;refGene_tx=XM_019836068.2;refGene_gn=KCNAB2;refGene_tx=XM_019836069.2;refGene_gn=KCNAB2;refGene_tx=XM_019836070.2;refGene_gn=KCNAB2;refGene_tx=XM_019836071.2
C1 3941893 3941953 refGene_gn=CHD5;refGene_tx=XM_023258012.1
C1 3942818 3942872
refGene_gn=CHD5;refGene_tx=XM_023258009.1;refGene_gn=CHD5;refGene_tx=XM_023258010.1;refGene_gn=CHD5;refGene_tx=XM_023258011.1;refGene_gn=CHD5;refGene_tx=XM_023258012.1;refGene_gn=CHD5;refGene_tx=XM_023258013.1
C1 3942973 3943088
refGene_gn=CHD5;refGene_tx=XM_023258009.1;refGene_gn=CHD5;refGene_tx=XM_023258010.1;refGene_gn=CHD5;refGene_tx=XM_023258011.1;refGene_gn=CHD5;refGene_tx=XM_023258013.1
C1 3943210 3943374
refGene_gn=CHD5;refGene_tx=XM_023258009.1;refGene_gn=CHD5;refGene_tx=XM_023258010.1;refGene_gn=CHD5;refGene_tx=XM_023258011.1;refGene_gn=CHD5;refGene_tx=XM_023258012.1;refGene_gn=CHD5;refGene_tx=XM_023258013.1
C1 3944130 3944326
refGene_gn=CHD5;refGene_tx=XM_023258009.1;refGene_gn=CHD5;refGene_tx=XM_023258010.1;refGene_gn=CHD5;refGene_tx=XM_023258011.1;refGene_gn=CHD5;refGene_tx=XM_023258012.1;refGene_gn=CHD5;refGene_tx=XM_023258013.1
C1 3944719 3944852
refGene_gn=CHD5;refGene_tx=XM_023258009.1;refGene_gn=CHD5;refGene_tx=XM_023258010.1;refGene_gn=CHD5;refGene_tx=XM_023258011.1;refGene_gn=CHD5;refGene_tx=XM_023258012.1;refGene_gn=CHD5;refGene_tx=XM_023258013.1
C1 3947039 3947148
refGene_gn=CHD5;refGene_tx=XM_023258009.1;refGene_gn=CHD5;refGene_tx=XM_023258010.1;refGene_gn=CHD5;refGene_tx=XM_023258011.1;refGene_gn=CHD5;refGene_tx=XM_023258012.1;refGene_gn=CHD5;refGene_tx=XM_023258013.1
C1 3947415 3947553
refGene_gn=CHD5;refGene_tx=XM_023258009.1;refGene_gn=CHD5;refGene_tx=XM_023258010.1;refGene_gn=CHD5;refGene_tx=XM_023258011.1;refGene_gn=CHD5;refGene_tx=XM_023258012.1;refGene_gn=CHD5;refGene_tx=XM_023258013.1
C1 3948338 3948428
refGene_gn=CHD5;refGene_tx=XM_023258009.1;refGene_gn=CHD5;refGene_tx=XM_023258010.1;refGene_gn=CHD5;refGene_tx=XM_023258011.1;refGene_gn=CHD5;refGene_tx=XM_023258012.1;refGene_gn=CHD5;refGene_tx=XM_023258013.1
C1 3955417 3955535
refGene_gn=CHD5;refGene_tx=XM_023258009.1;refGene_gn=CHD5;refGene_tx=XM_023258010.1;refGene_gn=CHD5;refGene_tx=XM_023258011.1;refGene_gn=CHD5;refGene_tx=XM_023258012.1;refGene_gn=CHD5;refGene_tx=XM_023258013.1
C1 3955791 3955871
refGene_gn=CHD5;refGene_tx=XM_023258009.1;refGene_gn=CHD5;refGene_tx=XM_023258010.1;refGene_gn=CHD5;refGene_tx=XM_023258011.1;refGene_gn=CHD5;refGene_tx=XM_023258012.1;refGene_gn=CHD5;refGene_tx=XM_023258013.1
C1 3957680 3957840
refGene_gn=CHD5;refGene_tx=XM_023258009.1;refGene_gn=CHD5;refGene_tx=XM_023258010.1;refGene_gn=CHD5;refGene_tx=XM_023258011.1;refGene_gn=CHD5;refGene_tx=XM_023258012.1;refGene_gn=CHD5;refGene_tx=XM_023258013.1
C1 3958222 3958367

refGene_gn=CHD5;refGene_tx=XM_023258009.1;refGene_gn=CHD5;refGene_tx=XM_023258010.1;refGene_gn=CHD5;refGene_tx=XM_023258011.1;refGene_gn=CHD5;refGene_tx=XM_023258012.1;refGene_gn=CHD5;refGene_tx=XM_023258013.1
C1 3958769 3958903
refGene_gn=CHD5;refGene_tx=XM_023258009.1;refGene_gn=CHD5;refGene_tx=XM_023258010.1;refGene_gn=CHD5;refGene_tx=XM_023258011.1;refGene_gn=CHD5;refGene_tx=XM_023258012.1;refGene_gn=CHD5;refGene_tx=XM_023258013.1
C1 3959186 3959275
refGene_gn=CHD5;refGene_tx=XM_023258009.1;refGene_gn=CHD5;refGene_tx=XM_023258010.1;refGene_gn=CHD5;refGene_tx=XM_023258011.1;refGene_gn=CHD5;refGene_tx=XM_023258012.1;refGene_gn=CHD5;refGene_tx=XM_023258013.1
C1 3959491 3959584
refGene_gn=CHD5;refGene_tx=XM_023258009.1;refGene_gn=CHD5;refGene_tx=XM_023258010.1;refGene_gn=CHD5;refGene_tx=XM_023258011.1;refGene_gn=CHD5;refGene_tx=XM_023258012.1;refGene_gn=CHD5;refGene_tx=XM_023258013.1
C1 3959956 3960131
refGene_gn=CHD5;refGene_tx=XM_023258009.1;refGene_gn=CHD5;refGene_tx=XM_023258010.1;refGene_gn=CHD5;refGene_tx=XM_023258011.1;refGene_gn=CHD5;refGene_tx=XM_023258012.1;refGene_gn=CHD5;refGene_tx=XM_023258013.1
C1 3960744 3960917
refGene_gn=CHD5;refGene_tx=XM_023258009.1;refGene_gn=CHD5;refGene_tx=XM_023258010.1;refGene_gn=CHD5;refGene_tx=XM_023258011.1;refGene_gn=CHD5;refGene_tx=XM_023258012.1;refGene_gn=CHD5;refGene_tx=XM_023258013.1
C1 3961254 3961365
refGene_gn=CHD5;refGene_tx=XM_023258009.1;refGene_gn=CHD5;refGene_tx=XM_023258010.1;refGene_gn=CHD5;refGene_tx=XM_023258011.1;refGene_gn=CHD5;refGene_tx=XM_023258012.1;refGene_gn=CHD5;refGene_tx=XM_023258013.1
C1 3961543 3961775
refGene_gn=CHD5;refGene_tx=XM_023258009.1;refGene_gn=CHD5;refGene_tx=XM_023258010.1;refGene_gn=CHD5;refGene_tx=XM_023258011.1;refGene_gn=CHD5;refGene_tx=XM_023258012.1;refGene_gn=CHD5;refGene_tx=XM_023258013.1
C1 3962832 3962957
refGene_gn=CHD5;refGene_tx=XM_023258009.1;refGene_gn=CHD5;refGene_tx=XM_023258010.1;refGene_gn=CHD5;refGene_tx=XM_023258011.1;refGene_gn=CHD5;refGene_tx=XM_023258012.1;refGene_gn=CHD5;refGene_tx=XM_023258013.1
C1 3964201 3964319
refGene_gn=CHD5;refGene_tx=XM_023258009.1;refGene_gn=CHD5;refGene_tx=XM_023258010.1;refGene_gn=CHD5;refGene_tx=XM_023258011.1;refGene_gn=CHD5;refGene_tx=XM_023258012.1;refGene_gn=CHD5;refGene_tx=XM_023258013.1
C1 3966563 3966695
refGene_gn=CHD5;refGene_tx=XM_023258009.1;refGene_gn=CHD5;refGene_tx=XM_023258010.1;refGene_gn=CHD5;refGene_tx=XM_023258011.1;refGene_gn=CHD5;refGene_tx=XM_023258012.1;refGene_gn=CHD5;refGene_tx=XM_023258013.1
C1 3967674 3967816
refGene_gn=CHD5;refGene_tx=XM_023258009.1;refGene_gn=CHD5;refGene_tx=XM_023258010.1;refGene_gn=CHD5;refGene_tx=XM_023258011.1;refGene_gn=CHD5;refGene_tx=XM_023258012.1;refGene_gn=CHD5;refGene_tx=XM_023258013.1
C1 3968239 3968413
refGene_gn=CHD5;refGene_tx=XM_023258009.1;refGene_gn=CHD5;refGene_tx=XM_023258010.1;refGene_gn=CHD5;refGene_tx=XM_023258011.1;refGene_gn=CHD

5;refGene_tx=XM_023258012.1;refGene_gn=CHD5;refGene_tx=XM_023258013.1
C1 3969281 3969403
refGene_gn=CHD5;refGene_tx=XM_023258009.1;refGene_gn=CHD5;refGene_tx=XM_023258010.1;refGene_gn=CHD5;refGene_tx=XM_023258011.1;refGene_gn=CHD5;refGene_tx=XM_023258012.1;refGene_gn=CHD5;refGene_tx=XM_023258013.1
C1 3969523 3969661
refGene_gn=CHD5;refGene_tx=XM_023258009.1;refGene_gn=CHD5;refGene_tx=XM_023258010.1;refGene_gn=CHD5;refGene_tx=XM_023258011.1;refGene_gn=CHD5;refGene_tx=XM_023258012.1;refGene_gn=CHD5;refGene_tx=XM_023258013.1
C1 3972680 3972881
refGene_gn=CHD5;refGene_tx=XM_023258009.1;refGene_gn=CHD5;refGene_tx=XM_023258010.1;refGene_gn=CHD5;refGene_tx=XM_023258011.1;refGene_gn=CHD5;refGene_tx=XM_023258012.1;refGene_gn=CHD5;refGene_tx=XM_023258013.1
C1 3972975 3973167
refGene_gn=CHD5;refGene_tx=XM_023258009.1;refGene_gn=CHD5;refGene_tx=XM_023258010.1;refGene_gn=CHD5;refGene_tx=XM_023258011.1;refGene_gn=CHD5;refGene_tx=XM_023258012.1;refGene_gn=CHD5;refGene_tx=XM_023258013.1
C1 3974228 3974334
refGene_gn=CHD5;refGene_tx=XM_023258009.1;refGene_gn=CHD5;refGene_tx=XM_023258010.1;refGene_gn=CHD5;refGene_tx=XM_023258011.1;refGene_gn=CHD5;refGene_tx=XM_023258012.1;refGene_gn=CHD5;refGene_tx=XM_023258013.1
C1 3974416 3974548
refGene_gn=CHD5;refGene_tx=XM_023258009.1;refGene_gn=CHD5;refGene_tx=XM_023258010.1;refGene_gn=CHD5;refGene_tx=XM_023258011.1;refGene_gn=CHD5;refGene_tx=XM_023258012.1;refGene_gn=CHD5;refGene_tx=XM_023258013.1
C1 3976883 3977098
refGene_gn=CHD5;refGene_tx=XM_023258009.1;refGene_gn=CHD5;refGene_tx=XM_023258010.1;refGene_gn=CHD5;refGene_tx=XM_023258011.1;refGene_gn=CHD5;refGene_tx=XM_023258012.1;refGene_gn=CHD5;refGene_tx=XM_023258013.1
C1 3977312 3977519
refGene_gn=CHD5;refGene_tx=XM_023258009.1;refGene_gn=CHD5;refGene_tx=XM_023258010.1;refGene_gn=CHD5;refGene_tx=XM_023258011.1;refGene_gn=CHD5;refGene_tx=XM_023258012.1;refGene_gn=CHD5;refGene_tx=XM_023258013.1
C1 3978640 3978862
refGene_gn=CHD5;refGene_tx=XM_023258009.1;refGene_gn=CHD5;refGene_tx=XM_023258010.1;refGene_gn=CHD5;refGene_tx=XM_023258011.1;refGene_gn=CHD5;refGene_tx=XM_023258012.1;refGene_gn=CHD5;refGene_tx=XM_023258013.1
C1 3979005 3979172
refGene_gn=CHD5;refGene_tx=XM_023258009.1;refGene_gn=CHD5;refGene_tx=XM_023258010.1;refGene_gn=CHD5;refGene_tx=XM_023258011.1;refGene_gn=CHD5;refGene_tx=XM_023258012.1;refGene_gn=CHD5;refGene_tx=XM_023258013.1
C1 3979697 3979821
refGene_gn=CHD5;refGene_tx=XM_023258009.1;refGene_gn=CHD5;refGene_tx=XM_023258010.1;refGene_gn=CHD5;refGene_tx=XM_023258011.1;refGene_gn=CHD5;refGene_tx=XM_023258012.1;refGene_gn=CHD5;refGene_tx=XM_023258013.1
C1 3981020 3981145
refGene_gn=CHD5;refGene_tx=XM_023258009.1;refGene_gn=CHD5;refGene_tx=XM_023258010.1;refGene_gn=CHD5;refGene_tx=XM_023258011.1;refGene_gn=CHD5;refGene_tx=XM_023258012.1;refGene_gn=CHD5;refGene_tx=XM_023258013.1
C1 3982817 3983056

refGene_gn=CHD5;refGene_tx=XM_023258009.1;refGene_gn=CHD5;refGene_tx=XM_023258010.1;refGene_gn=CHD5;refGene_tx=XM_023258011.1;refGene_gn=CHD5;refGene_tx=XM_023258012.1;refGene_gn=CHD5;refGene_tx=XM_023258013.1
C1 3983582 3983701
refGene_gn=CHD5;refGene_tx=XM_023258009.1;refGene_gn=CHD5;refGene_tx=XM_023258010.1;refGene_gn=CHD5;refGene_tx=XM_023258011.1;refGene_gn=CHD5;refGene_tx=XM_023258012.1;refGene_gn=CHD5;refGene_tx=XM_023258013.1
C1 3986239 3986419
refGene_gn=CHD5;refGene_tx=XM_023258009.1;refGene_gn=CHD5;refGene_tx=XM_023258010.1;refGene_gn=CHD5;refGene_tx=XM_023258011.1;refGene_gn=CHD5;refGene_tx=XM_023258012.1
C1 3991648 3991773
refGene_gn=CHD5;refGene_tx=XM_023258009.1;refGene_gn=CHD5;refGene_tx=XM_023258010.1;refGene_gn=CHD5;refGene_tx=XM_023258011.1;refGene_gn=CHD5;refGene_tx=XM_023258012.1;refGene_gn=CHD5;refGene_tx=XM_023258013.1
C1 4001935 4002014
refGene_gn=CHD5;refGene_tx=XM_023258009.1;refGene_gn=CHD5;refGene_tx=XM_023258010.1;refGene_gn=CHD5;refGene_tx=XM_023258011.1;refGene_gn=CHD5;refGene_tx=XM_023258012.1;refGene_gn=CHD5;refGene_tx=XM_023258013.1
C1 4008498 4008643 refGene_gn=RPL22;refGene_tx=XM_011284453.3
C1 4011591 4011716 refGene_gn=RPL22;refGene_tx=XM_011284453.3
C1 4014164 4014269 refGene_gn=RPL22;refGene_tx=XM_011284453.3
C1 4016076 4016088 refGene_gn=RPL22;refGene_tx=XM_011284453.3
C1 4020165 4020356
refGene_gn=RNF207;refGene_tx=XM_011284475.3;refGene_gn=RNF207;refGene_tx=XM_023258018.1
C1 4020976 4021109
refGene_gn=RNF207;refGene_tx=XM_011284475.3;refGene_gn=RNF207;refGene_tx=XM_023258018.1
C1 4022134 4022279
refGene_gn=RNF207;refGene_tx=XM_011284475.3;refGene_gn=RNF207;refGene_tx=XM_023258018.1
C1 4022377 4022459
refGene_gn=RNF207;refGene_tx=XM_011284475.3;refGene_gn=RNF207;refGene_tx=XM_023258018.1
C1 4022537 4022613
refGene_gn=RNF207;refGene_tx=XM_011284475.3;refGene_gn=RNF207;refGene_tx=XM_023258018.1
C1 4022687 4022813
refGene_gn=RNF207;refGene_tx=XM_011284475.3;refGene_gn=RNF207;refGene_tx=XM_023258018.1
C1 4023219 4023266
refGene_gn=RNF207;refGene_tx=XM_011284475.3;refGene_gn=RNF207;refGene_tx=XM_023258018.1
C1 4023519 4023592
refGene_gn=RNF207;refGene_tx=XM_011284475.3;refGene_gn=RNF207;refGene_tx=XM_023258018.1
C1 4023676 4023745
refGene_gn=RNF207;refGene_tx=XM_011284475.3;refGene_gn=RNF207;refGene_tx=XM_023258018.1

C1 4024203 4024272
refGene_gn=RNF207;refGene_tx=XM_011284475.3;refGene_gn=RNF207;refGene_tx=XM_023258018.1
C1 4024356 4024454
refGene_gn=RNF207;refGene_tx=XM_011284475.3;refGene_gn=RNF207;refGene_tx=XM_023258018.1
C1 4025107 4025294
refGene_gn=RNF207;refGene_tx=XM_011284475.3;refGene_gn=RNF207;refGene_tx=XM_023258018.1
C1 4025434 4025620
refGene_gn=RNF207;refGene_tx=XM_011284475.3;refGene_gn=RNF207;refGene_tx=XM_023258018.1
C1 4025879 4025938
refGene_gn=RNF207;refGene_tx=XM_011284475.3;refGene_gn=RNF207;refGene_tx=XM_023258018.1
C1 4026257 4026375
refGene_gn=RNF207;refGene_tx=XM_011284475.3;refGene_gn=RNF207;refGene_tx=XM_023258018.1
C1 4030896 4030977 refGene_gn=RNF207;refGene_tx=XM_011284475.3
C1 4031800 4031897 refGene_gn=RNF207;refGene_tx=XM_011284475.3
C1 4036596 4036779 refGene_gn=ICMT;refGene_tx=XM_023258019.1
C1 4039285 4039503 refGene_gn=ICMT;refGene_tx=XM_023258019.1
C1 4040281 4040451 refGene_gn=ICMT;refGene_tx=XM_023258019.1
C1 4041340 4041429 refGene_gn=ICMT;refGene_tx=XM_023258019.1
C1 4042305 4042500 refGene_gn=ICMT;refGene_tx=XM_023258019.1
C1 4050672 4050732 refGene_gn=HES3;refGene_tx=XM_023258021.1
C1 4050839 4050935
refGene_gn=HES3;refGene_tx=XM_023258021.1;refGene_gn=HES3;refGene_tx=XM_023258022.1
C1 4051036 4051118
refGene_gn=HES3;refGene_tx=XM_023258021.1;refGene_gn=HES3;refGene_tx=XM_023258022.1
C1 4051496 4051870
refGene_gn=HES3;refGene_tx=XM_023258021.1;refGene_gn=HES3;refGene_tx=XM_023258022.1
C1 4055547 4056213
refGene_gn=GPR153;refGene_tx=XM_011284473.3;refGene_gn=GPR153;refGene_tx=XM_023258020.1
C1 4056373 4056558
refGene_gn=GPR153;refGene_tx=XM_011284473.3;refGene_gn=GPR153;refGene_tx=XM_023258020.1
C1 4056926 4057119
refGene_gn=GPR153;refGene_tx=XM_011284473.3;refGene_gn=GPR153;refGene_tx=XM_023258020.1
C1 4058602 4059032
refGene_gn=GPR153;refGene_tx=XM_011284473.3;refGene_gn=GPR153;refGene_tx=XM_023258020.1
C1 4059472 4059828
refGene_gn=GPR153;refGene_tx=XM_011284473.3;refGene_gn=GPR153;refGene_tx=XM_023258020.1

C1 4067441 4067540
refGene_gn=ACOT7;refGene_tx=XM_006934247.4;refGene_gn=ACOT7;refGene_tx
=XM_023258023.1
C1 4083727 4083912
refGene_gn=ACOT7;refGene_tx=XM_006934247.4;refGene_gn=ACOT7;refGene_tx
=XM_023258023.1
C1 4094984 4095101
refGene_gn=ACOT7;refGene_tx=XM_006934247.4;refGene_gn=ACOT7;refGene_tx
=XM_023258023.1
C1 4111274 4111361
refGene_gn=ACOT7;refGene_tx=XM_006934247.4;refGene_gn=ACOT7;refGene_tx
=XM_023258023.1
C1 4119190 4119305
refGene_gn=ACOT7;refGene_tx=XM_006934247.4;refGene_gn=ACOT7;refGene_tx
=XM_023258023.1
C1 4124630 4124722
refGene_gn=ACOT7;refGene_tx=XM_006934247.4;refGene_gn=ACOT7;refGene_tx
=XM_023258023.1
C1 4129483 4129640
refGene_gn=ACOT7;refGene_tx=XM_006934247.4;refGene_gn=ACOT7;refGene_tx
=XM_023258023.1
C1 4135041 4135159
refGene_gn=ACOT7;refGene_tx=XM_006934247.4;refGene_gn=ACOT7;refGene_tx
=XM_023258023.1
C1 4152751 4152933 refGene_gn=ACOT7;refGene_tx=XM_006934247.4
C1 4167504 4167635 refGene_gn=ACOT7;refGene_tx=XM_023258023.1
C1 4189081 4189332 refGene_gn=HES2;refGene_tx=XM_023258024.1
C1 4189413 4189513 refGene_gn=HES2;refGene_tx=XM_023258024.1
C1 4189781 4189877 refGene_gn=HES2;refGene_tx=XM_023258024.1
C1 4189963 4190008 refGene_gn=HES2;refGene_tx=XM_023258024.1
C1 4195676 4195970
refGene_gn=ESPN;refGene_tx=XM_023258025.1;refGene_gn=ESPN;refGene_tx=X
M_023258026.1;refGene_gn=ESPN;refGene_tx=XM_023258029.1;refGene_gn=ESP
N;refGene_tx=XM_023258030.1;refGene_gn=ESPN;refGene_tx=XM_023258031.1
C1 4198884 4199078
refGene_gn=ESPN;refGene_tx=XM_023258025.1;refGene_gn=ESPN;refGene_tx=X
M_023258026.1;refGene_gn=ESPN;refGene_tx=XM_023258029.1;refGene_gn=ESP
N;refGene_tx=XM_023258030.1;refGene_gn=ESPN;refGene_tx=XM_023258031.1
C1 4210344 4210531
refGene_gn=ESPN;refGene_tx=XM_023258025.1;refGene_gn=ESPN;refGene_tx=X
M_023258026.1;refGene_gn=ESPN;refGene_tx=XM_023258029.1;refGene_gn=ESP
N;refGene_tx=XM_023258030.1;refGene_gn=ESPN;refGene_tx=XM_023258031.1
C1 4210666 4210849
refGene_gn=ESPN;refGene_tx=XM_023258025.1;refGene_gn=ESPN;refGene_tx=X
M_023258026.1;refGene_gn=ESPN;refGene_tx=XM_023258029.1;refGene_gn=ESP
N;refGene_tx=XM_023258030.1;refGene_gn=ESPN;refGene_tx=XM_023258031.1
C1 4211013 4211145
refGene_gn=ESPN;refGene_tx=XM_023258025.1;refGene_gn=ESPN;refGene_tx=X
M_023258026.1;refGene_gn=ESPN;refGene_tx=XM_023258029.1;refGene_gn=ESP
N;refGene_tx=XM_023258030.1;refGene_gn=ESPN;refGene_tx=XM_023258031.1

C1	4212959	4212968	refGene_gn=ESPN;refGene_tx=XM_023258027.1
C1	4213639	4213841
refGene_gn=ESPN;refGene_tx=XM_023258025.1;refGene_gn=ESPN;refGene_tx=XM_023258026.1;refGene_gn=ESPN;refGene_tx=XM_023258027.1;refGene_gn=ESPN;refGene_tx=XM_023258029.1;refGene_gn=ESPN;refGene_tx=XM_023258030.1;refGene_gn=ESPN;refGene_tx=XM_023258031.1
C1	4214739	4215008
refGene_gn=ESPN;refGene_tx=XM_023258025.1;refGene_gn=ESPN;refGene_tx=XM_023258026.1;refGene_gn=ESPN;refGene_tx=XM_023258027.1;refGene_gn=ESPN;refGene_tx=XM_023258029.1;refGene_gn=ESPN;refGene_tx=XM_023258030.1;refGene_gn=ESPN;refGene_tx=XM_023258031.1
C1	4217074	4217125	refGene_gn=ESPN;refGene_tx=XM_023258028.1
C1	4217641	4218092
refGene_gn=ESPN;refGene_tx=XM_023258025.1;refGene_gn=ESPN;refGene_tx=XM_023258026.1;refGene_gn=ESPN;refGene_tx=XM_023258027.1;refGene_gn=ESPN;refGene_tx=XM_023258028.1;refGene_gn=ESPN;refGene_tx=XM_023258029.1;refGene_gn=ESPN;refGene_tx=XM_023258030.1;refGene_gn=ESPN;refGene_tx=XM_023258031.1
C1	4219402	4219429
refGene_gn=ESPN;refGene_tx=XM_023258025.1;refGene_gn=ESPN;refGene_tx=XM_023258027.1;refGene_gn=ESPN;refGene_tx=XM_023258028.1;refGene_gn=ESPN;refGene_tx=XM_023258030.1
C1	4220329	4220369	refGene_gn=ESPN;refGene_tx=XM_023258032.1
C1	4220602	4220748
refGene_gn=ESPN;refGene_tx=XM_023258025.1;refGene_gn=ESPN;refGene_tx=XM_023258026.1;refGene_gn=ESPN;refGene_tx=XM_023258027.1;refGene_gn=ESPN;refGene_tx=XM_023258028.1;refGene_gn=ESPN;refGene_tx=XM_023258029.1;refGene_gn=ESPN;refGene_tx=XM_023258030.1;refGene_gn=ESPN;refGene_tx=XM_023258031.1;refGene_gn=ESPN;refGene_tx=XM_023258032.1
C1	4220832	4221096
refGene_gn=ESPN;refGene_tx=XM_023258025.1;refGene_gn=ESPN;refGene_tx=XM_023258026.1;refGene_gn=ESPN;refGene_tx=XM_023258027.1;refGene_gn=ESPN;refGene_tx=XM_023258028.1;refGene_gn=ESPN;refGene_tx=XM_023258029.1;refGene_gn=ESPN;refGene_tx=XM_023258030.1;refGene_gn=ESPN;refGene_tx=XM_023258031.1;refGene_gn=ESPN;refGene_tx=XM_023258032.1
C1	4222803	4224606
refGene_gn=ESPN;refGene_tx=XM_023258025.1;refGene_gn=ESPN;refGene_tx=XM_023258026.1;refGene_gn=ESPN;refGene_tx=XM_023258027.1;refGene_gn=ESPN;refGene_tx=XM_023258028.1;refGene_gn=ESPN;refGene_tx=XM_023258032.1
C1	4225557	4225637
refGene_gn=ESPN;refGene_tx=XM_023258029.1;refGene_gn=ESPN;refGene_tx=XM_023258030.1;refGene_gn=ESPN;refGene_tx=XM_023258031.1
C1	4225734	4225746
refGene_gn=ESPN;refGene_tx=XM_023258029.1;refGene_gn=ESPN;refGene_tx=XM_023258030.1;refGene_gn=ESPN;refGene_tx=XM_023258031.1
C1	4227035	4227183
refGene_gn=ESPN;refGene_tx=XM_023258029.1;refGene_gn=ESPN;refGene_tx=XM_023258030.1;refGene_gn=ESPN;refGene_tx=XM_023258031.1
C1	4228463	4228801
refGene_gn=TNFRSF25;refGene_tx=XM_023258042.1;refGene_gn=TNFRSF25;refG

ene_tx=XM_023258043.1
C1	4228992	4229167
refGene_gn=TNFRSF25;refGene_tx=XM_023258042.1;refGene_gn=TNFRSF25;refGene_tx=XM_023258043.1
C1	4229661	4229699
refGene_gn=TNFRSF25;refGene_tx=XM_023258042.1;refGene_gn=TNFRSF25;refGene_tx=XM_023258043.1
C1	4229908	4230016
refGene_gn=TNFRSF25;refGene_tx=XM_023258042.1;refGene_gn=TNFRSF25;refGene_tx=XM_023258043.1
C1	4230111	4230167
refGene_gn=TNFRSF25;refGene_tx=XM_023258042.1;refGene_gn=TNFRSF25;refGene_tx=XM_023258043.1
C1	4230909	4230988
refGene_gn=TNFRSF25;refGene_tx=XM_023258042.1;refGene_gn=TNFRSF25;refGene_tx=XM_023258043.1
C1	4231087	4231255
refGene_gn=TNFRSF25;refGene_tx=XM_023258042.1;refGene_gn=TNFRSF25;refGene_tx=XM_023258043.1
C1	4231622	4231757
refGene_gn=TNFRSF25;refGene_tx=XM_023258042.1;refGene_gn=TNFRSF25;refGene_tx=XM_023258043.1
C1	4231949	4232070
refGene_gn=TNFRSF25;refGene_tx=XM_023258042.1;refGene_gn=TNFRSF25;refGene_tx=XM_023258043.1
C1	4232601	4232643
refGene_gn=TNFRSF25;refGene_tx=XM_023258042.1;refGene_gn=TNFRSF25;refGene_tx=XM_023258043.1
C1	4233890	4233900
refGene_gn=PLEKHG5;refGene_tx=XM_023258033.1;refGene_gn=PLEKHG5;refGene_tx=XM_023258035.1;refGene_gn=PLEKHG5;refGene_tx=XM_023258036.1;refGene_gn=PLEKHG5;refGene_tx=XM_023258038.1;refGene_gn=PLEKHG5;refGene_tx=XM_023258039.1;refGene_gn=PLEKHG5;refGene_tx=XM_023258040.1;refGene_gn=PLEKHG5;refGene_tx=XM_023258041.1
C1	4234181	4234962
refGene_gn=PLEKHG5;refGene_tx=XM_023258033.1;refGene_gn=PLEKHG5;refGene_tx=XM_023258034.1;refGene_gn=PLEKHG5;refGene_tx=XM_023258035.1;refGene_gn=PLEKHG5;refGene_tx=XM_023258036.1;refGene_gn=PLEKHG5;refGene_tx=XM_023258037.1;refGene_gn=PLEKHG5;refGene_tx=XM_023258038.1;refGene_gn=PLEKHG5;refGene_tx=XM_023258039.1;refGene_gn=PLEKHG5;refGene_tx=XM_023258040.1;refGene_gn=PLEKHG5;refGene_tx=XM_023258041.1
C1	4235758	4235946
refGene_gn=PLEKHG5;refGene_tx=XM_023258033.1;refGene_gn=PLEKHG5;refGene_tx=XM_023258034.1;refGene_gn=PLEKHG5;refGene_tx=XM_023258035.1;refGene_gn=PLEKHG5;refGene_tx=XM_023258036.1;refGene_gn=PLEKHG5;refGene_tx=XM_023258037.1;refGene_gn=PLEKHG5;refGene_tx=XM_023258038.1;refGene_gn=PLEKHG5;refGene_tx=XM_023258039.1;refGene_gn=PLEKHG5;refGene_tx=XM_023258040.1;refGene_gn=PLEKHG5;refGene_tx=XM_023258041.1
C1	4236028	4236144
refGene_gn=PLEKHG5;refGene_tx=XM_023258033.1;refGene_gn=PLEKHG5;refGen

e_tx=XM_023258034.1;refGene_gn=PLEKHG5;refGene_tx=XM_023258035.1;refGene_gn=PLEKHG5;refGene_tx=XM_023258036.1;refGene_gn=PLEKHG5;refGene_tx=XM_023258037.1;refGene_gn=PLEKHG5;refGene_tx=XM_023258038.1;refGene_gn=PLEKHG5;refGene_tx=XM_023258039.1;refGene_gn=PLEKHG5;refGene_tx=XM_023258040.1;refGene_gn=PLEKHG5;refGene_tx=XM_023258041.1
C1 4236243 4236376
refGene_gn=PLEKHG5;refGene_tx=XM_023258033.1;refGene_gn=PLEKHG5;refGene_tx=XM_023258034.1;refGene_gn=PLEKHG5;refGene_tx=XM_023258035.1;refGene_gn=PLEKHG5;refGene_tx=XM_023258036.1;refGene_gn=PLEKHG5;refGene_tx=XM_023258037.1;refGene_gn=PLEKHG5;refGene_tx=XM_023258038.1;refGene_gn=PLEKHG5;refGene_tx=XM_023258039.1;refGene_gn=PLEKHG5;refGene_tx=XM_023258040.1;refGene_gn=PLEKHG5;refGene_tx=XM_023258041.1
C1 4238026 4238146
refGene_gn=PLEKHG5;refGene_tx=XM_023258033.1;refGene_gn=PLEKHG5;refGene_tx=XM_023258034.1;refGene_gn=PLEKHG5;refGene_tx=XM_023258035.1;refGene_gn=PLEKHG5;refGene_tx=XM_023258036.1;refGene_gn=PLEKHG5;refGene_tx=XM_023258037.1;refGene_gn=PLEKHG5;refGene_tx=XM_023258038.1;refGene_gn=PLEKHG5;refGene_tx=XM_023258039.1;refGene_gn=PLEKHG5;refGene_tx=XM_023258040.1;refGene_gn=PLEKHG5;refGene_tx=XM_023258041.1
C1 4238320 4238458
refGene_gn=PLEKHG5;refGene_tx=XM_023258033.1;refGene_gn=PLEKHG5;refGene_tx=XM_023258034.1;refGene_gn=PLEKHG5;refGene_tx=XM_023258035.1;refGene_gn=PLEKHG5;refGene_tx=XM_023258036.1;refGene_gn=PLEKHG5;refGene_tx=XM_023258037.1;refGene_gn=PLEKHG5;refGene_tx=XM_023258038.1;refGene_gn=PLEKHG5;refGene_tx=XM_023258039.1;refGene_gn=PLEKHG5;refGene_tx=XM_023258040.1;refGene_gn=PLEKHG5;refGene_tx=XM_023258041.1
C1 4238551 4238701
refGene_gn=PLEKHG5;refGene_tx=XM_023258033.1;refGene_gn=PLEKHG5;refGene_tx=XM_023258034.1;refGene_gn=PLEKHG5;refGene_tx=XM_023258035.1;refGene_gn=PLEKHG5;refGene_tx=XM_023258036.1;refGene_gn=PLEKHG5;refGene_tx=XM_023258037.1;refGene_gn=PLEKHG5;refGene_tx=XM_023258038.1;refGene_gn=PLEKHG5;refGene_tx=XM_023258039.1;refGene_gn=PLEKHG5;refGene_tx=XM_023258040.1;refGene_gn=PLEKHG5;refGene_tx=XM_023258041.1
C1 4238816 4238927
refGene_gn=PLEKHG5;refGene_tx=XM_023258033.1;refGene_gn=PLEKHG5;refGene_tx=XM_023258034.1;refGene_gn=PLEKHG5;refGene_tx=XM_023258035.1;refGene_gn=PLEKHG5;refGene_tx=XM_023258036.1;refGene_gn=PLEKHG5;refGene_tx=XM_023258037.1;refGene_gn=PLEKHG5;refGene_tx=XM_023258038.1;refGene_gn=PLEKHG5;refGene_tx=XM_023258039.1;refGene_gn=PLEKHG5;refGene_tx=XM_023258040.1;refGene_gn=PLEKHG5;refGene_tx=XM_023258041.1
C1 4239242 4239392
refGene_gn=PLEKHG5;refGene_tx=XM_023258033.1;refGene_gn=PLEKHG5;refGene_tx=XM_023258034.1;refGene_gn=PLEKHG5;refGene_tx=XM_023258035.1;refGene_gn=PLEKHG5;refGene_tx=XM_023258036.1;refGene_gn=PLEKHG5;refGene_tx=XM_023258037.1;refGene_gn=PLEKHG5;refGene_tx=XM_023258038.1;refGene_gn=PLEKHG5;refGene_tx=XM_023258039.1;refGene_gn=PLEKHG5;refGene_tx=XM_023258040.1;refGene_gn=PLEKHG5;refGene_tx=XM_023258041.1
C1 4239517 4239568
refGene_gn=PLEKHG5;refGene_tx=XM_023258033.1;refGene_gn=PLEKHG5;refGene_tx=XM_023258034.1;refGene_gn=PLEKHG5;refGene_tx=XM_023258035.1;refGe

ne_gn=PLEKHG5;refGene_tx=XM_023258036.1;refGene_gn=PLEKHG5;refGene_tx=XM_023258037.1;refGene_gn=PLEKHG5;refGene_tx=XM_023258038.1;refGene_gn=PLEKHG5;refGene_tx=XM_023258039.1;refGene_gn=PLEKHG5;refGene_tx=XM_023258040.1;refGene_gn=PLEKHG5;refGene_tx=XM_023258041.1
C1 4240184 4240280
refGene_gn=PLEKHG5;refGene_tx=XM_023258033.1;refGene_gn=PLEKHG5;refGene_tx=XM_023258034.1;refGene_gn=PLEKHG5;refGene_tx=XM_023258035.1;refGene_gn=PLEKHG5;refGene_tx=XM_023258036.1;refGene_gn=PLEKHG5;refGene_tx=XM_023258037.1;refGene_gn=PLEKHG5;refGene_tx=XM_023258038.1;refGene_gn=PLEKHG5;refGene_tx=XM_023258039.1;refGene_gn=PLEKHG5;refGene_tx=XM_023258040.1;refGene_gn=PLEKHG5;refGene_tx=XM_023258041.1
C1 4240568 4240763
refGene_gn=PLEKHG5;refGene_tx=XM_023258033.1;refGene_gn=PLEKHG5;refGene_tx=XM_023258034.1;refGene_gn=PLEKHG5;refGene_tx=XM_023258035.1;refGene_gn=PLEKHG5;refGene_tx=XM_023258036.1;refGene_gn=PLEKHG5;refGene_tx=XM_023258037.1;refGene_gn=PLEKHG5;refGene_tx=XM_023258038.1;refGene_gn=PLEKHG5;refGene_tx=XM_023258039.1;refGene_gn=PLEKHG5;refGene_tx=XM_023258040.1;refGene_gn=PLEKHG5;refGene_tx=XM_023258041.1
C1 4240840 4241035
refGene_gn=PLEKHG5;refGene_tx=XM_023258033.1;refGene_gn=PLEKHG5;refGene_tx=XM_023258034.1;refGene_gn=PLEKHG5;refGene_tx=XM_023258035.1;refGene_gn=PLEKHG5;refGene_tx=XM_023258036.1;refGene_gn=PLEKHG5;refGene_tx=XM_023258037.1;refGene_gn=PLEKHG5;refGene_tx=XM_023258038.1;refGene_gn=PLEKHG5;refGene_tx=XM_023258039.1;refGene_gn=PLEKHG5;refGene_tx=XM_023258040.1;refGene_gn=PLEKHG5;refGene_tx=XM_023258041.1
C1 4241497 4241643
refGene_gn=PLEKHG5;refGene_tx=XM_023258033.1;refGene_gn=PLEKHG5;refGene_tx=XM_023258034.1;refGene_gn=PLEKHG5;refGene_tx=XM_023258035.1;refGene_gn=PLEKHG5;refGene_tx=XM_023258036.1;refGene_gn=PLEKHG5;refGene_tx=XM_023258037.1;refGene_gn=PLEKHG5;refGene_tx=XM_023258038.1;refGene_gn=PLEKHG5;refGene_tx=XM_023258039.1;refGene_gn=PLEKHG5;refGene_tx=XM_023258040.1;refGene_gn=PLEKHG5;refGene_tx=XM_023258041.1
C1 4241924 4242061
refGene_gn=PLEKHG5;refGene_tx=XM_023258033.1;refGene_gn=PLEKHG5;refGene_tx=XM_023258034.1;refGene_gn=PLEKHG5;refGene_tx=XM_023258035.1;refGene_gn=PLEKHG5;refGene_tx=XM_023258036.1;refGene_gn=PLEKHG5;refGene_tx=XM_023258037.1;refGene_gn=PLEKHG5;refGene_tx=XM_023258038.1;refGene_gn=PLEKHG5;refGene_tx=XM_023258039.1;refGene_gn=PLEKHG5;refGene_tx=XM_023258040.1;refGene_gn=PLEKHG5;refGene_tx=XM_023258041.1
C1 4242483 4242575
refGene_gn=PLEKHG5;refGene_tx=XM_023258033.1;refGene_gn=PLEKHG5;refGene_tx=XM_023258034.1;refGene_gn=PLEKHG5;refGene_tx=XM_023258035.1;refGene_gn=PLEKHG5;refGene_tx=XM_023258036.1;refGene_gn=PLEKHG5;refGene_tx=XM_023258037.1;refGene_gn=PLEKHG5;refGene_tx=XM_023258038.1;refGene_gn=PLEKHG5;refGene_tx=XM_023258039.1;refGene_gn=PLEKHG5;refGene_tx=XM_023258040.1;refGene_gn=PLEKHG5;refGene_tx=XM_023258041.1
C1 4242941 4242999
refGene_gn=PLEKHG5;refGene_tx=XM_023258033.1;refGene_gn=PLEKHG5;refGene_tx=XM_023258034.1;refGene_gn=PLEKHG5;refGene_tx=XM_023258035.1;refGene_gn=PLEKHG5;refGene_tx=XM_023258036.1;refGene_gn=PLEKHG5;refGene_tx=

XM_023258037.1;refGene_gn=PLEKHG5;refGene_tx=XM_023258038.1;refGene_gn=PLEKHG5;refGene_tx=XM_023258039.1;refGene_gn=PLEKHG5;refGene_tx=XM_023258040.1;refGene_gn=PLEKHG5;refGene_tx=XM_023258041.1
C1 4243379 4243482
refGene_gn=PLEKHG5;refGene_tx=XM_023258033.1;refGene_gn=PLEKHG5;refGene_tx=XM_023258034.1;refGene_gn=PLEKHG5;refGene_tx=XM_023258035.1;refGene_gn=PLEKHG5;refGene_tx=XM_023258036.1;refGene_gn=PLEKHG5;refGene_tx=XM_023258037.1;refGene_gn=PLEKHG5;refGene_tx=XM_023258038.1;refGene_gn=PLEKHG5;refGene_tx=XM_023258039.1;refGene_gn=PLEKHG5;refGene_tx=XM_023258040.1;refGene_gn=PLEKHG5;refGene_tx=XM_023258041.1
C1 4244668 4244798
refGene_gn=PLEKHG5;refGene_tx=XM_023258033.1;refGene_gn=PLEKHG5;refGene_tx=XM_023258034.1;refGene_gn=PLEKHG5;refGene_tx=XM_023258035.1;refGene_gn=PLEKHG5;refGene_tx=XM_023258036.1;refGene_gn=PLEKHG5;refGene_tx=XM_023258037.1;refGene_gn=PLEKHG5;refGene_tx=XM_023258038.1;refGene_gn=PLEKHG5;refGene_tx=XM_023258039.1;refGene_gn=PLEKHG5;refGene_tx=XM_023258040.1;refGene_gn=PLEKHG5;refGene_tx=XM_023258041.1
C1 4250521 4250623
refGene_gn=PLEKHG5;refGene_tx=XM_023258036.1;refGene_gn=PLEKHG5;refGene_tx=XM_023258037.1
C1 4260126 4260203
refGene_gn=PLEKHG5;refGene_tx=XM_023258033.1;refGene_gn=PLEKHG5;refGene_tx=XM_023258034.1;refGene_gn=PLEKHG5;refGene_tx=XM_023258035.1;refGene_gn=PLEKHG5;refGene_tx=XM_023258038.1;refGene_gn=PLEKHG5;refGene_tx=XM_023258039.1
C1 4261063 4261067 refGene_gn=PLEKHG5;refGene_tx=XM_023258038.1
C1 4285195 4285345
refGene_gn=NOL9;refGene_tx=XM_023258044.1;refGene_gn=NOL9;refGene_tx=XM_023258045.1
C1 4286030 4286164
refGene_gn=NOL9;refGene_tx=XM_023258044.1;refGene_gn=NOL9;refGene_tx=XM_023258045.1
C1 4287670 4287848
refGene_gn=NOL9;refGene_tx=XM_023258044.1;refGene_gn=NOL9;refGene_tx=XM_023258045.1
C1 4289118 4289230
refGene_gn=NOL9;refGene_tx=XM_023258044.1;refGene_gn=NOL9;refGene_tx=XM_023258045.1
C1 4290214 4290512
refGene_gn=NOL9;refGene_tx=XM_023258044.1;refGene_gn=NOL9;refGene_tx=XM_023258045.1
C1 4290712 4290874
refGene_gn=NOL9;refGene_tx=XM_023258044.1;refGene_gn=NOL9;refGene_tx=XM_023258045.1
C1 4295535 4295633
refGene_gn=NOL9;refGene_tx=XM_023258044.1;refGene_gn=NOL9;refGene_tx=XM_023258045.1
C1 4296190 4296287
refGene_gn=NOL9;refGene_tx=XM_023258044.1;refGene_gn=NOL9;refGene_tx=XM_023258045.1

C1	4296409	4296545
refGene_gn=NOL9;refGene_tx=XM_023258044.1;refGene_gn=NOL9;refGene_tx=XM_023258045.1
C1	4299575	4299703
refGene_gn=NOL9;refGene_tx=XM_023258044.1;refGene_gn=NOL9;refGene_tx=XM_023258045.1
C1	4301738	4301958
refGene_gn=NOL9;refGene_tx=XM_023258044.1;refGene_gn=NOL9;refGene_tx=XM_023258045.1
C1	4303593	4303950
refGene_gn=NOL9;refGene_tx=XM_023258044.1;refGene_gn=NOL9;refGene_tx=XM_023258045.1
C1	4304901	4305092
refGene_gn=TAS1R1;refGene_tx=XM_003989461.4;refGene_gn=TAS1R1;refGene_tx=XM_019836077.2
C1	4306891	4307198
refGene_gn=TAS1R1;refGene_tx=XM_003989461.4;refGene_gn=TAS1R1;refGene_tx=XM_019836077.2
C1	4309513	4310275
refGene_gn=TAS1R1;refGene_tx=XM_003989461.4;refGene_gn=TAS1R1;refGene_tx=XM_019836077.2
C1	4310687	4310900	refGene_gn=TAS1R1;refGene_tx=XM_003989461.4
C1	4311211	4311332
refGene_gn=TAS1R1;refGene_tx=XM_003989461.4;refGene_gn=TAS1R1;refGene_tx=XM_019836077.2
C1	4312328	4313260
refGene_gn=TAS1R1;refGene_tx=XM_003989461.4;refGene_gn=TAS1R1;refGene_tx=XM_019836077.2
C1	4314479	4315169
refGene_gn=ZBTB48;refGene_tx=XM_011284447.2;refGene_gn=ZBTB48;refGene_tx=XM_011284448.2;refGene_gn=ZBTB48;refGene_tx=XM_011284449.2
C1	4315646	4315888
refGene_gn=ZBTB48;refGene_tx=XM_011284447.2;refGene_gn=ZBTB48;refGene_tx=XM_011284448.2;refGene_gn=ZBTB48;refGene_tx=XM_011284449.2
C1	4317911	4318023
refGene_gn=ZBTB48;refGene_tx=XM_011284447.2;refGene_gn=ZBTB48;refGene_tx=XM_011284448.2;refGene_gn=ZBTB48;refGene_tx=XM_011284449.2
C1	4318626	4318719
refGene_gn=ZBTB48;refGene_tx=XM_011284447.2;refGene_gn=ZBTB48;refGene_tx=XM_011284448.2;refGene_gn=ZBTB48;refGene_tx=XM_011284449.2
C1	4319109	4319196
refGene_gn=ZBTB48;refGene_tx=XM_011284447.2;refGene_gn=ZBTB48;refGene_tx=XM_011284448.2;refGene_gn=ZBTB48;refGene_tx=XM_011284449.2
C1	4319348	4319503
refGene_gn=ZBTB48;refGene_tx=XM_011284447.2;refGene_gn=ZBTB48;refGene_tx=XM_011284448.2;refGene_gn=ZBTB48;refGene_tx=XM_011284449.2
C1	4319940	4320077
refGene_gn=ZBTB48;refGene_tx=XM_011284447.2;refGene_gn=ZBTB48;refGene_tx=XM_011284448.2;refGene_gn=ZBTB48;refGene_tx=XM_011284449.2
C1	4320148	4320313

refGene_gn=ZBTB48;refGene_tx=XM_011284447.2;refGene_gn=ZBTB48;refGene_
tx=XM_011284448.2;refGene_gn=ZBTB48;refGene_tx=XM_011284449.2
C1 4320707 4320796
refGene_gn=ZBTB48;refGene_tx=XM_011284447.2;refGene_gn=ZBTB48;refGene_
tx=XM_011284448.2;refGene_gn=ZBTB48;refGene_tx=XM_011284449.2
C1 4320882 4321179
refGene_gn=ZBTB48;refGene_tx=XM_011284447.2;refGene_gn=ZBTB48;refGene_
tx=XM_011284448.2;refGene_gn=ZBTB48;refGene_tx=XM_011284449.2
C1 4326226 4326520 refGene_gn=KLHL21;refGene_tx=XM_006934332.4
C1 4327736 4327809 refGene_gn=KLHL21;refGene_tx=XM_006934332.4
C1 4331413 4331819 refGene_gn=KLHL21;refGene_tx=XM_006934332.4
C1 4333986 4335007 refGene_gn=KLHL21;refGene_tx=XM_006934332.4
C1 4342559 4342598 refGene_gn=PHF13;refGene_tx=XM_003989464.5
C1 4345007 4345109 refGene_gn=PHF13;refGene_tx=XM_003989464.5
C1 4346374 4346909 refGene_gn=PHF13;refGene_tx=XM_003989464.5
C1 4347424 4347651 refGene_gn=PHF13;refGene_tx=XM_003989464.5
C1 4351127 4351201
refGene_gn=THAP3;refGene_tx=XM_023258048.1;refGene_gn=THAP3;refGene_tx
=XM_023258049.1
C1 4354187 4354380
refGene_gn=THAP3;refGene_tx=XM_023258048.1;refGene_gn=THAP3;refGene_tx
=XM_023258049.1
C1 4359357 4359420
refGene_gn=THAP3;refGene_tx=XM_023258048.1;refGene_gn=THAP3;refGene_tx
=XM_023258049.1
C1 4359781 4359880
refGene_gn=THAP3;refGene_tx=XM_023258048.1;refGene_gn=THAP3;refGene_tx
=XM_023258049.1
C1 4360153 4360414
refGene_gn=THAP3;refGene_tx=XM_023258048.1;refGene_gn=THAP3;refGene_tx
=XM_023258049.1
C1 4363365 4363391
refGene_gn=DNAJC11;refGene_tx=XM_023258046.1;refGene_gn=DNAJC11;refGen
e_tx=XM_023258047.1
C1 4363785 4363915
refGene_gn=DNAJC11;refGene_tx=XM_023258046.1;refGene_gn=DNAJC11;refGen
e_tx=XM_023258047.1
C1 4364537 4364680
refGene_gn=DNAJC11;refGene_tx=XM_023258046.1;refGene_gn=DNAJC11;refGen
e_tx=XM_023258047.1
C1 4364813 4364871
refGene_gn=DNAJC11;refGene_tx=XM_023258046.1;refGene_gn=DNAJC11;refGen
e_tx=XM_023258047.1
C1 4365366 4365436
refGene_gn=DNAJC11;refGene_tx=XM_023258046.1;refGene_gn=DNAJC11;refGen
e_tx=XM_023258047.1
C1 4366485 4366641
refGene_gn=DNAJC11;refGene_tx=XM_023258046.1;refGene_gn=DNAJC11;refGen
e_tx=XM_023258047.1
C1 4373742 4373859

refGene_gn=DNAJC11;refGene_tx=XM_023258046.1;refGene_gn=DNAJC11;refGene_tx=XM_023258047.1
C1	4374112	4374198
refGene_gn=DNAJC11;refGene_tx=XM_023258046.1;refGene_gn=DNAJC11;refGene_tx=XM_023258047.1
C1	4374780	4374970
refGene_gn=DNAJC11;refGene_tx=XM_023258046.1;refGene_gn=DNAJC11;refGene_tx=XM_023258047.1
C1	4378077	4378151
refGene_gn=DNAJC11;refGene_tx=XM_023258046.1;refGene_gn=DNAJC11;refGene_tx=XM_023258047.1
C1	4379094	4379217
refGene_gn=DNAJC11;refGene_tx=XM_023258046.1;refGene_gn=DNAJC11;refGene_tx=XM_023258047.1
C1	4380193	4380322
refGene_gn=DNAJC11;refGene_tx=XM_023258046.1;refGene_gn=DNAJC11;refGene_tx=XM_023258047.1
C1	4397791	4397893
refGene_gn=DNAJC11;refGene_tx=XM_023258046.1;refGene_gn=DNAJC11;refGene_tx=XM_023258047.1
C1	4404518	4404592
refGene_gn=DNAJC11;refGene_tx=XM_023258046.1;refGene_gn=DNAJC11;refGene_tx=XM_023258047.1
C1	4405573	4405631	refGene_gn=DNAJC11;refGene_tx=XM_023258047.1
C1	4407701	4407831	refGene_gn=DNAJC11;refGene_tx=XM_023258046.1
C1	4433188	4433260	refGene_gn=DNAJC11;refGene_tx=XM_023258046.1
C1	4517471	4517561
refGene_gn=CAMTA1;refGene_tx=XM_011284441.3;refGene_gn=CAMTA1;refGene_tx=XM_011284442.3;refGene_gn=CAMTA1;refGene_tx=XM_011284444.3;refGene_gn=CAMTA1;refGene_tx=XM_023258051.1;refGene_gn=CAMTA1;refGene_tx=XM_023258052.1;refGene_gn=CAMTA1;refGene_tx=XM_023258053.1
C1	4522129	4522199
refGene_gn=CAMTA1;refGene_tx=XM_011284441.3;refGene_gn=CAMTA1;refGene_tx=XM_011284442.3;refGene_gn=CAMTA1;refGene_tx=XM_011284444.3;refGene_gn=CAMTA1;refGene_tx=XM_023258051.1;refGene_gn=CAMTA1;refGene_tx=XM_023258052.1;refGene_gn=CAMTA1;refGene_tx=XM_023258053.1
C1	4527241	4527360
refGene_gn=CAMTA1;refGene_tx=XM_011284441.3;refGene_gn=CAMTA1;refGene_tx=XM_011284442.3;refGene_gn=CAMTA1;refGene_tx=XM_011284444.3;refGene_gn=CAMTA1;refGene_tx=XM_023258051.1;refGene_gn=CAMTA1;refGene_tx=XM_023258052.1;refGene_gn=CAMTA1;refGene_tx=XM_023258053.1
C1	4782118	4782186
refGene_gn=CAMTA1;refGene_tx=XM_011284441.3;refGene_gn=CAMTA1;refGene_tx=XM_011284442.3;refGene_gn=CAMTA1;refGene_tx=XM_011284444.3;refGene_gn=CAMTA1;refGene_tx=XM_023258051.1;refGene_gn=CAMTA1;refGene_tx=XM_023258052.1;refGene_gn=CAMTA1;refGene_tx=XM_023258053.1
C1	4915326	4915462
refGene_gn=CAMTA1;refGene_tx=XM_011284441.3;refGene_gn=CAMTA1;refGene_tx=XM_011284442.3;refGene_gn=CAMTA1;refGene_tx=XM_011284444.3;refGene_gn=CAMTA1;refGene_tx=XM_023258051.1;refGene_gn=CAMTA1;refGene_tx=XM_02

3258052.1;refGene_gn=CAMTA1;refGene_tx=XM_023258053.1
C1	5096934	5097006
refGene_gn=CAMTA1;refGene_tx=XM_011284441.3;refGene_gn=CAMTA1;refGene_tx=XM_011284442.3;refGene_gn=CAMTA1;refGene_tx=XM_011284444.3;refGene_gn=CAMTA1;refGene_tx=XM_023258051.1;refGene_gn=CAMTA1;refGene_tx=XM_023258052.1;refGene_gn=CAMTA1;refGene_tx=XM_023258053.1
C1	5236942	5237096
refGene_gn=CAMTA1;refGene_tx=XM_011284441.3;refGene_gn=CAMTA1;refGene_tx=XM_011284442.3;refGene_gn=CAMTA1;refGene_tx=XM_011284444.3;refGene_gn=CAMTA1;refGene_tx=XM_023258051.1;refGene_gn=CAMTA1;refGene_tx=XM_023258052.1;refGene_gn=CAMTA1;refGene_tx=XM_023258053.1
C1	5252017	5252158
refGene_gn=CAMTA1;refGene_tx=XM_011284441.3;refGene_gn=CAMTA1;refGene_tx=XM_011284442.3;refGene_gn=CAMTA1;refGene_tx=XM_011284444.3;refGene_gn=CAMTA1;refGene_tx=XM_023258051.1;refGene_gn=CAMTA1;refGene_tx=XM_023258052.1;refGene_gn=CAMTA1;refGene_tx=XM_023258053.1
C1	5253674	5255521
refGene_gn=CAMTA1;refGene_tx=XM_011284441.3;refGene_gn=CAMTA1;refGene_tx=XM_011284442.3;refGene_gn=CAMTA1;refGene_tx=XM_011284444.3;refGene_gn=CAMTA1;refGene_tx=XM_023258051.1;refGene_gn=CAMTA1;refGene_tx=XM_023258052.1;refGene_gn=CAMTA1;refGene_tx=XM_023258053.1
C1	5259833	5259960
refGene_gn=CAMTA1;refGene_tx=XM_011284441.3;refGene_gn=CAMTA1;refGene_tx=XM_011284442.3;refGene_gn=CAMTA1;refGene_tx=XM_011284444.3;refGene_gn=CAMTA1;refGene_tx=XM_023258051.1;refGene_gn=CAMTA1;refGene_tx=XM_023258052.1;refGene_gn=CAMTA1;refGene_tx=XM_023258053.1
C1	5265327	5265462
refGene_gn=CAMTA1;refGene_tx=XM_011284441.3;refGene_gn=CAMTA1;refGene_tx=XM_011284442.3;refGene_gn=CAMTA1;refGene_tx=XM_011284444.3;refGene_gn=CAMTA1;refGene_tx=XM_023258051.1;refGene_gn=CAMTA1;refGene_tx=XM_023258052.1;refGene_gn=CAMTA1;refGene_tx=XM_023258053.1
C1	5317078	5317230
refGene_gn=CAMTA1;refGene_tx=XM_011284441.3;refGene_gn=CAMTA1;refGene_tx=XM_011284442.3;refGene_gn=CAMTA1;refGene_tx=XM_011284444.3;refGene_gn=CAMTA1;refGene_tx=XM_023258051.1;refGene_gn=CAMTA1;refGene_tx=XM_023258052.1;refGene_gn=CAMTA1;refGene_tx=XM_023258053.1
C1	5319559	5319756
refGene_gn=CAMTA1;refGene_tx=XM_011284441.3;refGene_gn=CAMTA1;refGene_tx=XM_011284442.3;refGene_gn=CAMTA1;refGene_tx=XM_011284444.3;refGene_gn=CAMTA1;refGene_tx=XM_023258051.1;refGene_gn=CAMTA1;refGene_tx=XM_023258052.1;refGene_gn=CAMTA1;refGene_tx=XM_023258053.1
C1	5320134	5320213
refGene_gn=CAMTA1;refGene_tx=XM_011284441.3;refGene_gn=CAMTA1;refGene_tx=XM_011284442.3;refGene_gn=CAMTA1;refGene_tx=XM_011284444.3;refGene_gn=CAMTA1;refGene_tx=XM_023258051.1;refGene_gn=CAMTA1;refGene_tx=XM_023258052.1;refGene_gn=CAMTA1;refGene_tx=XM_023258053.1
C1	5320395	5320711
refGene_gn=CAMTA1;refGene_tx=XM_011284441.3;refGene_gn=CAMTA1;refGene_tx=XM_011284442.3;refGene_gn=CAMTA1;refGene_tx=XM_011284444.3;refGene_gn=CAMTA1;refGene_tx=XM_023258051.1;refGene_gn=CAMTA1;refGene_tx=XM_02

3258052.1;refGene_gn=CAMTA1;refGene_tx=XM_023258053.1
C1 5321144 5321668
refGene_gn=CAMTA1;refGene_tx=XM_011284441.3;refGene_gn=CAMTA1;refGene_tx=XM_011284442.3;refGene_gn=CAMTA1;refGene_tx=XM_011284444.3;refGene_gn=CAMTA1;refGene_tx=XM_023258051.1;refGene_gn=CAMTA1;refGene_tx=XM_023258052.1;refGene_gn=CAMTA1;refGene_tx=XM_023258053.1
C1 5327206 5327394
refGene_gn=CAMTA1;refGene_tx=XM_011284441.3;refGene_gn=CAMTA1;refGene_tx=XM_011284442.3;refGene_gn=CAMTA1;refGene_tx=XM_011284444.3;refGene_gn=CAMTA1;refGene_tx=XM_023258051.1;refGene_gn=CAMTA1;refGene_tx=XM_023258052.1;refGene_gn=CAMTA1;refGene_tx=XM_023258053.1
C1 5327866 5328113
refGene_gn=CAMTA1;refGene_tx=XM_011284441.3;refGene_gn=CAMTA1;refGene_tx=XM_011284442.3;refGene_gn=CAMTA1;refGene_tx=XM_011284444.3;refGene_gn=CAMTA1;refGene_tx=XM_023258051.1;refGene_gn=CAMTA1;refGene_tx=XM_023258052.1;refGene_gn=CAMTA1;refGene_tx=XM_023258053.1
C1 5328939 5329011
refGene_gn=CAMTA1;refGene_tx=XM_011284441.3;refGene_gn=CAMTA1;refGene_tx=XM_011284442.3;refGene_gn=CAMTA1;refGene_tx=XM_011284444.3;refGene_gn=CAMTA1;refGene_tx=XM_023258051.1;refGene_gn=CAMTA1;refGene_tx=XM_023258052.1;refGene_gn=CAMTA1;refGene_tx=XM_023258053.1
C1 5330788 5330809
refGene_gn=CAMTA1;refGene_tx=XM_011284441.3;refGene_gn=CAMTA1;refGene_tx=XM_011284442.3;refGene_gn=CAMTA1;refGene_tx=XM_011284444.3;refGene_gn=CAMTA1;refGene_tx=XM_023258052.1
C1 5332256 5332450
refGene_gn=CAMTA1;refGene_tx=XM_011284441.3;refGene_gn=CAMTA1;refGene_tx=XM_011284442.3;refGene_gn=CAMTA1;refGene_tx=XM_011284444.3;refGene_gn=CAMTA1;refGene_tx=XM_023258051.1;refGene_gn=CAMTA1;refGene_tx=XM_023258052.1;refGene_gn=CAMTA1;refGene_tx=XM_023258053.1
C1 5333220 5333295
refGene_gn=CAMTA1;refGene_tx=XM_011284441.3;refGene_gn=CAMTA1;refGene_tx=XM_011284442.3;refGene_gn=CAMTA1;refGene_tx=XM_023258051.1;refGene_gn=CAMTA1;refGene_tx=XM_023258052.1;refGene_gn=CAMTA1;refGene_tx=XM_023258053.1
C1 5335885 5335916 refGene_gn=CAMTA1;refGene_tx=XM_011284442.3
C1 5345241 5345305
refGene_gn=CAMTA1;refGene_tx=XM_011284441.3;refGene_gn=CAMTA1;refGene_tx=XM_011284442.3;refGene_gn=CAMTA1;refGene_tx=XM_011284444.3;refGene_gn=CAMTA1;refGene_tx=XM_023258051.1;refGene_gn=CAMTA1;refGene_tx=XM_023258052.1;refGene_gn=CAMTA1;refGene_tx=XM_023258053.1
C1 5350328 5350330 refGene_gn=VAMP3;refGene_tx=XM_023258054.1
C1 5352753 5352835 refGene_gn=VAMP3;refGene_tx=XM_023258054.1
C1 5356010 5356169 refGene_gn=VAMP3;refGene_tx=XM_023258054.1
C1 5356955 5357004 refGene_gn=VAMP3;refGene_tx=XM_023258054.1
C1 5358061 5358081 refGene_gn=VAMP3;refGene_tx=XM_023258054.1
C1 5365760 5365939 refGene_gn=PER3;refGene_tx=XM_023260051.1
C1 5366364 5366510 refGene_gn=PER3;refGene_tx=XM_023260051.1
C1 5367848 5367964 refGene_gn=PER3;refGene_tx=XM_023260051.1
C1 5369170 5369372 refGene_gn=PER3;refGene_tx=XM_023260051.1

C1	5370564	5370655	refGene_gn=PER3;refGene_tx=XM_023260051.1
C1	5371884	5372002	refGene_gn=PER3;refGene_tx=XM_023260051.1
C1	5388004	5388064
refGene_gn=LOC101096081;refGene_tx=XM_023258055.1
C1	5388305	5388432
refGene_gn=LOC101096081;refGene_tx=XM_023258055.1;refGene_gn=LOC101096
081;refGene_tx=XM_023258056.1;refGene_gn=LOC101096081;refGene_tx=XM_02
3258057.1
C1	5399574	5399707
refGene_gn=LOC101096081;refGene_tx=XM_023258055.1;refGene_gn=LOC101096
081;refGene_tx=XM_023258056.1;refGene_gn=LOC101096081;refGene_tx=XM_02
3258057.1
C1	5400166	5400273
refGene_gn=LOC101096081;refGene_tx=XM_023258055.1;refGene_gn=LOC101096
081;refGene_tx=XM_023258056.1;refGene_gn=LOC101096081;refGene_tx=XM_02
3258057.1
C1	5400494	5400668
refGene_gn=LOC101096081;refGene_tx=XM_023258055.1;refGene_gn=LOC101096
081;refGene_tx=XM_023258056.1;refGene_gn=LOC101096081;refGene_tx=XM_02
3258057.1
C1	5405885	5406131
refGene_gn=LOC101096081;refGene_tx=XM_023258055.1;refGene_gn=LOC101096
081;refGene_tx=XM_023258056.1;refGene_gn=LOC101096081;refGene_tx=XM_02
3258057.1
C1	5406544	5407203
refGene_gn=LOC101096081;refGene_tx=XM_023258055.1;refGene_gn=LOC101096
081;refGene_tx=XM_023258056.1;refGene_gn=LOC101096081;refGene_tx=XM_02
3258057.1
C1	5408779	5408867
refGene_gn=LOC101096081;refGene_tx=XM_023258055.1;refGene_gn=LOC101096
081;refGene_tx=XM_023258056.1;refGene_gn=LOC101096081;refGene_tx=XM_02
3258057.1
C1	5419641	5419825
refGene_gn=LOC101096081;refGene_tx=XM_023258055.1;refGene_gn=LOC101096
081;refGene_tx=XM_023258056.1;refGene_gn=LOC101096081;refGene_tx=XM_02
3258057.1
C1	5420610	5420761
refGene_gn=LOC101096081;refGene_tx=XM_023258055.1;refGene_gn=LOC101096
081;refGene_tx=XM_023258056.1;refGene_gn=LOC101096081;refGene_tx=XM_02
3258057.1
C1	5424763	5424970
refGene_gn=LOC101096081;refGene_tx=XM_023258055.1;refGene_gn=LOC101096
081;refGene_tx=XM_023258056.1;refGene_gn=LOC101096081;refGene_tx=XM_02
3258057.1
C1	5425591	5425711
refGene_gn=UTS2;refGene_tx=XM_003989563.3;refGene_gn=UTS2;refGene_tx=X
M_019836082.1
C1	5426569	5426613
refGene_gn=UTS2;refGene_tx=XM_003989563.3;refGene_gn=UTS2;refGene_tx=X
M_019836082.1

```
C1	5428074	5428182	refGene_gn=UTS2;refGene_tx=XM_003989563.3;refGene_gn=UTS2;refGene_tx=XM_019836082.1
C1	5430784	5430884	refGene_gn=UTS2;refGene_tx=XM_003989563.3;refGene_gn=UTS2;refGene_tx=XM_019836082.1
C1	5478986	5479084	refGene_gn=TNFRSF9;refGene_tx=XM_023258059.1
C1	5483366	5483527	refGene_gn=TNFRSF9;refGene_tx=XM_023258058.1
C1	5483965	5484100	refGene_gn=TNFRSF9;refGene_tx=XM_023258058.1;refGene_gn=TNFRSF9;refGene_tx=XM_023258059.1
C1	5484630	5484761	refGene_gn=TNFRSF9;refGene_tx=XM_023258058.1;refGene_gn=TNFRSF9;refGene_tx=XM_023258059.1
C1	5487261	5487328	refGene_gn=TNFRSF9;refGene_tx=XM_023258058.1;refGene_gn=TNFRSF9;refGene_tx=XM_023258059.1
C1	5487769	5487907	refGene_gn=TNFRSF9;refGene_tx=XM_023258058.1;refGene_gn=TNFRSF9;refGene_tx=XM_023258059.1
C1	5488904	5489012	refGene_gn=TNFRSF9;refGene_tx=XM_023258058.1;refGene_gn=TNFRSF9;refGene_tx=XM_023258059.1
C1	5489746	5489861	refGene_gn=TNFRSF9;refGene_tx=XM_023258058.1;refGene_gn=TNFRSF9;refGene_tx=XM_023258059.1
C1	5507907	5507997	refGene_gn=PARK7;refGene_tx=XM_023258060.1;refGene_gn=PARK7;refGene_tx=XM_023258061.1
C1	5509600	5509702	refGene_gn=PARK7;refGene_tx=XM_023258060.1;refGene_gn=PARK7;refGene_tx=XM_023258061.1
C1	5512021	5512081	refGene_gn=PARK7;refGene_tx=XM_023258060.1;refGene_gn=PARK7;refGene_tx=XM_023258061.1
C1	5512982	5513052	refGene_gn=PARK7;refGene_tx=XM_023258060.1;refGene_gn=PARK7;refGene_tx=XM_023258061.1
C1	5516536	5516623	refGene_gn=PARK7;refGene_tx=XM_023258060.1;refGene_gn=PARK7;refGene_tx=XM_023258061.1
C1	5523811	5523972	refGene_gn=PARK7;refGene_tx=XM_023258060.1;refGene_gn=PARK7;refGene_tx=XM_023258061.1
C1	5551432	5552616	refGene_gn=ERRFI1;refGene_tx=XM_023258062.1
C1	5553506	5553583	refGene_gn=ERRFI1;refGene_tx=XM_023258062.1
C1	5553695	5553817	refGene_gn=ERRFI1;refGene_tx=XM_023258062.1
C1	5763627	5763738	refGene_gn=SLC45A1;refGene_tx=XM_023258068.1;refGene_gn=SLC45A1;refGene_tx=XM_023258069.1
```

C1 5768447 5768868
refGene_gn=SLC45A1;refGene_tx=XM_003989468.5;refGene_gn=SLC45A1;refGene_tx=XM_023258068.1;refGene_gn=SLC45A1;refGene_tx=XM_023258069.1
C1 5769453 5769546
refGene_gn=SLC45A1;refGene_tx=XM_003989468.5;refGene_gn=SLC45A1;refGene_tx=XM_023258068.1;refGene_gn=SLC45A1;refGene_tx=XM_023258069.1
C1 5770008 5770233
refGene_gn=SLC45A1;refGene_tx=XM_003989468.5;refGene_gn=SLC45A1;refGene_tx=XM_023258068.1;refGene_gn=SLC45A1;refGene_tx=XM_023258069.1
C1 5773206 5773946
refGene_gn=SLC45A1;refGene_tx=XM_003989468.5;refGene_gn=SLC45A1;refGene_tx=XM_023258068.1;refGene_gn=SLC45A1;refGene_tx=XM_023258069.1
C1 5776446 5776600
refGene_gn=SLC45A1;refGene_tx=XM_003989468.5;refGene_gn=SLC45A1;refGene_tx=XM_023258068.1;refGene_gn=SLC45A1;refGene_tx=XM_023258069.1
C1 5778683 5778860
refGene_gn=SLC45A1;refGene_tx=XM_003989468.5;refGene_gn=SLC45A1;refGene_tx=XM_023258068.1;refGene_gn=SLC45A1;refGene_tx=XM_023258069.1
C1 5779958 5780164
refGene_gn=SLC45A1;refGene_tx=XM_003989468.5;refGene_gn=SLC45A1;refGene_tx=XM_023258068.1;refGene_gn=SLC45A1;refGene_tx=XM_023258069.1
C1 5782592 5782859
refGene_gn=SLC45A1;refGene_tx=XM_003989468.5;refGene_gn=SLC45A1;refGene_tx=XM_023258068.1
C1 5785523 5785781 refGene_gn=SLC45A1;refGene_tx=XM_023258069.1
C1 5792591 5792625
refGene_gn=RERE;refGene_tx=XM_023258063.1;refGene_gn=RERE;refGene_tx=XM_023258064.1;refGene_gn=RERE;refGene_tx=XM_023258065.1;refGene_gn=RERE;refGene_tx=XM_023258066.1;refGene_gn=RERE;refGene_tx=XM_023258067.1
C1 5792978 5793159
refGene_gn=RERE;refGene_tx=XM_023258063.1;refGene_gn=RERE;refGene_tx=XM_023258064.1;refGene_gn=RERE;refGene_tx=XM_023258065.1;refGene_gn=RERE;refGene_tx=XM_023258066.1;refGene_gn=RERE;refGene_tx=XM_023258067.1
C1 5793645 5793792
refGene_gn=RERE;refGene_tx=XM_023258063.1;refGene_gn=RERE;refGene_tx=XM_023258064.1;refGene_gn=RERE;refGene_tx=XM_023258065.1;refGene_gn=RERE;refGene_tx=XM_023258066.1;refGene_gn=RERE;refGene_tx=XM_023258067.1
C1 5795418 5796142
refGene_gn=RERE;refGene_tx=XM_023258063.1;refGene_gn=RERE;refGene_tx=XM_023258064.1;refGene_gn=RERE;refGene_tx=XM_023258065.1;refGene_gn=RERE;refGene_tx=XM_023258066.1;refGene_gn=RERE;refGene_tx=XM_023258067.1
C1 5796458 5796681
refGene_gn=RERE;refGene_tx=XM_023258063.1;refGene_gn=RERE;refGene_tx=XM_023258064.1;refGene_gn=RERE;refGene_tx=XM_023258065.1;refGene_gn=RERE;refGene_tx=XM_023258066.1;refGene_gn=RERE;refGene_tx=XM_023258067.1
C1 5796790 5798112
refGene_gn=RERE;refGene_tx=XM_023258063.1;refGene_gn=RERE;refGene_tx=XM_023258064.1;refGene_gn=RERE;refGene_tx=XM_023258065.1;refGene_gn=RERE;refGene_tx=XM_023258066.1;refGene_gn=RERE;refGene_tx=XM_023258067.1
C1 5798389 5798503

refGene_gn=RERE;refGene_tx=XM_023258063.1;refGene_gn=RERE;refGene_tx=XM_023258064.1;refGene_gn=RERE;refGene_tx=XM_023258065.1;refGene_gn=RERE;refGene_tx=XM_023258066.1;refGene_gn=RERE;refGene_tx=XM_023258067.1
C1 5799004 5799166
refGene_gn=RERE;refGene_tx=XM_023258063.1;refGene_gn=RERE;refGene_tx=XM_023258064.1;refGene_gn=RERE;refGene_tx=XM_023258065.1;refGene_gn=RERE;refGene_tx=XM_023258066.1;refGene_gn=RERE;refGene_tx=XM_023258067.1
C1 5800379 5800579
refGene_gn=RERE;refGene_tx=XM_023258063.1;refGene_gn=RERE;refGene_tx=XM_023258064.1;refGene_gn=RERE;refGene_tx=XM_023258065.1;refGene_gn=RERE;refGene_tx=XM_023258066.1;refGene_gn=RERE;refGene_tx=XM_023258067.1
C1 5800911 5801004
refGene_gn=RERE;refGene_tx=XM_023258063.1;refGene_gn=RERE;refGene_tx=XM_023258064.1;refGene_gn=RERE;refGene_tx=XM_023258065.1;refGene_gn=RERE;refGene_tx=XM_023258066.1;refGene_gn=RERE;refGene_tx=XM_023258067.1
C1 5802127 5802290
refGene_gn=RERE;refGene_tx=XM_023258063.1;refGene_gn=RERE;refGene_tx=XM_023258064.1;refGene_gn=RERE;refGene_tx=XM_023258065.1;refGene_gn=RERE;refGene_tx=XM_023258066.1;refGene_gn=RERE;refGene_tx=XM_023258067.1
C1 5847532 5847613
refGene_gn=RERE;refGene_tx=XM_023258063.1;refGene_gn=RERE;refGene_tx=XM_023258064.1;refGene_gn=RERE;refGene_tx=XM_023258065.1;refGene_gn=RERE;refGene_tx=XM_023258066.1;refGene_gn=RERE;refGene_tx=XM_023258067.1
C1 5882744 5882843
refGene_gn=RERE;refGene_tx=XM_023258063.1;refGene_gn=RERE;refGene_tx=XM_023258064.1;refGene_gn=RERE;refGene_tx=XM_023258065.1;refGene_gn=RERE;refGene_tx=XM_023258066.1;refGene_gn=RERE;refGene_tx=XM_023258067.1
C1 5908123 5908223
refGene_gn=RERE;refGene_tx=XM_023258063.1;refGene_gn=RERE;refGene_tx=XM_023258064.1;refGene_gn=RERE;refGene_tx=XM_023258065.1;refGene_gn=RERE;refGene_tx=XM_023258066.1;refGene_gn=RERE;refGene_tx=XM_023258067.1
C1 5909621 5909746
refGene_gn=RERE;refGene_tx=XM_023258063.1;refGene_gn=RERE;refGene_tx=XM_023258064.1;refGene_gn=RERE;refGene_tx=XM_023258065.1;refGene_gn=RERE;refGene_tx=XM_023258066.1;refGene_gn=RERE;refGene_tx=XM_023258067.1
C1 5915978 5916027
refGene_gn=RERE;refGene_tx=XM_023258063.1;refGene_gn=RERE;refGene_tx=XM_023258064.1;refGene_gn=RERE;refGene_tx=XM_023258065.1;refGene_gn=RERE;refGene_tx=XM_023258066.1;refGene_gn=RERE;refGene_tx=XM_023258067.1
C1 5929432 5929458 refGene_gn=RERE;refGene_tx=XM_023258067.1
C1 5945218 5945323
refGene_gn=RERE;refGene_tx=XM_023258063.1;refGene_gn=RERE;refGene_tx=XM_023258064.1;refGene_gn=RERE;refGene_tx=XM_023258065.1;refGene_gn=RERE;refGene_tx=XM_023258066.1
C1 5958664 5958761
refGene_gn=RERE;refGene_tx=XM_023258063.1;refGene_gn=RERE;refGene_tx=XM_023258064.1;refGene_gn=RERE;refGene_tx=XM_023258065.1;refGene_gn=RERE;refGene_tx=XM_023258066.1
C1 5960116 5960222
refGene_gn=RERE;refGene_tx=XM_023258063.1;refGene_gn=RERE;refGene_tx=X

M_023258064.1;refGene_gn=RERE;refGene_tx=XM_023258065.1;refGene_gn=RERE;refGene_tx=XM_023258066.1
C1 6010357 6010483
refGene_gn=RERE;refGene_tx=XM_023258063.1;refGene_gn=RERE;refGene_tx=XM_023258064.1;refGene_gn=RERE;refGene_tx=XM_023258065.1
C1 6019660 6019731
refGene_gn=RERE;refGene_tx=XM_023258063.1;refGene_gn=RERE;refGene_tx=XM_023258064.1;refGene_gn=RERE;refGene_tx=XM_023258065.1;refGene_gn=RERE;refGene_tx=XM_023258066.1
C1 6051060 6051385
refGene_gn=RERE;refGene_tx=XM_023258063.1;refGene_gn=RERE;refGene_tx=XM_023258064.1;refGene_gn=RERE;refGene_tx=XM_023258065.1;refGene_gn=RERE;refGene_tx=XM_023258066.1
C1 6254856 6254926
refGene_gn=ENO1;refGene_tx=XM_003989469.4;refGene_gn=ENO1;refGene_tx=XM_006934255.3
C1 6256071 6256130
refGene_gn=ENO1;refGene_tx=XM_003989469.4;refGene_gn=ENO1;refGene_tx=XM_006934255.3
C1 6256304 6256413
refGene_gn=ENO1;refGene_tx=XM_003989469.4;refGene_gn=ENO1;refGene_tx=XM_006934255.3
C1 6256917 6257119
refGene_gn=ENO1;refGene_tx=XM_003989469.4;refGene_gn=ENO1;refGene_tx=XM_006934255.3
C1 6257913 6258111
refGene_gn=ENO1;refGene_tx=XM_003989469.4;refGene_gn=ENO1;refGene_tx=XM_006934255.3
C1 6259483 6259706
refGene_gn=ENO1;refGene_tx=XM_003989469.4;refGene_gn=ENO1;refGene_tx=XM_006934255.3
C1 6260120 6260254
refGene_gn=ENO1;refGene_tx=XM_003989469.4;refGene_gn=ENO1;refGene_tx=XM_006934255.3
C1 6260847 6260917
refGene_gn=ENO1;refGene_tx=XM_003989469.4;refGene_gn=ENO1;refGene_tx=XM_006934255.3
C1 6263248 6263307 refGene_gn=ENO1;refGene_tx=XM_003989469.4
C1 6263628 6263687 refGene_gn=ENO1;refGene_tx=XM_006934255.3
C1 6264732 6264828
refGene_gn=ENO1;refGene_tx=XM_003989469.4;refGene_gn=ENO1;refGene_tx=XM_006934255.3
C1 6267498 6267583
refGene_gn=ENO1;refGene_tx=XM_003989469.4;refGene_gn=ENO1;refGene_tx=XM_006934255.3
C1 6299502 6299581 refGene_gn=CA6;refGene_tx=XM_003989566.3
C1 6301476 6301656 refGene_gn=CA6;refGene_tx=XM_003989566.3
C1 6309456 6309605 refGene_gn=CA6;refGene_tx=XM_003989566.3
C1 6310501 6310594 refGene_gn=CA6;refGene_tx=XM_003989566.3
C1 6313995 6314065 refGene_gn=CA6;refGene_tx=XM_003989566.3

C1 6316411 6316569 refGene_gn=CA6;refGene_tx=XM_003989566.3
C1 6317482 6317606 refGene_gn=CA6;refGene_tx=XM_003989566.3
C1 6319673 6319774 refGene_gn=CA6;refGene_tx=XM_003989566.3
C1 6334496 6334700
refGene_gn=LOC101097571;refGene_tx=XM_019836092.1;refGene_gn=LOC101097571;refGene_tx=XM_019836094.1;refGene_gn=LOC101097571;refGene_tx=XM_023258070.1;refGene_gn=LOC101097571;refGene_tx=XM_023258071.1
C1 6334805 6334933
refGene_gn=LOC101097571;refGene_tx=XM_019836092.1;refGene_gn=LOC101097571;refGene_tx=XM_019836094.1;refGene_gn=LOC101097571;refGene_tx=XM_023258070.1;refGene_gn=LOC101097571;refGene_tx=XM_023258071.1
C1 6335408 6335484
refGene_gn=LOC101097571;refGene_tx=XM_019836092.1;refGene_gn=LOC101097571;refGene_tx=XM_019836094.1;refGene_gn=LOC101097571;refGene_tx=XM_023258070.1;refGene_gn=LOC101097571;refGene_tx=XM_023258071.1
C1 6336067 6336169
refGene_gn=LOC101097571;refGene_tx=XM_019836092.1;refGene_gn=LOC101097571;refGene_tx=XM_019836094.1;refGene_gn=LOC101097571;refGene_tx=XM_023258070.1;refGene_gn=LOC101097571;refGene_tx=XM_023258071.1
C1 6336624 6336735
refGene_gn=LOC101097571;refGene_tx=XM_019836092.1;refGene_gn=LOC101097571;refGene_tx=XM_019836094.1;refGene_gn=LOC101097571;refGene_tx=XM_023258070.1;refGene_gn=LOC101097571;refGene_tx=XM_023258071.1
C1 6336826 6337014
refGene_gn=LOC101097571;refGene_tx=XM_019836092.1;refGene_gn=LOC101097571;refGene_tx=XM_019836094.1;refGene_gn=LOC101097571;refGene_tx=XM_023258070.1;refGene_gn=LOC101097571;refGene_tx=XM_023258071.1
C1 6337107 6337233
refGene_gn=LOC101097571;refGene_tx=XM_019836092.1;refGene_gn=LOC101097571;refGene_tx=XM_019836094.1;refGene_gn=LOC101097571;refGene_tx=XM_023258070.1
C1 6339396 6339549
refGene_gn=LOC101097571;refGene_tx=XM_019836092.1;refGene_gn=LOC101097571;refGene_tx=XM_019836094.1;refGene_gn=LOC101097571;refGene_tx=XM_023258070.1;refGene_gn=LOC101097571;refGene_tx=XM_023258071.1
C1 6343322 6343447
refGene_gn=LOC101097571;refGene_tx=XM_019836092.1;refGene_gn=LOC101097571;refGene_tx=XM_019836094.1;refGene_gn=LOC101097571;refGene_tx=XM_023258070.1;refGene_gn=LOC101097571;refGene_tx=XM_023258071.1
C1 6345929 6346090
refGene_gn=LOC101097571;refGene_tx=XM_019836092.1;refGene_gn=LOC101097571;refGene_tx=XM_019836094.1;refGene_gn=LOC101097571;refGene_tx=XM_023258070.1;refGene_gn=LOC101097571;refGene_tx=XM_023258071.1
C1 6346610 6346709
refGene_gn=LOC101097571;refGene_tx=XM_019836092.1;refGene_gn=LOC101097571;refGene_tx=XM_023258070.1;refGene_gn=LOC101097571;refGene_tx=XM_023258071.1
C1 6354275 6354305
refGene_gn=LOC101097571;refGene_tx=XM_019836092.1;refGene_gn=LOC101097571;refGene_tx=XM_023258070.1;refGene_gn=LOC101097571;refGene_tx=XM_02

3258071.1
C1	6369278	6369494
refGene_gn=GPR157;refGene_tx=XM_006934339.2;refGene_gn=GPR157;refGene_tx=XM_023258072.1
C1	6369974	6370169
refGene_gn=GPR157;refGene_tx=XM_006934339.2;refGene_gn=GPR157;refGene_tx=XM_023258072.1
C1	6372763	6372977
refGene_gn=GPR157;refGene_tx=XM_006934339.2;refGene_gn=GPR157;refGene_tx=XM_023258072.1
C1	6385519	6385902
refGene_gn=GPR157;refGene_tx=XM_006934339.2;refGene_gn=GPR157;refGene_tx=XM_023258072.1
C1	6398822	6399621
refGene_gn=LOC102899466;refGene_tx=XM_006934257.4
C1	6399781	6399882
refGene_gn=LOC102899466;refGene_tx=XM_006934257.4
C1	6455993	6456043	refGene_gn=H6PD;refGene_tx=XM_023258074.1
C1	6456799	6457041	refGene_gn=H6PD;refGene_tx=XM_006934258.4
C1	6457047	6457079	refGene_gn=H6PD;refGene_tx=XM_023258073.1
C1	6457493	6457643
refGene_gn=H6PD;refGene_tx=XM_006934258.4;refGene_gn=H6PD;refGene_tx=XM_023258073.1;refGene_gn=H6PD;refGene_tx=XM_023258074.1
C1	6458333	6458338	refGene_gn=H6PD;refGene_tx=XM_023258075.1
C1	6463460	6463549	refGene_gn=H6PD;refGene_tx=XM_003989569.5
C1	6466317	6466954
refGene_gn=H6PD;refGene_tx=XM_003989569.5;refGene_gn=H6PD;refGene_tx=XM_006934258.4;refGene_gn=H6PD;refGene_tx=XM_019836100.2;refGene_gn=H6PD;refGene_tx=XM_019836101.2;refGene_gn=H6PD;refGene_tx=XM_023258073.1;refGene_gn=H6PD;refGene_tx=XM_023258074.1;refGene_gn=H6PD;refGene_tx=XM_023258075.1
C1	6468022	6468140
refGene_gn=H6PD;refGene_tx=XM_003989569.5;refGene_gn=H6PD;refGene_tx=XM_006934258.4;refGene_gn=H6PD;refGene_tx=XM_019836100.2;refGene_gn=H6PD;refGene_tx=XM_019836101.2;refGene_gn=H6PD;refGene_tx=XM_023258073.1;refGene_gn=H6PD;refGene_tx=XM_023258074.1;refGene_gn=H6PD;refGene_tx=XM_023258075.1
C1	6479904	6480174
refGene_gn=H6PD;refGene_tx=XM_003989569.5;refGene_gn=H6PD;refGene_tx=XM_006934258.4;refGene_gn=H6PD;refGene_tx=XM_019836100.2;refGene_gn=H6PD;refGene_tx=XM_019836101.2;refGene_gn=H6PD;refGene_tx=XM_023258073.1;refGene_gn=H6PD;refGene_tx=XM_023258074.1;refGene_gn=H6PD;refGene_tx=XM_023258075.1
C1	6481303	6482667
refGene_gn=H6PD;refGene_tx=XM_003989569.5;refGene_gn=H6PD;refGene_tx=XM_006934258.4;refGene_gn=H6PD;refGene_tx=XM_019836100.2;refGene_gn=H6PD;refGene_tx=XM_019836101.2;refGene_gn=H6PD;refGene_tx=XM_023258073.1;refGene_gn=H6PD;refGene_tx=XM_023258074.1;refGene_gn=H6PD;refGene_tx=XM_023258075.1
C1	6489943	6490048

refGene_gn=LOC111561966;refGene_tx=XM_023260053.1
C1	6506127	6506570
refGene_gn=LOC111561966;refGene_tx=XM_023260053.1
C1	6511530	6511675
refGene_gn=LOC111561966;refGene_tx=XM_023260053.1
C1	6512181	6512336
refGene_gn=LOC111561966;refGene_tx=XM_023260053.1
C1	6513007	6513091
refGene_gn=LOC111561966;refGene_tx=XM_023260053.1
C1	6527583	6527767
refGene_gn=LOC111561966;refGene_tx=XM_023260053.1
C1	6528721	6528920
refGene_gn=LOC111561966;refGene_tx=XM_023260053.1
C1	6529563	6529669
refGene_gn=LOC111561966;refGene_tx=XM_023260053.1
C1	6545618	6545822
refGene_gn=LOC111561966;refGene_tx=XM_023260053.1
C1	6546878	6547068
refGene_gn=LOC111561966;refGene_tx=XM_023260053.1
C1	6560970	6561664
refGene_gn=SPSB1;refGene_tx=XM_003989471.5;refGene_gn=SPSB1;refGene_tx
=XM_019836102.2;refGene_gn=SPSB1;refGene_tx=XM_023258076.1;refGene_gn=
SPSB1;refGene_tx=XM_023258077.1
C1	6571421	6571549
refGene_gn=SPSB1;refGene_tx=XM_003989471.5;refGene_gn=SPSB1;refGene_tx
=XM_019836102.2;refGene_gn=SPSB1;refGene_tx=XM_023258076.1;refGene_gn=
SPSB1;refGene_tx=XM_023258077.1
C1	6696182	6696238	refGene_gn=SLC25A33;refGene_tx=XM_023258078.1
C1	6710310	6710490	refGene_gn=SLC25A33;refGene_tx=XM_023258078.1
C1	6720155	6720232	refGene_gn=SLC25A33;refGene_tx=XM_023258079.1
C1	6721087	6721165
refGene_gn=SLC25A33;refGene_tx=XM_023258078.1;refGene_gn=SLC25A33;refG
ene_tx=XM_023258079.1
C1	6722954	6723055
refGene_gn=SLC25A33;refGene_tx=XM_023258078.1;refGene_gn=SLC25A33;refG
ene_tx=XM_023258079.1
C1	6725271	6725338
refGene_gn=SLC25A33;refGene_tx=XM_023258078.1;refGene_gn=SLC25A33;refG
ene_tx=XM_023258079.1
C1	6728338	6728619
refGene_gn=SLC25A33;refGene_tx=XM_023258078.1;refGene_gn=SLC25A33;refG
ene_tx=XM_023258079.1
C1	6729816	6730019
refGene_gn=SLC25A33;refGene_tx=XM_023258078.1;refGene_gn=SLC25A33;refG
ene_tx=XM_023258079.1
C1	6734581	6734694	refGene_gn=TMEM201;refGene_tx=XM_023258080.1
C1	6741775	6741896
refGene_gn=TMEM201;refGene_tx=XM_023258080.1;refGene_gn=TMEM201;refGen
e_tx=XM_023258081.1;refGene_gn=TMEM201;refGene_tx=XM_023258082.1
C1	6742390	6742585

refGene_gn=TMEM201;refGene_tx=XM_023258080.1;refGene_gn=TMEM201;refGene_tx=XM_023258081.1;refGene_gn=TMEM201;refGene_tx=XM_023258082.1
C1 6743774 6743951
refGene_gn=TMEM201;refGene_tx=XM_023258080.1;refGene_gn=TMEM201;refGene_tx=XM_023258081.1;refGene_gn=TMEM201;refGene_tx=XM_023258082.1
C1 6745473 6745823
refGene_gn=TMEM201;refGene_tx=XM_023258080.1;refGene_gn=TMEM201;refGene_tx=XM_023258081.1;refGene_gn=TMEM201;refGene_tx=XM_023258082.1
C1 6746417 6746621
refGene_gn=TMEM201;refGene_tx=XM_023258080.1;refGene_gn=TMEM201;refGene_tx=XM_023258081.1;refGene_gn=TMEM201;refGene_tx=XM_023258082.1
C1 6751917 6752150
refGene_gn=TMEM201;refGene_tx=XM_023258080.1;refGene_gn=TMEM201;refGene_tx=XM_023258081.1;refGene_gn=TMEM201;refGene_tx=XM_023258082.1
C1 6757223 6757295
refGene_gn=TMEM201;refGene_tx=XM_023258080.1;refGene_gn=TMEM201;refGene_tx=XM_023258081.1;refGene_gn=TMEM201;refGene_tx=XM_023258082.1
C1 6757844 6758144
refGene_gn=TMEM201;refGene_tx=XM_023258080.1;refGene_gn=TMEM201;refGene_tx=XM_023258081.1;refGene_gn=TMEM201;refGene_tx=XM_023258082.1
C1 6758974 6759112
refGene_gn=TMEM201;refGene_tx=XM_023258080.1;refGene_gn=TMEM201;refGene_tx=XM_023258081.1;refGene_gn=TMEM201;refGene_tx=XM_023258082.1
C1 6760515 6760613
refGene_gn=TMEM201;refGene_tx=XM_023258080.1;refGene_gn=TMEM201;refGene_tx=XM_023258081.1;refGene_gn=TMEM201;refGene_tx=XM_023258082.1
C1 6829580 6829721
refGene_gn=PIK3CD;refGene_tx=XM_006934259.2;refGene_gn=PIK3CD;refGene_tx=XM_006934260.2;refGene_gn=PIK3CD;refGene_tx=XM_006934262.4;refGene_gn=PIK3CD;refGene_tx=XM_019836105.1
C1 6832219 6832448
refGene_gn=PIK3CD;refGene_tx=XM_006934259.2;refGene_gn=PIK3CD;refGene_tx=XM_006934260.2;refGene_gn=PIK3CD;refGene_tx=XM_006934262.4;refGene_gn=PIK3CD;refGene_tx=XM_019836105.1
C1 6832733 6832963
refGene_gn=PIK3CD;refGene_tx=XM_006934259.2;refGene_gn=PIK3CD;refGene_tx=XM_006934260.2;refGene_gn=PIK3CD;refGene_tx=XM_006934262.4;refGene_gn=PIK3CD;refGene_tx=XM_019836105.1
C1 6833413 6833593
refGene_gn=PIK3CD;refGene_tx=XM_006934259.2;refGene_gn=PIK3CD;refGene_tx=XM_006934260.2;refGene_gn=PIK3CD;refGene_tx=XM_006934262.4;refGene_gn=PIK3CD;refGene_tx=XM_019836105.1
C1 6833864 6834014
refGene_gn=PIK3CD;refGene_tx=XM_006934259.2;refGene_gn=PIK3CD;refGene_tx=XM_006934260.2;refGene_gn=PIK3CD;refGene_tx=XM_006934262.4;refGene_gn=PIK3CD;refGene_tx=XM_006934263.2;refGene_gn=PIK3CD;refGene_tx=XM_019836105.1
C1 6834451 6834541
refGene_gn=PIK3CD;refGene_tx=XM_006934259.2;refGene_gn=PIK3CD;refGene_tx=XM_006934260.2;refGene_gn=PIK3CD;refGene_tx=XM_006934262.4;refGene_

gn=PIK3CD;refGene_tx=XM_006934263.2;refGene_gn=PIK3CD;refGene_tx=XM_019836105.1
C1 6835628 6835850
refGene_gn=PIK3CD;refGene_tx=XM_006934259.2;refGene_gn=PIK3CD;refGene_tx=XM_006934260.2;refGene_gn=PIK3CD;refGene_tx=XM_006934262.4;refGene_gn=PIK3CD;refGene_tx=XM_006934263.2;refGene_gn=PIK3CD;refGene_tx=XM_019836105.1
C1 6836489 6836586
refGene_gn=PIK3CD;refGene_tx=XM_006934259.2;refGene_gn=PIK3CD;refGene_tx=XM_006934260.2;refGene_gn=PIK3CD;refGene_tx=XM_006934262.4;refGene_gn=PIK3CD;refGene_tx=XM_006934263.2;refGene_gn=PIK3CD;refGene_tx=XM_019836105.1
C1 6836675 6836806
refGene_gn=PIK3CD;refGene_tx=XM_006934259.2;refGene_gn=PIK3CD;refGene_tx=XM_006934260.2;refGene_gn=PIK3CD;refGene_tx=XM_006934262.4;refGene_gn=PIK3CD;refGene_tx=XM_006934263.2;refGene_gn=PIK3CD;refGene_tx=XM_019836105.1
C1 6837286 6837337
refGene_gn=PIK3CD;refGene_tx=XM_006934259.2;refGene_gn=PIK3CD;refGene_tx=XM_006934260.2;refGene_gn=PIK3CD;refGene_tx=XM_006934262.4;refGene_gn=PIK3CD;refGene_tx=XM_006934263.2;refGene_gn=PIK3CD;refGene_tx=XM_019836105.1
C1 6837405 6837573
refGene_gn=PIK3CD;refGene_tx=XM_006934259.2;refGene_gn=PIK3CD;refGene_tx=XM_006934260.2;refGene_gn=PIK3CD;refGene_tx=XM_006934262.4;refGene_gn=PIK3CD;refGene_tx=XM_006934263.2;refGene_gn=PIK3CD;refGene_tx=XM_019836105.1
C1 6837667 6837789
refGene_gn=PIK3CD;refGene_tx=XM_006934259.2;refGene_gn=PIK3CD;refGene_tx=XM_006934260.2;refGene_gn=PIK3CD;refGene_tx=XM_006934262.4;refGene_gn=PIK3CD;refGene_tx=XM_006934263.2;refGene_gn=PIK3CD;refGene_tx=XM_019836105.1
C1 6837939 6838083
refGene_gn=PIK3CD;refGene_tx=XM_006934259.2;refGene_gn=PIK3CD;refGene_tx=XM_006934260.2;refGene_gn=PIK3CD;refGene_tx=XM_006934262.4;refGene_gn=PIK3CD;refGene_tx=XM_006934263.2;refGene_gn=PIK3CD;refGene_tx=XM_019836105.1
C1 6838175 6838275
refGene_gn=PIK3CD;refGene_tx=XM_006934259.2;refGene_gn=PIK3CD;refGene_tx=XM_006934260.2;refGene_gn=PIK3CD;refGene_tx=XM_006934262.4;refGene_gn=PIK3CD;refGene_tx=XM_006934263.2;refGene_gn=PIK3CD;refGene_tx=XM_019836105.1
C1 6838425 6838604
refGene_gn=PIK3CD;refGene_tx=XM_006934259.2;refGene_gn=PIK3CD;refGene_tx=XM_006934260.2;refGene_gn=PIK3CD;refGene_tx=XM_006934262.4;refGene_gn=PIK3CD;refGene_tx=XM_006934263.2;refGene_gn=PIK3CD;refGene_tx=XM_019836105.1
C1 6838695 6838808
refGene_gn=PIK3CD;refGene_tx=XM_006934259.2;refGene_gn=PIK3CD;refGene_tx=XM_006934260.2;refGene_gn=PIK3CD;refGene_tx=XM_006934262.4;refGene_

gn=PIK3CD;refGene_tx=XM_006934263.2;refGene_gn=PIK3CD;refGene_tx=XM_01
9836105.1
C1 6838959 6839038
refGene_gn=PIK3CD;refGene_tx=XM_006934259.2;refGene_gn=PIK3CD;refGene_
tx=XM_006934260.2;refGene_gn=PIK3CD;refGene_tx=XM_006934262.4;refGene_
gn=PIK3CD;refGene_tx=XM_006934263.2;refGene_gn=PIK3CD;refGene_tx=XM_01
9836105.1
C1 6839611 6839779
refGene_gn=PIK3CD;refGene_tx=XM_006934259.2;refGene_gn=PIK3CD;refGene_
tx=XM_006934260.2;refGene_gn=PIK3CD;refGene_tx=XM_006934262.4;refGene_
gn=PIK3CD;refGene_tx=XM_006934263.2;refGene_gn=PIK3CD;refGene_tx=XM_01
9836105.1
C1 6840381 6840505
refGene_gn=PIK3CD;refGene_tx=XM_006934259.2;refGene_gn=PIK3CD;refGene_
tx=XM_006934260.2;refGene_gn=PIK3CD;refGene_tx=XM_006934262.4;refGene_
gn=PIK3CD;refGene_tx=XM_006934263.2;refGene_gn=PIK3CD;refGene_tx=XM_01
9836105.1
C1 6840632 6840778
refGene_gn=PIK3CD;refGene_tx=XM_006934259.2;refGene_gn=PIK3CD;refGene_
tx=XM_006934260.2;refGene_gn=PIK3CD;refGene_tx=XM_006934262.4;refGene_
gn=PIK3CD;refGene_tx=XM_006934263.2;refGene_gn=PIK3CD;refGene_tx=XM_01
9836105.1
C1 6840960 6841093
refGene_gn=PIK3CD;refGene_tx=XM_006934259.2;refGene_gn=PIK3CD;refGene_
tx=XM_006934260.2;refGene_gn=PIK3CD;refGene_tx=XM_006934262.4;refGene_
gn=PIK3CD;refGene_tx=XM_006934263.2;refGene_gn=PIK3CD;refGene_tx=XM_01
9836105.1
C1 6841836 6841974
refGene_gn=PIK3CD;refGene_tx=XM_006934259.2;refGene_gn=PIK3CD;refGene_
tx=XM_006934260.2;refGene_gn=PIK3CD;refGene_tx=XM_006934262.4;refGene_
gn=PIK3CD;refGene_tx=XM_006934263.2;refGene_gn=PIK3CD;refGene_tx=XM_01
9836105.1
C1 6844922 6845120
refGene_gn=CLSTN1;refGene_tx=XM_019836106.1;refGene_gn=CLSTN1;refGene_
tx=XM_023258083.1;refGene_gn=CLSTN1;refGene_tx=XM_023258084.1;refGene_
gn=CLSTN1;refGene_tx=XM_023258085.1;refGene_gn=CLSTN1;refGene_tx=XM_02
3258086.1
C1 6845494 6845679
refGene_gn=CLSTN1;refGene_tx=XM_019836106.1;refGene_gn=CLSTN1;refGene_
tx=XM_023258083.1;refGene_gn=CLSTN1;refGene_tx=XM_023258084.1;refGene_
gn=CLSTN1;refGene_tx=XM_023258085.1;refGene_gn=CLSTN1;refGene_tx=XM_02
3258086.1
C1 6846042 6846178
refGene_gn=CLSTN1;refGene_tx=XM_019836106.1;refGene_gn=CLSTN1;refGene_
tx=XM_023258083.1;refGene_gn=CLSTN1;refGene_tx=XM_023258084.1;refGene_
gn=CLSTN1;refGene_tx=XM_023258085.1;refGene_gn=CLSTN1;refGene_tx=XM_02
3258086.1
C1 6847153 6847299
refGene_gn=CLSTN1;refGene_tx=XM_019836106.1;refGene_gn=CLSTN1;refGene_
tx=XM_023258083.1;refGene_gn=CLSTN1;refGene_tx=XM_023258084.1;refGene_

gn=CLSTN1;refGene_tx=XM_023258085.1;refGene_gn=CLSTN1;refGene_tx=XM_023258086.1
C1 6847639 6847810
refGene_gn=CLSTN1;refGene_tx=XM_019836106.1;refGene_gn=CLSTN1;refGene_tx=XM_023258083.1;refGene_gn=CLSTN1;refGene_tx=XM_023258084.1;refGene_gn=CLSTN1;refGene_tx=XM_023258085.1;refGene_gn=CLSTN1;refGene_tx=XM_023258086.1
C1 6848561 6848788
refGene_gn=CLSTN1;refGene_tx=XM_019836106.1;refGene_gn=CLSTN1;refGene_tx=XM_023258083.1;refGene_gn=CLSTN1;refGene_tx=XM_023258084.1;refGene_gn=CLSTN1;refGene_tx=XM_023258085.1;refGene_gn=CLSTN1;refGene_tx=XM_023258086.1
C1 6849094 6849243
refGene_gn=CLSTN1;refGene_tx=XM_019836106.1;refGene_gn=CLSTN1;refGene_tx=XM_023258083.1;refGene_gn=CLSTN1;refGene_tx=XM_023258084.1;refGene_gn=CLSTN1;refGene_tx=XM_023258085.1;refGene_gn=CLSTN1;refGene_tx=XM_023258086.1
C1 6850654 6850809
refGene_gn=CLSTN1;refGene_tx=XM_019836106.1;refGene_gn=CLSTN1;refGene_tx=XM_023258083.1;refGene_gn=CLSTN1;refGene_tx=XM_023258084.1;refGene_gn=CLSTN1;refGene_tx=XM_023258085.1;refGene_gn=CLSTN1;refGene_tx=XM_023258086.1
C1 6852233 6852290
refGene_gn=CLSTN1;refGene_tx=XM_019836106.1;refGene_gn=CLSTN1;refGene_tx=XM_023258083.1;refGene_gn=CLSTN1;refGene_tx=XM_023258084.1
C1 6855188 6855351
refGene_gn=CLSTN1;refGene_tx=XM_019836106.1;refGene_gn=CLSTN1;refGene_tx=XM_023258083.1;refGene_gn=CLSTN1;refGene_tx=XM_023258084.1;refGene_gn=CLSTN1;refGene_tx=XM_023258085.1;refGene_gn=CLSTN1;refGene_tx=XM_023258086.1
C1 6865833 6865988
refGene_gn=CLSTN1;refGene_tx=XM_019836106.1;refGene_gn=CLSTN1;refGene_tx=XM_023258083.1;refGene_gn=CLSTN1;refGene_tx=XM_023258084.1;refGene_gn=CLSTN1;refGene_tx=XM_023258085.1;refGene_gn=CLSTN1;refGene_tx=XM_023258086.1
C1 6866321 6866570
refGene_gn=CLSTN1;refGene_tx=XM_019836106.1;refGene_gn=CLSTN1;refGene_tx=XM_023258083.1;refGene_gn=CLSTN1;refGene_tx=XM_023258084.1;refGene_gn=CLSTN1;refGene_tx=XM_023258085.1;refGene_gn=CLSTN1;refGene_tx=XM_023258086.1
C1 6868868 6869054
refGene_gn=CLSTN1;refGene_tx=XM_019836106.1;refGene_gn=CLSTN1;refGene_tx=XM_023258083.1;refGene_gn=CLSTN1;refGene_tx=XM_023258084.1;refGene_gn=CLSTN1;refGene_tx=XM_023258085.1;refGene_gn=CLSTN1;refGene_tx=XM_023258086.1
C1 6869229 6869379
refGene_gn=CLSTN1;refGene_tx=XM_019836106.1;refGene_gn=CLSTN1;refGene_tx=XM_023258083.1;refGene_gn=CLSTN1;refGene_tx=XM_023258084.1;refGene_gn=CLSTN1;refGene_tx=XM_023258085.1;refGene_gn=CLSTN1;refGene_tx=XM_023258086.1

C1 6871228 6871437
refGene_gn=CLSTN1;refGene_tx=XM_019836106.1;refGene_gn=CLSTN1;refGene_tx=XM_023258083.1;refGene_gn=CLSTN1;refGene_tx=XM_023258084.1;refGene_gn=CLSTN1;refGene_tx=XM_023258085.1;refGene_gn=CLSTN1;refGene_tx=XM_023258086.1
C1 6872837 6873033
refGene_gn=CLSTN1;refGene_tx=XM_019836106.1;refGene_gn=CLSTN1;refGene_tx=XM_023258083.1;refGene_gn=CLSTN1;refGene_tx=XM_023258084.1;refGene_gn=CLSTN1;refGene_tx=XM_023258085.1;refGene_gn=CLSTN1;refGene_tx=XM_023258086.1
C1 6874224 6874254
refGene_gn=CLSTN1;refGene_tx=XM_019836106.1;refGene_gn=CLSTN1;refGene_tx=XM_023258083.1;refGene_gn=CLSTN1;refGene_tx=XM_023258085.1
C1 6886622 6886745
refGene_gn=CLSTN1;refGene_tx=XM_019836106.1;refGene_gn=CLSTN1;refGene_tx=XM_023258083.1;refGene_gn=CLSTN1;refGene_tx=XM_023258084.1;refGene_gn=CLSTN1;refGene_tx=XM_023258085.1;refGene_gn=CLSTN1;refGene_tx=XM_023258086.1
C1 6890216 6890226 refGene_gn=CLSTN1;refGene_tx=XM_019836106.1
C1 6931364 6931455
refGene_gn=CLSTN1;refGene_tx=XM_023258083.1;refGene_gn=CLSTN1;refGene_tx=XM_023258084.1;refGene_gn=CLSTN1;refGene_tx=XM_023258085.1;refGene_gn=CLSTN1;refGene_tx=XM_023258086.1
C1 6956894 6956953
refGene_gn=CTNNBIP1;refGene_tx=XM_003989473.5;refGene_gn=CTNNBIP1;refGene_tx=XM_023258087.1
C1 6977163 6977254
refGene_gn=CTNNBIP1;refGene_tx=XM_003989473.5;refGene_gn=CTNNBIP1;refGene_tx=XM_023258087.1
C1 6977841 6977937
refGene_gn=CTNNBIP1;refGene_tx=XM_003989473.5;refGene_gn=CTNNBIP1;refGene_tx=XM_023258087.1
C1 7026710 7026812
refGene_gn=LZIC;refGene_tx=XM_011284484.3;refGene_gn=LZIC;refGene_tx=XM_019836114.2;refGene_gn=LZIC;refGene_tx=XM_019836115.2
C1 7028075 7028157
refGene_gn=LZIC;refGene_tx=XM_011284484.3;refGene_gn=LZIC;refGene_tx=XM_019836115.2
C1 7028341 7028437
refGene_gn=LZIC;refGene_tx=XM_011284484.3;refGene_gn=LZIC;refGene_tx=XM_019836115.2
C1 7030519 7030618
refGene_gn=LZIC;refGene_tx=XM_011284484.3;refGene_gn=LZIC;refGene_tx=XM_019836114.2;refGene_gn=LZIC;refGene_tx=XM_019836115.2
C1 7030886 7031022
refGene_gn=LZIC;refGene_tx=XM_011284484.3;refGene_gn=LZIC;refGene_tx=XM_019836114.2;refGene_gn=LZIC;refGene_tx=XM_019836115.2
C1 7032044 7032153
refGene_gn=LZIC;refGene_tx=XM_011284484.3;refGene_gn=LZIC;refGene_tx=XM_019836114.2;refGene_gn=LZIC;refGene_tx=XM_019836115.2

C1 7038135 7038190 refGene_gn=LZIC;refGene_tx=XM_019836115.2
C1 7059366 7059481
refGene_gn=NMNAT1;refGene_tx=XM_003989474.5;refGene_gn=NMNAT1;refGene_tx=XM_006934265.4
C1 7061270 7061454
refGene_gn=NMNAT1;refGene_tx=XM_003989474.5;refGene_gn=NMNAT1;refGene_tx=XM_006934265.4
C1 7063961 7064104
refGene_gn=NMNAT1;refGene_tx=XM_003989474.5;refGene_gn=NMNAT1;refGene_tx=XM_006934265.4
C1 7064447 7064848
refGene_gn=NMNAT1;refGene_tx=XM_003989474.5;refGene_gn=NMNAT1;refGene_tx=XM_006934265.4
C1 7077400 7077473 refGene_gn=RBP7;refGene_tx=XM_023258088.1
C1 7090340 7090519 refGene_gn=RBP7;refGene_tx=XM_023258088.1
C1 7090641 7090743 refGene_gn=RBP7;refGene_tx=XM_023258088.1
C1 7093079 7093130 refGene_gn=RBP7;refGene_tx=XM_023258088.1
C1 7126429 7126453
refGene_gn=UBE4B;refGene_tx=XM_003989476.5;refGene_gn=UBE4B;refGene_tx=XM_006934267.4;refGene_gn=UBE4B;refGene_tx=XM_019836116.2;refGene_gn=UBE4B;refGene_tx=XM_019836117.2;refGene_gn=UBE4B;refGene_tx=XM_023258089.1;refGene_gn=UBE4B;refGene_tx=XM_023258090.1;refGene_gn=UBE4B;refGene_tx=XM_023258092.1
C1 7164691 7164878
refGene_gn=UBE4B;refGene_tx=XM_003989476.5;refGene_gn=UBE4B;refGene_tx=XM_006934267.4;refGene_gn=UBE4B;refGene_tx=XM_019836116.2;refGene_gn=UBE4B;refGene_tx=XM_019836117.2;refGene_gn=UBE4B;refGene_tx=XM_023258089.1;refGene_gn=UBE4B;refGene_tx=XM_023258090.1;refGene_gn=UBE4B;refGene_tx=XM_023258091.1;refGene_gn=UBE4B;refGene_tx=XM_023258092.1
C1 7181967 7182103
refGene_gn=UBE4B;refGene_tx=XM_003989476.5;refGene_gn=UBE4B;refGene_tx=XM_006934267.4;refGene_gn=UBE4B;refGene_tx=XM_019836116.2;refGene_gn=UBE4B;refGene_tx=XM_019836117.2;refGene_gn=UBE4B;refGene_tx=XM_023258089.1;refGene_gn=UBE4B;refGene_tx=XM_023258090.1;refGene_gn=UBE4B;refGene_tx=XM_023258091.1;refGene_gn=UBE4B;refGene_tx=XM_023258092.1
C1 7188069 7188157
refGene_gn=UBE4B;refGene_tx=XM_003989476.5;refGene_gn=UBE4B;refGene_tx=XM_006934267.4;refGene_gn=UBE4B;refGene_tx=XM_019836116.2;refGene_gn=UBE4B;refGene_tx=XM_019836117.2;refGene_gn=UBE4B;refGene_tx=XM_023258089.1;refGene_gn=UBE4B;refGene_tx=XM_023258090.1;refGene_gn=UBE4B;refGene_tx=XM_023258091.1;refGene_gn=UBE4B;refGene_tx=XM_023258092.1
C1 7189390 7189535
refGene_gn=UBE4B;refGene_tx=XM_003989476.5;refGene_gn=UBE4B;refGene_tx=XM_006934267.4;refGene_gn=UBE4B;refGene_tx=XM_019836116.2;refGene_gn=UBE4B;refGene_tx=XM_019836117.2;refGene_gn=UBE4B;refGene_tx=XM_023258089.1;refGene_gn=UBE4B;refGene_tx=XM_023258090.1;refGene_gn=UBE4B;refGene_tx=XM_023258091.1;refGene_gn=UBE4B;refGene_tx=XM_023258092.1
C1 7191151 7191377
refGene_gn=UBE4B;refGene_tx=XM_003989476.5;refGene_gn=UBE4B;refGene_tx=XM_006934267.4;refGene_gn=UBE4B;refGene_tx=XM_019836116.2;refGene_gn=

UBE4B;refGene_tx=XM_019836117.2;refGene_gn=UBE4B;refGene_tx=XM_0232580
89.1;refGene_gn=UBE4B;refGene_tx=XM_023258090.1;refGene_gn=UBE4B;refGe
ne_tx=XM_023258091.1;refGene_gn=UBE4B;refGene_tx=XM_023258092.1
C1 7191822 7192233
refGene_gn=UBE4B;refGene_tx=XM_006934267.4;refGene_gn=UBE4B;refGene_tx
=XM_019836116.2;refGene_gn=UBE4B;refGene_tx=XM_019836117.2;refGene_gn=
UBE4B;refGene_tx=XM_023258089.1;refGene_gn=UBE4B;refGene_tx=XM_0232580
90.1;refGene_gn=UBE4B;refGene_tx=XM_023258091.1
C1 7193025 7193175
refGene_gn=UBE4B;refGene_tx=XM_006934267.4;refGene_gn=UBE4B;refGene_tx
=XM_019836117.2;refGene_gn=UBE4B;refGene_tx=XM_023258089.1;refGene_gn=
UBE4B;refGene_tx=XM_023258091.1
C1 7200132 7200274
refGene_gn=UBE4B;refGene_tx=XM_003989476.5;refGene_gn=UBE4B;refGene_tx
=XM_006934267.4;refGene_gn=UBE4B;refGene_tx=XM_019836116.2;refGene_gn=
UBE4B;refGene_tx=XM_019836117.2;refGene_gn=UBE4B;refGene_tx=XM_0232580
89.1;refGene_gn=UBE4B;refGene_tx=XM_023258090.1;refGene_gn=UBE4B;refGe
ne_tx=XM_023258091.1;refGene_gn=UBE4B;refGene_tx=XM_023258092.1
C1 7201145 7201246
refGene_gn=UBE4B;refGene_tx=XM_003989476.5;refGene_gn=UBE4B;refGene_tx
=XM_006934267.4;refGene_gn=UBE4B;refGene_tx=XM_019836116.2;refGene_gn=
UBE4B;refGene_tx=XM_019836117.2;refGene_gn=UBE4B;refGene_tx=XM_0232580
89.1;refGene_gn=UBE4B;refGene_tx=XM_023258090.1;refGene_gn=UBE4B;refGe
ne_tx=XM_023258091.1;refGene_gn=UBE4B;refGene_tx=XM_023258092.1
C1 7202470 7202585
refGene_gn=UBE4B;refGene_tx=XM_003989476.5;refGene_gn=UBE4B;refGene_tx
=XM_006934267.4;refGene_gn=UBE4B;refGene_tx=XM_019836116.2;refGene_gn=
UBE4B;refGene_tx=XM_019836117.2;refGene_gn=UBE4B;refGene_tx=XM_0232580
89.1;refGene_gn=UBE4B;refGene_tx=XM_023258090.1;refGene_gn=UBE4B;refGe
ne_tx=XM_023258091.1;refGene_gn=UBE4B;refGene_tx=XM_023258092.1
C1 7206037 7206121
refGene_gn=UBE4B;refGene_tx=XM_003989476.5;refGene_gn=UBE4B;refGene_tx
=XM_006934267.4;refGene_gn=UBE4B;refGene_tx=XM_019836116.2;refGene_gn=
UBE4B;refGene_tx=XM_019836117.2;refGene_gn=UBE4B;refGene_tx=XM_0232580
89.1;refGene_gn=UBE4B;refGene_tx=XM_023258090.1;refGene_gn=UBE4B;refGe
ne_tx=XM_023258091.1;refGene_gn=UBE4B;refGene_tx=XM_023258092.1
C1 7208650 7208707
refGene_gn=UBE4B;refGene_tx=XM_003989476.5;refGene_gn=UBE4B;refGene_tx
=XM_006934267.4;refGene_gn=UBE4B;refGene_tx=XM_019836116.2;refGene_gn=
UBE4B;refGene_tx=XM_019836117.2;refGene_gn=UBE4B;refGene_tx=XM_0232580
89.1;refGene_gn=UBE4B;refGene_tx=XM_023258090.1;refGene_gn=UBE4B;refGe
ne_tx=XM_023258091.1;refGene_gn=UBE4B;refGene_tx=XM_023258092.1
C1 7209289 7209406
refGene_gn=UBE4B;refGene_tx=XM_003989476.5;refGene_gn=UBE4B;refGene_tx
=XM_006934267.4;refGene_gn=UBE4B;refGene_tx=XM_019836116.2;refGene_gn=
UBE4B;refGene_tx=XM_019836117.2;refGene_gn=UBE4B;refGene_tx=XM_0232580
89.1;refGene_gn=UBE4B;refGene_tx=XM_023258090.1;refGene_gn=UBE4B;refGe
ne_tx=XM_023258091.1;refGene_gn=UBE4B;refGene_tx=XM_023258092.1
C1 7209504 7209603
refGene_gn=UBE4B;refGene_tx=XM_003989476.5;refGene_gn=UBE4B;refGene_tx

=XM_006934267.4;refGene_gn=UBE4B;refGene_tx=XM_019836116.2;refGene_gn=
UBE4B;refGene_tx=XM_019836117.2;refGene_gn=UBE4B;refGene_tx=XM_0232580
89.1;refGene_gn=UBE4B;refGene_tx=XM_023258090.1;refGene_gn=UBE4B;refGe
ne_tx=XM_023258091.1;refGene_gn=UBE4B;refGene_tx=XM_023258092.1
C1 7210183 7210297
refGene_gn=UBE4B;refGene_tx=XM_003989476.5;refGene_gn=UBE4B;refGene_tx
=XM_006934267.4;refGene_gn=UBE4B;refGene_tx=XM_019836116.2;refGene_gn=
UBE4B;refGene_tx=XM_019836117.2;refGene_gn=UBE4B;refGene_tx=XM_0232580
89.1;refGene_gn=UBE4B;refGene_tx=XM_023258090.1;refGene_gn=UBE4B;refGe
ne_tx=XM_023258091.1;refGene_gn=UBE4B;refGene_tx=XM_023258092.1
C1 7211751 7211950
refGene_gn=UBE4B;refGene_tx=XM_003989476.5;refGene_gn=UBE4B;refGene_tx
=XM_006934267.4;refGene_gn=UBE4B;refGene_tx=XM_019836116.2;refGene_gn=
UBE4B;refGene_tx=XM_019836117.2;refGene_gn=UBE4B;refGene_tx=XM_0232580
89.1;refGene_gn=UBE4B;refGene_tx=XM_023258090.1;refGene_gn=UBE4B;refGe
ne_tx=XM_023258091.1;refGene_gn=UBE4B;refGene_tx=XM_023258092.1
C1 7214857 7214996
refGene_gn=UBE4B;refGene_tx=XM_003989476.5;refGene_gn=UBE4B;refGene_tx
=XM_006934267.4;refGene_gn=UBE4B;refGene_tx=XM_019836116.2;refGene_gn=
UBE4B;refGene_tx=XM_019836117.2;refGene_gn=UBE4B;refGene_tx=XM_0232580
89.1;refGene_gn=UBE4B;refGene_tx=XM_023258090.1;refGene_gn=UBE4B;refGe
ne_tx=XM_023258091.1;refGene_gn=UBE4B;refGene_tx=XM_023258092.1
C1 7219705 7219805
refGene_gn=UBE4B;refGene_tx=XM_003989476.5;refGene_gn=UBE4B;refGene_tx
=XM_006934267.4;refGene_gn=UBE4B;refGene_tx=XM_019836116.2;refGene_gn=
UBE4B;refGene_tx=XM_019836117.2;refGene_gn=UBE4B;refGene_tx=XM_0232580
89.1;refGene_gn=UBE4B;refGene_tx=XM_023258090.1;refGene_gn=UBE4B;refGe
ne_tx=XM_023258091.1;refGene_gn=UBE4B;refGene_tx=XM_023258092.1
C1 7221052 7221180
refGene_gn=UBE4B;refGene_tx=XM_003989476.5;refGene_gn=UBE4B;refGene_tx
=XM_006934267.4;refGene_gn=UBE4B;refGene_tx=XM_019836116.2;refGene_gn=
UBE4B;refGene_tx=XM_019836117.2;refGene_gn=UBE4B;refGene_tx=XM_0232580
89.1;refGene_gn=UBE4B;refGene_tx=XM_023258090.1;refGene_gn=UBE4B;refGe
ne_tx=XM_023258091.1;refGene_gn=UBE4B;refGene_tx=XM_023258092.1
C1 7221962 7222061
refGene_gn=UBE4B;refGene_tx=XM_003989476.5;refGene_gn=UBE4B;refGene_tx
=XM_006934267.4;refGene_gn=UBE4B;refGene_tx=XM_019836116.2;refGene_gn=
UBE4B;refGene_tx=XM_019836117.2;refGene_gn=UBE4B;refGene_tx=XM_0232580
89.1;refGene_gn=UBE4B;refGene_tx=XM_023258090.1;refGene_gn=UBE4B;refGe
ne_tx=XM_023258091.1;refGene_gn=UBE4B;refGene_tx=XM_023258092.1
C1 7222959 7223195
refGene_gn=UBE4B;refGene_tx=XM_003989476.5;refGene_gn=UBE4B;refGene_tx
=XM_006934267.4;refGene_gn=UBE4B;refGene_tx=XM_019836116.2;refGene_gn=
UBE4B;refGene_tx=XM_019836117.2;refGene_gn=UBE4B;refGene_tx=XM_0232580
89.1;refGene_gn=UBE4B;refGene_tx=XM_023258090.1;refGene_gn=UBE4B;refGe
ne_tx=XM_023258091.1;refGene_gn=UBE4B;refGene_tx=XM_023258092.1
C1 7230618 7230745
refGene_gn=UBE4B;refGene_tx=XM_003989476.5;refGene_gn=UBE4B;refGene_tx
=XM_006934267.4;refGene_gn=UBE4B;refGene_tx=XM_019836116.2;refGene_gn=
UBE4B;refGene_tx=XM_019836117.2;refGene_gn=UBE4B;refGene_tx=XM_0232580

89.1;refGene_gn=UBE4B;refGene_tx=XM_023258090.1;refGene_gn=UBE4B;refGene_tx=XM_023258091.1;refGene_gn=UBE4B;refGene_tx=XM_023258092.1
C1 7232663 7232808
refGene_gn=UBE4B;refGene_tx=XM_003989476.5;refGene_gn=UBE4B;refGene_tx=XM_006934267.4;refGene_gn=UBE4B;refGene_tx=XM_019836116.2;refGene_gn=UBE4B;refGene_tx=XM_019836117.2;refGene_gn=UBE4B;refGene_tx=XM_023258089.1;refGene_gn=UBE4B;refGene_tx=XM_023258090.1;refGene_gn=UBE4B;refGene_tx=XM_023258091.1;refGene_gn=UBE4B;refGene_tx=XM_023258092.1
C1 7237239 7237374
refGene_gn=UBE4B;refGene_tx=XM_003989476.5;refGene_gn=UBE4B;refGene_tx=XM_006934267.4;refGene_gn=UBE4B;refGene_tx=XM_019836116.2;refGene_gn=UBE4B;refGene_tx=XM_019836117.2;refGene_gn=UBE4B;refGene_tx=XM_023258089.1;refGene_gn=UBE4B;refGene_tx=XM_023258090.1;refGene_gn=UBE4B;refGene_tx=XM_023258091.1;refGene_gn=UBE4B;refGene_tx=XM_023258092.1
C1 7240109 7240301
refGene_gn=UBE4B;refGene_tx=XM_003989476.5;refGene_gn=UBE4B;refGene_tx=XM_006934267.4;refGene_gn=UBE4B;refGene_tx=XM_019836116.2;refGene_gn=UBE4B;refGene_tx=XM_019836117.2;refGene_gn=UBE4B;refGene_tx=XM_023258089.1;refGene_gn=UBE4B;refGene_tx=XM_023258090.1;refGene_gn=UBE4B;refGene_tx=XM_023258091.1;refGene_gn=UBE4B;refGene_tx=XM_023258092.1
C1 7240753 7240774 refGene_gn=UBE4B;refGene_tx=XM_019836117.2
C1 7246074 7246328
refGene_gn=UBE4B;refGene_tx=XM_003989476.5;refGene_gn=UBE4B;refGene_tx=XM_006934267.4;refGene_gn=UBE4B;refGene_tx=XM_019836116.2;refGene_gn=UBE4B;refGene_tx=XM_023258089.1;refGene_gn=UBE4B;refGene_tx=XM_023258090.1;refGene_gn=UBE4B;refGene_tx=XM_023258091.1;refGene_gn=UBE4B;refGene_tx=XM_023258092.1
C1 7246764 7246911
refGene_gn=UBE4B;refGene_tx=XM_003989476.5;refGene_gn=UBE4B;refGene_tx=XM_006934267.4;refGene_gn=UBE4B;refGene_tx=XM_019836116.2;refGene_gn=UBE4B;refGene_tx=XM_023258089.1;refGene_gn=UBE4B;refGene_tx=XM_023258090.1;refGene_gn=UBE4B;refGene_tx=XM_023258091.1;refGene_gn=UBE4B;refGene_tx=XM_023258092.1
C1 7247227 7247507
refGene_gn=UBE4B;refGene_tx=XM_003989476.5;refGene_gn=UBE4B;refGene_tx=XM_006934267.4;refGene_gn=UBE4B;refGene_tx=XM_019836116.2;refGene_gn=UBE4B;refGene_tx=XM_023258089.1;refGene_gn=UBE4B;refGene_tx=XM_023258090.1;refGene_gn=UBE4B;refGene_tx=XM_023258091.1;refGene_gn=UBE4B;refGene_tx=XM_023258092.1
C1 7295147 7295253
refGene_gn=KIF1B;refGene_tx=XM_019836118.2;refGene_gn=KIF1B;refGene_tx=XM_019836119.2;refGene_gn=KIF1B;refGene_tx=XM_023258093.1;refGene_gn=KIF1B;refGene_tx=XM_023258094.1;refGene_gn=KIF1B;refGene_tx=XM_023258095.1;refGene_gn=KIF1B;refGene_tx=XM_023258096.1;refGene_gn=KIF1B;refGene_tx=XM_023258097.1;refGene_gn=KIF1B;refGene_tx=XM_023258098.1;refGene_gn=KIF1B;refGene_tx=XM_023258099.1;refGene_gn=KIF1B;refGene_tx=XM_023258100.1;refGene_gn=KIF1B;refGene_tx=XM_023258101.1;refGene_gn=KIF1B;refGene_tx=XM_023258102.1;refGene_gn=KIF1B;refGene_tx=XM_023258103.1;refGene_gn=KIF1B;refGene_tx=XM_023258104.1;refGene_gn=KIF1B;refGene_tx=XM_023258105.1;refGene_gn=KIF1B;refGene_tx=XM_023258106.1;refGene_gn=K

IF1B;refGene_tx=XM_023258107.1;refGene_gn=KIF1B;refGene_tx=XM_02325810
8.1;refGene_gn=KIF1B;refGene_tx=XM_023258109.1;refGene_gn=KIF1B;refGen
e_tx=XM_023258110.1;refGene_gn=KIF1B;refGene_tx=XM_023258111.1;refGene
_gn=KIF1B;refGene_tx=XM_023258112.1;refGene_gn=KIF1B;refGene_tx=XM_023
258113.1;refGene_gn=KIF1B;refGene_tx=XM_023258114.1;refGene_gn=KIF1B;r
efGene_tx=XM_023258115.1;refGene_gn=KIF1B;refGene_tx=XM_023258116.1;re
fGene_gn=KIF1B;refGene_tx=XM_023258117.1
C1 7318138 7318215
refGene_gn=KIF1B;refGene_tx=XM_019836118.2;refGene_gn=KIF1B;refGene_tx
=XM_019836119.2;refGene_gn=KIF1B;refGene_tx=XM_023258093.1;refGene_gn=
KIF1B;refGene_tx=XM_023258094.1;refGene_gn=KIF1B;refGene_tx=XM_0232580
95.1;refGene_gn=KIF1B;refGene_tx=XM_023258096.1;refGene_gn=KIF1B;refGe
ne_tx=XM_023258097.1;refGene_gn=KIF1B;refGene_tx=XM_023258098.1;refGen
e_gn=KIF1B;refGene_tx=XM_023258099.1;refGene_gn=KIF1B;refGene_tx=XM_02
3258100.1;refGene_gn=KIF1B;refGene_tx=XM_023258101.1;refGene_gn=KIF1B;
refGene_tx=XM_023258102.1;refGene_gn=KIF1B;refGene_tx=XM_023258103.1;r
efGene_gn=KIF1B;refGene_tx=XM_023258104.1;refGene_gn=KIF1B;refGene_tx=
XM_023258105.1;refGene_gn=KIF1B;refGene_tx=XM_023258106.1;refGene_gn=K
IF1B;refGene_tx=XM_023258107.1;refGene_gn=KIF1B;refGene_tx=XM_02325810
8.1;refGene_gn=KIF1B;refGene_tx=XM_023258109.1;refGene_gn=KIF1B;refGen
e_tx=XM_023258110.1;refGene_gn=KIF1B;refGene_tx=XM_023258111.1;refGene
_gn=KIF1B;refGene_tx=XM_023258112.1;refGene_gn=KIF1B;refGene_tx=XM_023
258113.1;refGene_gn=KIF1B;refGene_tx=XM_023258114.1;refGene_gn=KIF1B;r
efGene_tx=XM_023258115.1;refGene_gn=KIF1B;refGene_tx=XM_023258116.1;re
fGene_gn=KIF1B;refGene_tx=XM_023258117.1
C1 7318979 7319159
refGene_gn=KIF1B;refGene_tx=XM_019836118.2;refGene_gn=KIF1B;refGene_tx
=XM_019836119.2;refGene_gn=KIF1B;refGene_tx=XM_023258093.1;refGene_gn=
KIF1B;refGene_tx=XM_023258094.1;refGene_gn=KIF1B;refGene_tx=XM_0232580
95.1;refGene_gn=KIF1B;refGene_tx=XM_023258096.1;refGene_gn=KIF1B;refGe
ne_tx=XM_023258097.1;refGene_gn=KIF1B;refGene_tx=XM_023258098.1;refGen
e_gn=KIF1B;refGene_tx=XM_023258099.1;refGene_gn=KIF1B;refGene_tx=XM_02
3258100.1;refGene_gn=KIF1B;refGene_tx=XM_023258101.1;refGene_gn=KIF1B;
refGene_tx=XM_023258102.1;refGene_gn=KIF1B;refGene_tx=XM_023258103.1;r
efGene_gn=KIF1B;refGene_tx=XM_023258104.1;refGene_gn=KIF1B;refGene_tx=
XM_023258105.1;refGene_gn=KIF1B;refGene_tx=XM_023258106.1;refGene_gn=K
IF1B;refGene_tx=XM_023258107.1;refGene_gn=KIF1B;refGene_tx=XM_02325810
8.1;refGene_gn=KIF1B;refGene_tx=XM_023258109.1;refGene_gn=KIF1B;refGen
e_tx=XM_023258110.1;refGene_gn=KIF1B;refGene_tx=XM_023258111.1;refGene
_gn=KIF1B;refGene_tx=XM_023258112.1;refGene_gn=KIF1B;refGene_tx=XM_023
258113.1;refGene_gn=KIF1B;refGene_tx=XM_023258114.1;refGene_gn=KIF1B;r
efGene_tx=XM_023258115.1;refGene_gn=KIF1B;refGene_tx=XM_023258116.1;re
fGene_gn=KIF1B;refGene_tx=XM_023258117.1
C1 7324293 7324359
refGene_gn=KIF1B;refGene_tx=XM_019836118.2;refGene_gn=KIF1B;refGene_tx
=XM_019836119.2;refGene_gn=KIF1B;refGene_tx=XM_023258093.1;refGene_gn=
KIF1B;refGene_tx=XM_023258094.1;refGene_gn=KIF1B;refGene_tx=XM_0232580
95.1;refGene_gn=KIF1B;refGene_tx=XM_023258096.1;refGene_gn=KIF1B;refGe
ne_tx=XM_023258097.1;refGene_gn=KIF1B;refGene_tx=XM_023258098.1;refGen
e_gn=KIF1B;refGene_tx=XM_023258099.1;refGene_gn=KIF1B;refGene_tx=XM_02

3258100.1;refGene_gn=KIF1B;refGene_tx=XM_023258101.1;refGene_gn=KIF1B;
refGene_tx=XM_023258102.1;refGene_gn=KIF1B;refGene_tx=XM_023258103.1;r
efGene_gn=KIF1B;refGene_tx=XM_023258104.1;refGene_gn=KIF1B;refGene_tx=
XM_023258105.1;refGene_gn=KIF1B;refGene_tx=XM_023258106.1;refGene_gn=K
IF1B;refGene_tx=XM_023258107.1;refGene_gn=KIF1B;refGene_tx=XM_02325810
8.1;refGene_gn=KIF1B;refGene_tx=XM_023258109.1;refGene_gn=KIF1B;refGen
e_tx=XM_023258110.1;refGene_gn=KIF1B;refGene_tx=XM_023258111.1;refGene
_gn=KIF1B;refGene_tx=XM_023258112.1;refGene_gn=KIF1B;refGene_tx=XM_023
258113.1;refGene_gn=KIF1B;refGene_tx=XM_023258114.1;refGene_gn=KIF1B;r
efGene_tx=XM_023258115.1;refGene_gn=KIF1B;refGene_tx=XM_023258116.1;re
fGene_gn=KIF1B;refGene_tx=XM_023258117.1
C1 7328244 7328423
refGene_gn=KIF1B;refGene_tx=XM_019836118.2;refGene_gn=KIF1B;refGene_tx
=XM_019836119.2;refGene_gn=KIF1B;refGene_tx=XM_023258093.1;refGene_gn=
KIF1B;refGene_tx=XM_023258094.1;refGene_gn=KIF1B;refGene_tx=XM_0232580
95.1;refGene_gn=KIF1B;refGene_tx=XM_023258096.1;refGene_gn=KIF1B;refGe
ne_tx=XM_023258097.1;refGene_gn=KIF1B;refGene_tx=XM_023258098.1;refGen
e_gn=KIF1B;refGene_tx=XM_023258099.1;refGene_gn=KIF1B;refGene_tx=XM_02
3258100.1;refGene_gn=KIF1B;refGene_tx=XM_023258101.1;refGene_gn=KIF1B;
refGene_tx=XM_023258102.1;refGene_gn=KIF1B;refGene_tx=XM_023258103.1;r
efGene_gn=KIF1B;refGene_tx=XM_023258104.1;refGene_gn=KIF1B;refGene_tx=
XM_023258105.1;refGene_gn=KIF1B;refGene_tx=XM_023258106.1;refGene_gn=K
IF1B;refGene_tx=XM_023258107.1;refGene_gn=KIF1B;refGene_tx=XM_02325810
8.1;refGene_gn=KIF1B;refGene_tx=XM_023258109.1;refGene_gn=KIF1B;refGen
e_tx=XM_023258110.1;refGene_gn=KIF1B;refGene_tx=XM_023258111.1;refGene
_gn=KIF1B;refGene_tx=XM_023258112.1;refGene_gn=KIF1B;refGene_tx=XM_023
258113.1;refGene_gn=KIF1B;refGene_tx=XM_023258114.1;refGene_gn=KIF1B;r
efGene_tx=XM_023258115.1;refGene_gn=KIF1B;refGene_tx=XM_023258116.1;re
fGene_gn=KIF1B;refGene_tx=XM_023258117.1
C1 7328987 7329099
refGene_gn=KIF1B;refGene_tx=XM_019836118.2;refGene_gn=KIF1B;refGene_tx
=XM_019836119.2;refGene_gn=KIF1B;refGene_tx=XM_023258093.1;refGene_gn=
KIF1B;refGene_tx=XM_023258094.1;refGene_gn=KIF1B;refGene_tx=XM_0232580
95.1;refGene_gn=KIF1B;refGene_tx=XM_023258096.1;refGene_gn=KIF1B;refGe
ne_tx=XM_023258097.1;refGene_gn=KIF1B;refGene_tx=XM_023258098.1;refGen
e_gn=KIF1B;refGene_tx=XM_023258099.1;refGene_gn=KIF1B;refGene_tx=XM_02
3258100.1;refGene_gn=KIF1B;refGene_tx=XM_023258101.1;refGene_gn=KIF1B;
refGene_tx=XM_023258102.1;refGene_gn=KIF1B;refGene_tx=XM_023258103.1;r
efGene_gn=KIF1B;refGene_tx=XM_023258104.1;refGene_gn=KIF1B;refGene_tx=
XM_023258105.1;refGene_gn=KIF1B;refGene_tx=XM_023258106.1;refGene_gn=K
IF1B;refGene_tx=XM_023258107.1;refGene_gn=KIF1B;refGene_tx=XM_02325810
8.1;refGene_gn=KIF1B;refGene_tx=XM_023258109.1;refGene_gn=KIF1B;refGen
e_tx=XM_023258110.1;refGene_gn=KIF1B;refGene_tx=XM_023258111.1;refGene
_gn=KIF1B;refGene_tx=XM_023258112.1;refGene_gn=KIF1B;refGene_tx=XM_023
258113.1;refGene_gn=KIF1B;refGene_tx=XM_023258114.1;refGene_gn=KIF1B;r
efGene_tx=XM_023258115.1;refGene_gn=KIF1B;refGene_tx=XM_023258116.1;re
fGene_gn=KIF1B;refGene_tx=XM_023258117.1
C1 7331631 7331709
refGene_gn=KIF1B;refGene_tx=XM_019836118.2;refGene_gn=KIF1B;refGene_tx
=XM_019836119.2;refGene_gn=KIF1B;refGene_tx=XM_023258093.1;refGene_gn=

KIF1B;refGene_tx=XM_023258094.1;refGene_gn=KIF1B;refGene_tx=XM_0232580
95.1;refGene_gn=KIF1B;refGene_tx=XM_023258096.1;refGene_gn=KIF1B;refGe
ne_tx=XM_023258097.1;refGene_gn=KIF1B;refGene_tx=XM_023258098.1;refGen
e_gn=KIF1B;refGene_tx=XM_023258099.1;refGene_gn=KIF1B;refGene_tx=XM_02
3258100.1;refGene_gn=KIF1B;refGene_tx=XM_023258101.1;refGene_gn=KIF1B;
refGene_tx=XM_023258102.1;refGene_gn=KIF1B;refGene_tx=XM_023258103.1;r
efGene_gn=KIF1B;refGene_tx=XM_023258104.1;refGene_gn=KIF1B;refGene_tx=
XM_023258105.1;refGene_gn=KIF1B;refGene_tx=XM_023258106.1;refGene_gn=K
IF1B;refGene_tx=XM_023258107.1;refGene_gn=KIF1B;refGene_tx=XM_02325810
8.1;refGene_gn=KIF1B;refGene_tx=XM_023258109.1;refGene_gn=KIF1B;refGen
e_tx=XM_023258110.1;refGene_gn=KIF1B;refGene_tx=XM_023258111.1;refGene
_gn=KIF1B;refGene_tx=XM_023258112.1;refGene_gn=KIF1B;refGene_tx=XM_023
258113.1;refGene_gn=KIF1B;refGene_tx=XM_023258114.1;refGene_gn=KIF1B;r
efGene_tx=XM_023258115.1;refGene_gn=KIF1B;refGene_tx=XM_023258116.1;re
fGene_gn=KIF1B;refGene_tx=XM_023258117.1
C1 7332385 7332451
refGene_gn=KIF1B;refGene_tx=XM_019836118.2;refGene_gn=KIF1B;refGene_tx
=XM_019836119.2;refGene_gn=KIF1B;refGene_tx=XM_023258093.1;refGene_gn=
KIF1B;refGene_tx=XM_023258094.1;refGene_gn=KIF1B;refGene_tx=XM_0232580
95.1;refGene_gn=KIF1B;refGene_tx=XM_023258096.1;refGene_gn=KIF1B;refGe
ne_tx=XM_023258097.1;refGene_gn=KIF1B;refGene_tx=XM_023258098.1;refGen
e_gn=KIF1B;refGene_tx=XM_023258099.1;refGene_gn=KIF1B;refGene_tx=XM_02
3258100.1;refGene_gn=KIF1B;refGene_tx=XM_023258101.1;refGene_gn=KIF1B;
refGene_tx=XM_023258102.1;refGene_gn=KIF1B;refGene_tx=XM_023258103.1;r
efGene_gn=KIF1B;refGene_tx=XM_023258104.1;refGene_gn=KIF1B;refGene_tx=
XM_023258105.1;refGene_gn=KIF1B;refGene_tx=XM_023258106.1;refGene_gn=K
IF1B;refGene_tx=XM_023258107.1;refGene_gn=KIF1B;refGene_tx=XM_02325810
8.1;refGene_gn=KIF1B;refGene_tx=XM_023258109.1;refGene_gn=KIF1B;refGen
e_tx=XM_023258110.1;refGene_gn=KIF1B;refGene_tx=XM_023258111.1;refGene
_gn=KIF1B;refGene_tx=XM_023258112.1;refGene_gn=KIF1B;refGene_tx=XM_023
258113.1;refGene_gn=KIF1B;refGene_tx=XM_023258114.1;refGene_gn=KIF1B;r
efGene_tx=XM_023258115.1;refGene_gn=KIF1B;refGene_tx=XM_023258116.1;re
fGene_gn=KIF1B;refGene_tx=XM_023258117.1
C1 7333207 7333225
refGene_gn=KIF1B;refGene_tx=XM_019836118.2;refGene_gn=KIF1B;refGene_tx
=XM_019836119.2;refGene_gn=KIF1B;refGene_tx=XM_023258093.1;refGene_gn=
KIF1B;refGene_tx=XM_023258095.1;refGene_gn=KIF1B;refGene_tx=XM_0232580
96.1;refGene_gn=KIF1B;refGene_tx=XM_023258097.1;refGene_gn=KIF1B;refGe
ne_tx=XM_023258098.1;refGene_gn=KIF1B;refGene_tx=XM_023258099.1;refGen
e_gn=KIF1B;refGene_tx=XM_023258100.1;refGene_gn=KIF1B;refGene_tx=XM_02
3258101.1;refGene_gn=KIF1B;refGene_tx=XM_023258102.1;refGene_gn=KIF1B;
refGene_tx=XM_023258103.1;refGene_gn=KIF1B;refGene_tx=XM_023258104.1;r
efGene_gn=KIF1B;refGene_tx=XM_023258105.1;refGene_gn=KIF1B;refGene_tx=
XM_023258106.1;refGene_gn=KIF1B;refGene_tx=XM_023258107.1;refGene_gn=K
IF1B;refGene_tx=XM_023258108.1;refGene_gn=KIF1B;refGene_tx=XM_02325810
9.1;refGene_gn=KIF1B;refGene_tx=XM_023258110.1;refGene_gn=KIF1B;refGen
e_tx=XM_023258111.1;refGene_gn=KIF1B;refGene_tx=XM_023258113.1;refGene
_gn=KIF1B;refGene_tx=XM_023258116.1;refGene_gn=KIF1B;refGene_tx=XM_023
258117.1
C1 7336998 7337074

refGene_gn=KIF1B;refGene_tx=XM_019836118.2;refGene_gn=KIF1B;refGene_tx
=XM_019836119.2;refGene_gn=KIF1B;refGene_tx=XM_023258093.1;refGene_gn=
KIF1B;refGene_tx=XM_023258094.1;refGene_gn=KIF1B;refGene_tx=XM_0232580
95.1;refGene_gn=KIF1B;refGene_tx=XM_023258096.1;refGene_gn=KIF1B;refGe
ne_tx=XM_023258097.1;refGene_gn=KIF1B;refGene_tx=XM_023258098.1;refGen
e_gn=KIF1B;refGene_tx=XM_023258099.1;refGene_gn=KIF1B;refGene_tx=XM_02
3258100.1;refGene_gn=KIF1B;refGene_tx=XM_023258101.1;refGene_gn=KIF1B;
refGene_tx=XM_023258102.1;refGene_gn=KIF1B;refGene_tx=XM_023258103.1;r
efGene_gn=KIF1B;refGene_tx=XM_023258104.1;refGene_gn=KIF1B;refGene_tx=
XM_023258105.1;refGene_gn=KIF1B;refGene_tx=XM_023258106.1;refGene_gn=K
IF1B;refGene_tx=XM_023258107.1;refGene_gn=KIF1B;refGene_tx=XM_02325810
8.1;refGene_gn=KIF1B;refGene_tx=XM_023258109.1;refGene_gn=KIF1B;refGen
e_tx=XM_023258110.1;refGene_gn=KIF1B;refGene_tx=XM_023258111.1;refGene
_gn=KIF1B;refGene_tx=XM_023258112.1;refGene_gn=KIF1B;refGene_tx=XM_023
258113.1;refGene_gn=KIF1B;refGene_tx=XM_023258114.1;refGene_gn=KIF1B;r
efGene_tx=XM_023258115.1;refGene_gn=KIF1B;refGene_tx=XM_023258116.1;re
fGene_gn=KIF1B;refGene_tx=XM_023258117.1
C1 7337801 7337880
refGene_gn=KIF1B;refGene_tx=XM_019836118.2;refGene_gn=KIF1B;refGene_tx
=XM_019836119.2;refGene_gn=KIF1B;refGene_tx=XM_023258093.1;refGene_gn=
KIF1B;refGene_tx=XM_023258094.1;refGene_gn=KIF1B;refGene_tx=XM_0232580
95.1;refGene_gn=KIF1B;refGene_tx=XM_023258096.1;refGene_gn=KIF1B;refGe
ne_tx=XM_023258097.1;refGene_gn=KIF1B;refGene_tx=XM_023258098.1;refGen
e_gn=KIF1B;refGene_tx=XM_023258099.1;refGene_gn=KIF1B;refGene_tx=XM_02
3258100.1;refGene_gn=KIF1B;refGene_tx=XM_023258101.1;refGene_gn=KIF1B;
refGene_tx=XM_023258102.1;refGene_gn=KIF1B;refGene_tx=XM_023258103.1;r
efGene_gn=KIF1B;refGene_tx=XM_023258104.1;refGene_gn=KIF1B;refGene_tx=
XM_023258105.1;refGene_gn=KIF1B;refGene_tx=XM_023258106.1;refGene_gn=K
IF1B;refGene_tx=XM_023258107.1;refGene_gn=KIF1B;refGene_tx=XM_02325810
8.1;refGene_gn=KIF1B;refGene_tx=XM_023258109.1;refGene_gn=KIF1B;refGen
e_tx=XM_023258110.1;refGene_gn=KIF1B;refGene_tx=XM_023258111.1;refGene
_gn=KIF1B;refGene_tx=XM_023258112.1;refGene_gn=KIF1B;refGene_tx=XM_023
258113.1;refGene_gn=KIF1B;refGene_tx=XM_023258114.1;refGene_gn=KIF1B;r
efGene_tx=XM_023258115.1;refGene_gn=KIF1B;refGene_tx=XM_023258116.1;re
fGene_gn=KIF1B;refGene_tx=XM_023258117.1
C1 7339237 7339380
refGene_gn=KIF1B;refGene_tx=XM_019836118.2;refGene_gn=KIF1B;refGene_tx
=XM_019836119.2;refGene_gn=KIF1B;refGene_tx=XM_023258093.1;refGene_gn=
KIF1B;refGene_tx=XM_023258094.1;refGene_gn=KIF1B;refGene_tx=XM_0232580
95.1;refGene_gn=KIF1B;refGene_tx=XM_023258096.1;refGene_gn=KIF1B;refGe
ne_tx=XM_023258097.1;refGene_gn=KIF1B;refGene_tx=XM_023258098.1;refGen
e_gn=KIF1B;refGene_tx=XM_023258099.1;refGene_gn=KIF1B;refGene_tx=XM_02
3258100.1;refGene_gn=KIF1B;refGene_tx=XM_023258101.1;refGene_gn=KIF1B;
refGene_tx=XM_023258102.1;refGene_gn=KIF1B;refGene_tx=XM_023258103.1;r
efGene_gn=KIF1B;refGene_tx=XM_023258104.1;refGene_gn=KIF1B;refGene_tx=
XM_023258105.1;refGene_gn=KIF1B;refGene_tx=XM_023258106.1;refGene_gn=K
IF1B;refGene_tx=XM_023258107.1;refGene_gn=KIF1B;refGene_tx=XM_02325810
8.1;refGene_gn=KIF1B;refGene_tx=XM_023258109.1;refGene_gn=KIF1B;refGen
e_tx=XM_023258110.1;refGene_gn=KIF1B;refGene_tx=XM_023258111.1;refGene
_gn=KIF1B;refGene_tx=XM_023258112.1;refGene_gn=KIF1B;refGene_tx=XM_023

258113.1;refGene_gn=KIF1B;refGene_tx=XM_023258114.1;refGene_gn=KIF1B;refGene_tx=XM_023258115.1;refGene_gn=KIF1B;refGene_tx=XM_023258116.1;refGene_gn=KIF1B;refGene_tx=XM_023258117.1
C1 7340819 7340861
refGene_gn=KIF1B;refGene_tx=XM_019836118.2;refGene_gn=KIF1B;refGene_tx=XM_019836119.2;refGene_gn=KIF1B;refGene_tx=XM_023258093.1;refGene_gn=KIF1B;refGene_tx=XM_023258094.1;refGene_gn=KIF1B;refGene_tx=XM_023258095.1;refGene_gn=KIF1B;refGene_tx=XM_023258096.1;refGene_gn=KIF1B;refGene_tx=XM_023258098.1;refGene_gn=KIF1B;refGene_tx=XM_023258099.1;refGene_gn=KIF1B;refGene_tx=XM_023258100.1;refGene_gn=KIF1B;refGene_tx=XM_023258102.1;refGene_gn=KIF1B;refGene_tx=XM_023258103.1;refGene_gn=KIF1B;refGene_tx=XM_023258104.1;refGene_gn=KIF1B;refGene_tx=XM_023258105.1;refGene_gn=KIF1B;refGene_tx=XM_023258106.1;refGene_gn=KIF1B;refGene_tx=XM_023258107.1;refGene_gn=KIF1B;refGene_tx=XM_023258108.1;refGene_gn=KIF1B;refGene_tx=XM_023258110.1;refGene_gn=KIF1B;refGene_tx=XM_023258117.1
C1 7343612 7343824
refGene_gn=KIF1B;refGene_tx=XM_019836118.2;refGene_gn=KIF1B;refGene_tx=XM_019836119.2;refGene_gn=KIF1B;refGene_tx=XM_023258093.1;refGene_gn=KIF1B;refGene_tx=XM_023258094.1;refGene_gn=KIF1B;refGene_tx=XM_023258095.1;refGene_gn=KIF1B;refGene_tx=XM_023258096.1;refGene_gn=KIF1B;refGene_tx=XM_023258097.1;refGene_gn=KIF1B;refGene_tx=XM_023258098.1;refGene_gn=KIF1B;refGene_tx=XM_023258099.1;refGene_gn=KIF1B;refGene_tx=XM_023258100.1;refGene_gn=KIF1B;refGene_tx=XM_023258101.1;refGene_gn=KIF1B;refGene_tx=XM_023258102.1;refGene_gn=KIF1B;refGene_tx=XM_023258103.1;refGene_gn=KIF1B;refGene_tx=XM_023258104.1;refGene_gn=KIF1B;refGene_tx=XM_023258105.1;refGene_gn=KIF1B;refGene_tx=XM_023258106.1;refGene_gn=KIF1B;refGene_tx=XM_023258107.1;refGene_gn=KIF1B;refGene_tx=XM_023258108.1;refGene_gn=KIF1B;refGene_tx=XM_023258109.1;refGene_gn=KIF1B;refGene_tx=XM_023258110.1;refGene_gn=KIF1B;refGene_tx=XM_023258111.1;refGene_gn=KIF1B;refGene_tx=XM_023258112.1;refGene_gn=KIF1B;refGene_tx=XM_023258113.1;refGene_gn=KIF1B;refGene_tx=XM_023258114.1;refGene_gn=KIF1B;refGene_tx=XM_023258115.1;refGene_gn=KIF1B;refGene_tx=XM_023258116.1;refGene_gn=KIF1B;refGene_tx=XM_023258117.1
C1 7350949 7351029
refGene_gn=KIF1B;refGene_tx=XM_019836118.2;refGene_gn=KIF1B;refGene_tx=XM_019836119.2;refGene_gn=KIF1B;refGene_tx=XM_023258093.1;refGene_gn=KIF1B;refGene_tx=XM_023258094.1;refGene_gn=KIF1B;refGene_tx=XM_023258095.1;refGene_gn=KIF1B;refGene_tx=XM_023258096.1;refGene_gn=KIF1B;refGene_tx=XM_023258097.1;refGene_gn=KIF1B;refGene_tx=XM_023258098.1;refGene_gn=KIF1B;refGene_tx=XM_023258099.1;refGene_gn=KIF1B;refGene_tx=XM_023258100.1;refGene_gn=KIF1B;refGene_tx=XM_023258101.1;refGene_gn=KIF1B;refGene_tx=XM_023258102.1;refGene_gn=KIF1B;refGene_tx=XM_023258103.1;refGene_gn=KIF1B;refGene_tx=XM_023258104.1;refGene_gn=KIF1B;refGene_tx=XM_023258105.1;refGene_gn=KIF1B;refGene_tx=XM_023258106.1;refGene_gn=KIF1B;refGene_tx=XM_023258107.1;refGene_gn=KIF1B;refGene_tx=XM_023258108.1;refGene_gn=KIF1B;refGene_tx=XM_023258109.1;refGene_gn=KIF1B;refGene_tx=XM_023258110.1;refGene_gn=KIF1B;refGene_tx=XM_023258111.1;refGene_gn=KIF1B;refGene_tx=XM_023258112.1;refGene_gn=KIF1B;refGene_tx=XM_023258113.1;refGene_gn=KIF1B;refGene_tx=XM_023258114.1;refGene_gn=KIF1B;r

efGene_tx=XM_023258115.1;refGene_gn=KIF1B;refGene_tx=XM_023258116.1;re
fGene_gn=KIF1B;refGene_tx=XM_023258117.1
C1 7352368 7352444
refGene_gn=KIF1B;refGene_tx=XM_019836118.2;refGene_gn=KIF1B;refGene_tx
=XM_019836119.2;refGene_gn=KIF1B;refGene_tx=XM_023258093.1;refGene_gn=
KIF1B;refGene_tx=XM_023258094.1;refGene_gn=KIF1B;refGene_tx=XM_0232580
95.1;refGene_gn=KIF1B;refGene_tx=XM_023258096.1;refGene_gn=KIF1B;refGe
ne_tx=XM_023258097.1;refGene_gn=KIF1B;refGene_tx=XM_023258098.1;refGen
e_gn=KIF1B;refGene_tx=XM_023258099.1;refGene_gn=KIF1B;refGene_tx=XM_02
3258100.1;refGene_gn=KIF1B;refGene_tx=XM_023258101.1;refGene_gn=KIF1B;
refGene_tx=XM_023258102.1;refGene_gn=KIF1B;refGene_tx=XM_023258103.1;r
efGene_gn=KIF1B;refGene_tx=XM_023258104.1;refGene_gn=KIF1B;refGene_tx=
XM_023258105.1;refGene_gn=KIF1B;refGene_tx=XM_023258106.1;refGene_gn=K
IF1B;refGene_tx=XM_023258107.1;refGene_gn=KIF1B;refGene_tx=XM_02325810
8.1;refGene_gn=KIF1B;refGene_tx=XM_023258109.1;refGene_gn=KIF1B;refGen
e_tx=XM_023258110.1;refGene_gn=KIF1B;refGene_tx=XM_023258111.1;refGene
_gn=KIF1B;refGene_tx=XM_023258112.1;refGene_gn=KIF1B;refGene_tx=XM_023
258113.1;refGene_gn=KIF1B;refGene_tx=XM_023258114.1;refGene_gn=KIF1B;r
efGene_tx=XM_023258115.1;refGene_gn=KIF1B;refGene_tx=XM_023258116.1;re
fGene_gn=KIF1B;refGene_tx=XM_023258117.1
C1 7352786 7352849
refGene_gn=KIF1B;refGene_tx=XM_019836118.2;refGene_gn=KIF1B;refGene_tx
=XM_019836119.2;refGene_gn=KIF1B;refGene_tx=XM_023258093.1;refGene_gn=
KIF1B;refGene_tx=XM_023258094.1;refGene_gn=KIF1B;refGene_tx=XM_0232580
95.1;refGene_gn=KIF1B;refGene_tx=XM_023258096.1;refGene_gn=KIF1B;refGe
ne_tx=XM_023258097.1;refGene_gn=KIF1B;refGene_tx=XM_023258099.1;refGen
e_gn=KIF1B;refGene_tx=XM_023258100.1;refGene_gn=KIF1B;refGene_tx=XM_02
3258101.1;refGene_gn=KIF1B;refGene_tx=XM_023258103.1;refGene_gn=KIF1B;
refGene_tx=XM_023258105.1;refGene_gn=KIF1B;refGene_tx=XM_023258106.1
C1 7354292 7354355
refGene_gn=KIF1B;refGene_tx=XM_019836118.2;refGene_gn=KIF1B;refGene_tx
=XM_019836119.2;refGene_gn=KIF1B;refGene_tx=XM_023258093.1;refGene_gn=
KIF1B;refGene_tx=XM_023258094.1;refGene_gn=KIF1B;refGene_tx=XM_0232580
95.1;refGene_gn=KIF1B;refGene_tx=XM_023258096.1;refGene_gn=KIF1B;refGe
ne_tx=XM_023258097.1;refGene_gn=KIF1B;refGene_tx=XM_023258098.1;refGen
e_gn=KIF1B;refGene_tx=XM_023258099.1;refGene_gn=KIF1B;refGene_tx=XM_02
3258100.1;refGene_gn=KIF1B;refGene_tx=XM_023258101.1;refGene_gn=KIF1B;
refGene_tx=XM_023258103.1;refGene_gn=KIF1B;refGene_tx=XM_023258105.1;r
efGene_gn=KIF1B;refGene_tx=XM_023258106.1
C1 7355491 7355571
refGene_gn=KIF1B;refGene_tx=XM_019836118.2;refGene_gn=KIF1B;refGene_tx
=XM_019836119.2;refGene_gn=KIF1B;refGene_tx=XM_023258093.1;refGene_gn=
KIF1B;refGene_tx=XM_023258094.1;refGene_gn=KIF1B;refGene_tx=XM_0232580
95.1;refGene_gn=KIF1B;refGene_tx=XM_023258096.1;refGene_gn=KIF1B;refGe
ne_tx=XM_023258097.1;refGene_gn=KIF1B;refGene_tx=XM_023258098.1;refGen
e_gn=KIF1B;refGene_tx=XM_023258099.1;refGene_gn=KIF1B;refGene_tx=XM_02
3258100.1;refGene_gn=KIF1B;refGene_tx=XM_023258101.1;refGene_gn=KIF1B;
refGene_tx=XM_023258102.1;refGene_gn=KIF1B;refGene_tx=XM_023258103.1;r
efGene_gn=KIF1B;refGene_tx=XM_023258104.1;refGene_gn=KIF1B;refGene_tx=
XM_023258105.1;refGene_gn=KIF1B;refGene_tx=XM_023258106.1;refGene_gn=K

IF1B;refGene_tx=XM_023258107.1;refGene_gn=KIF1B;refGene_tx=XM_02325810
8.1;refGene_gn=KIF1B;refGene_tx=XM_023258109.1;refGene_gn=KIF1B;refGen
e_tx=XM_023258110.1;refGene_gn=KIF1B;refGene_tx=XM_023258111.1;refGene
_gn=KIF1B;refGene_tx=XM_023258112.1;refGene_gn=KIF1B;refGene_tx=XM_023
258113.1;refGene_gn=KIF1B;refGene_tx=XM_023258114.1;refGene_gn=KIF1B;r
efGene_tx=XM_023258115.1;refGene_gn=KIF1B;refGene_tx=XM_023258116.1;re
fGene_gn=KIF1B;refGene_tx=XM_023258117.1
C1 7356069 7356176
refGene_gn=KIF1B;refGene_tx=XM_019836118.2;refGene_gn=KIF1B;refGene_tx
=XM_019836119.2;refGene_gn=KIF1B;refGene_tx=XM_023258093.1;refGene_gn=
KIF1B;refGene_tx=XM_023258094.1;refGene_gn=KIF1B;refGene_tx=XM_0232580
95.1;refGene_gn=KIF1B;refGene_tx=XM_023258096.1;refGene_gn=KIF1B;refGe
ne_tx=XM_023258097.1;refGene_gn=KIF1B;refGene_tx=XM_023258098.1;refGen
e_gn=KIF1B;refGene_tx=XM_023258099.1;refGene_gn=KIF1B;refGene_tx=XM_02
3258100.1;refGene_gn=KIF1B;refGene_tx=XM_023258101.1;refGene_gn=KIF1B;
refGene_tx=XM_023258102.1;refGene_gn=KIF1B;refGene_tx=XM_023258103.1;r
efGene_gn=KIF1B;refGene_tx=XM_023258104.1;refGene_gn=KIF1B;refGene_tx=
XM_023258105.1;refGene_gn=KIF1B;refGene_tx=XM_023258106.1;refGene_gn=K
IF1B;refGene_tx=XM_023258107.1;refGene_gn=KIF1B;refGene_tx=XM_02325810
8.1;refGene_gn=KIF1B;refGene_tx=XM_023258109.1;refGene_gn=KIF1B;refGen
e_tx=XM_023258110.1;refGene_gn=KIF1B;refGene_tx=XM_023258111.1;refGene
_gn=KIF1B;refGene_tx=XM_023258112.1;refGene_gn=KIF1B;refGene_tx=XM_023
258113.1;refGene_gn=KIF1B;refGene_tx=XM_023258114.1;refGene_gn=KIF1B;r
efGene_tx=XM_023258115.1;refGene_gn=KIF1B;refGene_tx=XM_023258116.1;re
fGene_gn=KIF1B;refGene_tx=XM_023258117.1
C1 7356947 7357031
refGene_gn=KIF1B;refGene_tx=XM_019836118.2;refGene_gn=KIF1B;refGene_tx
=XM_019836119.2;refGene_gn=KIF1B;refGene_tx=XM_023258093.1;refGene_gn=
KIF1B;refGene_tx=XM_023258094.1;refGene_gn=KIF1B;refGene_tx=XM_0232580
95.1;refGene_gn=KIF1B;refGene_tx=XM_023258096.1;refGene_gn=KIF1B;refGe
ne_tx=XM_023258097.1;refGene_gn=KIF1B;refGene_tx=XM_023258098.1;refGen
e_gn=KIF1B;refGene_tx=XM_023258099.1;refGene_gn=KIF1B;refGene_tx=XM_02
3258100.1;refGene_gn=KIF1B;refGene_tx=XM_023258101.1;refGene_gn=KIF1B;
refGene_tx=XM_023258102.1;refGene_gn=KIF1B;refGene_tx=XM_023258103.1;r
efGene_gn=KIF1B;refGene_tx=XM_023258104.1;refGene_gn=KIF1B;refGene_tx=
XM_023258105.1;refGene_gn=KIF1B;refGene_tx=XM_023258106.1;refGene_gn=K
IF1B;refGene_tx=XM_023258107.1;refGene_gn=KIF1B;refGene_tx=XM_02325810
8.1;refGene_gn=KIF1B;refGene_tx=XM_023258109.1;refGene_gn=KIF1B;refGen
e_tx=XM_023258110.1;refGene_gn=KIF1B;refGene_tx=XM_023258111.1;refGene
_gn=KIF1B;refGene_tx=XM_023258112.1;refGene_gn=KIF1B;refGene_tx=XM_023
258113.1;refGene_gn=KIF1B;refGene_tx=XM_023258114.1;refGene_gn=KIF1B;r
efGene_tx=XM_023258115.1;refGene_gn=KIF1B;refGene_tx=XM_023258116.1;re
fGene_gn=KIF1B;refGene_tx=XM_023258117.1
C1 7357263 7357444
refGene_gn=KIF1B;refGene_tx=XM_019836118.2;refGene_gn=KIF1B;refGene_tx
=XM_019836119.2;refGene_gn=KIF1B;refGene_tx=XM_023258093.1;refGene_gn=
KIF1B;refGene_tx=XM_023258094.1;refGene_gn=KIF1B;refGene_tx=XM_0232580
95.1;refGene_gn=KIF1B;refGene_tx=XM_023258096.1;refGene_gn=KIF1B;refGe
ne_tx=XM_023258097.1;refGene_gn=KIF1B;refGene_tx=XM_023258098.1;refGen
e_gn=KIF1B;refGene_tx=XM_023258099.1;refGene_gn=KIF1B;refGene_tx=XM_02

3258100.1;refGene_gn=KIF1B;refGene_tx=XM_023258101.1;refGene_gn=KIF1B;
refGene_tx=XM_023258102.1;refGene_gn=KIF1B;refGene_tx=XM_023258103.1;r
efGene_gn=KIF1B;refGene_tx=XM_023258104.1;refGene_gn=KIF1B;refGene_tx=
XM_023258105.1;refGene_gn=KIF1B;refGene_tx=XM_023258106.1;refGene_gn=K
IF1B;refGene_tx=XM_023258107.1;refGene_gn=KIF1B;refGene_tx=XM_02325810
8.1;refGene_gn=KIF1B;refGene_tx=XM_023258109.1;refGene_gn=KIF1B;refGen
e_tx=XM_023258110.1;refGene_gn=KIF1B;refGene_tx=XM_023258111.1;refGene
_gn=KIF1B;refGene_tx=XM_023258112.1;refGene_gn=KIF1B;refGene_tx=XM_023
258113.1;refGene_gn=KIF1B;refGene_tx=XM_023258114.1;refGene_gn=KIF1B;r
efGene_tx=XM_023258115.1;refGene_gn=KIF1B;refGene_tx=XM_023258116.1;re
fGene_gn=KIF1B;refGene_tx=XM_023258117.1
C1 7357573 7357646
refGene_gn=KIF1B;refGene_tx=XM_019836118.2;refGene_gn=KIF1B;refGene_tx
=XM_019836119.2;refGene_gn=KIF1B;refGene_tx=XM_023258093.1;refGene_gn=
KIF1B;refGene_tx=XM_023258094.1;refGene_gn=KIF1B;refGene_tx=XM_0232580
95.1;refGene_gn=KIF1B;refGene_tx=XM_023258096.1;refGene_gn=KIF1B;refGe
ne_tx=XM_023258097.1;refGene_gn=KIF1B;refGene_tx=XM_023258098.1;refGen
e_gn=KIF1B;refGene_tx=XM_023258099.1;refGene_gn=KIF1B;refGene_tx=XM_02
3258100.1;refGene_gn=KIF1B;refGene_tx=XM_023258101.1;refGene_gn=KIF1B;
refGene_tx=XM_023258102.1;refGene_gn=KIF1B;refGene_tx=XM_023258103.1;r
efGene_gn=KIF1B;refGene_tx=XM_023258104.1;refGene_gn=KIF1B;refGene_tx=
XM_023258105.1;refGene_gn=KIF1B;refGene_tx=XM_023258106.1;refGene_gn=K
IF1B;refGene_tx=XM_023258107.1;refGene_gn=KIF1B;refGene_tx=XM_02325810
8.1;refGene_gn=KIF1B;refGene_tx=XM_023258109.1;refGene_gn=KIF1B;refGen
e_tx=XM_023258110.1;refGene_gn=KIF1B;refGene_tx=XM_023258111.1;refGene
_gn=KIF1B;refGene_tx=XM_023258112.1;refGene_gn=KIF1B;refGene_tx=XM_023
258113.1;refGene_gn=KIF1B;refGene_tx=XM_023258114.1;refGene_gn=KIF1B;r
efGene_tx=XM_023258115.1;refGene_gn=KIF1B;refGene_tx=XM_023258116.1;re
fGene_gn=KIF1B;refGene_tx=XM_023258117.1
C1 7362940 7364425 refGene_gn=KIF1B;refGene_tx=XM_023258115.1
C1 7376774 7376868
refGene_gn=KIF1B;refGene_tx=XM_019836118.2;refGene_gn=KIF1B;refGene_tx
=XM_019836119.2;refGene_gn=KIF1B;refGene_tx=XM_023258093.1;refGene_gn=
KIF1B;refGene_tx=XM_023258094.1;refGene_gn=KIF1B;refGene_tx=XM_0232580
95.1;refGene_gn=KIF1B;refGene_tx=XM_023258096.1;refGene_gn=KIF1B;refGe
ne_tx=XM_023258097.1;refGene_gn=KIF1B;refGene_tx=XM_023258098.1;refGen
e_gn=KIF1B;refGene_tx=XM_023258099.1;refGene_gn=KIF1B;refGene_tx=XM_02
3258100.1;refGene_gn=KIF1B;refGene_tx=XM_023258101.1;refGene_gn=KIF1B;
refGene_tx=XM_023258102.1;refGene_gn=KIF1B;refGene_tx=XM_023258103.1;r
efGene_gn=KIF1B;refGene_tx=XM_023258104.1;refGene_gn=KIF1B;refGene_tx=
XM_023258105.1;refGene_gn=KIF1B;refGene_tx=XM_023258106.1;refGene_gn=K
IF1B;refGene_tx=XM_023258107.1;refGene_gn=KIF1B;refGene_tx=XM_02325810
8.1;refGene_gn=KIF1B;refGene_tx=XM_023258109.1;refGene_gn=KIF1B;refGen
e_tx=XM_023258110.1;refGene_gn=KIF1B;refGene_tx=XM_023258111.1;refGene
_gn=KIF1B;refGene_tx=XM_023258112.1;refGene_gn=KIF1B;refGene_tx=XM_023
258113.1;refGene_gn=KIF1B;refGene_tx=XM_023258114.1;refGene_gn=KIF1B;r
efGene_tx=XM_023258116.1;refGene_gn=KIF1B;refGene_tx=XM_023258117.1
C1 7377868 7378017
refGene_gn=KIF1B;refGene_tx=XM_019836118.2;refGene_gn=KIF1B;refGene_tx
=XM_019836119.2;refGene_gn=KIF1B;refGene_tx=XM_023258093.1;refGene_gn=

KIF1B;refGene_tx=XM_023258094.1;refGene_gn=KIF1B;refGene_tx=XM_0232580
95.1;refGene_gn=KIF1B;refGene_tx=XM_023258096.1;refGene_gn=KIF1B;refGe
ne_tx=XM_023258097.1;refGene_gn=KIF1B;refGene_tx=XM_023258098.1;refGen
e_gn=KIF1B;refGene_tx=XM_023258099.1;refGene_gn=KIF1B;refGene_tx=XM_02
3258100.1;refGene_gn=KIF1B;refGene_tx=XM_023258101.1;refGene_gn=KIF1B;
refGene_tx=XM_023258102.1;refGene_gn=KIF1B;refGene_tx=XM_023258103.1;r
efGene_gn=KIF1B;refGene_tx=XM_023258104.1;refGene_gn=KIF1B;refGene_tx=
XM_023258105.1;refGene_gn=KIF1B;refGene_tx=XM_023258106.1;refGene_gn=K
IF1B;refGene_tx=XM_023258107.1;refGene_gn=KIF1B;refGene_tx=XM_02325810
8.1;refGene_gn=KIF1B;refGene_tx=XM_023258109.1;refGene_gn=KIF1B;refGen
e_tx=XM_023258110.1;refGene_gn=KIF1B;refGene_tx=XM_023258111.1;refGene
_gn=KIF1B;refGene_tx=XM_023258112.1;refGene_gn=KIF1B;refGene_tx=XM_023
258113.1;refGene_gn=KIF1B;refGene_tx=XM_023258114.1;refGene_gn=KIF1B;r
efGene_tx=XM_023258116.1;refGene_gn=KIF1B;refGene_tx=XM_023258117.1
C1 7379718 7379897
refGene_gn=KIF1B;refGene_tx=XM_019836118.2;refGene_gn=KIF1B;refGene_tx
=XM_019836119.2;refGene_gn=KIF1B;refGene_tx=XM_023258093.1;refGene_gn=
KIF1B;refGene_tx=XM_023258094.1;refGene_gn=KIF1B;refGene_tx=XM_0232580
95.1;refGene_gn=KIF1B;refGene_tx=XM_023258096.1;refGene_gn=KIF1B;refGe
ne_tx=XM_023258097.1;refGene_gn=KIF1B;refGene_tx=XM_023258098.1;refGen
e_gn=KIF1B;refGene_tx=XM_023258099.1;refGene_gn=KIF1B;refGene_tx=XM_02
3258100.1;refGene_gn=KIF1B;refGene_tx=XM_023258101.1;refGene_gn=KIF1B;
refGene_tx=XM_023258102.1;refGene_gn=KIF1B;refGene_tx=XM_023258103.1;r
efGene_gn=KIF1B;refGene_tx=XM_023258104.1;refGene_gn=KIF1B;refGene_tx=
XM_023258105.1;refGene_gn=KIF1B;refGene_tx=XM_023258106.1;refGene_gn=K
IF1B;refGene_tx=XM_023258107.1;refGene_gn=KIF1B;refGene_tx=XM_02325810
8.1;refGene_gn=KIF1B;refGene_tx=XM_023258109.1;refGene_gn=KIF1B;refGen
e_tx=XM_023258110.1;refGene_gn=KIF1B;refGene_tx=XM_023258111.1;refGene
_gn=KIF1B;refGene_tx=XM_023258112.1;refGene_gn=KIF1B;refGene_tx=XM_023
258113.1;refGene_gn=KIF1B;refGene_tx=XM_023258114.1;refGene_gn=KIF1B;r
efGene_tx=XM_023258116.1;refGene_gn=KIF1B;refGene_tx=XM_023258117.1
C1 7380858 7380996
refGene_gn=KIF1B;refGene_tx=XM_019836118.2;refGene_gn=KIF1B;refGene_tx
=XM_019836119.2;refGene_gn=KIF1B;refGene_tx=XM_023258093.1;refGene_gn=
KIF1B;refGene_tx=XM_023258094.1;refGene_gn=KIF1B;refGene_tx=XM_0232580
95.1;refGene_gn=KIF1B;refGene_tx=XM_023258096.1;refGene_gn=KIF1B;refGe
ne_tx=XM_023258097.1;refGene_gn=KIF1B;refGene_tx=XM_023258098.1;refGen
e_gn=KIF1B;refGene_tx=XM_023258099.1;refGene_gn=KIF1B;refGene_tx=XM_02
3258100.1;refGene_gn=KIF1B;refGene_tx=XM_023258101.1;refGene_gn=KIF1B;
refGene_tx=XM_023258102.1;refGene_gn=KIF1B;refGene_tx=XM_023258103.1;r
efGene_gn=KIF1B;refGene_tx=XM_023258104.1;refGene_gn=KIF1B;refGene_tx=
XM_023258105.1;refGene_gn=KIF1B;refGene_tx=XM_023258106.1;refGene_gn=K
IF1B;refGene_tx=XM_023258107.1;refGene_gn=KIF1B;refGene_tx=XM_02325810
8.1;refGene_gn=KIF1B;refGene_tx=XM_023258109.1;refGene_gn=KIF1B;refGen
e_tx=XM_023258110.1;refGene_gn=KIF1B;refGene_tx=XM_023258111.1;refGene
_gn=KIF1B;refGene_tx=XM_023258112.1;refGene_gn=KIF1B;refGene_tx=XM_023
258113.1;refGene_gn=KIF1B;refGene_tx=XM_023258114.1;refGene_gn=KIF1B;r
efGene_tx=XM_023258116.1;refGene_gn=KIF1B;refGene_tx=XM_023258117.1
C1 7382386 7382728
refGene_gn=KIF1B;refGene_tx=XM_019836118.2;refGene_gn=KIF1B;refGene_tx

=XM_019836119.2;refGene_gn=KIF1B;refGene_tx=XM_023258093.1;refGene_gn=
KIF1B;refGene_tx=XM_023258094.1;refGene_gn=KIF1B;refGene_tx=XM_0232580
95.1;refGene_gn=KIF1B;refGene_tx=XM_023258096.1;refGene_gn=KIF1B;refGe
ne_tx=XM_023258097.1;refGene_gn=KIF1B;refGene_tx=XM_023258098.1;refGen
e_gn=KIF1B;refGene_tx=XM_023258099.1;refGene_gn=KIF1B;refGene_tx=XM_02
3258100.1;refGene_gn=KIF1B;refGene_tx=XM_023258101.1;refGene_gn=KIF1B;
refGene_tx=XM_023258102.1;refGene_gn=KIF1B;refGene_tx=XM_023258103.1;r
efGene_gn=KIF1B;refGene_tx=XM_023258104.1;refGene_gn=KIF1B;refGene_tx=
XM_023258105.1;refGene_gn=KIF1B;refGene_tx=XM_023258107.1;refGene_gn=K
IF1B;refGene_tx=XM_023258109.1;refGene_gn=KIF1B;refGene_tx=XM_02325811
0.1;refGene_gn=KIF1B;refGene_tx=XM_023258111.1;refGene_gn=KIF1B;refGen
e_tx=XM_023258112.1
C1 7387891 7388010
refGene_gn=KIF1B;refGene_tx=XM_019836118.2;refGene_gn=KIF1B;refGene_tx
=XM_019836119.2;refGene_gn=KIF1B;refGene_tx=XM_023258093.1;refGene_gn=
KIF1B;refGene_tx=XM_023258094.1;refGene_gn=KIF1B;refGene_tx=XM_0232580
95.1;refGene_gn=KIF1B;refGene_tx=XM_023258096.1;refGene_gn=KIF1B;refGe
ne_tx=XM_023258097.1;refGene_gn=KIF1B;refGene_tx=XM_023258098.1;refGen
e_gn=KIF1B;refGene_tx=XM_023258099.1;refGene_gn=KIF1B;refGene_tx=XM_02
3258100.1;refGene_gn=KIF1B;refGene_tx=XM_023258101.1;refGene_gn=KIF1B;
refGene_tx=XM_023258102.1;refGene_gn=KIF1B;refGene_tx=XM_023258103.1;r
efGene_gn=KIF1B;refGene_tx=XM_023258104.1;refGene_gn=KIF1B;refGene_tx=
XM_023258105.1;refGene_gn=KIF1B;refGene_tx=XM_023258106.1;refGene_gn=K
IF1B;refGene_tx=XM_023258107.1;refGene_gn=KIF1B;refGene_tx=XM_02325810
8.1;refGene_gn=KIF1B;refGene_tx=XM_023258109.1;refGene_gn=KIF1B;refGen
e_tx=XM_023258110.1;refGene_gn=KIF1B;refGene_tx=XM_023258111.1;refGene
_gn=KIF1B;refGene_tx=XM_023258112.1;refGene_gn=KIF1B;refGene_tx=XM_023
258113.1;refGene_gn=KIF1B;refGene_tx=XM_023258114.1;refGene_gn=KIF1B;r
efGene_tx=XM_023258116.1;refGene_gn=KIF1B;refGene_tx=XM_023258117.1
C1 7388931 7389017
refGene_gn=KIF1B;refGene_tx=XM_019836118.2;refGene_gn=KIF1B;refGene_tx
=XM_019836119.2;refGene_gn=KIF1B;refGene_tx=XM_023258093.1;refGene_gn=
KIF1B;refGene_tx=XM_023258094.1;refGene_gn=KIF1B;refGene_tx=XM_0232580
95.1;refGene_gn=KIF1B;refGene_tx=XM_023258096.1;refGene_gn=KIF1B;refGe
ne_tx=XM_023258097.1;refGene_gn=KIF1B;refGene_tx=XM_023258098.1;refGen
e_gn=KIF1B;refGene_tx=XM_023258099.1;refGene_gn=KIF1B;refGene_tx=XM_02
3258100.1;refGene_gn=KIF1B;refGene_tx=XM_023258101.1;refGene_gn=KIF1B;
refGene_tx=XM_023258102.1;refGene_gn=KIF1B;refGene_tx=XM_023258103.1;r
efGene_gn=KIF1B;refGene_tx=XM_023258104.1;refGene_gn=KIF1B;refGene_tx=
XM_023258105.1;refGene_gn=KIF1B;refGene_tx=XM_023258106.1;refGene_gn=K
IF1B;refGene_tx=XM_023258107.1;refGene_gn=KIF1B;refGene_tx=XM_02325810
8.1;refGene_gn=KIF1B;refGene_tx=XM_023258109.1;refGene_gn=KIF1B;refGen
e_tx=XM_023258110.1;refGene_gn=KIF1B;refGene_tx=XM_023258111.1;refGene
_gn=KIF1B;refGene_tx=XM_023258112.1;refGene_gn=KIF1B;refGene_tx=XM_023
258113.1;refGene_gn=KIF1B;refGene_tx=XM_023258114.1;refGene_gn=KIF1B;r
efGene_tx=XM_023258116.1;refGene_gn=KIF1B;refGene_tx=XM_023258117.1
C1 7389349 7389479
refGene_gn=KIF1B;refGene_tx=XM_019836118.2;refGene_gn=KIF1B;refGene_tx
=XM_019836119.2;refGene_gn=KIF1B;refGene_tx=XM_023258093.1;refGene_gn=
KIF1B;refGene_tx=XM_023258094.1;refGene_gn=KIF1B;refGene_tx=XM_0232580

95.1;refGene_gn=KIF1B;refGene_tx=XM_023258096.1;refGene_gn=KIF1B;refGe
ne_tx=XM_023258097.1;refGene_gn=KIF1B;refGene_tx=XM_023258098.1;refGen
e_gn=KIF1B;refGene_tx=XM_023258099.1;refGene_gn=KIF1B;refGene_tx=XM_02
3258100.1;refGene_gn=KIF1B;refGene_tx=XM_023258101.1;refGene_gn=KIF1B;
refGene_tx=XM_023258102.1;refGene_gn=KIF1B;refGene_tx=XM_023258103.1;r
efGene_gn=KIF1B;refGene_tx=XM_023258104.1;refGene_gn=KIF1B;refGene_tx=
XM_023258105.1;refGene_gn=KIF1B;refGene_tx=XM_023258106.1;refGene_gn=K
IF1B;refGene_tx=XM_023258107.1;refGene_gn=KIF1B;refGene_tx=XM_02325810
8.1;refGene_gn=KIF1B;refGene_tx=XM_023258109.1;refGene_gn=KIF1B;refGen
e_tx=XM_023258110.1;refGene_gn=KIF1B;refGene_tx=XM_023258111.1;refGene
_gn=KIF1B;refGene_tx=XM_023258112.1;refGene_gn=KIF1B;refGene_tx=XM_023
258113.1;refGene_gn=KIF1B;refGene_tx=XM_023258114.1;refGene_gn=KIF1B;r
efGene_tx=XM_023258116.1;refGene_gn=KIF1B;refGene_tx=XM_023258117.1
C1 7389655 7389818
refGene_gn=KIF1B;refGene_tx=XM_019836118.2;refGene_gn=KIF1B;refGene_tx
=XM_019836119.2;refGene_gn=KIF1B;refGene_tx=XM_023258093.1;refGene_gn=
KIF1B;refGene_tx=XM_023258094.1;refGene_gn=KIF1B;refGene_tx=XM_0232580
95.1;refGene_gn=KIF1B;refGene_tx=XM_023258096.1;refGene_gn=KIF1B;refGe
ne_tx=XM_023258097.1;refGene_gn=KIF1B;refGene_tx=XM_023258098.1;refGen
e_gn=KIF1B;refGene_tx=XM_023258099.1;refGene_gn=KIF1B;refGene_tx=XM_02
3258100.1;refGene_gn=KIF1B;refGene_tx=XM_023258101.1;refGene_gn=KIF1B;
refGene_tx=XM_023258102.1;refGene_gn=KIF1B;refGene_tx=XM_023258103.1;r
efGene_gn=KIF1B;refGene_tx=XM_023258104.1;refGene_gn=KIF1B;refGene_tx=
XM_023258105.1;refGene_gn=KIF1B;refGene_tx=XM_023258106.1;refGene_gn=K
IF1B;refGene_tx=XM_023258107.1;refGene_gn=KIF1B;refGene_tx=XM_02325810
8.1;refGene_gn=KIF1B;refGene_tx=XM_023258109.1;refGene_gn=KIF1B;refGen
e_tx=XM_023258110.1;refGene_gn=KIF1B;refGene_tx=XM_023258111.1;refGene
_gn=KIF1B;refGene_tx=XM_023258112.1;refGene_gn=KIF1B;refGene_tx=XM_023
258113.1;refGene_gn=KIF1B;refGene_tx=XM_023258114.1;refGene_gn=KIF1B;r
efGene_tx=XM_023258116.1;refGene_gn=KIF1B;refGene_tx=XM_023258117.1
C1 7392496 7392587
refGene_gn=KIF1B;refGene_tx=XM_019836118.2;refGene_gn=KIF1B;refGene_tx
=XM_019836119.2;refGene_gn=KIF1B;refGene_tx=XM_023258093.1;refGene_gn=
KIF1B;refGene_tx=XM_023258094.1;refGene_gn=KIF1B;refGene_tx=XM_0232580
95.1;refGene_gn=KIF1B;refGene_tx=XM_023258096.1;refGene_gn=KIF1B;refGe
ne_tx=XM_023258097.1;refGene_gn=KIF1B;refGene_tx=XM_023258098.1;refGen
e_gn=KIF1B;refGene_tx=XM_023258099.1;refGene_gn=KIF1B;refGene_tx=XM_02
3258100.1;refGene_gn=KIF1B;refGene_tx=XM_023258101.1;refGene_gn=KIF1B;
refGene_tx=XM_023258102.1;refGene_gn=KIF1B;refGene_tx=XM_023258103.1;r
efGene_gn=KIF1B;refGene_tx=XM_023258104.1;refGene_gn=KIF1B;refGene_tx=
XM_023258105.1;refGene_gn=KIF1B;refGene_tx=XM_023258106.1;refGene_gn=K
IF1B;refGene_tx=XM_023258107.1;refGene_gn=KIF1B;refGene_tx=XM_02325810
8.1;refGene_gn=KIF1B;refGene_tx=XM_023258109.1;refGene_gn=KIF1B;refGen
e_tx=XM_023258110.1;refGene_gn=KIF1B;refGene_tx=XM_023258111.1;refGene
_gn=KIF1B;refGene_tx=XM_023258112.1;refGene_gn=KIF1B;refGene_tx=XM_023
258113.1;refGene_gn=KIF1B;refGene_tx=XM_023258114.1;refGene_gn=KIF1B;r
efGene_tx=XM_023258116.1;refGene_gn=KIF1B;refGene_tx=XM_023258117.1
C1 7394349 7394468
refGene_gn=KIF1B;refGene_tx=XM_019836118.2;refGene_gn=KIF1B;refGene_tx
=XM_019836119.2;refGene_gn=KIF1B;refGene_tx=XM_023258093.1;refGene_gn=

KIF1B;refGene_tx=XM_023258094.1;refGene_gn=KIF1B;refGene_tx=XM_0232580
95.1;refGene_gn=KIF1B;refGene_tx=XM_023258096.1;refGene_gn=KIF1B;refGe
ne_tx=XM_023258097.1;refGene_gn=KIF1B;refGene_tx=XM_023258098.1;refGen
e_gn=KIF1B;refGene_tx=XM_023258099.1;refGene_gn=KIF1B;refGene_tx=XM_02
3258100.1;refGene_gn=KIF1B;refGene_tx=XM_023258101.1;refGene_gn=KIF1B;
refGene_tx=XM_023258102.1;refGene_gn=KIF1B;refGene_tx=XM_023258103.1;r
efGene_gn=KIF1B;refGene_tx=XM_023258104.1;refGene_gn=KIF1B;refGene_tx=
XM_023258105.1;refGene_gn=KIF1B;refGene_tx=XM_023258106.1;refGene_gn=K
IF1B;refGene_tx=XM_023258107.1;refGene_gn=KIF1B;refGene_tx=XM_02325810
8.1;refGene_gn=KIF1B;refGene_tx=XM_023258109.1;refGene_gn=KIF1B;refGen
e_tx=XM_023258110.1;refGene_gn=KIF1B;refGene_tx=XM_023258111.1;refGene
_gn=KIF1B;refGene_tx=XM_023258112.1;refGene_gn=KIF1B;refGene_tx=XM_023
258113.1;refGene_gn=KIF1B;refGene_tx=XM_023258114.1;refGene_gn=KIF1B;r
efGene_tx=XM_023258116.1;refGene_gn=KIF1B;refGene_tx=XM_023258117.1
C1 7395097 7395153
refGene_gn=KIF1B;refGene_tx=XM_019836118.2;refGene_gn=KIF1B;refGene_tx
=XM_019836119.2;refGene_gn=KIF1B;refGene_tx=XM_023258093.1;refGene_gn=
KIF1B;refGene_tx=XM_023258094.1;refGene_gn=KIF1B;refGene_tx=XM_0232580
95.1;refGene_gn=KIF1B;refGene_tx=XM_023258096.1;refGene_gn=KIF1B;refGe
ne_tx=XM_023258097.1;refGene_gn=KIF1B;refGene_tx=XM_023258098.1;refGen
e_gn=KIF1B;refGene_tx=XM_023258099.1;refGene_gn=KIF1B;refGene_tx=XM_02
3258100.1;refGene_gn=KIF1B;refGene_tx=XM_023258101.1;refGene_gn=KIF1B;
refGene_tx=XM_023258102.1;refGene_gn=KIF1B;refGene_tx=XM_023258103.1;r
efGene_gn=KIF1B;refGene_tx=XM_023258104.1;refGene_gn=KIF1B;refGene_tx=
XM_023258105.1;refGene_gn=KIF1B;refGene_tx=XM_023258106.1;refGene_gn=K
IF1B;refGene_tx=XM_023258107.1;refGene_gn=KIF1B;refGene_tx=XM_02325810
8.1;refGene_gn=KIF1B;refGene_tx=XM_023258109.1;refGene_gn=KIF1B;refGen
e_tx=XM_023258110.1;refGene_gn=KIF1B;refGene_tx=XM_023258111.1;refGene
_gn=KIF1B;refGene_tx=XM_023258112.1;refGene_gn=KIF1B;refGene_tx=XM_023
258113.1;refGene_gn=KIF1B;refGene_tx=XM_023258114.1;refGene_gn=KIF1B;r
efGene_tx=XM_023258116.1;refGene_gn=KIF1B;refGene_tx=XM_023258117.1
C1 7395586 7395787
refGene_gn=KIF1B;refGene_tx=XM_019836118.2;refGene_gn=KIF1B;refGene_tx
=XM_019836119.2;refGene_gn=KIF1B;refGene_tx=XM_023258093.1;refGene_gn=
KIF1B;refGene_tx=XM_023258094.1;refGene_gn=KIF1B;refGene_tx=XM_0232580
95.1;refGene_gn=KIF1B;refGene_tx=XM_023258096.1;refGene_gn=KIF1B;refGe
ne_tx=XM_023258097.1;refGene_gn=KIF1B;refGene_tx=XM_023258098.1;refGen
e_gn=KIF1B;refGene_tx=XM_023258099.1;refGene_gn=KIF1B;refGene_tx=XM_02
3258100.1;refGene_gn=KIF1B;refGene_tx=XM_023258101.1;refGene_gn=KIF1B;
refGene_tx=XM_023258102.1;refGene_gn=KIF1B;refGene_tx=XM_023258104.1;r
efGene_gn=KIF1B;refGene_tx=XM_023258106.1;refGene_gn=KIF1B;refGene_tx=
XM_023258108.1;refGene_gn=KIF1B;refGene_tx=XM_023258113.1
C1 7397248 7397357
refGene_gn=KIF1B;refGene_tx=XM_019836118.2;refGene_gn=KIF1B;refGene_tx
=XM_019836119.2;refGene_gn=KIF1B;refGene_tx=XM_023258093.1;refGene_gn=
KIF1B;refGene_tx=XM_023258094.1;refGene_gn=KIF1B;refGene_tx=XM_0232580
95.1;refGene_gn=KIF1B;refGene_tx=XM_023258096.1;refGene_gn=KIF1B;refGe
ne_tx=XM_023258097.1;refGene_gn=KIF1B;refGene_tx=XM_023258098.1;refGen
e_gn=KIF1B;refGene_tx=XM_023258099.1;refGene_gn=KIF1B;refGene_tx=XM_02
3258100.1;refGene_gn=KIF1B;refGene_tx=XM_023258101.1;refGene_gn=KIF1B;

refGene_tx=XM_023258102.1;refGene_gn=KIF1B;refGene_tx=XM_023258103.1;r
efGene_gn=KIF1B;refGene_tx=XM_023258104.1;refGene_gn=KIF1B;refGene_tx=
XM_023258105.1;refGene_gn=KIF1B;refGene_tx=XM_023258106.1;refGene_gn=K
IF1B;refGene_tx=XM_023258107.1;refGene_gn=KIF1B;refGene_tx=XM_02325810
8.1;refGene_gn=KIF1B;refGene_tx=XM_023258109.1;refGene_gn=KIF1B;refGen
e_tx=XM_023258110.1;refGene_gn=KIF1B;refGene_tx=XM_023258111.1;refGene
_gn=KIF1B;refGene_tx=XM_023258112.1;refGene_gn=KIF1B;refGene_tx=XM_023
258113.1;refGene_gn=KIF1B;refGene_tx=XM_023258114.1;refGene_gn=KIF1B;r
efGene_tx=XM_023258116.1;refGene_gn=KIF1B;refGene_tx=XM_023258117.1
C1 7398692 7398759
refGene_gn=KIF1B;refGene_tx=XM_019836118.2;refGene_gn=KIF1B;refGene_tx
=XM_019836119.2;refGene_gn=KIF1B;refGene_tx=XM_023258093.1;refGene_gn=
KIF1B;refGene_tx=XM_023258094.1;refGene_gn=KIF1B;refGene_tx=XM_0232580
95.1;refGene_gn=KIF1B;refGene_tx=XM_023258096.1;refGene_gn=KIF1B;refGe
ne_tx=XM_023258097.1;refGene_gn=KIF1B;refGene_tx=XM_023258098.1;refGen
e_gn=KIF1B;refGene_tx=XM_023258099.1;refGene_gn=KIF1B;refGene_tx=XM_02
3258100.1;refGene_gn=KIF1B;refGene_tx=XM_023258101.1;refGene_gn=KIF1B;
refGene_tx=XM_023258102.1;refGene_gn=KIF1B;refGene_tx=XM_023258103.1;r
efGene_gn=KIF1B;refGene_tx=XM_023258104.1;refGene_gn=KIF1B;refGene_tx=
XM_023258105.1;refGene_gn=KIF1B;refGene_tx=XM_023258106.1;refGene_gn=K
IF1B;refGene_tx=XM_023258107.1;refGene_gn=KIF1B;refGene_tx=XM_02325810
8.1;refGene_gn=KIF1B;refGene_tx=XM_023258109.1;refGene_gn=KIF1B;refGen
e_tx=XM_023258110.1;refGene_gn=KIF1B;refGene_tx=XM_023258111.1;refGene
_gn=KIF1B;refGene_tx=XM_023258112.1;refGene_gn=KIF1B;refGene_tx=XM_023
258113.1;refGene_gn=KIF1B;refGene_tx=XM_023258114.1;refGene_gn=KIF1B;r
efGene_tx=XM_023258116.1;refGene_gn=KIF1B;refGene_tx=XM_023258117.1
C1 7399698 7399783
refGene_gn=KIF1B;refGene_tx=XM_019836118.2;refGene_gn=KIF1B;refGene_tx
=XM_019836119.2;refGene_gn=KIF1B;refGene_tx=XM_023258093.1;refGene_gn=
KIF1B;refGene_tx=XM_023258094.1;refGene_gn=KIF1B;refGene_tx=XM_0232580
95.1;refGene_gn=KIF1B;refGene_tx=XM_023258096.1;refGene_gn=KIF1B;refGe
ne_tx=XM_023258097.1;refGene_gn=KIF1B;refGene_tx=XM_023258098.1;refGen
e_gn=KIF1B;refGene_tx=XM_023258099.1;refGene_gn=KIF1B;refGene_tx=XM_02
3258100.1;refGene_gn=KIF1B;refGene_tx=XM_023258101.1;refGene_gn=KIF1B;
refGene_tx=XM_023258102.1;refGene_gn=KIF1B;refGene_tx=XM_023258103.1;r
efGene_gn=KIF1B;refGene_tx=XM_023258104.1;refGene_gn=KIF1B;refGene_tx=
XM_023258105.1;refGene_gn=KIF1B;refGene_tx=XM_023258106.1;refGene_gn=K
IF1B;refGene_tx=XM_023258107.1;refGene_gn=KIF1B;refGene_tx=XM_02325810
8.1;refGene_gn=KIF1B;refGene_tx=XM_023258109.1;refGene_gn=KIF1B;refGen
e_tx=XM_023258110.1;refGene_gn=KIF1B;refGene_tx=XM_023258111.1;refGene
_gn=KIF1B;refGene_tx=XM_023258112.1;refGene_gn=KIF1B;refGene_tx=XM_023
258113.1;refGene_gn=KIF1B;refGene_tx=XM_023258114.1;refGene_gn=KIF1B;r
efGene_tx=XM_023258116.1;refGene_gn=KIF1B;refGene_tx=XM_023258117.1
C1 7404779 7404885
refGene_gn=KIF1B;refGene_tx=XM_019836118.2;refGene_gn=KIF1B;refGene_tx
=XM_019836119.2;refGene_gn=KIF1B;refGene_tx=XM_023258093.1;refGene_gn=
KIF1B;refGene_tx=XM_023258094.1;refGene_gn=KIF1B;refGene_tx=XM_0232580
95.1;refGene_gn=KIF1B;refGene_tx=XM_023258096.1;refGene_gn=KIF1B;refGe
ne_tx=XM_023258097.1;refGene_gn=KIF1B;refGene_tx=XM_023258098.1;refGen
e_gn=KIF1B;refGene_tx=XM_023258099.1;refGene_gn=KIF1B;refGene_tx=XM_02

3258100.1;refGene_gn=KIF1B;refGene_tx=XM_023258101.1;refGene_gn=KIF1B;
refGene_tx=XM_023258102.1;refGene_gn=KIF1B;refGene_tx=XM_023258103.1;r
efGene_gn=KIF1B;refGene_tx=XM_023258104.1;refGene_gn=KIF1B;refGene_tx=
XM_023258105.1;refGene_gn=KIF1B;refGene_tx=XM_023258106.1;refGene_gn=K
IF1B;refGene_tx=XM_023258107.1;refGene_gn=KIF1B;refGene_tx=XM_02325810
8.1;refGene_gn=KIF1B;refGene_tx=XM_023258109.1;refGene_gn=KIF1B;refGen
e_tx=XM_023258110.1;refGene_gn=KIF1B;refGene_tx=XM_023258111.1;refGene
_gn=KIF1B;refGene_tx=XM_023258112.1;refGene_gn=KIF1B;refGene_tx=XM_023
258113.1;refGene_gn=KIF1B;refGene_tx=XM_023258114.1;refGene_gn=KIF1B;r
efGene_tx=XM_023258116.1;refGene_gn=KIF1B;refGene_tx=XM_023258117.1
C1 7407780 7407807
refGene_gn=KIF1B;refGene_tx=XM_019836118.2;refGene_gn=KIF1B;refGene_tx
=XM_019836119.2;refGene_gn=KIF1B;refGene_tx=XM_023258093.1;refGene_gn=
KIF1B;refGene_tx=XM_023258094.1;refGene_gn=KIF1B;refGene_tx=XM_0232580
96.1;refGene_gn=KIF1B;refGene_tx=XM_023258097.1;refGene_gn=KIF1B;refGe
ne_tx=XM_023258098.1;refGene_gn=KIF1B;refGene_tx=XM_023258099.1;refGen
e_gn=KIF1B;refGene_tx=XM_023258100.1;refGene_gn=KIF1B;refGene_tx=XM_02
3258101.1;refGene_gn=KIF1B;refGene_tx=XM_023258102.1;refGene_gn=KIF1B;
refGene_tx=XM_023258103.1;refGene_gn=KIF1B;refGene_tx=XM_023258104.1;r
efGene_gn=KIF1B;refGene_tx=XM_023258106.1;refGene_gn=KIF1B;refGene_tx=
XM_023258108.1;refGene_gn=KIF1B;refGene_tx=XM_023258113.1;refGene_gn=K
IF1B;refGene_tx=XM_023258117.1
C1 7410687 7410802
refGene_gn=KIF1B;refGene_tx=XM_019836118.2;refGene_gn=KIF1B;refGene_tx
=XM_019836119.2;refGene_gn=KIF1B;refGene_tx=XM_023258093.1;refGene_gn=
KIF1B;refGene_tx=XM_023258094.1;refGene_gn=KIF1B;refGene_tx=XM_0232580
95.1;refGene_gn=KIF1B;refGene_tx=XM_023258096.1;refGene_gn=KIF1B;refGe
ne_tx=XM_023258097.1;refGene_gn=KIF1B;refGene_tx=XM_023258098.1;refGen
e_gn=KIF1B;refGene_tx=XM_023258099.1;refGene_gn=KIF1B;refGene_tx=XM_02
3258100.1;refGene_gn=KIF1B;refGene_tx=XM_023258101.1;refGene_gn=KIF1B;
refGene_tx=XM_023258102.1;refGene_gn=KIF1B;refGene_tx=XM_023258103.1;r
efGene_gn=KIF1B;refGene_tx=XM_023258104.1;refGene_gn=KIF1B;refGene_tx=
XM_023258105.1;refGene_gn=KIF1B;refGene_tx=XM_023258106.1;refGene_gn=K
IF1B;refGene_tx=XM_023258107.1;refGene_gn=KIF1B;refGene_tx=XM_02325810
8.1;refGene_gn=KIF1B;refGene_tx=XM_023258109.1;refGene_gn=KIF1B;refGen
e_tx=XM_023258110.1;refGene_gn=KIF1B;refGene_tx=XM_023258111.1;refGene
_gn=KIF1B;refGene_tx=XM_023258112.1;refGene_gn=KIF1B;refGene_tx=XM_023
258113.1;refGene_gn=KIF1B;refGene_tx=XM_023258114.1;refGene_gn=KIF1B;r
efGene_tx=XM_023258116.1;refGene_gn=KIF1B;refGene_tx=XM_023258117.1
C1 7411668 7411802
refGene_gn=KIF1B;refGene_tx=XM_019836118.2;refGene_gn=KIF1B;refGene_tx
=XM_019836119.2;refGene_gn=KIF1B;refGene_tx=XM_023258093.1;refGene_gn=
KIF1B;refGene_tx=XM_023258094.1;refGene_gn=KIF1B;refGene_tx=XM_0232580
95.1;refGene_gn=KIF1B;refGene_tx=XM_023258096.1;refGene_gn=KIF1B;refGe
ne_tx=XM_023258097.1;refGene_gn=KIF1B;refGene_tx=XM_023258098.1;refGen
e_gn=KIF1B;refGene_tx=XM_023258099.1;refGene_gn=KIF1B;refGene_tx=XM_02
3258100.1;refGene_gn=KIF1B;refGene_tx=XM_023258101.1;refGene_gn=KIF1B;
refGene_tx=XM_023258102.1;refGene_gn=KIF1B;refGene_tx=XM_023258103.1;r
efGene_gn=KIF1B;refGene_tx=XM_023258104.1;refGene_gn=KIF1B;refGene_tx=
XM_023258105.1;refGene_gn=KIF1B;refGene_tx=XM_023258106.1;refGene_gn=K

IF1B;refGene_tx=XM_023258107.1;refGene_gn=KIF1B;refGene_tx=XM_02325810
8.1;refGene_gn=KIF1B;refGene_tx=XM_023258109.1;refGene_gn=KIF1B;refGen
e_tx=XM_023258110.1;refGene_gn=KIF1B;refGene_tx=XM_023258111.1;refGene
_gn=KIF1B;refGene_tx=XM_023258112.1;refGene_gn=KIF1B;refGene_tx=XM_023
258113.1;refGene_gn=KIF1B;refGene_tx=XM_023258114.1;refGene_gn=KIF1B;r
efGene_tx=XM_023258116.1;refGene_gn=KIF1B;refGene_tx=XM_023258117.1
C1 7413013 7413075
refGene_gn=KIF1B;refGene_tx=XM_019836118.2;refGene_gn=KIF1B;refGene_tx
=XM_019836119.2;refGene_gn=KIF1B;refGene_tx=XM_023258093.1;refGene_gn=
KIF1B;refGene_tx=XM_023258094.1;refGene_gn=KIF1B;refGene_tx=XM_0232580
95.1;refGene_gn=KIF1B;refGene_tx=XM_023258096.1;refGene_gn=KIF1B;refGe
ne_tx=XM_023258097.1;refGene_gn=KIF1B;refGene_tx=XM_023258098.1;refGen
e_gn=KIF1B;refGene_tx=XM_023258099.1;refGene_gn=KIF1B;refGene_tx=XM_02
3258100.1;refGene_gn=KIF1B;refGene_tx=XM_023258101.1;refGene_gn=KIF1B;
refGene_tx=XM_023258102.1;refGene_gn=KIF1B;refGene_tx=XM_023258103.1;r
efGene_gn=KIF1B;refGene_tx=XM_023258104.1;refGene_gn=KIF1B;refGene_tx=
XM_023258105.1;refGene_gn=KIF1B;refGene_tx=XM_023258106.1;refGene_gn=K
IF1B;refGene_tx=XM_023258107.1;refGene_gn=KIF1B;refGene_tx=XM_02325810
8.1;refGene_gn=KIF1B;refGene_tx=XM_023258109.1;refGene_gn=KIF1B;refGen
e_tx=XM_023258110.1;refGene_gn=KIF1B;refGene_tx=XM_023258111.1;refGene
_gn=KIF1B;refGene_tx=XM_023258112.1;refGene_gn=KIF1B;refGene_tx=XM_023
258113.1;refGene_gn=KIF1B;refGene_tx=XM_023258114.1;refGene_gn=KIF1B;r
efGene_tx=XM_023258116.1;refGene_gn=KIF1B;refGene_tx=XM_023258117.1
C1 7414472 7414618
refGene_gn=KIF1B;refGene_tx=XM_019836118.2;refGene_gn=KIF1B;refGene_tx
=XM_019836119.2;refGene_gn=KIF1B;refGene_tx=XM_023258093.1;refGene_gn=
KIF1B;refGene_tx=XM_023258094.1;refGene_gn=KIF1B;refGene_tx=XM_0232580
95.1;refGene_gn=KIF1B;refGene_tx=XM_023258096.1;refGene_gn=KIF1B;refGe
ne_tx=XM_023258097.1;refGene_gn=KIF1B;refGene_tx=XM_023258098.1;refGen
e_gn=KIF1B;refGene_tx=XM_023258099.1;refGene_gn=KIF1B;refGene_tx=XM_02
3258100.1;refGene_gn=KIF1B;refGene_tx=XM_023258101.1;refGene_gn=KIF1B;
refGene_tx=XM_023258102.1;refGene_gn=KIF1B;refGene_tx=XM_023258103.1;r
efGene_gn=KIF1B;refGene_tx=XM_023258104.1;refGene_gn=KIF1B;refGene_tx=
XM_023258105.1;refGene_gn=KIF1B;refGene_tx=XM_023258106.1;refGene_gn=K
IF1B;refGene_tx=XM_023258107.1;refGene_gn=KIF1B;refGene_tx=XM_02325810
8.1;refGene_gn=KIF1B;refGene_tx=XM_023258109.1;refGene_gn=KIF1B;refGen
e_tx=XM_023258110.1;refGene_gn=KIF1B;refGene_tx=XM_023258111.1;refGene
_gn=KIF1B;refGene_tx=XM_023258112.1;refGene_gn=KIF1B;refGene_tx=XM_023
258113.1;refGene_gn=KIF1B;refGene_tx=XM_023258114.1;refGene_gn=KIF1B;r
efGene_tx=XM_023258116.1;refGene_gn=KIF1B;refGene_tx=XM_023258117.1
C1 7414804 7415044
refGene_gn=KIF1B;refGene_tx=XM_019836118.2;refGene_gn=KIF1B;refGene_tx
=XM_019836119.2;refGene_gn=KIF1B;refGene_tx=XM_023258093.1;refGene_gn=
KIF1B;refGene_tx=XM_023258094.1;refGene_gn=KIF1B;refGene_tx=XM_0232580
95.1;refGene_gn=KIF1B;refGene_tx=XM_023258096.1;refGene_gn=KIF1B;refGe
ne_tx=XM_023258097.1;refGene_gn=KIF1B;refGene_tx=XM_023258098.1;refGen
e_gn=KIF1B;refGene_tx=XM_023258099.1;refGene_gn=KIF1B;refGene_tx=XM_02
3258100.1;refGene_gn=KIF1B;refGene_tx=XM_023258101.1;refGene_gn=KIF1B;
refGene_tx=XM_023258102.1;refGene_gn=KIF1B;refGene_tx=XM_023258103.1;r
efGene_gn=KIF1B;refGene_tx=XM_023258104.1;refGene_gn=KIF1B;refGene_tx=

XM_023258105.1;refGene_gn=KIF1B;refGene_tx=XM_023258106.1;refGene_gn=KIF1B;refGene_tx=XM_023258107.1;refGene_gn=KIF1B;refGene_tx=XM_023258108.1;refGene_gn=KIF1B;refGene_tx=XM_023258109.1;refGene_gn=KIF1B;refGene_tx=XM_023258110.1;refGene_gn=KIF1B;refGene_tx=XM_023258111.1;refGene_gn=KIF1B;refGene_tx=XM_023258112.1;refGene_gn=KIF1B;refGene_tx=XM_023258113.1;refGene_gn=KIF1B;refGene_tx=XM_023258114.1;refGene_gn=KIF1B;refGene_tx=XM_023258116.1;refGene_gn=KIF1B;refGene_tx=XM_023258117.1
C1 7416154 7416226
refGene_gn=KIF1B;refGene_tx=XM_019836118.2;refGene_gn=KIF1B;refGene_tx=XM_019836119.2;refGene_gn=KIF1B;refGene_tx=XM_023258093.1;refGene_gn=KIF1B;refGene_tx=XM_023258094.1;refGene_gn=KIF1B;refGene_tx=XM_023258095.1;refGene_gn=KIF1B;refGene_tx=XM_023258096.1;refGene_gn=KIF1B;refGene_tx=XM_023258097.1;refGene_gn=KIF1B;refGene_tx=XM_023258098.1;refGene_gn=KIF1B;refGene_tx=XM_023258099.1;refGene_gn=KIF1B;refGene_tx=XM_023258100.1;refGene_gn=KIF1B;refGene_tx=XM_023258101.1;refGene_gn=KIF1B;refGene_tx=XM_023258102.1;refGene_gn=KIF1B;refGene_tx=XM_023258103.1;refGene_gn=KIF1B;refGene_tx=XM_023258104.1;refGene_gn=KIF1B;refGene_tx=XM_023258105.1;refGene_gn=KIF1B;refGene_tx=XM_023258106.1;refGene_gn=KIF1B;refGene_tx=XM_023258107.1;refGene_gn=KIF1B;refGene_tx=XM_023258108.1;refGene_gn=KIF1B;refGene_tx=XM_023258109.1;refGene_gn=KIF1B;refGene_tx=XM_023258110.1;refGene_gn=KIF1B;refGene_tx=XM_023258111.1;refGene_gn=KIF1B;refGene_tx=XM_023258112.1;refGene_gn=KIF1B;refGene_tx=XM_023258113.1;refGene_gn=KIF1B;refGene_tx=XM_023258114.1;refGene_gn=KIF1B;refGene_tx=XM_023258116.1;refGene_gn=KIF1B;refGene_tx=XM_023258117.1
C1 7418503 7418625
refGene_gn=KIF1B;refGene_tx=XM_019836118.2;refGene_gn=KIF1B;refGene_tx=XM_019836119.2;refGene_gn=KIF1B;refGene_tx=XM_023258093.1;refGene_gn=KIF1B;refGene_tx=XM_023258094.1;refGene_gn=KIF1B;refGene_tx=XM_023258095.1;refGene_gn=KIF1B;refGene_tx=XM_023258096.1;refGene_gn=KIF1B;refGene_tx=XM_023258097.1;refGene_gn=KIF1B;refGene_tx=XM_023258098.1;refGene_gn=KIF1B;refGene_tx=XM_023258099.1;refGene_gn=KIF1B;refGene_tx=XM_023258100.1;refGene_gn=KIF1B;refGene_tx=XM_023258101.1;refGene_gn=KIF1B;refGene_tx=XM_023258102.1;refGene_gn=KIF1B;refGene_tx=XM_023258103.1;refGene_gn=KIF1B;refGene_tx=XM_023258104.1;refGene_gn=KIF1B;refGene_tx=XM_023258105.1;refGene_gn=KIF1B;refGene_tx=XM_023258106.1;refGene_gn=KIF1B;refGene_tx=XM_023258107.1;refGene_gn=KIF1B;refGene_tx=XM_023258108.1;refGene_gn=KIF1B;refGene_tx=XM_023258109.1;refGene_gn=KIF1B;refGene_tx=XM_023258110.1;refGene_gn=KIF1B;refGene_tx=XM_023258111.1;refGene_gn=KIF1B;refGene_tx=XM_023258112.1;refGene_gn=KIF1B;refGene_tx=XM_023258113.1;refGene_gn=KIF1B;refGene_tx=XM_023258114.1;refGene_gn=KIF1B;refGene_tx=XM_023258116.1;refGene_gn=KIF1B;refGene_tx=XM_023258117.1
C1 7420668 7420818
refGene_gn=KIF1B;refGene_tx=XM_019836118.2;refGene_gn=KIF1B;refGene_tx=XM_019836119.2;refGene_gn=KIF1B;refGene_tx=XM_023258093.1;refGene_gn=KIF1B;refGene_tx=XM_023258094.1;refGene_gn=KIF1B;refGene_tx=XM_023258095.1;refGene_gn=KIF1B;refGene_tx=XM_023258096.1;refGene_gn=KIF1B;refGene_tx=XM_023258097.1;refGene_gn=KIF1B;refGene_tx=XM_023258098.1;refGene_gn=KIF1B;refGene_tx=XM_023258099.1;refGene_gn=KIF1B;refGene_tx=XM_023258100.1;refGene_gn=KIF1B;refGene_tx=XM_023258101.1;refGene_gn=KIF1B;refGene_tx=XM_023258102.1;refGene_gn=KIF1B;refGene_tx=XM_023258103.1;r

efGene_gn=KIF1B;refGene_tx=XM_023258104.1;refGene_gn=KIF1B;refGene_tx=
XM_023258105.1;refGene_gn=KIF1B;refGene_tx=XM_023258106.1;refGene_gn=K
IF1B;refGene_tx=XM_023258107.1;refGene_gn=KIF1B;refGene_tx=XM_02325810
8.1;refGene_gn=KIF1B;refGene_tx=XM_023258109.1;refGene_gn=KIF1B;refGen
e_tx=XM_023258110.1;refGene_gn=KIF1B;refGene_tx=XM_023258111.1;refGene
_gn=KIF1B;refGene_tx=XM_023258112.1;refGene_gn=KIF1B;refGene_tx=XM_023
258113.1;refGene_gn=KIF1B;refGene_tx=XM_023258114.1;refGene_gn=KIF1B;r
efGene_tx=XM_023258116.1;refGene_gn=KIF1B;refGene_tx=XM_023258117.1
C1 7421168 7421361
refGene_gn=KIF1B;refGene_tx=XM_019836118.2;refGene_gn=KIF1B;refGene_tx
=XM_019836119.2;refGene_gn=KIF1B;refGene_tx=XM_023258093.1;refGene_gn=
KIF1B;refGene_tx=XM_023258094.1;refGene_gn=KIF1B;refGene_tx=XM_0232580
95.1;refGene_gn=KIF1B;refGene_tx=XM_023258096.1;refGene_gn=KIF1B;refGe
ne_tx=XM_023258097.1;refGene_gn=KIF1B;refGene_tx=XM_023258098.1;refGen
e_gn=KIF1B;refGene_tx=XM_023258099.1;refGene_gn=KIF1B;refGene_tx=XM_02
3258100.1;refGene_gn=KIF1B;refGene_tx=XM_023258101.1;refGene_gn=KIF1B;
refGene_tx=XM_023258102.1;refGene_gn=KIF1B;refGene_tx=XM_023258103.1;r
efGene_gn=KIF1B;refGene_tx=XM_023258104.1;refGene_gn=KIF1B;refGene_tx=
XM_023258105.1;refGene_gn=KIF1B;refGene_tx=XM_023258106.1;refGene_gn=K
IF1B;refGene_tx=XM_023258107.1;refGene_gn=KIF1B;refGene_tx=XM_02325810
8.1;refGene_gn=KIF1B;refGene_tx=XM_023258109.1;refGene_gn=KIF1B;refGen
e_tx=XM_023258110.1;refGene_gn=KIF1B;refGene_tx=XM_023258111.1;refGene
_gn=KIF1B;refGene_tx=XM_023258112.1;refGene_gn=KIF1B;refGene_tx=XM_023
258113.1;refGene_gn=KIF1B;refGene_tx=XM_023258114.1;refGene_gn=KIF1B;r
efGene_tx=XM_023258116.1;refGene_gn=KIF1B;refGene_tx=XM_023258117.1
C1 7421569 7421688
refGene_gn=KIF1B;refGene_tx=XM_019836118.2;refGene_gn=KIF1B;refGene_tx
=XM_019836119.2;refGene_gn=KIF1B;refGene_tx=XM_023258093.1;refGene_gn=
KIF1B;refGene_tx=XM_023258094.1;refGene_gn=KIF1B;refGene_tx=XM_0232580
95.1;refGene_gn=KIF1B;refGene_tx=XM_023258096.1;refGene_gn=KIF1B;refGe
ne_tx=XM_023258097.1;refGene_gn=KIF1B;refGene_tx=XM_023258098.1;refGen
e_gn=KIF1B;refGene_tx=XM_023258099.1;refGene_gn=KIF1B;refGene_tx=XM_02
3258100.1;refGene_gn=KIF1B;refGene_tx=XM_023258101.1;refGene_gn=KIF1B;
refGene_tx=XM_023258102.1;refGene_gn=KIF1B;refGene_tx=XM_023258103.1;r
efGene_gn=KIF1B;refGene_tx=XM_023258104.1;refGene_gn=KIF1B;refGene_tx=
XM_023258105.1;refGene_gn=KIF1B;refGene_tx=XM_023258106.1;refGene_gn=K
IF1B;refGene_tx=XM_023258107.1;refGene_gn=KIF1B;refGene_tx=XM_02325810
8.1;refGene_gn=KIF1B;refGene_tx=XM_023258109.1;refGene_gn=KIF1B;refGen
e_tx=XM_023258110.1;refGene_gn=KIF1B;refGene_tx=XM_023258111.1;refGene
_gn=KIF1B;refGene_tx=XM_023258112.1;refGene_gn=KIF1B;refGene_tx=XM_023
258113.1;refGene_gn=KIF1B;refGene_tx=XM_023258114.1;refGene_gn=KIF1B;r
efGene_tx=XM_023258116.1;refGene_gn=KIF1B;refGene_tx=XM_023258117.1
C1 7422334 7422377
refGene_gn=KIF1B;refGene_tx=XM_019836118.2;refGene_gn=KIF1B;refGene_tx
=XM_019836119.2;refGene_gn=KIF1B;refGene_tx=XM_023258093.1;refGene_gn=
KIF1B;refGene_tx=XM_023258094.1;refGene_gn=KIF1B;refGene_tx=XM_0232580
95.1;refGene_gn=KIF1B;refGene_tx=XM_023258096.1;refGene_gn=KIF1B;refGe
ne_tx=XM_023258097.1;refGene_gn=KIF1B;refGene_tx=XM_023258098.1;refGen
e_gn=KIF1B;refGene_tx=XM_023258099.1;refGene_gn=KIF1B;refGene_tx=XM_02
3258100.1;refGene_gn=KIF1B;refGene_tx=XM_023258101.1;refGene_gn=KIF1B;

```
refGene_tx=XM_023258102.1;refGene_gn=KIF1B;refGene_tx=XM_023258103.1;r
efGene_gn=KIF1B;refGene_tx=XM_023258104.1;refGene_gn=KIF1B;refGene_tx=
XM_023258105.1;refGene_gn=KIF1B;refGene_tx=XM_023258106.1;refGene_gn=K
IF1B;refGene_tx=XM_023258107.1;refGene_gn=KIF1B;refGene_tx=XM_02325810
8.1;refGene_gn=KIF1B;refGene_tx=XM_023258109.1;refGene_gn=KIF1B;refGen
e_tx=XM_023258110.1;refGene_gn=KIF1B;refGene_tx=XM_023258111.1;refGene
_gn=KIF1B;refGene_tx=XM_023258112.1;refGene_gn=KIF1B;refGene_tx=XM_023
258113.1;refGene_gn=KIF1B;refGene_tx=XM_023258114.1;refGene_gn=KIF1B;r
efGene_tx=XM_023258116.1;refGene_gn=KIF1B;refGene_tx=XM_023258117.1
C1	7451328	7451336	refGene_gn=PGD;refGene_tx=XM_003989481.5
C1	7451930	7452006	refGene_gn=PGD;refGene_tx=XM_003989481.5
C1	7452555	7452735	refGene_gn=PGD;refGene_tx=XM_003989481.5
C1	7455183	7455249	refGene_gn=PGD;refGene_tx=XM_003989481.5
C1	7456895	7457014	refGene_gn=PGD;refGene_tx=XM_003989481.5
C1	7458970	7459040	refGene_gn=PGD;refGene_tx=XM_003989481.5
C1	7461271	7461406	refGene_gn=PGD;refGene_tx=XM_003989481.5
C1	7463322	7463512	refGene_gn=PGD;refGene_tx=XM_003989481.5
C1	7466403	7466534	refGene_gn=PGD;refGene_tx=XM_003989481.5
C1	7466736	7466870	refGene_gn=PGD;refGene_tx=XM_003989481.5
C1	7467233	7467333	refGene_gn=PGD;refGene_tx=XM_003989481.5
C1	7467488	7467611	refGene_gn=PGD;refGene_tx=XM_003989481.5
C1	7467702	7467822	refGene_gn=PGD;refGene_tx=XM_003989481.5
C1	7475472	7475523
refGene_gn=LOC101096495;refGene_tx=XM_003989482.5
C1	7478130	7478254
refGene_gn=LOC101096495;refGene_tx=XM_003989482.5
C1	7479121	7479155
refGene_gn=LOC101096495;refGene_tx=XM_003989482.5
C1	7482592	7482659
refGene_gn=LOC101096495;refGene_tx=XM_003989482.5
C1	7484631	7484781
refGene_gn=LOC101096495;refGene_tx=XM_003989482.5
C1	7498588	7498801
refGene_gn=DFFA;refGene_tx=XM_003989483.5;refGene_gn=DFFA;refGene_tx=X
M_006934272.4;refGene_gn=DFFA;refGene_tx=XM_019836144.2
C1	7499042	7499104
refGene_gn=DFFA;refGene_tx=XM_006934273.4;refGene_gn=DFFA;refGene_tx=X
M_019836144.2;refGene_gn=DFFA;refGene_tx=XM_023258118.1
C1	7499416	7499498
refGene_gn=DFFA;refGene_tx=XM_006934272.4;refGene_gn=DFFA;refGene_tx=X
M_006934273.4;refGene_gn=DFFA;refGene_tx=XM_023258118.1
C1	7499597	7499746
refGene_gn=DFFA;refGene_tx=XM_003989483.5;refGene_gn=DFFA;refGene_tx=X
M_006934272.4;refGene_gn=DFFA;refGene_tx=XM_006934273.4;refGene_gn=DFF
A;refGene_tx=XM_019836144.2;refGene_gn=DFFA;refGene_tx=XM_023258118.1
C1	7500186	7500376
refGene_gn=DFFA;refGene_tx=XM_003989483.5;refGene_gn=DFFA;refGene_tx=X
M_006934272.4;refGene_gn=DFFA;refGene_tx=XM_006934273.4;refGene_gn=DFF
A;refGene_tx=XM_019836144.2;refGene_gn=DFFA;refGene_tx=XM_023258118.1
C1	7502731	7502874
```

refGene_gn=DFFA;refGene_tx=XM_003989483.5;refGene_gn=DFFA;refGene_tx=XM_006934272.4;refGene_gn=DFFA;refGene_tx=XM_006934273.4;refGene_gn=DFFA;refGene_tx=XM_019836144.2;refGene_gn=DFFA;refGene_tx=XM_023258118.1
C1 7503333 7503495
refGene_gn=DFFA;refGene_tx=XM_003989483.5;refGene_gn=DFFA;refGene_tx=XM_006934272.4;refGene_gn=DFFA;refGene_tx=XM_006934273.4;refGene_gn=DFFA;refGene_tx=XM_019836144.2;refGene_gn=DFFA;refGene_tx=XM_023258118.1
C1 7505225 7505349
refGene_gn=DFFA;refGene_tx=XM_003989483.5;refGene_gn=DFFA;refGene_tx=XM_006934272.4;refGene_gn=DFFA;refGene_tx=XM_006934273.4;refGene_gn=DFFA;refGene_tx=XM_019836144.2;refGene_gn=DFFA;refGene_tx=XM_023258118.1
C1 7507247 7507283
refGene_gn=PEX14;refGene_tx=XM_003989484.4;refGene_gn=PEX14;refGene_tx=XM_019836145.2
C1 7531082 7531130
refGene_gn=PEX14;refGene_tx=XM_003989484.4;refGene_gn=PEX14;refGene_tx=XM_019836145.2
C1 7566056 7566141
refGene_gn=PEX14;refGene_tx=XM_003989484.4;refGene_gn=PEX14;refGene_tx=XM_019836145.2
C1 7625557 7625686
refGene_gn=PEX14;refGene_tx=XM_003989484.4;refGene_gn=PEX14;refGene_tx=XM_019836145.2
C1 7642636 7642722
refGene_gn=PEX14;refGene_tx=XM_003989484.4;refGene_gn=PEX14;refGene_tx=XM_019836145.2
C1 7645860 7645963 refGene_gn=PEX14;refGene_tx=XM_003989484.4
C1 7647136 7647234 refGene_gn=PEX14;refGene_tx=XM_003989484.4
C1 7651010 7651102
refGene_gn=PEX14;refGene_tx=XM_003989484.4;refGene_gn=PEX14;refGene_tx=XM_019836145.2
C1 7652910 7653355
refGene_gn=PEX14;refGene_tx=XM_003989484.4;refGene_gn=PEX14;refGene_tx=XM_019836145.2
C1 7662694 7663842
refGene_gn=CASZ1;refGene_tx=XM_023258119.1;refGene_gn=CASZ1;refGene_tx=XM_023258120.1;refGene_gn=CASZ1;refGene_tx=XM_023258121.1
C1 7665293 7665823 refGene_gn=CASZ1;refGene_tx=XM_023258122.1
C1 7666469 7666611
refGene_gn=CASZ1;refGene_tx=XM_023258119.1;refGene_gn=CASZ1;refGene_tx=XM_023258120.1;refGene_gn=CASZ1;refGene_tx=XM_023258121.1;refGene_gn=CASZ1;refGene_tx=XM_023258122.1
C1 7666732 7666884
refGene_gn=CASZ1;refGene_tx=XM_023258119.1;refGene_gn=CASZ1;refGene_tx=XM_023258120.1;refGene_gn=CASZ1;refGene_tx=XM_023258121.1;refGene_gn=CASZ1;refGene_tx=XM_023258122.1
C1 7668717 7668889
refGene_gn=CASZ1;refGene_tx=XM_023258119.1;refGene_gn=CASZ1;refGene_tx=XM_023258120.1;refGene_gn=CASZ1;refGene_tx=XM_023258121.1;refGene_gn=CASZ1;refGene_tx=XM_023258122.1

C1 7669663 7669862
refGene_gn=CASZ1;refGene_tx=XM_023258119.1;refGene_gn=CASZ1;refGene_tx=XM_023258120.1;refGene_gn=CASZ1;refGene_tx=XM_023258121.1;refGene_gn=CASZ1;refGene_tx=XM_023258122.1
C1 7671384 7671714
refGene_gn=CASZ1;refGene_tx=XM_023258119.1;refGene_gn=CASZ1;refGene_tx=XM_023258120.1;refGene_gn=CASZ1;refGene_tx=XM_023258121.1;refGene_gn=CASZ1;refGene_tx=XM_023258122.1
C1 7672666 7672789
refGene_gn=CASZ1;refGene_tx=XM_023258119.1;refGene_gn=CASZ1;refGene_tx=XM_023258120.1;refGene_gn=CASZ1;refGene_tx=XM_023258121.1;refGene_gn=CASZ1;refGene_tx=XM_023258122.1
C1 7672893 7673048
refGene_gn=CASZ1;refGene_tx=XM_023258119.1;refGene_gn=CASZ1;refGene_tx=XM_023258120.1;refGene_gn=CASZ1;refGene_tx=XM_023258121.1;refGene_gn=CASZ1;refGene_tx=XM_023258122.1
C1 7674325 7674389
refGene_gn=CASZ1;refGene_tx=XM_023258119.1;refGene_gn=CASZ1;refGene_tx=XM_023258120.1;refGene_gn=CASZ1;refGene_tx=XM_023258121.1;refGene_gn=CASZ1;refGene_tx=XM_023258122.1
C1 7674633 7674769
refGene_gn=CASZ1;refGene_tx=XM_023258119.1;refGene_gn=CASZ1;refGene_tx=XM_023258120.1;refGene_gn=CASZ1;refGene_tx=XM_023258121.1;refGene_gn=CASZ1;refGene_tx=XM_023258122.1
C1 7677121 7677969
refGene_gn=CASZ1;refGene_tx=XM_023258119.1;refGene_gn=CASZ1;refGene_tx=XM_023258120.1;refGene_gn=CASZ1;refGene_tx=XM_023258121.1;refGene_gn=CASZ1;refGene_tx=XM_023258122.1
C1 7678209 7678382
refGene_gn=CASZ1;refGene_tx=XM_023258119.1;refGene_gn=CASZ1;refGene_tx=XM_023258120.1;refGene_gn=CASZ1;refGene_tx=XM_023258121.1;refGene_gn=CASZ1;refGene_tx=XM_023258122.1
C1 7679316 7679481
refGene_gn=CASZ1;refGene_tx=XM_023258119.1;refGene_gn=CASZ1;refGene_tx=XM_023258120.1;refGene_gn=CASZ1;refGene_tx=XM_023258121.1;refGene_gn=CASZ1;refGene_tx=XM_023258122.1
C1 7680246 7680337
refGene_gn=CASZ1;refGene_tx=XM_023258119.1;refGene_gn=CASZ1;refGene_tx=XM_023258120.1;refGene_gn=CASZ1;refGene_tx=XM_023258121.1;refGene_gn=CASZ1;refGene_tx=XM_023258122.1
C1 7681973 7682042
refGene_gn=CASZ1;refGene_tx=XM_023258119.1;refGene_gn=CASZ1;refGene_tx=XM_023258120.1;refGene_gn=CASZ1;refGene_tx=XM_023258121.1;refGene_gn=CASZ1;refGene_tx=XM_023258122.1
C1 7683007 7683842
refGene_gn=CASZ1;refGene_tx=XM_023258119.1;refGene_gn=CASZ1;refGene_tx=XM_023258120.1;refGene_gn=CASZ1;refGene_tx=XM_023258121.1;refGene_gn=CASZ1;refGene_tx=XM_023258122.1
C1 7686929 7687406
refGene_gn=CASZ1;refGene_tx=XM_023258119.1;refGene_gn=CASZ1;refGene_tx

=XM_023258120.1;refGene_gn=CASZ1;refGene_tx=XM_023258121.1;refGene_gn=CASZ1;refGene_tx=XM_023258122.1
C1 7715114 7715153
refGene_gn=CASZ1;refGene_tx=XM_023258119.1;refGene_gn=CASZ1;refGene_tx=XM_023258120.1;refGene_gn=CASZ1;refGene_tx=XM_023258121.1;refGene_gn=CASZ1;refGene_tx=XM_023258122.1
C1 7715593 7715642
refGene_gn=CASZ1;refGene_tx=XM_023258119.1;refGene_gn=CASZ1;refGene_tx=XM_023258122.1
C1 7933780 7933915
refGene_gn=CC1H1orf127;refGene_tx=XM_023260054.1
C1 7934023 7934153
refGene_gn=CC1H1orf127;refGene_tx=XM_023260054.1
C1 7934600 7934669
refGene_gn=CC1H1orf127;refGene_tx=XM_023260054.1
C1 7934670 7934733
refGene_gn=CC1H1orf127;refGene_tx=XM_023260054.1
C1 7935360 7935582
refGene_gn=CC1H1orf127;refGene_tx=XM_023260054.1
C1 7938375 7938575
refGene_gn=CC1H1orf127;refGene_tx=XM_023260054.1
C1 7938773 7939168
refGene_gn=CC1H1orf127;refGene_tx=XM_023260054.1
C1 7940449 7940633
refGene_gn=CC1H1orf127;refGene_tx=XM_023260054.1
C1 7945082 7945667
refGene_gn=CC1H1orf127;refGene_tx=XM_023260054.1
C1 7946969 7947027
refGene_gn=CC1H1orf127;refGene_tx=XM_023260054.1
C1 7955125 7955227
refGene_gn=CC1H1orf127;refGene_tx=XM_023260054.1
C1 7957761 7957768
refGene_gn=CC1H1orf127;refGene_tx=XM_023260054.1
C1 7959442 7959535
refGene_gn=CC1H1orf127;refGene_tx=XM_023260054.1
C1 7963552 7963613
refGene_gn=CC1H1orf127;refGene_tx=XM_023260054.1
C1 7976955 7977193
refGene_gn=TARDBP;refGene_tx=XM_003989485.5;refGene_gn=TARDBP;refGene_tx=XM_023258123.1
C1 7979101 7979265
refGene_gn=TARDBP;refGene_tx=XM_003989485.5;refGene_gn=TARDBP;refGene_tx=XM_023258123.1
C1 7980413 7980554
refGene_gn=TARDBP;refGene_tx=XM_003989485.5;refGene_gn=TARDBP;refGene_tx=XM_023258123.1
C1 7981301 7981472
refGene_gn=TARDBP;refGene_tx=XM_003989485.5;refGene_gn=TARDBP;refGene_tx=XM_023258123.1
C1 7981850 7982381

refGene_gn=TARDBP;refGene_tx=XM_003989485.5;refGene_gn=TARDBP;refGene_tx=XM_023258123.1
C1 7982938 7982994 refGene_gn=TARDBP;refGene_tx=XM_023258123.1
C1 7986561 7987325
refGene_gn=MASP2;refGene_tx=XM_003989575.3;refGene_gn=MASP2;refGene_tx=XM_019836146.1
C1 7988557 7988664
refGene_gn=MASP2;refGene_tx=XM_003989575.3;refGene_gn=MASP2;refGene_tx=XM_019836146.1;refGene_gn=MASP2;refGene_tx=XM_019836147.1
C1 7989165 7989300
refGene_gn=MASP2;refGene_tx=XM_003989575.3;refGene_gn=MASP2;refGene_tx=XM_019836146.1;refGene_gn=MASP2;refGene_tx=XM_019836147.1
C1 7991018 7991097
refGene_gn=MASP2;refGene_tx=XM_003989575.3;refGene_gn=MASP2;refGene_tx=XM_019836146.1;refGene_gn=MASP2;refGene_tx=XM_019836147.1
C1 7992943 7993062
refGene_gn=MASP2;refGene_tx=XM_003989575.3;refGene_gn=MASP2;refGene_tx=XM_019836146.1;refGene_gn=MASP2;refGene_tx=XM_019836147.1
C1 7995958 7996106
refGene_gn=MASP2;refGene_tx=XM_003989575.3;refGene_gn=MASP2;refGene_tx=XM_019836146.1;refGene_gn=MASP2;refGene_tx=XM_019836147.1
C1 7996179 7996376
refGene_gn=MASP2;refGene_tx=XM_003989575.3;refGene_gn=MASP2;refGene_tx=XM_019836146.1;refGene_gn=MASP2;refGene_tx=XM_019836147.1
C1 7997638 7997652 refGene_gn=MASP2;refGene_tx=XM_019836149.1
C1 7998281 7998413
refGene_gn=MASP2;refGene_tx=XM_003989575.3;refGene_gn=MASP2;refGene_tx=XM_019836146.1;refGene_gn=MASP2;refGene_tx=XM_019836147.1;refGene_gn=MASP2;refGene_tx=XM_019836149.1
C1 7998812 7998894 refGene_gn=MASP2;refGene_tx=XM_019836146.1
C1 7999126 7999304
refGene_gn=MASP2;refGene_tx=XM_003989575.3;refGene_gn=MASP2;refGene_tx=XM_019836147.1;refGene_gn=MASP2;refGene_tx=XM_019836149.1
C1 7999498 7999730
refGene_gn=MASP2;refGene_tx=XM_003989575.3;refGene_gn=MASP2;refGene_tx=XM_019836147.1;refGene_gn=MASP2;refGene_tx=XM_019836149.1
C1 7999833 7999880
refGene_gn=MASP2;refGene_tx=XM_003989575.3;refGene_gn=MASP2;refGene_tx=XM_019836147.1;refGene_gn=MASP2;refGene_tx=XM_019836149.1
C1 8005209 8005230 refGene_gn=SRM;refGene_tx=XM_003989486.5
C1 8005336 8005459 refGene_gn=SRM;refGene_tx=XM_003989486.5
C1 8005542 8005688 refGene_gn=SRM;refGene_tx=XM_003989486.5
C1 8005780 8005864 refGene_gn=SRM;refGene_tx=XM_003989486.5
C1 8006409 8006563 refGene_gn=SRM;refGene_tx=XM_003989486.5
C1 8007387 8007480 refGene_gn=SRM;refGene_tx=XM_003989486.5
C1 8007997 8008118 refGene_gn=SRM;refGene_tx=XM_003989486.5
C1 8008718 8008885 refGene_gn=SRM;refGene_tx=XM_003989486.5
C1 8014055 8014086 refGene_gn=EXOSC10;refGene_tx=XM_003989487.4
C1 8014632 8014693
refGene_gn=EXOSC10;refGene_tx=XM_019836151.2;refGene_gn=EXOSC10;refGen

e_tx=XM_019836152.1
C1 8014741 8014818
refGene_gn=EXOSC10;refGene_tx=XM_003989487.4;refGene_gn=EXOSC10;refGene_tx=XM_019836151.2;refGene_gn=EXOSC10;refGene_tx=XM_019836152.1
C1 8015734 8015796
refGene_gn=EXOSC10;refGene_tx=XM_003989487.4;refGene_gn=EXOSC10;refGene_tx=XM_019836151.2;refGene_gn=EXOSC10;refGene_tx=XM_019836152.1
C1 8016562 8016734
refGene_gn=EXOSC10;refGene_tx=XM_003989487.4;refGene_gn=EXOSC10;refGene_tx=XM_019836151.2;refGene_gn=EXOSC10;refGene_tx=XM_019836152.1
C1 8017258 8017332
refGene_gn=EXOSC10;refGene_tx=XM_003989487.4;refGene_gn=EXOSC10;refGene_tx=XM_019836151.2;refGene_gn=EXOSC10;refGene_tx=XM_019836152.1
C1 8018455 8018543
refGene_gn=EXOSC10;refGene_tx=XM_003989487.4;refGene_gn=EXOSC10;refGene_tx=XM_019836151.2;refGene_gn=EXOSC10;refGene_tx=XM_019836152.1
C1 8020861 8020936
refGene_gn=EXOSC10;refGene_tx=XM_003989487.4;refGene_gn=EXOSC10;refGene_tx=XM_019836151.2;refGene_gn=EXOSC10;refGene_tx=XM_019836152.1
C1 8021256 8021352
refGene_gn=EXOSC10;refGene_tx=XM_003989487.4;refGene_gn=EXOSC10;refGene_tx=XM_019836151.2;refGene_gn=EXOSC10;refGene_tx=XM_019836152.1
C1 8023044 8023154
refGene_gn=EXOSC10;refGene_tx=XM_003989487.4;refGene_gn=EXOSC10;refGene_tx=XM_019836151.2;refGene_gn=EXOSC10;refGene_tx=XM_019836152.1
C1 8023240 8023319
refGene_gn=EXOSC10;refGene_tx=XM_003989487.4;refGene_gn=EXOSC10;refGene_tx=XM_019836151.2;refGene_gn=EXOSC10;refGene_tx=XM_019836152.1
C1 8023494 8023545
refGene_gn=EXOSC10;refGene_tx=XM_003989487.4;refGene_gn=EXOSC10;refGene_tx=XM_019836151.2;refGene_gn=EXOSC10;refGene_tx=XM_019836152.1
C1 8025312 8025424
refGene_gn=EXOSC10;refGene_tx=XM_003989487.4;refGene_gn=EXOSC10;refGene_tx=XM_019836151.2;refGene_gn=EXOSC10;refGene_tx=XM_019836152.1
C1 8026077 8026128
refGene_gn=EXOSC10;refGene_tx=XM_003989487.4;refGene_gn=EXOSC10;refGene_tx=XM_019836151.2;refGene_gn=EXOSC10;refGene_tx=XM_019836152.1
C1 8026310 8026459
refGene_gn=EXOSC10;refGene_tx=XM_003989487.4;refGene_gn=EXOSC10;refGene_tx=XM_019836151.2;refGene_gn=EXOSC10;refGene_tx=XM_019836152.1
C1 8026612 8026769
refGene_gn=EXOSC10;refGene_tx=XM_003989487.4;refGene_gn=EXOSC10;refGene_tx=XM_019836151.2;refGene_gn=EXOSC10;refGene_tx=XM_019836152.1
C1 8027790 8027981
refGene_gn=EXOSC10;refGene_tx=XM_003989487.4;refGene_gn=EXOSC10;refGene_tx=XM_019836151.2;refGene_gn=EXOSC10;refGene_tx=XM_019836152.1
C1 8028506 8028650
refGene_gn=EXOSC10;refGene_tx=XM_003989487.4;refGene_gn=EXOSC10;refGene_tx=XM_019836151.2;refGene_gn=EXOSC10;refGene_tx=XM_019836152.1
C1 8028851 8028962

refGene_gn=EXOSC10;refGene_tx=XM_003989487.4;refGene_gn=EXOSC10;refGene_tx=XM_019836151.2;refGene_gn=EXOSC10;refGene_tx=XM_019836152.1
C1 8029167 8029243
refGene_gn=EXOSC10;refGene_tx=XM_003989487.4;refGene_gn=EXOSC10;refGene_tx=XM_019836151.2;refGene_gn=EXOSC10;refGene_tx=XM_019836152.1
C1 8031065 8031180
refGene_gn=EXOSC10;refGene_tx=XM_003989487.4;refGene_gn=EXOSC10;refGene_tx=XM_019836151.2;refGene_gn=EXOSC10;refGene_tx=XM_019836152.1
C1 8031543 8031709
refGene_gn=EXOSC10;refGene_tx=XM_003989487.4;refGene_gn=EXOSC10;refGene_tx=XM_019836151.2;refGene_gn=EXOSC10;refGene_tx=XM_019836152.1
C1 8031998 8032103
refGene_gn=EXOSC10;refGene_tx=XM_003989487.4;refGene_gn=EXOSC10;refGene_tx=XM_019836151.2;refGene_gn=EXOSC10;refGene_tx=XM_019836152.1
C1 8033631 8033755
refGene_gn=EXOSC10;refGene_tx=XM_003989487.4;refGene_gn=EXOSC10;refGene_tx=XM_019836151.2;refGene_gn=EXOSC10;refGene_tx=XM_019836152.1
C1 8035932 8036069
refGene_gn=EXOSC10;refGene_tx=XM_003989487.4;refGene_gn=EXOSC10;refGene_tx=XM_019836151.2
C1 8038599 8038710
refGene_gn=EXOSC10;refGene_tx=XM_003989487.4;refGene_gn=EXOSC10;refGene_tx=XM_019836151.2
C1 8042474 8042490
refGene_gn=MTOR;refGene_tx=XM_003989489.3;refGene_gn=MTOR;refGene_tx=XM_019836150.2
C1 8043181 8043287
refGene_gn=MTOR;refGene_tx=XM_003989489.3;refGene_gn=MTOR;refGene_tx=XM_019836150.2
C1 8045559 8045640
refGene_gn=MTOR;refGene_tx=XM_003989489.3;refGene_gn=MTOR;refGene_tx=XM_019836150.2
C1 8045918 8045999
refGene_gn=MTOR;refGene_tx=XM_003989489.3;refGene_gn=MTOR;refGene_tx=XM_019836150.2
C1 8047993 8048059
refGene_gn=MTOR;refGene_tx=XM_003989489.3;refGene_gn=MTOR;refGene_tx=XM_019836150.2
C1 8049483 8049619
refGene_gn=MTOR;refGene_tx=XM_003989489.3;refGene_gn=MTOR;refGene_tx=XM_019836150.2
C1 8049999 8050074
refGene_gn=MTOR;refGene_tx=XM_003989489.3;refGene_gn=MTOR;refGene_tx=XM_019836150.2
C1 8050576 8050649
refGene_gn=MTOR;refGene_tx=XM_003989489.3;refGene_gn=MTOR;refGene_tx=XM_019836150.2
C1 8051772 8051855
refGene_gn=MTOR;refGene_tx=XM_003989489.3;refGene_gn=MTOR;refGene_tx=XM_019836150.2

C1 8053704 8053827
refGene_gn=MTOR;refGene_tx=XM_003989489.3;refGene_gn=MTOR;refGene_tx=XM_019836150.2
C1 8054462 8054610
refGene_gn=MTOR;refGene_tx=XM_003989489.3;refGene_gn=MTOR;refGene_tx=XM_019836150.2
C1 8055164 8055300
refGene_gn=MTOR;refGene_tx=XM_003989489.3;refGene_gn=MTOR;refGene_tx=XM_019836150.2
C1 8056828 8057003
refGene_gn=MTOR;refGene_tx=XM_003989489.3;refGene_gn=MTOR;refGene_tx=XM_019836150.2
C1 8057215 8057350
refGene_gn=MTOR;refGene_tx=XM_003989489.3;refGene_gn=MTOR;refGene_tx=XM_019836150.2
C1 8057784 8057967
refGene_gn=MTOR;refGene_tx=XM_003989489.3;refGene_gn=MTOR;refGene_tx=XM_019836150.2
C1 8058341 8058464
refGene_gn=MTOR;refGene_tx=XM_003989489.3;refGene_gn=MTOR;refGene_tx=XM_019836150.2
C1 8058753 8058852
refGene_gn=MTOR;refGene_tx=XM_003989489.3;refGene_gn=MTOR;refGene_tx=XM_019836150.2
C1 8059101 8059198
refGene_gn=MTOR;refGene_tx=XM_003989489.3;refGene_gn=MTOR;refGene_tx=XM_019836150.2
C1 8059850 8059951
refGene_gn=MTOR;refGene_tx=XM_003989489.3;refGene_gn=MTOR;refGene_tx=XM_019836150.2
C1 8060604 8060856
refGene_gn=MTOR;refGene_tx=XM_003989489.3;refGene_gn=MTOR;refGene_tx=XM_019836150.2
C1 8063074 8063192
refGene_gn=MTOR;refGene_tx=XM_003989489.3;refGene_gn=MTOR;refGene_tx=XM_019836150.2
C1 8064157 8064273
refGene_gn=MTOR;refGene_tx=XM_003989489.3;refGene_gn=MTOR;refGene_tx=XM_019836150.2
C1 8068348 8068480
refGene_gn=MTOR;refGene_tx=XM_003989489.3;refGene_gn=MTOR;refGene_tx=XM_019836150.2
C1 8068578 8068704
refGene_gn=MTOR;refGene_tx=XM_003989489.3;refGene_gn=MTOR;refGene_tx=XM_019836150.2
C1 8072045 8072153
refGene_gn=MTOR;refGene_tx=XM_003989489.3;refGene_gn=MTOR;refGene_tx=XM_019836150.2
C1 8072360 8072438
refGene_gn=MTOR;refGene_tx=XM_003989489.3;refGene_gn=MTOR;refGene_tx=X

```
M_019836150.2
C1	8073904	8074020	
refGene_gn=MTOR;refGene_tx=XM_003989489.3;refGene_gn=MTOR;refGene_tx=X
M_019836150.2
C1	8075853	8075954	
refGene_gn=MTOR;refGene_tx=XM_003989489.3;refGene_gn=MTOR;refGene_tx=X
M_019836150.2
C1	8082555	8082695	
refGene_gn=MTOR;refGene_tx=XM_003989489.3;refGene_gn=MTOR;refGene_tx=X
M_019836150.2
C1	8085660	8085736	
refGene_gn=MTOR;refGene_tx=XM_003989489.3;refGene_gn=MTOR;refGene_tx=X
M_019836150.2
C1	8103622	8103992	refGene_gn=ANGPTL7;refGene_tx=XM_003989490.4
C1	8106395	8106496	refGene_gn=ANGPTL7;refGene_tx=XM_003989490.4
C1	8107073	8107268	refGene_gn=ANGPTL7;refGene_tx=XM_003989490.4
C1	8107929	8108128	refGene_gn=ANGPTL7;refGene_tx=XM_003989490.4
C1	8108577	8108747	refGene_gn=ANGPTL7;refGene_tx=XM_003989490.4
C1	8116996	8117142	
refGene_gn=MTOR;refGene_tx=XM_003989489.3;refGene_gn=MTOR;refGene_tx=X
M_019836150.2
C1	8117248	8117411	
refGene_gn=MTOR;refGene_tx=XM_003989489.3;refGene_gn=MTOR;refGene_tx=X
M_019836150.2
C1	8120246	8120389	
refGene_gn=MTOR;refGene_tx=XM_003989489.3;refGene_gn=MTOR;refGene_tx=X
M_019836150.2
C1	8125839	8125986	
refGene_gn=MTOR;refGene_tx=XM_003989489.3;refGene_gn=MTOR;refGene_tx=X
M_019836150.2
C1	8126697	8126790	
refGene_gn=MTOR;refGene_tx=XM_003989489.3;refGene_gn=MTOR;refGene_tx=X
M_019836150.2
C1	8127912	8128075	
refGene_gn=MTOR;refGene_tx=XM_003989489.3;refGene_gn=MTOR;refGene_tx=X
M_019836150.2
C1	8128404	8128517	
refGene_gn=MTOR;refGene_tx=XM_003989489.3;refGene_gn=MTOR;refGene_tx=X
M_019836150.2
C1	8128996	8129164	
refGene_gn=MTOR;refGene_tx=XM_003989489.3;refGene_gn=MTOR;refGene_tx=X
M_019836150.2
C1	8131720	8131807	
refGene_gn=MTOR;refGene_tx=XM_003989489.3;refGene_gn=MTOR;refGene_tx=X
M_019836150.2
C1	8142442	8142693	
refGene_gn=MTOR;refGene_tx=XM_003989489.3;refGene_gn=MTOR;refGene_tx=X
M_019836150.2
C1	8144000	8144130	
refGene_gn=MTOR;refGene_tx=XM_003989489.3;refGene_gn=MTOR;refGene_tx=X
```

M_019836150.2
C1	8144371	8144506
refGene_gn=MTOR;refGene_tx=XM_003989489.3;refGene_gn=MTOR;refGene_tx=X
M_019836150.2
C1	8145187	8145280
refGene_gn=MTOR;refGene_tx=XM_003989489.3;refGene_gn=MTOR;refGene_tx=X
M_019836150.2
C1	8145396	8145486
refGene_gn=MTOR;refGene_tx=XM_003989489.3;refGene_gn=MTOR;refGene_tx=X
M_019836150.2
C1	8146096	8146219
refGene_gn=MTOR;refGene_tx=XM_003989489.3;refGene_gn=MTOR;refGene_tx=X
M_019836150.2
C1	8153461	8153667
refGene_gn=MTOR;refGene_tx=XM_003989489.3;refGene_gn=MTOR;refGene_tx=X
M_019836150.2
C1	8153971	8154187
refGene_gn=MTOR;refGene_tx=XM_003989489.3;refGene_gn=MTOR;refGene_tx=X
M_019836150.2
C1	8155266	8155511
refGene_gn=MTOR;refGene_tx=XM_003989489.3;refGene_gn=MTOR;refGene_tx=X
M_019836150.2
C1	8156224	8156353
refGene_gn=MTOR;refGene_tx=XM_003989489.3;refGene_gn=MTOR;refGene_tx=X
M_019836150.2
C1	8157023	8157210
refGene_gn=MTOR;refGene_tx=XM_003989489.3;refGene_gn=MTOR;refGene_tx=X
M_019836150.2
C1	8159966	8160075
refGene_gn=MTOR;refGene_tx=XM_003989489.3;refGene_gn=MTOR;refGene_tx=X
M_019836150.2
C1	8160157	8160433
refGene_gn=MTOR;refGene_tx=XM_003989489.3;refGene_gn=MTOR;refGene_tx=X
M_019836150.2
C1	8168059	8168194	refGene_gn=MTOR;refGene_tx=XM_003989489.3
C1	8169148	8169349	refGene_gn=MTOR;refGene_tx=XM_003989489.3
C1	8170842	8171075	refGene_gn=MTOR;refGene_tx=XM_003989489.3
C1	8171769	8171878	refGene_gn=MTOR;refGene_tx=XM_003989489.3
C1	8172653	8172815	refGene_gn=MTOR;refGene_tx=XM_003989489.3
C1	8179228	8179754
refGene_gn=LOC105259936;refGene_tx=XM_023258124.1
C1	8185148	8185348
refGene_gn=LOC105259936;refGene_tx=XM_023258124.1
C1	8185401	8185927
refGene_gn=LOC101099816;refGene_tx=XM_011284490.3;refGene_gn=LOC101099
816;refGene_tx=XM_019836153.2
C1	8192881	8193003
refGene_gn=LOC101099816;refGene_tx=XM_019836153.2
C1	8195265	8195753
refGene_gn=LOC101099816;refGene_tx=XM_011284490.3;refGene_gn=LOC101099

816;refGene_tx=XM_019836153.2
C1	8377477	8378573
refGene_gn=DISP3;refGene_tx=XM_011284494.3;refGene_gn=DISP3;refGene_tx=XM_019836154.2;refGene_gn=DISP3;refGene_tx=XM_019836155.2
C1	8379497	8379717
refGene_gn=DISP3;refGene_tx=XM_011284494.3;refGene_gn=DISP3;refGene_tx=XM_019836154.2;refGene_gn=DISP3;refGene_tx=XM_019836155.2
C1	8389581	8389718
refGene_gn=DISP3;refGene_tx=XM_011284494.3;refGene_gn=DISP3;refGene_tx=XM_019836154.2;refGene_gn=DISP3;refGene_tx=XM_019836155.2
C1	8390592	8390727
refGene_gn=DISP3;refGene_tx=XM_011284494.3;refGene_gn=DISP3;refGene_tx=XM_019836154.2
C1	8391031	8391192
refGene_gn=DISP3;refGene_tx=XM_011284494.3;refGene_gn=DISP3;refGene_tx=XM_019836154.2;refGene_gn=DISP3;refGene_tx=XM_019836155.2
C1	8392438	8392578
refGene_gn=DISP3;refGene_tx=XM_011284494.3;refGene_gn=DISP3;refGene_tx=XM_019836154.2;refGene_gn=DISP3;refGene_tx=XM_019836155.2
C1	8393532	8393681
refGene_gn=DISP3;refGene_tx=XM_011284494.3;refGene_gn=DISP3;refGene_tx=XM_019836154.2;refGene_gn=DISP3;refGene_tx=XM_019836155.2
C1	8393904	8394066
refGene_gn=DISP3;refGene_tx=XM_011284494.3;refGene_gn=DISP3;refGene_tx=XM_019836154.2;refGene_gn=DISP3;refGene_tx=XM_019836155.2
C1	8395099	8395261
refGene_gn=DISP3;refGene_tx=XM_011284494.3;refGene_gn=DISP3;refGene_tx=XM_019836154.2;refGene_gn=DISP3;refGene_tx=XM_019836155.2
C1	8397455	8397569
refGene_gn=DISP3;refGene_tx=XM_011284494.3;refGene_gn=DISP3;refGene_tx=XM_019836154.2;refGene_gn=DISP3;refGene_tx=XM_019836155.2
C1	8398340	8398477
refGene_gn=DISP3;refGene_tx=XM_011284494.3;refGene_gn=DISP3;refGene_tx=XM_019836154.2;refGene_gn=DISP3;refGene_tx=XM_019836155.2
C1	8399578	8399763
refGene_gn=DISP3;refGene_tx=XM_011284494.3;refGene_gn=DISP3;refGene_tx=XM_019836154.2;refGene_gn=DISP3;refGene_tx=XM_019836155.2
C1	8401558	8401689
refGene_gn=DISP3;refGene_tx=XM_011284494.3;refGene_gn=DISP3;refGene_tx=XM_019836154.2;refGene_gn=DISP3;refGene_tx=XM_019836155.2
C1	8401768	8401941
refGene_gn=DISP3;refGene_tx=XM_011284494.3;refGene_gn=DISP3;refGene_tx=XM_019836154.2;refGene_gn=DISP3;refGene_tx=XM_019836155.2
C1	8402890	8403017
refGene_gn=DISP3;refGene_tx=XM_011284494.3;refGene_gn=DISP3;refGene_tx=XM_019836154.2;refGene_gn=DISP3;refGene_tx=XM_019836155.2
C1	8403530	8403676
refGene_gn=DISP3;refGene_tx=XM_011284494.3;refGene_gn=DISP3;refGene_tx=XM_019836154.2;refGene_gn=DISP3;refGene_tx=XM_019836155.2
C1	8404928	8405088

refGene_gn=DISP3;refGene_tx=XM_011284494.3;refGene_gn=DISP3;refGene_tx=XM_019836154.2;refGene_gn=DISP3;refGene_tx=XM_019836155.2
C1 8405635 8405749
refGene_gn=DISP3;refGene_tx=XM_011284494.3;refGene_gn=DISP3;refGene_tx=XM_019836154.2;refGene_gn=DISP3;refGene_tx=XM_019836155.2
C1 8406210 8406377
refGene_gn=DISP3;refGene_tx=XM_011284494.3;refGene_gn=DISP3;refGene_tx=XM_019836154.2;refGene_gn=DISP3;refGene_tx=XM_019836155.2
C1 8407153 8407516
refGene_gn=DISP3;refGene_tx=XM_011284494.3;refGene_gn=DISP3;refGene_tx=XM_019836154.2;refGene_gn=DISP3;refGene_tx=XM_019836155.2
C1 8503266 8503401 refGene_gn=FBXO2;refGene_tx=XM_003989492.5
C1 8503642 8503778 refGene_gn=FBXO2;refGene_tx=XM_003989492.5
C1 8504271 8504367 refGene_gn=FBXO2;refGene_tx=XM_003989492.5
C1 8504448 8504578 refGene_gn=FBXO2;refGene_tx=XM_003989492.5
C1 8505006 8505060 refGene_gn=FBXO2;refGene_tx=XM_003989492.5
C1 8505062 8505383 refGene_gn=FBXO2;refGene_tx=XM_003989492.5
C1 8508569 8508591 refGene_gn=FBXO2;refGene_tx=XM_003989492.5
C1 8510149 8510414
refGene_gn=FBXO44;refGene_tx=XM_023258125.1;refGene_gn=FBXO44;refGene_tx=XM_023258126.1;refGene_gn=FBXO44;refGene_tx=XM_023258127.1
C1 8511913 8512040
refGene_gn=FBXO44;refGene_tx=XM_023258125.1;refGene_gn=FBXO44;refGene_tx=XM_023258126.1;refGene_gn=FBXO44;refGene_tx=XM_023258127.1
C1 8512195 8512291
refGene_gn=FBXO44;refGene_tx=XM_023258125.1;refGene_gn=FBXO44;refGene_tx=XM_023258126.1
C1 8512401 8512537
refGene_gn=FBXO44;refGene_tx=XM_023258125.1;refGene_gn=FBXO44;refGene_tx=XM_023258126.1;refGene_gn=FBXO44;refGene_tx=XM_023258127.1
C1 8513328 8513489
refGene_gn=FBXO44;refGene_tx=XM_023258125.1;refGene_gn=FBXO44;refGene_tx=XM_023258126.1;refGene_gn=FBXO44;refGene_tx=XM_023258127.1
C1 8517905 8518170 refGene_gn=FBXO6;refGene_tx=XM_011284602.2
C1 8520066 8520193 refGene_gn=FBXO6;refGene_tx=XM_011284602.2
C1 8520627 8520723 refGene_gn=FBXO6;refGene_tx=XM_011284602.2
C1 8521622 8521758 refGene_gn=FBXO6;refGene_tx=XM_011284602.2
C1 8522003 8522192 refGene_gn=FBXO6;refGene_tx=XM_011284602.2
C1 8522876 8522918
refGene_gn=MAD2L2;refGene_tx=XM_006934278.4;refGene_gn=MAD2L2;refGene_tx=XM_019836159.2
C1 8523159 8523252
refGene_gn=MAD2L2;refGene_tx=XM_006934278.4;refGene_gn=MAD2L2;refGene_tx=XM_019836159.2
C1 8523783 8523857
refGene_gn=MAD2L2;refGene_tx=XM_006934278.4;refGene_gn=MAD2L2;refGene_tx=XM_019836159.2
C1 8524185 8524280
refGene_gn=MAD2L2;refGene_tx=XM_006934278.4;refGene_gn=MAD2L2;refGene_tx=XM_019836159.2

C1	8525383	8525484
refGene_gn=MAD2L2;refGene_tx=XM_006934278.4;refGene_gn=MAD2L2;refGene_tx=XM_019836159.2
C1	8526043	8526115
refGene_gn=MAD2L2;refGene_tx=XM_006934278.4;refGene_gn=MAD2L2;refGene_tx=XM_019836159.2
C1	8527105	8527224
refGene_gn=MAD2L2;refGene_tx=XM_006934278.4;refGene_gn=MAD2L2;refGene_tx=XM_019836159.2
C1	8527321	8527361
refGene_gn=MAD2L2;refGene_tx=XM_006934278.4;refGene_gn=MAD2L2;refGene_tx=XM_019836159.2
C1	8547297	8547748	refGene_gn=DRAXIN;refGene_tx=XM_019836156.2
C1	8550039	8550230	refGene_gn=DRAXIN;refGene_tx=XM_019836156.2
C1	8550962	8551074	refGene_gn=DRAXIN;refGene_tx=XM_019836156.2
C1	8551452	8551542	refGene_gn=DRAXIN;refGene_tx=XM_019836156.2
C1	8552369	8552459	refGene_gn=DRAXIN;refGene_tx=XM_019836156.2
C1	8556210	8556323	refGene_gn=DRAXIN;refGene_tx=XM_019836156.2
C1	8573618	8573645
refGene_gn=AGTRAP;refGene_tx=XM_006934279.3;refGene_gn=AGTRAP;refGene_tx=XM_006934280.3
C1	8581545	8581580
refGene_gn=AGTRAP;refGene_tx=XM_006934279.3;refGene_gn=AGTRAP;refGene_tx=XM_006934280.3
C1	8583301	8583407
refGene_gn=AGTRAP;refGene_tx=XM_006934279.3;refGene_gn=AGTRAP;refGene_tx=XM_006934280.3
C1	8584301	8584497
refGene_gn=AGTRAP;refGene_tx=XM_006934279.3;refGene_gn=AGTRAP;refGene_tx=XM_006934280.3
C1	8585862	8585978
refGene_gn=AGTRAP;refGene_tx=XM_006934279.3;refGene_gn=AGTRAP;refGene_tx=XM_006934280.3
C1	8597896	8597978
refGene_gn=CC1H1orf167;refGene_tx=XM_023258128.1;refGene_gn=CC1H1orf167;refGene_tx=XM_023258129.1;refGene_gn=CC1H1orf167;refGene_tx=XM_023258130.1;refGene_gn=CC1H1orf167;refGene_tx=XM_023258131.1;refGene_gn=CC1H1orf167;refGene_tx=XM_023258132.1;refGene_gn=CC1H1orf167;refGene_tx=XM_023258133.1;refGene_gn=CC1H1orf167;refGene_tx=XM_023258134.1
C1	8598907	8600157
refGene_gn=CC1H1orf167;refGene_tx=XM_023258128.1;refGene_gn=CC1H1orf167;refGene_tx=XM_023258129.1;refGene_gn=CC1H1orf167;refGene_tx=XM_023258130.1;refGene_gn=CC1H1orf167;refGene_tx=XM_023258131.1;refGene_gn=CC1H1orf167;refGene_tx=XM_023258132.1;refGene_gn=CC1H1orf167;refGene_tx=XM_023258133.1;refGene_gn=CC1H1orf167;refGene_tx=XM_023258134.1
C1	8600354	8600398
refGene_gn=CC1H1orf167;refGene_tx=XM_023258128.1;refGene_gn=CC1H1orf167;refGene_tx=XM_023258129.1;refGene_gn=CC1H1orf167;refGene_tx=XM_023258130.1;refGene_gn=CC1H1orf167;refGene_tx=XM_023258131.1;refGene_gn=CC1H1orf167;refGene_tx=XM_023258132.1;refGene_gn=CC1H1orf167;refGene_tx=X

M_023258133.1;refGene_gn=CC1H1orf167;refGene_tx=XM_023258134.1
C1 8600573 8600763
refGene_gn=CC1H1orf167;refGene_tx=XM_023258128.1;refGene_gn=CC1H1orf16
7;refGene_tx=XM_023258129.1;refGene_gn=CC1H1orf167;refGene_tx=XM_02325
8130.1;refGene_gn=CC1H1orf167;refGene_tx=XM_023258131.1;refGene_gn=CC1
H1orf167;refGene_tx=XM_023258132.1;refGene_gn=CC1H1orf167;refGene_tx=X
M_023258133.1;refGene_gn=CC1H1orf167;refGene_tx=XM_023258134.1
C1 8601401 8601538
refGene_gn=CC1H1orf167;refGene_tx=XM_023258128.1;refGene_gn=CC1H1orf16
7;refGene_tx=XM_023258129.1;refGene_gn=CC1H1orf167;refGene_tx=XM_02325
8130.1;refGene_gn=CC1H1orf167;refGene_tx=XM_023258131.1;refGene_gn=CC1
H1orf167;refGene_tx=XM_023258132.1;refGene_gn=CC1H1orf167;refGene_tx=X
M_023258133.1;refGene_gn=CC1H1orf167;refGene_tx=XM_023258134.1
C1 8603067 8603162
refGene_gn=CC1H1orf167;refGene_tx=XM_023258128.1;refGene_gn=CC1H1orf16
7;refGene_tx=XM_023258129.1;refGene_gn=CC1H1orf167;refGene_tx=XM_02325
8130.1;refGene_gn=CC1H1orf167;refGene_tx=XM_023258131.1;refGene_gn=CC1
H1orf167;refGene_tx=XM_023258132.1;refGene_gn=CC1H1orf167;refGene_tx=X
M_023258133.1;refGene_gn=CC1H1orf167;refGene_tx=XM_023258134.1
C1 8603591 8603778
refGene_gn=CC1H1orf167;refGene_tx=XM_023258128.1;refGene_gn=CC1H1orf16
7;refGene_tx=XM_023258129.1;refGene_gn=CC1H1orf167;refGene_tx=XM_02325
8130.1;refGene_gn=CC1H1orf167;refGene_tx=XM_023258131.1;refGene_gn=CC1
H1orf167;refGene_tx=XM_023258132.1;refGene_gn=CC1H1orf167;refGene_tx=X
M_023258133.1;refGene_gn=CC1H1orf167;refGene_tx=XM_023258134.1
C1 8606685 8606861
refGene_gn=CC1H1orf167;refGene_tx=XM_023258128.1;refGene_gn=CC1H1orf16
7;refGene_tx=XM_023258129.1;refGene_gn=CC1H1orf167;refGene_tx=XM_02325
8130.1;refGene_gn=CC1H1orf167;refGene_tx=XM_023258131.1;refGene_gn=CC1
H1orf167;refGene_tx=XM_023258132.1;refGene_gn=CC1H1orf167;refGene_tx=X
M_023258133.1;refGene_gn=CC1H1orf167;refGene_tx=XM_023258134.1
C1 8607423 8607592
refGene_gn=CC1H1orf167;refGene_tx=XM_023258128.1;refGene_gn=CC1H1orf16
7;refGene_tx=XM_023258129.1;refGene_gn=CC1H1orf167;refGene_tx=XM_02325
8130.1;refGene_gn=CC1H1orf167;refGene_tx=XM_023258131.1;refGene_gn=CC1
H1orf167;refGene_tx=XM_023258132.1;refGene_gn=CC1H1orf167;refGene_tx=X
M_023258133.1;refGene_gn=CC1H1orf167;refGene_tx=XM_023258134.1
C1 8609201 8609388
refGene_gn=CC1H1orf167;refGene_tx=XM_023258128.1;refGene_gn=CC1H1orf16
7;refGene_tx=XM_023258129.1;refGene_gn=CC1H1orf167;refGene_tx=XM_02325
8130.1;refGene_gn=CC1H1orf167;refGene_tx=XM_023258131.1;refGene_gn=CC1
H1orf167;refGene_tx=XM_023258132.1;refGene_gn=CC1H1orf167;refGene_tx=X
M_023258133.1;refGene_gn=CC1H1orf167;refGene_tx=XM_023258134.1
C1 8609491 8609646
refGene_gn=CC1H1orf167;refGene_tx=XM_023258128.1;refGene_gn=CC1H1orf16
7;refGene_tx=XM_023258129.1;refGene_gn=CC1H1orf167;refGene_tx=XM_02325
8130.1;refGene_gn=CC1H1orf167;refGene_tx=XM_023258131.1;refGene_gn=CC1
H1orf167;refGene_tx=XM_023258132.1;refGene_gn=CC1H1orf167;refGene_tx=X
M_023258133.1;refGene_gn=CC1H1orf167;refGene_tx=XM_023258134.1
C1 8610372 8610581

refGene_gn=CC1H1orf167;refGene_tx=XM_023258128.1;refGene_gn=CC1H1orf16
7;refGene_tx=XM_023258129.1;refGene_gn=CC1H1orf167;refGene_tx=XM_02325
8130.1;refGene_gn=CC1H1orf167;refGene_tx=XM_023258131.1;refGene_gn=CC1
H1orf167;refGene_tx=XM_023258132.1;refGene_gn=CC1H1orf167;refGene_tx=X
M_023258133.1;refGene_gn=CC1H1orf167;refGene_tx=XM_023258134.1
C1 8611698 8611843
refGene_gn=CC1H1orf167;refGene_tx=XM_023258128.1;refGene_gn=CC1H1orf16
7;refGene_tx=XM_023258129.1;refGene_gn=CC1H1orf167;refGene_tx=XM_02325
8130.1;refGene_gn=CC1H1orf167;refGene_tx=XM_023258131.1;refGene_gn=CC1
H1orf167;refGene_tx=XM_023258132.1;refGene_gn=CC1H1orf167;refGene_tx=X
M_023258133.1;refGene_gn=CC1H1orf167;refGene_tx=XM_023258134.1
C1 8613243 8613465
refGene_gn=CC1H1orf167;refGene_tx=XM_023258128.1;refGene_gn=CC1H1orf16
7;refGene_tx=XM_023258129.1;refGene_gn=CC1H1orf167;refGene_tx=XM_02325
8130.1;refGene_gn=CC1H1orf167;refGene_tx=XM_023258131.1;refGene_gn=CC1
H1orf167;refGene_tx=XM_023258132.1;refGene_gn=CC1H1orf167;refGene_tx=X
M_023258133.1;refGene_gn=CC1H1orf167;refGene_tx=XM_023258134.1
C1 8614168 8614427
refGene_gn=CC1H1orf167;refGene_tx=XM_023258128.1;refGene_gn=CC1H1orf16
7;refGene_tx=XM_023258129.1;refGene_gn=CC1H1orf167;refGene_tx=XM_02325
8130.1;refGene_gn=CC1H1orf167;refGene_tx=XM_023258131.1;refGene_gn=CC1
H1orf167;refGene_tx=XM_023258132.1;refGene_gn=CC1H1orf167;refGene_tx=X
M_023258133.1;refGene_gn=CC1H1orf167;refGene_tx=XM_023258134.1
C1 8615529 8615589
refGene_gn=CC1H1orf167;refGene_tx=XM_023258134.1
C1 8617036 8617398
refGene_gn=CC1H1orf167;refGene_tx=XM_023258128.1;refGene_gn=CC1H1orf16
7;refGene_tx=XM_023258129.1;refGene_gn=CC1H1orf167;refGene_tx=XM_02325
8130.1;refGene_gn=CC1H1orf167;refGene_tx=XM_023258131.1;refGene_gn=CC1
H1orf167;refGene_tx=XM_023258132.1;refGene_gn=CC1H1orf167;refGene_tx=X
M_023258133.1
C1 8617594 8617730
refGene_gn=CC1H1orf167;refGene_tx=XM_023258128.1;refGene_gn=CC1H1orf16
7;refGene_tx=XM_023258129.1;refGene_gn=CC1H1orf167;refGene_tx=XM_02325
8130.1;refGene_gn=CC1H1orf167;refGene_tx=XM_023258131.1;refGene_gn=CC1
H1orf167;refGene_tx=XM_023258132.1;refGene_gn=CC1H1orf167;refGene_tx=X
M_023258133.1
C1 8618010 8618121
refGene_gn=CC1H1orf167;refGene_tx=XM_023258128.1;refGene_gn=CC1H1orf16
7;refGene_tx=XM_023258129.1;refGene_gn=CC1H1orf167;refGene_tx=XM_02325
8130.1;refGene_gn=CC1H1orf167;refGene_tx=XM_023258131.1;refGene_gn=CC1
H1orf167;refGene_tx=XM_023258132.1;refGene_gn=CC1H1orf167;refGene_tx=X
M_023258133.1
C1 8618691 8618904
refGene_gn=CC1H1orf167;refGene_tx=XM_023258128.1;refGene_gn=CC1H1orf16
7;refGene_tx=XM_023258129.1;refGene_gn=CC1H1orf167;refGene_tx=XM_02325
8130.1;refGene_gn=CC1H1orf167;refGene_tx=XM_023258131.1;refGene_gn=CC1
H1orf167;refGene_tx=XM_023258132.1;refGene_gn=CC1H1orf167;refGene_tx=X
M_023258133.1
C1 8620233 8620449

refGene_gn=MTHFR;refGene_tx=XM_003989579.5;refGene_gn=MTHFR;refGene_tx=XM_006934283.4;refGene_gn=MTHFR;refGene_tx=XM_006934284.4;refGene_gn=MTHFR;refGene_tx=XM_006934285.3;refGene_gn=MTHFR;refGene_tx=XM_006934286.4;refGene_gn=MTHFR;refGene_tx=XM_019836171.2;refGene_gn=MTHFR;refGene_tx=XM_019836172.2;refGene_gn=MTHFR;refGene_tx=XM_019836173.2;refGene_gn=MTHFR;refGene_tx=XM_019836174.2;refGene_gn=MTHFR;refGene_tx=XM_019836175.2
C1 8620747 8620867
refGene_gn=MTHFR;refGene_tx=XM_003989579.5;refGene_gn=MTHFR;refGene_tx=XM_006934283.4;refGene_gn=MTHFR;refGene_tx=XM_006934284.4;refGene_gn=MTHFR;refGene_tx=XM_006934285.3;refGene_gn=MTHFR;refGene_tx=XM_006934286.4;refGene_gn=MTHFR;refGene_tx=XM_019836171.2;refGene_gn=MTHFR;refGene_tx=XM_019836172.2;refGene_gn=MTHFR;refGene_tx=XM_019836173.2;refGene_gn=MTHFR;refGene_tx=XM_019836174.2;refGene_gn=MTHFR;refGene_tx=XM_019836175.2
C1 8621767 8621869
refGene_gn=MTHFR;refGene_tx=XM_003989579.5;refGene_gn=MTHFR;refGene_tx=XM_006934283.4;refGene_gn=MTHFR;refGene_tx=XM_006934284.4;refGene_gn=MTHFR;refGene_tx=XM_006934285.3;refGene_gn=MTHFR;refGene_tx=XM_006934286.4;refGene_gn=MTHFR;refGene_tx=XM_019836171.2;refGene_gn=MTHFR;refGene_tx=XM_019836172.2;refGene_gn=MTHFR;refGene_tx=XM_019836173.2;refGene_gn=MTHFR;refGene_tx=XM_019836174.2;refGene_gn=MTHFR;refGene_tx=XM_019836175.2
C1 8622706 8622889
refGene_gn=MTHFR;refGene_tx=XM_003989579.5;refGene_gn=MTHFR;refGene_tx=XM_006934283.4;refGene_gn=MTHFR;refGene_tx=XM_006934284.4;refGene_gn=MTHFR;refGene_tx=XM_006934285.3;refGene_gn=MTHFR;refGene_tx=XM_006934286.4;refGene_gn=MTHFR;refGene_tx=XM_019836171.2;refGene_gn=MTHFR;refGene_tx=XM_019836172.2;refGene_gn=MTHFR;refGene_tx=XM_019836173.2;refGene_gn=MTHFR;refGene_tx=XM_019836175.2
C1 8623139 8623320
refGene_gn=MTHFR;refGene_tx=XM_003989579.5;refGene_gn=MTHFR;refGene_tx=XM_006934283.4;refGene_gn=MTHFR;refGene_tx=XM_006934284.4;refGene_gn=MTHFR;refGene_tx=XM_006934285.3;refGene_gn=MTHFR;refGene_tx=XM_006934286.4;refGene_gn=MTHFR;refGene_tx=XM_019836171.2;refGene_gn=MTHFR;refGene_tx=XM_019836172.2;refGene_gn=MTHFR;refGene_tx=XM_019836173.2;refGene_gn=MTHFR;refGene_tx=XM_019836174.2;refGene_gn=MTHFR;refGene_tx=XM_019836175.2
C1 8623531 8623666
refGene_gn=MTHFR;refGene_tx=XM_003989579.5;refGene_gn=MTHFR;refGene_tx=XM_006934283.4;refGene_gn=MTHFR;refGene_tx=XM_006934284.4;refGene_gn=MTHFR;refGene_tx=XM_006934285.3;refGene_gn=MTHFR;refGene_tx=XM_006934286.4;refGene_gn=MTHFR;refGene_tx=XM_019836171.2;refGene_gn=MTHFR;refGene_tx=XM_019836172.2;refGene_gn=MTHFR;refGene_tx=XM_019836173.2;refGene_gn=MTHFR;refGene_tx=XM_019836174.2;refGene_gn=MTHFR;refGene_tx=XM_019836175.2
C1 8623946 8624197
refGene_gn=MTHFR;refGene_tx=XM_003989579.5;refGene_gn=MTHFR;refGene_tx=XM_006934283.4;refGene_gn=MTHFR;refGene_tx=XM_006934284.4;refGene_gn=MTHFR;refGene_tx=XM_006934285.3;refGene_gn=MTHFR;refGene_tx=XM_0069342

86.4;refGene_gn=MTHFR;refGene_tx=XM_019836171.2;refGene_gn=MTHFR;refGene_tx=XM_019836172.2;refGene_gn=MTHFR;refGene_tx=XM_019836173.2;refGene_gn=MTHFR;refGene_tx=XM_019836174.2
C1	8624907	8625101	refGene_gn=MTHFR;refGene_tx=XM_003989579.5;refGene_gn=MTHFR;refGene_tx=XM_006934283.4;refGene_gn=MTHFR;refGene_tx=XM_006934284.4;refGene_gn=MTHFR;refGene_tx=XM_006934285.3;refGene_gn=MTHFR;refGene_tx=XM_006934286.4;refGene_gn=MTHFR;refGene_tx=XM_019836171.2;refGene_gn=MTHFR;refGene_tx=XM_019836172.2;refGene_gn=MTHFR;refGene_tx=XM_019836173.2;refGene_gn=MTHFR;refGene_tx=XM_019836174.2
C1	8628062	8628173	refGene_gn=MTHFR;refGene_tx=XM_003989579.5;refGene_gn=MTHFR;refGene_tx=XM_006934283.4;refGene_gn=MTHFR;refGene_tx=XM_006934284.4;refGene_gn=MTHFR;refGene_tx=XM_006934285.3;refGene_gn=MTHFR;refGene_tx=XM_006934286.4;refGene_gn=MTHFR;refGene_tx=XM_019836171.2;refGene_gn=MTHFR;refGene_tx=XM_019836172.2;refGene_gn=MTHFR;refGene_tx=XM_019836173.2;refGene_gn=MTHFR;refGene_tx=XM_019836174.2;refGene_gn=MTHFR;refGene_tx=XM_019836175.2
C1	8629007	8629246	refGene_gn=MTHFR;refGene_tx=XM_003989579.5;refGene_gn=MTHFR;refGene_tx=XM_006934283.4;refGene_gn=MTHFR;refGene_tx=XM_006934284.4;refGene_gn=MTHFR;refGene_tx=XM_006934285.3;refGene_gn=MTHFR;refGene_tx=XM_006934286.4;refGene_gn=MTHFR;refGene_tx=XM_019836171.2;refGene_gn=MTHFR;refGene_tx=XM_019836172.2;refGene_gn=MTHFR;refGene_tx=XM_019836173.2;refGene_gn=MTHFR;refGene_tx=XM_019836174.2;refGene_gn=MTHFR;refGene_tx=XM_019836175.2
C1	8630410	8630731	refGene_gn=MTHFR;refGene_tx=XM_003989579.5;refGene_gn=MTHFR;refGene_tx=XM_006934283.4;refGene_gn=MTHFR;refGene_tx=XM_006934284.4;refGene_gn=MTHFR;refGene_tx=XM_006934285.3;refGene_gn=MTHFR;refGene_tx=XM_006934286.4;refGene_gn=MTHFR;refGene_tx=XM_019836171.2;refGene_gn=MTHFR;refGene_tx=XM_019836172.2;refGene_gn=MTHFR;refGene_tx=XM_019836173.2;refGene_gn=MTHFR;refGene_tx=XM_019836174.2;refGene_gn=MTHFR;refGene_tx=XM_019836175.2
C1	8630933	8631043	refGene_gn=MTHFR;refGene_tx=XM_003989579.5;refGene_gn=MTHFR;refGene_tx=XM_019836171.2;refGene_gn=MTHFR;refGene_tx=XM_019836172.2;refGene_gn=MTHFR;refGene_tx=XM_019836174.2;refGene_gn=MTHFR;refGene_tx=XM_019836175.2
C1	8633570	8633657	refGene_gn=CLCN6;refGene_tx=XM_003989496.4
C1	8634442	8634502	refGene_gn=CLCN6;refGene_tx=XM_003989496.4
C1	8640507	8640573	refGene_gn=CLCN6;refGene_tx=XM_003989496.4
C1	8640928	8640994	refGene_gn=CLCN6;refGene_tx=XM_003989496.4
C1	8643272	8643339	refGene_gn=CLCN6;refGene_tx=XM_003989496.4
C1	8645926	8646033	refGene_gn=CLCN6;refGene_tx=XM_003989496.4
C1	8646608	8646735	refGene_gn=CLCN6;refGene_tx=XM_003989496.4
C1	8647504	8647572	refGene_gn=CLCN6;refGene_tx=XM_003989496.4
C1	8648968	8649027	refGene_gn=CLCN6;refGene_tx=XM_003989496.4
C1	8649847	8649980	refGene_gn=CLCN6;refGene_tx=XM_003989496.4
C1	8650565	8650679	refGene_gn=CLCN6;refGene_tx=XM_003989496.4

C1 8650986 8651153 refGene_gn=CLCN6;refGene_tx=XM_003989496.4
C1 8651679 8651806 refGene_gn=CLCN6;refGene_tx=XM_003989496.4
C1 8654468 8654595 refGene_gn=CLCN6;refGene_tx=XM_003989496.4
C1 8654882 8655036 refGene_gn=CLCN6;refGene_tx=XM_003989496.4
C1 8655133 8655293 refGene_gn=CLCN6;refGene_tx=XM_003989496.4
C1 8655406 8655513 refGene_gn=CLCN6;refGene_tx=XM_003989496.4
C1 8656665 8656852 refGene_gn=CLCN6;refGene_tx=XM_003989496.4
C1 8657724 8657879 refGene_gn=CLCN6;refGene_tx=XM_003989496.4
C1 8658029 8658186 refGene_gn=CLCN6;refGene_tx=XM_003989496.4
C1 8659802 8659910 refGene_gn=CLCN6;refGene_tx=XM_003989496.4
C1 8660052 8660178 refGene_gn=CLCN6;refGene_tx=XM_003989496.4
C1 8662994 8663075 refGene_gn=CLCN6;refGene_tx=XM_003989496.4
C1 8669042 8669054 refGene_gn=NPPA;refGene_tx=XM_003989498.3
C1 8669562 8669889 refGene_gn=NPPA;refGene_tx=XM_003989498.3
C1 8669990 8670113 refGene_gn=NPPA;refGene_tx=XM_003989498.3
C1 8677263 8677280 refGene_gn=NPPB;refGene_tx=NM_001009244.1
C1 8677854 8678110 refGene_gn=NPPB;refGene_tx=NM_001009244.1
C1 8678346 8678472 refGene_gn=NPPB;refGene_tx=NM_001009244.1
C1 8726804 8726822
refGene_gn=KIAA2013;refGene_tx=XM_003989580.4;refGene_gn=KIAA2013;refG
ene_tx=XM_023258135.1
C1 8728361 8729215
refGene_gn=KIAA2013;refGene_tx=XM_003989580.4;refGene_gn=KIAA2013;refG
ene_tx=XM_023258135.1
C1 8732035 8733068
refGene_gn=KIAA2013;refGene_tx=XM_003989580.4;refGene_gn=KIAA2013;refG
ene_tx=XM_023258135.1
C1 8740255 8740331 refGene_gn=PLOD1;refGene_tx=XM_003989499.3
C1 8747970 8748062 refGene_gn=PLOD1;refGene_tx=XM_003989499.3
C1 8749165 8749299 refGene_gn=PLOD1;refGene_tx=XM_003989499.3
C1 8749681 8749845 refGene_gn=PLOD1;refGene_tx=XM_003989499.3
C1 8750895 8751008 refGene_gn=PLOD1;refGene_tx=XM_003989499.3
C1 8752649 8752713 refGene_gn=PLOD1;refGene_tx=XM_003989499.3
C1 8753939 8754037 refGene_gn=PLOD1;refGene_tx=XM_003989499.3
C1 8754889 8754991 refGene_gn=PLOD1;refGene_tx=XM_003989499.3
C1 8756284 8756416 refGene_gn=PLOD1;refGene_tx=XM_003989499.3
C1 8757865 8757987 refGene_gn=PLOD1;refGene_tx=XM_003989499.3
C1 8759482 8759587 refGene_gn=PLOD1;refGene_tx=XM_003989499.3
C1 8759970 8760096 refGene_gn=PLOD1;refGene_tx=XM_003989499.3
C1 8760372 8760514 refGene_gn=PLOD1;refGene_tx=XM_003989499.3
C1 8761354 8761468 refGene_gn=PLOD1;refGene_tx=XM_003989499.3
C1 8762166 8762232 refGene_gn=PLOD1;refGene_tx=XM_003989499.3
C1 8762828 8762933 refGene_gn=PLOD1;refGene_tx=XM_003989499.3
C1 8764281 8764428 refGene_gn=PLOD1;refGene_tx=XM_003989499.3
C1 8765205 8765331 refGene_gn=PLOD1;refGene_tx=XM_003989499.3
C1 8766876 8767032 refGene_gn=PLOD1;refGene_tx=XM_003989499.3
C1 8786521 8786696
refGene_gn=MFN2;refGene_tx=XM_003989500.5;refGene_gn=MFN2;refGene_tx=X
M_019836176.2;refGene_gn=MFN2;refGene_tx=XM_019836177.2;refGene_gn=MFN
2;refGene_tx=XM_019836178.2;refGene_gn=MFN2;refGene_tx=XM_019836179.2

C1 8790997 8791133
refGene_gn=MFN2;refGene_tx=XM_003989500.5;refGene_gn=MFN2;refGene_tx=XM_019836176.2;refGene_gn=MFN2;refGene_tx=XM_019836177.2;refGene_gn=MFN2;refGene_tx=XM_019836178.2;refGene_gn=MFN2;refGene_tx=XM_019836179.2
C1 8794570 8794733
refGene_gn=MFN2;refGene_tx=XM_003989500.5;refGene_gn=MFN2;refGene_tx=XM_019836176.2;refGene_gn=MFN2;refGene_tx=XM_019836177.2;refGene_gn=MFN2;refGene_tx=XM_019836178.2;refGene_gn=MFN2;refGene_tx=XM_019836179.2
C1 8795427 8795552
refGene_gn=MFN2;refGene_tx=XM_003989500.5;refGene_gn=MFN2;refGene_tx=XM_019836176.2;refGene_gn=MFN2;refGene_tx=XM_019836177.2;refGene_gn=MFN2;refGene_tx=XM_019836178.2;refGene_gn=MFN2;refGene_tx=XM_019836179.2
C1 8796699 8796808
refGene_gn=MFN2;refGene_tx=XM_003989500.5;refGene_gn=MFN2;refGene_tx=XM_019836176.2;refGene_gn=MFN2;refGene_tx=XM_019836177.2;refGene_gn=MFN2;refGene_tx=XM_019836178.2;refGene_gn=MFN2;refGene_tx=XM_019836179.2
C1 8796927 8797035
refGene_gn=MFN2;refGene_tx=XM_003989500.5;refGene_gn=MFN2;refGene_tx=XM_019836176.2;refGene_gn=MFN2;refGene_tx=XM_019836177.2;refGene_gn=MFN2;refGene_tx=XM_019836178.2;refGene_gn=MFN2;refGene_tx=XM_019836179.2
C1 8798079 8798233
refGene_gn=MFN2;refGene_tx=XM_003989500.5;refGene_gn=MFN2;refGene_tx=XM_019836176.2;refGene_gn=MFN2;refGene_tx=XM_019836177.2;refGene_gn=MFN2;refGene_tx=XM_019836178.2;refGene_gn=MFN2;refGene_tx=XM_019836179.2
C1 8798443 8798511
refGene_gn=MFN2;refGene_tx=XM_003989500.5;refGene_gn=MFN2;refGene_tx=XM_019836176.2;refGene_gn=MFN2;refGene_tx=XM_019836177.2;refGene_gn=MFN2;refGene_tx=XM_019836178.2;refGene_gn=MFN2;refGene_tx=XM_019836179.2
C1 8798657 8798779
refGene_gn=MFN2;refGene_tx=XM_003989500.5;refGene_gn=MFN2;refGene_tx=XM_019836176.2;refGene_gn=MFN2;refGene_tx=XM_019836177.2;refGene_gn=MFN2;refGene_tx=XM_019836178.2;refGene_gn=MFN2;refGene_tx=XM_019836179.2
C1 8800753 8800880
refGene_gn=MFN2;refGene_tx=XM_003989500.5;refGene_gn=MFN2;refGene_tx=XM_019836176.2;refGene_gn=MFN2;refGene_tx=XM_019836177.2;refGene_gn=MFN2;refGene_tx=XM_019836178.2;refGene_gn=MFN2;refGene_tx=XM_019836179.2
C1 8801267 8801372
refGene_gn=MFN2;refGene_tx=XM_003989500.5;refGene_gn=MFN2;refGene_tx=XM_019836176.2;refGene_gn=MFN2;refGene_tx=XM_019836177.2;refGene_gn=MFN2;refGene_tx=XM_019836178.2;refGene_gn=MFN2;refGene_tx=XM_019836179.2
C1 8801642 8801745
refGene_gn=MFN2;refGene_tx=XM_003989500.5;refGene_gn=MFN2;refGene_tx=XM_019836176.2;refGene_gn=MFN2;refGene_tx=XM_019836177.2;refGene_gn=MFN2;refGene_tx=XM_019836178.2;refGene_gn=MFN2;refGene_tx=XM_019836179.2
C1 8802196 8802453
refGene_gn=MFN2;refGene_tx=XM_003989500.5;refGene_gn=MFN2;refGene_tx=XM_019836176.2;refGene_gn=MFN2;refGene_tx=XM_019836177.2;refGene_gn=MFN2;refGene_tx=XM_019836178.2;refGene_gn=MFN2;refGene_tx=XM_019836179.2
C1 8803266 8803422
refGene_gn=MFN2;refGene_tx=XM_003989500.5;refGene_gn=MFN2;refGene_tx=X

M_019836176.2;refGene_gn=MFN2;refGene_tx=XM_019836177.2;refGene_gn=MFN2;refGene_tx=XM_019836178.2;refGene_gn=MFN2;refGene_tx=XM_019836179.2
C1 8804856 8805053 refGene_gn=MFN2;refGene_tx=XM_003989500.5;refGene_gn=MFN2;refGene_tx=XM_019836176.2;refGene_gn=MFN2;refGene_tx=XM_019836177.2;refGene_gn=MFN2;refGene_tx=XM_019836178.2;refGene_gn=MFN2;refGene_tx=XM_019836179.2
C1 8805853 8805988 refGene_gn=MFN2;refGene_tx=XM_003989500.5;refGene_gn=MFN2;refGene_tx=XM_019836176.2;refGene_gn=MFN2;refGene_tx=XM_019836177.2;refGene_gn=MFN2;refGene_tx=XM_019836178.2;refGene_gn=MFN2;refGene_tx=XM_019836179.2
C1 8807421 8807491 refGene_gn=MFN2;refGene_tx=XM_003989500.5;refGene_gn=MFN2;refGene_tx=XM_019836176.2;refGene_gn=MFN2;refGene_tx=XM_019836177.2;refGene_gn=MFN2;refGene_tx=XM_019836178.2;refGene_gn=MFN2;refGene_tx=XM_019836179.2
C1 8819674 8819803 refGene_gn=MIIP;refGene_tx=XM_006934292.3;refGene_gn=MIIP;refGene_tx=XM_019836181.2;refGene_gn=MIIP;refGene_tx=XM_019836182.2
C1 8820048 8820384 refGene_gn=MIIP;refGene_tx=XM_006934292.3;refGene_gn=MIIP;refGene_tx=XM_019836181.2
C1 8820796 8820881 refGene_gn=MIIP;refGene_tx=XM_006934292.3;refGene_gn=MIIP;refGene_tx=XM_019836181.2;refGene_gn=MIIP;refGene_tx=XM_019836182.2
C1 8824406 8824515 refGene_gn=MIIP;refGene_tx=XM_006934292.3;refGene_gn=MIIP;refGene_tx=XM_019836181.2;refGene_gn=MIIP;refGene_tx=XM_019836182.2
C1 8824596 8824649 refGene_gn=MIIP;refGene_tx=XM_006934292.3;refGene_gn=MIIP;refGene_tx=XM_019836181.2;refGene_gn=MIIP;refGene_tx=XM_019836182.2
C1 8825341 8825471 refGene_gn=MIIP;refGene_tx=XM_006934292.3;refGene_gn=MIIP;refGene_tx=XM_019836181.2;refGene_gn=MIIP;refGene_tx=XM_019836182.2
C1 8825655 8825752 refGene_gn=MIIP;refGene_tx=XM_006934292.3;refGene_gn=MIIP;refGene_tx=XM_019836181.2;refGene_gn=MIIP;refGene_tx=XM_019836182.2
C1 8826749 8826875 refGene_gn=MIIP;refGene_tx=XM_006934292.3;refGene_gn=MIIP;refGene_tx=XM_019836181.2;refGene_gn=MIIP;refGene_tx=XM_019836182.2
C1 8827130 8827208 refGene_gn=MIIP;refGene_tx=XM_006934292.3;refGene_gn=MIIP;refGene_tx=XM_019836181.2;refGene_gn=MIIP;refGene_tx=XM_019836182.2
C1 8861843 8861906 refGene_gn=TNFRSF8;refGene_tx=XM_023258136.1
C1 8875676 8875737 refGene_gn=TNFRSF8;refGene_tx=XM_023258136.1
C1 8885730 8885835 refGene_gn=TNFRSF8;refGene_tx=XM_023258136.1
C1 8894022 8894175 refGene_gn=TNFRSF8;refGene_tx=XM_023258136.1
C1 8897737 8897831 refGene_gn=TNFRSF8;refGene_tx=XM_023258136.1
C1 8898192 8898356 refGene_gn=TNFRSF8;refGene_tx=XM_023258136.1
C1 8900579 8900696 refGene_gn=TNFRSF8;refGene_tx=XM_023258136.1
C1 8903240 8903393 refGene_gn=TNFRSF8;refGene_tx=XM_023258136.1
C1 8910866 8910963 refGene_gn=TNFRSF8;refGene_tx=XM_023258136.1

C1	8911155	8911301	refGene_gn=TNFRSF8;refGene_tx=XM_023258136.1
C1	8913414	8913516	refGene_gn=TNFRSF8;refGene_tx=XM_023258136.1
C1	8913633	8913687	refGene_gn=TNFRSF8;refGene_tx=XM_023258136.1
C1	8920486	8920512	refGene_gn=TNFRSF8;refGene_tx=XM_023258136.1
C1	8923056	8923261	refGene_gn=TNFRSF8;refGene_tx=XM_023258136.1
C1	8926425	8926673	refGene_gn=TNFRSF8;refGene_tx=XM_023258136.1
C1	8949583	8949661	refGene_gn=TNFRSF1B;refGene_tx=XM_023258137.1
C1	8965532	8965632	refGene_gn=TNFRSF1B;refGene_tx=XM_023258137.1
C1	8967029	8967158	refGene_gn=TNFRSF1B;refGene_tx=XM_023258137.1
C1	8967828	8967978	refGene_gn=TNFRSF1B;refGene_tx=XM_023258137.1
C1	8968434	8968528	refGene_gn=TNFRSF1B;refGene_tx=XM_023258137.1
C1	8968931	8969170	refGene_gn=TNFRSF1B;refGene_tx=XM_023258137.1
C1	8969706	8969784	refGene_gn=TNFRSF1B;refGene_tx=XM_023258137.1
C1	8970738	8970773	refGene_gn=TNFRSF1B;refGene_tx=XM_023258137.1
C1	8976826	8977034	refGene_gn=TNFRSF1B;refGene_tx=XM_023258137.1
C1	8980169	8980450	refGene_gn=TNFRSF1B;refGene_tx=XM_023258137.1
C1	8992494	8992591
refGene_gn=VPS13D;refGene_tx=XM_023258138.1;refGene_gn=VPS13D;refGene_
tx=XM_023258139.1;refGene_gn=VPS13D;refGene_tx=XM_023258140.1;refGene_
gn=VPS13D;refGene_tx=XM_023258141.1;refGene_gn=VPS13D;refGene_tx=XM_02
3258142.1
C1	8998637	8998715
refGene_gn=VPS13D;refGene_tx=XM_023258138.1;refGene_gn=VPS13D;refGene_
tx=XM_023258139.1;refGene_gn=VPS13D;refGene_tx=XM_023258140.1;refGene_
gn=VPS13D;refGene_tx=XM_023258141.1;refGene_gn=VPS13D;refGene_tx=XM_02
3258142.1
C1	9001787	9001978
refGene_gn=VPS13D;refGene_tx=XM_023258138.1;refGene_gn=VPS13D;refGene_
tx=XM_023258139.1;refGene_gn=VPS13D;refGene_tx=XM_023258140.1;refGene_
gn=VPS13D;refGene_tx=XM_023258141.1;refGene_gn=VPS13D;refGene_tx=XM_02
3258142.1
C1	9002076	9002157
refGene_gn=VPS13D;refGene_tx=XM_023258138.1;refGene_gn=VPS13D;refGene_
tx=XM_023258139.1;refGene_gn=VPS13D;refGene_tx=XM_023258140.1;refGene_
gn=VPS13D;refGene_tx=XM_023258141.1;refGene_gn=VPS13D;refGene_tx=XM_02
3258142.1
C1	9005363	9005480
refGene_gn=VPS13D;refGene_tx=XM_023258138.1;refGene_gn=VPS13D;refGene_
tx=XM_023258139.1;refGene_gn=VPS13D;refGene_tx=XM_023258140.1;refGene_
gn=VPS13D;refGene_tx=XM_023258141.1;refGene_gn=VPS13D;refGene_tx=XM_02
3258142.1
C1	9007927	9008086
refGene_gn=VPS13D;refGene_tx=XM_019836189.2;refGene_gn=VPS13D;refGene_
tx=XM_023258138.1;refGene_gn=VPS13D;refGene_tx=XM_023258139.1;refGene_
gn=VPS13D;refGene_tx=XM_023258140.1;refGene_gn=VPS13D;refGene_tx=XM_02
3258141.1;refGene_gn=VPS13D;refGene_tx=XM_023258142.1
C1	9009761	9009932
refGene_gn=VPS13D;refGene_tx=XM_019836189.2;refGene_gn=VPS13D;refGene_
tx=XM_023258138.1;refGene_gn=VPS13D;refGene_tx=XM_023258139.1;refGene_
gn=VPS13D;refGene_tx=XM_023258140.1;refGene_gn=VPS13D;refGene_tx=XM_02

3258141.1;refGene_gn=VPS13D;refGene_tx=XM_023258142.1
C1 9010366 9010467
refGene_gn=VPS13D;refGene_tx=XM_019836189.2;refGene_gn=VPS13D;refGene_tx=XM_023258138.1;refGene_gn=VPS13D;refGene_tx=XM_023258139.1;refGene_gn=VPS13D;refGene_tx=XM_023258140.1;refGene_gn=VPS13D;refGene_tx=XM_023258141.1;refGene_gn=VPS13D;refGene_tx=XM_023258142.1
C1 9011513 9011682
refGene_gn=VPS13D;refGene_tx=XM_019836189.2;refGene_gn=VPS13D;refGene_tx=XM_023258138.1;refGene_gn=VPS13D;refGene_tx=XM_023258139.1;refGene_gn=VPS13D;refGene_tx=XM_023258140.1;refGene_gn=VPS13D;refGene_tx=XM_023258141.1;refGene_gn=VPS13D;refGene_tx=XM_023258142.1
C1 9014543 9014645
refGene_gn=VPS13D;refGene_tx=XM_019836189.2;refGene_gn=VPS13D;refGene_tx=XM_023258138.1;refGene_gn=VPS13D;refGene_tx=XM_023258139.1;refGene_gn=VPS13D;refGene_tx=XM_023258140.1;refGene_gn=VPS13D;refGene_tx=XM_023258141.1;refGene_gn=VPS13D;refGene_tx=XM_023258142.1
C1 9014800 9015002
refGene_gn=VPS13D;refGene_tx=XM_019836189.2;refGene_gn=VPS13D;refGene_tx=XM_023258138.1;refGene_gn=VPS13D;refGene_tx=XM_023258139.1;refGene_gn=VPS13D;refGene_tx=XM_023258140.1;refGene_gn=VPS13D;refGene_tx=XM_023258141.1;refGene_gn=VPS13D;refGene_tx=XM_023258142.1
C1 9015984 9016164
refGene_gn=VPS13D;refGene_tx=XM_019836189.2;refGene_gn=VPS13D;refGene_tx=XM_023258138.1;refGene_gn=VPS13D;refGene_tx=XM_023258139.1;refGene_gn=VPS13D;refGene_tx=XM_023258140.1;refGene_gn=VPS13D;refGene_tx=XM_023258141.1;refGene_gn=VPS13D;refGene_tx=XM_023258142.1
C1 9017252 9017383
refGene_gn=VPS13D;refGene_tx=XM_019836189.2;refGene_gn=VPS13D;refGene_tx=XM_023258138.1;refGene_gn=VPS13D;refGene_tx=XM_023258139.1;refGene_gn=VPS13D;refGene_tx=XM_023258140.1;refGene_gn=VPS13D;refGene_tx=XM_023258141.1;refGene_gn=VPS13D;refGene_tx=XM_023258142.1
C1 9018218 9018294
refGene_gn=VPS13D;refGene_tx=XM_019836189.2;refGene_gn=VPS13D;refGene_tx=XM_023258138.1;refGene_gn=VPS13D;refGene_tx=XM_023258139.1;refGene_gn=VPS13D;refGene_tx=XM_023258140.1;refGene_gn=VPS13D;refGene_tx=XM_023258141.1;refGene_gn=VPS13D;refGene_tx=XM_023258142.1
C1 9018706 9018877
refGene_gn=VPS13D;refGene_tx=XM_019836189.2;refGene_gn=VPS13D;refGene_tx=XM_023258138.1;refGene_gn=VPS13D;refGene_tx=XM_023258139.1;refGene_gn=VPS13D;refGene_tx=XM_023258140.1;refGene_gn=VPS13D;refGene_tx=XM_023258141.1;refGene_gn=VPS13D;refGene_tx=XM_023258142.1
C1 9019882 9020013
refGene_gn=VPS13D;refGene_tx=XM_019836189.2;refGene_gn=VPS13D;refGene_tx=XM_019836191.2;refGene_gn=VPS13D;refGene_tx=XM_023258138.1;refGene_gn=VPS13D;refGene_tx=XM_023258139.1;refGene_gn=VPS13D;refGene_tx=XM_023258140.1;refGene_gn=VPS13D;refGene_tx=XM_023258141.1;refGene_gn=VPS13D;refGene_tx=XM_023258142.1
C1 9021734 9021867
refGene_gn=VPS13D;refGene_tx=XM_019836189.2;refGene_gn=VPS13D;refGene_tx=XM_019836191.2;refGene_gn=VPS13D;refGene_tx=XM_023258138.1;refGene_

gn=VPS13D;refGene_tx=XM_023258139.1;refGene_gn=VPS13D;refGene_tx=XM_023258140.1;refGene_gn=VPS13D;refGene_tx=XM_023258141.1;refGene_gn=VPS13D;refGene_tx=XM_023258142.1
C1 9023903 9026117
refGene_gn=VPS13D;refGene_tx=XM_019836189.2;refGene_gn=VPS13D;refGene_tx=XM_019836191.2;refGene_gn=VPS13D;refGene_tx=XM_023258138.1;refGene_gn=VPS13D;refGene_tx=XM_023258139.1;refGene_gn=VPS13D;refGene_tx=XM_023258140.1;refGene_gn=VPS13D;refGene_tx=XM_023258141.1;refGene_gn=VPS13D;refGene_tx=XM_023258142.1
C1 9027879 9028031
refGene_gn=VPS13D;refGene_tx=XM_019836189.2;refGene_gn=VPS13D;refGene_tx=XM_019836191.2;refGene_gn=VPS13D;refGene_tx=XM_023258138.1;refGene_gn=VPS13D;refGene_tx=XM_023258139.1;refGene_gn=VPS13D;refGene_tx=XM_023258140.1;refGene_gn=VPS13D;refGene_tx=XM_023258141.1;refGene_gn=VPS13D;refGene_tx=XM_023258142.1
C1 9030736 9031768
refGene_gn=VPS13D;refGene_tx=XM_019836189.2;refGene_gn=VPS13D;refGene_tx=XM_019836191.2;refGene_gn=VPS13D;refGene_tx=XM_023258138.1;refGene_gn=VPS13D;refGene_tx=XM_023258139.1;refGene_gn=VPS13D;refGene_tx=XM_023258140.1;refGene_gn=VPS13D;refGene_tx=XM_023258141.1;refGene_gn=VPS13D;refGene_tx=XM_023258142.1
C1 9035584 9035675
refGene_gn=VPS13D;refGene_tx=XM_019836189.2;refGene_gn=VPS13D;refGene_tx=XM_019836191.2;refGene_gn=VPS13D;refGene_tx=XM_023258138.1;refGene_gn=VPS13D;refGene_tx=XM_023258139.1;refGene_gn=VPS13D;refGene_tx=XM_023258140.1;refGene_gn=VPS13D;refGene_tx=XM_023258141.1;refGene_gn=VPS13D;refGene_tx=XM_023258142.1
C1 9037888 9038015
refGene_gn=VPS13D;refGene_tx=XM_019836189.2;refGene_gn=VPS13D;refGene_tx=XM_019836191.2;refGene_gn=VPS13D;refGene_tx=XM_023258138.1;refGene_gn=VPS13D;refGene_tx=XM_023258139.1;refGene_gn=VPS13D;refGene_tx=XM_023258140.1;refGene_gn=VPS13D;refGene_tx=XM_023258141.1;refGene_gn=VPS13D;refGene_tx=XM_023258142.1
C1 9039212 9039393
refGene_gn=VPS13D;refGene_tx=XM_019836189.2;refGene_gn=VPS13D;refGene_tx=XM_019836191.2;refGene_gn=VPS13D;refGene_tx=XM_023258138.1;refGene_gn=VPS13D;refGene_tx=XM_023258139.1;refGene_gn=VPS13D;refGene_tx=XM_023258140.1;refGene_gn=VPS13D;refGene_tx=XM_023258141.1;refGene_gn=VPS13D;refGene_tx=XM_023258142.1
C1 9050759 9050942
refGene_gn=VPS13D;refGene_tx=XM_019836189.2;refGene_gn=VPS13D;refGene_tx=XM_019836191.2;refGene_gn=VPS13D;refGene_tx=XM_023258138.1;refGene_gn=VPS13D;refGene_tx=XM_023258139.1;refGene_gn=VPS13D;refGene_tx=XM_023258140.1;refGene_gn=VPS13D;refGene_tx=XM_023258141.1;refGene_gn=VPS13D;refGene_tx=XM_023258142.1
C1 9056237 9056460
refGene_gn=VPS13D;refGene_tx=XM_019836189.2;refGene_gn=VPS13D;refGene_tx=XM_019836191.2;refGene_gn=VPS13D;refGene_tx=XM_023258138.1;refGene_gn=VPS13D;refGene_tx=XM_023258139.1;refGene_gn=VPS13D;refGene_tx=XM_023258140.1;refGene_gn=VPS13D;refGene_tx=XM_023258141.1;refGene_gn=VPS13

D;refGene_tx=XM_023258142.1
C1 9060623 9060834
refGene_gn=VPS13D;refGene_tx=XM_019836189.2;refGene_gn=VPS13D;refGene_tx=XM_019836191.2;refGene_gn=VPS13D;refGene_tx=XM_023258138.1;refGene_gn=VPS13D;refGene_tx=XM_023258139.1;refGene_gn=VPS13D;refGene_tx=XM_023258140.1;refGene_gn=VPS13D;refGene_tx=XM_023258141.1;refGene_gn=VPS13D;refGene_tx=XM_023258142.1
C1 9062950 9063122
refGene_gn=VPS13D;refGene_tx=XM_019836189.2;refGene_gn=VPS13D;refGene_tx=XM_019836191.2;refGene_gn=VPS13D;refGene_tx=XM_023258138.1;refGene_gn=VPS13D;refGene_tx=XM_023258139.1;refGene_gn=VPS13D;refGene_tx=XM_023258140.1;refGene_gn=VPS13D;refGene_tx=XM_023258141.1;refGene_gn=VPS13D;refGene_tx=XM_023258142.1
C1 9063277 9063390
refGene_gn=VPS13D;refGene_tx=XM_019836189.2;refGene_gn=VPS13D;refGene_tx=XM_019836191.2;refGene_gn=VPS13D;refGene_tx=XM_023258138.1;refGene_gn=VPS13D;refGene_tx=XM_023258139.1;refGene_gn=VPS13D;refGene_tx=XM_023258140.1;refGene_gn=VPS13D;refGene_tx=XM_023258141.1;refGene_gn=VPS13D;refGene_tx=XM_023258142.1
C1 9065738 9065951
refGene_gn=VPS13D;refGene_tx=XM_019836189.2;refGene_gn=VPS13D;refGene_tx=XM_019836191.2;refGene_gn=VPS13D;refGene_tx=XM_023258138.1;refGene_gn=VPS13D;refGene_tx=XM_023258139.1;refGene_gn=VPS13D;refGene_tx=XM_023258140.1;refGene_gn=VPS13D;refGene_tx=XM_023258141.1;refGene_gn=VPS13D;refGene_tx=XM_023258142.1
C1 9069440 9069709
refGene_gn=VPS13D;refGene_tx=XM_019836189.2;refGene_gn=VPS13D;refGene_tx=XM_019836191.2;refGene_gn=VPS13D;refGene_tx=XM_023258138.1;refGene_gn=VPS13D;refGene_tx=XM_023258139.1;refGene_gn=VPS13D;refGene_tx=XM_023258140.1;refGene_gn=VPS13D;refGene_tx=XM_023258141.1;refGene_gn=VPS13D;refGene_tx=XM_023258142.1
C1 9071109 9071243
refGene_gn=VPS13D;refGene_tx=XM_019836189.2;refGene_gn=VPS13D;refGene_tx=XM_019836191.2;refGene_gn=VPS13D;refGene_tx=XM_023258138.1;refGene_gn=VPS13D;refGene_tx=XM_023258139.1;refGene_gn=VPS13D;refGene_tx=XM_023258140.1;refGene_gn=VPS13D;refGene_tx=XM_023258141.1;refGene_gn=VPS13D;refGene_tx=XM_023258142.1
C1 9073959 9074115
refGene_gn=VPS13D;refGene_tx=XM_019836189.2;refGene_gn=VPS13D;refGene_tx=XM_019836191.2;refGene_gn=VPS13D;refGene_tx=XM_023258138.1;refGene_gn=VPS13D;refGene_tx=XM_023258139.1;refGene_gn=VPS13D;refGene_tx=XM_023258140.1;refGene_gn=VPS13D;refGene_tx=XM_023258141.1;refGene_gn=VPS13D;refGene_tx=XM_023258142.1
C1 9075893 9076104
refGene_gn=VPS13D;refGene_tx=XM_019836189.2;refGene_gn=VPS13D;refGene_tx=XM_019836191.2;refGene_gn=VPS13D;refGene_tx=XM_023258138.1;refGene_gn=VPS13D;refGene_tx=XM_023258139.1;refGene_gn=VPS13D;refGene_tx=XM_023258140.1;refGene_gn=VPS13D;refGene_tx=XM_023258141.1;refGene_gn=VPS13D;refGene_tx=XM_023258142.1
C1 9076633 9076708

refGene_gn=VPS13D;refGene_tx=XM_019836189.2;refGene_gn=VPS13D;refGene_tx=XM_019836191.2;refGene_gn=VPS13D;refGene_tx=XM_023258138.1;refGene_gn=VPS13D;refGene_tx=XM_023258139.1;refGene_gn=VPS13D;refGene_tx=XM_023258140.1;refGene_gn=VPS13D;refGene_tx=XM_023258141.1;refGene_gn=VPS13D;refGene_tx=XM_023258142.1
C1 9079116 9079323
refGene_gn=VPS13D;refGene_tx=XM_019836189.2;refGene_gn=VPS13D;refGene_tx=XM_019836191.2;refGene_gn=VPS13D;refGene_tx=XM_023258138.1;refGene_gn=VPS13D;refGene_tx=XM_023258139.1;refGene_gn=VPS13D;refGene_tx=XM_023258140.1;refGene_gn=VPS13D;refGene_tx=XM_023258141.1;refGene_gn=VPS13D;refGene_tx=XM_023258142.1
C1 9080543 9080633
refGene_gn=VPS13D;refGene_tx=XM_019836189.2;refGene_gn=VPS13D;refGene_tx=XM_019836191.2;refGene_gn=VPS13D;refGene_tx=XM_023258138.1;refGene_gn=VPS13D;refGene_tx=XM_023258139.1;refGene_gn=VPS13D;refGene_tx=XM_023258140.1;refGene_gn=VPS13D;refGene_tx=XM_023258141.1;refGene_gn=VPS13D;refGene_tx=XM_023258142.1
C1 9082803 9082944
refGene_gn=VPS13D;refGene_tx=XM_019836189.2;refGene_gn=VPS13D;refGene_tx=XM_019836191.2;refGene_gn=VPS13D;refGene_tx=XM_023258138.1;refGene_gn=VPS13D;refGene_tx=XM_023258139.1;refGene_gn=VPS13D;refGene_tx=XM_023258140.1;refGene_gn=VPS13D;refGene_tx=XM_023258141.1;refGene_gn=VPS13D;refGene_tx=XM_023258142.1
C1 9085252 9085375
refGene_gn=VPS13D;refGene_tx=XM_019836189.2;refGene_gn=VPS13D;refGene_tx=XM_019836191.2;refGene_gn=VPS13D;refGene_tx=XM_023258138.1;refGene_gn=VPS13D;refGene_tx=XM_023258139.1;refGene_gn=VPS13D;refGene_tx=XM_023258140.1;refGene_gn=VPS13D;refGene_tx=XM_023258141.1;refGene_gn=VPS13D;refGene_tx=XM_023258142.1
C1 9087744 9087819
refGene_gn=VPS13D;refGene_tx=XM_019836189.2;refGene_gn=VPS13D;refGene_tx=XM_019836191.2;refGene_gn=VPS13D;refGene_tx=XM_023258138.1;refGene_gn=VPS13D;refGene_tx=XM_023258139.1;refGene_gn=VPS13D;refGene_tx=XM_023258142.1
C1 9091472 9091578
refGene_gn=VPS13D;refGene_tx=XM_019836189.2;refGene_gn=VPS13D;refGene_tx=XM_019836191.2;refGene_gn=VPS13D;refGene_tx=XM_023258138.1;refGene_gn=VPS13D;refGene_tx=XM_023258139.1;refGene_gn=VPS13D;refGene_tx=XM_023258140.1;refGene_gn=VPS13D;refGene_tx=XM_023258141.1;refGene_gn=VPS13D;refGene_tx=XM_023258142.1
C1 9092564 9092717
refGene_gn=VPS13D;refGene_tx=XM_019836189.2;refGene_gn=VPS13D;refGene_tx=XM_019836191.2;refGene_gn=VPS13D;refGene_tx=XM_023258138.1;refGene_gn=VPS13D;refGene_tx=XM_023258139.1;refGene_gn=VPS13D;refGene_tx=XM_023258140.1;refGene_gn=VPS13D;refGene_tx=XM_023258141.1;refGene_gn=VPS13D;refGene_tx=XM_023258142.1
C1 9095026 9095162
refGene_gn=VPS13D;refGene_tx=XM_019836189.2;refGene_gn=VPS13D;refGene_tx=XM_019836191.2;refGene_gn=VPS13D;refGene_tx=XM_023258138.1;refGene_gn=VPS13D;refGene_tx=XM_023258139.1;refGene_gn=VPS13D;refGene_tx=XM_02

3258140.1;refGene_gn=VPS13D;refGene_tx=XM_023258141.1;refGene_gn=VPS13D;refGene_tx=XM_023258142.1
C1 9096302 9096350
refGene_gn=VPS13D;refGene_tx=XM_019836189.2;refGene_gn=VPS13D;refGene_tx=XM_019836191.2;refGene_gn=VPS13D;refGene_tx=XM_023258138.1;refGene_gn=VPS13D;refGene_tx=XM_023258139.1;refGene_gn=VPS13D;refGene_tx=XM_023258140.1;refGene_gn=VPS13D;refGene_tx=XM_023258141.1;refGene_gn=VPS13D;refGene_tx=XM_023258142.1
C1 9098279 9098430
refGene_gn=VPS13D;refGene_tx=XM_019836189.2;refGene_gn=VPS13D;refGene_tx=XM_019836191.2;refGene_gn=VPS13D;refGene_tx=XM_023258138.1;refGene_gn=VPS13D;refGene_tx=XM_023258139.1;refGene_gn=VPS13D;refGene_tx=XM_023258140.1;refGene_gn=VPS13D;refGene_tx=XM_023258141.1;refGene_gn=VPS13D;refGene_tx=XM_023258142.1
C1 9098531 9098742
refGene_gn=VPS13D;refGene_tx=XM_019836189.2;refGene_gn=VPS13D;refGene_tx=XM_019836191.2;refGene_gn=VPS13D;refGene_tx=XM_023258138.1;refGene_gn=VPS13D;refGene_tx=XM_023258139.1;refGene_gn=VPS13D;refGene_tx=XM_023258140.1;refGene_gn=VPS13D;refGene_tx=XM_023258141.1;refGene_gn=VPS13D;refGene_tx=XM_023258142.1
C1 9103482 9103730
refGene_gn=VPS13D;refGene_tx=XM_019836189.2;refGene_gn=VPS13D;refGene_tx=XM_019836191.2;refGene_gn=VPS13D;refGene_tx=XM_023258138.1;refGene_gn=VPS13D;refGene_tx=XM_023258139.1;refGene_gn=VPS13D;refGene_tx=XM_023258140.1;refGene_gn=VPS13D;refGene_tx=XM_023258141.1;refGene_gn=VPS13D;refGene_tx=XM_023258142.1
C1 9105317 9105509
refGene_gn=VPS13D;refGene_tx=XM_019836189.2;refGene_gn=VPS13D;refGene_tx=XM_019836191.2;refGene_gn=VPS13D;refGene_tx=XM_023258138.1;refGene_gn=VPS13D;refGene_tx=XM_023258139.1;refGene_gn=VPS13D;refGene_tx=XM_023258140.1;refGene_gn=VPS13D;refGene_tx=XM_023258141.1;refGene_gn=VPS13D;refGene_tx=XM_023258142.1
C1 9105818 9105945
refGene_gn=VPS13D;refGene_tx=XM_019836189.2;refGene_gn=VPS13D;refGene_tx=XM_019836191.2;refGene_gn=VPS13D;refGene_tx=XM_023258138.1;refGene_gn=VPS13D;refGene_tx=XM_023258139.1;refGene_gn=VPS13D;refGene_tx=XM_023258140.1;refGene_gn=VPS13D;refGene_tx=XM_023258141.1;refGene_gn=VPS13D;refGene_tx=XM_023258142.1
C1 9107605 9107748
refGene_gn=VPS13D;refGene_tx=XM_019836189.2;refGene_gn=VPS13D;refGene_tx=XM_019836191.2;refGene_gn=VPS13D;refGene_tx=XM_023258138.1;refGene_gn=VPS13D;refGene_tx=XM_023258139.1;refGene_gn=VPS13D;refGene_tx=XM_023258140.1;refGene_gn=VPS13D;refGene_tx=XM_023258141.1;refGene_gn=VPS13D;refGene_tx=XM_023258142.1
C1 9111644 9111775
refGene_gn=VPS13D;refGene_tx=XM_019836189.2;refGene_gn=VPS13D;refGene_tx=XM_019836191.2;refGene_gn=VPS13D;refGene_tx=XM_023258138.1;refGene_gn=VPS13D;refGene_tx=XM_023258139.1;refGene_gn=VPS13D;refGene_tx=XM_023258140.1;refGene_gn=VPS13D;refGene_tx=XM_023258141.1;refGene_gn=VPS13D;refGene_tx=XM_023258142.1

C1 9112016 9112192
refGene_gn=VPS13D;refGene_tx=XM_019836189.2;refGene_gn=VPS13D;refGene_tx=XM_019836191.2;refGene_gn=VPS13D;refGene_tx=XM_023258138.1;refGene_gn=VPS13D;refGene_tx=XM_023258139.1;refGene_gn=VPS13D;refGene_tx=XM_023258140.1;refGene_gn=VPS13D;refGene_tx=XM_023258141.1;refGene_gn=VPS13D;refGene_tx=XM_023258142.1
C1 9120602 9120726
refGene_gn=VPS13D;refGene_tx=XM_019836189.2;refGene_gn=VPS13D;refGene_tx=XM_019836191.2;refGene_gn=VPS13D;refGene_tx=XM_023258138.1;refGene_gn=VPS13D;refGene_tx=XM_023258139.1;refGene_gn=VPS13D;refGene_tx=XM_023258140.1;refGene_gn=VPS13D;refGene_tx=XM_023258141.1;refGene_gn=VPS13D;refGene_tx=XM_023258142.1
C1 9121184 9121420
refGene_gn=VPS13D;refGene_tx=XM_019836189.2;refGene_gn=VPS13D;refGene_tx=XM_019836191.2;refGene_gn=VPS13D;refGene_tx=XM_023258138.1;refGene_gn=VPS13D;refGene_tx=XM_023258139.1;refGene_gn=VPS13D;refGene_tx=XM_023258140.1;refGene_gn=VPS13D;refGene_tx=XM_023258141.1;refGene_gn=VPS13D;refGene_tx=XM_023258142.1
C1 9124700 9124809
refGene_gn=VPS13D;refGene_tx=XM_019836189.2;refGene_gn=VPS13D;refGene_tx=XM_019836191.2;refGene_gn=VPS13D;refGene_tx=XM_023258138.1;refGene_gn=VPS13D;refGene_tx=XM_023258139.1;refGene_gn=VPS13D;refGene_tx=XM_023258140.1;refGene_gn=VPS13D;refGene_tx=XM_023258141.1;refGene_gn=VPS13D;refGene_tx=XM_023258142.1
C1 9130262 9130426
refGene_gn=VPS13D;refGene_tx=XM_019836189.2;refGene_gn=VPS13D;refGene_tx=XM_019836191.2;refGene_gn=VPS13D;refGene_tx=XM_023258138.1;refGene_gn=VPS13D;refGene_tx=XM_023258139.1;refGene_gn=VPS13D;refGene_tx=XM_023258140.1;refGene_gn=VPS13D;refGene_tx=XM_023258141.1;refGene_gn=VPS13D;refGene_tx=XM_023258142.1
C1 9131753 9131862
refGene_gn=VPS13D;refGene_tx=XM_019836189.2;refGene_gn=VPS13D;refGene_tx=XM_019836191.2;refGene_gn=VPS13D;refGene_tx=XM_023258138.1;refGene_gn=VPS13D;refGene_tx=XM_023258139.1;refGene_gn=VPS13D;refGene_tx=XM_023258140.1;refGene_gn=VPS13D;refGene_tx=XM_023258141.1;refGene_gn=VPS13D;refGene_tx=XM_023258142.1
C1 9133960 9134140
refGene_gn=VPS13D;refGene_tx=XM_019836189.2;refGene_gn=VPS13D;refGene_tx=XM_019836191.2;refGene_gn=VPS13D;refGene_tx=XM_023258138.1;refGene_gn=VPS13D;refGene_tx=XM_023258139.1;refGene_gn=VPS13D;refGene_tx=XM_023258140.1;refGene_gn=VPS13D;refGene_tx=XM_023258141.1;refGene_gn=VPS13D;refGene_tx=XM_023258142.1
C1 9136456 9136573
refGene_gn=VPS13D;refGene_tx=XM_019836189.2;refGene_gn=VPS13D;refGene_tx=XM_019836191.2;refGene_gn=VPS13D;refGene_tx=XM_023258138.1;refGene_gn=VPS13D;refGene_tx=XM_023258139.1;refGene_gn=VPS13D;refGene_tx=XM_023258140.1;refGene_gn=VPS13D;refGene_tx=XM_023258141.1;refGene_gn=VPS13D;refGene_tx=XM_023258142.1
C1 9137384 9137534
refGene_gn=VPS13D;refGene_tx=XM_019836189.2;refGene_gn=VPS13D;refGene_

tx=XM_019836191.2;refGene_gn=VPS13D;refGene_tx=XM_023258138.1;refGene_gn=VPS13D;refGene_tx=XM_023258139.1;refGene_gn=VPS13D;refGene_tx=XM_023258140.1;refGene_gn=VPS13D;refGene_tx=XM_023258141.1;refGene_gn=VPS13D;refGene_tx=XM_023258142.1
C1 9143052 9143202
refGene_gn=VPS13D;refGene_tx=XM_019836189.2;refGene_gn=VPS13D;refGene_tx=XM_019836191.2;refGene_gn=VPS13D;refGene_tx=XM_023258138.1;refGene_gn=VPS13D;refGene_tx=XM_023258139.1;refGene_gn=VPS13D;refGene_tx=XM_023258140.1;refGene_gn=VPS13D;refGene_tx=XM_023258141.1;refGene_gn=VPS13D;refGene_tx=XM_023258142.1
C1 9144098 9144195
refGene_gn=VPS13D;refGene_tx=XM_019836189.2;refGene_gn=VPS13D;refGene_tx=XM_019836191.2;refGene_gn=VPS13D;refGene_tx=XM_023258138.1;refGene_gn=VPS13D;refGene_tx=XM_023258139.1;refGene_gn=VPS13D;refGene_tx=XM_023258140.1;refGene_gn=VPS13D;refGene_tx=XM_023258141.1;refGene_gn=VPS13D;refGene_tx=XM_023258142.1
C1 9146181 9146330
refGene_gn=VPS13D;refGene_tx=XM_019836189.2;refGene_gn=VPS13D;refGene_tx=XM_019836191.2;refGene_gn=VPS13D;refGene_tx=XM_023258138.1;refGene_gn=VPS13D;refGene_tx=XM_023258139.1;refGene_gn=VPS13D;refGene_tx=XM_023258140.1;refGene_gn=VPS13D;refGene_tx=XM_023258141.1;refGene_gn=VPS13D;refGene_tx=XM_023258142.1
C1 9148531 9148597 refGene_gn=VPS13D;refGene_tx=XM_023258142.1
C1 9157063 9157198
refGene_gn=VPS13D;refGene_tx=XM_019836189.2;refGene_gn=VPS13D;refGene_tx=XM_019836191.2;refGene_gn=VPS13D;refGene_tx=XM_023258138.1;refGene_gn=VPS13D;refGene_tx=XM_023258139.1;refGene_gn=VPS13D;refGene_tx=XM_023258140.1;refGene_gn=VPS13D;refGene_tx=XM_023258141.1
C1 9158297 9158465
refGene_gn=VPS13D;refGene_tx=XM_019836189.2;refGene_gn=VPS13D;refGene_tx=XM_019836191.2;refGene_gn=VPS13D;refGene_tx=XM_023258138.1;refGene_gn=VPS13D;refGene_tx=XM_023258139.1;refGene_gn=VPS13D;refGene_tx=XM_023258140.1;refGene_gn=VPS13D;refGene_tx=XM_023258141.1
C1 9192534 9192667
refGene_gn=VPS13D;refGene_tx=XM_019836189.2;refGene_gn=VPS13D;refGene_tx=XM_019836191.2;refGene_gn=VPS13D;refGene_tx=XM_023258138.1;refGene_gn=VPS13D;refGene_tx=XM_023258139.1;refGene_gn=VPS13D;refGene_tx=XM_023258140.1;refGene_gn=VPS13D;refGene_tx=XM_023258141.1
C1 9195665 9195861
refGene_gn=VPS13D;refGene_tx=XM_019836189.2;refGene_gn=VPS13D;refGene_tx=XM_019836191.2;refGene_gn=VPS13D;refGene_tx=XM_023258138.1;refGene_gn=VPS13D;refGene_tx=XM_023258139.1;refGene_gn=VPS13D;refGene_tx=XM_023258140.1;refGene_gn=VPS13D;refGene_tx=XM_023258141.1
C1 9229594 9229726
refGene_gn=VPS13D;refGene_tx=XM_019836189.2;refGene_gn=VPS13D;refGene_tx=XM_019836191.2;refGene_gn=VPS13D;refGene_tx=XM_023258138.1;refGene_gn=VPS13D;refGene_tx=XM_023258139.1;refGene_gn=VPS13D;refGene_tx=XM_023258140.1;refGene_gn=VPS13D;refGene_tx=XM_023258141.1
C1 9240287 9240528
refGene_gn=VPS13D;refGene_tx=XM_019836189.2;refGene_gn=VPS13D;refGene_

tx=XM_019836191.2;refGene_gn=VPS13D;refGene_tx=XM_023258138.1;refGene_gn=VPS13D;refGene_tx=XM_023258139.1;refGene_gn=VPS13D;refGene_tx=XM_023258140.1;refGene_gn=VPS13D;refGene_tx=XM_023258141.1
C1	9242166	9242298	refGene_gn=VPS13D;refGene_tx=XM_019836189.2;refGene_gn=VPS13D;refGene_tx=XM_019836191.2;refGene_gn=VPS13D;refGene_tx=XM_023258138.1;refGene_gn=VPS13D;refGene_tx=XM_023258139.1;refGene_gn=VPS13D;refGene_tx=XM_023258140.1;refGene_gn=VPS13D;refGene_tx=XM_023258141.1
C1	9282639	9282724	refGene_gn=DHRS3;refGene_tx=XM_003989503.4;refGene_gn=DHRS3;refGene_tx=XM_006934295.3;refGene_gn=DHRS3;refGene_tx=XM_006934296.4;refGene_gn=DHRS3;refGene_tx=XM_011284598.3;refGene_gn=DHRS3;refGene_tx=XM_023258143.1
C1	9284903	9285029	refGene_gn=DHRS3;refGene_tx=XM_003989503.4;refGene_gn=DHRS3;refGene_tx=XM_006934295.3;refGene_gn=DHRS3;refGene_tx=XM_006934296.4;refGene_gn=DHRS3;refGene_tx=XM_011284598.3;refGene_gn=DHRS3;refGene_tx=XM_023258143.1
C1	9289515	9289754	refGene_gn=DHRS3;refGene_tx=XM_003989503.4;refGene_gn=DHRS3;refGene_tx=XM_006934295.3;refGene_gn=DHRS3;refGene_tx=XM_006934296.4;refGene_gn=DHRS3;refGene_tx=XM_011284598.3;refGene_gn=DHRS3;refGene_tx=XM_023258143.1
C1	9290118	9290238	refGene_gn=DHRS3;refGene_tx=XM_003989503.4;refGene_gn=DHRS3;refGene_tx=XM_006934295.3;refGene_gn=DHRS3;refGene_tx=XM_006934296.4;refGene_gn=DHRS3;refGene_tx=XM_011284598.3;refGene_gn=DHRS3;refGene_tx=XM_023258143.1
C1	9290932	9291076	refGene_gn=DHRS3;refGene_tx=XM_003989503.4;refGene_gn=DHRS3;refGene_tx=XM_006934295.3;refGene_gn=DHRS3;refGene_tx=XM_006934296.4;refGene_gn=DHRS3;refGene_tx=XM_011284598.3;refGene_gn=DHRS3;refGene_tx=XM_023258143.1
C1	9320860	9321055	refGene_gn=DHRS3;refGene_tx=XM_003989503.4;refGene_gn=DHRS3;refGene_tx=XM_006934295.3;refGene_gn=DHRS3;refGene_tx=XM_006934296.4;refGene_gn=DHRS3;refGene_tx=XM_011284598.3;refGene_gn=DHRS3;refGene_tx=XM_023258143.1
C1	9369240	9369399	refGene_gn=LOC101080646;refGene_tx=XM_023258146.1
C1	9372893	9372941	refGene_gn=LOC101080646;refGene_tx=XM_023258146.1
C1	9372942	9373114	refGene_gn=LOC101080646;refGene_tx=XM_023258146.1
C1	9377014	9377078	refGene_gn=LOC101080646;refGene_tx=XM_023258146.1
C1	9390347	9390622	refGene_gn=LOC101080646;refGene_tx=XM_023258146.1
C1	9390624	9391121	refGene_gn=LOC101080646;refGene_tx=XM_023258146.1

C1 9421039 9421081
refGene_gn=AADACL3;refGene_tx=XM_019836195.2;refGene_gn=AADACL3;refGene_tx=XM_023258144.1;refGene_gn=AADACL3;refGene_tx=XM_023258145.1
C1 9427469 9427637 refGene_gn=AADACL3;refGene_tx=XM_019836192.1
C1 9432031 9432248
refGene_gn=AADACL3;refGene_tx=XM_019836192.1;refGene_gn=AADACL3;refGene_tx=XM_019836195.2;refGene_gn=AADACL3;refGene_tx=XM_019836197.2;refGene_gn=AADACL3;refGene_tx=XM_023258144.1;refGene_gn=AADACL3;refGene_tx=XM_023258145.1
C1 9434066 9434130
refGene_gn=AADACL3;refGene_tx=XM_019836192.1;refGene_gn=AADACL3;refGene_tx=XM_019836195.2;refGene_gn=AADACL3;refGene_tx=XM_019836197.2;refGene_gn=AADACL3;refGene_tx=XM_023258144.1;refGene_gn=AADACL3;refGene_tx=XM_023258145.1
C1 9436323 9437098
refGene_gn=AADACL3;refGene_tx=XM_019836192.1;refGene_gn=AADACL3;refGene_tx=XM_019836195.2;refGene_gn=AADACL3;refGene_tx=XM_019836197.2;refGene_gn=AADACL3;refGene_tx=XM_023258144.1;refGene_gn=AADACL3;refGene_tx=XM_023258145.1
C1 9451676 9451790
refGene_gn=CC1H1orf158;refGene_tx=XM_003989586.4;refGene_gn=CC1H1orf158;refGene_tx=XM_019836201.2
C1 9458959 9459073
refGene_gn=CC1H1orf158;refGene_tx=XM_003989586.4;refGene_gn=CC1H1orf158;refGene_tx=XM_019836201.2
C1 9462370 9462548
refGene_gn=CC1H1orf158;refGene_tx=XM_003989586.4;refGene_gn=CC1H1orf158;refGene_tx=XM_019836201.2
C1 9463716 9463898
refGene_gn=CC1H1orf158;refGene_tx=XM_003989586.4;refGene_gn=CC1H1orf158;refGene_tx=XM_019836201.2
C1 9479478 9479765
refGene_gn=LOC101081397;refGene_tx=XM_023258147.1
C1 9480209 9480770
refGene_gn=LOC101081397;refGene_tx=XM_023258147.1
C1 9481967 9482526
refGene_gn=LOC101081397;refGene_tx=XM_023258147.1
C1 9529298 9529555 refGene_gn=LRRC38;refGene_tx=XM_011284635.3
C1 9562541 9563205 refGene_gn=LRRC38;refGene_tx=XM_011284635.3
C1 9607277 9607344
refGene_gn=PDPN;refGene_tx=XM_006934300.4;refGene_gn=PDPN;refGene_tx=XM_006934301.3
C1 9626442 9626579
refGene_gn=PDPN;refGene_tx=XM_006934300.4;refGene_gn=PDPN;refGene_tx=XM_006934301.3
C1 9628239 9628369
refGene_gn=PDPN;refGene_tx=XM_006934300.4;refGene_gn=PDPN;refGene_tx=XM_006934301.3
C1 9632963 9633002
refGene_gn=PDPN;refGene_tx=XM_006934300.4;refGene_gn=PDPN;refGene_tx=X

M_006934301.3
C1	9633559	9633671
refGene_gn=PDPN;refGene_tx=XM_006934300.4;refGene_gn=PDPN;refGene_tx=X
M_006934301.3
C1	9634929	9634936
refGene_gn=PDPN;refGene_tx=XM_006934300.4;refGene_gn=PDPN;refGene_tx=X
M_006934301.3
C1	9709221	9709230
refGene_gn=PRDM2;refGene_tx=XM_023260031.1;refGene_gn=PRDM2;refGene_tx
=XM_023260032.1;refGene_gn=PRDM2;refGene_tx=XM_023260033.1;refGene_gn=
PRDM2;refGene_tx=XM_023260037.1
C1	9726648	9726766
refGene_gn=PRDM2;refGene_tx=XM_023260031.1;refGene_gn=PRDM2;refGene_tx
=XM_023260032.1;refGene_gn=PRDM2;refGene_tx=XM_023260033.1;refGene_gn=
PRDM2;refGene_tx=XM_023260037.1
C1	9728616	9728720
refGene_gn=PRDM2;refGene_tx=XM_023260031.1;refGene_gn=PRDM2;refGene_tx
=XM_023260032.1;refGene_gn=PRDM2;refGene_tx=XM_023260033.1;refGene_gn=
PRDM2;refGene_tx=XM_023260036.1;refGene_gn=PRDM2;refGene_tx=XM_0232600
37.1
C1	9740762	9740915
refGene_gn=PRDM2;refGene_tx=XM_023260031.1;refGene_gn=PRDM2;refGene_tx
=XM_023260032.1;refGene_gn=PRDM2;refGene_tx=XM_023260033.1;refGene_gn=
PRDM2;refGene_tx=XM_023260034.1;refGene_gn=PRDM2;refGene_tx=XM_0232600
35.1;refGene_gn=PRDM2;refGene_tx=XM_023260036.1;refGene_gn=PRDM2;refGe
ne_tx=XM_023260037.1
C1	9747045	9747172
refGene_gn=PRDM2;refGene_tx=XM_023260031.1;refGene_gn=PRDM2;refGene_tx
=XM_023260032.1;refGene_gn=PRDM2;refGene_tx=XM_023260033.1;refGene_gn=
PRDM2;refGene_tx=XM_023260034.1;refGene_gn=PRDM2;refGene_tx=XM_0232600
35.1;refGene_gn=PRDM2;refGene_tx=XM_023260036.1;refGene_gn=PRDM2;refGe
ne_tx=XM_023260037.1
C1	9772152	9772290
refGene_gn=PRDM2;refGene_tx=XM_019836205.2;refGene_gn=PRDM2;refGene_tx
=XM_023260031.1;refGene_gn=PRDM2;refGene_tx=XM_023260032.1;refGene_gn=
PRDM2;refGene_tx=XM_023260033.1;refGene_gn=PRDM2;refGene_tx=XM_0232600
34.1;refGene_gn=PRDM2;refGene_tx=XM_023260035.1;refGene_gn=PRDM2;refGe
ne_tx=XM_023260036.1
C1	9778338	9782716
refGene_gn=PRDM2;refGene_tx=XM_019836205.2;refGene_gn=PRDM2;refGene_tx
=XM_023260031.1;refGene_gn=PRDM2;refGene_tx=XM_023260032.1;refGene_gn=
PRDM2;refGene_tx=XM_023260033.1;refGene_gn=PRDM2;refGene_tx=XM_0232600
34.1;refGene_gn=PRDM2;refGene_tx=XM_023260035.1;refGene_gn=PRDM2;refGe
ne_tx=XM_023260036.1
C1	9811978	9812122
refGene_gn=PRDM2;refGene_tx=XM_019836205.2;refGene_gn=PRDM2;refGene_tx
=XM_023260031.1;refGene_gn=PRDM2;refGene_tx=XM_023260032.1;refGene_gn=
PRDM2;refGene_tx=XM_023260033.1;refGene_gn=PRDM2;refGene_tx=XM_0232600
34.1;refGene_gn=PRDM2;refGene_tx=XM_023260035.1;refGene_gn=PRDM2;refGe
ne_tx=XM_023260036.1;refGene_gn=PRDM2;refGene_tx=XM_023260037.1

C1	9816709	9816735	refGene_gn=PRDM2;refGene_tx=XM_023260037.1
C1	9820895	9821168
refGene_gn=LOC111561930;refGene_tx=XM_023260044.1
C1	9861136	9861631
refGene_gn=LOC111561930;refGene_tx=XM_023260044.1
C1	9861826	9861917
refGene_gn=KAZN;refGene_tx=XM_023260038.1;refGene_gn=KAZN;refGene_tx=XM_023260039.1
C1	10159172	10159330
refGene_gn=KAZN;refGene_tx=XM_023260038.1;refGene_gn=KAZN;refGene_tx=XM_023260039.1
C1	10527000	10527003	refGene_gn=KAZN;refGene_tx=XM_023260040.1
C1	10527327	10527592
refGene_gn=KAZN;refGene_tx=XM_023260038.1;refGene_gn=KAZN;refGene_tx=XM_023260039.1;refGene_gn=KAZN;refGene_tx=XM_023260040.1
C1	10829546	10829754
refGene_gn=KAZN;refGene_tx=XM_023260042.1;refGene_gn=KAZN;refGene_tx=XM_023260043.1
C1	10834740	10835002	refGene_gn=KAZN;refGene_tx=XM_023260041.1
C1	10865285	10865477
refGene_gn=KAZN;refGene_tx=XM_023260038.1;refGene_gn=KAZN;refGene_tx=XM_023260039.1;refGene_gn=KAZN;refGene_tx=XM_023260040.1;refGene_gn=KAZN;refGene_tx=XM_023260041.1;refGene_gn=KAZN;refGene_tx=XM_023260042.1;refGene_gn=KAZN;refGene_tx=XM_023260043.1
C1	10922933	10923070
refGene_gn=KAZN;refGene_tx=XM_023260038.1;refGene_gn=KAZN;refGene_tx=XM_023260039.1;refGene_gn=KAZN;refGene_tx=XM_023260040.1;refGene_gn=KAZN;refGene_tx=XM_023260041.1;refGene_gn=KAZN;refGene_tx=XM_023260042.1;refGene_gn=KAZN;refGene_tx=XM_023260043.1
C1	10927856	10928027
refGene_gn=KAZN;refGene_tx=XM_023260038.1;refGene_gn=KAZN;refGene_tx=XM_023260039.1;refGene_gn=KAZN;refGene_tx=XM_023260040.1;refGene_gn=KAZN;refGene_tx=XM_023260041.1;refGene_gn=KAZN;refGene_tx=XM_023260042.1;refGene_gn=KAZN;refGene_tx=XM_023260043.1
C1	10935756	10935946
refGene_gn=KAZN;refGene_tx=XM_023260038.1;refGene_gn=KAZN;refGene_tx=XM_023260039.1;refGene_gn=KAZN;refGene_tx=XM_023260040.1;refGene_gn=KAZN;refGene_tx=XM_023260041.1;refGene_gn=KAZN;refGene_tx=XM_023260042.1;refGene_gn=KAZN;refGene_tx=XM_023260043.1
C1	10938080	10938211
refGene_gn=KAZN;refGene_tx=XM_023260038.1;refGene_gn=KAZN;refGene_tx=XM_023260039.1;refGene_gn=KAZN;refGene_tx=XM_023260040.1;refGene_gn=KAZN;refGene_tx=XM_023260041.1;refGene_gn=KAZN;refGene_tx=XM_023260042.1;refGene_gn=KAZN;refGene_tx=XM_023260043.1
C1	10941073	10941124
refGene_gn=KAZN;refGene_tx=XM_023260038.1;refGene_gn=KAZN;refGene_tx=XM_023260039.1;refGene_gn=KAZN;refGene_tx=XM_023260040.1;refGene_gn=KAZN;refGene_tx=XM_023260041.1;refGene_gn=KAZN;refGene_tx=XM_023260042.1;refGene_gn=KAZN;refGene_tx=XM_023260043.1
C1	10942505	10942673

refGene_gn=KAZN;refGene_tx=XM_023260038.1;refGene_gn=KAZN;refGene_tx=XM_023260039.1;refGene_gn=KAZN;refGene_tx=XM_023260040.1;refGene_gn=KAZN;refGene_tx=XM_023260041.1;refGene_gn=KAZN;refGene_tx=XM_023260042.1;refGene_gn=KAZN;refGene_tx=XM_023260043.1
C1 10964741 10964947 refGene_gn=KAZN;refGene_tx=XM_023260038.1;refGene_gn=KAZN;refGene_tx=XM_023260039.1;refGene_gn=KAZN;refGene_tx=XM_023260040.1;refGene_gn=KAZN;refGene_tx=XM_023260041.1;refGene_gn=KAZN;refGene_tx=XM_023260042.1
C1 10965462 10965581 refGene_gn=KAZN;refGene_tx=XM_023260038.1;refGene_gn=KAZN;refGene_tx=XM_023260039.1;refGene_gn=KAZN;refGene_tx=XM_023260040.1;refGene_gn=KAZN;refGene_tx=XM_023260041.1;refGene_gn=KAZN;refGene_tx=XM_023260042.1
C1 10973974 10974206 refGene_gn=KAZN;refGene_tx=XM_023260038.1;refGene_gn=KAZN;refGene_tx=XM_023260039.1;refGene_gn=KAZN;refGene_tx=XM_023260040.1;refGene_gn=KAZN;refGene_tx=XM_023260041.1;refGene_gn=KAZN;refGene_tx=XM_023260042.1
C1 10975521 10975623 refGene_gn=KAZN;refGene_tx=XM_023260038.1;refGene_gn=KAZN;refGene_tx=XM_023260039.1;refGene_gn=KAZN;refGene_tx=XM_023260040.1;refGene_gn=KAZN;refGene_tx=XM_023260041.1;refGene_gn=KAZN;refGene_tx=XM_023260042.1
C1 10976157 10976324 refGene_gn=KAZN;refGene_tx=XM_023260038.1;refGene_gn=KAZN;refGene_tx=XM_023260039.1;refGene_gn=KAZN;refGene_tx=XM_023260040.1;refGene_gn=KAZN;refGene_tx=XM_023260041.1;refGene_gn=KAZN;refGene_tx=XM_023260042.1
C1 10976662 10976903 refGene_gn=KAZN;refGene_tx=XM_023260039.1
C1 10990459 10990574 refGene_gn=KAZN;refGene_tx=XM_023260038.1;refGene_gn=KAZN;refGene_tx=XM_023260040.1;refGene_gn=KAZN;refGene_tx=XM_023260041.1;refGene_gn=KAZN;refGene_tx=XM_023260042.1
C1 10992410 10992575 refGene_gn=KAZN;refGene_tx=XM_023260038.1;refGene_gn=KAZN;refGene_tx=XM_023260040.1;refGene_gn=KAZN;refGene_tx=XM_023260041.1;refGene_gn=KAZN;refGene_tx=XM_023260042.1
C1 11080722 11081066 refGene_gn=TMEM51;refGene_tx=XM_019836214.2;refGene_gn=TMEM51;refGene_tx=XM_019836215.2;refGene_gn=TMEM51;refGene_tx=XM_019836216.2;refGene_gn=TMEM51;refGene_tx=XM_019836217.2;refGene_gn=TMEM51;refGene_tx=XM_023258148.1;refGene_gn=TMEM51;refGene_tx=XM_023258149.1;refGene_gn=TMEM51;refGene_tx=XM_023258150.1;refGene_gn=TMEM51;refGene_tx=XM_023258151.1;refGene_gn=TMEM51;refGene_tx=XM_023258152.1;refGene_gn=TMEM51;refGene_tx=XM_023258153.1
C1 11083620 11084032 refGene_gn=TMEM51;refGene_tx=XM_019836214.2;refGene_gn=TMEM51;refGene_tx=XM_019836215.2;refGene_gn=TMEM51;refGene_tx=XM_019836216.2;refGene_gn=TMEM51;refGene_tx=XM_019836217.2;refGene_gn=TMEM51;refGene_tx=XM_023258148.1;refGene_gn=TMEM51;refGene_tx=XM_023258149.1;refGene_gn=TMEM51;refGene_tx=XM_023258150.1;refGene_gn=TMEM51;refGene_tx=XM_023258151.1;refGene_gn=TMEM51;refGene_tx=XM_023258152.1;refGene_gn=TMEM51;refGene_tx=XM_023258153.1
C1 11105681 11105793 refGene_gn=FHAD1;refGene_tx=XM_023258154.1

C1 11105883 11105990 refGene_gn=FHAD1;refGene_tx=XM_023258154.1
C1 11126734 11126941 refGene_gn=FHAD1;refGene_tx=XM_023258154.1
C1 11134997 11135319 refGene_gn=FHAD1;refGene_tx=XM_023258154.1
C1 11144377 11144487 refGene_gn=FHAD1;refGene_tx=XM_023258154.1
C1 11147801 11148038 refGene_gn=FHAD1;refGene_tx=XM_023258154.1
C1 11152180 11152304 refGene_gn=FHAD1;refGene_tx=XM_023258154.1
C1 11156796 11156927 refGene_gn=FHAD1;refGene_tx=XM_023258154.1
C1 11159838 11159928 refGene_gn=FHAD1;refGene_tx=XM_023258154.1
C1 11160942 11161047 refGene_gn=FHAD1;refGene_tx=XM_023258154.1
C1 11164692 11164800 refGene_gn=FHAD1;refGene_tx=XM_023258154.1
C1 11167463 11167547 refGene_gn=FHAD1;refGene_tx=XM_023258154.1
C1 11168297 11168450 refGene_gn=FHAD1;refGene_tx=XM_023258154.1
C1 11169646 11169842 refGene_gn=FHAD1;refGene_tx=XM_023258154.1
C1 11178803 11178874 refGene_gn=FHAD1;refGene_tx=XM_023258154.1
C1 11181348 11181489 refGene_gn=FHAD1;refGene_tx=XM_023258154.1
C1 11186394 11186502 refGene_gn=FHAD1;refGene_tx=XM_023258154.1
C1 11186737 11186845 refGene_gn=FHAD1;refGene_tx=XM_023258154.1
C1 11190501 11190609 refGene_gn=FHAD1;refGene_tx=XM_023258154.1
C1 11193892 11194000 refGene_gn=FHAD1;refGene_tx=XM_023258154.1
C1 11199505 11199679 refGene_gn=FHAD1;refGene_tx=XM_023258154.1
C1 11201266 11201483 refGene_gn=FHAD1;refGene_tx=XM_023258154.1
C1 11202833 11202918 refGene_gn=FHAD1;refGene_tx=XM_023258154.1
C1 11203441 11203548 refGene_gn=FHAD1;refGene_tx=XM_023258154.1
C1 11205381 11205541 refGene_gn=FHAD1;refGene_tx=XM_023258154.1
C1 11207058 11207191 refGene_gn=FHAD1;refGene_tx=XM_023258154.1
C1 11210553 11210680 refGene_gn=FHAD1;refGene_tx=XM_023258154.1
C1 11211335 11211460 refGene_gn=FHAD1;refGene_tx=XM_023258154.1
C1 11214597 11214687 refGene_gn=FHAD1;refGene_tx=XM_023258154.1
C1 11215136 11215357 refGene_gn=FHAD1;refGene_tx=XM_023258154.1
C1 11215841 11216007 refGene_gn=FHAD1;refGene_tx=XM_023258154.1
C1 11226888 11227005 refGene_gn=FHAD1;refGene_tx=XM_023258154.1
C1 11240749 11241060 refGene_gn=EFHD2;refGene_tx=XM_023258155.1
C1 11252715 11252863
refGene_gn=EFHD2;refGene_tx=XM_023258155.1;refGene_gn=EFHD2;refGene_tx
=XM_023258156.1
C1 11254116 11254251
refGene_gn=EFHD2;refGene_tx=XM_023258155.1;refGene_gn=EFHD2;refGene_tx
=XM_023258156.1
C1 11255143 11255275
refGene_gn=EFHD2;refGene_tx=XM_023258155.1;refGene_gn=EFHD2;refGene_tx
=XM_023258156.1
C1 11261583 11261623 refGene_gn=CTRC;refGene_tx=XM_003989507.3
C1 11262780 11262872 refGene_gn=CTRC;refGene_tx=XM_003989507.3
C1 11262960 11263058 refGene_gn=CTRC;refGene_tx=XM_003989507.3
C1 11263886 11264012 refGene_gn=CTRC;refGene_tx=XM_003989507.3
C1 11264847 11264984 refGene_gn=CTRC;refGene_tx=XM_003989507.3
C1 11265743 11265889 refGene_gn=CTRC;refGene_tx=XM_003989507.3
C1 11266653 11266806 refGene_gn=CTRC;refGene_tx=XM_003989507.3
C1 11267391 11267406 refGene_gn=CTRC;refGene_tx=XM_003989507.3
C1 11271828 11271876 refGene_gn=CASP9;refGene_tx=XM_019836232.2

C1 11289622 11289712 refGene_gn=CASP9;refGene_tx=XM_011284592.3
C1 11290056 11290104 refGene_gn=CASP9;refGene_tx=XM_019836231.2
C1 11291734 11291844
refGene_gn=CASP9;refGene_tx=XM_011284592.3;refGene_gn=CASP9;refGene_tx=XM_019836231.2;refGene_gn=CASP9;refGene_tx=XM_019836232.2
C1 11292451 11292631
refGene_gn=CASP9;refGene_tx=XM_011284592.3;refGene_gn=CASP9;refGene_tx=XM_019836231.2;refGene_gn=CASP9;refGene_tx=XM_019836232.2
C1 11298949 11299097
refGene_gn=CASP9;refGene_tx=XM_011284592.3;refGene_gn=CASP9;refGene_tx=XM_019836231.2;refGene_gn=CASP9;refGene_tx=XM_019836232.2
C1 11300283 11300373
refGene_gn=CASP9;refGene_tx=XM_011284592.3;refGene_gn=CASP9;refGene_tx=XM_019836231.2;refGene_gn=CASP9;refGene_tx=XM_019836232.2
C1 11300811 11300988
refGene_gn=CASP9;refGene_tx=XM_011284592.3;refGene_gn=CASP9;refGene_tx=XM_019836231.2;refGene_gn=CASP9;refGene_tx=XM_019836232.2
C1 11301946 11301981
refGene_gn=CASP9;refGene_tx=XM_011284592.3;refGene_gn=CASP9;refGene_tx=XM_019836231.2;refGene_gn=CASP9;refGene_tx=XM_019836232.2
C1 11304520 11304914
refGene_gn=CASP9;refGene_tx=XM_011284592.3;refGene_gn=CASP9;refGene_tx=XM_019836231.2;refGene_gn=CASP9;refGene_tx=XM_019836232.2
C1 11307986 11308118
refGene_gn=CASP9;refGene_tx=XM_011284592.3;refGene_gn=CASP9;refGene_tx=XM_019836231.2;refGene_gn=CASP9;refGene_tx=XM_019836232.2
C1 11312221 11312388
refGene_gn=DNAJC16;refGene_tx=XM_023258157.1;refGene_gn=DNAJC16;refGene_tx=XM_023258158.1
C1 11316510 11316577
refGene_gn=DNAJC16;refGene_tx=XM_023258157.1;refGene_gn=DNAJC16;refGene_tx=XM_023258158.1
C1 11318289 11318626
refGene_gn=DNAJC16;refGene_tx=XM_023258157.1;refGene_gn=DNAJC16;refGene_tx=XM_023258158.1
C1 11328919 11329104
refGene_gn=DNAJC16;refGene_tx=XM_023258157.1;refGene_gn=DNAJC16;refGene_tx=XM_023258158.1
C1 11331154 11331259
refGene_gn=DNAJC16;refGene_tx=XM_023258157.1;refGene_gn=DNAJC16;refGene_tx=XM_023258158.1;refGene_gn=DNAJC16;refGene_tx=XM_023258159.1
C1 11332034 11332193
refGene_gn=DNAJC16;refGene_tx=XM_023258157.1;refGene_gn=DNAJC16;refGene_tx=XM_023258158.1;refGene_gn=DNAJC16;refGene_tx=XM_023258159.1
C1 11343609 11343740
refGene_gn=DNAJC16;refGene_tx=XM_023258157.1;refGene_gn=DNAJC16;refGene_tx=XM_023258158.1;refGene_gn=DNAJC16;refGene_tx=XM_023258159.1
C1 11345901 11346085
refGene_gn=DNAJC16;refGene_tx=XM_023258157.1;refGene_gn=DNAJC16;refGene_tx=XM_023258158.1;refGene_gn=DNAJC16;refGene_tx=XM_023258159.1

C1 11346850 11347033
refGene_gn=DNAJC16;refGene_tx=XM_023258157.1;refGene_gn=DNAJC16;refGene_tx=XM_023258158.1;refGene_gn=DNAJC16;refGene_tx=XM_023258159.1
C1 11347182 11347259
refGene_gn=DNAJC16;refGene_tx=XM_023258157.1;refGene_gn=DNAJC16;refGene_tx=XM_023258158.1;refGene_gn=DNAJC16;refGene_tx=XM_023258159.1
C1 11348528 11348609
refGene_gn=DNAJC16;refGene_tx=XM_023258157.1;refGene_gn=DNAJC16;refGene_tx=XM_023258158.1;refGene_gn=DNAJC16;refGene_tx=XM_023258159.1
C1 11348962 11349061
refGene_gn=DNAJC16;refGene_tx=XM_023258157.1;refGene_gn=DNAJC16;refGene_tx=XM_023258158.1;refGene_gn=DNAJC16;refGene_tx=XM_023258159.1
C1 11349646 11349817
refGene_gn=DNAJC16;refGene_tx=XM_023258157.1;refGene_gn=DNAJC16;refGene_tx=XM_023258158.1;refGene_gn=DNAJC16;refGene_tx=XM_023258159.1
C1 11350307 11350707
refGene_gn=DNAJC16;refGene_tx=XM_023258157.1;refGene_gn=DNAJC16;refGene_tx=XM_023258158.1;refGene_gn=DNAJC16;refGene_tx=XM_023258159.1
C1 11354395 11354469 refGene_gn=AGMAT;refGene_tx=XM_011284632.3
C1 11355141 11355226 refGene_gn=AGMAT;refGene_tx=XM_011284632.3
C1 11356082 11356139
refGene_gn=AGMAT;refGene_tx=XM_023258160.1;refGene_gn=AGMAT;refGene_tx=XM_023258161.1
C1 11357093 11357273
refGene_gn=AGMAT;refGene_tx=XM_011284632.3;refGene_gn=AGMAT;refGene_tx=XM_023258160.1;refGene_gn=AGMAT;refGene_tx=XM_023258161.1
C1 11357989 11358185
refGene_gn=AGMAT;refGene_tx=XM_011284632.3;refGene_gn=AGMAT;refGene_tx=XM_023258160.1;refGene_gn=AGMAT;refGene_tx=XM_023258161.1
C1 11359245 11359294
refGene_gn=AGMAT;refGene_tx=XM_011284632.3;refGene_gn=AGMAT;refGene_tx=XM_023258160.1;refGene_gn=AGMAT;refGene_tx=XM_023258161.1
C1 11361533 11361736
refGene_gn=AGMAT;refGene_tx=XM_011284632.3;refGene_gn=AGMAT;refGene_tx=XM_023258160.1;refGene_gn=AGMAT;refGene_tx=XM_023258161.1
C1 11362968 11363240
refGene_gn=AGMAT;refGene_tx=XM_011284632.3;refGene_gn=AGMAT;refGene_tx=XM_023258160.1;refGene_gn=AGMAT;refGene_tx=XM_023258161.1
C1 11386114 11386252
refGene_gn=LOC101083950;refGene_tx=XM_006934309.4;refGene_gn=LOC101083950;refGene_tx=XM_019836235.2;refGene_gn=LOC101083950;refGene_tx=XM_019836239.2;refGene_gn=LOC101083950;refGene_tx=XM_019836240.2
C1 11393880 11394010
refGene_gn=LOC101083950;refGene_tx=XM_006934309.4;refGene_gn=LOC101083950;refGene_tx=XM_019836235.2;refGene_gn=LOC101083950;refGene_tx=XM_019836239.2;refGene_gn=LOC101083950;refGene_tx=XM_019836240.2
C1 11397298 11397535
refGene_gn=LOC101083950;refGene_tx=XM_006934309.4;refGene_gn=LOC101083950;refGene_tx=XM_019836239.2;refGene_gn=LOC101083950;refGene_tx=XM_019836240.2

C1 11399353 11399480
refGene_gn=LOC101083950;refGene_tx=XM_006934309.4;refGene_gn=LOC101083950;refGene_tx=XM_019836235.2;refGene_gn=LOC101083950;refGene_tx=XM_019836239.2;refGene_gn=LOC101083950;refGene_tx=XM_019836240.2
C1 11403866 11403994
refGene_gn=LOC101083950;refGene_tx=XM_006934309.4;refGene_gn=LOC101083950;refGene_tx=XM_019836235.2;refGene_gn=LOC101083950;refGene_tx=XM_019836239.2;refGene_gn=LOC101083950;refGene_tx=XM_019836240.2
C1 11408066 11408195
refGene_gn=LOC101083950;refGene_tx=XM_006934309.4;refGene_gn=LOC101083950;refGene_tx=XM_019836235.2;refGene_gn=LOC101083950;refGene_tx=XM_019836239.2;refGene_gn=LOC101083950;refGene_tx=XM_019836240.2
C1 11415034 11415138
refGene_gn=LOC101083950;refGene_tx=XM_006934309.4;refGene_gn=LOC101083950;refGene_tx=XM_019836235.2;refGene_gn=LOC101083950;refGene_tx=XM_019836239.2;refGene_gn=LOC101083950;refGene_tx=XM_019836240.2
C1 11417299 11417489
refGene_gn=LOC101083950;refGene_tx=XM_006934309.4;refGene_gn=LOC101083950;refGene_tx=XM_019836235.2;refGene_gn=LOC101083950;refGene_tx=XM_019836239.2;refGene_gn=LOC101083950;refGene_tx=XM_019836240.2
C1 11417834 11417884
refGene_gn=LOC101083950;refGene_tx=XM_019836239.2
C1 11419819 11419836
refGene_gn=LOC101083950;refGene_tx=XM_019836240.2
C1 11422382 11424265
refGene_gn=LOC101083950;refGene_tx=XM_006934309.4;refGene_gn=LOC101083950;refGene_tx=XM_019836235.2;refGene_gn=LOC101083950;refGene_tx=XM_019836236.1;refGene_gn=LOC101083950;refGene_tx=XM_019836237.1;refGene_gn=LOC101083950;refGene_tx=XM_019836238.1
C1 11442362 11443070
refGene_gn=PLEKHM2;refGene_tx=XM_023258162.1;refGene_gn=PLEKHM2;refGene_tx=XM_023258163.1
C1 11462574 11462681
refGene_gn=PLEKHM2;refGene_tx=XM_023258162.1;refGene_gn=PLEKHM2;refGene_tx=XM_023258163.1
C1 11463006 11463116
refGene_gn=PLEKHM2;refGene_tx=XM_023258162.1;refGene_gn=PLEKHM2;refGene_tx=XM_023258163.1
C1 11463668 11463768
refGene_gn=PLEKHM2;refGene_tx=XM_023258162.1;refGene_gn=PLEKHM2;refGene_tx=XM_023258163.1
C1 11464305 11464393
refGene_gn=PLEKHM2;refGene_tx=XM_023258162.1;refGene_gn=PLEKHM2;refGene_tx=XM_023258163.1
C1 11465214 11465401
refGene_gn=PLEKHM2;refGene_tx=XM_023258162.1;refGene_gn=PLEKHM2;refGene_tx=XM_023258163.1
C1 11466674 11466734 refGene_gn=PLEKHM2;refGene_tx=XM_023258162.1
C1 11469871 11470100
refGene_gn=PLEKHM2;refGene_tx=XM_023258162.1;refGene_gn=PLEKHM2;refGen

e_tx=XM_023258163.1
C1 11471180 11472023
refGene_gn=PLEKHM2;refGene_tx=XM_023258162.1;refGene_gn=PLEKHM2;refGene_tx=XM_023258163.1
C1 11472286 11472356
refGene_gn=PLEKHM2;refGene_tx=XM_023258162.1;refGene_gn=PLEKHM2;refGene_tx=XM_023258163.1
C1 11472455 11472546
refGene_gn=PLEKHM2;refGene_tx=XM_023258162.1;refGene_gn=PLEKHM2;refGene_tx=XM_023258163.1
C1 11472867 11472932
refGene_gn=PLEKHM2;refGene_tx=XM_023258162.1;refGene_gn=PLEKHM2;refGene_tx=XM_023258163.1
C1 11473319 11473408
refGene_gn=PLEKHM2;refGene_tx=XM_023258162.1;refGene_gn=PLEKHM2;refGene_tx=XM_023258163.1
C1 11474015 11474148
refGene_gn=PLEKHM2;refGene_tx=XM_023258162.1;refGene_gn=PLEKHM2;refGene_tx=XM_023258163.1
C1 11474383 11474574
refGene_gn=PLEKHM2;refGene_tx=XM_023258162.1;refGene_gn=PLEKHM2;refGene_tx=XM_023258163.1
C1 11475022 11475088
refGene_gn=PLEKHM2;refGene_tx=XM_023258162.1;refGene_gn=PLEKHM2;refGene_tx=XM_023258163.1
C1 11475781 11475941
refGene_gn=PLEKHM2;refGene_tx=XM_023258162.1;refGene_gn=PLEKHM2;refGene_tx=XM_023258163.1
C1 11476290 11476470
refGene_gn=PLEKHM2;refGene_tx=XM_023258162.1;refGene_gn=PLEKHM2;refGene_tx=XM_023258163.1
C1 11476552 11476669
refGene_gn=PLEKHM2;refGene_tx=XM_023258162.1;refGene_gn=PLEKHM2;refGene_tx=XM_023258163.1
C1 11476962 11477097
refGene_gn=PLEKHM2;refGene_tx=XM_023258162.1;refGene_gn=PLEKHM2;refGene_tx=XM_023258163.1
C1 11479918 11480338 refGene_gn=SLC25A34;refGene_tx=XM_003989510.4
C1 11480882 11480948 refGene_gn=SLC25A34;refGene_tx=XM_003989510.4
C1 11481038 11481191 refGene_gn=SLC25A34;refGene_tx=XM_003989510.4
C1 11481591 11481726 refGene_gn=SLC25A34;refGene_tx=XM_003989510.4
C1 11482388 11482571 refGene_gn=SLC25A34;refGene_tx=XM_003989510.4
C1 11484576 11484664 refGene_gn=TMEM82;refGene_tx=XM_023258165.1
C1 11484819 11484892 refGene_gn=TMEM82;refGene_tx=XM_023258165.1
C1 11485079 11485254 refGene_gn=TMEM82;refGene_tx=XM_023258165.1
C1 11486109 11486527 refGene_gn=TMEM82;refGene_tx=XM_023258165.1
C1 11487697 11487885 refGene_gn=TMEM82;refGene_tx=XM_023258165.1
C1 11488126 11488222 refGene_gn=TMEM82;refGene_tx=XM_023258165.1
C1 11494092 11494126 refGene_gn=FBLIM1;refGene_tx=XM_011284589.3
C1 11495764 11495895 refGene_gn=FBLIM1;refGene_tx=XM_011284589.3

C1 11497453 11497610 refGene_gn=FBLIM1;refGene_tx=XM_011284589.3
C1 11502621 11502891
refGene_gn=FBLIM1;refGene_tx=XM_011284589.3;refGene_gn=FBLIM1;refGene_tx=XM_023258164.1
C1 11508525 11508695
refGene_gn=FBLIM1;refGene_tx=XM_011284589.3;refGene_gn=FBLIM1;refGene_tx=XM_023258164.1
C1 11509197 11509312
refGene_gn=FBLIM1;refGene_tx=XM_011284589.3;refGene_gn=FBLIM1;refGene_tx=XM_023258164.1
C1 11511620 11511790
refGene_gn=FBLIM1;refGene_tx=XM_011284589.3;refGene_gn=FBLIM1;refGene_tx=XM_023258164.1
C1 11515358 11515537
refGene_gn=FBLIM1;refGene_tx=XM_011284589.3;refGene_gn=FBLIM1;refGene_tx=XM_023258164.1
C1 11516298 11516416
refGene_gn=FBLIM1;refGene_tx=XM_011284589.3;refGene_gn=FBLIM1;refGene_tx=XM_023258164.1
C1 11519508 11519622
refGene_gn=FBLIM1;refGene_tx=XM_011284589.3;refGene_gn=FBLIM1;refGene_tx=XM_023258164.1
C1 11566225 11566308
refGene_gn=SPEN;refGene_tx=XM_023258166.1;refGene_gn=SPEN;refGene_tx=XM_023258167.1
C1 11595581 11595902
refGene_gn=SPEN;refGene_tx=XM_023258166.1;refGene_gn=SPEN;refGene_tx=XM_023258167.1
C1 11598989 11599466 refGene_gn=SPEN;refGene_tx=XM_023258166.1
C1 11630134 11630295
refGene_gn=SPEN;refGene_tx=XM_023258166.1;refGene_gn=SPEN;refGene_tx=XM_023258167.1
C1 11630947 11631148
refGene_gn=SPEN;refGene_tx=XM_023258166.1;refGene_gn=SPEN;refGene_tx=XM_023258167.1
C1 11634981 11635133
refGene_gn=SPEN;refGene_tx=XM_023258166.1;refGene_gn=SPEN;refGene_tx=XM_023258167.1
C1 11638301 11638427
refGene_gn=SPEN;refGene_tx=XM_023258166.1;refGene_gn=SPEN;refGene_tx=XM_023258167.1
C1 11638750 11638864
refGene_gn=SPEN;refGene_tx=XM_023258166.1;refGene_gn=SPEN;refGene_tx=XM_023258167.1
C1 11639904 11640018
refGene_gn=SPEN;refGene_tx=XM_023258166.1;refGene_gn=SPEN;refGene_tx=XM_023258167.1
C1 11641307 11641408
refGene_gn=SPEN;refGene_tx=XM_023258166.1;refGene_gn=SPEN;refGene_tx=XM_023258167.1

C1 11647159 11655350
refGene_gn=SPEN;refGene_tx=XM_023258166.1;refGene_gn=SPEN;refGene_tx=XM_023258167.1
C1 11655743 11656253
refGene_gn=SPEN;refGene_tx=XM_023258166.1;refGene_gn=SPEN;refGene_tx=XM_023258167.1
C1 11656418 11656613
refGene_gn=SPEN;refGene_tx=XM_023258166.1;refGene_gn=SPEN;refGene_tx=XM_023258167.1
C1 11656938 11657097
refGene_gn=SPEN;refGene_tx=XM_023258166.1;refGene_gn=SPEN;refGene_tx=XM_023258167.1
C1 11657549 11657681
refGene_gn=SPEN;refGene_tx=XM_023258166.1;refGene_gn=SPEN;refGene_tx=XM_023258167.1
C1 11659989 11660273
refGene_gn=ZBTB17;refGene_tx=XM_023258168.1;refGene_gn=ZBTB17;refGene_tx=XM_023258169.1;refGene_gn=ZBTB17;refGene_tx=XM_023258170.1;refGene_gn=ZBTB17;refGene_tx=XM_023258171.1
C1 11660356 11660446
refGene_gn=ZBTB17;refGene_tx=XM_023258168.1;refGene_gn=ZBTB17;refGene_tx=XM_023258169.1;refGene_gn=ZBTB17;refGene_tx=XM_023258170.1;refGene_gn=ZBTB17;refGene_tx=XM_023258171.1
C1 11660550 11660760
refGene_gn=ZBTB17;refGene_tx=XM_023258168.1;refGene_gn=ZBTB17;refGene_tx=XM_023258169.1;refGene_gn=ZBTB17;refGene_tx=XM_023258170.1;refGene_gn=ZBTB17;refGene_tx=XM_023258171.1
C1 11661098 11661229
refGene_gn=ZBTB17;refGene_tx=XM_023258168.1;refGene_gn=ZBTB17;refGene_tx=XM_023258169.1;refGene_gn=ZBTB17;refGene_tx=XM_023258170.1;refGene_gn=ZBTB17;refGene_tx=XM_023258171.1
C1 11661451 11661614
refGene_gn=ZBTB17;refGene_tx=XM_023258168.1;refGene_gn=ZBTB17;refGene_tx=XM_023258169.1;refGene_gn=ZBTB17;refGene_tx=XM_023258170.1;refGene_gn=ZBTB17;refGene_tx=XM_023258171.1
C1 11661716 11661833
refGene_gn=ZBTB17;refGene_tx=XM_023258168.1;refGene_gn=ZBTB17;refGene_tx=XM_023258169.1;refGene_gn=ZBTB17;refGene_tx=XM_023258170.1;refGene_gn=ZBTB17;refGene_tx=XM_023258171.1
C1 11661935 11662023
refGene_gn=ZBTB17;refGene_tx=XM_023258168.1;refGene_gn=ZBTB17;refGene_tx=XM_023258169.1;refGene_gn=ZBTB17;refGene_tx=XM_023258170.1;refGene_gn=ZBTB17;refGene_tx=XM_023258171.1
C1 11662342 11662643
refGene_gn=ZBTB17;refGene_tx=XM_023258168.1;refGene_gn=ZBTB17;refGene_tx=XM_023258169.1;refGene_gn=ZBTB17;refGene_tx=XM_023258170.1;refGene_gn=ZBTB17;refGene_tx=XM_023258171.1
C1 11662752 11662895
refGene_gn=ZBTB17;refGene_tx=XM_023258168.1;refGene_gn=ZBTB17;refGene_tx=XM_023258169.1;refGene_gn=ZBTB17;refGene_tx=XM_023258170.1;refGene_

gn=ZBTB17;refGene_tx=XM_023258171.1
C1 11662992 11663231
refGene_gn=ZBTB17;refGene_tx=XM_023258168.1;refGene_gn=ZBTB17;refGene_tx=XM_023258169.1;refGene_gn=ZBTB17;refGene_tx=XM_023258170.1;refGene_gn=ZBTB17;refGene_tx=XM_023258171.1
C1 11663723 11663849
refGene_gn=ZBTB17;refGene_tx=XM_023258168.1;refGene_gn=ZBTB17;refGene_tx=XM_023258169.1;refGene_gn=ZBTB17;refGene_tx=XM_023258170.1;refGene_gn=ZBTB17;refGene_tx=XM_023258171.1
C1 11664126 11664267
refGene_gn=ZBTB17;refGene_tx=XM_023258168.1;refGene_gn=ZBTB17;refGene_tx=XM_023258169.1;refGene_gn=ZBTB17;refGene_tx=XM_023258170.1;refGene_gn=ZBTB17;refGene_tx=XM_023258171.1
C1 11664882 11665071
refGene_gn=ZBTB17;refGene_tx=XM_023258168.1;refGene_gn=ZBTB17;refGene_tx=XM_023258169.1;refGene_gn=ZBTB17;refGene_tx=XM_023258170.1;refGene_gn=ZBTB17;refGene_tx=XM_023258171.1
C1 11666091 11666298
refGene_gn=ZBTB17;refGene_tx=XM_023258168.1;refGene_gn=ZBTB17;refGene_tx=XM_023258169.1;refGene_gn=ZBTB17;refGene_tx=XM_023258170.1
C1 11666748 11666797
refGene_gn=ZBTB17;refGene_tx=XM_023258168.1;refGene_gn=ZBTB17;refGene_tx=XM_023258169.1
C1 11702199 11702281 refGene_gn=SRARP;refGene_tx=XM_003989514.4
C1 11703962 11704390 refGene_gn=SRARP;refGene_tx=XM_003989514.4
C1 11713417 11713597 refGene_gn=HSPB7;refGene_tx=XM_003989515.5
C1 11714853 11714987 refGene_gn=HSPB7;refGene_tx=XM_003989515.5
C1 11715555 11715751 refGene_gn=HSPB7;refGene_tx=XM_003989515.5
C1 11720989 11721089 refGene_gn=CLCNKA;refGene_tx=XM_023258172.1
C1 11721806 11721935 refGene_gn=CLCNKA;refGene_tx=XM_023258172.1
C1 11722722 11722851 refGene_gn=CLCNKA;refGene_tx=XM_023258172.1
C1 11723452 11723592 refGene_gn=CLCNKA;refGene_tx=XM_023258172.1
C1 11723930 11724008 refGene_gn=CLCNKA;refGene_tx=XM_023258172.1
C1 11724199 11724278 refGene_gn=CLCNKA;refGene_tx=XM_023258172.1
C1 11724808 11724934 refGene_gn=CLCNKA;refGene_tx=XM_023258172.1
C1 11725249 11725334 refGene_gn=CLCNKA;refGene_tx=XM_023258172.1
C1 11725440 11725542 refGene_gn=CLCNKA;refGene_tx=XM_023258172.1
C1 11726135 11726220 refGene_gn=CLCNKA;refGene_tx=XM_023258172.1
C1 11726480 11726654 refGene_gn=CLCNKA;refGene_tx=XM_023258172.1
C1 11727104 11727174 refGene_gn=CLCNKA;refGene_tx=XM_023258172.1
C1 11727343 11727454 refGene_gn=CLCNKA;refGene_tx=XM_023258172.1
C1 11727796 11728010 refGene_gn=CLCNKA;refGene_tx=XM_023258172.1
C1 11728914 11728972 refGene_gn=CLCNKA;refGene_tx=XM_023258172.1
C1 11728973 11729049 refGene_gn=CLCNKA;refGene_tx=XM_023258172.1
C1 11730820 11730909 refGene_gn=CLCNKA;refGene_tx=XM_023258172.1
C1 11731045 11731126 refGene_gn=CLCNKA;refGene_tx=XM_023258172.1
C1 11731885 11731972 refGene_gn=CLCNKA;refGene_tx=XM_023258172.1
C1 11732325 11732373 refGene_gn=CLCNKA;refGene_tx=XM_023258172.1
C1 11733872 11734156
refGene_gn=FAM131C;refGene_tx=XM_019836244.2;refGene_gn=FAM131C;refGen

e_tx=XM_019836246.1;refGene_gn=FAM131C;refGene_tx=XM_023258173.1
C1 11734969 11735080
refGene_gn=FAM131C;refGene_tx=XM_019836244.2;refGene_gn=FAM131C;refGen
e_tx=XM_019836246.1;refGene_gn=FAM131C;refGene_tx=XM_023258173.1
C1 11735334 11735517
refGene_gn=FAM131C;refGene_tx=XM_019836244.2;refGene_gn=FAM131C;refGen
e_tx=XM_019836246.1;refGene_gn=FAM131C;refGene_tx=XM_023258173.1
C1 11737682 11737776
refGene_gn=FAM131C;refGene_tx=XM_019836244.2;refGene_gn=FAM131C;refGen
e_tx=XM_019836246.1;refGene_gn=FAM131C;refGene_tx=XM_023258173.1
C1 11738041 11738077
refGene_gn=FAM131C;refGene_tx=XM_019836244.2;refGene_gn=FAM131C;refGen
e_tx=XM_019836246.1;refGene_gn=FAM131C;refGene_tx=XM_023258173.1
C1 11739080 11739196
refGene_gn=FAM131C;refGene_tx=XM_019836244.2;refGene_gn=FAM131C;refGen
e_tx=XM_019836246.1;refGene_gn=FAM131C;refGene_tx=XM_023258173.1
C1 11751830 11751852 refGene_gn=FAM131C;refGene_tx=XM_023258173.1
C1 11788564 11788670 refGene_gn=EPHA2;refGene_tx=XM_023258174.1
C1 11793370 11793526 refGene_gn=EPHA2;refGene_tx=XM_023258174.1
C1 11794132 11794326 refGene_gn=EPHA2;refGene_tx=XM_023258174.1
C1 11794999 11795149 refGene_gn=EPHA2;refGene_tx=XM_023258174.1
C1 11795353 11795563 refGene_gn=EPHA2;refGene_tx=XM_023258174.1
C1 11795664 11795726 refGene_gn=EPHA2;refGene_tx=XM_023258174.1
C1 11795938 11796124 refGene_gn=EPHA2;refGene_tx=XM_023258174.1
C1 11796221 11796347 refGene_gn=EPHA2;refGene_tx=XM_023258174.1
C1 11796558 11796614 refGene_gn=EPHA2;refGene_tx=XM_023258174.1
C1 11797121 11797221 refGene_gn=EPHA2;refGene_tx=XM_023258174.1
C1 11797715 11797971 refGene_gn=EPHA2;refGene_tx=XM_023258174.1
C1 11798532 11798540 refGene_gn=EPHA2;refGene_tx=XM_023258174.1
C1 11798820 11799153 refGene_gn=EPHA2;refGene_tx=XM_023258174.1
C1 11799252 11799408 refGene_gn=EPHA2;refGene_tx=XM_023258174.1
C1 11807394 11808064 refGene_gn=EPHA2;refGene_tx=XM_023258174.1
C1 11809348 11809416 refGene_gn=EPHA2;refGene_tx=XM_023258174.1
C1 11813932 11814017 refGene_gn=EPHA2;refGene_tx=XM_023258174.1
C1 11847386 11847544 refGene_gn=ARHGEF19;refGene_tx=XM_023258177.1
C1 11847966 11848071 refGene_gn=ARHGEF19;refGene_tx=XM_023258177.1
C1 11849404 11849792
refGene_gn=ARHGEF19;refGene_tx=XM_023258175.1;refGene_gn=ARHGEF19;refG
ene_tx=XM_023258176.1;refGene_gn=ARHGEF19;refGene_tx=XM_023258177.1
C1 11849815 11849864 refGene_gn=ARHGEF19;refGene_tx=XM_023258179.1
C1 11850105 11850264
refGene_gn=ARHGEF19;refGene_tx=XM_023258175.1;refGene_gn=ARHGEF19;refG
ene_tx=XM_023258176.1;refGene_gn=ARHGEF19;refGene_tx=XM_023258177.1;re
fGene_gn=ARHGEF19;refGene_tx=XM_023258179.1
C1 11852227 11852388
refGene_gn=ARHGEF19;refGene_tx=XM_023258175.1;refGene_gn=ARHGEF19;refG
ene_tx=XM_023258176.1;refGene_gn=ARHGEF19;refGene_tx=XM_023258177.1;re
fGene_gn=ARHGEF19;refGene_tx=XM_023258179.1
C1 11852566 11852656
refGene_gn=ARHGEF19;refGene_tx=XM_023258175.1;refGene_gn=ARHGEF19;refG

ene_tx=XM_023258176.1;refGene_gn=ARHGEF19;refGene_tx=XM_023258177.1;refGene_gn=ARHGEF19;refGene_tx=XM_023258179.1
C1 11852824 11852899
refGene_gn=ARHGEF19;refGene_tx=XM_023258175.1;refGene_gn=ARHGEF19;refGene_tx=XM_023258176.1;refGene_gn=ARHGEF19;refGene_tx=XM_023258177.1;refGene_gn=ARHGEF19;refGene_tx=XM_023258179.1
C1 11853036 11853166
refGene_gn=ARHGEF19;refGene_tx=XM_023258175.1;refGene_gn=ARHGEF19;refGene_tx=XM_023258176.1;refGene_gn=ARHGEF19;refGene_tx=XM_023258177.1;refGene_gn=ARHGEF19;refGene_tx=XM_023258179.1
C1 11853465 11853618
refGene_gn=ARHGEF19;refGene_tx=XM_023258175.1;refGene_gn=ARHGEF19;refGene_tx=XM_023258176.1;refGene_gn=ARHGEF19;refGene_tx=XM_023258177.1;refGene_gn=ARHGEF19;refGene_tx=XM_023258179.1
C1 11853729 11853890
refGene_gn=ARHGEF19;refGene_tx=XM_023258175.1;refGene_gn=ARHGEF19;refGene_tx=XM_023258176.1;refGene_gn=ARHGEF19;refGene_tx=XM_023258177.1;refGene_gn=ARHGEF19;refGene_tx=XM_023258179.1
C1 11854674 11854937
refGene_gn=ARHGEF19;refGene_tx=XM_023258175.1;refGene_gn=ARHGEF19;refGene_tx=XM_023258176.1;refGene_gn=ARHGEF19;refGene_tx=XM_023258177.1;refGene_gn=ARHGEF19;refGene_tx=XM_023258179.1
C1 11855233 11855310
refGene_gn=ARHGEF19;refGene_tx=XM_023258175.1;refGene_gn=ARHGEF19;refGene_tx=XM_023258176.1;refGene_gn=ARHGEF19;refGene_tx=XM_023258177.1;refGene_gn=ARHGEF19;refGene_tx=XM_023258179.1
C1 11855386 11855495
refGene_gn=ARHGEF19;refGene_tx=XM_023258175.1;refGene_gn=ARHGEF19;refGene_tx=XM_023258176.1;refGene_gn=ARHGEF19;refGene_tx=XM_023258177.1;refGene_gn=ARHGEF19;refGene_tx=XM_023258179.1
C1 11855656 11855938
refGene_gn=ARHGEF19;refGene_tx=XM_023258175.1;refGene_gn=ARHGEF19;refGene_tx=XM_023258176.1;refGene_gn=ARHGEF19;refGene_tx=XM_023258177.1;refGene_gn=ARHGEF19;refGene_tx=XM_023258179.1
C1 11856354 11856769
refGene_gn=ARHGEF19;refGene_tx=XM_023258175.1;refGene_gn=ARHGEF19;refGene_tx=XM_023258176.1;refGene_gn=ARHGEF19;refGene_tx=XM_023258177.1;refGene_gn=ARHGEF19;refGene_tx=XM_023258179.1
C1 11874152 11874402 refGene_gn=RSG1;refGene_tx=XM_003989604.3
C1 11874636 11874773 refGene_gn=RSG1;refGene_tx=XM_003989604.3
C1 11875076 11875201 refGene_gn=RSG1;refGene_tx=XM_003989604.3
C1 11875751 11875904 refGene_gn=RSG1;refGene_tx=XM_003989604.3
C1 11877162 11877301 refGene_gn=RSG1;refGene_tx=XM_003989604.3
C1 11877719 11877722 refGene_gn=RSG1;refGene_tx=XM_003989604.3
C1 11891488 11892604
refGene_gn=FBXO42;refGene_tx=XM_011284587.3;refGene_gn=FBXO42;refGene_tx=XM_019836250.1
C1 11893486 11893603
refGene_gn=FBXO42;refGene_tx=XM_011284587.3;refGene_gn=FBXO42;refGene_tx=XM_019836250.1

C1 11894252 11894309
refGene_gn=FBXO42;refGene_tx=XM_011284587.3;refGene_gn=FBXO42;refGene_tx=XM_019836250.1
C1 11894795 11894892
refGene_gn=FBXO42;refGene_tx=XM_011284587.3;refGene_gn=FBXO42;refGene_tx=XM_019836250.1
C1 11896346 11896457
refGene_gn=FBXO42;refGene_tx=XM_011284587.3;refGene_gn=FBXO42;refGene_tx=XM_019836250.1
C1 11897630 11897784
refGene_gn=FBXO42;refGene_tx=XM_011284587.3;refGene_gn=FBXO42;refGene_tx=XM_019836250.1
C1 11913428 11913558 refGene_gn=FBXO42;refGene_tx=XM_019836250.1
C1 11920805 11920940 refGene_gn=FBXO42;refGene_tx=XM_011284587.3
C1 11932344 11932461 refGene_gn=FBXO42;refGene_tx=XM_011284587.3
C1 11935387 11935637 refGene_gn=FBXO42;refGene_tx=XM_011284587.3
C1 11996595 11996646
refGene_gn=SZRD1;refGene_tx=XM_023258188.1;refGene_gn=SZRD1;refGene_tx=XM_023258189.1
C1 12016263 12016313
refGene_gn=SZRD1;refGene_tx=XM_023258188.1;refGene_gn=SZRD1;refGene_tx=XM_023258189.1
C1 12018083 12018338
refGene_gn=SZRD1;refGene_tx=XM_023258188.1;refGene_gn=SZRD1;refGene_tx=XM_023258189.1
C1 12020470 12020573
refGene_gn=SZRD1;refGene_tx=XM_023258188.1;refGene_gn=SZRD1;refGene_tx=XM_023258189.1
C1 12024382 12024440 refGene_gn=SPATA21;refGene_tx=XM_023258186.1
C1 12024916 12025164
refGene_gn=SPATA21;refGene_tx=XM_023258180.1;refGene_gn=SPATA21;refGene_tx=XM_023258181.1;refGene_gn=SPATA21;refGene_tx=XM_023258182.1;refGene_gn=SPATA21;refGene_tx=XM_023258184.1;refGene_gn=SPATA21;refGene_tx=XM_023258185.1;refGene_gn=SPATA21;refGene_tx=XM_023258186.1
C1 12026044 12026220
refGene_gn=SPATA21;refGene_tx=XM_023258180.1;refGene_gn=SPATA21;refGene_tx=XM_023258181.1;refGene_gn=SPATA21;refGene_tx=XM_023258182.1;refGene_gn=SPATA21;refGene_tx=XM_023258184.1;refGene_gn=SPATA21;refGene_tx=XM_023258185.1;refGene_gn=SPATA21;refGene_tx=XM_023258186.1
C1 12039705 12039823
refGene_gn=SPATA21;refGene_tx=XM_023258180.1;refGene_gn=SPATA21;refGene_tx=XM_023258181.1;refGene_gn=SPATA21;refGene_tx=XM_023258182.1;refGene_gn=SPATA21;refGene_tx=XM_023258184.1;refGene_gn=SPATA21;refGene_tx=XM_023258185.1;refGene_gn=SPATA21;refGene_tx=XM_023258186.1
C1 12039938 12040010
refGene_gn=SPATA21;refGene_tx=XM_023258180.1;refGene_gn=SPATA21;refGene_tx=XM_023258181.1;refGene_gn=SPATA21;refGene_tx=XM_023258182.1;refGene_gn=SPATA21;refGene_tx=XM_023258184.1;refGene_gn=SPATA21;refGene_tx=XM_023258185.1;refGene_gn=SPATA21;refGene_tx=XM_023258186.1
C1 12042123 12042261

refGene_gn=SPATA21;refGene_tx=XM_023258180.1;refGene_gn=SPATA21;refGene_tx=XM_023258181.1;refGene_gn=SPATA21;refGene_tx=XM_023258182.1;refGene_gn=SPATA21;refGene_tx=XM_023258184.1;refGene_gn=SPATA21;refGene_tx=XM_023258185.1;refGene_gn=SPATA21;refGene_tx=XM_023258186.1
C1 12044122 12044211
refGene_gn=SPATA21;refGene_tx=XM_023258180.1;refGene_gn=SPATA21;refGene_tx=XM_023258181.1;refGene_gn=SPATA21;refGene_tx=XM_023258182.1;refGene_gn=SPATA21;refGene_tx=XM_023258184.1;refGene_gn=SPATA21;refGene_tx=XM_023258185.1;refGene_gn=SPATA21;refGene_tx=XM_023258186.1
C1 12044629 12045066
refGene_gn=SPATA21;refGene_tx=XM_023258180.1;refGene_gn=SPATA21;refGene_tx=XM_023258181.1;refGene_gn=SPATA21;refGene_tx=XM_023258182.1;refGene_gn=SPATA21;refGene_tx=XM_023258184.1;refGene_gn=SPATA21;refGene_tx=XM_023258185.1;refGene_gn=SPATA21;refGene_tx=XM_023258186.1
C1 12045699 12045748
refGene_gn=SPATA21;refGene_tx=XM_023258180.1;refGene_gn=SPATA21;refGene_tx=XM_023258181.1;refGene_gn=SPATA21;refGene_tx=XM_023258182.1;refGene_gn=SPATA21;refGene_tx=XM_023258184.1;refGene_gn=SPATA21;refGene_tx=XM_023258185.1;refGene_gn=SPATA21;refGene_tx=XM_023258186.1
C1 12046129 12046425
refGene_gn=SPATA21;refGene_tx=XM_023258180.1;refGene_gn=SPATA21;refGene_tx=XM_023258181.1;refGene_gn=SPATA21;refGene_tx=XM_023258182.1;refGene_gn=SPATA21;refGene_tx=XM_023258184.1;refGene_gn=SPATA21;refGene_tx=XM_023258185.1;refGene_gn=SPATA21;refGene_tx=XM_023258186.1
C1 12048705 12048925
refGene_gn=SPATA21;refGene_tx=XM_023258180.1;refGene_gn=SPATA21;refGene_tx=XM_023258181.1;refGene_gn=SPATA21;refGene_tx=XM_023258182.1;refGene_gn=SPATA21;refGene_tx=XM_023258184.1;refGene_gn=SPATA21;refGene_tx=XM_023258185.1;refGene_gn=SPATA21;refGene_tx=XM_023258186.1
C1 12050903 12051055
refGene_gn=SPATA21;refGene_tx=XM_023258180.1;refGene_gn=SPATA21;refGene_tx=XM_023258181.1;refGene_gn=SPATA21;refGene_tx=XM_023258182.1;refGene_gn=SPATA21;refGene_tx=XM_023258184.1;refGene_gn=SPATA21;refGene_tx=XM_023258185.1;refGene_gn=SPATA21;refGene_tx=XM_023258186.1
C1 12052661 12052790
refGene_gn=SPATA21;refGene_tx=XM_023258180.1;refGene_gn=SPATA21;refGene_tx=XM_023258181.1;refGene_gn=SPATA21;refGene_tx=XM_023258182.1;refGene_gn=SPATA21;refGene_tx=XM_023258184.1;refGene_gn=SPATA21;refGene_tx=XM_023258185.1;refGene_gn=SPATA21;refGene_tx=XM_023258186.1
C1 12062393 12062485
refGene_gn=NECAP2;refGene_tx=XM_003989520.5;refGene_gn=NECAP2;refGene_tx=XM_023258187.1
C1 12065037 12065138
refGene_gn=NECAP2;refGene_tx=XM_003989520.5;refGene_gn=NECAP2;refGene_tx=XM_023258187.1
C1 12067394 12067499
refGene_gn=NECAP2;refGene_tx=XM_003989520.5;refGene_gn=NECAP2;refGene_tx=XM_023258187.1
C1 12067585 12067667
refGene_gn=NECAP2;refGene_tx=XM_003989520.5;refGene_gn=NECAP2;refGene_

tx=XM_023258187.1
C1 12068571 12068680
refGene_gn=NECAP2;refGene_tx=XM_003989520.5;refGene_gn=NECAP2;refGene_
tx=XM_023258187.1
C1 12070584 12070762
refGene_gn=NECAP2;refGene_tx=XM_003989520.5;refGene_gn=NECAP2;refGene_
tx=XM_023258187.1
C1 12073402 12073490
refGene_gn=NECAP2;refGene_tx=XM_003989520.5;refGene_gn=NECAP2;refGene_
tx=XM_023258187.1
C1 12074212 12074261
refGene_gn=NECAP2;refGene_tx=XM_003989520.5;refGene_gn=NECAP2;refGene_
tx=XM_023258187.1
C1 12085634 12085694 refGene_gn=CROCC;refGene_tx=XM_023258193.1
C1 12086003 12086409
refGene_gn=CROCC;refGene_tx=XM_023258190.1;refGene_gn=CROCC;refGene_tx
=XM_023258191.1;refGene_gn=CROCC;refGene_tx=XM_023258192.1;refGene_gn=
CROCC;refGene_tx=XM_023258193.1
C1 12087612 12087767
refGene_gn=CROCC;refGene_tx=XM_023258190.1;refGene_gn=CROCC;refGene_tx
=XM_023258191.1;refGene_gn=CROCC;refGene_tx=XM_023258192.1;refGene_gn=
CROCC;refGene_tx=XM_023258193.1
C1 12092235 12092421
refGene_gn=CROCC;refGene_tx=XM_023258190.1;refGene_gn=CROCC;refGene_tx
=XM_023258191.1;refGene_gn=CROCC;refGene_tx=XM_023258193.1
C1 12092517 12092601
refGene_gn=CROCC;refGene_tx=XM_023258190.1;refGene_gn=CROCC;refGene_tx
=XM_023258191.1;refGene_gn=CROCC;refGene_tx=XM_023258192.1;refGene_gn=
CROCC;refGene_tx=XM_023258193.1
C1 12092703 12092765
refGene_gn=CROCC;refGene_tx=XM_023258190.1;refGene_gn=CROCC;refGene_tx
=XM_023258191.1;refGene_gn=CROCC;refGene_tx=XM_023258192.1;refGene_gn=
CROCC;refGene_tx=XM_023258193.1
C1 12092851 12093017
refGene_gn=CROCC;refGene_tx=XM_023258190.1;refGene_gn=CROCC;refGene_tx
=XM_023258191.1;refGene_gn=CROCC;refGene_tx=XM_023258192.1;refGene_gn=
CROCC;refGene_tx=XM_023258193.1;refGene_gn=CROCC;refGene_tx=XM_0232581
94.1
C1 12093630 12093737
refGene_gn=CROCC;refGene_tx=XM_023258190.1;refGene_gn=CROCC;refGene_tx
=XM_023258191.1;refGene_gn=CROCC;refGene_tx=XM_023258192.1;refGene_gn=
CROCC;refGene_tx=XM_023258193.1;refGene_gn=CROCC;refGene_tx=XM_0232581
94.1
C1 12097929 12098166
refGene_gn=CROCC;refGene_tx=XM_023258190.1;refGene_gn=CROCC;refGene_tx
=XM_023258191.1;refGene_gn=CROCC;refGene_tx=XM_023258192.1;refGene_gn=
CROCC;refGene_tx=XM_023258193.1;refGene_gn=CROCC;refGene_tx=XM_0232581
94.1
C1 12098868 12098965
refGene_gn=CROCC;refGene_tx=XM_023258190.1;refGene_gn=CROCC;refGene_tx

=XM_023258191.1;refGene_gn=CROCC;refGene_tx=XM_023258192.1;refGene_gn=CROCC;refGene_tx=XM_023258193.1;refGene_gn=CROCC;refGene_tx=XM_023258194.1
C1 12099632 12099716
refGene_gn=CROCC;refGene_tx=XM_023258190.1;refGene_gn=CROCC;refGene_tx=XM_023258191.1;refGene_gn=CROCC;refGene_tx=XM_023258192.1;refGene_gn=CROCC;refGene_tx=XM_023258193.1;refGene_gn=CROCC;refGene_tx=XM_023258194.1
C1 12100155 12100389
refGene_gn=CROCC;refGene_tx=XM_023258190.1;refGene_gn=CROCC;refGene_tx=XM_023258191.1;refGene_gn=CROCC;refGene_tx=XM_023258192.1;refGene_gn=CROCC;refGene_tx=XM_023258193.1;refGene_gn=CROCC;refGene_tx=XM_023258194.1
C1 12101132 12101332
refGene_gn=CROCC;refGene_tx=XM_023258190.1;refGene_gn=CROCC;refGene_tx=XM_023258191.1;refGene_gn=CROCC;refGene_tx=XM_023258192.1;refGene_gn=CROCC;refGene_tx=XM_023258193.1;refGene_gn=CROCC;refGene_tx=XM_023258194.1
C1 12103181 12103364
refGene_gn=CROCC;refGene_tx=XM_023258190.1;refGene_gn=CROCC;refGene_tx=XM_023258191.1;refGene_gn=CROCC;refGene_tx=XM_023258192.1;refGene_gn=CROCC;refGene_tx=XM_023258193.1;refGene_gn=CROCC;refGene_tx=XM_023258194.1
C1 12104692 12104837
refGene_gn=CROCC;refGene_tx=XM_023258190.1;refGene_gn=CROCC;refGene_tx=XM_023258191.1;refGene_gn=CROCC;refGene_tx=XM_023258192.1;refGene_gn=CROCC;refGene_tx=XM_023258193.1;refGene_gn=CROCC;refGene_tx=XM_023258194.1
C1 12105479 12105626
refGene_gn=CROCC;refGene_tx=XM_023258190.1;refGene_gn=CROCC;refGene_tx=XM_023258191.1;refGene_gn=CROCC;refGene_tx=XM_023258192.1;refGene_gn=CROCC;refGene_tx=XM_023258193.1;refGene_gn=CROCC;refGene_tx=XM_023258194.1
C1 12106070 12106301
refGene_gn=CROCC;refGene_tx=XM_023258190.1;refGene_gn=CROCC;refGene_tx=XM_023258191.1;refGene_gn=CROCC;refGene_tx=XM_023258192.1;refGene_gn=CROCC;refGene_tx=XM_023258193.1;refGene_gn=CROCC;refGene_tx=XM_023258194.1
C1 12107485 12107679
refGene_gn=CROCC;refGene_tx=XM_023258190.1;refGene_gn=CROCC;refGene_tx=XM_023258191.1;refGene_gn=CROCC;refGene_tx=XM_023258192.1;refGene_gn=CROCC;refGene_tx=XM_023258193.1;refGene_gn=CROCC;refGene_tx=XM_023258194.1
C1 12107964 12108092
refGene_gn=CROCC;refGene_tx=XM_023258190.1;refGene_gn=CROCC;refGene_tx=XM_023258191.1;refGene_gn=CROCC;refGene_tx=XM_023258192.1;refGene_gn=CROCC;refGene_tx=XM_023258193.1;refGene_gn=CROCC;refGene_tx=XM_023258194.1
C1 12109229 12109399
refGene_gn=CROCC;refGene_tx=XM_023258190.1;refGene_gn=CROCC;refGene_tx

=XM_023258191.1;refGene_gn=CROCC;refGene_tx=XM_023258192.1;refGene_gn=CROCC;refGene_tx=XM_023258193.1;refGene_gn=CROCC;refGene_tx=XM_023258194.1
C1 12111181 12111361
refGene_gn=CROCC;refGene_tx=XM_023258190.1;refGene_gn=CROCC;refGene_tx=XM_023258191.1;refGene_gn=CROCC;refGene_tx=XM_023258192.1;refGene_gn=CROCC;refGene_tx=XM_023258193.1;refGene_gn=CROCC;refGene_tx=XM_023258194.1
C1 12112101 12112236
refGene_gn=CROCC;refGene_tx=XM_023258190.1;refGene_gn=CROCC;refGene_tx=XM_023258191.1;refGene_gn=CROCC;refGene_tx=XM_023258192.1;refGene_gn=CROCC;refGene_tx=XM_023258193.1;refGene_gn=CROCC;refGene_tx=XM_023258194.1
C1 12112555 12112699
refGene_gn=CROCC;refGene_tx=XM_023258190.1;refGene_gn=CROCC;refGene_tx=XM_023258191.1;refGene_gn=CROCC;refGene_tx=XM_023258192.1;refGene_gn=CROCC;refGene_tx=XM_023258193.1;refGene_gn=CROCC;refGene_tx=XM_023258194.1
C1 12113119 12113358
refGene_gn=CROCC;refGene_tx=XM_023258190.1;refGene_gn=CROCC;refGene_tx=XM_023258191.1;refGene_gn=CROCC;refGene_tx=XM_023258192.1;refGene_gn=CROCC;refGene_tx=XM_023258193.1;refGene_gn=CROCC;refGene_tx=XM_023258194.1
C1 12114076 12114236
refGene_gn=CROCC;refGene_tx=XM_023258190.1;refGene_gn=CROCC;refGene_tx=XM_023258191.1;refGene_gn=CROCC;refGene_tx=XM_023258192.1;refGene_gn=CROCC;refGene_tx=XM_023258193.1;refGene_gn=CROCC;refGene_tx=XM_023258194.1
C1 12115741 12115909
refGene_gn=CROCC;refGene_tx=XM_023258190.1;refGene_gn=CROCC;refGene_tx=XM_023258191.1;refGene_gn=CROCC;refGene_tx=XM_023258192.1;refGene_gn=CROCC;refGene_tx=XM_023258193.1;refGene_gn=CROCC;refGene_tx=XM_023258194.1
C1 12117384 12117757
refGene_gn=CROCC;refGene_tx=XM_023258190.1;refGene_gn=CROCC;refGene_tx=XM_023258191.1;refGene_gn=CROCC;refGene_tx=XM_023258192.1;refGene_gn=CROCC;refGene_tx=XM_023258193.1;refGene_gn=CROCC;refGene_tx=XM_023258194.1
C1 12123712 12123882
refGene_gn=CROCC;refGene_tx=XM_023258190.1;refGene_gn=CROCC;refGene_tx=XM_023258191.1;refGene_gn=CROCC;refGene_tx=XM_023258192.1;refGene_gn=CROCC;refGene_tx=XM_023258193.1;refGene_gn=CROCC;refGene_tx=XM_023258194.1
C1 12123999 12124120
refGene_gn=CROCC;refGene_tx=XM_023258190.1;refGene_gn=CROCC;refGene_tx=XM_023258191.1;refGene_gn=CROCC;refGene_tx=XM_023258192.1;refGene_gn=CROCC;refGene_tx=XM_023258193.1;refGene_gn=CROCC;refGene_tx=XM_023258194.1
C1 12124423 12124587
refGene_gn=CROCC;refGene_tx=XM_023258190.1;refGene_gn=CROCC;refGene_tx

=XM_023258191.1;refGene_gn=CROCC;refGene_tx=XM_023258192.1;refGene_gn=CROCC;refGene_tx=XM_023258193.1;refGene_gn=CROCC;refGene_tx=XM_023258194.1
C1 12126215 12126431
refGene_gn=CROCC;refGene_tx=XM_023258190.1;refGene_gn=CROCC;refGene_tx=XM_023258191.1;refGene_gn=CROCC;refGene_tx=XM_023258192.1;refGene_gn=CROCC;refGene_tx=XM_023258193.1;refGene_gn=CROCC;refGene_tx=XM_023258194.1
C1 12127081 12127306
refGene_gn=CROCC;refGene_tx=XM_023258190.1;refGene_gn=CROCC;refGene_tx=XM_023258191.1;refGene_gn=CROCC;refGene_tx=XM_023258192.1;refGene_gn=CROCC;refGene_tx=XM_023258193.1;refGene_gn=CROCC;refGene_tx=XM_023258194.1
C1 12127685 12127835
refGene_gn=CROCC;refGene_tx=XM_023258190.1;refGene_gn=CROCC;refGene_tx=XM_023258191.1;refGene_gn=CROCC;refGene_tx=XM_023258192.1;refGene_gn=CROCC;refGene_tx=XM_023258193.1;refGene_gn=CROCC;refGene_tx=XM_023258194.1
C1 12128148 12128349
refGene_gn=CROCC;refGene_tx=XM_023258190.1;refGene_gn=CROCC;refGene_tx=XM_023258191.1;refGene_gn=CROCC;refGene_tx=XM_023258192.1;refGene_gn=CROCC;refGene_tx=XM_023258193.1;refGene_gn=CROCC;refGene_tx=XM_023258194.1
C1 12128551 12128683
refGene_gn=CROCC;refGene_tx=XM_023258190.1;refGene_gn=CROCC;refGene_tx=XM_023258191.1;refGene_gn=CROCC;refGene_tx=XM_023258192.1;refGene_gn=CROCC;refGene_tx=XM_023258193.1;refGene_gn=CROCC;refGene_tx=XM_023258194.1
C1 12129422 12129689
refGene_gn=CROCC;refGene_tx=XM_023258190.1;refGene_gn=CROCC;refGene_tx=XM_023258191.1;refGene_gn=CROCC;refGene_tx=XM_023258192.1;refGene_gn=CROCC;refGene_tx=XM_023258193.1;refGene_gn=CROCC;refGene_tx=XM_023258194.1
C1 12130309 12130396
refGene_gn=CROCC;refGene_tx=XM_023258190.1;refGene_gn=CROCC;refGene_tx=XM_023258191.1;refGene_gn=CROCC;refGene_tx=XM_023258193.1
C1 12132380 12132484 refGene_gn=MFAP2;refGene_tx=XM_003989522.5
C1 12132723 12132797 refGene_gn=MFAP2;refGene_tx=XM_003989522.5
C1 12132985 12133073 refGene_gn=MFAP2;refGene_tx=XM_003989522.5
C1 12133876 12133921 refGene_gn=MFAP2;refGene_tx=XM_003989522.5
C1 12134034 12134115 refGene_gn=MFAP2;refGene_tx=XM_003989522.5
C1 12134211 12134238 refGene_gn=MFAP2;refGene_tx=XM_003989522.5
C1 12134450 12134540 refGene_gn=MFAP2;refGene_tx=XM_003989522.5
C1 12135499 12135536 refGene_gn=MFAP2;refGene_tx=XM_003989522.5
C1 12141919 12142054
refGene_gn=ATP13A2;refGene_tx=XM_023258195.1;refGene_gn=ATP13A2;refGene_tx=XM_023258196.1;refGene_gn=ATP13A2;refGene_tx=XM_023258197.1;refGene_gn=ATP13A2;refGene_tx=XM_023258198.1;refGene_gn=ATP13A2;refGene_tx=XM_023258199.1;refGene_gn=ATP13A2;refGene_tx=XM_023258200.1
C1 12142146 12142316

refGene_gn=ATP13A2;refGene_tx=XM_023258195.1;refGene_gn=ATP13A2;refGene_tx=XM_023258196.1;refGene_gn=ATP13A2;refGene_tx=XM_023258197.1;refGene_gn=ATP13A2;refGene_tx=XM_023258198.1;refGene_gn=ATP13A2;refGene_tx=XM_023258199.1;refGene_gn=ATP13A2;refGene_tx=XM_023258200.1
C1 12142403 12142555
refGene_gn=ATP13A2;refGene_tx=XM_023258195.1;refGene_gn=ATP13A2;refGene_tx=XM_023258196.1;refGene_gn=ATP13A2;refGene_tx=XM_023258197.1;refGene_gn=ATP13A2;refGene_tx=XM_023258198.1;refGene_gn=ATP13A2;refGene_tx=XM_023258199.1;refGene_gn=ATP13A2;refGene_tx=XM_023258200.1
C1 12142637 12142861
refGene_gn=ATP13A2;refGene_tx=XM_023258195.1;refGene_gn=ATP13A2;refGene_tx=XM_023258196.1;refGene_gn=ATP13A2;refGene_tx=XM_023258197.1;refGene_gn=ATP13A2;refGene_tx=XM_023258198.1;refGene_gn=ATP13A2;refGene_tx=XM_023258199.1;refGene_gn=ATP13A2;refGene_tx=XM_023258200.1
C1 12143677 12143774
refGene_gn=ATP13A2;refGene_tx=XM_023258195.1;refGene_gn=ATP13A2;refGene_tx=XM_023258196.1;refGene_gn=ATP13A2;refGene_tx=XM_023258197.1;refGene_gn=ATP13A2;refGene_tx=XM_023258198.1;refGene_gn=ATP13A2;refGene_tx=XM_023258199.1;refGene_gn=ATP13A2;refGene_tx=XM_023258200.1
C1 12143862 12144015
refGene_gn=ATP13A2;refGene_tx=XM_023258195.1;refGene_gn=ATP13A2;refGene_tx=XM_023258196.1;refGene_gn=ATP13A2;refGene_tx=XM_023258197.1;refGene_gn=ATP13A2;refGene_tx=XM_023258198.1;refGene_gn=ATP13A2;refGene_tx=XM_023258199.1;refGene_gn=ATP13A2;refGene_tx=XM_023258200.1
C1 12144331 12144411
refGene_gn=ATP13A2;refGene_tx=XM_023258195.1;refGene_gn=ATP13A2;refGene_tx=XM_023258196.1;refGene_gn=ATP13A2;refGene_tx=XM_023258197.1;refGene_gn=ATP13A2;refGene_tx=XM_023258198.1;refGene_gn=ATP13A2;refGene_tx=XM_023258199.1;refGene_gn=ATP13A2;refGene_tx=XM_023258200.1
C1 12144537 12144654
refGene_gn=ATP13A2;refGene_tx=XM_023258195.1;refGene_gn=ATP13A2;refGene_tx=XM_023258196.1;refGene_gn=ATP13A2;refGene_tx=XM_023258197.1;refGene_gn=ATP13A2;refGene_tx=XM_023258199.1;refGene_gn=ATP13A2;refGene_tx=XM_023258200.1
C1 12144778 12144939
refGene_gn=ATP13A2;refGene_tx=XM_023258195.1;refGene_gn=ATP13A2;refGene_tx=XM_023258196.1;refGene_gn=ATP13A2;refGene_tx=XM_023258197.1;refGene_gn=ATP13A2;refGene_tx=XM_023258198.1;refGene_gn=ATP13A2;refGene_tx=XM_023258199.1;refGene_gn=ATP13A2;refGene_tx=XM_023258200.1
C1 12145678 12145803
refGene_gn=ATP13A2;refGene_tx=XM_023258195.1;refGene_gn=ATP13A2;refGene_tx=XM_023258196.1;refGene_gn=ATP13A2;refGene_tx=XM_023258197.1;refGene_gn=ATP13A2;refGene_tx=XM_023258198.1;refGene_gn=ATP13A2;refGene_tx=XM_023258199.1;refGene_gn=ATP13A2;refGene_tx=XM_023258200.1
C1 12145947 12146068
refGene_gn=ATP13A2;refGene_tx=XM_023258195.1;refGene_gn=ATP13A2;refGene_tx=XM_023258196.1;refGene_gn=ATP13A2;refGene_tx=XM_023258197.1;refGene_gn=ATP13A2;refGene_tx=XM_023258198.1;refGene_gn=ATP13A2;refGene_tx=XM_023258199.1;refGene_gn=ATP13A2;refGene_tx=XM_023258200.1
C1 12146171 12146331

refGene_gn=ATP13A2;refGene_tx=XM_023258195.1;refGene_gn=ATP13A2;refGene_tx=XM_023258196.1;refGene_gn=ATP13A2;refGene_tx=XM_023258197.1;refGene_gn=ATP13A2;refGene_tx=XM_023258198.1;refGene_gn=ATP13A2;refGene_tx=XM_023258199.1;refGene_gn=ATP13A2;refGene_tx=XM_023258200.1
C1 12146417 12146513
refGene_gn=ATP13A2;refGene_tx=XM_023258195.1;refGene_gn=ATP13A2;refGene_tx=XM_023258196.1;refGene_gn=ATP13A2;refGene_tx=XM_023258197.1;refGene_gn=ATP13A2;refGene_tx=XM_023258198.1;refGene_gn=ATP13A2;refGene_tx=XM_023258199.1;refGene_gn=ATP13A2;refGene_tx=XM_023258200.1
C1 12146965 12147172
refGene_gn=ATP13A2;refGene_tx=XM_023258195.1;refGene_gn=ATP13A2;refGene_tx=XM_023258196.1;refGene_gn=ATP13A2;refGene_tx=XM_023258197.1;refGene_gn=ATP13A2;refGene_tx=XM_023258198.1;refGene_gn=ATP13A2;refGene_tx=XM_023258199.1
C1 12148020 12148209
refGene_gn=ATP13A2;refGene_tx=XM_023258195.1;refGene_gn=ATP13A2;refGene_tx=XM_023258196.1;refGene_gn=ATP13A2;refGene_tx=XM_023258197.1;refGene_gn=ATP13A2;refGene_tx=XM_023258198.1;refGene_gn=ATP13A2;refGene_tx=XM_023258199.1;refGene_gn=ATP13A2;refGene_tx=XM_023258200.1
C1 12148293 12148340
refGene_gn=ATP13A2;refGene_tx=XM_023258195.1;refGene_gn=ATP13A2;refGene_tx=XM_023258196.1;refGene_gn=ATP13A2;refGene_tx=XM_023258197.1;refGene_gn=ATP13A2;refGene_tx=XM_023258198.1;refGene_gn=ATP13A2;refGene_tx=XM_023258199.1;refGene_gn=ATP13A2;refGene_tx=XM_023258200.1
C1 12148426 12148537
refGene_gn=ATP13A2;refGene_tx=XM_023258195.1;refGene_gn=ATP13A2;refGene_tx=XM_023258196.1;refGene_gn=ATP13A2;refGene_tx=XM_023258197.1;refGene_gn=ATP13A2;refGene_tx=XM_023258198.1;refGene_gn=ATP13A2;refGene_tx=XM_023258199.1;refGene_gn=ATP13A2;refGene_tx=XM_023258200.1
C1 12149012 12149168
refGene_gn=ATP13A2;refGene_tx=XM_023258195.1;refGene_gn=ATP13A2;refGene_tx=XM_023258196.1;refGene_gn=ATP13A2;refGene_tx=XM_023258197.1;refGene_gn=ATP13A2;refGene_tx=XM_023258198.1;refGene_gn=ATP13A2;refGene_tx=XM_023258199.1;refGene_gn=ATP13A2;refGene_tx=XM_023258200.1
C1 12150048 12150180
refGene_gn=ATP13A2;refGene_tx=XM_023258195.1;refGene_gn=ATP13A2;refGene_tx=XM_023258196.1;refGene_gn=ATP13A2;refGene_tx=XM_023258197.1;refGene_gn=ATP13A2;refGene_tx=XM_023258198.1;refGene_gn=ATP13A2;refGene_tx=XM_023258199.1;refGene_gn=ATP13A2;refGene_tx=XM_023258200.1
C1 12150258 12150325
refGene_gn=ATP13A2;refGene_tx=XM_023258195.1;refGene_gn=ATP13A2;refGene_tx=XM_023258196.1;refGene_gn=ATP13A2;refGene_tx=XM_023258197.1;refGene_gn=ATP13A2;refGene_tx=XM_023258198.1;refGene_gn=ATP13A2;refGene_tx=XM_023258199.1;refGene_gn=ATP13A2;refGene_tx=XM_023258200.1
C1 12150414 12150549
refGene_gn=ATP13A2;refGene_tx=XM_023258195.1;refGene_gn=ATP13A2;refGene_tx=XM_023258196.1;refGene_gn=ATP13A2;refGene_tx=XM_023258197.1;refGene_gn=ATP13A2;refGene_tx=XM_023258198.1;refGene_gn=ATP13A2;refGene_tx=XM_023258199.1;refGene_gn=ATP13A2;refGene_tx=XM_023258200.1
C1 12151315 12151385

refGene_gn=ATP13A2;refGene_tx=XM_023258195.1;refGene_gn=ATP13A2;refGene_tx=XM_023258196.1;refGene_gn=ATP13A2;refGene_tx=XM_023258197.1;refGene_gn=ATP13A2;refGene_tx=XM_023258198.1;refGene_gn=ATP13A2;refGene_tx=XM_023258199.1;refGene_gn=ATP13A2;refGene_tx=XM_023258200.1
C1 12151564 12151642
refGene_gn=ATP13A2;refGene_tx=XM_023258195.1;refGene_gn=ATP13A2;refGene_tx=XM_023258196.1;refGene_gn=ATP13A2;refGene_tx=XM_023258197.1;refGene_gn=ATP13A2;refGene_tx=XM_023258198.1;refGene_gn=ATP13A2;refGene_tx=XM_023258199.1;refGene_gn=ATP13A2;refGene_tx=XM_023258200.1
C1 12152825 12152905
refGene_gn=ATP13A2;refGene_tx=XM_023258195.1;refGene_gn=ATP13A2;refGene_tx=XM_023258196.1;refGene_gn=ATP13A2;refGene_tx=XM_023258197.1;refGene_gn=ATP13A2;refGene_tx=XM_023258198.1;refGene_gn=ATP13A2;refGene_tx=XM_023258199.1;refGene_gn=ATP13A2;refGene_tx=XM_023258200.1
C1 12153166 12153281
refGene_gn=ATP13A2;refGene_tx=XM_023258195.1;refGene_gn=ATP13A2;refGene_tx=XM_023258196.1;refGene_gn=ATP13A2;refGene_tx=XM_023258197.1;refGene_gn=ATP13A2;refGene_tx=XM_023258198.1;refGene_gn=ATP13A2;refGene_tx=XM_023258199.1;refGene_gn=ATP13A2;refGene_tx=XM_023258200.1
C1 12153449 12153508
refGene_gn=ATP13A2;refGene_tx=XM_023258195.1;refGene_gn=ATP13A2;refGene_tx=XM_023258196.1;refGene_gn=ATP13A2;refGene_tx=XM_023258197.1;refGene_gn=ATP13A2;refGene_tx=XM_023258198.1;refGene_gn=ATP13A2;refGene_tx=XM_023258199.1;refGene_gn=ATP13A2;refGene_tx=XM_023258200.1
C1 12153843 12154026
refGene_gn=ATP13A2;refGene_tx=XM_023258195.1;refGene_gn=ATP13A2;refGene_tx=XM_023258196.1;refGene_gn=ATP13A2;refGene_tx=XM_023258197.1;refGene_gn=ATP13A2;refGene_tx=XM_023258198.1;refGene_gn=ATP13A2;refGene_tx=XM_023258199.1;refGene_gn=ATP13A2;refGene_tx=XM_023258200.1
C1 12154127 12154222
refGene_gn=ATP13A2;refGene_tx=XM_023258195.1;refGene_gn=ATP13A2;refGene_tx=XM_023258196.1;refGene_gn=ATP13A2;refGene_tx=XM_023258197.1;refGene_gn=ATP13A2;refGene_tx=XM_023258198.1;refGene_gn=ATP13A2;refGene_tx=XM_023258199.1;refGene_gn=ATP13A2;refGene_tx=XM_023258200.1
C1 12160656 12160666
refGene_gn=ATP13A2;refGene_tx=XM_023258195.1;refGene_gn=ATP13A2;refGene_tx=XM_023258196.1;refGene_gn=ATP13A2;refGene_tx=XM_023258197.1;refGene_gn=ATP13A2;refGene_tx=XM_023258198.1;refGene_gn=ATP13A2;refGene_tx=XM_023258199.1;refGene_gn=ATP13A2;refGene_tx=XM_023258200.1
C1 12166638 12166761 refGene_gn=SDHB;refGene_tx=XM_019836254.2
C1 12167225 12167303 refGene_gn=SDHB;refGene_tx=XM_003989523.3
C1 12170791 12170914
refGene_gn=SDHB;refGene_tx=XM_003989523.3;refGene_gn=SDHB;refGene_tx=XM_019836254.2
C1 12172147 12172249
refGene_gn=SDHB;refGene_tx=XM_003989523.3;refGene_gn=SDHB;refGene_tx=XM_019836254.2
C1 12173362 12173479
refGene_gn=SDHB;refGene_tx=XM_003989523.3;refGene_gn=SDHB;refGene_tx=XM_019836254.2

C1 12174121 12174258
refGene_gn=SDHB;refGene_tx=XM_003989523.3;refGene_gn=SDHB;refGene_tx=XM_019836254.2
C1 12176732 12176818
refGene_gn=SDHB;refGene_tx=XM_003989523.3;refGene_gn=SDHB;refGene_tx=XM_019836254.2
C1 12187079 12187207
refGene_gn=SDHB;refGene_tx=XM_003989523.3;refGene_gn=SDHB;refGene_tx=XM_019836254.2
C1 12197240 12197315 refGene_gn=SDHB;refGene_tx=XM_003989523.3
C1 12211145 12211379 refGene_gn=PADI2;refGene_tx=XM_003989524.5
C1 12212025 12212154 refGene_gn=PADI2;refGene_tx=XM_003989524.5
C1 12212849 12212935 refGene_gn=PADI2;refGene_tx=XM_003989524.5
C1 12214649 12214743 refGene_gn=PADI2;refGene_tx=XM_003989524.5
C1 12215435 12215580 refGene_gn=PADI2;refGene_tx=XM_003989524.5
C1 12217381 12217533 refGene_gn=PADI2;refGene_tx=XM_003989524.5
C1 12220254 12220362 refGene_gn=PADI2;refGene_tx=XM_003989524.5
C1 12220877 12220989 refGene_gn=PADI2;refGene_tx=XM_003989524.5
C1 12221663 12221767 refGene_gn=PADI2;refGene_tx=XM_003989524.5
C1 12223511 12223690 refGene_gn=PADI2;refGene_tx=XM_003989524.5
C1 12229494 12229620 refGene_gn=PADI2;refGene_tx=XM_003989524.5
C1 12230077 12230195 refGene_gn=PADI2;refGene_tx=XM_003989524.5
C1 12232288 12232350 refGene_gn=PADI2;refGene_tx=XM_003989524.5
C1 12237897 12237970 refGene_gn=PADI2;refGene_tx=XM_003989524.5
C1 12238884 12239068 refGene_gn=PADI2;refGene_tx=XM_003989524.5
C1 12251910 12252002 refGene_gn=PADI2;refGene_tx=XM_003989524.5
C1 12324481 12324573 refGene_gn=PADI1;refGene_tx=XM_003989525.2
C1 12341537 12341718 refGene_gn=PADI1;refGene_tx=XM_003989525.2
C1 12342869 12342942 refGene_gn=PADI1;refGene_tx=XM_003989525.2
C1 12343568 12343630 refGene_gn=PADI1;refGene_tx=XM_003989525.2
C1 12344956 12345074 refGene_gn=PADI1;refGene_tx=XM_003989525.2
C1 12345219 12345345 refGene_gn=PADI1;refGene_tx=XM_003989525.2
C1 12347024 12347197 refGene_gn=PADI1;refGene_tx=XM_003989525.2
C1 12347346 12347450 refGene_gn=PADI1;refGene_tx=XM_003989525.2
C1 12348424 12348548 refGene_gn=PADI1;refGene_tx=XM_003989525.2
C1 12348923 12349031 refGene_gn=PADI1;refGene_tx=XM_003989525.2
C1 12350592 12350744 refGene_gn=PADI1;refGene_tx=XM_003989525.2
C1 12355703 12355848 refGene_gn=PADI1;refGene_tx=XM_003989525.2
C1 12356643 12356737 refGene_gn=PADI1;refGene_tx=XM_003989525.2
C1 12357734 12357814 refGene_gn=PADI1;refGene_tx=XM_003989525.2
C1 12358565 12358691 refGene_gn=PADI1;refGene_tx=XM_003989525.2
C1 12361522 12361756 refGene_gn=PADI1;refGene_tx=XM_003989525.2
C1 12365767 12365859 refGene_gn=PADI3;refGene_tx=XM_011284582.3
C1 12373348 12373529 refGene_gn=PADI3;refGene_tx=XM_011284582.3
C1 12376073 12376146 refGene_gn=PADI3;refGene_tx=XM_011284582.3
C1 12379579 12379641 refGene_gn=PADI3;refGene_tx=XM_011284582.3
C1 12380607 12380725 refGene_gn=PADI3;refGene_tx=XM_011284582.3
C1 12381354 12381480 refGene_gn=PADI3;refGene_tx=XM_011284582.3
C1 12383933 12384112 refGene_gn=PADI3;refGene_tx=XM_011284582.3
C1 12384337 12384441 refGene_gn=PADI3;refGene_tx=XM_011284582.3

C1 12384558 12384670 refGene_gn=PADI3;refGene_tx=XM_011284582.3
C1 12386437 12386545 refGene_gn=PADI3;refGene_tx=XM_011284582.3
C1 12387239 12387391 refGene_gn=PADI3;refGene_tx=XM_011284582.3
C1 12388906 12389051 refGene_gn=PADI3;refGene_tx=XM_011284582.3
C1 12389150 12389253 refGene_gn=PADI3;refGene_tx=XM_011284582.3
C1 12391634 12391714 refGene_gn=PADI3;refGene_tx=XM_011284582.3
C1 12393792 12393918 refGene_gn=PADI3;refGene_tx=XM_011284582.3
C1 12395600 12395834 refGene_gn=PADI3;refGene_tx=XM_011284582.3
C1 12418606 12418698
refGene_gn=PADI4;refGene_tx=XM_003989527.4;refGene_gn=PADI4;refGene_tx
=XM_019836256.2;refGene_gn=PADI4;refGene_tx=XM_023258201.1
C1 12429565 12429746
refGene_gn=PADI4;refGene_tx=XM_003989527.4;refGene_gn=PADI4;refGene_tx
=XM_019836256.2;refGene_gn=PADI4;refGene_tx=XM_023258201.1
C1 12432047 12432114
refGene_gn=PADI4;refGene_tx=XM_003989527.4;refGene_gn=PADI4;refGene_tx
=XM_019836256.2;refGene_gn=PADI4;refGene_tx=XM_023258201.1
C1 12433653 12433721
refGene_gn=PADI4;refGene_tx=XM_003989527.4;refGene_gn=PADI4;refGene_tx
=XM_019836256.2;refGene_gn=PADI4;refGene_tx=XM_023258201.1
C1 12435405 12435523
refGene_gn=PADI4;refGene_tx=XM_003989527.4;refGene_gn=PADI4;refGene_tx
=XM_019836256.2;refGene_gn=PADI4;refGene_tx=XM_023258201.1
C1 12437613 12437739
refGene_gn=PADI4;refGene_tx=XM_003989527.4;refGene_gn=PADI4;refGene_tx
=XM_019836256.2;refGene_gn=PADI4;refGene_tx=XM_023258201.1
C1 12439188 12439367
refGene_gn=PADI4;refGene_tx=XM_003989527.4;refGene_gn=PADI4;refGene_tx
=XM_019836256.2;refGene_gn=PADI4;refGene_tx=XM_023258201.1
C1 12439534 12439638
refGene_gn=PADI4;refGene_tx=XM_003989527.4;refGene_gn=PADI4;refGene_tx
=XM_019836256.2;refGene_gn=PADI4;refGene_tx=XM_023258201.1
C1 12440578 12440690
refGene_gn=PADI4;refGene_tx=XM_003989527.4;refGene_gn=PADI4;refGene_tx
=XM_019836256.2;refGene_gn=PADI4;refGene_tx=XM_023258201.1
C1 12442051 12442159
refGene_gn=PADI4;refGene_tx=XM_003989527.4;refGene_gn=PADI4;refGene_tx
=XM_019836256.2;refGene_gn=PADI4;refGene_tx=XM_023258201.1
C1 12447351 12447506
refGene_gn=PADI4;refGene_tx=XM_003989527.4;refGene_gn=PADI4;refGene_tx
=XM_019836256.2;refGene_gn=PADI4;refGene_tx=XM_023258201.1
C1 12448251 12448390
refGene_gn=PADI4;refGene_tx=XM_003989527.4;refGene_gn=PADI4;refGene_tx
=XM_019836256.2;refGene_gn=PADI4;refGene_tx=XM_023258201.1
C1 12448620 12448723
refGene_gn=PADI4;refGene_tx=XM_003989527.4;refGene_gn=PADI4;refGene_tx
=XM_019836256.2;refGene_gn=PADI4;refGene_tx=XM_023258201.1
C1 12449549 12449617
refGene_gn=PADI4;refGene_tx=XM_003989527.4;refGene_gn=PADI4;refGene_tx
=XM_019836256.2;refGene_gn=PADI4;refGene_tx=XM_023258201.1

C1 12450216 12450357
refGene_gn=PADI4;refGene_tx=XM_003989527.4;refGene_gn=PADI4;refGene_tx=XM_019836256.2;refGene_gn=PADI4;refGene_tx=XM_023258201.1
C1 12450531 12450639 refGene_gn=PADI4;refGene_tx=XM_019836256.2
C1 12451141 12451225 refGene_gn=PADI4;refGene_tx=XM_023258201.1
C1 12452217 12452451 refGene_gn=PADI4;refGene_tx=XM_003989527.4
C1 12457105 12457197 refGene_gn=PADI6;refGene_tx=NM_001079697.2
C1 12457897 12458081 refGene_gn=PADI6;refGene_tx=NM_001079697.2
C1 12459903 12459976 refGene_gn=PADI6;refGene_tx=NM_001079697.2
C1 12462468 12462536 refGene_gn=PADI6;refGene_tx=NM_001079697.2
C1 12462997 12463112 refGene_gn=PADI6;refGene_tx=NM_001079697.2
C1 12464148 12464274 refGene_gn=PADI6;refGene_tx=NM_001079697.2
C1 12468477 12468656 refGene_gn=PADI6;refGene_tx=NM_001079697.2
C1 12468917 12469021 refGene_gn=PADI6;refGene_tx=NM_001079697.2
C1 12472166 12472278 refGene_gn=PADI6;refGene_tx=NM_001079697.2
C1 12473666 12473774 refGene_gn=PADI6;refGene_tx=NM_001079697.2
C1 12473992 12474147 refGene_gn=PADI6;refGene_tx=NM_001079697.2
C1 12474933 12475090 refGene_gn=PADI6;refGene_tx=NM_001079697.2
C1 12475208 12475332 refGene_gn=PADI6;refGene_tx=NM_001079697.2
C1 12476390 12476461 refGene_gn=PADI6;refGene_tx=NM_001079697.2
C1 12477524 12477686 refGene_gn=PADI6;refGene_tx=NM_001079697.2
C1 12478515 12478761 refGene_gn=PADI6;refGene_tx=NM_001079697.2
C1 12482381 12482486
refGene_gn=RCC2;refGene_tx=XM_023258202.1;refGene_gn=RCC2;refGene_tx=XM_023258203.1;refGene_gn=RCC2;refGene_tx=XM_023258204.1
C1 12483148 12483226
refGene_gn=RCC2;refGene_tx=XM_023258202.1;refGene_gn=RCC2;refGene_tx=XM_023258203.1;refGene_gn=RCC2;refGene_tx=XM_023258204.1
C1 12484800 12484873
refGene_gn=RCC2;refGene_tx=XM_023258202.1;refGene_gn=RCC2;refGene_tx=XM_023258203.1;refGene_gn=RCC2;refGene_tx=XM_023258204.1
C1 12485761 12485867
refGene_gn=RCC2;refGene_tx=XM_023258202.1;refGene_gn=RCC2;refGene_tx=XM_023258203.1;refGene_gn=RCC2;refGene_tx=XM_023258204.1
C1 12486104 12486285
refGene_gn=RCC2;refGene_tx=XM_023258202.1;refGene_gn=RCC2;refGene_tx=XM_023258203.1;refGene_gn=RCC2;refGene_tx=XM_023258204.1
C1 12487703 12487870
refGene_gn=RCC2;refGene_tx=XM_023258202.1;refGene_gn=RCC2;refGene_tx=XM_023258203.1;refGene_gn=RCC2;refGene_tx=XM_023258204.1
C1 12490047 12490162
refGene_gn=RCC2;refGene_tx=XM_023258202.1;refGene_gn=RCC2;refGene_tx=XM_023258203.1;refGene_gn=RCC2;refGene_tx=XM_023258204.1
C1 12491396 12491485
refGene_gn=RCC2;refGene_tx=XM_023258202.1;refGene_gn=RCC2;refGene_tx=XM_023258203.1;refGene_gn=RCC2;refGene_tx=XM_023258204.1
C1 12491904 12492036
refGene_gn=RCC2;refGene_tx=XM_023258202.1;refGene_gn=RCC2;refGene_tx=XM_023258203.1;refGene_gn=RCC2;refGene_tx=XM_023258204.1
C1 12494781 12494925

refGene_gn=RCC2;refGene_tx=XM_023258202.1;refGene_gn=RCC2;refGene_tx=XM_023258203.1;refGene_gn=RCC2;refGene_tx=XM_023258204.1
C1 12498198 12498292 refGene_gn=RCC2;refGene_tx=XM_023258202.1;refGene_gn=RCC2;refGene_tx=XM_023258203.1;refGene_gn=RCC2;refGene_tx=XM_023258204.1
C1 12502554 12502608 refGene_gn=RCC2;refGene_tx=XM_023258204.1
C1 12503801 12504086 refGene_gn=RCC2;refGene_tx=XM_023258202.1;refGene_gn=RCC2;refGene_tx=XM_023258203.1
C1 12599572 12599609 refGene_gn=ARHGEF10L;refGene_tx=XM_023258205.1;refGene_gn=ARHGEF10L;refGene_tx=XM_023258206.1;refGene_gn=ARHGEF10L;refGene_tx=XM_023258207.1;refGene_gn=ARHGEF10L;refGene_tx=XM_023258208.1;refGene_gn=ARHGEF10L;refGene_tx=XM_023258209.1;refGene_gn=ARHGEF10L;refGene_tx=XM_023258210.1;refGene_gn=ARHGEF10L;refGene_tx=XM_023258211.1;refGene_gn=ARHGEF10L;refGene_tx=XM_023258212.1
C1 12604908 12605094 refGene_gn=ARHGEF10L;refGene_tx=XM_023258205.1;refGene_gn=ARHGEF10L;refGene_tx=XM_023258206.1;refGene_gn=ARHGEF10L;refGene_tx=XM_023258207.1;refGene_gn=ARHGEF10L;refGene_tx=XM_023258208.1;refGene_gn=ARHGEF10L;refGene_tx=XM_023258209.1;refGene_gn=ARHGEF10L;refGene_tx=XM_023258210.1;refGene_gn=ARHGEF10L;refGene_tx=XM_023258211.1;refGene_gn=ARHGEF10L;refGene_tx=XM_023258212.1
C1 12605845 12605897 refGene_gn=ARHGEF10L;refGene_tx=XM_023258205.1;refGene_gn=ARHGEF10L;refGene_tx=XM_023258206.1;refGene_gn=ARHGEF10L;refGene_tx=XM_023258207.1;refGene_gn=ARHGEF10L;refGene_tx=XM_023258208.1;refGene_gn=ARHGEF10L;refGene_tx=XM_023258209.1;refGene_gn=ARHGEF10L;refGene_tx=XM_023258210.1;refGene_gn=ARHGEF10L;refGene_tx=XM_023258211.1;refGene_gn=ARHGEF10L;refGene_tx=XM_023258212.1
C1 12618981 12619073 refGene_gn=ARHGEF10L;refGene_tx=XM_023258205.1;refGene_gn=ARHGEF10L;refGene_tx=XM_023258206.1;refGene_gn=ARHGEF10L;refGene_tx=XM_023258207.1;refGene_gn=ARHGEF10L;refGene_tx=XM_023258208.1;refGene_gn=ARHGEF10L;refGene_tx=XM_023258209.1;refGene_gn=ARHGEF10L;refGene_tx=XM_023258210.1;refGene_gn=ARHGEF10L;refGene_tx=XM_023258211.1;refGene_gn=ARHGEF10L;refGene_tx=XM_023258212.1;refGene_gn=ARHGEF10L;refGene_tx=XM_023258213.1
C1 12619357 12619442 refGene_gn=ARHGEF10L;refGene_tx=XM_023258213.1
C1 12621048 12621132 refGene_gn=ARHGEF10L;refGene_tx=XM_023258205.1;refGene_gn=ARHGEF10L;refGene_tx=XM_023258206.1;refGene_gn=ARHGEF10L;refGene_tx=XM_023258207.1;refGene_gn=ARHGEF10L;refGene_tx=XM_023258208.1;refGene_gn=ARHGEF10L;refGene_tx=XM_023258209.1;refGene_gn=ARHGEF10L;refGene_tx=XM_023258210.1;refGene_gn=ARHGEF10L;refGene_tx=XM_023258211.1;refGene_gn=ARHGEF10L;refGene_tx=XM_023258212.1;refGene_gn=ARHGEF10L;refGene_tx=XM_023258213.1
C1 12624605 12624775 refGene_gn=ARHGEF10L;refGene_tx=XM_023258205.1;refGene_gn=ARHGEF10L;re

fGene_tx=XM_023258206.1;refGene_gn=ARHGEF10L;refGene_tx=XM_023258207.1
;refGene_gn=ARHGEF10L;refGene_tx=XM_023258208.1;refGene_gn=ARHGEF10L;r
efGene_tx=XM_023258209.1;refGene_gn=ARHGEF10L;refGene_tx=XM_023258210.
1;refGene_gn=ARHGEF10L;refGene_tx=XM_023258211.1;refGene_gn=ARHGEF10L;
refGene_tx=XM_023258212.1;refGene_gn=ARHGEF10L;refGene_tx=XM_023258213
.1
C1 12629189 12629306
refGene_gn=ARHGEF10L;refGene_tx=XM_023258205.1;refGene_gn=ARHGEF10L;re
fGene_tx=XM_023258206.1;refGene_gn=ARHGEF10L;refGene_tx=XM_023258207.1
;refGene_gn=ARHGEF10L;refGene_tx=XM_023258208.1;refGene_gn=ARHGEF10L;r
efGene_tx=XM_023258209.1;refGene_gn=ARHGEF10L;refGene_tx=XM_023258213.
1
C1 12632155 12632264
refGene_gn=ARHGEF10L;refGene_tx=XM_019836261.2;refGene_gn=ARHGEF10L;re
fGene_tx=XM_023258205.1;refGene_gn=ARHGEF10L;refGene_tx=XM_023258206.1
;refGene_gn=ARHGEF10L;refGene_tx=XM_023258207.1;refGene_gn=ARHGEF10L;r
efGene_tx=XM_023258208.1;refGene_gn=ARHGEF10L;refGene_tx=XM_023258209.
1;refGene_gn=ARHGEF10L;refGene_tx=XM_023258210.1;refGene_gn=ARHGEF10L;
refGene_tx=XM_023258211.1;refGene_gn=ARHGEF10L;refGene_tx=XM_023258212
.1;refGene_gn=ARHGEF10L;refGene_tx=XM_023258213.1
C1 12633773 12633945
refGene_gn=ARHGEF10L;refGene_tx=XM_019836258.2;refGene_gn=ARHGEF10L;re
fGene_tx=XM_019836260.2;refGene_gn=ARHGEF10L;refGene_tx=XM_019836262.2
C1 12634388 12634560
refGene_gn=ARHGEF10L;refGene_tx=XM_019836259.2
C1 12634694 12634801
refGene_gn=ARHGEF10L;refGene_tx=XM_019836258.2;refGene_gn=ARHGEF10L;re
fGene_tx=XM_019836259.2;refGene_gn=ARHGEF10L;refGene_tx=XM_019836260.2
;refGene_gn=ARHGEF10L;refGene_tx=XM_019836261.2;refGene_gn=ARHGEF10L;r
efGene_tx=XM_019836262.2;refGene_gn=ARHGEF10L;refGene_tx=XM_023258205.
1;refGene_gn=ARHGEF10L;refGene_tx=XM_023258206.1;refGene_gn=ARHGEF10L;
refGene_tx=XM_023258207.1;refGene_gn=ARHGEF10L;refGene_tx=XM_023258208
.
1;refGene_gn=ARHGEF10L;refGene_tx=XM_023258209.1;refGene_gn=ARHGEF10L;
refGene_tx=XM_023258210.1;refGene_gn=ARHGEF10L;refGene_tx=XM_023258211
.
1;refGene_gn=ARHGEF10L;refGene_tx=XM_023258212.1;refGene_gn=ARHGEF10L;
refGene_tx=XM_023258213.1
C1 12636929 12637007
refGene_gn=ARHGEF10L;refGene_tx=XM_019836258.2;refGene_gn=ARHGEF10L;re
fGene_tx=XM_019836259.2;refGene_gn=ARHGEF10L;refGene_tx=XM_019836260.2
;refGene_gn=ARHGEF10L;refGene_tx=XM_019836261.2;refGene_gn=ARHGEF10L;r
efGene_tx=XM_019836262.2;refGene_gn=ARHGEF10L;refGene_tx=XM_023258205.
1;refGene_gn=ARHGEF10L;refGene_tx=XM_023258206.1;refGene_gn=ARHGEF10L;
refGene_tx=XM_023258207.1;refGene_gn=ARHGEF10L;refGene_tx=XM_023258208
.
1;refGene_gn=ARHGEF10L;refGene_tx=XM_023258209.1;refGene_gn=ARHGEF10L;
refGene_tx=XM_023258210.1;refGene_gn=ARHGEF10L;refGene_tx=XM_023258211
.
1;refGene_gn=ARHGEF10L;refGene_tx=XM_023258212.1;refGene_gn=ARHGEF10L;

refGene_tx=XM_023258213.1
C1 12638040 12638220
refGene_gn=ARHGEF10L;refGene_tx=XM_019836258.2;refGene_gn=ARHGEF10L;refGene_tx=XM_019836259.2;refGene_gn=ARHGEF10L;refGene_tx=XM_019836260.2;refGene_gn=ARHGEF10L;refGene_tx=XM_019836261.2;refGene_gn=ARHGEF10L;refGene_tx=XM_019836262.2;refGene_gn=ARHGEF10L;refGene_tx=XM_023258205.1;refGene_gn=ARHGEF10L;refGene_tx=XM_023258206.1;refGene_gn=ARHGEF10L;refGene_tx=XM_023258207.1;refGene_gn=ARHGEF10L;refGene_tx=XM_023258208.1;refGene_gn=ARHGEF10L;refGene_tx=XM_023258209.1;refGene_gn=ARHGEF10L;refGene_tx=XM_023258210.1;refGene_gn=ARHGEF10L;refGene_tx=XM_023258211.1;refGene_gn=ARHGEF10L;refGene_tx=XM_023258212.1;refGene_gn=ARHGEF10L;refGene_tx=XM_023258213.1
C1 12639287 12639404
refGene_gn=ARHGEF10L;refGene_tx=XM_019836258.2;refGene_gn=ARHGEF10L;refGene_tx=XM_019836259.2;refGene_gn=ARHGEF10L;refGene_tx=XM_019836260.2;refGene_gn=ARHGEF10L;refGene_tx=XM_019836261.2;refGene_gn=ARHGEF10L;refGene_tx=XM_019836262.2;refGene_gn=ARHGEF10L;refGene_tx=XM_023258205.1;refGene_gn=ARHGEF10L;refGene_tx=XM_023258206.1;refGene_gn=ARHGEF10L;refGene_tx=XM_023258207.1;refGene_gn=ARHGEF10L;refGene_tx=XM_023258208.1;refGene_gn=ARHGEF10L;refGene_tx=XM_023258209.1;refGene_gn=ARHGEF10L;refGene_tx=XM_023258210.1;refGene_gn=ARHGEF10L;refGene_tx=XM_023258211.1;refGene_gn=ARHGEF10L;refGene_tx=XM_023258212.1;refGene_gn=ARHGEF10L;refGene_tx=XM_023258213.1
C1 12639800 12639893
refGene_gn=ARHGEF10L;refGene_tx=XM_019836258.2;refGene_gn=ARHGEF10L;refGene_tx=XM_019836259.2;refGene_gn=ARHGEF10L;refGene_tx=XM_019836260.2;refGene_gn=ARHGEF10L;refGene_tx=XM_019836261.2;refGene_gn=ARHGEF10L;refGene_tx=XM_019836262.2;refGene_gn=ARHGEF10L;refGene_tx=XM_023258205.1;refGene_gn=ARHGEF10L;refGene_tx=XM_023258206.1;refGene_gn=ARHGEF10L;refGene_tx=XM_023258207.1;refGene_gn=ARHGEF10L;refGene_tx=XM_023258208.1;refGene_gn=ARHGEF10L;refGene_tx=XM_023258209.1;refGene_gn=ARHGEF10L;refGene_tx=XM_023258210.1;refGene_gn=ARHGEF10L;refGene_tx=XM_023258211.1;refGene_gn=ARHGEF10L;refGene_tx=XM_023258212.1;refGene_gn=ARHGEF10L;refGene_tx=XM_023258213.1
C1 12640293 12640377
refGene_gn=ARHGEF10L;refGene_tx=XM_019836258.2;refGene_gn=ARHGEF10L;refGene_tx=XM_019836259.2;refGene_gn=ARHGEF10L;refGene_tx=XM_019836260.2;refGene_gn=ARHGEF10L;refGene_tx=XM_019836261.2;refGene_gn=ARHGEF10L;refGene_tx=XM_023258205.1;refGene_gn=ARHGEF10L;refGene_tx=XM_023258206.1;refGene_gn=ARHGEF10L;refGene_tx=XM_023258207.1;refGene_gn=ARHGEF10L;refGene_tx=XM_023258210.1;refGene_gn=ARHGEF10L;refGene_tx=XM_023258213.1
C1 12640469 12640643
refGene_gn=ARHGEF10L;refGene_tx=XM_019836258.2;refGene_gn=ARHGEF10L;re

fGene_tx=XM_019836259.2;refGene_gn=ARHGEF10L;refGene_tx=XM_019836260.2
;refGene_gn=ARHGEF10L;refGene_tx=XM_019836261.2;refGene_gn=ARHGEF10L;r
efGene_tx=XM_019836262.2;refGene_gn=ARHGEF10L;refGene_tx=XM_023258205.
1;refGene_gn=ARHGEF10L;refGene_tx=XM_023258206.1;refGene_gn=ARHGEF10L;
refGene_tx=XM_023258207.1;refGene_gn=ARHGEF10L;refGene_tx=XM_023258208
.
1;refGene_gn=ARHGEF10L;refGene_tx=XM_023258209.1;refGene_gn=ARHGEF10L;
refGene_tx=XM_023258210.1;refGene_gn=ARHGEF10L;refGene_tx=XM_023258211
.
1;refGene_gn=ARHGEF10L;refGene_tx=XM_023258212.1;refGene_gn=ARHGEF10L;
refGene_tx=XM_023258213.1
C1 12644257 12644403
refGene_gn=ARHGEF10L;refGene_tx=XM_019836258.2;refGene_gn=ARHGEF10L;re
fGene_tx=XM_019836259.2;refGene_gn=ARHGEF10L;refGene_tx=XM_019836260.2
;refGene_gn=ARHGEF10L;refGene_tx=XM_019836261.2;refGene_gn=ARHGEF10L;r
efGene_tx=XM_019836262.2;refGene_gn=ARHGEF10L;refGene_tx=XM_023258205.
1;refGene_gn=ARHGEF10L;refGene_tx=XM_023258206.1;refGene_gn=ARHGEF10L;
refGene_tx=XM_023258207.1;refGene_gn=ARHGEF10L;refGene_tx=XM_023258208
.
1;refGene_gn=ARHGEF10L;refGene_tx=XM_023258209.1;refGene_gn=ARHGEF10L;
refGene_tx=XM_023258210.1;refGene_gn=ARHGEF10L;refGene_tx=XM_023258211
.
1;refGene_gn=ARHGEF10L;refGene_tx=XM_023258212.1;refGene_gn=ARHGEF10L;
refGene_tx=XM_023258213.1
C1 12646022 12646037
refGene_gn=ARHGEF10L;refGene_tx=XM_019836258.2;refGene_gn=ARHGEF10L;re
fGene_tx=XM_019836259.2;refGene_gn=ARHGEF10L;refGene_tx=XM_019836261.2
;refGene_gn=ARHGEF10L;refGene_tx=XM_019836262.2;refGene_gn=ARHGEF10L;r
efGene_tx=XM_023258205.1;refGene_gn=ARHGEF10L;refGene_tx=XM_023258206.
1;refGene_gn=ARHGEF10L;refGene_tx=XM_023258208.1;refGene_gn=ARHGEF10L;
refGene_tx=XM_023258210.1;refGene_gn=ARHGEF10L;refGene_tx=XM_023258211
.1;refGene_gn=ARHGEF10L;refGene_tx=XM_023258213.1
C1 12646264 12646446
refGene_gn=ARHGEF10L;refGene_tx=XM_019836258.2;refGene_gn=ARHGEF10L;re
fGene_tx=XM_019836259.2;refGene_gn=ARHGEF10L;refGene_tx=XM_019836260.2
;refGene_gn=ARHGEF10L;refGene_tx=XM_019836261.2;refGene_gn=ARHGEF10L;r
efGene_tx=XM_019836262.2;refGene_gn=ARHGEF10L;refGene_tx=XM_023258205.
1;refGene_gn=ARHGEF10L;refGene_tx=XM_023258206.1;refGene_gn=ARHGEF10L;
refGene_tx=XM_023258207.1;refGene_gn=ARHGEF10L;refGene_tx=XM_023258208
.
1;refGene_gn=ARHGEF10L;refGene_tx=XM_023258209.1;refGene_gn=ARHGEF10L;
refGene_tx=XM_023258210.1;refGene_gn=ARHGEF10L;refGene_tx=XM_023258211
.
1;refGene_gn=ARHGEF10L;refGene_tx=XM_023258212.1;refGene_gn=ARHGEF10L;
refGene_tx=XM_023258213.1
C1 12649875 12649991
refGene_gn=ARHGEF10L;refGene_tx=XM_019836258.2;refGene_gn=ARHGEF10L;re
fGene_tx=XM_019836259.2;refGene_gn=ARHGEF10L;refGene_tx=XM_019836260.2
;refGene_gn=ARHGEF10L;refGene_tx=XM_019836261.2;refGene_gn=ARHGEF10L;r
efGene_tx=XM_019836262.2;refGene_gn=ARHGEF10L;refGene_tx=XM_023258205.

1;refGene_gn=ARHGEF10L;refGene_tx=XM_023258206.1;refGene_gn=ARHGEF10L;
refGene_tx=XM_023258207.1;refGene_gn=ARHGEF10L;refGene_tx=XM_023258208
.
1;refGene_gn=ARHGEF10L;refGene_tx=XM_023258209.1;refGene_gn=ARHGEF10L;
refGene_tx=XM_023258210.1;refGene_gn=ARHGEF10L;refGene_tx=XM_023258211
.
1;refGene_gn=ARHGEF10L;refGene_tx=XM_023258212.1;refGene_gn=ARHGEF10L;
refGene_tx=XM_023258213.1
C1 12650548 12650676
refGene_gn=ARHGEF10L;refGene_tx=XM_019836258.2;refGene_gn=ARHGEF10L;re
fGene_tx=XM_019836259.2;refGene_gn=ARHGEF10L;refGene_tx=XM_019836260.2
;refGene_gn=ARHGEF10L;refGene_tx=XM_019836261.2;refGene_gn=ARHGEF10L;r
efGene_tx=XM_019836262.2;refGene_gn=ARHGEF10L;refGene_tx=XM_023258205.
1;refGene_gn=ARHGEF10L;refGene_tx=XM_023258206.1;refGene_gn=ARHGEF10L;
refGene_tx=XM_023258207.1;refGene_gn=ARHGEF10L;refGene_tx=XM_023258208
.
1;refGene_gn=ARHGEF10L;refGene_tx=XM_023258209.1;refGene_gn=ARHGEF10L;
refGene_tx=XM_023258210.1;refGene_gn=ARHGEF10L;refGene_tx=XM_023258211
.
1;refGene_gn=ARHGEF10L;refGene_tx=XM_023258212.1;refGene_gn=ARHGEF10L;
refGene_tx=XM_023258213.1
C1 12652056 12652157
refGene_gn=ARHGEF10L;refGene_tx=XM_019836258.2;refGene_gn=ARHGEF10L;re
fGene_tx=XM_019836259.2;refGene_gn=ARHGEF10L;refGene_tx=XM_019836260.2
;refGene_gn=ARHGEF10L;refGene_tx=XM_019836261.2;refGene_gn=ARHGEF10L;r
efGene_tx=XM_019836262.2;refGene_gn=ARHGEF10L;refGene_tx=XM_023258205.
1;refGene_gn=ARHGEF10L;refGene_tx=XM_023258206.1;refGene_gn=ARHGEF10L;
refGene_tx=XM_023258207.1;refGene_gn=ARHGEF10L;refGene_tx=XM_023258208
.
1;refGene_gn=ARHGEF10L;refGene_tx=XM_023258209.1;refGene_gn=ARHGEF10L;
refGene_tx=XM_023258210.1;refGene_gn=ARHGEF10L;refGene_tx=XM_023258211
.
1;refGene_gn=ARHGEF10L;refGene_tx=XM_023258212.1;refGene_gn=ARHGEF10L;
refGene_tx=XM_023258213.1
C1 12660037 12660159
refGene_gn=ARHGEF10L;refGene_tx=XM_019836258.2;refGene_gn=ARHGEF10L;re
fGene_tx=XM_019836259.2;refGene_gn=ARHGEF10L;refGene_tx=XM_019836260.2
;refGene_gn=ARHGEF10L;refGene_tx=XM_019836261.2;refGene_gn=ARHGEF10L;r
efGene_tx=XM_019836262.2;refGene_gn=ARHGEF10L;refGene_tx=XM_023258205.
1;refGene_gn=ARHGEF10L;refGene_tx=XM_023258206.1;refGene_gn=ARHGEF10L;
refGene_tx=XM_023258207.1;refGene_gn=ARHGEF10L;refGene_tx=XM_023258208
.
1;refGene_gn=ARHGEF10L;refGene_tx=XM_023258209.1;refGene_gn=ARHGEF10L;
refGene_tx=XM_023258210.1;refGene_gn=ARHGEF10L;refGene_tx=XM_023258211
.
1;refGene_gn=ARHGEF10L;refGene_tx=XM_023258212.1;refGene_gn=ARHGEF10L;
refGene_tx=XM_023258213.1
C1 12665612 12665699
refGene_gn=ARHGEF10L;refGene_tx=XM_019836258.2;refGene_gn=ARHGEF10L;re
fGene_tx=XM_019836259.2;refGene_gn=ARHGEF10L;refGene_tx=XM_019836260.2

;refGene_gn=ARHGEF10L;refGene_tx=XM_019836261.2;refGene_gn=ARHGEF10L;refGene_tx=XM_019836262.2;refGene_gn=ARHGEF10L;refGene_tx=XM_023258205.1;refGene_gn=ARHGEF10L;refGene_tx=XM_023258206.1;refGene_gn=ARHGEF10L;refGene_tx=XM_023258207.1;refGene_gn=ARHGEF10L;refGene_tx=XM_023258208.1;refGene_gn=ARHGEF10L;refGene_tx=XM_023258209.1;refGene_gn=ARHGEF10L;refGene_tx=XM_023258210.1;refGene_gn=ARHGEF10L;refGene_tx=XM_023258211.1;refGene_gn=ARHGEF10L;refGene_tx=XM_023258212.1;refGene_gn=ARHGEF10L;refGene_tx=XM_023258213.1
C1 12667162 12667383
refGene_gn=ARHGEF10L;refGene_tx=XM_019836258.2;refGene_gn=ARHGEF10L;refGene_tx=XM_019836259.2;refGene_gn=ARHGEF10L;refGene_tx=XM_019836260.2;refGene_gn=ARHGEF10L;refGene_tx=XM_019836261.2;refGene_gn=ARHGEF10L;refGene_tx=XM_019836262.2;refGene_gn=ARHGEF10L;refGene_tx=XM_023258205.1;refGene_gn=ARHGEF10L;refGene_tx=XM_023258206.1;refGene_gn=ARHGEF10L;refGene_tx=XM_023258207.1;refGene_gn=ARHGEF10L;refGene_tx=XM_023258208.1;refGene_gn=ARHGEF10L;refGene_tx=XM_023258209.1;refGene_gn=ARHGEF10L;refGene_tx=XM_023258210.1;refGene_gn=ARHGEF10L;refGene_tx=XM_023258211.1;refGene_gn=ARHGEF10L;refGene_tx=XM_023258212.1;refGene_gn=ARHGEF10L;refGene_tx=XM_023258213.1
C1 12668054 12668209
refGene_gn=ARHGEF10L;refGene_tx=XM_019836258.2;refGene_gn=ARHGEF10L;refGene_tx=XM_019836259.2;refGene_gn=ARHGEF10L;refGene_tx=XM_019836260.2;refGene_gn=ARHGEF10L;refGene_tx=XM_019836261.2;refGene_gn=ARHGEF10L;refGene_tx=XM_019836262.2;refGene_gn=ARHGEF10L;refGene_tx=XM_023258205.1;refGene_gn=ARHGEF10L;refGene_tx=XM_023258206.1;refGene_gn=ARHGEF10L;refGene_tx=XM_023258207.1;refGene_gn=ARHGEF10L;refGene_tx=XM_023258208.1;refGene_gn=ARHGEF10L;refGene_tx=XM_023258209.1;refGene_gn=ARHGEF10L;refGene_tx=XM_023258210.1;refGene_gn=ARHGEF10L;refGene_tx=XM_023258211.1;refGene_gn=ARHGEF10L;refGene_tx=XM_023258212.1;refGene_gn=ARHGEF10L;refGene_tx=XM_023258213.1
C1 12675792 12675941
refGene_gn=ARHGEF10L;refGene_tx=XM_019836258.2;refGene_gn=ARHGEF10L;refGene_tx=XM_019836259.2;refGene_gn=ARHGEF10L;refGene_tx=XM_019836260.2;refGene_gn=ARHGEF10L;refGene_tx=XM_019836261.2;refGene_gn=ARHGEF10L;refGene_tx=XM_019836262.2;refGene_gn=ARHGEF10L;refGene_tx=XM_023258205.1;refGene_gn=ARHGEF10L;refGene_tx=XM_023258206.1;refGene_gn=ARHGEF10L;refGene_tx=XM_023258207.1;refGene_gn=ARHGEF10L;refGene_tx=XM_023258208.1;refGene_gn=ARHGEF10L;refGene_tx=XM_023258209.1;refGene_gn=ARHGEF10L;refGene_tx=XM_023258210.1;refGene_gn=ARHGEF10L;refGene_tx=XM_023258211.1;refGene_gn=ARHGEF10L;refGene_tx=XM_023258212.1;refGene_gn=ARHGEF10L;refGene_tx=XM_023258213.1
C1 12677315 12678230

refGene_gn=LOC109502114;refGene_tx=XM_023258214.1
C1 12696675 12696850
refGene_gn=ARHGEF10L;refGene_tx=XM_019836258.2;refGene_gn=ARHGEF10L;refGene_tx=XM_019836259.2;refGene_gn=ARHGEF10L;refGene_tx=XM_019836260.2;refGene_gn=ARHGEF10L;refGene_tx=XM_019836261.2;refGene_gn=ARHGEF10L;refGene_tx=XM_019836262.2;refGene_gn=ARHGEF10L;refGene_tx=XM_023258205.1;refGene_gn=ARHGEF10L;refGene_tx=XM_023258206.1;refGene_gn=ARHGEF10L;refGene_tx=XM_023258207.1;refGene_gn=ARHGEF10L;refGene_tx=XM_023258208.1;refGene_gn=ARHGEF10L;refGene_tx=XM_023258209.1;refGene_gn=ARHGEF10L;refGene_tx=XM_023258210.1;refGene_gn=ARHGEF10L;refGene_tx=XM_023258211.1;refGene_gn=ARHGEF10L;refGene_tx=XM_023258212.1;refGene_gn=ARHGEF10L;refGene_tx=XM_023258213.1
C1 12703777 12703900
refGene_gn=ARHGEF10L;refGene_tx=XM_019836258.2;refGene_gn=ARHGEF10L;refGene_tx=XM_019836259.2;refGene_gn=ARHGEF10L;refGene_tx=XM_019836260.2;refGene_gn=ARHGEF10L;refGene_tx=XM_019836261.2;refGene_gn=ARHGEF10L;refGene_tx=XM_019836262.2;refGene_gn=ARHGEF10L;refGene_tx=XM_023258205.1;refGene_gn=ARHGEF10L;refGene_tx=XM_023258206.1;refGene_gn=ARHGEF10L;refGene_tx=XM_023258207.1;refGene_gn=ARHGEF10L;refGene_tx=XM_023258208.1;refGene_gn=ARHGEF10L;refGene_tx=XM_023258209.1;refGene_gn=ARHGEF10L;refGene_tx=XM_023258210.1;refGene_gn=ARHGEF10L;refGene_tx=XM_023258211.1;refGene_gn=ARHGEF10L;refGene_tx=XM_023258212.1;refGene_gn=ARHGEF10L;refGene_tx=XM_023258213.1
C1 12705078 12705611
refGene_gn=ARHGEF10L;refGene_tx=XM_019836258.2;refGene_gn=ARHGEF10L;refGene_tx=XM_019836259.2;refGene_gn=ARHGEF10L;refGene_tx=XM_019836260.2;refGene_gn=ARHGEF10L;refGene_tx=XM_019836261.2;refGene_gn=ARHGEF10L;refGene_tx=XM_019836262.2;refGene_gn=ARHGEF10L;refGene_tx=XM_023258205.1;refGene_gn=ARHGEF10L;refGene_tx=XM_023258206.1;refGene_gn=ARHGEF10L;refGene_tx=XM_023258207.1;refGene_gn=ARHGEF10L;refGene_tx=XM_023258208.1;refGene_gn=ARHGEF10L;refGene_tx=XM_023258209.1;refGene_gn=ARHGEF10L;refGene_tx=XM_023258210.1;refGene_gn=ARHGEF10L;refGene_tx=XM_023258211.1;refGene_gn=ARHGEF10L;refGene_tx=XM_023258212.1;refGene_gn=ARHGEF10L;refGene_tx=XM_023258213.1
C1 12812007 12812355 refGene_gn=ACTL8;refGene_tx=XM_003989528.4
C1 12814939 12815692 refGene_gn=ACTL8;refGene_tx=XM_003989528.4
C1 13051929 13051999 refGene_gn=IGSF21;refGene_tx=XM_023258215.1
C1 13153919 13154032 refGene_gn=IGSF21;refGene_tx=XM_023258215.1
C1 13210747 13210869 refGene_gn=IGSF21;refGene_tx=XM_023258215.1
C1 13252997 13253116 refGene_gn=IGSF21;refGene_tx=XM_023258215.1
C1 13276853 13276969 refGene_gn=IGSF21;refGene_tx=XM_023258215.1
C1 13279676 13280151 refGene_gn=IGSF21;refGene_tx=XM_023258215.1
C1 13287958 13288044 refGene_gn=IGSF21;refGene_tx=XM_023258215.1
C1 13288409 13288602 refGene_gn=IGSF21;refGene_tx=XM_023258215.1

C1 13288997 13289036 refGene_gn=IGSF21;refGene_tx=XM_023258215.1
C1 13289840 13289914 refGene_gn=IGSF21;refGene_tx=XM_023258215.1
C1 13356177 13356685 refGene_gn=KLHDC7A;refGene_tx=XM_019839042.2
C1 13356687 13358549 refGene_gn=KLHDC7A;refGene_tx=XM_019839042.2
C1 13498044 13498129
refGene_gn=PAX7;refGene_tx=XM_003989610.5;refGene_gn=PAX7;refGene_tx=XM_006934358.4;refGene_gn=PAX7;refGene_tx=XM_006934359.4
C1 13500767 13501003
refGene_gn=PAX7;refGene_tx=XM_003989610.5;refGene_gn=PAX7;refGene_tx=XM_006934358.4;refGene_gn=PAX7;refGene_tx=XM_006934359.4
C1 13501593 13501723
refGene_gn=PAX7;refGene_tx=XM_003989610.5;refGene_gn=PAX7;refGene_tx=XM_006934358.4;refGene_gn=PAX7;refGene_tx=XM_006934359.4
C1 13502639 13502774
refGene_gn=PAX7;refGene_tx=XM_003989610.5;refGene_gn=PAX7;refGene_tx=XM_006934358.4;refGene_gn=PAX7;refGene_tx=XM_006934359.4
C1 13549254 13549454
refGene_gn=PAX7;refGene_tx=XM_003989610.5;refGene_gn=PAX7;refGene_tx=XM_006934358.4;refGene_gn=PAX7;refGene_tx=XM_006934359.4
C1 13555626 13555792
refGene_gn=PAX7;refGene_tx=XM_003989610.5;refGene_gn=PAX7;refGene_tx=XM_006934358.4;refGene_gn=PAX7;refGene_tx=XM_006934359.4
C1 13557168 13557365
refGene_gn=PAX7;refGene_tx=XM_003989610.5;refGene_gn=PAX7;refGene_tx=XM_006934358.4;refGene_gn=PAX7;refGene_tx=XM_006934359.4
C1 13585389 13585636
refGene_gn=PAX7;refGene_tx=XM_003989610.5;refGene_gn=PAX7;refGene_tx=XM_006934358.4;refGene_gn=PAX7;refGene_tx=XM_006934359.4
C1 13593978 13594094
refGene_gn=PAX7;refGene_tx=XM_003989610.5;refGene_gn=PAX7;refGene_tx=XM_006934358.4;refGene_gn=PAX7;refGene_tx=XM_006934359.4
C1 13616014 13616220
refGene_gn=LOC111561980;refGene_tx=XM_023260057.1
C1 13616303 13616817
refGene_gn=LOC111561980;refGene_tx=XM_023260057.1
C1 13678084 13678195 refGene_gn=TAS1R2;refGene_tx=XM_019839044.1
C1 13678196 13678759 refGene_gn=TAS1R2;refGene_tx=XM_019839044.1
C1 13678761 13679015 refGene_gn=TAS1R2;refGene_tx=XM_019839044.1
C1 13680188 13680304 refGene_gn=TAS1R2;refGene_tx=XM_019839044.1
C1 13686888 13686977 refGene_gn=TAS1R2;refGene_tx=XM_019839044.1
C1 13686979 13687150 refGene_gn=TAS1R2;refGene_tx=XM_019839044.1
C1 13691780 13692306 refGene_gn=TAS1R2;refGene_tx=XM_019839044.1
C1 13692623 13692803 refGene_gn=TAS1R2;refGene_tx=XM_019839044.1
C1 13700371 13700484 refGene_gn=ALDH4A1;refGene_tx=XM_003989611.4
C1 13701142 13701261 refGene_gn=ALDH4A1;refGene_tx=XM_003989611.4
C1 13702041 13702163 refGene_gn=ALDH4A1;refGene_tx=XM_003989611.4
C1 13702838 13702991 refGene_gn=ALDH4A1;refGene_tx=XM_003989611.4
C1 13703825 13703873 refGene_gn=ALDH4A1;refGene_tx=XM_003989611.4
C1 13704045 13704242 refGene_gn=ALDH4A1;refGene_tx=XM_003989611.4
C1 13705576 13705650 refGene_gn=ALDH4A1;refGene_tx=XM_003989611.4

C1 13707982 13708170 refGene_gn=ALDH4A1;refGene_tx=XM_003989611.4
C1 13709209 13709284 refGene_gn=ALDH4A1;refGene_tx=XM_003989611.4
C1 13709362 13709512 refGene_gn=ALDH4A1;refGene_tx=XM_003989611.4
C1 13710982 13711138 refGene_gn=ALDH4A1;refGene_tx=XM_003989611.4
C1 13711948 13711996 refGene_gn=ALDH4A1;refGene_tx=XM_003989611.4
C1 13716571 13716664 refGene_gn=ALDH4A1;refGene_tx=XM_003989611.4
C1 13717237 13717331 refGene_gn=ALDH4A1;refGene_tx=XM_003989611.4
C1 13726144 13726206 refGene_gn=ALDH4A1;refGene_tx=XM_003989611.4
C1 13732652 13732758
refGene_gn=IFFO2;refGene_tx=XM_011284622.3;refGene_gn=IFFO2;refGene_tx
=XM_023258216.1
C1 13734606 13734737
refGene_gn=IFFO2;refGene_tx=XM_011284622.3;refGene_gn=IFFO2;refGene_tx
=XM_023258216.1
C1 13735663 13735756
refGene_gn=IFFO2;refGene_tx=XM_011284622.3;refGene_gn=IFFO2;refGene_tx
=XM_023258216.1
C1 13735868 13735989
refGene_gn=IFFO2;refGene_tx=XM_011284622.3;refGene_gn=IFFO2;refGene_tx
=XM_023258216.1
C1 13740093 13740233
refGene_gn=IFFO2;refGene_tx=XM_011284622.3;refGene_gn=IFFO2;refGene_tx
=XM_023258216.1
C1 13741407 13741548 refGene_gn=IFFO2;refGene_tx=XM_011284622.3
C1 13742766 13742862
refGene_gn=IFFO2;refGene_tx=XM_011284622.3;refGene_gn=IFFO2;refGene_tx
=XM_023258216.1
C1 13744359 13744420
refGene_gn=IFFO2;refGene_tx=XM_011284622.3;refGene_gn=IFFO2;refGene_tx
=XM_023258216.1
C1 13777030 13777695
refGene_gn=IFFO2;refGene_tx=XM_011284622.3;refGene_gn=IFFO2;refGene_tx
=XM_023258216.1
C1 13875645 13875710
refGene_gn=UBR4;refGene_tx=XM_011284581.3;refGene_gn=UBR4;refGene_tx=X
M_019836275.2;refGene_gn=UBR4;refGene_tx=XM_019836276.2;refGene_gn=UBR
4;refGene_tx=XM_023258217.1;refGene_gn=UBR4;refGene_tx=XM_023258218.1;
refGene_gn=UBR4;refGene_tx=XM_023258219.1;refGene_gn=UBR4;refGene_tx=X
M_023258220.1;refGene_gn=UBR4;refGene_tx=XM_023258221.1
C1 13877659 13877822
refGene_gn=UBR4;refGene_tx=XM_011284581.3;refGene_gn=UBR4;refGene_tx=X
M_019836275.2;refGene_gn=UBR4;refGene_tx=XM_019836276.2;refGene_gn=UBR
4;refGene_tx=XM_023258217.1;refGene_gn=UBR4;refGene_tx=XM_023258218.1;
refGene_gn=UBR4;refGene_tx=XM_023258219.1;refGene_gn=UBR4;refGene_tx=X
M_023258220.1;refGene_gn=UBR4;refGene_tx=XM_023258221.1
C1 13878511 13878602
refGene_gn=UBR4;refGene_tx=XM_011284581.3;refGene_gn=UBR4;refGene_tx=X
M_019836275.2;refGene_gn=UBR4;refGene_tx=XM_019836276.2;refGene_gn=UBR
4;refGene_tx=XM_023258217.1;refGene_gn=UBR4;refGene_tx=XM_023258218.1;
refGene_gn=UBR4;refGene_tx=XM_023258219.1;refGene_gn=UBR4;refGene_tx=X

M_023258220.1;refGene_gn=UBR4;refGene_tx=XM_023258221.1
C1 13881983 13882208
refGene_gn=UBR4;refGene_tx=XM_011284581.3;refGene_gn=UBR4;refGene_tx=XM_019836275.2;refGene_gn=UBR4;refGene_tx=XM_019836276.2;refGene_gn=UBR4;refGene_tx=XM_023258217.1;refGene_gn=UBR4;refGene_tx=XM_023258218.1;refGene_gn=UBR4;refGene_tx=XM_023258219.1;refGene_gn=UBR4;refGene_tx=XM_023258220.1;refGene_gn=UBR4;refGene_tx=XM_023258221.1
C1 13884992 13885187
refGene_gn=UBR4;refGene_tx=XM_011284581.3;refGene_gn=UBR4;refGene_tx=XM_019836275.2;refGene_gn=UBR4;refGene_tx=XM_019836276.2;refGene_gn=UBR4;refGene_tx=XM_023258217.1;refGene_gn=UBR4;refGene_tx=XM_023258218.1;refGene_gn=UBR4;refGene_tx=XM_023258219.1;refGene_gn=UBR4;refGene_tx=XM_023258220.1;refGene_gn=UBR4;refGene_tx=XM_023258221.1
C1 13886668 13886794
refGene_gn=UBR4;refGene_tx=XM_011284581.3;refGene_gn=UBR4;refGene_tx=XM_019836275.2;refGene_gn=UBR4;refGene_tx=XM_019836276.2;refGene_gn=UBR4;refGene_tx=XM_023258217.1;refGene_gn=UBR4;refGene_tx=XM_023258218.1;refGene_gn=UBR4;refGene_tx=XM_023258219.1;refGene_gn=UBR4;refGene_tx=XM_023258220.1;refGene_gn=UBR4;refGene_tx=XM_023258221.1
C1 13887214 13887357
refGene_gn=UBR4;refGene_tx=XM_011284581.3;refGene_gn=UBR4;refGene_tx=XM_019836275.2;refGene_gn=UBR4;refGene_tx=XM_019836276.2;refGene_gn=UBR4;refGene_tx=XM_023258217.1;refGene_gn=UBR4;refGene_tx=XM_023258218.1;refGene_gn=UBR4;refGene_tx=XM_023258219.1;refGene_gn=UBR4;refGene_tx=XM_023258220.1;refGene_gn=UBR4;refGene_tx=XM_023258221.1
C1 13888303 13888417
refGene_gn=UBR4;refGene_tx=XM_011284581.3;refGene_gn=UBR4;refGene_tx=XM_019836275.2;refGene_gn=UBR4;refGene_tx=XM_019836276.2;refGene_gn=UBR4;refGene_tx=XM_023258217.1;refGene_gn=UBR4;refGene_tx=XM_023258218.1;refGene_gn=UBR4;refGene_tx=XM_023258219.1;refGene_gn=UBR4;refGene_tx=XM_023258220.1;refGene_gn=UBR4;refGene_tx=XM_023258221.1
C1 13889290 13889509
refGene_gn=UBR4;refGene_tx=XM_011284581.3;refGene_gn=UBR4;refGene_tx=XM_019836275.2;refGene_gn=UBR4;refGene_tx=XM_019836276.2;refGene_gn=UBR4;refGene_tx=XM_023258217.1;refGene_gn=UBR4;refGene_tx=XM_023258218.1;refGene_gn=UBR4;refGene_tx=XM_023258219.1;refGene_gn=UBR4;refGene_tx=XM_023258220.1;refGene_gn=UBR4;refGene_tx=XM_023258221.1
C1 13893133 13893233
refGene_gn=UBR4;refGene_tx=XM_011284581.3;refGene_gn=UBR4;refGene_tx=XM_019836275.2;refGene_gn=UBR4;refGene_tx=XM_019836276.2;refGene_gn=UBR4;refGene_tx=XM_023258217.1;refGene_gn=UBR4;refGene_tx=XM_023258218.1;refGene_gn=UBR4;refGene_tx=XM_023258219.1;refGene_gn=UBR4;refGene_tx=XM_023258220.1;refGene_gn=UBR4;refGene_tx=XM_023258221.1
C1 13893608 13893782
refGene_gn=UBR4;refGene_tx=XM_011284581.3;refGene_gn=UBR4;refGene_tx=XM_019836275.2;refGene_gn=UBR4;refGene_tx=XM_019836276.2;refGene_gn=UBR4;refGene_tx=XM_023258217.1;refGene_gn=UBR4;refGene_tx=XM_023258218.1;refGene_gn=UBR4;refGene_tx=XM_023258219.1;refGene_gn=UBR4;refGene_tx=XM_023258220.1;refGene_gn=UBR4;refGene_tx=XM_023258221.1
C1 13894204 13894395

refGene_gn=UBR4;refGene_tx=XM_011284581.3;refGene_gn=UBR4;refGene_tx=XM_019836275.2;refGene_gn=UBR4;refGene_tx=XM_019836276.2;refGene_gn=UBR4;refGene_tx=XM_023258217.1;refGene_gn=UBR4;refGene_tx=XM_023258218.1;refGene_gn=UBR4;refGene_tx=XM_023258219.1;refGene_gn=UBR4;refGene_tx=XM_023258220.1;refGene_gn=UBR4;refGene_tx=XM_023258221.1
C1 13895136 13895256
refGene_gn=UBR4;refGene_tx=XM_011284581.3;refGene_gn=UBR4;refGene_tx=XM_019836275.2;refGene_gn=UBR4;refGene_tx=XM_019836276.2;refGene_gn=UBR4;refGene_tx=XM_023258217.1;refGene_gn=UBR4;refGene_tx=XM_023258218.1;refGene_gn=UBR4;refGene_tx=XM_023258219.1;refGene_gn=UBR4;refGene_tx=XM_023258220.1;refGene_gn=UBR4;refGene_tx=XM_023258221.1
C1 13895750 13895858
refGene_gn=UBR4;refGene_tx=XM_011284581.3;refGene_gn=UBR4;refGene_tx=XM_019836275.2;refGene_gn=UBR4;refGene_tx=XM_019836276.2;refGene_gn=UBR4;refGene_tx=XM_023258217.1;refGene_gn=UBR4;refGene_tx=XM_023258218.1;refGene_gn=UBR4;refGene_tx=XM_023258219.1;refGene_gn=UBR4;refGene_tx=XM_023258220.1;refGene_gn=UBR4;refGene_tx=XM_023258221.1
C1 13896867 13896995
refGene_gn=UBR4;refGene_tx=XM_011284581.3;refGene_gn=UBR4;refGene_tx=XM_019836275.2;refGene_gn=UBR4;refGene_tx=XM_019836276.2;refGene_gn=UBR4;refGene_tx=XM_023258217.1;refGene_gn=UBR4;refGene_tx=XM_023258218.1;refGene_gn=UBR4;refGene_tx=XM_023258219.1;refGene_gn=UBR4;refGene_tx=XM_023258220.1;refGene_gn=UBR4;refGene_tx=XM_023258221.1
C1 13897498 13897586
refGene_gn=UBR4;refGene_tx=XM_011284581.3;refGene_gn=UBR4;refGene_tx=XM_019836275.2;refGene_gn=UBR4;refGene_tx=XM_019836276.2;refGene_gn=UBR4;refGene_tx=XM_023258217.1;refGene_gn=UBR4;refGene_tx=XM_023258218.1;refGene_gn=UBR4;refGene_tx=XM_023258219.1;refGene_gn=UBR4;refGene_tx=XM_023258220.1;refGene_gn=UBR4;refGene_tx=XM_023258221.1
C1 13899727 13899808
refGene_gn=UBR4;refGene_tx=XM_011284581.3;refGene_gn=UBR4;refGene_tx=XM_019836275.2;refGene_gn=UBR4;refGene_tx=XM_019836276.2;refGene_gn=UBR4;refGene_tx=XM_023258217.1;refGene_gn=UBR4;refGene_tx=XM_023258218.1;refGene_gn=UBR4;refGene_tx=XM_023258219.1;refGene_gn=UBR4;refGene_tx=XM_023258220.1;refGene_gn=UBR4;refGene_tx=XM_023258221.1
C1 13900525 13900723
refGene_gn=UBR4;refGene_tx=XM_011284581.3;refGene_gn=UBR4;refGene_tx=XM_019836275.2;refGene_gn=UBR4;refGene_tx=XM_019836276.2;refGene_gn=UBR4;refGene_tx=XM_023258217.1;refGene_gn=UBR4;refGene_tx=XM_023258218.1;refGene_gn=UBR4;refGene_tx=XM_023258219.1;refGene_gn=UBR4;refGene_tx=XM_023258220.1;refGene_gn=UBR4;refGene_tx=XM_023258221.1
C1 13901611 13901733
refGene_gn=UBR4;refGene_tx=XM_011284581.3;refGene_gn=UBR4;refGene_tx=XM_019836275.2;refGene_gn=UBR4;refGene_tx=XM_019836276.2;refGene_gn=UBR4;refGene_tx=XM_023258217.1;refGene_gn=UBR4;refGene_tx=XM_023258218.1;refGene_gn=UBR4;refGene_tx=XM_023258219.1;refGene_gn=UBR4;refGene_tx=XM_023258220.1;refGene_gn=UBR4;refGene_tx=XM_023258221.1
C1 13903820 13903994
refGene_gn=UBR4;refGene_tx=XM_011284581.3;refGene_gn=UBR4;refGene_tx=XM_019836275.2;refGene_gn=UBR4;refGene_tx=XM_019836276.2;refGene_gn=UBR

4;refGene_tx=XM_023258217.1;refGene_gn=UBR4;refGene_tx=XM_023258218.1;
refGene_gn=UBR4;refGene_tx=XM_023258219.1;refGene_gn=UBR4;refGene_tx=X
M_023258220.1;refGene_gn=UBR4;refGene_tx=XM_023258221.1
C1 13904311 13904393
refGene_gn=UBR4;refGene_tx=XM_011284581.3;refGene_gn=UBR4;refGene_tx=X
M_019836275.2;refGene_gn=UBR4;refGene_tx=XM_019836276.2;refGene_gn=UBR
4;refGene_tx=XM_023258217.1;refGene_gn=UBR4;refGene_tx=XM_023258218.1;
refGene_gn=UBR4;refGene_tx=XM_023258219.1;refGene_gn=UBR4;refGene_tx=X
M_023258220.1;refGene_gn=UBR4;refGene_tx=XM_023258221.1
C1 13904767 13904909
refGene_gn=UBR4;refGene_tx=XM_011284581.3;refGene_gn=UBR4;refGene_tx=X
M_019836275.2;refGene_gn=UBR4;refGene_tx=XM_019836276.2;refGene_gn=UBR
4;refGene_tx=XM_023258217.1;refGene_gn=UBR4;refGene_tx=XM_023258218.1;
refGene_gn=UBR4;refGene_tx=XM_023258219.1;refGene_gn=UBR4;refGene_tx=X
M_023258220.1;refGene_gn=UBR4;refGene_tx=XM_023258221.1
C1 13905606 13905716
refGene_gn=UBR4;refGene_tx=XM_011284581.3;refGene_gn=UBR4;refGene_tx=X
M_019836275.2;refGene_gn=UBR4;refGene_tx=XM_019836276.2;refGene_gn=UBR
4;refGene_tx=XM_023258217.1;refGene_gn=UBR4;refGene_tx=XM_023258218.1;
refGene_gn=UBR4;refGene_tx=XM_023258219.1;refGene_gn=UBR4;refGene_tx=X
M_023258220.1;refGene_gn=UBR4;refGene_tx=XM_023258221.1
C1 13906671 13906829
refGene_gn=UBR4;refGene_tx=XM_011284581.3;refGene_gn=UBR4;refGene_tx=X
M_019836275.2;refGene_gn=UBR4;refGene_tx=XM_019836276.2;refGene_gn=UBR
4;refGene_tx=XM_023258217.1;refGene_gn=UBR4;refGene_tx=XM_023258218.1;
refGene_gn=UBR4;refGene_tx=XM_023258219.1;refGene_gn=UBR4;refGene_tx=X
M_023258220.1;refGene_gn=UBR4;refGene_tx=XM_023258221.1
C1 13906953 13907083
refGene_gn=UBR4;refGene_tx=XM_011284581.3;refGene_gn=UBR4;refGene_tx=X
M_019836275.2;refGene_gn=UBR4;refGene_tx=XM_019836276.2;refGene_gn=UBR
4;refGene_tx=XM_023258217.1;refGene_gn=UBR4;refGene_tx=XM_023258218.1;
refGene_gn=UBR4;refGene_tx=XM_023258219.1;refGene_gn=UBR4;refGene_tx=X
M_023258220.1;refGene_gn=UBR4;refGene_tx=XM_023258221.1
C1 13910505 13910633
refGene_gn=UBR4;refGene_tx=XM_011284581.3;refGene_gn=UBR4;refGene_tx=X
M_019836275.2;refGene_gn=UBR4;refGene_tx=XM_019836276.2;refGene_gn=UBR
4;refGene_tx=XM_023258217.1;refGene_gn=UBR4;refGene_tx=XM_023258218.1;
refGene_gn=UBR4;refGene_tx=XM_023258219.1;refGene_gn=UBR4;refGene_tx=X
M_023258220.1;refGene_gn=UBR4;refGene_tx=XM_023258221.1
C1 13910794 13910879
refGene_gn=UBR4;refGene_tx=XM_011284581.3;refGene_gn=UBR4;refGene_tx=X
M_019836275.2;refGene_gn=UBR4;refGene_tx=XM_019836276.2;refGene_gn=UBR
4;refGene_tx=XM_023258217.1;refGene_gn=UBR4;refGene_tx=XM_023258218.1;
refGene_gn=UBR4;refGene_tx=XM_023258219.1;refGene_gn=UBR4;refGene_tx=X
M_023258220.1;refGene_gn=UBR4;refGene_tx=XM_023258221.1
C1 13911186 13911277
refGene_gn=UBR4;refGene_tx=XM_011284581.3;refGene_gn=UBR4;refGene_tx=X
M_019836275.2;refGene_gn=UBR4;refGene_tx=XM_019836276.2;refGene_gn=UBR
4;refGene_tx=XM_023258217.1;refGene_gn=UBR4;refGene_tx=XM_023258218.1;
refGene_gn=UBR4;refGene_tx=XM_023258219.1;refGene_gn=UBR4;refGene_tx=X

M_023258220.1;refGene_gn=UBR4;refGene_tx=XM_023258221.1
C1 13912793 13913137
refGene_gn=UBR4;refGene_tx=XM_011284581.3;refGene_gn=UBR4;refGene_tx=XM_019836275.2;refGene_gn=UBR4;refGene_tx=XM_019836276.2;refGene_gn=UBR4;refGene_tx=XM_023258217.1;refGene_gn=UBR4;refGene_tx=XM_023258218.1;refGene_gn=UBR4;refGene_tx=XM_023258219.1;refGene_gn=UBR4;refGene_tx=XM_023258220.1;refGene_gn=UBR4;refGene_tx=XM_023258221.1
C1 13913931 13914060
refGene_gn=UBR4;refGene_tx=XM_011284581.3;refGene_gn=UBR4;refGene_tx=XM_019836275.2;refGene_gn=UBR4;refGene_tx=XM_019836276.2;refGene_gn=UBR4;refGene_tx=XM_023258217.1;refGene_gn=UBR4;refGene_tx=XM_023258218.1;refGene_gn=UBR4;refGene_tx=XM_023258219.1;refGene_gn=UBR4;refGene_tx=XM_023258220.1;refGene_gn=UBR4;refGene_tx=XM_023258221.1
C1 13914140 13914266
refGene_gn=UBR4;refGene_tx=XM_011284581.3;refGene_gn=UBR4;refGene_tx=XM_019836275.2;refGene_gn=UBR4;refGene_tx=XM_019836276.2;refGene_gn=UBR4;refGene_tx=XM_023258217.1;refGene_gn=UBR4;refGene_tx=XM_023258218.1;refGene_gn=UBR4;refGene_tx=XM_023258219.1;refGene_gn=UBR4;refGene_tx=XM_023258220.1;refGene_gn=UBR4;refGene_tx=XM_023258221.1
C1 13914764 13914903
refGene_gn=UBR4;refGene_tx=XM_011284581.3;refGene_gn=UBR4;refGene_tx=XM_019836275.2;refGene_gn=UBR4;refGene_tx=XM_019836276.2;refGene_gn=UBR4;refGene_tx=XM_023258217.1;refGene_gn=UBR4;refGene_tx=XM_023258218.1;refGene_gn=UBR4;refGene_tx=XM_023258219.1;refGene_gn=UBR4;refGene_tx=XM_023258220.1;refGene_gn=UBR4;refGene_tx=XM_023258221.1
C1 13915278 13915518
refGene_gn=UBR4;refGene_tx=XM_011284581.3;refGene_gn=UBR4;refGene_tx=XM_019836275.2;refGene_gn=UBR4;refGene_tx=XM_019836276.2;refGene_gn=UBR4;refGene_tx=XM_023258217.1;refGene_gn=UBR4;refGene_tx=XM_023258218.1;refGene_gn=UBR4;refGene_tx=XM_023258219.1;refGene_gn=UBR4;refGene_tx=XM_023258220.1;refGene_gn=UBR4;refGene_tx=XM_023258221.1
C1 13917169 13917363
refGene_gn=UBR4;refGene_tx=XM_011284581.3;refGene_gn=UBR4;refGene_tx=XM_019836275.2;refGene_gn=UBR4;refGene_tx=XM_019836276.2;refGene_gn=UBR4;refGene_tx=XM_023258217.1;refGene_gn=UBR4;refGene_tx=XM_023258218.1;refGene_gn=UBR4;refGene_tx=XM_023258219.1;refGene_gn=UBR4;refGene_tx=XM_023258220.1;refGene_gn=UBR4;refGene_tx=XM_023258221.1
C1 13917625 13917713
refGene_gn=UBR4;refGene_tx=XM_011284581.3;refGene_gn=UBR4;refGene_tx=XM_019836275.2;refGene_gn=UBR4;refGene_tx=XM_019836276.2;refGene_gn=UBR4;refGene_tx=XM_023258217.1;refGene_gn=UBR4;refGene_tx=XM_023258218.1;refGene_gn=UBR4;refGene_tx=XM_023258219.1;refGene_gn=UBR4;refGene_tx=XM_023258220.1;refGene_gn=UBR4;refGene_tx=XM_023258221.1
C1 13918413 13918499
refGene_gn=UBR4;refGene_tx=XM_011284581.3;refGene_gn=UBR4;refGene_tx=XM_019836275.2;refGene_gn=UBR4;refGene_tx=XM_019836276.2;refGene_gn=UBR4;refGene_tx=XM_023258217.1;refGene_gn=UBR4;refGene_tx=XM_023258218.1;refGene_gn=UBR4;refGene_tx=XM_023258219.1;refGene_gn=UBR4;refGene_tx=XM_023258220.1;refGene_gn=UBR4;refGene_tx=XM_023258221.1
C1 13919086 13919231

refGene_gn=UBR4;refGene_tx=XM_011284581.3;refGene_gn=UBR4;refGene_tx=XM_019836275.2;refGene_gn=UBR4;refGene_tx=XM_019836276.2;refGene_gn=UBR4;refGene_tx=XM_023258217.1;refGene_gn=UBR4;refGene_tx=XM_023258218.1;refGene_gn=UBR4;refGene_tx=XM_023258219.1;refGene_gn=UBR4;refGene_tx=XM_023258220.1;refGene_gn=UBR4;refGene_tx=XM_023258221.1
C1 13919681 13919850
refGene_gn=UBR4;refGene_tx=XM_011284581.3;refGene_gn=UBR4;refGene_tx=XM_019836275.2;refGene_gn=UBR4;refGene_tx=XM_019836276.2;refGene_gn=UBR4;refGene_tx=XM_023258217.1;refGene_gn=UBR4;refGene_tx=XM_023258218.1;refGene_gn=UBR4;refGene_tx=XM_023258219.1;refGene_gn=UBR4;refGene_tx=XM_023258220.1;refGene_gn=UBR4;refGene_tx=XM_023258221.1
C1 13920607 13920853
refGene_gn=UBR4;refGene_tx=XM_011284581.3;refGene_gn=UBR4;refGene_tx=XM_019836275.2;refGene_gn=UBR4;refGene_tx=XM_019836276.2;refGene_gn=UBR4;refGene_tx=XM_023258217.1;refGene_gn=UBR4;refGene_tx=XM_023258218.1;refGene_gn=UBR4;refGene_tx=XM_023258219.1;refGene_gn=UBR4;refGene_tx=XM_023258220.1;refGene_gn=UBR4;refGene_tx=XM_023258221.1
C1 13921364 13921443
refGene_gn=UBR4;refGene_tx=XM_011284581.3;refGene_gn=UBR4;refGene_tx=XM_019836275.2;refGene_gn=UBR4;refGene_tx=XM_019836276.2;refGene_gn=UBR4;refGene_tx=XM_023258217.1;refGene_gn=UBR4;refGene_tx=XM_023258218.1;refGene_gn=UBR4;refGene_tx=XM_023258219.1;refGene_gn=UBR4;refGene_tx=XM_023258220.1;refGene_gn=UBR4;refGene_tx=XM_023258221.1
C1 13922201 13922429
refGene_gn=UBR4;refGene_tx=XM_011284581.3;refGene_gn=UBR4;refGene_tx=XM_019836275.2;refGene_gn=UBR4;refGene_tx=XM_019836276.2;refGene_gn=UBR4;refGene_tx=XM_023258217.1;refGene_gn=UBR4;refGene_tx=XM_023258218.1;refGene_gn=UBR4;refGene_tx=XM_023258219.1;refGene_gn=UBR4;refGene_tx=XM_023258220.1;refGene_gn=UBR4;refGene_tx=XM_023258221.1
C1 13923645 13923795
refGene_gn=UBR4;refGene_tx=XM_011284581.3;refGene_gn=UBR4;refGene_tx=XM_019836275.2;refGene_gn=UBR4;refGene_tx=XM_019836276.2;refGene_gn=UBR4;refGene_tx=XM_023258217.1;refGene_gn=UBR4;refGene_tx=XM_023258218.1;refGene_gn=UBR4;refGene_tx=XM_023258219.1;refGene_gn=UBR4;refGene_tx=XM_023258220.1;refGene_gn=UBR4;refGene_tx=XM_023258221.1
C1 13925516 13925726
refGene_gn=UBR4;refGene_tx=XM_011284581.3;refGene_gn=UBR4;refGene_tx=XM_019836275.2;refGene_gn=UBR4;refGene_tx=XM_019836276.2;refGene_gn=UBR4;refGene_tx=XM_023258217.1;refGene_gn=UBR4;refGene_tx=XM_023258218.1;refGene_gn=UBR4;refGene_tx=XM_023258219.1;refGene_gn=UBR4;refGene_tx=XM_023258220.1;refGene_gn=UBR4;refGene_tx=XM_023258221.1
C1 13926630 13926747
refGene_gn=UBR4;refGene_tx=XM_011284581.3;refGene_gn=UBR4;refGene_tx=XM_019836275.2;refGene_gn=UBR4;refGene_tx=XM_019836276.2;refGene_gn=UBR4;refGene_tx=XM_023258217.1;refGene_gn=UBR4;refGene_tx=XM_023258218.1;refGene_gn=UBR4;refGene_tx=XM_023258219.1;refGene_gn=UBR4;refGene_tx=XM_023258220.1;refGene_gn=UBR4;refGene_tx=XM_023258221.1
C1 13927199 13927307
refGene_gn=UBR4;refGene_tx=XM_011284581.3;refGene_gn=UBR4;refGene_tx=XM_019836275.2;refGene_gn=UBR4;refGene_tx=XM_019836276.2;refGene_gn=UBR

4;refGene_tx=XM_023258217.1;refGene_gn=UBR4;refGene_tx=XM_023258218.1;
refGene_gn=UBR4;refGene_tx=XM_023258219.1;refGene_gn=UBR4;refGene_tx=X
M_023258220.1;refGene_gn=UBR4;refGene_tx=XM_023258221.1
C1 13927966 13928063
refGene_gn=UBR4;refGene_tx=XM_011284581.3;refGene_gn=UBR4;refGene_tx=X
M_019836275.2;refGene_gn=UBR4;refGene_tx=XM_019836276.2;refGene_gn=UBR
4;refGene_tx=XM_023258217.1;refGene_gn=UBR4;refGene_tx=XM_023258218.1;
refGene_gn=UBR4;refGene_tx=XM_023258219.1;refGene_gn=UBR4;refGene_tx=X
M_023258220.1;refGene_gn=UBR4;refGene_tx=XM_023258221.1
C1 13934689 13934864
refGene_gn=UBR4;refGene_tx=XM_011284581.3;refGene_gn=UBR4;refGene_tx=X
M_019836275.2;refGene_gn=UBR4;refGene_tx=XM_019836276.2;refGene_gn=UBR
4;refGene_tx=XM_023258217.1;refGene_gn=UBR4;refGene_tx=XM_023258218.1;
refGene_gn=UBR4;refGene_tx=XM_023258219.1;refGene_gn=UBR4;refGene_tx=X
M_023258220.1;refGene_gn=UBR4;refGene_tx=XM_023258221.1
C1 13935710 13935866
refGene_gn=UBR4;refGene_tx=XM_011284581.3;refGene_gn=UBR4;refGene_tx=X
M_019836275.2;refGene_gn=UBR4;refGene_tx=XM_019836276.2;refGene_gn=UBR
4;refGene_tx=XM_023258217.1;refGene_gn=UBR4;refGene_tx=XM_023258218.1;
refGene_gn=UBR4;refGene_tx=XM_023258219.1;refGene_gn=UBR4;refGene_tx=X
M_023258220.1;refGene_gn=UBR4;refGene_tx=XM_023258221.1
C1 13937403 13937508
refGene_gn=UBR4;refGene_tx=XM_011284581.3;refGene_gn=UBR4;refGene_tx=X
M_019836275.2;refGene_gn=UBR4;refGene_tx=XM_019836276.2;refGene_gn=UBR
4;refGene_tx=XM_023258217.1;refGene_gn=UBR4;refGene_tx=XM_023258218.1;
refGene_gn=UBR4;refGene_tx=XM_023258219.1;refGene_gn=UBR4;refGene_tx=X
M_023258220.1
C1 13937971 13938149
refGene_gn=UBR4;refGene_tx=XM_011284581.3;refGene_gn=UBR4;refGene_tx=X
M_019836275.2;refGene_gn=UBR4;refGene_tx=XM_019836276.2;refGene_gn=UBR
4;refGene_tx=XM_023258217.1;refGene_gn=UBR4;refGene_tx=XM_023258218.1;
refGene_gn=UBR4;refGene_tx=XM_023258219.1;refGene_gn=UBR4;refGene_tx=X
M_023258220.1;refGene_gn=UBR4;refGene_tx=XM_023258221.1
C1 13938270 13938401
refGene_gn=UBR4;refGene_tx=XM_011284581.3;refGene_gn=UBR4;refGene_tx=X
M_019836275.2;refGene_gn=UBR4;refGene_tx=XM_019836276.2;refGene_gn=UBR
4;refGene_tx=XM_023258217.1;refGene_gn=UBR4;refGene_tx=XM_023258218.1;
refGene_gn=UBR4;refGene_tx=XM_023258219.1;refGene_gn=UBR4;refGene_tx=X
M_023258220.1;refGene_gn=UBR4;refGene_tx=XM_023258221.1
C1 13940167 13940279
refGene_gn=UBR4;refGene_tx=XM_011284581.3;refGene_gn=UBR4;refGene_tx=X
M_019836275.2;refGene_gn=UBR4;refGene_tx=XM_019836276.2;refGene_gn=UBR
4;refGene_tx=XM_023258217.1;refGene_gn=UBR4;refGene_tx=XM_023258218.1;
refGene_gn=UBR4;refGene_tx=XM_023258219.1;refGene_gn=UBR4;refGene_tx=X
M_023258220.1;refGene_gn=UBR4;refGene_tx=XM_023258221.1
C1 13940950 13941072
refGene_gn=UBR4;refGene_tx=XM_011284581.3;refGene_gn=UBR4;refGene_tx=X
M_019836275.2;refGene_gn=UBR4;refGene_tx=XM_019836276.2;refGene_gn=UBR
4;refGene_tx=XM_023258217.1;refGene_gn=UBR4;refGene_tx=XM_023258218.1;
refGene_gn=UBR4;refGene_tx=XM_023258219.1;refGene_gn=UBR4;refGene_tx=X

M_023258220.1;refGene_gn=UBR4;refGene_tx=XM_023258221.1
C1 13941941 13942082
refGene_gn=UBR4;refGene_tx=XM_011284581.3;refGene_gn=UBR4;refGene_tx=XM_019836275.2;refGene_gn=UBR4;refGene_tx=XM_019836276.2;refGene_gn=UBR4;refGene_tx=XM_023258217.1;refGene_gn=UBR4;refGene_tx=XM_023258218.1;refGene_gn=UBR4;refGene_tx=XM_023258219.1;refGene_gn=UBR4;refGene_tx=XM_023258220.1;refGene_gn=UBR4;refGene_tx=XM_023258221.1
C1 13942960 13943135
refGene_gn=UBR4;refGene_tx=XM_011284581.3;refGene_gn=UBR4;refGene_tx=XM_019836275.2;refGene_gn=UBR4;refGene_tx=XM_019836276.2;refGene_gn=UBR4;refGene_tx=XM_023258217.1;refGene_gn=UBR4;refGene_tx=XM_023258218.1;refGene_gn=UBR4;refGene_tx=XM_023258219.1;refGene_gn=UBR4;refGene_tx=XM_023258220.1;refGene_gn=UBR4;refGene_tx=XM_023258221.1
C1 13944119 13944254
refGene_gn=UBR4;refGene_tx=XM_011284581.3;refGene_gn=UBR4;refGene_tx=XM_019836275.2;refGene_gn=UBR4;refGene_tx=XM_019836276.2;refGene_gn=UBR4;refGene_tx=XM_023258217.1;refGene_gn=UBR4;refGene_tx=XM_023258218.1;refGene_gn=UBR4;refGene_tx=XM_023258219.1;refGene_gn=UBR4;refGene_tx=XM_023258220.1;refGene_gn=UBR4;refGene_tx=XM_023258221.1
C1 13944691 13944755
refGene_gn=UBR4;refGene_tx=XM_011284581.3;refGene_gn=UBR4;refGene_tx=XM_019836275.2;refGene_gn=UBR4;refGene_tx=XM_019836276.2;refGene_gn=UBR4;refGene_tx=XM_023258217.1;refGene_gn=UBR4;refGene_tx=XM_023258218.1;refGene_gn=UBR4;refGene_tx=XM_023258219.1;refGene_gn=UBR4;refGene_tx=XM_023258220.1;refGene_gn=UBR4;refGene_tx=XM_023258221.1
C1 13945910 13945943
refGene_gn=UBR4;refGene_tx=XM_019836275.2;refGene_gn=UBR4;refGene_tx=XM_023258217.1;refGene_gn=UBR4;refGene_tx=XM_023258218.1;refGene_gn=UBR4;refGene_tx=XM_023258219.1;refGene_gn=UBR4;refGene_tx=XM_023258220.1;refGene_gn=UBR4;refGene_tx=XM_023258221.1
C1 13946697 13946914
refGene_gn=UBR4;refGene_tx=XM_011284581.3;refGene_gn=UBR4;refGene_tx=XM_019836275.2;refGene_gn=UBR4;refGene_tx=XM_019836276.2;refGene_gn=UBR4;refGene_tx=XM_023258217.1;refGene_gn=UBR4;refGene_tx=XM_023258218.1;refGene_gn=UBR4;refGene_tx=XM_023258219.1;refGene_gn=UBR4;refGene_tx=XM_023258220.1;refGene_gn=UBR4;refGene_tx=XM_023258221.1
C1 13947895 13948112
refGene_gn=UBR4;refGene_tx=XM_011284581.3;refGene_gn=UBR4;refGene_tx=XM_019836275.2;refGene_gn=UBR4;refGene_tx=XM_019836276.2;refGene_gn=UBR4;refGene_tx=XM_023258217.1;refGene_gn=UBR4;refGene_tx=XM_023258218.1;refGene_gn=UBR4;refGene_tx=XM_023258219.1;refGene_gn=UBR4;refGene_tx=XM_023258220.1;refGene_gn=UBR4;refGene_tx=XM_023258221.1
C1 13948553 13948717
refGene_gn=UBR4;refGene_tx=XM_011284581.3;refGene_gn=UBR4;refGene_tx=XM_019836275.2;refGene_gn=UBR4;refGene_tx=XM_019836276.2;refGene_gn=UBR4;refGene_tx=XM_023258217.1;refGene_gn=UBR4;refGene_tx=XM_023258218.1;refGene_gn=UBR4;refGene_tx=XM_023258219.1;refGene_gn=UBR4;refGene_tx=XM_023258220.1;refGene_gn=UBR4;refGene_tx=XM_023258221.1
C1 13949872 13950074
refGene_gn=UBR4;refGene_tx=XM_011284581.3;refGene_gn=UBR4;refGene_tx=X

M_019836275.2;refGene_gn=UBR4;refGene_tx=XM_019836276.2;refGene_gn=UBR4;refGene_tx=XM_023258217.1;refGene_gn=UBR4;refGene_tx=XM_023258218.1;refGene_gn=UBR4;refGene_tx=XM_023258219.1;refGene_gn=UBR4;refGene_tx=XM_023258220.1;refGene_gn=UBR4;refGene_tx=XM_023258221.1
C1 13950355 13950527
refGene_gn=UBR4;refGene_tx=XM_011284581.3;refGene_gn=UBR4;refGene_tx=XM_019836275.2;refGene_gn=UBR4;refGene_tx=XM_019836276.2;refGene_gn=UBR4;refGene_tx=XM_023258217.1;refGene_gn=UBR4;refGene_tx=XM_023258218.1;refGene_gn=UBR4;refGene_tx=XM_023258219.1;refGene_gn=UBR4;refGene_tx=XM_023258220.1;refGene_gn=UBR4;refGene_tx=XM_023258221.1
C1 13951499 13951657
refGene_gn=UBR4;refGene_tx=XM_011284581.3;refGene_gn=UBR4;refGene_tx=XM_019836275.2;refGene_gn=UBR4;refGene_tx=XM_019836276.2;refGene_gn=UBR4;refGene_tx=XM_023258217.1;refGene_gn=UBR4;refGene_tx=XM_023258218.1;refGene_gn=UBR4;refGene_tx=XM_023258219.1;refGene_gn=UBR4;refGene_tx=XM_023258220.1;refGene_gn=UBR4;refGene_tx=XM_023258221.1
C1 13951952 13952180
refGene_gn=UBR4;refGene_tx=XM_011284581.3;refGene_gn=UBR4;refGene_tx=XM_019836275.2;refGene_gn=UBR4;refGene_tx=XM_019836276.2;refGene_gn=UBR4;refGene_tx=XM_023258217.1;refGene_gn=UBR4;refGene_tx=XM_023258218.1;refGene_gn=UBR4;refGene_tx=XM_023258219.1;refGene_gn=UBR4;refGene_tx=XM_023258220.1;refGene_gn=UBR4;refGene_tx=XM_023258221.1
C1 13952627 13952780
refGene_gn=UBR4;refGene_tx=XM_011284581.3;refGene_gn=UBR4;refGene_tx=XM_019836275.2;refGene_gn=UBR4;refGene_tx=XM_019836276.2;refGene_gn=UBR4;refGene_tx=XM_023258217.1;refGene_gn=UBR4;refGene_tx=XM_023258218.1;refGene_gn=UBR4;refGene_tx=XM_023258219.1;refGene_gn=UBR4;refGene_tx=XM_023258220.1;refGene_gn=UBR4;refGene_tx=XM_023258221.1
C1 13953342 13953501
refGene_gn=UBR4;refGene_tx=XM_011284581.3;refGene_gn=UBR4;refGene_tx=XM_019836275.2;refGene_gn=UBR4;refGene_tx=XM_019836276.2;refGene_gn=UBR4;refGene_tx=XM_023258217.1;refGene_gn=UBR4;refGene_tx=XM_023258218.1;refGene_gn=UBR4;refGene_tx=XM_023258219.1;refGene_gn=UBR4;refGene_tx=XM_023258220.1;refGene_gn=UBR4;refGene_tx=XM_023258221.1
C1 13954444 13954627
refGene_gn=UBR4;refGene_tx=XM_011284581.3;refGene_gn=UBR4;refGene_tx=XM_019836275.2;refGene_gn=UBR4;refGene_tx=XM_019836276.2;refGene_gn=UBR4;refGene_tx=XM_023258217.1;refGene_gn=UBR4;refGene_tx=XM_023258218.1;refGene_gn=UBR4;refGene_tx=XM_023258219.1;refGene_gn=UBR4;refGene_tx=XM_023258220.1;refGene_gn=UBR4;refGene_tx=XM_023258221.1
C1 13956568 13956739
refGene_gn=UBR4;refGene_tx=XM_011284581.3;refGene_gn=UBR4;refGene_tx=XM_019836275.2;refGene_gn=UBR4;refGene_tx=XM_019836276.2;refGene_gn=UBR4;refGene_tx=XM_023258217.1;refGene_gn=UBR4;refGene_tx=XM_023258218.1;refGene_gn=UBR4;refGene_tx=XM_023258219.1;refGene_gn=UBR4;refGene_tx=XM_023258220.1;refGene_gn=UBR4;refGene_tx=XM_023258221.1
C1 13957401 13957629
refGene_gn=UBR4;refGene_tx=XM_011284581.3;refGene_gn=UBR4;refGene_tx=XM_019836275.2;refGene_gn=UBR4;refGene_tx=XM_019836276.2;refGene_gn=UBR4;refGene_tx=XM_023258217.1;refGene_gn=UBR4;refGene_tx=XM_023258218.1;

refGene_gn=UBR4;refGene_tx=XM_023258219.1;refGene_gn=UBR4;refGene_tx=XM_023258220.1;refGene_gn=UBR4;refGene_tx=XM_023258221.1
C1 13958085 13958233
refGene_gn=UBR4;refGene_tx=XM_011284581.3;refGene_gn=UBR4;refGene_tx=XM_019836275.2;refGene_gn=UBR4;refGene_tx=XM_019836276.2;refGene_gn=UBR4;refGene_tx=XM_023258217.1;refGene_gn=UBR4;refGene_tx=XM_023258218.1;refGene_gn=UBR4;refGene_tx=XM_023258219.1;refGene_gn=UBR4;refGene_tx=XM_023258220.1;refGene_gn=UBR4;refGene_tx=XM_023258221.1
C1 13958325 13958396
refGene_gn=UBR4;refGene_tx=XM_011284581.3;refGene_gn=UBR4;refGene_tx=XM_019836275.2;refGene_gn=UBR4;refGene_tx=XM_019836276.2;refGene_gn=UBR4;refGene_tx=XM_023258217.1;refGene_gn=UBR4;refGene_tx=XM_023258218.1;refGene_gn=UBR4;refGene_tx=XM_023258219.1;refGene_gn=UBR4;refGene_tx=XM_023258220.1;refGene_gn=UBR4;refGene_tx=XM_023258221.1
C1 13958955 13959147
refGene_gn=UBR4;refGene_tx=XM_011284581.3;refGene_gn=UBR4;refGene_tx=XM_019836275.2;refGene_gn=UBR4;refGene_tx=XM_019836276.2;refGene_gn=UBR4;refGene_tx=XM_023258217.1;refGene_gn=UBR4;refGene_tx=XM_023258218.1;refGene_gn=UBR4;refGene_tx=XM_023258219.1;refGene_gn=UBR4;refGene_tx=XM_023258220.1;refGene_gn=UBR4;refGene_tx=XM_023258221.1
C1 13960620 13960684
refGene_gn=UBR4;refGene_tx=XM_011284581.3;refGene_gn=UBR4;refGene_tx=XM_019836275.2;refGene_gn=UBR4;refGene_tx=XM_019836276.2;refGene_gn=UBR4;refGene_tx=XM_023258217.1;refGene_gn=UBR4;refGene_tx=XM_023258218.1;refGene_gn=UBR4;refGene_tx=XM_023258219.1;refGene_gn=UBR4;refGene_tx=XM_023258220.1;refGene_gn=UBR4;refGene_tx=XM_023258221.1
C1 13961400 13961589
refGene_gn=UBR4;refGene_tx=XM_011284581.3;refGene_gn=UBR4;refGene_tx=XM_019836275.2;refGene_gn=UBR4;refGene_tx=XM_019836276.2;refGene_gn=UBR4;refGene_tx=XM_023258217.1;refGene_gn=UBR4;refGene_tx=XM_023258218.1;refGene_gn=UBR4;refGene_tx=XM_023258219.1;refGene_gn=UBR4;refGene_tx=XM_023258220.1;refGene_gn=UBR4;refGene_tx=XM_023258221.1
C1 13961978 13962177
refGene_gn=UBR4;refGene_tx=XM_011284581.3;refGene_gn=UBR4;refGene_tx=XM_019836275.2;refGene_gn=UBR4;refGene_tx=XM_019836276.2;refGene_gn=UBR4;refGene_tx=XM_023258217.1;refGene_gn=UBR4;refGene_tx=XM_023258218.1;refGene_gn=UBR4;refGene_tx=XM_023258219.1;refGene_gn=UBR4;refGene_tx=XM_023258220.1;refGene_gn=UBR4;refGene_tx=XM_023258221.1
C1 13962419 13962520
refGene_gn=UBR4;refGene_tx=XM_011284581.3;refGene_gn=UBR4;refGene_tx=XM_019836275.2;refGene_gn=UBR4;refGene_tx=XM_019836276.2;refGene_gn=UBR4;refGene_tx=XM_023258217.1;refGene_gn=UBR4;refGene_tx=XM_023258218.1;refGene_gn=UBR4;refGene_tx=XM_023258219.1;refGene_gn=UBR4;refGene_tx=XM_023258220.1;refGene_gn=UBR4;refGene_tx=XM_023258221.1
C1 13962821 13962923
refGene_gn=UBR4;refGene_tx=XM_011284581.3;refGene_gn=UBR4;refGene_tx=XM_019836275.2;refGene_gn=UBR4;refGene_tx=XM_019836276.2;refGene_gn=UBR4;refGene_tx=XM_023258217.1;refGene_gn=UBR4;refGene_tx=XM_023258218.1;refGene_gn=UBR4;refGene_tx=XM_023258219.1;refGene_gn=UBR4;refGene_tx=XM_023258220.1;refGene_gn=UBR4;refGene_tx=XM_023258221.1

C1 13963932 13964142
refGene_gn=UBR4;refGene_tx=XM_011284581.3;refGene_gn=UBR4;refGene_tx=XM_019836275.2;refGene_gn=UBR4;refGene_tx=XM_019836276.2;refGene_gn=UBR4;refGene_tx=XM_023258217.1;refGene_gn=UBR4;refGene_tx=XM_023258218.1;refGene_gn=UBR4;refGene_tx=XM_023258219.1;refGene_gn=UBR4;refGene_tx=XM_023258220.1;refGene_gn=UBR4;refGene_tx=XM_023258221.1
C1 13964903 13965061
refGene_gn=UBR4;refGene_tx=XM_011284581.3;refGene_gn=UBR4;refGene_tx=XM_019836275.2;refGene_gn=UBR4;refGene_tx=XM_019836276.2;refGene_gn=UBR4;refGene_tx=XM_023258217.1;refGene_gn=UBR4;refGene_tx=XM_023258218.1;refGene_gn=UBR4;refGene_tx=XM_023258219.1;refGene_gn=UBR4;refGene_tx=XM_023258220.1;refGene_gn=UBR4;refGene_tx=XM_023258221.1
C1 13966048 13966146
refGene_gn=UBR4;refGene_tx=XM_011284581.3;refGene_gn=UBR4;refGene_tx=XM_019836275.2;refGene_gn=UBR4;refGene_tx=XM_019836276.2;refGene_gn=UBR4;refGene_tx=XM_023258217.1;refGene_gn=UBR4;refGene_tx=XM_023258218.1;refGene_gn=UBR4;refGene_tx=XM_023258219.1;refGene_gn=UBR4;refGene_tx=XM_023258220.1;refGene_gn=UBR4;refGene_tx=XM_023258221.1
C1 13967390 13967512
refGene_gn=UBR4;refGene_tx=XM_011284581.3;refGene_gn=UBR4;refGene_tx=XM_019836275.2;refGene_gn=UBR4;refGene_tx=XM_019836276.2;refGene_gn=UBR4;refGene_tx=XM_023258217.1;refGene_gn=UBR4;refGene_tx=XM_023258218.1;refGene_gn=UBR4;refGene_tx=XM_023258219.1;refGene_gn=UBR4;refGene_tx=XM_023258220.1;refGene_gn=UBR4;refGene_tx=XM_023258221.1
C1 13969494 13969724
refGene_gn=UBR4;refGene_tx=XM_011284581.3;refGene_gn=UBR4;refGene_tx=XM_019836275.2;refGene_gn=UBR4;refGene_tx=XM_019836276.2;refGene_gn=UBR4;refGene_tx=XM_023258217.1;refGene_gn=UBR4;refGene_tx=XM_023258218.1;refGene_gn=UBR4;refGene_tx=XM_023258219.1;refGene_gn=UBR4;refGene_tx=XM_023258220.1;refGene_gn=UBR4;refGene_tx=XM_023258221.1
C1 13969813 13969936
refGene_gn=UBR4;refGene_tx=XM_011284581.3;refGene_gn=UBR4;refGene_tx=XM_019836275.2;refGene_gn=UBR4;refGene_tx=XM_019836276.2;refGene_gn=UBR4;refGene_tx=XM_023258217.1;refGene_gn=UBR4;refGene_tx=XM_023258218.1;refGene_gn=UBR4;refGene_tx=XM_023258219.1;refGene_gn=UBR4;refGene_tx=XM_023258220.1;refGene_gn=UBR4;refGene_tx=XM_023258221.1
C1 13970088 13970271
refGene_gn=UBR4;refGene_tx=XM_011284581.3;refGene_gn=UBR4;refGene_tx=XM_019836275.2;refGene_gn=UBR4;refGene_tx=XM_019836276.2;refGene_gn=UBR4;refGene_tx=XM_023258217.1;refGene_gn=UBR4;refGene_tx=XM_023258218.1;refGene_gn=UBR4;refGene_tx=XM_023258219.1;refGene_gn=UBR4;refGene_tx=XM_023258220.1;refGene_gn=UBR4;refGene_tx=XM_023258221.1
C1 13971105 13971234
refGene_gn=UBR4;refGene_tx=XM_011284581.3;refGene_gn=UBR4;refGene_tx=XM_019836275.2;refGene_gn=UBR4;refGene_tx=XM_019836276.2;refGene_gn=UBR4;refGene_tx=XM_023258217.1;refGene_gn=UBR4;refGene_tx=XM_023258218.1;refGene_gn=UBR4;refGene_tx=XM_023258219.1;refGene_gn=UBR4;refGene_tx=XM_023258220.1;refGene_gn=UBR4;refGene_tx=XM_023258221.1
C1 13971696 13971776
refGene_gn=UBR4;refGene_tx=XM_011284581.3;refGene_gn=UBR4;refGene_tx=X

M_019836275.2;refGene_gn=UBR4;refGene_tx=XM_019836276.2;refGene_gn=UBR4;refGene_tx=XM_023258217.1;refGene_gn=UBR4;refGene_tx=XM_023258218.1;refGene_gn=UBR4;refGene_tx=XM_023258219.1;refGene_gn=UBR4;refGene_tx=XM_023258220.1;refGene_gn=UBR4;refGene_tx=XM_023258221.1
C1 13972700 13972836
refGene_gn=UBR4;refGene_tx=XM_011284581.3;refGene_gn=UBR4;refGene_tx=XM_019836275.2;refGene_gn=UBR4;refGene_tx=XM_019836276.2;refGene_gn=UBR4;refGene_tx=XM_023258217.1;refGene_gn=UBR4;refGene_tx=XM_023258218.1;refGene_gn=UBR4;refGene_tx=XM_023258219.1;refGene_gn=UBR4;refGene_tx=XM_023258220.1;refGene_gn=UBR4;refGene_tx=XM_023258221.1
C1 13973499 13973782
refGene_gn=UBR4;refGene_tx=XM_011284581.3;refGene_gn=UBR4;refGene_tx=XM_019836275.2;refGene_gn=UBR4;refGene_tx=XM_019836276.2;refGene_gn=UBR4;refGene_tx=XM_023258217.1;refGene_gn=UBR4;refGene_tx=XM_023258218.1;refGene_gn=UBR4;refGene_tx=XM_023258219.1;refGene_gn=UBR4;refGene_tx=XM_023258220.1;refGene_gn=UBR4;refGene_tx=XM_023258221.1
C1 13975131 13975301
refGene_gn=UBR4;refGene_tx=XM_011284581.3;refGene_gn=UBR4;refGene_tx=XM_019836275.2;refGene_gn=UBR4;refGene_tx=XM_019836276.2;refGene_gn=UBR4;refGene_tx=XM_023258217.1;refGene_gn=UBR4;refGene_tx=XM_023258218.1;refGene_gn=UBR4;refGene_tx=XM_023258219.1;refGene_gn=UBR4;refGene_tx=XM_023258220.1;refGene_gn=UBR4;refGene_tx=XM_023258221.1
C1 13982492 13982578
refGene_gn=UBR4;refGene_tx=XM_011284581.3;refGene_gn=UBR4;refGene_tx=XM_019836275.2;refGene_gn=UBR4;refGene_tx=XM_019836276.2;refGene_gn=UBR4;refGene_tx=XM_023258217.1;refGene_gn=UBR4;refGene_tx=XM_023258218.1;refGene_gn=UBR4;refGene_tx=XM_023258219.1;refGene_gn=UBR4;refGene_tx=XM_023258220.1;refGene_gn=UBR4;refGene_tx=XM_023258221.1
C1 13982698 13982858
refGene_gn=UBR4;refGene_tx=XM_011284581.3;refGene_gn=UBR4;refGene_tx=XM_019836275.2;refGene_gn=UBR4;refGene_tx=XM_019836276.2;refGene_gn=UBR4;refGene_tx=XM_023258217.1;refGene_gn=UBR4;refGene_tx=XM_023258218.1;refGene_gn=UBR4;refGene_tx=XM_023258219.1;refGene_gn=UBR4;refGene_tx=XM_023258220.1;refGene_gn=UBR4;refGene_tx=XM_023258221.1
C1 13983806 13983994
refGene_gn=UBR4;refGene_tx=XM_011284581.3;refGene_gn=UBR4;refGene_tx=XM_019836275.2;refGene_gn=UBR4;refGene_tx=XM_019836276.2;refGene_gn=UBR4;refGene_tx=XM_023258217.1;refGene_gn=UBR4;refGene_tx=XM_023258218.1;refGene_gn=UBR4;refGene_tx=XM_023258219.1;refGene_gn=UBR4;refGene_tx=XM_023258220.1;refGene_gn=UBR4;refGene_tx=XM_023258221.1
C1 13985043 13985161
refGene_gn=UBR4;refGene_tx=XM_011284581.3;refGene_gn=UBR4;refGene_tx=XM_019836275.2;refGene_gn=UBR4;refGene_tx=XM_019836276.2;refGene_gn=UBR4;refGene_tx=XM_023258217.1;refGene_gn=UBR4;refGene_tx=XM_023258218.1;refGene_gn=UBR4;refGene_tx=XM_023258219.1;refGene_gn=UBR4;refGene_tx=XM_023258220.1;refGene_gn=UBR4;refGene_tx=XM_023258221.1
C1 13986462 13986600
refGene_gn=UBR4;refGene_tx=XM_011284581.3;refGene_gn=UBR4;refGene_tx=XM_019836275.2;refGene_gn=UBR4;refGene_tx=XM_019836276.2;refGene_gn=UBR4;refGene_tx=XM_023258217.1;refGene_gn=UBR4;refGene_tx=XM_023258218.1;

refGene_gn=UBR4;refGene_tx=XM_023258219.1;refGene_gn=UBR4;refGene_tx=XM_023258220.1;refGene_gn=UBR4;refGene_tx=XM_023258221.1
C1 13986736 13986836
refGene_gn=UBR4;refGene_tx=XM_011284581.3;refGene_gn=UBR4;refGene_tx=XM_019836275.2;refGene_gn=UBR4;refGene_tx=XM_019836276.2;refGene_gn=UBR4;refGene_tx=XM_023258217.1;refGene_gn=UBR4;refGene_tx=XM_023258218.1;refGene_gn=UBR4;refGene_tx=XM_023258219.1;refGene_gn=UBR4;refGene_tx=XM_023258220.1;refGene_gn=UBR4;refGene_tx=XM_023258221.1
C1 13990946 13991137
refGene_gn=UBR4;refGene_tx=XM_011284581.3;refGene_gn=UBR4;refGene_tx=XM_019836275.2;refGene_gn=UBR4;refGene_tx=XM_019836276.2;refGene_gn=UBR4;refGene_tx=XM_023258217.1;refGene_gn=UBR4;refGene_tx=XM_023258218.1;refGene_gn=UBR4;refGene_tx=XM_023258219.1;refGene_gn=UBR4;refGene_tx=XM_023258220.1;refGene_gn=UBR4;refGene_tx=XM_023258221.1
C1 13991222 13991282
refGene_gn=UBR4;refGene_tx=XM_011284581.3;refGene_gn=UBR4;refGene_tx=XM_019836275.2;refGene_gn=UBR4;refGene_tx=XM_019836276.2;refGene_gn=UBR4;refGene_tx=XM_023258217.1;refGene_gn=UBR4;refGene_tx=XM_023258218.1;refGene_gn=UBR4;refGene_tx=XM_023258219.1;refGene_gn=UBR4;refGene_tx=XM_023258220.1;refGene_gn=UBR4;refGene_tx=XM_023258221.1
C1 13991880 13992005
refGene_gn=UBR4;refGene_tx=XM_011284581.3;refGene_gn=UBR4;refGene_tx=XM_019836275.2;refGene_gn=UBR4;refGene_tx=XM_019836276.2;refGene_gn=UBR4;refGene_tx=XM_023258217.1;refGene_gn=UBR4;refGene_tx=XM_023258218.1;refGene_gn=UBR4;refGene_tx=XM_023258219.1;refGene_gn=UBR4;refGene_tx=XM_023258220.1;refGene_gn=UBR4;refGene_tx=XM_023258221.1
C1 13994949 13995074
refGene_gn=UBR4;refGene_tx=XM_011284581.3;refGene_gn=UBR4;refGene_tx=XM_019836275.2;refGene_gn=UBR4;refGene_tx=XM_019836276.2;refGene_gn=UBR4;refGene_tx=XM_023258217.1;refGene_gn=UBR4;refGene_tx=XM_023258218.1;refGene_gn=UBR4;refGene_tx=XM_023258219.1;refGene_gn=UBR4;refGene_tx=XM_023258220.1;refGene_gn=UBR4;refGene_tx=XM_023258221.1
C1 13995231 13995373
refGene_gn=UBR4;refGene_tx=XM_011284581.3;refGene_gn=UBR4;refGene_tx=XM_019836275.2;refGene_gn=UBR4;refGene_tx=XM_019836276.2;refGene_gn=UBR4;refGene_tx=XM_023258217.1;refGene_gn=UBR4;refGene_tx=XM_023258218.1;refGene_gn=UBR4;refGene_tx=XM_023258219.1;refGene_gn=UBR4;refGene_tx=XM_023258220.1;refGene_gn=UBR4;refGene_tx=XM_023258221.1
C1 13995494 13995597
refGene_gn=UBR4;refGene_tx=XM_011284581.3;refGene_gn=UBR4;refGene_tx=XM_019836275.2;refGene_gn=UBR4;refGene_tx=XM_019836276.2;refGene_gn=UBR4;refGene_tx=XM_023258217.1;refGene_gn=UBR4;refGene_tx=XM_023258218.1;refGene_gn=UBR4;refGene_tx=XM_023258219.1;refGene_gn=UBR4;refGene_tx=XM_023258220.1;refGene_gn=UBR4;refGene_tx=XM_023258221.1
C1 13996049 13996189
refGene_gn=UBR4;refGene_tx=XM_011284581.3;refGene_gn=UBR4;refGene_tx=XM_019836275.2;refGene_gn=UBR4;refGene_tx=XM_019836276.2;refGene_gn=UBR4;refGene_tx=XM_023258217.1;refGene_gn=UBR4;refGene_tx=XM_023258218.1;refGene_gn=UBR4;refGene_tx=XM_023258219.1;refGene_gn=UBR4;refGene_tx=XM_023258220.1;refGene_gn=UBR4;refGene_tx=XM_023258221.1

C1 13996301 13996431
refGene_gn=UBR4;refGene_tx=XM_011284581.3;refGene_gn=UBR4;refGene_tx=XM_019836275.2;refGene_gn=UBR4;refGene_tx=XM_019836276.2;refGene_gn=UBR4;refGene_tx=XM_023258217.1;refGene_gn=UBR4;refGene_tx=XM_023258218.1;refGene_gn=UBR4;refGene_tx=XM_023258219.1;refGene_gn=UBR4;refGene_tx=XM_023258220.1;refGene_gn=UBR4;refGene_tx=XM_023258221.1
C1 13997197 13997301
refGene_gn=UBR4;refGene_tx=XM_011284581.3;refGene_gn=UBR4;refGene_tx=XM_019836275.2;refGene_gn=UBR4;refGene_tx=XM_019836276.2;refGene_gn=UBR4;refGene_tx=XM_023258217.1;refGene_gn=UBR4;refGene_tx=XM_023258218.1;refGene_gn=UBR4;refGene_tx=XM_023258219.1;refGene_gn=UBR4;refGene_tx=XM_023258220.1;refGene_gn=UBR4;refGene_tx=XM_023258221.1
C1 13998524 13998622
refGene_gn=UBR4;refGene_tx=XM_011284581.3;refGene_gn=UBR4;refGene_tx=XM_019836275.2;refGene_gn=UBR4;refGene_tx=XM_019836276.2;refGene_gn=UBR4;refGene_tx=XM_023258217.1;refGene_gn=UBR4;refGene_tx=XM_023258218.1;refGene_gn=UBR4;refGene_tx=XM_023258219.1;refGene_gn=UBR4;refGene_tx=XM_023258220.1;refGene_gn=UBR4;refGene_tx=XM_023258221.1
C1 14006354 14006524
refGene_gn=UBR4;refGene_tx=XM_011284581.3;refGene_gn=UBR4;refGene_tx=XM_019836275.2;refGene_gn=UBR4;refGene_tx=XM_019836276.2;refGene_gn=UBR4;refGene_tx=XM_023258217.1;refGene_gn=UBR4;refGene_tx=XM_023258218.1;refGene_gn=UBR4;refGene_tx=XM_023258219.1;refGene_gn=UBR4;refGene_tx=XM_023258220.1;refGene_gn=UBR4;refGene_tx=XM_023258221.1
C1 14015330 14015510
refGene_gn=EMC1;refGene_tx=XM_003989612.4;refGene_gn=EMC1;refGene_tx=XM_006934360.4;refGene_gn=EMC1;refGene_tx=XM_006934361.3;refGene_gn=EMC1;refGene_tx=XM_006934362.3
C1 14015665 14015795
refGene_gn=EMC1;refGene_tx=XM_003989612.4;refGene_gn=EMC1;refGene_tx=XM_006934360.4;refGene_gn=EMC1;refGene_tx=XM_006934361.3;refGene_gn=EMC1;refGene_tx=XM_006934362.3
C1 14016901 14016986
refGene_gn=EMC1;refGene_tx=XM_003989612.4;refGene_gn=EMC1;refGene_tx=XM_006934360.4;refGene_gn=EMC1;refGene_tx=XM_006934361.3;refGene_gn=EMC1;refGene_tx=XM_006934362.3
C1 14017877 14018088
refGene_gn=EMC1;refGene_tx=XM_003989612.4;refGene_gn=EMC1;refGene_tx=XM_006934360.4;refGene_gn=EMC1;refGene_tx=XM_006934361.3;refGene_gn=EMC1;refGene_tx=XM_006934362.3
C1 14018668 14018842
refGene_gn=EMC1;refGene_tx=XM_003989612.4;refGene_gn=EMC1;refGene_tx=XM_006934360.4;refGene_gn=EMC1;refGene_tx=XM_006934361.3;refGene_gn=EMC1;refGene_tx=XM_006934362.3
C1 14021950 14022088
refGene_gn=EMC1;refGene_tx=XM_003989612.4;refGene_gn=EMC1;refGene_tx=XM_006934360.4;refGene_gn=EMC1;refGene_tx=XM_006934361.3;refGene_gn=EMC1;refGene_tx=XM_006934362.3
C1 14026662 14026782
refGene_gn=EMC1;refGene_tx=XM_003989612.4;refGene_gn=EMC1;refGene_tx=X

M_006934360.4;refGene_gn=EMC1;refGene_tx=XM_006934361.3;refGene_gn=EMC
1;refGene_tx=XM_006934362.3
C1	14027112 14027274
refGene_gn=EMC1;refGene_tx=XM_003989612.4;refGene_gn=EMC1;refGene_tx=X
M_006934360.4;refGene_gn=EMC1;refGene_tx=XM_006934361.3;refGene_gn=EMC
1;refGene_tx=XM_006934362.3
C1	14027921 14028071
refGene_gn=EMC1;refGene_tx=XM_003989612.4;refGene_gn=EMC1;refGene_tx=X
M_006934360.4;refGene_gn=EMC1;refGene_tx=XM_006934361.3;refGene_gn=EMC
1;refGene_tx=XM_006934362.3
C1	14028234 14028434
refGene_gn=EMC1;refGene_tx=XM_003989612.4;refGene_gn=EMC1;refGene_tx=X
M_006934360.4;refGene_gn=EMC1;refGene_tx=XM_006934361.3;refGene_gn=EMC
1;refGene_tx=XM_006934362.3
C1	14029446 14029455
refGene_gn=EMC1;refGene_tx=XM_006934360.4;refGene_gn=EMC1;refGene_tx=X
M_006934361.3
C1	14030062 14030185
refGene_gn=EMC1;refGene_tx=XM_003989612.4;refGene_gn=EMC1;refGene_tx=X
M_006934360.4;refGene_gn=EMC1;refGene_tx=XM_006934361.3;refGene_gn=EMC
1;refGene_tx=XM_006934362.3
C1	14030521 14030618
refGene_gn=EMC1;refGene_tx=XM_003989612.4;refGene_gn=EMC1;refGene_tx=X
M_006934360.4;refGene_gn=EMC1;refGene_tx=XM_006934361.3;refGene_gn=EMC
1;refGene_tx=XM_006934362.3
C1	14031391 14031514
refGene_gn=EMC1;refGene_tx=XM_003989612.4;refGene_gn=EMC1;refGene_tx=X
M_006934360.4;refGene_gn=EMC1;refGene_tx=XM_006934361.3;refGene_gn=EMC
1;refGene_tx=XM_006934362.3
C1	14032161 14032221
refGene_gn=EMC1;refGene_tx=XM_003989612.4;refGene_gn=EMC1;refGene_tx=X
M_006934360.4;refGene_gn=EMC1;refGene_tx=XM_006934361.3;refGene_gn=EMC
1;refGene_tx=XM_006934362.3
C1	14032567 14032639
refGene_gn=EMC1;refGene_tx=XM_003989612.4;refGene_gn=EMC1;refGene_tx=X
M_006934360.4;refGene_gn=EMC1;refGene_tx=XM_006934361.3;refGene_gn=EMC
1;refGene_tx=XM_006934362.3
C1	14033133 14033301
refGene_gn=EMC1;refGene_tx=XM_003989612.4;refGene_gn=EMC1;refGene_tx=X
M_006934360.4;refGene_gn=EMC1;refGene_tx=XM_006934361.3;refGene_gn=EMC
1;refGene_tx=XM_006934362.3
C1	14033617 14033767
refGene_gn=EMC1;refGene_tx=XM_003989612.4;refGene_gn=EMC1;refGene_tx=X
M_006934360.4;refGene_gn=EMC1;refGene_tx=XM_006934361.3;refGene_gn=EMC
1;refGene_tx=XM_006934362.3
C1	14034324 14034451
refGene_gn=EMC1;refGene_tx=XM_003989612.4;refGene_gn=EMC1;refGene_tx=X
M_006934360.4;refGene_gn=EMC1;refGene_tx=XM_006934361.3;refGene_gn=EMC
1;refGene_tx=XM_006934362.3
C1	14036527 14036656

refGene_gn=EMC1;refGene_tx=XM_003989612.4;refGene_gn=EMC1;refGene_tx=XM_006934360.4;refGene_gn=EMC1;refGene_tx=XM_006934361.3;refGene_gn=EMC1;refGene_tx=XM_006934362.3
C1 14037511 14037605
refGene_gn=EMC1;refGene_tx=XM_003989612.4;refGene_gn=EMC1;refGene_tx=XM_006934360.4;refGene_gn=EMC1;refGene_tx=XM_006934361.3;refGene_gn=EMC1;refGene_tx=XM_006934362.3
C1 14037851 14037917
refGene_gn=EMC1;refGene_tx=XM_003989612.4;refGene_gn=EMC1;refGene_tx=XM_006934360.4;refGene_gn=EMC1;refGene_tx=XM_006934361.3;refGene_gn=EMC1;refGene_tx=XM_006934362.3
C1 14038508 14038633
refGene_gn=EMC1;refGene_tx=XM_003989612.4;refGene_gn=EMC1;refGene_tx=XM_006934360.4;refGene_gn=EMC1;refGene_tx=XM_006934361.3;refGene_gn=EMC1;refGene_tx=XM_006934362.3
C1 14041468 14041635
refGene_gn=EMC1;refGene_tx=XM_003989612.4;refGene_gn=EMC1;refGene_tx=XM_006934360.4;refGene_gn=EMC1;refGene_tx=XM_006934361.3;refGene_gn=EMC1;refGene_tx=XM_006934362.3
C1 14041860 14041888 refGene_gn=MRTO4;refGene_tx=XM_003989613.5
C1 14044246 14044305 refGene_gn=MRTO4;refGene_tx=XM_003989613.5
C1 14045760 14045864 refGene_gn=MRTO4;refGene_tx=XM_003989613.5
C1 14046783 14046865 refGene_gn=MRTO4;refGene_tx=XM_003989613.5
C1 14047043 14047111 refGene_gn=MRTO4;refGene_tx=XM_003989613.5
C1 14047376 14047528 refGene_gn=MRTO4;refGene_tx=XM_003989613.5
C1 14047924 14048001 refGene_gn=MRTO4;refGene_tx=XM_003989613.5
C1 14048111 14048258 refGene_gn=MRTO4;refGene_tx=XM_003989613.5
C1 14058500 14058662
refGene_gn=LOC101089785;refGene_tx=XM_003989614.5
C1 14059701 14059831
refGene_gn=LOC101089785;refGene_tx=XM_003989614.5
C1 14060444 14060544
refGene_gn=LOC101089785;refGene_tx=XM_003989614.5
C1 14060761 14060858
refGene_gn=LOC101089785;refGene_tx=XM_003989614.5
C1 14062633 14062738
refGene_gn=LOC101089785;refGene_tx=XM_003989614.5
C1 14062935 14063123
refGene_gn=LOC101089785;refGene_tx=XM_003989614.5
C1 14064949 14065124 refGene_gn=PQLC2;refGene_tx=XM_019836279.2
C1 14065314 14065651
refGene_gn=LOC101089785;refGene_tx=XM_003989614.5
C1 14070290 14070459
refGene_gn=PQLC2;refGene_tx=XM_003989615.5;refGene_gn=PQLC2;refGene_tx=XM_006934363.3;refGene_gn=PQLC2;refGene_tx=XM_006934364.3;refGene_gn=PQLC2;refGene_tx=XM_006934365.4;refGene_gn=PQLC2;refGene_tx=XM_019836279.2
C1 14077394 14077524
refGene_gn=PQLC2;refGene_tx=XM_003989615.5;refGene_gn=PQLC2;refGene_tx=XM_006934363.3;refGene_gn=PQLC2;refGene_tx=XM_006934364.3;refGene_gn=

PQLC2;refGene_tx=XM_006934365.4;refGene_gn=PQLC2;refGene_tx=XM_0198362
79.2
C1 14078342 14078430
refGene_gn=PQLC2;refGene_tx=XM_003989615.5;refGene_gn=PQLC2;refGene_tx
=XM_006934363.3;refGene_gn=PQLC2;refGene_tx=XM_006934364.3;refGene_gn=
PQLC2;refGene_tx=XM_006934365.4;refGene_gn=PQLC2;refGene_tx=XM_0198362
79.2
C1 14079021 14079164
refGene_gn=PQLC2;refGene_tx=XM_003989615.5;refGene_gn=PQLC2;refGene_tx
=XM_006934363.3;refGene_gn=PQLC2;refGene_tx=XM_006934364.3;refGene_gn=
PQLC2;refGene_tx=XM_006934365.4;refGene_gn=PQLC2;refGene_tx=XM_0198362
79.2
C1 14079339 14079432
refGene_gn=PQLC2;refGene_tx=XM_003989615.5;refGene_gn=PQLC2;refGene_tx
=XM_006934363.3;refGene_gn=PQLC2;refGene_tx=XM_006934364.3;refGene_gn=
PQLC2;refGene_tx=XM_006934365.4;refGene_gn=PQLC2;refGene_tx=XM_0198362
79.2
C1 14079925 14080111
refGene_gn=PQLC2;refGene_tx=XM_003989615.5;refGene_gn=PQLC2;refGene_tx
=XM_006934363.3;refGene_gn=PQLC2;refGene_tx=XM_006934364.3;refGene_gn=
PQLC2;refGene_tx=XM_006934365.4;refGene_gn=PQLC2;refGene_tx=XM_0198362
79.2
C1 14081010 14081082
refGene_gn=PQLC2;refGene_tx=XM_003989615.5;refGene_gn=PQLC2;refGene_tx
=XM_006934363.3;refGene_gn=PQLC2;refGene_tx=XM_006934364.3;refGene_gn=
PQLC2;refGene_tx=XM_006934365.4;refGene_gn=PQLC2;refGene_tx=XM_0198362
79.2
C1 14087500 14087588 refGene_gn=CAPZB;refGene_tx=XM_006934367.4
C1 14090341 14090444
refGene_gn=CAPZB;refGene_tx=XM_006934366.3;refGene_gn=CAPZB;refGene_tx
=XM_006934368.3;refGene_gn=CAPZB;refGene_tx=XM_023258222.1
C1 14091743 14091820
refGene_gn=CAPZB;refGene_tx=XM_006934366.3;refGene_gn=CAPZB;refGene_tx
=XM_006934367.4;refGene_gn=CAPZB;refGene_tx=XM_006934368.3;refGene_gn=
CAPZB;refGene_tx=XM_023258222.1
C1 14092566 14092632
refGene_gn=CAPZB;refGene_tx=XM_006934366.3;refGene_gn=CAPZB;refGene_tx
=XM_006934367.4;refGene_gn=CAPZB;refGene_tx=XM_006934368.3;refGene_gn=
CAPZB;refGene_tx=XM_023258222.1
C1 14103270 14103387
refGene_gn=CAPZB;refGene_tx=XM_006934366.3;refGene_gn=CAPZB;refGene_tx
=XM_006934367.4;refGene_gn=CAPZB;refGene_tx=XM_006934368.3;refGene_gn=
CAPZB;refGene_tx=XM_023258222.1
C1 14103894 14104036
refGene_gn=CAPZB;refGene_tx=XM_006934366.3;refGene_gn=CAPZB;refGene_tx
=XM_006934367.4;refGene_gn=CAPZB;refGene_tx=XM_006934368.3;refGene_gn=
CAPZB;refGene_tx=XM_023258222.1
C1 14122377 14122491
refGene_gn=CAPZB;refGene_tx=XM_006934366.3;refGene_gn=CAPZB;refGene_tx
=XM_006934367.4;refGene_gn=CAPZB;refGene_tx=XM_006934368.3;refGene_gn=

CAPZB;refGene_tx=XM_023258222.1
C1 14127660 14127782
refGene_gn=CAPZB;refGene_tx=XM_006934366.3;refGene_gn=CAPZB;refGene_tx
=XM_006934367.4;refGene_gn=CAPZB;refGene_tx=XM_006934368.3;refGene_gn=
CAPZB;refGene_tx=XM_023258222.1
C1 14159391 14159481
refGene_gn=CAPZB;refGene_tx=XM_006934366.3;refGene_gn=CAPZB;refGene_tx
=XM_006934367.4;refGene_gn=CAPZB;refGene_tx=XM_006934368.3;refGene_gn=
CAPZB;refGene_tx=XM_023258222.1
C1 14177382 14177391 refGene_gn=CAPZB;refGene_tx=XM_006934368.3
C1 14220779 14220869
refGene_gn=CAPZB;refGene_tx=XM_006934366.3;refGene_gn=CAPZB;refGene_tx
=XM_006934367.4
C1 14222048 14222051 refGene_gn=CAPZB;refGene_tx=XM_023258222.1
C1 14311418 14311482 refGene_gn=MINOS1;refGene_tx=XM_011284573.3
C1 14342555 14342603 refGene_gn=MINOS1;refGene_tx=XM_011284573.3
C1 14343667 14343777 refGene_gn=MINOS1;refGene_tx=XM_011284573.3
C1 14346308 14346323 refGene_gn=MINOS1;refGene_tx=XM_011284573.3
C1 14371230 14371400
refGene_gn=NBL1;refGene_tx=XM_003989619.5;refGene_gn=NBL1;refGene_tx=X
M_023258223.1
C1 14371528 14371640
refGene_gn=NBL1;refGene_tx=XM_003989619.5;refGene_gn=NBL1;refGene_tx=X
M_023258223.1
C1 14373014 14373263
refGene_gn=NBL1;refGene_tx=XM_003989619.5;refGene_gn=NBL1;refGene_tx=X
M_023258223.1
C1 14379538 14380252 refGene_gn=HTR6;refGene_tx=XM_023258232.1
C1 14390802 14390961 refGene_gn=HTR6;refGene_tx=XM_023258232.1
C1 14391161 14391611 refGene_gn=HTR6;refGene_tx=XM_023258232.1
C1 14394314 14394656
refGene_gn=TMCO4;refGene_tx=XM_023258224.1;refGene_gn=TMCO4;refGene_tx
=XM_023258225.1;refGene_gn=TMCO4;refGene_tx=XM_023258226.1
C1 14401752 14401870
refGene_gn=TMCO4;refGene_tx=XM_023258224.1;refGene_gn=TMCO4;refGene_tx
=XM_023258225.1;refGene_gn=TMCO4;refGene_tx=XM_023258226.1
C1 14408207 14408325
refGene_gn=TMCO4;refGene_tx=XM_023258224.1;refGene_gn=TMCO4;refGene_tx
=XM_023258225.1;refGene_gn=TMCO4;refGene_tx=XM_023258226.1
C1 14419255 14419575 refGene_gn=TMCO4;refGene_tx=XM_023258227.1
C1 14424023 14424280 refGene_gn=TMCO4;refGene_tx=XM_023258228.1
C1 14425888 14425932 refGene_gn=TMCO4;refGene_tx=XM_023258229.1
C1 14427452 14427487 refGene_gn=TMCO4;refGene_tx=XM_023258230.1
C1 14434987 14435072
refGene_gn=TMCO4;refGene_tx=XM_023258224.1;refGene_gn=TMCO4;refGene_tx
=XM_023258225.1;refGene_gn=TMCO4;refGene_tx=XM_023258226.1;refGene_gn=
TMCO4;refGene_tx=XM_023258227.1;refGene_gn=TMCO4;refGene_tx=XM_0232582
28.1;refGene_gn=TMCO4;refGene_tx=XM_023258229.1;refGene_gn=TMCO4;refGe
ne_tx=XM_023258230.1
C1 14438185 14438322

refGene_gn=TMCO4;refGene_tx=XM_023258224.1;refGene_gn=TMCO4;refGene_tx=XM_023258225.1;refGene_gn=TMCO4;refGene_tx=XM_023258226.1;refGene_gn=TMCO4;refGene_tx=XM_023258227.1;refGene_gn=TMCO4;refGene_tx=XM_023258228.1;refGene_gn=TMCO4;refGene_tx=XM_023258229.1;refGene_gn=TMCO4;refGene_tx=XM_023258230.1
C1 14439225 14439390
refGene_gn=TMCO4;refGene_tx=XM_023258224.1;refGene_gn=TMCO4;refGene_tx=XM_023258225.1;refGene_gn=TMCO4;refGene_tx=XM_023258226.1;refGene_gn=TMCO4;refGene_tx=XM_023258227.1;refGene_gn=TMCO4;refGene_tx=XM_023258228.1;refGene_gn=TMCO4;refGene_tx=XM_023258229.1;refGene_gn=TMCO4;refGene_tx=XM_023258230.1
C1 14442432 14442491 refGene_gn=TMCO4;refGene_tx=XM_023258231.1
C1 14444502 14444622
refGene_gn=TMCO4;refGene_tx=XM_023258224.1;refGene_gn=TMCO4;refGene_tx=XM_023258225.1;refGene_gn=TMCO4;refGene_tx=XM_023258226.1;refGene_gn=TMCO4;refGene_tx=XM_023258227.1;refGene_gn=TMCO4;refGene_tx=XM_023258228.1;refGene_gn=TMCO4;refGene_tx=XM_023258229.1;refGene_gn=TMCO4;refGene_tx=XM_023258230.1;refGene_gn=TMCO4;refGene_tx=XM_023258231.1
C1 14445413 14445557
refGene_gn=TMCO4;refGene_tx=XM_023258224.1;refGene_gn=TMCO4;refGene_tx=XM_023258225.1;refGene_gn=TMCO4;refGene_tx=XM_023258226.1;refGene_gn=TMCO4;refGene_tx=XM_023258227.1;refGene_gn=TMCO4;refGene_tx=XM_023258228.1;refGene_gn=TMCO4;refGene_tx=XM_023258229.1;refGene_gn=TMCO4;refGene_tx=XM_023258230.1;refGene_gn=TMCO4;refGene_tx=XM_023258231.1
C1 14445808 14445906
refGene_gn=TMCO4;refGene_tx=XM_023258224.1;refGene_gn=TMCO4;refGene_tx=XM_023258225.1;refGene_gn=TMCO4;refGene_tx=XM_023258226.1;refGene_gn=TMCO4;refGene_tx=XM_023258227.1;refGene_gn=TMCO4;refGene_tx=XM_023258228.1;refGene_gn=TMCO4;refGene_tx=XM_023258229.1;refGene_gn=TMCO4;refGene_tx=XM_023258230.1;refGene_gn=TMCO4;refGene_tx=XM_023258231.1
C1 14452480 14452613
refGene_gn=TMCO4;refGene_tx=XM_023258224.1;refGene_gn=TMCO4;refGene_tx=XM_023258225.1;refGene_gn=TMCO4;refGene_tx=XM_023258226.1;refGene_gn=TMCO4;refGene_tx=XM_023258227.1;refGene_gn=TMCO4;refGene_tx=XM_023258228.1;refGene_gn=TMCO4;refGene_tx=XM_023258229.1;refGene_gn=TMCO4;refGene_tx=XM_023258230.1;refGene_gn=TMCO4;refGene_tx=XM_023258231.1
C1 14466524 14466552
refGene_gn=TMCO4;refGene_tx=XM_023258224.1;refGene_gn=TMCO4;refGene_tx=XM_023258225.1;refGene_gn=TMCO4;refGene_tx=XM_023258226.1;refGene_gn=TMCO4;refGene_tx=XM_023258227.1;refGene_gn=TMCO4;refGene_tx=XM_023258228.1;refGene_gn=TMCO4;refGene_tx=XM_023258229.1;refGene_gn=TMCO4;refGene_tx=XM_023258230.1;refGene_gn=TMCO4;refGene_tx=XM_023258231.1
C1 14467390 14467565
refGene_gn=TMCO4;refGene_tx=XM_023258224.1;refGene_gn=TMCO4;refGene_tx=XM_023258225.1;refGene_gn=TMCO4;refGene_tx=XM_023258226.1;refGene_gn=TMCO4;refGene_tx=XM_023258227.1;refGene_gn=TMCO4;refGene_tx=XM_023258228.1;refGene_gn=TMCO4;refGene_tx=XM_023258229.1;refGene_gn=TMCO4;refGene_tx=XM_023258230.1;refGene_gn=TMCO4;refGene_tx=XM_023258231.1
C1 14473699 14473869
refGene_gn=TMCO4;refGene_tx=XM_023258224.1;refGene_gn=TMCO4;refGene_tx

=XM_023258225.1;refGene_gn=TMCO4;refGene_tx=XM_023258226.1;refGene_gn=TMCO4;refGene_tx=XM_023258227.1;refGene_gn=TMCO4;refGene_tx=XM_023258228.1;refGene_gn=TMCO4;refGene_tx=XM_023258229.1;refGene_gn=TMCO4;refGene_tx=XM_023258230.1;refGene_gn=TMCO4;refGene_tx=XM_023258231.1
C1 14504314 14504977 refGene_gn=RNF186;refGene_tx=XM_003989708.3
C1 14548217 14548438
refGene_gn=OTUD3;refGene_tx=XM_023258233.1;refGene_gn=OTUD3;refGene_tx=XM_023258234.1
C1 14556512 14556535 refGene_gn=OTUD3;refGene_tx=XM_019836285.2
C1 14556757 14556906
refGene_gn=OTUD3;refGene_tx=XM_019836285.2;refGene_gn=OTUD3;refGene_tx=XM_023258233.1;refGene_gn=OTUD3;refGene_tx=XM_023258234.1
C1 14561183 14561296
refGene_gn=OTUD3;refGene_tx=XM_019836285.2;refGene_gn=OTUD3;refGene_tx=XM_023258233.1;refGene_gn=OTUD3;refGene_tx=XM_023258234.1
C1 14565522 14565645
refGene_gn=OTUD3;refGene_tx=XM_019836285.2;refGene_gn=OTUD3;refGene_tx=XM_023258233.1;refGene_gn=OTUD3;refGene_tx=XM_023258234.1
C1 14569568 14569700
refGene_gn=OTUD3;refGene_tx=XM_019836285.2;refGene_gn=OTUD3;refGene_tx=XM_023258233.1;refGene_gn=OTUD3;refGene_tx=XM_023258234.1
C1 14570158 14570255
refGene_gn=OTUD3;refGene_tx=XM_019836285.2;refGene_gn=OTUD3;refGene_tx=XM_023258233.1;refGene_gn=OTUD3;refGene_tx=XM_023258234.1
C1 14571700 14571885
refGene_gn=OTUD3;refGene_tx=XM_019836285.2;refGene_gn=OTUD3;refGene_tx=XM_023258233.1;refGene_gn=OTUD3;refGene_tx=XM_023258234.1
C1 14572665 14572758 refGene_gn=OTUD3;refGene_tx=XM_023258234.1
C1 14572906 14573083
refGene_gn=OTUD3;refGene_tx=XM_019836285.2;refGene_gn=OTUD3;refGene_tx=XM_023258233.1
C1 14584709 14584852
refGene_gn=LOC101091384;refGene_tx=XM_003989621.5
C1 14586734 14586841
refGene_gn=LOC101091384;refGene_tx=XM_003989621.5
C1 14587007 14587146
refGene_gn=LOC101091384;refGene_tx=XM_003989621.5
C1 14587919 14587959
refGene_gn=LOC101091384;refGene_tx=XM_003989621.5
C1 14637861 14637911
refGene_gn=PLA2G5;refGene_tx=XM_006934373.3;refGene_gn=PLA2G5;refGene_tx=XM_019836287.2
C1 14651415 14651465
refGene_gn=PLA2G5;refGene_tx=XM_003989710.4;refGene_gn=PLA2G5;refGene_tx=XM_006934373.3;refGene_gn=PLA2G5;refGene_tx=XM_019836287.2;refGene_gn=PLA2G5;refGene_tx=XM_019836290.2
C1 14652366 14652379
refGene_gn=PLA2G5;refGene_tx=XM_019836293.2;refGene_gn=PLA2G5;refGene_tx=XM_023258235.1
C1 14652708 14652853

refGene_gn=PLA2G5;refGene_tx=XM_003989710.4;refGene_gn=PLA2G5;refGene_tx=XM_006934373.3;refGene_gn=PLA2G5;refGene_tx=XM_019836287.2;refGene_gn=PLA2G5;refGene_tx=XM_019836290.2;refGene_gn=PLA2G5;refGene_tx=XM_019836293.2;refGene_gn=PLA2G5;refGene_tx=XM_023258235.1
C1 14656132 14656239
refGene_gn=PLA2G5;refGene_tx=XM_003989710.4;refGene_gn=PLA2G5;refGene_tx=XM_006934373.3;refGene_gn=PLA2G5;refGene_tx=XM_019836287.2;refGene_gn=PLA2G5;refGene_tx=XM_019836290.2;refGene_gn=PLA2G5;refGene_tx=XM_019836293.2;refGene_gn=PLA2G5;refGene_tx=XM_023258235.1
C1 14656935 14657060
refGene_gn=PLA2G5;refGene_tx=XM_003989710.4;refGene_gn=PLA2G5;refGene_tx=XM_006934373.3;refGene_gn=PLA2G5;refGene_tx=XM_019836287.2;refGene_gn=PLA2G5;refGene_tx=XM_019836290.2;refGene_gn=PLA2G5;refGene_tx=XM_019836293.2;refGene_gn=PLA2G5;refGene_tx=XM_023258235.1
C1 14674864 14675010 refGene_gn=PLA2G2D;refGene_tx=XM_003989622.4
C1 14676442 14676549 refGene_gn=PLA2G2D;refGene_tx=XM_003989622.4
C1 14677231 14677376 refGene_gn=PLA2G2D;refGene_tx=XM_003989622.4
C1 14680800 14680840 refGene_gn=PLA2G2D;refGene_tx=XM_003989622.4
C1 14697356 14697472 refGene_gn=PLA2G2F;refGene_tx=XM_003989623.3
C1 14698102 14698155 refGene_gn=PLA2G2F;refGene_tx=XM_003989623.3
C1 14700744 14700889 refGene_gn=PLA2G2F;refGene_tx=XM_003989623.3
C1 14701792 14701902 refGene_gn=PLA2G2F;refGene_tx=XM_003989623.3
C1 14703934 14704146 refGene_gn=PLA2G2F;refGene_tx=XM_003989623.3
C1 14723277 14723444 refGene_gn=PLA2G2C;refGene_tx=XM_023258236.1
C1 14731282 14731386 refGene_gn=PLA2G2C;refGene_tx=XM_023258236.1
C1 14733032 14733177 refGene_gn=PLA2G2C;refGene_tx=XM_023258236.1
C1 14735216 14735256 refGene_gn=PLA2G2C;refGene_tx=XM_023258236.1
C1 14746827 14747676 refGene_gn=UBXN10;refGene_tx=XM_023258237.1
C1 14850763 14850902
refGene_gn=VWA5B1;refGene_tx=XM_019836295.2;refGene_gn=VWA5B1;refGene_tx=XM_023258238.1;refGene_gn=VWA5B1;refGene_tx=XM_023258239.1
C1 14852558 14852711
refGene_gn=VWA5B1;refGene_tx=XM_019836295.2;refGene_gn=VWA5B1;refGene_tx=XM_023258238.1;refGene_gn=VWA5B1;refGene_tx=XM_023258239.1
C1 14854344 14854615
refGene_gn=VWA5B1;refGene_tx=XM_019836295.2;refGene_gn=VWA5B1;refGene_tx=XM_023258238.1;refGene_gn=VWA5B1;refGene_tx=XM_023258239.1
C1 14858058 14858204
refGene_gn=VWA5B1;refGene_tx=XM_019836295.2;refGene_gn=VWA5B1;refGene_tx=XM_023258238.1;refGene_gn=VWA5B1;refGene_tx=XM_023258239.1
C1 14858983 14859115
refGene_gn=VWA5B1;refGene_tx=XM_019836295.2;refGene_gn=VWA5B1;refGene_tx=XM_023258238.1;refGene_gn=VWA5B1;refGene_tx=XM_023258239.1
C1 14859745 14859870
refGene_gn=VWA5B1;refGene_tx=XM_019836295.2;refGene_gn=VWA5B1;refGene_tx=XM_023258238.1;refGene_gn=VWA5B1;refGene_tx=XM_023258239.1
C1 14863497 14863674
refGene_gn=VWA5B1;refGene_tx=XM_019836295.2;refGene_gn=VWA5B1;refGene_tx=XM_023258238.1;refGene_gn=VWA5B1;refGene_tx=XM_023258239.1
C1 14868059 14868170

refGene_gn=VWA5B1;refGene_tx=XM_019836295.2;refGene_gn=VWA5B1;refGene_tx=XM_023258238.1;refGene_gn=VWA5B1;refGene_tx=XM_023258239.1
C1 14870047 14870250
refGene_gn=VWA5B1;refGene_tx=XM_019836295.2;refGene_gn=VWA5B1;refGene_tx=XM_023258238.1;refGene_gn=VWA5B1;refGene_tx=XM_023258239.1
C1 14870704 14870819
refGene_gn=VWA5B1;refGene_tx=XM_019836295.2;refGene_gn=VWA5B1;refGene_tx=XM_023258238.1;refGene_gn=VWA5B1;refGene_tx=XM_023258239.1
C1 14874360 14874546
refGene_gn=VWA5B1;refGene_tx=XM_019836295.2;refGene_gn=VWA5B1;refGene_tx=XM_023258238.1;refGene_gn=VWA5B1;refGene_tx=XM_023258239.1
C1 14876423 14876607
refGene_gn=VWA5B1;refGene_tx=XM_019836295.2;refGene_gn=VWA5B1;refGene_tx=XM_023258238.1;refGene_gn=VWA5B1;refGene_tx=XM_023258239.1
C1 14877553 14877741
refGene_gn=VWA5B1;refGene_tx=XM_019836295.2;refGene_gn=VWA5B1;refGene_tx=XM_023258238.1;refGene_gn=VWA5B1;refGene_tx=XM_023258239.1
C1 14881493 14881695
refGene_gn=VWA5B1;refGene_tx=XM_019836295.2;refGene_gn=VWA5B1;refGene_tx=XM_023258238.1;refGene_gn=VWA5B1;refGene_tx=XM_023258239.1
C1 14882165 14882480
refGene_gn=VWA5B1;refGene_tx=XM_019836295.2;refGene_gn=VWA5B1;refGene_tx=XM_023258238.1;refGene_gn=VWA5B1;refGene_tx=XM_023258239.1
C1 14884803 14884941
refGene_gn=VWA5B1;refGene_tx=XM_019836295.2;refGene_gn=VWA5B1;refGene_tx=XM_023258238.1;refGene_gn=VWA5B1;refGene_tx=XM_023258239.1
C1 14887384 14887516
refGene_gn=VWA5B1;refGene_tx=XM_019836295.2;refGene_gn=VWA5B1;refGene_tx=XM_023258238.1;refGene_gn=VWA5B1;refGene_tx=XM_023258239.1
C1 14889860 14889935
refGene_gn=VWA5B1;refGene_tx=XM_019836295.2;refGene_gn=VWA5B1;refGene_tx=XM_023258238.1;refGene_gn=VWA5B1;refGene_tx=XM_023258239.1
C1 14890795 14890865
refGene_gn=VWA5B1;refGene_tx=XM_019836295.2;refGene_gn=VWA5B1;refGene_tx=XM_023258238.1;refGene_gn=VWA5B1;refGene_tx=XM_023258239.1
C1 14891382 14891500
refGene_gn=VWA5B1;refGene_tx=XM_019836295.2;refGene_gn=VWA5B1;refGene_tx=XM_023258238.1;refGene_gn=VWA5B1;refGene_tx=XM_023258239.1
C1 14892365 14892872
refGene_gn=VWA5B1;refGene_tx=XM_019836295.2;refGene_gn=VWA5B1;refGene_tx=XM_023258238.1;refGene_gn=VWA5B1;refGene_tx=XM_023258239.1
C1 14999934 15000005 refGene_gn=CAMK2N1;refGene_tx=XM_003989712.3
C1 15001502 15001668 refGene_gn=CAMK2N1;refGene_tx=XM_003989712.3
C1 15015718 15016448 refGene_gn=MUL1;refGene_tx=XM_003989627.5
C1 15017902 15018023 refGene_gn=MUL1;refGene_tx=XM_003989627.5
C1 15018430 15018518 refGene_gn=MUL1;refGene_tx=XM_003989627.5
C1 15022393 15022513 refGene_gn=MUL1;refGene_tx=XM_003989627.5
C1 15056345 15057200 refGene_gn=FAM43B;refGene_tx=XM_023258240.1
C1 15057663 15057825 refGene_gn=FAM43B;refGene_tx=XM_023258240.1
C1 15092093 15092247 refGene_gn=CDA;refGene_tx=XM_023258241.1

C1 15104218 15104330 refGene_gn=CDA;refGene_tx=XM_023258241.1
C1 15109658 15109716 refGene_gn=CDA;refGene_tx=XM_023258241.1
C1 15113345 15113462 refGene_gn=CDA;refGene_tx=XM_023258241.1
C1 15124199 15124595
refGene_gn=PINK1;refGene_tx=XM_023258242.1;refGene_gn=PINK1;refGene_tx
=XM_023258243.1
C1 15135031 15135310
refGene_gn=PINK1;refGene_tx=XM_023258242.1;refGene_gn=PINK1;refGene_tx
=XM_023258243.1
C1 15136903 15137004
refGene_gn=PINK1;refGene_tx=XM_023258242.1;refGene_gn=PINK1;refGene_tx
=XM_023258243.1
C1 15139901 15140084
refGene_gn=PINK1;refGene_tx=XM_023258242.1;refGene_gn=PINK1;refGene_tx
=XM_023258243.1
C1 15140611 15140775
refGene_gn=PINK1;refGene_tx=XM_023258242.1;refGene_gn=PINK1;refGene_tx
=XM_023258243.1
C1 15141720 15141848
refGene_gn=PINK1;refGene_tx=XM_023258242.1;refGene_gn=PINK1;refGene_tx
=XM_023258243.1
C1 15142144 15142381
refGene_gn=PINK1;refGene_tx=XM_023258242.1;refGene_gn=PINK1;refGene_tx
=XM_023258243.1
C1 15143117 15143375 refGene_gn=PINK1;refGene_tx=XM_023258242.1
C1 15145133 15145283 refGene_gn=DDOST;refGene_tx=XM_003989628.4
C1 15145374 15145481 refGene_gn=DDOST;refGene_tx=XM_003989628.4
C1 15145588 15145709 refGene_gn=DDOST;refGene_tx=XM_003989628.4
C1 15146185 15146333 refGene_gn=DDOST;refGene_tx=XM_003989628.4
C1 15146699 15146848 refGene_gn=DDOST;refGene_tx=XM_003989628.4
C1 15147113 15147207 refGene_gn=DDOST;refGene_tx=XM_003989628.4
C1 15147489 15147584 refGene_gn=DDOST;refGene_tx=XM_003989628.4
C1 15147718 15147822 refGene_gn=DDOST;refGene_tx=XM_003989628.4
C1 15148614 15148701 refGene_gn=DDOST;refGene_tx=XM_003989628.4
C1 15150666 15150714 refGene_gn=PINK1;refGene_tx=XM_023258243.1
C1 15151914 15152025 refGene_gn=DDOST;refGene_tx=XM_003989628.4
C1 15152234 15152388 refGene_gn=DDOST;refGene_tx=XM_003989628.4
C1 15155639 15155818
refGene_gn=KIF17;refGene_tx=XM_006934375.3;refGene_gn=KIF17;refGene_tx
=XM_019836298.2;refGene_gn=KIF17;refGene_tx=XM_019836299.2;refGene_gn=
KIF17;refGene_tx=XM_019836300.2;refGene_gn=KIF17;refGene_tx=XM_0198363
01.2
C1 15156732 15156850
refGene_gn=KIF17;refGene_tx=XM_006934375.3;refGene_gn=KIF17;refGene_tx
=XM_019836298.2;refGene_gn=KIF17;refGene_tx=XM_019836299.2;refGene_gn=
KIF17;refGene_tx=XM_019836300.2;refGene_gn=KIF17;refGene_tx=XM_0198363
01.2
C1 15160398 15160469
refGene_gn=KIF17;refGene_tx=XM_006934375.3;refGene_gn=KIF17;refGene_tx
=XM_019836298.2;refGene_gn=KIF17;refGene_tx=XM_019836299.2;refGene_gn=

KIF17;refGene_tx=XM_019836300.2;refGene_gn=KIF17;refGene_tx=XM_0198363
01.2
C1	15161728	15161987
refGene_gn=KIF17;refGene_tx=XM_006934375.3;refGene_gn=KIF17;refGene_tx
=XM_019836298.2;refGene_gn=KIF17;refGene_tx=XM_019836299.2;refGene_gn=
KIF17;refGene_tx=XM_019836300.2;refGene_gn=KIF17;refGene_tx=XM_0198363
01.2
C1	15174377	15174609
refGene_gn=KIF17;refGene_tx=XM_006934375.3;refGene_gn=KIF17;refGene_tx
=XM_019836298.2;refGene_gn=KIF17;refGene_tx=XM_019836299.2;refGene_gn=
KIF17;refGene_tx=XM_019836300.2;refGene_gn=KIF17;refGene_tx=XM_0198363
01.2
C1	15176161	15176373
refGene_gn=KIF17;refGene_tx=XM_006934375.3;refGene_gn=KIF17;refGene_tx
=XM_019836298.2;refGene_gn=KIF17;refGene_tx=XM_019836299.2;refGene_gn=
KIF17;refGene_tx=XM_019836300.2;refGene_gn=KIF17;refGene_tx=XM_0198363
01.2
C1	15176997	15177096
refGene_gn=KIF17;refGene_tx=XM_006934375.3;refGene_gn=KIF17;refGene_tx
=XM_019836298.2;refGene_gn=KIF17;refGene_tx=XM_019836301.2
C1	15177477	15177564
refGene_gn=KIF17;refGene_tx=XM_006934375.3;refGene_gn=KIF17;refGene_tx
=XM_019836298.2;refGene_gn=KIF17;refGene_tx=XM_019836299.2;refGene_gn=
KIF17;refGene_tx=XM_019836301.2
C1	15178542	15179096
refGene_gn=KIF17;refGene_tx=XM_006934375.3;refGene_gn=KIF17;refGene_tx
=XM_019836298.2;refGene_gn=KIF17;refGene_tx=XM_019836299.2;refGene_gn=
KIF17;refGene_tx=XM_019836300.2;refGene_gn=KIF17;refGene_tx=XM_0198363
01.2
C1	15180402	15180550
refGene_gn=KIF17;refGene_tx=XM_006934375.3;refGene_gn=KIF17;refGene_tx
=XM_019836298.2;refGene_gn=KIF17;refGene_tx=XM_019836299.2;refGene_gn=
KIF17;refGene_tx=XM_019836300.2;refGene_gn=KIF17;refGene_tx=XM_0198363
01.2
C1	15184392	15184502
refGene_gn=KIF17;refGene_tx=XM_006934375.3;refGene_gn=KIF17;refGene_tx
=XM_019836298.2;refGene_gn=KIF17;refGene_tx=XM_019836299.2;refGene_gn=
KIF17;refGene_tx=XM_019836300.2;refGene_gn=KIF17;refGene_tx=XM_0198363
01.2
C1	15188870	15189323
refGene_gn=KIF17;refGene_tx=XM_006934375.3;refGene_gn=KIF17;refGene_tx
=XM_019836298.2;refGene_gn=KIF17;refGene_tx=XM_019836299.2;refGene_gn=
KIF17;refGene_tx=XM_019836300.2;refGene_gn=KIF17;refGene_tx=XM_0198363
01.2
C1	15192092	15192282
refGene_gn=KIF17;refGene_tx=XM_006934375.3;refGene_gn=KIF17;refGene_tx
=XM_019836298.2;refGene_gn=KIF17;refGene_tx=XM_019836299.2;refGene_gn=
KIF17;refGene_tx=XM_019836300.2;refGene_gn=KIF17;refGene_tx=XM_0198363
01.2
C1	15197133	15197235

refGene_gn=KIF17;refGene_tx=XM_006934375.3;refGene_gn=KIF17;refGene_tx=XM_019836298.2;refGene_gn=KIF17;refGene_tx=XM_019836299.2;refGene_gn=KIF17;refGene_tx=XM_019836300.2;refGene_gn=KIF17;refGene_tx=XM_019836301.2
C1 15197975 15198122
refGene_gn=KIF17;refGene_tx=XM_006934375.3;refGene_gn=KIF17;refGene_tx=XM_019836298.2;refGene_gn=KIF17;refGene_tx=XM_019836299.2;refGene_gn=KIF17;refGene_tx=XM_019836300.2;refGene_gn=KIF17;refGene_tx=XM_019836301.2
C1 15203991 15204222
refGene_gn=KIF17;refGene_tx=XM_006934375.3;refGene_gn=KIF17;refGene_tx=XM_019836298.2;refGene_gn=KIF17;refGene_tx=XM_019836299.2;refGene_gn=KIF17;refGene_tx=XM_019836300.2
C1 15207679 15207883
refGene_gn=SH2D5;refGene_tx=XM_003989629.4;refGene_gn=SH2D5;refGene_tx=XM_019836302.2;refGene_gn=SH2D5;refGene_tx=XM_019836306.2
C1 15208547 15208707
refGene_gn=SH2D5;refGene_tx=XM_003989629.4;refGene_gn=SH2D5;refGene_tx=XM_019836302.2;refGene_gn=SH2D5;refGene_tx=XM_019836306.2
C1 15209204 15209313
refGene_gn=SH2D5;refGene_tx=XM_003989629.4;refGene_gn=SH2D5;refGene_tx=XM_019836302.2;refGene_gn=SH2D5;refGene_tx=XM_019836306.2
C1 15209642 15209817
refGene_gn=SH2D5;refGene_tx=XM_003989629.4;refGene_gn=SH2D5;refGene_tx=XM_019836302.2;refGene_gn=SH2D5;refGene_tx=XM_019836306.2
C1 15209984 15210233
refGene_gn=SH2D5;refGene_tx=XM_003989629.4;refGene_gn=SH2D5;refGene_tx=XM_019836302.2;refGene_gn=SH2D5;refGene_tx=XM_019836306.2
C1 15211500 15211647
refGene_gn=SH2D5;refGene_tx=XM_003989629.4;refGene_gn=SH2D5;refGene_tx=XM_019836302.2;refGene_gn=SH2D5;refGene_tx=XM_019836306.2
C1 15212615 15212690
refGene_gn=SH2D5;refGene_tx=XM_003989629.4;refGene_gn=SH2D5;refGene_tx=XM_019836302.2
C1 15213177 15213261
refGene_gn=SH2D5;refGene_tx=XM_003989629.4;refGene_gn=SH2D5;refGene_tx=XM_019836302.2
C1 15213607 15213694
refGene_gn=SH2D5;refGene_tx=XM_003989629.4;refGene_gn=SH2D5;refGene_tx=XM_019836302.2;refGene_gn=SH2D5;refGene_tx=XM_019836306.2
C1 15229147 15229442
refGene_gn=HP1BP3;refGene_tx=XM_023258244.1;refGene_gn=HP1BP3;refGene_tx=XM_023258245.1;refGene_gn=HP1BP3;refGene_tx=XM_023258246.1;refGene_gn=HP1BP3;refGene_tx=XM_023258247.1;refGene_gn=HP1BP3;refGene_tx=XM_023258248.1
C1 15229901 15230015
refGene_gn=HP1BP3;refGene_tx=XM_023258244.1;refGene_gn=HP1BP3;refGene_tx=XM_023258245.1;refGene_gn=HP1BP3;refGene_tx=XM_023258246.1;refGene_gn=HP1BP3;refGene_tx=XM_023258247.1;refGene_gn=HP1BP3;refGene_tx=XM_023258248.1

C1 15231892 15232004
refGene_gn=HP1BP3;refGene_tx=XM_023258244.1;refGene_gn=HP1BP3;refGene_tx=XM_023258245.1;refGene_gn=HP1BP3;refGene_tx=XM_023258246.1;refGene_gn=HP1BP3;refGene_tx=XM_023258247.1;refGene_gn=HP1BP3;refGene_tx=XM_023258248.1
C1 15233777 15233937
refGene_gn=HP1BP3;refGene_tx=XM_023258244.1;refGene_gn=HP1BP3;refGene_tx=XM_023258245.1;refGene_gn=HP1BP3;refGene_tx=XM_023258246.1;refGene_gn=HP1BP3;refGene_tx=XM_023258247.1;refGene_gn=HP1BP3;refGene_tx=XM_023258248.1
C1 15239274 15239365
refGene_gn=HP1BP3;refGene_tx=XM_023258244.1;refGene_gn=HP1BP3;refGene_tx=XM_023258245.1;refGene_gn=HP1BP3;refGene_tx=XM_023258246.1;refGene_gn=HP1BP3;refGene_tx=XM_023258247.1;refGene_gn=HP1BP3;refGene_tx=XM_023258248.1
C1 15251094 15251249
refGene_gn=HP1BP3;refGene_tx=XM_023258244.1;refGene_gn=HP1BP3;refGene_tx=XM_023258245.1;refGene_gn=HP1BP3;refGene_tx=XM_023258246.1;refGene_gn=HP1BP3;refGene_tx=XM_023258247.1;refGene_gn=HP1BP3;refGene_tx=XM_023258248.1
C1 15253280 15253361
refGene_gn=HP1BP3;refGene_tx=XM_023258244.1;refGene_gn=HP1BP3;refGene_tx=XM_023258245.1;refGene_gn=HP1BP3;refGene_tx=XM_023258246.1;refGene_gn=HP1BP3;refGene_tx=XM_023258247.1;refGene_gn=HP1BP3;refGene_tx=XM_023258248.1
C1 15255163 15255307
refGene_gn=HP1BP3;refGene_tx=XM_023258244.1;refGene_gn=HP1BP3;refGene_tx=XM_023258245.1;refGene_gn=HP1BP3;refGene_tx=XM_023258246.1;refGene_gn=HP1BP3;refGene_tx=XM_023258247.1;refGene_gn=HP1BP3;refGene_tx=XM_023258248.1
C1 15258308 15258468
refGene_gn=HP1BP3;refGene_tx=XM_023258244.1;refGene_gn=HP1BP3;refGene_tx=XM_023258245.1;refGene_gn=HP1BP3;refGene_tx=XM_023258246.1;refGene_gn=HP1BP3;refGene_tx=XM_023258247.1;refGene_gn=HP1BP3;refGene_tx=XM_023258248.1
C1 15259783 15259937
refGene_gn=HP1BP3;refGene_tx=XM_023258244.1;refGene_gn=HP1BP3;refGene_tx=XM_023258245.1;refGene_gn=HP1BP3;refGene_tx=XM_023258246.1;refGene_gn=HP1BP3;refGene_tx=XM_023258247.1;refGene_gn=HP1BP3;refGene_tx=XM_023258248.1
C1 15262917 15263017
refGene_gn=HP1BP3;refGene_tx=XM_023258244.1;refGene_gn=HP1BP3;refGene_tx=XM_023258245.1;refGene_gn=HP1BP3;refGene_tx=XM_023258246.1;refGene_gn=HP1BP3;refGene_tx=XM_023258247.1;refGene_gn=HP1BP3;refGene_tx=XM_023258248.1
C1 15263467 15263563
refGene_gn=HP1BP3;refGene_tx=XM_023258244.1;refGene_gn=HP1BP3;refGene_tx=XM_023258245.1;refGene_gn=HP1BP3;refGene_tx=XM_023258246.1;refGene_gn=HP1BP3;refGene_tx=XM_023258247.1
C1 15284276 15284458

refGene_gn=EIF4G3;refGene_tx=XM_019836309.2;refGene_gn=EIF4G3;refGene_tx=XM_019836311.2;refGene_gn=EIF4G3;refGene_tx=XM_019836312.2;refGene_gn=EIF4G3;refGene_tx=XM_019836318.2;refGene_gn=EIF4G3;refGene_tx=XM_019836319.2;refGene_gn=EIF4G3;refGene_tx=XM_019836320.2;refGene_gn=EIF4G3;refGene_tx=XM_019836322.2;refGene_gn=EIF4G3;refGene_tx=XM_019836326.2;refGene_gn=EIF4G3;refGene_tx=XM_019836327.2;refGene_gn=EIF4G3;refGene_tx=XM_019836328.2;refGene_gn=EIF4G3;refGene_tx=XM_019836329.2;refGene_gn=EIF4G3;refGene_tx=XM_019836330.2;refGene_gn=EIF4G3;refGene_tx=XM_019836331.2;refGene_gn=EIF4G3;refGene_tx=XM_023258249.1;refGene_gn=EIF4G3;refGene_tx=XM_023258250.1;refGene_gn=EIF4G3;refGene_tx=XM_023258251.1;refGene_gn=EIF4G3;refGene_tx=XM_023258252.1;refGene_gn=EIF4G3;refGene_tx=XM_023258253.1;refGene_gn=EIF4G3;refGene_tx=XM_023258254.1;refGene_gn=EIF4G3;refGene_tx=XM_023258255.1;refGene_gn=EIF4G3;refGene_tx=XM_023258256.1;refGene_gn=EIF4G3;refGene_tx=XM_023258257.1;refGene_gn=EIF4G3;refGene_tx=XM_023258258.1;refGene_gn=EIF4G3;refGene_tx=XM_023258259.1;refGene_gn=EIF4G3;refGene_tx=XM_023258260.1
C1 15287208 15287355
refGene_gn=EIF4G3;refGene_tx=XM_019836309.2;refGene_gn=EIF4G3;refGene_tx=XM_019836311.2;refGene_gn=EIF4G3;refGene_tx=XM_019836312.2;refGene_gn=EIF4G3;refGene_tx=XM_019836318.2;refGene_gn=EIF4G3;refGene_tx=XM_019836319.2;refGene_gn=EIF4G3;refGene_tx=XM_019836320.2;refGene_gn=EIF4G3;refGene_tx=XM_019836322.2;refGene_gn=EIF4G3;refGene_tx=XM_019836326.2;refGene_gn=EIF4G3;refGene_tx=XM_019836327.2;refGene_gn=EIF4G3;refGene_tx=XM_019836328.2;refGene_gn=EIF4G3;refGene_tx=XM_019836329.2;refGene_gn=EIF4G3;refGene_tx=XM_019836330.2;refGene_gn=EIF4G3;refGene_tx=XM_019836331.2;refGene_gn=EIF4G3;refGene_tx=XM_023258249.1;refGene_gn=EIF4G3;refGene_tx=XM_023258250.1;refGene_gn=EIF4G3;refGene_tx=XM_023258251.1;refGene_gn=EIF4G3;refGene_tx=XM_023258252.1;refGene_gn=EIF4G3;refGene_tx=XM_023258253.1;refGene_gn=EIF4G3;refGene_tx=XM_023258254.1;refGene_gn=EIF4G3;refGene_tx=XM_023258255.1;refGene_gn=EIF4G3;refGene_tx=XM_023258256.1;refGene_gn=EIF4G3;refGene_tx=XM_023258257.1;refGene_gn=EIF4G3;refGene_tx=XM_023258258.1;refGene_gn=EIF4G3;refGene_tx=XM_023258259.1;refGene_gn=EIF4G3;refGene_tx=XM_023258260.1
C1 15288773 15288855
refGene_gn=EIF4G3;refGene_tx=XM_019836309.2;refGene_gn=EIF4G3;refGene_tx=XM_019836311.2;refGene_gn=EIF4G3;refGene_tx=XM_019836312.2;refGene_gn=EIF4G3;refGene_tx=XM_019836318.2;refGene_gn=EIF4G3;refGene_tx=XM_019836319.2;refGene_gn=EIF4G3;refGene_tx=XM_019836320.2;refGene_gn=EIF4G3;refGene_tx=XM_019836322.2;refGene_gn=EIF4G3;refGene_tx=XM_019836326.2;refGene_gn=EIF4G3;refGene_tx=XM_019836327.2;refGene_gn=EIF4G3;refGene_tx=XM_019836328.2;refGene_gn=EIF4G3;refGene_tx=XM_019836329.2;refGene_gn=EIF4G3;refGene_tx=XM_019836330.2;refGene_gn=EIF4G3;refGene_tx=XM_019836331.2;refGene_gn=EIF4G3;refGene_tx=XM_023258249.1;refGene_gn=EIF4G3;refGene_tx=XM_023258250.1;refGene_gn=EIF4G3;refGene_tx=XM_023258251.1;refGene_gn=EIF4G3;refGene_tx=XM_023258252.1;refGene_gn=EIF4G3;refGene_tx=XM_023258253.1;refGene_gn=EIF4G3;refGene_tx=XM_023258254.1;refGene_gn=EIF4G3;refGene_tx=XM_023258255.1;refGene_gn=EIF4G3;refGene_tx=XM_023258256.1;refGene_gn=EIF4G3;refGene_tx=XM_023258257.1;refGene_gn=EIF4G3;refGene_tx=XM_023258258.1;refGene_gn=EIF4G3;refGene_tx=XM_023258259.1;refGene_gn=EIF4G3;refGene_tx=XM_023258260.1

C1	15294187	15294334
refGene_gn=EIF4G3;refGene_tx=XM_019836309.2;refGene_gn=EIF4G3;refGene_tx=XM_019836311.2;refGene_gn=EIF4G3;refGene_tx=XM_019836312.2;refGene_gn=EIF4G3;refGene_tx=XM_019836318.2;refGene_gn=EIF4G3;refGene_tx=XM_019836319.2;refGene_gn=EIF4G3;refGene_tx=XM_019836320.2;refGene_gn=EIF4G3;refGene_tx=XM_019836322.2;refGene_gn=EIF4G3;refGene_tx=XM_019836326.2;refGene_gn=EIF4G3;refGene_tx=XM_019836327.2;refGene_gn=EIF4G3;refGene_tx=XM_019836328.2;refGene_gn=EIF4G3;refGene_tx=XM_019836329.2;refGene_gn=EIF4G3;refGene_tx=XM_019836330.2;refGene_gn=EIF4G3;refGene_tx=XM_019836331.2;refGene_gn=EIF4G3;refGene_tx=XM_023258249.1;refGene_gn=EIF4G3;refGene_tx=XM_023258250.1;refGene_gn=EIF4G3;refGene_tx=XM_023258251.1;refGene_gn=EIF4G3;refGene_tx=XM_023258252.1;refGene_gn=EIF4G3;refGene_tx=XM_023258253.1;refGene_gn=EIF4G3;refGene_tx=XM_023258254.1;refGene_gn=EIF4G3;refGene_tx=XM_023258255.1;refGene_gn=EIF4G3;refGene_tx=XM_023258256.1;refGene_gn=EIF4G3;refGene_tx=XM_023258257.1;refGene_gn=EIF4G3;refGene_tx=XM_023258258.1;refGene_gn=EIF4G3;refGene_tx=XM_023258259.1;refGene_gn=EIF4G3;refGene_tx=XM_023258260.1
C1	15303248	15303347
refGene_gn=EIF4G3;refGene_tx=XM_019836309.2;refGene_gn=EIF4G3;refGene_tx=XM_019836311.2;refGene_gn=EIF4G3;refGene_tx=XM_019836312.2;refGene_gn=EIF4G3;refGene_tx=XM_019836318.2;refGene_gn=EIF4G3;refGene_tx=XM_019836319.2;refGene_gn=EIF4G3;refGene_tx=XM_019836320.2;refGene_gn=EIF4G3;refGene_tx=XM_019836322.2;refGene_gn=EIF4G3;refGene_tx=XM_019836326.2;refGene_gn=EIF4G3;refGene_tx=XM_019836327.2;refGene_gn=EIF4G3;refGene_tx=XM_019836328.2;refGene_gn=EIF4G3;refGene_tx=XM_019836329.2;refGene_gn=EIF4G3;refGene_tx=XM_019836330.2;refGene_gn=EIF4G3;refGene_tx=XM_019836331.2;refGene_gn=EIF4G3;refGene_tx=XM_023258249.1;refGene_gn=EIF4G3;refGene_tx=XM_023258250.1;refGene_gn=EIF4G3;refGene_tx=XM_023258251.1;refGene_gn=EIF4G3;refGene_tx=XM_023258252.1;refGene_gn=EIF4G3;refGene_tx=XM_023258253.1;refGene_gn=EIF4G3;refGene_tx=XM_023258254.1;refGene_gn=EIF4G3;refGene_tx=XM_023258255.1;refGene_gn=EIF4G3;refGene_tx=XM_023258256.1;refGene_gn=EIF4G3;refGene_tx=XM_023258257.1;refGene_gn=EIF4G3;refGene_tx=XM_023258258.1;refGene_gn=EIF4G3;refGene_tx=XM_023258259.1;refGene_gn=EIF4G3;refGene_tx=XM_023258260.1
C1	15305819	15305901
refGene_gn=EIF4G3;refGene_tx=XM_019836309.2;refGene_gn=EIF4G3;refGene_tx=XM_019836311.2;refGene_gn=EIF4G3;refGene_tx=XM_019836312.2;refGene_gn=EIF4G3;refGene_tx=XM_019836318.2;refGene_gn=EIF4G3;refGene_tx=XM_019836319.2;refGene_gn=EIF4G3;refGene_tx=XM_019836320.2;refGene_gn=EIF4G3;refGene_tx=XM_019836322.2;refGene_gn=EIF4G3;refGene_tx=XM_019836326.2;refGene_gn=EIF4G3;refGene_tx=XM_019836327.2;refGene_gn=EIF4G3;refGene_tx=XM_019836328.2;refGene_gn=EIF4G3;refGene_tx=XM_019836329.2;refGene_gn=EIF4G3;refGene_tx=XM_019836330.2;refGene_gn=EIF4G3;refGene_tx=XM_019836331.2;refGene_gn=EIF4G3;refGene_tx=XM_023258249.1;refGene_gn=EIF4G3;refGene_tx=XM_023258250.1;refGene_gn=EIF4G3;refGene_tx=XM_023258251.1;refGene_gn=EIF4G3;refGene_tx=XM_023258252.1;refGene_gn=EIF4G3;refGene_tx=XM_023258253.1;refGene_gn=EIF4G3;refGene_tx=XM_023258254.1;refGene_gn=EIF4G3;refGene_tx=XM_023258255.1;refGene_gn=EIF4G3;refGene_tx=XM_023258256.1;refGene_gn=EIF4G3;refGene_tx=XM_023258257.1;refGene_gn=EIF4G3;refGene_tx=XM_023258258.1;refGene_gn=EIF4G3;refGene_tx=XM_023258

259.1;refGene_gn=EIF4G3;refGene_tx=XM_023258260.1
C1 15307660 15307786
refGene_gn=EIF4G3;refGene_tx=XM_019836309.2;refGene_gn=EIF4G3;refGene_tx=XM_019836311.2;refGene_gn=EIF4G3;refGene_tx=XM_019836312.2;refGene_gn=EIF4G3;refGene_tx=XM_019836318.2;refGene_gn=EIF4G3;refGene_tx=XM_019836319.2;refGene_gn=EIF4G3;refGene_tx=XM_019836320.2;refGene_gn=EIF4G3;refGene_tx=XM_019836322.2;refGene_gn=EIF4G3;refGene_tx=XM_019836326.2;refGene_gn=EIF4G3;refGene_tx=XM_019836327.2;refGene_gn=EIF4G3;refGene_tx=XM_019836328.2;refGene_gn=EIF4G3;refGene_tx=XM_019836329.2;refGene_gn=EIF4G3;refGene_tx=XM_019836330.2;refGene_gn=EIF4G3;refGene_tx=XM_019836331.2;refGene_gn=EIF4G3;refGene_tx=XM_023258249.1;refGene_gn=EIF4G3;refGene_tx=XM_023258250.1;refGene_gn=EIF4G3;refGene_tx=XM_023258251.1;refGene_gn=EIF4G3;refGene_tx=XM_023258252.1;refGene_gn=EIF4G3;refGene_tx=XM_023258253.1;refGene_gn=EIF4G3;refGene_tx=XM_023258254.1;refGene_gn=EIF4G3;refGene_tx=XM_023258255.1;refGene_gn=EIF4G3;refGene_tx=XM_023258256.1;refGene_gn=EIF4G3;refGene_tx=XM_023258257.1;refGene_gn=EIF4G3;refGene_tx=XM_023258258.1;refGene_gn=EIF4G3;refGene_tx=XM_023258259.1;refGene_gn=EIF4G3;refGene_tx=XM_023258260.1
C1 15314369 15314542
refGene_gn=EIF4G3;refGene_tx=XM_019836309.2;refGene_gn=EIF4G3;refGene_tx=XM_019836311.2;refGene_gn=EIF4G3;refGene_tx=XM_019836312.2;refGene_gn=EIF4G3;refGene_tx=XM_019836318.2;refGene_gn=EIF4G3;refGene_tx=XM_019836319.2;refGene_gn=EIF4G3;refGene_tx=XM_019836320.2;refGene_gn=EIF4G3;refGene_tx=XM_019836322.2;refGene_gn=EIF4G3;refGene_tx=XM_019836326.2;refGene_gn=EIF4G3;refGene_tx=XM_019836327.2;refGene_gn=EIF4G3;refGene_tx=XM_019836328.2;refGene_gn=EIF4G3;refGene_tx=XM_019836329.2;refGene_gn=EIF4G3;refGene_tx=XM_019836330.2;refGene_gn=EIF4G3;refGene_tx=XM_019836331.2;refGene_gn=EIF4G3;refGene_tx=XM_023258249.1;refGene_gn=EIF4G3;refGene_tx=XM_023258250.1;refGene_gn=EIF4G3;refGene_tx=XM_023258251.1;refGene_gn=EIF4G3;refGene_tx=XM_023258252.1;refGene_gn=EIF4G3;refGene_tx=XM_023258253.1;refGene_gn=EIF4G3;refGene_tx=XM_023258254.1;refGene_gn=EIF4G3;refGene_tx=XM_023258255.1;refGene_gn=EIF4G3;refGene_tx=XM_023258256.1;refGene_gn=EIF4G3;refGene_tx=XM_023258257.1;refGene_gn=EIF4G3;refGene_tx=XM_023258258.1;refGene_gn=EIF4G3;refGene_tx=XM_023258259.1;refGene_gn=EIF4G3;refGene_tx=XM_023258260.1
C1 15318489 15318608
refGene_gn=EIF4G3;refGene_tx=XM_019836309.2;refGene_gn=EIF4G3;refGene_tx=XM_019836311.2;refGene_gn=EIF4G3;refGene_tx=XM_019836312.2;refGene_gn=EIF4G3;refGene_tx=XM_019836318.2;refGene_gn=EIF4G3;refGene_tx=XM_019836319.2;refGene_gn=EIF4G3;refGene_tx=XM_019836320.2;refGene_gn=EIF4G3;refGene_tx=XM_019836322.2;refGene_gn=EIF4G3;refGene_tx=XM_019836326.2;refGene_gn=EIF4G3;refGene_tx=XM_019836327.2;refGene_gn=EIF4G3;refGene_tx=XM_019836328.2;refGene_gn=EIF4G3;refGene_tx=XM_019836329.2;refGene_gn=EIF4G3;refGene_tx=XM_019836330.2;refGene_gn=EIF4G3;refGene_tx=XM_019836331.2;refGene_gn=EIF4G3;refGene_tx=XM_023258249.1;refGene_gn=EIF4G3;refGene_tx=XM_023258250.1;refGene_gn=EIF4G3;refGene_tx=XM_023258251.1;refGene_gn=EIF4G3;refGene_tx=XM_023258252.1;refGene_gn=EIF4G3;refGene_tx=XM_023258253.1;refGene_gn=EIF4G3;refGene_tx=XM_023258254.1;refGene_gn=EIF4G3;refGene_tx=XM_023258255.1;refGene_gn=EIF4G3;refGene_tx=XM_023258256.1;refGene_gn=EIF4G3;refGene_tx=XM_023258257.1;refGene_gn=E

IF4G3;refGene_tx=XM_023258258.1;refGene_gn=EIF4G3;refGene_tx=XM_023258259.1;refGene_gn=EIF4G3;refGene_tx=XM_023258260.1
C1 15320314 15320535
refGene_gn=EIF4G3;refGene_tx=XM_019836309.2;refGene_gn=EIF4G3;refGene_tx=XM_019836311.2;refGene_gn=EIF4G3;refGene_tx=XM_019836312.2;refGene_gn=EIF4G3;refGene_tx=XM_019836318.2;refGene_gn=EIF4G3;refGene_tx=XM_019836319.2;refGene_gn=EIF4G3;refGene_tx=XM_019836320.2;refGene_gn=EIF4G3;refGene_tx=XM_019836322.2;refGene_gn=EIF4G3;refGene_tx=XM_019836326.2;refGene_gn=EIF4G3;refGene_tx=XM_019836327.2;refGene_gn=EIF4G3;refGene_tx=XM_019836328.2;refGene_gn=EIF4G3;refGene_tx=XM_019836329.2;refGene_gn=EIF4G3;refGene_tx=XM_019836330.2;refGene_gn=EIF4G3;refGene_tx=XM_019836331.2;refGene_gn=EIF4G3;refGene_tx=XM_023258249.1;refGene_gn=EIF4G3;refGene_tx=XM_023258250.1;refGene_gn=EIF4G3;refGene_tx=XM_023258251.1;refGene_gn=EIF4G3;refGene_tx=XM_023258252.1;refGene_gn=EIF4G3;refGene_tx=XM_023258253.1;refGene_gn=EIF4G3;refGene_tx=XM_023258254.1;refGene_gn=EIF4G3;refGene_tx=XM_023258255.1;refGene_gn=EIF4G3;refGene_tx=XM_023258256.1;refGene_gn=EIF4G3;refGene_tx=XM_023258257.1;refGene_gn=EIF4G3;refGene_tx=XM_023258258.1;refGene_gn=EIF4G3;refGene_tx=XM_023258259.1;refGene_gn=EIF4G3;refGene_tx=XM_023258260.1
C1 15322402 15322520
refGene_gn=EIF4G3;refGene_tx=XM_019836309.2;refGene_gn=EIF4G3;refGene_tx=XM_019836311.2;refGene_gn=EIF4G3;refGene_tx=XM_019836312.2;refGene_gn=EIF4G3;refGene_tx=XM_019836318.2;refGene_gn=EIF4G3;refGene_tx=XM_019836319.2;refGene_gn=EIF4G3;refGene_tx=XM_019836320.2;refGene_gn=EIF4G3;refGene_tx=XM_019836322.2;refGene_gn=EIF4G3;refGene_tx=XM_019836326.2;refGene_gn=EIF4G3;refGene_tx=XM_019836327.2;refGene_gn=EIF4G3;refGene_tx=XM_019836328.2;refGene_gn=EIF4G3;refGene_tx=XM_019836329.2;refGene_gn=EIF4G3;refGene_tx=XM_019836330.2;refGene_gn=EIF4G3;refGene_tx=XM_019836331.2;refGene_gn=EIF4G3;refGene_tx=XM_023258249.1;refGene_gn=EIF4G3;refGene_tx=XM_023258250.1;refGene_gn=EIF4G3;refGene_tx=XM_023258251.1;refGene_gn=EIF4G3;refGene_tx=XM_023258252.1;refGene_gn=EIF4G3;refGene_tx=XM_023258253.1;refGene_gn=EIF4G3;refGene_tx=XM_023258254.1;refGene_gn=EIF4G3;refGene_tx=XM_023258255.1;refGene_gn=EIF4G3;refGene_tx=XM_023258256.1;refGene_gn=EIF4G3;refGene_tx=XM_023258257.1;refGene_gn=EIF4G3;refGene_tx=XM_023258258.1;refGene_gn=EIF4G3;refGene_tx=XM_023258259.1;refGene_gn=EIF4G3;refGene_tx=XM_023258260.1
C1 15323868 15323962
refGene_gn=EIF4G3;refGene_tx=XM_019836309.2;refGene_gn=EIF4G3;refGene_tx=XM_019836311.2;refGene_gn=EIF4G3;refGene_tx=XM_019836312.2;refGene_gn=EIF4G3;refGene_tx=XM_019836318.2;refGene_gn=EIF4G3;refGene_tx=XM_019836319.2;refGene_gn=EIF4G3;refGene_tx=XM_019836320.2;refGene_gn=EIF4G3;refGene_tx=XM_019836322.2;refGene_gn=EIF4G3;refGene_tx=XM_019836326.2;refGene_gn=EIF4G3;refGene_tx=XM_019836327.2;refGene_gn=EIF4G3;refGene_tx=XM_019836328.2;refGene_gn=EIF4G3;refGene_tx=XM_019836329.2;refGene_gn=EIF4G3;refGene_tx=XM_019836330.2;refGene_gn=EIF4G3;refGene_tx=XM_019836331.2;refGene_gn=EIF4G3;refGene_tx=XM_023258249.1;refGene_gn=EIF4G3;refGene_tx=XM_023258250.1;refGene_gn=EIF4G3;refGene_tx=XM_023258251.1;refGene_gn=EIF4G3;refGene_tx=XM_023258252.1;refGene_gn=EIF4G3;refGene_tx=XM_023258253.1;refGene_gn=EIF4G3;refGene_tx=XM_023258254.1;refGene_gn=EIF4G3;refGene_tx=XM_023258255.1;refGene_gn=EIF4G3;refGene_tx=X

M_023258256.1;refGene_gn=EIF4G3;refGene_tx=XM_023258257.1;refGene_gn=EIF4G3;refGene_tx=XM_023258258.1;refGene_gn=EIF4G3;refGene_tx=XM_023258259.1;refGene_gn=EIF4G3;refGene_tx=XM_023258260.1
C1 15325971 15326066
refGene_gn=EIF4G3;refGene_tx=XM_019836309.2;refGene_gn=EIF4G3;refGene_tx=XM_019836311.2;refGene_gn=EIF4G3;refGene_tx=XM_019836312.2;refGene_gn=EIF4G3;refGene_tx=XM_019836318.2;refGene_gn=EIF4G3;refGene_tx=XM_019836319.2;refGene_gn=EIF4G3;refGene_tx=XM_019836320.2;refGene_gn=EIF4G3;refGene_tx=XM_019836322.2;refGene_gn=EIF4G3;refGene_tx=XM_019836326.2;refGene_gn=EIF4G3;refGene_tx=XM_019836327.2;refGene_gn=EIF4G3;refGene_tx=XM_019836328.2;refGene_gn=EIF4G3;refGene_tx=XM_019836329.2;refGene_gn=EIF4G3;refGene_tx=XM_019836330.2;refGene_gn=EIF4G3;refGene_tx=XM_019836331.2;refGene_gn=EIF4G3;refGene_tx=XM_023258249.1;refGene_gn=EIF4G3;refGene_tx=XM_023258250.1;refGene_gn=EIF4G3;refGene_tx=XM_023258251.1;refGene_gn=EIF4G3;refGene_tx=XM_023258252.1;refGene_gn=EIF4G3;refGene_tx=XM_023258253.1;refGene_gn=EIF4G3;refGene_tx=XM_023258254.1;refGene_gn=EIF4G3;refGene_tx=XM_023258255.1;refGene_gn=EIF4G3;refGene_tx=XM_023258256.1;refGene_gn=EIF4G3;refGene_tx=XM_023258257.1;refGene_gn=EIF4G3;refGene_tx=XM_023258258.1;refGene_gn=EIF4G3;refGene_tx=XM_023258259.1;refGene_gn=EIF4G3;refGene_tx=XM_023258260.1
C1 15329007 15329140
refGene_gn=EIF4G3;refGene_tx=XM_019836309.2;refGene_gn=EIF4G3;refGene_tx=XM_019836311.2;refGene_gn=EIF4G3;refGene_tx=XM_019836312.2;refGene_gn=EIF4G3;refGene_tx=XM_019836318.2;refGene_gn=EIF4G3;refGene_tx=XM_019836319.2;refGene_gn=EIF4G3;refGene_tx=XM_019836320.2;refGene_gn=EIF4G3;refGene_tx=XM_019836322.2;refGene_gn=EIF4G3;refGene_tx=XM_019836326.2;refGene_gn=EIF4G3;refGene_tx=XM_019836327.2;refGene_gn=EIF4G3;refGene_tx=XM_019836328.2;refGene_gn=EIF4G3;refGene_tx=XM_019836329.2;refGene_gn=EIF4G3;refGene_tx=XM_019836330.2;refGene_gn=EIF4G3;refGene_tx=XM_019836331.2;refGene_gn=EIF4G3;refGene_tx=XM_023258249.1;refGene_gn=EIF4G3;refGene_tx=XM_023258250.1;refGene_gn=EIF4G3;refGene_tx=XM_023258251.1;refGene_gn=EIF4G3;refGene_tx=XM_023258252.1;refGene_gn=EIF4G3;refGene_tx=XM_023258253.1;refGene_gn=EIF4G3;refGene_tx=XM_023258254.1;refGene_gn=EIF4G3;refGene_tx=XM_023258255.1;refGene_gn=EIF4G3;refGene_tx=XM_023258256.1;refGene_gn=EIF4G3;refGene_tx=XM_023258257.1;refGene_gn=EIF4G3;refGene_tx=XM_023258258.1;refGene_gn=EIF4G3;refGene_tx=XM_023258259.1;refGene_gn=EIF4G3;refGene_tx=XM_023258260.1
C1 15331537 15331642
refGene_gn=EIF4G3;refGene_tx=XM_019836309.2;refGene_gn=EIF4G3;refGene_tx=XM_019836311.2;refGene_gn=EIF4G3;refGene_tx=XM_019836312.2;refGene_gn=EIF4G3;refGene_tx=XM_019836318.2;refGene_gn=EIF4G3;refGene_tx=XM_019836319.2;refGene_gn=EIF4G3;refGene_tx=XM_019836320.2;refGene_gn=EIF4G3;refGene_tx=XM_019836322.2;refGene_gn=EIF4G3;refGene_tx=XM_019836326.2;refGene_gn=EIF4G3;refGene_tx=XM_019836327.2;refGene_gn=EIF4G3;refGene_tx=XM_019836328.2;refGene_gn=EIF4G3;refGene_tx=XM_019836329.2;refGene_gn=EIF4G3;refGene_tx=XM_019836330.2;refGene_gn=EIF4G3;refGene_tx=XM_019836331.2;refGene_gn=EIF4G3;refGene_tx=XM_023258249.1;refGene_gn=EIF4G3;refGene_tx=XM_023258250.1;refGene_gn=EIF4G3;refGene_tx=XM_023258251.1;refGene_gn=EIF4G3;refGene_tx=XM_023258252.1;refGene_gn=EIF4G3;refGene_tx=XM_023258253.1;refGene_gn=EIF4G3;refGene_tx=XM_023258254.1;refG

ene_gn=EIF4G3;refGene_tx=XM_023258255.1;refGene_gn=EIF4G3;refGene_tx=XM_023258256.1;refGene_gn=EIF4G3;refGene_tx=XM_023258257.1;refGene_gn=EIF4G3;refGene_tx=XM_023258258.1;refGene_gn=EIF4G3;refGene_tx=XM_023258259.1;refGene_gn=EIF4G3;refGene_tx=XM_023258260.1
C1 15333018 15333294
refGene_gn=EIF4G3;refGene_tx=XM_019836309.2;refGene_gn=EIF4G3;refGene_tx=XM_019836311.2;refGene_gn=EIF4G3;refGene_tx=XM_019836312.2;refGene_gn=EIF4G3;refGene_tx=XM_019836318.2;refGene_gn=EIF4G3;refGene_tx=XM_019836319.2;refGene_gn=EIF4G3;refGene_tx=XM_019836320.2;refGene_gn=EIF4G3;refGene_tx=XM_019836322.2;refGene_gn=EIF4G3;refGene_tx=XM_019836326.2;refGene_gn=EIF4G3;refGene_tx=XM_019836327.2;refGene_gn=EIF4G3;refGene_tx=XM_019836328.2;refGene_gn=EIF4G3;refGene_tx=XM_019836329.2;refGene_gn=EIF4G3;refGene_tx=XM_019836330.2;refGene_gn=EIF4G3;refGene_tx=XM_019836331.2;refGene_gn=EIF4G3;refGene_tx=XM_023258249.1;refGene_gn=EIF4G3;refGene_tx=XM_023258250.1;refGene_gn=EIF4G3;refGene_tx=XM_023258251.1;refGene_gn=EIF4G3;refGene_tx=XM_023258252.1;refGene_gn=EIF4G3;refGene_tx=XM_023258253.1;refGene_gn=EIF4G3;refGene_tx=XM_023258254.1;refGene_gn=EIF4G3;refGene_tx=XM_023258255.1;refGene_gn=EIF4G3;refGene_tx=XM_023258256.1;refGene_gn=EIF4G3;refGene_tx=XM_023258257.1;refGene_gn=EIF4G3;refGene_tx=XM_023258258.1;refGene_gn=EIF4G3;refGene_tx=XM_023258259.1;refGene_gn=EIF4G3;refGene_tx=XM_023258260.1
C1 15333702 15333849
refGene_gn=EIF4G3;refGene_tx=XM_019836309.2;refGene_gn=EIF4G3;refGene_tx=XM_019836311.2;refGene_gn=EIF4G3;refGene_tx=XM_019836312.2;refGene_gn=EIF4G3;refGene_tx=XM_019836318.2;refGene_gn=EIF4G3;refGene_tx=XM_019836319.2;refGene_gn=EIF4G3;refGene_tx=XM_019836320.2;refGene_gn=EIF4G3;refGene_tx=XM_019836322.2;refGene_gn=EIF4G3;refGene_tx=XM_019836326.2;refGene_gn=EIF4G3;refGene_tx=XM_019836327.2;refGene_gn=EIF4G3;refGene_tx=XM_019836328.2;refGene_gn=EIF4G3;refGene_tx=XM_019836329.2;refGene_gn=EIF4G3;refGene_tx=XM_019836330.2;refGene_gn=EIF4G3;refGene_tx=XM_019836331.2;refGene_gn=EIF4G3;refGene_tx=XM_023258249.1;refGene_gn=EIF4G3;refGene_tx=XM_023258250.1;refGene_gn=EIF4G3;refGene_tx=XM_023258251.1;refGene_gn=EIF4G3;refGene_tx=XM_023258252.1;refGene_gn=EIF4G3;refGene_tx=XM_023258253.1;refGene_gn=EIF4G3;refGene_tx=XM_023258254.1;refGene_gn=EIF4G3;refGene_tx=XM_023258255.1;refGene_gn=EIF4G3;refGene_tx=XM_023258256.1;refGene_gn=EIF4G3;refGene_tx=XM_023258257.1;refGene_gn=EIF4G3;refGene_tx=XM_023258258.1;refGene_gn=EIF4G3;refGene_tx=XM_023258259.1;refGene_gn=EIF4G3;refGene_tx=XM_023258260.1;refGene_gn=EIF4G3;refGene_tx=XM_023258261.1
C1 15350476 15350715
refGene_gn=EIF4G3;refGene_tx=XM_019836309.2;refGene_gn=EIF4G3;refGene_tx=XM_019836311.2;refGene_gn=EIF4G3;refGene_tx=XM_019836312.2;refGene_gn=EIF4G3;refGene_tx=XM_019836318.2;refGene_gn=EIF4G3;refGene_tx=XM_019836319.2;refGene_gn=EIF4G3;refGene_tx=XM_019836320.2;refGene_gn=EIF4G3;refGene_tx=XM_019836322.2;refGene_gn=EIF4G3;refGene_tx=XM_019836326.2;refGene_gn=EIF4G3;refGene_tx=XM_019836327.2;refGene_gn=EIF4G3;refGene_tx=XM_019836328.2;refGene_gn=EIF4G3;refGene_tx=XM_019836329.2;refGene_gn=EIF4G3;refGene_tx=XM_019836330.2;refGene_gn=EIF4G3;refGene_tx=XM_019836331.2;refGene_gn=EIF4G3;refGene_tx=XM_023258249.1;refGene_gn=EIF4G3;refGene_tx=XM_023258250.1;refGene_gn=EIF4G3;refGene_tx=XM_02325825

1.1;refGene_gn=EIF4G3;refGene_tx=XM_023258252.1;refGene_gn=EIF4G3;refGene_tx=XM_023258253.1;refGene_gn=EIF4G3;refGene_tx=XM_023258254.1;refGene_gn=EIF4G3;refGene_tx=XM_023258255.1;refGene_gn=EIF4G3;refGene_tx=XM_023258256.1;refGene_gn=EIF4G3;refGene_tx=XM_023258257.1;refGene_gn=EIF4G3;refGene_tx=XM_023258258.1;refGene_gn=EIF4G3;refGene_tx=XM_023258259.1;refGene_gn=EIF4G3;refGene_tx=XM_023258260.1;refGene_gn=EIF4G3;refGene_tx=XM_023258261.1
C1 15356686 15356857
refGene_gn=EIF4G3;refGene_tx=XM_019836309.2;refGene_gn=EIF4G3;refGene_tx=XM_019836311.2;refGene_gn=EIF4G3;refGene_tx=XM_019836312.2;refGene_gn=EIF4G3;refGene_tx=XM_019836318.2;refGene_gn=EIF4G3;refGene_tx=XM_019836319.2;refGene_gn=EIF4G3;refGene_tx=XM_019836320.2;refGene_gn=EIF4G3;refGene_tx=XM_019836322.2;refGene_gn=EIF4G3;refGene_tx=XM_019836326.2;refGene_gn=EIF4G3;refGene_tx=XM_019836327.2;refGene_gn=EIF4G3;refGene_tx=XM_019836328.2;refGene_gn=EIF4G3;refGene_tx=XM_019836329.2;refGene_gn=EIF4G3;refGene_tx=XM_019836330.2;refGene_gn=EIF4G3;refGene_tx=XM_019836331.2;refGene_gn=EIF4G3;refGene_tx=XM_023258249.1;refGene_gn=EIF4G3;refGene_tx=XM_023258250.1;refGene_gn=EIF4G3;refGene_tx=XM_023258251.1;refGene_gn=EIF4G3;refGene_tx=XM_023258252.1;refGene_gn=EIF4G3;refGene_tx=XM_023258253.1;refGene_gn=EIF4G3;refGene_tx=XM_023258254.1;refGene_gn=EIF4G3;refGene_tx=XM_023258255.1;refGene_gn=EIF4G3;refGene_tx=XM_023258256.1;refGene_gn=EIF4G3;refGene_tx=XM_023258257.1;refGene_gn=EIF4G3;refGene_tx=XM_023258258.1;refGene_gn=EIF4G3;refGene_tx=XM_023258259.1;refGene_gn=EIF4G3;refGene_tx=XM_023258260.1;refGene_gn=EIF4G3;refGene_tx=XM_023258261.1
C1 15362373 15362484
refGene_gn=EIF4G3;refGene_tx=XM_019836309.2;refGene_gn=EIF4G3;refGene_tx=XM_019836311.2;refGene_gn=EIF4G3;refGene_tx=XM_019836312.2;refGene_gn=EIF4G3;refGene_tx=XM_019836318.2;refGene_gn=EIF4G3;refGene_tx=XM_019836319.2;refGene_gn=EIF4G3;refGene_tx=XM_019836320.2;refGene_gn=EIF4G3;refGene_tx=XM_019836322.2;refGene_gn=EIF4G3;refGene_tx=XM_019836326.2;refGene_gn=EIF4G3;refGene_tx=XM_019836327.2;refGene_gn=EIF4G3;refGene_tx=XM_019836328.2;refGene_gn=EIF4G3;refGene_tx=XM_019836329.2;refGene_gn=EIF4G3;refGene_tx=XM_019836330.2;refGene_gn=EIF4G3;refGene_tx=XM_019836331.2;refGene_gn=EIF4G3;refGene_tx=XM_023258249.1;refGene_gn=EIF4G3;refGene_tx=XM_023258250.1;refGene_gn=EIF4G3;refGene_tx=XM_023258251.1;refGene_gn=EIF4G3;refGene_tx=XM_023258252.1;refGene_gn=EIF4G3;refGene_tx=XM_023258253.1;refGene_gn=EIF4G3;refGene_tx=XM_023258257.1;refGene_gn=EIF4G3;refGene_tx=XM_023258258.1;refGene_gn=EIF4G3;refGene_tx=XM_023258259.1;refGene_gn=EIF4G3;refGene_tx=XM_023258260.1;refGene_gn=EIF4G3;refGene_tx=XM_023258261.1
C1 15362869 15362989
refGene_gn=EIF4G3;refGene_tx=XM_019836309.2;refGene_gn=EIF4G3;refGene_tx=XM_019836311.2;refGene_gn=EIF4G3;refGene_tx=XM_019836312.2;refGene_gn=EIF4G3;refGene_tx=XM_019836318.2;refGene_gn=EIF4G3;refGene_tx=XM_019836319.2;refGene_gn=EIF4G3;refGene_tx=XM_019836320.2;refGene_gn=EIF4G3;refGene_tx=XM_019836322.2;refGene_gn=EIF4G3;refGene_tx=XM_019836326.2;refGene_gn=EIF4G3;refGene_tx=XM_019836327.2;refGene_gn=EIF4G3;refGene_tx=XM_019836328.2;refGene_gn=EIF4G3;refGene_tx=XM_019836329.2;refGene_gn=EIF4G3;refGene_tx=XM_019836330.2;refGene_gn=EIF4G3;refGene_tx=XM_

019836331.2;refGene_gn=EIF4G3;refGene_tx=XM_023258249.1;refGene_gn=EIF4G3;refGene_tx=XM_023258250.1;refGene_gn=EIF4G3;refGene_tx=XM_023258251.1;refGene_gn=EIF4G3;refGene_tx=XM_023258252.1;refGene_gn=EIF4G3;refGene_tx=XM_023258253.1;refGene_gn=EIF4G3;refGene_tx=XM_023258254.1;refGene_gn=EIF4G3;refGene_tx=XM_023258255.1;refGene_gn=EIF4G3;refGene_tx=XM_023258256.1;refGene_gn=EIF4G3;refGene_tx=XM_023258257.1;refGene_gn=EIF4G3;refGene_tx=XM_023258258.1;refGene_gn=EIF4G3;refGene_tx=XM_023258259.1;refGene_gn=EIF4G3;refGene_tx=XM_023258260.1;refGene_gn=EIF4G3;refGene_tx=XM_023258261.1
C1 15364966 15365100
refGene_gn=EIF4G3;refGene_tx=XM_019836309.2;refGene_gn=EIF4G3;refGene_tx=XM_019836311.2;refGene_gn=EIF4G3;refGene_tx=XM_019836312.2;refGene_gn=EIF4G3;refGene_tx=XM_019836318.2;refGene_gn=EIF4G3;refGene_tx=XM_019836319.2;refGene_gn=EIF4G3;refGene_tx=XM_019836320.2;refGene_gn=EIF4G3;refGene_tx=XM_019836322.2;refGene_gn=EIF4G3;refGene_tx=XM_019836326.2;refGene_gn=EIF4G3;refGene_tx=XM_019836327.2;refGene_gn=EIF4G3;refGene_tx=XM_019836328.2;refGene_gn=EIF4G3;refGene_tx=XM_019836329.2;refGene_gn=EIF4G3;refGene_tx=XM_019836330.2;refGene_gn=EIF4G3;refGene_tx=XM_019836331.2;refGene_gn=EIF4G3;refGene_tx=XM_023258249.1;refGene_gn=EIF4G3;refGene_tx=XM_023258250.1;refGene_gn=EIF4G3;refGene_tx=XM_023258251.1;refGene_gn=EIF4G3;refGene_tx=XM_023258252.1;refGene_gn=EIF4G3;refGene_tx=XM_023258253.1;refGene_gn=EIF4G3;refGene_tx=XM_023258254.1;refGene_gn=EIF4G3;refGene_tx=XM_023258255.1;refGene_gn=EIF4G3;refGene_tx=XM_023258256.1;refGene_gn=EIF4G3;refGene_tx=XM_023258257.1;refGene_gn=EIF4G3;refGene_tx=XM_023258258.1;refGene_gn=EIF4G3;refGene_tx=XM_023258259.1;refGene_gn=EIF4G3;refGene_tx=XM_023258260.1;refGene_gn=EIF4G3;refGene_tx=XM_023258261.1
C1 15369251 15369498
refGene_gn=EIF4G3;refGene_tx=XM_019836309.2;refGene_gn=EIF4G3;refGene_tx=XM_019836311.2;refGene_gn=EIF4G3;refGene_tx=XM_019836312.2;refGene_gn=EIF4G3;refGene_tx=XM_019836318.2;refGene_gn=EIF4G3;refGene_tx=XM_019836319.2;refGene_gn=EIF4G3;refGene_tx=XM_019836320.2;refGene_gn=EIF4G3;refGene_tx=XM_019836322.2;refGene_gn=EIF4G3;refGene_tx=XM_019836326.2;refGene_gn=EIF4G3;refGene_tx=XM_019836327.2;refGene_gn=EIF4G3;refGene_tx=XM_019836328.2;refGene_gn=EIF4G3;refGene_tx=XM_019836329.2;refGene_gn=EIF4G3;refGene_tx=XM_019836330.2;refGene_gn=EIF4G3;refGene_tx=XM_019836331.2;refGene_gn=EIF4G3;refGene_tx=XM_023258249.1;refGene_gn=EIF4G3;refGene_tx=XM_023258250.1;refGene_gn=EIF4G3;refGene_tx=XM_023258251.1;refGene_gn=EIF4G3;refGene_tx=XM_023258252.1;refGene_gn=EIF4G3;refGene_tx=XM_023258253.1;refGene_gn=EIF4G3;refGene_tx=XM_023258254.1;refGene_gn=EIF4G3;refGene_tx=XM_023258255.1;refGene_gn=EIF4G3;refGene_tx=XM_023258256.1;refGene_gn=EIF4G3;refGene_tx=XM_023258257.1;refGene_gn=EIF4G3;refGene_tx=XM_023258258.1;refGene_gn=EIF4G3;refGene_tx=XM_023258259.1;refGene_gn=EIF4G3;refGene_tx=XM_023258260.1;refGene_gn=EIF4G3;refGene_tx=XM_023258261.1
C1 15373518 15373607
refGene_gn=EIF4G3;refGene_tx=XM_019836309.2;refGene_gn=EIF4G3;refGene_tx=XM_019836311.2;refGene_gn=EIF4G3;refGene_tx=XM_019836312.2;refGene_gn=EIF4G3;refGene_tx=XM_019836318.2;refGene_gn=EIF4G3;refGene_tx=XM_019836319.2;refGene_gn=EIF4G3;refGene_tx=XM_019836320.2;refGene_gn=EIF4G

3;refGene_tx=XM_019836322.2;refGene_gn=EIF4G3;refGene_tx=XM_019836326.
2;refGene_gn=EIF4G3;refGene_tx=XM_019836327.2;refGene_gn=EIF4G3;refGen
e_tx=XM_019836328.2;refGene_gn=EIF4G3;refGene_tx=XM_019836329.2;refGen
e_gn=EIF4G3;refGene_tx=XM_019836330.2;refGene_gn=EIF4G3;refGene_tx=XM_
019836331.2;refGene_gn=EIF4G3;refGene_tx=XM_023258249.1;refGene_gn=EIF
4G3;refGene_tx=XM_023258250.1;refGene_gn=EIF4G3;refGene_tx=XM_02325825
1.1;refGene_gn=EIF4G3;refGene_tx=XM_023258252.1;refGene_gn=EIF4G3;refG
ene_tx=XM_023258253.1;refGene_gn=EIF4G3;refGene_tx=XM_023258254.1;refG
ene_gn=EIF4G3;refGene_tx=XM_023258255.1;refGene_gn=EIF4G3;refGene_tx=X
M_023258256.1;refGene_gn=EIF4G3;refGene_tx=XM_023258257.1;refGene_gn=E
IF4G3;refGene_tx=XM_023258258.1;refGene_gn=EIF4G3;refGene_tx=XM_023258
259.1;refGene_gn=EIF4G3;refGene_tx=XM_023258260.1;refGene_gn=EIF4G3;re
fGene_tx=XM_023258261.1
C1 15399243 15400083
refGene_gn=EIF4G3;refGene_tx=XM_019836309.2;refGene_gn=EIF4G3;refGene_
tx=XM_019836311.2;refGene_gn=EIF4G3;refGene_tx=XM_019836312.2;refGene_
gn=EIF4G3;refGene_tx=XM_019836318.2;refGene_gn=EIF4G3;refGene_tx=XM_01
9836319.2;refGene_gn=EIF4G3;refGene_tx=XM_019836320.2;refGene_gn=EIF4G
3;refGene_tx=XM_019836322.2;refGene_gn=EIF4G3;refGene_tx=XM_019836326.
2;refGene_gn=EIF4G3;refGene_tx=XM_019836327.2;refGene_gn=EIF4G3;refGen
e_tx=XM_019836328.2;refGene_gn=EIF4G3;refGene_tx=XM_019836329.2;refGen
e_gn=EIF4G3;refGene_tx=XM_019836330.2;refGene_gn=EIF4G3;refGene_tx=XM_
019836331.2;refGene_gn=EIF4G3;refGene_tx=XM_023258249.1;refGene_gn=EIF
4G3;refGene_tx=XM_023258250.1;refGene_gn=EIF4G3;refGene_tx=XM_02325825
1.1;refGene_gn=EIF4G3;refGene_tx=XM_023258252.1;refGene_gn=EIF4G3;refG
ene_tx=XM_023258253.1;refGene_gn=EIF4G3;refGene_tx=XM_023258254.1;refG
ene_gn=EIF4G3;refGene_tx=XM_023258255.1;refGene_gn=EIF4G3;refGene_tx=X
M_023258256.1;refGene_gn=EIF4G3;refGene_tx=XM_023258257.1;refGene_gn=E
IF4G3;refGene_tx=XM_023258258.1;refGene_gn=EIF4G3;refGene_tx=XM_023258
259.1;refGene_gn=EIF4G3;refGene_tx=XM_023258260.1;refGene_gn=EIF4G3;re
fGene_tx=XM_023258261.1
C1 15407135 15407244
refGene_gn=EIF4G3;refGene_tx=XM_019836309.2;refGene_gn=EIF4G3;refGene_
tx=XM_019836311.2;refGene_gn=EIF4G3;refGene_tx=XM_019836312.2;refGene_
gn=EIF4G3;refGene_tx=XM_019836318.2;refGene_gn=EIF4G3;refGene_tx=XM_01
9836319.2;refGene_gn=EIF4G3;refGene_tx=XM_019836320.2;refGene_gn=EIF4G
3;refGene_tx=XM_019836322.2;refGene_gn=EIF4G3;refGene_tx=XM_019836326.
2;refGene_gn=EIF4G3;refGene_tx=XM_019836327.2;refGene_gn=EIF4G3;refGen
e_tx=XM_019836328.2;refGene_gn=EIF4G3;refGene_tx=XM_019836329.2;refGen
e_gn=EIF4G3;refGene_tx=XM_019836330.2;refGene_gn=EIF4G3;refGene_tx=XM_
019836331.2;refGene_gn=EIF4G3;refGene_tx=XM_023258249.1;refGene_gn=EIF
4G3;refGene_tx=XM_023258250.1;refGene_gn=EIF4G3;refGene_tx=XM_02325825
1.1;refGene_gn=EIF4G3;refGene_tx=XM_023258252.1;refGene_gn=EIF4G3;refG
ene_tx=XM_023258253.1;refGene_gn=EIF4G3;refGene_tx=XM_023258254.1;refG
ene_gn=EIF4G3;refGene_tx=XM_023258255.1;refGene_gn=EIF4G3;refGene_tx=X
M_023258256.1;refGene_gn=EIF4G3;refGene_tx=XM_023258257.1;refGene_gn=E
IF4G3;refGene_tx=XM_023258258.1;refGene_gn=EIF4G3;refGene_tx=XM_023258
259.1;refGene_gn=EIF4G3;refGene_tx=XM_023258260.1;refGene_gn=EIF4G3;re
fGene_tx=XM_023258261.1
C1 15415133 15415256

refGene_gn=EIF4G3;refGene_tx=XM_019836309.2;refGene_gn=EIF4G3;refGene_tx=XM_019836311.2;refGene_gn=EIF4G3;refGene_tx=XM_019836312.2;refGene_gn=EIF4G3;refGene_tx=XM_019836318.2;refGene_gn=EIF4G3;refGene_tx=XM_019836319.2;refGene_gn=EIF4G3;refGene_tx=XM_019836320.2;refGene_gn=EIF4G3;refGene_tx=XM_019836322.2;refGene_gn=EIF4G3;refGene_tx=XM_019836326.2;refGene_gn=EIF4G3;refGene_tx=XM_019836327.2;refGene_gn=EIF4G3;refGene_tx=XM_019836328.2;refGene_gn=EIF4G3;refGene_tx=XM_019836329.2;refGene_gn=EIF4G3;refGene_tx=XM_019836330.2;refGene_gn=EIF4G3;refGene_tx=XM_019836331.2;refGene_gn=EIF4G3;refGene_tx=XM_023258249.1;refGene_gn=EIF4G3;refGene_tx=XM_023258250.1;refGene_gn=EIF4G3;refGene_tx=XM_023258251.1;refGene_gn=EIF4G3;refGene_tx=XM_023258252.1;refGene_gn=EIF4G3;refGene_tx=XM_023258253.1;refGene_gn=EIF4G3;refGene_tx=XM_023258254.1;refGene_gn=EIF4G3;refGene_tx=XM_023258255.1;refGene_gn=EIF4G3;refGene_tx=XM_023258256.1;refGene_gn=EIF4G3;refGene_tx=XM_023258257.1;refGene_gn=EIF4G3;refGene_tx=XM_023258258.1;refGene_gn=EIF4G3;refGene_tx=XM_023258259.1;refGene_gn=EIF4G3;refGene_tx=XM_023258260.1;refGene_gn=EIF4G3;refGene_tx=XM_023258261.1
C1 15417262 15417360
refGene_gn=EIF4G3;refGene_tx=XM_019836309.2;refGene_gn=EIF4G3;refGene_tx=XM_019836311.2;refGene_gn=EIF4G3;refGene_tx=XM_019836312.2;refGene_gn=EIF4G3;refGene_tx=XM_019836318.2;refGene_gn=EIF4G3;refGene_tx=XM_019836319.2;refGene_gn=EIF4G3;refGene_tx=XM_019836320.2;refGene_gn=EIF4G3;refGene_tx=XM_019836322.2;refGene_gn=EIF4G3;refGene_tx=XM_019836326.2;refGene_gn=EIF4G3;refGene_tx=XM_019836327.2;refGene_gn=EIF4G3;refGene_tx=XM_019836328.2;refGene_gn=EIF4G3;refGene_tx=XM_019836329.2;refGene_gn=EIF4G3;refGene_tx=XM_019836330.2;refGene_gn=EIF4G3;refGene_tx=XM_019836331.2;refGene_gn=EIF4G3;refGene_tx=XM_023258249.1;refGene_gn=EIF4G3;refGene_tx=XM_023258250.1;refGene_gn=EIF4G3;refGene_tx=XM_023258251.1;refGene_gn=EIF4G3;refGene_tx=XM_023258252.1;refGene_gn=EIF4G3;refGene_tx=XM_023258253.1;refGene_gn=EIF4G3;refGene_tx=XM_023258254.1;refGene_gn=EIF4G3;refGene_tx=XM_023258255.1;refGene_gn=EIF4G3;refGene_tx=XM_023258256.1;refGene_gn=EIF4G3;refGene_tx=XM_023258257.1;refGene_gn=EIF4G3;refGene_tx=XM_023258258.1;refGene_gn=EIF4G3;refGene_tx=XM_023258259.1;refGene_gn=EIF4G3;refGene_tx=XM_023258260.1;refGene_gn=EIF4G3;refGene_tx=XM_023258261.1
C1 15426184 15426299
refGene_gn=EIF4G3;refGene_tx=XM_019836309.2;refGene_gn=EIF4G3;refGene_tx=XM_019836311.2;refGene_gn=EIF4G3;refGene_tx=XM_019836312.2;refGene_gn=EIF4G3;refGene_tx=XM_019836318.2;refGene_gn=EIF4G3;refGene_tx=XM_019836319.2;refGene_gn=EIF4G3;refGene_tx=XM_019836320.2;refGene_gn=EIF4G3;refGene_tx=XM_019836322.2;refGene_gn=EIF4G3;refGene_tx=XM_019836326.2;refGene_gn=EIF4G3;refGene_tx=XM_019836327.2;refGene_gn=EIF4G3;refGene_tx=XM_019836328.2;refGene_gn=EIF4G3;refGene_tx=XM_019836329.2;refGene_gn=EIF4G3;refGene_tx=XM_019836330.2;refGene_gn=EIF4G3;refGene_tx=XM_019836331.2;refGene_gn=EIF4G3;refGene_tx=XM_023258249.1;refGene_gn=EIF4G3;refGene_tx=XM_023258250.1;refGene_gn=EIF4G3;refGene_tx=XM_023258251.1;refGene_gn=EIF4G3;refGene_tx=XM_023258252.1;refGene_gn=EIF4G3;refGene_tx=XM_023258253.1;refGene_gn=EIF4G3;refGene_tx=XM_023258254.1;refGene_gn=EIF4G3;refGene_tx=XM_023258255.1;refGene_gn=EIF4G3;refGene_tx=XM_023258256.1;refGene_gn=EIF4G3;refGene_tx=XM_023258257.1;refGene_gn=E

IF4G3;refGene_tx=XM_023258258.1;refGene_gn=EIF4G3;refGene_tx=XM_023258
259.1;refGene_gn=EIF4G3;refGene_tx=XM_023258260.1;refGene_gn=EIF4G3;re
fGene_tx=XM_023258261.1
C1 15426879 15427059
refGene_gn=EIF4G3;refGene_tx=XM_019836309.2;refGene_gn=EIF4G3;refGene_
tx=XM_019836311.2;refGene_gn=EIF4G3;refGene_tx=XM_019836312.2;refGene_
gn=EIF4G3;refGene_tx=XM_019836318.2;refGene_gn=EIF4G3;refGene_tx=XM_01
9836319.2;refGene_gn=EIF4G3;refGene_tx=XM_019836320.2;refGene_gn=EIF4G
3;refGene_tx=XM_019836322.2;refGene_gn=EIF4G3;refGene_tx=XM_019836326.
2;refGene_gn=EIF4G3;refGene_tx=XM_019836327.2;refGene_gn=EIF4G3;refGen
e_tx=XM_019836328.2;refGene_gn=EIF4G3;refGene_tx=XM_019836329.2;refGen
e_gn=EIF4G3;refGene_tx=XM_019836330.2;refGene_gn=EIF4G3;refGene_tx=XM_
019836331.2;refGene_gn=EIF4G3;refGene_tx=XM_023258249.1;refGene_gn=EIF
4G3;refGene_tx=XM_023258250.1;refGene_gn=EIF4G3;refGene_tx=XM_02325825
1.1;refGene_gn=EIF4G3;refGene_tx=XM_023258252.1;refGene_gn=EIF4G3;refG
ene_tx=XM_023258253.1;refGene_gn=EIF4G3;refGene_tx=XM_023258254.1;refG
ene_gn=EIF4G3;refGene_tx=XM_023258255.1;refGene_gn=EIF4G3;refGene_tx=X
M_023258256.1;refGene_gn=EIF4G3;refGene_tx=XM_023258257.1;refGene_gn=E
IF4G3;refGene_tx=XM_023258258.1;refGene_gn=EIF4G3;refGene_tx=XM_023258
259.1;refGene_gn=EIF4G3;refGene_tx=XM_023258260.1;refGene_gn=EIF4G3;re
fGene_tx=XM_023258261.1
C1 15427936 15427957
refGene_gn=EIF4G3;refGene_tx=XM_019836309.2;refGene_gn=EIF4G3;refGene_
tx=XM_019836311.2;refGene_gn=EIF4G3;refGene_tx=XM_019836318.2;refGene_
gn=EIF4G3;refGene_tx=XM_019836320.2;refGene_gn=EIF4G3;refGene_tx=XM_01
9836326.2;refGene_gn=EIF4G3;refGene_tx=XM_019836327.2;refGene_gn=EIF4G
3;refGene_tx=XM_019836329.2;refGene_gn=EIF4G3;refGene_tx=XM_019836330.
2;refGene_gn=EIF4G3;refGene_tx=XM_023258249.1;refGene_gn=EIF4G3;refGen
e_tx=XM_023258250.1;refGene_gn=EIF4G3;refGene_tx=XM_023258251.1;refGen
e_gn=EIF4G3;refGene_tx=XM_023258252.1;refGene_gn=EIF4G3;refGene_tx=XM_
023258253.1;refGene_gn=EIF4G3;refGene_tx=XM_023258254.1;refGene_gn=EIF
4G3;refGene_tx=XM_023258255.1;refGene_gn=EIF4G3;refGene_tx=XM_02325825
6.1;refGene_gn=EIF4G3;refGene_tx=XM_023258257.1;refGene_gn=EIF4G3;refG
ene_tx=XM_023258258.1;refGene_gn=EIF4G3;refGene_tx=XM_023258259.1;refG
ene_gn=EIF4G3;refGene_tx=XM_023258260.1;refGene_gn=EIF4G3;refGene_tx=X
M_023258261.1
C1 15438808 15438841
refGene_gn=EIF4G3;refGene_tx=XM_019836309.2;refGene_gn=EIF4G3;refGene_
tx=XM_019836311.2;refGene_gn=EIF4G3;refGene_tx=XM_019836312.2;refGene_
gn=EIF4G3;refGene_tx=XM_019836318.2;refGene_gn=EIF4G3;refGene_tx=XM_01
9836319.2;refGene_gn=EIF4G3;refGene_tx=XM_019836326.2;refGene_gn=EIF4G
3;refGene_tx=XM_019836327.2;refGene_gn=EIF4G3;refGene_tx=XM_019836328.
2;refGene_gn=EIF4G3;refGene_tx=XM_023258249.1;refGene_gn=EIF4G3;refGen
e_tx=XM_023258250.1;refGene_gn=EIF4G3;refGene_tx=XM_023258251.1;refGen
e_gn=EIF4G3;refGene_tx=XM_023258252.1;refGene_gn=EIF4G3;refGene_tx=XM_
023258253.1;refGene_gn=EIF4G3;refGene_tx=XM_023258254.1;refGene_gn=EIF
4G3;refGene_tx=XM_023258255.1;refGene_gn=EIF4G3;refGene_tx=XM_02325825
6.1;refGene_gn=EIF4G3;refGene_tx=XM_023258257.1;refGene_gn=EIF4G3;refG
ene_tx=XM_023258258.1;refGene_gn=EIF4G3;refGene_tx=XM_023258259.1;refG
ene_gn=EIF4G3;refGene_tx=XM_023258260.1;refGene_gn=EIF4G3;refGene_tx=X

M_023258261.1
C1 15442453 15442579
refGene_gn=EIF4G3;refGene_tx=XM_019836309.2;refGene_gn=EIF4G3;refGene_tx=XM_019836311.2;refGene_gn=EIF4G3;refGene_tx=XM_019836312.2;refGene_gn=EIF4G3;refGene_tx=XM_019836318.2;refGene_gn=EIF4G3;refGene_tx=XM_019836319.2;refGene_gn=EIF4G3;refGene_tx=XM_019836320.2;refGene_gn=EIF4G3;refGene_tx=XM_019836322.2;refGene_gn=EIF4G3;refGene_tx=XM_023258249.1;refGene_gn=EIF4G3;refGene_tx=XM_023258250.1;refGene_gn=EIF4G3;refGene_tx=XM_023258251.1;refGene_gn=EIF4G3;refGene_tx=XM_023258252.1;refGene_gn=EIF4G3;refGene_tx=XM_023258253.1;refGene_gn=EIF4G3;refGene_tx=XM_023258254.1;refGene_gn=EIF4G3;refGene_tx=XM_023258255.1;refGene_gn=EIF4G3;refGene_tx=XM_023258256.1;refGene_gn=EIF4G3;refGene_tx=XM_023258257.1;refGene_gn=EIF4G3;refGene_tx=XM_023258258.1;refGene_gn=EIF4G3;refGene_tx=XM_023258259.1;refGene_gn=EIF4G3;refGene_tx=XM_023258260.1;refGene_gn=EIF4G3;refGene_tx=XM_023258261.1
C1 15443966 15444062
refGene_gn=EIF4G3;refGene_tx=XM_019836309.2;refGene_gn=EIF4G3;refGene_tx=XM_019836311.2;refGene_gn=EIF4G3;refGene_tx=XM_019836312.2;refGene_gn=EIF4G3;refGene_tx=XM_019836318.2;refGene_gn=EIF4G3;refGene_tx=XM_019836319.2;refGene_gn=EIF4G3;refGene_tx=XM_019836320.2;refGene_gn=EIF4G3;refGene_tx=XM_019836322.2;refGene_gn=EIF4G3;refGene_tx=XM_019836326.2;refGene_gn=EIF4G3;refGene_tx=XM_019836327.2;refGene_gn=EIF4G3;refGene_tx=XM_019836328.2;refGene_gn=EIF4G3;refGene_tx=XM_019836329.2;refGene_gn=EIF4G3;refGene_tx=XM_019836330.2;refGene_gn=EIF4G3;refGene_tx=XM_019836331.2;refGene_gn=EIF4G3;refGene_tx=XM_023258249.1;refGene_gn=EIF4G3;refGene_tx=XM_023258250.1;refGene_gn=EIF4G3;refGene_tx=XM_023258251.1;refGene_gn=EIF4G3;refGene_tx=XM_023258252.1;refGene_gn=EIF4G3;refGene_tx=XM_023258253.1;refGene_gn=EIF4G3;refGene_tx=XM_023258254.1;refGene_gn=EIF4G3;refGene_tx=XM_023258255.1;refGene_gn=EIF4G3;refGene_tx=XM_023258256.1;refGene_gn=EIF4G3;refGene_tx=XM_023258257.1;refGene_gn=EIF4G3;refGene_tx=XM_023258258.1;refGene_gn=EIF4G3;refGene_tx=XM_023258259.1;refGene_gn=EIF4G3;refGene_tx=XM_023258260.1;refGene_gn=EIF4G3;refGene_tx=XM_023258261.1
C1 15459848 15459873
refGene_gn=EIF4G3;refGene_tx=XM_019836309.2;refGene_gn=EIF4G3;refGene_tx=XM_019836311.2;refGene_gn=EIF4G3;refGene_tx=XM_019836312.2
C1 15489013 15489142
refGene_gn=EIF4G3;refGene_tx=XM_019836319.2;refGene_gn=EIF4G3;refGene_tx=XM_019836320.2;refGene_gn=EIF4G3;refGene_tx=XM_019836322.2;refGene_gn=EIF4G3;refGene_tx=XM_019836326.2;refGene_gn=EIF4G3;refGene_tx=XM_019836328.2;refGene_gn=EIF4G3;refGene_tx=XM_019836329.2;refGene_gn=EIF4G3;refGene_tx=XM_019836331.2;refGene_gn=EIF4G3;refGene_tx=XM_023258249.1;refGene_gn=EIF4G3;refGene_tx=XM_023258250.1;refGene_gn=EIF4G3;refGene_tx=XM_023258251.1;refGene_gn=EIF4G3;refGene_tx=XM_023258252.1;refGene_gn=EIF4G3;refGene_tx=XM_023258253.1;refGene_gn=EIF4G3;refGene_tx=XM_023258254.1;refGene_gn=EIF4G3;refGene_tx=XM_023258255.1;refGene_gn=EIF4G3;refGene_tx=XM_023258256.1;refGene_gn=EIF4G3;refGene_tx=XM_023258261.1
C1 15502733 15502748
refGene_gn=EIF4G3;refGene_tx=XM_019836319.2;refGene_gn=EIF4G3;refGene_

tx=XM_019836320.2;refGene_gn=EIF4G3;refGene_tx=XM_019836322.2;refGene_gn=EIF4G3;refGene_tx=XM_019836326.2;refGene_gn=EIF4G3;refGene_tx=XM_019836328.2;refGene_gn=EIF4G3;refGene_tx=XM_019836329.2;refGene_gn=EIF4G3;refGene_tx=XM_019836331.2;refGene_gn=EIF4G3;refGene_tx=XM_023258249.1;refGene_gn=EIF4G3;refGene_tx=XM_023258250.1;refGene_gn=EIF4G3;refGene_tx=XM_023258251.1;refGene_gn=EIF4G3;refGene_tx=XM_023258252.1;refGene_gn=EIF4G3;refGene_tx=XM_023258253.1;refGene_gn=EIF4G3;refGene_tx=XM_023258254.1;refGene_gn=EIF4G3;refGene_tx=XM_023258255.1;refGene_gn=EIF4G3;refGene_tx=XM_023258256.1;refGene_gn=EIF4G3;refGene_tx=XM_023258261.1
C1 15626485 15626662
refGene_gn=ECE1;refGene_tx=XM_023258262.1;refGene_gn=ECE1;refGene_tx=XM_023258263.1
C1 15627320 15627416
refGene_gn=ECE1;refGene_tx=XM_023258262.1;refGene_gn=ECE1;refGene_tx=XM_023258263.1
C1 15630332 15630523
refGene_gn=ECE1;refGene_tx=XM_023258262.1;refGene_gn=ECE1;refGene_tx=XM_023258263.1
C1 15631557 15631625
refGene_gn=ECE1;refGene_tx=XM_023258262.1;refGene_gn=ECE1;refGene_tx=XM_023258263.1
C1 15632313 15632424
refGene_gn=ECE1;refGene_tx=XM_023258262.1;refGene_gn=ECE1;refGene_tx=XM_023258263.1
C1 15637478 15637582
refGene_gn=ECE1;refGene_tx=XM_023258262.1;refGene_gn=ECE1;refGene_tx=XM_023258263.1
C1 15639101 15639179
refGene_gn=ECE1;refGene_tx=XM_023258262.1;refGene_gn=ECE1;refGene_tx=XM_023258263.1
C1 15639673 15639772
refGene_gn=ECE1;refGene_tx=XM_023258262.1;refGene_gn=ECE1;refGene_tx=XM_023258263.1
C1 15641053 15641164
refGene_gn=ECE1;refGene_tx=XM_023258262.1;refGene_gn=ECE1;refGene_tx=XM_023258263.1
C1 15645981 15646096
refGene_gn=ECE1;refGene_tx=XM_023258262.1;refGene_gn=ECE1;refGene_tx=XM_023258263.1
C1 15647373 15647516
refGene_gn=ECE1;refGene_tx=XM_023258262.1;refGene_gn=ECE1;refGene_tx=XM_023258263.1
C1 15653673 15653865
refGene_gn=ECE1;refGene_tx=XM_023258262.1;refGene_gn=ECE1;refGene_tx=XM_023258263.1
C1 15654899 15654965
refGene_gn=ECE1;refGene_tx=XM_023258262.1;refGene_gn=ECE1;refGene_tx=XM_023258263.1
C1 15656058 15656205

refGene_gn=ECE1;refGene_tx=XM_023258262.1;refGene_gn=ECE1;refGene_tx=XM_023258263.1
C1 15657925 15658047
refGene_gn=ECE1;refGene_tx=XM_023258262.1;refGene_gn=ECE1;refGene_tx=XM_023258263.1
C1 15672826 15673039
refGene_gn=ECE1;refGene_tx=XM_023258262.1;refGene_gn=ECE1;refGene_tx=XM_023258263.1
C1 15677850 15677992
refGene_gn=ECE1;refGene_tx=XM_023258262.1;refGene_gn=ECE1;refGene_tx=XM_023258263.1
C1 15686183 15686270
refGene_gn=ECE1;refGene_tx=XM_023258262.1;refGene_gn=ECE1;refGene_tx=XM_023258263.1
C1 15686496 15686541 refGene_gn=ECE1;refGene_tx=XM_023258262.1
C1 15738260 15738263 refGene_gn=ECE1;refGene_tx=XM_023258263.1
C1 15842698 15842759
refGene_gn=ALPL;refGene_tx=NM_001042563.1;refGene_gn=ALPL;refGene_tx=XM_023257816.1;refGene_gn=ALPL;refGene_tx=XM_023257817.1
C1 15846658 15846778
refGene_gn=ALPL;refGene_tx=NM_001042563.1;refGene_gn=ALPL;refGene_tx=XM_023257816.1;refGene_gn=ALPL;refGene_tx=XM_023257817.1
C1 15847107 15847223
refGene_gn=ALPL;refGene_tx=NM_001042563.1;refGene_gn=ALPL;refGene_tx=XM_023257816.1;refGene_gn=ALPL;refGene_tx=XM_023257817.1
C1 15848352 15848527
refGene_gn=ALPL;refGene_tx=NM_001042563.1;refGene_gn=ALPL;refGene_tx=XM_023257816.1;refGene_gn=ALPL;refGene_tx=XM_023257817.1
C1 15849297 15849473
refGene_gn=ALPL;refGene_tx=NM_001042563.1;refGene_gn=ALPL;refGene_tx=XM_023257816.1;refGene_gn=ALPL;refGene_tx=XM_023257817.1
C1 15853140 15853284
refGene_gn=ALPL;refGene_tx=NM_001042563.1;refGene_gn=ALPL;refGene_tx=XM_023257816.1;refGene_gn=ALPL;refGene_tx=XM_023257817.1
C1 15855236 15855306
refGene_gn=ALPL;refGene_tx=NM_001042563.1;refGene_gn=ALPL;refGene_tx=XM_023257816.1;refGene_gn=ALPL;refGene_tx=XM_023257817.1
C1 15857569 15857704
refGene_gn=ALPL;refGene_tx=NM_001042563.1;refGene_gn=ALPL;refGene_tx=XM_023257816.1;refGene_gn=ALPL;refGene_tx=XM_023257817.1
C1 15859614 15859806
refGene_gn=ALPL;refGene_tx=NM_001042563.1;refGene_gn=ALPL;refGene_tx=XM_023257816.1;refGene_gn=ALPL;refGene_tx=XM_023257817.1
C1 15860251 15860371
refGene_gn=ALPL;refGene_tx=NM_001042563.1;refGene_gn=ALPL;refGene_tx=XM_023257816.1;refGene_gn=ALPL;refGene_tx=XM_023257817.1
C1 15861088 15861354
refGene_gn=ALPL;refGene_tx=NM_001042563.1;refGene_gn=ALPL;refGene_tx=XM_023257816.1;refGene_gn=ALPL;refGene_tx=XM_023257817.1
C1 15875649 15876012

refGene_gn=RAP1GAP;refGene_tx=XM_006934381.4;refGene_gn=RAP1GAP;refGene_tx=XM_019836348.2;refGene_gn=RAP1GAP;refGene_tx=XM_019836349.2;refGene_gn=RAP1GAP;refGene_tx=XM_019836350.2;refGene_gn=RAP1GAP;refGene_tx=XM_019836351.2;refGene_gn=RAP1GAP;refGene_tx=XM_019836355.2;refGene_gn=RAP1GAP;refGene_tx=XM_019836356.2;refGene_gn=RAP1GAP;refGene_tx=XM_019836357.2;refGene_gn=RAP1GAP;refGene_tx=XM_023258264.1;refGene_gn=RAP1GAP;refGene_tx=XM_023258265.1;refGene_gn=RAP1GAP;refGene_tx=XM_023258266.1;refGene_gn=RAP1GAP;refGene_tx=XM_023258267.1;refGene_gn=RAP1GAP;refGene_tx=XM_023258268.1
C1 15876458 15876467 refGene_gn=RAP1GAP;refGene_tx=XM_019836359.2
C1 15876744 15876879
refGene_gn=RAP1GAP;refGene_tx=XM_006934381.4;refGene_gn=RAP1GAP;refGene_tx=XM_019836348.2;refGene_gn=RAP1GAP;refGene_tx=XM_019836349.2;refGene_gn=RAP1GAP;refGene_tx=XM_019836350.2;refGene_gn=RAP1GAP;refGene_tx=XM_019836351.2;refGene_gn=RAP1GAP;refGene_tx=XM_019836355.2;refGene_gn=RAP1GAP;refGene_tx=XM_019836356.2;refGene_gn=RAP1GAP;refGene_tx=XM_019836357.2;refGene_gn=RAP1GAP;refGene_tx=XM_019836358.2;refGene_gn=RAP1GAP;refGene_tx=XM_019836359.2;refGene_gn=RAP1GAP;refGene_tx=XM_023258264.1;refGene_gn=RAP1GAP;refGene_tx=XM_023258265.1;refGene_gn=RAP1GAP;refGene_tx=XM_023258266.1;refGene_gn=RAP1GAP;refGene_tx=XM_023258267.1;refGene_gn=RAP1GAP;refGene_tx=XM_023258268.1
C1 15877193 15877296
refGene_gn=RAP1GAP;refGene_tx=XM_006934381.4;refGene_gn=RAP1GAP;refGene_tx=XM_019836348.2;refGene_gn=RAP1GAP;refGene_tx=XM_019836349.2;refGene_gn=RAP1GAP;refGene_tx=XM_019836350.2;refGene_gn=RAP1GAP;refGene_tx=XM_019836351.2;refGene_gn=RAP1GAP;refGene_tx=XM_019836355.2;refGene_gn=RAP1GAP;refGene_tx=XM_019836356.2;refGene_gn=RAP1GAP;refGene_tx=XM_019836357.2;refGene_gn=RAP1GAP;refGene_tx=XM_019836358.2;refGene_gn=RAP1GAP;refGene_tx=XM_019836359.2;refGene_gn=RAP1GAP;refGene_tx=XM_023258264.1;refGene_gn=RAP1GAP;refGene_tx=XM_023258265.1;refGene_gn=RAP1GAP;refGene_tx=XM_023258266.1;refGene_gn=RAP1GAP;refGene_tx=XM_023258267.1;refGene_gn=RAP1GAP;refGene_tx=XM_023258268.1
C1 15878292 15878416
refGene_gn=RAP1GAP;refGene_tx=XM_006934381.4;refGene_gn=RAP1GAP;refGene_tx=XM_019836348.2;refGene_gn=RAP1GAP;refGene_tx=XM_019836349.2;refGene_gn=RAP1GAP;refGene_tx=XM_019836350.2;refGene_gn=RAP1GAP;refGene_tx=XM_019836351.2;refGene_gn=RAP1GAP;refGene_tx=XM_019836355.2;refGene_gn=RAP1GAP;refGene_tx=XM_019836356.2;refGene_gn=RAP1GAP;refGene_tx=XM_019836357.2;refGene_gn=RAP1GAP;refGene_tx=XM_019836358.2;refGene_gn=RAP1GAP;refGene_tx=XM_019836359.2;refGene_gn=RAP1GAP;refGene_tx=XM_023258264.1;refGene_gn=RAP1GAP;refGene_tx=XM_023258265.1;refGene_gn=RAP1GAP;refGene_tx=XM_023258266.1;refGene_gn=RAP1GAP;refGene_tx=XM_023258267.1;refGene_gn=RAP1GAP;refGene_tx=XM_023258268.1
C1 15879923 15880079
refGene_gn=RAP1GAP;refGene_tx=XM_006934381.4;refGene_gn=RAP1GAP;refGene_tx=XM_019836348.2;refGene_gn=RAP1GAP;refGene_tx=XM_019836349.2;refGene_gn=RAP1GAP;refGene_tx=XM_019836350.2;refGene_gn=RAP1GAP;refGene_tx=XM_019836351.2;refGene_gn=RAP1GAP;refGene_tx=XM_019836355.2;refGene_gn=RAP1GAP;refGene_tx=XM_019836357.2;refGene_gn=RAP1GAP;refGene_tx=XM_019836358.2;refGene_gn=RAP1GAP;refGene_tx=XM_019836359.2;refGene_gn=RAP1

GAP;refGene_tx=XM_023258264.1;refGene_gn=RAP1GAP;refGene_tx=XM_0232582
65.1;refGene_gn=RAP1GAP;refGene_tx=XM_023258266.1;refGene_gn=RAP1GAP;r
efGene_tx=XM_023258267.1;refGene_gn=RAP1GAP;refGene_tx=XM_023258268.1
C1 15881003 15881113
refGene_gn=RAP1GAP;refGene_tx=XM_006934381.4;refGene_gn=RAP1GAP;refGen
e_tx=XM_019836348.2;refGene_gn=RAP1GAP;refGene_tx=XM_019836349.2;refGe
ne_gn=RAP1GAP;refGene_tx=XM_019836350.2;refGene_gn=RAP1GAP;refGene_tx=
XM_019836351.2;refGene_gn=RAP1GAP;refGene_tx=XM_019836355.2;refGene_gn
=RAP1GAP;refGene_tx=XM_019836356.2;refGene_gn=RAP1GAP;refGene_tx=XM_01
9836357.2;refGene_gn=RAP1GAP;refGene_tx=XM_019836358.2;refGene_gn=RAP1
GAP;refGene_tx=XM_019836359.2;refGene_gn=RAP1GAP;refGene_tx=XM_0232582
64.1;refGene_gn=RAP1GAP;refGene_tx=XM_023258265.1;refGene_gn=RAP1GAP;r
efGene_tx=XM_023258266.1;refGene_gn=RAP1GAP;refGene_tx=XM_023258267.1;
refGene_gn=RAP1GAP;refGene_tx=XM_023258268.1
C1 15882019 15882097
refGene_gn=RAP1GAP;refGene_tx=XM_006934381.4;refGene_gn=RAP1GAP;refGen
e_tx=XM_019836348.2;refGene_gn=RAP1GAP;refGene_tx=XM_019836349.2;refGe
ne_gn=RAP1GAP;refGene_tx=XM_019836351.2;refGene_gn=RAP1GAP;refGene_tx=
XM_019836356.2;refGene_gn=RAP1GAP;refGene_tx=XM_019836357.2;refGene_gn
=RAP1GAP;refGene_tx=XM_019836358.2;refGene_gn=RAP1GAP;refGene_tx=XM_01
9836359.2;refGene_gn=RAP1GAP;refGene_tx=XM_023258264.1;refGene_gn=RAP1
GAP;refGene_tx=XM_023258265.1;refGene_gn=RAP1GAP;refGene_tx=XM_0232582
66.1;refGene_gn=RAP1GAP;refGene_tx=XM_023258267.1;refGene_gn=RAP1GAP;r
efGene_tx=XM_023258268.1
C1 15883881 15884013
refGene_gn=RAP1GAP;refGene_tx=XM_006934381.4;refGene_gn=RAP1GAP;refGen
e_tx=XM_019836348.2;refGene_gn=RAP1GAP;refGene_tx=XM_019836349.2;refGe
ne_gn=RAP1GAP;refGene_tx=XM_019836350.2;refGene_gn=RAP1GAP;refGene_tx=
XM_019836351.2;refGene_gn=RAP1GAP;refGene_tx=XM_019836355.2;refGene_gn
=RAP1GAP;refGene_tx=XM_019836356.2;refGene_gn=RAP1GAP;refGene_tx=XM_01
9836357.2;refGene_gn=RAP1GAP;refGene_tx=XM_019836358.2;refGene_gn=RAP1
GAP;refGene_tx=XM_019836359.2;refGene_gn=RAP1GAP;refGene_tx=XM_0232582
64.1;refGene_gn=RAP1GAP;refGene_tx=XM_023258265.1;refGene_gn=RAP1GAP;r
efGene_tx=XM_023258266.1;refGene_gn=RAP1GAP;refGene_tx=XM_023258267.1;
refGene_gn=RAP1GAP;refGene_tx=XM_023258268.1
C1 15885891 15886029
refGene_gn=RAP1GAP;refGene_tx=XM_006934381.4;refGene_gn=RAP1GAP;refGen
e_tx=XM_019836348.2;refGene_gn=RAP1GAP;refGene_tx=XM_019836349.2;refGe
ne_gn=RAP1GAP;refGene_tx=XM_019836350.2;refGene_gn=RAP1GAP;refGene_tx=
XM_019836351.2;refGene_gn=RAP1GAP;refGene_tx=XM_019836355.2;refGene_gn
=RAP1GAP;refGene_tx=XM_019836356.2;refGene_gn=RAP1GAP;refGene_tx=XM_01
9836357.2;refGene_gn=RAP1GAP;refGene_tx=XM_019836358.2;refGene_gn=RAP1
GAP;refGene_tx=XM_019836359.2;refGene_gn=RAP1GAP;refGene_tx=XM_0232582
64.1;refGene_gn=RAP1GAP;refGene_tx=XM_023258265.1;refGene_gn=RAP1GAP;r
efGene_tx=XM_023258266.1;refGene_gn=RAP1GAP;refGene_tx=XM_023258267.1;
refGene_gn=RAP1GAP;refGene_tx=XM_023258268.1
C1 15886476 15886563
refGene_gn=RAP1GAP;refGene_tx=XM_006934381.4;refGene_gn=RAP1GAP;refGen
e_tx=XM_019836348.2;refGene_gn=RAP1GAP;refGene_tx=XM_019836349.2;refGe
ne_gn=RAP1GAP;refGene_tx=XM_019836350.2;refGene_gn=RAP1GAP;refGene_tx=

XM_019836351.2;refGene_gn=RAP1GAP;refGene_tx=XM_019836355.2;refGene_gn=RAP1GAP;refGene_tx=XM_019836356.2;refGene_gn=RAP1GAP;refGene_tx=XM_019836357.2;refGene_gn=RAP1GAP;refGene_tx=XM_019836358.2;refGene_gn=RAP1GAP;refGene_tx=XM_019836359.2;refGene_gn=RAP1GAP;refGene_tx=XM_023258264.1;refGene_gn=RAP1GAP;refGene_tx=XM_023258265.1;refGene_gn=RAP1GAP;refGene_tx=XM_023258266.1;refGene_gn=RAP1GAP;refGene_tx=XM_023258267.1;refGene_gn=RAP1GAP;refGene_tx=XM_023258268.1

C1 15887173 15887245

refGene_gn=RAP1GAP;refGene_tx=XM_006934381.4;refGene_gn=RAP1GAP;refGene_tx=XM_019836348.2;refGene_gn=RAP1GAP;refGene_tx=XM_019836349.2;refGene_gn=RAP1GAP;refGene_tx=XM_019836350.2;refGene_gn=RAP1GAP;refGene_tx=XM_019836351.2;refGene_gn=RAP1GAP;refGene_tx=XM_019836355.2;refGene_gn=RAP1GAP;refGene_tx=XM_019836356.2;refGene_gn=RAP1GAP;refGene_tx=XM_019836357.2;refGene_gn=RAP1GAP;refGene_tx=XM_019836358.2;refGene_gn=RAP1GAP;refGene_tx=XM_019836359.2;refGene_gn=RAP1GAP;refGene_tx=XM_023258264.1;refGene_gn=RAP1GAP;refGene_tx=XM_023258265.1;refGene_gn=RAP1GAP;refGene_tx=XM_023258266.1;refGene_gn=RAP1GAP;refGene_tx=XM_023258267.1;refGene_gn=RAP1GAP;refGene_tx=XM_023258268.1

C1 15887641 15887797

refGene_gn=RAP1GAP;refGene_tx=XM_006934381.4;refGene_gn=RAP1GAP;refGene_tx=XM_019836348.2;refGene_gn=RAP1GAP;refGene_tx=XM_019836349.2;refGene_gn=RAP1GAP;refGene_tx=XM_019836350.2;refGene_gn=RAP1GAP;refGene_tx=XM_019836351.2;refGene_gn=RAP1GAP;refGene_tx=XM_019836355.2;refGene_gn=RAP1GAP;refGene_tx=XM_019836356.2;refGene_gn=RAP1GAP;refGene_tx=XM_019836357.2;refGene_gn=RAP1GAP;refGene_tx=XM_019836358.2;refGene_gn=RAP1GAP;refGene_tx=XM_019836359.2;refGene_gn=RAP1GAP;refGene_tx=XM_023258264.1;refGene_gn=RAP1GAP;refGene_tx=XM_023258265.1;refGene_gn=RAP1GAP;refGene_tx=XM_023258266.1;refGene_gn=RAP1GAP;refGene_tx=XM_023258267.1;refGene_gn=RAP1GAP;refGene_tx=XM_023258268.1

C1 15888840 15888970

refGene_gn=RAP1GAP;refGene_tx=XM_006934381.4;refGene_gn=RAP1GAP;refGene_tx=XM_019836348.2;refGene_gn=RAP1GAP;refGene_tx=XM_019836349.2;refGene_gn=RAP1GAP;refGene_tx=XM_019836350.2;refGene_gn=RAP1GAP;refGene_tx=XM_019836351.2;refGene_gn=RAP1GAP;refGene_tx=XM_019836355.2;refGene_gn=RAP1GAP;refGene_tx=XM_019836356.2;refGene_gn=RAP1GAP;refGene_tx=XM_019836357.2;refGene_gn=RAP1GAP;refGene_tx=XM_019836358.2;refGene_gn=RAP1GAP;refGene_tx=XM_019836359.2;refGene_gn=RAP1GAP;refGene_tx=XM_023258264.1;refGene_gn=RAP1GAP;refGene_tx=XM_023258265.1;refGene_gn=RAP1GAP;refGene_tx=XM_023258266.1;refGene_gn=RAP1GAP;refGene_tx=XM_023258267.1;refGene_gn=RAP1GAP;refGene_tx=XM_023258268.1

C1 15889103 15889204

refGene_gn=RAP1GAP;refGene_tx=XM_006934381.4;refGene_gn=RAP1GAP;refGene_tx=XM_019836348.2;refGene_gn=RAP1GAP;refGene_tx=XM_019836349.2;refGene_gn=RAP1GAP;refGene_tx=XM_019836350.2;refGene_gn=RAP1GAP;refGene_tx=XM_019836351.2;refGene_gn=RAP1GAP;refGene_tx=XM_019836355.2;refGene_gn=RAP1GAP;refGene_tx=XM_019836356.2;refGene_gn=RAP1GAP;refGene_tx=XM_019836357.2;refGene_gn=RAP1GAP;refGene_tx=XM_019836358.2;refGene_gn=RAP1GAP;refGene_tx=XM_019836359.2;refGene_gn=RAP1GAP;refGene_tx=XM_023258264.1;refGene_gn=RAP1GAP;refGene_tx=XM_023258265.1;refGene_gn=RAP1GAP;refGene_tx=XM_023258266.1;refGene_gn=RAP1GAP;refGene_tx=XM_023258267.1;

refGene_gn=RAP1GAP;refGene_tx=XM_023258268.1
C1	15889410	15889494
refGene_gn=RAP1GAP;refGene_tx=XM_006934381.4;refGene_gn=RAP1GAP;refGene_tx=XM_019836348.2;refGene_gn=RAP1GAP;refGene_tx=XM_019836349.2;refGene_gn=RAP1GAP;refGene_tx=XM_019836350.2;refGene_gn=RAP1GAP;refGene_tx=XM_019836351.2;refGene_gn=RAP1GAP;refGene_tx=XM_019836355.2;refGene_gn=RAP1GAP;refGene_tx=XM_019836356.2;refGene_gn=RAP1GAP;refGene_tx=XM_019836357.2;refGene_gn=RAP1GAP;refGene_tx=XM_019836358.2;refGene_gn=RAP1GAP;refGene_tx=XM_019836359.2;refGene_gn=RAP1GAP;refGene_tx=XM_023258264.1;refGene_gn=RAP1GAP;refGene_tx=XM_023258265.1;refGene_gn=RAP1GAP;refGene_tx=XM_023258266.1;refGene_gn=RAP1GAP;refGene_tx=XM_023258267.1;refGene_gn=RAP1GAP;refGene_tx=XM_023258268.1
C1	15890575	15890629
refGene_gn=RAP1GAP;refGene_tx=XM_006934381.4;refGene_gn=RAP1GAP;refGene_tx=XM_019836348.2;refGene_gn=RAP1GAP;refGene_tx=XM_019836349.2;refGene_gn=RAP1GAP;refGene_tx=XM_019836350.2;refGene_gn=RAP1GAP;refGene_tx=XM_019836351.2;refGene_gn=RAP1GAP;refGene_tx=XM_019836355.2;refGene_gn=RAP1GAP;refGene_tx=XM_019836356.2;refGene_gn=RAP1GAP;refGene_tx=XM_019836357.2;refGene_gn=RAP1GAP;refGene_tx=XM_019836358.2;refGene_gn=RAP1GAP;refGene_tx=XM_019836359.2;refGene_gn=RAP1GAP;refGene_tx=XM_023258264.1;refGene_gn=RAP1GAP;refGene_tx=XM_023258265.1;refGene_gn=RAP1GAP;refGene_tx=XM_023258266.1;refGene_gn=RAP1GAP;refGene_tx=XM_023258267.1;refGene_gn=RAP1GAP;refGene_tx=XM_023258268.1
C1	15890987	15891066
refGene_gn=RAP1GAP;refGene_tx=XM_006934381.4;refGene_gn=RAP1GAP;refGene_tx=XM_019836348.2;refGene_gn=RAP1GAP;refGene_tx=XM_019836349.2;refGene_gn=RAP1GAP;refGene_tx=XM_019836350.2;refGene_gn=RAP1GAP;refGene_tx=XM_019836351.2;refGene_gn=RAP1GAP;refGene_tx=XM_019836355.2;refGene_gn=RAP1GAP;refGene_tx=XM_019836356.2;refGene_gn=RAP1GAP;refGene_tx=XM_019836357.2;refGene_gn=RAP1GAP;refGene_tx=XM_019836358.2;refGene_gn=RAP1GAP;refGene_tx=XM_019836359.2;refGene_gn=RAP1GAP;refGene_tx=XM_023258264.1;refGene_gn=RAP1GAP;refGene_tx=XM_023258265.1;refGene_gn=RAP1GAP;refGene_tx=XM_023258266.1;refGene_gn=RAP1GAP;refGene_tx=XM_023258267.1;refGene_gn=RAP1GAP;refGene_tx=XM_023258268.1
C1	15891327	15891431
refGene_gn=RAP1GAP;refGene_tx=XM_006934381.4;refGene_gn=RAP1GAP;refGene_tx=XM_019836348.2;refGene_gn=RAP1GAP;refGene_tx=XM_019836349.2;refGene_gn=RAP1GAP;refGene_tx=XM_019836350.2;refGene_gn=RAP1GAP;refGene_tx=XM_019836351.2;refGene_gn=RAP1GAP;refGene_tx=XM_019836355.2;refGene_gn=RAP1GAP;refGene_tx=XM_019836356.2;refGene_gn=RAP1GAP;refGene_tx=XM_019836357.2;refGene_gn=RAP1GAP;refGene_tx=XM_019836358.2;refGene_gn=RAP1GAP;refGene_tx=XM_019836359.2;refGene_gn=RAP1GAP;refGene_tx=XM_023258264.1;refGene_gn=RAP1GAP;refGene_tx=XM_023258265.1;refGene_gn=RAP1GAP;refGene_tx=XM_023258266.1;refGene_gn=RAP1GAP;refGene_tx=XM_023258267.1;refGene_gn=RAP1GAP;refGene_tx=XM_023258268.1
C1	15893551	15893737
refGene_gn=RAP1GAP;refGene_tx=XM_006934381.4;refGene_gn=RAP1GAP;refGene_tx=XM_019836348.2;refGene_gn=RAP1GAP;refGene_tx=XM_019836349.2;refGene_gn=RAP1GAP;refGene_tx=XM_019836350.2;refGene_gn=RAP1GAP;refGene_tx=XM_019836351.2;refGene_gn=RAP1GAP;refGene_tx=XM_019836355.2;refGene_gn

=RAP1GAP;refGene_tx=XM_019836356.2;refGene_gn=RAP1GAP;refGene_tx=XM_019836357.2;refGene_gn=RAP1GAP;refGene_tx=XM_019836358.2;refGene_gn=RAP1GAP;refGene_tx=XM_019836359.2;refGene_gn=RAP1GAP;refGene_tx=XM_023258264.1;refGene_gn=RAP1GAP;refGene_tx=XM_023258265.1;refGene_gn=RAP1GAP;refGene_tx=XM_023258266.1;refGene_gn=RAP1GAP;refGene_tx=XM_023258267.1;refGene_gn=RAP1GAP;refGene_tx=XM_023258268.1
C1 15894195 15894234
refGene_gn=RAP1GAP;refGene_tx=XM_006934381.4;refGene_gn=RAP1GAP;refGene_tx=XM_019836348.2;refGene_gn=RAP1GAP;refGene_tx=XM_019836349.2;refGene_gn=RAP1GAP;refGene_tx=XM_019836350.2;refGene_gn=RAP1GAP;refGene_tx=XM_019836351.2;refGene_gn=RAP1GAP;refGene_tx=XM_019836355.2;refGene_gn=RAP1GAP;refGene_tx=XM_019836356.2;refGene_gn=RAP1GAP;refGene_tx=XM_019836357.2;refGene_gn=RAP1GAP;refGene_tx=XM_019836358.2;refGene_gn=RAP1GAP;refGene_tx=XM_019836359.2;refGene_gn=RAP1GAP;refGene_tx=XM_023258264.1;refGene_gn=RAP1GAP;refGene_tx=XM_023258265.1;refGene_gn=RAP1GAP;refGene_tx=XM_023258266.1;refGene_gn=RAP1GAP;refGene_tx=XM_023258267.1;refGene_gn=RAP1GAP;refGene_tx=XM_023258268.1
C1 15895164 15895212
refGene_gn=RAP1GAP;refGene_tx=XM_006934381.4;refGene_gn=RAP1GAP;refGene_tx=XM_019836348.2;refGene_gn=RAP1GAP;refGene_tx=XM_019836349.2;refGene_gn=RAP1GAP;refGene_tx=XM_019836350.2;refGene_gn=RAP1GAP;refGene_tx=XM_019836351.2;refGene_gn=RAP1GAP;refGene_tx=XM_019836355.2;refGene_gn=RAP1GAP;refGene_tx=XM_019836356.2;refGene_gn=RAP1GAP;refGene_tx=XM_019836357.2;refGene_gn=RAP1GAP;refGene_tx=XM_019836358.2;refGene_gn=RAP1GAP;refGene_tx=XM_019836359.2;refGene_gn=RAP1GAP;refGene_tx=XM_023258264.1;refGene_gn=RAP1GAP;refGene_tx=XM_023258265.1;refGene_gn=RAP1GAP;refGene_tx=XM_023258266.1;refGene_gn=RAP1GAP;refGene_tx=XM_023258267.1;refGene_gn=RAP1GAP;refGene_tx=XM_023258268.1
C1 15896292 15896328
refGene_gn=RAP1GAP;refGene_tx=XM_006934381.4;refGene_gn=RAP1GAP;refGene_tx=XM_019836348.2;refGene_gn=RAP1GAP;refGene_tx=XM_019836349.2;refGene_gn=RAP1GAP;refGene_tx=XM_019836350.2;refGene_gn=RAP1GAP;refGene_tx=XM_019836351.2;refGene_gn=RAP1GAP;refGene_tx=XM_019836355.2;refGene_gn=RAP1GAP;refGene_tx=XM_019836356.2;refGene_gn=RAP1GAP;refGene_tx=XM_019836357.2;refGene_gn=RAP1GAP;refGene_tx=XM_019836358.2;refGene_gn=RAP1GAP;refGene_tx=XM_019836359.2;refGene_gn=RAP1GAP;refGene_tx=XM_023258264.1;refGene_gn=RAP1GAP;refGene_tx=XM_023258265.1;refGene_gn=RAP1GAP;refGene_tx=XM_023258266.1;refGene_gn=RAP1GAP;refGene_tx=XM_023258267.1;refGene_gn=RAP1GAP;refGene_tx=XM_023258268.1
C1 15898476 15898551 refGene_gn=RAP1GAP;refGene_tx=XM_019836351.2
C1 15901470 15901536
refGene_gn=RAP1GAP;refGene_tx=XM_023258264.1;refGene_gn=RAP1GAP;refGene_tx=XM_023258265.1;refGene_gn=RAP1GAP;refGene_tx=XM_023258266.1
C1 15902387 15902481
refGene_gn=RAP1GAP;refGene_tx=XM_006934381.4;refGene_gn=RAP1GAP;refGene_tx=XM_019836348.2;refGene_gn=RAP1GAP;refGene_tx=XM_019836349.2;refGene_gn=RAP1GAP;refGene_tx=XM_019836350.2;refGene_gn=RAP1GAP;refGene_tx=XM_019836355.2;refGene_gn=RAP1GAP;refGene_tx=XM_019836356.2;refGene_gn=RAP1GAP;refGene_tx=XM_019836358.2;refGene_gn=RAP1GAP;refGene_tx=XM_019836359.2

C1 15923106 15923142
refGene_gn=RAP1GAP;refGene_tx=XM_006934381.4;refGene_gn=RAP1GAP;refGene_tx=XM_019836348.2;refGene_gn=RAP1GAP;refGene_tx=XM_019836349.2;refGene_gn=RAP1GAP;refGene_tx=XM_019836350.2;refGene_gn=RAP1GAP;refGene_tx=XM_019836355.2;refGene_gn=RAP1GAP;refGene_tx=XM_019836356.2;refGene_gn=RAP1GAP;refGene_tx=XM_019836358.2;refGene_gn=RAP1GAP;refGene_tx=XM_019836359.2
C1 15925150 15925194
refGene_gn=RAP1GAP;refGene_tx=XM_006934381.4;refGene_gn=RAP1GAP;refGene_tx=XM_019836348.2;refGene_gn=RAP1GAP;refGene_tx=XM_019836349.2;refGene_gn=RAP1GAP;refGene_tx=XM_019836350.2;refGene_gn=RAP1GAP;refGene_tx=XM_019836355.2;refGene_gn=RAP1GAP;refGene_tx=XM_019836356.2;refGene_gn=RAP1GAP;refGene_tx=XM_019836358.2;refGene_gn=RAP1GAP;refGene_tx=XM_019836359.2
C1 15951410 15951433
refGene_gn=USP48;refGene_tx=XM_023258269.1;refGene_gn=USP48;refGene_tx=XM_023258270.1;refGene_gn=USP48;refGene_tx=XM_023258271.1;refGene_gn=USP48;refGene_tx=XM_023258272.1;refGene_gn=USP48;refGene_tx=XM_023258273.1;refGene_gn=USP48;refGene_tx=XM_023258274.1;refGene_gn=USP48;refGene_tx=XM_023258275.1;refGene_gn=USP48;refGene_tx=XM_023258276.1
C1 15952500 15952527
refGene_gn=USP48;refGene_tx=XM_023258269.1;refGene_gn=USP48;refGene_tx=XM_023258270.1;refGene_gn=USP48;refGene_tx=XM_023258271.1;refGene_gn=USP48;refGene_tx=XM_023258272.1;refGene_gn=USP48;refGene_tx=XM_023258273.1;refGene_gn=USP48;refGene_tx=XM_023258274.1;refGene_gn=USP48;refGene_tx=XM_023258275.1;refGene_gn=USP48;refGene_tx=XM_023258276.1
C1 15954981 15955030
refGene_gn=USP48;refGene_tx=XM_023258269.1;refGene_gn=USP48;refGene_tx=XM_023258270.1;refGene_gn=USP48;refGene_tx=XM_023258271.1;refGene_gn=USP48;refGene_tx=XM_023258272.1;refGene_gn=USP48;refGene_tx=XM_023258273.1;refGene_gn=USP48;refGene_tx=XM_023258274.1;refGene_gn=USP48;refGene_tx=XM_023258275.1;refGene_gn=USP48;refGene_tx=XM_023258276.1
C1 15956414 15956540
refGene_gn=USP48;refGene_tx=XM_023258269.1;refGene_gn=USP48;refGene_tx=XM_023258270.1;refGene_gn=USP48;refGene_tx=XM_023258271.1;refGene_gn=USP48;refGene_tx=XM_023258272.1;refGene_gn=USP48;refGene_tx=XM_023258273.1;refGene_gn=USP48;refGene_tx=XM_023258274.1;refGene_gn=USP48;refGene_tx=XM_023258275.1;refGene_gn=USP48;refGene_tx=XM_023258276.1
C1 15961312 15961468
refGene_gn=USP48;refGene_tx=XM_023258269.1;refGene_gn=USP48;refGene_tx=XM_023258270.1;refGene_gn=USP48;refGene_tx=XM_023258271.1;refGene_gn=USP48;refGene_tx=XM_023258272.1;refGene_gn=USP48;refGene_tx=XM_023258273.1;refGene_gn=USP48;refGene_tx=XM_023258274.1;refGene_gn=USP48;refGene_tx=XM_023258275.1;refGene_gn=USP48;refGene_tx=XM_023258276.1
C1 15966221 15966326
refGene_gn=USP48;refGene_tx=XM_023258269.1;refGene_gn=USP48;refGene_tx=XM_023258270.1;refGene_gn=USP48;refGene_tx=XM_023258271.1;refGene_gn=USP48;refGene_tx=XM_023258272.1;refGene_gn=USP48;refGene_tx=XM_023258273.1;refGene_gn=USP48;refGene_tx=XM_023258274.1;refGene_gn=USP48;refGene_tx=XM_023258275.1;refGene_gn=USP48;refGene_tx=XM_023258276.1

C1 15973063 15973170
refGene_gn=USP48;refGene_tx=XM_023258269.1;refGene_gn=USP48;refGene_tx=XM_023258270.1;refGene_gn=USP48;refGene_tx=XM_023258271.1;refGene_gn=USP48;refGene_tx=XM_023258272.1;refGene_gn=USP48;refGene_tx=XM_023258273.1;refGene_gn=USP48;refGene_tx=XM_023258274.1;refGene_gn=USP48;refGene_tx=XM_023258275.1;refGene_gn=USP48;refGene_tx=XM_023258276.1
C1 15973509 15973640
refGene_gn=USP48;refGene_tx=XM_023258269.1;refGene_gn=USP48;refGene_tx=XM_023258270.1;refGene_gn=USP48;refGene_tx=XM_023258271.1;refGene_gn=USP48;refGene_tx=XM_023258272.1;refGene_gn=USP48;refGene_tx=XM_023258273.1;refGene_gn=USP48;refGene_tx=XM_023258274.1;refGene_gn=USP48;refGene_tx=XM_023258275.1;refGene_gn=USP48;refGene_tx=XM_023258276.1
C1 15975483 15975594
refGene_gn=USP48;refGene_tx=XM_023258269.1;refGene_gn=USP48;refGene_tx=XM_023258270.1;refGene_gn=USP48;refGene_tx=XM_023258271.1;refGene_gn=USP48;refGene_tx=XM_023258272.1;refGene_gn=USP48;refGene_tx=XM_023258273.1;refGene_gn=USP48;refGene_tx=XM_023258274.1;refGene_gn=USP48;refGene_tx=XM_023258275.1;refGene_gn=USP48;refGene_tx=XM_023258276.1
C1 15976222 15976284
refGene_gn=USP48;refGene_tx=XM_023258269.1;refGene_gn=USP48;refGene_tx=XM_023258270.1;refGene_gn=USP48;refGene_tx=XM_023258271.1;refGene_gn=USP48;refGene_tx=XM_023258272.1;refGene_gn=USP48;refGene_tx=XM_023258273.1;refGene_gn=USP48;refGene_tx=XM_023258274.1;refGene_gn=USP48;refGene_tx=XM_023258275.1;refGene_gn=USP48;refGene_tx=XM_023258276.1
C1 15976561 15976684
refGene_gn=USP48;refGene_tx=XM_023258269.1;refGene_gn=USP48;refGene_tx=XM_023258270.1;refGene_gn=USP48;refGene_tx=XM_023258271.1;refGene_gn=USP48;refGene_tx=XM_023258272.1;refGene_gn=USP48;refGene_tx=XM_023258273.1;refGene_gn=USP48;refGene_tx=XM_023258274.1;refGene_gn=USP48;refGene_tx=XM_023258275.1;refGene_gn=USP48;refGene_tx=XM_023258276.1
C1 15976847 15976972
refGene_gn=USP48;refGene_tx=XM_023258269.1;refGene_gn=USP48;refGene_tx=XM_023258270.1;refGene_gn=USP48;refGene_tx=XM_023258271.1;refGene_gn=USP48;refGene_tx=XM_023258272.1;refGene_gn=USP48;refGene_tx=XM_023258273.1;refGene_gn=USP48;refGene_tx=XM_023258274.1;refGene_gn=USP48;refGene_tx=XM_023258275.1;refGene_gn=USP48;refGene_tx=XM_023258276.1
C1 15982944 15983016
refGene_gn=USP48;refGene_tx=XM_023258269.1;refGene_gn=USP48;refGene_tx=XM_023258270.1;refGene_gn=USP48;refGene_tx=XM_023258271.1;refGene_gn=USP48;refGene_tx=XM_023258272.1;refGene_gn=USP48;refGene_tx=XM_023258273.1;refGene_gn=USP48;refGene_tx=XM_023258274.1;refGene_gn=USP48;refGene_tx=XM_023258275.1;refGene_gn=USP48;refGene_tx=XM_023258276.1
C1 15984417 15984548
refGene_gn=USP48;refGene_tx=XM_023258269.1;refGene_gn=USP48;refGene_tx=XM_023258270.1;refGene_gn=USP48;refGene_tx=XM_023258271.1;refGene_gn=USP48;refGene_tx=XM_023258272.1;refGene_gn=USP48;refGene_tx=XM_023258273.1;refGene_gn=USP48;refGene_tx=XM_023258274.1;refGene_gn=USP48;refGene_tx=XM_023258275.1;refGene_gn=USP48;refGene_tx=XM_023258276.1
C1 15985482 15985597
refGene_gn=USP48;refGene_tx=XM_023258269.1;refGene_gn=USP48;refGene_tx

=XM_023258270.1;refGene_gn=USP48;refGene_tx=XM_023258271.1;refGene_gn=USP48;refGene_tx=XM_023258272.1;refGene_gn=USP48;refGene_tx=XM_023258273.1;refGene_gn=USP48;refGene_tx=XM_023258274.1;refGene_gn=USP48;refGene_tx=XM_023258275.1;refGene_gn=USP48;refGene_tx=XM_023258276.1
C1 15987225 15987423
refGene_gn=USP48;refGene_tx=XM_023258269.1;refGene_gn=USP48;refGene_tx=XM_023258270.1;refGene_gn=USP48;refGene_tx=XM_023258271.1;refGene_gn=USP48;refGene_tx=XM_023258272.1;refGene_gn=USP48;refGene_tx=XM_023258273.1;refGene_gn=USP48;refGene_tx=XM_023258274.1;refGene_gn=USP48;refGene_tx=XM_023258275.1;refGene_gn=USP48;refGene_tx=XM_023258276.1
C1 15988739 15988791
refGene_gn=USP48;refGene_tx=XM_023258269.1;refGene_gn=USP48;refGene_tx=XM_023258270.1;refGene_gn=USP48;refGene_tx=XM_023258273.1;refGene_gn=USP48;refGene_tx=XM_023258274.1;refGene_gn=USP48;refGene_tx=XM_023258275.1;refGene_gn=USP48;refGene_tx=XM_023258276.1
C1 15991614 15991768
refGene_gn=USP48;refGene_tx=XM_023258269.1;refGene_gn=USP48;refGene_tx=XM_023258270.1;refGene_gn=USP48;refGene_tx=XM_023258271.1;refGene_gn=USP48;refGene_tx=XM_023258272.1;refGene_gn=USP48;refGene_tx=XM_023258273.1;refGene_gn=USP48;refGene_tx=XM_023258274.1;refGene_gn=USP48;refGene_tx=XM_023258275.1;refGene_gn=USP48;refGene_tx=XM_023258276.1
C1 15993281 15993410
refGene_gn=USP48;refGene_tx=XM_023258269.1;refGene_gn=USP48;refGene_tx=XM_023258270.1;refGene_gn=USP48;refGene_tx=XM_023258271.1;refGene_gn=USP48;refGene_tx=XM_023258272.1;refGene_gn=USP48;refGene_tx=XM_023258274.1;refGene_gn=USP48;refGene_tx=XM_023258275.1;refGene_gn=USP48;refGene_tx=XM_023258276.1
C1 15996819 15996999
refGene_gn=USP48;refGene_tx=XM_023258269.1;refGene_gn=USP48;refGene_tx=XM_023258270.1;refGene_gn=USP48;refGene_tx=XM_023258271.1;refGene_gn=USP48;refGene_tx=XM_023258272.1;refGene_gn=USP48;refGene_tx=XM_023258273.1;refGene_gn=USP48;refGene_tx=XM_023258276.1
C1 16004070 16004153
refGene_gn=USP48;refGene_tx=XM_023258269.1;refGene_gn=USP48;refGene_tx=XM_023258270.1;refGene_gn=USP48;refGene_tx=XM_023258271.1;refGene_gn=USP48;refGene_tx=XM_023258272.1;refGene_gn=USP48;refGene_tx=XM_023258273.1;refGene_gn=USP48;refGene_tx=XM_023258274.1;refGene_gn=USP48;refGene_tx=XM_023258275.1;refGene_gn=USP48;refGene_tx=XM_023258276.1
C1 16004861 16004995
refGene_gn=USP48;refGene_tx=XM_023258269.1;refGene_gn=USP48;refGene_tx=XM_023258270.1;refGene_gn=USP48;refGene_tx=XM_023258271.1;refGene_gn=USP48;refGene_tx=XM_023258272.1;refGene_gn=USP48;refGene_tx=XM_023258273.1;refGene_gn=USP48;refGene_tx=XM_023258274.1;refGene_gn=USP48;refGene_tx=XM_023258275.1;refGene_gn=USP48;refGene_tx=XM_023258276.1
C1 16007758 16007867
refGene_gn=USP48;refGene_tx=XM_023258269.1;refGene_gn=USP48;refGene_tx=XM_023258270.1;refGene_gn=USP48;refGene_tx=XM_023258271.1;refGene_gn=USP48;refGene_tx=XM_023258272.1;refGene_gn=USP48;refGene_tx=XM_023258273.1;refGene_gn=USP48;refGene_tx=XM_023258274.1;refGene_gn=USP48;refGene_tx=XM_023258275.1;refGene_gn=USP48;refGene_tx=XM_023258276.1

C1 16008384 16008509
refGene_gn=USP48;refGene_tx=XM_023258269.1;refGene_gn=USP48;refGene_tx
=XM_023258270.1;refGene_gn=USP48;refGene_tx=XM_023258271.1;refGene_gn=
USP48;refGene_tx=XM_023258272.1;refGene_gn=USP48;refGene_tx=XM_0232582
73.1;refGene_gn=USP48;refGene_tx=XM_023258274.1;refGene_gn=USP48;refGe
ne_tx=XM_023258275.1;refGene_gn=USP48;refGene_tx=XM_023258276.1
C1 16009129 16009257
refGene_gn=USP48;refGene_tx=XM_023258269.1;refGene_gn=USP48;refGene_tx
=XM_023258270.1;refGene_gn=USP48;refGene_tx=XM_023258271.1;refGene_gn=
USP48;refGene_tx=XM_023258272.1;refGene_gn=USP48;refGene_tx=XM_0232582
73.1;refGene_gn=USP48;refGene_tx=XM_023258274.1;refGene_gn=USP48;refGe
ne_tx=XM_023258275.1;refGene_gn=USP48;refGene_tx=XM_023258276.1
C1 16013109 16013266
refGene_gn=USP48;refGene_tx=XM_023258269.1;refGene_gn=USP48;refGene_tx
=XM_023258270.1;refGene_gn=USP48;refGene_tx=XM_023258271.1;refGene_gn=
USP48;refGene_tx=XM_023258272.1;refGene_gn=USP48;refGene_tx=XM_0232582
73.1;refGene_gn=USP48;refGene_tx=XM_023258274.1;refGene_gn=USP48;refGe
ne_tx=XM_023258275.1;refGene_gn=USP48;refGene_tx=XM_023258276.1
C1 16014220 16014341
refGene_gn=USP48;refGene_tx=XM_023258269.1;refGene_gn=USP48;refGene_tx
=XM_023258270.1;refGene_gn=USP48;refGene_tx=XM_023258271.1;refGene_gn=
USP48;refGene_tx=XM_023258272.1;refGene_gn=USP48;refGene_tx=XM_0232582
73.1;refGene_gn=USP48;refGene_tx=XM_023258274.1;refGene_gn=USP48;refGe
ne_tx=XM_023258275.1
C1 16041391 16041525
refGene_gn=USP48;refGene_tx=XM_023258269.1;refGene_gn=USP48;refGene_tx
=XM_023258270.1;refGene_gn=USP48;refGene_tx=XM_023258271.1;refGene_gn=
USP48;refGene_tx=XM_023258272.1;refGene_gn=USP48;refGene_tx=XM_0232582
73.1;refGene_gn=USP48;refGene_tx=XM_023258274.1;refGene_gn=USP48;refGe
ne_tx=XM_023258275.1
C1 16060837 16060919 refGene_gn=LDLRAD2;refGene_tx=XM_011284560.3
C1 16063128 16063572 refGene_gn=LDLRAD2;refGene_tx=XM_011284560.3
C1 16066889 16067021 refGene_gn=LDLRAD2;refGene_tx=XM_011284560.3
C1 16069858 16070052 refGene_gn=LDLRAD2;refGene_tx=XM_011284560.3
C1 16071757 16071930 refGene_gn=HSPG2;refGene_tx=XM_023258277.1
C1 16072077 16072181 refGene_gn=HSPG2;refGene_tx=XM_023258277.1
C1 16072532 16072616 refGene_gn=HSPG2;refGene_tx=XM_023258277.1
C1 16072791 16072862 refGene_gn=HSPG2;refGene_tx=XM_023258277.1
C1 16073008 16073087 refGene_gn=HSPG2;refGene_tx=XM_023258277.1
C1 16073173 16073249 refGene_gn=HSPG2;refGene_tx=XM_023258277.1
C1 16075243 16075300 refGene_gn=HSPG2;refGene_tx=XM_023258277.1
C1 16075405 16075528 refGene_gn=HSPG2;refGene_tx=XM_023258277.1
C1 16075619 16075791 refGene_gn=HSPG2;refGene_tx=XM_023258277.1
C1 16076133 16076378 refGene_gn=HSPG2;refGene_tx=XM_023258277.1
C1 16076641 16076863 refGene_gn=HSPG2;refGene_tx=XM_023258277.1
C1 16077217 16077316 refGene_gn=HSPG2;refGene_tx=XM_023258277.1
C1 16077914 16078023 refGene_gn=HSPG2;refGene_tx=XM_023258277.1
C1 16078147 16078257 refGene_gn=HSPG2;refGene_tx=XM_023258277.1
C1 16078397 16078497 refGene_gn=HSPG2;refGene_tx=XM_023258277.1
C1 16078585 16078730 refGene_gn=HSPG2;refGene_tx=XM_023258277.1

| | | | |
|---|---|---|---|
| C1 | 16079396 | 16079508 | refGene_gn=HSPG2;refGene_tx=XM_023258277.1 |
| C1 | 16080164 | 16080281 | refGene_gn=HSPG2;refGene_tx=XM_023258277.1 |
| C1 | 16080363 | 16080511 | refGene_gn=HSPG2;refGene_tx=XM_023258277.1 |
| C1 | 16080709 | 16080819 | refGene_gn=HSPG2;refGene_tx=XM_023258277.1 |
| C1 | 16081340 | 16081607 | refGene_gn=HSPG2;refGene_tx=XM_023258277.1 |
| C1 | 16082496 | 16082594 | refGene_gn=HSPG2;refGene_tx=XM_023258277.1 |
| C1 | 16083122 | 16083327 | refGene_gn=HSPG2;refGene_tx=XM_023258277.1 |
| C1 | 16083492 | 16083753 | refGene_gn=HSPG2;refGene_tx=XM_023258277.1 |
| C1 | 16084055 | 16084235 | refGene_gn=HSPG2;refGene_tx=XM_023258277.1 |
| C1 | 16084489 | 16084685 | refGene_gn=HSPG2;refGene_tx=XM_023258277.1 |
| C1 | 16085246 | 16085428 | refGene_gn=HSPG2;refGene_tx=XM_023258277.1 |
| C1 | 16085679 | 16085814 | refGene_gn=HSPG2;refGene_tx=XM_023258277.1 |
| C1 | 16086196 | 16086337 | refGene_gn=HSPG2;refGene_tx=XM_023258277.1 |
| C1 | 16086453 | 16086595 | refGene_gn=HSPG2;refGene_tx=XM_023258277.1 |
| C1 | 16086974 | 16087126 | refGene_gn=HSPG2;refGene_tx=XM_023258277.1 |
| C1 | 16087463 | 16087605 | refGene_gn=HSPG2;refGene_tx=XM_023258277.1 |
| C1 | 16088022 | 16088174 | refGene_gn=HSPG2;refGene_tx=XM_023258277.1 |
| C1 | 16089462 | 16089610 | refGene_gn=HSPG2;refGene_tx=XM_023258277.1 |
| C1 | 16089838 | 16089990 | refGene_gn=HSPG2;refGene_tx=XM_023258277.1 |
| C1 | 16090535 | 16090674 | refGene_gn=HSPG2;refGene_tx=XM_023258277.1 |
| C1 | 16090921 | 16091073 | refGene_gn=HSPG2;refGene_tx=XM_023258277.1 |
| C1 | 16091190 | 16091326 | refGene_gn=HSPG2;refGene_tx=XM_023258277.1 |
| C1 | 16091943 | 16092095 | refGene_gn=HSPG2;refGene_tx=XM_023258277.1 |
| C1 | 16092184 | 16092323 | refGene_gn=HSPG2;refGene_tx=XM_023258277.1 |
| C1 | 16092754 | 16092906 | refGene_gn=HSPG2;refGene_tx=XM_023258277.1 |
| C1 | 16093049 | 16093185 | refGene_gn=HSPG2;refGene_tx=XM_023258277.1 |
| C1 | 16094080 | 16094232 | refGene_gn=HSPG2;refGene_tx=XM_023258277.1 |
| C1 | 16094328 | 16094464 | refGene_gn=HSPG2;refGene_tx=XM_023258277.1 |
| C1 | 16094642 | 16094794 | refGene_gn=HSPG2;refGene_tx=XM_023258277.1 |
| C1 | 16095207 | 16095355 | refGene_gn=HSPG2;refGene_tx=XM_023258277.1 |
| C1 | 16095438 | 16095608 | refGene_gn=HSPG2;refGene_tx=XM_023258277.1 |
| C1 | 16096321 | 16096472 | refGene_gn=HSPG2;refGene_tx=XM_023258277.1 |
| C1 | 16096718 | 16096867 | refGene_gn=HSPG2;refGene_tx=XM_023258277.1 |
| C1 | 16096953 | 16097089 | refGene_gn=HSPG2;refGene_tx=XM_023258277.1 |
| C1 | 16097407 | 16097550 | refGene_gn=HSPG2;refGene_tx=XM_023258277.1 |
| C1 | 16097630 | 16097784 | refGene_gn=HSPG2;refGene_tx=XM_023258277.1 |
| C1 | 16097916 | 16098050 | refGene_gn=HSPG2;refGene_tx=XM_023258277.1 |
| C1 | 16099413 | 16099594 | refGene_gn=HSPG2;refGene_tx=XM_023258277.1 |
| C1 | 16099698 | 16099799 | refGene_gn=HSPG2;refGene_tx=XM_023258277.1 |
| C1 | 16101697 | 16101808 | refGene_gn=HSPG2;refGene_tx=XM_023258277.1 |
| C1 | 16101959 | 16102127 | refGene_gn=HSPG2;refGene_tx=XM_023258277.1 |
| C1 | 16102313 | 16102372 | refGene_gn=HSPG2;refGene_tx=XM_023258277.1 |
| C1 | 16103352 | 16103439 | refGene_gn=HSPG2;refGene_tx=XM_023258277.1 |
| C1 | 16103563 | 16103691 | refGene_gn=HSPG2;refGene_tx=XM_023258277.1 |
| C1 | 16104255 | 16104369 | refGene_gn=HSPG2;refGene_tx=XM_023258277.1 |
| C1 | 16106555 | 16106786 | refGene_gn=HSPG2;refGene_tx=XM_023258277.1 |
| C1 | 16106997 | 16107078 | refGene_gn=HSPG2;refGene_tx=XM_023258277.1 |
| C1 | 16107417 | 16107510 | refGene_gn=HSPG2;refGene_tx=XM_023258277.1 |
| C1 | 16112663 | 16112855 | refGene_gn=HSPG2;refGene_tx=XM_023258277.1 |
| C1 | 16113254 | 16113395 | refGene_gn=HSPG2;refGene_tx=XM_023258277.1 |

C1 16113714 16113809 refGene_gn=HSPG2;refGene_tx=XM_023258277.1
C1 16114132 16114182 refGene_gn=HSPG2;refGene_tx=XM_023258277.1
C1 16114629 16114716 refGene_gn=HSPG2;refGene_tx=XM_023258277.1
C1 16115068 16115196 refGene_gn=HSPG2;refGene_tx=XM_023258277.1
C1 16115279 16115393 refGene_gn=HSPG2;refGene_tx=XM_023258277.1
C1 16115565 16115677 refGene_gn=HSPG2;refGene_tx=XM_023258277.1
C1 16116303 16116422 refGene_gn=HSPG2;refGene_tx=XM_023258277.1
C1 16116513 16116693 refGene_gn=HSPG2;refGene_tx=XM_023258277.1
C1 16116846 16117023 refGene_gn=HSPG2;refGene_tx=XM_023258277.1
C1 16117136 16117277 refGene_gn=HSPG2;refGene_tx=XM_023258277.1
C1 16118150 16118218 refGene_gn=HSPG2;refGene_tx=XM_023258277.1
C1 16118373 16118432 refGene_gn=HSPG2;refGene_tx=XM_023258277.1
C1 16118522 16118609 refGene_gn=HSPG2;refGene_tx=XM_023258277.1
C1 16119337 16119465 refGene_gn=HSPG2;refGene_tx=XM_023258277.1
C1 16120130 16120278 refGene_gn=HSPG2;refGene_tx=XM_023258277.1
C1 16120372 16120569 refGene_gn=HSPG2;refGene_tx=XM_023258277.1
C1 16120668 16120848 refGene_gn=HSPG2;refGene_tx=XM_023258277.1
C1 16121146 16121310 refGene_gn=HSPG2;refGene_tx=XM_023258277.1
C1 16123737 16123884 refGene_gn=HSPG2;refGene_tx=XM_023258277.1
C1 16123979 16124131 refGene_gn=HSPG2;refGene_tx=XM_023258277.1
C1 16124224 16124369 refGene_gn=HSPG2;refGene_tx=XM_023258277.1
C1 16124527 16124659 refGene_gn=HSPG2;refGene_tx=XM_023258277.1
C1 16126387 16126507 refGene_gn=HSPG2;refGene_tx=XM_023258277.1
C1 16126583 16126838 refGene_gn=HSPG2;refGene_tx=XM_023258277.1
C1 16127068 16127197 refGene_gn=HSPG2;refGene_tx=XM_023258277.1
C1 16128973 16129134 refGene_gn=HSPG2;refGene_tx=XM_023258277.1
C1 16129426 16129485 refGene_gn=HSPG2;refGene_tx=XM_023258277.1
C1 16129581 16129691 refGene_gn=HSPG2;refGene_tx=XM_023258277.1
C1 16135677 16135722 refGene_gn=HSPG2;refGene_tx=XM_023258277.1
C1 16136340 16136476 refGene_gn=HSPG2;refGene_tx=XM_023258277.1
C1 16170309 16170372 refGene_gn=HSPG2;refGene_tx=XM_023258277.1
C1 16181363 16182203
refGene_gn=LOC102901553;refGene_tx=XM_006934383.4;refGene_gn=LOC102901553;refGene_tx=XM_006934384.4;refGene_gn=LOC102901553;refGene_tx=XM_011284559.3
C1 16211761 16211804
refGene_gn=LOC101094275;refGene_tx=XM_023258278.1
C1 16212795 16212881
refGene_gn=LOC101094275;refGene_tx=XM_023258278.1
C1 16213775 16213873
refGene_gn=LOC101094275;refGene_tx=XM_023258278.1
C1 16213992 16214127
refGene_gn=LOC101094275;refGene_tx=XM_023258278.1
C1 16215288 16215425
refGene_gn=LOC101094275;refGene_tx=XM_023258278.1
C1 16215807 16215950
refGene_gn=LOC101094275;refGene_tx=XM_023258278.1
C1 16216556 16216709
refGene_gn=LOC101094275;refGene_tx=XM_023258278.1
C1 16217388 16217406

refGene_gn=LOC101094275;refGene_tx=XM_023258278.1
C1 16226380 16226833
refGene_gn=LOC101094519;refGene_tx=XM_003989634.5;refGene_gn=LOC101094519;refGene_tx=XM_006934385.4;refGene_gn=LOC101094519;refGene_tx=XM_006934386.3
C1 16271882 16271987
refGene_gn=CDC42;refGene_tx=XM_023258279.1;refGene_gn=CDC42;refGene_tx=XM_023258280.1
C1 16274310 16274383
refGene_gn=CDC42;refGene_tx=XM_023258279.1;refGene_gn=CDC42;refGene_tx=XM_023258280.1
C1 16290130 16290240
refGene_gn=CDC42;refGene_tx=XM_023258279.1;refGene_gn=CDC42;refGene_tx=XM_023258280.1
C1 16290364 16290562
refGene_gn=CDC42;refGene_tx=XM_023258279.1;refGene_gn=CDC42;refGene_tx=XM_023258280.1
C1 16294199 16294289 refGene_gn=CDC42;refGene_tx=XM_023258280.1
C1 16295695 16295785 refGene_gn=CDC42;refGene_tx=XM_023258279.1
C1 16321862 16322330
refGene_gn=WNT4;refGene_tx=XM_023258281.1;refGene_gn=WNT4;refGene_tx=XM_023258282.1;refGene_gn=WNT4;refGene_tx=XM_023258283.1;refGene_gn=WNT4;refGene_tx=XM_023258284.1
C1 16322985 16323128
refGene_gn=WNT4;refGene_tx=XM_023258281.1;refGene_gn=WNT4;refGene_tx=XM_023258282.1;refGene_gn=WNT4;refGene_tx=XM_023258283.1;refGene_gn=WNT4;refGene_tx=XM_023258284.1
C1 16323222 16323354
refGene_gn=WNT4;refGene_tx=XM_023258281.1;refGene_gn=WNT4;refGene_tx=XM_023258282.1;refGene_gn=WNT4;refGene_tx=XM_023258283.1;refGene_gn=WNT4;refGene_tx=XM_023258284.1
C1 16333762 16333998
refGene_gn=WNT4;refGene_tx=XM_023258281.1;refGene_gn=WNT4;refGene_tx=XM_023258282.1;refGene_gn=WNT4;refGene_tx=XM_023258283.1;refGene_gn=WNT4;refGene_tx=XM_023258284.1
C1 16348009 16348086
refGene_gn=WNT4;refGene_tx=XM_023258281.1;refGene_gn=WNT4;refGene_tx=XM_023258282.1
C1 16642886 16642923 refGene_gn=ZBTB40;refGene_tx=XM_019836360.2
C1 16644213 16644978
refGene_gn=ZBTB40;refGene_tx=XM_006934388.4;refGene_gn=ZBTB40;refGene_tx=XM_019836360.2;refGene_gn=ZBTB40;refGene_tx=XM_019836361.2;refGene_gn=ZBTB40;refGene_tx=XM_019836362.2;refGene_gn=ZBTB40;refGene_tx=XM_023258285.1;refGene_gn=ZBTB40;refGene_tx=XM_023258286.1
C1 16645681 16645815
refGene_gn=ZBTB40;refGene_tx=XM_006934388.4;refGene_gn=ZBTB40;refGene_tx=XM_019836360.2;refGene_gn=ZBTB40;refGene_tx=XM_019836361.2;refGene_gn=ZBTB40;refGene_tx=XM_019836362.2;refGene_gn=ZBTB40;refGene_tx=XM_023258285.1;refGene_gn=ZBTB40;refGene_tx=XM_023258286.1
C1 16653670 16653863

refGene_gn=ZBTB40;refGene_tx=XM_006934388.4;refGene_gn=ZBTB40;refGene_tx=XM_019836360.2;refGene_gn=ZBTB40;refGene_tx=XM_019836361.2;refGene_gn=ZBTB40;refGene_tx=XM_019836362.2;refGene_gn=ZBTB40;refGene_tx=XM_023258285.1;refGene_gn=ZBTB40;refGene_tx=XM_023258286.1
C1 16654472 16654615
refGene_gn=ZBTB40;refGene_tx=XM_006934388.4;refGene_gn=ZBTB40;refGene_tx=XM_019836360.2;refGene_gn=ZBTB40;refGene_tx=XM_019836361.2;refGene_gn=ZBTB40;refGene_tx=XM_019836362.2;refGene_gn=ZBTB40;refGene_tx=XM_023258285.1;refGene_gn=ZBTB40;refGene_tx=XM_023258286.1
C1 16658934 16659127
refGene_gn=ZBTB40;refGene_tx=XM_006934388.4;refGene_gn=ZBTB40;refGene_tx=XM_019836360.2;refGene_gn=ZBTB40;refGene_tx=XM_019836361.2;refGene_gn=ZBTB40;refGene_tx=XM_019836362.2;refGene_gn=ZBTB40;refGene_tx=XM_023258285.1;refGene_gn=ZBTB40;refGene_tx=XM_023258286.1
C1 16661543 16661680
refGene_gn=ZBTB40;refGene_tx=XM_006934388.4;refGene_gn=ZBTB40;refGene_tx=XM_019836360.2;refGene_gn=ZBTB40;refGene_tx=XM_019836361.2;refGene_gn=ZBTB40;refGene_tx=XM_019836362.2;refGene_gn=ZBTB40;refGene_tx=XM_023258285.1;refGene_gn=ZBTB40;refGene_tx=XM_023258286.1
C1 16662089 16662291
refGene_gn=ZBTB40;refGene_tx=XM_006934388.4;refGene_gn=ZBTB40;refGene_tx=XM_019836360.2;refGene_gn=ZBTB40;refGene_tx=XM_019836361.2;refGene_gn=ZBTB40;refGene_tx=XM_019836362.2;refGene_gn=ZBTB40;refGene_tx=XM_023258285.1;refGene_gn=ZBTB40;refGene_tx=XM_023258286.1
C1 16662645 16662779
refGene_gn=ZBTB40;refGene_tx=XM_006934388.4;refGene_gn=ZBTB40;refGene_tx=XM_019836360.2;refGene_gn=ZBTB40;refGene_tx=XM_019836361.2;refGene_gn=ZBTB40;refGene_tx=XM_019836362.2;refGene_gn=ZBTB40;refGene_tx=XM_023258285.1;refGene_gn=ZBTB40;refGene_tx=XM_023258286.1
C1 16664401 16664570
refGene_gn=ZBTB40;refGene_tx=XM_006934388.4;refGene_gn=ZBTB40;refGene_tx=XM_019836360.2;refGene_gn=ZBTB40;refGene_tx=XM_019836361.2;refGene_gn=ZBTB40;refGene_tx=XM_019836362.2;refGene_gn=ZBTB40;refGene_tx=XM_023258285.1;refGene_gn=ZBTB40;refGene_tx=XM_023258286.1
C1 16664888 16665356
refGene_gn=ZBTB40;refGene_tx=XM_006934388.4;refGene_gn=ZBTB40;refGene_tx=XM_019836360.2;refGene_gn=ZBTB40;refGene_tx=XM_019836361.2;refGene_gn=ZBTB40;refGene_tx=XM_019836362.2;refGene_gn=ZBTB40;refGene_tx=XM_023258285.1;refGene_gn=ZBTB40;refGene_tx=XM_023258286.1
C1 16666116 16666323
refGene_gn=ZBTB40;refGene_tx=XM_006934388.4;refGene_gn=ZBTB40;refGene_tx=XM_019836360.2;refGene_gn=ZBTB40;refGene_tx=XM_019836361.2;refGene_gn=ZBTB40;refGene_tx=XM_019836362.2;refGene_gn=ZBTB40;refGene_tx=XM_023258285.1;refGene_gn=ZBTB40;refGene_tx=XM_023258286.1
C1 16670106 16670271
refGene_gn=ZBTB40;refGene_tx=XM_006934388.4;refGene_gn=ZBTB40;refGene_tx=XM_019836360.2;refGene_gn=ZBTB40;refGene_tx=XM_019836361.2;refGene_gn=ZBTB40;refGene_tx=XM_019836362.2;refGene_gn=ZBTB40;refGene_tx=XM_023258285.1;refGene_gn=ZBTB40;refGene_tx=XM_023258286.1
C1 16672814 16673029

refGene_gn=ZBTB40;refGene_tx=XM_006934388.4;refGene_gn=ZBTB40;refGene_tx=XM_019836360.2;refGene_gn=ZBTB40;refGene_tx=XM_019836361.2;refGene_gn=ZBTB40;refGene_tx=XM_019836362.2;refGene_gn=ZBTB40;refGene_tx=XM_023258285.1;refGene_gn=ZBTB40;refGene_tx=XM_023258286.1
C1 16674196 16674359 refGene_gn=ZBTB40;refGene_tx=XM_006934388.4;refGene_gn=ZBTB40;refGene_tx=XM_019836360.2;refGene_gn=ZBTB40;refGene_tx=XM_019836361.2;refGene_gn=ZBTB40;refGene_tx=XM_019836362.2;refGene_gn=ZBTB40;refGene_tx=XM_023258285.1;refGene_gn=ZBTB40;refGene_tx=XM_023258286.1
C1 16675059 16675146 refGene_gn=ZBTB40;refGene_tx=XM_006934388.4;refGene_gn=ZBTB40;refGene_tx=XM_019836360.2;refGene_gn=ZBTB40;refGene_tx=XM_019836361.2;refGene_gn=ZBTB40;refGene_tx=XM_019836362.2;refGene_gn=ZBTB40;refGene_tx=XM_023258285.1;refGene_gn=ZBTB40;refGene_tx=XM_023258286.1
C1 16676334 16676552 refGene_gn=ZBTB40;refGene_tx=XM_006934388.4;refGene_gn=ZBTB40;refGene_tx=XM_019836360.2;refGene_gn=ZBTB40;refGene_tx=XM_019836361.2;refGene_gn=ZBTB40;refGene_tx=XM_019836362.2;refGene_gn=ZBTB40;refGene_tx=XM_023258285.1;refGene_gn=ZBTB40;refGene_tx=XM_023258286.1
C1 16678364 16678559 refGene_gn=ZBTB40;refGene_tx=XM_006934388.4;refGene_gn=ZBTB40;refGene_tx=XM_019836360.2;refGene_gn=ZBTB40;refGene_tx=XM_019836361.2;refGene_gn=ZBTB40;refGene_tx=XM_019836362.2;refGene_gn=ZBTB40;refGene_tx=XM_023258285.1;refGene_gn=ZBTB40;refGene_tx=XM_023258286.1
C1 16712093 16712187 refGene_gn=EPHA8;refGene_tx=XM_003989639.5
C1 16717758 16717823 refGene_gn=EPHA8;refGene_tx=XM_003989639.5
C1 16724636 16725300 refGene_gn=EPHA8;refGene_tx=XM_003989639.5
C1 16732474 16732630 refGene_gn=EPHA8;refGene_tx=XM_003989639.5
C1 16734727 16735063 refGene_gn=EPHA8;refGene_tx=XM_003989639.5
C1 16739281 16739406 refGene_gn=EPHA8;refGene_tx=XM_003989639.5
C1 16739483 16739646 refGene_gn=EPHA8;refGene_tx=XM_003989639.5
C1 16740873 16740967 refGene_gn=EPHA8;refGene_tx=XM_003989639.5
C1 16741939 16742004 refGene_gn=EPHA8;refGene_tx=XM_003989639.5
C1 16742811 16742976 refGene_gn=EPHA8;refGene_tx=XM_003989639.5
C1 16743166 16743352 refGene_gn=EPHA8;refGene_tx=XM_003989639.5
C1 16743659 16743721 refGene_gn=EPHA8;refGene_tx=XM_003989639.5
C1 16744197 16744407 refGene_gn=EPHA8;refGene_tx=XM_003989639.5
C1 16745296 16745446 refGene_gn=EPHA8;refGene_tx=XM_003989639.5
C1 16745538 16745578 refGene_gn=EPHA8;refGene_tx=XM_003989639.5
C1 16745580 16745728 refGene_gn=EPHA8;refGene_tx=XM_003989639.5
C1 16746185 16746359 refGene_gn=EPHA8;refGene_tx=XM_003989639.5
C1 16746501 16746616 refGene_gn=EPHA8;refGene_tx=XM_003989639.5
C1 16780980 16781143 refGene_gn=C1QA;refGene_tx=XM_006934389.4;refGene_gn=C1QA;refGene_tx=XM_006934390.4
C1 16782415 16782990 refGene_gn=C1QA;refGene_tx=XM_006934389.4;refGene_gn=C1QA;refGene_tx=XM_006934390.4
C1 16786973 16787154 refGene_gn=C1QC;refGene_tx=XM_019836365.2
C1 16790227 16790784 refGene_gn=C1QC;refGene_tx=XM_019836365.2

C1 16798289 16798479 refGene_gn=C1QB;refGene_tx=XM_003989642.5
C1 16799631 16800206 refGene_gn=C1QB;refGene_tx=XM_003989642.5
C1 16839067 16839128 refGene_gn=EPHB2;refGene_tx=XM_023258287.1
C1 16901674 16901703 refGene_gn=EPHB2;refGene_tx=XM_023258287.1
C1 16901704 16901740 refGene_gn=EPHB2;refGene_tx=XM_023258287.1
C1 16904751 16905436 refGene_gn=EPHB2;refGene_tx=XM_023258287.1
C1 16981786 16981942 refGene_gn=EPHB2;refGene_tx=XM_023258287.1
C1 16983113 16983449 refGene_gn=EPHB2;refGene_tx=XM_023258287.1
C1 17000955 17001080 refGene_gn=EPHB2;refGene_tx=XM_023258287.1
C1 17010300 17010463 refGene_gn=EPHB2;refGene_tx=XM_023258287.1
C1 17012565 17012674 refGene_gn=EPHB2;refGene_tx=XM_023258287.1
C1 17013423 17013491 refGene_gn=EPHB2;refGene_tx=XM_023258287.1
C1 17020688 17020811 refGene_gn=EPHB2;refGene_tx=XM_023258287.1
C1 17021348 17021596 refGene_gn=EPHB2;refGene_tx=XM_023258287.1
C1 17022573 17022789 refGene_gn=EPHB2;refGene_tx=XM_023258287.1
C1 17023679 17023829 refGene_gn=EPHB2;refGene_tx=XM_023258287.1
C1 17024793 17024987 refGene_gn=EPHB2;refGene_tx=XM_023258287.1
C1 17026564 17026720 refGene_gn=EPHB2;refGene_tx=XM_023258287.1
C1 17027228 17027337 refGene_gn=EPHB2;refGene_tx=XM_023258287.1
C1 17042429 17043408
refGene_gn=LACTBL1;refGene_tx=XM_023258288.1;refGene_gn=LACTBL1;refGene_tx=XM_023258289.1
C1 17044350 17044456
refGene_gn=LACTBL1;refGene_tx=XM_023258288.1;refGene_gn=LACTBL1;refGene_tx=XM_023258289.1
C1 17047894 17048133
refGene_gn=LACTBL1;refGene_tx=XM_023258288.1;refGene_gn=LACTBL1;refGene_tx=XM_023258289.1
C1 17048490 17048630 refGene_gn=LACTBL1;refGene_tx=XM_023258289.1
C1 17049078 17049236 refGene_gn=LACTBL1;refGene_tx=XM_023258288.1
C1 17051497 17051607 refGene_gn=LACTBL1;refGene_tx=XM_023258288.1
C1 17053959 17054095 refGene_gn=LACTBL1;refGene_tx=XM_023258288.1
C1 17059894 17059906 refGene_gn=LACTBL1;refGene_tx=XM_023258288.1
C1 17069128 17069425 refGene_gn=TEX46;refGene_tx=XM_006934393.3
C1 17071406 17071598 refGene_gn=TEX46;refGene_tx=XM_006934393.3
C1 17076940 17077291
refGene_gn=KDM1A;refGene_tx=XM_023258294.1;refGene_gn=KDM1A;refGene_tx=XM_023258295.1;refGene_gn=KDM1A;refGene_tx=XM_023258296.1;refGene_gn=KDM1A;refGene_tx=XM_023258297.1
C1 17085602 17085768
refGene_gn=KDM1A;refGene_tx=XM_023258294.1;refGene_gn=KDM1A;refGene_tx=XM_023258295.1;refGene_gn=KDM1A;refGene_tx=XM_023258296.1;refGene_gn=KDM1A;refGene_tx=XM_023258297.1
C1 17099426 17099486
refGene_gn=KDM1A;refGene_tx=XM_023258294.1;refGene_gn=KDM1A;refGene_tx=XM_023258295.1
C1 17104818 17104952
refGene_gn=KDM1A;refGene_tx=XM_023258294.1;refGene_gn=KDM1A;refGene_tx=XM_023258295.1;refGene_gn=KDM1A;refGene_tx=XM_023258296.1;refGene_gn=KDM1A;refGene_tx=XM_023258297.1

C1 17110280 17110359
refGene_gn=KDM1A;refGene_tx=XM_023258294.1;refGene_gn=KDM1A;refGene_tx
=XM_023258295.1;refGene_gn=KDM1A;refGene_tx=XM_023258296.1;refGene_gn=
KDM1A;refGene_tx=XM_023258297.1
C1 17110935 17111028
refGene_gn=KDM1A;refGene_tx=XM_023258294.1;refGene_gn=KDM1A;refGene_tx
=XM_023258295.1;refGene_gn=KDM1A;refGene_tx=XM_023258296.1;refGene_gn=
KDM1A;refGene_tx=XM_023258297.1
C1 17112019 17112126
refGene_gn=KDM1A;refGene_tx=XM_023258294.1;refGene_gn=KDM1A;refGene_tx
=XM_023258295.1;refGene_gn=KDM1A;refGene_tx=XM_023258296.1;refGene_gn=
KDM1A;refGene_tx=XM_023258297.1
C1 17113681 17113763
refGene_gn=KDM1A;refGene_tx=XM_023258294.1;refGene_gn=KDM1A;refGene_tx
=XM_023258295.1;refGene_gn=KDM1A;refGene_tx=XM_023258296.1;refGene_gn=
KDM1A;refGene_tx=XM_023258297.1
C1 17114467 17114562
refGene_gn=KDM1A;refGene_tx=XM_023258294.1;refGene_gn=KDM1A;refGene_tx
=XM_023258295.1;refGene_gn=KDM1A;refGene_tx=XM_023258296.1;refGene_gn=
KDM1A;refGene_tx=XM_023258297.1
C1 17122186 17122198
refGene_gn=KDM1A;refGene_tx=XM_023258294.1;refGene_gn=KDM1A;refGene_tx
=XM_023258296.1
C1 17124660 17124803
refGene_gn=KDM1A;refGene_tx=XM_023258294.1;refGene_gn=KDM1A;refGene_tx
=XM_023258295.1;refGene_gn=KDM1A;refGene_tx=XM_023258296.1;refGene_gn=
KDM1A;refGene_tx=XM_023258297.1
C1 17125175 17125266
refGene_gn=KDM1A;refGene_tx=XM_023258294.1;refGene_gn=KDM1A;refGene_tx
=XM_023258295.1;refGene_gn=KDM1A;refGene_tx=XM_023258296.1;refGene_gn=
KDM1A;refGene_tx=XM_023258297.1
C1 17126498 17126633
refGene_gn=KDM1A;refGene_tx=XM_023258294.1;refGene_gn=KDM1A;refGene_tx
=XM_023258295.1;refGene_gn=KDM1A;refGene_tx=XM_023258296.1;refGene_gn=
KDM1A;refGene_tx=XM_023258297.1
C1 17127805 17127879
refGene_gn=KDM1A;refGene_tx=XM_023258294.1;refGene_gn=KDM1A;refGene_tx
=XM_023258295.1;refGene_gn=KDM1A;refGene_tx=XM_023258296.1;refGene_gn=
KDM1A;refGene_tx=XM_023258297.1
C1 17128796 17128908
refGene_gn=KDM1A;refGene_tx=XM_023258294.1;refGene_gn=KDM1A;refGene_tx
=XM_023258295.1;refGene_gn=KDM1A;refGene_tx=XM_023258296.1;refGene_gn=
KDM1A;refGene_tx=XM_023258297.1
C1 17132064 17132197
refGene_gn=KDM1A;refGene_tx=XM_023258294.1;refGene_gn=KDM1A;refGene_tx
=XM_023258295.1;refGene_gn=KDM1A;refGene_tx=XM_023258296.1;refGene_gn=
KDM1A;refGene_tx=XM_023258297.1
C1 17133674 17133862
refGene_gn=KDM1A;refGene_tx=XM_023258294.1;refGene_gn=KDM1A;refGene_tx
=XM_023258295.1;refGene_gn=KDM1A;refGene_tx=XM_023258296.1;refGene_gn=

KDM1A;refGene_tx=XM_023258297.1
C1 17134282 17134397
refGene_gn=KDM1A;refGene_tx=XM_023258294.1;refGene_gn=KDM1A;refGene_tx=XM_023258295.1;refGene_gn=KDM1A;refGene_tx=XM_023258296.1;refGene_gn=KDM1A;refGene_tx=XM_023258297.1
C1 17135887 17136015
refGene_gn=KDM1A;refGene_tx=XM_023258294.1;refGene_gn=KDM1A;refGene_tx=XM_023258295.1;refGene_gn=KDM1A;refGene_tx=XM_023258296.1;refGene_gn=KDM1A;refGene_tx=XM_023258297.1
C1 17136882 17137029
refGene_gn=KDM1A;refGene_tx=XM_023258294.1;refGene_gn=KDM1A;refGene_tx=XM_023258295.1;refGene_gn=KDM1A;refGene_tx=XM_023258296.1;refGene_gn=KDM1A;refGene_tx=XM_023258297.1
C1 17137881 17138067
refGene_gn=KDM1A;refGene_tx=XM_023258294.1;refGene_gn=KDM1A;refGene_tx=XM_023258295.1;refGene_gn=KDM1A;refGene_tx=XM_023258296.1;refGene_gn=KDM1A;refGene_tx=XM_023258297.1
C1 17143554 17143710
refGene_gn=LUZP1;refGene_tx=XM_006934394.4;refGene_gn=LUZP1;refGene_tx=XM_019836367.2;refGene_gn=LUZP1;refGene_tx=XM_019836368.2;refGene_gn=LUZP1;refGene_tx=XM_019836369.2;refGene_gn=LUZP1;refGene_tx=XM_019836370.2;refGene_gn=LUZP1;refGene_tx=XM_019836373.2;refGene_gn=LUZP1;refGene_tx=XM_023258290.1;refGene_gn=LUZP1;refGene_tx=XM_023258291.1;refGene_gn=LUZP1;refGene_tx=XM_023258292.1;refGene_gn=LUZP1;refGene_tx=XM_023258293.1
C1 17145502 17148571
refGene_gn=LUZP1;refGene_tx=XM_006934394.4;refGene_gn=LUZP1;refGene_tx=XM_019836367.2;refGene_gn=LUZP1;refGene_tx=XM_019836368.2;refGene_gn=LUZP1;refGene_tx=XM_019836369.2;refGene_gn=LUZP1;refGene_tx=XM_019836370.2;refGene_gn=LUZP1;refGene_tx=XM_019836373.2;refGene_gn=LUZP1;refGene_tx=XM_023258290.1;refGene_gn=LUZP1;refGene_tx=XM_023258291.1;refGene_gn=LUZP1;refGene_tx=XM_023258292.1;refGene_gn=LUZP1;refGene_tx=XM_023258293.1
C1 17261896 17263030 refGene_gn=HTR1D;refGene_tx=XM_023258298.1
C1 17358044 17358657
refGene_gn=HNRNPR;refGene_tx=XM_011284547.3;refGene_gn=HNRNPR;refGene_tx=XM_011284548.3;refGene_gn=HNRNPR;refGene_tx=XM_011284550.3;refGene_gn=HNRNPR;refGene_tx=XM_019836377.2
C1 17358793 17358915
refGene_gn=HNRNPR;refGene_tx=XM_011284547.3;refGene_gn=HNRNPR;refGene_tx=XM_011284548.3;refGene_gn=HNRNPR;refGene_tx=XM_011284550.3;refGene_gn=HNRNPR;refGene_tx=XM_019836377.2
C1 17361522 17361672
refGene_gn=HNRNPR;refGene_tx=XM_011284547.3;refGene_gn=HNRNPR;refGene_tx=XM_011284548.3;refGene_gn=HNRNPR;refGene_tx=XM_011284550.3;refGene_gn=HNRNPR;refGene_tx=XM_019836377.2
C1 17365577 17365792
refGene_gn=HNRNPR;refGene_tx=XM_011284547.3;refGene_gn=HNRNPR;refGene_tx=XM_011284548.3;refGene_gn=HNRNPR;refGene_tx=XM_011284550.3;refGene_gn=HNRNPR;refGene_tx=XM_019836377.2

C1	17368435	17368571
refGene_gn=HNRNPR;refGene_tx=XM_011284547.3;refGene_gn=HNRNPR;refGene_tx=XM_011284548.3;refGene_gn=HNRNPR;refGene_tx=XM_011284550.3;refGene_gn=HNRNPR;refGene_tx=XM_019836377.2
C1	17370238	17370415
refGene_gn=HNRNPR;refGene_tx=XM_011284547.3;refGene_gn=HNRNPR;refGene_tx=XM_011284548.3;refGene_gn=HNRNPR;refGene_tx=XM_011284550.3;refGene_gn=HNRNPR;refGene_tx=XM_019836377.2
C1	17379910	17380024
refGene_gn=HNRNPR;refGene_tx=XM_011284547.3;refGene_gn=HNRNPR;refGene_tx=XM_011284548.3;refGene_gn=HNRNPR;refGene_tx=XM_011284550.3;refGene_gn=HNRNPR;refGene_tx=XM_019836377.2
C1	17383444	17383552
refGene_gn=HNRNPR;refGene_tx=XM_011284547.3;refGene_gn=HNRNPR;refGene_tx=XM_011284548.3;refGene_gn=HNRNPR;refGene_tx=XM_011284550.3;refGene_gn=HNRNPR;refGene_tx=XM_019836377.2
C1	17384176	17384295
refGene_gn=HNRNPR;refGene_tx=XM_011284547.3;refGene_gn=HNRNPR;refGene_tx=XM_011284548.3;refGene_gn=HNRNPR;refGene_tx=XM_019836377.2
C1	17386574	17386731
refGene_gn=HNRNPR;refGene_tx=XM_011284547.3;refGene_gn=HNRNPR;refGene_tx=XM_011284548.3;refGene_gn=HNRNPR;refGene_tx=XM_019836377.2
C1	17406821	17407070	refGene_gn=ZNF436;refGene_tx=XM_003989647.5
C1	17407071	17408075	refGene_gn=ZNF436;refGene_tx=XM_003989647.5
C1	17410557	17410684	refGene_gn=ZNF436;refGene_tx=XM_003989647.5
C1	17411483	17411516	refGene_gn=ZNF436;refGene_tx=XM_003989647.5
C1	17422062	17422071
refGene_gn=TCEA3;refGene_tx=XM_011284545.3;refGene_gn=TCEA3;refGene_tx=XM_011284546.3
C1	17424242	17424314
refGene_gn=TCEA3;refGene_tx=XM_011284545.3;refGene_gn=TCEA3;refGene_tx=XM_011284546.3
C1	17426027	17426174
refGene_gn=TCEA3;refGene_tx=XM_011284545.3;refGene_gn=TCEA3;refGene_tx=XM_011284546.3
C1	17430591	17430746
refGene_gn=TCEA3;refGene_tx=XM_011284545.3;refGene_gn=TCEA3;refGene_tx=XM_011284546.3
C1	17434093	17434150
refGene_gn=TCEA3;refGene_tx=XM_011284545.3;refGene_gn=TCEA3;refGene_tx=XM_011284546.3
C1	17434323	17434487
refGene_gn=TCEA3;refGene_tx=XM_011284545.3;refGene_gn=TCEA3;refGene_tx=XM_011284546.3
C1	17443819	17443882
refGene_gn=TCEA3;refGene_tx=XM_011284545.3;refGene_gn=TCEA3;refGene_tx=XM_011284546.3
C1	17452701	17452840
refGene_gn=TCEA3;refGene_tx=XM_011284545.3;refGene_gn=TCEA3;refGene_tx=XM_011284546.3

C1 17453317 17453423
refGene_gn=TCEA3;refGene_tx=XM_011284545.3;refGene_gn=TCEA3;refGene_tx
=XM_011284546.3
C1 17454171 17454234 refGene_gn=TCEA3;refGene_tx=XM_011284545.3
C1 17460117 17460186
refGene_gn=TCEA3;refGene_tx=XM_011284545.3;refGene_gn=TCEA3;refGene_tx
=XM_011284546.3
C1 17464655 17464699 refGene_gn=ASAP3;refGene_tx=XM_023258299.1
C1 17464701 17464729 refGene_gn=ASAP3;refGene_tx=XM_023258299.1
C1 17465916 17465976 refGene_gn=ASAP3;refGene_tx=XM_023258299.1
C1 17465977 17466008 refGene_gn=ASAP3;refGene_tx=XM_023258299.1
C1 17466522 17466751 refGene_gn=ASAP3;refGene_tx=XM_023258299.1
C1 17467627 17467823 refGene_gn=ASAP3;refGene_tx=XM_023258299.1
C1 17467958 17468066 refGene_gn=ASAP3;refGene_tx=XM_023258299.1
C1 17468160 17468228 refGene_gn=ASAP3;refGene_tx=XM_023258299.1
C1 17468824 17468940 refGene_gn=ASAP3;refGene_tx=XM_023258299.1
C1 17469095 17469181 refGene_gn=ASAP3;refGene_tx=XM_023258299.1
C1 17470123 17470301 refGene_gn=ASAP3;refGene_tx=XM_023258299.1
C1 17470474 17470569 refGene_gn=ASAP3;refGene_tx=XM_023258299.1
C1 17470795 17470929 refGene_gn=ASAP3;refGene_tx=XM_023258299.1
C1 17471068 17471259 refGene_gn=ASAP3;refGene_tx=XM_023258299.1
C1 17471384 17471433 refGene_gn=ASAP3;refGene_tx=XM_023258299.1
C1 17472583 17472671 refGene_gn=ASAP3;refGene_tx=XM_023258299.1
C1 17472985 17473055 refGene_gn=ASAP3;refGene_tx=XM_023258299.1
C1 17473923 17474033 refGene_gn=ASAP3;refGene_tx=XM_023258299.1
C1 17474209 17474296 refGene_gn=ASAP3;refGene_tx=XM_023258299.1
C1 17474477 17474553 refGene_gn=ASAP3;refGene_tx=XM_023258299.1
C1 17475021 17475107 refGene_gn=ASAP3;refGene_tx=XM_023258299.1
C1 17475308 17475420 refGene_gn=ASAP3;refGene_tx=XM_023258299.1
C1 17481944 17481994 refGene_gn=ASAP3;refGene_tx=XM_023258299.1
C1 17483407 17483482 refGene_gn=ASAP3;refGene_tx=XM_023258299.1
C1 17485875 17486021 refGene_gn=ASAP3;refGene_tx=XM_023258299.1
C1 17486120 17486193 refGene_gn=ASAP3;refGene_tx=XM_023258299.1
C1 17511682 17511811 refGene_gn=ASAP3;refGene_tx=XM_023258299.1
C1 17534235 17534519 refGene_gn=E2F2;refGene_tx=XM_003989649.4
C1 17540282 17540475 refGene_gn=E2F2;refGene_tx=XM_003989649.4
C1 17542306 17542421 refGene_gn=E2F2;refGene_tx=XM_003989649.4
C1 17543850 17544009 refGene_gn=E2F2;refGene_tx=XM_003989649.4
C1 17544754 17544974 refGene_gn=E2F2;refGene_tx=XM_003989649.4
C1 17547300 17547406 refGene_gn=E2F2;refGene_tx=XM_003989649.4
C1 17552940 17553192 refGene_gn=E2F2;refGene_tx=XM_003989649.4
C1 17572251 17572311 refGene_gn=ID3;refGene_tx=XM_019836380.2
C1 17572446 17572746 refGene_gn=ID3;refGene_tx=XM_019836380.2
C1 17655139 17655283
refGene_gn=LOC109502152;refGene_tx=XM_023258300.1
C1 17658317 17658497
refGene_gn=LOC109502152;refGene_tx=XM_023258300.1
C1 17658595 17658892
refGene_gn=LOC109502152;refGene_tx=XM_023258300.1
C1 17662308 17662459

refGene_gn=LOC109502152;refGene_tx=XM_023258300.1
C1 17662545 17662802
refGene_gn=LOC109502152;refGene_tx=XM_023258300.1
C1 17664518 17664764
refGene_gn=LOC109502152;refGene_tx=XM_023258300.1
C1 17669121 17669127 refGene_gn=RPL11;refGene_tx=XM_003989652.4
C1 17669900 17670051 refGene_gn=RPL11;refGene_tx=XM_003989652.4
C1 17671479 17671586 refGene_gn=RPL11;refGene_tx=XM_003989652.4
C1 17671818 17671950 refGene_gn=RPL11;refGene_tx=XM_003989652.4
C1 17672994 17673105 refGene_gn=RPL11;refGene_tx=XM_003989652.4
C1 17673535 17673565 refGene_gn=RPL11;refGene_tx=XM_003989652.4
C1 17703995 17704148 refGene_gn=ELOA;refGene_tx=XM_023258301.1
C1 17708944 17709001 refGene_gn=ELOA;refGene_tx=XM_023258301.1
C1 17709842 17709949 refGene_gn=ELOA;refGene_tx=XM_023258301.1
C1 17711119 17712305 refGene_gn=ELOA;refGene_tx=XM_023258301.1
C1 17712688 17712803 refGene_gn=ELOA;refGene_tx=XM_023258301.1
C1 17713556 17713712 refGene_gn=ELOA;refGene_tx=XM_023258301.1
C1 17713804 17713902 refGene_gn=ELOA;refGene_tx=XM_023258301.1
C1 17715072 17715253 refGene_gn=ELOA;refGene_tx=XM_023258301.1
C1 17715589 17715701 refGene_gn=ELOA;refGene_tx=XM_023258301.1
C1 17716414 17716590 refGene_gn=ELOA;refGene_tx=XM_023258301.1
C1 17718073 17718135 refGene_gn=ELOA;refGene_tx=XM_023258301.1
C1 17730079 17730277
refGene_gn=PITHD1;refGene_tx=XM_023258302.1;refGene_gn=PITHD1;refGene_
tx=XM_023258303.1
C1 17730978 17731022
refGene_gn=PITHD1;refGene_tx=XM_023258302.1;refGene_gn=PITHD1;refGene_
tx=XM_023258303.1
C1 17731380 17731458 refGene_gn=PITHD1;refGene_tx=XM_023258302.1
C1 17733999 17734104
refGene_gn=PITHD1;refGene_tx=XM_023258302.1;refGene_gn=PITHD1;refGene_
tx=XM_023258303.1
C1 17734802 17734911
refGene_gn=PITHD1;refGene_tx=XM_023258302.1;refGene_gn=PITHD1;refGene_
tx=XM_023258303.1
C1 17735668 17735770
refGene_gn=PITHD1;refGene_tx=XM_023258302.1;refGene_gn=PITHD1;refGene_
tx=XM_023258303.1
C1 17738733 17738857 refGene_gn=LYPLA2;refGene_tx=XM_003989653.5
C1 17740323 17740427 refGene_gn=LYPLA2;refGene_tx=XM_003989653.5
C1 17740699 17740731 refGene_gn=LYPLA2;refGene_tx=XM_003989653.5
C1 17740858 17740924 refGene_gn=LYPLA2;refGene_tx=XM_003989653.5
C1 17741380 17741428 refGene_gn=LYPLA2;refGene_tx=XM_003989653.5
C1 17741518 17741589 refGene_gn=LYPLA2;refGene_tx=XM_003989653.5
C1 17741734 17741808 refGene_gn=LYPLA2;refGene_tx=XM_003989653.5
C1 17741883 17741985 refGene_gn=LYPLA2;refGene_tx=XM_003989653.5
C1 17742087 17742261 refGene_gn=LYPLA2;refGene_tx=XM_003989653.5
C1 17742356 17742407 refGene_gn=LYPLA2;refGene_tx=XM_003989653.5
C1 17743594 17743653
refGene_gn=GALE;refGene_tx=XM_003989654.4;refGene_gn=GALE;refGene_tx=X

M_006934400.3;refGene_gn=GALE;refGene_tx=XM_006934401.3;refGene_gn=GALE;refGene_tx=XM_019836383.2
C1 17743902 17744017 refGene_gn=GALE;refGene_tx=XM_003989654.4;refGene_gn=GALE;refGene_tx=XM_006934400.3;refGene_gn=GALE;refGene_tx=XM_006934401.3;refGene_gn=GALE;refGene_tx=XM_019836383.2
C1 17744215 17744293 refGene_gn=GALE;refGene_tx=XM_003989654.4;refGene_gn=GALE;refGene_tx=XM_006934400.3;refGene_gn=GALE;refGene_tx=XM_006934401.3;refGene_gn=GALE;refGene_tx=XM_019836383.2
C1 17744392 17744478 refGene_gn=GALE;refGene_tx=XM_003989654.4;refGene_gn=GALE;refGene_tx=XM_006934400.3;refGene_gn=GALE;refGene_tx=XM_006934401.3;refGene_gn=GALE;refGene_tx=XM_019836383.2
C1 17744581 17744648 refGene_gn=GALE;refGene_tx=XM_003989654.4;refGene_gn=GALE;refGene_tx=XM_006934400.3;refGene_gn=GALE;refGene_tx=XM_006934401.3;refGene_gn=GALE;refGene_tx=XM_019836383.2
C1 17744744 17744858 refGene_gn=GALE;refGene_tx=XM_003989654.4;refGene_gn=GALE;refGene_tx=XM_006934400.3;refGene_gn=GALE;refGene_tx=XM_006934401.3;refGene_gn=GALE;refGene_tx=XM_019836383.2
C1 17745160 17745337 refGene_gn=GALE;refGene_tx=XM_003989654.4;refGene_gn=GALE;refGene_tx=XM_006934400.3;refGene_gn=GALE;refGene_tx=XM_006934401.3;refGene_gn=GALE;refGene_tx=XM_019836383.2
C1 17745602 17745716 refGene_gn=GALE;refGene_tx=XM_003989654.4;refGene_gn=GALE;refGene_tx=XM_006934400.3;refGene_gn=GALE;refGene_tx=XM_006934401.3;refGene_gn=GALE;refGene_tx=XM_019836383.2
C1 17745800 17745916 refGene_gn=GALE;refGene_tx=XM_003989654.4;refGene_gn=GALE;refGene_tx=XM_006934400.3;refGene_gn=GALE;refGene_tx=XM_006934401.3;refGene_gn=GALE;refGene_tx=XM_019836383.2
C1 17746109 17746230 refGene_gn=GALE;refGene_tx=XM_003989654.4;refGene_gn=GALE;refGene_tx=XM_006934400.3;refGene_gn=GALE;refGene_tx=XM_006934401.3;refGene_gn=GALE;refGene_tx=XM_019836383.2
C1 17749492 17749594 refGene_gn=HMGCL;refGene_tx=XM_003989655.5
C1 17750860 17750986 refGene_gn=HMGCL;refGene_tx=XM_003989655.5
C1 17752596 17752785 refGene_gn=HMGCL;refGene_tx=XM_003989655.5
C1 17754182 17754246 refGene_gn=HMGCL;refGene_tx=XM_003989655.5
C1 17756872 17757021 refGene_gn=HMGCL;refGene_tx=XM_003989655.5
C1 17758525 17758621 refGene_gn=HMGCL;refGene_tx=XM_003989655.5
C1 17759289 17759397 refGene_gn=HMGCL;refGene_tx=XM_003989655.5
C1 17762212 17762296 refGene_gn=HMGCL;refGene_tx=XM_003989655.5
C1 17767183 17767243 refGene_gn=HMGCL;refGene_tx=XM_003989655.5
C1 17787716 17787805 refGene_gn=FUCA1;refGene_tx=XM_003989657.5
C1 17787807 17787856 refGene_gn=FUCA1;refGene_tx=XM_003989657.5
C1 17788077 17788177 refGene_gn=FUCA1;refGene_tx=XM_003989657.5

C1	17789317	17789508	refGene_gn=FUCA1;refGene_tx=XM_003989657.5
C1	17791240	17791441	refGene_gn=FUCA1;refGene_tx=XM_003989657.5
C1	17793929	17794035	refGene_gn=FUCA1;refGene_tx=XM_003989657.5
C1	17795953	17796091	refGene_gn=FUCA1;refGene_tx=XM_003989657.5
C1	17798203	17798338	refGene_gn=FUCA1;refGene_tx=XM_003989657.5
C1	17800212	17800589	refGene_gn=FUCA1;refGene_tx=XM_003989657.5
C1	17805526	17806609
refGene_gn=CNR2;refGene_tx=XM_019836386.2;refGene_gn=CNR2;refGene_tx=XM_023258304.1
C1	17863134	17863554	refGene_gn=PNRC2;refGene_tx=XM_023258311.1
C1	17868775	17868836
refGene_gn=SRSF10;refGene_tx=XM_023258307.1;refGene_gn=SRSF10;refGene_tx=XM_023258308.1;refGene_gn=SRSF10;refGene_tx=XM_023258309.1;refGene_gn=SRSF10;refGene_tx=XM_023258310.1
C1	17872003	17872301
refGene_gn=SRSF10;refGene_tx=XM_023258305.1;refGene_gn=SRSF10;refGene_tx=XM_023258306.1;refGene_gn=SRSF10;refGene_tx=XM_023258310.1
C1	17872434	17872488
refGene_gn=SRSF10;refGene_tx=XM_023258305.1;refGene_gn=SRSF10;refGene_tx=XM_023258306.1;refGene_gn=SRSF10;refGene_tx=XM_023258307.1;refGene_gn=SRSF10;refGene_tx=XM_023258308.1;refGene_gn=SRSF10;refGene_tx=XM_023258310.1
C1	17872717	17872880
refGene_gn=SRSF10;refGene_tx=XM_023258305.1;refGene_gn=SRSF10;refGene_tx=XM_023258306.1;refGene_gn=SRSF10;refGene_tx=XM_023258307.1;refGene_gn=SRSF10;refGene_tx=XM_023258308.1;refGene_gn=SRSF10;refGene_tx=XM_023258309.1;refGene_gn=SRSF10;refGene_tx=XM_023258310.1
C1	17875204	17875308
refGene_gn=SRSF10;refGene_tx=XM_023258305.1;refGene_gn=SRSF10;refGene_tx=XM_023258306.1;refGene_gn=SRSF10;refGene_tx=XM_023258307.1;refGene_gn=SRSF10;refGene_tx=XM_023258308.1;refGene_gn=SRSF10;refGene_tx=XM_023258309.1;refGene_gn=SRSF10;refGene_tx=XM_023258310.1
C1	17878879	17878984
refGene_gn=SRSF10;refGene_tx=XM_023258305.1;refGene_gn=SRSF10;refGene_tx=XM_023258306.1;refGene_gn=SRSF10;refGene_tx=XM_023258307.1;refGene_gn=SRSF10;refGene_tx=XM_023258308.1;refGene_gn=SRSF10;refGene_tx=XM_023258309.1;refGene_gn=SRSF10;refGene_tx=XM_023258310.1
C1	17880443	17880508
refGene_gn=SRSF10;refGene_tx=XM_023258305.1;refGene_gn=SRSF10;refGene_tx=XM_023258306.1;refGene_gn=SRSF10;refGene_tx=XM_023258307.1;refGene_gn=SRSF10;refGene_tx=XM_023258308.1;refGene_gn=SRSF10;refGene_tx=XM_023258309.1;refGene_gn=SRSF10;refGene_tx=XM_023258310.1
C1	17947735	17948029
refGene_gn=MYOM3;refGene_tx=XM_003989727.5;refGene_gn=MYOM3;refGene_tx=XM_019836393.2;refGene_gn=MYOM3;refGene_tx=XM_023258312.1
C1	17949506	17949562
refGene_gn=MYOM3;refGene_tx=XM_003989727.5;refGene_gn=MYOM3;refGene_tx=XM_019836393.2;refGene_gn=MYOM3;refGene_tx=XM_023258312.1
C1	17950291	17950314
refGene_gn=MYOM3;refGene_tx=XM_003989727.5;refGene_gn=MYOM3;refGene_tx

=XM_019836393.2;refGene_gn=MYOM3;refGene_tx=XM_023258312.1
C1 17950503 17950540
refGene_gn=MYOM3;refGene_tx=XM_003989727.5;refGene_gn=MYOM3;refGene_tx=XM_019836393.2;refGene_gn=MYOM3;refGene_tx=XM_023258312.1
C1 17950775 17950939
refGene_gn=MYOM3;refGene_tx=XM_003989727.5;refGene_gn=MYOM3;refGene_tx=XM_019836393.2;refGene_gn=MYOM3;refGene_tx=XM_023258312.1
C1 17952122 17952231
refGene_gn=MYOM3;refGene_tx=XM_003989727.5;refGene_gn=MYOM3;refGene_tx=XM_019836393.2;refGene_gn=MYOM3;refGene_tx=XM_023258312.1
C1 17952473 17952512
refGene_gn=MYOM3;refGene_tx=XM_003989727.5;refGene_gn=MYOM3;refGene_tx=XM_019836393.2;refGene_gn=MYOM3;refGene_tx=XM_023258312.1
C1 17953047 17953135
refGene_gn=MYOM3;refGene_tx=XM_003989727.5;refGene_gn=MYOM3;refGene_tx=XM_019836393.2;refGene_gn=MYOM3;refGene_tx=XM_023258312.1
C1 17954885 17954996
refGene_gn=MYOM3;refGene_tx=XM_003989727.5;refGene_gn=MYOM3;refGene_tx=XM_019836393.2;refGene_gn=MYOM3;refGene_tx=XM_023258312.1
C1 17955788 17955856
refGene_gn=MYOM3;refGene_tx=XM_003989727.5;refGene_gn=MYOM3;refGene_tx=XM_019836393.2;refGene_gn=MYOM3;refGene_tx=XM_023258312.1
C1 17956267 17956327
refGene_gn=MYOM3;refGene_tx=XM_003989727.5;refGene_gn=MYOM3;refGene_tx=XM_019836393.2;refGene_gn=MYOM3;refGene_tx=XM_023258312.1
C1 17956440 17956585
refGene_gn=MYOM3;refGene_tx=XM_003989727.5;refGene_gn=MYOM3;refGene_tx=XM_019836393.2;refGene_gn=MYOM3;refGene_tx=XM_023258312.1
C1 17958933 17959070
refGene_gn=MYOM3;refGene_tx=XM_003989727.5;refGene_gn=MYOM3;refGene_tx=XM_019836393.2;refGene_gn=MYOM3;refGene_tx=XM_023258312.1
C1 17959758 17959803
refGene_gn=MYOM3;refGene_tx=XM_003989727.5;refGene_gn=MYOM3;refGene_tx=XM_019836393.2;refGene_gn=MYOM3;refGene_tx=XM_023258312.1
C1 17961026 17961136
refGene_gn=MYOM3;refGene_tx=XM_003989727.5;refGene_gn=MYOM3;refGene_tx=XM_019836393.2;refGene_gn=MYOM3;refGene_tx=XM_023258312.1
C1 17961807 17961964
refGene_gn=MYOM3;refGene_tx=XM_003989727.5;refGene_gn=MYOM3;refGene_tx=XM_019836393.2;refGene_gn=MYOM3;refGene_tx=XM_023258312.1
C1 17962965 17963080
refGene_gn=MYOM3;refGene_tx=XM_003989727.5;refGene_gn=MYOM3;refGene_tx=XM_019836393.2;refGene_gn=MYOM3;refGene_tx=XM_023258312.1
C1 17966267 17966446
refGene_gn=MYOM3;refGene_tx=XM_003989727.5;refGene_gn=MYOM3;refGene_tx=XM_019836393.2;refGene_gn=MYOM3;refGene_tx=XM_023258312.1
C1 17967454 17967581
refGene_gn=MYOM3;refGene_tx=XM_003989727.5;refGene_gn=MYOM3;refGene_tx=XM_019836393.2;refGene_gn=MYOM3;refGene_tx=XM_023258312.1
C1 17967841 17968029

refGene_gn=MYOM3;refGene_tx=XM_003989727.5;refGene_gn=MYOM3;refGene_tx=XM_019836393.2;refGene_gn=MYOM3;refGene_tx=XM_023258312.1
C1 17968435 17968557
refGene_gn=MYOM3;refGene_tx=XM_003989727.5;refGene_gn=MYOM3;refGene_tx=XM_019836393.2;refGene_gn=MYOM3;refGene_tx=XM_023258312.1
C1 17971650 17971825
refGene_gn=MYOM3;refGene_tx=XM_003989727.5;refGene_gn=MYOM3;refGene_tx=XM_019836393.2;refGene_gn=MYOM3;refGene_tx=XM_023258312.1
C1 17973198 17973382
refGene_gn=MYOM3;refGene_tx=XM_003989727.5;refGene_gn=MYOM3;refGene_tx=XM_019836393.2;refGene_gn=MYOM3;refGene_tx=XM_023258312.1
C1 17976036 17976164
refGene_gn=MYOM3;refGene_tx=XM_003989727.5;refGene_gn=MYOM3;refGene_tx=XM_019836393.2;refGene_gn=MYOM3;refGene_tx=XM_023258312.1
C1 17976548 17976602
refGene_gn=MYOM3;refGene_tx=XM_003989727.5;refGene_gn=MYOM3;refGene_tx=XM_019836393.2;refGene_gn=MYOM3;refGene_tx=XM_023258312.1
C1 17978396 17978596
refGene_gn=MYOM3;refGene_tx=XM_003989727.5;refGene_gn=MYOM3;refGene_tx=XM_019836393.2;refGene_gn=MYOM3;refGene_tx=XM_023258312.1
C1 17979152 17979294
refGene_gn=MYOM3;refGene_tx=XM_003989727.5;refGene_gn=MYOM3;refGene_tx=XM_019836393.2;refGene_gn=MYOM3;refGene_tx=XM_023258312.1
C1 17979945 17980107
refGene_gn=MYOM3;refGene_tx=XM_003989727.5;refGene_gn=MYOM3;refGene_tx=XM_019836393.2;refGene_gn=MYOM3;refGene_tx=XM_023258312.1
C1 17981257 17981398
refGene_gn=MYOM3;refGene_tx=XM_003989727.5;refGene_gn=MYOM3;refGene_tx=XM_019836393.2;refGene_gn=MYOM3;refGene_tx=XM_023258312.1
C1 17981860 17981905
refGene_gn=MYOM3;refGene_tx=XM_003989727.5;refGene_gn=MYOM3;refGene_tx=XM_019836393.2
C1 17983877 17983966
refGene_gn=MYOM3;refGene_tx=XM_003989727.5;refGene_gn=MYOM3;refGene_tx=XM_019836393.2
C1 17985576 17985672
refGene_gn=MYOM3;refGene_tx=XM_003989727.5;refGene_gn=MYOM3;refGene_tx=XM_019836393.2
C1 17989858 17990016 refGene_gn=MYOM3;refGene_tx=XM_003989727.5
C1 17990762 17990922 refGene_gn=MYOM3;refGene_tx=XM_003989727.5
C1 17991513 17991594 refGene_gn=MYOM3;refGene_tx=XM_003989727.5
C1 17993066 17993230 refGene_gn=MYOM3;refGene_tx=XM_003989727.5
C1 18002443 18003397 refGene_gn=IL22RA1;refGene_tx=XM_019836395.2
C1 18004404 18004526 refGene_gn=IL22RA1;refGene_tx=XM_019836395.2
C1 18008606 18008745 refGene_gn=IL22RA1;refGene_tx=XM_019836395.2
C1 18014224 18014400 refGene_gn=IL22RA1;refGene_tx=XM_019836395.2
C1 18016070 18016249 refGene_gn=IL22RA1;refGene_tx=XM_019836395.2
C1 18017153 18017286 refGene_gn=IL22RA1;refGene_tx=XM_019836395.2
C1 18020364 18020407 refGene_gn=IL22RA1;refGene_tx=XM_019836395.2
C1 18031629 18032400 refGene_gn=IFNLR1;refGene_tx=XM_023260058.1

C1 18033377 18033511 refGene_gn=IFNLR1;refGene_tx=XM_023260058.1
C1 18033823 18033983 refGene_gn=IFNLR1;refGene_tx=XM_023260058.1
C1 18036265 18036408 refGene_gn=IFNLR1;refGene_tx=XM_023260058.1
C1 18038825 18039010 refGene_gn=IFNLR1;refGene_tx=XM_023260058.1
C1 18046503 18046627 refGene_gn=IFNLR1;refGene_tx=XM_023260058.1
C1 18050038 18050357 refGene_gn=IFNLR1;refGene_tx=XM_023260058.1
C1 18157609 18157626
refGene_gn=GRHL3;refGene_tx=XM_003989661.5;refGene_gn=GRHL3;refGene_tx
=XM_019836398.2
C1 18169260 18169447
refGene_gn=GRHL3;refGene_tx=XM_003989661.5;refGene_gn=GRHL3;refGene_tx
=XM_019836398.2
C1 18172489 18172551
refGene_gn=GRHL3;refGene_tx=XM_003989661.5;refGene_gn=GRHL3;refGene_tx
=XM_019836398.2
C1 18173941 18174287
refGene_gn=GRHL3;refGene_tx=XM_003989661.5;refGene_gn=GRHL3;refGene_tx
=XM_019836398.2
C1 18174543 18174617
refGene_gn=GRHL3;refGene_tx=XM_003989661.5;refGene_gn=GRHL3;refGene_tx
=XM_019836398.2
C1 18175547 18175701
refGene_gn=GRHL3;refGene_tx=XM_003989661.5;refGene_gn=GRHL3;refGene_tx
=XM_019836398.2
C1 18175911 18176023
refGene_gn=GRHL3;refGene_tx=XM_003989661.5;refGene_gn=GRHL3;refGene_tx
=XM_019836398.2
C1 18177411 18177506
refGene_gn=GRHL3;refGene_tx=XM_003989661.5;refGene_gn=GRHL3;refGene_tx
=XM_019836398.2
C1 18179133 18179292
refGene_gn=GRHL3;refGene_tx=XM_003989661.5;refGene_gn=GRHL3;refGene_tx
=XM_019836398.2
C1 18179844 18179923
refGene_gn=GRHL3;refGene_tx=XM_003989661.5;refGene_gn=GRHL3;refGene_tx
=XM_019836398.2
C1 18180042 18180179
refGene_gn=GRHL3;refGene_tx=XM_003989661.5;refGene_gn=GRHL3;refGene_tx
=XM_019836398.2
C1 18182184 18182219
refGene_gn=GRHL3;refGene_tx=XM_003989661.5;refGene_gn=GRHL3;refGene_tx
=XM_019836398.2
C1 18183353 18183442
refGene_gn=GRHL3;refGene_tx=XM_003989661.5;refGene_gn=GRHL3;refGene_tx
=XM_019836398.2
C1 18184335 18184421
refGene_gn=GRHL3;refGene_tx=XM_003989661.5;refGene_gn=GRHL3;refGene_tx
=XM_019836398.2
C1 18186820 18186885
refGene_gn=GRHL3;refGene_tx=XM_003989661.5;refGene_gn=GRHL3;refGene_tx

=XM_019836398.2
C1	18190996	18191111
refGene_gn=GRHL3;refGene_tx=XM_003989661.5;refGene_gn=GRHL3;refGene_tx=XM_019836398.2
C1	18196871	18197043
refGene_gn=STPG1;refGene_tx=XM_006934404.3;refGene_gn=STPG1;refGene_tx=XM_006934405.3;refGene_gn=STPG1;refGene_tx=XM_006934406.3;refGene_gn=STPG1;refGene_tx=XM_011284540.3;refGene_gn=STPG1;refGene_tx=XM_019836399.2;refGene_gn=STPG1;refGene_tx=XM_019836400.2;refGene_gn=STPG1;refGene_tx=XM_019836401.2;refGene_gn=STPG1;refGene_tx=XM_019836402.2;refGene_gn=STPG1;refGene_tx=XM_019836403.2;refGene_gn=STPG1;refGene_tx=XM_019836404.2;refGene_gn=STPG1;refGene_tx=XM_019836405.2
C1	18199014	18199205
refGene_gn=STPG1;refGene_tx=XM_006934404.3;refGene_gn=STPG1;refGene_tx=XM_006934405.3;refGene_gn=STPG1;refGene_tx=XM_006934406.3;refGene_gn=STPG1;refGene_tx=XM_011284540.3;refGene_gn=STPG1;refGene_tx=XM_019836399.2;refGene_gn=STPG1;refGene_tx=XM_019836400.2;refGene_gn=STPG1;refGene_tx=XM_019836401.2;refGene_gn=STPG1;refGene_tx=XM_019836402.2;refGene_gn=STPG1;refGene_tx=XM_019836403.2;refGene_gn=STPG1;refGene_tx=XM_019836405.2
C1	18207900	18208066
refGene_gn=STPG1;refGene_tx=XM_006934404.3;refGene_gn=STPG1;refGene_tx=XM_006934405.3;refGene_gn=STPG1;refGene_tx=XM_006934406.3;refGene_gn=STPG1;refGene_tx=XM_011284540.3;refGene_gn=STPG1;refGene_tx=XM_019836399.2;refGene_gn=STPG1;refGene_tx=XM_019836400.2;refGene_gn=STPG1;refGene_tx=XM_019836401.2;refGene_gn=STPG1;refGene_tx=XM_019836402.2;refGene_gn=STPG1;refGene_tx=XM_019836403.2;refGene_gn=STPG1;refGene_tx=XM_019836404.2;refGene_gn=STPG1;refGene_tx=XM_019836405.2
C1	18215271	18215380
refGene_gn=STPG1;refGene_tx=XM_006934404.3;refGene_gn=STPG1;refGene_tx=XM_006934405.3;refGene_gn=STPG1;refGene_tx=XM_006934406.3;refGene_gn=STPG1;refGene_tx=XM_011284540.3;refGene_gn=STPG1;refGene_tx=XM_019836399.2;refGene_gn=STPG1;refGene_tx=XM_019836400.2;refGene_gn=STPG1;refGene_tx=XM_019836401.2;refGene_gn=STPG1;refGene_tx=XM_019836402.2;refGene_gn=STPG1;refGene_tx=XM_019836403.2;refGene_gn=STPG1;refGene_tx=XM_019836404.2;refGene_gn=STPG1;refGene_tx=XM_019836405.2
C1	18220148	18220319
refGene_gn=STPG1;refGene_tx=XM_006934404.3;refGene_gn=STPG1;refGene_tx=XM_006934405.3;refGene_gn=STPG1;refGene_tx=XM_011284540.3;refGene_gn=STPG1;refGene_tx=XM_019836399.2;refGene_gn=STPG1;refGene_tx=XM_019836400.2;refGene_gn=STPG1;refGene_tx=XM_019836401.2;refGene_gn=STPG1;refGene_tx=XM_019836402.2;refGene_gn=STPG1;refGene_tx=XM_019836403.2;refGene_gn=STPG1;refGene_tx=XM_019836404.2;refGene_gn=STPG1;refGene_tx=XM_019836405.2
C1	18224762	18224864
refGene_gn=STPG1;refGene_tx=XM_006934404.3;refGene_gn=STPG1;refGene_tx=XM_006934406.3;refGene_gn=STPG1;refGene_tx=XM_011284540.3;refGene_gn=STPG1;refGene_tx=XM_019836399.2;refGene_gn=STPG1;refGene_tx=XM_019836400.2;refGene_gn=STPG1;refGene_tx=XM_019836401.2;refGene_gn=STPG1;refGene_tx=XM_019836402.2;refGene_gn=STPG1;refGene_tx=XM_019836403.2;refGen

e_gn=STPG1;refGene_tx=XM_019836404.2;refGene_gn=STPG1;refGene_tx=XM_019836405.2
C1 18235482 18235604 refGene_gn=STPG1;refGene_tx=XM_006934404.3;refGene_gn=STPG1;refGene_tx=XM_006934405.3;refGene_gn=STPG1;refGene_tx=XM_006934406.3;refGene_gn=STPG1;refGene_tx=XM_011284540.3;refGene_gn=STPG1;refGene_tx=XM_019836399.2;refGene_gn=STPG1;refGene_tx=XM_019836400.2;refGene_gn=STPG1;refGene_tx=XM_019836401.2;refGene_gn=STPG1;refGene_tx=XM_019836402.2;refGene_gn=STPG1;refGene_tx=XM_019836403.2;refGene_gn=STPG1;refGene_tx=XM_019836404.2
C1 18249580 18249653 refGene_gn=STPG1;refGene_tx=XM_006934404.3;refGene_gn=STPG1;refGene_tx=XM_006934405.3;refGene_gn=STPG1;refGene_tx=XM_006934406.3;refGene_gn=STPG1;refGene_tx=XM_011284540.3;refGene_gn=STPG1;refGene_tx=XM_019836399.2;refGene_gn=STPG1;refGene_tx=XM_019836400.2;refGene_gn=STPG1;refGene_tx=XM_019836401.2;refGene_gn=STPG1;refGene_tx=XM_019836402.2;refGene_gn=STPG1;refGene_tx=XM_019836403.2;refGene_gn=STPG1;refGene_tx=XM_019836404.2
C1 18261360 18261456 refGene_gn=NIPAL3;refGene_tx=XM_023258313.1;refGene_gn=NIPAL3;refGene_tx=XM_023258314.1
C1 18281859 18281928 refGene_gn=NIPAL3;refGene_tx=XM_023258313.1;refGene_gn=NIPAL3;refGene_tx=XM_023258314.1
C1 18283397 18283569 refGene_gn=NIPAL3;refGene_tx=XM_023258313.1;refGene_gn=NIPAL3;refGene_tx=XM_023258314.1
C1 18286019 18286079 refGene_gn=NIPAL3;refGene_tx=XM_023258313.1;refGene_gn=NIPAL3;refGene_tx=XM_023258314.1
C1 18288649 18288795 refGene_gn=NIPAL3;refGene_tx=XM_023258313.1;refGene_gn=NIPAL3;refGene_tx=XM_023258314.1
C1 18292474 18292571 refGene_gn=NIPAL3;refGene_tx=XM_023258313.1;refGene_gn=NIPAL3;refGene_tx=XM_023258314.1
C1 18294642 18294778 refGene_gn=NIPAL3;refGene_tx=XM_023258313.1;refGene_gn=NIPAL3;refGene_tx=XM_023258314.1
C1 18296864 18296953 refGene_gn=NIPAL3;refGene_tx=XM_023258313.1;refGene_gn=NIPAL3;refGene_tx=XM_023258314.1
C1 18298383 18298447 refGene_gn=NIPAL3;refGene_tx=XM_023258313.1;refGene_gn=NIPAL3;refGene_tx=XM_023258314.1
C1 18301258 18301353 refGene_gn=NIPAL3;refGene_tx=XM_023258313.1;refGene_gn=NIPAL3;refGene_tx=XM_023258314.1
C1 18302780 18302797 refGene_gn=NIPAL3;refGene_tx=XM_023258314.1
C1 18305697 18305897 refGene_gn=NIPAL3;refGene_tx=XM_023258313.1

C1 18338470 18338665
refGene_gn=RCAN3;refGene_tx=XM_003989663.5;refGene_gn=RCAN3;refGene_tx
=XM_011284537.3
C1 18358436 18358439 refGene_gn=RCAN3;refGene_tx=XM_019836408.2
C1 18363546 18363603 refGene_gn=RCAN3;refGene_tx=XM_019836407.2
C1 18363718 18363892
refGene_gn=RCAN3;refGene_tx=XM_003989663.5;refGene_gn=RCAN3;refGene_tx
=XM_011284537.3;refGene_gn=RCAN3;refGene_tx=XM_019836407.2;refGene_gn=
RCAN3;refGene_tx=XM_019836408.2
C1 18365164 18365336
refGene_gn=RCAN3;refGene_tx=XM_003989663.5;refGene_gn=RCAN3;refGene_tx
=XM_011284537.3;refGene_gn=RCAN3;refGene_tx=XM_011284539.3;refGene_gn=
RCAN3;refGene_tx=XM_019836407.2;refGene_gn=RCAN3;refGene_tx=XM_0198364
08.2
C1 18367465 18367650
refGene_gn=RCAN3;refGene_tx=XM_003989663.5;refGene_gn=RCAN3;refGene_tx
=XM_011284537.3;refGene_gn=RCAN3;refGene_tx=XM_011284539.3;refGene_gn=
RCAN3;refGene_tx=XM_019836407.2;refGene_gn=RCAN3;refGene_tx=XM_0198364
08.2
C1 18370834 18370860 refGene_gn=RCAN3;refGene_tx=XM_011284537.3
C1 18423059 18423141 refGene_gn=NCMAP;refGene_tx=XM_003989664.5
C1 18426080 18426165 refGene_gn=NCMAP;refGene_tx=XM_003989664.5
C1 18429164 18429306 refGene_gn=NCMAP;refGene_tx=XM_003989664.5
C1 18453702 18453723 refGene_gn=SRRM1;refGene_tx=XM_019836411.2
C1 18456611 18456701 refGene_gn=SRRM1;refGene_tx=XM_019836411.2
C1 18456954 18457077 refGene_gn=SRRM1;refGene_tx=XM_019836411.2
C1 18459204 18459375 refGene_gn=SRRM1;refGene_tx=XM_019836411.2
C1 18459965 18460081 refGene_gn=SRRM1;refGene_tx=XM_019836411.2
C1 18461403 18461607 refGene_gn=SRRM1;refGene_tx=XM_019836411.2
C1 18462245 18462440 refGene_gn=SRRM1;refGene_tx=XM_019836411.2
C1 18462763 18462883 refGene_gn=SRRM1;refGene_tx=XM_019836411.2
C1 18464733 18465002 refGene_gn=SRRM1;refGene_tx=XM_019836411.2
C1 18470151 18470232 refGene_gn=SRRM1;refGene_tx=XM_019836411.2
C1 18471299 18471386 refGene_gn=SRRM1;refGene_tx=XM_019836411.2
C1 18473862 18474007 refGene_gn=SRRM1;refGene_tx=XM_019836411.2
C1 18474394 18474436 refGene_gn=SRRM1;refGene_tx=XM_019836411.2
C1 18478142 18478253 refGene_gn=SRRM1;refGene_tx=XM_019836411.2
C1 18480309 18480553 refGene_gn=SRRM1;refGene_tx=XM_019836411.2
C1 18480554 18480775 refGene_gn=SRRM1;refGene_tx=XM_019836411.2
C1 18481349 18481545 refGene_gn=SRRM1;refGene_tx=XM_019836411.2
C1 18483060 18483270 refGene_gn=SRRM1;refGene_tx=XM_019836411.2
C1 18484083 18484188 refGene_gn=SRRM1;refGene_tx=XM_019836411.2
C1 18531201 18531273 refGene_gn=CLIC4;refGene_tx=XM_023258315.1
C1 18567568 18567678 refGene_gn=CLIC4;refGene_tx=XM_023258315.1
C1 18578069 18578195 refGene_gn=CLIC4;refGene_tx=XM_023258315.1
C1 18585356 18585463 refGene_gn=CLIC4;refGene_tx=XM_023258315.1
C1 18594596 18594778 refGene_gn=CLIC4;refGene_tx=XM_023258315.1
C1 18595503 18595668 refGene_gn=CLIC4;refGene_tx=XM_023258315.1
C1 18644991 18645515 refGene_gn=RUNX3;refGene_tx=XM_023258316.1
C1 18650184 18650343 refGene_gn=RUNX3;refGene_tx=XM_023258316.1

C1	18661121	18661226	refGene_gn=RUNX3;refGene_tx=XM_023258316.1
C1	18668464	18668621	refGene_gn=RUNX3;refGene_tx=XM_023258316.1
C1	18670462	18670740	refGene_gn=RUNX3;refGene_tx=XM_023258316.1
C1	18700807	18700865	refGene_gn=RUNX3;refGene_tx=XM_023258316.1
C1	18933268	18933434	refGene_gn=SYF2;refGene_tx=XM_003989666.4
C1	18934701	18934800	refGene_gn=SYF2;refGene_tx=XM_003989666.4
C1	18935544	18935635	refGene_gn=SYF2;refGene_tx=XM_003989666.4
C1	18935991	18936109	refGene_gn=SYF2;refGene_tx=XM_003989666.4
C1	18936799	18936925	refGene_gn=SYF2;refGene_tx=XM_003989666.4
C1	18939338	18939452	refGene_gn=SYF2;refGene_tx=XM_003989666.4
C1	18939681	18939708	refGene_gn=SYF2;refGene_tx=XM_003989666.4
C1	18947809	18947833
refGene_gn=LOC101098728;refGene_tx=XM_006934408.3
C1	18947997	18948111
refGene_gn=LOC101098728;refGene_tx=XM_006934408.3
C1	18950957	18951083
refGene_gn=LOC101098728;refGene_tx=XM_006934408.3
C1	18951793	18951911
refGene_gn=LOC101098728;refGene_tx=XM_006934408.3
C1	18951999	18952090
refGene_gn=LOC101098728;refGene_tx=XM_006934408.3
C1	18952965	18953064
refGene_gn=LOC101098728;refGene_tx=XM_006934408.3
C1	18953579	18953745
refGene_gn=LOC101098728;refGene_tx=XM_006934408.3
C1	18958511	18958628	refGene_gn=RSRP1;refGene_tx=XM_023258317.1
C1	18959380	18959464	refGene_gn=RSRP1;refGene_tx=XM_023258317.1
C1	18960713	18960865	refGene_gn=RSRP1;refGene_tx=XM_023258317.1
C1	18961940	18962445	refGene_gn=RSRP1;refGene_tx=XM_023258317.1
C1	18986308	18986401
refGene_gn=TMEM50A;refGene_tx=XM_003989667.5;refGene_gn=TMEM50A;refGene_tx=XM_006934410.4
C1	18988615	18988728
refGene_gn=TMEM50A;refGene_tx=XM_003989667.5;refGene_gn=TMEM50A;refGene_tx=XM_006934410.4
C1	18991390	18991458
refGene_gn=TMEM50A;refGene_tx=XM_003989667.5;refGene_gn=TMEM50A;refGene_tx=XM_006934410.4
C1	18993240	18993333
refGene_gn=TMEM50A;refGene_tx=XM_003989667.5;refGene_gn=TMEM50A;refGene_tx=XM_006934410.4
C1	18996689	18996750
refGene_gn=TMEM50A;refGene_tx=XM_003989667.5;refGene_gn=TMEM50A;refGene_tx=XM_006934410.4
C1	19002214	19002260
refGene_gn=TMEM50A;refGene_tx=XM_003989667.5;refGene_gn=TMEM50A;refGene_tx=XM_006934410.4
C1	19004636	19004663	refGene_gn=RHCE;refGene_tx=XM_003989668.4
C1	19009177	19009251	refGene_gn=RHCE;refGene_tx=XM_003989668.4
C1	19012799	19013022

refGene_gn=RHCE;refGene_tx=XM_003989668.4;refGene_gn=RHCE;refGene_tx=XM_019836417.2
C1 19017826 19017960
refGene_gn=RHCE;refGene_tx=XM_003989668.4;refGene_gn=RHCE;refGene_tx=XM_019836417.2
C1 19020124 19020262
refGene_gn=RHCE;refGene_tx=XM_003989668.4;refGene_gn=RHCE;refGene_tx=XM_019836417.2
C1 19021944 19022114
refGene_gn=RHCE;refGene_tx=XM_003989668.4;refGene_gn=RHCE;refGene_tx=XM_019836417.2
C1 19022862 19023010
refGene_gn=RHCE;refGene_tx=XM_003989668.4;refGene_gn=RHCE;refGene_tx=XM_019836417.2
C1 19027106 19027257
refGene_gn=RHCE;refGene_tx=XM_003989668.4;refGene_gn=RHCE;refGene_tx=XM_019836417.2
C1 19030971 19031158
refGene_gn=RHCE;refGene_tx=XM_003989668.4;refGene_gn=RHCE;refGene_tx=XM_019836417.2
C1 19041286 19041473
refGene_gn=RHCE;refGene_tx=XM_003989668.4;refGene_gn=RHCE;refGene_tx=XM_019836417.2
C1 19050438 19050518 refGene_gn=MACO1;refGene_tx=XM_023258318.1
C1 19065556 19065698 refGene_gn=MACO1;refGene_tx=XM_023258318.1
C1 19067439 19067566 refGene_gn=MACO1;refGene_tx=XM_023258318.1
C1 19072219 19072343 refGene_gn=MACO1;refGene_tx=XM_023258318.1
C1 19073694 19073873 refGene_gn=MACO1;refGene_tx=XM_023258318.1
C1 19075270 19075772 refGene_gn=MACO1;refGene_tx=XM_023258318.1
C1 19099489 19099648 refGene_gn=MACO1;refGene_tx=XM_023258318.1
C1 19100904 19101087 refGene_gn=MACO1;refGene_tx=XM_023258318.1
C1 19103260 19103381 refGene_gn=MACO1;refGene_tx=XM_023258318.1
C1 19107592 19107767 refGene_gn=MACO1;refGene_tx=XM_023258318.1
C1 19113186 19113389 refGene_gn=MACO1;refGene_tx=XM_023258318.1
C1 19130973 19131037
refGene_gn=LOC109502159;refGene_tx=XM_023258320.1
C1 19134103 19134286
refGene_gn=LOC109502159;refGene_tx=XM_023258320.1
C1 19134648 19134791
refGene_gn=LOC109502159;refGene_tx=XM_023258320.1
C1 19137199 19137380
refGene_gn=LOC109502159;refGene_tx=XM_023258320.1
C1 19145551 19145607
refGene_gn=LOC109502159;refGene_tx=XM_023258320.1
C1 19148404 19148492 refGene_gn=LDLRAP1;refGene_tx=XM_023258319.1
C1 19153977 19154119
refGene_gn=LDLRAP1;refGene_tx=XM_006934412.4;refGene_gn=LDLRAP1;refGene_tx=XM_006934413.4
C1 19157543 19157686
refGene_gn=LDLRAP1;refGene_tx=XM_006934412.4;refGene_gn=LDLRAP1;refGen

e_tx=XM_006934413.4;refGene_gn=LDLRAP1;refGene_tx=XM_023258319.1
C1 19158528 19158641
refGene_gn=LDLRAP1;refGene_tx=XM_006934412.4;refGene_gn=LDLRAP1;refGene_tx=XM_006934413.4;refGene_gn=LDLRAP1;refGene_tx=XM_023258319.1
C1 19160592 19160707
refGene_gn=LDLRAP1;refGene_tx=XM_006934412.4;refGene_gn=LDLRAP1;refGene_tx=XM_006934413.4;refGene_gn=LDLRAP1;refGene_tx=XM_023258319.1
C1 19166191 19166264
refGene_gn=LDLRAP1;refGene_tx=XM_006934412.4;refGene_gn=LDLRAP1;refGene_tx=XM_006934413.4;refGene_gn=LDLRAP1;refGene_tx=XM_023258319.1
C1 19166658 19166745
refGene_gn=LDLRAP1;refGene_tx=XM_006934412.4;refGene_gn=LDLRAP1;refGene_tx=XM_006934413.4;refGene_gn=LDLRAP1;refGene_tx=XM_023258319.1
C1 19167289 19167420
refGene_gn=LDLRAP1;refGene_tx=XM_006934412.4;refGene_gn=LDLRAP1;refGene_tx=XM_006934413.4;refGene_gn=LDLRAP1;refGene_tx=XM_023258319.1
C1 19168874 19168909
refGene_gn=LDLRAP1;refGene_tx=XM_006934412.4;refGene_gn=LDLRAP1;refGene_tx=XM_006934413.4;refGene_gn=LDLRAP1;refGene_tx=XM_023258319.1
C1 19170496 19170641
refGene_gn=LDLRAP1;refGene_tx=XM_006934412.4;refGene_gn=LDLRAP1;refGene_tx=XM_006934413.4;refGene_gn=LDLRAP1;refGene_tx=XM_023258319.1
C1 19208699 19209239
refGene_gn=MAN1C1;refGene_tx=XM_011284525.3;refGene_gn=MAN1C1;refGene_tx=XM_011284526.3
C1 19264082 19264179
refGene_gn=MAN1C1;refGene_tx=XM_011284525.3;refGene_gn=MAN1C1;refGene_tx=XM_011284526.3
C1 19315532 19315648
refGene_gn=MAN1C1;refGene_tx=XM_011284525.3;refGene_gn=MAN1C1;refGene_tx=XM_011284526.3
C1 19317542 19317623
refGene_gn=MAN1C1;refGene_tx=XM_011284525.3;refGene_gn=MAN1C1;refGene_tx=XM_011284526.3
C1 19323317 19323412
refGene_gn=MAN1C1;refGene_tx=XM_011284525.3;refGene_gn=MAN1C1;refGene_tx=XM_011284526.3
C1 19329811 19329938
refGene_gn=MAN1C1;refGene_tx=XM_011284525.3;refGene_gn=MAN1C1;refGene_tx=XM_011284526.3
C1 19332945 19333039
refGene_gn=MAN1C1;refGene_tx=XM_011284525.3;refGene_gn=MAN1C1;refGene_tx=XM_011284526.3
C1 19338494 19338610
refGene_gn=MAN1C1;refGene_tx=XM_011284525.3;refGene_gn=MAN1C1;refGene_tx=XM_011284526.3
C1 19343948 19344168
refGene_gn=MAN1C1;refGene_tx=XM_011284525.3;refGene_gn=MAN1C1;refGene_tx=XM_011284526.3
C1 19346695 19346868

refGene_gn=MAN1C1;refGene_tx=XM_011284525.3;refGene_gn=MAN1C1;refGene_tx=XM_011284526.3
C1	19348156	19348272
refGene_gn=MAN1C1;refGene_tx=XM_011284525.3;refGene_gn=MAN1C1;refGene_tx=XM_011284526.3
C1	19349180	19349307
refGene_gn=MAN1C1;refGene_tx=XM_011284525.3;refGene_gn=MAN1C1;refGene_tx=XM_011284526.3
C1	19364702	19364888	refGene_gn=SELENON;refGene_tx=XM_023258321.1
C1	19365650	19365768	refGene_gn=SELENON;refGene_tx=XM_023258321.1
C1	19369581	19369712	refGene_gn=SELENON;refGene_tx=XM_023258321.1
C1	19372649	19372862	refGene_gn=SELENON;refGene_tx=XM_023258321.1
C1	19373106	19373231	refGene_gn=SELENON;refGene_tx=XM_023258321.1
C1	19373803	19373941	refGene_gn=SELENON;refGene_tx=XM_023258321.1
C1	19375562	19375644	refGene_gn=SELENON;refGene_tx=XM_023258321.1
C1	19376033	19376222	refGene_gn=SELENON;refGene_tx=XM_023258321.1
C1	19376543	19376649	refGene_gn=SELENON;refGene_tx=XM_023258321.1
C1	19377909	19378022	refGene_gn=SELENON;refGene_tx=XM_023258321.1
C1	19378115	19378217	refGene_gn=SELENON;refGene_tx=XM_023258321.1
C1	19379497	19379668	refGene_gn=SELENON;refGene_tx=XM_023258321.1
C1	19387275	19387299
refGene_gn=MTFR1L;refGene_tx=XM_003989669.5;refGene_gn=MTFR1L;refGene_tx=XM_003989671.4;refGene_gn=MTFR1L;refGene_tx=XM_006934414.4;refGene_gn=MTFR1L;refGene_tx=XM_019836424.2
C1	19387810	19387915
refGene_gn=MTFR1L;refGene_tx=XM_003989669.5;refGene_gn=MTFR1L;refGene_tx=XM_003989671.4;refGene_gn=MTFR1L;refGene_tx=XM_006934414.4;refGene_gn=MTFR1L;refGene_tx=XM_019836424.2
C1	19390678	19390788
refGene_gn=MTFR1L;refGene_tx=XM_003989669.5;refGene_gn=MTFR1L;refGene_tx=XM_003989671.4;refGene_gn=MTFR1L;refGene_tx=XM_006934414.4;refGene_gn=MTFR1L;refGene_tx=XM_019836424.2;refGene_gn=MTFR1L;refGene_tx=XM_019836425.2
C1	19391007	19391219
refGene_gn=MTFR1L;refGene_tx=XM_003989669.5;refGene_gn=MTFR1L;refGene_tx=XM_003989671.4;refGene_gn=MTFR1L;refGene_tx=XM_006934414.4;refGene_gn=MTFR1L;refGene_tx=XM_019836424.2;refGene_gn=MTFR1L;refGene_tx=XM_019836425.2
C1	19392425	19392747
refGene_gn=MTFR1L;refGene_tx=XM_003989669.5;refGene_gn=MTFR1L;refGene_tx=XM_003989671.4;refGene_gn=MTFR1L;refGene_tx=XM_006934414.4;refGene_gn=MTFR1L;refGene_tx=XM_019836424.2;refGene_gn=MTFR1L;refGene_tx=XM_019836425.2
C1	19401193	19401299
refGene_gn=MTFR1L;refGene_tx=XM_003989669.5;refGene_gn=MTFR1L;refGene_tx=XM_003989671.4;refGene_gn=MTFR1L;refGene_tx=XM_006934414.4;refGene_gn=MTFR1L;refGene_tx=XM_019836425.2
C1	19402727	19402732	refGene_gn=AUNIP;refGene_tx=XM_023258323.1
C1	19403303	19404157
refGene_gn=AUNIP;refGene_tx=XM_019836422.2;refGene_gn=AUNIP;refGene_tx

=XM_023258322.1;refGene_gn=AUNIP;refGene_tx=XM_023258323.1;refGene_gn=AUNIP;refGene_tx=XM_023258324.1
C1 19405637 19405776
refGene_gn=AUNIP;refGene_tx=XM_019836422.2;refGene_gn=AUNIP;refGene_tx=XM_023258322.1;refGene_gn=AUNIP;refGene_tx=XM_023258323.1;refGene_gn=AUNIP;refGene_tx=XM_023258324.1
C1 19417121 19417169 refGene_gn=AUNIP;refGene_tx=XM_019836422.2
C1 19425626 19425707
refGene_gn=AUNIP;refGene_tx=XM_023258322.1;refGene_gn=AUNIP;refGene_tx=XM_023258323.1
C1 19428937 19429019 refGene_gn=MTFR1L;refGene_tx=XM_019836424.2
C1 19429162 19430209
refGene_gn=PAQR7;refGene_tx=XM_023258325.1;refGene_gn=PAQR7;refGene_tx=XM_023258326.1;refGene_gn=PAQR7;refGene_tx=XM_023258327.1;refGene_gn=PAQR7;refGene_tx=XM_023258328.1
C1 19473620 19473692
refGene_gn=STMN1;refGene_tx=XM_003989674.5;refGene_gn=STMN1;refGene_tx=XM_006934418.4;refGene_gn=STMN1;refGene_tx=XM_023258330.1
C1 19474228 19474420
refGene_gn=STMN1;refGene_tx=XM_003989674.5;refGene_gn=STMN1;refGene_tx=XM_006934418.4;refGene_gn=STMN1;refGene_tx=XM_023258330.1
C1 19476509 19476682
refGene_gn=STMN1;refGene_tx=XM_003989674.5;refGene_gn=STMN1;refGene_tx=XM_006934418.4;refGene_gn=STMN1;refGene_tx=XM_023258330.1
C1 19477171 19477184
refGene_gn=STMN1;refGene_tx=XM_003989674.5;refGene_gn=STMN1;refGene_tx=XM_006934418.4;refGene_gn=STMN1;refGene_tx=XM_023258330.1
C1 19489534 19489684
refGene_gn=PAFAH2;refGene_tx=XM_019836426.2;refGene_gn=PAFAH2;refGene_tx=XM_019836427.2;refGene_gn=PAFAH2;refGene_tx=XM_023258329.1
C1 19502333 19502402 refGene_gn=PAFAH2;refGene_tx=XM_019836429.2
C1 19508855 19508974
refGene_gn=PAFAH2;refGene_tx=XM_019836426.2;refGene_gn=PAFAH2;refGene_tx=XM_019836427.2;refGene_gn=PAFAH2;refGene_tx=XM_019836429.2;refGene_gn=PAFAH2;refGene_tx=XM_023258329.1
C1 19523409 19523504
refGene_gn=PAFAH2;refGene_tx=XM_003989677.4;refGene_gn=PAFAH2;refGene_tx=XM_006934420.4
C1 19530909 19531064
refGene_gn=PAFAH2;refGene_tx=XM_003989677.4;refGene_gn=PAFAH2;refGene_tx=XM_019836426.2;refGene_gn=PAFAH2;refGene_tx=XM_019836427.2;refGene_gn=PAFAH2;refGene_tx=XM_019836429.2;refGene_gn=PAFAH2;refGene_tx=XM_023258329.1
C1 19533017 19533188
refGene_gn=PAFAH2;refGene_tx=XM_003989677.4;refGene_gn=PAFAH2;refGene_tx=XM_006934420.4;refGene_gn=PAFAH2;refGene_tx=XM_019836426.2;refGene_gn=PAFAH2;refGene_tx=XM_019836427.2;refGene_gn=PAFAH2;refGene_tx=XM_019836429.2;refGene_gn=PAFAH2;refGene_tx=XM_023258329.1
C1 19534803 19534895
refGene_gn=PAFAH2;refGene_tx=XM_003989677.4;refGene_gn=PAFAH2;refGene_

tx=XM_006934420.4;refGene_gn=PAFAH2;refGene_tx=XM_019836426.2;refGene_gn=PAFAH2;refGene_tx=XM_019836427.2;refGene_gn=PAFAH2;refGene_tx=XM_019836429.2;refGene_gn=PAFAH2;refGene_tx=XM_023258329.1
C1 19539369 19539483
refGene_gn=PAFAH2;refGene_tx=XM_003989677.4;refGene_gn=PAFAH2;refGene_tx=XM_006934420.4;refGene_gn=PAFAH2;refGene_tx=XM_019836426.2;refGene_gn=PAFAH2;refGene_tx=XM_019836427.2;refGene_gn=PAFAH2;refGene_tx=XM_019836429.2;refGene_gn=PAFAH2;refGene_tx=XM_023258329.1
C1 19540482 19540624
refGene_gn=PAFAH2;refGene_tx=XM_003989677.4;refGene_gn=PAFAH2;refGene_tx=XM_006934420.4;refGene_gn=PAFAH2;refGene_tx=XM_019836426.2;refGene_gn=PAFAH2;refGene_tx=XM_019836427.2;refGene_gn=PAFAH2;refGene_tx=XM_019836429.2;refGene_gn=PAFAH2;refGene_tx=XM_023258329.1
C1 19541263 19541332
refGene_gn=PAFAH2;refGene_tx=XM_003989677.4;refGene_gn=PAFAH2;refGene_tx=XM_006934420.4;refGene_gn=PAFAH2;refGene_tx=XM_019836426.2;refGene_gn=PAFAH2;refGene_tx=XM_019836427.2;refGene_gn=PAFAH2;refGene_tx=XM_019836429.2;refGene_gn=PAFAH2;refGene_tx=XM_023258329.1
C1 19547039 19547136
refGene_gn=PAFAH2;refGene_tx=XM_003989677.4;refGene_gn=PAFAH2;refGene_tx=XM_006934420.4;refGene_gn=PAFAH2;refGene_tx=XM_019836426.2;refGene_gn=PAFAH2;refGene_tx=XM_019836427.2;refGene_gn=PAFAH2;refGene_tx=XM_019836429.2
C1 19547701 19547855
refGene_gn=PAFAH2;refGene_tx=XM_003989677.4;refGene_gn=PAFAH2;refGene_tx=XM_006934420.4;refGene_gn=PAFAH2;refGene_tx=XM_019836426.2;refGene_gn=PAFAH2;refGene_tx=XM_019836427.2;refGene_gn=PAFAH2;refGene_tx=XM_019836429.2
C1 19548804 19548894
refGene_gn=PAFAH2;refGene_tx=XM_003989677.4;refGene_gn=PAFAH2;refGene_tx=XM_006934420.4;refGene_gn=PAFAH2;refGene_tx=XM_019836426.2;refGene_gn=PAFAH2;refGene_tx=XM_019836427.2;refGene_gn=PAFAH2;refGene_tx=XM_019836429.2
C1 19578479 19579252
refGene_gn=EXTL1;refGene_tx=XM_006934421.4;refGene_gn=EXTL1;refGene_tx=XM_006934422.4;refGene_gn=EXTL1;refGene_tx=XM_006934423.4
C1 19585574 19585665
refGene_gn=EXTL1;refGene_tx=XM_006934421.4;refGene_gn=EXTL1;refGene_tx=XM_006934422.4;refGene_gn=EXTL1;refGene_tx=XM_006934423.4
C1 19585953 19586061
refGene_gn=EXTL1;refGene_tx=XM_006934421.4;refGene_gn=EXTL1;refGene_tx=XM_006934422.4;refGene_gn=EXTL1;refGene_tx=XM_006934423.4
C1 19586887 19587007
refGene_gn=EXTL1;refGene_tx=XM_006934421.4;refGene_gn=EXTL1;refGene_tx=XM_006934422.4;refGene_gn=EXTL1;refGene_tx=XM_006934423.4
C1 19587591 19587724
refGene_gn=EXTL1;refGene_tx=XM_006934421.4;refGene_gn=EXTL1;refGene_tx=XM_006934422.4;refGene_gn=EXTL1;refGene_tx=XM_006934423.4
C1 19587947 19588054
refGene_gn=EXTL1;refGene_tx=XM_006934421.4;refGene_gn=EXTL1;refGene_tx

=XM_006934422.4;refGene_gn=EXTL1;refGene_tx=XM_006934423.4
C1	19589024	19589114
refGene_gn=EXTL1;refGene_tx=XM_006934421.4;refGene_gn=EXTL1;refGene_tx=XM_006934422.4
C1	19589942	19590029
refGene_gn=EXTL1;refGene_tx=XM_006934421.4;refGene_gn=EXTL1;refGene_tx=XM_006934422.4;refGene_gn=EXTL1;refGene_tx=XM_006934423.4
C1	19590418	19590579
refGene_gn=EXTL1;refGene_tx=XM_006934421.4;refGene_gn=EXTL1;refGene_tx=XM_006934422.4;refGene_gn=EXTL1;refGene_tx=XM_006934423.4
C1	19591627	19591799
refGene_gn=EXTL1;refGene_tx=XM_006934421.4;refGene_gn=EXTL1;refGene_tx=XM_006934422.4;refGene_gn=EXTL1;refGene_tx=XM_006934423.4
C1	19591969	19592140
refGene_gn=EXTL1;refGene_tx=XM_006934421.4;refGene_gn=EXTL1;refGene_tx=XM_006934422.4;refGene_gn=EXTL1;refGene_tx=XM_006934423.4
C1	19595671	19595817
refGene_gn=SLC30A2;refGene_tx=XM_011284520.3;refGene_gn=SLC30A2;refGene_tx=XM_011284521.3;refGene_gn=SLC30A2;refGene_tx=XM_019836431.2;refGene_gn=SLC30A2;refGene_tx=XM_019836432.2;refGene_gn=SLC30A2;refGene_tx=XM_019836433.2
C1	19596298	19596433
refGene_gn=SLC30A2;refGene_tx=XM_011284520.3;refGene_gn=SLC30A2;refGene_tx=XM_011284521.3;refGene_gn=SLC30A2;refGene_tx=XM_019836431.2;refGene_gn=SLC30A2;refGene_tx=XM_019836432.2;refGene_gn=SLC30A2;refGene_tx=XM_019836433.2
C1	19597293	19597342
refGene_gn=SLC30A2;refGene_tx=XM_019836431.2;refGene_gn=SLC30A2;refGene_tx=XM_019836433.2
C1	19597573	19597679
refGene_gn=SLC30A2;refGene_tx=XM_011284520.3;refGene_gn=SLC30A2;refGene_tx=XM_011284521.3;refGene_gn=SLC30A2;refGene_tx=XM_019836431.2;refGene_gn=SLC30A2;refGene_tx=XM_019836432.2;refGene_gn=SLC30A2;refGene_tx=XM_019836433.2
C1	19598413	19598570
refGene_gn=SLC30A2;refGene_tx=XM_011284520.3;refGene_gn=SLC30A2;refGene_tx=XM_011284521.3;refGene_gn=SLC30A2;refGene_tx=XM_019836431.2;refGene_gn=SLC30A2;refGene_tx=XM_019836432.2;refGene_gn=SLC30A2;refGene_tx=XM_019836433.2
C1	19599319	19599473
refGene_gn=SLC30A2;refGene_tx=XM_011284520.3;refGene_gn=SLC30A2;refGene_tx=XM_011284521.3;refGene_gn=SLC30A2;refGene_tx=XM_019836431.2;refGene_gn=SLC30A2;refGene_tx=XM_019836432.2;refGene_gn=SLC30A2;refGene_tx=XM_019836433.2
C1	19600184	19600331
refGene_gn=SLC30A2;refGene_tx=XM_011284520.3;refGene_gn=SLC30A2;refGene_tx=XM_019836431.2;refGene_gn=SLC30A2;refGene_tx=XM_019836432.2;refGene_gn=SLC30A2;refGene_tx=XM_019836433.2
C1	19601116	19601337
refGene_gn=SLC30A2;refGene_tx=XM_011284520.3;refGene_gn=SLC30A2;refGen

e_tx=XM_011284521.3;refGene_gn=SLC30A2;refGene_tx=XM_019836431.2;refGene_gn=SLC30A2;refGene_tx=XM_019836432.2;refGene_gn=SLC30A2;refGene_tx=XM_019836433.2
C1 19601993 19602043 refGene_gn=SLC30A2;refGene_tx=XM_011284520.3;refGene_gn=SLC30A2;refGene_tx=XM_011284521.3;refGene_gn=SLC30A2;refGene_tx=XM_019836431.2;refGene_gn=SLC30A2;refGene_tx=XM_019836432.2;refGene_gn=SLC30A2;refGene_tx=XM_019836433.2
C1 19616788 19616827 refGene_gn=TRIM63;refGene_tx=XM_003989679.5;refGene_gn=TRIM63;refGene_tx=XM_019836434.2
C1 19617779 19617851 refGene_gn=TRIM63;refGene_tx=XM_003989679.5;refGene_gn=TRIM63;refGene_tx=XM_019836434.2
C1 19620635 19620760 refGene_gn=TRIM63;refGene_tx=XM_003989679.5
C1 19621112 19621135 refGene_gn=TRIM63;refGene_tx=XM_003989679.5
C1 19621883 19622117 refGene_gn=TRIM63;refGene_tx=XM_003989679.5;refGene_gn=TRIM63;refGene_tx=XM_019836434.2
C1 19623178 19623274 refGene_gn=TRIM63;refGene_tx=XM_003989679.5;refGene_gn=TRIM63;refGene_tx=XM_019836434.2
C1 19624043 19624212 refGene_gn=TRIM63;refGene_tx=XM_003989679.5;refGene_gn=TRIM63;refGene_tx=XM_019836434.2
C1 19626999 19627172 refGene_gn=TRIM63;refGene_tx=XM_003989679.5;refGene_gn=TRIM63;refGene_tx=XM_019836434.2
C1 19628037 19628196 refGene_gn=TRIM63;refGene_tx=XM_003989679.5;refGene_gn=TRIM63;refGene_tx=XM_019836434.2
C1 19661424 19661709 refGene_gn=PDIK1L;refGene_tx=XM_003989680.4;refGene_gn=PDIK1L;refGene_tx=XM_006934425.3;refGene_gn=PDIK1L;refGene_tx=XM_023258331.1
C1 19667778 19668519 refGene_gn=PDIK1L;refGene_tx=XM_003989680.4;refGene_gn=PDIK1L;refGene_tx=XM_006934425.3;refGene_gn=PDIK1L;refGene_tx=XM_023258331.1
C1 19693142 19693967 refGene_gn=FAM110D;refGene_tx=XM_019836436.2;refGene_gn=FAM110D;refGene_tx=XM_023258332.1
C1 19695504 19695805 refGene_gn=LOC111561697;refGene_tx=XM_023258333.1;refGene_gn=LOC111561697;refGene_tx=XM_023258334.1
C1 19697268 19697366 refGene_gn=LOC111561697;refGene_tx=XM_023258333.1;refGene_gn=LOC111561697;refGene_tx=XM_023258334.1
C1 19697476 19697560 refGene_gn=LOC111561697;refGene_tx=XM_023258333.1;refGene_gn=LOC111561697;refGene_tx=XM_023258334.1
C1 19697868 19697916

refGene_gn=LOC111561697;refGene_tx=XM_023258334.1
C1 19698680 19698764
refGene_gn=LOC111561697;refGene_tx=XM_023258333.1
C1 19700249 19700449 refGene_gn=ZNF593;refGene_tx=XM_003989681.5
C1 19700644 19700707 refGene_gn=ZNF593;refGene_tx=XM_003989681.5
C1 19700801 19700943 refGene_gn=ZNF593;refGene_tx=XM_003989681.5
C1 19705578 19705621 refGene_gn=CNKSR1;refGene_tx=XM_023258335.1
C1 19705752 19705804 refGene_gn=CNKSR1;refGene_tx=XM_011284519.3
C1 19708157 19708315
refGene_gn=CNKSR1;refGene_tx=XM_011284519.3;refGene_gn=CNKSR1;refGene_
tx=XM_023258335.1
C1 19708421 19708603
refGene_gn=CNKSR1;refGene_tx=XM_011284519.3;refGene_gn=CNKSR1;refGene_
tx=XM_023258335.1
C1 19709161 19709246
refGene_gn=CNKSR1;refGene_tx=XM_011284519.3;refGene_gn=CNKSR1;refGene_
tx=XM_023258335.1
C1 19709661 19709703
refGene_gn=CNKSR1;refGene_tx=XM_011284519.3;refGene_gn=CNKSR1;refGene_
tx=XM_023258335.1
C1 19709783 19709888
refGene_gn=CNKSR1;refGene_tx=XM_011284519.3;refGene_gn=CNKSR1;refGene_
tx=XM_023258335.1
C1 19710449 19710509
refGene_gn=CNKSR1;refGene_tx=XM_011284519.3;refGene_gn=CNKSR1;refGene_
tx=XM_023258335.1
C1 19710610 19710679
refGene_gn=CNKSR1;refGene_tx=XM_011284519.3;refGene_gn=CNKSR1;refGene_
tx=XM_023258335.1
C1 19710978 19711080
refGene_gn=CNKSR1;refGene_tx=XM_011284519.3;refGene_gn=CNKSR1;refGene_
tx=XM_023258335.1
C1 19711164 19711235
refGene_gn=CNKSR1;refGene_tx=XM_011284519.3;refGene_gn=CNKSR1;refGene_
tx=XM_023258335.1
C1 19711314 19711388
refGene_gn=CNKSR1;refGene_tx=XM_011284519.3;refGene_gn=CNKSR1;refGene_
tx=XM_023258335.1
C1 19711537 19711644
refGene_gn=CNKSR1;refGene_tx=XM_011284519.3;refGene_gn=CNKSR1;refGene_
tx=XM_023258335.1
C1 19711721 19711752
refGene_gn=CNKSR1;refGene_tx=XM_011284519.3;refGene_gn=CNKSR1;refGene_
tx=XM_023258335.1
C1 19712344 19712517
refGene_gn=CNKSR1;refGene_tx=XM_011284519.3;refGene_gn=CNKSR1;refGene_
tx=XM_023258335.1
C1 19713366 19713440
refGene_gn=CNKSR1;refGene_tx=XM_011284519.3;refGene_gn=CNKSR1;refGene_
tx=XM_023258335.1

C1 19713593 19713665
refGene_gn=CNKSR1;refGene_tx=XM_011284519.3;refGene_gn=CNKSR1;refGene_tx=XM_023258335.1
C1 19714119 19714193
refGene_gn=CNKSR1;refGene_tx=XM_011284519.3;refGene_gn=CNKSR1;refGene_tx=XM_023258335.1
C1 19714345 19714407
refGene_gn=CNKSR1;refGene_tx=XM_011284519.3;refGene_gn=CNKSR1;refGene_tx=XM_023258335.1
C1 19714489 19714589
refGene_gn=CNKSR1;refGene_tx=XM_011284519.3;refGene_gn=CNKSR1;refGene_tx=XM_023258335.1
C1 19714683 19714865
refGene_gn=CNKSR1;refGene_tx=XM_011284519.3;refGene_gn=CNKSR1;refGene_tx=XM_023258335.1
C1 19715122 19715392
refGene_gn=CNKSR1;refGene_tx=XM_011284519.3;refGene_gn=CNKSR1;refGene_tx=XM_023258335.1
C1 19716537 19716606 refGene_gn=CATSPER4;refGene_tx=XM_003989684.4
C1 19716729 19716804 refGene_gn=CATSPER4;refGene_tx=XM_003989684.4
C1 19717279 19717423 refGene_gn=CATSPER4;refGene_tx=XM_003989684.4
C1 19721294 19721396 refGene_gn=CATSPER4;refGene_tx=XM_003989684.4
C1 19728922 19729020 refGene_gn=CATSPER4;refGene_tx=XM_003989684.4
C1 19729174 19729295 refGene_gn=CATSPER4;refGene_tx=XM_003989684.4
C1 19729505 19729639 refGene_gn=CATSPER4;refGene_tx=XM_003989684.4
C1 19731368 19731543 refGene_gn=CATSPER4;refGene_tx=XM_003989684.4
C1 19732522 19732734 refGene_gn=CATSPER4;refGene_tx=XM_003989684.4
C1 19733050 19733216 refGene_gn=CATSPER4;refGene_tx=XM_003989684.4
C1 19734153 19734207 refGene_gn=CATSPER4;refGene_tx=XM_003989684.4
C1 19762373 19762428
refGene_gn=CEP85;refGene_tx=XM_003989685.5;refGene_gn=CEP85;refGene_tx=XM_019836437.2;refGene_gn=CEP85;refGene_tx=XM_023258336.1;refGene_gn=CEP85;refGene_tx=XM_023258337.1
C1 19765440 19765593
refGene_gn=CEP85;refGene_tx=XM_003989685.5;refGene_gn=CEP85;refGene_tx=XM_019836437.2;refGene_gn=CEP85;refGene_tx=XM_023258336.1;refGene_gn=CEP85;refGene_tx=XM_023258337.1
C1 19771165 19771860
refGene_gn=CEP85;refGene_tx=XM_003989685.5;refGene_gn=CEP85;refGene_tx=XM_019836437.2;refGene_gn=CEP85;refGene_tx=XM_019836438.2;refGene_gn=CEP85;refGene_tx=XM_023258336.1;refGene_gn=CEP85;refGene_tx=XM_023258337.1
C1 19772689 19772823
refGene_gn=CEP85;refGene_tx=XM_003989685.5;refGene_gn=CEP85;refGene_tx=XM_019836437.2;refGene_gn=CEP85;refGene_tx=XM_019836438.2;refGene_gn=CEP85;refGene_tx=XM_023258336.1;refGene_gn=CEP85;refGene_tx=XM_023258337.1
C1 19773245 19773363
refGene_gn=CEP85;refGene_tx=XM_003989685.5;refGene_gn=CEP85;refGene_tx=XM_019836437.2;refGene_gn=CEP85;refGene_tx=XM_019836438.2;refGene_gn=

CEP85;refGene_tx=XM_023258336.1;refGene_gn=CEP85;refGene_tx=XM_0232583
37.1
C1 19774459 19774645
refGene_gn=CEP85;refGene_tx=XM_003989685.5;refGene_gn=CEP85;refGene_tx
=XM_019836437.2;refGene_gn=CEP85;refGene_tx=XM_019836438.2;refGene_gn=
CEP85;refGene_tx=XM_023258336.1;refGene_gn=CEP85;refGene_tx=XM_0232583
37.1
C1 19778335 19778488
refGene_gn=CEP85;refGene_tx=XM_003989685.5;refGene_gn=CEP85;refGene_tx
=XM_019836437.2;refGene_gn=CEP85;refGene_tx=XM_019836438.2;refGene_gn=
CEP85;refGene_tx=XM_023258336.1;refGene_gn=CEP85;refGene_tx=XM_0232583
37.1
C1 19778714 19778866
refGene_gn=CEP85;refGene_tx=XM_003989685.5;refGene_gn=CEP85;refGene_tx
=XM_019836437.2;refGene_gn=CEP85;refGene_tx=XM_019836438.2;refGene_gn=
CEP85;refGene_tx=XM_023258336.1;refGene_gn=CEP85;refGene_tx=XM_0232583
37.1
C1 19780006 19780121
refGene_gn=CEP85;refGene_tx=XM_003989685.5;refGene_gn=CEP85;refGene_tx
=XM_019836437.2;refGene_gn=CEP85;refGene_tx=XM_019836438.2;refGene_gn=
CEP85;refGene_tx=XM_023258336.1;refGene_gn=CEP85;refGene_tx=XM_0232583
37.1
C1 19781234 19781285
refGene_gn=CEP85;refGene_tx=XM_003989685.5;refGene_gn=CEP85;refGene_tx
=XM_019836437.2;refGene_gn=CEP85;refGene_tx=XM_019836438.2;refGene_gn=
CEP85;refGene_tx=XM_023258336.1;refGene_gn=CEP85;refGene_tx=XM_0232583
37.1
C1 19783652 19783760
refGene_gn=CEP85;refGene_tx=XM_003989685.5;refGene_gn=CEP85;refGene_tx
=XM_019836437.2;refGene_gn=CEP85;refGene_tx=XM_019836438.2;refGene_gn=
CEP85;refGene_tx=XM_023258336.1;refGene_gn=CEP85;refGene_tx=XM_0232583
37.1
C1 19784884 19785110
refGene_gn=CEP85;refGene_tx=XM_003989685.5;refGene_gn=CEP85;refGene_tx
=XM_019836437.2;refGene_gn=CEP85;refGene_tx=XM_019836438.2;refGene_gn=
CEP85;refGene_tx=XM_023258336.1;refGene_gn=CEP85;refGene_tx=XM_0232583
37.1
C1 19785484 19785645
refGene_gn=CEP85;refGene_tx=XM_003989685.5;refGene_gn=CEP85;refGene_tx
=XM_019836437.2;refGene_gn=CEP85;refGene_tx=XM_019836438.2;refGene_gn=
CEP85;refGene_tx=XM_023258336.1;refGene_gn=CEP85;refGene_tx=XM_0232583
37.1
C1 19788511 19788559 refGene_gn=SH3BGRL3;refGene_tx=XM_011284518.3
C1 19789179 19789347 refGene_gn=SH3BGRL3;refGene_tx=XM_011284518.3
C1 19789464 19789530 refGene_gn=SH3BGRL3;refGene_tx=XM_011284518.3
C1 19790637 19790797 refGene_gn=UBXN11;refGene_tx=XM_023258338.1
C1 19790882 19791023 refGene_gn=UBXN11;refGene_tx=XM_023258338.1
C1 19791097 19791171 refGene_gn=UBXN11;refGene_tx=XM_023258338.1
C1 19792354 19792458 refGene_gn=UBXN11;refGene_tx=XM_023258338.1
C1 19792589 19792707 refGene_gn=UBXN11;refGene_tx=XM_023258338.1

C1 19793667 19793745 refGene_gn=UBXN11;refGene_tx=XM_023258338.1
C1 19794032 19794247 refGene_gn=UBXN11;refGene_tx=XM_023258338.1
C1 19801063 19801190 refGene_gn=UBXN11;refGene_tx=XM_023258338.1
C1 19802175 19802257 refGene_gn=UBXN11;refGene_tx=XM_023258338.1
C1 19802728 19802778 refGene_gn=UBXN11;refGene_tx=XM_023258338.1
C1 19803275 19803376 refGene_gn=UBXN11;refGene_tx=XM_023258338.1
C1 19807001 19807100 refGene_gn=UBXN11;refGene_tx=XM_023258338.1
C1 19807697 19807726 refGene_gn=UBXN11;refGene_tx=XM_023258338.1
C1 19808501 19808572 refGene_gn=UBXN11;refGene_tx=XM_023258338.1
C1 19824426 19824515 refGene_gn=CRYBG2;refGene_tx=XM_003989742.5
C1 19824618 19824778 refGene_gn=CRYBG2;refGene_tx=XM_003989742.5
C1 19826170 19826329 refGene_gn=CRYBG2;refGene_tx=XM_003989742.5
C1 19828432 19828556 refGene_gn=CRYBG2;refGene_tx=XM_003989742.5
C1 19828908 19829048 refGene_gn=CRYBG2;refGene_tx=XM_003989742.5
C1 19830404 19830534 refGene_gn=CRYBG2;refGene_tx=XM_003989742.5
C1 19834197 19834310 refGene_gn=CRYBG2;refGene_tx=XM_003989742.5
C1 19834400 19834433 refGene_gn=CRYBG2;refGene_tx=XM_003989742.5
C1 19834668 19834795 refGene_gn=CRYBG2;refGene_tx=XM_003989742.5
C1 19834892 19835032 refGene_gn=CRYBG2;refGene_tx=XM_003989742.5
C1 19835315 19835442 refGene_gn=CRYBG2;refGene_tx=XM_003989742.5
C1 19835598 19835735 refGene_gn=CRYBG2;refGene_tx=XM_003989742.5
C1 19836085 19836121 refGene_gn=CRYBG2;refGene_tx=XM_003989742.5
C1 19836363 19836490 refGene_gn=CRYBG2;refGene_tx=XM_003989742.5
C1 19837539 19837679 refGene_gn=CRYBG2;refGene_tx=XM_003989742.5
C1 19839826 19839956 refGene_gn=CRYBG2;refGene_tx=XM_003989742.5
C1 19840121 19840234 refGene_gn=CRYBG2;refGene_tx=XM_003989742.5
C1 19840319 19840367 refGene_gn=CRYBG2;refGene_tx=XM_003989742.5
C1 19840801 19843424 refGene_gn=CRYBG2;refGene_tx=XM_003989742.5
C1 19849261 19849986 refGene_gn=CRYBG2;refGene_tx=XM_003989742.5
C1 19851547 19851617 refGene_gn=CRYBG2;refGene_tx=XM_003989742.5
C1 19853663 19853779 refGene_gn=CRYBG2;refGene_tx=XM_003989742.5
C1 19855997 19856280 refGene_gn=ZNF683;refGene_tx=XM_023258339.1
C1 19856281 19856577 refGene_gn=ZNF683;refGene_tx=XM_023258339.1
C1 19857621 19857750 refGene_gn=ZNF683;refGene_tx=XM_023258339.1
C1 19859020 19859721 refGene_gn=ZNF683;refGene_tx=XM_023258339.1
C1 19861399 19861604 refGene_gn=ZNF683;refGene_tx=XM_023258339.1
C1 19862267 19862381 refGene_gn=ZNF683;refGene_tx=XM_023258339.1
C1 19894837 19894868 refGene_gn=LIN28A;refGene_tx=NM_001173445.1
C1 19895333 19895518 refGene_gn=LIN28A;refGene_tx=NM_001173445.1
C1 19904902 19905087 refGene_gn=LIN28A;refGene_tx=NM_001173445.1
C1 19905762 19905982 refGene_gn=LIN28A;refGene_tx=NM_001173445.1
C1 19911097 19911160
refGene_gn=DHDDS;refGene_tx=XM_006934427.4;refGene_gn=DHDDS;refGene_tx
=XM_011284515.3
C1 19912844 19912961
refGene_gn=DHDDS;refGene_tx=XM_006934427.4;refGene_gn=DHDDS;refGene_tx
=XM_011284515.3
C1 19915788 19915931
refGene_gn=DHDDS;refGene_tx=XM_006934427.4;refGene_gn=DHDDS;refGene_tx
=XM_011284515.3

C1 19919538 19919655
refGene_gn=DHDDS;refGene_tx=XM_006934427.4;refGene_gn=DHDDS;refGene_tx=XM_011284515.3
C1 19920572 19920674
refGene_gn=DHDDS;refGene_tx=XM_006934427.4;refGene_gn=DHDDS;refGene_tx=XM_011284515.3
C1 19926799 19926914
refGene_gn=DHDDS;refGene_tx=XM_006934427.4;refGene_gn=DHDDS;refGene_tx=XM_011284515.3
C1 19928894 19929002
refGene_gn=DHDDS;refGene_tx=XM_006934427.4;refGene_gn=DHDDS;refGene_tx=XM_011284515.3
C1 19936464 19936704
refGene_gn=DHDDS;refGene_tx=XM_006934427.4;refGene_gn=DHDDS;refGene_tx=XM_011284515.3
C1 19940083 19940098 refGene_gn=HMGN2;refGene_tx=XM_023258341.1
C1 19941104 19941149 refGene_gn=HMGN2;refGene_tx=XM_023258341.1
C1 19941350 19941380 refGene_gn=HMGN2;refGene_tx=XM_023258341.1
C1 19941743 19941794 refGene_gn=HMGN2;refGene_tx=XM_023258341.1
C1 19942632 19942728 refGene_gn=HMGN2;refGene_tx=XM_023258341.1
C1 19943166 19943202 refGene_gn=HMGN2;refGene_tx=XM_023258341.1
C1 19970443 19970494
refGene_gn=RPS6KA1;refGene_tx=XM_006934428.4;refGene_gn=RPS6KA1;refGene_tx=XM_019836449.2
C1 19985317 19985380
refGene_gn=RPS6KA1;refGene_tx=XM_023258342.1;refGene_gn=RPS6KA1;refGene_tx=XM_023258344.1
C1 19991540 19991585
refGene_gn=RPS6KA1;refGene_tx=XM_006934428.4;refGene_gn=RPS6KA1;refGene_tx=XM_019836449.2;refGene_gn=RPS6KA1;refGene_tx=XM_023258342.1;refGene_gn=RPS6KA1;refGene_tx=XM_023258344.1
C1 19997124 19997184 refGene_gn=RPS6KA1;refGene_tx=XM_006934429.4
C1 19998862 19999022
refGene_gn=RPS6KA1;refGene_tx=XM_003989690.5;refGene_gn=RPS6KA1;refGene_tx=XM_019836447.2
C1 19999867 20000001
refGene_gn=RPS6KA1;refGene_tx=XM_003989690.5;refGene_gn=RPS6KA1;refGene_tx=XM_006934428.4;refGene_gn=RPS6KA1;refGene_tx=XM_006934429.4;refGene_gn=RPS6KA1;refGene_tx=XM_019836447.2;refGene_gn=RPS6KA1;refGene_tx=XM_023258342.1
C1 20000266 20000348
refGene_gn=RPS6KA1;refGene_tx=XM_003989690.5;refGene_gn=RPS6KA1;refGene_tx=XM_006934428.4;refGene_gn=RPS6KA1;refGene_tx=XM_006934429.4;refGene_gn=RPS6KA1;refGene_tx=XM_019836447.2;refGene_gn=RPS6KA1;refGene_tx=XM_019836449.2;refGene_gn=RPS6KA1;refGene_tx=XM_023258342.1;refGene_gn=RPS6KA1;refGene_tx=XM_023258344.1
C1 20002064 20002145
refGene_gn=RPS6KA1;refGene_tx=XM_003989690.5;refGene_gn=RPS6KA1;refGene_tx=XM_006934428.4;refGene_gn=RPS6KA1;refGene_tx=XM_006934429.4;refGene_gn=RPS6KA1;refGene_tx=XM_019836447.2;refGene_gn=RPS6KA1;refGene_tx=

XM_019836449.2;refGene_gn=RPS6KA1;refGene_tx=XM_023258342.1;refGene_gn=RPS6KA1;refGene_tx=XM_023258344.1
C1 20002337 20002417
refGene_gn=RPS6KA1;refGene_tx=XM_003989690.5;refGene_gn=RPS6KA1;refGene_tx=XM_006934428.4;refGene_gn=RPS6KA1;refGene_tx=XM_006934429.4;refGene_gn=RPS6KA1;refGene_tx=XM_019836447.2;refGene_gn=RPS6KA1;refGene_tx=XM_019836449.2;refGene_gn=RPS6KA1;refGene_tx=XM_023258342.1;refGene_gn=RPS6KA1;refGene_tx=XM_023258344.1
C1 20004262 20004369
refGene_gn=RPS6KA1;refGene_tx=XM_003989690.5;refGene_gn=RPS6KA1;refGene_tx=XM_006934428.4;refGene_gn=RPS6KA1;refGene_tx=XM_006934429.4;refGene_gn=RPS6KA1;refGene_tx=XM_019836447.2;refGene_gn=RPS6KA1;refGene_tx=XM_019836449.2;refGene_gn=RPS6KA1;refGene_tx=XM_023258342.1;refGene_gn=RPS6KA1;refGene_tx=XM_023258344.1
C1 20005098 20005136
refGene_gn=RPS6KA1;refGene_tx=XM_003989690.5;refGene_gn=RPS6KA1;refGene_tx=XM_006934428.4;refGene_gn=RPS6KA1;refGene_tx=XM_006934429.4;refGene_gn=RPS6KA1;refGene_tx=XM_019836447.2;refGene_gn=RPS6KA1;refGene_tx=XM_019836449.2;refGene_gn=RPS6KA1;refGene_tx=XM_023258342.1;refGene_gn=RPS6KA1;refGene_tx=XM_023258344.1
C1 20005426 20005569
refGene_gn=RPS6KA1;refGene_tx=XM_003989690.5;refGene_gn=RPS6KA1;refGene_tx=XM_006934428.4;refGene_gn=RPS6KA1;refGene_tx=XM_006934429.4;refGene_gn=RPS6KA1;refGene_tx=XM_019836447.2;refGene_gn=RPS6KA1;refGene_tx=XM_019836449.2;refGene_gn=RPS6KA1;refGene_tx=XM_023258342.1;refGene_gn=RPS6KA1;refGene_tx=XM_023258344.1
C1 20005977 20006048
refGene_gn=RPS6KA1;refGene_tx=XM_003989690.5;refGene_gn=RPS6KA1;refGene_tx=XM_006934428.4;refGene_gn=RPS6KA1;refGene_tx=XM_006934429.4;refGene_gn=RPS6KA1;refGene_tx=XM_019836447.2;refGene_gn=RPS6KA1;refGene_tx=XM_019836449.2;refGene_gn=RPS6KA1;refGene_tx=XM_023258342.1;refGene_gn=RPS6KA1;refGene_tx=XM_023258344.1
C1 20006363 20006452
refGene_gn=RPS6KA1;refGene_tx=XM_003989690.5;refGene_gn=RPS6KA1;refGene_tx=XM_006934428.4;refGene_gn=RPS6KA1;refGene_tx=XM_006934429.4;refGene_gn=RPS6KA1;refGene_tx=XM_019836447.2;refGene_gn=RPS6KA1;refGene_tx=XM_019836449.2;refGene_gn=RPS6KA1;refGene_tx=XM_023258342.1;refGene_gn=RPS6KA1;refGene_tx=XM_023258344.1
C1 20007466 20007531
refGene_gn=RPS6KA1;refGene_tx=XM_003989690.5;refGene_gn=RPS6KA1;refGene_tx=XM_006934428.4;refGene_gn=RPS6KA1;refGene_tx=XM_006934429.4;refGene_gn=RPS6KA1;refGene_tx=XM_019836447.2;refGene_gn=RPS6KA1;refGene_tx=XM_019836449.2;refGene_gn=RPS6KA1;refGene_tx=XM_023258342.1;refGene_gn=RPS6KA1;refGene_tx=XM_023258344.1
C1 20007792 20007895
refGene_gn=RPS6KA1;refGene_tx=XM_003989690.5;refGene_gn=RPS6KA1;refGene_tx=XM_006934428.4;refGene_gn=RPS6KA1;refGene_tx=XM_006934429.4;refGene_gn=RPS6KA1;refGene_tx=XM_019836447.2;refGene_gn=RPS6KA1;refGene_tx=XM_019836449.2;refGene_gn=RPS6KA1;refGene_tx=XM_023258342.1;refGene_gn=RPS6KA1;refGene_tx=XM_023258344.1

C1 20009351 20009482
refGene_gn=RPS6KA1;refGene_tx=XM_003989690.5;refGene_gn=RPS6KA1;refGene_tx=XM_006934428.4;refGene_gn=RPS6KA1;refGene_tx=XM_006934429.4;refGene_gn=RPS6KA1;refGene_tx=XM_019836447.2;refGene_gn=RPS6KA1;refGene_tx=XM_019836449.2;refGene_gn=RPS6KA1;refGene_tx=XM_023258342.1;refGene_gn=RPS6KA1;refGene_tx=XM_023258344.1
C1 20010490 20010616
refGene_gn=RPS6KA1;refGene_tx=XM_003989690.5;refGene_gn=RPS6KA1;refGene_tx=XM_006934428.4;refGene_gn=RPS6KA1;refGene_tx=XM_006934429.4;refGene_gn=RPS6KA1;refGene_tx=XM_019836447.2;refGene_gn=RPS6KA1;refGene_tx=XM_019836449.2;refGene_gn=RPS6KA1;refGene_tx=XM_023258342.1;refGene_gn=RPS6KA1;refGene_tx=XM_023258344.1
C1 20010840 20010930
refGene_gn=RPS6KA1;refGene_tx=XM_003989690.5;refGene_gn=RPS6KA1;refGene_tx=XM_006934428.4;refGene_gn=RPS6KA1;refGene_tx=XM_006934429.4;refGene_gn=RPS6KA1;refGene_tx=XM_019836447.2;refGene_gn=RPS6KA1;refGene_tx=XM_019836449.2;refGene_gn=RPS6KA1;refGene_tx=XM_023258342.1;refGene_gn=RPS6KA1;refGene_tx=XM_023258344.1
C1 20011322 20011481
refGene_gn=RPS6KA1;refGene_tx=XM_003989690.5;refGene_gn=RPS6KA1;refGene_tx=XM_006934428.4;refGene_gn=RPS6KA1;refGene_tx=XM_006934429.4;refGene_gn=RPS6KA1;refGene_tx=XM_019836447.2;refGene_gn=RPS6KA1;refGene_tx=XM_019836449.2;refGene_gn=RPS6KA1;refGene_tx=XM_023258342.1;refGene_gn=RPS6KA1;refGene_tx=XM_023258344.1
C1 20021233 20021395
refGene_gn=RPS6KA1;refGene_tx=XM_003989690.5;refGene_gn=RPS6KA1;refGene_tx=XM_006934428.4;refGene_gn=RPS6KA1;refGene_tx=XM_006934429.4;refGene_gn=RPS6KA1;refGene_tx=XM_019836447.2;refGene_gn=RPS6KA1;refGene_tx=XM_019836449.2;refGene_gn=RPS6KA1;refGene_tx=XM_023258342.1;refGene_gn=RPS6KA1;refGene_tx=XM_023258344.1
C1 20021888 20021965
refGene_gn=RPS6KA1;refGene_tx=XM_003989690.5;refGene_gn=RPS6KA1;refGene_tx=XM_006934428.4;refGene_gn=RPS6KA1;refGene_tx=XM_006934429.4;refGene_gn=RPS6KA1;refGene_tx=XM_019836447.2;refGene_gn=RPS6KA1;refGene_tx=XM_019836449.2;refGene_gn=RPS6KA1;refGene_tx=XM_023258342.1;refGene_gn=RPS6KA1;refGene_tx=XM_023258344.1
C1 20022225 20022343
refGene_gn=RPS6KA1;refGene_tx=XM_003989690.5;refGene_gn=RPS6KA1;refGene_tx=XM_006934428.4;refGene_gn=RPS6KA1;refGene_tx=XM_006934429.4;refGene_gn=RPS6KA1;refGene_tx=XM_019836447.2;refGene_gn=RPS6KA1;refGene_tx=XM_019836449.2;refGene_gn=RPS6KA1;refGene_tx=XM_023258342.1;refGene_gn=RPS6KA1;refGene_tx=XM_023258344.1
C1 20023258 20023396
refGene_gn=RPS6KA1;refGene_tx=XM_003989690.5;refGene_gn=RPS6KA1;refGene_tx=XM_006934428.4;refGene_gn=RPS6KA1;refGene_tx=XM_006934429.4;refGene_gn=RPS6KA1;refGene_tx=XM_019836447.2;refGene_gn=RPS6KA1;refGene_tx=XM_019836449.2;refGene_gn=RPS6KA1;refGene_tx=XM_023258342.1;refGene_gn=RPS6KA1;refGene_tx=XM_023258344.1
C1 20023734 20023857
refGene_gn=RPS6KA1;refGene_tx=XM_003989690.5;refGene_gn=RPS6KA1;refGen

e_tx=XM_006934428.4;refGene_gn=RPS6KA1;refGene_tx=XM_006934429.4;refGene_gn=RPS6KA1;refGene_tx=XM_019836447.2;refGene_gn=RPS6KA1;refGene_tx=XM_019836449.2;refGene_gn=RPS6KA1;refGene_tx=XM_023258342.1;refGene_gn=RPS6KA1;refGene_tx=XM_023258344.1
C1 20112299 20113439 refGene_gn=ARID1A;refGene_tx=XM_023258345.1;refGene_gn=ARID1A;refGene_tx=XM_023258346.1
C1 20118700 20118817 refGene_gn=ARID1A;refGene_tx=XM_019836451.2
C1 20137736 20137949 refGene_gn=ARID1A;refGene_tx=XM_019836451.2;refGene_gn=ARID1A;refGene_tx=XM_023258345.1;refGene_gn=ARID1A;refGene_tx=XM_023258346.1
C1 20139303 20139756 refGene_gn=ARID1A;refGene_tx=XM_019836451.2;refGene_gn=ARID1A;refGene_tx=XM_023258345.1;refGene_gn=ARID1A;refGene_tx=XM_023258346.1
C1 20140946 20141063 refGene_gn=ARID1A;refGene_tx=XM_019836451.2;refGene_gn=ARID1A;refGene_tx=XM_023258345.1;refGene_gn=ARID1A;refGene_tx=XM_023258346.1
C1 20164370 20164611 refGene_gn=ARID1A;refGene_tx=XM_019836451.2;refGene_gn=ARID1A;refGene_tx=XM_023258345.1;refGene_gn=ARID1A;refGene_tx=XM_023258346.1
C1 20164893 20164983 refGene_gn=ARID1A;refGene_tx=XM_019836451.2;refGene_gn=ARID1A;refGene_tx=XM_023258345.1;refGene_gn=ARID1A;refGene_tx=XM_023258346.1
C1 20165544 20165712 refGene_gn=ARID1A;refGene_tx=XM_019836451.2;refGene_gn=ARID1A;refGene_tx=XM_023258345.1;refGene_gn=ARID1A;refGene_tx=XM_023258346.1
C1 20166403 20166716 refGene_gn=ARID1A;refGene_tx=XM_019836451.2;refGene_gn=ARID1A;refGene_tx=XM_023258345.1;refGene_gn=ARID1A;refGene_tx=XM_023258346.1
C1 20168005 20168151 refGene_gn=ARID1A;refGene_tx=XM_019836451.2;refGene_gn=ARID1A;refGene_tx=XM_023258345.1;refGene_gn=ARID1A;refGene_tx=XM_023258346.1
C1 20168248 20168358 refGene_gn=ARID1A;refGene_tx=XM_019836451.2;refGene_gn=ARID1A;refGene_tx=XM_023258345.1;refGene_gn=ARID1A;refGene_tx=XM_023258346.1
C1 20169573 20169783 refGene_gn=ARID1A;refGene_tx=XM_019836451.2;refGene_gn=ARID1A;refGene_tx=XM_023258345.1;refGene_gn=ARID1A;refGene_tx=XM_023258346.1
C1 20172314 20172522 refGene_gn=ARID1A;refGene_tx=XM_019836451.2;refGene_gn=ARID1A;refGene_tx=XM_023258345.1;refGene_gn=ARID1A;refGene_tx=XM_023258346.1
C1 20173690 20173823 refGene_gn=ARID1A;refGene_tx=XM_019836451.2;refGene_gn=ARID1A;refGene_tx=XM_023258345.1;refGene_gn=ARID1A;refGene_tx=XM_023258346.1
C1 20173996 20174172 refGene_gn=ARID1A;refGene_tx=XM_019836451.2;refGene_gn=ARID1A;refGene_tx=XM_023258345.1;refGene_gn=ARID1A;refGene_tx=XM_023258346.1
C1 20174535 20174686 refGene_gn=ARID1A;refGene_tx=XM_019836451.2;refGene_gn=ARID1A;refGene_tx=XM_023258345.1;refGene_gn=ARID1A;refGene_tx=XM_023258346.1

C1 20174767 20174902 refGene_gn=ARID1A;refGene_tx=XM_019836451.2;refGene_gn=ARID1A;refGene_tx=XM_023258345.1;refGene_gn=ARID1A;refGene_tx=XM_023258346.1
C1 20174986 20175083 refGene_gn=ARID1A;refGene_tx=XM_019836451.2;refGene_gn=ARID1A;refGene_tx=XM_023258345.1;refGene_gn=ARID1A;refGene_tx=XM_023258346.1
C1 20175561 20176453 refGene_gn=ARID1A;refGene_tx=XM_019836451.2;refGene_gn=ARID1A;refGene_tx=XM_023258345.1;refGene_gn=ARID1A;refGene_tx=XM_023258346.1
C1 20176799 20176930 refGene_gn=ARID1A;refGene_tx=XM_019836451.2;refGene_gn=ARID1A;refGene_tx=XM_023258345.1;refGene_gn=ARID1A;refGene_tx=XM_023258346.1
C1 20178753 20180484 refGene_gn=ARID1A;refGene_tx=XM_019836451.2;refGene_gn=ARID1A;refGene_tx=XM_023258345.1;refGene_gn=ARID1A;refGene_tx=XM_023258346.1
C1 20191506 20191584 refGene_gn=PIGV;refGene_tx=XM_006934433.4;refGene_gn=PIGV;refGene_tx=XM_006934434.4;refGene_gn=PIGV;refGene_tx=XM_006934435.4;refGene_gn=PIGV;refGene_tx=XM_006934436.4;refGene_gn=PIGV;refGene_tx=XM_019836453.2
C1 20194743 20195865 refGene_gn=PIGV;refGene_tx=XM_006934433.4;refGene_gn=PIGV;refGene_tx=XM_006934434.4;refGene_gn=PIGV;refGene_tx=XM_006934435.4;refGene_gn=PIGV;refGene_tx=XM_006934436.4;refGene_gn=PIGV;refGene_tx=XM_019836453.2
C1 20197204 20197486 refGene_gn=PIGV;refGene_tx=XM_006934433.4;refGene_gn=PIGV;refGene_tx=XM_006934434.4;refGene_gn=PIGV;refGene_tx=XM_006934435.4;refGene_gn=PIGV;refGene_tx=XM_006934436.4;refGene_gn=PIGV;refGene_tx=XM_019836453.2
C1 20210604 20210927 refGene_gn=ZDHHC18;refGene_tx=XM_023258348.1
C1 20215139 20215300 refGene_gn=ZDHHC18;refGene_tx=XM_023258348.1
C1 20230844 20230994 refGene_gn=ZDHHC18;refGene_tx=XM_023258348.1
C1 20232566 20232704 refGene_gn=ZDHHC18;refGene_tx=XM_023258348.1
C1 20232823 20232872 refGene_gn=ZDHHC18;refGene_tx=XM_023258348.1
C1 20233412 20233515 refGene_gn=ZDHHC18;refGene_tx=XM_023258348.1
C1 20235124 20235237 refGene_gn=ZDHHC18;refGene_tx=XM_023258348.1
C1 20236359 20236477 refGene_gn=ZDHHC18;refGene_tx=XM_023258348.1
C1 20244270 20245017 refGene_gn=SFN;refGene_tx=XM_003989694.4
C1 20254068 20254141 refGene_gn=GPN2;refGene_tx=XM_003989695.5
C1 20258180 20258311 refGene_gn=GPN2;refGene_tx=XM_003989695.5
C1 20259311 20259472 refGene_gn=GPN2;refGene_tx=XM_003989695.5
C1 20261759 20261916 refGene_gn=GPN2;refGene_tx=XM_003989695.5
C1 20262492 20262903 refGene_gn=GPN2;refGene_tx=XM_003989695.5
C1 20263783 20264000 refGene_gn=GPATCH3;refGene_tx=XM_003989745.4
C1 20264507 20264635 refGene_gn=GPATCH3;refGene_tx=XM_003989745.4
C1 20264756 20264878 refGene_gn=GPATCH3;refGene_tx=XM_003989745.4
C1 20265705 20265765 refGene_gn=GPATCH3;refGene_tx=XM_003989745.4
C1 20266369 20266544 refGene_gn=GPATCH3;refGene_tx=XM_003989745.4
C1 20268103 20268528 refGene_gn=GPATCH3;refGene_tx=XM_003989745.4
C1 20270111 20270547 refGene_gn=GPATCH3;refGene_tx=XM_003989745.4
C1 20277807 20277848 refGene_gn=NR0B2;refGene_tx=XM_003989746.4
C1 20278530 20278800 refGene_gn=NR0B2;refGene_tx=XM_003989746.4

C1 20279602 20280143 refGene_gn=NR0B2;refGene_tx=XM_003989746.4
C1 20287611 20287692 refGene_gn=NUDC;refGene_tx=XM_003989696.3
C1 20290078 20290156
refGene_gn=NUDC;refGene_tx=XM_003989696.3;refGene_gn=NUDC;refGene_tx=XM_019836457.2
C1 20294936 20295140
refGene_gn=NUDC;refGene_tx=XM_003989696.3;refGene_gn=NUDC;refGene_tx=XM_019836457.2
C1 20295218 20295284
refGene_gn=NUDC;refGene_tx=XM_003989696.3;refGene_gn=NUDC;refGene_tx=XM_019836457.2
C1 20296053 20296173
refGene_gn=NUDC;refGene_tx=XM_003989696.3;refGene_gn=NUDC;refGene_tx=XM_019836457.2
C1 20296249 20296444
refGene_gn=NUDC;refGene_tx=XM_003989696.3;refGene_gn=NUDC;refGene_tx=XM_019836457.2
C1 20297609 20297693
refGene_gn=NUDC;refGene_tx=XM_003989696.3;refGene_gn=NUDC;refGene_tx=XM_019836457.2
C1 20297787 20297906
refGene_gn=NUDC;refGene_tx=XM_003989696.3;refGene_gn=NUDC;refGene_tx=XM_019836457.2
C1 20298324 20298376
refGene_gn=NUDC;refGene_tx=XM_003989696.3;refGene_gn=NUDC;refGene_tx=XM_019836457.2
C1 20300401 20300484
refGene_gn=KDF1;refGene_tx=XM_023258349.1;refGene_gn=KDF1;refGene_tx=XM_023258350.1
C1 20301041 20301116
refGene_gn=KDF1;refGene_tx=XM_023258349.1;refGene_gn=KDF1;refGene_tx=XM_023258350.1
C1 20301571 20302607
refGene_gn=KDF1;refGene_tx=XM_023258349.1;refGene_gn=KDF1;refGene_tx=XM_023258350.1
C1 20331645 20332329 refGene_gn=TRNP1;refGene_tx=XM_023258352.1
C1 20341850 20342870 refGene_gn=FAM46B;refGene_tx=XM_003989698.5
C1 20347580 20347844 refGene_gn=FAM46B;refGene_tx=XM_003989698.5
C1 20406456 20406794 refGene_gn=SLC9A1;refGene_tx=XM_003989699.4
C1 20407455 20407528 refGene_gn=SLC9A1;refGene_tx=XM_003989699.4
C1 20408745 20408847 refGene_gn=SLC9A1;refGene_tx=XM_003989699.4
C1 20408982 20409097 refGene_gn=SLC9A1;refGene_tx=XM_003989699.4
C1 20409334 20409508 refGene_gn=SLC9A1;refGene_tx=XM_003989699.4
C1 20409624 20409695 refGene_gn=SLC9A1;refGene_tx=XM_003989699.4
C1 20410174 20410264 refGene_gn=SLC9A1;refGene_tx=XM_003989699.4
C1 20411776 20411979 refGene_gn=SLC9A1;refGene_tx=XM_003989699.4
C1 20413194 20413412 refGene_gn=SLC9A1;refGene_tx=XM_003989699.4
C1 20414318 20414569 refGene_gn=SLC9A1;refGene_tx=XM_003989699.4
C1 20417827 20418288 refGene_gn=SLC9A1;refGene_tx=XM_003989699.4
C1 20451699 20452054 refGene_gn=SLC9A1;refGene_tx=XM_003989699.4

C1 20506586 20506634
refGene_gn=WDTC1;refGene_tx=XM_023258354.1;refGene_gn=WDTC1;refGene_tx
=XM_023258355.1;refGene_gn=WDTC1;refGene_tx=XM_023258356.1;refGene_gn=
WDTC1;refGene_tx=XM_023258357.1;refGene_gn=WDTC1;refGene_tx=XM_0232583
58.1;refGene_gn=WDTC1;refGene_tx=XM_023258359.1
C1 20508401 20508485
refGene_gn=WDTC1;refGene_tx=XM_023258354.1;refGene_gn=WDTC1;refGene_tx
=XM_023258355.1;refGene_gn=WDTC1;refGene_tx=XM_023258356.1;refGene_gn=
WDTC1;refGene_tx=XM_023258357.1;refGene_gn=WDTC1;refGene_tx=XM_0232583
58.1;refGene_gn=WDTC1;refGene_tx=XM_023258359.1
C1 20530809 20530872
refGene_gn=WDTC1;refGene_tx=XM_023258354.1;refGene_gn=WDTC1;refGene_tx
=XM_023258355.1;refGene_gn=WDTC1;refGene_tx=XM_023258356.1;refGene_gn=
WDTC1;refGene_tx=XM_023258359.1
C1 20534899 20534946
refGene_gn=WDTC1;refGene_tx=XM_023258354.1;refGene_gn=WDTC1;refGene_tx
=XM_023258355.1;refGene_gn=WDTC1;refGene_tx=XM_023258356.1;refGene_gn=
WDTC1;refGene_tx=XM_023258357.1;refGene_gn=WDTC1;refGene_tx=XM_0232583
58.1;refGene_gn=WDTC1;refGene_tx=XM_023258359.1
C1 20535345 20535457
refGene_gn=WDTC1;refGene_tx=XM_023258354.1;refGene_gn=WDTC1;refGene_tx
=XM_023258355.1;refGene_gn=WDTC1;refGene_tx=XM_023258356.1;refGene_gn=
WDTC1;refGene_tx=XM_023258357.1;refGene_gn=WDTC1;refGene_tx=XM_0232583
58.1;refGene_gn=WDTC1;refGene_tx=XM_023258359.1
C1 20538166 20538354
refGene_gn=WDTC1;refGene_tx=XM_023258354.1;refGene_gn=WDTC1;refGene_tx
=XM_023258355.1;refGene_gn=WDTC1;refGene_tx=XM_023258356.1;refGene_gn=
WDTC1;refGene_tx=XM_023258357.1;refGene_gn=WDTC1;refGene_tx=XM_0232583
58.1;refGene_gn=WDTC1;refGene_tx=XM_023258359.1
C1 20540591 20540774
refGene_gn=WDTC1;refGene_tx=XM_023258354.1;refGene_gn=WDTC1;refGene_tx
=XM_023258355.1;refGene_gn=WDTC1;refGene_tx=XM_023258356.1;refGene_gn=
WDTC1;refGene_tx=XM_023258357.1;refGene_gn=WDTC1;refGene_tx=XM_0232583
58.1;refGene_gn=WDTC1;refGene_tx=XM_023258359.1
C1 20541735 20541830
refGene_gn=WDTC1;refGene_tx=XM_023258354.1;refGene_gn=WDTC1;refGene_tx
=XM_023258355.1;refGene_gn=WDTC1;refGene_tx=XM_023258356.1;refGene_gn=
WDTC1;refGene_tx=XM_023258357.1;refGene_gn=WDTC1;refGene_tx=XM_0232583
58.1;refGene_gn=WDTC1;refGene_tx=XM_023258359.1
C1 20542239 20542355
refGene_gn=WDTC1;refGene_tx=XM_023258354.1;refGene_gn=WDTC1;refGene_tx
=XM_023258355.1;refGene_gn=WDTC1;refGene_tx=XM_023258356.1;refGene_gn=
WDTC1;refGene_tx=XM_023258357.1;refGene_gn=WDTC1;refGene_tx=XM_0232583
58.1;refGene_gn=WDTC1;refGene_tx=XM_023258359.1
C1 20543816 20543892
refGene_gn=WDTC1;refGene_tx=XM_023258354.1;refGene_gn=WDTC1;refGene_tx
=XM_023258355.1;refGene_gn=WDTC1;refGene_tx=XM_023258356.1;refGene_gn=
WDTC1;refGene_tx=XM_023258357.1;refGene_gn=WDTC1;refGene_tx=XM_0232583
58.1;refGene_gn=WDTC1;refGene_tx=XM_023258359.1
C1 20544503 20544612

refGene_gn=WDTC1;refGene_tx=XM_023258354.1;refGene_gn=WDTC1;refGene_tx=XM_023258355.1;refGene_gn=WDTC1;refGene_tx=XM_023258356.1;refGene_gn=WDTC1;refGene_tx=XM_023258357.1;refGene_gn=WDTC1;refGene_tx=XM_023258358.1;refGene_gn=WDTC1;refGene_tx=XM_023258359.1
C1 20545645 20545819
refGene_gn=WDTC1;refGene_tx=XM_023258354.1;refGene_gn=WDTC1;refGene_tx=XM_023258355.1;refGene_gn=WDTC1;refGene_tx=XM_023258356.1;refGene_gn=WDTC1;refGene_tx=XM_023258357.1;refGene_gn=WDTC1;refGene_tx=XM_023258358.1;refGene_gn=WDTC1;refGene_tx=XM_023258359.1
C1 20548876 20549112
refGene_gn=WDTC1;refGene_tx=XM_023258354.1;refGene_gn=WDTC1;refGene_tx=XM_023258355.1;refGene_gn=WDTC1;refGene_tx=XM_023258356.1;refGene_gn=WDTC1;refGene_tx=XM_023258357.1;refGene_gn=WDTC1;refGene_tx=XM_023258358.1;refGene_gn=WDTC1;refGene_tx=XM_023258359.1
C1 20550683 20550864
refGene_gn=WDTC1;refGene_tx=XM_023258354.1;refGene_gn=WDTC1;refGene_tx=XM_023258355.1;refGene_gn=WDTC1;refGene_tx=XM_023258356.1;refGene_gn=WDTC1;refGene_tx=XM_023258357.1;refGene_gn=WDTC1;refGene_tx=XM_023258358.1;refGene_gn=WDTC1;refGene_tx=XM_023258359.1
C1 20551814 20552007
refGene_gn=WDTC1;refGene_tx=XM_023258354.1;refGene_gn=WDTC1;refGene_tx=XM_023258355.1;refGene_gn=WDTC1;refGene_tx=XM_023258356.1;refGene_gn=WDTC1;refGene_tx=XM_023258357.1;refGene_gn=WDTC1;refGene_tx=XM_023258358.1;refGene_gn=WDTC1;refGene_tx=XM_023258359.1
C1 20553345 20553543
refGene_gn=WDTC1;refGene_tx=XM_023258354.1;refGene_gn=WDTC1;refGene_tx=XM_023258355.1;refGene_gn=WDTC1;refGene_tx=XM_023258356.1;refGene_gn=WDTC1;refGene_tx=XM_023258357.1;refGene_gn=WDTC1;refGene_tx=XM_023258358.1;refGene_gn=WDTC1;refGene_tx=XM_023258359.1
C1 20565416 20565610 refGene_gn=TMEM222;refGene_tx=XM_003989701.5
C1 20571454 20571539 refGene_gn=TMEM222;refGene_tx=XM_003989701.5
C1 20572772 20572804 refGene_gn=TMEM222;refGene_tx=XM_003989701.5
C1 20574610 20574707 refGene_gn=TMEM222;refGene_tx=XM_003989701.5
C1 20574791 20574922 refGene_gn=TMEM222;refGene_tx=XM_003989701.5
C1 20576058 20576146 refGene_gn=TMEM222;refGene_tx=XM_003989701.5
C1 20582406 20582597 refGene_gn=SYTL1;refGene_tx=XM_023260060.1
C1 20583569 20583718 refGene_gn=SYTL1;refGene_tx=XM_023260060.1
C1 20583910 20583989 refGene_gn=SYTL1;refGene_tx=XM_023260060.1
C1 20584070 20584115 refGene_gn=SYTL1;refGene_tx=XM_023260060.1
C1 20584208 20584245 refGene_gn=SYTL1;refGene_tx=XM_023260060.1
C1 20584532 20584605 refGene_gn=SYTL1;refGene_tx=XM_023260060.1
C1 20584939 20585037 refGene_gn=SYTL1;refGene_tx=XM_023260060.1
C1 20585174 20585288 refGene_gn=SYTL1;refGene_tx=XM_023260060.1
C1 20585460 20585627 refGene_gn=SYTL1;refGene_tx=XM_023260060.1
C1 20585863 20585960 refGene_gn=SYTL1;refGene_tx=XM_023260060.1
C1 20586297 20586456 refGene_gn=SYTL1;refGene_tx=XM_023260060.1
C1 20586721 20586800 refGene_gn=SYTL1;refGene_tx=XM_023260060.1
C1 20586907 20587007 refGene_gn=SYTL1;refGene_tx=XM_023260060.1
C1 20587779 20587985 refGene_gn=SYTL1;refGene_tx=XM_023260060.1
C1 20588192 20588332 refGene_gn=SYTL1;refGene_tx=XM_023260060.1

C1 20588811 20588902
refGene_gn=MAP3K6;refGene_tx=XM_011284512.3;refGene_gn=MAP3K6;refGene_tx=XM_011284513.3;refGene_gn=MAP3K6;refGene_tx=XM_019836459.2
C1 20589114 20589191
refGene_gn=MAP3K6;refGene_tx=XM_011284512.3;refGene_gn=MAP3K6;refGene_tx=XM_011284513.3;refGene_gn=MAP3K6;refGene_tx=XM_019836459.2
C1 20589487 20589561
refGene_gn=MAP3K6;refGene_tx=XM_011284512.3;refGene_gn=MAP3K6;refGene_tx=XM_011284513.3;refGene_gn=MAP3K6;refGene_tx=XM_019836459.2
C1 20589647 20589760
refGene_gn=MAP3K6;refGene_tx=XM_011284512.3;refGene_gn=MAP3K6;refGene_tx=XM_011284513.3;refGene_gn=MAP3K6;refGene_tx=XM_019836459.2
C1 20589854 20590026
refGene_gn=MAP3K6;refGene_tx=XM_011284512.3;refGene_gn=MAP3K6;refGene_tx=XM_011284513.3;refGene_gn=MAP3K6;refGene_tx=XM_019836459.2
C1 20590282 20590424
refGene_gn=MAP3K6;refGene_tx=XM_011284512.3;refGene_gn=MAP3K6;refGene_tx=XM_011284513.3;refGene_gn=MAP3K6;refGene_tx=XM_019836459.2
C1 20590703 20590880
refGene_gn=MAP3K6;refGene_tx=XM_011284512.3;refGene_gn=MAP3K6;refGene_tx=XM_011284513.3;refGene_gn=MAP3K6;refGene_tx=XM_019836459.2
C1 20591024 20591184
refGene_gn=MAP3K6;refGene_tx=XM_011284512.3;refGene_gn=MAP3K6;refGene_tx=XM_011284513.3;refGene_gn=MAP3K6;refGene_tx=XM_019836459.2
C1 20591485 20591624
refGene_gn=MAP3K6;refGene_tx=XM_011284512.3;refGene_gn=MAP3K6;refGene_tx=XM_011284513.3;refGene_gn=MAP3K6;refGene_tx=XM_019836459.2
C1 20591731 20591924
refGene_gn=MAP3K6;refGene_tx=XM_011284512.3;refGene_gn=MAP3K6;refGene_tx=XM_011284513.3;refGene_gn=MAP3K6;refGene_tx=XM_019836459.2
C1 20592027 20592185
refGene_gn=MAP3K6;refGene_tx=XM_011284512.3;refGene_gn=MAP3K6;refGene_tx=XM_011284513.3;refGene_gn=MAP3K6;refGene_tx=XM_019836459.2
C1 20592683 20592789
refGene_gn=MAP3K6;refGene_tx=XM_011284512.3;refGene_gn=MAP3K6;refGene_tx=XM_011284513.3;refGene_gn=MAP3K6;refGene_tx=XM_019836459.2
C1 20594259 20594396
refGene_gn=MAP3K6;refGene_tx=XM_011284512.3;refGene_gn=MAP3K6;refGene_tx=XM_011284513.3;refGene_gn=MAP3K6;refGene_tx=XM_019836459.2
C1 20594629 20594778
refGene_gn=MAP3K6;refGene_tx=XM_011284512.3;refGene_gn=MAP3K6;refGene_tx=XM_011284513.3;refGene_gn=MAP3K6;refGene_tx=XM_019836459.2
C1 20595129 20595263
refGene_gn=MAP3K6;refGene_tx=XM_011284512.3;refGene_gn=MAP3K6;refGene_tx=XM_011284513.3;refGene_gn=MAP3K6;refGene_tx=XM_019836459.2
C1 20595357 20595445
refGene_gn=MAP3K6;refGene_tx=XM_011284512.3;refGene_gn=MAP3K6;refGene_tx=XM_011284513.3;refGene_gn=MAP3K6;refGene_tx=XM_019836459.2
C1 20595597 20595693
refGene_gn=MAP3K6;refGene_tx=XM_011284512.3;refGene_gn=MAP3K6;refGene_

tx=XM_011284513.3;refGene_gn=MAP3K6;refGene_tx=XM_019836459.2
C1 20595783 20595833
refGene_gn=MAP3K6;refGene_tx=XM_011284512.3;refGene_gn=MAP3K6;refGene_tx=XM_011284513.3;refGene_gn=MAP3K6;refGene_tx=XM_019836459.2
C1 20595958 20596066
refGene_gn=MAP3K6;refGene_tx=XM_011284512.3;refGene_gn=MAP3K6;refGene_tx=XM_011284513.3;refGene_gn=MAP3K6;refGene_tx=XM_019836459.2
C1 20596145 20596308
refGene_gn=MAP3K6;refGene_tx=XM_011284512.3;refGene_gn=MAP3K6;refGene_tx=XM_011284513.3;refGene_gn=MAP3K6;refGene_tx=XM_019836459.2
C1 20596532 20596692
refGene_gn=MAP3K6;refGene_tx=XM_011284512.3;refGene_gn=MAP3K6;refGene_tx=XM_011284513.3;refGene_gn=MAP3K6;refGene_tx=XM_019836459.2
C1 20597315 20597428
refGene_gn=MAP3K6;refGene_tx=XM_011284512.3;refGene_gn=MAP3K6;refGene_tx=XM_011284513.3;refGene_gn=MAP3K6;refGene_tx=XM_019836459.2
C1 20597513 20597684
refGene_gn=MAP3K6;refGene_tx=XM_011284512.3;refGene_gn=MAP3K6;refGene_tx=XM_011284513.3;refGene_gn=MAP3K6;refGene_tx=XM_019836459.2
C1 20598113 20598220
refGene_gn=MAP3K6;refGene_tx=XM_011284512.3;refGene_gn=MAP3K6;refGene_tx=XM_011284513.3;refGene_gn=MAP3K6;refGene_tx=XM_019836459.2
C1 20598604 20598773
refGene_gn=MAP3K6;refGene_tx=XM_011284512.3;refGene_gn=MAP3K6;refGene_tx=XM_011284513.3;refGene_gn=MAP3K6;refGene_tx=XM_019836459.2
C1 20598888 20599079
refGene_gn=MAP3K6;refGene_tx=XM_011284512.3;refGene_gn=MAP3K6;refGene_tx=XM_011284513.3
C1 20599327 20599351
refGene_gn=MAP3K6;refGene_tx=XM_011284512.3;refGene_gn=MAP3K6;refGene_tx=XM_011284513.3;refGene_gn=MAP3K6;refGene_tx=XM_019836459.2
C1 20599437 20599577
refGene_gn=MAP3K6;refGene_tx=XM_011284512.3;refGene_gn=MAP3K6;refGene_tx=XM_011284513.3
C1 20600827 20601170
refGene_gn=MAP3K6;refGene_tx=XM_011284512.3;refGene_gn=MAP3K6;refGene_tx=XM_011284513.3
C1 20603157 20603399 refGene_gn=FCN3;refGene_tx=XM_003989748.4
C1 20604217 20604352 refGene_gn=FCN3;refGene_tx=XM_003989748.4
C1 20604460 20604476 refGene_gn=FCN3;refGene_tx=XM_003989748.4
C1 20604477 20604591 refGene_gn=FCN3;refGene_tx=XM_003989748.4
C1 20605855 20605983 refGene_gn=FCN3;refGene_tx=XM_003989748.4
C1 20606369 20606414 refGene_gn=FCN3;refGene_tx=XM_003989748.4
C1 20606751 20606846 refGene_gn=FCN3;refGene_tx=XM_003989748.4
C1 20607100 20607125 refGene_gn=FCN3;refGene_tx=XM_003989748.4
C1 20607127 20607179 refGene_gn=FCN3;refGene_tx=XM_003989748.4
C1 20610768 20610775
refGene_gn=CD164L2;refGene_tx=XM_006934460.4;refGene_gn=CD164L2;refGene_tx=XM_019836461.2;refGene_gn=CD164L2;refGene_tx=XM_023258361.1;refGene_gn=CD164L2;refGene_tx=XM_023258362.1

C1 20611324 20611521
refGene_gn=CD164L2;refGene_tx=XM_003989749.4;refGene_gn=CD164L2;refGene_tx=XM_006934460.4;refGene_gn=CD164L2;refGene_tx=XM_019836461.2;refGene_gn=CD164L2;refGene_tx=XM_023258361.1;refGene_gn=CD164L2;refGene_tx=XM_023258362.1
C1 20613025 20613070
refGene_gn=CD164L2;refGene_tx=XM_003989749.4;refGene_gn=CD164L2;refGene_tx=XM_006934460.4;refGene_gn=CD164L2;refGene_tx=XM_019836461.2;refGene_gn=CD164L2;refGene_tx=XM_023258361.1;refGene_gn=CD164L2;refGene_tx=XM_023258362.1
C1 20613676 20613748
refGene_gn=CD164L2;refGene_tx=XM_003989749.4;refGene_gn=CD164L2;refGene_tx=XM_006934460.4;refGene_gn=CD164L2;refGene_tx=XM_019836461.2;refGene_gn=CD164L2;refGene_tx=XM_023258361.1;refGene_gn=CD164L2;refGene_tx=XM_023258362.1
C1 20613848 20614016
refGene_gn=CD164L2;refGene_tx=XM_003989749.4;refGene_gn=CD164L2;refGene_tx=XM_006934460.4;refGene_gn=CD164L2;refGene_tx=XM_019836461.2;refGene_gn=CD164L2;refGene_tx=XM_023258361.1;refGene_gn=CD164L2;refGene_tx=XM_023258362.1
C1 20614479 20614567
refGene_gn=CD164L2;refGene_tx=XM_003989749.4;refGene_gn=CD164L2;refGene_tx=XM_006934460.4;refGene_gn=CD164L2;refGene_tx=XM_019836461.2
C1 20622183 20623176 refGene_gn=GPR3;refGene_tx=XM_023258363.1
C1 20635290 20635448
refGene_gn=WASF2;refGene_tx=XM_006934461.4;refGene_gn=WASF2;refGene_tx=XM_019836460.2
C1 20636868 20637383
refGene_gn=WASF2;refGene_tx=XM_006934461.4;refGene_gn=WASF2;refGene_tx=XM_019836460.2
C1 20639346 20639502
refGene_gn=WASF2;refGene_tx=XM_006934461.4;refGene_gn=WASF2;refGene_tx=XM_019836460.2
C1 20641592 20641723
refGene_gn=WASF2;refGene_tx=XM_006934461.4;refGene_gn=WASF2;refGene_tx=XM_019836460.2
C1 20642749 20642867
refGene_gn=WASF2;refGene_tx=XM_006934461.4;refGene_gn=WASF2;refGene_tx=XM_019836460.2
C1 20645019 20645173
refGene_gn=WASF2;refGene_tx=XM_006934461.4;refGene_gn=WASF2;refGene_tx=XM_019836460.2
C1 20645692 20645827
refGene_gn=WASF2;refGene_tx=XM_006934461.4;refGene_gn=WASF2;refGene_tx=XM_019836460.2
C1 20653115 20653245
refGene_gn=WASF2;refGene_tx=XM_006934461.4;refGene_gn=WASF2;refGene_tx=XM_019836460.2
C1 20750523 20755323
refGene_gn=AHDC1;refGene_tx=XM_006934462.4;refGene_gn=AHDC1;refGene_tx

=XM_006934463.4;refGene_gn=AHDC1;refGene_tx=XM_023258364.1;refGene_gn=AHDC1;refGene_tx=XM_023258365.1;refGene_gn=AHDC1;refGene_tx=XM_023258366.1;refGene_gn=AHDC1;refGene_tx=XM_023258367.1;refGene_gn=AHDC1;refGene_tx=XM_023258368.1
C1 20813151 20813360
refGene_gn=FGR;refGene_tx=XM_006934468.4;refGene_gn=FGR;refGene_tx=XM_011284510.3;refGene_gn=FGR;refGene_tx=XM_011284511.3;refGene_gn=FGR;refGene_tx=XM_019836464.2;refGene_gn=FGR;refGene_tx=XM_019836465.2;refGene_gn=FGR;refGene_tx=XM_023258369.1;refGene_gn=FGR;refGene_tx=XM_023258370.1
C1 20813470 20813602
refGene_gn=FGR;refGene_tx=XM_006934468.4;refGene_gn=FGR;refGene_tx=XM_011284510.3;refGene_gn=FGR;refGene_tx=XM_011284511.3;refGene_gn=FGR;refGene_tx=XM_019836464.2;refGene_gn=FGR;refGene_tx=XM_019836465.2;refGene_gn=FGR;refGene_tx=XM_023258369.1;refGene_gn=FGR;refGene_tx=XM_023258370.1
C1 20814631 20814785
refGene_gn=FGR;refGene_tx=XM_006934468.4;refGene_gn=FGR;refGene_tx=XM_011284510.3;refGene_gn=FGR;refGene_tx=XM_011284511.3;refGene_gn=FGR;refGene_tx=XM_019836464.2;refGene_gn=FGR;refGene_tx=XM_019836465.2;refGene_gn=FGR;refGene_tx=XM_023258369.1;refGene_gn=FGR;refGene_tx=XM_023258370.1
C1 20815031 20815108
refGene_gn=FGR;refGene_tx=XM_006934468.4;refGene_gn=FGR;refGene_tx=XM_011284510.3;refGene_gn=FGR;refGene_tx=XM_011284511.3;refGene_gn=FGR;refGene_tx=XM_019836464.2;refGene_gn=FGR;refGene_tx=XM_019836465.2;refGene_gn=FGR;refGene_tx=XM_023258369.1;refGene_gn=FGR;refGene_tx=XM_023258370.1
C1 20815707 20815887
refGene_gn=FGR;refGene_tx=XM_006934468.4;refGene_gn=FGR;refGene_tx=XM_011284510.3;refGene_gn=FGR;refGene_tx=XM_011284511.3;refGene_gn=FGR;refGene_tx=XM_019836464.2;refGene_gn=FGR;refGene_tx=XM_019836465.2;refGene_gn=FGR;refGene_tx=XM_023258369.1;refGene_gn=FGR;refGene_tx=XM_023258370.1
C1 20815972 20816128
refGene_gn=FGR;refGene_tx=XM_006934468.4;refGene_gn=FGR;refGene_tx=XM_011284510.3;refGene_gn=FGR;refGene_tx=XM_011284511.3;refGene_gn=FGR;refGene_tx=XM_019836464.2;refGene_gn=FGR;refGene_tx=XM_019836465.2;refGene_gn=FGR;refGene_tx=XM_023258369.1;refGene_gn=FGR;refGene_tx=XM_023258370.1
C1 20816830 20816980
refGene_gn=FGR;refGene_tx=XM_006934468.4;refGene_gn=FGR;refGene_tx=XM_011284510.3;refGene_gn=FGR;refGene_tx=XM_011284511.3;refGene_gn=FGR;refGene_tx=XM_019836464.2;refGene_gn=FGR;refGene_tx=XM_019836465.2;refGene_gn=FGR;refGene_tx=XM_023258369.1;refGene_gn=FGR;refGene_tx=XM_023258370.1
C1 20817227 20817331
refGene_gn=FGR;refGene_tx=XM_006934468.4;refGene_gn=FGR;refGene_tx=XM_011284510.3;refGene_gn=FGR;refGene_tx=XM_011284511.3;refGene_gn=FGR;refGene_tx=XM_019836464.2;refGene_gn=FGR;refGene_tx=XM_019836465.2;refGe

ne_gn=FGR;refGene_tx=XM_023258369.1;refGene_gn=FGR;refGene_tx=XM_02325
8370.1
C1 20820855 20820954
refGene_gn=FGR;refGene_tx=XM_006934468.4;refGene_gn=FGR;refGene_tx=XM_
011284510.3;refGene_gn=FGR;refGene_tx=XM_011284511.3;refGene_gn=FGR;re
fGene_tx=XM_019836464.2;refGene_gn=FGR;refGene_tx=XM_019836465.2;refGe
ne_gn=FGR;refGene_tx=XM_023258369.1;refGene_gn=FGR;refGene_tx=XM_02325
8370.1
C1 20821771 20821874
refGene_gn=FGR;refGene_tx=XM_006934468.4;refGene_gn=FGR;refGene_tx=XM_
011284510.3;refGene_gn=FGR;refGene_tx=XM_011284511.3;refGene_gn=FGR;re
fGene_tx=XM_019836464.2;refGene_gn=FGR;refGene_tx=XM_019836465.2;refGe
ne_gn=FGR;refGene_tx=XM_023258369.1;refGene_gn=FGR;refGene_tx=XM_02325
8370.1
C1 20822367 20822587
refGene_gn=FGR;refGene_tx=XM_006934468.4;refGene_gn=FGR;refGene_tx=XM_
011284510.3;refGene_gn=FGR;refGene_tx=XM_011284511.3;refGene_gn=FGR;re
fGene_tx=XM_019836464.2;refGene_gn=FGR;refGene_tx=XM_019836465.2;refGe
ne_gn=FGR;refGene_tx=XM_023258369.1;refGene_gn=FGR;refGene_tx=XM_02325
8370.1
C1 20841351 20842000
refGene_gn=LOC101084040;refGene_tx=XM_019836466.2
C1 20842002 20842085
refGene_gn=LOC101084040;refGene_tx=XM_019836466.2
C1 20858512 20858628
refGene_gn=IFI6;refGene_tx=XM_006934470.2;refGene_gn=IFI6;refGene_tx=X
M_019836467.2
C1 20860616 20860766
refGene_gn=IFI6;refGene_tx=XM_006934470.2;refGene_gn=IFI6;refGene_tx=X
M_019836467.2
C1 20860845 20860932
refGene_gn=IFI6;refGene_tx=XM_006934470.2;refGene_gn=IFI6;refGene_tx=X
M_019836467.2
C1 20861272 20861368
refGene_gn=IFI6;refGene_tx=XM_006934470.2;refGene_gn=IFI6;refGene_tx=X
M_019836467.2
C1 20862377 20862378 refGene_gn=IFI6;refGene_tx=XM_019836467.2
C1 20888598 20888679 refGene_gn=FAM76A;refGene_tx=XM_011284509.3
C1 20889989 20890054 refGene_gn=FAM76A;refGene_tx=XM_011284509.3
C1 20893338 20893393 refGene_gn=FAM76A;refGene_tx=XM_011284509.3
C1 20894136 20894289 refGene_gn=FAM76A;refGene_tx=XM_011284509.3
C1 20896985 20897143 refGene_gn=FAM76A;refGene_tx=XM_011284509.3
C1 20901360 20901447 refGene_gn=FAM76A;refGene_tx=XM_011284509.3
C1 20906934 20907070 refGene_gn=FAM76A;refGene_tx=XM_011284509.3
C1 20910003 20910105 refGene_gn=FAM76A;refGene_tx=XM_011284509.3
C1 20910853 20910940 refGene_gn=FAM76A;refGene_tx=XM_011284509.3
C1 20920521 20920639 refGene_gn=STX12;refGene_tx=XM_003989756.5
C1 20935570 20935640 refGene_gn=STX12;refGene_tx=XM_003989756.5
C1 20937158 20937258 refGene_gn=STX12;refGene_tx=XM_003989756.5
C1 20945941 20946079 refGene_gn=STX12;refGene_tx=XM_003989756.5

C1 20952034 20952078 refGene_gn=STX12;refGene_tx=XM_003989756.5
C1 20953395 20953501 refGene_gn=STX12;refGene_tx=XM_003989756.5
C1 20958278 20958351 refGene_gn=STX12;refGene_tx=XM_003989756.5
C1 20959567 20959650 refGene_gn=STX12;refGene_tx=XM_003989756.5
C1 20960936 20961029 refGene_gn=STX12;refGene_tx=XM_003989756.5
C1 20970574 20970630 refGene_gn=PPP1R8;refGene_tx=XM_003989757.5
C1 20972523 20972584 refGene_gn=PPP1R8;refGene_tx=XM_003989757.5
C1 20978413 20978567 refGene_gn=PPP1R8;refGene_tx=XM_003989757.5
C1 20979873 20980094
refGene_gn=PPP1R8;refGene_tx=XM_003989757.5;refGene_gn=PPP1R8;refGene_
tx=XM_006934471.3
C1 20981623 20981768
refGene_gn=PPP1R8;refGene_tx=XM_003989757.5;refGene_gn=PPP1R8;refGene_
tx=XM_006934471.3
C1 20987576 20987641
refGene_gn=PPP1R8;refGene_tx=XM_003989757.5;refGene_gn=PPP1R8;refGene_
tx=XM_006934471.3
C1 20989954 20990308
refGene_gn=PPP1R8;refGene_tx=XM_003989757.5;refGene_gn=PPP1R8;refGene_
tx=XM_006934471.3
C1 21008659 21008753 refGene_gn=THEMIS2;refGene_tx=XM_023258371.1
C1 21010918 21011059 refGene_gn=THEMIS2;refGene_tx=XM_023258371.1
C1 21013563 21013974 refGene_gn=THEMIS2;refGene_tx=XM_023258371.1
C1 21018207 21019274 refGene_gn=THEMIS2;refGene_tx=XM_023258371.1
C1 21021494 21021651 refGene_gn=THEMIS2;refGene_tx=XM_023258371.1
C1 21022083 21022139 refGene_gn=THEMIS2;refGene_tx=XM_023258371.1
C1 21026055 21026140 refGene_gn=RPA2;refGene_tx=XM_006934472.4
C1 21027335 21027430 refGene_gn=RPA2;refGene_tx=XM_006934472.4
C1 21027845 21027953 refGene_gn=RPA2;refGene_tx=XM_006934472.4
C1 21033857 21033974 refGene_gn=RPA2;refGene_tx=XM_006934472.4
C1 21034481 21034556 refGene_gn=RPA2;refGene_tx=XM_006934472.4
C1 21036724 21036838 refGene_gn=RPA2;refGene_tx=XM_006934472.4
C1 21036967 21037069 refGene_gn=RPA2;refGene_tx=XM_006934472.4
C1 21041243 21041350 refGene_gn=RPA2;refGene_tx=XM_006934472.4
C1 21041587 21041597 refGene_gn=RPA2;refGene_tx=XM_006934472.4
C1 21052512 21052573
refGene_gn=SMPDL3B;refGene_tx=XM_003989760.5;refGene_gn=SMPDL3B;refGen
e_tx=XM_019836469.2
C1 21061268 21061482 refGene_gn=SMPDL3B;refGene_tx=XM_003989760.5
C1 21066205 21066303 refGene_gn=SMPDL3B;refGene_tx=XM_003989760.5
C1 21070259 21070403
refGene_gn=SMPDL3B;refGene_tx=XM_003989760.5;refGene_gn=SMPDL3B;refGen
e_tx=XM_019836469.2
C1 21071369 21071542
refGene_gn=SMPDL3B;refGene_tx=XM_003989760.5;refGene_gn=SMPDL3B;refGen
e_tx=XM_019836469.2
C1 21072381 21072562
refGene_gn=SMPDL3B;refGene_tx=XM_003989760.5;refGene_gn=SMPDL3B;refGen
e_tx=XM_019836469.2
C1 21072647 21072781

refGene_gn=SMPDL3B;refGene_tx=XM_003989760.5;refGene_gn=SMPDL3B;refGene_tx=XM_019836469.2
C1	21075200	21075566	refGene_gn=SMPDL3B;refGene_tx=XM_003989760.5;refGene_gn=SMPDL3B;refGene_tx=XM_019836469.2
C1	21076429	21076722	refGene_gn=XKR8;refGene_tx=XM_023258372.1
C1	21078906	21079103	refGene_gn=XKR8;refGene_tx=XM_023258372.1
C1	21080684	21081394	refGene_gn=XKR8;refGene_tx=XM_023258372.1
C1	21088654	21088735	refGene_gn=EYA3;refGene_tx=XM_003989761.5;refGene_gn=EYA3;refGene_tx=XM_006934473.4;refGene_gn=EYA3;refGene_tx=XM_019836470.2
C1	21092424	21092525	refGene_gn=EYA3;refGene_tx=XM_003989761.5;refGene_gn=EYA3;refGene_tx=XM_006934473.4;refGene_gn=EYA3;refGene_tx=XM_019836470.2
C1	21100991	21101113	refGene_gn=EYA3;refGene_tx=XM_003989761.5;refGene_gn=EYA3;refGene_tx=XM_006934473.4;refGene_gn=EYA3;refGene_tx=XM_019836470.2
C1	21102140	21102255	refGene_gn=EYA3;refGene_tx=XM_003989761.5;refGene_gn=EYA3;refGene_tx=XM_006934473.4;refGene_gn=EYA3;refGene_tx=XM_019836470.2
C1	21105053	21105214	refGene_gn=EYA3;refGene_tx=XM_003989761.5;refGene_gn=EYA3;refGene_tx=XM_006934473.4;refGene_gn=EYA3;refGene_tx=XM_019836470.2
C1	21107341	21107400	refGene_gn=EYA3;refGene_tx=XM_003989761.5;refGene_gn=EYA3;refGene_tx=XM_006934473.4;refGene_gn=EYA3;refGene_tx=XM_019836470.2
C1	21109673	21109763	refGene_gn=EYA3;refGene_tx=XM_003989761.5;refGene_gn=EYA3;refGene_tx=XM_006934473.4;refGene_gn=EYA3;refGene_tx=XM_019836470.2
C1	21111413	21111497	refGene_gn=EYA3;refGene_tx=XM_003989761.5;refGene_gn=EYA3;refGene_tx=XM_006934473.4;refGene_gn=EYA3;refGene_tx=XM_019836470.2
C1	21116729	21116869	refGene_gn=EYA3;refGene_tx=XM_003989761.5;refGene_gn=EYA3;refGene_tx=XM_006934473.4;refGene_gn=EYA3;refGene_tx=XM_019836470.2
C1	21118805	21118989	refGene_gn=EYA3;refGene_tx=XM_003989761.5;refGene_gn=EYA3;refGene_tx=XM_006934473.4;refGene_gn=EYA3;refGene_tx=XM_019836470.2
C1	21122001	21122087	refGene_gn=EYA3;refGene_tx=XM_003989761.5;refGene_gn=EYA3;refGene_tx=XM_006934473.4;refGene_gn=EYA3;refGene_tx=XM_019836470.2
C1	21129645	21129783	refGene_gn=EYA3;refGene_tx=XM_006934473.4;refGene_gn=EYA3;refGene_tx=XM_019836470.2
C1	21137455	21137592	refGene_gn=EYA3;refGene_tx=XM_003989761.5;refGene_gn=EYA3;refGene_tx=XM_006934473.4;refGene_gn=EYA3;refGene_tx=XM_019836470.2
C1	21140066	21140133	refGene_gn=EYA3;refGene_tx=XM_003989761.5;refGene_gn=EYA3;refGene_tx=XM_006934473.4;refGene_gn=EYA3;refGene_tx=XM_019836470.2

C1 21145650 21145730
refGene_gn=EYA3;refGene_tx=XM_003989761.5;refGene_gn=EYA3;refGene_tx=XM_006934473.4;refGene_gn=EYA3;refGene_tx=XM_019836470.2
C1 21149855 21149899
refGene_gn=EYA3;refGene_tx=XM_003989761.5;refGene_gn=EYA3;refGene_tx=XM_006934473.4;refGene_gn=EYA3;refGene_tx=XM_019836470.2
C1 21155875 21155908
refGene_gn=EYA3;refGene_tx=XM_003989761.5;refGene_gn=EYA3;refGene_tx=XM_006934473.4;refGene_gn=EYA3;refGene_tx=XM_019836470.2
C1 21239085 21240114
refGene_gn=PTAFR;refGene_tx=XM_006934474.4;refGene_gn=PTAFR;refGene_tx=XM_019836495.2
C1 21274651 21274774 refGene_gn=DNAJC8;refGene_tx=XM_006934475.3
C1 21275841 21275917 refGene_gn=DNAJC8;refGene_tx=XM_006934475.3
C1 21276782 21276874 refGene_gn=DNAJC8;refGene_tx=XM_006934475.3
C1 21281528 21281600 refGene_gn=DNAJC8;refGene_tx=XM_006934475.3
C1 21283187 21283282 refGene_gn=DNAJC8;refGene_tx=XM_006934475.3
C1 21283803 21283870 refGene_gn=DNAJC8;refGene_tx=XM_006934475.3
C1 21287776 21287833 refGene_gn=DNAJC8;refGene_tx=XM_006934475.3
C1 21299053 21299155 refGene_gn=DNAJC8;refGene_tx=XM_006934475.3
C1 21301665 21301743 refGene_gn=DNAJC8;refGene_tx=XM_006934475.3
C1 21303883 21303970 refGene_gn=ATP5IF1;refGene_tx=XM_003989765.4
C1 21304059 21304145 refGene_gn=ATP5IF1;refGene_tx=XM_003989765.4
C1 21306030 21306184 refGene_gn=ATP5IF1;refGene_tx=XM_003989765.4
C1 21325724 21325820 refGene_gn=SESN2;refGene_tx=XM_011284508.3
C1 21332268 21332334 refGene_gn=SESN2;refGene_tx=XM_011284508.3
C1 21334183 21334381 refGene_gn=SESN2;refGene_tx=XM_011284508.3
C1 21334958 21335141 refGene_gn=SESN2;refGene_tx=XM_011284508.3
C1 21335255 21335468 refGene_gn=SESN2;refGene_tx=XM_011284508.3
C1 21336022 21336173 refGene_gn=SESN2;refGene_tx=XM_011284508.3
C1 21336691 21336810 refGene_gn=SESN2;refGene_tx=XM_011284508.3
C1 21337173 21337364 refGene_gn=SESN2;refGene_tx=XM_011284508.3
C1 21340092 21340237 refGene_gn=SESN2;refGene_tx=XM_011284508.3
C1 21340809 21340896 refGene_gn=SESN2;refGene_tx=XM_011284508.3
C1 21374857 21374930
refGene_gn=LOC101086663;refGene_tx=XM_003989766.5;refGene_gn=LOC101086663;refGene_tx=XM_019836480.2;refGene_gn=LOC101086663;refGene_tx=XM_019836481.2;refGene_gn=LOC101086663;refGene_tx=XM_019836482.2;refGene_gn=LOC101086663;refGene_tx=XM_023258373.1;refGene_gn=LOC101086663;refGene_tx=XM_023258374.1;refGene_gn=LOC101086663;refGene_tx=XM_023258375.1
C1 21381316 21381870
refGene_gn=LOC101086663;refGene_tx=XM_003989766.5
C1 21432594 21432648
refGene_gn=LOC101086663;refGene_tx=XM_011284503.3;refGene_gn=LOC101086663;refGene_tx=XM_011284505.3;refGene_gn=LOC101086663;refGene_tx=XM_019836480.2;refGene_gn=LOC101086663;refGene_tx=XM_019836481.2;refGene_gn=LOC101086663;refGene_tx=XM_019836482.2;refGene_gn=LOC101086663;refGene_tx=XM_023258373.1;refGene_gn=LOC101086663;refGene_tx=XM_023258374.1;refGene_gn=LOC101086663;refGene_tx=XM_023258375.1
C1 21463882 21463928

refGene_gn=LOC101086663;refGene_tx=XM_011284502.3
C1 21478033 21478057
refGene_gn=LOC101086663;refGene_tx=XM_011284503.3;refGene_gn=LOC101086663;refGene_tx=XM_019836480.2;refGene_gn=LOC101086663;refGene_tx=XM_019836481.2;refGene_gn=LOC101086663;refGene_tx=XM_023258373.1;refGene_gn=LOC101086663;refGene_tx=XM_023258374.1;refGene_gn=LOC101086663;refGene_tx=XM_023258375.1
C1 21479474 21479648
refGene_gn=LOC101086663;refGene_tx=XM_011284502.3;refGene_gn=LOC101086663;refGene_tx=XM_011284503.3;refGene_gn=LOC101086663;refGene_tx=XM_011284505.3;refGene_gn=LOC101086663;refGene_tx=XM_019836480.2;refGene_gn=LOC101086663;refGene_tx=XM_019836481.2;refGene_gn=LOC101086663;refGene_tx=XM_019836482.2;refGene_gn=LOC101086663;refGene_tx=XM_023258373.1;refGene_gn=LOC101086663;refGene_tx=XM_023258374.1;refGene_gn=LOC101086663;refGene_tx=XM_023258375.1;refGene_gn=LOC101086663;refGene_tx=XM_023258376.1;refGene_gn=LOC101086663;refGene_tx=XM_023258377.1
C1 21480071 21480152
refGene_gn=LOC101086663;refGene_tx=XM_011284502.3;refGene_gn=LOC101086663;refGene_tx=XM_011284503.3;refGene_gn=LOC101086663;refGene_tx=XM_011284505.3;refGene_gn=LOC101086663;refGene_tx=XM_019836480.2;refGene_gn=LOC101086663;refGene_tx=XM_019836481.2;refGene_gn=LOC101086663;refGene_tx=XM_019836482.2;refGene_gn=LOC101086663;refGene_tx=XM_023258373.1;refGene_gn=LOC101086663;refGene_tx=XM_023258374.1;refGene_gn=LOC101086663;refGene_tx=XM_023258375.1;refGene_gn=LOC101086663;refGene_tx=XM_023258376.1;refGene_gn=LOC101086663;refGene_tx=XM_023258377.1
C1 21484550 21484715
refGene_gn=LOC101086663;refGene_tx=XM_011284502.3;refGene_gn=LOC101086663;refGene_tx=XM_011284503.3;refGene_gn=LOC101086663;refGene_tx=XM_011284505.3;refGene_gn=LOC101086663;refGene_tx=XM_019836480.2;refGene_gn=LOC101086663;refGene_tx=XM_019836481.2;refGene_gn=LOC101086663;refGene_tx=XM_019836482.2;refGene_gn=LOC101086663;refGene_tx=XM_023258373.1;refGene_gn=LOC101086663;refGene_tx=XM_023258374.1;refGene_gn=LOC101086663;refGene_tx=XM_023258375.1;refGene_gn=LOC101086663;refGene_tx=XM_023258376.1;refGene_gn=LOC101086663;refGene_tx=XM_023258377.1
C1 21485249 21485645
refGene_gn=LOC101086663;refGene_tx=XM_011284502.3;refGene_gn=LOC101086663;refGene_tx=XM_011284503.3;refGene_gn=LOC101086663;refGene_tx=XM_011284505.3;refGene_gn=LOC101086663;refGene_tx=XM_019836480.2;refGene_gn=LOC101086663;refGene_tx=XM_019836481.2;refGene_gn=LOC101086663;refGene_tx=XM_019836482.2;refGene_gn=LOC101086663;refGene_tx=XM_023258373.1;refGene_gn=LOC101086663;refGene_tx=XM_023258374.1;refGene_gn=LOC101086663;refGene_tx=XM_023258375.1;refGene_gn=LOC101086663;refGene_tx=XM_023258376.1;refGene_gn=LOC101086663;refGene_tx=XM_023258377.1
C1 21490032 21490633
refGene_gn=LOC101086663;refGene_tx=XM_011284502.3;refGene_gn=LOC101086663;refGene_tx=XM_011284503.3;refGene_gn=LOC101086663;refGene_tx=XM_011284505.3;refGene_gn=LOC101086663;refGene_tx=XM_019836480.2;refGene_gn=LOC101086663;refGene_tx=XM_019836481.2;refGene_gn=LOC101086663;refGene_tx=XM_019836482.2;refGene_gn=LOC101086663;refGene_tx=XM_023258373.1;refGene_gn=LOC101086663;refGene_tx=XM_023258374.1;refGene_gn=LOC101086

663;refGene_tx=XM_023258375.1;refGene_gn=LOC101086663;refGene_tx=XM_02
3258376.1;refGene_gn=LOC101086663;refGene_tx=XM_023258377.1
C1 21491962 21492150
refGene_gn=LOC101086663;refGene_tx=XM_011284502.3;refGene_gn=LOC101086
663;refGene_tx=XM_011284503.3;refGene_gn=LOC101086663;refGene_tx=XM_01
1284505.3;refGene_gn=LOC101086663;refGene_tx=XM_019836480.2;refGene_gn
=LOC101086663;refGene_tx=XM_019836481.2;refGene_gn=LOC101086663;refGen
e_tx=XM_019836482.2;refGene_gn=LOC101086663;refGene_tx=XM_023258373.1;
refGene_gn=LOC101086663;refGene_tx=XM_023258374.1;refGene_gn=LOC101086
663;refGene_tx=XM_023258375.1;refGene_gn=LOC101086663;refGene_tx=XM_02
3258376.1;refGene_gn=LOC101086663;refGene_tx=XM_023258377.1
C1 21493479 21493633
refGene_gn=LOC101086663;refGene_tx=XM_011284502.3;refGene_gn=LOC101086
663;refGene_tx=XM_011284503.3;refGene_gn=LOC101086663;refGene_tx=XM_01
1284505.3;refGene_gn=LOC101086663;refGene_tx=XM_019836480.2;refGene_gn
=LOC101086663;refGene_tx=XM_019836481.2;refGene_gn=LOC101086663;refGen
e_tx=XM_019836482.2;refGene_gn=LOC101086663;refGene_tx=XM_023258373.1;
refGene_gn=LOC101086663;refGene_tx=XM_023258374.1;refGene_gn=LOC101086
663;refGene_tx=XM_023258375.1;refGene_gn=LOC101086663;refGene_tx=XM_02
3258376.1;refGene_gn=LOC101086663;refGene_tx=XM_023258377.1
C1 21498490 21498546
refGene_gn=LOC101086663;refGene_tx=XM_011284502.3;refGene_gn=LOC101086
663;refGene_tx=XM_011284503.3;refGene_gn=LOC101086663;refGene_tx=XM_01
1284505.3;refGene_gn=LOC101086663;refGene_tx=XM_019836480.2;refGene_gn
=LOC101086663;refGene_tx=XM_019836481.2;refGene_gn=LOC101086663;refGen
e_tx=XM_019836482.2;refGene_gn=LOC101086663;refGene_tx=XM_023258373.1;
refGene_gn=LOC101086663;refGene_tx=XM_023258374.1;refGene_gn=LOC101086
663;refGene_tx=XM_023258375.1;refGene_gn=LOC101086663;refGene_tx=XM_02
3258376.1;refGene_gn=LOC101086663;refGene_tx=XM_023258377.1
C1 21499888 21499950
refGene_gn=LOC101086663;refGene_tx=XM_011284502.3;refGene_gn=LOC101086
663;refGene_tx=XM_011284503.3;refGene_gn=LOC101086663;refGene_tx=XM_01
1284505.3;refGene_gn=LOC101086663;refGene_tx=XM_019836480.2;refGene_gn
=LOC101086663;refGene_tx=XM_019836481.2;refGene_gn=LOC101086663;refGen
e_tx=XM_019836482.2;refGene_gn=LOC101086663;refGene_tx=XM_023258373.1;
refGene_gn=LOC101086663;refGene_tx=XM_023258374.1;refGene_gn=LOC101086
663;refGene_tx=XM_023258375.1;refGene_gn=LOC101086663;refGene_tx=XM_02
3258376.1;refGene_gn=LOC101086663;refGene_tx=XM_023258377.1
C1 21502001 21502139
refGene_gn=LOC101086663;refGene_tx=XM_011284502.3;refGene_gn=LOC101086
663;refGene_tx=XM_011284503.3;refGene_gn=LOC101086663;refGene_tx=XM_01
1284505.3;refGene_gn=LOC101086663;refGene_tx=XM_019836480.2;refGene_gn
=LOC101086663;refGene_tx=XM_019836481.2;refGene_gn=LOC101086663;refGen
e_tx=XM_019836482.2;refGene_gn=LOC101086663;refGene_tx=XM_023258373.1;
refGene_gn=LOC101086663;refGene_tx=XM_023258374.1;refGene_gn=LOC101086
663;refGene_tx=XM_023258375.1;refGene_gn=LOC101086663;refGene_tx=XM_02
3258376.1;refGene_gn=LOC101086663;refGene_tx=XM_023258377.1
C1 21502840 21502917
refGene_gn=LOC101086663;refGene_tx=XM_011284502.3;refGene_gn=LOC101086
663;refGene_tx=XM_011284503.3;refGene_gn=LOC101086663;refGene_tx=XM_01

1284505.3;refGene_gn=LOC101086663;refGene_tx=XM_019836480.2;refGene_gn
=LOC101086663;refGene_tx=XM_019836481.2;refGene_gn=LOC101086663;refGen
e_tx=XM_019836482.2;refGene_gn=LOC101086663;refGene_tx=XM_023258373.1;
refGene_gn=LOC101086663;refGene_tx=XM_023258374.1;refGene_gn=LOC101086
663;refGene_tx=XM_023258375.1;refGene_gn=LOC101086663;refGene_tx=XM_02
3258376.1;refGene_gn=LOC101086663;refGene_tx=XM_023258377.1
C1 21506136 21506152
refGene_gn=LOC101086663;refGene_tx=XM_011284502.3;refGene_gn=LOC101086
663;refGene_tx=XM_011284503.3;refGene_gn=LOC101086663;refGene_tx=XM_01
1284505.3;refGene_gn=LOC101086663;refGene_tx=XM_019836480.2;refGene_gn
=LOC101086663;refGene_tx=XM_019836481.2;refGene_gn=LOC101086663;refGen
e_tx=XM_019836482.2;refGene_gn=LOC101086663;refGene_tx=XM_023258373.1;
refGene_gn=LOC101086663;refGene_tx=XM_023258374.1;refGene_gn=LOC101086
663;refGene_tx=XM_023258375.1;refGene_gn=LOC101086663;refGene_tx=XM_02
3258376.1;refGene_gn=LOC101086663;refGene_tx=XM_023258377.1
C1 21535510 21535583
refGene_gn=RCC1;refGene_tx=XM_003989769.4;refGene_gn=RCC1;refGene_tx=X
M_019836489.2;refGene_gn=RCC1;refGene_tx=XM_019836492.2;refGene_gn=RCC
1;refGene_tx=XM_019836494.2;refGene_gn=RCC1;refGene_tx=XM_023258378.1;
refGene_gn=RCC1;refGene_tx=XM_023258379.1;refGene_gn=RCC1;refGene_tx=X
M_023258380.1;refGene_gn=RCC1;refGene_tx=XM_023258381.1
C1 21536157 21536247
refGene_gn=RCC1;refGene_tx=XM_003989769.4;refGene_gn=RCC1;refGene_tx=X
M_019836489.2;refGene_gn=RCC1;refGene_tx=XM_023258378.1;refGene_gn=RCC
1;refGene_tx=XM_023258379.1
C1 21537137 21537325
refGene_gn=RCC1;refGene_tx=XM_003989769.4;refGene_gn=RCC1;refGene_tx=X
M_019836489.2;refGene_gn=RCC1;refGene_tx=XM_019836492.2;refGene_gn=RCC
1;refGene_tx=XM_019836494.2;refGene_gn=RCC1;refGene_tx=XM_023258378.1;
refGene_gn=RCC1;refGene_tx=XM_023258379.1;refGene_gn=RCC1;refGene_tx=X
M_023258380.1;refGene_gn=RCC1;refGene_tx=XM_023258381.1
C1 21537503 21537683
refGene_gn=RCC1;refGene_tx=XM_003989769.4;refGene_gn=RCC1;refGene_tx=X
M_019836489.2;refGene_gn=RCC1;refGene_tx=XM_019836492.2;refGene_gn=RCC
1;refGene_tx=XM_019836494.2;refGene_gn=RCC1;refGene_tx=XM_023258378.1;
refGene_gn=RCC1;refGene_tx=XM_023258379.1;refGene_gn=RCC1;refGene_tx=X
M_023258380.1;refGene_gn=RCC1;refGene_tx=XM_023258381.1
C1 21538511 21538608
refGene_gn=RCC1;refGene_tx=XM_003989769.4;refGene_gn=RCC1;refGene_tx=X
M_019836489.2;refGene_gn=RCC1;refGene_tx=XM_019836492.2;refGene_gn=RCC
1;refGene_tx=XM_019836494.2;refGene_gn=RCC1;refGene_tx=XM_023258378.1;
refGene_gn=RCC1;refGene_tx=XM_023258379.1;refGene_gn=RCC1;refGene_tx=X
M_023258380.1;refGene_gn=RCC1;refGene_tx=XM_023258381.1
C1 21538722 21538845
refGene_gn=RCC1;refGene_tx=XM_003989769.4;refGene_gn=RCC1;refGene_tx=X
M_019836489.2;refGene_gn=RCC1;refGene_tx=XM_019836492.2;refGene_gn=RCC
1;refGene_tx=XM_019836494.2;refGene_gn=RCC1;refGene_tx=XM_023258378.1;
refGene_gn=RCC1;refGene_tx=XM_023258379.1;refGene_gn=RCC1;refGene_tx=X
M_023258380.1;refGene_gn=RCC1;refGene_tx=XM_023258381.1
C1 21539336 21539492

refGene_gn=RCC1;refGene_tx=XM_003989769.4;refGene_gn=RCC1;refGene_tx=XM_019836489.2;refGene_gn=RCC1;refGene_tx=XM_019836492.2;refGene_gn=RCC1;refGene_tx=XM_019836494.2;refGene_gn=RCC1;refGene_tx=XM_023258378.1;refGene_gn=RCC1;refGene_tx=XM_023258379.1;refGene_gn=RCC1;refGene_tx=XM_023258380.1;refGene_gn=RCC1;refGene_tx=XM_023258381.1
C1 21539746 21539866
refGene_gn=RCC1;refGene_tx=XM_003989769.4;refGene_gn=RCC1;refGene_tx=XM_019836489.2;refGene_gn=RCC1;refGene_tx=XM_019836492.2;refGene_gn=RCC1;refGene_tx=XM_019836494.2;refGene_gn=RCC1;refGene_tx=XM_023258378.1;refGene_gn=RCC1;refGene_tx=XM_023258379.1;refGene_gn=RCC1;refGene_tx=XM_023258380.1;refGene_gn=RCC1;refGene_tx=XM_023258381.1
C1 21540186 21540339
refGene_gn=RCC1;refGene_tx=XM_003989769.4;refGene_gn=RCC1;refGene_tx=XM_019836489.2;refGene_gn=RCC1;refGene_tx=XM_019836492.2;refGene_gn=RCC1;refGene_tx=XM_019836494.2;refGene_gn=RCC1;refGene_tx=XM_023258378.1;refGene_gn=RCC1;refGene_tx=XM_023258379.1;refGene_gn=RCC1;refGene_tx=XM_023258380.1;refGene_gn=RCC1;refGene_tx=XM_023258381.1
C1 21541265 21541441
refGene_gn=RCC1;refGene_tx=XM_003989769.4;refGene_gn=RCC1;refGene_tx=XM_019836489.2;refGene_gn=RCC1;refGene_tx=XM_019836492.2;refGene_gn=RCC1;refGene_tx=XM_019836494.2;refGene_gn=RCC1;refGene_tx=XM_023258378.1;refGene_gn=RCC1;refGene_tx=XM_023258379.1;refGene_gn=RCC1;refGene_tx=XM_023258380.1;refGene_gn=RCC1;refGene_tx=XM_023258381.1
C1 21546635 21546662
refGene_gn=TRNAU1AP;refGene_tx=XM_003989770.5;refGene_gn=TRNAU1AP;refGene_tx=XM_006934485.4;refGene_gn=TRNAU1AP;refGene_tx=XM_019836499.2
C1 21547175 21547273
refGene_gn=TRNAU1AP;refGene_tx=XM_003989770.5;refGene_gn=TRNAU1AP;refGene_tx=XM_006934485.4;refGene_gn=TRNAU1AP;refGene_tx=XM_006934486.4;refGene_gn=TRNAU1AP;refGene_tx=XM_019836499.2
C1 21554485 21554585
refGene_gn=TRNAU1AP;refGene_tx=XM_003989770.5;refGene_gn=TRNAU1AP;refGene_tx=XM_006934485.4;refGene_gn=TRNAU1AP;refGene_tx=XM_006934486.4
C1 21555204 21555257
refGene_gn=TRNAU1AP;refGene_tx=XM_003989770.5;refGene_gn=TRNAU1AP;refGene_tx=XM_006934485.4;refGene_gn=TRNAU1AP;refGene_tx=XM_006934486.4
C1 21558340 21558472
refGene_gn=TRNAU1AP;refGene_tx=XM_003989770.5;refGene_gn=TRNAU1AP;refGene_tx=XM_006934485.4;refGene_gn=TRNAU1AP;refGene_tx=XM_006934486.4;refGene_gn=TRNAU1AP;refGene_tx=XM_019836499.2
C1 21559500 21559620
refGene_gn=TRNAU1AP;refGene_tx=XM_003989770.5;refGene_gn=TRNAU1AP;refGene_tx=XM_006934485.4;refGene_gn=TRNAU1AP;refGene_tx=XM_006934486.4;refGene_gn=TRNAU1AP;refGene_tx=XM_019836499.2
C1 21564192 21564355
refGene_gn=TRNAU1AP;refGene_tx=XM_003989770.5;refGene_gn=TRNAU1AP;refGene_tx=XM_006934485.4;refGene_gn=TRNAU1AP;refGene_tx=XM_006934486.4;refGene_gn=TRNAU1AP;refGene_tx=XM_019836499.2
C1 21564773 21564807
refGene_gn=TRNAU1AP;refGene_tx=XM_003989770.5;refGene_gn=TRNAU1AP;refG

ene_tx=XM_006934485.4;refGene_gn=TRNAU1AP;refGene_tx=XM_006934486.4;refGene_gn=TRNAU1AP;refGene_tx=XM_019836499.2
C1 21569730 21569799
refGene_gn=TRNAU1AP;refGene_tx=XM_006934485.4;refGene_gn=TRNAU1AP;refGene_tx=XM_006934486.4;refGene_gn=TRNAU1AP;refGene_tx=XM_019836499.2
C1 21572520 21572657
refGene_gn=TRNAU1AP;refGene_tx=XM_003989770.5;refGene_gn=TRNAU1AP;refGene_tx=XM_006934485.4;refGene_gn=TRNAU1AP;refGene_tx=XM_006934486.4;refGene_gn=TRNAU1AP;refGene_tx=XM_019836499.2
C1 21583140 21583397 refGene_gn=RAB42;refGene_tx=XM_023258382.1
C1 21583999 21584423 refGene_gn=RAB42;refGene_tx=XM_023258382.1
C1 21591582 21591618
refGene_gn=TAF12;refGene_tx=XM_003989771.4;refGene_gn=TAF12;refGene_tx=XM_006934487.4
C1 21593099 21593188
refGene_gn=TAF12;refGene_tx=XM_003989771.4;refGene_gn=TAF12;refGene_tx=XM_006934487.4
C1 21599122 21599237
refGene_gn=TAF12;refGene_tx=XM_003989771.4;refGene_gn=TAF12;refGene_tx=XM_006934487.4
C1 21604850 21604928
refGene_gn=TAF12;refGene_tx=XM_003989771.4;refGene_gn=TAF12;refGene_tx=XM_006934487.4
C1 21607973 21608225
refGene_gn=TAF12;refGene_tx=XM_003989771.4;refGene_gn=TAF12;refGene_tx=XM_006934487.4
C1 21608945 21609017 refGene_gn=TAF12;refGene_tx=XM_006934487.4
C1 21652010 21652138
refGene_gn=GMEB1;refGene_tx=XM_003989773.5;refGene_gn=GMEB1;refGene_tx=XM_003989774.5;refGene_gn=GMEB1;refGene_tx=XM_006934488.4;refGene_gn=GMEB1;refGene_tx=XM_006934489.4;refGene_gn=GMEB1;refGene_tx=XM_019836500.2;refGene_gn=GMEB1;refGene_tx=XM_023258383.1
C1 21659698 21659811
refGene_gn=GMEB1;refGene_tx=XM_003989773.5;refGene_gn=GMEB1;refGene_tx=XM_003989774.5;refGene_gn=GMEB1;refGene_tx=XM_006934488.4;refGene_gn=GMEB1;refGene_tx=XM_006934489.4;refGene_gn=GMEB1;refGene_tx=XM_019836500.2;refGene_gn=GMEB1;refGene_tx=XM_023258383.1
C1 21660520 21660645
refGene_gn=GMEB1;refGene_tx=XM_003989773.5;refGene_gn=GMEB1;refGene_tx=XM_003989774.5;refGene_gn=GMEB1;refGene_tx=XM_006934488.4;refGene_gn=GMEB1;refGene_tx=XM_006934489.4;refGene_gn=GMEB1;refGene_tx=XM_019836500.2;refGene_gn=GMEB1;refGene_tx=XM_023258383.1
C1 21661935 21662039
refGene_gn=GMEB1;refGene_tx=XM_003989773.5;refGene_gn=GMEB1;refGene_tx=XM_003989774.5;refGene_gn=GMEB1;refGene_tx=XM_006934488.4;refGene_gn=GMEB1;refGene_tx=XM_006934489.4;refGene_gn=GMEB1;refGene_tx=XM_019836500.2;refGene_gn=GMEB1;refGene_tx=XM_023258383.1
C1 21667217 21667375
refGene_gn=GMEB1;refGene_tx=XM_003989773.5;refGene_gn=GMEB1;refGene_tx=XM_003989774.5;refGene_gn=GMEB1;refGene_tx=XM_006934488.4;refGene_gn=

GMEB1;refGene_tx=XM_006934489.4;refGene_gn=GMEB1;refGene_tx=XM_019836500.2;refGene_gn=GMEB1;refGene_tx=XM_023258383.1
C1 21670225 21670357
refGene_gn=GMEB1;refGene_tx=XM_003989773.5;refGene_gn=GMEB1;refGene_tx=XM_003989774.5;refGene_gn=GMEB1;refGene_tx=XM_006934488.4;refGene_gn=GMEB1;refGene_tx=XM_006934489.4;refGene_gn=GMEB1;refGene_tx=XM_019836500.2;refGene_gn=GMEB1;refGene_tx=XM_023258383.1
C1 21671611 21671749
refGene_gn=GMEB1;refGene_tx=XM_003989773.5;refGene_gn=GMEB1;refGene_tx=XM_003989774.5;refGene_gn=GMEB1;refGene_tx=XM_006934488.4;refGene_gn=GMEB1;refGene_tx=XM_006934489.4;refGene_gn=GMEB1;refGene_tx=XM_019836500.2;refGene_gn=GMEB1;refGene_tx=XM_023258383.1
C1 21675721 21675844
refGene_gn=GMEB1;refGene_tx=XM_003989773.5;refGene_gn=GMEB1;refGene_tx=XM_003989774.5;refGene_gn=GMEB1;refGene_tx=XM_006934488.4;refGene_gn=GMEB1;refGene_tx=XM_006934489.4;refGene_gn=GMEB1;refGene_tx=XM_019836500.2;refGene_gn=GMEB1;refGene_tx=XM_023258383.1
C1 21679185 21679886
refGene_gn=GMEB1;refGene_tx=XM_003989773.5;refGene_gn=GMEB1;refGene_tx=XM_003989774.5;refGene_gn=GMEB1;refGene_tx=XM_006934488.4;refGene_gn=GMEB1;refGene_tx=XM_006934489.4;refGene_gn=GMEB1;refGene_tx=XM_019836500.2;refGene_gn=GMEB1;refGene_tx=XM_023258383.1
C1 21693818 21693964
refGene_gn=LOC109502189;refGene_tx=XM_019836503.1
C1 21694175 21694260
refGene_gn=LOC109502189;refGene_tx=XM_019836503.1
C1 21696645 21696672
refGene_gn=YTHDF2;refGene_tx=XM_023258384.1;refGene_gn=YTHDF2;refGene_tx=XM_023258385.1
C1 21697175 21697200
refGene_gn=YTHDF2;refGene_tx=XM_023258384.1;refGene_gn=YTHDF2;refGene_tx=XM_023258385.1
C1 21697813 21697893
refGene_gn=YTHDF2;refGene_tx=XM_023258384.1;refGene_gn=YTHDF2;refGene_tx=XM_023258385.1
C1 21699406 21699472 refGene_gn=YTHDF2;refGene_tx=XM_019836501.2
C1 21700866 21702453
refGene_gn=YTHDF2;refGene_tx=XM_019836501.2;refGene_gn=YTHDF2;refGene_tx=XM_023258384.1;refGene_gn=YTHDF2;refGene_tx=XM_023258385.1
C1 21729572 21729596
refGene_gn=YTHDF2;refGene_tx=XM_019836501.2;refGene_gn=YTHDF2;refGene_tx=XM_023258384.1;refGene_gn=YTHDF2;refGene_tx=XM_023258385.1
C1 21767629 21767868 refGene_gn=OPRD1;refGene_tx=XM_003989776.4
C1 21796109 21796459 refGene_gn=OPRD1;refGene_tx=XM_003989776.4
C1 21798818 21799360 refGene_gn=OPRD1;refGene_tx=XM_003989776.4
C1 21899740 21900211
refGene_gn=EPB41;refGene_tx=XM_019836510.2;refGene_gn=EPB41;refGene_tx=XM_023258386.1;refGene_gn=EPB41;refGene_tx=XM_023258387.1;refGene_gn=EPB41;refGene_tx=XM_023258388.1;refGene_gn=EPB41;refGene_tx=XM_023258389.1;refGene_gn=EPB41;refGene_tx=XM_023258390.1;refGene_gn=EPB41;refGe

ne_tx=XM_023258391.1;refGene_gn=EPB41;refGene_tx=XM_023258392.1;refGene_gn=EPB41;refGene_tx=XM_023258393.1;refGene_gn=EPB41;refGene_tx=XM_023258394.1;refGene_gn=EPB41;refGene_tx=XM_023258395.1;refGene_gn=EPB41;refGene_tx=XM_023258396.1;refGene_gn=EPB41;refGene_tx=XM_023258397.1
C1 21907589 21907802
refGene_gn=EPB41;refGene_tx=XM_019836510.2;refGene_gn=EPB41;refGene_tx=XM_019836513.2;refGene_gn=EPB41;refGene_tx=XM_023258386.1;refGene_gn=EPB41;refGene_tx=XM_023258387.1;refGene_gn=EPB41;refGene_tx=XM_023258388.1;refGene_gn=EPB41;refGene_tx=XM_023258389.1;refGene_gn=EPB41;refGene_tx=XM_023258390.1;refGene_gn=EPB41;refGene_tx=XM_023258391.1;refGene_gn=EPB41;refGene_tx=XM_023258392.1;refGene_gn=EPB41;refGene_tx=XM_023258393.1;refGene_gn=EPB41;refGene_tx=XM_023258394.1;refGene_gn=EPB41;refGene_tx=XM_023258395.1;refGene_gn=EPB41;refGene_tx=XM_023258396.1;refGene_gn=EPB41;refGene_tx=XM_023258397.1
C1 21910112 21910217
refGene_gn=EPB41;refGene_tx=XM_019836510.2;refGene_gn=EPB41;refGene_tx=XM_019836513.2;refGene_gn=EPB41;refGene_tx=XM_023258386.1;refGene_gn=EPB41;refGene_tx=XM_023258387.1;refGene_gn=EPB41;refGene_tx=XM_023258388.1;refGene_gn=EPB41;refGene_tx=XM_023258389.1;refGene_gn=EPB41;refGene_tx=XM_023258390.1;refGene_gn=EPB41;refGene_tx=XM_023258391.1;refGene_gn=EPB41;refGene_tx=XM_023258392.1;refGene_gn=EPB41;refGene_tx=XM_023258393.1;refGene_gn=EPB41;refGene_tx=XM_023258394.1;refGene_gn=EPB41;refGene_tx=XM_023258395.1;refGene_gn=EPB41;refGene_tx=XM_023258396.1;refGene_gn=EPB41;refGene_tx=XM_023258397.1
C1 21925514 21925557
refGene_gn=EPB41;refGene_tx=XM_019836510.2;refGene_gn=EPB41;refGene_tx=XM_019836513.2;refGene_gn=EPB41;refGene_tx=XM_023258386.1;refGene_gn=EPB41;refGene_tx=XM_023258387.1;refGene_gn=EPB41;refGene_tx=XM_023258388.1;refGene_gn=EPB41;refGene_tx=XM_023258389.1;refGene_gn=EPB41;refGene_tx=XM_023258390.1;refGene_gn=EPB41;refGene_tx=XM_023258391.1;refGene_gn=EPB41;refGene_tx=XM_023258392.1;refGene_gn=EPB41;refGene_tx=XM_023258393.1;refGene_gn=EPB41;refGene_tx=XM_023258394.1;refGene_gn=EPB41;refGene_tx=XM_023258395.1;refGene_gn=EPB41;refGene_tx=XM_023258396.1;refGene_gn=EPB41;refGene_tx=XM_023258397.1
C1 21933195 21933271
refGene_gn=EPB41;refGene_tx=XM_019836510.2;refGene_gn=EPB41;refGene_tx=XM_019836513.2;refGene_gn=EPB41;refGene_tx=XM_023258386.1;refGene_gn=EPB41;refGene_tx=XM_023258387.1;refGene_gn=EPB41;refGene_tx=XM_023258388.1;refGene_gn=EPB41;refGene_tx=XM_023258389.1;refGene_gn=EPB41;refGene_tx=XM_023258390.1;refGene_gn=EPB41;refGene_tx=XM_023258391.1;refGene_gn=EPB41;refGene_tx=XM_023258392.1;refGene_gn=EPB41;refGene_tx=XM_023258393.1;refGene_gn=EPB41;refGene_tx=XM_023258394.1;refGene_gn=EPB41;refGene_tx=XM_023258395.1;refGene_gn=EPB41;refGene_tx=XM_023258396.1;refGene_gn=EPB41;refGene_tx=XM_023258397.1
C1 21937397 21937616
refGene_gn=EPB41;refGene_tx=XM_019836510.2;refGene_gn=EPB41;refGene_tx=XM_019836513.2;refGene_gn=EPB41;refGene_tx=XM_023258386.1;refGene_gn=EPB41;refGene_tx=XM_023258387.1;refGene_gn=EPB41;refGene_tx=XM_023258388.1;refGene_gn=EPB41;refGene_tx=XM_023258389.1;refGene_gn=EPB41;refGene_tx=XM_023258390.1;refGene_gn=EPB41;refGene_tx=XM_023258391.1;refGen

e_gn=EPB41;refGene_tx=XM_023258392.1;refGene_gn=EPB41;refGene_tx=XM_023258393.1;refGene_gn=EPB41;refGene_tx=XM_023258394.1;refGene_gn=EPB41;refGene_tx=XM_023258395.1;refGene_gn=EPB41;refGene_tx=XM_023258396.1;refGene_gn=EPB41;refGene_tx=XM_023258397.1
C1 21945128 21945216
refGene_gn=EPB41;refGene_tx=XM_019836510.2;refGene_gn=EPB41;refGene_tx=XM_019836513.2;refGene_gn=EPB41;refGene_tx=XM_023258386.1;refGene_gn=EPB41;refGene_tx=XM_023258387.1;refGene_gn=EPB41;refGene_tx=XM_023258388.1;refGene_gn=EPB41;refGene_tx=XM_023258389.1;refGene_gn=EPB41;refGene_tx=XM_023258390.1;refGene_gn=EPB41;refGene_tx=XM_023258391.1;refGene_gn=EPB41;refGene_tx=XM_023258392.1;refGene_gn=EPB41;refGene_tx=XM_023258393.1;refGene_gn=EPB41;refGene_tx=XM_023258394.1;refGene_gn=EPB41;refGene_tx=XM_023258395.1;refGene_gn=EPB41;refGene_tx=XM_023258396.1;refGene_gn=EPB41;refGene_tx=XM_023258397.1
C1 21948180 21948333
refGene_gn=EPB41;refGene_tx=XM_019836510.2;refGene_gn=EPB41;refGene_tx=XM_019836513.2;refGene_gn=EPB41;refGene_tx=XM_023258386.1;refGene_gn=EPB41;refGene_tx=XM_023258387.1;refGene_gn=EPB41;refGene_tx=XM_023258388.1;refGene_gn=EPB41;refGene_tx=XM_023258389.1;refGene_gn=EPB41;refGene_tx=XM_023258390.1;refGene_gn=EPB41;refGene_tx=XM_023258391.1;refGene_gn=EPB41;refGene_tx=XM_023258392.1;refGene_gn=EPB41;refGene_tx=XM_023258393.1;refGene_gn=EPB41;refGene_tx=XM_023258394.1;refGene_gn=EPB41;refGene_tx=XM_023258395.1;refGene_gn=EPB41;refGene_tx=XM_023258396.1;refGene_gn=EPB41;refGene_tx=XM_023258397.1
C1 21951416 21951514
refGene_gn=EPB41;refGene_tx=XM_019836510.2;refGene_gn=EPB41;refGene_tx=XM_019836513.2;refGene_gn=EPB41;refGene_tx=XM_023258386.1;refGene_gn=EPB41;refGene_tx=XM_023258387.1;refGene_gn=EPB41;refGene_tx=XM_023258388.1;refGene_gn=EPB41;refGene_tx=XM_023258389.1;refGene_gn=EPB41;refGene_tx=XM_023258390.1;refGene_gn=EPB41;refGene_tx=XM_023258391.1;refGene_gn=EPB41;refGene_tx=XM_023258392.1;refGene_gn=EPB41;refGene_tx=XM_023258393.1;refGene_gn=EPB41;refGene_tx=XM_023258394.1;refGene_gn=EPB41;refGene_tx=XM_023258395.1;refGene_gn=EPB41;refGene_tx=XM_023258396.1;refGene_gn=EPB41;refGene_tx=XM_023258397.1
C1 21954078 21954251
refGene_gn=EPB41;refGene_tx=XM_019836510.2;refGene_gn=EPB41;refGene_tx=XM_019836513.2;refGene_gn=EPB41;refGene_tx=XM_023258386.1;refGene_gn=EPB41;refGene_tx=XM_023258387.1;refGene_gn=EPB41;refGene_tx=XM_023258388.1;refGene_gn=EPB41;refGene_tx=XM_023258389.1;refGene_gn=EPB41;refGene_tx=XM_023258390.1;refGene_gn=EPB41;refGene_tx=XM_023258391.1;refGene_gn=EPB41;refGene_tx=XM_023258392.1;refGene_gn=EPB41;refGene_tx=XM_023258393.1;refGene_gn=EPB41;refGene_tx=XM_023258394.1;refGene_gn=EPB41;refGene_tx=XM_023258395.1;refGene_gn=EPB41;refGene_tx=XM_023258396.1;refGene_gn=EPB41;refGene_tx=XM_023258397.1
C1 21964411 21964617
refGene_gn=EPB41;refGene_tx=XM_019836510.2;refGene_gn=EPB41;refGene_tx=XM_019836513.2;refGene_gn=EPB41;refGene_tx=XM_023258386.1;refGene_gn=EPB41;refGene_tx=XM_023258387.1;refGene_gn=EPB41;refGene_tx=XM_023258388.1;refGene_gn=EPB41;refGene_tx=XM_023258389.1;refGene_gn=EPB41;refGene_tx=XM_023258390.1;refGene_gn=EPB41;refGene_tx=XM_023258391.1;refGen

e_gn=EPB41;refGene_tx=XM_023258392.1;refGene_gn=EPB41;refGene_tx=XM_02
3258393.1;refGene_gn=EPB41;refGene_tx=XM_023258394.1;refGene_gn=EPB41;
refGene_tx=XM_023258395.1;refGene_gn=EPB41;refGene_tx=XM_023258396.1;r
efGene_gn=EPB41;refGene_tx=XM_023258397.1
C1 21968340 21968397
refGene_gn=EPB41;refGene_tx=XM_019836510.2;refGene_gn=EPB41;refGene_tx
=XM_019836513.2;refGene_gn=EPB41;refGene_tx=XM_023258386.1;refGene_gn=
EPB41;refGene_tx=XM_023258387.1;refGene_gn=EPB41;refGene_tx=XM_0232583
88.1;refGene_gn=EPB41;refGene_tx=XM_023258389.1;refGene_gn=EPB41;refGe
ne_tx=XM_023258392.1;refGene_gn=EPB41;refGene_tx=XM_023258393.1;refGen
e_gn=EPB41;refGene_tx=XM_023258394.1;refGene_gn=EPB41;refGene_tx=XM_02
3258395.1
C1 21968560 21968602
refGene_gn=EPB41;refGene_tx=XM_019836510.2;refGene_gn=EPB41;refGene_tx
=XM_019836513.2;refGene_gn=EPB41;refGene_tx=XM_023258386.1;refGene_gn=
EPB41;refGene_tx=XM_023258387.1;refGene_gn=EPB41;refGene_tx=XM_0232583
89.1;refGene_gn=EPB41;refGene_tx=XM_023258392.1;refGene_gn=EPB41;refGe
ne_tx=XM_023258393.1;refGene_gn=EPB41;refGene_tx=XM_023258394.1
C1 21969582 21969645
refGene_gn=EPB41;refGene_tx=XM_019836510.2;refGene_gn=EPB41;refGene_tx
=XM_019836513.2;refGene_gn=EPB41;refGene_tx=XM_023258386.1;refGene_gn=
EPB41;refGene_tx=XM_023258387.1;refGene_gn=EPB41;refGene_tx=XM_0232583
88.1;refGene_gn=EPB41;refGene_tx=XM_023258390.1;refGene_gn=EPB41;refGe
ne_tx=XM_023258392.1;refGene_gn=EPB41;refGene_tx=XM_023258393.1;refGen
e_gn=EPB41;refGene_tx=XM_023258394.1;refGene_gn=EPB41;refGene_tx=XM_02
3258395.1;refGene_gn=EPB41;refGene_tx=XM_023258396.1
C1 21974057 21974234
refGene_gn=EPB41;refGene_tx=XM_019836510.2;refGene_gn=EPB41;refGene_tx
=XM_019836513.2;refGene_gn=EPB41;refGene_tx=XM_023258386.1;refGene_gn=
EPB41;refGene_tx=XM_023258387.1;refGene_gn=EPB41;refGene_tx=XM_0232583
88.1;refGene_gn=EPB41;refGene_tx=XM_023258389.1;refGene_gn=EPB41;refGe
ne_tx=XM_023258390.1;refGene_gn=EPB41;refGene_tx=XM_023258391.1;refGen
e_gn=EPB41;refGene_tx=XM_023258392.1;refGene_gn=EPB41;refGene_tx=XM_02
3258393.1;refGene_gn=EPB41;refGene_tx=XM_023258394.1;refGene_gn=EPB41;
refGene_tx=XM_023258395.1;refGene_gn=EPB41;refGene_tx=XM_023258396.1;r
efGene_gn=EPB41;refGene_tx=XM_023258397.1
C1 21975975 21976026
refGene_gn=EPB41;refGene_tx=XM_023258392.1;refGene_gn=EPB41;refGene_tx
=XM_023258393.1
C1 21999093 21999543
refGene_gn=EPB41;refGene_tx=XM_019836510.2;refGene_gn=EPB41;refGene_tx
=XM_019836513.2;refGene_gn=EPB41;refGene_tx=XM_023258386.1;refGene_gn=
EPB41;refGene_tx=XM_023258387.1;refGene_gn=EPB41;refGene_tx=XM_0232583
88.1;refGene_gn=EPB41;refGene_tx=XM_023258389.1;refGene_gn=EPB41;refGe
ne_tx=XM_023258390.1;refGene_gn=EPB41;refGene_tx=XM_023258391.1
C1 22000763 22000892
refGene_gn=EPB41;refGene_tx=XM_019836510.2;refGene_gn=EPB41;refGene_tx
=XM_019836513.2;refGene_gn=EPB41;refGene_tx=XM_023258386.1;refGene_gn=
EPB41;refGene_tx=XM_023258387.1;refGene_gn=EPB41;refGene_tx=XM_0232583
88.1;refGene_gn=EPB41;refGene_tx=XM_023258389.1;refGene_gn=EPB41;refGe

ne_tx=XM_023258390.1;refGene_gn=EPB41;refGene_tx=XM_023258391.1;refGene_gn=EPB41;refGene_tx=XM_023258392.1;refGene_gn=EPB41;refGene_tx=XM_023258393.1;refGene_gn=EPB41;refGene_tx=XM_023258394.1;refGene_gn=EPB41;refGene_tx=XM_023258395.1;refGene_gn=EPB41;refGene_tx=XM_023258396.1;refGene_gn=EPB41;refGene_tx=XM_023258397.1
C1 22008324 22008426
refGene_gn=EPB41;refGene_tx=XM_019836510.2;refGene_gn=EPB41;refGene_tx=XM_019836513.2;refGene_gn=EPB41;refGene_tx=XM_023258386.1;refGene_gn=EPB41;refGene_tx=XM_023258387.1;refGene_gn=EPB41;refGene_tx=XM_023258388.1;refGene_gn=EPB41;refGene_tx=XM_023258389.1;refGene_gn=EPB41;refGene_tx=XM_023258390.1;refGene_gn=EPB41;refGene_tx=XM_023258391.1;refGene_gn=EPB41;refGene_tx=XM_023258392.1;refGene_gn=EPB41;refGene_tx=XM_023258393.1;refGene_gn=EPB41;refGene_tx=XM_023258394.1;refGene_gn=EPB41;refGene_tx=XM_023258395.1;refGene_gn=EPB41;refGene_tx=XM_023258396.1;refGene_gn=EPB41;refGene_tx=XM_023258397.1
C1 22009573 22009654
refGene_gn=EPB41;refGene_tx=XM_019836510.2;refGene_gn=EPB41;refGene_tx=XM_019836513.2;refGene_gn=EPB41;refGene_tx=XM_023258386.1;refGene_gn=EPB41;refGene_tx=XM_023258387.1;refGene_gn=EPB41;refGene_tx=XM_023258388.1;refGene_gn=EPB41;refGene_tx=XM_023258389.1;refGene_gn=EPB41;refGene_tx=XM_023258390.1;refGene_gn=EPB41;refGene_tx=XM_023258391.1;refGene_gn=EPB41;refGene_tx=XM_023258392.1;refGene_gn=EPB41;refGene_tx=XM_023258393.1;refGene_gn=EPB41;refGene_tx=XM_023258394.1;refGene_gn=EPB41;refGene_tx=XM_023258395.1;refGene_gn=EPB41;refGene_tx=XM_023258396.1;refGene_gn=EPB41;refGene_tx=XM_023258397.1
C1 22013030 22013129
refGene_gn=EPB41;refGene_tx=XM_019836510.2;refGene_gn=EPB41;refGene_tx=XM_019836513.2;refGene_gn=EPB41;refGene_tx=XM_023258386.1;refGene_gn=EPB41;refGene_tx=XM_023258387.1;refGene_gn=EPB41;refGene_tx=XM_023258388.1;refGene_gn=EPB41;refGene_tx=XM_023258389.1;refGene_gn=EPB41;refGene_tx=XM_023258390.1;refGene_gn=EPB41;refGene_tx=XM_023258391.1;refGene_gn=EPB41;refGene_tx=XM_023258392.1;refGene_gn=EPB41;refGene_tx=XM_023258393.1;refGene_gn=EPB41;refGene_tx=XM_023258394.1;refGene_gn=EPB41;refGene_tx=XM_023258395.1;refGene_gn=EPB41;refGene_tx=XM_023258396.1;refGene_gn=EPB41;refGene_tx=XM_023258397.1
C1 22017765 22018689
refGene_gn=TMEM200B;refGene_tx=XM_003989778.5;refGene_gn=TMEM200B;refGene_tx=XM_023258398.1
C1 22052041 22052873 refGene_gn=SRSF4;refGene_tx=XM_003989779.4
C1 22053114 22053204 refGene_gn=SRSF4;refGene_tx=XM_003989779.4
C1 22056384 22056599 refGene_gn=SRSF4;refGene_tx=XM_003989779.4
C1 22060484 22060597 refGene_gn=SRSF4;refGene_tx=XM_003989779.4
C1 22061503 22061646 refGene_gn=SRSF4;refGene_tx=XM_003989779.4
C1 22079464 22079571 refGene_gn=SRSF4;refGene_tx=XM_003989779.4
C1 22080079 22080162 refGene_gn=MECR;refGene_tx=XM_019836523.2
C1 22086496 22086657
refGene_gn=MECR;refGene_tx=XM_003989787.5;refGene_gn=MECR;refGene_tx=XM_019836524.2
C1 22087374 22087526 refGene_gn=MECR;refGene_tx=XM_019836522.2
C1 22088058 22088131

refGene_gn=MECR;refGene_tx=XM_003989787.5;refGene_gn=MECR;refGene_tx=XM_019836522.2;refGene_gn=MECR;refGene_tx=XM_019836523.2;refGene_gn=MECR;refGene_tx=XM_019836524.2
C1 22088287 22088348
refGene_gn=MECR;refGene_tx=XM_003989787.5;refGene_gn=MECR;refGene_tx=XM_019836522.2;refGene_gn=MECR;refGene_tx=XM_019836523.2;refGene_gn=MECR;refGene_tx=XM_019836524.2
C1 22091171 22091245
refGene_gn=MECR;refGene_tx=XM_003989787.5;refGene_gn=MECR;refGene_tx=XM_019836522.2;refGene_gn=MECR;refGene_tx=XM_019836523.2;refGene_gn=MECR;refGene_tx=XM_019836524.2
C1 22092597 22092700
refGene_gn=MECR;refGene_tx=XM_003989787.5;refGene_gn=MECR;refGene_tx=XM_019836522.2;refGene_gn=MECR;refGene_tx=XM_019836523.2;refGene_gn=MECR;refGene_tx=XM_019836524.2
C1 22093769 22093872
refGene_gn=MECR;refGene_tx=XM_003989787.5;refGene_gn=MECR;refGene_tx=XM_019836522.2;refGene_gn=MECR;refGene_tx=XM_019836523.2;refGene_gn=MECR;refGene_tx=XM_019836524.2
C1 22096488 22096632
refGene_gn=MECR;refGene_tx=XM_003989787.5;refGene_gn=MECR;refGene_tx=XM_019836522.2;refGene_gn=MECR;refGene_tx=XM_019836523.2;refGene_gn=MECR;refGene_tx=XM_019836524.2
C1 22103704 22103836
refGene_gn=MECR;refGene_tx=XM_003989787.5;refGene_gn=MECR;refGene_tx=XM_019836522.2;refGene_gn=MECR;refGene_tx=XM_019836523.2;refGene_gn=MECR;refGene_tx=XM_019836524.2
C1 22104296 22104394
refGene_gn=MECR;refGene_tx=XM_003989787.5;refGene_gn=MECR;refGene_tx=XM_019836522.2;refGene_gn=MECR;refGene_tx=XM_019836523.2;refGene_gn=MECR;refGene_tx=XM_019836524.2
C1 22117522 22117698
refGene_gn=MECR;refGene_tx=XM_003989787.5;refGene_gn=MECR;refGene_tx=XM_019836522.2;refGene_gn=MECR;refGene_tx=XM_019836523.2
C1 22124295 22124368 refGene_gn=PTPRU;refGene_tx=XM_023260061.1
C1 22139300 22139432 refGene_gn=PTPRU;refGene_tx=XM_023260061.1
C1 22142174 22142446 refGene_gn=PTPRU;refGene_tx=XM_023260061.1
C1 22142979 22143061 refGene_gn=PTPRU;refGene_tx=XM_023260061.1
C1 22143165 22143281 refGene_gn=PTPRU;refGene_tx=XM_023260061.1
C1 22143694 22143869 refGene_gn=PTPRU;refGene_tx=XM_023260061.1
C1 22144453 22144747 refGene_gn=PTPRU;refGene_tx=XM_023260061.1
C1 22156341 22156650 refGene_gn=PTPRU;refGene_tx=XM_023260061.1
C1 22161550 22161660 refGene_gn=PTPRU;refGene_tx=XM_023260061.1
C1 22161980 22162182 refGene_gn=PTPRU;refGene_tx=XM_023260061.1
C1 22162531 22162634 refGene_gn=PTPRU;refGene_tx=XM_023260061.1
C1 22165040 22165314 refGene_gn=PTPRU;refGene_tx=XM_023260061.1
C1 22166004 22166041 refGene_gn=PTPRU;refGene_tx=XM_023260061.1
C1 22166754 22166893 refGene_gn=PTPRU;refGene_tx=XM_023260061.1
C1 22171036 22171066 refGene_gn=PTPRU;refGene_tx=XM_023260061.1
C1 22174476 22174667 refGene_gn=PTPRU;refGene_tx=XM_023260061.1

C1	22185356	22185452	refGene_gn=PTPRU;refGene_tx=XM_023260061.1
C1	22185929	22186006	refGene_gn=PTPRU;refGene_tx=XM_023260061.1
C1	22192196	22192233	refGene_gn=PTPRU;refGene_tx=XM_023260061.1
C1	22192883	22192981	refGene_gn=PTPRU;refGene_tx=XM_023260061.1
C1	22193064	22193181	refGene_gn=PTPRU;refGene_tx=XM_023260061.1
C1	22194147	22194302	refGene_gn=PTPRU;refGene_tx=XM_023260061.1
C1	22196545	22196681	refGene_gn=PTPRU;refGene_tx=XM_023260061.1
C1	22197148	22197298	refGene_gn=PTPRU;refGene_tx=XM_023260061.1
C1	22198784	22198958	refGene_gn=PTPRU;refGene_tx=XM_023260061.1
C1	22199799	22199940	refGene_gn=PTPRU;refGene_tx=XM_023260061.1
C1	22201950	22202076	refGene_gn=PTPRU;refGene_tx=XM_023260061.1
C1	22202209	22202367	refGene_gn=PTPRU;refGene_tx=XM_023260061.1
C1	22203573	22203709	refGene_gn=PTPRU;refGene_tx=XM_023260061.1
C1	22203975	22204038	refGene_gn=PTPRU;refGene_tx=XM_023260061.1
C1	23169717	23169778	refGene_gn=MATN1;refGene_tx=XM_023260062.1
C1	23272556	23272674	refGene_gn=MATN1;refGene_tx=XM_023260062.1
C1	23273440	23273593	refGene_gn=MATN1;refGene_tx=XM_023260062.1
C1	23274397	23274814	refGene_gn=MATN1;refGene_tx=XM_023260062.1
C1	23275273	23275399	refGene_gn=MATN1;refGene_tx=XM_023260062.1
C1	23275890	23276113	refGene_gn=MATN1;refGene_tx=XM_023260062.1
C1	23278545	23278892	refGene_gn=MATN1;refGene_tx=XM_023260062.1
C1	23280514	23280608	refGene_gn=MATN1;refGene_tx=XM_023260062.1
C1	23292625	23292715	refGene_gn=LAPTM5;refGene_tx=XM_006934508.4
C1	23294017	23294110	refGene_gn=LAPTM5;refGene_tx=XM_006934508.4
C1	23296141	23296237	refGene_gn=LAPTM5;refGene_tx=XM_006934508.4
C1	23297043	23297166	refGene_gn=LAPTM5;refGene_tx=XM_006934508.4
C1	23297875	23298007	refGene_gn=LAPTM5;refGene_tx=XM_006934508.4
C1	23299828	23299905	refGene_gn=LAPTM5;refGene_tx=XM_006934508.4
C1	23300572	23300666	refGene_gn=LAPTM5;refGene_tx=XM_006934508.4
C1	23315775	23315862	refGene_gn=LAPTM5;refGene_tx=XM_006934508.4
C1	23416422	23416589	refGene_gn=SDC3;refGene_tx=XM_023260063.1
C1	23416984	23417276	refGene_gn=SDC3;refGene_tx=XM_023260063.1
C1	23419092	23419706	refGene_gn=SDC3;refGene_tx=XM_023260063.1
C1	23421665	23421783	refGene_gn=SDC3;refGene_tx=XM_023260063.1
C1	23447854	23447992	refGene_gn=SDC3;refGene_tx=XM_023260063.1
C1	23460574	23460706
refGene_gn=PUM1;refGene_tx=XM_011284760.3;refGene_gn=PUM1;refGene_tx=X
M_019836533.2;refGene_gn=PUM1;refGene_tx=XM_019836534.2;refGene_gn=PUM
1;refGene_tx=XM_019836535.2;refGene_gn=PUM1;refGene_tx=XM_019836536.2;
refGene_gn=PUM1;refGene_tx=XM_019836538.2;refGene_gn=PUM1;refGene_tx=X
M_019836539.2;refGene_gn=PUM1;refGene_tx=XM_019836540.2;refGene_gn=PUM
1;refGene_tx=XM_019836541.2;refGene_gn=PUM1;refGene_tx=XM_019836542.2;
refGene_gn=PUM1;refGene_tx=XM_019836543.2;refGene_gn=PUM1;refGene_tx=X
M_019836544.2;refGene_gn=PUM1;refGene_tx=XM_019836545.2;refGene_gn=PUM
1;refGene_tx=XM_019836546.2;refGene_gn=PUM1;refGene_tx=XM_019836548.2;
refGene_gn=PUM1;refGene_tx=XM_019836549.2;refGene_gn=PUM1;refGene_tx=X
M_019836550.2
C1	23463778	23463971
refGene_gn=PUM1;refGene_tx=XM_011284760.3;refGene_gn=PUM1;refGene_tx=X
M_019836533.2;refGene_gn=PUM1;refGene_tx=XM_019836534.2;refGene_gn=PUM

1;refGene_tx=XM_019836535.2;refGene_gn=PUM1;refGene_tx=XM_019836536.2;
refGene_gn=PUM1;refGene_tx=XM_019836538.2;refGene_gn=PUM1;refGene_tx=X
M_019836539.2;refGene_gn=PUM1;refGene_tx=XM_019836540.2;refGene_gn=PUM
1;refGene_tx=XM_019836541.2;refGene_gn=PUM1;refGene_tx=XM_019836542.2;
refGene_gn=PUM1;refGene_tx=XM_019836543.2;refGene_gn=PUM1;refGene_tx=X
M_019836544.2;refGene_gn=PUM1;refGene_tx=XM_019836545.2;refGene_gn=PUM
1;refGene_tx=XM_019836546.2;refGene_gn=PUM1;refGene_tx=XM_019836548.2;
refGene_gn=PUM1;refGene_tx=XM_019836549.2;refGene_gn=PUM1;refGene_tx=X
M_019836550.2
C1 23469311 23469433
refGene_gn=PUM1;refGene_tx=XM_011284760.3;refGene_gn=PUM1;refGene_tx=X
M_019836533.2;refGene_gn=PUM1;refGene_tx=XM_019836534.2;refGene_gn=PUM
1;refGene_tx=XM_019836535.2;refGene_gn=PUM1;refGene_tx=XM_019836536.2;
refGene_gn=PUM1;refGene_tx=XM_019836538.2;refGene_gn=PUM1;refGene_tx=X
M_019836539.2;refGene_gn=PUM1;refGene_tx=XM_019836540.2;refGene_gn=PUM
1;refGene_tx=XM_019836541.2;refGene_gn=PUM1;refGene_tx=XM_019836542.2;
refGene_gn=PUM1;refGene_tx=XM_019836543.2;refGene_gn=PUM1;refGene_tx=X
M_019836544.2;refGene_gn=PUM1;refGene_tx=XM_019836545.2;refGene_gn=PUM
1;refGene_tx=XM_019836546.2;refGene_gn=PUM1;refGene_tx=XM_019836548.2;
refGene_gn=PUM1;refGene_tx=XM_019836549.2;refGene_gn=PUM1;refGene_tx=X
M_019836550.2
C1 23470737 23470863
refGene_gn=PUM1;refGene_tx=XM_011284760.3;refGene_gn=PUM1;refGene_tx=X
M_019836533.2;refGene_gn=PUM1;refGene_tx=XM_019836534.2;refGene_gn=PUM
1;refGene_tx=XM_019836535.2;refGene_gn=PUM1;refGene_tx=XM_019836536.2;
refGene_gn=PUM1;refGene_tx=XM_019836538.2;refGene_gn=PUM1;refGene_tx=X
M_019836539.2;refGene_gn=PUM1;refGene_tx=XM_019836540.2;refGene_gn=PUM
1;refGene_tx=XM_019836541.2;refGene_gn=PUM1;refGene_tx=XM_019836542.2;
refGene_gn=PUM1;refGene_tx=XM_019836543.2;refGene_gn=PUM1;refGene_tx=X
M_019836544.2;refGene_gn=PUM1;refGene_tx=XM_019836545.2;refGene_gn=PUM
1;refGene_tx=XM_019836546.2;refGene_gn=PUM1;refGene_tx=XM_019836548.2;
refGene_gn=PUM1;refGene_tx=XM_019836549.2;refGene_gn=PUM1;refGene_tx=X
M_019836550.2
C1 23472991 23473129
refGene_gn=PUM1;refGene_tx=XM_011284760.3;refGene_gn=PUM1;refGene_tx=X
M_019836533.2;refGene_gn=PUM1;refGene_tx=XM_019836534.2;refGene_gn=PUM
1;refGene_tx=XM_019836535.2;refGene_gn=PUM1;refGene_tx=XM_019836536.2;
refGene_gn=PUM1;refGene_tx=XM_019836538.2;refGene_gn=PUM1;refGene_tx=X
M_019836539.2;refGene_gn=PUM1;refGene_tx=XM_019836540.2;refGene_gn=PUM
1;refGene_tx=XM_019836541.2;refGene_gn=PUM1;refGene_tx=XM_019836542.2;
refGene_gn=PUM1;refGene_tx=XM_019836543.2;refGene_gn=PUM1;refGene_tx=X
M_019836544.2;refGene_gn=PUM1;refGene_tx=XM_019836545.2;refGene_gn=PUM
1;refGene_tx=XM_019836546.2;refGene_gn=PUM1;refGene_tx=XM_019836548.2;
refGene_gn=PUM1;refGene_tx=XM_019836549.2;refGene_gn=PUM1;refGene_tx=X
M_019836550.2
C1 23476703 23476838
refGene_gn=PUM1;refGene_tx=XM_011284760.3;refGene_gn=PUM1;refGene_tx=X
M_019836533.2;refGene_gn=PUM1;refGene_tx=XM_019836534.2;refGene_gn=PUM
1;refGene_tx=XM_019836535.2;refGene_gn=PUM1;refGene_tx=XM_019836536.2;
refGene_gn=PUM1;refGene_tx=XM_019836538.2;refGene_gn=PUM1;refGene_tx=X

M_019836539.2;refGene_gn=PUM1;refGene_tx=XM_019836540.2;refGene_gn=PUM
1;refGene_tx=XM_019836541.2;refGene_gn=PUM1;refGene_tx=XM_019836542.2;
refGene_gn=PUM1;refGene_tx=XM_019836543.2;refGene_gn=PUM1;refGene_tx=X
M_019836544.2;refGene_gn=PUM1;refGene_tx=XM_019836545.2;refGene_gn=PUM
1;refGene_tx=XM_019836546.2;refGene_gn=PUM1;refGene_tx=XM_019836548.2;
refGene_gn=PUM1;refGene_tx=XM_019836549.2;refGene_gn=PUM1;refGene_tx=X
M_019836550.2
C1 23478887 23479017
refGene_gn=PUM1;refGene_tx=XM_011284760.3;refGene_gn=PUM1;refGene_tx=X
M_019836533.2;refGene_gn=PUM1;refGene_tx=XM_019836534.2;refGene_gn=PUM
1;refGene_tx=XM_019836535.2;refGene_gn=PUM1;refGene_tx=XM_019836536.2;
refGene_gn=PUM1;refGene_tx=XM_019836538.2;refGene_gn=PUM1;refGene_tx=X
M_019836539.2;refGene_gn=PUM1;refGene_tx=XM_019836540.2;refGene_gn=PUM
1;refGene_tx=XM_019836541.2;refGene_gn=PUM1;refGene_tx=XM_019836542.2;
refGene_gn=PUM1;refGene_tx=XM_019836543.2;refGene_gn=PUM1;refGene_tx=X
M_019836544.2;refGene_gn=PUM1;refGene_tx=XM_019836545.2;refGene_gn=PUM
1;refGene_tx=XM_019836546.2;refGene_gn=PUM1;refGene_tx=XM_019836548.2;
refGene_gn=PUM1;refGene_tx=XM_019836549.2;refGene_gn=PUM1;refGene_tx=X
M_019836550.2
C1 23480573 23480841
refGene_gn=PUM1;refGene_tx=XM_011284760.3;refGene_gn=PUM1;refGene_tx=X
M_019836533.2;refGene_gn=PUM1;refGene_tx=XM_019836534.2;refGene_gn=PUM
1;refGene_tx=XM_019836535.2;refGene_gn=PUM1;refGene_tx=XM_019836536.2;
refGene_gn=PUM1;refGene_tx=XM_019836538.2;refGene_gn=PUM1;refGene_tx=X
M_019836539.2;refGene_gn=PUM1;refGene_tx=XM_019836540.2;refGene_gn=PUM
1;refGene_tx=XM_019836541.2;refGene_gn=PUM1;refGene_tx=XM_019836542.2;
refGene_gn=PUM1;refGene_tx=XM_019836543.2;refGene_gn=PUM1;refGene_tx=X
M_019836544.2;refGene_gn=PUM1;refGene_tx=XM_019836545.2;refGene_gn=PUM
1;refGene_tx=XM_019836546.2;refGene_gn=PUM1;refGene_tx=XM_019836548.2;
refGene_gn=PUM1;refGene_tx=XM_019836549.2;refGene_gn=PUM1;refGene_tx=X
M_019836550.2
C1 23493291 23493528
refGene_gn=PUM1;refGene_tx=XM_011284760.3;refGene_gn=PUM1;refGene_tx=X
M_019836533.2;refGene_gn=PUM1;refGene_tx=XM_019836534.2;refGene_gn=PUM
1;refGene_tx=XM_019836535.2;refGene_gn=PUM1;refGene_tx=XM_019836536.2;
refGene_gn=PUM1;refGene_tx=XM_019836538.2;refGene_gn=PUM1;refGene_tx=X
M_019836539.2;refGene_gn=PUM1;refGene_tx=XM_019836542.2;refGene_gn=PUM
1;refGene_tx=XM_019836543.2;refGene_gn=PUM1;refGene_tx=XM_019836544.2;
refGene_gn=PUM1;refGene_tx=XM_019836545.2;refGene_gn=PUM1;refGene_tx=X
M_019836546.2;refGene_gn=PUM1;refGene_tx=XM_019836548.2;refGene_gn=PUM
1;refGene_tx=XM_019836549.2;refGene_gn=PUM1;refGene_tx=XM_019836550.2
C1 23494614 23494911
refGene_gn=PUM1;refGene_tx=XM_011284760.3;refGene_gn=PUM1;refGene_tx=X
M_019836533.2;refGene_gn=PUM1;refGene_tx=XM_019836534.2;refGene_gn=PUM
1;refGene_tx=XM_019836535.2;refGene_gn=PUM1;refGene_tx=XM_019836536.2;
refGene_gn=PUM1;refGene_tx=XM_019836538.2;refGene_gn=PUM1;refGene_tx=X
M_019836539.2;refGene_gn=PUM1;refGene_tx=XM_019836540.2;refGene_gn=PUM
1;refGene_tx=XM_019836541.2;refGene_gn=PUM1;refGene_tx=XM_019836542.2;
refGene_gn=PUM1;refGene_tx=XM_019836543.2;refGene_gn=PUM1;refGene_tx=X
M_019836544.2;refGene_gn=PUM1;refGene_tx=XM_019836545.2;refGene_gn=PUM

1;refGene_tx=XM_019836546.2;refGene_gn=PUM1;refGene_tx=XM_019836548.2;refGene_gn=PUM1;refGene_tx=XM_019836549.2;refGene_gn=PUM1;refGene_tx=XM_019836550.2

C1 23495815 23495959 refGene_gn=PUM1;refGene_tx=XM_011284760.3;refGene_gn=PUM1;refGene_tx=XM_019836533.2;refGene_gn=PUM1;refGene_tx=XM_019836534.2;refGene_gn=PUM1;refGene_tx=XM_019836535.2;refGene_gn=PUM1;refGene_tx=XM_019836536.2;refGene_gn=PUM1;refGene_tx=XM_019836538.2;refGene_gn=PUM1;refGene_tx=XM_019836539.2;refGene_gn=PUM1;refGene_tx=XM_019836540.2;refGene_gn=PUM1;refGene_tx=XM_019836541.2;refGene_gn=PUM1;refGene_tx=XM_019836542.2;refGene_gn=PUM1;refGene_tx=XM_019836543.2;refGene_gn=PUM1;refGene_tx=XM_019836544.2;refGene_gn=PUM1;refGene_tx=XM_019836545.2;refGene_gn=PUM1;refGene_tx=XM_019836546.2;refGene_gn=PUM1;refGene_tx=XM_019836548.2;refGene_gn=PUM1;refGene_tx=XM_019836549.2;refGene_gn=PUM1;refGene_tx=XM_019836550.2

C1 23497255 23497394 refGene_gn=PUM1;refGene_tx=XM_011284760.3;refGene_gn=PUM1;refGene_tx=XM_019836533.2;refGene_gn=PUM1;refGene_tx=XM_019836534.2;refGene_gn=PUM1;refGene_tx=XM_019836535.2;refGene_gn=PUM1;refGene_tx=XM_019836536.2;refGene_gn=PUM1;refGene_tx=XM_019836538.2;refGene_gn=PUM1;refGene_tx=XM_019836539.2;refGene_gn=PUM1;refGene_tx=XM_019836540.2;refGene_gn=PUM1;refGene_tx=XM_019836541.2;refGene_gn=PUM1;refGene_tx=XM_019836542.2;refGene_gn=PUM1;refGene_tx=XM_019836543.2;refGene_gn=PUM1;refGene_tx=XM_019836544.2;refGene_gn=PUM1;refGene_tx=XM_019836545.2;refGene_gn=PUM1;refGene_tx=XM_019836546.2;refGene_gn=PUM1;refGene_tx=XM_019836548.2;refGene_gn=PUM1;refGene_tx=XM_019836549.2;refGene_gn=PUM1;refGene_tx=XM_019836550.2

C1 23502245 23502397 refGene_gn=PUM1;refGene_tx=XM_011284760.3;refGene_gn=PUM1;refGene_tx=XM_019836533.2;refGene_gn=PUM1;refGene_tx=XM_019836534.2;refGene_gn=PUM1;refGene_tx=XM_019836535.2;refGene_gn=PUM1;refGene_tx=XM_019836536.2;refGene_gn=PUM1;refGene_tx=XM_019836538.2;refGene_gn=PUM1;refGene_tx=XM_019836539.2;refGene_gn=PUM1;refGene_tx=XM_019836540.2;refGene_gn=PUM1;refGene_tx=XM_019836541.2;refGene_gn=PUM1;refGene_tx=XM_019836542.2;refGene_gn=PUM1;refGene_tx=XM_019836543.2;refGene_gn=PUM1;refGene_tx=XM_019836544.2;refGene_gn=PUM1;refGene_tx=XM_019836545.2;refGene_gn=PUM1;refGene_tx=XM_019836546.2;refGene_gn=PUM1;refGene_tx=XM_019836548.2;refGene_gn=PUM1;refGene_tx=XM_019836549.2;refGene_gn=PUM1;refGene_tx=XM_019836550.2

C1 23506782 23506887 refGene_gn=PUM1;refGene_tx=XM_011284760.3;refGene_gn=PUM1;refGene_tx=XM_019836533.2;refGene_gn=PUM1;refGene_tx=XM_019836534.2;refGene_gn=PUM1;refGene_tx=XM_019836535.2;refGene_gn=PUM1;refGene_tx=XM_019836536.2;refGene_gn=PUM1;refGene_tx=XM_019836538.2;refGene_gn=PUM1;refGene_tx=XM_019836539.2;refGene_gn=PUM1;refGene_tx=XM_019836540.2;refGene_gn=PUM1;refGene_tx=XM_019836541.2;refGene_gn=PUM1;refGene_tx=XM_019836542.2;refGene_gn=PUM1;refGene_tx=XM_019836543.2;refGene_gn=PUM1;refGene_tx=XM_019836544.2;refGene_gn=PUM1;refGene_tx=XM_019836545.2;refGene_gn=PUM1;refGene_tx=XM_019836546.2;refGene_gn=PUM1;refGene_tx=XM_019836548.2;refGene_gn=PUM1;refGene_tx=XM_019836549.2;refGene_gn=PUM1;refGene_tx=X

M_019836550.2
C1 23508938 23509032
refGene_gn=PUM1;refGene_tx=XM_011284760.3;refGene_gn=PUM1;refGene_tx=XM_019836533.2;refGene_gn=PUM1;refGene_tx=XM_019836534.2;refGene_gn=PUM1;refGene_tx=XM_019836535.2;refGene_gn=PUM1;refGene_tx=XM_019836536.2;refGene_gn=PUM1;refGene_tx=XM_019836538.2;refGene_gn=PUM1;refGene_tx=XM_019836539.2;refGene_gn=PUM1;refGene_tx=XM_019836540.2;refGene_gn=PUM1;refGene_tx=XM_019836541.2;refGene_gn=PUM1;refGene_tx=XM_019836542.2;refGene_gn=PUM1;refGene_tx=XM_019836543.2;refGene_gn=PUM1;refGene_tx=XM_019836544.2;refGene_gn=PUM1;refGene_tx=XM_019836545.2;refGene_gn=PUM1;refGene_tx=XM_019836546.2;refGene_gn=PUM1;refGene_tx=XM_019836548.2;refGene_gn=PUM1;refGene_tx=XM_019836549.2;refGene_gn=PUM1;refGene_tx=XM_019836550.2
C1 23519660 23519931
refGene_gn=PUM1;refGene_tx=XM_011284760.3;refGene_gn=PUM1;refGene_tx=XM_019836533.2;refGene_gn=PUM1;refGene_tx=XM_019836534.2;refGene_gn=PUM1;refGene_tx=XM_019836535.2;refGene_gn=PUM1;refGene_tx=XM_019836536.2;refGene_gn=PUM1;refGene_tx=XM_019836538.2;refGene_gn=PUM1;refGene_tx=XM_019836539.2;refGene_gn=PUM1;refGene_tx=XM_019836540.2;refGene_gn=PUM1;refGene_tx=XM_019836541.2;refGene_gn=PUM1;refGene_tx=XM_019836542.2;refGene_gn=PUM1;refGene_tx=XM_019836543.2;refGene_gn=PUM1;refGene_tx=XM_019836544.2;refGene_gn=PUM1;refGene_tx=XM_019836545.2;refGene_gn=PUM1;refGene_tx=XM_019836550.2
C1 23521961 23522128
refGene_gn=PUM1;refGene_tx=XM_011284760.3;refGene_gn=PUM1;refGene_tx=XM_019836533.2;refGene_gn=PUM1;refGene_tx=XM_019836534.2;refGene_gn=PUM1;refGene_tx=XM_019836535.2;refGene_gn=PUM1;refGene_tx=XM_019836536.2;refGene_gn=PUM1;refGene_tx=XM_019836538.2;refGene_gn=PUM1;refGene_tx=XM_019836539.2;refGene_gn=PUM1;refGene_tx=XM_019836540.2;refGene_gn=PUM1;refGene_tx=XM_019836541.2;refGene_gn=PUM1;refGene_tx=XM_019836542.2;refGene_gn=PUM1;refGene_tx=XM_019836543.2;refGene_gn=PUM1;refGene_tx=XM_019836544.2;refGene_gn=PUM1;refGene_tx=XM_019836545.2;refGene_gn=PUM1;refGene_tx=XM_019836550.2
C1 23531732 23531911
refGene_gn=PUM1;refGene_tx=XM_011284760.3;refGene_gn=PUM1;refGene_tx=XM_019836533.2;refGene_gn=PUM1;refGene_tx=XM_019836534.2;refGene_gn=PUM1;refGene_tx=XM_019836535.2;refGene_gn=PUM1;refGene_tx=XM_019836536.2;refGene_gn=PUM1;refGene_tx=XM_019836538.2;refGene_gn=PUM1;refGene_tx=XM_019836539.2;refGene_gn=PUM1;refGene_tx=XM_019836540.2;refGene_gn=PUM1;refGene_tx=XM_019836541.2
C1 23532884 23532993
refGene_gn=PUM1;refGene_tx=XM_011284760.3;refGene_gn=PUM1;refGene_tx=XM_019836533.2;refGene_gn=PUM1;refGene_tx=XM_019836534.2;refGene_gn=PUM1;refGene_tx=XM_019836535.2;refGene_gn=PUM1;refGene_tx=XM_019836536.2;refGene_gn=PUM1;refGene_tx=XM_019836538.2;refGene_gn=PUM1;refGene_tx=XM_019836539.2;refGene_gn=PUM1;refGene_tx=XM_019836540.2;refGene_gn=PUM1;refGene_tx=XM_019836541.2;refGene_gn=PUM1;refGene_tx=XM_019836544.2;refGene_gn=PUM1;refGene_tx=XM_019836545.2
C1 23550716 23550785
refGene_gn=PUM1;refGene_tx=XM_011284760.3;refGene_gn=PUM1;refGene_tx=X

M_019836533.2;refGene_gn=PUM1;refGene_tx=XM_019836534.2;refGene_gn=PUM1;refGene_tx=XM_019836535.2;refGene_gn=PUM1;refGene_tx=XM_019836536.2;refGene_gn=PUM1;refGene_tx=XM_019836538.2;refGene_gn=PUM1;refGene_tx=XM_019836539.2;refGene_gn=PUM1;refGene_tx=XM_019836540.2;refGene_gn=PUM1;refGene_tx=XM_019836541.2;refGene_gn=PUM1;refGene_tx=XM_019836542.2;refGene_gn=PUM1;refGene_tx=XM_019836543.2;refGene_gn=PUM1;refGene_tx=XM_019836544.2;refGene_gn=PUM1;refGene_tx=XM_019836545.2;refGene_gn=PUM1;refGene_tx=XM_019836546.2;refGene_gn=PUM1;refGene_tx=XM_019836548.2;refGene_gn=PUM1;refGene_tx=XM_019836549.2;refGene_gn=PUM1;refGene_tx=XM_019836550.2
C1 23586504 23586878
refGene_gn=PUM1;refGene_tx=XM_011284760.3;refGene_gn=PUM1;refGene_tx=XM_019836533.2;refGene_gn=PUM1;refGene_tx=XM_019836534.2;refGene_gn=PUM1;refGene_tx=XM_019836535.2;refGene_gn=PUM1;refGene_tx=XM_019836536.2;refGene_gn=PUM1;refGene_tx=XM_019836538.2;refGene_gn=PUM1;refGene_tx=XM_019836539.2;refGene_gn=PUM1;refGene_tx=XM_019836540.2;refGene_gn=PUM1;refGene_tx=XM_019836541.2;refGene_gn=PUM1;refGene_tx=XM_019836542.2;refGene_gn=PUM1;refGene_tx=XM_019836543.2;refGene_gn=PUM1;refGene_tx=XM_019836546.2;refGene_gn=PUM1;refGene_tx=XM_019836548.2;refGene_gn=PUM1;refGene_tx=XM_019836549.2;refGene_gn=PUM1;refGene_tx=XM_019836550.2
C1 23591513 23591610
refGene_gn=PUM1;refGene_tx=XM_019836533.2;refGene_gn=PUM1;refGene_tx=XM_019836534.2;refGene_gn=PUM1;refGene_tx=XM_019836535.2;refGene_gn=PUM1;refGene_tx=XM_019836536.2;refGene_gn=PUM1;refGene_tx=XM_019836538.2;refGene_gn=PUM1;refGene_tx=XM_019836539.2;refGene_gn=PUM1;refGene_tx=XM_019836540.2;refGene_gn=PUM1;refGene_tx=XM_019836541.2;refGene_gn=PUM1;refGene_tx=XM_019836542.2;refGene_gn=PUM1;refGene_tx=XM_019836543.2;refGene_gn=PUM1;refGene_tx=XM_019836546.2;refGene_gn=PUM1;refGene_tx=XM_019836548.2;refGene_gn=PUM1;refGene_tx=XM_019836549.2
C1 23682100 23682110
refGene_gn=NKAIN1;refGene_tx=XM_023258399.1;refGene_gn=NKAIN1;refGene_tx=XM_023258400.1
C1 23682266 23682348
refGene_gn=NKAIN1;refGene_tx=XM_023258399.1;refGene_gn=NKAIN1;refGene_tx=XM_023258400.1
C1 23682978 23683039
refGene_gn=NKAIN1;refGene_tx=XM_023258399.1;refGene_gn=NKAIN1;refGene_tx=XM_023258400.1
C1 23683709 23683907 refGene_gn=NKAIN1;refGene_tx=XM_023258399.1
C1 23685321 23685402
refGene_gn=NKAIN1;refGene_tx=XM_023258399.1;refGene_gn=NKAIN1;refGene_tx=XM_023258400.1
C1 23686812 23686950
refGene_gn=NKAIN1;refGene_tx=XM_023258399.1;refGene_gn=NKAIN1;refGene_tx=XM_023258400.1
C1 23723500 23723554
refGene_gn=NKAIN1;refGene_tx=XM_023258399.1;refGene_gn=NKAIN1;refGene_tx=XM_023258400.1
C1 23739092 23739142 refGene_gn=SNRNP40;refGene_tx=XM_011284757.3
C1 23740133 23740237 refGene_gn=SNRNP40;refGene_tx=XM_011284757.3

C1 23744488 23744550 refGene_gn=SNRNP40;refGene_tx=XM_011284757.3
C1 23746253 23746336 refGene_gn=SNRNP40;refGene_tx=XM_011284757.3
C1 23748071 23748192 refGene_gn=SNRNP40;refGene_tx=XM_011284757.3
C1 23753516 23753639 refGene_gn=SNRNP40;refGene_tx=XM_011284757.3
C1 23766949 23767115 refGene_gn=SNRNP40;refGene_tx=XM_011284757.3
C1 23768301 23768395 refGene_gn=SNRNP40;refGene_tx=XM_011284757.3
C1 23769791 23769921 refGene_gn=SNRNP40;refGene_tx=XM_011284757.3
C1 23772126 23772270 refGene_gn=SNRNP40;refGene_tx=XM_011284757.3
C1 23772465 23772590 refGene_gn=ZCCHC17;refGene_tx=XM_019836552.2
C1 23781748 23781869
refGene_gn=ZCCHC17;refGene_tx=XM_019836552.2;refGene_gn=ZCCHC17;refGene_tx=XM_019836554.2
C1 23789439 23789497
refGene_gn=ZCCHC17;refGene_tx=XM_019836552.2;refGene_gn=ZCCHC17;refGene_tx=XM_019836554.2
C1 23803624 23803725
refGene_gn=ZCCHC17;refGene_tx=XM_019836552.2;refGene_gn=ZCCHC17;refGene_tx=XM_019836554.2;refGene_gn=ZCCHC17;refGene_tx=XM_019836555.2
C1 23805360 23805452
refGene_gn=ZCCHC17;refGene_tx=XM_019836552.2;refGene_gn=ZCCHC17;refGene_tx=XM_019836554.2;refGene_gn=ZCCHC17;refGene_tx=XM_019836555.2
C1 23813315 23813416
refGene_gn=ZCCHC17;refGene_tx=XM_019836552.2;refGene_gn=ZCCHC17;refGene_tx=XM_019836554.2;refGene_gn=ZCCHC17;refGene_tx=XM_019836555.2
C1 23815174 23815320
refGene_gn=ZCCHC17;refGene_tx=XM_019836552.2;refGene_gn=ZCCHC17;refGene_tx=XM_019836554.2;refGene_gn=ZCCHC17;refGene_tx=XM_019836555.2
C1 23833440 23833602
refGene_gn=ZCCHC17;refGene_tx=XM_019836552.2;refGene_gn=ZCCHC17;refGene_tx=XM_019836554.2;refGene_gn=ZCCHC17;refGene_tx=XM_019836555.2
C1 23835318 23835372
refGene_gn=FABP3;refGene_tx=XM_003989794.5;refGene_gn=FABP3;refGene_tx=XM_019836556.2
C1 23836687 23836789
refGene_gn=FABP3;refGene_tx=XM_003989794.5;refGene_gn=FABP3;refGene_tx=XM_019836556.2
C1 23838726 23838899
refGene_gn=FABP3;refGene_tx=XM_003989794.5;refGene_gn=FABP3;refGene_tx=XM_019836556.2
C1 23842258 23842331
refGene_gn=FABP3;refGene_tx=XM_003989794.5;refGene_gn=FABP3;refGene_tx=XM_019836556.2
C1 23871181 23871196 refGene_gn=SERINC2;refGene_tx=XM_019836557.2
C1 23871720 23871771 refGene_gn=SERINC2;refGene_tx=XM_019836557.2
C1 23874180 23874219 refGene_gn=SERINC2;refGene_tx=XM_023258401.1
C1 23881267 23881276 refGene_gn=SERINC2;refGene_tx=XM_006934521.3
C1 23883109 23883271
refGene_gn=SERINC2;refGene_tx=XM_006934521.3;refGene_gn=SERINC2;refGene_tx=XM_019836557.2;refGene_gn=SERINC2;refGene_tx=XM_023258401.1
C1 23884070 23884261

refGene_gn=SERINC2;refGene_tx=XM_006934521.3;refGene_gn=SERINC2;refGene_tx=XM_019836557.2;refGene_gn=SERINC2;refGene_tx=XM_023258401.1
C1 23884650 23884730
refGene_gn=SERINC2;refGene_tx=XM_006934521.3;refGene_gn=SERINC2;refGene_tx=XM_019836557.2;refGene_gn=SERINC2;refGene_tx=XM_023258401.1
C1 23885205 23885343
refGene_gn=SERINC2;refGene_tx=XM_006934521.3;refGene_gn=SERINC2;refGene_tx=XM_019836557.2;refGene_gn=SERINC2;refGene_tx=XM_023258401.1
C1 23886490 23886660
refGene_gn=SERINC2;refGene_tx=XM_006934521.3;refGene_gn=SERINC2;refGene_tx=XM_019836557.2;refGene_gn=SERINC2;refGene_tx=XM_023258401.1
C1 23888266 23888357
refGene_gn=SERINC2;refGene_tx=XM_006934521.3;refGene_gn=SERINC2;refGene_tx=XM_019836557.2;refGene_gn=SERINC2;refGene_tx=XM_023258401.1
C1 23888652 23888794
refGene_gn=SERINC2;refGene_tx=XM_006934521.3;refGene_gn=SERINC2;refGene_tx=XM_019836557.2;refGene_gn=SERINC2;refGene_tx=XM_023258401.1
C1 23893259 23893475
refGene_gn=SERINC2;refGene_tx=XM_006934521.3;refGene_gn=SERINC2;refGene_tx=XM_019836557.2;refGene_gn=SERINC2;refGene_tx=XM_023258401.1
C1 23894342 23894478
refGene_gn=SERINC2;refGene_tx=XM_006934521.3;refGene_gn=SERINC2;refGene_tx=XM_019836557.2;refGene_gn=SERINC2;refGene_tx=XM_023258401.1
C1 23928659 23928824
refGene_gn=LOC101101148;refGene_tx=XM_023257820.1
C1 23929808 23929982
refGene_gn=LOC101101148;refGene_tx=XM_023257820.1
C1 23930401 23930574
refGene_gn=LOC101101148;refGene_tx=XM_023257820.1
C1 23931245 23931380
refGene_gn=LOC101101148;refGene_tx=XM_023257820.1
C1 23932282 23932448
refGene_gn=LOC101101148;refGene_tx=XM_023257820.1
C1 23932789 23932937
refGene_gn=LOC101101148;refGene_tx=XM_023257820.1
C1 23934241 23934342
refGene_gn=LOC101101148;refGene_tx=XM_023257820.1
C1 23935139 23935357
refGene_gn=LOC101101148;refGene_tx=XM_023257820.1
C1 23936920 23937002
refGene_gn=LOC101101148;refGene_tx=XM_023257820.1
C1 23961109 23961545 refGene_gn=TINAGL1;refGene_tx=XM_023258403.1
C1 23962055 23962365
refGene_gn=TINAGL1;refGene_tx=XM_023258402.1;refGene_gn=TINAGL1;refGene_tx=XM_023258404.1
C1 23963353 23963417
refGene_gn=TINAGL1;refGene_tx=XM_023258402.1;refGene_gn=TINAGL1;refGene_tx=XM_023258403.1;refGene_gn=TINAGL1;refGene_tx=XM_023258404.1
C1 23967054 23967147
refGene_gn=TINAGL1;refGene_tx=XM_023258402.1;refGene_gn=TINAGL1;refGen

e_tx=XM_023258403.1;refGene_gn=TINAGL1;refGene_tx=XM_023258404.1
C1	23967340 23967455
refGene_gn=TINAGL1;refGene_tx=XM_023258402.1;refGene_gn=TINAGL1;refGen
e_tx=XM_023258403.1;refGene_gn=TINAGL1;refGene_tx=XM_023258404.1
C1	23968565 23968689
refGene_gn=TINAGL1;refGene_tx=XM_023258402.1;refGene_gn=TINAGL1;refGen
e_tx=XM_023258403.1;refGene_gn=TINAGL1;refGene_tx=XM_023258404.1
C1	23968782 23968933
refGene_gn=TINAGL1;refGene_tx=XM_023258402.1;refGene_gn=TINAGL1;refGen
e_tx=XM_023258403.1;refGene_gn=TINAGL1;refGene_tx=XM_023258404.1
C1	23969062 23969252
refGene_gn=TINAGL1;refGene_tx=XM_023258402.1;refGene_gn=TINAGL1;refGen
e_tx=XM_023258403.1;refGene_gn=TINAGL1;refGene_tx=XM_023258404.1
C1	23969353 23969399
refGene_gn=TINAGL1;refGene_tx=XM_023258402.1;refGene_gn=TINAGL1;refGen
e_tx=XM_023258403.1;refGene_gn=TINAGL1;refGene_tx=XM_023258404.1
C1	23969655 23969779
refGene_gn=TINAGL1;refGene_tx=XM_023258402.1;refGene_gn=TINAGL1;refGen
e_tx=XM_023258403.1;refGene_gn=TINAGL1;refGene_tx=XM_023258404.1
C1	23970553 23970599
refGene_gn=TINAGL1;refGene_tx=XM_023258402.1;refGene_gn=TINAGL1;refGen
e_tx=XM_023258403.1;refGene_gn=TINAGL1;refGene_tx=XM_023258404.1
C1	23970696 23970837
refGene_gn=TINAGL1;refGene_tx=XM_023258402.1;refGene_gn=TINAGL1;refGen
e_tx=XM_023258403.1;refGene_gn=TINAGL1;refGene_tx=XM_023258404.1
C1	24000747 24000952 refGene_gn=HCRTR1;refGene_tx=XM_003989799.3
C1	24001186 24001365 refGene_gn=HCRTR1;refGene_tx=XM_003989799.3
C1	24002589 24002833 refGene_gn=HCRTR1;refGene_tx=XM_003989799.3
C1	24003262 24003378 refGene_gn=HCRTR1;refGene_tx=XM_003989799.3
C1	24004531 24004758 refGene_gn=HCRTR1;refGene_tx=XM_003989799.3
C1	24005546 24005668 refGene_gn=HCRTR1;refGene_tx=XM_003989799.3
C1	24007344 24007535 refGene_gn=HCRTR1;refGene_tx=XM_003989799.3
C1	24011109 24011437
refGene_gn=PEF1;refGene_tx=XM_003989800.3;refGene_gn=PEF1;refGene_tx=X
M_006934523.3;refGene_gn=PEF1;refGene_tx=XM_019836562.1
C1	24012641 24012769
refGene_gn=PEF1;refGene_tx=XM_006934523.3;refGene_gn=PEF1;refGene_tx=X
M_019836562.1
C1	24013104 24013248
refGene_gn=PEF1;refGene_tx=XM_003989800.3;refGene_gn=PEF1;refGene_tx=X
M_006934523.3;refGene_gn=PEF1;refGene_tx=XM_019836562.1
C1	24013735 24013891
refGene_gn=PEF1;refGene_tx=XM_003989800.3;refGene_gn=PEF1;refGene_tx=X
M_006934523.3;refGene_gn=PEF1;refGene_tx=XM_019836562.1
C1	24016998 24017299
refGene_gn=PEF1;refGene_tx=XM_003989800.3;refGene_gn=PEF1;refGene_tx=X
M_006934523.3;refGene_gn=PEF1;refGene_tx=XM_019836562.1
C1	24026462 24026486
refGene_gn=PEF1;refGene_tx=XM_003989800.3;refGene_gn=PEF1;refGene_tx=X
M_006934523.3;refGene_gn=PEF1;refGene_tx=XM_019836562.1

| | | | |
|---|---|---|---|
| C1 | 24033128 | 24033331 | refGene_gn=COL16A1;refGene_tx=XM_006934524.4 |
| C1 | 24034033 | 24034111 | refGene_gn=COL16A1;refGene_tx=XM_006934524.4 |
| C1 | 24034311 | 24034488 | refGene_gn=COL16A1;refGene_tx=XM_006934524.4 |
| C1 | 24035184 | 24035251 | refGene_gn=COL16A1;refGene_tx=XM_006934524.4 |
| C1 | 24035623 | 24035812 | refGene_gn=COL16A1;refGene_tx=XM_006934524.4 |
| C1 | 24036737 | 24036782 | refGene_gn=COL16A1;refGene_tx=XM_006934524.4 |
| C1 | 24037164 | 24037200 | refGene_gn=COL16A1;refGene_tx=XM_006934524.4 |
| C1 | 24038679 | 24038769 | refGene_gn=COL16A1;refGene_tx=XM_006934524.4 |
| C1 | 24039115 | 24039166 | refGene_gn=COL16A1;refGene_tx=XM_006934524.4 |
| C1 | 24041287 | 24041341 | refGene_gn=COL16A1;refGene_tx=XM_006934524.4 |
| C1 | 24041772 | 24041808 | refGene_gn=COL16A1;refGene_tx=XM_006934524.4 |
| C1 | 24041809 | 24041827 | refGene_gn=COL16A1;refGene_tx=XM_006934524.4 |
| C1 | 24042130 | 24042175 | refGene_gn=COL16A1;refGene_tx=XM_006934524.4 |
| C1 | 24042390 | 24042435 | refGene_gn=COL16A1;refGene_tx=XM_006934524.4 |
| C1 | 24043046 | 24043100 | refGene_gn=COL16A1;refGene_tx=XM_006934524.4 |
| C1 | 24043289 | 24043361 | refGene_gn=COL16A1;refGene_tx=XM_006934524.4 |
| C1 | 24045662 | 24045725 | refGene_gn=COL16A1;refGene_tx=XM_006934524.4 |
| C1 | 24046072 | 24046108 | refGene_gn=COL16A1;refGene_tx=XM_006934524.4 |
| C1 | 24046445 | 24046499 | refGene_gn=COL16A1;refGene_tx=XM_006934524.4 |
| C1 | 24046613 | 24046658 | refGene_gn=COL16A1;refGene_tx=XM_006934524.4 |
| C1 | 24048269 | 24048323 | refGene_gn=COL16A1;refGene_tx=XM_006934524.4 |
| C1 | 24048852 | 24048906 | refGene_gn=COL16A1;refGene_tx=XM_006934524.4 |
| C1 | 24049486 | 24049540 | refGene_gn=COL16A1;refGene_tx=XM_006934524.4 |
| C1 | 24051305 | 24051350 | refGene_gn=COL16A1;refGene_tx=XM_006934524.4 |
| C1 | 24052221 | 24052266 | refGene_gn=COL16A1;refGene_tx=XM_006934524.4 |
| C1 | 24053066 | 24053153 | refGene_gn=COL16A1;refGene_tx=XM_006934524.4 |
| C1 | 24053244 | 24053286 | refGene_gn=COL16A1;refGene_tx=XM_006934524.4 |
| C1 | 24053370 | 24053487 | refGene_gn=COL16A1;refGene_tx=XM_006934524.4 |
| C1 | 24055401 | 24055434 | refGene_gn=COL16A1;refGene_tx=XM_006934524.4 |
| C1 | 24055674 | 24055728 | refGene_gn=COL16A1;refGene_tx=XM_006934524.4 |
| C1 | 24059021 | 24059075 | refGene_gn=COL16A1;refGene_tx=XM_006934524.4 |
| C1 | 24059207 | 24059255 | refGene_gn=COL16A1;refGene_tx=XM_006934524.4 |
| C1 | 24059440 | 24059500 | refGene_gn=COL16A1;refGene_tx=XM_006934524.4 |
| C1 | 24059955 | 24059982 | refGene_gn=COL16A1;refGene_tx=XM_006934524.4 |
| C1 | 24060073 | 24060118 | refGene_gn=COL16A1;refGene_tx=XM_006934524.4 |
| C1 | 24062193 | 24062262 | refGene_gn=COL16A1;refGene_tx=XM_006934524.4 |
| C1 | 24062433 | 24062487 | refGene_gn=COL16A1;refGene_tx=XM_006934524.4 |
| C1 | 24062577 | 24062613 | refGene_gn=COL16A1;refGene_tx=XM_006934524.4 |
| C1 | 24062939 | 24062981 | refGene_gn=COL16A1;refGene_tx=XM_006934524.4 |
| C1 | 24063181 | 24063235 | refGene_gn=COL16A1;refGene_tx=XM_006934524.4 |
| C1 | 24063342 | 24063465 | refGene_gn=COL16A1;refGene_tx=XM_006934524.4 |
| C1 | 24063785 | 24063893 | refGene_gn=COL16A1;refGene_tx=XM_006934524.4 |
| C1 | 24064095 | 24064131 | refGene_gn=COL16A1;refGene_tx=XM_006934524.4 |
| C1 | 24065134 | 24065266 | refGene_gn=COL16A1;refGene_tx=XM_006934524.4 |
| C1 | 24065568 | 24065613 | refGene_gn=COL16A1;refGene_tx=XM_006934524.4 |
| C1 | 24065783 | 24065819 | refGene_gn=COL16A1;refGene_tx=XM_006934524.4 |
| C1 | 24067624 | 24067669 | refGene_gn=COL16A1;refGene_tx=XM_006934524.4 |
| C1 | 24068014 | 24068125 | refGene_gn=COL16A1;refGene_tx=XM_006934524.4 |
| C1 | 24068202 | 24068238 | refGene_gn=COL16A1;refGene_tx=XM_006934524.4 |
| C1 | 24068464 | 24068500 | refGene_gn=COL16A1;refGene_tx=XM_006934524.4 |

C1 24068915 24069026 refGene_gn=COL16A1;refGene_tx=XM_006934524.4
C1 24069511 24069547 refGene_gn=COL16A1;refGene_tx=XM_006934524.4
C1 24069708 24069753 refGene_gn=COL16A1;refGene_tx=XM_006934524.4
C1 24070232 24070271 refGene_gn=COL16A1;refGene_tx=XM_006934524.4
C1 24070709 24070805 refGene_gn=COL16A1;refGene_tx=XM_006934524.4
C1 24070890 24070935 refGene_gn=COL16A1;refGene_tx=XM_006934524.4
C1 24071289 24071352 refGene_gn=COL16A1;refGene_tx=XM_006934524.4
C1 24071731 24071767 refGene_gn=COL16A1;refGene_tx=XM_006934524.4
C1 24071865 24071910 refGene_gn=COL16A1;refGene_tx=XM_006934524.4
C1 24072017 24072059 refGene_gn=COL16A1;refGene_tx=XM_006934524.4
C1 24072328 24072391 refGene_gn=COL16A1;refGene_tx=XM_006934524.4
C1 24073520 24073547 refGene_gn=COL16A1;refGene_tx=XM_006934524.4
C1 24074572 24074608 refGene_gn=COL16A1;refGene_tx=XM_006934524.4
C1 24075139 24075166 refGene_gn=COL16A1;refGene_tx=XM_006934524.4
C1 24075466 24075520 refGene_gn=COL16A1;refGene_tx=XM_006934524.4
C1 24076312 24076438 refGene_gn=COL16A1;refGene_tx=XM_006934524.4
C1 24076605 24076686 refGene_gn=COL16A1;refGene_tx=XM_006934524.4
C1 24077309 24077576 refGene_gn=COL16A1;refGene_tx=XM_006934524.4
C1 24077896 24078020 refGene_gn=COL16A1;refGene_tx=XM_006934524.4
C1 24078895 24079013 refGene_gn=COL16A1;refGene_tx=XM_006934524.4
C1 24079122 24079197 refGene_gn=COL16A1;refGene_tx=XM_006934524.4
C1 24081168 24081241 refGene_gn=COL16A1;refGene_tx=XM_006934524.4
C1 24105566 24105752
refGene_gn=ADGRB2;refGene_tx=XM_006934526.4;refGene_gn=ADGRB2;refGene_tx=XM_019836565.2;refGene_gn=ADGRB2;refGene_tx=XM_019836566.2;refGene_gn=ADGRB2;refGene_tx=XM_019836567.2;refGene_gn=ADGRB2;refGene_tx=XM_019836568.2;refGene_gn=ADGRB2;refGene_tx=XM_019836569.2;refGene_gn=ADGRB2;refGene_tx=XM_019836570.2;refGene_gn=ADGRB2;refGene_tx=XM_019836571.2;refGene_gn=ADGRB2;refGene_tx=XM_019836573.2;refGene_gn=ADGRB2;refGene_tx=XM_019836574.2;refGene_gn=ADGRB2;refGene_tx=XM_019836575.2;refGene_gn=ADGRB2;refGene_tx=XM_019836576.2
C1 24106113 24106170
refGene_gn=ADGRB2;refGene_tx=XM_006934526.4;refGene_gn=ADGRB2;refGene_tx=XM_019836565.2;refGene_gn=ADGRB2;refGene_tx=XM_019836566.2;refGene_gn=ADGRB2;refGene_tx=XM_019836569.2;refGene_gn=ADGRB2;refGene_tx=XM_019836570.2;refGene_gn=ADGRB2;refGene_tx=XM_019836571.2;refGene_gn=ADGRB2;refGene_tx=XM_019836573.2;refGene_gn=ADGRB2;refGene_tx=XM_019836574.2;refGene_gn=ADGRB2;refGene_tx=XM_019836575.2;refGene_gn=ADGRB2;refGene_tx=XM_019836576.2
C1 24106269 24106368
refGene_gn=ADGRB2;refGene_tx=XM_006934526.4;refGene_gn=ADGRB2;refGene_tx=XM_019836565.2;refGene_gn=ADGRB2;refGene_tx=XM_019836566.2;refGene_gn=ADGRB2;refGene_tx=XM_019836567.2;refGene_gn=ADGRB2;refGene_tx=XM_019836568.2;refGene_gn=ADGRB2;refGene_tx=XM_019836569.2;refGene_gn=ADGRB2;refGene_tx=XM_019836570.2;refGene_gn=ADGRB2;refGene_tx=XM_019836571.2;refGene_gn=ADGRB2;refGene_tx=XM_019836573.2;refGene_gn=ADGRB2;refGene_tx=XM_019836574.2;refGene_gn=ADGRB2;refGene_tx=XM_019836575.2;refGene_gn=ADGRB2;refGene_tx=XM_019836576.2
C1 24106727 24106763
refGene_gn=ADGRB2;refGene_tx=XM_006934526.4;refGene_gn=ADGRB2;refGene_

tx=XM_019836565.2;refGene_gn=ADGRB2;refGene_tx=XM_019836566.2;refGene_gn=ADGRB2;refGene_tx=XM_019836567.2;refGene_gn=ADGRB2;refGene_tx=XM_019836568.2;refGene_gn=ADGRB2;refGene_tx=XM_019836569.2;refGene_gn=ADGRB2;refGene_tx=XM_019836570.2;refGene_gn=ADGRB2;refGene_tx=XM_019836571.2;refGene_gn=ADGRB2;refGene_tx=XM_019836573.2;refGene_gn=ADGRB2;refGene_tx=XM_019836574.2;refGene_gn=ADGRB2;refGene_tx=XM_019836575.2;refGene_gn=ADGRB2;refGene_tx=XM_019836576.2
C1 24109104 24109724
refGene_gn=ADGRB2;refGene_tx=XM_006934526.4;refGene_gn=ADGRB2;refGene_tx=XM_019836565.2;refGene_gn=ADGRB2;refGene_tx=XM_019836566.2;refGene_gn=ADGRB2;refGene_tx=XM_019836567.2;refGene_gn=ADGRB2;refGene_tx=XM_019836568.2;refGene_gn=ADGRB2;refGene_tx=XM_019836569.2;refGene_gn=ADGRB2;refGene_tx=XM_019836570.2;refGene_gn=ADGRB2;refGene_tx=XM_019836571.2;refGene_gn=ADGRB2;refGene_tx=XM_019836573.2;refGene_gn=ADGRB2;refGene_tx=XM_019836574.2;refGene_gn=ADGRB2;refGene_tx=XM_019836575.2;refGene_gn=ADGRB2;refGene_tx=XM_019836576.2
C1 24110362 24110402
refGene_gn=ADGRB2;refGene_tx=XM_006934526.4;refGene_gn=ADGRB2;refGene_tx=XM_019836565.2;refGene_gn=ADGRB2;refGene_tx=XM_019836566.2;refGene_gn=ADGRB2;refGene_tx=XM_019836567.2;refGene_gn=ADGRB2;refGene_tx=XM_019836568.2;refGene_gn=ADGRB2;refGene_tx=XM_019836569.2;refGene_gn=ADGRB2;refGene_tx=XM_019836570.2;refGene_gn=ADGRB2;refGene_tx=XM_019836571.2;refGene_gn=ADGRB2;refGene_tx=XM_019836573.2;refGene_gn=ADGRB2;refGene_tx=XM_019836574.2;refGene_gn=ADGRB2;refGene_tx=XM_019836575.2;refGene_gn=ADGRB2;refGene_tx=XM_019836576.2
C1 24110775 24110871
refGene_gn=ADGRB2;refGene_tx=XM_006934526.4;refGene_gn=ADGRB2;refGene_tx=XM_019836565.2;refGene_gn=ADGRB2;refGene_tx=XM_019836566.2;refGene_gn=ADGRB2;refGene_tx=XM_019836567.2;refGene_gn=ADGRB2;refGene_tx=XM_019836568.2;refGene_gn=ADGRB2;refGene_tx=XM_019836569.2;refGene_gn=ADGRB2;refGene_tx=XM_019836570.2;refGene_gn=ADGRB2;refGene_tx=XM_019836571.2;refGene_gn=ADGRB2;refGene_tx=XM_019836573.2;refGene_gn=ADGRB2;refGene_tx=XM_019836574.2;refGene_gn=ADGRB2;refGene_tx=XM_019836575.2;refGene_gn=ADGRB2;refGene_tx=XM_019836576.2
C1 24111169 24111341
refGene_gn=ADGRB2;refGene_tx=XM_006934526.4;refGene_gn=ADGRB2;refGene_tx=XM_019836565.2;refGene_gn=ADGRB2;refGene_tx=XM_019836566.2;refGene_gn=ADGRB2;refGene_tx=XM_019836567.2;refGene_gn=ADGRB2;refGene_tx=XM_019836568.2;refGene_gn=ADGRB2;refGene_tx=XM_019836569.2;refGene_gn=ADGRB2;refGene_tx=XM_019836570.2;refGene_gn=ADGRB2;refGene_tx=XM_019836571.2;refGene_gn=ADGRB2;refGene_tx=XM_019836573.2;refGene_gn=ADGRB2;refGene_tx=XM_019836574.2;refGene_gn=ADGRB2;refGene_tx=XM_019836575.2;refGene_gn=ADGRB2;refGene_tx=XM_019836576.2
C1 24113389 24113488
refGene_gn=ADGRB2;refGene_tx=XM_006934526.4;refGene_gn=ADGRB2;refGene_tx=XM_019836565.2;refGene_gn=ADGRB2;refGene_tx=XM_019836566.2;refGene_gn=ADGRB2;refGene_tx=XM_019836567.2;refGene_gn=ADGRB2;refGene_tx=XM_019836568.2;refGene_gn=ADGRB2;refGene_tx=XM_019836570.2;refGene_gn=ADGRB2;refGene_tx=XM_019836571.2;refGene_gn=ADGRB2;refGene_tx=XM_019836573.2;refGene_gn=ADGRB2;refGene_tx=XM_019836574.2;refGene_gn=ADGRB2;refGen

e_tx=XM_019836575.2
C1 24113780 24113866
refGene_gn=ADGRB2;refGene_tx=XM_006934526.4;refGene_gn=ADGRB2;refGene_tx=XM_019836565.2;refGene_gn=ADGRB2;refGene_tx=XM_019836566.2;refGene_gn=ADGRB2;refGene_tx=XM_019836567.2;refGene_gn=ADGRB2;refGene_tx=XM_019836568.2;refGene_gn=ADGRB2;refGene_tx=XM_019836569.2;refGene_gn=ADGRB2;refGene_tx=XM_019836570.2;refGene_gn=ADGRB2;refGene_tx=XM_019836571.2;refGene_gn=ADGRB2;refGene_tx=XM_019836573.2;refGene_gn=ADGRB2;refGene_tx=XM_019836574.2;refGene_gn=ADGRB2;refGene_tx=XM_019836575.2;refGene_gn=ADGRB2;refGene_tx=XM_019836576.2
C1 24114006 24114073
refGene_gn=ADGRB2;refGene_tx=XM_006934526.4;refGene_gn=ADGRB2;refGene_tx=XM_019836565.2;refGene_gn=ADGRB2;refGene_tx=XM_019836566.2;refGene_gn=ADGRB2;refGene_tx=XM_019836567.2;refGene_gn=ADGRB2;refGene_tx=XM_019836568.2;refGene_gn=ADGRB2;refGene_tx=XM_019836569.2;refGene_gn=ADGRB2;refGene_tx=XM_019836570.2;refGene_gn=ADGRB2;refGene_tx=XM_019836571.2;refGene_gn=ADGRB2;refGene_tx=XM_019836573.2;refGene_gn=ADGRB2;refGene_tx=XM_019836574.2;refGene_gn=ADGRB2;refGene_tx=XM_019836575.2;refGene_gn=ADGRB2;refGene_tx=XM_019836576.2
C1 24114463 24114533
refGene_gn=ADGRB2;refGene_tx=XM_006934526.4;refGene_gn=ADGRB2;refGene_tx=XM_019836565.2;refGene_gn=ADGRB2;refGene_tx=XM_019836566.2;refGene_gn=ADGRB2;refGene_tx=XM_019836567.2;refGene_gn=ADGRB2;refGene_tx=XM_019836568.2;refGene_gn=ADGRB2;refGene_tx=XM_019836569.2;refGene_gn=ADGRB2;refGene_tx=XM_019836570.2;refGene_gn=ADGRB2;refGene_tx=XM_019836571.2;refGene_gn=ADGRB2;refGene_tx=XM_019836573.2;refGene_gn=ADGRB2;refGene_tx=XM_019836574.2;refGene_gn=ADGRB2;refGene_tx=XM_019836575.2;refGene_gn=ADGRB2;refGene_tx=XM_019836576.2
C1 24114724 24114875
refGene_gn=ADGRB2;refGene_tx=XM_006934526.4;refGene_gn=ADGRB2;refGene_tx=XM_019836565.2;refGene_gn=ADGRB2;refGene_tx=XM_019836566.2;refGene_gn=ADGRB2;refGene_tx=XM_019836567.2;refGene_gn=ADGRB2;refGene_tx=XM_019836568.2;refGene_gn=ADGRB2;refGene_tx=XM_019836569.2;refGene_gn=ADGRB2;refGene_tx=XM_019836570.2;refGene_gn=ADGRB2;refGene_tx=XM_019836571.2;refGene_gn=ADGRB2;refGene_tx=XM_019836573.2;refGene_gn=ADGRB2;refGene_tx=XM_019836574.2;refGene_gn=ADGRB2;refGene_tx=XM_019836575.2;refGene_gn=ADGRB2;refGene_tx=XM_019836576.2
C1 24115593 24115696
refGene_gn=ADGRB2;refGene_tx=XM_006934526.4;refGene_gn=ADGRB2;refGene_tx=XM_019836565.2;refGene_gn=ADGRB2;refGene_tx=XM_019836566.2;refGene_gn=ADGRB2;refGene_tx=XM_019836567.2;refGene_gn=ADGRB2;refGene_tx=XM_019836568.2;refGene_gn=ADGRB2;refGene_tx=XM_019836569.2;refGene_gn=ADGRB2;refGene_tx=XM_019836570.2;refGene_gn=ADGRB2;refGene_tx=XM_019836571.2;refGene_gn=ADGRB2;refGene_tx=XM_019836573.2;refGene_gn=ADGRB2;refGene_tx=XM_019836574.2;refGene_gn=ADGRB2;refGene_tx=XM_019836575.2;refGene_gn=ADGRB2;refGene_tx=XM_019836576.2
C1 24115797 24115901
refGene_gn=ADGRB2;refGene_tx=XM_006934526.4;refGene_gn=ADGRB2;refGene_tx=XM_019836565.2;refGene_gn=ADGRB2;refGene_tx=XM_019836566.2;refGene_gn=ADGRB2;refGene_tx=XM_019836567.2;refGene_gn=ADGRB2;refGene_tx=XM_01

9836568.2;refGene_gn=ADGRB2;refGene_tx=XM_019836569.2;refGene_gn=ADGRB2;refGene_tx=XM_019836570.2;refGene_gn=ADGRB2;refGene_tx=XM_019836571.2;refGene_gn=ADGRB2;refGene_tx=XM_019836573.2;refGene_gn=ADGRB2;refGene_tx=XM_019836574.2;refGene_gn=ADGRB2;refGene_tx=XM_019836575.2;refGene_gn=ADGRB2;refGene_tx=XM_019836576.2
C1 24116382 24116509
refGene_gn=ADGRB2;refGene_tx=XM_006934526.4;refGene_gn=ADGRB2;refGene_tx=XM_019836565.2;refGene_gn=ADGRB2;refGene_tx=XM_019836566.2;refGene_gn=ADGRB2;refGene_tx=XM_019836567.2;refGene_gn=ADGRB2;refGene_tx=XM_019836568.2;refGene_gn=ADGRB2;refGene_tx=XM_019836569.2;refGene_gn=ADGRB2;refGene_tx=XM_019836570.2;refGene_gn=ADGRB2;refGene_tx=XM_019836571.2;refGene_gn=ADGRB2;refGene_tx=XM_019836573.2;refGene_gn=ADGRB2;refGene_tx=XM_019836574.2;refGene_gn=ADGRB2;refGene_tx=XM_019836575.2;refGene_gn=ADGRB2;refGene_tx=XM_019836576.2
C1 24116772 24116816
refGene_gn=ADGRB2;refGene_tx=XM_006934526.4;refGene_gn=ADGRB2;refGene_tx=XM_019836565.2;refGene_gn=ADGRB2;refGene_tx=XM_019836566.2;refGene_gn=ADGRB2;refGene_tx=XM_019836567.2;refGene_gn=ADGRB2;refGene_tx=XM_019836568.2;refGene_gn=ADGRB2;refGene_tx=XM_019836569.2;refGene_gn=ADGRB2;refGene_tx=XM_019836570.2;refGene_gn=ADGRB2;refGene_tx=XM_019836571.2;refGene_gn=ADGRB2;refGene_tx=XM_019836573.2;refGene_gn=ADGRB2;refGene_tx=XM_019836574.2;refGene_gn=ADGRB2;refGene_tx=XM_019836575.2;refGene_gn=ADGRB2;refGene_tx=XM_019836576.2
C1 24116999 24117105
refGene_gn=ADGRB2;refGene_tx=XM_006934526.4;refGene_gn=ADGRB2;refGene_tx=XM_019836565.2;refGene_gn=ADGRB2;refGene_tx=XM_019836566.2;refGene_gn=ADGRB2;refGene_tx=XM_019836567.2;refGene_gn=ADGRB2;refGene_tx=XM_019836568.2;refGene_gn=ADGRB2;refGene_tx=XM_019836569.2;refGene_gn=ADGRB2;refGene_tx=XM_019836570.2;refGene_gn=ADGRB2;refGene_tx=XM_019836571.2;refGene_gn=ADGRB2;refGene_tx=XM_019836573.2;refGene_gn=ADGRB2;refGene_tx=XM_019836574.2;refGene_gn=ADGRB2;refGene_tx=XM_019836575.2;refGene_gn=ADGRB2;refGene_tx=XM_019836576.2
C1 24117433 24117764
refGene_gn=ADGRB2;refGene_tx=XM_006934526.4;refGene_gn=ADGRB2;refGene_tx=XM_019836565.2;refGene_gn=ADGRB2;refGene_tx=XM_019836566.2;refGene_gn=ADGRB2;refGene_tx=XM_019836567.2;refGene_gn=ADGRB2;refGene_tx=XM_019836568.2;refGene_gn=ADGRB2;refGene_tx=XM_019836569.2;refGene_gn=ADGRB2;refGene_tx=XM_019836570.2;refGene_gn=ADGRB2;refGene_tx=XM_019836571.2;refGene_gn=ADGRB2;refGene_tx=XM_019836573.2;refGene_gn=ADGRB2;refGene_tx=XM_019836574.2;refGene_gn=ADGRB2;refGene_tx=XM_019836575.2;refGene_gn=ADGRB2;refGene_tx=XM_019836576.2
C1 24118084 24118193
refGene_gn=ADGRB2;refGene_tx=XM_006934526.4;refGene_gn=ADGRB2;refGene_tx=XM_019836565.2;refGene_gn=ADGRB2;refGene_tx=XM_019836566.2;refGene_gn=ADGRB2;refGene_tx=XM_019836567.2;refGene_gn=ADGRB2;refGene_tx=XM_019836568.2;refGene_gn=ADGRB2;refGene_tx=XM_019836569.2;refGene_gn=ADGRB2;refGene_tx=XM_019836570.2;refGene_gn=ADGRB2;refGene_tx=XM_019836571.2;refGene_gn=ADGRB2;refGene_tx=XM_019836573.2;refGene_gn=ADGRB2;refGene_tx=XM_019836574.2;refGene_gn=ADGRB2;refGene_tx=XM_019836575.2;refGene_gn=ADGRB2;refGene_tx=XM_019836576.2

C1 24118270 24118339
refGene_gn=ADGRB2;refGene_tx=XM_006934526.4;refGene_gn=ADGRB2;refGene_tx=XM_019836565.2;refGene_gn=ADGRB2;refGene_tx=XM_019836566.2;refGene_gn=ADGRB2;refGene_tx=XM_019836567.2;refGene_gn=ADGRB2;refGene_tx=XM_019836568.2;refGene_gn=ADGRB2;refGene_tx=XM_019836569.2;refGene_gn=ADGRB2;refGene_tx=XM_019836570.2;refGene_gn=ADGRB2;refGene_tx=XM_019836571.2;refGene_gn=ADGRB2;refGene_tx=XM_019836573.2;refGene_gn=ADGRB2;refGene_tx=XM_019836574.2;refGene_gn=ADGRB2;refGene_tx=XM_019836575.2;refGene_gn=ADGRB2;refGene_tx=XM_019836576.2
C1 24118499 24118694
refGene_gn=ADGRB2;refGene_tx=XM_006934526.4;refGene_gn=ADGRB2;refGene_tx=XM_019836565.2;refGene_gn=ADGRB2;refGene_tx=XM_019836566.2;refGene_gn=ADGRB2;refGene_tx=XM_019836567.2;refGene_gn=ADGRB2;refGene_tx=XM_019836568.2;refGene_gn=ADGRB2;refGene_tx=XM_019836569.2;refGene_gn=ADGRB2;refGene_tx=XM_019836570.2;refGene_gn=ADGRB2;refGene_tx=XM_019836571.2;refGene_gn=ADGRB2;refGene_tx=XM_019836573.2;refGene_gn=ADGRB2;refGene_tx=XM_019836574.2;refGene_gn=ADGRB2;refGene_tx=XM_019836575.2;refGene_gn=ADGRB2;refGene_tx=XM_019836576.2
C1 24119483 24119590
refGene_gn=ADGRB2;refGene_tx=XM_006934526.4;refGene_gn=ADGRB2;refGene_tx=XM_019836565.2;refGene_gn=ADGRB2;refGene_tx=XM_019836566.2;refGene_gn=ADGRB2;refGene_tx=XM_019836567.2;refGene_gn=ADGRB2;refGene_tx=XM_019836568.2;refGene_gn=ADGRB2;refGene_tx=XM_019836569.2;refGene_gn=ADGRB2;refGene_tx=XM_019836570.2;refGene_gn=ADGRB2;refGene_tx=XM_019836571.2;refGene_gn=ADGRB2;refGene_tx=XM_019836573.2;refGene_gn=ADGRB2;refGene_tx=XM_019836574.2;refGene_gn=ADGRB2;refGene_tx=XM_019836575.2;refGene_gn=ADGRB2;refGene_tx=XM_019836576.2
C1 24119749 24119851
refGene_gn=ADGRB2;refGene_tx=XM_006934526.4;refGene_gn=ADGRB2;refGene_tx=XM_019836565.2;refGene_gn=ADGRB2;refGene_tx=XM_019836566.2;refGene_gn=ADGRB2;refGene_tx=XM_019836567.2;refGene_gn=ADGRB2;refGene_tx=XM_019836568.2;refGene_gn=ADGRB2;refGene_tx=XM_019836569.2;refGene_gn=ADGRB2;refGene_tx=XM_019836570.2;refGene_gn=ADGRB2;refGene_tx=XM_019836571.2;refGene_gn=ADGRB2;refGene_tx=XM_019836573.2;refGene_gn=ADGRB2;refGene_tx=XM_019836574.2;refGene_gn=ADGRB2;refGene_tx=XM_019836575.2;refGene_gn=ADGRB2;refGene_tx=XM_019836576.2
C1 24119928 24120096
refGene_gn=ADGRB2;refGene_tx=XM_006934526.4;refGene_gn=ADGRB2;refGene_tx=XM_019836565.2;refGene_gn=ADGRB2;refGene_tx=XM_019836566.2;refGene_gn=ADGRB2;refGene_tx=XM_019836567.2;refGene_gn=ADGRB2;refGene_tx=XM_019836568.2;refGene_gn=ADGRB2;refGene_tx=XM_019836569.2;refGene_gn=ADGRB2;refGene_tx=XM_019836570.2;refGene_gn=ADGRB2;refGene_tx=XM_019836571.2;refGene_gn=ADGRB2;refGene_tx=XM_019836573.2;refGene_gn=ADGRB2;refGene_tx=XM_019836574.2;refGene_gn=ADGRB2;refGene_tx=XM_019836575.2;refGene_gn=ADGRB2;refGene_tx=XM_019836576.2
C1 24120179 24120344
refGene_gn=ADGRB2;refGene_tx=XM_006934526.4;refGene_gn=ADGRB2;refGene_tx=XM_019836565.2;refGene_gn=ADGRB2;refGene_tx=XM_019836566.2;refGene_gn=ADGRB2;refGene_tx=XM_019836567.2;refGene_gn=ADGRB2;refGene_tx=XM_019836568.2;refGene_gn=ADGRB2;refGene_tx=XM_019836569.2;refGene_gn=ADGRB

2;refGene_tx=XM_019836570.2;refGene_gn=ADGRB2;refGene_tx=XM_019836571.
2;refGene_gn=ADGRB2;refGene_tx=XM_019836573.2;refGene_gn=ADGRB2;refGen
e_tx=XM_019836574.2;refGene_gn=ADGRB2;refGene_tx=XM_019836575.2;refGen
e_gn=ADGRB2;refGene_tx=XM_019836576.2
C1 24120967 24121132
refGene_gn=ADGRB2;refGene_tx=XM_006934526.4;refGene_gn=ADGRB2;refGene_
tx=XM_019836565.2;refGene_gn=ADGRB2;refGene_tx=XM_019836566.2;refGene_
gn=ADGRB2;refGene_tx=XM_019836567.2;refGene_gn=ADGRB2;refGene_tx=XM_01
9836568.2;refGene_gn=ADGRB2;refGene_tx=XM_019836569.2;refGene_gn=ADGRB
2;refGene_tx=XM_019836571.2;refGene_gn=ADGRB2;refGene_tx=XM_019836574.
2;refGene_gn=ADGRB2;refGene_tx=XM_019836575.2;refGene_gn=ADGRB2;refGen
e_tx=XM_019836576.2
C1 24122328 24122493
refGene_gn=ADGRB2;refGene_tx=XM_006934526.4;refGene_gn=ADGRB2;refGene_
tx=XM_019836565.2;refGene_gn=ADGRB2;refGene_tx=XM_019836566.2;refGene_
gn=ADGRB2;refGene_tx=XM_019836567.2;refGene_gn=ADGRB2;refGene_tx=XM_01
9836568.2;refGene_gn=ADGRB2;refGene_tx=XM_019836569.2;refGene_gn=ADGRB
2;refGene_tx=XM_019836570.2;refGene_gn=ADGRB2;refGene_tx=XM_019836574.
2;refGene_gn=ADGRB2;refGene_tx=XM_019836575.2;refGene_gn=ADGRB2;refGen
e_tx=XM_019836576.2
C1 24122748 24122832
refGene_gn=ADGRB2;refGene_tx=XM_006934526.4;refGene_gn=ADGRB2;refGene_
tx=XM_019836565.2;refGene_gn=ADGRB2;refGene_tx=XM_019836566.2;refGene_
gn=ADGRB2;refGene_tx=XM_019836567.2;refGene_gn=ADGRB2;refGene_tx=XM_01
9836568.2;refGene_gn=ADGRB2;refGene_tx=XM_019836569.2;refGene_gn=ADGRB
2;refGene_tx=XM_019836570.2;refGene_gn=ADGRB2;refGene_tx=XM_019836571.
2;refGene_gn=ADGRB2;refGene_tx=XM_019836573.2;refGene_gn=ADGRB2;refGen
e_tx=XM_019836574.2;refGene_gn=ADGRB2;refGene_tx=XM_019836575.2;refGen
e_gn=ADGRB2;refGene_tx=XM_019836576.2
C1 24133608 24134404
refGene_gn=ADGRB2;refGene_tx=XM_006934526.4;refGene_gn=ADGRB2;refGene_
tx=XM_019836565.2;refGene_gn=ADGRB2;refGene_tx=XM_019836566.2;refGene_
gn=ADGRB2;refGene_tx=XM_019836567.2;refGene_gn=ADGRB2;refGene_tx=XM_01
9836568.2;refGene_gn=ADGRB2;refGene_tx=XM_019836569.2;refGene_gn=ADGRB
2;refGene_tx=XM_019836570.2;refGene_gn=ADGRB2;refGene_tx=XM_019836571.
2;refGene_gn=ADGRB2;refGene_tx=XM_019836573.2;refGene_gn=ADGRB2;refGen
e_tx=XM_019836574.2;refGene_gn=ADGRB2;refGene_tx=XM_019836575.2;refGen
e_gn=ADGRB2;refGene_tx=XM_019836576.2
C1 24167796 24168461
refGene_gn=SPOCD1;refGene_tx=XM_023258406.1;refGene_gn=SPOCD1;refGene_
tx=XM_023258407.1
C1 24169453 24169640
refGene_gn=SPOCD1;refGene_tx=XM_023258406.1;refGene_gn=SPOCD1;refGene_
tx=XM_023258407.1
C1 24169926 24170016
refGene_gn=SPOCD1;refGene_tx=XM_023258406.1;refGene_gn=SPOCD1;refGene_
tx=XM_023258407.1
C1 24170964 24171100
refGene_gn=SPOCD1;refGene_tx=XM_023258406.1;refGene_gn=SPOCD1;refGene_
tx=XM_023258407.1

C1 24171409 24171560
refGene_gn=SPOCD1;refGene_tx=XM_023258406.1;refGene_gn=SPOCD1;refGene_tx=XM_023258407.1
C1 24171789 24171922
refGene_gn=SPOCD1;refGene_tx=XM_023258406.1;refGene_gn=SPOCD1;refGene_tx=XM_023258407.1
C1 24174044 24174173
refGene_gn=SPOCD1;refGene_tx=XM_023258406.1;refGene_gn=SPOCD1;refGene_tx=XM_023258407.1
C1 24175698 24175815
refGene_gn=SPOCD1;refGene_tx=XM_023258406.1;refGene_gn=SPOCD1;refGene_tx=XM_023258407.1
C1 24175934 24176094
refGene_gn=SPOCD1;refGene_tx=XM_023258406.1;refGene_gn=SPOCD1;refGene_tx=XM_023258407.1
C1 24176834 24176925
refGene_gn=SPOCD1;refGene_tx=XM_023258406.1;refGene_gn=SPOCD1;refGene_tx=XM_023258407.1
C1 24177196 24177257
refGene_gn=SPOCD1;refGene_tx=XM_023258406.1;refGene_gn=SPOCD1;refGene_tx=XM_023258407.1
C1 24177380 24177422
refGene_gn=SPOCD1;refGene_tx=XM_023258406.1;refGene_gn=SPOCD1;refGene_tx=XM_023258407.1
C1 24178084 24178261
refGene_gn=SPOCD1;refGene_tx=XM_023258406.1;refGene_gn=SPOCD1;refGene_tx=XM_023258407.1
C1 24179253 24179295 refGene_gn=SPOCD1;refGene_tx=XM_023258406.1
C1 24189533 24190874
refGene_gn=SPOCD1;refGene_tx=XM_023258406.1;refGene_gn=SPOCD1;refGene_tx=XM_023258407.1
C1 24254918 24255027 refGene_gn=PTP4A2;refGene_tx=XM_023258408.1
C1 24255848 24255923 refGene_gn=PTP4A2;refGene_tx=XM_023258408.1
C1 24258545 24258676 refGene_gn=PTP4A2;refGene_tx=XM_023258408.1
C1 24263290 24263356 refGene_gn=PTP4A2;refGene_tx=XM_023258408.1
C1 24263357 24263384 refGene_gn=PTP4A2;refGene_tx=XM_023258408.1
C1 24265621 24265631 refGene_gn=PTP4A2;refGene_tx=XM_023258408.1
C1 24267158 24267244 refGene_gn=PTP4A2;refGene_tx=XM_023258408.1
C1 24340078 24340460 refGene_gn=KHDRBS1;refGene_tx=XM_011284754.3
C1 24351280 24351405 refGene_gn=KHDRBS1;refGene_tx=XM_011284754.3
C1 24352713 24352830 refGene_gn=KHDRBS1;refGene_tx=XM_011284754.3
C1 24354862 24355009 refGene_gn=KHDRBS1;refGene_tx=XM_011284754.3
C1 24357675 24357809 refGene_gn=KHDRBS1;refGene_tx=XM_011284754.3
C1 24358770 24358972 refGene_gn=KHDRBS1;refGene_tx=XM_011284754.3
C1 24359481 24359549 refGene_gn=KHDRBS1;refGene_tx=XM_011284754.3
C1 24360304 24360363 refGene_gn=KHDRBS1;refGene_tx=XM_011284754.3
C1 24362688 24362786 refGene_gn=KHDRBS1;refGene_tx=XM_011284754.3
C1 24386594 24386598 refGene_gn=TMEM39B;refGene_tx=XM_003989808.4
C1 24386825 24386889 refGene_gn=TMEM39B;refGene_tx=XM_019836584.2
C1 24388258 24388385

refGene_gn=TMEM39B;refGene_tx=XM_003989808.4;refGene_gn=TMEM39B;refGene_tx=XM_019836584.2
C1 24388894 24389114
refGene_gn=TMEM39B;refGene_tx=XM_003989808.4;refGene_gn=TMEM39B;refGene_tx=XM_019836584.2
C1 24389726 24389810
refGene_gn=TMEM39B;refGene_tx=XM_003989808.4;refGene_gn=TMEM39B;refGene_tx=XM_019836584.2
C1 24390129 24390284
refGene_gn=TMEM39B;refGene_tx=XM_003989808.4;refGene_gn=TMEM39B;refGene_tx=XM_019836584.2
C1 24397366 24397703
refGene_gn=TMEM39B;refGene_tx=XM_003989808.4;refGene_gn=TMEM39B;refGene_tx=XM_019836584.2;refGene_gn=TMEM39B;refGene_tx=XM_019836586.2
C1 24398640 24398828
refGene_gn=TMEM39B;refGene_tx=XM_003989808.4;refGene_gn=TMEM39B;refGene_tx=XM_019836584.2;refGene_gn=TMEM39B;refGene_tx=XM_019836586.2
C1 24403094 24403215
refGene_gn=TMEM39B;refGene_tx=XM_003989808.4;refGene_gn=TMEM39B;refGene_tx=XM_019836584.2;refGene_gn=TMEM39B;refGene_tx=XM_019836586.2
C1 24404628 24404871
refGene_gn=TMEM39B;refGene_tx=XM_003989808.4;refGene_gn=TMEM39B;refGene_tx=XM_019836584.2;refGene_gn=TMEM39B;refGene_tx=XM_019836586.2
C1 24408481 24408485 refGene_gn=KPNA6;refGene_tx=XM_006934532.4
C1 24438422 24438441 refGene_gn=KPNA6;refGene_tx=XM_003989809.5
C1 24458076 24458210
refGene_gn=KPNA6;refGene_tx=XM_003989809.5;refGene_gn=KPNA6;refGene_tx=XM_006934532.4
C1 24459101 24459194
refGene_gn=KPNA6;refGene_tx=XM_003989809.5;refGene_gn=KPNA6;refGene_tx=XM_006934532.4
C1 24459592 24459692
refGene_gn=KPNA6;refGene_tx=XM_003989809.5;refGene_gn=KPNA6;refGene_tx=XM_006934532.4
C1 24460555 24460650
refGene_gn=KPNA6;refGene_tx=XM_003989809.5;refGene_gn=KPNA6;refGene_tx=XM_006934532.4
C1 24461619 24461751
refGene_gn=KPNA6;refGene_tx=XM_003989809.5;refGene_gn=KPNA6;refGene_tx=XM_006934532.4
C1 24462932 24463021
refGene_gn=KPNA6;refGene_tx=XM_003989809.5;refGene_gn=KPNA6;refGene_tx=XM_006934532.4
C1 24463912 24464012
refGene_gn=KPNA6;refGene_tx=XM_003989809.5;refGene_gn=KPNA6;refGene_tx=XM_006934532.4
C1 24464329 24464493
refGene_gn=KPNA6;refGene_tx=XM_003989809.5;refGene_gn=KPNA6;refGene_tx=XM_006934532.4
C1 24464679 24464758

refGene_gn=KPNA6;refGene_tx=XM_003989809.5;refGene_gn=KPNA6;refGene_tx=XM_006934532.4
C1 24466376 24466502 refGene_gn=KPNA6;refGene_tx=XM_003989809.5;refGene_gn=KPNA6;refGene_tx=XM_006934532.4
C1 24466794 24466922 refGene_gn=KPNA6;refGene_tx=XM_003989809.5;refGene_gn=KPNA6;refGene_tx=XM_006934532.4
C1 24467957 24468136 refGene_gn=KPNA6;refGene_tx=XM_003989809.5;refGene_gn=KPNA6;refGene_tx=XM_006934532.4
C1 24468750 24468938 refGene_gn=KPNA6;refGene_tx=XM_003989809.5;refGene_gn=KPNA6;refGene_tx=XM_006934532.4
C1 24477430 24477608 refGene_gn=TXLNA;refGene_tx=XM_011284753.2;refGene_gn=TXLNA;refGene_tx=XM_019836587.2
C1 24478224 24478563 refGene_gn=TXLNA;refGene_tx=XM_011284753.2;refGene_gn=TXLNA;refGene_tx=XM_019836587.2
C1 24480008 24480100 refGene_gn=TXLNA;refGene_tx=XM_011284753.2;refGene_gn=TXLNA;refGene_tx=XM_019836587.2
C1 24483237 24483408 refGene_gn=TXLNA;refGene_tx=XM_011284753.2;refGene_gn=TXLNA;refGene_tx=XM_019836587.2
C1 24484769 24484964 refGene_gn=TXLNA;refGene_tx=XM_011284753.2;refGene_gn=TXLNA;refGene_tx=XM_019836587.2
C1 24485586 24485706 refGene_gn=TXLNA;refGene_tx=XM_011284753.2;refGene_gn=TXLNA;refGene_tx=XM_019836587.2
C1 24485938 24486013 refGene_gn=TXLNA;refGene_tx=XM_011284753.2;refGene_gn=TXLNA;refGene_tx=XM_019836587.2
C1 24486765 24486858 refGene_gn=TXLNA;refGene_tx=XM_011284753.2;refGene_gn=TXLNA;refGene_tx=XM_019836587.2
C1 24487272 24487368 refGene_gn=TXLNA;refGene_tx=XM_011284753.2;refGene_gn=TXLNA;refGene_tx=XM_019836587.2
C1 24488085 24488412 refGene_gn=TXLNA;refGene_tx=XM_011284753.2;refGene_gn=TXLNA;refGene_tx=XM_019836587.2
C1 24495412 24495576 refGene_gn=CCDC28B;refGene_tx=XM_003989810.5
C1 24496698 24496865 refGene_gn=CCDC28B;refGene_tx=XM_003989810.5
C1 24496992 24497186 refGene_gn=CCDC28B;refGene_tx=XM_003989810.5
C1 24497388 24497411 refGene_gn=CCDC28B;refGene_tx=XM_003989810.5
C1 24498106 24498161 refGene_gn=CCDC28B;refGene_tx=XM_003989810.5
C1 24498593 24498635 refGene_gn=IQCC;refGene_tx=XM_003989917.4

C1	24499053	24499197	refGene_gn=IQCC;refGene_tx=XM_003989917.4
C1	24499427	24499692	refGene_gn=IQCC;refGene_tx=XM_003989917.4
C1	24499922	24500041	refGene_gn=IQCC;refGene_tx=XM_003989917.4
C1	24500161	24500986	refGene_gn=IQCC;refGene_tx=XM_003989917.4
C1	24501713	24501982	refGene_gn=DCDC2B;refGene_tx=XM_003989918.4
C1	24502868	24502920	refGene_gn=DCDC2B;refGene_tx=XM_003989918.4
C1	24503375	24503452	refGene_gn=DCDC2B;refGene_tx=XM_003989918.4
C1	24503691	24503823	refGene_gn=DCDC2B;refGene_tx=XM_003989918.4
C1	24504109	24504256	refGene_gn=DCDC2B;refGene_tx=XM_003989918.4
C1	24504366	24504406	refGene_gn=DCDC2B;refGene_tx=XM_003989918.4
C1	24505062	24505198	refGene_gn=DCDC2B;refGene_tx=XM_003989918.4
C1	24505604	24505708	refGene_gn=DCDC2B;refGene_tx=XM_003989918.4
C1	24505959	24506175	refGene_gn=DCDC2B;refGene_tx=XM_003989918.4
C1	24507027	24507122	refGene_gn=TMEM234;refGene_tx=XM_003989919.3
C1	24507393	24507486	refGene_gn=TMEM234;refGene_tx=XM_003989919.3
C1	24512444	24512511	refGene_gn=TMEM234;refGene_tx=XM_003989919.3
C1	24513292	24513444	refGene_gn=TMEM234;refGene_tx=XM_003989919.3
C1	24513722	24513738	refGene_gn=TMEM234;refGene_tx=XM_003989919.3
C1	24513856	24513859	refGene_gn=EIF3I;refGene_tx=XM_006934533.3
C1	24513964	24514057	refGene_gn=EIF3I;refGene_tx=XM_006934533.3
C1	24515209	24515297	refGene_gn=EIF3I;refGene_tx=XM_006934533.3
C1	24515607	24515673	refGene_gn=EIF3I;refGene_tx=XM_006934533.3
C1	24516780	24516930	refGene_gn=EIF3I;refGene_tx=XM_006934533.3
C1	24517019	24517147	refGene_gn=EIF3I;refGene_tx=XM_006934533.3
C1	24519848	24519959	refGene_gn=EIF3I;refGene_tx=XM_006934533.3
C1	24520078	24520168	refGene_gn=EIF3I;refGene_tx=XM_006934533.3
C1	24520473	24520547	refGene_gn=EIF3I;refGene_tx=XM_006934533.3
C1	24521014	24521107	refGene_gn=EIF3I;refGene_tx=XM_006934533.3
C1	24521238	24521320	refGene_gn=EIF3I;refGene_tx=XM_006934533.3
C1	24521666	24521828
refGene_gn=LOC101081743;refGene_tx=XM_006934534.4;refGene_gn=LOC101081743;refGene_tx=XM_019836588.2
C1	24521882	24522034
refGene_gn=LOC101081743;refGene_tx=XM_006934534.4;refGene_gn=LOC101081743;refGene_tx=XM_019836588.2
C1	24523317	24523538
refGene_gn=LOC101081743;refGene_tx=XM_006934534.4;refGene_gn=LOC101081743;refGene_tx=XM_019836588.2
C1	24523620	24523762
refGene_gn=LOC101081743;refGene_tx=XM_006934534.4;refGene_gn=LOC101081743;refGene_tx=XM_019836588.2
C1	24523936	24524098
refGene_gn=LOC101081743;refGene_tx=XM_006934534.4;refGene_gn=LOC101081743;refGene_tx=XM_019836588.2
C1	24525451	24525678
refGene_gn=LOC101081743;refGene_tx=XM_006934534.4;refGene_gn=LOC101081743;refGene_tx=XM_019836588.2
C1	24526182	24526356
refGene_gn=LOC101081743;refGene_tx=XM_006934534.4;refGene_gn=LOC101081743;refGene_tx=XM_019836588.2

C1 24526471 24526597
refGene_gn=LOC101081743;refGene_tx=XM_006934534.4;refGene_gn=LOC101081743;refGene_tx=XM_019836588.2
C1 24526837 24526946
refGene_gn=LOC101081743;refGene_tx=XM_006934534.4;refGene_gn=LOC101081743;refGene_tx=XM_019836588.2
C1 24527124 24527309
refGene_gn=LOC101081743;refGene_tx=XM_006934534.4;refGene_gn=LOC101081743;refGene_tx=XM_019836588.2
C1 24529515 24529776 refGene_gn=FAM167B;refGene_tx=XM_003989812.5
C1 24530450 24530681 refGene_gn=FAM167B;refGene_tx=XM_003989812.5
C1 24545719 24545747 refGene_gn=LCK;refGene_tx=XM_019836589.2
C1 24545844 24545954
refGene_gn=LCK;refGene_tx=XM_003989813.5;refGene_gn=LCK;refGene_tx=XM_019836589.2
C1 24546227 24546309
refGene_gn=LCK;refGene_tx=XM_003989813.5;refGene_gn=LCK;refGene_tx=XM_019836589.2
C1 24546504 24546595
refGene_gn=LCK;refGene_tx=XM_003989813.5;refGene_gn=LCK;refGene_tx=XM_019836589.2
C1 24546828 24546927
refGene_gn=LCK;refGene_tx=XM_003989813.5;refGene_gn=LCK;refGene_tx=XM_019836589.2
C1 24547060 24547164
refGene_gn=LCK;refGene_tx=XM_003989813.5;refGene_gn=LCK;refGene_tx=XM_019836589.2
C1 24547336 24547486
refGene_gn=LCK;refGene_tx=XM_003989813.5;refGene_gn=LCK;refGene_tx=XM_019836589.2
C1 24547776 24547929
refGene_gn=LCK;refGene_tx=XM_003989813.5;refGene_gn=LCK;refGene_tx=XM_019836589.2
C1 24548074 24548254
refGene_gn=LCK;refGene_tx=XM_003989813.5;refGene_gn=LCK;refGene_tx=XM_019836589.2
C1 24550397 24550474
refGene_gn=LCK;refGene_tx=XM_003989813.5;refGene_gn=LCK;refGene_tx=XM_019836589.2
C1 24550566 24550720
refGene_gn=LCK;refGene_tx=XM_003989813.5;refGene_gn=LCK;refGene_tx=XM_019836589.2
C1 24550803 24550935
refGene_gn=LCK;refGene_tx=XM_003989813.5;refGene_gn=LCK;refGene_tx=XM_019836589.2
C1 24553054 24553257
refGene_gn=LCK;refGene_tx=XM_003989813.5;refGene_gn=LCK;refGene_tx=XM_019836589.2
C1 24557686 24557735
refGene_gn=HDAC1;refGene_tx=XM_003989815.3;refGene_gn=HDAC1;refGene_tx

=XM_019836592.2
C1	24560958 24560986 refGene_gn=HDAC1;refGene_tx=XM_023258409.1
C1	24564338 24564451
refGene_gn=HDAC1;refGene_tx=XM_003989815.3;refGene_gn=HDAC1;refGene_tx
=XM_019836592.2;refGene_gn=HDAC1;refGene_tx=XM_023258409.1
C1	24575813 24575931
refGene_gn=HDAC1;refGene_tx=XM_003989815.3;refGene_gn=HDAC1;refGene_tx
=XM_019836592.2;refGene_gn=HDAC1;refGene_tx=XM_023258409.1;refGene_gn=
HDAC1;refGene_tx=XM_023258410.1
C1	24581700 24581775
refGene_gn=HDAC1;refGene_tx=XM_003989815.3;refGene_gn=HDAC1;refGene_tx
=XM_019836592.2;refGene_gn=HDAC1;refGene_tx=XM_023258409.1;refGene_gn=
HDAC1;refGene_tx=XM_023258410.1
C1	24583152 24583291
refGene_gn=HDAC1;refGene_tx=XM_003989815.3;refGene_gn=HDAC1;refGene_tx
=XM_019836592.2;refGene_gn=HDAC1;refGene_tx=XM_023258409.1;refGene_gn=
HDAC1;refGene_tx=XM_023258410.1
C1	24583754 24583896
refGene_gn=HDAC1;refGene_tx=XM_003989815.3;refGene_gn=HDAC1;refGene_tx
=XM_019836592.2;refGene_gn=HDAC1;refGene_tx=XM_023258409.1;refGene_gn=
HDAC1;refGene_tx=XM_023258410.1
C1	24584901 24584994
refGene_gn=HDAC1;refGene_tx=XM_003989815.3;refGene_gn=HDAC1;refGene_tx
=XM_019836592.2;refGene_gn=HDAC1;refGene_tx=XM_023258409.1;refGene_gn=
HDAC1;refGene_tx=XM_023258410.1
C1	24586138 24586247
refGene_gn=HDAC1;refGene_tx=XM_003989815.3;refGene_gn=HDAC1;refGene_tx
=XM_019836592.2;refGene_gn=HDAC1;refGene_tx=XM_023258409.1;refGene_gn=
HDAC1;refGene_tx=XM_023258410.1
C1	24586328 24586469
refGene_gn=HDAC1;refGene_tx=XM_003989815.3;refGene_gn=HDAC1;refGene_tx
=XM_019836592.2;refGene_gn=HDAC1;refGene_tx=XM_023258409.1;refGene_gn=
HDAC1;refGene_tx=XM_023258410.1
C1	24587005 24587114
refGene_gn=HDAC1;refGene_tx=XM_003989815.3;refGene_gn=HDAC1;refGene_tx
=XM_019836592.2;refGene_gn=HDAC1;refGene_tx=XM_023258409.1;refGene_gn=
HDAC1;refGene_tx=XM_023258410.1
C1	24587208 24587339
refGene_gn=HDAC1;refGene_tx=XM_003989815.3;refGene_gn=HDAC1;refGene_tx
=XM_019836592.2;refGene_gn=HDAC1;refGene_tx=XM_023258409.1;refGene_gn=
HDAC1;refGene_tx=XM_023258410.1
C1	24587614 24587767
refGene_gn=HDAC1;refGene_tx=XM_003989815.3;refGene_gn=HDAC1;refGene_tx
=XM_019836592.2;refGene_gn=HDAC1;refGene_tx=XM_023258409.1;refGene_gn=
HDAC1;refGene_tx=XM_023258410.1
C1	24588285 24588334 refGene_gn=HDAC1;refGene_tx=XM_003989815.3
C1	24588605 24588730
refGene_gn=HDAC1;refGene_tx=XM_003989815.3;refGene_gn=HDAC1;refGene_tx
=XM_019836592.2;refGene_gn=HDAC1;refGene_tx=XM_023258409.1;refGene_gn=
HDAC1;refGene_tx=XM_023258410.1

C1 24590200 24590710 refGene_gn=MARCKSL1;refGene_tx=XM_023258411.1
C1 24591571 24591658 refGene_gn=MARCKSL1;refGene_tx=XM_023258411.1
C1 24612784 24612887 refGene_gn=FAM229A;refGene_tx=XM_023258412.1
C1 24612928 24613001 refGene_gn=FAM229A;refGene_tx=XM_023258413.1
C1 24613095 24613242
refGene_gn=FAM229A;refGene_tx=XM_023258412.1;refGene_gn=FAM229A;refGen
e_tx=XM_023258413.1
C1 24613323 24613460
refGene_gn=FAM229A;refGene_tx=XM_023258412.1;refGene_gn=FAM229A;refGen
e_tx=XM_023258413.1
C1 24613800 24613813 refGene_gn=TSSK3;refGene_tx=XM_019836597.1
C1 24614074 24614219 refGene_gn=TSSK3;refGene_tx=XM_003989820.4
C1 24614947 24615609
refGene_gn=TSSK3;refGene_tx=XM_003989820.4;refGene_gn=TSSK3;refGene_tx
=XM_019836597.1
C1 24617756 24617789 refGene_gn=BSDC1;refGene_tx=XM_003989817.4
C1 24619246 24619350 refGene_gn=BSDC1;refGene_tx=XM_003989817.4
C1 24625644 24625819 refGene_gn=BSDC1;refGene_tx=XM_003989817.4
C1 24625820 24625976 refGene_gn=BSDC1;refGene_tx=XM_003989817.4
C1 24625977 24626126 refGene_gn=BSDC1;refGene_tx=XM_003989817.4
C1 24627892 24627971 refGene_gn=BSDC1;refGene_tx=XM_003989817.4
C1 24628132 24628201 refGene_gn=BSDC1;refGene_tx=XM_003989817.4
C1 24628619 24628735 refGene_gn=BSDC1;refGene_tx=XM_003989817.4
C1 24631364 24631419 refGene_gn=BSDC1;refGene_tx=XM_003989817.4
C1 24633066 24633234 refGene_gn=BSDC1;refGene_tx=XM_003989817.4
C1 24636172 24636289 refGene_gn=BSDC1;refGene_tx=XM_003989817.4
C1 24641126 24641187 refGene_gn=BSDC1;refGene_tx=XM_003989817.4
C1 24641453 24641464 refGene_gn=BSDC1;refGene_tx=XM_003989817.4
C1 24668992 24669968
refGene_gn=ZBTB8B;refGene_tx=XM_019836594.1;refGene_gn=ZBTB8B;refGene_
tx=XM_023258414.1;refGene_gn=ZBTB8B;refGene_tx=XM_023258415.1;refGene_
gn=ZBTB8B;refGene_tx=XM_023258416.1
C1 24680937 24681116
refGene_gn=ZBTB8B;refGene_tx=XM_019836594.1;refGene_gn=ZBTB8B;refGene_
tx=XM_023258414.1;refGene_gn=ZBTB8B;refGene_tx=XM_023258415.1;refGene_
gn=ZBTB8B;refGene_tx=XM_023258416.1
C1 24684561 24684879
refGene_gn=ZBTB8B;refGene_tx=XM_019836594.1;refGene_gn=ZBTB8B;refGene_
tx=XM_023258414.1;refGene_gn=ZBTB8B;refGene_tx=XM_023258415.1;refGene_
gn=ZBTB8B;refGene_tx=XM_023258416.1
C1 24785923 24786746
refGene_gn=ZBTB8A;refGene_tx=XM_023258418.1;refGene_gn=ZBTB8A;refGene_
tx=XM_023258419.1;refGene_gn=ZBTB8A;refGene_tx=XM_023258420.1
C1 24790257 24790427
refGene_gn=ZBTB8A;refGene_tx=XM_023258418.1;refGene_gn=ZBTB8A;refGene_
tx=XM_023258419.1;refGene_gn=ZBTB8A;refGene_tx=XM_023258420.1
C1 24796785 24797118
refGene_gn=ZBTB8A;refGene_tx=XM_023258418.1;refGene_gn=ZBTB8A;refGene_
tx=XM_023258419.1;refGene_gn=ZBTB8A;refGene_tx=XM_023258420.1
C1 24808121 24808203 refGene_gn=ZBTB8OS;refGene_tx=XM_019836608.2

C1 24813006 24813093
refGene_gn=ZBTB8OS;refGene_tx=XM_003989823.5;refGene_gn=ZBTB8OS;refGene_tx=XM_019836609.1
C1 24815787 24815824
refGene_gn=ZBTB8OS;refGene_tx=XM_003989823.5;refGene_gn=ZBTB8OS;refGene_tx=XM_019836609.1
C1 24817566 24817619
refGene_gn=ZBTB8OS;refGene_tx=XM_003989823.5;refGene_gn=ZBTB8OS;refGene_tx=XM_019836608.2;refGene_gn=ZBTB8OS;refGene_tx=XM_019836609.1
C1 24818599 24818682
refGene_gn=ZBTB8OS;refGene_tx=XM_003989823.5;refGene_gn=ZBTB8OS;refGene_tx=XM_019836608.2;refGene_gn=ZBTB8OS;refGene_tx=XM_019836609.1
C1 24818904 24819026
refGene_gn=ZBTB8OS;refGene_tx=XM_003989823.5;refGene_gn=ZBTB8OS;refGene_tx=XM_019836608.2;refGene_gn=ZBTB8OS;refGene_tx=XM_019836609.1
C1 24819555 24819580
refGene_gn=ZBTB8OS;refGene_tx=XM_003989823.5;refGene_gn=ZBTB8OS;refGene_tx=XM_019836608.2
C1 24829003 24829136
refGene_gn=ZBTB8OS;refGene_tx=XM_003989823.5;refGene_gn=ZBTB8OS;refGene_tx=XM_019836608.2
C1 24829886 24829902 refGene_gn=RBBP4;refGene_tx=XM_003989824.5
C1 24830481 24830629 refGene_gn=RBBP4;refGene_tx=XM_003989824.5
C1 24835063 24835209 refGene_gn=RBBP4;refGene_tx=XM_003989824.5
C1 24841608 24841782 refGene_gn=RBBP4;refGene_tx=XM_003989824.5
C1 24842130 24842246 refGene_gn=RBBP4;refGene_tx=XM_003989824.5
C1 24842363 24842524 refGene_gn=RBBP4;refGene_tx=XM_003989824.5
C1 24842620 24842747 refGene_gn=RBBP4;refGene_tx=XM_003989824.5
C1 24842884 24842962 refGene_gn=RBBP4;refGene_tx=XM_003989824.5
C1 24844752 24844829 refGene_gn=RBBP4;refGene_tx=XM_003989824.5
C1 24844935 24844993 refGene_gn=RBBP4;refGene_tx=XM_003989824.5
C1 24845083 24845194 refGene_gn=RBBP4;refGene_tx=XM_003989824.5
C1 24849600 24849666 refGene_gn=RBBP4;refGene_tx=XM_003989824.5
C1 24851719 24851795
refGene_gn=SYNC;refGene_tx=XM_023258421.1;refGene_gn=SYNC;refGene_tx=XM_023258422.1
C1 24853125 24853202 refGene_gn=SYNC;refGene_tx=XM_023258421.1
C1 24853380 24853505
refGene_gn=SYNC;refGene_tx=XM_023258421.1;refGene_gn=SYNC;refGene_tx=XM_023258422.1
C1 24859733 24860874
refGene_gn=SYNC;refGene_tx=XM_023258421.1;refGene_gn=SYNC;refGene_tx=XM_023258422.1
C1 24866382 24866435
refGene_gn=SYNC;refGene_tx=XM_023258421.1;refGene_gn=SYNC;refGene_tx=XM_023258422.1
C1 24923248 24923359 refGene_gn=YARS;refGene_tx=XM_023257821.1
C1 24925063 24925205 refGene_gn=YARS;refGene_tx=XM_023257821.1
C1 24925748 24925942 refGene_gn=YARS;refGene_tx=XM_023257821.1
C1 24926817 24926915 refGene_gn=YARS;refGene_tx=XM_023257821.1

C1	24927553	24927756	refGene_gn=YARS;refGene_tx=XM_023257821.1
C1	24934624	24934725	refGene_gn=YARS;refGene_tx=XM_023257821.1
C1	24940823	24940904	refGene_gn=YARS;refGene_tx=XM_023257821.1
C1	24943232	24943362	refGene_gn=YARS;refGene_tx=XM_023257821.1
C1	24948146	24948322	refGene_gn=YARS;refGene_tx=XM_023257821.1
C1	24948461	24948608	refGene_gn=YARS;refGene_tx=XM_023257821.1
C1	24951920	24952085	refGene_gn=YARS;refGene_tx=XM_023257821.1
C1	24959550	24960393
refGene_gn=S100PBP;refGene_tx=XM_006934539.4;refGene_gn=S100PBP;refGene_tx=XM_006934540.4;refGene_gn=S100PBP;refGene_tx=XM_006934541.4;refGene_gn=S100PBP;refGene_tx=XM_019836601.2;refGene_gn=S100PBP;refGene_tx=XM_019836602.2;refGene_gn=S100PBP;refGene_tx=XM_019836603.2;refGene_gn=S100PBP;refGene_tx=XM_019836604.2;refGene_gn=S100PBP;refGene_tx=XM_019836605.2;refGene_gn=S100PBP;refGene_tx=XM_019836606.2;refGene_gn=S100PBP;refGene_tx=XM_023258423.1
C1	24961021	24961110
refGene_gn=S100PBP;refGene_tx=XM_006934539.4;refGene_gn=S100PBP;refGene_tx=XM_006934540.4;refGene_gn=S100PBP;refGene_tx=XM_006934541.4;refGene_gn=S100PBP;refGene_tx=XM_019836601.2;refGene_gn=S100PBP;refGene_tx=XM_019836602.2;refGene_gn=S100PBP;refGene_tx=XM_019836603.2;refGene_gn=S100PBP;refGene_tx=XM_019836604.2;refGene_gn=S100PBP;refGene_tx=XM_019836605.2;refGene_gn=S100PBP;refGene_tx=XM_019836606.2;refGene_gn=S100PBP;refGene_tx=XM_023258423.1
C1	24963048	24963152
refGene_gn=S100PBP;refGene_tx=XM_006934539.4;refGene_gn=S100PBP;refGene_tx=XM_006934540.4;refGene_gn=S100PBP;refGene_tx=XM_006934541.4;refGene_gn=S100PBP;refGene_tx=XM_019836601.2;refGene_gn=S100PBP;refGene_tx=XM_019836602.2;refGene_gn=S100PBP;refGene_tx=XM_019836603.2;refGene_gn=S100PBP;refGene_tx=XM_019836604.2;refGene_gn=S100PBP;refGene_tx=XM_019836605.2;refGene_gn=S100PBP;refGene_tx=XM_019836606.2;refGene_gn=S100PBP;refGene_tx=XM_023258423.1
C1	24966771	24966776	refGene_gn=S100PBP;refGene_tx=XM_019836606.2
C1	24985212	24985300
refGene_gn=S100PBP;refGene_tx=XM_006934539.4;refGene_gn=S100PBP;refGene_tx=XM_006934540.4;refGene_gn=S100PBP;refGene_tx=XM_006934541.4;refGene_gn=S100PBP;refGene_tx=XM_019836601.2;refGene_gn=S100PBP;refGene_tx=XM_019836602.2;refGene_gn=S100PBP;refGene_tx=XM_019836603.2;refGene_gn=S100PBP;refGene_tx=XM_019836604.2;refGene_gn=S100PBP;refGene_tx=XM_019836605.2;refGene_gn=S100PBP;refGene_tx=XM_023258423.1
C1	24987122	24987264
refGene_gn=S100PBP;refGene_tx=XM_006934539.4;refGene_gn=S100PBP;refGene_tx=XM_006934540.4;refGene_gn=S100PBP;refGene_tx=XM_006934541.4;refGene_gn=S100PBP;refGene_tx=XM_019836601.2;refGene_gn=S100PBP;refGene_tx=XM_019836602.2;refGene_gn=S100PBP;refGene_tx=XM_019836603.2;refGene_gn=S100PBP;refGene_tx=XM_019836604.2;refGene_gn=S100PBP;refGene_tx=XM_019836605.2;refGene_gn=S100PBP;refGene_tx=XM_023258423.1
C1	24994313	24994352	refGene_gn=FNDC5;refGene_tx=XM_023258424.1
C1	24994905	24994911	refGene_gn=FNDC5;refGene_tx=XM_023258425.1
C1	24995220	24995354
refGene_gn=FNDC5;refGene_tx=XM_023258424.1;refGene_gn=FNDC5;refGene_tx

=XM_023258425.1
C1	24997396	24997486
refGene_gn=FNDC5;refGene_tx=XM_023258424.1;refGene_gn=FNDC5;refGene_tx
=XM_023258425.1
C1	24997831	24998030
refGene_gn=FNDC5;refGene_tx=XM_023258424.1;refGene_gn=FNDC5;refGene_tx
=XM_023258425.1
C1	24998529	24998645
refGene_gn=FNDC5;refGene_tx=XM_023258424.1;refGene_gn=FNDC5;refGene_tx
=XM_023258425.1
C1	25000273	25000346
refGene_gn=FNDC5;refGene_tx=XM_023258424.1;refGene_gn=FNDC5;refGene_tx
=XM_023258425.1
C1	25017222	25017600	refGene_gn=HPCA;refGene_tx=XM_006934543.4
C1	25021911	25022017	refGene_gn=HPCA;refGene_tx=XM_006934543.4
C1	25022138	25022236	refGene_gn=HPCA;refGene_tx=XM_006934543.4
C1	25023270	25023345	refGene_gn=TMEM54;refGene_tx=XM_003989829.4
C1	25023763	25023898	refGene_gn=TMEM54;refGene_tx=XM_003989829.4
C1	25024048	25024237	refGene_gn=TMEM54;refGene_tx=XM_003989829.4
C1	25024380	25024440	refGene_gn=TMEM54;refGene_tx=XM_003989829.4
C1	25026242	25026436	refGene_gn=TMEM54;refGene_tx=XM_003989829.4
C1	25029058	25029074	refGene_gn=TMEM54;refGene_tx=XM_003989829.4
C1	25053472	25053935
refGene_gn=RNF19B;refGene_tx=XM_023258426.1;refGene_gn=RNF19B;refGene_
tx=XM_023258427.1
C1	25053963	25053985	refGene_gn=RNF19B;refGene_tx=XM_023258428.1
C1	25054744	25054876
refGene_gn=RNF19B;refGene_tx=XM_023258426.1;refGene_gn=RNF19B;refGene_
tx=XM_023258427.1;refGene_gn=RNF19B;refGene_tx=XM_023258428.1
C1	25057234	25057442
refGene_gn=RNF19B;refGene_tx=XM_023258426.1;refGene_gn=RNF19B;refGene_
tx=XM_023258427.1;refGene_gn=RNF19B;refGene_tx=XM_023258428.1
C1	25058399	25058540
refGene_gn=RNF19B;refGene_tx=XM_023258426.1;refGene_gn=RNF19B;refGene_
tx=XM_023258427.1;refGene_gn=RNF19B;refGene_tx=XM_023258428.1
C1	25059596	25059711
refGene_gn=RNF19B;refGene_tx=XM_023258426.1;refGene_gn=RNF19B;refGene_
tx=XM_023258427.1;refGene_gn=RNF19B;refGene_tx=XM_023258428.1
C1	25060158	25060321
refGene_gn=RNF19B;refGene_tx=XM_023258426.1;refGene_gn=RNF19B;refGene_
tx=XM_023258427.1;refGene_gn=RNF19B;refGene_tx=XM_023258428.1
C1	25061757	25061902
refGene_gn=RNF19B;refGene_tx=XM_023258426.1;refGene_gn=RNF19B;refGene_
tx=XM_023258427.1;refGene_gn=RNF19B;refGene_tx=XM_023258428.1
C1	25063292	25063498
refGene_gn=RNF19B;refGene_tx=XM_023258426.1;refGene_gn=RNF19B;refGene_
tx=XM_023258427.1;refGene_gn=RNF19B;refGene_tx=XM_023258428.1
C1	25072806	25073438
refGene_gn=RNF19B;refGene_tx=XM_023258426.1;refGene_gn=RNF19B;refGene_
tx=XM_023258427.1;refGene_gn=RNF19B;refGene_tx=XM_023258428.1

C1 25112996 25113001 refGene_gn=AK2;refGene_tx=XM_006934544.4
C1 25115007 25115229
refGene_gn=AK2;refGene_tx=XM_003989831.5;refGene_gn=AK2;refGene_tx=XM_006934544.4
C1 25116391 25116464
refGene_gn=AK2;refGene_tx=XM_003989831.5;refGene_gn=AK2;refGene_tx=XM_006934544.4;refGene_gn=AK2;refGene_tx=XM_019836614.1
C1 25121345 25121440
refGene_gn=AK2;refGene_tx=XM_003989831.5;refGene_gn=AK2;refGene_tx=XM_006934544.4
C1 25121570 25121681
refGene_gn=AK2;refGene_tx=XM_003989831.5;refGene_gn=AK2;refGene_tx=XM_006934544.4;refGene_gn=AK2;refGene_tx=XM_019836614.1
C1 25123861 25123987
refGene_gn=AK2;refGene_tx=XM_003989831.5;refGene_gn=AK2;refGene_tx=XM_006934544.4;refGene_gn=AK2;refGene_tx=XM_019836614.1
C1 25132496 25132595
refGene_gn=AK2;refGene_tx=XM_003989831.5;refGene_gn=AK2;refGene_tx=XM_006934544.4;refGene_gn=AK2;refGene_tx=XM_019836614.1
C1 25165796 25165901
refGene_gn=AZIN2;refGene_tx=NM_001293721.1;refGene_gn=AZIN2;refGene_tx=XM_006934545.4;refGene_gn=AZIN2;refGene_tx=XM_019835927.2;refGene_gn=AZIN2;refGene_tx=XM_019835929.2;refGene_gn=AZIN2;refGene_tx=XM_019835931.2;refGene_gn=AZIN2;refGene_tx=XM_019835932.2
C1 25167405 25167579
refGene_gn=AZIN2;refGene_tx=NM_001293721.1;refGene_gn=AZIN2;refGene_tx=XM_006934545.4;refGene_gn=AZIN2;refGene_tx=XM_019835927.2;refGene_gn=AZIN2;refGene_tx=XM_019835929.2;refGene_gn=AZIN2;refGene_tx=XM_019835931.2;refGene_gn=AZIN2;refGene_tx=XM_019835932.2
C1 25171791 25172019
refGene_gn=AZIN2;refGene_tx=XM_006934545.4;refGene_gn=AZIN2;refGene_tx=XM_019835927.2;refGene_gn=AZIN2;refGene_tx=XM_019835929.2;refGene_gn=AZIN2;refGene_tx=XM_019835931.2;refGene_gn=AZIN2;refGene_tx=XM_019835932.2
C1 25176136 25176309
refGene_gn=AZIN2;refGene_tx=NM_001293721.1;refGene_gn=AZIN2;refGene_tx=XM_006934545.4;refGene_gn=AZIN2;refGene_tx=XM_019835927.2;refGene_gn=AZIN2;refGene_tx=XM_019835929.2;refGene_gn=AZIN2;refGene_tx=XM_019835931.2;refGene_gn=AZIN2;refGene_tx=XM_019835932.2
C1 25177316 25177445
refGene_gn=AZIN2;refGene_tx=NM_001293721.1;refGene_gn=AZIN2;refGene_tx=XM_006934545.4;refGene_gn=AZIN2;refGene_tx=XM_019835927.2;refGene_gn=AZIN2;refGene_tx=XM_019835929.2;refGene_gn=AZIN2;refGene_tx=XM_019835931.2;refGene_gn=AZIN2;refGene_tx=XM_019835932.2;refGene_gn=AZIN2;refGene_tx=XM_023257808.1
C1 25178263 25178429
refGene_gn=AZIN2;refGene_tx=NM_001293721.1;refGene_gn=AZIN2;refGene_tx=XM_006934545.4;refGene_gn=AZIN2;refGene_tx=XM_019835927.2;refGene_gn=AZIN2;refGene_tx=XM_019835929.2;refGene_gn=AZIN2;refGene_tx=XM_019835931.2;refGene_gn=AZIN2;refGene_tx=XM_019835932.2;refGene_gn=AZIN2;refGe

ne_tx=XM_023257808.1
C1 25179988 25180151
refGene_gn=AZIN2;refGene_tx=NM_001293721.1;refGene_gn=AZIN2;refGene_tx=XM_006934545.4;refGene_gn=AZIN2;refGene_tx=XM_019835927.2;refGene_gn=AZIN2;refGene_tx=XM_019835929.2;refGene_gn=AZIN2;refGene_tx=XM_019835931.2;refGene_gn=AZIN2;refGene_tx=XM_019835932.2;refGene_gn=AZIN2;refGene_tx=XM_023257808.1
C1 25181350 25181463
refGene_gn=AZIN2;refGene_tx=NM_001293721.1;refGene_gn=AZIN2;refGene_tx=XM_006934545.4;refGene_gn=AZIN2;refGene_tx=XM_019835927.2;refGene_gn=AZIN2;refGene_tx=XM_019835929.2;refGene_gn=AZIN2;refGene_tx=XM_019835931.2;refGene_gn=AZIN2;refGene_tx=XM_019835932.2;refGene_gn=AZIN2;refGene_tx=XM_023257808.1
C1 25182133 25182172 refGene_gn=AZIN2;refGene_tx=XM_019835932.2
C1 25192802 25192844 refGene_gn=AZIN2;refGene_tx=XM_019835931.2
C1 25194996 25195211
refGene_gn=AZIN2;refGene_tx=NM_001293721.1;refGene_gn=AZIN2;refGene_tx=XM_006934545.4;refGene_gn=AZIN2;refGene_tx=XM_019835927.2;refGene_gn=AZIN2;refGene_tx=XM_019835929.2;refGene_gn=AZIN2;refGene_tx=XM_023257808.1
C1 25197846 25198023
refGene_gn=AZIN2;refGene_tx=NM_001293721.1;refGene_gn=AZIN2;refGene_tx=XM_006934545.4;refGene_gn=AZIN2;refGene_tx=XM_019835927.2;refGene_gn=AZIN2;refGene_tx=XM_019835929.2;refGene_gn=AZIN2;refGene_tx=XM_023257808.1
C1 25218169 25218720
refGene_gn=TRIM62;refGene_tx=NM_001290247.1;refGene_gn=TRIM62;refGene_tx=XM_011284638.3;refGene_gn=TRIM62;refGene_tx=XM_019835933.2
C1 25228477 25228593
refGene_gn=TRIM62;refGene_tx=NM_001290247.1;refGene_gn=TRIM62;refGene_tx=XM_011284638.3;refGene_gn=TRIM62;refGene_tx=XM_019835933.2
C1 25229418 25229675
refGene_gn=TRIM62;refGene_tx=NM_001290247.1;refGene_gn=TRIM62;refGene_tx=XM_011284638.3;refGene_gn=TRIM62;refGene_tx=XM_019835933.2
C1 25234520 25234616
refGene_gn=TRIM62;refGene_tx=NM_001290247.1;refGene_gn=TRIM62;refGene_tx=XM_011284638.3;refGene_gn=TRIM62;refGene_tx=XM_019835933.2
C1 25241616 25241811 refGene_gn=TRIM62;refGene_tx=XM_019835933.2
C1 25246371 25246779
refGene_gn=TRIM62;refGene_tx=NM_001290247.1;refGene_gn=TRIM62;refGene_tx=XM_011284638.3
C1 25320787 25320825 refGene_gn=ZNF362;refGene_tx=XM_023258429.1
C1 25326159 25326223 refGene_gn=ZNF362;refGene_tx=XM_023258429.1
C1 25326411 25326658 refGene_gn=ZNF362;refGene_tx=XM_023258429.1
C1 25330858 25331180 refGene_gn=ZNF362;refGene_tx=XM_023258429.1
C1 25331944 25332169 refGene_gn=ZNF362;refGene_tx=XM_023258429.1
C1 25341012 25341091 refGene_gn=ZNF362;refGene_tx=XM_023258429.1
C1 25341203 25341362 refGene_gn=ZNF362;refGene_tx=XM_023258429.1
C1 25345056 25345173 refGene_gn=ZNF362;refGene_tx=XM_023258429.1
C1 25350830 25351518 refGene_gn=A3GALT2;refGene_tx=XM_023258430.1

C1 25352325 25352463 refGene_gn=A3GALT2;refGene_tx=XM_023258430.1
C1 25352744 25352834 refGene_gn=A3GALT2;refGene_tx=XM_023258430.1
C1 25353013 25353097 refGene_gn=A3GALT2;refGene_tx=XM_023258430.1
C1 25369476 25369499 refGene_gn=A3GALT2;refGene_tx=XM_023258430.1
C1 25373878 25374033
refGene_gn=PHC2;refGene_tx=XM_011284748.3;refGene_gn=PHC2;refGene_tx=XM_019836615.2;refGene_gn=PHC2;refGene_tx=XM_019836616.2;refGene_gn=PHC2;refGene_tx=XM_023258431.1;refGene_gn=PHC2;refGene_tx=XM_023258432.1;refGene_gn=PHC2;refGene_tx=XM_023258434.1;refGene_gn=PHC2;refGene_tx=XM_023258435.1
C1 25377065 25377342
refGene_gn=PHC2;refGene_tx=XM_011284748.3;refGene_gn=PHC2;refGene_tx=XM_019836615.2;refGene_gn=PHC2;refGene_tx=XM_019836616.2;refGene_gn=PHC2;refGene_tx=XM_023258431.1;refGene_gn=PHC2;refGene_tx=XM_023258432.1;refGene_gn=PHC2;refGene_tx=XM_023258434.1;refGene_gn=PHC2;refGene_tx=XM_023258435.1
C1 25378484 25378626
refGene_gn=PHC2;refGene_tx=XM_011284748.3;refGene_gn=PHC2;refGene_tx=XM_019836615.2;refGene_gn=PHC2;refGene_tx=XM_019836616.2;refGene_gn=PHC2;refGene_tx=XM_023258431.1;refGene_gn=PHC2;refGene_tx=XM_023258432.1;refGene_gn=PHC2;refGene_tx=XM_023258434.1;refGene_gn=PHC2;refGene_tx=XM_023258435.1
C1 25382562 25382677
refGene_gn=PHC2;refGene_tx=XM_011284748.3;refGene_gn=PHC2;refGene_tx=XM_019836615.2;refGene_gn=PHC2;refGene_tx=XM_019836616.2;refGene_gn=PHC2;refGene_tx=XM_023258431.1;refGene_gn=PHC2;refGene_tx=XM_023258432.1;refGene_gn=PHC2;refGene_tx=XM_023258434.1;refGene_gn=PHC2;refGene_tx=XM_023258435.1
C1 25383520 25383650
refGene_gn=PHC2;refGene_tx=XM_011284748.3;refGene_gn=PHC2;refGene_tx=XM_019836615.2;refGene_gn=PHC2;refGene_tx=XM_019836616.2;refGene_gn=PHC2;refGene_tx=XM_023258431.1;refGene_gn=PHC2;refGene_tx=XM_023258432.1;refGene_gn=PHC2;refGene_tx=XM_023258434.1;refGene_gn=PHC2;refGene_tx=XM_023258435.1
C1 25385377 25385580
refGene_gn=PHC2;refGene_tx=XM_011284748.3;refGene_gn=PHC2;refGene_tx=XM_019836615.2;refGene_gn=PHC2;refGene_tx=XM_019836616.2;refGene_gn=PHC2;refGene_tx=XM_023258431.1;refGene_gn=PHC2;refGene_tx=XM_023258432.1;refGene_gn=PHC2;refGene_tx=XM_023258434.1;refGene_gn=PHC2;refGene_tx=XM_023258435.1
C1 25403109 25403275
refGene_gn=PHC2;refGene_tx=XM_011284748.3;refGene_gn=PHC2;refGene_tx=XM_023258431.1;refGene_gn=PHC2;refGene_tx=XM_023258432.1;refGene_gn=PHC2;refGene_tx=XM_023258434.1
C1 25403526 25403927
refGene_gn=PHC2;refGene_tx=XM_011284748.3;refGene_gn=PHC2;refGene_tx=XM_023258431.1;refGene_gn=PHC2;refGene_tx=XM_023258432.1;refGene_gn=PHC2;refGene_tx=XM_023258434.1
C1 25417694 25418007
refGene_gn=PHC2;refGene_tx=XM_011284748.3;refGene_gn=PHC2;refGene_tx=X

M_023258431.1;refGene_gn=PHC2;refGene_tx=XM_023258432.1;refGene_gn=PHC2;refGene_tx=XM_023258434.1
C1 25419132 25419285
refGene_gn=PHC2;refGene_tx=XM_011284748.3;refGene_gn=PHC2;refGene_tx=XM_023258431.1;refGene_gn=PHC2;refGene_tx=XM_023258432.1
C1 25421164 25421329
refGene_gn=PHC2;refGene_tx=XM_011284748.3;refGene_gn=PHC2;refGene_tx=XM_023258431.1;refGene_gn=PHC2;refGene_tx=XM_023258432.1;refGene_gn=PHC2;refGene_tx=XM_023258434.1
C1 25421763 25421841
refGene_gn=PHC2;refGene_tx=XM_011284748.3;refGene_gn=PHC2;refGene_tx=XM_023258431.1;refGene_gn=PHC2;refGene_tx=XM_023258432.1;refGene_gn=PHC2;refGene_tx=XM_023258434.1
C1 25422870 25423029
refGene_gn=PHC2;refGene_tx=XM_011284748.3;refGene_gn=PHC2;refGene_tx=XM_023258431.1;refGene_gn=PHC2;refGene_tx=XM_023258432.1;refGene_gn=PHC2;refGene_tx=XM_023258434.1
C1 25425658 25425817
refGene_gn=PHC2;refGene_tx=XM_011284748.3;refGene_gn=PHC2;refGene_tx=XM_023258431.1;refGene_gn=PHC2;refGene_tx=XM_023258432.1;refGene_gn=PHC2;refGene_tx=XM_023258434.1
C1 25527187 25527598
refGene_gn=ZSCAN20;refGene_tx=XM_003989837.5;refGene_gn=ZSCAN20;refGene_tx=XM_006934549.4;refGene_gn=ZSCAN20;refGene_tx=XM_011284745.3;refGene_gn=ZSCAN20;refGene_tx=XM_011284746.3;refGene_gn=ZSCAN20;refGene_tx=XM_019836617.2;refGene_gn=ZSCAN20;refGene_tx=XM_019836618.2
C1 25532293 25532480
refGene_gn=ZSCAN20;refGene_tx=XM_003989837.5;refGene_gn=ZSCAN20;refGene_tx=XM_006934549.4;refGene_gn=ZSCAN20;refGene_tx=XM_006934552.4;refGene_gn=ZSCAN20;refGene_tx=XM_011284745.3;refGene_gn=ZSCAN20;refGene_tx=XM_011284746.3;refGene_gn=ZSCAN20;refGene_tx=XM_019836617.2;refGene_gn=ZSCAN20;refGene_tx=XM_019836618.2
C1 25532930 25533007
refGene_gn=ZSCAN20;refGene_tx=XM_003989837.5;refGene_gn=ZSCAN20;refGene_tx=XM_006934549.4;refGene_gn=ZSCAN20;refGene_tx=XM_006934552.4;refGene_gn=ZSCAN20;refGene_tx=XM_011284745.3;refGene_gn=ZSCAN20;refGene_tx=XM_011284746.3;refGene_gn=ZSCAN20;refGene_tx=XM_019836617.2;refGene_gn=ZSCAN20;refGene_tx=XM_019836618.2
C1 25533287 25533372
refGene_gn=ZSCAN20;refGene_tx=XM_003989837.5;refGene_gn=ZSCAN20;refGene_tx=XM_006934549.4;refGene_gn=ZSCAN20;refGene_tx=XM_006934552.4;refGene_gn=ZSCAN20;refGene_tx=XM_011284745.3;refGene_gn=ZSCAN20;refGene_tx=XM_011284746.3;refGene_gn=ZSCAN20;refGene_tx=XM_019836617.2;refGene_gn=ZSCAN20;refGene_tx=XM_019836618.2
C1 25536759 25537446
refGene_gn=ZSCAN20;refGene_tx=XM_003989837.5;refGene_gn=ZSCAN20;refGene_tx=XM_006934549.4;refGene_gn=ZSCAN20;refGene_tx=XM_006934552.4;refGene_gn=ZSCAN20;refGene_tx=XM_011284745.3;refGene_gn=ZSCAN20;refGene_tx=XM_011284746.3;refGene_gn=ZSCAN20;refGene_tx=XM_019836617.2;refGene_gn=ZSCAN20;refGene_tx=XM_019836618.2

C1	25539101	25539530
refGene_gn=ZSCAN20;refGene_tx=XM_003989837.5;refGene_gn=ZSCAN20;refGen
e_tx=XM_006934549.4;refGene_gn=ZSCAN20;refGene_tx=XM_006934552.4;refGe
ne_gn=ZSCAN20;refGene_tx=XM_011284745.3;refGene_gn=ZSCAN20;refGene_tx=
XM_011284746.3;refGene_gn=ZSCAN20;refGene_tx=XM_019836617.2;refGene_gn
=ZSCAN20;refGene_tx=XM_019836618.2
C1	25540313	25541569
refGene_gn=ZSCAN20;refGene_tx=XM_003989837.5;refGene_gn=ZSCAN20;refGen
e_tx=XM_006934549.4;refGene_gn=ZSCAN20;refGene_tx=XM_006934552.4;refGe
ne_gn=ZSCAN20;refGene_tx=XM_011284745.3;refGene_gn=ZSCAN20;refGene_tx=
XM_011284746.3;refGene_gn=ZSCAN20;refGene_tx=XM_019836617.2;refGene_gn
=ZSCAN20;refGene_tx=XM_019836618.2
C1	25561800	25562012
refGene_gn=CSMD2;refGene_tx=XM_023258437.1;refGene_gn=CSMD2;refGene_tx
=XM_023258438.1;refGene_gn=CSMD2;refGene_tx=XM_023258441.1
C1	25564300	25564516
refGene_gn=CSMD2;refGene_tx=XM_023258437.1;refGene_gn=CSMD2;refGene_tx
=XM_023258438.1;refGene_gn=CSMD2;refGene_tx=XM_023258439.1;refGene_gn=
CSMD2;refGene_tx=XM_023258440.1;refGene_gn=CSMD2;refGene_tx=XM_0232584
41.1
C1	25564724	25564863
refGene_gn=CSMD2;refGene_tx=XM_023258437.1;refGene_gn=CSMD2;refGene_tx
=XM_023258438.1;refGene_gn=CSMD2;refGene_tx=XM_023258439.1;refGene_gn=
CSMD2;refGene_tx=XM_023258440.1;refGene_gn=CSMD2;refGene_tx=XM_0232584
41.1
C1	25566240	25566328
refGene_gn=CSMD2;refGene_tx=XM_023258437.1;refGene_gn=CSMD2;refGene_tx
=XM_023258438.1;refGene_gn=CSMD2;refGene_tx=XM_023258439.1;refGene_gn=
CSMD2;refGene_tx=XM_023258440.1;refGene_gn=CSMD2;refGene_tx=XM_0232584
41.1
C1	25567713	25567826
refGene_gn=CSMD2;refGene_tx=XM_023258437.1;refGene_gn=CSMD2;refGene_tx
=XM_023258438.1;refGene_gn=CSMD2;refGene_tx=XM_023258439.1;refGene_gn=
CSMD2;refGene_tx=XM_023258440.1;refGene_gn=CSMD2;refGene_tx=XM_0232584
41.1
C1	25569551	25569713
refGene_gn=CSMD2;refGene_tx=XM_023258437.1;refGene_gn=CSMD2;refGene_tx
=XM_023258438.1;refGene_gn=CSMD2;refGene_tx=XM_023258439.1;refGene_gn=
CSMD2;refGene_tx=XM_023258440.1;refGene_gn=CSMD2;refGene_tx=XM_0232584
41.1
C1	25570407	25570473
refGene_gn=CSMD2;refGene_tx=XM_023258437.1;refGene_gn=CSMD2;refGene_tx
=XM_023258438.1;refGene_gn=CSMD2;refGene_tx=XM_023258439.1;refGene_gn=
CSMD2;refGene_tx=XM_023258440.1;refGene_gn=CSMD2;refGene_tx=XM_0232584
41.1
C1	25577174	25577354
refGene_gn=CSMD2;refGene_tx=XM_023258437.1;refGene_gn=CSMD2;refGene_tx
=XM_023258438.1;refGene_gn=CSMD2;refGene_tx=XM_023258439.1;refGene_gn=
CSMD2;refGene_tx=XM_023258440.1;refGene_gn=CSMD2;refGene_tx=XM_0232584
41.1

C1 25577924 25578036
refGene_gn=CSMD2;refGene_tx=XM_023258437.1;refGene_gn=CSMD2;refGene_tx=XM_023258438.1;refGene_gn=CSMD2;refGene_tx=XM_023258439.1;refGene_gn=CSMD2;refGene_tx=XM_023258440.1;refGene_gn=CSMD2;refGene_tx=XM_023258441.1
C1 25581608 25581682
refGene_gn=CSMD2;refGene_tx=XM_023258437.1;refGene_gn=CSMD2;refGene_tx=XM_023258438.1;refGene_gn=CSMD2;refGene_tx=XM_023258439.1;refGene_gn=CSMD2;refGene_tx=XM_023258440.1;refGene_gn=CSMD2;refGene_tx=XM_023258441.1
C1 25582019 25582193
refGene_gn=CSMD2;refGene_tx=XM_023258437.1;refGene_gn=CSMD2;refGene_tx=XM_023258438.1;refGene_gn=CSMD2;refGene_tx=XM_023258439.1;refGene_gn=CSMD2;refGene_tx=XM_023258440.1;refGene_gn=CSMD2;refGene_tx=XM_023258441.1
C1 25586976 25587150
refGene_gn=CSMD2;refGene_tx=XM_023258437.1;refGene_gn=CSMD2;refGene_tx=XM_023258438.1;refGene_gn=CSMD2;refGene_tx=XM_023258439.1;refGene_gn=CSMD2;refGene_tx=XM_023258440.1;refGene_gn=CSMD2;refGene_tx=XM_023258441.1
C1 25587584 25587764
refGene_gn=CSMD2;refGene_tx=XM_023258437.1;refGene_gn=CSMD2;refGene_tx=XM_023258438.1;refGene_gn=CSMD2;refGene_tx=XM_023258439.1;refGene_gn=CSMD2;refGene_tx=XM_023258440.1;refGene_gn=CSMD2;refGene_tx=XM_023258441.1
C1 25588967 25589144
refGene_gn=CSMD2;refGene_tx=XM_023258437.1;refGene_gn=CSMD2;refGene_tx=XM_023258438.1;refGene_gn=CSMD2;refGene_tx=XM_023258439.1;refGene_gn=CSMD2;refGene_tx=XM_023258440.1;refGene_gn=CSMD2;refGene_tx=XM_023258441.1
C1 25593634 25593817
refGene_gn=CSMD2;refGene_tx=XM_023258437.1;refGene_gn=CSMD2;refGene_tx=XM_023258438.1;refGene_gn=CSMD2;refGene_tx=XM_023258439.1;refGene_gn=CSMD2;refGene_tx=XM_023258440.1;refGene_gn=CSMD2;refGene_tx=XM_023258441.1;refGene_gn=CSMD2;refGene_tx=XM_023258442.1
C1 25598593 25598767
refGene_gn=CSMD2;refGene_tx=XM_023258437.1;refGene_gn=CSMD2;refGene_tx=XM_023258438.1;refGene_gn=CSMD2;refGene_tx=XM_023258439.1;refGene_gn=CSMD2;refGene_tx=XM_023258440.1;refGene_gn=CSMD2;refGene_tx=XM_023258441.1;refGene_gn=CSMD2;refGene_tx=XM_023258442.1
C1 25611976 25612165
refGene_gn=CSMD2;refGene_tx=XM_023258437.1;refGene_gn=CSMD2;refGene_tx=XM_023258438.1;refGene_gn=CSMD2;refGene_tx=XM_023258439.1;refGene_gn=CSMD2;refGene_tx=XM_023258440.1;refGene_gn=CSMD2;refGene_tx=XM_023258441.1;refGene_gn=CSMD2;refGene_tx=XM_023258442.1
C1 25615116 25615290
refGene_gn=CSMD2;refGene_tx=XM_023258437.1;refGene_gn=CSMD2;refGene_tx=XM_023258438.1;refGene_gn=CSMD2;refGene_tx=XM_023258439.1;refGene_gn=CSMD2;refGene_tx=XM_023258440.1;refGene_gn=CSMD2;refGene_tx=XM_023258441.1;refGene_gn=CSMD2;refGene_tx=XM_023258442.1

C1 25620830 25621004
refGene_gn=CSMD2;refGene_tx=XM_023258437.1;refGene_gn=CSMD2;refGene_tx=XM_023258438.1;refGene_gn=CSMD2;refGene_tx=XM_023258439.1;refGene_gn=CSMD2;refGene_tx=XM_023258440.1;refGene_gn=CSMD2;refGene_tx=XM_023258441.1;refGene_gn=CSMD2;refGene_tx=XM_023258442.1
C1 25622065 25622239
refGene_gn=CSMD2;refGene_tx=XM_023258437.1;refGene_gn=CSMD2;refGene_tx=XM_023258438.1;refGene_gn=CSMD2;refGene_tx=XM_023258439.1;refGene_gn=CSMD2;refGene_tx=XM_023258440.1;refGene_gn=CSMD2;refGene_tx=XM_023258441.1;refGene_gn=CSMD2;refGene_tx=XM_023258442.1
C1 25623913 25624108
refGene_gn=CSMD2;refGene_tx=XM_023258437.1;refGene_gn=CSMD2;refGene_tx=XM_023258438.1;refGene_gn=CSMD2;refGene_tx=XM_023258439.1;refGene_gn=CSMD2;refGene_tx=XM_023258440.1;refGene_gn=CSMD2;refGene_tx=XM_023258441.1;refGene_gn=CSMD2;refGene_tx=XM_023258442.1
C1 25625183 25625369
refGene_gn=CSMD2;refGene_tx=XM_023258437.1;refGene_gn=CSMD2;refGene_tx=XM_023258438.1;refGene_gn=CSMD2;refGene_tx=XM_023258439.1;refGene_gn=CSMD2;refGene_tx=XM_023258440.1;refGene_gn=CSMD2;refGene_tx=XM_023258441.1;refGene_gn=CSMD2;refGene_tx=XM_023258442.1
C1 25631236 25631425
refGene_gn=CSMD2;refGene_tx=XM_023258437.1;refGene_gn=CSMD2;refGene_tx=XM_023258438.1;refGene_gn=CSMD2;refGene_tx=XM_023258439.1;refGene_gn=CSMD2;refGene_tx=XM_023258440.1;refGene_gn=CSMD2;refGene_tx=XM_023258441.1;refGene_gn=CSMD2;refGene_tx=XM_023258442.1
C1 25633974 25634121
refGene_gn=CSMD2;refGene_tx=XM_023258437.1;refGene_gn=CSMD2;refGene_tx=XM_023258438.1;refGene_gn=CSMD2;refGene_tx=XM_023258439.1;refGene_gn=CSMD2;refGene_tx=XM_023258440.1;refGene_gn=CSMD2;refGene_tx=XM_023258441.1;refGene_gn=CSMD2;refGene_tx=XM_023258442.1
C1 25637021 25637210
refGene_gn=CSMD2;refGene_tx=XM_023258437.1;refGene_gn=CSMD2;refGene_tx=XM_023258438.1;refGene_gn=CSMD2;refGene_tx=XM_023258439.1;refGene_gn=CSMD2;refGene_tx=XM_023258440.1;refGene_gn=CSMD2;refGene_tx=XM_023258441.1;refGene_gn=CSMD2;refGene_tx=XM_023258442.1
C1 25647690 25647804
refGene_gn=CSMD2;refGene_tx=XM_023258437.1;refGene_gn=CSMD2;refGene_tx=XM_023258438.1;refGene_gn=CSMD2;refGene_tx=XM_023258439.1;refGene_gn=CSMD2;refGene_tx=XM_023258440.1;refGene_gn=CSMD2;refGene_tx=XM_023258441.1;refGene_gn=CSMD2;refGene_tx=XM_023258442.1
C1 25648529 25648610
refGene_gn=CSMD2;refGene_tx=XM_023258437.1;refGene_gn=CSMD2;refGene_tx=XM_023258438.1;refGene_gn=CSMD2;refGene_tx=XM_023258439.1;refGene_gn=CSMD2;refGene_tx=XM_023258440.1;refGene_gn=CSMD2;refGene_tx=XM_023258441.1;refGene_gn=CSMD2;refGene_tx=XM_023258442.1
C1 25656985 25657131
refGene_gn=CSMD2;refGene_tx=XM_023258437.1;refGene_gn=CSMD2;refGene_tx=XM_023258438.1;refGene_gn=CSMD2;refGene_tx=XM_023258439.1;refGene_gn=CSMD2;refGene_tx=XM_023258440.1;refGene_gn=CSMD2;refGene_tx=XM_023258441.1;refGene_gn=CSMD2;refGene_tx=XM_023258442.1

C1 25658516 25658694
refGene_gn=CSMD2;refGene_tx=XM_023258437.1;refGene_gn=CSMD2;refGene_tx=XM_023258438.1;refGene_gn=CSMD2;refGene_tx=XM_023258439.1;refGene_gn=CSMD2;refGene_tx=XM_023258440.1;refGene_gn=CSMD2;refGene_tx=XM_023258441.1;refGene_gn=CSMD2;refGene_tx=XM_023258442.1
C1 25660938 25661127
refGene_gn=CSMD2;refGene_tx=XM_023258437.1;refGene_gn=CSMD2;refGene_tx=XM_023258438.1;refGene_gn=CSMD2;refGene_tx=XM_023258439.1;refGene_gn=CSMD2;refGene_tx=XM_023258440.1;refGene_gn=CSMD2;refGene_tx=XM_023258441.1;refGene_gn=CSMD2;refGene_tx=XM_023258442.1
C1 25666428 25666638
refGene_gn=CSMD2;refGene_tx=XM_023258437.1;refGene_gn=CSMD2;refGene_tx=XM_023258438.1;refGene_gn=CSMD2;refGene_tx=XM_023258439.1;refGene_gn=CSMD2;refGene_tx=XM_023258440.1;refGene_gn=CSMD2;refGene_tx=XM_023258441.1;refGene_gn=CSMD2;refGene_tx=XM_023258442.1
C1 25669495 25669612
refGene_gn=CSMD2;refGene_tx=XM_023258437.1;refGene_gn=CSMD2;refGene_tx=XM_023258438.1;refGene_gn=CSMD2;refGene_tx=XM_023258439.1;refGene_gn=CSMD2;refGene_tx=XM_023258440.1;refGene_gn=CSMD2;refGene_tx=XM_023258441.1;refGene_gn=CSMD2;refGene_tx=XM_023258442.1
C1 25671610 25671680
refGene_gn=CSMD2;refGene_tx=XM_023258437.1;refGene_gn=CSMD2;refGene_tx=XM_023258438.1;refGene_gn=CSMD2;refGene_tx=XM_023258439.1;refGene_gn=CSMD2;refGene_tx=XM_023258440.1;refGene_gn=CSMD2;refGene_tx=XM_023258441.1;refGene_gn=CSMD2;refGene_tx=XM_023258442.1
C1 25672198 25672317
refGene_gn=CSMD2;refGene_tx=XM_023258437.1;refGene_gn=CSMD2;refGene_tx=XM_023258438.1;refGene_gn=CSMD2;refGene_tx=XM_023258439.1;refGene_gn=CSMD2;refGene_tx=XM_023258440.1;refGene_gn=CSMD2;refGene_tx=XM_023258441.1;refGene_gn=CSMD2;refGene_tx=XM_023258442.1
C1 25676893 25676998
refGene_gn=CSMD2;refGene_tx=XM_023258437.1;refGene_gn=CSMD2;refGene_tx=XM_023258438.1;refGene_gn=CSMD2;refGene_tx=XM_023258439.1;refGene_gn=CSMD2;refGene_tx=XM_023258440.1;refGene_gn=CSMD2;refGene_tx=XM_023258441.1;refGene_gn=CSMD2;refGene_tx=XM_023258442.1
C1 25678314 25678411
refGene_gn=CSMD2;refGene_tx=XM_023258437.1;refGene_gn=CSMD2;refGene_tx=XM_023258438.1;refGene_gn=CSMD2;refGene_tx=XM_023258439.1;refGene_gn=CSMD2;refGene_tx=XM_023258440.1;refGene_gn=CSMD2;refGene_tx=XM_023258441.1;refGene_gn=CSMD2;refGene_tx=XM_023258442.1
C1 25679363 25679488
refGene_gn=CSMD2;refGene_tx=XM_023258437.1;refGene_gn=CSMD2;refGene_tx=XM_023258438.1;refGene_gn=CSMD2;refGene_tx=XM_023258439.1;refGene_gn=CSMD2;refGene_tx=XM_023258440.1;refGene_gn=CSMD2;refGene_tx=XM_023258441.1;refGene_gn=CSMD2;refGene_tx=XM_023258442.1
C1 25679836 25680040
refGene_gn=CSMD2;refGene_tx=XM_023258437.1;refGene_gn=CSMD2;refGene_tx=XM_023258438.1;refGene_gn=CSMD2;refGene_tx=XM_023258439.1;refGene_gn=CSMD2;refGene_tx=XM_023258440.1;refGene_gn=CSMD2;refGene_tx=XM_023258441.1;refGene_gn=CSMD2;refGene_tx=XM_023258442.1

C1 25681209 25681305
refGene_gn=CSMD2;refGene_tx=XM_023258437.1;refGene_gn=CSMD2;refGene_tx=XM_023258438.1;refGene_gn=CSMD2;refGene_tx=XM_023258439.1;refGene_gn=CSMD2;refGene_tx=XM_023258440.1;refGene_gn=CSMD2;refGene_tx=XM_023258441.1;refGene_gn=CSMD2;refGene_tx=XM_023258442.1
C1 25688027 25688141
refGene_gn=CSMD2;refGene_tx=XM_023258437.1;refGene_gn=CSMD2;refGene_tx=XM_023258438.1;refGene_gn=CSMD2;refGene_tx=XM_023258439.1;refGene_gn=CSMD2;refGene_tx=XM_023258440.1;refGene_gn=CSMD2;refGene_tx=XM_023258441.1;refGene_gn=CSMD2;refGene_tx=XM_023258442.1
C1 25690850 25690967
refGene_gn=CSMD2;refGene_tx=XM_023258437.1;refGene_gn=CSMD2;refGene_tx=XM_023258438.1;refGene_gn=CSMD2;refGene_tx=XM_023258439.1;refGene_gn=CSMD2;refGene_tx=XM_023258440.1;refGene_gn=CSMD2;refGene_tx=XM_023258441.1;refGene_gn=CSMD2;refGene_tx=XM_023258442.1
C1 25691867 25692062
refGene_gn=CSMD2;refGene_tx=XM_023258437.1;refGene_gn=CSMD2;refGene_tx=XM_023258438.1;refGene_gn=CSMD2;refGene_tx=XM_023258439.1;refGene_gn=CSMD2;refGene_tx=XM_023258440.1;refGene_gn=CSMD2;refGene_tx=XM_023258441.1;refGene_gn=CSMD2;refGene_tx=XM_023258442.1
C1 25702758 25702946
refGene_gn=CSMD2;refGene_tx=XM_023258437.1;refGene_gn=CSMD2;refGene_tx=XM_023258438.1;refGene_gn=CSMD2;refGene_tx=XM_023258439.1;refGene_gn=CSMD2;refGene_tx=XM_023258440.1;refGene_gn=CSMD2;refGene_tx=XM_023258441.1;refGene_gn=CSMD2;refGene_tx=XM_023258442.1
C1 25706899 25707038
refGene_gn=CSMD2;refGene_tx=XM_023258437.1;refGene_gn=CSMD2;refGene_tx=XM_023258438.1;refGene_gn=CSMD2;refGene_tx=XM_023258439.1;refGene_gn=CSMD2;refGene_tx=XM_023258440.1;refGene_gn=CSMD2;refGene_tx=XM_023258441.1;refGene_gn=CSMD2;refGene_tx=XM_023258442.1
C1 25712450 25712642
refGene_gn=CSMD2;refGene_tx=XM_023258437.1;refGene_gn=CSMD2;refGene_tx=XM_023258438.1;refGene_gn=CSMD2;refGene_tx=XM_023258439.1;refGene_gn=CSMD2;refGene_tx=XM_023258440.1;refGene_gn=CSMD2;refGene_tx=XM_023258441.1;refGene_gn=CSMD2;refGene_tx=XM_023258442.1
C1 25717684 25717887
refGene_gn=CSMD2;refGene_tx=XM_023258437.1;refGene_gn=CSMD2;refGene_tx=XM_023258438.1;refGene_gn=CSMD2;refGene_tx=XM_023258439.1;refGene_gn=CSMD2;refGene_tx=XM_023258440.1;refGene_gn=CSMD2;refGene_tx=XM_023258441.1;refGene_gn=CSMD2;refGene_tx=XM_023258442.1
C1 25741064 25741191
refGene_gn=CSMD2;refGene_tx=XM_023258437.1;refGene_gn=CSMD2;refGene_tx=XM_023258438.1;refGene_gn=CSMD2;refGene_tx=XM_023258439.1;refGene_gn=CSMD2;refGene_tx=XM_023258440.1;refGene_gn=CSMD2;refGene_tx=XM_023258441.1;refGene_gn=CSMD2;refGene_tx=XM_023258442.1
C1 25746308 25746500
refGene_gn=CSMD2;refGene_tx=XM_023258437.1;refGene_gn=CSMD2;refGene_tx=XM_023258438.1;refGene_gn=CSMD2;refGene_tx=XM_023258439.1;refGene_gn=CSMD2;refGene_tx=XM_023258440.1;refGene_gn=CSMD2;refGene_tx=XM_023258441.1;refGene_gn=CSMD2;refGene_tx=XM_023258442.1

C1 25748279 25748436
refGene_gn=CSMD2;refGene_tx=XM_023258437.1;refGene_gn=CSMD2;refGene_tx=XM_023258438.1;refGene_gn=CSMD2;refGene_tx=XM_023258439.1;refGene_gn=CSMD2;refGene_tx=XM_023258440.1;refGene_gn=CSMD2;refGene_tx=XM_023258441.1;refGene_gn=CSMD2;refGene_tx=XM_023258442.1
C1 25758179 25758349
refGene_gn=CSMD2;refGene_tx=XM_023258437.1;refGene_gn=CSMD2;refGene_tx=XM_023258438.1;refGene_gn=CSMD2;refGene_tx=XM_023258439.1;refGene_gn=CSMD2;refGene_tx=XM_023258440.1;refGene_gn=CSMD2;refGene_tx=XM_023258441.1;refGene_gn=CSMD2;refGene_tx=XM_023258442.1
C1 25763132 25763321
refGene_gn=CSMD2;refGene_tx=XM_023258437.1;refGene_gn=CSMD2;refGene_tx=XM_023258438.1;refGene_gn=CSMD2;refGene_tx=XM_023258439.1;refGene_gn=CSMD2;refGene_tx=XM_023258440.1;refGene_gn=CSMD2;refGene_tx=XM_023258441.1;refGene_gn=CSMD2;refGene_tx=XM_023258442.1
C1 25764738 25764954
refGene_gn=CSMD2;refGene_tx=XM_023258437.1;refGene_gn=CSMD2;refGene_tx=XM_023258438.1;refGene_gn=CSMD2;refGene_tx=XM_023258439.1;refGene_gn=CSMD2;refGene_tx=XM_023258440.1;refGene_gn=CSMD2;refGene_tx=XM_023258441.1;refGene_gn=CSMD2;refGene_tx=XM_023258442.1
C1 25771433 25771550
refGene_gn=CSMD2;refGene_tx=XM_023258437.1;refGene_gn=CSMD2;refGene_tx=XM_023258438.1;refGene_gn=CSMD2;refGene_tx=XM_023258439.1;refGene_gn=CSMD2;refGene_tx=XM_023258440.1;refGene_gn=CSMD2;refGene_tx=XM_023258441.1;refGene_gn=CSMD2;refGene_tx=XM_023258442.1
C1 25771757 25771946
refGene_gn=CSMD2;refGene_tx=XM_023258437.1;refGene_gn=CSMD2;refGene_tx=XM_023258438.1;refGene_gn=CSMD2;refGene_tx=XM_023258439.1;refGene_gn=CSMD2;refGene_tx=XM_023258440.1;refGene_gn=CSMD2;refGene_tx=XM_023258441.1;refGene_gn=CSMD2;refGene_tx=XM_023258442.1
C1 25772721 25772909
refGene_gn=CSMD2;refGene_tx=XM_023258437.1;refGene_gn=CSMD2;refGene_tx=XM_023258438.1;refGene_gn=CSMD2;refGene_tx=XM_023258439.1;refGene_gn=CSMD2;refGene_tx=XM_023258440.1;refGene_gn=CSMD2;refGene_tx=XM_023258441.1;refGene_gn=CSMD2;refGene_tx=XM_023258442.1
C1 25773921 25774060
refGene_gn=CSMD2;refGene_tx=XM_023258437.1;refGene_gn=CSMD2;refGene_tx=XM_023258438.1;refGene_gn=CSMD2;refGene_tx=XM_023258439.1;refGene_gn=CSMD2;refGene_tx=XM_023258440.1;refGene_gn=CSMD2;refGene_tx=XM_023258441.1;refGene_gn=CSMD2;refGene_tx=XM_023258442.1
C1 25783631 25783826
refGene_gn=CSMD2;refGene_tx=XM_023258437.1;refGene_gn=CSMD2;refGene_tx=XM_023258438.1;refGene_gn=CSMD2;refGene_tx=XM_023258439.1;refGene_gn=CSMD2;refGene_tx=XM_023258440.1;refGene_gn=CSMD2;refGene_tx=XM_023258441.1;refGene_gn=CSMD2;refGene_tx=XM_023258442.1
C1 25787938 25788265
refGene_gn=CSMD2;refGene_tx=XM_023258437.1;refGene_gn=CSMD2;refGene_tx=XM_023258438.1;refGene_gn=CSMD2;refGene_tx=XM_023258439.1;refGene_gn=CSMD2;refGene_tx=XM_023258440.1;refGene_gn=CSMD2;refGene_tx=XM_023258441.1;refGene_gn=CSMD2;refGene_tx=XM_023258442.1

C1 25811665 25811848
refGene_gn=CSMD2;refGene_tx=XM_023258437.1;refGene_gn=CSMD2;refGene_tx=XM_023258438.1;refGene_gn=CSMD2;refGene_tx=XM_023258439.1;refGene_gn=CSMD2;refGene_tx=XM_023258440.1;refGene_gn=CSMD2;refGene_tx=XM_023258441.1;refGene_gn=CSMD2;refGene_tx=XM_023258442.1
C1 25829221 25829334
refGene_gn=CSMD2;refGene_tx=XM_023258437.1;refGene_gn=CSMD2;refGene_tx=XM_023258438.1;refGene_gn=CSMD2;refGene_tx=XM_023258439.1;refGene_gn=CSMD2;refGene_tx=XM_023258440.1;refGene_gn=CSMD2;refGene_tx=XM_023258441.1;refGene_gn=CSMD2;refGene_tx=XM_023258442.1
C1 25832454 25832558
refGene_gn=CSMD2;refGene_tx=XM_023258437.1;refGene_gn=CSMD2;refGene_tx=XM_023258438.1;refGene_gn=CSMD2;refGene_tx=XM_023258439.1;refGene_gn=CSMD2;refGene_tx=XM_023258440.1;refGene_gn=CSMD2;refGene_tx=XM_023258441.1;refGene_gn=CSMD2;refGene_tx=XM_023258442.1
C1 25849712 25849834
refGene_gn=CSMD2;refGene_tx=XM_023258437.1;refGene_gn=CSMD2;refGene_tx=XM_023258438.1;refGene_gn=CSMD2;refGene_tx=XM_023258439.1;refGene_gn=CSMD2;refGene_tx=XM_023258440.1;refGene_gn=CSMD2;refGene_tx=XM_023258441.1;refGene_gn=CSMD2;refGene_tx=XM_023258442.1
C1 25859157 25859282
refGene_gn=CSMD2;refGene_tx=XM_023258437.1;refGene_gn=CSMD2;refGene_tx=XM_023258438.1;refGene_gn=CSMD2;refGene_tx=XM_023258439.1;refGene_gn=CSMD2;refGene_tx=XM_023258440.1;refGene_gn=CSMD2;refGene_tx=XM_023258441.1;refGene_gn=CSMD2;refGene_tx=XM_023258442.1
C1 25859984 25860072
refGene_gn=CSMD2;refGene_tx=XM_023258437.1;refGene_gn=CSMD2;refGene_tx=XM_023258438.1;refGene_gn=CSMD2;refGene_tx=XM_023258439.1;refGene_gn=CSMD2;refGene_tx=XM_023258440.1;refGene_gn=CSMD2;refGene_tx=XM_023258441.1;refGene_gn=CSMD2;refGene_tx=XM_023258442.1
C1 25865108 25865186
refGene_gn=CSMD2;refGene_tx=XM_023258437.1;refGene_gn=CSMD2;refGene_tx=XM_023258438.1;refGene_gn=CSMD2;refGene_tx=XM_023258439.1;refGene_gn=CSMD2;refGene_tx=XM_023258440.1;refGene_gn=CSMD2;refGene_tx=XM_023258441.1;refGene_gn=CSMD2;refGene_tx=XM_023258442.1
C1 25878806 25878919
refGene_gn=CSMD2;refGene_tx=XM_023258437.1;refGene_gn=CSMD2;refGene_tx=XM_023258438.1;refGene_gn=CSMD2;refGene_tx=XM_023258439.1;refGene_gn=CSMD2;refGene_tx=XM_023258440.1;refGene_gn=CSMD2;refGene_tx=XM_023258442.1
C1 25897192 25897774 refGene_gn=HMGB4;refGene_tx=XM_003989838.5
C1 25948307 25948515
refGene_gn=CSMD2;refGene_tx=XM_023258437.1;refGene_gn=CSMD2;refGene_tx=XM_023258438.1;refGene_gn=CSMD2;refGene_tx=XM_023258439.1;refGene_gn=CSMD2;refGene_tx=XM_023258440.1;refGene_gn=CSMD2;refGene_tx=XM_023258442.1
C1 25961503 25961698
refGene_gn=CSMD2;refGene_tx=XM_023258437.1;refGene_gn=CSMD2;refGene_tx=XM_023258438.1;refGene_gn=CSMD2;refGene_tx=XM_023258439.1;refGene_gn=CSMD2;refGene_tx=XM_023258440.1;refGene_gn=CSMD2;refGene_tx=XM_0232584

42.1
C1 26056135 26056248
refGene_gn=CSMD2;refGene_tx=XM_023258437.1;refGene_gn=CSMD2;refGene_tx=XM_023258438.1;refGene_gn=CSMD2;refGene_tx=XM_023258439.1;refGene_gn=CSMD2;refGene_tx=XM_023258440.1;refGene_gn=CSMD2;refGene_tx=XM_023258442.1
C1 26102529 26102746
refGene_gn=CSMD2;refGene_tx=XM_023258437.1;refGene_gn=CSMD2;refGene_tx=XM_023258438.1;refGene_gn=CSMD2;refGene_tx=XM_023258439.1;refGene_gn=CSMD2;refGene_tx=XM_023258440.1;refGene_gn=CSMD2;refGene_tx=XM_023258442.1
C1 26163002 26163192
refGene_gn=CSMD2;refGene_tx=XM_023258437.1;refGene_gn=CSMD2;refGene_tx=XM_023258438.1;refGene_gn=CSMD2;refGene_tx=XM_023258439.1;refGene_gn=CSMD2;refGene_tx=XM_023258440.1;refGene_gn=CSMD2;refGene_tx=XM_023258442.1
C1 26176640 26176969
refGene_gn=CC1H1orf94;refGene_tx=XM_011284744.3;refGene_gn=CC1H1orf94;refGene_tx=XM_019836619.2;refGene_gn=CC1H1orf94;refGene_tx=XM_023258443.1
C1 26195814 26196503
refGene_gn=CC1H1orf94;refGene_tx=XM_011284744.3;refGene_gn=CC1H1orf94;refGene_tx=XM_019836619.2;refGene_gn=CC1H1orf94;refGene_tx=XM_023258443.1
C1 26199307 26199568
refGene_gn=CC1H1orf94;refGene_tx=XM_011284744.3;refGene_gn=CC1H1orf94;refGene_tx=XM_019836619.2;refGene_gn=CC1H1orf94;refGene_tx=XM_023258443.1
C1 26200600 26200776
refGene_gn=CC1H1orf94;refGene_tx=XM_011284744.3;refGene_gn=CC1H1orf94;refGene_tx=XM_019836619.2;refGene_gn=CC1H1orf94;refGene_tx=XM_023258443.1
C1 26202593 26202713
refGene_gn=CC1H1orf94;refGene_tx=XM_023258443.1
C1 26205029 26205107
refGene_gn=CC1H1orf94;refGene_tx=XM_011284744.3
C1 26208941 26209138
refGene_gn=CC1H1orf94;refGene_tx=XM_011284744.3;refGene_gn=CC1H1orf94;refGene_tx=XM_019836619.2
C1 26213035 26213111
refGene_gn=CC1H1orf94;refGene_tx=XM_011284744.3;refGene_gn=CC1H1orf94;refGene_tx=XM_019836619.2
C1 26731349 26732171
refGene_gn=GJB5;refGene_tx=XM_003989839.3;refGene_gn=GJB5;refGene_tx=XM_006934553.2
C1 26735314 26736115 refGene_gn=GJB4;refGene_tx=XM_003989840.4
C1 26758275 26759088
refGene_gn=GJB3;refGene_tx=XM_006934554.3;refGene_gn=GJB3;refGene_tx=XM_019836620.1;refGene_gn=GJB3;refGene_tx=XM_019836621.2
C1 26767598 26768600 refGene_gn=GJA4;refGene_tx=XM_003989842.3

C1 26821005 26821284
refGene_gn=SMIM12;refGene_tx=XM_003989843.3;refGene_gn=SMIM12;refGene_tx=XM_023258444.1
C1 26829787 26830006
refGene_gn=DLGAP3;refGene_tx=XM_011284742.3;refGene_gn=DLGAP3;refGene_tx=XM_023258445.1;refGene_gn=DLGAP3;refGene_tx=XM_023258446.1;refGene_gn=DLGAP3;refGene_tx=XM_023258447.1
C1 26830718 26830862
refGene_gn=DLGAP3;refGene_tx=XM_011284742.3;refGene_gn=DLGAP3;refGene_tx=XM_023258445.1;refGene_gn=DLGAP3;refGene_tx=XM_023258446.1;refGene_gn=DLGAP3;refGene_tx=XM_023258447.1
C1 26831099 26831191
refGene_gn=DLGAP3;refGene_tx=XM_011284742.3;refGene_gn=DLGAP3;refGene_tx=XM_023258445.1;refGene_gn=DLGAP3;refGene_tx=XM_023258446.1;refGene_gn=DLGAP3;refGene_tx=XM_023258447.1
C1 26832138 26832623
refGene_gn=DLGAP3;refGene_tx=XM_011284742.3;refGene_gn=DLGAP3;refGene_tx=XM_023258445.1;refGene_gn=DLGAP3;refGene_tx=XM_023258446.1;refGene_gn=DLGAP3;refGene_tx=XM_023258447.1
C1 26850468 26850554
refGene_gn=DLGAP3;refGene_tx=XM_011284742.3;refGene_gn=DLGAP3;refGene_tx=XM_023258445.1;refGene_gn=DLGAP3;refGene_tx=XM_023258446.1;refGene_gn=DLGAP3;refGene_tx=XM_023258447.1
C1 26850885 26851199
refGene_gn=DLGAP3;refGene_tx=XM_011284742.3;refGene_gn=DLGAP3;refGene_tx=XM_023258445.1;refGene_gn=DLGAP3;refGene_tx=XM_023258446.1;refGene_gn=DLGAP3;refGene_tx=XM_023258447.1
C1 26851482 26851696
refGene_gn=DLGAP3;refGene_tx=XM_011284742.3;refGene_gn=DLGAP3;refGene_tx=XM_023258445.1;refGene_gn=DLGAP3;refGene_tx=XM_023258446.1;refGene_gn=DLGAP3;refGene_tx=XM_023258447.1
C1 26862616 26862689
refGene_gn=DLGAP3;refGene_tx=XM_011284742.3;refGene_gn=DLGAP3;refGene_tx=XM_023258445.1;refGene_gn=DLGAP3;refGene_tx=XM_023258446.1;refGene_gn=DLGAP3;refGene_tx=XM_023258447.1
C1 26863002 26863208
refGene_gn=DLGAP3;refGene_tx=XM_011284742.3;refGene_gn=DLGAP3;refGene_tx=XM_023258445.1;refGene_gn=DLGAP3;refGene_tx=XM_023258446.1;refGene_gn=DLGAP3;refGene_tx=XM_023258447.1
C1 26867629 26868736
refGene_gn=DLGAP3;refGene_tx=XM_011284742.3;refGene_gn=DLGAP3;refGene_tx=XM_023258445.1;refGene_gn=DLGAP3;refGene_tx=XM_023258446.1;refGene_gn=DLGAP3;refGene_tx=XM_023258447.1
C1 26928617 26929625
refGene_gn=LOC111561626;refGene_tx=XM_023257822.1
C1 26934118 26934222
refGene_gn=TMEM35B;refGene_tx=XM_023258457.1;refGene_gn=TMEM35B;refGene_tx=XM_023258458.1
C1 26934498 26934680 refGene_gn=TMEM35B;refGene_tx=XM_023258458.1
C1 26934977 26935032

refGene_gn=TMEM35B;refGene_tx=XM_023258455.1;refGene_gn=TMEM35B;refGene_tx=XM_023258456.1;refGene_gn=TMEM35B;refGene_tx=XM_023258457.1;refGene_gn=TMEM35B;refGene_tx=XM_023258458.1
C1	26935134	26935315
refGene_gn=TMEM35B;refGene_tx=XM_023258455.1;refGene_gn=TMEM35B;refGene_tx=XM_023258456.1
C1	26936268	26936424	refGene_gn=TMEM35B;refGene_tx=XM_023258455.1
C1	26936720	26936828
refGene_gn=TMEM35B;refGene_tx=XM_023258456.1;refGene_gn=TMEM35B;refGene_tx=XM_023258457.1;refGene_gn=TMEM35B;refGene_tx=XM_023258458.1
C1	26938373	26940205
refGene_gn=ZMYM6;refGene_tx=XM_003989845.5;refGene_gn=ZMYM6;refGene_tx=XM_011284741.3;refGene_gn=ZMYM6;refGene_tx=XM_019836622.2;refGene_gn=ZMYM6;refGene_tx=XM_019836624.2;refGene_gn=ZMYM6;refGene_tx=XM_019836625.2;refGene_gn=ZMYM6;refGene_tx=XM_019836626.1;refGene_gn=ZMYM6;refGene_tx=XM_019836627.2;refGene_gn=ZMYM6;refGene_tx=XM_019836628.2;refGene_gn=ZMYM6;refGene_tx=XM_019836629.2;refGene_gn=ZMYM6;refGene_tx=XM_019836631.1;refGene_gn=ZMYM6;refGene_tx=XM_019836632.1;refGene_gn=ZMYM6;refGene_tx=XM_023258448.1;refGene_gn=ZMYM6;refGene_tx=XM_023258449.1;refGene_gn=ZMYM6;refGene_tx=XM_023258450.1
C1	26941917	26942062
refGene_gn=ZMYM6;refGene_tx=XM_003989845.5;refGene_gn=ZMYM6;refGene_tx=XM_011284741.3;refGene_gn=ZMYM6;refGene_tx=XM_019836622.2;refGene_gn=ZMYM6;refGene_tx=XM_019836624.2;refGene_gn=ZMYM6;refGene_tx=XM_019836625.2;refGene_gn=ZMYM6;refGene_tx=XM_019836626.1;refGene_gn=ZMYM6;refGene_tx=XM_019836627.2;refGene_gn=ZMYM6;refGene_tx=XM_019836628.2;refGene_gn=ZMYM6;refGene_tx=XM_019836629.2;refGene_gn=ZMYM6;refGene_tx=XM_019836631.1;refGene_gn=ZMYM6;refGene_tx=XM_019836632.1;refGene_gn=ZMYM6;refGene_tx=XM_023258448.1;refGene_gn=ZMYM6;refGene_tx=XM_023258449.1;refGene_gn=ZMYM6;refGene_tx=XM_023258450.1
C1	26953943	26953984
refGene_gn=ZMYM6;refGene_tx=XM_003989845.5;refGene_gn=ZMYM6;refGene_tx=XM_011284741.3;refGene_gn=ZMYM6;refGene_tx=XM_019836622.2;refGene_gn=ZMYM6;refGene_tx=XM_019836624.2;refGene_gn=ZMYM6;refGene_tx=XM_019836625.2;refGene_gn=ZMYM6;refGene_tx=XM_019836626.1;refGene_gn=ZMYM6;refGene_tx=XM_019836627.2;refGene_gn=ZMYM6;refGene_tx=XM_019836628.2;refGene_gn=ZMYM6;refGene_tx=XM_019836629.2;refGene_gn=ZMYM6;refGene_tx=XM_019836631.1;refGene_gn=ZMYM6;refGene_tx=XM_019836632.1;refGene_gn=ZMYM6;refGene_tx=XM_023258448.1;refGene_gn=ZMYM6;refGene_tx=XM_023258449.1;refGene_gn=ZMYM6;refGene_tx=XM_023258450.1
C1	26954429	26954570
refGene_gn=ZMYM6;refGene_tx=XM_003989845.5;refGene_gn=ZMYM6;refGene_tx=XM_011284741.3;refGene_gn=ZMYM6;refGene_tx=XM_019836622.2;refGene_gn=ZMYM6;refGene_tx=XM_019836624.2;refGene_gn=ZMYM6;refGene_tx=XM_019836625.2;refGene_gn=ZMYM6;refGene_tx=XM_019836626.1;refGene_gn=ZMYM6;refGene_tx=XM_019836627.2;refGene_gn=ZMYM6;refGene_tx=XM_019836628.2;refGene_gn=ZMYM6;refGene_tx=XM_019836629.2;refGene_gn=ZMYM6;refGene_tx=XM_019836631.1;refGene_gn=ZMYM6;refGene_tx=XM_019836632.1;refGene_gn=ZMYM6;refGene_tx=XM_023258448.1;refGene_gn=ZMYM6;refGene_tx=XM_023258449.1;refGene_gn=ZMYM6;refGene_tx=XM_023258450.1

C1	26956069	26956217
refGene_gn=ZMYM6;refGene_tx=XM_003989845.5;refGene_gn=ZMYM6;refGene_tx=XM_011284741.3;refGene_gn=ZMYM6;refGene_tx=XM_019836622.2;refGene_gn=ZMYM6;refGene_tx=XM_019836624.2;refGene_gn=ZMYM6;refGene_tx=XM_019836625.2;refGene_gn=ZMYM6;refGene_tx=XM_019836626.1;refGene_gn=ZMYM6;refGene_tx=XM_019836627.2;refGene_gn=ZMYM6;refGene_tx=XM_019836628.2;refGene_gn=ZMYM6;refGene_tx=XM_019836629.2;refGene_gn=ZMYM6;refGene_tx=XM_019836631.1;refGene_gn=ZMYM6;refGene_tx=XM_019836632.1;refGene_gn=ZMYM6;refGene_tx=XM_023258448.1;refGene_gn=ZMYM6;refGene_tx=XM_023258449.1;refGene_gn=ZMYM6;refGene_tx=XM_023258450.1
C1	26957553	26957726
refGene_gn=ZMYM6;refGene_tx=XM_003989845.5;refGene_gn=ZMYM6;refGene_tx=XM_011284741.3;refGene_gn=ZMYM6;refGene_tx=XM_019836622.2;refGene_gn=ZMYM6;refGene_tx=XM_019836624.2;refGene_gn=ZMYM6;refGene_tx=XM_019836625.2;refGene_gn=ZMYM6;refGene_tx=XM_019836626.1;refGene_gn=ZMYM6;refGene_tx=XM_019836627.2;refGene_gn=ZMYM6;refGene_tx=XM_019836628.2;refGene_gn=ZMYM6;refGene_tx=XM_019836629.2;refGene_gn=ZMYM6;refGene_tx=XM_019836631.1;refGene_gn=ZMYM6;refGene_tx=XM_019836632.1;refGene_gn=ZMYM6;refGene_tx=XM_023258448.1;refGene_gn=ZMYM6;refGene_tx=XM_023258449.1;refGene_gn=ZMYM6;refGene_tx=XM_023258450.1
C1	26958310	26958461
refGene_gn=ZMYM6;refGene_tx=XM_003989845.5;refGene_gn=ZMYM6;refGene_tx=XM_011284741.3;refGene_gn=ZMYM6;refGene_tx=XM_019836622.2;refGene_gn=ZMYM6;refGene_tx=XM_019836624.2;refGene_gn=ZMYM6;refGene_tx=XM_019836625.2;refGene_gn=ZMYM6;refGene_tx=XM_019836626.1;refGene_gn=ZMYM6;refGene_tx=XM_019836627.2;refGene_gn=ZMYM6;refGene_tx=XM_019836628.2;refGene_gn=ZMYM6;refGene_tx=XM_019836629.2;refGene_gn=ZMYM6;refGene_tx=XM_019836631.1;refGene_gn=ZMYM6;refGene_tx=XM_019836632.1;refGene_gn=ZMYM6;refGene_tx=XM_023258448.1;refGene_gn=ZMYM6;refGene_tx=XM_023258449.1;refGene_gn=ZMYM6;refGene_tx=XM_023258450.1
C1	26958627	26958903
refGene_gn=ZMYM6;refGene_tx=XM_003989845.5;refGene_gn=ZMYM6;refGene_tx=XM_011284741.3;refGene_gn=ZMYM6;refGene_tx=XM_019836622.2;refGene_gn=ZMYM6;refGene_tx=XM_019836624.2;refGene_gn=ZMYM6;refGene_tx=XM_019836625.2;refGene_gn=ZMYM6;refGene_tx=XM_019836626.1;refGene_gn=ZMYM6;refGene_tx=XM_019836627.2;refGene_gn=ZMYM6;refGene_tx=XM_019836628.2;refGene_gn=ZMYM6;refGene_tx=XM_019836629.2;refGene_gn=ZMYM6;refGene_tx=XM_019836631.1;refGene_gn=ZMYM6;refGene_tx=XM_019836632.1;refGene_gn=ZMYM6;refGene_tx=XM_023258448.1;refGene_gn=ZMYM6;refGene_tx=XM_023258449.1;refGene_gn=ZMYM6;refGene_tx=XM_023258450.1
C1	26959971	26960087
refGene_gn=ZMYM6;refGene_tx=XM_003989845.5;refGene_gn=ZMYM6;refGene_tx=XM_011284741.3;refGene_gn=ZMYM6;refGene_tx=XM_019836622.2;refGene_gn=ZMYM6;refGene_tx=XM_019836624.2;refGene_gn=ZMYM6;refGene_tx=XM_019836625.2;refGene_gn=ZMYM6;refGene_tx=XM_019836626.1;refGene_gn=ZMYM6;refGene_tx=XM_019836627.2;refGene_gn=ZMYM6;refGene_tx=XM_019836628.2;refGene_gn=ZMYM6;refGene_tx=XM_019836629.2;refGene_gn=ZMYM6;refGene_tx=XM_019836631.1;refGene_gn=ZMYM6;refGene_tx=XM_019836632.1;refGene_gn=ZMYM6;refGene_tx=XM_023258448.1;refGene_gn=ZMYM6;refGene_tx=XM_023258449.1;refGene_gn=ZMYM6;refGene_tx=XM_023258450.1

C1 26960512 26960663
refGene_gn=ZMYM6;refGene_tx=XM_003989845.5;refGene_gn=ZMYM6;refGene_tx=XM_011284741.3;refGene_gn=ZMYM6;refGene_tx=XM_019836622.2;refGene_gn=ZMYM6;refGene_tx=XM_019836624.2;refGene_gn=ZMYM6;refGene_tx=XM_019836625.2;refGene_gn=ZMYM6;refGene_tx=XM_019836626.1;refGene_gn=ZMYM6;refGene_tx=XM_019836627.2;refGene_gn=ZMYM6;refGene_tx=XM_019836628.2;refGene_gn=ZMYM6;refGene_tx=XM_019836629.2;refGene_gn=ZMYM6;refGene_tx=XM_019836631.1;refGene_gn=ZMYM6;refGene_tx=XM_019836632.1;refGene_gn=ZMYM6;refGene_tx=XM_023258448.1;refGene_gn=ZMYM6;refGene_tx=XM_023258449.1;refGene_gn=ZMYM6;refGene_tx=XM_023258450.1
C1 26963310 26963502
refGene_gn=ZMYM6;refGene_tx=XM_003989845.5;refGene_gn=ZMYM6;refGene_tx=XM_011284741.3;refGene_gn=ZMYM6;refGene_tx=XM_019836622.2;refGene_gn=ZMYM6;refGene_tx=XM_019836624.2;refGene_gn=ZMYM6;refGene_tx=XM_019836625.2;refGene_gn=ZMYM6;refGene_tx=XM_019836626.1;refGene_gn=ZMYM6;refGene_tx=XM_019836627.2;refGene_gn=ZMYM6;refGene_tx=XM_019836628.2;refGene_gn=ZMYM6;refGene_tx=XM_019836629.2;refGene_gn=ZMYM6;refGene_tx=XM_019836631.1;refGene_gn=ZMYM6;refGene_tx=XM_019836632.1;refGene_gn=ZMYM6;refGene_tx=XM_023258448.1;refGene_gn=ZMYM6;refGene_tx=XM_023258449.1
C1 26965288 26965309
refGene_gn=ZMYM6;refGene_tx=XM_023258449.1;refGene_gn=ZMYM6;refGene_tx=XM_023258450.1
C1 26966819 26966988
refGene_gn=ZMYM6;refGene_tx=XM_003989845.5;refGene_gn=ZMYM6;refGene_tx=XM_011284741.3;refGene_gn=ZMYM6;refGene_tx=XM_019836622.2;refGene_gn=ZMYM6;refGene_tx=XM_019836624.2;refGene_gn=ZMYM6;refGene_tx=XM_019836625.2;refGene_gn=ZMYM6;refGene_tx=XM_019836626.1;refGene_gn=ZMYM6;refGene_tx=XM_019836627.2;refGene_gn=ZMYM6;refGene_tx=XM_019836628.2;refGene_gn=ZMYM6;refGene_tx=XM_019836629.2;refGene_gn=ZMYM6;refGene_tx=XM_019836631.1;refGene_gn=ZMYM6;refGene_tx=XM_019836632.1;refGene_gn=ZMYM6;refGene_tx=XM_023258448.1
C1 26969939 26970189
refGene_gn=ZMYM6;refGene_tx=XM_003989845.5;refGene_gn=ZMYM6;refGene_tx=XM_011284741.3;refGene_gn=ZMYM6;refGene_tx=XM_019836622.2;refGene_gn=ZMYM6;refGene_tx=XM_019836624.2;refGene_gn=ZMYM6;refGene_tx=XM_019836625.2;refGene_gn=ZMYM6;refGene_tx=XM_019836626.1;refGene_gn=ZMYM6;refGene_tx=XM_019836627.2;refGene_gn=ZMYM6;refGene_tx=XM_019836628.2;refGene_gn=ZMYM6;refGene_tx=XM_019836629.2;refGene_gn=ZMYM6;refGene_tx=XM_023258448.1
C1 26971013 26971098
refGene_gn=ZMYM6;refGene_tx=XM_003989845.5;refGene_gn=ZMYM6;refGene_tx=XM_011284741.3;refGene_gn=ZMYM6;refGene_tx=XM_019836622.2;refGene_gn=ZMYM6;refGene_tx=XM_019836624.2;refGene_gn=ZMYM6;refGene_tx=XM_019836625.2;refGene_gn=ZMYM6;refGene_tx=XM_019836626.1;refGene_gn=ZMYM6;refGene_tx=XM_019836627.2;refGene_gn=ZMYM6;refGene_tx=XM_019836628.2;refGene_gn=ZMYM6;refGene_tx=XM_023258448.1
C1 26978424 26978436 refGene_gn=ZMYM6;refGene_tx=XM_019836628.2
C1 26979543 26979636
refGene_gn=ZMYM6;refGene_tx=XM_003989845.5;refGene_gn=ZMYM6;refGene_tx=XM_011284741.3;refGene_gn=ZMYM6;refGene_tx=XM_019836622.2;refGene_gn=

ZMYM6;refGene_tx=XM_019836624.2;refGene_gn=ZMYM6;refGene_tx=XM_0198366
25.2;refGene_gn=ZMYM6;refGene_tx=XM_019836626.1;refGene_gn=ZMYM6;refGe
ne_tx=XM_019836627.2;refGene_gn=ZMYM6;refGene_tx=XM_023258448.1
C1 27000759 27000793 refGene_gn=ZMYM1;refGene_tx=XM_023258459.1
C1 27009208 27009378 refGene_gn=ZMYM1;refGene_tx=XM_023258459.1
C1 27011472 27011545 refGene_gn=ZMYM1;refGene_tx=XM_023258459.1
C1 27011734 27011984 refGene_gn=ZMYM1;refGene_tx=XM_023258459.1
C1 27018407 27018576 refGene_gn=ZMYM1;refGene_tx=XM_023258459.1
C1 27018662 27018784 refGene_gn=ZMYM1;refGene_tx=XM_023258459.1
C1 27018785 27018867 refGene_gn=ZMYM1;refGene_tx=XM_023258459.1
C1 27021193 27021347 refGene_gn=ZMYM1;refGene_tx=XM_023258459.1
C1 27022228 27022366 refGene_gn=ZMYM1;refGene_tx=XM_023258459.1
C1 27022794 27022832 refGene_gn=ZMYM1;refGene_tx=XM_023258459.1
C1 27023575 27025876 refGene_gn=ZMYM1;refGene_tx=XM_023258459.1
C1 27059237 27059261 refGene_gn=SFPQ;refGene_tx=XM_023258454.1
C1 27065485 27065623
refGene_gn=SFPQ;refGene_tx=XM_023258452.1;refGene_gn=SFPQ;refGene_tx=X
M_023258453.1
C1 27068037 27068159
refGene_gn=SFPQ;refGene_tx=XM_023258452.1;refGene_gn=SFPQ;refGene_tx=X
M_023258453.1;refGene_gn=SFPQ;refGene_tx=XM_023258454.1
C1 27068239 27068288
refGene_gn=SFPQ;refGene_tx=XM_023258452.1;refGene_gn=SFPQ;refGene_tx=X
M_023258453.1;refGene_gn=SFPQ;refGene_tx=XM_023258454.1
C1 27068620 27068738
refGene_gn=SFPQ;refGene_tx=XM_023258452.1;refGene_gn=SFPQ;refGene_tx=X
M_023258453.1;refGene_gn=SFPQ;refGene_tx=XM_023258454.1
C1 27069453 27069538
refGene_gn=SFPQ;refGene_tx=XM_023258452.1;refGene_gn=SFPQ;refGene_tx=X
M_023258453.1;refGene_gn=SFPQ;refGene_tx=XM_023258454.1
C1 27069656 27069853
refGene_gn=SFPQ;refGene_tx=XM_023258452.1;refGene_gn=SFPQ;refGene_tx=X
M_023258453.1;refGene_gn=SFPQ;refGene_tx=XM_023258454.1
C1 27071108 27071204
refGene_gn=SFPQ;refGene_tx=XM_023258452.1;refGene_gn=SFPQ;refGene_tx=X
M_023258453.1;refGene_gn=SFPQ;refGene_tx=XM_023258454.1
C1 27071304 27071606
refGene_gn=SFPQ;refGene_tx=XM_023258452.1;refGene_gn=SFPQ;refGene_tx=X
M_023258453.1;refGene_gn=SFPQ;refGene_tx=XM_023258454.1
C1 27071945 27072134
refGene_gn=SFPQ;refGene_tx=XM_023258452.1;refGene_gn=SFPQ;refGene_tx=X
M_023258453.1;refGene_gn=SFPQ;refGene_tx=XM_023258454.1
C1 27072827 27073637
refGene_gn=SFPQ;refGene_tx=XM_023258452.1;refGene_gn=SFPQ;refGene_tx=X
M_023258453.1;refGene_gn=SFPQ;refGene_tx=XM_023258454.1
C1 27128605 27128644
refGene_gn=ZMYM4;refGene_tx=XM_011284739.3;refGene_gn=ZMYM4;refGene_tx
=XM_023258461.1
C1 27188803 27188849
refGene_gn=ZMYM4;refGene_tx=XM_011284739.3;refGene_gn=ZMYM4;refGene_tx

=XM_023258461.1
C1 27212370 27212895
refGene_gn=ZMYM4;refGene_tx=XM_011284739.3;refGene_gn=ZMYM4;refGene_tx=XM_019836636.2;refGene_gn=ZMYM4;refGene_tx=XM_023258461.1
C1 27213140 27213225 refGene_gn=ZMYM4;refGene_tx=XM_019836637.2
C1 27214628 27214690
refGene_gn=ZMYM4;refGene_tx=XM_011284739.3;refGene_gn=ZMYM4;refGene_tx=XM_019836636.2;refGene_gn=ZMYM4;refGene_tx=XM_019836637.2;refGene_gn=ZMYM4;refGene_tx=XM_023258461.1
C1 27215042 27215213
refGene_gn=ZMYM4;refGene_tx=XM_011284739.3;refGene_gn=ZMYM4;refGene_tx=XM_019836636.2;refGene_gn=ZMYM4;refGene_tx=XM_019836637.2;refGene_gn=ZMYM4;refGene_tx=XM_023258461.1
C1 27219559 27219622
refGene_gn=ZMYM4;refGene_tx=XM_019836636.2;refGene_gn=ZMYM4;refGene_tx=XM_019836637.2;refGene_gn=ZMYM4;refGene_tx=XM_023258461.1
C1 27222357 27222442
refGene_gn=ZMYM4;refGene_tx=XM_011284739.3;refGene_gn=ZMYM4;refGene_tx=XM_019836636.2;refGene_gn=ZMYM4;refGene_tx=XM_019836637.2;refGene_gn=ZMYM4;refGene_tx=XM_023258461.1
C1 27222677 27222933
refGene_gn=ZMYM4;refGene_tx=XM_011284739.3;refGene_gn=ZMYM4;refGene_tx=XM_019836636.2;refGene_gn=ZMYM4;refGene_tx=XM_019836637.2;refGene_gn=ZMYM4;refGene_tx=XM_023258461.1;refGene_gn=ZMYM4;refGene_tx=XM_023258462.1;refGene_gn=ZMYM4;refGene_tx=XM_023258463.1
C1 27227060 27227235
refGene_gn=ZMYM4;refGene_tx=XM_011284739.3;refGene_gn=ZMYM4;refGene_tx=XM_019836636.2;refGene_gn=ZMYM4;refGene_tx=XM_019836637.2;refGene_gn=ZMYM4;refGene_tx=XM_023258461.1;refGene_gn=ZMYM4;refGene_tx=XM_023258462.1;refGene_gn=ZMYM4;refGene_tx=XM_023258463.1
C1 27227529 27227742
refGene_gn=ZMYM4;refGene_tx=XM_011284739.3;refGene_gn=ZMYM4;refGene_tx=XM_019836636.2;refGene_gn=ZMYM4;refGene_tx=XM_019836637.2;refGene_gn=ZMYM4;refGene_tx=XM_023258461.1;refGene_gn=ZMYM4;refGene_tx=XM_023258462.1;refGene_gn=ZMYM4;refGene_tx=XM_023258463.1
C1 27230367 27230518
refGene_gn=ZMYM4;refGene_tx=XM_011284739.3;refGene_gn=ZMYM4;refGene_tx=XM_019836636.2;refGene_gn=ZMYM4;refGene_tx=XM_019836637.2;refGene_gn=ZMYM4;refGene_tx=XM_023258461.1;refGene_gn=ZMYM4;refGene_tx=XM_023258462.1;refGene_gn=ZMYM4;refGene_tx=XM_023258463.1
C1 27231014 27231130
refGene_gn=ZMYM4;refGene_tx=XM_011284739.3;refGene_gn=ZMYM4;refGene_tx=XM_019836636.2;refGene_gn=ZMYM4;refGene_tx=XM_019836637.2;refGene_gn=ZMYM4;refGene_tx=XM_023258461.1;refGene_gn=ZMYM4;refGene_tx=XM_023258462.1;refGene_gn=ZMYM4;refGene_tx=XM_023258463.1
C1 27232176 27232452
refGene_gn=ZMYM4;refGene_tx=XM_011284739.3;refGene_gn=ZMYM4;refGene_tx=XM_019836636.2;refGene_gn=ZMYM4;refGene_tx=XM_019836637.2;refGene_gn=ZMYM4;refGene_tx=XM_023258461.1;refGene_gn=ZMYM4;refGene_tx=XM_023258462.1;refGene_gn=ZMYM4;refGene_tx=XM_023258463.1

C1 27232626 27232777
refGene_gn=ZMYM4;refGene_tx=XM_011284739.3;refGene_gn=ZMYM4;refGene_tx=XM_019836636.2;refGene_gn=ZMYM4;refGene_tx=XM_019836637.2;refGene_gn=ZMYM4;refGene_tx=XM_023258461.1;refGene_gn=ZMYM4;refGene_tx=XM_023258462.1;refGene_gn=ZMYM4;refGene_tx=XM_023258463.1
C1 27233823 27233996
refGene_gn=ZMYM4;refGene_tx=XM_011284739.3;refGene_gn=ZMYM4;refGene_tx=XM_019836636.2;refGene_gn=ZMYM4;refGene_tx=XM_019836637.2;refGene_gn=ZMYM4;refGene_tx=XM_023258461.1;refGene_gn=ZMYM4;refGene_tx=XM_023258462.1;refGene_gn=ZMYM4;refGene_tx=XM_023258463.1
C1 27234781 27234932
refGene_gn=ZMYM4;refGene_tx=XM_011284739.3;refGene_gn=ZMYM4;refGene_tx=XM_019836636.2;refGene_gn=ZMYM4;refGene_tx=XM_019836637.2;refGene_gn=ZMYM4;refGene_tx=XM_023258461.1;refGene_gn=ZMYM4;refGene_tx=XM_023258462.1;refGene_gn=ZMYM4;refGene_tx=XM_023258463.1
C1 27235710 27235851
refGene_gn=ZMYM4;refGene_tx=XM_011284739.3;refGene_gn=ZMYM4;refGene_tx=XM_019836636.2;refGene_gn=ZMYM4;refGene_tx=XM_019836637.2;refGene_gn=ZMYM4;refGene_tx=XM_023258461.1;refGene_gn=ZMYM4;refGene_tx=XM_023258462.1;refGene_gn=ZMYM4;refGene_tx=XM_023258463.1
C1 27236173 27236211
refGene_gn=ZMYM4;refGene_tx=XM_011284739.3;refGene_gn=ZMYM4;refGene_tx=XM_019836636.2;refGene_gn=ZMYM4;refGene_tx=XM_019836637.2;refGene_gn=ZMYM4;refGene_tx=XM_023258461.1;refGene_gn=ZMYM4;refGene_tx=XM_023258462.1;refGene_gn=ZMYM4;refGene_tx=XM_023258463.1
C1 27237149 27237294
refGene_gn=ZMYM4;refGene_tx=XM_011284739.3;refGene_gn=ZMYM4;refGene_tx=XM_019836636.2;refGene_gn=ZMYM4;refGene_tx=XM_019836637.2;refGene_gn=ZMYM4;refGene_tx=XM_023258461.1;refGene_gn=ZMYM4;refGene_tx=XM_023258462.1;refGene_gn=ZMYM4;refGene_tx=XM_023258463.1
C1 27241150 27241269
refGene_gn=ZMYM4;refGene_tx=XM_011284739.3;refGene_gn=ZMYM4;refGene_tx=XM_019836636.2;refGene_gn=ZMYM4;refGene_tx=XM_019836637.2;refGene_gn=ZMYM4;refGene_tx=XM_023258461.1;refGene_gn=ZMYM4;refGene_tx=XM_023258462.1;refGene_gn=ZMYM4;refGene_tx=XM_023258463.1
C1 27242181 27242353
refGene_gn=ZMYM4;refGene_tx=XM_011284739.3;refGene_gn=ZMYM4;refGene_tx=XM_019836636.2;refGene_gn=ZMYM4;refGene_tx=XM_019836637.2;refGene_gn=ZMYM4;refGene_tx=XM_023258461.1;refGene_gn=ZMYM4;refGene_tx=XM_023258462.1;refGene_gn=ZMYM4;refGene_tx=XM_023258463.1
C1 27245392 27245446
refGene_gn=ZMYM4;refGene_tx=XM_011284739.3;refGene_gn=ZMYM4;refGene_tx=XM_019836636.2;refGene_gn=ZMYM4;refGene_tx=XM_019836637.2;refGene_gn=ZMYM4;refGene_tx=XM_023258461.1;refGene_gn=ZMYM4;refGene_tx=XM_023258462.1;refGene_gn=ZMYM4;refGene_tx=XM_023258463.1
C1 27245840 27246020
refGene_gn=ZMYM4;refGene_tx=XM_011284739.3;refGene_gn=ZMYM4;refGene_tx=XM_019836636.2;refGene_gn=ZMYM4;refGene_tx=XM_019836637.2;refGene_gn=ZMYM4;refGene_tx=XM_023258461.1;refGene_gn=ZMYM4;refGene_tx=XM_023258462.1;refGene_gn=ZMYM4;refGene_tx=XM_023258463.1

C1 27246356 27246451
refGene_gn=ZMYM4;refGene_tx=XM_011284739.3;refGene_gn=ZMYM4;refGene_tx=XM_019836636.2;refGene_gn=ZMYM4;refGene_tx=XM_019836637.2;refGene_gn=ZMYM4;refGene_tx=XM_023258461.1;refGene_gn=ZMYM4;refGene_tx=XM_023258462.1;refGene_gn=ZMYM4;refGene_tx=XM_023258463.1
C1 27253163 27253335
refGene_gn=ZMYM4;refGene_tx=XM_011284739.3;refGene_gn=ZMYM4;refGene_tx=XM_019836636.2;refGene_gn=ZMYM4;refGene_tx=XM_019836637.2;refGene_gn=ZMYM4;refGene_tx=XM_023258461.1;refGene_gn=ZMYM4;refGene_tx=XM_023258462.1;refGene_gn=ZMYM4;refGene_tx=XM_023258463.1
C1 27253512 27253608
refGene_gn=ZMYM4;refGene_tx=XM_011284739.3;refGene_gn=ZMYM4;refGene_tx=XM_019836636.2;refGene_gn=ZMYM4;refGene_tx=XM_019836637.2;refGene_gn=ZMYM4;refGene_tx=XM_023258461.1;refGene_gn=ZMYM4;refGene_tx=XM_023258462.1;refGene_gn=ZMYM4;refGene_tx=XM_023258463.1
C1 27255590 27255742
refGene_gn=ZMYM4;refGene_tx=XM_011284739.3;refGene_gn=ZMYM4;refGene_tx=XM_019836636.2;refGene_gn=ZMYM4;refGene_tx=XM_019836637.2;refGene_gn=ZMYM4;refGene_tx=XM_023258461.1;refGene_gn=ZMYM4;refGene_tx=XM_023258462.1;refGene_gn=ZMYM4;refGene_tx=XM_023258463.1
C1 27266085 27266197
refGene_gn=ZMYM4;refGene_tx=XM_011284739.3;refGene_gn=ZMYM4;refGene_tx=XM_019836636.2;refGene_gn=ZMYM4;refGene_tx=XM_019836637.2;refGene_gn=ZMYM4;refGene_tx=XM_023258461.1;refGene_gn=ZMYM4;refGene_tx=XM_023258462.1;refGene_gn=ZMYM4;refGene_tx=XM_023258463.1
C1 27267678 27267927
refGene_gn=ZMYM4;refGene_tx=XM_011284739.3;refGene_gn=ZMYM4;refGene_tx=XM_019836636.2;refGene_gn=ZMYM4;refGene_tx=XM_019836637.2;refGene_gn=ZMYM4;refGene_tx=XM_023258461.1;refGene_gn=ZMYM4;refGene_tx=XM_023258462.1;refGene_gn=ZMYM4;refGene_tx=XM_023258463.1
C1 27270493 27270623
refGene_gn=ZMYM4;refGene_tx=XM_011284739.3;refGene_gn=ZMYM4;refGene_tx=XM_019836636.2;refGene_gn=ZMYM4;refGene_tx=XM_019836637.2;refGene_gn=ZMYM4;refGene_tx=XM_023258461.1;refGene_gn=ZMYM4;refGene_tx=XM_023258462.1;refGene_gn=ZMYM4;refGene_tx=XM_023258463.1
C1 27271510 27271718
refGene_gn=ZMYM4;refGene_tx=XM_011284739.3;refGene_gn=ZMYM4;refGene_tx=XM_019836636.2;refGene_gn=ZMYM4;refGene_tx=XM_019836637.2;refGene_gn=ZMYM4;refGene_tx=XM_023258461.1;refGene_gn=ZMYM4;refGene_tx=XM_023258462.1;refGene_gn=ZMYM4;refGene_tx=XM_023258463.1
C1 27289878 27289931
refGene_gn=KIAA0319L;refGene_tx=XM_006934561.4
C1 27291580 27291768
refGene_gn=KIAA0319L;refGene_tx=XM_003989848.5
C1 27291986 27292186
refGene_gn=KIAA0319L;refGene_tx=XM_019836638.2;refGene_gn=KIAA0319L;refGene_tx=XM_019836639.2;refGene_gn=KIAA0319L;refGene_tx=XM_019836640.2;refGene_gn=KIAA0319L;refGene_tx=XM_023258465.1
C1 27296291 27296431
refGene_gn=KIAA0319L;refGene_tx=XM_006934560.4

C1	27296913	27297005
refGene_gn=KIAA0319L;refGene_tx=XM_003989848.5;refGene_gn=KIAA0319L;refGene_tx=XM_006934560.4;refGene_gn=KIAA0319L;refGene_tx=XM_006934561.4;refGene_gn=KIAA0319L;refGene_tx=XM_019836638.2;refGene_gn=KIAA0319L;refGene_tx=XM_019836639.2;refGene_gn=KIAA0319L;refGene_tx=XM_019836640.2;refGene_gn=KIAA0319L;refGene_tx=XM_023258465.1
C1	27299156	27299247
refGene_gn=KIAA0319L;refGene_tx=XM_003989848.5;refGene_gn=KIAA0319L;refGene_tx=XM_006934560.4;refGene_gn=KIAA0319L;refGene_tx=XM_006934561.4;refGene_gn=KIAA0319L;refGene_tx=XM_019836638.2;refGene_gn=KIAA0319L;refGene_tx=XM_019836639.2;refGene_gn=KIAA0319L;refGene_tx=XM_019836640.2;refGene_gn=KIAA0319L;refGene_tx=XM_023258465.1
C1	27299801	27299924
refGene_gn=KIAA0319L;refGene_tx=XM_003989848.5;refGene_gn=KIAA0319L;refGene_tx=XM_006934560.4;refGene_gn=KIAA0319L;refGene_tx=XM_006934561.4;refGene_gn=KIAA0319L;refGene_tx=XM_019836638.2;refGene_gn=KIAA0319L;refGene_tx=XM_019836639.2;refGene_gn=KIAA0319L;refGene_tx=XM_019836640.2;refGene_gn=KIAA0319L;refGene_tx=XM_023258465.1
C1	27300758	27300901
refGene_gn=KIAA0319L;refGene_tx=XM_003989848.5;refGene_gn=KIAA0319L;refGene_tx=XM_006934560.4;refGene_gn=KIAA0319L;refGene_tx=XM_006934561.4;refGene_gn=KIAA0319L;refGene_tx=XM_019836638.2;refGene_gn=KIAA0319L;refGene_tx=XM_019836639.2;refGene_gn=KIAA0319L;refGene_tx=XM_019836640.2;refGene_gn=KIAA0319L;refGene_tx=XM_023258465.1
C1	27305088	27305248
refGene_gn=KIAA0319L;refGene_tx=XM_003989848.5;refGene_gn=KIAA0319L;refGene_tx=XM_006934560.4;refGene_gn=KIAA0319L;refGene_tx=XM_006934561.4;refGene_gn=KIAA0319L;refGene_tx=XM_019836638.2;refGene_gn=KIAA0319L;refGene_tx=XM_019836639.2;refGene_gn=KIAA0319L;refGene_tx=XM_019836640.2;refGene_gn=KIAA0319L;refGene_tx=XM_023258465.1
C1	27306764	27306903
refGene_gn=KIAA0319L;refGene_tx=XM_003989848.5;refGene_gn=KIAA0319L;refGene_tx=XM_006934560.4;refGene_gn=KIAA0319L;refGene_tx=XM_006934561.4;refGene_gn=KIAA0319L;refGene_tx=XM_019836638.2;refGene_gn=KIAA0319L;refGene_tx=XM_019836639.2;refGene_gn=KIAA0319L;refGene_tx=XM_019836640.2;refGene_gn=KIAA0319L;refGene_tx=XM_023258465.1
C1	27307247	27307399
refGene_gn=KIAA0319L;refGene_tx=XM_003989848.5;refGene_gn=KIAA0319L;refGene_tx=XM_006934560.4;refGene_gn=KIAA0319L;refGene_tx=XM_006934561.4;refGene_gn=KIAA0319L;refGene_tx=XM_019836638.2;refGene_gn=KIAA0319L;refGene_tx=XM_019836639.2;refGene_gn=KIAA0319L;refGene_tx=XM_019836640.2;refGene_gn=KIAA0319L;refGene_tx=XM_023258465.1
C1	27309008	27309157
refGene_gn=KIAA0319L;refGene_tx=XM_003989848.5;refGene_gn=KIAA0319L;refGene_tx=XM_006934560.4;refGene_gn=KIAA0319L;refGene_tx=XM_006934561.4;refGene_gn=KIAA0319L;refGene_tx=XM_019836638.2;refGene_gn=KIAA0319L;refGene_tx=XM_019836639.2;refGene_gn=KIAA0319L;refGene_tx=XM_019836640.2;refGene_gn=KIAA0319L;refGene_tx=XM_023258465.1
C1	27310917	27311050
refGene_gn=KIAA0319L;refGene_tx=XM_003989848.5;refGene_gn=KIAA0319L;re

fGene_tx=XM_006934560.4;refGene_gn=KIAA0319L;refGene_tx=XM_006934561.4;refGene_gn=KIAA0319L;refGene_tx=XM_019836638.2;refGene_gn=KIAA0319L;refGene_tx=XM_019836639.2;refGene_gn=KIAA0319L;refGene_tx=XM_019836640.2;refGene_gn=KIAA0319L;refGene_tx=XM_023258465.1
C1 27311643 27311767
refGene_gn=KIAA0319L;refGene_tx=XM_003989848.5;refGene_gn=KIAA0319L;refGene_tx=XM_006934560.4;refGene_gn=KIAA0319L;refGene_tx=XM_006934561.4;refGene_gn=KIAA0319L;refGene_tx=XM_019836638.2;refGene_gn=KIAA0319L;refGene_tx=XM_019836639.2;refGene_gn=KIAA0319L;refGene_tx=XM_019836640.2;refGene_gn=KIAA0319L;refGene_tx=XM_023258465.1
C1 27313155 27313384
refGene_gn=KIAA0319L;refGene_tx=XM_003989848.5;refGene_gn=KIAA0319L;refGene_tx=XM_006934560.4;refGene_gn=KIAA0319L;refGene_tx=XM_006934561.4;refGene_gn=KIAA0319L;refGene_tx=XM_019836638.2;refGene_gn=KIAA0319L;refGene_tx=XM_019836639.2;refGene_gn=KIAA0319L;refGene_tx=XM_019836640.2;refGene_gn=KIAA0319L;refGene_tx=XM_023258465.1
C1 27316867 27317000
refGene_gn=KIAA0319L;refGene_tx=XM_003989848.5;refGene_gn=KIAA0319L;refGene_tx=XM_006934560.4;refGene_gn=KIAA0319L;refGene_tx=XM_006934561.4;refGene_gn=KIAA0319L;refGene_tx=XM_019836638.2;refGene_gn=KIAA0319L;refGene_tx=XM_019836639.2;refGene_gn=KIAA0319L;refGene_tx=XM_019836640.2;refGene_gn=KIAA0319L;refGene_tx=XM_023258465.1
C1 27318814 27318907
refGene_gn=KIAA0319L;refGene_tx=XM_003989848.5;refGene_gn=KIAA0319L;refGene_tx=XM_006934560.4;refGene_gn=KIAA0319L;refGene_tx=XM_006934561.4;refGene_gn=KIAA0319L;refGene_tx=XM_019836638.2;refGene_gn=KIAA0319L;refGene_tx=XM_019836639.2;refGene_gn=KIAA0319L;refGene_tx=XM_019836640.2;refGene_gn=KIAA0319L;refGene_tx=XM_023258465.1
C1 27320082 27320170
refGene_gn=KIAA0319L;refGene_tx=XM_003989848.5;refGene_gn=KIAA0319L;refGene_tx=XM_006934560.4;refGene_gn=KIAA0319L;refGene_tx=XM_006934561.4;refGene_gn=KIAA0319L;refGene_tx=XM_019836638.2;refGene_gn=KIAA0319L;refGene_tx=XM_019836639.2;refGene_gn=KIAA0319L;refGene_tx=XM_019836640.2;refGene_gn=KIAA0319L;refGene_tx=XM_023258465.1
C1 27320860 27320934
refGene_gn=LOC109502223;refGene_tx=XM_023258467.1
C1 27320936 27321096
refGene_gn=LOC109502223;refGene_tx=XM_023258467.1
C1 27321324 27321663
refGene_gn=LOC109502223;refGene_tx=XM_023258467.1
C1 27324022 27324120
refGene_gn=KIAA0319L;refGene_tx=XM_003989848.5;refGene_gn=KIAA0319L;refGene_tx=XM_006934560.4;refGene_gn=KIAA0319L;refGene_tx=XM_006934561.4;refGene_gn=KIAA0319L;refGene_tx=XM_019836638.2;refGene_gn=KIAA0319L;refGene_tx=XM_019836639.2;refGene_gn=KIAA0319L;refGene_tx=XM_019836640.2;refGene_gn=KIAA0319L;refGene_tx=XM_023258465.1
C1 27327039 27327141
refGene_gn=KIAA0319L;refGene_tx=XM_003989848.5;refGene_gn=KIAA0319L;refGene_tx=XM_006934560.4;refGene_gn=KIAA0319L;refGene_tx=XM_006934561.4;refGene_gn=KIAA0319L;refGene_tx=XM_019836638.2;refGene_gn=KIAA0319L;r

efGene_tx=XM_019836639.2;refGene_gn=KIAA0319L;refGene_tx=XM_019836640.2;refGene_gn=KIAA0319L;refGene_tx=XM_023258465.1
C1	27329532	27329779	refGene_gn=KIAA0319L;refGene_tx=XM_003989848.5;refGene_gn=KIAA0319L;refGene_tx=XM_006934560.4;refGene_gn=KIAA0319L;refGene_tx=XM_006934561.4;refGene_gn=KIAA0319L;refGene_tx=XM_019836638.2;refGene_gn=KIAA0319L;refGene_tx=XM_019836639.2;refGene_gn=KIAA0319L;refGene_tx=XM_019836640.2;refGene_gn=KIAA0319L;refGene_tx=XM_023258465.1
C1	27356637	27357167	refGene_gn=KIAA0319L;refGene_tx=XM_003989848.5;refGene_gn=KIAA0319L;refGene_tx=XM_006934560.4;refGene_gn=KIAA0319L;refGene_tx=XM_006934561.4;refGene_gn=KIAA0319L;refGene_tx=XM_019836638.2;refGene_gn=KIAA0319L;refGene_tx=XM_019836639.2;refGene_gn=KIAA0319L;refGene_tx=XM_019836640.2;refGene_gn=KIAA0319L;refGene_tx=XM_023258465.1
C1	27384074	27384216	refGene_gn=KIAA0319L;refGene_tx=XM_003989848.5;refGene_gn=KIAA0319L;refGene_tx=XM_006934560.4;refGene_gn=KIAA0319L;refGene_tx=XM_006934561.4;refGene_gn=KIAA0319L;refGene_tx=XM_019836638.2;refGene_gn=KIAA0319L;refGene_tx=XM_019836639.2;refGene_gn=KIAA0319L;refGene_tx=XM_023258465.1
C1	27387892	27387925	refGene_gn=NCDN;refGene_tx=XM_003989849.5
C1	27388806	27388947	refGene_gn=NCDN;refGene_tx=XM_003989849.5;refGene_gn=NCDN;refGene_tx=XM_011284737.3;refGene_gn=NCDN;refGene_tx=XM_023258466.1
C1	27390042	27391011	refGene_gn=NCDN;refGene_tx=XM_003989849.5;refGene_gn=NCDN;refGene_tx=XM_011284737.3;refGene_gn=NCDN;refGene_tx=XM_023258466.1
C1	27391806	27392048	refGene_gn=NCDN;refGene_tx=XM_003989849.5;refGene_gn=NCDN;refGene_tx=XM_011284737.3;refGene_gn=NCDN;refGene_tx=XM_023258466.1
C1	27392625	27392850	refGene_gn=NCDN;refGene_tx=XM_003989849.5;refGene_gn=NCDN;refGene_tx=XM_011284737.3;refGene_gn=NCDN;refGene_tx=XM_023258466.1
C1	27393236	27393379	refGene_gn=NCDN;refGene_tx=XM_003989849.5;refGene_gn=NCDN;refGene_tx=XM_011284737.3;refGene_gn=NCDN;refGene_tx=XM_023258466.1
C1	27394697	27395134	refGene_gn=NCDN;refGene_tx=XM_003989849.5;refGene_gn=NCDN;refGene_tx=XM_011284737.3;refGene_gn=NCDN;refGene_tx=XM_023258466.1
C1	27397718	27397791	refGene_gn=TFAP2E;refGene_tx=XM_019836641.2
C1	27402224	27402589	refGene_gn=TFAP2E;refGene_tx=XM_019836641.2
C1	27402777	27403257	refGene_gn=TFAP2E;refGene_tx=XM_019836641.2
C1	27403803	27403855	refGene_gn=TFAP2E;refGene_tx=XM_019836641.2
C1	27414488	27414711	refGene_gn=TFAP2E;refGene_tx=XM_019836641.2
C1	27416039	27416158	refGene_gn=TFAP2E;refGene_tx=XM_019836641.2
C1	27416696	27416838	refGene_gn=TFAP2E;refGene_tx=XM_019836641.2
C1	27418532	27418815	refGene_gn=TFAP2E;refGene_tx=XM_019836641.2
C1	27426228	27426336	refGene_gn=PSMB2;refGene_tx=XM_003989851.4;refGene_gn=PSMB2;refGene_tx=XM_023258468.1

C1 27428890 27428940
refGene_gn=PSMB2;refGene_tx=XM_003989851.4;refGene_gn=PSMB2;refGene_tx=XM_023258468.1
C1 27430425 27430522 refGene_gn=PSMB2;refGene_tx=XM_023258468.1
C1 27432165 27432328 refGene_gn=PSMB2;refGene_tx=XM_003989851.4
C1 27453527 27453598 refGene_gn=PSMB2;refGene_tx=XM_003989851.4
C1 27455974 27456097 refGene_gn=PSMB2;refGene_tx=XM_003989851.4
C1 27461189 27461280 refGene_gn=PSMB2;refGene_tx=XM_003989851.4
C1 27524616 27525303
refGene_gn=CC1H1orf216;refGene_tx=XM_019836643.1;refGene_gn=CC1H1orf216;refGene_tx=XM_023258469.1;refGene_gn=CC1H1orf216;refGene_tx=XM_023258470.1;refGene_gn=CC1H1orf216;refGene_tx=XM_023258471.1
C1 27543544 27543655
refGene_gn=CLSPN;refGene_tx=XM_011284735.3;refGene_gn=CLSPN;refGene_tx=XM_019836645.2;refGene_gn=CLSPN;refGene_tx=XM_019836646.2;refGene_gn=CLSPN;refGene_tx=XM_019836647.2
C1 27543963 27544125
refGene_gn=CLSPN;refGene_tx=XM_011284735.3;refGene_gn=CLSPN;refGene_tx=XM_019836645.2;refGene_gn=CLSPN;refGene_tx=XM_019836646.2;refGene_gn=CLSPN;refGene_tx=XM_019836647.2
C1 27544388 27544471
refGene_gn=CLSPN;refGene_tx=XM_011284735.3;refGene_gn=CLSPN;refGene_tx=XM_019836645.2;refGene_gn=CLSPN;refGene_tx=XM_019836646.2;refGene_gn=CLSPN;refGene_tx=XM_019836647.2
C1 27545193 27545296
refGene_gn=CLSPN;refGene_tx=XM_011284735.3;refGene_gn=CLSPN;refGene_tx=XM_019836645.2;refGene_gn=CLSPN;refGene_tx=XM_019836646.2;refGene_gn=CLSPN;refGene_tx=XM_019836647.2
C1 27545616 27545744
refGene_gn=CLSPN;refGene_tx=XM_011284735.3;refGene_gn=CLSPN;refGene_tx=XM_019836645.2;refGene_gn=CLSPN;refGene_tx=XM_019836646.2;refGene_gn=CLSPN;refGene_tx=XM_019836647.2
C1 27545987 27546109
refGene_gn=CLSPN;refGene_tx=XM_011284735.3;refGene_gn=CLSPN;refGene_tx=XM_019836645.2;refGene_gn=CLSPN;refGene_tx=XM_019836646.2;refGene_gn=CLSPN;refGene_tx=XM_019836647.2
C1 27546213 27546378
refGene_gn=CLSPN;refGene_tx=XM_011284735.3;refGene_gn=CLSPN;refGene_tx=XM_019836645.2;refGene_gn=CLSPN;refGene_tx=XM_019836646.2;refGene_gn=CLSPN;refGene_tx=XM_019836647.2
C1 27549149 27549250
refGene_gn=CLSPN;refGene_tx=XM_011284735.3;refGene_gn=CLSPN;refGene_tx=XM_019836645.2;refGene_gn=CLSPN;refGene_tx=XM_019836646.2;refGene_gn=CLSPN;refGene_tx=XM_019836647.2
C1 27549456 27549535
refGene_gn=CLSPN;refGene_tx=XM_011284735.3;refGene_gn=CLSPN;refGene_tx=XM_019836645.2;refGene_gn=CLSPN;refGene_tx=XM_019836646.2;refGene_gn=CLSPN;refGene_tx=XM_019836647.2
C1 27549781 27549787
refGene_gn=CLSPN;refGene_tx=XM_019836645.2;refGene_gn=CLSPN;refGene_tx

=XM_019836647.2
C1 27551145 27551257
refGene_gn=CLSPN;refGene_tx=XM_011284735.3;refGene_gn=CLSPN;refGene_tx=XM_019836645.2;refGene_gn=CLSPN;refGene_tx=XM_019836646.2;refGene_gn=CLSPN;refGene_tx=XM_019836647.2
C1 27552059 27552286
refGene_gn=CLSPN;refGene_tx=XM_011284735.3;refGene_gn=CLSPN;refGene_tx=XM_019836645.2;refGene_gn=CLSPN;refGene_tx=XM_019836646.2;refGene_gn=CLSPN;refGene_tx=XM_019836647.2
C1 27552869 27553024
refGene_gn=CLSPN;refGene_tx=XM_011284735.3;refGene_gn=CLSPN;refGene_tx=XM_019836645.2;refGene_gn=CLSPN;refGene_tx=XM_019836646.2;refGene_gn=CLSPN;refGene_tx=XM_019836647.2
C1 27553354 27553554
refGene_gn=CLSPN;refGene_tx=XM_011284735.3;refGene_gn=CLSPN;refGene_tx=XM_019836645.2;refGene_gn=CLSPN;refGene_tx=XM_019836646.2;refGene_gn=CLSPN;refGene_tx=XM_019836647.2
C1 27554124 27554189
refGene_gn=CLSPN;refGene_tx=XM_011284735.3;refGene_gn=CLSPN;refGene_tx=XM_019836645.2;refGene_gn=CLSPN;refGene_tx=XM_019836646.2;refGene_gn=CLSPN;refGene_tx=XM_019836647.2
C1 27554286 27554465
refGene_gn=CLSPN;refGene_tx=XM_011284735.3;refGene_gn=CLSPN;refGene_tx=XM_019836645.2;refGene_gn=CLSPN;refGene_tx=XM_019836646.2;refGene_gn=CLSPN;refGene_tx=XM_019836647.2
C1 27555411 27555632
refGene_gn=CLSPN;refGene_tx=XM_011284735.3;refGene_gn=CLSPN;refGene_tx=XM_019836645.2;refGene_gn=CLSPN;refGene_tx=XM_019836646.2;refGene_gn=CLSPN;refGene_tx=XM_019836647.2
C1 27558733 27558925
refGene_gn=CLSPN;refGene_tx=XM_011284735.3;refGene_gn=CLSPN;refGene_tx=XM_019836645.2;refGene_gn=CLSPN;refGene_tx=XM_019836646.2;refGene_gn=CLSPN;refGene_tx=XM_019836647.2
C1 27564505 27565083
refGene_gn=CLSPN;refGene_tx=XM_011284735.3;refGene_gn=CLSPN;refGene_tx=XM_019836645.2;refGene_gn=CLSPN;refGene_tx=XM_019836646.2;refGene_gn=CLSPN;refGene_tx=XM_019836647.2
C1 27565267 27565376
refGene_gn=CLSPN;refGene_tx=XM_011284735.3;refGene_gn=CLSPN;refGene_tx=XM_019836645.2;refGene_gn=CLSPN;refGene_tx=XM_019836646.2;refGene_gn=CLSPN;refGene_tx=XM_019836647.2
C1 27566159 27566232
refGene_gn=CLSPN;refGene_tx=XM_011284735.3;refGene_gn=CLSPN;refGene_tx=XM_019836645.2;refGene_gn=CLSPN;refGene_tx=XM_019836646.2;refGene_gn=CLSPN;refGene_tx=XM_019836647.2
C1 27567418 27567496
refGene_gn=CLSPN;refGene_tx=XM_011284735.3;refGene_gn=CLSPN;refGene_tx=XM_019836645.2;refGene_gn=CLSPN;refGene_tx=XM_019836646.2;refGene_gn=CLSPN;refGene_tx=XM_019836647.2
C1 27569325 27569487

refGene_gn=CLSPN;refGene_tx=XM_011284735.3;refGene_gn=CLSPN;refGene_tx=XM_019836645.2;refGene_gn=CLSPN;refGene_tx=XM_019836646.2;refGene_gn=CLSPN;refGene_tx=XM_019836647.2
C1 27570272 27570718
refGene_gn=CLSPN;refGene_tx=XM_011284735.3;refGene_gn=CLSPN;refGene_tx=XM_019836645.2;refGene_gn=CLSPN;refGene_tx=XM_019836646.2;refGene_gn=CLSPN;refGene_tx=XM_019836647.2
C1 27571232 27571341
refGene_gn=CLSPN;refGene_tx=XM_011284735.3;refGene_gn=CLSPN;refGene_tx=XM_019836645.2;refGene_gn=CLSPN;refGene_tx=XM_019836646.2;refGene_gn=CLSPN;refGene_tx=XM_019836647.2
C1 27574794 27574818
refGene_gn=CLSPN;refGene_tx=XM_011284735.3;refGene_gn=CLSPN;refGene_tx=XM_019836647.2
C1 27612473 27612492 refGene_gn=AGO4;refGene_tx=XM_023258472.1
C1 27612597 27612648
refGene_gn=CLSPN;refGene_tx=XM_019836645.2;refGene_gn=CLSPN;refGene_tx=XM_019836646.2
C1 27621700 27621866 refGene_gn=AGO4;refGene_tx=XM_023258472.1
C1 27625926 27626047 refGene_gn=AGO4;refGene_tx=XM_023258472.1
C1 27628158 27628340 refGene_gn=AGO4;refGene_tx=XM_023258472.1
C1 27628528 27628665 refGene_gn=AGO4;refGene_tx=XM_023258472.1
C1 27628836 27628971 refGene_gn=AGO4;refGene_tx=XM_023258472.1
C1 27629656 27629744 refGene_gn=AGO4;refGene_tx=XM_023258472.1
C1 27632781 27632929 refGene_gn=AGO4;refGene_tx=XM_023258472.1
C1 27633190 27633310 refGene_gn=AGO4;refGene_tx=XM_023258472.1
C1 27633433 27633562 refGene_gn=AGO4;refGene_tx=XM_023258472.1
C1 27633802 27633936 refGene_gn=AGO4;refGene_tx=XM_023258472.1
C1 27635038 27635223 refGene_gn=AGO4;refGene_tx=XM_023258472.1
C1 27636756 27636916 refGene_gn=AGO4;refGene_tx=XM_023258472.1
C1 27640113 27640429 refGene_gn=AGO4;refGene_tx=XM_023258472.1
C1 27640570 27640705 refGene_gn=AGO4;refGene_tx=XM_023258472.1
C1 27647200 27647302 refGene_gn=AGO4;refGene_tx=XM_023258472.1
C1 27647567 27647767 refGene_gn=AGO4;refGene_tx=XM_023258472.1
C1 27649895 27650004 refGene_gn=AGO4;refGene_tx=XM_023258472.1
C1 27680689 27680714
refGene_gn=LOC101086581;refGene_tx=XM_011284734.3;refGene_gn=LOC101086581;refGene_tx=XM_019836650.2
C1 27685840 27686024
refGene_gn=LOC101086581;refGene_tx=XM_011284734.3;refGene_gn=LOC101086581;refGene_tx=XM_019836650.2
C1 27689452 27689573
refGene_gn=LOC101086581;refGene_tx=XM_011284734.3;refGene_gn=LOC101086581;refGene_tx=XM_019836650.2
C1 27690008 27690190
refGene_gn=LOC101086581;refGene_tx=XM_011284734.3;refGene_gn=LOC101086581;refGene_tx=XM_019836650.2
C1 27690532 27690728
refGene_gn=LOC101086581;refGene_tx=XM_011284734.3;refGene_gn=LOC101086581;refGene_tx=XM_019836650.2;refGene_gn=LOC101086581;refGene_tx=XM_01

9836651.1
C1	27690952	27691087
refGene_gn=LOC101086581;refGene_tx=XM_011284734.3;refGene_gn=LOC101086581;refGene_tx=XM_019836650.2;refGene_gn=LOC101086581;refGene_tx=XM_019836651.1
C1	27691237	27691325
refGene_gn=LOC101086581;refGene_tx=XM_011284734.3;refGene_gn=LOC101086581;refGene_tx=XM_019836650.2;refGene_gn=LOC101086581;refGene_tx=XM_019836651.1
C1	27691933	27692081
refGene_gn=LOC101086581;refGene_tx=XM_011284734.3;refGene_gn=LOC101086581;refGene_tx=XM_019836650.2;refGene_gn=LOC101086581;refGene_tx=XM_019836651.1
C1	27695866	27695986
refGene_gn=LOC101086581;refGene_tx=XM_011284734.3;refGene_gn=LOC101086581;refGene_tx=XM_019836650.2;refGene_gn=LOC101086581;refGene_tx=XM_019836651.1
C1	27696352	27696484
refGene_gn=LOC101086581;refGene_tx=XM_011284734.3;refGene_gn=LOC101086581;refGene_tx=XM_019836650.2;refGene_gn=LOC101086581;refGene_tx=XM_019836651.1
C1	27696612	27696746
refGene_gn=LOC101086581;refGene_tx=XM_011284734.3;refGene_gn=LOC101086581;refGene_tx=XM_019836650.2;refGene_gn=LOC101086581;refGene_tx=XM_019836651.1
C1	27699826	27700011
refGene_gn=LOC101086581;refGene_tx=XM_011284734.3;refGene_gn=LOC101086581;refGene_tx=XM_019836650.2;refGene_gn=LOC101086581;refGene_tx=XM_019836651.1
C1	27706322	27706482
refGene_gn=LOC101086581;refGene_tx=XM_011284734.3;refGene_gn=LOC101086581;refGene_tx=XM_019836650.2;refGene_gn=LOC101086581;refGene_tx=XM_019836651.1
C1	27706663	27706754
refGene_gn=LOC101086581;refGene_tx=XM_011284734.3;refGene_gn=LOC101086581;refGene_tx=XM_019836650.2;refGene_gn=LOC101086581;refGene_tx=XM_019836651.1
C1	27707816	27708011
refGene_gn=LOC101086581;refGene_tx=XM_011284734.3;refGene_gn=LOC101086581;refGene_tx=XM_019836650.2;refGene_gn=LOC101086581;refGene_tx=XM_019836651.1
C1	27710186	27710321
refGene_gn=LOC101086581;refGene_tx=XM_011284734.3;refGene_gn=LOC101086581;refGene_tx=XM_019836650.2;refGene_gn=LOC101086581;refGene_tx=XM_019836651.1
C1	27710906	27711008
refGene_gn=LOC101086581;refGene_tx=XM_011284734.3;refGene_gn=LOC101086581;refGene_tx=XM_019836650.2;refGene_gn=LOC101086581;refGene_tx=XM_019836651.1
C1	27711335	27711535

refGene_gn=LOC101086581;refGene_tx=XM_011284734.3;refGene_gn=LOC101086
581;refGene_tx=XM_019836650.2;refGene_gn=LOC101086581;refGene_tx=XM_01
9836651.1
C1 27711769 27711878
refGene_gn=LOC101086581;refGene_tx=XM_011284734.3;refGene_gn=LOC101086
581;refGene_tx=XM_019836650.2;refGene_gn=LOC101086581;refGene_tx=XM_01
9836651.1
C1 27723048 27723067
refGene_gn=LOC101087079;refGene_tx=XM_003989853.5;refGene_gn=LOC101087
079;refGene_tx=XM_023258473.1
C1 27741901 27742073
refGene_gn=LOC101087079;refGene_tx=XM_003989853.5;refGene_gn=LOC101087
079;refGene_tx=XM_023258473.1
C1 27752300 27752350
refGene_gn=LOC101087079;refGene_tx=XM_019836652.2
C1 27759265 27759386
refGene_gn=LOC101087079;refGene_tx=XM_003989853.5;refGene_gn=LOC101087
079;refGene_tx=XM_019836652.2;refGene_gn=LOC101087079;refGene_tx=XM_02
3258473.1
C1 27763614 27763823
refGene_gn=LOC101087079;refGene_tx=XM_003989853.5;refGene_gn=LOC101087
079;refGene_tx=XM_019836652.2;refGene_gn=LOC101087079;refGene_tx=XM_02
3258473.1
C1 27764636 27764773
refGene_gn=LOC101087079;refGene_tx=XM_003989853.5;refGene_gn=LOC101087
079;refGene_tx=XM_019836652.2;refGene_gn=LOC101087079;refGene_tx=XM_02
3258473.1
C1 27774849 27774985
refGene_gn=LOC101087079;refGene_tx=XM_019836655.2
C1 27793317 27793452
refGene_gn=LOC101087079;refGene_tx=XM_003989853.5;refGene_gn=LOC101087
079;refGene_tx=XM_019836652.2;refGene_gn=LOC101087079;refGene_tx=XM_01
9836655.2;refGene_gn=LOC101087079;refGene_tx=XM_023258473.1
C1 27797726 27797814
refGene_gn=LOC101087079;refGene_tx=XM_003989853.5;refGene_gn=LOC101087
079;refGene_tx=XM_019836652.2;refGene_gn=LOC101087079;refGene_tx=XM_01
9836655.2;refGene_gn=LOC101087079;refGene_tx=XM_023258473.1
C1 27797945 27798093
refGene_gn=LOC101087079;refGene_tx=XM_003989853.5;refGene_gn=LOC101087
079;refGene_tx=XM_019836652.2;refGene_gn=LOC101087079;refGene_tx=XM_01
9836655.2;refGene_gn=LOC101087079;refGene_tx=XM_023258473.1
C1 27798708 27798828
refGene_gn=LOC101087079;refGene_tx=XM_003989853.5;refGene_gn=LOC101087
079;refGene_tx=XM_019836652.2;refGene_gn=LOC101087079;refGene_tx=XM_01
9836655.2;refGene_gn=LOC101087079;refGene_tx=XM_023258473.1
C1 27801729 27801852
refGene_gn=LOC101087079;refGene_tx=XM_003989853.5;refGene_gn=LOC101087
079;refGene_tx=XM_019836652.2;refGene_gn=LOC101087079;refGene_tx=XM_01
9836655.2;refGene_gn=LOC101087079;refGene_tx=XM_023258473.1
C1 27802174 27802308

refGene_gn=LOC101087079;refGene_tx=XM_003989853.5;refGene_gn=LOC101087079;refGene_tx=XM_019836652.2;refGene_gn=LOC101087079;refGene_tx=XM_019836655.2;refGene_gn=LOC101087079;refGene_tx=XM_023258473.1
C1 27804841 27805026
refGene_gn=LOC101087079;refGene_tx=XM_003989853.5;refGene_gn=LOC101087079;refGene_tx=XM_019836652.2;refGene_gn=LOC101087079;refGene_tx=XM_019836655.2;refGene_gn=LOC101087079;refGene_tx=XM_023258473.1
C1 27815514 27815674
refGene_gn=LOC101087079;refGene_tx=XM_003989853.5;refGene_gn=LOC101087079;refGene_tx=XM_019836652.2;refGene_gn=LOC101087079;refGene_tx=XM_019836655.2;refGene_gn=LOC101087079;refGene_tx=XM_023258473.1
C1 27817125 27817216
refGene_gn=LOC101087079;refGene_tx=XM_003989853.5;refGene_gn=LOC101087079;refGene_tx=XM_019836652.2;refGene_gn=LOC101087079;refGene_tx=XM_019836655.2;refGene_gn=LOC101087079;refGene_tx=XM_023258473.1
C1 27821644 27821839
refGene_gn=LOC101087079;refGene_tx=XM_003989853.5;refGene_gn=LOC101087079;refGene_tx=XM_019836652.2;refGene_gn=LOC101087079;refGene_tx=XM_019836655.2;refGene_gn=LOC101087079;refGene_tx=XM_023258473.1
C1 27822178 27822313
refGene_gn=LOC101087079;refGene_tx=XM_003989853.5;refGene_gn=LOC101087079;refGene_tx=XM_019836652.2;refGene_gn=LOC101087079;refGene_tx=XM_019836655.2;refGene_gn=LOC101087079;refGene_tx=XM_023258473.1
C1 27826853 27826955
refGene_gn=LOC101087079;refGene_tx=XM_003989853.5;refGene_gn=LOC101087079;refGene_tx=XM_019836652.2;refGene_gn=LOC101087079;refGene_tx=XM_019836655.2;refGene_gn=LOC101087079;refGene_tx=XM_023258473.1
C1 27834924 27835124
refGene_gn=LOC101087079;refGene_tx=XM_003989853.5;refGene_gn=LOC101087079;refGene_tx=XM_019836652.2;refGene_gn=LOC101087079;refGene_tx=XM_019836655.2;refGene_gn=LOC101087079;refGene_tx=XM_023258473.1
C1 27835645 27835754
refGene_gn=LOC101087079;refGene_tx=XM_003989853.5;refGene_gn=LOC101087079;refGene_tx=XM_019836652.2;refGene_gn=LOC101087079;refGene_tx=XM_019836655.2;refGene_gn=LOC101087079;refGene_tx=XM_023258473.1
C1 27858674 27858830
refGene_gn=TEKT2;refGene_tx=XM_011284728.3;refGene_gn=TEKT2;refGene_tx=XM_011284730.3;refGene_gn=TEKT2;refGene_tx=XM_011284731.3;refGene_gn=TEKT2;refGene_tx=XM_011284732.3;refGene_gn=TEKT2;refGene_tx=XM_011284733.3;refGene_gn=TEKT2;refGene_tx=XM_019836658.1
C1 27858908 27859034
refGene_gn=TEKT2;refGene_tx=XM_011284728.3;refGene_gn=TEKT2;refGene_tx=XM_011284730.3;refGene_gn=TEKT2;refGene_tx=XM_011284731.3;refGene_gn=TEKT2;refGene_tx=XM_011284732.3;refGene_gn=TEKT2;refGene_tx=XM_011284733.3;refGene_gn=TEKT2;refGene_tx=XM_019836658.1
C1 27859224 27859430
refGene_gn=TEKT2;refGene_tx=XM_011284728.3;refGene_gn=TEKT2;refGene_tx=XM_011284730.3;refGene_gn=TEKT2;refGene_tx=XM_011284731.3;refGene_gn=TEKT2;refGene_tx=XM_011284732.3;refGene_gn=TEKT2;refGene_tx=XM_011284733.3;refGene_gn=TEKT2;refGene_tx=XM_019836658.1

C1 27859959 27860103
refGene_gn=TEKT2;refGene_tx=XM_011284728.3;refGene_gn=TEKT2;refGene_tx=XM_011284730.3;refGene_gn=TEKT2;refGene_tx=XM_011284731.3;refGene_gn=TEKT2;refGene_tx=XM_011284732.3;refGene_gn=TEKT2;refGene_tx=XM_011284733.3;refGene_gn=TEKT2;refGene_tx=XM_019836658.1
C1 27860184 27860299
refGene_gn=TEKT2;refGene_tx=XM_011284728.3;refGene_gn=TEKT2;refGene_tx=XM_011284730.3;refGene_gn=TEKT2;refGene_tx=XM_011284731.3;refGene_gn=TEKT2;refGene_tx=XM_011284732.3;refGene_gn=TEKT2;refGene_tx=XM_011284733.3;refGene_gn=TEKT2;refGene_tx=XM_019836658.1
C1 27860454 27860562
refGene_gn=TEKT2;refGene_tx=XM_011284728.3;refGene_gn=TEKT2;refGene_tx=XM_011284730.3;refGene_gn=TEKT2;refGene_tx=XM_011284731.3;refGene_gn=TEKT2;refGene_tx=XM_011284732.3;refGene_gn=TEKT2;refGene_tx=XM_011284733.3;refGene_gn=TEKT2;refGene_tx=XM_019836658.1
C1 27860690 27860834
refGene_gn=TEKT2;refGene_tx=XM_011284728.3;refGene_gn=TEKT2;refGene_tx=XM_011284730.3;refGene_gn=TEKT2;refGene_tx=XM_011284731.3;refGene_gn=TEKT2;refGene_tx=XM_011284732.3;refGene_gn=TEKT2;refGene_tx=XM_011284733.3;refGene_gn=TEKT2;refGene_tx=XM_019836658.1
C1 27860997 27861077
refGene_gn=TEKT2;refGene_tx=XM_011284728.3;refGene_gn=TEKT2;refGene_tx=XM_011284730.3;refGene_gn=TEKT2;refGene_tx=XM_011284731.3;refGene_gn=TEKT2;refGene_tx=XM_011284732.3;refGene_gn=TEKT2;refGene_tx=XM_011284733.3;refGene_gn=TEKT2;refGene_tx=XM_019836658.1
C1 27861243 27861457
refGene_gn=TEKT2;refGene_tx=XM_011284728.3;refGene_gn=TEKT2;refGene_tx=XM_011284730.3;refGene_gn=TEKT2;refGene_tx=XM_011284731.3;refGene_gn=TEKT2;refGene_tx=XM_011284732.3;refGene_gn=TEKT2;refGene_tx=XM_011284733.3;refGene_gn=TEKT2;refGene_tx=XM_019836658.1
C1 27862264 27862487 refGene_gn=ADPRHL2;refGene_tx=XM_023258475.1
C1 27865937 27866034 refGene_gn=ADPRHL2;refGene_tx=XM_023258475.1
C1 27866320 27866528 refGene_gn=ADPRHL2;refGene_tx=XM_023258475.1
C1 27866631 27866816 refGene_gn=ADPRHL2;refGene_tx=XM_023258475.1
C1 27867824 27867925 refGene_gn=ADPRHL2;refGene_tx=XM_023258475.1
C1 27868556 27868849 refGene_gn=ADPRHL2;refGene_tx=XM_023258475.1
C1 27873009 27874928
refGene_gn=COL8A2;refGene_tx=XM_006934566.4;refGene_gn=COL8A2;refGene_tx=XM_006934567.4;refGene_gn=COL8A2;refGene_tx=XM_019836659.2;refGene_gn=COL8A2;refGene_tx=XM_023258474.1
C1 27875453 27875662
refGene_gn=COL8A2;refGene_tx=XM_006934566.4;refGene_gn=COL8A2;refGene_tx=XM_006934567.4;refGene_gn=COL8A2;refGene_tx=XM_019836659.2;refGene_gn=COL8A2;refGene_tx=XM_023258474.1
C1 27890534 27890551 refGene_gn=COL8A2;refGene_tx=XM_019836659.2
C1 27904683 27904803
refGene_gn=TRAPPC3;refGene_tx=XM_003989857.5;refGene_gn=TRAPPC3;refGene_tx=XM_006934568.4
C1 27905260 27905443
refGene_gn=TRAPPC3;refGene_tx=XM_003989857.5;refGene_gn=TRAPPC3;refGen

e_tx=XM_006934568.4
C1 27906491 27906591
refGene_gn=TRAPPC3;refGene_tx=XM_003989857.5;refGene_gn=TRAPPC3;refGen
e_tx=XM_006934568.4
C1 27906836 27906934
refGene_gn=TRAPPC3;refGene_tx=XM_003989857.5;refGene_gn=TRAPPC3;refGen
e_tx=XM_006934568.4
C1 27913768 27913810 refGene_gn=TRAPPC3;refGene_tx=XM_003989857.5
C1 27919422 27919468 refGene_gn=MAP7D1;refGene_tx=XM_023257823.1
C1 27933712 27934030 refGene_gn=MAP7D1;refGene_tx=XM_023257823.1
C1 27935025 27935427 refGene_gn=MAP7D1;refGene_tx=XM_023257823.1
C1 27935627 27935696 refGene_gn=MAP7D1;refGene_tx=XM_023257823.1
C1 27936781 27936945 refGene_gn=MAP7D1;refGene_tx=XM_023257823.1
C1 27937668 27937783 refGene_gn=MAP7D1;refGene_tx=XM_023257823.1
C1 27939225 27939336 refGene_gn=MAP7D1;refGene_tx=XM_023257823.1
C1 27940486 27940609 refGene_gn=MAP7D1;refGene_tx=XM_023257823.1
C1 27940705 27940872 refGene_gn=MAP7D1;refGene_tx=XM_023257823.1
C1 27940974 27941117 refGene_gn=MAP7D1;refGene_tx=XM_023257823.1
C1 27941664 27941993 refGene_gn=MAP7D1;refGene_tx=XM_023257823.1
C1 27942278 27942456 refGene_gn=MAP7D1;refGene_tx=XM_023257823.1
C1 27942543 27942682 refGene_gn=MAP7D1;refGene_tx=XM_023257823.1
C1 27942819 27942924 refGene_gn=MAP7D1;refGene_tx=XM_023257823.1
C1 27943196 27943250 refGene_gn=MAP7D1;refGene_tx=XM_023257823.1
C1 27943422 27943465 refGene_gn=MAP7D1;refGene_tx=XM_023257823.1
C1 27943567 27943658 refGene_gn=MAP7D1;refGene_tx=XM_023257823.1
C1 27943779 27943973 refGene_gn=MAP7D1;refGene_tx=XM_023257823.1
C1 27944141 27944152 refGene_gn=MAP7D1;refGene_tx=XM_023257823.1
C1 28030276 28030413 refGene_gn=THRAP3;refGene_tx=XM_003989860.5
C1 28034773 28035679 refGene_gn=THRAP3;refGene_tx=XM_003989860.5
C1 28037221 28037935 refGene_gn=THRAP3;refGene_tx=XM_003989860.5
C1 28038878 28039051 refGene_gn=THRAP3;refGene_tx=XM_003989860.5
C1 28039803 28039915 refGene_gn=THRAP3;refGene_tx=XM_003989860.5
C1 28040949 28041034 refGene_gn=THRAP3;refGene_tx=XM_003989860.5
C1 28043003 28043191 refGene_gn=THRAP3;refGene_tx=XM_003989860.5
C1 28045999 28046195 refGene_gn=THRAP3;refGene_tx=XM_003989860.5
C1 28046618 28046762 refGene_gn=THRAP3;refGene_tx=XM_003989860.5
C1 28048832 28049054 refGene_gn=THRAP3;refGene_tx=XM_003989860.5
C1 28051572 28051576
refGene_gn=SH3D21;refGene_tx=XM_023258476.1;refGene_gn=SH3D21;refGene_
tx=XM_023258477.1;refGene_gn=SH3D21;refGene_tx=XM_023258478.1;refGene_
gn=SH3D21;refGene_tx=XM_023258479.1
C1 28051729 28052042
refGene_gn=SH3D21;refGene_tx=XM_023258476.1;refGene_gn=SH3D21;refGene_
tx=XM_023258477.1;refGene_gn=SH3D21;refGene_tx=XM_023258478.1;refGene_
gn=SH3D21;refGene_tx=XM_023258479.1
C1 28052292 28052411
refGene_gn=SH3D21;refGene_tx=XM_023258476.1;refGene_gn=SH3D21;refGene_
tx=XM_023258477.1;refGene_gn=SH3D21;refGene_tx=XM_023258478.1;refGene_
gn=SH3D21;refGene_tx=XM_023258479.1
C1 28052652 28052743

refGene_gn=SH3D21;refGene_tx=XM_023258476.1;refGene_gn=SH3D21;refGene_tx=XM_023258478.1;refGene_gn=SH3D21;refGene_tx=XM_023258479.1
C1 28052917 28052973
refGene_gn=SH3D21;refGene_tx=XM_023258476.1;refGene_gn=SH3D21;refGene_tx=XM_023258477.1;refGene_gn=SH3D21;refGene_tx=XM_023258478.1;refGene_gn=SH3D21;refGene_tx=XM_023258479.1
C1 28053062 28053108
refGene_gn=SH3D21;refGene_tx=XM_023258476.1;refGene_gn=SH3D21;refGene_tx=XM_023258477.1;refGene_gn=SH3D21;refGene_tx=XM_023258478.1;refGene_gn=SH3D21;refGene_tx=XM_023258479.1
C1 28053257 28053358
refGene_gn=SH3D21;refGene_tx=XM_023258476.1;refGene_gn=SH3D21;refGene_tx=XM_023258477.1;refGene_gn=SH3D21;refGene_tx=XM_023258478.1;refGene_gn=SH3D21;refGene_tx=XM_023258479.1
C1 28053477 28053588
refGene_gn=SH3D21;refGene_tx=XM_023258476.1;refGene_gn=SH3D21;refGene_tx=XM_023258477.1;refGene_gn=SH3D21;refGene_tx=XM_023258478.1;refGene_gn=SH3D21;refGene_tx=XM_023258479.1
C1 28053726 28053769
refGene_gn=SH3D21;refGene_tx=XM_023258476.1;refGene_gn=SH3D21;refGene_tx=XM_023258477.1;refGene_gn=SH3D21;refGene_tx=XM_023258478.1;refGene_gn=SH3D21;refGene_tx=XM_023258479.1
C1 28062906 28063000
refGene_gn=SH3D21;refGene_tx=XM_023258476.1;refGene_gn=SH3D21;refGene_tx=XM_023258477.1;refGene_gn=SH3D21;refGene_tx=XM_023258478.1;refGene_gn=SH3D21;refGene_tx=XM_023258479.1
C1 28063087 28063140
refGene_gn=SH3D21;refGene_tx=XM_023258476.1;refGene_gn=SH3D21;refGene_tx=XM_023258477.1;refGene_gn=SH3D21;refGene_tx=XM_023258478.1;refGene_gn=SH3D21;refGene_tx=XM_023258479.1
C1 28063276 28063371
refGene_gn=SH3D21;refGene_tx=XM_023258476.1;refGene_gn=SH3D21;refGene_tx=XM_023258477.1;refGene_gn=SH3D21;refGene_tx=XM_023258478.1;refGene_gn=SH3D21;refGene_tx=XM_023258479.1
C1 28063529 28064377
refGene_gn=SH3D21;refGene_tx=XM_023258476.1;refGene_gn=SH3D21;refGene_tx=XM_023258477.1;refGene_gn=SH3D21;refGene_tx=XM_023258478.1;refGene_gn=SH3D21;refGene_tx=XM_023258479.1
C1 28064462 28064526
refGene_gn=SH3D21;refGene_tx=XM_023258476.1;refGene_gn=SH3D21;refGene_tx=XM_023258477.1;refGene_gn=SH3D21;refGene_tx=XM_023258478.1;refGene_gn=SH3D21;refGene_tx=XM_023258479.1
C1 28064604 28064676
refGene_gn=SH3D21;refGene_tx=XM_023258476.1;refGene_gn=SH3D21;refGene_tx=XM_023258477.1;refGene_gn=SH3D21;refGene_tx=XM_023258478.1;refGene_gn=SH3D21;refGene_tx=XM_023258479.1
C1 28075728 28075947
refGene_gn=STK40;refGene_tx=XM_006934571.3;refGene_gn=STK40;refGene_tx=XM_006934572.4;refGene_gn=STK40;refGene_tx=XM_006934573.4
C1 28077305 28077390

refGene_gn=STK40;refGene_tx=XM_006934571.3;refGene_gn=STK40;refGene_tx=XM_006934572.4;refGene_gn=STK40;refGene_tx=XM_006934573.4
C1 28077828 28077948
refGene_gn=STK40;refGene_tx=XM_006934571.3;refGene_gn=STK40;refGene_tx=XM_006934572.4;refGene_gn=STK40;refGene_tx=XM_006934573.4
C1 28078092 28078237
refGene_gn=STK40;refGene_tx=XM_006934571.3;refGene_gn=STK40;refGene_tx=XM_006934572.4;refGene_gn=STK40;refGene_tx=XM_006934573.4
C1 28081798 28081914
refGene_gn=STK40;refGene_tx=XM_006934571.3;refGene_gn=STK40;refGene_tx=XM_006934572.4;refGene_gn=STK40;refGene_tx=XM_006934573.4
C1 28087251 28087304
refGene_gn=STK40;refGene_tx=XM_006934571.3;refGene_gn=STK40;refGene_tx=XM_006934572.4;refGene_gn=STK40;refGene_tx=XM_006934573.4
C1 28088201 28088429
refGene_gn=STK40;refGene_tx=XM_006934571.3;refGene_gn=STK40;refGene_tx=XM_006934572.4;refGene_gn=STK40;refGene_tx=XM_006934573.4
C1 28090850 28090994
refGene_gn=STK40;refGene_tx=XM_006934571.3;refGene_gn=STK40;refGene_tx=XM_006934572.4;refGene_gn=STK40;refGene_tx=XM_006934573.4
C1 28091354 28091440
refGene_gn=STK40;refGene_tx=XM_006934571.3;refGene_gn=STK40;refGene_tx=XM_006934572.4;refGene_gn=STK40;refGene_tx=XM_006934573.4
C1 28093180 28093292
refGene_gn=STK40;refGene_tx=XM_006934571.3;refGene_gn=STK40;refGene_tx=XM_006934572.4;refGene_gn=STK40;refGene_tx=XM_006934573.4
C1 28120111 28120506
refGene_gn=LSM10;refGene_tx=XM_006934574.2;refGene_gn=LSM10;refGene_tx=XM_006934575.2;refGene_gn=LSM10;refGene_tx=XM_006934576.2
C1 28120897 28120928 refGene_gn=LSM10;refGene_tx=XM_006934574.2
C1 28131655 28131772
refGene_gn=OSCP1;refGene_tx=XM_023258480.1;refGene_gn=OSCP1;refGene_tx=XM_023258481.1;refGene_gn=OSCP1;refGene_tx=XM_023258482.1;refGene_gn=OSCP1;refGene_tx=XM_023258483.1;refGene_gn=OSCP1;refGene_tx=XM_023258484.1
C1 28132012 28132076
refGene_gn=OSCP1;refGene_tx=XM_023258480.1;refGene_gn=OSCP1;refGene_tx=XM_023258481.1;refGene_gn=OSCP1;refGene_tx=XM_023258482.1;refGene_gn=OSCP1;refGene_tx=XM_023258483.1;refGene_gn=OSCP1;refGene_tx=XM_023258484.1
C1 28133318 28133458
refGene_gn=OSCP1;refGene_tx=XM_023258480.1;refGene_gn=OSCP1;refGene_tx=XM_023258481.1;refGene_gn=OSCP1;refGene_tx=XM_023258482.1;refGene_gn=OSCP1;refGene_tx=XM_023258483.1;refGene_gn=OSCP1;refGene_tx=XM_023258484.1
C1 28134837 28134907
refGene_gn=OSCP1;refGene_tx=XM_023258480.1;refGene_gn=OSCP1;refGene_tx=XM_023258481.1;refGene_gn=OSCP1;refGene_tx=XM_023258482.1;refGene_gn=OSCP1;refGene_tx=XM_023258483.1;refGene_gn=OSCP1;refGene_tx=XM_023258484.1

C1	28135446	28135575
refGene_gn=OSCP1;refGene_tx=XM_023258480.1;refGene_gn=OSCP1;refGene_tx=XM_023258481.1;refGene_gn=OSCP1;refGene_tx=XM_023258482.1;refGene_gn=OSCP1;refGene_tx=XM_023258483.1;refGene_gn=OSCP1;refGene_tx=XM_023258484.1
C1	28136016	28136120
refGene_gn=OSCP1;refGene_tx=XM_023258480.1;refGene_gn=OSCP1;refGene_tx=XM_023258481.1;refGene_gn=OSCP1;refGene_tx=XM_023258482.1;refGene_gn=OSCP1;refGene_tx=XM_023258483.1;refGene_gn=OSCP1;refGene_tx=XM_023258484.1
C1	28142902	28143016
refGene_gn=OSCP1;refGene_tx=XM_023258480.1;refGene_gn=OSCP1;refGene_tx=XM_023258481.1;refGene_gn=OSCP1;refGene_tx=XM_023258482.1;refGene_gn=OSCP1;refGene_tx=XM_023258483.1;refGene_gn=OSCP1;refGene_tx=XM_023258484.1
C1	28143748	28143916
refGene_gn=OSCP1;refGene_tx=XM_023258480.1;refGene_gn=OSCP1;refGene_tx=XM_023258481.1;refGene_gn=OSCP1;refGene_tx=XM_023258482.1;refGene_gn=OSCP1;refGene_tx=XM_023258483.1;refGene_gn=OSCP1;refGene_tx=XM_023258484.1
C1	28148137	28148292
refGene_gn=OSCP1;refGene_tx=XM_023258480.1;refGene_gn=OSCP1;refGene_tx=XM_023258481.1;refGene_gn=OSCP1;refGene_tx=XM_023258482.1;refGene_gn=OSCP1;refGene_tx=XM_023258483.1;refGene_gn=OSCP1;refGene_tx=XM_023258484.1
C1	28153304	28153334
refGene_gn=OSCP1;refGene_tx=XM_023258480.1;refGene_gn=OSCP1;refGene_tx=XM_023258482.1
C1	28158292	28158404
refGene_gn=OSCP1;refGene_tx=XM_023258480.1;refGene_gn=OSCP1;refGene_tx=XM_023258481.1;refGene_gn=OSCP1;refGene_tx=XM_023258482.1;refGene_gn=OSCP1;refGene_tx=XM_023258483.1
C1	28162571	28162709	refGene_gn=MRPS15;refGene_tx=XM_006934579.4
C1	28162958	28163150	refGene_gn=MRPS15;refGene_tx=XM_006934579.4
C1	28164689	28164748	refGene_gn=MRPS15;refGene_tx=XM_006934579.4
C1	28166183	28166268	refGene_gn=MRPS15;refGene_tx=XM_006934579.4
C1	28166744	28166793	refGene_gn=MRPS15;refGene_tx=XM_006934579.4
C1	28167691	28167767	refGene_gn=MRPS15;refGene_tx=XM_006934579.4
C1	28168875	28168920	refGene_gn=MRPS15;refGene_tx=XM_006934579.4
C1	28169197	28169333	refGene_gn=MRPS15;refGene_tx=XM_006934579.4
C1	28171649	28172129
refGene_gn=CSF3R;refGene_tx=XM_003989865.5;refGene_gn=CSF3R;refGene_tx=XM_011284725.3
C1	28172546	28172628
refGene_gn=CSF3R;refGene_tx=XM_003989865.5;refGene_gn=CSF3R;refGene_tx=XM_011284725.3
C1	28172868	28172962
refGene_gn=CSF3R;refGene_tx=XM_003989865.5;refGene_gn=CSF3R;refGene_tx=XM_011284725.3
C1	28173198	28173339

refGene_gn=CSF3R;refGene_tx=XM_003989865.5;refGene_gn=CSF3R;refGene_tx=XM_011284725.3
C1 28173453 28173600
refGene_gn=CSF3R;refGene_tx=XM_003989865.5;refGene_gn=CSF3R;refGene_tx=XM_011284725.3
C1 28174247 28174349
refGene_gn=CSF3R;refGene_tx=XM_003989865.5;refGene_gn=CSF3R;refGene_tx=XM_011284725.3
C1 28174730 28174919
refGene_gn=CSF3R;refGene_tx=XM_003989865.5;refGene_gn=CSF3R;refGene_tx=XM_011284725.3
C1 28175894 28176102
refGene_gn=CSF3R;refGene_tx=XM_003989865.5;refGene_gn=CSF3R;refGene_tx=XM_011284725.3
C1 28176528 28176608
refGene_gn=CSF3R;refGene_tx=XM_003989865.5;refGene_gn=CSF3R;refGene_tx=XM_011284725.3
C1 28176700 28176854
refGene_gn=CSF3R;refGene_tx=XM_003989865.5;refGene_gn=CSF3R;refGene_tx=XM_011284725.3
C1 28176968 28177138
refGene_gn=CSF3R;refGene_tx=XM_003989865.5;refGene_gn=CSF3R;refGene_tx=XM_011284725.3
C1 28177979 28178167
refGene_gn=CSF3R;refGene_tx=XM_003989865.5;refGene_gn=CSF3R;refGene_tx=XM_011284725.3
C1 28178316 28178440
refGene_gn=CSF3R;refGene_tx=XM_003989865.5;refGene_gn=CSF3R;refGene_tx=XM_011284725.3
C1 28179364 28179661
refGene_gn=CSF3R;refGene_tx=XM_003989865.5;refGene_gn=CSF3R;refGene_tx=XM_011284725.3
C1 28181762 28181826
refGene_gn=CSF3R;refGene_tx=XM_003989865.5;refGene_gn=CSF3R;refGene_tx=XM_011284725.3
C1 28459936 28460131 refGene_gn=GRIK3;refGene_tx=XM_003989866.5
C1 28462737 28462988 refGene_gn=GRIK3;refGene_tx=XM_003989866.5
C1 28463511 28463734 refGene_gn=GRIK3;refGene_tx=XM_003989866.5
C1 28474361 28474579 refGene_gn=GRIK3;refGene_tx=XM_003989866.5
C1 28477043 28477162 refGene_gn=GRIK3;refGene_tx=XM_003989866.5
C1 28482173 28482397 refGene_gn=GRIK3;refGene_tx=XM_003989866.5
C1 28495702 28495816 refGene_gn=GRIK3;refGene_tx=XM_019836665.2
C1 28496813 28497017
refGene_gn=GRIK3;refGene_tx=XM_003989866.5;refGene_gn=GRIK3;refGene_tx=XM_019836665.2
C1 28504388 28504403 refGene_gn=GRIK3;refGene_tx=XM_019836667.2
C1 28505729 28505843
refGene_gn=GRIK3;refGene_tx=XM_003989866.5;refGene_gn=GRIK3;refGene_tx=XM_019836665.2;refGene_gn=GRIK3;refGene_tx=XM_019836667.2
C1 28508946 28509054

refGene_gn=GRIK3;refGene_tx=XM_003989866.5;refGene_gn=GRIK3;refGene_tx=XM_019836665.2;refGene_gn=GRIK3;refGene_tx=XM_019836667.2
C1 28514297 28514441
refGene_gn=GRIK3;refGene_tx=XM_003989866.5;refGene_gn=GRIK3;refGene_tx=XM_019836665.2;refGene_gn=GRIK3;refGene_tx=XM_019836667.2
C1 28515028 28515202
refGene_gn=GRIK3;refGene_tx=XM_003989866.5;refGene_gn=GRIK3;refGene_tx=XM_019836665.2;refGene_gn=GRIK3;refGene_tx=XM_019836667.2
C1 28524761 28524815
refGene_gn=GRIK3;refGene_tx=XM_003989866.5;refGene_gn=GRIK3;refGene_tx=XM_019836665.2;refGene_gn=GRIK3;refGene_tx=XM_019836667.2
C1 28527250 28527432
refGene_gn=GRIK3;refGene_tx=XM_003989866.5;refGene_gn=GRIK3;refGene_tx=XM_019836665.2;refGene_gn=GRIK3;refGene_tx=XM_019836667.2
C1 28534541 28534799
refGene_gn=GRIK3;refGene_tx=XM_003989866.5;refGene_gn=GRIK3;refGene_tx=XM_019836665.2;refGene_gn=GRIK3;refGene_tx=XM_019836667.2
C1 28546255 28546432
refGene_gn=GRIK3;refGene_tx=XM_003989866.5;refGene_gn=GRIK3;refGene_tx=XM_019836665.2;refGene_gn=GRIK3;refGene_tx=XM_019836667.2
C1 28683395 28683510
refGene_gn=GRIK3;refGene_tx=XM_003989866.5;refGene_gn=GRIK3;refGene_tx=XM_019836665.2;refGene_gn=GRIK3;refGene_tx=XM_019836667.2
C1 29039875 29040267 refGene_gn=ZC3H12A;refGene_tx=XM_023258487.1
C1 29044643 29044783 refGene_gn=ZC3H12A;refGene_tx=XM_023258487.1
C1 29045271 29045506 refGene_gn=ZC3H12A;refGene_tx=XM_023258487.1
C1 29045954 29046061 refGene_gn=ZC3H12A;refGene_tx=XM_023258487.1
C1 29046239 29047114 refGene_gn=ZC3H12A;refGene_tx=XM_023258487.1
C1 29057013 29057022
refGene_gn=MEAF6;refGene_tx=XM_023258488.1;refGene_gn=MEAF6;refGene_tx=XM_023258489.1
C1 29057622 29057656
refGene_gn=MEAF6;refGene_tx=XM_023258488.1;refGene_gn=MEAF6;refGene_tx=XM_023258489.1
C1 29058442 29058472 refGene_gn=MEAF6;refGene_tx=XM_023258488.1
C1 29062824 29062858 refGene_gn=MEAF6;refGene_tx=XM_023258490.1
C1 29063011 29063201
refGene_gn=MEAF6;refGene_tx=XM_023258488.1;refGene_gn=MEAF6;refGene_tx=XM_023258489.1;refGene_gn=MEAF6;refGene_tx=XM_023258490.1
C1 29078253 29078299
refGene_gn=MEAF6;refGene_tx=XM_023258488.1;refGene_gn=MEAF6;refGene_tx=XM_023258489.1;refGene_gn=MEAF6;refGene_tx=XM_023258490.1
C1 29078448 29078536
refGene_gn=MEAF6;refGene_tx=XM_023258488.1;refGene_gn=MEAF6;refGene_tx=XM_023258489.1;refGene_gn=MEAF6;refGene_tx=XM_023258490.1
C1 29081347 29081463
refGene_gn=MEAF6;refGene_tx=XM_023258488.1;refGene_gn=MEAF6;refGene_tx=XM_023258489.1;refGene_gn=MEAF6;refGene_tx=XM_023258490.1
C1 29082672 29082762
refGene_gn=MEAF6;refGene_tx=XM_023258488.1;refGene_gn=MEAF6;refGene_tx

=XM_023258489.1;refGene_gn=MEAF6;refGene_tx=XM_023258490.1
C1	29090408 29090673 refGene_gn=SNIP1;refGene_tx=XM_003989867.4
C1	29091609 29092208 refGene_gn=SNIP1;refGene_tx=XM_003989867.4
C1	29098581 29098684 refGene_gn=SNIP1;refGene_tx=XM_003989867.4
C1	29099941 29100177 refGene_gn=SNIP1;refGene_tx=XM_003989867.4
C1	29101590 29101615 refGene_gn=DNALI1;refGene_tx=XM_004001551.4
C1	29101617 29101736 refGene_gn=DNALI1;refGene_tx=XM_004001551.4
C1	29102283 29102429 refGene_gn=DNALI1;refGene_tx=XM_004001551.4
C1	29104508 29104678 refGene_gn=DNALI1;refGene_tx=XM_004001551.4
C1	29105865 29106044 refGene_gn=DNALI1;refGene_tx=XM_004001551.4
C1	29106386 29106551 refGene_gn=DNALI1;refGene_tx=XM_004001551.4
C1	29108419 29108455 refGene_gn=DNALI1;refGene_tx=XM_004001551.4
C1	29110158 29110311 refGene_gn=GNL2;refGene_tx=XM_003989868.5
C1	29110994 29111080 refGene_gn=GNL2;refGene_tx=XM_003989868.5
C1	29111898 29111975 refGene_gn=GNL2;refGene_tx=XM_003989868.5
C1	29112302 29112751 refGene_gn=GNL2;refGene_tx=XM_003989868.5
C1	29115910 29116024 refGene_gn=GNL2;refGene_tx=XM_003989868.5
C1	29116236 29116395 refGene_gn=GNL2;refGene_tx=XM_003989868.5
C1	29117181 29117286 refGene_gn=GNL2;refGene_tx=XM_003989868.5
C1	29118034 29118163 refGene_gn=GNL2;refGene_tx=XM_003989868.5
C1	29122050 29122164 refGene_gn=GNL2;refGene_tx=XM_003989868.5
C1	29122929 29123088 refGene_gn=GNL2;refGene_tx=XM_003989868.5
C1	29123943 29124010 refGene_gn=GNL2;refGene_tx=XM_003989868.5
C1	29128260 29128445 refGene_gn=GNL2;refGene_tx=XM_003989868.5
C1	29131007 29131147 refGene_gn=GNL2;refGene_tx=XM_003989868.5
C1	29132874 29132969 refGene_gn=GNL2;refGene_tx=XM_003989868.5
C1	29133958 29134043 refGene_gn=GNL2;refGene_tx=XM_003989868.5
C1	29135872 29135936 refGene_gn=GNL2;refGene_tx=XM_003989868.5
C1	29150511 29150678
refGene_gn=RSPO1;refGene_tx=XM_003989869.5;refGene_gn=RSPO1;refGene_tx
=XM_019836670.2;refGene_gn=RSPO1;refGene_tx=XM_019836671.2;refGene_gn=
RSPO1;refGene_tx=XM_019836672.2;refGene_gn=RSPO1;refGene_tx=XM_0198366
73.2;refGene_gn=RSPO1;refGene_tx=XM_023258492.1
C1	29151390 29151579
refGene_gn=RSPO1;refGene_tx=XM_003989869.5;refGene_gn=RSPO1;refGene_tx
=XM_019836670.2;refGene_gn=RSPO1;refGene_tx=XM_019836671.2;refGene_gn=
RSPO1;refGene_tx=XM_019836672.2;refGene_gn=RSPO1;refGene_tx=XM_0198366
73.2;refGene_gn=RSPO1;refGene_tx=XM_023258492.1
C1	29151867 29152017
refGene_gn=RSPO1;refGene_tx=XM_003989869.5;refGene_gn=RSPO1;refGene_tx
=XM_019836670.2;refGene_gn=RSPO1;refGene_tx=XM_019836671.2;refGene_gn=
RSPO1;refGene_tx=XM_019836672.2;refGene_gn=RSPO1;refGene_tx=XM_0198366
73.2;refGene_gn=RSPO1;refGene_tx=XM_023258492.1
C1	29154031 29154223
refGene_gn=RSPO1;refGene_tx=XM_003989869.5;refGene_gn=RSPO1;refGene_tx
=XM_019836670.2;refGene_gn=RSPO1;refGene_tx=XM_019836671.2;refGene_gn=
RSPO1;refGene_tx=XM_019836672.2;refGene_gn=RSPO1;refGene_tx=XM_0198366
73.2;refGene_gn=RSPO1;refGene_tx=XM_023258492.1
C1	29162436 29162530
refGene_gn=RSPO1;refGene_tx=XM_003989869.5;refGene_gn=RSPO1;refGene_tx

=XM_019836670.2;refGene_gn=RSPO1;refGene_tx=XM_019836671.2;refGene_gn=RSPO1;refGene_tx=XM_019836672.2;refGene_gn=RSPO1;refGene_tx=XM_019836673.2;refGene_gn=RSPO1;refGene_tx=XM_023258492.1
C1 29211578 29211616 refGene_gn=CC1H1orf109;refGene_tx=XM_003989871.5
C1 29211679 29211893 refGene_gn=CC1H1orf109;refGene_tx=XM_003989871.5;refGene_gn=CC1H1orf109;refGene_tx=XM_006934582.4
C1 29213901 29214081 refGene_gn=CC1H1orf109;refGene_tx=XM_003989871.5;refGene_gn=CC1H1orf109;refGene_tx=XM_006934582.4
C1 29215944 29216219 refGene_gn=CC1H1orf109;refGene_tx=XM_003989871.5;refGene_gn=CC1H1orf109;refGene_tx=XM_006934582.4
C1 29217128 29217225 refGene_gn=CDCA8;refGene_tx=XM_006934583.3
C1 29217357 29217486 refGene_gn=CDCA8;refGene_tx=XM_006934583.3
C1 29220003 29220044 refGene_gn=CDCA8;refGene_tx=XM_006934583.3
C1 29222973 29223046 refGene_gn=CDCA8;refGene_tx=XM_006934583.3
C1 29225000 29225083 refGene_gn=CDCA8;refGene_tx=XM_006934583.3
C1 29226432 29226497 refGene_gn=CDCA8;refGene_tx=XM_006934583.3
C1 29227065 29227161 refGene_gn=CDCA8;refGene_tx=XM_006934583.3
C1 29228285 29228412 refGene_gn=CDCA8;refGene_tx=XM_006934583.3
C1 29229181 29229268 refGene_gn=CDCA8;refGene_tx=XM_006934583.3
C1 29230396 29230441 refGene_gn=CDCA8;refGene_tx=XM_006934583.3
C1 29239028 29239143 refGene_gn=EPHA10;refGene_tx=XM_023258494.1
C1 29239301 29239457 refGene_gn=EPHA10;refGene_tx=XM_023258494.1
C1 29240144 29240338 refGene_gn=EPHA10;refGene_tx=XM_023258494.1
C1 29240530 29240680 refGene_gn=EPHA10;refGene_tx=XM_023258494.1
C1 29240980 29241184 refGene_gn=EPHA10;refGene_tx=XM_023258494.1
C1 29241410 29241472 refGene_gn=EPHA10;refGene_tx=XM_023258494.1
C1 29241913 29242099 refGene_gn=EPHA10;refGene_tx=XM_023258494.1
C1 29243175 29243301 refGene_gn=EPHA10;refGene_tx=XM_023258494.1
C1 29243417 29243479 refGene_gn=EPHA10;refGene_tx=XM_023258494.1
C1 29246921 29247030 refGene_gn=EPHA10;refGene_tx=XM_023258494.1
C1 29250620 29250792 refGene_gn=EPHA10;refGene_tx=XM_023258494.1
C1 29253341 29253475 refGene_gn=EPHA10;refGene_tx=XM_023258494.1
C1 29266940 29267291 refGene_gn=EPHA10;refGene_tx=XM_023258494.1
C1 29267809 29267965 refGene_gn=EPHA10;refGene_tx=XM_023258494.1
C1 29273959 29274638 refGene_gn=EPHA10;refGene_tx=XM_023258494.1
C1 29275354 29275419 refGene_gn=EPHA10;refGene_tx=XM_023258494.1
C1 29277327 29277433 refGene_gn=EPHA10;refGene_tx=XM_023258494.1
C1 29298438 29298988 refGene_gn=MANEAL;refGene_tx=XM_023258496.1
C1 29300251 29300361 refGene_gn=MANEAL;refGene_tx=XM_011284723.3;refGene_gn=MANEAL;refGene_tx=XM_023258496.1
C1 29301233 29301310 refGene_gn=MANEAL;refGene_tx=XM_023258496.1
C1 29302422 29303059 refGene_gn=MANEAL;refGene_tx=XM_011284723.3;refGene_gn=MANEAL;refGene_tx=XM_023258496.1
C1 29305533 29305606 refGene_gn=YRDC;refGene_tx=XM_023258497.1

C1 29305903 29306046 refGene_gn=YRDC;refGene_tx=XM_023258497.1
C1 29307460 29307580 refGene_gn=YRDC;refGene_tx=XM_023258497.1
C1 29307704 29307819 refGene_gn=YRDC;refGene_tx=XM_023258497.1
C1 29308416 29308808 refGene_gn=YRDC;refGene_tx=XM_023258497.1
C1 29309032 29309067
refGene_gn=CC1H1orf122;refGene_tx=XM_023258498.1
C1 29309159 29309361
refGene_gn=CC1H1orf122;refGene_tx=XM_023258498.1;refGene_gn=CC1H1orf12
2;refGene_tx=XM_023258499.1
C1 29309604 29309700
refGene_gn=CC1H1orf122;refGene_tx=XM_023258498.1;refGene_gn=CC1H1orf12
2;refGene_tx=XM_023258499.1
C1 29315536 29315964
refGene_gn=MTF1;refGene_tx=XM_003989875.5;refGene_gn=MTF1;refGene_tx=X
M_006934584.4;refGene_gn=MTF1;refGene_tx=XM_006934585.4;refGene_gn=MTF
1;refGene_tx=XM_019836682.2
C1 29317235 29317299
refGene_gn=MTF1;refGene_tx=XM_003989875.5;refGene_gn=MTF1;refGene_tx=X
M_006934584.4;refGene_gn=MTF1;refGene_tx=XM_006934585.4;refGene_gn=MTF
1;refGene_tx=XM_019836682.2
C1 29319603 29320187
refGene_gn=MTF1;refGene_tx=XM_003989875.5;refGene_gn=MTF1;refGene_tx=X
M_006934584.4;refGene_gn=MTF1;refGene_tx=XM_006934585.4;refGene_gn=MTF
1;refGene_tx=XM_019836682.2
C1 29320912 29321009
refGene_gn=MTF1;refGene_tx=XM_003989875.5;refGene_gn=MTF1;refGene_tx=X
M_006934584.4;refGene_gn=MTF1;refGene_tx=XM_006934585.4;refGene_gn=MTF
1;refGene_tx=XM_019836682.2
C1 29328360 29328435
refGene_gn=MTF1;refGene_tx=XM_003989875.5;refGene_gn=MTF1;refGene_tx=X
M_006934584.4;refGene_gn=MTF1;refGene_tx=XM_006934585.4;refGene_gn=MTF
1;refGene_tx=XM_019836682.2
C1 29332312 29332449
refGene_gn=MTF1;refGene_tx=XM_003989875.5;refGene_gn=MTF1;refGene_tx=X
M_006934584.4;refGene_gn=MTF1;refGene_tx=XM_006934585.4;refGene_gn=MTF
1;refGene_tx=XM_019836682.2
C1 29332933 29333007
refGene_gn=MTF1;refGene_tx=XM_003989875.5;refGene_gn=MTF1;refGene_tx=X
M_006934584.4;refGene_gn=MTF1;refGene_tx=XM_006934585.4;refGene_gn=MTF
1;refGene_tx=XM_019836682.2
C1 29335475 29335607
refGene_gn=MTF1;refGene_tx=XM_003989875.5;refGene_gn=MTF1;refGene_tx=X
M_006934584.4;refGene_gn=MTF1;refGene_tx=XM_006934585.4;refGene_gn=MTF
1;refGene_tx=XM_019836682.2
C1 29336368 29336607
refGene_gn=MTF1;refGene_tx=XM_003989875.5;refGene_gn=MTF1;refGene_tx=X
M_006934584.4;refGene_gn=MTF1;refGene_tx=XM_006934585.4;refGene_gn=MTF
1;refGene_tx=XM_019836682.2
C1 29354643 29355051
refGene_gn=MTF1;refGene_tx=XM_003989875.5;refGene_gn=MTF1;refGene_tx=X

M_006934584.4;refGene_gn=MTF1;refGene_tx=XM_006934585.4;refGene_gn=MTF1;refGene_tx=XM_019836682.2

| | | | |
|---|---|---|---|
| C1 | 29358639 | 29358755 | refGene_gn=INPP5B;refGene_tx=XM_023258500.1 |
| C1 | 29359005 | 29359117 | refGene_gn=INPP5B;refGene_tx=XM_023258500.1 |
| C1 | 29360421 | 29360549 | refGene_gn=INPP5B;refGene_tx=XM_023258500.1 |
| C1 | 29360940 | 29361025 | refGene_gn=INPP5B;refGene_tx=XM_023258500.1 |
| C1 | 29363803 | 29363917 | refGene_gn=INPP5B;refGene_tx=XM_023258500.1 |
| C1 | 29364685 | 29364921 | refGene_gn=INPP5B;refGene_tx=XM_023258500.1 |
| C1 | 29365755 | 29365918 | refGene_gn=INPP5B;refGene_tx=XM_023258500.1 |
| C1 | 29366875 | 29366986 | refGene_gn=INPP5B;refGene_tx=XM_023258500.1 |
| C1 | 29368282 | 29368418 | refGene_gn=INPP5B;refGene_tx=XM_023258500.1 |
| C1 | 29369455 | 29369569 | refGene_gn=INPP5B;refGene_tx=XM_023258500.1 |
| C1 | 29371565 | 29371673 | refGene_gn=INPP5B;refGene_tx=XM_023258500.1 |
| C1 | 29373172 | 29373360 | refGene_gn=INPP5B;refGene_tx=XM_023258500.1 |
| C1 | 29374240 | 29374357 | refGene_gn=INPP5B;refGene_tx=XM_023258500.1 |
| C1 | 29374717 | 29374832 | refGene_gn=INPP5B;refGene_tx=XM_023258500.1 |
| C1 | 29375623 | 29375725 | refGene_gn=INPP5B;refGene_tx=XM_023258500.1 |
| C1 | 29376858 | 29377026 | refGene_gn=INPP5B;refGene_tx=XM_023258500.1 |
| C1 | 29378734 | 29378831 | refGene_gn=INPP5B;refGene_tx=XM_023258500.1 |
| C1 | 29397423 | 29397564 | refGene_gn=INPP5B;refGene_tx=XM_023258500.1 |
| C1 | 29401413 | 29401524 | refGene_gn=INPP5B;refGene_tx=XM_023258500.1 |
| C1 | 29403471 | 29403501 | refGene_gn=INPP5B;refGene_tx=XM_023258500.1 |
| C1 | 29403617 | 29403715 | refGene_gn=INPP5B;refGene_tx=XM_023258500.1 |
| C1 | 29404535 | 29404630 | refGene_gn=INPP5B;refGene_tx=XM_023258500.1 |
| C1 | 29405016 | 29405073 | refGene_gn=INPP5B;refGene_tx=XM_023258500.1 |
| C1 | 29414010 | 29414088 | refGene_gn=SF3A3;refGene_tx=XM_011284720.3 |
| C1 | 29415082 | 29415138 | refGene_gn=SF3A3;refGene_tx=XM_011284720.3 |
| C1 | 29423535 | 29423626 | refGene_gn=SF3A3;refGene_tx=XM_011284720.3 |
| C1 | 29424810 | 29424921 | refGene_gn=SF3A3;refGene_tx=XM_011284720.3 |
| C1 | 29425011 | 29425176 | refGene_gn=SF3A3;refGene_tx=XM_011284720.3 |
| C1 | 29427348 | 29427418 | refGene_gn=SF3A3;refGene_tx=XM_011284720.3 |
| C1 | 29428624 | 29428732 | refGene_gn=SF3A3;refGene_tx=XM_011284720.3 |
| C1 | 29428892 | 29428960 | refGene_gn=SF3A3;refGene_tx=XM_011284720.3 |
| C1 | 29429611 | 29429680 | refGene_gn=SF3A3;refGene_tx=XM_011284720.3 |
| C1 | 29430184 | 29430323 | refGene_gn=SF3A3;refGene_tx=XM_011284720.3 |
| C1 | 29430998 | 29431081 | refGene_gn=SF3A3;refGene_tx=XM_011284720.3 |
| C1 | 29432523 | 29432615 | refGene_gn=SF3A3;refGene_tx=XM_011284720.3 |
| C1 | 29432977 | 29433050 | refGene_gn=SF3A3;refGene_tx=XM_011284720.3 |
| C1 | 29435539 | 29435645 | refGene_gn=SF3A3;refGene_tx=XM_011284720.3 |
| C1 | 29435750 | 29435803 | refGene_gn=SF3A3;refGene_tx=XM_011284720.3 |
| C1 | 29438255 | 29438303 | refGene_gn=SF3A3;refGene_tx=XM_011284720.3 |
| C1 | 29438590 | 29438686 | refGene_gn=SF3A3;refGene_tx=XM_011284720.3 |
| C1 | 29444675 | 29444830 | refGene_gn=FHL3;refGene_tx=XM_003989878.4 |
| C1 | 29444956 | 29445143 | refGene_gn=FHL3;refGene_tx=XM_003989878.4 |
| C1 | 29445236 | 29445406 | refGene_gn=FHL3;refGene_tx=XM_003989878.4 |
| C1 | 29446421 | 29446596 | refGene_gn=FHL3;refGene_tx=XM_003989878.4 |
| C1 | 29446684 | 29446840 | refGene_gn=FHL3;refGene_tx=XM_003989878.4 |
| C1 | 29457132 | 29457195 | refGene_gn=UTP11;refGene_tx=XM_003989879.5 |
| C1 | 29461727 | 29461789 | refGene_gn=UTP11;refGene_tx=XM_003989879.5 |
| C1 | 29463239 | 29463342 | refGene_gn=UTP11;refGene_tx=XM_003989879.5 |

C1	29463965	29464079	refGene_gn=UTP11;refGene_tx=XM_003989879.5
C1	29464563	29464657	refGene_gn=UTP11;refGene_tx=XM_003989879.5
C1	29464744	29464875	refGene_gn=UTP11;refGene_tx=XM_003989879.5
C1	29466971	29467082	refGene_gn=UTP11;refGene_tx=XM_003989879.5
C1	29467818	29467902	refGene_gn=UTP11;refGene_tx=XM_003989879.5
C1	29487938	29489388
refGene_gn=LOC111561628;refGene_tx=XM_023257824.1;refGene_gn=POU3F1;refGene_tx=XM_011279681.2
C1	29489509	29490179
refGene_gn=LOC111561628;refGene_tx=XM_023257824.1
C1	29537624	29537770
refGene_gn=LOC111561628;refGene_tx=XM_023257824.1
C1	29541953	29542001
refGene_gn=LOC111561628;refGene_tx=XM_023257824.1
C1	30175691	30175843	refGene_gn=RRAGC;refGene_tx=XM_023258503.1
C1	30180224	30180373	refGene_gn=RRAGC;refGene_tx=XM_023258503.1
C1	30182068	30182211	refGene_gn=RRAGC;refGene_tx=XM_023258503.1
C1	30183205	30183252	refGene_gn=RRAGC;refGene_tx=XM_023258503.1
C1	30183254	30183319	refGene_gn=RRAGC;refGene_tx=XM_023258503.1
C1	30185870	30186070	refGene_gn=RRAGC;refGene_tx=XM_023258503.1
C1	30187060	30187264	refGene_gn=RRAGC;refGene_tx=XM_023258503.1
C1	30189036	30189273	refGene_gn=RRAGC;refGene_tx=XM_023258503.1
C1	30193707	30193752
refGene_gn=MYCBP;refGene_tx=XM_003989880.4;refGene_gn=MYCBP;refGene_tx=XM_019836695.2
C1	30194993	30195123
refGene_gn=MYCBP;refGene_tx=XM_003989880.4;refGene_gn=MYCBP;refGene_tx=XM_019836695.2
C1	30195367	30195416	refGene_gn=MYCBP;refGene_tx=XM_003989880.4
C1	30200039	30200239
refGene_gn=MYCBP;refGene_tx=XM_003989880.4;refGene_gn=MYCBP;refGene_tx=XM_019836695.2
C1	30200307	30200391	refGene_gn=MYCBP;refGene_tx=XM_003989880.4
C1	30201643	30203182	refGene_gn=GJA9;refGene_tx=XM_004001469.5
C1	30209663	30209843
refGene_gn=RHBDL2;refGene_tx=XM_019836689.2;refGene_gn=RHBDL2;refGene_tx=XM_019836692.2;refGene_gn=RHBDL2;refGene_tx=XM_023258505.1;refGene_gn=RHBDL2;refGene_tx=XM_023258506.1
C1	30211056	30211118
refGene_gn=RHBDL2;refGene_tx=XM_019836689.2;refGene_gn=RHBDL2;refGene_tx=XM_019836692.2;refGene_gn=RHBDL2;refGene_tx=XM_023258505.1;refGene_gn=RHBDL2;refGene_tx=XM_023258506.1
C1	30215084	30215145
refGene_gn=RHBDL2;refGene_tx=XM_019836689.2;refGene_gn=RHBDL2;refGene_tx=XM_019836692.2;refGene_gn=RHBDL2;refGene_tx=XM_023258505.1;refGene_gn=RHBDL2;refGene_tx=XM_023258506.1
C1	30217849	30217950
refGene_gn=RHBDL2;refGene_tx=XM_019836689.2;refGene_gn=RHBDL2;refGene_tx=XM_019836692.2;refGene_gn=RHBDL2;refGene_tx=XM_023258505.1;refGene_gn=RHBDL2;refGene_tx=XM_023258506.1

C1 30230866 30230979
refGene_gn=RHBDL2;refGene_tx=XM_019836689.2;refGene_gn=RHBDL2;refGene_tx=XM_019836692.2;refGene_gn=RHBDL2;refGene_tx=XM_023258505.1;refGene_gn=RHBDL2;refGene_tx=XM_023258506.1
C1 30234512 30234661
refGene_gn=RHBDL2;refGene_tx=XM_019836689.2;refGene_gn=RHBDL2;refGene_tx=XM_019836692.2;refGene_gn=RHBDL2;refGene_tx=XM_023258505.1;refGene_gn=RHBDL2;refGene_tx=XM_023258506.1
C1 30238508 30238775
refGene_gn=RHBDL2;refGene_tx=XM_019836689.2;refGene_gn=RHBDL2;refGene_tx=XM_019836692.2;refGene_gn=RHBDL2;refGene_tx=XM_023258505.1;refGene_gn=RHBDL2;refGene_tx=XM_023258506.1
C1 30245303 30245351 refGene_gn=RHBDL2;refGene_tx=XM_019836689.2
C1 30289383 30289603 refGene_gn=AKIRIN1;refGene_tx=XM_023258507.1
C1 30294183 30294324 refGene_gn=AKIRIN1;refGene_tx=XM_023258507.1
C1 30296879 30297014 refGene_gn=AKIRIN1;refGene_tx=XM_023258507.1
C1 30298218 30298290 refGene_gn=AKIRIN1;refGene_tx=XM_023258507.1
C1 30298945 30298956 refGene_gn=AKIRIN1;refGene_tx=XM_023258507.1
C1 30312904 30313120 refGene_gn=NDUFS5;refGene_tx=XM_003989884.4
C1 30316737 30316842 refGene_gn=NDUFS5;refGene_tx=XM_003989884.4
C1 30353012 30353232
refGene_gn=MACF1;refGene_tx=XM_011284718.3;refGene_gn=MACF1;refGene_tx=XM_019836716.2;refGene_gn=MACF1;refGene_tx=XM_019836717.2;refGene_gn=MACF1;refGene_tx=XM_019836718.2;refGene_gn=MACF1;refGene_tx=XM_019836720.2
C1 30479613 30479675
refGene_gn=MACF1;refGene_tx=XM_011284718.3;refGene_gn=MACF1;refGene_tx=XM_019836716.2;refGene_gn=MACF1;refGene_tx=XM_019836717.2;refGene_gn=MACF1;refGene_tx=XM_019836718.2;refGene_gn=MACF1;refGene_tx=XM_019836720.2
C1 30491549 30491639
refGene_gn=MACF1;refGene_tx=XM_011284718.3;refGene_gn=MACF1;refGene_tx=XM_019836716.2;refGene_gn=MACF1;refGene_tx=XM_019836717.2;refGene_gn=MACF1;refGene_tx=XM_019836718.2;refGene_gn=MACF1;refGene_tx=XM_019836720.2
C1 30496083 30496161
refGene_gn=MACF1;refGene_tx=XM_011284718.3;refGene_gn=MACF1;refGene_tx=XM_019836716.2;refGene_gn=MACF1;refGene_tx=XM_019836717.2;refGene_gn=MACF1;refGene_tx=XM_019836718.2;refGene_gn=MACF1;refGene_tx=XM_019836720.2
C1 30499942 30500035
refGene_gn=MACF1;refGene_tx=XM_011284718.3;refGene_gn=MACF1;refGene_tx=XM_019836716.2;refGene_gn=MACF1;refGene_tx=XM_019836717.2;refGene_gn=MACF1;refGene_tx=XM_019836718.2;refGene_gn=MACF1;refGene_tx=XM_019836720.2
C1 30511312 30512335
refGene_gn=MACF1;refGene_tx=XM_019836714.2;refGene_gn=MACF1;refGene_tx=XM_019836715.2;refGene_gn=MACF1;refGene_tx=XM_023258509.1;refGene_gn=MACF1;refGene_tx=XM_023258510.1
C1 30524112 30524279

refGene_gn=MACF1;refGene_tx=XM_011284718.3;refGene_gn=MACF1;refGene_tx=XM_019836714.2;refGene_gn=MACF1;refGene_tx=XM_019836715.2;refGene_gn=MACF1;refGene_tx=XM_019836716.2;refGene_gn=MACF1;refGene_tx=XM_019836717.2;refGene_gn=MACF1;refGene_tx=XM_019836718.2;refGene_gn=MACF1;refGene_tx=XM_019836720.2;refGene_gn=MACF1;refGene_tx=XM_023258509.1;refGene_gn=MACF1;refGene_tx=XM_023258510.1
C1 30525072 30525185
refGene_gn=MACF1;refGene_tx=XM_011284718.3;refGene_gn=MACF1;refGene_tx=XM_019836714.2;refGene_gn=MACF1;refGene_tx=XM_019836715.2;refGene_gn=MACF1;refGene_tx=XM_019836716.2;refGene_gn=MACF1;refGene_tx=XM_019836717.2;refGene_gn=MACF1;refGene_tx=XM_019836718.2;refGene_gn=MACF1;refGene_tx=XM_019836720.2;refGene_gn=MACF1;refGene_tx=XM_023258509.1;refGene_gn=MACF1;refGene_tx=XM_023258510.1
C1 30525285 30525392
refGene_gn=MACF1;refGene_tx=XM_011284718.3;refGene_gn=MACF1;refGene_tx=XM_019836714.2;refGene_gn=MACF1;refGene_tx=XM_019836715.2;refGene_gn=MACF1;refGene_tx=XM_019836716.2;refGene_gn=MACF1;refGene_tx=XM_019836717.2;refGene_gn=MACF1;refGene_tx=XM_019836718.2;refGene_gn=MACF1;refGene_tx=XM_019836720.2;refGene_gn=MACF1;refGene_tx=XM_023258509.1;refGene_gn=MACF1;refGene_tx=XM_023258510.1
C1 30525806 30525926
refGene_gn=MACF1;refGene_tx=XM_011284718.3;refGene_gn=MACF1;refGene_tx=XM_019836714.2;refGene_gn=MACF1;refGene_tx=XM_019836715.2;refGene_gn=MACF1;refGene_tx=XM_019836716.2;refGene_gn=MACF1;refGene_tx=XM_019836717.2;refGene_gn=MACF1;refGene_tx=XM_019836718.2;refGene_gn=MACF1;refGene_tx=XM_019836720.2;refGene_gn=MACF1;refGene_tx=XM_023258509.1;refGene_gn=MACF1;refGene_tx=XM_023258510.1
C1 30526079 30526175
refGene_gn=MACF1;refGene_tx=XM_011284718.3;refGene_gn=MACF1;refGene_tx=XM_019836714.2;refGene_gn=MACF1;refGene_tx=XM_019836715.2;refGene_gn=MACF1;refGene_tx=XM_019836716.2;refGene_gn=MACF1;refGene_tx=XM_019836717.2;refGene_gn=MACF1;refGene_tx=XM_019836718.2;refGene_gn=MACF1;refGene_tx=XM_019836720.2;refGene_gn=MACF1;refGene_tx=XM_023258509.1;refGene_gn=MACF1;refGene_tx=XM_023258510.1
C1 30526825 30526953
refGene_gn=MACF1;refGene_tx=XM_011284718.3;refGene_gn=MACF1;refGene_tx=XM_019836714.2;refGene_gn=MACF1;refGene_tx=XM_019836715.2;refGene_gn=MACF1;refGene_tx=XM_019836716.2;refGene_gn=MACF1;refGene_tx=XM_019836717.2;refGene_gn=MACF1;refGene_tx=XM_019836718.2;refGene_gn=MACF1;refGene_tx=XM_019836720.2;refGene_gn=MACF1;refGene_tx=XM_023258509.1;refGene_gn=MACF1;refGene_tx=XM_023258510.1
C1 30527041 30527135
refGene_gn=MACF1;refGene_tx=XM_011284718.3;refGene_gn=MACF1;refGene_tx=XM_019836714.2;refGene_gn=MACF1;refGene_tx=XM_019836715.2;refGene_gn=MACF1;refGene_tx=XM_019836716.2;refGene_gn=MACF1;refGene_tx=XM_019836717.2;refGene_gn=MACF1;refGene_tx=XM_019836718.2;refGene_gn=MACF1;refGene_tx=XM_019836720.2;refGene_gn=MACF1;refGene_tx=XM_023258509.1;refGene_gn=MACF1;refGene_tx=XM_023258510.1
C1 30527297 30527452
refGene_gn=MACF1;refGene_tx=XM_011284718.3;refGene_gn=MACF1;refGene_tx

=XM_019836714.2;refGene_gn=MACF1;refGene_tx=XM_019836715.2;refGene_gn=MACF1;refGene_tx=XM_019836716.2;refGene_gn=MACF1;refGene_tx=XM_019836717.2;refGene_gn=MACF1;refGene_tx=XM_019836718.2;refGene_gn=MACF1;refGene_tx=XM_019836720.2;refGene_gn=MACF1;refGene_tx=XM_023258509.1;refGene_gn=MACF1;refGene_tx=XM_023258510.1
C1 30528236 30528513
refGene_gn=MACF1;refGene_tx=XM_011284718.3;refGene_gn=MACF1;refGene_tx=XM_019836714.2;refGene_gn=MACF1;refGene_tx=XM_019836715.2;refGene_gn=MACF1;refGene_tx=XM_019836716.2;refGene_gn=MACF1;refGene_tx=XM_019836717.2;refGene_gn=MACF1;refGene_tx=XM_019836718.2;refGene_gn=MACF1;refGene_tx=XM_019836720.2;refGene_gn=MACF1;refGene_tx=XM_023258509.1;refGene_gn=MACF1;refGene_tx=XM_023258510.1
C1 30529479 30529608
refGene_gn=MACF1;refGene_tx=XM_011284718.3;refGene_gn=MACF1;refGene_tx=XM_019836714.2;refGene_gn=MACF1;refGene_tx=XM_019836715.2;refGene_gn=MACF1;refGene_tx=XM_019836716.2;refGene_gn=MACF1;refGene_tx=XM_019836717.2;refGene_gn=MACF1;refGene_tx=XM_019836718.2;refGene_gn=MACF1;refGene_tx=XM_019836720.2;refGene_gn=MACF1;refGene_tx=XM_023258509.1;refGene_gn=MACF1;refGene_tx=XM_023258510.1
C1 30530852 30530930
refGene_gn=MACF1;refGene_tx=XM_011284718.3;refGene_gn=MACF1;refGene_tx=XM_019836714.2;refGene_gn=MACF1;refGene_tx=XM_019836715.2;refGene_gn=MACF1;refGene_tx=XM_019836716.2;refGene_gn=MACF1;refGene_tx=XM_019836717.2;refGene_gn=MACF1;refGene_tx=XM_019836718.2;refGene_gn=MACF1;refGene_tx=XM_019836720.2;refGene_gn=MACF1;refGene_tx=XM_023258509.1;refGene_gn=MACF1;refGene_tx=XM_023258510.1
C1 30531548 30531710
refGene_gn=MACF1;refGene_tx=XM_011284718.3;refGene_gn=MACF1;refGene_tx=XM_019836714.2;refGene_gn=MACF1;refGene_tx=XM_019836715.2;refGene_gn=MACF1;refGene_tx=XM_019836716.2;refGene_gn=MACF1;refGene_tx=XM_019836717.2;refGene_gn=MACF1;refGene_tx=XM_019836718.2;refGene_gn=MACF1;refGene_tx=XM_019836720.2;refGene_gn=MACF1;refGene_tx=XM_023258509.1;refGene_gn=MACF1;refGene_tx=XM_023258510.1
C1 30533524 30533629
refGene_gn=MACF1;refGene_tx=XM_011284718.3;refGene_gn=MACF1;refGene_tx=XM_019836714.2;refGene_gn=MACF1;refGene_tx=XM_019836715.2;refGene_gn=MACF1;refGene_tx=XM_019836716.2;refGene_gn=MACF1;refGene_tx=XM_019836717.2;refGene_gn=MACF1;refGene_tx=XM_019836718.2;refGene_gn=MACF1;refGene_tx=XM_019836720.2;refGene_gn=MACF1;refGene_tx=XM_023258509.1;refGene_gn=MACF1;refGene_tx=XM_023258510.1
C1 30534772 30534868
refGene_gn=MACF1;refGene_tx=XM_011284718.3;refGene_gn=MACF1;refGene_tx=XM_019836714.2;refGene_gn=MACF1;refGene_tx=XM_019836715.2;refGene_gn=MACF1;refGene_tx=XM_019836716.2;refGene_gn=MACF1;refGene_tx=XM_019836717.2;refGene_gn=MACF1;refGene_tx=XM_019836718.2;refGene_gn=MACF1;refGene_tx=XM_019836720.2;refGene_gn=MACF1;refGene_tx=XM_023258509.1;refGene_gn=MACF1;refGene_tx=XM_023258510.1
C1 30536662 30536788
refGene_gn=MACF1;refGene_tx=XM_011284718.3;refGene_gn=MACF1;refGene_tx=XM_019836714.2;refGene_gn=MACF1;refGene_tx=XM_019836715.2;refGene_gn=

MACF1;refGene_tx=XM_019836716.2;refGene_gn=MACF1;refGene_tx=XM_019836717.2;refGene_gn=MACF1;refGene_tx=XM_019836718.2;refGene_gn=MACF1;refGene_tx=XM_019836720.2;refGene_gn=MACF1;refGene_tx=XM_023258509.1;refGene_gn=MACF1;refGene_tx=XM_023258510.1
C1 30538320 30538326
refGene_gn=MACF1;refGene_tx=XM_019836714.2;refGene_gn=MACF1;refGene_tx=XM_023258509.1;refGene_gn=MACF1;refGene_tx=XM_023258510.1
C1 30539313 30539466
refGene_gn=MACF1;refGene_tx=XM_011284718.3;refGene_gn=MACF1;refGene_tx=XM_019836714.2;refGene_gn=MACF1;refGene_tx=XM_019836715.2;refGene_gn=MACF1;refGene_tx=XM_019836716.2;refGene_gn=MACF1;refGene_tx=XM_019836717.2;refGene_gn=MACF1;refGene_tx=XM_019836718.2;refGene_gn=MACF1;refGene_tx=XM_019836720.2;refGene_gn=MACF1;refGene_tx=XM_023258509.1;refGene_gn=MACF1;refGene_tx=XM_023258510.1
C1 30540766 30540921
refGene_gn=MACF1;refGene_tx=XM_011284718.3;refGene_gn=MACF1;refGene_tx=XM_019836714.2;refGene_gn=MACF1;refGene_tx=XM_019836715.2;refGene_gn=MACF1;refGene_tx=XM_019836716.2;refGene_gn=MACF1;refGene_tx=XM_019836717.2;refGene_gn=MACF1;refGene_tx=XM_019836718.2;refGene_gn=MACF1;refGene_tx=XM_019836720.2;refGene_gn=MACF1;refGene_tx=XM_023258509.1;refGene_gn=MACF1;refGene_tx=XM_023258510.1
C1 30542298 30542425
refGene_gn=MACF1;refGene_tx=XM_011284718.3;refGene_gn=MACF1;refGene_tx=XM_019836714.2;refGene_gn=MACF1;refGene_tx=XM_019836715.2;refGene_gn=MACF1;refGene_tx=XM_019836716.2;refGene_gn=MACF1;refGene_tx=XM_019836717.2;refGene_gn=MACF1;refGene_tx=XM_019836718.2;refGene_gn=MACF1;refGene_tx=XM_019836720.2;refGene_gn=MACF1;refGene_tx=XM_023258509.1;refGene_gn=MACF1;refGene_tx=XM_023258510.1
C1 30542959 30543143
refGene_gn=MACF1;refGene_tx=XM_011284718.3;refGene_gn=MACF1;refGene_tx=XM_019836714.2;refGene_gn=MACF1;refGene_tx=XM_019836715.2;refGene_gn=MACF1;refGene_tx=XM_019836716.2;refGene_gn=MACF1;refGene_tx=XM_019836717.2;refGene_gn=MACF1;refGene_tx=XM_019836718.2;refGene_gn=MACF1;refGene_tx=XM_019836720.2;refGene_gn=MACF1;refGene_tx=XM_023258509.1;refGene_gn=MACF1;refGene_tx=XM_023258510.1
C1 30543564 30543734
refGene_gn=MACF1;refGene_tx=XM_011284718.3;refGene_gn=MACF1;refGene_tx=XM_019836714.2;refGene_gn=MACF1;refGene_tx=XM_019836715.2;refGene_gn=MACF1;refGene_tx=XM_019836716.2;refGene_gn=MACF1;refGene_tx=XM_019836717.2;refGene_gn=MACF1;refGene_tx=XM_019836718.2;refGene_gn=MACF1;refGene_tx=XM_019836720.2;refGene_gn=MACF1;refGene_tx=XM_023258509.1;refGene_gn=MACF1;refGene_tx=XM_023258510.1
C1 30547727 30547906
refGene_gn=MACF1;refGene_tx=XM_011284718.3;refGene_gn=MACF1;refGene_tx=XM_019836714.2;refGene_gn=MACF1;refGene_tx=XM_019836715.2;refGene_gn=MACF1;refGene_tx=XM_019836716.2;refGene_gn=MACF1;refGene_tx=XM_019836717.2;refGene_gn=MACF1;refGene_tx=XM_019836718.2;refGene_gn=MACF1;refGene_tx=XM_019836720.2;refGene_gn=MACF1;refGene_tx=XM_023258509.1;refGene_gn=MACF1;refGene_tx=XM_023258510.1
C1 30548554 30548693

refGene_gn=MACF1;refGene_tx=XM_011284718.3;refGene_gn=MACF1;refGene_tx=XM_019836714.2;refGene_gn=MACF1;refGene_tx=XM_019836715.2;refGene_gn=MACF1;refGene_tx=XM_019836716.2;refGene_gn=MACF1;refGene_tx=XM_019836717.2;refGene_gn=MACF1;refGene_tx=XM_019836718.2;refGene_gn=MACF1;refGene_tx=XM_019836720.2;refGene_gn=MACF1;refGene_tx=XM_023258509.1;refGene_gn=MACF1;refGene_tx=XM_023258510.1
C1 30549212 30549406
refGene_gn=MACF1;refGene_tx=XM_011284718.3;refGene_gn=MACF1;refGene_tx=XM_019836714.2;refGene_gn=MACF1;refGene_tx=XM_019836715.2;refGene_gn=MACF1;refGene_tx=XM_019836716.2;refGene_gn=MACF1;refGene_tx=XM_019836717.2;refGene_gn=MACF1;refGene_tx=XM_019836718.2;refGene_gn=MACF1;refGene_tx=XM_019836720.2;refGene_gn=MACF1;refGene_tx=XM_023258509.1;refGene_gn=MACF1;refGene_tx=XM_023258510.1
C1 30550469 30550632
refGene_gn=MACF1;refGene_tx=XM_011284718.3;refGene_gn=MACF1;refGene_tx=XM_019836714.2;refGene_gn=MACF1;refGene_tx=XM_019836715.2;refGene_gn=MACF1;refGene_tx=XM_019836716.2;refGene_gn=MACF1;refGene_tx=XM_019836717.2;refGene_gn=MACF1;refGene_tx=XM_019836718.2;refGene_gn=MACF1;refGene_tx=XM_019836720.2;refGene_gn=MACF1;refGene_tx=XM_023258509.1;refGene_gn=MACF1;refGene_tx=XM_023258510.1
C1 30551514 30551598
refGene_gn=MACF1;refGene_tx=XM_011284718.3;refGene_gn=MACF1;refGene_tx=XM_019836714.2;refGene_gn=MACF1;refGene_tx=XM_019836715.2;refGene_gn=MACF1;refGene_tx=XM_019836716.2;refGene_gn=MACF1;refGene_tx=XM_019836717.2;refGene_gn=MACF1;refGene_tx=XM_019836718.2;refGene_gn=MACF1;refGene_tx=XM_019836720.2;refGene_gn=MACF1;refGene_tx=XM_023258509.1;refGene_gn=MACF1;refGene_tx=XM_023258510.1
C1 30554189 30554297
refGene_gn=MACF1;refGene_tx=XM_011284718.3;refGene_gn=MACF1;refGene_tx=XM_019836714.2;refGene_gn=MACF1;refGene_tx=XM_019836715.2;refGene_gn=MACF1;refGene_tx=XM_019836716.2;refGene_gn=MACF1;refGene_tx=XM_019836717.2;refGene_gn=MACF1;refGene_tx=XM_019836718.2;refGene_gn=MACF1;refGene_tx=XM_019836720.2;refGene_gn=MACF1;refGene_tx=XM_023258509.1;refGene_gn=MACF1;refGene_tx=XM_023258510.1
C1 30554490 30554589
refGene_gn=MACF1;refGene_tx=XM_011284718.3;refGene_gn=MACF1;refGene_tx=XM_019836714.2;refGene_gn=MACF1;refGene_tx=XM_019836715.2;refGene_gn=MACF1;refGene_tx=XM_019836716.2;refGene_gn=MACF1;refGene_tx=XM_019836717.2;refGene_gn=MACF1;refGene_tx=XM_019836718.2;refGene_gn=MACF1;refGene_tx=XM_019836720.2;refGene_gn=MACF1;refGene_tx=XM_023258509.1;refGene_gn=MACF1;refGene_tx=XM_023258510.1
C1 30556433 30556586
refGene_gn=MACF1;refGene_tx=XM_011284718.3;refGene_gn=MACF1;refGene_tx=XM_019836714.2;refGene_gn=MACF1;refGene_tx=XM_019836715.2;refGene_gn=MACF1;refGene_tx=XM_019836716.2;refGene_gn=MACF1;refGene_tx=XM_019836717.2;refGene_gn=MACF1;refGene_tx=XM_019836718.2;refGene_gn=MACF1;refGene_tx=XM_019836720.2;refGene_gn=MACF1;refGene_tx=XM_023258509.1;refGene_gn=MACF1;refGene_tx=XM_023258510.1
C1 30556891 30556980
refGene_gn=MACF1;refGene_tx=XM_011284718.3;refGene_gn=MACF1;refGene_tx

=XM_019836714.2;refGene_gn=MACF1;refGene_tx=XM_019836715.2;refGene_gn=MACF1;refGene_tx=XM_019836716.2;refGene_gn=MACF1;refGene_tx=XM_019836717.2;refGene_gn=MACF1;refGene_tx=XM_019836718.2;refGene_gn=MACF1;refGene_tx=XM_019836720.2;refGene_gn=MACF1;refGene_tx=XM_023258509.1;refGene_gn=MACF1;refGene_tx=XM_023258510.1
C1 30559059 30559195
refGene_gn=MACF1;refGene_tx=XM_011284718.3;refGene_gn=MACF1;refGene_tx=XM_019836714.2;refGene_gn=MACF1;refGene_tx=XM_019836715.2;refGene_gn=MACF1;refGene_tx=XM_019836716.2;refGene_gn=MACF1;refGene_tx=XM_019836717.2;refGene_gn=MACF1;refGene_tx=XM_019836718.2;refGene_gn=MACF1;refGene_tx=XM_019836720.2;refGene_gn=MACF1;refGene_tx=XM_023258509.1;refGene_gn=MACF1;refGene_tx=XM_023258510.1
C1 30575790 30575940
refGene_gn=MACF1;refGene_tx=XM_011284718.3;refGene_gn=MACF1;refGene_tx=XM_019836714.2;refGene_gn=MACF1;refGene_tx=XM_019836715.2;refGene_gn=MACF1;refGene_tx=XM_019836716.2;refGene_gn=MACF1;refGene_tx=XM_019836717.2;refGene_gn=MACF1;refGene_tx=XM_019836718.2;refGene_gn=MACF1;refGene_tx=XM_019836720.2;refGene_gn=MACF1;refGene_tx=XM_023258509.1;refGene_gn=MACF1;refGene_tx=XM_023258510.1
C1 30576980 30577214
refGene_gn=MACF1;refGene_tx=XM_011284718.3;refGene_gn=MACF1;refGene_tx=XM_019836714.2;refGene_gn=MACF1;refGene_tx=XM_019836715.2;refGene_gn=MACF1;refGene_tx=XM_019836716.2;refGene_gn=MACF1;refGene_tx=XM_019836717.2;refGene_gn=MACF1;refGene_tx=XM_019836718.2;refGene_gn=MACF1;refGene_tx=XM_019836720.2;refGene_gn=MACF1;refGene_tx=XM_023258509.1;refGene_gn=MACF1;refGene_tx=XM_023258510.1
C1 30578550 30578754
refGene_gn=MACF1;refGene_tx=XM_011284718.3;refGene_gn=MACF1;refGene_tx=XM_019836714.2;refGene_gn=MACF1;refGene_tx=XM_019836715.2;refGene_gn=MACF1;refGene_tx=XM_019836716.2;refGene_gn=MACF1;refGene_tx=XM_019836717.2;refGene_gn=MACF1;refGene_tx=XM_019836718.2;refGene_gn=MACF1;refGene_tx=XM_019836720.2;refGene_gn=MACF1;refGene_tx=XM_023258509.1;refGene_gn=MACF1;refGene_tx=XM_023258510.1
C1 30581123 30581642
refGene_gn=MACF1;refGene_tx=XM_011284718.3;refGene_gn=MACF1;refGene_tx=XM_019836714.2;refGene_gn=MACF1;refGene_tx=XM_019836715.2;refGene_gn=MACF1;refGene_tx=XM_019836716.2;refGene_gn=MACF1;refGene_tx=XM_019836717.2;refGene_gn=MACF1;refGene_tx=XM_019836718.2;refGene_gn=MACF1;refGene_tx=XM_019836720.2;refGene_gn=MACF1;refGene_tx=XM_023258509.1;refGene_gn=MACF1;refGene_tx=XM_023258510.1
C1 30582413 30582590
refGene_gn=MACF1;refGene_tx=XM_011284718.3;refGene_gn=MACF1;refGene_tx=XM_019836714.2;refGene_gn=MACF1;refGene_tx=XM_019836715.2;refGene_gn=MACF1;refGene_tx=XM_019836716.2;refGene_gn=MACF1;refGene_tx=XM_019836717.2;refGene_gn=MACF1;refGene_tx=XM_019836718.2;refGene_gn=MACF1;refGene_tx=XM_019836720.2;refGene_gn=MACF1;refGene_tx=XM_023258509.1;refGene_gn=MACF1;refGene_tx=XM_023258510.1
C1 30582856 30582980
refGene_gn=MACF1;refGene_tx=XM_011284718.3;refGene_gn=MACF1;refGene_tx=XM_019836714.2;refGene_gn=MACF1;refGene_tx=XM_019836715.2;refGene_gn=

MACF1;refGene_tx=XM_019836716.2;refGene_gn=MACF1;refGene_tx=XM_019836717.2;refGene_gn=MACF1;refGene_tx=XM_019836718.2;refGene_gn=MACF1;refGene_tx=XM_019836720.2;refGene_gn=MACF1;refGene_tx=XM_023258509.1;refGene_gn=MACF1;refGene_tx=XM_023258510.1
C1 30583791 30584000
refGene_gn=MACF1;refGene_tx=XM_011284718.3;refGene_gn=MACF1;refGene_tx=XM_019836714.2;refGene_gn=MACF1;refGene_tx=XM_019836715.2;refGene_gn=MACF1;refGene_tx=XM_019836716.2;refGene_gn=MACF1;refGene_tx=XM_019836717.2;refGene_gn=MACF1;refGene_tx=XM_019836718.2;refGene_gn=MACF1;refGene_tx=XM_019836720.2;refGene_gn=MACF1;refGene_tx=XM_023258509.1;refGene_gn=MACF1;refGene_tx=XM_023258510.1
C1 30584378 30584696
refGene_gn=MACF1;refGene_tx=XM_011284718.3;refGene_gn=MACF1;refGene_tx=XM_019836714.2;refGene_gn=MACF1;refGene_tx=XM_019836715.2;refGene_gn=MACF1;refGene_tx=XM_019836716.2;refGene_gn=MACF1;refGene_tx=XM_019836717.2;refGene_gn=MACF1;refGene_tx=XM_019836718.2;refGene_gn=MACF1;refGene_tx=XM_019836720.2;refGene_gn=MACF1;refGene_tx=XM_023258509.1;refGene_gn=MACF1;refGene_tx=XM_023258510.1
C1 30587320 30587487
refGene_gn=MACF1;refGene_tx=XM_011284718.3;refGene_gn=MACF1;refGene_tx=XM_019836714.2;refGene_gn=MACF1;refGene_tx=XM_019836715.2;refGene_gn=MACF1;refGene_tx=XM_019836716.2;refGene_gn=MACF1;refGene_tx=XM_019836717.2;refGene_gn=MACF1;refGene_tx=XM_019836718.2;refGene_gn=MACF1;refGene_tx=XM_019836720.2;refGene_gn=MACF1;refGene_tx=XM_023258509.1;refGene_gn=MACF1;refGene_tx=XM_023258510.1
C1 30588786 30588943
refGene_gn=MACF1;refGene_tx=XM_011284718.3;refGene_gn=MACF1;refGene_tx=XM_019836714.2;refGene_gn=MACF1;refGene_tx=XM_019836715.2;refGene_gn=MACF1;refGene_tx=XM_019836716.2;refGene_gn=MACF1;refGene_tx=XM_019836717.2;refGene_gn=MACF1;refGene_tx=XM_019836718.2;refGene_gn=MACF1;refGene_tx=XM_019836720.2;refGene_gn=MACF1;refGene_tx=XM_023258509.1;refGene_gn=MACF1;refGene_tx=XM_023258510.1
C1 30591240 30591358
refGene_gn=MACF1;refGene_tx=XM_011284718.3;refGene_gn=MACF1;refGene_tx=XM_019836714.2;refGene_gn=MACF1;refGene_tx=XM_019836715.2;refGene_gn=MACF1;refGene_tx=XM_019836716.2;refGene_gn=MACF1;refGene_tx=XM_019836717.2;refGene_gn=MACF1;refGene_tx=XM_019836718.2;refGene_gn=MACF1;refGene_tx=XM_019836720.2;refGene_gn=MACF1;refGene_tx=XM_023258509.1;refGene_gn=MACF1;refGene_tx=XM_023258510.1
C1 30594481 30594544
refGene_gn=MACF1;refGene_tx=XM_011284718.3;refGene_gn=MACF1;refGene_tx=XM_019836714.2;refGene_gn=MACF1;refGene_tx=XM_019836715.2;refGene_gn=MACF1;refGene_tx=XM_019836716.2;refGene_gn=MACF1;refGene_tx=XM_019836718.2;refGene_gn=MACF1;refGene_tx=XM_019836720.2;refGene_gn=MACF1;refGene_tx=XM_023258509.1
C1 30595167 30595409
refGene_gn=MACF1;refGene_tx=XM_011284718.3;refGene_gn=MACF1;refGene_tx=XM_019836714.2;refGene_gn=MACF1;refGene_tx=XM_019836715.2;refGene_gn=MACF1;refGene_tx=XM_019836716.2;refGene_gn=MACF1;refGene_tx=XM_019836717.2;refGene_gn=MACF1;refGene_tx=XM_019836718.2;refGene_gn=MACF1;refGe

ne_tx=XM_019836720.2;refGene_gn=MACF1;refGene_tx=XM_023258509.1;refGene_gn=MACF1;refGene_tx=XM_023258510.1
C1 30596222 30596352
refGene_gn=MACF1;refGene_tx=XM_011284718.3;refGene_gn=MACF1;refGene_tx=XM_019836714.2;refGene_gn=MACF1;refGene_tx=XM_019836715.2;refGene_gn=MACF1;refGene_tx=XM_019836716.2;refGene_gn=MACF1;refGene_tx=XM_019836717.2;refGene_gn=MACF1;refGene_tx=XM_019836718.2;refGene_gn=MACF1;refGene_tx=XM_019836720.2;refGene_gn=MACF1;refGene_tx=XM_023258509.1;refGene_gn=MACF1;refGene_tx=XM_023258510.1
C1 30597049 30597249
refGene_gn=MACF1;refGene_tx=XM_011284718.3;refGene_gn=MACF1;refGene_tx=XM_019836714.2;refGene_gn=MACF1;refGene_tx=XM_019836715.2;refGene_gn=MACF1;refGene_tx=XM_019836716.2;refGene_gn=MACF1;refGene_tx=XM_019836717.2;refGene_gn=MACF1;refGene_tx=XM_019836718.2;refGene_gn=MACF1;refGene_tx=XM_019836720.2;refGene_gn=MACF1;refGene_tx=XM_023258509.1;refGene_gn=MACF1;refGene_tx=XM_023258510.1
C1 30601826 30602322
refGene_gn=MACF1;refGene_tx=XM_011284718.3;refGene_gn=MACF1;refGene_tx=XM_019836714.2;refGene_gn=MACF1;refGene_tx=XM_019836715.2;refGene_gn=MACF1;refGene_tx=XM_019836716.2;refGene_gn=MACF1;refGene_tx=XM_019836717.2;refGene_gn=MACF1;refGene_tx=XM_019836718.2;refGene_gn=MACF1;refGene_tx=XM_019836720.2;refGene_gn=MACF1;refGene_tx=XM_023258509.1;refGene_gn=MACF1;refGene_tx=XM_023258510.1
C1 30603003 30604475
refGene_gn=MACF1;refGene_tx=XM_011284718.3;refGene_gn=MACF1;refGene_tx=XM_019836714.2;refGene_gn=MACF1;refGene_tx=XM_019836715.2;refGene_gn=MACF1;refGene_tx=XM_019836716.2;refGene_gn=MACF1;refGene_tx=XM_019836717.2;refGene_gn=MACF1;refGene_tx=XM_019836718.2;refGene_gn=MACF1;refGene_tx=XM_019836720.2;refGene_gn=MACF1;refGene_tx=XM_023258509.1;refGene_gn=MACF1;refGene_tx=XM_023258510.1
C1 30621418 30625084 refGene_gn=MACF1;refGene_tx=XM_023258512.1
C1 30631423 30631585
refGene_gn=MACF1;refGene_tx=XM_011284718.3;refGene_gn=MACF1;refGene_tx=XM_019836714.2;refGene_gn=MACF1;refGene_tx=XM_019836715.2;refGene_gn=MACF1;refGene_tx=XM_019836716.2;refGene_gn=MACF1;refGene_tx=XM_019836717.2;refGene_gn=MACF1;refGene_tx=XM_019836718.2;refGene_gn=MACF1;refGene_tx=XM_019836720.2;refGene_gn=MACF1;refGene_tx=XM_023258509.1;refGene_gn=MACF1;refGene_tx=XM_023258510.1;refGene_gn=MACF1;refGene_tx=XM_023258512.1
C1 30631780 30631951
refGene_gn=MACF1;refGene_tx=XM_011284718.3;refGene_gn=MACF1;refGene_tx=XM_019836714.2;refGene_gn=MACF1;refGene_tx=XM_019836715.2;refGene_gn=MACF1;refGene_tx=XM_019836716.2;refGene_gn=MACF1;refGene_tx=XM_019836717.2;refGene_gn=MACF1;refGene_tx=XM_019836718.2;refGene_gn=MACF1;refGene_tx=XM_019836720.2;refGene_gn=MACF1;refGene_tx=XM_023258509.1;refGene_gn=MACF1;refGene_tx=XM_023258510.1;refGene_gn=MACF1;refGene_tx=XM_023258512.1
C1 30632584 30632751
refGene_gn=MACF1;refGene_tx=XM_011284718.3;refGene_gn=MACF1;refGene_tx=XM_019836714.2;refGene_gn=MACF1;refGene_tx=XM_019836715.2;refGene_gn=

MACF1;refGene_tx=XM_019836716.2;refGene_gn=MACF1;refGene_tx=XM_0198367
17.2;refGene_gn=MACF1;refGene_tx=XM_019836718.2;refGene_gn=MACF1;refGe
ne_tx=XM_019836720.2;refGene_gn=MACF1;refGene_tx=XM_023258509.1;refGen
e_gn=MACF1;refGene_tx=XM_023258510.1;refGene_gn=MACF1;refGene_tx=XM_02
3258512.1
C1 30636925 30637085
refGene_gn=MACF1;refGene_tx=XM_011284718.3;refGene_gn=MACF1;refGene_tx
=XM_019836714.2;refGene_gn=MACF1;refGene_tx=XM_019836715.2;refGene_gn=
MACF1;refGene_tx=XM_019836716.2;refGene_gn=MACF1;refGene_tx=XM_0198367
17.2;refGene_gn=MACF1;refGene_tx=XM_019836718.2;refGene_gn=MACF1;refGe
ne_tx=XM_019836720.2;refGene_gn=MACF1;refGene_tx=XM_023258509.1;refGen
e_gn=MACF1;refGene_tx=XM_023258510.1;refGene_gn=MACF1;refGene_tx=XM_02
3258512.1
C1 30637441 30637768
refGene_gn=MACF1;refGene_tx=XM_011284718.3;refGene_gn=MACF1;refGene_tx
=XM_019836714.2;refGene_gn=MACF1;refGene_tx=XM_019836715.2;refGene_gn=
MACF1;refGene_tx=XM_019836716.2;refGene_gn=MACF1;refGene_tx=XM_0198367
17.2;refGene_gn=MACF1;refGene_tx=XM_023258509.1;refGene_gn=MACF1;refGe
ne_tx=XM_023258510.1;refGene_gn=MACF1;refGene_tx=XM_023258512.1
C1 30638735 30638820
refGene_gn=MACF1;refGene_tx=XM_011284718.3;refGene_gn=MACF1;refGene_tx
=XM_019836714.2;refGene_gn=MACF1;refGene_tx=XM_019836715.2;refGene_gn=
MACF1;refGene_tx=XM_019836716.2;refGene_gn=MACF1;refGene_tx=XM_0198367
17.2;refGene_gn=MACF1;refGene_tx=XM_019836718.2;refGene_gn=MACF1;refGe
ne_tx=XM_019836720.2;refGene_gn=MACF1;refGene_tx=XM_023258509.1;refGen
e_gn=MACF1;refGene_tx=XM_023258510.1;refGene_gn=MACF1;refGene_tx=XM_02
3258512.1
C1 30639330 30639572
refGene_gn=MACF1;refGene_tx=XM_011284718.3;refGene_gn=MACF1;refGene_tx
=XM_019836714.2;refGene_gn=MACF1;refGene_tx=XM_019836715.2;refGene_gn=
MACF1;refGene_tx=XM_019836716.2;refGene_gn=MACF1;refGene_tx=XM_0198367
17.2;refGene_gn=MACF1;refGene_tx=XM_019836718.2;refGene_gn=MACF1;refGe
ne_tx=XM_019836720.2;refGene_gn=MACF1;refGene_tx=XM_023258509.1;refGen
e_gn=MACF1;refGene_tx=XM_023258510.1;refGene_gn=MACF1;refGene_tx=XM_02
3258512.1
C1 30640273 30640480
refGene_gn=MACF1;refGene_tx=XM_011284718.3;refGene_gn=MACF1;refGene_tx
=XM_019836714.2;refGene_gn=MACF1;refGene_tx=XM_019836715.2;refGene_gn=
MACF1;refGene_tx=XM_019836716.2;refGene_gn=MACF1;refGene_tx=XM_0198367
17.2;refGene_gn=MACF1;refGene_tx=XM_019836718.2;refGene_gn=MACF1;refGe
ne_tx=XM_019836720.2;refGene_gn=MACF1;refGene_tx=XM_023258509.1;refGen
e_gn=MACF1;refGene_tx=XM_023258510.1;refGene_gn=MACF1;refGene_tx=XM_02
3258512.1
C1 30642012 30642132
refGene_gn=MACF1;refGene_tx=XM_011284718.3;refGene_gn=MACF1;refGene_tx
=XM_019836714.2;refGene_gn=MACF1;refGene_tx=XM_019836715.2;refGene_gn=
MACF1;refGene_tx=XM_019836716.2;refGene_gn=MACF1;refGene_tx=XM_0198367
17.2;refGene_gn=MACF1;refGene_tx=XM_019836718.2;refGene_gn=MACF1;refGe
ne_tx=XM_019836720.2;refGene_gn=MACF1;refGene_tx=XM_023258509.1;refGen
e_gn=MACF1;refGene_tx=XM_023258510.1;refGene_gn=MACF1;refGene_tx=XM_02

3258512.1
C1 30642566 30642674
refGene_gn=MACF1;refGene_tx=XM_011284718.3;refGene_gn=MACF1;refGene_tx=XM_019836714.2;refGene_gn=MACF1;refGene_tx=XM_019836715.2;refGene_gn=MACF1;refGene_tx=XM_019836716.2;refGene_gn=MACF1;refGene_tx=XM_019836717.2;refGene_gn=MACF1;refGene_tx=XM_019836718.2;refGene_gn=MACF1;refGene_tx=XM_019836720.2;refGene_gn=MACF1;refGene_tx=XM_023258509.1;refGene_gn=MACF1;refGene_tx=XM_023258510.1;refGene_gn=MACF1;refGene_tx=XM_023258512.1
C1 30643571 30643790
refGene_gn=MACF1;refGene_tx=XM_011284718.3;refGene_gn=MACF1;refGene_tx=XM_019836714.2;refGene_gn=MACF1;refGene_tx=XM_019836715.2;refGene_gn=MACF1;refGene_tx=XM_019836716.2;refGene_gn=MACF1;refGene_tx=XM_019836717.2;refGene_gn=MACF1;refGene_tx=XM_019836718.2;refGene_gn=MACF1;refGene_tx=XM_019836720.2;refGene_gn=MACF1;refGene_tx=XM_023258509.1;refGene_gn=MACF1;refGene_tx=XM_023258510.1;refGene_gn=MACF1;refGene_tx=XM_023258512.1
C1 30644759 30644963
refGene_gn=MACF1;refGene_tx=XM_011284718.3;refGene_gn=MACF1;refGene_tx=XM_019836714.2;refGene_gn=MACF1;refGene_tx=XM_019836715.2;refGene_gn=MACF1;refGene_tx=XM_019836716.2;refGene_gn=MACF1;refGene_tx=XM_019836717.2;refGene_gn=MACF1;refGene_tx=XM_019836718.2;refGene_gn=MACF1;refGene_tx=XM_019836720.2;refGene_gn=MACF1;refGene_tx=XM_023258509.1;refGene_gn=MACF1;refGene_tx=XM_023258510.1;refGene_gn=MACF1;refGene_tx=XM_023258512.1
C1 30645713 30645722
refGene_gn=MACF1;refGene_tx=XM_011284718.3;refGene_gn=MACF1;refGene_tx=XM_019836714.2;refGene_gn=MACF1;refGene_tx=XM_019836715.2;refGene_gn=MACF1;refGene_tx=XM_019836716.2;refGene_gn=MACF1;refGene_tx=XM_019836717.2;refGene_gn=MACF1;refGene_tx=XM_023258510.1;refGene_gn=MACF1;refGene_tx=XM_023258512.1
C1 30646684 30646916
refGene_gn=MACF1;refGene_tx=XM_011284718.3;refGene_gn=MACF1;refGene_tx=XM_019836714.2;refGene_gn=MACF1;refGene_tx=XM_019836715.2;refGene_gn=MACF1;refGene_tx=XM_019836716.2;refGene_gn=MACF1;refGene_tx=XM_019836717.2;refGene_gn=MACF1;refGene_tx=XM_019836718.2;refGene_gn=MACF1;refGene_tx=XM_019836720.2;refGene_gn=MACF1;refGene_tx=XM_023258509.1;refGene_gn=MACF1;refGene_tx=XM_023258510.1;refGene_gn=MACF1;refGene_tx=XM_023258512.1
C1 30647536 30647763
refGene_gn=MACF1;refGene_tx=XM_011284718.3;refGene_gn=MACF1;refGene_tx=XM_019836714.2;refGene_gn=MACF1;refGene_tx=XM_019836715.2;refGene_gn=MACF1;refGene_tx=XM_019836716.2;refGene_gn=MACF1;refGene_tx=XM_019836717.2;refGene_gn=MACF1;refGene_tx=XM_019836718.2;refGene_gn=MACF1;refGene_tx=XM_019836720.2;refGene_gn=MACF1;refGene_tx=XM_023258509.1;refGene_gn=MACF1;refGene_tx=XM_023258510.1;refGene_gn=MACF1;refGene_tx=XM_023258512.1
C1 30649098 30649221
refGene_gn=MACF1;refGene_tx=XM_011284718.3;refGene_gn=MACF1;refGene_tx=XM_019836714.2;refGene_gn=MACF1;refGene_tx=XM_019836715.2;refGene_gn=

MACF1;refGene_tx=XM_019836716.2;refGene_gn=MACF1;refGene_tx=XM_0198367
17.2;refGene_gn=MACF1;refGene_tx=XM_019836718.2;refGene_gn=MACF1;refGe
ne_tx=XM_019836720.2;refGene_gn=MACF1;refGene_tx=XM_023258509.1;refGen
e_gn=MACF1;refGene_tx=XM_023258510.1;refGene_gn=MACF1;refGene_tx=XM_02
3258512.1
C1 30649313 30649415
refGene_gn=MACF1;refGene_tx=XM_011284718.3;refGene_gn=MACF1;refGene_tx
=XM_019836714.2;refGene_gn=MACF1;refGene_tx=XM_019836715.2;refGene_gn=
MACF1;refGene_tx=XM_019836716.2;refGene_gn=MACF1;refGene_tx=XM_0198367
17.2;refGene_gn=MACF1;refGene_tx=XM_019836718.2;refGene_gn=MACF1;refGe
ne_tx=XM_019836720.2;refGene_gn=MACF1;refGene_tx=XM_023258509.1;refGen
e_gn=MACF1;refGene_tx=XM_023258510.1;refGene_gn=MACF1;refGene_tx=XM_02
3258512.1
C1 30650030 30650132
refGene_gn=MACF1;refGene_tx=XM_011284718.3;refGene_gn=MACF1;refGene_tx
=XM_019836714.2;refGene_gn=MACF1;refGene_tx=XM_019836715.2;refGene_gn=
MACF1;refGene_tx=XM_019836716.2;refGene_gn=MACF1;refGene_tx=XM_0198367
17.2;refGene_gn=MACF1;refGene_tx=XM_019836718.2;refGene_gn=MACF1;refGe
ne_tx=XM_019836720.2;refGene_gn=MACF1;refGene_tx=XM_023258509.1;refGen
e_gn=MACF1;refGene_tx=XM_023258510.1;refGene_gn=MACF1;refGene_tx=XM_02
3258512.1
C1 30650224 30650398
refGene_gn=MACF1;refGene_tx=XM_011284718.3;refGene_gn=MACF1;refGene_tx
=XM_019836714.2;refGene_gn=MACF1;refGene_tx=XM_019836715.2;refGene_gn=
MACF1;refGene_tx=XM_019836716.2;refGene_gn=MACF1;refGene_tx=XM_0198367
17.2;refGene_gn=MACF1;refGene_tx=XM_019836718.2;refGene_gn=MACF1;refGe
ne_tx=XM_019836720.2;refGene_gn=MACF1;refGene_tx=XM_023258509.1;refGen
e_gn=MACF1;refGene_tx=XM_023258510.1;refGene_gn=MACF1;refGene_tx=XM_02
3258512.1
C1 30650489 30650645
refGene_gn=MACF1;refGene_tx=XM_011284718.3;refGene_gn=MACF1;refGene_tx
=XM_019836714.2;refGene_gn=MACF1;refGene_tx=XM_019836715.2;refGene_gn=
MACF1;refGene_tx=XM_019836716.2;refGene_gn=MACF1;refGene_tx=XM_0198367
17.2;refGene_gn=MACF1;refGene_tx=XM_019836718.2;refGene_gn=MACF1;refGe
ne_tx=XM_019836720.2;refGene_gn=MACF1;refGene_tx=XM_023258509.1;refGen
e_gn=MACF1;refGene_tx=XM_023258510.1;refGene_gn=MACF1;refGene_tx=XM_02
3258512.1
C1 30650789 30650987
refGene_gn=MACF1;refGene_tx=XM_011284718.3;refGene_gn=MACF1;refGene_tx
=XM_019836714.2;refGene_gn=MACF1;refGene_tx=XM_019836715.2;refGene_gn=
MACF1;refGene_tx=XM_019836716.2;refGene_gn=MACF1;refGene_tx=XM_0198367
17.2;refGene_gn=MACF1;refGene_tx=XM_019836718.2;refGene_gn=MACF1;refGe
ne_tx=XM_019836720.2;refGene_gn=MACF1;refGene_tx=XM_023258509.1;refGen
e_gn=MACF1;refGene_tx=XM_023258510.1;refGene_gn=MACF1;refGene_tx=XM_02
3258512.1
C1 30651845 30651974
refGene_gn=MACF1;refGene_tx=XM_011284718.3;refGene_gn=MACF1;refGene_tx
=XM_019836714.2;refGene_gn=MACF1;refGene_tx=XM_019836715.2;refGene_gn=
MACF1;refGene_tx=XM_019836716.2;refGene_gn=MACF1;refGene_tx=XM_0198367
17.2;refGene_gn=MACF1;refGene_tx=XM_019836718.2;refGene_gn=MACF1;refGe

ne_tx=XM_019836720.2;refGene_gn=MACF1;refGene_tx=XM_023258509.1;refGene_gn=MACF1;refGene_tx=XM_023258510.1;refGene_gn=MACF1;refGene_tx=XM_023258512.1
C1 30652650 30652824
refGene_gn=MACF1;refGene_tx=XM_011284718.3;refGene_gn=MACF1;refGene_tx=XM_019836714.2;refGene_gn=MACF1;refGene_tx=XM_019836715.2;refGene_gn=MACF1;refGene_tx=XM_019836716.2;refGene_gn=MACF1;refGene_tx=XM_019836717.2;refGene_gn=MACF1;refGene_tx=XM_019836718.2;refGene_gn=MACF1;refGene_tx=XM_019836720.2;refGene_gn=MACF1;refGene_tx=XM_023258509.1;refGene_gn=MACF1;refGene_tx=XM_023258510.1;refGene_gn=MACF1;refGene_tx=XM_023258512.1
C1 30655149 30655305
refGene_gn=MACF1;refGene_tx=XM_011284718.3;refGene_gn=MACF1;refGene_tx=XM_019836714.2;refGene_gn=MACF1;refGene_tx=XM_019836715.2;refGene_gn=MACF1;refGene_tx=XM_019836716.2;refGene_gn=MACF1;refGene_tx=XM_019836717.2;refGene_gn=MACF1;refGene_tx=XM_019836718.2;refGene_gn=MACF1;refGene_tx=XM_019836720.2;refGene_gn=MACF1;refGene_tx=XM_023258509.1;refGene_gn=MACF1;refGene_tx=XM_023258510.1;refGene_gn=MACF1;refGene_tx=XM_023258512.1
C1 30655409 30655616
refGene_gn=MACF1;refGene_tx=XM_011284718.3;refGene_gn=MACF1;refGene_tx=XM_019836714.2;refGene_gn=MACF1;refGene_tx=XM_019836715.2;refGene_gn=MACF1;refGene_tx=XM_019836716.2;refGene_gn=MACF1;refGene_tx=XM_019836717.2;refGene_gn=MACF1;refGene_tx=XM_019836718.2;refGene_gn=MACF1;refGene_tx=XM_019836720.2;refGene_gn=MACF1;refGene_tx=XM_023258509.1;refGene_gn=MACF1;refGene_tx=XM_023258510.1;refGene_gn=MACF1;refGene_tx=XM_023258512.1
C1 30655739 30655859
refGene_gn=MACF1;refGene_tx=XM_011284718.3;refGene_gn=MACF1;refGene_tx=XM_019836714.2;refGene_gn=MACF1;refGene_tx=XM_019836715.2;refGene_gn=MACF1;refGene_tx=XM_019836716.2;refGene_gn=MACF1;refGene_tx=XM_019836717.2;refGene_gn=MACF1;refGene_tx=XM_019836718.2;refGene_gn=MACF1;refGene_tx=XM_019836720.2;refGene_gn=MACF1;refGene_tx=XM_023258509.1;refGene_gn=MACF1;refGene_tx=XM_023258510.1;refGene_gn=MACF1;refGene_tx=XM_023258512.1
C1 30656734 30656904
refGene_gn=MACF1;refGene_tx=XM_011284718.3;refGene_gn=MACF1;refGene_tx=XM_019836714.2;refGene_gn=MACF1;refGene_tx=XM_019836715.2;refGene_gn=MACF1;refGene_tx=XM_019836716.2;refGene_gn=MACF1;refGene_tx=XM_019836717.2;refGene_gn=MACF1;refGene_tx=XM_019836718.2;refGene_gn=MACF1;refGene_tx=XM_019836720.2;refGene_gn=MACF1;refGene_tx=XM_023258509.1;refGene_gn=MACF1;refGene_tx=XM_023258510.1;refGene_gn=MACF1;refGene_tx=XM_023258512.1
C1 30657966 30658126
refGene_gn=MACF1;refGene_tx=XM_011284718.3;refGene_gn=MACF1;refGene_tx=XM_019836714.2;refGene_gn=MACF1;refGene_tx=XM_019836715.2;refGene_gn=MACF1;refGene_tx=XM_019836716.2;refGene_gn=MACF1;refGene_tx=XM_019836717.2;refGene_gn=MACF1;refGene_tx=XM_019836718.2;refGene_gn=MACF1;refGene_tx=XM_019836720.2;refGene_gn=MACF1;refGene_tx=XM_023258509.1;refGene_gn=MACF1;refGene_tx=XM_023258510.1;refGene_gn=MACF1;refGene_tx=XM_02

3258512.1
C1 30658918 30659113
refGene_gn=MACF1;refGene_tx=XM_011284718.3;refGene_gn=MACF1;refGene_tx=XM_019836714.2;refGene_gn=MACF1;refGene_tx=XM_019836715.2;refGene_gn=MACF1;refGene_tx=XM_019836716.2;refGene_gn=MACF1;refGene_tx=XM_019836717.2;refGene_gn=MACF1;refGene_tx=XM_019836718.2;refGene_gn=MACF1;refGene_tx=XM_019836720.2;refGene_gn=MACF1;refGene_tx=XM_023258509.1;refGene_gn=MACF1;refGene_tx=XM_023258510.1;refGene_gn=MACF1;refGene_tx=XM_023258512.1
C1 30659450 30659579
refGene_gn=MACF1;refGene_tx=XM_011284718.3;refGene_gn=MACF1;refGene_tx=XM_019836714.2;refGene_gn=MACF1;refGene_tx=XM_019836715.2;refGene_gn=MACF1;refGene_tx=XM_019836716.2;refGene_gn=MACF1;refGene_tx=XM_019836717.2;refGene_gn=MACF1;refGene_tx=XM_019836718.2;refGene_gn=MACF1;refGene_tx=XM_019836720.2;refGene_gn=MACF1;refGene_tx=XM_023258509.1;refGene_gn=MACF1;refGene_tx=XM_023258510.1;refGene_gn=MACF1;refGene_tx=XM_023258512.1
C1 30660703 30660847
refGene_gn=MACF1;refGene_tx=XM_011284718.3;refGene_gn=MACF1;refGene_tx=XM_019836714.2;refGene_gn=MACF1;refGene_tx=XM_019836715.2;refGene_gn=MACF1;refGene_tx=XM_019836716.2;refGene_gn=MACF1;refGene_tx=XM_019836717.2;refGene_gn=MACF1;refGene_tx=XM_019836718.2;refGene_gn=MACF1;refGene_tx=XM_019836720.2;refGene_gn=MACF1;refGene_tx=XM_023258509.1;refGene_gn=MACF1;refGene_tx=XM_023258510.1;refGene_gn=MACF1;refGene_tx=XM_023258512.1
C1 30661327 30661516
refGene_gn=MACF1;refGene_tx=XM_011284718.3;refGene_gn=MACF1;refGene_tx=XM_019836714.2;refGene_gn=MACF1;refGene_tx=XM_019836715.2;refGene_gn=MACF1;refGene_tx=XM_019836716.2;refGene_gn=MACF1;refGene_tx=XM_019836717.2;refGene_gn=MACF1;refGene_tx=XM_019836718.2;refGene_gn=MACF1;refGene_tx=XM_019836720.2;refGene_gn=MACF1;refGene_tx=XM_023258509.1;refGene_gn=MACF1;refGene_tx=XM_023258510.1;refGene_gn=MACF1;refGene_tx=XM_023258512.1
C1 30662238 30662359
refGene_gn=MACF1;refGene_tx=XM_011284718.3;refGene_gn=MACF1;refGene_tx=XM_019836714.2;refGene_gn=MACF1;refGene_tx=XM_019836715.2;refGene_gn=MACF1;refGene_tx=XM_019836716.2;refGene_gn=MACF1;refGene_tx=XM_019836717.2;refGene_gn=MACF1;refGene_tx=XM_019836718.2;refGene_gn=MACF1;refGene_tx=XM_019836720.2;refGene_gn=MACF1;refGene_tx=XM_023258509.1;refGene_gn=MACF1;refGene_tx=XM_023258510.1;refGene_gn=MACF1;refGene_tx=XM_023258512.1
C1 30663064 30663228
refGene_gn=MACF1;refGene_tx=XM_011284718.3;refGene_gn=MACF1;refGene_tx=XM_019836714.2;refGene_gn=MACF1;refGene_tx=XM_019836715.2;refGene_gn=MACF1;refGene_tx=XM_019836716.2;refGene_gn=MACF1;refGene_tx=XM_019836717.2;refGene_gn=MACF1;refGene_tx=XM_019836718.2;refGene_gn=MACF1;refGene_tx=XM_019836720.2;refGene_gn=MACF1;refGene_tx=XM_023258509.1;refGene_gn=MACF1;refGene_tx=XM_023258510.1;refGene_gn=MACF1;refGene_tx=XM_023258512.1
C1 30664612 30664775

refGene_gn=MACF1;refGene_tx=XM_011284718.3;refGene_gn=MACF1;refGene_tx=XM_019836714.2;refGene_gn=MACF1;refGene_tx=XM_019836715.2;refGene_gn=MACF1;refGene_tx=XM_019836716.2;refGene_gn=MACF1;refGene_tx=XM_019836717.2;refGene_gn=MACF1;refGene_tx=XM_019836718.2;refGene_gn=MACF1;refGene_tx=XM_019836720.2;refGene_gn=MACF1;refGene_tx=XM_023258509.1;refGene_gn=MACF1;refGene_tx=XM_023258510.1;refGene_gn=MACF1;refGene_tx=XM_023258512.1
C1 30666031 30666186
refGene_gn=MACF1;refGene_tx=XM_011284718.3;refGene_gn=MACF1;refGene_tx=XM_019836714.2;refGene_gn=MACF1;refGene_tx=XM_019836715.2;refGene_gn=MACF1;refGene_tx=XM_019836716.2;refGene_gn=MACF1;refGene_tx=XM_019836717.2;refGene_gn=MACF1;refGene_tx=XM_019836718.2;refGene_gn=MACF1;refGene_tx=XM_019836720.2;refGene_gn=MACF1;refGene_tx=XM_023258509.1;refGene_gn=MACF1;refGene_tx=XM_023258510.1;refGene_gn=MACF1;refGene_tx=XM_023258512.1
C1 30667043 30667118
refGene_gn=MACF1;refGene_tx=XM_011284718.3;refGene_gn=MACF1;refGene_tx=XM_019836714.2;refGene_gn=MACF1;refGene_tx=XM_019836715.2;refGene_gn=MACF1;refGene_tx=XM_019836716.2;refGene_gn=MACF1;refGene_tx=XM_019836717.2;refGene_gn=MACF1;refGene_tx=XM_019836718.2;refGene_gn=MACF1;refGene_tx=XM_019836720.2;refGene_gn=MACF1;refGene_tx=XM_023258509.1;refGene_gn=MACF1;refGene_tx=XM_023258510.1;refGene_gn=MACF1;refGene_tx=XM_023258512.1
C1 30668244 30668262
refGene_gn=MACF1;refGene_tx=XM_019836714.2;refGene_gn=MACF1;refGene_tx=XM_019836715.2;refGene_gn=MACF1;refGene_tx=XM_019836716.2;refGene_gn=MACF1;refGene_tx=XM_019836718.2;refGene_gn=MACF1;refGene_tx=XM_019836720.2;refGene_gn=MACF1;refGene_tx=XM_023258509.1;refGene_gn=MACF1;refGene_tx=XM_023258510.1;refGene_gn=MACF1;refGene_tx=XM_023258512.1
C1 30671271 30671389
refGene_gn=MACF1;refGene_tx=XM_011284718.3;refGene_gn=MACF1;refGene_tx=XM_019836714.2;refGene_gn=MACF1;refGene_tx=XM_019836715.2;refGene_gn=MACF1;refGene_tx=XM_019836716.2;refGene_gn=MACF1;refGene_tx=XM_019836717.2;refGene_gn=MACF1;refGene_tx=XM_019836718.2;refGene_gn=MACF1;refGene_tx=XM_019836720.2;refGene_gn=MACF1;refGene_tx=XM_023258509.1;refGene_gn=MACF1;refGene_tx=XM_023258510.1;refGene_gn=MACF1;refGene_tx=XM_023258512.1
C1 30672181 30672250
refGene_gn=MACF1;refGene_tx=XM_019836714.2;refGene_gn=MACF1;refGene_tx=XM_019836715.2;refGene_gn=MACF1;refGene_tx=XM_019836716.2;refGene_gn=MACF1;refGene_tx=XM_019836718.2;refGene_gn=MACF1;refGene_tx=XM_023258509.1;refGene_gn=MACF1;refGene_tx=XM_023258510.1;refGene_gn=MACF1;refGene_tx=XM_023258512.1
C1 30678446 30678658
refGene_gn=MACF1;refGene_tx=XM_011284718.3;refGene_gn=MACF1;refGene_tx=XM_019836714.2;refGene_gn=MACF1;refGene_tx=XM_019836715.2;refGene_gn=MACF1;refGene_tx=XM_019836716.2;refGene_gn=MACF1;refGene_tx=XM_019836717.2;refGene_gn=MACF1;refGene_tx=XM_019836718.2;refGene_gn=MACF1;refGene_tx=XM_019836720.2;refGene_gn=MACF1;refGene_tx=XM_023258509.1;refGene_gn=MACF1;refGene_tx=XM_023258510.1;refGene_gn=MACF1;refGene_tx=XM_02

3258512.1
C1	30679181	30679292
refGene_gn=MACF1;refGene_tx=XM_019836714.2;refGene_gn=MACF1;refGene_tx=XM_019836715.2;refGene_gn=MACF1;refGene_tx=XM_019836716.2;refGene_gn=MACF1;refGene_tx=XM_019836718.2;refGene_gn=MACF1;refGene_tx=XM_023258509.1;refGene_gn=MACF1;refGene_tx=XM_023258510.1;refGene_gn=MACF1;refGene_tx=XM_023258512.1
C1	30681767	30681897
refGene_gn=MACF1;refGene_tx=XM_011284718.3;refGene_gn=MACF1;refGene_tx=XM_019836714.2;refGene_gn=MACF1;refGene_tx=XM_019836715.2;refGene_gn=MACF1;refGene_tx=XM_019836716.2;refGene_gn=MACF1;refGene_tx=XM_019836717.2;refGene_gn=MACF1;refGene_tx=XM_019836718.2;refGene_gn=MACF1;refGene_tx=XM_019836720.2;refGene_gn=MACF1;refGene_tx=XM_023258509.1;refGene_gn=MACF1;refGene_tx=XM_023258510.1;refGene_gn=MACF1;refGene_tx=XM_023258512.1
C1	30682742	30682999
refGene_gn=MACF1;refGene_tx=XM_011284718.3;refGene_gn=MACF1;refGene_tx=XM_019836714.2;refGene_gn=MACF1;refGene_tx=XM_019836715.2;refGene_gn=MACF1;refGene_tx=XM_019836716.2;refGene_gn=MACF1;refGene_tx=XM_019836717.2;refGene_gn=MACF1;refGene_tx=XM_019836718.2;refGene_gn=MACF1;refGene_tx=XM_019836720.2;refGene_gn=MACF1;refGene_tx=XM_023258509.1;refGene_gn=MACF1;refGene_tx=XM_023258510.1;refGene_gn=MACF1;refGene_tx=XM_023258512.1
C1	30688452	30688786	refGene_gn=BMP8A;refGene_tx=XM_023257825.1
C1	30700678	30700868	refGene_gn=BMP8A;refGene_tx=XM_023257825.1
C1	30701291	30701440	refGene_gn=BMP8A;refGene_tx=XM_023257825.1
C1	30707812	30707917	refGene_gn=BMP8A;refGene_tx=XM_023257825.1
C1	30707919	30708365	refGene_gn=BMP8A;refGene_tx=XM_023257825.1
C1	30709320	30709483	refGene_gn=BMP8A;refGene_tx=XM_023257825.1
C1	30721385	30721575	refGene_gn=BMP8A;refGene_tx=XM_023257825.1
C1	30721998	30722147	refGene_gn=BMP8A;refGene_tx=XM_023257825.1
C1	30724625	30725303
refGene_gn=LOC101088562;refGene_tx=XM_011284765.3
C1	30725304	30725898
refGene_gn=LOC101088562;refGene_tx=XM_011284765.3
C1	30725899	30726190
refGene_gn=LOC101088562;refGene_tx=XM_011284765.3
C1	30728511	30728706	refGene_gn=BMP8A;refGene_tx=XM_023257825.1
C1	30729464	30729544	refGene_gn=BMP8A;refGene_tx=XM_023257825.1
C1	30730106	30730217	refGene_gn=BMP8A;refGene_tx=XM_023257825.1
C1	30737544	30737694	refGene_gn=BMP8A;refGene_tx=XM_023257825.1
C1	30742733	30742775
refGene_gn=LOC105260819;refGene_tx=XM_019836723.2;refGene_gn=LOC105260819;refGene_tx=XM_023258513.1;refGene_gn=LOC105260819;refGene_tx=XM_023258514.1
C1	30742932	30743098
refGene_gn=LOC105260819;refGene_tx=XM_019836723.2;refGene_gn=LOC105260819;refGene_tx=XM_023258513.1;refGene_gn=LOC105260819;refGene_tx=XM_023258514.1
C1	30751713	30751837

refGene_gn=LOC105260819;refGene_tx=XM_019836723.2;refGene_gn=LOC105260
819;refGene_tx=XM_023258513.1;refGene_gn=LOC105260819;refGene_tx=XM_02
3258514.1
C1 30760909 30761071
refGene_gn=LOC105260819;refGene_tx=XM_019836723.2;refGene_gn=LOC105260
819;refGene_tx=XM_023258513.1;refGene_gn=LOC105260819;refGene_tx=XM_02
3258514.1
C1 30763071 30763089 refGene_gn=PABPC4;refGene_tx=XM_006934591.3
C1 30764105 30764195
refGene_gn=PABPC4;refGene_tx=XM_003989886.4;refGene_gn=PABPC4;refGene_
tx=XM_003989887.4;refGene_gn=PABPC4;refGene_tx=XM_003989888.4;refGene_
gn=PABPC4;refGene_tx=XM_006934591.3;refGene_gn=PABPC4;refGene_tx=XM_00
6934592.3;refGene_gn=PABPC4;refGene_tx=XM_006934593.3;refGene_gn=PABPC
4;refGene_tx=XM_019836725.2;refGene_gn=PABPC4;refGene_tx=XM_019836726.
2
C1 30764415 30764546
refGene_gn=PABPC4;refGene_tx=XM_003989886.4;refGene_gn=PABPC4;refGene_
tx=XM_003989887.4;refGene_gn=PABPC4;refGene_tx=XM_003989888.4;refGene_
gn=PABPC4;refGene_tx=XM_006934591.3;refGene_gn=PABPC4;refGene_tx=XM_00
6934592.3;refGene_gn=PABPC4;refGene_tx=XM_006934593.3;refGene_gn=PABPC
4;refGene_tx=XM_019836725.2;refGene_gn=PABPC4;refGene_tx=XM_019836726.
2
C1 30764631 30764725
refGene_gn=PABPC4;refGene_tx=XM_003989886.4;refGene_gn=PABPC4;refGene_
tx=XM_003989887.4;refGene_gn=PABPC4;refGene_tx=XM_003989888.4;refGene_
gn=PABPC4;refGene_tx=XM_006934591.3;refGene_gn=PABPC4;refGene_tx=XM_00
6934592.3;refGene_gn=PABPC4;refGene_tx=XM_006934593.3;refGene_gn=PABPC
4;refGene_tx=XM_019836725.2;refGene_gn=PABPC4;refGene_tx=XM_019836726.
2
C1 30766111 30766239
refGene_gn=PABPC4;refGene_tx=XM_003989886.4;refGene_gn=PABPC4;refGene_
tx=XM_003989887.4;refGene_gn=PABPC4;refGene_tx=XM_003989888.4;refGene_
gn=PABPC4;refGene_tx=XM_006934591.3;refGene_gn=PABPC4;refGene_tx=XM_00
6934592.3;refGene_gn=PABPC4;refGene_tx=XM_006934593.3;refGene_gn=PABPC
4;refGene_tx=XM_019836725.2;refGene_gn=PABPC4;refGene_tx=XM_019836726.
2
C1 30766333 30766420
refGene_gn=PABPC4;refGene_tx=XM_003989886.4;refGene_gn=PABPC4;refGene_
tx=XM_003989887.4;refGene_gn=PABPC4;refGene_tx=XM_006934591.3;refGene_
gn=PABPC4;refGene_tx=XM_006934592.3;refGene_gn=PABPC4;refGene_tx=XM_01
9836725.2
C1 30766923 30767043
refGene_gn=PABPC4;refGene_tx=XM_003989886.4;refGene_gn=PABPC4;refGene_
tx=XM_003989887.4;refGene_gn=PABPC4;refGene_tx=XM_003989888.4;refGene_
gn=PABPC4;refGene_tx=XM_006934591.3;refGene_gn=PABPC4;refGene_tx=XM_00
6934592.3;refGene_gn=PABPC4;refGene_tx=XM_006934593.3;refGene_gn=PABPC
4;refGene_tx=XM_019836725.2;refGene_gn=PABPC4;refGene_tx=XM_019836726.
2
C1 30767186 30767274
refGene_gn=PABPC4;refGene_tx=XM_003989886.4;refGene_gn=PABPC4;refGene_

tx=XM_003989887.4;refGene_gn=PABPC4;refGene_tx=XM_003989888.4;refGene_
gn=PABPC4;refGene_tx=XM_006934591.3;refGene_gn=PABPC4;refGene_tx=XM_00
6934592.3;refGene_gn=PABPC4;refGene_tx=XM_006934593.3;refGene_gn=PABPC
4;refGene_tx=XM_019836725.2;refGene_gn=PABPC4;refGene_tx=XM_019836726.
2
C1 30767647 30767920
refGene_gn=PABPC4;refGene_tx=XM_003989886.4;refGene_gn=PABPC4;refGene_
tx=XM_003989887.4;refGene_gn=PABPC4;refGene_tx=XM_003989888.4;refGene_
gn=PABPC4;refGene_tx=XM_006934591.3;refGene_gn=PABPC4;refGene_tx=XM_00
6934592.3;refGene_gn=PABPC4;refGene_tx=XM_006934593.3;refGene_gn=PABPC
4;refGene_tx=XM_019836725.2;refGene_gn=PABPC4;refGene_tx=XM_019836726.
2
C1 30769887 30769983
refGene_gn=PABPC4;refGene_tx=XM_003989886.4;refGene_gn=PABPC4;refGene_
tx=XM_003989887.4;refGene_gn=PABPC4;refGene_tx=XM_003989888.4;refGene_
gn=PABPC4;refGene_tx=XM_006934591.3;refGene_gn=PABPC4;refGene_tx=XM_00
6934592.3;refGene_gn=PABPC4;refGene_tx=XM_006934593.3;refGene_gn=PABPC
4;refGene_tx=XM_019836725.2;refGene_gn=PABPC4;refGene_tx=XM_019836726.
2
C1 30770574 30770712
refGene_gn=PABPC4;refGene_tx=XM_003989886.4;refGene_gn=PABPC4;refGene_
tx=XM_003989887.4;refGene_gn=PABPC4;refGene_tx=XM_003989888.4;refGene_
gn=PABPC4;refGene_tx=XM_006934591.3;refGene_gn=PABPC4;refGene_tx=XM_00
6934592.3;refGene_gn=PABPC4;refGene_tx=XM_006934593.3;refGene_gn=PABPC
4;refGene_tx=XM_019836725.2;refGene_gn=PABPC4;refGene_tx=XM_019836726.
2;refGene_gn=PABPC4;refGene_tx=XM_019836727.2
C1 30771379 30771474
refGene_gn=PABPC4;refGene_tx=XM_003989886.4;refGene_gn=PABPC4;refGene_
tx=XM_003989887.4;refGene_gn=PABPC4;refGene_tx=XM_003989888.4;refGene_
gn=PABPC4;refGene_tx=XM_006934591.3;refGene_gn=PABPC4;refGene_tx=XM_00
6934592.3;refGene_gn=PABPC4;refGene_tx=XM_006934593.3;refGene_gn=PABPC
4;refGene_tx=XM_019836725.2;refGene_gn=PABPC4;refGene_tx=XM_019836726.
2
C1 30771827 30771967
refGene_gn=PABPC4;refGene_tx=XM_003989886.4;refGene_gn=PABPC4;refGene_
tx=XM_003989887.4;refGene_gn=PABPC4;refGene_tx=XM_003989888.4;refGene_
gn=PABPC4;refGene_tx=XM_006934591.3;refGene_gn=PABPC4;refGene_tx=XM_00
6934592.3;refGene_gn=PABPC4;refGene_tx=XM_006934593.3;refGene_gn=PABPC
4;refGene_tx=XM_019836725.2;refGene_gn=PABPC4;refGene_tx=XM_019836726.
2;refGene_gn=PABPC4;refGene_tx=XM_019836727.2
C1 30773212 30773328
refGene_gn=PABPC4;refGene_tx=XM_003989886.4;refGene_gn=PABPC4;refGene_
tx=XM_003989887.4;refGene_gn=PABPC4;refGene_tx=XM_003989888.4;refGene_
gn=PABPC4;refGene_tx=XM_006934591.3;refGene_gn=PABPC4;refGene_tx=XM_00
6934592.3;refGene_gn=PABPC4;refGene_tx=XM_006934593.3;refGene_gn=PABPC
4;refGene_tx=XM_019836725.2;refGene_gn=PABPC4;refGene_tx=XM_019836726.
2;refGene_gn=PABPC4;refGene_tx=XM_019836727.2
C1 30774072 30774266
refGene_gn=PABPC4;refGene_tx=XM_003989886.4;refGene_gn=PABPC4;refGene_
tx=XM_003989887.4;refGene_gn=PABPC4;refGene_tx=XM_003989888.4;refGene_

gn=PABPC4;refGene_tx=XM_006934591.3;refGene_gn=PABPC4;refGene_tx=XM_006934592.3;refGene_gn=PABPC4;refGene_tx=XM_006934593.3;refGene_gn=PABPC4;refGene_tx=XM_019836725.2;refGene_gn=PABPC4;refGene_tx=XM_019836726.2;refGene_gn=PABPC4;refGene_tx=XM_019836727.2
C1 30777039 30777232
refGene_gn=PABPC4;refGene_tx=XM_003989886.4;refGene_gn=PABPC4;refGene_tx=XM_003989887.4;refGene_gn=PABPC4;refGene_tx=XM_003989888.4;refGene_gn=PABPC4;refGene_tx=XM_006934591.3;refGene_gn=PABPC4;refGene_tx=XM_006934592.3;refGene_gn=PABPC4;refGene_tx=XM_006934593.3;refGene_gn=PABPC4;refGene_tx=XM_019836725.2;refGene_gn=PABPC4;refGene_tx=XM_019836726.2;refGene_gn=PABPC4;refGene_tx=XM_019836727.2
C1 30812375 30813049 refGene_gn=HEYL;refGene_tx=XM_011284715.3
C1 30816328 30816410 refGene_gn=HEYL;refGene_tx=XM_011284715.3
C1 30817601 30817685 refGene_gn=HEYL;refGene_tx=XM_011284715.3
C1 30818694 30818761 refGene_gn=HEYL;refGene_tx=XM_011284715.3
C1 30825748 30825828 refGene_gn=HEYL;refGene_tx=XM_011284715.3
C1 30844968 30845334 refGene_gn=NT5C1A;refGene_tx=XM_003989889.5
C1 30846870 30847055 refGene_gn=NT5C1A;refGene_tx=XM_003989889.5
C1 30848958 30849081 refGene_gn=NT5C1A;refGene_tx=XM_003989889.5
C1 30850276 30850406 refGene_gn=NT5C1A;refGene_tx=XM_003989889.5
C1 30850866 30851034 refGene_gn=NT5C1A;refGene_tx=XM_003989889.5
C1 30857763 30857898 refGene_gn=NT5C1A;refGene_tx=XM_003989889.5
C1 30866660 30866858 refGene_gn=HPCAL4;refGene_tx=XM_003989890.5
C1 30868187 30868403 refGene_gn=HPCAL4;refGene_tx=XM_003989890.5
C1 30868709 30868871 refGene_gn=HPCAL4;refGene_tx=XM_003989890.5
C1 30900182 30900213 refGene_gn=PPIE;refGene_tx=XM_003989891.4
C1 30901141 30901240 refGene_gn=PPIE;refGene_tx=XM_003989891.4
C1 30902746 30902790 refGene_gn=PPIE;refGene_tx=XM_003989891.4
C1 30903240 30903267
refGene_gn=PPIE;refGene_tx=XM_003989891.4;refGene_gn=PPIE;refGene_tx=XM_019836728.2;refGene_gn=PPIE;refGene_tx=XM_019836729.2
C1 30904419 30904501
refGene_gn=PPIE;refGene_tx=XM_003989891.4;refGene_gn=PPIE;refGene_tx=XM_019836728.2;refGene_gn=PPIE;refGene_tx=XM_019836729.2;refGene_gn=PPIE;refGene_tx=XM_019836730.2
C1 30905033 30905134
refGene_gn=PPIE;refGene_tx=XM_003989891.4;refGene_gn=PPIE;refGene_tx=XM_019836728.2;refGene_gn=PPIE;refGene_tx=XM_019836729.2;refGene_gn=PPIE;refGene_tx=XM_019836730.2
C1 30907696 30907820
refGene_gn=PPIE;refGene_tx=XM_003989891.4;refGene_gn=PPIE;refGene_tx=XM_019836728.2;refGene_gn=PPIE;refGene_tx=XM_019836729.2;refGene_gn=PPIE;refGene_tx=XM_019836730.2
C1 30911993 30912179
refGene_gn=PPIE;refGene_tx=XM_003989891.4;refGene_gn=PPIE;refGene_tx=XM_019836728.2;refGene_gn=PPIE;refGene_tx=XM_019836729.2;refGene_gn=PPIE;refGene_tx=XM_019836730.2
C1 30913139 30913282
refGene_gn=PPIE;refGene_tx=XM_003989891.4;refGene_gn=PPIE;refGene_tx=XM_019836728.2;refGene_gn=PPIE;refGene_tx=XM_019836729.2;refGene_gn=PPI

E;refGene_tx=XM_019836730.2
C1 30913463 30913532
refGene_gn=PPIE;refGene_tx=XM_003989891.4;refGene_gn=PPIE;refGene_tx=XM_019836728.2;refGene_gn=PPIE;refGene_tx=XM_019836729.2;refGene_gn=PPIE;refGene_tx=XM_019836730.2
C1 30919744 30919894 refGene_gn=BMP8B;refGene_tx=XM_023258517.1
C1 30923000 30923303
refGene_gn=BMP8B;refGene_tx=XM_023258516.1;refGene_gn=BMP8B;refGene_tx=XM_023258517.1
C1 30923865 30923945
refGene_gn=BMP8B;refGene_tx=XM_023258516.1;refGene_gn=BMP8B;refGene_tx=XM_023258517.1
C1 30924703 30924898
refGene_gn=BMP8B;refGene_tx=XM_023258516.1;refGene_gn=BMP8B;refGene_tx=XM_023258517.1
C1 30927244 30928536 refGene_gn=OXCT2;refGene_tx=XM_023258515.1
C1 30928538 30928821 refGene_gn=OXCT2;refGene_tx=XM_023258515.1
C1 30931287 30931436
refGene_gn=BMP8B;refGene_tx=XM_023258516.1;refGene_gn=BMP8B;refGene_tx=XM_023258517.1
C1 30931859 30932049
refGene_gn=BMP8B;refGene_tx=XM_023258516.1;refGene_gn=BMP8B;refGene_tx=XM_023258517.1
C1 30948345 30948679
refGene_gn=BMP8B;refGene_tx=XM_023258516.1;refGene_gn=BMP8B;refGene_tx=XM_023258517.1
C1 31022733 31022903
refGene_gn=TRIT1;refGene_tx=XM_003989893.5;refGene_gn=TRIT1;refGene_tx=XM_006934595.4;refGene_gn=TRIT1;refGene_tx=XM_006934596.4;refGene_gn=TRIT1;refGene_tx=XM_006934597.4;refGene_gn=TRIT1;refGene_tx=XM_019836731.2;refGene_gn=TRIT1;refGene_tx=XM_019836732.2
C1 31025377 31025495
refGene_gn=TRIT1;refGene_tx=XM_003989893.5;refGene_gn=TRIT1;refGene_tx=XM_006934595.4;refGene_gn=TRIT1;refGene_tx=XM_006934597.4;refGene_gn=TRIT1;refGene_tx=XM_019836731.2;refGene_gn=TRIT1;refGene_tx=XM_019836732.2
C1 31025831 31025941
refGene_gn=TRIT1;refGene_tx=XM_003989893.5;refGene_gn=TRIT1;refGene_tx=XM_006934595.4;refGene_gn=TRIT1;refGene_tx=XM_006934597.4;refGene_gn=TRIT1;refGene_tx=XM_019836731.2;refGene_gn=TRIT1;refGene_tx=XM_019836732.2
C1 31027957 31027986 refGene_gn=TRIT1;refGene_tx=XM_019836733.2
C1 31028736 31028814
refGene_gn=TRIT1;refGene_tx=XM_003989893.5;refGene_gn=TRIT1;refGene_tx=XM_006934595.4;refGene_gn=TRIT1;refGene_tx=XM_006934596.4;refGene_gn=TRIT1;refGene_tx=XM_006934597.4;refGene_gn=TRIT1;refGene_tx=XM_019836731.2;refGene_gn=TRIT1;refGene_tx=XM_019836732.2;refGene_gn=TRIT1;refGene_tx=XM_019836733.2
C1 31029061 31029174
refGene_gn=TRIT1;refGene_tx=XM_003989893.5;refGene_gn=TRIT1;refGene_tx

=XM_006934595.4;refGene_gn=TRIT1;refGene_tx=XM_006934596.4;refGene_gn=
TRIT1;refGene_tx=XM_006934597.4;refGene_gn=TRIT1;refGene_tx=XM_0198367
31.2;refGene_gn=TRIT1;refGene_tx=XM_019836732.2;refGene_gn=TRIT1;refGe
ne_tx=XM_019836733.2
C1 31029544 31029656
refGene_gn=TRIT1;refGene_tx=XM_003989893.5;refGene_gn=TRIT1;refGene_tx
=XM_006934595.4;refGene_gn=TRIT1;refGene_tx=XM_006934596.4;refGene_gn=
TRIT1;refGene_tx=XM_006934597.4;refGene_gn=TRIT1;refGene_tx=XM_0198367
31.2;refGene_gn=TRIT1;refGene_tx=XM_019836732.2;refGene_gn=TRIT1;refGe
ne_tx=XM_019836733.2
C1 31030672 31030807
refGene_gn=TRIT1;refGene_tx=XM_006934595.4;refGene_gn=TRIT1;refGene_tx
=XM_006934596.4;refGene_gn=TRIT1;refGene_tx=XM_006934597.4;refGene_gn=
TRIT1;refGene_tx=XM_019836731.2;refGene_gn=TRIT1;refGene_tx=XM_0198367
32.2;refGene_gn=TRIT1;refGene_tx=XM_019836733.2
C1 31031391 31031534
refGene_gn=TRIT1;refGene_tx=XM_003989893.5;refGene_gn=TRIT1;refGene_tx
=XM_006934595.4;refGene_gn=TRIT1;refGene_tx=XM_006934596.4;refGene_gn=
TRIT1;refGene_tx=XM_006934597.4;refGene_gn=TRIT1;refGene_tx=XM_0198367
31.2;refGene_gn=TRIT1;refGene_tx=XM_019836732.2;refGene_gn=TRIT1;refGe
ne_tx=XM_019836733.2
C1 31034164 31034310
refGene_gn=TRIT1;refGene_tx=XM_003989893.5;refGene_gn=TRIT1;refGene_tx
=XM_006934595.4;refGene_gn=TRIT1;refGene_tx=XM_006934596.4;refGene_gn=
TRIT1;refGene_tx=XM_006934597.4;refGene_gn=TRIT1;refGene_tx=XM_0198367
31.2;refGene_gn=TRIT1;refGene_tx=XM_019836732.2;refGene_gn=TRIT1;refGe
ne_tx=XM_019836733.2
C1 31035027 31035126
refGene_gn=TRIT1;refGene_tx=XM_003989893.5;refGene_gn=TRIT1;refGene_tx
=XM_006934595.4;refGene_gn=TRIT1;refGene_tx=XM_006934596.4;refGene_gn=
TRIT1;refGene_tx=XM_019836731.2;refGene_gn=TRIT1;refGene_tx=XM_0198367
32.2;refGene_gn=TRIT1;refGene_tx=XM_019836733.2
C1 31039049 31039190
refGene_gn=TRIT1;refGene_tx=XM_003989893.5;refGene_gn=TRIT1;refGene_tx
=XM_006934595.4;refGene_gn=TRIT1;refGene_tx=XM_006934596.4;refGene_gn=
TRIT1;refGene_tx=XM_019836731.2;refGene_gn=TRIT1;refGene_tx=XM_0198367
32.2;refGene_gn=TRIT1;refGene_tx=XM_019836733.2
C1 31063472 31063538 refGene_gn=TRIT1;refGene_tx=XM_019836732.2
C1 31064390 31064564
refGene_gn=TRIT1;refGene_tx=XM_003989893.5;refGene_gn=TRIT1;refGene_tx
=XM_006934595.4;refGene_gn=TRIT1;refGene_tx=XM_006934596.4;refGene_gn=
TRIT1;refGene_tx=XM_006934597.4;refGene_gn=TRIT1;refGene_tx=XM_0198367
31.2;refGene_gn=TRIT1;refGene_tx=XM_019836733.2
C1 31076864 31077463
refGene_gn=MYCL;refGene_tx=XM_023258520.1;refGene_gn=MYCL;refGene_tx=X
M_023258521.1
C1 31080405 31080910
refGene_gn=MYCL;refGene_tx=XM_023258520.1;refGene_gn=MYCL;refGene_tx=X
M_023258521.1
C1 31081247 31081328 refGene_gn=MYCL;refGene_tx=XM_023258520.1

C1 31115586 31115679
refGene_gn=MFSD2A;refGene_tx=XM_023258518.1;refGene_gn=MFSD2A;refGene_tx=XM_023258519.1
C1 31117245 31117377
refGene_gn=MFSD2A;refGene_tx=XM_023258518.1;refGene_gn=MFSD2A;refGene_tx=XM_023258519.1
C1 31118834 31118959
refGene_gn=MFSD2A;refGene_tx=XM_023258518.1;refGene_gn=MFSD2A;refGene_tx=XM_023258519.1
C1 31123000 31123124
refGene_gn=MFSD2A;refGene_tx=XM_023258518.1;refGene_gn=MFSD2A;refGene_tx=XM_023258519.1
C1 31123258 31123337
refGene_gn=MFSD2A;refGene_tx=XM_023258518.1;refGene_gn=MFSD2A;refGene_tx=XM_023258519.1
C1 31123693 31123851
refGene_gn=MFSD2A;refGene_tx=XM_023258518.1;refGene_gn=MFSD2A;refGene_tx=XM_023258519.1
C1 31124246 31124337
refGene_gn=MFSD2A;refGene_tx=XM_023258518.1;refGene_gn=MFSD2A;refGene_tx=XM_023258519.1
C1 31124456 31124578
refGene_gn=MFSD2A;refGene_tx=XM_023258518.1;refGene_gn=MFSD2A;refGene_tx=XM_023258519.1
C1 31124812 31124896
refGene_gn=MFSD2A;refGene_tx=XM_023258518.1;refGene_gn=MFSD2A;refGene_tx=XM_023258519.1
C1 31125323 31125407
refGene_gn=MFSD2A;refGene_tx=XM_023258518.1;refGene_gn=MFSD2A;refGene_tx=XM_023258519.1
C1 31125505 31125618
refGene_gn=MFSD2A;refGene_tx=XM_023258518.1;refGene_gn=MFSD2A;refGene_tx=XM_023258519.1
C1 31126026 31126170
refGene_gn=MFSD2A;refGene_tx=XM_023258518.1;refGene_gn=MFSD2A;refGene_tx=XM_023258519.1
C1 31126261 31126438
refGene_gn=MFSD2A;refGene_tx=XM_023258518.1;refGene_gn=MFSD2A;refGene_tx=XM_023258519.1
C1 31127308 31127372
refGene_gn=MFSD2A;refGene_tx=XM_023258518.1;refGene_gn=MFSD2A;refGene_tx=XM_023258519.1
C1 31192950 31193062
refGene_gn=CAP1;refGene_tx=XM_011284711.3;refGene_gn=CAP1;refGene_tx=XM_019836734.2;refGene_gn=CAP1;refGene_tx=XM_019836735.2;refGene_gn=CAP1;refGene_tx=XM_019836736.2;refGene_gn=CAP1;refGene_tx=XM_019836737.2
C1 31193668 31193772
refGene_gn=CAP1;refGene_tx=XM_011284711.3;refGene_gn=CAP1;refGene_tx=XM_019836734.2;refGene_gn=CAP1;refGene_tx=XM_019836735.2;refGene_gn=CAP1;refGene_tx=XM_019836736.2;refGene_gn=CAP1;refGene_tx=XM_019836737.2

C1 31195195 31195273
refGene_gn=CAP1;refGene_tx=XM_011284711.3;refGene_gn=CAP1;refGene_tx=XM_019836734.2;refGene_gn=CAP1;refGene_tx=XM_019836735.2;refGene_gn=CAP1;refGene_tx=XM_019836736.2;refGene_gn=CAP1;refGene_tx=XM_019836737.2
C1 31197265 31197409
refGene_gn=CAP1;refGene_tx=XM_011284711.3;refGene_gn=CAP1;refGene_tx=XM_019836734.2;refGene_gn=CAP1;refGene_tx=XM_019836735.2;refGene_gn=CAP1;refGene_tx=XM_019836736.2;refGene_gn=CAP1;refGene_tx=XM_019836737.2
C1 31197510 31197596
refGene_gn=CAP1;refGene_tx=XM_011284711.3;refGene_gn=CAP1;refGene_tx=XM_019836734.2;refGene_gn=CAP1;refGene_tx=XM_019836735.2;refGene_gn=CAP1;refGene_tx=XM_019836736.2;refGene_gn=CAP1;refGene_tx=XM_019836737.2
C1 31198930 31199036
refGene_gn=CAP1;refGene_tx=XM_011284711.3;refGene_gn=CAP1;refGene_tx=XM_019836734.2;refGene_gn=CAP1;refGene_tx=XM_019836735.2;refGene_gn=CAP1;refGene_tx=XM_019836736.2;refGene_gn=CAP1;refGene_tx=XM_019836737.2
C1 31200202 31200380
refGene_gn=CAP1;refGene_tx=XM_011284711.3;refGene_gn=CAP1;refGene_tx=XM_019836734.2;refGene_gn=CAP1;refGene_tx=XM_019836735.2;refGene_gn=CAP1;refGene_tx=XM_019836736.2;refGene_gn=CAP1;refGene_tx=XM_019836737.2
C1 31202664 31202849
refGene_gn=CAP1;refGene_tx=XM_011284711.3;refGene_gn=CAP1;refGene_tx=XM_019836734.2;refGene_gn=CAP1;refGene_tx=XM_019836735.2;refGene_gn=CAP1;refGene_tx=XM_019836736.2;refGene_gn=CAP1;refGene_tx=XM_019836737.2
C1 31203291 31203415
refGene_gn=CAP1;refGene_tx=XM_011284711.3;refGene_gn=CAP1;refGene_tx=XM_019836734.2;refGene_gn=CAP1;refGene_tx=XM_019836735.2;refGene_gn=CAP1;refGene_tx=XM_019836736.2;refGene_gn=CAP1;refGene_tx=XM_019836737.2
C1 31203539 31203622
refGene_gn=CAP1;refGene_tx=XM_011284711.3;refGene_gn=CAP1;refGene_tx=XM_019836734.2;refGene_gn=CAP1;refGene_tx=XM_019836735.2;refGene_gn=CAP1;refGene_tx=XM_019836736.2;refGene_gn=CAP1;refGene_tx=XM_019836737.2
C1 31203946 31204090
refGene_gn=CAP1;refGene_tx=XM_011284711.3;refGene_gn=CAP1;refGene_tx=XM_019836734.2;refGene_gn=CAP1;refGene_tx=XM_019836735.2;refGene_gn=CAP1;refGene_tx=XM_019836736.2;refGene_gn=CAP1;refGene_tx=XM_019836737.2
C1 31204687 31204771
refGene_gn=CAP1;refGene_tx=XM_011284711.3;refGene_gn=CAP1;refGene_tx=XM_019836734.2;refGene_gn=CAP1;refGene_tx=XM_019836735.2;refGene_gn=CAP1;refGene_tx=XM_019836736.2;refGene_gn=CAP1;refGene_tx=XM_019836737.2
C1 31207474 31207597 refGene_gn=PPT1;refGene_tx=XM_003989895.5
C1 31208801 31208873 refGene_gn=PPT1;refGene_tx=XM_003989895.5
C1 31210150 31210249 refGene_gn=PPT1;refGene_tx=XM_003989895.5
C1 31211503 31211594 refGene_gn=PPT1;refGene_tx=XM_003989895.5
C1 31225662 31225765 refGene_gn=PPT1;refGene_tx=XM_003989895.5
C1 31227569 31227640 refGene_gn=PPT1;refGene_tx=XM_003989895.5
C1 31227950 31228078 refGene_gn=PPT1;refGene_tx=XM_003989895.5
C1 31228300 31228410 refGene_gn=PPT1;refGene_tx=XM_003989895.5
C1 31232653 31232777 refGene_gn=PPT1;refGene_tx=XM_003989895.5
C1 31276864 31277101 refGene_gn=RLF;refGene_tx=XM_003989897.3

C1	31310615	31310770	refGene_gn=RLF;refGene_tx=XM_003989897.3
C1	31311848	31311930	refGene_gn=RLF;refGene_tx=XM_003989897.3
C1	31315466	31315599	refGene_gn=RLF;refGene_tx=XM_003989897.3
C1	31322554	31322757	refGene_gn=RLF;refGene_tx=XM_003989897.3
C1	31340593	31340730	refGene_gn=RLF;refGene_tx=XM_003989897.3
C1	31351257	31351399	refGene_gn=RLF;refGene_tx=XM_003989897.3
C1	31358204	31362848	refGene_gn=RLF;refGene_tx=XM_003989897.3
C1	31370418	31370661	refGene_gn=TMCO2;refGene_tx=XM_003989946.5
C1	31373458	31373773	refGene_gn=TMCO2;refGene_tx=XM_003989946.5
C1	31380499	31380622
refGene_gn=ZMPSTE24;refGene_tx=XM_003989898.5;refGene_gn=ZMPSTE24;refGene_tx=XM_019836740.2;refGene_gn=ZMPSTE24;refGene_tx=XM_019836741.2
C1	31383016	31383163
refGene_gn=ZMPSTE24;refGene_tx=XM_003989898.5;refGene_gn=ZMPSTE24;refGene_tx=XM_019836741.2
C1	31390045	31390132
refGene_gn=ZMPSTE24;refGene_tx=XM_003989898.5;refGene_gn=ZMPSTE24;refGene_tx=XM_019836740.2;refGene_gn=ZMPSTE24;refGene_tx=XM_019836741.2
C1	31391770	31391887
refGene_gn=ZMPSTE24;refGene_tx=XM_003989898.5;refGene_gn=ZMPSTE24;refGene_tx=XM_019836740.2;refGene_gn=ZMPSTE24;refGene_tx=XM_019836741.2
C1	31392691	31392844
refGene_gn=ZMPSTE24;refGene_tx=XM_003989898.5;refGene_gn=ZMPSTE24;refGene_tx=XM_019836740.2;refGene_gn=ZMPSTE24;refGene_tx=XM_019836741.2
C1	31394373	31394515
refGene_gn=ZMPSTE24;refGene_tx=XM_003989898.5;refGene_gn=ZMPSTE24;refGene_tx=XM_019836740.2;refGene_gn=ZMPSTE24;refGene_tx=XM_019836741.2
C1	31400543	31400728
refGene_gn=ZMPSTE24;refGene_tx=XM_003989898.5;refGene_gn=ZMPSTE24;refGene_tx=XM_019836740.2;refGene_gn=ZMPSTE24;refGene_tx=XM_019836741.2
C1	31411988	31412093
refGene_gn=ZMPSTE24;refGene_tx=XM_003989898.5;refGene_gn=ZMPSTE24;refGene_tx=XM_019836740.2;refGene_gn=ZMPSTE24;refGene_tx=XM_019836741.2
C1	31414952	31415096
refGene_gn=ZMPSTE24;refGene_tx=XM_003989898.5;refGene_gn=ZMPSTE24;refGene_tx=XM_019836740.2;refGene_gn=ZMPSTE24;refGene_tx=XM_019836741.2
C1	31418572	31418797
refGene_gn=ZMPSTE24;refGene_tx=XM_003989898.5;refGene_gn=ZMPSTE24;refGene_tx=XM_019836740.2
C1	31420361	31420403	refGene_gn=ZMPSTE24;refGene_tx=XM_019836741.2
C1	31425942	31426142	refGene_gn=COL9A2;refGene_tx=XM_023258522.1
C1	31426597	31426675	refGene_gn=COL9A2;refGene_tx=XM_023258522.1
C1	31427500	31427689	refGene_gn=COL9A2;refGene_tx=XM_023258522.1
C1	31428012	31428067	refGene_gn=COL9A2;refGene_tx=XM_023258522.1
C1	31428300	31428447	refGene_gn=COL9A2;refGene_tx=XM_023258522.1
C1	31428661	31428694	refGene_gn=COL9A2;refGene_tx=XM_023258522.1
C1	31428796	31428841	refGene_gn=COL9A2;refGene_tx=XM_023258522.1
C1	31428945	31428981	refGene_gn=COL9A2;refGene_tx=XM_023258522.1
C1	31429243	31429315	refGene_gn=COL9A2;refGene_tx=XM_023258522.1
C1	31429404	31429458	refGene_gn=COL9A2;refGene_tx=XM_023258522.1

C1 31429774 31429828 refGene_gn=COL9A2;refGene_tx=XM_023258522.1
C1 31430435 31430489 refGene_gn=COL9A2;refGene_tx=XM_023258522.1
C1 31430848 31430893 refGene_gn=COL9A2;refGene_tx=XM_023258522.1
C1 31431860 31431914 refGene_gn=COL9A2;refGene_tx=XM_023258522.1
C1 31432147 31432201 refGene_gn=COL9A2;refGene_tx=XM_023258522.1
C1 31432615 31432669 refGene_gn=COL9A2;refGene_tx=XM_023258522.1
C1 31434268 31434322 refGene_gn=COL9A2;refGene_tx=XM_023258522.1
C1 31434440 31434494 refGene_gn=COL9A2;refGene_tx=XM_023258522.1
C1 31434591 31434645 refGene_gn=COL9A2;refGene_tx=XM_023258522.1
C1 31434991 31435045 refGene_gn=COL9A2;refGene_tx=XM_023258522.1
C1 31435348 31435402 refGene_gn=COL9A2;refGene_tx=XM_023258522.1
C1 31435484 31435541 refGene_gn=COL9A2;refGene_tx=XM_023258522.1
C1 31435726 31435774 refGene_gn=COL9A2;refGene_tx=XM_023258522.1
C1 31435880 31435934 refGene_gn=COL9A2;refGene_tx=XM_023258522.1
C1 31436283 31436337 refGene_gn=COL9A2;refGene_tx=XM_023258522.1
C1 31436658 31436682 refGene_gn=COL9A2;refGene_tx=XM_023258522.1
C1 31436787 31436823 refGene_gn=COL9A2;refGene_tx=XM_023258522.1
C1 31436935 31436989 refGene_gn=COL9A2;refGene_tx=XM_023258522.1
C1 31437962 31438025 refGene_gn=COL9A2;refGene_tx=XM_023258522.1
C1 31438111 31438147 refGene_gn=COL9A2;refGene_tx=XM_023258522.1
C1 31439387 31439462 refGene_gn=COL9A2;refGene_tx=XM_023258522.1
C1 31440674 31440746 refGene_gn=COL9A2;refGene_tx=XM_023258522.1
C1 31474228 31474331 refGene_gn=SMAP2;refGene_tx=XM_023258523.1
C1 31506430 31506564 refGene_gn=SMAP2;refGene_tx=XM_023258523.1
C1 31507839 31507925 refGene_gn=SMAP2;refGene_tx=XM_023258523.1
C1 31508897 31508976 refGene_gn=SMAP2;refGene_tx=XM_023258523.1
C1 31512383 31512470 refGene_gn=SMAP2;refGene_tx=XM_023258523.1
C1 31513502 31513584 refGene_gn=SMAP2;refGene_tx=XM_023258523.1
C1 31514829 31514939 refGene_gn=SMAP2;refGene_tx=XM_023258523.1
C1 31515746 31515912 refGene_gn=SMAP2;refGene_tx=XM_023258523.1
C1 31516370 31516687 refGene_gn=SMAP2;refGene_tx=XM_023258523.1
C1 31521458 31521584 refGene_gn=SMAP2;refGene_tx=XM_023258523.1
C1 31547304 31547431
refGene_gn=ZFP69B;refGene_tx=XM_006934599.3;refGene_gn=ZFP69B;refGene_
tx=XM_019836751.2
C1 31550376 31550462
refGene_gn=ZFP69B;refGene_tx=XM_006934599.3;refGene_gn=ZFP69B;refGene_
tx=XM_019836751.2
C1 31552176 31552303
refGene_gn=ZFP69B;refGene_tx=XM_006934599.3;refGene_gn=ZFP69B;refGene_
tx=XM_019836751.2;refGene_gn=ZFP69B;refGene_tx=XM_019836752.1
C1 31552557 31552653
refGene_gn=ZFP69B;refGene_tx=XM_006934599.3;refGene_gn=ZFP69B;refGene_
tx=XM_019836752.1
C1 31556721 31557887
refGene_gn=ZFP69B;refGene_tx=XM_006934599.3;refGene_gn=ZFP69B;refGene_
tx=XM_019836751.2;refGene_gn=ZFP69B;refGene_tx=XM_019836752.1
C1 31567947 31568074
refGene_gn=ZFP69;refGene_tx=XM_003989900.4;refGene_gn=ZFP69;refGene_tx
=XM_006934600.3;refGene_gn=ZFP69;refGene_tx=XM_011284710.2;refGene_gn=

ZFP69;refGene_tx=XM_023258524.1
C1 31568088 31568347
refGene_gn=ZFP69;refGene_tx=XM_019836746.1;refGene_gn=ZFP69;refGene_tx=XM_019836747.1;refGene_gn=ZFP69;refGene_tx=XM_019836748.1
C1 31569338 31569424
refGene_gn=ZFP69;refGene_tx=XM_003989900.4;refGene_gn=ZFP69;refGene_tx=XM_006934600.3;refGene_gn=ZFP69;refGene_tx=XM_011284710.2;refGene_gn=ZFP69;refGene_tx=XM_019836746.1;refGene_gn=ZFP69;refGene_tx=XM_019836747.1;refGene_gn=ZFP69;refGene_tx=XM_019836748.1;refGene_gn=ZFP69;refGene_tx=XM_023258524.1
C1 31574955 31575082
refGene_gn=ZFP69;refGene_tx=XM_003989900.4;refGene_gn=ZFP69;refGene_tx=XM_006934600.3;refGene_gn=ZFP69;refGene_tx=XM_011284710.2;refGene_gn=ZFP69;refGene_tx=XM_019836746.1;refGene_gn=ZFP69;refGene_tx=XM_019836747.1;refGene_gn=ZFP69;refGene_tx=XM_019836748.1;refGene_gn=ZFP69;refGene_tx=XM_019836750.2;refGene_gn=ZFP69;refGene_tx=XM_023258524.1
C1 31575404 31575503
refGene_gn=ZFP69;refGene_tx=XM_003989900.4;refGene_gn=ZFP69;refGene_tx=XM_006934600.3;refGene_gn=ZFP69;refGene_tx=XM_011284710.2;refGene_gn=ZFP69;refGene_tx=XM_019836746.1;refGene_gn=ZFP69;refGene_tx=XM_019836747.1;refGene_gn=ZFP69;refGene_tx=XM_019836750.2;refGene_gn=ZFP69;refGene_tx=XM_023258524.1
C1 31580467 31581600
refGene_gn=ZFP69;refGene_tx=XM_003989900.4;refGene_gn=ZFP69;refGene_tx=XM_006934600.3;refGene_gn=ZFP69;refGene_tx=XM_011284710.2;refGene_gn=ZFP69;refGene_tx=XM_019836746.1;refGene_gn=ZFP69;refGene_tx=XM_019836747.1;refGene_gn=ZFP69;refGene_tx=XM_019836748.1;refGene_gn=ZFP69;refGene_tx=XM_019836750.2;refGene_gn=ZFP69;refGene_tx=XM_023258524.1
C1 31595873 31596995
refGene_gn=EXO5;refGene_tx=XM_019836756.2;refGene_gn=EXO5;refGene_tx=XM_019836757.2
C1 31608801 31608816 refGene_gn=ZNF684;refGene_tx=XM_006934602.4
C1 31614236 31614363 refGene_gn=ZNF684;refGene_tx=XM_006934602.4
C1 31614693 31614789 refGene_gn=ZNF684;refGene_tx=XM_006934602.4
C1 31620545 31620931 refGene_gn=ZNF684;refGene_tx=XM_006934602.4
C1 31620932 31621451 refGene_gn=ZNF684;refGene_tx=XM_006934602.4
C1 31666384 31666408
refGene_gn=LOC101091386;refGene_tx=XM_019836758.2
C1 31667066 31667193
refGene_gn=LOC101091386;refGene_tx=XM_019836758.2
C1 31671655 31672263
refGene_gn=LOC101091386;refGene_tx=XM_019836758.2
C1 31749287 31749500
refGene_gn=RIMS3;refGene_tx=XM_003989902.5;refGene_gn=RIMS3;refGene_tx=XM_019836759.2;refGene_gn=RIMS3;refGene_tx=XM_019836761.2;refGene_gn=RIMS3;refGene_tx=XM_023258525.1
C1 31750703 31750805 refGene_gn=RIMS3;refGene_tx=XM_023258526.1
C1 31751060 31751200
refGene_gn=RIMS3;refGene_tx=XM_003989902.5;refGene_gn=RIMS3;refGene_tx=XM_019836759.2;refGene_gn=RIMS3;refGene_tx=XM_019836761.2;refGene_gn=

RIMS3;refGene_tx=XM_023258525.1;refGene_gn=RIMS3;refGene_tx=XM_0232585
26.1
C1	31751531	31751633
refGene_gn=RIMS3;refGene_tx=XM_003989902.5;refGene_gn=RIMS3;refGene_tx
=XM_019836759.2;refGene_gn=RIMS3;refGene_tx=XM_019836761.2;refGene_gn=
RIMS3;refGene_tx=XM_023258525.1;refGene_gn=RIMS3;refGene_tx=XM_0232585
26.1
C1	31753823	31753936
refGene_gn=RIMS3;refGene_tx=XM_003989902.5;refGene_gn=RIMS3;refGene_tx
=XM_019836759.2;refGene_gn=RIMS3;refGene_tx=XM_019836761.2;refGene_gn=
RIMS3;refGene_tx=XM_023258525.1;refGene_gn=RIMS3;refGene_tx=XM_0232585
26.1
C1	31756322	31756464
refGene_gn=RIMS3;refGene_tx=XM_003989902.5;refGene_gn=RIMS3;refGene_tx
=XM_019836759.2;refGene_gn=RIMS3;refGene_tx=XM_019836761.2;refGene_gn=
RIMS3;refGene_tx=XM_023258525.1;refGene_gn=RIMS3;refGene_tx=XM_0232585
26.1
C1	31761857	31762074
refGene_gn=RIMS3;refGene_tx=XM_003989902.5;refGene_gn=RIMS3;refGene_tx
=XM_019836759.2;refGene_gn=RIMS3;refGene_tx=XM_019836761.2;refGene_gn=
RIMS3;refGene_tx=XM_023258525.1;refGene_gn=RIMS3;refGene_tx=XM_0232585
26.1
C1	31839851	31839956
refGene_gn=NFYC;refGene_tx=XM_019836763.2;refGene_gn=NFYC;refGene_tx=X
M_019836764.2;refGene_gn=NFYC;refGene_tx=XM_023258527.1;refGene_gn=NFY
C;refGene_tx=XM_023258528.1;refGene_gn=NFYC;refGene_tx=XM_023258529.1
C1	31847385	31847457
refGene_gn=NFYC;refGene_tx=XM_019836763.2;refGene_gn=NFYC;refGene_tx=X
M_019836764.2;refGene_gn=NFYC;refGene_tx=XM_023258527.1;refGene_gn=NFY
C;refGene_tx=XM_023258528.1;refGene_gn=NFYC;refGene_tx=XM_023258529.1
C1	31849186	31849300
refGene_gn=NFYC;refGene_tx=XM_019836763.2;refGene_gn=NFYC;refGene_tx=X
M_019836764.2;refGene_gn=NFYC;refGene_tx=XM_023258528.1
C1	31852173	31852269
refGene_gn=NFYC;refGene_tx=XM_019836763.2;refGene_gn=NFYC;refGene_tx=X
M_019836764.2;refGene_gn=NFYC;refGene_tx=XM_023258527.1;refGene_gn=NFY
C;refGene_tx=XM_023258528.1;refGene_gn=NFYC;refGene_tx=XM_023258529.1
C1	31857262	31857436
refGene_gn=NFYC;refGene_tx=XM_019836763.2;refGene_gn=NFYC;refGene_tx=X
M_019836764.2;refGene_gn=NFYC;refGene_tx=XM_023258527.1;refGene_gn=NFY
C;refGene_tx=XM_023258528.1;refGene_gn=NFYC;refGene_tx=XM_023258529.1
C1	31860936	31861095
refGene_gn=NFYC;refGene_tx=XM_019836763.2;refGene_gn=NFYC;refGene_tx=X
M_019836764.2;refGene_gn=NFYC;refGene_tx=XM_023258527.1;refGene_gn=NFY
C;refGene_tx=XM_023258528.1;refGene_gn=NFYC;refGene_tx=XM_023258529.1
C1	31864320	31864428
refGene_gn=NFYC;refGene_tx=XM_019836763.2;refGene_gn=NFYC;refGene_tx=X
M_019836764.2;refGene_gn=NFYC;refGene_tx=XM_023258527.1;refGene_gn=NFY
C;refGene_tx=XM_023258528.1;refGene_gn=NFYC;refGene_tx=XM_023258529.1
C1	31866883	31866943

refGene_gn=NFYC;refGene_tx=XM_019836763.2;refGene_gn=NFYC;refGene_tx=XM_019836764.2;refGene_gn=NFYC;refGene_tx=XM_023258527.1;refGene_gn=NFYC;refGene_tx=XM_023258528.1;refGene_gn=NFYC;refGene_tx=XM_023258529.1
C1 31867952 31868384
refGene_gn=NFYC;refGene_tx=XM_019836763.2;refGene_gn=NFYC;refGene_tx=XM_019836764.2;refGene_gn=NFYC;refGene_tx=XM_023258527.1;refGene_gn=NFYC;refGene_tx=XM_023258528.1;refGene_gn=NFYC;refGene_tx=XM_023258529.1
C1 31880000 31880314
refGene_gn=KCNQ4;refGene_tx=XM_023258530.1;refGene_gn=KCNQ4;refGene_tx=XM_023258531.1
C1 31910186 31910277
refGene_gn=KCNQ4;refGene_tx=XM_023258530.1;refGene_gn=KCNQ4;refGene_tx=XM_023258531.1
C1 31910943 31911070
refGene_gn=KCNQ4;refGene_tx=XM_023258530.1;refGene_gn=KCNQ4;refGene_tx=XM_023258531.1;refGene_gn=KCNQ4;refGene_tx=XM_023258532.1
C1 31911362 31911538
refGene_gn=KCNQ4;refGene_tx=XM_023258530.1;refGene_gn=KCNQ4;refGene_tx=XM_023258531.1;refGene_gn=KCNQ4;refGene_tx=XM_023258532.1
C1 31912157 31912283
refGene_gn=KCNQ4;refGene_tx=XM_023258530.1;refGene_gn=KCNQ4;refGene_tx=XM_023258531.1;refGene_gn=KCNQ4;refGene_tx=XM_023258532.1
C1 31912670 31912781
refGene_gn=KCNQ4;refGene_tx=XM_023258530.1;refGene_gn=KCNQ4;refGene_tx=XM_023258531.1;refGene_gn=KCNQ4;refGene_tx=XM_023258532.1
C1 31912971 31913067
refGene_gn=KCNQ4;refGene_tx=XM_023258530.1;refGene_gn=KCNQ4;refGene_tx=XM_023258531.1;refGene_gn=KCNQ4;refGene_tx=XM_023258532.1
C1 31915953 31916042
refGene_gn=KCNQ4;refGene_tx=XM_023258530.1;refGene_gn=KCNQ4;refGene_tx=XM_023258531.1;refGene_gn=KCNQ4;refGene_tx=XM_023258532.1
C1 31917710 31917872
refGene_gn=KCNQ4;refGene_tx=XM_023258530.1;refGene_gn=KCNQ4;refGene_tx=XM_023258532.1
C1 31924757 31924978
refGene_gn=KCNQ4;refGene_tx=XM_023258530.1;refGene_gn=KCNQ4;refGene_tx=XM_023258531.1;refGene_gn=KCNQ4;refGene_tx=XM_023258532.1
C1 31926136 31926236
refGene_gn=KCNQ4;refGene_tx=XM_023258530.1;refGene_gn=KCNQ4;refGene_tx=XM_023258531.1;refGene_gn=KCNQ4;refGene_tx=XM_023258532.1
C1 31928568 31928700
refGene_gn=KCNQ4;refGene_tx=XM_023258530.1;refGene_gn=KCNQ4;refGene_tx=XM_023258531.1;refGene_gn=KCNQ4;refGene_tx=XM_023258532.1
C1 31930953 31931083
refGene_gn=KCNQ4;refGene_tx=XM_023258530.1;refGene_gn=KCNQ4;refGene_tx=XM_023258531.1;refGene_gn=KCNQ4;refGene_tx=XM_023258532.1
C1 31931673 31931886
refGene_gn=KCNQ4;refGene_tx=XM_023258530.1;refGene_gn=KCNQ4;refGene_tx=XM_023258531.1;refGene_gn=KCNQ4;refGene_tx=XM_023258532.1
C1 31954421 31954976 refGene_gn=CITED4;refGene_tx=XM_023258533.1

C1 32048672 32048838
refGene_gn=CTPS1;refGene_tx=XM_023258534.1;refGene_gn=CTPS1;refGene_tx=XM_023258535.1;refGene_gn=CTPS1;refGene_tx=XM_023258536.1
C1 32049712 32049883
refGene_gn=CTPS1;refGene_tx=XM_023258534.1;refGene_gn=CTPS1;refGene_tx=XM_023258535.1;refGene_gn=CTPS1;refGene_tx=XM_023258536.1
C1 32052386 32052487
refGene_gn=CTPS1;refGene_tx=XM_023258534.1;refGene_gn=CTPS1;refGene_tx=XM_023258535.1;refGene_gn=CTPS1;refGene_tx=XM_023258536.1
C1 32053063 32053180
refGene_gn=CTPS1;refGene_tx=XM_023258534.1;refGene_gn=CTPS1;refGene_tx=XM_023258535.1;refGene_gn=CTPS1;refGene_tx=XM_023258536.1
C1 32054286 32054370
refGene_gn=CTPS1;refGene_tx=XM_023258534.1;refGene_gn=CTPS1;refGene_tx=XM_023258535.1;refGene_gn=CTPS1;refGene_tx=XM_023258536.1
C1 32055541 32055622
refGene_gn=CTPS1;refGene_tx=XM_023258534.1;refGene_gn=CTPS1;refGene_tx=XM_023258535.1;refGene_gn=CTPS1;refGene_tx=XM_023258536.1
C1 32060229 32060381 refGene_gn=CTPS1;refGene_tx=XM_023258536.1
C1 32061500 32061652
refGene_gn=CTPS1;refGene_tx=XM_023258534.1;refGene_gn=CTPS1;refGene_tx=XM_023258535.1
C1 32063608 32063741
refGene_gn=CTPS1;refGene_tx=XM_023258534.1;refGene_gn=CTPS1;refGene_tx=XM_023258535.1;refGene_gn=CTPS1;refGene_tx=XM_023258536.1
C1 32066205 32066294
refGene_gn=CTPS1;refGene_tx=XM_023258534.1;refGene_gn=CTPS1;refGene_tx=XM_023258535.1;refGene_gn=CTPS1;refGene_tx=XM_023258536.1
C1 32067532 32067627
refGene_gn=CTPS1;refGene_tx=XM_023258534.1;refGene_gn=CTPS1;refGene_tx=XM_023258535.1;refGene_gn=CTPS1;refGene_tx=XM_023258536.1
C1 32068303 32068366
refGene_gn=CTPS1;refGene_tx=XM_023258534.1;refGene_gn=CTPS1;refGene_tx=XM_023258535.1;refGene_gn=CTPS1;refGene_tx=XM_023258536.1
C1 32070241 32070285
refGene_gn=CTPS1;refGene_tx=XM_023258534.1;refGene_gn=CTPS1;refGene_tx=XM_023258535.1;refGene_gn=CTPS1;refGene_tx=XM_023258536.1
C1 32071564 32071661
refGene_gn=CTPS1;refGene_tx=XM_023258534.1;refGene_gn=CTPS1;refGene_tx=XM_023258535.1;refGene_gn=CTPS1;refGene_tx=XM_023258536.1
C1 32072694 32072750
refGene_gn=CTPS1;refGene_tx=XM_023258534.1;refGene_gn=CTPS1;refGene_tx=XM_023258535.1;refGene_gn=CTPS1;refGene_tx=XM_023258536.1
C1 32072850 32072947
refGene_gn=CTPS1;refGene_tx=XM_023258534.1;refGene_gn=CTPS1;refGene_tx=XM_023258535.1;refGene_gn=CTPS1;refGene_tx=XM_023258536.1
C1 32073427 32073572
refGene_gn=CTPS1;refGene_tx=XM_023258534.1;refGene_gn=CTPS1;refGene_tx=XM_023258535.1;refGene_gn=CTPS1;refGene_tx=XM_023258536.1
C1 32074052 32074137

refGene_gn=CTPS1;refGene_tx=XM_023258534.1;refGene_gn=CTPS1;refGene_tx=XM_023258535.1;refGene_gn=CTPS1;refGene_tx=XM_023258536.1
C1 32080269 32080392 refGene_gn=SLFNL1;refGene_tx=XM_019836768.2
C1 32081066 32081210 refGene_gn=SLFNL1;refGene_tx=XM_019836768.2
C1 32081429 32081897 refGene_gn=SLFNL1;refGene_tx=XM_019836768.2
C1 32083549 32084002 refGene_gn=SLFNL1;refGene_tx=XM_019836768.2
C1 32091160 32091286
refGene_gn=SCMH1;refGene_tx=XM_011284702.3;refGene_gn=SCMH1;refGene_tx=XM_011284706.3;refGene_gn=SCMH1;refGene_tx=XM_023258537.1;refGene_gn=SCMH1;refGene_tx=XM_023258538.1;refGene_gn=SCMH1;refGene_tx=XM_023258539.1;refGene_gn=SCMH1;refGene_tx=XM_023258540.1;refGene_gn=SCMH1;refGene_tx=XM_023258541.1;refGene_gn=SCMH1;refGene_tx=XM_023258542.1;refGene_gn=SCMH1;refGene_tx=XM_023258543.1;refGene_gn=SCMH1;refGene_tx=XM_023258544.1;refGene_gn=SCMH1;refGene_tx=XM_023258545.1;refGene_gn=SCMH1;refGene_tx=XM_023258546.1;refGene_gn=SCMH1;refGene_tx=XM_023258547.1;refGene_gn=SCMH1;refGene_tx=XM_023258548.1;refGene_gn=SCMH1;refGene_tx=XM_023258550.1;refGene_gn=SCMH1;refGene_tx=XM_023258552.1;refGene_gn=SCMH1;refGene_tx=XM_023258553.1;refGene_gn=SCMH1;refGene_tx=XM_023258554.1;refGene_gn=SCMH1;refGene_tx=XM_023258555.1;refGene_gn=SCMH1;refGene_tx=XM_023258556.1;refGene_gn=SCMH1;refGene_tx=XM_023258557.1;refGene_gn=SCMH1;refGene_tx=XM_023258559.1;refGene_gn=SCMH1;refGene_tx=XM_023258560.1
C1 32092332 32092559
refGene_gn=SCMH1;refGene_tx=XM_011284702.3;refGene_gn=SCMH1;refGene_tx=XM_011284706.3;refGene_gn=SCMH1;refGene_tx=XM_023258537.1;refGene_gn=SCMH1;refGene_tx=XM_023258538.1;refGene_gn=SCMH1;refGene_tx=XM_023258539.1;refGene_gn=SCMH1;refGene_tx=XM_023258540.1;refGene_gn=SCMH1;refGene_tx=XM_023258541.1;refGene_gn=SCMH1;refGene_tx=XM_023258542.1;refGene_gn=SCMH1;refGene_tx=XM_023258543.1;refGene_gn=SCMH1;refGene_tx=XM_023258544.1;refGene_gn=SCMH1;refGene_tx=XM_023258545.1;refGene_gn=SCMH1;refGene_tx=XM_023258546.1;refGene_gn=SCMH1;refGene_tx=XM_023258547.1;refGene_gn=SCMH1;refGene_tx=XM_023258548.1;refGene_gn=SCMH1;refGene_tx=XM_023258549.1;refGene_gn=SCMH1;refGene_tx=XM_023258550.1;refGene_gn=SCMH1;refGene_tx=XM_023258552.1;refGene_gn=SCMH1;refGene_tx=XM_023258553.1;refGene_gn=SCMH1;refGene_tx=XM_023258554.1;refGene_gn=SCMH1;refGene_tx=XM_023258555.1;refGene_gn=SCMH1;refGene_tx=XM_023258556.1;refGene_gn=SCMH1;refGene_tx=XM_023258557.1;refGene_gn=SCMH1;refGene_tx=XM_023258559.1;refGene_gn=SCMH1;refGene_tx=XM_023258560.1
C1 32099326 32099521
refGene_gn=SCMH1;refGene_tx=XM_011284702.3;refGene_gn=SCMH1;refGene_tx=XM_011284706.3;refGene_gn=SCMH1;refGene_tx=XM_023258537.1;refGene_gn=SCMH1;refGene_tx=XM_023258538.1;refGene_gn=SCMH1;refGene_tx=XM_023258539.1;refGene_gn=SCMH1;refGene_tx=XM_023258540.1;refGene_gn=SCMH1;refGene_tx=XM_023258541.1;refGene_gn=SCMH1;refGene_tx=XM_023258542.1;refGene_gn=SCMH1;refGene_tx=XM_023258543.1;refGene_gn=SCMH1;refGene_tx=XM_023258544.1;refGene_gn=SCMH1;refGene_tx=XM_023258545.1;refGene_gn=SCMH1;refGene_tx=XM_023258546.1;refGene_gn=SCMH1;refGene_tx=XM_023258547.1;refGene_gn=SCMH1;refGene_tx=XM_023258548.1;refGene_gn=SCMH1;refGene_tx=XM_023258549.1;refGene_gn=SCMH1;refGene_tx=XM_023258550.1;refGene_gn=SCMH1;refGene_tx=XM_023258552.1;refGene_gn=SCMH1;refGene_tx=XM_02325855

3.1;refGene_gn=SCMH1;refGene_tx=XM_023258554.1;refGene_gn=SCMH1;refGen
e_tx=XM_023258555.1;refGene_gn=SCMH1;refGene_tx=XM_023258556.1;refGene
_gn=SCMH1;refGene_tx=XM_023258557.1;refGene_gn=SCMH1;refGene_tx=XM_023
258558.1;refGene_gn=SCMH1;refGene_tx=XM_023258559.1;refGene_gn=SCMH1;r
efGene_tx=XM_023258560.1
C1 32110207 32110405
refGene_gn=SCMH1;refGene_tx=XM_011284702.3;refGene_gn=SCMH1;refGene_tx
=XM_011284706.3;refGene_gn=SCMH1;refGene_tx=XM_023258537.1;refGene_gn=
SCMH1;refGene_tx=XM_023258538.1;refGene_gn=SCMH1;refGene_tx=XM_0232585
39.1;refGene_gn=SCMH1;refGene_tx=XM_023258540.1;refGene_gn=SCMH1;refGe
ne_tx=XM_023258541.1;refGene_gn=SCMH1;refGene_tx=XM_023258542.1;refGen
e_gn=SCMH1;refGene_tx=XM_023258543.1;refGene_gn=SCMH1;refGene_tx=XM_02
3258544.1;refGene_gn=SCMH1;refGene_tx=XM_023258545.1;refGene_gn=SCMH1;
refGene_tx=XM_023258546.1;refGene_gn=SCMH1;refGene_tx=XM_023258547.1;r
efGene_gn=SCMH1;refGene_tx=XM_023258548.1;refGene_gn=SCMH1;refGene_tx=
XM_023258549.1;refGene_gn=SCMH1;refGene_tx=XM_023258550.1;refGene_gn=S
CMH1;refGene_tx=XM_023258552.1;refGene_gn=SCMH1;refGene_tx=XM_02325855
3.1;refGene_gn=SCMH1;refGene_tx=XM_023258554.1;refGene_gn=SCMH1;refGen
e_tx=XM_023258555.1;refGene_gn=SCMH1;refGene_tx=XM_023258556.1;refGene
_gn=SCMH1;refGene_tx=XM_023258557.1;refGene_gn=SCMH1;refGene_tx=XM_023
258558.1;refGene_gn=SCMH1;refGene_tx=XM_023258559.1;refGene_gn=SCMH1;r
efGene_tx=XM_023258560.1
C1 32110646 32110747
refGene_gn=SCMH1;refGene_tx=XM_023258537.1;refGene_gn=SCMH1;refGene_tx
=XM_023258538.1;refGene_gn=SCMH1;refGene_tx=XM_023258539.1;refGene_gn=
SCMH1;refGene_tx=XM_023258540.1;refGene_gn=SCMH1;refGene_tx=XM_0232585
41.1;refGene_gn=SCMH1;refGene_tx=XM_023258542.1;refGene_gn=SCMH1;refGe
ne_tx=XM_023258543.1;refGene_gn=SCMH1;refGene_tx=XM_023258545.1;refGen
e_gn=SCMH1;refGene_tx=XM_023258546.1;refGene_gn=SCMH1;refGene_tx=XM_02
3258547.1;refGene_gn=SCMH1;refGene_tx=XM_023258549.1;refGene_gn=SCMH1;
refGene_tx=XM_023258550.1;refGene_gn=SCMH1;refGene_tx=XM_023258552.1;r
efGene_gn=SCMH1;refGene_tx=XM_023258553.1;refGene_gn=SCMH1;refGene_tx=
XM_023258554.1;refGene_gn=SCMH1;refGene_tx=XM_023258555.1;refGene_gn=S
CMH1;refGene_tx=XM_023258556.1;refGene_gn=SCMH1;refGene_tx=XM_02325855
8.1;refGene_gn=SCMH1;refGene_tx=XM_023258559.1;refGene_gn=SCMH1;refGen
e_tx=XM_023258560.1
C1 32112127 32112328
refGene_gn=SCMH1;refGene_tx=XM_011284702.3;refGene_gn=SCMH1;refGene_tx
=XM_011284706.3;refGene_gn=SCMH1;refGene_tx=XM_023258537.1;refGene_gn=
SCMH1;refGene_tx=XM_023258538.1;refGene_gn=SCMH1;refGene_tx=XM_0232585
39.1;refGene_gn=SCMH1;refGene_tx=XM_023258540.1;refGene_gn=SCMH1;refGe
ne_tx=XM_023258541.1;refGene_gn=SCMH1;refGene_tx=XM_023258542.1;refGen
e_gn=SCMH1;refGene_tx=XM_023258543.1;refGene_gn=SCMH1;refGene_tx=XM_02
3258544.1;refGene_gn=SCMH1;refGene_tx=XM_023258545.1;refGene_gn=SCMH1;
refGene_tx=XM_023258546.1;refGene_gn=SCMH1;refGene_tx=XM_023258547.1;r
efGene_gn=SCMH1;refGene_tx=XM_023258548.1;refGene_gn=SCMH1;refGene_tx=
XM_023258549.1;refGene_gn=SCMH1;refGene_tx=XM_023258550.1;refGene_gn=S
CMH1;refGene_tx=XM_023258552.1;refGene_gn=SCMH1;refGene_tx=XM_02325855
3.1;refGene_gn=SCMH1;refGene_tx=XM_023258554.1;refGene_gn=SCMH1;refGen
e_tx=XM_023258555.1;refGene_gn=SCMH1;refGene_tx=XM_023258556.1;refGene

_gn=SCMH1;refGene_tx=XM_023258557.1;refGene_gn=SCMH1;refGene_tx=XM_023258558.1;refGene_gn=SCMH1;refGene_tx=XM_023258559.1;refGene_gn=SCMH1;refGene_tx=XM_023258560.1
C1 32129739 32129866
refGene_gn=SCMH1;refGene_tx=XM_011284702.3;refGene_gn=SCMH1;refGene_tx=XM_011284706.3;refGene_gn=SCMH1;refGene_tx=XM_023258537.1;refGene_gn=SCMH1;refGene_tx=XM_023258538.1;refGene_gn=SCMH1;refGene_tx=XM_023258539.1;refGene_gn=SCMH1;refGene_tx=XM_023258540.1;refGene_gn=SCMH1;refGene_tx=XM_023258541.1;refGene_gn=SCMH1;refGene_tx=XM_023258542.1;refGene_gn=SCMH1;refGene_tx=XM_023258543.1;refGene_gn=SCMH1;refGene_tx=XM_023258544.1;refGene_gn=SCMH1;refGene_tx=XM_023258545.1;refGene_gn=SCMH1;refGene_tx=XM_023258546.1;refGene_gn=SCMH1;refGene_tx=XM_023258547.1;refGene_gn=SCMH1;refGene_tx=XM_023258548.1;refGene_gn=SCMH1;refGene_tx=XM_023258549.1;refGene_gn=SCMH1;refGene_tx=XM_023258550.1;refGene_gn=SCMH1;refGene_tx=XM_023258552.1;refGene_gn=SCMH1;refGene_tx=XM_023258553.1;refGene_gn=SCMH1;refGene_tx=XM_023258555.1;refGene_gn=SCMH1;refGene_tx=XM_023258556.1;refGene_gn=SCMH1;refGene_tx=XM_023258557.1;refGene_gn=SCMH1;refGene_tx=XM_023258558.1;refGene_gn=SCMH1;refGene_tx=XM_023258559.1;refGene_gn=SCMH1;refGene_tx=XM_023258560.1
C1 32133401 32133634
refGene_gn=SCMH1;refGene_tx=XM_011284702.3;refGene_gn=SCMH1;refGene_tx=XM_011284706.3;refGene_gn=SCMH1;refGene_tx=XM_023258537.1;refGene_gn=SCMH1;refGene_tx=XM_023258538.1;refGene_gn=SCMH1;refGene_tx=XM_023258539.1;refGene_gn=SCMH1;refGene_tx=XM_023258540.1;refGene_gn=SCMH1;refGene_tx=XM_023258541.1;refGene_gn=SCMH1;refGene_tx=XM_023258542.1;refGene_gn=SCMH1;refGene_tx=XM_023258543.1;refGene_gn=SCMH1;refGene_tx=XM_023258544.1;refGene_gn=SCMH1;refGene_tx=XM_023258545.1;refGene_gn=SCMH1;refGene_tx=XM_023258546.1;refGene_gn=SCMH1;refGene_tx=XM_023258547.1;refGene_gn=SCMH1;refGene_tx=XM_023258548.1;refGene_gn=SCMH1;refGene_tx=XM_023258549.1;refGene_gn=SCMH1;refGene_tx=XM_023258550.1;refGene_gn=SCMH1;refGene_tx=XM_023258552.1;refGene_gn=SCMH1;refGene_tx=XM_023258553.1;refGene_gn=SCMH1;refGene_tx=XM_023258555.1;refGene_gn=SCMH1;refGene_tx=XM_023258557.1;refGene_gn=SCMH1;refGene_tx=XM_023258558.1;refGene_gn=SCMH1;refGene_tx=XM_023258559.1;refGene_gn=SCMH1;refGene_tx=XM_023258560.1
C1 32172430 32172674
refGene_gn=SCMH1;refGene_tx=XM_011284702.3;refGene_gn=SCMH1;refGene_tx=XM_011284706.3;refGene_gn=SCMH1;refGene_tx=XM_023258537.1;refGene_gn=SCMH1;refGene_tx=XM_023258538.1;refGene_gn=SCMH1;refGene_tx=XM_023258539.1;refGene_gn=SCMH1;refGene_tx=XM_023258540.1;refGene_gn=SCMH1;refGene_tx=XM_023258541.1;refGene_gn=SCMH1;refGene_tx=XM_023258542.1;refGene_gn=SCMH1;refGene_tx=XM_023258543.1;refGene_gn=SCMH1;refGene_tx=XM_023258544.1;refGene_gn=SCMH1;refGene_tx=XM_023258545.1;refGene_gn=SCMH1;refGene_tx=XM_023258546.1;refGene_gn=SCMH1;refGene_tx=XM_023258547.1;refGene_gn=SCMH1;refGene_tx=XM_023258548.1;refGene_gn=SCMH1;refGene_tx=XM_023258549.1;refGene_gn=SCMH1;refGene_tx=XM_023258550.1;refGene_gn=SCMH1;refGene_tx=XM_023258553.1;refGene_gn=SCMH1;refGene_tx=XM_023258554.1;refGene_gn=SCMH1;refGene_tx=XM_023258555.1;refGene_gn=SCMH1;refGene_tx=XM_023258558.1;refGene_gn=SCMH1;refGene_tx=XM_023258559.1;refGene_gn=SCMH1;refGene_tx=XM_023258560.1

C1 32177771 32177860
refGene_gn=SCMH1;refGene_tx=XM_011284702.3;refGene_gn=SCMH1;refGene_tx=XM_011284706.3;refGene_gn=SCMH1;refGene_tx=XM_023258537.1;refGene_gn=SCMH1;refGene_tx=XM_023258538.1;refGene_gn=SCMH1;refGene_tx=XM_023258539.1;refGene_gn=SCMH1;refGene_tx=XM_023258540.1;refGene_gn=SCMH1;refGene_tx=XM_023258541.1;refGene_gn=SCMH1;refGene_tx=XM_023258542.1;refGene_gn=SCMH1;refGene_tx=XM_023258543.1;refGene_gn=SCMH1;refGene_tx=XM_023258544.1;refGene_gn=SCMH1;refGene_tx=XM_023258545.1;refGene_gn=SCMH1;refGene_tx=XM_023258546.1;refGene_gn=SCMH1;refGene_tx=XM_023258547.1;refGene_gn=SCMH1;refGene_tx=XM_023258548.1;refGene_gn=SCMH1;refGene_tx=XM_023258549.1;refGene_gn=SCMH1;refGene_tx=XM_023258550.1;refGene_gn=SCMH1;refGene_tx=XM_023258553.1;refGene_gn=SCMH1;refGene_tx=XM_023258554.1;refGene_gn=SCMH1;refGene_tx=XM_023258555.1;refGene_gn=SCMH1;refGene_tx=XM_023258556.1;refGene_gn=SCMH1;refGene_tx=XM_023258558.1;refGene_gn=SCMH1;refGene_tx=XM_023258559.1;refGene_gn=SCMH1;refGene_tx=XM_023258560.1
C1 32185018 32185064
refGene_gn=SCMH1;refGene_tx=XM_011284706.3;refGene_gn=SCMH1;refGene_tx=XM_023258553.1
C1 32196553 32196788
refGene_gn=SCMH1;refGene_tx=XM_011284702.3;refGene_gn=SCMH1;refGene_tx=XM_011284706.3;refGene_gn=SCMH1;refGene_tx=XM_023258537.1;refGene_gn=SCMH1;refGene_tx=XM_023258538.1;refGene_gn=SCMH1;refGene_tx=XM_023258539.1;refGene_gn=SCMH1;refGene_tx=XM_023258540.1;refGene_gn=SCMH1;refGene_tx=XM_023258541.1;refGene_gn=SCMH1;refGene_tx=XM_023258542.1;refGene_gn=SCMH1;refGene_tx=XM_023258543.1;refGene_gn=SCMH1;refGene_tx=XM_023258544.1;refGene_gn=SCMH1;refGene_tx=XM_023258545.1;refGene_gn=SCMH1;refGene_tx=XM_023258546.1;refGene_gn=SCMH1;refGene_tx=XM_023258547.1;refGene_gn=SCMH1;refGene_tx=XM_023258548.1;refGene_gn=SCMH1;refGene_tx=XM_023258549.1;refGene_gn=SCMH1;refGene_tx=XM_023258550.1;refGene_gn=SCMH1;refGene_tx=XM_023258552.1;refGene_gn=SCMH1;refGene_tx=XM_023258553.1;refGene_gn=SCMH1;refGene_tx=XM_023258554.1;refGene_gn=SCMH1;refGene_tx=XM_023258555.1;refGene_gn=SCMH1;refGene_tx=XM_023258556.1;refGene_gn=SCMH1;refGene_tx=XM_023258557.1;refGene_gn=SCMH1;refGene_tx=XM_023258558.1;refGene_gn=SCMH1;refGene_tx=XM_023258559.1;refGene_gn=SCMH1;refGene_tx=XM_023258560.1
C1 32209909 32209980
refGene_gn=SCMH1;refGene_tx=XM_011284702.3;refGene_gn=SCMH1;refGene_tx=XM_023258537.1;refGene_gn=SCMH1;refGene_tx=XM_023258538.1;refGene_gn=SCMH1;refGene_tx=XM_023258539.1;refGene_gn=SCMH1;refGene_tx=XM_023258540.1;refGene_gn=SCMH1;refGene_tx=XM_023258541.1;refGene_gn=SCMH1;refGene_tx=XM_023258542.1;refGene_gn=SCMH1;refGene_tx=XM_023258543.1;refGene_gn=SCMH1;refGene_tx=XM_023258544.1;refGene_gn=SCMH1;refGene_tx=XM_023258545.1;refGene_gn=SCMH1;refGene_tx=XM_023258546.1;refGene_gn=SCMH1;refGene_tx=XM_023258547.1;refGene_gn=SCMH1;refGene_tx=XM_023258548.1;refGene_gn=SCMH1;refGene_tx=XM_023258549.1;refGene_gn=SCMH1;refGene_tx=XM_023258552.1;refGene_gn=SCMH1;refGene_tx=XM_023258554.1;refGene_gn=SCMH1;refGene_tx=XM_023258555.1;refGene_gn=SCMH1;refGene_tx=XM_023258556.1;refGene_gn=SCMH1;refGene_tx=XM_023258557.1;refGene_gn=SCMH1;refGene_tx=XM_023258558.1

C1 32210848 32210924
refGene_gn=SCMH1;refGene_tx=XM_011284702.3;refGene_gn=SCMH1;refGene_tx=XM_023258546.1;refGene_gn=SCMH1;refGene_tx=XM_023258547.1
C1 32219964 32219988
refGene_gn=SCMH1;refGene_tx=XM_023258537.1;refGene_gn=SCMH1;refGene_tx=XM_023258538.1;refGene_gn=SCMH1;refGene_tx=XM_023258539.1;refGene_gn=SCMH1;refGene_tx=XM_023258540.1;refGene_gn=SCMH1;refGene_tx=XM_023258541.1;refGene_gn=SCMH1;refGene_tx=XM_023258542.1;refGene_gn=SCMH1;refGene_tx=XM_023258543.1;refGene_gn=SCMH1;refGene_tx=XM_023258544.1;refGene_gn=SCMH1;refGene_tx=XM_023258548.1;refGene_gn=SCMH1;refGene_tx=XM_023258549.1;refGene_gn=SCMH1;refGene_tx=XM_023258552.1;refGene_gn=SCMH1;refGene_tx=XM_023258554.1;refGene_gn=SCMH1;refGene_tx=XM_023258555.1;refGene_gn=SCMH1;refGene_tx=XM_023258556.1;refGene_gn=SCMH1;refGene_tx=XM_023258557.1;refGene_gn=SCMH1;refGene_tx=XM_023258558.1
C1 32220448 32220517
refGene_gn=SCMH1;refGene_tx=XM_023258537.1;refGene_gn=SCMH1;refGene_tx=XM_023258538.1;refGene_gn=SCMH1;refGene_tx=XM_023258539.1;refGene_gn=SCMH1;refGene_tx=XM_023258540.1;refGene_gn=SCMH1;refGene_tx=XM_023258541.1;refGene_gn=SCMH1;refGene_tx=XM_023258542.1;refGene_gn=SCMH1;refGene_tx=XM_023258543.1;refGene_gn=SCMH1;refGene_tx=XM_023258544.1;refGene_gn=SCMH1;refGene_tx=XM_023258545.1;refGene_gn=SCMH1;refGene_tx=XM_023258548.1;refGene_gn=SCMH1;refGene_tx=XM_023258549.1;refGene_gn=SCMH1;refGene_tx=XM_023258552.1;refGene_gn=SCMH1;refGene_tx=XM_023258554.1;refGene_gn=SCMH1;refGene_tx=XM_023258555.1;refGene_gn=SCMH1;refGene_tx=XM_023258556.1;refGene_gn=SCMH1;refGene_tx=XM_023258557.1;refGene_gn=SCMH1;refGene_tx=XM_023258558.1
C1 32229857 32229870
refGene_gn=SCMH1;refGene_tx=XM_023258541.1;refGene_gn=SCMH1;refGene_tx=XM_023258542.1;refGene_gn=SCMH1;refGene_tx=XM_023258543.1;refGene_gn=SCMH1;refGene_tx=XM_023258545.1
C1 32238177 32238258
refGene_gn=SCMH1;refGene_tx=XM_023258537.1;refGene_gn=SCMH1;refGene_tx=XM_023258538.1;refGene_gn=SCMH1;refGene_tx=XM_023258539.1;refGene_gn=SCMH1;refGene_tx=XM_023258540.1;refGene_gn=SCMH1;refGene_tx=XM_023258544.1;refGene_gn=SCMH1;refGene_tx=XM_023258548.1;refGene_gn=SCMH1;refGene_tx=XM_023258549.1;refGene_gn=SCMH1;refGene_tx=XM_023258552.1;refGene_gn=SCMH1;refGene_tx=XM_023258554.1;refGene_gn=SCMH1;refGene_tx=XM_023258555.1;refGene_gn=SCMH1;refGene_tx=XM_023258556.1;refGene_gn=SCMH1;refGene_tx=XM_023258557.1;refGene_gn=SCMH1;refGene_tx=XM_023258558.1
C1 32243147 32243166
refGene_gn=SCMH1;refGene_tx=XM_023258537.1;refGene_gn=SCMH1;refGene_tx=XM_023258538.1;refGene_gn=SCMH1;refGene_tx=XM_023258540.1;refGene_gn=SCMH1;refGene_tx=XM_023258544.1;refGene_gn=SCMH1;refGene_tx=XM_023258548.1;refGene_gn=SCMH1;refGene_tx=XM_023258549.1;refGene_gn=SCMH1;refGene_tx=XM_023258552.1;refGene_gn=SCMH1;refGene_tx=XM_023258554.1;refGene_gn=SCMH1;refGene_tx=XM_023258556.1;refGene_gn=SCMH1;refGene_tx=XM_023258557.1;refGene_gn=SCMH1;refGene_tx=XM_023258558.1
C1 32388366 32388780 refGene_gn=FOXO6;refGene_tx=XM_023258561.1
C1 32407219 32408479 refGene_gn=FOXO6;refGene_tx=XM_023258561.1
C1 32508026 32508120 refGene_gn=EDN2;refGene_tx=NM_001009387.1

C1 32509677 32509776 refGene_gn=EDN2;refGene_tx=NM_001009387.1
C1 32510760 32510883 refGene_gn=EDN2;refGene_tx=NM_001009387.1
C1 32512362 32512519 refGene_gn=EDN2;refGene_tx=NM_001009387.1
C1 32512887 32512951 refGene_gn=EDN2;refGene_tx=NM_001009387.1
C1 32539287 32540100 refGene_gn=HIVEP3;refGene_tx=XM_023258562.1
C1 32541078 32541992 refGene_gn=HIVEP3;refGene_tx=XM_023258562.1
C1 32546572 32546659 refGene_gn=HIVEP3;refGene_tx=XM_023258562.1
C1 32553003 32553179 refGene_gn=HIVEP3;refGene_tx=XM_023258562.1
C1 32588982 32589128 refGene_gn=HIVEP3;refGene_tx=XM_023258562.1
C1 32593086 32596611 refGene_gn=HIVEP3;refGene_tx=XM_023258562.1
C1 32596613 32598146 refGene_gn=HIVEP3;refGene_tx=XM_023258562.1
C1 33086716 33086806 refGene_gn=GUCA2B;refGene_tx=XM_003989954.3
C1 33087925 33088112 refGene_gn=GUCA2B;refGene_tx=XM_003989954.3
C1 33088846 33088908 refGene_gn=GUCA2B;refGene_tx=XM_003989954.3
C1 33092576 33092641 refGene_gn=GUCA2A;refGene_tx=XM_003989955.3
C1 33093033 33093229 refGene_gn=GUCA2A;refGene_tx=XM_003989955.3
C1 33093721 33093796 refGene_gn=GUCA2A;refGene_tx=XM_003989955.3
C1 33106541 33106657
refGene_gn=FOXJ3;refGene_tx=XM_003989956.5;refGene_gn=FOXJ3;refGene_tx=XM_019836783.1;refGene_gn=FOXJ3;refGene_tx=XM_019836784.1;refGene_gn=FOXJ3;refGene_tx=XM_019836785.1;refGene_gn=FOXJ3;refGene_tx=XM_019836786.1;refGene_gn=FOXJ3;refGene_tx=XM_019836787.2;refGene_gn=FOXJ3;refGene_tx=XM_019836789.1;refGene_gn=FOXJ3;refGene_tx=XM_023258563.1;refGene_gn=FOXJ3;refGene_tx=XM_023258564.1
C1 33108686 33108794
refGene_gn=FOXJ3;refGene_tx=XM_003989956.5;refGene_gn=FOXJ3;refGene_tx=XM_019836783.1;refGene_gn=FOXJ3;refGene_tx=XM_019836784.1;refGene_gn=FOXJ3;refGene_tx=XM_019836785.1;refGene_gn=FOXJ3;refGene_tx=XM_019836786.1;refGene_gn=FOXJ3;refGene_tx=XM_019836787.2;refGene_gn=FOXJ3;refGene_tx=XM_019836789.1;refGene_gn=FOXJ3;refGene_tx=XM_023258563.1;refGene_gn=FOXJ3;refGene_tx=XM_023258564.1
C1 33122557 33122749
refGene_gn=FOXJ3;refGene_tx=XM_003989956.5;refGene_gn=FOXJ3;refGene_tx=XM_019836783.1;refGene_gn=FOXJ3;refGene_tx=XM_019836784.1;refGene_gn=FOXJ3;refGene_tx=XM_019836785.1;refGene_gn=FOXJ3;refGene_tx=XM_019836786.1;refGene_gn=FOXJ3;refGene_tx=XM_019836787.2;refGene_gn=FOXJ3;refGene_tx=XM_019836789.1;refGene_gn=FOXJ3;refGene_tx=XM_023258563.1;refGene_gn=FOXJ3;refGene_tx=XM_023258564.1
C1 33123376 33123478
refGene_gn=FOXJ3;refGene_tx=XM_003989956.5;refGene_gn=FOXJ3;refGene_tx=XM_019836783.1;refGene_gn=FOXJ3;refGene_tx=XM_019836784.1;refGene_gn=FOXJ3;refGene_tx=XM_019836785.1;refGene_gn=FOXJ3;refGene_tx=XM_019836786.1;refGene_gn=FOXJ3;refGene_tx=XM_019836787.2;refGene_gn=FOXJ3;refGene_tx=XM_019836789.1;refGene_gn=FOXJ3;refGene_tx=XM_023258563.1;refGene_gn=FOXJ3;refGene_tx=XM_023258564.1
C1 33127230 33127647
refGene_gn=FOXJ3;refGene_tx=XM_003989956.5;refGene_gn=FOXJ3;refGene_tx=XM_019836783.1;refGene_gn=FOXJ3;refGene_tx=XM_019836784.1;refGene_gn=FOXJ3;refGene_tx=XM_019836785.1;refGene_gn=FOXJ3;refGene_tx=XM_019836786.1;refGene_gn=FOXJ3;refGene_tx=XM_019836787.2;refGene_gn=FOXJ3;refGe

ne_tx=XM_019836789.1;refGene_gn=FOXJ3;refGene_tx=XM_023258563.1;refGene_gn=FOXJ3;refGene_tx=XM_023258564.1
C1	33132023	33132198
refGene_gn=FOXJ3;refGene_tx=XM_003989956.5;refGene_gn=FOXJ3;refGene_tx=XM_019836783.1;refGene_gn=FOXJ3;refGene_tx=XM_019836784.1;refGene_gn=FOXJ3;refGene_tx=XM_019836785.1;refGene_gn=FOXJ3;refGene_tx=XM_019836786.1;refGene_gn=FOXJ3;refGene_tx=XM_019836787.2;refGene_gn=FOXJ3;refGene_tx=XM_019836789.1;refGene_gn=FOXJ3;refGene_tx=XM_023258563.1;refGene_gn=FOXJ3;refGene_tx=XM_023258564.1
C1	33135375	33135504
refGene_gn=FOXJ3;refGene_tx=XM_003989956.5;refGene_gn=FOXJ3;refGene_tx=XM_019836783.1;refGene_gn=FOXJ3;refGene_tx=XM_019836784.1;refGene_gn=FOXJ3;refGene_tx=XM_019836785.1;refGene_gn=FOXJ3;refGene_tx=XM_019836786.1;refGene_gn=FOXJ3;refGene_tx=XM_019836787.2;refGene_gn=FOXJ3;refGene_tx=XM_019836789.1;refGene_gn=FOXJ3;refGene_tx=XM_023258563.1;refGene_gn=FOXJ3;refGene_tx=XM_023258564.1
C1	33147734	33147836
refGene_gn=FOXJ3;refGene_tx=XM_003989956.5;refGene_gn=FOXJ3;refGene_tx=XM_019836783.1;refGene_gn=FOXJ3;refGene_tx=XM_019836784.1;refGene_gn=FOXJ3;refGene_tx=XM_019836785.1;refGene_gn=FOXJ3;refGene_tx=XM_019836787.2;refGene_gn=FOXJ3;refGene_tx=XM_019836789.1;refGene_gn=FOXJ3;refGene_tx=XM_023258563.1;refGene_gn=FOXJ3;refGene_tx=XM_023258564.1
C1	33164880	33164964
refGene_gn=FOXJ3;refGene_tx=XM_003989956.5;refGene_gn=FOXJ3;refGene_tx=XM_019836783.1;refGene_gn=FOXJ3;refGene_tx=XM_019836784.1;refGene_gn=FOXJ3;refGene_tx=XM_019836785.1;refGene_gn=FOXJ3;refGene_tx=XM_019836786.1;refGene_gn=FOXJ3;refGene_tx=XM_019836787.2;refGene_gn=FOXJ3;refGene_tx=XM_019836789.1;refGene_gn=FOXJ3;refGene_tx=XM_023258563.1;refGene_gn=FOXJ3;refGene_tx=XM_023258564.1
C1	33178003	33178135	refGene_gn=FOXJ3;refGene_tx=XM_019836787.2
C1	33188658	33188733
refGene_gn=FOXJ3;refGene_tx=XM_003989956.5;refGene_gn=FOXJ3;refGene_tx=XM_019836783.1;refGene_gn=FOXJ3;refGene_tx=XM_019836784.1;refGene_gn=FOXJ3;refGene_tx=XM_019836785.1;refGene_gn=FOXJ3;refGene_tx=XM_019836786.1;refGene_gn=FOXJ3;refGene_tx=XM_019836789.1;refGene_gn=FOXJ3;refGene_tx=XM_023258563.1;refGene_gn=FOXJ3;refGene_tx=XM_023258564.1
C1	33201708	33202033
refGene_gn=FOXJ3;refGene_tx=XM_003989956.5;refGene_gn=FOXJ3;refGene_tx=XM_019836783.1;refGene_gn=FOXJ3;refGene_tx=XM_019836784.1;refGene_gn=FOXJ3;refGene_tx=XM_019836785.1;refGene_gn=FOXJ3;refGene_tx=XM_019836786.1;refGene_gn=FOXJ3;refGene_tx=XM_019836789.1;refGene_gn=FOXJ3;refGene_tx=XM_023258563.1;refGene_gn=FOXJ3;refGene_tx=XM_023258564.1
C1	33229253	33229314
refGene_gn=FOXJ3;refGene_tx=XM_003989956.5;refGene_gn=FOXJ3;refGene_tx=XM_019836783.1;refGene_gn=FOXJ3;refGene_tx=XM_019836784.1;refGene_gn=FOXJ3;refGene_tx=XM_019836786.1;refGene_gn=FOXJ3;refGene_tx=XM_019836789.1;refGene_gn=FOXJ3;refGene_tx=XM_023258563.1;refGene_gn=FOXJ3;refGene_tx=XM_023258564.1
C1	33235154	33235179
refGene_gn=FOXJ3;refGene_tx=XM_019836783.1;refGene_gn=FOXJ3;refGene_tx

=XM_019836786.1
C1 33305308 33305471 refGene_gn=RIMKLA;refGene_tx=XM_023258565.1
C1 33320275 33320312
refGene_gn=RIMKLA;refGene_tx=XM_019836790.2;refGene_gn=RIMKLA;refGene_tx=XM_023258566.1;refGene_gn=RIMKLA;refGene_tx=XM_023258568.1
C1 33321813 33322044
refGene_gn=RIMKLA;refGene_tx=XM_019836790.2;refGene_gn=RIMKLA;refGene_tx=XM_023258565.1;refGene_gn=RIMKLA;refGene_tx=XM_023258566.1;refGene_gn=RIMKLA;refGene_tx=XM_023258568.1
C1 33326117 33326204
refGene_gn=RIMKLA;refGene_tx=XM_019836790.2;refGene_gn=RIMKLA;refGene_tx=XM_023258565.1;refGene_gn=RIMKLA;refGene_tx=XM_023258566.1;refGene_gn=RIMKLA;refGene_tx=XM_023258568.1
C1 33331576 33331780
refGene_gn=RIMKLA;refGene_tx=XM_019836790.2;refGene_gn=RIMKLA;refGene_tx=XM_023258565.1;refGene_gn=RIMKLA;refGene_tx=XM_023258566.1;refGene_gn=RIMKLA;refGene_tx=XM_023258568.1
C1 33334504 33334995
refGene_gn=RIMKLA;refGene_tx=XM_019836790.2;refGene_gn=RIMKLA;refGene_tx=XM_023258565.1;refGene_gn=RIMKLA;refGene_tx=XM_023258566.1;refGene_gn=RIMKLA;refGene_tx=XM_023258568.1
C1 33351060 33351186
refGene_gn=ZMYND12;refGene_tx=XM_011284691.3;refGene_gn=ZMYND12;refGene_tx=XM_011284692.3;refGene_gn=ZMYND12;refGene_tx=XM_019836794.2
C1 33352789 33352935
refGene_gn=ZMYND12;refGene_tx=XM_011284691.3;refGene_gn=ZMYND12;refGene_tx=XM_011284692.3;refGene_gn=ZMYND12;refGene_tx=XM_019836794.2
C1 33354893 33355005
refGene_gn=ZMYND12;refGene_tx=XM_011284691.3;refGene_gn=ZMYND12;refGene_tx=XM_011284692.3;refGene_gn=ZMYND12;refGene_tx=XM_019836794.2
C1 33355237 33355360
refGene_gn=ZMYND12;refGene_tx=XM_011284691.3;refGene_gn=ZMYND12;refGene_tx=XM_011284692.3;refGene_gn=ZMYND12;refGene_tx=XM_019836794.2
C1 33357664 33357834
refGene_gn=ZMYND12;refGene_tx=XM_011284691.3;refGene_gn=ZMYND12;refGene_tx=XM_011284692.3;refGene_gn=ZMYND12;refGene_tx=XM_019836794.2
C1 33366323 33366495
refGene_gn=ZMYND12;refGene_tx=XM_011284691.3;refGene_gn=ZMYND12;refGene_tx=XM_011284692.3;refGene_gn=ZMYND12;refGene_tx=XM_019836794.2
C1 33367606 33367748
refGene_gn=ZMYND12;refGene_tx=XM_011284691.3;refGene_gn=ZMYND12;refGene_tx=XM_011284692.3;refGene_gn=ZMYND12;refGene_tx=XM_019836794.2
C1 33378657 33378767
refGene_gn=ZMYND12;refGene_tx=XM_011284691.3;refGene_gn=ZMYND12;refGene_tx=XM_011284692.3;refGene_gn=ZMYND12;refGene_tx=XM_019836794.2
C1 33379363 33379871
refGene_gn=PPCS;refGene_tx=XM_003989958.4;refGene_gn=PPCS;refGene_tx=XM_019836796.2
C1 33380042 33380146
refGene_gn=PPCS;refGene_tx=XM_003989958.4;refGene_gn=PPCS;refGene_tx=X

M_006934649.3;refGene_gn=PPCS;refGene_tx=XM_006934650.3
C1 33380695 33380746 refGene_gn=PPCS;refGene_tx=XM_019836797.2
C1 33381852 33382176
refGene_gn=PPCS;refGene_tx=XM_003989958.4;refGene_gn=PPCS;refGene_tx=XM_006934649.3;refGene_gn=PPCS;refGene_tx=XM_006934650.3;refGene_gn=PPCS;refGene_tx=XM_006934651.3;refGene_gn=PPCS;refGene_tx=XM_019836796.2;refGene_gn=PPCS;refGene_tx=XM_019836797.2
C1 33393950 33393959
refGene_gn=CCDC30;refGene_tx=XM_019836798.2;refGene_gn=CCDC30;refGene_tx=XM_019836799.2;refGene_gn=CCDC30;refGene_tx=XM_019836800.2;refGene_gn=CCDC30;refGene_tx=XM_019836802.2;refGene_gn=CCDC30;refGene_tx=XM_019836803.2;refGene_gn=CCDC30;refGene_tx=XM_019836807.2
C1 33394712 33394769
refGene_gn=CCDC30;refGene_tx=XM_019836798.2;refGene_gn=CCDC30;refGene_tx=XM_019836799.2;refGene_gn=CCDC30;refGene_tx=XM_019836800.2;refGene_gn=CCDC30;refGene_tx=XM_019836803.2;refGene_gn=CCDC30;refGene_tx=XM_019836807.2
C1 33396105 33396226
refGene_gn=CCDC30;refGene_tx=XM_019836798.2;refGene_gn=CCDC30;refGene_tx=XM_019836799.2;refGene_gn=CCDC30;refGene_tx=XM_019836800.2;refGene_gn=CCDC30;refGene_tx=XM_019836802.2;refGene_gn=CCDC30;refGene_tx=XM_019836803.2;refGene_gn=CCDC30;refGene_tx=XM_019836804.2;refGene_gn=CCDC30;refGene_tx=XM_019836805.2;refGene_gn=CCDC30;refGene_tx=XM_019836807.2
C1 33399892 33400086
refGene_gn=CCDC30;refGene_tx=XM_019836798.2;refGene_gn=CCDC30;refGene_tx=XM_019836799.2;refGene_gn=CCDC30;refGene_tx=XM_019836800.2;refGene_gn=CCDC30;refGene_tx=XM_019836802.2;refGene_gn=CCDC30;refGene_tx=XM_019836803.2;refGene_gn=CCDC30;refGene_tx=XM_019836807.2
C1 33403315 33403405
refGene_gn=CCDC30;refGene_tx=XM_019836798.2;refGene_gn=CCDC30;refGene_tx=XM_019836799.2;refGene_gn=CCDC30;refGene_tx=XM_019836800.2;refGene_gn=CCDC30;refGene_tx=XM_019836802.2;refGene_gn=CCDC30;refGene_tx=XM_019836803.2;refGene_gn=CCDC30;refGene_tx=XM_019836804.2;refGene_gn=CCDC30;refGene_tx=XM_019836805.2;refGene_gn=CCDC30;refGene_tx=XM_019836807.2
C1 33405218 33405372
refGene_gn=CCDC30;refGene_tx=XM_019836798.2;refGene_gn=CCDC30;refGene_tx=XM_019836799.2;refGene_gn=CCDC30;refGene_tx=XM_019836800.2;refGene_gn=CCDC30;refGene_tx=XM_019836802.2;refGene_gn=CCDC30;refGene_tx=XM_019836803.2;refGene_gn=CCDC30;refGene_tx=XM_019836804.2;refGene_gn=CCDC30;refGene_tx=XM_019836805.2;refGene_gn=CCDC30;refGene_tx=XM_019836807.2
C1 33415504 33415576
refGene_gn=CCDC30;refGene_tx=XM_019836798.2;refGene_gn=CCDC30;refGene_tx=XM_019836799.2;refGene_gn=CCDC30;refGene_tx=XM_019836800.2;refGene_gn=CCDC30;refGene_tx=XM_019836802.2;refGene_gn=CCDC30;refGene_tx=XM_019836803.2;refGene_gn=CCDC30;refGene_tx=XM_019836804.2;refGene_gn=CCDC30;refGene_tx=XM_019836805.2;refGene_gn=CCDC30;refGene_tx=XM_019836807.2

C1 33421971 33422087
refGene_gn=CCDC30;refGene_tx=XM_019836798.2;refGene_gn=CCDC30;refGene_
tx=XM_019836799.2;refGene_gn=CCDC30;refGene_tx=XM_019836800.2;refGene_
gn=CCDC30;refGene_tx=XM_019836802.2;refGene_gn=CCDC30;refGene_tx=XM_01
9836803.2;refGene_gn=CCDC30;refGene_tx=XM_019836804.2;refGene_gn=CCDC3
0;refGene_tx=XM_019836805.2;refGene_gn=CCDC30;refGene_tx=XM_019836807.
2
C1 33424658 33424757
refGene_gn=CCDC30;refGene_tx=XM_019836798.2;refGene_gn=CCDC30;refGene_
tx=XM_019836799.2;refGene_gn=CCDC30;refGene_tx=XM_019836800.2;refGene_
gn=CCDC30;refGene_tx=XM_019836802.2;refGene_gn=CCDC30;refGene_tx=XM_01
9836803.2;refGene_gn=CCDC30;refGene_tx=XM_019836804.2;refGene_gn=CCDC3
0;refGene_tx=XM_019836805.2;refGene_gn=CCDC30;refGene_tx=XM_019836807.
2
C1 33442348 33442474
refGene_gn=CCDC30;refGene_tx=XM_011284688.3;refGene_gn=CCDC30;refGene_
tx=XM_019836798.2;refGene_gn=CCDC30;refGene_tx=XM_019836799.2;refGene_
gn=CCDC30;refGene_tx=XM_019836800.2;refGene_gn=CCDC30;refGene_tx=XM_01
9836802.2;refGene_gn=CCDC30;refGene_tx=XM_019836803.2;refGene_gn=CCDC3
0;refGene_tx=XM_019836804.2;refGene_gn=CCDC30;refGene_tx=XM_019836805.
2
C1 33443824 33443962
refGene_gn=CCDC30;refGene_tx=XM_011284688.3;refGene_gn=CCDC30;refGene_
tx=XM_019836798.2;refGene_gn=CCDC30;refGene_tx=XM_019836799.2;refGene_
gn=CCDC30;refGene_tx=XM_019836800.2;refGene_gn=CCDC30;refGene_tx=XM_01
9836802.2;refGene_gn=CCDC30;refGene_tx=XM_019836803.2;refGene_gn=CCDC3
0;refGene_tx=XM_019836804.2;refGene_gn=CCDC30;refGene_tx=XM_019836805.
2
C1 33445934 33446104
refGene_gn=CCDC30;refGene_tx=XM_011284688.3;refGene_gn=CCDC30;refGene_
tx=XM_019836798.2;refGene_gn=CCDC30;refGene_tx=XM_019836799.2;refGene_
gn=CCDC30;refGene_tx=XM_019836800.2;refGene_gn=CCDC30;refGene_tx=XM_01
9836802.2;refGene_gn=CCDC30;refGene_tx=XM_019836803.2;refGene_gn=CCDC3
0;refGene_tx=XM_019836804.2;refGene_gn=CCDC30;refGene_tx=XM_019836805.
2
C1 33452005 33452055 refGene_gn=CCDC30;refGene_tx=XM_023258571.1
C1 33458384 33458628
refGene_gn=CCDC30;refGene_tx=XM_011284688.3;refGene_gn=CCDC30;refGene_
tx=XM_019836798.2;refGene_gn=CCDC30;refGene_tx=XM_019836799.2;refGene_
gn=CCDC30;refGene_tx=XM_019836800.2;refGene_gn=CCDC30;refGene_tx=XM_01
9836802.2;refGene_gn=CCDC30;refGene_tx=XM_019836803.2;refGene_gn=CCDC3
0;refGene_tx=XM_019836804.2;refGene_gn=CCDC30;refGene_tx=XM_019836805.
2;refGene_gn=CCDC30;refGene_tx=XM_023258571.1
C1 33463323 33463503
refGene_gn=CCDC30;refGene_tx=XM_011284688.3;refGene_gn=CCDC30;refGene_
tx=XM_019836798.2;refGene_gn=CCDC30;refGene_tx=XM_019836799.2;refGene_
gn=CCDC30;refGene_tx=XM_019836800.2;refGene_gn=CCDC30;refGene_tx=XM_01
9836802.2;refGene_gn=CCDC30;refGene_tx=XM_019836803.2;refGene_gn=CCDC3
0;refGene_tx=XM_019836804.2;refGene_gn=CCDC30;refGene_tx=XM_019836805.
2;refGene_gn=CCDC30;refGene_tx=XM_019836807.2;refGene_gn=CCDC30;refGen

e_tx=XM_023258571.1
C1 33472161 33472371
refGene_gn=CCDC30;refGene_tx=XM_011284688.3;refGene_gn=CCDC30;refGene_tx=XM_019836798.2;refGene_gn=CCDC30;refGene_tx=XM_019836799.2;refGene_gn=CCDC30;refGene_tx=XM_019836800.2;refGene_gn=CCDC30;refGene_tx=XM_019836802.2;refGene_gn=CCDC30;refGene_tx=XM_019836803.2;refGene_gn=CCDC30;refGene_tx=XM_019836804.2;refGene_gn=CCDC30;refGene_tx=XM_019836805.2;refGene_gn=CCDC30;refGene_tx=XM_019836807.2;refGene_gn=CCDC30;refGene_tx=XM_023258571.1
C1 33476005 33476160
refGene_gn=CCDC30;refGene_tx=XM_011284688.3;refGene_gn=CCDC30;refGene_tx=XM_019836798.2;refGene_gn=CCDC30;refGene_tx=XM_019836799.2;refGene_gn=CCDC30;refGene_tx=XM_019836800.2;refGene_gn=CCDC30;refGene_tx=XM_019836802.2;refGene_gn=CCDC30;refGene_tx=XM_019836803.2;refGene_gn=CCDC30;refGene_tx=XM_019836804.2;refGene_gn=CCDC30;refGene_tx=XM_019836805.2;refGene_gn=CCDC30;refGene_tx=XM_019836807.2;refGene_gn=CCDC30;refGene_tx=XM_023258571.1
C1 33485787 33485953
refGene_gn=CCDC30;refGene_tx=XM_011284688.3;refGene_gn=CCDC30;refGene_tx=XM_019836798.2;refGene_gn=CCDC30;refGene_tx=XM_019836799.2;refGene_gn=CCDC30;refGene_tx=XM_019836800.2;refGene_gn=CCDC30;refGene_tx=XM_019836802.2;refGene_gn=CCDC30;refGene_tx=XM_019836803.2;refGene_gn=CCDC30;refGene_tx=XM_019836804.2;refGene_gn=CCDC30;refGene_tx=XM_019836805.2;refGene_gn=CCDC30;refGene_tx=XM_019836807.2;refGene_gn=CCDC30;refGene_tx=XM_023258571.1
C1 33486883 33486996
refGene_gn=CCDC30;refGene_tx=XM_011284688.3;refGene_gn=CCDC30;refGene_tx=XM_019836798.2;refGene_gn=CCDC30;refGene_tx=XM_019836799.2;refGene_gn=CCDC30;refGene_tx=XM_019836800.2;refGene_gn=CCDC30;refGene_tx=XM_019836802.2;refGene_gn=CCDC30;refGene_tx=XM_019836803.2;refGene_gn=CCDC30;refGene_tx=XM_019836804.2;refGene_gn=CCDC30;refGene_tx=XM_019836805.2;refGene_gn=CCDC30;refGene_tx=XM_019836807.2;refGene_gn=CCDC30;refGene_tx=XM_023258571.1
C1 33502544 33502686
refGene_gn=CCDC30;refGene_tx=XM_011284688.3;refGene_gn=CCDC30;refGene_tx=XM_019836798.2;refGene_gn=CCDC30;refGene_tx=XM_019836799.2;refGene_gn=CCDC30;refGene_tx=XM_019836800.2;refGene_gn=CCDC30;refGene_tx=XM_019836802.2;refGene_gn=CCDC30;refGene_tx=XM_019836803.2;refGene_gn=CCDC30;refGene_tx=XM_019836804.2;refGene_gn=CCDC30;refGene_tx=XM_019836805.2;refGene_gn=CCDC30;refGene_tx=XM_019836807.2;refGene_gn=CCDC30;refGene_tx=XM_023258571.1
C1 33506048 33506185
refGene_gn=CCDC30;refGene_tx=XM_011284688.3;refGene_gn=CCDC30;refGene_tx=XM_019836798.2;refGene_gn=CCDC30;refGene_tx=XM_019836799.2;refGene_gn=CCDC30;refGene_tx=XM_019836800.2;refGene_gn=CCDC30;refGene_tx=XM_019836802.2;refGene_gn=CCDC30;refGene_tx=XM_019836803.2;refGene_gn=CCDC30;refGene_tx=XM_019836804.2;refGene_gn=CCDC30;refGene_tx=XM_019836805.2;refGene_gn=CCDC30;refGene_tx=XM_019836807.2;refGene_gn=CCDC30;refGene_tx=XM_023258571.1
C1 33509066 33509181

refGene_gn=CCDC30;refGene_tx=XM_011284688.3;refGene_gn=CCDC30;refGene_tx=XM_019836798.2;refGene_gn=CCDC30;refGene_tx=XM_019836799.2;refGene_gn=CCDC30;refGene_tx=XM_019836800.2;refGene_gn=CCDC30;refGene_tx=XM_019836802.2;refGene_gn=CCDC30;refGene_tx=XM_019836803.2;refGene_gn=CCDC30;refGene_tx=XM_019836804.2;refGene_gn=CCDC30;refGene_tx=XM_019836805.2;refGene_gn=CCDC30;refGene_tx=XM_019836807.2;refGene_gn=CCDC30;refGene_tx=XM_023258571.1
C1 33515442 33515510
refGene_gn=CCDC30;refGene_tx=XM_011284688.3;refGene_gn=CCDC30;refGene_tx=XM_019836798.2;refGene_gn=CCDC30;refGene_tx=XM_019836799.2;refGene_gn=CCDC30;refGene_tx=XM_019836800.2;refGene_gn=CCDC30;refGene_tx=XM_019836802.2;refGene_gn=CCDC30;refGene_tx=XM_019836804.2;refGene_gn=CCDC30;refGene_tx=XM_019836805.2;refGene_gn=CCDC30;refGene_tx=XM_019836807.2;refGene_gn=CCDC30;refGene_tx=XM_023258571.1
C1 33515867 33516066
refGene_gn=CCDC30;refGene_tx=XM_011284688.3;refGene_gn=CCDC30;refGene_tx=XM_019836798.2;refGene_gn=CCDC30;refGene_tx=XM_019836799.2;refGene_gn=CCDC30;refGene_tx=XM_019836800.2;refGene_gn=CCDC30;refGene_tx=XM_019836802.2;refGene_gn=CCDC30;refGene_tx=XM_019836803.2;refGene_gn=CCDC30;refGene_tx=XM_019836804.2;refGene_gn=CCDC30;refGene_tx=XM_019836805.2;refGene_gn=CCDC30;refGene_tx=XM_019836807.2;refGene_gn=CCDC30;refGene_tx=XM_023258571.1
C1 33518249 33518315
refGene_gn=PPIH;refGene_tx=NM_001278861.1;refGene_gn=PPIH;refGene_tx=XM_006934646.4;refGene_gn=PPIH;refGene_tx=XM_006934647.4;refGene_gn=PPIH;refGene_tx=XM_019835942.2
C1 33518656 33518721
refGene_gn=PPIH;refGene_tx=NM_001278861.1;refGene_gn=PPIH;refGene_tx=XM_006934646.4;refGene_gn=PPIH;refGene_tx=XM_006934647.4;refGene_gn=PPIH;refGene_tx=XM_019835942.2
C1 33519042 33519066
refGene_gn=PPIH;refGene_tx=NM_001278861.1;refGene_gn=PPIH;refGene_tx=XM_006934646.4;refGene_gn=PPIH;refGene_tx=XM_006934647.4;refGene_gn=PPIH;refGene_tx=XM_019835942.2
C1 33519336 33519381
refGene_gn=PPIH;refGene_tx=NM_001278861.1;refGene_gn=PPIH;refGene_tx=XM_006934646.4;refGene_gn=PPIH;refGene_tx=XM_006934647.4;refGene_gn=PPIH;refGene_tx=XM_019835942.2
C1 33519650 33519693
refGene_gn=PPIH;refGene_tx=NM_001278861.1;refGene_gn=PPIH;refGene_tx=XM_006934646.4;refGene_gn=PPIH;refGene_tx=XM_006934647.4;refGene_gn=PPIH;refGene_tx=XM_019835942.2
C1 33523211 33523304
refGene_gn=PPIH;refGene_tx=NM_001278861.1;refGene_gn=PPIH;refGene_tx=XM_006934646.4;refGene_gn=PPIH;refGene_tx=XM_006934647.4;refGene_gn=PPIH;refGene_tx=XM_019835942.2
C1 33526653 33526741
refGene_gn=PPIH;refGene_tx=NM_001278861.1;refGene_gn=PPIH;refGene_tx=XM_006934646.4;refGene_gn=PPIH;refGene_tx=XM_006934647.4;refGene_gn=PPIH;refGene_tx=XM_019835942.2

C1 33527217 33527258
refGene_gn=PPIH;refGene_tx=NM_001278861.1;refGene_gn=PPIH;refGene_tx=XM_006934646.4;refGene_gn=PPIH;refGene_tx=XM_006934647.4;refGene_gn=PPIH;refGene_tx=XM_019835942.2
C1 33528083 33528152
refGene_gn=PPIH;refGene_tx=NM_001278861.1;refGene_gn=PPIH;refGene_tx=XM_006934646.4;refGene_gn=PPIH;refGene_tx=XM_019835942.2
C1 33534213 33534234 refGene_gn=PPIH;refGene_tx=XM_006934647.4
C1 33539231 33539397
refGene_gn=YBX1;refGene_tx=XM_023258572.1;refGene_gn=YBX1;refGene_tx=XM_023258573.1
C1 33540093 33540157
refGene_gn=YBX1;refGene_tx=XM_023258572.1;refGene_gn=YBX1;refGene_tx=XM_023258573.1
C1 33549769 33549803
refGene_gn=YBX1;refGene_tx=XM_023258572.1;refGene_gn=YBX1;refGene_tx=XM_023258573.1
C1 33552630 33552720 refGene_gn=YBX1;refGene_tx=XM_023258572.1
C1 33553016 33553319
refGene_gn=YBX1;refGene_tx=XM_023258572.1;refGene_gn=YBX1;refGene_tx=XM_023258573.1
C1 33553594 33553677
refGene_gn=YBX1;refGene_tx=XM_023258572.1;refGene_gn=YBX1;refGene_tx=XM_023258573.1
C1 33557688 33557923
refGene_gn=YBX1;refGene_tx=XM_023258572.1;refGene_gn=YBX1;refGene_tx=XM_023258573.1
C1 33579232 33579281 refGene_gn=CLDN19;refGene_tx=XM_003989960.5
C1 33580032 33580185 refGene_gn=CLDN19;refGene_tx=XM_003989960.5
C1 33581958 33582043 refGene_gn=CLDN19;refGene_tx=XM_003989960.5
C1 33582152 33582317 refGene_gn=CLDN19;refGene_tx=XM_003989960.5
C1 33583341 33583564 refGene_gn=CLDN19;refGene_tx=XM_003989960.5
C1 33589549 33589684 refGene_gn=P3H1;refGene_tx=XM_003989962.4
C1 33590100 33590241 refGene_gn=P3H1;refGene_tx=XM_003989962.4
C1 33590519 33590595 refGene_gn=P3H1;refGene_tx=XM_003989962.4
C1 33590976 33591094 refGene_gn=P3H1;refGene_tx=XM_003989962.4
C1 33593412 33593563 refGene_gn=P3H1;refGene_tx=XM_003989962.4
C1 33593982 33594078 refGene_gn=P3H1;refGene_tx=XM_003989962.4
C1 33594203 33594331 refGene_gn=P3H1;refGene_tx=XM_003989962.4
C1 33596129 33596251 refGene_gn=P3H1;refGene_tx=XM_003989962.4
C1 33596425 33596478 refGene_gn=P3H1;refGene_tx=XM_003989962.4
C1 33596775 33596865 refGene_gn=P3H1;refGene_tx=XM_003989962.4
C1 33602453 33602593 refGene_gn=P3H1;refGene_tx=XM_003989962.4
C1 33603864 33603996 refGene_gn=P3H1;refGene_tx=XM_003989962.4
C1 33604201 33604391 refGene_gn=P3H1;refGene_tx=XM_003989962.4
C1 33606205 33606358 refGene_gn=P3H1;refGene_tx=XM_003989962.4
C1 33609292 33609757 refGene_gn=P3H1;refGene_tx=XM_003989962.4
C1 33628549 33628627
refGene_gn=TMEM269;refGene_tx=XM_023258574.1;refGene_gn=TMEM269;refGene_tx=XM_023258576.1;refGene_gn=TMEM269;refGene_tx=XM_023258577.1

C1 33629672 33629781
refGene_gn=TMEM269;refGene_tx=XM_023258574.1;refGene_gn=TMEM269;refGene_tx=XM_023258576.1;refGene_gn=TMEM269;refGene_tx=XM_023258577.1
C1 33632685 33632830
refGene_gn=TMEM269;refGene_tx=XM_023258574.1;refGene_gn=TMEM269;refGene_tx=XM_023258576.1;refGene_gn=TMEM269;refGene_tx=XM_023258577.1
C1 33637344 33637443
refGene_gn=TMEM269;refGene_tx=XM_023258574.1;refGene_gn=TMEM269;refGene_tx=XM_023258576.1;refGene_gn=TMEM269;refGene_tx=XM_023258577.1
C1 33638123 33638266
refGene_gn=TMEM269;refGene_tx=XM_023258574.1;refGene_gn=TMEM269;refGene_tx=XM_023258576.1;refGene_gn=TMEM269;refGene_tx=XM_023258577.1
C1 33638895 33639096
refGene_gn=TMEM269;refGene_tx=XM_023258574.1;refGene_gn=TMEM269;refGene_tx=XM_023258576.1;refGene_gn=TMEM269;refGene_tx=XM_023258577.1
C1 33644025 33644162
refGene_gn=TMEM269;refGene_tx=XM_023258574.1;refGene_gn=TMEM269;refGene_tx=XM_023258576.1;refGene_gn=TMEM269;refGene_tx=XM_023258577.1
C1 33652052 33652139
refGene_gn=SVBP;refGene_tx=XM_023258578.1;refGene_gn=SVBP;refGene_tx=XM_023258579.1
C1 33657478 33657592
refGene_gn=SVBP;refGene_tx=XM_023258578.1;refGene_gn=SVBP;refGene_tx=XM_023258579.1
C1 33670751 33670836
refGene_gn=ERMAP;refGene_tx=XM_019836813.2;refGene_gn=ERMAP;refGene_tx=XM_019836815.2
C1 33671075 33671414
refGene_gn=ERMAP;refGene_tx=XM_019836813.2;refGene_gn=ERMAP;refGene_tx=XM_019836815.2
C1 33671687 33671966
refGene_gn=ERMAP;refGene_tx=XM_019836813.2;refGene_gn=ERMAP;refGene_tx=XM_019836815.2
C1 33674279 33674396
refGene_gn=ERMAP;refGene_tx=XM_019836813.2;refGene_gn=ERMAP;refGene_tx=XM_019836815.2
C1 33674945 33674978
refGene_gn=ERMAP;refGene_tx=XM_019836813.2;refGene_gn=ERMAP;refGene_tx=XM_019836815.2
C1 33675991 33676024
refGene_gn=ERMAP;refGene_tx=XM_019836813.2;refGene_gn=ERMAP;refGene_tx=XM_019836815.2
C1 33677797 33677818
refGene_gn=ERMAP;refGene_tx=XM_019836813.2;refGene_gn=ERMAP;refGene_tx=XM_019836815.2
C1 33678617 33678638
refGene_gn=ERMAP;refGene_tx=XM_019836813.2;refGene_gn=ERMAP;refGene_tx=XM_019836815.2
C1 33678736 33678763
refGene_gn=ERMAP;refGene_tx=XM_019836813.2;refGene_gn=ERMAP;refGene_tx

=XM_019836815.2
C1	33678856	33678883	refGene_gn=ERMAP;refGene_tx=XM_019836813.2
C1	33681040	33681765
refGene_gn=ERMAP;refGene_tx=XM_019836813.2;refGene_gn=ERMAP;refGene_tx
=XM_019836815.2
C1	33689319	33690180
refGene_gn=ZNF691;refGene_tx=XM_011284681.3;refGene_gn=ZNF691;refGene_
tx=XM_019836818.1
C1	33728809	33729010	refGene_gn=SLC2A1;refGene_tx=XM_003989965.5
C1	33729398	33729602	refGene_gn=SLC2A1;refGene_tx=XM_003989965.5
C1	33730811	33730913	refGene_gn=SLC2A1;refGene_tx=XM_003989965.5
C1	33731370	33731475	refGene_gn=SLC2A1;refGene_tx=XM_003989965.5
C1	33731719	33731907	refGene_gn=SLC2A1;refGene_tx=XM_003989965.5
C1	33732000	33732163	refGene_gn=SLC2A1;refGene_tx=XM_003989965.5
C1	33732773	33733014	refGene_gn=SLC2A1;refGene_tx=XM_003989965.5
C1	33733162	33733323	refGene_gn=SLC2A1;refGene_tx=XM_003989965.5
C1	33744244	33744340	refGene_gn=SLC2A1;refGene_tx=XM_003989965.5
C1	33756885	33756903	refGene_gn=SLC2A1;refGene_tx=XM_003989965.5
C1	33877321	33878153
refGene_gn=LOC111561629;refGene_tx=XM_023257826.1
C1	33878716	33878853
refGene_gn=LOC111561629;refGene_tx=XM_023257826.1
C1	33879059	33879128
refGene_gn=LOC111561629;refGene_tx=XM_023257826.1
C1	33879647	33879849
refGene_gn=LOC111561629;refGene_tx=XM_023257826.1
C1	33880868	33880981
refGene_gn=LOC111561629;refGene_tx=XM_023257826.1
C1	33881468	33881828
refGene_gn=LOC111561629;refGene_tx=XM_023257826.1
C1	33907566	33908514
refGene_gn=LOC111561721;refGene_tx=XM_023258580.1
C1	33938687	33939683
refGene_gn=LOC101101149;refGene_tx=XM_019836826.1
C1	33954306	33955254
refGene_gn=LOC101095757;refGene_tx=XM_003989967.2
C1	33973119	33974058
refGene_gn=LOC111556297;refGene_tx=XM_023258581.1
C1	33984391	33985339
refGene_gn=LOC101101398;refGene_tx=XM_003989986.2
C1	34012756	34012886
refGene_gn=FAM183A;refGene_tx=XM_011284679.3;refGene_gn=FAM183A;refGen
e_tx=XM_019836831.2;refGene_gn=FAM183A;refGene_tx=XM_023258583.1;refGe
ne_gn=FAM183A;refGene_tx=XM_023258584.1
C1	34016146	34016206
refGene_gn=FAM183A;refGene_tx=XM_011284679.3;refGene_gn=FAM183A;refGen
e_tx=XM_019836831.2;refGene_gn=FAM183A;refGene_tx=XM_023258583.1;refGe
ne_gn=FAM183A;refGene_tx=XM_023258584.1
C1	34018090	34018194
refGene_gn=FAM183A;refGene_tx=XM_011284679.3;refGene_gn=FAM183A;refGen

e_tx=XM_019836831.2;refGene_gn=FAM183A;refGene_tx=XM_023258583.1;refGene_gn=FAM183A;refGene_tx=XM_023258584.1
C1 34025601 34025685 refGene_gn=FAM183A;refGene_tx=XM_019836831.2
C1 34027838 34027949
refGene_gn=FAM183A;refGene_tx=XM_011284679.3;refGene_gn=FAM183A;refGene_tx=XM_023258583.1;refGene_gn=FAM183A;refGene_tx=XM_023258584.1
C1 34033298 34033349 refGene_gn=EBNA1BP2;refGene_tx=XM_003989970.4
C1 34033484 34033647 refGene_gn=EBNA1BP2;refGene_tx=XM_003989970.4
C1 34035547 34035641 refGene_gn=EBNA1BP2;refGene_tx=XM_003989970.4
C1 34035875 34035951 refGene_gn=EBNA1BP2;refGene_tx=XM_003989970.4
C1 34037622 34037712 refGene_gn=EBNA1BP2;refGene_tx=XM_003989970.4
C1 34038986 34039110 refGene_gn=EBNA1BP2;refGene_tx=XM_003989970.4
C1 34039976 34040146 refGene_gn=EBNA1BP2;refGene_tx=XM_003989970.4
C1 34040345 34040429 refGene_gn=EBNA1BP2;refGene_tx=XM_003989970.4
C1 34040512 34040919
refGene_gn=CFAP57;refGene_tx=XM_011284678.3;refGene_gn=CFAP57;refGene_tx=XM_019836828.2;refGene_gn=CFAP57;refGene_tx=XM_023258585.1;refGene_gn=EBNA1BP2;refGene_tx=XM_003989970.4
C1 34041163 34041340
refGene_gn=CFAP57;refGene_tx=XM_011284678.3;refGene_gn=CFAP57;refGene_tx=XM_019836828.2;refGene_gn=CFAP57;refGene_tx=XM_023258585.1
C1 34045382 34045741
refGene_gn=CFAP57;refGene_tx=XM_011284678.3;refGene_gn=CFAP57;refGene_tx=XM_019836828.2;refGene_gn=CFAP57;refGene_tx=XM_023258585.1
C1 34047302 34047589
refGene_gn=CFAP57;refGene_tx=XM_011284678.3;refGene_gn=CFAP57;refGene_tx=XM_019836828.2;refGene_gn=CFAP57;refGene_tx=XM_023258585.1
C1 34049451 34049659
refGene_gn=CFAP57;refGene_tx=XM_011284678.3;refGene_gn=CFAP57;refGene_tx=XM_019836828.2;refGene_gn=CFAP57;refGene_tx=XM_023258585.1
C1 34052723 34052876
refGene_gn=CFAP57;refGene_tx=XM_011284678.3;refGene_gn=CFAP57;refGene_tx=XM_019836828.2;refGene_gn=CFAP57;refGene_tx=XM_023258585.1
C1 34061290 34061430
refGene_gn=CFAP57;refGene_tx=XM_011284678.3;refGene_gn=CFAP57;refGene_tx=XM_019836828.2;refGene_gn=CFAP57;refGene_tx=XM_023258585.1
C1 34062198 34062364
refGene_gn=CFAP57;refGene_tx=XM_011284678.3;refGene_gn=CFAP57;refGene_tx=XM_019836828.2;refGene_gn=CFAP57;refGene_tx=XM_023258585.1
C1 34063091 34063205
refGene_gn=CFAP57;refGene_tx=XM_011284678.3;refGene_gn=CFAP57;refGene_tx=XM_019836828.2;refGene_gn=CFAP57;refGene_tx=XM_023258585.1
C1 34068619 34068832
refGene_gn=CFAP57;refGene_tx=XM_011284678.3;refGene_gn=CFAP57;refGene_tx=XM_019836828.2;refGene_gn=CFAP57;refGene_tx=XM_023258585.1
C1 34070634 34070808
refGene_gn=CFAP57;refGene_tx=XM_011284678.3;refGene_gn=CFAP57;refGene_tx=XM_019836828.2;refGene_gn=CFAP57;refGene_tx=XM_023258585.1
C1 34074757 34074919
refGene_gn=CFAP57;refGene_tx=XM_011284678.3;refGene_gn=CFAP57;refGene_

tx=XM_019836828.2;refGene_gn=CFAP57;refGene_tx=XM_023258585.1
C1	34078916	34079072	refGene_gn=CFAP57;refGene_tx=XM_011284678.3;refGene_gn=CFAP57;refGene_tx=XM_019836828.2;refGene_gn=CFAP57;refGene_tx=XM_023258585.1
C1	34080240	34080334	refGene_gn=CFAP57;refGene_tx=XM_011284678.3;refGene_gn=CFAP57;refGene_tx=XM_019836828.2;refGene_gn=CFAP57;refGene_tx=XM_023258585.1
C1	34080957	34081148	refGene_gn=CFAP57;refGene_tx=XM_011284678.3;refGene_gn=CFAP57;refGene_tx=XM_019836828.2;refGene_gn=CFAP57;refGene_tx=XM_023258585.1
C1	34084754	34084928	refGene_gn=CFAP57;refGene_tx=XM_011284678.3;refGene_gn=CFAP57;refGene_tx=XM_019836828.2;refGene_gn=CFAP57;refGene_tx=XM_023258585.1
C1	34085836	34085995	refGene_gn=CFAP57;refGene_tx=XM_011284678.3;refGene_gn=CFAP57;refGene_tx=XM_019836828.2;refGene_gn=CFAP57;refGene_tx=XM_023258585.1
C1	34088158	34088302	refGene_gn=CFAP57;refGene_tx=XM_011284678.3;refGene_gn=CFAP57;refGene_tx=XM_019836828.2;refGene_gn=CFAP57;refGene_tx=XM_023258585.1
C1	34092763	34092880	refGene_gn=CFAP57;refGene_tx=XM_011284678.3;refGene_gn=CFAP57;refGene_tx=XM_019836828.2;refGene_gn=CFAP57;refGene_tx=XM_023258585.1
C1	34094229	34094364	refGene_gn=CFAP57;refGene_tx=XM_011284678.3;refGene_gn=CFAP57;refGene_tx=XM_019836828.2;refGene_gn=CFAP57;refGene_tx=XM_023258585.1
C1	34094446	34094590	refGene_gn=CFAP57;refGene_tx=XM_011284678.3;refGene_gn=CFAP57;refGene_tx=XM_019836828.2;refGene_gn=CFAP57;refGene_tx=XM_023258585.1
C1	34101790	34101923	refGene_gn=CFAP57;refGene_tx=XM_011284678.3;refGene_gn=CFAP57;refGene_tx=XM_019836828.2;refGene_gn=CFAP57;refGene_tx=XM_023258585.1
C1	34111844	34112059	refGene_gn=CFAP57;refGene_tx=XM_011284678.3;refGene_gn=CFAP57;refGene_tx=XM_019836828.2;refGene_gn=CFAP57;refGene_tx=XM_023258585.1
C1	34123399	34123795	refGene_gn=TMEM125;refGene_tx=XM_019836832.2
C1	34123796	34124060	refGene_gn=TMEM125;refGene_tx=XM_019836832.2
C1	34134853	34135173	refGene_gn=CC1H1orf210;refGene_tx=XM_003989988.5;refGene_gn=CC1H1orf210;refGene_tx=XM_019836833.2;refGene_gn=CC1H1orf210;refGene_tx=XM_019836834.2;refGene_gn=CC1H1orf210;refGene_tx=XM_019836835.2;refGene_gn=CC1H1orf210;refGene_tx=XM_019836836.2;refGene_gn=CC1H1orf210;refGene_tx=XM_019836837.2
C1	34135294	34135313	refGene_gn=CC1H1orf210;refGene_tx=XM_003989988.5;refGene_gn=CC1H1orf210;refGene_tx=XM_019836833.2;refGene_gn=CC1H1orf210;refGene_tx=XM_019836834.2;refGene_gn=CC1H1orf210;refGene_tx=XM_019836835.2;refGene_gn=CC1H1orf210;refGene_tx=XM_019836836.2;refGene_gn=CC1H1orf210;refGene_tx=XM_019836837.2
C1	34160304	34160362	refGene_gn=TIE1;refGene_tx=XM_023258586.1
C1	34162572	34162884	refGene_gn=TIE1;refGene_tx=XM_023258586.1

C1 34162972 34163083 refGene_gn=TIE1;refGene_tx=XM_023258586.1
C1 34164456 34164612 refGene_gn=TIE1;refGene_tx=XM_023258586.1
C1 34164756 34164888 refGene_gn=TIE1;refGene_tx=XM_023258586.1
C1 34165040 34165181 refGene_gn=TIE1;refGene_tx=XM_023258586.1
C1 34165369 34165498 refGene_gn=TIE1;refGene_tx=XM_023258586.1
C1 34166283 34166429 refGene_gn=TIE1;refGene_tx=XM_023258586.1
C1 34166668 34166813 refGene_gn=TIE1;refGene_tx=XM_023258586.1
C1 34169156 34169315 refGene_gn=TIE1;refGene_tx=XM_023258586.1
C1 34169460 34169598 refGene_gn=TIE1;refGene_tx=XM_023258586.1
C1 34169798 34170095 refGene_gn=TIE1;refGene_tx=XM_023258586.1
C1 34170670 34170961 refGene_gn=TIE1;refGene_tx=XM_023258586.1
C1 34171208 34171399 refGene_gn=TIE1;refGene_tx=XM_023258586.1
C1 34173913 34174124 refGene_gn=TIE1;refGene_tx=XM_023258586.1
C1 34174286 34174397 refGene_gn=TIE1;refGene_tx=XM_023258586.1
C1 34174576 34174767 refGene_gn=TIE1;refGene_tx=XM_023258586.1
C1 34175739 34175853 refGene_gn=TIE1;refGene_tx=XM_023258586.1
C1 34175955 34176026 refGene_gn=TIE1;refGene_tx=XM_023258586.1
C1 34177194 34177235 refGene_gn=TIE1;refGene_tx=XM_023258586.1
C1 34177324 34177421 refGene_gn=TIE1;refGene_tx=XM_023258586.1
C1 34177505 34177605 refGene_gn=TIE1;refGene_tx=XM_023258586.1
C1 34179854 34179926 refGene_gn=TIE1;refGene_tx=XM_023258586.1
C1 34191364 34191437 refGene_gn=MPL;refGene_tx=XM_019836840.2
C1 34192946 34193369
refGene_gn=MPL;refGene_tx=XM_003989971.4;refGene_gn=MPL;refGene_tx=XM_023258587.1;refGene_gn=MPL;refGene_tx=XM_023258588.1
C1 34193518 34193651
refGene_gn=MPL;refGene_tx=XM_003989971.4;refGene_gn=MPL;refGene_tx=XM_019836840.2;refGene_gn=MPL;refGene_tx=XM_023258587.1;refGene_gn=MPL;refGene_tx=XM_023258588.1
C1 34193957 34194136
refGene_gn=MPL;refGene_tx=XM_003989971.4;refGene_gn=MPL;refGene_tx=XM_019836840.2;refGene_gn=MPL;refGene_tx=XM_023258587.1;refGene_gn=MPL;refGene_tx=XM_023258588.1
C1 34194686 34194985
refGene_gn=MPL;refGene_tx=XM_003989971.4;refGene_gn=MPL;refGene_tx=XM_019836840.2;refGene_gn=MPL;refGene_tx=XM_023258587.1;refGene_gn=MPL;refGene_tx=XM_023258588.1
C1 34195403 34195566
refGene_gn=MPL;refGene_tx=XM_003989971.4;refGene_gn=MPL;refGene_tx=XM_019836840.2;refGene_gn=MPL;refGene_tx=XM_023258587.1;refGene_gn=MPL;refGene_tx=XM_023258588.1
C1 34195820 34195947
refGene_gn=MPL;refGene_tx=XM_003989971.4;refGene_gn=MPL;refGene_tx=XM_019836840.2;refGene_gn=MPL;refGene_tx=XM_023258587.1;refGene_gn=MPL;refGene_tx=XM_023258588.1
C1 34200249 34200428
refGene_gn=MPL;refGene_tx=XM_003989971.4;refGene_gn=MPL;refGene_tx=XM_019836840.2;refGene_gn=MPL;refGene_tx=XM_023258587.1;refGene_gn=MPL;refGene_tx=XM_023258588.1
C1 34200590 34200733

refGene_gn=MPL;refGene_tx=XM_003989971.4;refGene_gn=MPL;refGene_tx=XM_019836840.2;refGene_gn=MPL;refGene_tx=XM_023258587.1;refGene_gn=MPL;refGene_tx=XM_023258588.1
C1 34202361 34202521
refGene_gn=MPL;refGene_tx=XM_003989971.4;refGene_gn=MPL;refGene_tx=XM_019836840.2;refGene_gn=MPL;refGene_tx=XM_023258587.1;refGene_gn=MPL;refGene_tx=XM_023258588.1
C1 34202685 34202782
refGene_gn=MPL;refGene_tx=XM_003989971.4;refGene_gn=MPL;refGene_tx=XM_019836840.2;refGene_gn=MPL;refGene_tx=XM_023258587.1;refGene_gn=MPL;refGene_tx=XM_023258588.1
C1 34204889 34204977
refGene_gn=MPL;refGene_tx=XM_003989971.4;refGene_gn=MPL;refGene_tx=XM_019836840.2;refGene_gn=MPL;refGene_tx=XM_023258587.1
C1 34205189 34205585
refGene_gn=MPL;refGene_tx=XM_003989971.4;refGene_gn=MPL;refGene_tx=XM_019836840.2;refGene_gn=MPL;refGene_tx=XM_023258587.1;refGene_gn=MPL;refGene_tx=XM_023258588.1
C1 34210520 34210650 refGene_gn=CDC20;refGene_tx=XM_006934655.4
C1 34210821 34211010
refGene_gn=CDC20;refGene_tx=XM_003989972.3;refGene_gn=CDC20;refGene_tx=XM_006934655.4
C1 34211122 34211271
refGene_gn=CDC20;refGene_tx=XM_003989972.3;refGene_gn=CDC20;refGene_tx=XM_006934655.4
C1 34211366 34211463
refGene_gn=CDC20;refGene_tx=XM_003989972.3;refGene_gn=CDC20;refGene_tx=XM_006934655.4
C1 34211645 34211774
refGene_gn=CDC20;refGene_tx=XM_003989972.3;refGene_gn=CDC20;refGene_tx=XM_006934655.4
C1 34211873 34212070
refGene_gn=CDC20;refGene_tx=XM_003989972.3;refGene_gn=CDC20;refGene_tx=XM_006934655.4
C1 34212183 34212278
refGene_gn=CDC20;refGene_tx=XM_003989972.3;refGene_gn=CDC20;refGene_tx=XM_006934655.4
C1 34212426 34212655
refGene_gn=CDC20;refGene_tx=XM_003989972.3;refGene_gn=CDC20;refGene_tx=XM_006934655.4
C1 34212805 34212931
refGene_gn=CDC20;refGene_tx=XM_003989972.3;refGene_gn=CDC20;refGene_tx=XM_006934655.4
C1 34213851 34213969
refGene_gn=CDC20;refGene_tx=XM_003989972.3;refGene_gn=CDC20;refGene_tx=XM_006934655.4
C1 34214337 34214516
refGene_gn=CDC20;refGene_tx=XM_003989972.3;refGene_gn=CDC20;refGene_tx=XM_006934655.4
C1 34215243 34215465

refGene_gn=ELOVL1;refGene_tx=XM_003989973.4;refGene_gn=ELOVL1;refGene_tx=XM_006934658.3
C1 34215616 34215753
refGene_gn=ELOVL1;refGene_tx=XM_003989973.4;refGene_gn=ELOVL1;refGene_tx=XM_006934658.3
C1 34215842 34215948
refGene_gn=ELOVL1;refGene_tx=XM_003989973.4;refGene_gn=ELOVL1;refGene_tx=XM_006934658.3
C1 34216034 34216091
refGene_gn=ELOVL1;refGene_tx=XM_003989973.4;refGene_gn=ELOVL1;refGene_tx=XM_006934658.3
C1 34216235 34216316
refGene_gn=ELOVL1;refGene_tx=XM_003989973.4;refGene_gn=ELOVL1;refGene_tx=XM_006934658.3
C1 34216490 34216681
refGene_gn=ELOVL1;refGene_tx=XM_003989973.4;refGene_gn=ELOVL1;refGene_tx=XM_006934658.3
C1 34216862 34216908
refGene_gn=ELOVL1;refGene_tx=XM_003989973.4;refGene_gn=ELOVL1;refGene_tx=XM_006934658.3
C1 34223732 34223796
refGene_gn=MED8;refGene_tx=XM_003989976.4;refGene_gn=MED8;refGene_tx=XM_019836843.2
C1 34229663 34229670 refGene_gn=MED8;refGene_tx=XM_019836844.2
C1 34235780 34235845
refGene_gn=MED8;refGene_tx=XM_003989976.4;refGene_gn=MED8;refGene_tx=XM_006934660.4;refGene_gn=MED8;refGene_tx=XM_019836843.2;refGene_gn=MED8;refGene_tx=XM_019836844.2
C1 34236629 34236878
refGene_gn=MED8;refGene_tx=XM_003989976.4;refGene_gn=MED8;refGene_tx=XM_006934660.4;refGene_gn=MED8;refGene_tx=XM_019836843.2;refGene_gn=MED8;refGene_tx=XM_019836844.2
C1 34237233 34237315
refGene_gn=MED8;refGene_tx=XM_003989976.4;refGene_gn=MED8;refGene_tx=XM_006934660.4;refGene_gn=MED8;refGene_tx=XM_019836843.2;refGene_gn=MED8;refGene_tx=XM_019836844.2
C1 34237510 34237651
refGene_gn=MED8;refGene_tx=XM_003989976.4;refGene_gn=MED8;refGene_tx=XM_006934660.4;refGene_gn=MED8;refGene_tx=XM_019836843.2;refGene_gn=MED8;refGene_tx=XM_019836844.2
C1 34238177 34238322
refGene_gn=MED8;refGene_tx=XM_003989976.4;refGene_gn=MED8;refGene_tx=XM_006934660.4;refGene_gn=MED8;refGene_tx=XM_019836843.2;refGene_gn=MED8;refGene_tx=XM_019836844.2
C1 34239006 34239173
refGene_gn=MED8;refGene_tx=XM_003989976.4;refGene_gn=MED8;refGene_tx=XM_006934660.4;refGene_gn=MED8;refGene_tx=XM_019836843.2;refGene_gn=MED8;refGene_tx=XM_019836844.2
C1 34240438 34240444 refGene_gn=MED8;refGene_tx=XM_003989976.4
C1 34240648 34240675

refGene_gn=SZT2;refGene_tx=XM_003989977.4;refGene_gn=SZT2;refGene_tx=XM_019836838.2;refGene_gn=SZT2;refGene_tx=XM_019836839.1
C1 34250480 34250606
refGene_gn=SZT2;refGene_tx=XM_003989977.4;refGene_gn=SZT2;refGene_tx=XM_019836838.2;refGene_gn=SZT2;refGene_tx=XM_019836839.1
C1 34251148 34251322
refGene_gn=SZT2;refGene_tx=XM_003989977.4;refGene_gn=SZT2;refGene_tx=XM_019836838.2;refGene_gn=SZT2;refGene_tx=XM_019836839.1
C1 34251621 34251792
refGene_gn=SZT2;refGene_tx=XM_003989977.4;refGene_gn=SZT2;refGene_tx=XM_019836838.2;refGene_gn=SZT2;refGene_tx=XM_019836839.1
C1 34255646 34255778
refGene_gn=SZT2;refGene_tx=XM_003989977.4;refGene_gn=SZT2;refGene_tx=XM_019836838.2;refGene_gn=SZT2;refGene_tx=XM_019836839.1
C1 34256515 34256657
refGene_gn=SZT2;refGene_tx=XM_003989977.4;refGene_gn=SZT2;refGene_tx=XM_019836838.2;refGene_gn=SZT2;refGene_tx=XM_019836839.1
C1 34257053 34257160
refGene_gn=SZT2;refGene_tx=XM_003989977.4;refGene_gn=SZT2;refGene_tx=XM_019836838.2;refGene_gn=SZT2;refGene_tx=XM_019836839.1
C1 34260660 34260871
refGene_gn=SZT2;refGene_tx=XM_003989977.4;refGene_gn=SZT2;refGene_tx=XM_019836838.2;refGene_gn=SZT2;refGene_tx=XM_019836839.1
C1 34261102 34261273
refGene_gn=SZT2;refGene_tx=XM_003989977.4;refGene_gn=SZT2;refGene_tx=XM_019836838.2;refGene_gn=SZT2;refGene_tx=XM_019836839.1
C1 34261687 34261922
refGene_gn=SZT2;refGene_tx=XM_003989977.4;refGene_gn=SZT2;refGene_tx=XM_019836838.2;refGene_gn=SZT2;refGene_tx=XM_019836839.1
C1 34262121 34262251
refGene_gn=SZT2;refGene_tx=XM_003989977.4;refGene_gn=SZT2;refGene_tx=XM_019836838.2;refGene_gn=SZT2;refGene_tx=XM_019836839.1
C1 34263002 34263145
refGene_gn=SZT2;refGene_tx=XM_003989977.4;refGene_gn=SZT2;refGene_tx=XM_019836838.2;refGene_gn=SZT2;refGene_tx=XM_019836839.1
C1 34263373 34263526
refGene_gn=SZT2;refGene_tx=XM_003989977.4;refGene_gn=SZT2;refGene_tx=XM_019836838.2;refGene_gn=SZT2;refGene_tx=XM_019836839.1
C1 34263664 34263779
refGene_gn=SZT2;refGene_tx=XM_003989977.4;refGene_gn=SZT2;refGene_tx=XM_019836838.2;refGene_gn=SZT2;refGene_tx=XM_019836839.1
C1 34263993 34264211
refGene_gn=SZT2;refGene_tx=XM_003989977.4;refGene_gn=SZT2;refGene_tx=XM_019836838.2;refGene_gn=SZT2;refGene_tx=XM_019836839.1
C1 34264307 34264523
refGene_gn=SZT2;refGene_tx=XM_003989977.4;refGene_gn=SZT2;refGene_tx=XM_019836838.2;refGene_gn=SZT2;refGene_tx=XM_019836839.1
C1 34264905 34264984
refGene_gn=SZT2;refGene_tx=XM_003989977.4;refGene_gn=SZT2;refGene_tx=XM_019836838.2;refGene_gn=SZT2;refGene_tx=XM_019836839.1

C1 34265228 34265323
refGene_gn=SZT2;refGene_tx=XM_003989977.4;refGene_gn=SZT2;refGene_tx=XM_019836838.2;refGene_gn=SZT2;refGene_tx=XM_019836839.1
C1 34265776 34265945
refGene_gn=SZT2;refGene_tx=XM_003989977.4;refGene_gn=SZT2;refGene_tx=XM_019836838.2;refGene_gn=SZT2;refGene_tx=XM_019836839.1
C1 34266137 34266252
refGene_gn=SZT2;refGene_tx=XM_003989977.4;refGene_gn=SZT2;refGene_tx=XM_019836838.2;refGene_gn=SZT2;refGene_tx=XM_019836839.1
C1 34266366 34266480
refGene_gn=SZT2;refGene_tx=XM_003989977.4;refGene_gn=SZT2;refGene_tx=XM_019836838.2;refGene_gn=SZT2;refGene_tx=XM_019836839.1
C1 34266644 34266815
refGene_gn=SZT2;refGene_tx=XM_003989977.4;refGene_gn=SZT2;refGene_tx=XM_019836838.2;refGene_gn=SZT2;refGene_tx=XM_019836839.1
C1 34266981 34267076
refGene_gn=SZT2;refGene_tx=XM_003989977.4;refGene_gn=SZT2;refGene_tx=XM_019836838.2;refGene_gn=SZT2;refGene_tx=XM_019836839.1
C1 34267325 34267449
refGene_gn=SZT2;refGene_tx=XM_003989977.4;refGene_gn=SZT2;refGene_tx=XM_019836838.2;refGene_gn=SZT2;refGene_tx=XM_019836839.1
C1 34267552 34267717
refGene_gn=SZT2;refGene_tx=XM_003989977.4;refGene_gn=SZT2;refGene_tx=XM_019836838.2;refGene_gn=SZT2;refGene_tx=XM_019836839.1
C1 34267806 34268011
refGene_gn=SZT2;refGene_tx=XM_003989977.4;refGene_gn=SZT2;refGene_tx=XM_019836838.2;refGene_gn=SZT2;refGene_tx=XM_019836839.1
C1 34268297 34268413
refGene_gn=SZT2;refGene_tx=XM_003989977.4;refGene_gn=SZT2;refGene_tx=XM_019836838.2;refGene_gn=SZT2;refGene_tx=XM_019836839.1
C1 34268523 34268770
refGene_gn=SZT2;refGene_tx=XM_003989977.4;refGene_gn=SZT2;refGene_tx=XM_019836838.2;refGene_gn=SZT2;refGene_tx=XM_019836839.1
C1 34269658 34269800
refGene_gn=SZT2;refGene_tx=XM_003989977.4;refGene_gn=SZT2;refGene_tx=XM_019836838.2;refGene_gn=SZT2;refGene_tx=XM_019836839.1
C1 34269964 34270057
refGene_gn=SZT2;refGene_tx=XM_003989977.4;refGene_gn=SZT2;refGene_tx=XM_019836838.2;refGene_gn=SZT2;refGene_tx=XM_019836839.1
C1 34270208 34270287
refGene_gn=SZT2;refGene_tx=XM_003989977.4;refGene_gn=SZT2;refGene_tx=XM_019836838.2;refGene_gn=SZT2;refGene_tx=XM_019836839.1
C1 34270392 34270683
refGene_gn=SZT2;refGene_tx=XM_003989977.4;refGene_gn=SZT2;refGene_tx=XM_019836838.2;refGene_gn=SZT2;refGene_tx=XM_019836839.1
C1 34270835 34270977
refGene_gn=SZT2;refGene_tx=XM_003989977.4;refGene_gn=SZT2;refGene_tx=XM_019836838.2;refGene_gn=SZT2;refGene_tx=XM_019836839.1
C1 34271164 34271272
refGene_gn=SZT2;refGene_tx=XM_003989977.4;refGene_gn=SZT2;refGene_tx=X

M_019836838.2;refGene_gn=SZT2;refGene_tx=XM_019836839.1
C1 34271359 34271423
refGene_gn=SZT2;refGene_tx=XM_003989977.4;refGene_gn=SZT2;refGene_tx=X
M_019836838.2;refGene_gn=SZT2;refGene_tx=XM_019836839.1
C1 34271645 34271831
refGene_gn=SZT2;refGene_tx=XM_003989977.4;refGene_gn=SZT2;refGene_tx=X
M_019836838.2;refGene_gn=SZT2;refGene_tx=XM_019836839.1
C1 34272177 34272345
refGene_gn=SZT2;refGene_tx=XM_003989977.4;refGene_gn=SZT2;refGene_tx=X
M_019836838.2;refGene_gn=SZT2;refGene_tx=XM_019836839.1
C1 34272436 34272515
refGene_gn=SZT2;refGene_tx=XM_003989977.4;refGene_gn=SZT2;refGene_tx=X
M_019836838.2;refGene_gn=SZT2;refGene_tx=XM_019836839.1
C1 34272649 34272721
refGene_gn=SZT2;refGene_tx=XM_003989977.4;refGene_gn=SZT2;refGene_tx=X
M_019836838.2;refGene_gn=SZT2;refGene_tx=XM_019836839.1
C1 34272903 34273105
refGene_gn=SZT2;refGene_tx=XM_003989977.4;refGene_gn=SZT2;refGene_tx=X
M_019836838.2;refGene_gn=SZT2;refGene_tx=XM_019836839.1
C1 34273899 34273999
refGene_gn=SZT2;refGene_tx=XM_003989977.4;refGene_gn=SZT2;refGene_tx=X
M_019836838.2;refGene_gn=SZT2;refGene_tx=XM_019836839.1
C1 34274679 34274809
refGene_gn=SZT2;refGene_tx=XM_003989977.4;refGene_gn=SZT2;refGene_tx=X
M_019836838.2;refGene_gn=SZT2;refGene_tx=XM_019836839.1
C1 34276574 34276727
refGene_gn=SZT2;refGene_tx=XM_003989977.4;refGene_gn=SZT2;refGene_tx=X
M_019836838.2;refGene_gn=SZT2;refGene_tx=XM_019836839.1
C1 34276809 34276912
refGene_gn=SZT2;refGene_tx=XM_003989977.4;refGene_gn=SZT2;refGene_tx=X
M_019836838.2;refGene_gn=SZT2;refGene_tx=XM_019836839.1
C1 34277001 34277107
refGene_gn=SZT2;refGene_tx=XM_003989977.4;refGene_gn=SZT2;refGene_tx=X
M_019836838.2;refGene_gn=SZT2;refGene_tx=XM_019836839.1
C1 34277192 34277304
refGene_gn=SZT2;refGene_tx=XM_003989977.4;refGene_gn=SZT2;refGene_tx=X
M_019836838.2;refGene_gn=SZT2;refGene_tx=XM_019836839.1
C1 34277934 34278053
refGene_gn=SZT2;refGene_tx=XM_003989977.4;refGene_gn=SZT2;refGene_tx=X
M_019836838.2;refGene_gn=SZT2;refGene_tx=XM_019836839.1
C1 34278157 34278322
refGene_gn=SZT2;refGene_tx=XM_003989977.4;refGene_gn=SZT2;refGene_tx=X
M_019836838.2;refGene_gn=SZT2;refGene_tx=XM_019836839.1
C1 34278574 34278659
refGene_gn=SZT2;refGene_tx=XM_003989977.4;refGene_gn=SZT2;refGene_tx=X
M_019836838.2;refGene_gn=SZT2;refGene_tx=XM_019836839.1
C1 34278817 34278982
refGene_gn=SZT2;refGene_tx=XM_003989977.4;refGene_gn=SZT2;refGene_tx=X
M_019836838.2;refGene_gn=SZT2;refGene_tx=XM_019836839.1
C1 34279070 34279238

refGene_gn=SZT2;refGene_tx=XM_003989977.4;refGene_gn=SZT2;refGene_tx=XM_019836838.2;refGene_gn=SZT2;refGene_tx=XM_019836839.1
C1 34279596 34279736
refGene_gn=SZT2;refGene_tx=XM_003989977.4;refGene_gn=SZT2;refGene_tx=XM_019836838.2;refGene_gn=SZT2;refGene_tx=XM_019836839.1
C1 34280162 34280329
refGene_gn=SZT2;refGene_tx=XM_003989977.4;refGene_gn=SZT2;refGene_tx=XM_019836838.2;refGene_gn=SZT2;refGene_tx=XM_019836839.1
C1 34280456 34280554
refGene_gn=SZT2;refGene_tx=XM_003989977.4;refGene_gn=SZT2;refGene_tx=XM_019836838.2;refGene_gn=SZT2;refGene_tx=XM_019836839.1
C1 34280645 34280778
refGene_gn=SZT2;refGene_tx=XM_003989977.4;refGene_gn=SZT2;refGene_tx=XM_019836838.2;refGene_gn=SZT2;refGene_tx=XM_019836839.1
C1 34280889 34281020
refGene_gn=SZT2;refGene_tx=XM_003989977.4;refGene_gn=SZT2;refGene_tx=XM_019836838.2;refGene_gn=SZT2;refGene_tx=XM_019836839.1
C1 34281123 34281224
refGene_gn=SZT2;refGene_tx=XM_003989977.4;refGene_gn=SZT2;refGene_tx=XM_019836838.2;refGene_gn=SZT2;refGene_tx=XM_019836839.1
C1 34281297 34281474
refGene_gn=SZT2;refGene_tx=XM_003989977.4;refGene_gn=SZT2;refGene_tx=XM_019836838.2;refGene_gn=SZT2;refGene_tx=XM_019836839.1
C1 34281677 34281945
refGene_gn=SZT2;refGene_tx=XM_003989977.4;refGene_gn=SZT2;refGene_tx=XM_019836838.2;refGene_gn=SZT2;refGene_tx=XM_019836839.1
C1 34282026 34282106
refGene_gn=SZT2;refGene_tx=XM_003989977.4;refGene_gn=SZT2;refGene_tx=XM_019836838.2;refGene_gn=SZT2;refGene_tx=XM_019836839.1
C1 34282193 34282319
refGene_gn=SZT2;refGene_tx=XM_003989977.4;refGene_gn=SZT2;refGene_tx=XM_019836838.2;refGene_gn=SZT2;refGene_tx=XM_019836839.1
C1 34282443 34282643
refGene_gn=SZT2;refGene_tx=XM_003989977.4;refGene_gn=SZT2;refGene_tx=XM_019836838.2
C1 34284758 34284849
refGene_gn=SZT2;refGene_tx=XM_003989977.4;refGene_gn=SZT2;refGene_tx=XM_019836838.2;refGene_gn=SZT2;refGene_tx=XM_019836839.1
C1 34284994 34285075
refGene_gn=SZT2;refGene_tx=XM_003989977.4;refGene_gn=SZT2;refGene_tx=XM_019836838.2
C1 34285157 34285232
refGene_gn=SZT2;refGene_tx=XM_003989977.4;refGene_gn=SZT2;refGene_tx=XM_019836838.2
C1 34286768 34286771 refGene_gn=SZT2;refGene_tx=XM_019836838.2
C1 34287693 34287907 refGene_gn=SZT2;refGene_tx=XM_003989977.4
C1 34288312 34288466 refGene_gn=SZT2;refGene_tx=XM_003989977.4
C1 34288616 34288739 refGene_gn=SZT2;refGene_tx=XM_003989977.4
C1 34288848 34289254 refGene_gn=SZT2;refGene_tx=XM_003989977.4
C1 34289383 34289500 refGene_gn=SZT2;refGene_tx=XM_003989977.4

C1 34290816 34290885 refGene_gn=SZT2;refGene_tx=XM_003989977.4
C1 34291021 34291165 refGene_gn=SZT2;refGene_tx=XM_003989977.4
C1 34291974 34292074
refGene_gn=HYI;refGene_tx=XM_003989979.4;refGene_gn=HYI;refGene_tx=XM_023258590.1
C1 34292168 34292303
refGene_gn=HYI;refGene_tx=XM_003989979.4;refGene_gn=HYI;refGene_tx=XM_023258590.1
C1 34292437 34292507
refGene_gn=HYI;refGene_tx=XM_003989979.4;refGene_gn=HYI;refGene_tx=XM_023258590.1
C1 34292620 34292670
refGene_gn=HYI;refGene_tx=XM_003989979.4;refGene_gn=HYI;refGene_tx=XM_023258590.1
C1 34292760 34292839
refGene_gn=HYI;refGene_tx=XM_003989979.4;refGene_gn=HYI;refGene_tx=XM_023258590.1
C1 34293013 34293128
refGene_gn=HYI;refGene_tx=XM_003989979.4;refGene_gn=HYI;refGene_tx=XM_023258590.1
C1 34294150 34294262
refGene_gn=HYI;refGene_tx=XM_003989979.4;refGene_gn=HYI;refGene_tx=XM_023258590.1
C1 34294358 34294557
refGene_gn=HYI;refGene_tx=XM_003989979.4;refGene_gn=HYI;refGene_tx=XM_023258590.1
C1 34360234 34360325
refGene_gn=PTPRF;refGene_tx=XM_023258591.1;refGene_gn=PTPRF;refGene_tx=XM_023258592.1;refGene_gn=PTPRF;refGene_tx=XM_023258593.1;refGene_gn=PTPRF;refGene_tx=XM_023258595.1;refGene_gn=PTPRF;refGene_tx=XM_023258596.1;refGene_gn=PTPRF;refGene_tx=XM_023258597.1;refGene_gn=PTPRF;refGene_tx=XM_023258598.1;refGene_gn=PTPRF;refGene_tx=XM_023258599.1;refGene_gn=PTPRF;refGene_tx=XM_023258600.1;refGene_gn=PTPRF;refGene_tx=XM_023258601.1;refGene_gn=PTPRF;refGene_tx=XM_023258602.1;refGene_gn=PTPRF;refGene_tx=XM_023258603.1;refGene_gn=PTPRF;refGene_tx=XM_023258604.1;refGene_gn=PTPRF;refGene_tx=XM_023258605.1;refGene_gn=PTPRF;refGene_tx=XM_023258606.1;refGene_gn=PTPRF;refGene_tx=XM_023258607.1;refGene_gn=PTPRF;refGene_tx=XM_023258608.1;refGene_gn=PTPRF;refGene_tx=XM_023258609.1;refGene_gn=PTPRF;refGene_tx=XM_023258610.1;refGene_gn=PTPRF;refGene_tx=XM_023258611.1;refGene_gn=PTPRF;refGene_tx=XM_023258612.1;refGene_gn=PTPRF;refGene_tx=XM_023258613.1;refGene_gn=PTPRF;refGene_tx=XM_023258614.1
C1 34368973 34369119
refGene_gn=PTPRF;refGene_tx=XM_023258591.1;refGene_gn=PTPRF;refGene_tx=XM_023258592.1;refGene_gn=PTPRF;refGene_tx=XM_023258593.1;refGene_gn=PTPRF;refGene_tx=XM_023258595.1;refGene_gn=PTPRF;refGene_tx=XM_023258596.1;refGene_gn=PTPRF;refGene_tx=XM_023258597.1;refGene_gn=PTPRF;refGene_tx=XM_023258598.1;refGene_gn=PTPRF;refGene_tx=XM_023258599.1;refGene_gn=PTPRF;refGene_tx=XM_023258600.1;refGene_gn=PTPRF;refGene_tx=XM_023258601.1;refGene_gn=PTPRF;refGene_tx=XM_023258602.1;refGene_gn=PTPRF;

refGene_tx=XM_023258603.1;refGene_gn=PTPRF;refGene_tx=XM_023258604.1;refGene_gn=PTPRF;refGene_tx=XM_023258605.1;refGene_gn=PTPRF;refGene_tx=XM_023258606.1;refGene_gn=PTPRF;refGene_tx=XM_023258607.1;refGene_gn=PTPRF;refGene_tx=XM_023258608.1;refGene_gn=PTPRF;refGene_tx=XM_023258609.1;refGene_gn=PTPRF;refGene_tx=XM_023258610.1;refGene_gn=PTPRF;refGene_tx=XM_023258611.1;refGene_gn=PTPRF;refGene_tx=XM_023258612.1;refGene_gn=PTPRF;refGene_tx=XM_023258613.1;refGene_gn=PTPRF;refGene_tx=XM_023258614.1
C1 34369286 34369428
refGene_gn=PTPRF;refGene_tx=XM_023258591.1;refGene_gn=PTPRF;refGene_tx=XM_023258592.1;refGene_gn=PTPRF;refGene_tx=XM_023258593.1;refGene_gn=PTPRF;refGene_tx=XM_023258595.1;refGene_gn=PTPRF;refGene_tx=XM_023258596.1;refGene_gn=PTPRF;refGene_tx=XM_023258597.1;refGene_gn=PTPRF;refGene_tx=XM_023258598.1;refGene_gn=PTPRF;refGene_tx=XM_023258599.1;refGene_gn=PTPRF;refGene_tx=XM_023258600.1;refGene_gn=PTPRF;refGene_tx=XM_023258601.1;refGene_gn=PTPRF;refGene_tx=XM_023258602.1;refGene_gn=PTPRF;refGene_tx=XM_023258603.1;refGene_gn=PTPRF;refGene_tx=XM_023258604.1;refGene_gn=PTPRF;refGene_tx=XM_023258605.1;refGene_gn=PTPRF;refGene_tx=XM_023258606.1;refGene_gn=PTPRF;refGene_tx=XM_023258607.1;refGene_gn=PTPRF;refGene_tx=XM_023258608.1;refGene_gn=PTPRF;refGene_tx=XM_023258609.1;refGene_gn=PTPRF;refGene_tx=XM_023258610.1;refGene_gn=PTPRF;refGene_tx=XM_023258611.1;refGene_gn=PTPRF;refGene_tx=XM_023258612.1;refGene_gn=PTPRF;refGene_tx=XM_023258613.1;refGene_gn=PTPRF;refGene_tx=XM_023258614.1
C1 34382879 34383068
refGene_gn=PTPRF;refGene_tx=XM_023258591.1;refGene_gn=PTPRF;refGene_tx=XM_023258592.1;refGene_gn=PTPRF;refGene_tx=XM_023258593.1;refGene_gn=PTPRF;refGene_tx=XM_023258595.1;refGene_gn=PTPRF;refGene_tx=XM_023258596.1;refGene_gn=PTPRF;refGene_tx=XM_023258597.1;refGene_gn=PTPRF;refGene_tx=XM_023258598.1;refGene_gn=PTPRF;refGene_tx=XM_023258599.1;refGene_gn=PTPRF;refGene_tx=XM_023258600.1;refGene_gn=PTPRF;refGene_tx=XM_023258601.1;refGene_gn=PTPRF;refGene_tx=XM_023258602.1;refGene_gn=PTPRF;refGene_tx=XM_023258603.1;refGene_gn=PTPRF;refGene_tx=XM_023258604.1;refGene_gn=PTPRF;refGene_tx=XM_023258605.1;refGene_gn=PTPRF;refGene_tx=XM_023258606.1;refGene_gn=PTPRF;refGene_tx=XM_023258607.1;refGene_gn=PTPRF;refGene_tx=XM_023258608.1;refGene_gn=PTPRF;refGene_tx=XM_023258609.1;refGene_gn=PTPRF;refGene_tx=XM_023258610.1;refGene_gn=PTPRF;refGene_tx=XM_023258611.1;refGene_gn=PTPRF;refGene_tx=XM_023258612.1;refGene_gn=PTPRF;refGene_tx=XM_023258613.1;refGene_gn=PTPRF;refGene_tx=XM_023258614.1
C1 34389026 34389044
refGene_gn=PTPRF;refGene_tx=XM_023258591.1;refGene_gn=PTPRF;refGene_tx=XM_023258592.1;refGene_gn=PTPRF;refGene_tx=XM_023258593.1;refGene_gn=PTPRF;refGene_tx=XM_023258596.1;refGene_gn=PTPRF;refGene_tx=XM_023258598.1;refGene_gn=PTPRF;refGene_tx=XM_023258599.1;refGene_gn=PTPRF;refGene_tx=XM_023258600.1;refGene_gn=PTPRF;refGene_tx=XM_023258603.1;refGene_gn=PTPRF;refGene_tx=XM_023258604.1;refGene_gn=PTPRF;refGene_tx=XM_023258607.1;refGene_gn=PTPRF;refGene_tx=XM_023258608.1;refGene_gn=PTPRF;refGene_tx=XM_023258609.1;refGene_gn=PTPRF;refGene_tx=XM_023258610.1;refGene_gn=PTPRF;refGene_tx=XM_023258611.1

C1	34391944	34392055
refGene_gn=PTPRF;refGene_tx=XM_023258591.1;refGene_gn=PTPRF;refGene_tx=XM_023258592.1;refGene_gn=PTPRF;refGene_tx=XM_023258593.1;refGene_gn=PTPRF;refGene_tx=XM_023258595.1;refGene_gn=PTPRF;refGene_tx=XM_023258596.1;refGene_gn=PTPRF;refGene_tx=XM_023258597.1;refGene_gn=PTPRF;refGene_tx=XM_023258598.1;refGene_gn=PTPRF;refGene_tx=XM_023258599.1;refGene_gn=PTPRF;refGene_tx=XM_023258600.1;refGene_gn=PTPRF;refGene_tx=XM_023258601.1;refGene_gn=PTPRF;refGene_tx=XM_023258602.1;refGene_gn=PTPRF;refGene_tx=XM_023258603.1;refGene_gn=PTPRF;refGene_tx=XM_023258604.1;refGene_gn=PTPRF;refGene_tx=XM_023258605.1;refGene_gn=PTPRF;refGene_tx=XM_023258606.1;refGene_gn=PTPRF;refGene_tx=XM_023258607.1;refGene_gn=PTPRF;refGene_tx=XM_023258608.1;refGene_gn=PTPRF;refGene_tx=XM_023258609.1;refGene_gn=PTPRF;refGene_tx=XM_023258610.1;refGene_gn=PTPRF;refGene_tx=XM_023258611.1;refGene_gn=PTPRF;refGene_tx=XM_023258612.1;refGene_gn=PTPRF;refGene_tx=XM_023258613.1;refGene_gn=PTPRF;refGene_tx=XM_023258614.1
C1	34395822	34395834
refGene_gn=PTPRF;refGene_tx=XM_023258591.1;refGene_gn=PTPRF;refGene_tx=XM_023258592.1;refGene_gn=PTPRF;refGene_tx=XM_023258595.1;refGene_gn=PTPRF;refGene_tx=XM_023258596.1;refGene_gn=PTPRF;refGene_tx=XM_023258598.1;refGene_gn=PTPRF;refGene_tx=XM_023258599.1;refGene_gn=PTPRF;refGene_tx=XM_023258600.1;refGene_gn=PTPRF;refGene_tx=XM_023258603.1;refGene_gn=PTPRF;refGene_tx=XM_023258604.1;refGene_gn=PTPRF;refGene_tx=XM_023258605.1;refGene_gn=PTPRF;refGene_tx=XM_023258607.1;refGene_gn=PTPRF;refGene_tx=XM_023258608.1;refGene_gn=PTPRF;refGene_tx=XM_023258609.1;refGene_gn=PTPRF;refGene_tx=XM_023258610.1;refGene_gn=PTPRF;refGene_tx=XM_023258614.1
C1	34401473	34401743
refGene_gn=PTPRF;refGene_tx=XM_023258591.1;refGene_gn=PTPRF;refGene_tx=XM_023258592.1;refGene_gn=PTPRF;refGene_tx=XM_023258593.1;refGene_gn=PTPRF;refGene_tx=XM_023258595.1;refGene_gn=PTPRF;refGene_tx=XM_023258596.1;refGene_gn=PTPRF;refGene_tx=XM_023258597.1;refGene_gn=PTPRF;refGene_tx=XM_023258598.1;refGene_gn=PTPRF;refGene_tx=XM_023258599.1;refGene_gn=PTPRF;refGene_tx=XM_023258600.1;refGene_gn=PTPRF;refGene_tx=XM_023258601.1;refGene_gn=PTPRF;refGene_tx=XM_023258602.1;refGene_gn=PTPRF;refGene_tx=XM_023258603.1;refGene_gn=PTPRF;refGene_tx=XM_023258604.1;refGene_gn=PTPRF;refGene_tx=XM_023258605.1;refGene_gn=PTPRF;refGene_tx=XM_023258606.1;refGene_gn=PTPRF;refGene_tx=XM_023258607.1;refGene_gn=PTPRF;refGene_tx=XM_023258608.1;refGene_gn=PTPRF;refGene_tx=XM_023258609.1;refGene_gn=PTPRF;refGene_tx=XM_023258610.1;refGene_gn=PTPRF;refGene_tx=XM_023258611.1;refGene_gn=PTPRF;refGene_tx=XM_023258612.1;refGene_gn=PTPRF;refGene_tx=XM_023258613.1;refGene_gn=PTPRF;refGene_tx=XM_023258614.1
C1	34403220	34403802
refGene_gn=PTPRF;refGene_tx=XM_023258591.1;refGene_gn=PTPRF;refGene_tx=XM_023258592.1;refGene_gn=PTPRF;refGene_tx=XM_023258593.1;refGene_gn=PTPRF;refGene_tx=XM_023258595.1;refGene_gn=PTPRF;refGene_tx=XM_023258596.1;refGene_gn=PTPRF;refGene_tx=XM_023258597.1;refGene_gn=PTPRF;refGene_tx=XM_023258598.1;refGene_gn=PTPRF;refGene_tx=XM_023258599.1;refGene_gn=PTPRF;refGene_tx=XM_023258600.1;refGene_gn=PTPRF;refGene_tx=XM_02

3258601.1;refGene_gn=PTPRF;refGene_tx=XM_023258602.1;refGene_gn=PTPRF;refGene_tx=XM_023258603.1;refGene_gn=PTPRF;refGene_tx=XM_023258604.1;refGene_gn=PTPRF;refGene_tx=XM_023258605.1;refGene_gn=PTPRF;refGene_tx=XM_023258606.1;refGene_gn=PTPRF;refGene_tx=XM_023258607.1;refGene_gn=PTPRF;refGene_tx=XM_023258608.1;refGene_gn=PTPRF;refGene_tx=XM_023258609.1;refGene_gn=PTPRF;refGene_tx=XM_023258610.1;refGene_gn=PTPRF;refGene_tx=XM_023258611.1;refGene_gn=PTPRF;refGene_tx=XM_023258612.1;refGene_gn=PTPRF;refGene_tx=XM_023258613.1;refGene_gn=PTPRF;refGene_tx=XM_023258614.1
C1 34404014 34404151
refGene_gn=PTPRF;refGene_tx=XM_023258591.1;refGene_gn=PTPRF;refGene_tx=XM_023258592.1;refGene_gn=PTPRF;refGene_tx=XM_023258593.1;refGene_gn=PTPRF;refGene_tx=XM_023258595.1;refGene_gn=PTPRF;refGene_tx=XM_023258596.1;refGene_gn=PTPRF;refGene_tx=XM_023258597.1;refGene_gn=PTPRF;refGene_tx=XM_023258598.1;refGene_gn=PTPRF;refGene_tx=XM_023258599.1;refGene_gn=PTPRF;refGene_tx=XM_023258600.1;refGene_gn=PTPRF;refGene_tx=XM_023258601.1;refGene_gn=PTPRF;refGene_tx=XM_023258602.1;refGene_gn=PTPRF;refGene_tx=XM_023258603.1;refGene_gn=PTPRF;refGene_tx=XM_023258604.1;refGene_gn=PTPRF;refGene_tx=XM_023258605.1;refGene_gn=PTPRF;refGene_tx=XM_023258606.1;refGene_gn=PTPRF;refGene_tx=XM_023258607.1;refGene_gn=PTPRF;refGene_tx=XM_023258608.1;refGene_gn=PTPRF;refGene_tx=XM_023258609.1;refGene_gn=PTPRF;refGene_tx=XM_023258610.1;refGene_gn=PTPRF;refGene_tx=XM_023258611.1;refGene_gn=PTPRF;refGene_tx=XM_023258612.1;refGene_gn=PTPRF;refGene_tx=XM_023258613.1;refGene_gn=PTPRF;refGene_tx=XM_023258614.1
C1 34404645 34404790
refGene_gn=PTPRF;refGene_tx=XM_023258591.1;refGene_gn=PTPRF;refGene_tx=XM_023258592.1;refGene_gn=PTPRF;refGene_tx=XM_023258593.1;refGene_gn=PTPRF;refGene_tx=XM_023258595.1;refGene_gn=PTPRF;refGene_tx=XM_023258596.1;refGene_gn=PTPRF;refGene_tx=XM_023258597.1;refGene_gn=PTPRF;refGene_tx=XM_023258598.1;refGene_gn=PTPRF;refGene_tx=XM_023258599.1;refGene_gn=PTPRF;refGene_tx=XM_023258600.1;refGene_gn=PTPRF;refGene_tx=XM_023258601.1;refGene_gn=PTPRF;refGene_tx=XM_023258602.1;refGene_gn=PTPRF;refGene_tx=XM_023258603.1;refGene_gn=PTPRF;refGene_tx=XM_023258604.1;refGene_gn=PTPRF;refGene_tx=XM_023258605.1;refGene_gn=PTPRF;refGene_tx=XM_023258606.1;refGene_gn=PTPRF;refGene_tx=XM_023258607.1;refGene_gn=PTPRF;refGene_tx=XM_023258608.1;refGene_gn=PTPRF;refGene_tx=XM_023258609.1;refGene_gn=PTPRF;refGene_tx=XM_023258610.1;refGene_gn=PTPRF;refGene_tx=XM_023258611.1;refGene_gn=PTPRF;refGene_tx=XM_023258612.1;refGene_gn=PTPRF;refGene_tx=XM_023258613.1;refGene_gn=PTPRF;refGene_tx=XM_023258614.1
C1 34408255 34408561
refGene_gn=PTPRF;refGene_tx=XM_023258591.1;refGene_gn=PTPRF;refGene_tx=XM_023258592.1;refGene_gn=PTPRF;refGene_tx=XM_023258593.1;refGene_gn=PTPRF;refGene_tx=XM_023258595.1;refGene_gn=PTPRF;refGene_tx=XM_023258596.1;refGene_gn=PTPRF;refGene_tx=XM_023258597.1;refGene_gn=PTPRF;refGene_tx=XM_023258598.1;refGene_gn=PTPRF;refGene_tx=XM_023258599.1;refGene_gn=PTPRF;refGene_tx=XM_023258600.1;refGene_gn=PTPRF;refGene_tx=XM_023258601.1;refGene_gn=PTPRF;refGene_tx=XM_023258602.1;refGene_gn=PTPRF;refGene_tx=XM_023258604.1;refGene_gn=PTPRF;refGene_tx=XM_023258605.1;r

efGene_gn=PTPRF;refGene_tx=XM_023258607.1;refGene_gn=PTPRF;refGene_tx=XM_023258608.1;refGene_gn=PTPRF;refGene_tx=XM_023258614.1
C1 34409203 34409397
refGene_gn=PTPRF;refGene_tx=XM_023258591.1;refGene_gn=PTPRF;refGene_tx=XM_023258592.1;refGene_gn=PTPRF;refGene_tx=XM_023258593.1;refGene_gn=PTPRF;refGene_tx=XM_023258595.1;refGene_gn=PTPRF;refGene_tx=XM_023258596.1;refGene_gn=PTPRF;refGene_tx=XM_023258597.1;refGene_gn=PTPRF;refGene_tx=XM_023258598.1;refGene_gn=PTPRF;refGene_tx=XM_023258599.1;refGene_gn=PTPRF;refGene_tx=XM_023258600.1;refGene_gn=PTPRF;refGene_tx=XM_023258601.1;refGene_gn=PTPRF;refGene_tx=XM_023258602.1;refGene_gn=PTPRF;refGene_tx=XM_023258603.1;refGene_gn=PTPRF;refGene_tx=XM_023258604.1;refGene_gn=PTPRF;refGene_tx=XM_023258605.1;refGene_gn=PTPRF;refGene_tx=XM_023258606.1;refGene_gn=PTPRF;refGene_tx=XM_023258607.1;refGene_gn=PTPRF;refGene_tx=XM_023258608.1;refGene_gn=PTPRF;refGene_tx=XM_023258609.1;refGene_gn=PTPRF;refGene_tx=XM_023258610.1;refGene_gn=PTPRF;refGene_tx=XM_023258611.1;refGene_gn=PTPRF;refGene_tx=XM_023258612.1;refGene_gn=PTPRF;refGene_tx=XM_023258613.1;refGene_gn=PTPRF;refGene_tx=XM_023258614.1
C1 34412649 34412676
refGene_gn=PTPRF;refGene_tx=XM_023258591.1;refGene_gn=PTPRF;refGene_tx=XM_023258592.1;refGene_gn=PTPRF;refGene_tx=XM_023258593.1;refGene_gn=PTPRF;refGene_tx=XM_023258595.1;refGene_gn=PTPRF;refGene_tx=XM_023258597.1;refGene_gn=PTPRF;refGene_tx=XM_023258598.1;refGene_gn=PTPRF;refGene_tx=XM_023258599.1;refGene_gn=PTPRF;refGene_tx=XM_023258601.1;refGene_gn=PTPRF;refGene_tx=XM_023258603.1;refGene_gn=PTPRF;refGene_tx=XM_023258604.1;refGene_gn=PTPRF;refGene_tx=XM_023258605.1;refGene_gn=PTPRF;refGene_tx=XM_023258607.1;refGene_gn=PTPRF;refGene_tx=XM_023258608.1;refGene_gn=PTPRF;refGene_tx=XM_023258609.1;refGene_gn=PTPRF;refGene_tx=XM_023258610.1;refGene_gn=PTPRF;refGene_tx=XM_023258611.1;refGene_gn=PTPRF;refGene_tx=XM_023258612.1;refGene_gn=PTPRF;refGene_tx=XM_023258614.1
C1 34414037 34414155
refGene_gn=PTPRF;refGene_tx=XM_023258591.1;refGene_gn=PTPRF;refGene_tx=XM_023258592.1;refGene_gn=PTPRF;refGene_tx=XM_023258593.1;refGene_gn=PTPRF;refGene_tx=XM_023258595.1;refGene_gn=PTPRF;refGene_tx=XM_023258596.1;refGene_gn=PTPRF;refGene_tx=XM_023258597.1;refGene_gn=PTPRF;refGene_tx=XM_023258598.1;refGene_gn=PTPRF;refGene_tx=XM_023258599.1;refGene_gn=PTPRF;refGene_tx=XM_023258600.1;refGene_gn=PTPRF;refGene_tx=XM_023258601.1;refGene_gn=PTPRF;refGene_tx=XM_023258602.1;refGene_gn=PTPRF;refGene_tx=XM_023258603.1;refGene_gn=PTPRF;refGene_tx=XM_023258604.1;refGene_gn=PTPRF;refGene_tx=XM_023258605.1;refGene_gn=PTPRF;refGene_tx=XM_023258606.1;refGene_gn=PTPRF;refGene_tx=XM_023258607.1;refGene_gn=PTPRF;refGene_tx=XM_023258608.1;refGene_gn=PTPRF;refGene_tx=XM_023258609.1;refGene_gn=PTPRF;refGene_tx=XM_023258610.1;refGene_gn=PTPRF;refGene_tx=XM_023258611.1;refGene_gn=PTPRF;refGene_tx=XM_023258612.1;refGene_gn=PTPRF;refGene_tx=XM_023258613.1;refGene_gn=PTPRF;refGene_tx=XM_023258614.1
C1 34414234 34414813
refGene_gn=PTPRF;refGene_tx=XM_023258591.1;refGene_gn=PTPRF;refGene_tx=XM_023258592.1;refGene_gn=PTPRF;refGene_tx=XM_023258593.1;refGene_gn=

PTPRF;refGene_tx=XM_023258595.1;refGene_gn=PTPRF;refGene_tx=XM_0232585
96.1;refGene_gn=PTPRF;refGene_tx=XM_023258597.1;refGene_gn=PTPRF;refGe
ne_tx=XM_023258598.1;refGene_gn=PTPRF;refGene_tx=XM_023258599.1;refGen
e_gn=PTPRF;refGene_tx=XM_023258600.1;refGene_gn=PTPRF;refGene_tx=XM_02
3258601.1;refGene_gn=PTPRF;refGene_tx=XM_023258602.1;refGene_gn=PTPRF;
refGene_tx=XM_023258603.1;refGene_gn=PTPRF;refGene_tx=XM_023258604.1;r
efGene_gn=PTPRF;refGene_tx=XM_023258605.1;refGene_gn=PTPRF;refGene_tx=
XM_023258606.1
C1 34415449 34415547
refGene_gn=PTPRF;refGene_tx=XM_023258591.1;refGene_gn=PTPRF;refGene_tx
=XM_023258592.1;refGene_gn=PTPRF;refGene_tx=XM_023258593.1;refGene_gn=
PTPRF;refGene_tx=XM_023258595.1;refGene_gn=PTPRF;refGene_tx=XM_0232585
96.1;refGene_gn=PTPRF;refGene_tx=XM_023258597.1;refGene_gn=PTPRF;refGe
ne_tx=XM_023258598.1;refGene_gn=PTPRF;refGene_tx=XM_023258599.1;refGen
e_gn=PTPRF;refGene_tx=XM_023258600.1;refGene_gn=PTPRF;refGene_tx=XM_02
3258601.1;refGene_gn=PTPRF;refGene_tx=XM_023258602.1;refGene_gn=PTPRF;
refGene_tx=XM_023258603.1;refGene_gn=PTPRF;refGene_tx=XM_023258604.1;r
efGene_gn=PTPRF;refGene_tx=XM_023258605.1;refGene_gn=PTPRF;refGene_tx=
XM_023258606.1;refGene_gn=PTPRF;refGene_tx=XM_023258607.1;refGene_gn=P
TPRF;refGene_tx=XM_023258608.1;refGene_gn=PTPRF;refGene_tx=XM_02325860
9.1;refGene_gn=PTPRF;refGene_tx=XM_023258610.1;refGene_gn=PTPRF;refGen
e_tx=XM_023258611.1;refGene_gn=PTPRF;refGene_tx=XM_023258612.1;refGene
_gn=PTPRF;refGene_tx=XM_023258613.1;refGene_gn=PTPRF;refGene_tx=XM_023
258614.1
C1 34415736 34415990
refGene_gn=PTPRF;refGene_tx=XM_023258591.1;refGene_gn=PTPRF;refGene_tx
=XM_023258592.1;refGene_gn=PTPRF;refGene_tx=XM_023258593.1;refGene_gn=
PTPRF;refGene_tx=XM_023258595.1;refGene_gn=PTPRF;refGene_tx=XM_0232585
96.1;refGene_gn=PTPRF;refGene_tx=XM_023258597.1;refGene_gn=PTPRF;refGe
ne_tx=XM_023258598.1;refGene_gn=PTPRF;refGene_tx=XM_023258599.1;refGen
e_gn=PTPRF;refGene_tx=XM_023258600.1;refGene_gn=PTPRF;refGene_tx=XM_02
3258601.1;refGene_gn=PTPRF;refGene_tx=XM_023258602.1;refGene_gn=PTPRF;
refGene_tx=XM_023258603.1;refGene_gn=PTPRF;refGene_tx=XM_023258604.1;r
efGene_gn=PTPRF;refGene_tx=XM_023258605.1;refGene_gn=PTPRF;refGene_tx=
XM_023258606.1;refGene_gn=PTPRF;refGene_tx=XM_023258607.1;refGene_gn=P
TPRF;refGene_tx=XM_023258608.1;refGene_gn=PTPRF;refGene_tx=XM_02325860
9.1;refGene_gn=PTPRF;refGene_tx=XM_023258610.1;refGene_gn=PTPRF;refGen
e_tx=XM_023258611.1;refGene_gn=PTPRF;refGene_tx=XM_023258612.1;refGene
_gn=PTPRF;refGene_tx=XM_023258613.1;refGene_gn=PTPRF;refGene_tx=XM_023
258614.1
C1 34416104 34416198
refGene_gn=PTPRF;refGene_tx=XM_023258591.1;refGene_gn=PTPRF;refGene_tx
=XM_023258592.1;refGene_gn=PTPRF;refGene_tx=XM_023258593.1;refGene_gn=
PTPRF;refGene_tx=XM_023258595.1;refGene_gn=PTPRF;refGene_tx=XM_0232585
96.1;refGene_gn=PTPRF;refGene_tx=XM_023258597.1;refGene_gn=PTPRF;refGe
ne_tx=XM_023258598.1;refGene_gn=PTPRF;refGene_tx=XM_023258599.1;refGen
e_gn=PTPRF;refGene_tx=XM_023258600.1;refGene_gn=PTPRF;refGene_tx=XM_02
3258601.1;refGene_gn=PTPRF;refGene_tx=XM_023258602.1;refGene_gn=PTPRF;
refGene_tx=XM_023258603.1;refGene_gn=PTPRF;refGene_tx=XM_023258604.1;r
efGene_gn=PTPRF;refGene_tx=XM_023258605.1;refGene_gn=PTPRF;refGene_tx=

XM_023258606.1;refGene_gn=PTPRF;refGene_tx=XM_023258607.1;refGene_gn=PTPRF;refGene_tx=XM_023258608.1;refGene_gn=PTPRF;refGene_tx=XM_023258609.1;refGene_gn=PTPRF;refGene_tx=XM_023258610.1;refGene_gn=PTPRF;refGene_tx=XM_023258611.1;refGene_gn=PTPRF;refGene_tx=XM_023258612.1;refGene_gn=PTPRF;refGene_tx=XM_023258613.1;refGene_gn=PTPRF;refGene_tx=XM_023258614.1

C1 34416781 34416997
refGene_gn=PTPRF;refGene_tx=XM_023258591.1;refGene_gn=PTPRF;refGene_tx=XM_023258592.1;refGene_gn=PTPRF;refGene_tx=XM_023258593.1;refGene_gn=PTPRF;refGene_tx=XM_023258595.1;refGene_gn=PTPRF;refGene_tx=XM_023258596.1;refGene_gn=PTPRF;refGene_tx=XM_023258597.1;refGene_gn=PTPRF;refGene_tx=XM_023258598.1;refGene_gn=PTPRF;refGene_tx=XM_023258599.1;refGene_gn=PTPRF;refGene_tx=XM_023258600.1;refGene_gn=PTPRF;refGene_tx=XM_023258601.1;refGene_gn=PTPRF;refGene_tx=XM_023258602.1;refGene_gn=PTPRF;refGene_tx=XM_023258603.1;refGene_gn=PTPRF;refGene_tx=XM_023258604.1;refGene_gn=PTPRF;refGene_tx=XM_023258605.1;refGene_gn=PTPRF;refGene_tx=XM_023258606.1;refGene_gn=PTPRF;refGene_tx=XM_023258607.1;refGene_gn=PTPRF;refGene_tx=XM_023258608.1;refGene_gn=PTPRF;refGene_tx=XM_023258609.1;refGene_gn=PTPRF;refGene_tx=XM_023258610.1;refGene_gn=PTPRF;refGene_tx=XM_023258611.1;refGene_gn=PTPRF;refGene_tx=XM_023258612.1;refGene_gn=PTPRF;refGene_tx=XM_023258613.1;refGene_gn=PTPRF;refGene_tx=XM_023258614.1

C1 34417313 34417468
refGene_gn=PTPRF;refGene_tx=XM_023258591.1;refGene_gn=PTPRF;refGene_tx=XM_023258592.1;refGene_gn=PTPRF;refGene_tx=XM_023258593.1;refGene_gn=PTPRF;refGene_tx=XM_023258595.1;refGene_gn=PTPRF;refGene_tx=XM_023258596.1;refGene_gn=PTPRF;refGene_tx=XM_023258597.1;refGene_gn=PTPRF;refGene_tx=XM_023258598.1;refGene_gn=PTPRF;refGene_tx=XM_023258599.1;refGene_gn=PTPRF;refGene_tx=XM_023258600.1;refGene_gn=PTPRF;refGene_tx=XM_023258601.1;refGene_gn=PTPRF;refGene_tx=XM_023258602.1;refGene_gn=PTPRF;refGene_tx=XM_023258603.1;refGene_gn=PTPRF;refGene_tx=XM_023258604.1;refGene_gn=PTPRF;refGene_tx=XM_023258605.1;refGene_gn=PTPRF;refGene_tx=XM_023258606.1;refGene_gn=PTPRF;refGene_tx=XM_023258607.1;refGene_gn=PTPRF;refGene_tx=XM_023258608.1;refGene_gn=PTPRF;refGene_tx=XM_023258609.1;refGene_gn=PTPRF;refGene_tx=XM_023258610.1;refGene_gn=PTPRF;refGene_tx=XM_023258611.1;refGene_gn=PTPRF;refGene_tx=XM_023258612.1;refGene_gn=PTPRF;refGene_tx=XM_023258613.1;refGene_gn=PTPRF;refGene_tx=XM_023258614.1

C1 34419133 34419145
refGene_gn=PTPRF;refGene_tx=XM_023258591.1;refGene_gn=PTPRF;refGene_tx=XM_023258593.1;refGene_gn=PTPRF;refGene_tx=XM_023258595.1;refGene_gn=PTPRF;refGene_tx=XM_023258596.1;refGene_gn=PTPRF;refGene_tx=XM_023258597.1;refGene_gn=PTPRF;refGene_tx=XM_023258598.1;refGene_gn=PTPRF;refGene_tx=XM_023258603.1;refGene_gn=PTPRF;refGene_tx=XM_023258604.1;refGene_gn=PTPRF;refGene_tx=XM_023258605.1;refGene_gn=PTPRF;refGene_tx=XM_023258607.1;refGene_gn=PTPRF;refGene_tx=XM_023258609.1

C1 34420138 34420254
refGene_gn=PTPRF;refGene_tx=XM_023258591.1;refGene_gn=PTPRF;refGene_tx=XM_023258592.1;refGene_gn=PTPRF;refGene_tx=XM_023258593.1;refGene_gn=PTPRF;refGene_tx=XM_023258595.1;refGene_gn=PTPRF;refGene_tx=XM_0232585

96.1;refGene_gn=PTPRF;refGene_tx=XM_023258597.1;refGene_gn=PTPRF;refGene_tx=XM_023258598.1;refGene_gn=PTPRF;refGene_tx=XM_023258599.1;refGene_gn=PTPRF;refGene_tx=XM_023258600.1;refGene_gn=PTPRF;refGene_tx=XM_023258601.1;refGene_gn=PTPRF;refGene_tx=XM_023258602.1;refGene_gn=PTPRF;refGene_tx=XM_023258603.1;refGene_gn=PTPRF;refGene_tx=XM_023258604.1;refGene_gn=PTPRF;refGene_tx=XM_023258605.1;refGene_gn=PTPRF;refGene_tx=XM_023258606.1;refGene_gn=PTPRF;refGene_tx=XM_023258607.1;refGene_gn=PTPRF;refGene_tx=XM_023258608.1;refGene_gn=PTPRF;refGene_tx=XM_023258609.1;refGene_gn=PTPRF;refGene_tx=XM_023258610.1;refGene_gn=PTPRF;refGene_tx=XM_023258611.1;refGene_gn=PTPRF;refGene_tx=XM_023258612.1;refGene_gn=PTPRF;refGene_tx=XM_023258613.1;refGene_gn=PTPRF;refGene_tx=XM_023258614.1

C1 34424016 34424049

refGene_gn=PTPRF;refGene_tx=XM_023258591.1;refGene_gn=PTPRF;refGene_tx=XM_023258592.1;refGene_gn=PTPRF;refGene_tx=XM_023258593.1;refGene_gn=PTPRF;refGene_tx=XM_023258595.1;refGene_gn=PTPRF;refGene_tx=XM_023258596.1;refGene_gn=PTPRF;refGene_tx=XM_023258597.1;refGene_gn=PTPRF;refGene_tx=XM_023258603.1;refGene_gn=PTPRF;refGene_tx=XM_023258604.1;refGene_gn=PTPRF;refGene_tx=XM_023258605.1;refGene_gn=PTPRF;refGene_tx=XM_023258607.1;refGene_gn=PTPRF;refGene_tx=XM_023258608.1;refGene_gn=PTPRF;refGene_tx=XM_023258609.1;refGene_gn=PTPRF;refGene_tx=XM_023258610.1;refGene_gn=PTPRF;refGene_tx=XM_023258611.1;refGene_gn=PTPRF;refGene_tx=XM_023258612.1;refGene_gn=PTPRF;refGene_tx=XM_023258614.1

C1 34424848 34424946

refGene_gn=PTPRF;refGene_tx=XM_023258591.1;refGene_gn=PTPRF;refGene_tx=XM_023258592.1;refGene_gn=PTPRF;refGene_tx=XM_023258593.1;refGene_gn=PTPRF;refGene_tx=XM_023258595.1;refGene_gn=PTPRF;refGene_tx=XM_023258596.1;refGene_gn=PTPRF;refGene_tx=XM_023258597.1;refGene_gn=PTPRF;refGene_tx=XM_023258598.1;refGene_gn=PTPRF;refGene_tx=XM_023258599.1;refGene_gn=PTPRF;refGene_tx=XM_023258600.1;refGene_gn=PTPRF;refGene_tx=XM_023258601.1;refGene_gn=PTPRF;refGene_tx=XM_023258602.1;refGene_gn=PTPRF;refGene_tx=XM_023258603.1;refGene_gn=PTPRF;refGene_tx=XM_023258604.1;refGene_gn=PTPRF;refGene_tx=XM_023258605.1;refGene_gn=PTPRF;refGene_tx=XM_023258606.1;refGene_gn=PTPRF;refGene_tx=XM_023258607.1;refGene_gn=PTPRF;refGene_tx=XM_023258608.1;refGene_gn=PTPRF;refGene_tx=XM_023258609.1;refGene_gn=PTPRF;refGene_tx=XM_023258610.1;refGene_gn=PTPRF;refGene_tx=XM_023258611.1;refGene_gn=PTPRF;refGene_tx=XM_023258612.1;refGene_gn=PTPRF;refGene_tx=XM_023258613.1;refGene_gn=PTPRF;refGene_tx=XM_023258614.1

C1 34427834 34427958

refGene_gn=PTPRF;refGene_tx=XM_023258591.1;refGene_gn=PTPRF;refGene_tx=XM_023258592.1;refGene_gn=PTPRF;refGene_tx=XM_023258593.1;refGene_gn=PTPRF;refGene_tx=XM_023258595.1;refGene_gn=PTPRF;refGene_tx=XM_023258596.1;refGene_gn=PTPRF;refGene_tx=XM_023258597.1;refGene_gn=PTPRF;refGene_tx=XM_023258598.1;refGene_gn=PTPRF;refGene_tx=XM_023258599.1;refGene_gn=PTPRF;refGene_tx=XM_023258600.1;refGene_gn=PTPRF;refGene_tx=XM_023258601.1;refGene_gn=PTPRF;refGene_tx=XM_023258602.1;refGene_gn=PTPRF;refGene_tx=XM_023258603.1;refGene_gn=PTPRF;refGene_tx=XM_023258604.1;refGene_gn=PTPRF;refGene_tx=XM_023258605.1;refGene_gn=PTPRF;refGene_tx=XM_023258606.1;refGene_gn=PTPRF;refGene_tx=XM_023258607.1;refGene_gn=P

TPRF;refGene_tx=XM_023258608.1;refGene_gn=PTPRF;refGene_tx=XM_02325860
9.1;refGene_gn=PTPRF;refGene_tx=XM_023258610.1;refGene_gn=PTPRF;refGen
e_tx=XM_023258611.1;refGene_gn=PTPRF;refGene_tx=XM_023258612.1;refGene
_gn=PTPRF;refGene_tx=XM_023258613.1;refGene_gn=PTPRF;refGene_tx=XM_023
258614.1
C1 34428127 34428303
refGene_gn=PTPRF;refGene_tx=XM_023258591.1;refGene_gn=PTPRF;refGene_tx
=XM_023258592.1;refGene_gn=PTPRF;refGene_tx=XM_023258593.1;refGene_gn=
PTPRF;refGene_tx=XM_023258595.1;refGene_gn=PTPRF;refGene_tx=XM_0232585
96.1;refGene_gn=PTPRF;refGene_tx=XM_023258597.1;refGene_gn=PTPRF;refGe
ne_tx=XM_023258598.1;refGene_gn=PTPRF;refGene_tx=XM_023258599.1;refGen
e_gn=PTPRF;refGene_tx=XM_023258600.1;refGene_gn=PTPRF;refGene_tx=XM_02
3258601.1;refGene_gn=PTPRF;refGene_tx=XM_023258602.1;refGene_gn=PTPRF;
refGene_tx=XM_023258603.1;refGene_gn=PTPRF;refGene_tx=XM_023258604.1;r
efGene_gn=PTPRF;refGene_tx=XM_023258605.1;refGene_gn=PTPRF;refGene_tx=
XM_023258606.1;refGene_gn=PTPRF;refGene_tx=XM_023258607.1;refGene_gn=P
TPRF;refGene_tx=XM_023258608.1;refGene_gn=PTPRF;refGene_tx=XM_02325860
9.1;refGene_gn=PTPRF;refGene_tx=XM_023258610.1;refGene_gn=PTPRF;refGen
e_tx=XM_023258611.1;refGene_gn=PTPRF;refGene_tx=XM_023258612.1;refGene
_gn=PTPRF;refGene_tx=XM_023258613.1;refGene_gn=PTPRF;refGene_tx=XM_023
258614.1
C1 34429027 34429147
refGene_gn=PTPRF;refGene_tx=XM_023258591.1;refGene_gn=PTPRF;refGene_tx
=XM_023258592.1;refGene_gn=PTPRF;refGene_tx=XM_023258593.1;refGene_gn=
PTPRF;refGene_tx=XM_023258595.1;refGene_gn=PTPRF;refGene_tx=XM_0232585
96.1;refGene_gn=PTPRF;refGene_tx=XM_023258597.1;refGene_gn=PTPRF;refGe
ne_tx=XM_023258598.1;refGene_gn=PTPRF;refGene_tx=XM_023258599.1;refGen
e_gn=PTPRF;refGene_tx=XM_023258600.1;refGene_gn=PTPRF;refGene_tx=XM_02
3258601.1;refGene_gn=PTPRF;refGene_tx=XM_023258602.1;refGene_gn=PTPRF;
refGene_tx=XM_023258603.1;refGene_gn=PTPRF;refGene_tx=XM_023258604.1;r
efGene_gn=PTPRF;refGene_tx=XM_023258605.1;refGene_gn=PTPRF;refGene_tx=
XM_023258606.1;refGene_gn=PTPRF;refGene_tx=XM_023258607.1;refGene_gn=P
TPRF;refGene_tx=XM_023258608.1;refGene_gn=PTPRF;refGene_tx=XM_02325860
9.1;refGene_gn=PTPRF;refGene_tx=XM_023258610.1;refGene_gn=PTPRF;refGen
e_tx=XM_023258611.1;refGene_gn=PTPRF;refGene_tx=XM_023258612.1;refGene
_gn=PTPRF;refGene_tx=XM_023258613.1;refGene_gn=PTPRF;refGene_tx=XM_023
258614.1
C1 34429429 34429584
refGene_gn=PTPRF;refGene_tx=XM_023258591.1;refGene_gn=PTPRF;refGene_tx
=XM_023258592.1;refGene_gn=PTPRF;refGene_tx=XM_023258593.1;refGene_gn=
PTPRF;refGene_tx=XM_023258595.1;refGene_gn=PTPRF;refGene_tx=XM_0232585
96.1;refGene_gn=PTPRF;refGene_tx=XM_023258597.1;refGene_gn=PTPRF;refGe
ne_tx=XM_023258598.1;refGene_gn=PTPRF;refGene_tx=XM_023258599.1;refGen
e_gn=PTPRF;refGene_tx=XM_023258600.1;refGene_gn=PTPRF;refGene_tx=XM_02
3258601.1;refGene_gn=PTPRF;refGene_tx=XM_023258602.1;refGene_gn=PTPRF;
refGene_tx=XM_023258603.1;refGene_gn=PTPRF;refGene_tx=XM_023258604.1;r
efGene_gn=PTPRF;refGene_tx=XM_023258605.1;refGene_gn=PTPRF;refGene_tx=
XM_023258606.1;refGene_gn=PTPRF;refGene_tx=XM_023258607.1;refGene_gn=P
TPRF;refGene_tx=XM_023258608.1;refGene_gn=PTPRF;refGene_tx=XM_02325860
9.1;refGene_gn=PTPRF;refGene_tx=XM_023258610.1;refGene_gn=PTPRF;refGen

e_tx=XM_023258611.1;refGene_gn=PTPRF;refGene_tx=XM_023258612.1;refGene_gn=PTPRF;refGene_tx=XM_023258613.1;refGene_gn=PTPRF;refGene_tx=XM_023258614.1
C1 34429665 34429951
refGene_gn=PTPRF;refGene_tx=XM_023258591.1;refGene_gn=PTPRF;refGene_tx=XM_023258592.1;refGene_gn=PTPRF;refGene_tx=XM_023258593.1;refGene_gn=PTPRF;refGene_tx=XM_023258595.1;refGene_gn=PTPRF;refGene_tx=XM_023258596.1;refGene_gn=PTPRF;refGene_tx=XM_023258597.1;refGene_gn=PTPRF;refGene_tx=XM_023258598.1;refGene_gn=PTPRF;refGene_tx=XM_023258599.1;refGene_gn=PTPRF;refGene_tx=XM_023258600.1;refGene_gn=PTPRF;refGene_tx=XM_023258601.1;refGene_gn=PTPRF;refGene_tx=XM_023258602.1;refGene_gn=PTPRF;refGene_tx=XM_023258603.1;refGene_gn=PTPRF;refGene_tx=XM_023258604.1;refGene_gn=PTPRF;refGene_tx=XM_023258605.1;refGene_gn=PTPRF;refGene_tx=XM_023258606.1;refGene_gn=PTPRF;refGene_tx=XM_023258607.1;refGene_gn=PTPRF;refGene_tx=XM_023258608.1;refGene_gn=PTPRF;refGene_tx=XM_023258609.1;refGene_gn=PTPRF;refGene_tx=XM_023258610.1;refGene_gn=PTPRF;refGene_tx=XM_023258611.1;refGene_gn=PTPRF;refGene_tx=XM_023258612.1;refGene_gn=PTPRF;refGene_tx=XM_023258613.1;refGene_gn=PTPRF;refGene_tx=XM_023258614.1
C1 34430031 34430210
refGene_gn=PTPRF;refGene_tx=XM_023258591.1;refGene_gn=PTPRF;refGene_tx=XM_023258592.1;refGene_gn=PTPRF;refGene_tx=XM_023258593.1;refGene_gn=PTPRF;refGene_tx=XM_023258595.1;refGene_gn=PTPRF;refGene_tx=XM_023258596.1;refGene_gn=PTPRF;refGene_tx=XM_023258597.1;refGene_gn=PTPRF;refGene_tx=XM_023258598.1;refGene_gn=PTPRF;refGene_tx=XM_023258599.1;refGene_gn=PTPRF;refGene_tx=XM_023258600.1;refGene_gn=PTPRF;refGene_tx=XM_023258601.1;refGene_gn=PTPRF;refGene_tx=XM_023258602.1;refGene_gn=PTPRF;refGene_tx=XM_023258603.1;refGene_gn=PTPRF;refGene_tx=XM_023258604.1;refGene_gn=PTPRF;refGene_tx=XM_023258605.1;refGene_gn=PTPRF;refGene_tx=XM_023258606.1;refGene_gn=PTPRF;refGene_tx=XM_023258607.1;refGene_gn=PTPRF;refGene_tx=XM_023258608.1;refGene_gn=PTPRF;refGene_tx=XM_023258609.1;refGene_gn=PTPRF;refGene_tx=XM_023258610.1;refGene_gn=PTPRF;refGene_tx=XM_023258611.1;refGene_gn=PTPRF;refGene_tx=XM_023258612.1;refGene_gn=PTPRF;refGene_tx=XM_023258613.1;refGene_gn=PTPRF;refGene_tx=XM_023258614.1
C1 34430427 34430554
refGene_gn=PTPRF;refGene_tx=XM_023258591.1;refGene_gn=PTPRF;refGene_tx=XM_023258592.1;refGene_gn=PTPRF;refGene_tx=XM_023258593.1;refGene_gn=PTPRF;refGene_tx=XM_023258595.1;refGene_gn=PTPRF;refGene_tx=XM_023258596.1;refGene_gn=PTPRF;refGene_tx=XM_023258597.1;refGene_gn=PTPRF;refGene_tx=XM_023258598.1;refGene_gn=PTPRF;refGene_tx=XM_023258599.1;refGene_gn=PTPRF;refGene_tx=XM_023258600.1;refGene_gn=PTPRF;refGene_tx=XM_023258601.1;refGene_gn=PTPRF;refGene_tx=XM_023258602.1;refGene_gn=PTPRF;refGene_tx=XM_023258603.1;refGene_gn=PTPRF;refGene_tx=XM_023258604.1;refGene_gn=PTPRF;refGene_tx=XM_023258605.1;refGene_gn=PTPRF;refGene_tx=XM_023258606.1;refGene_gn=PTPRF;refGene_tx=XM_023258607.1;refGene_gn=PTPRF;refGene_tx=XM_023258608.1;refGene_gn=PTPRF;refGene_tx=XM_023258609.1;refGene_gn=PTPRF;refGene_tx=XM_023258610.1;refGene_gn=PTPRF;refGene_tx=XM_023258611.1;refGene_gn=PTPRF;refGene_tx=XM_023258612.1;refGene_gn=PTPRF;refGene_tx=XM_023258613.1;refGene_gn=PTPRF;refGene_tx=XM_023

258614.1
C1	34430764	34430890
refGene_gn=PTPRF;refGene_tx=XM_023258591.1;refGene_gn=PTPRF;refGene_tx=XM_023258592.1;refGene_gn=PTPRF;refGene_tx=XM_023258593.1;refGene_gn=PTPRF;refGene_tx=XM_023258595.1;refGene_gn=PTPRF;refGene_tx=XM_023258596.1;refGene_gn=PTPRF;refGene_tx=XM_023258597.1;refGene_gn=PTPRF;refGene_tx=XM_023258598.1;refGene_gn=PTPRF;refGene_tx=XM_023258599.1;refGene_gn=PTPRF;refGene_tx=XM_023258600.1;refGene_gn=PTPRF;refGene_tx=XM_023258601.1;refGene_gn=PTPRF;refGene_tx=XM_023258602.1;refGene_gn=PTPRF;refGene_tx=XM_023258603.1;refGene_gn=PTPRF;refGene_tx=XM_023258604.1;refGene_gn=PTPRF;refGene_tx=XM_023258605.1;refGene_gn=PTPRF;refGene_tx=XM_023258606.1;refGene_gn=PTPRF;refGene_tx=XM_023258607.1;refGene_gn=PTPRF;refGene_tx=XM_023258608.1;refGene_gn=PTPRF;refGene_tx=XM_023258609.1;refGene_gn=PTPRF;refGene_tx=XM_023258610.1;refGene_gn=PTPRF;refGene_tx=XM_023258611.1;refGene_gn=PTPRF;refGene_tx=XM_023258612.1;refGene_gn=PTPRF;refGene_tx=XM_023258613.1;refGene_gn=PTPRF;refGene_tx=XM_023258614.1
C1	34431167	34431322
refGene_gn=PTPRF;refGene_tx=XM_023258591.1;refGene_gn=PTPRF;refGene_tx=XM_023258592.1;refGene_gn=PTPRF;refGene_tx=XM_023258593.1;refGene_gn=PTPRF;refGene_tx=XM_023258595.1;refGene_gn=PTPRF;refGene_tx=XM_023258596.1;refGene_gn=PTPRF;refGene_tx=XM_023258597.1;refGene_gn=PTPRF;refGene_tx=XM_023258598.1;refGene_gn=PTPRF;refGene_tx=XM_023258599.1;refGene_gn=PTPRF;refGene_tx=XM_023258600.1;refGene_gn=PTPRF;refGene_tx=XM_023258601.1;refGene_gn=PTPRF;refGene_tx=XM_023258602.1;refGene_gn=PTPRF;refGene_tx=XM_023258603.1;refGene_gn=PTPRF;refGene_tx=XM_023258604.1;refGene_gn=PTPRF;refGene_tx=XM_023258605.1;refGene_gn=PTPRF;refGene_tx=XM_023258606.1;refGene_gn=PTPRF;refGene_tx=XM_023258607.1;refGene_gn=PTPRF;refGene_tx=XM_023258608.1;refGene_gn=PTPRF;refGene_tx=XM_023258609.1;refGene_gn=PTPRF;refGene_tx=XM_023258610.1;refGene_gn=PTPRF;refGene_tx=XM_023258611.1;refGene_gn=PTPRF;refGene_tx=XM_023258612.1;refGene_gn=PTPRF;refGene_tx=XM_023258613.1;refGene_gn=PTPRF;refGene_tx=XM_023258614.1
C1	34431410	34431546
refGene_gn=PTPRF;refGene_tx=XM_023258591.1;refGene_gn=PTPRF;refGene_tx=XM_023258592.1;refGene_gn=PTPRF;refGene_tx=XM_023258593.1;refGene_gn=PTPRF;refGene_tx=XM_023258595.1;refGene_gn=PTPRF;refGene_tx=XM_023258596.1;refGene_gn=PTPRF;refGene_tx=XM_023258597.1;refGene_gn=PTPRF;refGene_tx=XM_023258598.1;refGene_gn=PTPRF;refGene_tx=XM_023258599.1;refGene_gn=PTPRF;refGene_tx=XM_023258600.1;refGene_gn=PTPRF;refGene_tx=XM_023258601.1;refGene_gn=PTPRF;refGene_tx=XM_023258602.1;refGene_gn=PTPRF;refGene_tx=XM_023258603.1;refGene_gn=PTPRF;refGene_tx=XM_023258604.1;refGene_gn=PTPRF;refGene_tx=XM_023258605.1;refGene_gn=PTPRF;refGene_tx=XM_023258606.1;refGene_gn=PTPRF;refGene_tx=XM_023258607.1;refGene_gn=PTPRF;refGene_tx=XM_023258608.1;refGene_gn=PTPRF;refGene_tx=XM_023258609.1;refGene_gn=PTPRF;refGene_tx=XM_023258610.1;refGene_gn=PTPRF;refGene_tx=XM_023258611.1;refGene_gn=PTPRF;refGene_tx=XM_023258612.1;refGene_gn=PTPRF;refGene_tx=XM_023258613.1;refGene_gn=PTPRF;refGene_tx=XM_023258614.1
C1	34431974	34432043

refGene_gn=PTPRF;refGene_tx=XM_023258591.1;refGene_gn=PTPRF;refGene_tx
=XM_023258592.1;refGene_gn=PTPRF;refGene_tx=XM_023258593.1;refGene_gn=
PTPRF;refGene_tx=XM_023258595.1;refGene_gn=PTPRF;refGene_tx=XM_0232585
96.1;refGene_gn=PTPRF;refGene_tx=XM_023258597.1;refGene_gn=PTPRF;refGe
ne_tx=XM_023258598.1;refGene_gn=PTPRF;refGene_tx=XM_023258599.1;refGen
e_gn=PTPRF;refGene_tx=XM_023258600.1;refGene_gn=PTPRF;refGene_tx=XM_02
3258601.1;refGene_gn=PTPRF;refGene_tx=XM_023258602.1;refGene_gn=PTPRF;
refGene_tx=XM_023258603.1;refGene_gn=PTPRF;refGene_tx=XM_023258604.1;r
efGene_gn=PTPRF;refGene_tx=XM_023258605.1;refGene_gn=PTPRF;refGene_tx=
XM_023258606.1;refGene_gn=PTPRF;refGene_tx=XM_023258607.1;refGene_gn=P
TPRF;refGene_tx=XM_023258608.1;refGene_gn=PTPRF;refGene_tx=XM_02325860
9.1;refGene_gn=PTPRF;refGene_tx=XM_023258610.1;refGene_gn=PTPRF;refGen
e_tx=XM_023258611.1;refGene_gn=PTPRF;refGene_tx=XM_023258612.1;refGene
_gn=PTPRF;refGene_tx=XM_023258613.1;refGene_gn=PTPRF;refGene_tx=XM_023
258614.1
C1 34456614 34456752
refGene_gn=KDM4A;refGene_tx=XM_003989992.3;refGene_gn=KDM4A;refGene_tx
=XM_006934673.2;refGene_gn=KDM4A;refGene_tx=XM_019836867.2
C1 34457380 34457556
refGene_gn=KDM4A;refGene_tx=XM_003989992.3;refGene_gn=KDM4A;refGene_tx
=XM_006934673.2;refGene_gn=KDM4A;refGene_tx=XM_019836867.2
C1 34460187 34460302
refGene_gn=KDM4A;refGene_tx=XM_003989992.3;refGene_gn=KDM4A;refGene_tx
=XM_006934673.2;refGene_gn=KDM4A;refGene_tx=XM_019836867.2
C1 34461805 34461999
refGene_gn=KDM4A;refGene_tx=XM_003989992.3;refGene_gn=KDM4A;refGene_tx
=XM_006934673.2;refGene_gn=KDM4A;refGene_tx=XM_019836867.2
C1 34464296 34464346
refGene_gn=KDM4A;refGene_tx=XM_003989992.3;refGene_gn=KDM4A;refGene_tx
=XM_006934673.2;refGene_gn=KDM4A;refGene_tx=XM_019836867.2
C1 34465082 34465186
refGene_gn=KDM4A;refGene_tx=XM_003989992.3;refGene_gn=KDM4A;refGene_tx
=XM_006934673.2;refGene_gn=KDM4A;refGene_tx=XM_019836867.2
C1 34465579 34465717
refGene_gn=KDM4A;refGene_tx=XM_003989992.3;refGene_gn=KDM4A;refGene_tx
=XM_006934673.2;refGene_gn=KDM4A;refGene_tx=XM_019836867.2
C1 34466411 34466662
refGene_gn=KDM4A;refGene_tx=XM_003989992.3;refGene_gn=KDM4A;refGene_tx
=XM_006934673.2;refGene_gn=KDM4A;refGene_tx=XM_019836867.2
C1 34467365 34467565
refGene_gn=KDM4A;refGene_tx=XM_003989992.3;refGene_gn=KDM4A;refGene_tx
=XM_006934673.2;refGene_gn=KDM4A;refGene_tx=XM_019836867.2
C1 34468891 34469268
refGene_gn=KDM4A;refGene_tx=XM_003989992.3;refGene_gn=KDM4A;refGene_tx
=XM_006934673.2;refGene_gn=KDM4A;refGene_tx=XM_019836867.2
C1 34481678 34481799
refGene_gn=KDM4A;refGene_tx=XM_003989992.3;refGene_gn=KDM4A;refGene_tx
=XM_006934673.2;refGene_gn=KDM4A;refGene_tx=XM_019836867.2
C1 34486695 34486877
refGene_gn=KDM4A;refGene_tx=XM_003989992.3;refGene_gn=KDM4A;refGene_tx

=XM_006934673.2;refGene_gn=KDM4A;refGene_tx=XM_019836867.2
C1 34488400 34488605
refGene_gn=KDM4A;refGene_tx=XM_003989992.3;refGene_gn=KDM4A;refGene_tx=XM_006934673.2;refGene_gn=KDM4A;refGene_tx=XM_019836867.2
C1 34489087 34489164
refGene_gn=KDM4A;refGene_tx=XM_003989992.3;refGene_gn=KDM4A;refGene_tx=XM_006934673.2;refGene_gn=KDM4A;refGene_tx=XM_019836867.2
C1 34489848 34489904
refGene_gn=KDM4A;refGene_tx=XM_003989992.3;refGene_gn=KDM4A;refGene_tx=XM_006934673.2;refGene_gn=KDM4A;refGene_tx=XM_019836867.2
C1 34491544 34491653
refGene_gn=KDM4A;refGene_tx=XM_003989992.3;refGene_gn=KDM4A;refGene_tx=XM_006934673.2;refGene_gn=KDM4A;refGene_tx=XM_019836867.2
C1 34491972 34492158
refGene_gn=KDM4A;refGene_tx=XM_003989992.3;refGene_gn=KDM4A;refGene_tx=XM_006934673.2;refGene_gn=KDM4A;refGene_tx=XM_019836867.2
C1 34495596 34495767
refGene_gn=KDM4A;refGene_tx=XM_003989992.3;refGene_gn=KDM4A;refGene_tx=XM_006934673.2;refGene_gn=KDM4A;refGene_tx=XM_019836867.2
C1 34498185 34498305
refGene_gn=KDM4A;refGene_tx=XM_003989992.3;refGene_gn=KDM4A;refGene_tx=XM_006934673.2;refGene_gn=KDM4A;refGene_tx=XM_019836867.2
C1 34498587 34498680
refGene_gn=KDM4A;refGene_tx=XM_003989992.3;refGene_gn=KDM4A;refGene_tx=XM_006934673.2;refGene_gn=KDM4A;refGene_tx=XM_019836867.2
C1 34498812 34498953
refGene_gn=KDM4A;refGene_tx=XM_003989992.3;refGene_gn=KDM4A;refGene_tx=XM_006934673.2;refGene_gn=KDM4A;refGene_tx=XM_019836867.2
C1 34529022 34529137
refGene_gn=ST3GAL3;refGene_tx=XM_003989993.5;refGene_gn=ST3GAL3;refGene_tx=XM_006934674.4;refGene_gn=ST3GAL3;refGene_tx=XM_011284672.3;refGene_gn=ST3GAL3;refGene_tx=XM_011284674.3
C1 34540325 34540628
refGene_gn=LOC111561722;refGene_tx=XM_023258615.1
C1 34588000 34588048
refGene_gn=ST3GAL3;refGene_tx=XM_003989993.5;refGene_gn=ST3GAL3;refGene_tx=XM_006934674.4
C1 34606436 34606479
refGene_gn=ST3GAL3;refGene_tx=XM_003989993.5;refGene_gn=ST3GAL3;refGene_tx=XM_006934674.4;refGene_gn=ST3GAL3;refGene_tx=XM_011284672.3;refGene_gn=ST3GAL3;refGene_tx=XM_011284674.3
C1 34627674 34627767
refGene_gn=ST3GAL3;refGene_tx=XM_003989993.5;refGene_gn=ST3GAL3;refGene_tx=XM_006934674.4;refGene_gn=ST3GAL3;refGene_tx=XM_011284672.3;refGene_gn=ST3GAL3;refGene_tx=XM_011284674.3
C1 34677986 34678081
refGene_gn=ST3GAL3;refGene_tx=XM_003989993.5;refGene_gn=ST3GAL3;refGene_tx=XM_006934674.4;refGene_gn=ST3GAL3;refGene_tx=XM_011284672.3;refGene_gn=ST3GAL3;refGene_tx=XM_011284674.3
C1 34681238 34681302

refGene_gn=ST3GAL3;refGene_tx=XM_003989993.5;refGene_gn=ST3GAL3;refGene_tx=XM_006934674.4;refGene_gn=ST3GAL3;refGene_tx=XM_011284672.3;refGene_gn=ST3GAL3;refGene_tx=XM_011284674.3
C1 34682106 34682202 refGene_gn=ST3GAL3;refGene_tx=XM_003989993.5;refGene_gn=ST3GAL3;refGene_tx=XM_006934674.4;refGene_gn=ST3GAL3;refGene_tx=XM_011284672.3;refGene_gn=ST3GAL3;refGene_tx=XM_011284674.3
C1 34682490 34682677 refGene_gn=ST3GAL3;refGene_tx=XM_003989993.5;refGene_gn=ST3GAL3;refGene_tx=XM_006934674.4;refGene_gn=ST3GAL3;refGene_tx=XM_011284672.3;refGene_gn=ST3GAL3;refGene_tx=XM_011284674.3
C1 34690838 34690943 refGene_gn=ST3GAL3;refGene_tx=XM_006934674.4;refGene_gn=ST3GAL3;refGene_tx=XM_011284672.3
C1 34699254 34699401 refGene_gn=ST3GAL3;refGene_tx=XM_003989993.5;refGene_gn=ST3GAL3;refGene_tx=XM_006934674.4;refGene_gn=ST3GAL3;refGene_tx=XM_011284672.3;refGene_gn=ST3GAL3;refGene_tx=XM_011284674.3
C1 34699597 34699744 refGene_gn=ST3GAL3;refGene_tx=XM_003989993.5;refGene_gn=ST3GAL3;refGene_tx=XM_006934674.4;refGene_gn=ST3GAL3;refGene_tx=XM_011284672.3;refGene_gn=ST3GAL3;refGene_tx=XM_011284674.3
C1 34706837 34706927 refGene_gn=ST3GAL3;refGene_tx=XM_003989993.5;refGene_gn=ST3GAL3;refGene_tx=XM_006934674.4;refGene_gn=ST3GAL3;refGene_tx=XM_011284672.3;refGene_gn=ST3GAL3;refGene_tx=XM_011284674.3
C1 34712627 34712730 refGene_gn=ARTN;refGene_tx=XM_023258616.1
C1 34712732 34712853 refGene_gn=ARTN;refGene_tx=XM_023258616.1
C1 34712854 34712880 refGene_gn=ARTN;refGene_tx=XM_023258616.1
C1 34712951 34713421 refGene_gn=ARTN;refGene_tx=XM_023258616.1
C1 34718739 34718823 refGene_gn=IPO13;refGene_tx=XM_006934675.4;refGene_gn=IPO13;refGene_tx=XM_019836869.2
C1 34720542 34721279 refGene_gn=IPO13;refGene_tx=XM_006934675.4;refGene_gn=IPO13;refGene_tx=XM_019836869.2
C1 34726444 34726585 refGene_gn=IPO13;refGene_tx=XM_006934675.4;refGene_gn=IPO13;refGene_tx=XM_019836869.2
C1 34726694 34726836 refGene_gn=IPO13;refGene_tx=XM_006934675.4;refGene_gn=IPO13;refGene_tx=XM_019836869.2
C1 34726944 34727111 refGene_gn=IPO13;refGene_tx=XM_006934675.4;refGene_gn=IPO13;refGene_tx=XM_019836869.2
C1 34727318 34727439 refGene_gn=IPO13;refGene_tx=XM_006934675.4;refGene_gn=IPO13;refGene_tx=XM_019836869.2
C1 34727525 34727673 refGene_gn=IPO13;refGene_tx=XM_006934675.4;refGene_gn=IPO13;refGene_tx

=XM_019836869.2
C1 34728093 34728275
refGene_gn=IPO13;refGene_tx=XM_006934675.4;refGene_gn=IPO13;refGene_tx
=XM_019836869.2
C1 34728355 34728382
refGene_gn=IPO13;refGene_tx=XM_006934675.4;refGene_gn=IPO13;refGene_tx
=XM_019836869.2
C1 34728563 34728710
refGene_gn=IPO13;refGene_tx=XM_006934675.4;refGene_gn=IPO13;refGene_tx
=XM_019836869.2
C1 34728887 34729031
refGene_gn=IPO13;refGene_tx=XM_006934675.4;refGene_gn=IPO13;refGene_tx
=XM_019836869.2
C1 34730357 34730438
refGene_gn=IPO13;refGene_tx=XM_006934675.4;refGene_gn=IPO13;refGene_tx
=XM_019836869.2
C1 34731236 34731374
refGene_gn=IPO13;refGene_tx=XM_006934675.4;refGene_gn=IPO13;refGene_tx
=XM_019836869.2
C1 34731565 34731662
refGene_gn=IPO13;refGene_tx=XM_006934675.4;refGene_gn=IPO13;refGene_tx
=XM_019836869.2
C1 34734633 34734686
refGene_gn=IPO13;refGene_tx=XM_006934675.4;refGene_gn=IPO13;refGene_tx
=XM_019836869.2
C1 34736929 34736996
refGene_gn=IPO13;refGene_tx=XM_006934675.4;refGene_gn=IPO13;refGene_tx
=XM_019836869.2
C1 34737084 34737143
refGene_gn=IPO13;refGene_tx=XM_006934675.4;refGene_gn=IPO13;refGene_tx
=XM_019836869.2
C1 34737304 34737394
refGene_gn=IPO13;refGene_tx=XM_006934675.4;refGene_gn=IPO13;refGene_tx
=XM_019836869.2
C1 34737690 34737872
refGene_gn=IPO13;refGene_tx=XM_006934675.4;refGene_gn=IPO13;refGene_tx
=XM_019836869.2
C1 34737960 34738057
refGene_gn=IPO13;refGene_tx=XM_006934675.4;refGene_gn=IPO13;refGene_tx
=XM_019836869.2
C1 34740438 34740735
refGene_gn=DPH2;refGene_tx=XM_003990005.5;refGene_gn=DPH2;refGene_tx=X
M_006934676.4;refGene_gn=DPH2;refGene_tx=XM_019836870.2
C1 34740948 34741061
refGene_gn=DPH2;refGene_tx=XM_003990005.5;refGene_gn=DPH2;refGene_tx=X
M_006934676.4;refGene_gn=DPH2;refGene_tx=XM_019836870.2;refGene_gn=DPH
2;refGene_tx=XM_019836871.1
C1 34741316 34741540
refGene_gn=DPH2;refGene_tx=XM_003990005.5;refGene_gn=DPH2;refGene_tx=X
M_006934676.4;refGene_gn=DPH2;refGene_tx=XM_019836870.2

C1 34741718 34742402
refGene_gn=DPH2;refGene_tx=XM_003990005.5;refGene_gn=DPH2;refGene_tx=XM_006934676.4;refGene_gn=DPH2;refGene_tx=XM_019836870.2;refGene_gn=DPH2;refGene_tx=XM_019836871.1
C1 34742482 34742659
refGene_gn=DPH2;refGene_tx=XM_003990005.5;refGene_gn=DPH2;refGene_tx=XM_006934676.4;refGene_gn=DPH2;refGene_tx=XM_019836870.2;refGene_gn=DPH2;refGene_tx=XM_019836871.1
C1 34742740 34742865
refGene_gn=DPH2;refGene_tx=XM_003990005.5;refGene_gn=DPH2;refGene_tx=XM_006934676.4;refGene_gn=DPH2;refGene_tx=XM_019836870.2;refGene_gn=DPH2;refGene_tx=XM_019836871.1
C1 34745818 34745885 refGene_gn=ATP6V0B;refGene_tx=XM_003990006.4
C1 34746648 34746697 refGene_gn=ATP6V0B;refGene_tx=XM_003990006.4
C1 34746931 34747015
refGene_gn=ATP6V0B;refGene_tx=XM_003990006.4;refGene_gn=ATP6V0B;refGene_tx=XM_019836872.2
C1 34747142 34747220
refGene_gn=ATP6V0B;refGene_tx=XM_003990006.4;refGene_gn=ATP6V0B;refGene_tx=XM_019836872.2
C1 34747427 34747497
refGene_gn=ATP6V0B;refGene_tx=XM_003990006.4;refGene_gn=ATP6V0B;refGene_tx=XM_019836872.2
C1 34747618 34747670
refGene_gn=ATP6V0B;refGene_tx=XM_003990006.4;refGene_gn=ATP6V0B;refGene_tx=XM_019836872.2
C1 34747886 34748077
refGene_gn=ATP6V0B;refGene_tx=XM_003990006.4;refGene_gn=ATP6V0B;refGene_tx=XM_019836872.2
C1 34748779 34748806
refGene_gn=ATP6V0B;refGene_tx=XM_003990006.4;refGene_gn=ATP6V0B;refGene_tx=XM_019836872.2
C1 34750771 34750806 refGene_gn=B4GALT2;refGene_tx=XM_003990008.5
C1 34751887 34752243
refGene_gn=B4GALT2;refGene_tx=XM_003990008.5;refGene_gn=B4GALT2;refGene_tx=XM_006934677.4;refGene_gn=B4GALT2;refGene_tx=XM_023258617.1
C1 34752453 34752689
refGene_gn=B4GALT2;refGene_tx=XM_003990008.5;refGene_gn=B4GALT2;refGene_tx=XM_006934677.4;refGene_gn=B4GALT2;refGene_tx=XM_023258617.1
C1 34755517 34755708
refGene_gn=B4GALT2;refGene_tx=XM_003990008.5;refGene_gn=B4GALT2;refGene_tx=XM_006934677.4;refGene_gn=B4GALT2;refGene_tx=XM_023258617.1
C1 34755878 34756001
refGene_gn=B4GALT2;refGene_tx=XM_003990008.5;refGene_gn=B4GALT2;refGene_tx=XM_006934677.4;refGene_gn=B4GALT2;refGene_tx=XM_023258617.1
C1 34756112 34756217
refGene_gn=B4GALT2;refGene_tx=XM_003990008.5;refGene_gn=B4GALT2;refGene_tx=XM_006934677.4;refGene_gn=B4GALT2;refGene_tx=XM_023258617.1
C1 34758797 34758948
refGene_gn=B4GALT2;refGene_tx=XM_003990008.5;refGene_gn=B4GALT2;refGen

e_tx=XM_006934677.4;refGene_gn=B4GALT2;refGene_tx=XM_023258617.1
C1 34760460 34760586
refGene_gn=CCDC24;refGene_tx=XM_006934682.3;refGene_gn=CCDC24;refGene_tx=XM_019836874.2
C1 34760806 34760982
refGene_gn=CCDC24;refGene_tx=XM_006934682.3;refGene_gn=CCDC24;refGene_tx=XM_019836874.2;refGene_gn=CCDC24;refGene_tx=XM_019836876.1
C1 34761119 34761236
refGene_gn=CCDC24;refGene_tx=XM_006934682.3;refGene_gn=CCDC24;refGene_tx=XM_019836874.2;refGene_gn=CCDC24;refGene_tx=XM_019836876.1
C1 34761754 34761832
refGene_gn=CCDC24;refGene_tx=XM_006934682.3;refGene_gn=CCDC24;refGene_tx=XM_019836874.2;refGene_gn=CCDC24;refGene_tx=XM_019836876.1
C1 34762384 34762439
refGene_gn=CCDC24;refGene_tx=XM_006934682.3;refGene_gn=CCDC24;refGene_tx=XM_019836874.2;refGene_gn=CCDC24;refGene_tx=XM_019836876.1
C1 34762898 34762968
refGene_gn=CCDC24;refGene_tx=XM_006934682.3;refGene_gn=CCDC24;refGene_tx=XM_019836874.2;refGene_gn=CCDC24;refGene_tx=XM_019836876.1
C1 34763078 34763157
refGene_gn=CCDC24;refGene_tx=XM_006934682.3;refGene_gn=CCDC24;refGene_tx=XM_019836874.2;refGene_gn=CCDC24;refGene_tx=XM_019836876.1
C1 34763239 34763474
refGene_gn=CCDC24;refGene_tx=XM_006934682.3;refGene_gn=CCDC24;refGene_tx=XM_019836874.2;refGene_gn=CCDC24;refGene_tx=XM_019836876.1
C1 34764876 34765071 refGene_gn=SLC6A9;refGene_tx=XM_019836873.2
C1 34765185 34765356 refGene_gn=SLC6A9;refGene_tx=XM_019836873.2
C1 34768104 34768205 refGene_gn=SLC6A9;refGene_tx=XM_019836873.2
C1 34768294 34768394 refGene_gn=SLC6A9;refGene_tx=XM_019836873.2
C1 34768502 34768637 refGene_gn=SLC6A9;refGene_tx=XM_019836873.2
C1 34768727 34768965 refGene_gn=SLC6A9;refGene_tx=XM_019836873.2
C1 34769583 34769687 refGene_gn=SLC6A9;refGene_tx=XM_019836873.2
C1 34769778 34769913 refGene_gn=SLC6A9;refGene_tx=XM_019836873.2
C1 34770138 34770271 refGene_gn=SLC6A9;refGene_tx=XM_019836873.2
C1 34776435 34776706 refGene_gn=SLC6A9;refGene_tx=XM_019836873.2
C1 34777091 34777223 refGene_gn=SLC6A9;refGene_tx=XM_019836873.2
C1 34777833 34777990 refGene_gn=SLC6A9;refGene_tx=XM_019836873.2
C1 34784127 34784193 refGene_gn=SLC6A9;refGene_tx=XM_019836873.2
C1 34784195 34784219 refGene_gn=SLC6A9;refGene_tx=XM_019836873.2
C1 34878127 34878211 refGene_gn=KLF17;refGene_tx=XM_003990011.4
C1 34887169 34888025
refGene_gn=KLF17;refGene_tx=XM_003990011.4;refGene_gn=KLF17;refGene_tx=XM_011284667.3;refGene_gn=KLF17;refGene_tx=XM_019836878.2
C1 34888376 34888621
refGene_gn=KLF17;refGene_tx=XM_003990011.4;refGene_gn=KLF17;refGene_tx=XM_011284667.3;refGene_gn=KLF17;refGene_tx=XM_019836878.2
C1 34903924 34904824
refGene_gn=LOC111561630;refGene_tx=XM_023257827.1
C1 34904947 34905055
refGene_gn=LOC111561630;refGene_tx=XM_023257827.1

C1	34951540	34951645	refGene_gn=DMAP1;refGene_tx=XM_006934685.3
C1	34952195	34952287
refGene_gn=DMAP1;refGene_tx=XM_006934685.3;refGene_gn=DMAP1;refGene_tx=XM_023258618.1
C1	34952552	34952748
refGene_gn=DMAP1;refGene_tx=XM_006934685.3;refGene_gn=DMAP1;refGene_tx=XM_023258618.1
C1	34957509	34957668
refGene_gn=DMAP1;refGene_tx=XM_006934685.3;refGene_gn=DMAP1;refGene_tx=XM_023258618.1
C1	34957796	34957964
refGene_gn=DMAP1;refGene_tx=XM_006934685.3;refGene_gn=DMAP1;refGene_tx=XM_023258618.1
C1	34958220	34958406
refGene_gn=DMAP1;refGene_tx=XM_006934685.3;refGene_gn=DMAP1;refGene_tx=XM_023258618.1
C1	34958535	34958607
refGene_gn=DMAP1;refGene_tx=XM_006934685.3;refGene_gn=DMAP1;refGene_tx=XM_023258618.1
C1	34958910	34958983
refGene_gn=DMAP1;refGene_tx=XM_006934685.3;refGene_gn=DMAP1;refGene_tx=XM_023258618.1
C1	34959120	34959416
refGene_gn=DMAP1;refGene_tx=XM_006934685.3;refGene_gn=DMAP1;refGene_tx=XM_023258618.1
C1	34959660	34959720
refGene_gn=DMAP1;refGene_tx=XM_006934685.3;refGene_gn=DMAP1;refGene_tx=XM_023258618.1
C1	34960630	34960713
refGene_gn=ERI3;refGene_tx=XM_003990014.2;refGene_gn=ERI3;refGene_tx=XM_006934686.1;refGene_gn=ERI3;refGene_tx=XM_011284664.3;refGene_gn=ERI3;refGene_tx=XM_011284665.3;refGene_gn=ERI3;refGene_tx=XM_011284666.3
C1	34971125	34971130	refGene_gn=ERI3;refGene_tx=XM_019836885.1
C1	34973455	34973508	refGene_gn=ERI3;refGene_tx=XM_023258622.1
C1	34986494	34986594
refGene_gn=ERI3;refGene_tx=XM_003990014.2;refGene_gn=ERI3;refGene_tx=XM_006934686.1;refGene_gn=ERI3;refGene_tx=XM_011284664.3;refGene_gn=ERI3;refGene_tx=XM_011284665.3;refGene_gn=ERI3;refGene_tx=XM_011284666.3;refGene_gn=ERI3;refGene_tx=XM_019836885.1;refGene_gn=ERI3;refGene_tx=XM_019836887.1;refGene_gn=ERI3;refGene_tx=XM_023258622.1
C1	35022434	35022507
refGene_gn=ERI3;refGene_tx=XM_003990014.2;refGene_gn=ERI3;refGene_tx=XM_006934686.1;refGene_gn=ERI3;refGene_tx=XM_011284664.3;refGene_gn=ERI3;refGene_tx=XM_011284665.3;refGene_gn=ERI3;refGene_tx=XM_011284666.3;refGene_gn=ERI3;refGene_tx=XM_019836885.1;refGene_gn=ERI3;refGene_tx=XM_023258622.1
C1	35043775	35043867
refGene_gn=ERI3;refGene_tx=XM_003990014.2;refGene_gn=ERI3;refGene_tx=XM_006934686.1;refGene_gn=ERI3;refGene_tx=XM_011284664.3;refGene_gn=ERI3;refGene_tx=XM_011284665.3;refGene_gn=ERI3;refGene_tx=XM_011284666.3;

refGene_gn=ERI3;refGene_tx=XM_019836885.1;refGene_gn=ERI3;refGene_tx=XM_019836887.1;refGene_gn=ERI3;refGene_tx=XM_023258622.1
C1 35048211 35048271
refGene_gn=ERI3;refGene_tx=XM_003990014.2;refGene_gn=ERI3;refGene_tx=XM_006934686.1;refGene_gn=ERI3;refGene_tx=XM_011284664.3;refGene_gn=ERI3;refGene_tx=XM_011284665.3;refGene_gn=ERI3;refGene_tx=XM_011284666.3;refGene_gn=ERI3;refGene_tx=XM_019836885.1;refGene_gn=ERI3;refGene_tx=XM_019836887.1;refGene_gn=ERI3;refGene_tx=XM_023258622.1
C1 35054809 35054926
refGene_gn=ERI3;refGene_tx=XM_003990014.2;refGene_gn=ERI3;refGene_tx=XM_006934686.1;refGene_gn=ERI3;refGene_tx=XM_011284664.3;refGene_gn=ERI3;refGene_tx=XM_011284665.3;refGene_gn=ERI3;refGene_tx=XM_011284666.3;refGene_gn=ERI3;refGene_tx=XM_019836885.1;refGene_gn=ERI3;refGene_tx=XM_019836887.1;refGene_gn=ERI3;refGene_tx=XM_023258622.1
C1 35074028 35074306
refGene_gn=ERI3;refGene_tx=XM_003990014.2;refGene_gn=ERI3;refGene_tx=XM_006934686.1;refGene_gn=ERI3;refGene_tx=XM_011284664.3;refGene_gn=ERI3;refGene_tx=XM_011284665.3;refGene_gn=ERI3;refGene_tx=XM_019836885.1;refGene_gn=ERI3;refGene_tx=XM_019836887.1;refGene_gn=ERI3;refGene_tx=XM_023258622.1
C1 35086160 35086236
refGene_gn=ERI3;refGene_tx=XM_003990014.2;refGene_gn=ERI3;refGene_tx=XM_006934686.1;refGene_gn=ERI3;refGene_tx=XM_019836885.1;refGene_gn=ERI3;refGene_tx=XM_019836887.1;refGene_gn=ERI3;refGene_tx=XM_023258622.1
C1 35088253 35088388
refGene_gn=ERI3;refGene_tx=XM_003990014.2;refGene_gn=ERI3;refGene_tx=XM_006934686.1;refGene_gn=ERI3;refGene_tx=XM_019836885.1;refGene_gn=ERI3;refGene_tx=XM_019836887.1;refGene_gn=ERI3;refGene_tx=XM_023258622.1
C1 35145068 35145693
refGene_gn=RNF220;refGene_tx=XM_019836881.2;refGene_gn=RNF220;refGene_tx=XM_019836883.2
C1 35336151 35336284
refGene_gn=RNF220;refGene_tx=XM_019836881.2;refGene_gn=RNF220;refGene_tx=XM_019836883.2
C1 35344161 35344207
refGene_gn=RNF220;refGene_tx=XM_019836881.2;refGene_gn=RNF220;refGene_tx=XM_019836883.2
C1 35347756 35347858
refGene_gn=RNF220;refGene_tx=XM_019836881.2;refGene_gn=RNF220;refGene_tx=XM_019836883.2
C1 35352596 35352639
refGene_gn=RNF220;refGene_tx=XM_019836881.2;refGene_gn=RNF220;refGene_tx=XM_019836883.2
C1 35355746 35355868
refGene_gn=RNF220;refGene_tx=XM_019836881.2;refGene_gn=RNF220;refGene_tx=XM_019836883.2
C1 35356238 35356371
refGene_gn=RNF220;refGene_tx=XM_019836881.2;refGene_gn=RNF220;refGene_tx=XM_019836883.2
C1 35364526 35364623

refGene_gn=RNF220;refGene_tx=XM_019836881.2;refGene_gn=RNF220;refGene_tx=XM_019836883.2
C1 35364824 35364911
refGene_gn=RNF220;refGene_tx=XM_019836881.2;refGene_gn=RNF220;refGene_tx=XM_019836883.2
C1 35365049 35365105
refGene_gn=RNF220;refGene_tx=XM_019836881.2;refGene_gn=RNF220;refGene_tx=XM_019836883.2
C1 35365241 35365320
refGene_gn=RNF220;refGene_tx=XM_019836881.2;refGene_gn=RNF220;refGene_tx=XM_019836883.2
C1 35369561 35369670
refGene_gn=RNF220;refGene_tx=XM_019836881.2;refGene_gn=RNF220;refGene_tx=XM_019836883.2
C1 35369817 35369892
refGene_gn=RNF220;refGene_tx=XM_019836881.2;refGene_gn=RNF220;refGene_tx=XM_019836883.2
C1 35370639 35370711
refGene_gn=RNF220;refGene_tx=XM_019836881.2;refGene_gn=RNF220;refGene_tx=XM_019836883.2
C1 35374037 35374691
refGene_gn=TMEM53;refGene_tx=XM_003990016.5;refGene_gn=TMEM53;refGene_tx=XM_011284662.3
C1 35378512 35378620 refGene_gn=TMEM53;refGene_tx=XM_011284662.3
C1 35378901 35379023 refGene_gn=TMEM53;refGene_tx=XM_003990016.5
C1 35394326 35394387 refGene_gn=TMEM53;refGene_tx=XM_003990016.5
C1 35408208 35408414
refGene_gn=CC1H1orf228;refGene_tx=XM_023258619.1;refGene_gn=CC1H1orf228;refGene_tx=XM_023258620.1;refGene_gn=CC1H1orf228;refGene_tx=XM_023258621.1
C1 35413465 35413534
refGene_gn=CC1H1orf228;refGene_tx=XM_023258619.1
C1 35414231 35414398
refGene_gn=CC1H1orf228;refGene_tx=XM_023258619.1;refGene_gn=CC1H1orf228;refGene_tx=XM_023258620.1;refGene_gn=CC1H1orf228;refGene_tx=XM_023258621.1
C1 35415439 35415636
refGene_gn=CC1H1orf228;refGene_tx=XM_023258619.1;refGene_gn=CC1H1orf228;refGene_tx=XM_023258620.1;refGene_gn=CC1H1orf228;refGene_tx=XM_023258621.1
C1 35419336 35419421
refGene_gn=CC1H1orf228;refGene_tx=XM_023258619.1;refGene_gn=CC1H1orf228;refGene_tx=XM_023258620.1;refGene_gn=CC1H1orf228;refGene_tx=XM_023258621.1
C1 35443205 35443328
refGene_gn=CC1H1orf228;refGene_tx=XM_023258619.1;refGene_gn=CC1H1orf228;refGene_tx=XM_023258620.1;refGene_gn=CC1H1orf228;refGene_tx=XM_023258621.1
C1 35443415 35443488
refGene_gn=CC1H1orf228;refGene_tx=XM_023258619.1;refGene_gn=CC1H1orf22

8;refGene_tx=XM_023258620.1;refGene_gn=CC1H1orf228;refGene_tx=XM_023258621.1
C1 35443620 35443750 refGene_gn=CC1H1orf228;refGene_tx=XM_023258619.1;refGene_gn=CC1H1orf228;refGene_tx=XM_023258620.1;refGene_gn=CC1H1orf228;refGene_tx=XM_023258621.1
C1 35443839 35443917 refGene_gn=CC1H1orf228;refGene_tx=XM_023258619.1;refGene_gn=CC1H1orf228;refGene_tx=XM_023258620.1;refGene_gn=CC1H1orf228;refGene_tx=XM_023258621.1
C1 35444222 35444304 refGene_gn=CC1H1orf228;refGene_tx=XM_023258619.1;refGene_gn=CC1H1orf228;refGene_tx=XM_023258620.1;refGene_gn=CC1H1orf228;refGene_tx=XM_023258621.1
C1 35444380 35444499 refGene_gn=CC1H1orf228;refGene_tx=XM_023258619.1;refGene_gn=CC1H1orf228;refGene_tx=XM_023258620.1;refGene_gn=CC1H1orf228;refGene_tx=XM_023258621.1
C1 35463273 35463337 refGene_gn=KIF2C;refGene_tx=XM_003990019.4;refGene_gn=KIF2C;refGene_tx=XM_011284660.3
C1 35464211 35464306 refGene_gn=KIF2C;refGene_tx=XM_003990019.4;refGene_gn=KIF2C;refGene_tx=XM_011284660.3
C1 35467196 35467199 refGene_gn=KIF2C;refGene_tx=XM_003990020.4
C1 35467361 35467463 refGene_gn=KIF2C;refGene_tx=XM_003990019.4;refGene_gn=KIF2C;refGene_tx=XM_003990020.4;refGene_gn=KIF2C;refGene_tx=XM_011284660.3
C1 35467635 35467684 refGene_gn=KIF2C;refGene_tx=XM_003990019.4;refGene_gn=KIF2C;refGene_tx=XM_003990020.4;refGene_gn=KIF2C;refGene_tx=XM_011284660.3
C1 35469299 35469422 refGene_gn=KIF2C;refGene_tx=XM_003990019.4;refGene_gn=KIF2C;refGene_tx=XM_003990020.4;refGene_gn=KIF2C;refGene_tx=XM_011284660.3
C1 35470563 35470686 refGene_gn=KIF2C;refGene_tx=XM_003990019.4;refGene_gn=KIF2C;refGene_tx=XM_003990020.4
C1 35471234 35471335 refGene_gn=KIF2C;refGene_tx=XM_003990019.4;refGene_gn=KIF2C;refGene_tx=XM_003990020.4;refGene_gn=KIF2C;refGene_tx=XM_011284660.3
C1 35472009 35472105 refGene_gn=KIF2C;refGene_tx=XM_003990019.4;refGene_gn=KIF2C;refGene_tx=XM_003990020.4;refGene_gn=KIF2C;refGene_tx=XM_011284660.3
C1 35472516 35472571 refGene_gn=KIF2C;refGene_tx=XM_003990019.4;refGene_gn=KIF2C;refGene_tx=XM_003990020.4;refGene_gn=KIF2C;refGene_tx=XM_011284660.3
C1 35472660 35472823 refGene_gn=KIF2C;refGene_tx=XM_003990019.4;refGene_gn=KIF2C;refGene_tx=XM_003990020.4;refGene_gn=KIF2C;refGene_tx=XM_011284660.3
C1 35475354 35475445

refGene_gn=KIF2C;refGene_tx=XM_003990019.4;refGene_gn=KIF2C;refGene_tx=XM_003990020.4;refGene_gn=KIF2C;refGene_tx=XM_011284660.3
C1 35475703 35475767
refGene_gn=KIF2C;refGene_tx=XM_003990019.4;refGene_gn=KIF2C;refGene_tx=XM_003990020.4;refGene_gn=KIF2C;refGene_tx=XM_011284660.3
C1 35475845 35475937
refGene_gn=KIF2C;refGene_tx=XM_003990019.4;refGene_gn=KIF2C;refGene_tx=XM_003990020.4;refGene_gn=KIF2C;refGene_tx=XM_011284660.3
C1 35476732 35476875
refGene_gn=KIF2C;refGene_tx=XM_003990019.4;refGene_gn=KIF2C;refGene_tx=XM_003990020.4;refGene_gn=KIF2C;refGene_tx=XM_011284660.3
C1 35477656 35477861
refGene_gn=KIF2C;refGene_tx=XM_003990019.4;refGene_gn=KIF2C;refGene_tx=XM_003990020.4;refGene_gn=KIF2C;refGene_tx=XM_011284660.3
C1 35477980 35478091
refGene_gn=KIF2C;refGene_tx=XM_003990019.4;refGene_gn=KIF2C;refGene_tx=XM_003990020.4;refGene_gn=KIF2C;refGene_tx=XM_011284660.3
C1 35479138 35479206
refGene_gn=KIF2C;refGene_tx=XM_003990019.4;refGene_gn=KIF2C;refGene_tx=XM_003990020.4;refGene_gn=KIF2C;refGene_tx=XM_011284660.3
C1 35479482 35479588
refGene_gn=KIF2C;refGene_tx=XM_003990019.4;refGene_gn=KIF2C;refGene_tx=XM_003990020.4;refGene_gn=KIF2C;refGene_tx=XM_011284660.3
C1 35479680 35479794
refGene_gn=KIF2C;refGene_tx=XM_003990019.4;refGene_gn=KIF2C;refGene_tx=XM_003990020.4;refGene_gn=KIF2C;refGene_tx=XM_011284660.3
C1 35481812 35481936
refGene_gn=KIF2C;refGene_tx=XM_003990019.4;refGene_gn=KIF2C;refGene_tx=XM_003990020.4;refGene_gn=KIF2C;refGene_tx=XM_011284660.3
C1 35482031 35482114
refGene_gn=KIF2C;refGene_tx=XM_003990019.4;refGene_gn=KIF2C;refGene_tx=XM_003990020.4;refGene_gn=KIF2C;refGene_tx=XM_011284660.3
C1 35488289 35488293 refGene_gn=RPS8;refGene_tx=XM_003990022.3
C1 35488690 35488797 refGene_gn=RPS8;refGene_tx=XM_003990022.3
C1 35489273 35489373 refGene_gn=RPS8;refGene_tx=XM_003990022.3
C1 35489770 35489946 refGene_gn=RPS8;refGene_tx=XM_003990022.3
C1 35490180 35490310 refGene_gn=RPS8;refGene_tx=XM_003990022.3
C1 35490816 35490926 refGene_gn=RPS8;refGene_tx=XM_003990022.3
C1 35495982 35496256 refGene_gn=BEST4;refGene_tx=XM_023260048.1
C1 35496404 35496559 refGene_gn=BEST4;refGene_tx=XM_023260048.1
C1 35496663 35496744 refGene_gn=BEST4;refGene_tx=XM_023260048.1
C1 35496847 35497045 refGene_gn=BEST4;refGene_tx=XM_023260048.1
C1 35497335 35497413 refGene_gn=BEST4;refGene_tx=XM_023260048.1
C1 35497857 35498012 refGene_gn=BEST4;refGene_tx=XM_023260048.1
C1 35498188 35498422 refGene_gn=BEST4;refGene_tx=XM_023260048.1
C1 35499004 35499099 refGene_gn=BEST4;refGene_tx=XM_023260048.1
C1 35499184 35499336 refGene_gn=BEST4;refGene_tx=XM_023260048.1
C1 35508905 35509115 refGene_gn=PLK3;refGene_tx=XM_003990024.3
C1 35509324 35509432 refGene_gn=PLK3;refGene_tx=XM_003990024.3
C1 35509521 35509638 refGene_gn=PLK3;refGene_tx=XM_003990024.3

C1 35509778 35509908 refGene_gn=PLK3;refGene_tx=XM_003990024.3
C1 35509989 35510077 refGene_gn=PLK3;refGene_tx=XM_003990024.3
C1 35510894 35510990 refGene_gn=PLK3;refGene_tx=XM_003990024.3
C1 35511081 35511280 refGene_gn=PLK3;refGene_tx=XM_003990024.3
C1 35511389 35511513 refGene_gn=PLK3;refGene_tx=XM_003990024.3
C1 35511697 35511789 refGene_gn=PLK3;refGene_tx=XM_003990024.3
C1 35512029 35512123 refGene_gn=PLK3;refGene_tx=XM_003990024.3
C1 35512266 35512350 refGene_gn=PLK3;refGene_tx=XM_003990024.3
C1 35512443 35512606 refGene_gn=PLK3;refGene_tx=XM_003990024.3
C1 35512746 35512876 refGene_gn=PLK3;refGene_tx=XM_003990024.3
C1 35513046 35513160 refGene_gn=PLK3;refGene_tx=XM_003990024.3
C1 35513270 35513462 refGene_gn=PLK3;refGene_tx=XM_003990024.3
C1 35513812 35514472 refGene_gn=TCTEX1D4;refGene_tx=XM_023258623.1
C1 35516665 35516751
refGene_gn=BTBD19;refGene_tx=XM_003990025.4;refGene_gn=BTBD19;refGene_tx=XM_019836889.2
C1 35517795 35517821 refGene_gn=BTBD19;refGene_tx=XM_019836888.2
C1 35518034 35518248
refGene_gn=BTBD19;refGene_tx=XM_003990025.4;refGene_gn=BTBD19;refGene_tx=XM_019836888.2;refGene_gn=BTBD19;refGene_tx=XM_019836889.2
C1 35518333 35518387
refGene_gn=BTBD19;refGene_tx=XM_003990025.4;refGene_gn=BTBD19;refGene_tx=XM_019836888.2;refGene_gn=BTBD19;refGene_tx=XM_019836889.2
C1 35519548 35519608
refGene_gn=BTBD19;refGene_tx=XM_003990025.4;refGene_gn=BTBD19;refGene_tx=XM_019836888.2;refGene_gn=BTBD19;refGene_tx=XM_019836889.2
C1 35519709 35519778
refGene_gn=BTBD19;refGene_tx=XM_003990025.4;refGene_gn=BTBD19;refGene_tx=XM_019836888.2;refGene_gn=BTBD19;refGene_tx=XM_019836889.2
C1 35519855 35519987
refGene_gn=BTBD19;refGene_tx=XM_003990025.4;refGene_gn=BTBD19;refGene_tx=XM_019836888.2;refGene_gn=BTBD19;refGene_tx=XM_019836889.2
C1 35520112 35520238
refGene_gn=BTBD19;refGene_tx=XM_003990025.4;refGene_gn=BTBD19;refGene_tx=XM_019836888.2
C1 35520376 35520511
refGene_gn=BTBD19;refGene_tx=XM_003990025.4;refGene_gn=BTBD19;refGene_tx=XM_019836888.2;refGene_gn=BTBD19;refGene_tx=XM_019836889.2
C1 35529076 35529343
refGene_gn=PTCH2;refGene_tx=XM_003990026.5;refGene_gn=PTCH2;refGene_tx=XM_019836891.2;refGene_gn=PTCH2;refGene_tx=XM_019836892.2;refGene_gn=PTCH2;refGene_tx=XM_019836893.2;refGene_gn=PTCH2;refGene_tx=XM_019836894.2;refGene_gn=PTCH2;refGene_tx=XM_019836896.2;refGene_gn=PTCH2;refGene_tx=XM_019836897.2
C1 35529397 35529472 refGene_gn=PTCH2;refGene_tx=XM_019836895.2
C1 35529756 35529856
refGene_gn=PTCH2;refGene_tx=XM_003990026.5;refGene_gn=PTCH2;refGene_tx=XM_019836891.2;refGene_gn=PTCH2;refGene_tx=XM_019836892.2;refGene_gn=PTCH2;refGene_tx=XM_019836893.2;refGene_gn=PTCH2;refGene_tx=XM_019836894.2;refGene_gn=PTCH2;refGene_tx=XM_019836895.2;refGene_gn=PTCH2;refGe

ne_tx=XM_019836896.2;refGene_gn=PTCH2;refGene_tx=XM_019836897.2
C1 35529940 35530083
refGene_gn=PTCH2;refGene_tx=XM_003990026.5;refGene_gn=PTCH2;refGene_tx=XM_019836891.2;refGene_gn=PTCH2;refGene_tx=XM_019836892.2;refGene_gn=PTCH2;refGene_tx=XM_019836893.2;refGene_gn=PTCH2;refGene_tx=XM_019836894.2;refGene_gn=PTCH2;refGene_tx=XM_019836895.2;refGene_gn=PTCH2;refGene_tx=XM_019836896.2;refGene_gn=PTCH2;refGene_tx=XM_019836897.2
C1 35531324 35531462
refGene_gn=PTCH2;refGene_tx=XM_003990026.5;refGene_gn=PTCH2;refGene_tx=XM_019836891.2;refGene_gn=PTCH2;refGene_tx=XM_019836892.2;refGene_gn=PTCH2;refGene_tx=XM_019836893.2;refGene_gn=PTCH2;refGene_tx=XM_019836894.2;refGene_gn=PTCH2;refGene_tx=XM_019836895.2;refGene_gn=PTCH2;refGene_tx=XM_019836896.2;refGene_gn=PTCH2;refGene_tx=XM_019836897.2
C1 35531571 35531852
refGene_gn=PTCH2;refGene_tx=XM_003990026.5;refGene_gn=PTCH2;refGene_tx=XM_019836891.2;refGene_gn=PTCH2;refGene_tx=XM_019836892.2;refGene_gn=PTCH2;refGene_tx=XM_019836893.2;refGene_gn=PTCH2;refGene_tx=XM_019836894.2;refGene_gn=PTCH2;refGene_tx=XM_019836895.2;refGene_gn=PTCH2;refGene_tx=XM_019836896.2;refGene_gn=PTCH2;refGene_tx=XM_019836897.2
C1 35531982 35532392
refGene_gn=PTCH2;refGene_tx=XM_003990026.5;refGene_gn=PTCH2;refGene_tx=XM_019836891.2;refGene_gn=PTCH2;refGene_tx=XM_019836892.2;refGene_gn=PTCH2;refGene_tx=XM_019836893.2;refGene_gn=PTCH2;refGene_tx=XM_019836894.2;refGene_gn=PTCH2;refGene_tx=XM_019836895.2;refGene_gn=PTCH2;refGene_tx=XM_019836896.2;refGene_gn=PTCH2;refGene_tx=XM_019836897.2;refGene_gn=PTCH2;refGene_tx=XM_023258624.1;refGene_gn=PTCH2;refGene_tx=XM_023258625.1;refGene_gn=PTCH2;refGene_tx=XM_023258626.1
C1 35532483 35532796
refGene_gn=PTCH2;refGene_tx=XM_003990026.5;refGene_gn=PTCH2;refGene_tx=XM_019836891.2;refGene_gn=PTCH2;refGene_tx=XM_019836892.2;refGene_gn=PTCH2;refGene_tx=XM_019836893.2;refGene_gn=PTCH2;refGene_tx=XM_019836894.2;refGene_gn=PTCH2;refGene_tx=XM_019836895.2;refGene_gn=PTCH2;refGene_tx=XM_019836896.2;refGene_gn=PTCH2;refGene_tx=XM_019836897.2;refGene_gn=PTCH2;refGene_tx=XM_023258624.1;refGene_gn=PTCH2;refGene_tx=XM_023258625.1;refGene_gn=PTCH2;refGene_tx=XM_023258626.1
C1 35532941 35533523
refGene_gn=PTCH2;refGene_tx=XM_003990026.5;refGene_gn=PTCH2;refGene_tx=XM_019836891.2;refGene_gn=PTCH2;refGene_tx=XM_019836892.2;refGene_gn=PTCH2;refGene_tx=XM_019836893.2;refGene_gn=PTCH2;refGene_tx=XM_019836894.2;refGene_gn=PTCH2;refGene_tx=XM_019836895.2;refGene_gn=PTCH2;refGene_tx=XM_019836896.2;refGene_gn=PTCH2;refGene_tx=XM_019836897.2;refGene_gn=PTCH2;refGene_tx=XM_023258624.1;refGene_gn=PTCH2;refGene_tx=XM_023258625.1;refGene_gn=PTCH2;refGene_tx=XM_023258626.1;refGene_gn=PTCH2;refGene_tx=XM_023258627.1
C1 35533616 35533742
refGene_gn=PTCH2;refGene_tx=XM_003990026.5;refGene_gn=PTCH2;refGene_tx=XM_019836891.2;refGene_gn=PTCH2;refGene_tx=XM_019836892.2;refGene_gn=PTCH2;refGene_tx=XM_019836893.2;refGene_gn=PTCH2;refGene_tx=XM_019836895.2;refGene_gn=PTCH2;refGene_tx=XM_019836896.2;refGene_gn=PTCH2;refGene_tx=XM_019836897.2;refGene_gn=PTCH2;refGene_tx=XM_023258624.1;refGen

e_gn=PTCH2;refGene_tx=XM_023258625.1;refGene_gn=PTCH2;refGene_tx=XM_023258626.1;refGene_gn=PTCH2;refGene_tx=XM_023258627.1
C1 35533772 35533802 refGene_gn=PTCH2;refGene_tx=XM_023258628.1
C1 35534080 35534173
refGene_gn=PTCH2;refGene_tx=XM_003990026.5;refGene_gn=PTCH2;refGene_tx=XM_019836891.2;refGene_gn=PTCH2;refGene_tx=XM_019836892.2;refGene_gn=PTCH2;refGene_tx=XM_019836893.2;refGene_gn=PTCH2;refGene_tx=XM_019836894.2;refGene_gn=PTCH2;refGene_tx=XM_019836895.2;refGene_gn=PTCH2;refGene_tx=XM_019836896.2;refGene_gn=PTCH2;refGene_tx=XM_019836897.2;refGene_gn=PTCH2;refGene_tx=XM_023258624.1;refGene_gn=PTCH2;refGene_tx=XM_023258625.1;refGene_gn=PTCH2;refGene_tx=XM_023258626.1;refGene_gn=PTCH2;refGene_tx=XM_023258627.1;refGene_gn=PTCH2;refGene_tx=XM_023258628.1
C1 35534257 35534413
refGene_gn=PTCH2;refGene_tx=XM_003990026.5;refGene_gn=PTCH2;refGene_tx=XM_019836891.2;refGene_gn=PTCH2;refGene_tx=XM_019836892.2;refGene_gn=PTCH2;refGene_tx=XM_019836893.2;refGene_gn=PTCH2;refGene_tx=XM_019836894.2;refGene_gn=PTCH2;refGene_tx=XM_019836895.2;refGene_gn=PTCH2;refGene_tx=XM_019836896.2;refGene_gn=PTCH2;refGene_tx=XM_019836897.2;refGene_gn=PTCH2;refGene_tx=XM_023258624.1;refGene_gn=PTCH2;refGene_tx=XM_023258625.1;refGene_gn=PTCH2;refGene_tx=XM_023258626.1;refGene_gn=PTCH2;refGene_tx=XM_023258627.1;refGene_gn=PTCH2;refGene_tx=XM_023258628.1
C1 35534503 35534635
refGene_gn=PTCH2;refGene_tx=XM_003990026.5;refGene_gn=PTCH2;refGene_tx=XM_019836891.2;refGene_gn=PTCH2;refGene_tx=XM_019836892.2;refGene_gn=PTCH2;refGene_tx=XM_019836893.2;refGene_gn=PTCH2;refGene_tx=XM_019836894.2;refGene_gn=PTCH2;refGene_tx=XM_019836895.2;refGene_gn=PTCH2;refGene_tx=XM_019836896.2;refGene_gn=PTCH2;refGene_tx=XM_019836897.2;refGene_gn=PTCH2;refGene_tx=XM_023258624.1;refGene_gn=PTCH2;refGene_tx=XM_023258625.1;refGene_gn=PTCH2;refGene_tx=XM_023258626.1;refGene_gn=PTCH2;refGene_tx=XM_023258627.1;refGene_gn=PTCH2;refGene_tx=XM_023258628.1
C1 35534714 35534862
refGene_gn=PTCH2;refGene_tx=XM_003990026.5;refGene_gn=PTCH2;refGene_tx=XM_019836891.2;refGene_gn=PTCH2;refGene_tx=XM_019836892.2;refGene_gn=PTCH2;refGene_tx=XM_019836893.2;refGene_gn=PTCH2;refGene_tx=XM_019836894.2;refGene_gn=PTCH2;refGene_tx=XM_019836895.2;refGene_gn=PTCH2;refGene_tx=XM_019836896.2;refGene_gn=PTCH2;refGene_tx=XM_019836897.2;refGene_gn=PTCH2;refGene_tx=XM_023258624.1;refGene_gn=PTCH2;refGene_tx=XM_023258625.1;refGene_gn=PTCH2;refGene_tx=XM_023258626.1;refGene_gn=PTCH2;refGene_tx=XM_023258627.1;refGene_gn=PTCH2;refGene_tx=XM_023258628.1
C1 35535010 35535132
refGene_gn=PTCH2;refGene_tx=XM_003990026.5;refGene_gn=PTCH2;refGene_tx=XM_019836891.2;refGene_gn=PTCH2;refGene_tx=XM_019836892.2;refGene_gn=PTCH2;refGene_tx=XM_019836893.2;refGene_gn=PTCH2;refGene_tx=XM_019836894.2;refGene_gn=PTCH2;refGene_tx=XM_019836895.2;refGene_gn=PTCH2;refGene_tx=XM_019836896.2;refGene_gn=PTCH2;refGene_tx=XM_019836897.2;refGene_gn=PTCH2;refGene_tx=XM_023258624.1;refGene_gn=PTCH2;refGene_tx=XM_023258625.1;refGene_gn=PTCH2;refGene_tx=XM_023258626.1;refGene_gn=PTCH2;refGene_tx=XM_023258627.1;refGene_gn=PTCH2;refGene_tx=XM_023258628.1
C1 35535723 35535919
refGene_gn=PTCH2;refGene_tx=XM_003990026.5;refGene_gn=PTCH2;refGene_tx

=XM_019836891.2;refGene_gn=PTCH2;refGene_tx=XM_019836892.2;refGene_gn=PTCH2;refGene_tx=XM_019836893.2;refGene_gn=PTCH2;refGene_tx=XM_019836894.2;refGene_gn=PTCH2;refGene_tx=XM_019836895.2;refGene_gn=PTCH2;refGene_tx=XM_019836896.2;refGene_gn=PTCH2;refGene_tx=XM_019836897.2;refGene_gn=PTCH2;refGene_tx=XM_023258624.1;refGene_gn=PTCH2;refGene_tx=XM_023258625.1;refGene_gn=PTCH2;refGene_tx=XM_023258626.1;refGene_gn=PTCH2;refGene_tx=XM_023258627.1;refGene_gn=PTCH2;refGene_tx=XM_023258628.1
C1 35536542 35536634
refGene_gn=PTCH2;refGene_tx=XM_003990026.5;refGene_gn=PTCH2;refGene_tx=XM_019836891.2;refGene_gn=PTCH2;refGene_tx=XM_019836892.2;refGene_gn=PTCH2;refGene_tx=XM_019836893.2;refGene_gn=PTCH2;refGene_tx=XM_019836894.2;refGene_gn=PTCH2;refGene_tx=XM_019836895.2;refGene_gn=PTCH2;refGene_tx=XM_019836896.2;refGene_gn=PTCH2;refGene_tx=XM_019836897.2;refGene_gn=PTCH2;refGene_tx=XM_023258624.1;refGene_gn=PTCH2;refGene_tx=XM_023258625.1;refGene_gn=PTCH2;refGene_tx=XM_023258626.1;refGene_gn=PTCH2;refGene_tx=XM_023258627.1;refGene_gn=PTCH2;refGene_tx=XM_023258628.1
C1 35536824 35536894
refGene_gn=PTCH2;refGene_tx=XM_003990026.5;refGene_gn=PTCH2;refGene_tx=XM_019836891.2;refGene_gn=PTCH2;refGene_tx=XM_019836892.2;refGene_gn=PTCH2;refGene_tx=XM_019836893.2;refGene_gn=PTCH2;refGene_tx=XM_019836894.2;refGene_gn=PTCH2;refGene_tx=XM_019836895.2;refGene_gn=PTCH2;refGene_tx=XM_019836896.2;refGene_gn=PTCH2;refGene_tx=XM_019836897.2;refGene_gn=PTCH2;refGene_tx=XM_023258624.1;refGene_gn=PTCH2;refGene_tx=XM_023258625.1;refGene_gn=PTCH2;refGene_tx=XM_023258626.1;refGene_gn=PTCH2;refGene_tx=XM_023258627.1;refGene_gn=PTCH2;refGene_tx=XM_023258628.1
C1 35537000 35537190
refGene_gn=PTCH2;refGene_tx=XM_003990026.5;refGene_gn=PTCH2;refGene_tx=XM_019836891.2;refGene_gn=PTCH2;refGene_tx=XM_019836892.2;refGene_gn=PTCH2;refGene_tx=XM_019836893.2;refGene_gn=PTCH2;refGene_tx=XM_019836894.2;refGene_gn=PTCH2;refGene_tx=XM_019836895.2;refGene_gn=PTCH2;refGene_tx=XM_019836896.2;refGene_gn=PTCH2;refGene_tx=XM_019836897.2;refGene_gn=PTCH2;refGene_tx=XM_023258624.1;refGene_gn=PTCH2;refGene_tx=XM_023258625.1;refGene_gn=PTCH2;refGene_tx=XM_023258626.1;refGene_gn=PTCH2;refGene_tx=XM_023258627.1;refGene_gn=PTCH2;refGene_tx=XM_023258628.1
C1 35543063 35543256
refGene_gn=PTCH2;refGene_tx=XM_003990026.5;refGene_gn=PTCH2;refGene_tx=XM_019836891.2;refGene_gn=PTCH2;refGene_tx=XM_019836892.2;refGene_gn=PTCH2;refGene_tx=XM_019836893.2;refGene_gn=PTCH2;refGene_tx=XM_019836894.2;refGene_gn=PTCH2;refGene_tx=XM_019836895.2;refGene_gn=PTCH2;refGene_tx=XM_023258624.1;refGene_gn=PTCH2;refGene_tx=XM_023258625.1;refGene_gn=PTCH2;refGene_tx=XM_023258626.1;refGene_gn=PTCH2;refGene_tx=XM_023258627.1;refGene_gn=PTCH2;refGene_tx=XM_023258628.1
C1 35543702 35543774
refGene_gn=PTCH2;refGene_tx=XM_003990026.5;refGene_gn=PTCH2;refGene_tx=XM_019836891.2;refGene_gn=PTCH2;refGene_tx=XM_019836892.2;refGene_gn=PTCH2;refGene_tx=XM_019836893.2;refGene_gn=PTCH2;refGene_tx=XM_019836894.2;refGene_gn=PTCH2;refGene_tx=XM_019836895.2;refGene_gn=PTCH2;refGene_tx=XM_023258624.1;refGene_gn=PTCH2;refGene_tx=XM_023258625.1;refGene_gn=PTCH2;refGene_tx=XM_023258626.1;refGene_gn=PTCH2;refGene_tx=XM_023258627.1;refGene_gn=PTCH2;refGene_tx=XM_023258628.1

C1	35549855	35549908
refGene_gn=EIF2B3;refGene_tx=XM_011284653.3;refGene_gn=EIF2B3;refGene_tx=XM_011284654.3;refGene_gn=EIF2B3;refGene_tx=XM_011284655.3
C1	35556795	35556899
refGene_gn=EIF2B3;refGene_tx=XM_011284653.3;refGene_gn=EIF2B3;refGene_tx=XM_011284654.3;refGene_gn=EIF2B3;refGene_tx=XM_011284655.3
C1	35572790	35572939
refGene_gn=EIF2B3;refGene_tx=XM_011284653.3;refGene_gn=EIF2B3;refGene_tx=XM_011284654.3;refGene_gn=EIF2B3;refGene_tx=XM_011284655.3
C1	35573444	35573522
refGene_gn=EIF2B3;refGene_tx=XM_011284653.3;refGene_gn=EIF2B3;refGene_tx=XM_011284654.3;refGene_gn=EIF2B3;refGene_tx=XM_011284655.3
C1	35576010	35576201
refGene_gn=EIF2B3;refGene_tx=XM_011284653.3;refGene_gn=EIF2B3;refGene_tx=XM_011284654.3;refGene_gn=EIF2B3;refGene_tx=XM_011284655.3
C1	35577935	35578063
refGene_gn=EIF2B3;refGene_tx=XM_011284653.3;refGene_gn=EIF2B3;refGene_tx=XM_011284654.3;refGene_gn=EIF2B3;refGene_tx=XM_011284655.3
C1	35594431	35594521
refGene_gn=EIF2B3;refGene_tx=XM_011284653.3;refGene_gn=EIF2B3;refGene_tx=XM_011284655.3
C1	35626095	35626207
refGene_gn=EIF2B3;refGene_tx=XM_011284653.3;refGene_gn=EIF2B3;refGene_tx=XM_011284654.3;refGene_gn=EIF2B3;refGene_tx=XM_011284655.3
C1	35639255	35639415
refGene_gn=EIF2B3;refGene_tx=XM_011284653.3;refGene_gn=EIF2B3;refGene_tx=XM_011284654.3;refGene_gn=EIF2B3;refGene_tx=XM_011284655.3
C1	35682493	35682639
refGene_gn=EIF2B3;refGene_tx=XM_011284653.3;refGene_gn=EIF2B3;refGene_tx=XM_011284654.3;refGene_gn=EIF2B3;refGene_tx=XM_011284655.3
C1	35686370	35686518
refGene_gn=EIF2B3;refGene_tx=XM_011284653.3;refGene_gn=EIF2B3;refGene_tx=XM_011284654.3;refGene_gn=EIF2B3;refGene_tx=XM_011284655.3
C1	35699610	35699698	refGene_gn=HECTD3;refGene_tx=XM_019836898.2
C1	35700289	35700425
refGene_gn=HECTD3;refGene_tx=XM_003990073.5;refGene_gn=HECTD3;refGene_tx=XM_011284652.3
C1	35700571	35700643
refGene_gn=HECTD3;refGene_tx=XM_003990073.5;refGene_gn=HECTD3;refGene_tx=XM_011284652.3;refGene_gn=HECTD3;refGene_tx=XM_019836898.2
C1	35700784	35700866
refGene_gn=HECTD3;refGene_tx=XM_003990073.5;refGene_gn=HECTD3;refGene_tx=XM_011284652.3;refGene_gn=HECTD3;refGene_tx=XM_019836898.2
C1	35700989	35701064
refGene_gn=HECTD3;refGene_tx=XM_003990073.5;refGene_gn=HECTD3;refGene_tx=XM_011284652.3;refGene_gn=HECTD3;refGene_tx=XM_019836898.2
C1	35701151	35701281
refGene_gn=HECTD3;refGene_tx=XM_003990073.5;refGene_gn=HECTD3;refGene_tx=XM_011284652.3;refGene_gn=HECTD3;refGene_tx=XM_019836898.2
C1	35701572	35701779

refGene_gn=HECTD3;refGene_tx=XM_003990073.5;refGene_gn=HECTD3;refGene_tx=XM_011284652.3;refGene_gn=HECTD3;refGene_tx=XM_019836898.2
C1 35702753 35702843
refGene_gn=HECTD3;refGene_tx=XM_003990073.5;refGene_gn=HECTD3;refGene_tx=XM_011284652.3;refGene_gn=HECTD3;refGene_tx=XM_019836898.2
C1 35702958 35703078
refGene_gn=HECTD3;refGene_tx=XM_003990073.5;refGene_gn=HECTD3;refGene_tx=XM_011284652.3;refGene_gn=HECTD3;refGene_tx=XM_019836898.2
C1 35703338 35703442
refGene_gn=HECTD3;refGene_tx=XM_003990073.5;refGene_gn=HECTD3;refGene_tx=XM_011284652.3;refGene_gn=HECTD3;refGene_tx=XM_019836898.2
C1 35703597 35703662
refGene_gn=HECTD3;refGene_tx=XM_003990073.5;refGene_gn=HECTD3;refGene_tx=XM_011284652.3;refGene_gn=HECTD3;refGene_tx=XM_019836898.2
C1 35703839 35703892
refGene_gn=HECTD3;refGene_tx=XM_003990073.5;refGene_gn=HECTD3;refGene_tx=XM_011284652.3;refGene_gn=HECTD3;refGene_tx=XM_019836898.2
C1 35704043 35704226
refGene_gn=HECTD3;refGene_tx=XM_003990073.5;refGene_gn=HECTD3;refGene_tx=XM_011284652.3;refGene_gn=HECTD3;refGene_tx=XM_019836898.2
C1 35704900 35704982
refGene_gn=HECTD3;refGene_tx=XM_003990073.5;refGene_gn=HECTD3;refGene_tx=XM_011284652.3;refGene_gn=HECTD3;refGene_tx=XM_019836898.2
C1 35705209 35705375
refGene_gn=HECTD3;refGene_tx=XM_003990073.5;refGene_gn=HECTD3;refGene_tx=XM_011284652.3;refGene_gn=HECTD3;refGene_tx=XM_019836898.2
C1 35705808 35705891
refGene_gn=HECTD3;refGene_tx=XM_003990073.5;refGene_gn=HECTD3;refGene_tx=XM_011284652.3;refGene_gn=HECTD3;refGene_tx=XM_019836898.2
C1 35706038 35706152
refGene_gn=HECTD3;refGene_tx=XM_003990073.5;refGene_gn=HECTD3;refGene_tx=XM_011284652.3;refGene_gn=HECTD3;refGene_tx=XM_019836898.2
C1 35706238 35706354
refGene_gn=HECTD3;refGene_tx=XM_003990073.5;refGene_gn=HECTD3;refGene_tx=XM_011284652.3;refGene_gn=HECTD3;refGene_tx=XM_019836898.2
C1 35706631 35706767
refGene_gn=HECTD3;refGene_tx=XM_003990073.5;refGene_gn=HECTD3;refGene_tx=XM_011284652.3;refGene_gn=HECTD3;refGene_tx=XM_019836898.2
C1 35706846 35706939
refGene_gn=HECTD3;refGene_tx=XM_003990073.5;refGene_gn=HECTD3;refGene_tx=XM_011284652.3;refGene_gn=HECTD3;refGene_tx=XM_019836898.2
C1 35707189 35707350
refGene_gn=HECTD3;refGene_tx=XM_003990073.5;refGene_gn=HECTD3;refGene_tx=XM_011284652.3;refGene_gn=HECTD3;refGene_tx=XM_019836898.2
C1 35707531 35707900
refGene_gn=HECTD3;refGene_tx=XM_003990073.5;refGene_gn=HECTD3;refGene_tx=XM_011284652.3;refGene_gn=HECTD3;refGene_tx=XM_019836898.2
C1 35710206 35710214 refGene_gn=UROD;refGene_tx=XM_006934688.3
C1 35710489 35710509
refGene_gn=UROD;refGene_tx=XM_003990027.4;refGene_gn=UROD;refGene_tx=X

M_019836899.2;refGene_gn=UROD;refGene_tx=XM_019836900.2
C1 35711134 35711247
refGene_gn=UROD;refGene_tx=XM_003990027.4;refGene_gn=UROD;refGene_tx=XM_006934688.3;refGene_gn=UROD;refGene_tx=XM_019836899.2;refGene_gn=UROD;refGene_tx=XM_019836900.2;refGene_gn=UROD;refGene_tx=XM_019836901.2;refGene_gn=UROD;refGene_tx=XM_019836902.2
C1 35711366 35711446
refGene_gn=UROD;refGene_tx=XM_003990027.4;refGene_gn=UROD;refGene_tx=XM_006934688.3;refGene_gn=UROD;refGene_tx=XM_019836899.2;refGene_gn=UROD;refGene_tx=XM_019836900.2
C1 35711522 35711585
refGene_gn=UROD;refGene_tx=XM_003990027.4;refGene_gn=UROD;refGene_tx=XM_006934688.3;refGene_gn=UROD;refGene_tx=XM_019836899.2
C1 35711795 35711993
refGene_gn=UROD;refGene_tx=XM_003990027.4;refGene_gn=UROD;refGene_tx=XM_006934688.3;refGene_gn=UROD;refGene_tx=XM_019836899.2;refGene_gn=UROD;refGene_tx=XM_019836900.2;refGene_gn=UROD;refGene_tx=XM_019836901.2;refGene_gn=UROD;refGene_tx=XM_019836902.2
C1 35712118 35712280
refGene_gn=UROD;refGene_tx=XM_003990027.4;refGene_gn=UROD;refGene_tx=XM_006934688.3;refGene_gn=UROD;refGene_tx=XM_019836899.2;refGene_gn=UROD;refGene_tx=XM_019836900.2;refGene_gn=UROD;refGene_tx=XM_019836901.2;refGene_gn=UROD;refGene_tx=XM_019836902.2
C1 35712644 35712782
refGene_gn=UROD;refGene_tx=XM_003990027.4;refGene_gn=UROD;refGene_tx=XM_006934688.3;refGene_gn=UROD;refGene_tx=XM_019836899.2;refGene_gn=UROD;refGene_tx=XM_019836900.2;refGene_gn=UROD;refGene_tx=XM_019836901.2;refGene_gn=UROD;refGene_tx=XM_019836902.2
C1 35712944 35713045
refGene_gn=UROD;refGene_tx=XM_003990027.4;refGene_gn=UROD;refGene_tx=XM_006934688.3;refGene_gn=UROD;refGene_tx=XM_019836899.2;refGene_gn=UROD;refGene_tx=XM_019836900.2;refGene_gn=UROD;refGene_tx=XM_019836901.2;refGene_gn=UROD;refGene_tx=XM_019836902.2
C1 35713126 35713193
refGene_gn=UROD;refGene_tx=XM_003990027.4;refGene_gn=UROD;refGene_tx=XM_006934688.3;refGene_gn=UROD;refGene_tx=XM_019836899.2;refGene_gn=UROD;refGene_tx=XM_019836900.2;refGene_gn=UROD;refGene_tx=XM_019836901.2;refGene_gn=UROD;refGene_tx=XM_019836902.2
C1 35713529 35713691
refGene_gn=UROD;refGene_tx=XM_003990027.4;refGene_gn=UROD;refGene_tx=XM_006934688.3;refGene_gn=UROD;refGene_tx=XM_019836899.2;refGene_gn=UROD;refGene_tx=XM_019836900.2;refGene_gn=UROD;refGene_tx=XM_019836901.2;refGene_gn=UROD;refGene_tx=XM_019836902.2
C1 35716894 35717757 refGene_gn=ZSWIM5;refGene_tx=XM_019836903.2
C1 35718473 35718555 refGene_gn=ZSWIM5;refGene_tx=XM_019836903.2
C1 35719048 35719212 refGene_gn=ZSWIM5;refGene_tx=XM_019836903.2
C1 35746588 35746746 refGene_gn=ZSWIM5;refGene_tx=XM_019836903.2
C1 35747811 35747947 refGene_gn=ZSWIM5;refGene_tx=XM_019836903.2
C1 35748150 35748411 refGene_gn=ZSWIM5;refGene_tx=XM_019836903.2
C1 35749833 35749971 refGene_gn=ZSWIM5;refGene_tx=XM_019836903.2

C1	35750702	35750849	refGene_gn=ZSWIM5;refGene_tx=XM_019836903.2
C1	35752484	35752661	refGene_gn=ZSWIM5;refGene_tx=XM_019836903.2
C1	35758064	35758244	refGene_gn=ZSWIM5;refGene_tx=XM_019836903.2
C1	35770448	35770599	refGene_gn=ZSWIM5;refGene_tx=XM_019836903.2
C1	35771879	35772028	refGene_gn=ZSWIM5;refGene_tx=XM_019836903.2
C1	35790926	35791283	refGene_gn=ZSWIM5;refGene_tx=XM_019836903.2
C1	35965421	35966016	refGene_gn=ZSWIM5;refGene_tx=XM_019836903.2
C1	36046180	36047299	refGene_gn=HPDL;refGene_tx=XM_003990028.5
C1	36048142	36048274
refGene_gn=MUTYH;refGene_tx=XM_003990029.5;refGene_gn=MUTYH;refGene_tx=XM_003990030.4;refGene_gn=MUTYH;refGene_tx=XM_011284649.2;refGene_gn=MUTYH;refGene_tx=XM_011284650.3;refGene_gn=MUTYH;refGene_tx=XM_019836904.1;refGene_gn=MUTYH;refGene_tx=XM_023258629.1
C1	36048971	36049013
refGene_gn=MUTYH;refGene_tx=XM_003990029.5;refGene_gn=MUTYH;refGene_tx=XM_003990030.4;refGene_gn=MUTYH;refGene_tx=XM_011284649.2;refGene_gn=MUTYH;refGene_tx=XM_011284650.3;refGene_gn=MUTYH;refGene_tx=XM_019836904.1;refGene_gn=MUTYH;refGene_tx=XM_023258629.1
C1	36049202	36049355
refGene_gn=MUTYH;refGene_tx=XM_003990029.5;refGene_gn=MUTYH;refGene_tx=XM_003990030.4;refGene_gn=MUTYH;refGene_tx=XM_011284649.2;refGene_gn=MUTYH;refGene_tx=XM_011284650.3;refGene_gn=MUTYH;refGene_tx=XM_019836904.1;refGene_gn=MUTYH;refGene_tx=XM_023258629.1
C1	36049439	36049576
refGene_gn=MUTYH;refGene_tx=XM_003990029.5;refGene_gn=MUTYH;refGene_tx=XM_003990030.4;refGene_gn=MUTYH;refGene_tx=XM_011284649.2;refGene_gn=MUTYH;refGene_tx=XM_011284650.3;refGene_gn=MUTYH;refGene_tx=XM_019836904.1;refGene_gn=MUTYH;refGene_tx=XM_023258629.1
C1	36049679	36049871
refGene_gn=MUTYH;refGene_tx=XM_003990029.5;refGene_gn=MUTYH;refGene_tx=XM_003990030.4;refGene_gn=MUTYH;refGene_tx=XM_011284649.2;refGene_gn=MUTYH;refGene_tx=XM_011284650.3;refGene_gn=MUTYH;refGene_tx=XM_019836904.1;refGene_gn=MUTYH;refGene_tx=XM_023258629.1
C1	36050047	36050111
refGene_gn=MUTYH;refGene_tx=XM_003990029.5;refGene_gn=MUTYH;refGene_tx=XM_003990030.4;refGene_gn=MUTYH;refGene_tx=XM_011284649.2;refGene_gn=MUTYH;refGene_tx=XM_011284650.3;refGene_gn=MUTYH;refGene_tx=XM_019836904.1;refGene_gn=MUTYH;refGene_tx=XM_023258629.1
C1	36050191	36050336
refGene_gn=MUTYH;refGene_tx=XM_003990029.5;refGene_gn=MUTYH;refGene_tx=XM_003990030.4;refGene_gn=MUTYH;refGene_tx=XM_011284649.2;refGene_gn=MUTYH;refGene_tx=XM_011284650.3;refGene_gn=MUTYH;refGene_tx=XM_019836904.1;refGene_gn=MUTYH;refGene_tx=XM_023258629.1
C1	36050415	36050513
refGene_gn=MUTYH;refGene_tx=XM_003990029.5;refGene_gn=MUTYH;refGene_tx=XM_003990030.4;refGene_gn=MUTYH;refGene_tx=XM_011284649.2;refGene_gn=MUTYH;refGene_tx=XM_011284650.3;refGene_gn=MUTYH;refGene_tx=XM_019836904.1;refGene_gn=MUTYH;refGene_tx=XM_023258629.1
C1	36050598	36050712
refGene_gn=MUTYH;refGene_tx=XM_003990029.5;refGene_gn=MUTYH;refGene_tx

=XM_003990030.4;refGene_gn=MUTYH;refGene_tx=XM_011284649.2;refGene_gn=MUTYH;refGene_tx=XM_011284650.3;refGene_gn=MUTYH;refGene_tx=XM_019836904.1;refGene_gn=MUTYH;refGene_tx=XM_023258629.1
C1 36050787 36050859
refGene_gn=MUTYH;refGene_tx=XM_003990029.5;refGene_gn=MUTYH;refGene_tx=XM_003990030.4;refGene_gn=MUTYH;refGene_tx=XM_011284649.2;refGene_gn=MUTYH;refGene_tx=XM_011284650.3;refGene_gn=MUTYH;refGene_tx=XM_019836904.1;refGene_gn=MUTYH;refGene_tx=XM_023258629.1
C1 36050934 36050976
refGene_gn=MUTYH;refGene_tx=XM_003990029.5;refGene_gn=MUTYH;refGene_tx=XM_003990030.4;refGene_gn=MUTYH;refGene_tx=XM_011284649.2;refGene_gn=MUTYH;refGene_tx=XM_011284650.3;refGene_gn=MUTYH;refGene_tx=XM_019836904.1;refGene_gn=MUTYH;refGene_tx=XM_023258629.1
C1 36051114 36051188
refGene_gn=MUTYH;refGene_tx=XM_003990029.5;refGene_gn=MUTYH;refGene_tx=XM_003990030.4;refGene_gn=MUTYH;refGene_tx=XM_011284649.2;refGene_gn=MUTYH;refGene_tx=XM_011284650.3;refGene_gn=MUTYH;refGene_tx=XM_019836904.1
C1 36051310 36051350
refGene_gn=MUTYH;refGene_tx=XM_003990029.5;refGene_gn=MUTYH;refGene_tx=XM_003990030.4;refGene_gn=MUTYH;refGene_tx=XM_011284649.2;refGene_gn=MUTYH;refGene_tx=XM_011284650.3;refGene_gn=MUTYH;refGene_tx=XM_019836904.1;refGene_gn=MUTYH;refGene_tx=XM_023258629.1
C1 36051440 36051622
refGene_gn=MUTYH;refGene_tx=XM_003990029.5;refGene_gn=MUTYH;refGene_tx=XM_003990030.4;refGene_gn=MUTYH;refGene_tx=XM_011284649.2;refGene_gn=MUTYH;refGene_tx=XM_011284650.3;refGene_gn=MUTYH;refGene_tx=XM_019836904.1
C1 36052139 36052260
refGene_gn=MUTYH;refGene_tx=XM_003990029.5;refGene_gn=MUTYH;refGene_tx=XM_003990030.4;refGene_gn=MUTYH;refGene_tx=XM_011284649.2;refGene_gn=MUTYH;refGene_tx=XM_011284650.3;refGene_gn=MUTYH;refGene_tx=XM_019836904.1
C1 36052375 36052480
refGene_gn=MUTYH;refGene_tx=XM_003990029.5;refGene_gn=MUTYH;refGene_tx=XM_011284649.2;refGene_gn=MUTYH;refGene_tx=XM_011284650.3;refGene_gn=MUTYH;refGene_tx=XM_019836904.1
C1 36052562 36052610
refGene_gn=MUTYH;refGene_tx=XM_003990029.5;refGene_gn=MUTYH;refGene_tx=XM_011284649.2
C1 36055800 36055852
refGene_gn=TOE1;refGene_tx=XM_003990033.4;refGene_gn=TOE1;refGene_tx=XM_019836905.2
C1 36056525 36056668
refGene_gn=TOE1;refGene_tx=XM_003990033.4;refGene_gn=TOE1;refGene_tx=XM_019836905.2
C1 36056759 36056800
refGene_gn=TOE1;refGene_tx=XM_003990033.4;refGene_gn=TOE1;refGene_tx=XM_019836905.2
C1 36056933 36057030

refGene_gn=TOE1;refGene_tx=XM_003990033.4;refGene_gn=TOE1;refGene_tx=XM_019836905.2
C1 36057157 36057316 refGene_gn=TOE1;refGene_tx=XM_003990033.4
C1 36057570 36057830
refGene_gn=TOE1;refGene_tx=XM_003990033.4;refGene_gn=TOE1;refGene_tx=XM_019836905.2
C1 36058027 36058187
refGene_gn=TOE1;refGene_tx=XM_003990033.4;refGene_gn=TOE1;refGene_tx=XM_019836905.2
C1 36058275 36058896
refGene_gn=TOE1;refGene_tx=XM_003990033.4;refGene_gn=TOE1;refGene_tx=XM_019836905.2
C1 36060061 36060786 refGene_gn=TESK2;refGene_tx=XM_003990031.4
C1 36061054 36061172 refGene_gn=TESK2;refGene_tx=XM_003990031.4
C1 36061881 36061968 refGene_gn=TESK2;refGene_tx=XM_003990031.4
C1 36062174 36062258 refGene_gn=TESK2;refGene_tx=XM_003990031.4
C1 36062509 36062594 refGene_gn=TESK2;refGene_tx=XM_003990031.4
C1 36062833 36062916 refGene_gn=TESK2;refGene_tx=XM_003990031.4
C1 36071596 36071743 refGene_gn=TESK2;refGene_tx=XM_003990031.4
C1 36099396 36099445 refGene_gn=TESK2;refGene_tx=XM_003990031.4
C1 36127904 36128026 refGene_gn=TESK2;refGene_tx=XM_003990031.4
C1 36167452 36167674 refGene_gn=TESK2;refGene_tx=XM_003990031.4
C1 36199154 36199262
refGene_gn=LOC101080469;refGene_tx=XM_006934691.4;refGene_gn=LOC101080469;refGene_tx=XM_006934692.4;refGene_gn=LOC101080469;refGene_tx=XM_011284646.3;refGene_gn=LOC101080469;refGene_tx=XM_011284647.3;refGene_gn=LOC101080469;refGene_tx=XM_011284648.3;refGene_gn=LOC101080469;refGene_tx=XM_019836906.2;refGene_gn=LOC101080469;refGene_tx=XM_023258630.1;refGene_gn=LOC101080469;refGene_tx=XM_023258631.1
C1 36199604 36199631
refGene_gn=LOC101080469;refGene_tx=XM_006934693.3
C1 36200350 36200504
refGene_gn=LOC101080469;refGene_tx=XM_006934691.4;refGene_gn=LOC101080469;refGene_tx=XM_006934692.4;refGene_gn=LOC101080469;refGene_tx=XM_006934693.3;refGene_gn=LOC101080469;refGene_tx=XM_011284646.3;refGene_gn=LOC101080469;refGene_tx=XM_011284647.3;refGene_gn=LOC101080469;refGene_tx=XM_011284648.3;refGene_gn=LOC101080469;refGene_tx=XM_019836906.2;refGene_gn=LOC101080469;refGene_tx=XM_019836907.2;refGene_gn=LOC101080469;refGene_tx=XM_019836908.2;refGene_gn=LOC101080469;refGene_tx=XM_023258630.1
C1 36200703 36200771
refGene_gn=LOC101080469;refGene_tx=XM_006934691.4;refGene_gn=LOC101080469;refGene_tx=XM_006934692.4;refGene_gn=LOC101080469;refGene_tx=XM_006934693.3;refGene_gn=LOC101080469;refGene_tx=XM_011284646.3;refGene_gn=LOC101080469;refGene_tx=XM_011284647.3;refGene_gn=LOC101080469;refGene_tx=XM_011284648.3;refGene_gn=LOC101080469;refGene_tx=XM_019836906.2;refGene_gn=LOC101080469;refGene_tx=XM_019836907.2;refGene_gn=LOC101080469;refGene_tx=XM_019836908.2;refGene_gn=LOC101080469;refGene_tx=XM_023258630.1;refGene_gn=LOC101080469;refGene_tx=XM_023258631.1
C1 36200884 36201071

refGene_gn=LOC101080469;refGene_tx=XM_006934691.4;refGene_gn=LOC101080469;refGene_tx=XM_006934692.4;refGene_gn=LOC101080469;refGene_tx=XM_006934693.3;refGene_gn=LOC101080469;refGene_tx=XM_011284646.3;refGene_gn=LOC101080469;refGene_tx=XM_011284647.3;refGene_gn=LOC101080469;refGene_tx=XM_011284648.3;refGene_gn=LOC101080469;refGene_tx=XM_019836906.2;refGene_gn=LOC101080469;refGene_tx=XM_019836907.2;refGene_gn=LOC101080469;refGene_tx=XM_019836908.2;refGene_gn=LOC101080469;refGene_tx=XM_023258630.1;refGene_gn=LOC101080469;refGene_tx=XM_023258631.1
C1 36201235 36201317
refGene_gn=LOC101080469;refGene_tx=XM_006934691.4;refGene_gn=LOC101080469;refGene_tx=XM_006934692.4;refGene_gn=LOC101080469;refGene_tx=XM_006934693.3;refGene_gn=LOC101080469;refGene_tx=XM_011284646.3;refGene_gn=LOC101080469;refGene_tx=XM_019836906.2;refGene_gn=LOC101080469;refGene_tx=XM_019836907.2;refGene_gn=LOC101080469;refGene_tx=XM_019836908.2;refGene_gn=LOC101080469;refGene_tx=XM_023258630.1;refGene_gn=LOC101080469;refGene_tx=XM_023258631.1
C1 36201408 36201599
refGene_gn=LOC101080469;refGene_tx=XM_006934691.4;refGene_gn=LOC101080469;refGene_tx=XM_006934692.4;refGene_gn=LOC101080469;refGene_tx=XM_006934693.3;refGene_gn=LOC101080469;refGene_tx=XM_011284646.3;refGene_gn=LOC101080469;refGene_tx=XM_019836906.2;refGene_gn=LOC101080469;refGene_tx=XM_019836907.2;refGene_gn=LOC101080469;refGene_tx=XM_019836908.2;refGene_gn=LOC101080469;refGene_tx=XM_023258630.1;refGene_gn=LOC101080469;refGene_tx=XM_023258631.1
C1 36201768 36201802
refGene_gn=LOC101080469;refGene_tx=XM_006934691.4;refGene_gn=LOC101080469;refGene_tx=XM_006934692.4;refGene_gn=LOC101080469;refGene_tx=XM_006934693.3;refGene_gn=LOC101080469;refGene_tx=XM_011284646.3;refGene_gn=LOC101080469;refGene_tx=XM_011284647.3;refGene_gn=LOC101080469;refGene_tx=XM_011284648.3;refGene_gn=LOC101080469;refGene_tx=XM_019836906.2;refGene_gn=LOC101080469;refGene_tx=XM_019836907.2;refGene_gn=LOC101080469;refGene_tx=XM_019836908.2;refGene_gn=LOC101080469;refGene_tx=XM_023258630.1;refGene_gn=LOC101080469;refGene_tx=XM_023258631.1
C1 36201890 36201968
refGene_gn=LOC101080469;refGene_tx=XM_006934691.4;refGene_gn=LOC101080469;refGene_tx=XM_006934692.4;refGene_gn=LOC101080469;refGene_tx=XM_006934693.3;refGene_gn=LOC101080469;refGene_tx=XM_011284646.3;refGene_gn=LOC101080469;refGene_tx=XM_011284647.3;refGene_gn=LOC101080469;refGene_tx=XM_011284648.3;refGene_gn=LOC101080469;refGene_tx=XM_019836906.2;refGene_gn=LOC101080469;refGene_tx=XM_019836907.2;refGene_gn=LOC101080469;refGene_tx=XM_019836908.2;refGene_gn=LOC101080469;refGene_tx=XM_023258630.1;refGene_gn=LOC101080469;refGene_tx=XM_023258631.1
C1 36202661 36202742
refGene_gn=MMACHC;refGene_tx=XM_003990034.4;refGene_gn=MMACHC;refGene_tx=XM_011284645.3
C1 36204693 36204888
refGene_gn=MMACHC;refGene_tx=XM_003990034.4;refGene_gn=MMACHC;refGene_tx=XM_011284645.3
C1 36205615 36205768 refGene_gn=MMACHC;refGene_tx=XM_003990034.4
C1 36206254 36206674

refGene_gn=MMACHC;refGene_tx=XM_003990034.4;refGene_gn=MMACHC;refGene_tx=XM_011284645.3
C1 36208111 36208197 refGene_gn=PRDX1;refGene_tx=XM_003990035.4
C1 36215499 36215630 refGene_gn=PRDX1;refGene_tx=XM_003990035.4
C1 36216383 36216506 refGene_gn=PRDX1;refGene_tx=XM_003990035.4
C1 36216888 36217042 refGene_gn=PRDX1;refGene_tx=XM_003990035.4
C1 36219892 36219998 refGene_gn=PRDX1;refGene_tx=XM_003990035.4
C1 36262676 36262760
refGene_gn=AKR1A1;refGene_tx=XM_006934694.4;refGene_gn=AKR1A1;refGene_tx=XM_023258632.1
C1 36264489 36264609
refGene_gn=AKR1A1;refGene_tx=XM_006934694.4;refGene_gn=AKR1A1;refGene_tx=XM_023258632.1
C1 36264774 36264926
refGene_gn=AKR1A1;refGene_tx=XM_006934694.4;refGene_gn=AKR1A1;refGene_tx=XM_023258632.1
C1 36265554 36265750
refGene_gn=AKR1A1;refGene_tx=XM_006934694.4;refGene_gn=AKR1A1;refGene_tx=XM_023258632.1
C1 36266908 36267108
refGene_gn=AKR1A1;refGene_tx=XM_006934694.4;refGene_gn=AKR1A1;refGene_tx=XM_023258632.1
C1 36267344 36267417
refGene_gn=AKR1A1;refGene_tx=XM_006934694.4;refGene_gn=AKR1A1;refGene_tx=XM_023258632.1
C1 36267558 36267645
refGene_gn=AKR1A1;refGene_tx=XM_006934694.4;refGene_gn=AKR1A1;refGene_tx=XM_023258632.1
C1 36268293 36268359
refGene_gn=AKR1A1;refGene_tx=XM_006934694.4;refGene_gn=AKR1A1;refGene_tx=XM_023258632.1
C1 36279728 36279787
refGene_gn=NASP;refGene_tx=XM_006934695.4;refGene_gn=NASP;refGene_tx=XM_006934697.4;refGene_gn=NASP;refGene_tx=XM_006934699.4;refGene_gn=NASP;refGene_tx=XM_019836911.2
C1 36304985 36305036
refGene_gn=NASP;refGene_tx=XM_006934695.4;refGene_gn=NASP;refGene_tx=XM_006934697.4;refGene_gn=NASP;refGene_tx=XM_006934699.4;refGene_gn=NASP;refGene_tx=XM_019836911.2
C1 36306594 36306705
refGene_gn=NASP;refGene_tx=XM_006934695.4;refGene_gn=NASP;refGene_tx=XM_006934697.4;refGene_gn=NASP;refGene_tx=XM_006934699.4;refGene_gn=NASP;refGene_tx=XM_019836911.2
C1 36309276 36309357
refGene_gn=NASP;refGene_tx=XM_006934695.4;refGene_gn=NASP;refGene_tx=XM_006934697.4;refGene_gn=NASP;refGene_tx=XM_006934699.4;refGene_gn=NASP;refGene_tx=XM_019836911.2
C1 36310298 36310408
refGene_gn=NASP;refGene_tx=XM_006934695.4;refGene_gn=NASP;refGene_tx=XM_006934697.4;refGene_gn=NASP;refGene_tx=XM_006934699.4;refGene_gn=NAS

P;refGene_tx=XM_019836911.2
C1 36311110 36312130
refGene_gn=NASP;refGene_tx=XM_006934695.4;refGene_gn=NASP;refGene_tx=XM_006934697.4;refGene_gn=NASP;refGene_tx=XM_019836911.2
C1 36317309 36317389
refGene_gn=NASP;refGene_tx=XM_006934695.4;refGene_gn=NASP;refGene_tx=XM_006934697.4;refGene_gn=NASP;refGene_tx=XM_006934699.4;refGene_gn=NASP;refGene_tx=XM_019836911.2
C1 36317561 36317647
refGene_gn=NASP;refGene_tx=XM_006934695.4;refGene_gn=NASP;refGene_tx=XM_006934697.4;refGene_gn=NASP;refGene_tx=XM_006934699.4;refGene_gn=NASP;refGene_tx=XM_019836911.2
C1 36317766 36317840
refGene_gn=NASP;refGene_tx=XM_006934695.4;refGene_gn=NASP;refGene_tx=XM_006934697.4;refGene_gn=NASP;refGene_tx=XM_006934699.4;refGene_gn=NASP;refGene_tx=XM_019836911.2
C1 36318436 36318625
refGene_gn=NASP;refGene_tx=XM_006934695.4;refGene_gn=NASP;refGene_tx=XM_006934697.4;refGene_gn=NASP;refGene_tx=XM_006934699.4;refGene_gn=NASP;refGene_tx=XM_019836911.2
C1 36318744 36318911
refGene_gn=NASP;refGene_tx=XM_006934695.4;refGene_gn=NASP;refGene_tx=XM_006934697.4;refGene_gn=NASP;refGene_tx=XM_006934699.4;refGene_gn=NASP;refGene_tx=XM_019836911.2
C1 36319630 36319687
refGene_gn=NASP;refGene_tx=XM_006934695.4;refGene_gn=NASP;refGene_tx=XM_006934697.4;refGene_gn=NASP;refGene_tx=XM_006934699.4;refGene_gn=NASP;refGene_tx=XM_019836911.2
C1 36319774 36319852
refGene_gn=NASP;refGene_tx=XM_006934695.4;refGene_gn=NASP;refGene_tx=XM_006934697.4;refGene_gn=NASP;refGene_tx=XM_006934699.4;refGene_gn=NASP;refGene_tx=XM_019836911.2
C1 36320531 36320660
refGene_gn=NASP;refGene_tx=XM_006934695.4;refGene_gn=NASP;refGene_tx=XM_006934697.4;refGene_gn=NASP;refGene_tx=XM_006934699.4;refGene_gn=NASP;refGene_tx=XM_019836911.2
C1 36321156 36321237
refGene_gn=NASP;refGene_tx=XM_006934695.4;refGene_gn=NASP;refGene_tx=XM_006934697.4;refGene_gn=NASP;refGene_tx=XM_006934699.4;refGene_gn=NASP;refGene_tx=XM_019836911.2
C1 36323094 36323229 refGene_gn=CCDC17;refGene_tx=XM_023258633.1
C1 36323478 36323589 refGene_gn=CCDC17;refGene_tx=XM_023258633.1
C1 36323658 36323869 refGene_gn=CCDC17;refGene_tx=XM_023258633.1
C1 36324011 36324215 refGene_gn=CCDC17;refGene_tx=XM_023258633.1
C1 36324370 36324468 refGene_gn=CCDC17;refGene_tx=XM_023258633.1
C1 36324580 36324726 refGene_gn=CCDC17;refGene_tx=XM_023258633.1
C1 36324932 36325070 refGene_gn=CCDC17;refGene_tx=XM_023258633.1
C1 36325353 36325472 refGene_gn=CCDC17;refGene_tx=XM_023258633.1
C1 36325563 36325647 refGene_gn=CCDC17;refGene_tx=XM_023258633.1
C1 36325868 36326031 refGene_gn=CCDC17;refGene_tx=XM_023258633.1

C1	36326131	36326354	refGene_gn=CCDC17;refGene_tx=XM_023258633.1
C1	36326459	36326540	refGene_gn=CCDC17;refGene_tx=XM_023258633.1
C1	36326542	36326758	refGene_gn=CCDC17;refGene_tx=XM_023258633.1
C1	36329561	36329714
refGene_gn=GPBP1L1;refGene_tx=XM_011284642.3;refGene_gn=GPBP1L1;refGene_tx=XM_011284643.3;refGene_gn=GPBP1L1;refGene_tx=XM_011284644.3;refGene_gn=GPBP1L1;refGene_tx=XM_019836913.2
C1	36330633	36330736
refGene_gn=GPBP1L1;refGene_tx=XM_011284642.3;refGene_gn=GPBP1L1;refGene_tx=XM_011284643.3;refGene_gn=GPBP1L1;refGene_tx=XM_011284644.3;refGene_gn=GPBP1L1;refGene_tx=XM_019836913.2
C1	36330910	36330977	refGene_gn=GPBP1L1;refGene_tx=XM_019836914.1
C1	36331684	36331809
refGene_gn=GPBP1L1;refGene_tx=XM_011284642.3;refGene_gn=GPBP1L1;refGene_tx=XM_011284643.3;refGene_gn=GPBP1L1;refGene_tx=XM_011284644.3;refGene_gn=GPBP1L1;refGene_tx=XM_019836913.2;refGene_gn=GPBP1L1;refGene_tx=XM_019836914.1
C1	36333331	36333490
refGene_gn=GPBP1L1;refGene_tx=XM_011284642.3;refGene_gn=GPBP1L1;refGene_tx=XM_011284643.3;refGene_gn=GPBP1L1;refGene_tx=XM_011284644.3;refGene_gn=GPBP1L1;refGene_tx=XM_019836913.2;refGene_gn=GPBP1L1;refGene_tx=XM_019836914.1
C1	36333945	36334086
refGene_gn=GPBP1L1;refGene_tx=XM_011284642.3;refGene_gn=GPBP1L1;refGene_tx=XM_011284643.3;refGene_gn=GPBP1L1;refGene_tx=XM_011284644.3;refGene_gn=GPBP1L1;refGene_tx=XM_019836913.2;refGene_gn=GPBP1L1;refGene_tx=XM_019836914.1
C1	36342135	36342329
refGene_gn=GPBP1L1;refGene_tx=XM_011284642.3;refGene_gn=GPBP1L1;refGene_tx=XM_011284643.3;refGene_gn=GPBP1L1;refGene_tx=XM_011284644.3;refGene_gn=GPBP1L1;refGene_tx=XM_019836913.2;refGene_gn=GPBP1L1;refGene_tx=XM_019836914.1
C1	36343581	36343654
refGene_gn=GPBP1L1;refGene_tx=XM_011284642.3;refGene_gn=GPBP1L1;refGene_tx=XM_011284643.3;refGene_gn=GPBP1L1;refGene_tx=XM_011284644.3;refGene_gn=GPBP1L1;refGene_tx=XM_019836913.2;refGene_gn=GPBP1L1;refGene_tx=XM_019836914.1
C1	36361111	36361398
refGene_gn=GPBP1L1;refGene_tx=XM_011284642.3;refGene_gn=GPBP1L1;refGene_tx=XM_011284643.3;refGene_gn=GPBP1L1;refGene_tx=XM_011284644.3;refGene_gn=GPBP1L1;refGene_tx=XM_019836913.2;refGene_gn=GPBP1L1;refGene_tx=XM_019836914.1
C1	36361973	36362103
refGene_gn=GPBP1L1;refGene_tx=XM_011284642.3;refGene_gn=GPBP1L1;refGene_tx=XM_011284643.3;refGene_gn=GPBP1L1;refGene_tx=XM_011284644.3;refGene_gn=GPBP1L1;refGene_tx=XM_019836913.2;refGene_gn=GPBP1L1;refGene_tx=XM_019836914.1
C1	36364689	36364749
refGene_gn=GPBP1L1;refGene_tx=XM_011284642.3;refGene_gn=GPBP1L1;refGene_tx=XM_011284643.3;refGene_gn=GPBP1L1;refGene_tx=XM_011284644.3;refGe

ne_gn=GPBP1L1;refGene_tx=XM_019836913.2;refGene_gn=GPBP1L1;refGene_tx=XM_019836914.1
C1 36396722 36397100 refGene_gn=TMEM69;refGene_tx=XM_019836915.1
C1 36400387 36400524
refGene_gn=TMEM69;refGene_tx=XM_006934706.4;refGene_gn=TMEM69;refGene_tx=XM_011284641.3;refGene_gn=TMEM69;refGene_tx=XM_019836915.1
C1 36401408 36401502 refGene_gn=TMEM69;refGene_tx=XM_019836915.1
C1 36401667 36402378
refGene_gn=TMEM69;refGene_tx=XM_006934706.4;refGene_gn=TMEM69;refGene_tx=XM_011284641.3
C1 36406771 36406912
refGene_gn=IPP;refGene_tx=XM_003990042.5;refGene_gn=IPP;refGene_tx=XM_006934701.3;refGene_gn=IPP;refGene_tx=XM_006934702.3;refGene_gn=IPP;refGene_tx=XM_023258634.1
C1 36420034 36420255
refGene_gn=IPP;refGene_tx=XM_003990042.5;refGene_gn=IPP;refGene_tx=XM_006934701.3;refGene_gn=IPP;refGene_tx=XM_006934702.3;refGene_gn=IPP;refGene_tx=XM_019836912.1;refGene_gn=IPP;refGene_tx=XM_023258634.1
C1 36422738 36422861
refGene_gn=IPP;refGene_tx=XM_003990042.5;refGene_gn=IPP;refGene_tx=XM_006934701.3;refGene_gn=IPP;refGene_tx=XM_006934702.3;refGene_gn=IPP;refGene_tx=XM_019836912.1;refGene_gn=IPP;refGene_tx=XM_023258634.1
C1 36423408 36423546
refGene_gn=IPP;refGene_tx=XM_003990042.5;refGene_gn=IPP;refGene_tx=XM_006934701.3;refGene_gn=IPP;refGene_tx=XM_006934702.3;refGene_gn=IPP;refGene_tx=XM_019836912.1;refGene_gn=IPP;refGene_tx=XM_023258634.1
C1 36429459 36429627
refGene_gn=IPP;refGene_tx=XM_003990042.5;refGene_gn=IPP;refGene_tx=XM_006934701.3;refGene_gn=IPP;refGene_tx=XM_019836912.1;refGene_gn=IPP;refGene_tx=XM_023258634.1
C1 36431044 36431200
refGene_gn=IPP;refGene_tx=XM_003990042.5;refGene_gn=IPP;refGene_tx=XM_006934701.3;refGene_gn=IPP;refGene_tx=XM_006934702.3;refGene_gn=IPP;refGene_tx=XM_019836912.1;refGene_gn=IPP;refGene_tx=XM_023258634.1
C1 36438983 36439415
refGene_gn=IPP;refGene_tx=XM_003990042.5;refGene_gn=IPP;refGene_tx=XM_006934701.3;refGene_gn=IPP;refGene_tx=XM_006934702.3;refGene_gn=IPP;refGene_tx=XM_019836912.1;refGene_gn=IPP;refGene_tx=XM_023258634.1
C1 36443998 36444290
refGene_gn=IPP;refGene_tx=XM_003990042.5;refGene_gn=IPP;refGene_tx=XM_006934701.3;refGene_gn=IPP;refGene_tx=XM_006934702.3;refGene_gn=IPP;refGene_tx=XM_019836912.1;refGene_gn=IPP;refGene_tx=XM_023258634.1
C1 36461288 36461436
refGene_gn=LOC101093034;refGene_tx=XM_003990043.3
C1 36461562 36461732
refGene_gn=LOC101093034;refGene_tx=XM_003990043.3
C1 36526201 36526387
refGene_gn=MAST2;refGene_tx=XM_023258635.1;refGene_gn=MAST2;refGene_tx=XM_023258636.1;refGene_gn=MAST2;refGene_tx=XM_023258637.1;refGene_gn=MAST2;refGene_tx=XM_023258638.1;refGene_gn=MAST2;refGene_tx=XM_0232586

39.1
C1 36559422 36559570
refGene_gn=MAST2;refGene_tx=XM_023258635.1;refGene_gn=MAST2;refGene_tx=XM_023258636.1;refGene_gn=MAST2;refGene_tx=XM_023258637.1;refGene_gn=MAST2;refGene_tx=XM_023258638.1;refGene_gn=MAST2;refGene_tx=XM_023258639.1
C1 36565661 36565804
refGene_gn=MAST2;refGene_tx=XM_023258635.1;refGene_gn=MAST2;refGene_tx=XM_023258636.1;refGene_gn=MAST2;refGene_tx=XM_023258637.1;refGene_gn=MAST2;refGene_tx=XM_023258638.1;refGene_gn=MAST2;refGene_tx=XM_023258639.1
C1 36623880 36623912
refGene_gn=MAST2;refGene_tx=XM_023258635.1;refGene_gn=MAST2;refGene_tx=XM_023258636.1;refGene_gn=MAST2;refGene_tx=XM_023258637.1;refGene_gn=MAST2;refGene_tx=XM_023258638.1;refGene_gn=MAST2;refGene_tx=XM_023258639.1
C1 36657508 36657558 refGene_gn=MAST2;refGene_tx=XM_023258640.1
C1 36660282 36660477
refGene_gn=MAST2;refGene_tx=XM_023258635.1;refGene_gn=MAST2;refGene_tx=XM_023258636.1;refGene_gn=MAST2;refGene_tx=XM_023258637.1;refGene_gn=MAST2;refGene_tx=XM_023258640.1
C1 36697128 36697220
refGene_gn=MAST2;refGene_tx=XM_023258635.1;refGene_gn=MAST2;refGene_tx=XM_023258636.1;refGene_gn=MAST2;refGene_tx=XM_023258637.1;refGene_gn=MAST2;refGene_tx=XM_023258638.1;refGene_gn=MAST2;refGene_tx=XM_023258639.1;refGene_gn=MAST2;refGene_tx=XM_023258640.1
C1 36719394 36719470
refGene_gn=MAST2;refGene_tx=XM_023258635.1;refGene_gn=MAST2;refGene_tx=XM_023258636.1;refGene_gn=MAST2;refGene_tx=XM_023258637.1;refGene_gn=MAST2;refGene_tx=XM_023258638.1;refGene_gn=MAST2;refGene_tx=XM_023258639.1;refGene_gn=MAST2;refGene_tx=XM_023258640.1
C1 36722037 36722058
refGene_gn=MAST2;refGene_tx=XM_023258635.1;refGene_gn=MAST2;refGene_tx=XM_023258637.1;refGene_gn=MAST2;refGene_tx=XM_023258638.1;refGene_gn=MAST2;refGene_tx=XM_023258640.1
C1 36723855 36723934
refGene_gn=MAST2;refGene_tx=XM_023258635.1;refGene_gn=MAST2;refGene_tx=XM_023258636.1;refGene_gn=MAST2;refGene_tx=XM_023258637.1;refGene_gn=MAST2;refGene_tx=XM_023258638.1;refGene_gn=MAST2;refGene_tx=XM_023258639.1;refGene_gn=MAST2;refGene_tx=XM_023258640.1
C1 36725688 36725843
refGene_gn=MAST2;refGene_tx=XM_023258635.1;refGene_gn=MAST2;refGene_tx=XM_023258636.1;refGene_gn=MAST2;refGene_tx=XM_023258637.1;refGene_gn=MAST2;refGene_tx=XM_023258638.1;refGene_gn=MAST2;refGene_tx=XM_023258639.1;refGene_gn=MAST2;refGene_tx=XM_023258640.1
C1 36728195 36728271
refGene_gn=MAST2;refGene_tx=XM_023258635.1;refGene_gn=MAST2;refGene_tx=XM_023258636.1;refGene_gn=MAST2;refGene_tx=XM_023258637.1;refGene_gn=MAST2;refGene_tx=XM_023258638.1;refGene_gn=MAST2;refGene_tx=XM_023258639.1;refGene_gn=MAST2;refGene_tx=XM_023258640.1

C1 36731620 36731830
refGene_gn=MAST2;refGene_tx=XM_023258635.1;refGene_gn=MAST2;refGene_tx=XM_023258636.1;refGene_gn=MAST2;refGene_tx=XM_023258637.1;refGene_gn=MAST2;refGene_tx=XM_023258638.1;refGene_gn=MAST2;refGene_tx=XM_023258639.1;refGene_gn=MAST2;refGene_tx=XM_023258640.1
C1 36737230 36737332
refGene_gn=MAST2;refGene_tx=XM_023258635.1;refGene_gn=MAST2;refGene_tx=XM_023258636.1;refGene_gn=MAST2;refGene_tx=XM_023258637.1;refGene_gn=MAST2;refGene_tx=XM_023258638.1;refGene_gn=MAST2;refGene_tx=XM_023258639.1;refGene_gn=MAST2;refGene_tx=XM_023258640.1
C1 36739511 36739644
refGene_gn=MAST2;refGene_tx=XM_023258635.1;refGene_gn=MAST2;refGene_tx=XM_023258636.1;refGene_gn=MAST2;refGene_tx=XM_023258637.1;refGene_gn=MAST2;refGene_tx=XM_023258638.1;refGene_gn=MAST2;refGene_tx=XM_023258639.1;refGene_gn=MAST2;refGene_tx=XM_023258640.1
C1 36740412 36740474
refGene_gn=MAST2;refGene_tx=XM_023258635.1;refGene_gn=MAST2;refGene_tx=XM_023258636.1;refGene_gn=MAST2;refGene_tx=XM_023258637.1;refGene_gn=MAST2;refGene_tx=XM_023258638.1;refGene_gn=MAST2;refGene_tx=XM_023258639.1;refGene_gn=MAST2;refGene_tx=XM_023258640.1
C1 36740744 36740827
refGene_gn=MAST2;refGene_tx=XM_023258635.1;refGene_gn=MAST2;refGene_tx=XM_023258636.1;refGene_gn=MAST2;refGene_tx=XM_023258637.1;refGene_gn=MAST2;refGene_tx=XM_023258638.1;refGene_gn=MAST2;refGene_tx=XM_023258639.1;refGene_gn=MAST2;refGene_tx=XM_023258640.1
C1 36741297 36741506
refGene_gn=MAST2;refGene_tx=XM_023258635.1;refGene_gn=MAST2;refGene_tx=XM_023258636.1;refGene_gn=MAST2;refGene_tx=XM_023258637.1;refGene_gn=MAST2;refGene_tx=XM_023258638.1;refGene_gn=MAST2;refGene_tx=XM_023258639.1;refGene_gn=MAST2;refGene_tx=XM_023258640.1
C1 36742554 36742693
refGene_gn=MAST2;refGene_tx=XM_023258635.1;refGene_gn=MAST2;refGene_tx=XM_023258636.1;refGene_gn=MAST2;refGene_tx=XM_023258637.1;refGene_gn=MAST2;refGene_tx=XM_023258638.1;refGene_gn=MAST2;refGene_tx=XM_023258639.1;refGene_gn=MAST2;refGene_tx=XM_023258640.1
C1 36745193 36745326
refGene_gn=MAST2;refGene_tx=XM_023258635.1;refGene_gn=MAST2;refGene_tx=XM_023258636.1;refGene_gn=MAST2;refGene_tx=XM_023258637.1;refGene_gn=MAST2;refGene_tx=XM_023258638.1;refGene_gn=MAST2;refGene_tx=XM_023258639.1;refGene_gn=MAST2;refGene_tx=XM_023258640.1
C1 36746100 36746266
refGene_gn=MAST2;refGene_tx=XM_023258635.1;refGene_gn=MAST2;refGene_tx=XM_023258636.1;refGene_gn=MAST2;refGene_tx=XM_023258637.1;refGene_gn=MAST2;refGene_tx=XM_023258638.1;refGene_gn=MAST2;refGene_tx=XM_023258639.1;refGene_gn=MAST2;refGene_tx=XM_023258640.1
C1 36746796 36746898
refGene_gn=MAST2;refGene_tx=XM_023258635.1;refGene_gn=MAST2;refGene_tx=XM_023258636.1;refGene_gn=MAST2;refGene_tx=XM_023258637.1;refGene_gn=MAST2;refGene_tx=XM_023258638.1;refGene_gn=MAST2;refGene_tx=XM_023258639.1;refGene_gn=MAST2;refGene_tx=XM_023258640.1

C1 36747160 36747283
refGene_gn=MAST2;refGene_tx=XM_023258635.1;refGene_gn=MAST2;refGene_tx=XM_023258636.1;refGene_gn=MAST2;refGene_tx=XM_023258637.1;refGene_gn=MAST2;refGene_tx=XM_023258638.1;refGene_gn=MAST2;refGene_tx=XM_023258639.1;refGene_gn=MAST2;refGene_tx=XM_023258640.1
C1 36747460 36747570
refGene_gn=MAST2;refGene_tx=XM_023258635.1;refGene_gn=MAST2;refGene_tx=XM_023258636.1;refGene_gn=MAST2;refGene_tx=XM_023258637.1;refGene_gn=MAST2;refGene_tx=XM_023258638.1;refGene_gn=MAST2;refGene_tx=XM_023258639.1;refGene_gn=MAST2;refGene_tx=XM_023258640.1
C1 36747923 36748078
refGene_gn=MAST2;refGene_tx=XM_023258635.1;refGene_gn=MAST2;refGene_tx=XM_023258636.1;refGene_gn=MAST2;refGene_tx=XM_023258637.1;refGene_gn=MAST2;refGene_tx=XM_023258638.1;refGene_gn=MAST2;refGene_tx=XM_023258639.1;refGene_gn=MAST2;refGene_tx=XM_023258640.1
C1 36748325 36748609
refGene_gn=MAST2;refGene_tx=XM_023258635.1;refGene_gn=MAST2;refGene_tx=XM_023258636.1;refGene_gn=MAST2;refGene_tx=XM_023258637.1;refGene_gn=MAST2;refGene_tx=XM_023258638.1;refGene_gn=MAST2;refGene_tx=XM_023258639.1;refGene_gn=MAST2;refGene_tx=XM_023258640.1
C1 36748717 36748915
refGene_gn=MAST2;refGene_tx=XM_023258635.1;refGene_gn=MAST2;refGene_tx=XM_023258636.1;refGene_gn=MAST2;refGene_tx=XM_023258637.1;refGene_gn=MAST2;refGene_tx=XM_023258638.1;refGene_gn=MAST2;refGene_tx=XM_023258639.1;refGene_gn=MAST2;refGene_tx=XM_023258640.1
C1 36749537 36749761
refGene_gn=MAST2;refGene_tx=XM_023258635.1;refGene_gn=MAST2;refGene_tx=XM_023258636.1;refGene_gn=MAST2;refGene_tx=XM_023258637.1;refGene_gn=MAST2;refGene_tx=XM_023258638.1;refGene_gn=MAST2;refGene_tx=XM_023258639.1;refGene_gn=MAST2;refGene_tx=XM_023258640.1
C1 36749959 36750082
refGene_gn=MAST2;refGene_tx=XM_023258635.1;refGene_gn=MAST2;refGene_tx=XM_023258636.1;refGene_gn=MAST2;refGene_tx=XM_023258637.1;refGene_gn=MAST2;refGene_tx=XM_023258638.1;refGene_gn=MAST2;refGene_tx=XM_023258639.1;refGene_gn=MAST2;refGene_tx=XM_023258640.1
C1 36750508 36750645
refGene_gn=MAST2;refGene_tx=XM_023258635.1;refGene_gn=MAST2;refGene_tx=XM_023258636.1;refGene_gn=MAST2;refGene_tx=XM_023258637.1;refGene_gn=MAST2;refGene_tx=XM_023258638.1;refGene_gn=MAST2;refGene_tx=XM_023258639.1;refGene_gn=MAST2;refGene_tx=XM_023258640.1
C1 36750762 36750956
refGene_gn=MAST2;refGene_tx=XM_023258635.1;refGene_gn=MAST2;refGene_tx=XM_023258636.1;refGene_gn=MAST2;refGene_tx=XM_023258637.1;refGene_gn=MAST2;refGene_tx=XM_023258638.1;refGene_gn=MAST2;refGene_tx=XM_023258639.1;refGene_gn=MAST2;refGene_tx=XM_023258640.1
C1 36751265 36752791
refGene_gn=MAST2;refGene_tx=XM_023258635.1;refGene_gn=MAST2;refGene_tx=XM_023258636.1;refGene_gn=MAST2;refGene_tx=XM_023258637.1;refGene_gn=MAST2;refGene_tx=XM_023258638.1;refGene_gn=MAST2;refGene_tx=XM_023258639.1;refGene_gn=MAST2;refGene_tx=XM_023258640.1

C1 36760554 36760753
refGene_gn=PIK3R3;refGene_tx=XM_011284796.3;refGene_gn=PIK3R3;refGene_tx=XM_019836920.2
C1 36769243 36769414
refGene_gn=PIK3R3;refGene_tx=XM_011284796.3;refGene_gn=PIK3R3;refGene_tx=XM_019836920.2
C1 36769834 36769909
refGene_gn=PIK3R3;refGene_tx=XM_011284796.3;refGene_gn=PIK3R3;refGene_tx=XM_019836920.2
C1 36772851 36773028
refGene_gn=PIK3R3;refGene_tx=XM_011284796.3;refGene_gn=PIK3R3;refGene_tx=XM_019836920.2
C1 36775471 36775614
refGene_gn=PIK3R3;refGene_tx=XM_011284796.3;refGene_gn=PIK3R3;refGene_tx=XM_019836920.2
C1 36790162 36790288
refGene_gn=PIK3R3;refGene_tx=XM_011284796.3;refGene_gn=PIK3R3;refGene_tx=XM_019836920.2
C1 36790739 36790920
refGene_gn=PIK3R3;refGene_tx=XM_011284796.3;refGene_gn=PIK3R3;refGene_tx=XM_019836920.2
C1 36805063 36805162 refGene_gn=PIK3R3;refGene_tx=XM_011284796.3
C1 36807319 36807428 refGene_gn=PIK3R3;refGene_tx=XM_011284796.3
C1 36905630 36905736 refGene_gn=PIK3R3;refGene_tx=XM_011284796.3
C1 36905854 36905996
refGene_gn=LOC109503010;refGene_tx=XM_019839075.2
C1 36906082 36906185
refGene_gn=LOC109503010;refGene_tx=XM_019839075.2
C1 36906711 36906965
refGene_gn=LOC109503010;refGene_tx=XM_019839075.2
C1 36910153 36910249
refGene_gn=LOC109503010;refGene_tx=XM_019839075.2
C1 36910883 36910983
refGene_gn=LOC109503010;refGene_tx=XM_019839075.2
C1 36913539 36913654
refGene_gn=LOC109503010;refGene_tx=XM_019839075.2
C1 36949017 36949074 refGene_gn=TSPAN1;refGene_tx=XM_019836921.2
C1 36951529 36951736 refGene_gn=TSPAN1;refGene_tx=XM_019836921.2
C1 36951944 36952019 refGene_gn=TSPAN1;refGene_tx=XM_019836921.2
C1 36952134 36952233 refGene_gn=TSPAN1;refGene_tx=XM_019836921.2
C1 36952312 36952468 refGene_gn=TSPAN1;refGene_tx=XM_019836921.2
C1 36952568 36952652 refGene_gn=TSPAN1;refGene_tx=XM_019836921.2
C1 36952838 36952886 refGene_gn=TSPAN1;refGene_tx=XM_019836921.2
C1 36956846 36956934
refGene_gn=POMGNT1;refGene_tx=XM_019836923.2;refGene_gn=POMGNT1;refGene_tx=XM_023258641.1;refGene_gn=POMGNT1;refGene_tx=XM_023258642.1
C1 36957033 36957143
refGene_gn=POMGNT1;refGene_tx=XM_019836923.2;refGene_gn=POMGNT1;refGene_tx=XM_023258641.1;refGene_gn=POMGNT1;refGene_tx=XM_023258642.1
C1 36957407 36957543

refGene_gn=POMGNT1;refGene_tx=XM_019836923.2;refGene_gn=POMGNT1;refGene_tx=XM_023258641.1;refGene_gn=POMGNT1;refGene_tx=XM_023258642.1
C1 36957774 36957819
refGene_gn=POMGNT1;refGene_tx=XM_019836923.2;refGene_gn=POMGNT1;refGene_tx=XM_023258641.1;refGene_gn=POMGNT1;refGene_tx=XM_023258642.1
C1 36958013 36958078
refGene_gn=POMGNT1;refGene_tx=XM_019836923.2;refGene_gn=POMGNT1;refGene_tx=XM_023258641.1;refGene_gn=POMGNT1;refGene_tx=XM_023258642.1
C1 36959300 36959426
refGene_gn=POMGNT1;refGene_tx=XM_019836923.2;refGene_gn=POMGNT1;refGene_tx=XM_023258641.1;refGene_gn=POMGNT1;refGene_tx=XM_023258642.1
C1 36959510 36959639
refGene_gn=POMGNT1;refGene_tx=XM_019836923.2;refGene_gn=POMGNT1;refGene_tx=XM_023258641.1;refGene_gn=POMGNT1;refGene_tx=XM_023258642.1
C1 36959727 36959800
refGene_gn=POMGNT1;refGene_tx=XM_019836923.2;refGene_gn=POMGNT1;refGene_tx=XM_023258641.1;refGene_gn=POMGNT1;refGene_tx=XM_023258642.1
C1 36960139 36960198
refGene_gn=POMGNT1;refGene_tx=XM_019836923.2;refGene_gn=POMGNT1;refGene_tx=XM_023258641.1;refGene_gn=POMGNT1;refGene_tx=XM_023258642.1
C1 36960410 36960452
refGene_gn=POMGNT1;refGene_tx=XM_019836923.2;refGene_gn=POMGNT1;refGene_tx=XM_023258641.1;refGene_gn=POMGNT1;refGene_tx=XM_023258642.1
C1 36960541 36960625
refGene_gn=POMGNT1;refGene_tx=XM_019836923.2;refGene_gn=POMGNT1;refGene_tx=XM_023258641.1;refGene_gn=POMGNT1;refGene_tx=XM_023258642.1
C1 36960805 36960881
refGene_gn=POMGNT1;refGene_tx=XM_019836923.2;refGene_gn=POMGNT1;refGene_tx=XM_023258641.1;refGene_gn=POMGNT1;refGene_tx=XM_023258642.1
C1 36961107 36961178
refGene_gn=POMGNT1;refGene_tx=XM_019836923.2;refGene_gn=POMGNT1;refGene_tx=XM_023258641.1;refGene_gn=POMGNT1;refGene_tx=XM_023258642.1
C1 36961551 36961679
refGene_gn=POMGNT1;refGene_tx=XM_019836923.2;refGene_gn=POMGNT1;refGene_tx=XM_023258641.1;refGene_gn=POMGNT1;refGene_tx=XM_023258642.1
C1 36961846 36961945
refGene_gn=POMGNT1;refGene_tx=XM_019836923.2;refGene_gn=POMGNT1;refGene_tx=XM_023258641.1;refGene_gn=POMGNT1;refGene_tx=XM_023258642.1
C1 36962138 36962256
refGene_gn=POMGNT1;refGene_tx=XM_019836923.2;refGene_gn=POMGNT1;refGene_tx=XM_023258641.1;refGene_gn=POMGNT1;refGene_tx=XM_023258642.1
C1 36963080 36963194
refGene_gn=POMGNT1;refGene_tx=XM_019836923.2;refGene_gn=POMGNT1;refGene_tx=XM_023258641.1;refGene_gn=POMGNT1;refGene_tx=XM_023258642.1
C1 36963283 36963349
refGene_gn=POMGNT1;refGene_tx=XM_023258641.1;refGene_gn=POMGNT1;refGene_tx=XM_023258642.1
C1 36963997 36964116
refGene_gn=POMGNT1;refGene_tx=XM_023258641.1;refGene_gn=POMGNT1;refGene_tx=XM_023258642.1

C1 36964267 36964382
refGene_gn=POMGNT1;refGene_tx=XM_023258641.1;refGene_gn=POMGNT1;refGen
e_tx=XM_023258642.1
C1 36965006 36965126
refGene_gn=POMGNT1;refGene_tx=XM_023258641.1;refGene_gn=POMGNT1;refGen
e_tx=XM_023258642.1
C1 36969249 36969447 refGene_gn=LURAP1;refGene_tx=XM_003990047.5
C1 36978916 36979438 refGene_gn=LURAP1;refGene_tx=XM_003990047.5
C1 36999466 36999469
refGene_gn=RAD54L;refGene_tx=XM_019836925.2;refGene_gn=RAD54L;refGene_
tx=XM_023258643.1
C1 36999574 36999661
refGene_gn=RAD54L;refGene_tx=XM_019836925.2;refGene_gn=RAD54L;refGene_
tx=XM_023258643.1
C1 37000917 37001037
refGene_gn=RAD54L;refGene_tx=XM_019836925.2;refGene_gn=RAD54L;refGene_
tx=XM_023258643.1
C1 37006996 37007057
refGene_gn=RAD54L;refGene_tx=XM_019836925.2;refGene_gn=RAD54L;refGene_
tx=XM_023258643.1
C1 37007778 37007914
refGene_gn=RAD54L;refGene_tx=XM_019836925.2;refGene_gn=RAD54L;refGene_
tx=XM_023258643.1
C1 37008365 37008435
refGene_gn=RAD54L;refGene_tx=XM_019836925.2;refGene_gn=RAD54L;refGene_
tx=XM_023258643.1
C1 37008534 37008823
refGene_gn=RAD54L;refGene_tx=XM_019836925.2;refGene_gn=RAD54L;refGene_
tx=XM_023258643.1
C1 37009028 37009153
refGene_gn=RAD54L;refGene_tx=XM_019836925.2;refGene_gn=RAD54L;refGene_
tx=XM_023258643.1
C1 37014531 37014682
refGene_gn=RAD54L;refGene_tx=XM_019836925.2;refGene_gn=RAD54L;refGene_
tx=XM_023258643.1
C1 37017826 37017953
refGene_gn=RAD54L;refGene_tx=XM_019836925.2;refGene_gn=RAD54L;refGene_
tx=XM_023258643.1
C1 37019163 37019238
refGene_gn=RAD54L;refGene_tx=XM_019836925.2;refGene_gn=RAD54L;refGene_
tx=XM_023258643.1
C1 37019369 37019500
refGene_gn=RAD54L;refGene_tx=XM_019836925.2;refGene_gn=RAD54L;refGene_
tx=XM_023258643.1
C1 37020033 37020144
refGene_gn=RAD54L;refGene_tx=XM_019836925.2;refGene_gn=RAD54L;refGene_
tx=XM_023258643.1
C1 37020295 37020419
refGene_gn=RAD54L;refGene_tx=XM_019836925.2;refGene_gn=RAD54L;refGene_
tx=XM_023258643.1

C1	37020777	37020856
refGene_gn=RAD54L;refGene_tx=XM_019836925.2;refGene_gn=RAD54L;refGene_tx=XM_023258643.1
C1	37021181	37021361
refGene_gn=RAD54L;refGene_tx=XM_019836925.2;refGene_gn=RAD54L;refGene_tx=XM_023258643.1
C1	37023734	37023770	refGene_gn=RAD54L;refGene_tx=XM_023258643.1
C1	37024095	37024259	refGene_gn=RAD54L;refGene_tx=XM_019836925.2
C1	37024355	37024566	refGene_gn=RAD54L;refGene_tx=XM_019836925.2
C1	37025162	37025382
refGene_gn=LRRC41;refGene_tx=XM_011284798.3;refGene_gn=LRRC41;refGene_tx=XM_011284800.3
C1	37025476	37025552
refGene_gn=LRRC41;refGene_tx=XM_011284798.3;refGene_gn=LRRC41;refGene_tx=XM_011284800.3
C1	37025772	37025895
refGene_gn=LRRC41;refGene_tx=XM_011284798.3;refGene_gn=LRRC41;refGene_tx=XM_011284800.3
C1	37026433	37026532
refGene_gn=LRRC41;refGene_tx=XM_011284798.3;refGene_gn=LRRC41;refGene_tx=XM_011284800.3
C1	37026621	37026786
refGene_gn=LRRC41;refGene_tx=XM_011284798.3;refGene_gn=LRRC41;refGene_tx=XM_011284800.3
C1	37027302	37027563
refGene_gn=LRRC41;refGene_tx=XM_011284798.3;refGene_gn=LRRC41;refGene_tx=XM_011284800.3
C1	37030257	37031401
refGene_gn=LRRC41;refGene_tx=XM_011284798.3;refGene_gn=LRRC41;refGene_tx=XM_011284800.3
C1	37038854	37038925
refGene_gn=LRRC41;refGene_tx=XM_011284798.3;refGene_gn=LRRC41;refGene_tx=XM_011284800.3
C1	37039829	37039916	refGene_gn=LRRC41;refGene_tx=XM_011284798.3
C1	37043986	37044185	refGene_gn=LRRC41;refGene_tx=XM_011284798.3
C1	37044628	37044682
refGene_gn=LOC101095032;refGene_tx=XM_003990050.5
C1	37048492	37048519
refGene_gn=LOC101095032;refGene_tx=XM_003990050.5
C1	37049092	37049155
refGene_gn=LOC101095032;refGene_tx=XM_019836926.1
C1	37049260	37049422
refGene_gn=LOC101095032;refGene_tx=XM_003990050.5;refGene_gn=LOC101095032;refGene_tx=XM_019836926.1
C1	37052504	37052537
refGene_gn=LOC101095032;refGene_tx=XM_003990050.5;refGene_gn=LOC101095032;refGene_tx=XM_019836926.1
C1	37064947	37065040	refGene_gn=NSUN4;refGene_tx=XM_003990051.5
C1	37065292	37068355
refGene_gn=LOC111561731;refGene_tx=XM_023258644.1

C1 37069457 37069801
refGene_gn=NSUN4;refGene_tx=XM_003990051.5;refGene_gn=NSUN4;refGene_tx=XM_011284802.3;refGene_gn=NSUN4;refGene_tx=XM_019836927.2;refGene_gn=NSUN4;refGene_tx=XM_019836928.2
C1 37074001 37074156
refGene_gn=NSUN4;refGene_tx=XM_003990051.5;refGene_gn=NSUN4;refGene_tx=XM_011284802.3;refGene_gn=NSUN4;refGene_tx=XM_019836927.2;refGene_gn=NSUN4;refGene_tx=XM_019836928.2
C1 37077593 37077754
refGene_gn=NSUN4;refGene_tx=XM_003990051.5;refGene_gn=NSUN4;refGene_tx=XM_011284802.3;refGene_gn=NSUN4;refGene_tx=XM_019836927.2;refGene_gn=NSUN4;refGene_tx=XM_019836928.2
C1 37078928 37079053
refGene_gn=NSUN4;refGene_tx=XM_003990051.5;refGene_gn=NSUN4;refGene_tx=XM_011284802.3;refGene_gn=NSUN4;refGene_tx=XM_019836927.2;refGene_gn=NSUN4;refGene_tx=XM_019836928.2
C1 37079640 37079917
refGene_gn=NSUN4;refGene_tx=XM_003990051.5;refGene_gn=NSUN4;refGene_tx=XM_011284802.3;refGene_gn=NSUN4;refGene_tx=XM_019836927.2;refGene_gn=NSUN4;refGene_tx=XM_019836928.2
C1 37094427 37094622 refGene_gn=FAAH;refGene_tx=XM_003990052.5
C1 37100874 37100988 refGene_gn=FAAH;refGene_tx=XM_003990052.5
C1 37103421 37103556 refGene_gn=FAAH;refGene_tx=XM_003990052.5
C1 37103785 37103919 refGene_gn=FAAH;refGene_tx=XM_003990052.5
C1 37103996 37104203 refGene_gn=FAAH;refGene_tx=XM_003990052.5
C1 37104454 37104495 refGene_gn=FAAH;refGene_tx=XM_003990052.5
C1 37104717 37104842 refGene_gn=FAAH;refGene_tx=XM_003990052.5
C1 37105701 37105827 refGene_gn=FAAH;refGene_tx=XM_003990052.5
C1 37106368 37106466 refGene_gn=FAAH;refGene_tx=XM_003990052.5
C1 37107937 37108037 refGene_gn=FAAH;refGene_tx=XM_003990052.5
C1 37108327 37108368 refGene_gn=FAAH;refGene_tx=XM_003990052.5
C1 37109141 37109181 refGene_gn=FAAH;refGene_tx=XM_003990052.5
C1 37109656 37109765 refGene_gn=FAAH;refGene_tx=XM_003990052.5
C1 37110331 37110477 refGene_gn=FAAH;refGene_tx=XM_003990052.5
C1 37110857 37110986 refGene_gn=FAAH;refGene_tx=XM_003990052.5
C1 37119795 37119975
refGene_gn=LOC101081996;refGene_tx=XM_023257828.1
C1 37126226 37126340
refGene_gn=LOC101081996;refGene_tx=XM_023257828.1
C1 37126581 37126731
refGene_gn=LOC101081996;refGene_tx=XM_023257828.1
C1 37127053 37127187
refGene_gn=LOC101081996;refGene_tx=XM_023257828.1
C1 37127773 37127980
refGene_gn=LOC101081996;refGene_tx=XM_023257828.1
C1 37129228 37129269
refGene_gn=LOC101081996;refGene_tx=XM_023257828.1
C1 37130047 37130172
refGene_gn=LOC101081996;refGene_tx=XM_023257828.1
C1 37131366 37131492

refGene_gn=LOC101081996;refGene_tx=XM_023257828.1
C1	37132199	37132297
refGene_gn=LOC101081996;refGene_tx=XM_023257828.1
C1	37132623	37132723
refGene_gn=LOC101081996;refGene_tx=XM_023257828.1
C1	37134178	37134287
refGene_gn=LOC101081996;refGene_tx=XM_023257828.1
C1	37136550	37136686
refGene_gn=LOC101081996;refGene_tx=XM_023257828.1
C1	37154408	37154529
refGene_gn=LOC101081996;refGene_tx=XM_023257828.1
C1	37154749	37154805
refGene_gn=LOC101081996;refGene_tx=XM_023257828.1
C1	37156501	37156662
refGene_gn=LOC101081996;refGene_tx=XM_023257828.1
C1	37160796	37161140
refGene_gn=LOC101081996;refGene_tx=XM_023257828.1
C1	37198223	37198377	refGene_gn=DMBX1;refGene_tx=XM_003990053.3
C1	37201325	37201504	refGene_gn=DMBX1;refGene_tx=XM_003990053.3
C1	37201762	37202111	refGene_gn=DMBX1;refGene_tx=XM_003990053.3
C1	37203048	37203491	refGene_gn=DMBX1;refGene_tx=XM_003990053.3
C1	37234769	37234849	refGene_gn=KNCN;refGene_tx=XM_019836932.2
C1	37235662	37235737	refGene_gn=KNCN;refGene_tx=XM_019836932.2
C1	37236448	37236517	refGene_gn=KNCN;refGene_tx=XM_019836932.2
C1	37237605	37237756	refGene_gn=KNCN;refGene_tx=XM_019836932.2
C1	37244600	37244829
refGene_gn=MKNK1;refGene_tx=XM_006934712.3;refGene_gn=MKNK1;refGene_tx=XM_011284803.2;refGene_gn=MKNK1;refGene_tx=XM_011284805.3;refGene_gn=MKNK1;refGene_tx=XM_011284806.2;refGene_gn=MKNK1;refGene_tx=XM_011284807.3;refGene_gn=MKNK1;refGene_tx=XM_019836930.1;refGene_gn=MKNK1;refGene_tx=XM_019836931.1
C1	37246158	37246202
refGene_gn=MKNK1;refGene_tx=XM_011284803.2;refGene_gn=MKNK1;refGene_tx=XM_011284805.3;refGene_gn=MKNK1;refGene_tx=XM_011284806.2;refGene_gn=MKNK1;refGene_tx=XM_011284807.3;refGene_gn=MKNK1;refGene_tx=XM_019836930.1;refGene_gn=MKNK1;refGene_tx=XM_019836931.1
C1	37247628	37247793
refGene_gn=MKNK1;refGene_tx=XM_006934712.3;refGene_gn=MKNK1;refGene_tx=XM_011284803.2;refGene_gn=MKNK1;refGene_tx=XM_011284805.3;refGene_gn=MKNK1;refGene_tx=XM_011284806.2;refGene_gn=MKNK1;refGene_tx=XM_011284807.3;refGene_gn=MKNK1;refGene_tx=XM_019836930.1;refGene_gn=MKNK1;refGene_tx=XM_019836931.1
C1	37248502	37248697
refGene_gn=MKNK1;refGene_tx=XM_006934712.3;refGene_gn=MKNK1;refGene_tx=XM_011284803.2;refGene_gn=MKNK1;refGene_tx=XM_011284805.3;refGene_gn=MKNK1;refGene_tx=XM_011284806.2;refGene_gn=MKNK1;refGene_tx=XM_011284807.3;refGene_gn=MKNK1;refGene_tx=XM_019836930.1;refGene_gn=MKNK1;refGene_tx=XM_019836931.1
C1	37250236	37250332
refGene_gn=MKNK1;refGene_tx=XM_006934712.3;refGene_gn=MKNK1;refGene_tx

=XM_011284803.2;refGene_gn=MKNK1;refGene_tx=XM_011284805.3;refGene_gn=MKNK1;refGene_tx=XM_011284806.2;refGene_gn=MKNK1;refGene_tx=XM_011284807.3;refGene_gn=MKNK1;refGene_tx=XM_019836930.1;refGene_gn=MKNK1;refGene_tx=XM_019836931.1
C1 37253283 37253339
refGene_gn=MKNK1;refGene_tx=XM_006934712.3;refGene_gn=MKNK1;refGene_tx=XM_011284803.2;refGene_gn=MKNK1;refGene_tx=XM_011284805.3;refGene_gn=MKNK1;refGene_tx=XM_011284806.2;refGene_gn=MKNK1;refGene_tx=XM_011284807.3;refGene_gn=MKNK1;refGene_tx=XM_019836930.1;refGene_gn=MKNK1;refGene_tx=XM_019836931.1
C1 37256557 37256662
refGene_gn=MKNK1;refGene_tx=XM_006934712.3;refGene_gn=MKNK1;refGene_tx=XM_011284803.2;refGene_gn=MKNK1;refGene_tx=XM_011284805.3;refGene_gn=MKNK1;refGene_tx=XM_011284806.2;refGene_gn=MKNK1;refGene_tx=XM_011284807.3;refGene_gn=MKNK1;refGene_tx=XM_019836930.1;refGene_gn=MKNK1;refGene_tx=XM_019836931.1
C1 37258797 37258871
refGene_gn=MKNK1;refGene_tx=XM_006934712.3;refGene_gn=MKNK1;refGene_tx=XM_011284803.2;refGene_gn=MKNK1;refGene_tx=XM_011284805.3;refGene_gn=MKNK1;refGene_tx=XM_011284806.2;refGene_gn=MKNK1;refGene_tx=XM_011284807.3;refGene_gn=MKNK1;refGene_tx=XM_019836930.1;refGene_gn=MKNK1;refGene_tx=XM_019836931.1
C1 37260417 37260497
refGene_gn=MKNK1;refGene_tx=XM_006934712.3;refGene_gn=MKNK1;refGene_tx=XM_011284803.2;refGene_gn=MKNK1;refGene_tx=XM_011284805.3;refGene_gn=MKNK1;refGene_tx=XM_011284806.2;refGene_gn=MKNK1;refGene_tx=XM_011284807.3;refGene_gn=MKNK1;refGene_tx=XM_019836930.1;refGene_gn=MKNK1;refGene_tx=XM_019836931.1
C1 37263953 37264051
refGene_gn=MKNK1;refGene_tx=XM_006934712.3;refGene_gn=MKNK1;refGene_tx=XM_011284803.2;refGene_gn=MKNK1;refGene_tx=XM_011284805.3;refGene_gn=MKNK1;refGene_tx=XM_011284806.2;refGene_gn=MKNK1;refGene_tx=XM_011284807.3;refGene_gn=MKNK1;refGene_tx=XM_019836930.1;refGene_gn=MKNK1;refGene_tx=XM_019836931.1
C1 37266106 37266208
refGene_gn=MKNK1;refGene_tx=XM_006934712.3;refGene_gn=MKNK1;refGene_tx=XM_011284803.2;refGene_gn=MKNK1;refGene_tx=XM_011284805.3;refGene_gn=MKNK1;refGene_tx=XM_011284806.2;refGene_gn=MKNK1;refGene_tx=XM_011284807.3;refGene_gn=MKNK1;refGene_tx=XM_019836930.1;refGene_gn=MKNK1;refGene_tx=XM_019836931.1
C1 37268613 37268629
refGene_gn=MKNK1;refGene_tx=XM_006934712.3;refGene_gn=MKNK1;refGene_tx=XM_011284803.2;refGene_gn=MKNK1;refGene_tx=XM_011284805.3;refGene_gn=MKNK1;refGene_tx=XM_011284806.2
C1 37289853 37289883
refGene_gn=MOB3C;refGene_tx=XM_006934713.4;refGene_gn=MOB3C;refGene_tx=XM_006934716.3;refGene_gn=MOB3C;refGene_tx=XM_006934717.3;refGene_gn=MOB3C;refGene_tx=XM_019836933.2
C1 37290198 37290401
refGene_gn=MOB3C;refGene_tx=XM_006934713.4;refGene_gn=MOB3C;refGene_tx

=XM_006934716.3;refGene_gn=MOB3C;refGene_tx=XM_006934717.3;refGene_gn=MOB3C;refGene_tx=XM_019836933.2
C1 37292514 37292932
refGene_gn=MOB3C;refGene_tx=XM_006934713.4;refGene_gn=MOB3C;refGene_tx=XM_006934716.3;refGene_gn=MOB3C;refGene_tx=XM_006934717.3;refGene_gn=MOB3C;refGene_tx=XM_019836933.2
C1 37313671 37313866 refGene_gn=ATPAF1;refGene_tx=XM_023258645.1
C1 37319974 37320082 refGene_gn=ATPAF1;refGene_tx=XM_023258645.1
C1 37321997 37322093 refGene_gn=ATPAF1;refGene_tx=XM_023258645.1
C1 37325889 37325937 refGene_gn=ATPAF1;refGene_tx=XM_023258645.1
C1 37327114 37327165 refGene_gn=ATPAF1;refGene_tx=XM_023258645.1
C1 37330956 37331019 refGene_gn=ATPAF1;refGene_tx=XM_023258645.1
C1 37331656 37331707 refGene_gn=ATPAF1;refGene_tx=XM_023258645.1
C1 37335942 37336051 refGene_gn=ATPAF1;refGene_tx=XM_023258645.1
C1 37338871 37339194 refGene_gn=ATPAF1;refGene_tx=XM_023258645.1
C1 37343110 37343114 refGene_gn=TEX38;refGene_tx=XM_006934719.4
C1 37343168 37343266
refGene_gn=TEX38;refGene_tx=XM_003990055.3;refGene_gn=TEX38;refGene_tx=XM_019836935.2
C1 37343428 37343432 refGene_gn=TEX38;refGene_tx=XM_006934720.3
C1 37343836 37344444
refGene_gn=TEX38;refGene_tx=XM_003990055.3;refGene_gn=TEX38;refGene_tx=XM_006934719.4;refGene_gn=TEX38;refGene_tx=XM_006934720.3;refGene_gn=TEX38;refGene_tx=XM_011284812.3;refGene_gn=TEX38;refGene_tx=XM_019836935.2;refGene_gn=TEX38;refGene_tx=XM_019836936.2
C1 37385243 37385267
refGene_gn=LOC101096660;refGene_tx=XM_011284815.3
C1 37401537 37401567
refGene_gn=LOC101096660;refGene_tx=XM_019836940.2
C1 37427625 37427805
refGene_gn=LOC101096660;refGene_tx=XM_011284813.3;refGene_gn=LOC101096660;refGene_tx=XM_019836938.2
C1 37439549 37439691
refGene_gn=LOC101096660;refGene_tx=XM_011284813.3;refGene_gn=LOC101096660;refGene_tx=XM_011284815.3;refGene_gn=LOC101096660;refGene_tx=XM_019836938.2;refGene_gn=LOC101096660;refGene_tx=XM_019836940.2
C1 37439874 37439919
refGene_gn=LOC101096660;refGene_tx=XM_011284813.3;refGene_gn=LOC101096660;refGene_tx=XM_011284815.3;refGene_gn=LOC101096660;refGene_tx=XM_019836938.2;refGene_gn=LOC101096660;refGene_tx=XM_019836940.2
C1 37440462 37440590
refGene_gn=LOC101096660;refGene_tx=XM_011284813.3;refGene_gn=LOC101096660;refGene_tx=XM_011284815.3;refGene_gn=LOC101096660;refGene_tx=XM_019836938.2;refGene_gn=LOC101096660;refGene_tx=XM_019836940.2
C1 37441395 37441520
refGene_gn=LOC101096660;refGene_tx=XM_011284813.3;refGene_gn=LOC101096660;refGene_tx=XM_011284815.3;refGene_gn=LOC101096660;refGene_tx=XM_019836938.2;refGene_gn=LOC101096660;refGene_tx=XM_019836940.2
C1 37441836 37441988
refGene_gn=LOC101096660;refGene_tx=XM_011284813.3;refGene_gn=LOC101096

660;refGene_tx=XM_011284815.3;refGene_gn=LOC101096660;refGene_tx=XM_019836938.2;refGene_gn=LOC101096660;refGene_tx=XM_019836940.2
C1 37442141 37442248
refGene_gn=LOC101096660;refGene_tx=XM_011284813.3;refGene_gn=LOC101096660;refGene_tx=XM_011284815.3;refGene_gn=LOC101096660;refGene_tx=XM_019836938.2;refGene_gn=LOC101096660;refGene_tx=XM_019836940.2
C1 37442949 37443140
refGene_gn=LOC101096660;refGene_tx=XM_011284813.3;refGene_gn=LOC101096660;refGene_tx=XM_011284815.3;refGene_gn=LOC101096660;refGene_tx=XM_019836938.2;refGene_gn=LOC101096660;refGene_tx=XM_019836940.2
C1 37445335 37445469
refGene_gn=LOC101096660;refGene_tx=XM_011284813.3;refGene_gn=LOC101096660;refGene_tx=XM_011284815.3;refGene_gn=LOC101096660;refGene_tx=XM_019836938.2;refGene_gn=LOC101096660;refGene_tx=XM_019836940.2
C1 37446267 37446332
refGene_gn=LOC101096660;refGene_tx=XM_011284813.3;refGene_gn=LOC101096660;refGene_tx=XM_011284815.3;refGene_gn=LOC101096660;refGene_tx=XM_019836938.2;refGene_gn=LOC101096660;refGene_tx=XM_019836940.2
C1 37446432 37446515
refGene_gn=LOC101096660;refGene_tx=XM_011284813.3;refGene_gn=LOC101096660;refGene_tx=XM_011284815.3;refGene_gn=LOC101096660;refGene_tx=XM_019836938.2;refGene_gn=LOC101096660;refGene_tx=XM_019836940.2
C1 37446682 37446854
refGene_gn=LOC101096660;refGene_tx=XM_019836938.2
C1 37446943 37447127
refGene_gn=LOC101096660;refGene_tx=XM_011284813.3;refGene_gn=LOC101096660;refGene_tx=XM_011284815.3;refGene_gn=LOC101096660;refGene_tx=XM_019836940.2
C1 37514797 37514966
refGene_gn=LOC101094994;refGene_tx=XM_023258648.1;refGene_gn=LOC101094994;refGene_tx=XM_023258649.1
C1 37516369 37516446
refGene_gn=LOC101094994;refGene_tx=XM_023258648.1;refGene_gn=LOC101094994;refGene_tx=XM_023258649.1
C1 37516594 37516659
refGene_gn=LOC101094994;refGene_tx=XM_023258648.1;refGene_gn=LOC101094994;refGene_tx=XM_023258649.1
C1 37517944 37518078
refGene_gn=LOC101094994;refGene_tx=XM_023258648.1;refGene_gn=LOC101094994;refGene_tx=XM_023258649.1
C1 37518174 37518365
refGene_gn=LOC101094994;refGene_tx=XM_023258648.1;refGene_gn=LOC101094994;refGene_tx=XM_023258649.1
C1 37518451 37518558
refGene_gn=LOC101094994;refGene_tx=XM_023258648.1;refGene_gn=LOC101094994;refGene_tx=XM_023258649.1
C1 37518995 37519150
refGene_gn=LOC101094994;refGene_tx=XM_023258648.1;refGene_gn=LOC101094994;refGene_tx=XM_023258649.1
C1 37519505 37519630

refGene_gn=LOC101094994;refGene_tx=XM_023258648.1;refGene_gn=LOC101094994;refGene_tx=XM_023258649.1
C1 37520780 37520908
refGene_gn=LOC101094994;refGene_tx=XM_023258648.1;refGene_gn=LOC101094994;refGene_tx=XM_023258649.1
C1 37521661 37521706
refGene_gn=LOC101094994;refGene_tx=XM_023258648.1;refGene_gn=LOC101094994;refGene_tx=XM_023258649.1
C1 37522540 37522682
refGene_gn=LOC101094994;refGene_tx=XM_023258648.1;refGene_gn=LOC101094994;refGene_tx=XM_023258649.1
C1 37524666 37524675
refGene_gn=LOC101094994;refGene_tx=XM_023258649.1
C1 37525263 37525458
refGene_gn=LOC101094994;refGene_tx=XM_023258648.1
C1 37554380 37554557
refGene_gn=LOC101096906;refGene_tx=XM_023258650.1
C1 37562785 37562927
refGene_gn=LOC101096906;refGene_tx=XM_023258650.1;refGene_gn=LOC101096906;refGene_tx=XM_023258651.1
C1 37564378 37564423
refGene_gn=LOC101096906;refGene_tx=XM_023258650.1;refGene_gn=LOC101096906;refGene_tx=XM_023258651.1
C1 37565808 37565936
refGene_gn=LOC101096906;refGene_tx=XM_023258650.1;refGene_gn=LOC101096906;refGene_tx=XM_023258651.1
C1 37567887 37568015
refGene_gn=LOC101096906;refGene_tx=XM_023258650.1;refGene_gn=LOC101096906;refGene_tx=XM_023258651.1
C1 37568098 37568253
refGene_gn=LOC101096906;refGene_tx=XM_023258650.1;refGene_gn=LOC101096906;refGene_tx=XM_023258651.1
C1 37575347 37575454
refGene_gn=LOC101096906;refGene_tx=XM_023258650.1;refGene_gn=LOC101096906;refGene_tx=XM_023258651.1
C1 37576021 37576212
refGene_gn=LOC101096906;refGene_tx=XM_023258650.1;refGene_gn=LOC101096906;refGene_tx=XM_023258651.1
C1 37579393 37579527
refGene_gn=LOC101096906;refGene_tx=XM_023258650.1;refGene_gn=LOC101096906;refGene_tx=XM_023258651.1
C1 37580822 37580887
refGene_gn=LOC101096906;refGene_tx=XM_023258650.1;refGene_gn=LOC101096906;refGene_tx=XM_023258651.1
C1 37589105 37589188
refGene_gn=LOC101096906;refGene_tx=XM_023258650.1;refGene_gn=LOC101096906;refGene_tx=XM_023258651.1
C1 37590004 37590173
refGene_gn=LOC101096906;refGene_tx=XM_023258650.1;refGene_gn=LOC101096906;refGene_tx=XM_023258651.1

C1 37621983 37622056
refGene_gn=PDZK1IP1;refGene_tx=XM_019836947.1;refGene_gn=PDZK1IP1;refGene_tx=XM_019836948.1
C1 37622679 37622775
refGene_gn=PDZK1IP1;refGene_tx=XM_019836947.1;refGene_gn=PDZK1IP1;refGene_tx=XM_019836948.1
C1 37624652 37624761
refGene_gn=PDZK1IP1;refGene_tx=XM_019836947.1;refGene_gn=PDZK1IP1;refGene_tx=XM_019836948.1
C1 37627891 37627958
refGene_gn=PDZK1IP1;refGene_tx=XM_019836947.1;refGene_gn=PDZK1IP1;refGene_tx=XM_019836948.1
C1 37648172 37648718
refGene_gn=TAL1;refGene_tx=XM_003990060.5;refGene_gn=TAL1;refGene_tx=XM_023258653.1;refGene_gn=TAL1;refGene_tx=XM_023258654.1;refGene_gn=TAL1;refGene_tx=XM_023258655.1;refGene_gn=TAL1;refGene_tx=XM_023258656.1;refGene_gn=TAL1;refGene_tx=XM_023258657.1;refGene_gn=TAL1;refGene_tx=XM_023258658.1
C1 37652413 37652508
refGene_gn=TAL1;refGene_tx=XM_003990060.5;refGene_gn=TAL1;refGene_tx=XM_023258653.1;refGene_gn=TAL1;refGene_tx=XM_023258654.1;refGene_gn=TAL1;refGene_tx=XM_023258655.1;refGene_gn=TAL1;refGene_tx=XM_023258656.1;refGene_gn=TAL1;refGene_tx=XM_023258657.1;refGene_gn=TAL1;refGene_tx=XM_023258658.1
C1 37653408 37653458 refGene_gn=TAL1;refGene_tx=XM_003990060.5
C1 37653879 37654316
refGene_gn=TAL1;refGene_tx=XM_023258653.1;refGene_gn=TAL1;refGene_tx=XM_023258654.1;refGene_gn=TAL1;refGene_tx=XM_023258655.1;refGene_gn=TAL1;refGene_tx=XM_023258656.1;refGene_gn=TAL1;refGene_tx=XM_023258657.1;refGene_gn=TAL1;refGene_tx=XM_023258658.1
C1 37675815 37676602
refGene_gn=STIL;refGene_tx=XM_011284819.3;refGene_gn=STIL;refGene_tx=XM_019836949.2;refGene_gn=STIL;refGene_tx=XM_019836954.2;refGene_gn=STIL;refGene_tx=XM_019836957.2;refGene_gn=STIL;refGene_tx=XM_019836958.2;refGene_gn=STIL;refGene_tx=XM_019836959.2;refGene_gn=STIL;refGene_tx=XM_019836961.2;refGene_gn=STIL;refGene_tx=XM_019836963.2;refGene_gn=STIL;refGene_tx=XM_023258659.1;refGene_gn=STIL;refGene_tx=XM_023258661.1;refGene_gn=STIL;refGene_tx=XM_023258662.1
C1 37685577 37685828
refGene_gn=STIL;refGene_tx=XM_011284819.3;refGene_gn=STIL;refGene_tx=XM_019836949.2;refGene_gn=STIL;refGene_tx=XM_019836954.2;refGene_gn=STIL;refGene_tx=XM_019836957.2;refGene_gn=STIL;refGene_tx=XM_019836958.2;refGene_gn=STIL;refGene_tx=XM_019836959.2;refGene_gn=STIL;refGene_tx=XM_019836961.2;refGene_gn=STIL;refGene_tx=XM_019836963.2;refGene_gn=STIL;refGene_tx=XM_023258659.1;refGene_gn=STIL;refGene_tx=XM_023258661.1;refGene_gn=STIL;refGene_tx=XM_023258662.1
C1 37687301 37687515
refGene_gn=STIL;refGene_tx=XM_011284819.3;refGene_gn=STIL;refGene_tx=XM_019836949.2;refGene_gn=STIL;refGene_tx=XM_019836954.2;refGene_gn=STIL;refGene_tx=XM_019836957.2;refGene_gn=STIL;refGene_tx=XM_019836958.2;

refGene_gn=STIL;refGene_tx=XM_019836959.2;refGene_gn=STIL;refGene_tx=XM_019836961.2;refGene_gn=STIL;refGene_tx=XM_019836963.2;refGene_gn=STIL;refGene_tx=XM_023258659.1;refGene_gn=STIL;refGene_tx=XM_023258661.1;refGene_gn=STIL;refGene_tx=XM_023258662.1
C1 37700796 37701028
refGene_gn=STIL;refGene_tx=XM_011284819.3;refGene_gn=STIL;refGene_tx=XM_019836949.2;refGene_gn=STIL;refGene_tx=XM_019836954.2;refGene_gn=STIL;refGene_tx=XM_019836957.2;refGene_gn=STIL;refGene_tx=XM_019836958.2;refGene_gn=STIL;refGene_tx=XM_019836959.2;refGene_gn=STIL;refGene_tx=XM_019836961.2;refGene_gn=STIL;refGene_tx=XM_019836963.2;refGene_gn=STIL;refGene_tx=XM_023258659.1;refGene_gn=STIL;refGene_tx=XM_023258661.1;refGene_gn=STIL;refGene_tx=XM_023258662.1
C1 37704172 37704338
refGene_gn=STIL;refGene_tx=XM_011284819.3;refGene_gn=STIL;refGene_tx=XM_019836949.2;refGene_gn=STIL;refGene_tx=XM_019836954.2;refGene_gn=STIL;refGene_tx=XM_019836957.2;refGene_gn=STIL;refGene_tx=XM_019836958.2;refGene_gn=STIL;refGene_tx=XM_019836959.2;refGene_gn=STIL;refGene_tx=XM_019836961.2;refGene_gn=STIL;refGene_tx=XM_019836963.2;refGene_gn=STIL;refGene_tx=XM_023258659.1;refGene_gn=STIL;refGene_tx=XM_023258661.1;refGene_gn=STIL;refGene_tx=XM_023258662.1
C1 37709602 37710541
refGene_gn=STIL;refGene_tx=XM_011284819.3;refGene_gn=STIL;refGene_tx=XM_019836949.2;refGene_gn=STIL;refGene_tx=XM_019836954.2;refGene_gn=STIL;refGene_tx=XM_019836957.2;refGene_gn=STIL;refGene_tx=XM_019836958.2;refGene_gn=STIL;refGene_tx=XM_019836959.2;refGene_gn=STIL;refGene_tx=XM_019836961.2;refGene_gn=STIL;refGene_tx=XM_019836963.2;refGene_gn=STIL;refGene_tx=XM_023258659.1;refGene_gn=STIL;refGene_tx=XM_023258661.1
C1 37712773 37712888
refGene_gn=STIL;refGene_tx=XM_011284819.3;refGene_gn=STIL;refGene_tx=XM_019836949.2;refGene_gn=STIL;refGene_tx=XM_019836954.2;refGene_gn=STIL;refGene_tx=XM_019836957.2;refGene_gn=STIL;refGene_tx=XM_019836958.2;refGene_gn=STIL;refGene_tx=XM_019836959.2;refGene_gn=STIL;refGene_tx=XM_019836961.2;refGene_gn=STIL;refGene_tx=XM_019836963.2;refGene_gn=STIL;refGene_tx=XM_023258659.1;refGene_gn=STIL;refGene_tx=XM_023258661.1;refGene_gn=STIL;refGene_tx=XM_023258662.1
C1 37717104 37717214
refGene_gn=STIL;refGene_tx=XM_011284819.3;refGene_gn=STIL;refGene_tx=XM_019836949.2;refGene_gn=STIL;refGene_tx=XM_019836954.2;refGene_gn=STIL;refGene_tx=XM_019836957.2;refGene_gn=STIL;refGene_tx=XM_019836958.2;refGene_gn=STIL;refGene_tx=XM_019836959.2;refGene_gn=STIL;refGene_tx=XM_019836961.2;refGene_gn=STIL;refGene_tx=XM_019836963.2;refGene_gn=STIL;refGene_tx=XM_023258659.1;refGene_gn=STIL;refGene_tx=XM_023258661.1;refGene_gn=STIL;refGene_tx=XM_023258662.1
C1 37720324 37720478
refGene_gn=STIL;refGene_tx=XM_011284819.3;refGene_gn=STIL;refGene_tx=XM_019836949.2;refGene_gn=STIL;refGene_tx=XM_019836954.2;refGene_gn=STIL;refGene_tx=XM_019836957.2;refGene_gn=STIL;refGene_tx=XM_019836958.2;refGene_gn=STIL;refGene_tx=XM_019836959.2;refGene_gn=STIL;refGene_tx=XM_019836961.2;refGene_gn=STIL;refGene_tx=XM_019836963.2;refGene_gn=STIL;refGene_tx=XM_023258659.1;refGene_gn=STIL;refGene_tx=XM_023258661.1;

refGene_gn=STIL;refGene_tx=XM_023258662.1
C1 37723785 37723872
refGene_gn=STIL;refGene_tx=XM_011284819.3;refGene_gn=STIL;refGene_tx=XM_019836949.2;refGene_gn=STIL;refGene_tx=XM_019836954.2;refGene_gn=STIL;refGene_tx=XM_019836957.2;refGene_gn=STIL;refGene_tx=XM_019836958.2;refGene_gn=STIL;refGene_tx=XM_019836959.2;refGene_gn=STIL;refGene_tx=XM_019836961.2;refGene_gn=STIL;refGene_tx=XM_019836963.2;refGene_gn=STIL;refGene_tx=XM_023258659.1;refGene_gn=STIL;refGene_tx=XM_023258661.1;refGene_gn=STIL;refGene_tx=XM_023258662.1
C1 37725176 37725260
refGene_gn=STIL;refGene_tx=XM_011284819.3;refGene_gn=STIL;refGene_tx=XM_019836949.2;refGene_gn=STIL;refGene_tx=XM_019836954.2;refGene_gn=STIL;refGene_tx=XM_019836957.2;refGene_gn=STIL;refGene_tx=XM_019836958.2;refGene_gn=STIL;refGene_tx=XM_019836959.2;refGene_gn=STIL;refGene_tx=XM_019836961.2;refGene_gn=STIL;refGene_tx=XM_019836963.2;refGene_gn=STIL;refGene_tx=XM_023258659.1;refGene_gn=STIL;refGene_tx=XM_023258661.1;refGene_gn=STIL;refGene_tx=XM_023258662.1
C1 37728064 37728312
refGene_gn=STIL;refGene_tx=XM_011284819.3;refGene_gn=STIL;refGene_tx=XM_019836949.2;refGene_gn=STIL;refGene_tx=XM_019836954.2;refGene_gn=STIL;refGene_tx=XM_019836957.2;refGene_gn=STIL;refGene_tx=XM_019836958.2;refGene_gn=STIL;refGene_tx=XM_019836959.2;refGene_gn=STIL;refGene_tx=XM_019836961.2;refGene_gn=STIL;refGene_tx=XM_019836963.2;refGene_gn=STIL;refGene_tx=XM_023258659.1;refGene_gn=STIL;refGene_tx=XM_023258661.1;refGene_gn=STIL;refGene_tx=XM_023258662.1
C1 37729667 37729855
refGene_gn=STIL;refGene_tx=XM_011284819.3;refGene_gn=STIL;refGene_tx=XM_019836949.2;refGene_gn=STIL;refGene_tx=XM_019836954.2;refGene_gn=STIL;refGene_tx=XM_019836957.2;refGene_gn=STIL;refGene_tx=XM_019836958.2;refGene_gn=STIL;refGene_tx=XM_019836959.2;refGene_gn=STIL;refGene_tx=XM_019836961.2;refGene_gn=STIL;refGene_tx=XM_019836963.2;refGene_gn=STIL;refGene_tx=XM_023258659.1;refGene_gn=STIL;refGene_tx=XM_023258661.1;refGene_gn=STIL;refGene_tx=XM_023258662.1
C1 37730171 37730284
refGene_gn=STIL;refGene_tx=XM_011284819.3;refGene_gn=STIL;refGene_tx=XM_019836949.2;refGene_gn=STIL;refGene_tx=XM_019836954.2;refGene_gn=STIL;refGene_tx=XM_019836957.2;refGene_gn=STIL;refGene_tx=XM_019836958.2;refGene_gn=STIL;refGene_tx=XM_019836959.2;refGene_gn=STIL;refGene_tx=XM_019836961.2;refGene_gn=STIL;refGene_tx=XM_019836963.2;refGene_gn=STIL;refGene_tx=XM_023258659.1;refGene_gn=STIL;refGene_tx=XM_023258661.1;refGene_gn=STIL;refGene_tx=XM_023258662.1
C1 37732623 37732731
refGene_gn=STIL;refGene_tx=XM_011284819.3;refGene_gn=STIL;refGene_tx=XM_019836949.2;refGene_gn=STIL;refGene_tx=XM_019836954.2;refGene_gn=STIL;refGene_tx=XM_019836957.2;refGene_gn=STIL;refGene_tx=XM_019836958.2;refGene_gn=STIL;refGene_tx=XM_019836959.2;refGene_gn=STIL;refGene_tx=XM_019836961.2;refGene_gn=STIL;refGene_tx=XM_019836963.2;refGene_gn=STIL;refGene_tx=XM_023258659.1;refGene_gn=STIL;refGene_tx=XM_023258661.1;refGene_gn=STIL;refGene_tx=XM_023258662.1
C1 37736994 37737038

refGene_gn=STIL;refGene_tx=XM_011284819.3;refGene_gn=STIL;refGene_tx=XM_019836949.2;refGene_gn=STIL;refGene_tx=XM_019836954.2;refGene_gn=STIL;refGene_tx=XM_019836957.2;refGene_gn=STIL;refGene_tx=XM_019836958.2;refGene_gn=STIL;refGene_tx=XM_019836959.2;refGene_gn=STIL;refGene_tx=XM_019836961.2;refGene_gn=STIL;refGene_tx=XM_019836963.2;refGene_gn=STIL;refGene_tx=XM_023258659.1;refGene_gn=STIL;refGene_tx=XM_023258661.1;refGene_gn=STIL;refGene_tx=XM_023258662.1
C1 37752649 37752820 refGene_gn=CMPK1;refGene_tx=XM_023258663.1
C1 37778539 37778686 refGene_gn=CMPK1;refGene_tx=XM_023258663.1
C1 37782861 37783014 refGene_gn=CMPK1;refGene_tx=XM_023258663.1
C1 37784262 37784339 refGene_gn=CMPK1;refGene_tx=XM_023258663.1
C1 37784583 37784680 refGene_gn=CMPK1;refGene_tx=XM_023258663.1
C1 37785422 37785464 refGene_gn=CMPK1;refGene_tx=XM_023258663.1
C1 37823166 37824120 refGene_gn=FOXE3;refGene_tx=XM_023258665.1
C1 37844302 37845802 refGene_gn=FOXD2;refGene_tx=XM_023258664.1
C1 38142456 38142676 refGene_gn=TRABD2B;refGene_tx=XM_003990063.5
C1 38151243 38151513 refGene_gn=TRABD2B;refGene_tx=XM_003990063.5
C1 38154625 38154716 refGene_gn=TRABD2B;refGene_tx=XM_003990063.5
C1 38170222 38170397 refGene_gn=TRABD2B;refGene_tx=XM_003990063.5
C1 38176766 38176913 refGene_gn=TRABD2B;refGene_tx=XM_003990063.5
C1 38351874 38352435 refGene_gn=TRABD2B;refGene_tx=XM_003990063.5
C1 38354483 38354585 refGene_gn=TRABD2B;refGene_tx=XM_003990063.5
C1 38480642 38480783 refGene_gn=SLC5A9;refGene_tx=XM_003990064.5
C1 38482696 38482768 refGene_gn=SLC5A9;refGene_tx=XM_003990064.5
C1 38485909 38486014 refGene_gn=SLC5A9;refGene_tx=XM_003990064.5
C1 38486362 38486527 refGene_gn=SLC5A9;refGene_tx=XM_003990064.5
C1 38487720 38487826 refGene_gn=SLC5A9;refGene_tx=XM_003990064.5
C1 38488415 38488496 refGene_gn=SLC5A9;refGene_tx=XM_003990064.5
C1 38488793 38488999 refGene_gn=SLC5A9;refGene_tx=XM_003990064.5
C1 38489179 38489315 refGene_gn=SLC5A9;refGene_tx=XM_003990064.5
C1 38489768 38489876 refGene_gn=SLC5A9;refGene_tx=XM_003990064.5
C1 38492008 38492159 refGene_gn=SLC5A9;refGene_tx=XM_003990064.5
C1 38493920 38494089 refGene_gn=SLC5A9;refGene_tx=XM_003990064.5
C1 38495472 38495688 refGene_gn=SLC5A9;refGene_tx=XM_003990064.5
C1 38498905 38499065 refGene_gn=SLC5A9;refGene_tx=XM_003990064.5
C1 38502594 38502657 refGene_gn=SLC5A9;refGene_tx=XM_003990064.5
C1 38502659 38502802 refGene_gn=SLC5A9;refGene_tx=XM_003990064.5
C1 38531687 38531868 refGene_gn=SPATA6;refGene_tx=XM_023258667.1
C1 38545698 38545790 refGene_gn=SPATA6;refGene_tx=XM_023258667.1
C1 38575562 38575614 refGene_gn=SPATA6;refGene_tx=XM_023258667.1
C1 38580130 38580315 refGene_gn=SPATA6;refGene_tx=XM_023258667.1
C1 38595965 38596006 refGene_gn=SPATA6;refGene_tx=XM_023258667.1
C1 38604195 38604283 refGene_gn=SPATA6;refGene_tx=XM_023258667.1
C1 38608649 38608943 refGene_gn=SPATA6;refGene_tx=XM_023258667.1
C1 38613430 38613511 refGene_gn=SPATA6;refGene_tx=XM_023258667.1
C1 38623769 38623894 refGene_gn=SPATA6;refGene_tx=XM_023258667.1
C1 38656200 38656249 refGene_gn=SPATA6;refGene_tx=XM_023258667.1
C1 38657524 38657662 refGene_gn=SPATA6;refGene_tx=XM_023258667.1
C1 38680678 38680921 refGene_gn=SPATA6;refGene_tx=XM_023258667.1
C1 38713976 38714129

refGene_gn=AGBL4;refGene_tx=XM_023258671.1;refGene_gn=AGBL4;refGene_tx
=XM_023258674.1
C1 38725024 38725145 refGene_gn=AGBL4;refGene_tx=XM_023258670.1
C1 38729620 38729711 refGene_gn=AGBL4;refGene_tx=XM_023258672.1
C1 38731099 38731196
refGene_gn=AGBL4;refGene_tx=XM_023258670.1;refGene_gn=AGBL4;refGene_tx
=XM_023258671.1;refGene_gn=AGBL4;refGene_tx=XM_023258672.1;refGene_gn=
AGBL4;refGene_tx=XM_023258674.1
C1 38772890 38773053
refGene_gn=AGBL4;refGene_tx=XM_023258670.1;refGene_gn=AGBL4;refGene_tx
=XM_023258671.1;refGene_gn=AGBL4;refGene_tx=XM_023258672.1;refGene_gn=
AGBL4;refGene_tx=XM_023258674.1
C1 38775506 38775659
refGene_gn=AGBL4;refGene_tx=XM_023258670.1;refGene_gn=AGBL4;refGene_tx
=XM_023258671.1;refGene_gn=AGBL4;refGene_tx=XM_023258672.1;refGene_gn=
AGBL4;refGene_tx=XM_023258674.1
C1 38819046 38819158
refGene_gn=AGBL4;refGene_tx=XM_023258670.1;refGene_gn=AGBL4;refGene_tx
=XM_023258671.1;refGene_gn=AGBL4;refGene_tx=XM_023258672.1;refGene_gn=
AGBL4;refGene_tx=XM_023258674.1
C1 38839343 38839458
refGene_gn=AGBL4;refGene_tx=XM_023258670.1;refGene_gn=AGBL4;refGene_tx
=XM_023258671.1;refGene_gn=AGBL4;refGene_tx=XM_023258672.1;refGene_gn=
AGBL4;refGene_tx=XM_023258674.1
C1 38849373 38849463
refGene_gn=AGBL4;refGene_tx=XM_023258670.1;refGene_gn=AGBL4;refGene_tx
=XM_023258671.1;refGene_gn=AGBL4;refGene_tx=XM_023258672.1;refGene_gn=
AGBL4;refGene_tx=XM_023258674.1
C1 38921942 38922100
refGene_gn=AGBL4;refGene_tx=XM_023258668.1;refGene_gn=AGBL4;refGene_tx
=XM_023258669.1;refGene_gn=AGBL4;refGene_tx=XM_023258675.1
C1 38926081 38926223
refGene_gn=AGBL4;refGene_tx=XM_023258668.1;refGene_gn=AGBL4;refGene_tx
=XM_023258675.1
C1 38929860 38930074
refGene_gn=AGBL4;refGene_tx=XM_023258668.1;refGene_gn=AGBL4;refGene_tx
=XM_023258669.1;refGene_gn=AGBL4;refGene_tx=XM_023258675.1
C1 38935876 38936025
refGene_gn=AGBL4;refGene_tx=XM_023258668.1;refGene_gn=AGBL4;refGene_tx
=XM_023258669.1;refGene_gn=AGBL4;refGene_tx=XM_023258675.1
C1 38951157 38951542
refGene_gn=AGBL4;refGene_tx=XM_023258668.1;refGene_gn=AGBL4;refGene_tx
=XM_023258669.1;refGene_gn=AGBL4;refGene_tx=XM_023258675.1
C1 38953692 38953826
refGene_gn=AGBL4;refGene_tx=XM_023258668.1;refGene_gn=AGBL4;refGene_tx
=XM_023258669.1;refGene_gn=AGBL4;refGene_tx=XM_023258675.1
C1 39048666 39048706
refGene_gn=AGBL4;refGene_tx=XM_023258668.1;refGene_gn=AGBL4;refGene_tx
=XM_023258669.1;refGene_gn=AGBL4;refGene_tx=XM_023258670.1;refGene_gn=
AGBL4;refGene_tx=XM_023258671.1;refGene_gn=AGBL4;refGene_tx=XM_0232586

72.1;refGene_gn=AGBL4;refGene_tx=XM_023258674.1;refGene_gn=AGBL4;refGene_tx=XM_023258675.1
C1 39228851 39229068
refGene_gn=AGBL4;refGene_tx=XM_023258668.1;refGene_gn=AGBL4;refGene_tx=XM_023258669.1;refGene_gn=AGBL4;refGene_tx=XM_023258670.1;refGene_gn=AGBL4;refGene_tx=XM_023258671.1;refGene_gn=AGBL4;refGene_tx=XM_023258672.1;refGene_gn=AGBL4;refGene_tx=XM_023258674.1;refGene_gn=AGBL4;refGene_tx=XM_023258675.1
C1 39439563 39439658
refGene_gn=AGBL4;refGene_tx=XM_023258668.1;refGene_gn=AGBL4;refGene_tx=XM_023258669.1;refGene_gn=AGBL4;refGene_tx=XM_023258670.1;refGene_gn=AGBL4;refGene_tx=XM_023258671.1;refGene_gn=AGBL4;refGene_tx=XM_023258672.1;refGene_gn=AGBL4;refGene_tx=XM_023258674.1;refGene_gn=AGBL4;refGene_tx=XM_023258675.1
C1 39889202 39889327
refGene_gn=AGBL4;refGene_tx=XM_023258668.1;refGene_gn=AGBL4;refGene_tx=XM_023258669.1;refGene_gn=AGBL4;refGene_tx=XM_023258670.1;refGene_gn=AGBL4;refGene_tx=XM_023258671.1;refGene_gn=AGBL4;refGene_tx=XM_023258672.1;refGene_gn=AGBL4;refGene_tx=XM_023258674.1
C1 40036460 40036583
refGene_gn=AGBL4;refGene_tx=XM_023258668.1;refGene_gn=AGBL4;refGene_tx=XM_023258669.1;refGene_gn=AGBL4;refGene_tx=XM_023258670.1;refGene_gn=AGBL4;refGene_tx=XM_023258671.1;refGene_gn=AGBL4;refGene_tx=XM_023258672.1;refGene_gn=AGBL4;refGene_tx=XM_023258674.1;refGene_gn=AGBL4;refGene_tx=XM_023258675.1
C1 40175855 40175889
refGene_gn=AGBL4;refGene_tx=XM_023258668.1;refGene_gn=AGBL4;refGene_tx=XM_023258669.1;refGene_gn=AGBL4;refGene_tx=XM_023258670.1;refGene_gn=AGBL4;refGene_tx=XM_023258671.1;refGene_gn=AGBL4;refGene_tx=XM_023258672.1;refGene_gn=AGBL4;refGene_tx=XM_023258674.1
C1 40205578 40205596 refGene_gn=ELAVL4;refGene_tx=XM_023258677.1
C1 40258728 40258776
refGene_gn=ELAVL4;refGene_tx=XM_019836975.2;refGene_gn=ELAVL4;refGene_tx=XM_019836976.2;refGene_gn=ELAVL4;refGene_tx=XM_019836977.2
C1 40262062 40262071 refGene_gn=ELAVL4;refGene_tx=XM_011285003.3
C1 40262835 40262859 refGene_gn=ELAVL4;refGene_tx=XM_011285002.3
C1 40300135 40300376
refGene_gn=ELAVL4;refGene_tx=XM_011285002.3;refGene_gn=ELAVL4;refGene_tx=XM_011285003.3;refGene_gn=ELAVL4;refGene_tx=XM_019836975.2;refGene_gn=ELAVL4;refGene_tx=XM_019836976.2;refGene_gn=ELAVL4;refGene_tx=XM_019836977.2;refGene_gn=ELAVL4;refGene_tx=XM_023258677.1
C1 40326950 40327054
refGene_gn=ELAVL4;refGene_tx=XM_011285002.3;refGene_gn=ELAVL4;refGene_tx=XM_011285003.3;refGene_gn=ELAVL4;refGene_tx=XM_019836975.2;refGene_gn=ELAVL4;refGene_tx=XM_019836976.2;refGene_gn=ELAVL4;refGene_tx=XM_019836977.2;refGene_gn=ELAVL4;refGene_tx=XM_023258677.1
C1 40342545 40342699
refGene_gn=ELAVL4;refGene_tx=XM_011285002.3;refGene_gn=ELAVL4;refGene_tx=XM_011285003.3;refGene_gn=ELAVL4;refGene_tx=XM_019836975.2;refGene_gn=ELAVL4;refGene_tx=XM_019836976.2;refGene_gn=ELAVL4;refGene_tx=XM_01

9836977.2;refGene_gn=ELAVL4;refGene_tx=XM_023258677.1
C1 40344377 40344603
refGene_gn=ELAVL4;refGene_tx=XM_011285002.3;refGene_gn=ELAVL4;refGene_tx=XM_011285003.3;refGene_gn=ELAVL4;refGene_tx=XM_019836975.2;refGene_gn=ELAVL4;refGene_tx=XM_019836976.2;refGene_gn=ELAVL4;refGene_tx=XM_019836977.2;refGene_gn=ELAVL4;refGene_tx=XM_023258677.1
C1 40346227 40346266
refGene_gn=ELAVL4;refGene_tx=XM_011285002.3;refGene_gn=ELAVL4;refGene_tx=XM_011285003.3;refGene_gn=ELAVL4;refGene_tx=XM_019836975.2;refGene_gn=ELAVL4;refGene_tx=XM_019836976.2;refGene_gn=ELAVL4;refGene_tx=XM_023258677.1
C1 40349465 40349835
refGene_gn=ELAVL4;refGene_tx=XM_011285002.3;refGene_gn=ELAVL4;refGene_tx=XM_011285003.3;refGene_gn=ELAVL4;refGene_tx=XM_019836975.2;refGene_gn=ELAVL4;refGene_tx=XM_019836976.2;refGene_gn=ELAVL4;refGene_tx=XM_019836977.2;refGene_gn=ELAVL4;refGene_tx=XM_023258677.1
C1 40547732 40548784 refGene_gn=DMRTA2;refGene_tx=XM_011285032.3
C1 40550044 40550600 refGene_gn=DMRTA2;refGene_tx=XM_011285032.3
C1 40569402 40569486
refGene_gn=FAF1;refGene_tx=XM_019836980.2;refGene_gn=FAF1;refGene_tx=XM_019836981.2;refGene_gn=FAF1;refGene_tx=XM_023258678.1
C1 40609646 40609862
refGene_gn=FAF1;refGene_tx=XM_019836980.2;refGene_gn=FAF1;refGene_tx=XM_019836981.2;refGene_gn=FAF1;refGene_tx=XM_023258678.1
C1 40624270 40624348
refGene_gn=FAF1;refGene_tx=XM_019836980.2;refGene_gn=FAF1;refGene_tx=XM_019836981.2;refGene_gn=FAF1;refGene_tx=XM_023258678.1
C1 40625326 40625407
refGene_gn=FAF1;refGene_tx=XM_019836980.2;refGene_gn=FAF1;refGene_tx=XM_019836981.2;refGene_gn=FAF1;refGene_tx=XM_023258678.1
C1 40673232 40673321
refGene_gn=FAF1;refGene_tx=XM_019836980.2;refGene_gn=FAF1;refGene_tx=XM_019836981.2;refGene_gn=FAF1;refGene_tx=XM_023258678.1
C1 40678514 40678651
refGene_gn=FAF1;refGene_tx=XM_019836980.2;refGene_gn=FAF1;refGene_tx=XM_019836981.2;refGene_gn=FAF1;refGene_tx=XM_023258678.1
C1 40709812 40709967
refGene_gn=FAF1;refGene_tx=XM_019836980.2;refGene_gn=FAF1;refGene_tx=XM_019836981.2;refGene_gn=FAF1;refGene_tx=XM_023258678.1
C1 40725392 40725474
refGene_gn=FAF1;refGene_tx=XM_019836980.2;refGene_gn=FAF1;refGene_tx=XM_019836981.2;refGene_gn=FAF1;refGene_tx=XM_023258678.1
C1 40726388 40726452
refGene_gn=FAF1;refGene_tx=XM_019836980.2;refGene_gn=FAF1;refGene_tx=XM_019836981.2;refGene_gn=FAF1;refGene_tx=XM_023258678.1
C1 40727404 40727531
refGene_gn=FAF1;refGene_tx=XM_019836980.2;refGene_gn=FAF1;refGene_tx=XM_019836981.2;refGene_gn=FAF1;refGene_tx=XM_023258678.1
C1 40737074 40737170
refGene_gn=FAF1;refGene_tx=XM_019836980.2;refGene_gn=FAF1;refGene_tx=X

M_019836981.2;refGene_gn=FAF1;refGene_tx=XM_023258678.1
C1 40791277 40791364
refGene_gn=FAF1;refGene_tx=XM_019836980.2;refGene_gn=FAF1;refGene_tx=XM_019836981.2;refGene_gn=FAF1;refGene_tx=XM_023258678.1
C1 40836116 40836222
refGene_gn=FAF1;refGene_tx=XM_019836980.2;refGene_gn=FAF1;refGene_tx=XM_019836981.2;refGene_gn=FAF1;refGene_tx=XM_023258678.1
C1 40862626 40862718
refGene_gn=FAF1;refGene_tx=XM_019836980.2;refGene_gn=FAF1;refGene_tx=XM_019836981.2;refGene_gn=FAF1;refGene_tx=XM_023258678.1
C1 40875990 40876082
refGene_gn=FAF1;refGene_tx=XM_019836980.2;refGene_gn=FAF1;refGene_tx=XM_019836981.2;refGene_gn=FAF1;refGene_tx=XM_023258678.1
C1 40918272 40918478
refGene_gn=FAF1;refGene_tx=XM_019836980.2;refGene_gn=FAF1;refGene_tx=XM_019836981.2;refGene_gn=FAF1;refGene_tx=XM_023258678.1
C1 40928939 40928986
refGene_gn=FAF1;refGene_tx=XM_019836980.2;refGene_gn=FAF1;refGene_tx=XM_023258678.1
C1 40935746 40935803 refGene_gn=FAF1;refGene_tx=XM_019836980.2
C1 41017927 41017996 refGene_gn=FAF1;refGene_tx=XM_023258678.1
C1 41093138 41093183 refGene_gn=FAF1;refGene_tx=XM_023258678.1
C1 41101560 41101689
refGene_gn=CDKN2C;refGene_tx=XM_006934733.3;refGene_gn=CDKN2C;refGene_tx=XM_006934734.3
C1 41105130 41105508
refGene_gn=CDKN2C;refGene_tx=XM_006934733.3;refGene_gn=CDKN2C;refGene_tx=XM_006934734.3
C1 41269785 41269846
refGene_gn=CC1H1orf185;refGene_tx=XM_003990089.5
C1 41279025 41279131
refGene_gn=CC1H1orf185;refGene_tx=XM_003990089.5
C1 41291772 41291908
refGene_gn=CC1H1orf185;refGene_tx=XM_003990089.5
C1 41302524 41302561
refGene_gn=CC1H1orf185;refGene_tx=XM_003990089.5
C1 41304624 41305019
refGene_gn=CC1H1orf185;refGene_tx=XM_003990089.5
C1 41357368 41357491 refGene_gn=RNF11;refGene_tx=XM_003990090.5
C1 41393140 41393310 refGene_gn=RNF11;refGene_tx=XM_003990090.5
C1 41394813 41394985 refGene_gn=RNF11;refGene_tx=XM_003990090.5
C1 41404043 41404167
refGene_gn=TTC39A;refGene_tx=XM_019836987.2;refGene_gn=TTC39A;refGene_tx=XM_019836988.2;refGene_gn=TTC39A;refGene_tx=XM_019836989.2;refGene_gn=TTC39A;refGene_tx=XM_019836990.2;refGene_gn=TTC39A;refGene_tx=XM_019836995.1;refGene_gn=TTC39A;refGene_tx=XM_019836996.1;refGene_gn=TTC39A;refGene_tx=XM_023258680.1;refGene_gn=TTC39A;refGene_tx=XM_023258681.1;refGene_gn=TTC39A;refGene_tx=XM_023258682.1;refGene_gn=TTC39A;refGene_tx=XM_023258683.1;refGene_gn=TTC39A;refGene_tx=XM_023258684.1;refGene_gn=TTC39A;refGene_tx=XM_023258686.1;refGene_gn=TTC39A;refGene_tx=XM_

023258687.1
C1	41404696	41404813
refGene_gn=TTC39A;refGene_tx=XM_019836987.2;refGene_gn=TTC39A;refGene_tx=XM_019836988.2;refGene_gn=TTC39A;refGene_tx=XM_019836989.2;refGene_gn=TTC39A;refGene_tx=XM_019836990.2;refGene_gn=TTC39A;refGene_tx=XM_019836995.1;refGene_gn=TTC39A;refGene_tx=XM_019836996.1;refGene_gn=TTC39A;refGene_tx=XM_023258680.1;refGene_gn=TTC39A;refGene_tx=XM_023258681.1;refGene_gn=TTC39A;refGene_tx=XM_023258682.1;refGene_gn=TTC39A;refGene_tx=XM_023258683.1;refGene_gn=TTC39A;refGene_tx=XM_023258684.1;refGene_gn=TTC39A;refGene_tx=XM_023258686.1
C1	41405907	41406022
refGene_gn=TTC39A;refGene_tx=XM_019836987.2;refGene_gn=TTC39A;refGene_tx=XM_019836988.2;refGene_gn=TTC39A;refGene_tx=XM_019836989.2;refGene_gn=TTC39A;refGene_tx=XM_019836990.2;refGene_gn=TTC39A;refGene_tx=XM_019836995.1;refGene_gn=TTC39A;refGene_tx=XM_019836996.1;refGene_gn=TTC39A;refGene_tx=XM_023258680.1;refGene_gn=TTC39A;refGene_tx=XM_023258681.1;refGene_gn=TTC39A;refGene_tx=XM_023258682.1;refGene_gn=TTC39A;refGene_tx=XM_023258683.1;refGene_gn=TTC39A;refGene_tx=XM_023258684.1;refGene_gn=TTC39A;refGene_tx=XM_023258686.1;refGene_gn=TTC39A;refGene_tx=XM_023258687.1
C1	41406426	41406598
refGene_gn=TTC39A;refGene_tx=XM_019836987.2;refGene_gn=TTC39A;refGene_tx=XM_019836988.2;refGene_gn=TTC39A;refGene_tx=XM_019836989.2;refGene_gn=TTC39A;refGene_tx=XM_019836990.2;refGene_gn=TTC39A;refGene_tx=XM_019836995.1;refGene_gn=TTC39A;refGene_tx=XM_019836996.1;refGene_gn=TTC39A;refGene_tx=XM_023258680.1;refGene_gn=TTC39A;refGene_tx=XM_023258681.1;refGene_gn=TTC39A;refGene_tx=XM_023258682.1;refGene_gn=TTC39A;refGene_tx=XM_023258683.1;refGene_gn=TTC39A;refGene_tx=XM_023258684.1;refGene_gn=TTC39A;refGene_tx=XM_023258686.1;refGene_gn=TTC39A;refGene_tx=XM_023258687.1
C1	41408599	41408720
refGene_gn=TTC39A;refGene_tx=XM_019836987.2;refGene_gn=TTC39A;refGene_tx=XM_019836988.2;refGene_gn=TTC39A;refGene_tx=XM_019836989.2;refGene_gn=TTC39A;refGene_tx=XM_019836990.2;refGene_gn=TTC39A;refGene_tx=XM_019836995.1;refGene_gn=TTC39A;refGene_tx=XM_019836996.1;refGene_gn=TTC39A;refGene_tx=XM_023258680.1;refGene_gn=TTC39A;refGene_tx=XM_023258681.1;refGene_gn=TTC39A;refGene_tx=XM_023258682.1;refGene_gn=TTC39A;refGene_tx=XM_023258683.1;refGene_gn=TTC39A;refGene_tx=XM_023258684.1;refGene_gn=TTC39A;refGene_tx=XM_023258686.1;refGene_gn=TTC39A;refGene_tx=XM_023258687.1
C1	41410896	41410988
refGene_gn=TTC39A;refGene_tx=XM_019836987.2;refGene_gn=TTC39A;refGene_tx=XM_019836988.2;refGene_gn=TTC39A;refGene_tx=XM_019836989.2;refGene_gn=TTC39A;refGene_tx=XM_019836990.2;refGene_gn=TTC39A;refGene_tx=XM_019836995.1;refGene_gn=TTC39A;refGene_tx=XM_023258680.1;refGene_gn=TTC39A;refGene_tx=XM_023258681.1;refGene_gn=TTC39A;refGene_tx=XM_023258682.1;refGene_gn=TTC39A;refGene_tx=XM_023258683.1;refGene_gn=TTC39A;refGene_tx=XM_023258684.1;refGene_gn=TTC39A;refGene_tx=XM_023258686.1;refGene_gn=TTC39A;refGene_tx=XM_023258687.1
C1	41415588	41415750

refGene_gn=TTC39A;refGene_tx=XM_019836987.2;refGene_gn=TTC39A;refGene_tx=XM_019836988.2;refGene_gn=TTC39A;refGene_tx=XM_019836989.2;refGene_gn=TTC39A;refGene_tx=XM_019836990.2;refGene_gn=TTC39A;refGene_tx=XM_019836995.1;refGene_gn=TTC39A;refGene_tx=XM_019836996.1;refGene_gn=TTC39A;refGene_tx=XM_023258680.1;refGene_gn=TTC39A;refGene_tx=XM_023258681.1;refGene_gn=TTC39A;refGene_tx=XM_023258682.1;refGene_gn=TTC39A;refGene_tx=XM_023258683.1;refGene_gn=TTC39A;refGene_tx=XM_023258684.1;refGene_gn=TTC39A;refGene_tx=XM_023258686.1;refGene_gn=TTC39A;refGene_tx=XM_023258687.1
C1 41416636 41416696
refGene_gn=TTC39A;refGene_tx=XM_019836987.2;refGene_gn=TTC39A;refGene_tx=XM_019836988.2;refGene_gn=TTC39A;refGene_tx=XM_019836989.2;refGene_gn=TTC39A;refGene_tx=XM_019836990.2;refGene_gn=TTC39A;refGene_tx=XM_019836995.1;refGene_gn=TTC39A;refGene_tx=XM_023258680.1;refGene_gn=TTC39A;refGene_tx=XM_023258681.1;refGene_gn=TTC39A;refGene_tx=XM_023258682.1;refGene_gn=TTC39A;refGene_tx=XM_023258683.1;refGene_gn=TTC39A;refGene_tx=XM_023258684.1;refGene_gn=TTC39A;refGene_tx=XM_023258686.1;refGene_gn=TTC39A;refGene_tx=XM_023258687.1
C1 41416779 41416847
refGene_gn=TTC39A;refGene_tx=XM_019836987.2;refGene_gn=TTC39A;refGene_tx=XM_019836988.2;refGene_gn=TTC39A;refGene_tx=XM_019836989.2;refGene_gn=TTC39A;refGene_tx=XM_019836990.2;refGene_gn=TTC39A;refGene_tx=XM_019836995.1;refGene_gn=TTC39A;refGene_tx=XM_023258680.1;refGene_gn=TTC39A;refGene_tx=XM_023258681.1;refGene_gn=TTC39A;refGene_tx=XM_023258682.1;refGene_gn=TTC39A;refGene_tx=XM_023258683.1;refGene_gn=TTC39A;refGene_tx=XM_023258684.1;refGene_gn=TTC39A;refGene_tx=XM_023258686.1;refGene_gn=TTC39A;refGene_tx=XM_023258687.1
C1 41417367 41417476
refGene_gn=TTC39A;refGene_tx=XM_019836987.2;refGene_gn=TTC39A;refGene_tx=XM_019836988.2;refGene_gn=TTC39A;refGene_tx=XM_019836989.2;refGene_gn=TTC39A;refGene_tx=XM_019836990.2;refGene_gn=TTC39A;refGene_tx=XM_019836995.1;refGene_gn=TTC39A;refGene_tx=XM_023258680.1;refGene_gn=TTC39A;refGene_tx=XM_023258681.1;refGene_gn=TTC39A;refGene_tx=XM_023258682.1;refGene_gn=TTC39A;refGene_tx=XM_023258683.1;refGene_gn=TTC39A;refGene_tx=XM_023258684.1;refGene_gn=TTC39A;refGene_tx=XM_023258686.1;refGene_gn=TTC39A;refGene_tx=XM_023258687.1
C1 41419379 41419445
refGene_gn=TTC39A;refGene_tx=XM_019836987.2;refGene_gn=TTC39A;refGene_tx=XM_019836988.2;refGene_gn=TTC39A;refGene_tx=XM_019836989.2;refGene_gn=TTC39A;refGene_tx=XM_019836990.2;refGene_gn=TTC39A;refGene_tx=XM_019836995.1;refGene_gn=TTC39A;refGene_tx=XM_023258680.1;refGene_gn=TTC39A;refGene_tx=XM_023258681.1;refGene_gn=TTC39A;refGene_tx=XM_023258682.1;refGene_gn=TTC39A;refGene_tx=XM_023258683.1;refGene_gn=TTC39A;refGene_tx=XM_023258684.1;refGene_gn=TTC39A;refGene_tx=XM_023258686.1;refGene_gn=TTC39A;refGene_tx=XM_023258687.1
C1 41420738 41420838
refGene_gn=TTC39A;refGene_tx=XM_019836987.2;refGene_gn=TTC39A;refGene_tx=XM_019836988.2;refGene_gn=TTC39A;refGene_tx=XM_019836989.2;refGene_gn=TTC39A;refGene_tx=XM_019836990.2;refGene_gn=TTC39A;refGene_tx=XM_019836995.1;refGene_gn=TTC39A;refGene_tx=XM_023258680.1;refGene_gn=TTC39

A;refGene_tx=XM_023258681.1;refGene_gn=TTC39A;refGene_tx=XM_023258682.1;refGene_gn=TTC39A;refGene_tx=XM_023258683.1;refGene_gn=TTC39A;refGene_tx=XM_023258684.1;refGene_gn=TTC39A;refGene_tx=XM_023258686.1;refGene_gn=TTC39A;refGene_tx=XM_023258687.1
C1 41422856 41422921
refGene_gn=TTC39A;refGene_tx=XM_019836987.2;refGene_gn=TTC39A;refGene_tx=XM_019836988.2;refGene_gn=TTC39A;refGene_tx=XM_019836989.2;refGene_gn=TTC39A;refGene_tx=XM_019836990.2;refGene_gn=TTC39A;refGene_tx=XM_019836995.1;refGene_gn=TTC39A;refGene_tx=XM_023258680.1;refGene_gn=TTC39A;refGene_tx=XM_023258681.1;refGene_gn=TTC39A;refGene_tx=XM_023258682.1;refGene_gn=TTC39A;refGene_tx=XM_023258683.1;refGene_gn=TTC39A;refGene_tx=XM_023258684.1;refGene_gn=TTC39A;refGene_tx=XM_023258686.1;refGene_gn=TTC39A;refGene_tx=XM_023258687.1
C1 41424571 41424639
refGene_gn=TTC39A;refGene_tx=XM_019836987.2;refGene_gn=TTC39A;refGene_tx=XM_019836988.2;refGene_gn=TTC39A;refGene_tx=XM_019836989.2;refGene_gn=TTC39A;refGene_tx=XM_019836990.2;refGene_gn=TTC39A;refGene_tx=XM_023258680.1;refGene_gn=TTC39A;refGene_tx=XM_023258681.1;refGene_gn=TTC39A;refGene_tx=XM_023258682.1;refGene_gn=TTC39A;refGene_tx=XM_023258683.1;refGene_gn=TTC39A;refGene_tx=XM_023258684.1;refGene_gn=TTC39A;refGene_tx=XM_023258686.1;refGene_gn=TTC39A;refGene_tx=XM_023258687.1
C1 41425804 41425881
refGene_gn=TTC39A;refGene_tx=XM_019836987.2;refGene_gn=TTC39A;refGene_tx=XM_019836988.2;refGene_gn=TTC39A;refGene_tx=XM_019836989.2;refGene_gn=TTC39A;refGene_tx=XM_019836990.2;refGene_gn=TTC39A;refGene_tx=XM_023258680.1;refGene_gn=TTC39A;refGene_tx=XM_023258681.1;refGene_gn=TTC39A;refGene_tx=XM_023258682.1;refGene_gn=TTC39A;refGene_tx=XM_023258683.1;refGene_gn=TTC39A;refGene_tx=XM_023258684.1;refGene_gn=TTC39A;refGene_tx=XM_023258686.1;refGene_gn=TTC39A;refGene_tx=XM_023258687.1
C1 41426697 41426829
refGene_gn=TTC39A;refGene_tx=XM_019836987.2;refGene_gn=TTC39A;refGene_tx=XM_019836988.2;refGene_gn=TTC39A;refGene_tx=XM_019836989.2;refGene_gn=TTC39A;refGene_tx=XM_019836990.2;refGene_gn=TTC39A;refGene_tx=XM_023258680.1;refGene_gn=TTC39A;refGene_tx=XM_023258681.1;refGene_gn=TTC39A;refGene_tx=XM_023258682.1;refGene_gn=TTC39A;refGene_tx=XM_023258683.1;refGene_gn=TTC39A;refGene_tx=XM_023258684.1;refGene_gn=TTC39A;refGene_tx=XM_023258686.1;refGene_gn=TTC39A;refGene_tx=XM_023258687.1
C1 41435432 41435540
refGene_gn=TTC39A;refGene_tx=XM_019836987.2;refGene_gn=TTC39A;refGene_tx=XM_019836988.2;refGene_gn=TTC39A;refGene_tx=XM_019836989.2;refGene_gn=TTC39A;refGene_tx=XM_019836990.2;refGene_gn=TTC39A;refGene_tx=XM_023258680.1;refGene_gn=TTC39A;refGene_tx=XM_023258681.1;refGene_gn=TTC39A;refGene_tx=XM_023258682.1;refGene_gn=TTC39A;refGene_tx=XM_023258683.1;refGene_gn=TTC39A;refGene_tx=XM_023258684.1;refGene_gn=TTC39A;refGene_tx=XM_023258686.1;refGene_gn=TTC39A;refGene_tx=XM_023258687.1
C1 41445432 41445470
refGene_gn=TTC39A;refGene_tx=XM_023258680.1;refGene_gn=TTC39A;refGene_tx=XM_023258681.1;refGene_gn=TTC39A;refGene_tx=XM_023258682.1;refGene_gn=TTC39A;refGene_tx=XM_023258683.1;refGene_gn=TTC39A;refGene_tx=XM_023258687.1

C1 41458810 41459088
refGene_gn=TTC39A;refGene_tx=XM_023258684.1;refGene_gn=TTC39A;refGene_tx=XM_023258686.1
C1 41470465 41470612
refGene_gn=EPS15;refGene_tx=XM_003990092.5;refGene_gn=EPS15;refGene_tx=XM_006934738.3
C1 41472958 41473152 refGene_gn=EPS15;refGene_tx=XM_003990092.5
C1 41477604 41477767
refGene_gn=EPS15;refGene_tx=XM_003990092.5;refGene_gn=EPS15;refGene_tx=XM_006934738.3
C1 41480517 41480594
refGene_gn=EPS15;refGene_tx=XM_003990092.5;refGene_gn=EPS15;refGene_tx=XM_006934738.3
C1 41496736 41496806
refGene_gn=EPS15;refGene_tx=XM_003990092.5;refGene_gn=EPS15;refGene_tx=XM_006934738.3
C1 41502335 41502472
refGene_gn=EPS15;refGene_tx=XM_003990092.5;refGene_gn=EPS15;refGene_tx=XM_006934738.3
C1 41503374 41503410
refGene_gn=EPS15;refGene_tx=XM_003990092.5;refGene_gn=EPS15;refGene_tx=XM_006934738.3
C1 41506047 41506138
refGene_gn=EPS15;refGene_tx=XM_003990092.5;refGene_gn=EPS15;refGene_tx=XM_006934738.3
C1 41506365 41506482
refGene_gn=EPS15;refGene_tx=XM_003990092.5;refGene_gn=EPS15;refGene_tx=XM_006934738.3
C1 41508303 41508507
refGene_gn=EPS15;refGene_tx=XM_003990092.5;refGene_gn=EPS15;refGene_tx=XM_006934738.3
C1 41514787 41514985
refGene_gn=EPS15;refGene_tx=XM_003990092.5;refGene_gn=EPS15;refGene_tx=XM_006934738.3
C1 41515395 41515557
refGene_gn=EPS15;refGene_tx=XM_003990092.5;refGene_gn=EPS15;refGene_tx=XM_006934738.3
C1 41527752 41527825
refGene_gn=EPS15;refGene_tx=XM_003990092.5;refGene_gn=EPS15;refGene_tx=XM_006934738.3
C1 41545016 41545102
refGene_gn=EPS15;refGene_tx=XM_003990092.5;refGene_gn=EPS15;refGene_tx=XM_006934738.3
C1 41549368 41549525
refGene_gn=EPS15;refGene_tx=XM_003990092.5;refGene_gn=EPS15;refGene_tx=XM_006934738.3
C1 41551348 41551494
refGene_gn=EPS15;refGene_tx=XM_003990092.5;refGene_gn=EPS15;refGene_tx=XM_006934738.3
C1 41552509 41552599

refGene_gn=EPS15;refGene_tx=XM_003990092.5;refGene_gn=EPS15;refGene_tx
=XM_006934738.3
C1 41566457 41566517
refGene_gn=EPS15;refGene_tx=XM_003990092.5;refGene_gn=EPS15;refGene_tx
=XM_006934738.3
C1 41569196 41569322
refGene_gn=EPS15;refGene_tx=XM_003990092.5;refGene_gn=EPS15;refGene_tx
=XM_006934738.3
C1 41570508 41570574
refGene_gn=EPS15;refGene_tx=XM_003990092.5;refGene_gn=EPS15;refGene_tx
=XM_006934738.3
C1 41573030 41573126
refGene_gn=EPS15;refGene_tx=XM_003990092.5;refGene_gn=EPS15;refGene_tx
=XM_006934738.3
C1 41576093 41576141
refGene_gn=EPS15;refGene_tx=XM_003990092.5;refGene_gn=EPS15;refGene_tx
=XM_006934738.3
C1 41577023 41577113
refGene_gn=EPS15;refGene_tx=XM_003990092.5;refGene_gn=EPS15;refGene_tx
=XM_006934738.3
C1 41580158 41580200
refGene_gn=EPS15;refGene_tx=XM_003990092.5;refGene_gn=EPS15;refGene_tx
=XM_006934738.3
C1 41615143 41615176
refGene_gn=EPS15;refGene_tx=XM_003990092.5;refGene_gn=EPS15;refGene_tx
=XM_006934738.3
C1 41706020 41706131
refGene_gn=OSBPL9;refGene_tx=XM_011284993.3;refGene_gn=OSBPL9;refGene_
tx=XM_011284994.3;refGene_gn=OSBPL9;refGene_tx=XM_011284995.3;refGene_
gn=OSBPL9;refGene_tx=XM_011284996.3
C1 41748230 41748281
refGene_gn=OSBPL9;refGene_tx=XM_011284993.3;refGene_gn=OSBPL9;refGene_
tx=XM_011284994.3;refGene_gn=OSBPL9;refGene_tx=XM_011284995.3;refGene_
gn=OSBPL9;refGene_tx=XM_011284996.3;refGene_gn=OSBPL9;refGene_tx=XM_01
1284997.2;refGene_gn=OSBPL9;refGene_tx=XM_019836999.1
C1 41781862 41781941
refGene_gn=OSBPL9;refGene_tx=XM_011284993.3;refGene_gn=OSBPL9;refGene_
tx=XM_011284994.3;refGene_gn=OSBPL9;refGene_tx=XM_011284995.3;refGene_
gn=OSBPL9;refGene_tx=XM_011284996.3
C1 41819478 41819555
refGene_gn=OSBPL9;refGene_tx=XM_011284993.3;refGene_gn=OSBPL9;refGene_
tx=XM_011284994.3;refGene_gn=OSBPL9;refGene_tx=XM_011284995.3;refGene_
gn=OSBPL9;refGene_tx=XM_011284996.3;refGene_gn=OSBPL9;refGene_tx=XM_01
1284997.2;refGene_gn=OSBPL9;refGene_tx=XM_019836999.1
C1 41834659 41834686 refGene_gn=OSBPL9;refGene_tx=XM_011284998.3
C1 41844643 41844712
refGene_gn=OSBPL9;refGene_tx=XM_011284993.3;refGene_gn=OSBPL9;refGene_
tx=XM_011284994.3;refGene_gn=OSBPL9;refGene_tx=XM_011284997.2
C1 41850143 41850239
refGene_gn=OSBPL9;refGene_tx=XM_011284993.3;refGene_gn=OSBPL9;refGene_

tx=XM_011284994.3;refGene_gn=OSBPL9;refGene_tx=XM_011284995.3;refGene_
gn=OSBPL9;refGene_tx=XM_011284996.3;refGene_gn=OSBPL9;refGene_tx=XM_01
1284997.2;refGene_gn=OSBPL9;refGene_tx=XM_011284998.3;refGene_gn=OSBPL
9;refGene_tx=XM_019836999.1
C1 41851222 41851270
refGene_gn=OSBPL9;refGene_tx=XM_011284993.3;refGene_gn=OSBPL9;refGene_
tx=XM_011284994.3;refGene_gn=OSBPL9;refGene_tx=XM_011284995.3;refGene_
gn=OSBPL9;refGene_tx=XM_011284996.3;refGene_gn=OSBPL9;refGene_tx=XM_01
1284997.2;refGene_gn=OSBPL9;refGene_tx=XM_011284998.3;refGene_gn=OSBPL
9;refGene_tx=XM_019836999.1
C1 41852315 41852345
refGene_gn=OSBPL9;refGene_tx=XM_011284993.3;refGene_gn=OSBPL9;refGene_
tx=XM_011284994.3;refGene_gn=OSBPL9;refGene_tx=XM_011284995.3;refGene_
gn=OSBPL9;refGene_tx=XM_011284996.3;refGene_gn=OSBPL9;refGene_tx=XM_01
1284997.2;refGene_gn=OSBPL9;refGene_tx=XM_011284998.3;refGene_gn=OSBPL
9;refGene_tx=XM_019836999.1
C1 41853665 41853716
refGene_gn=OSBPL9;refGene_tx=XM_011284993.3;refGene_gn=OSBPL9;refGene_
tx=XM_011284994.3;refGene_gn=OSBPL9;refGene_tx=XM_011284995.3;refGene_
gn=OSBPL9;refGene_tx=XM_011284996.3;refGene_gn=OSBPL9;refGene_tx=XM_01
1284997.2;refGene_gn=OSBPL9;refGene_tx=XM_011284998.3;refGene_gn=OSBPL
9;refGene_tx=XM_019836999.1;refGene_gn=OSBPL9;refGene_tx=XM_019837000.
1
C1 41863808 41863847
refGene_gn=OSBPL9;refGene_tx=XM_011284993.3;refGene_gn=OSBPL9;refGene_
tx=XM_011284995.3;refGene_gn=OSBPL9;refGene_tx=XM_011284997.2;refGene_
gn=OSBPL9;refGene_tx=XM_011284998.3;refGene_gn=OSBPL9;refGene_tx=XM_01
9836999.1;refGene_gn=OSBPL9;refGene_tx=XM_019837000.1
C1 41867568 41867659
refGene_gn=OSBPL9;refGene_tx=XM_011284993.3;refGene_gn=OSBPL9;refGene_
tx=XM_011284994.3;refGene_gn=OSBPL9;refGene_tx=XM_011284995.3;refGene_
gn=OSBPL9;refGene_tx=XM_011284996.3;refGene_gn=OSBPL9;refGene_tx=XM_01
1284997.2;refGene_gn=OSBPL9;refGene_tx=XM_011284998.3;refGene_gn=OSBPL
9;refGene_tx=XM_019836999.1;refGene_gn=OSBPL9;refGene_tx=XM_019837000.
1
C1 41870638 41870743
refGene_gn=OSBPL9;refGene_tx=XM_011284993.3;refGene_gn=OSBPL9;refGene_
tx=XM_011284994.3;refGene_gn=OSBPL9;refGene_tx=XM_011284995.3;refGene_
gn=OSBPL9;refGene_tx=XM_011284996.3;refGene_gn=OSBPL9;refGene_tx=XM_01
1284997.2;refGene_gn=OSBPL9;refGene_tx=XM_011284998.3;refGene_gn=OSBPL
9;refGene_tx=XM_019836999.1;refGene_gn=OSBPL9;refGene_tx=XM_019837000.
1
C1 41876064 41876224
refGene_gn=OSBPL9;refGene_tx=XM_011284993.3;refGene_gn=OSBPL9;refGene_
tx=XM_011284994.3;refGene_gn=OSBPL9;refGene_tx=XM_011284995.3;refGene_
gn=OSBPL9;refGene_tx=XM_011284996.3;refGene_gn=OSBPL9;refGene_tx=XM_01
1284997.2;refGene_gn=OSBPL9;refGene_tx=XM_011284998.3;refGene_gn=OSBPL
9;refGene_tx=XM_019836999.1;refGene_gn=OSBPL9;refGene_tx=XM_019837000.
1
C1 41882062 41882175

refGene_gn=OSBPL9;refGene_tx=XM_011284993.3;refGene_gn=OSBPL9;refGene_tx=XM_011284994.3;refGene_gn=OSBPL9;refGene_tx=XM_011284995.3;refGene_gn=OSBPL9;refGene_tx=XM_011284996.3;refGene_gn=OSBPL9;refGene_tx=XM_011284997.2;refGene_gn=OSBPL9;refGene_tx=XM_011284998.3;refGene_gn=OSBPL9;refGene_tx=XM_019836999.1;refGene_gn=OSBPL9;refGene_tx=XM_019837000.1
C1 41882297 41882416
refGene_gn=OSBPL9;refGene_tx=XM_011284993.3;refGene_gn=OSBPL9;refGene_tx=XM_011284994.3;refGene_gn=OSBPL9;refGene_tx=XM_011284995.3;refGene_gn=OSBPL9;refGene_tx=XM_011284996.3;refGene_gn=OSBPL9;refGene_tx=XM_011284997.2;refGene_gn=OSBPL9;refGene_tx=XM_011284998.3;refGene_gn=OSBPL9;refGene_tx=XM_019836999.1;refGene_gn=OSBPL9;refGene_tx=XM_019837000.1
C1 41887321 41887407
refGene_gn=OSBPL9;refGene_tx=XM_011284993.3;refGene_gn=OSBPL9;refGene_tx=XM_011284994.3;refGene_gn=OSBPL9;refGene_tx=XM_011284995.3;refGene_gn=OSBPL9;refGene_tx=XM_011284996.3;refGene_gn=OSBPL9;refGene_tx=XM_011284997.2;refGene_gn=OSBPL9;refGene_tx=XM_011284998.3;refGene_gn=OSBPL9;refGene_tx=XM_019836999.1;refGene_gn=OSBPL9;refGene_tx=XM_019837000.1
C1 41888626 41888798
refGene_gn=OSBPL9;refGene_tx=XM_011284993.3;refGene_gn=OSBPL9;refGene_tx=XM_011284994.3;refGene_gn=OSBPL9;refGene_tx=XM_011284995.3;refGene_gn=OSBPL9;refGene_tx=XM_011284996.3;refGene_gn=OSBPL9;refGene_tx=XM_011284997.2;refGene_gn=OSBPL9;refGene_tx=XM_011284998.3;refGene_gn=OSBPL9;refGene_tx=XM_019836999.1;refGene_gn=OSBPL9;refGene_tx=XM_019837000.1
C1 41890025 41890110
refGene_gn=OSBPL9;refGene_tx=XM_011284993.3;refGene_gn=OSBPL9;refGene_tx=XM_011284994.3;refGene_gn=OSBPL9;refGene_tx=XM_011284995.3;refGene_gn=OSBPL9;refGene_tx=XM_011284996.3;refGene_gn=OSBPL9;refGene_tx=XM_011284997.2;refGene_gn=OSBPL9;refGene_tx=XM_011284998.3;refGene_gn=OSBPL9;refGene_tx=XM_019836999.1;refGene_gn=OSBPL9;refGene_tx=XM_019837000.1
C1 41890827 41890938
refGene_gn=OSBPL9;refGene_tx=XM_011284993.3;refGene_gn=OSBPL9;refGene_tx=XM_011284994.3;refGene_gn=OSBPL9;refGene_tx=XM_011284995.3;refGene_gn=OSBPL9;refGene_tx=XM_011284996.3;refGene_gn=OSBPL9;refGene_tx=XM_011284997.2;refGene_gn=OSBPL9;refGene_tx=XM_011284998.3;refGene_gn=OSBPL9;refGene_tx=XM_019836999.1;refGene_gn=OSBPL9;refGene_tx=XM_019837000.1
C1 41891211 41891275
refGene_gn=OSBPL9;refGene_tx=XM_011284993.3;refGene_gn=OSBPL9;refGene_tx=XM_011284994.3;refGene_gn=OSBPL9;refGene_tx=XM_011284995.3;refGene_gn=OSBPL9;refGene_tx=XM_011284996.3;refGene_gn=OSBPL9;refGene_tx=XM_011284997.2;refGene_gn=OSBPL9;refGene_tx=XM_011284998.3;refGene_gn=OSBPL9;refGene_tx=XM_019836999.1;refGene_gn=OSBPL9;refGene_tx=XM_019837000.1
C1 41891396 41891537
refGene_gn=OSBPL9;refGene_tx=XM_011284993.3;refGene_gn=OSBPL9;refGene_

tx=XM_011284994.3;refGene_gn=OSBPL9;refGene_tx=XM_011284995.3;refGene_gn=OSBPL9;refGene_tx=XM_011284996.3;refGene_gn=OSBPL9;refGene_tx=XM_011284997.2;refGene_gn=OSBPL9;refGene_tx=XM_011284998.3;refGene_gn=OSBPL9;refGene_tx=XM_019836999.1;refGene_gn=OSBPL9;refGene_tx=XM_019837000.1
C1 41893971 41894050
refGene_gn=OSBPL9;refGene_tx=XM_011284993.3;refGene_gn=OSBPL9;refGene_tx=XM_011284994.3;refGene_gn=OSBPL9;refGene_tx=XM_011284995.3;refGene_gn=OSBPL9;refGene_tx=XM_011284996.3;refGene_gn=OSBPL9;refGene_tx=XM_011284997.2;refGene_gn=OSBPL9;refGene_tx=XM_011284998.3;refGene_gn=OSBPL9;refGene_tx=XM_019836999.1;refGene_gn=OSBPL9;refGene_tx=XM_019837000.1
C1 41894931 41895023
refGene_gn=OSBPL9;refGene_tx=XM_011284993.3;refGene_gn=OSBPL9;refGene_tx=XM_011284994.3;refGene_gn=OSBPL9;refGene_tx=XM_011284995.3;refGene_gn=OSBPL9;refGene_tx=XM_011284996.3;refGene_gn=OSBPL9;refGene_tx=XM_011284997.2;refGene_gn=OSBPL9;refGene_tx=XM_011284998.3;refGene_gn=OSBPL9;refGene_tx=XM_019836999.1;refGene_gn=OSBPL9;refGene_tx=XM_019837000.1
C1 41896739 41896875
refGene_gn=OSBPL9;refGene_tx=XM_011284993.3;refGene_gn=OSBPL9;refGene_tx=XM_011284994.3;refGene_gn=OSBPL9;refGene_tx=XM_011284995.3;refGene_gn=OSBPL9;refGene_tx=XM_011284996.3;refGene_gn=OSBPL9;refGene_tx=XM_011284997.2;refGene_gn=OSBPL9;refGene_tx=XM_011284998.3;refGene_gn=OSBPL9;refGene_tx=XM_019836999.1;refGene_gn=OSBPL9;refGene_tx=XM_019837000.1
C1 41897132 41897207
refGene_gn=OSBPL9;refGene_tx=XM_011284993.3;refGene_gn=OSBPL9;refGene_tx=XM_011284994.3;refGene_gn=OSBPL9;refGene_tx=XM_011284995.3;refGene_gn=OSBPL9;refGene_tx=XM_011284996.3;refGene_gn=OSBPL9;refGene_tx=XM_011284997.2;refGene_gn=OSBPL9;refGene_tx=XM_011284998.3;refGene_gn=OSBPL9;refGene_tx=XM_019836999.1;refGene_gn=OSBPL9;refGene_tx=XM_019837000.1
C1 41898559 41898757
refGene_gn=NRDC;refGene_tx=XM_006934739.4;refGene_gn=NRDC;refGene_tx=XM_006934740.4
C1 41898914 41899004
refGene_gn=NRDC;refGene_tx=XM_006934739.4;refGene_gn=NRDC;refGene_tx=XM_006934740.4
C1 41900106 41900223
refGene_gn=NRDC;refGene_tx=XM_006934739.4;refGene_gn=NRDC;refGene_tx=XM_006934740.4
C1 41900507 41900598
refGene_gn=NRDC;refGene_tx=XM_006934739.4;refGene_gn=NRDC;refGene_tx=XM_006934740.4
C1 41901165 41901249
refGene_gn=NRDC;refGene_tx=XM_006934739.4;refGene_gn=NRDC;refGene_tx=XM_006934740.4
C1 41901681 41901734
refGene_gn=NRDC;refGene_tx=XM_006934739.4;refGene_gn=NRDC;refGene_tx=X

M_006934740.4
C1 41902026 41902074
refGene_gn=NRDC;refGene_tx=XM_006934739.4;refGene_gn=NRDC;refGene_tx=X
M_006934740.4
C1 41904299 41904438
refGene_gn=NRDC;refGene_tx=XM_006934739.4;refGene_gn=NRDC;refGene_tx=X
M_006934740.4
C1 41904691 41904723
refGene_gn=NRDC;refGene_tx=XM_006934739.4;refGene_gn=NRDC;refGene_tx=X
M_006934740.4
C1 41906540 41906703
refGene_gn=NRDC;refGene_tx=XM_006934739.4;refGene_gn=NRDC;refGene_tx=X
M_006934740.4
C1 41908517 41908645
refGene_gn=NRDC;refGene_tx=XM_006934739.4;refGene_gn=NRDC;refGene_tx=X
M_006934740.4
C1 41911419 41911570
refGene_gn=NRDC;refGene_tx=XM_006934739.4;refGene_gn=NRDC;refGene_tx=X
M_006934740.4
C1 41912732 41912784
refGene_gn=NRDC;refGene_tx=XM_006934739.4;refGene_gn=NRDC;refGene_tx=X
M_006934740.4
C1 41914062 41914182
refGene_gn=NRDC;refGene_tx=XM_006934739.4;refGene_gn=NRDC;refGene_tx=X
M_006934740.4
C1 41915908 41915995
refGene_gn=NRDC;refGene_tx=XM_006934739.4;refGene_gn=NRDC;refGene_tx=X
M_006934740.4
C1 41916543 41916667
refGene_gn=NRDC;refGene_tx=XM_006934739.4;refGene_gn=NRDC;refGene_tx=X
M_006934740.4
C1 41918697 41918802
refGene_gn=NRDC;refGene_tx=XM_006934739.4;refGene_gn=NRDC;refGene_tx=X
M_006934740.4
C1 41923794 41923849
refGene_gn=NRDC;refGene_tx=XM_006934739.4;refGene_gn=NRDC;refGene_tx=X
M_006934740.4
C1 41924139 41924198
refGene_gn=NRDC;refGene_tx=XM_006934739.4;refGene_gn=NRDC;refGene_tx=X
M_006934740.4
C1 41924274 41924395
refGene_gn=NRDC;refGene_tx=XM_006934739.4;refGene_gn=NRDC;refGene_tx=X
M_006934740.4
C1 41925703 41925781
refGene_gn=NRDC;refGene_tx=XM_006934739.4;refGene_gn=NRDC;refGene_tx=X
M_006934740.4
C1 41928428 41928498
refGene_gn=NRDC;refGene_tx=XM_006934739.4;refGene_gn=NRDC;refGene_tx=X
M_006934740.4
C1 41929836 41929910

refGene_gn=NRDC;refGene_tx=XM_006934739.4;refGene_gn=NRDC;refGene_tx=XM_006934740.4
C1 41931831 41931889
refGene_gn=NRDC;refGene_tx=XM_006934739.4;refGene_gn=NRDC;refGene_tx=XM_006934740.4
C1 41933696 41933819
refGene_gn=NRDC;refGene_tx=XM_006934739.4;refGene_gn=NRDC;refGene_tx=XM_006934740.4
C1 41935044 41935140
refGene_gn=NRDC;refGene_tx=XM_006934739.4;refGene_gn=NRDC;refGene_tx=XM_006934740.4
C1 41940136 41940210
refGene_gn=NRDC;refGene_tx=XM_006934739.4;refGene_gn=NRDC;refGene_tx=XM_006934740.4
C1 41954090 41954244
refGene_gn=NRDC;refGene_tx=XM_006934739.4;refGene_gn=NRDC;refGene_tx=XM_006934740.4
C1 41955307 41955389
refGene_gn=NRDC;refGene_tx=XM_006934739.4;refGene_gn=NRDC;refGene_tx=XM_006934740.4
C1 41955544 41955614 refGene_gn=NRDC;refGene_tx=XM_006934739.4
C1 41956518 41956649 refGene_gn=NRDC;refGene_tx=XM_006934739.4
C1 41959479 41959789
refGene_gn=NRDC;refGene_tx=XM_006934739.4;refGene_gn=NRDC;refGene_tx=XM_006934740.4
C1 41999868 42000212
refGene_gn=NRDC;refGene_tx=XM_006934739.4;refGene_gn=NRDC;refGene_tx=XM_006934740.4
C1 42040095 42040283
refGene_gn=RAB3B;refGene_tx=XM_011284992.2;refGene_gn=RAB3B;refGene_tx=XM_019837002.2
C1 42053111 42053236
refGene_gn=RAB3B;refGene_tx=XM_011284992.2;refGene_gn=RAB3B;refGene_tx=XM_019837002.2
C1 42056150 42056269
refGene_gn=RAB3B;refGene_tx=XM_011284992.2;refGene_gn=RAB3B;refGene_tx=XM_019837002.2
C1 42095106 42095334
refGene_gn=RAB3B;refGene_tx=XM_011284992.2;refGene_gn=RAB3B;refGene_tx=XM_019837002.2
C1 42132407 42132487 refGene_gn=TXNDC12;refGene_tx=XM_003990099.4
C1 42134426 42134510 refGene_gn=TXNDC12;refGene_tx=XM_003990099.4
C1 42135475 42135545 refGene_gn=TXNDC12;refGene_tx=XM_003990099.4
C1 42136418 42136492 refGene_gn=TXNDC12;refGene_tx=XM_003990099.4
C1 42137606 42137659 refGene_gn=TXNDC12;refGene_tx=XM_003990099.4
C1 42140914 42141979 refGene_gn=KTI12;refGene_tx=XM_003990100.5
C1 42145081 42145142 refGene_gn=TXNDC12;refGene_tx=XM_003990099.4
C1 42158818 42158915 refGene_gn=TXNDC12;refGene_tx=XM_003990099.4
C1 42163604 42163658
refGene_gn=BTF3L4;refGene_tx=XM_011284990.3;refGene_gn=BTF3L4;refGene_

tx=XM_019837005.2;refGene_gn=BTF3L4;refGene_tx=XM_019837006.2
C1 42166966 42167080
refGene_gn=BTF3L4;refGene_tx=XM_011284990.3;refGene_gn=BTF3L4;refGene_
tx=XM_019837005.2;refGene_gn=BTF3L4;refGene_tx=XM_019837006.2
C1 42179553 42179755
refGene_gn=BTF3L4;refGene_tx=XM_011284990.3;refGene_gn=BTF3L4;refGene_
tx=XM_019837005.2;refGene_gn=BTF3L4;refGene_tx=XM_019837006.2
C1 42180589 42180649
refGene_gn=BTF3L4;refGene_tx=XM_011284990.3;refGene_gn=BTF3L4;refGene_
tx=XM_019837005.2;refGene_gn=BTF3L4;refGene_tx=XM_019837006.2
C1 42181350 42181397
refGene_gn=BTF3L4;refGene_tx=XM_011284990.3;refGene_gn=BTF3L4;refGene_
tx=XM_019837005.2;refGene_gn=BTF3L4;refGene_tx=XM_019837006.2
C1 42314480 42314550
refGene_gn=ZFYVE9;refGene_tx=XM_011284987.3;refGene_gn=ZFYVE9;refGene_
tx=XM_011284988.3;refGene_gn=ZFYVE9;refGene_tx=XM_011284989.3
C1 42317243 42319348
refGene_gn=ZFYVE9;refGene_tx=XM_011284987.3;refGene_gn=ZFYVE9;refGene_
tx=XM_011284988.3;refGene_gn=ZFYVE9;refGene_tx=XM_011284989.3
C1 42349267 42349367
refGene_gn=ZFYVE9;refGene_tx=XM_011284987.3;refGene_gn=ZFYVE9;refGene_
tx=XM_011284988.3;refGene_gn=ZFYVE9;refGene_tx=XM_011284989.3
C1 42351827 42352004
refGene_gn=ZFYVE9;refGene_tx=XM_011284987.3;refGene_gn=ZFYVE9;refGene_
tx=XM_011284988.3
C1 42353626 42353796
refGene_gn=ZFYVE9;refGene_tx=XM_011284987.3;refGene_gn=ZFYVE9;refGene_
tx=XM_011284988.3;refGene_gn=ZFYVE9;refGene_tx=XM_011284989.3
C1 42356715 42356836
refGene_gn=ZFYVE9;refGene_tx=XM_011284987.3;refGene_gn=ZFYVE9;refGene_
tx=XM_011284988.3;refGene_gn=ZFYVE9;refGene_tx=XM_011284989.3
C1 42362657 42362780
refGene_gn=ZFYVE9;refGene_tx=XM_011284987.3;refGene_gn=ZFYVE9;refGene_
tx=XM_011284988.3;refGene_gn=ZFYVE9;refGene_tx=XM_011284989.3
C1 42364692 42364848
refGene_gn=ZFYVE9;refGene_tx=XM_011284987.3;refGene_gn=ZFYVE9;refGene_
tx=XM_011284988.3;refGene_gn=ZFYVE9;refGene_tx=XM_011284989.3
C1 42380914 42381139
refGene_gn=ZFYVE9;refGene_tx=XM_011284987.3;refGene_gn=ZFYVE9;refGene_
tx=XM_011284988.3;refGene_gn=ZFYVE9;refGene_tx=XM_011284989.3
C1 42382673 42382756
refGene_gn=ZFYVE9;refGene_tx=XM_011284987.3;refGene_gn=ZFYVE9;refGene_
tx=XM_011284988.3;refGene_gn=ZFYVE9;refGene_tx=XM_011284989.3
C1 42391826 42391931
refGene_gn=ZFYVE9;refGene_tx=XM_011284987.3;refGene_gn=ZFYVE9;refGene_
tx=XM_011284988.3;refGene_gn=ZFYVE9;refGene_tx=XM_011284989.3
C1 42424318 42424469
refGene_gn=ZFYVE9;refGene_tx=XM_011284987.3;refGene_gn=ZFYVE9;refGene_
tx=XM_011284988.3;refGene_gn=ZFYVE9;refGene_tx=XM_011284989.3
C1 42426945 42427026

refGene_gn=ZFYVE9;refGene_tx=XM_011284987.3;refGene_gn=ZFYVE9;refGene_tx=XM_011284988.3;refGene_gn=ZFYVE9;refGene_tx=XM_011284989.3
C1 42429727 42429890
refGene_gn=ZFYVE9;refGene_tx=XM_011284987.3;refGene_gn=ZFYVE9;refGene_tx=XM_011284988.3;refGene_gn=ZFYVE9;refGene_tx=XM_011284989.3
C1 42432943 42433049
refGene_gn=ZFYVE9;refGene_tx=XM_011284987.3;refGene_gn=ZFYVE9;refGene_tx=XM_011284988.3;refGene_gn=ZFYVE9;refGene_tx=XM_011284989.3
C1 42435698 42435875
refGene_gn=ZFYVE9;refGene_tx=XM_011284987.3;refGene_gn=ZFYVE9;refGene_tx=XM_011284988.3;refGene_gn=ZFYVE9;refGene_tx=XM_011284989.3
C1 42437512 42437674
refGene_gn=ZFYVE9;refGene_tx=XM_011284987.3;refGene_gn=ZFYVE9;refGene_tx=XM_011284988.3;refGene_gn=ZFYVE9;refGene_tx=XM_011284989.3
C1 42444033 42444162
refGene_gn=CC2D1B;refGene_tx=XM_006934741.4;refGene_gn=CC2D1B;refGene_tx=XM_019837008.2;refGene_gn=CC2D1B;refGene_tx=XM_019837009.2;refGene_gn=CC2D1B;refGene_tx=XM_019837010.2
C1 42445373 42445464
refGene_gn=CC2D1B;refGene_tx=XM_006934741.4;refGene_gn=CC2D1B;refGene_tx=XM_019837008.2;refGene_gn=CC2D1B;refGene_tx=XM_019837009.2;refGene_gn=CC2D1B;refGene_tx=XM_019837010.2
C1 42445628 42445728
refGene_gn=CC2D1B;refGene_tx=XM_006934741.4;refGene_gn=CC2D1B;refGene_tx=XM_019837008.2;refGene_gn=CC2D1B;refGene_tx=XM_019837009.2;refGene_gn=CC2D1B;refGene_tx=XM_019837010.2
C1 42446134 42446186
refGene_gn=CC2D1B;refGene_tx=XM_006934741.4;refGene_gn=CC2D1B;refGene_tx=XM_019837008.2;refGene_gn=CC2D1B;refGene_tx=XM_019837009.2;refGene_gn=CC2D1B;refGene_tx=XM_019837010.2
C1 42446415 42446474
refGene_gn=CC2D1B;refGene_tx=XM_006934741.4;refGene_gn=CC2D1B;refGene_tx=XM_019837008.2;refGene_gn=CC2D1B;refGene_tx=XM_019837009.2;refGene_gn=CC2D1B;refGene_tx=XM_019837010.2
C1 42446619 42446693
refGene_gn=CC2D1B;refGene_tx=XM_006934741.4;refGene_gn=CC2D1B;refGene_tx=XM_019837008.2;refGene_gn=CC2D1B;refGene_tx=XM_019837009.2;refGene_gn=CC2D1B;refGene_tx=XM_019837010.2
C1 42447068 42447185
refGene_gn=CC2D1B;refGene_tx=XM_006934741.4;refGene_gn=CC2D1B;refGene_tx=XM_019837008.2;refGene_gn=CC2D1B;refGene_tx=XM_019837009.2
C1 42447273 42447332
refGene_gn=CC2D1B;refGene_tx=XM_006934741.4;refGene_gn=CC2D1B;refGene_tx=XM_019837008.2;refGene_gn=CC2D1B;refGene_tx=XM_019837009.2;refGene_gn=CC2D1B;refGene_tx=XM_019837010.2
C1 42447836 42447962
refGene_gn=CC2D1B;refGene_tx=XM_006934741.4;refGene_gn=CC2D1B;refGene_tx=XM_019837008.2;refGene_gn=CC2D1B;refGene_tx=XM_019837009.2;refGene_gn=CC2D1B;refGene_tx=XM_019837010.2
C1 42448357 42448533

refGene_gn=CC2D1B;refGene_tx=XM_006934741.4;refGene_gn=CC2D1B;refGene_tx=XM_019837008.2;refGene_gn=CC2D1B;refGene_tx=XM_019837009.2;refGene_gn=CC2D1B;refGene_tx=XM_019837010.2
C1 42448617 42448753
refGene_gn=CC2D1B;refGene_tx=XM_006934741.4;refGene_gn=CC2D1B;refGene_tx=XM_019837008.2;refGene_gn=CC2D1B;refGene_tx=XM_019837009.2;refGene_gn=CC2D1B;refGene_tx=XM_019837010.2
C1 42449036 42449173
refGene_gn=CC2D1B;refGene_tx=XM_006934741.4;refGene_gn=CC2D1B;refGene_tx=XM_019837008.2;refGene_gn=CC2D1B;refGene_tx=XM_019837009.2;refGene_gn=CC2D1B;refGene_tx=XM_019837010.2
C1 42449375 42449448
refGene_gn=CC2D1B;refGene_tx=XM_006934741.4;refGene_gn=CC2D1B;refGene_tx=XM_019837008.2;refGene_gn=CC2D1B;refGene_tx=XM_019837009.2;refGene_gn=CC2D1B;refGene_tx=XM_019837010.2
C1 42449661 42449792
refGene_gn=CC2D1B;refGene_tx=XM_006934741.4;refGene_gn=CC2D1B;refGene_tx=XM_019837008.2;refGene_gn=CC2D1B;refGene_tx=XM_019837009.2;refGene_gn=CC2D1B;refGene_tx=XM_019837010.2
C1 42449874 42449985
refGene_gn=CC2D1B;refGene_tx=XM_006934741.4;refGene_gn=CC2D1B;refGene_tx=XM_019837008.2;refGene_gn=CC2D1B;refGene_tx=XM_019837009.2;refGene_gn=CC2D1B;refGene_tx=XM_019837010.2
C1 42450095 42450171
refGene_gn=CC2D1B;refGene_tx=XM_006934741.4;refGene_gn=CC2D1B;refGene_tx=XM_019837008.2;refGene_gn=CC2D1B;refGene_tx=XM_019837009.2;refGene_gn=CC2D1B;refGene_tx=XM_019837010.2
C1 42450332 42450514
refGene_gn=CC2D1B;refGene_tx=XM_006934741.4;refGene_gn=CC2D1B;refGene_tx=XM_019837008.2;refGene_gn=CC2D1B;refGene_tx=XM_019837009.2;refGene_gn=CC2D1B;refGene_tx=XM_019837010.2
C1 42450725 42450885
refGene_gn=CC2D1B;refGene_tx=XM_006934741.4;refGene_gn=CC2D1B;refGene_tx=XM_019837008.2;refGene_gn=CC2D1B;refGene_tx=XM_019837009.2;refGene_gn=CC2D1B;refGene_tx=XM_019837010.2
C1 42451059 42451185
refGene_gn=CC2D1B;refGene_tx=XM_006934741.4;refGene_gn=CC2D1B;refGene_tx=XM_019837008.2;refGene_gn=CC2D1B;refGene_tx=XM_019837009.2;refGene_gn=CC2D1B;refGene_tx=XM_019837010.2
C1 42451613 42451772
refGene_gn=CC2D1B;refGene_tx=XM_006934741.4;refGene_gn=CC2D1B;refGene_tx=XM_019837008.2;refGene_gn=CC2D1B;refGene_tx=XM_019837009.2;refGene_gn=CC2D1B;refGene_tx=XM_019837010.2
C1 42452134 42452241
refGene_gn=CC2D1B;refGene_tx=XM_006934741.4;refGene_gn=CC2D1B;refGene_tx=XM_019837008.2;refGene_gn=CC2D1B;refGene_tx=XM_019837009.2;refGene_gn=CC2D1B;refGene_tx=XM_019837010.2
C1 42453031 42453176
refGene_gn=CC2D1B;refGene_tx=XM_006934741.4;refGene_gn=CC2D1B;refGene_tx=XM_019837008.2;refGene_gn=CC2D1B;refGene_tx=XM_019837009.2;refGene_

gn=CC2D1B;refGene_tx=XM_019837010.2
C1 42454461 42454527
refGene_gn=CC2D1B;refGene_tx=XM_006934741.4;refGene_gn=CC2D1B;refGene_tx=XM_019837008.2;refGene_gn=CC2D1B;refGene_tx=XM_019837009.2;refGene_gn=CC2D1B;refGene_tx=XM_019837010.2
C1 42458881 42459073 refGene_gn=ORC1;refGene_tx=XM_023258689.1
C1 42460788 42460876 refGene_gn=ORC1;refGene_tx=XM_023258689.1
C1 42462103 42462273 refGene_gn=ORC1;refGene_tx=XM_023258689.1
C1 42466329 42466449 refGene_gn=ORC1;refGene_tx=XM_023258689.1
C1 42467622 42467772 refGene_gn=ORC1;refGene_tx=XM_023258689.1
C1 42468059 42468167 refGene_gn=ORC1;refGene_tx=XM_023258689.1
C1 42468767 42468939 refGene_gn=ORC1;refGene_tx=XM_023258689.1
C1 42469346 42469448 refGene_gn=ORC1;refGene_tx=XM_023258689.1
C1 42470292 42470390 refGene_gn=ORC1;refGene_tx=XM_023258689.1
C1 42472827 42473023 refGene_gn=ORC1;refGene_tx=XM_023258689.1
C1 42473603 42473708 refGene_gn=ORC1;refGene_tx=XM_023258689.1
C1 42474803 42475072 refGene_gn=ORC1;refGene_tx=XM_023258689.1
C1 42475073 42475171 refGene_gn=ORC1;refGene_tx=XM_023258689.1
C1 42479992 42480311 refGene_gn=ORC1;refGene_tx=XM_023258689.1
C1 42481738 42481917 refGene_gn=ORC1;refGene_tx=XM_023258689.1
C1 42483919 42484047 refGene_gn=ORC1;refGene_tx=XM_023258689.1
C1 42484683 42484778 refGene_gn=ORC1;refGene_tx=XM_023258689.1
C1 42491497 42491627
refGene_gn=PRPF38A;refGene_tx=XM_003990104.5;refGene_gn=PRPF38A;refGene_tx=XM_006934744.4
C1 42493925 42494085
refGene_gn=PRPF38A;refGene_tx=XM_003990104.5;refGene_gn=PRPF38A;refGene_tx=XM_006934744.4
C1 42495794 42495916
refGene_gn=PRPF38A;refGene_tx=XM_003990104.5;refGene_gn=PRPF38A;refGene_tx=XM_006934744.4
C1 42497659 42497745
refGene_gn=PRPF38A;refGene_tx=XM_003990104.5;refGene_gn=PRPF38A;refGene_tx=XM_006934744.4
C1 42500619 42500730
refGene_gn=PRPF38A;refGene_tx=XM_003990104.5;refGene_gn=PRPF38A;refGene_tx=XM_006934744.4
C1 42503536 42503649
refGene_gn=PRPF38A;refGene_tx=XM_003990104.5;refGene_gn=PRPF38A;refGene_tx=XM_006934744.4
C1 42504255 42504282
refGene_gn=PRPF38A;refGene_tx=XM_003990104.5;refGene_gn=PRPF38A;refGene_tx=XM_006934744.4
C1 42504378 42504488
refGene_gn=PRPF38A;refGene_tx=XM_003990104.5;refGene_gn=PRPF38A;refGene_tx=XM_006934744.4
C1 42504966 42505015
refGene_gn=PRPF38A;refGene_tx=XM_003990104.5;refGene_gn=PRPF38A;refGene_tx=XM_006934744.4
C1 42505829 42505872

refGene_gn=PRPF38A;refGene_tx=XM_003990104.5;refGene_gn=PRPF38A;refGene_tx=XM_006934744.4
C1 42514809 42514877 refGene_gn=ZCCHC11;refGene_tx=XM_023258690.1
C1 42516235 42516394 refGene_gn=ZCCHC11;refGene_tx=XM_023258690.1
C1 42521202 42521650 refGene_gn=ZCCHC11;refGene_tx=XM_023258690.1
C1 42524633 42524734 refGene_gn=ZCCHC11;refGene_tx=XM_023258690.1
C1 42525484 42525708 refGene_gn=ZCCHC11;refGene_tx=XM_023258690.1
C1 42526626 42526742 refGene_gn=ZCCHC11;refGene_tx=XM_023258690.1
C1 42534285 42534368 refGene_gn=ZCCHC11;refGene_tx=XM_023258690.1
C1 42534457 42534505 refGene_gn=ZCCHC11;refGene_tx=XM_023258690.1
C1 42534767 42534945 refGene_gn=ZCCHC11;refGene_tx=XM_023258690.1
C1 42535079 42535157 refGene_gn=ZCCHC11;refGene_tx=XM_023258690.1
C1 42540232 42540346 refGene_gn=ZCCHC11;refGene_tx=XM_023258690.1
C1 42543270 42543360 refGene_gn=ZCCHC11;refGene_tx=XM_023258690.1
C1 42543643 42543747 refGene_gn=ZCCHC11;refGene_tx=XM_023258690.1
C1 42543841 42543899 refGene_gn=ZCCHC11;refGene_tx=XM_023258690.1
C1 42546818 42546922 refGene_gn=ZCCHC11;refGene_tx=XM_023258690.1
C1 42548862 42548949 refGene_gn=ZCCHC11;refGene_tx=XM_023258690.1
C1 42561771 42561923 refGene_gn=ZCCHC11;refGene_tx=XM_023258690.1
C1 42565622 42566323 refGene_gn=ZCCHC11;refGene_tx=XM_023258690.1
C1 42569239 42569414 refGene_gn=ZCCHC11;refGene_tx=XM_023258690.1
C1 42573359 42573572 refGene_gn=ZCCHC11;refGene_tx=XM_023258690.1
C1 42573741 42573861 refGene_gn=ZCCHC11;refGene_tx=XM_023258690.1
C1 42583073 42583200 refGene_gn=ZCCHC11;refGene_tx=XM_023258690.1
C1 42585037 42585107 refGene_gn=ZCCHC11;refGene_tx=XM_023258690.1
C1 42587509 42587561 refGene_gn=ZCCHC11;refGene_tx=XM_023258690.1
C1 42588646 42588735 refGene_gn=ZCCHC11;refGene_tx=XM_023258690.1
C1 42590694 42590872 refGene_gn=ZCCHC11;refGene_tx=XM_023258690.1
C1 42608152 42608269 refGene_gn=ZCCHC11;refGene_tx=XM_023258690.1
C1 42613762 42613926 refGene_gn=ZCCHC11;refGene_tx=XM_023258690.1
C1 42622300 42623060 refGene_gn=ZCCHC11;refGene_tx=XM_023258690.1
C1 42662344 42662635 refGene_gn=LOC101091387;refGene_tx=XM_003990105.1
C1 42707381 42707516 refGene_gn=GPX7;refGene_tx=XM_003990106.4
C1 42712696 42712958 refGene_gn=GPX7;refGene_tx=XM_003990106.4
C1 42714504 42714668 refGene_gn=GPX7;refGene_tx=XM_003990106.4
C1 42727787 42727969 refGene_gn=SHISAL2A;refGene_tx=XM_011285030.3
C1 42733377 42733517 refGene_gn=SHISAL2A;refGene_tx=XM_011285030.3
C1 42743152 42743406 refGene_gn=SHISAL2A;refGene_tx=XM_011285030.3
C1 42769243 42769692 refGene_gn=LOC101091887;refGene_tx=XM_003990107.4
C1 42782652 42782793 refGene_gn=LOC101091887;refGene_tx=XM_003990107.4
C1 42786229 42786335 refGene_gn=LOC101091887;refGene_tx=XM_003990107.4
C1 42818186 42818189 refGene_gn=ZYG11B;refGene_tx=XM_006934745.4
C1 42844362 42844528 refGene_gn=ZYG11B;refGene_tx=XM_006934745.4
C1 42855446 42856201 refGene_gn=ZYG11B;refGene_tx=XM_006934745.4
C1 42864626 42864767 refGene_gn=ZYG11B;refGene_tx=XM_006934745.4
C1 42867414 42867591 refGene_gn=ZYG11B;refGene_tx=XM_006934745.4

C1 42868945 42869010 refGene_gn=ZYG11B;refGene_tx=XM_006934745.4
C1 42872823 42872923 refGene_gn=ZYG11B;refGene_tx=XM_006934745.4
C1 42873423 42873474 refGene_gn=ZYG11B;refGene_tx=XM_006934745.4
C1 42875928 42876090 refGene_gn=ZYG11B;refGene_tx=XM_006934745.4
C1 42876195 42876243 refGene_gn=ZYG11B;refGene_tx=XM_006934745.4
C1 42879933 42880131 refGene_gn=ZYG11B;refGene_tx=XM_006934745.4
C1 42880264 42880317 refGene_gn=ZYG11B;refGene_tx=XM_006934745.4
C1 42886523 42886621 refGene_gn=ZYG11B;refGene_tx=XM_006934745.4
C1 42906905 42907096 refGene_gn=ZYG11B;refGene_tx=XM_006934745.4
C1 42920886 42920976 refGene_gn=ZYG11A;refGene_tx=XM_003990190.4
C1 42935261 42935427 refGene_gn=ZYG11A;refGene_tx=XM_003990190.4
C1 42951226 42951978 refGene_gn=ZYG11A;refGene_tx=XM_003990190.4
C1 42967372 42967513 refGene_gn=ZYG11A;refGene_tx=XM_003990190.4
C1 42970051 42970228 refGene_gn=ZYG11A;refGene_tx=XM_003990190.4
C1 42972340 42972405 refGene_gn=ZYG11A;refGene_tx=XM_003990190.4
C1 42978617 42978717 refGene_gn=ZYG11A;refGene_tx=XM_003990190.4
C1 42978813 42978867 refGene_gn=ZYG11A;refGene_tx=XM_003990190.4
C1 42983578 42983740 refGene_gn=ZYG11A;refGene_tx=XM_003990190.4
C1 42983815 42983860 refGene_gn=ZYG11A;refGene_tx=XM_003990190.4
C1 42988836 42989031 refGene_gn=ZYG11A;refGene_tx=XM_003990190.4
C1 42992448 42992510 refGene_gn=ZYG11A;refGene_tx=XM_003990190.4
C1 42992902 42993000 refGene_gn=ZYG11A;refGene_tx=XM_003990190.4
C1 42997383 42997547 refGene_gn=ZYG11A;refGene_tx=XM_003990190.4
C1 43000371 43000449
refGene_gn=ECHDC2;refGene_tx=XM_003990108.4;refGene_gn=ECHDC2;refGene_tx=XM_006934746.3;refGene_gn=ECHDC2;refGene_tx=XM_006934747.3;refGene_gn=ECHDC2;refGene_tx=XM_006934748.4;refGene_gn=ECHDC2;refGene_tx=XM_006934749.4;refGene_gn=ECHDC2;refGene_tx=XM_019837018.1;refGene_gn=ECHDC2;refGene_tx=XM_019837019.2
C1 43001324 43001372
refGene_gn=ECHDC2;refGene_tx=XM_003990108.4;refGene_gn=ECHDC2;refGene_tx=XM_006934746.3;refGene_gn=ECHDC2;refGene_tx=XM_006934747.3;refGene_gn=ECHDC2;refGene_tx=XM_006934748.4;refGene_gn=ECHDC2;refGene_tx=XM_006934749.4;refGene_gn=ECHDC2;refGene_tx=XM_019837018.1;refGene_gn=ECHDC2;refGene_tx=XM_019837019.2
C1 43002869 43002920
refGene_gn=ECHDC2;refGene_tx=XM_003990108.4;refGene_gn=ECHDC2;refGene_tx=XM_006934746.3;refGene_gn=ECHDC2;refGene_tx=XM_006934747.3;refGene_gn=ECHDC2;refGene_tx=XM_006934748.4;refGene_gn=ECHDC2;refGene_tx=XM_006934749.4;refGene_gn=ECHDC2;refGene_tx=XM_019837019.2
C1 43006633 43006821
refGene_gn=ECHDC2;refGene_tx=XM_003990108.4;refGene_gn=ECHDC2;refGene_tx=XM_006934746.3;refGene_gn=ECHDC2;refGene_tx=XM_006934747.3;refGene_gn=ECHDC2;refGene_tx=XM_006934748.4;refGene_gn=ECHDC2;refGene_tx=XM_006934749.4;refGene_gn=ECHDC2;refGene_tx=XM_019837018.1;refGene_gn=ECHDC2;refGene_tx=XM_019837019.2
C1 43007038 43007098
refGene_gn=ECHDC2;refGene_tx=XM_003990108.4;refGene_gn=ECHDC2;refGene_tx=XM_006934746.3;refGene_gn=ECHDC2;refGene_tx=XM_006934747.3;refGene_gn=ECHDC2;refGene_tx=XM_006934748.4;refGene_gn=ECHDC2;refGene_tx=XM_01

9837018.1
C1 43008529 43008622
refGene_gn=ECHDC2;refGene_tx=XM_003990108.4;refGene_gn=ECHDC2;refGene_tx=XM_006934746.3;refGene_gn=ECHDC2;refGene_tx=XM_006934747.3;refGene_gn=ECHDC2;refGene_tx=XM_006934748.4;refGene_gn=ECHDC2;refGene_tx=XM_006934749.4;refGene_gn=ECHDC2;refGene_tx=XM_019837018.1;refGene_gn=ECHDC2;refGene_tx=XM_019837019.2
C1 43009454 43009541
refGene_gn=ECHDC2;refGene_tx=XM_003990108.4;refGene_gn=ECHDC2;refGene_tx=XM_006934746.3;refGene_gn=ECHDC2;refGene_tx=XM_006934747.3;refGene_gn=ECHDC2;refGene_tx=XM_006934748.4;refGene_gn=ECHDC2;refGene_tx=XM_006934749.4;refGene_gn=ECHDC2;refGene_tx=XM_019837018.1;refGene_gn=ECHDC2;refGene_tx=XM_019837019.2
C1 43012460 43012548
refGene_gn=ECHDC2;refGene_tx=XM_003990108.4;refGene_gn=ECHDC2;refGene_tx=XM_006934746.3;refGene_gn=ECHDC2;refGene_tx=XM_006934747.3;refGene_gn=ECHDC2;refGene_tx=XM_006934748.4;refGene_gn=ECHDC2;refGene_tx=XM_006934749.4;refGene_gn=ECHDC2;refGene_tx=XM_019837018.1;refGene_gn=ECHDC2;refGene_tx=XM_019837019.2
C1 43012628 43012696
refGene_gn=ECHDC2;refGene_tx=XM_003990108.4;refGene_gn=ECHDC2;refGene_tx=XM_006934746.3;refGene_gn=ECHDC2;refGene_tx=XM_006934747.3;refGene_gn=ECHDC2;refGene_tx=XM_006934748.4;refGene_gn=ECHDC2;refGene_tx=XM_006934749.4;refGene_gn=ECHDC2;refGene_tx=XM_019837018.1;refGene_gn=ECHDC2;refGene_tx=XM_019837019.2
C1 43018280 43018413
refGene_gn=ECHDC2;refGene_tx=XM_003990108.4;refGene_gn=ECHDC2;refGene_tx=XM_006934746.3;refGene_gn=ECHDC2;refGene_tx=XM_006934747.3;refGene_gn=ECHDC2;refGene_tx=XM_006934748.4;refGene_gn=ECHDC2;refGene_tx=XM_006934749.4;refGene_gn=ECHDC2;refGene_tx=XM_019837018.1;refGene_gn=ECHDC2;refGene_tx=XM_019837019.2
C1 43022683 43022752 refGene_gn=SCP2;refGene_tx=XM_011284984.3
C1 43030507 43030565 refGene_gn=SCP2;refGene_tx=XM_011284984.3
C1 43031188 43031260 refGene_gn=SCP2;refGene_tx=XM_011284984.3
C1 43035034 43035166 refGene_gn=SCP2;refGene_tx=XM_011284984.3
C1 43038079 43038156 refGene_gn=SCP2;refGene_tx=XM_011284984.3
C1 43050109 43050236 refGene_gn=SCP2;refGene_tx=XM_011284984.3
C1 43062152 43062216 refGene_gn=SCP2;refGene_tx=XM_011284984.3
C1 43067766 43067853 refGene_gn=SCP2;refGene_tx=XM_011284984.3
C1 43069227 43069378 refGene_gn=SCP2;refGene_tx=XM_011284984.3
C1 43070995 43071143 refGene_gn=SCP2;refGene_tx=XM_011284984.3
C1 43080711 43080819 refGene_gn=SCP2;refGene_tx=XM_011284984.3
C1 43110475 43110629 refGene_gn=SCP2;refGene_tx=XM_011284984.3
C1 43133504 43133598 refGene_gn=SCP2;refGene_tx=XM_011284984.3
C1 43139000 43139130 refGene_gn=SCP2;refGene_tx=XM_011284984.3
C1 43150763 43150843 refGene_gn=SCP2;refGene_tx=XM_011284984.3
C1 43154038 43154134 refGene_gn=SCP2;refGene_tx=XM_011284984.3
C1 43165494 43165584
refGene_gn=PODN;refGene_tx=XM_023258691.1;refGene_gn=PODN;refGene_tx=XM_023258692.1

C1 43172500 43172842
refGene_gn=PODN;refGene_tx=XM_006934751.4;refGene_gn=PODN;refGene_tx=XM_023258691.1;refGene_gn=PODN;refGene_tx=XM_023258692.1;refGene_gn=PODN;refGene_tx=XM_023258693.1
C1 43174081 43174175
refGene_gn=PODN;refGene_tx=XM_006934751.4;refGene_gn=PODN;refGene_tx=XM_023258691.1;refGene_gn=PODN;refGene_tx=XM_023258692.1;refGene_gn=PODN;refGene_tx=XM_023258693.1
C1 43175731 43175796
refGene_gn=PODN;refGene_tx=XM_006934751.4;refGene_gn=PODN;refGene_tx=XM_023258691.1;refGene_gn=PODN;refGene_tx=XM_023258692.1;refGene_gn=PODN;refGene_tx=XM_023258693.1
C1 43177371 43177481
refGene_gn=PODN;refGene_tx=XM_006934751.4;refGene_gn=PODN;refGene_tx=XM_023258691.1;refGene_gn=PODN;refGene_tx=XM_023258692.1;refGene_gn=PODN;refGene_tx=XM_023258693.1
C1 43178611 43178768
refGene_gn=PODN;refGene_tx=XM_006934751.4;refGene_gn=PODN;refGene_tx=XM_023258691.1;refGene_gn=PODN;refGene_tx=XM_023258692.1;refGene_gn=PODN;refGene_tx=XM_023258693.1
C1 43179068 43179184
refGene_gn=PODN;refGene_tx=XM_006934751.4;refGene_gn=PODN;refGene_tx=XM_023258691.1;refGene_gn=PODN;refGene_tx=XM_023258692.1;refGene_gn=PODN;refGene_tx=XM_023258693.1
C1 43179958 43180616
refGene_gn=PODN;refGene_tx=XM_006934751.4;refGene_gn=PODN;refGene_tx=XM_023258691.1;refGene_gn=PODN;refGene_tx=XM_023258692.1;refGene_gn=PODN;refGene_tx=XM_023258693.1
C1 43183834 43183983
refGene_gn=PODN;refGene_tx=XM_006934751.4;refGene_gn=PODN;refGene_tx=XM_023258691.1;refGene_gn=PODN;refGene_tx=XM_023258692.1;refGene_gn=PODN;refGene_tx=XM_023258693.1
C1 43185126 43185310
refGene_gn=PODN;refGene_tx=XM_006934751.4;refGene_gn=PODN;refGene_tx=XM_023258691.1;refGene_gn=PODN;refGene_tx=XM_023258692.1;refGene_gn=PODN;refGene_tx=XM_023258693.1
C1 43191365 43191603
refGene_gn=SLC1A7;refGene_tx=XM_023258694.1;refGene_gn=SLC1A7;refGene_tx=XM_023258695.1;refGene_gn=SLC1A7;refGene_tx=XM_023258696.1;refGene_gn=SLC1A7;refGene_tx=XM_023258697.1;refGene_gn=SLC1A7;refGene_tx=XM_023258698.1;refGene_gn=SLC1A7;refGene_tx=XM_023258699.1;refGene_gn=SLC1A7;refGene_tx=XM_023258700.1;refGene_gn=SLC1A7;refGene_tx=XM_023258702.1;refGene_gn=SLC1A7;refGene_tx=XM_023258704.1
C1 43192147 43192250
refGene_gn=SLC1A7;refGene_tx=XM_023258694.1;refGene_gn=SLC1A7;refGene_tx=XM_023258695.1;refGene_gn=SLC1A7;refGene_tx=XM_023258696.1;refGene_gn=SLC1A7;refGene_tx=XM_023258697.1;refGene_gn=SLC1A7;refGene_tx=XM_023258698.1;refGene_gn=SLC1A7;refGene_tx=XM_023258699.1;refGene_gn=SLC1A7;refGene_tx=XM_023258700.1;refGene_gn=SLC1A7;refGene_tx=XM_023258702.1;refGene_gn=SLC1A7;refGene_tx=XM_023258704.1

C1 43192695 43192927
refGene_gn=SLC1A7;refGene_tx=XM_023258694.1;refGene_gn=SLC1A7;refGene_tx=XM_023258695.1;refGene_gn=SLC1A7;refGene_tx=XM_023258696.1;refGene_gn=SLC1A7;refGene_tx=XM_023258697.1;refGene_gn=SLC1A7;refGene_tx=XM_023258698.1;refGene_gn=SLC1A7;refGene_tx=XM_023258699.1;refGene_gn=SLC1A7;refGene_tx=XM_023258700.1;refGene_gn=SLC1A7;refGene_tx=XM_023258701.1;refGene_gn=SLC1A7;refGene_tx=XM_023258702.1;refGene_gn=SLC1A7;refGene_tx=XM_023258704.1
C1 43192956 43193056 refGene_gn=SLC1A7;refGene_tx=XM_023258703.1
C1 43193630 43193825
refGene_gn=SLC1A7;refGene_tx=XM_023258694.1;refGene_gn=SLC1A7;refGene_tx=XM_023258695.1;refGene_gn=SLC1A7;refGene_tx=XM_023258696.1;refGene_gn=SLC1A7;refGene_tx=XM_023258697.1;refGene_gn=SLC1A7;refGene_tx=XM_023258698.1;refGene_gn=SLC1A7;refGene_tx=XM_023258699.1;refGene_gn=SLC1A7;refGene_tx=XM_023258700.1;refGene_gn=SLC1A7;refGene_tx=XM_023258701.1;refGene_gn=SLC1A7;refGene_tx=XM_023258702.1;refGene_gn=SLC1A7;refGene_tx=XM_023258703.1;refGene_gn=SLC1A7;refGene_tx=XM_023258704.1
C1 43195776 43196010
refGene_gn=SLC1A7;refGene_tx=XM_023258694.1;refGene_gn=SLC1A7;refGene_tx=XM_023258695.1;refGene_gn=SLC1A7;refGene_tx=XM_023258696.1;refGene_gn=SLC1A7;refGene_tx=XM_023258697.1;refGene_gn=SLC1A7;refGene_tx=XM_023258698.1;refGene_gn=SLC1A7;refGene_tx=XM_023258699.1;refGene_gn=SLC1A7;refGene_tx=XM_023258700.1;refGene_gn=SLC1A7;refGene_tx=XM_023258701.1;refGene_gn=SLC1A7;refGene_tx=XM_023258702.1;refGene_gn=SLC1A7;refGene_tx=XM_023258703.1;refGene_gn=SLC1A7;refGene_tx=XM_023258704.1
C1 43196700 43196800
refGene_gn=SLC1A7;refGene_tx=XM_023258694.1;refGene_gn=SLC1A7;refGene_tx=XM_023258695.1;refGene_gn=SLC1A7;refGene_tx=XM_023258696.1;refGene_gn=SLC1A7;refGene_tx=XM_023258697.1;refGene_gn=SLC1A7;refGene_tx=XM_023258698.1;refGene_gn=SLC1A7;refGene_tx=XM_023258699.1;refGene_gn=SLC1A7;refGene_tx=XM_023258700.1;refGene_gn=SLC1A7;refGene_tx=XM_023258701.1;refGene_gn=SLC1A7;refGene_tx=XM_023258702.1;refGene_gn=SLC1A7;refGene_tx=XM_023258703.1;refGene_gn=SLC1A7;refGene_tx=XM_023258704.1
C1 43200403 43200626
refGene_gn=SLC1A7;refGene_tx=XM_023258694.1;refGene_gn=SLC1A7;refGene_tx=XM_023258695.1;refGene_gn=SLC1A7;refGene_tx=XM_023258696.1;refGene_gn=SLC1A7;refGene_tx=XM_023258697.1;refGene_gn=SLC1A7;refGene_tx=XM_023258698.1;refGene_gn=SLC1A7;refGene_tx=XM_023258699.1;refGene_gn=SLC1A7;refGene_tx=XM_023258700.1;refGene_gn=SLC1A7;refGene_tx=XM_023258701.1;refGene_gn=SLC1A7;refGene_tx=XM_023258702.1;refGene_gn=SLC1A7;refGene_tx=XM_023258703.1;refGene_gn=SLC1A7;refGene_tx=XM_023258704.1
C1 43202832 43202875
refGene_gn=SLC1A7;refGene_tx=XM_023258694.1;refGene_gn=SLC1A7;refGene_tx=XM_023258695.1;refGene_gn=SLC1A7;refGene_tx=XM_023258696.1;refGene_gn=SLC1A7;refGene_tx=XM_023258697.1;refGene_gn=SLC1A7;refGene_tx=XM_023258698.1;refGene_gn=SLC1A7;refGene_tx=XM_023258699.1;refGene_gn=SLC1A7;refGene_tx=XM_023258700.1;refGene_gn=SLC1A7;refGene_tx=XM_023258701.1;refGene_gn=SLC1A7;refGene_tx=XM_023258702.1;refGene_gn=SLC1A7;refGene_tx=XM_023258703.1;refGene_gn=SLC1A7;refGene_tx=XM_023258704.1
C1 43211081 43211297

refGene_gn=SLC1A7;refGene_tx=XM_023258694.1;refGene_gn=SLC1A7;refGene_
tx=XM_023258695.1;refGene_gn=SLC1A7;refGene_tx=XM_023258696.1;refGene_
gn=SLC1A7;refGene_tx=XM_023258697.1;refGene_gn=SLC1A7;refGene_tx=XM_02
3258698.1;refGene_gn=SLC1A7;refGene_tx=XM_023258699.1;refGene_gn=SLC1A
7;refGene_tx=XM_023258700.1;refGene_gn=SLC1A7;refGene_tx=XM_023258701.
1;refGene_gn=SLC1A7;refGene_tx=XM_023258702.1;refGene_gn=SLC1A7;refGen
e_tx=XM_023258703.1;refGene_gn=SLC1A7;refGene_tx=XM_023258704.1
C1 43227295 43227373
refGene_gn=SLC1A7;refGene_tx=XM_023258694.1;refGene_gn=SLC1A7;refGene_
tx=XM_023258695.1;refGene_gn=SLC1A7;refGene_tx=XM_023258697.1;refGene_
gn=SLC1A7;refGene_tx=XM_023258700.1;refGene_gn=SLC1A7;refGene_tx=XM_02
3258701.1;refGene_gn=SLC1A7;refGene_tx=XM_023258702.1;refGene_gn=SLC1A
7;refGene_tx=XM_023258703.1
C1 43229876 43229956
refGene_gn=SLC1A7;refGene_tx=XM_023258694.1;refGene_gn=SLC1A7;refGene_
tx=XM_023258695.1;refGene_gn=SLC1A7;refGene_tx=XM_023258696.1;refGene_
gn=SLC1A7;refGene_tx=XM_023258697.1;refGene_gn=SLC1A7;refGene_tx=XM_02
3258698.1;refGene_gn=SLC1A7;refGene_tx=XM_023258699.1;refGene_gn=SLC1A
7;refGene_tx=XM_023258700.1;refGene_gn=SLC1A7;refGene_tx=XM_023258701.
1;refGene_gn=SLC1A7;refGene_tx=XM_023258702.1;refGene_gn=SLC1A7;refGen
e_tx=XM_023258703.1;refGene_gn=SLC1A7;refGene_tx=XM_023258704.1
C1 43230702 43230807
refGene_gn=SLC1A7;refGene_tx=XM_023258694.1;refGene_gn=SLC1A7;refGene_
tx=XM_023258696.1;refGene_gn=SLC1A7;refGene_tx=XM_023258700.1;refGene_
gn=SLC1A7;refGene_tx=XM_023258701.1;refGene_gn=SLC1A7;refGene_tx=XM_02
3258703.1
C1 43235704 43235922
refGene_gn=SLC1A7;refGene_tx=XM_023258694.1;refGene_gn=SLC1A7;refGene_
tx=XM_023258695.1;refGene_gn=SLC1A7;refGene_tx=XM_023258696.1;refGene_
gn=SLC1A7;refGene_tx=XM_023258697.1;refGene_gn=SLC1A7;refGene_tx=XM_02
3258698.1;refGene_gn=SLC1A7;refGene_tx=XM_023258699.1;refGene_gn=SLC1A
7;refGene_tx=XM_023258700.1;refGene_gn=SLC1A7;refGene_tx=XM_023258701.
1;refGene_gn=SLC1A7;refGene_tx=XM_023258703.1
C1 43277383 43277756
refGene_gn=CPT2;refGene_tx=XM_003990112.4;refGene_gn=CPT2;refGene_tx=X
M_019837021.1;refGene_gn=CPT2;refGene_tx=XM_019837022.1
C1 43287981 43288073
refGene_gn=CPT2;refGene_tx=XM_003990112.4;refGene_gn=CPT2;refGene_tx=X
M_006934754.2;refGene_gn=CPT2;refGene_tx=XM_019837022.1;refGene_gn=CPT
2;refGene_tx=XM_019837024.2
C1 43289828 43289935 refGene_gn=CPT2;refGene_tx=XM_003990112.4
C1 43292526 43292611 refGene_gn=CPT2;refGene_tx=XM_006934753.4
C1 43293294 43294599
refGene_gn=CPT2;refGene_tx=XM_003990112.4;refGene_gn=CPT2;refGene_tx=X
M_006934753.4;refGene_gn=CPT2;refGene_tx=XM_006934754.2;refGene_gn=CPT
2;refGene_tx=XM_019837021.1;refGene_gn=CPT2;refGene_tx=XM_019837022.1;
refGene_gn=CPT2;refGene_tx=XM_019837024.2
C1 43295574 43295906
refGene_gn=CPT2;refGene_tx=XM_003990112.4;refGene_gn=CPT2;refGene_tx=X
M_006934753.4;refGene_gn=CPT2;refGene_tx=XM_006934754.2;refGene_gn=CPT

2;refGene_tx=XM_019837021.1;refGene_gn=CPT2;refGene_tx=XM_019837022.1;refGene_gn=CPT2;refGene_tx=XM_019837024.2
C1 43305197 43305275
refGene_gn=CC1H1orf123;refGene_tx=XM_003990113.3;refGene_gn=CC1H1orf123;refGene_tx=XM_006934755.2
C1 43306140 43306206
refGene_gn=CC1H1orf123;refGene_tx=XM_003990113.3;refGene_gn=CC1H1orf123;refGene_tx=XM_006934755.2
C1 43306868 43306946
refGene_gn=CC1H1orf123;refGene_tx=XM_003990113.3;refGene_gn=CC1H1orf123;refGene_tx=XM_006934755.2
C1 43308709 43308741
refGene_gn=CC1H1orf123;refGene_tx=XM_003990113.3;refGene_gn=CC1H1orf123;refGene_tx=XM_006934755.2
C1 43308948 43309030
refGene_gn=CC1H1orf123;refGene_tx=XM_003990113.3;refGene_gn=CC1H1orf123;refGene_tx=XM_006934755.2
C1 43309397 43309454
refGene_gn=CC1H1orf123;refGene_tx=XM_003990113.3;refGene_gn=CC1H1orf123;refGene_tx=XM_006934755.2
C1 43310582 43310666
refGene_gn=CC1H1orf123;refGene_tx=XM_003990113.3
C1 43310854 43310860
refGene_gn=CC1H1orf123;refGene_tx=XM_003990113.3
C1 43312780 43312880 refGene_gn=MAGOH;refGene_tx=XM_003990114.3
C1 43313991 43314074 refGene_gn=MAGOH;refGene_tx=XM_003990114.3
C1 43318717 43318828 refGene_gn=MAGOH;refGene_tx=XM_003990114.3
C1 43319213 43319272 refGene_gn=MAGOH;refGene_tx=XM_003990114.3
C1 43321408 43321496 refGene_gn=MAGOH;refGene_tx=XM_003990114.3
C1 43330744 43330783
refGene_gn=LRP8;refGene_tx=XM_023258706.1;refGene_gn=LRP8;refGene_tx=XM_023258707.1;refGene_gn=LRP8;refGene_tx=XM_023258708.1;refGene_gn=LRP8;refGene_tx=XM_023258709.1;refGene_gn=LRP8;refGene_tx=XM_023258710.1;refGene_gn=LRP8;refGene_tx=XM_023258711.1;refGene_gn=LRP8;refGene_tx=XM_023258712.1;refGene_gn=LRP8;refGene_tx=XM_023258713.1
C1 43333055 43333232
refGene_gn=LRP8;refGene_tx=XM_023258706.1;refGene_gn=LRP8;refGene_tx=XM_023258707.1;refGene_gn=LRP8;refGene_tx=XM_023258708.1;refGene_gn=LRP8;refGene_tx=XM_023258711.1;refGene_gn=LRP8;refGene_tx=XM_023258712.1
C1 43334278 43334451
refGene_gn=LRP8;refGene_tx=XM_023258706.1;refGene_gn=LRP8;refGene_tx=XM_023258707.1;refGene_gn=LRP8;refGene_tx=XM_023258708.1;refGene_gn=LRP8;refGene_tx=XM_023258709.1;refGene_gn=LRP8;refGene_tx=XM_023258710.1;refGene_gn=LRP8;refGene_tx=XM_023258711.1;refGene_gn=LRP8;refGene_tx=XM_023258712.1;refGene_gn=LRP8;refGene_tx=XM_023258713.1
C1 43338016 43338085
refGene_gn=LRP8;refGene_tx=XM_023258706.1;refGene_gn=LRP8;refGene_tx=XM_023258707.1;refGene_gn=LRP8;refGene_tx=XM_023258708.1;refGene_gn=LRP8;refGene_tx=XM_023258709.1;refGene_gn=LRP8;refGene_tx=XM_023258710.1;refGene_gn=LRP8;refGene_tx=XM_023258711.1;refGene_gn=LRP8;refGene_tx=X

M_023258712.1;refGene_gn=LRP8;refGene_tx=XM_023258713.1
C1 43339992 43340217
refGene_gn=LRP8;refGene_tx=XM_023258706.1;refGene_gn=LRP8;refGene_tx=XM_023258707.1;refGene_gn=LRP8;refGene_tx=XM_023258708.1;refGene_gn=LRP8;refGene_tx=XM_023258709.1;refGene_gn=LRP8;refGene_tx=XM_023258710.1;refGene_gn=LRP8;refGene_tx=XM_023258711.1;refGene_gn=LRP8;refGene_tx=XM_023258712.1;refGene_gn=LRP8;refGene_tx=XM_023258713.1
C1 43341089 43341242
refGene_gn=LRP8;refGene_tx=XM_023258706.1;refGene_gn=LRP8;refGene_tx=XM_023258707.1;refGene_gn=LRP8;refGene_tx=XM_023258708.1;refGene_gn=LRP8;refGene_tx=XM_023258709.1;refGene_gn=LRP8;refGene_tx=XM_023258710.1;refGene_gn=LRP8;refGene_tx=XM_023258711.1;refGene_gn=LRP8;refGene_tx=XM_023258712.1;refGene_gn=LRP8;refGene_tx=XM_023258713.1
C1 43342711 43342853
refGene_gn=LRP8;refGene_tx=XM_023258706.1;refGene_gn=LRP8;refGene_tx=XM_023258707.1;refGene_gn=LRP8;refGene_tx=XM_023258708.1;refGene_gn=LRP8;refGene_tx=XM_023258709.1;refGene_gn=LRP8;refGene_tx=XM_023258710.1;refGene_gn=LRP8;refGene_tx=XM_023258711.1;refGene_gn=LRP8;refGene_tx=XM_023258712.1;refGene_gn=LRP8;refGene_tx=XM_023258713.1
C1 43344259 43344399
refGene_gn=LRP8;refGene_tx=XM_023258706.1;refGene_gn=LRP8;refGene_tx=XM_023258707.1;refGene_gn=LRP8;refGene_tx=XM_023258708.1;refGene_gn=LRP8;refGene_tx=XM_023258709.1;refGene_gn=LRP8;refGene_tx=XM_023258710.1;refGene_gn=LRP8;refGene_tx=XM_023258711.1;refGene_gn=LRP8;refGene_tx=XM_023258712.1;refGene_gn=LRP8;refGene_tx=XM_023258713.1
C1 43344652 43344771
refGene_gn=LRP8;refGene_tx=XM_023258706.1;refGene_gn=LRP8;refGene_tx=XM_023258707.1;refGene_gn=LRP8;refGene_tx=XM_023258708.1;refGene_gn=LRP8;refGene_tx=XM_023258709.1;refGene_gn=LRP8;refGene_tx=XM_023258710.1;refGene_gn=LRP8;refGene_tx=XM_023258711.1;refGene_gn=LRP8;refGene_tx=XM_023258712.1;refGene_gn=LRP8;refGene_tx=XM_023258713.1
C1 43347571 43347799
refGene_gn=LRP8;refGene_tx=XM_023258706.1;refGene_gn=LRP8;refGene_tx=XM_023258707.1;refGene_gn=LRP8;refGene_tx=XM_023258708.1;refGene_gn=LRP8;refGene_tx=XM_023258709.1;refGene_gn=LRP8;refGene_tx=XM_023258710.1;refGene_gn=LRP8;refGene_tx=XM_023258711.1;refGene_gn=LRP8;refGene_tx=XM_023258712.1;refGene_gn=LRP8;refGene_tx=XM_023258713.1
C1 43350086 43350261
refGene_gn=LRP8;refGene_tx=XM_023258706.1;refGene_gn=LRP8;refGene_tx=XM_023258707.1;refGene_gn=LRP8;refGene_tx=XM_023258708.1;refGene_gn=LRP8;refGene_tx=XM_023258709.1;refGene_gn=LRP8;refGene_tx=XM_023258710.1;refGene_gn=LRP8;refGene_tx=XM_023258711.1;refGene_gn=LRP8;refGene_tx=XM_023258712.1;refGene_gn=LRP8;refGene_tx=XM_023258713.1
C1 43353316 43353442
refGene_gn=LRP8;refGene_tx=XM_023258706.1;refGene_gn=LRP8;refGene_tx=XM_023258707.1;refGene_gn=LRP8;refGene_tx=XM_023258708.1;refGene_gn=LRP8;refGene_tx=XM_023258709.1;refGene_gn=LRP8;refGene_tx=XM_023258710.1;refGene_gn=LRP8;refGene_tx=XM_023258711.1;refGene_gn=LRP8;refGene_tx=XM_023258712.1;refGene_gn=LRP8;refGene_tx=XM_023258713.1
C1 43353530 43353650

refGene_gn=LRP8;refGene_tx=XM_023258706.1;refGene_gn=LRP8;refGene_tx=XM_023258707.1;refGene_gn=LRP8;refGene_tx=XM_023258708.1;refGene_gn=LRP8;refGene_tx=XM_023258709.1;refGene_gn=LRP8;refGene_tx=XM_023258710.1;refGene_gn=LRP8;refGene_tx=XM_023258711.1;refGene_gn=LRP8;refGene_tx=XM_023258712.1;refGene_gn=LRP8;refGene_tx=XM_023258713.1
C1 43355128 43355167 refGene_gn=LRP8;refGene_tx=XM_023258707.1
C1 43357184 43357307
refGene_gn=LRP8;refGene_tx=XM_023258706.1;refGene_gn=LRP8;refGene_tx=XM_023258709.1;refGene_gn=LRP8;refGene_tx=XM_023258711.1
C1 43358105 43358228
refGene_gn=LRP8;refGene_tx=XM_023258706.1;refGene_gn=LRP8;refGene_tx=XM_023258707.1;refGene_gn=LRP8;refGene_tx=XM_023258708.1;refGene_gn=LRP8;refGene_tx=XM_023258709.1;refGene_gn=LRP8;refGene_tx=XM_023258710.1;refGene_gn=LRP8;refGene_tx=XM_023258711.1;refGene_gn=LRP8;refGene_tx=XM_023258712.1;refGene_gn=LRP8;refGene_tx=XM_023258713.1
C1 43359153 43359549
refGene_gn=LRP8;refGene_tx=XM_023258706.1;refGene_gn=LRP8;refGene_tx=XM_023258707.1;refGene_gn=LRP8;refGene_tx=XM_023258708.1;refGene_gn=LRP8;refGene_tx=XM_023258709.1;refGene_gn=LRP8;refGene_tx=XM_023258710.1
C1 43362672 43362801
refGene_gn=LRP8;refGene_tx=XM_023258706.1;refGene_gn=LRP8;refGene_tx=XM_023258707.1;refGene_gn=LRP8;refGene_tx=XM_023258708.1;refGene_gn=LRP8;refGene_tx=XM_023258709.1;refGene_gn=LRP8;refGene_tx=XM_023258710.1;refGene_gn=LRP8;refGene_tx=XM_023258711.1;refGene_gn=LRP8;refGene_tx=XM_023258712.1;refGene_gn=LRP8;refGene_tx=XM_023258713.1
C1 43370469 43370592
refGene_gn=LRP8;refGene_tx=XM_023258706.1;refGene_gn=LRP8;refGene_tx=XM_023258707.1;refGene_gn=LRP8;refGene_tx=XM_023258708.1;refGene_gn=LRP8;refGene_tx=XM_023258709.1;refGene_gn=LRP8;refGene_tx=XM_023258710.1;refGene_gn=LRP8;refGene_tx=XM_023258711.1;refGene_gn=LRP8;refGene_tx=XM_023258712.1;refGene_gn=LRP8;refGene_tx=XM_023258713.1
C1 43405324 43405444
refGene_gn=LRP8;refGene_tx=XM_023258706.1;refGene_gn=LRP8;refGene_tx=XM_023258707.1;refGene_gn=LRP8;refGene_tx=XM_023258708.1;refGene_gn=LRP8;refGene_tx=XM_023258709.1;refGene_gn=LRP8;refGene_tx=XM_023258710.1;refGene_gn=LRP8;refGene_tx=XM_023258711.1;refGene_gn=LRP8;refGene_tx=XM_023258712.1;refGene_gn=LRP8;refGene_tx=XM_023258713.1
C1 43406218 43406327
refGene_gn=LRP8;refGene_tx=XM_023258706.1;refGene_gn=LRP8;refGene_tx=XM_023258707.1;refGene_gn=LRP8;refGene_tx=XM_023258708.1;refGene_gn=LRP8;refGene_tx=XM_023258709.1;refGene_gn=LRP8;refGene_tx=XM_023258710.1;refGene_gn=LRP8;refGene_tx=XM_023258711.1;refGene_gn=LRP8;refGene_tx=XM_023258712.1;refGene_gn=LRP8;refGene_tx=XM_023258713.1
C1 43514150 43514748 refGene_gn=DMRTB1;refGene_tx=XM_023258714.1
C1 43516355 43516561 refGene_gn=DMRTB1;refGene_tx=XM_023258714.1
C1 43519941 43520131 refGene_gn=DMRTB1;refGene_tx=XM_023258714.1
C1 43521494 43521661 refGene_gn=DMRTB1;refGene_tx=XM_023258714.1
C1 43559431 43559589
refGene_gn=GLIS1;refGene_tx=XM_023258715.1;refGene_gn=GLIS1;refGene_tx=XM_023258716.1

C1 43561872 43562040
refGene_gn=GLIS1;refGene_tx=XM_023258715.1;refGene_gn=GLIS1;refGene_tx=XM_023258716.1
C1 43562659 43562838
refGene_gn=GLIS1;refGene_tx=XM_023258715.1;refGene_gn=GLIS1;refGene_tx=XM_023258716.1
C1 43567724 43567881
refGene_gn=GLIS1;refGene_tx=XM_023258715.1;refGene_gn=GLIS1;refGene_tx=XM_023258716.1
C1 43572480 43572607
refGene_gn=GLIS1;refGene_tx=XM_023258715.1;refGene_gn=GLIS1;refGene_tx=XM_023258716.1
C1 43576341 43576452
refGene_gn=GLIS1;refGene_tx=XM_023258715.1;refGene_gn=GLIS1;refGene_tx=XM_023258716.1
C1 43580976 43581138
refGene_gn=GLIS1;refGene_tx=XM_023258715.1;refGene_gn=GLIS1;refGene_tx=XM_023258716.1
C1 43642192 43643075
refGene_gn=GLIS1;refGene_tx=XM_023258715.1;refGene_gn=GLIS1;refGene_tx=XM_023258716.1
C1 43649580 43649758
refGene_gn=GLIS1;refGene_tx=XM_023258715.1;refGene_gn=GLIS1;refGene_tx=XM_023258716.1
C1 43787446 43787705
refGene_gn=GLIS1;refGene_tx=XM_023258715.1;refGene_gn=GLIS1;refGene_tx=XM_023258716.1
C1 43810380 43810444 refGene_gn=NDC1;refGene_tx=XM_006934756.2
C1 43813318 43813479 refGene_gn=NDC1;refGene_tx=XM_006934756.2
C1 43823135 43823236 refGene_gn=NDC1;refGene_tx=XM_006934756.2
C1 43824018 43824082 refGene_gn=NDC1;refGene_tx=XM_006934756.2
C1 43825839 43825890 refGene_gn=NDC1;refGene_tx=XM_006934756.2
C1 43828212 43828328 refGene_gn=NDC1;refGene_tx=XM_006934756.2
C1 43828432 43828672 refGene_gn=NDC1;refGene_tx=XM_006934756.2
C1 43831046 43831202 refGene_gn=NDC1;refGene_tx=XM_006934756.2
C1 43833065 43833147 refGene_gn=NDC1;refGene_tx=XM_006934756.2
C1 43835368 43835461 refGene_gn=NDC1;refGene_tx=XM_006934756.2
C1 43836244 43836380 refGene_gn=NDC1;refGene_tx=XM_006934756.2
C1 43839267 43839319 refGene_gn=NDC1;refGene_tx=XM_006934756.2
C1 43841699 43841808 refGene_gn=NDC1;refGene_tx=XM_006934756.2
C1 43846646 43846785 refGene_gn=NDC1;refGene_tx=XM_006934756.2
C1 43847945 43848120 refGene_gn=NDC1;refGene_tx=XM_006934756.2
C1 43850771 43850873 refGene_gn=NDC1;refGene_tx=XM_006934756.2
C1 43853128 43853249 refGene_gn=NDC1;refGene_tx=XM_006934756.2
C1 43854536 43854899 refGene_gn=NDC1;refGene_tx=XM_006934756.2
C1 43875052 43875142
refGene_gn=YIPF1;refGene_tx=XM_006934757.4;refGene_gn=YIPF1;refGene_tx=XM_006934758.3
C1 43879718 43879901
refGene_gn=YIPF1;refGene_tx=XM_006934757.4;refGene_gn=YIPF1;refGene_tx

=XM_006934758.3
C1 43880259 43880426
refGene_gn=YIPF1;refGene_tx=XM_006934757.4;refGene_gn=YIPF1;refGene_tx
=XM_006934758.3
C1 43883647 43883764
refGene_gn=YIPF1;refGene_tx=XM_006934757.4;refGene_gn=YIPF1;refGene_tx
=XM_006934758.3
C1 43890286 43890374
refGene_gn=YIPF1;refGene_tx=XM_006934757.4;refGene_gn=YIPF1;refGene_tx
=XM_006934758.3
C1 43890569 43890650
refGene_gn=YIPF1;refGene_tx=XM_006934757.4;refGene_gn=YIPF1;refGene_tx
=XM_006934758.3
C1 43893776 43893940
refGene_gn=YIPF1;refGene_tx=XM_006934757.4;refGene_gn=YIPF1;refGene_tx
=XM_006934758.3
C1 43899321 43899352
refGene_gn=YIPF1;refGene_tx=XM_006934757.4;refGene_gn=YIPF1;refGene_tx
=XM_006934758.3
C1 43924397 43924719 refGene_gn=DIO1;refGene_tx=NM_001009267.1
C1 43933310 43933454 refGene_gn=DIO1;refGene_tx=NM_001009267.1
C1 43939166 43939366 refGene_gn=DIO1;refGene_tx=NM_001009267.1
C1 43941347 43941415 refGene_gn=DIO1;refGene_tx=NM_001009267.1
C1 43942205 43942228
refGene_gn=HSPB11;refGene_tx=XM_019837038.2;refGene_gn=HSPB11;refGene_
tx=XM_019837042.2;refGene_gn=HSPB11;refGene_tx=XM_023258718.1;refGene_
gn=HSPB11;refGene_tx=XM_023258719.1;refGene_gn=HSPB11;refGene_tx=XM_02
3258722.1;refGene_gn=HSPB11;refGene_tx=XM_023258723.1;refGene_gn=HSPB1
1;refGene_tx=XM_023258724.1;refGene_gn=HSPB11;refGene_tx=XM_023258725.
1
C1 43943841 43943959
refGene_gn=HSPB11;refGene_tx=XM_019837038.2;refGene_gn=HSPB11;refGene_
tx=XM_019837042.2;refGene_gn=HSPB11;refGene_tx=XM_023258718.1;refGene_
gn=HSPB11;refGene_tx=XM_023258719.1;refGene_gn=HSPB11;refGene_tx=XM_02
3258722.1;refGene_gn=HSPB11;refGene_tx=XM_023258723.1;refGene_gn=HSPB1
1;refGene_tx=XM_023258724.1;refGene_gn=HSPB11;refGene_tx=XM_023258725.
1
C1 43954237 43954351 refGene_gn=HSPB11;refGene_tx=XM_023258720.1
C1 43956146 43956197 refGene_gn=HSPB11;refGene_tx=XM_023258721.1
C1 43956502 43956546
refGene_gn=HSPB11;refGene_tx=XM_019837038.2;refGene_gn=HSPB11;refGene_
tx=XM_019837042.2;refGene_gn=HSPB11;refGene_tx=XM_023258718.1;refGene_
gn=HSPB11;refGene_tx=XM_023258719.1;refGene_gn=HSPB11;refGene_tx=XM_02
3258720.1;refGene_gn=HSPB11;refGene_tx=XM_023258721.1;refGene_gn=HSPB1
1;refGene_tx=XM_023258722.1;refGene_gn=HSPB11;refGene_tx=XM_023258723.
1;refGene_gn=HSPB11;refGene_tx=XM_023258724.1;refGene_gn=HSPB11;refGen
e_tx=XM_023258725.1
C1 43960468 43960537
refGene_gn=HSPB11;refGene_tx=XM_019837038.2;refGene_gn=HSPB11;refGene_
tx=XM_019837042.2;refGene_gn=HSPB11;refGene_tx=XM_023258718.1;refGene_

gn=HSPB11;refGene_tx=XM_023258719.1;refGene_gn=HSPB11;refGene_tx=XM_023258720.1;refGene_gn=HSPB11;refGene_tx=XM_023258721.1;refGene_gn=HSPB11;refGene_tx=XM_023258722.1;refGene_gn=HSPB11;refGene_tx=XM_023258723.1;refGene_gn=HSPB11;refGene_tx=XM_023258724.1;refGene_gn=HSPB11;refGene_tx=XM_023258725.1
C1 43962425 43962532
refGene_gn=HSPB11;refGene_tx=XM_019837038.2;refGene_gn=HSPB11;refGene_tx=XM_019837042.2;refGene_gn=HSPB11;refGene_tx=XM_023258718.1;refGene_gn=HSPB11;refGene_tx=XM_023258719.1;refGene_gn=HSPB11;refGene_tx=XM_023258720.1;refGene_gn=HSPB11;refGene_tx=XM_023258721.1;refGene_gn=HSPB11;refGene_tx=XM_023258722.1;refGene_gn=HSPB11;refGene_tx=XM_023258723.1;refGene_gn=HSPB11;refGene_tx=XM_023258724.1;refGene_gn=HSPB11;refGene_tx=XM_023258725.1
C1 43974589 43974702
refGene_gn=HSPB11;refGene_tx=XM_019837038.2;refGene_gn=HSPB11;refGene_tx=XM_019837042.2;refGene_gn=HSPB11;refGene_tx=XM_023258718.1;refGene_gn=HSPB11;refGene_tx=XM_023258719.1;refGene_gn=HSPB11;refGene_tx=XM_023258720.1;refGene_gn=HSPB11;refGene_tx=XM_023258721.1;refGene_gn=HSPB11;refGene_tx=XM_023258722.1;refGene_gn=HSPB11;refGene_tx=XM_023258723.1;refGene_gn=HSPB11;refGene_tx=XM_023258724.1;refGene_gn=HSPB11;refGene_tx=XM_023258725.1
C1 43976259 43976329 refGene_gn=HSPB11;refGene_tx=XM_023258719.1
C1 43984464 43984568 refGene_gn=HSPB11;refGene_tx=XM_023258719.1
C1 43990587 43991060 refGene_gn=LRRC42;refGene_tx=XM_023258717.1
C1 43996369 43996501 refGene_gn=LRRC42;refGene_tx=XM_023258717.1
C1 43997671 43997790 refGene_gn=LRRC42;refGene_tx=XM_023258717.1
C1 43999583 43999672 refGene_gn=LRRC42;refGene_tx=XM_023258717.1
C1 43999841 43999955 refGene_gn=LRRC42;refGene_tx=XM_023258717.1
C1 44003466 44003551 refGene_gn=LRRC42;refGene_tx=XM_023258717.1
C1 44005267 44005542 refGene_gn=LRRC42;refGene_tx=XM_023258717.1
C1 44022790 44022939 refGene_gn=LDLRAD1;refGene_tx=XM_023260049.1
C1 44024149 44024278
refGene_gn=LDLRAD1;refGene_tx=XM_011284981.3;refGene_gn=LDLRAD1;refGene_tx=XM_023260049.1
C1 44026021 44026159
refGene_gn=LDLRAD1;refGene_tx=XM_011284981.3;refGene_gn=LDLRAD1;refGene_tx=XM_023260049.1
C1 44028206 44028332 refGene_gn=LDLRAD1;refGene_tx=XM_023260049.1
C1 44028645 44028648 refGene_gn=LDLRAD1;refGene_tx=XM_023260049.1
C1 44030906 44030961
refGene_gn=LDLRAD1;refGene_tx=XM_011284981.3;refGene_gn=LDLRAD1;refGene_tx=XM_023260049.1
C1 44031650 44031722
refGene_gn=LDLRAD1;refGene_tx=XM_011284981.3;refGene_gn=LDLRAD1;refGene_tx=XM_023260049.1
C1 44041485 44041641
refGene_gn=TMEM59;refGene_tx=XM_019837052.2;refGene_gn=TMEM59;refGene_tx=XM_019837053.2;refGene_gn=TMEM59;refGene_tx=XM_019837054.2;refGene_gn=TMEM59;refGene_tx=XM_019837055.2;refGene_gn=TMEM59;refGene_tx=XM_019837056.2;refGene_gn=TMEM59;refGene_tx=XM_023258727.1;refGene_gn=TMEM5

9;refGene_tx=XM_023258728.1
C1 44051014 44051123
refGene_gn=TMEM59;refGene_tx=XM_019837052.2;refGene_gn=TMEM59;refGene_tx=XM_019837053.2;refGene_gn=TMEM59;refGene_tx=XM_019837054.2;refGene_gn=TMEM59;refGene_tx=XM_019837055.2;refGene_gn=TMEM59;refGene_tx=XM_019837056.2;refGene_gn=TMEM59;refGene_tx=XM_023258727.1;refGene_gn=TMEM59;refGene_tx=XM_023258728.1
C1 44054486 44054571
refGene_gn=TMEM59;refGene_tx=XM_019837052.2;refGene_gn=TMEM59;refGene_tx=XM_019837053.2;refGene_gn=TMEM59;refGene_tx=XM_019837054.2;refGene_gn=TMEM59;refGene_tx=XM_019837055.2;refGene_gn=TMEM59;refGene_tx=XM_019837056.2;refGene_gn=TMEM59;refGene_tx=XM_023258727.1;refGene_gn=TMEM59;refGene_tx=XM_023258728.1
C1 44055382 44055464
refGene_gn=TMEM59;refGene_tx=XM_019837052.2;refGene_gn=TMEM59;refGene_tx=XM_019837053.2;refGene_gn=TMEM59;refGene_tx=XM_019837054.2;refGene_gn=TMEM59;refGene_tx=XM_019837055.2;refGene_gn=TMEM59;refGene_tx=XM_019837056.2;refGene_gn=TMEM59;refGene_tx=XM_023258727.1;refGene_gn=TMEM59;refGene_tx=XM_023258728.1
C1 44057016 44057169
refGene_gn=TMEM59;refGene_tx=XM_019837052.2;refGene_gn=TMEM59;refGene_tx=XM_019837053.2;refGene_gn=TMEM59;refGene_tx=XM_019837054.2;refGene_gn=TMEM59;refGene_tx=XM_019837055.2;refGene_gn=TMEM59;refGene_tx=XM_019837056.2;refGene_gn=TMEM59;refGene_tx=XM_023258727.1;refGene_gn=TMEM59;refGene_tx=XM_023258728.1
C1 44059867 44059962
refGene_gn=TMEM59;refGene_tx=XM_019837052.2;refGene_gn=TMEM59;refGene_tx=XM_019837053.2;refGene_gn=TMEM59;refGene_tx=XM_019837054.2;refGene_gn=TMEM59;refGene_tx=XM_019837055.2;refGene_gn=TMEM59;refGene_tx=XM_019837056.2;refGene_gn=TMEM59;refGene_tx=XM_023258727.1;refGene_gn=TMEM59;refGene_tx=XM_023258728.1
C1 44061946 44062052
refGene_gn=TMEM59;refGene_tx=XM_019837052.2;refGene_gn=TMEM59;refGene_tx=XM_019837053.2;refGene_gn=TMEM59;refGene_tx=XM_019837054.2;refGene_gn=TMEM59;refGene_tx=XM_019837055.2;refGene_gn=TMEM59;refGene_tx=XM_019837056.2;refGene_gn=TMEM59;refGene_tx=XM_023258727.1;refGene_gn=TMEM59;refGene_tx=XM_023258728.1
C1 44063047 44063080
refGene_gn=TMEM59;refGene_tx=XM_019837052.2;refGene_gn=TMEM59;refGene_tx=XM_019837053.2
C1 44067092 44067281
refGene_gn=TMEM59;refGene_tx=XM_019837052.2;refGene_gn=TMEM59;refGene_tx=XM_019837053.2;refGene_gn=TMEM59;refGene_tx=XM_019837054.2;refGene_gn=TMEM59;refGene_tx=XM_019837055.2
C1 44067528 44067534 refGene_gn=TMEM59;refGene_tx=XM_023258727.1
C1 44068504 44068606
refGene_gn=TCEANC2;refGene_tx=XM_003990126.5;refGene_gn=TCEANC2;refGene_tx=XM_019837057.2
C1 44078494 44078636
refGene_gn=TCEANC2;refGene_tx=XM_003990126.5;refGene_gn=TCEANC2;refGen

e_tx=XM_019837057.2
C1 44093957 44094151
refGene_gn=TCEANC2;refGene_tx=XM_003990126.5;refGene_gn=TCEANC2;refGen
e_tx=XM_019837057.2
C1 44099765 44099853 refGene_gn=TCEANC2;refGene_tx=XM_019837057.2
C1 44102488 44102677
refGene_gn=TCEANC2;refGene_tx=XM_003990126.5;refGene_gn=TCEANC2;refGen
e_tx=XM_019837057.2
C1 44137694 44138021 refGene_gn=CDCP2;refGene_tx=XM_023258729.1
C1 44140472 44140651 refGene_gn=CDCP2;refGene_tx=XM_023258729.1
C1 44143572 44143926 refGene_gn=CDCP2;refGene_tx=XM_023258729.1
C1 44144821 44145157 refGene_gn=CDCP2;refGene_tx=XM_023258729.1
C1 44148138 44148486 refGene_gn=CDCP2;refGene_tx=XM_023258729.1
C1 44151415 44151494 refGene_gn=CDCP2;refGene_tx=XM_023258729.1
C1 44167457 44167661
refGene_gn=LOC101096748;refGene_tx=XM_023258726.1
C1 44171129 44171333
refGene_gn=LOC101096748;refGene_tx=XM_023258726.1
C1 44174818 44174923
refGene_gn=LOC101096748;refGene_tx=XM_023258726.1
C1 44178374 44178462
refGene_gn=LOC101096748;refGene_tx=XM_023258726.1
C1 44181119 44181268
refGene_gn=LOC101096748;refGene_tx=XM_023258726.1
C1 44182181 44182376
refGene_gn=LOC101096748;refGene_tx=XM_023258726.1
C1 44185650 44185996 refGene_gn=MRPL37;refGene_tx=XM_003990128.4
C1 44188788 44188972 refGene_gn=MRPL37;refGene_tx=XM_003990128.4
C1 44189067 44189183 refGene_gn=MRPL37;refGene_tx=XM_003990128.4
C1 44192142 44192328 refGene_gn=MRPL37;refGene_tx=XM_003990128.4
C1 44194632 44194790 refGene_gn=MRPL37;refGene_tx=XM_003990128.4
C1 44197777 44197981 refGene_gn=MRPL37;refGene_tx=XM_003990128.4
C1 44199763 44199841 refGene_gn=MRPL37;refGene_tx=XM_003990128.4
C1 44205288 44205318
refGene_gn=SSBP3;refGene_tx=XM_011284976.3;refGene_gn=SSBP3;refGene_tx
=XM_011284977.3;refGene_gn=SSBP3;refGene_tx=XM_019837059.2;refGene_gn=
SSBP3;refGene_tx=XM_019837060.2;refGene_gn=SSBP3;refGene_tx=XM_0198370
61.1
C1 44206100 44206202
refGene_gn=SSBP3;refGene_tx=XM_011284976.3;refGene_gn=SSBP3;refGene_tx
=XM_011284977.3;refGene_gn=SSBP3;refGene_tx=XM_019837059.2;refGene_gn=
SSBP3;refGene_tx=XM_019837060.2;refGene_gn=SSBP3;refGene_tx=XM_0198370
61.1
C1 44206338 44206367
refGene_gn=SSBP3;refGene_tx=XM_011284976.3;refGene_gn=SSBP3;refGene_tx
=XM_011284977.3;refGene_gn=SSBP3;refGene_tx=XM_019837059.2;refGene_gn=
SSBP3;refGene_tx=XM_019837060.2;refGene_gn=SSBP3;refGene_tx=XM_0198370
61.1
C1 44206614 44206693
refGene_gn=SSBP3;refGene_tx=XM_011284976.3;refGene_gn=SSBP3;refGene_tx

=XM_011284977.3;refGene_gn=SSBP3;refGene_tx=XM_019837059.2;refGene_gn=SSBP3;refGene_tx=XM_019837060.2;refGene_gn=SSBP3;refGene_tx=XM_019837061.1
C1 44213832 44213903
refGene_gn=SSBP3;refGene_tx=XM_011284976.3;refGene_gn=SSBP3;refGene_tx=XM_011284977.3;refGene_gn=SSBP3;refGene_tx=XM_019837059.2;refGene_gn=SSBP3;refGene_tx=XM_019837060.2;refGene_gn=SSBP3;refGene_tx=XM_019837061.1
C1 44215273 44215328
refGene_gn=SSBP3;refGene_tx=XM_011284976.3;refGene_gn=SSBP3;refGene_tx=XM_011284977.3;refGene_gn=SSBP3;refGene_tx=XM_019837059.2;refGene_gn=SSBP3;refGene_tx=XM_019837060.2;refGene_gn=SSBP3;refGene_tx=XM_019837061.1
C1 44215848 44215884
refGene_gn=SSBP3;refGene_tx=XM_011284976.3;refGene_gn=SSBP3;refGene_tx=XM_011284977.3;refGene_gn=SSBP3;refGene_tx=XM_019837059.2;refGene_gn=SSBP3;refGene_tx=XM_019837060.2;refGene_gn=SSBP3;refGene_tx=XM_019837061.1
C1 44216540 44216589
refGene_gn=SSBP3;refGene_tx=XM_011284976.3;refGene_gn=SSBP3;refGene_tx=XM_011284977.3;refGene_gn=SSBP3;refGene_tx=XM_019837059.2;refGene_gn=SSBP3;refGene_tx=XM_019837060.2;refGene_gn=SSBP3;refGene_tx=XM_019837061.1
C1 44217364 44217453
refGene_gn=SSBP3;refGene_tx=XM_011284976.3;refGene_gn=SSBP3;refGene_tx=XM_011284977.3;refGene_gn=SSBP3;refGene_tx=XM_019837059.2;refGene_gn=SSBP3;refGene_tx=XM_019837060.2;refGene_gn=SSBP3;refGene_tx=XM_019837061.1
C1 44224686 44224763
refGene_gn=SSBP3;refGene_tx=XM_011284976.3;refGene_gn=SSBP3;refGene_tx=XM_011284977.3;refGene_gn=SSBP3;refGene_tx=XM_019837059.2;refGene_gn=SSBP3;refGene_tx=XM_019837060.2;refGene_gn=SSBP3;refGene_tx=XM_019837061.1
C1 44224853 44224920
refGene_gn=SSBP3;refGene_tx=XM_011284976.3;refGene_gn=SSBP3;refGene_tx=XM_011284977.3;refGene_gn=SSBP3;refGene_tx=XM_019837059.2;refGene_gn=SSBP3;refGene_tx=XM_019837060.2;refGene_gn=SSBP3;refGene_tx=XM_019837061.1
C1 44229132 44229192
refGene_gn=SSBP3;refGene_tx=XM_011284976.3;refGene_gn=SSBP3;refGene_tx=XM_011284977.3;refGene_gn=SSBP3;refGene_tx=XM_019837059.2;refGene_gn=SSBP3;refGene_tx=XM_019837060.2;refGene_gn=SSBP3;refGene_tx=XM_019837061.1
C1 44230107 44230188
refGene_gn=SSBP3;refGene_tx=XM_011284976.3;refGene_gn=SSBP3;refGene_tx=XM_019837059.2;refGene_gn=SSBP3;refGene_tx=XM_019837061.1
C1 44252128 44252218
refGene_gn=SSBP3;refGene_tx=XM_011284976.3;refGene_gn=SSBP3;refGene_tx=XM_011284977.3;refGene_gn=SSBP3;refGene_tx=XM_019837059.2;refGene_gn=SSBP3;refGene_tx=XM_019837060.2;refGene_gn=SSBP3;refGene_tx=XM_0198370

61.1
C1 44321965 44322106 refGene_gn=SSBP3;refGene_tx=XM_019837061.1
C1 44365033 44365118
refGene_gn=SSBP3;refGene_tx=XM_011284976.3;refGene_gn=SSBP3;refGene_tx=XM_011284977.3;refGene_gn=SSBP3;refGene_tx=XM_019837059.2;refGene_gn=SSBP3;refGene_tx=XM_019837060.2
C1 44367720 44367782
refGene_gn=SSBP3;refGene_tx=XM_011284976.3;refGene_gn=SSBP3;refGene_tx=XM_011284977.3;refGene_gn=SSBP3;refGene_tx=XM_019837059.2;refGene_gn=SSBP3;refGene_tx=XM_019837060.2
C1 44368006 44368079
refGene_gn=SSBP3;refGene_tx=XM_011284976.3;refGene_gn=SSBP3;refGene_tx=XM_011284977.3;refGene_gn=SSBP3;refGene_tx=XM_019837059.2;refGene_gn=SSBP3;refGene_tx=XM_019837060.2
C1 44369087 44369143
refGene_gn=SSBP3;refGene_tx=XM_011284976.3;refGene_gn=SSBP3;refGene_tx=XM_011284977.3;refGene_gn=SSBP3;refGene_tx=XM_019837059.2;refGene_gn=SSBP3;refGene_tx=XM_019837060.2
C1 44474715 44474748
refGene_gn=ACOT11;refGene_tx=XM_019837065.2;refGene_gn=ACOT11;refGene_tx=XM_023258730.1
C1 44497038 44497249
refGene_gn=ACOT11;refGene_tx=XM_019837065.2;refGene_gn=ACOT11;refGene_tx=XM_019837066.2;refGene_gn=ACOT11;refGene_tx=XM_023258730.1;refGene_gn=ACOT11;refGene_tx=XM_023258731.1
C1 44498172 44498242
refGene_gn=ACOT11;refGene_tx=XM_019837065.2;refGene_gn=ACOT11;refGene_tx=XM_019837066.2;refGene_gn=ACOT11;refGene_tx=XM_023258730.1;refGene_gn=ACOT11;refGene_tx=XM_023258731.1
C1 44503170 44503240
refGene_gn=ACOT11;refGene_tx=XM_019837065.2;refGene_gn=ACOT11;refGene_tx=XM_019837066.2;refGene_gn=ACOT11;refGene_tx=XM_023258730.1;refGene_gn=ACOT11;refGene_tx=XM_023258731.1
C1 44504617 44504716
refGene_gn=ACOT11;refGene_tx=XM_019837065.2;refGene_gn=ACOT11;refGene_tx=XM_019837066.2;refGene_gn=ACOT11;refGene_tx=XM_023258730.1;refGene_gn=ACOT11;refGene_tx=XM_023258731.1
C1 44505201 44505337
refGene_gn=ACOT11;refGene_tx=XM_019837065.2;refGene_gn=ACOT11;refGene_tx=XM_019837066.2;refGene_gn=ACOT11;refGene_tx=XM_023258730.1;refGene_gn=ACOT11;refGene_tx=XM_023258731.1
C1 44507176 44507333
refGene_gn=ACOT11;refGene_tx=XM_019837065.2;refGene_gn=ACOT11;refGene_tx=XM_019837066.2;refGene_gn=ACOT11;refGene_tx=XM_023258730.1;refGene_gn=ACOT11;refGene_tx=XM_023258731.1
C1 44508753 44508873
refGene_gn=ACOT11;refGene_tx=XM_019837065.2;refGene_gn=ACOT11;refGene_tx=XM_019837066.2;refGene_gn=ACOT11;refGene_tx=XM_023258730.1;refGene_gn=ACOT11;refGene_tx=XM_023258731.1
C1 44509902 44510047

refGene_gn=ACOT11;refGene_tx=XM_019837065.2;refGene_gn=ACOT11;refGene_tx=XM_019837066.2;refGene_gn=ACOT11;refGene_tx=XM_023258730.1;refGene_gn=ACOT11;refGene_tx=XM_023258731.1
C1 44511352 44511408
refGene_gn=ACOT11;refGene_tx=XM_019837065.2;refGene_gn=ACOT11;refGene_tx=XM_019837066.2;refGene_gn=ACOT11;refGene_tx=XM_023258730.1;refGene_gn=ACOT11;refGene_tx=XM_023258731.1
C1 44512444 44512511
refGene_gn=ACOT11;refGene_tx=XM_019837065.2;refGene_gn=ACOT11;refGene_tx=XM_019837066.2;refGene_gn=ACOT11;refGene_tx=XM_023258730.1;refGene_gn=ACOT11;refGene_tx=XM_023258731.1
C1 44512878 44512962
refGene_gn=ACOT11;refGene_tx=XM_019837065.2;refGene_gn=ACOT11;refGene_tx=XM_019837066.2;refGene_gn=ACOT11;refGene_tx=XM_023258730.1;refGene_gn=ACOT11;refGene_tx=XM_023258731.1
C1 44513620 44513754
refGene_gn=ACOT11;refGene_tx=XM_019837065.2;refGene_gn=ACOT11;refGene_tx=XM_019837066.2;refGene_gn=ACOT11;refGene_tx=XM_023258730.1;refGene_gn=ACOT11;refGene_tx=XM_023258731.1
C1 44515616 44515748
refGene_gn=ACOT11;refGene_tx=XM_019837065.2;refGene_gn=ACOT11;refGene_tx=XM_019837066.2;refGene_gn=ACOT11;refGene_tx=XM_023258730.1;refGene_gn=ACOT11;refGene_tx=XM_023258731.1
C1 44516444 44516571
refGene_gn=ACOT11;refGene_tx=XM_019837065.2;refGene_gn=ACOT11;refGene_tx=XM_019837066.2;refGene_gn=ACOT11;refGene_tx=XM_023258730.1;refGene_gn=ACOT11;refGene_tx=XM_023258731.1
C1 44517417 44517573
refGene_gn=ACOT11;refGene_tx=XM_019837065.2;refGene_gn=ACOT11;refGene_tx=XM_019837066.2;refGene_gn=ACOT11;refGene_tx=XM_023258730.1;refGene_gn=ACOT11;refGene_tx=XM_023258731.1
C1 44517742 44518416 refGene_gn=FAM151A;refGene_tx=XM_011284975.3
C1 44518843 44518987 refGene_gn=FAM151A;refGene_tx=XM_011284975.3
C1 44519745 44519885 refGene_gn=FAM151A;refGene_tx=XM_011284975.3
C1 44520615 44520840 refGene_gn=FAM151A;refGene_tx=XM_011284975.3
C1 44522261 44522421 refGene_gn=FAM151A;refGene_tx=XM_011284975.3
C1 44523438 44523591 refGene_gn=FAM151A;refGene_tx=XM_011284975.3
C1 44528766 44528913 refGene_gn=FAM151A;refGene_tx=XM_011284975.3
C1 44529878 44530005 refGene_gn=FAM151A;refGene_tx=XM_011284975.3
C1 44548213 44549444
refGene_gn=MROH7;refGene_tx=XM_011284974.3;refGene_gn=MROH7;refGene_tx=XM_019837068.2;refGene_gn=MROH7;refGene_tx=XM_019837069.2;refGene_gn=MROH7;refGene_tx=XM_019837070.2;refGene_gn=MROH7;refGene_tx=XM_019837071.2;refGene_gn=MROH7;refGene_tx=XM_019837072.2;refGene_gn=MROH7;refGene_tx=XM_019837073.2
C1 44563184 44563258
refGene_gn=MROH7;refGene_tx=XM_011284974.3;refGene_gn=MROH7;refGene_tx=XM_019837068.2;refGene_gn=MROH7;refGene_tx=XM_019837069.2;refGene_gn=MROH7;refGene_tx=XM_019837070.2;refGene_gn=MROH7;refGene_tx=XM_019837071.2;refGene_gn=MROH7;refGene_tx=XM_019837072.2;refGene_gn=MROH7;refGe

ne_tx=XM_019837073.2
C1 44566447 44566531
refGene_gn=MROH7;refGene_tx=XM_011284974.3;refGene_gn=MROH7;refGene_tx=XM_019837068.2;refGene_gn=MROH7;refGene_tx=XM_019837069.2;refGene_gn=MROH7;refGene_tx=XM_019837070.2;refGene_gn=MROH7;refGene_tx=XM_019837071.2;refGene_gn=MROH7;refGene_tx=XM_019837072.2;refGene_gn=MROH7;refGene_tx=XM_019837073.2
C1 44567467 44567547
refGene_gn=MROH7;refGene_tx=XM_011284974.3;refGene_gn=MROH7;refGene_tx=XM_019837068.2;refGene_gn=MROH7;refGene_tx=XM_019837069.2;refGene_gn=MROH7;refGene_tx=XM_019837070.2;refGene_gn=MROH7;refGene_tx=XM_019837071.2;refGene_gn=MROH7;refGene_tx=XM_019837072.2;refGene_gn=MROH7;refGene_tx=XM_019837073.2
C1 44567842 44567972
refGene_gn=MROH7;refGene_tx=XM_011284974.3;refGene_gn=MROH7;refGene_tx=XM_019837068.2;refGene_gn=MROH7;refGene_tx=XM_019837069.2;refGene_gn=MROH7;refGene_tx=XM_019837070.2;refGene_gn=MROH7;refGene_tx=XM_019837071.2;refGene_gn=MROH7;refGene_tx=XM_019837072.2;refGene_gn=MROH7;refGene_tx=XM_019837073.2
C1 44569894 44569990
refGene_gn=MROH7;refGene_tx=XM_011284974.3;refGene_gn=MROH7;refGene_tx=XM_019837068.2;refGene_gn=MROH7;refGene_tx=XM_019837069.2;refGene_gn=MROH7;refGene_tx=XM_019837070.2;refGene_gn=MROH7;refGene_tx=XM_019837071.2;refGene_gn=MROH7;refGene_tx=XM_019837072.2;refGene_gn=MROH7;refGene_tx=XM_019837073.2
C1 44570479 44570584
refGene_gn=MROH7;refGene_tx=XM_011284974.3;refGene_gn=MROH7;refGene_tx=XM_019837068.2;refGene_gn=MROH7;refGene_tx=XM_019837069.2;refGene_gn=MROH7;refGene_tx=XM_019837070.2;refGene_gn=MROH7;refGene_tx=XM_019837071.2;refGene_gn=MROH7;refGene_tx=XM_019837072.2;refGene_gn=MROH7;refGene_tx=XM_019837073.2
C1 44570794 44570930
refGene_gn=MROH7;refGene_tx=XM_011284974.3;refGene_gn=MROH7;refGene_tx=XM_019837068.2;refGene_gn=MROH7;refGene_tx=XM_019837069.2;refGene_gn=MROH7;refGene_tx=XM_019837070.2;refGene_gn=MROH7;refGene_tx=XM_019837071.2;refGene_gn=MROH7;refGene_tx=XM_019837072.2;refGene_gn=MROH7;refGene_tx=XM_019837073.2
C1 44572930 44573043
refGene_gn=MROH7;refGene_tx=XM_011284974.3;refGene_gn=MROH7;refGene_tx=XM_019837068.2;refGene_gn=MROH7;refGene_tx=XM_019837069.2;refGene_gn=MROH7;refGene_tx=XM_019837070.2;refGene_gn=MROH7;refGene_tx=XM_019837071.2;refGene_gn=MROH7;refGene_tx=XM_019837072.2;refGene_gn=MROH7;refGene_tx=XM_019837073.2
C1 44573381 44573558
refGene_gn=MROH7;refGene_tx=XM_011284974.3;refGene_gn=MROH7;refGene_tx=XM_019837068.2;refGene_gn=MROH7;refGene_tx=XM_019837069.2;refGene_gn=MROH7;refGene_tx=XM_019837070.2;refGene_gn=MROH7;refGene_tx=XM_019837071.2;refGene_gn=MROH7;refGene_tx=XM_019837072.2;refGene_gn=MROH7;refGene_tx=XM_019837073.2
C1 44573994 44574265

refGene_gn=MROH7;refGene_tx=XM_011284974.3;refGene_gn=MROH7;refGene_tx
=XM_019837068.2;refGene_gn=MROH7;refGene_tx=XM_019837069.2;refGene_gn=
MROH7;refGene_tx=XM_019837070.2;refGene_gn=MROH7;refGene_tx=XM_0198370
71.2;refGene_gn=MROH7;refGene_tx=XM_019837072.2;refGene_gn=MROH7;refGe
ne_tx=XM_019837073.2
C1 44576320 44576459
refGene_gn=MROH7;refGene_tx=XM_011284974.3;refGene_gn=MROH7;refGene_tx
=XM_019837068.2;refGene_gn=MROH7;refGene_tx=XM_019837069.2;refGene_gn=
MROH7;refGene_tx=XM_019837070.2;refGene_gn=MROH7;refGene_tx=XM_0198370
71.2;refGene_gn=MROH7;refGene_tx=XM_019837072.2;refGene_gn=MROH7;refGe
ne_tx=XM_019837073.2
C1 44579892 44580095
refGene_gn=MROH7;refGene_tx=XM_011284974.3;refGene_gn=MROH7;refGene_tx
=XM_019837068.2;refGene_gn=MROH7;refGene_tx=XM_019837069.2;refGene_gn=
MROH7;refGene_tx=XM_019837070.2;refGene_gn=MROH7;refGene_tx=XM_0198370
71.2;refGene_gn=MROH7;refGene_tx=XM_019837072.2;refGene_gn=MROH7;refGe
ne_tx=XM_019837073.2
C1 44583433 44583571
refGene_gn=MROH7;refGene_tx=XM_011284974.3;refGene_gn=MROH7;refGene_tx
=XM_019837068.2;refGene_gn=MROH7;refGene_tx=XM_019837069.2;refGene_gn=
MROH7;refGene_tx=XM_019837070.2;refGene_gn=MROH7;refGene_tx=XM_0198370
71.2;refGene_gn=MROH7;refGene_tx=XM_019837072.2;refGene_gn=MROH7;refGe
ne_tx=XM_019837073.2
C1 44584689 44584804
refGene_gn=MROH7;refGene_tx=XM_011284974.3;refGene_gn=MROH7;refGene_tx
=XM_019837068.2;refGene_gn=MROH7;refGene_tx=XM_019837069.2;refGene_gn=
MROH7;refGene_tx=XM_019837070.2;refGene_gn=MROH7;refGene_tx=XM_0198370
71.2;refGene_gn=MROH7;refGene_tx=XM_019837072.2;refGene_gn=MROH7;refGe
ne_tx=XM_019837073.2
C1 44588184 44588325
refGene_gn=MROH7;refGene_tx=XM_011284974.3;refGene_gn=MROH7;refGene_tx
=XM_019837068.2;refGene_gn=MROH7;refGene_tx=XM_019837069.2;refGene_gn=
MROH7;refGene_tx=XM_019837070.2;refGene_gn=MROH7;refGene_tx=XM_0198370
71.2;refGene_gn=MROH7;refGene_tx=XM_019837072.2;refGene_gn=MROH7;refGe
ne_tx=XM_019837073.2
C1 44589482 44589662
refGene_gn=MROH7;refGene_tx=XM_011284974.3;refGene_gn=MROH7;refGene_tx
=XM_019837068.2;refGene_gn=MROH7;refGene_tx=XM_019837069.2;refGene_gn=
MROH7;refGene_tx=XM_019837070.2;refGene_gn=MROH7;refGene_tx=XM_0198370
71.2;refGene_gn=MROH7;refGene_tx=XM_019837072.2;refGene_gn=MROH7;refGe
ne_tx=XM_019837073.2
C1 44590197 44590410
refGene_gn=MROH7;refGene_tx=XM_011284974.3;refGene_gn=MROH7;refGene_tx
=XM_019837068.2;refGene_gn=MROH7;refGene_tx=XM_019837069.2;refGene_gn=
MROH7;refGene_tx=XM_019837070.2;refGene_gn=MROH7;refGene_tx=XM_0198370
71.2;refGene_gn=MROH7;refGene_tx=XM_019837072.2;refGene_gn=MROH7;refGe
ne_tx=XM_019837073.2
C1 44590976 44591099
refGene_gn=MROH7;refGene_tx=XM_011284974.3;refGene_gn=MROH7;refGene_tx
=XM_019837068.2;refGene_gn=MROH7;refGene_tx=XM_019837069.2;refGene_gn=

MROH7;refGene_tx=XM_019837070.2;refGene_gn=MROH7;refGene_tx=XM_0198370
71.2;refGene_gn=MROH7;refGene_tx=XM_019837072.2;refGene_gn=MROH7;refGe
ne_tx=XM_019837073.2
C1 44594403 44594506
refGene_gn=MROH7;refGene_tx=XM_011284974.3;refGene_gn=MROH7;refGene_tx
=XM_019837068.2;refGene_gn=MROH7;refGene_tx=XM_019837069.2;refGene_gn=
MROH7;refGene_tx=XM_019837070.2;refGene_gn=MROH7;refGene_tx=XM_0198370
71.2;refGene_gn=MROH7;refGene_tx=XM_019837072.2
C1 44596604 44596667
refGene_gn=MROH7;refGene_tx=XM_011284974.3;refGene_gn=MROH7;refGene_tx
=XM_019837068.2;refGene_gn=MROH7;refGene_tx=XM_019837069.2;refGene_gn=
MROH7;refGene_tx=XM_019837070.2;refGene_gn=MROH7;refGene_tx=XM_0198370
71.2;refGene_gn=MROH7;refGene_tx=XM_019837072.2;refGene_gn=MROH7;refGe
ne_tx=XM_019837073.2
C1 44597198 44597443
refGene_gn=MROH7;refGene_tx=XM_011284974.3;refGene_gn=MROH7;refGene_tx
=XM_019837068.2;refGene_gn=MROH7;refGene_tx=XM_019837069.2;refGene_gn=
MROH7;refGene_tx=XM_019837070.2;refGene_gn=MROH7;refGene_tx=XM_0198370
71.2;refGene_gn=MROH7;refGene_tx=XM_019837072.2
C1 44601793 44601910 refGene_gn=TTC4;refGene_tx=XM_003990132.4
C1 44602533 44602651 refGene_gn=TTC4;refGene_tx=XM_003990132.4
C1 44603399 44603561 refGene_gn=TTC4;refGene_tx=XM_003990132.4
C1 44608995 44609073 refGene_gn=TTC4;refGene_tx=XM_003990132.4
C1 44610517 44610642 refGene_gn=TTC4;refGene_tx=XM_003990132.4
C1 44618280 44618367 refGene_gn=TTC4;refGene_tx=XM_003990132.4
C1 44620377 44620595 refGene_gn=TTC4;refGene_tx=XM_003990132.4
C1 44621450 44621532 refGene_gn=TTC4;refGene_tx=XM_003990132.4
C1 44629467 44629550 refGene_gn=TTC4;refGene_tx=XM_003990132.4
C1 44632181 44632284 refGene_gn=TTC4;refGene_tx=XM_003990132.4
C1 44641070 44642526 refGene_gn=PARS2;refGene_tx=XM_006934768.3
C1 44644158 44644160 refGene_gn=PARS2;refGene_tx=XM_006934768.3
C1 44659879 44660416 refGene_gn=TTC22;refGene_tx=XM_023258733.1
C1 44660980 44661133 refGene_gn=TTC22;refGene_tx=XM_023258733.1
C1 44662747 44662909 refGene_gn=TTC22;refGene_tx=XM_023258733.1
C1 44663873 44663992 refGene_gn=TTC22;refGene_tx=XM_023258733.1
C1 44664337 44664453 refGene_gn=TTC22;refGene_tx=XM_023258733.1
C1 44664651 44664707 refGene_gn=TTC22;refGene_tx=XM_023258733.1
C1 44678055 44678622 refGene_gn=TTC22;refGene_tx=XM_023258733.1
C1 44683704 44683754
refGene_gn=LEXM;refGene_tx=XM_003990196.4;refGene_gn=LEXM;refGene_tx=X
M_023258734.1
C1 44684755 44684843
refGene_gn=LEXM;refGene_tx=XM_003990196.4;refGene_gn=LEXM;refGene_tx=X
M_023258734.1
C1 44685295 44685460
refGene_gn=LEXM;refGene_tx=XM_003990196.4;refGene_gn=LEXM;refGene_tx=X
M_023258734.1
C1 44685578 44685701
refGene_gn=LEXM;refGene_tx=XM_003990196.4;refGene_gn=LEXM;refGene_tx=X
M_023258734.1

C1	44690237	44690369
refGene_gn=LEXM;refGene_tx=XM_003990196.4;refGene_gn=LEXM;refGene_tx=XM_023258734.1
C1	44690524	44690668
refGene_gn=LEXM;refGene_tx=XM_003990196.4;refGene_gn=LEXM;refGene_tx=XM_023258734.1
C1	44692288	44692450
refGene_gn=LEXM;refGene_tx=XM_003990196.4;refGene_gn=LEXM;refGene_tx=XM_023258734.1
C1	44693413	44693539
refGene_gn=LEXM;refGene_tx=XM_003990196.4;refGene_gn=LEXM;refGene_tx=XM_023258734.1
C1	44695748	44695882
refGene_gn=LEXM;refGene_tx=XM_003990196.4;refGene_gn=LEXM;refGene_tx=XM_023258734.1
C1	44696107	44696111	refGene_gn=LEXM;refGene_tx=XM_023258734.1
C1	44713646	44713719	refGene_gn=LEXM;refGene_tx=XM_003990196.4
C1	44724005	44724159	refGene_gn=DHCR24;refGene_tx=XM_023258735.1
C1	44725122	44725301	refGene_gn=DHCR24;refGene_tx=XM_023258735.1
C1	44725626	44725824	refGene_gn=DHCR24;refGene_tx=XM_023258735.1
C1	44731587	44731731	refGene_gn=DHCR24;refGene_tx=XM_023258735.1
C1	44735510	44735774	refGene_gn=DHCR24;refGene_tx=XM_023258735.1
C1	44737442	44737561	refGene_gn=DHCR24;refGene_tx=XM_023258735.1
C1	44738243	44738349	refGene_gn=DHCR24;refGene_tx=XM_023258735.1
C1	44746301	44746457	refGene_gn=DHCR24;refGene_tx=XM_023258735.1
C1	44749743	44749974	refGene_gn=DHCR24;refGene_tx=XM_023258735.1
C1	44797501	44797516	refGene_gn=TMEM61;refGene_tx=XM_006934770.4
C1	44801602	44801952	refGene_gn=TMEM61;refGene_tx=XM_006934770.4
C1	44807631	44807950	refGene_gn=TMEM61;refGene_tx=XM_006934770.4
C1	44814450	44814627	refGene_gn=BSND;refGene_tx=XM_019837077.2
C1	44821627	44821722	refGene_gn=BSND;refGene_tx=XM_019837077.2
C1	44823417	44823687	refGene_gn=BSND;refGene_tx=XM_019837077.2
C1	44824608	44825023	refGene_gn=BSND;refGene_tx=XM_019837077.2
C1	44844107	44844170
refGene_gn=USP24;refGene_tx=XM_023258737.1;refGene_gn=USP24;refGene_tx=XM_023258738.1;refGene_gn=USP24;refGene_tx=XM_023258739.1
C1	44846430	44846541
refGene_gn=USP24;refGene_tx=XM_023258737.1;refGene_gn=USP24;refGene_tx=XM_023258738.1;refGene_gn=USP24;refGene_tx=XM_023258739.1
C1	44846887	44846974
refGene_gn=USP24;refGene_tx=XM_023258737.1;refGene_gn=USP24;refGene_tx=XM_023258738.1;refGene_gn=USP24;refGene_tx=XM_023258739.1
C1	44847589	44847665
refGene_gn=USP24;refGene_tx=XM_023258737.1;refGene_gn=USP24;refGene_tx=XM_023258738.1;refGene_gn=USP24;refGene_tx=XM_023258739.1
C1	44849695	44849774
refGene_gn=USP24;refGene_tx=XM_023258737.1;refGene_gn=USP24;refGene_tx=XM_023258738.1;refGene_gn=USP24;refGene_tx=XM_023258739.1
C1	44850768	44850835
refGene_gn=USP24;refGene_tx=XM_023258737.1;refGene_gn=USP24;refGene_tx

=XM_023258738.1;refGene_gn=USP24;refGene_tx=XM_023258739.1
C1 44853486 44853552
refGene_gn=USP24;refGene_tx=XM_023258737.1;refGene_gn=USP24;refGene_tx
=XM_023258738.1;refGene_gn=USP24;refGene_tx=XM_023258739.1
C1 44855433 44855547
refGene_gn=USP24;refGene_tx=XM_023258737.1;refGene_gn=USP24;refGene_tx
=XM_023258738.1;refGene_gn=USP24;refGene_tx=XM_023258739.1
C1 44856466 44856588
refGene_gn=USP24;refGene_tx=XM_023258737.1;refGene_gn=USP24;refGene_tx
=XM_023258738.1;refGene_gn=USP24;refGene_tx=XM_023258739.1
C1 44858144 44858247
refGene_gn=USP24;refGene_tx=XM_023258737.1;refGene_gn=USP24;refGene_tx
=XM_023258738.1;refGene_gn=USP24;refGene_tx=XM_023258739.1
C1 44860275 44860368
refGene_gn=USP24;refGene_tx=XM_023258737.1;refGene_gn=USP24;refGene_tx
=XM_023258738.1;refGene_gn=USP24;refGene_tx=XM_023258739.1
C1 44861035 44861152
refGene_gn=USP24;refGene_tx=XM_023258737.1;refGene_gn=USP24;refGene_tx
=XM_023258738.1;refGene_gn=USP24;refGene_tx=XM_023258739.1
C1 44861663 44861696 refGene_gn=USP24;refGene_tx=XM_023258740.1
C1 44863756 44863853
refGene_gn=USP24;refGene_tx=XM_023258737.1;refGene_gn=USP24;refGene_tx
=XM_023258738.1;refGene_gn=USP24;refGene_tx=XM_023258739.1;refGene_gn=
USP24;refGene_tx=XM_023258740.1
C1 44868654 44868768
refGene_gn=USP24;refGene_tx=XM_023258737.1;refGene_gn=USP24;refGene_tx
=XM_023258738.1;refGene_gn=USP24;refGene_tx=XM_023258739.1;refGene_gn=
USP24;refGene_tx=XM_023258740.1
C1 44870340 44870444
refGene_gn=USP24;refGene_tx=XM_023258737.1;refGene_gn=USP24;refGene_tx
=XM_023258738.1;refGene_gn=USP24;refGene_tx=XM_023258739.1;refGene_gn=
USP24;refGene_tx=XM_023258740.1
C1 44871097 44871193
refGene_gn=USP24;refGene_tx=XM_023258737.1;refGene_gn=USP24;refGene_tx
=XM_023258738.1;refGene_gn=USP24;refGene_tx=XM_023258740.1
C1 44872409 44872560
refGene_gn=USP24;refGene_tx=XM_023258737.1;refGene_gn=USP24;refGene_tx
=XM_023258738.1;refGene_gn=USP24;refGene_tx=XM_023258739.1;refGene_gn=
USP24;refGene_tx=XM_023258740.1
C1 44873238 44873380
refGene_gn=USP24;refGene_tx=XM_023258737.1;refGene_gn=USP24;refGene_tx
=XM_023258738.1;refGene_gn=USP24;refGene_tx=XM_023258739.1;refGene_gn=
USP24;refGene_tx=XM_023258740.1
C1 44874683 44874808
refGene_gn=USP24;refGene_tx=XM_023258737.1;refGene_gn=USP24;refGene_tx
=XM_023258738.1;refGene_gn=USP24;refGene_tx=XM_023258739.1;refGene_gn=
USP24;refGene_tx=XM_023258740.1
C1 44875222 44875443
refGene_gn=USP24;refGene_tx=XM_023258737.1;refGene_gn=USP24;refGene_tx
=XM_023258738.1;refGene_gn=USP24;refGene_tx=XM_023258739.1;refGene_gn=

USP24;refGene_tx=XM_023258740.1
C1 44875766 44875886
refGene_gn=USP24;refGene_tx=XM_023258737.1;refGene_gn=USP24;refGene_tx=XM_023258738.1;refGene_gn=USP24;refGene_tx=XM_023258739.1;refGene_gn=USP24;refGene_tx=XM_023258740.1
C1 44876107 44876249
refGene_gn=USP24;refGene_tx=XM_023258737.1;refGene_gn=USP24;refGene_tx=XM_023258738.1;refGene_gn=USP24;refGene_tx=XM_023258739.1;refGene_gn=USP24;refGene_tx=XM_023258740.1
C1 44878034 44878117
refGene_gn=USP24;refGene_tx=XM_023258737.1;refGene_gn=USP24;refGene_tx=XM_023258738.1;refGene_gn=USP24;refGene_tx=XM_023258739.1;refGene_gn=USP24;refGene_tx=XM_023258740.1
C1 44879009 44879108
refGene_gn=USP24;refGene_tx=XM_023258737.1;refGene_gn=USP24;refGene_tx=XM_023258738.1;refGene_gn=USP24;refGene_tx=XM_023258739.1;refGene_gn=USP24;refGene_tx=XM_023258740.1
C1 44880614 44880740
refGene_gn=USP24;refGene_tx=XM_023258737.1;refGene_gn=USP24;refGene_tx=XM_023258738.1;refGene_gn=USP24;refGene_tx=XM_023258739.1;refGene_gn=USP24;refGene_tx=XM_023258740.1
C1 44881999 44882119
refGene_gn=USP24;refGene_tx=XM_023258737.1;refGene_gn=USP24;refGene_tx=XM_023258738.1;refGene_gn=USP24;refGene_tx=XM_023258739.1;refGene_gn=USP24;refGene_tx=XM_023258740.1
C1 44884230 44884375
refGene_gn=USP24;refGene_tx=XM_023258737.1;refGene_gn=USP24;refGene_tx=XM_023258738.1;refGene_gn=USP24;refGene_tx=XM_023258739.1;refGene_gn=USP24;refGene_tx=XM_023258740.1
C1 44886574 44886692
refGene_gn=USP24;refGene_tx=XM_023258737.1;refGene_gn=USP24;refGene_tx=XM_023258738.1;refGene_gn=USP24;refGene_tx=XM_023258739.1;refGene_gn=USP24;refGene_tx=XM_023258740.1
C1 44887681 44887873
refGene_gn=USP24;refGene_tx=XM_023258737.1;refGene_gn=USP24;refGene_tx=XM_023258738.1;refGene_gn=USP24;refGene_tx=XM_023258739.1;refGene_gn=USP24;refGene_tx=XM_023258740.1
C1 44890315 44890377
refGene_gn=USP24;refGene_tx=XM_023258737.1;refGene_gn=USP24;refGene_tx=XM_023258738.1;refGene_gn=USP24;refGene_tx=XM_023258739.1;refGene_gn=USP24;refGene_tx=XM_023258740.1
C1 44893656 44893817
refGene_gn=USP24;refGene_tx=XM_023258737.1;refGene_gn=USP24;refGene_tx=XM_023258738.1;refGene_gn=USP24;refGene_tx=XM_023258739.1;refGene_gn=USP24;refGene_tx=XM_023258740.1
C1 44894480 44894551
refGene_gn=USP24;refGene_tx=XM_023258737.1;refGene_gn=USP24;refGene_tx=XM_023258738.1;refGene_gn=USP24;refGene_tx=XM_023258739.1;refGene_gn=USP24;refGene_tx=XM_023258740.1
C1 44896764 44896920

refGene_gn=USP24;refGene_tx=XM_023258737.1;refGene_gn=USP24;refGene_tx=XM_023258738.1;refGene_gn=USP24;refGene_tx=XM_023258739.1;refGene_gn=USP24;refGene_tx=XM_023258740.1
C1 44897784 44897944
refGene_gn=USP24;refGene_tx=XM_023258737.1;refGene_gn=USP24;refGene_tx=XM_023258738.1;refGene_gn=USP24;refGene_tx=XM_023258739.1;refGene_gn=USP24;refGene_tx=XM_023258740.1
C1 44898581 44898810
refGene_gn=USP24;refGene_tx=XM_023258737.1;refGene_gn=USP24;refGene_tx=XM_023258738.1;refGene_gn=USP24;refGene_tx=XM_023258739.1;refGene_gn=USP24;refGene_tx=XM_023258740.1
C1 44898925 44899021
refGene_gn=USP24;refGene_tx=XM_023258737.1;refGene_gn=USP24;refGene_tx=XM_023258738.1;refGene_gn=USP24;refGene_tx=XM_023258739.1;refGene_gn=USP24;refGene_tx=XM_023258740.1
C1 44902007 44902105
refGene_gn=USP24;refGene_tx=XM_023258737.1;refGene_gn=USP24;refGene_tx=XM_023258738.1;refGene_gn=USP24;refGene_tx=XM_023258739.1;refGene_gn=USP24;refGene_tx=XM_023258740.1
C1 44905920 44906076
refGene_gn=USP24;refGene_tx=XM_023258737.1;refGene_gn=USP24;refGene_tx=XM_023258738.1;refGene_gn=USP24;refGene_tx=XM_023258739.1;refGene_gn=USP24;refGene_tx=XM_023258740.1
C1 44907105 44907199
refGene_gn=USP24;refGene_tx=XM_023258737.1;refGene_gn=USP24;refGene_tx=XM_023258738.1;refGene_gn=USP24;refGene_tx=XM_023258739.1;refGene_gn=USP24;refGene_tx=XM_023258740.1
C1 44907348 44907434
refGene_gn=USP24;refGene_tx=XM_023258737.1;refGene_gn=USP24;refGene_tx=XM_023258738.1;refGene_gn=USP24;refGene_tx=XM_023258739.1;refGene_gn=USP24;refGene_tx=XM_023258740.1
C1 44911911 44912085
refGene_gn=USP24;refGene_tx=XM_023258737.1;refGene_gn=USP24;refGene_tx=XM_023258738.1;refGene_gn=USP24;refGene_tx=XM_023258739.1;refGene_gn=USP24;refGene_tx=XM_023258740.1
C1 44912173 44912272
refGene_gn=USP24;refGene_tx=XM_023258737.1;refGene_gn=USP24;refGene_tx=XM_023258738.1;refGene_gn=USP24;refGene_tx=XM_023258739.1;refGene_gn=USP24;refGene_tx=XM_023258740.1
C1 44912995 44913106
refGene_gn=USP24;refGene_tx=XM_023258737.1;refGene_gn=USP24;refGene_tx=XM_023258738.1;refGene_gn=USP24;refGene_tx=XM_023258739.1;refGene_gn=USP24;refGene_tx=XM_023258740.1
C1 44913230 44913297
refGene_gn=USP24;refGene_tx=XM_023258737.1;refGene_gn=USP24;refGene_tx=XM_023258738.1;refGene_gn=USP24;refGene_tx=XM_023258739.1;refGene_gn=USP24;refGene_tx=XM_023258740.1
C1 44916900 44917016
refGene_gn=USP24;refGene_tx=XM_023258737.1;refGene_gn=USP24;refGene_tx=XM_023258738.1;refGene_gn=USP24;refGene_tx=XM_023258739.1;refGene_gn=

USP24;refGene_tx=XM_023258740.1
C1 44918027 44918081
refGene_gn=USP24;refGene_tx=XM_023258737.1;refGene_gn=USP24;refGene_tx
=XM_023258738.1;refGene_gn=USP24;refGene_tx=XM_023258739.1;refGene_gn=
USP24;refGene_tx=XM_023258740.1
C1 44918272 44918413
refGene_gn=USP24;refGene_tx=XM_023258737.1;refGene_gn=USP24;refGene_tx
=XM_023258738.1;refGene_gn=USP24;refGene_tx=XM_023258739.1;refGene_gn=
USP24;refGene_tx=XM_023258740.1
C1 44919641 44919718
refGene_gn=USP24;refGene_tx=XM_023258737.1;refGene_gn=USP24;refGene_tx
=XM_023258738.1;refGene_gn=USP24;refGene_tx=XM_023258739.1;refGene_gn=
USP24;refGene_tx=XM_023258740.1
C1 44920839 44920951
refGene_gn=USP24;refGene_tx=XM_023258737.1;refGene_gn=USP24;refGene_tx
=XM_023258738.1;refGene_gn=USP24;refGene_tx=XM_023258739.1;refGene_gn=
USP24;refGene_tx=XM_023258740.1
C1 44921804 44921936
refGene_gn=USP24;refGene_tx=XM_023258737.1;refGene_gn=USP24;refGene_tx
=XM_023258738.1;refGene_gn=USP24;refGene_tx=XM_023258739.1;refGene_gn=
USP24;refGene_tx=XM_023258740.1
C1 44922558 44922708
refGene_gn=USP24;refGene_tx=XM_023258737.1;refGene_gn=USP24;refGene_tx
=XM_023258738.1;refGene_gn=USP24;refGene_tx=XM_023258739.1;refGene_gn=
USP24;refGene_tx=XM_023258740.1
C1 44923629 44923737
refGene_gn=USP24;refGene_tx=XM_023258737.1;refGene_gn=USP24;refGene_tx
=XM_023258738.1;refGene_gn=USP24;refGene_tx=XM_023258739.1;refGene_gn=
USP24;refGene_tx=XM_023258740.1
C1 44929046 44929094
refGene_gn=USP24;refGene_tx=XM_023258737.1;refGene_gn=USP24;refGene_tx
=XM_023258738.1;refGene_gn=USP24;refGene_tx=XM_023258739.1;refGene_gn=
USP24;refGene_tx=XM_023258740.1
C1 44929292 44929454
refGene_gn=USP24;refGene_tx=XM_023258737.1;refGene_gn=USP24;refGene_tx
=XM_023258738.1;refGene_gn=USP24;refGene_tx=XM_023258739.1;refGene_gn=
USP24;refGene_tx=XM_023258740.1
C1 44929544 44929640
refGene_gn=USP24;refGene_tx=XM_023258737.1;refGene_gn=USP24;refGene_tx
=XM_023258738.1;refGene_gn=USP24;refGene_tx=XM_023258739.1;refGene_gn=
USP24;refGene_tx=XM_023258740.1
C1 44929825 44929933
refGene_gn=USP24;refGene_tx=XM_023258737.1;refGene_gn=USP24;refGene_tx
=XM_023258738.1;refGene_gn=USP24;refGene_tx=XM_023258739.1;refGene_gn=
USP24;refGene_tx=XM_023258740.1
C1 44932028 44932132
refGene_gn=USP24;refGene_tx=XM_023258737.1;refGene_gn=USP24;refGene_tx
=XM_023258738.1;refGene_gn=USP24;refGene_tx=XM_023258739.1;refGene_gn=
USP24;refGene_tx=XM_023258740.1
C1 44932340 44932455

refGene_gn=USP24;refGene_tx=XM_023258737.1;refGene_gn=USP24;refGene_tx=XM_023258738.1;refGene_gn=USP24;refGene_tx=XM_023258739.1;refGene_gn=USP24;refGene_tx=XM_023258740.1
C1 44933165 44933324
refGene_gn=USP24;refGene_tx=XM_023258737.1;refGene_gn=USP24;refGene_tx=XM_023258738.1;refGene_gn=USP24;refGene_tx=XM_023258739.1;refGene_gn=USP24;refGene_tx=XM_023258740.1
C1 44933907 44933982
refGene_gn=USP24;refGene_tx=XM_023258737.1;refGene_gn=USP24;refGene_tx=XM_023258738.1;refGene_gn=USP24;refGene_tx=XM_023258739.1;refGene_gn=USP24;refGene_tx=XM_023258740.1
C1 44936449 44936515
refGene_gn=USP24;refGene_tx=XM_023258737.1;refGene_gn=USP24;refGene_tx=XM_023258738.1;refGene_gn=USP24;refGene_tx=XM_023258739.1;refGene_gn=USP24;refGene_tx=XM_023258740.1
C1 44940937 44941003
refGene_gn=USP24;refGene_tx=XM_023258737.1;refGene_gn=USP24;refGene_tx=XM_023258738.1;refGene_gn=USP24;refGene_tx=XM_023258739.1;refGene_gn=USP24;refGene_tx=XM_023258740.1
C1 44941341 44941377
refGene_gn=USP24;refGene_tx=XM_023258737.1;refGene_gn=USP24;refGene_tx=XM_023258738.1;refGene_gn=USP24;refGene_tx=XM_023258739.1;refGene_gn=USP24;refGene_tx=XM_023258740.1
C1 44947229 44947352
refGene_gn=USP24;refGene_tx=XM_023258737.1;refGene_gn=USP24;refGene_tx=XM_023258738.1;refGene_gn=USP24;refGene_tx=XM_023258739.1;refGene_gn=USP24;refGene_tx=XM_023258740.1
C1 44947999 44948143
refGene_gn=USP24;refGene_tx=XM_023258737.1;refGene_gn=USP24;refGene_tx=XM_023258738.1;refGene_gn=USP24;refGene_tx=XM_023258739.1;refGene_gn=USP24;refGene_tx=XM_023258740.1
C1 44952021 44952089
refGene_gn=USP24;refGene_tx=XM_023258737.1;refGene_gn=USP24;refGene_tx=XM_023258738.1;refGene_gn=USP24;refGene_tx=XM_023258739.1;refGene_gn=USP24;refGene_tx=XM_023258740.1
C1 44953491 44953657
refGene_gn=USP24;refGene_tx=XM_023258737.1;refGene_gn=USP24;refGene_tx=XM_023258738.1;refGene_gn=USP24;refGene_tx=XM_023258739.1;refGene_gn=USP24;refGene_tx=XM_023258740.1
C1 44983298 44983619
refGene_gn=USP24;refGene_tx=XM_023258737.1;refGene_gn=USP24;refGene_tx=XM_023258738.1;refGene_gn=USP24;refGene_tx=XM_023258739.1;refGene_gn=USP24;refGene_tx=XM_023258740.1
C1 45006989 45007020
refGene_gn=LOC102901879;refGene_tx=XM_023257829.1
C1 45007222 45007560
refGene_gn=LOC102901879;refGene_tx=XM_023257829.1
C1 45031062 45031168
refGene_gn=LOC102901879;refGene_tx=XM_023257829.1
C1 45032097 45032253

refGene_gn=LOC102901879;refGene_tx=XM_023257829.1
C1 45032744 45032847
refGene_gn=LOC102901879;refGene_tx=XM_023257829.1
C1 45034901 45034983
refGene_gn=LOC102901879;refGene_tx=XM_023257829.1
C1 46049067 46049193
refGene_gn=PLPP3;refGene_tx=XM_003990199.5;refGene_gn=PLPP3;refGene_tx=XM_019837081.2;refGene_gn=PLPP3;refGene_tx=XM_023258741.1;refGene_gn=PLPP3;refGene_tx=XM_023258742.1
C1 46062005 46062182
refGene_gn=PLPP3;refGene_tx=XM_003990199.5;refGene_gn=PLPP3;refGene_tx=XM_019837081.2;refGene_gn=PLPP3;refGene_tx=XM_023258741.1
C1 46072917 46072975
refGene_gn=PLPP3;refGene_tx=XM_003990199.5;refGene_gn=PLPP3;refGene_tx=XM_019837081.2;refGene_gn=PLPP3;refGene_tx=XM_023258741.1;refGene_gn=PLPP3;refGene_tx=XM_023258742.1
C1 46073406 46073687
refGene_gn=PLPP3;refGene_tx=XM_003990199.5;refGene_gn=PLPP3;refGene_tx=XM_019837081.2;refGene_gn=PLPP3;refGene_tx=XM_023258741.1;refGene_gn=PLPP3;refGene_tx=XM_023258742.1
C1 46089287 46089445
refGene_gn=PLPP3;refGene_tx=XM_003990199.5;refGene_gn=PLPP3;refGene_tx=XM_019837081.2;refGene_gn=PLPP3;refGene_tx=XM_023258741.1;refGene_gn=PLPP3;refGene_tx=XM_023258742.1
C1 46099105 46099108 refGene_gn=PLPP3;refGene_tx=XM_003990199.5
C1 46107145 46107146 refGene_gn=PLPP3;refGene_tx=XM_003990199.5
C1 46122924 46123030 refGene_gn=PLPP3;refGene_tx=XM_019837081.2
C1 46136634 46136773
refGene_gn=PLPP3;refGene_tx=XM_023258741.1;refGene_gn=PLPP3;refGene_tx=XM_023258742.1
C1 46186927 46187021
refGene_gn=PRKAA2;refGene_tx=XM_023258743.1;refGene_gn=PRKAA2;refGene_tx=XM_023258744.1
C1 46221895 46222037
refGene_gn=PRKAA2;refGene_tx=XM_023258743.1;refGene_gn=PRKAA2;refGene_tx=XM_023258744.1
C1 46236536 46236630
refGene_gn=PRKAA2;refGene_tx=XM_023258743.1;refGene_gn=PRKAA2;refGene_tx=XM_023258744.1;refGene_gn=PRKAA2;refGene_tx=XM_023258745.1;refGene_gn=PRKAA2;refGene_tx=XM_023258746.1
C1 46237324 46237469
refGene_gn=PRKAA2;refGene_tx=XM_023258743.1;refGene_gn=PRKAA2;refGene_tx=XM_023258744.1;refGene_gn=PRKAA2;refGene_tx=XM_023258745.1;refGene_gn=PRKAA2;refGene_tx=XM_023258746.1
C1 46247641 46247729
refGene_gn=PRKAA2;refGene_tx=XM_023258743.1;refGene_gn=PRKAA2;refGene_tx=XM_023258744.1;refGene_gn=PRKAA2;refGene_tx=XM_023258745.1;refGene_gn=PRKAA2;refGene_tx=XM_023258746.1
C1 46249730 46249955
refGene_gn=PRKAA2;refGene_tx=XM_023258743.1;refGene_gn=PRKAA2;refGene_

tx=XM_023258745.1;refGene_gn=PRKAA2;refGene_tx=XM_023258746.1
C1 46262138 46262643
refGene_gn=PRKAA2;refGene_tx=XM_023258743.1;refGene_gn=PRKAA2;refGene_tx=XM_023258744.1;refGene_gn=PRKAA2;refGene_tx=XM_023258745.1;refGene_gn=PRKAA2;refGene_tx=XM_023258746.1
C1 46264083 46264210
refGene_gn=PRKAA2;refGene_tx=XM_023258743.1;refGene_gn=PRKAA2;refGene_tx=XM_023258744.1;refGene_gn=PRKAA2;refGene_tx=XM_023258745.1;refGene_gn=PRKAA2;refGene_tx=XM_023258746.1
C1 46265480 46265719
refGene_gn=PRKAA2;refGene_tx=XM_023258743.1;refGene_gn=PRKAA2;refGene_tx=XM_023258744.1;refGene_gn=PRKAA2;refGene_tx=XM_023258745.1;refGene_gn=PRKAA2;refGene_tx=XM_023258746.1
C1 46277062 46277084 refGene_gn=FYB2;refGene_tx=XM_023257830.1
C1 46277323 46277357 refGene_gn=FYB2;refGene_tx=XM_023257830.1
C1 46277462 46277619 refGene_gn=FYB2;refGene_tx=XM_023257830.1
C1 46280327 46280409 refGene_gn=FYB2;refGene_tx=XM_023257830.1
C1 46282700 46282787 refGene_gn=FYB2;refGene_tx=XM_023257830.1
C1 46295111 46295172 refGene_gn=FYB2;refGene_tx=XM_023257830.1
C1 46296617 46296646 refGene_gn=FYB2;refGene_tx=XM_023257830.1
C1 46301578 46301677 refGene_gn=FYB2;refGene_tx=XM_023257830.1
C1 46302711 46302790 refGene_gn=FYB2;refGene_tx=XM_023257830.1
C1 46303860 46303901 refGene_gn=FYB2;refGene_tx=XM_023257830.1
C1 46303991 46304106 refGene_gn=FYB2;refGene_tx=XM_023257830.1
C1 46313971 46314134 refGene_gn=FYB2;refGene_tx=XM_023257830.1
C1 46316021 46316177 refGene_gn=FYB2;refGene_tx=XM_023257830.1
C1 46316767 46316856 refGene_gn=FYB2;refGene_tx=XM_023257830.1
C1 46323877 46323912 refGene_gn=FYB2;refGene_tx=XM_023257830.1
C1 46333322 46333423 refGene_gn=FYB2;refGene_tx=XM_023257830.1
C1 46364380 46364414 refGene_gn=FYB2;refGene_tx=XM_023257830.1
C1 46367592 46367754 refGene_gn=FYB2;refGene_tx=XM_023257830.1
C1 46371348 46372090 refGene_gn=FYB2;refGene_tx=XM_023257830.1
C1 46406735 46407068 refGene_gn=FYB2;refGene_tx=XM_023257830.1
C1 46439540 46439617 refGene_gn=C8A;refGene_tx=XM_019837087.2
C1 46461441 46461544 refGene_gn=C8A;refGene_tx=XM_019837087.2
C1 46466992 46467137 refGene_gn=C8A;refGene_tx=XM_019837087.2
C1 46468110 46468258 refGene_gn=C8A;refGene_tx=XM_019837087.2
C1 46471720 46471910 refGene_gn=C8A;refGene_tx=XM_019837087.2
C1 46474941 46475145 refGene_gn=C8A;refGene_tx=XM_019837087.2
C1 46477293 46477534 refGene_gn=C8A;refGene_tx=XM_019837087.2
C1 46498414 46498546 refGene_gn=C8A;refGene_tx=XM_019837087.2
C1 46499584 46499742 refGene_gn=C8A;refGene_tx=XM_019837087.2
C1 46503112 46503123 refGene_gn=C8B;refGene_tx=XM_019837086.2
C1 46504094 46504317 refGene_gn=C8A;refGene_tx=XM_019837087.2
C1 46508061 46508213 refGene_gn=C8A;refGene_tx=XM_019837087.2
C1 46520067 46520219 refGene_gn=C8B;refGene_tx=XM_003990136.4
C1 46522545 46522614
refGene_gn=C8B;refGene_tx=XM_003990136.4;refGene_gn=C8B;refGene_tx=XM_019837086.2
C1 46524662 46524816

refGene_gn=C8B;refGene_tx=XM_003990136.4;refGene_gn=C8B;refGene_tx=XM_019837086.2
C1 46531862 46532026
refGene_gn=C8B;refGene_tx=XM_003990136.4;refGene_gn=C8B;refGene_tx=XM_019837086.2
C1 46534955 46535084
refGene_gn=C8B;refGene_tx=XM_003990136.4;refGene_gn=C8B;refGene_tx=XM_019837086.2
C1 46536958 46537199
refGene_gn=C8B;refGene_tx=XM_003990136.4;refGene_gn=C8B;refGene_tx=XM_019837086.2
C1 46540698 46540896
refGene_gn=C8B;refGene_tx=XM_003990136.4;refGene_gn=C8B;refGene_tx=XM_019837086.2
C1 46543262 46543395
refGene_gn=C8B;refGene_tx=XM_003990136.4;refGene_gn=C8B;refGene_tx=XM_019837086.2
C1 46545678 46545820
refGene_gn=C8B;refGene_tx=XM_003990136.4;refGene_gn=C8B;refGene_tx=XM_019837086.2
C1 46547394 46547536
refGene_gn=C8B;refGene_tx=XM_003990136.4;refGene_gn=C8B;refGene_tx=XM_019837086.2
C1 46550798 46550955
refGene_gn=C8B;refGene_tx=XM_003990136.4;refGene_gn=C8B;refGene_tx=XM_019837086.2
C1 46557240 46557332
refGene_gn=C8B;refGene_tx=XM_003990136.4;refGene_gn=C8B;refGene_tx=XM_019837086.2
C1 46591595 46591640 refGene_gn=DAB1;refGene_tx=XM_023258754.1
C1 46602428 46602524
refGene_gn=DAB1;refGene_tx=XM_011284970.3;refGene_gn=DAB1;refGene_tx=XM_019837088.2;refGene_gn=DAB1;refGene_tx=XM_019837092.2;refGene_gn=DAB1;refGene_tx=XM_019837093.2;refGene_gn=DAB1;refGene_tx=XM_023258747.1;refGene_gn=DAB1;refGene_tx=XM_023258748.1;refGene_gn=DAB1;refGene_tx=XM_023258749.1;refGene_gn=DAB1;refGene_tx=XM_023258750.1;refGene_gn=DAB1;refGene_tx=XM_023258751.1;refGene_gn=DAB1;refGene_tx=XM_023258752.1;refGene_gn=DAB1;refGene_tx=XM_023258753.1;refGene_gn=DAB1;refGene_tx=XM_023258755.1;refGene_gn=DAB1;refGene_tx=XM_023258756.1;refGene_gn=DAB1;refGene_tx=XM_023258757.1;refGene_gn=DAB1;refGene_tx=XM_023258758.1;refGene_gn=DAB1;refGene_tx=XM_023258759.1;refGene_gn=DAB1;refGene_tx=XM_023258760.1;refGene_gn=DAB1;refGene_tx=XM_023258761.1;refGene_gn=DAB1;refGene_tx=XM_023258762.1
C1 46602863 46602991
refGene_gn=DAB1;refGene_tx=XM_011284970.3;refGene_gn=DAB1;refGene_tx=XM_019837088.2;refGene_gn=DAB1;refGene_tx=XM_019837092.2;refGene_gn=DAB1;refGene_tx=XM_019837093.2;refGene_gn=DAB1;refGene_tx=XM_023258747.1;refGene_gn=DAB1;refGene_tx=XM_023258748.1;refGene_gn=DAB1;refGene_tx=XM_023258749.1;refGene_gn=DAB1;refGene_tx=XM_023258750.1;refGene_gn=DAB1;refGene_tx=XM_023258751.1;refGene_gn=DAB1;refGene_tx=XM_023258752.1;

refGene_gn=DAB1;refGene_tx=XM_023258753.1;refGene_gn=DAB1;refGene_tx=XM_023258754.1;refGene_gn=DAB1;refGene_tx=XM_023258755.1;refGene_gn=DAB1;refGene_tx=XM_023258756.1;refGene_gn=DAB1;refGene_tx=XM_023258757.1;refGene_gn=DAB1;refGene_tx=XM_023258758.1;refGene_gn=DAB1;refGene_tx=XM_023258759.1;refGene_gn=DAB1;refGene_tx=XM_023258760.1;refGene_gn=DAB1;refGene_tx=XM_023258761.1;refGene_gn=DAB1;refGene_tx=XM_023258762.1
C1 46606754 46607303
refGene_gn=DAB1;refGene_tx=XM_011284970.3;refGene_gn=DAB1;refGene_tx=XM_019837088.2;refGene_gn=DAB1;refGene_tx=XM_019837092.2;refGene_gn=DAB1;refGene_tx=XM_019837093.2;refGene_gn=DAB1;refGene_tx=XM_023258747.1;refGene_gn=DAB1;refGene_tx=XM_023258748.1;refGene_gn=DAB1;refGene_tx=XM_023258749.1;refGene_gn=DAB1;refGene_tx=XM_023258750.1;refGene_gn=DAB1;refGene_tx=XM_023258751.1;refGene_gn=DAB1;refGene_tx=XM_023258752.1;refGene_gn=DAB1;refGene_tx=XM_023258753.1;refGene_gn=DAB1;refGene_tx=XM_023258754.1;refGene_gn=DAB1;refGene_tx=XM_023258755.1;refGene_gn=DAB1;refGene_tx=XM_023258756.1;refGene_gn=DAB1;refGene_tx=XM_023258757.1;refGene_gn=DAB1;refGene_tx=XM_023258758.1;refGene_gn=DAB1;refGene_tx=XM_023258759.1;refGene_gn=DAB1;refGene_tx=XM_023258760.1;refGene_gn=DAB1;refGene_tx=XM_023258761.1
C1 46614922 46615031
refGene_gn=DAB1;refGene_tx=XM_011284970.3;refGene_gn=DAB1;refGene_tx=XM_019837088.2;refGene_gn=DAB1;refGene_tx=XM_019837092.2;refGene_gn=DAB1;refGene_tx=XM_019837093.2;refGene_gn=DAB1;refGene_tx=XM_023258747.1;refGene_gn=DAB1;refGene_tx=XM_023258748.1;refGene_gn=DAB1;refGene_tx=XM_023258749.1;refGene_gn=DAB1;refGene_tx=XM_023258750.1;refGene_gn=DAB1;refGene_tx=XM_023258751.1;refGene_gn=DAB1;refGene_tx=XM_023258752.1;refGene_gn=DAB1;refGene_tx=XM_023258753.1;refGene_gn=DAB1;refGene_tx=XM_023258754.1;refGene_gn=DAB1;refGene_tx=XM_023258755.1;refGene_gn=DAB1;refGene_tx=XM_023258756.1;refGene_gn=DAB1;refGene_tx=XM_023258757.1;refGene_gn=DAB1;refGene_tx=XM_023258758.1;refGene_gn=DAB1;refGene_tx=XM_023258759.1;refGene_gn=DAB1;refGene_tx=XM_023258760.1;refGene_gn=DAB1;refGene_tx=XM_023258761.1;refGene_gn=DAB1;refGene_tx=XM_023258762.1
C1 46617366 46617429
refGene_gn=DAB1;refGene_tx=XM_011284970.3;refGene_gn=DAB1;refGene_tx=XM_019837088.2;refGene_gn=DAB1;refGene_tx=XM_019837092.2;refGene_gn=DAB1;refGene_tx=XM_019837093.2;refGene_gn=DAB1;refGene_tx=XM_023258747.1;refGene_gn=DAB1;refGene_tx=XM_023258748.1;refGene_gn=DAB1;refGene_tx=XM_023258749.1;refGene_gn=DAB1;refGene_tx=XM_023258750.1;refGene_gn=DAB1;refGene_tx=XM_023258751.1;refGene_gn=DAB1;refGene_tx=XM_023258752.1;refGene_gn=DAB1;refGene_tx=XM_023258753.1;refGene_gn=DAB1;refGene_tx=XM_023258754.1;refGene_gn=DAB1;refGene_tx=XM_023258755.1;refGene_gn=DAB1;refGene_tx=XM_023258756.1;refGene_gn=DAB1;refGene_tx=XM_023258757.1;refGene_gn=DAB1;refGene_tx=XM_023258758.1;refGene_gn=DAB1;refGene_tx=XM_023258759.1;refGene_gn=DAB1;refGene_tx=XM_023258760.1;refGene_gn=DAB1;refGene_tx=XM_023258761.1;refGene_gn=DAB1;refGene_tx=XM_023258762.1
C1 46622119 46622167
refGene_gn=DAB1;refGene_tx=XM_011284970.3;refGene_gn=DAB1;refGene_tx=XM_019837088.2;refGene_gn=DAB1;refGene_tx=XM_019837092.2;refGene_gn=DAB1;refGene_tx=XM_019837093.2;refGene_gn=DAB1;refGene_tx=XM_023258747.1;refGene_gn=DAB1;refGene_tx=XM_023258748.1;refGene_gn=DAB1;refGene_tx=X

M_023258749.1;refGene_gn=DAB1;refGene_tx=XM_023258750.1;refGene_gn=DAB1;refGene_tx=XM_023258751.1;refGene_gn=DAB1;refGene_tx=XM_023258752.1;refGene_gn=DAB1;refGene_tx=XM_023258754.1;refGene_gn=DAB1;refGene_tx=XM_023258756.1;refGene_gn=DAB1;refGene_tx=XM_023258758.1;refGene_gn=DAB1;refGene_tx=XM_023258762.1
C1 46622258 46622309
refGene_gn=DAB1;refGene_tx=XM_011284970.3;refGene_gn=DAB1;refGene_tx=XM_019837088.2;refGene_gn=DAB1;refGene_tx=XM_019837092.2;refGene_gn=DAB1;refGene_tx=XM_019837093.2;refGene_gn=DAB1;refGene_tx=XM_023258747.1;refGene_gn=DAB1;refGene_tx=XM_023258748.1;refGene_gn=DAB1;refGene_tx=XM_023258749.1;refGene_gn=DAB1;refGene_tx=XM_023258750.1;refGene_gn=DAB1;refGene_tx=XM_023258751.1;refGene_gn=DAB1;refGene_tx=XM_023258752.1;refGene_gn=DAB1;refGene_tx=XM_023258753.1;refGene_gn=DAB1;refGene_tx=XM_023258754.1;refGene_gn=DAB1;refGene_tx=XM_023258756.1;refGene_gn=DAB1;refGene_tx=XM_023258758.1;refGene_gn=DAB1;refGene_tx=XM_023258759.1;refGene_gn=DAB1;refGene_tx=XM_023258762.1
C1 46647112 46647172
refGene_gn=DAB1;refGene_tx=XM_011284970.3;refGene_gn=DAB1;refGene_tx=XM_019837088.2;refGene_gn=DAB1;refGene_tx=XM_019837092.2;refGene_gn=DAB1;refGene_tx=XM_019837093.2;refGene_gn=DAB1;refGene_tx=XM_023258747.1;refGene_gn=DAB1;refGene_tx=XM_023258748.1;refGene_gn=DAB1;refGene_tx=XM_023258749.1;refGene_gn=DAB1;refGene_tx=XM_023258750.1;refGene_gn=DAB1;refGene_tx=XM_023258751.1;refGene_gn=DAB1;refGene_tx=XM_023258752.1;refGene_gn=DAB1;refGene_tx=XM_023258753.1;refGene_gn=DAB1;refGene_tx=XM_023258754.1;refGene_gn=DAB1;refGene_tx=XM_023258755.1;refGene_gn=DAB1;refGene_tx=XM_023258756.1;refGene_gn=DAB1;refGene_tx=XM_023258757.1;refGene_gn=DAB1;refGene_tx=XM_023258758.1;refGene_gn=DAB1;refGene_tx=XM_023258759.1;refGene_gn=DAB1;refGene_tx=XM_023258760.1;refGene_gn=DAB1;refGene_tx=XM_023258761.1;refGene_gn=DAB1;refGene_tx=XM_023258762.1
C1 46654002 46654068
refGene_gn=DAB1;refGene_tx=XM_011284970.3;refGene_gn=DAB1;refGene_tx=XM_019837088.2;refGene_gn=DAB1;refGene_tx=XM_019837092.2;refGene_gn=DAB1;refGene_tx=XM_019837093.2;refGene_gn=DAB1;refGene_tx=XM_023258747.1;refGene_gn=DAB1;refGene_tx=XM_023258748.1;refGene_gn=DAB1;refGene_tx=XM_023258749.1;refGene_gn=DAB1;refGene_tx=XM_023258750.1;refGene_gn=DAB1;refGene_tx=XM_023258751.1;refGene_gn=DAB1;refGene_tx=XM_023258752.1;refGene_gn=DAB1;refGene_tx=XM_023258753.1;refGene_gn=DAB1;refGene_tx=XM_023258754.1;refGene_gn=DAB1;refGene_tx=XM_023258755.1;refGene_gn=DAB1;refGene_tx=XM_023258757.1;refGene_gn=DAB1;refGene_tx=XM_023258762.1
C1 46655613 46655652
refGene_gn=DAB1;refGene_tx=XM_011284970.3;refGene_gn=DAB1;refGene_tx=XM_019837088.2;refGene_gn=DAB1;refGene_tx=XM_019837092.2;refGene_gn=DAB1;refGene_tx=XM_019837093.2;refGene_gn=DAB1;refGene_tx=XM_023258747.1;refGene_gn=DAB1;refGene_tx=XM_023258748.1;refGene_gn=DAB1;refGene_tx=XM_023258749.1;refGene_gn=DAB1;refGene_tx=XM_023258750.1;refGene_gn=DAB1;refGene_tx=XM_023258751.1;refGene_gn=DAB1;refGene_tx=XM_023258753.1;refGene_gn=DAB1;refGene_tx=XM_023258754.1;refGene_gn=DAB1;refGene_tx=XM_023258755.1;refGene_gn=DAB1;refGene_tx=XM_023258757.1;refGene_gn=DAB1;refGene_tx=XM_023258762.1
C1 46656133 46656253

refGene_gn=DAB1;refGene_tx=XM_011284970.3;refGene_gn=DAB1;refGene_tx=XM_019837088.2;refGene_gn=DAB1;refGene_tx=XM_019837092.2;refGene_gn=DAB1;refGene_tx=XM_019837093.2;refGene_gn=DAB1;refGene_tx=XM_023258747.1;refGene_gn=DAB1;refGene_tx=XM_023258748.1;refGene_gn=DAB1;refGene_tx=XM_023258749.1;refGene_gn=DAB1;refGene_tx=XM_023258750.1;refGene_gn=DAB1;refGene_tx=XM_023258751.1;refGene_gn=DAB1;refGene_tx=XM_023258752.1;refGene_gn=DAB1;refGene_tx=XM_023258753.1;refGene_gn=DAB1;refGene_tx=XM_023258754.1;refGene_gn=DAB1;refGene_tx=XM_023258755.1;refGene_gn=DAB1;refGene_tx=XM_023258756.1;refGene_gn=DAB1;refGene_tx=XM_023258757.1;refGene_gn=DAB1;refGene_tx=XM_023258758.1;refGene_gn=DAB1;refGene_tx=XM_023258759.1;refGene_gn=DAB1;refGene_tx=XM_023258760.1;refGene_gn=DAB1;refGene_tx=XM_023258761.1;refGene_gn=DAB1;refGene_tx=XM_023258762.1
C1 46656859 46656991
refGene_gn=DAB1;refGene_tx=XM_011284970.3;refGene_gn=DAB1;refGene_tx=XM_019837088.2;refGene_gn=DAB1;refGene_tx=XM_019837092.2;refGene_gn=DAB1;refGene_tx=XM_019837093.2;refGene_gn=DAB1;refGene_tx=XM_023258747.1;refGene_gn=DAB1;refGene_tx=XM_023258748.1;refGene_gn=DAB1;refGene_tx=XM_023258749.1;refGene_gn=DAB1;refGene_tx=XM_023258750.1;refGene_gn=DAB1;refGene_tx=XM_023258751.1;refGene_gn=DAB1;refGene_tx=XM_023258752.1;refGene_gn=DAB1;refGene_tx=XM_023258753.1;refGene_gn=DAB1;refGene_tx=XM_023258754.1;refGene_gn=DAB1;refGene_tx=XM_023258755.1;refGene_gn=DAB1;refGene_tx=XM_023258756.1;refGene_gn=DAB1;refGene_tx=XM_023258757.1;refGene_gn=DAB1;refGene_tx=XM_023258758.1;refGene_gn=DAB1;refGene_tx=XM_023258759.1;refGene_gn=DAB1;refGene_tx=XM_023258760.1;refGene_gn=DAB1;refGene_tx=XM_023258761.1;refGene_gn=DAB1;refGene_tx=XM_023258762.1
C1 46726118 46726217
refGene_gn=DAB1;refGene_tx=XM_011284970.3;refGene_gn=DAB1;refGene_tx=XM_019837088.2;refGene_gn=DAB1;refGene_tx=XM_019837092.2;refGene_gn=DAB1;refGene_tx=XM_019837093.2;refGene_gn=DAB1;refGene_tx=XM_023258747.1;refGene_gn=DAB1;refGene_tx=XM_023258748.1;refGene_gn=DAB1;refGene_tx=XM_023258749.1;refGene_gn=DAB1;refGene_tx=XM_023258750.1;refGene_gn=DAB1;refGene_tx=XM_023258751.1;refGene_gn=DAB1;refGene_tx=XM_023258752.1;refGene_gn=DAB1;refGene_tx=XM_023258753.1;refGene_gn=DAB1;refGene_tx=XM_023258754.1;refGene_gn=DAB1;refGene_tx=XM_023258755.1;refGene_gn=DAB1;refGene_tx=XM_023258756.1;refGene_gn=DAB1;refGene_tx=XM_023258757.1;refGene_gn=DAB1;refGene_tx=XM_023258758.1;refGene_gn=DAB1;refGene_tx=XM_023258759.1;refGene_gn=DAB1;refGene_tx=XM_023258760.1;refGene_gn=DAB1;refGene_tx=XM_023258761.1;refGene_gn=DAB1;refGene_tx=XM_023258762.1
C1 46734765 46734905
refGene_gn=DAB1;refGene_tx=XM_011284970.3;refGene_gn=DAB1;refGene_tx=XM_019837088.2;refGene_gn=DAB1;refGene_tx=XM_019837092.2;refGene_gn=DAB1;refGene_tx=XM_019837093.2;refGene_gn=DAB1;refGene_tx=XM_023258747.1;refGene_gn=DAB1;refGene_tx=XM_023258748.1;refGene_gn=DAB1;refGene_tx=XM_023258749.1;refGene_gn=DAB1;refGene_tx=XM_023258750.1;refGene_gn=DAB1;refGene_tx=XM_023258751.1;refGene_gn=DAB1;refGene_tx=XM_023258752.1;refGene_gn=DAB1;refGene_tx=XM_023258753.1;refGene_gn=DAB1;refGene_tx=XM_023258754.1;refGene_gn=DAB1;refGene_tx=XM_023258755.1;refGene_gn=DAB1;refGene_tx=XM_023258756.1;refGene_gn=DAB1;refGene_tx=XM_023258757.1;refGene_gn=DAB1;refGene_tx=XM_023258758.1;refGene_gn=DAB1;refGene_tx=XM_023258759.1;refGene_gn=DAB1;refGene_tx=XM_023258760.1;refGene_gn=DAB

1;refGene_tx=XM_023258761.1;refGene_gn=DAB1;refGene_tx=XM_023258762.1
C1 46869255 46869322
refGene_gn=DAB1;refGene_tx=XM_011284970.3;refGene_gn=DAB1;refGene_tx=XM_019837088.2;refGene_gn=DAB1;refGene_tx=XM_019837092.2;refGene_gn=DAB1;refGene_tx=XM_019837093.2;refGene_gn=DAB1;refGene_tx=XM_023258747.1;refGene_gn=DAB1;refGene_tx=XM_023258748.1;refGene_gn=DAB1;refGene_tx=XM_023258749.1;refGene_gn=DAB1;refGene_tx=XM_023258750.1;refGene_gn=DAB1;refGene_tx=XM_023258751.1;refGene_gn=DAB1;refGene_tx=XM_023258752.1;refGene_gn=DAB1;refGene_tx=XM_023258753.1;refGene_gn=DAB1;refGene_tx=XM_023258754.1;refGene_gn=DAB1;refGene_tx=XM_023258755.1;refGene_gn=DAB1;refGene_tx=XM_023258756.1;refGene_gn=DAB1;refGene_tx=XM_023258757.1;refGene_gn=DAB1;refGene_tx=XM_023258758.1;refGene_gn=DAB1;refGene_tx=XM_023258759.1;refGene_gn=DAB1;refGene_tx=XM_023258760.1;refGene_gn=DAB1;refGene_tx=XM_023258761.1;refGene_gn=DAB1;refGene_tx=XM_023258762.1
C1 47918310 47918403 refGene_gn=OMA1;refGene_tx=XM_019837106.2
C1 47921798 47922011
refGene_gn=OMA1;refGene_tx=XM_003990140.4;refGene_gn=OMA1;refGene_tx=XM_006934772.3;refGene_gn=OMA1;refGene_tx=XM_019837105.1;refGene_gn=OMA1;refGene_tx=XM_019837107.1;refGene_gn=OMA1;refGene_tx=XM_019837108.1
C1 47939322 47939472
refGene_gn=OMA1;refGene_tx=XM_003990140.4;refGene_gn=OMA1;refGene_tx=XM_006934772.3;refGene_gn=OMA1;refGene_tx=XM_019837105.1;refGene_gn=OMA1;refGene_tx=XM_019837106.2;refGene_gn=OMA1;refGene_tx=XM_019837107.1;refGene_gn=OMA1;refGene_tx=XM_019837108.1
C1 47975836 47975911
refGene_gn=OMA1;refGene_tx=XM_003990140.4;refGene_gn=OMA1;refGene_tx=XM_006934772.3;refGene_gn=OMA1;refGene_tx=XM_019837105.1;refGene_gn=OMA1;refGene_tx=XM_019837106.2;refGene_gn=OMA1;refGene_tx=XM_019837107.1;refGene_gn=OMA1;refGene_tx=XM_019837108.1
C1 47980359 47980488
refGene_gn=OMA1;refGene_tx=XM_003990140.4;refGene_gn=OMA1;refGene_tx=XM_006934772.3;refGene_gn=OMA1;refGene_tx=XM_019837105.1;refGene_gn=OMA1;refGene_tx=XM_019837106.2;refGene_gn=OMA1;refGene_tx=XM_019837107.1;refGene_gn=OMA1;refGene_tx=XM_019837108.1
C1 47983065 47983173
refGene_gn=OMA1;refGene_tx=XM_003990140.4;refGene_gn=OMA1;refGene_tx=XM_006934772.3;refGene_gn=OMA1;refGene_tx=XM_019837105.1;refGene_gn=OMA1;refGene_tx=XM_019837106.2;refGene_gn=OMA1;refGene_tx=XM_019837107.1;refGene_gn=OMA1;refGene_tx=XM_019837108.1
C1 47983273 47983447
refGene_gn=OMA1;refGene_tx=XM_003990140.4;refGene_gn=OMA1;refGene_tx=XM_006934772.3;refGene_gn=OMA1;refGene_tx=XM_019837105.1;refGene_gn=OMA1;refGene_tx=XM_019837106.2;refGene_gn=OMA1;refGene_tx=XM_019837107.1;refGene_gn=OMA1;refGene_tx=XM_019837108.1
C1 47985996 47986225
refGene_gn=OMA1;refGene_tx=XM_003990140.4;refGene_gn=OMA1;refGene_tx=XM_006934772.3;refGene_gn=OMA1;refGene_tx=XM_019837105.1;refGene_gn=OMA1;refGene_tx=XM_019837106.2;refGene_gn=OMA1;refGene_tx=XM_019837107.1;refGene_gn=OMA1;refGene_tx=XM_019837108.1
C1 47986511 47986516 refGene_gn=OMA1;refGene_tx=XM_019837108.1

C1	47988618	47989124
refGene_gn=OMA1;refGene_tx=XM_003990140.4;refGene_gn=OMA1;refGene_tx=XM_006934772.3;refGene_gn=OMA1;refGene_tx=XM_019837105.1;refGene_gn=OMA1;refGene_tx=XM_019837106.2;refGene_gn=OMA1;refGene_tx=XM_019837107.1
C1	48022397	48023372	refGene_gn=TACSTD2;refGene_tx=XM_011285026.3
C1	48083399	48083567
refGene_gn=MYSM1;refGene_tx=XM_011284968.2;refGene_gn=MYSM1;refGene_tx=XM_011284969.2;refGene_gn=MYSM1;refGene_tx=XM_019837110.1;refGene_gn=MYSM1;refGene_tx=XM_019837111.1;refGene_gn=MYSM1;refGene_tx=XM_019837112.1;refGene_gn=MYSM1;refGene_tx=XM_019837113.1
C1	48084614	48084672
refGene_gn=MYSM1;refGene_tx=XM_011284968.2;refGene_gn=MYSM1;refGene_tx=XM_011284969.2;refGene_gn=MYSM1;refGene_tx=XM_019837110.1;refGene_gn=MYSM1;refGene_tx=XM_019837111.1;refGene_gn=MYSM1;refGene_tx=XM_019837112.1;refGene_gn=MYSM1;refGene_tx=XM_019837113.1
C1	48084850	48084956
refGene_gn=MYSM1;refGene_tx=XM_011284968.2;refGene_gn=MYSM1;refGene_tx=XM_011284969.2;refGene_gn=MYSM1;refGene_tx=XM_019837110.1;refGene_gn=MYSM1;refGene_tx=XM_019837111.1;refGene_gn=MYSM1;refGene_tx=XM_019837112.1;refGene_gn=MYSM1;refGene_tx=XM_019837113.1
C1	48086606	48086739
refGene_gn=MYSM1;refGene_tx=XM_011284968.2;refGene_gn=MYSM1;refGene_tx=XM_011284969.2;refGene_gn=MYSM1;refGene_tx=XM_019837110.1;refGene_gn=MYSM1;refGene_tx=XM_019837111.1;refGene_gn=MYSM1;refGene_tx=XM_019837112.1;refGene_gn=MYSM1;refGene_tx=XM_019837113.1
C1	48087673	48087862
refGene_gn=MYSM1;refGene_tx=XM_011284968.2;refGene_gn=MYSM1;refGene_tx=XM_011284969.2;refGene_gn=MYSM1;refGene_tx=XM_019837110.1;refGene_gn=MYSM1;refGene_tx=XM_019837111.1;refGene_gn=MYSM1;refGene_tx=XM_019837112.1;refGene_gn=MYSM1;refGene_tx=XM_019837113.1
C1	48088488	48088563
refGene_gn=MYSM1;refGene_tx=XM_011284968.2;refGene_gn=MYSM1;refGene_tx=XM_011284969.2;refGene_gn=MYSM1;refGene_tx=XM_019837110.1;refGene_gn=MYSM1;refGene_tx=XM_019837111.1;refGene_gn=MYSM1;refGene_tx=XM_019837112.1;refGene_gn=MYSM1;refGene_tx=XM_019837113.1
C1	48089256	48089307
refGene_gn=MYSM1;refGene_tx=XM_011284968.2;refGene_gn=MYSM1;refGene_tx=XM_011284969.2;refGene_gn=MYSM1;refGene_tx=XM_019837110.1;refGene_gn=MYSM1;refGene_tx=XM_019837111.1;refGene_gn=MYSM1;refGene_tx=XM_019837112.1;refGene_gn=MYSM1;refGene_tx=XM_019837113.1
C1	48089656	48089711
refGene_gn=MYSM1;refGene_tx=XM_011284968.2;refGene_gn=MYSM1;refGene_tx=XM_011284969.2;refGene_gn=MYSM1;refGene_tx=XM_019837110.1;refGene_gn=MYSM1;refGene_tx=XM_019837111.1;refGene_gn=MYSM1;refGene_tx=XM_019837112.1;refGene_gn=MYSM1;refGene_tx=XM_019837113.1
C1	48093177	48093263
refGene_gn=MYSM1;refGene_tx=XM_011284968.2;refGene_gn=MYSM1;refGene_tx=XM_011284969.2;refGene_gn=MYSM1;refGene_tx=XM_019837110.1;refGene_gn=MYSM1;refGene_tx=XM_019837111.1;refGene_gn=MYSM1;refGene_tx=XM_019837112.1;refGene_gn=MYSM1;refGene_tx=XM_019837113.1

C1 48094964 48095042
refGene_gn=MYSM1;refGene_tx=XM_011284968.2;refGene_gn=MYSM1;refGene_tx=XM_011284969.2;refGene_gn=MYSM1;refGene_tx=XM_019837110.1;refGene_gn=MYSM1;refGene_tx=XM_019837111.1;refGene_gn=MYSM1;refGene_tx=XM_019837112.1;refGene_gn=MYSM1;refGene_tx=XM_019837113.1
C1 48096489 48096593
refGene_gn=MYSM1;refGene_tx=XM_011284968.2;refGene_gn=MYSM1;refGene_tx=XM_011284969.2;refGene_gn=MYSM1;refGene_tx=XM_019837110.1;refGene_gn=MYSM1;refGene_tx=XM_019837111.1;refGene_gn=MYSM1;refGene_tx=XM_019837112.1;refGene_gn=MYSM1;refGene_tx=XM_019837113.1
C1 48097692 48097823
refGene_gn=MYSM1;refGene_tx=XM_011284968.2;refGene_gn=MYSM1;refGene_tx=XM_011284969.2;refGene_gn=MYSM1;refGene_tx=XM_019837110.1;refGene_gn=MYSM1;refGene_tx=XM_019837111.1;refGene_gn=MYSM1;refGene_tx=XM_019837112.1;refGene_gn=MYSM1;refGene_tx=XM_019837113.1
C1 48101874 48102647
refGene_gn=MYSM1;refGene_tx=XM_011284968.2;refGene_gn=MYSM1;refGene_tx=XM_011284969.2;refGene_gn=MYSM1;refGene_tx=XM_019837110.1;refGene_gn=MYSM1;refGene_tx=XM_019837111.1;refGene_gn=MYSM1;refGene_tx=XM_019837112.1;refGene_gn=MYSM1;refGene_tx=XM_019837113.1
C1 48104431 48104530
refGene_gn=MYSM1;refGene_tx=XM_011284968.2;refGene_gn=MYSM1;refGene_tx=XM_011284969.2;refGene_gn=MYSM1;refGene_tx=XM_019837110.1;refGene_gn=MYSM1;refGene_tx=XM_019837111.1;refGene_gn=MYSM1;refGene_tx=XM_019837112.1;refGene_gn=MYSM1;refGene_tx=XM_019837113.1
C1 48107724 48107889
refGene_gn=MYSM1;refGene_tx=XM_011284968.2;refGene_gn=MYSM1;refGene_tx=XM_011284969.2;refGene_gn=MYSM1;refGene_tx=XM_019837110.1;refGene_gn=MYSM1;refGene_tx=XM_019837111.1;refGene_gn=MYSM1;refGene_tx=XM_019837112.1
C1 48108919 48108943
refGene_gn=MYSM1;refGene_tx=XM_011284968.2;refGene_gn=MYSM1;refGene_tx=XM_011284969.2;refGene_gn=MYSM1;refGene_tx=XM_019837110.1;refGene_gn=MYSM1;refGene_tx=XM_019837111.1
C1 48109035 48109113
refGene_gn=MYSM1;refGene_tx=XM_011284968.2;refGene_gn=MYSM1;refGene_tx=XM_011284969.2;refGene_gn=MYSM1;refGene_tx=XM_019837110.1
C1 48110932 48111003
refGene_gn=MYSM1;refGene_tx=XM_011284968.2;refGene_gn=MYSM1;refGene_tx=XM_011284969.2;refGene_gn=MYSM1;refGene_tx=XM_019837110.1;refGene_gn=MYSM1;refGene_tx=XM_019837111.1
C1 48112726 48112802
refGene_gn=MYSM1;refGene_tx=XM_011284968.2;refGene_gn=MYSM1;refGene_tx=XM_011284969.2;refGene_gn=MYSM1;refGene_tx=XM_019837110.1;refGene_gn=MYSM1;refGene_tx=XM_019837111.1;refGene_gn=MYSM1;refGene_tx=XM_019837113.1
C1 48117134 48117217
refGene_gn=MYSM1;refGene_tx=XM_011284968.2;refGene_gn=MYSM1;refGene_tx=XM_011284969.2;refGene_gn=MYSM1;refGene_tx=XM_019837110.1;refGene_gn=MYSM1;refGene_tx=XM_019837111.1;refGene_gn=MYSM1;refGene_tx=XM_0198371

13.1
C1 48184939 48185944 refGene_gn=JUN;refGene_tx=XM_011284967.3
C1 48663785 48663986
refGene_gn=FGGY;refGene_tx=XM_011284961.2;refGene_gn=FGGY;refGene_tx=XM_011284962.2;refGene_gn=FGGY;refGene_tx=XM_011284964.2;refGene_gn=FGGY;refGene_tx=XM_019837115.2;refGene_gn=FGGY;refGene_tx=XM_019837116.1;refGene_gn=FGGY;refGene_tx=XM_019837117.2;refGene_gn=FGGY;refGene_tx=XM_019837118.1;refGene_gn=FGGY;refGene_tx=XM_019837120.2
C1 48679123 48679235
refGene_gn=FGGY;refGene_tx=XM_011284961.2;refGene_gn=FGGY;refGene_tx=XM_011284962.2;refGene_gn=FGGY;refGene_tx=XM_019837115.2;refGene_gn=FGGY;refGene_tx=XM_019837116.1;refGene_gn=FGGY;refGene_tx=XM_019837117.2;refGene_gn=FGGY;refGene_tx=XM_019837118.1
C1 48686013 48686165
refGene_gn=FGGY;refGene_tx=XM_011284961.2;refGene_gn=FGGY;refGene_tx=XM_011284962.2;refGene_gn=FGGY;refGene_tx=XM_011284965.3;refGene_gn=FGGY;refGene_tx=XM_011284966.3;refGene_gn=FGGY;refGene_tx=XM_019837115.2;refGene_gn=FGGY;refGene_tx=XM_019837116.1;refGene_gn=FGGY;refGene_tx=XM_019837117.2;refGene_gn=FGGY;refGene_tx=XM_019837118.1;refGene_gn=FGGY;refGene_tx=XM_019837120.2;refGene_gn=FGGY;refGene_tx=XM_023258763.1;refGene_gn=FGGY;refGene_tx=XM_023258764.1
C1 48711522 48711611
refGene_gn=FGGY;refGene_tx=XM_011284961.2;refGene_gn=FGGY;refGene_tx=XM_011284962.2;refGene_gn=FGGY;refGene_tx=XM_011284964.2;refGene_gn=FGGY;refGene_tx=XM_011284965.3;refGene_gn=FGGY;refGene_tx=XM_011284966.3;refGene_gn=FGGY;refGene_tx=XM_019837115.2;refGene_gn=FGGY;refGene_tx=XM_019837116.1;refGene_gn=FGGY;refGene_tx=XM_019837117.2;refGene_gn=FGGY;refGene_tx=XM_019837118.1;refGene_gn=FGGY;refGene_tx=XM_019837120.2;refGene_gn=FGGY;refGene_tx=XM_023258763.1;refGene_gn=FGGY;refGene_tx=XM_023258764.1
C1 48796520 48796636
refGene_gn=FGGY;refGene_tx=XM_011284961.2;refGene_gn=FGGY;refGene_tx=XM_011284962.2;refGene_gn=FGGY;refGene_tx=XM_011284964.2;refGene_gn=FGGY;refGene_tx=XM_011284965.3;refGene_gn=FGGY;refGene_tx=XM_011284966.3;refGene_gn=FGGY;refGene_tx=XM_019837115.2;refGene_gn=FGGY;refGene_tx=XM_019837116.1;refGene_gn=FGGY;refGene_tx=XM_019837117.2;refGene_gn=FGGY;refGene_tx=XM_019837118.1;refGene_gn=FGGY;refGene_tx=XM_019837120.2;refGene_gn=FGGY;refGene_tx=XM_023258763.1;refGene_gn=FGGY;refGene_tx=XM_023258764.1
C1 48842849 48842978
refGene_gn=FGGY;refGene_tx=XM_011284961.2;refGene_gn=FGGY;refGene_tx=XM_011284962.2;refGene_gn=FGGY;refGene_tx=XM_011284964.2;refGene_gn=FGGY;refGene_tx=XM_011284965.3;refGene_gn=FGGY;refGene_tx=XM_011284966.3;refGene_gn=FGGY;refGene_tx=XM_019837115.2;refGene_gn=FGGY;refGene_tx=XM_019837116.1;refGene_gn=FGGY;refGene_tx=XM_019837117.2;refGene_gn=FGGY;refGene_tx=XM_019837120.2;refGene_gn=FGGY;refGene_tx=XM_023258763.1;refGene_gn=FGGY;refGene_tx=XM_023258764.1
C1 48878703 48878807
refGene_gn=FGGY;refGene_tx=XM_011284961.2;refGene_gn=FGGY;refGene_tx=XM_011284962.2;refGene_gn=FGGY;refGene_tx=XM_011284964.2;refGene_gn=FGG

Y;refGene_tx=XM_011284965.3;refGene_gn=FGGY;refGene_tx=XM_011284966.3;
refGene_gn=FGGY;refGene_tx=XM_019837115.2;refGene_gn=FGGY;refGene_tx=X
M_019837116.1;refGene_gn=FGGY;refGene_tx=XM_019837117.2;refGene_gn=FGG
Y;refGene_tx=XM_019837118.1;refGene_gn=FGGY;refGene_tx=XM_019837120.2;
refGene_gn=FGGY;refGene_tx=XM_023258763.1;refGene_gn=FGGY;refGene_tx=X
M_023258764.1
C1 48920895 48921003
refGene_gn=FGGY;refGene_tx=XM_011284961.2;refGene_gn=FGGY;refGene_tx=X
M_011284962.2;refGene_gn=FGGY;refGene_tx=XM_011284964.2;refGene_gn=FGG
Y;refGene_tx=XM_011284965.3;refGene_gn=FGGY;refGene_tx=XM_011284966.3;
refGene_gn=FGGY;refGene_tx=XM_019837115.2;refGene_gn=FGGY;refGene_tx=X
M_019837116.1;refGene_gn=FGGY;refGene_tx=XM_019837117.2;refGene_gn=FGG
Y;refGene_tx=XM_019837118.1;refGene_gn=FGGY;refGene_tx=XM_019837120.2;
refGene_gn=FGGY;refGene_tx=XM_023258763.1;refGene_gn=FGGY;refGene_tx=X
M_023258764.1
C1 48942828 48942890
refGene_gn=FGGY;refGene_tx=XM_011284961.2;refGene_gn=FGGY;refGene_tx=X
M_011284962.2;refGene_gn=FGGY;refGene_tx=XM_011284964.2;refGene_gn=FGG
Y;refGene_tx=XM_011284965.3;refGene_gn=FGGY;refGene_tx=XM_011284966.3;
refGene_gn=FGGY;refGene_tx=XM_019837115.2;refGene_gn=FGGY;refGene_tx=X
M_019837116.1;refGene_gn=FGGY;refGene_tx=XM_019837117.2;refGene_gn=FGG
Y;refGene_tx=XM_019837118.1;refGene_gn=FGGY;refGene_tx=XM_019837120.2;
refGene_gn=FGGY;refGene_tx=XM_023258763.1;refGene_gn=FGGY;refGene_tx=X
M_023258764.1
C1 48954617 48954765
refGene_gn=FGGY;refGene_tx=XM_011284961.2;refGene_gn=FGGY;refGene_tx=X
M_011284962.2;refGene_gn=FGGY;refGene_tx=XM_011284964.2;refGene_gn=FGG
Y;refGene_tx=XM_011284965.3;refGene_gn=FGGY;refGene_tx=XM_011284966.3;
refGene_gn=FGGY;refGene_tx=XM_019837115.2;refGene_gn=FGGY;refGene_tx=X
M_019837116.1;refGene_gn=FGGY;refGene_tx=XM_019837117.2;refGene_gn=FGG
Y;refGene_tx=XM_019837118.1;refGene_gn=FGGY;refGene_tx=XM_019837120.2;
refGene_gn=FGGY;refGene_tx=XM_023258763.1;refGene_gn=FGGY;refGene_tx=X
M_023258764.1
C1 48969966 48970041
refGene_gn=FGGY;refGene_tx=XM_011284961.2;refGene_gn=FGGY;refGene_tx=X
M_011284964.2;refGene_gn=FGGY;refGene_tx=XM_011284965.3;refGene_gn=FGG
Y;refGene_tx=XM_011284966.3;refGene_gn=FGGY;refGene_tx=XM_019837115.2;
refGene_gn=FGGY;refGene_tx=XM_019837116.1;refGene_gn=FGGY;refGene_tx=X
M_019837117.2;refGene_gn=FGGY;refGene_tx=XM_019837118.1;refGene_gn=FGG
Y;refGene_tx=XM_019837120.2;refGene_gn=FGGY;refGene_tx=XM_023258763.1;
refGene_gn=FGGY;refGene_tx=XM_023258764.1
C1 48975521 48975642
refGene_gn=FGGY;refGene_tx=XM_011284961.2;refGene_gn=FGGY;refGene_tx=X
M_011284962.2;refGene_gn=FGGY;refGene_tx=XM_011284964.2;refGene_gn=FGG
Y;refGene_tx=XM_011284965.3;refGene_gn=FGGY;refGene_tx=XM_011284966.3;
refGene_gn=FGGY;refGene_tx=XM_019837115.2;refGene_gn=FGGY;refGene_tx=X
M_019837116.1;refGene_gn=FGGY;refGene_tx=XM_019837117.2;refGene_gn=FGG
Y;refGene_tx=XM_019837118.1;refGene_gn=FGGY;refGene_tx=XM_019837120.2;
refGene_gn=FGGY;refGene_tx=XM_023258763.1;refGene_gn=FGGY;refGene_tx=X
M_023258764.1

C1 48981672 48981767
refGene_gn=FGGY;refGene_tx=XM_011284961.2;refGene_gn=FGGY;refGene_tx=X
M_011284962.2;refGene_gn=FGGY;refGene_tx=XM_011284964.2;refGene_gn=FGG
Y;refGene_tx=XM_011284965.3;refGene_gn=FGGY;refGene_tx=XM_011284966.3;
refGene_gn=FGGY;refGene_tx=XM_019837115.2;refGene_gn=FGGY;refGene_tx=X
M_019837116.1;refGene_gn=FGGY;refGene_tx=XM_019837117.2;refGene_gn=FGG
Y;refGene_tx=XM_019837118.1;refGene_gn=FGGY;refGene_tx=XM_019837120.2;
refGene_gn=FGGY;refGene_tx=XM_023258763.1;refGene_gn=FGGY;refGene_tx=X
M_023258764.1
C1 49055177 49055239
refGene_gn=FGGY;refGene_tx=XM_011284961.2;refGene_gn=FGGY;refGene_tx=X
M_011284962.2;refGene_gn=FGGY;refGene_tx=XM_011284964.2;refGene_gn=FGG
Y;refGene_tx=XM_011284965.3;refGene_gn=FGGY;refGene_tx=XM_011284966.3;
refGene_gn=FGGY;refGene_tx=XM_019837115.2;refGene_gn=FGGY;refGene_tx=X
M_019837116.1;refGene_gn=FGGY;refGene_tx=XM_019837117.2;refGene_gn=FGG
Y;refGene_tx=XM_019837118.1;refGene_gn=FGGY;refGene_tx=XM_019837120.2;
refGene_gn=FGGY;refGene_tx=XM_023258763.1;refGene_gn=FGGY;refGene_tx=X
M_023258764.1
C1 49060702 49060784
refGene_gn=FGGY;refGene_tx=XM_011284961.2;refGene_gn=FGGY;refGene_tx=X
M_011284962.2;refGene_gn=FGGY;refGene_tx=XM_011284964.2;refGene_gn=FGG
Y;refGene_tx=XM_011284965.3;refGene_gn=FGGY;refGene_tx=XM_011284966.3;
refGene_gn=FGGY;refGene_tx=XM_019837115.2;refGene_gn=FGGY;refGene_tx=X
M_019837116.1;refGene_gn=FGGY;refGene_tx=XM_019837117.2;refGene_gn=FGG
Y;refGene_tx=XM_019837118.1;refGene_gn=FGGY;refGene_tx=XM_019837120.2;
refGene_gn=FGGY;refGene_tx=XM_023258763.1;refGene_gn=FGGY;refGene_tx=X
M_023258764.1
C1 49106010 49106073 refGene_gn=HOOK1;refGene_tx=XM_019839080.2
C1 49114484 49114570 refGene_gn=HOOK1;refGene_tx=XM_019839080.2
C1 49121344 49121417 refGene_gn=HOOK1;refGene_tx=XM_019839080.2
C1 49125156 49125207 refGene_gn=HOOK1;refGene_tx=XM_019839080.2
C1 49126614 49126747 refGene_gn=HOOK1;refGene_tx=XM_019839080.2
C1 49136853 49136921 refGene_gn=HOOK1;refGene_tx=XM_019839080.2
C1 49138544 49138607 refGene_gn=HOOK1;refGene_tx=XM_019839080.2
C1 49140469 49140553 refGene_gn=HOOK1;refGene_tx=XM_019839080.2
C1 49145978 49146145 refGene_gn=HOOK1;refGene_tx=XM_019839080.2
C1 49152800 49152941 refGene_gn=HOOK1;refGene_tx=XM_019839080.2
C1 49154642 49154844 refGene_gn=HOOK1;refGene_tx=XM_019839080.2
C1 49155396 49155507 refGene_gn=HOOK1;refGene_tx=XM_019839080.2
C1 49158172 49158260 refGene_gn=HOOK1;refGene_tx=XM_019839080.2
C1 49158339 49158400 refGene_gn=HOOK1;refGene_tx=XM_019839080.2
C1 49160359 49160500 refGene_gn=HOOK1;refGene_tx=XM_019839080.2
C1 49162693 49162787 refGene_gn=HOOK1;refGene_tx=XM_019839080.2
C1 49164457 49164492 refGene_gn=HOOK1;refGene_tx=XM_019839080.2
C1 49165027 49165110 refGene_gn=HOOK1;refGene_tx=XM_019839080.2
C1 49165736 49165837 refGene_gn=HOOK1;refGene_tx=XM_019839080.2
C1 49169358 49169460 refGene_gn=HOOK1;refGene_tx=XM_019839080.2
C1 49171805 49171874 refGene_gn=HOOK1;refGene_tx=XM_019839080.2
C1 49173595 49173766 refGene_gn=HOOK1;refGene_tx=XM_019839080.2
C1 49187500 49187679

refGene_gn=LOC101097746;refGene_tx=XM_019839081.2
C1 49191487 49191626
refGene_gn=LOC101097746;refGene_tx=XM_019839081.2
C1 49194450 49194638
refGene_gn=LOC101097746;refGene_tx=XM_019839081.2
C1 49197372 49197514
refGene_gn=LOC101097746;refGene_tx=XM_019839081.2
C1 49199566 49199743
refGene_gn=LOC101097746;refGene_tx=XM_019839081.2
C1 49200624 49200785
refGene_gn=LOC101097746;refGene_tx=XM_019839081.2
C1 49201219 49201369
refGene_gn=LOC101097746;refGene_tx=XM_019839081.2
C1 49206344 49206507
refGene_gn=LOC101097746;refGene_tx=XM_019839081.2
C1 49218544 49218754
refGene_gn=LOC101097746;refGene_tx=XM_019839081.2
C1 50293528 50293531 refGene_gn=NFIA;refGene_tx=XM_019837122.2
C1 50298772 50298799
refGene_gn=NFIA;refGene_tx=XM_003990146.5;refGene_gn=NFIA;refGene_tx=XM_003990147.5;refGene_gn=NFIA;refGene_tx=XM_006934781.4;refGene_gn=NFIA;refGene_tx=XM_006934782.4;refGene_gn=NFIA;refGene_tx=XM_006934783.4
C1 50304260 50304792
refGene_gn=NFIA;refGene_tx=XM_003990146.5;refGene_gn=NFIA;refGene_tx=XM_003990147.5;refGene_gn=NFIA;refGene_tx=XM_006934781.4;refGene_gn=NFIA;refGene_tx=XM_006934782.4;refGene_gn=NFIA;refGene_tx=XM_006934783.4;
refGene_gn=NFIA;refGene_tx=XM_019837122.2
C1 50487594 50487660
refGene_gn=NFIA;refGene_tx=XM_003990146.5;refGene_gn=NFIA;refGene_tx=XM_003990147.5;refGene_gn=NFIA;refGene_tx=XM_006934781.4;refGene_gn=NFIA;refGene_tx=XM_006934782.4;refGene_gn=NFIA;refGene_tx=XM_006934783.4;
refGene_gn=NFIA;refGene_tx=XM_019837122.2
C1 50541244 50541319
refGene_gn=NFIA;refGene_tx=XM_003990146.5;refGene_gn=NFIA;refGene_tx=XM_003990147.5;refGene_gn=NFIA;refGene_tx=XM_006934781.4;refGene_gn=NFIA;refGene_tx=XM_006934782.4;refGene_gn=NFIA;refGene_tx=XM_006934783.4;
refGene_gn=NFIA;refGene_tx=XM_019837122.2
C1 50558578 50558696
refGene_gn=NFIA;refGene_tx=XM_003990146.5;refGene_gn=NFIA;refGene_tx=XM_003990147.5;refGene_gn=NFIA;refGene_tx=XM_006934781.4;refGene_gn=NFIA;refGene_tx=XM_006934782.4;refGene_gn=NFIA;refGene_tx=XM_006934783.4;
refGene_gn=NFIA;refGene_tx=XM_019837122.2
C1 50565411 50565539
refGene_gn=NFIA;refGene_tx=XM_003990146.5;refGene_gn=NFIA;refGene_tx=XM_003990147.5;refGene_gn=NFIA;refGene_tx=XM_006934781.4;refGene_gn=NFIA;refGene_tx=XM_006934782.4;refGene_gn=NFIA;refGene_tx=XM_006934783.4;
refGene_gn=NFIA;refGene_tx=XM_019837122.2
C1 50589655 50589784
refGene_gn=NFIA;refGene_tx=XM_003990146.5;refGene_gn=NFIA;refGene_tx=XM_003990147.5;refGene_gn=NFIA;refGene_tx=XM_006934781.4;refGene_gn=NFI

A;refGene_tx=XM_006934782.4;refGene_gn=NFIA;refGene_tx=XM_006934783.4;
refGene_gn=NFIA;refGene_tx=XM_019837122.2
C1	50592060	50592114
refGene_gn=NFIA;refGene_tx=XM_006934781.4;refGene_gn=NFIA;refGene_tx=X
M_006934782.4;refGene_gn=NFIA;refGene_tx=XM_006934783.4;refGene_gn=NFI
A;refGene_tx=XM_019837122.2
C1	50610781	50610960
refGene_gn=NFIA;refGene_tx=XM_003990146.5;refGene_gn=NFIA;refGene_tx=X
M_003990147.5;refGene_gn=NFIA;refGene_tx=XM_006934781.4;refGene_gn=NFI
A;refGene_tx=XM_006934782.4;refGene_gn=NFIA;refGene_tx=XM_006934783.4;
refGene_gn=NFIA;refGene_tx=XM_019837122.2
C1	50613261	50613427
refGene_gn=NFIA;refGene_tx=XM_003990146.5;refGene_gn=NFIA;refGene_tx=X
M_003990147.5;refGene_gn=NFIA;refGene_tx=XM_006934781.4;refGene_gn=NFI
A;refGene_tx=XM_006934782.4;refGene_gn=NFIA;refGene_tx=XM_006934783.4;
refGene_gn=NFIA;refGene_tx=XM_019837122.2
C1	50632078	50632170
refGene_gn=NFIA;refGene_tx=XM_003990146.5;refGene_gn=NFIA;refGene_tx=X
M_006934781.4;refGene_gn=NFIA;refGene_tx=XM_006934782.4;refGene_gn=NFI
A;refGene_tx=XM_019837122.2
C1	50663190	50663267
refGene_gn=NFIA;refGene_tx=XM_003990146.5;refGene_gn=NFIA;refGene_tx=X
M_003990147.5;refGene_gn=NFIA;refGene_tx=XM_006934781.4;refGene_gn=NFI
A;refGene_tx=XM_006934782.4;refGene_gn=NFIA;refGene_tx=XM_006934783.4;
refGene_gn=NFIA;refGene_tx=XM_019837122.2
C1	50858475	50858586
refGene_gn=TM2D1;refGene_tx=XM_023258765.1;refGene_gn=TM2D1;refGene_tx
=XM_023258766.1
C1	50871766	50871840
refGene_gn=TM2D1;refGene_tx=XM_023258765.1;refGene_gn=TM2D1;refGene_tx
=XM_023258766.1
C1	50882433	50882525
refGene_gn=TM2D1;refGene_tx=XM_023258765.1;refGene_gn=TM2D1;refGene_tx
=XM_023258766.1
C1	50887994	50888103
refGene_gn=TM2D1;refGene_tx=XM_023258765.1;refGene_gn=TM2D1;refGene_tx
=XM_023258766.1
C1	50902269	50902343
refGene_gn=TM2D1;refGene_tx=XM_023258765.1;refGene_gn=TM2D1;refGene_tx
=XM_023258766.1
C1	50904344	50904511
refGene_gn=TM2D1;refGene_tx=XM_023258765.1;refGene_gn=TM2D1;refGene_tx
=XM_023258766.1
C1	50928596	50928618	refGene_gn=PATJ;refGene_tx=XM_023258767.1
C1	50928716	50928883	refGene_gn=PATJ;refGene_tx=XM_023258767.1
C1	50931599	50931794	refGene_gn=PATJ;refGene_tx=XM_023258767.1
C1	50933336	50933476	refGene_gn=PATJ;refGene_tx=XM_023258767.1
C1	50938813	50939009	refGene_gn=PATJ;refGene_tx=XM_023258767.1
C1	50941159	50941288	refGene_gn=PATJ;refGene_tx=XM_023258767.1
C1	50953989	50954208	refGene_gn=PATJ;refGene_tx=XM_023258767.1

C1	50957249	50957349	refGene_gn=PATJ;refGene_tx=XM_023258767.1
C1	50959740	50959832	refGene_gn=PATJ;refGene_tx=XM_023258767.1
C1	50961719	50961861	refGene_gn=PATJ;refGene_tx=XM_023258767.1
C1	50965306	50965438	refGene_gn=PATJ;refGene_tx=XM_023258767.1
C1	50968770	50968847	refGene_gn=PATJ;refGene_tx=XM_023258767.1
C1	50970901	50970958	refGene_gn=PATJ;refGene_tx=XM_023258767.1
C1	50982454	50982589	refGene_gn=PATJ;refGene_tx=XM_023258767.1
C1	50989108	50989270	refGene_gn=PATJ;refGene_tx=XM_023258767.1
C1	50992779	50992911	refGene_gn=PATJ;refGene_tx=XM_023258767.1
C1	51015983	51016193	refGene_gn=PATJ;refGene_tx=XM_023258767.1
C1	51023716	51023836	refGene_gn=PATJ;refGene_tx=XM_023258767.1
C1	51025638	51026034	refGene_gn=PATJ;refGene_tx=XM_023258767.1
C1	51032742	51032866	refGene_gn=PATJ;refGene_tx=XM_023258767.1
C1	51038233	51038408	refGene_gn=PATJ;refGene_tx=XM_023258767.1
C1	51052140	51052212	refGene_gn=PATJ;refGene_tx=XM_023258767.1
C1	51054254	51054432	refGene_gn=PATJ;refGene_tx=XM_023258767.1
C1	51059779	51059890	refGene_gn=PATJ;refGene_tx=XM_023258767.1
C1	51064403	51064481	refGene_gn=PATJ;refGene_tx=XM_023258767.1
C1	51076535	51076632	refGene_gn=PATJ;refGene_tx=XM_023258767.1
C1	51130338	51130535	refGene_gn=PATJ;refGene_tx=XM_023258767.1
C1	51154604	51154696	refGene_gn=PATJ;refGene_tx=XM_023258767.1
C1	51170936	51171009	refGene_gn=PATJ;refGene_tx=XM_023258767.1
C1	51185325	51185403	refGene_gn=PATJ;refGene_tx=XM_023258767.1
C1	51202773	51202845	refGene_gn=PATJ;refGene_tx=XM_023258767.1
C1	51207517	51207554	refGene_gn=PATJ;refGene_tx=XM_023258767.1
C1	51207556	51207634	refGene_gn=PATJ;refGene_tx=XM_023258767.1
C1	51211980	51212114	refGene_gn=PATJ;refGene_tx=XM_023258767.1
C1	51232175	51232259	refGene_gn=PATJ;refGene_tx=XM_023258767.1
C1	51238022	51238216	refGene_gn=PATJ;refGene_tx=XM_023258767.1
C1	51241363	51241511	refGene_gn=PATJ;refGene_tx=XM_023258767.1
C1	51242614	51242701	refGene_gn=PATJ;refGene_tx=XM_023258767.1
C1	51247085	51247200	refGene_gn=PATJ;refGene_tx=XM_023258767.1
C1	51248249	51248287	refGene_gn=PATJ;refGene_tx=XM_023258767.1
C1	51252451	51252574	refGene_gn=PATJ;refGene_tx=XM_023258767.1
C1	51253303	51253408	refGene_gn=PATJ;refGene_tx=XM_023258767.1
C1	51270042	51270149	refGene_gn=PATJ;refGene_tx=XM_023258767.1
C1	51273566	51273690	refGene_gn=PATJ;refGene_tx=XM_023258767.1
C1	51278200	51278347	refGene_gn=PATJ;refGene_tx=XM_023258767.1
C1	51310381	51311368	refGene_gn=L1TD1;refGene_tx=XM_011284959.3
C1	51313810	51315430	refGene_gn=L1TD1;refGene_tx=XM_011284959.3
C1	51348108	51348213
refGene_gn=KANK4;refGene_tx=XM_003990209.5;refGene_gn=KANK4;refGene_tx=XM_023258768.1;refGene_gn=KANK4;refGene_tx=XM_023258769.1;refGene_gn=KANK4;refGene_tx=XM_023258770.1;refGene_gn=KANK4;refGene_tx=XM_023258771.1
C1	51352489	51352690
refGene_gn=KANK4;refGene_tx=XM_003990209.5;refGene_gn=KANK4;refGene_tx=XM_023258768.1;refGene_gn=KANK4;refGene_tx=XM_023258769.1;refGene_gn=KANK4;refGene_tx=XM_023258770.1;refGene_gn=KANK4;refGene_tx=XM_023258771.1

C1 51356258 51356401
refGene_gn=KANK4;refGene_tx=XM_003990209.5;refGene_gn=KANK4;refGene_tx=XM_023258768.1;refGene_gn=KANK4;refGene_tx=XM_023258769.1;refGene_gn=KANK4;refGene_tx=XM_023258770.1;refGene_gn=KANK4;refGene_tx=XM_023258771.1
C1 51360389 51360609
refGene_gn=KANK4;refGene_tx=XM_003990209.5;refGene_gn=KANK4;refGene_tx=XM_023258768.1;refGene_gn=KANK4;refGene_tx=XM_023258769.1;refGene_gn=KANK4;refGene_tx=XM_023258770.1;refGene_gn=KANK4;refGene_tx=XM_023258771.1
C1 51362795 51362883
refGene_gn=KANK4;refGene_tx=XM_003990209.5;refGene_gn=KANK4;refGene_tx=XM_023258768.1;refGene_gn=KANK4;refGene_tx=XM_023258769.1;refGene_gn=KANK4;refGene_tx=XM_023258770.1;refGene_gn=KANK4;refGene_tx=XM_023258771.1
C1 51364284 51364503
refGene_gn=KANK4;refGene_tx=XM_003990209.5;refGene_gn=KANK4;refGene_tx=XM_023258768.1;refGene_gn=KANK4;refGene_tx=XM_023258769.1;refGene_gn=KANK4;refGene_tx=XM_023258770.1;refGene_gn=KANK4;refGene_tx=XM_023258771.1
C1 51367454 51367566
refGene_gn=KANK4;refGene_tx=XM_003990209.5;refGene_gn=KANK4;refGene_tx=XM_023258768.1;refGene_gn=KANK4;refGene_tx=XM_023258769.1;refGene_gn=KANK4;refGene_tx=XM_023258770.1;refGene_gn=KANK4;refGene_tx=XM_023258771.1
C1 51368868 51370806
refGene_gn=KANK4;refGene_tx=XM_003990209.5;refGene_gn=KANK4;refGene_tx=XM_023258768.1;refGene_gn=KANK4;refGene_tx=XM_023258769.1;refGene_gn=KANK4;refGene_tx=XM_023258770.1;refGene_gn=KANK4;refGene_tx=XM_023258771.1
C1 51377308 51377324
refGene_gn=KANK4;refGene_tx=XM_003990209.5;refGene_gn=KANK4;refGene_tx=XM_023258768.1;refGene_gn=KANK4;refGene_tx=XM_023258769.1;refGene_gn=KANK4;refGene_tx=XM_023258770.1;refGene_gn=KANK4;refGene_tx=XM_023258771.1
C1 51535984 51536154
refGene_gn=USP1;refGene_tx=XM_011284956.3;refGene_gn=USP1;refGene_tx=XM_019837132.2;refGene_gn=USP1;refGene_tx=XM_023258773.1
C1 51537617 51537738
refGene_gn=USP1;refGene_tx=XM_011284956.3;refGene_gn=USP1;refGene_tx=XM_019837132.2;refGene_gn=USP1;refGene_tx=XM_023258773.1
C1 51538488 51538593
refGene_gn=USP1;refGene_tx=XM_011284956.3;refGene_gn=USP1;refGene_tx=XM_019837132.2;refGene_gn=USP1;refGene_tx=XM_023258773.1
C1 51539126 51539287
refGene_gn=USP1;refGene_tx=XM_011284956.3;refGene_gn=USP1;refGene_tx=XM_019837132.2;refGene_gn=USP1;refGene_tx=XM_023258773.1
C1 51540458 51541150
refGene_gn=USP1;refGene_tx=XM_011284956.3;refGene_gn=USP1;refGene_tx=XM_019837132.2;refGene_gn=USP1;refGene_tx=XM_023258773.1

C1 51541840 51542011
refGene_gn=USP1;refGene_tx=XM_011284956.3;refGene_gn=USP1;refGene_tx=XM_019837132.2;refGene_gn=USP1;refGene_tx=XM_023258773.1
C1 51542681 51542883
refGene_gn=USP1;refGene_tx=XM_011284956.3;refGene_gn=USP1;refGene_tx=XM_019837132.2;refGene_gn=USP1;refGene_tx=XM_023258773.1
C1 51543916 51544649
refGene_gn=USP1;refGene_tx=XM_011284956.3;refGene_gn=USP1;refGene_tx=XM_019837132.2;refGene_gn=USP1;refGene_tx=XM_023258773.1
C1 51547956 51547999 refGene_gn=DOCK7;refGene_tx=XM_023258772.1
C1 51551070 51551238 refGene_gn=DOCK7;refGene_tx=XM_023258772.1
C1 51561609 51561716 refGene_gn=DOCK7;refGene_tx=XM_023258772.1
C1 51562852 51562996 refGene_gn=DOCK7;refGene_tx=XM_023258772.1
C1 51563337 51563574 refGene_gn=DOCK7;refGene_tx=XM_023258772.1
C1 51563691 51563781 refGene_gn=DOCK7;refGene_tx=XM_023258772.1
C1 51565589 51565715 refGene_gn=DOCK7;refGene_tx=XM_023258772.1
C1 51573881 51573896 refGene_gn=DOCK7;refGene_tx=XM_023258772.1
C1 51575444 51575576 refGene_gn=DOCK7;refGene_tx=XM_023258772.1
C1 51578452 51578596 refGene_gn=DOCK7;refGene_tx=XM_023258772.1
C1 51580236 51580429 refGene_gn=DOCK7;refGene_tx=XM_023258772.1
C1 51581725 51581826 refGene_gn=DOCK7;refGene_tx=XM_023258772.1
C1 51582515 51582674 refGene_gn=DOCK7;refGene_tx=XM_023258772.1
C1 51585606 51585759 refGene_gn=DOCK7;refGene_tx=XM_023258772.1
C1 51587080 51587215 refGene_gn=DOCK7;refGene_tx=XM_023258772.1
C1 51587333 51587430 refGene_gn=DOCK7;refGene_tx=XM_023258772.1
C1 51590467 51590564 refGene_gn=DOCK7;refGene_tx=XM_023258772.1
C1 51594459 51594595 refGene_gn=DOCK7;refGene_tx=XM_023258772.1
C1 51594705 51594750 refGene_gn=DOCK7;refGene_tx=XM_023258772.1
C1 51594751 51594889 refGene_gn=DOCK7;refGene_tx=XM_023258772.1
C1 51605192 51605347 refGene_gn=DOCK7;refGene_tx=XM_023258772.1
C1 51606328 51606498 refGene_gn=DOCK7;refGene_tx=XM_023258772.1
C1 51611218 51611358 refGene_gn=DOCK7;refGene_tx=XM_023258772.1
C1 51613071 51613242 refGene_gn=DOCK7;refGene_tx=XM_023258772.1
C1 51615268 51615382 refGene_gn=DOCK7;refGene_tx=XM_023258772.1
C1 51615475 51615616 refGene_gn=DOCK7;refGene_tx=XM_023258772.1
C1 51617248 51617344 refGene_gn=DOCK7;refGene_tx=XM_023258772.1
C1 51617929 51617965 refGene_gn=DOCK7;refGene_tx=XM_023258772.1
C1 51618288 51618378 refGene_gn=DOCK7;refGene_tx=XM_023258772.1
C1 51619566 51619659 refGene_gn=DOCK7;refGene_tx=XM_023258772.1
C1 51627977 51628147 refGene_gn=DOCK7;refGene_tx=XM_023258772.1
C1 51630068 51630233 refGene_gn=DOCK7;refGene_tx=XM_023258772.1
C1 51633462 51633694 refGene_gn=DOCK7;refGene_tx=XM_023258772.1
C1 51638213 51638300 refGene_gn=DOCK7;refGene_tx=XM_023258772.1
C1 51648692 51648794 refGene_gn=DOCK7;refGene_tx=XM_023258772.1
C1 51649690 51649829 refGene_gn=DOCK7;refGene_tx=XM_023258772.1
C1 51653802 51653873 refGene_gn=DOCK7;refGene_tx=XM_023258772.1
C1 51657217 51657335 refGene_gn=DOCK7;refGene_tx=XM_023258772.1
C1 51679136 51679631
refGene_gn=ANGPTL3;refGene_tx=XM_019837134.1;refGene_gn=ANGPTL3;refGene_tx=XM_023258774.1

C1 51680299 51680410
refGene_gn=ANGPTL3;refGene_tx=XM_019837134.1;refGene_gn=ANGPTL3;refGen
e_tx=XM_023258774.1
C1 51683084 51683199
refGene_gn=ANGPTL3;refGene_tx=XM_019837134.1;refGene_gn=ANGPTL3;refGen
e_tx=XM_023258774.1
C1 51684638 51684794
refGene_gn=ANGPTL3;refGene_tx=XM_019837134.1;refGene_gn=ANGPTL3;refGen
e_tx=XM_023258774.1
C1 51685176 51685272
refGene_gn=ANGPTL3;refGene_tx=XM_019837134.1;refGene_gn=ANGPTL3;refGen
e_tx=XM_023258774.1
C1 51687031 51687298
refGene_gn=ANGPTL3;refGene_tx=XM_019837134.1;refGene_gn=ANGPTL3;refGen
e_tx=XM_023258774.1
C1 51687705 51687890
refGene_gn=ANGPTL3;refGene_tx=XM_019837134.1;refGene_gn=ANGPTL3;refGen
e_tx=XM_023258774.1
C1 51705294 51705457 refGene_gn=DOCK7;refGene_tx=XM_023258772.1
C1 51706148 51706242 refGene_gn=DOCK7;refGene_tx=XM_023258772.1
C1 51710176 51710319 refGene_gn=DOCK7;refGene_tx=XM_023258772.1
C1 51716218 51716384 refGene_gn=DOCK7;refGene_tx=XM_023258772.1
C1 51718421 51718502 refGene_gn=DOCK7;refGene_tx=XM_023258772.1
C1 51729297 51729447 refGene_gn=DOCK7;refGene_tx=XM_023258772.1
C1 51730872 51730939 refGene_gn=DOCK7;refGene_tx=XM_023258772.1
C1 51739189 51739275 refGene_gn=DOCK7;refGene_tx=XM_023258772.1
C1 51739619 51739832 refGene_gn=DOCK7;refGene_tx=XM_023258772.1
C1 51739923 51740053 refGene_gn=DOCK7;refGene_tx=XM_023258772.1
C1 51744642 51744711 refGene_gn=DOCK7;refGene_tx=XM_023258772.1
C1 51744899 51745075 refGene_gn=DOCK7;refGene_tx=XM_023258772.1
C1 51756179 51756285 refGene_gn=DOCK7;refGene_tx=XM_023258772.1
C1 51781360 51781917 refGene_gn=DOCK7;refGene_tx=XM_023258772.1
C1 51868153 51868229
refGene_gn=ATG4C;refGene_tx=XM_011284952.2;refGene_gn=ATG4C;refGene_tx
=XM_011284954.2;refGene_gn=ATG4C;refGene_tx=XM_019837135.2;refGene_gn=
ATG4C;refGene_tx=XM_019837136.2;refGene_gn=ATG4C;refGene_tx=XM_0198371
37.1;refGene_gn=ATG4C;refGene_tx=XM_019837138.1
C1 51884115 51884199
refGene_gn=ATG4C;refGene_tx=XM_011284952.2;refGene_gn=ATG4C;refGene_tx
=XM_011284954.2;refGene_gn=ATG4C;refGene_tx=XM_019837135.2;refGene_gn=
ATG4C;refGene_tx=XM_019837136.2;refGene_gn=ATG4C;refGene_tx=XM_0198371
37.1;refGene_gn=ATG4C;refGene_tx=XM_019837138.1
C1 51892197 51892431
refGene_gn=ATG4C;refGene_tx=XM_011284952.2;refGene_gn=ATG4C;refGene_tx
=XM_011284954.2;refGene_gn=ATG4C;refGene_tx=XM_019837135.2;refGene_gn=
ATG4C;refGene_tx=XM_019837136.2;refGene_gn=ATG4C;refGene_tx=XM_0198371
37.1;refGene_gn=ATG4C;refGene_tx=XM_019837138.1
C1 51895251 51895582
refGene_gn=ATG4C;refGene_tx=XM_011284952.2;refGene_gn=ATG4C;refGene_tx
=XM_011284954.2;refGene_gn=ATG4C;refGene_tx=XM_019837135.2;refGene_gn=

ATG4C;refGene_tx=XM_019837136.2;refGene_gn=ATG4C;refGene_tx=XM_0198371
37.1;refGene_gn=ATG4C;refGene_tx=XM_019837138.1
C1 51897603 51897674
refGene_gn=ATG4C;refGene_tx=XM_011284952.2;refGene_gn=ATG4C;refGene_tx
=XM_011284954.2;refGene_gn=ATG4C;refGene_tx=XM_019837135.2;refGene_gn=
ATG4C;refGene_tx=XM_019837136.2;refGene_gn=ATG4C;refGene_tx=XM_0198371
37.1;refGene_gn=ATG4C;refGene_tx=XM_019837138.1
C1 51900284 51900421
refGene_gn=ATG4C;refGene_tx=XM_011284952.2;refGene_gn=ATG4C;refGene_tx
=XM_011284954.2;refGene_gn=ATG4C;refGene_tx=XM_019837135.2;refGene_gn=
ATG4C;refGene_tx=XM_019837136.2;refGene_gn=ATG4C;refGene_tx=XM_0198371
37.1
C1 51901669 51901707 refGene_gn=ATG4C;refGene_tx=XM_019837138.1
C1 51905990 51906069
refGene_gn=ATG4C;refGene_tx=XM_011284952.2;refGene_gn=ATG4C;refGene_tx
=XM_011284954.2;refGene_gn=ATG4C;refGene_tx=XM_019837135.2;refGene_gn=
ATG4C;refGene_tx=XM_019837136.2;refGene_gn=ATG4C;refGene_tx=XM_0198371
37.1;refGene_gn=ATG4C;refGene_tx=XM_019837138.1
C1 51906724 51906801
refGene_gn=ATG4C;refGene_tx=XM_011284952.2;refGene_gn=ATG4C;refGene_tx
=XM_011284954.2;refGene_gn=ATG4C;refGene_tx=XM_019837135.2;refGene_gn=
ATG4C;refGene_tx=XM_019837136.2;refGene_gn=ATG4C;refGene_tx=XM_0198371
37.1;refGene_gn=ATG4C;refGene_tx=XM_019837138.1
C1 51914352 51914472
refGene_gn=ATG4C;refGene_tx=XM_011284952.2;refGene_gn=ATG4C;refGene_tx
=XM_019837135.2;refGene_gn=ATG4C;refGene_tx=XM_019837136.2;refGene_gn=
ATG4C;refGene_tx=XM_019837137.1;refGene_gn=ATG4C;refGene_tx=XM_0198371
38.1
C1 51935039 51935207
refGene_gn=ATG4C;refGene_tx=XM_011284952.2;refGene_gn=ATG4C;refGene_tx
=XM_011284954.2;refGene_gn=ATG4C;refGene_tx=XM_019837135.2;refGene_gn=
ATG4C;refGene_tx=XM_019837136.2;refGene_gn=ATG4C;refGene_tx=XM_0198371
37.1;refGene_gn=ATG4C;refGene_tx=XM_019837138.1
C1 52386580 52388026 refGene_gn=FOXD3;refGene_tx=XM_023258775.1
C1 52431070 52431152
refGene_gn=ALG6;refGene_tx=XM_006934795.3;refGene_gn=ALG6;refGene_tx=X
M_006934796.4;refGene_gn=ALG6;refGene_tx=XM_006934797.4;refGene_gn=ALG
6;refGene_tx=XM_019837142.2;refGene_gn=ALG6;refGene_tx=XM_019837143.2
C1 52443237 52443262 refGene_gn=ALG6;refGene_tx=XM_019837144.2
C1 52444019 52444104
refGene_gn=ALG6;refGene_tx=XM_006934795.3;refGene_gn=ALG6;refGene_tx=X
M_006934796.4;refGene_gn=ALG6;refGene_tx=XM_006934797.4;refGene_gn=ALG
6;refGene_tx=XM_019837142.2;refGene_gn=ALG6;refGene_tx=XM_019837143.2;
refGene_gn=ALG6;refGene_tx=XM_019837144.2
C1 52456117 52456207
refGene_gn=ALG6;refGene_tx=XM_006934795.3;refGene_gn=ALG6;refGene_tx=X
M_006934796.4;refGene_gn=ALG6;refGene_tx=XM_006934797.4;refGene_gn=ALG
6;refGene_tx=XM_019837142.2;refGene_gn=ALG6;refGene_tx=XM_019837143.2;
refGene_gn=ALG6;refGene_tx=XM_019837144.2
C1 52457784 52457873

refGene_gn=ALG6;refGene_tx=XM_006934795.3;refGene_gn=ALG6;refGene_tx=XM_006934796.4;refGene_gn=ALG6;refGene_tx=XM_006934797.4;refGene_gn=ALG6;refGene_tx=XM_019837142.2;refGene_gn=ALG6;refGene_tx=XM_019837143.2;refGene_gn=ALG6;refGene_tx=XM_019837144.2
C1 52460391 52460474
refGene_gn=ALG6;refGene_tx=XM_006934795.3;refGene_gn=ALG6;refGene_tx=XM_006934796.4;refGene_gn=ALG6;refGene_tx=XM_006934797.4;refGene_gn=ALG6;refGene_tx=XM_019837142.2;refGene_gn=ALG6;refGene_tx=XM_019837143.2;refGene_gn=ALG6;refGene_tx=XM_019837144.2
C1 52461310 52461375
refGene_gn=ALG6;refGene_tx=XM_006934795.3;refGene_gn=ALG6;refGene_tx=XM_006934796.4;refGene_gn=ALG6;refGene_tx=XM_006934797.4;refGene_gn=ALG6;refGene_tx=XM_019837142.2;refGene_gn=ALG6;refGene_tx=XM_019837143.2;refGene_gn=ALG6;refGene_tx=XM_019837144.2
C1 52464836 52465022
refGene_gn=ALG6;refGene_tx=XM_006934795.3;refGene_gn=ALG6;refGene_tx=XM_006934796.4;refGene_gn=ALG6;refGene_tx=XM_006934797.4;refGene_gn=ALG6;refGene_tx=XM_019837142.2;refGene_gn=ALG6;refGene_tx=XM_019837143.2;refGene_gn=ALG6;refGene_tx=XM_019837144.2
C1 52465600 52465736
refGene_gn=ALG6;refGene_tx=XM_006934795.3;refGene_gn=ALG6;refGene_tx=XM_006934796.4;refGene_gn=ALG6;refGene_tx=XM_006934797.4;refGene_gn=ALG6;refGene_tx=XM_019837142.2;refGene_gn=ALG6;refGene_tx=XM_019837143.2;refGene_gn=ALG6;refGene_tx=XM_019837144.2
C1 52468453 52468539
refGene_gn=ALG6;refGene_tx=XM_006934795.3;refGene_gn=ALG6;refGene_tx=XM_006934796.4;refGene_gn=ALG6;refGene_tx=XM_006934797.4;refGene_gn=ALG6;refGene_tx=XM_019837142.2;refGene_gn=ALG6;refGene_tx=XM_019837143.2;refGene_gn=ALG6;refGene_tx=XM_019837144.2
C1 52470645 52470730
refGene_gn=ALG6;refGene_tx=XM_006934795.3;refGene_gn=ALG6;refGene_tx=XM_006934796.4;refGene_gn=ALG6;refGene_tx=XM_006934797.4;refGene_gn=ALG6;refGene_tx=XM_019837142.2;refGene_gn=ALG6;refGene_tx=XM_019837143.2;refGene_gn=ALG6;refGene_tx=XM_019837144.2
C1 52474416 52474487
refGene_gn=ALG6;refGene_tx=XM_006934795.3;refGene_gn=ALG6;refGene_tx=XM_006934796.4;refGene_gn=ALG6;refGene_tx=XM_006934797.4;refGene_gn=ALG6;refGene_tx=XM_019837142.2;refGene_gn=ALG6;refGene_tx=XM_019837143.2;refGene_gn=ALG6;refGene_tx=XM_019837144.2
C1 52482620 52482689
refGene_gn=ALG6;refGene_tx=XM_006934795.3;refGene_gn=ALG6;refGene_tx=XM_006934796.4;refGene_gn=ALG6;refGene_tx=XM_006934797.4;refGene_gn=ALG6;refGene_tx=XM_019837142.2;refGene_gn=ALG6;refGene_tx=XM_019837143.2;refGene_gn=ALG6;refGene_tx=XM_019837144.2
C1 52482817 52483016
refGene_gn=ALG6;refGene_tx=XM_006934795.3;refGene_gn=ALG6;refGene_tx=XM_006934796.4;refGene_gn=ALG6;refGene_tx=XM_006934797.4;refGene_gn=ALG6;refGene_tx=XM_019837142.2;refGene_gn=ALG6;refGene_tx=XM_019837143.2;refGene_gn=ALG6;refGene_tx=XM_019837144.2
C1 52490148 52490346

refGene_gn=ALG6;refGene_tx=XM_006934795.3;refGene_gn=ALG6;refGene_tx=XM_006934796.4;refGene_gn=ALG6;refGene_tx=XM_006934797.4;refGene_gn=ALG6;refGene_tx=XM_019837142.2;refGene_gn=ALG6;refGene_tx=XM_019837143.2;refGene_gn=ALG6;refGene_tx=XM_019837144.2
C1 52500032 52500082 refGene_gn=ITGB3BP;refGene_tx=XM_023257831.1
C1 52506588 52506648 refGene_gn=ITGB3BP;refGene_tx=XM_023257831.1
C1 52507095 52507192 refGene_gn=ITGB3BP;refGene_tx=XM_023257831.1
C1 52507634 52507713 refGene_gn=ITGB3BP;refGene_tx=XM_023257831.1
C1 52521628 52521698 refGene_gn=ITGB3BP;refGene_tx=XM_023257831.1
C1 52531443 52531579 refGene_gn=ITGB3BP;refGene_tx=XM_023257831.1
C1 52542278 52542321 refGene_gn=ITGB3BP;refGene_tx=XM_023257831.1
C1 52562997 52563387 refGene_gn=ITGB3BP;refGene_tx=XM_023257831.1
C1 52563957 52563962 refGene_gn=ITGB3BP;refGene_tx=XM_023257831.1
C1 52566502 52566689 refGene_gn=EFCAB7;refGene_tx=XM_003990155.5;refGene_gn=EFCAB7;refGene_tx=XM_019837141.2
C1 52571262 52571474 refGene_gn=EFCAB7;refGene_tx=XM_003990155.5;refGene_gn=EFCAB7;refGene_tx=XM_019837141.2
C1 52572113 52572200 refGene_gn=EFCAB7;refGene_tx=XM_003990155.5;refGene_gn=EFCAB7;refGene_tx=XM_019837141.2
C1 52572763 52572959 refGene_gn=EFCAB7;refGene_tx=XM_003990155.5;refGene_gn=EFCAB7;refGene_tx=XM_019837141.2
C1 52573410 52573532 refGene_gn=EFCAB7;refGene_tx=XM_003990155.5;refGene_gn=EFCAB7;refGene_tx=XM_019837141.2
C1 52589888 52590030 refGene_gn=EFCAB7;refGene_tx=XM_003990155.5
C1 52592276 52592386 refGene_gn=EFCAB7;refGene_tx=XM_003990155.5
C1 52594690 52594848 refGene_gn=EFCAB7;refGene_tx=XM_003990155.5;refGene_gn=EFCAB7;refGene_tx=XM_019837141.2
C1 52595747 52595881 refGene_gn=EFCAB7;refGene_tx=XM_003990155.5;refGene_gn=EFCAB7;refGene_tx=XM_019837141.2
C1 52600073 52600222 refGene_gn=EFCAB7;refGene_tx=XM_003990155.5;refGene_gn=EFCAB7;refGene_tx=XM_019837141.2
C1 52605760 52605970 refGene_gn=EFCAB7;refGene_tx=XM_003990155.5;refGene_gn=EFCAB7;refGene_tx=XM_019837141.2
C1 52608716 52608824 refGene_gn=EFCAB7;refGene_tx=XM_003990155.5;refGene_gn=EFCAB7;refGene_tx=XM_019837141.2
C1 52609802 52609877 refGene_gn=EFCAB7;refGene_tx=XM_003990155.5;refGene_gn=EFCAB7;refGene_tx=XM_019837141.2
C1 52633208 52633454 refGene_gn=PGM1;refGene_tx=XM_023258776.1
C1 52661384 52661684 refGene_gn=PGM1;refGene_tx=XM_019837155.2

C1	52667055	52667218
refGene_gn=PGM1;refGene_tx=XM_019837155.2;refGene_gn=PGM1;refGene_tx=XM_023258776.1
C1	52667564	52667711
refGene_gn=PGM1;refGene_tx=XM_019837155.2;refGene_gn=PGM1;refGene_tx=XM_023258776.1
C1	52669180	52669306
refGene_gn=PGM1;refGene_tx=XM_019837155.2;refGene_gn=PGM1;refGene_tx=XM_023258776.1
C1	52672411	52672602
refGene_gn=PGM1;refGene_tx=XM_019837155.2;refGene_gn=PGM1;refGene_tx=XM_023258776.1
C1	52673892	52674047
refGene_gn=PGM1;refGene_tx=XM_019837155.2;refGene_gn=PGM1;refGene_tx=XM_023258776.1
C1	52676827	52676943
refGene_gn=PGM1;refGene_tx=XM_019837155.2;refGene_gn=PGM1;refGene_tx=XM_023258776.1
C1	52686439	52686575
refGene_gn=PGM1;refGene_tx=XM_019837155.2;refGene_gn=PGM1;refGene_tx=XM_023258776.1
C1	52690101	52690285
refGene_gn=PGM1;refGene_tx=XM_019837155.2;refGene_gn=PGM1;refGene_tx=XM_023258776.1
C1	52692776	52692911
refGene_gn=PGM1;refGene_tx=XM_019837155.2;refGene_gn=PGM1;refGene_tx=XM_023258776.1
C1	52695355	52695445
refGene_gn=PGM1;refGene_tx=XM_019837155.2;refGene_gn=PGM1;refGene_tx=XM_023258776.1
C1	52807379	52807470	refGene_gn=ROR1;refGene_tx=XM_023257832.1
C1	53034672	53034744	refGene_gn=ROR1;refGene_tx=XM_023257832.1
C1	53076259	53076547	refGene_gn=ROR1;refGene_tx=XM_023257832.1
C1	53077225	53077256	refGene_gn=ROR1;refGene_tx=XM_023257832.1
C1	53166390	53166518	refGene_gn=ROR1;refGene_tx=XM_023257832.1
C1	53168444	53168762	refGene_gn=ROR1;refGene_tx=XM_023257832.1
C1	53170778	53171024	refGene_gn=ROR1;refGene_tx=XM_023257832.1
C1	53185783	53185995	refGene_gn=ROR1;refGene_tx=XM_023257832.1
C1	53201621	53203049	refGene_gn=ROR1;refGene_tx=XM_023257832.1
C1	53218387	53218453
refGene_gn=UBE2U;refGene_tx=XM_003990157.4;refGene_gn=UBE2U;refGene_tx=XM_019837157.2;refGene_gn=UBE2U;refGene_tx=XM_019837158.2;refGene_gn=UBE2U;refGene_tx=XM_019837160.2
C1	53220175	53220257
refGene_gn=UBE2U;refGene_tx=XM_003990157.4;refGene_gn=UBE2U;refGene_tx=XM_019837157.2;refGene_gn=UBE2U;refGene_tx=XM_019837158.2;refGene_gn=UBE2U;refGene_tx=XM_019837160.2
C1	53221272	53221365
refGene_gn=UBE2U;refGene_tx=XM_003990157.4;refGene_gn=UBE2U;refGene_tx=XM_019837157.2;refGene_gn=UBE2U;refGene_tx=XM_019837158.2;refGene_gn=

UBE2U;refGene_tx=XM_019837159.1;refGene_gn=UBE2U;refGene_tx=XM_0198371
60.2
C1	53225560	53225658
refGene_gn=UBE2U;refGene_tx=XM_003990157.4;refGene_gn=UBE2U;refGene_tx
=XM_019837157.2;refGene_gn=UBE2U;refGene_tx=XM_019837158.2;refGene_gn=
UBE2U;refGene_tx=XM_019837160.2
C1	53230914	53231035
refGene_gn=UBE2U;refGene_tx=XM_003990157.4;refGene_gn=UBE2U;refGene_tx
=XM_019837157.2;refGene_gn=UBE2U;refGene_tx=XM_019837158.2;refGene_gn=
UBE2U;refGene_tx=XM_019837159.1;refGene_gn=UBE2U;refGene_tx=XM_0198371
60.2
C1	53234607	53234656
refGene_gn=UBE2U;refGene_tx=XM_003990157.4;refGene_gn=UBE2U;refGene_tx
=XM_019837157.2;refGene_gn=UBE2U;refGene_tx=XM_019837158.2;refGene_gn=
UBE2U;refGene_tx=XM_019837159.1;refGene_gn=UBE2U;refGene_tx=XM_0198371
60.2
C1	53243744	53243838
refGene_gn=UBE2U;refGene_tx=XM_003990157.4;refGene_gn=UBE2U;refGene_tx
=XM_019837157.2;refGene_gn=UBE2U;refGene_tx=XM_019837158.2;refGene_gn=
UBE2U;refGene_tx=XM_019837159.1;refGene_gn=UBE2U;refGene_tx=XM_0198371
60.2
C1	53244015	53244039	refGene_gn=UBE2U;refGene_tx=XM_019837160.2
C1	53251309	53251395
refGene_gn=UBE2U;refGene_tx=XM_003990157.4;refGene_gn=UBE2U;refGene_tx
=XM_019837157.2;refGene_gn=UBE2U;refGene_tx=XM_019837158.2;refGene_gn=
UBE2U;refGene_tx=XM_019837159.1
C1	53262352	53262447
refGene_gn=UBE2U;refGene_tx=XM_003990157.4;refGene_gn=UBE2U;refGene_tx
=XM_019837159.1
C1	53275602	53275790
refGene_gn=UBE2U;refGene_tx=XM_003990157.4;refGene_gn=UBE2U;refGene_tx
=XM_019837159.1
C1	53462362	53462608	refGene_gn=CACHD1;refGene_tx=XM_023258777.1
C1	53551681	53551744
refGene_gn=CACHD1;refGene_tx=XM_023258777.1;refGene_gn=CACHD1;refGene_
tx=XM_023258778.1;refGene_gn=CACHD1;refGene_tx=XM_023258779.1
C1	53578370	53578519	refGene_gn=CACHD1;refGene_tx=XM_023258777.1
C1	53595746	53595802	refGene_gn=CACHD1;refGene_tx=XM_019837162.2
C1	53599807	53599914
refGene_gn=CACHD1;refGene_tx=XM_019837162.2;refGene_gn=CACHD1;refGene_
tx=XM_023258777.1;refGene_gn=CACHD1;refGene_tx=XM_023258778.1;refGene_
gn=CACHD1;refGene_tx=XM_023258779.1
C1	53617986	53618113
refGene_gn=CACHD1;refGene_tx=XM_019837162.2;refGene_gn=CACHD1;refGene_
tx=XM_023258777.1;refGene_gn=CACHD1;refGene_tx=XM_023258778.1;refGene_
gn=CACHD1;refGene_tx=XM_023258779.1
C1	53620998	53621143
refGene_gn=CACHD1;refGene_tx=XM_019837162.2;refGene_gn=CACHD1;refGene_
tx=XM_023258777.1;refGene_gn=CACHD1;refGene_tx=XM_023258778.1;refGene_
gn=CACHD1;refGene_tx=XM_023258779.1

C1 53622033 53622250
refGene_gn=CACHD1;refGene_tx=XM_019837162.2;refGene_gn=CACHD1;refGene_tx=XM_023258777.1;refGene_gn=CACHD1;refGene_tx=XM_023258778.1;refGene_gn=CACHD1;refGene_tx=XM_023258779.1
C1 53628977 53629127
refGene_gn=CACHD1;refGene_tx=XM_019837162.2;refGene_gn=CACHD1;refGene_tx=XM_023258777.1;refGene_gn=CACHD1;refGene_tx=XM_023258778.1;refGene_gn=CACHD1;refGene_tx=XM_023258779.1
C1 53633265 53633499
refGene_gn=CACHD1;refGene_tx=XM_019837162.2;refGene_gn=CACHD1;refGene_tx=XM_023258777.1;refGene_gn=CACHD1;refGene_tx=XM_023258778.1;refGene_gn=CACHD1;refGene_tx=XM_023258779.1
C1 53637742 53637892
refGene_gn=CACHD1;refGene_tx=XM_019837162.2;refGene_gn=CACHD1;refGene_tx=XM_023258777.1;refGene_gn=CACHD1;refGene_tx=XM_023258778.1;refGene_gn=CACHD1;refGene_tx=XM_023258779.1
C1 53638772 53638896
refGene_gn=CACHD1;refGene_tx=XM_019837162.2;refGene_gn=CACHD1;refGene_tx=XM_023258777.1;refGene_gn=CACHD1;refGene_tx=XM_023258778.1;refGene_gn=CACHD1;refGene_tx=XM_023258779.1
C1 53639711 53639829
refGene_gn=CACHD1;refGene_tx=XM_019837162.2;refGene_gn=CACHD1;refGene_tx=XM_023258777.1;refGene_gn=CACHD1;refGene_tx=XM_023258778.1;refGene_gn=CACHD1;refGene_tx=XM_023258779.1
C1 53644491 53644660
refGene_gn=CACHD1;refGene_tx=XM_019837162.2;refGene_gn=CACHD1;refGene_tx=XM_023258777.1;refGene_gn=CACHD1;refGene_tx=XM_023258778.1;refGene_gn=CACHD1;refGene_tx=XM_023258779.1
C1 53647821 53647964
refGene_gn=CACHD1;refGene_tx=XM_019837162.2;refGene_gn=CACHD1;refGene_tx=XM_023258777.1;refGene_gn=CACHD1;refGene_tx=XM_023258778.1;refGene_gn=CACHD1;refGene_tx=XM_023258779.1
C1 53648652 53648834
refGene_gn=CACHD1;refGene_tx=XM_019837162.2;refGene_gn=CACHD1;refGene_tx=XM_023258777.1;refGene_gn=CACHD1;refGene_tx=XM_023258778.1;refGene_gn=CACHD1;refGene_tx=XM_023258779.1
C1 53649307 53649418
refGene_gn=CACHD1;refGene_tx=XM_019837162.2;refGene_gn=CACHD1;refGene_tx=XM_023258777.1;refGene_gn=CACHD1;refGene_tx=XM_023258778.1;refGene_gn=CACHD1;refGene_tx=XM_023258779.1
C1 53654853 53654976
refGene_gn=CACHD1;refGene_tx=XM_019837162.2;refGene_gn=CACHD1;refGene_tx=XM_023258777.1;refGene_gn=CACHD1;refGene_tx=XM_023258778.1;refGene_gn=CACHD1;refGene_tx=XM_023258779.1
C1 53656362 53656462
refGene_gn=CACHD1;refGene_tx=XM_019837162.2;refGene_gn=CACHD1;refGene_tx=XM_023258777.1;refGene_gn=CACHD1;refGene_tx=XM_023258778.1;refGene_gn=CACHD1;refGene_tx=XM_023258779.1
C1 53656554 53656671
refGene_gn=CACHD1;refGene_tx=XM_019837162.2;refGene_gn=CACHD1;refGene_

tx=XM_023258777.1;refGene_gn=CACHD1;refGene_tx=XM_023258778.1;refGene_gn=CACHD1;refGene_tx=XM_023258779.1
C1 53659493 53659654
refGene_gn=CACHD1;refGene_tx=XM_019837162.2;refGene_gn=CACHD1;refGene_tx=XM_023258777.1;refGene_gn=CACHD1;refGene_tx=XM_023258778.1;refGene_gn=CACHD1;refGene_tx=XM_023258779.1
C1 53660006 53660093
refGene_gn=CACHD1;refGene_tx=XM_019837162.2;refGene_gn=CACHD1;refGene_tx=XM_023258777.1;refGene_gn=CACHD1;refGene_tx=XM_023258778.1;refGene_gn=CACHD1;refGene_tx=XM_023258779.1
C1 53660934 53661051
refGene_gn=CACHD1;refGene_tx=XM_019837162.2;refGene_gn=CACHD1;refGene_tx=XM_023258777.1;refGene_gn=CACHD1;refGene_tx=XM_023258778.1;refGene_gn=CACHD1;refGene_tx=XM_023258779.1
C1 53662477 53662629
refGene_gn=CACHD1;refGene_tx=XM_019837162.2;refGene_gn=CACHD1;refGene_tx=XM_023258777.1;refGene_gn=CACHD1;refGene_tx=XM_023258778.1;refGene_gn=CACHD1;refGene_tx=XM_023258779.1
C1 53663949 53664111
refGene_gn=CACHD1;refGene_tx=XM_019837162.2;refGene_gn=CACHD1;refGene_tx=XM_023258777.1;refGene_gn=CACHD1;refGene_tx=XM_023258778.1;refGene_gn=CACHD1;refGene_tx=XM_023258779.1
C1 53665399 53665477
refGene_gn=CACHD1;refGene_tx=XM_019837162.2;refGene_gn=CACHD1;refGene_tx=XM_023258777.1;refGene_gn=CACHD1;refGene_tx=XM_023258778.1;refGene_gn=CACHD1;refGene_tx=XM_023258779.1
C1 53666130 53666232
refGene_gn=CACHD1;refGene_tx=XM_019837162.2;refGene_gn=CACHD1;refGene_tx=XM_023258777.1;refGene_gn=CACHD1;refGene_tx=XM_023258778.1;refGene_gn=CACHD1;refGene_tx=XM_023258779.1
C1 53674269 53674508
refGene_gn=CACHD1;refGene_tx=XM_019837162.2;refGene_gn=CACHD1;refGene_tx=XM_023258777.1;refGene_gn=CACHD1;refGene_tx=XM_023258778.1;refGene_gn=CACHD1;refGene_tx=XM_023258779.1
C1 53727903 53728131
refGene_gn=RAVER2;refGene_tx=XM_023258780.1;refGene_gn=RAVER2;refGene_tx=XM_023258781.1
C1 53747960 53748027
refGene_gn=RAVER2;refGene_tx=XM_023258780.1;refGene_gn=RAVER2;refGene_tx=XM_023258781.1
C1 53752670 53753140
refGene_gn=RAVER2;refGene_tx=XM_023258780.1;refGene_gn=RAVER2;refGene_tx=XM_023258781.1
C1 53756131 53756323
refGene_gn=RAVER2;refGene_tx=XM_023258780.1;refGene_gn=RAVER2;refGene_tx=XM_023258781.1
C1 53768238 53768365
refGene_gn=RAVER2;refGene_tx=XM_023258780.1;refGene_gn=RAVER2;refGene_tx=XM_023258781.1
C1 53775864 53775950

refGene_gn=RAVER2;refGene_tx=XM_023258780.1;refGene_gn=RAVER2;refGene_tx=XM_023258781.1
C1 53777584 53777686
refGene_gn=RAVER2;refGene_tx=XM_023258780.1;refGene_gn=RAVER2;refGene_tx=XM_023258781.1
C1 53777843 53777955
refGene_gn=RAVER2;refGene_tx=XM_023258780.1;refGene_gn=RAVER2;refGene_tx=XM_023258781.1
C1 53780307 53780576
refGene_gn=RAVER2;refGene_tx=XM_023258780.1;refGene_gn=RAVER2;refGene_tx=XM_023258781.1
C1 53788233 53788345
refGene_gn=RAVER2;refGene_tx=XM_023258780.1;refGene_gn=RAVER2;refGene_tx=XM_023258781.1
C1 53789945 53790082
refGene_gn=RAVER2;refGene_tx=XM_023258780.1;refGene_gn=RAVER2;refGene_tx=XM_023258781.1
C1 53817514 53817661
refGene_gn=RAVER2;refGene_tx=XM_023258780.1;refGene_gn=RAVER2;refGene_tx=XM_023258781.1
C1 53820794 53820902
refGene_gn=JAK1;refGene_tx=XM_006934801.3;refGene_gn=JAK1;refGene_tx=XM_019837163.2;refGene_gn=JAK1;refGene_tx=XM_019837164.2;refGene_gn=JAK1;refGene_tx=XM_019837165.2
C1 53821712 53821823
refGene_gn=JAK1;refGene_tx=XM_006934801.3;refGene_gn=JAK1;refGene_tx=XM_019837163.2;refGene_gn=JAK1;refGene_tx=XM_019837164.2;refGene_gn=JAK1;refGene_tx=XM_019837165.2
C1 53822355 53822473
refGene_gn=JAK1;refGene_tx=XM_006934801.3;refGene_gn=JAK1;refGene_tx=XM_019837163.2;refGene_gn=JAK1;refGene_tx=XM_019837164.2;refGene_gn=JAK1;refGene_tx=XM_019837165.2
C1 53824197 53824370
refGene_gn=JAK1;refGene_tx=XM_006934801.3;refGene_gn=JAK1;refGene_tx=XM_019837163.2;refGene_gn=JAK1;refGene_tx=XM_019837164.2;refGene_gn=JAK1;refGene_tx=XM_019837165.2
C1 53824734 53824859
refGene_gn=JAK1;refGene_tx=XM_006934801.3;refGene_gn=JAK1;refGene_tx=XM_019837163.2;refGene_gn=JAK1;refGene_tx=XM_019837164.2;refGene_gn=JAK1;refGene_tx=XM_019837165.2
C1 53825309 53825502
refGene_gn=JAK1;refGene_tx=XM_006934801.3;refGene_gn=JAK1;refGene_tx=XM_019837163.2;refGene_gn=JAK1;refGene_tx=XM_019837164.2;refGene_gn=JAK1;refGene_tx=XM_019837165.2
C1 53826590 53826685
refGene_gn=JAK1;refGene_tx=XM_006934801.3;refGene_gn=JAK1;refGene_tx=XM_019837163.2;refGene_gn=JAK1;refGene_tx=XM_019837164.2;refGene_gn=JAK1;refGene_tx=XM_019837165.2
C1 53826809 53826960
refGene_gn=JAK1;refGene_tx=XM_006934801.3;refGene_gn=JAK1;refGene_tx=X

M_019837163.2;refGene_gn=JAK1;refGene_tx=XM_019837164.2;refGene_gn=JAK1;refGene_tx=XM_019837165.2
C1 53829833 53829985
refGene_gn=JAK1;refGene_tx=XM_006934801.3;refGene_gn=JAK1;refGene_tx=XM_019837163.2;refGene_gn=JAK1;refGene_tx=XM_019837164.2;refGene_gn=JAK1;refGene_tx=XM_019837165.2
C1 53831860 53831996
refGene_gn=JAK1;refGene_tx=XM_006934801.3;refGene_gn=JAK1;refGene_tx=XM_019837163.2;refGene_gn=JAK1;refGene_tx=XM_019837164.2;refGene_gn=JAK1;refGene_tx=XM_019837165.2
C1 53832613 53832741
refGene_gn=JAK1;refGene_tx=XM_006934801.3;refGene_gn=JAK1;refGene_tx=XM_019837163.2;refGene_gn=JAK1;refGene_tx=XM_019837164.2;refGene_gn=JAK1;refGene_tx=XM_019837165.2
C1 53833160 53833248
refGene_gn=JAK1;refGene_tx=XM_006934801.3;refGene_gn=JAK1;refGene_tx=XM_019837163.2;refGene_gn=JAK1;refGene_tx=XM_019837164.2;refGene_gn=JAK1;refGene_tx=XM_019837165.2
C1 53833983 53834127
refGene_gn=JAK1;refGene_tx=XM_006934801.3;refGene_gn=JAK1;refGene_tx=XM_019837163.2;refGene_gn=JAK1;refGene_tx=XM_019837164.2;refGene_gn=JAK1;refGene_tx=XM_019837165.2
C1 53836980 53837087
refGene_gn=JAK1;refGene_tx=XM_006934801.3;refGene_gn=JAK1;refGene_tx=XM_019837163.2;refGene_gn=JAK1;refGene_tx=XM_019837164.2;refGene_gn=JAK1;refGene_tx=XM_019837165.2
C1 53842578 53842765
refGene_gn=JAK1;refGene_tx=XM_006934801.3;refGene_gn=JAK1;refGene_tx=XM_019837163.2;refGene_gn=JAK1;refGene_tx=XM_019837164.2;refGene_gn=JAK1;refGene_tx=XM_019837165.2
C1 53844751 53844875
refGene_gn=JAK1;refGene_tx=XM_006934801.3;refGene_gn=JAK1;refGene_tx=XM_019837163.2;refGene_gn=JAK1;refGene_tx=XM_019837164.2;refGene_gn=JAK1;refGene_tx=XM_019837165.2
C1 53845365 53845523
refGene_gn=JAK1;refGene_tx=XM_006934801.3;refGene_gn=JAK1;refGene_tx=XM_019837163.2;refGene_gn=JAK1;refGene_tx=XM_019837164.2;refGene_gn=JAK1;refGene_tx=XM_019837165.2
C1 53849426 53849612
refGene_gn=JAK1;refGene_tx=XM_006934801.3;refGene_gn=JAK1;refGene_tx=XM_019837163.2;refGene_gn=JAK1;refGene_tx=XM_019837164.2;refGene_gn=JAK1;refGene_tx=XM_019837165.2
C1 53851149 53851489
refGene_gn=JAK1;refGene_tx=XM_006934801.3;refGene_gn=JAK1;refGene_tx=XM_019837163.2;refGene_gn=JAK1;refGene_tx=XM_019837164.2;refGene_gn=JAK1;refGene_tx=XM_019837165.2
C1 53853212 53853376
refGene_gn=JAK1;refGene_tx=XM_006934801.3;refGene_gn=JAK1;refGene_tx=XM_019837163.2;refGene_gn=JAK1;refGene_tx=XM_019837164.2;refGene_gn=JAK1;refGene_tx=XM_019837165.2

C1 53856900 53857054
refGene_gn=JAK1;refGene_tx=XM_006934801.3;refGene_gn=JAK1;refGene_tx=XM_019837163.2;refGene_gn=JAK1;refGene_tx=XM_019837164.2;refGene_gn=JAK1;refGene_tx=XM_019837165.2
C1 53862720 53862844
refGene_gn=JAK1;refGene_tx=XM_006934801.3;refGene_gn=JAK1;refGene_tx=XM_019837163.2;refGene_gn=JAK1;refGene_tx=XM_019837164.2;refGene_gn=JAK1;refGene_tx=XM_019837165.2
C1 53866235 53866434
refGene_gn=JAK1;refGene_tx=XM_006934801.3;refGene_gn=JAK1;refGene_tx=XM_019837163.2;refGene_gn=JAK1;refGene_tx=XM_019837164.2;refGene_gn=JAK1;refGene_tx=XM_019837165.2
C1 53869081 53869087
refGene_gn=JAK1;refGene_tx=XM_006934801.3;refGene_gn=JAK1;refGene_tx=XM_019837163.2;refGene_gn=JAK1;refGene_tx=XM_019837164.2;refGene_gn=JAK1;refGene_tx=XM_019837165.2
C1 54129312 54129457
refGene_gn=AK4;refGene_tx=XM_023258782.1;refGene_gn=AK4;refGene_tx=XM_023258783.1;refGene_gn=AK4;refGene_tx=XM_023258784.1
C1 54169665 54169785
refGene_gn=AK4;refGene_tx=XM_019837166.1;refGene_gn=AK4;refGene_tx=XM_019837167.2;refGene_gn=AK4;refGene_tx=XM_023258782.1;refGene_gn=AK4;refGene_tx=XM_023258783.1;refGene_gn=AK4;refGene_tx=XM_023258785.1
C1 54190165 54190338
refGene_gn=AK4;refGene_tx=XM_019837166.1;refGene_gn=AK4;refGene_tx=XM_019837167.2;refGene_gn=AK4;refGene_tx=XM_023258782.1;refGene_gn=AK4;refGene_tx=XM_023258783.1;refGene_gn=AK4;refGene_tx=XM_023258784.1;refGene_gn=AK4;refGene_tx=XM_023258785.1
C1 54193602 54193721
refGene_gn=AK4;refGene_tx=XM_019837166.1;refGene_gn=AK4;refGene_tx=XM_019837167.2;refGene_gn=AK4;refGene_tx=XM_023258782.1;refGene_gn=AK4;refGene_tx=XM_023258783.1;refGene_gn=AK4;refGene_tx=XM_023258784.1;refGene_gn=AK4;refGene_tx=XM_023258785.1
C1 54195856 54195971
refGene_gn=AK4;refGene_tx=XM_019837166.1;refGene_gn=AK4;refGene_tx=XM_019837167.2;refGene_gn=AK4;refGene_tx=XM_023258782.1;refGene_gn=AK4;refGene_tx=XM_023258783.1;refGene_gn=AK4;refGene_tx=XM_023258784.1;refGene_gn=AK4;refGene_tx=XM_023258785.1
C1 54228046 54228161 refGene_gn=DNAJC6;refGene_tx=XM_023258786.1
C1 54238411 54238433 refGene_gn=DNAJC6;refGene_tx=XM_011284948.3
C1 54277346 54277539 refGene_gn=DNAJC6;refGene_tx=XM_003990160.5
C1 54330352 54330503
refGene_gn=DNAJC6;refGene_tx=XM_003990160.5;refGene_gn=DNAJC6;refGene_tx=XM_011284948.3;refGene_gn=DNAJC6;refGene_tx=XM_011284949.3;refGene_gn=DNAJC6;refGene_tx=XM_023258786.1
C1 54331612 54331662
refGene_gn=DNAJC6;refGene_tx=XM_003990160.5;refGene_gn=DNAJC6;refGene_tx=XM_011284948.3;refGene_gn=DNAJC6;refGene_tx=XM_011284949.3;refGene_gn=DNAJC6;refGene_tx=XM_023258786.1
C1 54331761 54331910

refGene_gn=DNAJC6;refGene_tx=XM_003990160.5;refGene_gn=DNAJC6;refGene_tx=XM_011284948.3;refGene_gn=DNAJC6;refGene_tx=XM_011284949.3;refGene_gn=DNAJC6;refGene_tx=XM_023258786.1
C1 54343543 54343666
refGene_gn=DNAJC6;refGene_tx=XM_003990160.5;refGene_gn=DNAJC6;refGene_tx=XM_011284948.3;refGene_gn=DNAJC6;refGene_tx=XM_011284949.3;refGene_gn=DNAJC6;refGene_tx=XM_023258786.1
C1 54348235 54348369
refGene_gn=DNAJC6;refGene_tx=XM_003990160.5;refGene_gn=DNAJC6;refGene_tx=XM_011284948.3;refGene_gn=DNAJC6;refGene_tx=XM_011284949.3;refGene_gn=DNAJC6;refGene_tx=XM_023258786.1
C1 54349108 54349303
refGene_gn=DNAJC6;refGene_tx=XM_003990160.5;refGene_gn=DNAJC6;refGene_tx=XM_011284948.3;refGene_gn=DNAJC6;refGene_tx=XM_011284949.3;refGene_gn=DNAJC6;refGene_tx=XM_023258786.1
C1 54350109 54350227
refGene_gn=DNAJC6;refGene_tx=XM_003990160.5;refGene_gn=DNAJC6;refGene_tx=XM_011284948.3;refGene_gn=DNAJC6;refGene_tx=XM_011284949.3;refGene_gn=DNAJC6;refGene_tx=XM_023258786.1
C1 54351966 54352046
refGene_gn=DNAJC6;refGene_tx=XM_003990160.5;refGene_gn=DNAJC6;refGene_tx=XM_011284948.3;refGene_gn=DNAJC6;refGene_tx=XM_011284949.3;refGene_gn=DNAJC6;refGene_tx=XM_023258786.1
C1 54352901 54353095
refGene_gn=DNAJC6;refGene_tx=XM_003990160.5;refGene_gn=DNAJC6;refGene_tx=XM_011284948.3;refGene_gn=DNAJC6;refGene_tx=XM_011284949.3;refGene_gn=DNAJC6;refGene_tx=XM_023258786.1
C1 54353200 54353281
refGene_gn=DNAJC6;refGene_tx=XM_003990160.5;refGene_gn=DNAJC6;refGene_tx=XM_011284948.3;refGene_gn=DNAJC6;refGene_tx=XM_011284949.3;refGene_gn=DNAJC6;refGene_tx=XM_023258786.1
C1 54355634 54356069
refGene_gn=DNAJC6;refGene_tx=XM_003990160.5;refGene_gn=DNAJC6;refGene_tx=XM_011284948.3;refGene_gn=DNAJC6;refGene_tx=XM_011284949.3;refGene_gn=DNAJC6;refGene_tx=XM_023258786.1
C1 54358467 54358602
refGene_gn=DNAJC6;refGene_tx=XM_003990160.5;refGene_gn=DNAJC6;refGene_tx=XM_011284948.3;refGene_gn=DNAJC6;refGene_tx=XM_011284949.3;refGene_gn=DNAJC6;refGene_tx=XM_023258786.1
C1 54363352 54363421
refGene_gn=DNAJC6;refGene_tx=XM_003990160.5;refGene_gn=DNAJC6;refGene_tx=XM_011284948.3;refGene_gn=DNAJC6;refGene_tx=XM_011284949.3;refGene_gn=DNAJC6;refGene_tx=XM_023258786.1
C1 54364899 54365019
refGene_gn=DNAJC6;refGene_tx=XM_003990160.5;refGene_gn=DNAJC6;refGene_tx=XM_011284948.3;refGene_gn=DNAJC6;refGene_tx=XM_011284949.3;refGene_gn=DNAJC6;refGene_tx=XM_023258786.1
C1 54371204 54371468
refGene_gn=DNAJC6;refGene_tx=XM_003990160.5;refGene_gn=DNAJC6;refGene_tx=XM_011284948.3;refGene_gn=DNAJC6;refGene_tx=XM_011284949.3;refGene_

gn=DNAJC6;refGene_tx=XM_023258786.1
C1	54373654	54373797
refGene_gn=DNAJC6;refGene_tx=XM_003990160.5;refGene_gn=DNAJC6;refGene_
tx=XM_011284948.3;refGene_gn=DNAJC6;refGene_tx=XM_011284949.3;refGene_
gn=DNAJC6;refGene_tx=XM_023258786.1
C1	54375344	54375521
refGene_gn=DNAJC6;refGene_tx=XM_003990160.5;refGene_gn=DNAJC6;refGene_
tx=XM_011284948.3;refGene_gn=DNAJC6;refGene_tx=XM_011284949.3;refGene_
gn=DNAJC6;refGene_tx=XM_023258786.1
C1	54376452	54376554
refGene_gn=DNAJC6;refGene_tx=XM_003990160.5;refGene_gn=DNAJC6;refGene_
tx=XM_011284948.3;refGene_gn=DNAJC6;refGene_tx=XM_011284949.3;refGene_
gn=DNAJC6;refGene_tx=XM_023258786.1
C1	54383461	54383471	refGene_gn=LEPROT;refGene_tx=XM_019837173.2
C1	54383851	54383867	refGene_gn=LEPROT;refGene_tx=XM_003990161.5
C1	54385267	54385301	refGene_gn=LEPROT;refGene_tx=XM_019837172.2
C1	54388335	54388411
refGene_gn=LEPROT;refGene_tx=XM_003990161.5;refGene_gn=LEPROT;refGene_
tx=XM_019837172.2;refGene_gn=LEPROT;refGene_tx=XM_019837173.2
C1	54394024	54394211
refGene_gn=LEPROT;refGene_tx=XM_003990161.5;refGene_gn=LEPROT;refGene_
tx=XM_019837172.2;refGene_gn=LEPROT;refGene_tx=XM_019837173.2
C1	54395865	54395982
refGene_gn=LEPROT;refGene_tx=XM_003990161.5;refGene_gn=LEPROT;refGene_
tx=XM_019837172.2;refGene_gn=LEPROT;refGene_tx=XM_019837173.2
C1	54501808	54501848	refGene_gn=LEPR;refGene_tx=XM_023258787.1
C1	54506265	54506595	refGene_gn=LEPR;refGene_tx=XM_023258787.1
C1	54509368	54509492	refGene_gn=LEPR;refGene_tx=XM_023258787.1
C1	54526922	54527131	refGene_gn=LEPR;refGene_tx=XM_023258787.1
C1	54531788	54531934	refGene_gn=LEPR;refGene_tx=XM_023258787.1
C1	54534200	54534345	refGene_gn=LEPR;refGene_tx=XM_023258787.1
C1	54541790	54542081	refGene_gn=LEPR;refGene_tx=XM_023258787.1
C1	54542238	54542258	refGene_gn=LEPR;refGene_tx=XM_023258787.1
C1	54542259	54542357	refGene_gn=LEPR;refGene_tx=XM_023258787.1
C1	54545053	54545253	refGene_gn=LEPR;refGene_tx=XM_023258787.1
C1	54547064	54547213	refGene_gn=LEPR;refGene_tx=XM_023258787.1
C1	54548456	54548616	refGene_gn=LEPR;refGene_tx=XM_023258787.1
C1	54548735	54548818	refGene_gn=LEPR;refGene_tx=XM_023258787.1
C1	54549989	54550206	refGene_gn=LEPR;refGene_tx=XM_023258787.1
C1	54552345	54552528	refGene_gn=LEPR;refGene_tx=XM_023258787.1
C1	54553944	54554040	refGene_gn=LEPR;refGene_tx=XM_023258787.1
C1	54555677	54555783	refGene_gn=LEPR;refGene_tx=XM_023258787.1
C1	54556965	54557041	refGene_gn=LEPR;refGene_tx=XM_023258787.1
C1	54571640	54572468	refGene_gn=LEPR;refGene_tx=XM_023258787.1
C1	54781284	54781326	refGene_gn=PDE4B;refGene_tx=XM_023258788.1
C1	54788257	54788496	refGene_gn=PDE4B;refGene_tx=XM_023258788.1
C1	55085532	55085727	refGene_gn=PDE4B;refGene_tx=XM_023258788.1
C1	55094505	55094542	refGene_gn=PDE4B;refGene_tx=XM_023258788.1
C1	55094664	55094735	refGene_gn=PDE4B;refGene_tx=XM_023258788.1
C1	55105495	55105548	refGene_gn=PDE4B;refGene_tx=XM_023258788.1

C1	55170754	55170867	refGene_gn=PDE4B;refGene_tx=XM_023258788.1
C1	55194228	55194322	refGene_gn=PDE4B;refGene_tx=XM_023258788.1
C1	55200163	55200342	refGene_gn=PDE4B;refGene_tx=XM_023258788.1
C1	55201601	55201700	refGene_gn=PDE4B;refGene_tx=XM_023258788.1
C1	55201836	55202001	refGene_gn=PDE4B;refGene_tx=XM_023258788.1
C1	55203839	55203939	refGene_gn=PDE4B;refGene_tx=XM_023258788.1
C1	55206036	55206191	refGene_gn=PDE4B;refGene_tx=XM_023258788.1
C1	55206302	55206425	refGene_gn=PDE4B;refGene_tx=XM_023258788.1
C1	55207130	55207313	refGene_gn=PDE4B;refGene_tx=XM_023258788.1
C1	55210282	55210327	refGene_gn=PDE4B;refGene_tx=XM_023258788.1
C1	55210328	55210649	refGene_gn=PDE4B;refGene_tx=XM_023258788.1
C1	55354531	55354541
refGene_gn=SGIP1;refGene_tx=XM_003990163.5;refGene_gn=SGIP1;refGene_tx=XM_019837178.2;refGene_gn=SGIP1;refGene_tx=XM_019837179.2;refGene_gn=SGIP1;refGene_tx=XM_019837180.2;refGene_gn=SGIP1;refGene_tx=XM_019837181.2;refGene_gn=SGIP1;refGene_tx=XM_019837182.2;refGene_gn=SGIP1;refGene_tx=XM_019837183.2;refGene_gn=SGIP1;refGene_tx=XM_019837184.2;refGene_gn=SGIP1;refGene_tx=XM_019837185.2;refGene_gn=SGIP1;refGene_tx=XM_019837186.2;refGene_gn=SGIP1;refGene_tx=XM_019837187.2;refGene_gn=SGIP1;refGene_tx=XM_019837188.2;refGene_gn=SGIP1;refGene_tx=XM_019837189.2;refGene_gn=SGIP1;refGene_tx=XM_019837190.2;refGene_gn=SGIP1;refGene_tx=XM_019837191.2;refGene_gn=SGIP1;refGene_tx=XM_019837193.2;refGene_gn=SGIP1;refGene_tx=XM_019837194.2;refGene_gn=SGIP1;refGene_tx=XM_019837195.2;refGene_gn=SGIP1;refGene_tx=XM_019837196.2;refGene_gn=SGIP1;refGene_tx=XM_019837198.2;refGene_gn=SGIP1;refGene_tx=XM_019837199.2;refGene_gn=SGIP1;refGene_tx=XM_019837200.2;refGene_gn=SGIP1;refGene_tx=XM_019837201.2;refGene_gn=SGIP1;refGene_tx=XM_023258789.1;refGene_gn=SGIP1;refGene_tx=XM_023258792.1;refGene_gn=SGIP1;refGene_tx=XM_023258794.1
C1	55438344	55438408
refGene_gn=SGIP1;refGene_tx=XM_003990163.5;refGene_gn=SGIP1;refGene_tx=XM_019837178.2;refGene_gn=SGIP1;refGene_tx=XM_019837179.2;refGene_gn=SGIP1;refGene_tx=XM_019837180.2;refGene_gn=SGIP1;refGene_tx=XM_019837181.2;refGene_gn=SGIP1;refGene_tx=XM_019837182.2;refGene_gn=SGIP1;refGene_tx=XM_019837183.2;refGene_gn=SGIP1;refGene_tx=XM_019837184.2;refGene_gn=SGIP1;refGene_tx=XM_019837185.2;refGene_gn=SGIP1;refGene_tx=XM_019837186.2;refGene_gn=SGIP1;refGene_tx=XM_019837187.2;refGene_gn=SGIP1;refGene_tx=XM_019837188.2;refGene_gn=SGIP1;refGene_tx=XM_019837189.2;refGene_gn=SGIP1;refGene_tx=XM_019837190.2;refGene_gn=SGIP1;refGene_tx=XM_019837191.2;refGene_gn=SGIP1;refGene_tx=XM_019837193.2;refGene_gn=SGIP1;refGene_tx=XM_019837194.2;refGene_gn=SGIP1;refGene_tx=XM_019837195.2;refGene_gn=SGIP1;refGene_tx=XM_019837196.2;refGene_gn=SGIP1;refGene_tx=XM_019837198.2;refGene_gn=SGIP1;refGene_tx=XM_019837199.2;refGene_gn=SGIP1;refGene_tx=XM_019837200.2;refGene_gn=SGIP1;refGene_tx=XM_019837201.2;refGene_gn=SGIP1;refGene_tx=XM_023258789.1;refGene_gn=SGIP1;refGene_tx=XM_023258792.1;refGene_gn=SGIP1;refGene_tx=XM_023258794.1
C1	55442914	55442939
refGene_gn=SGIP1;refGene_tx=XM_003990163.5;refGene_gn=SGIP1;refGene_tx=XM_019837178.2;refGene_gn=SGIP1;refGene_tx=XM_019837179.2;refGene_gn=SGIP1;refGene_tx=XM_019837180.2;refGene_gn=SGIP1;refGene_tx=XM_019837181.2;refGene_gn=SGIP1;refGene_tx=XM_019837182.2;refGene_gn=SGIP1;refGe

ne_tx=XM_019837183.2;refGene_gn=SGIP1;refGene_tx=XM_019837184.2;refGen
e_gn=SGIP1;refGene_tx=XM_019837185.2;refGene_gn=SGIP1;refGene_tx=XM_01
9837186.2;refGene_gn=SGIP1;refGene_tx=XM_019837187.2;refGene_gn=SGIP1;
refGene_tx=XM_019837188.2;refGene_gn=SGIP1;refGene_tx=XM_019837189.2;r
efGene_gn=SGIP1;refGene_tx=XM_019837190.2;refGene_gn=SGIP1;refGene_tx=
XM_019837191.2;refGene_gn=SGIP1;refGene_tx=XM_019837192.2;refGene_gn=S
GIP1;refGene_tx=XM_019837193.2;refGene_gn=SGIP1;refGene_tx=XM_01983719
4.2;refGene_gn=SGIP1;refGene_tx=XM_019837195.2;refGene_gn=SGIP1;refGen
e_tx=XM_019837196.2;refGene_gn=SGIP1;refGene_tx=XM_019837198.2;refGene
_gn=SGIP1;refGene_tx=XM_019837199.2;refGene_gn=SGIP1;refGene_tx=XM_019
837200.2;refGene_gn=SGIP1;refGene_tx=XM_019837201.2;refGene_gn=SGIP1;r
efGene_tx=XM_023258789.1;refGene_gn=SGIP1;refGene_tx=XM_023258792.1;re
fGene_gn=SGIP1;refGene_tx=XM_023258794.1
C1 55443943 55444027
refGene_gn=SGIP1;refGene_tx=XM_019837178.2;refGene_gn=SGIP1;refGene_tx
=XM_019837179.2;refGene_gn=SGIP1;refGene_tx=XM_019837180.2;refGene_gn=
SGIP1;refGene_tx=XM_019837181.2;refGene_gn=SGIP1;refGene_tx=XM_0198371
82.2;refGene_gn=SGIP1;refGene_tx=XM_019837183.2;refGene_gn=SGIP1;refGe
ne_tx=XM_019837184.2;refGene_gn=SGIP1;refGene_tx=XM_019837185.2;refGen
e_gn=SGIP1;refGene_tx=XM_019837186.2;refGene_gn=SGIP1;refGene_tx=XM_01
9837187.2;refGene_gn=SGIP1;refGene_tx=XM_019837188.2;refGene_gn=SGIP1;
refGene_tx=XM_019837189.2;refGene_gn=SGIP1;refGene_tx=XM_019837193.2;r
efGene_gn=SGIP1;refGene_tx=XM_019837198.2;refGene_gn=SGIP1;refGene_tx=
XM_023258789.1;refGene_gn=SGIP1;refGene_tx=XM_023258792.1
C1 55445808 55445932
refGene_gn=SGIP1;refGene_tx=XM_003990163.5;refGene_gn=SGIP1;refGene_tx
=XM_019837178.2;refGene_gn=SGIP1;refGene_tx=XM_019837179.2;refGene_gn=
SGIP1;refGene_tx=XM_019837180.2;refGene_gn=SGIP1;refGene_tx=XM_0198371
81.2;refGene_gn=SGIP1;refGene_tx=XM_019837182.2;refGene_gn=SGIP1;refGe
ne_tx=XM_019837183.2;refGene_gn=SGIP1;refGene_tx=XM_019837184.2;refGen
e_gn=SGIP1;refGene_tx=XM_019837185.2;refGene_gn=SGIP1;refGene_tx=XM_01
9837186.2;refGene_gn=SGIP1;refGene_tx=XM_019837187.2;refGene_gn=SGIP1;
refGene_tx=XM_019837190.2;refGene_gn=SGIP1;refGene_tx=XM_019837191.2;r
efGene_gn=SGIP1;refGene_tx=XM_019837192.2;refGene_gn=SGIP1;refGene_tx=
XM_019837193.2;refGene_gn=SGIP1;refGene_tx=XM_019837195.2;refGene_gn=S
GIP1;refGene_tx=XM_019837198.2;refGene_gn=SGIP1;refGene_tx=XM_01983719
9.2;refGene_gn=SGIP1;refGene_tx=XM_019837200.2;refGene_gn=SGIP1;refGen
e_tx=XM_023258789.1
C1 55449569 55449626
refGene_gn=SGIP1;refGene_tx=XM_003990163.5;refGene_gn=SGIP1;refGene_tx
=XM_019837178.2;refGene_gn=SGIP1;refGene_tx=XM_019837179.2;refGene_gn=
SGIP1;refGene_tx=XM_019837180.2;refGene_gn=SGIP1;refGene_tx=XM_0198371
81.2;refGene_gn=SGIP1;refGene_tx=XM_019837182.2;refGene_gn=SGIP1;refGe
ne_tx=XM_019837183.2;refGene_gn=SGIP1;refGene_tx=XM_019837184.2;refGen
e_gn=SGIP1;refGene_tx=XM_019837185.2;refGene_gn=SGIP1;refGene_tx=XM_01
9837186.2;refGene_gn=SGIP1;refGene_tx=XM_019837187.2;refGene_gn=SGIP1;
refGene_tx=XM_019837188.2;refGene_gn=SGIP1;refGene_tx=XM_019837189.2;r
efGene_gn=SGIP1;refGene_tx=XM_019837190.2;refGene_gn=SGIP1;refGene_tx=
XM_019837191.2;refGene_gn=SGIP1;refGene_tx=XM_019837192.2;refGene_gn=S
GIP1;refGene_tx=XM_019837193.2;refGene_gn=SGIP1;refGene_tx=XM_01983719

4.2;refGene_gn=SGIP1;refGene_tx=XM_019837195.2;refGene_gn=SGIP1;refGene_tx=XM_019837196.2;refGene_gn=SGIP1;refGene_tx=XM_019837198.2;refGene_gn=SGIP1;refGene_tx=XM_019837199.2;refGene_gn=SGIP1;refGene_tx=XM_019837200.2;refGene_gn=SGIP1;refGene_tx=XM_019837201.2;refGene_gn=SGIP1;refGene_tx=XM_023258789.1;refGene_gn=SGIP1;refGene_tx=XM_023258792.1;refGene_gn=SGIP1;refGene_tx=XM_023258794.1
C1 55453986 55454041
refGene_gn=SGIP1;refGene_tx=XM_003990163.5;refGene_gn=SGIP1;refGene_tx=XM_019837178.2;refGene_gn=SGIP1;refGene_tx=XM_019837179.2;refGene_gn=SGIP1;refGene_tx=XM_019837180.2;refGene_gn=SGIP1;refGene_tx=XM_019837181.2;refGene_gn=SGIP1;refGene_tx=XM_019837182.2;refGene_gn=SGIP1;refGene_tx=XM_019837183.2;refGene_gn=SGIP1;refGene_tx=XM_019837184.2;refGene_gn=SGIP1;refGene_tx=XM_019837185.2;refGene_gn=SGIP1;refGene_tx=XM_019837186.2;refGene_gn=SGIP1;refGene_tx=XM_019837187.2;refGene_gn=SGIP1;refGene_tx=XM_019837188.2;refGene_gn=SGIP1;refGene_tx=XM_019837189.2;refGene_gn=SGIP1;refGene_tx=XM_019837190.2;refGene_gn=SGIP1;refGene_tx=XM_019837191.2;refGene_gn=SGIP1;refGene_tx=XM_019837192.2;refGene_gn=SGIP1;refGene_tx=XM_019837193.2;refGene_gn=SGIP1;refGene_tx=XM_019837194.2;refGene_gn=SGIP1;refGene_tx=XM_019837195.2;refGene_gn=SGIP1;refGene_tx=XM_019837196.2;refGene_gn=SGIP1;refGene_tx=XM_019837198.2;refGene_gn=SGIP1;refGene_tx=XM_019837199.2;refGene_gn=SGIP1;refGene_tx=XM_019837200.2;refGene_gn=SGIP1;refGene_tx=XM_019837201.2;refGene_gn=SGIP1;refGene_tx=XM_023258789.1;refGene_gn=SGIP1;refGene_tx=XM_023258792.1;refGene_gn=SGIP1;refGene_tx=XM_023258794.1
C1 55454800 55454976
refGene_gn=SGIP1;refGene_tx=XM_003990163.5;refGene_gn=SGIP1;refGene_tx=XM_019837178.2;refGene_gn=SGIP1;refGene_tx=XM_019837179.2;refGene_gn=SGIP1;refGene_tx=XM_019837180.2;refGene_gn=SGIP1;refGene_tx=XM_019837181.2;refGene_gn=SGIP1;refGene_tx=XM_019837182.2;refGene_gn=SGIP1;refGene_tx=XM_019837183.2;refGene_gn=SGIP1;refGene_tx=XM_019837184.2;refGene_gn=SGIP1;refGene_tx=XM_019837185.2;refGene_gn=SGIP1;refGene_tx=XM_019837186.2;refGene_gn=SGIP1;refGene_tx=XM_019837187.2;refGene_gn=SGIP1;refGene_tx=XM_019837188.2;refGene_gn=SGIP1;refGene_tx=XM_019837189.2;refGene_gn=SGIP1;refGene_tx=XM_019837190.2;refGene_gn=SGIP1;refGene_tx=XM_019837191.2;refGene_gn=SGIP1;refGene_tx=XM_019837192.2;refGene_gn=SGIP1;refGene_tx=XM_019837193.2;refGene_gn=SGIP1;refGene_tx=XM_019837194.2;refGene_gn=SGIP1;refGene_tx=XM_019837195.2;refGene_gn=SGIP1;refGene_tx=XM_019837196.2;refGene_gn=SGIP1;refGene_tx=XM_019837198.2;refGene_gn=SGIP1;refGene_tx=XM_019837199.2;refGene_gn=SGIP1;refGene_tx=XM_019837200.2;refGene_gn=SGIP1;refGene_tx=XM_019837201.2;refGene_gn=SGIP1;refGene_tx=XM_023258789.1;refGene_gn=SGIP1;refGene_tx=XM_023258792.1;refGene_gn=SGIP1;refGene_tx=XM_023258794.1
C1 55468550 55468562
refGene_gn=SGIP1;refGene_tx=XM_003990163.5;refGene_gn=SGIP1;refGene_tx=XM_019837178.2;refGene_gn=SGIP1;refGene_tx=XM_019837179.2;refGene_gn=SGIP1;refGene_tx=XM_019837180.2;refGene_gn=SGIP1;refGene_tx=XM_019837181.2;refGene_gn=SGIP1;refGene_tx=XM_019837182.2;refGene_gn=SGIP1;refGene_tx=XM_019837184.2;refGene_gn=SGIP1;refGene_tx=XM_019837186.2;refGene_gn=SGIP1;refGene_tx=XM_019837187.2;refGene_gn=SGIP1;refGene_tx=XM_019837188.2;refGene_gn=SGIP1;refGene_tx=XM_019837189.2;refGene_gn=SGIP1;

refGene_tx=XM_019837190.2;refGene_gn=SGIP1;refGene_tx=XM_019837191.2;r
efGene_gn=SGIP1;refGene_tx=XM_019837192.2;refGene_gn=SGIP1;refGene_tx=
XM_019837193.2;refGene_gn=SGIP1;refGene_tx=XM_019837194.2;refGene_gn=S
GIP1;refGene_tx=XM_019837195.2;refGene_gn=SGIP1;refGene_tx=XM_01983719
6.2;refGene_gn=SGIP1;refGene_tx=XM_019837198.2;refGene_gn=SGIP1;refGen
e_tx=XM_019837199.2;refGene_gn=SGIP1;refGene_tx=XM_019837200.2;refGene
_gn=SGIP1;refGene_tx=XM_023258789.1;refGene_gn=SGIP1;refGene_tx=XM_023
258792.1;refGene_gn=SGIP1;refGene_tx=XM_023258794.1
C1 55473677 55473689
refGene_gn=SGIP1;refGene_tx=XM_003990163.5;refGene_gn=SGIP1;refGene_tx
=XM_019837178.2;refGene_gn=SGIP1;refGene_tx=XM_019837179.2;refGene_gn=
SGIP1;refGene_tx=XM_019837180.2;refGene_gn=SGIP1;refGene_tx=XM_0198371
81.2;refGene_gn=SGIP1;refGene_tx=XM_019837182.2;refGene_gn=SGIP1;refGe
ne_tx=XM_019837183.2;refGene_gn=SGIP1;refGene_tx=XM_019837184.2;refGen
e_gn=SGIP1;refGene_tx=XM_019837186.2;refGene_gn=SGIP1;refGene_tx=XM_01
9837187.2;refGene_gn=SGIP1;refGene_tx=XM_019837188.2;refGene_gn=SGIP1;
refGene_tx=XM_019837189.2;refGene_gn=SGIP1;refGene_tx=XM_019837190.2;r
efGene_gn=SGIP1;refGene_tx=XM_019837191.2;refGene_gn=SGIP1;refGene_tx=
XM_019837192.2;refGene_gn=SGIP1;refGene_tx=XM_019837193.2;refGene_gn=S
GIP1;refGene_tx=XM_019837194.2;refGene_gn=SGIP1;refGene_tx=XM_01983719
5.2;refGene_gn=SGIP1;refGene_tx=XM_019837196.2;refGene_gn=SGIP1;refGen
e_tx=XM_019837198.2;refGene_gn=SGIP1;refGene_tx=XM_019837199.2;refGene
_gn=SGIP1;refGene_tx=XM_019837200.2;refGene_gn=SGIP1;refGene_tx=XM_023
258789.1;refGene_gn=SGIP1;refGene_tx=XM_023258792.1;refGene_gn=SGIP1;r
efGene_tx=XM_023258794.1
C1 55476847 55476872
refGene_gn=SGIP1;refGene_tx=XM_003990163.5;refGene_gn=SGIP1;refGene_tx
=XM_019837178.2;refGene_gn=SGIP1;refGene_tx=XM_019837179.2;refGene_gn=
SGIP1;refGene_tx=XM_019837180.2;refGene_gn=SGIP1;refGene_tx=XM_0198371
81.2;refGene_gn=SGIP1;refGene_tx=XM_019837182.2;refGene_gn=SGIP1;refGe
ne_tx=XM_019837183.2;refGene_gn=SGIP1;refGene_tx=XM_019837184.2;refGen
e_gn=SGIP1;refGene_tx=XM_019837185.2;refGene_gn=SGIP1;refGene_tx=XM_01
9837186.2;refGene_gn=SGIP1;refGene_tx=XM_019837187.2;refGene_gn=SGIP1;
refGene_tx=XM_019837188.2;refGene_gn=SGIP1;refGene_tx=XM_019837189.2;r
efGene_gn=SGIP1;refGene_tx=XM_019837190.2;refGene_gn=SGIP1;refGene_tx=
XM_019837191.2;refGene_gn=SGIP1;refGene_tx=XM_019837192.2;refGene_gn=S
GIP1;refGene_tx=XM_019837193.2;refGene_gn=SGIP1;refGene_tx=XM_01983719
4.2;refGene_gn=SGIP1;refGene_tx=XM_019837195.2;refGene_gn=SGIP1;refGen
e_tx=XM_019837196.2;refGene_gn=SGIP1;refGene_tx=XM_019837198.2;refGene
_gn=SGIP1;refGene_tx=XM_019837199.2;refGene_gn=SGIP1;refGene_tx=XM_019
837200.2;refGene_gn=SGIP1;refGene_tx=XM_019837201.2;refGene_gn=SGIP1;r
efGene_tx=XM_023258789.1;refGene_gn=SGIP1;refGene_tx=XM_023258792.1;re
fGene_gn=SGIP1;refGene_tx=XM_023258794.1
C1 55477813 55477865
refGene_gn=SGIP1;refGene_tx=XM_003990163.5;refGene_gn=SGIP1;refGene_tx
=XM_019837178.2;refGene_gn=SGIP1;refGene_tx=XM_019837179.2;refGene_gn=
SGIP1;refGene_tx=XM_019837180.2;refGene_gn=SGIP1;refGene_tx=XM_0198371
81.2;refGene_gn=SGIP1;refGene_tx=XM_019837182.2;refGene_gn=SGIP1;refGe
ne_tx=XM_019837183.2;refGene_gn=SGIP1;refGene_tx=XM_019837184.2;refGen
e_gn=SGIP1;refGene_tx=XM_019837185.2;refGene_gn=SGIP1;refGene_tx=XM_01

9837186.2;refGene_gn=SGIP1;refGene_tx=XM_019837187.2;refGene_gn=SGIP1;
refGene_tx=XM_019837188.2;refGene_gn=SGIP1;refGene_tx=XM_019837189.2;r
efGene_gn=SGIP1;refGene_tx=XM_019837190.2;refGene_gn=SGIP1;refGene_tx=
XM_019837191.2;refGene_gn=SGIP1;refGene_tx=XM_019837192.2;refGene_gn=S
GIP1;refGene_tx=XM_019837193.2;refGene_gn=SGIP1;refGene_tx=XM_01983719
4.2;refGene_gn=SGIP1;refGene_tx=XM_019837195.2;refGene_gn=SGIP1;refGen
e_tx=XM_019837196.2;refGene_gn=SGIP1;refGene_tx=XM_019837198.2;refGene
_gn=SGIP1;refGene_tx=XM_019837199.2;refGene_gn=SGIP1;refGene_tx=XM_019
837200.2;refGene_gn=SGIP1;refGene_tx=XM_019837201.2;refGene_gn=SGIP1;r
efGene_tx=XM_023258789.1;refGene_gn=SGIP1;refGene_tx=XM_023258792.1;re
fGene_gn=SGIP1;refGene_tx=XM_023258794.1
C1 55479170 55479256
refGene_gn=SGIP1;refGene_tx=XM_003990163.5;refGene_gn=SGIP1;refGene_tx
=XM_019837178.2;refGene_gn=SGIP1;refGene_tx=XM_019837179.2;refGene_gn=
SGIP1;refGene_tx=XM_019837180.2;refGene_gn=SGIP1;refGene_tx=XM_0198371
81.2;refGene_gn=SGIP1;refGene_tx=XM_019837182.2;refGene_gn=SGIP1;refGe
ne_tx=XM_019837183.2;refGene_gn=SGIP1;refGene_tx=XM_019837184.2;refGen
e_gn=SGIP1;refGene_tx=XM_019837185.2;refGene_gn=SGIP1;refGene_tx=XM_01
9837186.2;refGene_gn=SGIP1;refGene_tx=XM_019837187.2;refGene_gn=SGIP1;
refGene_tx=XM_019837188.2;refGene_gn=SGIP1;refGene_tx=XM_019837189.2;r
efGene_gn=SGIP1;refGene_tx=XM_019837190.2;refGene_gn=SGIP1;refGene_tx=
XM_019837191.2;refGene_gn=SGIP1;refGene_tx=XM_019837192.2;refGene_gn=S
GIP1;refGene_tx=XM_019837193.2;refGene_gn=SGIP1;refGene_tx=XM_01983719
4.2;refGene_gn=SGIP1;refGene_tx=XM_019837195.2;refGene_gn=SGIP1;refGen
e_tx=XM_019837196.2;refGene_gn=SGIP1;refGene_tx=XM_019837198.2;refGene
_gn=SGIP1;refGene_tx=XM_019837199.2;refGene_gn=SGIP1;refGene_tx=XM_019
837200.2;refGene_gn=SGIP1;refGene_tx=XM_019837201.2;refGene_gn=SGIP1;r
efGene_tx=XM_023258789.1;refGene_gn=SGIP1;refGene_tx=XM_023258792.1;re
fGene_gn=SGIP1;refGene_tx=XM_023258794.1
C1 55482519 55482612
refGene_gn=SGIP1;refGene_tx=XM_003990163.5;refGene_gn=SGIP1;refGene_tx
=XM_019837178.2;refGene_gn=SGIP1;refGene_tx=XM_019837179.2;refGene_gn=
SGIP1;refGene_tx=XM_019837180.2;refGene_gn=SGIP1;refGene_tx=XM_0198371
81.2;refGene_gn=SGIP1;refGene_tx=XM_019837182.2;refGene_gn=SGIP1;refGe
ne_tx=XM_019837183.2;refGene_gn=SGIP1;refGene_tx=XM_019837184.2;refGen
e_gn=SGIP1;refGene_tx=XM_019837185.2;refGene_gn=SGIP1;refGene_tx=XM_01
9837186.2;refGene_gn=SGIP1;refGene_tx=XM_019837187.2;refGene_gn=SGIP1;
refGene_tx=XM_019837188.2;refGene_gn=SGIP1;refGene_tx=XM_019837189.2;r
efGene_gn=SGIP1;refGene_tx=XM_019837190.2;refGene_gn=SGIP1;refGene_tx=
XM_019837191.2;refGene_gn=SGIP1;refGene_tx=XM_019837192.2;refGene_gn=S
GIP1;refGene_tx=XM_019837193.2;refGene_gn=SGIP1;refGene_tx=XM_01983719
4.2;refGene_gn=SGIP1;refGene_tx=XM_019837195.2;refGene_gn=SGIP1;refGen
e_tx=XM_019837196.2;refGene_gn=SGIP1;refGene_tx=XM_019837198.2;refGene
_gn=SGIP1;refGene_tx=XM_019837199.2;refGene_gn=SGIP1;refGene_tx=XM_019
837200.2;refGene_gn=SGIP1;refGene_tx=XM_019837201.2;refGene_gn=SGIP1;r
efGene_tx=XM_023258789.1;refGene_gn=SGIP1;refGene_tx=XM_023258792.1;re
fGene_gn=SGIP1;refGene_tx=XM_023258794.1
C1 55485789 55485864
refGene_gn=SGIP1;refGene_tx=XM_003990163.5;refGene_gn=SGIP1;refGene_tx
=XM_019837178.2;refGene_gn=SGIP1;refGene_tx=XM_019837179.2;refGene_gn=

SGIP1;refGene_tx=XM_019837180.2;refGene_gn=SGIP1;refGene_tx=XM_0198371
81.2;refGene_gn=SGIP1;refGene_tx=XM_019837182.2;refGene_gn=SGIP1;refGe
ne_tx=XM_019837183.2;refGene_gn=SGIP1;refGene_tx=XM_019837184.2;refGen
e_gn=SGIP1;refGene_tx=XM_019837185.2;refGene_gn=SGIP1;refGene_tx=XM_01
9837186.2;refGene_gn=SGIP1;refGene_tx=XM_019837187.2;refGene_gn=SGIP1;
refGene_tx=XM_019837188.2;refGene_gn=SGIP1;refGene_tx=XM_019837189.2;r
efGene_gn=SGIP1;refGene_tx=XM_019837190.2;refGene_gn=SGIP1;refGene_tx=
XM_019837191.2;refGene_gn=SGIP1;refGene_tx=XM_019837192.2;refGene_gn=S
GIP1;refGene_tx=XM_019837193.2;refGene_gn=SGIP1;refGene_tx=XM_01983719
4.2;refGene_gn=SGIP1;refGene_tx=XM_019837195.2;refGene_gn=SGIP1;refGen
e_tx=XM_019837196.2;refGene_gn=SGIP1;refGene_tx=XM_019837198.2;refGene
_gn=SGIP1;refGene_tx=XM_019837199.2;refGene_gn=SGIP1;refGene_tx=XM_019
837200.2;refGene_gn=SGIP1;refGene_tx=XM_019837201.2;refGene_gn=SGIP1;r
efGene_tx=XM_023258789.1;refGene_gn=SGIP1;refGene_tx=XM_023258792.1;re
fGene_gn=SGIP1;refGene_tx=XM_023258794.1
C1 55487922 55488423
refGene_gn=SGIP1;refGene_tx=XM_003990163.5;refGene_gn=SGIP1;refGene_tx
=XM_019837178.2;refGene_gn=SGIP1;refGene_tx=XM_019837179.2;refGene_gn=
SGIP1;refGene_tx=XM_019837180.2;refGene_gn=SGIP1;refGene_tx=XM_0198371
81.2;refGene_gn=SGIP1;refGene_tx=XM_019837182.2;refGene_gn=SGIP1;refGe
ne_tx=XM_019837183.2;refGene_gn=SGIP1;refGene_tx=XM_019837184.2;refGen
e_gn=SGIP1;refGene_tx=XM_019837185.2;refGene_gn=SGIP1;refGene_tx=XM_01
9837186.2;refGene_gn=SGIP1;refGene_tx=XM_019837187.2;refGene_gn=SGIP1;
refGene_tx=XM_019837188.2;refGene_gn=SGIP1;refGene_tx=XM_019837189.2;r
efGene_gn=SGIP1;refGene_tx=XM_019837190.2;refGene_gn=SGIP1;refGene_tx=
XM_019837191.2;refGene_gn=SGIP1;refGene_tx=XM_019837192.2;refGene_gn=S
GIP1;refGene_tx=XM_019837193.2;refGene_gn=SGIP1;refGene_tx=XM_01983719
4.2;refGene_gn=SGIP1;refGene_tx=XM_019837195.2;refGene_gn=SGIP1;refGen
e_tx=XM_019837196.2;refGene_gn=SGIP1;refGene_tx=XM_023258789.1;refGene
_gn=SGIP1;refGene_tx=XM_023258792.1
C1 55495039 55495167
refGene_gn=SGIP1;refGene_tx=XM_003990163.5;refGene_gn=SGIP1;refGene_tx
=XM_019837178.2;refGene_gn=SGIP1;refGene_tx=XM_019837179.2;refGene_gn=
SGIP1;refGene_tx=XM_019837180.2;refGene_gn=SGIP1;refGene_tx=XM_0198371
81.2;refGene_gn=SGIP1;refGene_tx=XM_019837182.2;refGene_gn=SGIP1;refGe
ne_tx=XM_019837183.2;refGene_gn=SGIP1;refGene_tx=XM_019837184.2;refGen
e_gn=SGIP1;refGene_tx=XM_019837185.2;refGene_gn=SGIP1;refGene_tx=XM_01
9837186.2;refGene_gn=SGIP1;refGene_tx=XM_019837187.2;refGene_gn=SGIP1;
refGene_tx=XM_019837188.2;refGene_gn=SGIP1;refGene_tx=XM_019837189.2;r
efGene_gn=SGIP1;refGene_tx=XM_019837190.2;refGene_gn=SGIP1;refGene_tx=
XM_019837191.2;refGene_gn=SGIP1;refGene_tx=XM_019837192.2;refGene_gn=S
GIP1;refGene_tx=XM_019837193.2;refGene_gn=SGIP1;refGene_tx=XM_01983719
4.2;refGene_gn=SGIP1;refGene_tx=XM_019837195.2;refGene_gn=SGIP1;refGen
e_tx=XM_019837196.2;refGene_gn=SGIP1;refGene_tx=XM_019837198.2;refGene
_gn=SGIP1;refGene_tx=XM_019837199.2;refGene_gn=SGIP1;refGene_tx=XM_019
837200.2;refGene_gn=SGIP1;refGene_tx=XM_019837201.2;refGene_gn=SGIP1;r
efGene_tx=XM_023258789.1;refGene_gn=SGIP1;refGene_tx=XM_023258792.1;re
fGene_gn=SGIP1;refGene_tx=XM_023258794.1
C1 55496144 55496271
refGene_gn=SGIP1;refGene_tx=XM_003990163.5;refGene_gn=SGIP1;refGene_tx

=XM_019837178.2;refGene_gn=SGIP1;refGene_tx=XM_019837179.2;refGene_gn=SGIP1;refGene_tx=XM_019837180.2;refGene_gn=SGIP1;refGene_tx=XM_019837181.2;refGene_gn=SGIP1;refGene_tx=XM_019837182.2;refGene_gn=SGIP1;refGene_tx=XM_019837183.2;refGene_gn=SGIP1;refGene_tx=XM_019837184.2;refGene_gn=SGIP1;refGene_tx=XM_019837185.2;refGene_gn=SGIP1;refGene_tx=XM_019837186.2;refGene_gn=SGIP1;refGene_tx=XM_019837187.2;refGene_gn=SGIP1;refGene_tx=XM_019837188.2;refGene_gn=SGIP1;refGene_tx=XM_019837189.2;refGene_gn=SGIP1;refGene_tx=XM_019837190.2;refGene_gn=SGIP1;refGene_tx=XM_019837191.2;refGene_gn=SGIP1;refGene_tx=XM_019837192.2;refGene_gn=SGIP1;refGene_tx=XM_019837193.2;refGene_gn=SGIP1;refGene_tx=XM_019837194.2;refGene_gn=SGIP1;refGene_tx=XM_019837195.2;refGene_gn=SGIP1;refGene_tx=XM_019837196.2;refGene_gn=SGIP1;refGene_tx=XM_019837198.2;refGene_gn=SGIP1;refGene_tx=XM_019837199.2;refGene_gn=SGIP1;refGene_tx=XM_019837200.2;refGene_gn=SGIP1;refGene_tx=XM_019837201.2;refGene_gn=SGIP1;refGene_tx=XM_023258789.1;refGene_gn=SGIP1;refGene_tx=XM_023258792.1;refGene_gn=SGIP1;refGene_tx=XM_023258794.1
C1 55501532 55501598
refGene_gn=SGIP1;refGene_tx=XM_019837178.2;refGene_gn=SGIP1;refGene_tx=XM_019837179.2;refGene_gn=SGIP1;refGene_tx=XM_019837180.2;refGene_gn=SGIP1;refGene_tx=XM_019837181.2;refGene_gn=SGIP1;refGene_tx=XM_019837182.2;refGene_gn=SGIP1;refGene_tx=XM_019837183.2;refGene_gn=SGIP1;refGene_tx=XM_019837184.2;refGene_gn=SGIP1;refGene_tx=XM_019837185.2;refGene_gn=SGIP1;refGene_tx=XM_019837186.2;refGene_gn=SGIP1;refGene_tx=XM_019837188.2;refGene_gn=SGIP1;refGene_tx=XM_019837189.2;refGene_gn=SGIP1;refGene_tx=XM_019837190.2;refGene_gn=SGIP1;refGene_tx=XM_019837191.2;refGene_gn=SGIP1;refGene_tx=XM_019837192.2;refGene_gn=SGIP1;refGene_tx=XM_019837194.2;refGene_gn=SGIP1;refGene_tx=XM_019837198.2;refGene_gn=SGIP1;refGene_tx=XM_019837201.2;refGene_gn=SGIP1;refGene_tx=XM_023258789.1
C1 55502517 55502577
refGene_gn=SGIP1;refGene_tx=XM_003990163.5;refGene_gn=SGIP1;refGene_tx=XM_019837178.2;refGene_gn=SGIP1;refGene_tx=XM_019837179.2;refGene_gn=SGIP1;refGene_tx=XM_019837180.2;refGene_gn=SGIP1;refGene_tx=XM_019837181.2;refGene_gn=SGIP1;refGene_tx=XM_019837182.2;refGene_gn=SGIP1;refGene_tx=XM_019837183.2;refGene_gn=SGIP1;refGene_tx=XM_019837184.2;refGene_gn=SGIP1;refGene_tx=XM_019837185.2;refGene_gn=SGIP1;refGene_tx=XM_019837187.2;refGene_gn=SGIP1;refGene_tx=XM_019837188.2;refGene_gn=SGIP1;refGene_tx=XM_019837189.2;refGene_gn=SGIP1;refGene_tx=XM_019837190.2;refGene_gn=SGIP1;refGene_tx=XM_019837191.2;refGene_gn=SGIP1;refGene_tx=XM_019837192.2;refGene_gn=SGIP1;refGene_tx=XM_019837194.2;refGene_gn=SGIP1;refGene_tx=XM_019837196.2;refGene_gn=SGIP1;refGene_tx=XM_019837198.2;refGene_gn=SGIP1;refGene_tx=XM_019837199.2;refGene_gn=SGIP1;refGene_tx=XM_023258789.1;refGene_gn=SGIP1;refGene_tx=XM_023258792.1;refGene_gn=SGIP1;refGene_tx=XM_023258794.1
C1 55529827 55529939
refGene_gn=SGIP1;refGene_tx=XM_003990163.5;refGene_gn=SGIP1;refGene_tx=XM_019837178.2;refGene_gn=SGIP1;refGene_tx=XM_019837179.2;refGene_gn=SGIP1;refGene_tx=XM_019837180.2;refGene_gn=SGIP1;refGene_tx=XM_019837181.2;refGene_gn=SGIP1;refGene_tx=XM_019837182.2;refGene_gn=SGIP1;refGene_tx=XM_019837183.2;refGene_gn=SGIP1;refGene_tx=XM_019837184.2;refGen

e_gn=SGIP1;refGene_tx=XM_019837185.2;refGene_gn=SGIP1;refGene_tx=XM_019837186.2;refGene_gn=SGIP1;refGene_tx=XM_019837187.2;refGene_gn=SGIP1;refGene_tx=XM_019837188.2;refGene_gn=SGIP1;refGene_tx=XM_019837189.2;refGene_gn=SGIP1;refGene_tx=XM_019837190.2;refGene_gn=SGIP1;refGene_tx=XM_019837191.2;refGene_gn=SGIP1;refGene_tx=XM_019837192.2;refGene_gn=SGIP1;refGene_tx=XM_019837193.2;refGene_gn=SGIP1;refGene_tx=XM_019837194.2;refGene_gn=SGIP1;refGene_tx=XM_019837195.2;refGene_gn=SGIP1;refGene_tx=XM_019837196.2;refGene_gn=SGIP1;refGene_tx=XM_019837198.2;refGene_gn=SGIP1;refGene_tx=XM_019837199.2;refGene_gn=SGIP1;refGene_tx=XM_019837200.2;refGene_gn=SGIP1;refGene_tx=XM_019837201.2;refGene_gn=SGIP1;refGene_tx=XM_023258789.1;refGene_gn=SGIP1;refGene_tx=XM_023258792.1;refGene_gn=SGIP1;refGene_tx=XM_023258794.1
C1 55538443 55538599
refGene_gn=SGIP1;refGene_tx=XM_003990163.5;refGene_gn=SGIP1;refGene_tx=XM_019837178.2;refGene_gn=SGIP1;refGene_tx=XM_019837179.2;refGene_gn=SGIP1;refGene_tx=XM_019837180.2;refGene_gn=SGIP1;refGene_tx=XM_019837181.2;refGene_gn=SGIP1;refGene_tx=XM_019837182.2;refGene_gn=SGIP1;refGene_tx=XM_019837183.2;refGene_gn=SGIP1;refGene_tx=XM_019837184.2;refGene_gn=SGIP1;refGene_tx=XM_019837185.2;refGene_gn=SGIP1;refGene_tx=XM_019837186.2;refGene_gn=SGIP1;refGene_tx=XM_019837187.2;refGene_gn=SGIP1;refGene_tx=XM_019837188.2;refGene_gn=SGIP1;refGene_tx=XM_019837189.2;refGene_gn=SGIP1;refGene_tx=XM_019837190.2;refGene_gn=SGIP1;refGene_tx=XM_019837191.2;refGene_gn=SGIP1;refGene_tx=XM_019837192.2;refGene_gn=SGIP1;refGene_tx=XM_019837193.2;refGene_gn=SGIP1;refGene_tx=XM_019837194.2;refGene_gn=SGIP1;refGene_tx=XM_019837195.2;refGene_gn=SGIP1;refGene_tx=XM_019837196.2;refGene_gn=SGIP1;refGene_tx=XM_019837198.2;refGene_gn=SGIP1;refGene_tx=XM_019837199.2;refGene_gn=SGIP1;refGene_tx=XM_019837200.2;refGene_gn=SGIP1;refGene_tx=XM_019837201.2;refGene_gn=SGIP1;refGene_tx=XM_023258789.1;refGene_gn=SGIP1;refGene_tx=XM_023258792.1;refGene_gn=SGIP1;refGene_tx=XM_023258794.1
C1 55542893 55543026
refGene_gn=SGIP1;refGene_tx=XM_003990163.5;refGene_gn=SGIP1;refGene_tx=XM_019837178.2;refGene_gn=SGIP1;refGene_tx=XM_019837179.2;refGene_gn=SGIP1;refGene_tx=XM_019837180.2;refGene_gn=SGIP1;refGene_tx=XM_019837181.2;refGene_gn=SGIP1;refGene_tx=XM_019837182.2;refGene_gn=SGIP1;refGene_tx=XM_019837183.2;refGene_gn=SGIP1;refGene_tx=XM_019837184.2;refGene_gn=SGIP1;refGene_tx=XM_019837185.2;refGene_gn=SGIP1;refGene_tx=XM_019837186.2;refGene_gn=SGIP1;refGene_tx=XM_019837187.2;refGene_gn=SGIP1;refGene_tx=XM_019837188.2;refGene_gn=SGIP1;refGene_tx=XM_019837189.2;refGene_gn=SGIP1;refGene_tx=XM_019837190.2;refGene_gn=SGIP1;refGene_tx=XM_019837191.2;refGene_gn=SGIP1;refGene_tx=XM_019837192.2;refGene_gn=SGIP1;refGene_tx=XM_019837193.2;refGene_gn=SGIP1;refGene_tx=XM_019837194.2;refGene_gn=SGIP1;refGene_tx=XM_019837195.2;refGene_gn=SGIP1;refGene_tx=XM_019837196.2;refGene_gn=SGIP1;refGene_tx=XM_019837198.2;refGene_gn=SGIP1;refGene_tx=XM_019837199.2;refGene_gn=SGIP1;refGene_tx=XM_019837200.2;refGene_gn=SGIP1;refGene_tx=XM_019837201.2;refGene_gn=SGIP1;refGene_tx=XM_023258789.1;refGene_gn=SGIP1;refGene_tx=XM_023258792.1;refGene_gn=SGIP1;refGene_tx=XM_023258794.1
C1 55547254 55547457
refGene_gn=SGIP1;refGene_tx=XM_003990163.5;refGene_gn=SGIP1;refGene_tx

=XM_019837178.2;refGene_gn=SGIP1;refGene_tx=XM_019837179.2;refGene_gn=SGIP1;refGene_tx=XM_019837180.2;refGene_gn=SGIP1;refGene_tx=XM_019837181.2;refGene_gn=SGIP1;refGene_tx=XM_019837182.2;refGene_gn=SGIP1;refGene_tx=XM_019837183.2;refGene_gn=SGIP1;refGene_tx=XM_019837184.2;refGene_gn=SGIP1;refGene_tx=XM_019837185.2;refGene_gn=SGIP1;refGene_tx=XM_019837186.2;refGene_gn=SGIP1;refGene_tx=XM_019837187.2;refGene_gn=SGIP1;refGene_tx=XM_019837188.2;refGene_gn=SGIP1;refGene_tx=XM_019837189.2;refGene_gn=SGIP1;refGene_tx=XM_019837190.2;refGene_gn=SGIP1;refGene_tx=XM_019837191.2;refGene_gn=SGIP1;refGene_tx=XM_019837192.2;refGene_gn=SGIP1;refGene_tx=XM_019837193.2;refGene_gn=SGIP1;refGene_tx=XM_019837194.2;refGene_gn=SGIP1;refGene_tx=XM_019837195.2;refGene_gn=SGIP1;refGene_tx=XM_019837196.2;refGene_gn=SGIP1;refGene_tx=XM_019837198.2;refGene_gn=SGIP1;refGene_tx=XM_019837199.2;refGene_gn=SGIP1;refGene_tx=XM_019837200.2;refGene_gn=SGIP1;refGene_tx=XM_019837201.2;refGene_gn=SGIP1;refGene_tx=XM_023258789.1;refGene_gn=SGIP1;refGene_tx=XM_023258792.1;refGene_gn=SGIP1;refGene_tx=XM_023258794.1
C1 55548635 55548700
refGene_gn=SGIP1;refGene_tx=XM_003990163.5;refGene_gn=SGIP1;refGene_tx=XM_019837178.2;refGene_gn=SGIP1;refGene_tx=XM_019837179.2;refGene_gn=SGIP1;refGene_tx=XM_019837180.2;refGene_gn=SGIP1;refGene_tx=XM_019837181.2;refGene_gn=SGIP1;refGene_tx=XM_019837182.2;refGene_gn=SGIP1;refGene_tx=XM_019837183.2;refGene_gn=SGIP1;refGene_tx=XM_019837184.2;refGene_gn=SGIP1;refGene_tx=XM_019837185.2;refGene_gn=SGIP1;refGene_tx=XM_019837186.2;refGene_gn=SGIP1;refGene_tx=XM_019837187.2;refGene_gn=SGIP1;refGene_tx=XM_019837188.2;refGene_gn=SGIP1;refGene_tx=XM_019837189.2;refGene_gn=SGIP1;refGene_tx=XM_019837190.2;refGene_gn=SGIP1;refGene_tx=XM_019837191.2;refGene_gn=SGIP1;refGene_tx=XM_019837192.2;refGene_gn=SGIP1;refGene_tx=XM_019837193.2;refGene_gn=SGIP1;refGene_tx=XM_019837194.2;refGene_gn=SGIP1;refGene_tx=XM_019837195.2;refGene_gn=SGIP1;refGene_tx=XM_019837196.2;refGene_gn=SGIP1;refGene_tx=XM_019837198.2;refGene_gn=SGIP1;refGene_tx=XM_019837199.2;refGene_gn=SGIP1;refGene_tx=XM_019837200.2;refGene_gn=SGIP1;refGene_tx=XM_019837201.2;refGene_gn=SGIP1;refGene_tx=XM_023258789.1;refGene_gn=SGIP1;refGene_tx=XM_023258792.1;refGene_gn=SGIP1;refGene_tx=XM_023258794.1
C1 55549277 55549442
refGene_gn=SGIP1;refGene_tx=XM_003990163.5;refGene_gn=SGIP1;refGene_tx=XM_019837178.2;refGene_gn=SGIP1;refGene_tx=XM_019837179.2;refGene_gn=SGIP1;refGene_tx=XM_019837180.2;refGene_gn=SGIP1;refGene_tx=XM_019837181.2;refGene_gn=SGIP1;refGene_tx=XM_019837182.2;refGene_gn=SGIP1;refGene_tx=XM_019837183.2;refGene_gn=SGIP1;refGene_tx=XM_019837184.2;refGene_gn=SGIP1;refGene_tx=XM_019837185.2;refGene_gn=SGIP1;refGene_tx=XM_019837186.2;refGene_gn=SGIP1;refGene_tx=XM_019837187.2;refGene_gn=SGIP1;refGene_tx=XM_019837188.2;refGene_gn=SGIP1;refGene_tx=XM_019837189.2;refGene_gn=SGIP1;refGene_tx=XM_019837190.2;refGene_gn=SGIP1;refGene_tx=XM_019837191.2;refGene_gn=SGIP1;refGene_tx=XM_019837192.2;refGene_gn=SGIP1;refGene_tx=XM_019837193.2;refGene_gn=SGIP1;refGene_tx=XM_019837194.2;refGene_gn=SGIP1;refGene_tx=XM_019837195.2;refGene_gn=SGIP1;refGene_tx=XM_019837196.2;refGene_gn=SGIP1;refGene_tx=XM_019837198.2;refGene_gn=SGIP1;refGene_tx=XM_019837199.2;refGene_gn=SGIP1;refGene_tx=XM_019837200.2;refGene_gn=SGIP1;refGene_tx=XM_019837201.2;refGene_gn=SGIP1;r

efGene_tx=XM_023258789.1;refGene_gn=SGIP1;refGene_tx=XM_023258792.1;refGene_gn=SGIP1;refGene_tx=XM_023258794.1
C1	55550493	55550516
refGene_gn=SGIP1;refGene_tx=XM_003990163.5;refGene_gn=SGIP1;refGene_tx=XM_019837178.2;refGene_gn=SGIP1;refGene_tx=XM_019837179.2;refGene_gn=SGIP1;refGene_tx=XM_019837180.2;refGene_gn=SGIP1;refGene_tx=XM_019837181.2;refGene_gn=SGIP1;refGene_tx=XM_019837182.2;refGene_gn=SGIP1;refGene_tx=XM_019837183.2;refGene_gn=SGIP1;refGene_tx=XM_019837184.2;refGene_gn=SGIP1;refGene_tx=XM_019837185.2;refGene_gn=SGIP1;refGene_tx=XM_019837186.2;refGene_gn=SGIP1;refGene_tx=XM_019837187.2;refGene_gn=SGIP1;refGene_tx=XM_019837188.2;refGene_gn=SGIP1;refGene_tx=XM_019837189.2;refGene_gn=SGIP1;refGene_tx=XM_019837190.2;refGene_gn=SGIP1;refGene_tx=XM_019837191.2;refGene_gn=SGIP1;refGene_tx=XM_019837192.2;refGene_gn=SGIP1;refGene_tx=XM_019837193.2;refGene_gn=SGIP1;refGene_tx=XM_019837194.2;refGene_gn=SGIP1;refGene_tx=XM_019837195.2;refGene_gn=SGIP1;refGene_tx=XM_019837196.2;refGene_gn=SGIP1;refGene_tx=XM_019837198.2;refGene_gn=SGIP1;refGene_tx=XM_019837199.2;refGene_gn=SGIP1;refGene_tx=XM_019837200.2;refGene_gn=SGIP1;refGene_tx=XM_019837201.2;refGene_gn=SGIP1;refGene_tx=XM_023258789.1;refGene_gn=SGIP1;refGene_tx=XM_023258792.1;refGene_gn=SGIP1;refGene_tx=XM_023258794.1
C1	55561755	55561874	refGene_gn=TCTEX1D1;refGene_tx=XM_003990165.4
C1	55580353	55580445	refGene_gn=TCTEX1D1;refGene_tx=XM_003990165.4
C1	55587991	55588116	refGene_gn=TCTEX1D1;refGene_tx=XM_003990165.4
C1	55588995	55589199	refGene_gn=TCTEX1D1;refGene_tx=XM_003990165.4
C1	55602787	55602899	refGene_gn=INSL5;refGene_tx=XM_023258795.1
C1	55602900	55603163	refGene_gn=INSL5;refGene_tx=XM_023258795.1
C1	55614240	55614291	refGene_gn=WDR78;refGene_tx=XM_003990166.4
C1	55624743	55624900	refGene_gn=WDR78;refGene_tx=XM_003990166.4
C1	55627880	55628107	refGene_gn=WDR78;refGene_tx=XM_003990166.4
C1	55628742	55628841	refGene_gn=WDR78;refGene_tx=XM_003990166.4
C1	55633947	55634069	refGene_gn=WDR78;refGene_tx=XM_003990166.4
C1	55634360	55634518	refGene_gn=WDR78;refGene_tx=XM_003990166.4
C1	55636440	55636592	refGene_gn=WDR78;refGene_tx=XM_003990166.4
C1	55637843	55637930	refGene_gn=WDR78;refGene_tx=XM_003990166.4
C1	55640914	55641123	refGene_gn=WDR78;refGene_tx=XM_003990166.4
C1	55645050	55645245	refGene_gn=WDR78;refGene_tx=XM_003990166.4
C1	55652997	55653153	refGene_gn=WDR78;refGene_tx=XM_003990166.4
C1	55661582	55661722	refGene_gn=WDR78;refGene_tx=XM_003990166.4
C1	55665351	55665508	refGene_gn=WDR78;refGene_tx=XM_003990166.4
C1	55667149	55667262	refGene_gn=WDR78;refGene_tx=XM_003990166.4
C1	55669095	55669280	refGene_gn=WDR78;refGene_tx=XM_003990166.4
C1	55678831	55679009	refGene_gn=WDR78;refGene_tx=XM_003990166.4
C1	55700865	55700973	refGene_gn=WDR78;refGene_tx=XM_003990166.4
C1	55700974	55701298
refGene_gn=MIER1;refGene_tx=XM_003990168.4;refGene_gn=MIER1;refGene_tx=XM_019837205.1;refGene_gn=MIER1;refGene_tx=XM_023258796.1;refGene_gn=MIER1;refGene_tx=XM_023258798.1;refGene_gn=MIER1;refGene_tx=XM_023258800.1;refGene_gn=MIER1;refGene_tx=XM_023258801.1;refGene_gn=WDR78;refGene_tx=XM_003990166.4
C1	55702398	55702499

refGene_gn=MIER1;refGene_tx=XM_003990168.4;refGene_gn=MIER1;refGene_tx=XM_019837205.1;refGene_gn=MIER1;refGene_tx=XM_023258796.1;refGene_gn=MIER1;refGene_tx=XM_023258798.1;refGene_gn=MIER1;refGene_tx=XM_023258800.1;refGene_gn=MIER1;refGene_tx=XM_023258801.1;refGene_gn=MIER1;refGene_tx=XM_023258806.1
C1 55705236 55705310
refGene_gn=MIER1;refGene_tx=XM_006934805.3;refGene_gn=MIER1;refGene_tx=XM_019837205.1;refGene_gn=MIER1;refGene_tx=XM_023258796.1;refGene_gn=MIER1;refGene_tx=XM_023258798.1;refGene_gn=MIER1;refGene_tx=XM_023258799.1;refGene_gn=MIER1;refGene_tx=XM_023258800.1;refGene_gn=MIER1;refGene_tx=XM_023258801.1;refGene_gn=MIER1;refGene_tx=XM_023258802.1;refGene_gn=MIER1;refGene_tx=XM_023258806.1
C1 55706758 55706767 refGene_gn=MIER1;refGene_tx=XM_023258803.1
C1 55716110 55716135
refGene_gn=MIER1;refGene_tx=XM_003990168.4;refGene_gn=MIER1;refGene_tx=XM_006934805.3;refGene_gn=MIER1;refGene_tx=XM_019837205.1;refGene_gn=MIER1;refGene_tx=XM_023258796.1;refGene_gn=MIER1;refGene_tx=XM_023258798.1;refGene_gn=MIER1;refGene_tx=XM_023258799.1;refGene_gn=MIER1;refGene_tx=XM_023258800.1;refGene_gn=MIER1;refGene_tx=XM_023258801.1;refGene_gn=MIER1;refGene_tx=XM_023258802.1;refGene_gn=MIER1;refGene_tx=XM_023258803.1;refGene_gn=MIER1;refGene_tx=XM_023258806.1
C1 55716255 55716292
refGene_gn=MIER1;refGene_tx=XM_019837205.1;refGene_gn=MIER1;refGene_tx=XM_023258796.1;refGene_gn=MIER1;refGene_tx=XM_023258798.1;refGene_gn=MIER1;refGene_tx=XM_023258799.1;refGene_gn=MIER1;refGene_tx=XM_023258800.1;refGene_gn=MIER1;refGene_tx=XM_023258801.1;refGene_gn=MIER1;refGene_tx=XM_023258804.1;refGene_gn=MIER1;refGene_tx=XM_023258806.1
C1 55720930 55721076
refGene_gn=MIER1;refGene_tx=XM_003990168.4;refGene_gn=MIER1;refGene_tx=XM_006934805.3;refGene_gn=MIER1;refGene_tx=XM_019837205.1;refGene_gn=MIER1;refGene_tx=XM_023258796.1;refGene_gn=MIER1;refGene_tx=XM_023258798.1;refGene_gn=MIER1;refGene_tx=XM_023258799.1;refGene_gn=MIER1;refGene_tx=XM_023258800.1;refGene_gn=MIER1;refGene_tx=XM_023258801.1;refGene_gn=MIER1;refGene_tx=XM_023258802.1;refGene_gn=MIER1;refGene_tx=XM_023258803.1;refGene_gn=MIER1;refGene_tx=XM_023258804.1;refGene_gn=MIER1;refGene_tx=XM_023258805.1;refGene_gn=MIER1;refGene_tx=XM_023258806.1
C1 55733312 55733474
refGene_gn=MIER1;refGene_tx=XM_003990168.4;refGene_gn=MIER1;refGene_tx=XM_006934805.3;refGene_gn=MIER1;refGene_tx=XM_019837205.1;refGene_gn=MIER1;refGene_tx=XM_023258796.1;refGene_gn=MIER1;refGene_tx=XM_023258798.1;refGene_gn=MIER1;refGene_tx=XM_023258799.1;refGene_gn=MIER1;refGene_tx=XM_023258800.1;refGene_gn=MIER1;refGene_tx=XM_023258801.1;refGene_gn=MIER1;refGene_tx=XM_023258802.1;refGene_gn=MIER1;refGene_tx=XM_023258803.1;refGene_gn=MIER1;refGene_tx=XM_023258804.1;refGene_gn=MIER1;refGene_tx=XM_023258805.1;refGene_gn=MIER1;refGene_tx=XM_023258806.1
C1 55734075 55734208
refGene_gn=MIER1;refGene_tx=XM_003990168.4;refGene_gn=MIER1;refGene_tx=XM_006934805.3;refGene_gn=MIER1;refGene_tx=XM_019837205.1;refGene_gn=MIER1;refGene_tx=XM_023258796.1;refGene_gn=MIER1;refGene_tx=XM_023258798.1;refGene_gn=MIER1;refGene_tx=XM_023258799.1;refGene_gn=MIER1;refGe

ne_tx=XM_023258800.1;refGene_gn=MIER1;refGene_tx=XM_023258801.1;refGen
e_gn=MIER1;refGene_tx=XM_023258802.1;refGene_gn=MIER1;refGene_tx=XM_02
3258803.1;refGene_gn=MIER1;refGene_tx=XM_023258804.1;refGene_gn=MIER1;
refGene_tx=XM_023258805.1;refGene_gn=MIER1;refGene_tx=XM_023258806.1
C1 55736128 55736193
refGene_gn=MIER1;refGene_tx=XM_003990168.4;refGene_gn=MIER1;refGene_tx
=XM_006934805.3;refGene_gn=MIER1;refGene_tx=XM_019837205.1;refGene_gn=
MIER1;refGene_tx=XM_023258796.1;refGene_gn=MIER1;refGene_tx=XM_0232587
98.1;refGene_gn=MIER1;refGene_tx=XM_023258799.1;refGene_gn=MIER1;refGe
ne_tx=XM_023258800.1;refGene_gn=MIER1;refGene_tx=XM_023258801.1;refGen
e_gn=MIER1;refGene_tx=XM_023258802.1;refGene_gn=MIER1;refGene_tx=XM_02
3258803.1;refGene_gn=MIER1;refGene_tx=XM_023258804.1;refGene_gn=MIER1;
refGene_tx=XM_023258805.1;refGene_gn=MIER1;refGene_tx=XM_023258806.1
C1 55739944 55740017
refGene_gn=MIER1;refGene_tx=XM_003990168.4;refGene_gn=MIER1;refGene_tx
=XM_006934805.3;refGene_gn=MIER1;refGene_tx=XM_019837205.1;refGene_gn=
MIER1;refGene_tx=XM_023258796.1;refGene_gn=MIER1;refGene_tx=XM_0232587
98.1;refGene_gn=MIER1;refGene_tx=XM_023258799.1;refGene_gn=MIER1;refGe
ne_tx=XM_023258800.1;refGene_gn=MIER1;refGene_tx=XM_023258801.1;refGen
e_gn=MIER1;refGene_tx=XM_023258802.1;refGene_gn=MIER1;refGene_tx=XM_02
3258803.1;refGene_gn=MIER1;refGene_tx=XM_023258804.1;refGene_gn=MIER1;
refGene_tx=XM_023258805.1;refGene_gn=MIER1;refGene_tx=XM_023258806.1
C1 55746884 55747036
refGene_gn=MIER1;refGene_tx=XM_003990168.4;refGene_gn=MIER1;refGene_tx
=XM_006934805.3;refGene_gn=MIER1;refGene_tx=XM_019837205.1;refGene_gn=
MIER1;refGene_tx=XM_023258796.1;refGene_gn=MIER1;refGene_tx=XM_0232587
98.1;refGene_gn=MIER1;refGene_tx=XM_023258799.1;refGene_gn=MIER1;refGe
ne_tx=XM_023258800.1;refGene_gn=MIER1;refGene_tx=XM_023258801.1;refGen
e_gn=MIER1;refGene_tx=XM_023258802.1;refGene_gn=MIER1;refGene_tx=XM_02
3258803.1;refGene_gn=MIER1;refGene_tx=XM_023258804.1;refGene_gn=MIER1;
refGene_tx=XM_023258805.1;refGene_gn=MIER1;refGene_tx=XM_023258806.1
C1 55748183 55748265
refGene_gn=MIER1;refGene_tx=XM_003990168.4;refGene_gn=MIER1;refGene_tx
=XM_006934805.3;refGene_gn=MIER1;refGene_tx=XM_019837205.1;refGene_gn=
MIER1;refGene_tx=XM_023258796.1;refGene_gn=MIER1;refGene_tx=XM_0232587
98.1;refGene_gn=MIER1;refGene_tx=XM_023258799.1;refGene_gn=MIER1;refGe
ne_tx=XM_023258800.1;refGene_gn=MIER1;refGene_tx=XM_023258801.1;refGen
e_gn=MIER1;refGene_tx=XM_023258802.1;refGene_gn=MIER1;refGene_tx=XM_02
3258803.1;refGene_gn=MIER1;refGene_tx=XM_023258804.1;refGene_gn=MIER1;
refGene_tx=XM_023258805.1;refGene_gn=MIER1;refGene_tx=XM_023258806.1
C1 55749607 55749702
refGene_gn=MIER1;refGene_tx=XM_003990168.4;refGene_gn=MIER1;refGene_tx
=XM_006934805.3;refGene_gn=MIER1;refGene_tx=XM_019837205.1;refGene_gn=
MIER1;refGene_tx=XM_023258796.1;refGene_gn=MIER1;refGene_tx=XM_0232587
98.1;refGene_gn=MIER1;refGene_tx=XM_023258799.1;refGene_gn=MIER1;refGe
ne_tx=XM_023258800.1;refGene_gn=MIER1;refGene_tx=XM_023258801.1;refGen
e_gn=MIER1;refGene_tx=XM_023258802.1;refGene_gn=MIER1;refGene_tx=XM_02
3258803.1;refGene_gn=MIER1;refGene_tx=XM_023258804.1;refGene_gn=MIER1;
refGene_tx=XM_023258805.1;refGene_gn=MIER1;refGene_tx=XM_023258806.1
C1 55753106 55753234

refGene_gn=MIER1;refGene_tx=XM_003990168.4;refGene_gn=MIER1;refGene_tx=XM_006934805.3;refGene_gn=MIER1;refGene_tx=XM_019837205.1;refGene_gn=MIER1;refGene_tx=XM_023258796.1;refGene_gn=MIER1;refGene_tx=XM_023258798.1;refGene_gn=MIER1;refGene_tx=XM_023258799.1;refGene_gn=MIER1;refGene_tx=XM_023258800.1;refGene_gn=MIER1;refGene_tx=XM_023258801.1;refGene_gn=MIER1;refGene_tx=XM_023258802.1;refGene_gn=MIER1;refGene_tx=XM_023258803.1;refGene_gn=MIER1;refGene_tx=XM_023258804.1;refGene_gn=MIER1;refGene_tx=XM_023258805.1;refGene_gn=MIER1;refGene_tx=XM_023258806.1
C1 55754721 55754779 refGene_gn=MIER1;refGene_tx=XM_023258800.1
C1 55754829 55754866 refGene_gn=MIER1;refGene_tx=XM_023258801.1
C1 55757519 55757659
refGene_gn=MIER1;refGene_tx=XM_003990168.4;refGene_gn=MIER1;refGene_tx=XM_006934805.3;refGene_gn=MIER1;refGene_tx=XM_019837205.1;refGene_gn=MIER1;refGene_tx=XM_023258796.1;refGene_gn=MIER1;refGene_tx=XM_023258798.1;refGene_gn=MIER1;refGene_tx=XM_023258799.1;refGene_gn=MIER1;refGene_tx=XM_023258802.1;refGene_gn=MIER1;refGene_tx=XM_023258803.1;refGene_gn=MIER1;refGene_tx=XM_023258804.1;refGene_gn=MIER1;refGene_tx=XM_023258805.1;refGene_gn=MIER1;refGene_tx=XM_023258806.1
C1 55760277 55760606
refGene_gn=MIER1;refGene_tx=XM_003990168.4;refGene_gn=MIER1;refGene_tx=XM_006934805.3;refGene_gn=MIER1;refGene_tx=XM_019837205.1;refGene_gn=MIER1;refGene_tx=XM_023258796.1;refGene_gn=MIER1;refGene_tx=XM_023258798.1;refGene_gn=MIER1;refGene_tx=XM_023258799.1;refGene_gn=MIER1;refGene_tx=XM_023258802.1;refGene_gn=MIER1;refGene_tx=XM_023258803.1;refGene_gn=MIER1;refGene_tx=XM_023258804.1;refGene_gn=MIER1;refGene_tx=XM_023258805.1;refGene_gn=MIER1;refGene_tx=XM_023258806.1
C1 55777681 55777790 refGene_gn=SLC35D1;refGene_tx=XM_023258808.1
C1 55791601 55791684 refGene_gn=SLC35D1;refGene_tx=XM_023258808.1
C1 55795136 55795215 refGene_gn=SLC35D1;refGene_tx=XM_023258808.1
C1 55796324 55796392 refGene_gn=SLC35D1;refGene_tx=XM_023258808.1
C1 55812637 55812730 refGene_gn=SLC35D1;refGene_tx=XM_023258808.1
C1 55816287 55816390 refGene_gn=SLC35D1;refGene_tx=XM_023258808.1
C1 55818559 55818628 refGene_gn=SLC35D1;refGene_tx=XM_023258808.1
C1 55819496 55819568 refGene_gn=SLC35D1;refGene_tx=XM_023258808.1
C1 55821183 55821251 refGene_gn=SLC35D1;refGene_tx=XM_023258808.1
C1 55821956 55822043 refGene_gn=SLC35D1;refGene_tx=XM_023258808.1
C1 55822140 55822174 refGene_gn=SLC35D1;refGene_tx=XM_023258808.1
C1 55822986 55823189 refGene_gn=SLC35D1;refGene_tx=XM_023258808.1
C1 55848404 55848977
refGene_gn=CC1H1orf141;refGene_tx=XM_011284942.3
C1 55850701 55850891
refGene_gn=CC1H1orf141;refGene_tx=XM_011284942.3
C1 55852183 55852253
refGene_gn=CC1H1orf141;refGene_tx=XM_011284942.3
C1 55860805 55860911
refGene_gn=CC1H1orf141;refGene_tx=XM_011284942.3
C1 55866671 55866739
refGene_gn=CC1H1orf141;refGene_tx=XM_011284942.3
C1 55872198 55872311
refGene_gn=CC1H1orf141;refGene_tx=XM_011284942.3

C1	55881409	55881567
refGene_gn=CC1H1orf141;refGene_tx=XM_011284942.3
C1	55884883	55884958
refGene_gn=CC1H1orf141;refGene_tx=XM_011284942.3
C1	55937698	55937768	refGene_gn=IL23R;refGene_tx=XM_023258809.1
C1	55940352	55940649	refGene_gn=IL23R;refGene_tx=XM_023258809.1
C1	55947754	55947878	refGene_gn=IL23R;refGene_tx=XM_023258809.1
C1	55961857	55962018	refGene_gn=IL23R;refGene_tx=XM_023258809.1
C1	55966817	55966963	refGene_gn=IL23R;refGene_tx=XM_023258809.1
C1	55972757	55972914	refGene_gn=IL23R;refGene_tx=XM_023258809.1
C1	55986078	55986168	refGene_gn=IL23R;refGene_tx=XM_023258809.1
C1	55988825	55988931	refGene_gn=IL23R;refGene_tx=XM_023258809.1
C1	55989865	55989956	refGene_gn=IL23R;refGene_tx=XM_023258809.1
C1	55993500	55993887	refGene_gn=IL23R;refGene_tx=XM_023258809.1
C1	55993888	55994155	refGene_gn=IL23R;refGene_tx=XM_023258809.1
C1	56045418	56045494
refGene_gn=IL12RB2;refGene_tx=XM_023258810.1;refGene_gn=IL12RB2;refGen
e_tx=XM_023258811.1
C1	56047241	56047529
refGene_gn=IL12RB2;refGene_tx=XM_023258810.1;refGene_gn=IL12RB2;refGen
e_tx=XM_023258811.1
C1	56056986	56057101
refGene_gn=IL12RB2;refGene_tx=XM_023258810.1;refGene_gn=IL12RB2;refGen
e_tx=XM_023258811.1
C1	56058141	56058326
refGene_gn=IL12RB2;refGene_tx=XM_023258810.1;refGene_gn=IL12RB2;refGen
e_tx=XM_023258811.1
C1	56059750	56059893
refGene_gn=IL12RB2;refGene_tx=XM_023258810.1;refGene_gn=IL12RB2;refGen
e_tx=XM_023258811.1
C1	56061054	56061205
refGene_gn=IL12RB2;refGene_tx=XM_023258810.1;refGene_gn=IL12RB2;refGen
e_tx=XM_023258811.1
C1	56070681	56070761
refGene_gn=IL12RB2;refGene_tx=XM_023258810.1;refGene_gn=IL12RB2;refGen
e_tx=XM_023258811.1
C1	56081817	56082037
refGene_gn=IL12RB2;refGene_tx=XM_023258810.1;refGene_gn=IL12RB2;refGen
e_tx=XM_023258811.1
C1	56092329	56092530
refGene_gn=IL12RB2;refGene_tx=XM_023258810.1;refGene_gn=IL12RB2;refGen
e_tx=XM_023258811.1
C1	56096721	56096820
refGene_gn=IL12RB2;refGene_tx=XM_023258810.1;refGene_gn=IL12RB2;refGen
e_tx=XM_023258811.1
C1	56096910	56097069
refGene_gn=IL12RB2;refGene_tx=XM_023258810.1;refGene_gn=IL12RB2;refGen
e_tx=XM_023258811.1
C1	56101884	56102022
refGene_gn=IL12RB2;refGene_tx=XM_023258810.1;refGene_gn=IL12RB2;refGen

e_tx=XM_023258811.1
C1 56104781 56104872
refGene_gn=IL12RB2;refGene_tx=XM_023258810.1;refGene_gn=IL12RB2;refGen
e_tx=XM_023258811.1
C1 56107338 56107438 refGene_gn=IL12RB2;refGene_tx=XM_023258810.1
C1 56112629 56113169
refGene_gn=IL12RB2;refGene_tx=XM_023258810.1;refGene_gn=IL12RB2;refGen
e_tx=XM_023258811.1
C1 56125420 56125477
refGene_gn=SERBP1;refGene_tx=XM_003990173.5;refGene_gn=SERBP1;refGene_
tx=XM_003990174.5;refGene_gn=SERBP1;refGene_tx=XM_003990175.5;refGene_
gn=SERBP1;refGene_tx=XM_003990176.5
C1 56126848 56127022
refGene_gn=SERBP1;refGene_tx=XM_003990173.5;refGene_gn=SERBP1;refGene_
tx=XM_003990174.5;refGene_gn=SERBP1;refGene_tx=XM_003990175.5;refGene_
gn=SERBP1;refGene_tx=XM_003990176.5
C1 56129977 56130155
refGene_gn=SERBP1;refGene_tx=XM_003990173.5;refGene_gn=SERBP1;refGene_
tx=XM_003990174.5;refGene_gn=SERBP1;refGene_tx=XM_003990175.5;refGene_
gn=SERBP1;refGene_tx=XM_003990176.5
C1 56132861 56132981
refGene_gn=SERBP1;refGene_tx=XM_003990173.5;refGene_gn=SERBP1;refGene_
tx=XM_003990174.5;refGene_gn=SERBP1;refGene_tx=XM_003990175.5;refGene_
gn=SERBP1;refGene_tx=XM_003990176.5
C1 56133563 56133653
refGene_gn=SERBP1;refGene_tx=XM_003990173.5;refGene_gn=SERBP1;refGene_
tx=XM_003990174.5;refGene_gn=SERBP1;refGene_tx=XM_003990175.5;refGene_
gn=SERBP1;refGene_tx=XM_003990176.5
C1 56133765 56133906
refGene_gn=SERBP1;refGene_tx=XM_003990173.5;refGene_gn=SERBP1;refGene_
tx=XM_003990174.5;refGene_gn=SERBP1;refGene_tx=XM_003990175.5;refGene_
gn=SERBP1;refGene_tx=XM_003990176.5
C1 56134552 56134703
refGene_gn=SERBP1;refGene_tx=XM_003990173.5;refGene_gn=SERBP1;refGene_
tx=XM_003990174.5;refGene_gn=SERBP1;refGene_tx=XM_003990175.5;refGene_
gn=SERBP1;refGene_tx=XM_003990176.5
C1 56138347 56138660
refGene_gn=SERBP1;refGene_tx=XM_003990173.5;refGene_gn=SERBP1;refGene_
tx=XM_003990174.5;refGene_gn=SERBP1;refGene_tx=XM_003990175.5;refGene_
gn=SERBP1;refGene_tx=XM_003990176.5
C1 56359923 56359967
refGene_gn=GADD45A;refGene_tx=NM_001009199.1;refGene_gn=GADD45A;refGen
e_tx=XM_023257813.1
C1 56360500 56360602 refGene_gn=GADD45A;refGene_tx=XM_023257813.1
C1 56360833 56361071 refGene_gn=GADD45A;refGene_tx=XM_023257813.1
C1 56362302 56362416
refGene_gn=GADD45A;refGene_tx=NM_001009199.1;refGene_gn=GADD45A;refGen
e_tx=XM_023257813.1
C1 56387196 56387322
refGene_gn=GNG12;refGene_tx=XM_019837214.2;refGene_gn=GNG12;refGene_tx

=XM_019837215.2
C1 56389010 56389103
refGene_gn=GNG12;refGene_tx=XM_019837214.2;refGene_gn=GNG12;refGene_tx
=XM_019837215.2
C1 56724861 56724971 refGene_gn=WLS;refGene_tx=XM_003990178.5
C1 56728039 56728077 refGene_gn=WLS;refGene_tx=XM_019837216.2
C1 56735071 56735225
refGene_gn=WLS;refGene_tx=XM_003990178.5;refGene_gn=WLS;refGene_tx=XM_
019837216.2
C1 56741758 56741842
refGene_gn=WLS;refGene_tx=XM_003990178.5;refGene_gn=WLS;refGene_tx=XM_
019837216.2
C1 56743001 56743145
refGene_gn=WLS;refGene_tx=XM_003990178.5;refGene_gn=WLS;refGene_tx=XM_
019837216.2
C1 56745402 56745466
refGene_gn=WLS;refGene_tx=XM_003990178.5;refGene_gn=WLS;refGene_tx=XM_
019837216.2
C1 56745823 56745921
refGene_gn=WLS;refGene_tx=XM_003990178.5;refGene_gn=WLS;refGene_tx=XM_
019837216.2
C1 56747237 56747406
refGene_gn=WLS;refGene_tx=XM_003990178.5;refGene_gn=WLS;refGene_tx=XM_
019837216.2
C1 56750287 56750424
refGene_gn=WLS;refGene_tx=XM_003990178.5;refGene_gn=WLS;refGene_tx=XM_
019837216.2
C1 56751732 56751894
refGene_gn=WLS;refGene_tx=XM_003990178.5;refGene_gn=WLS;refGene_tx=XM_
019837216.2
C1 56756840 56756965
refGene_gn=WLS;refGene_tx=XM_003990178.5;refGene_gn=WLS;refGene_tx=XM_
019837216.2
C1 56782220 56782493
refGene_gn=WLS;refGene_tx=XM_003990178.5;refGene_gn=WLS;refGene_tx=XM_
019837216.2
C1 56824964 56825070
refGene_gn=WLS;refGene_tx=XM_003990178.5;refGene_gn=WLS;refGene_tx=XM_
019837216.2
C1 56998701 56998853
refGene_gn=RPE65;refGene_tx=XM_003990180.5;refGene_gn=RPE65;refGene_tx
=XM_019837218.1
C1 56999950 57000062
refGene_gn=RPE65;refGene_tx=XM_003990180.5;refGene_gn=RPE65;refGene_tx
=XM_019837218.1
C1 57000169 57000264
refGene_gn=RPE65;refGene_tx=XM_003990180.5;refGene_gn=RPE65;refGene_tx
=XM_019837218.1
C1 57000345 57000460
refGene_gn=RPE65;refGene_tx=XM_003990180.5;refGene_gn=RPE65;refGene_tx

=XM_019837218.1
C1	57008134	57008264
refGene_gn=RPE65;refGene_tx=XM_003990180.5;refGene_gn=RPE65;refGene_tx=XM_019837218.1
C1	57008973	57009113
refGene_gn=RPE65;refGene_tx=XM_003990180.5;refGene_gn=RPE65;refGene_tx=XM_019837218.1
C1	57009220	57009353
refGene_gn=RPE65;refGene_tx=XM_003990180.5;refGene_gn=RPE65;refGene_tx=XM_019837218.1
C1	57009590	57009672
refGene_gn=RPE65;refGene_tx=XM_003990180.5;refGene_gn=RPE65;refGene_tx=XM_019837218.1
C1	57010887	57011035
refGene_gn=RPE65;refGene_tx=XM_003990180.5;refGene_gn=RPE65;refGene_tx=XM_019837218.1
C1	57013794	57013936
refGene_gn=RPE65;refGene_tx=XM_003990180.5;refGene_gn=RPE65;refGene_tx=XM_019837218.1
C1	57014038	57014146
refGene_gn=RPE65;refGene_tx=XM_003990180.5;refGene_gn=RPE65;refGene_tx=XM_019837218.1
C1	57015899	57016050	refGene_gn=RPE65;refGene_tx=XM_003990180.5
C1	57017665	57017748	refGene_gn=RPE65;refGene_tx=XM_003990180.5
C1	57018819	57018830	refGene_gn=RPE65;refGene_tx=XM_003990180.5
C1	57046788	57046929
refGene_gn=DEPDC1;refGene_tx=XM_003990181.5;refGene_gn=DEPDC1;refGene_tx=XM_019837217.2
C1	57047637	57047823
refGene_gn=DEPDC1;refGene_tx=XM_003990181.5;refGene_gn=DEPDC1;refGene_tx=XM_019837217.2
C1	57049237	57049414
refGene_gn=DEPDC1;refGene_tx=XM_003990181.5;refGene_gn=DEPDC1;refGene_tx=XM_019837217.2
C1	57051127	57051300
refGene_gn=DEPDC1;refGene_tx=XM_003990181.5;refGene_gn=DEPDC1;refGene_tx=XM_019837217.2
C1	57051758	57052610	refGene_gn=DEPDC1;refGene_tx=XM_003990181.5
C1	57053895	57054033
refGene_gn=DEPDC1;refGene_tx=XM_003990181.5;refGene_gn=DEPDC1;refGene_tx=XM_019837217.2
C1	57057661	57057709
refGene_gn=DEPDC1;refGene_tx=XM_003990181.5;refGene_gn=DEPDC1;refGene_tx=XM_019837217.2
C1	57059082	57059213
refGene_gn=DEPDC1;refGene_tx=XM_003990181.5;refGene_gn=DEPDC1;refGene_tx=XM_019837217.2
C1	57059640	57059759
refGene_gn=DEPDC1;refGene_tx=XM_003990181.5;refGene_gn=DEPDC1;refGene_tx=XM_019837217.2

C1 57060210 57060367
refGene_gn=DEPDC1;refGene_tx=XM_003990181.5;refGene_gn=DEPDC1;refGene_tx=XM_019837217.2
C1 57063595 57063861
refGene_gn=DEPDC1;refGene_tx=XM_003990181.5;refGene_gn=DEPDC1;refGene_tx=XM_019837217.2
C1 57065952 57066000
refGene_gn=DEPDC1;refGene_tx=XM_003990181.5;refGene_gn=DEPDC1;refGene_tx=XM_019837217.2
C1 57920259 57920261
refGene_gn=LRRC7;refGene_tx=XM_011284940.3;refGene_gn=LRRC7;refGene_tx=XM_019837219.2;refGene_gn=LRRC7;refGene_tx=XM_019837221.2;refGene_gn=LRRC7;refGene_tx=XM_019837222.2;refGene_gn=LRRC7;refGene_tx=XM_019837223.2;refGene_gn=LRRC7;refGene_tx=XM_019837224.2;refGene_gn=LRRC7;refGene_tx=XM_019837225.2
C1 58024393 58024491
refGene_gn=LRRC7;refGene_tx=XM_011284940.3;refGene_gn=LRRC7;refGene_tx=XM_019837219.2;refGene_gn=LRRC7;refGene_tx=XM_019837221.2;refGene_gn=LRRC7;refGene_tx=XM_019837222.2;refGene_gn=LRRC7;refGene_tx=XM_019837223.2;refGene_gn=LRRC7;refGene_tx=XM_019837224.2;refGene_gn=LRRC7;refGene_tx=XM_019837225.2
C1 58063205 58063206 refGene_gn=LRRC7;refGene_tx=XM_019837220.2
C1 58116321 58116524
refGene_gn=LRRC7;refGene_tx=XM_011284940.3;refGene_gn=LRRC7;refGene_tx=XM_019837219.2;refGene_gn=LRRC7;refGene_tx=XM_019837220.2;refGene_gn=LRRC7;refGene_tx=XM_019837221.2;refGene_gn=LRRC7;refGene_tx=XM_019837222.2;refGene_gn=LRRC7;refGene_tx=XM_019837223.2;refGene_gn=LRRC7;refGene_tx=XM_019837224.2;refGene_gn=LRRC7;refGene_tx=XM_019837225.2
C1 58162944 58163062
refGene_gn=LRRC7;refGene_tx=XM_011284940.3;refGene_gn=LRRC7;refGene_tx=XM_019837219.2;refGene_gn=LRRC7;refGene_tx=XM_019837220.2;refGene_gn=LRRC7;refGene_tx=XM_019837221.2;refGene_gn=LRRC7;refGene_tx=XM_019837222.2;refGene_gn=LRRC7;refGene_tx=XM_019837223.2;refGene_gn=LRRC7;refGene_tx=XM_019837224.2;refGene_gn=LRRC7;refGene_tx=XM_019837225.2
C1 58185969 58186048
refGene_gn=LRRC7;refGene_tx=XM_011284940.3;refGene_gn=LRRC7;refGene_tx=XM_019837219.2;refGene_gn=LRRC7;refGene_tx=XM_019837220.2;refGene_gn=LRRC7;refGene_tx=XM_019837221.2;refGene_gn=LRRC7;refGene_tx=XM_019837222.2;refGene_gn=LRRC7;refGene_tx=XM_019837223.2;refGene_gn=LRRC7;refGene_tx=XM_019837224.2;refGene_gn=LRRC7;refGene_tx=XM_019837225.2
C1 58195154 58195244
refGene_gn=LRRC7;refGene_tx=XM_011284940.3;refGene_gn=LRRC7;refGene_tx=XM_019837219.2;refGene_gn=LRRC7;refGene_tx=XM_019837220.2;refGene_gn=LRRC7;refGene_tx=XM_019837221.2;refGene_gn=LRRC7;refGene_tx=XM_019837222.2;refGene_gn=LRRC7;refGene_tx=XM_019837223.2;refGene_gn=LRRC7;refGene_tx=XM_019837224.2;refGene_gn=LRRC7;refGene_tx=XM_019837225.2
C1 58201031 58201088
refGene_gn=LRRC7;refGene_tx=XM_011284940.3;refGene_gn=LRRC7;refGene_tx=XM_019837219.2;refGene_gn=LRRC7;refGene_tx=XM_019837220.2;refGene_gn=LRRC7;refGene_tx=XM_019837221.2;refGene_gn=LRRC7;refGene_tx=XM_0198372

22.2;refGene_gn=LRRC7;refGene_tx=XM_019837223.2;refGene_gn=LRRC7;refGene_tx=XM_019837224.2;refGene_gn=LRRC7;refGene_tx=XM_019837225.2
C1 58266789 58266853
refGene_gn=LRRC7;refGene_tx=XM_011284940.3;refGene_gn=LRRC7;refGene_tx=XM_019837219.2;refGene_gn=LRRC7;refGene_tx=XM_019837220.2;refGene_gn=LRRC7;refGene_tx=XM_019837221.2;refGene_gn=LRRC7;refGene_tx=XM_019837222.2;refGene_gn=LRRC7;refGene_tx=XM_019837223.2;refGene_gn=LRRC7;refGene_tx=XM_019837224.2;refGene_gn=LRRC7;refGene_tx=XM_019837225.2
C1 58322318 58322393
refGene_gn=LRRC7;refGene_tx=XM_011284940.3;refGene_gn=LRRC7;refGene_tx=XM_019837219.2;refGene_gn=LRRC7;refGene_tx=XM_019837220.2;refGene_gn=LRRC7;refGene_tx=XM_019837221.2;refGene_gn=LRRC7;refGene_tx=XM_019837222.2;refGene_gn=LRRC7;refGene_tx=XM_019837223.2;refGene_gn=LRRC7;refGene_tx=XM_019837224.2;refGene_gn=LRRC7;refGene_tx=XM_019837225.2
C1 58328365 58328510
refGene_gn=LRRC7;refGene_tx=XM_011284940.3;refGene_gn=LRRC7;refGene_tx=XM_019837219.2;refGene_gn=LRRC7;refGene_tx=XM_019837220.2;refGene_gn=LRRC7;refGene_tx=XM_019837221.2;refGene_gn=LRRC7;refGene_tx=XM_019837222.2;refGene_gn=LRRC7;refGene_tx=XM_019837223.2;refGene_gn=LRRC7;refGene_tx=XM_019837224.2;refGene_gn=LRRC7;refGene_tx=XM_019837225.2
C1 58333571 58333644
refGene_gn=LRRC7;refGene_tx=XM_011284940.3;refGene_gn=LRRC7;refGene_tx=XM_019837219.2;refGene_gn=LRRC7;refGene_tx=XM_019837220.2;refGene_gn=LRRC7;refGene_tx=XM_019837221.2;refGene_gn=LRRC7;refGene_tx=XM_019837222.2;refGene_gn=LRRC7;refGene_tx=XM_019837223.2;refGene_gn=LRRC7;refGene_tx=XM_019837224.2;refGene_gn=LRRC7;refGene_tx=XM_019837225.2
C1 58352125 58352255
refGene_gn=LRRC7;refGene_tx=XM_011284940.3;refGene_gn=LRRC7;refGene_tx=XM_019837219.2;refGene_gn=LRRC7;refGene_tx=XM_019837220.2;refGene_gn=LRRC7;refGene_tx=XM_019837221.2;refGene_gn=LRRC7;refGene_tx=XM_019837222.2;refGene_gn=LRRC7;refGene_tx=XM_019837223.2;refGene_gn=LRRC7;refGene_tx=XM_019837224.2;refGene_gn=LRRC7;refGene_tx=XM_019837225.2
C1 58353262 58353378
refGene_gn=LRRC7;refGene_tx=XM_011284940.3;refGene_gn=LRRC7;refGene_tx=XM_019837219.2;refGene_gn=LRRC7;refGene_tx=XM_019837220.2;refGene_gn=LRRC7;refGene_tx=XM_019837221.2;refGene_gn=LRRC7;refGene_tx=XM_019837222.2;refGene_gn=LRRC7;refGene_tx=XM_019837223.2;refGene_gn=LRRC7;refGene_tx=XM_019837224.2;refGene_gn=LRRC7;refGene_tx=XM_019837225.2
C1 58358146 58358216
refGene_gn=LRRC7;refGene_tx=XM_011284940.3;refGene_gn=LRRC7;refGene_tx=XM_019837219.2;refGene_gn=LRRC7;refGene_tx=XM_019837220.2;refGene_gn=LRRC7;refGene_tx=XM_019837221.2;refGene_gn=LRRC7;refGene_tx=XM_019837222.2;refGene_gn=LRRC7;refGene_tx=XM_019837223.2;refGene_gn=LRRC7;refGene_tx=XM_019837224.2;refGene_gn=LRRC7;refGene_tx=XM_019837225.2
C1 58360160 58360260
refGene_gn=LRRC7;refGene_tx=XM_011284940.3;refGene_gn=LRRC7;refGene_tx=XM_019837219.2;refGene_gn=LRRC7;refGene_tx=XM_019837220.2;refGene_gn=LRRC7;refGene_tx=XM_019837221.2;refGene_gn=LRRC7;refGene_tx=XM_019837222.2;refGene_gn=LRRC7;refGene_tx=XM_019837223.2;refGene_gn=LRRC7;refGene_tx=XM_019837224.2;refGene_gn=LRRC7;refGene_tx=XM_019837225.2

C1 58362192 58362317
refGene_gn=LRRC7;refGene_tx=XM_011284940.3;refGene_gn=LRRC7;refGene_tx
=XM_019837219.2;refGene_gn=LRRC7;refGene_tx=XM_019837220.2;refGene_gn=
LRRC7;refGene_tx=XM_019837221.2;refGene_gn=LRRC7;refGene_tx=XM_0198372
22.2;refGene_gn=LRRC7;refGene_tx=XM_019837223.2;refGene_gn=LRRC7;refGe
ne_tx=XM_019837224.2;refGene_gn=LRRC7;refGene_tx=XM_019837225.2
C1 58364341 58364590
refGene_gn=LRRC7;refGene_tx=XM_011284940.3;refGene_gn=LRRC7;refGene_tx
=XM_019837219.2;refGene_gn=LRRC7;refGene_tx=XM_019837220.2;refGene_gn=
LRRC7;refGene_tx=XM_019837221.2;refGene_gn=LRRC7;refGene_tx=XM_0198372
23.2;refGene_gn=LRRC7;refGene_tx=XM_019837225.2
C1 58369498 58369699
refGene_gn=LRRC7;refGene_tx=XM_011284940.3;refGene_gn=LRRC7;refGene_tx
=XM_019837219.2;refGene_gn=LRRC7;refGene_tx=XM_019837220.2;refGene_gn=
LRRC7;refGene_tx=XM_019837221.2;refGene_gn=LRRC7;refGene_tx=XM_0198372
22.2;refGene_gn=LRRC7;refGene_tx=XM_019837223.2;refGene_gn=LRRC7;refGe
ne_tx=XM_019837225.2
C1 58377582 58377694
refGene_gn=LRRC7;refGene_tx=XM_011284940.3;refGene_gn=LRRC7;refGene_tx
=XM_019837219.2;refGene_gn=LRRC7;refGene_tx=XM_019837220.2;refGene_gn=
LRRC7;refGene_tx=XM_019837221.2;refGene_gn=LRRC7;refGene_tx=XM_0198372
22.2;refGene_gn=LRRC7;refGene_tx=XM_019837223.2;refGene_gn=LRRC7;refGe
ne_tx=XM_019837224.2;refGene_gn=LRRC7;refGene_tx=XM_019837225.2
C1 58377906 58378087
refGene_gn=LRRC7;refGene_tx=XM_011284940.3;refGene_gn=LRRC7;refGene_tx
=XM_019837219.2;refGene_gn=LRRC7;refGene_tx=XM_019837220.2;refGene_gn=
LRRC7;refGene_tx=XM_019837221.2;refGene_gn=LRRC7;refGene_tx=XM_0198372
22.2;refGene_gn=LRRC7;refGene_tx=XM_019837223.2;refGene_gn=LRRC7;refGe
ne_tx=XM_019837224.2;refGene_gn=LRRC7;refGene_tx=XM_019837225.2
C1 58379915 58381596
refGene_gn=LRRC7;refGene_tx=XM_011284940.3;refGene_gn=LRRC7;refGene_tx
=XM_019837219.2;refGene_gn=LRRC7;refGene_tx=XM_019837220.2;refGene_gn=
LRRC7;refGene_tx=XM_019837221.2;refGene_gn=LRRC7;refGene_tx=XM_0198372
22.2;refGene_gn=LRRC7;refGene_tx=XM_019837223.2;refGene_gn=LRRC7;refGe
ne_tx=XM_019837224.2;refGene_gn=LRRC7;refGene_tx=XM_019837225.2
C1 58382947 58383015 refGene_gn=LRRC7;refGene_tx=XM_019837225.2
C1 58385545 58385686
refGene_gn=LRRC7;refGene_tx=XM_019837219.2;refGene_gn=LRRC7;refGene_tx
=XM_019837220.2;refGene_gn=LRRC7;refGene_tx=XM_019837221.2;refGene_gn=
LRRC7;refGene_tx=XM_019837222.2;refGene_gn=LRRC7;refGene_tx=XM_0198372
24.2;refGene_gn=LRRC7;refGene_tx=XM_019837225.2
C1 58394766 58394886
refGene_gn=LRRC7;refGene_tx=XM_011284940.3;refGene_gn=LRRC7;refGene_tx
=XM_019837219.2;refGene_gn=LRRC7;refGene_tx=XM_019837220.2;refGene_gn=
LRRC7;refGene_tx=XM_019837221.2;refGene_gn=LRRC7;refGene_tx=XM_0198372
22.2;refGene_gn=LRRC7;refGene_tx=XM_019837224.2
C1 58425331 58425553
refGene_gn=LRRC7;refGene_tx=XM_011284940.3;refGene_gn=LRRC7;refGene_tx
=XM_019837219.2;refGene_gn=LRRC7;refGene_tx=XM_019837220.2;refGene_gn=
LRRC7;refGene_tx=XM_019837221.2;refGene_gn=LRRC7;refGene_tx=XM_0198372

22.2;refGene_gn=LRRC7;refGene_tx=XM_019837223.2;refGene_gn=LRRC7;refGene_tx=XM_019837224.2
C1 58438186 58438279
refGene_gn=LRRC7;refGene_tx=XM_011284940.3;refGene_gn=LRRC7;refGene_tx=XM_019837219.2;refGene_gn=LRRC7;refGene_tx=XM_019837220.2;refGene_gn=LRRC7;refGene_tx=XM_019837221.2;refGene_gn=LRRC7;refGene_tx=XM_019837222.2;refGene_gn=LRRC7;refGene_tx=XM_019837223.2;refGene_gn=LRRC7;refGene_tx=XM_019837224.2
C1 58452674 58452749
refGene_gn=LRRC7;refGene_tx=XM_011284940.3;refGene_gn=LRRC7;refGene_tx=XM_019837219.2;refGene_gn=LRRC7;refGene_tx=XM_019837220.2;refGene_gn=LRRC7;refGene_tx=XM_019837221.2;refGene_gn=LRRC7;refGene_tx=XM_019837222.2;refGene_gn=LRRC7;refGene_tx=XM_019837223.2;refGene_gn=LRRC7;refGene_tx=XM_019837224.2
C1 58468792 58468900
refGene_gn=LRRC7;refGene_tx=XM_011284940.3;refGene_gn=LRRC7;refGene_tx=XM_019837219.2;refGene_gn=LRRC7;refGene_tx=XM_019837220.2;refGene_gn=LRRC7;refGene_tx=XM_019837222.2;refGene_gn=LRRC7;refGene_tx=XM_019837223.2;refGene_gn=LRRC7;refGene_tx=XM_019837224.2
C1 58489642 58489748
refGene_gn=LRRC40;refGene_tx=XM_011284937.3;refGene_gn=LRRC40;refGene_tx=XM_011284938.3;refGene_gn=LRRC40;refGene_tx=XM_019837226.2
C1 58491956 58492142
refGene_gn=LRRC40;refGene_tx=XM_011284937.3;refGene_gn=LRRC40;refGene_tx=XM_011284938.3;refGene_gn=LRRC40;refGene_tx=XM_019837226.2
C1 58494718 58494796
refGene_gn=LRRC40;refGene_tx=XM_011284937.3;refGene_gn=LRRC40;refGene_tx=XM_011284938.3;refGene_gn=LRRC40;refGene_tx=XM_019837226.2
C1 58496845 58496956
refGene_gn=LRRC40;refGene_tx=XM_011284937.3;refGene_gn=LRRC40;refGene_tx=XM_011284938.3
C1 58500627 58500735
refGene_gn=LRRC40;refGene_tx=XM_011284937.3;refGene_gn=LRRC40;refGene_tx=XM_011284938.3;refGene_gn=LRRC40;refGene_tx=XM_019837226.2
C1 58503394 58503503
refGene_gn=LRRC40;refGene_tx=XM_011284937.3;refGene_gn=LRRC40;refGene_tx=XM_011284938.3;refGene_gn=LRRC40;refGene_tx=XM_019837226.2
C1 58509980 58510026
refGene_gn=LRRC40;refGene_tx=XM_011284937.3;refGene_gn=LRRC40;refGene_tx=XM_011284938.3;refGene_gn=LRRC40;refGene_tx=XM_019837226.2
C1 58510142 58510230
refGene_gn=LRRC40;refGene_tx=XM_011284937.3;refGene_gn=LRRC40;refGene_tx=XM_011284938.3;refGene_gn=LRRC40;refGene_tx=XM_019837226.2
C1 58512217 58512390
refGene_gn=LRRC40;refGene_tx=XM_011284937.3;refGene_gn=LRRC40;refGene_tx=XM_011284938.3;refGene_gn=LRRC40;refGene_tx=XM_019837226.2
C1 58513631 58513774
refGene_gn=LRRC40;refGene_tx=XM_011284937.3;refGene_gn=LRRC40;refGene_tx=XM_011284938.3;refGene_gn=LRRC40;refGene_tx=XM_019837226.2
C1 58517440 58517564

refGene_gn=LRRC40;refGene_tx=XM_011284937.3;refGene_gn=LRRC40;refGene_tx=XM_011284938.3;refGene_gn=LRRC40;refGene_tx=XM_019837226.2
C1 58520765 58520895
refGene_gn=LRRC40;refGene_tx=XM_011284937.3;refGene_gn=LRRC40;refGene_tx=XM_011284938.3;refGene_gn=LRRC40;refGene_tx=XM_019837226.2
C1 58522659 58522733
refGene_gn=LRRC40;refGene_tx=XM_011284937.3;refGene_gn=LRRC40;refGene_tx=XM_011284938.3;refGene_gn=LRRC40;refGene_tx=XM_019837226.2
C1 58524137 58524319
refGene_gn=LRRC40;refGene_tx=XM_011284937.3;refGene_gn=LRRC40;refGene_tx=XM_011284938.3;refGene_gn=LRRC40;refGene_tx=XM_019837226.2
C1 58535637 58535788
refGene_gn=LRRC40;refGene_tx=XM_011284937.3;refGene_gn=LRRC40;refGene_tx=XM_011284938.3;refGene_gn=LRRC40;refGene_tx=XM_019837226.2
C1 58551707 58551907
refGene_gn=SRSF11;refGene_tx=XM_023258816.1;refGene_gn=SRSF11;refGene_tx=XM_023258818.1;refGene_gn=SRSF11;refGene_tx=XM_023258819.1;refGene_gn=SRSF11;refGene_tx=XM_023258820.1;refGene_gn=SRSF11;refGene_tx=XM_023258821.1;refGene_gn=SRSF11;refGene_tx=XM_023258822.1
C1 58558541 58558675
refGene_gn=SRSF11;refGene_tx=XM_023258816.1;refGene_gn=SRSF11;refGene_tx=XM_023258818.1;refGene_gn=SRSF11;refGene_tx=XM_023258819.1;refGene_gn=SRSF11;refGene_tx=XM_023258820.1;refGene_gn=SRSF11;refGene_tx=XM_023258821.1;refGene_gn=SRSF11;refGene_tx=XM_023258822.1
C1 58562430 58562540
refGene_gn=SRSF11;refGene_tx=XM_023258816.1;refGene_gn=SRSF11;refGene_tx=XM_023258818.1;refGene_gn=SRSF11;refGene_tx=XM_023258819.1;refGene_gn=SRSF11;refGene_tx=XM_023258820.1;refGene_gn=SRSF11;refGene_tx=XM_023258821.1;refGene_gn=SRSF11;refGene_tx=XM_023258822.1
C1 58564474 58564567
refGene_gn=SRSF11;refGene_tx=XM_023258816.1;refGene_gn=SRSF11;refGene_tx=XM_023258818.1;refGene_gn=SRSF11;refGene_tx=XM_023258819.1;refGene_gn=SRSF11;refGene_tx=XM_023258820.1;refGene_gn=SRSF11;refGene_tx=XM_023258821.1;refGene_gn=SRSF11;refGene_tx=XM_023258822.1
C1 58565302 58565352
refGene_gn=SRSF11;refGene_tx=XM_023258816.1;refGene_gn=SRSF11;refGene_tx=XM_023258818.1;refGene_gn=SRSF11;refGene_tx=XM_023258819.1;refGene_gn=SRSF11;refGene_tx=XM_023258820.1;refGene_gn=SRSF11;refGene_tx=XM_023258821.1;refGene_gn=SRSF11;refGene_tx=XM_023258822.1
C1 58566758 58566886
refGene_gn=SRSF11;refGene_tx=XM_023258816.1;refGene_gn=SRSF11;refGene_tx=XM_023258818.1;refGene_gn=SRSF11;refGene_tx=XM_023258819.1;refGene_gn=SRSF11;refGene_tx=XM_023258820.1;refGene_gn=SRSF11;refGene_tx=XM_023258821.1;refGene_gn=SRSF11;refGene_tx=XM_023258822.1
C1 58568699 58568781
refGene_gn=SRSF11;refGene_tx=XM_023258816.1;refGene_gn=SRSF11;refGene_tx=XM_023258818.1;refGene_gn=SRSF11;refGene_tx=XM_023258819.1;refGene_gn=SRSF11;refGene_tx=XM_023258820.1;refGene_gn=SRSF11;refGene_tx=XM_023258821.1;refGene_gn=SRSF11;refGene_tx=XM_023258822.1
C1 58572822 58572954

refGene_gn=SRSF11;refGene_tx=XM_023258816.1;refGene_gn=SRSF11;refGene_tx=XM_023258818.1;refGene_gn=SRSF11;refGene_tx=XM_023258819.1;refGene_gn=SRSF11;refGene_tx=XM_023258820.1;refGene_gn=SRSF11;refGene_tx=XM_023258821.1;refGene_gn=SRSF11;refGene_tx=XM_023258822.1
C1 58575418 58575508
refGene_gn=SRSF11;refGene_tx=XM_023258816.1;refGene_gn=SRSF11;refGene_tx=XM_023258818.1;refGene_gn=SRSF11;refGene_tx=XM_023258819.1;refGene_gn=SRSF11;refGene_tx=XM_023258820.1;refGene_gn=SRSF11;refGene_tx=XM_023258821.1;refGene_gn=SRSF11;refGene_tx=XM_023258822.1
C1 58577639 58577756
refGene_gn=SRSF11;refGene_tx=XM_023258816.1;refGene_gn=SRSF11;refGene_tx=XM_023258818.1;refGene_gn=SRSF11;refGene_tx=XM_023258819.1;refGene_gn=SRSF11;refGene_tx=XM_023258820.1;refGene_gn=SRSF11;refGene_tx=XM_023258821.1;refGene_gn=SRSF11;refGene_tx=XM_023258822.1
C1 58578151 58578290
refGene_gn=SRSF11;refGene_tx=XM_023258816.1;refGene_gn=SRSF11;refGene_tx=XM_023258818.1;refGene_gn=SRSF11;refGene_tx=XM_023258819.1;refGene_gn=SRSF11;refGene_tx=XM_023258820.1;refGene_gn=SRSF11;refGene_tx=XM_023258821.1;refGene_gn=SRSF11;refGene_tx=XM_023258822.1
C1 58578395 58578593
refGene_gn=SRSF11;refGene_tx=XM_023258816.1;refGene_gn=SRSF11;refGene_tx=XM_023258818.1;refGene_gn=SRSF11;refGene_tx=XM_023258819.1;refGene_gn=SRSF11;refGene_tx=XM_023258820.1;refGene_gn=SRSF11;refGene_tx=XM_023258821.1;refGene_gn=SRSF11;refGene_tx=XM_023258822.1
C1 58587071 58587202
refGene_gn=ANKRD13C;refGene_tx=XM_011284936.3;refGene_gn=ANKRD13C;refGene_tx=XM_019837227.2
C1 58592160 58592261
refGene_gn=ANKRD13C;refGene_tx=XM_011284936.3;refGene_gn=ANKRD13C;refGene_tx=XM_019837227.2
C1 58597376 58597475 refGene_gn=ANKRD13C;refGene_tx=XM_011284936.3
C1 58599155 58599235
refGene_gn=ANKRD13C;refGene_tx=XM_011284936.3;refGene_gn=ANKRD13C;refGene_tx=XM_019837227.2
C1 58609424 58609586
refGene_gn=ANKRD13C;refGene_tx=XM_011284936.3;refGene_gn=ANKRD13C;refGene_tx=XM_019837227.2
C1 58612696 58612828
refGene_gn=ANKRD13C;refGene_tx=XM_011284936.3;refGene_gn=ANKRD13C;refGene_tx=XM_019837227.2
C1 58618589 58618734
refGene_gn=ANKRD13C;refGene_tx=XM_011284936.3;refGene_gn=ANKRD13C;refGene_tx=XM_019837227.2
C1 58623866 58623933
refGene_gn=ANKRD13C;refGene_tx=XM_011284936.3;refGene_gn=ANKRD13C;refGene_tx=XM_019837227.2
C1 58627946 58627992
refGene_gn=ANKRD13C;refGene_tx=XM_011284936.3;refGene_gn=ANKRD13C;refGene_tx=XM_019837227.2
C1 58629181 58629267

refGene_gn=ANKRD13C;refGene_tx=XM_011284936.3;refGene_gn=ANKRD13C;refGene_tx=XM_019837227.2
C1 58634961 58635066
refGene_gn=ANKRD13C;refGene_tx=XM_011284936.3;refGene_gn=ANKRD13C;refGene_tx=XM_019837227.2
C1 58652048 58652090
refGene_gn=ANKRD13C;refGene_tx=XM_011284936.3;refGene_gn=ANKRD13C;refGene_tx=XM_019837227.2
C1 58674860 58675305
refGene_gn=ANKRD13C;refGene_tx=XM_011284936.3;refGene_gn=ANKRD13C;refGene_tx=XM_019837227.2
C1 58709926 58710094 refGene_gn=CTH;refGene_tx=XM_003990218.4
C1 58713118 58713200 refGene_gn=CTH;refGene_tx=XM_003990218.4
C1 58715701 58715797 refGene_gn=CTH;refGene_tx=XM_003990218.4
C1 58717529 58717639 refGene_gn=CTH;refGene_tx=XM_003990218.4
C1 58719667 58719799 refGene_gn=CTH;refGene_tx=XM_003990218.4
C1 58723481 58723539 refGene_gn=CTH;refGene_tx=XM_003990218.4
C1 58723997 58724075 refGene_gn=CTH;refGene_tx=XM_003990218.4
C1 58725629 58725782 refGene_gn=CTH;refGene_tx=XM_003990218.4
C1 58726879 58727001 refGene_gn=CTH;refGene_tx=XM_003990218.4
C1 58728440 58728493 refGene_gn=CTH;refGene_tx=XM_003990218.4
C1 58732556 58732695 refGene_gn=CTH;refGene_tx=XM_003990218.4
C1 58733159 58733186 refGene_gn=CTH;refGene_tx=XM_003990218.4
C1 59048693 59048715 refGene_gn=PTGER3;refGene_tx=XM_019837230.1
C1 59118447 59118539 refGene_gn=PTGER3;refGene_tx=XM_019837230.1
C1 59161270 59161450 refGene_gn=PTGER3;refGene_tx=XM_019837230.1
C1 59192439 59192553 refGene_gn=PTGER3;refGene_tx=XM_019837230.1
C1 59192554 59193334 refGene_gn=PTGER3;refGene_tx=XM_019837230.1
C1 59207314 59207378 refGene_gn=ZRANB2;refGene_tx=XM_003990222.5
C1 59207967 59208001 refGene_gn=ZRANB2;refGene_tx=XM_011284930.3
C1 59209059 59209218
refGene_gn=ZRANB2;refGene_tx=XM_003990222.5;refGene_gn=ZRANB2;refGene_tx=XM_011284930.3
C1 59211130 59211217
refGene_gn=ZRANB2;refGene_tx=XM_003990222.5;refGene_gn=ZRANB2;refGene_tx=XM_011284930.3
C1 59212928 59213098
refGene_gn=ZRANB2;refGene_tx=XM_003990222.5;refGene_gn=ZRANB2;refGene_tx=XM_011284930.3
C1 59214199 59214334
refGene_gn=ZRANB2;refGene_tx=XM_003990222.5;refGene_gn=ZRANB2;refGene_tx=XM_011284930.3
C1 59214626 59214703
refGene_gn=ZRANB2;refGene_tx=XM_003990222.5;refGene_gn=ZRANB2;refGene_tx=XM_011284930.3
C1 59219046 59219129
refGene_gn=ZRANB2;refGene_tx=XM_003990222.5;refGene_gn=ZRANB2;refGene_tx=XM_011284930.3
C1 59220220 59220329
refGene_gn=ZRANB2;refGene_tx=XM_003990222.5;refGene_gn=ZRANB2;refGene_

tx=XM_011284930.3
C1 59220420 59220473
refGene_gn=ZRANB2;refGene_tx=XM_003990222.5;refGene_gn=ZRANB2;refGene_
tx=XM_011284930.3
C1 59222869 59222925
refGene_gn=ZRANB2;refGene_tx=XM_003990222.5;refGene_gn=ZRANB2;refGene_
tx=XM_011284930.3
C1 59479471 59479596 refGene_gn=NEGR1;refGene_tx=XM_003990224.4
C1 59635240 59635341 refGene_gn=NEGR1;refGene_tx=XM_019837231.2
C1 59649334 59649486
refGene_gn=NEGR1;refGene_tx=XM_003990224.4;refGene_gn=NEGR1;refGene_tx
=XM_019837231.2
C1 59665365 59665486
refGene_gn=NEGR1;refGene_tx=XM_003990224.4;refGene_gn=NEGR1;refGene_tx
=XM_019837231.2
C1 59753901 59754033
refGene_gn=NEGR1;refGene_tx=XM_003990224.4;refGene_gn=NEGR1;refGene_tx
=XM_019837231.2
C1 59818540 59818666
refGene_gn=NEGR1;refGene_tx=XM_003990224.4;refGene_gn=NEGR1;refGene_tx
=XM_019837231.2
C1 59959466 59959699
refGene_gn=NEGR1;refGene_tx=XM_003990224.4;refGene_gn=NEGR1;refGene_tx
=XM_019837231.2
C1 60319702 60319878
refGene_gn=NEGR1;refGene_tx=XM_003990224.4;refGene_gn=NEGR1;refGene_tx
=XM_019837231.2
C1 62035389 62035546 refGene_gn=LRRIQ3;refGene_tx=XM_003990225.4
C1 62044862 62045592 refGene_gn=LRRIQ3;refGene_tx=XM_003990225.4
C1 62100725 62100855 refGene_gn=LRRIQ3;refGene_tx=XM_003990225.4
C1 62125253 62125407 refGene_gn=LRRIQ3;refGene_tx=XM_003990225.4
C1 62199392 62199526 refGene_gn=LRRIQ3;refGene_tx=XM_003990225.4
C1 62225349 62225673 refGene_gn=LRRIQ3;refGene_tx=XM_003990225.4
C1 62228407 62228656 refGene_gn=LRRIQ3;refGene_tx=XM_003990225.4
C1 62255460 62255542
refGene_gn=FPGT;refGene_tx=XM_003990227.4;refGene_gn=FPGT;refGene_tx=X
M_019837237.2
C1 62256269 62256327 refGene_gn=FPGT;refGene_tx=XM_006934837.2
C1 62257011 62257179
refGene_gn=FPGT;refGene_tx=XM_003990227.4;refGene_gn=FPGT;refGene_tx=X
M_006934837.2;refGene_gn=FPGT;refGene_tx=XM_019837237.2
C1 62258679 62258772
refGene_gn=FPGT;refGene_tx=XM_003990227.4;refGene_gn=FPGT;refGene_tx=X
M_006934837.2
C1 62260530 62261987
refGene_gn=FPGT;refGene_tx=XM_003990227.4;refGene_gn=FPGT;refGene_tx=X
M_006934837.2;refGene_gn=FPGT;refGene_tx=XM_019837237.2
C1 62289558 62289598
refGene_gn=LOC101085560;refGene_tx=XM_003990226.4;refGene_gn=LOC101085
560;refGene_tx=XM_023258825.1

C1	62290185	62290294
refGene_gn=LOC101085560;refGene_tx=XM_003990226.4;refGene_gn=LOC101085560;refGene_tx=XM_023258825.1
C1	62300754	62300840
refGene_gn=LOC101085560;refGene_tx=XM_003990226.4;refGene_gn=LOC101085560;refGene_tx=XM_023258825.1
C1	62301962	62302060
refGene_gn=LOC101085560;refGene_tx=XM_003990226.4;refGene_gn=LOC101085560;refGene_tx=XM_023258825.1
C1	62322244	62322355
refGene_gn=LOC101085560;refGene_tx=XM_003990226.4;refGene_gn=LOC101085560;refGene_tx=XM_023258825.1
C1	62374027	62374126
refGene_gn=LOC101085560;refGene_tx=XM_003990226.4;refGene_gn=LOC101085560;refGene_tx=XM_023258825.1
C1	62380421	62380560
refGene_gn=LOC101085560;refGene_tx=XM_003990226.4;refGene_gn=LOC101085560;refGene_tx=XM_023258825.1
C1	62387235	62387380
refGene_gn=LOC101085560;refGene_tx=XM_003990226.4;refGene_gn=LOC101085560;refGene_tx=XM_023258825.1
C1	62387466	62387571
refGene_gn=LOC101085560;refGene_tx=XM_003990226.4;refGene_gn=LOC101085560;refGene_tx=XM_023258825.1
C1	62390871	62390966
refGene_gn=LOC101085560;refGene_tx=XM_003990226.4;refGene_gn=LOC101085560;refGene_tx=XM_023258825.1
C1	62391804	62391954
refGene_gn=LOC101085560;refGene_tx=XM_003990226.4;refGene_gn=LOC101085560;refGene_tx=XM_023258825.1
C1	62405081	62405168
refGene_gn=LOC101085560;refGene_tx=XM_003990226.4;refGene_gn=LOC101085560;refGene_tx=XM_023258825.1
C1	62406255	62406312
refGene_gn=LOC101085560;refGene_tx=XM_003990226.4;refGene_gn=LOC101085560;refGene_tx=XM_023258825.1
C1	62407284	62407377
refGene_gn=LOC101085560;refGene_tx=XM_003990226.4;refGene_gn=LOC101085560;refGene_tx=XM_023258825.1
C1	62407461	62407519
refGene_gn=LOC101085560;refGene_tx=XM_003990226.4;refGene_gn=LOC101085560;refGene_tx=XM_023258825.1
C1	62407645	62407840
refGene_gn=LOC101085560;refGene_tx=XM_003990226.4;refGene_gn=LOC101085560;refGene_tx=XM_023258825.1
C1	62408588	62408693
refGene_gn=LOC101085560;refGene_tx=XM_003990226.4;refGene_gn=LOC101085560;refGene_tx=XM_023258825.1
C1	62473962	62474015
refGene_gn=LOC101085560;refGene_tx=XM_003990226.4;refGene_gn=LOC101085

560;refGene_tx=XM_023258825.1
C1	62474335	62474388
refGene_gn=LOC101085560;refGene_tx=XM_003990226.4;refGene_gn=LOC101085560;refGene_tx=XM_023258825.1
C1	62478215	62478348
refGene_gn=LOC101085560;refGene_tx=XM_003990226.4;refGene_gn=LOC101085560;refGene_tx=XM_023258825.1
C1	62491033	62491143
refGene_gn=LOC101085560;refGene_tx=XM_003990226.4;refGene_gn=LOC101085560;refGene_tx=XM_023258825.1
C1	62518226	62518286
refGene_gn=LOC101085560;refGene_tx=XM_003990226.4
C1	62521345	62521515
refGene_gn=LOC101085560;refGene_tx=XM_003990226.4;refGene_gn=LOC101085560;refGene_tx=XM_023258825.1
C1	62570988	62571068
refGene_gn=LOC101085560;refGene_tx=XM_003990226.4;refGene_gn=LOC101085560;refGene_tx=XM_023258825.1
C1	62574272	62574349
refGene_gn=LOC101085560;refGene_tx=XM_003990226.4;refGene_gn=LOC101085560;refGene_tx=XM_023258825.1
C1	62595604	62597994
refGene_gn=ERICH3;refGene_tx=XM_006934846.2;refGene_gn=ERICH3;refGene_tx=XM_019837238.1;refGene_gn=ERICH3;refGene_tx=XM_019837240.2;refGene_gn=ERICH3;refGene_tx=XM_019837241.1;refGene_gn=ERICH3;refGene_tx=XM_019837242.1;refGene_gn=ERICH3;refGene_tx=XM_019837243.1;refGene_gn=ERICH3;refGene_tx=XM_019837244.1;refGene_gn=ERICH3;refGene_tx=XM_019837245.1;refGene_gn=ERICH3;refGene_tx=XM_019837246.1;refGene_gn=ERICH3;refGene_tx=XM_019837247.2;refGene_gn=ERICH3;refGene_tx=XM_019837248.1;refGene_gn=ERICH3;refGene_tx=XM_019837249.2;refGene_gn=ERICH3;refGene_tx=XM_023258828.1
C1	62601683	62601809
refGene_gn=ERICH3;refGene_tx=XM_006934846.2;refGene_gn=ERICH3;refGene_tx=XM_019837238.1;refGene_gn=ERICH3;refGene_tx=XM_019837240.2;refGene_gn=ERICH3;refGene_tx=XM_019837241.1;refGene_gn=ERICH3;refGene_tx=XM_019837242.1;refGene_gn=ERICH3;refGene_tx=XM_019837243.1;refGene_gn=ERICH3;refGene_tx=XM_019837244.1;refGene_gn=ERICH3;refGene_tx=XM_019837245.1;refGene_gn=ERICH3;refGene_tx=XM_019837246.1;refGene_gn=ERICH3;refGene_tx=XM_019837247.2;refGene_gn=ERICH3;refGene_tx=XM_019837248.1;refGene_gn=ERICH3;refGene_tx=XM_019837249.2;refGene_gn=ERICH3;refGene_tx=XM_023258828.1
C1	62612103	62612553
refGene_gn=ERICH3;refGene_tx=XM_006934846.2;refGene_gn=ERICH3;refGene_tx=XM_019837238.1;refGene_gn=ERICH3;refGene_tx=XM_019837240.2;refGene_gn=ERICH3;refGene_tx=XM_019837241.1;refGene_gn=ERICH3;refGene_tx=XM_019837242.1;refGene_gn=ERICH3;refGene_tx=XM_019837243.1;refGene_gn=ERICH3;refGene_tx=XM_019837244.1;refGene_gn=ERICH3;refGene_tx=XM_019837245.1;refGene_gn=ERICH3;refGene_tx=XM_019837246.1;refGene_gn=ERICH3;refGene_tx=XM_019837247.2;refGene_gn=ERICH3;refGene_tx=XM_019837248.1;refGene_gn=ERICH3;refGene_tx=XM_019837249.2;refGene_gn=ERICH3;refGene_tx=XM_

023258828.1
C1 62623333 62623570
refGene_gn=ERICH3;refGene_tx=XM_006934846.2;refGene_gn=ERICH3;refGene_tx=XM_019837238.1;refGene_gn=ERICH3;refGene_tx=XM_019837240.2;refGene_gn=ERICH3;refGene_tx=XM_019837241.1;refGene_gn=ERICH3;refGene_tx=XM_019837242.1;refGene_gn=ERICH3;refGene_tx=XM_019837243.1;refGene_gn=ERICH3;refGene_tx=XM_019837244.1;refGene_gn=ERICH3;refGene_tx=XM_019837245.1;refGene_gn=ERICH3;refGene_tx=XM_019837246.1;refGene_gn=ERICH3;refGene_tx=XM_019837247.2;refGene_gn=ERICH3;refGene_tx=XM_019837248.1;refGene_gn=ERICH3;refGene_tx=XM_019837249.2;refGene_gn=ERICH3;refGene_tx=XM_023258828.1
C1 62628736 62629035
refGene_gn=ERICH3;refGene_tx=XM_006934846.2;refGene_gn=ERICH3;refGene_tx=XM_019837238.1;refGene_gn=ERICH3;refGene_tx=XM_019837240.2;refGene_gn=ERICH3;refGene_tx=XM_019837241.1;refGene_gn=ERICH3;refGene_tx=XM_019837242.1;refGene_gn=ERICH3;refGene_tx=XM_019837243.1;refGene_gn=ERICH3;refGene_tx=XM_019837244.1;refGene_gn=ERICH3;refGene_tx=XM_019837245.1;refGene_gn=ERICH3;refGene_tx=XM_019837246.1;refGene_gn=ERICH3;refGene_tx=XM_019837247.2;refGene_gn=ERICH3;refGene_tx=XM_019837248.1;refGene_gn=ERICH3;refGene_tx=XM_019837249.2;refGene_gn=ERICH3;refGene_tx=XM_023258828.1
C1 62635286 62635473
refGene_gn=ERICH3;refGene_tx=XM_006934846.2;refGene_gn=ERICH3;refGene_tx=XM_019837238.1;refGene_gn=ERICH3;refGene_tx=XM_019837240.2;refGene_gn=ERICH3;refGene_tx=XM_019837241.1;refGene_gn=ERICH3;refGene_tx=XM_019837242.1;refGene_gn=ERICH3;refGene_tx=XM_019837243.1;refGene_gn=ERICH3;refGene_tx=XM_019837244.1;refGene_gn=ERICH3;refGene_tx=XM_019837245.1;refGene_gn=ERICH3;refGene_tx=XM_019837246.1;refGene_gn=ERICH3;refGene_tx=XM_019837247.2;refGene_gn=ERICH3;refGene_tx=XM_019837248.1;refGene_gn=ERICH3;refGene_tx=XM_019837249.2;refGene_gn=ERICH3;refGene_tx=XM_023258828.1
C1 62643401 62643582
refGene_gn=ERICH3;refGene_tx=XM_006934846.2;refGene_gn=ERICH3;refGene_tx=XM_019837238.1;refGene_gn=ERICH3;refGene_tx=XM_019837240.2;refGene_gn=ERICH3;refGene_tx=XM_019837241.1;refGene_gn=ERICH3;refGene_tx=XM_019837242.1;refGene_gn=ERICH3;refGene_tx=XM_019837243.1;refGene_gn=ERICH3;refGene_tx=XM_019837244.1;refGene_gn=ERICH3;refGene_tx=XM_019837245.1;refGene_gn=ERICH3;refGene_tx=XM_019837246.1;refGene_gn=ERICH3;refGene_tx=XM_019837247.2;refGene_gn=ERICH3;refGene_tx=XM_019837248.1;refGene_gn=ERICH3;refGene_tx=XM_019837249.2;refGene_gn=ERICH3;refGene_tx=XM_023258828.1
C1 62656626 62656845
refGene_gn=ERICH3;refGene_tx=XM_006934846.2;refGene_gn=ERICH3;refGene_tx=XM_019837238.1;refGene_gn=ERICH3;refGene_tx=XM_019837240.2;refGene_gn=ERICH3;refGene_tx=XM_019837241.1;refGene_gn=ERICH3;refGene_tx=XM_019837242.1;refGene_gn=ERICH3;refGene_tx=XM_019837243.1;refGene_gn=ERICH3;refGene_tx=XM_019837244.1;refGene_gn=ERICH3;refGene_tx=XM_019837245.1;refGene_gn=ERICH3;refGene_tx=XM_019837246.1;refGene_gn=ERICH3;refGene_tx=XM_019837247.2;refGene_gn=ERICH3;refGene_tx=XM_019837248.1;refGene_gn=ERICH3;refGene_tx=XM_019837249.2;refGene_gn=ERICH3;refGene_tx=XM_

023258828.1
C1	62659679 62659694 refGene_gn=ERICH3;refGene_tx=XM_019837247.2
C1	62661144 62661303
refGene_gn=ERICH3;refGene_tx=XM_006934846.2;refGene_gn=ERICH3;refGene_tx=XM_019837238.1;refGene_gn=ERICH3;refGene_tx=XM_019837240.2;refGene_gn=ERICH3;refGene_tx=XM_019837241.1;refGene_gn=ERICH3;refGene_tx=XM_019837242.1;refGene_gn=ERICH3;refGene_tx=XM_019837243.1;refGene_gn=ERICH3;refGene_tx=XM_019837244.1;refGene_gn=ERICH3;refGene_tx=XM_019837245.1;refGene_gn=ERICH3;refGene_tx=XM_019837246.1
C1	62666112 62666241
refGene_gn=ERICH3;refGene_tx=XM_006934846.2;refGene_gn=ERICH3;refGene_tx=XM_019837238.1;refGene_gn=ERICH3;refGene_tx=XM_019837240.2
C1	62668120 62668192
refGene_gn=ERICH3;refGene_tx=XM_006934846.2;refGene_gn=ERICH3;refGene_tx=XM_019837238.1;refGene_gn=ERICH3;refGene_tx=XM_019837241.1
C1	62671979 62672105
refGene_gn=ERICH3;refGene_tx=XM_006934846.2;refGene_gn=ERICH3;refGene_tx=XM_019837238.1;refGene_gn=ERICH3;refGene_tx=XM_019837240.2;refGene_gn=ERICH3;refGene_tx=XM_019837241.1;refGene_gn=ERICH3;refGene_tx=XM_019837242.1
C1	62674545 62674639
refGene_gn=ERICH3;refGene_tx=XM_006934846.2;refGene_gn=ERICH3;refGene_tx=XM_019837238.1;refGene_gn=ERICH3;refGene_tx=XM_019837240.2;refGene_gn=ERICH3;refGene_tx=XM_019837241.1;refGene_gn=ERICH3;refGene_tx=XM_019837242.1
C1	62699707 62699730 refGene_gn=ERICH3;refGene_tx=XM_006934846.2
C1	62700170 62700274
refGene_gn=ERICH3;refGene_tx=XM_019837238.1;refGene_gn=ERICH3;refGene_tx=XM_019837240.2;refGene_gn=ERICH3;refGene_tx=XM_019837241.1;refGene_gn=ERICH3;refGene_tx=XM_019837242.1
C1	62721843 62722026 refGene_gn=CRYZ;refGene_tx=XM_023258829.1
C1	62732698 62732863 refGene_gn=CRYZ;refGene_tx=XM_006934838.4
C1	62733301 62733397
refGene_gn=CRYZ;refGene_tx=XM_006934838.4;refGene_gn=CRYZ;refGene_tx=XM_023258829.1
C1	62733497 62733599
refGene_gn=CRYZ;refGene_tx=XM_006934838.4;refGene_gn=CRYZ;refGene_tx=XM_023258829.1
C1	62736793 62736943
refGene_gn=CRYZ;refGene_tx=XM_006934838.4;refGene_gn=CRYZ;refGene_tx=XM_023258829.1
C1	62741169 62741221
refGene_gn=CRYZ;refGene_tx=XM_006934838.4;refGene_gn=CRYZ;refGene_tx=XM_023258829.1
C1	62746059 62746223
refGene_gn=CRYZ;refGene_tx=XM_006934838.4;refGene_gn=CRYZ;refGene_tx=XM_023258829.1
C1	62750285 62750438
refGene_gn=CRYZ;refGene_tx=XM_006934838.4;refGene_gn=CRYZ;refGene_tx=XM_023258829.1

C1 62751880 62751991
refGene_gn=CRYZ;refGene_tx=XM_006934838.4;refGene_gn=CRYZ;refGene_tx=X
M_023258829.1
C1 62757620 62757794
refGene_gn=TYW3;refGene_tx=XM_003990229.4;refGene_gn=TYW3;refGene_tx=X
M_006934839.3;refGene_gn=TYW3;refGene_tx=XM_006934840.3;refGene_gn=TYW
3;refGene_tx=XM_019837258.1
C1 62760318 62760399
refGene_gn=TYW3;refGene_tx=XM_003990229.4;refGene_gn=TYW3;refGene_tx=X
M_006934839.3;refGene_gn=TYW3;refGene_tx=XM_006934840.3
C1 62764216 62764315
refGene_gn=TYW3;refGene_tx=XM_003990229.4;refGene_gn=TYW3;refGene_tx=X
M_006934839.3;refGene_gn=TYW3;refGene_tx=XM_006934840.3;refGene_gn=TYW
3;refGene_tx=XM_019837258.1
C1 62764545 62764617
refGene_gn=TYW3;refGene_tx=XM_003990229.4;refGene_gn=TYW3;refGene_tx=X
M_006934840.3;refGene_gn=TYW3;refGene_tx=XM_019837258.1
C1 62766818 62766952
refGene_gn=TYW3;refGene_tx=XM_003990229.4;refGene_gn=TYW3;refGene_tx=X
M_006934839.3;refGene_gn=TYW3;refGene_tx=XM_006934840.3;refGene_gn=TYW
3;refGene_tx=XM_019837258.1
C1 62772712 62772791 refGene_gn=TYW3;refGene_tx=XM_006934840.3
C1 62775759 62775973
refGene_gn=TYW3;refGene_tx=XM_003990229.4;refGene_gn=TYW3;refGene_tx=X
M_006934839.3;refGene_gn=TYW3;refGene_tx=XM_019837258.1
C1 63078672 63078752 refGene_gn=LHX8;refGene_tx=XM_023258831.1
C1 63079151 63079308 refGene_gn=LHX8;refGene_tx=XM_023258831.1
C1 63083158 63083280 refGene_gn=LHX8;refGene_tx=XM_023258831.1
C1 63085167 63085388 refGene_gn=LHX8;refGene_tx=XM_023258831.1
C1 63085897 63086001 refGene_gn=LHX8;refGene_tx=XM_023258831.1
C1 63090710 63090806 refGene_gn=LHX8;refGene_tx=XM_023258831.1
C1 63097154 63097338 refGene_gn=LHX8;refGene_tx=XM_023258831.1
C1 63101173 63101250 refGene_gn=LHX8;refGene_tx=XM_023258831.1
C1 63160213 63160320 refGene_gn=SLC44A5;refGene_tx=XM_023258835.1
C1 63163335 63163448
refGene_gn=SLC44A5;refGene_tx=XM_023258832.1;refGene_gn=SLC44A5;refGen
e_tx=XM_023258833.1;refGene_gn=SLC44A5;refGene_tx=XM_023258834.1;refGe
ne_gn=SLC44A5;refGene_tx=XM_023258837.1;refGene_gn=SLC44A5;refGene_tx=
XM_023258838.1
C1 63165700 63165785
refGene_gn=SLC44A5;refGene_tx=XM_023258832.1;refGene_gn=SLC44A5;refGen
e_tx=XM_023258833.1;refGene_gn=SLC44A5;refGene_tx=XM_023258834.1;refGe
ne_gn=SLC44A5;refGene_tx=XM_023258835.1;refGene_gn=SLC44A5;refGene_tx=
XM_023258837.1;refGene_gn=SLC44A5;refGene_tx=XM_023258838.1
C1 63167121 63167210
refGene_gn=SLC44A5;refGene_tx=XM_023258832.1;refGene_gn=SLC44A5;refGen
e_tx=XM_023258833.1;refGene_gn=SLC44A5;refGene_tx=XM_023258834.1;refGe
ne_gn=SLC44A5;refGene_tx=XM_023258835.1;refGene_gn=SLC44A5;refGene_tx=
XM_023258837.1;refGene_gn=SLC44A5;refGene_tx=XM_023258838.1
C1 63167319 63167390

refGene_gn=SLC44A5;refGene_tx=XM_023258832.1;refGene_gn=SLC44A5;refGene_tx=XM_023258833.1;refGene_gn=SLC44A5;refGene_tx=XM_023258834.1;refGene_gn=SLC44A5;refGene_tx=XM_023258835.1;refGene_gn=SLC44A5;refGene_tx=XM_023258837.1;refGene_gn=SLC44A5;refGene_tx=XM_023258838.1
C1 63168252 63168326
refGene_gn=SLC44A5;refGene_tx=XM_023258832.1;refGene_gn=SLC44A5;refGene_tx=XM_023258833.1;refGene_gn=SLC44A5;refGene_tx=XM_023258834.1;refGene_gn=SLC44A5;refGene_tx=XM_023258835.1;refGene_gn=SLC44A5;refGene_tx=XM_023258837.1;refGene_gn=SLC44A5;refGene_tx=XM_023258838.1
C1 63169381 63169485
refGene_gn=SLC44A5;refGene_tx=XM_023258832.1;refGene_gn=SLC44A5;refGene_tx=XM_023258833.1;refGene_gn=SLC44A5;refGene_tx=XM_023258834.1;refGene_gn=SLC44A5;refGene_tx=XM_023258835.1;refGene_gn=SLC44A5;refGene_tx=XM_023258837.1;refGene_gn=SLC44A5;refGene_tx=XM_023258838.1
C1 63176401 63176496
refGene_gn=SLC44A5;refGene_tx=XM_023258832.1;refGene_gn=SLC44A5;refGene_tx=XM_023258833.1;refGene_gn=SLC44A5;refGene_tx=XM_023258834.1;refGene_gn=SLC44A5;refGene_tx=XM_023258835.1;refGene_gn=SLC44A5;refGene_tx=XM_023258837.1;refGene_gn=SLC44A5;refGene_tx=XM_023258838.1
C1 63177150 63177413
refGene_gn=SLC44A5;refGene_tx=XM_023258832.1;refGene_gn=SLC44A5;refGene_tx=XM_023258833.1;refGene_gn=SLC44A5;refGene_tx=XM_023258834.1;refGene_gn=SLC44A5;refGene_tx=XM_023258835.1;refGene_gn=SLC44A5;refGene_tx=XM_023258837.1;refGene_gn=SLC44A5;refGene_tx=XM_023258838.1
C1 63177895 63177983
refGene_gn=SLC44A5;refGene_tx=XM_023258832.1;refGene_gn=SLC44A5;refGene_tx=XM_023258833.1;refGene_gn=SLC44A5;refGene_tx=XM_023258834.1;refGene_gn=SLC44A5;refGene_tx=XM_023258835.1;refGene_gn=SLC44A5;refGene_tx=XM_023258837.1;refGene_gn=SLC44A5;refGene_tx=XM_023258838.1
C1 63178465 63178558
refGene_gn=SLC44A5;refGene_tx=XM_023258832.1;refGene_gn=SLC44A5;refGene_tx=XM_023258833.1;refGene_gn=SLC44A5;refGene_tx=XM_023258834.1;refGene_gn=SLC44A5;refGene_tx=XM_023258835.1;refGene_gn=SLC44A5;refGene_tx=XM_023258837.1;refGene_gn=SLC44A5;refGene_tx=XM_023258838.1
C1 63180979 63181079
refGene_gn=SLC44A5;refGene_tx=XM_023258832.1;refGene_gn=SLC44A5;refGene_tx=XM_023258833.1;refGene_gn=SLC44A5;refGene_tx=XM_023258834.1;refGene_gn=SLC44A5;refGene_tx=XM_023258835.1;refGene_gn=SLC44A5;refGene_tx=XM_023258837.1;refGene_gn=SLC44A5;refGene_tx=XM_023258838.1
C1 63184570 63184702
refGene_gn=SLC44A5;refGene_tx=XM_023258832.1;refGene_gn=SLC44A5;refGene_tx=XM_023258833.1;refGene_gn=SLC44A5;refGene_tx=XM_023258834.1;refGene_gn=SLC44A5;refGene_tx=XM_023258835.1;refGene_gn=SLC44A5;refGene_tx=XM_023258837.1;refGene_gn=SLC44A5;refGene_tx=XM_023258838.1
C1 63193655 63193768
refGene_gn=SLC44A5;refGene_tx=XM_023258832.1;refGene_gn=SLC44A5;refGene_tx=XM_023258833.1;refGene_gn=SLC44A5;refGene_tx=XM_023258834.1;refGene_gn=SLC44A5;refGene_tx=XM_023258835.1;refGene_gn=SLC44A5;refGene_tx=XM_023258837.1;refGene_gn=SLC44A5;refGene_tx=XM_023258838.1
C1 63203211 63203295

refGene_gn=SLC44A5;refGene_tx=XM_023258832.1;refGene_gn=SLC44A5;refGene_tx=XM_023258833.1;refGene_gn=SLC44A5;refGene_tx=XM_023258834.1;refGene_gn=SLC44A5;refGene_tx=XM_023258835.1;refGene_gn=SLC44A5;refGene_tx=XM_023258837.1;refGene_gn=SLC44A5;refGene_tx=XM_023258838.1
C1 63204709 63204833
refGene_gn=SLC44A5;refGene_tx=XM_023258832.1;refGene_gn=SLC44A5;refGene_tx=XM_023258833.1;refGene_gn=SLC44A5;refGene_tx=XM_023258834.1;refGene_gn=SLC44A5;refGene_tx=XM_023258835.1;refGene_gn=SLC44A5;refGene_tx=XM_023258837.1;refGene_gn=SLC44A5;refGene_tx=XM_023258838.1
C1 63208091 63208152
refGene_gn=SLC44A5;refGene_tx=XM_023258832.1;refGene_gn=SLC44A5;refGene_tx=XM_023258833.1;refGene_gn=SLC44A5;refGene_tx=XM_023258834.1;refGene_gn=SLC44A5;refGene_tx=XM_023258835.1;refGene_gn=SLC44A5;refGene_tx=XM_023258837.1;refGene_gn=SLC44A5;refGene_tx=XM_023258838.1
C1 63208984 63209113
refGene_gn=SLC44A5;refGene_tx=XM_023258832.1;refGene_gn=SLC44A5;refGene_tx=XM_023258833.1;refGene_gn=SLC44A5;refGene_tx=XM_023258834.1;refGene_gn=SLC44A5;refGene_tx=XM_023258835.1;refGene_gn=SLC44A5;refGene_tx=XM_023258837.1;refGene_gn=SLC44A5;refGene_tx=XM_023258838.1
C1 63215734 63215819
refGene_gn=SLC44A5;refGene_tx=XM_023258832.1;refGene_gn=SLC44A5;refGene_tx=XM_023258833.1;refGene_gn=SLC44A5;refGene_tx=XM_023258834.1;refGene_gn=SLC44A5;refGene_tx=XM_023258835.1;refGene_gn=SLC44A5;refGene_tx=XM_023258837.1;refGene_gn=SLC44A5;refGene_tx=XM_023258838.1
C1 63241720 63241805
refGene_gn=SLC44A5;refGene_tx=XM_023258832.1;refGene_gn=SLC44A5;refGene_tx=XM_023258833.1;refGene_gn=SLC44A5;refGene_tx=XM_023258834.1;refGene_gn=SLC44A5;refGene_tx=XM_023258835.1;refGene_gn=SLC44A5;refGene_tx=XM_023258837.1;refGene_gn=SLC44A5;refGene_tx=XM_023258838.1
C1 63264321 63264395
refGene_gn=SLC44A5;refGene_tx=XM_023258832.1;refGene_gn=SLC44A5;refGene_tx=XM_023258833.1;refGene_gn=SLC44A5;refGene_tx=XM_023258834.1;refGene_gn=SLC44A5;refGene_tx=XM_023258835.1;refGene_gn=SLC44A5;refGene_tx=XM_023258837.1;refGene_gn=SLC44A5;refGene_tx=XM_023258838.1
C1 63314376 63314431
refGene_gn=SLC44A5;refGene_tx=XM_023258832.1;refGene_gn=SLC44A5;refGene_tx=XM_023258833.1;refGene_gn=SLC44A5;refGene_tx=XM_023258834.1;refGene_gn=SLC44A5;refGene_tx=XM_023258835.1;refGene_gn=SLC44A5;refGene_tx=XM_023258837.1;refGene_gn=SLC44A5;refGene_tx=XM_023258838.1
C1 63349546 63349577 refGene_gn=SLC44A5;refGene_tx=XM_023258837.1
C1 63353012 63353054
refGene_gn=SLC44A5;refGene_tx=XM_023258832.1;refGene_gn=SLC44A5;refGene_tx=XM_023258833.1;refGene_gn=SLC44A5;refGene_tx=XM_023258834.1;refGene_gn=SLC44A5;refGene_tx=XM_023258835.1;refGene_gn=SLC44A5;refGene_tx=XM_023258837.1
C1 63474822 63474835
refGene_gn=SLC44A5;refGene_tx=XM_023258832.1;refGene_gn=SLC44A5;refGene_tx=XM_023258833.1;refGene_gn=SLC44A5;refGene_tx=XM_023258834.1;refGene_gn=SLC44A5;refGene_tx=XM_023258835.1;refGene_gn=SLC44A5;refGene_tx=XM_023258838.1

C1 63581281 63581311 refGene_gn=ACADM;refGene_tx=XM_011284929.3
C1 63585284 63585372 refGene_gn=ACADM;refGene_tx=XM_011284929.3
C1 63588697 63588795 refGene_gn=ACADM;refGene_tx=XM_011284929.3
C1 63588916 63588986 refGene_gn=ACADM;refGene_tx=XM_011284929.3
C1 63590754 63590855 refGene_gn=ACADM;refGene_tx=XM_011284929.3
C1 63591478 63591559 refGene_gn=ACADM;refGene_tx=XM_011284929.3
C1 63598926 63599057 refGene_gn=ACADM;refGene_tx=XM_011284929.3
C1 63602576 63602685 refGene_gn=ACADM;refGene_tx=XM_011284929.3
C1 63606841 63606982 refGene_gn=ACADM;refGene_tx=XM_011284929.3
C1 63607561 63607657 refGene_gn=ACADM;refGene_tx=XM_011284929.3
C1 63611426 63611675 refGene_gn=ACADM;refGene_tx=XM_011284929.3
C1 63613142 63613214 refGene_gn=ACADM;refGene_tx=XM_011284929.3
C1 63623491 63623725
refGene_gn=RABGGTB;refGene_tx=XM_003990237.5;refGene_gn=RABGGTB;refGen
e_tx=XM_006934848.4
C1 63624801 63624909
refGene_gn=RABGGTB;refGene_tx=XM_003990237.5;refGene_gn=RABGGTB;refGen
e_tx=XM_006934848.4
C1 63626132 63626330 refGene_gn=RABGGTB;refGene_tx=XM_003990237.5
C1 63626895 63627001
refGene_gn=RABGGTB;refGene_tx=XM_003990237.5;refGene_gn=RABGGTB;refGen
e_tx=XM_006934848.4
C1 63627843 63627896
refGene_gn=RABGGTB;refGene_tx=XM_003990237.5;refGene_gn=RABGGTB;refGen
e_tx=XM_006934848.4
C1 63627981 63628092
refGene_gn=RABGGTB;refGene_tx=XM_003990237.5;refGene_gn=RABGGTB;refGen
e_tx=XM_006934848.4
C1 63628761 63628887
refGene_gn=RABGGTB;refGene_tx=XM_003990237.5;refGene_gn=RABGGTB;refGen
e_tx=XM_006934848.4
C1 63630676 63630826
refGene_gn=RABGGTB;refGene_tx=XM_003990237.5;refGene_gn=RABGGTB;refGen
e_tx=XM_006934848.4
C1 63631403 63631544
refGene_gn=RABGGTB;refGene_tx=XM_003990237.5;refGene_gn=RABGGTB;refGen
e_tx=XM_006934848.4
C1 63633665 63633906 refGene_gn=MSH4;refGene_tx=XM_011284928.3
C1 63635551 63635734 refGene_gn=MSH4;refGene_tx=XM_011284928.3
C1 63641294 63641455 refGene_gn=MSH4;refGene_tx=XM_011284928.3
C1 63642788 63642899 refGene_gn=MSH4;refGene_tx=XM_011284928.3
C1 63647122 63647238 refGene_gn=MSH4;refGene_tx=XM_011284928.3
C1 63648108 63648282 refGene_gn=MSH4;refGene_tx=XM_011284928.3
C1 63652571 63652744 refGene_gn=MSH4;refGene_tx=XM_011284928.3
C1 63658777 63658845 refGene_gn=MSH4;refGene_tx=XM_011284928.3
C1 63686803 63686878 refGene_gn=MSH4;refGene_tx=XM_011284928.3
C1 63691350 63691415 refGene_gn=MSH4;refGene_tx=XM_011284928.3
C1 63692775 63692945 refGene_gn=MSH4;refGene_tx=XM_011284928.3
C1 63697506 63697643 refGene_gn=MSH4;refGene_tx=XM_011284928.3
C1 63698617 63698721 refGene_gn=MSH4;refGene_tx=XM_011284928.3

C1 63700264 63700389 refGene_gn=MSH4;refGene_tx=XM_011284928.3
C1 63702271 63702472 refGene_gn=MSH4;refGene_tx=XM_011284928.3
C1 63705632 63705751 refGene_gn=MSH4;refGene_tx=XM_011284928.3
C1 63707078 63707207 refGene_gn=MSH4;refGene_tx=XM_011284928.3
C1 63713669 63713844 refGene_gn=MSH4;refGene_tx=XM_011284928.3
C1 63715514 63715603 refGene_gn=MSH4;refGene_tx=XM_011284928.3
C1 63720216 63720408 refGene_gn=MSH4;refGene_tx=XM_011284928.3
C1 63726191 63726398 refGene_gn=ASB17;refGene_tx=XM_003990239.4
C1 63728359 63728639 refGene_gn=ASB17;refGene_tx=XM_003990239.4
C1 63746678 63747079 refGene_gn=ASB17;refGene_tx=XM_003990239.4
C1 63899877 63899895
refGene_gn=ST6GALNAC3;refGene_tx=XM_019837261.1
C1 64036799 64036964
refGene_gn=ST6GALNAC3;refGene_tx=XM_006934850.4;refGene_gn=ST6GALNAC3;
refGene_tx=XM_019837262.2;refGene_gn=ST6GALNAC3;refGene_tx=XM_01983726
3.2
C1 64134363 64134558
refGene_gn=ST6GALNAC3;refGene_tx=XM_006934850.4;refGene_gn=ST6GALNAC3;
refGene_tx=XM_019837261.1;refGene_gn=ST6GALNAC3;refGene_tx=XM_01983726
2.2;refGene_gn=ST6GALNAC3;refGene_tx=XM_019837263.2
C1 64237771 64238181
refGene_gn=ST6GALNAC3;refGene_tx=XM_006934850.4;refGene_gn=ST6GALNAC3;
refGene_tx=XM_019837261.1;refGene_gn=ST6GALNAC3;refGene_tx=XM_01983726
2.2;refGene_gn=ST6GALNAC3;refGene_tx=XM_019837263.2
C1 64346247 64346257
refGene_gn=ST6GALNAC3;refGene_tx=XM_019837263.2
C1 64432210 64432318
refGene_gn=ST6GALNAC3;refGene_tx=XM_006934850.4;refGene_gn=ST6GALNAC3;
refGene_tx=XM_019837261.1;refGene_gn=ST6GALNAC3;refGene_tx=XM_01983726
2.2
C1 64433397 64433584
refGene_gn=ST6GALNAC3;refGene_tx=XM_006934850.4;refGene_gn=ST6GALNAC3;
refGene_tx=XM_019837261.1;refGene_gn=ST6GALNAC3;refGene_tx=XM_01983726
2.2
C1 64645643 64645658
refGene_gn=ST6GALNAC5;refGene_tx=XM_019837264.2
C1 64646499 64646724
refGene_gn=ST6GALNAC5;refGene_tx=XM_019837264.2
C1 64799652 64800062
refGene_gn=ST6GALNAC5;refGene_tx=XM_019837264.2
C1 64805589 64805697
refGene_gn=ST6GALNAC5;refGene_tx=XM_019837264.2
C1 64817544 64817776
refGene_gn=ST6GALNAC5;refGene_tx=XM_019837264.2
C1 64845185 64845302 refGene_gn=PIGK;refGene_tx=XM_003990242.4
C1 64882333 64882418 refGene_gn=PIGK;refGene_tx=XM_003990242.4
C1 64913461 64913634 refGene_gn=PIGK;refGene_tx=XM_003990242.4
C1 64918773 64918884 refGene_gn=PIGK;refGene_tx=XM_003990242.4
C1 64919666 64919784 refGene_gn=PIGK;refGene_tx=XM_003990242.4
C1 64921850 64921947 refGene_gn=PIGK;refGene_tx=XM_003990242.4

C1 64923178 64923290 refGene_gn=PIGK;refGene_tx=XM_003990242.4
C1 64925027 64925163 refGene_gn=PIGK;refGene_tx=XM_003990242.4
C1 64954719 64954811 refGene_gn=PIGK;refGene_tx=XM_003990242.4
C1 64960859 64960913 refGene_gn=PIGK;refGene_tx=XM_003990242.4
C1 64964190 64964283 refGene_gn=PIGK;refGene_tx=XM_003990242.4
C1 65018312 65018372
refGene_gn=AK5;refGene_tx=XM_003990243.5;refGene_gn=AK5;refGene_tx=XM_011284927.3;refGene_gn=AK5;refGene_tx=XM_019837266.2
C1 65022681 65022868
refGene_gn=AK5;refGene_tx=XM_003990243.5;refGene_gn=AK5;refGene_tx=XM_011284927.3;refGene_gn=AK5;refGene_tx=XM_019837266.2;refGene_gn=AK5;refGene_tx=XM_019837267.1
C1 65028458 65028626
refGene_gn=AK5;refGene_tx=XM_003990243.5;refGene_gn=AK5;refGene_tx=XM_011284927.3;refGene_gn=AK5;refGene_tx=XM_019837266.2;refGene_gn=AK5;refGene_tx=XM_019837267.1
C1 65033165 65033335
refGene_gn=AK5;refGene_tx=XM_003990243.5;refGene_gn=AK5;refGene_tx=XM_011284927.3;refGene_gn=AK5;refGene_tx=XM_019837266.2;refGene_gn=AK5;refGene_tx=XM_019837267.1
C1 65033437 65033551
refGene_gn=AK5;refGene_tx=XM_003990243.5;refGene_gn=AK5;refGene_tx=XM_011284927.3;refGene_gn=AK5;refGene_tx=XM_019837266.2;refGene_gn=AK5;refGene_tx=XM_019837267.1
C1 65084186 65084378
refGene_gn=AK5;refGene_tx=XM_003990243.5;refGene_gn=AK5;refGene_tx=XM_011284927.3;refGene_gn=AK5;refGene_tx=XM_019837266.2;refGene_gn=AK5;refGene_tx=XM_019837267.1
C1 65153168 65153259
refGene_gn=AK5;refGene_tx=XM_003990243.5;refGene_gn=AK5;refGene_tx=XM_011284927.3;refGene_gn=AK5;refGene_tx=XM_019837266.2;refGene_gn=AK5;refGene_tx=XM_019837267.1
C1 65162154 65162231
refGene_gn=AK5;refGene_tx=XM_003990243.5;refGene_gn=AK5;refGene_tx=XM_011284927.3;refGene_gn=AK5;refGene_tx=XM_019837266.2;refGene_gn=AK5;refGene_tx=XM_019837267.1
C1 65206234 65206277
refGene_gn=AK5;refGene_tx=XM_003990243.5;refGene_gn=AK5;refGene_tx=XM_011284927.3;refGene_gn=AK5;refGene_tx=XM_019837266.2;refGene_gn=AK5;refGene_tx=XM_019837267.1
C1 65211368 65211413
refGene_gn=AK5;refGene_tx=XM_003990243.5;refGene_gn=AK5;refGene_tx=XM_011284927.3;refGene_gn=AK5;refGene_tx=XM_019837267.1
C1 65238891 65239055
refGene_gn=AK5;refGene_tx=XM_003990243.5;refGene_gn=AK5;refGene_tx=XM_011284927.3;refGene_gn=AK5;refGene_tx=XM_019837266.2;refGene_gn=AK5;refGene_tx=XM_019837267.1
C1 65240965 65241082
refGene_gn=AK5;refGene_tx=XM_003990243.5;refGene_gn=AK5;refGene_tx=XM_011284927.3;refGene_gn=AK5;refGene_tx=XM_019837266.2;refGene_gn=AK5;re

fGene_tx=XM_019837267.1
C1 65259737 65259929
refGene_gn=AK5;refGene_tx=XM_003990243.5;refGene_gn=AK5;refGene_tx=XM_011284927.3;refGene_gn=AK5;refGene_tx=XM_019837266.2;refGene_gn=AK5;refGene_tx=XM_019837267.1
C1 65279879 65279948
refGene_gn=AK5;refGene_tx=XM_003990243.5;refGene_gn=AK5;refGene_tx=XM_011284927.3;refGene_gn=AK5;refGene_tx=XM_019837266.2;refGene_gn=AK5;refGene_tx=XM_019837267.1
C1 65286663 65286808
refGene_gn=ZZZ3;refGene_tx=XM_011284924.2;refGene_gn=ZZZ3;refGene_tx=XM_011284926.2;refGene_gn=ZZZ3;refGene_tx=XM_019837265.1
C1 65287112 65287213
refGene_gn=ZZZ3;refGene_tx=XM_011284924.2;refGene_gn=ZZZ3;refGene_tx=XM_011284926.2;refGene_gn=ZZZ3;refGene_tx=XM_019837265.1
C1 65290437 65290572
refGene_gn=ZZZ3;refGene_tx=XM_011284924.2;refGene_gn=ZZZ3;refGene_tx=XM_011284926.2;refGene_gn=ZZZ3;refGene_tx=XM_019837265.1
C1 65296116 65296269
refGene_gn=ZZZ3;refGene_tx=XM_011284924.2;refGene_gn=ZZZ3;refGene_tx=XM_011284926.2;refGene_gn=ZZZ3;refGene_tx=XM_019837265.1
C1 65298666 65298762
refGene_gn=ZZZ3;refGene_tx=XM_011284924.2;refGene_gn=ZZZ3;refGene_tx=XM_011284926.2;refGene_gn=ZZZ3;refGene_tx=XM_019837265.1
C1 65299410 65299512
refGene_gn=ZZZ3;refGene_tx=XM_011284924.2;refGene_gn=ZZZ3;refGene_tx=XM_011284926.2;refGene_gn=ZZZ3;refGene_tx=XM_019837265.1
C1 65300558 65300630
refGene_gn=ZZZ3;refGene_tx=XM_011284924.2;refGene_gn=ZZZ3;refGene_tx=XM_011284926.2;refGene_gn=ZZZ3;refGene_tx=XM_019837265.1
C1 65301318 65301434
refGene_gn=ZZZ3;refGene_tx=XM_011284924.2;refGene_gn=ZZZ3;refGene_tx=XM_011284926.2;refGene_gn=ZZZ3;refGene_tx=XM_019837265.1
C1 65301518 65301666
refGene_gn=ZZZ3;refGene_tx=XM_011284924.2;refGene_gn=ZZZ3;refGene_tx=XM_011284926.2;refGene_gn=ZZZ3;refGene_tx=XM_019837265.1
C1 65305212 65305351
refGene_gn=ZZZ3;refGene_tx=XM_011284924.2;refGene_gn=ZZZ3;refGene_tx=XM_011284926.2;refGene_gn=ZZZ3;refGene_tx=XM_019837265.1
C1 65341389 65342894
refGene_gn=ZZZ3;refGene_tx=XM_011284924.2;refGene_gn=ZZZ3;refGene_tx=XM_011284926.2;refGene_gn=ZZZ3;refGene_tx=XM_019837265.1
C1 65399930 65400088
refGene_gn=USP33;refGene_tx=XM_006934853.4;refGene_gn=USP33;refGene_tx=XM_011284923.3;refGene_gn=USP33;refGene_tx=XM_019837268.2;refGene_gn=USP33;refGene_tx=XM_019837269.2
C1 65400460 65400529
refGene_gn=USP33;refGene_tx=XM_006934853.4;refGene_gn=USP33;refGene_tx=XM_011284923.3;refGene_gn=USP33;refGene_tx=XM_019837268.2;refGene_gn=USP33;refGene_tx=XM_019837269.2

C1 65402252 65402355
refGene_gn=USP33;refGene_tx=XM_006934853.4;refGene_gn=USP33;refGene_tx=XM_011284923.3;refGene_gn=USP33;refGene_tx=XM_019837268.2;refGene_gn=USP33;refGene_tx=XM_019837269.2
C1 65412322 65412431
refGene_gn=USP33;refGene_tx=XM_006934853.4;refGene_gn=USP33;refGene_tx=XM_011284923.3;refGene_gn=USP33;refGene_tx=XM_019837268.2;refGene_gn=USP33;refGene_tx=XM_019837269.2
C1 65413576 65413658
refGene_gn=USP33;refGene_tx=XM_006934853.4;refGene_gn=USP33;refGene_tx=XM_011284923.3;refGene_gn=USP33;refGene_tx=XM_019837268.2;refGene_gn=USP33;refGene_tx=XM_019837269.2
C1 65414775 65414945
refGene_gn=USP33;refGene_tx=XM_006934853.4;refGene_gn=USP33;refGene_tx=XM_011284923.3;refGene_gn=USP33;refGene_tx=XM_019837268.2;refGene_gn=USP33;refGene_tx=XM_019837269.2
C1 65415619 65415746
refGene_gn=USP33;refGene_tx=XM_006934853.4;refGene_gn=USP33;refGene_tx=XM_011284923.3;refGene_gn=USP33;refGene_tx=XM_019837268.2;refGene_gn=USP33;refGene_tx=XM_019837269.2
C1 65417080 65417261
refGene_gn=USP33;refGene_tx=XM_006934853.4;refGene_gn=USP33;refGene_tx=XM_011284923.3;refGene_gn=USP33;refGene_tx=XM_019837268.2;refGene_gn=USP33;refGene_tx=XM_019837269.2
C1 65417919 65417965
refGene_gn=USP33;refGene_tx=XM_006934853.4;refGene_gn=USP33;refGene_tx=XM_011284923.3;refGene_gn=USP33;refGene_tx=XM_019837268.2;refGene_gn=USP33;refGene_tx=XM_019837269.2
C1 65420472 65420506
refGene_gn=USP33;refGene_tx=XM_006934853.4;refGene_gn=USP33;refGene_tx=XM_011284923.3;refGene_gn=USP33;refGene_tx=XM_019837268.2;refGene_gn=USP33;refGene_tx=XM_019837269.2
C1 65421126 65421221
refGene_gn=USP33;refGene_tx=XM_006934853.4;refGene_gn=USP33;refGene_tx=XM_011284923.3;refGene_gn=USP33;refGene_tx=XM_019837268.2;refGene_gn=USP33;refGene_tx=XM_019837269.2
C1 65421328 65421501
refGene_gn=USP33;refGene_tx=XM_006934853.4;refGene_gn=USP33;refGene_tx=XM_011284923.3;refGene_gn=USP33;refGene_tx=XM_019837268.2;refGene_gn=USP33;refGene_tx=XM_019837269.2
C1 65422832 65422948
refGene_gn=USP33;refGene_tx=XM_006934853.4;refGene_gn=USP33;refGene_tx=XM_011284923.3;refGene_gn=USP33;refGene_tx=XM_019837268.2;refGene_gn=USP33;refGene_tx=XM_019837269.2
C1 65424669 65424810
refGene_gn=USP33;refGene_tx=XM_006934853.4;refGene_gn=USP33;refGene_tx=XM_011284923.3;refGene_gn=USP33;refGene_tx=XM_019837268.2
C1 65427409 65427827
refGene_gn=USP33;refGene_tx=XM_006934853.4;refGene_gn=USP33;refGene_tx=XM_011284923.3;refGene_gn=USP33;refGene_tx=XM_019837268.2;refGene_gn=

USP33;refGene_tx=XM_019837269.2
C1	65429289	65429368
refGene_gn=USP33;refGene_tx=XM_006934853.4;refGene_gn=USP33;refGene_tx=XM_011284923.3;refGene_gn=USP33;refGene_tx=XM_019837268.2;refGene_gn=USP33;refGene_tx=XM_019837269.2
C1	65433138	65433252
refGene_gn=USP33;refGene_tx=XM_006934853.4;refGene_gn=USP33;refGene_tx=XM_011284923.3;refGene_gn=USP33;refGene_tx=XM_019837268.2;refGene_gn=USP33;refGene_tx=XM_019837269.2
C1	65436195	65436265
refGene_gn=USP33;refGene_tx=XM_006934853.4;refGene_gn=USP33;refGene_tx=XM_011284923.3;refGene_gn=USP33;refGene_tx=XM_019837268.2;refGene_gn=USP33;refGene_tx=XM_019837269.2
C1	65437181	65437284
refGene_gn=USP33;refGene_tx=XM_006934853.4;refGene_gn=USP33;refGene_tx=XM_011284923.3;refGene_gn=USP33;refGene_tx=XM_019837268.2;refGene_gn=USP33;refGene_tx=XM_019837269.2
C1	65440789	65440942
refGene_gn=USP33;refGene_tx=XM_006934853.4;refGene_gn=USP33;refGene_tx=XM_011284923.3;refGene_gn=USP33;refGene_tx=XM_019837268.2;refGene_gn=USP33;refGene_tx=XM_019837269.2
C1	65442913	65442976
refGene_gn=USP33;refGene_tx=XM_006934853.4;refGene_gn=USP33;refGene_tx=XM_011284923.3;refGene_gn=USP33;refGene_tx=XM_019837268.2;refGene_gn=USP33;refGene_tx=XM_019837269.2
C1	65444331	65444385
refGene_gn=USP33;refGene_tx=XM_006934853.4;refGene_gn=USP33;refGene_tx=XM_011284923.3;refGene_gn=USP33;refGene_tx=XM_019837268.2;refGene_gn=USP33;refGene_tx=XM_019837269.2
C1	65445077	65445158
refGene_gn=USP33;refGene_tx=XM_006934853.4;refGene_gn=USP33;refGene_tx=XM_011284923.3;refGene_gn=USP33;refGene_tx=XM_019837268.2;refGene_gn=USP33;refGene_tx=XM_019837269.2
C1	65476774	65476873
refGene_gn=MIGA1;refGene_tx=XM_011284922.3;refGene_gn=MIGA1;refGene_tx=XM_019837271.2
C1	65496672	65496850
refGene_gn=MIGA1;refGene_tx=XM_011284922.3;refGene_gn=MIGA1;refGene_tx=XM_019837270.2;refGene_gn=MIGA1;refGene_tx=XM_019837271.2
C1	65498569	65498706
refGene_gn=MIGA1;refGene_tx=XM_011284922.3;refGene_gn=MIGA1;refGene_tx=XM_019837270.2;refGene_gn=MIGA1;refGene_tx=XM_019837271.2
C1	65501377	65501504
refGene_gn=MIGA1;refGene_tx=XM_011284922.3;refGene_gn=MIGA1;refGene_tx=XM_019837270.2;refGene_gn=MIGA1;refGene_tx=XM_019837271.2
C1	65504647	65504781
refGene_gn=MIGA1;refGene_tx=XM_011284922.3;refGene_gn=MIGA1;refGene_tx=XM_019837270.2
C1	65505745	65505869
refGene_gn=MIGA1;refGene_tx=XM_011284922.3;refGene_gn=MIGA1;refGene_tx

=XM_019837270.2
C1	65540877	65540978
refGene_gn=MIGA1;refGene_tx=XM_011284922.3;refGene_gn=MIGA1;refGene_tx=XM_019837270.2;refGene_gn=MIGA1;refGene_tx=XM_019837271.2
C1	65561476	65561595
refGene_gn=MIGA1;refGene_tx=XM_011284922.3;refGene_gn=MIGA1;refGene_tx=XM_019837270.2;refGene_gn=MIGA1;refGene_tx=XM_019837271.2
C1	65561851	65561924
refGene_gn=MIGA1;refGene_tx=XM_011284922.3;refGene_gn=MIGA1;refGene_tx=XM_019837270.2;refGene_gn=MIGA1;refGene_tx=XM_019837271.2
C1	65562611	65562698
refGene_gn=MIGA1;refGene_tx=XM_011284922.3;refGene_gn=MIGA1;refGene_tx=XM_019837270.2;refGene_gn=MIGA1;refGene_tx=XM_019837271.2
C1	65564986	65565085
refGene_gn=MIGA1;refGene_tx=XM_011284922.3;refGene_gn=MIGA1;refGene_tx=XM_019837270.2;refGene_gn=MIGA1;refGene_tx=XM_019837271.2
C1	65567156	65567291
refGene_gn=MIGA1;refGene_tx=XM_011284922.3;refGene_gn=MIGA1;refGene_tx=XM_019837270.2;refGene_gn=MIGA1;refGene_tx=XM_019837271.2
C1	65568958	65569012
refGene_gn=MIGA1;refGene_tx=XM_011284922.3;refGene_gn=MIGA1;refGene_tx=XM_019837270.2;refGene_gn=MIGA1;refGene_tx=XM_019837271.2
C1	65570828	65570945
refGene_gn=MIGA1;refGene_tx=XM_011284922.3;refGene_gn=MIGA1;refGene_tx=XM_019837270.2;refGene_gn=MIGA1;refGene_tx=XM_019837271.2
C1	65574102	65574321
refGene_gn=MIGA1;refGene_tx=XM_011284922.3;refGene_gn=MIGA1;refGene_tx=XM_019837270.2;refGene_gn=MIGA1;refGene_tx=XM_019837271.2
C1	65610185	65610212
refGene_gn=NEXN;refGene_tx=XM_023258840.1;refGene_gn=NEXN;refGene_tx=XM_023258841.1;refGene_gn=NEXN;refGene_tx=XM_023258842.1;refGene_gn=NEXN;refGene_tx=XM_023258843.1;refGene_gn=NEXN;refGene_tx=XM_023258844.1
C1	65611624	65611816
refGene_gn=NEXN;refGene_tx=XM_023258840.1;refGene_gn=NEXN;refGene_tx=XM_023258841.1;refGene_gn=NEXN;refGene_tx=XM_023258842.1
C1	65612008	65612087
refGene_gn=NEXN;refGene_tx=XM_023258840.1;refGene_gn=NEXN;refGene_tx=XM_023258841.1;refGene_gn=NEXN;refGene_tx=XM_023258842.1;refGene_gn=NEXN;refGene_tx=XM_023258843.1;refGene_gn=NEXN;refGene_tx=XM_023258844.1
C1	65612171	65612320
refGene_gn=NEXN;refGene_tx=XM_023258840.1;refGene_gn=NEXN;refGene_tx=XM_023258841.1;refGene_gn=NEXN;refGene_tx=XM_023258842.1;refGene_gn=NEXN;refGene_tx=XM_023258843.1;refGene_gn=NEXN;refGene_tx=XM_023258844.1
C1	65618397	65618439
refGene_gn=NEXN;refGene_tx=XM_023258840.1;refGene_gn=NEXN;refGene_tx=XM_023258841.1;refGene_gn=NEXN;refGene_tx=XM_023258843.1;refGene_gn=NEXN;refGene_tx=XM_023258844.1
C1	65619665	65619863
refGene_gn=NEXN;refGene_tx=XM_023258840.1;refGene_gn=NEXN;refGene_tx=XM_023258841.1;refGene_gn=NEXN;refGene_tx=XM_023258842.1;refGene_gn=NEX

N;refGene_tx=XM_023258843.1;refGene_gn=NEXN;refGene_tx=XM_023258844.1
C1 65619980 65620157
refGene_gn=NEXN;refGene_tx=XM_023258840.1;refGene_gn=NEXN;refGene_tx=XM_023258841.1;refGene_gn=NEXN;refGene_tx=XM_023258842.1;refGene_gn=NEXN;refGene_tx=XM_023258843.1;refGene_gn=NEXN;refGene_tx=XM_023258844.1
C1 65621625 65621814
refGene_gn=NEXN;refGene_tx=XM_023258840.1;refGene_gn=NEXN;refGene_tx=XM_023258841.1;refGene_gn=NEXN;refGene_tx=XM_023258842.1;refGene_gn=NEXN;refGene_tx=XM_023258843.1;refGene_gn=NEXN;refGene_tx=XM_023258844.1
C1 65625449 65625647
refGene_gn=NEXN;refGene_tx=XM_023258840.1;refGene_gn=NEXN;refGene_tx=XM_023258841.1;refGene_gn=NEXN;refGene_tx=XM_023258842.1;refGene_gn=NEXN;refGene_tx=XM_023258843.1;refGene_gn=NEXN;refGene_tx=XM_023258844.1
C1 65629707 65629929
refGene_gn=NEXN;refGene_tx=XM_023258840.1;refGene_gn=NEXN;refGene_tx=XM_023258841.1;refGene_gn=NEXN;refGene_tx=XM_023258842.1;refGene_gn=NEXN;refGene_tx=XM_023258843.1;refGene_gn=NEXN;refGene_tx=XM_023258844.1
C1 65634603 65634789
refGene_gn=NEXN;refGene_tx=XM_023258840.1;refGene_gn=NEXN;refGene_tx=XM_023258841.1;refGene_gn=NEXN;refGene_tx=XM_023258842.1;refGene_gn=NEXN;refGene_tx=XM_023258843.1;refGene_gn=NEXN;refGene_tx=XM_023258844.1
C1 65635237 65635609
refGene_gn=NEXN;refGene_tx=XM_023258840.1;refGene_gn=NEXN;refGene_tx=XM_023258841.1;refGene_gn=NEXN;refGene_tx=XM_023258842.1;refGene_gn=NEXN;refGene_tx=XM_023258843.1;refGene_gn=NEXN;refGene_tx=XM_023258844.1
C1 65635688 65635702
refGene_gn=NEXN;refGene_tx=XM_023258840.1;refGene_gn=NEXN;refGene_tx=XM_023258842.1;refGene_gn=NEXN;refGene_tx=XM_023258843.1
C1 65640124 65640130 refGene_gn=FUBP1;refGene_tx=XM_023258839.1
C1 65641350 65641359 refGene_gn=FUBP1;refGene_tx=XM_011284916.3
C1 65641457 65641499
refGene_gn=FUBP1;refGene_tx=XM_019837273.1;refGene_gn=FUBP1;refGene_tx=XM_019837274.1;refGene_gn=FUBP1;refGene_tx=XM_019837275.1;refGene_gn=FUBP1;refGene_tx=XM_019837276.1;refGene_gn=FUBP1;refGene_tx=XM_019837277.1;refGene_gn=FUBP1;refGene_tx=XM_019837278.1;refGene_gn=FUBP1;refGene_tx=XM_019837279.1;refGene_gn=FUBP1;refGene_tx=XM_019837280.1;refGene_gn=FUBP1;refGene_tx=XM_019837281.1;refGene_gn=FUBP1;refGene_tx=XM_019837282.1
C1 65641742 65641888
refGene_gn=FUBP1;refGene_tx=XM_011284916.3;refGene_gn=FUBP1;refGene_tx=XM_019837273.1;refGene_gn=FUBP1;refGene_tx=XM_019837274.1;refGene_gn=FUBP1;refGene_tx=XM_019837275.1;refGene_gn=FUBP1;refGene_tx=XM_019837276.1;refGene_gn=FUBP1;refGene_tx=XM_019837277.1;refGene_gn=FUBP1;refGene_tx=XM_019837278.1;refGene_gn=FUBP1;refGene_tx=XM_019837279.1;refGene_gn=FUBP1;refGene_tx=XM_019837280.1;refGene_gn=FUBP1;refGene_tx=XM_019837281.1;refGene_gn=FUBP1;refGene_tx=XM_019837282.1;refGene_gn=FUBP1;refGene_tx=XM_023258839.1
C1 65641968 65642028
refGene_gn=FUBP1;refGene_tx=XM_011284916.3;refGene_gn=FUBP1;refGene_tx=XM_019837273.1;refGene_gn=FUBP1;refGene_tx=XM_019837274.1;refGene_gn=

FUBP1;refGene_tx=XM_019837275.1;refGene_gn=FUBP1;refGene_tx=XM_019837277.1;refGene_gn=FUBP1;refGene_tx=XM_019837278.1;refGene_gn=FUBP1;refGene_tx=XM_023258839.1
C1 65648062 65648137
refGene_gn=FUBP1;refGene_tx=XM_011284916.3;refGene_gn=FUBP1;refGene_tx=XM_019837273.1;refGene_gn=FUBP1;refGene_tx=XM_019837274.1;refGene_gn=FUBP1;refGene_tx=XM_019837275.1;refGene_gn=FUBP1;refGene_tx=XM_019837276.1;refGene_gn=FUBP1;refGene_tx=XM_019837277.1;refGene_gn=FUBP1;refGene_tx=XM_019837278.1;refGene_gn=FUBP1;refGene_tx=XM_019837279.1;refGene_gn=FUBP1;refGene_tx=XM_019837280.1;refGene_gn=FUBP1;refGene_tx=XM_019837281.1;refGene_gn=FUBP1;refGene_tx=XM_019837282.1;refGene_gn=FUBP1;refGene_tx=XM_023258839.1
C1 65650097 65650235
refGene_gn=FUBP1;refGene_tx=XM_011284916.3;refGene_gn=FUBP1;refGene_tx=XM_019837273.1;refGene_gn=FUBP1;refGene_tx=XM_019837274.1;refGene_gn=FUBP1;refGene_tx=XM_019837275.1;refGene_gn=FUBP1;refGene_tx=XM_019837276.1;refGene_gn=FUBP1;refGene_tx=XM_019837277.1;refGene_gn=FUBP1;refGene_tx=XM_019837278.1;refGene_gn=FUBP1;refGene_tx=XM_019837279.1;refGene_gn=FUBP1;refGene_tx=XM_019837280.1;refGene_gn=FUBP1;refGene_tx=XM_019837281.1;refGene_gn=FUBP1;refGene_tx=XM_019837282.1;refGene_gn=FUBP1;refGene_tx=XM_023258839.1
C1 65653313 65653393
refGene_gn=FUBP1;refGene_tx=XM_011284916.3;refGene_gn=FUBP1;refGene_tx=XM_019837273.1;refGene_gn=FUBP1;refGene_tx=XM_019837274.1;refGene_gn=FUBP1;refGene_tx=XM_019837275.1;refGene_gn=FUBP1;refGene_tx=XM_019837276.1;refGene_gn=FUBP1;refGene_tx=XM_019837277.1;refGene_gn=FUBP1;refGene_tx=XM_019837278.1;refGene_gn=FUBP1;refGene_tx=XM_019837279.1;refGene_gn=FUBP1;refGene_tx=XM_019837280.1;refGene_gn=FUBP1;refGene_tx=XM_019837281.1;refGene_gn=FUBP1;refGene_tx=XM_019837282.1;refGene_gn=FUBP1;refGene_tx=XM_023258839.1
C1 65653480 65653632
refGene_gn=FUBP1;refGene_tx=XM_011284916.3;refGene_gn=FUBP1;refGene_tx=XM_019837273.1;refGene_gn=FUBP1;refGene_tx=XM_019837274.1;refGene_gn=FUBP1;refGene_tx=XM_019837275.1;refGene_gn=FUBP1;refGene_tx=XM_019837276.1;refGene_gn=FUBP1;refGene_tx=XM_019837277.1;refGene_gn=FUBP1;refGene_tx=XM_019837278.1;refGene_gn=FUBP1;refGene_tx=XM_019837279.1;refGene_gn=FUBP1;refGene_tx=XM_019837280.1;refGene_gn=FUBP1;refGene_tx=XM_019837281.1;refGene_gn=FUBP1;refGene_tx=XM_019837282.1;refGene_gn=FUBP1;refGene_tx=XM_023258839.1
C1 65657018 65657179
refGene_gn=FUBP1;refGene_tx=XM_011284916.3;refGene_gn=FUBP1;refGene_tx=XM_019837273.1;refGene_gn=FUBP1;refGene_tx=XM_019837274.1;refGene_gn=FUBP1;refGene_tx=XM_019837275.1;refGene_gn=FUBP1;refGene_tx=XM_019837276.1;refGene_gn=FUBP1;refGene_tx=XM_019837277.1;refGene_gn=FUBP1;refGene_tx=XM_019837278.1;refGene_gn=FUBP1;refGene_tx=XM_019837279.1;refGene_gn=FUBP1;refGene_tx=XM_019837280.1;refGene_gn=FUBP1;refGene_tx=XM_019837281.1;refGene_gn=FUBP1;refGene_tx=XM_019837282.1;refGene_gn=FUBP1;refGene_tx=XM_023258839.1
C1 65657818 65657960
refGene_gn=FUBP1;refGene_tx=XM_011284916.3;refGene_gn=FUBP1;refGene_tx

=XM_019837273.1;refGene_gn=FUBP1;refGene_tx=XM_019837274.1;refGene_gn=FUBP1;refGene_tx=XM_019837275.1;refGene_gn=FUBP1;refGene_tx=XM_019837276.1;refGene_gn=FUBP1;refGene_tx=XM_019837277.1;refGene_gn=FUBP1;refGene_tx=XM_019837278.1;refGene_gn=FUBP1;refGene_tx=XM_019837279.1;refGene_gn=FUBP1;refGene_tx=XM_019837280.1;refGene_gn=FUBP1;refGene_tx=XM_019837281.1;refGene_gn=FUBP1;refGene_tx=XM_019837282.1;refGene_gn=FUBP1;refGene_tx=XM_023258839.1
C1 65658450 65658551
refGene_gn=FUBP1;refGene_tx=XM_011284916.3;refGene_gn=FUBP1;refGene_tx=XM_019837273.1;refGene_gn=FUBP1;refGene_tx=XM_019837274.1;refGene_gn=FUBP1;refGene_tx=XM_019837275.1;refGene_gn=FUBP1;refGene_tx=XM_019837276.1;refGene_gn=FUBP1;refGene_tx=XM_019837277.1;refGene_gn=FUBP1;refGene_tx=XM_019837278.1;refGene_gn=FUBP1;refGene_tx=XM_019837279.1;refGene_gn=FUBP1;refGene_tx=XM_019837280.1;refGene_gn=FUBP1;refGene_tx=XM_019837281.1;refGene_gn=FUBP1;refGene_tx=XM_019837282.1;refGene_gn=FUBP1;refGene_tx=XM_023258839.1
C1 65658647 65658750
refGene_gn=FUBP1;refGene_tx=XM_011284916.3;refGene_gn=FUBP1;refGene_tx=XM_019837273.1;refGene_gn=FUBP1;refGene_tx=XM_019837274.1;refGene_gn=FUBP1;refGene_tx=XM_019837275.1;refGene_gn=FUBP1;refGene_tx=XM_019837276.1;refGene_gn=FUBP1;refGene_tx=XM_019837277.1;refGene_gn=FUBP1;refGene_tx=XM_019837278.1;refGene_gn=FUBP1;refGene_tx=XM_019837279.1;refGene_gn=FUBP1;refGene_tx=XM_019837280.1;refGene_gn=FUBP1;refGene_tx=XM_019837281.1;refGene_gn=FUBP1;refGene_tx=XM_019837282.1;refGene_gn=FUBP1;refGene_tx=XM_023258839.1
C1 65659007 65659109
refGene_gn=FUBP1;refGene_tx=XM_011284916.3;refGene_gn=FUBP1;refGene_tx=XM_019837273.1;refGene_gn=FUBP1;refGene_tx=XM_019837274.1;refGene_gn=FUBP1;refGene_tx=XM_019837275.1;refGene_gn=FUBP1;refGene_tx=XM_019837276.1;refGene_gn=FUBP1;refGene_tx=XM_019837277.1;refGene_gn=FUBP1;refGene_tx=XM_019837278.1;refGene_gn=FUBP1;refGene_tx=XM_019837279.1;refGene_gn=FUBP1;refGene_tx=XM_019837280.1;refGene_gn=FUBP1;refGene_tx=XM_019837281.1;refGene_gn=FUBP1;refGene_tx=XM_019837282.1;refGene_gn=FUBP1;refGene_tx=XM_023258839.1
C1 65659237 65659336
refGene_gn=FUBP1;refGene_tx=XM_011284916.3;refGene_gn=FUBP1;refGene_tx=XM_019837273.1;refGene_gn=FUBP1;refGene_tx=XM_019837274.1;refGene_gn=FUBP1;refGene_tx=XM_019837275.1;refGene_gn=FUBP1;refGene_tx=XM_019837276.1;refGene_gn=FUBP1;refGene_tx=XM_019837277.1;refGene_gn=FUBP1;refGene_tx=XM_019837278.1;refGene_gn=FUBP1;refGene_tx=XM_019837279.1;refGene_gn=FUBP1;refGene_tx=XM_019837280.1;refGene_gn=FUBP1;refGene_tx=XM_019837281.1;refGene_gn=FUBP1;refGene_tx=XM_019837282.1;refGene_gn=FUBP1;refGene_tx=XM_023258839.1
C1 65659431 65659594
refGene_gn=FUBP1;refGene_tx=XM_011284916.3;refGene_gn=FUBP1;refGene_tx=XM_019837273.1;refGene_gn=FUBP1;refGene_tx=XM_019837274.1;refGene_gn=FUBP1;refGene_tx=XM_019837275.1;refGene_gn=FUBP1;refGene_tx=XM_019837276.1;refGene_gn=FUBP1;refGene_tx=XM_019837277.1;refGene_gn=FUBP1;refGene_tx=XM_019837278.1;refGene_gn=FUBP1;refGene_tx=XM_019837279.1;refGene_gn=FUBP1;refGene_tx=XM_019837280.1;refGene_gn=FUBP1;refGene_tx=XM_01

9837281.1;refGene_gn=FUBP1;refGene_tx=XM_019837282.1;refGene_gn=FUBP1;
refGene_tx=XM_023258839.1
C1 65661525 65661583
refGene_gn=FUBP1;refGene_tx=XM_011284916.3;refGene_gn=FUBP1;refGene_tx
=XM_019837273.1;refGene_gn=FUBP1;refGene_tx=XM_019837274.1;refGene_gn=
FUBP1;refGene_tx=XM_019837275.1;refGene_gn=FUBP1;refGene_tx=XM_0198372
76.1;refGene_gn=FUBP1;refGene_tx=XM_019837277.1;refGene_gn=FUBP1;refGe
ne_tx=XM_019837278.1;refGene_gn=FUBP1;refGene_tx=XM_019837279.1;refGen
e_gn=FUBP1;refGene_tx=XM_019837280.1;refGene_gn=FUBP1;refGene_tx=XM_01
9837281.1;refGene_gn=FUBP1;refGene_tx=XM_019837282.1;refGene_gn=FUBP1;
refGene_tx=XM_023258839.1
C1 65661709 65661781
refGene_gn=FUBP1;refGene_tx=XM_011284916.3;refGene_gn=FUBP1;refGene_tx
=XM_019837273.1;refGene_gn=FUBP1;refGene_tx=XM_019837274.1;refGene_gn=
FUBP1;refGene_tx=XM_019837275.1;refGene_gn=FUBP1;refGene_tx=XM_0198372
76.1;refGene_gn=FUBP1;refGene_tx=XM_019837277.1;refGene_gn=FUBP1;refGe
ne_tx=XM_019837278.1;refGene_gn=FUBP1;refGene_tx=XM_019837279.1;refGen
e_gn=FUBP1;refGene_tx=XM_019837280.1;refGene_gn=FUBP1;refGene_tx=XM_01
9837281.1;refGene_gn=FUBP1;refGene_tx=XM_019837282.1;refGene_gn=FUBP1;
refGene_tx=XM_023258839.1
C1 65661867 65661920
refGene_gn=FUBP1;refGene_tx=XM_011284916.3;refGene_gn=FUBP1;refGene_tx
=XM_019837273.1;refGene_gn=FUBP1;refGene_tx=XM_019837274.1;refGene_gn=
FUBP1;refGene_tx=XM_019837275.1;refGene_gn=FUBP1;refGene_tx=XM_0198372
76.1;refGene_gn=FUBP1;refGene_tx=XM_019837277.1;refGene_gn=FUBP1;refGe
ne_tx=XM_019837278.1;refGene_gn=FUBP1;refGene_tx=XM_019837279.1;refGen
e_gn=FUBP1;refGene_tx=XM_019837280.1;refGene_gn=FUBP1;refGene_tx=XM_01
9837281.1;refGene_gn=FUBP1;refGene_tx=XM_019837282.1;refGene_gn=FUBP1;
refGene_tx=XM_023258839.1
C1 65662414 65662454
refGene_gn=FUBP1;refGene_tx=XM_011284916.3;refGene_gn=FUBP1;refGene_tx
=XM_019837273.1;refGene_gn=FUBP1;refGene_tx=XM_019837274.1;refGene_gn=
FUBP1;refGene_tx=XM_019837275.1;refGene_gn=FUBP1;refGene_tx=XM_0198372
76.1;refGene_gn=FUBP1;refGene_tx=XM_019837277.1;refGene_gn=FUBP1;refGe
ne_tx=XM_019837278.1;refGene_gn=FUBP1;refGene_tx=XM_019837279.1;refGen
e_gn=FUBP1;refGene_tx=XM_019837280.1;refGene_gn=FUBP1;refGene_tx=XM_01
9837281.1;refGene_gn=FUBP1;refGene_tx=XM_019837282.1;refGene_gn=FUBP1;
refGene_tx=XM_023258839.1
C1 65662924 65662963
refGene_gn=FUBP1;refGene_tx=XM_011284916.3;refGene_gn=FUBP1;refGene_tx
=XM_019837273.1;refGene_gn=FUBP1;refGene_tx=XM_019837274.1;refGene_gn=
FUBP1;refGene_tx=XM_019837275.1;refGene_gn=FUBP1;refGene_tx=XM_0198372
76.1;refGene_gn=FUBP1;refGene_tx=XM_019837277.1;refGene_gn=FUBP1;refGe
ne_tx=XM_019837278.1;refGene_gn=FUBP1;refGene_tx=XM_019837279.1;refGen
e_gn=FUBP1;refGene_tx=XM_019837280.1;refGene_gn=FUBP1;refGene_tx=XM_01
9837281.1;refGene_gn=FUBP1;refGene_tx=XM_019837282.1;refGene_gn=FUBP1;
refGene_tx=XM_023258839.1
C1 65663779 65663842
refGene_gn=FUBP1;refGene_tx=XM_011284916.3;refGene_gn=FUBP1;refGene_tx
=XM_019837273.1;refGene_gn=FUBP1;refGene_tx=XM_019837274.1;refGene_gn=

FUBP1;refGene_tx=XM_019837275.1;refGene_gn=FUBP1;refGene_tx=XM_0198372
76.1;refGene_gn=FUBP1;refGene_tx=XM_019837279.1;refGene_gn=FUBP1;refGe
ne_tx=XM_023258839.1
C1 65664700 65664791
refGene_gn=FUBP1;refGene_tx=XM_011284916.3;refGene_gn=FUBP1;refGene_tx
=XM_019837273.1;refGene_gn=FUBP1;refGene_tx=XM_019837274.1;refGene_gn=
FUBP1;refGene_tx=XM_019837275.1;refGene_gn=FUBP1;refGene_tx=XM_0198372
76.1;refGene_gn=FUBP1;refGene_tx=XM_019837277.1;refGene_gn=FUBP1;refGe
ne_tx=XM_019837278.1;refGene_gn=FUBP1;refGene_tx=XM_019837279.1;refGen
e_gn=FUBP1;refGene_tx=XM_019837280.1;refGene_gn=FUBP1;refGene_tx=XM_01
9837281.1;refGene_gn=FUBP1;refGene_tx=XM_019837282.1;refGene_gn=FUBP1;
refGene_tx=XM_023258839.1
C1 65670463 65670583
refGene_gn=FUBP1;refGene_tx=XM_011284916.3;refGene_gn=FUBP1;refGene_tx
=XM_019837273.1;refGene_gn=FUBP1;refGene_tx=XM_019837274.1;refGene_gn=
FUBP1;refGene_tx=XM_019837275.1;refGene_gn=FUBP1;refGene_tx=XM_0198372
76.1;refGene_gn=FUBP1;refGene_tx=XM_019837277.1;refGene_gn=FUBP1;refGe
ne_tx=XM_019837278.1;refGene_gn=FUBP1;refGene_tx=XM_019837279.1;refGen
e_gn=FUBP1;refGene_tx=XM_019837280.1;refGene_gn=FUBP1;refGene_tx=XM_01
9837281.1;refGene_gn=FUBP1;refGene_tx=XM_019837282.1;refGene_gn=FUBP1;
refGene_tx=XM_023258839.1
C1 65710331 65710542
refGene_gn=DNAJB4;refGene_tx=XM_006934864.4;refGene_gn=DNAJB4;refGene_
tx=XM_011284910.2;refGene_gn=DNAJB4;refGene_tx=XM_019837283.1;refGene_
gn=DNAJB4;refGene_tx=XM_019837284.2;refGene_gn=DNAJB4;refGene_tx=XM_02
3258845.1
C1 65716600 65717169
refGene_gn=DNAJB4;refGene_tx=XM_006934864.4;refGene_gn=DNAJB4;refGene_
tx=XM_011284910.2;refGene_gn=DNAJB4;refGene_tx=XM_019837283.1;refGene_
gn=DNAJB4;refGene_tx=XM_019837284.2;refGene_gn=DNAJB4;refGene_tx=XM_01
9837286.1;refGene_gn=DNAJB4;refGene_tx=XM_023258845.1
C1 65718895 65719129
refGene_gn=DNAJB4;refGene_tx=XM_006934864.4;refGene_gn=DNAJB4;refGene_
tx=XM_011284910.2;refGene_gn=DNAJB4;refGene_tx=XM_019837283.1;refGene_
gn=DNAJB4;refGene_tx=XM_019837284.2;refGene_gn=DNAJB4;refGene_tx=XM_01
9837286.1;refGene_gn=DNAJB4;refGene_tx=XM_023258845.1
C1 65744424 65744664
refGene_gn=GIPC2;refGene_tx=XM_003990249.4;refGene_gn=GIPC2;refGene_tx
=XM_023258846.1
C1 65744789 65744846 refGene_gn=GIPC2;refGene_tx=XM_019837287.2
C1 65770990 65771176
refGene_gn=GIPC2;refGene_tx=XM_003990249.4;refGene_gn=GIPC2;refGene_tx
=XM_019837287.2;refGene_gn=GIPC2;refGene_tx=XM_023258846.1
C1 65785530 65785711
refGene_gn=GIPC2;refGene_tx=XM_003990249.4;refGene_gn=GIPC2;refGene_tx
=XM_019837287.2;refGene_gn=GIPC2;refGene_tx=XM_023258846.1
C1 65806809 65806916
refGene_gn=GIPC2;refGene_tx=XM_003990249.4;refGene_gn=GIPC2;refGene_tx
=XM_019837287.2;refGene_gn=GIPC2;refGene_tx=XM_023258846.1
C1 65813179 65813261

refGene_gn=GIPC2;refGene_tx=XM_003990249.4;refGene_gn=GIPC2;refGene_tx=XM_019837287.2;refGene_gn=GIPC2;refGene_tx=XM_023258846.1
C1 65825474 65825620 refGene_gn=GIPC2;refGene_tx=XM_003990249.4;refGene_gn=GIPC2;refGene_tx=XM_019837287.2
C1 65942823 65942918 refGene_gn=GIPC2;refGene_tx=XM_023258846.1
C1 66169771 66170569 refGene_gn=PTGFR;refGene_tx=NM_001009861.1;refGene_gn=PTGFR;refGene_tx=XM_006934847.4
C1 66216343 66216646 refGene_gn=PTGFR;refGene_tx=NM_001009861.1;refGene_gn=PTGFR;refGene_tx=XM_006934847.4
C1 66279090 66279568 refGene_gn=IFI44L;refGene_tx=XM_006934865.3
C1 66280095 66280144 refGene_gn=IFI44L;refGene_tx=XM_006934865.3
C1 66281103 66281299 refGene_gn=IFI44L;refGene_tx=XM_006934865.3
C1 66284744 66284897 refGene_gn=IFI44L;refGene_tx=XM_006934865.3
C1 66287628 66287800 refGene_gn=IFI44L;refGene_tx=XM_006934865.3
C1 66292385 66292486 refGene_gn=IFI44L;refGene_tx=XM_006934865.3
C1 66292803 66292974 refGene_gn=IFI44L;refGene_tx=XM_006934865.3
C1 66293678 66293810 refGene_gn=IFI44L;refGene_tx=XM_006934865.3
C1 66299699 66300153 refGene_gn=IFI44;refGene_tx=XM_003990313.5
C1 66305056 66305093 refGene_gn=IFI44;refGene_tx=XM_003990313.5
C1 66305801 66305997 refGene_gn=IFI44;refGene_tx=XM_003990313.5
C1 66306385 66306535 refGene_gn=IFI44;refGene_tx=XM_003990313.5
C1 66311453 66311625 refGene_gn=IFI44;refGene_tx=XM_003990313.5
C1 66312680 66312781 refGene_gn=IFI44;refGene_tx=XM_003990313.5
C1 66315000 66315192 refGene_gn=IFI44;refGene_tx=XM_003990313.5
C1 66493096 66493159 refGene_gn=ADGRL4;refGene_tx=XM_003990250.5;refGene_gn=ADGRL4;refGene_tx=XM_019837288.2
C1 66493515 66493684 refGene_gn=ADGRL4;refGene_tx=XM_003990250.5;refGene_gn=ADGRL4;refGene_tx=XM_019837288.2;refGene_gn=ADGRL4;refGene_tx=XM_019837289.2
C1 66495275 66495367 refGene_gn=ADGRL4;refGene_tx=XM_003990250.5;refGene_gn=ADGRL4;refGene_tx=XM_019837288.2;refGene_gn=ADGRL4;refGene_tx=XM_019837291.2
C1 66504482 66504506 refGene_gn=ADGRL4;refGene_tx=XM_019837290.2
C1 66520996 66521063 refGene_gn=ADGRL4;refGene_tx=XM_003990250.5;refGene_gn=ADGRL4;refGene_tx=XM_019837288.2;refGene_gn=ADGRL4;refGene_tx=XM_019837289.2;refGene_gn=ADGRL4;refGene_tx=XM_019837290.2
C1 66521185 66521406 refGene_gn=ADGRL4;refGene_tx=XM_003990250.5;refGene_gn=ADGRL4;refGene_tx=XM_019837288.2;refGene_gn=ADGRL4;refGene_tx=XM_019837289.2;refGene_gn=ADGRL4;refGene_tx=XM_019837290.2;refGene_gn=ADGRL4;refGene_tx=XM_019837291.2
C1 66523201 66523405 refGene_gn=ADGRL4;refGene_tx=XM_003990250.5;refGene_gn=ADGRL4;refGene_tx=XM_019837288.2;refGene_gn=ADGRL4;refGene_tx=XM_019837289.2;refGene_gn=ADGRL4;refGene_tx=XM_019837290.2;refGene_gn=ADGRL4;refGene_tx=XM_01

9837291.2
C1 66524816 66524990
refGene_gn=ADGRL4;refGene_tx=XM_003990250.5;refGene_gn=ADGRL4;refGene_tx=XM_019837288.2;refGene_gn=ADGRL4;refGene_tx=XM_019837289.2;refGene_gn=ADGRL4;refGene_tx=XM_019837290.2;refGene_gn=ADGRL4;refGene_tx=XM_019837291.2
C1 66525505 66525559
refGene_gn=ADGRL4;refGene_tx=XM_019837288.2;refGene_gn=ADGRL4;refGene_tx=XM_019837289.2;refGene_gn=ADGRL4;refGene_tx=XM_019837290.2;refGene_gn=ADGRL4;refGene_tx=XM_019837291.2
C1 66527431 66527637
refGene_gn=ADGRL4;refGene_tx=XM_003990250.5;refGene_gn=ADGRL4;refGene_tx=XM_019837288.2;refGene_gn=ADGRL4;refGene_tx=XM_019837289.2;refGene_gn=ADGRL4;refGene_tx=XM_019837290.2;refGene_gn=ADGRL4;refGene_tx=XM_019837291.2
C1 66532407 66532524
refGene_gn=ADGRL4;refGene_tx=XM_003990250.5;refGene_gn=ADGRL4;refGene_tx=XM_019837288.2;refGene_gn=ADGRL4;refGene_tx=XM_019837289.2;refGene_gn=ADGRL4;refGene_tx=XM_019837290.2;refGene_gn=ADGRL4;refGene_tx=XM_019837291.2
C1 66533810 66533994
refGene_gn=ADGRL4;refGene_tx=XM_003990250.5;refGene_gn=ADGRL4;refGene_tx=XM_019837288.2;refGene_gn=ADGRL4;refGene_tx=XM_019837289.2;refGene_gn=ADGRL4;refGene_tx=XM_019837290.2;refGene_gn=ADGRL4;refGene_tx=XM_019837291.2
C1 66534109 66534289
refGene_gn=ADGRL4;refGene_tx=XM_003990250.5;refGene_gn=ADGRL4;refGene_tx=XM_019837288.2;refGene_gn=ADGRL4;refGene_tx=XM_019837289.2;refGene_gn=ADGRL4;refGene_tx=XM_019837290.2;refGene_gn=ADGRL4;refGene_tx=XM_019837291.2
C1 66535294 66535365
refGene_gn=ADGRL4;refGene_tx=XM_003990250.5;refGene_gn=ADGRL4;refGene_tx=XM_019837288.2;refGene_gn=ADGRL4;refGene_tx=XM_019837289.2;refGene_gn=ADGRL4;refGene_tx=XM_019837290.2;refGene_gn=ADGRL4;refGene_tx=XM_019837291.2
C1 66535891 66536041
refGene_gn=ADGRL4;refGene_tx=XM_003990250.5;refGene_gn=ADGRL4;refGene_tx=XM_019837288.2;refGene_gn=ADGRL4;refGene_tx=XM_019837289.2;refGene_gn=ADGRL4;refGene_tx=XM_019837290.2;refGene_gn=ADGRL4;refGene_tx=XM_019837291.2
C1 66542778 66542931
refGene_gn=ADGRL4;refGene_tx=XM_003990250.5;refGene_gn=ADGRL4;refGene_tx=XM_019837288.2;refGene_gn=ADGRL4;refGene_tx=XM_019837289.2;refGene_gn=ADGRL4;refGene_tx=XM_019837290.2;refGene_gn=ADGRL4;refGene_tx=XM_019837291.2
C1 66600559 66600709
refGene_gn=ADGRL4;refGene_tx=XM_003990250.5;refGene_gn=ADGRL4;refGene_tx=XM_019837288.2;refGene_gn=ADGRL4;refGene_tx=XM_019837289.2;refGene_gn=ADGRL4;refGene_tx=XM_019837290.2;refGene_gn=ADGRL4;refGene_tx=XM_019837291.2

C1	66602096 66602346
refGene_gn=ADGRL4;refGene_tx=XM_003990250.5;refGene_gn=ADGRL4;refGene_tx=XM_019837288.2;refGene_gn=ADGRL4;refGene_tx=XM_019837289.2;refGene_gn=ADGRL4;refGene_tx=XM_019837290.2;refGene_gn=ADGRL4;refGene_tx=XM_019837291.2
C1	69025611 69025684
refGene_gn=ADGRL2;refGene_tx=XM_019837292.2;refGene_gn=ADGRL2;refGene_tx=XM_019837293.1;refGene_gn=ADGRL2;refGene_tx=XM_019837294.1;refGene_gn=ADGRL2;refGene_tx=XM_019837295.2;refGene_gn=ADGRL2;refGene_tx=XM_019837296.2;refGene_gn=ADGRL2;refGene_tx=XM_019837297.2;refGene_gn=ADGRL2;refGene_tx=XM_019837298.2;refGene_gn=ADGRL2;refGene_tx=XM_019837299.2;refGene_gn=ADGRL2;refGene_tx=XM_019837300.2;refGene_gn=ADGRL2;refGene_tx=XM_019837301.2;refGene_gn=ADGRL2;refGene_tx=XM_019837302.2;refGene_gn=ADGRL2;refGene_tx=XM_019837303.2;refGene_gn=ADGRL2;refGene_tx=XM_019837304.2;refGene_gn=ADGRL2;refGene_tx=XM_019837305.2;refGene_gn=ADGRL2;refGene_tx=XM_019837306.2;refGene_gn=ADGRL2;refGene_tx=XM_019837307.2;refGene_gn=ADGRL2;refGene_tx=XM_019837308.2;refGene_gn=ADGRL2;refGene_tx=XM_019837309.2;refGene_gn=ADGRL2;refGene_tx=XM_023258848.1;refGene_gn=ADGRL2;refGene_tx=XM_023258849.1;refGene_gn=ADGRL2;refGene_tx=XM_023258850.1;refGene_gn=ADGRL2;refGene_tx=XM_023258852.1
C1	69093567 69093781
refGene_gn=ADGRL2;refGene_tx=XM_019837292.2;refGene_gn=ADGRL2;refGene_tx=XM_019837293.1;refGene_gn=ADGRL2;refGene_tx=XM_019837294.1;refGene_gn=ADGRL2;refGene_tx=XM_019837295.2;refGene_gn=ADGRL2;refGene_tx=XM_019837296.2;refGene_gn=ADGRL2;refGene_tx=XM_019837297.2;refGene_gn=ADGRL2;refGene_tx=XM_019837298.2;refGene_gn=ADGRL2;refGene_tx=XM_019837299.2;refGene_gn=ADGRL2;refGene_tx=XM_019837300.2;refGene_gn=ADGRL2;refGene_tx=XM_019837301.2;refGene_gn=ADGRL2;refGene_tx=XM_019837302.2;refGene_gn=ADGRL2;refGene_tx=XM_019837303.2;refGene_gn=ADGRL2;refGene_tx=XM_019837304.2;refGene_gn=ADGRL2;refGene_tx=XM_019837305.2;refGene_gn=ADGRL2;refGene_tx=XM_019837306.2;refGene_gn=ADGRL2;refGene_tx=XM_019837307.2;refGene_gn=ADGRL2;refGene_tx=XM_019837308.2;refGene_gn=ADGRL2;refGene_tx=XM_019837309.2;refGene_gn=ADGRL2;refGene_tx=XM_023258848.1;refGene_gn=ADGRL2;refGene_tx=XM_023258849.1;refGene_gn=ADGRL2;refGene_tx=XM_023258850.1;refGene_gn=ADGRL2;refGene_tx=XM_023258852.1
C1	69121422 69121532
refGene_gn=ADGRL2;refGene_tx=XM_019837292.2;refGene_gn=ADGRL2;refGene_tx=XM_019837293.1;refGene_gn=ADGRL2;refGene_tx=XM_019837294.1;refGene_gn=ADGRL2;refGene_tx=XM_019837295.2;refGene_gn=ADGRL2;refGene_tx=XM_019837296.2;refGene_gn=ADGRL2;refGene_tx=XM_019837297.2;refGene_gn=ADGRL2;refGene_tx=XM_019837298.2;refGene_gn=ADGRL2;refGene_tx=XM_019837299.2;refGene_gn=ADGRL2;refGene_tx=XM_019837300.2;refGene_gn=ADGRL2;refGene_tx=XM_019837301.2;refGene_gn=ADGRL2;refGene_tx=XM_019837302.2;refGene_gn=ADGRL2;refGene_tx=XM_019837303.2;refGene_gn=ADGRL2;refGene_tx=XM_019837304.2;refGene_gn=ADGRL2;refGene_tx=XM_019837305.2;refGene_gn=ADGRL2;refGene_tx=XM_019837306.2;refGene_gn=ADGRL2;refGene_tx=XM_019837307.2;refGene_gn=ADGRL2;refGene_tx=XM_019837308.2;refGene_gn=ADGRL2;refGene_tx=XM_019837309.2;refGene_gn=ADGRL2;refGene_tx=XM_023258848.1;refGene_gn=ADGRL2;refGene_tx=XM_023258849.1;refGene_gn=ADGRL2;refGene_tx=XM_023258850.1;refGene_gn=ADGRL2;refGene_tx=XM_023258852.1

C1 69126718 69126733
refGene_gn=ADGRL2;refGene_tx=XM_019837292.2;refGene_gn=ADGRL2;refGene_tx=XM_019837293.1;refGene_gn=ADGRL2;refGene_tx=XM_019837294.1;refGene_gn=ADGRL2;refGene_tx=XM_019837295.2;refGene_gn=ADGRL2;refGene_tx=XM_019837296.2;refGene_gn=ADGRL2;refGene_tx=XM_019837297.2;refGene_gn=ADGRL2;refGene_tx=XM_019837298.2;refGene_gn=ADGRL2;refGene_tx=XM_019837299.2;refGene_gn=ADGRL2;refGene_tx=XM_019837301.2;refGene_gn=ADGRL2;refGene_tx=XM_019837303.2;refGene_gn=ADGRL2;refGene_tx=XM_019837304.2;refGene_gn=ADGRL2;refGene_tx=XM_019837305.2;refGene_gn=ADGRL2;refGene_tx=XM_019837307.2;refGene_gn=ADGRL2;refGene_tx=XM_023258848.1;refGene_gn=ADGRL2;refGene_tx=XM_023258849.1;refGene_gn=ADGRL2;refGene_tx=XM_023258850.1
C1 69127659 69128460
refGene_gn=ADGRL2;refGene_tx=XM_019837292.2;refGene_gn=ADGRL2;refGene_tx=XM_019837293.1;refGene_gn=ADGRL2;refGene_tx=XM_019837294.1;refGene_gn=ADGRL2;refGene_tx=XM_019837295.2;refGene_gn=ADGRL2;refGene_tx=XM_019837296.2;refGene_gn=ADGRL2;refGene_tx=XM_019837297.2;refGene_gn=ADGRL2;refGene_tx=XM_019837298.2;refGene_gn=ADGRL2;refGene_tx=XM_019837299.2;refGene_gn=ADGRL2;refGene_tx=XM_019837300.2;refGene_gn=ADGRL2;refGene_tx=XM_019837301.2;refGene_gn=ADGRL2;refGene_tx=XM_019837302.2;refGene_gn=ADGRL2;refGene_tx=XM_019837303.2;refGene_gn=ADGRL2;refGene_tx=XM_019837304.2;refGene_gn=ADGRL2;refGene_tx=XM_019837305.2;refGene_gn=ADGRL2;refGene_tx=XM_019837306.2;refGene_gn=ADGRL2;refGene_tx=XM_019837307.2;refGene_gn=ADGRL2;refGene_tx=XM_019837308.2;refGene_gn=ADGRL2;refGene_tx=XM_019837309.2;refGene_gn=ADGRL2;refGene_tx=XM_023258848.1;refGene_gn=ADGRL2;refGene_tx=XM_023258849.1;refGene_gn=ADGRL2;refGene_tx=XM_023258850.1;refGene_gn=ADGRL2;refGene_tx=XM_023258852.1
C1 69134818 69135112
refGene_gn=ADGRL2;refGene_tx=XM_019837292.2;refGene_gn=ADGRL2;refGene_tx=XM_019837293.1;refGene_gn=ADGRL2;refGene_tx=XM_019837294.1;refGene_gn=ADGRL2;refGene_tx=XM_019837295.2;refGene_gn=ADGRL2;refGene_tx=XM_019837296.2;refGene_gn=ADGRL2;refGene_tx=XM_019837297.2;refGene_gn=ADGRL2;refGene_tx=XM_019837298.2;refGene_gn=ADGRL2;refGene_tx=XM_019837299.2;refGene_gn=ADGRL2;refGene_tx=XM_019837300.2;refGene_gn=ADGRL2;refGene_tx=XM_019837301.2;refGene_gn=ADGRL2;refGene_tx=XM_019837302.2;refGene_gn=ADGRL2;refGene_tx=XM_019837303.2;refGene_gn=ADGRL2;refGene_tx=XM_019837304.2;refGene_gn=ADGRL2;refGene_tx=XM_019837305.2;refGene_gn=ADGRL2;refGene_tx=XM_019837306.2;refGene_gn=ADGRL2;refGene_tx=XM_019837307.2;refGene_gn=ADGRL2;refGene_tx=XM_019837308.2;refGene_gn=ADGRL2;refGene_tx=XM_019837309.2;refGene_gn=ADGRL2;refGene_tx=XM_023258848.1;refGene_gn=ADGRL2;refGene_tx=XM_023258849.1;refGene_gn=ADGRL2;refGene_tx=XM_023258850.1;refGene_gn=ADGRL2;refGene_tx=XM_023258852.1
C1 69135659 69135763
refGene_gn=ADGRL2;refGene_tx=XM_019837292.2;refGene_gn=ADGRL2;refGene_tx=XM_019837293.1;refGene_gn=ADGRL2;refGene_tx=XM_019837294.1;refGene_gn=ADGRL2;refGene_tx=XM_019837295.2;refGene_gn=ADGRL2;refGene_tx=XM_019837296.2;refGene_gn=ADGRL2;refGene_tx=XM_019837297.2;refGene_gn=ADGRL2;refGene_tx=XM_019837298.2;refGene_gn=ADGRL2;refGene_tx=XM_019837299.2;refGene_gn=ADGRL2;refGene_tx=XM_019837300.2;refGene_gn=ADGRL2;refGene_tx=XM_019837301.2;refGene_gn=ADGRL2;refGene_tx=XM_019837302.2;refGen

e_gn=ADGRL2;refGene_tx=XM_019837303.2;refGene_gn=ADGRL2;refGene_tx=XM_019837304.2;refGene_gn=ADGRL2;refGene_tx=XM_019837305.2;refGene_gn=ADGRL2;refGene_tx=XM_019837306.2;refGene_gn=ADGRL2;refGene_tx=XM_019837307.2;refGene_gn=ADGRL2;refGene_tx=XM_019837308.2;refGene_gn=ADGRL2;refGene_tx=XM_019837309.2;refGene_gn=ADGRL2;refGene_tx=XM_023258848.1;refGene_gn=ADGRL2;refGene_tx=XM_023258849.1;refGene_gn=ADGRL2;refGene_tx=XM_023258850.1;refGene_gn=ADGRL2;refGene_tx=XM_023258852.1
C1 69136879 69137065
refGene_gn=ADGRL2;refGene_tx=XM_019837292.2;refGene_gn=ADGRL2;refGene_tx=XM_019837293.1;refGene_gn=ADGRL2;refGene_tx=XM_019837294.1;refGene_gn=ADGRL2;refGene_tx=XM_019837295.2;refGene_gn=ADGRL2;refGene_tx=XM_019837296.2;refGene_gn=ADGRL2;refGene_tx=XM_019837297.2;refGene_gn=ADGRL2;refGene_tx=XM_019837298.2;refGene_gn=ADGRL2;refGene_tx=XM_019837299.2;refGene_gn=ADGRL2;refGene_tx=XM_019837300.2;refGene_gn=ADGRL2;refGene_tx=XM_019837301.2;refGene_gn=ADGRL2;refGene_tx=XM_019837302.2;refGene_gn=ADGRL2;refGene_tx=XM_019837303.2;refGene_gn=ADGRL2;refGene_tx=XM_019837304.2;refGene_gn=ADGRL2;refGene_tx=XM_019837305.2;refGene_gn=ADGRL2;refGene_tx=XM_019837306.2;refGene_gn=ADGRL2;refGene_tx=XM_019837307.2;refGene_gn=ADGRL2;refGene_tx=XM_019837308.2;refGene_gn=ADGRL2;refGene_tx=XM_019837309.2;refGene_gn=ADGRL2;refGene_tx=XM_023258848.1;refGene_gn=ADGRL2;refGene_tx=XM_023258849.1;refGene_gn=ADGRL2;refGene_tx=XM_023258850.1;refGene_gn=ADGRL2;refGene_tx=XM_023258852.1
C1 69137897 69137936
refGene_gn=ADGRL2;refGene_tx=XM_019837292.2;refGene_gn=ADGRL2;refGene_tx=XM_019837293.1;refGene_gn=ADGRL2;refGene_tx=XM_019837294.1;refGene_gn=ADGRL2;refGene_tx=XM_019837295.2;refGene_gn=ADGRL2;refGene_tx=XM_019837296.2;refGene_gn=ADGRL2;refGene_tx=XM_019837297.2;refGene_gn=ADGRL2;refGene_tx=XM_019837299.2;refGene_gn=ADGRL2;refGene_tx=XM_019837300.2;refGene_gn=ADGRL2;refGene_tx=XM_019837303.2;refGene_gn=ADGRL2;refGene_tx=XM_019837305.2;refGene_gn=ADGRL2;refGene_tx=XM_019837306.2;refGene_gn=ADGRL2;refGene_tx=XM_023258848.1;refGene_gn=ADGRL2;refGene_tx=XM_023258849.1;refGene_gn=ADGRL2;refGene_tx=XM_023258850.1
C1 69140773 69140957
refGene_gn=ADGRL2;refGene_tx=XM_019837292.2;refGene_gn=ADGRL2;refGene_tx=XM_019837293.1;refGene_gn=ADGRL2;refGene_tx=XM_019837294.1;refGene_gn=ADGRL2;refGene_tx=XM_019837295.2;refGene_gn=ADGRL2;refGene_tx=XM_019837296.2;refGene_gn=ADGRL2;refGene_tx=XM_019837297.2;refGene_gn=ADGRL2;refGene_tx=XM_019837298.2;refGene_gn=ADGRL2;refGene_tx=XM_019837299.2;refGene_gn=ADGRL2;refGene_tx=XM_019837300.2;refGene_gn=ADGRL2;refGene_tx=XM_019837301.2;refGene_gn=ADGRL2;refGene_tx=XM_019837302.2;refGene_gn=ADGRL2;refGene_tx=XM_019837303.2;refGene_gn=ADGRL2;refGene_tx=XM_019837304.2;refGene_gn=ADGRL2;refGene_tx=XM_019837305.2;refGene_gn=ADGRL2;refGene_tx=XM_019837306.2;refGene_gn=ADGRL2;refGene_tx=XM_019837307.2;refGene_gn=ADGRL2;refGene_tx=XM_019837308.2;refGene_gn=ADGRL2;refGene_tx=XM_019837309.2;refGene_gn=ADGRL2;refGene_tx=XM_023258848.1;refGene_gn=ADGRL2;refGene_tx=XM_023258849.1;refGene_gn=ADGRL2;refGene_tx=XM_023258850.1;refGene_gn=ADGRL2;refGene_tx=XM_023258852.1
C1 69150370 69150496
refGene_gn=ADGRL2;refGene_tx=XM_019837292.2;refGene_gn=ADGRL2;refGene_tx=XM_019837293.1;refGene_gn=ADGRL2;refGene_tx=XM_019837294.1;refGene_

gn=ADGRL2;refGene_tx=XM_019837295.2;refGene_gn=ADGRL2;refGene_tx=XM_01
9837296.2;refGene_gn=ADGRL2;refGene_tx=XM_019837297.2;refGene_gn=ADGRL
2;refGene_tx=XM_019837298.2;refGene_gn=ADGRL2;refGene_tx=XM_019837299.
2;refGene_gn=ADGRL2;refGene_tx=XM_019837300.2;refGene_gn=ADGRL2;refGen
e_tx=XM_019837301.2;refGene_gn=ADGRL2;refGene_tx=XM_019837302.2;refGen
e_gn=ADGRL2;refGene_tx=XM_019837303.2;refGene_gn=ADGRL2;refGene_tx=XM_
019837304.2;refGene_gn=ADGRL2;refGene_tx=XM_019837305.2;refGene_gn=ADG
RL2;refGene_tx=XM_019837306.2;refGene_gn=ADGRL2;refGene_tx=XM_01983730
7.2;refGene_gn=ADGRL2;refGene_tx=XM_019837308.2;refGene_gn=ADGRL2;refG
ene_tx=XM_019837309.2;refGene_gn=ADGRL2;refGene_tx=XM_023258848.1;refG
ene_gn=ADGRL2;refGene_tx=XM_023258849.1;refGene_gn=ADGRL2;refGene_tx=X
M_023258850.1;refGene_gn=ADGRL2;refGene_tx=XM_023258852.1
C1 69150767 69150973
refGene_gn=ADGRL2;refGene_tx=XM_019837292.2;refGene_gn=ADGRL2;refGene_
tx=XM_019837293.1;refGene_gn=ADGRL2;refGene_tx=XM_019837294.1;refGene_
gn=ADGRL2;refGene_tx=XM_019837295.2;refGene_gn=ADGRL2;refGene_tx=XM_01
9837296.2;refGene_gn=ADGRL2;refGene_tx=XM_019837297.2;refGene_gn=ADGRL
2;refGene_tx=XM_019837298.2;refGene_gn=ADGRL2;refGene_tx=XM_019837299.
2;refGene_gn=ADGRL2;refGene_tx=XM_019837300.2;refGene_gn=ADGRL2;refGen
e_tx=XM_019837301.2;refGene_gn=ADGRL2;refGene_tx=XM_019837302.2;refGen
e_gn=ADGRL2;refGene_tx=XM_019837303.2;refGene_gn=ADGRL2;refGene_tx=XM_
019837304.2;refGene_gn=ADGRL2;refGene_tx=XM_019837305.2;refGene_gn=ADG
RL2;refGene_tx=XM_019837306.2;refGene_gn=ADGRL2;refGene_tx=XM_01983730
7.2;refGene_gn=ADGRL2;refGene_tx=XM_019837308.2;refGene_gn=ADGRL2;refG
ene_tx=XM_019837309.2;refGene_gn=ADGRL2;refGene_tx=XM_023258848.1;refG
ene_gn=ADGRL2;refGene_tx=XM_023258849.1;refGene_gn=ADGRL2;refGene_tx=X
M_023258850.1;refGene_gn=ADGRL2;refGene_tx=XM_023258852.1
C1 69152068 69152242
refGene_gn=ADGRL2;refGene_tx=XM_019837292.2;refGene_gn=ADGRL2;refGene_
tx=XM_019837293.1;refGene_gn=ADGRL2;refGene_tx=XM_019837294.1;refGene_
gn=ADGRL2;refGene_tx=XM_019837295.2;refGene_gn=ADGRL2;refGene_tx=XM_01
9837296.2;refGene_gn=ADGRL2;refGene_tx=XM_019837297.2;refGene_gn=ADGRL
2;refGene_tx=XM_019837298.2;refGene_gn=ADGRL2;refGene_tx=XM_019837299.
2;refGene_gn=ADGRL2;refGene_tx=XM_019837300.2;refGene_gn=ADGRL2;refGen
e_tx=XM_019837301.2;refGene_gn=ADGRL2;refGene_tx=XM_019837302.2;refGen
e_gn=ADGRL2;refGene_tx=XM_019837303.2;refGene_gn=ADGRL2;refGene_tx=XM_
019837304.2;refGene_gn=ADGRL2;refGene_tx=XM_019837305.2;refGene_gn=ADG
RL2;refGene_tx=XM_019837306.2;refGene_gn=ADGRL2;refGene_tx=XM_01983730
7.2;refGene_gn=ADGRL2;refGene_tx=XM_019837308.2;refGene_gn=ADGRL2;refG
ene_tx=XM_019837309.2;refGene_gn=ADGRL2;refGene_tx=XM_023258848.1;refG
ene_gn=ADGRL2;refGene_tx=XM_023258849.1;refGene_gn=ADGRL2;refGene_tx=X
M_023258850.1;refGene_gn=ADGRL2;refGene_tx=XM_023258852.1
C1 69153219 69153429
refGene_gn=ADGRL2;refGene_tx=XM_019837292.2;refGene_gn=ADGRL2;refGene_
tx=XM_019837293.1;refGene_gn=ADGRL2;refGene_tx=XM_019837294.1;refGene_
gn=ADGRL2;refGene_tx=XM_019837295.2;refGene_gn=ADGRL2;refGene_tx=XM_01
9837296.2;refGene_gn=ADGRL2;refGene_tx=XM_019837297.2;refGene_gn=ADGRL
2;refGene_tx=XM_019837298.2;refGene_gn=ADGRL2;refGene_tx=XM_019837299.
2;refGene_gn=ADGRL2;refGene_tx=XM_019837300.2;refGene_gn=ADGRL2;refGen
e_tx=XM_019837301.2;refGene_gn=ADGRL2;refGene_tx=XM_019837302.2;refGen

e_gn=ADGRL2;refGene_tx=XM_019837303.2;refGene_gn=ADGRL2;refGene_tx=XM_019837304.2;refGene_gn=ADGRL2;refGene_tx=XM_019837305.2;refGene_gn=ADGRL2;refGene_tx=XM_019837306.2;refGene_gn=ADGRL2;refGene_tx=XM_019837307.2;refGene_gn=ADGRL2;refGene_tx=XM_019837308.2;refGene_gn=ADGRL2;refGene_tx=XM_019837309.2;refGene_gn=ADGRL2;refGene_tx=XM_023258848.1;refGene_gn=ADGRL2;refGene_tx=XM_023258849.1;refGene_gn=ADGRL2;refGene_tx=XM_023258850.1;refGene_gn=ADGRL2;refGene_tx=XM_023258852.1
C1 69154367 69154588
refGene_gn=ADGRL2;refGene_tx=XM_019837292.2;refGene_gn=ADGRL2;refGene_tx=XM_019837293.1;refGene_gn=ADGRL2;refGene_tx=XM_019837294.1;refGene_gn=ADGRL2;refGene_tx=XM_019837295.2;refGene_gn=ADGRL2;refGene_tx=XM_019837296.2;refGene_gn=ADGRL2;refGene_tx=XM_019837297.2;refGene_gn=ADGRL2;refGene_tx=XM_019837298.2;refGene_gn=ADGRL2;refGene_tx=XM_019837299.2;refGene_gn=ADGRL2;refGene_tx=XM_019837300.2;refGene_gn=ADGRL2;refGene_tx=XM_019837301.2;refGene_gn=ADGRL2;refGene_tx=XM_019837302.2;refGene_gn=ADGRL2;refGene_tx=XM_019837303.2;refGene_gn=ADGRL2;refGene_tx=XM_019837304.2;refGene_gn=ADGRL2;refGene_tx=XM_019837305.2;refGene_gn=ADGRL2;refGene_tx=XM_019837306.2;refGene_gn=ADGRL2;refGene_tx=XM_019837307.2;refGene_gn=ADGRL2;refGene_tx=XM_019837308.2;refGene_gn=ADGRL2;refGene_tx=XM_019837309.2;refGene_gn=ADGRL2;refGene_tx=XM_023258848.1;refGene_gn=ADGRL2;refGene_tx=XM_023258849.1;refGene_gn=ADGRL2;refGene_tx=XM_023258850.1;refGene_gn=ADGRL2;refGene_tx=XM_023258852.1
C1 69155921 69155988
refGene_gn=ADGRL2;refGene_tx=XM_019837292.2;refGene_gn=ADGRL2;refGene_tx=XM_019837293.1;refGene_gn=ADGRL2;refGene_tx=XM_019837294.1;refGene_gn=ADGRL2;refGene_tx=XM_019837295.2;refGene_gn=ADGRL2;refGene_tx=XM_019837296.2;refGene_gn=ADGRL2;refGene_tx=XM_019837297.2;refGene_gn=ADGRL2;refGene_tx=XM_019837298.2;refGene_gn=ADGRL2;refGene_tx=XM_019837299.2;refGene_gn=ADGRL2;refGene_tx=XM_019837300.2;refGene_gn=ADGRL2;refGene_tx=XM_019837301.2;refGene_gn=ADGRL2;refGene_tx=XM_019837302.2;refGene_gn=ADGRL2;refGene_tx=XM_019837303.2;refGene_gn=ADGRL2;refGene_tx=XM_019837304.2;refGene_gn=ADGRL2;refGene_tx=XM_019837305.2;refGene_gn=ADGRL2;refGene_tx=XM_019837306.2;refGene_gn=ADGRL2;refGene_tx=XM_019837307.2;refGene_gn=ADGRL2;refGene_tx=XM_019837308.2;refGene_gn=ADGRL2;refGene_tx=XM_019837309.2;refGene_gn=ADGRL2;refGene_tx=XM_023258848.1;refGene_gn=ADGRL2;refGene_tx=XM_023258849.1;refGene_gn=ADGRL2;refGene_tx=XM_023258850.1;refGene_gn=ADGRL2;refGene_tx=XM_023258852.1
C1 69164231 69164323
refGene_gn=ADGRL2;refGene_tx=XM_019837292.2;refGene_gn=ADGRL2;refGene_tx=XM_019837293.1;refGene_gn=ADGRL2;refGene_tx=XM_019837294.1;refGene_gn=ADGRL2;refGene_tx=XM_019837295.2;refGene_gn=ADGRL2;refGene_tx=XM_019837296.2;refGene_gn=ADGRL2;refGene_tx=XM_019837297.2;refGene_gn=ADGRL2;refGene_tx=XM_019837298.2;refGene_gn=ADGRL2;refGene_tx=XM_019837299.2;refGene_gn=ADGRL2;refGene_tx=XM_019837300.2;refGene_gn=ADGRL2;refGene_tx=XM_019837301.2;refGene_gn=ADGRL2;refGene_tx=XM_019837302.2;refGene_gn=ADGRL2;refGene_tx=XM_019837303.2;refGene_gn=ADGRL2;refGene_tx=XM_019837304.2;refGene_gn=ADGRL2;refGene_tx=XM_019837305.2;refGene_gn=ADGRL2;refGene_tx=XM_019837306.2;refGene_gn=ADGRL2;refGene_tx=XM_019837307.2;refGene_gn=ADGRL2;refGene_tx=XM_019837308.2;refGene_gn=ADGRL2;refGene_tx=XM_019837309.2;refGene_gn=ADGRL2;refGene_tx=XM_023258848.1;refG

ene_gn=ADGRL2;refGene_tx=XM_023258849.1;refGene_gn=ADGRL2;refGene_tx=XM_023258850.1;refGene_gn=ADGRL2;refGene_tx=XM_023258852.1
C1 69165149 69165194
refGene_gn=ADGRL2;refGene_tx=XM_019837292.2;refGene_gn=ADGRL2;refGene_tx=XM_019837293.1;refGene_gn=ADGRL2;refGene_tx=XM_019837294.1;refGene_gn=ADGRL2;refGene_tx=XM_019837295.2;refGene_gn=ADGRL2;refGene_tx=XM_019837296.2;refGene_gn=ADGRL2;refGene_tx=XM_019837297.2;refGene_gn=ADGRL2;refGene_tx=XM_019837298.2;refGene_gn=ADGRL2;refGene_tx=XM_019837303.2;refGene_gn=ADGRL2;refGene_tx=XM_019837304.2;refGene_gn=ADGRL2;refGene_tx=XM_023258848.1;refGene_gn=ADGRL2;refGene_tx=XM_023258849.1;refGene_gn=ADGRL2;refGene_tx=XM_023258850.1
C1 69165478 69165505
refGene_gn=ADGRL2;refGene_tx=XM_019837292.2;refGene_gn=ADGRL2;refGene_tx=XM_019837293.1;refGene_gn=ADGRL2;refGene_tx=XM_019837294.1;refGene_gn=ADGRL2;refGene_tx=XM_019837295.2;refGene_gn=ADGRL2;refGene_tx=XM_019837296.2;refGene_gn=ADGRL2;refGene_tx=XM_023258848.1;refGene_gn=ADGRL2;refGene_tx=XM_023258849.1;refGene_gn=ADGRL2;refGene_tx=XM_023258850.1
C1 69166164 69166333
refGene_gn=ADGRL2;refGene_tx=XM_019837292.2;refGene_gn=ADGRL2;refGene_tx=XM_019837293.1;refGene_gn=ADGRL2;refGene_tx=XM_019837294.1;refGene_gn=ADGRL2;refGene_tx=XM_019837295.2;refGene_gn=ADGRL2;refGene_tx=XM_019837296.2;refGene_gn=ADGRL2;refGene_tx=XM_019837297.2;refGene_gn=ADGRL2;refGene_tx=XM_019837298.2;refGene_gn=ADGRL2;refGene_tx=XM_019837299.2;refGene_gn=ADGRL2;refGene_tx=XM_019837300.2;refGene_gn=ADGRL2;refGene_tx=XM_019837301.2;refGene_gn=ADGRL2;refGene_tx=XM_019837302.2;refGene_gn=ADGRL2;refGene_tx=XM_019837303.2;refGene_gn=ADGRL2;refGene_tx=XM_019837304.2;refGene_gn=ADGRL2;refGene_tx=XM_019837305.2;refGene_gn=ADGRL2;refGene_tx=XM_019837306.2;refGene_gn=ADGRL2;refGene_tx=XM_019837307.2;refGene_gn=ADGRL2;refGene_tx=XM_019837308.2;refGene_gn=ADGRL2;refGene_tx=XM_019837309.2;refGene_gn=ADGRL2;refGene_tx=XM_023258848.1;refGene_gn=ADGRL2;refGene_tx=XM_023258849.1;refGene_gn=ADGRL2;refGene_tx=XM_023258850.1;refGene_gn=ADGRL2;refGene_tx=XM_023258852.1
C1 69168951 69169080
refGene_gn=ADGRL2;refGene_tx=XM_019837292.2;refGene_gn=ADGRL2;refGene_tx=XM_019837293.1;refGene_gn=ADGRL2;refGene_tx=XM_019837294.1;refGene_gn=ADGRL2;refGene_tx=XM_019837295.2;refGene_gn=ADGRL2;refGene_tx=XM_019837296.2;refGene_gn=ADGRL2;refGene_tx=XM_019837297.2;refGene_gn=ADGRL2;refGene_tx=XM_019837298.2;refGene_gn=ADGRL2;refGene_tx=XM_019837299.2;refGene_gn=ADGRL2;refGene_tx=XM_019837300.2;refGene_gn=ADGRL2;refGene_tx=XM_019837301.2;refGene_gn=ADGRL2;refGene_tx=XM_019837302.2;refGene_gn=ADGRL2;refGene_tx=XM_019837303.2;refGene_gn=ADGRL2;refGene_tx=XM_019837304.2;refGene_gn=ADGRL2;refGene_tx=XM_019837305.2;refGene_gn=ADGRL2;refGene_tx=XM_019837306.2;refGene_gn=ADGRL2;refGene_tx=XM_019837307.2;refGene_gn=ADGRL2;refGene_tx=XM_019837308.2;refGene_gn=ADGRL2;refGene_tx=XM_019837309.2;refGene_gn=ADGRL2;refGene_tx=XM_023258848.1;refGene_gn=ADGRL2;refGene_tx=XM_023258849.1;refGene_gn=ADGRL2;refGene_tx=XM_023258850.1;refGene_gn=ADGRL2;refGene_tx=XM_023258852.1
C1 69169638 69169735
refGene_gn=ADGRL2;refGene_tx=XM_019837292.2;refGene_gn=ADGRL2;refGene_

tx=XM_019837293.1;refGene_gn=ADGRL2;refGene_tx=XM_019837294.1;refGene_gn=ADGRL2;refGene_tx=XM_019837295.2;refGene_gn=ADGRL2;refGene_tx=XM_019837296.2;refGene_gn=ADGRL2;refGene_tx=XM_019837297.2;refGene_gn=ADGRL2;refGene_tx=XM_019837298.2;refGene_gn=ADGRL2;refGene_tx=XM_019837299.2;refGene_gn=ADGRL2;refGene_tx=XM_019837300.2;refGene_gn=ADGRL2;refGene_tx=XM_019837301.2;refGene_gn=ADGRL2;refGene_tx=XM_019837302.2;refGene_gn=ADGRL2;refGene_tx=XM_019837303.2;refGene_gn=ADGRL2;refGene_tx=XM_019837304.2;refGene_gn=ADGRL2;refGene_tx=XM_019837305.2;refGene_gn=ADGRL2;refGene_tx=XM_019837306.2;refGene_gn=ADGRL2;refGene_tx=XM_019837307.2;refGene_gn=ADGRL2;refGene_tx=XM_019837308.2;refGene_gn=ADGRL2;refGene_tx=XM_019837309.2;refGene_gn=ADGRL2;refGene_tx=XM_023258848.1;refGene_gn=ADGRL2;refGene_tx=XM_023258849.1;refGene_gn=ADGRL2;refGene_tx=XM_023258850.1;refGene_gn=ADGRL2;refGene_tx=XM_023258852.1
C1 69171444 69171573
refGene_gn=ADGRL2;refGene_tx=XM_019837292.2;refGene_gn=ADGRL2;refGene_tx=XM_019837293.1;refGene_gn=ADGRL2;refGene_tx=XM_019837294.1;refGene_gn=ADGRL2;refGene_tx=XM_019837295.2;refGene_gn=ADGRL2;refGene_tx=XM_019837296.2;refGene_gn=ADGRL2;refGene_tx=XM_019837297.2;refGene_gn=ADGRL2;refGene_tx=XM_019837298.2;refGene_gn=ADGRL2;refGene_tx=XM_019837299.2;refGene_gn=ADGRL2;refGene_tx=XM_019837300.2;refGene_gn=ADGRL2;refGene_tx=XM_019837301.2;refGene_gn=ADGRL2;refGene_tx=XM_019837302.2;refGene_gn=ADGRL2;refGene_tx=XM_023258848.1;refGene_gn=ADGRL2;refGene_tx=XM_023258849.1;refGene_gn=ADGRL2;refGene_tx=XM_023258850.1
C1 69171805 69171887
refGene_gn=ADGRL2;refGene_tx=XM_019837309.2;refGene_gn=ADGRL2;refGene_tx=XM_023258852.1
C1 69174932 69175687
refGene_gn=ADGRL2;refGene_tx=XM_019837292.2;refGene_gn=ADGRL2;refGene_tx=XM_019837293.1;refGene_gn=ADGRL2;refGene_tx=XM_019837294.1;refGene_gn=ADGRL2;refGene_tx=XM_019837295.2;refGene_gn=ADGRL2;refGene_tx=XM_019837296.2;refGene_gn=ADGRL2;refGene_tx=XM_019837297.2;refGene_gn=ADGRL2;refGene_tx=XM_019837298.2;refGene_gn=ADGRL2;refGene_tx=XM_019837299.2;refGene_gn=ADGRL2;refGene_tx=XM_019837300.2;refGene_gn=ADGRL2;refGene_tx=XM_019837301.2;refGene_gn=ADGRL2;refGene_tx=XM_019837302.2;refGene_gn=ADGRL2;refGene_tx=XM_019837303.2;refGene_gn=ADGRL2;refGene_tx=XM_019837304.2;refGene_gn=ADGRL2;refGene_tx=XM_019837305.2;refGene_gn=ADGRL2;refGene_tx=XM_019837306.2;refGene_gn=ADGRL2;refGene_tx=XM_019837307.2;refGene_gn=ADGRL2;refGene_tx=XM_019837308.2;refGene_gn=ADGRL2;refGene_tx=XM_019837309.2;refGene_gn=ADGRL2;refGene_tx=XM_023258848.1;refGene_gn=ADGRL2;refGene_tx=XM_023258849.1;refGene_gn=ADGRL2;refGene_tx=XM_023258850.1
C1 70932558 70932679
refGene_gn=TTLL7;refGene_tx=XM_011284903.3;refGene_gn=TTLL7;refGene_tx=XM_019837310.2
C1 70943439 70943613
refGene_gn=TTLL7;refGene_tx=XM_011284903.3;refGene_gn=TTLL7;refGene_tx=XM_019837310.2
C1 70957741 70957902
refGene_gn=TTLL7;refGene_tx=XM_011284903.3;refGene_gn=TTLL7;refGene_tx=XM_019837310.2

C1 70964369 70964408 refGene_gn=TTLL7;refGene_tx=XM_011284905.3
C1 70971034 70971115
refGene_gn=TTLL7;refGene_tx=XM_011284903.3;refGene_gn=TTLL7;refGene_tx=XM_011284905.3;refGene_gn=TTLL7;refGene_tx=XM_019837310.2
C1 70973198 70973333
refGene_gn=TTLL7;refGene_tx=XM_011284903.3;refGene_gn=TTLL7;refGene_tx=XM_011284905.3;refGene_gn=TTLL7;refGene_tx=XM_019837310.2
C1 70974325 70974531
refGene_gn=TTLL7;refGene_tx=XM_011284903.3;refGene_gn=TTLL7;refGene_tx=XM_011284905.3;refGene_gn=TTLL7;refGene_tx=XM_019837310.2
C1 70978299 70978498
refGene_gn=TTLL7;refGene_tx=XM_011284903.3;refGene_gn=TTLL7;refGene_tx=XM_011284905.3;refGene_gn=TTLL7;refGene_tx=XM_019837310.2
C1 70982973 70983060
refGene_gn=TTLL7;refGene_tx=XM_011284903.3;refGene_gn=TTLL7;refGene_tx=XM_011284905.3;refGene_gn=TTLL7;refGene_tx=XM_019837310.2
C1 70989755 70989891
refGene_gn=TTLL7;refGene_tx=XM_011284903.3;refGene_gn=TTLL7;refGene_tx=XM_011284905.3;refGene_gn=TTLL7;refGene_tx=XM_019837310.2
C1 70990912 70990986
refGene_gn=TTLL7;refGene_tx=XM_011284903.3;refGene_gn=TTLL7;refGene_tx=XM_011284905.3;refGene_gn=TTLL7;refGene_tx=XM_019837310.2
C1 70991071 70991219
refGene_gn=TTLL7;refGene_tx=XM_011284903.3;refGene_gn=TTLL7;refGene_tx=XM_011284905.3;refGene_gn=TTLL7;refGene_tx=XM_019837310.2
C1 70993447 70993542
refGene_gn=TTLL7;refGene_tx=XM_011284903.3;refGene_gn=TTLL7;refGene_tx=XM_011284905.3;refGene_gn=TTLL7;refGene_tx=XM_019837310.2
C1 70998682 70998841
refGene_gn=TTLL7;refGene_tx=XM_011284903.3;refGene_gn=TTLL7;refGene_tx=XM_011284905.3;refGene_gn=TTLL7;refGene_tx=XM_019837310.2
C1 71004210 71004375
refGene_gn=TTLL7;refGene_tx=XM_011284903.3;refGene_gn=TTLL7;refGene_tx=XM_011284905.3;refGene_gn=TTLL7;refGene_tx=XM_019837310.2
C1 71009591 71009808
refGene_gn=TTLL7;refGene_tx=XM_011284903.3;refGene_gn=TTLL7;refGene_tx=XM_011284905.3;refGene_gn=TTLL7;refGene_tx=XM_019837310.2
C1 71015301 71015460
refGene_gn=TTLL7;refGene_tx=XM_011284903.3;refGene_gn=TTLL7;refGene_tx=XM_011284905.3;refGene_gn=TTLL7;refGene_tx=XM_019837310.2
C1 71019400 71019468
refGene_gn=TTLL7;refGene_tx=XM_011284903.3;refGene_gn=TTLL7;refGene_tx=XM_011284905.3;refGene_gn=TTLL7;refGene_tx=XM_019837310.2
C1 71020407 71020529
refGene_gn=TTLL7;refGene_tx=XM_011284903.3;refGene_gn=TTLL7;refGene_tx=XM_011284905.3;refGene_gn=TTLL7;refGene_tx=XM_019837310.2
C1 71025040 71025172
refGene_gn=TTLL7;refGene_tx=XM_011284903.3;refGene_gn=TTLL7;refGene_tx=XM_011284905.3;refGene_gn=TTLL7;refGene_tx=XM_019837310.2
C1 71025384 71025409

refGene_gn=TTLL7;refGene_tx=XM_011284903.3;refGene_gn=TTLL7;refGene_tx=XM_011284905.3
C1 71126245 71126291
refGene_gn=PRKACB;refGene_tx=XM_023258853.1;refGene_gn=PRKACB;refGene_tx=XM_023258854.1
C1 71182089 71182273 refGene_gn=PRKACB;refGene_tx=XM_003990255.5
C1 71197923 71197933
refGene_gn=PRKACB;refGene_tx=XM_011284900.3;refGene_gn=PRKACB;refGene_tx=XM_011284902.3;refGene_gn=PRKACB;refGene_tx=XM_023258855.1
C1 71198520 71198527
refGene_gn=PRKACB;refGene_tx=XM_011284901.3;refGene_gn=PRKACB;refGene_tx=XM_019837312.2;refGene_gn=PRKACB;refGene_tx=XM_019837313.2
C1 71206995 71207019
refGene_gn=PRKACB;refGene_tx=XM_011284900.3;refGene_gn=PRKACB;refGene_tx=XM_011284901.3
C1 71208965 71208989
refGene_gn=PRKACB;refGene_tx=XM_011284900.3;refGene_gn=PRKACB;refGene_tx=XM_011284901.3;refGene_gn=PRKACB;refGene_tx=XM_019837312.2;refGene_gn=PRKACB;refGene_tx=XM_023258853.1;refGene_gn=PRKACB;refGene_tx=XM_023258855.1
C1 71213262 71213324
refGene_gn=PRKACB;refGene_tx=XM_003990255.5;refGene_gn=PRKACB;refGene_tx=XM_011284900.3;refGene_gn=PRKACB;refGene_tx=XM_011284901.3;refGene_gn=PRKACB;refGene_tx=XM_011284902.3;refGene_gn=PRKACB;refGene_tx=XM_019837312.2;refGene_gn=PRKACB;refGene_tx=XM_019837313.2;refGene_gn=PRKACB;refGene_tx=XM_023258853.1;refGene_gn=PRKACB;refGene_tx=XM_023258854.1;refGene_gn=PRKACB;refGene_tx=XM_023258855.1
C1 71215522 71215651
refGene_gn=PRKACB;refGene_tx=XM_003990255.5;refGene_gn=PRKACB;refGene_tx=XM_011284900.3;refGene_gn=PRKACB;refGene_tx=XM_011284901.3;refGene_gn=PRKACB;refGene_tx=XM_011284902.3;refGene_gn=PRKACB;refGene_tx=XM_019837312.2;refGene_gn=PRKACB;refGene_tx=XM_019837313.2;refGene_gn=PRKACB;refGene_tx=XM_023258853.1;refGene_gn=PRKACB;refGene_tx=XM_023258854.1;refGene_gn=PRKACB;refGene_tx=XM_023258855.1
C1 71223411 71223510
refGene_gn=PRKACB;refGene_tx=XM_003990255.5;refGene_gn=PRKACB;refGene_tx=XM_011284900.3;refGene_gn=PRKACB;refGene_tx=XM_011284901.3;refGene_gn=PRKACB;refGene_tx=XM_011284902.3;refGene_gn=PRKACB;refGene_tx=XM_019837312.2;refGene_gn=PRKACB;refGene_tx=XM_019837313.2;refGene_gn=PRKACB;refGene_tx=XM_023258853.1;refGene_gn=PRKACB;refGene_tx=XM_023258854.1;refGene_gn=PRKACB;refGene_tx=XM_023258855.1
C1 71224515 71224598
refGene_gn=PRKACB;refGene_tx=XM_003990255.5;refGene_gn=PRKACB;refGene_tx=XM_011284900.3;refGene_gn=PRKACB;refGene_tx=XM_011284901.3;refGene_gn=PRKACB;refGene_tx=XM_011284902.3;refGene_gn=PRKACB;refGene_tx=XM_019837312.2;refGene_gn=PRKACB;refGene_tx=XM_019837313.2;refGene_gn=PRKACB;refGene_tx=XM_023258853.1;refGene_gn=PRKACB;refGene_tx=XM_023258854.1;refGene_gn=PRKACB;refGene_tx=XM_023258855.1
C1 71227780 71227907
refGene_gn=PRKACB;refGene_tx=XM_003990255.5;refGene_gn=PRKACB;refGene_

tx=XM_011284900.3;refGene_gn=PRKACB;refGene_tx=XM_011284901.3;refGene_gn=PRKACB;refGene_tx=XM_011284902.3;refGene_gn=PRKACB;refGene_tx=XM_019837312.2;refGene_gn=PRKACB;refGene_tx=XM_019837313.2;refGene_gn=PRKACB;refGene_tx=XM_023258853.1;refGene_gn=PRKACB;refGene_tx=XM_023258854.1;refGene_gn=PRKACB;refGene_tx=XM_023258855.1
C1 71228868 71228964
refGene_gn=PRKACB;refGene_tx=XM_003990255.5;refGene_gn=PRKACB;refGene_tx=XM_011284900.3;refGene_gn=PRKACB;refGene_tx=XM_011284901.3;refGene_gn=PRKACB;refGene_tx=XM_011284902.3;refGene_gn=PRKACB;refGene_tx=XM_019837312.2;refGene_gn=PRKACB;refGene_tx=XM_019837313.2;refGene_gn=PRKACB;refGene_tx=XM_023258853.1;refGene_gn=PRKACB;refGene_tx=XM_023258854.1;refGene_gn=PRKACB;refGene_tx=XM_023258855.1
C1 71232471 71232594
refGene_gn=PRKACB;refGene_tx=XM_003990255.5;refGene_gn=PRKACB;refGene_tx=XM_011284900.3;refGene_gn=PRKACB;refGene_tx=XM_011284901.3;refGene_gn=PRKACB;refGene_tx=XM_011284902.3;refGene_gn=PRKACB;refGene_tx=XM_019837312.2;refGene_gn=PRKACB;refGene_tx=XM_019837313.2;refGene_gn=PRKACB;refGene_tx=XM_023258853.1;refGene_gn=PRKACB;refGene_tx=XM_023258854.1;refGene_gn=PRKACB;refGene_tx=XM_023258855.1
C1 71244619 71244784
refGene_gn=PRKACB;refGene_tx=XM_003990255.5;refGene_gn=PRKACB;refGene_tx=XM_011284900.3;refGene_gn=PRKACB;refGene_tx=XM_011284901.3;refGene_gn=PRKACB;refGene_tx=XM_011284902.3;refGene_gn=PRKACB;refGene_tx=XM_019837312.2;refGene_gn=PRKACB;refGene_tx=XM_019837313.2;refGene_gn=PRKACB;refGene_tx=XM_023258853.1;refGene_gn=PRKACB;refGene_tx=XM_023258854.1;refGene_gn=PRKACB;refGene_tx=XM_023258855.1
C1 71248289 71248415
refGene_gn=PRKACB;refGene_tx=XM_003990255.5;refGene_gn=PRKACB;refGene_tx=XM_011284900.3;refGene_gn=PRKACB;refGene_tx=XM_011284901.3;refGene_gn=PRKACB;refGene_tx=XM_011284902.3;refGene_gn=PRKACB;refGene_tx=XM_019837312.2;refGene_gn=PRKACB;refGene_tx=XM_019837313.2;refGene_gn=PRKACB;refGene_tx=XM_023258853.1;refGene_gn=PRKACB;refGene_tx=XM_023258854.1;refGene_gn=PRKACB;refGene_tx=XM_023258855.1
C1 71308752 71308879 refGene_gn=SAMD13;refGene_tx=XM_019839087.2
C1 71313605 71313690 refGene_gn=SAMD13;refGene_tx=XM_019839087.2
C1 71331464 71331576 refGene_gn=SAMD13;refGene_tx=XM_019839087.2
C1 71347542 71347686 refGene_gn=SAMD13;refGene_tx=XM_019839087.2
C1 71359546 71359621
refGene_gn=LOC101096750;refGene_tx=XM_003990261.5;refGene_gn=LOC101096750;refGene_tx=XM_019837322.2
C1 71361746 71361830
refGene_gn=LOC101096750;refGene_tx=XM_003990261.5;refGene_gn=LOC101096750;refGene_tx=XM_019837322.2
C1 71363997 71364121
refGene_gn=LOC101096750;refGene_tx=XM_003990261.5;refGene_gn=LOC101096750;refGene_tx=XM_019837322.2
C1 71365829 71366016
refGene_gn=LOC101096750;refGene_tx=XM_003990261.5;refGene_gn=LOC101096750;refGene_tx=XM_019837322.2
C1 71376850 71376929

refGene_gn=LOC101096750;refGene_tx=XM_003990261.5;refGene_gn=LOC101096750;refGene_tx=XM_019837322.2
C1 71377028 71377145
refGene_gn=LOC101096750;refGene_tx=XM_003990261.5;refGene_gn=LOC101096750;refGene_tx=XM_019837322.2
C1 71379926 71380145
refGene_gn=LOC101096750;refGene_tx=XM_003990261.5;refGene_gn=LOC101096750;refGene_tx=XM_019837322.2
C1 71390212 71390320
refGene_gn=LOC101096750;refGene_tx=XM_003990261.5
C1 71391028 71391141 refGene_gn=DNASE2B;refGene_tx=XM_019837317.2
C1 71394256 71394434
refGene_gn=DNASE2B;refGene_tx=XM_019837316.2;refGene_gn=DNASE2B;refGene_tx=XM_019837317.2;refGene_gn=DNASE2B;refGene_tx=XM_019837318.2;refGene_gn=DNASE2B;refGene_tx=XM_023258856.1
C1 71397790 71397872
refGene_gn=DNASE2B;refGene_tx=XM_019837316.2;refGene_gn=DNASE2B;refGene_tx=XM_019837317.2;refGene_gn=DNASE2B;refGene_tx=XM_019837318.2;refGene_gn=DNASE2B;refGene_tx=XM_023258856.1
C1 71400396 71400558
refGene_gn=DNASE2B;refGene_tx=XM_019837316.2;refGene_gn=DNASE2B;refGene_tx=XM_019837317.2;refGene_gn=DNASE2B;refGene_tx=XM_019837318.2;refGene_gn=DNASE2B;refGene_tx=XM_023258856.1
C1 71402107 71402305
refGene_gn=DNASE2B;refGene_tx=XM_006934875.4;refGene_gn=DNASE2B;refGene_tx=XM_019837316.2;refGene_gn=DNASE2B;refGene_tx=XM_019837317.2;refGene_gn=DNASE2B;refGene_tx=XM_019837318.2;refGene_gn=DNASE2B;refGene_tx=XM_019837319.2;refGene_gn=DNASE2B;refGene_tx=XM_023258856.1
C1 71406016 71406357
refGene_gn=DNASE2B;refGene_tx=XM_006934875.4;refGene_gn=DNASE2B;refGene_tx=XM_019837316.2;refGene_gn=DNASE2B;refGene_tx=XM_019837317.2;refGene_gn=DNASE2B;refGene_tx=XM_019837318.2;refGene_gn=DNASE2B;refGene_tx=XM_019837319.2;refGene_gn=DNASE2B;refGene_tx=XM_023258856.1
C1 71439134 71439185
refGene_gn=LOC101096750;refGene_tx=XM_019837322.2
C1 71457164 71457392 refGene_gn=RPF1;refGene_tx=XM_003990263.5
C1 71458891 71458948 refGene_gn=RPF1;refGene_tx=XM_003990263.5
C1 71461849 71461930 refGene_gn=RPF1;refGene_tx=XM_003990263.5
C1 71467759 71467855 refGene_gn=RPF1;refGene_tx=XM_003990263.5
C1 71468447 71468601 refGene_gn=RPF1;refGene_tx=XM_003990263.5
C1 71475494 71475577 refGene_gn=RPF1;refGene_tx=XM_003990263.5
C1 71475989 71476171 refGene_gn=RPF1;refGene_tx=XM_003990263.5
C1 71476359 71476486 refGene_gn=RPF1;refGene_tx=XM_003990263.5
C1 71477052 71477094 refGene_gn=RPF1;refGene_tx=XM_003990263.5
C1 71482347 71482473 refGene_gn=GNG5;refGene_tx=XM_023258859.1
C1 71486253 71486334 refGene_gn=GNG5;refGene_tx=XM_023258859.1
C1 71495499 71495601
refGene_gn=SPATA1;refGene_tx=XM_011284899.3;refGene_gn=SPATA1;refGene_tx=XM_019837323.2;refGene_gn=SPATA1;refGene_tx=XM_019837324.2
C1 71506014 71506050 refGene_gn=SPATA1;refGene_tx=XM_023258858.1

C1 71510535 71510642
refGene_gn=SPATA1;refGene_tx=XM_011284899.3;refGene_gn=SPATA1;refGene_tx=XM_019837323.2;refGene_gn=SPATA1;refGene_tx=XM_019837324.2;refGene_gn=SPATA1;refGene_tx=XM_023258858.1
C1 71512323 71512441
refGene_gn=SPATA1;refGene_tx=XM_011284899.3;refGene_gn=SPATA1;refGene_tx=XM_019837323.2;refGene_gn=SPATA1;refGene_tx=XM_019837324.2;refGene_gn=SPATA1;refGene_tx=XM_023258858.1
C1 71515514 71515568
refGene_gn=SPATA1;refGene_tx=XM_011284899.3;refGene_gn=SPATA1;refGene_tx=XM_019837323.2;refGene_gn=SPATA1;refGene_tx=XM_019837324.2;refGene_gn=SPATA1;refGene_tx=XM_023258858.1
C1 71515663 71515892
refGene_gn=SPATA1;refGene_tx=XM_011284899.3;refGene_gn=SPATA1;refGene_tx=XM_019837323.2;refGene_gn=SPATA1;refGene_tx=XM_019837324.2;refGene_gn=SPATA1;refGene_tx=XM_023258858.1
C1 71526218 71526330
refGene_gn=SPATA1;refGene_tx=XM_011284899.3;refGene_gn=SPATA1;refGene_tx=XM_019837323.2;refGene_gn=SPATA1;refGene_tx=XM_019837324.2;refGene_gn=SPATA1;refGene_tx=XM_023258858.1
C1 71526994 71527055
refGene_gn=SPATA1;refGene_tx=XM_011284899.3;refGene_gn=SPATA1;refGene_tx=XM_019837323.2;refGene_gn=SPATA1;refGene_tx=XM_019837324.2;refGene_gn=SPATA1;refGene_tx=XM_023258858.1
C1 71531525 71531637
refGene_gn=SPATA1;refGene_tx=XM_011284899.3;refGene_gn=SPATA1;refGene_tx=XM_019837323.2;refGene_gn=SPATA1;refGene_tx=XM_019837324.2;refGene_gn=SPATA1;refGene_tx=XM_023258858.1
C1 71532223 71532349
refGene_gn=SPATA1;refGene_tx=XM_011284899.3;refGene_gn=SPATA1;refGene_tx=XM_019837323.2;refGene_gn=SPATA1;refGene_tx=XM_019837324.2;refGene_gn=SPATA1;refGene_tx=XM_023258858.1
C1 71535200 71535379
refGene_gn=SPATA1;refGene_tx=XM_011284899.3;refGene_gn=SPATA1;refGene_tx=XM_019837323.2;refGene_gn=SPATA1;refGene_tx=XM_019837324.2;refGene_gn=SPATA1;refGene_tx=XM_023258858.1
C1 71536851 71536950
refGene_gn=SPATA1;refGene_tx=XM_011284899.3;refGene_gn=SPATA1;refGene_tx=XM_019837323.2;refGene_gn=SPATA1;refGene_tx=XM_019837324.2;refGene_gn=SPATA1;refGene_tx=XM_023258858.1
C1 71537689 71537777
refGene_gn=SPATA1;refGene_tx=XM_011284899.3;refGene_gn=SPATA1;refGene_tx=XM_019837323.2;refGene_gn=SPATA1;refGene_tx=XM_019837324.2;refGene_gn=SPATA1;refGene_tx=XM_023258858.1
C1 71539062 71539254 refGene_gn=CTBS;refGene_tx=XM_023258860.1
C1 71539710 71539733
refGene_gn=SPATA1;refGene_tx=XM_011284899.3;refGene_gn=SPATA1;refGene_tx=XM_019837323.2;refGene_gn=SPATA1;refGene_tx=XM_019837324.2;refGene_gn=SPATA1;refGene_tx=XM_023258858.1
C1 71541646 71541808 refGene_gn=CTBS;refGene_tx=XM_023258860.1

C1 71542080 71542178 refGene_gn=CTBS;refGene_tx=XM_023258860.1
C1 71543587 71543756 refGene_gn=CTBS;refGene_tx=XM_023258860.1
C1 71561195 71561404 refGene_gn=CTBS;refGene_tx=XM_023258860.1
C1 71561875 71562014 refGene_gn=CTBS;refGene_tx=XM_023258860.1
C1 71581926 71582094 refGene_gn=CTBS;refGene_tx=XM_023258860.1
C1 71590998 71591089
refGene_gn=LOC109502425;refGene_tx=XM_019837326.1
C1 71593436 71593452
refGene_gn=LOC109502425;refGene_tx=XM_019837327.1
C1 71599458 71599798
refGene_gn=LOC109502425;refGene_tx=XM_019837326.1;refGene_gn=LOC109502425;refGene_tx=XM_019837327.1;refGene_gn=LOC109502425;refGene_tx=XM_019837330.1;refGene_gn=LOC109502425;refGene_tx=XM_019837331.1;refGene_gn=LOC109502425;refGene_tx=XM_019837332.1;refGene_gn=LOC109502425;refGene_tx=XM_019837333.1;refGene_gn=LOC109502425;refGene_tx=XM_019837336.1
C1 71600798 71600958
refGene_gn=LOC109502425;refGene_tx=XM_019837326.1;refGene_gn=LOC109502425;refGene_tx=XM_019837327.1;refGene_gn=LOC109502425;refGene_tx=XM_019837330.1;refGene_gn=LOC109502425;refGene_tx=XM_019837331.1;refGene_gn=LOC109502425;refGene_tx=XM_019837332.1;refGene_gn=LOC109502425;refGene_tx=XM_019837333.1;refGene_gn=LOC109502425;refGene_tx=XM_019837336.1
C1 71605509 71605718
refGene_gn=LOC109502425;refGene_tx=XM_019837326.1;refGene_gn=LOC109502425;refGene_tx=XM_019837327.1;refGene_gn=LOC109502425;refGene_tx=XM_019837330.1;refGene_gn=LOC109502425;refGene_tx=XM_019837331.1;refGene_gn=LOC109502425;refGene_tx=XM_019837332.1;refGene_gn=LOC109502425;refGene_tx=XM_019837333.1;refGene_gn=LOC109502425;refGene_tx=XM_019837336.1
C1 71628111 71628286
refGene_gn=SSX2IP;refGene_tx=XM_011284896.3;refGene_gn=SSX2IP;refGene_tx=XM_019837338.2;refGene_gn=SSX2IP;refGene_tx=XM_023258861.1;refGene_gn=SSX2IP;refGene_tx=XM_023258862.1;refGene_gn=SSX2IP;refGene_tx=XM_023258863.1
C1 71631435 71631601
refGene_gn=SSX2IP;refGene_tx=XM_011284896.3;refGene_gn=SSX2IP;refGene_tx=XM_019837338.2;refGene_gn=SSX2IP;refGene_tx=XM_023258861.1;refGene_gn=SSX2IP;refGene_tx=XM_023258862.1;refGene_gn=SSX2IP;refGene_tx=XM_023258863.1
C1 71632163 71632278
refGene_gn=SSX2IP;refGene_tx=XM_011284896.3;refGene_gn=SSX2IP;refGene_tx=XM_019837338.2;refGene_gn=SSX2IP;refGene_tx=XM_023258861.1;refGene_gn=SSX2IP;refGene_tx=XM_023258862.1;refGene_gn=SSX2IP;refGene_tx=XM_023258863.1
C1 71635087 71635261
refGene_gn=SSX2IP;refGene_tx=XM_011284896.3;refGene_gn=SSX2IP;refGene_tx=XM_019837338.2;refGene_gn=SSX2IP;refGene_tx=XM_023258861.1;refGene_gn=SSX2IP;refGene_tx=XM_023258862.1;refGene_gn=SSX2IP;refGene_tx=XM_023258863.1
C1 71635588 71635725
refGene_gn=SSX2IP;refGene_tx=XM_011284896.3;refGene_gn=SSX2IP;refGene_tx=XM_019837338.2;refGene_gn=SSX2IP;refGene_tx=XM_023258861.1;refGene_

gn=SSX2IP;refGene_tx=XM_023258862.1;refGene_gn=SSX2IP;refGene_tx=XM_02
3258863.1
C1 71637491 71637642
refGene_gn=SSX2IP;refGene_tx=XM_011284896.3;refGene_gn=SSX2IP;refGene_
tx=XM_019837338.2;refGene_gn=SSX2IP;refGene_tx=XM_023258861.1;refGene_
gn=SSX2IP;refGene_tx=XM_023258862.1;refGene_gn=SSX2IP;refGene_tx=XM_02
3258863.1
C1 71641031 71641221
refGene_gn=SSX2IP;refGene_tx=XM_011284896.3;refGene_gn=SSX2IP;refGene_
tx=XM_019837338.2;refGene_gn=SSX2IP;refGene_tx=XM_023258861.1;refGene_
gn=SSX2IP;refGene_tx=XM_023258862.1;refGene_gn=SSX2IP;refGene_tx=XM_02
3258863.1
C1 71641291 71641367
refGene_gn=SSX2IP;refGene_tx=XM_011284896.3;refGene_gn=SSX2IP;refGene_
tx=XM_019837338.2;refGene_gn=SSX2IP;refGene_tx=XM_023258861.1;refGene_
gn=SSX2IP;refGene_tx=XM_023258862.1;refGene_gn=SSX2IP;refGene_tx=XM_02
3258863.1
C1 71643177 71643313
refGene_gn=SSX2IP;refGene_tx=XM_011284896.3;refGene_gn=SSX2IP;refGene_
tx=XM_019837338.2;refGene_gn=SSX2IP;refGene_tx=XM_023258861.1;refGene_
gn=SSX2IP;refGene_tx=XM_023258862.1;refGene_gn=SSX2IP;refGene_tx=XM_02
3258863.1
C1 71644913 71645024
refGene_gn=SSX2IP;refGene_tx=XM_011284896.3;refGene_gn=SSX2IP;refGene_
tx=XM_019837338.2;refGene_gn=SSX2IP;refGene_tx=XM_023258861.1;refGene_
gn=SSX2IP;refGene_tx=XM_023258862.1;refGene_gn=SSX2IP;refGene_tx=XM_02
3258863.1
C1 71647531 71647744
refGene_gn=SSX2IP;refGene_tx=XM_011284896.3;refGene_gn=SSX2IP;refGene_
tx=XM_019837338.2;refGene_gn=SSX2IP;refGene_tx=XM_023258861.1;refGene_
gn=SSX2IP;refGene_tx=XM_023258862.1;refGene_gn=SSX2IP;refGene_tx=XM_02
3258863.1
C1 71648212 71648382
refGene_gn=SSX2IP;refGene_tx=XM_011284896.3;refGene_gn=SSX2IP;refGene_
tx=XM_019837338.2;refGene_gn=SSX2IP;refGene_tx=XM_023258861.1;refGene_
gn=SSX2IP;refGene_tx=XM_023258862.1;refGene_gn=SSX2IP;refGene_tx=XM_02
3258863.1
C1 71649287 71649330
refGene_gn=SSX2IP;refGene_tx=XM_011284896.3;refGene_gn=SSX2IP;refGene_
tx=XM_019837338.2;refGene_gn=SSX2IP;refGene_tx=XM_023258861.1;refGene_
gn=SSX2IP;refGene_tx=XM_023258862.1;refGene_gn=SSX2IP;refGene_tx=XM_02
3258863.1
C1 71789383 71789712
refGene_gn=LPAR3;refGene_tx=XM_019837345.2;refGene_gn=LPAR3;refGene_tx
=XM_019837346.2;refGene_gn=LPAR3;refGene_tx=XM_023258864.1
C1 71837854 71838590
refGene_gn=LPAR3;refGene_tx=XM_019837345.2;refGene_gn=LPAR3;refGene_tx
=XM_019837346.2;refGene_gn=LPAR3;refGene_tx=XM_023258864.1
C1 71893955 71893992
refGene_gn=MCOLN2;refGene_tx=XM_006934876.3;refGene_gn=MCOLN2;refGene_

tx=XM_019837340.1;refGene_gn=MCOLN2;refGene_tx=XM_019837341.1;refGene_gn=MCOLN2;refGene_tx=XM_019837342.2;refGene_gn=MCOLN2;refGene_tx=XM_019837343.1
C1 71896412 71896534
refGene_gn=MCOLN2;refGene_tx=XM_006934876.3;refGene_gn=MCOLN2;refGene_tx=XM_019837340.1;refGene_gn=MCOLN2;refGene_tx=XM_019837341.1;refGene_gn=MCOLN2;refGene_tx=XM_019837342.2;refGene_gn=MCOLN2;refGene_tx=XM_019837343.1
C1 71898552 71898759
refGene_gn=MCOLN2;refGene_tx=XM_006934876.3;refGene_gn=MCOLN2;refGene_tx=XM_019837340.1;refGene_gn=MCOLN2;refGene_tx=XM_019837341.1;refGene_gn=MCOLN2;refGene_tx=XM_019837342.2;refGene_gn=MCOLN2;refGene_tx=XM_019837343.1
C1 71906634 71906757
refGene_gn=MCOLN2;refGene_tx=XM_006934876.3;refGene_gn=MCOLN2;refGene_tx=XM_019837340.1;refGene_gn=MCOLN2;refGene_tx=XM_019837341.1;refGene_gn=MCOLN2;refGene_tx=XM_019837342.2;refGene_gn=MCOLN2;refGene_tx=XM_019837343.1
C1 71906859 71906961
refGene_gn=MCOLN2;refGene_tx=XM_006934876.3;refGene_gn=MCOLN2;refGene_tx=XM_019837340.1;refGene_gn=MCOLN2;refGene_tx=XM_019837341.1;refGene_gn=MCOLN2;refGene_tx=XM_019837342.2;refGene_gn=MCOLN2;refGene_tx=XM_019837343.1
C1 71908601 71908751
refGene_gn=MCOLN2;refGene_tx=XM_006934876.3;refGene_gn=MCOLN2;refGene_tx=XM_019837340.1;refGene_gn=MCOLN2;refGene_tx=XM_019837341.1;refGene_gn=MCOLN2;refGene_tx=XM_019837342.2;refGene_gn=MCOLN2;refGene_tx=XM_019837343.1
C1 71910186 71910299
refGene_gn=MCOLN2;refGene_tx=XM_006934876.3;refGene_gn=MCOLN2;refGene_tx=XM_019837340.1;refGene_gn=MCOLN2;refGene_tx=XM_019837341.1;refGene_gn=MCOLN2;refGene_tx=XM_019837342.2;refGene_gn=MCOLN2;refGene_tx=XM_019837343.1
C1 71916740 71916840
refGene_gn=MCOLN2;refGene_tx=XM_006934876.3;refGene_gn=MCOLN2;refGene_tx=XM_019837340.1;refGene_gn=MCOLN2;refGene_tx=XM_019837341.1;refGene_gn=MCOLN2;refGene_tx=XM_019837342.2;refGene_gn=MCOLN2;refGene_tx=XM_019837343.1
C1 71919036 71919133
refGene_gn=MCOLN2;refGene_tx=XM_006934876.3;refGene_gn=MCOLN2;refGene_tx=XM_019837340.1;refGene_gn=MCOLN2;refGene_tx=XM_019837341.1;refGene_gn=MCOLN2;refGene_tx=XM_019837342.2;refGene_gn=MCOLN2;refGene_tx=XM_019837343.1
C1 71919238 71919314
refGene_gn=MCOLN2;refGene_tx=XM_006934876.3;refGene_gn=MCOLN2;refGene_tx=XM_019837340.1;refGene_gn=MCOLN2;refGene_tx=XM_019837341.1;refGene_gn=MCOLN2;refGene_tx=XM_019837342.2;refGene_gn=MCOLN2;refGene_tx=XM_019837343.1
C1 71923879 71924033
refGene_gn=MCOLN2;refGene_tx=XM_006934876.3;refGene_gn=MCOLN2;refGene_

tx=XM_019837340.1;refGene_gn=MCOLN2;refGene_tx=XM_019837341.1;refGene_gn=MCOLN2;refGene_tx=XM_019837342.2;refGene_gn=MCOLN2;refGene_tx=XM_019837343.1
C1 71926053 71926227 refGene_gn=MCOLN2;refGene_tx=XM_006934876.3;refGene_gn=MCOLN2;refGene_tx=XM_019837340.1;refGene_gn=MCOLN2;refGene_tx=XM_019837341.1;refGene_gn=MCOLN2;refGene_tx=XM_019837342.2;refGene_gn=MCOLN2;refGene_tx=XM_019837343.1
C1 71932171 71932331 refGene_gn=MCOLN2;refGene_tx=XM_006934876.3;refGene_gn=MCOLN2;refGene_tx=XM_019837340.1;refGene_gn=MCOLN2;refGene_tx=XM_019837341.1;refGene_gn=MCOLN2;refGene_tx=XM_019837342.2;refGene_gn=MCOLN2;refGene_tx=XM_019837343.1
C1 71942984 71943028 refGene_gn=MCOLN2;refGene_tx=XM_019837341.1
C1 71950336 71950413 refGene_gn=MCOLN2;refGene_tx=XM_019837340.1
C1 71954035 71954169 refGene_gn=MCOLN3;refGene_tx=XM_023258865.1
C1 71955660 71955871 refGene_gn=MCOLN3;refGene_tx=XM_023258865.1
C1 71956946 71957069 refGene_gn=MCOLN3;refGene_tx=XM_023258865.1
C1 71957179 71957281 refGene_gn=MCOLN3;refGene_tx=XM_023258865.1
C1 71961092 71961242 refGene_gn=MCOLN3;refGene_tx=XM_023258865.1
C1 71961324 71961437 refGene_gn=MCOLN3;refGene_tx=XM_023258865.1
C1 71965534 71965634 refGene_gn=MCOLN3;refGene_tx=XM_023258865.1
C1 71970049 71970146 refGene_gn=MCOLN3;refGene_tx=XM_023258865.1
C1 71970225 71970310 refGene_gn=MCOLN3;refGene_tx=XM_023258865.1
C1 71971718 71971872 refGene_gn=MCOLN3;refGene_tx=XM_023258865.1
C1 71976891 71977059 refGene_gn=MCOLN3;refGene_tx=XM_023258865.1
C1 71981655 71981883 refGene_gn=MCOLN3;refGene_tx=XM_023258865.1
C1 72009644 72009726 refGene_gn=WDR63;refGene_tx=XM_006934877.4;refGene_gn=WDR63;refGene_tx=XM_019837349.1;refGene_gn=WDR63;refGene_tx=XM_019837350.2;refGene_gn=WDR63;refGene_tx=XM_019837351.2
C1 72010118 72010157 refGene_gn=WDR63;refGene_tx=XM_006934877.4;refGene_gn=WDR63;refGene_tx=XM_019837349.1;refGene_gn=WDR63;refGene_tx=XM_019837350.2;refGene_gn=WDR63;refGene_tx=XM_019837351.2
C1 72016608 72016790 refGene_gn=WDR63;refGene_tx=XM_006934877.4;refGene_gn=WDR63;refGene_tx=XM_019837349.1;refGene_gn=WDR63;refGene_tx=XM_019837350.2;refGene_gn=WDR63;refGene_tx=XM_019837351.2
C1 72018059 72018164 refGene_gn=WDR63;refGene_tx=XM_006934877.4;refGene_gn=WDR63;refGene_tx=XM_019837349.1;refGene_gn=WDR63;refGene_tx=XM_019837350.2;refGene_gn=WDR63;refGene_tx=XM_019837351.2
C1 72018730 72018880 refGene_gn=WDR63;refGene_tx=XM_006934877.4;refGene_gn=WDR63;refGene_tx=XM_019837349.1;refGene_gn=WDR63;refGene_tx=XM_019837350.2;refGene_gn=WDR63;refGene_tx=XM_019837351.2
C1 72019954 72020154 refGene_gn=WDR63;refGene_tx=XM_006934877.4;refGene_gn=WDR63;refGene_tx=XM_019837349.1;refGene_gn=WDR63;refGene_tx=XM_019837350.2;refGene_gn=

WDR63;refGene_tx=XM_019837351.2
C1 72026114 72026231
refGene_gn=WDR63;refGene_tx=XM_006934877.4;refGene_gn=WDR63;refGene_tx=XM_019837349.1;refGene_gn=WDR63;refGene_tx=XM_019837350.2;refGene_gn=WDR63;refGene_tx=XM_019837351.2
C1 72032565 72032756
refGene_gn=WDR63;refGene_tx=XM_006934877.4;refGene_gn=WDR63;refGene_tx=XM_019837349.1;refGene_gn=WDR63;refGene_tx=XM_019837350.2;refGene_gn=WDR63;refGene_tx=XM_019837351.2
C1 72033718 72033843
refGene_gn=WDR63;refGene_tx=XM_006934877.4;refGene_gn=WDR63;refGene_tx=XM_019837349.1;refGene_gn=WDR63;refGene_tx=XM_019837350.2;refGene_gn=WDR63;refGene_tx=XM_019837351.2
C1 72035988 72036078
refGene_gn=WDR63;refGene_tx=XM_006934877.4;refGene_gn=WDR63;refGene_tx=XM_019837349.1;refGene_gn=WDR63;refGene_tx=XM_019837350.2;refGene_gn=WDR63;refGene_tx=XM_019837351.2
C1 72037859 72037946
refGene_gn=WDR63;refGene_tx=XM_006934877.4;refGene_gn=WDR63;refGene_tx=XM_019837349.1;refGene_gn=WDR63;refGene_tx=XM_019837350.2;refGene_gn=WDR63;refGene_tx=XM_019837351.2
C1 72038810 72038939
refGene_gn=WDR63;refGene_tx=XM_006934877.4;refGene_gn=WDR63;refGene_tx=XM_019837349.1;refGene_gn=WDR63;refGene_tx=XM_019837350.2;refGene_gn=WDR63;refGene_tx=XM_019837351.2
C1 72042218 72042292
refGene_gn=WDR63;refGene_tx=XM_006934877.4;refGene_gn=WDR63;refGene_tx=XM_019837349.1;refGene_gn=WDR63;refGene_tx=XM_019837350.2;refGene_gn=WDR63;refGene_tx=XM_019837351.2
C1 72051441 72051586
refGene_gn=WDR63;refGene_tx=XM_006934877.4;refGene_gn=WDR63;refGene_tx=XM_019837349.1;refGene_gn=WDR63;refGene_tx=XM_019837350.2;refGene_gn=WDR63;refGene_tx=XM_019837351.2
C1 72052957 72053045
refGene_gn=WDR63;refGene_tx=XM_006934877.4;refGene_gn=WDR63;refGene_tx=XM_019837349.1;refGene_gn=WDR63;refGene_tx=XM_019837350.2;refGene_gn=WDR63;refGene_tx=XM_019837351.2
C1 72061281 72061412
refGene_gn=WDR63;refGene_tx=XM_006934877.4;refGene_gn=WDR63;refGene_tx=XM_019837349.1;refGene_gn=WDR63;refGene_tx=XM_019837350.2;refGene_gn=WDR63;refGene_tx=XM_019837351.2
C1 72065292 72065356
refGene_gn=WDR63;refGene_tx=XM_006934877.4;refGene_gn=WDR63;refGene_tx=XM_019837349.1;refGene_gn=WDR63;refGene_tx=XM_019837350.2;refGene_gn=WDR63;refGene_tx=XM_019837351.2
C1 72071000 72071131
refGene_gn=WDR63;refGene_tx=XM_006934877.4;refGene_gn=WDR63;refGene_tx=XM_019837349.1;refGene_gn=WDR63;refGene_tx=XM_019837350.2;refGene_gn=WDR63;refGene_tx=XM_019837351.2
C1 72073477 72073682

refGene_gn=WDR63;refGene_tx=XM_006934877.4;refGene_gn=WDR63;refGene_tx=XM_019837349.1;refGene_gn=WDR63;refGene_tx=XM_019837350.2;refGene_gn=WDR63;refGene_tx=XM_019837351.2
C1 72075530 72075622 refGene_gn=WDR63;refGene_tx=XM_006934877.4;refGene_gn=WDR63;refGene_tx=XM_019837349.1;refGene_gn=WDR63;refGene_tx=XM_019837350.2;refGene_gn=WDR63;refGene_tx=XM_019837351.2
C1 72076955 72077078 refGene_gn=WDR63;refGene_tx=XM_006934877.4;refGene_gn=WDR63;refGene_tx=XM_019837349.1;refGene_gn=WDR63;refGene_tx=XM_019837350.2;refGene_gn=WDR63;refGene_tx=XM_019837351.2
C1 72080841 72080985 refGene_gn=WDR63;refGene_tx=XM_006934877.4;refGene_gn=WDR63;refGene_tx=XM_019837349.1;refGene_gn=WDR63;refGene_tx=XM_019837350.2;refGene_gn=WDR63;refGene_tx=XM_019837351.2
C1 72102548 72103048 refGene_gn=SYDE2;refGene_tx=XM_023258866.1
C1 72103653 72103664 refGene_gn=SYDE2;refGene_tx=XM_023258867.1
C1 72109300 72109529 refGene_gn=SYDE2;refGene_tx=XM_023258866.1;refGene_gn=SYDE2;refGene_tx=XM_023258867.1
C1 72112591 72112773 refGene_gn=SYDE2;refGene_tx=XM_023258866.1;refGene_gn=SYDE2;refGene_tx=XM_023258867.1
C1 72128372 72128499 refGene_gn=SYDE2;refGene_tx=XM_023258866.1;refGene_gn=SYDE2;refGene_tx=XM_023258867.1
C1 72132628 72133731 refGene_gn=SYDE2;refGene_tx=XM_023258866.1;refGene_gn=SYDE2;refGene_tx=XM_023258867.1
C1 72140877 72141561 refGene_gn=SYDE2;refGene_tx=XM_023258866.1;refGene_gn=SYDE2;refGene_tx=XM_023258867.1
C1 72151474 72152789 refGene_gn=SYDE2;refGene_tx=XM_023258866.1;refGene_gn=SYDE2;refGene_tx=XM_023258867.1
C1 72193718 72193792 refGene_gn=CC1H1orf52;refGene_tx=XM_003990270.5
C1 72201542 72201741 refGene_gn=CC1H1orf52;refGene_tx=XM_003990270.5
C1 72202383 72202659 refGene_gn=CC1H1orf52;refGene_tx=XM_003990270.5
C1 72208829 72209185 refGene_gn=BCL10;refGene_tx=XM_003990271.5
C1 72211879 72212168 refGene_gn=BCL10;refGene_tx=XM_003990271.5
C1 72219295 72219352 refGene_gn=BCL10;refGene_tx=XM_003990271.5
C1 72254952 72255069 refGene_gn=DDAH1;refGene_tx=XM_006934878.3;refGene_gn=DDAH1;refGene_tx=XM_019837353.1;refGene_gn=DDAH1;refGene_tx=XM_019837354.2;refGene_gn=DDAH1;refGene_tx=XM_023258868.1;refGene_gn=DDAH1;refGene_tx=XM_023258869.1
C1 72257330 72257474

refGene_gn=DDAH1;refGene_tx=XM_006934878.3;refGene_gn=DDAH1;refGene_tx=XM_019837353.1;refGene_gn=DDAH1;refGene_tx=XM_019837354.2;refGene_gn=DDAH1;refGene_tx=XM_023258868.1;refGene_gn=DDAH1;refGene_tx=XM_023258869.1
C1 72284760 72284880
refGene_gn=DDAH1;refGene_tx=XM_006934878.3;refGene_gn=DDAH1;refGene_tx=XM_019837353.1;refGene_gn=DDAH1;refGene_tx=XM_019837354.2;refGene_gn=DDAH1;refGene_tx=XM_023258868.1;refGene_gn=DDAH1;refGene_tx=XM_023258869.1
C1 72285705 72285779
refGene_gn=DDAH1;refGene_tx=XM_006934878.3;refGene_gn=DDAH1;refGene_tx=XM_019837353.1;refGene_gn=DDAH1;refGene_tx=XM_019837354.2;refGene_gn=DDAH1;refGene_tx=XM_023258868.1;refGene_gn=DDAH1;refGene_tx=XM_023258869.1
C1 72292269 72292369
refGene_gn=DDAH1;refGene_tx=XM_006934878.3;refGene_gn=DDAH1;refGene_tx=XM_019837353.1;refGene_gn=DDAH1;refGene_tx=XM_019837354.2;refGene_gn=DDAH1;refGene_tx=XM_023258868.1;refGene_gn=DDAH1;refGene_tx=XM_023258869.1
C1 72318493 72318544
refGene_gn=DDAH1;refGene_tx=XM_019837353.1;refGene_gn=DDAH1;refGene_tx=XM_023258868.1
C1 72328662 72328665 refGene_gn=DDAH1;refGene_tx=XM_019837354.2
C1 72389777 72390080
refGene_gn=DDAH1;refGene_tx=XM_023258868.1;refGene_gn=DDAH1;refGene_tx=XM_023258869.1
C1 72496691 72497140
refGene_gn=LOC111561763;refGene_tx=XM_023258870.1
C1 72499187 72499407
refGene_gn=LOC111561763;refGene_tx=XM_023258870.1
C1 72499897 72499960 refGene_gn=CYR61;refGene_tx=XM_003990273.4
C1 72500260 72500474 refGene_gn=CYR61;refGene_tx=XM_003990273.4
C1 72500823 72501180 refGene_gn=CYR61;refGene_tx=XM_003990273.4
C1 72501311 72501523 refGene_gn=CYR61;refGene_tx=XM_003990273.4
C1 72501640 72501943 refGene_gn=CYR61;refGene_tx=XM_003990273.4
C1 72510401 72510418 refGene_gn=ZNHIT6;refGene_tx=XM_011284895.3
C1 72549136 72549210 refGene_gn=ZNHIT6;refGene_tx=XM_019837352.2
C1 72557305 72557346 refGene_gn=ZNHIT6;refGene_tx=XM_011284894.3
C1 72561173 72561298
refGene_gn=ZNHIT6;refGene_tx=XM_011284894.3;refGene_gn=ZNHIT6;refGene_tx=XM_011284895.3;refGene_gn=ZNHIT6;refGene_tx=XM_019837352.2
C1 72578225 72578303
refGene_gn=ZNHIT6;refGene_tx=XM_011284894.3;refGene_gn=ZNHIT6;refGene_tx=XM_011284895.3;refGene_gn=ZNHIT6;refGene_tx=XM_019837352.2
C1 72579688 72579769
refGene_gn=ZNHIT6;refGene_tx=XM_011284894.3;refGene_gn=ZNHIT6;refGene_tx=XM_011284895.3;refGene_gn=ZNHIT6;refGene_tx=XM_019837352.2
C1 72580997 72581066
refGene_gn=ZNHIT6;refGene_tx=XM_011284894.3;refGene_gn=ZNHIT6;refGene_tx=XM_011284895.3;refGene_gn=ZNHIT6;refGene_tx=XM_019837352.2

C1 72601260 72601364
refGene_gn=ZNHIT6;refGene_tx=XM_011284894.3;refGene_gn=ZNHIT6;refGene_tx=XM_011284895.3;refGene_gn=ZNHIT6;refGene_tx=XM_019837352.2
C1 72610442 72610528
refGene_gn=ZNHIT6;refGene_tx=XM_011284894.3;refGene_gn=ZNHIT6;refGene_tx=XM_011284895.3;refGene_gn=ZNHIT6;refGene_tx=XM_019837352.2
C1 72610612 72610719
refGene_gn=ZNHIT6;refGene_tx=XM_011284894.3;refGene_gn=ZNHIT6;refGene_tx=XM_011284895.3;refGene_gn=ZNHIT6;refGene_tx=XM_019837352.2
C1 72610805 72610871
refGene_gn=ZNHIT6;refGene_tx=XM_011284894.3;refGene_gn=ZNHIT6;refGene_tx=XM_011284895.3;refGene_gn=ZNHIT6;refGene_tx=XM_019837352.2
C1 72612004 72612666
refGene_gn=ZNHIT6;refGene_tx=XM_011284894.3;refGene_gn=ZNHIT6;refGene_tx=XM_011284895.3;refGene_gn=ZNHIT6;refGene_tx=XM_019837352.2
C1 72630991 72631138
refGene_gn=COL24A1;refGene_tx=XM_019837358.2;refGene_gn=COL24A1;refGene_tx=XM_019837359.2;refGene_gn=COL24A1;refGene_tx=XM_019837363.1;refGene_gn=COL24A1;refGene_tx=XM_023258872.1;refGene_gn=COL24A1;refGene_tx=XM_023258873.1
C1 72634305 72634521
refGene_gn=COL24A1;refGene_tx=XM_019837358.2;refGene_gn=COL24A1;refGene_tx=XM_019837363.1;refGene_gn=COL24A1;refGene_tx=XM_023258872.1;refGene_gn=COL24A1;refGene_tx=XM_023258873.1
C1 72637114 72637224
refGene_gn=COL24A1;refGene_tx=XM_019837358.2;refGene_gn=COL24A1;refGene_tx=XM_019837359.2;refGene_gn=COL24A1;refGene_tx=XM_019837363.1;refGene_gn=COL24A1;refGene_tx=XM_023258872.1;refGene_gn=COL24A1;refGene_tx=XM_023258873.1
C1 72644628 72644797
refGene_gn=COL24A1;refGene_tx=XM_019837358.2;refGene_gn=COL24A1;refGene_tx=XM_019837359.2;refGene_gn=COL24A1;refGene_tx=XM_019837363.1;refGene_gn=COL24A1;refGene_tx=XM_023258872.1;refGene_gn=COL24A1;refGene_tx=XM_023258873.1
C1 72645433 72645499
refGene_gn=COL24A1;refGene_tx=XM_019837358.2;refGene_gn=COL24A1;refGene_tx=XM_019837359.2;refGene_gn=COL24A1;refGene_tx=XM_019837363.1;refGene_gn=COL24A1;refGene_tx=XM_023258872.1;refGene_gn=COL24A1;refGene_tx=XM_023258873.1
C1 72659384 72659411
refGene_gn=COL24A1;refGene_tx=XM_019837358.2;refGene_gn=COL24A1;refGene_tx=XM_019837359.2;refGene_gn=COL24A1;refGene_tx=XM_019837363.1;refGene_gn=COL24A1;refGene_tx=XM_023258872.1;refGene_gn=COL24A1;refGene_tx=XM_023258873.1
C1 72659515 72659551
refGene_gn=COL24A1;refGene_tx=XM_019837358.2;refGene_gn=COL24A1;refGene_tx=XM_019837359.2;refGene_gn=COL24A1;refGene_tx=XM_019837363.1;refGene_gn=COL24A1;refGene_tx=XM_023258872.1;refGene_gn=COL24A1;refGene_tx=XM_023258873.1
C1 72667615 72667651

refGene_gn=COL24A1;refGene_tx=XM_019837358.2;refGene_gn=COL24A1;refGene_tx=XM_019837359.2;refGene_gn=COL24A1;refGene_tx=XM_019837363.1;refGene_gn=COL24A1;refGene_tx=XM_023258872.1;refGene_gn=COL24A1;refGene_tx=XM_023258873.1
C1 72673082 72673136
refGene_gn=COL24A1;refGene_tx=XM_019837358.2;refGene_gn=COL24A1;refGene_tx=XM_019837359.2;refGene_gn=COL24A1;refGene_tx=XM_019837363.1;refGene_gn=COL24A1;refGene_tx=XM_023258872.1;refGene_gn=COL24A1;refGene_tx=XM_023258873.1
C1 72676564 72676627
refGene_gn=COL24A1;refGene_tx=XM_019837358.2;refGene_gn=COL24A1;refGene_tx=XM_019837359.2;refGene_gn=COL24A1;refGene_tx=XM_019837363.1;refGene_gn=COL24A1;refGene_tx=XM_023258872.1;refGene_gn=COL24A1;refGene_tx=XM_023258873.1
C1 72677192 72677246
refGene_gn=COL24A1;refGene_tx=XM_019837358.2;refGene_gn=COL24A1;refGene_tx=XM_019837359.2;refGene_gn=COL24A1;refGene_tx=XM_019837363.1;refGene_gn=COL24A1;refGene_tx=XM_023258872.1;refGene_gn=COL24A1;refGene_tx=XM_023258873.1
C1 72677340 72677448
refGene_gn=COL24A1;refGene_tx=XM_019837358.2;refGene_gn=COL24A1;refGene_tx=XM_019837359.2;refGene_gn=COL24A1;refGene_tx=XM_019837363.1;refGene_gn=COL24A1;refGene_tx=XM_023258872.1;refGene_gn=COL24A1;refGene_tx=XM_023258873.1
C1 72679229 72679337
refGene_gn=COL24A1;refGene_tx=XM_019837358.2;refGene_gn=COL24A1;refGene_tx=XM_019837359.2;refGene_gn=COL24A1;refGene_tx=XM_019837363.1;refGene_gn=COL24A1;refGene_tx=XM_023258872.1;refGene_gn=COL24A1;refGene_tx=XM_023258873.1
C1 72694469 72694577
refGene_gn=COL24A1;refGene_tx=XM_019837358.2;refGene_gn=COL24A1;refGene_tx=XM_019837359.2;refGene_gn=COL24A1;refGene_tx=XM_019837363.1;refGene_gn=COL24A1;refGene_tx=XM_023258872.1;refGene_gn=COL24A1;refGene_tx=XM_023258873.1
C1 72695705 72695759
refGene_gn=COL24A1;refGene_tx=XM_019837358.2;refGene_gn=COL24A1;refGene_tx=XM_019837359.2;refGene_gn=COL24A1;refGene_tx=XM_019837363.1;refGene_gn=COL24A1;refGene_tx=XM_023258872.1;refGene_gn=COL24A1;refGene_tx=XM_023258873.1
C1 72702253 72702307
refGene_gn=COL24A1;refGene_tx=XM_019837358.2;refGene_gn=COL24A1;refGene_tx=XM_019837359.2;refGene_gn=COL24A1;refGene_tx=XM_019837363.1;refGene_gn=COL24A1;refGene_tx=XM_023258872.1;refGene_gn=COL24A1;refGene_tx=XM_023258873.1
C1 72702417 72702471
refGene_gn=COL24A1;refGene_tx=XM_019837358.2;refGene_gn=COL24A1;refGene_tx=XM_019837359.2;refGene_gn=COL24A1;refGene_tx=XM_019837363.1;refGene_gn=COL24A1;refGene_tx=XM_023258872.1;refGene_gn=COL24A1;refGene_tx=XM_023258873.1
C1 72715263 72715317

refGene_gn=COL24A1;refGene_tx=XM_019837358.2;refGene_gn=COL24A1;refGene_tx=XM_019837359.2;refGene_gn=COL24A1;refGene_tx=XM_019837363.1;refGene_gn=COL24A1;refGene_tx=XM_023258872.1;refGene_gn=COL24A1;refGene_tx=XM_023258873.1
C1 72724003 72724060
refGene_gn=COL24A1;refGene_tx=XM_019837358.2;refGene_gn=COL24A1;refGene_tx=XM_019837359.2;refGene_gn=COL24A1;refGene_tx=XM_019837363.1;refGene_gn=COL24A1;refGene_tx=XM_023258872.1;refGene_gn=COL24A1;refGene_tx=XM_023258873.1
C1 72724700 72724754
refGene_gn=COL24A1;refGene_tx=XM_019837358.2;refGene_gn=COL24A1;refGene_tx=XM_019837359.2;refGene_gn=COL24A1;refGene_tx=XM_019837363.1;refGene_gn=COL24A1;refGene_tx=XM_023258872.1;refGene_gn=COL24A1;refGene_tx=XM_023258873.1
C1 72725025 72725079
refGene_gn=COL24A1;refGene_tx=XM_019837358.2;refGene_gn=COL24A1;refGene_tx=XM_019837359.2;refGene_gn=COL24A1;refGene_tx=XM_019837363.1;refGene_gn=COL24A1;refGene_tx=XM_023258872.1;refGene_gn=COL24A1;refGene_tx=XM_023258873.1
C1 72728612 72728720
refGene_gn=COL24A1;refGene_tx=XM_019837358.2;refGene_gn=COL24A1;refGene_tx=XM_019837359.2;refGene_gn=COL24A1;refGene_tx=XM_019837363.1;refGene_gn=COL24A1;refGene_tx=XM_023258872.1;refGene_gn=COL24A1;refGene_tx=XM_023258873.1
C1 72730589 72730643
refGene_gn=COL24A1;refGene_tx=XM_019837358.2;refGene_gn=COL24A1;refGene_tx=XM_019837359.2;refGene_gn=COL24A1;refGene_tx=XM_019837363.1;refGene_gn=COL24A1;refGene_tx=XM_023258872.1;refGene_gn=COL24A1;refGene_tx=XM_023258873.1
C1 72752693 72752801
refGene_gn=COL24A1;refGene_tx=XM_019837358.2;refGene_gn=COL24A1;refGene_tx=XM_019837359.2;refGene_gn=COL24A1;refGene_tx=XM_019837363.1;refGene_gn=COL24A1;refGene_tx=XM_023258872.1;refGene_gn=COL24A1;refGene_tx=XM_023258873.1
C1 72752960 72753014
refGene_gn=COL24A1;refGene_tx=XM_019837358.2;refGene_gn=COL24A1;refGene_tx=XM_019837359.2;refGene_gn=COL24A1;refGene_tx=XM_019837363.1;refGene_gn=COL24A1;refGene_tx=XM_023258872.1;refGene_gn=COL24A1;refGene_tx=XM_023258873.1
C1 72756618 72756672
refGene_gn=COL24A1;refGene_tx=XM_019837358.2;refGene_gn=COL24A1;refGene_tx=XM_019837359.2;refGene_gn=COL24A1;refGene_tx=XM_019837363.1;refGene_gn=COL24A1;refGene_tx=XM_023258872.1;refGene_gn=COL24A1;refGene_tx=XM_023258873.1
C1 72757116 72757170
refGene_gn=COL24A1;refGene_tx=XM_019837358.2;refGene_gn=COL24A1;refGene_tx=XM_019837359.2;refGene_gn=COL24A1;refGene_tx=XM_019837363.1;refGene_gn=COL24A1;refGene_tx=XM_023258872.1;refGene_gn=COL24A1;refGene_tx=XM_023258873.1
C1 72760403 72760457

refGene_gn=COL24A1;refGene_tx=XM_019837358.2;refGene_gn=COL24A1;refGene_tx=XM_019837359.2;refGene_gn=COL24A1;refGene_tx=XM_019837363.1;refGene_gn=COL24A1;refGene_tx=XM_023258872.1;refGene_gn=COL24A1;refGene_tx=XM_023258873.1
C1 72770268 72770322
refGene_gn=COL24A1;refGene_tx=XM_019837358.2;refGene_gn=COL24A1;refGene_tx=XM_019837359.2;refGene_gn=COL24A1;refGene_tx=XM_019837363.1;refGene_gn=COL24A1;refGene_tx=XM_023258872.1;refGene_gn=COL24A1;refGene_tx=XM_023258873.1
C1 72785803 72785848
refGene_gn=COL24A1;refGene_tx=XM_019837358.2;refGene_gn=COL24A1;refGene_tx=XM_019837359.2;refGene_gn=COL24A1;refGene_tx=XM_019837363.1;refGene_gn=COL24A1;refGene_tx=XM_023258872.1;refGene_gn=COL24A1;refGene_tx=XM_023258873.1
C1 72785968 72786013
refGene_gn=COL24A1;refGene_tx=XM_019837358.2;refGene_gn=COL24A1;refGene_tx=XM_019837359.2;refGene_gn=COL24A1;refGene_tx=XM_019837363.1;refGene_gn=COL24A1;refGene_tx=XM_023258872.1;refGene_gn=COL24A1;refGene_tx=XM_023258873.1
C1 72786267 72786321
refGene_gn=COL24A1;refGene_tx=XM_019837358.2;refGene_gn=COL24A1;refGene_tx=XM_019837359.2;refGene_gn=COL24A1;refGene_tx=XM_019837363.1;refGene_gn=COL24A1;refGene_tx=XM_023258872.1;refGene_gn=COL24A1;refGene_tx=XM_023258873.1
C1 72794794 72794848
refGene_gn=COL24A1;refGene_tx=XM_019837358.2;refGene_gn=COL24A1;refGene_tx=XM_019837359.2;refGene_gn=COL24A1;refGene_tx=XM_019837363.1;refGene_gn=COL24A1;refGene_tx=XM_023258872.1;refGene_gn=COL24A1;refGene_tx=XM_023258873.1
C1 72796134 72796188
refGene_gn=COL24A1;refGene_tx=XM_019837358.2;refGene_gn=COL24A1;refGene_tx=XM_019837359.2;refGene_gn=COL24A1;refGene_tx=XM_019837363.1;refGene_gn=COL24A1;refGene_tx=XM_023258872.1;refGene_gn=COL24A1;refGene_tx=XM_023258873.1
C1 72797866 72797920
refGene_gn=COL24A1;refGene_tx=XM_019837358.2;refGene_gn=COL24A1;refGene_tx=XM_019837359.2;refGene_gn=COL24A1;refGene_tx=XM_019837363.1;refGene_gn=COL24A1;refGene_tx=XM_023258872.1;refGene_gn=COL24A1;refGene_tx=XM_023258873.1
C1 72802589 72802643
refGene_gn=COL24A1;refGene_tx=XM_019837358.2;refGene_gn=COL24A1;refGene_tx=XM_019837359.2;refGene_gn=COL24A1;refGene_tx=XM_019837363.1;refGene_gn=COL24A1;refGene_tx=XM_023258872.1;refGene_gn=COL24A1;refGene_tx=XM_023258873.1
C1 72856121 72856166
refGene_gn=COL24A1;refGene_tx=XM_019837358.2;refGene_gn=COL24A1;refGene_tx=XM_019837359.2;refGene_gn=COL24A1;refGene_tx=XM_019837363.1;refGene_gn=COL24A1;refGene_tx=XM_023258872.1;refGene_gn=COL24A1;refGene_tx=XM_023258873.1
C1 72860036 72860090

refGene_gn=COL24A1;refGene_tx=XM_019837358.2;refGene_gn=COL24A1;refGene_tx=XM_019837359.2;refGene_gn=COL24A1;refGene_tx=XM_019837363.1;refGene_gn=COL24A1;refGene_tx=XM_023258872.1;refGene_gn=COL24A1;refGene_tx=XM_023258873.1
C1 72864388 72864433
refGene_gn=COL24A1;refGene_tx=XM_019837358.2;refGene_gn=COL24A1;refGene_tx=XM_019837359.2;refGene_gn=COL24A1;refGene_tx=XM_019837363.1;refGene_gn=COL24A1;refGene_tx=XM_023258872.1;refGene_gn=COL24A1;refGene_tx=XM_023258873.1
C1 72865862 72865916
refGene_gn=COL24A1;refGene_tx=XM_019837358.2;refGene_gn=COL24A1;refGene_tx=XM_019837359.2;refGene_gn=COL24A1;refGene_tx=XM_019837363.1;refGene_gn=COL24A1;refGene_tx=XM_023258872.1;refGene_gn=COL24A1;refGene_tx=XM_023258873.1
C1 72891007 72891061
refGene_gn=COL24A1;refGene_tx=XM_019837358.2;refGene_gn=COL24A1;refGene_tx=XM_019837359.2;refGene_gn=COL24A1;refGene_tx=XM_019837363.1;refGene_gn=COL24A1;refGene_tx=XM_023258872.1;refGene_gn=COL24A1;refGene_tx=XM_023258873.1
C1 72924107 72924161
refGene_gn=COL24A1;refGene_tx=XM_019837358.2;refGene_gn=COL24A1;refGene_tx=XM_019837359.2;refGene_gn=COL24A1;refGene_tx=XM_019837363.1;refGene_gn=COL24A1;refGene_tx=XM_023258872.1;refGene_gn=COL24A1;refGene_tx=XM_023258873.1
C1 72929009 72929063
refGene_gn=COL24A1;refGene_tx=XM_019837358.2;refGene_gn=COL24A1;refGene_tx=XM_019837359.2;refGene_gn=COL24A1;refGene_tx=XM_019837363.1;refGene_gn=COL24A1;refGene_tx=XM_023258872.1;refGene_gn=COL24A1;refGene_tx=XM_023258873.1
C1 72929327 72929381
refGene_gn=COL24A1;refGene_tx=XM_019837358.2;refGene_gn=COL24A1;refGene_tx=XM_019837359.2;refGene_gn=COL24A1;refGene_tx=XM_019837363.1;refGene_gn=COL24A1;refGene_tx=XM_023258872.1;refGene_gn=COL24A1;refGene_tx=XM_023258873.1
C1 72929629 72929674
refGene_gn=COL24A1;refGene_tx=XM_019837358.2;refGene_gn=COL24A1;refGene_tx=XM_019837359.2;refGene_gn=COL24A1;refGene_tx=XM_019837363.1;refGene_gn=COL24A1;refGene_tx=XM_023258872.1;refGene_gn=COL24A1;refGene_tx=XM_023258873.1
C1 72929751 72929805
refGene_gn=COL24A1;refGene_tx=XM_019837358.2;refGene_gn=COL24A1;refGene_tx=XM_019837359.2;refGene_gn=COL24A1;refGene_tx=XM_019837363.1;refGene_gn=COL24A1;refGene_tx=XM_023258872.1;refGene_gn=COL24A1;refGene_tx=XM_023258873.1
C1 72936085 72936130
refGene_gn=COL24A1;refGene_tx=XM_019837358.2;refGene_gn=COL24A1;refGene_tx=XM_019837359.2;refGene_gn=COL24A1;refGene_tx=XM_019837363.1;refGene_gn=COL24A1;refGene_tx=XM_023258872.1;refGene_gn=COL24A1;refGene_tx=XM_023258873.1
C1 72938573 72938627

refGene_gn=COL24A1;refGene_tx=XM_019837358.2;refGene_gn=COL24A1;refGene_tx=XM_019837359.2;refGene_gn=COL24A1;refGene_tx=XM_019837363.1;refGene_gn=COL24A1;refGene_tx=XM_023258872.1;refGene_gn=COL24A1;refGene_tx=XM_023258873.1
C1 72952729 72952774
refGene_gn=COL24A1;refGene_tx=XM_019837358.2;refGene_gn=COL24A1;refGene_tx=XM_019837359.2;refGene_gn=COL24A1;refGene_tx=XM_019837363.1;refGene_gn=COL24A1;refGene_tx=XM_023258872.1;refGene_gn=COL24A1;refGene_tx=XM_023258873.1
C1 72955942 72955996
refGene_gn=COL24A1;refGene_tx=XM_019837358.2;refGene_gn=COL24A1;refGene_tx=XM_019837359.2;refGene_gn=COL24A1;refGene_tx=XM_019837363.1;refGene_gn=COL24A1;refGene_tx=XM_023258872.1;refGene_gn=COL24A1;refGene_tx=XM_023258873.1
C1 72960361 72960406
refGene_gn=COL24A1;refGene_tx=XM_019837358.2;refGene_gn=COL24A1;refGene_tx=XM_019837359.2;refGene_gn=COL24A1;refGene_tx=XM_019837363.1;refGene_gn=COL24A1;refGene_tx=XM_023258872.1;refGene_gn=COL24A1;refGene_tx=XM_023258873.1
C1 72961822 72961876
refGene_gn=COL24A1;refGene_tx=XM_019837358.2;refGene_gn=COL24A1;refGene_tx=XM_019837359.2;refGene_gn=COL24A1;refGene_tx=XM_019837363.1;refGene_gn=COL24A1;refGene_tx=XM_023258872.1;refGene_gn=COL24A1;refGene_tx=XM_023258873.1
C1 72969293 72969338
refGene_gn=COL24A1;refGene_tx=XM_019837358.2;refGene_gn=COL24A1;refGene_tx=XM_019837359.2;refGene_gn=COL24A1;refGene_tx=XM_019837363.1;refGene_gn=COL24A1;refGene_tx=XM_023258872.1;refGene_gn=COL24A1;refGene_tx=XM_023258873.1
C1 72981575 72981629
refGene_gn=COL24A1;refGene_tx=XM_019837358.2;refGene_gn=COL24A1;refGene_tx=XM_019837359.2;refGene_gn=COL24A1;refGene_tx=XM_019837363.1;refGene_gn=COL24A1;refGene_tx=XM_023258872.1;refGene_gn=COL24A1;refGene_tx=XM_023258873.1
C1 72986108 72986162
refGene_gn=COL24A1;refGene_tx=XM_019837358.2;refGene_gn=COL24A1;refGene_tx=XM_019837359.2;refGene_gn=COL24A1;refGene_tx=XM_019837363.1;refGene_gn=COL24A1;refGene_tx=XM_023258872.1;refGene_gn=COL24A1;refGene_tx=XM_023258873.1
C1 73011233 73011287
refGene_gn=COL24A1;refGene_tx=XM_019837358.2;refGene_gn=COL24A1;refGene_tx=XM_019837359.2;refGene_gn=COL24A1;refGene_tx=XM_019837363.1;refGene_gn=COL24A1;refGene_tx=XM_023258872.1;refGene_gn=COL24A1;refGene_tx=XM_023258873.1
C1 73014116 73014170
refGene_gn=COL24A1;refGene_tx=XM_019837358.2;refGene_gn=COL24A1;refGene_tx=XM_019837359.2;refGene_gn=COL24A1;refGene_tx=XM_019837363.1;refGene_gn=COL24A1;refGene_tx=XM_023258872.1;refGene_gn=COL24A1;refGene_tx=XM_023258873.1
C1 73022281 73023633

refGene_gn=COL24A1;refGene_tx=XM_019837358.2;refGene_gn=COL24A1;refGene_tx=XM_019837359.2;refGene_gn=COL24A1;refGene_tx=XM_019837363.1;refGene_gn=COL24A1;refGene_tx=XM_023258872.1;refGene_gn=COL24A1;refGene_tx=XM_023258873.1
C1 73035289 73035354
refGene_gn=COL24A1;refGene_tx=XM_019837358.2;refGene_gn=COL24A1;refGene_tx=XM_019837359.2;refGene_gn=COL24A1;refGene_tx=XM_023258872.1;refGene_gn=COL24A1;refGene_tx=XM_023258873.1
C1 73044932 73044988
refGene_gn=COL24A1;refGene_tx=XM_019837358.2;refGene_gn=COL24A1;refGene_tx=XM_019837359.2;refGene_gn=COL24A1;refGene_tx=XM_023258872.1;refGene_gn=COL24A1;refGene_tx=XM_023258873.1
C1 73215025 73215154
refGene_gn=ODF2L;refGene_tx=XM_019837366.1;refGene_gn=ODF2L;refGene_tx=XM_019837367.1;refGene_gn=ODF2L;refGene_tx=XM_019837372.1
C1 73217663 73217789
refGene_gn=ODF2L;refGene_tx=XM_019837366.1;refGene_gn=ODF2L;refGene_tx=XM_019837367.1;refGene_gn=ODF2L;refGene_tx=XM_019837372.1
C1 73220565 73220728
refGene_gn=ODF2L;refGene_tx=XM_019837366.1;refGene_gn=ODF2L;refGene_tx=XM_019837367.1;refGene_gn=ODF2L;refGene_tx=XM_019837372.1
C1 73220810 73220896
refGene_gn=ODF2L;refGene_tx=XM_019837366.1;refGene_gn=ODF2L;refGene_tx=XM_019837367.1;refGene_gn=ODF2L;refGene_tx=XM_019837372.1
C1 73222329 73222488
refGene_gn=ODF2L;refGene_tx=XM_019837366.1;refGene_gn=ODF2L;refGene_tx=XM_019837367.1
C1 73224772 73224877
refGene_gn=ODF2L;refGene_tx=XM_019837366.1;refGene_gn=ODF2L;refGene_tx=XM_019837367.1;refGene_gn=ODF2L;refGene_tx=XM_019837372.1
C1 73226909 73227020
refGene_gn=ODF2L;refGene_tx=XM_019837366.1;refGene_gn=ODF2L;refGene_tx=XM_019837367.1;refGene_gn=ODF2L;refGene_tx=XM_019837372.1
C1 73231966 73232102
refGene_gn=ODF2L;refGene_tx=XM_019837366.1;refGene_gn=ODF2L;refGene_tx=XM_019837367.1;refGene_gn=ODF2L;refGene_tx=XM_019837372.1
C1 73233279 73233389
refGene_gn=ODF2L;refGene_tx=XM_019837366.1;refGene_gn=ODF2L;refGene_tx=XM_019837367.1;refGene_gn=ODF2L;refGene_tx=XM_019837372.1
C1 73235877 73236063
refGene_gn=ODF2L;refGene_tx=XM_019837366.1;refGene_gn=ODF2L;refGene_tx=XM_019837367.1;refGene_gn=ODF2L;refGene_tx=XM_019837372.1
C1 73240110 73240227
refGene_gn=ODF2L;refGene_tx=XM_019837366.1;refGene_gn=ODF2L;refGene_tx=XM_019837367.1;refGene_gn=ODF2L;refGene_tx=XM_019837372.1
C1 73240967 73241039
refGene_gn=ODF2L;refGene_tx=XM_019837366.1;refGene_gn=ODF2L;refGene_tx=XM_019837367.1;refGene_gn=ODF2L;refGene_tx=XM_019837372.1
C1 73241162 73241222
refGene_gn=ODF2L;refGene_tx=XM_019837366.1;refGene_gn=ODF2L;refGene_tx

=XM_019837367.1;refGene_gn=ODF2L;refGene_tx=XM_019837372.1
C1	73242934	73243060
refGene_gn=ODF2L;refGene_tx=XM_019837366.1;refGene_gn=ODF2L;refGene_tx
=XM_019837367.1;refGene_gn=ODF2L;refGene_tx=XM_019837372.1
C1	73243705	73243838
refGene_gn=ODF2L;refGene_tx=XM_019837366.1;refGene_gn=ODF2L;refGene_tx
=XM_019837367.1;refGene_gn=ODF2L;refGene_tx=XM_019837372.1
C1	73245892	73246005
refGene_gn=ODF2L;refGene_tx=XM_019837366.1;refGene_gn=ODF2L;refGene_tx
=XM_019837367.1;refGene_gn=ODF2L;refGene_tx=XM_019837372.1
C1	73278865	73279051
refGene_gn=CLCA2;refGene_tx=XM_003990276.4;refGene_gn=CLCA2;refGene_tx
=XM_019837374.2;refGene_gn=CLCA2;refGene_tx=XM_019837376.2
C1	73280026	73280119 refGene_gn=CLCA2;refGene_tx=XM_019837375.1
C1	73280146	73280284
refGene_gn=CLCA2;refGene_tx=XM_003990276.4;refGene_gn=CLCA2;refGene_tx
=XM_019837374.2;refGene_gn=CLCA2;refGene_tx=XM_019837376.2
C1	73283225	73283376
refGene_gn=CLCA2;refGene_tx=XM_003990276.4;refGene_gn=CLCA2;refGene_tx
=XM_019837374.2;refGene_gn=CLCA2;refGene_tx=XM_019837375.1;refGene_gn=
CLCA2;refGene_tx=XM_019837376.2
C1	73286729	73286838
refGene_gn=CLCA2;refGene_tx=XM_003990276.4;refGene_gn=CLCA2;refGene_tx
=XM_019837374.2;refGene_gn=CLCA2;refGene_tx=XM_019837375.1;refGene_gn=
CLCA2;refGene_tx=XM_019837376.2
C1	73287945	73288105
refGene_gn=CLCA2;refGene_tx=XM_003990276.4;refGene_gn=CLCA2;refGene_tx
=XM_019837374.2;refGene_gn=CLCA2;refGene_tx=XM_019837375.1;refGene_gn=
CLCA2;refGene_tx=XM_019837376.2
C1	73290231	73290459
refGene_gn=CLCA2;refGene_tx=XM_003990276.4;refGene_gn=CLCA2;refGene_tx
=XM_019837374.2;refGene_gn=CLCA2;refGene_tx=XM_019837375.1;refGene_gn=
CLCA2;refGene_tx=XM_019837376.2
C1	73293188	73293419
refGene_gn=CLCA2;refGene_tx=XM_003990276.4;refGene_gn=CLCA2;refGene_tx
=XM_019837374.2;refGene_gn=CLCA2;refGene_tx=XM_019837375.1;refGene_gn=
CLCA2;refGene_tx=XM_019837376.2
C1	73294214	73294392
refGene_gn=CLCA2;refGene_tx=XM_003990276.4;refGene_gn=CLCA2;refGene_tx
=XM_019837374.2;refGene_gn=CLCA2;refGene_tx=XM_019837375.1;refGene_gn=
CLCA2;refGene_tx=XM_019837376.2
C1	73295255	73295362
refGene_gn=CLCA2;refGene_tx=XM_003990276.4;refGene_gn=CLCA2;refGene_tx
=XM_019837374.2;refGene_gn=CLCA2;refGene_tx=XM_019837375.1;refGene_gn=
CLCA2;refGene_tx=XM_019837376.2
C1	73297358	73297583
refGene_gn=CLCA2;refGene_tx=XM_003990276.4;refGene_gn=CLCA2;refGene_tx
=XM_019837374.2;refGene_gn=CLCA2;refGene_tx=XM_019837375.1;refGene_gn=
CLCA2;refGene_tx=XM_019837376.2
C1	73300344	73300615

refGene_gn=CLCA2;refGene_tx=XM_003990276.4;refGene_gn=CLCA2;refGene_tx=XM_019837374.2;refGene_gn=CLCA2;refGene_tx=XM_019837375.1;refGene_gn=CLCA2;refGene_tx=XM_019837376.2
C1 73303917 73304088 refGene_gn=CLCA2;refGene_tx=XM_003990276.4;refGene_gn=CLCA2;refGene_tx=XM_019837375.1;refGene_gn=CLCA2;refGene_tx=XM_019837376.2
C1 73307879 73308113 refGene_gn=CLCA2;refGene_tx=XM_003990276.4;refGene_gn=CLCA2;refGene_tx=XM_019837374.2;refGene_gn=CLCA2;refGene_tx=XM_019837375.1;refGene_gn=CLCA2;refGene_tx=XM_019837376.2
C1 73312139 73312218 refGene_gn=CLCA2;refGene_tx=XM_019837376.2
C1 73315590 73316036 refGene_gn=CLCA2;refGene_tx=XM_003990276.4;refGene_gn=CLCA2;refGene_tx=XM_019837374.2;refGene_gn=CLCA2;refGene_tx=XM_019837375.1;refGene_gn=CLCA2;refGene_tx=XM_019837376.2
C1 73339689 73339896 refGene_gn=CLCA1;refGene_tx=XM_003990277.4
C1 73344661 73344802 refGene_gn=CLCA1;refGene_tx=XM_003990277.4
C1 73344967 73345115 refGene_gn=CLCA1;refGene_tx=XM_003990277.4
C1 73347650 73347756 refGene_gn=CLCA1;refGene_tx=XM_003990277.4
C1 73353371 73353549 refGene_gn=CLCA1;refGene_tx=XM_003990277.4
C1 73356398 73356617 refGene_gn=CLCA1;refGene_tx=XM_003990277.4
C1 73357189 73357417 refGene_gn=CLCA1;refGene_tx=XM_003990277.4
C1 73360031 73360206 refGene_gn=CLCA1;refGene_tx=XM_003990277.4
C1 73362051 73362158 refGene_gn=CLCA1;refGene_tx=XM_003990277.4
C1 73364187 73364403 refGene_gn=CLCA1;refGene_tx=XM_003990277.4
C1 73365272 73365534 refGene_gn=CLCA1;refGene_tx=XM_003990277.4
C1 73366562 73366733 refGene_gn=CLCA1;refGene_tx=XM_003990277.4
C1 73369212 73369452 refGene_gn=CLCA1;refGene_tx=XM_003990277.4
C1 73370300 73370689 refGene_gn=CLCA1;refGene_tx=XM_003990277.4
C1 73411172 73411328 refGene_gn=CLCA4;refGene_tx=XM_011285015.2
C1 73415317 73415458 refGene_gn=CLCA4;refGene_tx=XM_011285015.2
C1 73415598 73415746 refGene_gn=CLCA4;refGene_tx=XM_011285015.2
C1 73418611 73418720 refGene_gn=CLCA4;refGene_tx=XM_011285015.2
C1 73420258 73420436 refGene_gn=CLCA4;refGene_tx=XM_011285015.2
C1 73420768 73420987 refGene_gn=CLCA4;refGene_tx=XM_011285015.2
C1 73423254 73423482 refGene_gn=CLCA4;refGene_tx=XM_011285015.2
C1 73425567 73425745 refGene_gn=CLCA4;refGene_tx=XM_011285015.2
C1 73427064 73427171 refGene_gn=CLCA4;refGene_tx=XM_011285015.2
C1 73429852 73430068 refGene_gn=CLCA4;refGene_tx=XM_011285015.2
C1 73430661 73430929 refGene_gn=CLCA4;refGene_tx=XM_011285015.2
C1 73432757 73432928 refGene_gn=CLCA4;refGene_tx=XM_011285015.2
C1 73433872 73434106 refGene_gn=CLCA4;refGene_tx=XM_011285015.2
C1 73434456 73434848 refGene_gn=CLCA4;refGene_tx=XM_011285015.2
C1 73490183 73490342 refGene_gn=LOC101093450;refGene_tx=XM_003990321.2
C1 73491340 73491481 refGene_gn=LOC101093450;refGene_tx=XM_003990321.2
C1 73491678 73491826 refGene_gn=LOC101093450;refGene_tx=XM_003990321.2
C1 73492183 73492292

refGene_gn=LOC101093450;refGene_tx=XM_003990321.2
C1 73494528 73494709
refGene_gn=LOC101093450;refGene_tx=XM_003990321.2
C1 73498495 73498723
refGene_gn=LOC101093450;refGene_tx=XM_003990321.2
C1 73499548 73499773
refGene_gn=LOC101093450;refGene_tx=XM_003990321.2
C1 73502186 73502364
refGene_gn=LOC101093450;refGene_tx=XM_003990321.2
C1 73503214 73503315
refGene_gn=LOC101093450;refGene_tx=XM_003990321.2
C1 73503671 73503896
refGene_gn=LOC101093450;refGene_tx=XM_003990321.2
C1 73504181 73504452
refGene_gn=LOC101093450;refGene_tx=XM_003990321.2
C1 73505213 73505384
refGene_gn=LOC101093450;refGene_tx=XM_003990321.2
C1 73505919 73506159
refGene_gn=LOC101093450;refGene_tx=XM_003990321.2
C1 73510004 73510345
refGene_gn=LOC101093450;refGene_tx=XM_003990321.2
C1 73545973 73546045
refGene_gn=SH3GLB1;refGene_tx=XM_011284890.2;refGene_gn=SH3GLB1;refGene_tx=XM_011284891.2;refGene_gn=SH3GLB1;refGene_tx=XM_011284892.2;refGene_gn=SH3GLB1;refGene_tx=XM_019837377.2;refGene_gn=SH3GLB1;refGene_tx=XM_023258874.1;refGene_gn=SH3GLB1;refGene_tx=XM_023258875.1;refGene_gn=SH3GLB1;refGene_tx=XM_023258876.1;refGene_gn=SH3GLB1;refGene_tx=XM_023258877.1;refGene_gn=SH3GLB1;refGene_tx=XM_023258878.1
C1 73556658 73556800
refGene_gn=SH3GLB1;refGene_tx=XM_011284890.2;refGene_gn=SH3GLB1;refGene_tx=XM_011284891.2;refGene_gn=SH3GLB1;refGene_tx=XM_011284892.2;refGene_gn=SH3GLB1;refGene_tx=XM_019837377.2;refGene_gn=SH3GLB1;refGene_tx=XM_023258874.1;refGene_gn=SH3GLB1;refGene_tx=XM_023258875.1;refGene_gn=SH3GLB1;refGene_tx=XM_023258876.1;refGene_gn=SH3GLB1;refGene_tx=XM_023258877.1;refGene_gn=SH3GLB1;refGene_tx=XM_023258878.1
C1 73559453 73559582
refGene_gn=SH3GLB1;refGene_tx=XM_011284890.2;refGene_gn=SH3GLB1;refGene_tx=XM_011284891.2;refGene_gn=SH3GLB1;refGene_tx=XM_011284892.2;refGene_gn=SH3GLB1;refGene_tx=XM_019837377.2;refGene_gn=SH3GLB1;refGene_tx=XM_023258874.1;refGene_gn=SH3GLB1;refGene_tx=XM_023258875.1;refGene_gn=SH3GLB1;refGene_tx=XM_023258876.1;refGene_gn=SH3GLB1;refGene_tx=XM_023258877.1;refGene_gn=SH3GLB1;refGene_tx=XM_023258878.1
C1 73561711 73561845
refGene_gn=SH3GLB1;refGene_tx=XM_011284890.2;refGene_gn=SH3GLB1;refGene_tx=XM_011284891.2;refGene_gn=SH3GLB1;refGene_tx=XM_011284892.2;refGene_gn=SH3GLB1;refGene_tx=XM_019837377.2;refGene_gn=SH3GLB1;refGene_tx=XM_023258874.1;refGene_gn=SH3GLB1;refGene_tx=XM_023258875.1;refGene_gn=SH3GLB1;refGene_tx=XM_023258876.1;refGene_gn=SH3GLB1;refGene_tx=XM_023258877.1;refGene_gn=SH3GLB1;refGene_tx=XM_023258878.1
C1 73563194 73563287

refGene_gn=SH3GLB1;refGene_tx=XM_011284890.2;refGene_gn=SH3GLB1;refGene_tx=XM_011284891.2;refGene_gn=SH3GLB1;refGene_tx=XM_011284892.2;refGene_gn=SH3GLB1;refGene_tx=XM_019837377.2;refGene_gn=SH3GLB1;refGene_tx=XM_023258874.1;refGene_gn=SH3GLB1;refGene_tx=XM_023258875.1;refGene_gn=SH3GLB1;refGene_tx=XM_023258876.1;refGene_gn=SH3GLB1;refGene_tx=XM_023258877.1;refGene_gn=SH3GLB1;refGene_tx=XM_023258878.1
C1 73566840 73566927
refGene_gn=SH3GLB1;refGene_tx=XM_011284890.2;refGene_gn=SH3GLB1;refGene_tx=XM_011284891.2;refGene_gn=SH3GLB1;refGene_tx=XM_019837377.2;refGene_gn=SH3GLB1;refGene_tx=XM_023258874.1;refGene_gn=SH3GLB1;refGene_tx=XM_023258875.1;refGene_gn=SH3GLB1;refGene_tx=XM_023258876.1;refGene_gn=SH3GLB1;refGene_tx=XM_023258877.1
C1 73568535 73568559
refGene_gn=SH3GLB1;refGene_tx=XM_011284890.2;refGene_gn=SH3GLB1;refGene_tx=XM_011284891.2;refGene_gn=SH3GLB1;refGene_tx=XM_023258874.1;refGene_gn=SH3GLB1;refGene_tx=XM_023258876.1
C1 73572657 73572747
refGene_gn=SH3GLB1;refGene_tx=XM_011284890.2;refGene_gn=SH3GLB1;refGene_tx=XM_011284891.2;refGene_gn=SH3GLB1;refGene_tx=XM_011284892.2;refGene_gn=SH3GLB1;refGene_tx=XM_019837377.2;refGene_gn=SH3GLB1;refGene_tx=XM_023258874.1;refGene_gn=SH3GLB1;refGene_tx=XM_023258875.1;refGene_gn=SH3GLB1;refGene_tx=XM_023258876.1;refGene_gn=SH3GLB1;refGene_tx=XM_023258877.1;refGene_gn=SH3GLB1;refGene_tx=XM_023258878.1
C1 73573175 73573279
refGene_gn=SH3GLB1;refGene_tx=XM_011284890.2;refGene_gn=SH3GLB1;refGene_tx=XM_011284891.2;refGene_gn=SH3GLB1;refGene_tx=XM_011284892.2;refGene_gn=SH3GLB1;refGene_tx=XM_019837377.2;refGene_gn=SH3GLB1;refGene_tx=XM_023258874.1;refGene_gn=SH3GLB1;refGene_tx=XM_023258875.1;refGene_gn=SH3GLB1;refGene_tx=XM_023258876.1;refGene_gn=SH3GLB1;refGene_tx=XM_023258877.1;refGene_gn=SH3GLB1;refGene_tx=XM_023258878.1
C1 73579712 73579938
refGene_gn=SH3GLB1;refGene_tx=XM_011284890.2;refGene_gn=SH3GLB1;refGene_tx=XM_011284891.2;refGene_gn=SH3GLB1;refGene_tx=XM_011284892.2;refGene_gn=SH3GLB1;refGene_tx=XM_019837377.2;refGene_gn=SH3GLB1;refGene_tx=XM_023258874.1;refGene_gn=SH3GLB1;refGene_tx=XM_023258875.1;refGene_gn=SH3GLB1;refGene_tx=XM_023258876.1;refGene_gn=SH3GLB1;refGene_tx=XM_023258877.1;refGene_gn=SH3GLB1;refGene_tx=XM_023258878.1
C1 73580546 73580654
refGene_gn=SH3GLB1;refGene_tx=XM_011284890.2;refGene_gn=SH3GLB1;refGene_tx=XM_011284891.2;refGene_gn=SH3GLB1;refGene_tx=XM_011284892.2;refGene_gn=SH3GLB1;refGene_tx=XM_019837377.2;refGene_gn=SH3GLB1;refGene_tx=XM_023258874.1;refGene_gn=SH3GLB1;refGene_tx=XM_023258875.1;refGene_gn=SH3GLB1;refGene_tx=XM_023258876.1;refGene_gn=SH3GLB1;refGene_tx=XM_023258877.1;refGene_gn=SH3GLB1;refGene_tx=XM_023258878.1
C1 73667127 73667259
refGene_gn=SELENOF;refGene_tx=XM_003990279.5;refGene_gn=SELENOF;refGene_tx=XM_019837379.2
C1 73669585 73669635 refGene_gn=SELENOF;refGene_tx=XM_003990279.5
C1 73693043 73693107
refGene_gn=SELENOF;refGene_tx=XM_003990279.5;refGene_gn=SELENOF;refGen

e_tx=XM_019837379.2
C1 73715438 73715606
refGene_gn=SELENOF;refGene_tx=XM_003990279.5;refGene_gn=SELENOF;refGen
e_tx=XM_019837379.2
C1 73724032 73724116
refGene_gn=SELENOF;refGene_tx=XM_003990279.5;refGene_gn=SELENOF;refGen
e_tx=XM_019837379.2
C1 73725072 73725196
refGene_gn=HS2ST1;refGene_tx=XM_003990280.5;refGene_gn=HS2ST1;refGene_
tx=XM_006934882.4
C1 73873866 73874105
refGene_gn=HS2ST1;refGene_tx=XM_003990280.5;refGene_gn=HS2ST1;refGene_
tx=XM_006934882.4;refGene_gn=HS2ST1;refGene_tx=XM_019837380.2
C1 73884733 73884819
refGene_gn=HS2ST1;refGene_tx=XM_003990280.5;refGene_gn=HS2ST1;refGene_
tx=XM_006934882.4;refGene_gn=HS2ST1;refGene_tx=XM_019837380.2
C1 73893990 73894129
refGene_gn=HS2ST1;refGene_tx=XM_003990280.5;refGene_gn=HS2ST1;refGene_
tx=XM_006934882.4;refGene_gn=HS2ST1;refGene_tx=XM_019837380.2
C1 73904691 73904789
refGene_gn=HS2ST1;refGene_tx=XM_003990280.5;refGene_gn=HS2ST1;refGene_
tx=XM_006934882.4;refGene_gn=HS2ST1;refGene_tx=XM_019837380.2
C1 73906490 73906648
refGene_gn=HS2ST1;refGene_tx=XM_003990280.5;refGene_gn=HS2ST1;refGene_
tx=XM_006934882.4;refGene_gn=HS2ST1;refGene_tx=XM_019837380.2
C1 73907647 73907877
refGene_gn=HS2ST1;refGene_tx=XM_003990280.5;refGene_gn=HS2ST1;refGene_
tx=XM_006934882.4;refGene_gn=HS2ST1;refGene_tx=XM_019837380.2
C1 74135900 74136136
refGene_gn=LMO4;refGene_tx=XM_019837382.2;refGene_gn=LMO4;refGene_tx=X
M_019837383.2;refGene_gn=LMO4;refGene_tx=XM_019837384.2;refGene_gn=LMO
4;refGene_tx=XM_019837385.2
C1 74143496 74143593
refGene_gn=LMO4;refGene_tx=XM_019837382.2;refGene_gn=LMO4;refGene_tx=X
M_019837383.2;refGene_gn=LMO4;refGene_tx=XM_019837384.2;refGene_gn=LMO
4;refGene_tx=XM_019837385.2
C1 74143990 74144146
refGene_gn=LMO4;refGene_tx=XM_019837382.2;refGene_gn=LMO4;refGene_tx=X
M_019837383.2;refGene_gn=LMO4;refGene_tx=XM_019837384.2;refGene_gn=LMO
4;refGene_tx=XM_019837385.2
C1 74148822 74148831 refGene_gn=LMO4;refGene_tx=XM_019837382.2
C1 74183775 74183787
refGene_gn=LMO4;refGene_tx=XM_019837383.2;refGene_gn=LMO4;refGene_tx=X
M_019837384.2;refGene_gn=LMO4;refGene_tx=XM_019837385.2
C1 75401210 75401258
refGene_gn=PKN2;refGene_tx=XM_011284886.3;refGene_gn=PKN2;refGene_tx=X
M_011284888.3
C1 75450194 75450495
refGene_gn=PKN2;refGene_tx=XM_011284886.3;refGene_gn=PKN2;refGene_tx=X
M_011284888.3

C1	75467922	75468077
refGene_gn=PKN2;refGene_tx=XM_011284886.3;refGene_gn=PKN2;refGene_tx=XM_011284888.3
C1	75475483	75475601
refGene_gn=PKN2;refGene_tx=XM_011284886.3;refGene_gn=PKN2;refGene_tx=XM_011284888.3
C1	75475910	75476056
refGene_gn=PKN2;refGene_tx=XM_011284886.3;refGene_gn=PKN2;refGene_tx=XM_011284888.3
C1	75476147	75476361
refGene_gn=PKN2;refGene_tx=XM_011284886.3;refGene_gn=PKN2;refGene_tx=XM_011284888.3
C1	75485777	75485963
refGene_gn=PKN2;refGene_tx=XM_011284886.3;refGene_gn=PKN2;refGene_tx=XM_011284888.3
C1	75486886	75486996
refGene_gn=PKN2;refGene_tx=XM_011284886.3;refGene_gn=PKN2;refGene_tx=XM_011284888.3
C1	75503212	75503356
refGene_gn=PKN2;refGene_tx=XM_011284886.3;refGene_gn=PKN2;refGene_tx=XM_011284888.3
C1	75503674	75503750
refGene_gn=PKN2;refGene_tx=XM_011284886.3;refGene_gn=PKN2;refGene_tx=XM_011284888.3
C1	75504348	75504523
refGene_gn=PKN2;refGene_tx=XM_011284886.3;refGene_gn=PKN2;refGene_tx=XM_011284888.3
C1	75504761	75504888
refGene_gn=PKN2;refGene_tx=XM_011284886.3;refGene_gn=PKN2;refGene_tx=XM_011284888.3
C1	75505368	75505499
refGene_gn=PKN2;refGene_tx=XM_011284886.3;refGene_gn=PKN2;refGene_tx=XM_011284888.3
C1	75505581	75505657
refGene_gn=PKN2;refGene_tx=XM_011284886.3;refGene_gn=PKN2;refGene_tx=XM_011284888.3
C1	75505736	75505828
refGene_gn=PKN2;refGene_tx=XM_011284886.3;refGene_gn=PKN2;refGene_tx=XM_011284888.3
C1	75512283	75512460
refGene_gn=PKN2;refGene_tx=XM_011284886.3;refGene_gn=PKN2;refGene_tx=XM_011284888.3
C1	75517205	75517268
refGene_gn=PKN2;refGene_tx=XM_011284886.3;refGene_gn=PKN2;refGene_tx=XM_011284888.3
C1	75518446	75518523
refGene_gn=PKN2;refGene_tx=XM_011284886.3;refGene_gn=PKN2;refGene_tx=XM_011284888.3
C1	75523625	75523768
refGene_gn=PKN2;refGene_tx=XM_011284886.3;refGene_gn=PKN2;refGene_tx=X

M_011284888.3
C1	75525143	75525170	refGene_gn=PKN2;refGene_tx=XM_011284888.3
C1	75527025	75527133	refGene_gn=PKN2;refGene_tx=XM_011284886.3
C1	75527504	75527585	refGene_gn=PKN2;refGene_tx=XM_011284886.3
C1	75527670	75527874	refGene_gn=PKN2;refGene_tx=XM_011284886.3
C1	75560828	75560962	refGene_gn=GTF2B;refGene_tx=XM_019837386.1
C1	75562917	75563199	refGene_gn=GTF2B;refGene_tx=XM_019837386.1
C1	75565430	75565560	refGene_gn=GTF2B;refGene_tx=XM_019837386.1
C1	75565692	75565839	refGene_gn=GTF2B;refGene_tx=XM_019837386.1
C1	75569704	75569838	refGene_gn=GTF2B;refGene_tx=XM_019837386.1
C1	75587797	75587904	refGene_gn=GTF2B;refGene_tx=XM_019837386.1
C1	75593044	75593061	refGene_gn=GTF2B;refGene_tx=XM_019837386.1
C1	75614698	75614761
refGene_gn=KYAT3;refGene_tx=XM_003990286.4;refGene_gn=KYAT3;refGene_tx=XM_006934884.4;refGene_gn=KYAT3;refGene_tx=XM_006934885.4;refGene_gn=KYAT3;refGene_tx=XM_019837387.2
C1	75620717	75620804
refGene_gn=KYAT3;refGene_tx=XM_003990286.4;refGene_gn=KYAT3;refGene_tx=XM_006934884.4;refGene_gn=KYAT3;refGene_tx=XM_006934885.4;refGene_gn=KYAT3;refGene_tx=XM_019837387.2
C1	75621078	75621155
refGene_gn=KYAT3;refGene_tx=XM_003990286.4;refGene_gn=KYAT3;refGene_tx=XM_006934884.4;refGene_gn=KYAT3;refGene_tx=XM_006934885.4;refGene_gn=KYAT3;refGene_tx=XM_019837387.2
C1	75624963	75625150
refGene_gn=KYAT3;refGene_tx=XM_003990286.4;refGene_gn=KYAT3;refGene_tx=XM_006934884.4;refGene_gn=KYAT3;refGene_tx=XM_006934885.4;refGene_gn=KYAT3;refGene_tx=XM_019837387.2
C1	75630782	75630872
refGene_gn=KYAT3;refGene_tx=XM_003990286.4;refGene_gn=KYAT3;refGene_tx=XM_006934884.4;refGene_gn=KYAT3;refGene_tx=XM_006934885.4;refGene_gn=KYAT3;refGene_tx=XM_019837387.2
C1	75632314	75632391
refGene_gn=KYAT3;refGene_tx=XM_003990286.4;refGene_gn=KYAT3;refGene_tx=XM_006934884.4;refGene_gn=KYAT3;refGene_tx=XM_006934885.4;refGene_gn=KYAT3;refGene_tx=XM_019837387.2
C1	75639705	75639826
refGene_gn=KYAT3;refGene_tx=XM_003990286.4;refGene_gn=KYAT3;refGene_tx=XM_006934884.4;refGene_gn=KYAT3;refGene_tx=XM_006934885.4;refGene_gn=KYAT3;refGene_tx=XM_019837387.2
C1	75639925	75640051
refGene_gn=KYAT3;refGene_tx=XM_003990286.4;refGene_gn=KYAT3;refGene_tx=XM_006934884.4;refGene_gn=KYAT3;refGene_tx=XM_006934885.4;refGene_gn=KYAT3;refGene_tx=XM_019837387.2
C1	75641287	75641374
refGene_gn=KYAT3;refGene_tx=XM_003990286.4;refGene_gn=KYAT3;refGene_tx=XM_006934884.4;refGene_gn=KYAT3;refGene_tx=XM_006934885.4;refGene_gn=KYAT3;refGene_tx=XM_019837387.2
C1	75645598	75645748
refGene_gn=KYAT3;refGene_tx=XM_003990286.4;refGene_gn=KYAT3;refGene_tx

=XM_006934884.4;refGene_gn=KYAT3;refGene_tx=XM_006934885.4;refGene_gn=KYAT3;refGene_tx=XM_019837387.2
C1 75649365 75649510
refGene_gn=KYAT3;refGene_tx=XM_003990286.4;refGene_gn=KYAT3;refGene_tx=XM_006934884.4;refGene_gn=KYAT3;refGene_tx=XM_006934885.4;refGene_gn=KYAT3;refGene_tx=XM_019837387.2
C1 75650291 75650350
refGene_gn=KYAT3;refGene_tx=XM_003990286.4;refGene_gn=KYAT3;refGene_tx=XM_006934884.4;refGene_gn=KYAT3;refGene_tx=XM_006934885.4;refGene_gn=KYAT3;refGene_tx=XM_019837387.2
C1 75684013 75684112
refGene_gn=KYAT3;refGene_tx=XM_003990286.4;refGene_gn=KYAT3;refGene_tx=XM_006934884.4;refGene_gn=KYAT3;refGene_tx=XM_006934885.4;refGene_gn=KYAT3;refGene_tx=XM_019837387.2
C1 75719535 75719652
refGene_gn=LOC101082424;refGene_tx=XM_003990287.3
C1 75720234 75720428
refGene_gn=LOC101082424;refGene_tx=XM_003990287.3
C1 75720666 75720769
refGene_gn=LOC101082424;refGene_tx=XM_003990287.3
C1 75721554 75721767
refGene_gn=LOC101082424;refGene_tx=XM_003990287.3
C1 75723321 75723602
refGene_gn=LOC101082424;refGene_tx=XM_003990287.3
C1 75725082 75725325
refGene_gn=LOC101082424;refGene_tx=XM_003990287.3
C1 75725911 75726108
refGene_gn=LOC101082424;refGene_tx=XM_003990287.3
C1 75726626 75726736
refGene_gn=LOC101082424;refGene_tx=XM_003990287.3
C1 75728140 75728268
refGene_gn=LOC101082424;refGene_tx=XM_003990287.3
C1 75729567 75729757
refGene_gn=LOC101082424;refGene_tx=XM_003990287.3
C1 75772765 75772813
refGene_gn=LOC101093701;refGene_tx=XM_006934887.3;refGene_gn=LOC101093701;refGene_tx=XM_011284884.2;refGene_gn=LOC101093701;refGene_tx=XM_019837390.2;refGene_gn=LOC101093701;refGene_tx=XM_023258879.1
C1 75783013 75783207
refGene_gn=LOC101093701;refGene_tx=XM_006934887.3;refGene_gn=LOC101093701;refGene_tx=XM_011284884.2;refGene_gn=LOC101093701;refGene_tx=XM_023258879.1
C1 75783832 75784001
refGene_gn=LOC101093701;refGene_tx=XM_006934887.3;refGene_gn=LOC101093701;refGene_tx=XM_011284884.2;refGene_gn=LOC101093701;refGene_tx=XM_023258879.1
C1 75784862 75785056
refGene_gn=LOC101093701;refGene_tx=XM_006934887.3;refGene_gn=LOC101093701;refGene_tx=XM_011284884.2;refGene_gn=LOC101093701;refGene_tx=XM_019837390.2;refGene_gn=LOC101093701;refGene_tx=XM_023258879.1

C1	75785381	75785484
refGene_gn=LOC101093701;refGene_tx=XM_006934887.3;refGene_gn=LOC101093701;refGene_tx=XM_011284884.2;refGene_gn=LOC101093701;refGene_tx=XM_019837390.2;refGene_gn=LOC101093701;refGene_tx=XM_023258879.1
C1	75786568	75786781
refGene_gn=LOC101093701;refGene_tx=XM_006934887.3;refGene_gn=LOC101093701;refGene_tx=XM_011284884.2;refGene_gn=LOC101093701;refGene_tx=XM_019837390.2;refGene_gn=LOC101093701;refGene_tx=XM_023258879.1
C1	75787706	75787987
refGene_gn=LOC101093701;refGene_tx=XM_006934887.3;refGene_gn=LOC101093701;refGene_tx=XM_011284884.2;refGene_gn=LOC101093701;refGene_tx=XM_019837390.2;refGene_gn=LOC101093701;refGene_tx=XM_023258879.1
C1	75789040	75789286
refGene_gn=LOC101093701;refGene_tx=XM_006934887.3;refGene_gn=LOC101093701;refGene_tx=XM_011284884.2;refGene_gn=LOC101093701;refGene_tx=XM_019837390.2;refGene_gn=LOC101093701;refGene_tx=XM_023258879.1
C1	75791296	75791493
refGene_gn=LOC101093701;refGene_tx=XM_006934887.3;refGene_gn=LOC101093701;refGene_tx=XM_011284884.2;refGene_gn=LOC101093701;refGene_tx=XM_019837390.2;refGene_gn=LOC101093701;refGene_tx=XM_023258879.1
C1	75791702	75791812
refGene_gn=LOC101093701;refGene_tx=XM_006934887.3;refGene_gn=LOC101093701;refGene_tx=XM_011284884.2;refGene_gn=LOC101093701;refGene_tx=XM_019837390.2;refGene_gn=LOC101093701;refGene_tx=XM_023258879.1
C1	75799824	75799952
refGene_gn=LOC101093701;refGene_tx=XM_006934887.3;refGene_gn=LOC101093701;refGene_tx=XM_011284884.2;refGene_gn=LOC101093701;refGene_tx=XM_019837390.2;refGene_gn=LOC101093701;refGene_tx=XM_023258879.1
C1	75801676	75801872
refGene_gn=LOC101093701;refGene_tx=XM_006934887.3;refGene_gn=LOC101093701;refGene_tx=XM_011284884.2;refGene_gn=LOC101093701;refGene_tx=XM_019837390.2;refGene_gn=LOC101093701;refGene_tx=XM_023258879.1
C1	75803058	75803061
refGene_gn=LOC101093701;refGene_tx=XM_006934887.3;refGene_gn=LOC101093701;refGene_tx=XM_011284884.2;refGene_gn=LOC101093701;refGene_tx=XM_019837390.2
C1	75826588	75826807
refGene_gn=LOC101093951;refGene_tx=XM_003990323.5
C1	75827678	75827872
refGene_gn=LOC101093951;refGene_tx=XM_003990323.5
C1	75828211	75828314
refGene_gn=LOC101093951;refGene_tx=XM_003990323.5
C1	75829270	75829483
refGene_gn=LOC101093951;refGene_tx=XM_003990323.5
C1	75830929	75831210
refGene_gn=LOC101093951;refGene_tx=XM_003990323.5
C1	75832475	75832721
refGene_gn=LOC101093951;refGene_tx=XM_003990323.5
C1	75834506	75834700
refGene_gn=LOC101093951;refGene_tx=XM_003990323.5

C1 75834887 75834997
refGene_gn=LOC101093951;refGene_tx=XM_003990323.5
C1 75836948 75837076
refGene_gn=LOC101093951;refGene_tx=XM_003990323.5
C1 75839108 75839298
refGene_gn=LOC101093951;refGene_tx=XM_003990323.5
C1 75882408 75882525
refGene_gn=LOC101094195;refGene_tx=XM_003990324.3;refGene_gn=LOC101094195;refGene_tx=XM_019837391.1
C1 75883963 75884145
refGene_gn=LOC101094195;refGene_tx=XM_003990324.3;refGene_gn=LOC101094195;refGene_tx=XM_019837391.1
C1 75884421 75884524
refGene_gn=LOC101094195;refGene_tx=XM_003990324.3;refGene_gn=LOC101094195;refGene_tx=XM_019837391.1
C1 75885292 75885505
refGene_gn=LOC101094195;refGene_tx=XM_003990324.3;refGene_gn=LOC101094195;refGene_tx=XM_019837391.1
C1 75886530 75886811
refGene_gn=LOC101094195;refGene_tx=XM_003990324.3;refGene_gn=LOC101094195;refGene_tx=XM_019837391.1
C1 75887830 75888073
refGene_gn=LOC101094195;refGene_tx=XM_003990324.3;refGene_gn=LOC101094195;refGene_tx=XM_019837391.1
C1 75888418 75888615
refGene_gn=LOC101094195;refGene_tx=XM_003990324.3;refGene_gn=LOC101094195;refGene_tx=XM_019837391.1
C1 75888875 75888985
refGene_gn=LOC101094195;refGene_tx=XM_003990324.3;refGene_gn=LOC101094195;refGene_tx=XM_019837391.1
C1 75891595 75891723
refGene_gn=LOC101094195;refGene_tx=XM_003990324.3;refGene_gn=LOC101094195;refGene_tx=XM_019837391.1
C1 75892484 75892674
refGene_gn=LOC101094195;refGene_tx=XM_003990324.3;refGene_gn=LOC101094195;refGene_tx=XM_019837391.1
C1 75926732 75926906
refGene_gn=LOC102901600;refGene_tx=XM_006934890.4
C1 75927835 75928026
refGene_gn=LOC102901600;refGene_tx=XM_006934890.4
C1 75928367 75928452
refGene_gn=LOC102901600;refGene_tx=XM_006934890.4
C1 75929492 75929582
refGene_gn=LOC102901600;refGene_tx=XM_006934890.4
C1 76023373 76023601
refGene_gn=LOC101082677;refGene_tx=XM_003990288.4
C1 76024595 76024789
refGene_gn=LOC101082677;refGene_tx=XM_003990288.4
C1 76025111 76025214
refGene_gn=LOC101082677;refGene_tx=XM_003990288.4

C1 76026947 76027160
refGene_gn=LOC101082677;refGene_tx=XM_003990288.4
C1 76027863 76028144
refGene_gn=LOC101082677;refGene_tx=XM_003990288.4
C1 76029027 76029273
refGene_gn=LOC101082677;refGene_tx=XM_003990288.4
C1 76031119 76031316
refGene_gn=LOC101082677;refGene_tx=XM_003990288.4
C1 76031517 76031627
refGene_gn=LOC101082677;refGene_tx=XM_003990288.4
C1 76033225 76033353
refGene_gn=LOC101082677;refGene_tx=XM_003990288.4
C1 76034820 76035010
refGene_gn=LOC101082677;refGene_tx=XM_003990288.4
C1 76171339 76173478
refGene_gn=LRRC8B;refGene_tx=XM_023258880.1;refGene_gn=LRRC8B;refGene_tx=XM_023258881.1;refGene_gn=LRRC8B;refGene_tx=XM_023258882.1;refGene_gn=LRRC8B;refGene_tx=XM_023258883.1;refGene_gn=LRRC8B;refGene_tx=XM_023258884.1;refGene_gn=LRRC8B;refGene_tx=XM_023258885.1;refGene_gn=LRRC8B;refGene_tx=XM_023258886.1;refGene_gn=LRRC8B;refGene_tx=XM_023258887.1;refGene_gn=LRRC8B;refGene_tx=XM_023258888.1;refGene_gn=LRRC8B;refGene_tx=XM_023258889.1
C1 76178989 76179439
refGene_gn=LRRC8B;refGene_tx=XM_023258880.1;refGene_gn=LRRC8B;refGene_tx=XM_023258881.1;refGene_gn=LRRC8B;refGene_tx=XM_023258882.1;refGene_gn=LRRC8B;refGene_tx=XM_023258883.1;refGene_gn=LRRC8B;refGene_tx=XM_023258884.1;refGene_gn=LRRC8B;refGene_tx=XM_023258885.1;refGene_gn=LRRC8B;refGene_tx=XM_023258886.1
C1 76180586 76180607 refGene_gn=LRRC8B;refGene_tx=XM_023258889.1
C1 76182239 76182512
refGene_gn=LRRC8B;refGene_tx=XM_023258887.1;refGene_gn=LRRC8B;refGene_tx=XM_023258888.1
C1 76188360 76188407 refGene_gn=LRRC8B;refGene_tx=XM_023258888.1
C1 76273357 76273495 refGene_gn=LRRC8C;refGene_tx=XM_023258891.1
C1 76302838 76305112 refGene_gn=LRRC8C;refGene_tx=XM_023258891.1
C1 76515872 76518449
refGene_gn=LRRC8D;refGene_tx=XM_019837396.2;refGene_gn=LRRC8D;refGene_tx=XM_023258892.1
C1 76580106 76580122
refGene_gn=ZNF326;refGene_tx=XM_006934897.4;refGene_gn=ZNF326;refGene_tx=XM_023258895.1
C1 76580201 76580658 refGene_gn=ZNF326;refGene_tx=XM_023258893.1
C1 76582618 76582663
refGene_gn=ZNF326;refGene_tx=XM_006934897.4;refGene_gn=ZNF326;refGene_tx=XM_023258893.1;refGene_gn=ZNF326;refGene_tx=XM_023258895.1
C1 76591343 76591379
refGene_gn=ZNF326;refGene_tx=XM_006934897.4;refGene_gn=ZNF326;refGene_tx=XM_023258893.1;refGene_gn=ZNF326;refGene_tx=XM_023258894.1;refGene_gn=ZNF326;refGene_tx=XM_023258895.1
C1 76591474 76591590

refGene_gn=ZNF326;refGene_tx=XM_006934897.4;refGene_gn=ZNF326;refGene_tx=XM_023258893.1;refGene_gn=ZNF326;refGene_tx=XM_023258894.1;refGene_gn=ZNF326;refGene_tx=XM_023258895.1
C1 76593726 76594132
refGene_gn=ZNF326;refGene_tx=XM_006934897.4;refGene_gn=ZNF326;refGene_tx=XM_023258893.1;refGene_gn=ZNF326;refGene_tx=XM_023258894.1
C1 76596504 76596703
refGene_gn=ZNF326;refGene_tx=XM_006934897.4;refGene_gn=ZNF326;refGene_tx=XM_019837400.2;refGene_gn=ZNF326;refGene_tx=XM_019837401.2;refGene_gn=ZNF326;refGene_tx=XM_023258893.1;refGene_gn=ZNF326;refGene_tx=XM_023258894.1
C1 76603138 76603250
refGene_gn=ZNF326;refGene_tx=XM_006934897.4;refGene_gn=ZNF326;refGene_tx=XM_019837400.2;refGene_gn=ZNF326;refGene_tx=XM_019837401.2;refGene_gn=ZNF326;refGene_tx=XM_023258893.1;refGene_gn=ZNF326;refGene_tx=XM_023258894.1
C1 76608612 76608760
refGene_gn=ZNF326;refGene_tx=XM_006934897.4;refGene_gn=ZNF326;refGene_tx=XM_019837400.2;refGene_gn=ZNF326;refGene_tx=XM_019837401.2;refGene_gn=ZNF326;refGene_tx=XM_023258893.1;refGene_gn=ZNF326;refGene_tx=XM_023258894.1
C1 76610063 76610163
refGene_gn=ZNF326;refGene_tx=XM_006934897.4;refGene_gn=ZNF326;refGene_tx=XM_019837400.2;refGene_gn=ZNF326;refGene_tx=XM_019837401.2;refGene_gn=ZNF326;refGene_tx=XM_023258893.1;refGene_gn=ZNF326;refGene_tx=XM_023258894.1
C1 76611658 76612147
refGene_gn=LOC101084379;refGene_tx=XM_019837402.2
C1 76612269 76612686
refGene_gn=LOC101084379;refGene_tx=XM_019837402.2
C1 76614368 76614499
refGene_gn=ZNF326;refGene_tx=XM_006934897.4;refGene_gn=ZNF326;refGene_tx=XM_019837400.2;refGene_gn=ZNF326;refGene_tx=XM_019837401.2;refGene_gn=ZNF326;refGene_tx=XM_023258893.1;refGene_gn=ZNF326;refGene_tx=XM_023258894.1
C1 76616292 76616388
refGene_gn=ZNF326;refGene_tx=XM_006934897.4;refGene_gn=ZNF326;refGene_tx=XM_019837400.2;refGene_gn=ZNF326;refGene_tx=XM_019837401.2;refGene_gn=ZNF326;refGene_tx=XM_023258893.1;refGene_gn=ZNF326;refGene_tx=XM_023258894.1
C1 76618291 76618660
refGene_gn=ZNF326;refGene_tx=XM_006934897.4;refGene_gn=ZNF326;refGene_tx=XM_019837400.2;refGene_gn=ZNF326;refGene_tx=XM_019837401.2;refGene_gn=ZNF326;refGene_tx=XM_023258893.1;refGene_gn=ZNF326;refGene_tx=XM_023258894.1
C1 77275968 77276281 refGene_gn=BARHL2;refGene_tx=XM_011284878.3
C1 77278042 77278268 refGene_gn=BARHL2;refGene_tx=XM_011284878.3
C1 77280113 77280735 refGene_gn=BARHL2;refGene_tx=XM_011284878.3
C1 77281053 77281307
refGene_gn=LOC111561634;refGene_tx=XM_023257834.1

C1	77281381 77281682
refGene_gn=LOC111561634;refGene_tx=XM_023257834.1
C1	77462249 77462442
refGene_gn=ZNF644;refGene_tx=XM_006934898.4;refGene_gn=ZNF644;refGene_tx=XM_019837404.2;refGene_gn=ZNF644;refGene_tx=XM_019837405.2;refGene_gn=ZNF644;refGene_tx=XM_019837406.2;refGene_gn=ZNF644;refGene_tx=XM_019837407.2;refGene_gn=ZNF644;refGene_tx=XM_019837408.2;refGene_gn=ZNF644;refGene_tx=XM_019837409.2;refGene_gn=ZNF644;refGene_tx=XM_019837410.1;refGene_gn=ZNF644;refGene_tx=XM_019837411.2;refGene_gn=ZNF644;refGene_tx=XM_019837413.2;refGene_gn=ZNF644;refGene_tx=XM_019837414.2;refGene_gn=ZNF644;refGene_tx=XM_019837415.2;refGene_gn=ZNF644;refGene_tx=XM_019837416.2;refGene_gn=ZNF644;refGene_tx=XM_023258896.1;refGene_gn=ZNF644;refGene_tx=XM_023258897.1
C1	77463246 77463349
refGene_gn=ZNF644;refGene_tx=XM_006934898.4;refGene_gn=ZNF644;refGene_tx=XM_019837404.2;refGene_gn=ZNF644;refGene_tx=XM_019837405.2;refGene_gn=ZNF644;refGene_tx=XM_019837406.2;refGene_gn=ZNF644;refGene_tx=XM_019837407.2;refGene_gn=ZNF644;refGene_tx=XM_019837408.2;refGene_gn=ZNF644;refGene_tx=XM_019837409.2;refGene_gn=ZNF644;refGene_tx=XM_019837410.1;refGene_gn=ZNF644;refGene_tx=XM_019837411.2;refGene_gn=ZNF644;refGene_tx=XM_019837413.2;refGene_gn=ZNF644;refGene_tx=XM_019837414.2;refGene_gn=ZNF644;refGene_tx=XM_019837415.2;refGene_gn=ZNF644;refGene_tx=XM_019837416.2;refGene_gn=ZNF644;refGene_tx=XM_023258896.1;refGene_gn=ZNF644;refGene_tx=XM_023258897.1
C1	77465387 77465437 refGene_gn=ZNF644;refGene_tx=XM_019837412.2
C1	77478191 77478314
refGene_gn=ZNF644;refGene_tx=XM_006934898.4;refGene_gn=ZNF644;refGene_tx=XM_019837404.2;refGene_gn=ZNF644;refGene_tx=XM_019837405.2;refGene_gn=ZNF644;refGene_tx=XM_019837406.2;refGene_gn=ZNF644;refGene_tx=XM_019837407.2;refGene_gn=ZNF644;refGene_tx=XM_019837408.2;refGene_gn=ZNF644;refGene_tx=XM_019837409.2;refGene_gn=ZNF644;refGene_tx=XM_019837410.1;refGene_gn=ZNF644;refGene_tx=XM_023258896.1;refGene_gn=ZNF644;refGene_tx=XM_023258897.1
C1	77480206 77480812
refGene_gn=ZNF644;refGene_tx=XM_006934898.4;refGene_gn=ZNF644;refGene_tx=XM_019837404.2;refGene_gn=ZNF644;refGene_tx=XM_019837405.2;refGene_gn=ZNF644;refGene_tx=XM_019837406.2;refGene_gn=ZNF644;refGene_tx=XM_019837407.2;refGene_gn=ZNF644;refGene_tx=XM_019837408.2;refGene_gn=ZNF644;refGene_tx=XM_019837409.2;refGene_gn=ZNF644;refGene_tx=XM_019837410.1;refGene_gn=ZNF644;refGene_tx=XM_019837411.2;refGene_gn=ZNF644;refGene_tx=XM_019837412.2;refGene_gn=ZNF644;refGene_tx=XM_023258896.1;refGene_gn=ZNF644;refGene_tx=XM_023258897.1
C1	77480997 77484035
refGene_gn=ZNF644;refGene_tx=XM_006934898.4;refGene_gn=ZNF644;refGene_tx=XM_019837404.2;refGene_gn=ZNF644;refGene_tx=XM_019837405.2;refGene_gn=ZNF644;refGene_tx=XM_019837406.2;refGene_gn=ZNF644;refGene_tx=XM_019837407.2;refGene_gn=ZNF644;refGene_tx=XM_019837408.2;refGene_gn=ZNF644;refGene_tx=XM_019837409.2;refGene_gn=ZNF644;refGene_tx=XM_019837410.1;refGene_gn=ZNF644;refGene_tx=XM_019837411.2;refGene_gn=ZNF644;refGene_tx=XM_019837412.2;refGene_gn=ZNF644;refGene_tx=XM_023258896.1;refGen

e_gn=ZNF644;refGene_tx=XM_023258897.1
C1 77535223 77535267
refGene_gn=ZNF644;refGene_tx=XM_006934898.4;refGene_gn=ZNF644;refGene_tx=XM_019837404.2;refGene_gn=ZNF644;refGene_tx=XM_019837405.2;refGene_gn=ZNF644;refGene_tx=XM_019837406.2;refGene_gn=ZNF644;refGene_tx=XM_019837407.2;refGene_gn=ZNF644;refGene_tx=XM_019837408.2;refGene_gn=ZNF644;refGene_tx=XM_019837409.2;refGene_gn=ZNF644;refGene_tx=XM_019837410.1;refGene_gn=ZNF644;refGene_tx=XM_019837411.2;refGene_gn=ZNF644;refGene_tx=XM_019837412.2;refGene_gn=ZNF644;refGene_tx=XM_019837413.2;refGene_gn=ZNF644;refGene_tx=XM_019837414.2;refGene_gn=ZNF644;refGene_tx=XM_019837415.2;refGene_gn=ZNF644;refGene_tx=XM_019837416.2;refGene_gn=ZNF644;refGene_tx=XM_023258896.1;refGene_gn=ZNF644;refGene_tx=XM_023258897.1
C1 77711255 77711448
refGene_gn=HFM1;refGene_tx=XM_023258898.1;refGene_gn=HFM1;refGene_tx=XM_023258900.1
C1 77714728 77714798
refGene_gn=HFM1;refGene_tx=XM_006934913.4;refGene_gn=HFM1;refGene_tx=XM_023258899.1
C1 77715474 77715629
refGene_gn=HFM1;refGene_tx=XM_006934913.4;refGene_gn=HFM1;refGene_tx=XM_023258898.1;refGene_gn=HFM1;refGene_tx=XM_023258899.1;refGene_gn=HFM1;refGene_tx=XM_023258900.1
C1 77715721 77715833
refGene_gn=HFM1;refGene_tx=XM_006934913.4;refGene_gn=HFM1;refGene_tx=XM_023258898.1;refGene_gn=HFM1;refGene_tx=XM_023258899.1;refGene_gn=HFM1;refGene_tx=XM_023258900.1
C1 77722461 77722552
refGene_gn=HFM1;refGene_tx=XM_006934913.4;refGene_gn=HFM1;refGene_tx=XM_023258898.1;refGene_gn=HFM1;refGene_tx=XM_023258899.1;refGene_gn=HFM1;refGene_tx=XM_023258900.1
C1 77723928 77724174
refGene_gn=HFM1;refGene_tx=XM_006934913.4;refGene_gn=HFM1;refGene_tx=XM_023258898.1;refGene_gn=HFM1;refGene_tx=XM_023258899.1;refGene_gn=HFM1;refGene_tx=XM_023258900.1;refGene_gn=HFM1;refGene_tx=XM_023258901.1
C1 77730371 77730472
refGene_gn=HFM1;refGene_tx=XM_006934913.4;refGene_gn=HFM1;refGene_tx=XM_023258898.1;refGene_gn=HFM1;refGene_tx=XM_023258899.1;refGene_gn=HFM1;refGene_tx=XM_023258900.1
C1 77731451 77731531
refGene_gn=HFM1;refGene_tx=XM_006934913.4;refGene_gn=HFM1;refGene_tx=XM_023258898.1;refGene_gn=HFM1;refGene_tx=XM_023258899.1;refGene_gn=HFM1;refGene_tx=XM_023258900.1
C1 77733681 77733797
refGene_gn=HFM1;refGene_tx=XM_006934913.4;refGene_gn=HFM1;refGene_tx=XM_023258898.1;refGene_gn=HFM1;refGene_tx=XM_023258899.1;refGene_gn=HFM1;refGene_tx=XM_023258900.1;refGene_gn=HFM1;refGene_tx=XM_023258901.1
C1 77734019 77734100
refGene_gn=HFM1;refGene_tx=XM_006934913.4;refGene_gn=HFM1;refGene_tx=XM_023258898.1;refGene_gn=HFM1;refGene_tx=XM_023258899.1;refGene_gn=HFM

1;refGene_tx=XM_023258900.1;refGene_gn=HFM1;refGene_tx=XM_023258901.1
C1 77750697 77750844
refGene_gn=HFM1;refGene_tx=XM_006934913.4;refGene_gn=HFM1;refGene_tx=XM_023258898.1;refGene_gn=HFM1;refGene_tx=XM_023258899.1;refGene_gn=HFM1;refGene_tx=XM_023258901.1
C1 77751445 77751549
refGene_gn=HFM1;refGene_tx=XM_006934913.4;refGene_gn=HFM1;refGene_tx=XM_023258898.1;refGene_gn=HFM1;refGene_tx=XM_023258899.1;refGene_gn=HFM1;refGene_tx=XM_023258900.1;refGene_gn=HFM1;refGene_tx=XM_023258901.1
C1 77753192 77753350
refGene_gn=HFM1;refGene_tx=XM_006934913.4;refGene_gn=HFM1;refGene_tx=XM_023258898.1;refGene_gn=HFM1;refGene_tx=XM_023258899.1;refGene_gn=HFM1;refGene_tx=XM_023258900.1;refGene_gn=HFM1;refGene_tx=XM_023258901.1
C1 77753505 77753589
refGene_gn=HFM1;refGene_tx=XM_006934913.4;refGene_gn=HFM1;refGene_tx=XM_023258898.1;refGene_gn=HFM1;refGene_tx=XM_023258899.1;refGene_gn=HFM1;refGene_tx=XM_023258900.1;refGene_gn=HFM1;refGene_tx=XM_023258901.1
C1 77754720 77754806
refGene_gn=HFM1;refGene_tx=XM_006934913.4;refGene_gn=HFM1;refGene_tx=XM_023258898.1;refGene_gn=HFM1;refGene_tx=XM_023258899.1;refGene_gn=HFM1;refGene_tx=XM_023258900.1;refGene_gn=HFM1;refGene_tx=XM_023258901.1
C1 77757281 77757413
refGene_gn=HFM1;refGene_tx=XM_006934913.4;refGene_gn=HFM1;refGene_tx=XM_023258898.1;refGene_gn=HFM1;refGene_tx=XM_023258899.1;refGene_gn=HFM1;refGene_tx=XM_023258900.1;refGene_gn=HFM1;refGene_tx=XM_023258901.1
C1 77757496 77757594
refGene_gn=HFM1;refGene_tx=XM_006934913.4;refGene_gn=HFM1;refGene_tx=XM_023258898.1;refGene_gn=HFM1;refGene_tx=XM_023258899.1;refGene_gn=HFM1;refGene_tx=XM_023258900.1;refGene_gn=HFM1;refGene_tx=XM_023258901.1
C1 77758193 77758241
refGene_gn=HFM1;refGene_tx=XM_006934913.4;refGene_gn=HFM1;refGene_tx=XM_023258898.1;refGene_gn=HFM1;refGene_tx=XM_023258899.1;refGene_gn=HFM1;refGene_tx=XM_023258900.1;refGene_gn=HFM1;refGene_tx=XM_023258901.1
C1 77758332 77758439
refGene_gn=HFM1;refGene_tx=XM_006934913.4;refGene_gn=HFM1;refGene_tx=XM_023258898.1;refGene_gn=HFM1;refGene_tx=XM_023258899.1;refGene_gn=HFM1;refGene_tx=XM_023258900.1;refGene_gn=HFM1;refGene_tx=XM_023258901.1
C1 77760582 77760674
refGene_gn=HFM1;refGene_tx=XM_006934913.4;refGene_gn=HFM1;refGene_tx=XM_023258898.1;refGene_gn=HFM1;refGene_tx=XM_023258899.1;refGene_gn=HFM1;refGene_tx=XM_023258900.1;refGene_gn=HFM1;refGene_tx=XM_023258901.1
C1 77762530 77762611
refGene_gn=HFM1;refGene_tx=XM_006934913.4;refGene_gn=HFM1;refGene_tx=XM_023258898.1;refGene_gn=HFM1;refGene_tx=XM_023258899.1;refGene_gn=HFM1;refGene_tx=XM_023258900.1;refGene_gn=HFM1;refGene_tx=XM_023258901.1
C1 77765787 77765835
refGene_gn=HFM1;refGene_tx=XM_006934913.4;refGene_gn=HFM1;refGene_tx=XM_023258898.1;refGene_gn=HFM1;refGene_tx=XM_023258899.1;refGene_gn=HFM1;refGene_tx=XM_023258900.1;refGene_gn=HFM1;refGene_tx=XM_023258901.1
C1 77766805 77766939

refGene_gn=HFM1;refGene_tx=XM_006934913.4;refGene_gn=HFM1;refGene_tx=XM_023258898.1;refGene_gn=HFM1;refGene_tx=XM_023258899.1;refGene_gn=HFM1;refGene_tx=XM_023258900.1;refGene_gn=HFM1;refGene_tx=XM_023258901.1
C1 77767570 77767665
refGene_gn=HFM1;refGene_tx=XM_006934913.4;refGene_gn=HFM1;refGene_tx=XM_023258898.1;refGene_gn=HFM1;refGene_tx=XM_023258899.1;refGene_gn=HFM1;refGene_tx=XM_023258900.1;refGene_gn=HFM1;refGene_tx=XM_023258901.1
C1 77768549 77768695
refGene_gn=HFM1;refGene_tx=XM_006934913.4;refGene_gn=HFM1;refGene_tx=XM_023258898.1;refGene_gn=HFM1;refGene_tx=XM_023258899.1;refGene_gn=HFM1;refGene_tx=XM_023258900.1;refGene_gn=HFM1;refGene_tx=XM_023258901.1
C1 77769091 77769193
refGene_gn=HFM1;refGene_tx=XM_006934913.4;refGene_gn=HFM1;refGene_tx=XM_023258898.1;refGene_gn=HFM1;refGene_tx=XM_023258899.1;refGene_gn=HFM1;refGene_tx=XM_023258900.1;refGene_gn=HFM1;refGene_tx=XM_023258901.1
C1 77769300 77769344
refGene_gn=HFM1;refGene_tx=XM_006934913.4;refGene_gn=HFM1;refGene_tx=XM_023258898.1;refGene_gn=HFM1;refGene_tx=XM_023258899.1;refGene_gn=HFM1;refGene_tx=XM_023258900.1;refGene_gn=HFM1;refGene_tx=XM_023258901.1
C1 77774041 77774130
refGene_gn=HFM1;refGene_tx=XM_006934913.4;refGene_gn=HFM1;refGene_tx=XM_023258898.1;refGene_gn=HFM1;refGene_tx=XM_023258899.1;refGene_gn=HFM1;refGene_tx=XM_023258900.1;refGene_gn=HFM1;refGene_tx=XM_023258901.1
C1 77774212 77774413
refGene_gn=HFM1;refGene_tx=XM_006934913.4;refGene_gn=HFM1;refGene_tx=XM_023258898.1;refGene_gn=HFM1;refGene_tx=XM_023258899.1;refGene_gn=HFM1;refGene_tx=XM_023258900.1;refGene_gn=HFM1;refGene_tx=XM_023258901.1
C1 77776876 77777035
refGene_gn=HFM1;refGene_tx=XM_006934913.4;refGene_gn=HFM1;refGene_tx=XM_023258898.1;refGene_gn=HFM1;refGene_tx=XM_023258899.1;refGene_gn=HFM1;refGene_tx=XM_023258900.1;refGene_gn=HFM1;refGene_tx=XM_023258901.1
C1 77777245 77777323
refGene_gn=HFM1;refGene_tx=XM_006934913.4;refGene_gn=HFM1;refGene_tx=XM_023258898.1;refGene_gn=HFM1;refGene_tx=XM_023258899.1;refGene_gn=HFM1;refGene_tx=XM_023258900.1;refGene_gn=HFM1;refGene_tx=XM_023258901.1
C1 77777908 77778060
refGene_gn=HFM1;refGene_tx=XM_006934913.4;refGene_gn=HFM1;refGene_tx=XM_023258898.1;refGene_gn=HFM1;refGene_tx=XM_023258899.1;refGene_gn=HFM1;refGene_tx=XM_023258900.1;refGene_gn=HFM1;refGene_tx=XM_023258901.1
C1 77778717 77778850
refGene_gn=HFM1;refGene_tx=XM_006934913.4;refGene_gn=HFM1;refGene_tx=XM_023258898.1;refGene_gn=HFM1;refGene_tx=XM_023258899.1;refGene_gn=HFM1;refGene_tx=XM_023258900.1;refGene_gn=HFM1;refGene_tx=XM_023258901.1
C1 77779318 77779389
refGene_gn=HFM1;refGene_tx=XM_006934913.4;refGene_gn=HFM1;refGene_tx=XM_023258898.1;refGene_gn=HFM1;refGene_tx=XM_023258899.1;refGene_gn=HFM1;refGene_tx=XM_023258900.1;refGene_gn=HFM1;refGene_tx=XM_023258901.1
C1 77785426 77785474
refGene_gn=HFM1;refGene_tx=XM_006934913.4;refGene_gn=HFM1;refGene_tx=XM_023258898.1;refGene_gn=HFM1;refGene_tx=XM_023258899.1;refGene_gn=HFM

1;refGene_tx=XM_023258900.1;refGene_gn=HFM1;refGene_tx=XM_023258901.1
C1 77785850 77786116
refGene_gn=HFM1;refGene_tx=XM_006934913.4;refGene_gn=HFM1;refGene_tx=XM_023258898.1;refGene_gn=HFM1;refGene_tx=XM_023258899.1;refGene_gn=HFM1;refGene_tx=XM_023258900.1;refGene_gn=HFM1;refGene_tx=XM_023258901.1
C1 77794972 77795285
refGene_gn=HFM1;refGene_tx=XM_006934913.4;refGene_gn=HFM1;refGene_tx=XM_023258898.1;refGene_gn=HFM1;refGene_tx=XM_023258899.1;refGene_gn=HFM1;refGene_tx=XM_023258900.1;refGene_gn=HFM1;refGene_tx=XM_023258901.1
C1 77796487 77796600
refGene_gn=HFM1;refGene_tx=XM_006934913.4;refGene_gn=HFM1;refGene_tx=XM_023258898.1;refGene_gn=HFM1;refGene_tx=XM_023258899.1;refGene_gn=HFM1;refGene_tx=XM_023258900.1;refGene_gn=HFM1;refGene_tx=XM_023258901.1
C1 77800613 77800684
refGene_gn=HFM1;refGene_tx=XM_006934913.4;refGene_gn=HFM1;refGene_tx=XM_023258898.1;refGene_gn=HFM1;refGene_tx=XM_023258899.1;refGene_gn=HFM1;refGene_tx=XM_023258900.1;refGene_gn=HFM1;refGene_tx=XM_023258901.1
C1 77879255 77879370
refGene_gn=CDC7;refGene_tx=XM_003990296.5;refGene_gn=CDC7;refGene_tx=XM_003990297.5;refGene_gn=CDC7;refGene_tx=XM_019837421.1;refGene_gn=CDC7;refGene_tx=XM_019837422.2
C1 77887526 77887610
refGene_gn=CDC7;refGene_tx=XM_003990296.5;refGene_gn=CDC7;refGene_tx=XM_003990297.5;refGene_gn=CDC7;refGene_tx=XM_019837421.1;refGene_gn=CDC7;refGene_tx=XM_019837422.2
C1 77889047 77889183
refGene_gn=CDC7;refGene_tx=XM_003990296.5;refGene_gn=CDC7;refGene_tx=XM_003990297.5;refGene_gn=CDC7;refGene_tx=XM_019837421.1;refGene_gn=CDC7;refGene_tx=XM_019837422.2
C1 77892352 77892446
refGene_gn=CDC7;refGene_tx=XM_003990296.5;refGene_gn=CDC7;refGene_tx=XM_003990297.5;refGene_gn=CDC7;refGene_tx=XM_019837421.1;refGene_gn=CDC7;refGene_tx=XM_019837422.2
C1 77892531 77892674
refGene_gn=CDC7;refGene_tx=XM_003990296.5;refGene_gn=CDC7;refGene_tx=XM_003990297.5;refGene_gn=CDC7;refGene_tx=XM_019837421.1;refGene_gn=CDC7;refGene_tx=XM_019837422.2
C1 77894033 77894283
refGene_gn=CDC7;refGene_tx=XM_003990296.5;refGene_gn=CDC7;refGene_tx=XM_003990297.5;refGene_gn=CDC7;refGene_tx=XM_019837421.1;refGene_gn=CDC7;refGene_tx=XM_019837422.2
C1 77894885 77894981
refGene_gn=CDC7;refGene_tx=XM_003990296.5;refGene_gn=CDC7;refGene_tx=XM_003990297.5;refGene_gn=CDC7;refGene_tx=XM_019837421.1
C1 77895754 77895933
refGene_gn=CDC7;refGene_tx=XM_003990296.5;refGene_gn=CDC7;refGene_tx=XM_003990297.5;refGene_gn=CDC7;refGene_tx=XM_019837421.1;refGene_gn=CDC7;refGene_tx=XM_019837422.2
C1 77896787 77896870
refGene_gn=CDC7;refGene_tx=XM_003990296.5;refGene_gn=CDC7;refGene_tx=X

M_003990297.5;refGene_gn=CDC7;refGene_tx=XM_019837421.1;refGene_gn=CDC7;refGene_tx=XM_019837422.2
C1 77899550 77899700 refGene_gn=CDC7;refGene_tx=XM_003990296.5;refGene_gn=CDC7;refGene_tx=XM_003990297.5;refGene_gn=CDC7;refGene_tx=XM_019837421.1;refGene_gn=CDC7;refGene_tx=XM_019837422.2
C1 77903313 77903708 refGene_gn=CDC7;refGene_tx=XM_003990296.5;refGene_gn=CDC7;refGene_tx=XM_003990297.5;refGene_gn=CDC7;refGene_tx=XM_019837421.1;refGene_gn=CDC7;refGene_tx=XM_019837422.2
C1 78033012 78033131 refGene_gn=TGFBR3;refGene_tx=XM_003990327.5;refGene_gn=TGFBR3;refGene_tx=XM_006934902.4;refGene_gn=TGFBR3;refGene_tx=XM_023258903.1
C1 78033267 78033308 refGene_gn=TGFBR3;refGene_tx=XM_019837423.2
C1 78044791 78044899 refGene_gn=TGFBR3;refGene_tx=XM_003990327.5;refGene_gn=TGFBR3;refGene_tx=XM_006934902.4;refGene_gn=TGFBR3;refGene_tx=XM_019837423.2;refGene_gn=TGFBR3;refGene_tx=XM_023258903.1
C1 78047133 78047175 refGene_gn=TGFBR3;refGene_tx=XM_003990327.5;refGene_gn=TGFBR3;refGene_tx=XM_006934902.4;refGene_gn=TGFBR3;refGene_tx=XM_019837423.2;refGene_gn=TGFBR3;refGene_tx=XM_023258903.1
C1 78057555 78057676 refGene_gn=TGFBR3;refGene_tx=XM_003990327.5;refGene_gn=TGFBR3;refGene_tx=XM_006934902.4;refGene_gn=TGFBR3;refGene_tx=XM_019837423.2;refGene_gn=TGFBR3;refGene_tx=XM_023258903.1
C1 78060847 78061147 refGene_gn=TGFBR3;refGene_tx=XM_003990327.5;refGene_gn=TGFBR3;refGene_tx=XM_006934902.4;refGene_gn=TGFBR3;refGene_tx=XM_019837423.2;refGene_gn=TGFBR3;refGene_tx=XM_023258903.1
C1 78066052 78066211 refGene_gn=TGFBR3;refGene_tx=XM_003990327.5;refGene_gn=TGFBR3;refGene_tx=XM_006934902.4;refGene_gn=TGFBR3;refGene_tx=XM_019837423.2;refGene_gn=TGFBR3;refGene_tx=XM_023258903.1
C1 78066394 78066535 refGene_gn=TGFBR3;refGene_tx=XM_003990327.5;refGene_gn=TGFBR3;refGene_tx=XM_006934902.4;refGene_gn=TGFBR3;refGene_tx=XM_019837423.2;refGene_gn=TGFBR3;refGene_tx=XM_023258903.1
C1 78069142 78069295 refGene_gn=TGFBR3;refGene_tx=XM_003990327.5;refGene_gn=TGFBR3;refGene_tx=XM_006934902.4;refGene_gn=TGFBR3;refGene_tx=XM_019837423.2;refGene_gn=TGFBR3;refGene_tx=XM_023258903.1
C1 78069700 78070035 refGene_gn=TGFBR3;refGene_tx=XM_003990327.5;refGene_gn=TGFBR3;refGene_tx=XM_006934902.4;refGene_gn=TGFBR3;refGene_tx=XM_019837423.2;refGene_gn=TGFBR3;refGene_tx=XM_023258903.1
C1 78071951 78072141 refGene_gn=TGFBR3;refGene_tx=XM_003990327.5;refGene_gn=TGFBR3;refGene_tx=XM_006934902.4;refGene_gn=TGFBR3;refGene_tx=XM_019837423.2;refGene_gn=TGFBR3;refGene_tx=XM_023258903.1

C1	78076882 78077030
refGene_gn=TGFBR3;refGene_tx=XM_003990327.5;refGene_gn=TGFBR3;refGene_tx=XM_006934902.4;refGene_gn=TGFBR3;refGene_tx=XM_019837423.2;refGene_gn=TGFBR3;refGene_tx=XM_023258903.1
C1	78078789 78078958
refGene_gn=TGFBR3;refGene_tx=XM_003990327.5;refGene_gn=TGFBR3;refGene_tx=XM_006934902.4;refGene_gn=TGFBR3;refGene_tx=XM_019837423.2;refGene_gn=TGFBR3;refGene_tx=XM_023258903.1
C1	78083679 78083863
refGene_gn=TGFBR3;refGene_tx=XM_003990327.5;refGene_gn=TGFBR3;refGene_tx=XM_006934902.4;refGene_gn=TGFBR3;refGene_tx=XM_019837423.2;refGene_gn=TGFBR3;refGene_tx=XM_023258903.1
C1	78106728 78106866
refGene_gn=TGFBR3;refGene_tx=XM_003990327.5;refGene_gn=TGFBR3;refGene_tx=XM_006934902.4;refGene_gn=TGFBR3;refGene_tx=XM_019837423.2;refGene_gn=TGFBR3;refGene_tx=XM_023258903.1
C1	78150853 78151038
refGene_gn=TGFBR3;refGene_tx=XM_003990327.5;refGene_gn=TGFBR3;refGene_tx=XM_006934902.4;refGene_gn=TGFBR3;refGene_tx=XM_019837423.2;refGene_gn=TGFBR3;refGene_tx=XM_023258903.1
C1	78214344 78214432
refGene_gn=TGFBR3;refGene_tx=XM_003990327.5;refGene_gn=TGFBR3;refGene_tx=XM_006934902.4;refGene_gn=TGFBR3;refGene_tx=XM_019837423.2;refGene_gn=TGFBR3;refGene_tx=XM_023258903.1
C1	78310399 78310588
refGene_gn=BRDT;refGene_tx=XM_019837424.2;refGene_gn=BRDT;refGene_tx=XM_019837425.2;refGene_gn=BRDT;refGene_tx=XM_019837426.2;refGene_gn=BRDT;refGene_tx=XM_019837427.2;refGene_gn=BRDT;refGene_tx=XM_019837428.2;refGene_gn=BRDT;refGene_tx=XM_019837429.2;refGene_gn=BRDT;refGene_tx=XM_019837430.2
C1	78311376 78311514
refGene_gn=BRDT;refGene_tx=XM_019837424.2;refGene_gn=BRDT;refGene_tx=XM_019837425.2;refGene_gn=BRDT;refGene_tx=XM_019837426.2;refGene_gn=BRDT;refGene_tx=XM_019837427.2;refGene_gn=BRDT;refGene_tx=XM_019837428.2;refGene_gn=BRDT;refGene_tx=XM_019837429.2;refGene_gn=BRDT;refGene_tx=XM_019837430.2
C1	78315832 78315947
refGene_gn=BRDT;refGene_tx=XM_019837424.2;refGene_gn=BRDT;refGene_tx=XM_019837425.2;refGene_gn=BRDT;refGene_tx=XM_019837426.2;refGene_gn=BRDT;refGene_tx=XM_019837427.2;refGene_gn=BRDT;refGene_tx=XM_019837428.2;refGene_gn=BRDT;refGene_tx=XM_019837429.2;refGene_gn=BRDT;refGene_tx=XM_019837430.2
C1	78331575 78331748
refGene_gn=BRDT;refGene_tx=XM_019837424.2;refGene_gn=BRDT;refGene_tx=XM_019837425.2;refGene_gn=BRDT;refGene_tx=XM_019837426.2;refGene_gn=BRDT;refGene_tx=XM_019837427.2;refGene_gn=BRDT;refGene_tx=XM_019837428.2;refGene_gn=BRDT;refGene_tx=XM_019837429.2;refGene_gn=BRDT;refGene_tx=XM_019837430.2
C1	78332361 78332715
refGene_gn=BRDT;refGene_tx=XM_019837424.2;refGene_gn=BRDT;refGene_tx=X

M_019837425.2;refGene_gn=BRDT;refGene_tx=XM_019837426.2;refGene_gn=BRDT;refGene_tx=XM_019837427.2;refGene_gn=BRDT;refGene_tx=XM_019837428.2;refGene_gn=BRDT;refGene_tx=XM_019837429.2;refGene_gn=BRDT;refGene_tx=XM_019837430.2
C1 78337944 78338073
refGene_gn=BRDT;refGene_tx=XM_019837424.2;refGene_gn=BRDT;refGene_tx=XM_019837425.2;refGene_gn=BRDT;refGene_tx=XM_019837426.2;refGene_gn=BRDT;refGene_tx=XM_019837427.2;refGene_gn=BRDT;refGene_tx=XM_019837428.2;refGene_gn=BRDT;refGene_tx=XM_019837429.2;refGene_gn=BRDT;refGene_tx=XM_019837430.2
C1 78343030 78343219
refGene_gn=BRDT;refGene_tx=XM_019837424.2;refGene_gn=BRDT;refGene_tx=XM_019837425.2;refGene_gn=BRDT;refGene_tx=XM_019837426.2;refGene_gn=BRDT;refGene_tx=XM_019837427.2;refGene_gn=BRDT;refGene_tx=XM_019837428.2;refGene_gn=BRDT;refGene_tx=XM_019837429.2;refGene_gn=BRDT;refGene_tx=XM_019837430.2
C1 78343810 78343980
refGene_gn=BRDT;refGene_tx=XM_019837424.2;refGene_gn=BRDT;refGene_tx=XM_019837425.2;refGene_gn=BRDT;refGene_tx=XM_019837426.2;refGene_gn=BRDT;refGene_tx=XM_019837427.2;refGene_gn=BRDT;refGene_tx=XM_019837428.2;refGene_gn=BRDT;refGene_tx=XM_019837429.2;refGene_gn=BRDT;refGene_tx=XM_019837430.2
C1 78344062 78344352
refGene_gn=BRDT;refGene_tx=XM_019837424.2;refGene_gn=BRDT;refGene_tx=XM_019837425.2;refGene_gn=BRDT;refGene_tx=XM_019837426.2;refGene_gn=BRDT;refGene_tx=XM_019837427.2;refGene_gn=BRDT;refGene_tx=XM_019837428.2;refGene_gn=BRDT;refGene_tx=XM_019837429.2;refGene_gn=BRDT;refGene_tx=XM_019837430.2
C1 78344438 78344552
refGene_gn=BRDT;refGene_tx=XM_019837424.2;refGene_gn=BRDT;refGene_tx=XM_019837425.2;refGene_gn=BRDT;refGene_tx=XM_019837426.2;refGene_gn=BRDT;refGene_tx=XM_019837427.2;refGene_gn=BRDT;refGene_tx=XM_019837428.2;refGene_gn=BRDT;refGene_tx=XM_019837429.2;refGene_gn=BRDT;refGene_tx=XM_019837430.2
C1 78344628 78344784
refGene_gn=BRDT;refGene_tx=XM_019837424.2;refGene_gn=BRDT;refGene_tx=XM_019837425.2;refGene_gn=BRDT;refGene_tx=XM_019837426.2;refGene_gn=BRDT;refGene_tx=XM_019837427.2;refGene_gn=BRDT;refGene_tx=XM_019837428.2;refGene_gn=BRDT;refGene_tx=XM_019837429.2;refGene_gn=BRDT;refGene_tx=XM_019837430.2
C1 78352445 78352504
refGene_gn=BRDT;refGene_tx=XM_019837424.2;refGene_gn=BRDT;refGene_tx=XM_019837425.2;refGene_gn=BRDT;refGene_tx=XM_019837426.2;refGene_gn=BRDT;refGene_tx=XM_019837427.2;refGene_gn=BRDT;refGene_tx=XM_019837428.2;refGene_gn=BRDT;refGene_tx=XM_019837429.2;refGene_gn=BRDT;refGene_tx=XM_019837430.2
C1 78354840 78355012
refGene_gn=BRDT;refGene_tx=XM_019837424.2;refGene_gn=BRDT;refGene_tx=XM_019837425.2;refGene_gn=BRDT;refGene_tx=XM_019837426.2;refGene_gn=BRDT;refGene_tx=XM_019837427.2;refGene_gn=BRDT;refGene_tx=XM_019837428.2;

refGene_gn=BRDT;refGene_tx=XM_019837429.2;refGene_gn=BRDT;refGene_tx=XM_019837430.2
C1 78362051 78362152
refGene_gn=BRDT;refGene_tx=XM_019837424.2;refGene_gn=BRDT;refGene_tx=XM_019837425.2;refGene_gn=BRDT;refGene_tx=XM_019837426.2;refGene_gn=BRDT;refGene_tx=XM_019837427.2;refGene_gn=BRDT;refGene_tx=XM_019837428.2;refGene_gn=BRDT;refGene_tx=XM_019837429.2;refGene_gn=BRDT;refGene_tx=XM_019837430.2
C1 78380993 78381199
refGene_gn=BRDT;refGene_tx=XM_019837424.2;refGene_gn=BRDT;refGene_tx=XM_019837425.2;refGene_gn=BRDT;refGene_tx=XM_019837426.2;refGene_gn=BRDT;refGene_tx=XM_019837427.2;refGene_gn=BRDT;refGene_tx=XM_019837428.2;refGene_gn=BRDT;refGene_tx=XM_019837429.2;refGene_gn=BRDT;refGene_tx=XM_019837430.2
C1 78381838 78382013
refGene_gn=BRDT;refGene_tx=XM_019837424.2;refGene_gn=BRDT;refGene_tx=XM_019837425.2;refGene_gn=BRDT;refGene_tx=XM_019837426.2;refGene_gn=BRDT;refGene_tx=XM_019837427.2;refGene_gn=BRDT;refGene_tx=XM_019837428.2;refGene_gn=BRDT;refGene_tx=XM_019837429.2;refGene_gn=BRDT;refGene_tx=XM_019837430.2
C1 78387013 78387082
refGene_gn=BRDT;refGene_tx=XM_019837424.2;refGene_gn=BRDT;refGene_tx=XM_019837425.2;refGene_gn=BRDT;refGene_tx=XM_019837426.2;refGene_gn=BRDT;refGene_tx=XM_019837427.2;refGene_gn=BRDT;refGene_tx=XM_019837428.2;refGene_gn=BRDT;refGene_tx=XM_019837429.2
C1 78403647 78403878 refGene_gn=EPHX4;refGene_tx=XM_019839090.2
C1 78406134 78406220 refGene_gn=EPHX4;refGene_tx=XM_019839090.2
C1 78420488 78420646 refGene_gn=EPHX4;refGene_tx=XM_019839090.2
C1 78422413 78422542 refGene_gn=EPHX4;refGene_tx=XM_019839090.2
C1 78426366 78426470 refGene_gn=EPHX4;refGene_tx=XM_019839090.2
C1 78430543 78430692 refGene_gn=EPHX4;refGene_tx=XM_019839090.2
C1 78446259 78446491 refGene_gn=EPHX4;refGene_tx=XM_019839090.2
C1 78463325 78463474
refGene_gn=LOC101086076;refGene_tx=XM_019837431.2;refGene_gn=LOC101086076;refGene_tx=XM_019837433.2;refGene_gn=LOC101086076;refGene_tx=XM_019837434.1;refGene_gn=LOC101086076;refGene_tx=XM_019837435.1
C1 78468431 78468629
refGene_gn=LOC101086076;refGene_tx=XM_019837431.2;refGene_gn=LOC101086076;refGene_tx=XM_019837432.2;refGene_gn=LOC101086076;refGene_tx=XM_019837433.2;refGene_gn=LOC101086076;refGene_tx=XM_019837434.1;refGene_gn=LOC101086076;refGene_tx=XM_019837435.1
C1 78487742 78487963
refGene_gn=LOC101086076;refGene_tx=XM_019837431.2;refGene_gn=LOC101086076;refGene_tx=XM_019837432.2;refGene_gn=LOC101086076;refGene_tx=XM_019837433.2;refGene_gn=LOC101086076;refGene_tx=XM_019837434.1;refGene_gn=LOC101086076;refGene_tx=XM_019837435.1
C1 78490872 78490990
refGene_gn=LOC101086076;refGene_tx=XM_019837431.2;refGene_gn=LOC101086076;refGene_tx=XM_019837432.2;refGene_gn=LOC101086076;refGene_tx=XM_019837433.2;refGene_gn=LOC101086076;refGene_tx=XM_019837434.1;refGene_gn

=LOC101086076;refGene_tx=XM_019837435.1
C1 78524482 78524572
refGene_gn=LOC101086076;refGene_tx=XM_019837431.2;refGene_gn=LOC101086076;refGene_tx=XM_019837432.2;refGene_gn=LOC101086076;refGene_tx=XM_019837433.2;refGene_gn=LOC101086076;refGene_tx=XM_019837434.1;refGene_gn=LOC101086076;refGene_tx=XM_019837435.1
C1 78532966 78533047
refGene_gn=LOC101086076;refGene_tx=XM_019837431.2;refGene_gn=LOC101086076;refGene_tx=XM_019837432.2;refGene_gn=LOC101086076;refGene_tx=XM_019837433.2;refGene_gn=LOC101086076;refGene_tx=XM_019837434.1;refGene_gn=LOC101086076;refGene_tx=XM_019837435.1
C1 78534263 78534360
refGene_gn=LOC101086076;refGene_tx=XM_019837431.2;refGene_gn=LOC101086076;refGene_tx=XM_019837432.2;refGene_gn=LOC101086076;refGene_tx=XM_019837433.2;refGene_gn=LOC101086076;refGene_tx=XM_019837434.1;refGene_gn=LOC101086076;refGene_tx=XM_019837435.1
C1 78541539 78541628
refGene_gn=LOC101086076;refGene_tx=XM_019837431.2;refGene_gn=LOC101086076;refGene_tx=XM_019837432.2;refGene_gn=LOC101086076;refGene_tx=XM_019837433.2;refGene_gn=LOC101086076;refGene_tx=XM_019837434.1;refGene_gn=LOC101086076;refGene_tx=XM_019837435.1
C1 78542068 78542171
refGene_gn=LOC101086076;refGene_tx=XM_019837431.2;refGene_gn=LOC101086076;refGene_tx=XM_019837432.2;refGene_gn=LOC101086076;refGene_tx=XM_019837433.2;refGene_gn=LOC101086076;refGene_tx=XM_019837434.1;refGene_gn=LOC101086076;refGene_tx=XM_019837435.1
C1 78552230 78552413
refGene_gn=LOC101086076;refGene_tx=XM_003990300.5;refGene_gn=LOC101086076;refGene_tx=XM_019837431.2;refGene_gn=LOC101086076;refGene_tx=XM_019837432.2;refGene_gn=LOC101086076;refGene_tx=XM_019837433.2;refGene_gn=LOC101086076;refGene_tx=XM_019837434.1;refGene_gn=LOC101086076;refGene_tx=XM_019837435.1
C1 78553115 78553253
refGene_gn=LOC101086076;refGene_tx=XM_003990300.5;refGene_gn=LOC101086076;refGene_tx=XM_019837431.2;refGene_gn=LOC101086076;refGene_tx=XM_019837432.2;refGene_gn=LOC101086076;refGene_tx=XM_019837433.2;refGene_gn=LOC101086076;refGene_tx=XM_019837434.1
C1 78554139 78554269
refGene_gn=LOC101086076;refGene_tx=XM_003990300.5;refGene_gn=LOC101086076;refGene_tx=XM_019837431.2;refGene_gn=LOC101086076;refGene_tx=XM_019837432.2;refGene_gn=LOC101086076;refGene_tx=XM_019837433.2;refGene_gn=LOC101086076;refGene_tx=XM_019837434.1
C1 78556645 78556707
refGene_gn=LOC101086076;refGene_tx=XM_003990300.5;refGene_gn=LOC101086076;refGene_tx=XM_019837431.2;refGene_gn=LOC101086076;refGene_tx=XM_019837432.2;refGene_gn=LOC101086076;refGene_tx=XM_019837433.2;refGene_gn=LOC101086076;refGene_tx=XM_019837434.1
C1 78560754 78561466
refGene_gn=LOC101086076;refGene_tx=XM_003990300.5;refGene_gn=LOC101086076;refGene_tx=XM_019837431.2;refGene_gn=LOC101086076;refGene_tx=XM_01

9837432.2;refGene_gn=LOC101086076;refGene_tx=XM_019837433.2;refGene_gn=LOC101086076;refGene_tx=XM_019837434.1
C1 78561722 78561807
refGene_gn=LOC101086076;refGene_tx=XM_003990300.5;refGene_gn=LOC101086076;refGene_tx=XM_019837431.2;refGene_gn=LOC101086076;refGene_tx=XM_019837432.2;refGene_gn=LOC101086076;refGene_tx=XM_019837433.2;refGene_gn=LOC101086076;refGene_tx=XM_019837434.1
C1 78562672 78562812
refGene_gn=LOC101086076;refGene_tx=XM_003990300.5;refGene_gn=LOC101086076;refGene_tx=XM_019837431.2;refGene_gn=LOC101086076;refGene_tx=XM_019837432.2;refGene_gn=LOC101086076;refGene_tx=XM_019837433.2
C1 78565107 78567423
refGene_gn=LOC101086076;refGene_tx=XM_003990300.5;refGene_gn=LOC101086076;refGene_tx=XM_019837431.2;refGene_gn=LOC101086076;refGene_tx=XM_019837432.2;refGene_gn=LOC101086076;refGene_tx=XM_019837433.2;refGene_gn=LOC101086076;refGene_tx=XM_019837434.1
C1 78568709 78569176
refGene_gn=LOC101086076;refGene_tx=XM_003990300.5;refGene_gn=LOC101086076;refGene_tx=XM_019837431.2;refGene_gn=LOC101086076;refGene_tx=XM_019837432.2
C1 78580088 78580114
refGene_gn=LOC101086076;refGene_tx=XM_019837433.2
C1 78635130 78635199
refGene_gn=CC1H1orf146;refGene_tx=XM_003990301.5
C1 78644129 78644220
refGene_gn=CC1H1orf146;refGene_tx=XM_003990301.5
C1 78645681 78645850
refGene_gn=CC1H1orf146;refGene_tx=XM_003990301.5
C1 78646239 78646318
refGene_gn=CC1H1orf146;refGene_tx=XM_003990301.5
C1 78647202 78647337
refGene_gn=CC1H1orf146;refGene_tx=XM_003990301.5
C1 78648478 78648595
refGene_gn=GLMN;refGene_tx=XM_003990302.3;refGene_gn=GLMN;refGene_tx=XM_006934904.2;refGene_gn=GLMN;refGene_tx=XM_019837439.2
C1 78648701 78648784
refGene_gn=GLMN;refGene_tx=XM_003990302.3;refGene_gn=GLMN;refGene_tx=XM_006934904.2;refGene_gn=GLMN;refGene_tx=XM_019837439.2
C1 78649395 78649519
refGene_gn=GLMN;refGene_tx=XM_003990302.3;refGene_gn=GLMN;refGene_tx=XM_006934904.2;refGene_gn=GLMN;refGene_tx=XM_019837439.2
C1 78656098 78656162
refGene_gn=GLMN;refGene_tx=XM_003990302.3;refGene_gn=GLMN;refGene_tx=XM_006934904.2;refGene_gn=GLMN;refGene_tx=XM_019837439.2
C1 78657103 78657213
refGene_gn=GLMN;refGene_tx=XM_003990302.3;refGene_gn=GLMN;refGene_tx=XM_006934904.2;refGene_gn=GLMN;refGene_tx=XM_019837439.2
C1 78657691 78657776
refGene_gn=GLMN;refGene_tx=XM_003990302.3;refGene_gn=GLMN;refGene_tx=XM_006934904.2;refGene_gn=GLMN;refGene_tx=XM_019837439.2

C1 78658980 78659054
refGene_gn=GLMN;refGene_tx=XM_003990302.3;refGene_gn=GLMN;refGene_tx=XM_006934904.2;refGene_gn=GLMN;refGene_tx=XM_019837439.2
C1 78659753 78659795
refGene_gn=GLMN;refGene_tx=XM_003990302.3;refGene_gn=GLMN;refGene_tx=XM_006934904.2;refGene_gn=GLMN;refGene_tx=XM_019837439.2
C1 78660694 78660784
refGene_gn=GLMN;refGene_tx=XM_003990302.3;refGene_gn=GLMN;refGene_tx=XM_006934904.2;refGene_gn=GLMN;refGene_tx=XM_019837439.2
C1 78660870 78660901
refGene_gn=GLMN;refGene_tx=XM_003990302.3;refGene_gn=GLMN;refGene_tx=XM_006934904.2;refGene_gn=GLMN;refGene_tx=XM_019837439.2
C1 78662386 78662440
refGene_gn=GLMN;refGene_tx=XM_003990302.3;refGene_gn=GLMN;refGene_tx=XM_019837439.2
C1 78665026 78665214
refGene_gn=GLMN;refGene_tx=XM_003990302.3;refGene_gn=GLMN;refGene_tx=XM_006934904.2;refGene_gn=GLMN;refGene_tx=XM_019837439.2
C1 78672387 78672490
refGene_gn=GLMN;refGene_tx=XM_003990302.3;refGene_gn=GLMN;refGene_tx=XM_006934904.2;refGene_gn=GLMN;refGene_tx=XM_019837439.2
C1 78674838 78675085
refGene_gn=GLMN;refGene_tx=XM_003990302.3;refGene_gn=GLMN;refGene_tx=XM_006934904.2;refGene_gn=GLMN;refGene_tx=XM_019837439.2
C1 78676259 78676368
refGene_gn=GLMN;refGene_tx=XM_003990302.3;refGene_gn=GLMN;refGene_tx=XM_006934904.2;refGene_gn=GLMN;refGene_tx=XM_019837439.2
C1 78677510 78677627
refGene_gn=GLMN;refGene_tx=XM_003990302.3;refGene_gn=GLMN;refGene_tx=XM_006934904.2;refGene_gn=GLMN;refGene_tx=XM_019837439.2
C1 78683075 78683201
refGene_gn=GLMN;refGene_tx=XM_003990302.3;refGene_gn=GLMN;refGene_tx=XM_006934904.2;refGene_gn=GLMN;refGene_tx=XM_019837439.2
C1 78683640 78683679
refGene_gn=GLMN;refGene_tx=XM_003990302.3;refGene_gn=GLMN;refGene_tx=XM_006934904.2;refGene_gn=GLMN;refGene_tx=XM_019837439.2
C1 78684690 78684763 refGene_gn=RPAP2;refGene_tx=XM_011284871.2
C1 78685532 78685578 refGene_gn=RPAP2;refGene_tx=XM_011284871.2
C1 78687920 78688035 refGene_gn=RPAP2;refGene_tx=XM_011284871.2
C1 78697535 78697634 refGene_gn=RPAP2;refGene_tx=XM_011284871.2
C1 78697957 78698023 refGene_gn=RPAP2;refGene_tx=XM_011284871.2
C1 78699920 78700009 refGene_gn=RPAP2;refGene_tx=XM_011284871.2
C1 78720949 78720985 refGene_gn=RPAP2;refGene_tx=XM_011284871.2
C1 78723406 78724307 refGene_gn=RPAP2;refGene_tx=XM_011284871.2
C1 78724308 78724347 refGene_gn=RPAP2;refGene_tx=XM_011284871.2
C1 78730279 78730362 refGene_gn=RPAP2;refGene_tx=XM_011284871.2
C1 78733534 78733615 refGene_gn=RPAP2;refGene_tx=XM_011284871.2
C1 78744459 78744528 refGene_gn=RPAP2;refGene_tx=XM_011284871.2
C1 78773047 78773189 refGene_gn=RPAP2;refGene_tx=XM_011284871.2
C1 78865179 78865358 refGene_gn=GFI1;refGene_tx=NM_001112709.1

C1 78867663 78867829 refGene_gn=GFI1;refGene_tx=NM_001112709.1
C1 78868881 78869019 refGene_gn=GFI1;refGene_tx=NM_001112709.1
C1 78869123 78869611 refGene_gn=GFI1;refGene_tx=NM_001112709.1
C1 78871421 78871604 refGene_gn=GFI1;refGene_tx=NM_001112709.1
C1 78871929 78872044 refGene_gn=GFI1;refGene_tx=NM_001112709.1
C1 78904321 78904633
refGene_gn=EVI5;refGene_tx=XM_023258904.1;refGene_gn=EVI5;refGene_tx=XM_023258905.1;refGene_gn=EVI5;refGene_tx=XM_023258906.1;refGene_gn=EVI5;refGene_tx=XM_023258907.1;refGene_gn=EVI5;refGene_tx=XM_023258908.1;refGene_gn=EVI5;refGene_tx=XM_023258909.1;refGene_gn=EVI5;refGene_tx=XM_023258910.1
C1 78916258 78916271
refGene_gn=EVI5;refGene_tx=XM_023258909.1;refGene_gn=EVI5;refGene_tx=XM_023258910.1
C1 78936569 78936665
refGene_gn=EVI5;refGene_tx=XM_023258904.1;refGene_gn=EVI5;refGene_tx=XM_023258905.1;refGene_gn=EVI5;refGene_tx=XM_023258906.1;refGene_gn=EVI5;refGene_tx=XM_023258907.1;refGene_gn=EVI5;refGene_tx=XM_023258908.1;refGene_gn=EVI5;refGene_tx=XM_023258909.1;refGene_gn=EVI5;refGene_tx=XM_023258910.1
C1 78972660 78972756
refGene_gn=EVI5;refGene_tx=XM_023258904.1;refGene_gn=EVI5;refGene_tx=XM_023258905.1;refGene_gn=EVI5;refGene_tx=XM_023258906.1;refGene_gn=EVI5;refGene_tx=XM_023258907.1;refGene_gn=EVI5;refGene_tx=XM_023258908.1;refGene_gn=EVI5;refGene_tx=XM_023258909.1;refGene_gn=EVI5;refGene_tx=XM_023258910.1
C1 78980199 78980346
refGene_gn=EVI5;refGene_tx=XM_023258904.1;refGene_gn=EVI5;refGene_tx=XM_023258905.1;refGene_gn=EVI5;refGene_tx=XM_023258906.1;refGene_gn=EVI5;refGene_tx=XM_023258907.1;refGene_gn=EVI5;refGene_tx=XM_023258908.1;refGene_gn=EVI5;refGene_tx=XM_023258909.1;refGene_gn=EVI5;refGene_tx=XM_023258910.1
C1 78990265 78990424
refGene_gn=EVI5;refGene_tx=XM_023258904.1;refGene_gn=EVI5;refGene_tx=XM_023258905.1;refGene_gn=EVI5;refGene_tx=XM_023258906.1;refGene_gn=EVI5;refGene_tx=XM_023258907.1;refGene_gn=EVI5;refGene_tx=XM_023258908.1;refGene_gn=EVI5;refGene_tx=XM_023258909.1;refGene_gn=EVI5;refGene_tx=XM_023258910.1
C1 78991768 78991909
refGene_gn=EVI5;refGene_tx=XM_023258904.1;refGene_gn=EVI5;refGene_tx=XM_023258905.1;refGene_gn=EVI5;refGene_tx=XM_023258906.1;refGene_gn=EVI5;refGene_tx=XM_023258907.1;refGene_gn=EVI5;refGene_tx=XM_023258908.1;refGene_gn=EVI5;refGene_tx=XM_023258909.1;refGene_gn=EVI5;refGene_tx=XM_023258910.1
C1 78994834 78994969
refGene_gn=EVI5;refGene_tx=XM_023258904.1;refGene_gn=EVI5;refGene_tx=XM_023258905.1;refGene_gn=EVI5;refGene_tx=XM_023258906.1;refGene_gn=EVI5;refGene_tx=XM_023258907.1;refGene_gn=EVI5;refGene_tx=XM_023258908.1;refGene_gn=EVI5;refGene_tx=XM_023258909.1;refGene_gn=EVI5;refGene_tx=XM_023258910.1

C1 79018680 79018827
refGene_gn=EVI5;refGene_tx=XM_023258904.1;refGene_gn=EVI5;refGene_tx=XM_023258905.1;refGene_gn=EVI5;refGene_tx=XM_023258906.1;refGene_gn=EVI5;refGene_tx=XM_023258907.1;refGene_gn=EVI5;refGene_tx=XM_023258909.1;refGene_gn=EVI5;refGene_tx=XM_023258910.1
C1 79019394 79019427
refGene_gn=EVI5;refGene_tx=XM_023258904.1;refGene_gn=EVI5;refGene_tx=XM_023258905.1;refGene_gn=EVI5;refGene_tx=XM_023258906.1;refGene_gn=EVI5;refGene_tx=XM_023258907.1;refGene_gn=EVI5;refGene_tx=XM_023258909.1;refGene_gn=EVI5;refGene_tx=XM_023258910.1
C1 79021282 79021336
refGene_gn=EVI5;refGene_tx=XM_023258904.1;refGene_gn=EVI5;refGene_tx=XM_023258905.1;refGene_gn=EVI5;refGene_tx=XM_023258906.1;refGene_gn=EVI5;refGene_tx=XM_023258907.1;refGene_gn=EVI5;refGene_tx=XM_023258908.1;refGene_gn=EVI5;refGene_tx=XM_023258909.1;refGene_gn=EVI5;refGene_tx=XM_023258910.1
C1 79030596 79030657
refGene_gn=EVI5;refGene_tx=XM_023258904.1;refGene_gn=EVI5;refGene_tx=XM_023258905.1;refGene_gn=EVI5;refGene_tx=XM_023258906.1;refGene_gn=EVI5;refGene_tx=XM_023258907.1;refGene_gn=EVI5;refGene_tx=XM_023258908.1;refGene_gn=EVI5;refGene_tx=XM_023258909.1;refGene_gn=EVI5;refGene_tx=XM_023258910.1
C1 79047570 79047668
refGene_gn=EVI5;refGene_tx=XM_023258904.1;refGene_gn=EVI5;refGene_tx=XM_023258905.1;refGene_gn=EVI5;refGene_tx=XM_023258906.1;refGene_gn=EVI5;refGene_tx=XM_023258907.1;refGene_gn=EVI5;refGene_tx=XM_023258908.1;refGene_gn=EVI5;refGene_tx=XM_023258909.1;refGene_gn=EVI5;refGene_tx=XM_023258910.1
C1 79047772 79047862
refGene_gn=EVI5;refGene_tx=XM_023258904.1;refGene_gn=EVI5;refGene_tx=XM_023258905.1;refGene_gn=EVI5;refGene_tx=XM_023258906.1;refGene_gn=EVI5;refGene_tx=XM_023258907.1;refGene_gn=EVI5;refGene_tx=XM_023258908.1;refGene_gn=EVI5;refGene_tx=XM_023258909.1;refGene_gn=EVI5;refGene_tx=XM_023258910.1
C1 79049441 79049585
refGene_gn=EVI5;refGene_tx=XM_023258904.1;refGene_gn=EVI5;refGene_tx=XM_023258905.1;refGene_gn=EVI5;refGene_tx=XM_023258906.1;refGene_gn=EVI5;refGene_tx=XM_023258907.1;refGene_gn=EVI5;refGene_tx=XM_023258908.1;refGene_gn=EVI5;refGene_tx=XM_023258909.1;refGene_gn=EVI5;refGene_tx=XM_023258910.1
C1 79050934 79051060
refGene_gn=EVI5;refGene_tx=XM_023258904.1;refGene_gn=EVI5;refGene_tx=XM_023258905.1;refGene_gn=EVI5;refGene_tx=XM_023258906.1;refGene_gn=EVI5;refGene_tx=XM_023258907.1;refGene_gn=EVI5;refGene_tx=XM_023258908.1;refGene_gn=EVI5;refGene_tx=XM_023258909.1;refGene_gn=EVI5;refGene_tx=XM_023258910.1
C1 79057014 79057089
refGene_gn=EVI5;refGene_tx=XM_023258904.1;refGene_gn=EVI5;refGene_tx=XM_023258905.1;refGene_gn=EVI5;refGene_tx=XM_023258906.1;refGene_gn=EVI5;refGene_tx=XM_023258907.1;refGene_gn=EVI5;refGene_tx=XM_023258908.1;

refGene_gn=EVI5;refGene_tx=XM_023258909.1;refGene_gn=EVI5;refGene_tx=XM_023258910.1
C1 79057835 79058060
refGene_gn=EVI5;refGene_tx=XM_023258904.1;refGene_gn=EVI5;refGene_tx=XM_023258905.1;refGene_gn=EVI5;refGene_tx=XM_023258906.1;refGene_gn=EVI5;refGene_tx=XM_023258907.1;refGene_gn=EVI5;refGene_tx=XM_023258908.1;refGene_gn=EVI5;refGene_tx=XM_023258909.1;refGene_gn=EVI5;refGene_tx=XM_023258910.1
C1 79058714 79058904
refGene_gn=EVI5;refGene_tx=XM_023258904.1;refGene_gn=EVI5;refGene_tx=XM_023258905.1;refGene_gn=EVI5;refGene_tx=XM_023258906.1;refGene_gn=EVI5;refGene_tx=XM_023258907.1;refGene_gn=EVI5;refGene_tx=XM_023258908.1;refGene_gn=EVI5;refGene_tx=XM_023258909.1;refGene_gn=EVI5;refGene_tx=XM_023258910.1
C1 79071591 79071623 refGene_gn=EVI5;refGene_tx=XM_023258907.1
C1 79079933 79080082
refGene_gn=EVI5;refGene_tx=XM_023258904.1;refGene_gn=EVI5;refGene_tx=XM_023258905.1;refGene_gn=EVI5;refGene_tx=XM_023258906.1;refGene_gn=EVI5;refGene_tx=XM_023258908.1;refGene_gn=EVI5;refGene_tx=XM_023258909.1;refGene_gn=EVI5;refGene_tx=XM_023258910.1
C1 79081611 79081968 refGene_gn=TCEB2;refGene_tx=NM_001246228.1
C1 79175692 79175695
refGene_gn=RPL5;refGene_tx=XM_019837448.1;refGene_gn=RPL5;refGene_tx=XM_019837449.1
C1 79176673 79176743
refGene_gn=RPL5;refGene_tx=XM_019837448.1;refGene_gn=RPL5;refGene_tx=XM_019837449.1
C1 79176826 79176942
refGene_gn=RPL5;refGene_tx=XM_019837448.1;refGene_gn=RPL5;refGene_tx=XM_019837449.1
C1 79178075 79178210
refGene_gn=RPL5;refGene_tx=XM_019837448.1;refGene_gn=RPL5;refGene_tx=XM_019837449.1
C1 79179546 79179749
refGene_gn=RPL5;refGene_tx=XM_019837448.1;refGene_gn=RPL5;refGene_tx=XM_019837449.1
C1 79180850 79181028
refGene_gn=RPL5;refGene_tx=XM_019837448.1;refGene_gn=RPL5;refGene_tx=XM_019837449.1
C1 79183641 79183730
refGene_gn=RPL5;refGene_tx=XM_019837448.1;refGene_gn=RPL5;refGene_tx=XM_019837449.1
C1 79184163 79184263
refGene_gn=RPL5;refGene_tx=XM_019837448.1;refGene_gn=RPL5;refGene_tx=XM_019837449.1
C1 79185766 79186579 refGene_gn=FAM69A;refGene_tx=XM_023258911.1
C1 79187842 79188019 refGene_gn=FAM69A;refGene_tx=XM_023258911.1
C1 79192490 79192598 refGene_gn=FAM69A;refGene_tx=XM_023258911.1
C1 79217635 79217770 refGene_gn=FAM69A;refGene_tx=XM_023258911.1
C1 79290435 79290489 refGene_gn=FAM69A;refGene_tx=XM_023258911.1

C1 79371882 79371887 refGene_gn=MTF2;refGene_tx=XM_003990305.4
C1 79422988 79423187 refGene_gn=MTF2;refGene_tx=XM_003990305.4
C1 79423312 79423394 refGene_gn=MTF2;refGene_tx=XM_003990305.4
C1 79428002 79428098
refGene_gn=MTF2;refGene_tx=XM_003990305.4;refGene_gn=MTF2;refGene_tx=XM_006934906.3;refGene_gn=MTF2;refGene_tx=XM_006934907.2
C1 79428332 79428433
refGene_gn=MTF2;refGene_tx=XM_003990305.4;refGene_gn=MTF2;refGene_tx=XM_006934906.3;refGene_gn=MTF2;refGene_tx=XM_006934907.2
C1 79428823 79428972
refGene_gn=MTF2;refGene_tx=XM_003990305.4;refGene_gn=MTF2;refGene_tx=XM_006934906.3;refGene_gn=MTF2;refGene_tx=XM_006934907.2
C1 79431046 79431142
refGene_gn=MTF2;refGene_tx=XM_003990305.4;refGene_gn=MTF2;refGene_tx=XM_006934906.3;refGene_gn=MTF2;refGene_tx=XM_006934907.2
C1 79432486 79432555
refGene_gn=MTF2;refGene_tx=XM_003990305.4;refGene_gn=MTF2;refGene_tx=XM_006934906.3;refGene_gn=MTF2;refGene_tx=XM_006934907.2
C1 79433847 79433971
refGene_gn=MTF2;refGene_tx=XM_003990305.4;refGene_gn=MTF2;refGene_tx=XM_006934906.3;refGene_gn=MTF2;refGene_tx=XM_006934907.2
C1 79441281 79441349
refGene_gn=MTF2;refGene_tx=XM_003990305.4;refGene_gn=MTF2;refGene_tx=XM_006934906.3;refGene_gn=MTF2;refGene_tx=XM_006934907.2
C1 79442997 79443168
refGene_gn=MTF2;refGene_tx=XM_003990305.4;refGene_gn=MTF2;refGene_tx=XM_006934906.3;refGene_gn=MTF2;refGene_tx=XM_006934907.2
C1 79446978 79447084
refGene_gn=MTF2;refGene_tx=XM_003990305.4;refGene_gn=MTF2;refGene_tx=XM_006934906.3;refGene_gn=MTF2;refGene_tx=XM_006934907.2
C1 79447206 79447259
refGene_gn=MTF2;refGene_tx=XM_003990305.4;refGene_gn=MTF2;refGene_tx=XM_006934906.3;refGene_gn=MTF2;refGene_tx=XM_006934907.2
C1 79447370 79447475
refGene_gn=MTF2;refGene_tx=XM_003990305.4;refGene_gn=MTF2;refGene_tx=XM_006934906.3;refGene_gn=MTF2;refGene_tx=XM_006934907.2
C1 79449092 79449450
refGene_gn=MTF2;refGene_tx=XM_003990305.4;refGene_gn=MTF2;refGene_tx=XM_006934906.3;refGene_gn=MTF2;refGene_tx=XM_006934907.2
C1 79468146 79468365
refGene_gn=TMED5;refGene_tx=XM_003990333.5;refGene_gn=TMED5;refGene_tx=XM_003990334.5
C1 79468956 79469010 refGene_gn=TMED5;refGene_tx=XM_003990334.5
C1 79469094 79469095 refGene_gn=TMED5;refGene_tx=XM_003990334.5
C1 79471095 79471279
refGene_gn=TMED5;refGene_tx=XM_003990333.5;refGene_gn=TMED5;refGene_tx=XM_003990334.5
C1 79478185 79478283
refGene_gn=TMED5;refGene_tx=XM_003990333.5;refGene_gn=TMED5;refGene_tx=XM_003990334.5

C1 79484428 79484617
refGene_gn=TMED5;refGene_tx=XM_003990333.5;refGene_gn=TMED5;refGene_tx=XM_003990334.5
C1 79487285 79487419
refGene_gn=CCDC18;refGene_tx=XM_019837450.2;refGene_gn=CCDC18;refGene_tx=XM_019837451.2;refGene_gn=CCDC18;refGene_tx=XM_019837452.2;refGene_gn=CCDC18;refGene_tx=XM_019837455.2;refGene_gn=CCDC18;refGene_tx=XM_019837456.2;refGene_gn=CCDC18;refGene_tx=XM_019837457.2;refGene_gn=CCDC18;refGene_tx=XM_019837458.2
C1 79488170 79488345
refGene_gn=CCDC18;refGene_tx=XM_019837450.2;refGene_gn=CCDC18;refGene_tx=XM_019837451.2;refGene_gn=CCDC18;refGene_tx=XM_019837452.2;refGene_gn=CCDC18;refGene_tx=XM_019837455.2;refGene_gn=CCDC18;refGene_tx=XM_019837456.2;refGene_gn=CCDC18;refGene_tx=XM_019837457.2;refGene_gn=CCDC18;refGene_tx=XM_019837458.2
C1 79491539 79491698
refGene_gn=CCDC18;refGene_tx=XM_019837450.2;refGene_gn=CCDC18;refGene_tx=XM_019837451.2;refGene_gn=CCDC18;refGene_tx=XM_019837452.2;refGene_gn=CCDC18;refGene_tx=XM_019837455.2;refGene_gn=CCDC18;refGene_tx=XM_019837456.2;refGene_gn=CCDC18;refGene_tx=XM_019837457.2;refGene_gn=CCDC18;refGene_tx=XM_019837458.2;refGene_gn=CCDC18;refGene_tx=XM_019837459.2
C1 79498150 79498257
refGene_gn=CCDC18;refGene_tx=XM_019837450.2;refGene_gn=CCDC18;refGene_tx=XM_019837451.2;refGene_gn=CCDC18;refGene_tx=XM_019837452.2;refGene_gn=CCDC18;refGene_tx=XM_019837455.2;refGene_gn=CCDC18;refGene_tx=XM_019837456.2;refGene_gn=CCDC18;refGene_tx=XM_019837457.2;refGene_gn=CCDC18;refGene_tx=XM_019837458.2;refGene_gn=CCDC18;refGene_tx=XM_019837459.2
C1 79499436 79499565
refGene_gn=CCDC18;refGene_tx=XM_019837450.2;refGene_gn=CCDC18;refGene_tx=XM_019837451.2;refGene_gn=CCDC18;refGene_tx=XM_019837452.2;refGene_gn=CCDC18;refGene_tx=XM_019837455.2;refGene_gn=CCDC18;refGene_tx=XM_019837456.2;refGene_gn=CCDC18;refGene_tx=XM_019837457.2;refGene_gn=CCDC18;refGene_tx=XM_019837458.2;refGene_gn=CCDC18;refGene_tx=XM_019837459.2
C1 79511930 79512027
refGene_gn=CCDC18;refGene_tx=XM_019837450.2;refGene_gn=CCDC18;refGene_tx=XM_019837451.2;refGene_gn=CCDC18;refGene_tx=XM_019837452.2;refGene_gn=CCDC18;refGene_tx=XM_019837455.2;refGene_gn=CCDC18;refGene_tx=XM_019837456.2;refGene_gn=CCDC18;refGene_tx=XM_019837457.2;refGene_gn=CCDC18;refGene_tx=XM_019837458.2;refGene_gn=CCDC18;refGene_tx=XM_019837459.2
C1 79513816 79513938
refGene_gn=CCDC18;refGene_tx=XM_019837450.2;refGene_gn=CCDC18;refGene_tx=XM_019837451.2;refGene_gn=CCDC18;refGene_tx=XM_019837452.2;refGene_gn=CCDC18;refGene_tx=XM_019837455.2;refGene_gn=CCDC18;refGene_tx=XM_019837456.2;refGene_gn=CCDC18;refGene_tx=XM_019837457.2;refGene_gn=CCDC18;refGene_tx=XM_019837458.2;refGene_gn=CCDC18;refGene_tx=XM_019837459.2

C1 79514890 79515182
refGene_gn=CCDC18;refGene_tx=XM_019837450.2;refGene_gn=CCDC18;refGene_tx=XM_019837451.2;refGene_gn=CCDC18;refGene_tx=XM_019837452.2;refGene_gn=CCDC18;refGene_tx=XM_019837455.2;refGene_gn=CCDC18;refGene_tx=XM_019837456.2;refGene_gn=CCDC18;refGene_tx=XM_019837457.2;refGene_gn=CCDC18;refGene_tx=XM_019837458.2;refGene_gn=CCDC18;refGene_tx=XM_019837459.2
C1 79517154 79517279
refGene_gn=CCDC18;refGene_tx=XM_019837450.2;refGene_gn=CCDC18;refGene_tx=XM_019837451.2;refGene_gn=CCDC18;refGene_tx=XM_019837452.2;refGene_gn=CCDC18;refGene_tx=XM_019837455.2;refGene_gn=CCDC18;refGene_tx=XM_019837456.2;refGene_gn=CCDC18;refGene_tx=XM_019837457.2;refGene_gn=CCDC18;refGene_tx=XM_019837458.2;refGene_gn=CCDC18;refGene_tx=XM_019837459.2
C1 79518120 79518281
refGene_gn=CCDC18;refGene_tx=XM_019837450.2;refGene_gn=CCDC18;refGene_tx=XM_019837451.2;refGene_gn=CCDC18;refGene_tx=XM_019837452.2;refGene_gn=CCDC18;refGene_tx=XM_019837455.2;refGene_gn=CCDC18;refGene_tx=XM_019837456.2;refGene_gn=CCDC18;refGene_tx=XM_019837457.2;refGene_gn=CCDC18;refGene_tx=XM_019837458.2;refGene_gn=CCDC18;refGene_tx=XM_019837459.2
C1 79524663 79524887
refGene_gn=CCDC18;refGene_tx=XM_019837450.2;refGene_gn=CCDC18;refGene_tx=XM_019837451.2;refGene_gn=CCDC18;refGene_tx=XM_019837452.2;refGene_gn=CCDC18;refGene_tx=XM_019837455.2;refGene_gn=CCDC18;refGene_tx=XM_019837456.2;refGene_gn=CCDC18;refGene_tx=XM_019837457.2;refGene_gn=CCDC18;refGene_tx=XM_019837458.2;refGene_gn=CCDC18;refGene_tx=XM_019837459.2
C1 79528090 79528201
refGene_gn=CCDC18;refGene_tx=XM_019837450.2;refGene_gn=CCDC18;refGene_tx=XM_019837451.2;refGene_gn=CCDC18;refGene_tx=XM_019837452.2;refGene_gn=CCDC18;refGene_tx=XM_019837455.2;refGene_gn=CCDC18;refGene_tx=XM_019837456.2;refGene_gn=CCDC18;refGene_tx=XM_019837457.2;refGene_gn=CCDC18;refGene_tx=XM_019837458.2;refGene_gn=CCDC18;refGene_tx=XM_019837459.2
C1 79528791 79528923
refGene_gn=CCDC18;refGene_tx=XM_019837450.2;refGene_gn=CCDC18;refGene_tx=XM_019837451.2;refGene_gn=CCDC18;refGene_tx=XM_019837452.2;refGene_gn=CCDC18;refGene_tx=XM_019837455.2;refGene_gn=CCDC18;refGene_tx=XM_019837456.2;refGene_gn=CCDC18;refGene_tx=XM_019837457.2;refGene_gn=CCDC18;refGene_tx=XM_019837458.2;refGene_gn=CCDC18;refGene_tx=XM_019837459.2
C1 79536952 79537087
refGene_gn=CCDC18;refGene_tx=XM_019837450.2;refGene_gn=CCDC18;refGene_tx=XM_019837451.2;refGene_gn=CCDC18;refGene_tx=XM_019837452.2;refGene_gn=CCDC18;refGene_tx=XM_019837455.2;refGene_gn=CCDC18;refGene_tx=XM_019837456.2;refGene_gn=CCDC18;refGene_tx=XM_019837457.2;refGene_gn=CCDC18;refGene_tx=XM_019837458.2;refGene_gn=CCDC18;refGene_tx=XM_019837459.2
C1 79537251 79537329

refGene_gn=CCDC18;refGene_tx=XM_019837450.2;refGene_gn=CCDC18;refGene_tx=XM_019837451.2;refGene_gn=CCDC18;refGene_tx=XM_019837452.2;refGene_gn=CCDC18;refGene_tx=XM_019837455.2;refGene_gn=CCDC18;refGene_tx=XM_019837456.2;refGene_gn=CCDC18;refGene_tx=XM_019837457.2;refGene_gn=CCDC18;refGene_tx=XM_019837458.2;refGene_gn=CCDC18;refGene_tx=XM_019837459.2
C1 79541509 79541626
refGene_gn=CCDC18;refGene_tx=XM_019837450.2;refGene_gn=CCDC18;refGene_tx=XM_019837451.2;refGene_gn=CCDC18;refGene_tx=XM_019837452.2;refGene_gn=CCDC18;refGene_tx=XM_019837455.2;refGene_gn=CCDC18;refGene_tx=XM_019837456.2;refGene_gn=CCDC18;refGene_tx=XM_019837457.2;refGene_gn=CCDC18;refGene_tx=XM_019837458.2;refGene_gn=CCDC18;refGene_tx=XM_019837459.2
C1 79548003 79548171
refGene_gn=CCDC18;refGene_tx=XM_019837450.2;refGene_gn=CCDC18;refGene_tx=XM_019837451.2;refGene_gn=CCDC18;refGene_tx=XM_019837452.2;refGene_gn=CCDC18;refGene_tx=XM_019837455.2;refGene_gn=CCDC18;refGene_tx=XM_019837456.2;refGene_gn=CCDC18;refGene_tx=XM_019837457.2;refGene_gn=CCDC18;refGene_tx=XM_019837458.2;refGene_gn=CCDC18;refGene_tx=XM_019837459.2
C1 79550084 79550227
refGene_gn=CCDC18;refGene_tx=XM_019837450.2;refGene_gn=CCDC18;refGene_tx=XM_019837451.2;refGene_gn=CCDC18;refGene_tx=XM_019837452.2;refGene_gn=CCDC18;refGene_tx=XM_019837455.2;refGene_gn=CCDC18;refGene_tx=XM_019837456.2;refGene_gn=CCDC18;refGene_tx=XM_019837457.2;refGene_gn=CCDC18;refGene_tx=XM_019837458.2;refGene_gn=CCDC18;refGene_tx=XM_019837459.2
C1 79554919 79555083
refGene_gn=CCDC18;refGene_tx=XM_019837450.2;refGene_gn=CCDC18;refGene_tx=XM_019837451.2;refGene_gn=CCDC18;refGene_tx=XM_019837452.2;refGene_gn=CCDC18;refGene_tx=XM_019837455.2;refGene_gn=CCDC18;refGene_tx=XM_019837456.2;refGene_gn=CCDC18;refGene_tx=XM_019837457.2;refGene_gn=CCDC18;refGene_tx=XM_019837458.2;refGene_gn=CCDC18;refGene_tx=XM_019837459.2
C1 79555303 79555517
refGene_gn=CCDC18;refGene_tx=XM_019837450.2;refGene_gn=CCDC18;refGene_tx=XM_019837451.2;refGene_gn=CCDC18;refGene_tx=XM_019837452.2;refGene_gn=CCDC18;refGene_tx=XM_019837455.2;refGene_gn=CCDC18;refGene_tx=XM_019837456.2;refGene_gn=CCDC18;refGene_tx=XM_019837457.2;refGene_gn=CCDC18;refGene_tx=XM_019837458.2;refGene_gn=CCDC18;refGene_tx=XM_019837459.2
C1 79562219 79562319
refGene_gn=CCDC18;refGene_tx=XM_019837450.2;refGene_gn=CCDC18;refGene_tx=XM_019837451.2;refGene_gn=CCDC18;refGene_tx=XM_019837452.2;refGene_gn=CCDC18;refGene_tx=XM_019837455.2;refGene_gn=CCDC18;refGene_tx=XM_019837456.2;refGene_gn=CCDC18;refGene_tx=XM_019837457.2;refGene_gn=CCDC18;refGene_tx=XM_019837458.2;refGene_gn=CCDC18;refGene_tx=XM_019837459.2
C1 79567014 79567131
refGene_gn=CCDC18;refGene_tx=XM_019837450.2;refGene_gn=CCDC18;refGene_

tx=XM_019837451.2;refGene_gn=CCDC18;refGene_tx=XM_019837452.2;refGene_gn=CCDC18;refGene_tx=XM_019837455.2;refGene_gn=CCDC18;refGene_tx=XM_019837456.2;refGene_gn=CCDC18;refGene_tx=XM_019837457.2;refGene_gn=CCDC18;refGene_tx=XM_019837458.2;refGene_gn=CCDC18;refGene_tx=XM_019837459.2
C1 79568113 79568257 refGene_gn=CCDC18;refGene_tx=XM_019837450.2;refGene_gn=CCDC18;refGene_tx=XM_019837451.2;refGene_gn=CCDC18;refGene_tx=XM_019837452.2;refGene_gn=CCDC18;refGene_tx=XM_019837455.2;refGene_gn=CCDC18;refGene_tx=XM_019837456.2;refGene_gn=CCDC18;refGene_tx=XM_019837457.2;refGene_gn=CCDC18;refGene_tx=XM_019837458.2;refGene_gn=CCDC18;refGene_tx=XM_019837459.2
C1 79569743 79569947 refGene_gn=CCDC18;refGene_tx=XM_019837450.2;refGene_gn=CCDC18;refGene_tx=XM_019837451.2;refGene_gn=CCDC18;refGene_tx=XM_019837452.2;refGene_gn=CCDC18;refGene_tx=XM_019837455.2;refGene_gn=CCDC18;refGene_tx=XM_019837456.2;refGene_gn=CCDC18;refGene_tx=XM_019837457.2;refGene_gn=CCDC18;refGene_tx=XM_019837458.2;refGene_gn=CCDC18;refGene_tx=XM_019837459.2
C1 79573166 79573304 refGene_gn=CCDC18;refGene_tx=XM_019837450.2;refGene_gn=CCDC18;refGene_tx=XM_019837451.2;refGene_gn=CCDC18;refGene_tx=XM_019837452.2;refGene_gn=CCDC18;refGene_tx=XM_019837455.2;refGene_gn=CCDC18;refGene_tx=XM_019837456.2;refGene_gn=CCDC18;refGene_tx=XM_019837457.2;refGene_gn=CCDC18;refGene_tx=XM_019837458.2;refGene_gn=CCDC18;refGene_tx=XM_019837459.2
C1 79575659 79575860 refGene_gn=CCDC18;refGene_tx=XM_019837450.2;refGene_gn=CCDC18;refGene_tx=XM_019837451.2;refGene_gn=CCDC18;refGene_tx=XM_019837452.2;refGene_gn=CCDC18;refGene_tx=XM_019837455.2;refGene_gn=CCDC18;refGene_tx=XM_019837456.2;refGene_gn=CCDC18;refGene_tx=XM_019837457.2;refGene_gn=CCDC18;refGene_tx=XM_019837458.2;refGene_gn=CCDC18;refGene_tx=XM_019837459.2
C1 79584760 79585228 refGene_gn=CCDC18;refGene_tx=XM_019837450.2;refGene_gn=CCDC18;refGene_tx=XM_019837451.2;refGene_gn=CCDC18;refGene_tx=XM_019837452.2;refGene_gn=CCDC18;refGene_tx=XM_019837455.2;refGene_gn=CCDC18;refGene_tx=XM_019837456.2;refGene_gn=CCDC18;refGene_tx=XM_019837457.2;refGene_gn=CCDC18;refGene_tx=XM_019837458.2;refGene_gn=CCDC18;refGene_tx=XM_019837459.2
C1 79596418 79596433 refGene_gn=CCDC18;refGene_tx=XM_019837457.2;refGene_gn=CCDC18;refGene_tx=XM_019837458.2
C1 79602988 79603186 refGene_gn=CCDC18;refGene_tx=XM_019837450.2;refGene_gn=CCDC18;refGene_tx=XM_019837451.2;refGene_gn=CCDC18;refGene_tx=XM_019837452.2;refGene_gn=CCDC18;refGene_tx=XM_019837455.2;refGene_gn=CCDC18;refGene_tx=XM_019837456.2;refGene_gn=CCDC18;refGene_tx=XM_019837459.2
C1 79694856 79695076 refGene_gn=DR1;refGene_tx=XM_003990336.5
C1 79703849 79704013 refGene_gn=DR1;refGene_tx=XM_003990336.5

C1 79712109 79712256 refGene_gn=DR1;refGene_tx=XM_003990336.5
C1 79765977 79766001
refGene_gn=FNBP1L;refGene_tx=XM_006934916.4;refGene_gn=FNBP1L;refGene_tx=XM_023258912.1;refGene_gn=FNBP1L;refGene_tx=XM_023258915.1
C1 79829679 79829795
refGene_gn=FNBP1L;refGene_tx=XM_006934916.4;refGene_gn=FNBP1L;refGene_tx=XM_023258912.1;refGene_gn=FNBP1L;refGene_tx=XM_023258915.1
C1 79853971 79854025
refGene_gn=FNBP1L;refGene_tx=XM_006934916.4;refGene_gn=FNBP1L;refGene_tx=XM_023258912.1;refGene_gn=FNBP1L;refGene_tx=XM_023258915.1
C1 79855235 79855383
refGene_gn=FNBP1L;refGene_tx=XM_006934916.4;refGene_gn=FNBP1L;refGene_tx=XM_023258912.1;refGene_gn=FNBP1L;refGene_tx=XM_023258914.1;refGene_gn=FNBP1L;refGene_tx=XM_023258915.1
C1 79856153 79856216
refGene_gn=FNBP1L;refGene_tx=XM_006934916.4;refGene_gn=FNBP1L;refGene_tx=XM_023258912.1;refGene_gn=FNBP1L;refGene_tx=XM_023258914.1;refGene_gn=FNBP1L;refGene_tx=XM_023258915.1
C1 79860617 79860722
refGene_gn=FNBP1L;refGene_tx=XM_006934916.4;refGene_gn=FNBP1L;refGene_tx=XM_023258912.1;refGene_gn=FNBP1L;refGene_tx=XM_023258914.1;refGene_gn=FNBP1L;refGene_tx=XM_023258915.1
C1 79864101 79864230
refGene_gn=FNBP1L;refGene_tx=XM_006934916.4;refGene_gn=FNBP1L;refGene_tx=XM_023258912.1;refGene_gn=FNBP1L;refGene_tx=XM_023258914.1;refGene_gn=FNBP1L;refGene_tx=XM_023258915.1
C1 79866268 79866415
refGene_gn=FNBP1L;refGene_tx=XM_006934916.4;refGene_gn=FNBP1L;refGene_tx=XM_023258912.1;refGene_gn=FNBP1L;refGene_tx=XM_023258914.1;refGene_gn=FNBP1L;refGene_tx=XM_023258915.1
C1 79868339 79868543
refGene_gn=FNBP1L;refGene_tx=XM_006934916.4;refGene_gn=FNBP1L;refGene_tx=XM_023258912.1;refGene_gn=FNBP1L;refGene_tx=XM_023258914.1;refGene_gn=FNBP1L;refGene_tx=XM_023258915.1
C1 79869967 79870126 refGene_gn=FNBP1L;refGene_tx=XM_023258915.1
C1 79876012 79876027 refGene_gn=FNBP1L;refGene_tx=XM_023258915.1
C1 79879017 79879127
refGene_gn=FNBP1L;refGene_tx=XM_006934916.4;refGene_gn=FNBP1L;refGene_tx=XM_023258912.1;refGene_gn=FNBP1L;refGene_tx=XM_023258914.1;refGene_gn=FNBP1L;refGene_tx=XM_023258915.1
C1 79882571 79882704
refGene_gn=FNBP1L;refGene_tx=XM_006934916.4;refGene_gn=FNBP1L;refGene_tx=XM_023258912.1;refGene_gn=FNBP1L;refGene_tx=XM_023258914.1;refGene_gn=FNBP1L;refGene_tx=XM_023258915.1
C1 79883080 79883175
refGene_gn=FNBP1L;refGene_tx=XM_006934916.4;refGene_gn=FNBP1L;refGene_tx=XM_023258912.1;refGene_gn=FNBP1L;refGene_tx=XM_023258914.1;refGene_gn=FNBP1L;refGene_tx=XM_023258915.1
C1 79885279 79885428
refGene_gn=FNBP1L;refGene_tx=XM_006934916.4;refGene_gn=FNBP1L;refGene_

tx=XM_023258912.1;refGene_gn=FNBP1L;refGene_tx=XM_023258914.1;refGene_gn=FNBP1L;refGene_tx=XM_023258915.1
C1 79886845 79887024
refGene_gn=FNBP1L;refGene_tx=XM_006934916.4;refGene_gn=FNBP1L;refGene_tx=XM_023258912.1;refGene_gn=FNBP1L;refGene_tx=XM_023258914.1;refGene_gn=FNBP1L;refGene_tx=XM_023258915.1
C1 79888279 79888287 refGene_gn=FNBP1L;refGene_tx=XM_006934916.4
C1 79890855 79890881
refGene_gn=FNBP1L;refGene_tx=XM_023258912.1;refGene_gn=FNBP1L;refGene_tx=XM_023258914.1
C1 79897199 79897378
refGene_gn=BCAR3;refGene_tx=XM_003990338.5;refGene_gn=BCAR3;refGene_tx=XM_011284867.2;refGene_gn=BCAR3;refGene_tx=XM_019837461.1;refGene_gn=BCAR3;refGene_tx=XM_023258916.1;refGene_gn=BCAR3;refGene_tx=XM_023258917.1
C1 79900406 79900619
refGene_gn=BCAR3;refGene_tx=XM_003990338.5;refGene_gn=BCAR3;refGene_tx=XM_011284867.2;refGene_gn=BCAR3;refGene_tx=XM_019837461.1;refGene_gn=BCAR3;refGene_tx=XM_023258916.1;refGene_gn=BCAR3;refGene_tx=XM_023258917.1
C1 79908054 79908166
refGene_gn=BCAR3;refGene_tx=XM_003990338.5;refGene_gn=BCAR3;refGene_tx=XM_011284867.2;refGene_gn=BCAR3;refGene_tx=XM_019837461.1;refGene_gn=BCAR3;refGene_tx=XM_023258916.1;refGene_gn=BCAR3;refGene_tx=XM_023258917.1
C1 79910803 79910975
refGene_gn=BCAR3;refGene_tx=XM_003990338.5;refGene_gn=BCAR3;refGene_tx=XM_011284867.2;refGene_gn=BCAR3;refGene_tx=XM_019837461.1;refGene_gn=BCAR3;refGene_tx=XM_023258916.1;refGene_gn=BCAR3;refGene_tx=XM_023258917.1
C1 79915378 79915494
refGene_gn=BCAR3;refGene_tx=XM_003990338.5;refGene_gn=BCAR3;refGene_tx=XM_011284867.2;refGene_gn=BCAR3;refGene_tx=XM_019837461.1;refGene_gn=BCAR3;refGene_tx=XM_023258916.1;refGene_gn=BCAR3;refGene_tx=XM_023258917.1
C1 79921094 79921747
refGene_gn=BCAR3;refGene_tx=XM_003990338.5;refGene_gn=BCAR3;refGene_tx=XM_011284867.2;refGene_gn=BCAR3;refGene_tx=XM_019837461.1;refGene_gn=BCAR3;refGene_tx=XM_023258916.1;refGene_gn=BCAR3;refGene_tx=XM_023258917.1
C1 79922829 79922933
refGene_gn=BCAR3;refGene_tx=XM_003990338.5;refGene_gn=BCAR3;refGene_tx=XM_011284867.2;refGene_gn=BCAR3;refGene_tx=XM_019837461.1;refGene_gn=BCAR3;refGene_tx=XM_023258916.1;refGene_gn=BCAR3;refGene_tx=XM_023258917.1
C1 79926644 79927087
refGene_gn=BCAR3;refGene_tx=XM_003990338.5;refGene_gn=BCAR3;refGene_tx=XM_011284867.2;refGene_gn=BCAR3;refGene_tx=XM_023258916.1;refGene_gn=BCAR3;refGene_tx=XM_023258917.1
C1 79929595 79929724

refGene_gn=BCAR3;refGene_tx=XM_003990338.5;refGene_gn=BCAR3;refGene_tx=XM_011284867.2;refGene_gn=BCAR3;refGene_tx=XM_023258916.1;refGene_gn=BCAR3;refGene_tx=XM_023258917.1
C1 79951851 79951935 refGene_gn=BCAR3;refGene_tx=XM_003990338.5
C1 79977645 79977685
refGene_gn=BCAR3;refGene_tx=XM_011284867.2;refGene_gn=BCAR3;refGene_tx=XM_023258916.1;refGene_gn=BCAR3;refGene_tx=XM_023258917.1
C1 80005020 80005337
refGene_gn=BCAR3;refGene_tx=XM_011284867.2;refGene_gn=BCAR3;refGene_tx=XM_023258916.1;refGene_gn=BCAR3;refGene_tx=XM_023258917.1
C1 80170571 80170665 refGene_gn=DNTTIP2;refGene_tx=XM_011284866.3
C1 80178752 80178862 refGene_gn=DNTTIP2;refGene_tx=XM_011284866.3
C1 80180413 80180578 refGene_gn=DNTTIP2;refGene_tx=XM_011284866.3
C1 80181986 80182082 refGene_gn=DNTTIP2;refGene_tx=XM_011284866.3
C1 80184839 80184978 refGene_gn=DNTTIP2;refGene_tx=XM_011284866.3
C1 80186793 80188433 refGene_gn=DNTTIP2;refGene_tx=XM_011284866.3
C1 80189441 80189513 refGene_gn=DNTTIP2;refGene_tx=XM_011284866.3
C1 80196482 80196505 refGene_gn=GCLM;refGene_tx=XM_023258920.1
C1 80198119 80198289 refGene_gn=GCLM;refGene_tx=XM_023258919.1
C1 80202774 80202889
refGene_gn=GCLM;refGene_tx=XM_023258919.1;refGene_gn=GCLM;refGene_tx=XM_023258920.1
C1 80204508 80204711
refGene_gn=GCLM;refGene_tx=XM_023258919.1;refGene_gn=GCLM;refGene_tx=XM_023258920.1
C1 80205937 80205997
refGene_gn=GCLM;refGene_tx=XM_023258919.1;refGene_gn=GCLM;refGene_tx=XM_023258920.1
C1 80209117 80209202
refGene_gn=GCLM;refGene_tx=XM_023258919.1;refGene_gn=GCLM;refGene_tx=XM_023258920.1
C1 80210540 80210606
refGene_gn=GCLM;refGene_tx=XM_023258919.1;refGene_gn=GCLM;refGene_tx=XM_023258920.1
C1 80215816 80215942
refGene_gn=GCLM;refGene_tx=XM_023258919.1;refGene_gn=GCLM;refGene_tx=XM_023258920.1
C1 80247085 80247091 refGene_gn=ABCA4;refGene_tx=XM_019837464.2
C1 80251134 80251386
refGene_gn=ABCA4;refGene_tx=XM_019837462.1;refGene_gn=ABCA4;refGene_tx=XM_019837463.1;refGene_gn=ABCA4;refGene_tx=XM_019837464.2;refGene_gn=ABCA4;refGene_tx=XM_019837465.2
C1 80252680 80252930
refGene_gn=ABCA4;refGene_tx=XM_019837462.1;refGene_gn=ABCA4;refGene_tx=XM_019837463.1;refGene_gn=ABCA4;refGene_tx=XM_019837464.2;refGene_gn=ABCA4;refGene_tx=XM_019837465.2
C1 80254928 80255021
refGene_gn=ABCA4;refGene_tx=XM_019837462.1;refGene_gn=ABCA4;refGene_tx=XM_019837463.1;refGene_gn=ABCA4;refGene_tx=XM_019837464.2;refGene_gn=ABCA4;refGene_tx=XM_019837465.2

C1 80255106 80255210
refGene_gn=ABCA4;refGene_tx=XM_019837462.1;refGene_gn=ABCA4;refGene_tx=XM_019837463.1;refGene_gn=ABCA4;refGene_tx=XM_019837464.2;refGene_gn=ABCA4;refGene_tx=XM_019837465.2
C1 80256002 80256137
refGene_gn=ABCA4;refGene_tx=XM_019837462.1;refGene_gn=ABCA4;refGene_tx=XM_019837463.1;refGene_gn=ABCA4;refGene_tx=XM_019837464.2;refGene_gn=ABCA4;refGene_tx=XM_019837465.2
C1 80260232 80260374
refGene_gn=ABCA4;refGene_tx=XM_019837462.1;refGene_gn=ABCA4;refGene_tx=XM_019837463.1;refGene_gn=ABCA4;refGene_tx=XM_019837464.2;refGene_gn=ABCA4;refGene_tx=XM_019837465.2
C1 80262784 80262891
refGene_gn=ABCA4;refGene_tx=XM_019837462.1;refGene_gn=ABCA4;refGene_tx=XM_019837463.1;refGene_gn=ABCA4;refGene_tx=XM_019837464.2;refGene_gn=ABCA4;refGene_tx=XM_019837465.2
C1 80263955 80264018
refGene_gn=ABCA4;refGene_tx=XM_019837462.1;refGene_gn=ABCA4;refGene_tx=XM_019837463.1;refGene_gn=ABCA4;refGene_tx=XM_019837464.2;refGene_gn=ABCA4;refGene_tx=XM_019837465.2
C1 80264476 80264597
refGene_gn=ABCA4;refGene_tx=XM_019837462.1;refGene_gn=ABCA4;refGene_tx=XM_019837463.1;refGene_gn=ABCA4;refGene_tx=XM_019837464.2;refGene_gn=ABCA4;refGene_tx=XM_019837465.2
C1 80267009 80267139
refGene_gn=ABCA4;refGene_tx=XM_019837462.1;refGene_gn=ABCA4;refGene_tx=XM_019837463.1;refGene_gn=ABCA4;refGene_tx=XM_019837464.2;refGene_gn=ABCA4;refGene_tx=XM_019837465.2
C1 80267502 80267626
refGene_gn=ABCA4;refGene_tx=XM_019837462.1;refGene_gn=ABCA4;refGene_tx=XM_019837463.1;refGene_gn=ABCA4;refGene_tx=XM_019837464.2;refGene_gn=ABCA4;refGene_tx=XM_019837465.2
C1 80270664 80270812
refGene_gn=ABCA4;refGene_tx=XM_019837462.1;refGene_gn=ABCA4;refGene_tx=XM_019837463.1;refGene_gn=ABCA4;refGene_tx=XM_019837464.2;refGene_gn=ABCA4;refGene_tx=XM_019837465.2
C1 80271832 80271948
refGene_gn=ABCA4;refGene_tx=XM_019837462.1;refGene_gn=ABCA4;refGene_tx=XM_019837463.1;refGene_gn=ABCA4;refGene_tx=XM_019837464.2;refGene_gn=ABCA4;refGene_tx=XM_019837465.2
C1 80275851 80276029
refGene_gn=ABCA4;refGene_tx=XM_019837462.1;refGene_gn=ABCA4;refGene_tx=XM_019837463.1;refGene_gn=ABCA4;refGene_tx=XM_019837464.2;refGene_gn=ABCA4;refGene_tx=XM_019837465.2
C1 80277949 80278119
refGene_gn=ABCA4;refGene_tx=XM_019837462.1;refGene_gn=ABCA4;refGene_tx=XM_019837463.1;refGene_gn=ABCA4;refGene_tx=XM_019837464.2;refGene_gn=ABCA4;refGene_tx=XM_019837465.2
C1 80278350 80278425
refGene_gn=ABCA4;refGene_tx=XM_019837462.1;refGene_gn=ABCA4;refGene_tx

=XM_019837463.1;refGene_gn=ABCA4;refGene_tx=XM_019837464.2;refGene_gn=ABCA4;refGene_tx=XM_019837465.2
C1 80278544 80278650
refGene_gn=ABCA4;refGene_tx=XM_019837462.1;refGene_gn=ABCA4;refGene_tx=XM_019837463.1;refGene_gn=ABCA4;refGene_tx=XM_019837464.2;refGene_gn=ABCA4;refGene_tx=XM_019837465.2
C1 80280081 80280114
refGene_gn=ABCA4;refGene_tx=XM_019837462.1;refGene_gn=ABCA4;refGene_tx=XM_019837463.1;refGene_gn=ABCA4;refGene_tx=XM_019837464.2;refGene_gn=ABCA4;refGene_tx=XM_019837465.2
C1 80281600 80281695
refGene_gn=ABCA4;refGene_tx=XM_019837462.1;refGene_gn=ABCA4;refGene_tx=XM_019837463.1;refGene_gn=ABCA4;refGene_tx=XM_019837464.2;refGene_gn=ABCA4;refGene_tx=XM_019837465.2
C1 80285686 80285873
refGene_gn=ABCA4;refGene_tx=XM_019837462.1;refGene_gn=ABCA4;refGene_tx=XM_019837463.1;refGene_gn=ABCA4;refGene_tx=XM_019837464.2;refGene_gn=ABCA4;refGene_tx=XM_019837465.2
C1 80286709 80286808
refGene_gn=ABCA4;refGene_tx=XM_019837462.1;refGene_gn=ABCA4;refGene_tx=XM_019837463.1;refGene_gn=ABCA4;refGene_tx=XM_019837464.2;refGene_gn=ABCA4;refGene_tx=XM_019837465.2
C1 80287290 80287415
refGene_gn=ABCA4;refGene_tx=XM_019837462.1;refGene_gn=ABCA4;refGene_tx=XM_019837463.1;refGene_gn=ABCA4;refGene_tx=XM_019837464.2;refGene_gn=ABCA4;refGene_tx=XM_019837465.2
C1 80289778 80290026
refGene_gn=ABCA4;refGene_tx=XM_019837462.1;refGene_gn=ABCA4;refGene_tx=XM_019837463.1;refGene_gn=ABCA4;refGene_tx=XM_019837464.2;refGene_gn=ABCA4;refGene_tx=XM_019837465.2
C1 80294318 80294367
refGene_gn=ABCA4;refGene_tx=XM_019837462.1;refGene_gn=ABCA4;refGene_tx=XM_019837463.1;refGene_gn=ABCA4;refGene_tx=XM_019837464.2;refGene_gn=ABCA4;refGene_tx=XM_019837465.2
C1 80294728 80294934
refGene_gn=ABCA4;refGene_tx=XM_019837462.1;refGene_gn=ABCA4;refGene_tx=XM_019837463.1;refGene_gn=ABCA4;refGene_tx=XM_019837464.2;refGene_gn=ABCA4;refGene_tx=XM_019837465.2
C1 80297737 80297822
refGene_gn=ABCA4;refGene_tx=XM_019837462.1;refGene_gn=ABCA4;refGene_tx=XM_019837463.1;refGene_gn=ABCA4;refGene_tx=XM_019837464.2;refGene_gn=ABCA4;refGene_tx=XM_019837465.2
C1 80299146 80299340
refGene_gn=ABCA4;refGene_tx=XM_019837462.1;refGene_gn=ABCA4;refGene_tx=XM_019837463.1;refGene_gn=ABCA4;refGene_tx=XM_019837464.2;refGene_gn=ABCA4;refGene_tx=XM_019837465.2
C1 80300327 80300465
refGene_gn=ABCA4;refGene_tx=XM_019837462.1;refGene_gn=ABCA4;refGene_tx=XM_019837463.1;refGene_gn=ABCA4;refGene_tx=XM_019837464.2;refGene_gn=ABCA4;refGene_tx=XM_019837465.2

C1 80300918 80301058
refGene_gn=ABCA4;refGene_tx=XM_019837462.1;refGene_gn=ABCA4;refGene_tx=XM_019837463.1;refGene_gn=ABCA4;refGene_tx=XM_019837464.2;refGene_gn=ABCA4;refGene_tx=XM_019837465.2
C1 80302249 80302381
refGene_gn=ABCA4;refGene_tx=XM_019837462.1;refGene_gn=ABCA4;refGene_tx=XM_019837463.1;refGene_gn=ABCA4;refGene_tx=XM_019837464.2;refGene_gn=ABCA4;refGene_tx=XM_019837465.2
C1 80304847 80305022
refGene_gn=ABCA4;refGene_tx=XM_019837462.1;refGene_gn=ABCA4;refGene_tx=XM_019837463.1;refGene_gn=ABCA4;refGene_tx=XM_019837464.2;refGene_gn=ABCA4;refGene_tx=XM_019837465.2
C1 80306991 80307081
refGene_gn=ABCA4;refGene_tx=XM_019837462.1;refGene_gn=ABCA4;refGene_tx=XM_019837464.2;refGene_gn=ABCA4;refGene_tx=XM_019837465.2
C1 80309731 80309797
refGene_gn=ABCA4;refGene_tx=XM_019837462.1;refGene_gn=ABCA4;refGene_tx=XM_019837463.1;refGene_gn=ABCA4;refGene_tx=XM_019837464.2;refGene_gn=ABCA4;refGene_tx=XM_019837465.2
C1 80313453 80313658
refGene_gn=ABCA4;refGene_tx=XM_019837462.1;refGene_gn=ABCA4;refGene_tx=XM_019837463.1;refGene_gn=ABCA4;refGene_tx=XM_019837464.2;refGene_gn=ABCA4;refGene_tx=XM_019837465.2
C1 80315789 80316011
refGene_gn=ABCA4;refGene_tx=XM_019837462.1;refGene_gn=ABCA4;refGene_tx=XM_019837463.1;refGene_gn=ABCA4;refGene_tx=XM_019837464.2;refGene_gn=ABCA4;refGene_tx=XM_019837465.2
C1 80319136 80319359
refGene_gn=ABCA4;refGene_tx=XM_019837462.1;refGene_gn=ABCA4;refGene_tx=XM_019837463.1;refGene_gn=ABCA4;refGene_tx=XM_019837464.2;refGene_gn=ABCA4;refGene_tx=XM_019837465.2
C1 80321327 80321504
refGene_gn=ABCA4;refGene_tx=XM_019837462.1;refGene_gn=ABCA4;refGene_tx=XM_019837463.1;refGene_gn=ABCA4;refGene_tx=XM_019837464.2;refGene_gn=ABCA4;refGene_tx=XM_019837465.2
C1 80322136 80322342
refGene_gn=ABCA4;refGene_tx=XM_019837462.1;refGene_gn=ABCA4;refGene_tx=XM_019837463.1;refGene_gn=ABCA4;refGene_tx=XM_019837464.2;refGene_gn=ABCA4;refGene_tx=XM_019837465.2
C1 80337689 80337887
refGene_gn=ABCA4;refGene_tx=XM_019837462.1;refGene_gn=ABCA4;refGene_tx=XM_019837463.1;refGene_gn=ABCA4;refGene_tx=XM_019837464.2;refGene_gn=ABCA4;refGene_tx=XM_019837465.2
C1 80338572 80338689
refGene_gn=ABCA4;refGene_tx=XM_019837462.1;refGene_gn=ABCA4;refGene_tx=XM_019837463.1;refGene_gn=ABCA4;refGene_tx=XM_019837464.2;refGene_gn=ABCA4;refGene_tx=XM_019837465.2
C1 80339327 80339467
refGene_gn=ABCA4;refGene_tx=XM_019837462.1;refGene_gn=ABCA4;refGene_tx=XM_019837463.1;refGene_gn=ABCA4;refGene_tx=XM_019837464.2;refGene_gn=

ABCA4;refGene_tx=XM_019837465.2
C1 80340477 80340718
refGene_gn=ABCA4;refGene_tx=XM_019837462.1;refGene_gn=ABCA4;refGene_tx=XM_019837463.1;refGene_gn=ABCA4;refGene_tx=XM_019837464.2;refGene_gn=ABCA4;refGene_tx=XM_019837465.2
C1 80343002 80343092
refGene_gn=ABCA4;refGene_tx=XM_019837462.1;refGene_gn=ABCA4;refGene_tx=XM_019837463.1;refGene_gn=ABCA4;refGene_tx=XM_019837464.2;refGene_gn=ABCA4;refGene_tx=XM_019837465.2
C1 80352183 80352186 refGene_gn=ABCA4;refGene_tx=XM_019837465.2
C1 80356347 80356545
refGene_gn=ABCA4;refGene_tx=XM_019837462.1;refGene_gn=ABCA4;refGene_tx=XM_019837463.1;refGene_gn=ABCA4;refGene_tx=XM_019837464.2
C1 80361046 80361174
refGene_gn=ABCA4;refGene_tx=XM_019837462.1;refGene_gn=ABCA4;refGene_tx=XM_019837463.1;refGene_gn=ABCA4;refGene_tx=XM_019837464.2
C1 80366729 80366869
refGene_gn=ABCA4;refGene_tx=XM_019837462.1;refGene_gn=ABCA4;refGene_tx=XM_019837463.1;refGene_gn=ABCA4;refGene_tx=XM_019837464.2
C1 80368746 80368888
refGene_gn=ABCA4;refGene_tx=XM_019837462.1;refGene_gn=ABCA4;refGene_tx=XM_019837463.1;refGene_gn=ABCA4;refGene_tx=XM_019837464.2
C1 80369923 80370017
refGene_gn=ABCA4;refGene_tx=XM_019837462.1;refGene_gn=ABCA4;refGene_tx=XM_019837463.1;refGene_gn=ABCA4;refGene_tx=XM_019837464.2
C1 80378645 80378711
refGene_gn=ABCA4;refGene_tx=XM_019837462.1;refGene_gn=ABCA4;refGene_tx=XM_019837463.1;refGene_gn=ABCA4;refGene_tx=XM_019837464.2
C1 80413874 80414764 refGene_gn=ARHGAP29;refGene_tx=XM_023258922.1
C1 80416789 80416898 refGene_gn=ARHGAP29;refGene_tx=XM_023258922.1
C1 80417020 80417351 refGene_gn=ARHGAP29;refGene_tx=XM_023258922.1
C1 80418571 80418804 refGene_gn=ARHGAP29;refGene_tx=XM_023258922.1
C1 80423101 80423239 refGene_gn=ARHGAP29;refGene_tx=XM_023258922.1
C1 80423508 80423697 refGene_gn=ARHGAP29;refGene_tx=XM_023258922.1
C1 80423975 80424115 refGene_gn=ARHGAP29;refGene_tx=XM_023258922.1
C1 80425356 80425455 refGene_gn=ARHGAP29;refGene_tx=XM_023258922.1
C1 80427670 80427775 refGene_gn=ARHGAP29;refGene_tx=XM_023258922.1
C1 80428267 80428407 refGene_gn=ARHGAP29;refGene_tx=XM_023258922.1
C1 80428920 80429078 refGene_gn=ARHGAP29;refGene_tx=XM_023258922.1
C1 80440291 80440372 refGene_gn=ARHGAP29;refGene_tx=XM_023258922.1
C1 80440373 80440430 refGene_gn=ARHGAP29;refGene_tx=XM_023258922.1
C1 80441145 80441334 refGene_gn=ARHGAP29;refGene_tx=XM_023258922.1
C1 80441966 80442047 refGene_gn=ARHGAP29;refGene_tx=XM_023258922.1
C1 80442139 80442250 refGene_gn=ARHGAP29;refGene_tx=XM_023258922.1
C1 80442915 80442980 refGene_gn=ARHGAP29;refGene_tx=XM_023258922.1
C1 80443856 80443994 refGene_gn=ARHGAP29;refGene_tx=XM_023258922.1
C1 80444623 80444672 refGene_gn=ARHGAP29;refGene_tx=XM_023258922.1
C1 80447746 80447819 refGene_gn=ARHGAP29;refGene_tx=XM_023258922.1
C1 80448171 80448268 refGene_gn=ARHGAP29;refGene_tx=XM_023258922.1
C1 80459900 80460035 refGene_gn=ARHGAP29;refGene_tx=XM_023258922.1

C1	80481112 80481125 refGene_gn=ARHGAP29;refGene_tx=XM_023258922.1
C1	80492418 80493226
refGene_gn=LOC111561635;refGene_tx=XM_023257835.1
C1	80494603 80494737
refGene_gn=LOC111561635;refGene_tx=XM_023257835.1
C1	80572802 80572958
refGene_gn=LOC102901205;refGene_tx=XM_011284863.2
C1	80604379 80604381
refGene_gn=ABCD3;refGene_tx=XM_023258924.1;refGene_gn=ABCD3;refGene_tx
=XM_023258925.1;refGene_gn=ABCD3;refGene_tx=XM_023258926.1;refGene_gn=
ABCD3;refGene_tx=XM_023258927.1
C1	80605377 80605406 refGene_gn=ABCD3;refGene_tx=XM_023258923.1
C1	80606057 80606207
refGene_gn=ABCD3;refGene_tx=XM_023258923.1;refGene_gn=ABCD3;refGene_tx
=XM_023258924.1;refGene_gn=ABCD3;refGene_tx=XM_023258925.1;refGene_gn=
ABCD3;refGene_tx=XM_023258926.1;refGene_gn=ABCD3;refGene_tx=XM_0232589
27.1
C1	80643882 80643992 refGene_gn=ABCD3;refGene_tx=XM_003990341.5
C1	80668302 80668339
refGene_gn=ABCD3;refGene_tx=XM_003990341.5;refGene_gn=ABCD3;refGene_tx
=XM_023258923.1;refGene_gn=ABCD3;refGene_tx=XM_023258924.1;refGene_gn=
ABCD3;refGene_tx=XM_023258925.1;refGene_gn=ABCD3;refGene_tx=XM_0232589
26.1;refGene_gn=ABCD3;refGene_tx=XM_023258927.1
C1	80675140 80675239
refGene_gn=ABCD3;refGene_tx=XM_003990341.5;refGene_gn=ABCD3;refGene_tx
=XM_023258923.1;refGene_gn=ABCD3;refGene_tx=XM_023258924.1;refGene_gn=
ABCD3;refGene_tx=XM_023258925.1;refGene_gn=ABCD3;refGene_tx=XM_0232589
26.1;refGene_gn=ABCD3;refGene_tx=XM_023258927.1
C1	80679024 80679113
refGene_gn=ABCD3;refGene_tx=XM_003990341.5;refGene_gn=ABCD3;refGene_tx
=XM_023258923.1;refGene_gn=ABCD3;refGene_tx=XM_023258924.1;refGene_gn=
ABCD3;refGene_tx=XM_023258925.1;refGene_gn=ABCD3;refGene_tx=XM_0232589
26.1;refGene_gn=ABCD3;refGene_tx=XM_023258927.1
C1	80686110 80686180
refGene_gn=ABCD3;refGene_tx=XM_003990341.5;refGene_gn=ABCD3;refGene_tx
=XM_006934921.3;refGene_gn=ABCD3;refGene_tx=XM_023258923.1;refGene_gn=
ABCD3;refGene_tx=XM_023258924.1;refGene_gn=ABCD3;refGene_tx=XM_0232589
25.1;refGene_gn=ABCD3;refGene_tx=XM_023258926.1;refGene_gn=ABCD3;refGe
ne_tx=XM_023258927.1
C1	80687508 80687606
refGene_gn=ABCD3;refGene_tx=XM_003990341.5;refGene_gn=ABCD3;refGene_tx
=XM_006934921.3;refGene_gn=ABCD3;refGene_tx=XM_023258923.1;refGene_gn=
ABCD3;refGene_tx=XM_023258924.1;refGene_gn=ABCD3;refGene_tx=XM_0232589
25.1;refGene_gn=ABCD3;refGene_tx=XM_023258926.1;refGene_gn=ABCD3;refGe
ne_tx=XM_023258927.1
C1	80687950 80688074
refGene_gn=ABCD3;refGene_tx=XM_003990341.5;refGene_gn=ABCD3;refGene_tx
=XM_006934921.3;refGene_gn=ABCD3;refGene_tx=XM_023258923.1;refGene_gn=
ABCD3;refGene_tx=XM_023258924.1;refGene_gn=ABCD3;refGene_tx=XM_0232589
25.1;refGene_gn=ABCD3;refGene_tx=XM_023258926.1;refGene_gn=ABCD3;refGe

ne_tx=XM_023258927.1
C1 80688608 80688665
refGene_gn=ABCD3;refGene_tx=XM_003990341.5;refGene_gn=ABCD3;refGene_tx=XM_006934921.3;refGene_gn=ABCD3;refGene_tx=XM_023258923.1;refGene_gn=ABCD3;refGene_tx=XM_023258924.1;refGene_gn=ABCD3;refGene_tx=XM_023258925.1;refGene_gn=ABCD3;refGene_tx=XM_023258926.1;refGene_gn=ABCD3;refGene_tx=XM_023258927.1
C1 80690617 80690760
refGene_gn=ABCD3;refGene_tx=XM_003990341.5;refGene_gn=ABCD3;refGene_tx=XM_006934921.3;refGene_gn=ABCD3;refGene_tx=XM_023258923.1;refGene_gn=ABCD3;refGene_tx=XM_023258924.1;refGene_gn=ABCD3;refGene_tx=XM_023258925.1;refGene_gn=ABCD3;refGene_tx=XM_023258926.1;refGene_gn=ABCD3;refGene_tx=XM_023258927.1
C1 80694570 80694640
refGene_gn=ABCD3;refGene_tx=XM_003990341.5;refGene_gn=ABCD3;refGene_tx=XM_006934921.3;refGene_gn=ABCD3;refGene_tx=XM_023258923.1;refGene_gn=ABCD3;refGene_tx=XM_023258924.1;refGene_gn=ABCD3;refGene_tx=XM_023258925.1;refGene_gn=ABCD3;refGene_tx=XM_023258926.1;refGene_gn=ABCD3;refGene_tx=XM_023258927.1
C1 80696699 80696769
refGene_gn=ABCD3;refGene_tx=XM_003990341.5;refGene_gn=ABCD3;refGene_tx=XM_006934921.3;refGene_gn=ABCD3;refGene_tx=XM_023258923.1;refGene_gn=ABCD3;refGene_tx=XM_023258924.1;refGene_gn=ABCD3;refGene_tx=XM_023258925.1;refGene_gn=ABCD3;refGene_tx=XM_023258926.1;refGene_gn=ABCD3;refGene_tx=XM_023258927.1
C1 80696858 80696956
refGene_gn=ABCD3;refGene_tx=XM_003990341.5;refGene_gn=ABCD3;refGene_tx=XM_006934921.3;refGene_gn=ABCD3;refGene_tx=XM_023258923.1;refGene_gn=ABCD3;refGene_tx=XM_023258924.1;refGene_gn=ABCD3;refGene_tx=XM_023258925.1;refGene_gn=ABCD3;refGene_tx=XM_023258926.1;refGene_gn=ABCD3;refGene_tx=XM_023258927.1
C1 80697068 80697160
refGene_gn=ABCD3;refGene_tx=XM_003990341.5;refGene_gn=ABCD3;refGene_tx=XM_006934921.3;refGene_gn=ABCD3;refGene_tx=XM_023258923.1;refGene_gn=ABCD3;refGene_tx=XM_023258924.1;refGene_gn=ABCD3;refGene_tx=XM_023258925.1;refGene_gn=ABCD3;refGene_tx=XM_023258926.1;refGene_gn=ABCD3;refGene_tx=XM_023258927.1
C1 80698740 80698832
refGene_gn=ABCD3;refGene_tx=XM_003990341.5;refGene_gn=ABCD3;refGene_tx=XM_006934921.3;refGene_gn=ABCD3;refGene_tx=XM_023258923.1;refGene_gn=ABCD3;refGene_tx=XM_023258924.1;refGene_gn=ABCD3;refGene_tx=XM_023258925.1;refGene_gn=ABCD3;refGene_tx=XM_023258926.1;refGene_gn=ABCD3;refGene_tx=XM_023258927.1
C1 80698913 80698989
refGene_gn=ABCD3;refGene_tx=XM_003990341.5;refGene_gn=ABCD3;refGene_tx=XM_006934921.3;refGene_gn=ABCD3;refGene_tx=XM_023258923.1;refGene_gn=ABCD3;refGene_tx=XM_023258924.1;refGene_gn=ABCD3;refGene_tx=XM_023258925.1;refGene_gn=ABCD3;refGene_tx=XM_023258926.1;refGene_gn=ABCD3;refGene_tx=XM_023258927.1
C1 80699115 80699179

refGene_gn=ABCD3;refGene_tx=XM_003990341.5;refGene_gn=ABCD3;refGene_tx=XM_006934921.3;refGene_gn=ABCD3;refGene_tx=XM_023258923.1;refGene_gn=ABCD3;refGene_tx=XM_023258924.1;refGene_gn=ABCD3;refGene_tx=XM_023258925.1;refGene_gn=ABCD3;refGene_tx=XM_023258926.1;refGene_gn=ABCD3;refGene_tx=XM_023258927.1
C1 80704831 80704909
refGene_gn=ABCD3;refGene_tx=XM_003990341.5;refGene_gn=ABCD3;refGene_tx=XM_006934921.3;refGene_gn=ABCD3;refGene_tx=XM_023258923.1;refGene_gn=ABCD3;refGene_tx=XM_023258924.1;refGene_gn=ABCD3;refGene_tx=XM_023258925.1;refGene_gn=ABCD3;refGene_tx=XM_023258926.1;refGene_gn=ABCD3;refGene_tx=XM_023258927.1
C1 80705015 80705081
refGene_gn=ABCD3;refGene_tx=XM_003990341.5;refGene_gn=ABCD3;refGene_tx=XM_006934921.3;refGene_gn=ABCD3;refGene_tx=XM_023258923.1;refGene_gn=ABCD3;refGene_tx=XM_023258924.1;refGene_gn=ABCD3;refGene_tx=XM_023258925.1;refGene_gn=ABCD3;refGene_tx=XM_023258926.1;refGene_gn=ABCD3;refGene_tx=XM_023258927.1
C1 80705177 80705267
refGene_gn=ABCD3;refGene_tx=XM_003990341.5;refGene_gn=ABCD3;refGene_tx=XM_006934921.3;refGene_gn=ABCD3;refGene_tx=XM_023258923.1;refGene_gn=ABCD3;refGene_tx=XM_023258924.1;refGene_gn=ABCD3;refGene_tx=XM_023258925.1;refGene_gn=ABCD3;refGene_tx=XM_023258926.1;refGene_gn=ABCD3;refGene_tx=XM_023258927.1
C1 80705740 80705860
refGene_gn=ABCD3;refGene_tx=XM_003990341.5;refGene_gn=ABCD3;refGene_tx=XM_006934921.3;refGene_gn=ABCD3;refGene_tx=XM_023258923.1;refGene_gn=ABCD3;refGene_tx=XM_023258924.1;refGene_gn=ABCD3;refGene_tx=XM_023258925.1;refGene_gn=ABCD3;refGene_tx=XM_023258926.1;refGene_gn=ABCD3;refGene_tx=XM_023258927.1
C1 80712098 80712203
refGene_gn=ABCD3;refGene_tx=XM_003990341.5;refGene_gn=ABCD3;refGene_tx=XM_006934921.3;refGene_gn=ABCD3;refGene_tx=XM_023258923.1;refGene_gn=ABCD3;refGene_tx=XM_023258924.1;refGene_gn=ABCD3;refGene_tx=XM_023258925.1;refGene_gn=ABCD3;refGene_tx=XM_023258926.1;refGene_gn=ABCD3;refGene_tx=XM_023258927.1
C1 80715190 80715247
refGene_gn=ABCD3;refGene_tx=XM_003990341.5;refGene_gn=ABCD3;refGene_tx=XM_006934921.3;refGene_gn=ABCD3;refGene_tx=XM_023258923.1;refGene_gn=ABCD3;refGene_tx=XM_023258924.1;refGene_gn=ABCD3;refGene_tx=XM_023258925.1;refGene_gn=ABCD3;refGene_tx=XM_023258926.1;refGene_gn=ABCD3;refGene_tx=XM_023258927.1
C1 80717212 80717290
refGene_gn=ABCD3;refGene_tx=XM_003990341.5;refGene_gn=ABCD3;refGene_tx=XM_006934921.3;refGene_gn=ABCD3;refGene_tx=XM_023258923.1;refGene_gn=ABCD3;refGene_tx=XM_023258924.1;refGene_gn=ABCD3;refGene_tx=XM_023258925.1;refGene_gn=ABCD3;refGene_tx=XM_023258926.1;refGene_gn=ABCD3;refGene_tx=XM_023258927.1
C1 80743002 80743139 refGene_gn=F3;refGene_tx=XM_006934929.4
C1 80744513 80744661 refGene_gn=F3;refGene_tx=XM_006934929.4
C1 80745214 80745393 refGene_gn=F3;refGene_tx=XM_006934929.4

C1 80747309 80747500 refGene_gn=F3;refGene_tx=XM_006934929.4
C1 80751633 80751745 refGene_gn=F3;refGene_tx=XM_006934929.4
C1 80752867 80752967 refGene_gn=F3;refGene_tx=XM_006934929.4
C1 80979838 80979865
refGene_gn=SLC44A3;refGene_tx=XM_019837471.2;refGene_gn=SLC44A3;refGene_tx=XM_019837473.2;refGene_gn=SLC44A3;refGene_tx=XM_019837474.2;refGene_gn=SLC44A3;refGene_tx=XM_019837477.2
C1 80980376 80980484
refGene_gn=SLC44A3;refGene_tx=XM_019837471.2;refGene_gn=SLC44A3;refGene_tx=XM_019837473.2;refGene_gn=SLC44A3;refGene_tx=XM_019837474.2;refGene_gn=SLC44A3;refGene_tx=XM_019837477.2
C1 80984046 80984189
refGene_gn=SLC44A3;refGene_tx=XM_019837471.2;refGene_gn=SLC44A3;refGene_tx=XM_019837473.2;refGene_gn=SLC44A3;refGene_tx=XM_019837474.2;refGene_gn=SLC44A3;refGene_tx=XM_019837477.2
C1 80986326 80986463
refGene_gn=SLC44A3;refGene_tx=XM_019837471.2;refGene_gn=SLC44A3;refGene_tx=XM_019837473.2;refGene_gn=SLC44A3;refGene_tx=XM_019837474.2;refGene_gn=SLC44A3;refGene_tx=XM_019837477.2
C1 80987388 80987482
refGene_gn=SLC44A3;refGene_tx=XM_019837471.2;refGene_gn=SLC44A3;refGene_tx=XM_019837473.2;refGene_gn=SLC44A3;refGene_tx=XM_019837474.2;refGene_gn=SLC44A3;refGene_tx=XM_019837477.2
C1 81011615 81011776
refGene_gn=SLC44A3;refGene_tx=XM_019837471.2;refGene_gn=SLC44A3;refGene_tx=XM_019837473.2;refGene_gn=SLC44A3;refGene_tx=XM_019837474.2;refGene_gn=SLC44A3;refGene_tx=XM_019837477.2
C1 81013660 81013750
refGene_gn=SLC44A3;refGene_tx=XM_019837471.2;refGene_gn=SLC44A3;refGene_tx=XM_019837473.2;refGene_gn=SLC44A3;refGene_tx=XM_019837474.2;refGene_gn=SLC44A3;refGene_tx=XM_019837477.2
C1 81014969 81015094
refGene_gn=SLC44A3;refGene_tx=XM_019837471.2;refGene_gn=SLC44A3;refGene_tx=XM_019837473.2;refGene_gn=SLC44A3;refGene_tx=XM_019837474.2;refGene_gn=SLC44A3;refGene_tx=XM_019837477.2
C1 81019613 81019800
refGene_gn=SLC44A3;refGene_tx=XM_019837471.2;refGene_gn=SLC44A3;refGene_tx=XM_019837473.2;refGene_gn=SLC44A3;refGene_tx=XM_019837474.2;refGene_gn=SLC44A3;refGene_tx=XM_019837477.2
C1 81031000 81031166
refGene_gn=SLC44A3;refGene_tx=XM_019837471.2;refGene_gn=SLC44A3;refGene_tx=XM_019837473.2;refGene_gn=SLC44A3;refGene_tx=XM_019837474.2;refGene_gn=SLC44A3;refGene_tx=XM_019837477.2
C1 81040492 81040649
refGene_gn=SLC44A3;refGene_tx=XM_019837471.2;refGene_gn=SLC44A3;refGene_tx=XM_019837473.2;refGene_gn=SLC44A3;refGene_tx=XM_019837474.2;refGene_gn=SLC44A3;refGene_tx=XM_019837477.2
C1 81043861 81043945
refGene_gn=SLC44A3;refGene_tx=XM_019837471.2;refGene_gn=SLC44A3;refGene_tx=XM_019837473.2;refGene_gn=SLC44A3;refGene_tx=XM_019837474.2;refGe

ne_gn=SLC44A3;refGene_tx=XM_019837477.2
C1 81067413 81067587
refGene_gn=SLC44A3;refGene_tx=XM_019837471.2;refGene_gn=SLC44A3;refGene_tx=XM_019837473.2;refGene_gn=SLC44A3;refGene_tx=XM_019837474.2
C1 81072662 81072800
refGene_gn=SLC44A3;refGene_tx=XM_019837471.2;refGene_gn=SLC44A3;refGene_tx=XM_019837473.2;refGene_gn=SLC44A3;refGene_tx=XM_019837474.2;refGene_gn=SLC44A3;refGene_tx=XM_019837477.2
C1 81075094 81075331
refGene_gn=SLC44A3;refGene_tx=XM_019837471.2;refGene_gn=SLC44A3;refGene_tx=XM_019837473.2;refGene_gn=SLC44A3;refGene_tx=XM_019837474.2;refGene_gn=SLC44A3;refGene_tx=XM_019837477.2
C1 81076720 81076822
refGene_gn=SLC44A3;refGene_tx=XM_019837471.2;refGene_gn=SLC44A3;refGene_tx=XM_019837473.2;refGene_gn=SLC44A3;refGene_tx=XM_019837474.2;refGene_gn=SLC44A3;refGene_tx=XM_019837477.2
C1 81078134 81078476 refGene_gn=CNN3;refGene_tx=XM_011284861.3
C1 81079798 81079945 refGene_gn=CNN3;refGene_tx=XM_011284861.3
C1 81081638 81081755 refGene_gn=CNN3;refGene_tx=XM_011284861.3
C1 81082130 81082268 refGene_gn=CNN3;refGene_tx=XM_011284861.3
C1 81082992 81083059 refGene_gn=CNN3;refGene_tx=XM_011284861.3
C1 81083366 81083488 refGene_gn=CNN3;refGene_tx=XM_011284861.3
C1 81104204 81104261 refGene_gn=CNN3;refGene_tx=XM_011284861.3
C1 81166665 81166896 refGene_gn=ALG14;refGene_tx=XM_003990347.4
C1 81188751 81188874 refGene_gn=ALG14;refGene_tx=XM_019837480.2
C1 81235989 81236121
refGene_gn=ALG14;refGene_tx=XM_003990347.4;refGene_gn=ALG14;refGene_tx=XM_019837480.2
C1 81267969 81268121
refGene_gn=ALG14;refGene_tx=XM_003990347.4;refGene_gn=ALG14;refGene_tx=XM_019837480.2
C1 81273904 81274043
refGene_gn=ALG14;refGene_tx=XM_003990347.4;refGene_gn=ALG14;refGene_tx=XM_019837480.2
C1 81329019 81329174
refGene_gn=LOC101080312;refGene_tx=XM_011284847.3;refGene_gn=LOC101080312;refGene_tx=XM_019837482.2;refGene_gn=LOC101080312;refGene_tx=XM_023258929.1
C1 81333643 81333733
refGene_gn=LOC101080312;refGene_tx=XM_011284847.3;refGene_gn=LOC101080312;refGene_tx=XM_019837482.2;refGene_gn=LOC101080312;refGene_tx=XM_023258929.1
C1 81335404 81335463
refGene_gn=LOC101080312;refGene_tx=XM_011284847.3;refGene_gn=LOC101080312;refGene_tx=XM_019837482.2;refGene_gn=LOC101080312;refGene_tx=XM_023258929.1
C1 81337811 81337906
refGene_gn=LOC101080312;refGene_tx=XM_011284847.3;refGene_gn=LOC101080312;refGene_tx=XM_019837482.2;refGene_gn=LOC101080312;refGene_tx=XM_023258929.1

C1 81358521 81358595
refGene_gn=LOC101080312;refGene_tx=XM_011284847.3;refGene_gn=LOC101080312;refGene_tx=XM_019837482.2;refGene_gn=LOC101080312;refGene_tx=XM_023258929.1
C1 81373905 81374224
refGene_gn=LOC101080312;refGene_tx=XM_011284847.3;refGene_gn=LOC101080312;refGene_tx=XM_019837482.2;refGene_gn=LOC101080312;refGene_tx=XM_023258929.1
C1 81411468 81411517 refGene_gn=RWDD3;refGene_tx=XM_006934930.3
C1 81423064 81423149
refGene_gn=RWDD3;refGene_tx=XM_023258930.1;refGene_gn=RWDD3;refGene_tx=XM_023258931.1
C1 81430010 81430498
refGene_gn=RWDD3;refGene_tx=XM_006934930.3;refGene_gn=RWDD3;refGene_tx=XM_023258930.1;refGene_gn=RWDD3;refGene_tx=XM_023258931.1
C1 81432452 81432568
refGene_gn=RWDD3;refGene_tx=XM_006934930.3;refGene_gn=RWDD3;refGene_tx=XM_023258930.1;refGene_gn=RWDD3;refGene_tx=XM_023258931.1
C1 81432666 81432781
refGene_gn=RWDD3;refGene_tx=XM_006934930.3;refGene_gn=RWDD3;refGene_tx=XM_023258930.1
C1 82609166 82609174
refGene_gn=PTBP2;refGene_tx=XM_006934931.4;refGene_gn=PTBP2;refGene_tx=XM_006934932.4;refGene_gn=PTBP2;refGene_tx=XM_006934933.4;refGene_gn=PTBP2;refGene_tx=XM_011284845.3
C1 82610874 82610905
refGene_gn=PTBP2;refGene_tx=XM_006934931.4;refGene_gn=PTBP2;refGene_tx=XM_006934932.4;refGene_gn=PTBP2;refGene_tx=XM_006934933.4;refGene_gn=PTBP2;refGene_tx=XM_011284845.3
C1 82637456 82637532
refGene_gn=PTBP2;refGene_tx=XM_006934931.4;refGene_gn=PTBP2;refGene_tx=XM_006934932.4;refGene_gn=PTBP2;refGene_tx=XM_006934933.4;refGene_gn=PTBP2;refGene_tx=XM_006934935.4;refGene_gn=PTBP2;refGene_tx=XM_011284845.3;refGene_gn=PTBP2;refGene_tx=XM_019837483.2
C1 82654633 82654806
refGene_gn=PTBP2;refGene_tx=XM_006934931.4;refGene_gn=PTBP2;refGene_tx=XM_006934932.4;refGene_gn=PTBP2;refGene_tx=XM_006934933.4;refGene_gn=PTBP2;refGene_tx=XM_006934935.4;refGene_gn=PTBP2;refGene_tx=XM_011284845.3;refGene_gn=PTBP2;refGene_tx=XM_019837483.2
C1 82655670 82655814
refGene_gn=PTBP2;refGene_tx=XM_006934931.4;refGene_gn=PTBP2;refGene_tx=XM_006934932.4;refGene_gn=PTBP2;refGene_tx=XM_006934933.4;refGene_gn=PTBP2;refGene_tx=XM_006934935.4;refGene_gn=PTBP2;refGene_tx=XM_011284845.3;refGene_gn=PTBP2;refGene_tx=XM_019837483.2
C1 82661749 82661914
refGene_gn=PTBP2;refGene_tx=XM_006934931.4;refGene_gn=PTBP2;refGene_tx=XM_006934932.4;refGene_gn=PTBP2;refGene_tx=XM_006934933.4;refGene_gn=PTBP2;refGene_tx=XM_006934935.4;refGene_gn=PTBP2;refGene_tx=XM_011284845.3;refGene_gn=PTBP2;refGene_tx=XM_019837483.2
C1 82662006 82662117

refGene_gn=PTBP2;refGene_tx=XM_006934931.4;refGene_gn=PTBP2;refGene_tx=XM_006934932.4;refGene_gn=PTBP2;refGene_tx=XM_006934933.4;refGene_gn=PTBP2;refGene_tx=XM_006934935.4;refGene_gn=PTBP2;refGene_tx=XM_011284845.3;refGene_gn=PTBP2;refGene_tx=XM_019837483.2
C1 82667873 82668069
refGene_gn=PTBP2;refGene_tx=XM_006934931.4;refGene_gn=PTBP2;refGene_tx=XM_006934932.4;refGene_gn=PTBP2;refGene_tx=XM_006934933.4;refGene_gn=PTBP2;refGene_tx=XM_006934935.4;refGene_gn=PTBP2;refGene_tx=XM_011284845.3;refGene_gn=PTBP2;refGene_tx=XM_019837483.2
C1 82682533 82682688
refGene_gn=PTBP2;refGene_tx=XM_006934931.4;refGene_gn=PTBP2;refGene_tx=XM_006934932.4;refGene_gn=PTBP2;refGene_tx=XM_006934933.4;refGene_gn=PTBP2;refGene_tx=XM_006934935.4;refGene_gn=PTBP2;refGene_tx=XM_011284845.3;refGene_gn=PTBP2;refGene_tx=XM_019837483.2
C1 82684157 82684191
refGene_gn=PTBP2;refGene_tx=XM_006934931.4;refGene_gn=PTBP2;refGene_tx=XM_006934932.4;refGene_gn=PTBP2;refGene_tx=XM_006934933.4;refGene_gn=PTBP2;refGene_tx=XM_006934935.4;refGene_gn=PTBP2;refGene_tx=XM_011284845.3;refGene_gn=PTBP2;refGene_tx=XM_019837483.2
C1 82684608 82684701
refGene_gn=PTBP2;refGene_tx=XM_006934931.4;refGene_gn=PTBP2;refGene_tx=XM_006934932.4;refGene_gn=PTBP2;refGene_tx=XM_006934933.4;refGene_gn=PTBP2;refGene_tx=XM_006934935.4;refGene_gn=PTBP2;refGene_tx=XM_011284845.3;refGene_gn=PTBP2;refGene_tx=XM_019837483.2
C1 82691010 82691227
refGene_gn=PTBP2;refGene_tx=XM_006934931.4;refGene_gn=PTBP2;refGene_tx=XM_006934932.4;refGene_gn=PTBP2;refGene_tx=XM_006934933.4;refGene_gn=PTBP2;refGene_tx=XM_006934935.4;refGene_gn=PTBP2;refGene_tx=XM_011284845.3;refGene_gn=PTBP2;refGene_tx=XM_019837483.2
C1 82691327 82691405
refGene_gn=PTBP2;refGene_tx=XM_006934931.4;refGene_gn=PTBP2;refGene_tx=XM_006934932.4;refGene_gn=PTBP2;refGene_tx=XM_006934933.4;refGene_gn=PTBP2;refGene_tx=XM_006934935.4;refGene_gn=PTBP2;refGene_tx=XM_011284845.3;refGene_gn=PTBP2;refGene_tx=XM_019837483.2
C1 82691580 82691713
refGene_gn=PTBP2;refGene_tx=XM_006934931.4;refGene_gn=PTBP2;refGene_tx=XM_006934932.4;refGene_gn=PTBP2;refGene_tx=XM_006934933.4;refGene_gn=PTBP2;refGene_tx=XM_006934935.4;refGene_gn=PTBP2;refGene_tx=XM_011284845.3;refGene_gn=PTBP2;refGene_tx=XM_019837483.2
C1 83023470 83023644 refGene_gn=DPYD;refGene_tx=XM_011284844.3
C1 83026180 83026321 refGene_gn=DPYD;refGene_tx=XM_011284844.3
C1 83052014 83052158 refGene_gn=DPYD;refGene_tx=XM_011284844.3
C1 83149878 83150058 refGene_gn=DPYD;refGene_tx=XM_011284844.3
C1 83181157 83181300 refGene_gn=DPYD;refGene_tx=XM_011284844.3
C1 83247040 83247160 refGene_gn=DPYD;refGene_tx=XM_011284844.3
C1 83248001 83248122 refGene_gn=DPYD;refGene_tx=XM_011284844.3
C1 83299283 83299367 refGene_gn=DPYD;refGene_tx=XM_011284844.3
C1 83308301 83308370 refGene_gn=DPYD;refGene_tx=XM_011284844.3
C1 83365671 83365836 refGene_gn=DPYD;refGene_tx=XM_011284844.3
C1 83421853 83422069 refGene_gn=DPYD;refGene_tx=XM_011284844.3

C1 83447433 83447618 refGene_gn=DPYD;refGene_tx=XM_011284844.3
C1 83471285 83471496 refGene_gn=DPYD;refGene_tx=XM_011284844.3
C1 83494108 83494278 refGene_gn=DPYD;refGene_tx=XM_011284844.3
C1 83496074 83496182 refGene_gn=DPYD;refGene_tx=XM_011284844.3
C1 83598719 83598807 refGene_gn=DPYD;refGene_tx=XM_011284844.3
C1 83607900 83607982 refGene_gn=DPYD;refGene_tx=XM_011284844.3
C1 83616520 83616717 refGene_gn=DPYD;refGene_tx=XM_011284844.3
C1 83657451 83657613 refGene_gn=DPYD;refGene_tx=XM_011284844.3
C1 83680599 83680687 refGene_gn=DPYD;refGene_tx=XM_011284844.3
C1 83768625 83768708 refGene_gn=DPYD;refGene_tx=XM_011284844.3
C1 83853805 83853916 refGene_gn=DPYD;refGene_tx=XM_011284844.3
C1 83893208 83893247 refGene_gn=DPYD;refGene_tx=XM_011284844.3
C1 84464865 84464892 refGene_gn=SNX7;refGene_tx=XM_019837492.2
C1 84610967 84611147
refGene_gn=SNX7;refGene_tx=XM_011284841.3;refGene_gn=SNX7;refGene_tx=XM_011284842.3;refGene_gn=SNX7;refGene_tx=XM_023258932.1
C1 84637353 84637536
refGene_gn=SNX7;refGene_tx=XM_011284841.3;refGene_gn=SNX7;refGene_tx=XM_011284842.3;refGene_gn=SNX7;refGene_tx=XM_019837492.2;refGene_gn=SNX7;refGene_tx=XM_019837493.2;refGene_gn=SNX7;refGene_tx=XM_023258932.1
C1 84643447 84643558
refGene_gn=SNX7;refGene_tx=XM_011284841.3;refGene_gn=SNX7;refGene_tx=XM_011284842.3;refGene_gn=SNX7;refGene_tx=XM_019837492.2;refGene_gn=SNX7;refGene_tx=XM_019837493.2;refGene_gn=SNX7;refGene_tx=XM_023258932.1
C1 84644081 84644246
refGene_gn=SNX7;refGene_tx=XM_011284841.3;refGene_gn=SNX7;refGene_tx=XM_011284842.3;refGene_gn=SNX7;refGene_tx=XM_019837492.2;refGene_gn=SNX7;refGene_tx=XM_019837493.2;refGene_gn=SNX7;refGene_tx=XM_023258932.1
C1 84646302 84646501
refGene_gn=SNX7;refGene_tx=XM_011284841.3;refGene_gn=SNX7;refGene_tx=XM_011284842.3;refGene_gn=SNX7;refGene_tx=XM_019837492.2;refGene_gn=SNX7;refGene_tx=XM_019837493.2;refGene_gn=SNX7;refGene_tx=XM_023258932.1
C1 84652210 84652410
refGene_gn=SNX7;refGene_tx=XM_011284841.3;refGene_gn=SNX7;refGene_tx=XM_011284842.3;refGene_gn=SNX7;refGene_tx=XM_019837492.2;refGene_gn=SNX7;refGene_tx=XM_019837493.2;refGene_gn=SNX7;refGene_tx=XM_023258932.1
C1 84655169 84655256
refGene_gn=SNX7;refGene_tx=XM_011284841.3;refGene_gn=SNX7;refGene_tx=XM_011284842.3;refGene_gn=SNX7;refGene_tx=XM_019837492.2;refGene_gn=SNX7;refGene_tx=XM_019837493.2;refGene_gn=SNX7;refGene_tx=XM_023258932.1
C1 84692409 84692562
refGene_gn=SNX7;refGene_tx=XM_011284841.3;refGene_gn=SNX7;refGene_tx=XM_011284842.3;refGene_gn=SNX7;refGene_tx=XM_019837492.2;refGene_gn=SNX7;refGene_tx=XM_019837493.2;refGene_gn=SNX7;refGene_tx=XM_023258932.1
C1 84720096 84720207
refGene_gn=SNX7;refGene_tx=XM_011284841.3;refGene_gn=SNX7;refGene_tx=XM_019837492.2;refGene_gn=SNX7;refGene_tx=XM_019837493.2
C1 84720268 84720355 refGene_gn=SNX7;refGene_tx=XM_011284842.3
C1 84737143 84737194 refGene_gn=SNX7;refGene_tx=XM_023258932.1
C1 84823122 84823155

refGene_gn=PLPPR5;refGene_tx=XM_023258933.1;refGene_gn=PLPPR5;refGene_tx=XM_023258934.1
C1 84852596 84852731
refGene_gn=PLPPR5;refGene_tx=XM_023258933.1;refGene_gn=PLPPR5;refGene_tx=XM_023258934.1
C1 84863732 84863909
refGene_gn=PLPPR5;refGene_tx=XM_023258933.1;refGene_gn=PLPPR5;refGene_tx=XM_023258934.1
C1 84902122 84902373
refGene_gn=PLPPR5;refGene_tx=XM_023258933.1;refGene_gn=PLPPR5;refGene_tx=XM_023258934.1
C1 84906635 84906768
refGene_gn=PLPPR5;refGene_tx=XM_023258933.1;refGene_gn=PLPPR5;refGene_tx=XM_023258934.1
C1 84949194 84949431
refGene_gn=PLPPR5;refGene_tx=XM_023258933.1;refGene_gn=PLPPR5;refGene_tx=XM_023258934.1
C1 85169100 85169322 refGene_gn=PLPPR4;refGene_tx=NM_001128199.1
C1 85192274 85192460 refGene_gn=PLPPR4;refGene_tx=NM_001128199.1
C1 85199870 85200000 refGene_gn=PLPPR4;refGene_tx=NM_001128199.1
C1 85201244 85201356 refGene_gn=PLPPR4;refGene_tx=XM_019835926.2
C1 85202063 85202259
refGene_gn=PLPPR4;refGene_tx=NM_001128199.1;refGene_gn=PLPPR4;refGene_tx=XM_019835926.2
C1 85203885 85203943
refGene_gn=PLPPR4;refGene_tx=NM_001128199.1;refGene_gn=PLPPR4;refGene_tx=XM_019835926.2
C1 85204708 85204882
refGene_gn=PLPPR4;refGene_tx=NM_001128199.1;refGene_gn=PLPPR4;refGene_tx=XM_019835926.2
C1 85207840 85209166
refGene_gn=PLPPR4;refGene_tx=NM_001128199.1;refGene_gn=PLPPR4;refGene_tx=XM_019835926.2
C1 85494195 85494240 refGene_gn=PALMD;refGene_tx=XM_003990357.5
C1 85511514 85511595 refGene_gn=PALMD;refGene_tx=XM_003990357.5
C1 85517049 85517174 refGene_gn=PALMD;refGene_tx=XM_003990357.5
C1 85534603 85534718 refGene_gn=PALMD;refGene_tx=XM_003990357.5
C1 85534858 85534892 refGene_gn=PALMD;refGene_tx=XM_003990357.5
C1 85535010 85535124 refGene_gn=PALMD;refGene_tx=XM_003990357.5
C1 85536726 85537824 refGene_gn=PALMD;refGene_tx=XM_003990357.5
C1 85542456 85542500 refGene_gn=PALMD;refGene_tx=XM_003990357.5
C1 85556101 85556194
refGene_gn=FRRS1;refGene_tx=XM_019837487.1;refGene_gn=FRRS1;refGene_tx=XM_019837488.1;refGene_gn=FRRS1;refGene_tx=XM_023258935.1;refGene_gn=FRRS1;refGene_tx=XM_023258936.1
C1 85556275 85556337
refGene_gn=FRRS1;refGene_tx=XM_019837487.1;refGene_gn=FRRS1;refGene_tx=XM_019837488.1;refGene_gn=FRRS1;refGene_tx=XM_023258935.1;refGene_gn=FRRS1;refGene_tx=XM_023258936.1
C1 85557684 85557828

refGene_gn=FRRS1;refGene_tx=XM_019837487.1;refGene_gn=FRRS1;refGene_tx=XM_019837488.1;refGene_gn=FRRS1;refGene_tx=XM_023258935.1;refGene_gn=FRRS1;refGene_tx=XM_023258936.1
C1 85558636 85558695
refGene_gn=FRRS1;refGene_tx=XM_019837487.1;refGene_gn=FRRS1;refGene_tx=XM_019837488.1;refGene_gn=FRRS1;refGene_tx=XM_023258935.1;refGene_gn=FRRS1;refGene_tx=XM_023258936.1
C1 85559171 85559269
refGene_gn=FRRS1;refGene_tx=XM_019837487.1;refGene_gn=FRRS1;refGene_tx=XM_019837488.1;refGene_gn=FRRS1;refGene_tx=XM_023258935.1;refGene_gn=FRRS1;refGene_tx=XM_023258936.1
C1 85561768 85561855
refGene_gn=FRRS1;refGene_tx=XM_019837487.1;refGene_gn=FRRS1;refGene_tx=XM_019837488.1;refGene_gn=FRRS1;refGene_tx=XM_023258935.1;refGene_gn=FRRS1;refGene_tx=XM_023258936.1
C1 85563459 85563572
refGene_gn=FRRS1;refGene_tx=XM_019837487.1;refGene_gn=FRRS1;refGene_tx=XM_019837488.1;refGene_gn=FRRS1;refGene_tx=XM_023258935.1;refGene_gn=FRRS1;refGene_tx=XM_023258936.1
C1 85565149 85565263
refGene_gn=FRRS1;refGene_tx=XM_019837487.1;refGene_gn=FRRS1;refGene_tx=XM_019837488.1;refGene_gn=FRRS1;refGene_tx=XM_023258935.1;refGene_gn=FRRS1;refGene_tx=XM_023258936.1
C1 85573730 85573878
refGene_gn=FRRS1;refGene_tx=XM_019837487.1;refGene_gn=FRRS1;refGene_tx=XM_019837488.1;refGene_gn=FRRS1;refGene_tx=XM_023258935.1;refGene_gn=FRRS1;refGene_tx=XM_023258936.1
C1 85575198 85575297
refGene_gn=FRRS1;refGene_tx=XM_019837487.1;refGene_gn=FRRS1;refGene_tx=XM_019837488.1;refGene_gn=FRRS1;refGene_tx=XM_023258935.1;refGene_gn=FRRS1;refGene_tx=XM_023258936.1
C1 85582683 85582866
refGene_gn=FRRS1;refGene_tx=XM_019837487.1;refGene_gn=FRRS1;refGene_tx=XM_019837488.1;refGene_gn=FRRS1;refGene_tx=XM_023258935.1;refGene_gn=FRRS1;refGene_tx=XM_023258936.1
C1 85585801 85585949
refGene_gn=FRRS1;refGene_tx=XM_019837487.1;refGene_gn=FRRS1;refGene_tx=XM_019837488.1;refGene_gn=FRRS1;refGene_tx=XM_023258935.1;refGene_gn=FRRS1;refGene_tx=XM_023258936.1
C1 85587262 85587357
refGene_gn=FRRS1;refGene_tx=XM_019837487.1;refGene_gn=FRRS1;refGene_tx=XM_019837488.1;refGene_gn=FRRS1;refGene_tx=XM_023258935.1;refGene_gn=FRRS1;refGene_tx=XM_023258936.1
C1 85588443 85588580
refGene_gn=FRRS1;refGene_tx=XM_019837487.1;refGene_gn=FRRS1;refGene_tx=XM_019837488.1;refGene_gn=FRRS1;refGene_tx=XM_023258935.1;refGene_gn=FRRS1;refGene_tx=XM_023258936.1
C1 85589656 85589852
refGene_gn=FRRS1;refGene_tx=XM_019837487.1;refGene_gn=FRRS1;refGene_tx=XM_019837488.1;refGene_gn=FRRS1;refGene_tx=XM_023258935.1;refGene_gn=

FRRS1;refGene_tx=XM_023258936.1
C1 85659895 85659977
refGene_gn=AGL;refGene_tx=XM_003990359.5;refGene_gn=AGL;refGene_tx=XM_011284840.2;refGene_gn=AGL;refGene_tx=XM_019837485.1
C1 85675603 85675814
refGene_gn=AGL;refGene_tx=XM_003990359.5;refGene_gn=AGL;refGene_tx=XM_011284840.2;refGene_gn=AGL;refGene_tx=XM_019837485.1
C1 85676444 85676611
refGene_gn=AGL;refGene_tx=XM_003990359.5;refGene_gn=AGL;refGene_tx=XM_011284840.2;refGene_gn=AGL;refGene_tx=XM_019837485.1
C1 85677452 85677656
refGene_gn=AGL;refGene_tx=XM_003990359.5;refGene_gn=AGL;refGene_tx=XM_011284840.2;refGene_gn=AGL;refGene_tx=XM_019837485.1
C1 85686344 85686526
refGene_gn=AGL;refGene_tx=XM_003990359.5;refGene_gn=AGL;refGene_tx=XM_011284840.2;refGene_gn=AGL;refGene_tx=XM_019837485.1
C1 85686888 85687000
refGene_gn=AGL;refGene_tx=XM_003990359.5;refGene_gn=AGL;refGene_tx=XM_011284840.2;refGene_gn=AGL;refGene_tx=XM_019837485.1
C1 85691931 85692058
refGene_gn=AGL;refGene_tx=XM_003990359.5;refGene_gn=AGL;refGene_tx=XM_011284840.2;refGene_gn=AGL;refGene_tx=XM_019837485.1
C1 85692402 85692505
refGene_gn=AGL;refGene_tx=XM_003990359.5;refGene_gn=AGL;refGene_tx=XM_011284840.2;refGene_gn=AGL;refGene_tx=XM_019837485.1
C1 85692600 85692698
refGene_gn=AGL;refGene_tx=XM_003990359.5;refGene_gn=AGL;refGene_tx=XM_011284840.2;refGene_gn=AGL;refGene_tx=XM_019837485.1
C1 85693590 85693730
refGene_gn=AGL;refGene_tx=XM_003990359.5;refGene_gn=AGL;refGene_tx=XM_011284840.2;refGene_gn=AGL;refGene_tx=XM_019837485.1
C1 85694777 85694965
refGene_gn=AGL;refGene_tx=XM_003990359.5;refGene_gn=AGL;refGene_tx=XM_011284840.2;refGene_gn=AGL;refGene_tx=XM_019837485.1
C1 85696549 85696673
refGene_gn=AGL;refGene_tx=XM_003990359.5;refGene_gn=AGL;refGene_tx=XM_011284840.2;refGene_gn=AGL;refGene_tx=XM_019837485.1
C1 85697485 85697649
refGene_gn=AGL;refGene_tx=XM_003990359.5;refGene_gn=AGL;refGene_tx=XM_011284840.2;refGene_gn=AGL;refGene_tx=XM_019837485.1
C1 85698222 85698324
refGene_gn=AGL;refGene_tx=XM_003990359.5;refGene_gn=AGL;refGene_tx=XM_011284840.2;refGene_gn=AGL;refGene_tx=XM_019837485.1
C1 85698443 85698599
refGene_gn=AGL;refGene_tx=XM_003990359.5;refGene_gn=AGL;refGene_tx=XM_011284840.2;refGene_gn=AGL;refGene_tx=XM_019837485.1
C1 85698696 85698847
refGene_gn=AGL;refGene_tx=XM_003990359.5;refGene_gn=AGL;refGene_tx=XM_011284840.2;refGene_gn=AGL;refGene_tx=XM_019837485.1
C1 85701485 85701610

refGene_gn=AGL;refGene_tx=XM_003990359.5;refGene_gn=AGL;refGene_tx=XM_011284840.2;refGene_gn=AGL;refGene_tx=XM_019837485.1
C1 85701699 85701812
refGene_gn=AGL;refGene_tx=XM_003990359.5;refGene_gn=AGL;refGene_tx=XM_011284840.2;refGene_gn=AGL;refGene_tx=XM_019837485.1
C1 85702148 85702283
refGene_gn=AGL;refGene_tx=XM_003990359.5;refGene_gn=AGL;refGene_tx=XM_011284840.2;refGene_gn=AGL;refGene_tx=XM_019837485.1
C1 85705534 85705665
refGene_gn=AGL;refGene_tx=XM_003990359.5;refGene_gn=AGL;refGene_tx=XM_011284840.2;refGene_gn=AGL;refGene_tx=XM_019837485.1
C1 85709549 85709686
refGene_gn=AGL;refGene_tx=XM_003990359.5;refGene_gn=AGL;refGene_tx=XM_011284840.2;refGene_gn=AGL;refGene_tx=XM_019837485.1
C1 85709934 85710068
refGene_gn=AGL;refGene_tx=XM_003990359.5;refGene_gn=AGL;refGene_tx=XM_011284840.2;refGene_gn=AGL;refGene_tx=XM_019837485.1
C1 85711354 85711530
refGene_gn=AGL;refGene_tx=XM_003990359.5;refGene_gn=AGL;refGene_tx=XM_011284840.2;refGene_gn=AGL;refGene_tx=XM_019837485.1
C1 85715557 85715660
refGene_gn=AGL;refGene_tx=XM_003990359.5;refGene_gn=AGL;refGene_tx=XM_011284840.2;refGene_gn=AGL;refGene_tx=XM_019837485.1
C1 85718919 85719145
refGene_gn=AGL;refGene_tx=XM_003990359.5;refGene_gn=AGL;refGene_tx=XM_011284840.2;refGene_gn=AGL;refGene_tx=XM_019837485.1
C1 85720378 85720490
refGene_gn=AGL;refGene_tx=XM_003990359.5;refGene_gn=AGL;refGene_tx=XM_011284840.2;refGene_gn=AGL;refGene_tx=XM_019837485.1
C1 85727657 85727793
refGene_gn=AGL;refGene_tx=XM_003990359.5;refGene_gn=AGL;refGene_tx=XM_011284840.2;refGene_gn=AGL;refGene_tx=XM_019837485.1
C1 85729033 85729146
refGene_gn=AGL;refGene_tx=XM_003990359.5;refGene_gn=AGL;refGene_tx=XM_011284840.2;refGene_gn=AGL;refGene_tx=XM_019837485.1
C1 85731245 85731457
refGene_gn=AGL;refGene_tx=XM_003990359.5;refGene_gn=AGL;refGene_tx=XM_011284840.2;refGene_gn=AGL;refGene_tx=XM_019837485.1
C1 85732652 85732750
refGene_gn=AGL;refGene_tx=XM_003990359.5;refGene_gn=AGL;refGene_tx=XM_011284840.2;refGene_gn=AGL;refGene_tx=XM_019837485.1
C1 85735512 85735600
refGene_gn=AGL;refGene_tx=XM_003990359.5;refGene_gn=AGL;refGene_tx=XM_011284840.2;refGene_gn=AGL;refGene_tx=XM_019837485.1
C1 85735715 85735849
refGene_gn=AGL;refGene_tx=XM_003990359.5;refGene_gn=AGL;refGene_tx=XM_011284840.2;refGene_gn=AGL;refGene_tx=XM_019837485.1
C1 85741261 85741379
refGene_gn=AGL;refGene_tx=XM_003990359.5;refGene_gn=AGL;refGene_tx=XM_011284840.2;refGene_gn=AGL;refGene_tx=XM_019837485.1

C1 85765701 85765719 refGene_gn=SLC35A3;refGene_tx=XM_019837500.2
C1 85794719 85794924
refGene_gn=SLC35A3;refGene_tx=XM_003990360.4;refGene_gn=SLC35A3;refGene_tx=XM_019837500.2;refGene_gn=SLC35A3;refGene_tx=XM_019837501.2;refGene_gn=SLC35A3;refGene_tx=XM_019837502.2
C1 85801240 85801395
refGene_gn=SLC35A3;refGene_tx=XM_003990360.4;refGene_gn=SLC35A3;refGene_tx=XM_019837500.2;refGene_gn=SLC35A3;refGene_tx=XM_019837501.2;refGene_gn=SLC35A3;refGene_tx=XM_019837502.2
C1 85801472 85801595
refGene_gn=SLC35A3;refGene_tx=XM_003990360.4;refGene_gn=SLC35A3;refGene_tx=XM_019837500.2;refGene_gn=SLC35A3;refGene_tx=XM_019837501.2;refGene_gn=SLC35A3;refGene_tx=XM_019837502.2
C1 85805536 85805708
refGene_gn=SLC35A3;refGene_tx=XM_003990360.4;refGene_gn=SLC35A3;refGene_tx=XM_019837500.2;refGene_gn=SLC35A3;refGene_tx=XM_019837501.2;refGene_gn=SLC35A3;refGene_tx=XM_019837502.2
C1 85808637 85808756
refGene_gn=SLC35A3;refGene_tx=XM_003990360.4;refGene_gn=SLC35A3;refGene_tx=XM_019837500.2;refGene_gn=SLC35A3;refGene_tx=XM_019837501.2;refGene_gn=SLC35A3;refGene_tx=XM_019837502.2
C1 85811185 85811319
refGene_gn=SLC35A3;refGene_tx=XM_003990360.4;refGene_gn=SLC35A3;refGene_tx=XM_019837500.2;refGene_gn=SLC35A3;refGene_tx=XM_019837501.2;refGene_gn=SLC35A3;refGene_tx=XM_019837502.2
C1 85816381 85816472
refGene_gn=SLC35A3;refGene_tx=XM_003990360.4;refGene_gn=SLC35A3;refGene_tx=XM_019837500.2;refGene_gn=SLC35A3;refGene_tx=XM_019837501.2;refGene_gn=SLC35A3;refGene_tx=XM_019837502.2
C1 85832347 85832431 refGene_gn=MFSD14A;refGene_tx=XM_011284838.3
C1 85848147 85848243 refGene_gn=MFSD14A;refGene_tx=XM_011284838.3
C1 85856233 85856296 refGene_gn=MFSD14A;refGene_tx=XM_011284838.3
C1 85857050 85857178 refGene_gn=MFSD14A;refGene_tx=XM_011284838.3
C1 85859177 85859292 refGene_gn=MFSD14A;refGene_tx=XM_011284838.3
C1 85863575 85863794 refGene_gn=MFSD14A;refGene_tx=XM_011284838.3
C1 85864068 85864182 refGene_gn=MFSD14A;refGene_tx=XM_011284838.3
C1 85864942 85865014 refGene_gn=MFSD14A;refGene_tx=XM_011284838.3
C1 85871379 85871492 refGene_gn=MFSD14A;refGene_tx=XM_011284838.3
C1 85872607 85872705 refGene_gn=MFSD14A;refGene_tx=XM_011284838.3
C1 85874066 85874230 refGene_gn=MFSD14A;refGene_tx=XM_011284838.3
C1 85875275 85875482 refGene_gn=MFSD14A;refGene_tx=XM_011284838.3
C1 85878739 85878846 refGene_gn=SASS6;refGene_tx=XM_003990361.5
C1 85878945 85879040 refGene_gn=SASS6;refGene_tx=XM_003990361.5
C1 85881482 85881580 refGene_gn=SASS6;refGene_tx=XM_003990361.5
C1 85900731 85900860 refGene_gn=SASS6;refGene_tx=XM_003990361.5
C1 85903147 85903284 refGene_gn=SASS6;refGene_tx=XM_003990361.5
C1 85904214 85904296 refGene_gn=SASS6;refGene_tx=XM_003990361.5
C1 85904911 85905091 refGene_gn=SASS6;refGene_tx=XM_003990361.5
C1 85905166 85905256 refGene_gn=SASS6;refGene_tx=XM_003990361.5
C1 85905362 85905557 refGene_gn=SASS6;refGene_tx=XM_003990361.5

C1 85908051 85908243 refGene_gn=SASS6;refGene_tx=XM_003990361.5
C1 85919752 85919872 refGene_gn=SASS6;refGene_tx=XM_003990361.5
C1 85920737 85920803 refGene_gn=SASS6;refGene_tx=XM_003990361.5
C1 85921490 85921643 refGene_gn=SASS6;refGene_tx=XM_003990361.5
C1 85921644 85921663 refGene_gn=SASS6;refGene_tx=XM_003990361.5
C1 85922626 85922731 refGene_gn=SASS6;refGene_tx=XM_003990361.5
C1 85923232 85923312 refGene_gn=SASS6;refGene_tx=XM_003990361.5
C1 85925123 85925184 refGene_gn=SASS6;refGene_tx=XM_003990361.5
C1 85935333 85935398 refGene_gn=SASS6;refGene_tx=XM_003990361.5
C1 85935738 85935885
refGene_gn=TRMT13;refGene_tx=XM_003990362.4;refGene_gn=TRMT13;refGene_
tx=XM_023258937.1
C1 85938325 85938372
refGene_gn=TRMT13;refGene_tx=XM_003990362.4;refGene_gn=TRMT13;refGene_
tx=XM_023258937.1
C1 85938460 85938527
refGene_gn=TRMT13;refGene_tx=XM_003990362.4;refGene_gn=TRMT13;refGene_
tx=XM_023258937.1
C1 85940459 85940522
refGene_gn=TRMT13;refGene_tx=XM_003990362.4;refGene_gn=TRMT13;refGene_
tx=XM_023258937.1
C1 85940714 85940784
refGene_gn=TRMT13;refGene_tx=XM_003990362.4;refGene_gn=TRMT13;refGene_
tx=XM_023258937.1
C1 85940939 85941046 refGene_gn=TRMT13;refGene_tx=XM_003990362.4
C1 85941394 85941562
refGene_gn=TRMT13;refGene_tx=XM_003990362.4;refGene_gn=TRMT13;refGene_
tx=XM_023258937.1
C1 85943914 85943987
refGene_gn=TRMT13;refGene_tx=XM_003990362.4;refGene_gn=TRMT13;refGene_
tx=XM_023258937.1
C1 85949187 85949262
refGene_gn=TRMT13;refGene_tx=XM_003990362.4;refGene_gn=TRMT13;refGene_
tx=XM_023258937.1
C1 85952026 85952459
refGene_gn=TRMT13;refGene_tx=XM_003990362.4;refGene_gn=TRMT13;refGene_
tx=XM_023258937.1
C1 85952762 85952958
refGene_gn=TRMT13;refGene_tx=XM_003990362.4;refGene_gn=TRMT13;refGene_
tx=XM_023258937.1
C1 85953181 85953237
refGene_gn=LRRC39;refGene_tx=XM_003990363.4;refGene_gn=LRRC39;refGene_
tx=XM_019837506.2
C1 85956266 85956406
refGene_gn=LRRC39;refGene_tx=XM_003990363.4;refGene_gn=LRRC39;refGene_
tx=XM_019837506.2
C1 85957637 85957790
refGene_gn=LRRC39;refGene_tx=XM_003990363.4;refGene_gn=LRRC39;refGene_
tx=XM_019837506.2
C1 85958505 85958651

refGene_gn=LRRC39;refGene_tx=XM_003990363.4;refGene_gn=LRRC39;refGene_tx=XM_019837506.2
C1 85961405 85961542
refGene_gn=LRRC39;refGene_tx=XM_003990363.4;refGene_gn=LRRC39;refGene_tx=XM_019837506.2
C1 85962114 85962271
refGene_gn=LRRC39;refGene_tx=XM_003990363.4;refGene_gn=LRRC39;refGene_tx=XM_019837506.2
C1 85963807 85963913
refGene_gn=LRRC39;refGene_tx=XM_003990363.4;refGene_gn=LRRC39;refGene_tx=XM_019837506.2
C1 85969986 85970099
refGene_gn=LRRC39;refGene_tx=XM_003990363.4;refGene_gn=LRRC39;refGene_tx=XM_019837506.2
C1 85993377 85993545 refGene_gn=DBT;refGene_tx=XM_003990364.5
C1 85998138 85998210 refGene_gn=DBT;refGene_tx=XM_003990364.5
C1 85998368 85998560 refGene_gn=DBT;refGene_tx=XM_003990364.5
C1 85999794 85999872 refGene_gn=DBT;refGene_tx=XM_003990364.5
C1 86003788 86003955 refGene_gn=DBT;refGene_tx=XM_003990364.5
C1 86004592 86004809 refGene_gn=DBT;refGene_tx=XM_003990364.5
C1 86009805 86009927 refGene_gn=DBT;refGene_tx=XM_003990364.5
C1 86019668 86019850 refGene_gn=DBT;refGene_tx=XM_003990364.5
C1 86036459 86036535 refGene_gn=DBT;refGene_tx=XM_003990364.5
C1 86040538 86040662 refGene_gn=DBT;refGene_tx=XM_003990364.5
C1 86046063 86046114 refGene_gn=DBT;refGene_tx=XM_003990364.5
C1 86061857 86061902 refGene_gn=RTCA;refGene_tx=XM_023258938.1
C1 86062010 86062111 refGene_gn=RTCA;refGene_tx=XM_023258938.1
C1 86063879 86064023
refGene_gn=RTCA;refGene_tx=XM_023258938.1;refGene_gn=RTCA;refGene_tx=XM_023258939.1
C1 86067536 86067660
refGene_gn=RTCA;refGene_tx=XM_023258938.1;refGene_gn=RTCA;refGene_tx=XM_023258939.1
C1 86070145 86070204
refGene_gn=RTCA;refGene_tx=XM_023258938.1;refGene_gn=RTCA;refGene_tx=XM_023258939.1
C1 86071735 86071877
refGene_gn=RTCA;refGene_tx=XM_023258938.1;refGene_gn=RTCA;refGene_tx=XM_023258939.1
C1 86072485 86072610
refGene_gn=RTCA;refGene_tx=XM_023258938.1;refGene_gn=RTCA;refGene_tx=XM_023258939.1
C1 86074891 86074950
refGene_gn=RTCA;refGene_tx=XM_023258938.1;refGene_gn=RTCA;refGene_tx=XM_023258939.1
C1 86079693 86079788
refGene_gn=RTCA;refGene_tx=XM_023258938.1;refGene_gn=RTCA;refGene_tx=XM_023258939.1
C1 86082320 86082425
refGene_gn=RTCA;refGene_tx=XM_023258938.1;refGene_gn=RTCA;refGene_tx=X

M_023258939.1
C1 86085349 86085451
refGene_gn=RTCA;refGene_tx=XM_023258938.1;refGene_gn=RTCA;refGene_tx=XM_023258939.1
C1 86125805 86125899 refGene_gn=CDC14A;refGene_tx=XM_023257836.1
C1 86127835 86127926 refGene_gn=CDC14A;refGene_tx=XM_023257836.1
C1 86148055 86148131 refGene_gn=CDC14A;refGene_tx=XM_023257836.1
C1 86154191 86154236 refGene_gn=CDC14A;refGene_tx=XM_023257836.1
C1 86161844 86161948 refGene_gn=CDC14A;refGene_tx=XM_023257836.1
C1 86209041 86209225 refGene_gn=CDC14A;refGene_tx=XM_023257836.1
C1 86212484 86212547 refGene_gn=CDC14A;refGene_tx=XM_023257836.1
C1 86227946 86228034 refGene_gn=CDC14A;refGene_tx=XM_023257836.1
C1 86235837 86236068 refGene_gn=CDC14A;refGene_tx=XM_023257836.1
C1 86245072 86245211 refGene_gn=CDC14A;refGene_tx=XM_023257836.1
C1 86275980 86276140 refGene_gn=CDC14A;refGene_tx=XM_023257836.1
C1 86293066 86293182 refGene_gn=CDC14A;refGene_tx=XM_023257836.1
C1 86295469 86295517 refGene_gn=CDC14A;refGene_tx=XM_023257836.1
C1 86297879 86298002 refGene_gn=CDC14A;refGene_tx=XM_023257836.1
C1 86298694 86299148 refGene_gn=CDC14A;refGene_tx=XM_023257836.1
C1 86340882 86342037 refGene_gn=GPR88;refGene_tx=XM_019839093.2
C1 86473205 86473296
refGene_gn=VCAM1;refGene_tx=XM_003990367.4;refGene_gn=VCAM1;refGene_tx=XM_011284836.3
C1 86473825 86474101
refGene_gn=VCAM1;refGene_tx=XM_003990367.4;refGene_gn=VCAM1;refGene_tx=XM_011284836.3
C1 86476550 86476871
refGene_gn=VCAM1;refGene_tx=XM_003990367.4;refGene_gn=VCAM1;refGene_tx=XM_011284836.3
C1 86478099 86478366
refGene_gn=VCAM1;refGene_tx=XM_003990367.4;refGene_gn=VCAM1;refGene_tx=XM_011284836.3
C1 86481721 86481997
refGene_gn=VCAM1;refGene_tx=XM_003990367.4;refGene_gn=VCAM1;refGene_tx=XM_011284836.3
C1 86483848 86484169
refGene_gn=VCAM1;refGene_tx=XM_003990367.4;refGene_gn=VCAM1;refGene_tx=XM_011284836.3
C1 86485042 86485309
refGene_gn=VCAM1;refGene_tx=XM_003990367.4;refGene_gn=VCAM1;refGene_tx=XM_011284836.3
C1 86486813 86487080 refGene_gn=VCAM1;refGene_tx=XM_003990367.4
C1 86490317 86490478
refGene_gn=VCAM1;refGene_tx=XM_003990367.4;refGene_gn=VCAM1;refGene_tx=XM_011284836.3
C1 86595741 86596230
refGene_gn=EXTL2;refGene_tx=XM_006934945.4;refGene_gn=EXTL2;refGene_tx=XM_006934946.4;refGene_gn=EXTL2;refGene_tx=XM_006934947.4;refGene_gn=EXTL2;refGene_tx=XM_011284833.3
C1 86599129 86599200

refGene_gn=EXTL2;refGene_tx=XM_006934945.4;refGene_gn=EXTL2;refGene_tx=XM_006934946.4;refGene_gn=EXTL2;refGene_tx=XM_006934947.4;refGene_gn=EXTL2;refGene_tx=XM_011284833.3
C1 86599844 86600272 refGene_gn=EXTL2;refGene_tx=XM_006934945.4;refGene_gn=EXTL2;refGene_tx=XM_006934946.4;refGene_gn=EXTL2;refGene_tx=XM_006934947.4;refGene_gn=EXTL2;refGene_tx=XM_011284833.3
C1 86617830 86617910 refGene_gn=SLC30A7;refGene_tx=XM_003990369.4
C1 86618131 86618233 refGene_gn=SLC30A7;refGene_tx=XM_003990369.4
C1 86631195 86631309 refGene_gn=SLC30A7;refGene_tx=XM_003990369.4
C1 86635589 86635677 refGene_gn=SLC30A7;refGene_tx=XM_003990369.4
C1 86636331 86636458 refGene_gn=SLC30A7;refGene_tx=XM_003990369.4
C1 86645225 86645369 refGene_gn=SLC30A7;refGene_tx=XM_003990369.4
C1 86650021 86650072 refGene_gn=SLC30A7;refGene_tx=XM_003990369.4
C1 86653998 86654134 refGene_gn=SLC30A7;refGene_tx=XM_003990369.4
C1 86683037 86683128 refGene_gn=SLC30A7;refGene_tx=XM_003990369.4
C1 86689791 86689941 refGene_gn=SLC30A7;refGene_tx=XM_003990369.4
C1 86698283 86698331 refGene_gn=SLC30A7;refGene_tx=XM_003990369.4
C1 86716134 86716340 refGene_gn=DPH5;refGene_tx=XM_019837520.2;refGene_gn=DPH5;refGene_tx=XM_019837521.2
C1 86719005 86719109 refGene_gn=DPH5;refGene_tx=XM_019837520.2;refGene_gn=DPH5;refGene_tx=XM_019837521.2
C1 86720556 86720596 refGene_gn=DPH5;refGene_tx=XM_019837520.2;refGene_gn=DPH5;refGene_tx=XM_019837521.2
C1 86723361 86723482 refGene_gn=DPH5;refGene_tx=XM_019837520.2;refGene_gn=DPH5;refGene_tx=XM_019837521.2
C1 86734885 86734994 refGene_gn=DPH5;refGene_tx=XM_019837520.2;refGene_gn=DPH5;refGene_tx=XM_019837521.2
C1 86742565 86742690 refGene_gn=DPH5;refGene_tx=XM_019837520.2;refGene_gn=DPH5;refGene_tx=XM_019837521.2
C1 86750780 86750915 refGene_gn=DPH5;refGene_tx=XM_019837520.2;refGene_gn=DPH5;refGene_tx=XM_019837521.2
C1 86963821 86964967 refGene_gn=S1PR1;refGene_tx=XM_003990374.5;refGene_gn=S1PR1;refGene_tx=XM_006934949.4;refGene_gn=S1PR1;refGene_tx=XM_006934950.4
C1 87425832 87426510 refGene_gn=OLFM3;refGene_tx=XM_003990375.5;refGene_gn=OLFM3;refGene_tx=XM_011284827.3
C1 87427593 87427700 refGene_gn=OLFM3;refGene_tx=XM_003990375.5;refGene_gn=OLFM3;refGene_tx=XM_011284827.3
C1 87446725 87446945 refGene_gn=OLFM3;refGene_tx=XM_003990375.5;refGene_gn=OLFM3;refGene_tx

=XM_011284827.3
C1	87451983	87452139
refGene_gn=OLFM3;refGene_tx=XM_003990375.5;refGene_gn=OLFM3;refGene_tx=XM_011284827.3
C1	87459062	87459209
refGene_gn=OLFM3;refGene_tx=XM_003990375.5;refGene_gn=OLFM3;refGene_tx=XM_011284827.3
C1	87469216	87469345	refGene_gn=OLFM3;refGene_tx=XM_003990375.5
C1	87613788	87613857	refGene_gn=OLFM3;refGene_tx=XM_011284827.3
C1	88300558	88300705
refGene_gn=COL11A1;refGene_tx=XM_023258940.1;refGene_gn=COL11A1;refGene_tx=XM_023258941.1;refGene_gn=COL11A1;refGene_tx=XM_023258942.1;refGene_gn=COL11A1;refGene_tx=XM_023258943.1;refGene_gn=COL11A1;refGene_tx=XM_023258944.1;refGene_gn=COL11A1;refGene_tx=XM_023258945.1;refGene_gn=COL11A1;refGene_tx=XM_023258946.1;refGene_gn=COL11A1;refGene_tx=XM_023258947.1;refGene_gn=COL11A1;refGene_tx=XM_023258948.1
C1	88301932	88302166
refGene_gn=COL11A1;refGene_tx=XM_023258940.1;refGene_gn=COL11A1;refGene_tx=XM_023258941.1;refGene_gn=COL11A1;refGene_tx=XM_023258942.1;refGene_gn=COL11A1;refGene_tx=XM_023258943.1;refGene_gn=COL11A1;refGene_tx=XM_023258944.1;refGene_gn=COL11A1;refGene_tx=XM_023258945.1;refGene_gn=COL11A1;refGene_tx=XM_023258946.1;refGene_gn=COL11A1;refGene_tx=XM_023258947.1;refGene_gn=COL11A1;refGene_tx=XM_023258948.1
C1	88304215	88304284
refGene_gn=COL11A1;refGene_tx=XM_023258940.1;refGene_gn=COL11A1;refGene_tx=XM_023258941.1;refGene_gn=COL11A1;refGene_tx=XM_023258942.1;refGene_gn=COL11A1;refGene_tx=XM_023258943.1;refGene_gn=COL11A1;refGene_tx=XM_023258944.1;refGene_gn=COL11A1;refGene_tx=XM_023258945.1;refGene_gn=COL11A1;refGene_tx=XM_023258946.1;refGene_gn=COL11A1;refGene_tx=XM_023258947.1;refGene_gn=COL11A1;refGene_tx=XM_023258948.1
C1	88305682	88305795
refGene_gn=COL11A1;refGene_tx=XM_023258940.1;refGene_gn=COL11A1;refGene_tx=XM_023258941.1;refGene_gn=COL11A1;refGene_tx=XM_023258942.1;refGene_gn=COL11A1;refGene_tx=XM_023258943.1;refGene_gn=COL11A1;refGene_tx=XM_023258944.1;refGene_gn=COL11A1;refGene_tx=XM_023258945.1;refGene_gn=COL11A1;refGene_tx=XM_023258946.1;refGene_gn=COL11A1;refGene_tx=XM_023258947.1;refGene_gn=COL11A1;refGene_tx=XM_023258948.1
C1	88308526	88308776
refGene_gn=COL11A1;refGene_tx=XM_023258940.1;refGene_gn=COL11A1;refGene_tx=XM_023258941.1;refGene_gn=COL11A1;refGene_tx=XM_023258942.1;refGene_gn=COL11A1;refGene_tx=XM_023258943.1;refGene_gn=COL11A1;refGene_tx=XM_023258944.1;refGene_gn=COL11A1;refGene_tx=XM_023258945.1;refGene_gn=COL11A1;refGene_tx=XM_023258946.1;refGene_gn=COL11A1;refGene_tx=XM_023258947.1;refGene_gn=COL11A1;refGene_tx=XM_023258948.1
C1	88310556	88310610
refGene_gn=COL11A1;refGene_tx=XM_023258940.1;refGene_gn=COL11A1;refGene_tx=XM_023258941.1;refGene_gn=COL11A1;refGene_tx=XM_023258942.1;refGene_gn=COL11A1;refGene_tx=XM_023258943.1;refGene_gn=COL11A1;refGene_tx=XM_023258944.1;refGene_gn=COL11A1;refGene_tx=XM_023258945.1;refGene_gn=COL11A1;refGene_tx=XM_023258946.1;refGene_gn=COL11A1;refGene_tx=XM_02

3258947.1;refGene_gn=COL11A1;refGene_tx=XM_023258948.1
C1 88310702 88310738
refGene_gn=COL11A1;refGene_tx=XM_023258940.1;refGene_gn=COL11A1;refGene_tx=XM_023258941.1;refGene_gn=COL11A1;refGene_tx=XM_023258942.1;refGene_gn=COL11A1;refGene_tx=XM_023258943.1;refGene_gn=COL11A1;refGene_tx=XM_023258944.1;refGene_gn=COL11A1;refGene_tx=XM_023258945.1;refGene_gn=COL11A1;refGene_tx=XM_023258946.1;refGene_gn=COL11A1;refGene_tx=XM_023258947.1;refGene_gn=COL11A1;refGene_tx=XM_023258948.1
C1 88310839 88310893
refGene_gn=COL11A1;refGene_tx=XM_023258940.1;refGene_gn=COL11A1;refGene_tx=XM_023258941.1;refGene_gn=COL11A1;refGene_tx=XM_023258942.1;refGene_gn=COL11A1;refGene_tx=XM_023258943.1;refGene_gn=COL11A1;refGene_tx=XM_023258944.1;refGene_gn=COL11A1;refGene_tx=XM_023258945.1;refGene_gn=COL11A1;refGene_tx=XM_023258946.1;refGene_gn=COL11A1;refGene_tx=XM_023258947.1;refGene_gn=COL11A1;refGene_tx=XM_023258948.1
C1 88311370 88311478
refGene_gn=COL11A1;refGene_tx=XM_023258940.1;refGene_gn=COL11A1;refGene_tx=XM_023258941.1;refGene_gn=COL11A1;refGene_tx=XM_023258942.1;refGene_gn=COL11A1;refGene_tx=XM_023258943.1;refGene_gn=COL11A1;refGene_tx=XM_023258944.1;refGene_gn=COL11A1;refGene_tx=XM_023258945.1;refGene_gn=COL11A1;refGene_tx=XM_023258946.1;refGene_gn=COL11A1;refGene_tx=XM_023258947.1;refGene_gn=COL11A1;refGene_tx=XM_023258948.1
C1 88312366 88312420
refGene_gn=COL11A1;refGene_tx=XM_023258940.1;refGene_gn=COL11A1;refGene_tx=XM_023258941.1;refGene_gn=COL11A1;refGene_tx=XM_023258942.1;refGene_gn=COL11A1;refGene_tx=XM_023258943.1;refGene_gn=COL11A1;refGene_tx=XM_023258944.1;refGene_gn=COL11A1;refGene_tx=XM_023258945.1;refGene_gn=COL11A1;refGene_tx=XM_023258946.1;refGene_gn=COL11A1;refGene_tx=XM_023258947.1;refGene_gn=COL11A1;refGene_tx=XM_023258948.1
C1 88321715 88321769
refGene_gn=COL11A1;refGene_tx=XM_023258940.1;refGene_gn=COL11A1;refGene_tx=XM_023258941.1;refGene_gn=COL11A1;refGene_tx=XM_023258942.1;refGene_gn=COL11A1;refGene_tx=XM_023258943.1;refGene_gn=COL11A1;refGene_tx=XM_023258944.1;refGene_gn=COL11A1;refGene_tx=XM_023258945.1;refGene_gn=COL11A1;refGene_tx=XM_023258946.1;refGene_gn=COL11A1;refGene_tx=XM_023258947.1;refGene_gn=COL11A1;refGene_tx=XM_023258948.1
C1 88322228 88322336
refGene_gn=COL11A1;refGene_tx=XM_023258940.1;refGene_gn=COL11A1;refGene_tx=XM_023258941.1;refGene_gn=COL11A1;refGene_tx=XM_023258942.1;refGene_gn=COL11A1;refGene_tx=XM_023258943.1;refGene_gn=COL11A1;refGene_tx=XM_023258944.1;refGene_gn=COL11A1;refGene_tx=XM_023258945.1;refGene_gn=COL11A1;refGene_tx=XM_023258946.1;refGene_gn=COL11A1;refGene_tx=XM_023258947.1;refGene_gn=COL11A1;refGene_tx=XM_023258948.1
C1 88322461 88322515
refGene_gn=COL11A1;refGene_tx=XM_023258940.1;refGene_gn=COL11A1;refGene_tx=XM_023258941.1;refGene_gn=COL11A1;refGene_tx=XM_023258942.1;refGene_gn=COL11A1;refGene_tx=XM_023258943.1;refGene_gn=COL11A1;refGene_tx=XM_023258944.1;refGene_gn=COL11A1;refGene_tx=XM_023258945.1;refGene_gn=COL11A1;refGene_tx=XM_023258946.1;refGene_gn=COL11A1;refGene_tx=XM_023258947.1;refGene_gn=COL11A1;refGene_tx=XM_023258948.1

C1 88328466 88328520
refGene_gn=COL11A1;refGene_tx=XM_023258940.1;refGene_gn=COL11A1;refGene_tx=XM_023258941.1;refGene_gn=COL11A1;refGene_tx=XM_023258942.1;refGene_gn=COL11A1;refGene_tx=XM_023258943.1;refGene_gn=COL11A1;refGene_tx=XM_023258944.1;refGene_gn=COL11A1;refGene_tx=XM_023258945.1;refGene_gn=COL11A1;refGene_tx=XM_023258946.1;refGene_gn=COL11A1;refGene_tx=XM_023258947.1;refGene_gn=COL11A1;refGene_tx=XM_023258948.1
C1 88329546 88329600
refGene_gn=COL11A1;refGene_tx=XM_023258940.1;refGene_gn=COL11A1;refGene_tx=XM_023258941.1;refGene_gn=COL11A1;refGene_tx=XM_023258942.1;refGene_gn=COL11A1;refGene_tx=XM_023258943.1;refGene_gn=COL11A1;refGene_tx=XM_023258944.1;refGene_gn=COL11A1;refGene_tx=XM_023258945.1;refGene_gn=COL11A1;refGene_tx=XM_023258946.1;refGene_gn=COL11A1;refGene_tx=XM_023258947.1;refGene_gn=COL11A1;refGene_tx=XM_023258948.1
C1 88330231 88330285
refGene_gn=COL11A1;refGene_tx=XM_023258940.1;refGene_gn=COL11A1;refGene_tx=XM_023258941.1;refGene_gn=COL11A1;refGene_tx=XM_023258942.1;refGene_gn=COL11A1;refGene_tx=XM_023258943.1;refGene_gn=COL11A1;refGene_tx=XM_023258944.1;refGene_gn=COL11A1;refGene_tx=XM_023258945.1;refGene_gn=COL11A1;refGene_tx=XM_023258946.1;refGene_gn=COL11A1;refGene_tx=XM_023258947.1;refGene_gn=COL11A1;refGene_tx=XM_023258948.1
C1 88330579 88330687
refGene_gn=COL11A1;refGene_tx=XM_023258940.1;refGene_gn=COL11A1;refGene_tx=XM_023258941.1;refGene_gn=COL11A1;refGene_tx=XM_023258942.1;refGene_gn=COL11A1;refGene_tx=XM_023258943.1;refGene_gn=COL11A1;refGene_tx=XM_023258944.1;refGene_gn=COL11A1;refGene_tx=XM_023258945.1;refGene_gn=COL11A1;refGene_tx=XM_023258946.1;refGene_gn=COL11A1;refGene_tx=XM_023258947.1;refGene_gn=COL11A1;refGene_tx=XM_023258948.1
C1 88331540 88331594
refGene_gn=COL11A1;refGene_tx=XM_023258940.1;refGene_gn=COL11A1;refGene_tx=XM_023258941.1;refGene_gn=COL11A1;refGene_tx=XM_023258942.1;refGene_gn=COL11A1;refGene_tx=XM_023258943.1;refGene_gn=COL11A1;refGene_tx=XM_023258944.1;refGene_gn=COL11A1;refGene_tx=XM_023258945.1;refGene_gn=COL11A1;refGene_tx=XM_023258946.1;refGene_gn=COL11A1;refGene_tx=XM_023258947.1;refGene_gn=COL11A1;refGene_tx=XM_023258948.1
C1 88336760 88336814
refGene_gn=COL11A1;refGene_tx=XM_023258940.1;refGene_gn=COL11A1;refGene_tx=XM_023258941.1;refGene_gn=COL11A1;refGene_tx=XM_023258942.1;refGene_gn=COL11A1;refGene_tx=XM_023258943.1;refGene_gn=COL11A1;refGene_tx=XM_023258944.1;refGene_gn=COL11A1;refGene_tx=XM_023258945.1;refGene_gn=COL11A1;refGene_tx=XM_023258946.1;refGene_gn=COL11A1;refGene_tx=XM_023258947.1;refGene_gn=COL11A1;refGene_tx=XM_023258948.1
C1 88338420 88338474
refGene_gn=COL11A1;refGene_tx=XM_023258940.1;refGene_gn=COL11A1;refGene_tx=XM_023258941.1;refGene_gn=COL11A1;refGene_tx=XM_023258942.1;refGene_gn=COL11A1;refGene_tx=XM_023258943.1;refGene_gn=COL11A1;refGene_tx=XM_023258944.1;refGene_gn=COL11A1;refGene_tx=XM_023258945.1;refGene_gn=COL11A1;refGene_tx=XM_023258946.1;refGene_gn=COL11A1;refGene_tx=XM_023258947.1;refGene_gn=COL11A1;refGene_tx=XM_023258948.1
C1 88340073 88340127

refGene_gn=COL11A1;refGene_tx=XM_023258940.1;refGene_gn=COL11A1;refGen
e_tx=XM_023258941.1;refGene_gn=COL11A1;refGene_tx=XM_023258942.1;refGe
ne_gn=COL11A1;refGene_tx=XM_023258943.1;refGene_gn=COL11A1;refGene_tx=
XM_023258944.1;refGene_gn=COL11A1;refGene_tx=XM_023258945.1;refGene_gn
=COL11A1;refGene_tx=XM_023258946.1;refGene_gn=COL11A1;refGene_tx=XM_02
3258947.1;refGene_gn=COL11A1;refGene_tx=XM_023258948.1
C1 88344459 88344567
refGene_gn=COL11A1;refGene_tx=XM_023258940.1;refGene_gn=COL11A1;refGen
e_tx=XM_023258941.1;refGene_gn=COL11A1;refGene_tx=XM_023258942.1;refGe
ne_gn=COL11A1;refGene_tx=XM_023258943.1;refGene_gn=COL11A1;refGene_tx=
XM_023258944.1;refGene_gn=COL11A1;refGene_tx=XM_023258945.1;refGene_gn
=COL11A1;refGene_tx=XM_023258946.1;refGene_gn=COL11A1;refGene_tx=XM_02
3258947.1;refGene_gn=COL11A1;refGene_tx=XM_023258948.1
C1 88345059 88345113
refGene_gn=COL11A1;refGene_tx=XM_023258940.1;refGene_gn=COL11A1;refGen
e_tx=XM_023258941.1;refGene_gn=COL11A1;refGene_tx=XM_023258942.1;refGe
ne_gn=COL11A1;refGene_tx=XM_023258943.1;refGene_gn=COL11A1;refGene_tx=
XM_023258944.1;refGene_gn=COL11A1;refGene_tx=XM_023258945.1;refGene_gn
=COL11A1;refGene_tx=XM_023258946.1;refGene_gn=COL11A1;refGene_tx=XM_02
3258947.1;refGene_gn=COL11A1;refGene_tx=XM_023258948.1
C1 88346261 88346315
refGene_gn=COL11A1;refGene_tx=XM_023258940.1;refGene_gn=COL11A1;refGen
e_tx=XM_023258941.1;refGene_gn=COL11A1;refGene_tx=XM_023258942.1;refGe
ne_gn=COL11A1;refGene_tx=XM_023258943.1;refGene_gn=COL11A1;refGene_tx=
XM_023258944.1;refGene_gn=COL11A1;refGene_tx=XM_023258945.1;refGene_gn
=COL11A1;refGene_tx=XM_023258946.1;refGene_gn=COL11A1;refGene_tx=XM_02
3258947.1;refGene_gn=COL11A1;refGene_tx=XM_023258948.1
C1 88347962 88348070
refGene_gn=COL11A1;refGene_tx=XM_023258940.1;refGene_gn=COL11A1;refGen
e_tx=XM_023258941.1;refGene_gn=COL11A1;refGene_tx=XM_023258942.1;refGe
ne_gn=COL11A1;refGene_tx=XM_023258943.1;refGene_gn=COL11A1;refGene_tx=
XM_023258944.1;refGene_gn=COL11A1;refGene_tx=XM_023258945.1;refGene_gn
=COL11A1;refGene_tx=XM_023258946.1;refGene_gn=COL11A1;refGene_tx=XM_02
3258947.1;refGene_gn=COL11A1;refGene_tx=XM_023258948.1
C1 88352464 88352572
refGene_gn=COL11A1;refGene_tx=XM_023258940.1;refGene_gn=COL11A1;refGen
e_tx=XM_023258941.1;refGene_gn=COL11A1;refGene_tx=XM_023258942.1;refGe
ne_gn=COL11A1;refGene_tx=XM_023258943.1;refGene_gn=COL11A1;refGene_tx=
XM_023258944.1;refGene_gn=COL11A1;refGene_tx=XM_023258945.1;refGene_gn
=COL11A1;refGene_tx=XM_023258946.1;refGene_gn=COL11A1;refGene_tx=XM_02
3258947.1;refGene_gn=COL11A1;refGene_tx=XM_023258948.1
C1 88367254 88367308
refGene_gn=COL11A1;refGene_tx=XM_023258940.1;refGene_gn=COL11A1;refGen
e_tx=XM_023258941.1;refGene_gn=COL11A1;refGene_tx=XM_023258942.1;refGe
ne_gn=COL11A1;refGene_tx=XM_023258943.1;refGene_gn=COL11A1;refGene_tx=
XM_023258944.1;refGene_gn=COL11A1;refGene_tx=XM_023258945.1;refGene_gn
=COL11A1;refGene_tx=XM_023258946.1;refGene_gn=COL11A1;refGene_tx=XM_02
3258947.1;refGene_gn=COL11A1;refGene_tx=XM_023258948.1
C1 88367565 88367655
refGene_gn=COL11A1;refGene_tx=XM_023258940.1;refGene_gn=COL11A1;refGen

e_tx=XM_023258941.1;refGene_gn=COL11A1;refGene_tx=XM_023258942.1;refGene_gn=COL11A1;refGene_tx=XM_023258943.1;refGene_gn=COL11A1;refGene_tx=XM_023258944.1;refGene_gn=COL11A1;refGene_tx=XM_023258945.1;refGene_gn=COL11A1;refGene_tx=XM_023258946.1;refGene_gn=COL11A1;refGene_tx=XM_023258947.1;refGene_gn=COL11A1;refGene_tx=XM_023258948.1
C1 88368260 88368368
refGene_gn=COL11A1;refGene_tx=XM_023258940.1;refGene_gn=COL11A1;refGene_tx=XM_023258941.1;refGene_gn=COL11A1;refGene_tx=XM_023258942.1;refGene_gn=COL11A1;refGene_tx=XM_023258943.1;refGene_gn=COL11A1;refGene_tx=XM_023258944.1;refGene_gn=COL11A1;refGene_tx=XM_023258945.1;refGene_gn=COL11A1;refGene_tx=XM_023258946.1;refGene_gn=COL11A1;refGene_tx=XM_023258947.1;refGene_gn=COL11A1;refGene_tx=XM_023258948.1
C1 88371053 88371107
refGene_gn=COL11A1;refGene_tx=XM_023258940.1;refGene_gn=COL11A1;refGene_tx=XM_023258941.1;refGene_gn=COL11A1;refGene_tx=XM_023258942.1;refGene_gn=COL11A1;refGene_tx=XM_023258943.1;refGene_gn=COL11A1;refGene_tx=XM_023258944.1;refGene_gn=COL11A1;refGene_tx=XM_023258945.1;refGene_gn=COL11A1;refGene_tx=XM_023258946.1;refGene_gn=COL11A1;refGene_tx=XM_023258947.1;refGene_gn=COL11A1;refGene_tx=XM_023258948.1
C1 88376064 88376118
refGene_gn=COL11A1;refGene_tx=XM_023258940.1;refGene_gn=COL11A1;refGene_tx=XM_023258941.1;refGene_gn=COL11A1;refGene_tx=XM_023258942.1;refGene_gn=COL11A1;refGene_tx=XM_023258943.1;refGene_gn=COL11A1;refGene_tx=XM_023258944.1;refGene_gn=COL11A1;refGene_tx=XM_023258945.1;refGene_gn=COL11A1;refGene_tx=XM_023258946.1;refGene_gn=COL11A1;refGene_tx=XM_023258947.1;refGene_gn=COL11A1;refGene_tx=XM_023258948.1
C1 88382278 88382332
refGene_gn=COL11A1;refGene_tx=XM_023258940.1;refGene_gn=COL11A1;refGene_tx=XM_023258941.1;refGene_gn=COL11A1;refGene_tx=XM_023258942.1;refGene_gn=COL11A1;refGene_tx=XM_023258943.1;refGene_gn=COL11A1;refGene_tx=XM_023258944.1;refGene_gn=COL11A1;refGene_tx=XM_023258945.1;refGene_gn=COL11A1;refGene_tx=XM_023258946.1;refGene_gn=COL11A1;refGene_tx=XM_023258947.1;refGene_gn=COL11A1;refGene_tx=XM_023258948.1
C1 88386288 88386333
refGene_gn=COL11A1;refGene_tx=XM_023258940.1;refGene_gn=COL11A1;refGene_tx=XM_023258941.1;refGene_gn=COL11A1;refGene_tx=XM_023258942.1;refGene_gn=COL11A1;refGene_tx=XM_023258943.1;refGene_gn=COL11A1;refGene_tx=XM_023258944.1;refGene_gn=COL11A1;refGene_tx=XM_023258945.1;refGene_gn=COL11A1;refGene_tx=XM_023258946.1;refGene_gn=COL11A1;refGene_tx=XM_023258947.1;refGene_gn=COL11A1;refGene_tx=XM_023258948.1
C1 88386846 88386900
refGene_gn=COL11A1;refGene_tx=XM_023258940.1;refGene_gn=COL11A1;refGene_tx=XM_023258941.1;refGene_gn=COL11A1;refGene_tx=XM_023258942.1;refGene_gn=COL11A1;refGene_tx=XM_023258943.1;refGene_gn=COL11A1;refGene_tx=XM_023258944.1;refGene_gn=COL11A1;refGene_tx=XM_023258945.1;refGene_gn=COL11A1;refGene_tx=XM_023258946.1;refGene_gn=COL11A1;refGene_tx=XM_023258947.1;refGene_gn=COL11A1;refGene_tx=XM_023258948.1
C1 88387040 88387085
refGene_gn=COL11A1;refGene_tx=XM_023258940.1;refGene_gn=COL11A1;refGene_tx=XM_023258941.1;refGene_gn=COL11A1;refGene_tx=XM_023258942.1;refGe

ne_gn=COL11A1;refGene_tx=XM_023258943.1;refGene_gn=COL11A1;refGene_tx=XM_023258944.1;refGene_gn=COL11A1;refGene_tx=XM_023258945.1;refGene_gn=COL11A1;refGene_tx=XM_023258946.1;refGene_gn=COL11A1;refGene_tx=XM_023258947.1;refGene_gn=COL11A1;refGene_tx=XM_023258948.1
C1 88387218 88387272
refGene_gn=COL11A1;refGene_tx=XM_023258940.1;refGene_gn=COL11A1;refGene_tx=XM_023258941.1;refGene_gn=COL11A1;refGene_tx=XM_023258942.1;refGene_gn=COL11A1;refGene_tx=XM_023258943.1;refGene_gn=COL11A1;refGene_tx=XM_023258944.1;refGene_gn=COL11A1;refGene_tx=XM_023258945.1;refGene_gn=COL11A1;refGene_tx=XM_023258946.1;refGene_gn=COL11A1;refGene_tx=XM_023258947.1;refGene_gn=COL11A1;refGene_tx=XM_023258948.1
C1 88391836 88391890
refGene_gn=COL11A1;refGene_tx=XM_023258940.1;refGene_gn=COL11A1;refGene_tx=XM_023258941.1;refGene_gn=COL11A1;refGene_tx=XM_023258942.1;refGene_gn=COL11A1;refGene_tx=XM_023258943.1;refGene_gn=COL11A1;refGene_tx=XM_023258944.1;refGene_gn=COL11A1;refGene_tx=XM_023258945.1;refGene_gn=COL11A1;refGene_tx=XM_023258946.1;refGene_gn=COL11A1;refGene_tx=XM_023258947.1;refGene_gn=COL11A1;refGene_tx=XM_023258948.1
C1 88394856 88394964
refGene_gn=COL11A1;refGene_tx=XM_023258940.1;refGene_gn=COL11A1;refGene_tx=XM_023258941.1;refGene_gn=COL11A1;refGene_tx=XM_023258942.1;refGene_gn=COL11A1;refGene_tx=XM_023258943.1;refGene_gn=COL11A1;refGene_tx=XM_023258944.1;refGene_gn=COL11A1;refGene_tx=XM_023258945.1;refGene_gn=COL11A1;refGene_tx=XM_023258946.1;refGene_gn=COL11A1;refGene_tx=XM_023258947.1;refGene_gn=COL11A1;refGene_tx=XM_023258948.1
C1 88397022 88397076
refGene_gn=COL11A1;refGene_tx=XM_023258940.1;refGene_gn=COL11A1;refGene_tx=XM_023258941.1;refGene_gn=COL11A1;refGene_tx=XM_023258942.1;refGene_gn=COL11A1;refGene_tx=XM_023258943.1;refGene_gn=COL11A1;refGene_tx=XM_023258944.1;refGene_gn=COL11A1;refGene_tx=XM_023258945.1;refGene_gn=COL11A1;refGene_tx=XM_023258946.1;refGene_gn=COL11A1;refGene_tx=XM_023258947.1;refGene_gn=COL11A1;refGene_tx=XM_023258948.1
C1 88400977 88401022
refGene_gn=COL11A1;refGene_tx=XM_023258940.1;refGene_gn=COL11A1;refGene_tx=XM_023258941.1;refGene_gn=COL11A1;refGene_tx=XM_023258942.1;refGene_gn=COL11A1;refGene_tx=XM_023258943.1;refGene_gn=COL11A1;refGene_tx=XM_023258944.1;refGene_gn=COL11A1;refGene_tx=XM_023258945.1;refGene_gn=COL11A1;refGene_tx=XM_023258946.1;refGene_gn=COL11A1;refGene_tx=XM_023258947.1;refGene_gn=COL11A1;refGene_tx=XM_023258948.1
C1 88401102 88401156
refGene_gn=COL11A1;refGene_tx=XM_023258940.1;refGene_gn=COL11A1;refGene_tx=XM_023258941.1;refGene_gn=COL11A1;refGene_tx=XM_023258942.1;refGene_gn=COL11A1;refGene_tx=XM_023258943.1;refGene_gn=COL11A1;refGene_tx=XM_023258944.1;refGene_gn=COL11A1;refGene_tx=XM_023258945.1;refGene_gn=COL11A1;refGene_tx=XM_023258946.1;refGene_gn=COL11A1;refGene_tx=XM_023258947.1;refGene_gn=COL11A1;refGene_tx=XM_023258948.1
C1 88402198 88402243
refGene_gn=COL11A1;refGene_tx=XM_023258940.1;refGene_gn=COL11A1;refGene_tx=XM_023258941.1;refGene_gn=COL11A1;refGene_tx=XM_023258942.1;refGene_gn=COL11A1;refGene_tx=XM_023258943.1;refGene_gn=COL11A1;refGene_tx=

XM_023258944.1;refGene_gn=COL11A1;refGene_tx=XM_023258945.1;refGene_gn=COL11A1;refGene_tx=XM_023258946.1;refGene_gn=COL11A1;refGene_tx=XM_023258947.1;refGene_gn=COL11A1;refGene_tx=XM_023258948.1
C1 88403383 88403437
refGene_gn=COL11A1;refGene_tx=XM_023258940.1;refGene_gn=COL11A1;refGene_tx=XM_023258941.1;refGene_gn=COL11A1;refGene_tx=XM_023258942.1;refGene_gn=COL11A1;refGene_tx=XM_023258943.1;refGene_gn=COL11A1;refGene_tx=XM_023258944.1;refGene_gn=COL11A1;refGene_tx=XM_023258945.1;refGene_gn=COL11A1;refGene_tx=XM_023258946.1;refGene_gn=COL11A1;refGene_tx=XM_023258947.1;refGene_gn=COL11A1;refGene_tx=XM_023258948.1
C1 88409008 88409053
refGene_gn=COL11A1;refGene_tx=XM_023258940.1;refGene_gn=COL11A1;refGene_tx=XM_023258941.1;refGene_gn=COL11A1;refGene_tx=XM_023258942.1;refGene_gn=COL11A1;refGene_tx=XM_023258943.1;refGene_gn=COL11A1;refGene_tx=XM_023258944.1;refGene_gn=COL11A1;refGene_tx=XM_023258945.1;refGene_gn=COL11A1;refGene_tx=XM_023258946.1;refGene_gn=COL11A1;refGene_tx=XM_023258947.1;refGene_gn=COL11A1;refGene_tx=XM_023258948.1
C1 88409510 88409564
refGene_gn=COL11A1;refGene_tx=XM_023258940.1;refGene_gn=COL11A1;refGene_tx=XM_023258941.1;refGene_gn=COL11A1;refGene_tx=XM_023258942.1;refGene_gn=COL11A1;refGene_tx=XM_023258943.1;refGene_gn=COL11A1;refGene_tx=XM_023258944.1;refGene_gn=COL11A1;refGene_tx=XM_023258945.1;refGene_gn=COL11A1;refGene_tx=XM_023258946.1;refGene_gn=COL11A1;refGene_tx=XM_023258947.1;refGene_gn=COL11A1;refGene_tx=XM_023258948.1
C1 88409817 88409862
refGene_gn=COL11A1;refGene_tx=XM_023258940.1;refGene_gn=COL11A1;refGene_tx=XM_023258941.1;refGene_gn=COL11A1;refGene_tx=XM_023258942.1;refGene_gn=COL11A1;refGene_tx=XM_023258943.1;refGene_gn=COL11A1;refGene_tx=XM_023258944.1;refGene_gn=COL11A1;refGene_tx=XM_023258945.1;refGene_gn=COL11A1;refGene_tx=XM_023258946.1;refGene_gn=COL11A1;refGene_tx=XM_023258947.1;refGene_gn=COL11A1;refGene_tx=XM_023258948.1
C1 88410328 88410382
refGene_gn=COL11A1;refGene_tx=XM_023258940.1;refGene_gn=COL11A1;refGene_tx=XM_023258941.1;refGene_gn=COL11A1;refGene_tx=XM_023258942.1;refGene_gn=COL11A1;refGene_tx=XM_023258943.1;refGene_gn=COL11A1;refGene_tx=XM_023258944.1;refGene_gn=COL11A1;refGene_tx=XM_023258945.1;refGene_gn=COL11A1;refGene_tx=XM_023258946.1;refGene_gn=COL11A1;refGene_tx=XM_023258947.1;refGene_gn=COL11A1;refGene_tx=XM_023258948.1
C1 88411556 88411601
refGene_gn=COL11A1;refGene_tx=XM_023258940.1;refGene_gn=COL11A1;refGene_tx=XM_023258941.1;refGene_gn=COL11A1;refGene_tx=XM_023258942.1;refGene_gn=COL11A1;refGene_tx=XM_023258943.1;refGene_gn=COL11A1;refGene_tx=XM_023258944.1;refGene_gn=COL11A1;refGene_tx=XM_023258945.1;refGene_gn=COL11A1;refGene_tx=XM_023258946.1;refGene_gn=COL11A1;refGene_tx=XM_023258947.1;refGene_gn=COL11A1;refGene_tx=XM_023258948.1
C1 88411720 88411774
refGene_gn=COL11A1;refGene_tx=XM_023258940.1;refGene_gn=COL11A1;refGene_tx=XM_023258941.1;refGene_gn=COL11A1;refGene_tx=XM_023258942.1;refGene_gn=COL11A1;refGene_tx=XM_023258943.1;refGene_gn=COL11A1;refGene_tx=XM_023258944.1;refGene_gn=COL11A1;refGene_tx=XM_023258945.1;refGene_gn

=COL11A1;refGene_tx=XM_023258946.1;refGene_gn=COL11A1;refGene_tx=XM_023258947.1;refGene_gn=COL11A1;refGene_tx=XM_023258948.1
C1 88412953 88413007
refGene_gn=COL11A1;refGene_tx=XM_023258940.1;refGene_gn=COL11A1;refGene_tx=XM_023258941.1;refGene_gn=COL11A1;refGene_tx=XM_023258942.1;refGene_gn=COL11A1;refGene_tx=XM_023258943.1;refGene_gn=COL11A1;refGene_tx=XM_023258944.1;refGene_gn=COL11A1;refGene_tx=XM_023258945.1;refGene_gn=COL11A1;refGene_tx=XM_023258946.1;refGene_gn=COL11A1;refGene_tx=XM_023258947.1;refGene_gn=COL11A1;refGene_tx=XM_023258948.1
C1 88413182 88413236
refGene_gn=COL11A1;refGene_tx=XM_023258940.1;refGene_gn=COL11A1;refGene_tx=XM_023258941.1;refGene_gn=COL11A1;refGene_tx=XM_023258942.1;refGene_gn=COL11A1;refGene_tx=XM_023258943.1;refGene_gn=COL11A1;refGene_tx=XM_023258944.1;refGene_gn=COL11A1;refGene_tx=XM_023258945.1;refGene_gn=COL11A1;refGene_tx=XM_023258946.1;refGene_gn=COL11A1;refGene_tx=XM_023258947.1;refGene_gn=COL11A1;refGene_tx=XM_023258948.1
C1 88413369 88413423
refGene_gn=COL11A1;refGene_tx=XM_023258940.1;refGene_gn=COL11A1;refGene_tx=XM_023258941.1;refGene_gn=COL11A1;refGene_tx=XM_023258942.1;refGene_gn=COL11A1;refGene_tx=XM_023258943.1;refGene_gn=COL11A1;refGene_tx=XM_023258944.1;refGene_gn=COL11A1;refGene_tx=XM_023258945.1;refGene_gn=COL11A1;refGene_tx=XM_023258946.1;refGene_gn=COL11A1;refGene_tx=XM_023258947.1;refGene_gn=COL11A1;refGene_tx=XM_023258948.1
C1 88417162 88417216
refGene_gn=COL11A1;refGene_tx=XM_023258940.1;refGene_gn=COL11A1;refGene_tx=XM_023258941.1;refGene_gn=COL11A1;refGene_tx=XM_023258942.1;refGene_gn=COL11A1;refGene_tx=XM_023258943.1;refGene_gn=COL11A1;refGene_tx=XM_023258944.1;refGene_gn=COL11A1;refGene_tx=XM_023258945.1;refGene_gn=COL11A1;refGene_tx=XM_023258946.1;refGene_gn=COL11A1;refGene_tx=XM_023258947.1;refGene_gn=COL11A1;refGene_tx=XM_023258948.1
C1 88420990 88421047
refGene_gn=COL11A1;refGene_tx=XM_023258940.1;refGene_gn=COL11A1;refGene_tx=XM_023258941.1;refGene_gn=COL11A1;refGene_tx=XM_023258942.1;refGene_gn=COL11A1;refGene_tx=XM_023258943.1;refGene_gn=COL11A1;refGene_tx=XM_023258944.1;refGene_gn=COL11A1;refGene_tx=XM_023258945.1;refGene_gn=COL11A1;refGene_tx=XM_023258946.1;refGene_gn=COL11A1;refGene_tx=XM_023258947.1;refGene_gn=COL11A1;refGene_tx=XM_023258948.1
C1 88423199 88423283
refGene_gn=COL11A1;refGene_tx=XM_023258940.1;refGene_gn=COL11A1;refGene_tx=XM_023258941.1;refGene_gn=COL11A1;refGene_tx=XM_023258942.1;refGene_gn=COL11A1;refGene_tx=XM_023258943.1;refGene_gn=COL11A1;refGene_tx=XM_023258944.1;refGene_gn=COL11A1;refGene_tx=XM_023258945.1;refGene_gn=COL11A1;refGene_tx=XM_023258946.1;refGene_gn=COL11A1;refGene_tx=XM_023258947.1;refGene_gn=COL11A1;refGene_tx=XM_023258948.1
C1 88424865 88424940
refGene_gn=COL11A1;refGene_tx=XM_023258940.1;refGene_gn=COL11A1;refGene_tx=XM_023258941.1;refGene_gn=COL11A1;refGene_tx=XM_023258942.1;refGene_gn=COL11A1;refGene_tx=XM_023258943.1;refGene_gn=COL11A1;refGene_tx=XM_023258944.1;refGene_gn=COL11A1;refGene_tx=XM_023258945.1;refGene_gn=COL11A1;refGene_tx=XM_023258946.1;refGene_gn=COL11A1;refGene_tx=XM_02

3258947.1;refGene_gn=COL11A1;refGene_tx=XM_023258948.1
C1 88425604 88425667
refGene_gn=COL11A1;refGene_tx=XM_023258940.1;refGene_gn=COL11A1;refGene_tx=XM_023258941.1;refGene_gn=COL11A1;refGene_tx=XM_023258942.1;refGene_gn=COL11A1;refGene_tx=XM_023258943.1;refGene_gn=COL11A1;refGene_tx=XM_023258944.1;refGene_gn=COL11A1;refGene_tx=XM_023258945.1;refGene_gn=COL11A1;refGene_tx=XM_023258946.1;refGene_gn=COL11A1;refGene_tx=XM_023258947.1;refGene_gn=COL11A1;refGene_tx=XM_023258948.1
C1 88426636 88426678
refGene_gn=COL11A1;refGene_tx=XM_023258940.1;refGene_gn=COL11A1;refGene_tx=XM_023258941.1;refGene_gn=COL11A1;refGene_tx=XM_023258942.1;refGene_gn=COL11A1;refGene_tx=XM_023258943.1;refGene_gn=COL11A1;refGene_tx=XM_023258944.1;refGene_gn=COL11A1;refGene_tx=XM_023258945.1;refGene_gn=COL11A1;refGene_tx=XM_023258946.1;refGene_gn=COL11A1;refGene_tx=XM_023258947.1;refGene_gn=COL11A1;refGene_tx=XM_023258948.1
C1 88428180 88428243
refGene_gn=COL11A1;refGene_tx=XM_023258940.1;refGene_gn=COL11A1;refGene_tx=XM_023258941.1;refGene_gn=COL11A1;refGene_tx=XM_023258942.1;refGene_gn=COL11A1;refGene_tx=XM_023258943.1;refGene_gn=COL11A1;refGene_tx=XM_023258944.1;refGene_gn=COL11A1;refGene_tx=XM_023258945.1;refGene_gn=COL11A1;refGene_tx=XM_023258946.1;refGene_gn=COL11A1;refGene_tx=XM_023258947.1;refGene_gn=COL11A1;refGene_tx=XM_023258948.1
C1 88429007 88429262
refGene_gn=COL11A1;refGene_tx=XM_023258940.1;refGene_gn=COL11A1;refGene_tx=XM_023258941.1;refGene_gn=COL11A1;refGene_tx=XM_023258942.1;refGene_gn=COL11A1;refGene_tx=XM_023258943.1;refGene_gn=COL11A1;refGene_tx=XM_023258946.1
C1 88431597 88431690
refGene_gn=COL11A1;refGene_tx=XM_023258940.1;refGene_gn=COL11A1;refGene_tx=XM_023258941.1;refGene_gn=COL11A1;refGene_tx=XM_023258942.1;refGene_gn=COL11A1;refGene_tx=XM_023258943.1;refGene_gn=COL11A1;refGene_tx=XM_023258944.1;refGene_gn=COL11A1;refGene_tx=XM_023258945.1;refGene_gn=COL11A1;refGene_tx=XM_023258946.1;refGene_gn=COL11A1;refGene_tx=XM_023258947.1;refGene_gn=COL11A1;refGene_tx=XM_023258948.1
C1 88431879 88432032
refGene_gn=COL11A1;refGene_tx=XM_023258940.1;refGene_gn=COL11A1;refGene_tx=XM_023258941.1;refGene_gn=COL11A1;refGene_tx=XM_023258942.1;refGene_gn=COL11A1;refGene_tx=XM_023258944.1;refGene_gn=COL11A1;refGene_tx=XM_023258945.1
C1 88432296 88432413
refGene_gn=COL11A1;refGene_tx=XM_023258940.1;refGene_gn=COL11A1;refGene_tx=XM_023258941.1;refGene_gn=COL11A1;refGene_tx=XM_023258943.1;refGene_gn=COL11A1;refGene_tx=XM_023258944.1;refGene_gn=COL11A1;refGene_tx=XM_023258945.1;refGene_gn=COL11A1;refGene_tx=XM_023258947.1
C1 88437793 88437922
refGene_gn=COL11A1;refGene_tx=XM_023258940.1;refGene_gn=COL11A1;refGene_tx=XM_023258941.1;refGene_gn=COL11A1;refGene_tx=XM_023258942.1;refGene_gn=COL11A1;refGene_tx=XM_023258943.1;refGene_gn=COL11A1;refGene_tx=XM_023258944.1;refGene_gn=COL11A1;refGene_tx=XM_023258945.1;refGene_gn=COL11A1;refGene_tx=XM_023258946.1;refGene_gn=COL11A1;refGene_tx=XM_02

3258947.1;refGene_gn=COL11A1;refGene_tx=XM_023258948.1
C1 88476364 88476527
refGene_gn=COL11A1;refGene_tx=XM_023258940.1;refGene_gn=COL11A1;refGene_tx=XM_023258941.1;refGene_gn=COL11A1;refGene_tx=XM_023258942.1;refGene_gn=COL11A1;refGene_tx=XM_023258943.1;refGene_gn=COL11A1;refGene_tx=XM_023258944.1;refGene_gn=COL11A1;refGene_tx=XM_023258945.1;refGene_gn=COL11A1;refGene_tx=XM_023258946.1;refGene_gn=COL11A1;refGene_tx=XM_023258947.1;refGene_gn=COL11A1;refGene_tx=XM_023258948.1
C1 88479572 88479786
refGene_gn=COL11A1;refGene_tx=XM_023258940.1;refGene_gn=COL11A1;refGene_tx=XM_023258941.1;refGene_gn=COL11A1;refGene_tx=XM_023258942.1;refGene_gn=COL11A1;refGene_tx=XM_023258943.1;refGene_gn=COL11A1;refGene_tx=XM_023258944.1;refGene_gn=COL11A1;refGene_tx=XM_023258945.1;refGene_gn=COL11A1;refGene_tx=XM_023258946.1;refGene_gn=COL11A1;refGene_tx=XM_023258947.1;refGene_gn=COL11A1;refGene_tx=XM_023258948.1
C1 88486023 88486191
refGene_gn=COL11A1;refGene_tx=XM_023258940.1;refGene_gn=COL11A1;refGene_tx=XM_023258941.1;refGene_gn=COL11A1;refGene_tx=XM_023258942.1;refGene_gn=COL11A1;refGene_tx=XM_023258943.1;refGene_gn=COL11A1;refGene_tx=XM_023258944.1;refGene_gn=COL11A1;refGene_tx=XM_023258945.1;refGene_gn=COL11A1;refGene_tx=XM_023258946.1;refGene_gn=COL11A1;refGene_tx=XM_023258947.1;refGene_gn=COL11A1;refGene_tx=XM_023258948.1
C1 88513877 88513983
refGene_gn=COL11A1;refGene_tx=XM_023258940.1;refGene_gn=COL11A1;refGene_tx=XM_023258941.1;refGene_gn=COL11A1;refGene_tx=XM_023258942.1;refGene_gn=COL11A1;refGene_tx=XM_023258943.1;refGene_gn=COL11A1;refGene_tx=XM_023258944.1;refGene_gn=COL11A1;refGene_tx=XM_023258945.1;refGene_gn=COL11A1;refGene_tx=XM_023258946.1;refGene_gn=COL11A1;refGene_tx=XM_023258947.1;refGene_gn=COL11A1;refGene_tx=XM_023258948.1
C1 88894541 88894736
refGene_gn=RNPC3;refGene_tx=XM_003990378.5;refGene_gn=RNPC3;refGene_tx=XM_019837522.2;refGene_gn=RNPC3;refGene_tx=XM_019837523.2;refGene_gn=RNPC3;refGene_tx=XM_019837524.2;refGene_gn=RNPC3;refGene_tx=XM_019837526.2
C1 88896392 88896440
refGene_gn=RNPC3;refGene_tx=XM_003990378.5;refGene_gn=RNPC3;refGene_tx=XM_019837522.2;refGene_gn=RNPC3;refGene_tx=XM_019837523.2;refGene_gn=RNPC3;refGene_tx=XM_019837524.2;refGene_gn=RNPC3;refGene_tx=XM_019837526.2
C1 88900194 88900313
refGene_gn=RNPC3;refGene_tx=XM_003990378.5;refGene_gn=RNPC3;refGene_tx=XM_019837522.2;refGene_gn=RNPC3;refGene_tx=XM_019837523.2;refGene_gn=RNPC3;refGene_tx=XM_019837524.2;refGene_gn=RNPC3;refGene_tx=XM_019837526.2
C1 88901531 88901615
refGene_gn=RNPC3;refGene_tx=XM_003990378.5;refGene_gn=RNPC3;refGene_tx=XM_019837522.2;refGene_gn=RNPC3;refGene_tx=XM_019837523.2;refGene_gn=RNPC3;refGene_tx=XM_019837524.2;refGene_gn=RNPC3;refGene_tx=XM_019837526.2
C1 88902269 88902381

refGene_gn=RNPC3;refGene_tx=XM_003990378.5;refGene_gn=RNPC3;refGene_tx=XM_019837522.2;refGene_gn=RNPC3;refGene_tx=XM_019837523.2;refGene_gn=RNPC3;refGene_tx=XM_019837524.2;refGene_gn=RNPC3;refGene_tx=XM_019837526.2
C1 88903697 88903766 refGene_gn=RNPC3;refGene_tx=XM_003990378.5;refGene_gn=RNPC3;refGene_tx=XM_019837522.2;refGene_gn=RNPC3;refGene_tx=XM_019837523.2;refGene_gn=RNPC3;refGene_tx=XM_019837524.2;refGene_gn=RNPC3;refGene_tx=XM_019837526.2
C1 88905434 88905577 refGene_gn=RNPC3;refGene_tx=XM_003990378.5;refGene_gn=RNPC3;refGene_tx=XM_019837522.2;refGene_gn=RNPC3;refGene_tx=XM_019837523.2;refGene_gn=RNPC3;refGene_tx=XM_019837524.2;refGene_gn=RNPC3;refGene_tx=XM_019837526.2
C1 88909021 88909144 refGene_gn=RNPC3;refGene_tx=XM_003990378.5;refGene_gn=RNPC3;refGene_tx=XM_019837522.2;refGene_gn=RNPC3;refGene_tx=XM_019837523.2;refGene_gn=RNPC3;refGene_tx=XM_019837524.2;refGene_gn=RNPC3;refGene_tx=XM_019837526.2
C1 88910168 88910320 refGene_gn=RNPC3;refGene_tx=XM_003990378.5;refGene_gn=RNPC3;refGene_tx=XM_019837522.2;refGene_gn=RNPC3;refGene_tx=XM_019837523.2;refGene_gn=RNPC3;refGene_tx=XM_019837524.2;refGene_gn=RNPC3;refGene_tx=XM_019837526.2
C1 88914025 88914187 refGene_gn=RNPC3;refGene_tx=XM_003990378.5;refGene_gn=RNPC3;refGene_tx=XM_019837522.2;refGene_gn=RNPC3;refGene_tx=XM_019837523.2;refGene_gn=RNPC3;refGene_tx=XM_019837524.2;refGene_gn=RNPC3;refGene_tx=XM_019837526.2
C1 88915477 88915572 refGene_gn=RNPC3;refGene_tx=XM_003990378.5;refGene_gn=RNPC3;refGene_tx=XM_019837522.2;refGene_gn=RNPC3;refGene_tx=XM_019837523.2;refGene_gn=RNPC3;refGene_tx=XM_019837524.2;refGene_gn=RNPC3;refGene_tx=XM_019837526.2
C1 88915660 88915719 refGene_gn=RNPC3;refGene_tx=XM_003990378.5;refGene_gn=RNPC3;refGene_tx=XM_019837522.2;refGene_gn=RNPC3;refGene_tx=XM_019837523.2;refGene_gn=RNPC3;refGene_tx=XM_019837524.2;refGene_gn=RNPC3;refGene_tx=XM_019837526.2
C1 88917848 88917981 refGene_gn=RNPC3;refGene_tx=XM_003990378.5;refGene_gn=RNPC3;refGene_tx=XM_019837522.2;refGene_gn=RNPC3;refGene_tx=XM_019837523.2;refGene_gn=RNPC3;refGene_tx=XM_019837524.2;refGene_gn=RNPC3;refGene_tx=XM_019837526.2
C1 88918892 88918952 refGene_gn=RNPC3;refGene_tx=XM_003990378.5;refGene_gn=RNPC3;refGene_tx=XM_019837522.2;refGene_gn=RNPC3;refGene_tx=XM_019837523.2;refGene_gn=RNPC3;refGene_tx=XM_019837524.2;refGene_gn=RNPC3;refGene_tx=XM_019837526.2
C1 88970870 88971038 refGene_gn=AMY2B;refGene_tx=XM_003990379.3

C1 88971365 88971512 refGene_gn=AMY2B;refGene_tx=XM_003990379.3
C1 88972326 88972524 refGene_gn=AMY2B;refGene_tx=XM_003990379.3
C1 88972938 88973169 refGene_gn=AMY2B;refGene_tx=XM_003990379.3
C1 88973944 88974078 refGene_gn=AMY2B;refGene_tx=XM_003990379.3
C1 88974870 88974993 refGene_gn=AMY2B;refGene_tx=XM_003990379.3
C1 88975095 88975195 refGene_gn=AMY2B;refGene_tx=XM_003990379.3
C1 88976698 88976817 refGene_gn=AMY2B;refGene_tx=XM_003990379.3
C1 88976924 88977050 refGene_gn=AMY2B;refGene_tx=XM_003990379.3
C1 88978664 88978854 refGene_gn=AMY2B;refGene_tx=XM_003990379.3
C1 89832775 89833199
refGene_gn=LOC109503023;refGene_tx=XM_023260050.1
C1 89833956 89834066
refGene_gn=LOC109503023;refGene_tx=XM_023260050.1
C1 89838920 89838995
refGene_gn=LOC109503023;refGene_tx=XM_023260050.1
C1 89839868 89839937
refGene_gn=LOC109503023;refGene_tx=XM_023260050.1
C1 91639602 91640730 refGene_gn=PRMT6;refGene_tx=XM_019837527.1
C1 91727422 91727668
refGene_gn=NTNG1;refGene_tx=XM_011285033.3;refGene_gn=NTNG1;refGene_tx=XM_011285034.3;refGene_gn=NTNG1;refGene_tx=XM_019837529.2;refGene_gn=NTNG1;refGene_tx=XM_019837532.2;refGene_gn=NTNG1;refGene_tx=XM_019837534.2;refGene_gn=NTNG1;refGene_tx=XM_019837535.2;refGene_gn=NTNG1;refGene_tx=XM_023258949.1;refGene_gn=NTNG1;refGene_tx=XM_023258950.1
C1 91902844 91903485
refGene_gn=NTNG1;refGene_tx=XM_011285033.3;refGene_gn=NTNG1;refGene_tx=XM_011285034.3;refGene_gn=NTNG1;refGene_tx=XM_019837529.2;refGene_gn=NTNG1;refGene_tx=XM_019837532.2;refGene_gn=NTNG1;refGene_tx=XM_019837534.2;refGene_gn=NTNG1;refGene_tx=XM_019837535.2;refGene_gn=NTNG1;refGene_tx=XM_023258949.1;refGene_gn=NTNG1;refGene_tx=XM_023258950.1;refGene_gn=NTNG1;refGene_tx=XM_023258951.1
C1 91917547 91918273
refGene_gn=LOC101093877;refGene_tx=XM_003990381.5
C1 91969813 91969986
refGene_gn=NTNG1;refGene_tx=XM_011285033.3;refGene_gn=NTNG1;refGene_tx=XM_011285034.3;refGene_gn=NTNG1;refGene_tx=XM_019837529.2;refGene_gn=NTNG1;refGene_tx=XM_019837532.2;refGene_gn=NTNG1;refGene_tx=XM_019837534.2;refGene_gn=NTNG1;refGene_tx=XM_019837535.2;refGene_gn=NTNG1;refGene_tx=XM_023258949.1;refGene_gn=NTNG1;refGene_tx=XM_023258950.1;refGene_gn=NTNG1;refGene_tx=XM_023258951.1
C1 91983642 91983669
refGene_gn=NTNG1;refGene_tx=XM_011285033.3;refGene_gn=NTNG1;refGene_tx=XM_011285034.3;refGene_gn=NTNG1;refGene_tx=XM_019837529.2;refGene_gn=NTNG1;refGene_tx=XM_019837532.2;refGene_gn=NTNG1;refGene_tx=XM_019837534.2;refGene_gn=NTNG1;refGene_tx=XM_019837535.2;refGene_gn=NTNG1;refGene_tx=XM_023258949.1;refGene_gn=NTNG1;refGene_tx=XM_023258950.1;refGene_gn=NTNG1;refGene_tx=XM_023258951.1
C1 91994197 91994263
refGene_gn=NTNG1;refGene_tx=XM_011285034.3;refGene_gn=NTNG1;refGene_tx=XM_019837535.2

C1 91996596 91996656 refGene_gn=NTNG1;refGene_tx=XM_011285034.3
C1 92004465 92004633
refGene_gn=NTNG1;refGene_tx=XM_019837529.2;refGene_gn=NTNG1;refGene_tx=XM_019837532.2;refGene_gn=NTNG1;refGene_tx=XM_023258949.1;refGene_gn=NTNG1;refGene_tx=XM_023258950.1;refGene_gn=NTNG1;refGene_tx=XM_023258951.1
C1 92010070 92010205
refGene_gn=NTNG1;refGene_tx=XM_019837529.2;refGene_gn=NTNG1;refGene_tx=XM_019837534.2;refGene_gn=NTNG1;refGene_tx=XM_023258949.1;refGene_gn=NTNG1;refGene_tx=XM_023258950.1;refGene_gn=NTNG1;refGene_tx=XM_023258951.1
C1 92054053 92054283
refGene_gn=NTNG1;refGene_tx=XM_011285033.3;refGene_gn=NTNG1;refGene_tx=XM_011285034.3;refGene_gn=NTNG1;refGene_tx=XM_019837529.2;refGene_gn=NTNG1;refGene_tx=XM_019837532.2;refGene_gn=NTNG1;refGene_tx=XM_019837534.2;refGene_gn=NTNG1;refGene_tx=XM_019837535.2;refGene_gn=NTNG1;refGene_tx=XM_023258949.1;refGene_gn=NTNG1;refGene_tx=XM_023258950.1;refGene_gn=NTNG1;refGene_tx=XM_023258951.1
C1 92163897 92163939 refGene_gn=VAV3;refGene_tx=XM_011285035.3
C1 92164593 92164745 refGene_gn=VAV3;refGene_tx=XM_011285035.3
C1 92183784 92183914 refGene_gn=VAV3;refGene_tx=XM_011285035.3
C1 92189719 92189807 refGene_gn=VAV3;refGene_tx=XM_011285035.3
C1 92190363 92190480 refGene_gn=VAV3;refGene_tx=XM_011285035.3
C1 92196864 92196899 refGene_gn=VAV3;refGene_tx=XM_011285035.3
C1 92202656 92202722 refGene_gn=VAV3;refGene_tx=XM_011285035.3
C1 92226770 92226907 refGene_gn=VAV3;refGene_tx=XM_011285035.3
C1 92267957 92268003 refGene_gn=VAV3;refGene_tx=XM_011285035.3
C1 92274097 92274123 refGene_gn=VAV3;refGene_tx=XM_011285035.3
C1 92285543 92285644 refGene_gn=VAV3;refGene_tx=XM_011285035.3
C1 92285950 92286052 refGene_gn=VAV3;refGene_tx=XM_011285035.3
C1 92321340 92321450 refGene_gn=VAV3;refGene_tx=XM_011285035.3
C1 92321834 92321967 refGene_gn=VAV3;refGene_tx=XM_011285035.3
C1 92323464 92323550 refGene_gn=VAV3;refGene_tx=XM_011285035.3
C1 92327910 92327997 refGene_gn=VAV3;refGene_tx=XM_011285035.3
C1 92329763 92329832 refGene_gn=VAV3;refGene_tx=XM_011285035.3
C1 92332920 92333016 refGene_gn=VAV3;refGene_tx=XM_011285035.3
C1 92337003 92337103 refGene_gn=VAV3;refGene_tx=XM_011285035.3
C1 92338530 92338634 refGene_gn=VAV3;refGene_tx=XM_011285035.3
C1 92340716 92340785 refGene_gn=VAV3;refGene_tx=XM_011285035.3
C1 92342969 92343062 refGene_gn=VAV3;refGene_tx=XM_011285035.3
C1 92345054 92345163 refGene_gn=VAV3;refGene_tx=XM_011285035.3
C1 92351823 92351889 refGene_gn=VAV3;refGene_tx=XM_011285035.3
C1 92354129 92354188 refGene_gn=VAV3;refGene_tx=XM_011285035.3
C1 92432279 92432396 refGene_gn=VAV3;refGene_tx=XM_011285035.3
C1 92516040 92516244 refGene_gn=VAV3;refGene_tx=XM_011285035.3
C1 92697311 92697496 refGene_gn=SLC25A24;refGene_tx=XM_003990389.5
C1 92700172 92700323 refGene_gn=SLC25A24;refGene_tx=XM_003990389.5
C1 92705013 92705181 refGene_gn=SLC25A24;refGene_tx=XM_003990389.5
C1 92706870 92706978 refGene_gn=SLC25A24;refGene_tx=XM_003990389.5
C1 92713910 92714063 refGene_gn=SLC25A24;refGene_tx=XM_003990389.5

C1 92716391 92716550 refGene_gn=SLC25A24;refGene_tx=XM_003990389.5
C1 92723336 92723448 refGene_gn=SLC25A24;refGene_tx=XM_003990389.5
C1 92733725 92733813 refGene_gn=SLC25A24;refGene_tx=XM_003990389.5
C1 92738013 92738140 refGene_gn=SLC25A24;refGene_tx=XM_003990389.5
C1 92748296 92748479 refGene_gn=SLC25A24;refGene_tx=XM_003990389.5
C1 92773896 92774075
refGene_gn=LOC101095594;refGene_tx=XM_011285036.3
C1 92775255 92775406
refGene_gn=LOC101095594;refGene_tx=XM_011285036.3
C1 92777711 92777879
refGene_gn=LOC101095594;refGene_tx=XM_011285036.3
C1 92780907 92781015
refGene_gn=LOC101095594;refGene_tx=XM_011285036.3
C1 92787223 92787376
refGene_gn=LOC101095594;refGene_tx=XM_011285036.3
C1 92789998 92790157
refGene_gn=LOC101095594;refGene_tx=XM_011285036.3
C1 92797467 92797579
refGene_gn=LOC101095594;refGene_tx=XM_011285036.3
C1 92798838 92798866
refGene_gn=LOC101095594;refGene_tx=XM_011285036.3
C1 92798868 92798925
refGene_gn=LOC101095594;refGene_tx=XM_011285036.3
C1 92804881 92805014
refGene_gn=LOC101095594;refGene_tx=XM_011285036.3
C1 92822613 92822976
refGene_gn=LOC101095594;refGene_tx=XM_011285036.3
C1 92930939 92931076 refGene_gn=FAM102B;refGene_tx=XM_003990390.5
C1 92979046 92979167 refGene_gn=FAM102B;refGene_tx=XM_003990390.5
C1 92990046 92990076 refGene_gn=FAM102B;refGene_tx=XM_003990390.5
C1 92995848 92995965 refGene_gn=FAM102B;refGene_tx=XM_003990390.5
C1 93000139 93000186 refGene_gn=FAM102B;refGene_tx=XM_003990390.5
C1 93009256 93009360 refGene_gn=FAM102B;refGene_tx=XM_003990390.5
C1 93012303 93012399 refGene_gn=FAM102B;refGene_tx=XM_003990390.5
C1 93012539 93012729 refGene_gn=FAM102B;refGene_tx=XM_003990390.5
C1 93012829 93012957 refGene_gn=FAM102B;refGene_tx=XM_003990390.5
C1 93014695 93014765 refGene_gn=FAM102B;refGene_tx=XM_003990390.5
C1 93021442 93021485 refGene_gn=FAM102B;refGene_tx=XM_003990390.5
C1 93033620 93034046
refGene_gn=HENMT1;refGene_tx=XM_003990434.4;refGene_gn=HENMT1;refGene_
tx=XM_023258952.1
C1 93035192 93035391
refGene_gn=HENMT1;refGene_tx=XM_003990434.4;refGene_gn=HENMT1;refGene_
tx=XM_023258952.1
C1 93036046 93036226
refGene_gn=HENMT1;refGene_tx=XM_003990434.4;refGene_gn=HENMT1;refGene_
tx=XM_023258952.1
C1 93037239 93037374
refGene_gn=HENMT1;refGene_tx=XM_003990434.4;refGene_gn=HENMT1;refGene_
tx=XM_023258952.1

C1 93037970 93038086
refGene_gn=HENMT1;refGene_tx=XM_003990434.4;refGene_gn=HENMT1;refGene_tx=XM_023258952.1
C1 93040379 93040490
refGene_gn=HENMT1;refGene_tx=XM_003990434.4;refGene_gn=HENMT1;refGene_tx=XM_023258952.1
C1 93041743 93041767
refGene_gn=HENMT1;refGene_tx=XM_003990434.4;refGene_gn=HENMT1;refGene_tx=XM_023258952.1
C1 93072756 93073035 refGene_gn=PRPF38B;refGene_tx=XM_003990391.4
C1 93075892 93075961 refGene_gn=PRPF38B;refGene_tx=XM_003990391.4
C1 93076243 93076395 refGene_gn=PRPF38B;refGene_tx=XM_003990391.4
C1 93076475 93076536 refGene_gn=PRPF38B;refGene_tx=XM_003990391.4
C1 93078740 93078964 refGene_gn=PRPF38B;refGene_tx=XM_003990391.4
C1 93079256 93080118 refGene_gn=PRPF38B;refGene_tx=XM_003990391.4
C1 93095401 93095464 refGene_gn=FNDC7;refGene_tx=XM_006934989.3
C1 93095994 93096013 refGene_gn=FNDC7;refGene_tx=XM_006934989.3
C1 93100116 93100371 refGene_gn=FNDC7;refGene_tx=XM_006934989.3
C1 93101798 93102059 refGene_gn=FNDC7;refGene_tx=XM_006934989.3
C1 93105985 93106243 refGene_gn=FNDC7;refGene_tx=XM_006934989.3
C1 93108661 93108916 refGene_gn=FNDC7;refGene_tx=XM_006934989.3
C1 93110430 93110688 refGene_gn=FNDC7;refGene_tx=XM_006934989.3
C1 93111266 93111521 refGene_gn=FNDC7;refGene_tx=XM_006934989.3
C1 93112717 93112972 refGene_gn=FNDC7;refGene_tx=XM_006934989.3
C1 93115966 93116227 refGene_gn=FNDC7;refGene_tx=XM_006934989.3
C1 93120577 93120607 refGene_gn=FNDC7;refGene_tx=XM_006934989.3
C1 93124299 93124331 refGene_gn=FNDC7;refGene_tx=XM_006934989.3
C1 93130252 93130301
refGene_gn=STXBP3;refGene_tx=XM_006934955.4;refGene_gn=STXBP3;refGene_tx=XM_023258953.1;refGene_gn=STXBP3;refGene_tx=XM_023258954.1
C1 93143369 93143419
refGene_gn=STXBP3;refGene_tx=XM_006934955.4;refGene_gn=STXBP3;refGene_tx=XM_023258953.1;refGene_gn=STXBP3;refGene_tx=XM_023258954.1
C1 93144002 93144084
refGene_gn=STXBP3;refGene_tx=XM_006934955.4;refGene_gn=STXBP3;refGene_tx=XM_023258953.1;refGene_gn=STXBP3;refGene_tx=XM_023258954.1
C1 93148159 93148236
refGene_gn=STXBP3;refGene_tx=XM_006934955.4;refGene_gn=STXBP3;refGene_tx=XM_023258953.1;refGene_gn=STXBP3;refGene_tx=XM_023258954.1
C1 93150674 93150753
refGene_gn=STXBP3;refGene_tx=XM_006934955.4;refGene_gn=STXBP3;refGene_tx=XM_023258953.1;refGene_gn=STXBP3;refGene_tx=XM_023258954.1
C1 93152071 93152172
refGene_gn=STXBP3;refGene_tx=XM_006934955.4;refGene_gn=STXBP3;refGene_tx=XM_023258953.1;refGene_gn=STXBP3;refGene_tx=XM_023258954.1
C1 93156122 93156277
refGene_gn=STXBP3;refGene_tx=XM_006934955.4;refGene_gn=STXBP3;refGene_tx=XM_023258953.1;refGene_gn=STXBP3;refGene_tx=XM_023258954.1
C1 93160788 93160879
refGene_gn=STXBP3;refGene_tx=XM_006934955.4;refGene_gn=STXBP3;refGene_

tx=XM_023258953.1;refGene_gn=STXBP3;refGene_tx=XM_023258954.1
C1 93163699 93163824
refGene_gn=STXBP3;refGene_tx=XM_006934955.4;refGene_gn=STXBP3;refGene_tx=XM_023258953.1;refGene_gn=STXBP3;refGene_tx=XM_023258954.1
C1 93166300 93166396
refGene_gn=STXBP3;refGene_tx=XM_006934955.4;refGene_gn=STXBP3;refGene_tx=XM_023258953.1;refGene_gn=STXBP3;refGene_tx=XM_023258954.1
C1 93166530 93166588
refGene_gn=STXBP3;refGene_tx=XM_006934955.4;refGene_gn=STXBP3;refGene_tx=XM_023258953.1;refGene_gn=STXBP3;refGene_tx=XM_023258954.1
C1 93171278 93171344
refGene_gn=STXBP3;refGene_tx=XM_006934955.4;refGene_gn=STXBP3;refGene_tx=XM_023258953.1;refGene_gn=STXBP3;refGene_tx=XM_023258954.1
C1 93172108 93172189
refGene_gn=STXBP3;refGene_tx=XM_006934955.4;refGene_gn=STXBP3;refGene_tx=XM_023258953.1;refGene_gn=STXBP3;refGene_tx=XM_023258954.1
C1 93174804 93174943
refGene_gn=STXBP3;refGene_tx=XM_006934955.4;refGene_gn=STXBP3;refGene_tx=XM_023258953.1;refGene_gn=STXBP3;refGene_tx=XM_023258954.1
C1 93175234 93175341
refGene_gn=STXBP3;refGene_tx=XM_006934955.4;refGene_gn=STXBP3;refGene_tx=XM_023258953.1;refGene_gn=STXBP3;refGene_tx=XM_023258954.1
C1 93175884 93175977
refGene_gn=STXBP3;refGene_tx=XM_006934955.4;refGene_gn=STXBP3;refGene_tx=XM_023258953.1;refGene_gn=STXBP3;refGene_tx=XM_023258954.1
C1 93177733 93177772 refGene_gn=STXBP3;refGene_tx=XM_023258953.1
C1 93178359 93178445 refGene_gn=STXBP3;refGene_tx=XM_006934955.4
C1 93181840 93181989
refGene_gn=STXBP3;refGene_tx=XM_006934955.4;refGene_gn=STXBP3;refGene_tx=XM_023258954.1
C1 93182771 93182866 refGene_gn=STXBP3;refGene_tx=XM_006934955.4
C1 93190324 93190456
refGene_gn=AKNAD1;refGene_tx=XM_006934956.4;refGene_gn=AKNAD1;refGene_tx=XM_019837537.2;refGene_gn=AKNAD1;refGene_tx=XM_019837538.2
C1 93191199 93191335
refGene_gn=AKNAD1;refGene_tx=XM_006934956.4;refGene_gn=AKNAD1;refGene_tx=XM_019837537.2;refGene_gn=AKNAD1;refGene_tx=XM_019837538.2;refGene_gn=AKNAD1;refGene_tx=XM_019837539.2;refGene_gn=AKNAD1;refGene_tx=XM_019837540.2
C1 93193789 93193871
refGene_gn=AKNAD1;refGene_tx=XM_006934956.4;refGene_gn=AKNAD1;refGene_tx=XM_019837537.2;refGene_gn=AKNAD1;refGene_tx=XM_019837538.2;refGene_gn=AKNAD1;refGene_tx=XM_019837539.2;refGene_gn=AKNAD1;refGene_tx=XM_019837540.2
C1 93197804 93197912
refGene_gn=AKNAD1;refGene_tx=XM_006934956.4;refGene_gn=AKNAD1;refGene_tx=XM_019837537.2;refGene_gn=AKNAD1;refGene_tx=XM_019837538.2;refGene_gn=AKNAD1;refGene_tx=XM_019837539.2;refGene_gn=AKNAD1;refGene_tx=XM_019837540.2
C1 93198003 93198126

refGene_gn=AKNAD1;refGene_tx=XM_006934956.4;refGene_gn=AKNAD1;refGene_tx=XM_019837537.2;refGene_gn=AKNAD1;refGene_tx=XM_019837538.2;refGene_gn=AKNAD1;refGene_tx=XM_019837539.2;refGene_gn=AKNAD1;refGene_tx=XM_019837540.2
C1	93201683	93201781
refGene_gn=AKNAD1;refGene_tx=XM_006934956.4;refGene_gn=AKNAD1;refGene_tx=XM_019837537.2;refGene_gn=AKNAD1;refGene_tx=XM_019837538.2;refGene_gn=AKNAD1;refGene_tx=XM_019837539.2;refGene_gn=AKNAD1;refGene_tx=XM_019837540.2
C1	93204968	93205054
refGene_gn=AKNAD1;refGene_tx=XM_006934956.4;refGene_gn=AKNAD1;refGene_tx=XM_019837537.2;refGene_gn=AKNAD1;refGene_tx=XM_019837538.2;refGene_gn=AKNAD1;refGene_tx=XM_019837539.2;refGene_gn=AKNAD1;refGene_tx=XM_019837540.2
C1	93211000	93211157
refGene_gn=AKNAD1;refGene_tx=XM_006934956.4;refGene_gn=AKNAD1;refGene_tx=XM_019837537.2;refGene_gn=AKNAD1;refGene_tx=XM_019837538.2;refGene_gn=AKNAD1;refGene_tx=XM_019837539.2;refGene_gn=AKNAD1;refGene_tx=XM_019837540.2
C1	93218615	93218746
refGene_gn=AKNAD1;refGene_tx=XM_006934956.4;refGene_gn=AKNAD1;refGene_tx=XM_019837537.2;refGene_gn=AKNAD1;refGene_tx=XM_019837538.2;refGene_gn=AKNAD1;refGene_tx=XM_019837539.2;refGene_gn=AKNAD1;refGene_tx=XM_019837540.2
C1	93224123	93224186
refGene_gn=AKNAD1;refGene_tx=XM_006934956.4;refGene_gn=AKNAD1;refGene_tx=XM_019837537.2;refGene_gn=AKNAD1;refGene_tx=XM_019837538.2;refGene_gn=AKNAD1;refGene_tx=XM_019837539.2;refGene_gn=AKNAD1;refGene_tx=XM_019837540.2
C1	93224279	93224428
refGene_gn=AKNAD1;refGene_tx=XM_006934956.4;refGene_gn=AKNAD1;refGene_tx=XM_019837537.2;refGene_gn=AKNAD1;refGene_tx=XM_019837538.2;refGene_gn=AKNAD1;refGene_tx=XM_019837539.2;refGene_gn=AKNAD1;refGene_tx=XM_019837540.2
C1	93225074	93225114
refGene_gn=AKNAD1;refGene_tx=XM_006934956.4;refGene_gn=AKNAD1;refGene_tx=XM_019837537.2;refGene_gn=AKNAD1;refGene_tx=XM_019837538.2;refGene_gn=AKNAD1;refGene_tx=XM_019837539.2;refGene_gn=AKNAD1;refGene_tx=XM_019837540.2
C1	93227239	93228223
refGene_gn=AKNAD1;refGene_tx=XM_006934956.4;refGene_gn=AKNAD1;refGene_tx=XM_019837537.2;refGene_gn=AKNAD1;refGene_tx=XM_019837538.2;refGene_gn=AKNAD1;refGene_tx=XM_019837539.2;refGene_gn=AKNAD1;refGene_tx=XM_019837540.2
C1	93254143	93254457
refGene_gn=GPSM2;refGene_tx=XM_023258955.1;refGene_gn=GPSM2;refGene_tx=XM_023258956.1;refGene_gn=GPSM2;refGene_tx=XM_023258957.1
C1	93266232	93266288
refGene_gn=GPSM2;refGene_tx=XM_023258958.1;refGene_gn=GPSM2;refGene_tx=XM_023258959.1

C1 93274650 93274872
refGene_gn=GPSM2;refGene_tx=XM_023258955.1;refGene_gn=GPSM2;refGene_tx=XM_023258956.1;refGene_gn=GPSM2;refGene_tx=XM_023258957.1;refGene_gn=GPSM2;refGene_tx=XM_023258958.1;refGene_gn=GPSM2;refGene_tx=XM_023258959.1
C1 93275254 93275390
refGene_gn=GPSM2;refGene_tx=XM_023258955.1;refGene_gn=GPSM2;refGene_tx=XM_023258956.1;refGene_gn=GPSM2;refGene_tx=XM_023258957.1;refGene_gn=GPSM2;refGene_tx=XM_023258958.1;refGene_gn=GPSM2;refGene_tx=XM_023258959.1
C1 93275720 93275863
refGene_gn=GPSM2;refGene_tx=XM_023258955.1;refGene_gn=GPSM2;refGene_tx=XM_023258956.1;refGene_gn=GPSM2;refGene_tx=XM_023258957.1;refGene_gn=GPSM2;refGene_tx=XM_023258958.1;refGene_gn=GPSM2;refGene_tx=XM_023258959.1
C1 93276403 93276527
refGene_gn=GPSM2;refGene_tx=XM_023258955.1;refGene_gn=GPSM2;refGene_tx=XM_023258956.1;refGene_gn=GPSM2;refGene_tx=XM_023258957.1;refGene_gn=GPSM2;refGene_tx=XM_023258958.1;refGene_gn=GPSM2;refGene_tx=XM_023258959.1
C1 93276635 93276751
refGene_gn=GPSM2;refGene_tx=XM_023258955.1;refGene_gn=GPSM2;refGene_tx=XM_023258956.1;refGene_gn=GPSM2;refGene_tx=XM_023258957.1;refGene_gn=GPSM2;refGene_tx=XM_023258958.1;refGene_gn=GPSM2;refGene_tx=XM_023258959.1
C1 93279510 93279666
refGene_gn=GPSM2;refGene_tx=XM_023258955.1;refGene_gn=GPSM2;refGene_tx=XM_023258956.1;refGene_gn=GPSM2;refGene_tx=XM_023258957.1;refGene_gn=GPSM2;refGene_tx=XM_023258958.1;refGene_gn=GPSM2;refGene_tx=XM_023258959.1
C1 93280786 93280895
refGene_gn=GPSM2;refGene_tx=XM_023258955.1;refGene_gn=GPSM2;refGene_tx=XM_023258956.1;refGene_gn=GPSM2;refGene_tx=XM_023258957.1;refGene_gn=GPSM2;refGene_tx=XM_023258958.1;refGene_gn=GPSM2;refGene_tx=XM_023258959.1
C1 93282072 93282202
refGene_gn=GPSM2;refGene_tx=XM_023258955.1;refGene_gn=GPSM2;refGene_tx=XM_023258956.1;refGene_gn=GPSM2;refGene_tx=XM_023258957.1;refGene_gn=GPSM2;refGene_tx=XM_023258958.1;refGene_gn=GPSM2;refGene_tx=XM_023258959.1
C1 93284709 93284780
refGene_gn=GPSM2;refGene_tx=XM_023258955.1;refGene_gn=GPSM2;refGene_tx=XM_023258956.1;refGene_gn=GPSM2;refGene_tx=XM_023258957.1;refGene_gn=GPSM2;refGene_tx=XM_023258958.1;refGene_gn=GPSM2;refGene_tx=XM_023258959.1
C1 93288798 93288975
refGene_gn=GPSM2;refGene_tx=XM_023258955.1;refGene_gn=GPSM2;refGene_tx=XM_023258956.1;refGene_gn=GPSM2;refGene_tx=XM_023258957.1;refGene_gn=GPSM2;refGene_tx=XM_023258958.1;refGene_gn=GPSM2;refGene_tx=XM_023258959.1

C1 93291762 93291922
refGene_gn=GPSM2;refGene_tx=XM_023258955.1;refGene_gn=GPSM2;refGene_tx
=XM_023258956.1;refGene_gn=GPSM2;refGene_tx=XM_023258957.1;refGene_gn=
GPSM2;refGene_tx=XM_023258958.1;refGene_gn=GPSM2;refGene_tx=XM_0232589
59.1
C1 93292775 93292993
refGene_gn=GPSM2;refGene_tx=XM_023258955.1;refGene_gn=GPSM2;refGene_tx
=XM_023258956.1;refGene_gn=GPSM2;refGene_tx=XM_023258957.1;refGene_gn=
GPSM2;refGene_tx=XM_023258958.1;refGene_gn=GPSM2;refGene_tx=XM_0232589
59.1
C1 93300042 93300081 refGene_gn=GPSM2;refGene_tx=XM_023258957.1
C1 93301047 93301275
refGene_gn=GPSM2;refGene_tx=XM_023258955.1;refGene_gn=GPSM2;refGene_tx
=XM_023258956.1;refGene_gn=GPSM2;refGene_tx=XM_023258958.1;refGene_gn=
GPSM2;refGene_tx=XM_023258959.1
C1 93304394 93304652
refGene_gn=CLCC1;refGene_tx=XM_003990438.5;refGene_gn=CLCC1;refGene_tx
=XM_006934960.3;refGene_gn=CLCC1;refGene_tx=XM_006934961.2;refGene_gn=
CLCC1;refGene_tx=XM_006934962.2;refGene_gn=CLCC1;refGene_tx=XM_0198375
44.2
C1 93306061 93306403
refGene_gn=CLCC1;refGene_tx=XM_003990438.5;refGene_gn=CLCC1;refGene_tx
=XM_006934960.3;refGene_gn=CLCC1;refGene_tx=XM_006934961.2;refGene_gn=
CLCC1;refGene_tx=XM_006934962.2;refGene_gn=CLCC1;refGene_tx=XM_0198375
44.2
C1 93308906 93309053
refGene_gn=CLCC1;refGene_tx=XM_003990438.5;refGene_gn=CLCC1;refGene_tx
=XM_006934960.3;refGene_gn=CLCC1;refGene_tx=XM_006934961.2;refGene_gn=
CLCC1;refGene_tx=XM_006934962.2;refGene_gn=CLCC1;refGene_tx=XM_0198375
44.2
C1 93309312 93309410
refGene_gn=CLCC1;refGene_tx=XM_003990438.5;refGene_gn=CLCC1;refGene_tx
=XM_006934960.3;refGene_gn=CLCC1;refGene_tx=XM_006934961.2;refGene_gn=
CLCC1;refGene_tx=XM_006934962.2;refGene_gn=CLCC1;refGene_tx=XM_0198375
44.2
C1 93310901 93310995
refGene_gn=CLCC1;refGene_tx=XM_003990438.5;refGene_gn=CLCC1;refGene_tx
=XM_006934960.3;refGene_gn=CLCC1;refGene_tx=XM_006934961.2;refGene_gn=
CLCC1;refGene_tx=XM_006934962.2;refGene_gn=CLCC1;refGene_tx=XM_0198375
44.2
C1 93312606 93312747
refGene_gn=CLCC1;refGene_tx=XM_003990438.5;refGene_gn=CLCC1;refGene_tx
=XM_006934960.3;refGene_gn=CLCC1;refGene_tx=XM_006934961.2;refGene_gn=
CLCC1;refGene_tx=XM_006934962.2;refGene_gn=CLCC1;refGene_tx=XM_0198375
44.2
C1 93312988 93313210
refGene_gn=CLCC1;refGene_tx=XM_003990438.5;refGene_gn=CLCC1;refGene_tx
=XM_006934960.3;refGene_gn=CLCC1;refGene_tx=XM_006934961.2;refGene_gn=
CLCC1;refGene_tx=XM_006934962.2;refGene_gn=CLCC1;refGene_tx=XM_0198375
44.2

C1 93315658 93315766
refGene_gn=CLCC1;refGene_tx=XM_003990438.5;refGene_gn=CLCC1;refGene_tx=XM_006934960.3;refGene_gn=CLCC1;refGene_tx=XM_006934961.2;refGene_gn=CLCC1;refGene_tx=XM_006934962.2;refGene_gn=CLCC1;refGene_tx=XM_019837544.2
C1 93319073 93319175
refGene_gn=CLCC1;refGene_tx=XM_003990438.5;refGene_gn=CLCC1;refGene_tx=XM_006934960.3;refGene_gn=CLCC1;refGene_tx=XM_006934961.2;refGene_gn=CLCC1;refGene_tx=XM_006934962.2
C1 93319587 93319716
refGene_gn=CLCC1;refGene_tx=XM_003990438.5;refGene_gn=CLCC1;refGene_tx=XM_006934960.3;refGene_gn=CLCC1;refGene_tx=XM_006934961.2;refGene_gn=CLCC1;refGene_tx=XM_006934962.2;refGene_gn=CLCC1;refGene_tx=XM_019837544.2
C1 93338321 93338464
refGene_gn=WDR47;refGene_tx=XM_019837545.2;refGene_gn=WDR47;refGene_tx=XM_019837546.2;refGene_gn=WDR47;refGene_tx=XM_019837547.2;refGene_gn=WDR47;refGene_tx=XM_019837548.2;refGene_gn=WDR47;refGene_tx=XM_019837549.2;refGene_gn=WDR47;refGene_tx=XM_019837550.2;refGene_gn=WDR47;refGene_tx=XM_019837551.2;refGene_gn=WDR47;refGene_tx=XM_019837552.2;refGene_gn=WDR47;refGene_tx=XM_019837553.2;refGene_gn=WDR47;refGene_tx=XM_019837554.1
C1 93341760 93341979
refGene_gn=WDR47;refGene_tx=XM_019837545.2;refGene_gn=WDR47;refGene_tx=XM_019837546.2;refGene_gn=WDR47;refGene_tx=XM_019837547.2;refGene_gn=WDR47;refGene_tx=XM_019837548.2;refGene_gn=WDR47;refGene_tx=XM_019837549.2;refGene_gn=WDR47;refGene_tx=XM_019837550.2;refGene_gn=WDR47;refGene_tx=XM_019837551.2;refGene_gn=WDR47;refGene_tx=XM_019837552.2;refGene_gn=WDR47;refGene_tx=XM_019837553.2;refGene_gn=WDR47;refGene_tx=XM_019837554.1
C1 93351175 93351307
refGene_gn=WDR47;refGene_tx=XM_019837545.2;refGene_gn=WDR47;refGene_tx=XM_019837546.2;refGene_gn=WDR47;refGene_tx=XM_019837547.2;refGene_gn=WDR47;refGene_tx=XM_019837548.2;refGene_gn=WDR47;refGene_tx=XM_019837549.2;refGene_gn=WDR47;refGene_tx=XM_019837550.2;refGene_gn=WDR47;refGene_tx=XM_019837551.2;refGene_gn=WDR47;refGene_tx=XM_019837552.2;refGene_gn=WDR47;refGene_tx=XM_019837553.2;refGene_gn=WDR47;refGene_tx=XM_019837554.1
C1 93351987 93352164
refGene_gn=WDR47;refGene_tx=XM_019837545.2;refGene_gn=WDR47;refGene_tx=XM_019837546.2;refGene_gn=WDR47;refGene_tx=XM_019837547.2;refGene_gn=WDR47;refGene_tx=XM_019837548.2;refGene_gn=WDR47;refGene_tx=XM_019837549.2;refGene_gn=WDR47;refGene_tx=XM_019837550.2;refGene_gn=WDR47;refGene_tx=XM_019837551.2;refGene_gn=WDR47;refGene_tx=XM_019837552.2;refGene_gn=WDR47;refGene_tx=XM_019837553.2;refGene_gn=WDR47;refGene_tx=XM_019837554.1
C1 93352929 93353099
refGene_gn=WDR47;refGene_tx=XM_019837545.2;refGene_gn=WDR47;refGene_tx=XM_019837546.2;refGene_gn=WDR47;refGene_tx=XM_019837547.2;refGene_gn=WDR47;refGene_tx=XM_019837548.2;refGene_gn=WDR47;refGene_tx=XM_0198375

49.2;refGene_gn=WDR47;refGene_tx=XM_019837550.2;refGene_gn=WDR47;refGene_tx=XM_019837551.2;refGene_gn=WDR47;refGene_tx=XM_019837552.2;refGene_gn=WDR47;refGene_tx=XM_019837553.2;refGene_gn=WDR47;refGene_tx=XM_019837554.1
C1 93355646 93355804
refGene_gn=WDR47;refGene_tx=XM_019837545.2;refGene_gn=WDR47;refGene_tx=XM_019837546.2;refGene_gn=WDR47;refGene_tx=XM_019837547.2;refGene_gn=WDR47;refGene_tx=XM_019837548.2;refGene_gn=WDR47;refGene_tx=XM_019837549.2;refGene_gn=WDR47;refGene_tx=XM_019837550.2;refGene_gn=WDR47;refGene_tx=XM_019837551.2;refGene_gn=WDR47;refGene_tx=XM_019837552.2;refGene_gn=WDR47;refGene_tx=XM_019837553.2;refGene_gn=WDR47;refGene_tx=XM_019837554.1
C1 93358054 93358130
refGene_gn=WDR47;refGene_tx=XM_019837545.2;refGene_gn=WDR47;refGene_tx=XM_019837546.2;refGene_gn=WDR47;refGene_tx=XM_019837547.2;refGene_gn=WDR47;refGene_tx=XM_019837548.2;refGene_gn=WDR47;refGene_tx=XM_019837549.2;refGene_gn=WDR47;refGene_tx=XM_019837550.2;refGene_gn=WDR47;refGene_tx=XM_019837551.2;refGene_gn=WDR47;refGene_tx=XM_019837552.2;refGene_gn=WDR47;refGene_tx=XM_019837553.2;refGene_gn=WDR47;refGene_tx=XM_019837554.1
C1 93360572 93360833
refGene_gn=WDR47;refGene_tx=XM_019837545.2;refGene_gn=WDR47;refGene_tx=XM_019837546.2;refGene_gn=WDR47;refGene_tx=XM_019837547.2;refGene_gn=WDR47;refGene_tx=XM_019837548.2;refGene_gn=WDR47;refGene_tx=XM_019837549.2;refGene_gn=WDR47;refGene_tx=XM_019837550.2;refGene_gn=WDR47;refGene_tx=XM_019837551.2;refGene_gn=WDR47;refGene_tx=XM_019837552.2;refGene_gn=WDR47;refGene_tx=XM_019837553.2;refGene_gn=WDR47;refGene_tx=XM_019837554.1
C1 93364760 93364939
refGene_gn=WDR47;refGene_tx=XM_019837545.2;refGene_gn=WDR47;refGene_tx=XM_019837546.2;refGene_gn=WDR47;refGene_tx=XM_019837547.2;refGene_gn=WDR47;refGene_tx=XM_019837548.2;refGene_gn=WDR47;refGene_tx=XM_019837549.2;refGene_gn=WDR47;refGene_tx=XM_019837550.2;refGene_gn=WDR47;refGene_tx=XM_019837551.2;refGene_gn=WDR47;refGene_tx=XM_019837552.2;refGene_gn=WDR47;refGene_tx=XM_019837553.2;refGene_gn=WDR47;refGene_tx=XM_019837554.1
C1 93374163 93374290
refGene_gn=WDR47;refGene_tx=XM_019837545.2;refGene_gn=WDR47;refGene_tx=XM_019837546.2;refGene_gn=WDR47;refGene_tx=XM_019837547.2;refGene_gn=WDR47;refGene_tx=XM_019837548.2;refGene_gn=WDR47;refGene_tx=XM_019837549.2;refGene_gn=WDR47;refGene_tx=XM_019837550.2;refGene_gn=WDR47;refGene_tx=XM_019837551.2;refGene_gn=WDR47;refGene_tx=XM_019837552.2;refGene_gn=WDR47;refGene_tx=XM_019837553.2;refGene_gn=WDR47;refGene_tx=XM_019837554.1
C1 93379281 93380084
refGene_gn=WDR47;refGene_tx=XM_019837545.2;refGene_gn=WDR47;refGene_tx=XM_019837546.2;refGene_gn=WDR47;refGene_tx=XM_019837547.2;refGene_gn=WDR47;refGene_tx=XM_019837548.2;refGene_gn=WDR47;refGene_tx=XM_019837549.2;refGene_gn=WDR47;refGene_tx=XM_019837550.2;refGene_gn=WDR47;refGene_tx=XM_019837551.2;refGene_gn=WDR47;refGene_tx=XM_019837552.2;refGen

e_gn=WDR47;refGene_tx=XM_019837553.2;refGene_gn=WDR47;refGene_tx=XM_019837554.1
C1 93381537 93381600 refGene_gn=WDR47;refGene_tx=XM_019837554.1
C1 93382510 93382616
refGene_gn=WDR47;refGene_tx=XM_019837545.2;refGene_gn=WDR47;refGene_tx=XM_019837546.2;refGene_gn=WDR47;refGene_tx=XM_019837547.2;refGene_gn=WDR47;refGene_tx=XM_019837548.2;refGene_gn=WDR47;refGene_tx=XM_019837549.2;refGene_gn=WDR47;refGene_tx=XM_019837550.2;refGene_gn=WDR47;refGene_tx=XM_019837551.2;refGene_gn=WDR47;refGene_tx=XM_019837552.2;refGene_gn=WDR47;refGene_tx=XM_019837553.2
C1 93387496 93387580
refGene_gn=WDR47;refGene_tx=XM_019837545.2;refGene_gn=WDR47;refGene_tx=XM_019837546.2;refGene_gn=WDR47;refGene_tx=XM_019837547.2;refGene_gn=WDR47;refGene_tx=XM_019837548.2;refGene_gn=WDR47;refGene_tx=XM_019837549.2;refGene_gn=WDR47;refGene_tx=XM_019837550.2;refGene_gn=WDR47;refGene_tx=XM_019837551.2;refGene_gn=WDR47;refGene_tx=XM_019837552.2;refGene_gn=WDR47;refGene_tx=XM_019837553.2
C1 93389785 93389943
refGene_gn=WDR47;refGene_tx=XM_019837545.2;refGene_gn=WDR47;refGene_tx=XM_019837546.2;refGene_gn=WDR47;refGene_tx=XM_019837547.2;refGene_gn=WDR47;refGene_tx=XM_019837548.2;refGene_gn=WDR47;refGene_tx=XM_019837549.2;refGene_gn=WDR47;refGene_tx=XM_019837550.2;refGene_gn=WDR47;refGene_tx=XM_019837551.2;refGene_gn=WDR47;refGene_tx=XM_019837552.2;refGene_gn=WDR47;refGene_tx=XM_019837553.2
C1 93420005 93420176 refGene_gn=TAF13;refGene_tx=XM_003990394.5
C1 93422199 93422297 refGene_gn=TAF13;refGene_tx=XM_003990394.5
C1 93426329 93426408 refGene_gn=TAF13;refGene_tx=XM_003990394.5
C1 93427506 93427533 refGene_gn=TAF13;refGene_tx=XM_003990394.5
C1 93438422 93438432 refGene_gn=TMEM167B;refGene_tx=XM_003990395.5
C1 93440575 93440707 refGene_gn=TMEM167B;refGene_tx=XM_003990395.5
C1 93441246 93441329 refGene_gn=TMEM167B;refGene_tx=XM_003990395.5
C1 93446871 93446942
refGene_gn=CC1H1orf194;refGene_tx=XM_006934966.4;refGene_gn=CC1H1orf194;refGene_tx=XM_019837559.2
C1 93447384 93447515
refGene_gn=CC1H1orf194;refGene_tx=XM_006934966.4;refGene_gn=CC1H1orf194;refGene_tx=XM_019837559.2;refGene_gn=CC1H1orf194;refGene_tx=XM_019837560.2
C1 93447879 93447995
refGene_gn=CC1H1orf194;refGene_tx=XM_006934966.4;refGene_gn=CC1H1orf194;refGene_tx=XM_019837559.2;refGene_gn=CC1H1orf194;refGene_tx=XM_019837560.2
C1 93448517 93448634
refGene_gn=CC1H1orf194;refGene_tx=XM_006934966.4;refGene_gn=CC1H1orf194;refGene_tx=XM_019837559.2;refGene_gn=CC1H1orf194;refGene_tx=XM_019837560.2
C1 93456768 93456849
refGene_gn=CC1H1orf194;refGene_tx=XM_006934966.4
C1 93457426 93457729
refGene_gn=CC1H1orf194;refGene_tx=XM_019837559.2;refGene_gn=CC1H1orf19

4;refGene_tx=XM_019837560.2
C1 93458012 93458165
refGene_gn=KIAA1324;refGene_tx=XM_006934965.4;refGene_gn=KIAA1324;refGene_tx=XM_019837556.2;refGene_gn=KIAA1324;refGene_tx=XM_019837557.2
C1 93495991 93496112
refGene_gn=KIAA1324;refGene_tx=XM_006934965.4;refGene_gn=KIAA1324;refGene_tx=XM_019837556.2;refGene_gn=KIAA1324;refGene_tx=XM_019837557.2
C1 93498674 93498867
refGene_gn=KIAA1324;refGene_tx=XM_006934965.4;refGene_gn=KIAA1324;refGene_tx=XM_019837556.2;refGene_gn=KIAA1324;refGene_tx=XM_019837557.2;refGene_gn=KIAA1324;refGene_tx=XM_023258960.1
C1 93503934 93504082
refGene_gn=KIAA1324;refGene_tx=XM_006934965.4;refGene_gn=KIAA1324;refGene_tx=XM_019837556.2;refGene_gn=KIAA1324;refGene_tx=XM_019837557.2;refGene_gn=KIAA1324;refGene_tx=XM_023258960.1
C1 93504784 93504865
refGene_gn=KIAA1324;refGene_tx=XM_006934965.4;refGene_gn=KIAA1324;refGene_tx=XM_019837556.2;refGene_gn=KIAA1324;refGene_tx=XM_019837557.2;refGene_gn=KIAA1324;refGene_tx=XM_023258960.1
C1 93505447 93505553
refGene_gn=KIAA1324;refGene_tx=XM_006934965.4;refGene_gn=KIAA1324;refGene_tx=XM_019837556.2;refGene_gn=KIAA1324;refGene_tx=XM_019837557.2;refGene_gn=KIAA1324;refGene_tx=XM_023258960.1
C1 93505694 93505844
refGene_gn=KIAA1324;refGene_tx=XM_006934965.4;refGene_gn=KIAA1324;refGene_tx=XM_019837556.2;refGene_gn=KIAA1324;refGene_tx=XM_019837557.2;refGene_gn=KIAA1324;refGene_tx=XM_023258960.1
C1 93512060 93512149
refGene_gn=KIAA1324;refGene_tx=XM_006934965.4;refGene_gn=KIAA1324;refGene_tx=XM_019837556.2;refGene_gn=KIAA1324;refGene_tx=XM_019837557.2;refGene_gn=KIAA1324;refGene_tx=XM_023258960.1
C1 93516245 93516423
refGene_gn=KIAA1324;refGene_tx=XM_006934965.4;refGene_gn=KIAA1324;refGene_tx=XM_019837556.2;refGene_gn=KIAA1324;refGene_tx=XM_019837557.2;refGene_gn=KIAA1324;refGene_tx=XM_023258960.1
C1 93517119 93517248
refGene_gn=KIAA1324;refGene_tx=XM_006934965.4;refGene_gn=KIAA1324;refGene_tx=XM_019837556.2;refGene_gn=KIAA1324;refGene_tx=XM_019837557.2;refGene_gn=KIAA1324;refGene_tx=XM_023258960.1
C1 93517643 93517734
refGene_gn=KIAA1324;refGene_tx=XM_006934965.4;refGene_gn=KIAA1324;refGene_tx=XM_019837556.2;refGene_gn=KIAA1324;refGene_tx=XM_019837557.2;refGene_gn=KIAA1324;refGene_tx=XM_023258960.1
C1 93519738 93519844
refGene_gn=KIAA1324;refGene_tx=XM_006934965.4;refGene_gn=KIAA1324;refGene_tx=XM_019837556.2;refGene_gn=KIAA1324;refGene_tx=XM_019837557.2;refGene_gn=KIAA1324;refGene_tx=XM_023258960.1
C1 93520112 93520250
refGene_gn=KIAA1324;refGene_tx=XM_006934965.4;refGene_gn=KIAA1324;refGene_tx=XM_019837556.2;refGene_gn=KIAA1324;refGene_tx=XM_019837557.2;re

fGene_gn=KIAA1324;refGene_tx=XM_023258960.1
C1 93520770 93521034
refGene_gn=KIAA1324;refGene_tx=XM_006934965.4;refGene_gn=KIAA1324;refGene_tx=XM_019837556.2;refGene_gn=KIAA1324;refGene_tx=XM_019837557.2;refGene_gn=KIAA1324;refGene_tx=XM_023258960.1
C1 93522333 93522507
refGene_gn=KIAA1324;refGene_tx=XM_006934965.4;refGene_gn=KIAA1324;refGene_tx=XM_019837556.2;refGene_gn=KIAA1324;refGene_tx=XM_019837557.2;refGene_gn=KIAA1324;refGene_tx=XM_023258960.1
C1 93527932 93528113
refGene_gn=KIAA1324;refGene_tx=XM_006934965.4;refGene_gn=KIAA1324;refGene_tx=XM_019837556.2;refGene_gn=KIAA1324;refGene_tx=XM_019837557.2;refGene_gn=KIAA1324;refGene_tx=XM_023258960.1
C1 93528418 93528515
refGene_gn=KIAA1324;refGene_tx=XM_006934965.4;refGene_gn=KIAA1324;refGene_tx=XM_019837556.2;refGene_gn=KIAA1324;refGene_tx=XM_019837557.2;refGene_gn=KIAA1324;refGene_tx=XM_023258960.1
C1 93528984 93529086
refGene_gn=KIAA1324;refGene_tx=XM_006934965.4;refGene_gn=KIAA1324;refGene_tx=XM_019837556.2;refGene_gn=KIAA1324;refGene_tx=XM_019837557.2;refGene_gn=KIAA1324;refGene_tx=XM_023258960.1
C1 93530415 93530542
refGene_gn=KIAA1324;refGene_tx=XM_006934965.4;refGene_gn=KIAA1324;refGene_tx=XM_019837556.2;refGene_gn=KIAA1324;refGene_tx=XM_019837557.2;refGene_gn=KIAA1324;refGene_tx=XM_023258960.1
C1 93530619 93530798
refGene_gn=KIAA1324;refGene_tx=XM_006934965.4;refGene_gn=KIAA1324;refGene_tx=XM_019837556.2;refGene_gn=KIAA1324;refGene_tx=XM_019837557.2;refGene_gn=KIAA1324;refGene_tx=XM_023258960.1
C1 93531316 93531482
refGene_gn=KIAA1324;refGene_tx=XM_006934965.4;refGene_gn=KIAA1324;refGene_tx=XM_019837556.2;refGene_gn=KIAA1324;refGene_tx=XM_019837557.2;refGene_gn=KIAA1324;refGene_tx=XM_023258960.1
C1 93531725 93531781
refGene_gn=KIAA1324;refGene_tx=XM_019837556.2;refGene_gn=KIAA1324;refGene_tx=XM_023258960.1
C1 93532613 93532713
refGene_gn=KIAA1324;refGene_tx=XM_006934965.4;refGene_gn=KIAA1324;refGene_tx=XM_019837556.2;refGene_gn=KIAA1324;refGene_tx=XM_019837557.2;refGene_gn=KIAA1324;refGene_tx=XM_023258960.1
C1 93540373 93540509 refGene_gn=SARS;refGene_tx=XM_006934990.3
C1 93547351 93547422 refGene_gn=SARS;refGene_tx=XM_006934990.3
C1 93552570 93552648 refGene_gn=SARS;refGene_tx=XM_006934990.3
C1 93553882 93554041 refGene_gn=SARS;refGene_tx=XM_006934990.3
C1 93555014 93555158 refGene_gn=SARS;refGene_tx=XM_006934990.3
C1 93555924 93556080 refGene_gn=SARS;refGene_tx=XM_006934990.3
C1 93558961 93559183 refGene_gn=SARS;refGene_tx=XM_006934990.3
C1 93559740 93559870 refGene_gn=SARS;refGene_tx=XM_006934990.3
C1 93560139 93560297 refGene_gn=SARS;refGene_tx=XM_006934990.3
C1 93561482 93561612 refGene_gn=SARS;refGene_tx=XM_006934990.3

C1 93561953 93562111 refGene_gn=SARS;refGene_tx=XM_006934990.3
C1 93572571 93575902
refGene_gn=CELSR2;refGene_tx=XM_003990400.5;refGene_gn=CELSR2;refGene_tx=XM_011285046.3;refGene_gn=CELSR2;refGene_tx=XM_011285047.3;refGene_gn=CELSR2;refGene_tx=XM_019837561.2
C1 93580251 93580899
refGene_gn=CELSR2;refGene_tx=XM_003990400.5;refGene_gn=CELSR2;refGene_tx=XM_011285046.3;refGene_gn=CELSR2;refGene_tx=XM_011285047.3;refGene_gn=CELSR2;refGene_tx=XM_019837561.2
C1 93582815 93583038
refGene_gn=CELSR2;refGene_tx=XM_003990400.5;refGene_gn=CELSR2;refGene_tx=XM_011285046.3;refGene_gn=CELSR2;refGene_tx=XM_011285047.3;refGene_gn=CELSR2;refGene_tx=XM_019837561.2
C1 93583452 93583568
refGene_gn=CELSR2;refGene_tx=XM_003990400.5;refGene_gn=CELSR2;refGene_tx=XM_011285046.3;refGene_gn=CELSR2;refGene_tx=XM_011285047.3;refGene_gn=CELSR2;refGene_tx=XM_019837561.2
C1 93583744 93583833
refGene_gn=CELSR2;refGene_tx=XM_003990400.5;refGene_gn=CELSR2;refGene_tx=XM_011285046.3;refGene_gn=CELSR2;refGene_tx=XM_011285047.3;refGene_gn=CELSR2;refGene_tx=XM_019837561.2
C1 93584216 93584374
refGene_gn=CELSR2;refGene_tx=XM_003990400.5;refGene_gn=CELSR2;refGene_tx=XM_011285046.3;refGene_gn=CELSR2;refGene_tx=XM_011285047.3;refGene_gn=CELSR2;refGene_tx=XM_019837561.2
C1 93584812 93584976
refGene_gn=CELSR2;refGene_tx=XM_003990400.5;refGene_gn=CELSR2;refGene_tx=XM_011285046.3;refGene_gn=CELSR2;refGene_tx=XM_011285047.3;refGene_gn=CELSR2;refGene_tx=XM_019837561.2
C1 93585129 93585255
refGene_gn=CELSR2;refGene_tx=XM_003990400.5;refGene_gn=CELSR2;refGene_tx=XM_011285046.3;refGene_gn=CELSR2;refGene_tx=XM_011285047.3;refGene_gn=CELSR2;refGene_tx=XM_019837561.2
C1 93585620 93585787
refGene_gn=CELSR2;refGene_tx=XM_003990400.5;refGene_gn=CELSR2;refGene_tx=XM_011285046.3;refGene_gn=CELSR2;refGene_tx=XM_011285047.3;refGene_gn=CELSR2;refGene_tx=XM_019837561.2
C1 93586067 93586355
refGene_gn=CELSR2;refGene_tx=XM_003990400.5;refGene_gn=CELSR2;refGene_tx=XM_011285046.3;refGene_gn=CELSR2;refGene_tx=XM_011285047.3;refGene_gn=CELSR2;refGene_tx=XM_019837561.2
C1 93586438 93586613
refGene_gn=CELSR2;refGene_tx=XM_003990400.5;refGene_gn=CELSR2;refGene_tx=XM_011285046.3;refGene_gn=CELSR2;refGene_tx=XM_011285047.3;refGene_gn=CELSR2;refGene_tx=XM_019837561.2
C1 93586847 93586989
refGene_gn=CELSR2;refGene_tx=XM_003990400.5;refGene_gn=CELSR2;refGene_tx=XM_011285046.3;refGene_gn=CELSR2;refGene_tx=XM_011285047.3;refGene_gn=CELSR2;refGene_tx=XM_019837561.2
C1 93587147 93587268

refGene_gn=CELSR2;refGene_tx=XM_003990400.5;refGene_gn=CELSR2;refGene_tx=XM_011285046.3;refGene_gn=CELSR2;refGene_tx=XM_011285047.3;refGene_gn=CELSR2;refGene_tx=XM_019837561.2
C1 93587667 93587851
refGene_gn=CELSR2;refGene_tx=XM_003990400.5;refGene_gn=CELSR2;refGene_tx=XM_011285046.3;refGene_gn=CELSR2;refGene_tx=XM_011285047.3;refGene_gn=CELSR2;refGene_tx=XM_019837561.2
C1 93588027 93588129
refGene_gn=CELSR2;refGene_tx=XM_003990400.5;refGene_gn=CELSR2;refGene_tx=XM_011285046.3;refGene_gn=CELSR2;refGene_tx=XM_011285047.3;refGene_gn=CELSR2;refGene_tx=XM_019837561.2
C1 93588767 93588862
refGene_gn=CELSR2;refGene_tx=XM_003990400.5;refGene_gn=CELSR2;refGene_tx=XM_011285046.3;refGene_gn=CELSR2;refGene_tx=XM_011285047.3;refGene_gn=CELSR2;refGene_tx=XM_019837561.2
C1 93589078 93589288
refGene_gn=CELSR2;refGene_tx=XM_003990400.5;refGene_gn=CELSR2;refGene_tx=XM_011285046.3;refGene_gn=CELSR2;refGene_tx=XM_011285047.3;refGene_gn=CELSR2;refGene_tx=XM_019837561.2
C1 93589771 93589955
refGene_gn=CELSR2;refGene_tx=XM_003990400.5;refGene_gn=CELSR2;refGene_tx=XM_011285046.3;refGene_gn=CELSR2;refGene_tx=XM_011285047.3;refGene_gn=CELSR2;refGene_tx=XM_019837561.2
C1 93590055 93590184
refGene_gn=CELSR2;refGene_tx=XM_003990400.5;refGene_gn=CELSR2;refGene_tx=XM_011285046.3;refGene_gn=CELSR2;refGene_tx=XM_011285047.3;refGene_gn=CELSR2;refGene_tx=XM_019837561.2
C1 93590270 93590451
refGene_gn=CELSR2;refGene_tx=XM_003990400.5;refGene_gn=CELSR2;refGene_tx=XM_011285046.3;refGene_gn=CELSR2;refGene_tx=XM_011285047.3;refGene_gn=CELSR2;refGene_tx=XM_019837561.2
C1 93590611 93590779
refGene_gn=CELSR2;refGene_tx=XM_003990400.5;refGene_gn=CELSR2;refGene_tx=XM_011285046.3;refGene_gn=CELSR2;refGene_tx=XM_011285047.3;refGene_gn=CELSR2;refGene_tx=XM_019837561.2
C1 93590877 93591004
refGene_gn=CELSR2;refGene_tx=XM_003990400.5;refGene_gn=CELSR2;refGene_tx=XM_011285046.3;refGene_gn=CELSR2;refGene_tx=XM_011285047.3;refGene_gn=CELSR2;refGene_tx=XM_019837561.2
C1 93591118 93591319
refGene_gn=CELSR2;refGene_tx=XM_003990400.5;refGene_gn=CELSR2;refGene_tx=XM_011285046.3;refGene_gn=CELSR2;refGene_tx=XM_011285047.3;refGene_gn=CELSR2;refGene_tx=XM_019837561.2
C1 93591681 93591856
refGene_gn=CELSR2;refGene_tx=XM_003990400.5;refGene_gn=CELSR2;refGene_tx=XM_011285046.3;refGene_gn=CELSR2;refGene_tx=XM_011285047.3;refGene_gn=CELSR2;refGene_tx=XM_019837561.2
C1 93592176 93592289
refGene_gn=CELSR2;refGene_tx=XM_003990400.5;refGene_gn=CELSR2;refGene_tx=XM_011285046.3;refGene_gn=CELSR2;refGene_tx=XM_011285047.3;refGene_

gn=CELSR2;refGene_tx=XM_019837561.2
C1 93592419 93592499
refGene_gn=CELSR2;refGene_tx=XM_003990400.5;refGene_gn=CELSR2;refGene_tx=XM_011285046.3;refGene_gn=CELSR2;refGene_tx=XM_011285047.3;refGene_gn=CELSR2;refGene_tx=XM_019837561.2
C1 93592586 93592713
refGene_gn=CELSR2;refGene_tx=XM_003990400.5;refGene_gn=CELSR2;refGene_tx=XM_011285046.3;refGene_gn=CELSR2;refGene_tx=XM_011285047.3;refGene_gn=CELSR2;refGene_tx=XM_019837561.2
C1 93592803 93592926
refGene_gn=CELSR2;refGene_tx=XM_003990400.5;refGene_gn=CELSR2;refGene_tx=XM_011285046.3;refGene_gn=CELSR2;refGene_tx=XM_011285047.3;refGene_gn=CELSR2;refGene_tx=XM_019837561.2
C1 93593549 93593677
refGene_gn=CELSR2;refGene_tx=XM_003990400.5;refGene_gn=CELSR2;refGene_tx=XM_011285046.3;refGene_gn=CELSR2;refGene_tx=XM_011285047.3;refGene_gn=CELSR2;refGene_tx=XM_019837561.2
C1 93594124 93594213
refGene_gn=CELSR2;refGene_tx=XM_003990400.5;refGene_gn=CELSR2;refGene_tx=XM_011285046.3;refGene_gn=CELSR2;refGene_tx=XM_011285047.3;refGene_gn=CELSR2;refGene_tx=XM_019837561.2
C1 93594318 93594513
refGene_gn=CELSR2;refGene_tx=XM_003990400.5;refGene_gn=CELSR2;refGene_tx=XM_011285046.3;refGene_gn=CELSR2;refGene_tx=XM_011285047.3;refGene_gn=CELSR2;refGene_tx=XM_019837561.2
C1 93594641 93594812
refGene_gn=CELSR2;refGene_tx=XM_003990400.5;refGene_gn=CELSR2;refGene_tx=XM_011285046.3;refGene_gn=CELSR2;refGene_tx=XM_011285047.3
C1 93594960 93595195
refGene_gn=CELSR2;refGene_tx=XM_003990400.5;refGene_gn=CELSR2;refGene_tx=XM_011285046.3;refGene_gn=CELSR2;refGene_tx=XM_011285047.3;refGene_gn=CELSR2;refGene_tx=XM_019837561.2
C1 93595515 93595543 refGene_gn=CELSR2;refGene_tx=XM_003990400.5
C1 93595658 93595680 refGene_gn=CELSR2;refGene_tx=XM_011285047.3
C1 93598287 93598573
refGene_gn=CELSR2;refGene_tx=XM_011285046.3;refGene_gn=CELSR2;refGene_tx=XM_019837561.2
C1 93600400 93600404
refGene_gn=PSRC1;refGene_tx=XM_006934967.4;refGene_gn=PSRC1;refGene_tx=XM_019837571.2
C1 93600833 93600931
refGene_gn=PSRC1;refGene_tx=XM_006934967.4;refGene_gn=PSRC1;refGene_tx=XM_019837571.2;refGene_gn=PSRC1;refGene_tx=XM_019837572.2
C1 93601237 93601486
refGene_gn=PSRC1;refGene_tx=XM_006934967.4;refGene_gn=PSRC1;refGene_tx=XM_019837571.2;refGene_gn=PSRC1;refGene_tx=XM_019837572.2
C1 93601576 93601691
refGene_gn=PSRC1;refGene_tx=XM_006934967.4;refGene_gn=PSRC1;refGene_tx=XM_019837571.2;refGene_gn=PSRC1;refGene_tx=XM_019837572.2
C1 93602044 93602486

refGene_gn=PSRC1;refGene_tx=XM_006934967.4;refGene_gn=PSRC1;refGene_tx=XM_019837571.2;refGene_gn=PSRC1;refGene_tx=XM_019837572.2
C1 93602942 93603000
refGene_gn=PSRC1;refGene_tx=XM_006934967.4;refGene_gn=PSRC1;refGene_tx=XM_019837571.2;refGene_gn=PSRC1;refGene_tx=XM_019837572.2
C1 93603091 93603149
refGene_gn=PSRC1;refGene_tx=XM_006934967.4;refGene_gn=PSRC1;refGene_tx=XM_019837571.2;refGene_gn=PSRC1;refGene_tx=XM_019837572.2
C1 93603621 93603654
refGene_gn=PSRC1;refGene_tx=XM_019837571.2;refGene_gn=PSRC1;refGene_tx=XM_019837572.2
C1 93609140 93609151
refGene_gn=MYBPHL;refGene_tx=XM_019837564.2;refGene_gn=MYBPHL;refGene_tx=XM_019837565.2;refGene_gn=MYBPHL;refGene_tx=XM_019837566.2;refGene_gn=MYBPHL;refGene_tx=XM_019837567.1;refGene_gn=MYBPHL;refGene_tx=XM_019837568.1;refGene_gn=MYBPHL;refGene_tx=XM_019837569.1
C1 93610308 93610495
refGene_gn=MYBPHL;refGene_tx=XM_019837564.2;refGene_gn=MYBPHL;refGene_tx=XM_019837565.2;refGene_gn=MYBPHL;refGene_tx=XM_019837566.2;refGene_gn=MYBPHL;refGene_tx=XM_019837567.1;refGene_gn=MYBPHL;refGene_tx=XM_019837568.1;refGene_gn=MYBPHL;refGene_tx=XM_019837569.1
C1 93611588 93611725
refGene_gn=MYBPHL;refGene_tx=XM_019837564.2;refGene_gn=MYBPHL;refGene_tx=XM_019837565.2;refGene_gn=MYBPHL;refGene_tx=XM_019837566.2;refGene_gn=MYBPHL;refGene_tx=XM_019837567.1;refGene_gn=MYBPHL;refGene_tx=XM_019837568.1;refGene_gn=MYBPHL;refGene_tx=XM_019837569.1
C1 93611827 93611987
refGene_gn=MYBPHL;refGene_tx=XM_019837564.2;refGene_gn=MYBPHL;refGene_tx=XM_019837565.2;refGene_gn=MYBPHL;refGene_tx=XM_019837566.2;refGene_gn=MYBPHL;refGene_tx=XM_019837567.1;refGene_gn=MYBPHL;refGene_tx=XM_019837568.1;refGene_gn=MYBPHL;refGene_tx=XM_019837569.1
C1 93612130 93612270
refGene_gn=MYBPHL;refGene_tx=XM_019837564.2;refGene_gn=MYBPHL;refGene_tx=XM_019837565.2;refGene_gn=MYBPHL;refGene_tx=XM_019837566.2;refGene_gn=MYBPHL;refGene_tx=XM_019837567.1;refGene_gn=MYBPHL;refGene_tx=XM_019837568.1;refGene_gn=MYBPHL;refGene_tx=XM_019837569.1
C1 93612483 93612679
refGene_gn=MYBPHL;refGene_tx=XM_019837564.2;refGene_gn=MYBPHL;refGene_tx=XM_019837565.2;refGene_gn=MYBPHL;refGene_tx=XM_019837566.2;refGene_gn=MYBPHL;refGene_tx=XM_019837567.1;refGene_gn=MYBPHL;refGene_tx=XM_019837568.1;refGene_gn=MYBPHL;refGene_tx=XM_019837569.1
C1 93613238 93613327
refGene_gn=MYBPHL;refGene_tx=XM_019837564.2;refGene_gn=MYBPHL;refGene_tx=XM_019837565.2;refGene_gn=MYBPHL;refGene_tx=XM_019837566.2;refGene_gn=MYBPHL;refGene_tx=XM_019837567.1;refGene_gn=MYBPHL;refGene_tx=XM_019837568.1;refGene_gn=MYBPHL;refGene_tx=XM_019837569.1
C1 93620219 93620502
refGene_gn=MYBPHL;refGene_tx=XM_019837564.2;refGene_gn=MYBPHL;refGene_tx=XM_019837565.2
C1 93621875 93622128

refGene_gn=MYBPHL;refGene_tx=XM_019837566.2;refGene_gn=MYBPHL;refGene_tx=XM_019837567.1;refGene_gn=MYBPHL;refGene_tx=XM_019837568.1;refGene_gn=MYBPHL;refGene_tx=XM_019837569.1
C1 93622540 93622562
refGene_gn=MYBPHL;refGene_tx=XM_019837567.1;refGene_gn=MYBPHL;refGene_tx=XM_019837568.1;refGene_gn=MYBPHL;refGene_tx=XM_019837569.1
C1 93629131 93629146
refGene_gn=SORT1;refGene_tx=XM_023258961.1;refGene_gn=SORT1;refGene_tx=XM_023258962.1
C1 93629362 93629486
refGene_gn=SORT1;refGene_tx=XM_023258961.1;refGene_gn=SORT1;refGene_tx=XM_023258962.1
C1 93629761 93629868
refGene_gn=SORT1;refGene_tx=XM_023258961.1;refGene_gn=SORT1;refGene_tx=XM_023258962.1
C1 93631297 93631406
refGene_gn=SORT1;refGene_tx=XM_023258961.1;refGene_gn=SORT1;refGene_tx=XM_023258962.1
C1 93632010 93632127
refGene_gn=SORT1;refGene_tx=XM_023258961.1;refGene_gn=SORT1;refGene_tx=XM_023258962.1
C1 93635791 93635981
refGene_gn=SORT1;refGene_tx=XM_023258961.1;refGene_gn=SORT1;refGene_tx=XM_023258962.1
C1 93638219 93638410
refGene_gn=SORT1;refGene_tx=XM_023258961.1;refGene_gn=SORT1;refGene_tx=XM_023258962.1
C1 93639248 93639417
refGene_gn=SORT1;refGene_tx=XM_023258961.1;refGene_gn=SORT1;refGene_tx=XM_023258962.1
C1 93639755 93639858
refGene_gn=SORT1;refGene_tx=XM_023258961.1;refGene_gn=SORT1;refGene_tx=XM_023258962.1
C1 93642726 93642833
refGene_gn=SORT1;refGene_tx=XM_023258961.1;refGene_gn=SORT1;refGene_tx=XM_023258962.1
C1 93644480 93644636
refGene_gn=SORT1;refGene_tx=XM_023258961.1;refGene_gn=SORT1;refGene_tx=XM_023258962.1
C1 93645616 93645761
refGene_gn=SORT1;refGene_tx=XM_023258961.1;refGene_gn=SORT1;refGene_tx=XM_023258962.1
C1 93648885 93649016
refGene_gn=SORT1;refGene_tx=XM_023258961.1;refGene_gn=SORT1;refGene_tx=XM_023258962.1
C1 93649894 93649944
refGene_gn=SORT1;refGene_tx=XM_023258961.1;refGene_gn=SORT1;refGene_tx=XM_023258962.1
C1 93654234 93654308
refGene_gn=SORT1;refGene_tx=XM_023258961.1;refGene_gn=SORT1;refGene_tx

=XM_023258962.1
C1	93657987 93658152
refGene_gn=SORT1;refGene_tx=XM_023258961.1;refGene_gn=SORT1;refGene_tx=XM_023258962.1
C1	93658685 93658788
refGene_gn=SORT1;refGene_tx=XM_023258961.1;refGene_gn=SORT1;refGene_tx=XM_023258962.1
C1	93668210 93668284
refGene_gn=SORT1;refGene_tx=XM_023258961.1;refGene_gn=SORT1;refGene_tx=XM_023258962.1
C1	93669722 93669782
refGene_gn=SORT1;refGene_tx=XM_023258961.1;refGene_gn=SORT1;refGene_tx=XM_023258962.1
C1	93692955 93693267
refGene_gn=SORT1;refGene_tx=XM_023258961.1;refGene_gn=SORT1;refGene_tx=XM_023258962.1
C1	93697429 93697507 refGene_gn=PSMA5;refGene_tx=XM_003990402.3
C1	93707762 93707849 refGene_gn=PSMA5;refGene_tx=XM_003990402.3
C1	93708864 93708967 refGene_gn=PSMA5;refGene_tx=XM_003990402.3
C1	93709850 93709909 refGene_gn=PSMA5;refGene_tx=XM_003990402.3
C1	93710048 93710156 refGene_gn=PSMA5;refGene_tx=XM_003990402.3
C1	93711064 93711132 refGene_gn=PSMA5;refGene_tx=XM_003990402.3
C1	93713564 93713691 refGene_gn=PSMA5;refGene_tx=XM_003990402.3
C1	93720037 93720104 refGene_gn=PSMA5;refGene_tx=XM_003990402.3
C1	93724395 93724424 refGene_gn=PSMA5;refGene_tx=XM_003990402.3
C1	93757128 93757182 refGene_gn=SYPL2;refGene_tx=XM_023258963.1
C1	93757342 93757417 refGene_gn=SYPL2;refGene_tx=XM_023258963.1
C1	93764243 93764368 refGene_gn=SYPL2;refGene_tx=XM_023258963.1
C1	93765454 93765656 refGene_gn=SYPL2;refGene_tx=XM_023258963.1
C1	93766802 93766994 refGene_gn=SYPL2;refGene_tx=XM_023258963.1
C1	93768357 93768510 refGene_gn=SYPL2;refGene_tx=XM_023258963.1
C1	93772852 93772979
refGene_gn=ATXN7L2;refGene_tx=XM_011285161.3;refGene_gn=ATXN7L2;refGene_tx=XM_019837574.2;refGene_gn=ATXN7L2;refGene_tx=XM_023258964.1
C1	93774892 93774958
refGene_gn=ATXN7L2;refGene_tx=XM_011285161.3;refGene_gn=ATXN7L2;refGene_tx=XM_019837574.2;refGene_gn=ATXN7L2;refGene_tx=XM_023258964.1
C1	93775120 93775225
refGene_gn=ATXN7L2;refGene_tx=XM_011285161.3;refGene_gn=ATXN7L2;refGene_tx=XM_019837574.2;refGene_gn=ATXN7L2;refGene_tx=XM_023258964.1
C1	93775629 93775837
refGene_gn=ATXN7L2;refGene_tx=XM_011285161.3;refGene_gn=ATXN7L2;refGene_tx=XM_019837574.2;refGene_gn=ATXN7L2;refGene_tx=XM_023258964.1
C1	93776140 93776427
refGene_gn=ATXN7L2;refGene_tx=XM_011285161.3;refGene_gn=ATXN7L2;refGene_tx=XM_019837574.2;refGene_gn=ATXN7L2;refGene_tx=XM_023258964.1
C1	93777019 93777102
refGene_gn=ATXN7L2;refGene_tx=XM_011285161.3;refGene_gn=ATXN7L2;refGene_tx=XM_019837574.2;refGene_gn=ATXN7L2;refGene_tx=XM_023258964.1
C1	93777458 93777709

refGene_gn=ATXN7L2;refGene_tx=XM_011285161.3;refGene_gn=ATXN7L2;refGene_tx=XM_019837574.2;refGene_gn=ATXN7L2;refGene_tx=XM_023258964.1
C1 93778508 93778707
refGene_gn=ATXN7L2;refGene_tx=XM_011285161.3;refGene_gn=ATXN7L2;refGene_tx=XM_019837574.2;refGene_gn=ATXN7L2;refGene_tx=XM_023258964.1
C1 93778850 93778972
refGene_gn=ATXN7L2;refGene_tx=XM_011285161.3;refGene_gn=ATXN7L2;refGene_tx=XM_019837574.2;refGene_gn=ATXN7L2;refGene_tx=XM_023258964.1
C1 93779462 93780354
refGene_gn=ATXN7L2;refGene_tx=XM_011285161.3;refGene_gn=ATXN7L2;refGene_tx=XM_019837574.2;refGene_gn=ATXN7L2;refGene_tx=XM_023258964.1
C1 93781276 93781291 refGene_gn=ATXN7L2;refGene_tx=XM_011285161.3
C1 93782849 93782997
refGene_gn=LOC101099231;refGene_tx=XM_003990404.5;refGene_gn=LOC101099231;refGene_tx=XM_003990406.5
C1 93783531 93783569
refGene_gn=LOC101099231;refGene_tx=XM_003990404.5
C1 93784166 93784670
refGene_gn=LOC101099231;refGene_tx=XM_003990404.5;refGene_gn=LOC101099231;refGene_tx=XM_003990406.5
C1 93795432 93796914
refGene_gn=AMIGO1;refGene_tx=XM_006934973.4;refGene_gn=AMIGO1;refGene_tx=XM_006934974.4
C1 93828165 93829521 refGene_gn=GPR61;refGene_tx=XM_019837575.1
C1 93833848 93833966 refGene_gn=GNAI3;refGene_tx=XM_003990409.5
C1 93854579 93854622 refGene_gn=GNAI3;refGene_tx=XM_003990409.5
C1 93854734 93854876 refGene_gn=GNAI3;refGene_tx=XM_003990409.5
C1 93859712 93859870 refGene_gn=GNAI3;refGene_tx=XM_003990409.5
C1 93862100 93862229 refGene_gn=GNAI3;refGene_tx=XM_003990409.5
C1 93865331 93865461 refGene_gn=GNAI3;refGene_tx=XM_003990409.5
C1 93865877 93866031 refGene_gn=GNAI3;refGene_tx=XM_003990409.5
C1 93870382 93870573 refGene_gn=GNAI3;refGene_tx=XM_003990409.5
C1 93880328 93880519
refGene_gn=GNAT2;refGene_tx=XM_003990410.4;refGene_gn=GNAT2;refGene_tx=XM_019837577.1;refGene_gn=GNAT2;refGene_tx=XM_019837578.1
C1 93881529 93881683
refGene_gn=GNAT2;refGene_tx=XM_003990410.4;refGene_gn=GNAT2;refGene_tx=XM_019837577.1;refGene_gn=GNAT2;refGene_tx=XM_019837578.1
C1 93883361 93883491
refGene_gn=GNAT2;refGene_tx=XM_003990410.4;refGene_gn=GNAT2;refGene_tx=XM_019837577.1;refGene_gn=GNAT2;refGene_tx=XM_019837578.1
C1 93883722 93883851
refGene_gn=GNAT2;refGene_tx=XM_003990410.4;refGene_gn=GNAT2;refGene_tx=XM_019837577.1;refGene_gn=GNAT2;refGene_tx=XM_019837578.1
C1 93886019 93886177
refGene_gn=GNAT2;refGene_tx=XM_003990410.4;refGene_gn=GNAT2;refGene_tx=XM_019837577.1;refGene_gn=GNAT2;refGene_tx=XM_019837578.1
C1 93886896 93887259
refGene_gn=GNAT2;refGene_tx=XM_003990410.4;refGene_gn=GNAT2;refGene_tx=XM_019837577.1;refGene_gn=GNAT2;refGene_tx=XM_019837578.1

C1	93887325	93887368
refGene_gn=GNAT2;refGene_tx=XM_003990410.4;refGene_gn=GNAT2;refGene_tx=XM_019837578.1
C1	93888437	93888459	refGene_gn=GNAT2;refGene_tx=XM_019837578.1
C1	93889055	93889173	refGene_gn=GNAT2;refGene_tx=XM_003990410.4
C1	93896177	93896187	refGene_gn=AMPD2;refGene_tx=XM_011285052.3
C1	93897014	93897170
refGene_gn=AMPD2;refGene_tx=XM_011285054.2;refGene_gn=AMPD2;refGene_tx=XM_011285055.3
C1	93900923	93901054
refGene_gn=AMPD2;refGene_tx=XM_011285052.3;refGene_gn=AMPD2;refGene_tx=XM_011285053.2;refGene_gn=AMPD2;refGene_tx=XM_011285054.2
C1	93901289	93901420
refGene_gn=AMPD2;refGene_tx=XM_011285052.3;refGene_gn=AMPD2;refGene_tx=XM_011285053.2;refGene_gn=AMPD2;refGene_tx=XM_011285054.2;refGene_gn=AMPD2;refGene_tx=XM_011285055.3
C1	93901970	93902039
refGene_gn=AMPD2;refGene_tx=XM_011285052.3;refGene_gn=AMPD2;refGene_tx=XM_011285053.2;refGene_gn=AMPD2;refGene_tx=XM_011285054.2;refGene_gn=AMPD2;refGene_tx=XM_011285055.3
C1	93902137	93902249
refGene_gn=AMPD2;refGene_tx=XM_011285052.3;refGene_gn=AMPD2;refGene_tx=XM_011285053.2;refGene_gn=AMPD2;refGene_tx=XM_011285054.2;refGene_gn=AMPD2;refGene_tx=XM_011285055.3
C1	93902498	93902685
refGene_gn=AMPD2;refGene_tx=XM_011285052.3;refGene_gn=AMPD2;refGene_tx=XM_011285053.2;refGene_gn=AMPD2;refGene_tx=XM_011285054.2;refGene_gn=AMPD2;refGene_tx=XM_011285055.3
C1	93902920	93903062
refGene_gn=AMPD2;refGene_tx=XM_011285052.3;refGene_gn=AMPD2;refGene_tx=XM_011285053.2;refGene_gn=AMPD2;refGene_tx=XM_011285054.2;refGene_gn=AMPD2;refGene_tx=XM_011285055.3
C1	93903177	93903267
refGene_gn=AMPD2;refGene_tx=XM_011285052.3;refGene_gn=AMPD2;refGene_tx=XM_011285053.2;refGene_gn=AMPD2;refGene_tx=XM_011285054.2;refGene_gn=AMPD2;refGene_tx=XM_011285055.3
C1	93903504	93903634
refGene_gn=AMPD2;refGene_tx=XM_011285052.3;refGene_gn=AMPD2;refGene_tx=XM_011285053.2;refGene_gn=AMPD2;refGene_tx=XM_011285054.2;refGene_gn=AMPD2;refGene_tx=XM_011285055.3
C1	93903791	93903986
refGene_gn=AMPD2;refGene_tx=XM_011285052.3;refGene_gn=AMPD2;refGene_tx=XM_011285053.2;refGene_gn=AMPD2;refGene_tx=XM_011285054.2;refGene_gn=AMPD2;refGene_tx=XM_011285055.3
C1	93904073	93904205
refGene_gn=AMPD2;refGene_tx=XM_011285052.3;refGene_gn=AMPD2;refGene_tx=XM_011285053.2;refGene_gn=AMPD2;refGene_tx=XM_011285054.2;refGene_gn=AMPD2;refGene_tx=XM_011285055.3
C1	93904411	93904575
refGene_gn=AMPD2;refGene_tx=XM_011285052.3;refGene_gn=AMPD2;refGene_tx

=XM_011285053.2;refGene_gn=AMPD2;refGene_tx=XM_011285054.2;refGene_gn=AMPD2;refGene_tx=XM_011285055.3
C1 93904859 93904986
refGene_gn=AMPD2;refGene_tx=XM_011285052.3;refGene_gn=AMPD2;refGene_tx=XM_011285053.2;refGene_gn=AMPD2;refGene_tx=XM_011285054.2;refGene_gn=AMPD2;refGene_tx=XM_011285055.3
C1 93905077 93905241
refGene_gn=AMPD2;refGene_tx=XM_011285052.3;refGene_gn=AMPD2;refGene_tx=XM_011285053.2;refGene_gn=AMPD2;refGene_tx=XM_011285054.2;refGene_gn=AMPD2;refGene_tx=XM_011285055.3
C1 93905499 93905620
refGene_gn=AMPD2;refGene_tx=XM_011285052.3;refGene_gn=AMPD2;refGene_tx=XM_011285053.2;refGene_gn=AMPD2;refGene_tx=XM_011285054.2;refGene_gn=AMPD2;refGene_tx=XM_011285055.3
C1 93905913 93906087
refGene_gn=AMPD2;refGene_tx=XM_011285052.3;refGene_gn=AMPD2;refGene_tx=XM_011285053.2;refGene_gn=AMPD2;refGene_tx=XM_011285054.2;refGene_gn=AMPD2;refGene_tx=XM_011285055.3
C1 93906324 93906435
refGene_gn=AMPD2;refGene_tx=XM_011285052.3;refGene_gn=AMPD2;refGene_tx=XM_011285053.2;refGene_gn=AMPD2;refGene_tx=XM_011285054.2;refGene_gn=AMPD2;refGene_tx=XM_011285055.3
C1 93906583 93906793
refGene_gn=AMPD2;refGene_tx=XM_011285052.3;refGene_gn=AMPD2;refGene_tx=XM_011285053.2;refGene_gn=AMPD2;refGene_tx=XM_011285054.2;refGene_gn=AMPD2;refGene_tx=XM_011285055.3
C1 93947677 93947738
refGene_gn=LOC101100824;refGene_tx=XM_023257837.1
C1 93948491 93948635
refGene_gn=LOC101100824;refGene_tx=XM_023257837.1
C1 93949080 93949145
refGene_gn=LOC101100824;refGene_tx=XM_023257837.1
C1 93949418 93949485
refGene_gn=LOC101100824;refGene_tx=XM_023257837.1
C1 93949587 93949691
refGene_gn=LOC101100824;refGene_tx=XM_023257837.1
C1 93950492 93950588
refGene_gn=LOC101100824;refGene_tx=XM_023257837.1
C1 93950672 93950783
refGene_gn=LOC101100824;refGene_tx=XM_023257837.1
C1 93953088 93953178
refGene_gn=LOC101100824;refGene_tx=XM_023257837.1
C1 93970476 93970575 refGene_gn=GSTM3;refGene_tx=XM_003990413.5
C1 93970885 93970996 refGene_gn=GSTM3;refGene_tx=XM_003990413.5
C1 93971084 93971180 refGene_gn=GSTM3;refGene_tx=XM_003990413.5
C1 93971501 93971602 refGene_gn=GSTM3;refGene_tx=XM_003990413.5
C1 93971695 93971777 refGene_gn=GSTM3;refGene_tx=XM_003990413.5
C1 93972724 93972789 refGene_gn=GSTM3;refGene_tx=XM_003990413.5
C1 93973150 93973226 refGene_gn=GSTM3;refGene_tx=XM_003990413.5
C1 93973622 93973673 refGene_gn=GSTM3;refGene_tx=XM_003990413.5

C1 93982338 93982350
refGene_gn=EPS8L3;refGene_tx=XM_019837582.1;refGene_gn=EPS8L3;refGene_tx=XM_019837583.2
C1 93982612 93982745
refGene_gn=EPS8L3;refGene_tx=XM_019837582.1;refGene_gn=EPS8L3;refGene_tx=XM_019837583.2
C1 93983232 93983306
refGene_gn=EPS8L3;refGene_tx=XM_019837582.1;refGene_gn=EPS8L3;refGene_tx=XM_019837583.2
C1 93983602 93983722
refGene_gn=EPS8L3;refGene_tx=XM_019837582.1;refGene_gn=EPS8L3;refGene_tx=XM_019837583.2
C1 93983927 93984114
refGene_gn=EPS8L3;refGene_tx=XM_019837582.1;refGene_gn=EPS8L3;refGene_tx=XM_019837583.2
C1 93984624 93984659
refGene_gn=EPS8L3;refGene_tx=XM_019837582.1;refGene_gn=EPS8L3;refGene_tx=XM_019837583.2
C1 93984917 93984999
refGene_gn=EPS8L3;refGene_tx=XM_019837582.1;refGene_gn=EPS8L3;refGene_tx=XM_019837583.2
C1 93988068 93988217
refGene_gn=EPS8L3;refGene_tx=XM_019837582.1;refGene_gn=EPS8L3;refGene_tx=XM_019837583.2
C1 93988574 93988649
refGene_gn=EPS8L3;refGene_tx=XM_019837582.1;refGene_gn=EPS8L3;refGene_tx=XM_019837583.2
C1 93988904 93988966
refGene_gn=EPS8L3;refGene_tx=XM_019837582.1;refGene_gn=EPS8L3;refGene_tx=XM_019837583.2
C1 93989054 93989175
refGene_gn=EPS8L3;refGene_tx=XM_019837582.1;refGene_gn=EPS8L3;refGene_tx=XM_019837583.2
C1 93989436 93989552
refGene_gn=EPS8L3;refGene_tx=XM_019837582.1;refGene_gn=EPS8L3;refGene_tx=XM_019837583.2
C1 93989639 93989773
refGene_gn=EPS8L3;refGene_tx=XM_019837582.1;refGene_gn=EPS8L3;refGene_tx=XM_019837583.2
C1 93990176 93990232
refGene_gn=EPS8L3;refGene_tx=XM_019837582.1;refGene_gn=EPS8L3;refGene_tx=XM_019837583.2
C1 93990348 93990498
refGene_gn=EPS8L3;refGene_tx=XM_019837582.1;refGene_gn=EPS8L3;refGene_tx=XM_019837583.2
C1 93990786 93990945
refGene_gn=EPS8L3;refGene_tx=XM_019837582.1;refGene_gn=EPS8L3;refGene_tx=XM_019837583.2
C1 93992291 93992356
refGene_gn=EPS8L3;refGene_tx=XM_019837582.1;refGene_gn=EPS8L3;refGene_

tx=XM_019837583.2
C1 93992507 93992538
refGene_gn=EPS8L3;refGene_tx=XM_019837582.1;refGene_gn=EPS8L3;refGene_tx=XM_019837583.2
C1 94122197 94122236
refGene_gn=CSF1;refGene_tx=XM_023258965.1;refGene_gn=CSF1;refGene_tx=XM_023258966.1;refGene_gn=CSF1;refGene_tx=XM_023258967.1;refGene_gn=CSF1;refGene_tx=XM_023258968.1
C1 94125444 94125567
refGene_gn=CSF1;refGene_tx=XM_023258965.1;refGene_gn=CSF1;refGene_tx=XM_023258966.1;refGene_gn=CSF1;refGene_tx=XM_023258967.1;refGene_gn=CSF1;refGene_tx=XM_023258968.1
C1 94126805 94126868
refGene_gn=CSF1;refGene_tx=XM_023258965.1;refGene_gn=CSF1;refGene_tx=XM_023258966.1;refGene_gn=CSF1;refGene_tx=XM_023258967.1;refGene_gn=CSF1;refGene_tx=XM_023258968.1
C1 94128445 94128616
refGene_gn=CSF1;refGene_tx=XM_023258965.1;refGene_gn=CSF1;refGene_tx=XM_023258966.1;refGene_gn=CSF1;refGene_tx=XM_023258967.1;refGene_gn=CSF1;refGene_tx=XM_023258968.1
C1 94132381 94132529
refGene_gn=CSF1;refGene_tx=XM_023258965.1;refGene_gn=CSF1;refGene_tx=XM_023258966.1;refGene_gn=CSF1;refGene_tx=XM_023258967.1;refGene_gn=CSF1;refGene_tx=XM_023258968.1
C1 94133720 94134787
refGene_gn=CSF1;refGene_tx=XM_023258965.1;refGene_gn=CSF1;refGene_tx=XM_023258966.1;refGene_gn=CSF1;refGene_tx=XM_023258967.1;refGene_gn=CSF1;refGene_tx=XM_023258968.1
C1 94135370 94135423
refGene_gn=CSF1;refGene_tx=XM_023258965.1;refGene_gn=CSF1;refGene_tx=XM_023258967.1;refGene_gn=CSF1;refGene_tx=XM_023258968.1
C1 94135758 94135804
refGene_gn=CSF1;refGene_tx=XM_023258965.1;refGene_gn=CSF1;refGene_tx=XM_023258966.1;refGene_gn=CSF1;refGene_tx=XM_023258967.1;refGene_gn=CSF1;refGene_tx=XM_023258968.1
C1 94190098 94190218
refGene_gn=AHCYL1;refGene_tx=XM_011285057.3;refGene_gn=AHCYL1;refGene_tx=XM_011285058.3
C1 94213705 94213817
refGene_gn=AHCYL1;refGene_tx=XM_011285057.3;refGene_gn=AHCYL1;refGene_tx=XM_011285058.3
C1 94216314 94216458
refGene_gn=AHCYL1;refGene_tx=XM_011285057.3;refGene_gn=AHCYL1;refGene_tx=XM_011285058.3
C1 94217461 94217562
refGene_gn=AHCYL1;refGene_tx=XM_011285057.3;refGene_gn=AHCYL1;refGene_tx=XM_011285058.3
C1 94217972 94218075
refGene_gn=AHCYL1;refGene_tx=XM_011285057.3;refGene_gn=AHCYL1;refGene_tx=XM_011285058.3

C1 94219448 94219543
refGene_gn=AHCYL1;refGene_tx=XM_011285057.3;refGene_gn=AHCYL1;refGene_tx=XM_011285058.3
C1 94220112 94220219
refGene_gn=AHCYL1;refGene_tx=XM_011285057.3;refGene_gn=AHCYL1;refGene_tx=XM_011285058.3
C1 94221037 94221154
refGene_gn=AHCYL1;refGene_tx=XM_011285057.3;refGene_gn=AHCYL1;refGene_tx=XM_011285058.3
C1 94221394 94221458
refGene_gn=AHCYL1;refGene_tx=XM_011285057.3;refGene_gn=AHCYL1;refGene_tx=XM_011285058.3
C1 94222264 94222353
refGene_gn=AHCYL1;refGene_tx=XM_011285057.3;refGene_gn=AHCYL1;refGene_tx=XM_011285058.3
C1 94222718 94222789
refGene_gn=AHCYL1;refGene_tx=XM_011285057.3;refGene_gn=AHCYL1;refGene_tx=XM_011285058.3
C1 94223155 94223250
refGene_gn=AHCYL1;refGene_tx=XM_011285057.3;refGene_gn=AHCYL1;refGene_tx=XM_011285058.3
C1 94223337 94223436
refGene_gn=AHCYL1;refGene_tx=XM_011285057.3;refGene_gn=AHCYL1;refGene_tx=XM_011285058.3
C1 94223830 94223899
refGene_gn=AHCYL1;refGene_tx=XM_011285057.3;refGene_gn=AHCYL1;refGene_tx=XM_011285058.3
C1 94224343 94224422
refGene_gn=AHCYL1;refGene_tx=XM_011285057.3;refGene_gn=AHCYL1;refGene_tx=XM_011285058.3
C1 94225767 94225892
refGene_gn=AHCYL1;refGene_tx=XM_011285057.3;refGene_gn=AHCYL1;refGene_tx=XM_011285058.3
C1 94226410 94226417 refGene_gn=AHCYL1;refGene_tx=XM_011285057.3
C1 94240833 94241013 refGene_gn=STRIP1;refGene_tx=XM_003990418.4
C1 94244100 94244170 refGene_gn=STRIP1;refGene_tx=XM_003990418.4
C1 94244904 94244979 refGene_gn=STRIP1;refGene_tx=XM_003990418.4
C1 94245411 94245546 refGene_gn=STRIP1;refGene_tx=XM_003990418.4
C1 94245625 94245746 refGene_gn=STRIP1;refGene_tx=XM_003990418.4
C1 94246010 94246079 refGene_gn=STRIP1;refGene_tx=XM_003990418.4
C1 94247019 94247126 refGene_gn=STRIP1;refGene_tx=XM_003990418.4
C1 94247239 94247367 refGene_gn=STRIP1;refGene_tx=XM_003990418.4
C1 94248473 94248656 refGene_gn=STRIP1;refGene_tx=XM_003990418.4
C1 94249012 94249230 refGene_gn=STRIP1;refGene_tx=XM_003990418.4
C1 94250468 94250534 refGene_gn=STRIP1;refGene_tx=XM_003990418.4
C1 94250657 94250721 refGene_gn=STRIP1;refGene_tx=XM_003990418.4
C1 94252017 94252089 refGene_gn=STRIP1;refGene_tx=XM_003990418.4
C1 94252574 94252649 refGene_gn=STRIP1;refGene_tx=XM_003990418.4
C1 94252859 94252957 refGene_gn=STRIP1;refGene_tx=XM_003990418.4
C1 94254185 94254312 refGene_gn=STRIP1;refGene_tx=XM_003990418.4

C1 94254518 94254619 refGene_gn=STRIP1;refGene_tx=XM_003990418.4
C1 94255340 94255407 refGene_gn=STRIP1;refGene_tx=XM_003990418.4
C1 94255956 94256061 refGene_gn=STRIP1;refGene_tx=XM_003990418.4
C1 94256634 94256839 refGene_gn=STRIP1;refGene_tx=XM_003990418.4
C1 94258226 94258474 refGene_gn=STRIP1;refGene_tx=XM_003990418.4
C1 94265323 94265632 refGene_gn=ALX3;refGene_tx=XM_023258969.1
C1 94266023 94266152 refGene_gn=ALX3;refGene_tx=XM_023258969.1
C1 94269087 94269404 refGene_gn=ALX3;refGene_tx=XM_023258969.1
C1 94274615 94274892 refGene_gn=ALX3;refGene_tx=XM_023258969.1
C1 94316703 94317234 refGene_gn=UBL4B;refGene_tx=XM_006934982.4
C1 94369365 94369651
refGene_gn=SLC6A17;refGene_tx=XM_011285060.3;refGene_gn=SLC6A17;refGene_tx=XM_019837588.1;refGene_gn=SLC6A17;refGene_tx=XM_019837589.1
C1 94374175 94374333
refGene_gn=SLC6A17;refGene_tx=XM_011285060.3;refGene_gn=SLC6A17;refGene_tx=XM_019837588.1;refGene_gn=SLC6A17;refGene_tx=XM_019837589.1
C1 94376064 94376191
refGene_gn=SLC6A17;refGene_tx=XM_011285060.3;refGene_gn=SLC6A17;refGene_tx=XM_019837588.1;refGene_gn=SLC6A17;refGene_tx=XM_019837589.1
C1 94376958 94377140
refGene_gn=SLC6A17;refGene_tx=XM_011285060.3;refGene_gn=SLC6A17;refGene_tx=XM_019837588.1;refGene_gn=SLC6A17;refGene_tx=XM_019837589.1
C1 94378824 94378935
refGene_gn=SLC6A17;refGene_tx=XM_011285060.3;refGene_gn=SLC6A17;refGene_tx=XM_019837588.1;refGene_gn=SLC6A17;refGene_tx=XM_019837589.1
C1 94394349 94394591
refGene_gn=SLC6A17;refGene_tx=XM_011285060.3;refGene_gn=SLC6A17;refGene_tx=XM_019837588.1;refGene_gn=SLC6A17;refGene_tx=XM_019837589.1
C1 94394864 94395057
refGene_gn=SLC6A17;refGene_tx=XM_011285060.3;refGene_gn=SLC6A17;refGene_tx=XM_019837588.1;refGene_gn=SLC6A17;refGene_tx=XM_019837589.1
C1 94396895 94397088
refGene_gn=SLC6A17;refGene_tx=XM_011285060.3;refGene_gn=SLC6A17;refGene_tx=XM_019837588.1;refGene_gn=SLC6A17;refGene_tx=XM_019837589.1
C1 94397565 94397725
refGene_gn=SLC6A17;refGene_tx=XM_011285060.3;refGene_gn=SLC6A17;refGene_tx=XM_019837588.1;refGene_gn=SLC6A17;refGene_tx=XM_019837589.1
C1 94399343 94399506
refGene_gn=SLC6A17;refGene_tx=XM_011285060.3;refGene_gn=SLC6A17;refGene_tx=XM_019837588.1;refGene_gn=SLC6A17;refGene_tx=XM_019837589.1
C1 94399988 94400357
refGene_gn=SLC6A17;refGene_tx=XM_011285060.3;refGene_gn=SLC6A17;refGene_tx=XM_019837588.1;refGene_gn=SLC6A17;refGene_tx=XM_019837589.1
C1 94409872 94410334
refGene_gn=LOC111561791;refGene_tx=XM_023258970.1;refGene_gn=LOC111561791;refGene_tx=XM_023258971.1
C1 94411471 94411910
refGene_gn=LOC111561791;refGene_tx=XM_023258970.1;refGene_gn=LOC111561791;refGene_tx=XM_023258971.1
C1 94412572 94413250

refGene_gn=KCNC4;refGene_tx=XM_003990420.5;refGene_gn=KCNC4;refGene_tx=XM_003990421.5;refGene_gn=KCNC4;refGene_tx=XM_006934983.4;refGene_gn=KCNC4;refGene_tx=XM_006934985.4
C1 94424443 94425380
refGene_gn=KCNC4;refGene_tx=XM_003990420.5;refGene_gn=KCNC4;refGene_tx=XM_003990421.5;refGene_gn=KCNC4;refGene_tx=XM_006934983.4;refGene_gn=KCNC4;refGene_tx=XM_006934985.4
C1 94427430 94427634
refGene_gn=KCNC4;refGene_tx=XM_003990420.5;refGene_gn=KCNC4;refGene_tx=XM_003990421.5;refGene_gn=KCNC4;refGene_tx=XM_006934983.4;refGene_gn=KCNC4;refGene_tx=XM_006934985.4
C1 94430073 94430243 refGene_gn=KCNC4;refGene_tx=XM_006934983.4
C1 94433620 94433676 refGene_gn=KCNC4;refGene_tx=XM_003990421.5
C1 94433682 94433744 refGene_gn=KCNC4;refGene_tx=XM_003990420.5
C1 94456695 94456715 refGene_gn=KCNC4;refGene_tx=XM_006934985.4
C1 94530777 94533643 refGene_gn=RBM15;refGene_tx=XM_003990422.4
C1 94537732 94537779 refGene_gn=RBM15;refGene_tx=XM_003990422.4
C1 94553127 94553180 refGene_gn=SLC16A4;refGene_tx=XM_019837593.2
C1 94553182 94553230 refGene_gn=SLC16A4;refGene_tx=XM_019837593.2
C1 94562558 94562652 refGene_gn=SLC16A4;refGene_tx=XM_019837593.2
C1 94563662 94563874 refGene_gn=SLC16A4;refGene_tx=XM_019837593.2
C1 94566474 94566996 refGene_gn=SLC16A4;refGene_tx=XM_019837593.2
C1 94568357 94568519 refGene_gn=SLC16A4;refGene_tx=XM_019837593.2
C1 94568683 94568827 refGene_gn=SLC16A4;refGene_tx=XM_019837593.2
C1 94569630 94569763 refGene_gn=SLC16A4;refGene_tx=XM_019837593.2
C1 94576372 94576459 refGene_gn=SLC16A4;refGene_tx=XM_019837593.2
C1 94593409 94593470 refGene_gn=LAMTOR5;refGene_tx=XM_003990426.4
C1 94594137 94594255 refGene_gn=LAMTOR5;refGene_tx=XM_003990426.4
C1 94596863 94596925 refGene_gn=LAMTOR5;refGene_tx=XM_003990426.4
C1 94598091 94598126 refGene_gn=LAMTOR5;refGene_tx=XM_003990426.4
C1 94625044 94625263 refGene_gn=PROK1;refGene_tx=XM_019837594.2
C1 94627985 94628057 refGene_gn=PROK1;refGene_tx=XM_003990427.3
C1 94630571 94630697
refGene_gn=PROK1;refGene_tx=XM_003990427.3;refGene_gn=PROK1;refGene_tx=XM_019837594.2
C1 94632292 94632412
refGene_gn=PROK1;refGene_tx=XM_003990427.3;refGene_gn=PROK1;refGene_tx=XM_019837594.2
C1 94645806 94645856
refGene_gn=LOC101083355;refGene_tx=XM_019839172.1
C1 94653049 94653111
refGene_gn=LOC101083355;refGene_tx=XM_003990428.2
C1 94655886 94656037
refGene_gn=LOC101083355;refGene_tx=XM_003990428.2;refGene_gn=LOC101083355;refGene_tx=XM_019839172.1
C1 94656877 94656995
refGene_gn=LOC101083355;refGene_tx=XM_003990428.2;refGene_gn=LOC101083355;refGene_tx=XM_019839172.1
C1 94657813 94657932
refGene_gn=LOC101083355;refGene_tx=XM_003990428.2;refGene_gn=LOC101083

355;refGene_tx=XM_019839172.1
C1 94659096 94659296
refGene_gn=LOC101083355;refGene_tx=XM_003990428.2;refGene_gn=LOC101083355;refGene_tx=XM_019839172.1
C1 94660930 94661044
refGene_gn=LOC101083355;refGene_tx=XM_003990428.2;refGene_gn=LOC101083355;refGene_tx=XM_019839172.1
C1 94661798 94661943
refGene_gn=LOC101083355;refGene_tx=XM_003990428.2;refGene_gn=LOC101083355;refGene_tx=XM_019839172.1
C1 94662449 94662548
refGene_gn=LOC101083355;refGene_tx=XM_003990428.2;refGene_gn=LOC101083355;refGene_tx=XM_019839172.1
C1 94663316 94663454
refGene_gn=LOC101083355;refGene_tx=XM_003990428.2;refGene_gn=LOC101083355;refGene_tx=XM_019839172.1
C1 94686177 94687713 refGene_gn=KCNA10;refGene_tx=XM_003990429.2
C1 94771165 94772665
refGene_gn=KCNA2;refGene_tx=XM_011285061.3;refGene_gn=KCNA2;refGene_tx=XM_019837595.2;refGene_gn=KCNA2;refGene_tx=XM_019837597.2;refGene_gn=KCNA2;refGene_tx=XM_019837598.2;refGene_gn=KCNA2;refGene_tx=XM_023258972.1
C1 94839318 94841061 refGene_gn=KCNA3;refGene_tx=XM_019839096.2
C1 95023932 95024886
refGene_gn=LOC101092952;refGene_tx=XM_003990451.1
C1 95076392 95076455 refGene_gn=CD53;refGene_tx=XM_003990431.3
C1 95077302 95077479 refGene_gn=CD53;refGene_tx=XM_003990431.3
C1 95078942 95079017 refGene_gn=CD53;refGene_tx=XM_003990431.3
C1 95079529 95079625 refGene_gn=CD53;refGene_tx=XM_003990431.3
C1 95080136 95080217 refGene_gn=CD53;refGene_tx=XM_003990431.3
C1 95081300 95081384 refGene_gn=CD53;refGene_tx=XM_003990431.3
C1 95082201 95082273 refGene_gn=CD53;refGene_tx=XM_003990431.3
C1 95103104 95103542
refGene_gn=LRIF1;refGene_tx=XM_003990432.5;refGene_gn=LRIF1;refGene_tx=XM_006934987.4;refGene_gn=LRIF1;refGene_tx=XM_019837600.2
C1 95104395 95104668
refGene_gn=LRIF1;refGene_tx=XM_003990432.5;refGene_gn=LRIF1;refGene_tx=XM_006934987.4;refGene_gn=LRIF1;refGene_tx=XM_019837600.2
C1 95105805 95107315
refGene_gn=LRIF1;refGene_tx=XM_003990432.5;refGene_gn=LRIF1;refGene_tx=XM_019837600.2
C1 95119121 95119189 refGene_gn=LRIF1;refGene_tx=XM_003990432.5
C1 95254095 95254203
refGene_gn=DRAM2;refGene_tx=XM_019837601.1;refGene_gn=DRAM2;refGene_tx=XM_023258974.1;refGene_gn=DRAM2;refGene_tx=XM_023258975.1
C1 95254736 95254829
refGene_gn=DRAM2;refGene_tx=XM_019837601.1;refGene_gn=DRAM2;refGene_tx=XM_023258974.1;refGene_gn=DRAM2;refGene_tx=XM_023258975.1
C1 95256553 95256636
refGene_gn=DRAM2;refGene_tx=XM_019837601.1;refGene_gn=DRAM2;refGene_tx

=XM_023258974.1;refGene_gn=DRAM2;refGene_tx=XM_023258975.1
C1	95257137	95257315
refGene_gn=DRAM2;refGene_tx=XM_019837601.1;refGene_gn=DRAM2;refGene_tx=XM_023258974.1;refGene_gn=DRAM2;refGene_tx=XM_023258975.1
C1	95264441	95264581
refGene_gn=DRAM2;refGene_tx=XM_019837601.1;refGene_gn=DRAM2;refGene_tx=XM_023258974.1;refGene_gn=DRAM2;refGene_tx=XM_023258975.1
C1	95265823	95265891
refGene_gn=DRAM2;refGene_tx=XM_019837601.1;refGene_gn=DRAM2;refGene_tx=XM_023258974.1;refGene_gn=DRAM2;refGene_tx=XM_023258975.1
C1	95269715	95269846
refGene_gn=DRAM2;refGene_tx=XM_019837601.1;refGene_gn=DRAM2;refGene_tx=XM_023258974.1
C1	95283864	95284203
refGene_gn=CEPT1;refGene_tx=XM_003990453.5;refGene_gn=CEPT1;refGene_tx=XM_006934996.4;refGene_gn=CEPT1;refGene_tx=XM_019837604.1
C1	95289137	95289285
refGene_gn=CEPT1;refGene_tx=XM_003990453.5;refGene_gn=CEPT1;refGene_tx=XM_006934996.4;refGene_gn=CEPT1;refGene_tx=XM_019837604.1
C1	95291303	95291445
refGene_gn=CEPT1;refGene_tx=XM_003990453.5;refGene_gn=CEPT1;refGene_tx=XM_006934996.4;refGene_gn=CEPT1;refGene_tx=XM_019837604.1
C1	95297443	95297528	refGene_gn=CEPT1;refGene_tx=XM_006934996.4
C1	95304362	95304447
refGene_gn=CEPT1;refGene_tx=XM_003990453.5;refGene_gn=CEPT1;refGene_tx=XM_019837604.1
C1	95310985	95311117
refGene_gn=CEPT1;refGene_tx=XM_003990453.5;refGene_gn=CEPT1;refGene_tx=XM_006934996.4;refGene_gn=CEPT1;refGene_tx=XM_019837604.1
C1	95311578	95311737
refGene_gn=CEPT1;refGene_tx=XM_003990453.5;refGene_gn=CEPT1;refGene_tx=XM_006934996.4;refGene_gn=CEPT1;refGene_tx=XM_019837604.1
C1	95312285	95312411
refGene_gn=CEPT1;refGene_tx=XM_003990453.5;refGene_gn=CEPT1;refGene_tx=XM_006934996.4;refGene_gn=CEPT1;refGene_tx=XM_019837604.1
C1	95313006	95313126
refGene_gn=CEPT1;refGene_tx=XM_003990453.5;refGene_gn=CEPT1;refGene_tx=XM_006934996.4;refGene_gn=CEPT1;refGene_tx=XM_019837604.1
C1	95316755	95316832
refGene_gn=DENND2D;refGene_tx=XM_003990454.5;refGene_gn=DENND2D;refGene_tx=XM_011285063.3;refGene_gn=DENND2D;refGene_tx=XM_023258976.1;refGene_gn=DENND2D;refGene_tx=XM_023258977.1
C1	95317255	95317498
refGene_gn=DENND2D;refGene_tx=XM_003990454.5;refGene_gn=DENND2D;refGene_tx=XM_011285063.3;refGene_gn=DENND2D;refGene_tx=XM_023258976.1;refGene_gn=DENND2D;refGene_tx=XM_023258977.1
C1	95317800	95317885
refGene_gn=DENND2D;refGene_tx=XM_003990454.5;refGene_gn=DENND2D;refGene_tx=XM_011285063.3;refGene_gn=DENND2D;refGene_tx=XM_023258976.1;refGene_gn=DENND2D;refGene_tx=XM_023258977.1

C1 95318275 95318317
refGene_gn=DENND2D;refGene_tx=XM_003990454.5;refGene_gn=DENND2D;refGene_tx=XM_011285063.3;refGene_gn=DENND2D;refGene_tx=XM_023258976.1;refGene_gn=DENND2D;refGene_tx=XM_023258977.1
C1 95321574 95321752
refGene_gn=DENND2D;refGene_tx=XM_003990454.5;refGene_gn=DENND2D;refGene_tx=XM_011285063.3;refGene_gn=DENND2D;refGene_tx=XM_023258976.1;refGene_gn=DENND2D;refGene_tx=XM_023258977.1
C1 95325866 95326015
refGene_gn=DENND2D;refGene_tx=XM_003990454.5;refGene_gn=DENND2D;refGene_tx=XM_011285063.3;refGene_gn=DENND2D;refGene_tx=XM_023258976.1;refGene_gn=DENND2D;refGene_tx=XM_023258977.1
C1 95327095 95327236
refGene_gn=DENND2D;refGene_tx=XM_003990454.5;refGene_gn=DENND2D;refGene_tx=XM_011285063.3;refGene_gn=DENND2D;refGene_tx=XM_023258976.1;refGene_gn=DENND2D;refGene_tx=XM_023258977.1
C1 95328280 95328358
refGene_gn=DENND2D;refGene_tx=XM_003990454.5;refGene_gn=DENND2D;refGene_tx=XM_011285063.3;refGene_gn=DENND2D;refGene_tx=XM_023258976.1;refGene_gn=DENND2D;refGene_tx=XM_023258977.1
C1 95329079 95329149
refGene_gn=DENND2D;refGene_tx=XM_003990454.5;refGene_gn=DENND2D;refGene_tx=XM_011285063.3;refGene_gn=DENND2D;refGene_tx=XM_023258976.1;refGene_gn=DENND2D;refGene_tx=XM_023258977.1
C1 95329844 95329957
refGene_gn=DENND2D;refGene_tx=XM_003990454.5;refGene_gn=DENND2D;refGene_tx=XM_011285063.3;refGene_gn=DENND2D;refGene_tx=XM_023258976.1;refGene_gn=DENND2D;refGene_tx=XM_023258977.1
C1 95330808 95330984
refGene_gn=DENND2D;refGene_tx=XM_003990454.5;refGene_gn=DENND2D;refGene_tx=XM_011285063.3;refGene_gn=DENND2D;refGene_tx=XM_023258976.1;refGene_gn=DENND2D;refGene_tx=XM_023258977.1
C1 95331585 95331952
refGene_gn=DENND2D;refGene_tx=XM_003990454.5;refGene_gn=DENND2D;refGene_tx=XM_011285063.3
C1 95335571 95335629
refGene_gn=DENND2D;refGene_tx=XM_023258976.1;refGene_gn=DENND2D;refGene_tx=XM_023258977.1
C1 95397039 95397195 refGene_gn=CHIA;refGene_tx=XM_023258978.1
C1 95399692 95399817 refGene_gn=CHIA;refGene_tx=XM_023258978.1
C1 95399998 95400122 refGene_gn=CHIA;refGene_tx=XM_023258978.1
C1 95400404 95400590 refGene_gn=CHIA;refGene_tx=XM_023258978.1
C1 95401326 95401446 refGene_gn=CHIA;refGene_tx=XM_023258978.1
C1 95401532 95401674 refGene_gn=CHIA;refGene_tx=XM_023258978.1
C1 95402471 95402752 refGene_gn=CHIA;refGene_tx=XM_023258978.1
C1 95406527 95406702 refGene_gn=PIFO;refGene_tx=XM_023258979.1
C1 95413226 95413245 refGene_gn=PIFO;refGene_tx=XM_003990455.4
C1 95413474 95413617
refGene_gn=PIFO;refGene_tx=XM_003990455.4;refGene_gn=PIFO;refGene_tx=XM_023258979.1

C1 95414219 95414318
refGene_gn=PIFO;refGene_tx=XM_003990455.4;refGene_gn=PIFO;refGene_tx=XM_023258979.1
C1 95415073 95415204
refGene_gn=PIFO;refGene_tx=XM_003990455.4;refGene_gn=PIFO;refGene_tx=XM_023258979.1
C1 95416636 95416754
refGene_gn=PIFO;refGene_tx=XM_003990455.4;refGene_gn=PIFO;refGene_tx=XM_023258979.1
C1 95418329 95418398
refGene_gn=PIFO;refGene_tx=XM_003990455.4;refGene_gn=PIFO;refGene_tx=XM_023258979.1
C1 95422666 95422819
refGene_gn=LOC111556254;refGene_tx=XM_023257838.1
C1 95423026 95423125
refGene_gn=LOC111556254;refGene_tx=XM_023257838.1
C1 95424025 95424173
refGene_gn=LOC111556254;refGene_tx=XM_023257838.1
C1 95425936 95426059
refGene_gn=LOC111556254;refGene_tx=XM_023257838.1
C1 95427627 95427827
refGene_gn=LOC111556254;refGene_tx=XM_023257838.1
C1 95428499 95428618
refGene_gn=LOC111556254;refGene_tx=XM_023257838.1
C1 95429150 95429268
refGene_gn=LOC111556254;refGene_tx=XM_023257838.1
C1 95430619 95430823
refGene_gn=LOC111556254;refGene_tx=XM_023257838.1
C1 95462791 95462944
refGene_gn=OVGP1;refGene_tx=XM_023257811.1;refGene_gn=OVGP1;refGene_tx=XM_023257812.1
C1 95463173 95463272
refGene_gn=OVGP1;refGene_tx=XM_023257811.1;refGene_gn=OVGP1;refGene_tx=XM_023257812.1
C1 95464132 95464280
refGene_gn=OVGP1;refGene_tx=XM_023257811.1;refGene_gn=OVGP1;refGene_tx=XM_023257812.1
C1 95465845 95465968
refGene_gn=OVGP1;refGene_tx=XM_023257811.1;refGene_gn=OVGP1;refGene_tx=XM_023257812.1
C1 95467179 95467379
refGene_gn=OVGP1;refGene_tx=XM_023257811.1;refGene_gn=OVGP1;refGene_tx=XM_023257812.1
C1 95468053 95468172
refGene_gn=OVGP1;refGene_tx=XM_023257811.1;refGene_gn=OVGP1;refGene_tx=XM_023257812.1
C1 95468677 95468795
refGene_gn=OVGP1;refGene_tx=XM_023257811.1;refGene_gn=OVGP1;refGene_tx=XM_023257812.1
C1 95469984 95470141

refGene_gn=OVGP1;refGene_tx=XM_023257811.1;refGene_gn=OVGP1;refGene_tx=XM_023257812.1
C1 95470690 95470746 refGene_gn=OVGP1;refGene_tx=XM_023257812.1
C1 95475903 95476424
refGene_gn=OVGP1;refGene_tx=NM_001278838.1;refGene_gn=OVGP1;refGene_tx=XM_019835944.2
C1 95477372 95477508
refGene_gn=OVGP1;refGene_tx=NM_001278838.1;refGene_gn=OVGP1;refGene_tx=XM_019835944.2
C1 95481923 95482040
refGene_gn=OVGP1;refGene_tx=NM_001278838.1;refGene_gn=OVGP1;refGene_tx=XM_019835944.2;refGene_gn=OVGP1;refGene_tx=XM_023257811.1
C1 95483629 95483815
refGene_gn=OVGP1;refGene_tx=NM_001278838.1;refGene_gn=OVGP1;refGene_tx=XM_019835944.2
C1 95483911 95484020
refGene_gn=OVGP1;refGene_tx=NM_001278838.1;refGene_gn=OVGP1;refGene_tx=XM_019835944.2
C1 95485292 95485417
refGene_gn=OVGP1;refGene_tx=NM_001278838.1;refGene_gn=OVGP1;refGene_tx=XM_019835944.2
C1 95485898 95486064
refGene_gn=OVGP1;refGene_tx=NM_001278838.1;refGene_gn=OVGP1;refGene_tx=XM_019835944.2
C1 95486287 95486344 refGene_gn=OVGP1;refGene_tx=NM_001278838.1
C1 95487578 95487783 refGene_gn=OVGP1;refGene_tx=NM_001278838.1
C1 95488196 95488226 refGene_gn=OVGP1;refGene_tx=NM_001278838.1
C1 95488832 95488857 refGene_gn=OVGP1;refGene_tx=NM_001278838.1
C1 95493988 95494148 refGene_gn=WDR77;refGene_tx=XM_003990457.5
C1 95494739 95494808 refGene_gn=WDR77;refGene_tx=XM_003990457.5
C1 95495357 95495466 refGene_gn=WDR77;refGene_tx=XM_003990457.5
C1 95496033 95496105 refGene_gn=WDR77;refGene_tx=XM_003990457.5
C1 95496578 95496633 refGene_gn=WDR77;refGene_tx=XM_003990457.5
C1 95496767 95496838 refGene_gn=WDR77;refGene_tx=XM_003990457.5
C1 95499707 95499757 refGene_gn=WDR77;refGene_tx=XM_003990457.5
C1 95500038 95500180 refGene_gn=WDR77;refGene_tx=XM_003990457.5
C1 95501180 95501366 refGene_gn=WDR77;refGene_tx=XM_003990457.5
C1 95501620 95501735 refGene_gn=WDR77;refGene_tx=XM_003990457.5
C1 95502091 95502131 refGene_gn=ATP5PB;refGene_tx=XM_011285065.2
C1 95502410 95502447 refGene_gn=ATP5PB;refGene_tx=XM_011285065.2
C1 95506376 95506522 refGene_gn=ATP5PB;refGene_tx=XM_011285065.2
C1 95508663 95508827 refGene_gn=ATP5PB;refGene_tx=XM_011285065.2
C1 95509622 95509748 refGene_gn=ATP5PB;refGene_tx=XM_011285065.2
C1 95514997 95515177 refGene_gn=ATP5PB;refGene_tx=XM_011285065.2
C1 95516469 95516547 refGene_gn=ATP5PB;refGene_tx=XM_011285065.2
C1 95532643 95532680
refGene_gn=CC1H1orf162;refGene_tx=XM_006934997.3;refGene_gn=CC1H1orf162;refGene_tx=XM_019837608.2;refGene_gn=CC1H1orf162;refGene_tx=XM_019837612.1;refGene_gn=CC1H1orf162;refGene_tx=XM_019837613.2
C1 95533859 95533920

refGene_gn=CC1H1orf162;refGene_tx=XM_006934997.3;refGene_gn=CC1H1orf162;refGene_tx=XM_019837608.2;refGene_gn=CC1H1orf162;refGene_tx=XM_019837612.1;refGene_gn=CC1H1orf162;refGene_tx=XM_019837613.2
C1 95534409 95534504
refGene_gn=CC1H1orf162;refGene_tx=XM_006934997.3;refGene_gn=CC1H1orf162;refGene_tx=XM_019837608.2;refGene_gn=CC1H1orf162;refGene_tx=XM_019837612.1;refGene_gn=CC1H1orf162;refGene_tx=XM_019837613.2
C1 95534764 95534836
refGene_gn=CC1H1orf162;refGene_tx=XM_006934997.3;refGene_gn=CC1H1orf162;refGene_tx=XM_019837608.2;refGene_gn=CC1H1orf162;refGene_tx=XM_019837612.1;refGene_gn=CC1H1orf162;refGene_tx=XM_019837613.2
C1 95535002 95535161
refGene_gn=CC1H1orf162;refGene_tx=XM_006934997.3;refGene_gn=CC1H1orf162;refGene_tx=XM_019837608.2;refGene_gn=CC1H1orf162;refGene_tx=XM_019837612.1;refGene_gn=CC1H1orf162;refGene_tx=XM_019837613.2
C1 95542164 95542209 refGene_gn=TMIGD3;refGene_tx=XM_023257839.1
C1 95542210 95542236 refGene_gn=TMIGD3;refGene_tx=XM_023257839.1
C1 95544192 95544292 refGene_gn=TMIGD3;refGene_tx=XM_023257839.1
C1 95545033 95545101 refGene_gn=TMIGD3;refGene_tx=XM_023257839.1
C1 95546653 95546889 refGene_gn=TMIGD3;refGene_tx=XM_023257839.1
C1 95547891 95547975 refGene_gn=TMIGD3;refGene_tx=XM_023257839.1
C1 95550019 95550084 refGene_gn=TMIGD3;refGene_tx=XM_023257839.1
C1 95555042 95555649 refGene_gn=ADORA3;refGene_tx=XM_006935002.4
C1 95557892 95558242 refGene_gn=ADORA3;refGene_tx=XM_006935002.4
C1 95678835 95678849 refGene_gn=RAP1A;refGene_tx=XM_019837615.1
C1 95728041 95728125
refGene_gn=RAP1A;refGene_tx=XM_006934999.3;refGene_gn=RAP1A;refGene_tx=XM_019837615.1
C1 95732367 95732436
refGene_gn=RAP1A;refGene_tx=XM_006934999.3;refGene_gn=RAP1A;refGene_tx=XM_019837615.1
C1 95735305 95735362
refGene_gn=RAP1A;refGene_tx=XM_006934999.3;refGene_gn=RAP1A;refGene_tx=XM_019837615.1
C1 95737669 95737810
refGene_gn=RAP1A;refGene_tx=XM_006934999.3;refGene_gn=RAP1A;refGene_tx=XM_019837615.1
C1 95739419 95739563
refGene_gn=RAP1A;refGene_tx=XM_006934999.3;refGene_gn=RAP1A;refGene_tx=XM_019837615.1
C1 95744319 95744406
refGene_gn=RAP1A;refGene_tx=XM_006934999.3;refGene_gn=RAP1A;refGene_tx=XM_019837615.1
C1 95761757 95762558
refGene_gn=FAM212B;refGene_tx=XM_003990461.5;refGene_gn=FAM212B;refGene_tx=XM_023258981.1
C1 95773183 95773219 refGene_gn=FAM212B;refGene_tx=XM_003990461.5
C1 95788859 95789160 refGene_gn=DDX20;refGene_tx=XM_023258980.1
C1 95789530 95789625 refGene_gn=DDX20;refGene_tx=XM_023258980.1
C1 95791225 95791394 refGene_gn=DDX20;refGene_tx=XM_023258980.1

C1 95792006 95792121 refGene_gn=DDX20;refGene_tx=XM_023258980.1
C1 95792243 95792386 refGene_gn=DDX20;refGene_tx=XM_023258980.1
C1 95792530 95792669 refGene_gn=DDX20;refGene_tx=XM_023258980.1
C1 95792768 95792827 refGene_gn=DDX20;refGene_tx=XM_023258980.1
C1 95794305 95794388 refGene_gn=DDX20;refGene_tx=XM_023258980.1
C1 95794957 95795063 refGene_gn=DDX20;refGene_tx=XM_023258980.1
C1 95795185 95795287 refGene_gn=DDX20;refGene_tx=XM_023258980.1
C1 95797860 95799020 refGene_gn=DDX20;refGene_tx=XM_023258980.1
C1 95808983 95809185
refGene_gn=KCND3;refGene_tx=XM_003990463.5;refGene_gn=KCND3;refGene_tx=XM_023258982.1;refGene_gn=KCND3;refGene_tx=XM_023258983.1
C1 95809987 95810235
refGene_gn=KCND3;refGene_tx=XM_003990463.5;refGene_gn=KCND3;refGene_tx=XM_023258982.1;refGene_gn=KCND3;refGene_tx=XM_023258983.1
C1 95811411 95811468
refGene_gn=KCND3;refGene_tx=XM_003990463.5;refGene_gn=KCND3;refGene_tx=XM_023258982.1
C1 95812921 95813011
refGene_gn=KCND3;refGene_tx=XM_003990463.5;refGene_gn=KCND3;refGene_tx=XM_023258982.1;refGene_gn=KCND3;refGene_tx=XM_023258983.1
C1 95813342 95813444
refGene_gn=KCND3;refGene_tx=XM_003990463.5;refGene_gn=KCND3;refGene_tx=XM_023258982.1;refGene_gn=KCND3;refGene_tx=XM_023258983.1
C1 95819705 95819868
refGene_gn=KCND3;refGene_tx=XM_003990463.5;refGene_gn=KCND3;refGene_tx=XM_023258982.1;refGene_gn=KCND3;refGene_tx=XM_023258983.1
C1 96010452 96011558
refGene_gn=KCND3;refGene_tx=XM_003990463.5;refGene_gn=KCND3;refGene_tx=XM_023258982.1;refGene_gn=KCND3;refGene_tx=XM_023258983.1
C1 96375665 96375764
refGene_gn=CTTNBP2NL;refGene_tx=XM_003990464.5;refGene_gn=CTTNBP2NL;refGene_tx=XM_006935000.4
C1 96401717 96401948
refGene_gn=CTTNBP2NL;refGene_tx=XM_003990464.5;refGene_gn=CTTNBP2NL;refGene_tx=XM_006935000.4
C1 96406930 96407038
refGene_gn=CTTNBP2NL;refGene_tx=XM_003990464.5;refGene_gn=CTTNBP2NL;refGene_tx=XM_006935000.4
C1 96408225 96409707
refGene_gn=CTTNBP2NL;refGene_tx=XM_003990464.5;refGene_gn=CTTNBP2NL;refGene_tx=XM_006935000.4
C1 96443899 96444087 refGene_gn=WNT2B;refGene_tx=XM_011285068.3
C1 96449815 96450036 refGene_gn=WNT2B;refGene_tx=XM_011285068.3
C1 96451080 96451358 refGene_gn=WNT2B;refGene_tx=XM_011285068.3
C1 96452014 96452279 refGene_gn=WNT2B;refGene_tx=XM_011285068.3
C1 96453321 96453384
refGene_gn=ST7L;refGene_tx=XM_011285067.3;refGene_gn=ST7L;refGene_tx=XM_023258984.1;refGene_gn=ST7L;refGene_tx=XM_023258985.1;refGene_gn=ST7L;refGene_tx=XM_023258986.1;refGene_gn=ST7L;refGene_tx=XM_023258987.1
C1 96455510 96455740 refGene_gn=WNT2B;refGene_tx=XM_011285068.3

C1 96475170 96475310
refGene_gn=ST7L;refGene_tx=XM_011285067.3;refGene_gn=ST7L;refGene_tx=XM_023258984.1;refGene_gn=ST7L;refGene_tx=XM_023258985.1;refGene_gn=ST7L;refGene_tx=XM_023258986.1;refGene_gn=ST7L;refGene_tx=XM_023258987.1
C1 96480991 96481084
refGene_gn=ST7L;refGene_tx=XM_011285067.3;refGene_gn=ST7L;refGene_tx=XM_023258984.1;refGene_gn=ST7L;refGene_tx=XM_023258985.1;refGene_gn=ST7L;refGene_tx=XM_023258986.1;refGene_gn=ST7L;refGene_tx=XM_023258987.1
C1 96485176 96485327
refGene_gn=ST7L;refGene_tx=XM_011285067.3;refGene_gn=ST7L;refGene_tx=XM_023258984.1;refGene_gn=ST7L;refGene_tx=XM_023258985.1;refGene_gn=ST7L;refGene_tx=XM_023258986.1;refGene_gn=ST7L;refGene_tx=XM_023258987.1
C1 96501880 96501983
refGene_gn=ST7L;refGene_tx=XM_011285067.3;refGene_gn=ST7L;refGene_tx=XM_023258984.1;refGene_gn=ST7L;refGene_tx=XM_023258985.1;refGene_gn=ST7L;refGene_tx=XM_023258986.1;refGene_gn=ST7L;refGene_tx=XM_023258987.1
C1 96503188 96503261
refGene_gn=ST7L;refGene_tx=XM_011285067.3;refGene_gn=ST7L;refGene_tx=XM_023258984.1;refGene_gn=ST7L;refGene_tx=XM_023258985.1;refGene_gn=ST7L;refGene_tx=XM_023258986.1;refGene_gn=ST7L;refGene_tx=XM_023258987.1
C1 96505656 96505771
refGene_gn=ST7L;refGene_tx=XM_011285067.3;refGene_gn=ST7L;refGene_tx=XM_023258984.1;refGene_gn=ST7L;refGene_tx=XM_023258985.1;refGene_gn=ST7L;refGene_tx=XM_023258986.1;refGene_gn=ST7L;refGene_tx=XM_023258987.1
C1 96506040 96506138
refGene_gn=ST7L;refGene_tx=XM_011285067.3;refGene_gn=ST7L;refGene_tx=XM_023258984.1;refGene_gn=ST7L;refGene_tx=XM_023258985.1;refGene_gn=ST7L;refGene_tx=XM_023258986.1;refGene_gn=ST7L;refGene_tx=XM_023258987.1
C1 96507634 96507789
refGene_gn=ST7L;refGene_tx=XM_011285067.3;refGene_gn=ST7L;refGene_tx=XM_023258984.1;refGene_gn=ST7L;refGene_tx=XM_023258985.1;refGene_gn=ST7L;refGene_tx=XM_023258986.1;refGene_gn=ST7L;refGene_tx=XM_023258987.1
C1 96517141 96517220
refGene_gn=ST7L;refGene_tx=XM_011285067.3;refGene_gn=ST7L;refGene_tx=XM_023258984.1;refGene_gn=ST7L;refGene_tx=XM_023258985.1;refGene_gn=ST7L;refGene_tx=XM_023258986.1;refGene_gn=ST7L;refGene_tx=XM_023258987.1
C1 96523181 96523297
refGene_gn=ST7L;refGene_tx=XM_011285067.3;refGene_gn=ST7L;refGene_tx=XM_023258984.1;refGene_gn=ST7L;refGene_tx=XM_023258985.1;refGene_gn=ST7L;refGene_tx=XM_023258986.1;refGene_gn=ST7L;refGene_tx=XM_023258987.1
C1 96526656 96526711
refGene_gn=ST7L;refGene_tx=XM_011285067.3;refGene_gn=ST7L;refGene_tx=XM_023258984.1;refGene_gn=ST7L;refGene_tx=XM_023258985.1;refGene_gn=ST7L;refGene_tx=XM_023258986.1;refGene_gn=ST7L;refGene_tx=XM_023258987.1
C1 96528760 96528920
refGene_gn=ST7L;refGene_tx=XM_011285067.3;refGene_gn=ST7L;refGene_tx=XM_023258984.1;refGene_gn=ST7L;refGene_tx=XM_023258985.1;refGene_gn=ST7L;refGene_tx=XM_023258986.1;refGene_gn=ST7L;refGene_tx=XM_023258987.1
C1 96532774 96532857
refGene_gn=ST7L;refGene_tx=XM_011285067.3;refGene_gn=ST7L;refGene_tx=X

M_023258984.1
C1 96535025 96535233
refGene_gn=ST7L;refGene_tx=XM_011285067.3;refGene_gn=ST7L;refGene_tx=X
M_023258984.1
C1 96535950 96535989 refGene_gn=CAPZA1;refGene_tx=XM_003990472.5
C1 96536671 96536707 refGene_gn=CAPZA1;refGene_tx=XM_019837627.2
C1 96558583 96558647
refGene_gn=CAPZA1;refGene_tx=XM_003990472.5;refGene_gn=CAPZA1;refGene_
tx=XM_019837627.2
C1 96561402 96561454
refGene_gn=CAPZA1;refGene_tx=XM_003990472.5;refGene_gn=CAPZA1;refGene_
tx=XM_019837627.2
C1 96565189 96565253
refGene_gn=CAPZA1;refGene_tx=XM_003990472.5;refGene_gn=CAPZA1;refGene_
tx=XM_019837627.2
C1 96570581 96570788
refGene_gn=CAPZA1;refGene_tx=XM_003990472.5;refGene_gn=CAPZA1;refGene_
tx=XM_019837627.2
C1 96573444 96573524
refGene_gn=CAPZA1;refGene_tx=XM_003990472.5;refGene_gn=CAPZA1;refGene_
tx=XM_019837627.2
C1 96574142 96574221
refGene_gn=CAPZA1;refGene_tx=XM_003990472.5;refGene_gn=CAPZA1;refGene_
tx=XM_019837627.2
C1 96581557 96581629
refGene_gn=CAPZA1;refGene_tx=XM_003990472.5;refGene_gn=CAPZA1;refGene_
tx=XM_019837627.2
C1 96583878 96583941
refGene_gn=CAPZA1;refGene_tx=XM_003990472.5;refGene_gn=CAPZA1;refGene_
tx=XM_019837627.2
C1 96584326 96584467
refGene_gn=CAPZA1;refGene_tx=XM_003990472.5;refGene_gn=CAPZA1;refGene_
tx=XM_019837627.2
C1 96590107 96590244
refGene_gn=MOV10;refGene_tx=XM_023258988.1;refGene_gn=MOV10;refGene_tx
=XM_023258989.1;refGene_gn=MOV10;refGene_tx=XM_023258990.1;refGene_gn=
MOV10;refGene_tx=XM_023258991.1
C1 96603246 96603450
refGene_gn=MOV10;refGene_tx=XM_023258988.1;refGene_gn=MOV10;refGene_tx
=XM_023258989.1;refGene_gn=MOV10;refGene_tx=XM_023258990.1;refGene_gn=
MOV10;refGene_tx=XM_023258991.1;refGene_gn=MOV10;refGene_tx=XM_0232589
92.1;refGene_gn=MOV10;refGene_tx=XM_023258993.1;refGene_gn=MOV10;refGe
ne_tx=XM_023258994.1
C1 96603724 96603960
refGene_gn=MOV10;refGene_tx=XM_023258988.1;refGene_gn=MOV10;refGene_tx
=XM_023258989.1;refGene_gn=MOV10;refGene_tx=XM_023258990.1;refGene_gn=
MOV10;refGene_tx=XM_023258991.1;refGene_gn=MOV10;refGene_tx=XM_0232589
92.1;refGene_gn=MOV10;refGene_tx=XM_023258993.1;refGene_gn=MOV10;refGe
ne_tx=XM_023258994.1
C1 96604129 96604388

refGene_gn=MOV10;refGene_tx=XM_023258988.1;refGene_gn=MOV10;refGene_tx=XM_023258989.1;refGene_gn=MOV10;refGene_tx=XM_023258990.1;refGene_gn=MOV10;refGene_tx=XM_023258991.1;refGene_gn=MOV10;refGene_tx=XM_023258992.1;refGene_gn=MOV10;refGene_tx=XM_023258993.1;refGene_gn=MOV10;refGene_tx=XM_023258994.1
C1 96605993 96606128
refGene_gn=MOV10;refGene_tx=XM_023258988.1;refGene_gn=MOV10;refGene_tx=XM_023258989.1;refGene_gn=MOV10;refGene_tx=XM_023258990.1;refGene_gn=MOV10;refGene_tx=XM_023258991.1;refGene_gn=MOV10;refGene_tx=XM_023258992.1;refGene_gn=MOV10;refGene_tx=XM_023258993.1;refGene_gn=MOV10;refGene_tx=XM_023258994.1
C1 96606793 96606962
refGene_gn=MOV10;refGene_tx=XM_023258988.1;refGene_gn=MOV10;refGene_tx=XM_023258989.1;refGene_gn=MOV10;refGene_tx=XM_023258990.1;refGene_gn=MOV10;refGene_tx=XM_023258991.1;refGene_gn=MOV10;refGene_tx=XM_023258992.1;refGene_gn=MOV10;refGene_tx=XM_023258993.1;refGene_gn=MOV10;refGene_tx=XM_023258994.1
C1 96607653 96607808
refGene_gn=MOV10;refGene_tx=XM_023258988.1;refGene_gn=MOV10;refGene_tx=XM_023258989.1;refGene_gn=MOV10;refGene_tx=XM_023258990.1;refGene_gn=MOV10;refGene_tx=XM_023258991.1;refGene_gn=MOV10;refGene_tx=XM_023258992.1;refGene_gn=MOV10;refGene_tx=XM_023258993.1;refGene_gn=MOV10;refGene_tx=XM_023258994.1
C1 96608303 96608480
refGene_gn=MOV10;refGene_tx=XM_023258988.1;refGene_gn=MOV10;refGene_tx=XM_023258989.1;refGene_gn=MOV10;refGene_tx=XM_023258990.1;refGene_gn=MOV10;refGene_tx=XM_023258991.1;refGene_gn=MOV10;refGene_tx=XM_023258992.1;refGene_gn=MOV10;refGene_tx=XM_023258993.1;refGene_gn=MOV10;refGene_tx=XM_023258994.1
C1 96608593 96608741
refGene_gn=MOV10;refGene_tx=XM_023258988.1;refGene_gn=MOV10;refGene_tx=XM_023258989.1;refGene_gn=MOV10;refGene_tx=XM_023258990.1;refGene_gn=MOV10;refGene_tx=XM_023258991.1;refGene_gn=MOV10;refGene_tx=XM_023258992.1;refGene_gn=MOV10;refGene_tx=XM_023258993.1;refGene_gn=MOV10;refGene_tx=XM_023258994.1
C1 96609157 96609316
refGene_gn=MOV10;refGene_tx=XM_023258988.1;refGene_gn=MOV10;refGene_tx=XM_023258989.1;refGene_gn=MOV10;refGene_tx=XM_023258990.1;refGene_gn=MOV10;refGene_tx=XM_023258991.1;refGene_gn=MOV10;refGene_tx=XM_023258992.1;refGene_gn=MOV10;refGene_tx=XM_023258993.1;refGene_gn=MOV10;refGene_tx=XM_023258994.1
C1 96609574 96609678
refGene_gn=MOV10;refGene_tx=XM_023258988.1;refGene_gn=MOV10;refGene_tx=XM_023258989.1;refGene_gn=MOV10;refGene_tx=XM_023258990.1;refGene_gn=MOV10;refGene_tx=XM_023258991.1;refGene_gn=MOV10;refGene_tx=XM_023258992.1;refGene_gn=MOV10;refGene_tx=XM_023258993.1;refGene_gn=MOV10;refGene_tx=XM_023258994.1
C1 96609846 96609944
refGene_gn=MOV10;refGene_tx=XM_023258988.1;refGene_gn=MOV10;refGene_tx=XM_023258989.1;refGene_gn=MOV10;refGene_tx=XM_023258990.1;refGene_gn=

MOV10;refGene_tx=XM_023258991.1;refGene_gn=MOV10;refGene_tx=XM_0232589
92.1;refGene_gn=MOV10;refGene_tx=XM_023258993.1;refGene_gn=MOV10;refGe
ne_tx=XM_023258994.1
C1 96610038 96610255
refGene_gn=MOV10;refGene_tx=XM_023258988.1;refGene_gn=MOV10;refGene_tx
=XM_023258989.1;refGene_gn=MOV10;refGene_tx=XM_023258990.1;refGene_gn=
MOV10;refGene_tx=XM_023258991.1;refGene_gn=MOV10;refGene_tx=XM_0232589
92.1;refGene_gn=MOV10;refGene_tx=XM_023258993.1;refGene_gn=MOV10;refGe
ne_tx=XM_023258994.1
C1 96611100 96611218
refGene_gn=MOV10;refGene_tx=XM_023258988.1;refGene_gn=MOV10;refGene_tx
=XM_023258989.1;refGene_gn=MOV10;refGene_tx=XM_023258990.1;refGene_gn=
MOV10;refGene_tx=XM_023258991.1;refGene_gn=MOV10;refGene_tx=XM_0232589
92.1;refGene_gn=MOV10;refGene_tx=XM_023258993.1;refGene_gn=MOV10;refGe
ne_tx=XM_023258994.1
C1 96611415 96611607
refGene_gn=MOV10;refGene_tx=XM_023258988.1;refGene_gn=MOV10;refGene_tx
=XM_023258989.1;refGene_gn=MOV10;refGene_tx=XM_023258990.1;refGene_gn=
MOV10;refGene_tx=XM_023258991.1;refGene_gn=MOV10;refGene_tx=XM_0232589
92.1;refGene_gn=MOV10;refGene_tx=XM_023258993.1;refGene_gn=MOV10;refGe
ne_tx=XM_023258994.1
C1 96611844 96611919
refGene_gn=MOV10;refGene_tx=XM_023258988.1;refGene_gn=MOV10;refGene_tx
=XM_023258989.1;refGene_gn=MOV10;refGene_tx=XM_023258990.1;refGene_gn=
MOV10;refGene_tx=XM_023258991.1;refGene_gn=MOV10;refGene_tx=XM_0232589
92.1;refGene_gn=MOV10;refGene_tx=XM_023258993.1;refGene_gn=MOV10;refGe
ne_tx=XM_023258994.1
C1 96612804 96612930
refGene_gn=MOV10;refGene_tx=XM_023258988.1;refGene_gn=MOV10;refGene_tx
=XM_023258989.1;refGene_gn=MOV10;refGene_tx=XM_023258990.1;refGene_gn=
MOV10;refGene_tx=XM_023258991.1;refGene_gn=MOV10;refGene_tx=XM_0232589
92.1;refGene_gn=MOV10;refGene_tx=XM_023258993.1;refGene_gn=MOV10;refGe
ne_tx=XM_023258994.1
C1 96613011 96613100
refGene_gn=MOV10;refGene_tx=XM_023258988.1;refGene_gn=MOV10;refGene_tx
=XM_023258989.1;refGene_gn=MOV10;refGene_tx=XM_023258990.1;refGene_gn=
MOV10;refGene_tx=XM_023258991.1;refGene_gn=MOV10;refGene_tx=XM_0232589
92.1;refGene_gn=MOV10;refGene_tx=XM_023258993.1;refGene_gn=MOV10;refGe
ne_tx=XM_023258994.1
C1 96613370 96613492
refGene_gn=MOV10;refGene_tx=XM_023258988.1;refGene_gn=MOV10;refGene_tx
=XM_023258989.1;refGene_gn=MOV10;refGene_tx=XM_023258990.1;refGene_gn=
MOV10;refGene_tx=XM_023258992.1;refGene_gn=MOV10;refGene_tx=XM_0232589
93.1;refGene_gn=MOV10;refGene_tx=XM_023258994.1
C1 96613566 96613658
refGene_gn=MOV10;refGene_tx=XM_023258988.1;refGene_gn=MOV10;refGene_tx
=XM_023258989.1;refGene_gn=MOV10;refGene_tx=XM_023258990.1;refGene_gn=
MOV10;refGene_tx=XM_023258991.1;refGene_gn=MOV10;refGene_tx=XM_0232589
92.1;refGene_gn=MOV10;refGene_tx=XM_023258993.1;refGene_gn=MOV10;refGe
ne_tx=XM_023258994.1

C1 96614815 96614989
refGene_gn=RHOC;refGene_tx=XM_011285072.3;refGene_gn=RHOC;refGene_tx=X
M_019837628.2
C1 96615742 96615873
refGene_gn=RHOC;refGene_tx=XM_011285072.3;refGene_gn=RHOC;refGene_tx=X
M_019837628.2
C1 96616211 96616332
refGene_gn=RHOC;refGene_tx=XM_011285072.3;refGene_gn=RHOC;refGene_tx=X
M_019837628.2
C1 96616865 96617021
refGene_gn=RHOC;refGene_tx=XM_011285072.3;refGene_gn=RHOC;refGene_tx=X
M_019837628.2
C1 96623058 96623206
refGene_gn=PPM1J;refGene_tx=XM_003990474.4;refGene_gn=PPM1J;refGene_tx
=XM_006935007.3
C1 96623350 96623502
refGene_gn=PPM1J;refGene_tx=XM_003990474.4;refGene_gn=PPM1J;refGene_tx
=XM_006935007.3
C1 96623618 96623726
refGene_gn=PPM1J;refGene_tx=XM_003990474.4;refGene_gn=PPM1J;refGene_tx
=XM_006935007.3
C1 96623878 96623942
refGene_gn=PPM1J;refGene_tx=XM_003990474.4;refGene_gn=PPM1J;refGene_tx
=XM_006935007.3
C1 96624146 96624265
refGene_gn=PPM1J;refGene_tx=XM_003990474.4;refGene_gn=PPM1J;refGene_tx
=XM_006935007.3
C1 96624813 96624898
refGene_gn=PPM1J;refGene_tx=XM_003990474.4;refGene_gn=PPM1J;refGene_tx
=XM_006935007.3
C1 96625183 96625296
refGene_gn=PPM1J;refGene_tx=XM_003990474.4;refGene_gn=PPM1J;refGene_tx
=XM_006935007.3
C1 96625581 96625869
refGene_gn=PPM1J;refGene_tx=XM_003990474.4;refGene_gn=PPM1J;refGene_tx
=XM_006935007.3
C1 96626372 96626487
refGene_gn=PPM1J;refGene_tx=XM_003990474.4;refGene_gn=PPM1J;refGene_tx
=XM_006935007.3
C1 96627833 96628159
refGene_gn=PPM1J;refGene_tx=XM_003990474.4;refGene_gn=PPM1J;refGene_tx
=XM_006935007.3
C1 96633600 96633782 refGene_gn=FAM19A3;refGene_tx=XM_023258995.1
C1 96635110 96635226 refGene_gn=FAM19A3;refGene_tx=XM_023258995.1
C1 96635908 96636058 refGene_gn=FAM19A3;refGene_tx=XM_023258995.1
C1 96636844 96636900 refGene_gn=FAM19A3;refGene_tx=XM_023258995.1
C1 96636902 96637042 refGene_gn=FAM19A3;refGene_tx=XM_023258995.1
C1 96638150 96638170 refGene_gn=FAM19A3;refGene_tx=XM_023258995.1
C1 96638172 96638330 refGene_gn=FAM19A3;refGene_tx=XM_023258995.1
C1 96697961 96698167

refGene_gn=LOC101081401;refGene_tx=XM_003990493.3;refGene_gn=LOC101081401;refGene_tx=XM_019837630.1
C1 96700390 96701257
refGene_gn=LOC101081401;refGene_tx=XM_003990493.3;refGene_gn=LOC101081401;refGene_tx=XM_019837630.1
C1 96704172 96704316
refGene_gn=LOC101081401;refGene_tx=XM_003990493.3;refGene_gn=LOC101081401;refGene_tx=XM_019837630.1
C1 96704865 96704891
refGene_gn=LOC101081401;refGene_tx=XM_019837630.1
C1 96705362 96705579
refGene_gn=LOC101081401;refGene_tx=XM_003990493.3;refGene_gn=LOC101081401;refGene_tx=XM_019837630.1
C1 96797411 96797683
refGene_gn=LOC101099065;refGene_tx=XM_023258996.1
C1 96800473 96801340
refGene_gn=LOC101099065;refGene_tx=XM_023258996.1
C1 96806445 96806589
refGene_gn=LOC101099065;refGene_tx=XM_023258996.1
C1 96811138 96811355
refGene_gn=LOC101099065;refGene_tx=XM_023258996.1
C1 96943236 96943475 refGene_gn=LRIG2;refGene_tx=XM_003990477.5
C1 96975109 96975175 refGene_gn=LRIG2;refGene_tx=XM_003990477.5
C1 96976987 96977013 refGene_gn=LRIG2;refGene_tx=XM_003990477.5
C1 96977015 96977061 refGene_gn=LRIG2;refGene_tx=XM_003990477.5
C1 96977231 96977366 refGene_gn=LRIG2;refGene_tx=XM_003990477.5
C1 96977868 96978012 refGene_gn=LRIG2;refGene_tx=XM_003990477.5
C1 96978129 96978273 refGene_gn=LRIG2;refGene_tx=XM_003990477.5
C1 96979013 96979157 refGene_gn=LRIG2;refGene_tx=XM_003990477.5
C1 96979362 96979506 refGene_gn=LRIG2;refGene_tx=XM_003990477.5
C1 96981856 96981937 refGene_gn=LRIG2;refGene_tx=XM_003990477.5
C1 96983083 96983155 refGene_gn=LRIG2;refGene_tx=XM_003990477.5
C1 96983292 96983361 refGene_gn=LRIG2;refGene_tx=XM_003990477.5
C1 96987835 96987999 refGene_gn=LRIG2;refGene_tx=XM_003990477.5
C1 96993170 96993491 refGene_gn=LRIG2;refGene_tx=XM_003990477.5
C1 96994792 96995074 refGene_gn=LRIG2;refGene_tx=XM_003990477.5
C1 96996773 96997223 refGene_gn=LRIG2;refGene_tx=XM_003990477.5
C1 96998263 96998413 refGene_gn=LRIG2;refGene_tx=XM_003990477.5
C1 97000056 97000347 refGene_gn=LRIG2;refGene_tx=XM_003990477.5
C1 97002544 97002771 refGene_gn=LRIG2;refGene_tx=XM_003990477.5
C1 97234276 97234592 refGene_gn=MAGI3;refGene_tx=XM_011285166.3
C1 97381057 97381174 refGene_gn=MAGI3;refGene_tx=XM_011285166.3
C1 97395974 97396094 refGene_gn=MAGI3;refGene_tx=XM_011285166.3
C1 97403135 97403345 refGene_gn=MAGI3;refGene_tx=XM_011285166.3
C1 97409552 97409727 refGene_gn=MAGI3;refGene_tx=XM_011285166.3
C1 97413094 97413174 refGene_gn=MAGI3;refGene_tx=XM_011285166.3
C1 97431758 97431816 refGene_gn=MAGI3;refGene_tx=XM_011285166.3
C1 97435981 97436076 refGene_gn=MAGI3;refGene_tx=XM_011285166.3
C1 97441423 97441612 refGene_gn=MAGI3;refGene_tx=XM_011285166.3
C1 97444114 97444720 refGene_gn=MAGI3;refGene_tx=XM_011285166.3

C1 97445949 97445981 refGene_gn=MAGI3;refGene_tx=XM_011285166.3
C1 97447997 97448154 refGene_gn=MAGI3;refGene_tx=XM_011285166.3
C1 97450260 97450352 refGene_gn=MAGI3;refGene_tx=XM_011285166.3
C1 97452040 97452233 refGene_gn=MAGI3;refGene_tx=XM_011285166.3
C1 97454546 97454735 refGene_gn=MAGI3;refGene_tx=XM_011285166.3
C1 97459736 97459922 refGene_gn=MAGI3;refGene_tx=XM_011285166.3
C1 97470501 97470604 refGene_gn=MAGI3;refGene_tx=XM_011285166.3
C1 97471373 97471500 refGene_gn=MAGI3;refGene_tx=XM_011285166.3
C1 97472344 97472488 refGene_gn=MAGI3;refGene_tx=XM_011285166.3
C1 97478012 97478151 refGene_gn=MAGI3;refGene_tx=XM_011285166.3
C1 97479675 97480022 refGene_gn=MAGI3;refGene_tx=XM_011285166.3
C1 97480023 97480176 refGene_gn=MAGI3;refGene_tx=XM_011285166.3
C1 97480177 97480798 refGene_gn=MAGI3;refGene_tx=XM_011285166.3
C1 97512012 97512033
refGene_gn=PHTF1;refGene_tx=XM_023258997.1;refGene_gn=PHTF1;refGene_tx=XM_023258998.1;refGene_gn=PHTF1;refGene_tx=XM_023258999.1;refGene_gn=PHTF1;refGene_tx=XM_023259000.1;refGene_gn=PHTF1;refGene_tx=XM_023259001.1;refGene_gn=PHTF1;refGene_tx=XM_023259002.1;refGene_gn=PHTF1;refGene_tx=XM_023259003.1;refGene_gn=PHTF1;refGene_tx=XM_023259004.1
C1 97512580 97512706
refGene_gn=PHTF1;refGene_tx=XM_023258997.1;refGene_gn=PHTF1;refGene_tx=XM_023258998.1;refGene_gn=PHTF1;refGene_tx=XM_023258999.1;refGene_gn=PHTF1;refGene_tx=XM_023259000.1;refGene_gn=PHTF1;refGene_tx=XM_023259001.1;refGene_gn=PHTF1;refGene_tx=XM_023259002.1;refGene_gn=PHTF1;refGene_tx=XM_023259003.1;refGene_gn=PHTF1;refGene_tx=XM_023259004.1
C1 97513142 97513238
refGene_gn=PHTF1;refGene_tx=XM_023258997.1;refGene_gn=PHTF1;refGene_tx=XM_023258998.1;refGene_gn=PHTF1;refGene_tx=XM_023258999.1;refGene_gn=PHTF1;refGene_tx=XM_023259000.1;refGene_gn=PHTF1;refGene_tx=XM_023259001.1;refGene_gn=PHTF1;refGene_tx=XM_023259002.1;refGene_gn=PHTF1;refGene_tx=XM_023259003.1;refGene_gn=PHTF1;refGene_tx=XM_023259004.1
C1 97514458 97514614
refGene_gn=PHTF1;refGene_tx=XM_023258997.1;refGene_gn=PHTF1;refGene_tx=XM_023258998.1;refGene_gn=PHTF1;refGene_tx=XM_023258999.1;refGene_gn=PHTF1;refGene_tx=XM_023259000.1;refGene_gn=PHTF1;refGene_tx=XM_023259001.1;refGene_gn=PHTF1;refGene_tx=XM_023259002.1;refGene_gn=PHTF1;refGene_tx=XM_023259003.1;refGene_gn=PHTF1;refGene_tx=XM_023259004.1
C1 97517979 97518066
refGene_gn=PHTF1;refGene_tx=XM_023258997.1;refGene_gn=PHTF1;refGene_tx=XM_023258998.1;refGene_gn=PHTF1;refGene_tx=XM_023258999.1;refGene_gn=PHTF1;refGene_tx=XM_023259000.1;refGene_gn=PHTF1;refGene_tx=XM_023259001.1;refGene_gn=PHTF1;refGene_tx=XM_023259002.1;refGene_gn=PHTF1;refGene_tx=XM_023259003.1;refGene_gn=PHTF1;refGene_tx=XM_023259004.1
C1 97518579 97518711
refGene_gn=PHTF1;refGene_tx=XM_023258997.1;refGene_gn=PHTF1;refGene_tx=XM_023258998.1;refGene_gn=PHTF1;refGene_tx=XM_023258999.1;refGene_gn=PHTF1;refGene_tx=XM_023259000.1;refGene_gn=PHTF1;refGene_tx=XM_023259001.1;refGene_gn=PHTF1;refGene_tx=XM_023259002.1;refGene_gn=PHTF1;refGene_tx=XM_023259003.1;refGene_gn=PHTF1;refGene_tx=XM_023259004.1
C1 97519474 97519747

refGene_gn=PHTF1;refGene_tx=XM_023258997.1;refGene_gn=PHTF1;refGene_tx=XM_023258998.1;refGene_gn=PHTF1;refGene_tx=XM_023258999.1;refGene_gn=PHTF1;refGene_tx=XM_023259000.1;refGene_gn=PHTF1;refGene_tx=XM_023259001.1;refGene_gn=PHTF1;refGene_tx=XM_023259002.1;refGene_gn=PHTF1;refGene_tx=XM_023259003.1;refGene_gn=PHTF1;refGene_tx=XM_023259004.1
C1 97520388 97520517
refGene_gn=PHTF1;refGene_tx=XM_023258997.1;refGene_gn=PHTF1;refGene_tx=XM_023258998.1;refGene_gn=PHTF1;refGene_tx=XM_023258999.1;refGene_gn=PHTF1;refGene_tx=XM_023259000.1;refGene_gn=PHTF1;refGene_tx=XM_023259001.1;refGene_gn=PHTF1;refGene_tx=XM_023259002.1;refGene_gn=PHTF1;refGene_tx=XM_023259003.1;refGene_gn=PHTF1;refGene_tx=XM_023259004.1
C1 97523064 97523286
refGene_gn=PHTF1;refGene_tx=XM_023258997.1;refGene_gn=PHTF1;refGene_tx=XM_023258998.1;refGene_gn=PHTF1;refGene_tx=XM_023258999.1;refGene_gn=PHTF1;refGene_tx=XM_023259000.1;refGene_gn=PHTF1;refGene_tx=XM_023259001.1;refGene_gn=PHTF1;refGene_tx=XM_023259002.1;refGene_gn=PHTF1;refGene_tx=XM_023259003.1;refGene_gn=PHTF1;refGene_tx=XM_023259004.1
C1 97524289 97524379
refGene_gn=PHTF1;refGene_tx=XM_023258997.1;refGene_gn=PHTF1;refGene_tx=XM_023258998.1;refGene_gn=PHTF1;refGene_tx=XM_023258999.1;refGene_gn=PHTF1;refGene_tx=XM_023259000.1;refGene_gn=PHTF1;refGene_tx=XM_023259001.1;refGene_gn=PHTF1;refGene_tx=XM_023259002.1;refGene_gn=PHTF1;refGene_tx=XM_023259003.1;refGene_gn=PHTF1;refGene_tx=XM_023259004.1
C1 97524484 97524658
refGene_gn=PHTF1;refGene_tx=XM_023258997.1;refGene_gn=PHTF1;refGene_tx=XM_023258998.1;refGene_gn=PHTF1;refGene_tx=XM_023258999.1;refGene_gn=PHTF1;refGene_tx=XM_023259000.1;refGene_gn=PHTF1;refGene_tx=XM_023259001.1;refGene_gn=PHTF1;refGene_tx=XM_023259002.1;refGene_gn=PHTF1;refGene_tx=XM_023259003.1;refGene_gn=PHTF1;refGene_tx=XM_023259004.1
C1 97525524 97525684
refGene_gn=PHTF1;refGene_tx=XM_023258997.1;refGene_gn=PHTF1;refGene_tx=XM_023258998.1;refGene_gn=PHTF1;refGene_tx=XM_023258999.1;refGene_gn=PHTF1;refGene_tx=XM_023259000.1;refGene_gn=PHTF1;refGene_tx=XM_023259001.1;refGene_gn=PHTF1;refGene_tx=XM_023259002.1;refGene_gn=PHTF1;refGene_tx=XM_023259003.1;refGene_gn=PHTF1;refGene_tx=XM_023259004.1
C1 97527845 97527980
refGene_gn=PHTF1;refGene_tx=XM_023258997.1;refGene_gn=PHTF1;refGene_tx=XM_023258998.1;refGene_gn=PHTF1;refGene_tx=XM_023258999.1;refGene_gn=PHTF1;refGene_tx=XM_023259000.1;refGene_gn=PHTF1;refGene_tx=XM_023259001.1;refGene_gn=PHTF1;refGene_tx=XM_023259002.1;refGene_gn=PHTF1;refGene_tx=XM_023259003.1;refGene_gn=PHTF1;refGene_tx=XM_023259004.1
C1 97529202 97529359
refGene_gn=PHTF1;refGene_tx=XM_023258997.1;refGene_gn=PHTF1;refGene_tx=XM_023258998.1;refGene_gn=PHTF1;refGene_tx=XM_023258999.1;refGene_gn=PHTF1;refGene_tx=XM_023259000.1;refGene_gn=PHTF1;refGene_tx=XM_023259001.1;refGene_gn=PHTF1;refGene_tx=XM_023259002.1;refGene_gn=PHTF1;refGene_tx=XM_023259003.1;refGene_gn=PHTF1;refGene_tx=XM_023259004.1
C1 97541548 97541626
refGene_gn=PHTF1;refGene_tx=XM_023258997.1;refGene_gn=PHTF1;refGene_tx=XM_023258998.1;refGene_gn=PHTF1;refGene_tx=XM_023259001.1;refGene_gn=

PHTF1;refGene_tx=XM_023259002.1;refGene_gn=PHTF1;refGene_tx=XM_0232590
03.1
C1 97541874 97542033
refGene_gn=PHTF1;refGene_tx=XM_023258997.1;refGene_gn=PHTF1;refGene_tx
=XM_023258998.1;refGene_gn=PHTF1;refGene_tx=XM_023258999.1;refGene_gn=
PHTF1;refGene_tx=XM_023259000.1;refGene_gn=PHTF1;refGene_tx=XM_0232590
01.1;refGene_gn=PHTF1;refGene_tx=XM_023259002.1;refGene_gn=PHTF1;refGe
ne_tx=XM_023259003.1;refGene_gn=PHTF1;refGene_tx=XM_023259004.1
C1 97542737 97542783
refGene_gn=PHTF1;refGene_tx=XM_023259002.1;refGene_gn=PHTF1;refGene_tx
=XM_023259004.1
C1 97543775 97543845
refGene_gn=PHTF1;refGene_tx=XM_023258997.1;refGene_gn=PHTF1;refGene_tx
=XM_023258998.1;refGene_gn=PHTF1;refGene_tx=XM_023258999.1;refGene_gn=
PHTF1;refGene_tx=XM_023259000.1;refGene_gn=PHTF1;refGene_tx=XM_0232590
01.1;refGene_gn=PHTF1;refGene_tx=XM_023259003.1
C1 97552468 97552498
refGene_gn=PHTF1;refGene_tx=XM_023258997.1;refGene_gn=PHTF1;refGene_tx
=XM_023258999.1;refGene_gn=PHTF1;refGene_tx=XM_023259001.1
C1 97572521 97572578
refGene_gn=PHTF1;refGene_tx=XM_023258997.1;refGene_gn=PHTF1;refGene_tx
=XM_023258998.1;refGene_gn=PHTF1;refGene_tx=XM_023258999.1;refGene_gn=
PHTF1;refGene_tx=XM_023259000.1
C1 97573234 97573279
refGene_gn=PHTF1;refGene_tx=XM_023258997.1;refGene_gn=PHTF1;refGene_tx
=XM_023258998.1;refGene_gn=PHTF1;refGene_tx=XM_023258999.1;refGene_gn=
PHTF1;refGene_tx=XM_023259000.1
C1 97580361 97580832 refGene_gn=RSBN1;refGene_tx=XM_023259005.1
C1 97581466 97581575 refGene_gn=RSBN1;refGene_tx=XM_023259005.1
C1 97582825 97582993 refGene_gn=RSBN1;refGene_tx=XM_023259005.1
C1 97590234 97590377 refGene_gn=RSBN1;refGene_tx=XM_023259005.1
C1 97590685 97590823 refGene_gn=RSBN1;refGene_tx=XM_023259005.1
C1 97611538 97612212 refGene_gn=RSBN1;refGene_tx=XM_023259005.1
C1 97622974 97623680 refGene_gn=RSBN1;refGene_tx=XM_023259005.1
C1 97625928 97625993
refGene_gn=PTPN22;refGene_tx=XM_003990480.4;refGene_gn=PTPN22;refGene_
tx=XM_019837643.2;refGene_gn=PTPN22;refGene_tx=XM_019837646.2
C1 97631627 97631705
refGene_gn=PTPN22;refGene_tx=XM_003990480.4;refGene_gn=PTPN22;refGene_
tx=XM_019837643.2;refGene_gn=PTPN22;refGene_tx=XM_019837646.2
C1 97636810 97636841
refGene_gn=PTPN22;refGene_tx=XM_003990480.4;refGene_gn=PTPN22;refGene_
tx=XM_019837643.2;refGene_gn=PTPN22;refGene_tx=XM_019837646.2
C1 97638036 97638152
refGene_gn=PTPN22;refGene_tx=XM_003990480.4;refGene_gn=PTPN22;refGene_
tx=XM_019837643.2;refGene_gn=PTPN22;refGene_tx=XM_019837646.2
C1 97638402 97638486
refGene_gn=PTPN22;refGene_tx=XM_003990480.4;refGene_gn=PTPN22;refGene_
tx=XM_019837643.2;refGene_gn=PTPN22;refGene_tx=XM_019837646.2
C1 97642780 97642808

refGene_gn=PTPN22;refGene_tx=XM_003990480.4;refGene_gn=PTPN22;refGene_tx=XM_019837643.2;refGene_gn=PTPN22;refGene_tx=XM_019837646.2
C1 97645521 97645652
refGene_gn=PTPN22;refGene_tx=XM_003990480.4;refGene_gn=PTPN22;refGene_tx=XM_019837643.2;refGene_gn=PTPN22;refGene_tx=XM_019837646.2
C1 97645807 97645891
refGene_gn=PTPN22;refGene_tx=XM_003990480.4;refGene_gn=PTPN22;refGene_tx=XM_019837643.2;refGene_gn=PTPN22;refGene_tx=XM_019837646.2
C1 97648061 97648867
refGene_gn=PTPN22;refGene_tx=XM_003990480.4;refGene_gn=PTPN22;refGene_tx=XM_019837643.2;refGene_gn=PTPN22;refGene_tx=XM_019837646.2
C1 97649005 97649079
refGene_gn=PTPN22;refGene_tx=XM_003990480.4;refGene_gn=PTPN22;refGene_tx=XM_019837643.2;refGene_gn=PTPN22;refGene_tx=XM_019837646.2
C1 97664182 97664269
refGene_gn=PTPN22;refGene_tx=XM_003990480.4;refGene_gn=PTPN22;refGene_tx=XM_019837643.2;refGene_gn=PTPN22;refGene_tx=XM_019837646.2
C1 97666842 97666920
refGene_gn=PTPN22;refGene_tx=XM_003990480.4;refGene_gn=PTPN22;refGene_tx=XM_019837643.2;refGene_gn=PTPN22;refGene_tx=XM_019837646.2
C1 97667946 97668013
refGene_gn=PTPN22;refGene_tx=XM_003990480.4;refGene_gn=PTPN22;refGene_tx=XM_019837643.2;refGene_gn=PTPN22;refGene_tx=XM_019837646.2
C1 97668520 97668663
refGene_gn=PTPN22;refGene_tx=XM_003990480.4;refGene_gn=PTPN22;refGene_tx=XM_019837643.2;refGene_gn=PTPN22;refGene_tx=XM_019837646.2
C1 97669711 97669771
refGene_gn=PTPN22;refGene_tx=XM_003990480.4;refGene_gn=PTPN22;refGene_tx=XM_019837643.2;refGene_gn=PTPN22;refGene_tx=XM_019837646.2
C1 97669887 97669959
refGene_gn=PTPN22;refGene_tx=XM_003990480.4;refGene_gn=PTPN22;refGene_tx=XM_019837643.2;refGene_gn=PTPN22;refGene_tx=XM_019837646.2
C1 97671755 97671794
refGene_gn=PTPN22;refGene_tx=XM_003990480.4;refGene_gn=PTPN22;refGene_tx=XM_019837643.2
C1 97672229 97672325
refGene_gn=PTPN22;refGene_tx=XM_003990480.4;refGene_gn=PTPN22;refGene_tx=XM_019837643.2;refGene_gn=PTPN22;refGene_tx=XM_019837646.2
C1 97672483 97672560
refGene_gn=PTPN22;refGene_tx=XM_003990480.4;refGene_gn=PTPN22;refGene_tx=XM_019837643.2;refGene_gn=PTPN22;refGene_tx=XM_019837646.2
C1 97672841 97672950
refGene_gn=PTPN22;refGene_tx=XM_003990480.4;refGene_gn=PTPN22;refGene_tx=XM_019837643.2;refGene_gn=PTPN22;refGene_tx=XM_019837646.2
C1 97685320 97685407
refGene_gn=PTPN22;refGene_tx=XM_003990480.4;refGene_gn=PTPN22;refGene_tx=XM_019837643.2;refGene_gn=PTPN22;refGene_tx=XM_019837646.2
C1 97694924 97694942
refGene_gn=BCL2L15;refGene_tx=XM_003990482.4;refGene_gn=BCL2L15;refGene_tx=XM_019837650.1

C1 97695546 97695771
refGene_gn=BCL2L15;refGene_tx=XM_003990482.4;refGene_gn=BCL2L15;refGene_tx=XM_019837650.1
C1 97698825 97698947
refGene_gn=BCL2L15;refGene_tx=XM_003990482.4;refGene_gn=BCL2L15;refGene_tx=XM_019837650.1
C1 97699238 97699365
refGene_gn=BCL2L15;refGene_tx=XM_003990482.4;refGene_gn=BCL2L15;refGene_tx=XM_019837650.1
C1 97705332 97705760
refGene_gn=AP4B1;refGene_tx=XM_003990483.5;refGene_gn=AP4B1;refGene_tx=XM_011285076.3;refGene_gn=AP4B1;refGene_tx=XM_019837647.1;refGene_gn=AP4B1;refGene_tx=XM_019837649.2
C1 97706022 97706304
refGene_gn=AP4B1;refGene_tx=XM_003990483.5;refGene_gn=AP4B1;refGene_tx=XM_011285076.3;refGene_gn=AP4B1;refGene_tx=XM_019837647.1;refGene_gn=AP4B1;refGene_tx=XM_019837649.2
C1 97706526 97706734
refGene_gn=AP4B1;refGene_tx=XM_003990483.5;refGene_gn=AP4B1;refGene_tx=XM_011285076.3;refGene_gn=AP4B1;refGene_tx=XM_019837647.1;refGene_gn=AP4B1;refGene_tx=XM_019837649.2
C1 97707421 97707525
refGene_gn=AP4B1;refGene_tx=XM_003990483.5;refGene_gn=AP4B1;refGene_tx=XM_011285076.3;refGene_gn=AP4B1;refGene_tx=XM_019837647.1;refGene_gn=AP4B1;refGene_tx=XM_019837649.2
C1 97708283 97708367
refGene_gn=AP4B1;refGene_tx=XM_003990483.5;refGene_gn=AP4B1;refGene_tx=XM_011285076.3;refGene_gn=AP4B1;refGene_tx=XM_019837647.1;refGene_gn=AP4B1;refGene_tx=XM_019837649.2
C1 97709369 97709866
refGene_gn=AP4B1;refGene_tx=XM_003990483.5;refGene_gn=AP4B1;refGene_tx=XM_011285076.3;refGene_gn=AP4B1;refGene_tx=XM_019837647.1;refGene_gn=AP4B1;refGene_tx=XM_019837649.2
C1 97710388 97710536
refGene_gn=AP4B1;refGene_tx=XM_003990483.5;refGene_gn=AP4B1;refGene_tx=XM_011285076.3;refGene_gn=AP4B1;refGene_tx=XM_019837647.1
C1 97711124 97711255
refGene_gn=AP4B1;refGene_tx=XM_003990483.5;refGene_gn=AP4B1;refGene_tx=XM_011285076.3;refGene_gn=AP4B1;refGene_tx=XM_019837647.1
C1 97711720 97711945
refGene_gn=AP4B1;refGene_tx=XM_003990483.5;refGene_gn=AP4B1;refGene_tx=XM_011285076.3;refGene_gn=AP4B1;refGene_tx=XM_019837647.1
C1 97713882 97713995
refGene_gn=AP4B1;refGene_tx=XM_003990483.5;refGene_gn=AP4B1;refGene_tx=XM_011285076.3;refGene_gn=AP4B1;refGene_tx=XM_019837649.2
C1 97714890 97715079 refGene_gn=DCLRE1B;refGene_tx=XM_003990496.5
C1 97715764 97715930 refGene_gn=DCLRE1B;refGene_tx=XM_003990496.5
C1 97716468 97716651 refGene_gn=DCLRE1B;refGene_tx=XM_003990496.5
C1 97718244 97719332 refGene_gn=DCLRE1B;refGene_tx=XM_003990496.5
C1 97747863 97748939

refGene_gn=HIPK1;refGene_tx=XM_011285077.3;refGene_gn=HIPK1;refGene_tx=XM_023259006.1;refGene_gn=HIPK1;refGene_tx=XM_023259007.1;refGene_gn=HIPK1;refGene_tx=XM_023259008.1;refGene_gn=HIPK1;refGene_tx=XM_023259009.1;refGene_gn=HIPK1;refGene_tx=XM_023259010.1;refGene_gn=HIPK1;refGene_tx=XM_023259011.1
C1 97757661 97757785
refGene_gn=HIPK1;refGene_tx=XM_011285077.3;refGene_gn=HIPK1;refGene_tx=XM_023259006.1;refGene_gn=HIPK1;refGene_tx=XM_023259007.1;refGene_gn=HIPK1;refGene_tx=XM_023259008.1;refGene_gn=HIPK1;refGene_tx=XM_023259009.1;refGene_gn=HIPK1;refGene_tx=XM_023259010.1;refGene_gn=HIPK1;refGene_tx=XM_023259011.1
C1 97767413 97767533
refGene_gn=HIPK1;refGene_tx=XM_011285077.3;refGene_gn=HIPK1;refGene_tx=XM_023259006.1;refGene_gn=HIPK1;refGene_tx=XM_023259007.1;refGene_gn=HIPK1;refGene_tx=XM_023259008.1;refGene_gn=HIPK1;refGene_tx=XM_023259009.1;refGene_gn=HIPK1;refGene_tx=XM_023259010.1;refGene_gn=HIPK1;refGene_tx=XM_023259011.1
C1 97768582 97768669
refGene_gn=HIPK1;refGene_tx=XM_011285077.3;refGene_gn=HIPK1;refGene_tx=XM_023259006.1;refGene_gn=HIPK1;refGene_tx=XM_023259007.1;refGene_gn=HIPK1;refGene_tx=XM_023259008.1;refGene_gn=HIPK1;refGene_tx=XM_023259009.1;refGene_gn=HIPK1;refGene_tx=XM_023259010.1;refGene_gn=HIPK1;refGene_tx=XM_023259011.1
C1 97769363 97769548
refGene_gn=HIPK1;refGene_tx=XM_011285077.3;refGene_gn=HIPK1;refGene_tx=XM_023259006.1;refGene_gn=HIPK1;refGene_tx=XM_023259007.1;refGene_gn=HIPK1;refGene_tx=XM_023259008.1;refGene_gn=HIPK1;refGene_tx=XM_023259009.1;refGene_gn=HIPK1;refGene_tx=XM_023259010.1;refGene_gn=HIPK1;refGene_tx=XM_023259011.1
C1 97769865 97770028
refGene_gn=HIPK1;refGene_tx=XM_011285077.3;refGene_gn=HIPK1;refGene_tx=XM_023259006.1;refGene_gn=HIPK1;refGene_tx=XM_023259007.1;refGene_gn=HIPK1;refGene_tx=XM_023259008.1;refGene_gn=HIPK1;refGene_tx=XM_023259009.1;refGene_gn=HIPK1;refGene_tx=XM_023259010.1;refGene_gn=HIPK1;refGene_tx=XM_023259011.1
C1 97770952 97771178
refGene_gn=HIPK1;refGene_tx=XM_011285077.3;refGene_gn=HIPK1;refGene_tx=XM_023259006.1;refGene_gn=HIPK1;refGene_tx=XM_023259007.1;refGene_gn=HIPK1;refGene_tx=XM_023259008.1;refGene_gn=HIPK1;refGene_tx=XM_023259009.1;refGene_gn=HIPK1;refGene_tx=XM_023259010.1;refGene_gn=HIPK1;refGene_tx=XM_023259011.1
C1 97775281 97775403
refGene_gn=HIPK1;refGene_tx=XM_011285077.3;refGene_gn=HIPK1;refGene_tx=XM_023259006.1;refGene_gn=HIPK1;refGene_tx=XM_023259007.1;refGene_gn=HIPK1;refGene_tx=XM_023259008.1;refGene_gn=HIPK1;refGene_tx=XM_023259009.1;refGene_gn=HIPK1;refGene_tx=XM_023259010.1;refGene_gn=HIPK1;refGene_tx=XM_023259011.1
C1 97776319 97776454
refGene_gn=HIPK1;refGene_tx=XM_011285077.3;refGene_gn=HIPK1;refGene_tx=XM_023259006.1;refGene_gn=HIPK1;refGene_tx=XM_023259007.1;refGene_gn=

HIPK1;refGene_tx=XM_023259008.1;refGene_gn=HIPK1;refGene_tx=XM_023259010.1;refGene_gn=HIPK1;refGene_tx=XM_023259011.1
C1 97779091 97779234
refGene_gn=HIPK1;refGene_tx=XM_011285077.3;refGene_gn=HIPK1;refGene_tx=XM_023259006.1;refGene_gn=HIPK1;refGene_tx=XM_023259007.1;refGene_gn=HIPK1;refGene_tx=XM_023259008.1;refGene_gn=HIPK1;refGene_tx=XM_023259009.1;refGene_gn=HIPK1;refGene_tx=XM_023259010.1;refGene_gn=HIPK1;refGene_tx=XM_023259011.1
C1 97779632 97779815
refGene_gn=HIPK1;refGene_tx=XM_011285077.3;refGene_gn=HIPK1;refGene_tx=XM_023259006.1;refGene_gn=HIPK1;refGene_tx=XM_023259007.1;refGene_gn=HIPK1;refGene_tx=XM_023259008.1;refGene_gn=HIPK1;refGene_tx=XM_023259009.1;refGene_gn=HIPK1;refGene_tx=XM_023259010.1;refGene_gn=HIPK1;refGene_tx=XM_023259011.1
C1 97780297 97780504
refGene_gn=HIPK1;refGene_tx=XM_011285077.3;refGene_gn=HIPK1;refGene_tx=XM_023259006.1;refGene_gn=HIPK1;refGene_tx=XM_023259007.1;refGene_gn=HIPK1;refGene_tx=XM_023259008.1;refGene_gn=HIPK1;refGene_tx=XM_023259009.1;refGene_gn=HIPK1;refGene_tx=XM_023259010.1;refGene_gn=HIPK1;refGene_tx=XM_023259011.1
C1 97782054 97782296
refGene_gn=HIPK1;refGene_tx=XM_011285077.3;refGene_gn=HIPK1;refGene_tx=XM_023259006.1;refGene_gn=HIPK1;refGene_tx=XM_023259007.1;refGene_gn=HIPK1;refGene_tx=XM_023259008.1;refGene_gn=HIPK1;refGene_tx=XM_023259009.1;refGene_gn=HIPK1;refGene_tx=XM_023259010.1;refGene_gn=HIPK1;refGene_tx=XM_023259011.1
C1 97783921 97784052
refGene_gn=HIPK1;refGene_tx=XM_011285077.3;refGene_gn=HIPK1;refGene_tx=XM_023259006.1;refGene_gn=HIPK1;refGene_tx=XM_023259007.1;refGene_gn=HIPK1;refGene_tx=XM_023259008.1;refGene_gn=HIPK1;refGene_tx=XM_023259009.1;refGene_gn=HIPK1;refGene_tx=XM_023259010.1;refGene_gn=HIPK1;refGene_tx=XM_023259011.1
C1 97784713 97784818
refGene_gn=HIPK1;refGene_tx=XM_023259010.1;refGene_gn=HIPK1;refGene_tx=XM_023259011.1
C1 97785135 97785624
refGene_gn=HIPK1;refGene_tx=XM_011285077.3;refGene_gn=HIPK1;refGene_tx=XM_023259006.1;refGene_gn=HIPK1;refGene_tx=XM_023259007.1;refGene_gn=HIPK1;refGene_tx=XM_023259008.1;refGene_gn=HIPK1;refGene_tx=XM_023259009.1
C1 97792020 97792134 refGene_gn=OLFML3;refGene_tx=XM_003990487.4
C1 97792723 97793009 refGene_gn=OLFML3;refGene_tx=XM_003990487.4
C1 97793333 97794154 refGene_gn=OLFML3;refGene_tx=XM_003990487.4
C1 97875472 97875671
refGene_gn=SYT6;refGene_tx=XM_019837655.2;refGene_gn=SYT6;refGene_tx=XM_019837656.2;refGene_gn=SYT6;refGene_tx=XM_019837657.2;refGene_gn=SYT6;refGene_tx=XM_019837661.1;refGene_gn=SYT6;refGene_tx=XM_023259012.1;refGene_gn=SYT6;refGene_tx=XM_023259013.1
C1 97877301 97877368 refGene_gn=SYT6;refGene_tx=XM_019837658.2
C1 97881740 97881917

refGene_gn=SYT6;refGene_tx=XM_019837655.2;refGene_gn=SYT6;refGene_tx=XM_019837656.2;refGene_gn=SYT6;refGene_tx=XM_019837660.1;refGene_gn=SYT6;refGene_tx=XM_019837661.1;refGene_gn=SYT6;refGene_tx=XM_023259012.1;refGene_gn=SYT6;refGene_tx=XM_023259013.1
C1 97883109 97883281
refGene_gn=SYT6;refGene_tx=XM_019837655.2;refGene_gn=SYT6;refGene_tx=XM_019837656.2;refGene_gn=SYT6;refGene_tx=XM_019837657.2;refGene_gn=SYT6;refGene_tx=XM_019837658.2;refGene_gn=SYT6;refGene_tx=XM_019837660.1;refGene_gn=SYT6;refGene_tx=XM_019837661.1;refGene_gn=SYT6;refGene_tx=XM_023259012.1;refGene_gn=SYT6;refGene_tx=XM_023259013.1
C1 97887909 97888030
refGene_gn=SYT6;refGene_tx=XM_019837655.2;refGene_gn=SYT6;refGene_tx=XM_019837656.2;refGene_gn=SYT6;refGene_tx=XM_019837657.2;refGene_gn=SYT6;refGene_tx=XM_019837658.2;refGene_gn=SYT6;refGene_tx=XM_019837660.1;refGene_gn=SYT6;refGene_tx=XM_019837661.1;refGene_gn=SYT6;refGene_tx=XM_023259012.1;refGene_gn=SYT6;refGene_tx=XM_023259013.1
C1 97920742 97921301
refGene_gn=SYT6;refGene_tx=XM_019837655.2;refGene_gn=SYT6;refGene_tx=XM_019837656.2;refGene_gn=SYT6;refGene_tx=XM_019837657.2;refGene_gn=SYT6;refGene_tx=XM_019837658.2;refGene_gn=SYT6;refGene_tx=XM_019837660.1;refGene_gn=SYT6;refGene_tx=XM_019837661.1;refGene_gn=SYT6;refGene_tx=XM_023259012.1;refGene_gn=SYT6;refGene_tx=XM_023259013.1
C1 97922829 97923178
refGene_gn=SYT6;refGene_tx=XM_019837655.2;refGene_gn=SYT6;refGene_tx=XM_019837656.2;refGene_gn=SYT6;refGene_tx=XM_019837657.2;refGene_gn=SYT6;refGene_tx=XM_019837658.2;refGene_gn=SYT6;refGene_tx=XM_019837660.1;refGene_gn=SYT6;refGene_tx=XM_019837661.1;refGene_gn=SYT6;refGene_tx=XM_023259012.1;refGene_gn=SYT6;refGene_tx=XM_023259013.1
C1 97934203 97934366
refGene_gn=SYT6;refGene_tx=XM_019837655.2;refGene_gn=SYT6;refGene_tx=XM_019837656.2;refGene_gn=SYT6;refGene_tx=XM_019837657.2;refGene_gn=SYT6;refGene_tx=XM_019837658.2;refGene_gn=SYT6;refGene_tx=XM_019837660.1
C1 98120768 98120981
refGene_gn=TRIM33;refGene_tx=XM_006944178.4;refGene_gn=TRIM33;refGene_tx=XM_023259014.1;refGene_gn=TRIM33;refGene_tx=XM_023259015.1
C1 98121060 98121111
refGene_gn=TRIM33;refGene_tx=XM_006944178.4;refGene_gn=TRIM33;refGene_tx=XM_023259015.1
C1 98122548 98122701
refGene_gn=TRIM33;refGene_tx=XM_006944178.4;refGene_gn=TRIM33;refGene_tx=XM_023259014.1;refGene_gn=TRIM33;refGene_tx=XM_023259015.1
C1 98125002 98125077
refGene_gn=TRIM33;refGene_tx=XM_006944178.4;refGene_gn=TRIM33;refGene_tx=XM_023259014.1
C1 98125503 98125627
refGene_gn=TRIM33;refGene_tx=XM_006944178.4;refGene_gn=TRIM33;refGene_tx=XM_023259014.1;refGene_gn=TRIM33;refGene_tx=XM_023259015.1
C1 98127784 98128134
refGene_gn=TRIM33;refGene_tx=XM_006944178.4;refGene_gn=TRIM33;refGene_tx=XM_023259014.1;refGene_gn=TRIM33;refGene_tx=XM_023259015.1

C1 98129429 98129589
refGene_gn=TRIM33;refGene_tx=XM_006944178.4;refGene_gn=TRIM33;refGene_tx=XM_023259014.1;refGene_gn=TRIM33;refGene_tx=XM_023259015.1
C1 98132027 98132091
refGene_gn=TRIM33;refGene_tx=XM_006944178.4;refGene_gn=TRIM33;refGene_tx=XM_023259014.1;refGene_gn=TRIM33;refGene_tx=XM_023259015.1
C1 98133865 98133998
refGene_gn=TRIM33;refGene_tx=XM_006944178.4;refGene_gn=TRIM33;refGene_tx=XM_023259014.1;refGene_gn=TRIM33;refGene_tx=XM_023259015.1
C1 98140603 98140804
refGene_gn=TRIM33;refGene_tx=XM_006944178.4;refGene_gn=TRIM33;refGene_tx=XM_023259014.1;refGene_gn=TRIM33;refGene_tx=XM_023259015.1
C1 98144758 98144923
refGene_gn=TRIM33;refGene_tx=XM_006944178.4;refGene_gn=TRIM33;refGene_tx=XM_023259014.1;refGene_gn=TRIM33;refGene_tx=XM_023259015.1
C1 98145574 98145849
refGene_gn=TRIM33;refGene_tx=XM_006944178.4;refGene_gn=TRIM33;refGene_tx=XM_023259014.1;refGene_gn=TRIM33;refGene_tx=XM_023259015.1
C1 98147240 98147358
refGene_gn=TRIM33;refGene_tx=XM_006944178.4;refGene_gn=TRIM33;refGene_tx=XM_023259014.1;refGene_gn=TRIM33;refGene_tx=XM_023259015.1
C1 98147796 98147943
refGene_gn=TRIM33;refGene_tx=XM_006944178.4;refGene_gn=TRIM33;refGene_tx=XM_023259014.1;refGene_gn=TRIM33;refGene_tx=XM_023259015.1
C1 98150231 98150346
refGene_gn=TRIM33;refGene_tx=XM_006944178.4;refGene_gn=TRIM33;refGene_tx=XM_023259014.1;refGene_gn=TRIM33;refGene_tx=XM_023259015.1
C1 98152996 98153113
refGene_gn=TRIM33;refGene_tx=XM_006944178.4;refGene_gn=TRIM33;refGene_tx=XM_023259014.1;refGene_gn=TRIM33;refGene_tx=XM_023259015.1
C1 98185422 98185555
refGene_gn=TRIM33;refGene_tx=XM_006944178.4;refGene_gn=TRIM33;refGene_tx=XM_023259014.1;refGene_gn=TRIM33;refGene_tx=XM_023259015.1
C1 98185722 98185867
refGene_gn=TRIM33;refGene_tx=XM_006944178.4;refGene_gn=TRIM33;refGene_tx=XM_023259014.1;refGene_gn=TRIM33;refGene_tx=XM_023259015.1
C1 98186225 98186344
refGene_gn=TRIM33;refGene_tx=XM_006944178.4;refGene_gn=TRIM33;refGene_tx=XM_023259014.1;refGene_gn=TRIM33;refGene_tx=XM_023259015.1
C1 98246443 98246969
refGene_gn=TRIM33;refGene_tx=XM_006944178.4;refGene_gn=TRIM33;refGene_tx=XM_023259014.1;refGene_gn=TRIM33;refGene_tx=XM_023259015.1
C1 98294440 98294567 refGene_gn=BCAS2;refGene_tx=XM_004001122.5
C1 98295576 98295657 refGene_gn=BCAS2;refGene_tx=XM_004001122.5
C1 98297031 98297082 refGene_gn=BCAS2;refGene_tx=XM_004001122.5
C1 98298529 98298691 refGene_gn=BCAS2;refGene_tx=XM_004001122.5
C1 98300562 98300633 refGene_gn=BCAS2;refGene_tx=XM_004001122.5
C1 98303316 98303409 refGene_gn=BCAS2;refGene_tx=XM_004001122.5
C1 98303517 98303679 refGene_gn=BCAS2;refGene_tx=XM_004001122.5
C1 98306517 98306549

refGene_gn=DENND2C;refGene_tx=XM_023259016.1;refGene_gn=DENND2C;refGene_tx=XM_023259017.1;refGene_gn=DENND2C;refGene_tx=XM_023259018.1
C1 98307998 98308085
refGene_gn=DENND2C;refGene_tx=XM_023259016.1;refGene_gn=DENND2C;refGene_tx=XM_023259017.1;refGene_gn=DENND2C;refGene_tx=XM_023259018.1
C1 98308324 98308564
refGene_gn=DENND2C;refGene_tx=XM_023259016.1;refGene_gn=DENND2C;refGene_tx=XM_023259017.1;refGene_gn=DENND2C;refGene_tx=XM_023259018.1
C1 98313731 98313837
refGene_gn=DENND2C;refGene_tx=XM_023259016.1;refGene_gn=DENND2C;refGene_tx=XM_023259017.1;refGene_gn=DENND2C;refGene_tx=XM_023259018.1
C1 98314736 98314778
refGene_gn=DENND2C;refGene_tx=XM_023259016.1;refGene_gn=DENND2C;refGene_tx=XM_023259017.1;refGene_gn=DENND2C;refGene_tx=XM_023259018.1
C1 98320507 98320685
refGene_gn=DENND2C;refGene_tx=XM_023259016.1;refGene_gn=DENND2C;refGene_tx=XM_023259017.1;refGene_gn=DENND2C;refGene_tx=XM_023259018.1
C1 98321399 98321548
refGene_gn=DENND2C;refGene_tx=XM_023259016.1;refGene_gn=DENND2C;refGene_tx=XM_023259017.1;refGene_gn=DENND2C;refGene_tx=XM_023259018.1
C1 98322170 98322311
refGene_gn=DENND2C;refGene_tx=XM_023259016.1;refGene_gn=DENND2C;refGene_tx=XM_023259017.1;refGene_gn=DENND2C;refGene_tx=XM_023259018.1
C1 98322909 98322987
refGene_gn=DENND2C;refGene_tx=XM_023259016.1;refGene_gn=DENND2C;refGene_tx=XM_023259017.1;refGene_gn=DENND2C;refGene_tx=XM_023259018.1
C1 98323617 98323687
refGene_gn=DENND2C;refGene_tx=XM_023259016.1;refGene_gn=DENND2C;refGene_tx=XM_023259017.1;refGene_gn=DENND2C;refGene_tx=XM_023259018.1
C1 98325645 98325755
refGene_gn=DENND2C;refGene_tx=XM_023259016.1;refGene_gn=DENND2C;refGene_tx=XM_023259017.1;refGene_gn=DENND2C;refGene_tx=XM_023259018.1
C1 98329022 98329210
refGene_gn=DENND2C;refGene_tx=XM_023259016.1;refGene_gn=DENND2C;refGene_tx=XM_023259017.1;refGene_gn=DENND2C;refGene_tx=XM_023259018.1
C1 98331945 98331990
refGene_gn=DENND2C;refGene_tx=XM_023259016.1;refGene_gn=DENND2C;refGene_tx=XM_023259017.1;refGene_gn=DENND2C;refGene_tx=XM_023259018.1
C1 98337574 98337671
refGene_gn=DENND2C;refGene_tx=XM_023259016.1;refGene_gn=DENND2C;refGene_tx=XM_023259017.1;refGene_gn=DENND2C;refGene_tx=XM_023259018.1
C1 98342515 98342686
refGene_gn=DENND2C;refGene_tx=XM_023259016.1;refGene_gn=DENND2C;refGene_tx=XM_023259017.1;refGene_gn=DENND2C;refGene_tx=XM_023259018.1
C1 98343327 98343440
refGene_gn=DENND2C;refGene_tx=XM_023259016.1;refGene_gn=DENND2C;refGene_tx=XM_023259017.1;refGene_gn=DENND2C;refGene_tx=XM_023259018.1
C1 98344034 98344171
refGene_gn=DENND2C;refGene_tx=XM_023259016.1;refGene_gn=DENND2C;refGene_tx=XM_023259017.1;refGene_gn=DENND2C;refGene_tx=XM_023259018.1

C1 98344694 98345515
refGene_gn=DENND2C;refGene_tx=XM_023259016.1;refGene_gn=DENND2C;refGene_tx=XM_023259017.1;refGene_gn=DENND2C;refGene_tx=XM_023259018.1
C1 98397009 98397168 refGene_gn=AMPD1;refGene_tx=XM_004001120.5
C1 98397526 98397637 refGene_gn=AMPD1;refGene_tx=XM_004001120.5
C1 98397858 98398032 refGene_gn=AMPD1;refGene_tx=XM_004001120.5
C1 98398693 98398814 refGene_gn=AMPD1;refGene_tx=XM_004001120.5
C1 98399491 98399655 refGene_gn=AMPD1;refGene_tx=XM_004001120.5
C1 98399837 98399964 refGene_gn=AMPD1;refGene_tx=XM_004001120.5
C1 98401890 98402054 refGene_gn=AMPD1;refGene_tx=XM_004001120.5
C1 98402697 98402829 refGene_gn=AMPD1;refGene_tx=XM_004001120.5
C1 98403123 98403318 refGene_gn=AMPD1;refGene_tx=XM_004001120.5
C1 98404550 98404680 refGene_gn=AMPD1;refGene_tx=XM_004001120.5
C1 98405197 98405417 refGene_gn=AMPD1;refGene_tx=XM_004001120.5
C1 98409368 98409534 refGene_gn=AMPD1;refGene_tx=XM_004001120.5
C1 98411882 98412048 refGene_gn=AMPD1;refGene_tx=XM_004001120.5
C1 98413138 98413319 refGene_gn=AMPD1;refGene_tx=XM_004001120.5
C1 98416744 98416756 refGene_gn=AMPD1;refGene_tx=XM_004001120.5
C1 98418075 98418160 refGene_gn=AMPD1;refGene_tx=XM_004001120.5
C1 98418161 98418197 refGene_gn=AMPD1;refGene_tx=XM_004001120.5
C1 98434137 98434257
refGene_gn=NRAS;refGene_tx=XM_019837665.1;refGene_gn=NRAS;refGene_tx=XM_019837666.1
C1 98434757 98434917
refGene_gn=NRAS;refGene_tx=XM_019837665.1;refGene_gn=NRAS;refGene_tx=XM_019837666.1
C1 98437309 98437488
refGene_gn=NRAS;refGene_tx=XM_019837665.1;refGene_gn=NRAS;refGene_tx=XM_019837666.1
C1 98439678 98439789
refGene_gn=NRAS;refGene_tx=XM_019837665.1;refGene_gn=NRAS;refGene_tx=XM_019837666.1
C1 98441842 98441890
refGene_gn=CSDE1;refGene_tx=XM_011285081.2;refGene_gn=CSDE1;refGene_tx=XM_011285082.3;refGene_gn=CSDE1;refGene_tx=XM_011285083.2;refGene_gn=CSDE1;refGene_tx=XM_011285084.2
C1 98442258 98442391
refGene_gn=CSDE1;refGene_tx=XM_011285081.2;refGene_gn=CSDE1;refGene_tx=XM_011285082.3;refGene_gn=CSDE1;refGene_tx=XM_011285083.2;refGene_gn=CSDE1;refGene_tx=XM_011285084.2
C1 98443368 98443532
refGene_gn=CSDE1;refGene_tx=XM_011285081.2;refGene_gn=CSDE1;refGene_tx=XM_011285082.3;refGene_gn=CSDE1;refGene_tx=XM_011285083.2;refGene_gn=CSDE1;refGene_tx=XM_011285084.2
C1 98444243 98444422
refGene_gn=CSDE1;refGene_tx=XM_011285081.2;refGene_gn=CSDE1;refGene_tx=XM_011285082.3;refGene_gn=CSDE1;refGene_tx=XM_011285083.2;refGene_gn=CSDE1;refGene_tx=XM_011285084.2
C1 98446966 98447086
refGene_gn=CSDE1;refGene_tx=XM_011285081.2;refGene_gn=CSDE1;refGene_tx

=XM_011285082.3;refGene_gn=CSDE1;refGene_tx=XM_011285083.2;refGene_gn=CSDE1;refGene_tx=XM_011285084.2
C1	98448431	98448544
refGene_gn=CSDE1;refGene_tx=XM_011285081.2;refGene_gn=CSDE1;refGene_tx=XM_011285082.3;refGene_gn=CSDE1;refGene_tx=XM_011285083.2;refGene_gn=CSDE1;refGene_tx=XM_011285084.2
C1	98449101	98449277
refGene_gn=CSDE1;refGene_tx=XM_011285081.2;refGene_gn=CSDE1;refGene_tx=XM_011285082.3;refGene_gn=CSDE1;refGene_tx=XM_011285083.2;refGene_gn=CSDE1;refGene_tx=XM_011285084.2
C1	98449869	98449977
refGene_gn=CSDE1;refGene_tx=XM_011285081.2;refGene_gn=CSDE1;refGene_tx=XM_011285082.3;refGene_gn=CSDE1;refGene_tx=XM_011285083.2;refGene_gn=CSDE1;refGene_tx=XM_011285084.2
C1	98452303	98452468
refGene_gn=CSDE1;refGene_tx=XM_011285081.2;refGene_gn=CSDE1;refGene_tx=XM_011285082.3;refGene_gn=CSDE1;refGene_tx=XM_011285083.2;refGene_gn=CSDE1;refGene_tx=XM_011285084.2
C1	98452551	98452692
refGene_gn=CSDE1;refGene_tx=XM_011285081.2;refGene_gn=CSDE1;refGene_tx=XM_011285082.3;refGene_gn=CSDE1;refGene_tx=XM_011285083.2;refGene_gn=CSDE1;refGene_tx=XM_011285084.2
C1	98454316	98454529
refGene_gn=CSDE1;refGene_tx=XM_011285081.2;refGene_gn=CSDE1;refGene_tx=XM_011285082.3;refGene_gn=CSDE1;refGene_tx=XM_011285083.2;refGene_gn=CSDE1;refGene_tx=XM_011285084.2
C1	98455386	98455512
refGene_gn=CSDE1;refGene_tx=XM_011285081.2;refGene_gn=CSDE1;refGene_tx=XM_011285082.3;refGene_gn=CSDE1;refGene_tx=XM_011285083.2;refGene_gn=CSDE1;refGene_tx=XM_011285084.2
C1	98455621	98455750
refGene_gn=CSDE1;refGene_tx=XM_011285081.2;refGene_gn=CSDE1;refGene_tx=XM_011285082.3;refGene_gn=CSDE1;refGene_tx=XM_011285083.2;refGene_gn=CSDE1;refGene_tx=XM_011285084.2
C1	98456098	98456180
refGene_gn=CSDE1;refGene_tx=XM_011285081.2;refGene_gn=CSDE1;refGene_tx=XM_011285082.3;refGene_gn=CSDE1;refGene_tx=XM_011285083.2;refGene_gn=CSDE1;refGene_tx=XM_011285084.2
C1	98459160	98459258
refGene_gn=CSDE1;refGene_tx=XM_011285081.2;refGene_gn=CSDE1;refGene_tx=XM_011285082.3;refGene_gn=CSDE1;refGene_tx=XM_011285083.2;refGene_gn=CSDE1;refGene_tx=XM_011285084.2
C1	98459877	98459970
refGene_gn=CSDE1;refGene_tx=XM_011285081.2;refGene_gn=CSDE1;refGene_tx=XM_011285082.3;refGene_gn=CSDE1;refGene_tx=XM_011285084.2
C1	98460370	98460480
refGene_gn=CSDE1;refGene_tx=XM_011285081.2;refGene_gn=CSDE1;refGene_tx=XM_011285082.3;refGene_gn=CSDE1;refGene_tx=XM_011285083.2;refGene_gn=CSDE1;refGene_tx=XM_011285084.2
C1	98461748	98461947

refGene_gn=CSDE1;refGene_tx=XM_011285081.2;refGene_gn=CSDE1;refGene_tx=XM_011285082.3;refGene_gn=CSDE1;refGene_tx=XM_011285083.2;refGene_gn=CSDE1;refGene_tx=XM_011285084.2
C1 98463613 98463751
refGene_gn=CSDE1;refGene_tx=XM_011285081.2;refGene_gn=CSDE1;refGene_tx=XM_011285082.3;refGene_gn=CSDE1;refGene_tx=XM_011285083.2
C1 98491892 98491994 refGene_gn=SIKE1;refGene_tx=XM_004001118.5
C1 98493835 98493949 refGene_gn=SIKE1;refGene_tx=XM_004001118.5
C1 98495572 98495715 refGene_gn=SIKE1;refGene_tx=XM_004001118.5
C1 98496578 98496684 refGene_gn=SIKE1;refGene_tx=XM_004001118.5
C1 98496921 98497080 refGene_gn=SIKE1;refGene_tx=XM_004001118.5
C1 98536107 98536174
refGene_gn=LOC101098124;refGene_tx=XM_019837668.2
C1 98538289 98538652
refGene_gn=LOC101098124;refGene_tx=XM_019837668.2
C1 98540258 98540399
refGene_gn=LOC101098124;refGene_tx=XM_019837668.2
C1 98542274 98542414
refGene_gn=LOC101098124;refGene_tx=XM_019837668.2
C1 98542519 98542624
refGene_gn=LOC101098124;refGene_tx=XM_019837668.2
C1 98542787 98542887
refGene_gn=LOC101098124;refGene_tx=XM_019837668.2
C1 98544237 98544360
refGene_gn=LOC101098124;refGene_tx=XM_019837668.2
C1 98544765 98544891
refGene_gn=LOC101098124;refGene_tx=XM_019837668.2
C1 98545042 98545156
refGene_gn=LOC101098124;refGene_tx=XM_019837668.2
C1 98547538 98547786
refGene_gn=LOC101098124;refGene_tx=XM_019837668.2
C1 98550867 98551053
refGene_gn=SYCP1;refGene_tx=XM_023259019.1;refGene_gn=SYCP1;refGene_tx=XM_023259020.1;refGene_gn=SYCP1;refGene_tx=XM_023259021.1
C1 98554271 98554356
refGene_gn=SYCP1;refGene_tx=XM_023259019.1;refGene_gn=SYCP1;refGene_tx=XM_023259020.1;refGene_gn=SYCP1;refGene_tx=XM_023259021.1
C1 98554534 98554578
refGene_gn=SYCP1;refGene_tx=XM_023259019.1;refGene_gn=SYCP1;refGene_tx=XM_023259020.1;refGene_gn=SYCP1;refGene_tx=XM_023259021.1
C1 98554772 98554829
refGene_gn=SYCP1;refGene_tx=XM_023259019.1;refGene_gn=SYCP1;refGene_tx=XM_023259020.1;refGene_gn=SYCP1;refGene_tx=XM_023259021.1
C1 98556164 98556329
refGene_gn=SYCP1;refGene_tx=XM_023259019.1;refGene_gn=SYCP1;refGene_tx=XM_023259020.1;refGene_gn=SYCP1;refGene_tx=XM_023259021.1
C1 98557014 98557082
refGene_gn=SYCP1;refGene_tx=XM_023259019.1;refGene_gn=SYCP1;refGene_tx=XM_023259020.1;refGene_gn=SYCP1;refGene_tx=XM_023259021.1
C1 98565531 98565605

refGene_gn=SYCP1;refGene_tx=XM_023259019.1;refGene_gn=SYCP1;refGene_tx=XM_023259020.1;refGene_gn=SYCP1;refGene_tx=XM_023259021.1;refGene_gn=SYCP1;refGene_tx=XM_023259022.1
C1 98575605 98575664
refGene_gn=SYCP1;refGene_tx=XM_023259019.1;refGene_gn=SYCP1;refGene_tx=XM_023259020.1;refGene_gn=SYCP1;refGene_tx=XM_023259021.1;refGene_gn=SYCP1;refGene_tx=XM_023259022.1
C1 98577102 98577172
refGene_gn=SYCP1;refGene_tx=XM_023259019.1;refGene_gn=SYCP1;refGene_tx=XM_023259020.1;refGene_gn=SYCP1;refGene_tx=XM_023259021.1;refGene_gn=SYCP1;refGene_tx=XM_023259022.1
C1 98578136 98578210
refGene_gn=SYCP1;refGene_tx=XM_023259019.1;refGene_gn=SYCP1;refGene_tx=XM_023259020.1;refGene_gn=SYCP1;refGene_tx=XM_023259021.1;refGene_gn=SYCP1;refGene_tx=XM_023259022.1
C1 98580568 98580677
refGene_gn=SYCP1;refGene_tx=XM_023259019.1;refGene_gn=SYCP1;refGene_tx=XM_023259020.1;refGene_gn=SYCP1;refGene_tx=XM_023259021.1;refGene_gn=SYCP1;refGene_tx=XM_023259022.1
C1 98587748 98587843
refGene_gn=SYCP1;refGene_tx=XM_023259019.1;refGene_gn=SYCP1;refGene_tx=XM_023259020.1;refGene_gn=SYCP1;refGene_tx=XM_023259021.1;refGene_gn=SYCP1;refGene_tx=XM_023259022.1
C1 98588313 98588498
refGene_gn=SYCP1;refGene_tx=XM_023259019.1;refGene_gn=SYCP1;refGene_tx=XM_023259020.1;refGene_gn=SYCP1;refGene_tx=XM_023259021.1;refGene_gn=SYCP1;refGene_tx=XM_023259022.1
C1 98589943 98590011
refGene_gn=SYCP1;refGene_tx=XM_023259019.1;refGene_gn=SYCP1;refGene_tx=XM_023259020.1;refGene_gn=SYCP1;refGene_tx=XM_023259021.1;refGene_gn=SYCP1;refGene_tx=XM_023259022.1
C1 98596696 98596758
refGene_gn=SYCP1;refGene_tx=XM_023259019.1;refGene_gn=SYCP1;refGene_tx=XM_023259020.1;refGene_gn=SYCP1;refGene_tx=XM_023259021.1;refGene_gn=SYCP1;refGene_tx=XM_023259022.1
C1 98603962 98604067
refGene_gn=SYCP1;refGene_tx=XM_023259019.1;refGene_gn=SYCP1;refGene_tx=XM_023259020.1;refGene_gn=SYCP1;refGene_tx=XM_023259021.1;refGene_gn=SYCP1;refGene_tx=XM_023259022.1
C1 98604881 98604985
refGene_gn=SYCP1;refGene_tx=XM_023259019.1;refGene_gn=SYCP1;refGene_tx=XM_023259020.1;refGene_gn=SYCP1;refGene_tx=XM_023259021.1;refGene_gn=SYCP1;refGene_tx=XM_023259022.1
C1 98606251 98606369
refGene_gn=SYCP1;refGene_tx=XM_023259019.1;refGene_gn=SYCP1;refGene_tx=XM_023259020.1;refGene_gn=SYCP1;refGene_tx=XM_023259021.1;refGene_gn=SYCP1;refGene_tx=XM_023259022.1
C1 98607164 98607235
refGene_gn=SYCP1;refGene_tx=XM_023259019.1;refGene_gn=SYCP1;refGene_tx=XM_023259020.1;refGene_gn=SYCP1;refGene_tx=XM_023259021.1;refGene_gn=

SYCP1;refGene_tx=XM_023259022.1
C1 98611769 98611851
refGene_gn=SYCP1;refGene_tx=XM_023259019.1;refGene_gn=SYCP1;refGene_tx=XM_023259020.1;refGene_gn=SYCP1;refGene_tx=XM_023259021.1;refGene_gn=SYCP1;refGene_tx=XM_023259022.1
C1 98614878 98614941
refGene_gn=SYCP1;refGene_tx=XM_023259019.1;refGene_gn=SYCP1;refGene_tx=XM_023259021.1;refGene_gn=SYCP1;refGene_tx=XM_023259022.1
C1 98617395 98617458
refGene_gn=SYCP1;refGene_tx=XM_023259019.1;refGene_gn=SYCP1;refGene_tx=XM_023259020.1;refGene_gn=SYCP1;refGene_tx=XM_023259021.1;refGene_gn=SYCP1;refGene_tx=XM_023259022.1
C1 98617619 98617682
refGene_gn=SYCP1;refGene_tx=XM_023259019.1;refGene_gn=SYCP1;refGene_tx=XM_023259020.1;refGene_gn=SYCP1;refGene_tx=XM_023259021.1;refGene_gn=SYCP1;refGene_tx=XM_023259022.1
C1 98625886 98626003
refGene_gn=SYCP1;refGene_tx=XM_023259019.1;refGene_gn=SYCP1;refGene_tx=XM_023259020.1;refGene_gn=SYCP1;refGene_tx=XM_023259021.1;refGene_gn=SYCP1;refGene_tx=XM_023259022.1
C1 98626422 98626533
refGene_gn=SYCP1;refGene_tx=XM_023259019.1;refGene_gn=SYCP1;refGene_tx=XM_023259020.1;refGene_gn=SYCP1;refGene_tx=XM_023259021.1;refGene_gn=SYCP1;refGene_tx=XM_023259022.1
C1 98627802 98627895
refGene_gn=SYCP1;refGene_tx=XM_023259019.1;refGene_gn=SYCP1;refGene_tx=XM_023259020.1;refGene_gn=SYCP1;refGene_tx=XM_023259021.1;refGene_gn=SYCP1;refGene_tx=XM_023259022.1
C1 98628420 98628495
refGene_gn=SYCP1;refGene_tx=XM_023259019.1;refGene_gn=SYCP1;refGene_tx=XM_023259020.1;refGene_gn=SYCP1;refGene_tx=XM_023259021.1;refGene_gn=SYCP1;refGene_tx=XM_023259022.1
C1 98649319 98649379
refGene_gn=SYCP1;refGene_tx=XM_023259019.1;refGene_gn=SYCP1;refGene_tx=XM_023259020.1;refGene_gn=SYCP1;refGene_tx=XM_023259021.1;refGene_gn=SYCP1;refGene_tx=XM_023259022.1
C1 98655424 98655604
refGene_gn=SYCP1;refGene_tx=XM_023259019.1;refGene_gn=SYCP1;refGene_tx=XM_023259020.1;refGene_gn=SYCP1;refGene_tx=XM_023259021.1;refGene_gn=SYCP1;refGene_tx=XM_023259022.1
C1 98661840 98661984
refGene_gn=SYCP1;refGene_tx=XM_023259019.1;refGene_gn=SYCP1;refGene_tx=XM_023259020.1;refGene_gn=SYCP1;refGene_tx=XM_023259021.1;refGene_gn=SYCP1;refGene_tx=XM_023259022.1
C1 98669629 98669719
refGene_gn=SYCP1;refGene_tx=XM_023259019.1;refGene_gn=SYCP1;refGene_tx=XM_023259020.1;refGene_gn=SYCP1;refGene_tx=XM_023259021.1;refGene_gn=SYCP1;refGene_tx=XM_023259022.1
C1 98669816 98669954
refGene_gn=SYCP1;refGene_tx=XM_023259019.1;refGene_gn=SYCP1;refGene_tx

=XM_023259020.1;refGene_gn=SYCP1;refGene_tx=XM_023259021.1;refGene_gn=SYCP1;refGene_tx=XM_023259022.1
C1	98712371	98712533
refGene_gn=TSHB;refGene_tx=NM_001048015.1;refGene_gn=TSHB;refGene_tx=XM_019835959.2
C1	98712951	98713206
refGene_gn=TSHB;refGene_tx=NM_001048015.1;refGene_gn=TSHB;refGene_tx=XM_019835959.2
C1	98730021	98730087	refGene_gn=TSPAN2;refGene_tx=XM_023259023.1
C1	98732584	98732668	refGene_gn=TSPAN2;refGene_tx=XM_023259023.1
C1	98753572	98753644	refGene_gn=TSPAN2;refGene_tx=XM_023259023.1
C1	98754902	98755001	refGene_gn=TSPAN2;refGene_tx=XM_023259023.1
C1	98756257	98756332	refGene_gn=TSPAN2;refGene_tx=XM_023259023.1
C1	98757619	98757717	refGene_gn=TSPAN2;refGene_tx=XM_023259023.1
C1	98769694	98769797	refGene_gn=TSPAN2;refGene_tx=XM_023259023.1
C1	98785644	98785713	refGene_gn=TSPAN2;refGene_tx=XM_023259023.1
C1	98947101	98947827	refGene_gn=NGF;refGene_tx=XM_004001117.4
C1	99236539	99236610	refGene_gn=VANGL1;refGene_tx=XM_023259024.1
C1	99243357	99243490	refGene_gn=VANGL1;refGene_tx=XM_023259024.1
C1	99247608	99248216	refGene_gn=VANGL1;refGene_tx=XM_023259024.1
C1	99267488	99267622	refGene_gn=VANGL1;refGene_tx=XM_023259024.1
C1	99269135	99269268	refGene_gn=VANGL1;refGene_tx=XM_023259024.1
C1	99270160	99270395	refGene_gn=VANGL1;refGene_tx=XM_023259024.1
C1	99274566	99274692	refGene_gn=VANGL1;refGene_tx=XM_023259024.1
C1	99274693	99274828	refGene_gn=VANGL1;refGene_tx=XM_023259024.1
C1	99289269	99289485	refGene_gn=CASQ2;refGene_tx=XM_011285085.3
C1	99291021	99291096	refGene_gn=CASQ2;refGene_tx=XM_011285085.3
C1	99293498	99293599	refGene_gn=CASQ2;refGene_tx=XM_011285085.3
C1	99303718	99303773	refGene_gn=CASQ2;refGene_tx=XM_011285085.3
C1	99310948	99310994	refGene_gn=CASQ2;refGene_tx=XM_011285085.3
C1	99312448	99312579	refGene_gn=CASQ2;refGene_tx=XM_011285085.3
C1	99318239	99318313	refGene_gn=CASQ2;refGene_tx=XM_011285085.3
C1	99323662	99323774	refGene_gn=CASQ2;refGene_tx=XM_011285085.3
C1	99325147	99325248	refGene_gn=CASQ2;refGene_tx=XM_011285085.3
C1	99327978	99328063	refGene_gn=CASQ2;refGene_tx=XM_011285085.3
C1	99351199	99351433	refGene_gn=CASQ2;refGene_tx=XM_011285085.3
C1	99416696	99417104
refGene_gn=NHLH2;refGene_tx=XM_023259025.1;refGene_gn=NHLH2;refGene_tx=XM_023259026.1
C1	99517364	99517451
refGene_gn=SLC22A15;refGene_tx=XM_023259027.1;refGene_gn=SLC22A15;refGene_tx=XM_023259028.1
C1	99532974	99533187
refGene_gn=SLC22A15;refGene_tx=XM_023259027.1;refGene_gn=SLC22A15;refGene_tx=XM_023259028.1
C1	99558694	99558827
refGene_gn=SLC22A15;refGene_tx=XM_023259027.1;refGene_gn=SLC22A15;refGene_tx=XM_023259028.1
C1	99559483	99559648
refGene_gn=SLC22A15;refGene_tx=XM_023259027.1;refGene_gn=SLC22A15;refG

ene_tx=XM_023259028.1
C1 99563013 99563143
refGene_gn=SLC22A15;refGene_tx=XM_023259027.1;refGene_gn=SLC22A15;refGene_tx=XM_023259028.1
C1 99568015 99568231
refGene_gn=SLC22A15;refGene_tx=XM_023259027.1;refGene_gn=SLC22A15;refGene_tx=XM_023259028.1
C1 99575636 99575777
refGene_gn=SLC22A15;refGene_tx=XM_023259027.1;refGene_gn=SLC22A15;refGene_tx=XM_023259028.1
C1 99577864 99577950
refGene_gn=SLC22A15;refGene_tx=XM_023259027.1;refGene_gn=SLC22A15;refGene_tx=XM_023259028.1
C1 99592350 99592471
refGene_gn=SLC22A15;refGene_tx=XM_023259027.1;refGene_gn=SLC22A15;refGene_tx=XM_023259028.1
C1 99594640 99594713 refGene_gn=SLC22A15;refGene_tx=XM_023259027.1
C1 99597938 99598127
refGene_gn=SLC22A15;refGene_tx=XM_023259027.1;refGene_gn=SLC22A15;refGene_tx=XM_023259028.1
C1 99598518 99598608 refGene_gn=SLC22A15;refGene_tx=XM_023259027.1
C1 99655500 99655548 refGene_gn=MAB21L3;refGene_tx=XM_023259032.1
C1 99663790 99663877
refGene_gn=MAB21L3;refGene_tx=XM_023259029.1;refGene_gn=MAB21L3;refGene_tx=XM_023259030.1;refGene_gn=MAB21L3;refGene_tx=XM_023259031.1
C1 99665869 99666010
refGene_gn=MAB21L3;refGene_tx=XM_019837673.2;refGene_gn=MAB21L3;refGene_tx=XM_019837680.1;refGene_gn=MAB21L3;refGene_tx=XM_019837684.1;refGene_gn=MAB21L3;refGene_tx=XM_019837685.1;refGene_gn=MAB21L3;refGene_tx=XM_023259029.1;refGene_gn=MAB21L3;refGene_tx=XM_023259030.1;refGene_gn=MAB21L3;refGene_tx=XM_023259031.1;refGene_gn=MAB21L3;refGene_tx=XM_023259032.1
C1 99668814 99669106
refGene_gn=MAB21L3;refGene_tx=XM_019837673.2;refGene_gn=MAB21L3;refGene_tx=XM_019837680.1;refGene_gn=MAB21L3;refGene_tx=XM_019837684.1;refGene_gn=MAB21L3;refGene_tx=XM_019837685.1;refGene_gn=MAB21L3;refGene_tx=XM_023259029.1;refGene_gn=MAB21L3;refGene_tx=XM_023259030.1;refGene_gn=MAB21L3;refGene_tx=XM_023259031.1;refGene_gn=MAB21L3;refGene_tx=XM_023259032.1
C1 99672574 99672753
refGene_gn=MAB21L3;refGene_tx=XM_019837673.2;refGene_gn=MAB21L3;refGene_tx=XM_019837680.1;refGene_gn=MAB21L3;refGene_tx=XM_019837684.1;refGene_gn=MAB21L3;refGene_tx=XM_019837685.1;refGene_gn=MAB21L3;refGene_tx=XM_023259029.1;refGene_gn=MAB21L3;refGene_tx=XM_023259030.1;refGene_gn=MAB21L3;refGene_tx=XM_023259031.1;refGene_gn=MAB21L3;refGene_tx=XM_023259032.1
C1 99673219 99673414
refGene_gn=MAB21L3;refGene_tx=XM_019837673.2;refGene_gn=MAB21L3;refGene_tx=XM_019837680.1;refGene_gn=MAB21L3;refGene_tx=XM_019837684.1;refGene_gn=MAB21L3;refGene_tx=XM_019837685.1;refGene_gn=MAB21L3;refGene_tx=

XM_023259029.1;refGene_gn=MAB21L3;refGene_tx=XM_023259030.1;refGene_gn=MAB21L3;refGene_tx=XM_023259031.1;refGene_gn=MAB21L3;refGene_tx=XM_023259032.1
C1 99678358 99678592
refGene_gn=MAB21L3;refGene_tx=XM_019837673.2;refGene_gn=MAB21L3;refGene_tx=XM_023259029.1;refGene_gn=MAB21L3;refGene_tx=XM_023259030.1;refGene_gn=MAB21L3;refGene_tx=XM_023259031.1;refGene_gn=MAB21L3;refGene_tx=XM_023259032.1
C1 99681495 99681501 refGene_gn=MAB21L3;refGene_tx=XM_019837684.1
C1 99681544 99681661 refGene_gn=MAB21L3;refGene_tx=XM_019837680.1
C1 99687130 99687136 refGene_gn=MAB21L3;refGene_tx=XM_019837685.1
C1 99702555 99702650
refGene_gn=LOC109502552;refGene_tx=XM_023259034.1
C1 99705086 99705408
refGene_gn=LOC109502552;refGene_tx=XM_023259033.1;refGene_gn=LOC109502552;refGene_tx=XM_023259034.1
C1 99706255 99706500
refGene_gn=LOC109502552;refGene_tx=XM_023259033.1;refGene_gn=LOC109502552;refGene_tx=XM_023259034.1
C1 99706929 99707058
refGene_gn=LOC109502552;refGene_tx=XM_023259033.1;refGene_gn=LOC109502552;refGene_tx=XM_023259034.1
C1 99904046 99904058 refGene_gn=ATP1A1;refGene_tx=XM_011285086.3
C1 99914841 99914946 refGene_gn=ATP1A1;refGene_tx=XM_011285086.3
C1 99915638 99915698 refGene_gn=ATP1A1;refGene_tx=XM_011285086.3
C1 99917636 99917840 refGene_gn=ATP1A1;refGene_tx=XM_011285086.3
C1 99918504 99918618 refGene_gn=ATP1A1;refGene_tx=XM_011285086.3
C1 99919023 99919158 refGene_gn=ATP1A1;refGene_tx=XM_011285086.3
C1 99919245 99919363 refGene_gn=ATP1A1;refGene_tx=XM_011285086.3
C1 99919807 99920076 refGene_gn=ATP1A1;refGene_tx=XM_011285086.3
C1 99920567 99920766 refGene_gn=ATP1A1;refGene_tx=XM_011285086.3
C1 99921046 99921156 refGene_gn=ATP1A1;refGene_tx=XM_011285086.3
C1 99923298 99923433 refGene_gn=ATP1A1;refGene_tx=XM_011285086.3
C1 99923981 99924174 refGene_gn=ATP1A1;refGene_tx=XM_011285086.3
C1 99925514 99925690 refGene_gn=ATP1A1;refGene_tx=XM_011285086.3
C1 99926706 99926843 refGene_gn=ATP1A1;refGene_tx=XM_011285086.3
C1 99927655 99927806 refGene_gn=ATP1A1;refGene_tx=XM_011285086.3
C1 99928351 99928520 refGene_gn=ATP1A1;refGene_tx=XM_011285086.3
C1 99928640 99928795 refGene_gn=ATP1A1;refGene_tx=XM_011285086.3
C1 99929110 99929234 refGene_gn=ATP1A1;refGene_tx=XM_011285086.3
C1 99930151 99930297 refGene_gn=ATP1A1;refGene_tx=XM_011285086.3
C1 99930417 99930548 refGene_gn=ATP1A1;refGene_tx=XM_011285086.3
C1 99930824 99930926 refGene_gn=ATP1A1;refGene_tx=XM_011285086.3
C1 99932946 99933038 refGene_gn=ATP1A1;refGene_tx=XM_011285086.3
C1 99933473 99933502 refGene_gn=ATP1A1;refGene_tx=XM_011285086.3
C1 100061578 100061588
refGene_gn=CD58;refGene_tx=XM_023259036.1
C1 100064580 100064641
refGene_gn=CD58;refGene_tx=XM_023259035.1
C1 100065884 100065921

refGene_gn=CD58;refGene_tx=XM_023259035.1;refGene_gn=CD58;refGene_tx=XM_023259036.1
C1 100067962 100068046
refGene_gn=CD58;refGene_tx=XM_023259035.1;refGene_gn=CD58;refGene_tx=XM_023259036.1
C1 100075892 100076156
refGene_gn=CD58;refGene_tx=XM_023259035.1;refGene_gn=CD58;refGene_tx=XM_023259036.1
C1 100084806 100085103
refGene_gn=CD58;refGene_tx=XM_023259035.1;refGene_gn=CD58;refGene_tx=XM_023259036.1
C1 100102070 100102140
refGene_gn=CD58;refGene_tx=XM_023259035.1;refGene_gn=CD58;refGene_tx=XM_023259036.1
C1 100109139 100109390
refGene_gn=IGSF3;refGene_tx=XM_019837688.2;refGene_gn=IGSF3;refGene_tx=XM_019837689.2;refGene_gn=IGSF3;refGene_tx=XM_023259037.1;refGene_gn=IGSF3;refGene_tx=XM_023259038.1;refGene_gn=IGSF3;refGene_tx=XM_023259039.1
C1 100111160 100111631
refGene_gn=IGSF3;refGene_tx=XM_019837688.2;refGene_gn=IGSF3;refGene_tx=XM_019837689.2;refGene_gn=IGSF3;refGene_tx=XM_023259037.1;refGene_gn=IGSF3;refGene_tx=XM_023259038.1;refGene_gn=IGSF3;refGene_tx=XM_023259039.1
C1 100114989 100115397
refGene_gn=IGSF3;refGene_tx=XM_019837688.2;refGene_gn=IGSF3;refGene_tx=XM_019837689.2;refGene_gn=IGSF3;refGene_tx=XM_023259037.1;refGene_gn=IGSF3;refGene_tx=XM_023259038.1;refGene_gn=IGSF3;refGene_tx=XM_023259039.1
C1 100119533 100119944
refGene_gn=IGSF3;refGene_tx=XM_019837688.2;refGene_gn=IGSF3;refGene_tx=XM_019837689.2;refGene_gn=IGSF3;refGene_tx=XM_023259037.1;refGene_gn=IGSF3;refGene_tx=XM_023259038.1;refGene_gn=IGSF3;refGene_tx=XM_023259039.1
C1 100131544 100131949
refGene_gn=IGSF3;refGene_tx=XM_019837688.2;refGene_gn=IGSF3;refGene_tx=XM_019837689.2;refGene_gn=IGSF3;refGene_tx=XM_023259037.1;refGene_gn=IGSF3;refGene_tx=XM_023259038.1;refGene_gn=IGSF3;refGene_tx=XM_023259039.1
C1 100135246 100135648
refGene_gn=IGSF3;refGene_tx=XM_019837688.2;refGene_gn=IGSF3;refGene_tx=XM_019837689.2;refGene_gn=IGSF3;refGene_tx=XM_023259037.1;refGene_gn=IGSF3;refGene_tx=XM_023259038.1;refGene_gn=IGSF3;refGene_tx=XM_023259039.1
C1 100138046 100138106
refGene_gn=IGSF3;refGene_tx=XM_019837688.2;refGene_gn=IGSF3;refGene_tx=XM_019837689.2;refGene_gn=IGSF3;refGene_tx=XM_023259037.1;refGene_gn=IGSF3;refGene_tx=XM_023259038.1
C1 100139517 100139907
refGene_gn=IGSF3;refGene_tx=XM_019837688.2;refGene_gn=IGSF3;refGene_tx

=XM_019837689.2;refGene_gn=IGSF3;refGene_tx=XM_023259037.1;refGene_gn=IGSF3;refGene_tx=XM_023259038.1;refGene_gn=IGSF3;refGene_tx=XM_023259039.1
C1 100143639 100144050
refGene_gn=IGSF3;refGene_tx=XM_019837688.2;refGene_gn=IGSF3;refGene_tx=XM_019837689.2;refGene_gn=IGSF3;refGene_tx=XM_023259037.1;refGene_gn=IGSF3;refGene_tx=XM_023259038.1;refGene_gn=IGSF3;refGene_tx=XM_023259039.1
C1 100146127 100146505
refGene_gn=IGSF3;refGene_tx=XM_019837688.2;refGene_gn=IGSF3;refGene_tx=XM_019837689.2;refGene_gn=IGSF3;refGene_tx=XM_023259037.1;refGene_gn=IGSF3;refGene_tx=XM_023259038.1;refGene_gn=IGSF3;refGene_tx=XM_023259039.1
C1 100199095 100199138
refGene_gn=IGSF3;refGene_tx=XM_019837688.2;refGene_gn=IGSF3;refGene_tx=XM_019837689.2;refGene_gn=IGSF3;refGene_tx=XM_023259037.1;refGene_gn=IGSF3;refGene_tx=XM_023259038.1;refGene_gn=IGSF3;refGene_tx=XM_023259039.1
C1 100270284 100270336
refGene_gn=CD2;refGene_tx=NM_001009841.1
C1 100270456 100270759
refGene_gn=CD2;refGene_tx=NM_001009841.1
C1 100274816 100275050
refGene_gn=CD2;refGene_tx=NM_001009841.1
C1 100278871 100278994
refGene_gn=CD2;refGene_tx=NM_001009841.1
C1 100282324 100282623
refGene_gn=CD2;refGene_tx=NM_001009841.1
C1 100405002 100405399
refGene_gn=PTGFRN;refGene_tx=XM_023259040.1
C1 100447724 100448093
refGene_gn=PTGFRN;refGene_tx=XM_023259040.1
C1 100450948 100451362
refGene_gn=PTGFRN;refGene_tx=XM_023259040.1
C1 100455627 100456008
refGene_gn=PTGFRN;refGene_tx=XM_023259040.1
C1 100468414 100468840
refGene_gn=PTGFRN;refGene_tx=XM_023259040.1
C1 100473471 100473891
refGene_gn=PTGFRN;refGene_tx=XM_023259040.1
C1 100482508 100482616
refGene_gn=PTGFRN;refGene_tx=XM_023259040.1
C1 100487127 100487433
refGene_gn=PTGFRN;refGene_tx=XM_023259040.1
C1 100488941 100489108
refGene_gn=PTGFRN;refGene_tx=XM_023259040.1
C1 100511823 100512232
refGene_gn=CD101;refGene_tx=XM_023259041.1
C1 100513546 100513963
refGene_gn=CD101;refGene_tx=XM_023259041.1

C1 100518996 100519383
refGene_gn=CD101;refGene_tx=XM_023259041.1
C1 100522474 100522858
refGene_gn=CD101;refGene_tx=XM_023259041.1
C1 100523549 100523954
refGene_gn=CD101;refGene_tx=XM_023259041.1
C1 100526267 100526678
refGene_gn=CD101;refGene_tx=XM_023259041.1
C1 100529720 100530116
refGene_gn=CD101;refGene_tx=XM_023259041.1
C1 100538454 100538684
refGene_gn=CD101;refGene_tx=XM_023259041.1
C1 100558194 100558222
refGene_gn=TTF2;refGene_tx=XM_006935022.4
C1 100558307 100558410
refGene_gn=TTF2;refGene_tx=XM_006935022.4
C1 100561203 100561290
refGene_gn=TTF2;refGene_tx=XM_006935022.4
C1 100571961 100572028
refGene_gn=TTF2;refGene_tx=XM_006935022.4
C1 100572880 100573840
refGene_gn=TTF2;refGene_tx=XM_006935022.4
C1 100574397 100574512
refGene_gn=TTF2;refGene_tx=XM_006935022.4
C1 100574855 100575038
refGene_gn=TTF2;refGene_tx=XM_006935022.4
C1 100576554 100576682
refGene_gn=TTF2;refGene_tx=XM_006935022.4
C1 100577878 100577960
refGene_gn=TTF2;refGene_tx=XM_006935022.4
C1 100580228 100580318
refGene_gn=TTF2;refGene_tx=XM_006935022.4
C1 100580320 100580347
refGene_gn=TTF2;refGene_tx=XM_006935022.4
C1 100581926 100582077
refGene_gn=TTF2;refGene_tx=XM_006935022.4
C1 100583859 100583968
refGene_gn=TTF2;refGene_tx=XM_006935022.4
C1 100586228 100586413
refGene_gn=TTF2;refGene_tx=XM_006935022.4
C1 100587376 100587530
refGene_gn=TTF2;refGene_tx=XM_006935022.4
C1 100587793 100587885
refGene_gn=TTF2;refGene_tx=XM_006935022.4
C1 100588742 100588828
refGene_gn=TTF2;refGene_tx=XM_006935022.4
C1 100589445 100589582
refGene_gn=TTF2;refGene_tx=XM_006935022.4
C1 100590549 100590720
refGene_gn=TTF2;refGene_tx=XM_006935022.4

C1 100593716 100593775
refGene_gn=TTF2;refGene_tx=XM_006935022.4
C1 100595274 100595425
refGene_gn=TTF2;refGene_tx=XM_006935022.4
C1 100596114 100596197
refGene_gn=TTF2;refGene_tx=XM_006935022.4
C1 100597925 100598000
refGene_gn=TTF2;refGene_tx=XM_006935022.4
C1 100599938 100600083
refGene_gn=TTF2;refGene_tx=XM_006935022.4
C1 100610989 100611138
refGene_gn=TRIM45;refGene_tx=XM_003990508.5
C1 100611996 100612123
refGene_gn=TRIM45;refGene_tx=XM_003990508.5
C1 100614146 100614261
refGene_gn=TRIM45;refGene_tx=XM_003990508.5
C1 100614840 100614970
refGene_gn=TRIM45;refGene_tx=XM_003990508.5
C1 100616242 100616976
refGene_gn=TRIM45;refGene_tx=XM_003990508.5
C1 100618635 100619123
refGene_gn=TRIM45;refGene_tx=XM_003990508.5
C1 100644560 100644685
refGene_gn=VTCN1;refGene_tx=XM_006935023.3
C1 100644992 100645108
refGene_gn=VTCN1;refGene_tx=XM_019837693.2
C1 100654503 100654782
refGene_gn=VTCN1;refGene_tx=XM_006935023.3;refGene_gn=VTCN1;refGene_tx=XM_019837693.2
C1 100657642 100657990
refGene_gn=VTCN1;refGene_tx=XM_006935023.3;refGene_gn=VTCN1;refGene_tx=XM_019837693.2
C1 100671359 100671424
refGene_gn=VTCN1;refGene_tx=XM_006935023.3;refGene_gn=VTCN1;refGene_tx=XM_019837693.2
C1 100705112 100705144
refGene_gn=VTCN1;refGene_tx=XM_006935023.3;refGene_gn=VTCN1;refGene_tx=XM_019837693.2
C1 100822711 100823013
refGene_gn=MAN1A2;refGene_tx=XM_003990510.5;refGene_gn=MAN1A2;refGene_tx=XM_019837695.2
C1 100861200 100861456
refGene_gn=MAN1A2;refGene_tx=XM_003990510.5;refGene_gn=MAN1A2;refGene_tx=XM_019837695.2
C1 100868250 100868347
refGene_gn=MAN1A2;refGene_tx=XM_003990510.5;refGene_gn=MAN1A2;refGene_tx=XM_019837695.2
C1 100879210 100879329
refGene_gn=MAN1A2;refGene_tx=XM_003990510.5;refGene_gn=MAN1A2;refGene_tx=XM_019837695.2

C1 100884900 100884981
refGene_gn=MAN1A2;refGene_tx=XM_003990510.5;refGene_gn=MAN1A2;refGene_tx=XM_019837695.2
C1 100902622 100902717
refGene_gn=MAN1A2;refGene_tx=XM_003990510.5;refGene_gn=MAN1A2;refGene_tx=XM_019837695.2
C1 100926746 100926870
refGene_gn=MAN1A2;refGene_tx=XM_003990510.5;refGene_gn=MAN1A2;refGene_tx=XM_019837695.2
C1 100931132 100931226
refGene_gn=MAN1A2;refGene_tx=XM_003990510.5;refGene_gn=MAN1A2;refGene_tx=XM_019837695.2
C1 100956191 100956307
refGene_gn=MAN1A2;refGene_tx=XM_003990510.5;refGene_gn=MAN1A2;refGene_tx=XM_019837695.2
C1 100959511 100959731
refGene_gn=MAN1A2;refGene_tx=XM_003990510.5;refGene_gn=MAN1A2;refGene_tx=XM_019837695.2
C1 100965055 100965228
refGene_gn=MAN1A2;refGene_tx=XM_003990510.5;refGene_gn=MAN1A2;refGene_tx=XM_019837695.2
C1 100969000 100969116
refGene_gn=MAN1A2;refGene_tx=XM_003990510.5;refGene_gn=MAN1A2;refGene_tx=XM_019837695.2
C1 100982156 100982208
refGene_gn=MAN1A2;refGene_tx=XM_019837695.2
C1 101001274 101001407
refGene_gn=MAN1A2;refGene_tx=XM_003990510.5;refGene_gn=MAN1A2;refGene_tx=XM_019837695.2
C1 101093400 101094576
refGene_gn=FAM46C;refGene_tx=XM_006935024.3;refGene_gn=FAM46C;refGene_tx=XM_011285088.3;refGene_gn=FAM46C;refGene_tx=XM_011285089.3;refGene_gn=FAM46C;refGene_tx=XM_023259042.1
C1 101286188 101286236
refGene_gn=GDAP2;refGene_tx=XM_006935025.4
C1 101290118 101290262
refGene_gn=GDAP2;refGene_tx=XM_006935025.4
C1 101301344 101301399
refGene_gn=GDAP2;refGene_tx=XM_006935025.4
C1 101302078 101302218
refGene_gn=GDAP2;refGene_tx=XM_006935025.4
C1 101305381 101305458
refGene_gn=GDAP2;refGene_tx=XM_006935025.4
C1 101306184 101306261
refGene_gn=GDAP2;refGene_tx=XM_006935025.4
C1 101315871 101316028
refGene_gn=GDAP2;refGene_tx=XM_006935025.4
C1 101318280 101318440
refGene_gn=GDAP2;refGene_tx=XM_006935025.4
C1 101319812 101319889

refGene_gn=GDAP2;refGene_tx=XM_006935025.4
C1	101325842	101325931
refGene_gn=GDAP2;refGene_tx=XM_006935025.4
C1	101326330	101326484
refGene_gn=GDAP2;refGene_tx=XM_006935025.4
C1	101335306	101335446
refGene_gn=GDAP2;refGene_tx=XM_006935025.4
C1	101336723	101336899
refGene_gn=GDAP2;refGene_tx=XM_006935025.4
C1	101346188	101346211
refGene_gn=WDR3;refGene_tx=XM_011285091.3;refGene_gn=WDR3;refGene_tx=XM_023259043.1
C1	101348324	101348526
refGene_gn=WDR3;refGene_tx=XM_011285091.3;refGene_gn=WDR3;refGene_tx=XM_011285092.2;refGene_gn=WDR3;refGene_tx=XM_023259043.1
C1	101349470	101349680
refGene_gn=WDR3;refGene_tx=XM_011285091.3;refGene_gn=WDR3;refGene_tx=XM_011285092.2;refGene_gn=WDR3;refGene_tx=XM_023259043.1
C1	101352492	101352611
refGene_gn=WDR3;refGene_tx=XM_011285091.3;refGene_gn=WDR3;refGene_tx=XM_011285092.2;refGene_gn=WDR3;refGene_tx=XM_023259043.1
C1	101354713	101354792
refGene_gn=WDR3;refGene_tx=XM_011285091.3;refGene_gn=WDR3;refGene_tx=XM_011285092.2;refGene_gn=WDR3;refGene_tx=XM_023259043.1
C1	101355677	101355773
refGene_gn=WDR3;refGene_tx=XM_011285091.3;refGene_gn=WDR3;refGene_tx=XM_011285092.2;refGene_gn=WDR3;refGene_tx=XM_023259043.1
C1	101356704	101356818
refGene_gn=WDR3;refGene_tx=XM_011285091.3;refGene_gn=WDR3;refGene_tx=XM_011285092.2;refGene_gn=WDR3;refGene_tx=XM_023259043.1
C1	101357024	101357126
refGene_gn=WDR3;refGene_tx=XM_011285091.3;refGene_gn=WDR3;refGene_tx=XM_011285092.2;refGene_gn=WDR3;refGene_tx=XM_023259043.1
C1	101357570	101357668
refGene_gn=WDR3;refGene_tx=XM_011285091.3;refGene_gn=WDR3;refGene_tx=XM_011285092.2;refGene_gn=WDR3;refGene_tx=XM_023259043.1
C1	101358530	101358638
refGene_gn=WDR3;refGene_tx=XM_011285091.3;refGene_gn=WDR3;refGene_tx=XM_011285092.2;refGene_gn=WDR3;refGene_tx=XM_023259043.1
C1	101359161	101359392
refGene_gn=WDR3;refGene_tx=XM_011285091.3;refGene_gn=WDR3;refGene_tx=XM_011285092.2;refGene_gn=WDR3;refGene_tx=XM_023259043.1
C1	101361739	101361833
refGene_gn=WDR3;refGene_tx=XM_011285091.3;refGene_gn=WDR3;refGene_tx=XM_011285092.2;refGene_gn=WDR3;refGene_tx=XM_023259043.1
C1	101363665	101363767
refGene_gn=WDR3;refGene_tx=XM_011285091.3;refGene_gn=WDR3;refGene_tx=XM_011285092.2;refGene_gn=WDR3;refGene_tx=XM_023259043.1
C1	101365335	101365421
refGene_gn=WDR3;refGene_tx=XM_011285091.3;refGene_gn=WDR3;refGene_tx=X

M_011285092.2;refGene_gn=WDR3;refGene_tx=XM_023259043.1
C1	101365808	101365944
refGene_gn=WDR3;refGene_tx=XM_011285091.3;refGene_gn=WDR3;refGene_tx=XM_011285092.2;refGene_gn=WDR3;refGene_tx=XM_023259043.1
C1	101366621	101366678
refGene_gn=WDR3;refGene_tx=XM_011285091.3;refGene_gn=WDR3;refGene_tx=XM_011285092.2;refGene_gn=WDR3;refGene_tx=XM_023259043.1
C1	101367949	101368050
refGene_gn=WDR3;refGene_tx=XM_011285091.3;refGene_gn=WDR3;refGene_tx=XM_011285092.2;refGene_gn=WDR3;refGene_tx=XM_023259043.1
C1	101368264	101368376
refGene_gn=WDR3;refGene_tx=XM_011285091.3;refGene_gn=WDR3;refGene_tx=XM_011285092.2;refGene_gn=WDR3;refGene_tx=XM_023259043.1
C1	101368834	101368969
refGene_gn=WDR3;refGene_tx=XM_011285091.3;refGene_gn=WDR3;refGene_tx=XM_011285092.2;refGene_gn=WDR3;refGene_tx=XM_023259043.1
C1	101369246	101369297
refGene_gn=WDR3;refGene_tx=XM_011285091.3;refGene_gn=WDR3;refGene_tx=XM_011285092.2;refGene_gn=WDR3;refGene_tx=XM_023259043.1
C1	101370301	101370367
refGene_gn=WDR3;refGene_tx=XM_011285091.3;refGene_gn=WDR3;refGene_tx=XM_011285092.2;refGene_gn=WDR3;refGene_tx=XM_023259043.1
C1	101370864	101370957
refGene_gn=WDR3;refGene_tx=XM_011285091.3;refGene_gn=WDR3;refGene_tx=XM_011285092.2;refGene_gn=WDR3;refGene_tx=XM_023259043.1
C1	101371446	101371494
refGene_gn=WDR3;refGene_tx=XM_011285091.3;refGene_gn=WDR3;refGene_tx=XM_011285092.2;refGene_gn=WDR3;refGene_tx=XM_023259043.1
C1	101372393	101372437
refGene_gn=WDR3;refGene_tx=XM_011285091.3;refGene_gn=WDR3;refGene_tx=XM_011285092.2;refGene_gn=WDR3;refGene_tx=XM_023259043.1
C1	101374087	101374216
refGene_gn=WDR3;refGene_tx=XM_011285091.3;refGene_gn=WDR3;refGene_tx=XM_011285092.2;refGene_gn=WDR3;refGene_tx=XM_023259043.1
C1	101375974	101376068
refGene_gn=WDR3;refGene_tx=XM_011285091.3;refGene_gn=WDR3;refGene_tx=XM_011285092.2;refGene_gn=WDR3;refGene_tx=XM_023259043.1
C1	101376367	101376523
refGene_gn=WDR3;refGene_tx=XM_011285091.3;refGene_gn=WDR3;refGene_tx=XM_011285092.2;refGene_gn=WDR3;refGene_tx=XM_023259043.1
C1	101394369	101394509
refGene_gn=SPAG17;refGene_tx=XM_023257840.1
C1	101396942	101397087
refGene_gn=SPAG17;refGene_tx=XM_023257840.1
C1	101400014	101400075
refGene_gn=SPAG17;refGene_tx=XM_023257840.1
C1	101401993	101402178
refGene_gn=SPAG17;refGene_tx=XM_023257840.1
C1	101403646	101403783
refGene_gn=SPAG17;refGene_tx=XM_023257840.1

C1 101412119 101412251
refGene_gn=SPAG17;refGene_tx=XM_023257840.1
C1 101414395 101414498
refGene_gn=SPAG17;refGene_tx=XM_023257840.1
C1 101415202 101415302
refGene_gn=SPAG17;refGene_tx=XM_023257840.1
C1 101417993 101418041
refGene_gn=SPAG17;refGene_tx=XM_023257840.1
C1 101418258 101418358
refGene_gn=SPAG17;refGene_tx=XM_023257840.1
C1 101420498 101420544
refGene_gn=SPAG17;refGene_tx=XM_023257840.1
C1 101421061 101421175
refGene_gn=SPAG17;refGene_tx=XM_023257840.1
C1 101422130 101422313
refGene_gn=SPAG17;refGene_tx=XM_023257840.1
C1 101424110 101424238
refGene_gn=SPAG17;refGene_tx=XM_023257840.1
C1 101425961 101426092
refGene_gn=SPAG17;refGene_tx=XM_023257840.1
C1 101426195 101426341
refGene_gn=SPAG17;refGene_tx=XM_023257840.1
C1 101427998 101428039
refGene_gn=SPAG17;refGene_tx=XM_023257840.1
C1 101428041 101428113
refGene_gn=SPAG17;refGene_tx=XM_023257840.1
C1 101428115 101428166
refGene_gn=SPAG17;refGene_tx=XM_023257840.1
C1 101428168 101428187
refGene_gn=SPAG17;refGene_tx=XM_023257840.1
C1 101437191 101437346
refGene_gn=SPAG17;refGene_tx=XM_023257840.1
C1 101439084 101439229
refGene_gn=SPAG17;refGene_tx=XM_023257840.1
C1 101441081 101441299
refGene_gn=SPAG17;refGene_tx=XM_023257840.1
C1 101449374 101449505
refGene_gn=SPAG17;refGene_tx=XM_023257840.1
C1 101449546 101449560
refGene_gn=SPAG17;refGene_tx=XM_023257840.1
C1 101451887 101451999
refGene_gn=SPAG17;refGene_tx=XM_023257840.1
C1 101452000 101452064
refGene_gn=SPAG17;refGene_tx=XM_023257840.1
C1 101455182 101455221
refGene_gn=SPAG17;refGene_tx=XM_023257840.1
C1 101455223 101455299
refGene_gn=SPAG17;refGene_tx=XM_023257840.1
C1 101457446 101457559
refGene_gn=SPAG17;refGene_tx=XM_023257840.1

C1 101457561 101457621
refGene_gn=SPAG17;refGene_tx=XM_023257840.1
C1 101468665 101468779
refGene_gn=SPAG17;refGene_tx=XM_023257840.1
C1 101470530 101470689
refGene_gn=SPAG17;refGene_tx=XM_023257840.1
C1 101472793 101472905
refGene_gn=SPAG17;refGene_tx=XM_023257840.1
C1 101473916 101474156
refGene_gn=SPAG17;refGene_tx=XM_023257840.1
C1 101482202 101482312
refGene_gn=SPAG17;refGene_tx=XM_023257840.1
C1 101484069 101484248
refGene_gn=SPAG17;refGene_tx=XM_023257840.1
C1 101487824 101487979
refGene_gn=SPAG17;refGene_tx=XM_023257840.1
C1 101494866 101494980
refGene_gn=SPAG17;refGene_tx=XM_023257840.1
C1 101495580 101495642
refGene_gn=SPAG17;refGene_tx=XM_023257840.1
C1 101502462 101502681
refGene_gn=SPAG17;refGene_tx=XM_023257840.1
C1 101502776 101503004
refGene_gn=SPAG17;refGene_tx=XM_023257840.1
C1 101506245 101506396
refGene_gn=SPAG17;refGene_tx=XM_023257840.1
C1 101507202 101507316
refGene_gn=SPAG17;refGene_tx=XM_023257840.1
C1 101507404 101507542
refGene_gn=SPAG17;refGene_tx=XM_023257840.1
C1 101517995 101518108
refGene_gn=SPAG17;refGene_tx=XM_023257840.1
C1 101518313 101518386
refGene_gn=SPAG17;refGene_tx=XM_023257840.1
C1 101519505 101519667
refGene_gn=SPAG17;refGene_tx=XM_023257840.1
C1 101523606 101523788
refGene_gn=SPAG17;refGene_tx=XM_023257840.1
C1 101526051 101526246
refGene_gn=SPAG17;refGene_tx=XM_023257840.1
C1 101527795 101527982
refGene_gn=SPAG17;refGene_tx=XM_023257840.1
C1 101539297 101539429
refGene_gn=SPAG17;refGene_tx=XM_023257840.1
C1 101571228 101571312
refGene_gn=SPAG17;refGene_tx=XM_023257840.1
C1 101571925 101572066
refGene_gn=SPAG17;refGene_tx=XM_023257840.1
C1 101606934 101607021
refGene_gn=SPAG17;refGene_tx=XM_023257840.1

C1	102199250	102200035
refGene_gn=TBX15;refGene_tx=XM_003990514.5;refGene_gn=TBX15;refGene_tx=XM_006935028.4
C1	102210130	102210228
refGene_gn=TBX15;refGene_tx=XM_003990514.5;refGene_gn=TBX15;refGene_tx=XM_006935028.4
C1	102225550	102225615
refGene_gn=TBX15;refGene_tx=XM_003990514.5;refGene_gn=TBX15;refGene_tx=XM_006935028.4
C1	102235508	102235676
refGene_gn=TBX15;refGene_tx=XM_003990514.5;refGene_gn=TBX15;refGene_tx=XM_006935028.4
C1	102236695	102236867
refGene_gn=TBX15;refGene_tx=XM_003990514.5;refGene_gn=TBX15;refGene_tx=XM_006935028.4
C1	102238627	102238729
refGene_gn=TBX15;refGene_tx=XM_003990514.5;refGene_gn=TBX15;refGene_tx=XM_006935028.4
C1	102244084	102244298
refGene_gn=TBX15;refGene_tx=XM_003990514.5;refGene_gn=TBX15;refGene_tx=XM_006935028.4
C1	102301894	102302099
refGene_gn=TBX15;refGene_tx=XM_003990514.5
C1	102339004	102339453
refGene_gn=WARS2;refGene_tx=XM_003990515.4
C1	102342380	102342499
refGene_gn=WARS2;refGene_tx=XM_003990515.4
C1	102354959	102355045
refGene_gn=WARS2;refGene_tx=XM_003990515.4
C1	102359057	102359138
refGene_gn=WARS2;refGene_tx=XM_003990515.4
C1	102384741	102384999
refGene_gn=WARS2;refGene_tx=XM_003990515.4
C1	102437525	102437615
refGene_gn=WARS2;refGene_tx=XM_003990515.4
C1	102638542	102638673
refGene_gn=HAO2;refGene_tx=XM_003990516.4;refGene_gn=HAO2;refGene_tx=XM_019837704.2;refGene_gn=HAO2;refGene_tx=XM_019837705.2;refGene_gn=HAO2;refGene_tx=XM_019837706.2
C1	102640374	102640550
refGene_gn=HAO2;refGene_tx=XM_003990516.4;refGene_gn=HAO2;refGene_tx=XM_019837704.2;refGene_gn=HAO2;refGene_tx=XM_019837705.2;refGene_gn=HAO2;refGene_tx=XM_019837706.2
C1	102642471	102642755
refGene_gn=HAO2;refGene_tx=XM_003990516.4;refGene_gn=HAO2;refGene_tx=XM_019837704.2;refGene_gn=HAO2;refGene_tx=XM_019837705.2;refGene_gn=HAO2;refGene_tx=XM_019837706.2
C1	102644599	102644809
refGene_gn=HAO2;refGene_tx=XM_003990516.4;refGene_gn=HAO2;refGene_tx=XM_019837704.2;refGene_gn=HAO2;refGene_tx=XM_019837705.2

C1 102649425 102649584
refGene_gn=HAO2;refGene_tx=XM_003990516.4;refGene_gn=HAO2;refGene_tx=XM_019837704.2;refGene_gn=HAO2;refGene_tx=XM_019837705.2;refGene_gn=HAO2;refGene_tx=XM_019837706.2
C1 102650191 102650261
refGene_gn=HAO2;refGene_tx=XM_003990516.4;refGene_gn=HAO2;refGene_tx=XM_019837704.2;refGene_gn=HAO2;refGene_tx=XM_019837705.2;refGene_gn=HAO2;refGene_tx=XM_019837706.2
C1 102655657 102655713
refGene_gn=HAO2;refGene_tx=XM_003990516.4;refGene_gn=HAO2;refGene_tx=XM_019837704.2;refGene_gn=HAO2;refGene_tx=XM_019837705.2;refGene_gn=HAO2;refGene_tx=XM_019837706.2
C1 102683942 102684087
refGene_gn=HSD3B2;refGene_tx=NM_001246194.1
C1 102687949 102688114
refGene_gn=HSD3B2;refGene_tx=NM_001246194.1
C1 102690361 102691173
refGene_gn=HSD3B2;refGene_tx=NM_001246194.1
C1 102707994 102709418
refGene_gn=ZNF697;refGene_tx=XM_011285093.3;refGene_gn=ZNF697;refGene_tx=XM_019837707.2;refGene_gn=ZNF697;refGene_tx=XM_023259044.1
C1 102709645 102709712
refGene_gn=ZNF697;refGene_tx=XM_019837707.2
C1 102710797 102711041
refGene_gn=ZNF697;refGene_tx=XM_011285093.3;refGene_gn=ZNF697;refGene_tx=XM_023259044.1
C1 102782786 102782924
refGene_gn=PHGDH;refGene_tx=XM_003990517.5
C1 102791403 102791555
refGene_gn=PHGDH;refGene_tx=XM_003990517.5
C1 102793283 102793349
refGene_gn=PHGDH;refGene_tx=XM_003990517.5
C1 102798783 102798838
refGene_gn=PHGDH;refGene_tx=XM_003990517.5
C1 102798936 102799035
refGene_gn=PHGDH;refGene_tx=XM_003990517.5
C1 102806746 102806879
refGene_gn=PHGDH;refGene_tx=XM_003990517.5
C1 102807377 102807526
refGene_gn=PHGDH;refGene_tx=XM_003990517.5
C1 102808940 102809093
refGene_gn=PHGDH;refGene_tx=XM_003990517.5
C1 102812622 102812755
refGene_gn=PHGDH;refGene_tx=XM_003990517.5
C1 102813387 102813518
refGene_gn=PHGDH;refGene_tx=XM_003990517.5
C1 102814580 102814818
refGene_gn=PHGDH;refGene_tx=XM_003990517.5
C1 102815737 102815892
refGene_gn=PHGDH;refGene_tx=XM_003990517.5

C1	102822274	102822381
refGene_gn=HMGCS2;refGene_tx=XM_006935029.4;refGene_gn=HMGCS2;refGene_tx=XM_019837709.2
C1	102823845	102823971
refGene_gn=HMGCS2;refGene_tx=XM_006935029.4;refGene_gn=HMGCS2;refGene_tx=XM_019837709.2
C1	102824575	102824682
refGene_gn=HMGCS2;refGene_tx=XM_006935029.4;refGene_gn=HMGCS2;refGene_tx=XM_019837709.2
C1	102825971	102826142
refGene_gn=HMGCS2;refGene_tx=XM_006935029.4;refGene_gn=HMGCS2;refGene_tx=XM_019837709.2
C1	102827960	102828126
refGene_gn=HMGCS2;refGene_tx=XM_006935029.4;refGene_gn=HMGCS2;refGene_tx=XM_019837709.2
C1	102829518	102829683
refGene_gn=HMGCS2;refGene_tx=XM_006935029.4
C1	102830270	102830396
refGene_gn=HMGCS2;refGene_tx=XM_006935029.4;refGene_gn=HMGCS2;refGene_tx=XM_019837709.2
C1	102833860	102834315
refGene_gn=HMGCS2;refGene_tx=XM_006935029.4;refGene_gn=HMGCS2;refGene_tx=XM_019837709.2
C1	102838616	102838720
refGene_gn=HMGCS2;refGene_tx=XM_006935029.4;refGene_gn=HMGCS2;refGene_tx=XM_019837709.2
C1	102865300	102865368
refGene_gn=REG4;refGene_tx=XM_023259045.1
C1	102869604	102869710
refGene_gn=REG4;refGene_tx=XM_023259045.1
C1	102871429	102871567
refGene_gn=REG4;refGene_tx=XM_023259045.1
C1	102874528	102874626
refGene_gn=REG4;refGene_tx=XM_023259045.1
C1	102877890	102877957
refGene_gn=REG4;refGene_tx=XM_023259045.1
C1	102929759	102931148
refGene_gn=NOTCH2;refGene_tx=XM_011285094.3;refGene_gn=NOTCH2;refGene_tx=XM_019837710.2
C1	102932408	102932506
refGene_gn=NOTCH2;refGene_tx=XM_011285094.3;refGene_gn=NOTCH2;refGene_tx=XM_019837710.2
C1	102933146	102933294
refGene_gn=NOTCH2;refGene_tx=XM_011285094.3;refGene_gn=NOTCH2;refGene_tx=XM_019837710.2
C1	102934070	102934372
refGene_gn=NOTCH2;refGene_tx=XM_011285094.3;refGene_gn=NOTCH2;refGene_tx=XM_019837710.2
C1	102934992	102935161
refGene_gn=NOTCH2;refGene_tx=XM_011285094.3;refGene_gn=NOTCH2;refGene_

tx=XM_019837710.2
C1	102936783	102936880
refGene_gn=NOTCH2;refGene_tx=XM_011285094.3;refGene_gn=NOTCH2;refGene_tx=XM_019837710.2
C1	102937311	102937522
refGene_gn=NOTCH2;refGene_tx=XM_011285094.3;refGene_gn=NOTCH2;refGene_tx=XM_019837710.2
C1	102937721	102937864
refGene_gn=NOTCH2;refGene_tx=XM_011285094.3;refGene_gn=NOTCH2;refGene_tx=XM_019837710.2
C1	102938610	102938958
refGene_gn=NOTCH2;refGene_tx=XM_011285094.3;refGene_gn=NOTCH2;refGene_tx=XM_019837710.2
C1	102940330	102940836
refGene_gn=NOTCH2;refGene_tx=XM_011285094.3;refGene_gn=NOTCH2;refGene_tx=XM_019837710.2
C1	102941746	102941859
refGene_gn=NOTCH2;refGene_tx=XM_011285094.3;refGene_gn=NOTCH2;refGene_tx=XM_019837710.2
C1	102944753	102944990
refGene_gn=NOTCH2;refGene_tx=XM_011285094.3;refGene_gn=NOTCH2;refGene_tx=XM_019837710.2
C1	102950052	102950185
refGene_gn=NOTCH2;refGene_tx=XM_011285094.3;refGene_gn=NOTCH2;refGene_tx=XM_019837710.2
C1	102951669	102951854
refGene_gn=NOTCH2;refGene_tx=XM_011285094.3;refGene_gn=NOTCH2;refGene_tx=XM_019837710.2
C1	102952207	102952361
refGene_gn=NOTCH2;refGene_tx=XM_011285094.3;refGene_gn=NOTCH2;refGene_tx=XM_019837710.2
C1	102953816	102954018
refGene_gn=NOTCH2;refGene_tx=XM_011285094.3;refGene_gn=NOTCH2;refGene_tx=XM_019837710.2
C1	102954789	102955018
refGene_gn=NOTCH2;refGene_tx=XM_011285094.3;refGene_gn=NOTCH2;refGene_tx=XM_019837710.2
C1	102963423	102963576
refGene_gn=NOTCH2;refGene_tx=XM_011285094.3;refGene_gn=NOTCH2;refGene_tx=XM_019837710.2
C1	102963976	102964096
refGene_gn=NOTCH2;refGene_tx=XM_011285094.3;refGene_gn=NOTCH2;refGene_tx=XM_019837710.2
C1	102966743	102966857
refGene_gn=NOTCH2;refGene_tx=XM_011285094.3;refGene_gn=NOTCH2;refGene_tx=XM_019837710.2
C1	102968613	102968759
refGene_gn=NOTCH2;refGene_tx=XM_011285094.3;refGene_gn=NOTCH2;refGene_tx=XM_019837710.2
C1	102970118	102970311

refGene_gn=NOTCH2;refGene_tx=XM_011285094.3;refGene_gn=NOTCH2;refGene_tx=XM_019837710.2
C1 102977004 102977115
refGene_gn=NOTCH2;refGene_tx=XM_011285094.3;refGene_gn=NOTCH2;refGene_tx=XM_019837710.2
C1 102980003 102980237
refGene_gn=NOTCH2;refGene_tx=XM_011285094.3;refGene_gn=NOTCH2;refGene_tx=XM_019837710.2
C1 102981916 102982030
refGene_gn=NOTCH2;refGene_tx=XM_011285094.3;refGene_gn=NOTCH2;refGene_tx=XM_019837710.2
C1 102982811 102982925
refGene_gn=NOTCH2;refGene_tx=XM_011285094.3;refGene_gn=NOTCH2;refGene_tx=XM_019837710.2
C1 102984079 102984268
refGene_gn=NOTCH2;refGene_tx=XM_011285094.3;refGene_gn=NOTCH2;refGene_tx=XM_019837710.2
C1 102984734 102984890
refGene_gn=NOTCH2;refGene_tx=XM_011285094.3;refGene_gn=NOTCH2;refGene_tx=XM_019837710.2
C1 102986241 102986475
refGene_gn=NOTCH2;refGene_tx=XM_011285094.3;refGene_gn=NOTCH2;refGene_tx=XM_019837710.2
C1 103007341 103007464
refGene_gn=NOTCH2;refGene_tx=XM_011285094.3;refGene_gn=NOTCH2;refGene_tx=XM_019837710.2
C1 103018047 103018383
refGene_gn=NOTCH2;refGene_tx=XM_011285094.3;refGene_gn=NOTCH2;refGene_tx=XM_019837710.2
C1 103022031 103022291
refGene_gn=NOTCH2;refGene_tx=XM_011285094.3;refGene_gn=NOTCH2;refGene_tx=XM_019837710.2
C1 103050230 103050312
refGene_gn=NOTCH2;refGene_tx=XM_011285094.3;refGene_gn=NOTCH2;refGene_tx=XM_019837710.2
C1 103077966 103078027
refGene_gn=NOTCH2;refGene_tx=XM_019837710.2
C1 103091409 103091482
refGene_gn=NOTCH2;refGene_tx=XM_011285094.3
C1 103169215 103169370
refGene_gn=SEC22B;refGene_tx=XM_003990519.5
C1 103172904 103173051
refGene_gn=SEC22B;refGene_tx=XM_003990519.5
C1 103175181 103175342
refGene_gn=SEC22B;refGene_tx=XM_003990519.5
C1 103178395 103178505
refGene_gn=SEC22B;refGene_tx=XM_003990519.5
C1 103187042 103187117
refGene_gn=SEC22B;refGene_tx=XM_003990519.5
C1 103201272 103201511

refGene_gn=LOC101093537;refGene_tx=XM_023259046.1;refGene_gn=LOC101093
537;refGene_tx=XM_023259047.1;refGene_gn=LOC101093537;refGene_tx=XM_02
3259048.1;refGene_gn=LOC101093537;refGene_tx=XM_023259049.1;refGene_gn
=LOC101093537;refGene_tx=XM_023259050.1;refGene_gn=LOC101093537;refGen
e_tx=XM_023259051.1;refGene_gn=LOC101093537;refGene_tx=XM_023259052.1;
refGene_gn=LOC101093537;refGene_tx=XM_023259053.1;refGene_gn=LOC101093
537;refGene_tx=XM_023259054.1;refGene_gn=LOC101093537;refGene_tx=XM_02
3259055.1;refGene_gn=LOC101093537;refGene_tx=XM_023259056.1;refGene_gn
=LOC101093537;refGene_tx=XM_023259058.1;refGene_gn=LOC101093537;refGen
e_tx=XM_023259059.1;refGene_gn=LOC101093537;refGene_tx=XM_023259060.1
C1 103229783 103229803
refGene_gn=LOC101093537;refGene_tx=XM_019837713.2;refGene_gn=LOC101093
537;refGene_tx=XM_023259057.1
C1 103248902 103248958
refGene_gn=LOC101093537;refGene_tx=XM_019837713.2;refGene_gn=LOC101093
537;refGene_tx=XM_023259046.1;refGene_gn=LOC101093537;refGene_tx=XM_02
3259047.1;refGene_gn=LOC101093537;refGene_tx=XM_023259048.1;refGene_gn
=LOC101093537;refGene_tx=XM_023259049.1;refGene_gn=LOC101093537;refGen
e_tx=XM_023259050.1;refGene_gn=LOC101093537;refGene_tx=XM_023259051.1;
refGene_gn=LOC101093537;refGene_tx=XM_023259052.1;refGene_gn=LOC101093
537;refGene_tx=XM_023259053.1;refGene_gn=LOC101093537;refGene_tx=XM_02
3259054.1;refGene_gn=LOC101093537;refGene_tx=XM_023259055.1;refGene_gn
=LOC101093537;refGene_tx=XM_023259056.1;refGene_gn=LOC101093537;refGen
e_tx=XM_023259057.1;refGene_gn=LOC101093537;refGene_tx=XM_023259058.1;
refGene_gn=LOC101093537;refGene_tx=XM_023259059.1;refGene_gn=LOC101093
537;refGene_tx=XM_023259060.1
C1 103253983 103254135
refGene_gn=LOC101093537;refGene_tx=XM_019837713.2;refGene_gn=LOC101093
537;refGene_tx=XM_023259046.1;refGene_gn=LOC101093537;refGene_tx=XM_02
3259047.1;refGene_gn=LOC101093537;refGene_tx=XM_023259048.1;refGene_gn
=LOC101093537;refGene_tx=XM_023259049.1;refGene_gn=LOC101093537;refGen
e_tx=XM_023259050.1;refGene_gn=LOC101093537;refGene_tx=XM_023259051.1;
refGene_gn=LOC101093537;refGene_tx=XM_023259052.1;refGene_gn=LOC101093
537;refGene_tx=XM_023259053.1;refGene_gn=LOC101093537;refGene_tx=XM_02
3259054.1;refGene_gn=LOC101093537;refGene_tx=XM_023259055.1;refGene_gn
=LOC101093537;refGene_tx=XM_023259056.1;refGene_gn=LOC101093537;refGen
e_tx=XM_023259057.1;refGene_gn=LOC101093537;refGene_tx=XM_023259058.1;
refGene_gn=LOC101093537;refGene_tx=XM_023259059.1;refGene_gn=LOC101093
537;refGene_tx=XM_023259060.1
C1 103277155 103277266
refGene_gn=LOC101093537;refGene_tx=XM_019837713.2;refGene_gn=LOC101093
537;refGene_tx=XM_023259046.1;refGene_gn=LOC101093537;refGene_tx=XM_02
3259047.1;refGene_gn=LOC101093537;refGene_tx=XM_023259048.1;refGene_gn
=LOC101093537;refGene_tx=XM_023259049.1;refGene_gn=LOC101093537;refGen
e_tx=XM_023259050.1;refGene_gn=LOC101093537;refGene_tx=XM_023259051.1;
refGene_gn=LOC101093537;refGene_tx=XM_023259052.1;refGene_gn=LOC101093
537;refGene_tx=XM_023259053.1;refGene_gn=LOC101093537;refGene_tx=XM_02
3259054.1;refGene_gn=LOC101093537;refGene_tx=XM_023259055.1;refGene_gn
=LOC101093537;refGene_tx=XM_023259056.1;refGene_gn=LOC101093537;refGen
e_tx=XM_023259057.1;refGene_gn=LOC101093537;refGene_tx=XM_023259058.1;

refGene_gn=LOC101093537;refGene_tx=XM_023259059.1;refGene_gn=LOC101093
537;refGene_tx=XM_023259060.1
C1	103312033	103312110
refGene_gn=LOC101093537;refGene_tx=XM_019837713.2;refGene_gn=LOC101093
537;refGene_tx=XM_023259046.1;refGene_gn=LOC101093537;refGene_tx=XM_02
3259047.1;refGene_gn=LOC101093537;refGene_tx=XM_023259048.1;refGene_gn
=LOC101093537;refGene_tx=XM_023259049.1;refGene_gn=LOC101093537;refGen
e_tx=XM_023259050.1;refGene_gn=LOC101093537;refGene_tx=XM_023259051.1;
refGene_gn=LOC101093537;refGene_tx=XM_023259052.1;refGene_gn=LOC101093
537;refGene_tx=XM_023259053.1;refGene_gn=LOC101093537;refGene_tx=XM_02
3259054.1;refGene_gn=LOC101093537;refGene_tx=XM_023259055.1;refGene_gn
=LOC101093537;refGene_tx=XM_023259056.1;refGene_gn=LOC101093537;refGen
e_tx=XM_023259057.1;refGene_gn=LOC101093537;refGene_tx=XM_023259058.1;
refGene_gn=LOC101093537;refGene_tx=XM_023259059.1;refGene_gn=LOC101093
537;refGene_tx=XM_023259060.1
C1	103314712	103314836
refGene_gn=LOC101093537;refGene_tx=XM_019837713.2;refGene_gn=LOC101093
537;refGene_tx=XM_023259046.1;refGene_gn=LOC101093537;refGene_tx=XM_02
3259047.1;refGene_gn=LOC101093537;refGene_tx=XM_023259048.1;refGene_gn
=LOC101093537;refGene_tx=XM_023259049.1;refGene_gn=LOC101093537;refGen
e_tx=XM_023259050.1;refGene_gn=LOC101093537;refGene_tx=XM_023259051.1;
refGene_gn=LOC101093537;refGene_tx=XM_023259052.1;refGene_gn=LOC101093
537;refGene_tx=XM_023259053.1;refGene_gn=LOC101093537;refGene_tx=XM_02
3259054.1;refGene_gn=LOC101093537;refGene_tx=XM_023259055.1;refGene_gn
=LOC101093537;refGene_tx=XM_023259056.1;refGene_gn=LOC101093537;refGen
e_tx=XM_023259057.1;refGene_gn=LOC101093537;refGene_tx=XM_023259058.1;
refGene_gn=LOC101093537;refGene_tx=XM_023259059.1;refGene_gn=LOC101093
537;refGene_tx=XM_023259060.1
C1	103315023	103315199
refGene_gn=LOC101093537;refGene_tx=XM_019837713.2;refGene_gn=LOC101093
537;refGene_tx=XM_023259046.1;refGene_gn=LOC101093537;refGene_tx=XM_02
3259047.1;refGene_gn=LOC101093537;refGene_tx=XM_023259048.1;refGene_gn
=LOC101093537;refGene_tx=XM_023259049.1;refGene_gn=LOC101093537;refGen
e_tx=XM_023259050.1;refGene_gn=LOC101093537;refGene_tx=XM_023259051.1;
refGene_gn=LOC101093537;refGene_tx=XM_023259052.1;refGene_gn=LOC101093
537;refGene_tx=XM_023259053.1;refGene_gn=LOC101093537;refGene_tx=XM_02
3259054.1;refGene_gn=LOC101093537;refGene_tx=XM_023259055.1;refGene_gn
=LOC101093537;refGene_tx=XM_023259056.1;refGene_gn=LOC101093537;refGen
e_tx=XM_023259057.1;refGene_gn=LOC101093537;refGene_tx=XM_023259058.1;
refGene_gn=LOC101093537;refGene_tx=XM_023259059.1;refGene_gn=LOC101093
537;refGene_tx=XM_023259060.1
C1	103319344	103319462
refGene_gn=LOC101093537;refGene_tx=XM_019837713.2;refGene_gn=LOC101093
537;refGene_tx=XM_023259046.1;refGene_gn=LOC101093537;refGene_tx=XM_02
3259047.1;refGene_gn=LOC101093537;refGene_tx=XM_023259048.1;refGene_gn
=LOC101093537;refGene_tx=XM_023259049.1;refGene_gn=LOC101093537;refGen
e_tx=XM_023259050.1;refGene_gn=LOC101093537;refGene_tx=XM_023259051.1;
refGene_gn=LOC101093537;refGene_tx=XM_023259052.1;refGene_gn=LOC101093
537;refGene_tx=XM_023259053.1;refGene_gn=LOC101093537;refGene_tx=XM_02
3259054.1;refGene_gn=LOC101093537;refGene_tx=XM_023259055.1;refGene_gn

=LOC101093537;refGene_tx=XM_023259056.1;refGene_gn=LOC101093537;refGen
e_tx=XM_023259057.1;refGene_gn=LOC101093537;refGene_tx=XM_023259058.1;
refGene_gn=LOC101093537;refGene_tx=XM_023259059.1;refGene_gn=LOC101093
537;refGene_tx=XM_023259060.1
C1 103341094 103342219
refGene_gn=LOC101093537;refGene_tx=XM_006935036.4;refGene_gn=LOC101093
537;refGene_tx=XM_006935041.4;refGene_gn=LOC101093537;refGene_tx=XM_01
9837717.2
C1 103348622 103348754
refGene_gn=LOC101093537;refGene_tx=XM_006935036.4;refGene_gn=LOC101093
537;refGene_tx=XM_006935041.4;refGene_gn=LOC101093537;refGene_tx=XM_01
9837713.2;refGene_gn=LOC101093537;refGene_tx=XM_019837717.2;refGene_gn
=LOC101093537;refGene_tx=XM_023259046.1;refGene_gn=LOC101093537;refGen
e_tx=XM_023259047.1;refGene_gn=LOC101093537;refGene_tx=XM_023259048.1;
refGene_gn=LOC101093537;refGene_tx=XM_023259049.1;refGene_gn=LOC101093
537;refGene_tx=XM_023259050.1;refGene_gn=LOC101093537;refGene_tx=XM_02
3259051.1;refGene_gn=LOC101093537;refGene_tx=XM_023259052.1;refGene_gn
=LOC101093537;refGene_tx=XM_023259053.1;refGene_gn=LOC101093537;refGen
e_tx=XM_023259054.1;refGene_gn=LOC101093537;refGene_tx=XM_023259055.1;
refGene_gn=LOC101093537;refGene_tx=XM_023259056.1;refGene_gn=LOC101093
537;refGene_tx=XM_023259057.1;refGene_gn=LOC101093537;refGene_tx=XM_02
3259058.1;refGene_gn=LOC101093537;refGene_tx=XM_023259059.1;refGene_gn
=LOC101093537;refGene_tx=XM_023259060.1
C1 103349850 103349970
refGene_gn=LOC101093537;refGene_tx=XM_006935036.4;refGene_gn=LOC101093
537;refGene_tx=XM_006935041.4;refGene_gn=LOC101093537;refGene_tx=XM_01
9837713.2;refGene_gn=LOC101093537;refGene_tx=XM_019837717.2;refGene_gn
=LOC101093537;refGene_tx=XM_023259046.1;refGene_gn=LOC101093537;refGen
e_tx=XM_023259047.1;refGene_gn=LOC101093537;refGene_tx=XM_023259048.1;
refGene_gn=LOC101093537;refGene_tx=XM_023259049.1;refGene_gn=LOC101093
537;refGene_tx=XM_023259050.1;refGene_gn=LOC101093537;refGene_tx=XM_02
3259051.1;refGene_gn=LOC101093537;refGene_tx=XM_023259052.1;refGene_gn
=LOC101093537;refGene_tx=XM_023259053.1;refGene_gn=LOC101093537;refGen
e_tx=XM_023259054.1;refGene_gn=LOC101093537;refGene_tx=XM_023259055.1;
refGene_gn=LOC101093537;refGene_tx=XM_023259056.1;refGene_gn=LOC101093
537;refGene_tx=XM_023259057.1;refGene_gn=LOC101093537;refGene_tx=XM_02
3259058.1;refGene_gn=LOC101093537;refGene_tx=XM_023259059.1;refGene_gn
=LOC101093537;refGene_tx=XM_023259060.1
C1 103350190 103350283
refGene_gn=LOC101093537;refGene_tx=XM_006935036.4;refGene_gn=LOC101093
537;refGene_tx=XM_006935041.4;refGene_gn=LOC101093537;refGene_tx=XM_01
9837713.2;refGene_gn=LOC101093537;refGene_tx=XM_019837717.2;refGene_gn
=LOC101093537;refGene_tx=XM_023259046.1;refGene_gn=LOC101093537;refGen
e_tx=XM_023259047.1;refGene_gn=LOC101093537;refGene_tx=XM_023259048.1;
refGene_gn=LOC101093537;refGene_tx=XM_023259049.1;refGene_gn=LOC101093
537;refGene_tx=XM_023259050.1;refGene_gn=LOC101093537;refGene_tx=XM_02
3259051.1;refGene_gn=LOC101093537;refGene_tx=XM_023259052.1;refGene_gn
=LOC101093537;refGene_tx=XM_023259053.1;refGene_gn=LOC101093537;refGen
e_tx=XM_023259054.1;refGene_gn=LOC101093537;refGene_tx=XM_023259055.1;
refGene_gn=LOC101093537;refGene_tx=XM_023259056.1;refGene_gn=LOC101093

537;refGene_tx=XM_023259057.1;refGene_gn=LOC101093537;refGene_tx=XM_02
3259058.1;refGene_gn=LOC101093537;refGene_tx=XM_023259059.1;refGene_gn
=LOC101093537;refGene_tx=XM_023259060.1
C1 103350425 103350638
refGene_gn=LOC101093537;refGene_tx=XM_006935036.4;refGene_gn=LOC101093
537;refGene_tx=XM_006935041.4;refGene_gn=LOC101093537;refGene_tx=XM_01
9837713.2;refGene_gn=LOC101093537;refGene_tx=XM_019837717.2;refGene_gn
=LOC101093537;refGene_tx=XM_023259046.1;refGene_gn=LOC101093537;refGen
e_tx=XM_023259047.1;refGene_gn=LOC101093537;refGene_tx=XM_023259048.1;
refGene_gn=LOC101093537;refGene_tx=XM_023259049.1;refGene_gn=LOC101093
537;refGene_tx=XM_023259050.1;refGene_gn=LOC101093537;refGene_tx=XM_02
3259051.1;refGene_gn=LOC101093537;refGene_tx=XM_023259052.1;refGene_gn
=LOC101093537;refGene_tx=XM_023259053.1;refGene_gn=LOC101093537;refGen
e_tx=XM_023259054.1;refGene_gn=LOC101093537;refGene_tx=XM_023259055.1;
refGene_gn=LOC101093537;refGene_tx=XM_023259056.1;refGene_gn=LOC101093
537;refGene_tx=XM_023259057.1;refGene_gn=LOC101093537;refGene_tx=XM_02
3259058.1;refGene_gn=LOC101093537;refGene_tx=XM_023259059.1;refGene_gn
=LOC101093537;refGene_tx=XM_023259060.1
C1 103351952 103352111
refGene_gn=LOC101093537;refGene_tx=XM_006935036.4;refGene_gn=LOC101093
537;refGene_tx=XM_006935041.4;refGene_gn=LOC101093537;refGene_tx=XM_01
9837713.2;refGene_gn=LOC101093537;refGene_tx=XM_019837717.2;refGene_gn
=LOC101093537;refGene_tx=XM_023259046.1;refGene_gn=LOC101093537;refGen
e_tx=XM_023259047.1;refGene_gn=LOC101093537;refGene_tx=XM_023259048.1;
refGene_gn=LOC101093537;refGene_tx=XM_023259049.1;refGene_gn=LOC101093
537;refGene_tx=XM_023259050.1;refGene_gn=LOC101093537;refGene_tx=XM_02
3259051.1;refGene_gn=LOC101093537;refGene_tx=XM_023259052.1;refGene_gn
=LOC101093537;refGene_tx=XM_023259053.1;refGene_gn=LOC101093537;refGen
e_tx=XM_023259054.1;refGene_gn=LOC101093537;refGene_tx=XM_023259055.1;
refGene_gn=LOC101093537;refGene_tx=XM_023259056.1;refGene_gn=LOC101093
537;refGene_tx=XM_023259057.1;refGene_gn=LOC101093537;refGene_tx=XM_02
3259058.1;refGene_gn=LOC101093537;refGene_tx=XM_023259059.1;refGene_gn
=LOC101093537;refGene_tx=XM_023259060.1
C1 103352986 103353100
refGene_gn=LOC101093537;refGene_tx=XM_006935036.4;refGene_gn=LOC101093
537;refGene_tx=XM_006935041.4;refGene_gn=LOC101093537;refGene_tx=XM_01
9837713.2;refGene_gn=LOC101093537;refGene_tx=XM_019837717.2;refGene_gn
=LOC101093537;refGene_tx=XM_023259046.1;refGene_gn=LOC101093537;refGen
e_tx=XM_023259047.1;refGene_gn=LOC101093537;refGene_tx=XM_023259048.1;
refGene_gn=LOC101093537;refGene_tx=XM_023259049.1;refGene_gn=LOC101093
537;refGene_tx=XM_023259050.1;refGene_gn=LOC101093537;refGene_tx=XM_02
3259051.1;refGene_gn=LOC101093537;refGene_tx=XM_023259052.1;refGene_gn
=LOC101093537;refGene_tx=XM_023259053.1;refGene_gn=LOC101093537;refGen
e_tx=XM_023259054.1;refGene_gn=LOC101093537;refGene_tx=XM_023259055.1;
refGene_gn=LOC101093537;refGene_tx=XM_023259056.1;refGene_gn=LOC101093
537;refGene_tx=XM_023259057.1;refGene_gn=LOC101093537;refGene_tx=XM_02
3259058.1;refGene_gn=LOC101093537;refGene_tx=XM_023259059.1;refGene_gn
=LOC101093537;refGene_tx=XM_023259060.1
C1 103353507 103353645
refGene_gn=LOC101093537;refGene_tx=XM_006935036.4;refGene_gn=LOC101093

537;refGene_tx=XM_006935041.4;refGene_gn=LOC101093537;refGene_tx=XM_01
9837713.2;refGene_gn=LOC101093537;refGene_tx=XM_019837717.2;refGene_gn
=LOC101093537;refGene_tx=XM_023259046.1;refGene_gn=LOC101093537;refGen
e_tx=XM_023259047.1;refGene_gn=LOC101093537;refGene_tx=XM_023259048.1;
refGene_gn=LOC101093537;refGene_tx=XM_023259049.1;refGene_gn=LOC101093
537;refGene_tx=XM_023259050.1;refGene_gn=LOC101093537;refGene_tx=XM_02
3259051.1;refGene_gn=LOC101093537;refGene_tx=XM_023259052.1;refGene_gn
=LOC101093537;refGene_tx=XM_023259053.1;refGene_gn=LOC101093537;refGen
e_tx=XM_023259054.1;refGene_gn=LOC101093537;refGene_tx=XM_023259055.1;
refGene_gn=LOC101093537;refGene_tx=XM_023259056.1;refGene_gn=LOC101093
537;refGene_tx=XM_023259057.1;refGene_gn=LOC101093537;refGene_tx=XM_02
3259058.1;refGene_gn=LOC101093537;refGene_tx=XM_023259059.1;refGene_gn
=LOC101093537;refGene_tx=XM_023259060.1
C1 103354586 103354766
refGene_gn=LOC101093537;refGene_tx=XM_006935036.4;refGene_gn=LOC101093
537;refGene_tx=XM_006935041.4;refGene_gn=LOC101093537;refGene_tx=XM_01
9837713.2;refGene_gn=LOC101093537;refGene_tx=XM_019837717.2;refGene_gn
=LOC101093537;refGene_tx=XM_023259046.1;refGene_gn=LOC101093537;refGen
e_tx=XM_023259047.1;refGene_gn=LOC101093537;refGene_tx=XM_023259048.1;
refGene_gn=LOC101093537;refGene_tx=XM_023259049.1;refGene_gn=LOC101093
537;refGene_tx=XM_023259050.1;refGene_gn=LOC101093537;refGene_tx=XM_02
3259051.1;refGene_gn=LOC101093537;refGene_tx=XM_023259052.1;refGene_gn
=LOC101093537;refGene_tx=XM_023259053.1;refGene_gn=LOC101093537;refGen
e_tx=XM_023259054.1;refGene_gn=LOC101093537;refGene_tx=XM_023259055.1;
refGene_gn=LOC101093537;refGene_tx=XM_023259056.1;refGene_gn=LOC101093
537;refGene_tx=XM_023259057.1;refGene_gn=LOC101093537;refGene_tx=XM_02
3259058.1;refGene_gn=LOC101093537;refGene_tx=XM_023259059.1;refGene_gn
=LOC101093537;refGene_tx=XM_023259060.1
C1 103355670 103355865
refGene_gn=LOC101093537;refGene_tx=XM_006935036.4;refGene_gn=LOC101093
537;refGene_tx=XM_006935041.4;refGene_gn=LOC101093537;refGene_tx=XM_01
9837713.2;refGene_gn=LOC101093537;refGene_tx=XM_019837717.2;refGene_gn
=LOC101093537;refGene_tx=XM_023259046.1;refGene_gn=LOC101093537;refGen
e_tx=XM_023259047.1;refGene_gn=LOC101093537;refGene_tx=XM_023259048.1;
refGene_gn=LOC101093537;refGene_tx=XM_023259049.1;refGene_gn=LOC101093
537;refGene_tx=XM_023259050.1;refGene_gn=LOC101093537;refGene_tx=XM_02
3259051.1;refGene_gn=LOC101093537;refGene_tx=XM_023259052.1;refGene_gn
=LOC101093537;refGene_tx=XM_023259053.1;refGene_gn=LOC101093537;refGen
e_tx=XM_023259054.1;refGene_gn=LOC101093537;refGene_tx=XM_023259055.1;
refGene_gn=LOC101093537;refGene_tx=XM_023259056.1;refGene_gn=LOC101093
537;refGene_tx=XM_023259057.1;refGene_gn=LOC101093537;refGene_tx=XM_02
3259058.1;refGene_gn=LOC101093537;refGene_tx=XM_023259059.1;refGene_gn
=LOC101093537;refGene_tx=XM_023259060.1
C1 103358657 103358825
refGene_gn=LOC101093537;refGene_tx=XM_006935036.4;refGene_gn=LOC101093
537;refGene_tx=XM_006935041.4;refGene_gn=LOC101093537;refGene_tx=XM_01
9837713.2;refGene_gn=LOC101093537;refGene_tx=XM_019837717.2;refGene_gn
=LOC101093537;refGene_tx=XM_023259046.1;refGene_gn=LOC101093537;refGen
e_tx=XM_023259047.1;refGene_gn=LOC101093537;refGene_tx=XM_023259048.1;
refGene_gn=LOC101093537;refGene_tx=XM_023259049.1;refGene_gn=LOC101093

537;refGene_tx=XM_023259050.1;refGene_gn=LOC101093537;refGene_tx=XM_02
3259051.1;refGene_gn=LOC101093537;refGene_tx=XM_023259052.1;refGene_gn
=LOC101093537;refGene_tx=XM_023259053.1;refGene_gn=LOC101093537;refGen
e_tx=XM_023259054.1;refGene_gn=LOC101093537;refGene_tx=XM_023259055.1;
refGene_gn=LOC101093537;refGene_tx=XM_023259056.1;refGene_gn=LOC101093
537;refGene_tx=XM_023259057.1;refGene_gn=LOC101093537;refGene_tx=XM_02
3259058.1;refGene_gn=LOC101093537;refGene_tx=XM_023259059.1;refGene_gn
=LOC101093537;refGene_tx=XM_023259060.1
C1 103358981 103359063
refGene_gn=LOC101093537;refGene_tx=XM_006935036.4;refGene_gn=LOC101093
537;refGene_tx=XM_006935041.4;refGene_gn=LOC101093537;refGene_tx=XM_01
9837713.2;refGene_gn=LOC101093537;refGene_tx=XM_019837717.2;refGene_gn
=LOC101093537;refGene_tx=XM_023259046.1;refGene_gn=LOC101093537;refGen
e_tx=XM_023259047.1;refGene_gn=LOC101093537;refGene_tx=XM_023259048.1;
refGene_gn=LOC101093537;refGene_tx=XM_023259049.1;refGene_gn=LOC101093
537;refGene_tx=XM_023259050.1;refGene_gn=LOC101093537;refGene_tx=XM_02
3259051.1;refGene_gn=LOC101093537;refGene_tx=XM_023259052.1;refGene_gn
=LOC101093537;refGene_tx=XM_023259053.1;refGene_gn=LOC101093537;refGen
e_tx=XM_023259054.1;refGene_gn=LOC101093537;refGene_tx=XM_023259055.1;
refGene_gn=LOC101093537;refGene_tx=XM_023259056.1;refGene_gn=LOC101093
537;refGene_tx=XM_023259057.1;refGene_gn=LOC101093537;refGene_tx=XM_02
3259058.1;refGene_gn=LOC101093537;refGene_tx=XM_023259059.1;refGene_gn
=LOC101093537;refGene_tx=XM_023259060.1
C1 103362473 103362565
refGene_gn=LOC101093537;refGene_tx=XM_006935036.4;refGene_gn=LOC101093
537;refGene_tx=XM_006935041.4;refGene_gn=LOC101093537;refGene_tx=XM_01
9837713.2;refGene_gn=LOC101093537;refGene_tx=XM_019837717.2;refGene_gn
=LOC101093537;refGene_tx=XM_023259046.1;refGene_gn=LOC101093537;refGen
e_tx=XM_023259047.1;refGene_gn=LOC101093537;refGene_tx=XM_023259048.1;
refGene_gn=LOC101093537;refGene_tx=XM_023259049.1;refGene_gn=LOC101093
537;refGene_tx=XM_023259050.1;refGene_gn=LOC101093537;refGene_tx=XM_02
3259051.1;refGene_gn=LOC101093537;refGene_tx=XM_023259052.1;refGene_gn
=LOC101093537;refGene_tx=XM_023259053.1;refGene_gn=LOC101093537;refGen
e_tx=XM_023259054.1;refGene_gn=LOC101093537;refGene_tx=XM_023259055.1;
refGene_gn=LOC101093537;refGene_tx=XM_023259056.1;refGene_gn=LOC101093
537;refGene_tx=XM_023259057.1;refGene_gn=LOC101093537;refGene_tx=XM_02
3259058.1;refGene_gn=LOC101093537;refGene_tx=XM_023259059.1;refGene_gn
=LOC101093537;refGene_tx=XM_023259060.1
C1 103365660 103365777
refGene_gn=LOC101093537;refGene_tx=XM_006935036.4;refGene_gn=LOC101093
537;refGene_tx=XM_006935041.4;refGene_gn=LOC101093537;refGene_tx=XM_01
9837713.2;refGene_gn=LOC101093537;refGene_tx=XM_019837717.2;refGene_gn
=LOC101093537;refGene_tx=XM_023259046.1;refGene_gn=LOC101093537;refGen
e_tx=XM_023259047.1;refGene_gn=LOC101093537;refGene_tx=XM_023259048.1;
refGene_gn=LOC101093537;refGene_tx=XM_023259049.1;refGene_gn=LOC101093
537;refGene_tx=XM_023259050.1;refGene_gn=LOC101093537;refGene_tx=XM_02
3259051.1;refGene_gn=LOC101093537;refGene_tx=XM_023259052.1;refGene_gn
=LOC101093537;refGene_tx=XM_023259053.1;refGene_gn=LOC101093537;refGen
e_tx=XM_023259054.1;refGene_gn=LOC101093537;refGene_tx=XM_023259055.1;
refGene_gn=LOC101093537;refGene_tx=XM_023259056.1;refGene_gn=LOC101093

537;refGene_tx=XM_023259057.1;refGene_gn=LOC101093537;refGene_tx=XM_02
3259058.1;refGene_gn=LOC101093537;refGene_tx=XM_023259059.1;refGene_gn
=LOC101093537;refGene_tx=XM_023259060.1
C1	103366105	103366243
refGene_gn=LOC101093537;refGene_tx=XM_006935036.4;refGene_gn=LOC101093
537;refGene_tx=XM_006935041.4;refGene_gn=LOC101093537;refGene_tx=XM_01
9837713.2;refGene_gn=LOC101093537;refGene_tx=XM_019837717.2;refGene_gn
=LOC101093537;refGene_tx=XM_023259046.1;refGene_gn=LOC101093537;refGen
e_tx=XM_023259047.1;refGene_gn=LOC101093537;refGene_tx=XM_023259048.1;
refGene_gn=LOC101093537;refGene_tx=XM_023259049.1;refGene_gn=LOC101093
537;refGene_tx=XM_023259050.1;refGene_gn=LOC101093537;refGene_tx=XM_02
3259051.1;refGene_gn=LOC101093537;refGene_tx=XM_023259052.1;refGene_gn
=LOC101093537;refGene_tx=XM_023259053.1;refGene_gn=LOC101093537;refGen
e_tx=XM_023259054.1;refGene_gn=LOC101093537;refGene_tx=XM_023259055.1;
refGene_gn=LOC101093537;refGene_tx=XM_023259056.1;refGene_gn=LOC101093
537;refGene_tx=XM_023259057.1;refGene_gn=LOC101093537;refGene_tx=XM_02
3259058.1;refGene_gn=LOC101093537;refGene_tx=XM_023259059.1;refGene_gn
=LOC101093537;refGene_tx=XM_023259060.1
C1	103367285	103367398
refGene_gn=LOC101093537;refGene_tx=XM_006935036.4;refGene_gn=LOC101093
537;refGene_tx=XM_006935041.4;refGene_gn=LOC101093537;refGene_tx=XM_01
9837713.2;refGene_gn=LOC101093537;refGene_tx=XM_019837717.2;refGene_gn
=LOC101093537;refGene_tx=XM_023259046.1;refGene_gn=LOC101093537;refGen
e_tx=XM_023259047.1;refGene_gn=LOC101093537;refGene_tx=XM_023259048.1;
refGene_gn=LOC101093537;refGene_tx=XM_023259049.1;refGene_gn=LOC101093
537;refGene_tx=XM_023259050.1;refGene_gn=LOC101093537;refGene_tx=XM_02
3259051.1;refGene_gn=LOC101093537;refGene_tx=XM_023259052.1;refGene_gn
=LOC101093537;refGene_tx=XM_023259053.1;refGene_gn=LOC101093537;refGen
e_tx=XM_023259054.1;refGene_gn=LOC101093537;refGene_tx=XM_023259055.1;
refGene_gn=LOC101093537;refGene_tx=XM_023259056.1;refGene_gn=LOC101093
537;refGene_tx=XM_023259057.1;refGene_gn=LOC101093537;refGene_tx=XM_02
3259058.1;refGene_gn=LOC101093537;refGene_tx=XM_023259059.1;refGene_gn
=LOC101093537;refGene_tx=XM_023259060.1
C1	103369177	103369305
refGene_gn=LOC101093537;refGene_tx=XM_006935036.4;refGene_gn=LOC101093
537;refGene_tx=XM_006935041.4;refGene_gn=LOC101093537;refGene_tx=XM_01
9837713.2;refGene_gn=LOC101093537;refGene_tx=XM_019837717.2;refGene_gn
=LOC101093537;refGene_tx=XM_023259046.1;refGene_gn=LOC101093537;refGen
e_tx=XM_023259047.1;refGene_gn=LOC101093537;refGene_tx=XM_023259048.1;
refGene_gn=LOC101093537;refGene_tx=XM_023259049.1;refGene_gn=LOC101093
537;refGene_tx=XM_023259050.1;refGene_gn=LOC101093537;refGene_tx=XM_02
3259051.1;refGene_gn=LOC101093537;refGene_tx=XM_023259052.1;refGene_gn
=LOC101093537;refGene_tx=XM_023259053.1;refGene_gn=LOC101093537;refGen
e_tx=XM_023259054.1;refGene_gn=LOC101093537;refGene_tx=XM_023259055.1;
refGene_gn=LOC101093537;refGene_tx=XM_023259056.1;refGene_gn=LOC101093
537;refGene_tx=XM_023259057.1;refGene_gn=LOC101093537;refGene_tx=XM_02
3259058.1;refGene_gn=LOC101093537;refGene_tx=XM_023259059.1;refGene_gn
=LOC101093537;refGene_tx=XM_023259060.1
C1	103379764	103379853
refGene_gn=LOC101093537;refGene_tx=XM_006935036.4;refGene_gn=LOC101093

537;refGene_tx=XM_006935041.4;refGene_gn=LOC101093537;refGene_tx=XM_019837713.2;refGene_gn=LOC101093537;refGene_tx=XM_019837717.2;refGene_gn=LOC101093537;refGene_tx=XM_023259046.1;refGene_gn=LOC101093537;refGene_tx=XM_023259047.1;refGene_gn=LOC101093537;refGene_tx=XM_023259048.1;refGene_gn=LOC101093537;refGene_tx=XM_023259049.1;refGene_gn=LOC101093537;refGene_tx=XM_023259050.1;refGene_gn=LOC101093537;refGene_tx=XM_023259051.1;refGene_gn=LOC101093537;refGene_tx=XM_023259052.1;refGene_gn=LOC101093537;refGene_tx=XM_023259053.1;refGene_gn=LOC101093537;refGene_tx=XM_023259054.1;refGene_gn=LOC101093537;refGene_tx=XM_023259055.1;refGene_gn=LOC101093537;refGene_tx=XM_023259056.1;refGene_gn=LOC101093537;refGene_tx=XM_023259057.1;refGene_gn=LOC101093537;refGene_tx=XM_023259058.1;refGene_gn=LOC101093537;refGene_tx=XM_023259059.1;refGene_gn=LOC101093537;refGene_tx=XM_023259060.1

C1 103380140 103380146
refGene_gn=LOC101093537;refGene_tx=XM_019837717.2

C1 103384929 103385162
refGene_gn=LOC101093537;refGene_tx=XM_006935036.4;refGene_gn=LOC101093537;refGene_tx=XM_006935041.4;refGene_gn=LOC101093537;refGene_tx=XM_019837713.2;refGene_gn=LOC101093537;refGene_tx=XM_023259046.1;refGene_gn=LOC101093537;refGene_tx=XM_023259047.1;refGene_gn=LOC101093537;refGene_tx=XM_023259048.1;refGene_gn=LOC101093537;refGene_tx=XM_023259049.1;refGene_gn=LOC101093537;refGene_tx=XM_023259050.1;refGene_gn=LOC101093537;refGene_tx=XM_023259051.1;refGene_gn=LOC101093537;refGene_tx=XM_023259052.1;refGene_gn=LOC101093537;refGene_tx=XM_023259053.1;refGene_gn=LOC101093537;refGene_tx=XM_023259054.1;refGene_gn=LOC101093537;refGene_tx=XM_023259055.1;refGene_gn=LOC101093537;refGene_tx=XM_023259056.1;refGene_gn=LOC101093537;refGene_tx=XM_023259057.1;refGene_gn=LOC101093537;refGene_tx=XM_023259058.1;refGene_gn=LOC101093537;refGene_tx=XM_023259059.1;refGene_gn=LOC101093537;refGene_tx=XM_023259060.1

C1 103388367 103388803
refGene_gn=LOC101093537;refGene_tx=XM_006935036.4;refGene_gn=LOC101093537;refGene_tx=XM_006935041.4;refGene_gn=LOC101093537;refGene_tx=XM_019837713.2;refGene_gn=LOC101093537;refGene_tx=XM_023259046.1;refGene_gn=LOC101093537;refGene_tx=XM_023259047.1;refGene_gn=LOC101093537;refGene_tx=XM_023259048.1;refGene_gn=LOC101093537;refGene_tx=XM_023259049.1;refGene_gn=LOC101093537;refGene_tx=XM_023259050.1;refGene_gn=LOC101093537;refGene_tx=XM_023259051.1;refGene_gn=LOC101093537;refGene_tx=XM_023259052.1;refGene_gn=LOC101093537;refGene_tx=XM_023259053.1;refGene_gn=LOC101093537;refGene_tx=XM_023259054.1;refGene_gn=LOC101093537;refGene_tx=XM_023259055.1;refGene_gn=LOC101093537;refGene_tx=XM_023259056.1;refGene_gn=LOC101093537;refGene_tx=XM_023259057.1;refGene_gn=LOC101093537;refGene_tx=XM_023259058.1;refGene_gn=LOC101093537;refGene_tx=XM_023259059.1;refGene_gn=LOC101093537;refGene_tx=XM_023259060.1

C1 103389628 103389821
refGene_gn=LOC101093537;refGene_tx=XM_006935036.4;refGene_gn=LOC101093537;refGene_tx=XM_006935041.4;refGene_gn=LOC101093537;refGene_tx=XM_019837713.2;refGene_gn=LOC101093537;refGene_tx=XM_023259046.1;refGene_gn=LOC101093537;refGene_tx=XM_023259047.1;refGene_gn=LOC101093537;refGene_tx=XM_023259048.1;refGene_gn=LOC101093537;refGene_tx=XM_023259049.1;refGene_gn=LOC101093537;refGene_tx=XM_023259050.1;refGene_gn=LOC101093

537;refGene_tx=XM_023259051.1;refGene_gn=LOC101093537;refGene_tx=XM_02
3259052.1;refGene_gn=LOC101093537;refGene_tx=XM_023259053.1;refGene_gn
=LOC101093537;refGene_tx=XM_023259054.1;refGene_gn=LOC101093537;refGen
e_tx=XM_023259055.1;refGene_gn=LOC101093537;refGene_tx=XM_023259056.1;
refGene_gn=LOC101093537;refGene_tx=XM_023259057.1;refGene_gn=LOC101093
537;refGene_tx=XM_023259058.1;refGene_gn=LOC101093537;refGene_tx=XM_02
3259059.1;refGene_gn=LOC101093537;refGene_tx=XM_023259060.1
C1 103391152 103391269
refGene_gn=LOC101093537;refGene_tx=XM_006935036.4;refGene_gn=LOC101093
537;refGene_tx=XM_006935041.4;refGene_gn=LOC101093537;refGene_tx=XM_01
9837713.2;refGene_gn=LOC101093537;refGene_tx=XM_023259046.1;refGene_gn
=LOC101093537;refGene_tx=XM_023259047.1;refGene_gn=LOC101093537;refGen
e_tx=XM_023259048.1;refGene_gn=LOC101093537;refGene_tx=XM_023259049.1;
refGene_gn=LOC101093537;refGene_tx=XM_023259050.1;refGene_gn=LOC101093
537;refGene_tx=XM_023259051.1;refGene_gn=LOC101093537;refGene_tx=XM_02
3259052.1;refGene_gn=LOC101093537;refGene_tx=XM_023259053.1;refGene_gn
=LOC101093537;refGene_tx=XM_023259054.1;refGene_gn=LOC101093537;refGen
e_tx=XM_023259055.1;refGene_gn=LOC101093537;refGene_tx=XM_023259056.1;
refGene_gn=LOC101093537;refGene_tx=XM_023259057.1;refGene_gn=LOC101093
537;refGene_tx=XM_023259058.1;refGene_gn=LOC101093537;refGene_tx=XM_02
3259059.1;refGene_gn=LOC101093537;refGene_tx=XM_023259060.1
C1 103391869 103392344
refGene_gn=LOC101093537;refGene_tx=XM_006935036.4;refGene_gn=LOC101093
537;refGene_tx=XM_006935041.4;refGene_gn=LOC101093537;refGene_tx=XM_01
9837713.2;refGene_gn=LOC101093537;refGene_tx=XM_023259046.1;refGene_gn
=LOC101093537;refGene_tx=XM_023259047.1;refGene_gn=LOC101093537;refGen
e_tx=XM_023259048.1;refGene_gn=LOC101093537;refGene_tx=XM_023259049.1;
refGene_gn=LOC101093537;refGene_tx=XM_023259050.1;refGene_gn=LOC101093
537;refGene_tx=XM_023259051.1;refGene_gn=LOC101093537;refGene_tx=XM_02
3259052.1;refGene_gn=LOC101093537;refGene_tx=XM_023259053.1;refGene_gn
=LOC101093537;refGene_tx=XM_023259054.1;refGene_gn=LOC101093537;refGen
e_tx=XM_023259055.1;refGene_gn=LOC101093537;refGene_tx=XM_023259056.1;
refGene_gn=LOC101093537;refGene_tx=XM_023259057.1;refGene_gn=LOC101093
537;refGene_tx=XM_023259058.1;refGene_gn=LOC101093537;refGene_tx=XM_02
3259059.1;refGene_gn=LOC101093537;refGene_tx=XM_023259060.1
C1 103394448 103394684
refGene_gn=LOC101093537;refGene_tx=XM_006935036.4;refGene_gn=LOC101093
537;refGene_tx=XM_006935041.4;refGene_gn=LOC101093537;refGene_tx=XM_01
9837713.2;refGene_gn=LOC101093537;refGene_tx=XM_023259046.1;refGene_gn
=LOC101093537;refGene_tx=XM_023259047.1;refGene_gn=LOC101093537;refGen
e_tx=XM_023259048.1;refGene_gn=LOC101093537;refGene_tx=XM_023259049.1;
refGene_gn=LOC101093537;refGene_tx=XM_023259050.1;refGene_gn=LOC101093
537;refGene_tx=XM_023259051.1;refGene_gn=LOC101093537;refGene_tx=XM_02
3259052.1;refGene_gn=LOC101093537;refGene_tx=XM_023259053.1;refGene_gn
=LOC101093537;refGene_tx=XM_023259054.1;refGene_gn=LOC101093537;refGen
e_tx=XM_023259055.1;refGene_gn=LOC101093537;refGene_tx=XM_023259056.1;
refGene_gn=LOC101093537;refGene_tx=XM_023259057.1;refGene_gn=LOC101093
537;refGene_tx=XM_023259058.1;refGene_gn=LOC101093537;refGene_tx=XM_02
3259059.1;refGene_gn=LOC101093537;refGene_tx=XM_023259060.1
C1 103395671 103395723

refGene_gn=LOC101093537;refGene_tx=XM_006935036.4;refGene_gn=LOC101093
537;refGene_tx=XM_006935041.4;refGene_gn=LOC101093537;refGene_tx=XM_01
9837713.2;refGene_gn=LOC101093537;refGene_tx=XM_023259046.1;refGene_gn
=LOC101093537;refGene_tx=XM_023259047.1;refGene_gn=LOC101093537;refGen
e_tx=XM_023259048.1;refGene_gn=LOC101093537;refGene_tx=XM_023259049.1;
refGene_gn=LOC101093537;refGene_tx=XM_023259050.1;refGene_gn=LOC101093
537;refGene_tx=XM_023259051.1;refGene_gn=LOC101093537;refGene_tx=XM_02
3259052.1;refGene_gn=LOC101093537;refGene_tx=XM_023259053.1;refGene_gn
=LOC101093537;refGene_tx=XM_023259054.1;refGene_gn=LOC101093537;refGen
e_tx=XM_023259055.1;refGene_gn=LOC101093537;refGene_tx=XM_023259056.1;
refGene_gn=LOC101093537;refGene_tx=XM_023259057.1;refGene_gn=LOC101093
537;refGene_tx=XM_023259058.1;refGene_gn=LOC101093537;refGene_tx=XM_02
3259059.1;refGene_gn=LOC101093537;refGene_tx=XM_023259060.1
C1 103396769 103396993
refGene_gn=LOC101093537;refGene_tx=XM_006935036.4;refGene_gn=LOC101093
537;refGene_tx=XM_006935041.4;refGene_gn=LOC101093537;refGene_tx=XM_01
9837713.2;refGene_gn=LOC101093537;refGene_tx=XM_023259046.1;refGene_gn
=LOC101093537;refGene_tx=XM_023259047.1;refGene_gn=LOC101093537;refGen
e_tx=XM_023259048.1;refGene_gn=LOC101093537;refGene_tx=XM_023259049.1;
refGene_gn=LOC101093537;refGene_tx=XM_023259050.1;refGene_gn=LOC101093
537;refGene_tx=XM_023259051.1;refGene_gn=LOC101093537;refGene_tx=XM_02
3259052.1;refGene_gn=LOC101093537;refGene_tx=XM_023259053.1;refGene_gn
=LOC101093537;refGene_tx=XM_023259054.1;refGene_gn=LOC101093537;refGen
e_tx=XM_023259055.1;refGene_gn=LOC101093537;refGene_tx=XM_023259056.1;
refGene_gn=LOC101093537;refGene_tx=XM_023259057.1;refGene_gn=LOC101093
537;refGene_tx=XM_023259058.1;refGene_gn=LOC101093537;refGene_tx=XM_02
3259059.1;refGene_gn=LOC101093537;refGene_tx=XM_023259060.1
C1 103397638 103397791
refGene_gn=LOC101093537;refGene_tx=XM_006935036.4;refGene_gn=LOC101093
537;refGene_tx=XM_006935041.4;refGene_gn=LOC101093537;refGene_tx=XM_01
9837713.2;refGene_gn=LOC101093537;refGene_tx=XM_023259046.1;refGene_gn
=LOC101093537;refGene_tx=XM_023259047.1;refGene_gn=LOC101093537;refGen
e_tx=XM_023259048.1;refGene_gn=LOC101093537;refGene_tx=XM_023259049.1;
refGene_gn=LOC101093537;refGene_tx=XM_023259051.1;refGene_gn=LOC101093
537;refGene_tx=XM_023259052.1;refGene_gn=LOC101093537;refGene_tx=XM_02
3259053.1;refGene_gn=LOC101093537;refGene_tx=XM_023259054.1;refGene_gn
=LOC101093537;refGene_tx=XM_023259055.1;refGene_gn=LOC101093537;refGen
e_tx=XM_023259056.1;refGene_gn=LOC101093537;refGene_tx=XM_023259057.1;
refGene_gn=LOC101093537;refGene_tx=XM_023259058.1;refGene_gn=LOC101093
537;refGene_tx=XM_023259060.1
C1 103399683 103399869
refGene_gn=LOC101093537;refGene_tx=XM_006935036.4;refGene_gn=LOC101093
537;refGene_tx=XM_006935041.4;refGene_gn=LOC101093537;refGene_tx=XM_01
9837713.2;refGene_gn=LOC101093537;refGene_tx=XM_023259046.1;refGene_gn
=LOC101093537;refGene_tx=XM_023259047.1;refGene_gn=LOC101093537;refGen
e_tx=XM_023259048.1;refGene_gn=LOC101093537;refGene_tx=XM_023259049.1;
refGene_gn=LOC101093537;refGene_tx=XM_023259050.1;refGene_gn=LOC101093
537;refGene_tx=XM_023259051.1;refGene_gn=LOC101093537;refGene_tx=XM_02
3259053.1;refGene_gn=LOC101093537;refGene_tx=XM_023259054.1;refGene_gn
=LOC101093537;refGene_tx=XM_023259055.1;refGene_gn=LOC101093537;refGen

e_tx=XM_023259056.1;refGene_gn=LOC101093537;refGene_tx=XM_023259057.1
C1 103403170 103403422
refGene_gn=LOC101093537;refGene_tx=XM_006935036.4;refGene_gn=LOC101093537;refGene_tx=XM_006935041.4;refGene_gn=LOC101093537;refGene_tx=XM_019837713.2;refGene_gn=LOC101093537;refGene_tx=XM_023259046.1;refGene_gn=LOC101093537;refGene_tx=XM_023259047.1;refGene_gn=LOC101093537;refGene_tx=XM_023259048.1;refGene_gn=LOC101093537;refGene_tx=XM_023259049.1;refGene_gn=LOC101093537;refGene_tx=XM_023259050.1;refGene_gn=LOC101093537;refGene_tx=XM_023259051.1;refGene_gn=LOC101093537;refGene_tx=XM_023259052.1;refGene_gn=LOC101093537;refGene_tx=XM_023259053.1;refGene_gn=LOC101093537;refGene_tx=XM_023259054.1;refGene_gn=LOC101093537;refGene_tx=XM_023259055.1;refGene_gn=LOC101093537;refGene_tx=XM_023259056.1;refGene_gn=LOC101093537;refGene_tx=XM_023259057.1;refGene_gn=LOC101093537;refGene_tx=XM_023259058.1;refGene_gn=LOC101093537;refGene_tx=XM_023259059.1;refGene_gn=LOC101093537;refGene_tx=XM_023259060.1
C1 103405071 103405203
refGene_gn=LOC101093537;refGene_tx=XM_006935036.4;refGene_gn=LOC101093537;refGene_tx=XM_006935041.4;refGene_gn=LOC101093537;refGene_tx=XM_019837713.2;refGene_gn=LOC101093537;refGene_tx=XM_023259046.1;refGene_gn=LOC101093537;refGene_tx=XM_023259047.1;refGene_gn=LOC101093537;refGene_tx=XM_023259048.1;refGene_gn=LOC101093537;refGene_tx=XM_023259049.1;refGene_gn=LOC101093537;refGene_tx=XM_023259050.1;refGene_gn=LOC101093537;refGene_tx=XM_023259051.1;refGene_gn=LOC101093537;refGene_tx=XM_023259052.1;refGene_gn=LOC101093537;refGene_tx=XM_023259053.1;refGene_gn=LOC101093537;refGene_tx=XM_023259054.1;refGene_gn=LOC101093537;refGene_tx=XM_023259055.1;refGene_gn=LOC101093537;refGene_tx=XM_023259056.1;refGene_gn=LOC101093537;refGene_tx=XM_023259057.1;refGene_gn=LOC101093537;refGene_tx=XM_023259058.1;refGene_gn=LOC101093537;refGene_tx=XM_023259059.1;refGene_gn=LOC101093537;refGene_tx=XM_023259060.1
C1 103405893 103406013
refGene_gn=LOC101093537;refGene_tx=XM_006935036.4;refGene_gn=LOC101093537;refGene_tx=XM_006935041.4;refGene_gn=LOC101093537;refGene_tx=XM_019837713.2;refGene_gn=LOC101093537;refGene_tx=XM_023259046.1;refGene_gn=LOC101093537;refGene_tx=XM_023259047.1;refGene_gn=LOC101093537;refGene_tx=XM_023259048.1;refGene_gn=LOC101093537;refGene_tx=XM_023259049.1;refGene_gn=LOC101093537;refGene_tx=XM_023259050.1;refGene_gn=LOC101093537;refGene_tx=XM_023259051.1;refGene_gn=LOC101093537;refGene_tx=XM_023259052.1;refGene_gn=LOC101093537;refGene_tx=XM_023259053.1;refGene_gn=LOC101093537;refGene_tx=XM_023259054.1;refGene_gn=LOC101093537;refGene_tx=XM_023259055.1;refGene_gn=LOC101093537;refGene_tx=XM_023259056.1;refGene_gn=LOC101093537;refGene_tx=XM_023259057.1;refGene_gn=LOC101093537;refGene_tx=XM_023259058.1;refGene_gn=LOC101093537;refGene_tx=XM_023259059.1;refGene_gn=LOC101093537;refGene_tx=XM_023259060.1
C1 103407391 103407581
refGene_gn=LOC101093537;refGene_tx=XM_006935036.4;refGene_gn=LOC101093537;refGene_tx=XM_006935041.4;refGene_gn=LOC101093537;refGene_tx=XM_019837713.2;refGene_gn=LOC101093537;refGene_tx=XM_023259046.1;refGene_gn=LOC101093537;refGene_tx=XM_023259047.1;refGene_gn=LOC101093537;refGene_tx=XM_023259048.1;refGene_gn=LOC101093537;refGene_tx=XM_023259049.1;refGene_gn=LOC101093537;refGene_tx=XM_023259050.1;refGene_gn=LOC101093

537;refGene_tx=XM_023259051.1;refGene_gn=LOC101093537;refGene_tx=XM_023259052.1;refGene_gn=LOC101093537;refGene_tx=XM_023259053.1;refGene_gn=LOC101093537;refGene_tx=XM_023259054.1;refGene_gn=LOC101093537;refGene_tx=XM_023259055.1;refGene_gn=LOC101093537;refGene_tx=XM_023259056.1;refGene_gn=LOC101093537;refGene_tx=XM_023259057.1;refGene_gn=LOC101093537;refGene_tx=XM_023259058.1;refGene_gn=LOC101093537;refGene_tx=XM_023259059.1;refGene_gn=LOC101093537;refGene_tx=XM_023259060.1
C1 103408148 103408273
refGene_gn=LOC101093537;refGene_tx=XM_006935036.4;refGene_gn=LOC101093537;refGene_tx=XM_006935041.4;refGene_gn=LOC101093537;refGene_tx=XM_019837713.2;refGene_gn=LOC101093537;refGene_tx=XM_023259046.1;refGene_gn=LOC101093537;refGene_tx=XM_023259047.1;refGene_gn=LOC101093537;refGene_tx=XM_023259048.1;refGene_gn=LOC101093537;refGene_tx=XM_023259049.1;refGene_gn=LOC101093537;refGene_tx=XM_023259050.1;refGene_gn=LOC101093537;refGene_tx=XM_023259051.1;refGene_gn=LOC101093537;refGene_tx=XM_023259052.1;refGene_gn=LOC101093537;refGene_tx=XM_023259053.1;refGene_gn=LOC101093537;refGene_tx=XM_023259054.1;refGene_gn=LOC101093537;refGene_tx=XM_023259055.1;refGene_gn=LOC101093537;refGene_tx=XM_023259056.1;refGene_gn=LOC101093537;refGene_tx=XM_023259057.1;refGene_gn=LOC101093537;refGene_tx=XM_023259058.1;refGene_gn=LOC101093537;refGene_tx=XM_023259059.1;refGene_gn=LOC101093537;refGene_tx=XM_023259060.1
C1 103412121 103412343
refGene_gn=LOC101093537;refGene_tx=XM_006935036.4;refGene_gn=LOC101093537;refGene_tx=XM_006935041.4;refGene_gn=LOC101093537;refGene_tx=XM_019837713.2;refGene_gn=LOC101093537;refGene_tx=XM_023259046.1;refGene_gn=LOC101093537;refGene_tx=XM_023259047.1;refGene_gn=LOC101093537;refGene_tx=XM_023259048.1;refGene_gn=LOC101093537;refGene_tx=XM_023259049.1;refGene_gn=LOC101093537;refGene_tx=XM_023259050.1;refGene_gn=LOC101093537;refGene_tx=XM_023259051.1;refGene_gn=LOC101093537;refGene_tx=XM_023259052.1;refGene_gn=LOC101093537;refGene_tx=XM_023259053.1;refGene_gn=LOC101093537;refGene_tx=XM_023259054.1;refGene_gn=LOC101093537;refGene_tx=XM_023259055.1;refGene_gn=LOC101093537;refGene_tx=XM_023259056.1;refGene_gn=LOC101093537;refGene_tx=XM_023259057.1;refGene_gn=LOC101093537;refGene_tx=XM_023259058.1;refGene_gn=LOC101093537;refGene_tx=XM_023259059.1;refGene_gn=LOC101093537;refGene_tx=XM_023259060.1
C1 103414466 103414583
refGene_gn=LOC101093537;refGene_tx=XM_006935036.4;refGene_gn=LOC101093537;refGene_tx=XM_006935041.4;refGene_gn=LOC101093537;refGene_tx=XM_019837713.2;refGene_gn=LOC101093537;refGene_tx=XM_023259046.1;refGene_gn=LOC101093537;refGene_tx=XM_023259047.1;refGene_gn=LOC101093537;refGene_tx=XM_023259048.1;refGene_gn=LOC101093537;refGene_tx=XM_023259049.1;refGene_gn=LOC101093537;refGene_tx=XM_023259050.1;refGene_gn=LOC101093537;refGene_tx=XM_023259051.1;refGene_gn=LOC101093537;refGene_tx=XM_023259052.1;refGene_gn=LOC101093537;refGene_tx=XM_023259053.1;refGene_gn=LOC101093537;refGene_tx=XM_023259054.1;refGene_gn=LOC101093537;refGene_tx=XM_023259055.1;refGene_gn=LOC101093537;refGene_tx=XM_023259056.1;refGene_gn=LOC101093537;refGene_tx=XM_023259057.1;refGene_gn=LOC101093537;refGene_tx=XM_023259058.1;refGene_gn=LOC101093537;refGene_tx=XM_023259059.1;refGene_gn=LOC101093537;refGene_tx=XM_023259060.1
C1 103415388 103415615

refGene_gn=LOC101093537;refGene_tx=XM_006935036.4;refGene_gn=LOC101093
537;refGene_tx=XM_006935041.4;refGene_gn=LOC101093537;refGene_tx=XM_01
9837713.2;refGene_gn=LOC101093537;refGene_tx=XM_023259046.1;refGene_gn
=LOC101093537;refGene_tx=XM_023259047.1;refGene_gn=LOC101093537;refGen
e_tx=XM_023259048.1;refGene_gn=LOC101093537;refGene_tx=XM_023259049.1;
refGene_gn=LOC101093537;refGene_tx=XM_023259050.1;refGene_gn=LOC101093
537;refGene_tx=XM_023259051.1;refGene_gn=LOC101093537;refGene_tx=XM_02
3259052.1;refGene_gn=LOC101093537;refGene_tx=XM_023259053.1;refGene_gn
=LOC101093537;refGene_tx=XM_023259054.1;refGene_gn=LOC101093537;refGen
e_tx=XM_023259055.1;refGene_gn=LOC101093537;refGene_tx=XM_023259056.1;
refGene_gn=LOC101093537;refGene_tx=XM_023259057.1;refGene_gn=LOC101093
537;refGene_tx=XM_023259058.1;refGene_gn=LOC101093537;refGene_tx=XM_02
3259059.1;refGene_gn=LOC101093537;refGene_tx=XM_023259060.1
C1 103416743 103416887
refGene_gn=LOC101093537;refGene_tx=XM_006935036.4;refGene_gn=LOC101093
537;refGene_tx=XM_006935041.4;refGene_gn=LOC101093537;refGene_tx=XM_01
9837713.2;refGene_gn=LOC101093537;refGene_tx=XM_023259046.1;refGene_gn
=LOC101093537;refGene_tx=XM_023259047.1;refGene_gn=LOC101093537;refGen
e_tx=XM_023259048.1;refGene_gn=LOC101093537;refGene_tx=XM_023259049.1;
refGene_gn=LOC101093537;refGene_tx=XM_023259050.1;refGene_gn=LOC101093
537;refGene_tx=XM_023259051.1;refGene_gn=LOC101093537;refGene_tx=XM_02
3259052.1;refGene_gn=LOC101093537;refGene_tx=XM_023259053.1;refGene_gn
=LOC101093537;refGene_tx=XM_023259054.1;refGene_gn=LOC101093537;refGen
e_tx=XM_023259055.1;refGene_gn=LOC101093537;refGene_tx=XM_023259056.1;
refGene_gn=LOC101093537;refGene_tx=XM_023259057.1;refGene_gn=LOC101093
537;refGene_tx=XM_023259058.1;refGene_gn=LOC101093537;refGene_tx=XM_02
3259059.1;refGene_gn=LOC101093537;refGene_tx=XM_023259060.1
C1 103417949 103418088
refGene_gn=LOC101093537;refGene_tx=XM_006935036.4;refGene_gn=LOC101093
537;refGene_tx=XM_006935041.4;refGene_gn=LOC101093537;refGene_tx=XM_01
9837713.2;refGene_gn=LOC101093537;refGene_tx=XM_023259046.1;refGene_gn
=LOC101093537;refGene_tx=XM_023259047.1;refGene_gn=LOC101093537;refGen
e_tx=XM_023259048.1;refGene_gn=LOC101093537;refGene_tx=XM_023259049.1;
refGene_gn=LOC101093537;refGene_tx=XM_023259050.1;refGene_gn=LOC101093
537;refGene_tx=XM_023259051.1;refGene_gn=LOC101093537;refGene_tx=XM_02
3259052.1;refGene_gn=LOC101093537;refGene_tx=XM_023259053.1;refGene_gn
=LOC101093537;refGene_tx=XM_023259054.1;refGene_gn=LOC101093537;refGen
e_tx=XM_023259055.1;refGene_gn=LOC101093537;refGene_tx=XM_023259056.1;
refGene_gn=LOC101093537;refGene_tx=XM_023259057.1;refGene_gn=LOC101093
537;refGene_tx=XM_023259058.1;refGene_gn=LOC101093537;refGene_tx=XM_02
3259059.1;refGene_gn=LOC101093537;refGene_tx=XM_023259060.1
C1 103418383 103418430
refGene_gn=LOC101093537;refGene_tx=XM_006935036.4;refGene_gn=LOC101093
537;refGene_tx=XM_006935041.4;refGene_gn=LOC101093537;refGene_tx=XM_01
9837713.2;refGene_gn=LOC101093537;refGene_tx=XM_023259046.1;refGene_gn
=LOC101093537;refGene_tx=XM_023259047.1;refGene_gn=LOC101093537;refGen
e_tx=XM_023259048.1;refGene_gn=LOC101093537;refGene_tx=XM_023259049.1;
refGene_gn=LOC101093537;refGene_tx=XM_023259050.1;refGene_gn=LOC101093
537;refGene_tx=XM_023259051.1;refGene_gn=LOC101093537;refGene_tx=XM_02
3259052.1;refGene_gn=LOC101093537;refGene_tx=XM_023259053.1;refGene_gn

=LOC101093537;refGene_tx=XM_023259054.1;refGene_gn=LOC101093537;refGene_tx=XM_023259055.1;refGene_gn=LOC101093537;refGene_tx=XM_023259056.1;refGene_gn=LOC101093537;refGene_tx=XM_023259057.1;refGene_gn=LOC101093537;refGene_tx=XM_023259058.1;refGene_gn=LOC101093537;refGene_tx=XM_023259059.1;refGene_gn=LOC101093537;refGene_tx=XM_023259060.1
C1 103418671 103418698
refGene_gn=LOC101093537;refGene_tx=XM_006935036.4;refGene_gn=LOC101093537;refGene_tx=XM_019837713.2;refGene_gn=LOC101093537;refGene_tx=XM_023259046.1;refGene_gn=LOC101093537;refGene_tx=XM_023259048.1;refGene_gn=LOC101093537;refGene_tx=XM_023259050.1;refGene_gn=LOC101093537;refGene_tx=XM_023259051.1;refGene_gn=LOC101093537;refGene_tx=XM_023259052.1;refGene_gn=LOC101093537;refGene_tx=XM_023259054.1;refGene_gn=LOC101093537;refGene_tx=XM_023259056.1;refGene_gn=LOC101093537;refGene_tx=XM_023259057.1;refGene_gn=LOC101093537;refGene_tx=XM_023259059.1
C1 103419286 103419347
refGene_gn=LOC101093537;refGene_tx=XM_006935036.4;refGene_gn=LOC101093537;refGene_tx=XM_019837713.2;refGene_gn=LOC101093537;refGene_tx=XM_023259046.1;refGene_gn=LOC101093537;refGene_tx=XM_023259047.1;refGene_gn=LOC101093537;refGene_tx=XM_023259050.1;refGene_gn=LOC101093537;refGene_tx=XM_023259051.1;refGene_gn=LOC101093537;refGene_tx=XM_023259052.1;refGene_gn=LOC101093537;refGene_tx=XM_023259053.1;refGene_gn=LOC101093537;refGene_tx=XM_023259056.1;refGene_gn=LOC101093537;refGene_tx=XM_023259057.1;refGene_gn=LOC101093537;refGene_tx=XM_023259059.1
C1 103419986 103420237
refGene_gn=LOC101093537;refGene_tx=XM_006935036.4;refGene_gn=LOC101093537;refGene_tx=XM_006935041.4;refGene_gn=LOC101093537;refGene_tx=XM_019837713.2;refGene_gn=LOC101093537;refGene_tx=XM_023259046.1;refGene_gn=LOC101093537;refGene_tx=XM_023259047.1;refGene_gn=LOC101093537;refGene_tx=XM_023259048.1;refGene_gn=LOC101093537;refGene_tx=XM_023259049.1;refGene_gn=LOC101093537;refGene_tx=XM_023259050.1;refGene_gn=LOC101093537;refGene_tx=XM_023259051.1;refGene_gn=LOC101093537;refGene_tx=XM_023259052.1;refGene_gn=LOC101093537;refGene_tx=XM_023259053.1;refGene_gn=LOC101093537;refGene_tx=XM_023259054.1;refGene_gn=LOC101093537;refGene_tx=XM_023259055.1;refGene_gn=LOC101093537;refGene_tx=XM_023259057.1;refGene_gn=LOC101093537;refGene_tx=XM_023259058.1;refGene_gn=LOC101093537;refGene_tx=XM_023259059.1;refGene_gn=LOC101093537;refGene_tx=XM_023259060.1
C1 103442657 103442880
refGene_gn=LOC111561809;refGene_tx=XM_023259064.1
C1 103444493 103444733
refGene_gn=LOC111561809;refGene_tx=XM_023259064.1
C1 103445795 103446001
refGene_gn=LOC111561809;refGene_tx=XM_023259064.1
C1 103473283 103473310
refGene_gn=LOC101095725;refGene_tx=XM_023259061.1;refGene_gn=LOC101095725;refGene_tx=XM_023259062.1;refGene_gn=LOC101095725;refGene_tx=XM_023259063.1
C1 103474470 103474719
refGene_gn=LOC101095725;refGene_tx=XM_023259061.1;refGene_gn=LOC101095725;refGene_tx=XM_023259062.1;refGene_gn=LOC101095725;refGene_tx=XM_02

3259063.1
C1	103480550	103480598
refGene_gn=LOC101095725;refGene_tx=XM_023259061.1;refGene_gn=LOC101095725;refGene_tx=XM_023259062.1;refGene_gn=LOC101095725;refGene_tx=XM_023259063.1
C1	103480698	103480909
refGene_gn=LOC101095725;refGene_tx=XM_023259061.1;refGene_gn=LOC101095725;refGene_tx=XM_023259062.1;refGene_gn=LOC101095725;refGene_tx=XM_023259063.1
C1	103483192	103483289
refGene_gn=LOC101095725;refGene_tx=XM_023259061.1;refGene_gn=LOC101095725;refGene_tx=XM_023259062.1;refGene_gn=LOC101095725;refGene_tx=XM_023259063.1
C1	103487708	103487923
refGene_gn=LOC101095725;refGene_tx=XM_023259061.1;refGene_gn=LOC101095725;refGene_tx=XM_023259062.1;refGene_gn=LOC101095725;refGene_tx=XM_023259063.1
C1	103488946	103489001
refGene_gn=LOC101095725;refGene_tx=XM_023259061.1;refGene_gn=LOC101095725;refGene_tx=XM_023259062.1;refGene_gn=LOC101095725;refGene_tx=XM_023259063.1
C1	103489836	103490051
refGene_gn=LOC101095725;refGene_tx=XM_023259061.1;refGene_gn=LOC101095725;refGene_tx=XM_023259062.1;refGene_gn=LOC101095725;refGene_tx=XM_023259063.1
C1	103491041	103491093
refGene_gn=LOC101095725;refGene_tx=XM_023259061.1;refGene_gn=LOC101095725;refGene_tx=XM_023259062.1;refGene_gn=LOC101095725;refGene_tx=XM_023259063.1
C1	103491881	103492057
refGene_gn=LOC101095725;refGene_tx=XM_023259061.1;refGene_gn=LOC101095725;refGene_tx=XM_023259062.1;refGene_gn=LOC101095725;refGene_tx=XM_023259063.1
C1	103493114	103493166
refGene_gn=LOC101095725;refGene_tx=XM_023259061.1;refGene_gn=LOC101095725;refGene_tx=XM_023259062.1;refGene_gn=LOC101095725;refGene_tx=XM_023259063.1
C1	103493988	103494164
refGene_gn=LOC101095725;refGene_tx=XM_023259061.1;refGene_gn=LOC101095725;refGene_tx=XM_023259062.1;refGene_gn=LOC101095725;refGene_tx=XM_023259063.1
C1	103495224	103495276
refGene_gn=LOC101095725;refGene_tx=XM_023259061.1;refGene_gn=LOC101095725;refGene_tx=XM_023259062.1
C1	103496062	103496244
refGene_gn=LOC101095725;refGene_tx=XM_023259061.1;refGene_gn=LOC101095725;refGene_tx=XM_023259062.1
C1	103497859	103498099
refGene_gn=LOC101095725;refGene_tx=XM_023259061.1;refGene_gn=LOC101095725;refGene_tx=XM_023259062.1;refGene_gn=LOC101095725;refGene_tx=XM_02

3259063.1
C1	103499160	103499366
refGene_gn=LOC101095725;refGene_tx=XM_023259061.1;refGene_gn=LOC101095725;refGene_tx=XM_023259063.1
C1	103502518	103502541
refGene_gn=LOC101095725;refGene_tx=XM_023259062.1
C1	103575246	103575324
refGene_gn=PRKAB2;refGene_tx=XM_019837733.2;refGene_gn=PRKAB2;refGene_tx=XM_023259066.1;refGene_gn=PRKAB2;refGene_tx=XM_023259067.1
C1	103576465	103576502
refGene_gn=PRKAB2;refGene_tx=XM_019837733.2
C1	103577402	103577471
refGene_gn=PRKAB2;refGene_tx=XM_019837733.2;refGene_gn=PRKAB2;refGene_tx=XM_023259066.1;refGene_gn=PRKAB2;refGene_tx=XM_023259067.1
C1	103577994	103578128
refGene_gn=PRKAB2;refGene_tx=XM_019837733.2;refGene_gn=PRKAB2;refGene_tx=XM_023259066.1;refGene_gn=PRKAB2;refGene_tx=XM_023259067.1
C1	103582196	103582317
refGene_gn=PRKAB2;refGene_tx=XM_019837733.2;refGene_gn=PRKAB2;refGene_tx=XM_023259066.1;refGene_gn=PRKAB2;refGene_tx=XM_023259067.1
C1	103582549	103582643
refGene_gn=PRKAB2;refGene_tx=XM_019837733.2;refGene_gn=PRKAB2;refGene_tx=XM_023259066.1;refGene_gn=PRKAB2;refGene_tx=XM_023259067.1
C1	103583489	103583656
refGene_gn=PRKAB2;refGene_tx=XM_019837733.2;refGene_gn=PRKAB2;refGene_tx=XM_023259066.1;refGene_gn=PRKAB2;refGene_tx=XM_023259067.1
C1	103587504	103587660
refGene_gn=PRKAB2;refGene_tx=XM_019837733.2;refGene_gn=PRKAB2;refGene_tx=XM_023259066.1;refGene_gn=PRKAB2;refGene_tx=XM_023259067.1
C1	103602576	103602804
refGene_gn=FMO5;refGene_tx=XM_023259065.1
C1	103602805	103602923
refGene_gn=FMO5;refGene_tx=XM_023259065.1
C1	103605575	103605648
refGene_gn=FMO5;refGene_tx=XM_023259065.1
C1	103607477	103607830
refGene_gn=FMO5;refGene_tx=XM_023259065.1
C1	103613407	103613607
refGene_gn=FMO5;refGene_tx=XM_023259065.1
C1	103616484	103616627
refGene_gn=FMO5;refGene_tx=XM_023259065.1
C1	103617663	103617826
refGene_gn=FMO5;refGene_tx=XM_023259065.1
C1	103619338	103619527
refGene_gn=FMO5;refGene_tx=XM_023259065.1
C1	103634976	103635111
refGene_gn=FMO5;refGene_tx=XM_023259065.1
C1	103656174	103656307
refGene_gn=CHD1L;refGene_tx=XM_006935047.4
C1	103668770	103668883

refGene_gn=CHD1L;refGene_tx=XM_006935047.4;refGene_gn=CHD1L;refGene_tx=XM_019837726.2
C1 103673057 103673164 refGene_gn=CHD1L;refGene_tx=XM_006935047.4;refGene_gn=CHD1L;refGene_tx=XM_019837726.2;refGene_gn=CHD1L;refGene_tx=XM_019837727.2
C1 103673710 103673825 refGene_gn=CHD1L;refGene_tx=XM_006935047.4;refGene_gn=CHD1L;refGene_tx=XM_019837726.2;refGene_gn=CHD1L;refGene_tx=XM_019837727.2
C1 103674429 103674461 refGene_gn=CHD1L;refGene_tx=XM_006935047.4;refGene_gn=CHD1L;refGene_tx=XM_019837726.2;refGene_gn=CHD1L;refGene_tx=XM_019837727.2
C1 103677356 103677438 refGene_gn=CHD1L;refGene_tx=XM_006935047.4;refGene_gn=CHD1L;refGene_tx=XM_019837726.2;refGene_gn=CHD1L;refGene_tx=XM_019837727.2
C1 103680411 103680574 refGene_gn=CHD1L;refGene_tx=XM_006935047.4;refGene_gn=CHD1L;refGene_tx=XM_019837726.2;refGene_gn=CHD1L;refGene_tx=XM_019837727.2
C1 103682240 103682396 refGene_gn=CHD1L;refGene_tx=XM_006935047.4;refGene_gn=CHD1L;refGene_tx=XM_019837726.2;refGene_gn=CHD1L;refGene_tx=XM_019837727.2
C1 103683131 103683224 refGene_gn=CHD1L;refGene_tx=XM_006935047.4;refGene_gn=CHD1L;refGene_tx=XM_019837726.2;refGene_gn=CHD1L;refGene_tx=XM_019837727.2
C1 103684540 103684637 refGene_gn=CHD1L;refGene_tx=XM_006935047.4;refGene_gn=CHD1L;refGene_tx=XM_019837726.2;refGene_gn=CHD1L;refGene_tx=XM_019837727.2
C1 103686347 103686421 refGene_gn=CHD1L;refGene_tx=XM_006935047.4;refGene_gn=CHD1L;refGene_tx=XM_019837726.2;refGene_gn=CHD1L;refGene_tx=XM_019837727.2
C1 103688036 103688147 refGene_gn=CHD1L;refGene_tx=XM_006935047.4;refGene_gn=CHD1L;refGene_tx=XM_019837726.2;refGene_gn=CHD1L;refGene_tx=XM_019837727.2
C1 103690859 103690974 refGene_gn=CHD1L;refGene_tx=XM_006935047.4;refGene_gn=CHD1L;refGene_tx=XM_019837726.2;refGene_gn=CHD1L;refGene_tx=XM_019837727.2
C1 103691601 103691755 refGene_gn=CHD1L;refGene_tx=XM_006935047.4;refGene_gn=CHD1L;refGene_tx=XM_019837726.2;refGene_gn=CHD1L;refGene_tx=XM_019837727.2
C1 103693692 103693858 refGene_gn=CHD1L;refGene_tx=XM_006935047.4;refGene_gn=CHD1L;refGene_tx=XM_019837726.2;refGene_gn=CHD1L;refGene_tx=XM_019837727.2
C1 103697672 103697821 refGene_gn=CHD1L;refGene_tx=XM_006935047.4;refGene_gn=CHD1L;refGene_tx=XM_019837726.2;refGene_gn=CHD1L;refGene_tx=XM_019837727.2
C1 103698785 103698949 refGene_gn=CHD1L;refGene_tx=XM_006935047.4;refGene_gn=CHD1L;refGene_tx=XM_019837726.2;refGene_gn=CHD1L;refGene_tx=XM_019837727.2
C1 103699484 103699693 refGene_gn=CHD1L;refGene_tx=XM_006935047.4;refGene_gn=CHD1L;refGene_tx=XM_019837726.2;refGene_gn=CHD1L;refGene_tx=XM_019837727.2

C1	103700728	103700827
refGene_gn=CHD1L;refGene_tx=XM_006935047.4;refGene_gn=CHD1L;refGene_tx
=XM_019837726.2;refGene_gn=CHD1L;refGene_tx=XM_019837727.2
C1	103710845	103710916
refGene_gn=CHD1L;refGene_tx=XM_006935047.4;refGene_gn=CHD1L;refGene_tx
=XM_019837726.2;refGene_gn=CHD1L;refGene_tx=XM_019837727.2
C1	103712513	103712628
refGene_gn=CHD1L;refGene_tx=XM_006935047.4;refGene_gn=CHD1L;refGene_tx
=XM_019837726.2;refGene_gn=CHD1L;refGene_tx=XM_019837727.2
C1	103713365	103713474
refGene_gn=CHD1L;refGene_tx=XM_006935047.4;refGene_gn=CHD1L;refGene_tx
=XM_019837726.2;refGene_gn=CHD1L;refGene_tx=XM_019837727.2
C1	103714563	103714639
refGene_gn=CHD1L;refGene_tx=XM_006935047.4;refGene_gn=CHD1L;refGene_tx
=XM_019837726.2;refGene_gn=CHD1L;refGene_tx=XM_019837727.2
C1	103867246	103868176
refGene_gn=LOC101094778;refGene_tx=XM_006935049.4
C1	103894834	103895794
refGene_gn=LOC101095034;refGene_tx=XM_003990531.1
C1	104093929	104093982
refGene_gn=BCL9;refGene_tx=XM_004001337.5;refGene_gn=BCL9;refGene_tx=X
M_011285100.3;refGene_gn=BCL9;refGene_tx=XM_019837734.2;refGene_gn=BCL
9;refGene_tx=XM_019837735.2;refGene_gn=BCL9;refGene_tx=XM_023259068.1
C1	104095202	104095519
refGene_gn=BCL9;refGene_tx=XM_004001337.5;refGene_gn=BCL9;refGene_tx=X
M_011285100.3;refGene_gn=BCL9;refGene_tx=XM_019837734.2;refGene_gn=BCL
9;refGene_tx=XM_019837735.2;refGene_gn=BCL9;refGene_tx=XM_023259068.1
C1	104096740	104096930
refGene_gn=BCL9;refGene_tx=XM_004001337.5;refGene_gn=BCL9;refGene_tx=X
M_011285100.3;refGene_gn=BCL9;refGene_tx=XM_019837734.2;refGene_gn=BCL
9;refGene_tx=XM_019837735.2;refGene_gn=BCL9;refGene_tx=XM_019837736.2;
refGene_gn=BCL9;refGene_tx=XM_023259068.1
C1	104097817	104097971
refGene_gn=BCL9;refGene_tx=XM_004001337.5;refGene_gn=BCL9;refGene_tx=X
M_011285100.3;refGene_gn=BCL9;refGene_tx=XM_019837734.2;refGene_gn=BCL
9;refGene_tx=XM_019837735.2;refGene_gn=BCL9;refGene_tx=XM_019837736.2;
refGene_gn=BCL9;refGene_tx=XM_023259068.1
C1	104100750	104102992
refGene_gn=BCL9;refGene_tx=XM_004001337.5;refGene_gn=BCL9;refGene_tx=X
M_011285100.3;refGene_gn=BCL9;refGene_tx=XM_019837734.2;refGene_gn=BCL
9;refGene_tx=XM_019837735.2;refGene_gn=BCL9;refGene_tx=XM_019837736.2;
refGene_gn=BCL9;refGene_tx=XM_023259068.1
C1	104104258	104104519
refGene_gn=BCL9;refGene_tx=XM_004001337.5;refGene_gn=BCL9;refGene_tx=X
M_011285100.3;refGene_gn=BCL9;refGene_tx=XM_019837734.2;refGene_gn=BCL
9;refGene_tx=XM_019837735.2;refGene_gn=BCL9;refGene_tx=XM_019837736.2;
refGene_gn=BCL9;refGene_tx=XM_023259068.1
C1	104105621	104106739
refGene_gn=BCL9;refGene_tx=XM_004001337.5;refGene_gn=BCL9;refGene_tx=X
M_011285100.3;refGene_gn=BCL9;refGene_tx=XM_019837734.2;refGene_gn=BCL

9;refGene_tx=XM_019837735.2;refGene_gn=BCL9;refGene_tx=XM_019837736.2;
refGene_gn=BCL9;refGene_tx=XM_023259068.1
C1 104129655 104129799
refGene_gn=ACP6;refGene_tx=XM_006935048.4
C1 104130436 104130602
refGene_gn=ACP6;refGene_tx=XM_006935048.4
C1 104132257 104132353
refGene_gn=ACP6;refGene_tx=XM_006935048.4
C1 104134778 104134879
refGene_gn=ACP6;refGene_tx=XM_006935048.4
C1 104135987 104136120
refGene_gn=ACP6;refGene_tx=XM_006935048.4
C1 104136990 104137078
refGene_gn=ACP6;refGene_tx=XM_006935048.4
C1 104140001 104140081
refGene_gn=ACP6;refGene_tx=XM_006935048.4
C1 104140416 104140547
refGene_gn=ACP6;refGene_tx=XM_006935048.4
C1 104140662 104140791
refGene_gn=ACP6;refGene_tx=XM_006935048.4
C1 104148656 104148872
refGene_gn=ACP6;refGene_tx=XM_006935048.4
C1 104212254 104213331
refGene_gn=GJA5;refGene_tx=XM_003990532.5
C1 104337707 104339030
refGene_gn=GJA8;refGene_tx=XM_019837741.2
C1 104351017 104351059
refGene_gn=GPR89A;refGene_tx=XM_003990534.5
C1 104359628 104359688
refGene_gn=GPR89A;refGene_tx=XM_003990534.5
C1 104360646 104360750
refGene_gn=GPR89A;refGene_tx=XM_003990534.5
C1 104365296 104365403
refGene_gn=GPR89A;refGene_tx=XM_003990534.5
C1 104365838 104365940
refGene_gn=GPR89A;refGene_tx=XM_003990534.5;refGene_gn=GPR89A;refGene_
tx=XM_023259072.1
C1 104372200 104372321
refGene_gn=GPR89A;refGene_tx=XM_003990534.5;refGene_gn=GPR89A;refGene_
tx=XM_023259072.1
C1 104373237 104373318
refGene_gn=GPR89A;refGene_tx=XM_003990534.5;refGene_gn=GPR89A;refGene_
tx=XM_023259072.1
C1 104390118 104390228
refGene_gn=GPR89A;refGene_tx=XM_003990534.5;refGene_gn=GPR89A;refGene_
tx=XM_023259072.1
C1 104392539 104392628
refGene_gn=GPR89A;refGene_tx=XM_003990534.5;refGene_gn=GPR89A;refGene_
tx=XM_023259072.1
C1 104393543 104393636

refGene_gn=GPR89A;refGene_tx=XM_003990534.5;refGene_gn=GPR89A;refGene_tx=XM_023259072.1
C1	104406711	104406807
refGene_gn=GPR89A;refGene_tx=XM_003990534.5;refGene_gn=GPR89A;refGene_tx=XM_023259072.1
C1	104408910	104409000
refGene_gn=GPR89A;refGene_tx=XM_003990534.5;refGene_gn=GPR89A;refGene_tx=XM_023259072.1
C1	104410110	104410176
refGene_gn=GPR89A;refGene_tx=XM_003990534.5;refGene_gn=GPR89A;refGene_tx=XM_023259072.1
C1	104410309	104410516
refGene_gn=GPR89A;refGene_tx=XM_003990534.5;refGene_gn=GPR89A;refGene_tx=XM_023259072.1
C1	104411915	104411969
refGene_gn=PDZK1;refGene_tx=XM_006935051.4;refGene_gn=PDZK1;refGene_tx=XM_006935053.4;refGene_gn=PDZK1;refGene_tx=XM_006935054.4;refGene_gn=PDZK1;refGene_tx=XM_019837738.2;refGene_gn=PDZK1;refGene_tx=XM_019837739.2;refGene_gn=PDZK1;refGene_tx=XM_023259071.1
C1	104413277	104413568
refGene_gn=PDZK1;refGene_tx=XM_006935051.4;refGene_gn=PDZK1;refGene_tx=XM_006935053.4;refGene_gn=PDZK1;refGene_tx=XM_006935054.4;refGene_gn=PDZK1;refGene_tx=XM_019837738.2;refGene_gn=PDZK1;refGene_tx=XM_019837739.2;refGene_gn=PDZK1;refGene_tx=XM_023259071.1
C1	104423160	104423379
refGene_gn=PDZK1;refGene_tx=XM_006935051.4;refGene_gn=PDZK1;refGene_tx=XM_006935053.4;refGene_gn=PDZK1;refGene_tx=XM_006935054.4;refGene_gn=PDZK1;refGene_tx=XM_019837738.2;refGene_gn=PDZK1;refGene_tx=XM_019837739.2
C1	104429715	104429912
refGene_gn=PDZK1;refGene_tx=XM_006935051.4;refGene_gn=PDZK1;refGene_tx=XM_006935053.4;refGene_gn=PDZK1;refGene_tx=XM_006935054.4;refGene_gn=PDZK1;refGene_tx=XM_019837738.2;refGene_gn=PDZK1;refGene_tx=XM_019837739.2;refGene_gn=PDZK1;refGene_tx=XM_023259071.1
C1	104432632	104432828
refGene_gn=PDZK1;refGene_tx=XM_006935051.4;refGene_gn=PDZK1;refGene_tx=XM_006935053.4;refGene_gn=PDZK1;refGene_tx=XM_006935054.4;refGene_gn=PDZK1;refGene_tx=XM_019837738.2;refGene_gn=PDZK1;refGene_tx=XM_019837739.2;refGene_gn=PDZK1;refGene_tx=XM_023259071.1
C1	104433651	104433788
refGene_gn=PDZK1;refGene_tx=XM_006935051.4;refGene_gn=PDZK1;refGene_tx=XM_006935053.4;refGene_gn=PDZK1;refGene_tx=XM_006935054.4;refGene_gn=PDZK1;refGene_tx=XM_019837738.2;refGene_gn=PDZK1;refGene_tx=XM_019837739.2;refGene_gn=PDZK1;refGene_tx=XM_023259071.1
C1	104435978	104436228
refGene_gn=PDZK1;refGene_tx=XM_006935051.4;refGene_gn=PDZK1;refGene_tx=XM_006935053.4;refGene_gn=PDZK1;refGene_tx=XM_006935054.4;refGene_gn=PDZK1;refGene_tx=XM_019837738.2;refGene_gn=PDZK1;refGene_tx=XM_019837739.2;refGene_gn=PDZK1;refGene_tx=XM_023259071.1
C1	104436895	104437105

refGene_gn=PDZK1;refGene_tx=XM_006935051.4;refGene_gn=PDZK1;refGene_tx=XM_006935053.4;refGene_gn=PDZK1;refGene_tx=XM_006935054.4;refGene_gn=PDZK1;refGene_tx=XM_019837738.2;refGene_gn=PDZK1;refGene_tx=XM_019837739.2;refGene_gn=PDZK1;refGene_tx=XM_023259071.1
C1 104469129 104469177
refGene_gn=CD160;refGene_tx=XM_003990577.5
C1 104478597 104478793
refGene_gn=CD160;refGene_tx=XM_003990577.5;refGene_gn=CD160;refGene_tx=XM_019837742.2;refGene_gn=CD160;refGene_tx=XM_019837743.2
C1 104479467 104479794
refGene_gn=CD160;refGene_tx=XM_003990577.5;refGene_gn=CD160;refGene_tx=XM_019837742.2;refGene_gn=CD160;refGene_tx=XM_019837743.2
C1 104484247 104484382
refGene_gn=CD160;refGene_tx=XM_003990577.5;refGene_gn=CD160;refGene_tx=XM_019837742.2;refGene_gn=CD160;refGene_tx=XM_019837743.2
C1 104486257 104486265
refGene_gn=CD160;refGene_tx=XM_003990577.5;refGene_gn=CD160;refGene_tx=XM_019837742.2;refGene_gn=CD160;refGene_tx=XM_019837743.2
C1 104494328 104494460
refGene_gn=RNF115;refGene_tx=XM_003990538.5;refGene_gn=RNF115;refGene_tx=XM_006935057.3;refGene_gn=RNF115;refGene_tx=XM_023259077.1
C1 104495139 104495255
refGene_gn=RNF115;refGene_tx=XM_003990538.5;refGene_gn=RNF115;refGene_tx=XM_006935057.3;refGene_gn=RNF115;refGene_tx=XM_023259077.1
C1 104497706 104497800
refGene_gn=RNF115;refGene_tx=XM_003990538.5;refGene_gn=RNF115;refGene_tx=XM_006935057.3;refGene_gn=RNF115;refGene_tx=XM_023259077.1
C1 104498743 104498816
refGene_gn=RNF115;refGene_tx=XM_003990538.5;refGene_gn=RNF115;refGene_tx=XM_006935057.3;refGene_gn=RNF115;refGene_tx=XM_023259077.1
C1 104499975 104500047
refGene_gn=RNF115;refGene_tx=XM_003990538.5;refGene_gn=RNF115;refGene_tx=XM_006935057.3;refGene_gn=RNF115;refGene_tx=XM_023259077.1
C1 104528942 104529154
refGene_gn=RNF115;refGene_tx=XM_003990538.5;refGene_gn=RNF115;refGene_tx=XM_006935057.3;refGene_gn=RNF115;refGene_tx=XM_023259077.1
C1 104547575 104547633
refGene_gn=RNF115;refGene_tx=XM_003990538.5;refGene_gn=RNF115;refGene_tx=XM_006935057.3;refGene_gn=RNF115;refGene_tx=XM_023259077.1
C1 104550519 104550571
refGene_gn=RNF115;refGene_tx=XM_006935057.3
C1 104551466 104551525
refGene_gn=RNF115;refGene_tx=XM_003990538.5;refGene_gn=RNF115;refGene_tx=XM_006935057.3;refGene_gn=RNF115;refGene_tx=XM_023259077.1
C1 104577566 104577698
refGene_gn=RNF115;refGene_tx=XM_023259077.1
C1 104579119 104579221
refGene_gn=RNF115;refGene_tx=XM_003990538.5
C1 104580768 104580915
refGene_gn=POLR3C;refGene_tx=XM_003990539.5;refGene_gn=POLR3C;refGene_

tx=XM_006935056.4
C1	104581441	104581697
refGene_gn=POLR3C;refGene_tx=XM_003990539.5;refGene_gn=POLR3C;refGene_tx=XM_006935056.4
C1	104581807	104581993
refGene_gn=POLR3C;refGene_tx=XM_003990539.5;refGene_gn=POLR3C;refGene_tx=XM_006935056.4
C1	104583652	104583741
refGene_gn=POLR3C;refGene_tx=XM_003990539.5;refGene_gn=POLR3C;refGene_tx=XM_006935056.4
C1	104584289	104584394
refGene_gn=POLR3C;refGene_tx=XM_003990539.5;refGene_gn=POLR3C;refGene_tx=XM_006935056.4
C1	104584527	104584620
refGene_gn=POLR3C;refGene_tx=XM_003990539.5
C1	104586270	104586351
refGene_gn=POLR3C;refGene_tx=XM_003990539.5;refGene_gn=POLR3C;refGene_tx=XM_006935056.4
C1	104586603	104586655
refGene_gn=POLR3C;refGene_tx=XM_003990539.5;refGene_gn=POLR3C;refGene_tx=XM_006935056.4
C1	104587006	104587067
refGene_gn=POLR3C;refGene_tx=XM_003990539.5;refGene_gn=POLR3C;refGene_tx=XM_006935056.4
C1	104587508	104587659
refGene_gn=POLR3C;refGene_tx=XM_003990539.5;refGene_gn=POLR3C;refGene_tx=XM_006935056.4
C1	104588379	104588481
refGene_gn=POLR3C;refGene_tx=XM_003990539.5;refGene_gn=POLR3C;refGene_tx=XM_006935056.4
C1	104588611	104588661
refGene_gn=POLR3C;refGene_tx=XM_003990539.5;refGene_gn=POLR3C;refGene_tx=XM_006935056.4
C1	104589077	104589227
refGene_gn=POLR3C;refGene_tx=XM_003990539.5;refGene_gn=POLR3C;refGene_tx=XM_006935056.4
C1	104589653	104589735
refGene_gn=POLR3C;refGene_tx=XM_003990539.5;refGene_gn=POLR3C;refGene_tx=XM_006935056.4
C1	104591590	104591782
refGene_gn=NUDT17;refGene_tx=XM_006935058.4;refGene_gn=NUDT17;refGene_tx=XM_006935059.4;refGene_gn=NUDT17;refGene_tx=XM_006935060.4;refGene_gn=NUDT17;refGene_tx=XM_019837747.2
C1	104592001	104592185
refGene_gn=NUDT17;refGene_tx=XM_006935058.4;refGene_gn=NUDT17;refGene_tx=XM_006935059.4;refGene_gn=NUDT17;refGene_tx=XM_006935060.4;refGene_gn=NUDT17;refGene_tx=XM_019837747.2
C1	104592431	104592457
refGene_gn=NUDT17;refGene_tx=XM_006935058.4;refGene_gn=NUDT17;refGene_tx=XM_006935059.4;refGene_gn=NUDT17;refGene_tx=XM_006935060.4;refGene_

gn=NUDT17;refGene_tx=XM_019837747.2
C1 104592585 104592678
refGene_gn=NUDT17;refGene_tx=XM_006935058.4;refGene_gn=NUDT17;refGene_tx=XM_006935059.4;refGene_gn=NUDT17;refGene_tx=XM_006935060.4;refGene_gn=NUDT17;refGene_tx=XM_019837747.2
C1 104592923 104593022
refGene_gn=NUDT17;refGene_tx=XM_006935058.4;refGene_gn=NUDT17;refGene_tx=XM_006935059.4;refGene_gn=NUDT17;refGene_tx=XM_006935060.4;refGene_gn=NUDT17;refGene_tx=XM_019837747.2
C1 104593236 104593373
refGene_gn=NUDT17;refGene_tx=XM_006935058.4;refGene_gn=NUDT17;refGene_tx=XM_006935059.4;refGene_gn=NUDT17;refGene_tx=XM_006935060.4;refGene_gn=NUDT17;refGene_tx=XM_019837747.2
C1 104593669 104593904
refGene_gn=NUDT17;refGene_tx=XM_006935058.4;refGene_gn=NUDT17;refGene_tx=XM_006935059.4;refGene_gn=NUDT17;refGene_tx=XM_006935060.4;refGene_gn=NUDT17;refGene_tx=XM_019837747.2
C1 104593992 104594090
refGene_gn=NUDT17;refGene_tx=XM_019837747.2
C1 104595037 104595304
refGene_gn=PIAS3;refGene_tx=XM_011285104.3;refGene_gn=PIAS3;refGene_tx=XM_023259074.1;refGene_gn=PIAS3;refGene_tx=XM_023259075.1
C1 104595874 104595912
refGene_gn=PIAS3;refGene_tx=XM_011285104.3;refGene_gn=PIAS3;refGene_tx=XM_023259074.1;refGene_gn=PIAS3;refGene_tx=XM_023259075.1
C1 104596098 104596232
refGene_gn=PIAS3;refGene_tx=XM_011285104.3;refGene_gn=PIAS3;refGene_tx=XM_023259074.1;refGene_gn=PIAS3;refGene_tx=XM_023259075.1
C1 104596398 104596567
refGene_gn=NUDT17;refGene_tx=XM_006935058.4;refGene_gn=NUDT17;refGene_tx=XM_006935059.4;refGene_gn=NUDT17;refGene_tx=XM_006935060.4;refGene_gn=PIAS3;refGene_tx=XM_011285104.3;refGene_gn=PIAS3;refGene_tx=XM_023259074.1;refGene_gn=PIAS3;refGene_tx=XM_023259075.1
C1 104596647 104596781
refGene_gn=PIAS3;refGene_tx=XM_011285104.3;refGene_gn=PIAS3;refGene_tx=XM_023259074.1;refGene_gn=PIAS3;refGene_tx=XM_023259075.1
C1 104597876 104598037
refGene_gn=PIAS3;refGene_tx=XM_011285104.3;refGene_gn=PIAS3;refGene_tx=XM_023259074.1;refGene_gn=PIAS3;refGene_tx=XM_023259075.1
C1 104598167 104598241
refGene_gn=PIAS3;refGene_tx=XM_011285104.3;refGene_gn=PIAS3;refGene_tx=XM_023259074.1;refGene_gn=PIAS3;refGene_tx=XM_023259075.1
C1 104598830 104598936
refGene_gn=PIAS3;refGene_tx=XM_011285104.3;refGene_gn=PIAS3;refGene_tx=XM_023259074.1;refGene_gn=PIAS3;refGene_tx=XM_023259075.1
C1 104599180 104599315
refGene_gn=PIAS3;refGene_tx=XM_011285104.3;refGene_gn=PIAS3;refGene_tx=XM_023259074.1;refGene_gn=PIAS3;refGene_tx=XM_023259075.1
C1 104599817 104599908
refGene_gn=PIAS3;refGene_tx=XM_011285104.3;refGene_gn=PIAS3;refGene_tx

=XM_023259074.1;refGene_gn=PIAS3;refGene_tx=XM_023259075.1
C1	104600155	104600206
refGene_gn=PIAS3;refGene_tx=XM_011285104.3;refGene_gn=PIAS3;refGene_tx=XM_023259074.1;refGene_gn=PIAS3;refGene_tx=XM_023259075.1
C1	104600428	104600513
refGene_gn=PIAS3;refGene_tx=XM_011285104.3;refGene_gn=PIAS3;refGene_tx=XM_023259074.1;refGene_gn=PIAS3;refGene_tx=XM_023259075.1
C1	104600671	104601089
refGene_gn=PIAS3;refGene_tx=XM_011285104.3;refGene_gn=PIAS3;refGene_tx=XM_023259074.1;refGene_gn=PIAS3;refGene_tx=XM_023259075.1
C1	104601310	104601406
refGene_gn=PIAS3;refGene_tx=XM_023259074.1
C1	104603077	104603101
refGene_gn=PIAS3;refGene_tx=XM_011285104.3
C1	104608743	104608806
refGene_gn=ANKRD35;refGene_tx=XM_019837748.2;refGene_gn=ANKRD35;refGene_tx=XM_023259078.1;refGene_gn=ANKRD35;refGene_tx=XM_023259079.1
C1	104609593	104609659
refGene_gn=ANKRD35;refGene_tx=XM_019837748.2;refGene_gn=ANKRD35;refGene_tx=XM_023259078.1;refGene_gn=ANKRD35;refGene_tx=XM_023259079.1
C1	104609940	104610030
refGene_gn=ANKRD35;refGene_tx=XM_019837748.2;refGene_gn=ANKRD35;refGene_tx=XM_023259078.1;refGene_gn=ANKRD35;refGene_tx=XM_023259079.1
C1	104611891	104613895
refGene_gn=ANKRD35;refGene_tx=XM_019837748.2;refGene_gn=ANKRD35;refGene_tx=XM_023259078.1;refGene_gn=ANKRD35;refGene_tx=XM_023259079.1
C1	104614072	104614110
refGene_gn=ANKRD35;refGene_tx=XM_019837748.2;refGene_gn=ANKRD35;refGene_tx=XM_023259078.1;refGene_gn=ANKRD35;refGene_tx=XM_023259079.1
C1	104614758	104614946
refGene_gn=ANKRD35;refGene_tx=XM_019837748.2;refGene_gn=ANKRD35;refGene_tx=XM_023259078.1;refGene_gn=ANKRD35;refGene_tx=XM_023259079.1
C1	104615540	104615647
refGene_gn=ANKRD35;refGene_tx=XM_019837748.2;refGene_gn=ANKRD35;refGene_tx=XM_023259078.1;refGene_gn=ANKRD35;refGene_tx=XM_023259079.1
C1	104615983	104616054
refGene_gn=ANKRD35;refGene_tx=XM_019837748.2;refGene_gn=ANKRD35;refGene_tx=XM_023259078.1;refGene_gn=ANKRD35;refGene_tx=XM_023259079.1
C1	104616243	104616301
refGene_gn=ANKRD35;refGene_tx=XM_019837748.2;refGene_gn=ANKRD35;refGene_tx=XM_023259078.1;refGene_gn=ANKRD35;refGene_tx=XM_023259079.1
C1	104617108	104617173
refGene_gn=ANKRD35;refGene_tx=XM_019837748.2;refGene_gn=ANKRD35;refGene_tx=XM_023259078.1;refGene_gn=ANKRD35;refGene_tx=XM_023259079.1
C1	104617561	104617650
refGene_gn=ANKRD35;refGene_tx=XM_019837748.2;refGene_gn=ANKRD35;refGene_tx=XM_023259078.1;refGene_gn=ANKRD35;refGene_tx=XM_023259079.1
C1	104618422	104618643
refGene_gn=ANKRD35;refGene_tx=XM_019837748.2;refGene_gn=ANKRD35;refGene_tx=XM_023259078.1;refGene_gn=ANKRD35;refGene_tx=XM_023259079.1

C1 104623816 104623855
refGene_gn=ANKRD35;refGene_tx=XM_019837748.2
C1 104638488 104638554
refGene_gn=ITGA10;refGene_tx=XM_003990541.5;refGene_gn=ITGA10;refGene_tx=XM_019837749.2;refGene_gn=ITGA10;refGene_tx=XM_019837750.2
C1 104638850 104638964
refGene_gn=ITGA10;refGene_tx=XM_003990541.5;refGene_gn=ITGA10;refGene_tx=XM_019837749.2;refGene_gn=ITGA10;refGene_tx=XM_019837750.2
C1 104639243 104639339
refGene_gn=ITGA10;refGene_tx=XM_003990541.5;refGene_gn=ITGA10;refGene_tx=XM_019837749.2;refGene_gn=ITGA10;refGene_tx=XM_019837750.2
C1 104640626 104640740
refGene_gn=ITGA10;refGene_tx=XM_003990541.5;refGene_gn=ITGA10;refGene_tx=XM_019837749.2;refGene_gn=ITGA10;refGene_tx=XM_019837750.2
C1 104640892 104640973
refGene_gn=ITGA10;refGene_tx=XM_003990541.5;refGene_gn=ITGA10;refGene_tx=XM_019837749.2;refGene_gn=ITGA10;refGene_tx=XM_019837750.2
C1 104641231 104641345
refGene_gn=ITGA10;refGene_tx=XM_003990541.5;refGene_gn=ITGA10;refGene_tx=XM_019837749.2;refGene_gn=ITGA10;refGene_tx=XM_019837750.2
C1 104641511 104641590
refGene_gn=ITGA10;refGene_tx=XM_003990541.5;refGene_gn=ITGA10;refGene_tx=XM_019837749.2;refGene_gn=ITGA10;refGene_tx=XM_019837750.2
C1 104642008 104642098
refGene_gn=ITGA10;refGene_tx=XM_003990541.5;refGene_gn=ITGA10;refGene_tx=XM_019837749.2;refGene_gn=ITGA10;refGene_tx=XM_019837750.2
C1 104642243 104642320
refGene_gn=ITGA10;refGene_tx=XM_003990541.5;refGene_gn=ITGA10;refGene_tx=XM_019837749.2;refGene_gn=ITGA10;refGene_tx=XM_019837750.2
C1 104642479 104642572
refGene_gn=ITGA10;refGene_tx=XM_003990541.5;refGene_gn=ITGA10;refGene_tx=XM_019837749.2;refGene_gn=ITGA10;refGene_tx=XM_019837750.2
C1 104642741 104642883
refGene_gn=ITGA10;refGene_tx=XM_003990541.5;refGene_gn=ITGA10;refGene_tx=XM_019837749.2;refGene_gn=ITGA10;refGene_tx=XM_019837750.2
C1 104643038 104643124
refGene_gn=ITGA10;refGene_tx=XM_003990541.5;refGene_gn=ITGA10;refGene_tx=XM_019837749.2;refGene_gn=ITGA10;refGene_tx=XM_019837750.2
C1 104643324 104643438
refGene_gn=ITGA10;refGene_tx=XM_003990541.5;refGene_gn=ITGA10;refGene_tx=XM_019837749.2;refGene_gn=ITGA10;refGene_tx=XM_019837750.2
C1 104644734 104644877
refGene_gn=ITGA10;refGene_tx=XM_003990541.5;refGene_gn=ITGA10;refGene_tx=XM_019837749.2;refGene_gn=ITGA10;refGene_tx=XM_019837750.2
C1 104644969 104645136
refGene_gn=ITGA10;refGene_tx=XM_003990541.5;refGene_gn=ITGA10;refGene_tx=XM_019837749.2;refGene_gn=ITGA10;refGene_tx=XM_019837750.2
C1 104645677 104645808
refGene_gn=ITGA10;refGene_tx=XM_003990541.5;refGene_gn=ITGA10;refGene_tx=XM_019837749.2;refGene_gn=ITGA10;refGene_tx=XM_019837750.2

C1 104646105 104646309
refGene_gn=ITGA10;refGene_tx=XM_003990541.5;refGene_gn=ITGA10;refGene_tx=XM_019837749.2;refGene_gn=ITGA10;refGene_tx=XM_019837750.2
C1 104646437 104646581
refGene_gn=ITGA10;refGene_tx=XM_003990541.5;refGene_gn=ITGA10;refGene_tx=XM_019837749.2;refGene_gn=ITGA10;refGene_tx=XM_019837750.2
C1 104646808 104646957
refGene_gn=ITGA10;refGene_tx=XM_003990541.5;refGene_gn=ITGA10;refGene_tx=XM_019837749.2;refGene_gn=ITGA10;refGene_tx=XM_019837750.2
C1 104647141 104647286
refGene_gn=ITGA10;refGene_tx=XM_003990541.5;refGene_gn=ITGA10;refGene_tx=XM_019837749.2;refGene_gn=ITGA10;refGene_tx=XM_019837750.2
C1 104647514 104647588
refGene_gn=ITGA10;refGene_tx=XM_003990541.5;refGene_gn=ITGA10;refGene_tx=XM_019837749.2;refGene_gn=ITGA10;refGene_tx=XM_019837750.2
C1 104647720 104647886
refGene_gn=ITGA10;refGene_tx=XM_003990541.5;refGene_gn=ITGA10;refGene_tx=XM_019837749.2;refGene_gn=ITGA10;refGene_tx=XM_019837750.2
C1 104648090 104648241
refGene_gn=ITGA10;refGene_tx=XM_003990541.5;refGene_gn=ITGA10;refGene_tx=XM_019837749.2;refGene_gn=ITGA10;refGene_tx=XM_019837750.2
C1 104648912 104649061
refGene_gn=ITGA10;refGene_tx=XM_003990541.5;refGene_gn=ITGA10;refGene_tx=XM_019837749.2;refGene_gn=ITGA10;refGene_tx=XM_019837750.2
C1 104649227 104649355
refGene_gn=ITGA10;refGene_tx=XM_003990541.5;refGene_gn=ITGA10;refGene_tx=XM_019837749.2;refGene_gn=ITGA10;refGene_tx=XM_019837750.2
C1 104650341 104650456
refGene_gn=ITGA10;refGene_tx=XM_003990541.5;refGene_gn=ITGA10;refGene_tx=XM_019837749.2;refGene_gn=ITGA10;refGene_tx=XM_019837750.2
C1 104650680 104650772
refGene_gn=ITGA10;refGene_tx=XM_003990541.5;refGene_gn=ITGA10;refGene_tx=XM_019837749.2;refGene_gn=ITGA10;refGene_tx=XM_019837750.2
C1 104650979 104651089
refGene_gn=ITGA10;refGene_tx=XM_003990541.5;refGene_gn=ITGA10;refGene_tx=XM_019837749.2;refGene_gn=ITGA10;refGene_tx=XM_019837750.2
C1 104651277 104651389
refGene_gn=ITGA10;refGene_tx=XM_003990541.5;refGene_gn=ITGA10;refGene_tx=XM_019837749.2
C1 104653018 104653148
refGene_gn=ITGA10;refGene_tx=XM_003990541.5;refGene_gn=ITGA10;refGene_tx=XM_019837749.2
C1 104654733 104655139
refGene_gn=PEX11B;refGene_tx=XM_003990545.5;refGene_gn=PEX11B;refGene_tx=XM_003990546.5
C1 104657713 104657915
refGene_gn=PEX11B;refGene_tx=XM_003990545.5;refGene_gn=PEX11B;refGene_tx=XM_003990546.5
C1 104658314 104658430
refGene_gn=PEX11B;refGene_tx=XM_003990545.5;refGene_gn=PEX11B;refGene_

tx=XM_003990546.5
C1	104659045	104659059
refGene_gn=PEX11B;refGene_tx=XM_003990546.5
C1	104659230	104659286
refGene_gn=PEX11B;refGene_tx=XM_003990545.5
C1	104665812	104665858
refGene_gn=RBM8A;refGene_tx=XM_003990544.5;refGene_gn=RBM8A;refGene_tx
=XM_006935061.4
C1	104665958	104666095
refGene_gn=RBM8A;refGene_tx=XM_003990544.5;refGene_gn=RBM8A;refGene_tx
=XM_006935061.4
C1	104666385	104666522
refGene_gn=RBM8A;refGene_tx=XM_003990544.5;refGene_gn=RBM8A;refGene_tx
=XM_006935061.4
C1	104666706	104666784
refGene_gn=RBM8A;refGene_tx=XM_003990544.5;refGene_gn=RBM8A;refGene_tx
=XM_006935061.4
C1	104666909	104666969
refGene_gn=RBM8A;refGene_tx=XM_003990544.5;refGene_gn=RBM8A;refGene_tx
=XM_006935061.4
C1	104667245	104667312
refGene_gn=RBM8A;refGene_tx=XM_003990544.5;refGene_gn=RBM8A;refGene_tx
=XM_006935061.4
C1	104672185	104672428
refGene_gn=LIX1L;refGene_tx=XM_023259082.1
C1	104672764	104672842
refGene_gn=LIX1L;refGene_tx=XM_023259082.1
C1	104673387	104673483
refGene_gn=LIX1L;refGene_tx=XM_023259082.1
C1	104677564	104677705
refGene_gn=LIX1L;refGene_tx=XM_023259082.1
C1	104680192	104680314
refGene_gn=LIX1L;refGene_tx=XM_023259082.1
C1	104680316	104680355
refGene_gn=LIX1L;refGene_tx=XM_023259082.1
C1	104689813	104690105
refGene_gn=LIX1L;refGene_tx=XM_023259082.1
C1	104692501	104694022
refGene_gn=ANKRD34A;refGene_tx=XM_011285106.2;refGene_gn=ANKRD34A;refG
ene_tx=XM_011285107.2;refGene_gn=ANKRD34A;refGene_tx=XM_019837751.1;re
fGene_gn=ANKRD34A;refGene_tx=XM_019837752.1;refGene_gn=ANKRD34A;refGen
e_tx=XM_019837753.1;refGene_gn=ANKRD34A;refGene_tx=XM_019837754.1
C1	104702691	104702817
refGene_gn=POLR3GL;refGene_tx=XM_003990548.5;refGene_gn=POLR3GL;refGen
e_tx=XM_019837755.2
C1	104703137	104703267
refGene_gn=POLR3GL;refGene_tx=XM_003990548.5;refGene_gn=POLR3GL;refGen
e_tx=XM_019837755.2
C1	104703973	104704042
refGene_gn=POLR3GL;refGene_tx=XM_003990548.5

C1	104704385	104704442
refGene_gn=POLR3GL;refGene_tx=XM_003990548.5;refGene_gn=POLR3GL;refGene_tx=XM_019837755.2
C1	104704679	104704753
refGene_gn=POLR3GL;refGene_tx=XM_003990548.5;refGene_gn=POLR3GL;refGene_tx=XM_019837755.2
C1	104704884	104704998
refGene_gn=POLR3GL;refGene_tx=XM_003990548.5;refGene_gn=POLR3GL;refGene_tx=XM_019837755.2
C1	104705304	104705391
refGene_gn=POLR3GL;refGene_tx=XM_003990548.5;refGene_gn=POLR3GL;refGene_tx=XM_019837755.2
C1	104718284	104718320
refGene_gn=TXNIP;refGene_tx=XM_003990549.5
C1	104718471	104718623
refGene_gn=TXNIP;refGene_tx=XM_003990549.5
C1	104718735	104718892
refGene_gn=TXNIP;refGene_tx=XM_003990549.5
C1	104718988	104719245
refGene_gn=TXNIP;refGene_tx=XM_003990549.5
C1	104719387	104719490
refGene_gn=TXNIP;refGene_tx=XM_003990549.5
C1	104719700	104719848
refGene_gn=TXNIP;refGene_tx=XM_003990549.5
C1	104719970	104720043
refGene_gn=TXNIP;refGene_tx=XM_003990549.5
C1	104720548	104720798
refGene_gn=TXNIP;refGene_tx=XM_003990549.5
C1	104733778	104734405
refGene_gn=HFE2;refGene_tx=XM_011285108.3;refGene_gn=HFE2;refGene_tx=XM_023259083.1;refGene_gn=HFE2;refGene_tx=XM_023259084.1;refGene_gn=HFE2;refGene_tx=XM_023259085.1
C1	104734848	104735469
refGene_gn=HFE2;refGene_tx=XM_011285108.3;refGene_gn=HFE2;refGene_tx=XM_023259083.1;refGene_gn=HFE2;refGene_tx=XM_023259084.1;refGene_gn=HFE2;refGene_tx=XM_023259085.1
C1	104735839	104735921
refGene_gn=HFE2;refGene_tx=XM_023259083.1;refGene_gn=HFE2;refGene_tx=XM_023259084.1;refGene_gn=HFE2;refGene_tx=XM_023259085.1
C1	104811659	104811784
refGene_gn=LOC111561813;refGene_tx=XM_023259086.1;refGene_gn=LOC111561813;refGene_tx=XM_023259087.1;refGene_gn=LOC111561813;refGene_tx=XM_023259088.1;refGene_gn=LOC111561813;refGene_tx=XM_023259089.1;refGene_gn=LOC111561813;refGene_tx=XM_023259090.1;refGene_gn=LOC111561813;refGene_tx=XM_023259091.1;refGene_gn=LOC111561813;refGene_tx=XM_023259092.1;refGene_gn=LOC111561813;refGene_tx=XM_023259093.1
C1	104919893	104919904
refGene_gn=LOC101087418;refGene_tx=XM_006935063.4;refGene_gn=LOC101087418;refGene_tx=XM_006935065.4;refGene_gn=LOC101087418;refGene_tx=XM_019837756.2;refGene_gn=LOC101087418;refGene_tx=XM_019837757.2;refGene_gn

=LOC101087418;refGene_tx=XM_019837759.2
C1	104925307	104925427
refGene_gn=LOC101087418;refGene_tx=XM_006935063.4;refGene_gn=LOC101087418;refGene_tx=XM_006935065.4;refGene_gn=LOC101087418;refGene_tx=XM_019837756.2;refGene_gn=LOC101087418;refGene_tx=XM_019837757.2;refGene_gn=LOC101087418;refGene_tx=XM_019837759.2
C1	104926208	104926364
refGene_gn=LOC101087418;refGene_tx=XM_006935063.4;refGene_gn=LOC101087418;refGene_tx=XM_019837756.2;refGene_gn=LOC101087418;refGene_tx=XM_019837757.2;refGene_gn=LOC101087418;refGene_tx=XM_019837759.2
C1	104927866	104928030
refGene_gn=LOC101087418;refGene_tx=XM_006935063.4;refGene_gn=LOC101087418;refGene_tx=XM_006935065.4;refGene_gn=LOC101087418;refGene_tx=XM_019837756.2;refGene_gn=LOC101087418;refGene_tx=XM_019837757.2;refGene_gn=LOC101087418;refGene_tx=XM_019837759.2
C1	104930315	104930367
refGene_gn=LOC101087418;refGene_tx=XM_006935063.4;refGene_gn=LOC101087418;refGene_tx=XM_006935065.4;refGene_gn=LOC101087418;refGene_tx=XM_019837756.2;refGene_gn=LOC101087418;refGene_tx=XM_019837757.2;refGene_gn=LOC101087418;refGene_tx=XM_019837759.2
C1	104933877	104934113
refGene_gn=LOC101087418;refGene_tx=XM_006935063.4;refGene_gn=LOC101087418;refGene_tx=XM_006935065.4;refGene_gn=LOC101087418;refGene_tx=XM_019837756.2;refGene_gn=LOC101087418;refGene_tx=XM_019837757.2;refGene_gn=LOC101087418;refGene_tx=XM_019837759.2
C1	104935975	104936075
refGene_gn=LOC101087418;refGene_tx=XM_006935063.4;refGene_gn=LOC101087418;refGene_tx=XM_006935065.4;refGene_gn=LOC101087418;refGene_tx=XM_019837756.2;refGene_gn=LOC101087418;refGene_tx=XM_019837757.2;refGene_gn=LOC101087418;refGene_tx=XM_019837759.2
C1	104937983	104938161
refGene_gn=LOC101087418;refGene_tx=XM_006935063.4;refGene_gn=LOC101087418;refGene_tx=XM_006935065.4;refGene_gn=LOC101087418;refGene_tx=XM_019837756.2;refGene_gn=LOC101087418;refGene_tx=XM_019837757.2;refGene_gn=LOC101087418;refGene_tx=XM_019837759.2
C1	104955666	104955796
refGene_gn=LOC111561813;refGene_tx=XM_023259086.1;refGene_gn=LOC111561813;refGene_tx=XM_023259087.1;refGene_gn=LOC111561813;refGene_tx=XM_023259088.1;refGene_gn=LOC111561813;refGene_tx=XM_023259089.1;refGene_gn=LOC111561813;refGene_tx=XM_023259090.1;refGene_gn=LOC111561813;refGene_tx=XM_023259091.1;refGene_gn=LOC111561813;refGene_tx=XM_023259092.1;refGene_gn=LOC111561813;refGene_tx=XM_023259093.1
C1	104955904	104956378
refGene_gn=LOC111561813;refGene_tx=XM_023259086.1;refGene_gn=LOC111561813;refGene_tx=XM_023259087.1;refGene_gn=LOC111561813;refGene_tx=XM_023259088.1;refGene_gn=LOC111561813;refGene_tx=XM_023259089.1;refGene_gn=LOC111561813;refGene_tx=XM_023259090.1;refGene_gn=LOC111561813;refGene_tx=XM_023259091.1;refGene_gn=LOC111561813;refGene_tx=XM_023259092.1
C1	104958013	104958033
refGene_gn=LOC111561813;refGene_tx=XM_023259092.1

C1	104981858	104982050
refGene_gn=LOC111561813;refGene_tx=XM_023259093.1
C1	105016438	105016530
refGene_gn=LOC101100497;refGene_tx=XM_023259094.1;refGene_gn=LOC101100497;refGene_tx=XM_023259095.1;refGene_gn=LOC101100497;refGene_tx=XM_023259096.1;refGene_gn=LOC101100497;refGene_tx=XM_023259097.1;refGene_gn=LOC101100497;refGene_tx=XM_023259098.1
C1	105016843	105016918
refGene_gn=LOC101100497;refGene_tx=XM_023259094.1;refGene_gn=LOC101100497;refGene_tx=XM_023259095.1;refGene_gn=LOC101100497;refGene_tx=XM_023259096.1;refGene_gn=LOC101100497;refGene_tx=XM_023259098.1
C1	105017042	105017068
refGene_gn=LOC101100497;refGene_tx=XM_023259099.1
C1	105017816	105017948
refGene_gn=LOC101100497;refGene_tx=XM_023259094.1;refGene_gn=LOC101100497;refGene_tx=XM_023259095.1;refGene_gn=LOC101100497;refGene_tx=XM_023259096.1;refGene_gn=LOC101100497;refGene_tx=XM_023259097.1;refGene_gn=LOC101100497;refGene_tx=XM_023259098.1;refGene_gn=LOC101100497;refGene_tx=XM_023259099.1
C1	105019241	105019475
refGene_gn=LOC101100497;refGene_tx=XM_023259094.1;refGene_gn=LOC101100497;refGene_tx=XM_023259095.1;refGene_gn=LOC101100497;refGene_tx=XM_023259096.1;refGene_gn=LOC101100497;refGene_tx=XM_023259097.1;refGene_gn=LOC101100497;refGene_tx=XM_023259098.1;refGene_gn=LOC101100497;refGene_tx=XM_023259099.1
C1	105023190	105023489
refGene_gn=LOC101100497;refGene_tx=XM_023259100.1
C1	105023920	105024108
refGene_gn=LOC101100497;refGene_tx=XM_023259094.1;refGene_gn=LOC101100497;refGene_tx=XM_023259095.1;refGene_gn=LOC101100497;refGene_tx=XM_023259096.1;refGene_gn=LOC101100497;refGene_tx=XM_023259097.1;refGene_gn=LOC101100497;refGene_tx=XM_023259098.1;refGene_gn=LOC101100497;refGene_tx=XM_023259099.1;refGene_gn=LOC101100497;refGene_tx=XM_023259100.1
C1	105025145	105025197
refGene_gn=LOC101100497;refGene_tx=XM_023259094.1;refGene_gn=LOC101100497;refGene_tx=XM_023259095.1;refGene_gn=LOC101100497;refGene_tx=XM_023259096.1;refGene_gn=LOC101100497;refGene_tx=XM_023259097.1;refGene_gn=LOC101100497;refGene_tx=XM_023259098.1;refGene_gn=LOC101100497;refGene_tx=XM_023259099.1;refGene_gn=LOC101100497;refGene_tx=XM_023259100.1
C1	105026293	105026529
refGene_gn=LOC101100497;refGene_tx=XM_023259094.1;refGene_gn=LOC101100497;refGene_tx=XM_023259095.1;refGene_gn=LOC101100497;refGene_tx=XM_023259096.1;refGene_gn=LOC101100497;refGene_tx=XM_023259097.1;refGene_gn=LOC101100497;refGene_tx=XM_023259098.1;refGene_gn=LOC101100497;refGene_tx=XM_023259099.1;refGene_gn=LOC101100497;refGene_tx=XM_023259100.1
C1	105028704	105028794
refGene_gn=LOC101100497;refGene_tx=XM_023259094.1;refGene_gn=LOC101100497;refGene_tx=XM_023259095.1;refGene_gn=LOC101100497;refGene_tx=XM_023259096.1;refGene_gn=LOC101100497;refGene_tx=XM_023259097.1;refGene_gn=LOC101100497;refGene_tx=XM_023259099.1;refGene_gn=LOC101100497;refGen

e_tx=XM_023259100.1
C1	105036948	105037067
refGene_gn=LOC101100497;refGene_tx=XM_023259094.1;refGene_gn=LOC101100497;refGene_tx=XM_023259095.1;refGene_gn=LOC101100497;refGene_tx=XM_023259096.1;refGene_gn=LOC101100497;refGene_tx=XM_023259097.1;refGene_gn=LOC101100497;refGene_tx=XM_023259098.1;refGene_gn=LOC101100497;refGene_tx=XM_023259099.1;refGene_gn=LOC101100497;refGene_tx=XM_023259100.1
C1	105084605	105084636
refGene_gn=LOC101087936;refGene_tx=XM_023259101.1
C1	105085040	105085061
refGene_gn=LOC101087936;refGene_tx=XM_023259101.1
C1	105085792	105086047
refGene_gn=LOC101087936;refGene_tx=XM_023259101.1
C1	105091193	105091445
refGene_gn=LOC101087936;refGene_tx=XM_023259101.1
C1	105092863	105093148
refGene_gn=LOC101087936;refGene_tx=XM_023259101.1
C1	105094144	105094347
refGene_gn=LOC101087936;refGene_tx=XM_023259101.1
C1	105094349	105094427
refGene_gn=LOC101087936;refGene_tx=XM_023259101.1
C1	105116764	105117145
refGene_gn=LOC101100989;refGene_tx=XM_006935069.4
C1	105127254	105127566
refGene_gn=LOC101088423;refGene_tx=XM_006935068.4
C1	105143870	105144149
refGene_gn=LOC101080314;refGene_tx=XM_019839103.2
C1	105144151	105144259
refGene_gn=LOC101080314;refGene_tx=XM_019839103.2
C1	105153966	105154347
refGene_gn=LOC101088681;refGene_tx=XM_019837760.2
C1	105154702	105155095
refGene_gn=LOC101101494;refGene_tx=XM_003990554.5
C1	105155993	105156569
refGene_gn=LOC101101243;refGene_tx=XM_023259102.1
C1	105180811	105181192
refGene_gn=LOC101080561;refGene_tx=XM_003990555.5
C1	105181519	105181852
refGene_gn=LOC101080814;refGene_tx=XM_003990556.5
C1	105182066	105182459
refGene_gn=LOC101081066;refGene_tx=XM_003990557.5
C1	105193216	105193630
refGene_gn=BOLA1;refGene_tx=XM_003990558.4;refGene_gn=BOLA1;refGene_tx=XM_006935070.2
C1	105198432	105198616
refGene_gn=SV2A;refGene_tx=XM_003990559.5
C1	105199016	105199176
refGene_gn=SV2A;refGene_tx=XM_003990559.5
C1	105199595	105199697
refGene_gn=BOLA1;refGene_tx=XM_006935070.2

C1 105199779 105199986
refGene_gn=SV2A;refGene_tx=XM_003990559.5
C1 105200569 105200584
refGene_gn=BOLA1;refGene_tx=XM_006935070.2
C1 105200883 105201017
refGene_gn=SV2A;refGene_tx=XM_003990559.5
C1 105201209 105201374
refGene_gn=SV2A;refGene_tx=XM_003990559.5
C1 105201676 105201765
refGene_gn=SV2A;refGene_tx=XM_003990559.5
C1 105201935 105202046
refGene_gn=SV2A;refGene_tx=XM_003990559.5
C1 105202285 105202375
refGene_gn=SV2A;refGene_tx=XM_003990559.5
C1 105203011 105203145
refGene_gn=SV2A;refGene_tx=XM_003990559.5
C1 105203256 105203408
refGene_gn=SV2A;refGene_tx=XM_003990559.5
C1 105204238 105204419
refGene_gn=SV2A;refGene_tx=XM_003990559.5
C1 105205711 105206333
refGene_gn=SV2A;refGene_tx=XM_003990559.5
C1 105220620 105220808
refGene_gn=SF3B4;refGene_tx=XM_004001496.4;refGene_gn=SF3B4;refGene_tx
=XM_019837764.1
C1 105221017 105221191
refGene_gn=SF3B4;refGene_tx=XM_004001496.4;refGene_gn=SF3B4;refGene_tx
=XM_019837764.1
C1 105222868 105223075
refGene_gn=SF3B4;refGene_tx=XM_004001496.4;refGene_gn=SF3B4;refGene_tx
=XM_019837764.1
C1 105223399 105223942
refGene_gn=SF3B4;refGene_tx=XM_004001496.4;refGene_gn=SF3B4;refGene_tx
=XM_019837764.1
C1 105224409 105224538
refGene_gn=SF3B4;refGene_tx=XM_004001496.4;refGene_gn=SF3B4;refGene_tx
=XM_019837764.1
C1 105225010 105225044
refGene_gn=SF3B4;refGene_tx=XM_004001496.4;refGene_gn=SF3B4;refGene_tx
=XM_019837764.1
C1 105226685 105226832
refGene_gn=MTMR11;refGene_tx=XM_006935071.4;refGene_gn=MTMR11;refGene_
tx=XM_011285113.3;refGene_gn=MTMR11;refGene_tx=XM_019837761.2;refGene_
gn=MTMR11;refGene_tx=XM_019837762.2;refGene_gn=MTMR11;refGene_tx=XM_01
9837763.1
C1 105227300 105227594
refGene_gn=MTMR11;refGene_tx=XM_006935071.4;refGene_gn=MTMR11;refGene_
tx=XM_011285113.3;refGene_gn=MTMR11;refGene_tx=XM_019837761.2;refGene_
gn=MTMR11;refGene_tx=XM_019837762.2;refGene_gn=MTMR11;refGene_tx=XM_01
9837763.1

C1	105228092	105228275
refGene_gn=MTMR11;refGene_tx=XM_006935071.4;refGene_gn=MTMR11;refGene_tx=XM_011285113.3;refGene_gn=MTMR11;refGene_tx=XM_019837761.2;refGene_gn=MTMR11;refGene_tx=XM_019837762.2;refGene_gn=MTMR11;refGene_tx=XM_019837763.1
C1	105228526	105228700
refGene_gn=MTMR11;refGene_tx=XM_006935071.4;refGene_gn=MTMR11;refGene_tx=XM_011285113.3;refGene_gn=MTMR11;refGene_tx=XM_019837761.2;refGene_gn=MTMR11;refGene_tx=XM_019837762.2;refGene_gn=MTMR11;refGene_tx=XM_019837763.1
C1	105228997	105229164
refGene_gn=MTMR11;refGene_tx=XM_006935071.4;refGene_gn=MTMR11;refGene_tx=XM_011285113.3;refGene_gn=MTMR11;refGene_tx=XM_019837761.2;refGene_gn=MTMR11;refGene_tx=XM_019837762.2;refGene_gn=MTMR11;refGene_tx=XM_019837763.1
C1	105229711	105229782
refGene_gn=MTMR11;refGene_tx=XM_006935071.4;refGene_gn=MTMR11;refGene_tx=XM_011285113.3;refGene_gn=MTMR11;refGene_tx=XM_019837761.2;refGene_gn=MTMR11;refGene_tx=XM_019837762.2;refGene_gn=MTMR11;refGene_tx=XM_019837763.1
C1	105230027	105230094
refGene_gn=MTMR11;refGene_tx=XM_006935071.4;refGene_gn=MTMR11;refGene_tx=XM_011285113.3;refGene_gn=MTMR11;refGene_tx=XM_019837761.2;refGene_gn=MTMR11;refGene_tx=XM_019837762.2;refGene_gn=MTMR11;refGene_tx=XM_019837763.1
C1	105230485	105230610
refGene_gn=MTMR11;refGene_tx=XM_006935071.4;refGene_gn=MTMR11;refGene_tx=XM_011285113.3;refGene_gn=MTMR11;refGene_tx=XM_019837761.2;refGene_gn=MTMR11;refGene_tx=XM_019837762.2;refGene_gn=MTMR11;refGene_tx=XM_019837763.1
C1	105230695	105230784
refGene_gn=MTMR11;refGene_tx=XM_006935071.4;refGene_gn=MTMR11;refGene_tx=XM_011285113.3;refGene_gn=MTMR11;refGene_tx=XM_019837761.2;refGene_gn=MTMR11;refGene_tx=XM_019837762.2;refGene_gn=MTMR11;refGene_tx=XM_019837763.1
C1	105230939	105231027
refGene_gn=MTMR11;refGene_tx=XM_006935071.4;refGene_gn=MTMR11;refGene_tx=XM_011285113.3;refGene_gn=MTMR11;refGene_tx=XM_019837761.2;refGene_gn=MTMR11;refGene_tx=XM_019837762.2;refGene_gn=MTMR11;refGene_tx=XM_019837763.1
C1	105231276	105231412
refGene_gn=MTMR11;refGene_tx=XM_006935071.4;refGene_gn=MTMR11;refGene_tx=XM_011285113.3;refGene_gn=MTMR11;refGene_tx=XM_019837761.2;refGene_gn=MTMR11;refGene_tx=XM_019837762.2;refGene_gn=MTMR11;refGene_tx=XM_019837763.1
C1	105231543	105231622
refGene_gn=MTMR11;refGene_tx=XM_006935071.4;refGene_gn=MTMR11;refGene_tx=XM_011285113.3;refGene_gn=MTMR11;refGene_tx=XM_019837761.2;refGene_gn=MTMR11;refGene_tx=XM_019837762.2;refGene_gn=MTMR11;refGene_tx=XM_019837763.1

C1	105232092	105232235
refGene_gn=MTMR11;refGene_tx=XM_006935071.4;refGene_gn=MTMR11;refGene_tx=XM_011285113.3;refGene_gn=MTMR11;refGene_tx=XM_019837761.2;refGene_gn=MTMR11;refGene_tx=XM_019837762.2;refGene_gn=MTMR11;refGene_tx=XM_019837763.1
C1	105232392	105232453
refGene_gn=MTMR11;refGene_tx=XM_006935071.4;refGene_gn=MTMR11;refGene_tx=XM_011285113.3;refGene_gn=MTMR11;refGene_tx=XM_019837761.2;refGene_gn=MTMR11;refGene_tx=XM_019837762.2
C1	105232676	105232798
refGene_gn=MTMR11;refGene_tx=XM_006935071.4;refGene_gn=MTMR11;refGene_tx=XM_011285113.3;refGene_gn=MTMR11;refGene_tx=XM_019837761.2;refGene_gn=MTMR11;refGene_tx=XM_019837762.2;refGene_gn=MTMR11;refGene_tx=XM_019837763.1
C1	105233215	105233306
refGene_gn=MTMR11;refGene_tx=XM_006935071.4;refGene_gn=MTMR11;refGene_tx=XM_011285113.3;refGene_gn=MTMR11;refGene_tx=XM_019837761.2;refGene_gn=MTMR11;refGene_tx=XM_019837762.2
C1	105233666	105233732
refGene_gn=MTMR11;refGene_tx=XM_006935071.4;refGene_gn=MTMR11;refGene_tx=XM_011285113.3;refGene_gn=MTMR11;refGene_tx=XM_019837761.2;refGene_gn=MTMR11;refGene_tx=XM_019837762.2
C1	105240018	105241227
refGene_gn=OTUD7B;refGene_tx=XM_011285114.2;refGene_gn=OTUD7B;refGene_tx=XM_023259104.1
C1	105242209	105242294
refGene_gn=OTUD7B;refGene_tx=XM_011285114.2;refGene_gn=OTUD7B;refGene_tx=XM_023259104.1
C1	105244087	105244202
refGene_gn=OTUD7B;refGene_tx=XM_011285114.2;refGene_gn=OTUD7B;refGene_tx=XM_023259104.1
C1	105244793	105244943
refGene_gn=OTUD7B;refGene_tx=XM_011285114.2;refGene_gn=OTUD7B;refGene_tx=XM_023259104.1
C1	105245247	105245375
refGene_gn=OTUD7B;refGene_tx=XM_011285114.2;refGene_gn=OTUD7B;refGene_tx=XM_023259104.1
C1	105247284	105247397
refGene_gn=OTUD7B;refGene_tx=XM_011285114.2;refGene_gn=OTUD7B;refGene_tx=XM_023259104.1
C1	105251380	105251508
refGene_gn=OTUD7B;refGene_tx=XM_011285114.2;refGene_gn=OTUD7B;refGene_tx=XM_023259104.1
C1	105252314	105252416
refGene_gn=OTUD7B;refGene_tx=XM_011285114.2;refGene_gn=OTUD7B;refGene_tx=XM_023259104.1
C1	105253872	105254100
refGene_gn=OTUD7B;refGene_tx=XM_011285114.2;refGene_gn=OTUD7B;refGene_tx=XM_023259104.1
C1	105255915	105256101

refGene_gn=OTUD7B;refGene_tx=XM_011285114.2;refGene_gn=OTUD7B;refGene_
tx=XM_023259104.1
C1 105260820 105260905
refGene_gn=OTUD7B;refGene_tx=XM_011285114.2;refGene_gn=OTUD7B;refGene_
tx=XM_023259104.1
C1 105346833 105346926
refGene_gn=VPS45;refGene_tx=XM_003990560.5;refGene_gn=VPS45;refGene_tx
=XM_006935074.4;refGene_gn=VPS45;refGene_tx=XM_019837765.2
C1 105347561 105347696
refGene_gn=VPS45;refGene_tx=XM_003990560.5;refGene_gn=VPS45;refGene_tx
=XM_006935074.4;refGene_gn=VPS45;refGene_tx=XM_019837765.2
C1 105350848 105350909
refGene_gn=VPS45;refGene_tx=XM_003990560.5;refGene_gn=VPS45;refGene_tx
=XM_006935074.4;refGene_gn=VPS45;refGene_tx=XM_019837765.2
C1 105353835 105353915
refGene_gn=VPS45;refGene_tx=XM_003990560.5;refGene_gn=VPS45;refGene_tx
=XM_006935074.4;refGene_gn=VPS45;refGene_tx=XM_019837765.2
C1 105355213 105355282
refGene_gn=VPS45;refGene_tx=XM_003990560.5;refGene_gn=VPS45;refGene_tx
=XM_006935074.4;refGene_gn=VPS45;refGene_tx=XM_019837765.2
C1 105355399 105355537
refGene_gn=VPS45;refGene_tx=XM_003990560.5;refGene_gn=VPS45;refGene_tx
=XM_006935074.4;refGene_gn=VPS45;refGene_tx=XM_019837765.2
C1 105355968 105356079
refGene_gn=VPS45;refGene_tx=XM_003990560.5;refGene_gn=VPS45;refGene_tx
=XM_006935074.4;refGene_gn=VPS45;refGene_tx=XM_019837765.2
C1 105358363 105358498
refGene_gn=VPS45;refGene_tx=XM_003990560.5;refGene_gn=VPS45;refGene_tx
=XM_006935074.4;refGene_gn=VPS45;refGene_tx=XM_019837765.2
C1 105359072 105359186
refGene_gn=VPS45;refGene_tx=XM_003990560.5;refGene_gn=VPS45;refGene_tx
=XM_006935074.4;refGene_gn=VPS45;refGene_tx=XM_019837765.2
C1 105359582 105359750
refGene_gn=VPS45;refGene_tx=XM_003990560.5;refGene_gn=VPS45;refGene_tx
=XM_006935074.4;refGene_gn=VPS45;refGene_tx=XM_019837765.2
C1 105368756 105368915
refGene_gn=VPS45;refGene_tx=XM_003990560.5;refGene_gn=VPS45;refGene_tx
=XM_006935074.4;refGene_gn=VPS45;refGene_tx=XM_019837765.2
C1 105369118 105369226
refGene_gn=VPS45;refGene_tx=XM_003990560.5;refGene_gn=VPS45;refGene_tx
=XM_006935074.4;refGene_gn=VPS45;refGene_tx=XM_019837765.2
C1 105370040 105370162
refGene_gn=VPS45;refGene_tx=XM_003990560.5;refGene_gn=VPS45;refGene_tx
=XM_006935074.4;refGene_gn=VPS45;refGene_tx=XM_019837765.2
C1 105386761 105386893
refGene_gn=VPS45;refGene_tx=XM_003990560.5;refGene_gn=VPS45;refGene_tx
=XM_006935074.4;refGene_gn=VPS45;refGene_tx=XM_019837765.2
C1 105396711 105396760
refGene_gn=VPS45;refGene_tx=XM_019837765.2
C1 105400989 105401008

refGene_gn=VPS45;refGene_tx=XM_006935074.4
C1 105411738 105411826
refGene_gn=VPS45;refGene_tx=XM_003990560.5
C1 105417972 105417999
refGene_gn=PLEKHO1;refGene_tx=XM_023259107.1
C1 105418574 105418721
refGene_gn=PLEKHO1;refGene_tx=XM_023259107.1
C1 105422905 105423049
refGene_gn=PLEKHO1;refGene_tx=XM_023259107.1
C1 105423773 105423878
refGene_gn=PLEKHO1;refGene_tx=XM_023259107.1
C1 105424223 105424349
refGene_gn=PLEKHO1;refGene_tx=XM_019837766.2;refGene_gn=PLEKHO1;refGen
e_tx=XM_019837767.1;refGene_gn=PLEKHO1;refGene_tx=XM_023259106.1;refGe
ne_gn=PLEKHO1;refGene_tx=XM_023259107.1
C1 105425449 105426154
refGene_gn=PLEKHO1;refGene_tx=XM_019837766.2;refGene_gn=PLEKHO1;refGen
e_tx=XM_019837767.1;refGene_gn=PLEKHO1;refGene_tx=XM_023259106.1;refGe
ne_gn=PLEKHO1;refGene_tx=XM_023259107.1
C1 105479747 105479818
refGene_gn=ANP32E;refGene_tx=XM_011285116.3
C1 105481079 105481134
refGene_gn=ANP32E;refGene_tx=XM_011285116.3
C1 105487158 105487343
refGene_gn=ANP32E;refGene_tx=XM_011285116.3;refGene_gn=ANP32E;refGene_
tx=XM_023259110.1
C1 105489071 105489237
refGene_gn=ANP32E;refGene_tx=XM_011285116.3
C1 105490002 105490125
refGene_gn=ANP32E;refGene_tx=XM_011285116.3;refGene_gn=ANP32E;refGene_
tx=XM_023259110.1
C1 105490596 105490746
refGene_gn=ANP32E;refGene_tx=XM_011285116.3;refGene_gn=ANP32E;refGene_
tx=XM_023259110.1
C1 105493827 105493881
refGene_gn=ANP32E;refGene_tx=XM_011285116.3;refGene_gn=ANP32E;refGene_
tx=XM_023259110.1
C1 105514021 105514076
refGene_gn=CA14;refGene_tx=XM_019837770.2;refGene_gn=CA14;refGene_tx=X
M_023259113.1;refGene_gn=CA14;refGene_tx=XM_023259114.1
C1 105516104 105516125
refGene_gn=CA14;refGene_tx=XM_019837770.2;refGene_gn=CA14;refGene_tx=X
M_023259113.1;refGene_gn=CA14;refGene_tx=XM_023259114.1
C1 105516195 105516202
refGene_gn=CA14;refGene_tx=XM_019837771.1
C1 105516995 105517175
refGene_gn=CA14;refGene_tx=XM_019837770.2;refGene_gn=CA14;refGene_tx=X
M_019837771.1;refGene_gn=CA14;refGene_tx=XM_023259113.1;refGene_gn=CA1
4;refGene_tx=XM_023259114.1
C1 105517547 105517690

```
refGene_gn=CA14;refGene_tx=XM_019837770.2;refGene_gn=CA14;refGene_tx=X
M_019837771.1;refGene_gn=CA14;refGene_tx=XM_019837773.1;refGene_gn=CA1
4;refGene_tx=XM_023259113.1;refGene_gn=CA14;refGene_tx=XM_023259114.1
C1	105517954	105518050
refGene_gn=CA14;refGene_tx=XM_019837770.2;refGene_gn=CA14;refGene_tx=X
M_019837771.1;refGene_gn=CA14;refGene_tx=XM_019837773.1;refGene_gn=CA1
4;refGene_tx=XM_023259113.1;refGene_gn=CA14;refGene_tx=XM_023259114.1
C1	105518142	105518296
refGene_gn=CA14;refGene_tx=XM_019837770.2;refGene_gn=CA14;refGene_tx=X
M_019837771.1;refGene_gn=CA14;refGene_tx=XM_019837773.1;refGene_gn=CA1
4;refGene_tx=XM_023259113.1;refGene_gn=CA14;refGene_tx=XM_023259114.1
C1	105518458	105518616
refGene_gn=CA14;refGene_tx=XM_019837770.2;refGene_gn=CA14;refGene_tx=X
M_019837771.1;refGene_gn=CA14;refGene_tx=XM_019837773.1;refGene_gn=CA1
4;refGene_tx=XM_023259113.1;refGene_gn=CA14;refGene_tx=XM_023259114.1
C1	105518950	105519071
refGene_gn=CA14;refGene_tx=XM_019837770.2;refGene_gn=CA14;refGene_tx=X
M_019837771.1;refGene_gn=CA14;refGene_tx=XM_019837773.1;refGene_gn=CA1
4;refGene_tx=XM_023259113.1;refGene_gn=CA14;refGene_tx=XM_023259114.1
C1	105519311	105519332
refGene_gn=CA14;refGene_tx=XM_019837770.2;refGene_gn=CA14;refGene_tx=X
M_019837771.1;refGene_gn=CA14;refGene_tx=XM_019837773.1;refGene_gn=CA1
4;refGene_tx=XM_023259113.1;refGene_gn=CA14;refGene_tx=XM_023259114.1
C1	105519449	105519534
refGene_gn=CA14;refGene_tx=XM_019837770.2;refGene_gn=CA14;refGene_tx=X
M_019837771.1;refGene_gn=CA14;refGene_tx=XM_019837773.1;refGene_gn=CA1
4;refGene_tx=XM_023259113.1;refGene_gn=CA14;refGene_tx=XM_023259114.1
C1	105519653	105519717
refGene_gn=CA14;refGene_tx=XM_019837770.2;refGene_gn=CA14;refGene_tx=X
M_019837771.1;refGene_gn=CA14;refGene_tx=XM_019837773.1;refGene_gn=CA1
4;refGene_tx=XM_023259113.1
C1	105521228	105521293
refGene_gn=APH1A;refGene_tx=XM_003990565.5
C1	105521630	105521754
refGene_gn=APH1A;refGene_tx=XM_003990565.5
C1	105522161	105522289
refGene_gn=APH1A;refGene_tx=XM_003990565.5
C1	105522442	105522565
refGene_gn=APH1A;refGene_tx=XM_003990565.5
C1	105522799	105522873
refGene_gn=APH1A;refGene_tx=XM_003990565.5
C1	105523050	105523221
refGene_gn=APH1A;refGene_tx=XM_003990565.5
C1	105523814	105523927
refGene_gn=APH1A;refGene_tx=XM_003990565.5
C1	105526479	105526525
refGene_gn=CC1H1orf54;refGene_tx=XM_006935076.4
C1	105527741	105527825
refGene_gn=CC1H1orf54;refGene_tx=XM_006935076.4
C1	105528265	105528324
```

refGene_gn=CC1H1orf54;refGene_tx=XM_006935076.4
C1 105528722 105528764
refGene_gn=CC1H1orf54;refGene_tx=XM_006935076.4
C1 105528765 105528837
refGene_gn=CC1H1orf54;refGene_tx=XM_006935076.4
C1 105531198 105531297
refGene_gn=CC1H1orf54;refGene_tx=XM_006935076.4
C1 105532728 105532734
refGene_gn=CIART;refGene_tx=XM_019837768.2
C1 105534298 105534748
refGene_gn=CIART;refGene_tx=XM_003990568.4;refGene_gn=CIART;refGene_tx=XM_006935079.3;refGene_gn=CIART;refGene_tx=XM_019837768.2;refGene_gn=CIART;refGene_tx=XM_023259111.1;refGene_gn=CIART;refGene_tx=XM_023259112.1
C1 105534893 105534969
refGene_gn=CIART;refGene_tx=XM_003990568.4;refGene_gn=CIART;refGene_tx=XM_006935079.3;refGene_gn=CIART;refGene_tx=XM_019837768.2;refGene_gn=CIART;refGene_tx=XM_023259111.1;refGene_gn=CIART;refGene_tx=XM_023259112.1
C1 105535224 105535303
refGene_gn=CIART;refGene_tx=XM_003990568.4;refGene_gn=CIART;refGene_tx=XM_006935079.3;refGene_gn=CIART;refGene_tx=XM_019837768.2;refGene_gn=CIART;refGene_tx=XM_023259111.1;refGene_gn=CIART;refGene_tx=XM_023259112.1
C1 105535415 105535527
refGene_gn=CIART;refGene_tx=XM_003990568.4;refGene_gn=CIART;refGene_tx=XM_006935079.3;refGene_gn=CIART;refGene_tx=XM_019837768.2;refGene_gn=CIART;refGene_tx=XM_023259111.1;refGene_gn=CIART;refGene_tx=XM_023259112.1
C1 105538352 105538877
refGene_gn=CIART;refGene_tx=XM_003990568.4;refGene_gn=CIART;refGene_tx=XM_006935079.3;refGene_gn=CIART;refGene_tx=XM_019837768.2;refGene_gn=CIART;refGene_tx=XM_023259111.1;refGene_gn=CIART;refGene_tx=XM_023259112.1
C1 105544622 105544705
refGene_gn=MRPS21;refGene_tx=XM_003990569.5;refGene_gn=MRPS21;refGene_tx=XM_006935080.3
C1 105557379 105557560
refGene_gn=MRPS21;refGene_tx=XM_003990569.5;refGene_gn=MRPS21;refGene_tx=XM_006935080.3
C1 105570111 105570256
refGene_gn=PRPF3;refGene_tx=XM_003990570.4;refGene_gn=PRPF3;refGene_tx=XM_006935082.3;refGene_gn=PRPF3;refGene_tx=XM_006935083.3;refGene_gn=PRPF3;refGene_tx=XM_019837776.2
C1 105571708 105571839
refGene_gn=PRPF3;refGene_tx=XM_003990570.4;refGene_gn=PRPF3;refGene_tx=XM_006935082.3;refGene_gn=PRPF3;refGene_tx=XM_006935083.3;refGene_gn=PRPF3;refGene_tx=XM_019837776.2
C1 105573821 105573968
refGene_gn=PRPF3;refGene_tx=XM_003990570.4;refGene_gn=PRPF3;refGene_tx

=XM_006935082.3;refGene_gn=PRPF3;refGene_tx=XM_006935083.3;refGene_gn=
PRPF3;refGene_tx=XM_006935084.2;refGene_gn=PRPF3;refGene_tx=XM_0198377
76.2
C1	105576339	105576423
refGene_gn=PRPF3;refGene_tx=XM_003990570.4;refGene_gn=PRPF3;refGene_tx
=XM_006935082.3;refGene_gn=PRPF3;refGene_tx=XM_006935083.3;refGene_gn=
PRPF3;refGene_tx=XM_006935084.2;refGene_gn=PRPF3;refGene_tx=XM_0198377
76.2
C1	105576623	105576844
refGene_gn=PRPF3;refGene_tx=XM_003990570.4;refGene_gn=PRPF3;refGene_tx
=XM_006935082.3;refGene_gn=PRPF3;refGene_tx=XM_006935083.3;refGene_gn=
PRPF3;refGene_tx=XM_006935084.2;refGene_gn=PRPF3;refGene_tx=XM_0198377
76.2
C1	105577498	105577805
refGene_gn=PRPF3;refGene_tx=XM_003990570.4;refGene_gn=PRPF3;refGene_tx
=XM_006935082.3;refGene_gn=PRPF3;refGene_tx=XM_006935083.3;refGene_gn=
PRPF3;refGene_tx=XM_006935084.2;refGene_gn=PRPF3;refGene_tx=XM_0198377
76.2
C1	105578920	105579087
refGene_gn=PRPF3;refGene_tx=XM_003990570.4;refGene_gn=PRPF3;refGene_tx
=XM_006935082.3;refGene_gn=PRPF3;refGene_tx=XM_006935083.3;refGene_gn=
PRPF3;refGene_tx=XM_006935084.2;refGene_gn=PRPF3;refGene_tx=XM_0198377
76.2
C1	105579824	105579904
refGene_gn=PRPF3;refGene_tx=XM_003990570.4;refGene_gn=PRPF3;refGene_tx
=XM_006935082.3;refGene_gn=PRPF3;refGene_tx=XM_006935083.3;refGene_gn=
PRPF3;refGene_tx=XM_006935084.2;refGene_gn=PRPF3;refGene_tx=XM_0198377
76.2
C1	105583526	105583670
refGene_gn=PRPF3;refGene_tx=XM_003990570.4;refGene_gn=PRPF3;refGene_tx
=XM_006935082.3;refGene_gn=PRPF3;refGene_tx=XM_006935083.3;refGene_gn=
PRPF3;refGene_tx=XM_006935084.2;refGene_gn=PRPF3;refGene_tx=XM_0198377
76.2
C1	105584688	105584788
refGene_gn=PRPF3;refGene_tx=XM_003990570.4;refGene_gn=PRPF3;refGene_tx
=XM_006935082.3;refGene_gn=PRPF3;refGene_tx=XM_006935083.3;refGene_gn=
PRPF3;refGene_tx=XM_006935084.2;refGene_gn=PRPF3;refGene_tx=XM_0198377
76.2
C1	105584961	105585075
refGene_gn=PRPF3;refGene_tx=XM_003990570.4;refGene_gn=PRPF3;refGene_tx
=XM_006935082.3;refGene_gn=PRPF3;refGene_tx=XM_006935083.3;refGene_gn=
PRPF3;refGene_tx=XM_006935084.2;refGene_gn=PRPF3;refGene_tx=XM_0198377
76.2
C1	105586120	105586239
refGene_gn=PRPF3;refGene_tx=XM_003990570.4;refGene_gn=PRPF3;refGene_tx
=XM_006935082.3;refGene_gn=PRPF3;refGene_tx=XM_006935083.3;refGene_gn=
PRPF3;refGene_tx=XM_006935084.2;refGene_gn=PRPF3;refGene_tx=XM_0198377
76.2
C1	105586523	105586607
refGene_gn=PRPF3;refGene_tx=XM_003990570.4;refGene_gn=PRPF3;refGene_tx

=XM_006935082.3;refGene_gn=PRPF3;refGene_tx=XM_006935083.3;refGene_gn=PRPF3;refGene_tx=XM_006935084.2;refGene_gn=PRPF3;refGene_tx=XM_019837776.2
C1 105587929 105587991
refGene_gn=PRPF3;refGene_tx=XM_003990570.4;refGene_gn=PRPF3;refGene_tx=XM_006935082.3;refGene_gn=PRPF3;refGene_tx=XM_006935084.2;refGene_gn=PRPF3;refGene_tx=XM_019837776.2
C1 105591021 105591168
refGene_gn=PRPF3;refGene_tx=XM_003990570.4;refGene_gn=PRPF3;refGene_tx=XM_006935082.3;refGene_gn=PRPF3;refGene_tx=XM_006935083.3;refGene_gn=PRPF3;refGene_tx=XM_006935084.2;refGene_gn=PRPF3;refGene_tx=XM_019837776.2
C1 105600990 105601195
refGene_gn=RPRD2;refGene_tx=XM_003990572.4;refGene_gn=RPRD2;refGene_tx=XM_003990573.4;refGene_gn=RPRD2;refGene_tx=XM_011285119.3
C1 105651635 105651765
refGene_gn=RPRD2;refGene_tx=XM_003990572.4;refGene_gn=RPRD2;refGene_tx=XM_003990573.4;refGene_gn=RPRD2;refGene_tx=XM_011285119.3
C1 105670233 105670334
refGene_gn=RPRD2;refGene_tx=XM_003990572.4;refGene_gn=RPRD2;refGene_tx=XM_003990573.4;refGene_gn=RPRD2;refGene_tx=XM_011285119.3;refGene_gn=RPRD2;refGene_tx=XM_019837777.2;refGene_gn=RPRD2;refGene_tx=XM_023259116.1;refGene_gn=RPRD2;refGene_tx=XM_023259117.1;refGene_gn=RPRD2;refGene_tx=XM_023259118.1;refGene_gn=RPRD2;refGene_tx=XM_023259119.1;refGene_gn=RPRD2;refGene_tx=XM_023259120.1
C1 105671207 105671285
refGene_gn=RPRD2;refGene_tx=XM_003990572.4;refGene_gn=RPRD2;refGene_tx=XM_019837777.2;refGene_gn=RPRD2;refGene_tx=XM_023259116.1;refGene_gn=RPRD2;refGene_tx=XM_023259117.1;refGene_gn=RPRD2;refGene_tx=XM_023259118.1;refGene_gn=RPRD2;refGene_tx=XM_023259120.1
C1 105672821 105672874
refGene_gn=RPRD2;refGene_tx=XM_003990572.4;refGene_gn=RPRD2;refGene_tx=XM_003990573.4;refGene_gn=RPRD2;refGene_tx=XM_011285119.3;refGene_gn=RPRD2;refGene_tx=XM_019837777.2;refGene_gn=RPRD2;refGene_tx=XM_023259116.1;refGene_gn=RPRD2;refGene_tx=XM_023259117.1;refGene_gn=RPRD2;refGene_tx=XM_023259118.1;refGene_gn=RPRD2;refGene_tx=XM_023259119.1;refGene_gn=RPRD2;refGene_tx=XM_023259120.1
C1 105673470 105673597
refGene_gn=RPRD2;refGene_tx=XM_003990572.4;refGene_gn=RPRD2;refGene_tx=XM_003990573.4;refGene_gn=RPRD2;refGene_tx=XM_011285119.3;refGene_gn=RPRD2;refGene_tx=XM_019837777.2;refGene_gn=RPRD2;refGene_tx=XM_023259116.1;refGene_gn=RPRD2;refGene_tx=XM_023259117.1;refGene_gn=RPRD2;refGene_tx=XM_023259118.1;refGene_gn=RPRD2;refGene_tx=XM_023259119.1;refGene_gn=RPRD2;refGene_tx=XM_023259120.1
C1 105675060 105675236
refGene_gn=RPRD2;refGene_tx=XM_003990572.4;refGene_gn=RPRD2;refGene_tx=XM_003990573.4;refGene_gn=RPRD2;refGene_tx=XM_011285119.3;refGene_gn=RPRD2;refGene_tx=XM_019837777.2;refGene_gn=RPRD2;refGene_tx=XM_023259116.1;refGene_gn=RPRD2;refGene_tx=XM_023259117.1;refGene_gn=RPRD2;refGene_tx=XM_023259118.1;refGene_gn=RPRD2;refGene_tx=XM_023259119.1;refGen

e_gn=RPRD2;refGene_tx=XM_023259120.1
C1 105688705 105688988
refGene_gn=RPRD2;refGene_tx=XM_003990572.4;refGene_gn=RPRD2;refGene_tx=XM_003990573.4;refGene_gn=RPRD2;refGene_tx=XM_011285119.3;refGene_gn=RPRD2;refGene_tx=XM_019837777.2;refGene_gn=RPRD2;refGene_tx=XM_023259116.1;refGene_gn=RPRD2;refGene_tx=XM_023259117.1;refGene_gn=RPRD2;refGene_tx=XM_023259118.1;refGene_gn=RPRD2;refGene_tx=XM_023259119.1;refGene_gn=RPRD2;refGene_tx=XM_023259120.1
C1 105691315 105691576
refGene_gn=RPRD2;refGene_tx=XM_003990572.4;refGene_gn=RPRD2;refGene_tx=XM_003990573.4;refGene_gn=RPRD2;refGene_tx=XM_011285119.3;refGene_gn=RPRD2;refGene_tx=XM_019837777.2;refGene_gn=RPRD2;refGene_tx=XM_023259116.1;refGene_gn=RPRD2;refGene_tx=XM_023259117.1;refGene_gn=RPRD2;refGene_tx=XM_023259118.1;refGene_gn=RPRD2;refGene_tx=XM_023259119.1;refGene_gn=RPRD2;refGene_tx=XM_023259120.1
C1 105693560 105693761
refGene_gn=RPRD2;refGene_tx=XM_003990572.4;refGene_gn=RPRD2;refGene_tx=XM_003990573.4;refGene_gn=RPRD2;refGene_tx=XM_011285119.3;refGene_gn=RPRD2;refGene_tx=XM_019837777.2;refGene_gn=RPRD2;refGene_tx=XM_023259116.1;refGene_gn=RPRD2;refGene_tx=XM_023259117.1;refGene_gn=RPRD2;refGene_tx=XM_023259118.1;refGene_gn=RPRD2;refGene_tx=XM_023259119.1;refGene_gn=RPRD2;refGene_tx=XM_023259120.1
C1 105697051 105699825
refGene_gn=RPRD2;refGene_tx=XM_003990572.4;refGene_gn=RPRD2;refGene_tx=XM_003990573.4;refGene_gn=RPRD2;refGene_tx=XM_011285119.3;refGene_gn=RPRD2;refGene_tx=XM_019837777.2;refGene_gn=RPRD2;refGene_tx=XM_023259116.1;refGene_gn=RPRD2;refGene_tx=XM_023259117.1;refGene_gn=RPRD2;refGene_tx=XM_023259118.1;refGene_gn=RPRD2;refGene_tx=XM_023259119.1;refGene_gn=RPRD2;refGene_tx=XM_023259120.1
C1 105702037 105702044
refGene_gn=RPRD2;refGene_tx=XM_023259118.1
C1 105703299 105703339
refGene_gn=RPRD2;refGene_tx=XM_019837777.2;refGene_gn=RPRD2;refGene_tx=XM_023259116.1;refGene_gn=RPRD2;refGene_tx=XM_023259119.1;refGene_gn=RPRD2;refGene_tx=XM_023259120.1
C1 105707407 105707417
refGene_gn=RPRD2;refGene_tx=XM_023259117.1
C1 105709308 105709374
refGene_gn=TARS2;refGene_tx=XM_003990574.5;refGene_gn=TARS2;refGene_tx=XM_019837782.2;refGene_gn=TARS2;refGene_tx=XM_019837783.2;refGene_gn=TARS2;refGene_tx=XM_019837784.2;refGene_gn=TARS2;refGene_tx=XM_019837785.2;refGene_gn=TARS2;refGene_tx=XM_019837786.2;refGene_gn=TARS2;refGene_tx=XM_019837787.2;refGene_gn=TARS2;refGene_tx=XM_019837788.2;refGene_gn=TARS2;refGene_tx=XM_019837789.2;refGene_gn=TARS2;refGene_tx=XM_019837790.2
C1 105709682 105709879
refGene_gn=TARS2;refGene_tx=XM_003990574.5;refGene_gn=TARS2;refGene_tx=XM_019837782.2;refGene_gn=TARS2;refGene_tx=XM_019837783.2;refGene_gn=TARS2;refGene_tx=XM_019837784.2;refGene_gn=TARS2;refGene_tx=XM_019837785.2;refGene_gn=TARS2;refGene_tx=XM_019837786.2;refGene_gn=TARS2;refGe

ne_tx=XM_019837787.2;refGene_gn=TARS2;refGene_tx=XM_019837788.2;refGene_gn=TARS2;refGene_tx=XM_019837789.2;refGene_gn=TARS2;refGene_tx=XM_019837790.2
C1 105710208 105710332
refGene_gn=TARS2;refGene_tx=XM_003990574.5;refGene_gn=TARS2;refGene_tx=XM_019837782.2;refGene_gn=TARS2;refGene_tx=XM_019837783.2;refGene_gn=TARS2;refGene_tx=XM_019837784.2;refGene_gn=TARS2;refGene_tx=XM_019837785.2;refGene_gn=TARS2;refGene_tx=XM_019837786.2;refGene_gn=TARS2;refGene_tx=XM_019837787.2;refGene_gn=TARS2;refGene_tx=XM_019837788.2;refGene_gn=TARS2;refGene_tx=XM_019837789.2;refGene_gn=TARS2;refGene_tx=XM_019837790.2
C1 105711728 105711853
refGene_gn=TARS2;refGene_tx=XM_003990574.5;refGene_gn=TARS2;refGene_tx=XM_019837782.2;refGene_gn=TARS2;refGene_tx=XM_019837783.2;refGene_gn=TARS2;refGene_tx=XM_019837784.2;refGene_gn=TARS2;refGene_tx=XM_019837785.2;refGene_gn=TARS2;refGene_tx=XM_019837786.2;refGene_gn=TARS2;refGene_tx=XM_019837787.2;refGene_gn=TARS2;refGene_tx=XM_019837788.2;refGene_gn=TARS2;refGene_tx=XM_019837789.2;refGene_gn=TARS2;refGene_tx=XM_019837790.2
C1 105712980 105713098
refGene_gn=TARS2;refGene_tx=XM_003990574.5;refGene_gn=TARS2;refGene_tx=XM_019837782.2;refGene_gn=TARS2;refGene_tx=XM_019837783.2;refGene_gn=TARS2;refGene_tx=XM_019837784.2;refGene_gn=TARS2;refGene_tx=XM_019837785.2;refGene_gn=TARS2;refGene_tx=XM_019837786.2;refGene_gn=TARS2;refGene_tx=XM_019837787.2;refGene_gn=TARS2;refGene_tx=XM_019837788.2;refGene_gn=TARS2;refGene_tx=XM_019837789.2;refGene_gn=TARS2;refGene_tx=XM_019837790.2
C1 105713470 105713535
refGene_gn=TARS2;refGene_tx=XM_003990574.5;refGene_gn=TARS2;refGene_tx=XM_019837782.2;refGene_gn=TARS2;refGene_tx=XM_019837784.2;refGene_gn=TARS2;refGene_tx=XM_019837785.2;refGene_gn=TARS2;refGene_tx=XM_019837786.2;refGene_gn=TARS2;refGene_tx=XM_019837787.2;refGene_gn=TARS2;refGene_tx=XM_019837788.2;refGene_gn=TARS2;refGene_tx=XM_019837789.2;refGene_gn=TARS2;refGene_tx=XM_019837790.2
C1 105713793 105713872
refGene_gn=TARS2;refGene_tx=XM_003990574.5;refGene_gn=TARS2;refGene_tx=XM_019837782.2;refGene_gn=TARS2;refGene_tx=XM_019837783.2;refGene_gn=TARS2;refGene_tx=XM_019837784.2;refGene_gn=TARS2;refGene_tx=XM_019837785.2;refGene_gn=TARS2;refGene_tx=XM_019837786.2;refGene_gn=TARS2;refGene_tx=XM_019837787.2;refGene_gn=TARS2;refGene_tx=XM_019837788.2;refGene_gn=TARS2;refGene_tx=XM_019837789.2;refGene_gn=TARS2;refGene_tx=XM_019837790.2
C1 105715005 105715152
refGene_gn=TARS2;refGene_tx=XM_003990574.5;refGene_gn=TARS2;refGene_tx=XM_019837782.2;refGene_gn=TARS2;refGene_tx=XM_019837783.2;refGene_gn=TARS2;refGene_tx=XM_019837784.2;refGene_gn=TARS2;refGene_tx=XM_019837785.2;refGene_gn=TARS2;refGene_tx=XM_019837786.2;refGene_gn=TARS2;refGene_tx=XM_019837787.2
C1 105715288 105715410
refGene_gn=TARS2;refGene_tx=XM_003990574.5;refGene_gn=TARS2;refGene_tx

=XM_019837782.2;refGene_gn=TARS2;refGene_tx=XM_019837783.2;refGene_gn=TARS2;refGene_tx=XM_019837784.2;refGene_gn=TARS2;refGene_tx=XM_019837786.2
C1 105715732 105715950
refGene_gn=TARS2;refGene_tx=XM_003990574.5;refGene_gn=TARS2;refGene_tx=XM_019837782.2;refGene_gn=TARS2;refGene_tx=XM_019837783.2;refGene_gn=TARS2;refGene_tx=XM_019837784.2;refGene_gn=TARS2;refGene_tx=XM_019837785.2;refGene_gn=TARS2;refGene_tx=XM_019837786.2;refGene_gn=TARS2;refGene_tx=XM_019837787.2;refGene_gn=TARS2;refGene_tx=XM_019837788.2;refGene_gn=TARS2;refGene_tx=XM_019837789.2;refGene_gn=TARS2;refGene_tx=XM_019837790.2
C1 105716433 105716596
refGene_gn=TARS2;refGene_tx=XM_003990574.5;refGene_gn=TARS2;refGene_tx=XM_019837782.2;refGene_gn=TARS2;refGene_tx=XM_019837783.2;refGene_gn=TARS2;refGene_tx=XM_019837784.2;refGene_gn=TARS2;refGene_tx=XM_019837785.2;refGene_gn=TARS2;refGene_tx=XM_019837786.2;refGene_gn=TARS2;refGene_tx=XM_019837787.2;refGene_gn=TARS2;refGene_tx=XM_019837788.2;refGene_gn=TARS2;refGene_tx=XM_019837789.2;refGene_gn=TARS2;refGene_tx=XM_019837790.2
C1 105716820 105716958
refGene_gn=TARS2;refGene_tx=XM_003990574.5;refGene_gn=TARS2;refGene_tx=XM_019837782.2;refGene_gn=TARS2;refGene_tx=XM_019837783.2;refGene_gn=TARS2;refGene_tx=XM_019837784.2;refGene_gn=TARS2;refGene_tx=XM_019837785.2;refGene_gn=TARS2;refGene_tx=XM_019837786.2;refGene_gn=TARS2;refGene_tx=XM_019837787.2;refGene_gn=TARS2;refGene_tx=XM_019837788.2;refGene_gn=TARS2;refGene_tx=XM_019837789.2;refGene_gn=TARS2;refGene_tx=XM_019837790.2
C1 105717127 105717205
refGene_gn=TARS2;refGene_tx=XM_003990574.5;refGene_gn=TARS2;refGene_tx=XM_019837782.2;refGene_gn=TARS2;refGene_tx=XM_019837783.2;refGene_gn=TARS2;refGene_tx=XM_019837784.2;refGene_gn=TARS2;refGene_tx=XM_019837785.2;refGene_gn=TARS2;refGene_tx=XM_019837786.2;refGene_gn=TARS2;refGene_tx=XM_019837787.2;refGene_gn=TARS2;refGene_tx=XM_019837788.2;refGene_gn=TARS2;refGene_tx=XM_019837789.2;refGene_gn=TARS2;refGene_tx=XM_019837790.2
C1 105719162 105719284
refGene_gn=TARS2;refGene_tx=XM_003990574.5;refGene_gn=TARS2;refGene_tx=XM_019837782.2;refGene_gn=TARS2;refGene_tx=XM_019837783.2;refGene_gn=TARS2;refGene_tx=XM_019837784.2;refGene_gn=TARS2;refGene_tx=XM_019837785.2;refGene_gn=TARS2;refGene_tx=XM_019837786.2;refGene_gn=TARS2;refGene_tx=XM_019837787.2;refGene_gn=TARS2;refGene_tx=XM_019837788.2;refGene_gn=TARS2;refGene_tx=XM_019837789.2;refGene_gn=TARS2;refGene_tx=XM_019837790.2
C1 105719488 105719590
refGene_gn=TARS2;refGene_tx=XM_003990574.5;refGene_gn=TARS2;refGene_tx=XM_019837782.2;refGene_gn=TARS2;refGene_tx=XM_019837783.2;refGene_gn=TARS2;refGene_tx=XM_019837784.2;refGene_gn=TARS2;refGene_tx=XM_019837785.2;refGene_gn=TARS2;refGene_tx=XM_019837786.2;refGene_gn=TARS2;refGene_tx=XM_019837787.2;refGene_gn=TARS2;refGene_tx=XM_019837788.2;refGene_gn=TARS2;refGene_tx=XM_019837789.2;refGene_gn=TARS2;refGene_tx=XM_01

9837790.2
C1	105719765	105719838
refGene_gn=TARS2;refGene_tx=XM_003990574.5;refGene_gn=TARS2;refGene_tx=XM_019837782.2;refGene_gn=TARS2;refGene_tx=XM_019837783.2;refGene_gn=TARS2;refGene_tx=XM_019837784.2;refGene_gn=TARS2;refGene_tx=XM_019837785.2;refGene_gn=TARS2;refGene_tx=XM_019837786.2;refGene_gn=TARS2;refGene_tx=XM_019837787.2;refGene_gn=TARS2;refGene_tx=XM_019837788.2;refGene_gn=TARS2;refGene_tx=XM_019837789.2;refGene_gn=TARS2;refGene_tx=XM_019837790.2
C1	105720425	105720540
refGene_gn=TARS2;refGene_tx=XM_003990574.5;refGene_gn=TARS2;refGene_tx=XM_019837782.2;refGene_gn=TARS2;refGene_tx=XM_019837783.2;refGene_gn=TARS2;refGene_tx=XM_019837784.2;refGene_gn=TARS2;refGene_tx=XM_019837785.2;refGene_gn=TARS2;refGene_tx=XM_019837786.2;refGene_gn=TARS2;refGene_tx=XM_019837787.2;refGene_gn=TARS2;refGene_tx=XM_019837788.2;refGene_gn=TARS2;refGene_tx=XM_019837789.2;refGene_gn=TARS2;refGene_tx=XM_019837790.2
C1	105723850	105723999
refGene_gn=TARS2;refGene_tx=XM_003990574.5;refGene_gn=TARS2;refGene_tx=XM_019837782.2;refGene_gn=TARS2;refGene_tx=XM_019837783.2;refGene_gn=TARS2;refGene_tx=XM_019837784.2;refGene_gn=TARS2;refGene_tx=XM_019837785.2;refGene_gn=TARS2;refGene_tx=XM_019837786.2;refGene_gn=TARS2;refGene_tx=XM_019837787.2;refGene_gn=TARS2;refGene_tx=XM_019837788.2;refGene_gn=TARS2;refGene_tx=XM_019837789.2;refGene_gn=TARS2;refGene_tx=XM_019837790.2
C1	105724826	105724899
refGene_gn=ECM1;refGene_tx=XM_006935088.4;refGene_gn=ECM1;refGene_tx=XM_011285121.3;refGene_gn=ECM1;refGene_tx=XM_023259122.1;refGene_gn=ECM1;refGene_tx=XM_023259123.1;refGene_gn=ECM1;refGene_tx=XM_023259124.1
C1	105726125	105726188
refGene_gn=ECM1;refGene_tx=XM_006935088.4;refGene_gn=ECM1;refGene_tx=XM_011285121.3;refGene_gn=ECM1;refGene_tx=XM_023259122.1;refGene_gn=ECM1;refGene_tx=XM_023259123.1;refGene_gn=ECM1;refGene_tx=XM_023259124.1
C1	105726273	105726633
refGene_gn=ECM1;refGene_tx=XM_006935088.4;refGene_gn=ECM1;refGene_tx=XM_011285121.3;refGene_gn=ECM1;refGene_tx=XM_023259122.1;refGene_gn=ECM1;refGene_tx=XM_023259123.1;refGene_gn=ECM1;refGene_tx=XM_023259124.1
C1	105726730	105726811
refGene_gn=ECM1;refGene_tx=XM_006935088.4;refGene_gn=ECM1;refGene_tx=XM_011285121.3;refGene_gn=ECM1;refGene_tx=XM_023259122.1;refGene_gn=ECM1;refGene_tx=XM_023259123.1;refGene_gn=ECM1;refGene_tx=XM_023259124.1
C1	105727111	105727168
refGene_gn=ECM1;refGene_tx=XM_006935088.4;refGene_gn=ECM1;refGene_tx=XM_023259122.1;refGene_gn=ECM1;refGene_tx=XM_023259124.1
C1	105727486	105727803
refGene_gn=ECM1;refGene_tx=XM_006935088.4;refGene_gn=ECM1;refGene_tx=XM_011285121.3;refGene_gn=ECM1;refGene_tx=XM_023259122.1;refGene_gn=ECM1;refGene_tx=XM_023259123.1;refGene_gn=ECM1;refGene_tx=XM_023259124.1
C1	105728083	105728458
refGene_gn=ECM1;refGene_tx=XM_006935088.4;refGene_gn=ECM1;refGene_tx=X

M_011285121.3;refGene_gn=ECM1;refGene_tx=XM_023259122.1;refGene_gn=ECM1;refGene_tx=XM_023259123.1
C1 105728954 105729175
refGene_gn=ECM1;refGene_tx=XM_006935088.4;refGene_gn=ECM1;refGene_tx=XM_011285121.3;refGene_gn=ECM1;refGene_tx=XM_023259122.1;refGene_gn=ECM1;refGene_tx=XM_023259123.1;refGene_gn=ECM1;refGene_tx=XM_023259124.1
C1 105729341 105729429
refGene_gn=ECM1;refGene_tx=XM_006935088.4;refGene_gn=ECM1;refGene_tx=XM_011285121.3;refGene_gn=ECM1;refGene_tx=XM_023259122.1;refGene_gn=ECM1;refGene_tx=XM_023259123.1;refGene_gn=ECM1;refGene_tx=XM_023259124.1
C1 105730091 105730322
refGene_gn=ECM1;refGene_tx=XM_006935088.4;refGene_gn=ECM1;refGene_tx=XM_011285121.3;refGene_gn=ECM1;refGene_tx=XM_023259122.1;refGene_gn=ECM1;refGene_tx=XM_023259123.1;refGene_gn=ECM1;refGene_tx=XM_023259124.1
C1 105761076 105761096
refGene_gn=ADAMTSL4;refGene_tx=XM_023259121.1
C1 105761279 105761337
refGene_gn=ADAMTSL4;refGene_tx=XM_023259121.1
C1 105761616 105761969
refGene_gn=ADAMTSL4;refGene_tx=XM_023259121.1
C1 105762178 105762836
refGene_gn=ADAMTSL4;refGene_tx=XM_019837781.2;refGene_gn=ADAMTSL4;refGene_tx=XM_023259121.1
C1 105763085 105763188
refGene_gn=ADAMTSL4;refGene_tx=XM_019837781.2;refGene_gn=ADAMTSL4;refGene_tx=XM_023259121.1
C1 105763908 105764045
refGene_gn=ADAMTSL4;refGene_tx=XM_019837781.2;refGene_gn=ADAMTSL4;refGene_tx=XM_023259121.1
C1 105764406 105764611
refGene_gn=ADAMTSL4;refGene_tx=XM_019837781.2;refGene_gn=ADAMTSL4;refGene_tx=XM_023259121.1
C1 105764843 105765016
refGene_gn=ADAMTSL4;refGene_tx=XM_019837781.2;refGene_gn=ADAMTSL4;refGene_tx=XM_023259121.1
C1 105765169 105765281
refGene_gn=ADAMTSL4;refGene_tx=XM_019837781.2;refGene_gn=ADAMTSL4;refGene_tx=XM_023259121.1
C1 105765384 105765570
refGene_gn=ADAMTSL4;refGene_tx=XM_019837781.2;refGene_gn=ADAMTSL4;refGene_tx=XM_023259121.1
C1 105765765 105765895
refGene_gn=ADAMTSL4;refGene_tx=XM_019837781.2;refGene_gn=ADAMTSL4;refGene_tx=XM_023259121.1
C1 105766236 105766441
refGene_gn=ADAMTSL4;refGene_tx=XM_019837781.2;refGene_gn=ADAMTSL4;refGene_tx=XM_023259121.1
C1 105766782 105766959
refGene_gn=ADAMTSL4;refGene_tx=XM_019837781.2;refGene_gn=ADAMTSL4;refGene_tx=XM_023259121.1

C1	105767249	105767456
refGene_gn=ADAMTSL4;refGene_tx=XM_019837781.2;refGene_gn=ADAMTSL4;refGene_tx=XM_023259121.1
C1	105767568	105767748
refGene_gn=ADAMTSL4;refGene_tx=XM_019837781.2;refGene_gn=ADAMTSL4;refGene_tx=XM_023259121.1
C1	105768020	105768165
refGene_gn=ADAMTSL4;refGene_tx=XM_019837781.2;refGene_gn=ADAMTSL4;refGene_tx=XM_023259121.1
C1	105768317	105768454
refGene_gn=ADAMTSL4;refGene_tx=XM_019837781.2;refGene_gn=ADAMTSL4;refGene_tx=XM_023259121.1
C1	105780131	105780248
refGene_gn=MCL1;refGene_tx=NM_001009374.1
C1	105780773	105781021
refGene_gn=MCL1;refGene_tx=NM_001009374.1
C1	105781375	105782063
refGene_gn=MCL1;refGene_tx=NM_001009374.1
C1	105812964	105812980
refGene_gn=ENSA;refGene_tx=XM_003990575.5;refGene_gn=ENSA;refGene_tx=XM_019837793.2
C1	105816023	105816190
refGene_gn=ENSA;refGene_tx=XM_003990575.5;refGene_gn=ENSA;refGene_tx=XM_019837793.2
C1	105817475	105817601
refGene_gn=ENSA;refGene_tx=XM_003990575.5
C1	105822320	105822377
refGene_gn=ENSA;refGene_tx=XM_003990575.5;refGene_gn=ENSA;refGene_tx=XM_019837793.2
C1	105832080	105832508
refGene_gn=GOLPH3L;refGene_tx=XM_011285122.3;refGene_gn=GOLPH3L;refGene_tx=XM_019837803.2
C1	105843426	105843541
refGene_gn=GOLPH3L;refGene_tx=XM_011285122.3;refGene_gn=GOLPH3L;refGene_tx=XM_019837803.2
C1	105845163	105845295
refGene_gn=GOLPH3L;refGene_tx=XM_011285122.3;refGene_gn=GOLPH3L;refGene_tx=XM_019837803.2
C1	105864877	105865060
refGene_gn=GOLPH3L;refGene_tx=XM_011285122.3;refGene_gn=GOLPH3L;refGene_tx=XM_019837803.2
C1	105868705	105868786
refGene_gn=HORMAD1;refGene_tx=XM_011285124.3;refGene_gn=HORMAD1;refGene_tx=XM_011285125.3;refGene_gn=HORMAD1;refGene_tx=XM_019837797.2;refGene_gn=HORMAD1;refGene_tx=XM_019837798.2;refGene_gn=HORMAD1;refGene_tx=XM_019837799.2
C1	105870805	105870877
refGene_gn=HORMAD1;refGene_tx=XM_011285124.3;refGene_gn=HORMAD1;refGene_tx=XM_011285125.3;refGene_gn=HORMAD1;refGene_tx=XM_019837797.2;refGene_gn=HORMAD1;refGene_tx=XM_019837798.2;refGene_gn=HORMAD1;refGene_tx=

XM_019837799.2
C1	105874317	105874401
refGene_gn=HORMAD1;refGene_tx=XM_011285124.3;refGene_gn=HORMAD1;refGene_tx=XM_011285125.3;refGene_gn=HORMAD1;refGene_tx=XM_019837797.2;refGene_gn=HORMAD1;refGene_tx=XM_019837798.2;refGene_gn=HORMAD1;refGene_tx=XM_019837799.2
C1	105875121	105875198
refGene_gn=HORMAD1;refGene_tx=XM_011285124.3;refGene_gn=HORMAD1;refGene_tx=XM_011285125.3;refGene_gn=HORMAD1;refGene_tx=XM_019837797.2;refGene_gn=HORMAD1;refGene_tx=XM_019837798.2;refGene_gn=HORMAD1;refGene_tx=XM_019837799.2
C1	105875281	105875348
refGene_gn=HORMAD1;refGene_tx=XM_011285124.3;refGene_gn=HORMAD1;refGene_tx=XM_011285125.3;refGene_gn=HORMAD1;refGene_tx=XM_019837797.2;refGene_gn=HORMAD1;refGene_tx=XM_019837798.2;refGene_gn=HORMAD1;refGene_tx=XM_019837799.2
C1	105877155	105877415
refGene_gn=HORMAD1;refGene_tx=XM_011285124.3;refGene_gn=HORMAD1;refGene_tx=XM_011285125.3;refGene_gn=HORMAD1;refGene_tx=XM_019837797.2;refGene_gn=HORMAD1;refGene_tx=XM_019837798.2;refGene_gn=HORMAD1;refGene_tx=XM_019837799.2
C1	105878822	105878974
refGene_gn=HORMAD1;refGene_tx=XM_011285124.3;refGene_gn=HORMAD1;refGene_tx=XM_011285125.3;refGene_gn=HORMAD1;refGene_tx=XM_019837797.2;refGene_gn=HORMAD1;refGene_tx=XM_019837798.2;refGene_gn=HORMAD1;refGene_tx=XM_019837799.2
C1	105879487	105879555
refGene_gn=HORMAD1;refGene_tx=XM_011285124.3;refGene_gn=HORMAD1;refGene_tx=XM_011285125.3;refGene_gn=HORMAD1;refGene_tx=XM_019837797.2;refGene_gn=HORMAD1;refGene_tx=XM_019837798.2;refGene_gn=HORMAD1;refGene_tx=XM_019837799.2
C1	105881071	105881098
refGene_gn=HORMAD1;refGene_tx=XM_011285124.3;refGene_gn=HORMAD1;refGene_tx=XM_011285125.3;refGene_gn=HORMAD1;refGene_tx=XM_019837797.2;refGene_gn=HORMAD1;refGene_tx=XM_019837798.2;refGene_gn=HORMAD1;refGene_tx=XM_019837799.2
C1	105881371	105881392
refGene_gn=HORMAD1;refGene_tx=XM_011285124.3;refGene_gn=HORMAD1;refGene_tx=XM_019837797.2;refGene_gn=HORMAD1;refGene_tx=XM_019837798.2;refGene_gn=HORMAD1;refGene_tx=XM_019837799.2
C1	105883857	105883894
refGene_gn=HORMAD1;refGene_tx=XM_011285124.3;refGene_gn=HORMAD1;refGene_tx=XM_011285125.3;refGene_gn=HORMAD1;refGene_tx=XM_019837797.2;refGene_gn=HORMAD1;refGene_tx=XM_019837798.2;refGene_gn=HORMAD1;refGene_tx=XM_019837799.2
C1	105884661	105884725
refGene_gn=HORMAD1;refGene_tx=XM_011285124.3;refGene_gn=HORMAD1;refGene_tx=XM_011285125.3;refGene_gn=HORMAD1;refGene_tx=XM_019837797.2;refGene_gn=HORMAD1;refGene_tx=XM_019837798.2;refGene_gn=HORMAD1;refGene_tx=XM_019837799.2

C1 105886320 105886465
refGene_gn=HORMAD1;refGene_tx=XM_011285124.3;refGene_gn=HORMAD1;refGen
e_tx=XM_011285125.3;refGene_gn=HORMAD1;refGene_tx=XM_019837797.2;refGe
ne_gn=HORMAD1;refGene_tx=XM_019837798.2;refGene_gn=HORMAD1;refGene_tx=
XM_019837799.2
C1 105887148 105887211
refGene_gn=HORMAD1;refGene_tx=XM_011285124.3;refGene_gn=HORMAD1;refGen
e_tx=XM_011285125.3;refGene_gn=HORMAD1;refGene_tx=XM_019837797.2;refGe
ne_gn=HORMAD1;refGene_tx=XM_019837798.2;refGene_gn=HORMAD1;refGene_tx=
XM_019837799.2
C1 105887824 105887845
refGene_gn=HORMAD1;refGene_tx=XM_011285124.3;refGene_gn=HORMAD1;refGen
e_tx=XM_011285125.3
C1 105896603 105896625
refGene_gn=CTSS;refGene_tx=XM_011285127.2
C1 105896734 105896738
refGene_gn=CTSS;refGene_tx=XM_011285128.2
C1 105900811 105900911
refGene_gn=CTSS;refGene_tx=XM_011285126.2
C1 105914836 105914939
refGene_gn=CTSS;refGene_tx=XM_011285126.2;refGene_gn=CTSS;refGene_tx=X
M_011285127.2;refGene_gn=CTSS;refGene_tx=XM_011285128.2
C1 105915797 105915963
refGene_gn=CTSS;refGene_tx=XM_011285126.2;refGene_gn=CTSS;refGene_tx=X
M_011285127.2;refGene_gn=CTSS;refGene_tx=XM_011285128.2
C1 105917179 105917407
refGene_gn=CTSS;refGene_tx=XM_011285126.2;refGene_gn=CTSS;refGene_tx=X
M_011285127.2;refGene_gn=CTSS;refGene_tx=XM_011285128.2
C1 105919774 105919924
refGene_gn=CTSS;refGene_tx=XM_011285126.2;refGene_gn=CTSS;refGene_tx=X
M_011285127.2;refGene_gn=CTSS;refGene_tx=XM_011285128.2
C1 105920777 105920900
refGene_gn=CTSS;refGene_tx=XM_011285126.2;refGene_gn=CTSS;refGene_tx=X
M_011285127.2;refGene_gn=CTSS;refGene_tx=XM_011285128.2
C1 105922740 105922867
refGene_gn=CTSS;refGene_tx=XM_011285126.2;refGene_gn=CTSS;refGene_tx=X
M_011285127.2;refGene_gn=CTSS;refGene_tx=XM_011285128.2
C1 105923810 105923845
refGene_gn=CTSS;refGene_tx=XM_011285126.2;refGene_gn=CTSS;refGene_tx=X
M_011285127.2;refGene_gn=CTSS;refGene_tx=XM_011285128.2
C1 105944162 105944262
refGene_gn=CTSK;refGene_tx=XM_019837802.2
C1 105945393 105945499
refGene_gn=CTSK;refGene_tx=XM_019837802.2
C1 105945762 105945850
refGene_gn=CTSK;refGene_tx=XM_019837802.2
C1 105945851 105945929
refGene_gn=CTSK;refGene_tx=XM_019837802.2
C1 105949904 105950123
refGene_gn=CTSK;refGene_tx=XM_019837802.2

C1	105951112	105951268
refGene_gn=CTSK;refGene_tx=XM_019837802.2
C1	105951353	105951476
refGene_gn=CTSK;refGene_tx=XM_019837802.2
C1	105951945	105952068
refGene_gn=CTSK;refGene_tx=XM_019837802.2
C1	105956964	105957054
refGene_gn=ARNT;refGene_tx=XM_023259125.1;refGene_gn=ARNT;refGene_tx=XM_023259126.1;refGene_gn=ARNT;refGene_tx=XM_023259127.1;refGene_gn=ARNT;refGene_tx=XM_023259128.1
C1	105957766	105957933
refGene_gn=ARNT;refGene_tx=XM_023259125.1;refGene_gn=ARNT;refGene_tx=XM_023259126.1;refGene_gn=ARNT;refGene_tx=XM_023259127.1;refGene_gn=ARNT;refGene_tx=XM_023259128.1
C1	105958531	105958694
refGene_gn=ARNT;refGene_tx=XM_023259125.1;refGene_gn=ARNT;refGene_tx=XM_023259126.1;refGene_gn=ARNT;refGene_tx=XM_023259127.1;refGene_gn=ARNT;refGene_tx=XM_023259128.1
C1	105959902	105960050
refGene_gn=ARNT;refGene_tx=XM_023259125.1;refGene_gn=ARNT;refGene_tx=XM_023259126.1;refGene_gn=ARNT;refGene_tx=XM_023259127.1;refGene_gn=ARNT;refGene_tx=XM_023259128.1
C1	105960433	105960536
refGene_gn=ARNT;refGene_tx=XM_023259125.1;refGene_gn=ARNT;refGene_tx=XM_023259126.1;refGene_gn=ARNT;refGene_tx=XM_023259127.1;refGene_gn=ARNT;refGene_tx=XM_023259128.1
C1	105960712	105960836
refGene_gn=ARNT;refGene_tx=XM_023259125.1;refGene_gn=ARNT;refGene_tx=XM_023259126.1;refGene_gn=ARNT;refGene_tx=XM_023259127.1;refGene_gn=ARNT;refGene_tx=XM_023259128.1
C1	105960979	105961052
refGene_gn=ARNT;refGene_tx=XM_023259125.1;refGene_gn=ARNT;refGene_tx=XM_023259126.1;refGene_gn=ARNT;refGene_tx=XM_023259127.1;refGene_gn=ARNT;refGene_tx=XM_023259128.1
C1	105961250	105961361
refGene_gn=ARNT;refGene_tx=XM_023259125.1;refGene_gn=ARNT;refGene_tx=XM_023259126.1;refGene_gn=ARNT;refGene_tx=XM_023259127.1;refGene_gn=ARNT;refGene_tx=XM_023259128.1
C1	105963850	105964002
refGene_gn=ARNT;refGene_tx=XM_023259125.1;refGene_gn=ARNT;refGene_tx=XM_023259126.1;refGene_gn=ARNT;refGene_tx=XM_023259127.1;refGene_gn=ARNT;refGene_tx=XM_023259128.1
C1	105967595	105967670
refGene_gn=ARNT;refGene_tx=XM_023259125.1;refGene_gn=ARNT;refGene_tx=XM_023259126.1;refGene_gn=ARNT;refGene_tx=XM_023259127.1;refGene_gn=ARNT;refGene_tx=XM_023259128.1
C1	105970262	105970397
refGene_gn=ARNT;refGene_tx=XM_023259125.1;refGene_gn=ARNT;refGene_tx=XM_023259126.1;refGene_gn=ARNT;refGene_tx=XM_023259127.1;refGene_gn=ARNT;refGene_tx=XM_023259128.1

C1	105975486	105975563
refGene_gn=ARNT;refGene_tx=XM_023259125.1;refGene_gn=ARNT;refGene_tx=XM_023259126.1;refGene_gn=ARNT;refGene_tx=XM_023259127.1;refGene_gn=ARNT;refGene_tx=XM_023259128.1
C1	105977127	105977213
refGene_gn=ARNT;refGene_tx=XM_023259125.1;refGene_gn=ARNT;refGene_tx=XM_023259126.1;refGene_gn=ARNT;refGene_tx=XM_023259127.1;refGene_gn=ARNT;refGene_tx=XM_023259128.1
C1	105977627	105977693
refGene_gn=ARNT;refGene_tx=XM_023259125.1;refGene_gn=ARNT;refGene_tx=XM_023259126.1;refGene_gn=ARNT;refGene_tx=XM_023259127.1;refGene_gn=ARNT;refGene_tx=XM_023259128.1
C1	105979372	105979475
refGene_gn=ARNT;refGene_tx=XM_023259125.1;refGene_gn=ARNT;refGene_tx=XM_023259126.1;refGene_gn=ARNT;refGene_tx=XM_023259127.1;refGene_gn=ARNT;refGene_tx=XM_023259128.1
C1	105980635	105980849
refGene_gn=ARNT;refGene_tx=XM_023259125.1;refGene_gn=ARNT;refGene_tx=XM_023259126.1;refGene_gn=ARNT;refGene_tx=XM_023259127.1;refGene_gn=ARNT;refGene_tx=XM_023259128.1
C1	105984938	105985152
refGene_gn=ARNT;refGene_tx=XM_023259125.1;refGene_gn=ARNT;refGene_tx=XM_023259126.1;refGene_gn=ARNT;refGene_tx=XM_023259127.1;refGene_gn=ARNT;refGene_tx=XM_023259128.1
C1	105987630	105987675
refGene_gn=ARNT;refGene_tx=XM_023259125.1;refGene_gn=ARNT;refGene_tx=XM_023259126.1;refGene_gn=ARNT;refGene_tx=XM_023259127.1;refGene_gn=ARNT;refGene_tx=XM_023259128.1
C1	105991380	105991425
refGene_gn=ARNT;refGene_tx=XM_023259125.1;refGene_gn=ARNT;refGene_tx=XM_023259126.1;refGene_gn=ARNT;refGene_tx=XM_023259127.1;refGene_gn=ARNT;refGene_tx=XM_023259128.1
C1	105998500	105998545
refGene_gn=ARNT;refGene_tx=XM_023259125.1;refGene_gn=ARNT;refGene_tx=XM_023259126.1;refGene_gn=ARNT;refGene_tx=XM_023259127.1;refGene_gn=ARNT;refGene_tx=XM_023259128.1
C1	106006402	106006514
refGene_gn=ARNT;refGene_tx=XM_023259125.1;refGene_gn=ARNT;refGene_tx=XM_023259126.1;refGene_gn=ARNT;refGene_tx=XM_023259127.1;refGene_gn=ARNT;refGene_tx=XM_023259128.1
C1	106030846	106030871
refGene_gn=ARNT;refGene_tx=XM_023259125.1;refGene_gn=ARNT;refGene_tx=XM_023259126.1;refGene_gn=ARNT;refGene_tx=XM_023259127.1;refGene_gn=ARNT;refGene_tx=XM_023259128.1
C1	106039451	106039814
refGene_gn=LOC101093878;refGene_tx=XM_006935099.2
C1	106068071	106068331
refGene_gn=SETDB1;refGene_tx=XM_006935100.4
C1	106069468	106069620
refGene_gn=SETDB1;refGene_tx=XM_006935100.4

C1	106083172	106083207
refGene_gn=SETDB1;refGene_tx=XM_006935100.4
C1	106083874	106083974
refGene_gn=SETDB1;refGene_tx=XM_006935100.4
C1	106085042	106085168
refGene_gn=SETDB1;refGene_tx=XM_006935100.4
C1	106085335	106085537
refGene_gn=SETDB1;refGene_tx=XM_006935100.4
C1	106086030	106086104
refGene_gn=SETDB1;refGene_tx=XM_006935100.4
C1	106087293	106087484
refGene_gn=SETDB1;refGene_tx=XM_006935100.4
C1	106088196	106088326
refGene_gn=SETDB1;refGene_tx=XM_006935100.4
C1	106088974	106089090
refGene_gn=SETDB1;refGene_tx=XM_006935100.4
C1	106089092	106089151
refGene_gn=SETDB1;refGene_tx=XM_006935100.4
C1	106089234	106089393
refGene_gn=SETDB1;refGene_tx=XM_006935100.4
C1	106090007	106090581
refGene_gn=SETDB1;refGene_tx=XM_006935100.4
C1	106090582	106090635
refGene_gn=SETDB1;refGene_tx=XM_006935100.4
C1	106091073	106091190
refGene_gn=SETDB1;refGene_tx=XM_006935100.4
C1	106096110	106096280
refGene_gn=SETDB1;refGene_tx=XM_006935100.4
C1	106096838	106097467
refGene_gn=SETDB1;refGene_tx=XM_006935100.4
C1	106098348	106098377
refGene_gn=SETDB1;refGene_tx=XM_006935100.4
C1	106099030	106099163
refGene_gn=SETDB1;refGene_tx=XM_006935100.4
C1	106099450	106099616
refGene_gn=SETDB1;refGene_tx=XM_006935100.4
C1	106099987	106100199
refGene_gn=SETDB1;refGene_tx=XM_006935100.4
C1	106100680	106100769
refGene_gn=SETDB1;refGene_tx=XM_006935100.4
C1	106100883	106101001
refGene_gn=SETDB1;refGene_tx=XM_006935100.4
C1	106102533	106102674
refGene_gn=CERS2;refGene_tx=XM_003990597.5;refGene_gn=CERS2;refGene_tx=XM_019837804.2;refGene_gn=CERS2;refGene_tx=XM_023259130.1;refGene_gn=CERS2;refGene_tx=XM_023259131.1
C1	106102852	106103006
refGene_gn=CERS2;refGene_tx=XM_003990597.5;refGene_gn=CERS2;refGene_tx=XM_019837804.2;refGene_gn=CERS2;refGene_tx=XM_023259130.1;refGene_gn=CERS2;refGene_tx=XM_023259131.1

C1 106103113 106103220
refGene_gn=CERS2;refGene_tx=XM_003990597.5;refGene_gn=CERS2;refGene_tx
=XM_019837804.2;refGene_gn=CERS2;refGene_tx=XM_023259130.1;refGene_gn=
CERS2;refGene_tx=XM_023259131.1
C1 106103432 106103561
refGene_gn=CERS2;refGene_tx=XM_003990597.5;refGene_gn=CERS2;refGene_tx
=XM_019837804.2;refGene_gn=CERS2;refGene_tx=XM_023259130.1;refGene_gn=
CERS2;refGene_tx=XM_023259131.1
C1 106103750 106103843
refGene_gn=CERS2;refGene_tx=XM_003990597.5;refGene_gn=CERS2;refGene_tx
=XM_019837804.2;refGene_gn=CERS2;refGene_tx=XM_023259130.1;refGene_gn=
CERS2;refGene_tx=XM_023259131.1
C1 106104062 106104113
refGene_gn=CERS2;refGene_tx=XM_003990597.5;refGene_gn=CERS2;refGene_tx
=XM_019837804.2;refGene_gn=CERS2;refGene_tx=XM_023259130.1;refGene_gn=
CERS2;refGene_tx=XM_023259131.1
C1 106104219 106104277
refGene_gn=CERS2;refGene_tx=XM_003990597.5;refGene_gn=CERS2;refGene_tx
=XM_019837804.2;refGene_gn=CERS2;refGene_tx=XM_023259130.1;refGene_gn=
CERS2;refGene_tx=XM_023259131.1
C1 106104490 106104609
refGene_gn=CERS2;refGene_tx=XM_003990597.5;refGene_gn=CERS2;refGene_tx
=XM_019837804.2;refGene_gn=CERS2;refGene_tx=XM_023259130.1;refGene_gn=
CERS2;refGene_tx=XM_023259131.1
C1 106104808 106104926
refGene_gn=CERS2;refGene_tx=XM_003990597.5;refGene_gn=CERS2;refGene_tx
=XM_019837804.2;refGene_gn=CERS2;refGene_tx=XM_023259130.1;refGene_gn=
CERS2;refGene_tx=XM_023259131.1
C1 106105414 106105587
refGene_gn=CERS2;refGene_tx=XM_003990597.5;refGene_gn=CERS2;refGene_tx
=XM_019837804.2;refGene_gn=CERS2;refGene_tx=XM_023259130.1;refGene_gn=
CERS2;refGene_tx=XM_023259131.1
C1 106117057 106117132
refGene_gn=ANXA9;refGene_tx=XM_003990598.4;refGene_gn=ANXA9;refGene_tx
=XM_019837815.2;refGene_gn=ANXA9;refGene_tx=XM_019837816.2;refGene_gn=
ANXA9;refGene_tx=XM_019837817.2;refGene_gn=ANXA9;refGene_tx=XM_0198378
19.2;refGene_gn=ANXA9;refGene_tx=XM_019837820.2
C1 106117314 106117411
refGene_gn=ANXA9;refGene_tx=XM_003990598.4;refGene_gn=ANXA9;refGene_tx
=XM_019837815.2;refGene_gn=ANXA9;refGene_tx=XM_019837816.2;refGene_gn=
ANXA9;refGene_tx=XM_019837817.2;refGene_gn=ANXA9;refGene_tx=XM_0198378
19.2;refGene_gn=ANXA9;refGene_tx=XM_019837820.2
C1 106117983 106118078
refGene_gn=ANXA9;refGene_tx=XM_003990598.4;refGene_gn=ANXA9;refGene_tx
=XM_019837815.2;refGene_gn=ANXA9;refGene_tx=XM_019837816.2;refGene_gn=
ANXA9;refGene_tx=XM_019837817.2;refGene_gn=ANXA9;refGene_tx=XM_0198378
19.2;refGene_gn=ANXA9;refGene_tx=XM_019837820.2
C1 106118281 106118395
refGene_gn=ANXA9;refGene_tx=XM_003990598.4;refGene_gn=ANXA9;refGene_tx
=XM_019837815.2;refGene_gn=ANXA9;refGene_tx=XM_019837816.2;refGene_gn=

ANXA9;refGene_tx=XM_019837817.2;refGene_gn=ANXA9;refGene_tx=XM_0198378
19.2;refGene_gn=ANXA9;refGene_tx=XM_019837820.2
C1	106118615	106118706
refGene_gn=ANXA9;refGene_tx=XM_003990598.4;refGene_gn=ANXA9;refGene_tx
=XM_019837815.2;refGene_gn=ANXA9;refGene_tx=XM_019837816.2;refGene_gn=
ANXA9;refGene_tx=XM_019837817.2;refGene_gn=ANXA9;refGene_tx=XM_0198378
19.2;refGene_gn=ANXA9;refGene_tx=XM_019837820.2
C1	106119668	106119748
refGene_gn=ANXA9;refGene_tx=XM_003990598.4;refGene_gn=ANXA9;refGene_tx
=XM_019837815.2;refGene_gn=ANXA9;refGene_tx=XM_019837816.2;refGene_gn=
ANXA9;refGene_tx=XM_019837817.2;refGene_gn=ANXA9;refGene_tx=XM_0198378
19.2;refGene_gn=ANXA9;refGene_tx=XM_019837820.2
C1	106119932	106119992
refGene_gn=ANXA9;refGene_tx=XM_003990598.4;refGene_gn=ANXA9;refGene_tx
=XM_019837815.2;refGene_gn=ANXA9;refGene_tx=XM_019837816.2;refGene_gn=
ANXA9;refGene_tx=XM_019837817.2;refGene_gn=ANXA9;refGene_tx=XM_0198378
19.2;refGene_gn=ANXA9;refGene_tx=XM_019837820.2
C1	106120531	106120616
refGene_gn=ANXA9;refGene_tx=XM_003990598.4;refGene_gn=ANXA9;refGene_tx
=XM_019837815.2;refGene_gn=ANXA9;refGene_tx=XM_019837816.2;refGene_gn=
ANXA9;refGene_tx=XM_019837817.2;refGene_gn=ANXA9;refGene_tx=XM_0198378
19.2;refGene_gn=ANXA9;refGene_tx=XM_019837820.2
C1	106120758	106120854
refGene_gn=ANXA9;refGene_tx=XM_003990598.4;refGene_gn=ANXA9;refGene_tx
=XM_019837815.2;refGene_gn=ANXA9;refGene_tx=XM_019837816.2;refGene_gn=
ANXA9;refGene_tx=XM_019837817.2;refGene_gn=ANXA9;refGene_tx=XM_0198378
19.2;refGene_gn=ANXA9;refGene_tx=XM_019837820.2
C1	106120937	106120996
refGene_gn=ANXA9;refGene_tx=XM_003990598.4;refGene_gn=ANXA9;refGene_tx
=XM_019837815.2;refGene_gn=ANXA9;refGene_tx=XM_019837816.2;refGene_gn=
ANXA9;refGene_tx=XM_019837817.2;refGene_gn=ANXA9;refGene_tx=XM_0198378
19.2;refGene_gn=ANXA9;refGene_tx=XM_019837820.2
C1	106124630	106124753
refGene_gn=ANXA9;refGene_tx=XM_003990598.4;refGene_gn=ANXA9;refGene_tx
=XM_019837815.2;refGene_gn=ANXA9;refGene_tx=XM_019837816.2;refGene_gn=
ANXA9;refGene_tx=XM_019837817.2;refGene_gn=ANXA9;refGene_tx=XM_0198378
19.2;refGene_gn=ANXA9;refGene_tx=XM_019837820.2
C1	106124989	106125052
refGene_gn=ANXA9;refGene_tx=XM_003990598.4;refGene_gn=ANXA9;refGene_tx
=XM_019837815.2;refGene_gn=ANXA9;refGene_tx=XM_019837816.2;refGene_gn=
ANXA9;refGene_tx=XM_019837817.2;refGene_gn=ANXA9;refGene_tx=XM_0198378
19.2;refGene_gn=ANXA9;refGene_tx=XM_019837820.2
C1	106126442	106126523
refGene_gn=MINDY1;refGene_tx=XM_003990599.5;refGene_gn=MINDY1;refGene_
tx=XM_006935101.4
C1	106126859	106127003
refGene_gn=MINDY1;refGene_tx=XM_003990599.5;refGene_gn=MINDY1;refGene_
tx=XM_006935101.4
C1	106127537	106127729
refGene_gn=MINDY1;refGene_tx=XM_003990599.5;refGene_gn=MINDY1;refGene_

tx=XM_006935101.4
C1	106128512	106128655
refGene_gn=MINDY1;refGene_tx=XM_003990599.5;refGene_gn=MINDY1;refGene_tx=XM_006935101.4
C1	106129274	106129377
refGene_gn=MINDY1;refGene_tx=XM_003990599.5;refGene_gn=MINDY1;refGene_tx=XM_006935101.4
C1	106129719	106129878
refGene_gn=MINDY1;refGene_tx=XM_003990599.5;refGene_gn=MINDY1;refGene_tx=XM_006935101.4
C1	106130316	106130381
refGene_gn=MINDY1;refGene_tx=XM_003990599.5;refGene_gn=MINDY1;refGene_tx=XM_006935101.4
C1	106130797	106130855
refGene_gn=MINDY1;refGene_tx=XM_003990599.5;refGene_gn=MINDY1;refGene_tx=XM_006935101.4
C1	106131508	106131955
refGene_gn=MINDY1;refGene_tx=XM_003990599.5;refGene_gn=MINDY1;refGene_tx=XM_006935101.4
C1	106137667	106137706
refGene_gn=PRUNE1;refGene_tx=XM_003990600.5
C1	106145164	106145257
refGene_gn=PRUNE1;refGene_tx=XM_003990600.5;refGene_gn=PRUNE1;refGene_tx=XM_006935102.4;refGene_gn=PRUNE1;refGene_tx=XM_019837805.2
C1	106145791	106145994
refGene_gn=PRUNE1;refGene_tx=XM_003990600.5;refGene_gn=PRUNE1;refGene_tx=XM_006935102.4;refGene_gn=PRUNE1;refGene_tx=XM_011285136.2;refGene_gn=PRUNE1;refGene_tx=XM_019837805.2
C1	106149020	106149205
refGene_gn=PRUNE1;refGene_tx=XM_003990600.5;refGene_gn=PRUNE1;refGene_tx=XM_006935102.4;refGene_gn=PRUNE1;refGene_tx=XM_019837805.2
C1	106149799	106149958
refGene_gn=PRUNE1;refGene_tx=XM_003990600.5;refGene_gn=PRUNE1;refGene_tx=XM_006935102.4;refGene_gn=PRUNE1;refGene_tx=XM_019837805.2
C1	106150587	106150682
refGene_gn=PRUNE1;refGene_tx=XM_003990600.5;refGene_gn=PRUNE1;refGene_tx=XM_006935102.4;refGene_gn=PRUNE1;refGene_tx=XM_011285136.2;refGene_gn=PRUNE1;refGene_tx=XM_019837805.2
C1	106152534	106152693
refGene_gn=PRUNE1;refGene_tx=XM_003990600.5;refGene_gn=PRUNE1;refGene_tx=XM_006935102.4;refGene_gn=PRUNE1;refGene_tx=XM_011285136.2;refGene_gn=PRUNE1;refGene_tx=XM_019837805.2
C1	106154833	106155262
refGene_gn=PRUNE1;refGene_tx=XM_003990600.5;refGene_gn=PRUNE1;refGene_tx=XM_006935102.4;refGene_gn=PRUNE1;refGene_tx=XM_011285136.2;refGene_gn=PRUNE1;refGene_tx=XM_019837805.2
C1	106157638	106157682
refGene_gn=BNIPL;refGene_tx=XM_006935104.4;refGene_gn=BNIPL;refGene_tx=XM_019837808.2;refGene_gn=BNIPL;refGene_tx=XM_019837809.2;refGene_gn=BNIPL;refGene_tx=XM_019837811.2;refGene_gn=BNIPL;refGene_tx=XM_0198378

13.2;refGene_gn=BNIPL;refGene_tx=XM_019837814.2
C1	106158951	106159125
refGene_gn=BNIPL;refGene_tx=XM_006935104.4;refGene_gn=BNIPL;refGene_tx=XM_019837808.2;refGene_gn=BNIPL;refGene_tx=XM_019837809.2;refGene_gn=BNIPL;refGene_tx=XM_019837811.2;refGene_gn=BNIPL;refGene_tx=XM_019837813.2
C1	106159285	106159350
refGene_gn=BNIPL;refGene_tx=XM_006935104.4;refGene_gn=BNIPL;refGene_tx=XM_019837808.2;refGene_gn=BNIPL;refGene_tx=XM_019837809.2;refGene_gn=BNIPL;refGene_tx=XM_019837811.2;refGene_gn=BNIPL;refGene_tx=XM_019837813.2;refGene_gn=BNIPL;refGene_tx=XM_019837814.2
C1	106159786	106160011
refGene_gn=BNIPL;refGene_tx=XM_006935104.4;refGene_gn=BNIPL;refGene_tx=XM_019837808.2;refGene_gn=BNIPL;refGene_tx=XM_019837809.2;refGene_gn=BNIPL;refGene_tx=XM_019837811.2;refGene_gn=BNIPL;refGene_tx=XM_019837813.2;refGene_gn=BNIPL;refGene_tx=XM_019837814.2
C1	106162136	106162319
refGene_gn=BNIPL;refGene_tx=XM_006935104.4;refGene_gn=BNIPL;refGene_tx=XM_019837808.2;refGene_gn=BNIPL;refGene_tx=XM_019837809.2;refGene_gn=BNIPL;refGene_tx=XM_019837811.2;refGene_gn=BNIPL;refGene_tx=XM_019837813.2;refGene_gn=BNIPL;refGene_tx=XM_019837814.2
C1	106162509	106162612
refGene_gn=BNIPL;refGene_tx=XM_006935104.4;refGene_gn=BNIPL;refGene_tx=XM_019837808.2;refGene_gn=BNIPL;refGene_tx=XM_019837809.2;refGene_gn=BNIPL;refGene_tx=XM_019837811.2;refGene_gn=BNIPL;refGene_tx=XM_019837813.2;refGene_gn=BNIPL;refGene_tx=XM_019837814.2
C1	106162760	106162907
refGene_gn=BNIPL;refGene_tx=XM_006935104.4;refGene_gn=BNIPL;refGene_tx=XM_019837808.2;refGene_gn=BNIPL;refGene_tx=XM_019837809.2;refGene_gn=BNIPL;refGene_tx=XM_019837811.2;refGene_gn=BNIPL;refGene_tx=XM_019837813.2;refGene_gn=BNIPL;refGene_tx=XM_019837814.2
C1	106163026	106163126
refGene_gn=BNIPL;refGene_tx=XM_019837813.2
C1	106164010	106164097
refGene_gn=BNIPL;refGene_tx=XM_006935104.4;refGene_gn=BNIPL;refGene_tx=XM_019837808.2;refGene_gn=BNIPL;refGene_tx=XM_019837809.2;refGene_gn=BNIPL;refGene_tx=XM_019837811.2;refGene_gn=BNIPL;refGene_tx=XM_019837814.2
C1	106164276	106164375
refGene_gn=BNIPL;refGene_tx=XM_006935104.4;refGene_gn=BNIPL;refGene_tx=XM_019837808.2;refGene_gn=BNIPL;refGene_tx=XM_019837809.2;refGene_gn=BNIPL;refGene_tx=XM_019837811.2;refGene_gn=BNIPL;refGene_tx=XM_019837814.2
C1	106164744	106164781
refGene_gn=BNIPL;refGene_tx=XM_006935104.4;refGene_gn=BNIPL;refGene_tx=XM_019837808.2;refGene_gn=BNIPL;refGene_tx=XM_019837809.2;refGene_gn=BNIPL;refGene_tx=XM_019837811.2;refGene_gn=BNIPL;refGene_tx=XM_019837814.2
C1	106165671	106166502
refGene_gn=CC1H1orf56;refGene_tx=XM_023259139.1

C1 106166503 106166604
refGene_gn=CC1H1orf56;refGene_tx=XM_023259139.1
C1 106166635 106166723
refGene_gn=CC1H1orf56;refGene_tx=XM_023259139.1
C1 106167555 106167576
refGene_gn=CC1H1orf56;refGene_tx=XM_023259139.1
C1 106170727 106170802
refGene_gn=CDC42SE1;refGene_tx=XM_023259138.1
C1 106171507 106171618
refGene_gn=CDC42SE1;refGene_tx=XM_023259138.1
C1 106172127 106172181
refGene_gn=CDC42SE1;refGene_tx=XM_023259138.1
C1 106180990 106181263
refGene_gn=MLLT11;refGene_tx=XM_003990604.5;refGene_gn=MLLT11;refGene_tx=XM_019837821.2;refGene_gn=MLLT11;refGene_tx=XM_023259135.1;refGene_gn=MLLT11;refGene_tx=XM_023259136.1;refGene_gn=MLLT11;refGene_tx=XM_023259137.1
C1 106189646 106189754
refGene_gn=GABPB2;refGene_tx=XM_006935105.4;refGene_gn=GABPB2;refGene_tx=XM_006935106.4;refGene_gn=GABPB2;refGene_tx=XM_006935107.4;refGene_gn=GABPB2;refGene_tx=XM_019837806.2;refGene_gn=GABPB2;refGene_tx=XM_019837807.2
C1 106191640 106191808
refGene_gn=GABPB2;refGene_tx=XM_006935105.4;refGene_gn=GABPB2;refGene_tx=XM_006935106.4;refGene_gn=GABPB2;refGene_tx=XM_019837806.2
C1 106192348 106192543
refGene_gn=GABPB2;refGene_tx=XM_006935105.4;refGene_gn=GABPB2;refGene_tx=XM_006935106.4;refGene_gn=GABPB2;refGene_tx=XM_006935107.4;refGene_gn=GABPB2;refGene_tx=XM_019837806.2;refGene_gn=GABPB2;refGene_tx=XM_019837807.2
C1 106195711 106195850
refGene_gn=GABPB2;refGene_tx=XM_006935105.4;refGene_gn=GABPB2;refGene_tx=XM_006935106.4;refGene_gn=GABPB2;refGene_tx=XM_006935107.4;refGene_gn=GABPB2;refGene_tx=XM_019837806.2;refGene_gn=GABPB2;refGene_tx=XM_019837807.2
C1 106204427 106204544
refGene_gn=GABPB2;refGene_tx=XM_006935105.4;refGene_gn=GABPB2;refGene_tx=XM_006935107.4;refGene_gn=GABPB2;refGene_tx=XM_019837806.2
C1 106208094 106208280
refGene_gn=GABPB2;refGene_tx=XM_006935105.4;refGene_gn=GABPB2;refGene_tx=XM_006935106.4;refGene_gn=GABPB2;refGene_tx=XM_006935107.4;refGene_gn=GABPB2;refGene_tx=XM_019837806.2;refGene_gn=GABPB2;refGene_tx=XM_019837807.2
C1 106213768 106213902
refGene_gn=GABPB2;refGene_tx=XM_006935105.4;refGene_gn=GABPB2;refGene_tx=XM_006935106.4;refGene_gn=GABPB2;refGene_tx=XM_006935107.4;refGene_gn=GABPB2;refGene_tx=XM_019837806.2;refGene_gn=GABPB2;refGene_tx=XM_019837807.2
C1 106214291 106214594
refGene_gn=GABPB2;refGene_tx=XM_006935105.4;refGene_gn=GABPB2;refGene_

tx=XM_006935106.4;refGene_gn=GABPB2;refGene_tx=XM_006935107.4;refGene_gn=GABPB2;refGene_tx=XM_019837806.2;refGene_gn=GABPB2;refGene_tx=XM_019837807.2
C1 106226800 106227846
refGene_gn=SEMA6C;refGene_tx=XM_023259140.1;refGene_gn=SEMA6C;refGene_tx=XM_023259141.1;refGene_gn=SEMA6C;refGene_tx=XM_023259142.1;refGene_gn=SEMA6C;refGene_tx=XM_023259143.1;refGene_gn=SEMA6C;refGene_tx=XM_023259144.1;refGene_gn=SEMA6C;refGene_tx=XM_023259145.1;refGene_gn=SEMA6C;refGene_tx=XM_023259146.1;refGene_gn=SEMA6C;refGene_tx=XM_023259147.1;refGene_gn=SEMA6C;refGene_tx=XM_023259148.1;refGene_gn=SEMA6C;refGene_tx=XM_023259149.1;refGene_gn=SEMA6C;refGene_tx=XM_023259151.1;refGene_gn=SEMA6C;refGene_tx=XM_023259152.1;refGene_gn=SEMA6C;refGene_tx=XM_023259153.1
C1 106228254 106228350
refGene_gn=SEMA6C;refGene_tx=XM_023259140.1;refGene_gn=SEMA6C;refGene_tx=XM_023259141.1;refGene_gn=SEMA6C;refGene_tx=XM_023259142.1;refGene_gn=SEMA6C;refGene_tx=XM_023259143.1;refGene_gn=SEMA6C;refGene_tx=XM_023259144.1;refGene_gn=SEMA6C;refGene_tx=XM_023259145.1;refGene_gn=SEMA6C;refGene_tx=XM_023259148.1;refGene_gn=SEMA6C;refGene_tx=XM_023259149.1;refGene_gn=SEMA6C;refGene_tx=XM_023259152.1;refGene_gn=SEMA6C;refGene_tx=XM_023259153.1
C1 106228722 106228767
refGene_gn=SEMA6C;refGene_tx=XM_023259140.1;refGene_gn=SEMA6C;refGene_tx=XM_023259141.1;refGene_gn=SEMA6C;refGene_tx=XM_023259142.1;refGene_gn=SEMA6C;refGene_tx=XM_023259143.1;refGene_gn=SEMA6C;refGene_tx=XM_023259144.1;refGene_gn=SEMA6C;refGene_tx=XM_023259145.1;refGene_gn=SEMA6C;refGene_tx=XM_023259146.1;refGene_gn=SEMA6C;refGene_tx=XM_023259147.1;refGene_gn=SEMA6C;refGene_tx=XM_023259148.1;refGene_gn=SEMA6C;refGene_tx=XM_023259149.1;refGene_gn=SEMA6C;refGene_tx=XM_023259151.1;refGene_gn=SEMA6C;refGene_tx=XM_023259152.1;refGene_gn=SEMA6C;refGene_tx=XM_023259153.1
C1 106228938 106228994
refGene_gn=SEMA6C;refGene_tx=XM_023259140.1;refGene_gn=SEMA6C;refGene_tx=XM_023259141.1;refGene_gn=SEMA6C;refGene_tx=XM_023259142.1;refGene_gn=SEMA6C;refGene_tx=XM_023259143.1;refGene_gn=SEMA6C;refGene_tx=XM_023259144.1;refGene_gn=SEMA6C;refGene_tx=XM_023259145.1;refGene_gn=SEMA6C;refGene_tx=XM_023259146.1;refGene_gn=SEMA6C;refGene_tx=XM_023259147.1;refGene_gn=SEMA6C;refGene_tx=XM_023259148.1;refGene_gn=SEMA6C;refGene_tx=XM_023259149.1;refGene_gn=SEMA6C;refGene_tx=XM_023259151.1;refGene_gn=SEMA6C;refGene_tx=XM_023259152.1;refGene_gn=SEMA6C;refGene_tx=XM_023259153.1
C1 106229131 106229209
refGene_gn=SEMA6C;refGene_tx=XM_023259140.1;refGene_gn=SEMA6C;refGene_tx=XM_023259141.1;refGene_gn=SEMA6C;refGene_tx=XM_023259142.1;refGene_gn=SEMA6C;refGene_tx=XM_023259143.1;refGene_gn=SEMA6C;refGene_tx=XM_023259144.1;refGene_gn=SEMA6C;refGene_tx=XM_023259145.1;refGene_gn=SEMA6C;refGene_tx=XM_023259146.1;refGene_gn=SEMA6C;refGene_tx=XM_023259147.1;refGene_gn=SEMA6C;refGene_tx=XM_023259148.1;refGene_gn=SEMA6C;refGene_tx=XM_023259149.1;refGene_gn=SEMA6C;refGene_tx=XM_023259151.1;refGene_gn=SEMA6C;refGene_tx=XM_023259152.1;refGene_gn=SEMA6C;refGene_tx=XM_

023259153.1
C1 106229456 106229603
refGene_gn=SEMA6C;refGene_tx=XM_023259140.1;refGene_gn=SEMA6C;refGene_tx=XM_023259141.1;refGene_gn=SEMA6C;refGene_tx=XM_023259142.1;refGene_gn=SEMA6C;refGene_tx=XM_023259143.1;refGene_gn=SEMA6C;refGene_tx=XM_023259144.1;refGene_gn=SEMA6C;refGene_tx=XM_023259145.1;refGene_gn=SEMA6C;refGene_tx=XM_023259146.1;refGene_gn=SEMA6C;refGene_tx=XM_023259147.1;refGene_gn=SEMA6C;refGene_tx=XM_023259148.1;refGene_gn=SEMA6C;refGene_tx=XM_023259149.1;refGene_gn=SEMA6C;refGene_tx=XM_023259151.1;refGene_gn=SEMA6C;refGene_tx=XM_023259152.1;refGene_gn=SEMA6C;refGene_tx=XM_023259153.1
C1 106229899 106230073
refGene_gn=SEMA6C;refGene_tx=XM_023259140.1;refGene_gn=SEMA6C;refGene_tx=XM_023259141.1;refGene_gn=SEMA6C;refGene_tx=XM_023259142.1;refGene_gn=SEMA6C;refGene_tx=XM_023259143.1;refGene_gn=SEMA6C;refGene_tx=XM_023259144.1;refGene_gn=SEMA6C;refGene_tx=XM_023259145.1;refGene_gn=SEMA6C;refGene_tx=XM_023259146.1;refGene_gn=SEMA6C;refGene_tx=XM_023259147.1;refGene_gn=SEMA6C;refGene_tx=XM_023259148.1;refGene_gn=SEMA6C;refGene_tx=XM_023259149.1;refGene_gn=SEMA6C;refGene_tx=XM_023259151.1;refGene_gn=SEMA6C;refGene_tx=XM_023259152.1;refGene_gn=SEMA6C;refGene_tx=XM_023259153.1
C1 106230311 106230464
refGene_gn=SEMA6C;refGene_tx=XM_023259140.1;refGene_gn=SEMA6C;refGene_tx=XM_023259141.1;refGene_gn=SEMA6C;refGene_tx=XM_023259142.1;refGene_gn=SEMA6C;refGene_tx=XM_023259143.1;refGene_gn=SEMA6C;refGene_tx=XM_023259144.1;refGene_gn=SEMA6C;refGene_tx=XM_023259145.1;refGene_gn=SEMA6C;refGene_tx=XM_023259146.1;refGene_gn=SEMA6C;refGene_tx=XM_023259147.1;refGene_gn=SEMA6C;refGene_tx=XM_023259148.1;refGene_gn=SEMA6C;refGene_tx=XM_023259149.1;refGene_gn=SEMA6C;refGene_tx=XM_023259151.1;refGene_gn=SEMA6C;refGene_tx=XM_023259152.1;refGene_gn=SEMA6C;refGene_tx=XM_023259153.1
C1 106230741 106230873
refGene_gn=SEMA6C;refGene_tx=XM_023259140.1;refGene_gn=SEMA6C;refGene_tx=XM_023259141.1;refGene_gn=SEMA6C;refGene_tx=XM_023259142.1;refGene_gn=SEMA6C;refGene_tx=XM_023259143.1;refGene_gn=SEMA6C;refGene_tx=XM_023259144.1;refGene_gn=SEMA6C;refGene_tx=XM_023259145.1;refGene_gn=SEMA6C;refGene_tx=XM_023259146.1;refGene_gn=SEMA6C;refGene_tx=XM_023259147.1;refGene_gn=SEMA6C;refGene_tx=XM_023259148.1;refGene_gn=SEMA6C;refGene_tx=XM_023259149.1;refGene_gn=SEMA6C;refGene_tx=XM_023259151.1;refGene_gn=SEMA6C;refGene_tx=XM_023259152.1;refGene_gn=SEMA6C;refGene_tx=XM_023259153.1
C1 106231160 106231378
refGene_gn=SEMA6C;refGene_tx=XM_023259140.1;refGene_gn=SEMA6C;refGene_tx=XM_023259141.1;refGene_gn=SEMA6C;refGene_tx=XM_023259142.1;refGene_gn=SEMA6C;refGene_tx=XM_023259143.1;refGene_gn=SEMA6C;refGene_tx=XM_023259144.1;refGene_gn=SEMA6C;refGene_tx=XM_023259145.1;refGene_gn=SEMA6C;refGene_tx=XM_023259146.1;refGene_gn=SEMA6C;refGene_tx=XM_023259147.1;refGene_gn=SEMA6C;refGene_tx=XM_023259148.1;refGene_gn=SEMA6C;refGene_tx=XM_023259149.1;refGene_gn=SEMA6C;refGene_tx=XM_023259151.1;refGene_gn=SEMA6C;refGene_tx=XM_023259152.1;refGene_gn=SEMA6C;refGene_tx=XM_

023259153.1
C1 106231710 106231799
refGene_gn=SEMA6C;refGene_tx=XM_023259140.1;refGene_gn=SEMA6C;refGene_tx=XM_023259141.1;refGene_gn=SEMA6C;refGene_tx=XM_023259142.1;refGene_gn=SEMA6C;refGene_tx=XM_023259143.1;refGene_gn=SEMA6C;refGene_tx=XM_023259144.1;refGene_gn=SEMA6C;refGene_tx=XM_023259145.1;refGene_gn=SEMA6C;refGene_tx=XM_023259146.1;refGene_gn=SEMA6C;refGene_tx=XM_023259147.1;refGene_gn=SEMA6C;refGene_tx=XM_023259148.1;refGene_gn=SEMA6C;refGene_tx=XM_023259149.1;refGene_gn=SEMA6C;refGene_tx=XM_023259151.1;refGene_gn=SEMA6C;refGene_tx=XM_023259152.1;refGene_gn=SEMA6C;refGene_tx=XM_023259153.1
C1 106231979 106232099
refGene_gn=SEMA6C;refGene_tx=XM_023259140.1;refGene_gn=SEMA6C;refGene_tx=XM_023259141.1;refGene_gn=SEMA6C;refGene_tx=XM_023259142.1;refGene_gn=SEMA6C;refGene_tx=XM_023259143.1;refGene_gn=SEMA6C;refGene_tx=XM_023259144.1;refGene_gn=SEMA6C;refGene_tx=XM_023259145.1;refGene_gn=SEMA6C;refGene_tx=XM_023259146.1;refGene_gn=SEMA6C;refGene_tx=XM_023259147.1;refGene_gn=SEMA6C;refGene_tx=XM_023259148.1;refGene_gn=SEMA6C;refGene_tx=XM_023259149.1;refGene_gn=SEMA6C;refGene_tx=XM_023259151.1;refGene_gn=SEMA6C;refGene_tx=XM_023259152.1;refGene_gn=SEMA6C;refGene_tx=XM_023259153.1
C1 106232306 106232397
refGene_gn=SEMA6C;refGene_tx=XM_023259140.1;refGene_gn=SEMA6C;refGene_tx=XM_023259141.1;refGene_gn=SEMA6C;refGene_tx=XM_023259142.1;refGene_gn=SEMA6C;refGene_tx=XM_023259143.1;refGene_gn=SEMA6C;refGene_tx=XM_023259144.1;refGene_gn=SEMA6C;refGene_tx=XM_023259145.1;refGene_gn=SEMA6C;refGene_tx=XM_023259146.1;refGene_gn=SEMA6C;refGene_tx=XM_023259147.1;refGene_gn=SEMA6C;refGene_tx=XM_023259148.1;refGene_gn=SEMA6C;refGene_tx=XM_023259149.1;refGene_gn=SEMA6C;refGene_tx=XM_023259151.1;refGene_gn=SEMA6C;refGene_tx=XM_023259152.1;refGene_gn=SEMA6C;refGene_tx=XM_023259153.1
C1 106232602 106232704
refGene_gn=SEMA6C;refGene_tx=XM_023259140.1;refGene_gn=SEMA6C;refGene_tx=XM_023259141.1;refGene_gn=SEMA6C;refGene_tx=XM_023259142.1;refGene_gn=SEMA6C;refGene_tx=XM_023259143.1;refGene_gn=SEMA6C;refGene_tx=XM_023259144.1;refGene_gn=SEMA6C;refGene_tx=XM_023259145.1;refGene_gn=SEMA6C;refGene_tx=XM_023259146.1;refGene_gn=SEMA6C;refGene_tx=XM_023259147.1;refGene_gn=SEMA6C;refGene_tx=XM_023259148.1;refGene_gn=SEMA6C;refGene_tx=XM_023259149.1;refGene_gn=SEMA6C;refGene_tx=XM_023259151.1;refGene_gn=SEMA6C;refGene_tx=XM_023259152.1
C1 106233059 106233116
refGene_gn=SEMA6C;refGene_tx=XM_023259140.1;refGene_gn=SEMA6C;refGene_tx=XM_023259141.1;refGene_gn=SEMA6C;refGene_tx=XM_023259142.1;refGene_gn=SEMA6C;refGene_tx=XM_023259143.1;refGene_gn=SEMA6C;refGene_tx=XM_023259144.1;refGene_gn=SEMA6C;refGene_tx=XM_023259145.1;refGene_gn=SEMA6C;refGene_tx=XM_023259146.1;refGene_gn=SEMA6C;refGene_tx=XM_023259147.1;refGene_gn=SEMA6C;refGene_tx=XM_023259148.1;refGene_gn=SEMA6C;refGene_tx=XM_023259149.1;refGene_gn=SEMA6C;refGene_tx=XM_023259151.1;refGene_gn=SEMA6C;refGene_tx=XM_023259152.1
C1 106233274 106233338

refGene_gn=SEMA6C;refGene_tx=XM_023259140.1;refGene_gn=SEMA6C;refGene_tx=XM_023259141.1;refGene_gn=SEMA6C;refGene_tx=XM_023259142.1;refGene_gn=SEMA6C;refGene_tx=XM_023259143.1;refGene_gn=SEMA6C;refGene_tx=XM_023259144.1;refGene_gn=SEMA6C;refGene_tx=XM_023259145.1;refGene_gn=SEMA6C;refGene_tx=XM_023259146.1;refGene_gn=SEMA6C;refGene_tx=XM_023259147.1;refGene_gn=SEMA6C;refGene_tx=XM_023259148.1;refGene_gn=SEMA6C;refGene_tx=XM_023259149.1;refGene_gn=SEMA6C;refGene_tx=XM_023259151.1
C1 106233465 106233588
refGene_gn=SEMA6C;refGene_tx=XM_023259148.1;refGene_gn=SEMA6C;refGene_tx=XM_023259149.1
C1 106233625 106233740
refGene_gn=SEMA6C;refGene_tx=XM_023259140.1;refGene_gn=SEMA6C;refGene_tx=XM_023259141.1;refGene_gn=SEMA6C;refGene_tx=XM_023259142.1;refGene_gn=SEMA6C;refGene_tx=XM_023259143.1;refGene_gn=SEMA6C;refGene_tx=XM_023259144.1;refGene_gn=SEMA6C;refGene_tx=XM_023259145.1;refGene_gn=SEMA6C;refGene_tx=XM_023259146.1;refGene_gn=SEMA6C;refGene_tx=XM_023259147.1;refGene_gn=SEMA6C;refGene_tx=XM_023259151.1
C1 106235500 106235612
refGene_gn=SEMA6C;refGene_tx=XM_023259140.1;refGene_gn=SEMA6C;refGene_tx=XM_023259141.1;refGene_gn=SEMA6C;refGene_tx=XM_023259142.1;refGene_gn=SEMA6C;refGene_tx=XM_023259143.1;refGene_gn=SEMA6C;refGene_tx=XM_023259144.1;refGene_gn=SEMA6C;refGene_tx=XM_023259145.1;refGene_gn=SEMA6C;refGene_tx=XM_023259146.1;refGene_gn=SEMA6C;refGene_tx=XM_023259147.1;refGene_gn=SEMA6C;refGene_tx=XM_023259151.1
C1 106252927 106253482
refGene_gn=TNFAIP8L2;refGene_tx=XM_003990607.5
C1 106255122 106255261
refGene_gn=LYSMD1;refGene_tx=XM_003990608.5
C1 106255674 106256039
refGene_gn=LYSMD1;refGene_tx=XM_003990608.5
C1 106260788 106260968
refGene_gn=LYSMD1;refGene_tx=XM_003990608.5
C1 106261875 106261926
refGene_gn=SCNM1;refGene_tx=XM_003990609.5
C1 106262161 106262232
refGene_gn=SCNM1;refGene_tx=XM_003990609.5;refGene_gn=SCNM1;refGene_tx=XM_019837827.2;refGene_gn=SCNM1;refGene_tx=XM_023259155.1
C1 106262419 106262508
refGene_gn=SCNM1;refGene_tx=XM_003990609.5;refGene_gn=SCNM1;refGene_tx=XM_019837827.2;refGene_gn=SCNM1;refGene_tx=XM_023259155.1
C1 106262607 106262705
refGene_gn=SCNM1;refGene_tx=XM_003990609.5;refGene_gn=SCNM1;refGene_tx=XM_019837827.2;refGene_gn=SCNM1;refGene_tx=XM_023259155.1
C1 106262808 106262897
refGene_gn=SCNM1;refGene_tx=XM_003990609.5;refGene_gn=SCNM1;refGene_tx=XM_019837827.2;refGene_gn=SCNM1;refGene_tx=XM_023259155.1
C1 106263769 106263964
refGene_gn=SCNM1;refGene_tx=XM_003990609.5;refGene_gn=SCNM1;refGene_tx=XM_019837827.2;refGene_gn=SCNM1;refGene_tx=XM_023259155.1
C1 106264113 106264213

refGene_gn=SCNM1;refGene_tx=XM_003990609.5;refGene_gn=SCNM1;refGene_tx=XM_019837827.2;refGene_gn=SCNM1;refGene_tx=XM_023259155.1
C1 106264889 106264912
refGene_gn=TMOD4;refGene_tx=XM_003990610.5;refGene_gn=TMOD4;refGene_tx=XM_006935111.3
C1 106265299 106265444
refGene_gn=TMOD4;refGene_tx=XM_003990610.5;refGene_gn=TMOD4;refGene_tx=XM_006935111.3
C1 106265700 106265844
refGene_gn=TMOD4;refGene_tx=XM_003990610.5;refGene_gn=TMOD4;refGene_tx=XM_006935111.3
C1 106266258 106266366
refGene_gn=TMOD4;refGene_tx=XM_003990610.5;refGene_gn=TMOD4;refGene_tx=XM_006935111.3
C1 106266462 106266593
refGene_gn=TMOD4;refGene_tx=XM_003990610.5;refGene_gn=TMOD4;refGene_tx=XM_006935111.3
C1 106266773 106266863
refGene_gn=TMOD4;refGene_tx=XM_003990610.5;refGene_gn=TMOD4;refGene_tx=XM_006935111.3
C1 106267642 106267759
refGene_gn=TMOD4;refGene_tx=XM_003990610.5;refGene_gn=TMOD4;refGene_tx=XM_006935111.3
C1 106268078 106268235
refGene_gn=TMOD4;refGene_tx=XM_003990610.5;refGene_gn=TMOD4;refGene_tx=XM_006935111.3
C1 106268446 106268569
refGene_gn=TMOD4;refGene_tx=XM_003990610.5;refGene_gn=TMOD4;refGene_tx=XM_006935111.3
C1 106270328 106270716
refGene_gn=VPS72;refGene_tx=XM_003990611.5;refGene_gn=VPS72;refGene_tx=XM_019837826.2
C1 106271030 106271175
refGene_gn=VPS72;refGene_tx=XM_003990611.5
C1 106276336 106276513
refGene_gn=VPS72;refGene_tx=XM_003990611.5;refGene_gn=VPS72;refGene_tx=XM_019837826.2
C1 106276904 106277019
refGene_gn=VPS72;refGene_tx=XM_003990611.5;refGene_gn=VPS72;refGene_tx=XM_019837826.2
C1 106277199 106277352
refGene_gn=VPS72;refGene_tx=XM_003990611.5;refGene_gn=VPS72;refGene_tx=XM_019837826.2
C1 106281453 106281570
refGene_gn=VPS72;refGene_tx=XM_003990611.5;refGene_gn=VPS72;refGene_tx=XM_019837826.2
C1 106285563 106285645
refGene_gn=PIP5K1A;refGene_tx=XM_006935116.4
C1 106295095 106295134
refGene_gn=PIP5K1A;refGene_tx=XM_006935116.4

C1 106302683 106302715
refGene_gn=PIP5K1A;refGene_tx=XM_006935116.4
C1 106305043 106305124
refGene_gn=PIP5K1A;refGene_tx=XM_006935116.4
C1 106307015 106307146
refGene_gn=PIP5K1A;refGene_tx=XM_006935116.4
C1 106307587 106307705
refGene_gn=PIP5K1A;refGene_tx=XM_006935116.4
C1 106307918 106308071
refGene_gn=PIP5K1A;refGene_tx=XM_006935116.4
C1 106308795 106309095
refGene_gn=PIP5K1A;refGene_tx=XM_006935116.4
C1 106313505 106313711
refGene_gn=PIP5K1A;refGene_tx=XM_006935116.4
C1 106314494 106314543
refGene_gn=PIP5K1A;refGene_tx=XM_006935116.4
C1 106314544 106314579
refGene_gn=PIP5K1A;refGene_tx=XM_006935116.4
C1 106315403 106315452
refGene_gn=PIP5K1A;refGene_tx=XM_006935116.4
C1 106315649 106315734
refGene_gn=PIP5K1A;refGene_tx=XM_006935116.4
C1 106316804 106316951
refGene_gn=PIP5K1A;refGene_tx=XM_006935116.4
C1 106317153 106317283
refGene_gn=PIP5K1A;refGene_tx=XM_006935116.4
C1 106320319 106320368
refGene_gn=PIP5K1A;refGene_tx=XM_006935116.4
C1 106320652 106320655
refGene_gn=PIP5K1A;refGene_tx=XM_006935116.4
C1 106332265 106332291
refGene_gn=PSMD4;refGene_tx=XM_011285139.3
C1 106334224 106334259
refGene_gn=PSMD4;refGene_tx=XM_019837825.2
C1 106337016 106337157
refGene_gn=PSMD4;refGene_tx=XM_011285139.3;refGene_gn=PSMD4;refGene_tx=XM_019837825.2
C1 106338118 106338233
refGene_gn=PSMD4;refGene_tx=XM_011285139.3;refGene_gn=PSMD4;refGene_tx=XM_019837825.2
C1 106338638 106338725
refGene_gn=PSMD4;refGene_tx=XM_011285139.3;refGene_gn=PSMD4;refGene_tx=XM_019837825.2
C1 106338954 106339023
refGene_gn=PSMD4;refGene_tx=XM_011285139.3;refGene_gn=PSMD4;refGene_tx=XM_019837825.2
C1 106339193 106339409
refGene_gn=PSMD4;refGene_tx=XM_011285139.3;refGene_gn=PSMD4;refGene_tx=XM_019837825.2
C1 106339798 106339907

refGene_gn=PSMD4;refGene_tx=XM_011285139.3;refGene_gn=PSMD4;refGene_tx=XM_019837825.2
C1 106340075 106340207
refGene_gn=PSMD4;refGene_tx=XM_011285139.3;refGene_gn=PSMD4;refGene_tx=XM_019837825.2
C1 106340292 106340360
refGene_gn=PSMD4;refGene_tx=XM_011285139.3;refGene_gn=PSMD4;refGene_tx=XM_019837825.2
C1 106340943 106341114
refGene_gn=PSMD4;refGene_tx=XM_011285139.3;refGene_gn=PSMD4;refGene_tx=XM_019837825.2
C1 106376536 106378642
refGene_gn=ZNF687;refGene_tx=XM_003990616.4;refGene_gn=ZNF687;refGene_tx=XM_006935118.4;refGene_gn=ZNF687;refGene_tx=XM_006935119.4;refGene_gn=ZNF687;refGene_tx=XM_019837829.2;refGene_gn=ZNF687;refGene_tx=XM_019837830.2;refGene_gn=ZNF687;refGene_tx=XM_019837831.2
C1 106378779 106378958
refGene_gn=ZNF687;refGene_tx=XM_003990616.4;refGene_gn=ZNF687;refGene_tx=XM_006935118.4;refGene_gn=ZNF687;refGene_tx=XM_006935119.4;refGene_gn=ZNF687;refGene_tx=XM_019837829.2;refGene_gn=ZNF687;refGene_tx=XM_019837830.2;refGene_gn=ZNF687;refGene_tx=XM_019837831.2
C1 106379333 106379510
refGene_gn=ZNF687;refGene_tx=XM_003990616.4;refGene_gn=ZNF687;refGene_tx=XM_006935118.4;refGene_gn=ZNF687;refGene_tx=XM_006935119.4;refGene_gn=ZNF687;refGene_tx=XM_019837829.2;refGene_gn=ZNF687;refGene_tx=XM_019837830.2;refGene_gn=ZNF687;refGene_tx=XM_019837831.2
C1 106379598 106379761
refGene_gn=ZNF687;refGene_tx=XM_003990616.4;refGene_gn=ZNF687;refGene_tx=XM_006935118.4;refGene_gn=ZNF687;refGene_tx=XM_006935119.4;refGene_gn=ZNF687;refGene_tx=XM_019837829.2;refGene_gn=ZNF687;refGene_tx=XM_019837830.2;refGene_gn=ZNF687;refGene_tx=XM_019837831.2
C1 106379932 106380268
refGene_gn=ZNF687;refGene_tx=XM_003990616.4;refGene_gn=ZNF687;refGene_tx=XM_006935118.4;refGene_gn=ZNF687;refGene_tx=XM_006935119.4;refGene_gn=ZNF687;refGene_tx=XM_019837829.2;refGene_gn=ZNF687;refGene_tx=XM_019837830.2;refGene_gn=ZNF687;refGene_tx=XM_019837831.2
C1 106380365 106380478
refGene_gn=ZNF687;refGene_tx=XM_003990616.4;refGene_gn=ZNF687;refGene_tx=XM_006935118.4;refGene_gn=ZNF687;refGene_tx=XM_006935119.4;refGene_gn=ZNF687;refGene_tx=XM_019837829.2;refGene_gn=ZNF687;refGene_tx=XM_019837830.2;refGene_gn=ZNF687;refGene_tx=XM_019837831.2
C1 106380744 106380886
refGene_gn=ZNF687;refGene_tx=XM_003990616.4;refGene_gn=ZNF687;refGene_tx=XM_006935118.4;refGene_gn=ZNF687;refGene_tx=XM_006935119.4;refGene_gn=ZNF687;refGene_tx=XM_019837829.2;refGene_gn=ZNF687;refGene_tx=XM_019837830.2;refGene_gn=ZNF687;refGene_tx=XM_019837831.2
C1 106381031 106381526
refGene_gn=ZNF687;refGene_tx=XM_003990616.4;refGene_gn=ZNF687;refGene_tx=XM_006935118.4;refGene_gn=ZNF687;refGene_tx=XM_006935119.4;refGene_gn=ZNF687;refGene_tx=XM_019837829.2;refGene_gn=ZNF687;refGene_tx=XM_01

9837830.2;refGene_gn=ZNF687;refGene_tx=XM_019837831.2
C1	106382991	106383173
refGene_gn=PI4KB;refGene_tx=XM_006935120.4;refGene_gn=PI4KB;refGene_tx=XM_006935121.4;refGene_gn=PI4KB;refGene_tx=XM_006935122.4;refGene_gn=PI4KB;refGene_tx=XM_006935123.4;refGene_gn=PI4KB;refGene_tx=XM_023259156.1
C1	106383477	106383540
refGene_gn=PI4KB;refGene_tx=XM_006935120.4;refGene_gn=PI4KB;refGene_tx=XM_006935121.4;refGene_gn=PI4KB;refGene_tx=XM_023259156.1
C1	106384209	106384330
refGene_gn=PI4KB;refGene_tx=XM_006935120.4;refGene_gn=PI4KB;refGene_tx=XM_006935121.4;refGene_gn=PI4KB;refGene_tx=XM_006935122.4;refGene_gn=PI4KB;refGene_tx=XM_006935123.4;refGene_gn=PI4KB;refGene_tx=XM_023259156.1
C1	106384583	106384716
refGene_gn=PI4KB;refGene_tx=XM_006935120.4;refGene_gn=PI4KB;refGene_tx=XM_006935121.4;refGene_gn=PI4KB;refGene_tx=XM_006935122.4;refGene_gn=PI4KB;refGene_tx=XM_006935123.4;refGene_gn=PI4KB;refGene_tx=XM_023259156.1
C1	106387273	106387539
refGene_gn=PI4KB;refGene_tx=XM_006935120.4;refGene_gn=PI4KB;refGene_tx=XM_006935121.4;refGene_gn=PI4KB;refGene_tx=XM_006935122.4;refGene_gn=PI4KB;refGene_tx=XM_006935123.4;refGene_gn=PI4KB;refGene_tx=XM_023259156.1
C1	106388198	106388322
refGene_gn=PI4KB;refGene_tx=XM_006935120.4;refGene_gn=PI4KB;refGene_tx=XM_006935121.4;refGene_gn=PI4KB;refGene_tx=XM_006935122.4;refGene_gn=PI4KB;refGene_tx=XM_006935123.4;refGene_gn=PI4KB;refGene_tx=XM_023259156.1
C1	106388537	106388642
refGene_gn=PI4KB;refGene_tx=XM_006935120.4;refGene_gn=PI4KB;refGene_tx=XM_006935121.4;refGene_gn=PI4KB;refGene_tx=XM_006935122.4;refGene_gn=PI4KB;refGene_tx=XM_006935123.4
C1	106389192	106389302
refGene_gn=PI4KB;refGene_tx=XM_006935120.4;refGene_gn=PI4KB;refGene_tx=XM_006935121.4;refGene_gn=PI4KB;refGene_tx=XM_006935122.4;refGene_gn=PI4KB;refGene_tx=XM_006935123.4;refGene_gn=PI4KB;refGene_tx=XM_023259156.1
C1	106390949	106391177
refGene_gn=PI4KB;refGene_tx=XM_006935120.4;refGene_gn=PI4KB;refGene_tx=XM_006935121.4;refGene_gn=PI4KB;refGene_tx=XM_006935122.4;refGene_gn=PI4KB;refGene_tx=XM_006935123.4;refGene_gn=PI4KB;refGene_tx=XM_023259156.1
C1	106392211	106392439
refGene_gn=PI4KB;refGene_tx=XM_006935120.4;refGene_gn=PI4KB;refGene_tx=XM_006935121.4;refGene_gn=PI4KB;refGene_tx=XM_006935122.4;refGene_gn=PI4KB;refGene_tx=XM_006935123.4;refGene_gn=PI4KB;refGene_tx=XM_023259156.1
C1	106394827	106394872
refGene_gn=PI4KB;refGene_tx=XM_006935120.4;refGene_gn=PI4KB;refGene_tx

=XM_006935122.4;refGene_gn=PI4KB;refGene_tx=XM_023259156.1
C1	106399410	106400319
refGene_gn=PI4KB;refGene_tx=XM_006935120.4;refGene_gn=PI4KB;refGene_tx=XM_006935121.4;refGene_gn=PI4KB;refGene_tx=XM_006935122.4;refGene_gn=PI4KB;refGene_tx=XM_006935123.4;refGene_gn=PI4KB;refGene_tx=XM_023259156.1
C1	106417811	106418450
refGene_gn=RFX5;refGene_tx=XM_019837833.2
C1	106418452	106418824
refGene_gn=RFX5;refGene_tx=XM_019837833.2
C1	106418969	106419070
refGene_gn=RFX5;refGene_tx=XM_019837833.2
C1	106419310	106419512
refGene_gn=RFX5;refGene_tx=XM_019837833.2
C1	106419770	106419852
refGene_gn=RFX5;refGene_tx=XM_019837833.2
C1	106419995	106420115
refGene_gn=RFX5;refGene_tx=XM_019837833.2
C1	106420322	106420442
refGene_gn=RFX5;refGene_tx=XM_019837833.2
C1	106420694	106420777
refGene_gn=RFX5;refGene_tx=XM_019837833.2
C1	106421209	106421243
refGene_gn=RFX5;refGene_tx=XM_019837833.2
C1	106421473	106421589
refGene_gn=RFX5;refGene_tx=XM_019837833.2
C1	106432492	106432655
refGene_gn=SELENBP1;refGene_tx=XM_003990619.3;refGene_gn=SELENBP1;refGene_tx=XM_019837840.2;refGene_gn=SELENBP1;refGene_tx=XM_019837842.2
C1	106432860	106432979
refGene_gn=SELENBP1;refGene_tx=XM_003990619.3;refGene_gn=SELENBP1;refGene_tx=XM_019837840.2;refGene_gn=SELENBP1;refGene_tx=XM_019837842.2
C1	106433150	106433243
refGene_gn=SELENBP1;refGene_tx=XM_003990619.3;refGene_gn=SELENBP1;refGene_tx=XM_019837840.2;refGene_gn=SELENBP1;refGene_tx=XM_019837842.2
C1	106433510	106433632
refGene_gn=SELENBP1;refGene_tx=XM_003990619.3;refGene_gn=SELENBP1;refGene_tx=XM_019837840.2;refGene_gn=SELENBP1;refGene_tx=XM_019837842.2
C1	106433726	106433805
refGene_gn=SELENBP1;refGene_tx=XM_003990619.3;refGene_gn=SELENBP1;refGene_tx=XM_019837840.2;refGene_gn=SELENBP1;refGene_tx=XM_019837842.2
C1	106434229	106434408
refGene_gn=SELENBP1;refGene_tx=XM_003990619.3;refGene_gn=SELENBP1;refGene_tx=XM_019837840.2;refGene_gn=SELENBP1;refGene_tx=XM_019837842.2
C1	106434685	106434868
refGene_gn=SELENBP1;refGene_tx=XM_003990619.3;refGene_gn=SELENBP1;refGene_tx=XM_019837840.2;refGene_gn=SELENBP1;refGene_tx=XM_019837842.2
C1	106436017	106436138
refGene_gn=SELENBP1;refGene_tx=XM_003990619.3;refGene_gn=SELENBP1;refGene_tx=XM_019837840.2;refGene_gn=SELENBP1;refGene_tx=XM_019837842.2

C1	106436794	106436980
refGene_gn=SELENBP1;refGene_tx=XM_003990619.3;refGene_gn=SELENBP1;refGene_tx=XM_019837840.2;refGene_gn=SELENBP1;refGene_tx=XM_019837842.2
C1	106437237	106437350
refGene_gn=SELENBP1;refGene_tx=XM_003990619.3;refGene_gn=SELENBP1;refGene_tx=XM_019837840.2;refGene_gn=SELENBP1;refGene_tx=XM_019837842.2
C1	106437533	106437590
refGene_gn=SELENBP1;refGene_tx=XM_003990619.3;refGene_gn=SELENBP1;refGene_tx=XM_019837840.2;refGene_gn=SELENBP1;refGene_tx=XM_019837842.2
C1	106437674	106437762
refGene_gn=SELENBP1;refGene_tx=XM_019837840.2
C1	106439987	106439991
refGene_gn=SELENBP1;refGene_tx=XM_003990619.3
C1	106449609	106449640
refGene_gn=SELENBP1;refGene_tx=XM_019837842.2
C1	106483990	106484130
refGene_gn=PSMB4;refGene_tx=XM_003990621.4
C1	106484380	106484587
refGene_gn=PSMB4;refGene_tx=XM_003990621.4
C1	106484863	106485010
refGene_gn=PSMB4;refGene_tx=XM_003990621.4
C1	106485190	106485272
refGene_gn=PSMB4;refGene_tx=XM_003990621.4
C1	106485658	106485775
refGene_gn=PSMB4;refGene_tx=XM_003990621.4
C1	106485983	106486072
refGene_gn=PSMB4;refGene_tx=XM_003990621.4
C1	106486282	106486295
refGene_gn=PSMB4;refGene_tx=XM_003990621.4
C1	106489899	106491571
refGene_gn=POGZ;refGene_tx=XM_011285146.3;refGene_gn=POGZ;refGene_tx=XM_023259157.1;refGene_gn=POGZ;refGene_tx=XM_023259158.1;refGene_gn=POGZ;refGene_tx=XM_023259159.1;refGene_gn=POGZ;refGene_tx=XM_023259160.1
C1	106491704	106491729
refGene_gn=POGZ;refGene_tx=XM_011285146.3;refGene_gn=POGZ;refGene_tx=XM_023259157.1;refGene_gn=POGZ;refGene_tx=XM_023259158.1;refGene_gn=POGZ;refGene_tx=XM_023259159.1;refGene_gn=POGZ;refGene_tx=XM_023259160.1
C1	106491947	106492060
refGene_gn=POGZ;refGene_tx=XM_011285146.3;refGene_gn=POGZ;refGene_tx=XM_023259157.1;refGene_gn=POGZ;refGene_tx=XM_023259158.1;refGene_gn=POGZ;refGene_tx=XM_023259159.1;refGene_gn=POGZ;refGene_tx=XM_023259160.1
C1	106492248	106492305
refGene_gn=POGZ;refGene_tx=XM_011285146.3;refGene_gn=POGZ;refGene_tx=XM_023259157.1;refGene_gn=POGZ;refGene_tx=XM_023259158.1;refGene_gn=POGZ;refGene_tx=XM_023259159.1;refGene_gn=POGZ;refGene_tx=XM_023259160.1
C1	106492995	106493136
refGene_gn=POGZ;refGene_tx=XM_011285146.3;refGene_gn=POGZ;refGene_tx=XM_023259157.1;refGene_gn=POGZ;refGene_tx=XM_023259158.1;refGene_gn=POGZ;refGene_tx=XM_023259159.1;refGene_gn=POGZ;refGene_tx=XM_023259160.1
C1	106493292	106493465

refGene_gn=POGZ;refGene_tx=XM_011285146.3;refGene_gn=POGZ;refGene_tx=XM_023259157.1;refGene_gn=POGZ;refGene_tx=XM_023259158.1;refGene_gn=POGZ;refGene_tx=XM_023259159.1;refGene_gn=POGZ;refGene_tx=XM_023259160.1
C1 106493579 106493714
refGene_gn=POGZ;refGene_tx=XM_011285146.3;refGene_gn=POGZ;refGene_tx=XM_023259157.1;refGene_gn=POGZ;refGene_tx=XM_023259158.1;refGene_gn=POGZ;refGene_tx=XM_023259159.1;refGene_gn=POGZ;refGene_tx=XM_023259160.1
C1 106496582 106496729
refGene_gn=POGZ;refGene_tx=XM_011285146.3;refGene_gn=POGZ;refGene_tx=XM_023259157.1;refGene_gn=POGZ;refGene_tx=XM_023259159.1;refGene_gn=POGZ;refGene_tx=XM_023259160.1
C1 106497265 106497366
refGene_gn=POGZ;refGene_tx=XM_011285146.3;refGene_gn=POGZ;refGene_tx=XM_023259157.1;refGene_gn=POGZ;refGene_tx=XM_023259158.1;refGene_gn=POGZ;refGene_tx=XM_023259159.1;refGene_gn=POGZ;refGene_tx=XM_023259160.1
C1 106503721 106503876
refGene_gn=POGZ;refGene_tx=XM_011285146.3;refGene_gn=POGZ;refGene_tx=XM_023259157.1;refGene_gn=POGZ;refGene_tx=XM_023259158.1;refGene_gn=POGZ;refGene_tx=XM_023259159.1;refGene_gn=POGZ;refGene_tx=XM_023259160.1
C1 106504247 106504585
refGene_gn=POGZ;refGene_tx=XM_011285146.3;refGene_gn=POGZ;refGene_tx=XM_023259157.1;refGene_gn=POGZ;refGene_tx=XM_023259158.1;refGene_gn=POGZ;refGene_tx=XM_023259159.1;refGene_gn=POGZ;refGene_tx=XM_023259160.1
C1 106505269 106505376
refGene_gn=POGZ;refGene_tx=XM_011285146.3;refGene_gn=POGZ;refGene_tx=XM_023259157.1;refGene_gn=POGZ;refGene_tx=XM_023259158.1;refGene_gn=POGZ;refGene_tx=XM_023259159.1;refGene_gn=POGZ;refGene_tx=XM_023259160.1
C1 106507523 106507742
refGene_gn=POGZ;refGene_tx=XM_011285146.3;refGene_gn=POGZ;refGene_tx=XM_023259157.1;refGene_gn=POGZ;refGene_tx=XM_023259158.1;refGene_gn=POGZ;refGene_tx=XM_023259159.1;refGene_gn=POGZ;refGene_tx=XM_023259160.1
C1 106507822 106508113
refGene_gn=POGZ;refGene_tx=XM_011285146.3;refGene_gn=POGZ;refGene_tx=XM_023259157.1;refGene_gn=POGZ;refGene_tx=XM_023259158.1;refGene_gn=POGZ;refGene_tx=XM_023259159.1;refGene_gn=POGZ;refGene_tx=XM_023259160.1
C1 106509340 106509449
refGene_gn=POGZ;refGene_tx=XM_011285146.3;refGene_gn=POGZ;refGene_tx=XM_023259157.1;refGene_gn=POGZ;refGene_tx=XM_023259158.1;refGene_gn=POGZ;refGene_tx=XM_023259159.1;refGene_gn=POGZ;refGene_tx=XM_023259160.1
C1 106510352 106510528
refGene_gn=POGZ;refGene_tx=XM_011285146.3;refGene_gn=POGZ;refGene_tx=XM_023259157.1;refGene_gn=POGZ;refGene_tx=XM_023259158.1;refGene_gn=POGZ;refGene_tx=XM_023259159.1;refGene_gn=POGZ;refGene_tx=XM_023259160.1
C1 106518033 106518192
refGene_gn=POGZ;refGene_tx=XM_011285146.3;refGene_gn=POGZ;refGene_tx=XM_023259157.1;refGene_gn=POGZ;refGene_tx=XM_023259158.1
C1 106519213 106519337
refGene_gn=POGZ;refGene_tx=XM_011285146.3;refGene_gn=POGZ;refGene_tx=XM_023259157.1;refGene_gn=POGZ;refGene_tx=XM_023259158.1;refGene_gn=POGZ;refGene_tx=XM_023259159.1;refGene_gn=POGZ;refGene_tx=XM_023259160.1

C1 106585196 106586060
refGene_gn=CGN;refGene_tx=XM_023259162.1;refGene_gn=CGN;refGene_tx=XM_023259163.1
C1 106586742 106586843
refGene_gn=CGN;refGene_tx=XM_023259162.1;refGene_gn=CGN;refGene_tx=XM_023259163.1
C1 106586996 106587066
refGene_gn=CGN;refGene_tx=XM_023259162.1;refGene_gn=CGN;refGene_tx=XM_023259163.1
C1 106587180 106587273
refGene_gn=CGN;refGene_tx=XM_023259162.1
C1 106589161 106589289
refGene_gn=CGN;refGene_tx=XM_023259162.1;refGene_gn=CGN;refGene_tx=XM_023259163.1
C1 106589887 106590020
refGene_gn=CGN;refGene_tx=XM_023259162.1;refGene_gn=CGN;refGene_tx=XM_023259163.1
C1 106590343 106590556
refGene_gn=CGN;refGene_tx=XM_023259162.1;refGene_gn=CGN;refGene_tx=XM_023259163.1
C1 106591003 106591152
refGene_gn=CGN;refGene_tx=XM_023259162.1;refGene_gn=CGN;refGene_tx=XM_023259163.1
C1 106591362 106591495
refGene_gn=CGN;refGene_tx=XM_023259162.1;refGene_gn=CGN;refGene_tx=XM_023259163.1
C1 106594492 106594693
refGene_gn=CGN;refGene_tx=XM_023259162.1;refGene_gn=CGN;refGene_tx=XM_023259163.1
C1 106595023 106595230
refGene_gn=CGN;refGene_tx=XM_023259162.1;refGene_gn=CGN;refGene_tx=XM_023259163.1
C1 106595588 106595846
refGene_gn=CGN;refGene_tx=XM_023259162.1;refGene_gn=CGN;refGene_tx=XM_023259163.1
C1 106597046 106597217
refGene_gn=CGN;refGene_tx=XM_023259162.1;refGene_gn=CGN;refGene_tx=XM_023259163.1
C1 106598492 106598654
refGene_gn=CGN;refGene_tx=XM_023259162.1;refGene_gn=CGN;refGene_tx=XM_023259163.1
C1 106599857 106599947
refGene_gn=CGN;refGene_tx=XM_023259162.1;refGene_gn=CGN;refGene_tx=XM_023259163.1
C1 106600399 106600483
refGene_gn=CGN;refGene_tx=XM_023259162.1;refGene_gn=CGN;refGene_tx=XM_023259163.1
C1 106600593 106600712
refGene_gn=CGN;refGene_tx=XM_023259162.1;refGene_gn=CGN;refGene_tx=XM_023259163.1

C1	106601015	106601124
refGene_gn=CGN;refGene_tx=XM_023259162.1;refGene_gn=CGN;refGene_tx=XM_023259163.1
C1	106601512	106601676
refGene_gn=CGN;refGene_tx=XM_023259162.1;refGene_gn=CGN;refGene_tx=XM_023259163.1
C1	106601986	106602125
refGene_gn=CGN;refGene_tx=XM_023259162.1;refGene_gn=CGN;refGene_tx=XM_023259163.1
C1	106604384	106604561
refGene_gn=TUFT1;refGene_tx=XM_019837848.2;refGene_gn=TUFT1;refGene_tx=XM_023259164.1;refGene_gn=TUFT1;refGene_tx=XM_023259165.1;refGene_gn=TUFT1;refGene_tx=XM_023259166.1;refGene_gn=TUFT1;refGene_tx=XM_023259167.1
C1	106604814	106604874
refGene_gn=TUFT1;refGene_tx=XM_003990627.5;refGene_gn=TUFT1;refGene_tx=XM_003990628.5;refGene_gn=TUFT1;refGene_tx=XM_006935138.4;refGene_gn=TUFT1;refGene_tx=XM_006935140.4
C1	106626003	106626078
refGene_gn=TUFT1;refGene_tx=XM_003990627.5;refGene_gn=TUFT1;refGene_tx=XM_006935138.4;refGene_gn=TUFT1;refGene_tx=XM_019837848.2;refGene_gn=TUFT1;refGene_tx=XM_023259164.1;refGene_gn=TUFT1;refGene_tx=XM_023259165.1;refGene_gn=TUFT1;refGene_tx=XM_023259166.1;refGene_gn=TUFT1;refGene_tx=XM_023259167.1
C1	106626472	106626574
refGene_gn=TUFT1;refGene_tx=XM_003990627.5;refGene_gn=TUFT1;refGene_tx=XM_003990628.5;refGene_gn=TUFT1;refGene_tx=XM_006935138.4;refGene_gn=TUFT1;refGene_tx=XM_006935140.4;refGene_gn=TUFT1;refGene_tx=XM_019837848.2;refGene_gn=TUFT1;refGene_tx=XM_023259164.1;refGene_gn=TUFT1;refGene_tx=XM_023259165.1;refGene_gn=TUFT1;refGene_tx=XM_023259166.1;refGene_gn=TUFT1;refGene_tx=XM_023259167.1
C1	106629937	106630024
refGene_gn=TUFT1;refGene_tx=XM_003990627.5;refGene_gn=TUFT1;refGene_tx=XM_003990628.5;refGene_gn=TUFT1;refGene_tx=XM_006935138.4;refGene_gn=TUFT1;refGene_tx=XM_006935140.4;refGene_gn=TUFT1;refGene_tx=XM_019837848.2;refGene_gn=TUFT1;refGene_tx=XM_023259164.1;refGene_gn=TUFT1;refGene_tx=XM_023259165.1;refGene_gn=TUFT1;refGene_tx=XM_023259166.1;refGene_gn=TUFT1;refGene_tx=XM_023259167.1
C1	106630565	106630655
refGene_gn=TUFT1;refGene_tx=XM_003990627.5;refGene_gn=TUFT1;refGene_tx=XM_003990628.5;refGene_gn=TUFT1;refGene_tx=XM_006935138.4;refGene_gn=TUFT1;refGene_tx=XM_006935140.4;refGene_gn=TUFT1;refGene_tx=XM_019837848.2;refGene_gn=TUFT1;refGene_tx=XM_023259164.1;refGene_gn=TUFT1;refGene_tx=XM_023259165.1
C1	106632178	106632253
refGene_gn=TUFT1;refGene_tx=XM_003990627.5;refGene_gn=TUFT1;refGene_tx=XM_003990628.5;refGene_gn=TUFT1;refGene_tx=XM_006935138.4;refGene_gn=TUFT1;refGene_tx=XM_006935140.4;refGene_gn=TUFT1;refGene_tx=XM_019837848.2;refGene_gn=TUFT1;refGene_tx=XM_023259164.1;refGene_gn=TUFT1;refGene_tx=XM_023259165.1;refGene_gn=TUFT1;refGene_tx=XM_023259166.1;refGen

e_gn=TUFT1;refGene_tx=XM_023259167.1
C1 106634743 106634857
refGene_gn=TUFT1;refGene_tx=XM_003990627.5;refGene_gn=TUFT1;refGene_tx=XM_003990628.5;refGene_gn=TUFT1;refGene_tx=XM_006935138.4;refGene_gn=TUFT1;refGene_tx=XM_006935140.4;refGene_gn=TUFT1;refGene_tx=XM_019837848.2;refGene_gn=TUFT1;refGene_tx=XM_023259164.1;refGene_gn=TUFT1;refGene_tx=XM_023259165.1;refGene_gn=TUFT1;refGene_tx=XM_023259166.1;refGene_gn=TUFT1;refGene_tx=XM_023259167.1
C1 106638451 106638580
refGene_gn=TUFT1;refGene_tx=XM_003990627.5;refGene_gn=TUFT1;refGene_tx=XM_003990628.5;refGene_gn=TUFT1;refGene_tx=XM_006935138.4;refGene_gn=TUFT1;refGene_tx=XM_006935140.4;refGene_gn=TUFT1;refGene_tx=XM_019837848.2;refGene_gn=TUFT1;refGene_tx=XM_023259164.1;refGene_gn=TUFT1;refGene_tx=XM_023259165.1;refGene_gn=TUFT1;refGene_tx=XM_023259166.1;refGene_gn=TUFT1;refGene_tx=XM_023259167.1
C1 106639164 106639259
refGene_gn=TUFT1;refGene_tx=XM_003990627.5;refGene_gn=TUFT1;refGene_tx=XM_003990628.5;refGene_gn=TUFT1;refGene_tx=XM_006935138.4;refGene_gn=TUFT1;refGene_tx=XM_006935140.4;refGene_gn=TUFT1;refGene_tx=XM_019837848.2;refGene_gn=TUFT1;refGene_tx=XM_023259164.1;refGene_gn=TUFT1;refGene_tx=XM_023259165.1;refGene_gn=TUFT1;refGene_tx=XM_023259166.1;refGene_gn=TUFT1;refGene_tx=XM_023259167.1
C1 106640989 106641095
refGene_gn=TUFT1;refGene_tx=XM_003990627.5;refGene_gn=TUFT1;refGene_tx=XM_003990628.5;refGene_gn=TUFT1;refGene_tx=XM_006935138.4;refGene_gn=TUFT1;refGene_tx=XM_006935140.4;refGene_gn=TUFT1;refGene_tx=XM_019837848.2;refGene_gn=TUFT1;refGene_tx=XM_023259164.1;refGene_gn=TUFT1;refGene_tx=XM_023259165.1;refGene_gn=TUFT1;refGene_tx=XM_023259166.1;refGene_gn=TUFT1;refGene_tx=XM_023259167.1
C1 106643593 106643677
refGene_gn=TUFT1;refGene_tx=XM_003990627.5;refGene_gn=TUFT1;refGene_tx=XM_003990628.5;refGene_gn=TUFT1;refGene_tx=XM_006935138.4;refGene_gn=TUFT1;refGene_tx=XM_006935140.4;refGene_gn=TUFT1;refGene_tx=XM_019837848.2;refGene_gn=TUFT1;refGene_tx=XM_023259164.1;refGene_gn=TUFT1;refGene_tx=XM_023259165.1;refGene_gn=TUFT1;refGene_tx=XM_023259166.1;refGene_gn=TUFT1;refGene_tx=XM_023259167.1
C1 106644715 106644816
refGene_gn=TUFT1;refGene_tx=XM_003990627.5;refGene_gn=TUFT1;refGene_tx=XM_003990628.5;refGene_gn=TUFT1;refGene_tx=XM_006935138.4;refGene_gn=TUFT1;refGene_tx=XM_006935140.4;refGene_gn=TUFT1;refGene_tx=XM_019837848.2;refGene_gn=TUFT1;refGene_tx=XM_023259164.1;refGene_gn=TUFT1;refGene_tx=XM_023259165.1;refGene_gn=TUFT1;refGene_tx=XM_023259166.1;refGene_gn=TUFT1;refGene_tx=XM_023259167.1
C1 106645386 106645453
refGene_gn=TUFT1;refGene_tx=XM_003990627.5;refGene_gn=TUFT1;refGene_tx=XM_003990628.5;refGene_gn=TUFT1;refGene_tx=XM_006935138.4;refGene_gn=TUFT1;refGene_tx=XM_006935140.4;refGene_gn=TUFT1;refGene_tx=XM_019837848.2;refGene_gn=TUFT1;refGene_tx=XM_023259164.1;refGene_gn=TUFT1;refGene_tx=XM_023259165.1;refGene_gn=TUFT1;refGene_tx=XM_023259166.1;refGene_gn=TUFT1;refGene_tx=XM_023259167.1

C1 106666546 106666857
refGene_gn=SNX27;refGene_tx=XM_023259168.1;refGene_gn=SNX27;refGene_tx
=XM_023259169.1
C1 106693006 106693238
refGene_gn=SNX27;refGene_tx=XM_023259168.1;refGene_gn=SNX27;refGene_tx
=XM_023259169.1
C1 106710060 106710253
refGene_gn=SNX27;refGene_tx=XM_023259168.1;refGene_gn=SNX27;refGene_tx
=XM_023259169.1
C1 106711859 106711924
refGene_gn=SNX27;refGene_tx=XM_023259168.1;refGene_gn=SNX27;refGene_tx
=XM_023259169.1
C1 106714583 106714688
refGene_gn=SNX27;refGene_tx=XM_023259168.1;refGene_gn=SNX27;refGene_tx
=XM_023259169.1
C1 106718312 106718391
refGene_gn=SNX27;refGene_tx=XM_023259168.1;refGene_gn=SNX27;refGene_tx
=XM_023259169.1
C1 106719963 106720127
refGene_gn=SNX27;refGene_tx=XM_023259168.1;refGene_gn=SNX27;refGene_tx
=XM_023259169.1
C1 106722724 106722814
refGene_gn=SNX27;refGene_tx=XM_023259168.1;refGene_gn=SNX27;refGene_tx
=XM_023259169.1
C1 106730468 106730618
refGene_gn=SNX27;refGene_tx=XM_023259168.1;refGene_gn=SNX27;refGene_tx
=XM_023259169.1
C1 106730918 106731047
refGene_gn=SNX27;refGene_tx=XM_023259168.1;refGene_gn=SNX27;refGene_tx
=XM_023259169.1
C1 106731411 106731471
refGene_gn=SNX27;refGene_tx=XM_023259168.1;refGene_gn=SNX27;refGene_tx
=XM_023259169.1
C1 106731574 106731583
refGene_gn=SNX27;refGene_tx=XM_023259169.1
C1 106732347 106732395
refGene_gn=SNX27;refGene_tx=XM_023259168.1
C1 106744320 106744448
refGene_gn=CELF3;refGene_tx=XM_006935142.4;refGene_gn=CELF3;refGene_tx
=XM_019837851.2;refGene_gn=CELF3;refGene_tx=XM_019837852.2;refGene_gn=
CELF3;refGene_tx=XM_019837853.2;refGene_gn=CELF3;refGene_tx=XM_0232591
70.1;refGene_gn=CELF3;refGene_tx=XM_023259171.1
C1 106745112 106745256
refGene_gn=CELF3;refGene_tx=XM_006935142.4;refGene_gn=CELF3;refGene_tx
=XM_019837851.2;refGene_gn=CELF3;refGene_tx=XM_019837852.2;refGene_gn=
CELF3;refGene_tx=XM_019837853.2;refGene_gn=CELF3;refGene_tx=XM_0232591
70.1;refGene_gn=CELF3;refGene_tx=XM_023259171.1
C1 106745503 106745635
refGene_gn=CELF3;refGene_tx=XM_006935142.4;refGene_gn=CELF3;refGene_tx
=XM_019837851.2;refGene_gn=CELF3;refGene_tx=XM_019837852.2;refGene_gn=

CELF3;refGene_tx=XM_019837853.2;refGene_gn=CELF3;refGene_tx=XM_0232591
70.1;refGene_gn=CELF3;refGene_tx=XM_023259171.1
C1 106745910 106745976
refGene_gn=CELF3;refGene_tx=XM_006935142.4;refGene_gn=CELF3;refGene_tx
=XM_019837851.2;refGene_gn=CELF3;refGene_tx=XM_019837852.2;refGene_gn=
CELF3;refGene_tx=XM_019837853.2;refGene_gn=CELF3;refGene_tx=XM_0232591
70.1;refGene_gn=CELF3;refGene_tx=XM_023259171.1
C1 106746380 106746530
refGene_gn=CELF3;refGene_tx=XM_006935142.4;refGene_gn=CELF3;refGene_tx
=XM_019837851.2;refGene_gn=CELF3;refGene_tx=XM_019837852.2;refGene_gn=
CELF3;refGene_tx=XM_019837853.2;refGene_gn=CELF3;refGene_tx=XM_0232591
70.1;refGene_gn=CELF3;refGene_tx=XM_023259171.1
C1 106746822 106746964
refGene_gn=CELF3;refGene_tx=XM_006935142.4;refGene_gn=CELF3;refGene_tx
=XM_019837851.2;refGene_gn=CELF3;refGene_tx=XM_019837852.2;refGene_gn=
CELF3;refGene_tx=XM_019837853.2;refGene_gn=CELF3;refGene_tx=XM_0232591
70.1;refGene_gn=CELF3;refGene_tx=XM_023259171.1
C1 106747124 106747268
refGene_gn=CELF3;refGene_tx=XM_006935142.4;refGene_gn=CELF3;refGene_tx
=XM_019837851.2;refGene_gn=CELF3;refGene_tx=XM_019837852.2;refGene_gn=
CELF3;refGene_tx=XM_019837853.2;refGene_gn=CELF3;refGene_tx=XM_0232591
70.1;refGene_gn=CELF3;refGene_tx=XM_023259171.1
C1 106748145 106748225
refGene_gn=CELF3;refGene_tx=XM_006935142.4;refGene_gn=CELF3;refGene_tx
=XM_019837851.2;refGene_gn=CELF3;refGene_tx=XM_019837852.2;refGene_gn=
CELF3;refGene_tx=XM_019837853.2;refGene_gn=CELF3;refGene_tx=XM_0232591
70.1;refGene_gn=CELF3;refGene_tx=XM_023259171.1
C1 106748367 106748496
refGene_gn=CELF3;refGene_tx=XM_006935142.4;refGene_gn=CELF3;refGene_tx
=XM_019837851.2;refGene_gn=CELF3;refGene_tx=XM_019837852.2;refGene_gn=
CELF3;refGene_tx=XM_019837853.2;refGene_gn=CELF3;refGene_tx=XM_0232591
70.1;refGene_gn=CELF3;refGene_tx=XM_023259171.1
C1 106748932 106748981
refGene_gn=CELF3;refGene_tx=XM_006935142.4;refGene_gn=CELF3;refGene_tx
=XM_019837851.2;refGene_gn=CELF3;refGene_tx=XM_019837852.2;refGene_gn=
CELF3;refGene_tx=XM_019837853.2;refGene_gn=CELF3;refGene_tx=XM_0232591
70.1;refGene_gn=CELF3;refGene_tx=XM_023259171.1
C1 106754124 106754207
refGene_gn=CELF3;refGene_tx=XM_006935142.4;refGene_gn=CELF3;refGene_tx
=XM_019837851.2;refGene_gn=CELF3;refGene_tx=XM_019837852.2
C1 106755421 106755566
refGene_gn=CELF3;refGene_tx=XM_006935142.4;refGene_gn=CELF3;refGene_tx
=XM_019837851.2;refGene_gn=CELF3;refGene_tx=XM_019837852.2
C1 106761655 106761739
refGene_gn=RIIAD1;refGene_tx=XM_003990631.4
C1 106762233 106762310
refGene_gn=RIIAD1;refGene_tx=XM_003990631.4
C1 106766237 106766284
refGene_gn=RIIAD1;refGene_tx=XM_003990631.4
C1 106767620 106767691

refGene_gn=RIIAD1;refGene_tx=XM_003990631.4
C1	106776759	106776864
refGene_gn=MRPL9;refGene_tx=XM_023259173.1
C1	106787963	106788121
refGene_gn=MRPL9;refGene_tx=XM_006935152.4;refGene_gn=MRPL9;refGene_tx=XM_019837856.2
C1	106788205	106788289
refGene_gn=MRPL9;refGene_tx=XM_006935152.4;refGene_gn=MRPL9;refGene_tx=XM_019837856.2;refGene_gn=MRPL9;refGene_tx=XM_023259173.1
C1	106788799	106788901
refGene_gn=MRPL9;refGene_tx=XM_006935152.4;refGene_gn=MRPL9;refGene_tx=XM_019837856.2;refGene_gn=MRPL9;refGene_tx=XM_023259173.1
C1	106789903	106789954
refGene_gn=MRPL9;refGene_tx=XM_006935152.4;refGene_gn=MRPL9;refGene_tx=XM_019837856.2;refGene_gn=MRPL9;refGene_tx=XM_023259173.1
C1	106790158	106790283
refGene_gn=MRPL9;refGene_tx=XM_006935152.4;refGene_gn=MRPL9;refGene_tx=XM_019837856.2;refGene_gn=MRPL9;refGene_tx=XM_023259173.1
C1	106790740	106790897
refGene_gn=MRPL9;refGene_tx=XM_006935152.4;refGene_gn=MRPL9;refGene_tx=XM_019837856.2;refGene_gn=MRPL9;refGene_tx=XM_023259173.1
C1	106791070	106791187
refGene_gn=MRPL9;refGene_tx=XM_006935152.4;refGene_gn=MRPL9;refGene_tx=XM_019837856.2;refGene_gn=MRPL9;refGene_tx=XM_023259173.1
C1	106793942	106794106
refGene_gn=OAZ3;refGene_tx=NM_001290358.1
C1	106794390	106794451
refGene_gn=OAZ3;refGene_tx=NM_001290358.1
C1	106794452	106794527
refGene_gn=OAZ3;refGene_tx=NM_001290358.1
C1	106795396	106795572
refGene_gn=OAZ3;refGene_tx=NM_001290358.1
C1	106796072	106796158
refGene_gn=OAZ3;refGene_tx=NM_001290358.1
C1	106796882	106797025
refGene_gn=OAZ3;refGene_tx=NM_001290358.1
C1	106799457	106799510
refGene_gn=TDRKH;refGene_tx=XM_003990634.5;refGene_gn=TDRKH;refGene_tx=XM_006935153.4;refGene_gn=TDRKH;refGene_tx=XM_006935155.4;refGene_gn=TDRKH;refGene_tx=XM_019837855.2;refGene_gn=TDRKH;refGene_tx=XM_023259172.1
C1	106799687	106799784
refGene_gn=TDRKH;refGene_tx=XM_003990634.5;refGene_gn=TDRKH;refGene_tx=XM_006935153.4;refGene_gn=TDRKH;refGene_tx=XM_006935155.4;refGene_gn=TDRKH;refGene_tx=XM_019837855.2;refGene_gn=TDRKH;refGene_tx=XM_023259172.1
C1	106800009	106800111
refGene_gn=TDRKH;refGene_tx=XM_003990634.5;refGene_gn=TDRKH;refGene_tx=XM_006935153.4;refGene_gn=TDRKH;refGene_tx=XM_006935155.4;refGene_gn=TDRKH;refGene_tx=XM_019837855.2;refGene_gn=TDRKH;refGene_tx=XM_0232591

72.1
C1 106800558 106800710
refGene_gn=TDRKH;refGene_tx=XM_003990634.5;refGene_gn=TDRKH;refGene_tx=XM_006935153.4;refGene_gn=TDRKH;refGene_tx=XM_006935155.4;refGene_gn=TDRKH;refGene_tx=XM_019837855.2;refGene_gn=TDRKH;refGene_tx=XM_023259172.1
C1 106800969 106801034
refGene_gn=TDRKH;refGene_tx=XM_003990634.5;refGene_gn=TDRKH;refGene_tx=XM_006935153.4;refGene_gn=TDRKH;refGene_tx=XM_006935155.4;refGene_gn=TDRKH;refGene_tx=XM_019837855.2;refGene_gn=TDRKH;refGene_tx=XM_023259172.1
C1 106801252 106801425
refGene_gn=TDRKH;refGene_tx=XM_003990634.5;refGene_gn=TDRKH;refGene_tx=XM_006935153.4;refGene_gn=TDRKH;refGene_tx=XM_006935155.4;refGene_gn=TDRKH;refGene_tx=XM_019837855.2;refGene_gn=TDRKH;refGene_tx=XM_023259172.1
C1 106801607 106801768
refGene_gn=TDRKH;refGene_tx=XM_003990634.5;refGene_gn=TDRKH;refGene_tx=XM_006935153.4;refGene_gn=TDRKH;refGene_tx=XM_006935155.4;refGene_gn=TDRKH;refGene_tx=XM_019837855.2;refGene_gn=TDRKH;refGene_tx=XM_023259172.1
C1 106803059 106803384
refGene_gn=TDRKH;refGene_tx=XM_003990634.5;refGene_gn=TDRKH;refGene_tx=XM_006935153.4;refGene_gn=TDRKH;refGene_tx=XM_006935155.4;refGene_gn=TDRKH;refGene_tx=XM_019837855.2;refGene_gn=TDRKH;refGene_tx=XM_023259172.1
C1 106803488 106803628
refGene_gn=TDRKH;refGene_tx=XM_003990634.5;refGene_gn=TDRKH;refGene_tx=XM_006935153.4;refGene_gn=TDRKH;refGene_tx=XM_006935155.4;refGene_gn=TDRKH;refGene_tx=XM_019837855.2;refGene_gn=TDRKH;refGene_tx=XM_023259172.1
C1 106804348 106804538
refGene_gn=TDRKH;refGene_tx=XM_003990634.5;refGene_gn=TDRKH;refGene_tx=XM_006935153.4;refGene_gn=TDRKH;refGene_tx=XM_006935155.4;refGene_gn=TDRKH;refGene_tx=XM_019837855.2;refGene_gn=TDRKH;refGene_tx=XM_023259172.1
C1 106806095 106806202
refGene_gn=TDRKH;refGene_tx=XM_003990634.5;refGene_gn=TDRKH;refGene_tx=XM_006935153.4;refGene_gn=TDRKH;refGene_tx=XM_006935155.4;refGene_gn=TDRKH;refGene_tx=XM_019837855.2;refGene_gn=TDRKH;refGene_tx=XM_023259172.1
C1 106807155 106807279
refGene_gn=TDRKH;refGene_tx=XM_003990634.5;refGene_gn=TDRKH;refGene_tx=XM_006935153.4;refGene_gn=TDRKH;refGene_tx=XM_006935155.4;refGene_gn=TDRKH;refGene_tx=XM_019837855.2;refGene_gn=TDRKH;refGene_tx=XM_023259172.1
C1 106823561 106825356
refGene_gn=LINGO4;refGene_tx=XM_003990635.4;refGene_gn=LINGO4;refGene_tx=XM_019837854.2
C1 106825876 106826011

refGene_gn=LINGO4;refGene_tx=XM_019837854.2
C1 106827842 106827907
refGene_gn=LINGO4;refGene_tx=XM_003990635.4;refGene_gn=LINGO4;refGene_tx=XM_019837854.2
C1 106829993 106830155
refGene_gn=RORC;refGene_tx=XM_023257841.1
C1 106833272 106833382
refGene_gn=RORC;refGene_tx=XM_023257841.1
C1 106835215 106835326
refGene_gn=RORC;refGene_tx=XM_023257841.1
C1 106835508 106835616
refGene_gn=RORC;refGene_tx=XM_023257841.1
C1 106835710 106835843
refGene_gn=RORC;refGene_tx=XM_023257841.1
C1 106836824 106836946
refGene_gn=RORC;refGene_tx=XM_023257841.1
C1 106837170 106837683
refGene_gn=RORC;refGene_tx=XM_023257841.1
C1 106838887 106839002
refGene_gn=RORC;refGene_tx=XM_023257841.1
C1 106839003 106839030
refGene_gn=RORC;refGene_tx=XM_023257841.1
C1 106839353 106839380
refGene_gn=RORC;refGene_tx=XM_023257841.1
C1 106839381 106839440
refGene_gn=RORC;refGene_tx=XM_023257841.1
C1 106850311 106850341
refGene_gn=RORC;refGene_tx=XM_023257841.1
C1 106850722 106850747
refGene_gn=RORC;refGene_tx=XM_023257841.1
C1 106859946 106860975
refGene_gn=C2CD4D;refGene_tx=XM_023259174.1
C1 106870666 106870775
refGene_gn=THEM5;refGene_tx=XM_003990639.4;refGene_gn=THEM5;refGene_tx=XM_006935156.2
C1 106871075 106871200
refGene_gn=THEM5;refGene_tx=XM_003990639.4
C1 106871528 106871639
refGene_gn=THEM5;refGene_tx=XM_003990639.4;refGene_gn=THEM5;refGene_tx=XM_006935156.2
C1 106875715 106875854
refGene_gn=THEM5;refGene_tx=XM_003990639.4;refGene_gn=THEM5;refGene_tx=XM_006935156.2
C1 106877061 106877263
refGene_gn=THEM5;refGene_tx=XM_003990639.4;refGene_gn=THEM5;refGene_tx=XM_006935156.2
C1 106878676 106878799
refGene_gn=THEM5;refGene_tx=XM_003990639.4;refGene_gn=THEM5;refGene_tx=XM_006935156.2
C1 106901860 106901901

refGene_gn=THEM4;refGene_tx=XM_011285154.3;refGene_gn=THEM4;refGene_tx=XM_023259175.1
C1 106905121 106905246
refGene_gn=THEM4;refGene_tx=XM_011285154.3;refGene_gn=THEM4;refGene_tx=XM_023259175.1
C1 106911163 106911274
refGene_gn=THEM4;refGene_tx=XM_011285154.3;refGene_gn=THEM4;refGene_tx=XM_023259175.1
C1 106911872 106912029
refGene_gn=THEM4;refGene_tx=XM_011285154.3;refGene_gn=THEM4;refGene_tx=XM_023259175.1
C1 106918225 106918382
refGene_gn=THEM4;refGene_tx=XM_011285154.3;refGene_gn=THEM4;refGene_tx=XM_023259175.1
C1 106981086 106981248
refGene_gn=S100A10;refGene_tx=XM_023259176.1
C1 106984242 106984374
refGene_gn=S100A10;refGene_tx=XM_023259176.1
C1 106989391 106989396
refGene_gn=LOC102899606;refGene_tx=XM_023257842.1
C1 106990275 106990769
refGene_gn=LOC102899606;refGene_tx=XM_023257842.1
C1 106990968 106991088
refGene_gn=LOC102899606;refGene_tx=XM_023257842.1
C1 106991806 106991817
refGene_gn=LOC102899606;refGene_tx=XM_023257842.1
C1 107017326 107017482
refGene_gn=S100A11;refGene_tx=XM_003990641.4
C1 107018410 107018563
refGene_gn=S100A11;refGene_tx=XM_003990641.4
C1 107022507 107022510
refGene_gn=S100A11;refGene_tx=XM_003990641.4
C1 107099604 107101917
refGene_gn=TCHHL1;refGene_tx=XM_019837858.2
C1 107102587 107102725
refGene_gn=TCHHL1;refGene_tx=XM_019837858.2
C1 107117203 107117785
refGene_gn=TCHH;refGene_tx=XM_023257843.1
C1 107117871 107118646
refGene_gn=TCHH;refGene_tx=XM_023257843.1
C1 107118677 107121925
refGene_gn=TCHH;refGene_tx=XM_023257843.1
C1 107122659 107122797
refGene_gn=TCHH;refGene_tx=XM_023257843.1
C1 107158629 107160320
refGene_gn=RPTN;refGene_tx=XM_023259178.1
C1 107160680 107161713
refGene_gn=RPTN;refGene_tx=XM_023259178.1
C1 107162391 107162529
refGene_gn=RPTN;refGene_tx=XM_023259178.1

C1 107221153 107223285
refGene_gn=HRNR;refGene_tx=XM_023259177.1
C1 107223744 107224203
refGene_gn=HRNR;refGene_tx=XM_023259177.1
C1 107224205 107226192
refGene_gn=HRNR;refGene_tx=XM_023259177.1
C1 107227862 107228000
refGene_gn=HRNR;refGene_tx=XM_023259177.1
C1 107250598 107250627
refGene_gn=FLG;refGene_tx=XM_023257844.1
C1 107251331 107251389
refGene_gn=FLG;refGene_tx=XM_023257844.1
C1 107257566 107258585
refGene_gn=FLG;refGene_tx=XM_023257844.1
C1 107258723 107258909
refGene_gn=FLG;refGene_tx=XM_023257844.1
C1 107261241 107264592
refGene_gn=FLG;refGene_tx=XM_023257844.1
C1 107265570 107266413
refGene_gn=FLG;refGene_tx=XM_023257844.1
C1 107267014 107267152
refGene_gn=FLG;refGene_tx=XM_023257844.1
C1 107326506 107326903
refGene_gn=LOC109502610;refGene_tx=XM_023257846.1
C1 107326991 107328124
refGene_gn=LOC109502610;refGene_tx=XM_023257846.1
C1 107330395 107331318
refGene_gn=LOC109502610;refGene_tx=XM_023257846.1
C1 107331577 107331996
refGene_gn=LOC109502610;refGene_tx=XM_023257846.1
C1 107332220 107332903
refGene_gn=LOC109502610;refGene_tx=XM_023257846.1
C1 107333079 107333457
refGene_gn=LOC109502610;refGene_tx=XM_023257846.1
C1 107333888 107333913
refGene_gn=LOC101086332;refGene_tx=XM_023257847.1
C1 107334023 107334204
refGene_gn=LOC101086332;refGene_tx=XM_023257847.1
C1 107334809 107336868
refGene_gn=LOC101086332;refGene_tx=XM_023257847.1
C1 107337913 107338051
refGene_gn=LOC101086332;refGene_tx=XM_023257847.1
C1 107405010 107406498
refGene_gn=CRNN;refGene_tx=XM_019837864.1
C1 107407604 107407742
refGene_gn=CRNN;refGene_tx=XM_019837864.1
C1 107496572 107496959
refGene_gn=LCE5A;refGene_tx=XM_019839175.2
C1 107500639 107500975
refGene_gn=CRCT1;refGene_tx=XM_006935165.3

C1 107551957 107552248
refGene_gn=LCE3C;refGene_tx=XM_003990655.3
C1 107559072 107559363
refGene_gn=LOC109503120;refGene_tx=XM_019839176.2
C1 107608584 107609337
refGene_gn=CC1H1orf68;refGene_tx=XM_003990652.2
C1 107639396 107639432
refGene_gn=KPRP;refGene_tx=XM_019837867.2
C1 107639708 107641496
refGene_gn=KPRP;refGene_tx=XM_003990656.3;refGene_gn=KPRP;refGene_tx=XM_019837867.2;refGene_gn=KPRP;refGene_tx=XM_019837868.2
C1 107712125 107712378
refGene_gn=LCE6A;refGene_tx=XM_023257848.1
C1 107728015 107728041
refGene_gn=LCE6A;refGene_tx=XM_023257848.1
C1 107782387 107783721
refGene_gn=IVL;refGene_tx=XM_011285157.3;refGene_gn=IVL;refGene_tx=XM_023259179.1
C1 107816247 107816329
refGene_gn=LOC109503125;refGene_tx=XM_019839178.2
C1 107817551 107817844
refGene_gn=LOC109503125;refGene_tx=XM_019839178.2
C1 107841951 107842182
refGene_gn=SPRR4;refGene_tx=XM_011285160.1
C1 107860211 107860481
refGene_gn=LOC101087594;refGene_tx=XM_003990654.3;refGene_gn=LOC101087594;refGene_tx=XM_019837869.1
C1 110117697 110117994
refGene_gn=LELP1;refGene_tx=XM_006935171.3
C1 110128422 110128749
refGene_gn=PRR9;refGene_tx=XM_006935172.3
C1 110182774 110183794
refGene_gn=LOR;refGene_tx=XM_023259180.1;refGene_gn=LOR;refGene_tx=XM_023259181.1;refGene_gn=LOR;refGene_tx=XM_023259182.1;refGene_gn=LOR;refGene_tx=XM_023259183.1
C1 110210999 110211178
refGene_gn=PGLYRP3;refGene_tx=XM_019837872.2;refGene_gn=PGLYRP3;refGene_tx=XM_019837874.2;refGene_gn=PGLYRP3;refGene_tx=XM_023259184.1;refGene_gn=PGLYRP3;refGene_tx=XM_023259185.1;refGene_gn=PGLYRP3;refGene_tx=XM_023259186.1
C1 110211433 110211552
refGene_gn=PGLYRP3;refGene_tx=XM_019837872.2;refGene_gn=PGLYRP3;refGene_tx=XM_019837874.2;refGene_gn=PGLYRP3;refGene_tx=XM_023259184.1;refGene_gn=PGLYRP3;refGene_tx=XM_023259185.1;refGene_gn=PGLYRP3;refGene_tx=XM_023259186.1
C1 110214964 110215163
refGene_gn=PGLYRP3;refGene_tx=XM_019837872.2;refGene_gn=PGLYRP3;refGene_tx=XM_019837874.2;refGene_gn=PGLYRP3;refGene_tx=XM_023259184.1;refGene_gn=PGLYRP3;refGene_tx=XM_023259185.1;refGene_gn=PGLYRP3;refGene_tx=XM_023259186.1

C1 110216401 110216554
refGene_gn=PGLYRP3;refGene_tx=XM_019837872.2;refGene_gn=PGLYRP3;refGen
e_tx=XM_019837874.2;refGene_gn=PGLYRP3;refGene_tx=XM_023259184.1;refGe
ne_gn=PGLYRP3;refGene_tx=XM_023259185.1;refGene_gn=PGLYRP3;refGene_tx=
XM_023259186.1
C1 110217705 110217824
refGene_gn=PGLYRP3;refGene_tx=XM_019837872.2;refGene_gn=PGLYRP3;refGen
e_tx=XM_019837874.2;refGene_gn=PGLYRP3;refGene_tx=XM_023259184.1;refGe
ne_gn=PGLYRP3;refGene_tx=XM_023259185.1;refGene_gn=PGLYRP3;refGene_tx=
XM_023259186.1
C1 110220365 110220567
refGene_gn=PGLYRP3;refGene_tx=XM_019837872.2;refGene_gn=PGLYRP3;refGen
e_tx=XM_019837874.2;refGene_gn=PGLYRP3;refGene_tx=XM_023259184.1;refGe
ne_gn=PGLYRP3;refGene_tx=XM_023259185.1;refGene_gn=PGLYRP3;refGene_tx=
XM_023259186.1
C1 110221394 110221502
refGene_gn=PGLYRP3;refGene_tx=XM_019837872.2;refGene_gn=PGLYRP3;refGen
e_tx=XM_019837874.2;refGene_gn=PGLYRP3;refGene_tx=XM_023259184.1;refGe
ne_gn=PGLYRP3;refGene_tx=XM_023259185.1;refGene_gn=PGLYRP3;refGene_tx=
XM_023259186.1
C1 110222651 110222700
refGene_gn=PGLYRP3;refGene_tx=XM_019837872.2;refGene_gn=PGLYRP3;refGen
e_tx=XM_019837874.2;refGene_gn=PGLYRP3;refGene_tx=XM_023259184.1;refGe
ne_gn=PGLYRP3;refGene_tx=XM_023259185.1;refGene_gn=PGLYRP3;refGene_tx=
XM_023259186.1
C1 110448314 110448365
refGene_gn=LOC101093953;refGene_tx=XM_023257849.1
C1 110449116 110449253
refGene_gn=LOC101093953;refGene_tx=XM_023257849.1
C1 110449518 110449628
refGene_gn=LOC101093953;refGene_tx=XM_023257849.1
C1 110460915 110461037
refGene_gn=LOC101093953;refGene_tx=XM_023257849.1
C1 110483915 110484233
refGene_gn=LOC101094197;refGene_tx=XM_011285190.3
C1 110487543 110487710
refGene_gn=LOC101094197;refGene_tx=XM_011285190.3
C1 110489281 110489556
refGene_gn=LOC101094197;refGene_tx=XM_011285190.3
C1 110491523 110491686
refGene_gn=LOC101094197;refGene_tx=XM_011285190.3
C1 110492503 110492627
refGene_gn=LOC101094197;refGene_tx=XM_011285190.3
C1 110516915 110516997
refGene_gn=LOC101094439;refGene_tx=XM_011285204.1
C1 110518042 110518139
refGene_gn=LOC101094439;refGene_tx=XM_011285204.1
C1 110518140 110518961
refGene_gn=LOC101094439;refGene_tx=XM_011285204.1
C1 110518963 110519079

refGene_gn=LOC101094439;refGene_tx=XM_011285204.1
C1	110543653	110546032
refGene_gn=AMER3;refGene_tx=XM_023259187.1
C1	110606155	110606266
refGene_gn=LOC111561637;refGene_tx=XM_023257850.1
C1	110606347	110606599
refGene_gn=LOC111561637;refGene_tx=XM_023257850.1
C1	110698520	110700725
refGene_gn=LOC111561637;refGene_tx=XM_023257850.1
C1	110700820	110702227
refGene_gn=ARHGEF4;refGene_tx=XM_011285191.3;refGene_gn=ARHGEF4;refGene_tx=XM_019837885.2;refGene_gn=ARHGEF4;refGene_tx=XM_023259188.1;refGene_gn=ARHGEF4;refGene_tx=XM_023259189.1;refGene_gn=ARHGEF4;refGene_tx=XM_023259190.1;refGene_gn=ARHGEF4;refGene_tx=XM_023259191.1
C1	110728443	110728752
refGene_gn=ARHGEF4;refGene_tx=XM_011285191.3;refGene_gn=ARHGEF4;refGene_tx=XM_019837885.2;refGene_gn=ARHGEF4;refGene_tx=XM_023259188.1;refGene_gn=ARHGEF4;refGene_tx=XM_023259189.1;refGene_gn=ARHGEF4;refGene_tx=XM_023259191.1
C1	110768611	110768738
refGene_gn=ARHGEF4;refGene_tx=XM_011285191.3;refGene_gn=ARHGEF4;refGene_tx=XM_019837885.2;refGene_gn=ARHGEF4;refGene_tx=XM_023259188.1;refGene_gn=ARHGEF4;refGene_tx=XM_023259189.1;refGene_gn=ARHGEF4;refGene_tx=XM_023259190.1;refGene_gn=ARHGEF4;refGene_tx=XM_023259191.1
C1	110786978	110787369
refGene_gn=ARHGEF4;refGene_tx=XM_023259192.1
C1	110856546	110856719
refGene_gn=ARHGEF4;refGene_tx=XM_011285191.3;refGene_gn=ARHGEF4;refGene_tx=XM_019837885.2;refGene_gn=ARHGEF4;refGene_tx=XM_023259188.1;refGene_gn=ARHGEF4;refGene_tx=XM_023259189.1;refGene_gn=ARHGEF4;refGene_tx=XM_023259190.1;refGene_gn=ARHGEF4;refGene_tx=XM_023259191.1;refGene_gn=ARHGEF4;refGene_tx=XM_023259192.1
C1	110868619	110868799
refGene_gn=ARHGEF4;refGene_tx=XM_011285191.3;refGene_gn=ARHGEF4;refGene_tx=XM_019837885.2;refGene_gn=ARHGEF4;refGene_tx=XM_023259188.1;refGene_gn=ARHGEF4;refGene_tx=XM_023259189.1;refGene_gn=ARHGEF4;refGene_tx=XM_023259190.1;refGene_gn=ARHGEF4;refGene_tx=XM_023259191.1;refGene_gn=ARHGEF4;refGene_tx=XM_023259192.1
C1	110869832	110870009
refGene_gn=ARHGEF4;refGene_tx=XM_011285191.3;refGene_gn=ARHGEF4;refGene_tx=XM_019837885.2;refGene_gn=ARHGEF4;refGene_tx=XM_023259188.1;refGene_gn=ARHGEF4;refGene_tx=XM_023259189.1;refGene_gn=ARHGEF4;refGene_tx=XM_023259190.1;refGene_gn=ARHGEF4;refGene_tx=XM_023259191.1;refGene_gn=ARHGEF4;refGene_tx=XM_023259192.1
C1	110870082	110870262
refGene_gn=ARHGEF4;refGene_tx=XM_011285191.3;refGene_gn=ARHGEF4;refGene_tx=XM_019837885.2;refGene_gn=ARHGEF4;refGene_tx=XM_023259190.1;refGene_gn=ARHGEF4;refGene_tx=XM_023259191.1;refGene_gn=ARHGEF4;refGene_tx=XM_023259192.1
C1	110872447	110872680

refGene_gn=ARHGEF4;refGene_tx=XM_011285191.3;refGene_gn=ARHGEF4;refGen
e_tx=XM_019837885.2;refGene_gn=ARHGEF4;refGene_tx=XM_023259188.1;refGe
ne_gn=ARHGEF4;refGene_tx=XM_023259189.1;refGene_gn=ARHGEF4;refGene_tx=
XM_023259190.1;refGene_gn=ARHGEF4;refGene_tx=XM_023259191.1;refGene_gn
=ARHGEF4;refGene_tx=XM_023259192.1
C1 110872970 110873100
refGene_gn=ARHGEF4;refGene_tx=XM_011285191.3;refGene_gn=ARHGEF4;refGen
e_tx=XM_019837885.2;refGene_gn=ARHGEF4;refGene_tx=XM_023259188.1;refGe
ne_gn=ARHGEF4;refGene_tx=XM_023259189.1;refGene_gn=ARHGEF4;refGene_tx=
XM_023259190.1;refGene_gn=ARHGEF4;refGene_tx=XM_023259191.1;refGene_gn
=ARHGEF4;refGene_tx=XM_023259192.1
C1 110874549 110874681
refGene_gn=ARHGEF4;refGene_tx=XM_011285191.3;refGene_gn=ARHGEF4;refGen
e_tx=XM_019837885.2;refGene_gn=ARHGEF4;refGene_tx=XM_023259188.1;refGe
ne_gn=ARHGEF4;refGene_tx=XM_023259189.1;refGene_gn=ARHGEF4;refGene_tx=
XM_023259190.1;refGene_gn=ARHGEF4;refGene_tx=XM_023259191.1;refGene_gn
=ARHGEF4;refGene_tx=XM_023259192.1
C1 110875448 110875692
refGene_gn=ARHGEF4;refGene_tx=XM_011285191.3;refGene_gn=ARHGEF4;refGen
e_tx=XM_019837885.2;refGene_gn=ARHGEF4;refGene_tx=XM_023259188.1;refGe
ne_gn=ARHGEF4;refGene_tx=XM_023259189.1;refGene_gn=ARHGEF4;refGene_tx=
XM_023259190.1;refGene_gn=ARHGEF4;refGene_tx=XM_023259192.1
C1 110876413 110876491
refGene_gn=ARHGEF4;refGene_tx=XM_011285191.3;refGene_gn=ARHGEF4;refGen
e_tx=XM_019837885.2;refGene_gn=ARHGEF4;refGene_tx=XM_023259188.1;refGe
ne_gn=ARHGEF4;refGene_tx=XM_023259189.1;refGene_gn=ARHGEF4;refGene_tx=
XM_023259190.1;refGene_gn=ARHGEF4;refGene_tx=XM_023259191.1;refGene_gn
=ARHGEF4;refGene_tx=XM_023259192.1
C1 110877059 110877211
refGene_gn=ARHGEF4;refGene_tx=XM_011285191.3;refGene_gn=ARHGEF4;refGen
e_tx=XM_019837885.2;refGene_gn=ARHGEF4;refGene_tx=XM_023259188.1;refGe
ne_gn=ARHGEF4;refGene_tx=XM_023259189.1;refGene_gn=ARHGEF4;refGene_tx=
XM_023259190.1;refGene_gn=ARHGEF4;refGene_tx=XM_023259192.1
C1 110884018 110884131
refGene_gn=FAM168B;refGene_tx=XM_023259193.1;refGene_gn=FAM168B;refGen
e_tx=XM_023259194.1;refGene_gn=FAM168B;refGene_tx=XM_023259195.1
C1 110886025 110886203
refGene_gn=FAM168B;refGene_tx=XM_023259193.1;refGene_gn=FAM168B;refGen
e_tx=XM_023259194.1;refGene_gn=FAM168B;refGene_tx=XM_023259195.1
C1 110886338 110886481
refGene_gn=FAM168B;refGene_tx=XM_023259193.1;refGene_gn=FAM168B;refGen
e_tx=XM_023259194.1;refGene_gn=FAM168B;refGene_tx=XM_023259195.1
C1 110900457 110900541
refGene_gn=FAM168B;refGene_tx=XM_023259193.1;refGene_gn=FAM168B;refGen
e_tx=XM_023259194.1;refGene_gn=FAM168B;refGene_tx=XM_023259195.1
C1 110909321 110909391
refGene_gn=FAM168B;refGene_tx=XM_023259193.1;refGene_gn=FAM168B;refGen
e_tx=XM_023259194.1;refGene_gn=FAM168B;refGene_tx=XM_023259195.1
C1 110939638 110939675
refGene_gn=PLEKHB2;refGene_tx=XM_003990664.5;refGene_gn=PLEKHB2;refGen

e_tx=XM_019837894.2
C1 110945216 110945369
refGene_gn=PLEKHB2;refGene_tx=XM_003990664.5;refGene_gn=PLEKHB2;refGene_tx=XM_019837894.2;refGene_gn=PLEKHB2;refGene_tx=XM_019837895.2;refGene_gn=PLEKHB2;refGene_tx=XM_023259196.1
C1 110946648 110946751
refGene_gn=PLEKHB2;refGene_tx=XM_003990664.5;refGene_gn=PLEKHB2;refGene_tx=XM_019837894.2;refGene_gn=PLEKHB2;refGene_tx=XM_019837895.2;refGene_gn=PLEKHB2;refGene_tx=XM_023259196.1
C1 110952537 110952577
refGene_gn=PLEKHB2;refGene_tx=XM_003990664.5;refGene_gn=PLEKHB2;refGene_tx=XM_019837894.2;refGene_gn=PLEKHB2;refGene_tx=XM_019837895.2;refGene_gn=PLEKHB2;refGene_tx=XM_023259196.1
C1 110954072 110954162
refGene_gn=PLEKHB2;refGene_tx=XM_003990664.5;refGene_gn=PLEKHB2;refGene_tx=XM_019837894.2;refGene_gn=PLEKHB2;refGene_tx=XM_019837895.2;refGene_gn=PLEKHB2;refGene_tx=XM_023259196.1
C1 110962291 110962400
refGene_gn=PLEKHB2;refGene_tx=XM_003990664.5;refGene_gn=PLEKHB2;refGene_tx=XM_019837894.2;refGene_gn=PLEKHB2;refGene_tx=XM_019837895.2;refGene_gn=PLEKHB2;refGene_tx=XM_023259196.1
C1 110968221 110968358
refGene_gn=PLEKHB2;refGene_tx=XM_003990664.5;refGene_gn=PLEKHB2;refGene_tx=XM_019837894.2;refGene_gn=PLEKHB2;refGene_tx=XM_019837895.2;refGene_gn=PLEKHB2;refGene_tx=XM_023259196.1
C1 112936969 112937496
refGene_gn=HS6ST1;refGene_tx=XM_023259197.1
C1 112979769 112980478
refGene_gn=HS6ST1;refGene_tx=XM_023259197.1
C1 113033692 113033718
refGene_gn=UGGT1;refGene_tx=XM_023259198.1
C1 113035908 113036074
refGene_gn=UGGT1;refGene_tx=XM_023259198.1
C1 113036557 113036674
refGene_gn=UGGT1;refGene_tx=XM_023259198.1
C1 113047630 113047745
refGene_gn=UGGT1;refGene_tx=XM_023259198.1
C1 113048961 113049122
refGene_gn=UGGT1;refGene_tx=XM_023259198.1
C1 113053787 113053970
refGene_gn=UGGT1;refGene_tx=XM_023259198.1
C1 113055129 113055214
refGene_gn=UGGT1;refGene_tx=XM_023259198.1
C1 113056449 113056554
refGene_gn=UGGT1;refGene_tx=XM_023259198.1
C1 113057083 113057172
refGene_gn=UGGT1;refGene_tx=XM_023259198.1
C1 113058251 113058336
refGene_gn=UGGT1;refGene_tx=XM_023259198.1
C1 113060090 113060176

refGene_gn=UGGT1;refGene_tx=XM_023259198.1
C1	113060581	113060740
refGene_gn=UGGT1;refGene_tx=XM_023259198.1
C1	113064711	113064901
refGene_gn=UGGT1;refGene_tx=XM_023259198.1
C1	113066371	113066451
refGene_gn=UGGT1;refGene_tx=XM_023259198.1
C1	113067817	113067920
refGene_gn=UGGT1;refGene_tx=XM_023259198.1
C1	113071141	113071237
refGene_gn=UGGT1;refGene_tx=XM_023259198.1
C1	113074187	113074319
refGene_gn=UGGT1;refGene_tx=XM_023259198.1
C1	113075128	113075271
refGene_gn=UGGT1;refGene_tx=XM_023259198.1
C1	113076253	113076460
refGene_gn=UGGT1;refGene_tx=XM_023259198.1
C1	113077794	113077889
refGene_gn=UGGT1;refGene_tx=XM_023259198.1
C1	113078384	113078408
refGene_gn=UGGT1;refGene_tx=XM_023259198.1
C1	113079391	113079490
refGene_gn=UGGT1;refGene_tx=XM_023259198.1
C1	113081967	113082088
refGene_gn=UGGT1;refGene_tx=XM_023259198.1
C1	113095732	113095897
refGene_gn=UGGT1;refGene_tx=XM_023259198.1
C1	113104766	113104898
refGene_gn=UGGT1;refGene_tx=XM_023259198.1
C1	113107289	113107425
refGene_gn=UGGT1;refGene_tx=XM_023259198.1
C1	113111440	113111526
refGene_gn=UGGT1;refGene_tx=XM_023259198.1
C1	113112604	113112724
refGene_gn=UGGT1;refGene_tx=XM_023259198.1
C1	113116445	113116596
refGene_gn=UGGT1;refGene_tx=XM_023259198.1
C1	113118407	113118499
refGene_gn=UGGT1;refGene_tx=XM_023259198.1
C1	113122742	113122803
refGene_gn=UGGT1;refGene_tx=XM_023259198.1
C1	113123900	113124000
refGene_gn=UGGT1;refGene_tx=XM_023259198.1
C1	113124709	113124810
refGene_gn=UGGT1;refGene_tx=XM_023259198.1
C1	113127395	113127474
refGene_gn=UGGT1;refGene_tx=XM_023259198.1
C1	113128633	113128730
refGene_gn=UGGT1;refGene_tx=XM_023259198.1
C1	113130565	113130740

refGene_gn=UGGT1;refGene_tx=XM_023259198.1
C1 113133158 113133271
refGene_gn=UGGT1;refGene_tx=XM_023259198.1
C1 113135817 113135948
refGene_gn=UGGT1;refGene_tx=XM_023259198.1
C1 113137447 113137530
refGene_gn=UGGT1;refGene_tx=XM_023259198.1
C1 113142801 113142937
refGene_gn=UGGT1;refGene_tx=XM_023259198.1
C1 113148298 113148356
refGene_gn=UGGT1;refGene_tx=XM_023259198.1
C1 113218701 113218773
refGene_gn=SAP130;refGene_tx=XM_003990672.4;refGene_gn=SAP130;refGene_tx=XM_003990673.4;refGene_gn=SAP130;refGene_tx=XM_019837899.2;refGene_gn=SAP130;refGene_tx=XM_019837901.2;refGene_gn=SAP130;refGene_tx=XM_019837902.2;refGene_gn=SAP130;refGene_tx=XM_019837903.2;refGene_gn=SAP130;refGene_tx=XM_019837905.2;refGene_gn=SAP130;refGene_tx=XM_019837906.2
C1 113219573 113219691
refGene_gn=SAP130;refGene_tx=XM_003990672.4;refGene_gn=SAP130;refGene_tx=XM_003990673.4;refGene_gn=SAP130;refGene_tx=XM_019837899.2;refGene_gn=SAP130;refGene_tx=XM_019837900.2;refGene_gn=SAP130;refGene_tx=XM_019837901.2;refGene_gn=SAP130;refGene_tx=XM_019837902.2;refGene_gn=SAP130;refGene_tx=XM_019837903.2;refGene_gn=SAP130;refGene_tx=XM_019837905.2;refGene_gn=SAP130;refGene_tx=XM_019837906.2
C1 113227684 113227920
refGene_gn=SAP130;refGene_tx=XM_003990672.4;refGene_gn=SAP130;refGene_tx=XM_003990673.4;refGene_gn=SAP130;refGene_tx=XM_019837899.2;refGene_gn=SAP130;refGene_tx=XM_019837900.2;refGene_gn=SAP130;refGene_tx=XM_019837901.2;refGene_gn=SAP130;refGene_tx=XM_019837902.2;refGene_gn=SAP130;refGene_tx=XM_019837903.2;refGene_gn=SAP130;refGene_tx=XM_019837904.2;refGene_gn=SAP130;refGene_tx=XM_019837905.2;refGene_gn=SAP130;refGene_tx=XM_019837906.2
C1 113228638 113228797
refGene_gn=SAP130;refGene_tx=XM_003990672.4;refGene_gn=SAP130;refGene_tx=XM_003990673.4;refGene_gn=SAP130;refGene_tx=XM_019837899.2;refGene_gn=SAP130;refGene_tx=XM_019837900.2;refGene_gn=SAP130;refGene_tx=XM_019837901.2;refGene_gn=SAP130;refGene_tx=XM_019837902.2;refGene_gn=SAP130;refGene_tx=XM_019837904.2;refGene_gn=SAP130;refGene_tx=XM_019837905.2
C1 113230071 113230183
refGene_gn=SAP130;refGene_tx=XM_003990672.4;refGene_gn=SAP130;refGene_tx=XM_003990673.4;refGene_gn=SAP130;refGene_tx=XM_019837899.2;refGene_gn=SAP130;refGene_tx=XM_019837900.2;refGene_gn=SAP130;refGene_tx=XM_019837901.2;refGene_gn=SAP130;refGene_tx=XM_019837902.2;refGene_gn=SAP130;refGene_tx=XM_019837903.2;refGene_gn=SAP130;refGene_tx=XM_019837904.2;refGene_gn=SAP130;refGene_tx=XM_019837905.2;refGene_gn=SAP130;refGene_tx=XM_019837906.2
C1 113231243 113231368
refGene_gn=SAP130;refGene_tx=XM_003990672.4;refGene_gn=SAP130;refGene_

tx=XM_003990673.4;refGene_gn=SAP130;refGene_tx=XM_019837899.2;refGene_gn=SAP130;refGene_tx=XM_019837900.2;refGene_gn=SAP130;refGene_tx=XM_019837901.2;refGene_gn=SAP130;refGene_tx=XM_019837902.2;refGene_gn=SAP130;refGene_tx=XM_019837903.2;refGene_gn=SAP130;refGene_tx=XM_019837904.2;refGene_gn=SAP130;refGene_tx=XM_019837905.2;refGene_gn=SAP130;refGene_tx=XM_019837906.2
C1 113233024 113233149
refGene_gn=SAP130;refGene_tx=XM_003990672.4;refGene_gn=SAP130;refGene_tx=XM_003990673.4;refGene_gn=SAP130;refGene_tx=XM_019837899.2;refGene_gn=SAP130;refGene_tx=XM_019837900.2;refGene_gn=SAP130;refGene_tx=XM_019837901.2;refGene_gn=SAP130;refGene_tx=XM_019837902.2;refGene_gn=SAP130;refGene_tx=XM_019837903.2;refGene_gn=SAP130;refGene_tx=XM_019837904.2;refGene_gn=SAP130;refGene_tx=XM_019837905.2;refGene_gn=SAP130;refGene_tx=XM_019837906.2
C1 113246677 113246825
refGene_gn=SAP130;refGene_tx=XM_003990672.4;refGene_gn=SAP130;refGene_tx=XM_003990673.4;refGene_gn=SAP130;refGene_tx=XM_019837899.2;refGene_gn=SAP130;refGene_tx=XM_019837900.2;refGene_gn=SAP130;refGene_tx=XM_019837901.2;refGene_gn=SAP130;refGene_tx=XM_019837902.2;refGene_gn=SAP130;refGene_tx=XM_019837903.2;refGene_gn=SAP130;refGene_tx=XM_019837904.2;refGene_gn=SAP130;refGene_tx=XM_019837905.2;refGene_gn=SAP130;refGene_tx=XM_019837906.2
C1 113246987 113247078
refGene_gn=SAP130;refGene_tx=XM_003990672.4;refGene_gn=SAP130;refGene_tx=XM_003990673.4;refGene_gn=SAP130;refGene_tx=XM_019837899.2;refGene_gn=SAP130;refGene_tx=XM_019837900.2;refGene_gn=SAP130;refGene_tx=XM_019837901.2;refGene_gn=SAP130;refGene_tx=XM_019837902.2;refGene_gn=SAP130;refGene_tx=XM_019837903.2;refGene_gn=SAP130;refGene_tx=XM_019837904.2;refGene_gn=SAP130;refGene_tx=XM_019837905.2;refGene_gn=SAP130;refGene_tx=XM_019837906.2
C1 113247283 113247388
refGene_gn=SAP130;refGene_tx=XM_003990672.4;refGene_gn=SAP130;refGene_tx=XM_003990673.4;refGene_gn=SAP130;refGene_tx=XM_019837899.2;refGene_gn=SAP130;refGene_tx=XM_019837900.2;refGene_gn=SAP130;refGene_tx=XM_019837901.2;refGene_gn=SAP130;refGene_tx=XM_019837902.2;refGene_gn=SAP130;refGene_tx=XM_019837903.2;refGene_gn=SAP130;refGene_tx=XM_019837904.2;refGene_gn=SAP130;refGene_tx=XM_019837905.2;refGene_gn=SAP130;refGene_tx=XM_019837906.2
C1 113251953 113252095
refGene_gn=SAP130;refGene_tx=XM_003990672.4;refGene_gn=SAP130;refGene_tx=XM_003990673.4;refGene_gn=SAP130;refGene_tx=XM_019837899.2;refGene_gn=SAP130;refGene_tx=XM_019837900.2;refGene_gn=SAP130;refGene_tx=XM_019837901.2;refGene_gn=SAP130;refGene_tx=XM_019837902.2;refGene_gn=SAP130;refGene_tx=XM_019837903.2;refGene_gn=SAP130;refGene_tx=XM_019837904.2;refGene_gn=SAP130;refGene_tx=XM_019837906.2
C1 113254615 113254737
refGene_gn=SAP130;refGene_tx=XM_003990672.4;refGene_gn=SAP130;refGene_tx=XM_003990673.4;refGene_gn=SAP130;refGene_tx=XM_019837899.2;refGene_gn=SAP130;refGene_tx=XM_019837900.2;refGene_gn=SAP130;refGene_tx=XM_019837901.2;refGene_gn=SAP130;refGene_tx=XM_019837902.2;refGene_gn=SAP13

0;refGene_tx=XM_019837903.2;refGene_gn=SAP130;refGene_tx=XM_019837904.2;refGene_gn=SAP130;refGene_tx=XM_019837906.2
C1 113260093 113260396
refGene_gn=SAP130;refGene_tx=XM_003990672.4;refGene_gn=SAP130;refGene_tx=XM_003990673.4;refGene_gn=SAP130;refGene_tx=XM_019837899.2;refGene_gn=SAP130;refGene_tx=XM_019837900.2;refGene_gn=SAP130;refGene_tx=XM_019837901.2;refGene_gn=SAP130;refGene_tx=XM_019837902.2;refGene_gn=SAP130;refGene_tx=XM_019837903.2;refGene_gn=SAP130;refGene_tx=XM_019837904.2;refGene_gn=SAP130;refGene_tx=XM_019837905.2;refGene_gn=SAP130;refGene_tx=XM_019837906.2
C1 113261302 113261480
refGene_gn=SAP130;refGene_tx=XM_003990672.4;refGene_gn=SAP130;refGene_tx=XM_003990673.4;refGene_gn=SAP130;refGene_tx=XM_019837899.2;refGene_gn=SAP130;refGene_tx=XM_019837900.2;refGene_gn=SAP130;refGene_tx=XM_019837901.2;refGene_gn=SAP130;refGene_tx=XM_019837902.2;refGene_gn=SAP130;refGene_tx=XM_019837903.2;refGene_gn=SAP130;refGene_tx=XM_019837904.2;refGene_gn=SAP130;refGene_tx=XM_019837905.2;refGene_gn=SAP130;refGene_tx=XM_019837906.2
C1 113268201 113268306
refGene_gn=SAP130;refGene_tx=XM_003990673.4;refGene_gn=SAP130;refGene_tx=XM_019837899.2;refGene_gn=SAP130;refGene_tx=XM_019837900.2;refGene_gn=SAP130;refGene_tx=XM_019837902.2;refGene_gn=SAP130;refGene_tx=XM_019837903.2;refGene_gn=SAP130;refGene_tx=XM_019837904.2;refGene_gn=SAP130;refGene_tx=XM_019837905.2;refGene_gn=SAP130;refGene_tx=XM_019837906.2
C1 113281050 113281409
refGene_gn=SAP130;refGene_tx=XM_003990672.4;refGene_gn=SAP130;refGene_tx=XM_003990673.4;refGene_gn=SAP130;refGene_tx=XM_019837899.2;refGene_gn=SAP130;refGene_tx=XM_019837900.2;refGene_gn=SAP130;refGene_tx=XM_019837901.2;refGene_gn=SAP130;refGene_tx=XM_019837902.2;refGene_gn=SAP130;refGene_tx=XM_019837903.2;refGene_gn=SAP130;refGene_tx=XM_019837904.2;refGene_gn=SAP130;refGene_tx=XM_019837905.2;refGene_gn=SAP130;refGene_tx=XM_019837906.2
C1 113284797 113285046
refGene_gn=SAP130;refGene_tx=XM_003990672.4;refGene_gn=SAP130;refGene_tx=XM_003990673.4;refGene_gn=SAP130;refGene_tx=XM_019837899.2;refGene_gn=SAP130;refGene_tx=XM_019837900.2;refGene_gn=SAP130;refGene_tx=XM_019837901.2;refGene_gn=SAP130;refGene_tx=XM_019837902.2;refGene_gn=SAP130;refGene_tx=XM_019837903.2;refGene_gn=SAP130;refGene_tx=XM_019837904.2;refGene_gn=SAP130;refGene_tx=XM_019837905.2;refGene_gn=SAP130;refGene_tx=XM_019837906.2
C1 113285211 113285337
refGene_gn=SAP130;refGene_tx=XM_003990672.4;refGene_gn=SAP130;refGene_tx=XM_003990673.4;refGene_gn=SAP130;refGene_tx=XM_019837899.2;refGene_gn=SAP130;refGene_tx=XM_019837900.2;refGene_gn=SAP130;refGene_tx=XM_019837901.2;refGene_gn=SAP130;refGene_tx=XM_019837902.2;refGene_gn=SAP130;refGene_tx=XM_019837903.2;refGene_gn=SAP130;refGene_tx=XM_019837904.2;refGene_gn=SAP130;refGene_tx=XM_019837905.2;refGene_gn=SAP130;refGene_tx=XM_019837906.2
C1 113296334 113296438

refGene_gn=SAP130;refGene_tx=XM_003990672.4;refGene_gn=SAP130;refGene_tx=XM_003990673.4;refGene_gn=SAP130;refGene_tx=XM_019837899.2;refGene_gn=SAP130;refGene_tx=XM_019837900.2;refGene_gn=SAP130;refGene_tx=XM_019837901.2;refGene_gn=SAP130;refGene_tx=XM_019837902.2;refGene_gn=SAP130;refGene_tx=XM_019837903.2;refGene_gn=SAP130;refGene_tx=XM_019837904.2;refGene_gn=SAP130;refGene_tx=XM_019837905.2;refGene_gn=SAP130;refGene_tx=XM_019837906.2
C1 113297543 113297657
refGene_gn=SAP130;refGene_tx=XM_003990672.4;refGene_gn=SAP130;refGene_tx=XM_003990673.4;refGene_gn=SAP130;refGene_tx=XM_019837899.2;refGene_gn=SAP130;refGene_tx=XM_019837900.2;refGene_gn=SAP130;refGene_tx=XM_019837901.2;refGene_gn=SAP130;refGene_tx=XM_019837902.2;refGene_gn=SAP130;refGene_tx=XM_019837903.2;refGene_gn=SAP130;refGene_tx=XM_019837904.2;refGene_gn=SAP130;refGene_tx=XM_019837905.2;refGene_gn=SAP130;refGene_tx=XM_019837906.2
C1 113297961 113298120
refGene_gn=SAP130;refGene_tx=XM_003990672.4;refGene_gn=SAP130;refGene_tx=XM_003990673.4;refGene_gn=SAP130;refGene_tx=XM_019837899.2;refGene_gn=SAP130;refGene_tx=XM_019837900.2;refGene_gn=SAP130;refGene_tx=XM_019837901.2;refGene_gn=SAP130;refGene_tx=XM_019837902.2;refGene_gn=SAP130;refGene_tx=XM_019837903.2;refGene_gn=SAP130;refGene_tx=XM_019837904.2;refGene_gn=SAP130;refGene_tx=XM_019837905.2;refGene_gn=SAP130;refGene_tx=XM_019837906.2
C1 113353211 113353618
refGene_gn=AMMECR1L;refGene_tx=XM_003990674.4;refGene_gn=AMMECR1L;refGene_tx=XM_023259199.1
C1 113355988 113356099
refGene_gn=AMMECR1L;refGene_tx=XM_003990674.4;refGene_gn=AMMECR1L;refGene_tx=XM_023259199.1
C1 113356409 113356524
refGene_gn=AMMECR1L;refGene_tx=XM_003990674.4;refGene_gn=AMMECR1L;refGene_tx=XM_023259199.1
C1 113357191 113357282
refGene_gn=AMMECR1L;refGene_tx=XM_003990674.4;refGene_gn=AMMECR1L;refGene_tx=XM_023259199.1
C1 113359259 113359356
refGene_gn=AMMECR1L;refGene_tx=XM_003990674.4;refGene_gn=AMMECR1L;refGene_tx=XM_023259199.1
C1 113361042 113361154
refGene_gn=AMMECR1L;refGene_tx=XM_003990674.4;refGene_gn=AMMECR1L;refGene_tx=XM_023259199.1
C1 113369316 113369389
refGene_gn=POLR2D;refGene_tx=XM_003990675.3
C1 113376508 113376689
refGene_gn=POLR2D;refGene_tx=XM_003990675.3
C1 113381606 113381702
refGene_gn=POLR2D;refGene_tx=XM_003990675.3
C1 113382668 113382747
refGene_gn=POLR2D;refGene_tx=XM_003990675.3
C1 113441310 113441514

refGene_gn=WDR33;refGene_tx=XM_003990676.4;refGene_gn=WDR33;refGene_tx=XM_006935188.3;refGene_gn=WDR33;refGene_tx=XM_006935189.3;refGene_gn=WDR33;refGene_tx=XM_006935190.4;refGene_gn=WDR33;refGene_tx=XM_006935191.3;refGene_gn=WDR33;refGene_tx=XM_019837907.2;refGene_gn=WDR33;refGene_tx=XM_019837908.2;refGene_gn=WDR33;refGene_tx=XM_019837909.2;refGene_gn=WDR33;refGene_tx=XM_019837910.2;refGene_gn=WDR33;refGene_tx=XM_019837911.2;refGene_gn=WDR33;refGene_tx=XM_019837912.2
C1 113443357 113443426
refGene_gn=WDR33;refGene_tx=XM_003990676.4;refGene_gn=WDR33;refGene_tx=XM_006935188.3;refGene_gn=WDR33;refGene_tx=XM_006935189.3;refGene_gn=WDR33;refGene_tx=XM_006935190.4;refGene_gn=WDR33;refGene_tx=XM_006935191.3;refGene_gn=WDR33;refGene_tx=XM_019837907.2;refGene_gn=WDR33;refGene_tx=XM_019837908.2;refGene_gn=WDR33;refGene_tx=XM_019837909.2;refGene_gn=WDR33;refGene_tx=XM_019837910.2;refGene_gn=WDR33;refGene_tx=XM_019837911.2;refGene_gn=WDR33;refGene_tx=XM_019837912.2
C1 113444012 113444117
refGene_gn=WDR33;refGene_tx=XM_003990676.4;refGene_gn=WDR33;refGene_tx=XM_006935188.3;refGene_gn=WDR33;refGene_tx=XM_006935189.3;refGene_gn=WDR33;refGene_tx=XM_006935190.4;refGene_gn=WDR33;refGene_tx=XM_006935191.3;refGene_gn=WDR33;refGene_tx=XM_019837907.2;refGene_gn=WDR33;refGene_tx=XM_019837908.2;refGene_gn=WDR33;refGene_tx=XM_019837909.2;refGene_gn=WDR33;refGene_tx=XM_019837910.2;refGene_gn=WDR33;refGene_tx=XM_019837911.2;refGene_gn=WDR33;refGene_tx=XM_019837912.2
C1 113447947 113448043
refGene_gn=WDR33;refGene_tx=XM_003990676.4;refGene_gn=WDR33;refGene_tx=XM_006935188.3;refGene_gn=WDR33;refGene_tx=XM_006935189.3;refGene_gn=WDR33;refGene_tx=XM_006935190.4;refGene_gn=WDR33;refGene_tx=XM_006935191.3;refGene_gn=WDR33;refGene_tx=XM_019837907.2;refGene_gn=WDR33;refGene_tx=XM_019837908.2;refGene_gn=WDR33;refGene_tx=XM_019837909.2;refGene_gn=WDR33;refGene_tx=XM_019837910.2;refGene_gn=WDR33;refGene_tx=XM_019837911.2;refGene_gn=WDR33;refGene_tx=XM_019837912.2
C1 113448229 113448381
refGene_gn=WDR33;refGene_tx=XM_003990676.4;refGene_gn=WDR33;refGene_tx=XM_006935188.3;refGene_gn=WDR33;refGene_tx=XM_006935189.3;refGene_gn=WDR33;refGene_tx=XM_006935190.4;refGene_gn=WDR33;refGene_tx=XM_006935191.3;refGene_gn=WDR33;refGene_tx=XM_019837907.2;refGene_gn=WDR33;refGene_tx=XM_019837908.2;refGene_gn=WDR33;refGene_tx=XM_019837909.2;refGene_gn=WDR33;refGene_tx=XM_019837910.2;refGene_gn=WDR33;refGene_tx=XM_019837911.2;refGene_gn=WDR33;refGene_tx=XM_019837912.2
C1 113450095 113450193
refGene_gn=WDR33;refGene_tx=XM_003990676.4;refGene_gn=WDR33;refGene_tx=XM_006935188.3;refGene_gn=WDR33;refGene_tx=XM_006935189.3;refGene_gn=WDR33;refGene_tx=XM_006935190.4;refGene_gn=WDR33;refGene_tx=XM_006935191.3;refGene_gn=WDR33;refGene_tx=XM_019837907.2;refGene_gn=WDR33;refGene_tx=XM_019837908.2;refGene_gn=WDR33;refGene_tx=XM_019837909.2;refGene_gn=WDR33;refGene_tx=XM_019837910.2;refGene_gn=WDR33;refGene_tx=XM_019837911.2;refGene_gn=WDR33;refGene_tx=XM_019837912.2
C1 113453062 113453097
refGene_gn=WDR33;refGene_tx=XM_019837911.2
C1 113462367 113462429

refGene_gn=WDR33;refGene_tx=XM_006935190.4
C1	113469428	113469439
refGene_gn=WDR33;refGene_tx=XM_006935191.3
C1	113488285	113488412
refGene_gn=WDR33;refGene_tx=XM_003990676.4;refGene_gn=WDR33;refGene_tx=XM_006935188.3;refGene_gn=WDR33;refGene_tx=XM_006935189.3;refGene_gn=WDR33;refGene_tx=XM_019837907.2;refGene_gn=WDR33;refGene_tx=XM_019837908.2;refGene_gn=WDR33;refGene_tx=XM_019837909.2;refGene_gn=WDR33;refGene_tx=XM_019837910.2
C1	113490914	113491069
refGene_gn=WDR33;refGene_tx=XM_003990676.4;refGene_gn=WDR33;refGene_tx=XM_006935188.3;refGene_gn=WDR33;refGene_tx=XM_006935189.3;refGene_gn=WDR33;refGene_tx=XM_019837907.2;refGene_gn=WDR33;refGene_tx=XM_019837908.2;refGene_gn=WDR33;refGene_tx=XM_019837909.2;refGene_gn=WDR33;refGene_tx=XM_019837910.2
C1	113491157	113491236
refGene_gn=WDR33;refGene_tx=XM_003990676.4;refGene_gn=WDR33;refGene_tx=XM_006935188.3;refGene_gn=WDR33;refGene_tx=XM_006935189.3;refGene_gn=WDR33;refGene_tx=XM_019837907.2;refGene_gn=WDR33;refGene_tx=XM_019837908.2;refGene_gn=WDR33;refGene_tx=XM_019837909.2;refGene_gn=WDR33;refGene_tx=XM_019837910.2
C1	113491708	113491819
refGene_gn=WDR33;refGene_tx=XM_003990676.4;refGene_gn=WDR33;refGene_tx=XM_006935188.3;refGene_gn=WDR33;refGene_tx=XM_006935189.3;refGene_gn=WDR33;refGene_tx=XM_019837907.2;refGene_gn=WDR33;refGene_tx=XM_019837908.2;refGene_gn=WDR33;refGene_tx=XM_019837909.2;refGene_gn=WDR33;refGene_tx=XM_019837910.2
C1	113492589	113492684
refGene_gn=WDR33;refGene_tx=XM_003990676.4;refGene_gn=WDR33;refGene_tx=XM_006935188.3;refGene_gn=WDR33;refGene_tx=XM_006935189.3;refGene_gn=WDR33;refGene_tx=XM_019837907.2;refGene_gn=WDR33;refGene_tx=XM_019837908.2;refGene_gn=WDR33;refGene_tx=XM_019837909.2;refGene_gn=WDR33;refGene_tx=XM_019837910.2
C1	113492884	113492971
refGene_gn=WDR33;refGene_tx=XM_003990676.4;refGene_gn=WDR33;refGene_tx=XM_006935188.3;refGene_gn=WDR33;refGene_tx=XM_006935189.3;refGene_gn=WDR33;refGene_tx=XM_019837907.2;refGene_gn=WDR33;refGene_tx=XM_019837908.2;refGene_gn=WDR33;refGene_tx=XM_019837909.2;refGene_gn=WDR33;refGene_tx=XM_019837910.2
C1	113493439	113493579
refGene_gn=WDR33;refGene_tx=XM_003990676.4;refGene_gn=WDR33;refGene_tx=XM_006935188.3;refGene_gn=WDR33;refGene_tx=XM_006935189.3;refGene_gn=WDR33;refGene_tx=XM_019837907.2;refGene_gn=WDR33;refGene_tx=XM_019837908.2;refGene_gn=WDR33;refGene_tx=XM_019837909.2;refGene_gn=WDR33;refGene_tx=XM_019837910.2
C1	113494173	113494326
refGene_gn=WDR33;refGene_tx=XM_003990676.4;refGene_gn=WDR33;refGene_tx=XM_006935188.3;refGene_gn=WDR33;refGene_tx=XM_006935189.3;refGene_gn=WDR33;refGene_tx=XM_019837907.2;refGene_gn=WDR33;refGene_tx=XM_019837908.2;refGene_gn=WDR33;refGene_tx=XM_019837909.2;refGene_gn=WDR33;refGe

ne_tx=XM_019837910.2
C1	113495231	113496302
refGene_gn=WDR33;refGene_tx=XM_003990676.4;refGene_gn=WDR33;refGene_tx=XM_006935188.3;refGene_gn=WDR33;refGene_tx=XM_006935189.3;refGene_gn=WDR33;refGene_tx=XM_019837907.2;refGene_gn=WDR33;refGene_tx=XM_019837908.2;refGene_gn=WDR33;refGene_tx=XM_019837909.2;refGene_gn=WDR33;refGene_tx=XM_019837910.2
C1	113498912	113499021
refGene_gn=WDR33;refGene_tx=XM_003990676.4;refGene_gn=WDR33;refGene_tx=XM_006935189.3;refGene_gn=WDR33;refGene_tx=XM_019837907.2;refGene_gn=WDR33;refGene_tx=XM_019837908.2;refGene_gn=WDR33;refGene_tx=XM_019837910.2
C1	113500293	113500400
refGene_gn=WDR33;refGene_tx=XM_006935189.3
C1	113502084	113502523
refGene_gn=WDR33;refGene_tx=XM_003990676.4;refGene_gn=WDR33;refGene_tx=XM_006935188.3;refGene_gn=WDR33;refGene_tx=XM_019837907.2;refGene_gn=WDR33;refGene_tx=XM_019837908.2;refGene_gn=WDR33;refGene_tx=XM_019837909.2
C1	113506704	113506868
refGene_gn=WDR33;refGene_tx=XM_003990676.4;refGene_gn=WDR33;refGene_tx=XM_006935188.3;refGene_gn=WDR33;refGene_tx=XM_019837907.2;refGene_gn=WDR33;refGene_tx=XM_019837908.2;refGene_gn=WDR33;refGene_tx=XM_019837909.2;refGene_gn=WDR33;refGene_tx=XM_019837910.2
C1	113506980	113507073
refGene_gn=WDR33;refGene_tx=XM_003990676.4;refGene_gn=WDR33;refGene_tx=XM_006935188.3;refGene_gn=WDR33;refGene_tx=XM_019837907.2;refGene_gn=WDR33;refGene_tx=XM_019837908.2;refGene_gn=WDR33;refGene_tx=XM_019837909.2;refGene_gn=WDR33;refGene_tx=XM_019837910.2
C1	113507957	113508173
refGene_gn=WDR33;refGene_tx=XM_003990676.4;refGene_gn=WDR33;refGene_tx=XM_006935188.3;refGene_gn=WDR33;refGene_tx=XM_019837907.2;refGene_gn=WDR33;refGene_tx=XM_019837910.2
C1	113510366	113510596
refGene_gn=WDR33;refGene_tx=XM_003990676.4;refGene_gn=WDR33;refGene_tx=XM_006935188.3;refGene_gn=WDR33;refGene_tx=XM_019837907.2;refGene_gn=WDR33;refGene_tx=XM_019837908.2;refGene_gn=WDR33;refGene_tx=XM_019837909.2;refGene_gn=WDR33;refGene_tx=XM_019837910.2
C1	113514163	113514820
refGene_gn=SFT2D3;refGene_tx=XM_023259203.1
C1	113545005	113545016
refGene_gn=LIMS2;refGene_tx=XM_023259204.1;refGene_gn=LIMS2;refGene_tx=XM_023259206.1
C1	113557345	113557431
refGene_gn=LIMS2;refGene_tx=XM_023259205.1
C1	113563446	113563606
refGene_gn=LIMS2;refGene_tx=XM_023259204.1;refGene_gn=LIMS2;refGene_tx=XM_023259205.1
C1	113566147	113566214
refGene_gn=LIMS2;refGene_tx=XM_023259204.1;refGene_gn=LIMS2;refGene_tx

=XM_023259205.1
C1 113566472 113566593
refGene_gn=LIMS2;refGene_tx=XM_023259204.1;refGene_gn=LIMS2;refGene_tx
=XM_023259205.1
C1 113570594 113571614
refGene_gn=GPR17;refGene_tx=XM_006935193.3;refGene_gn=GPR17;refGene_tx
=XM_019837913.2
C1 113578789 113578939
refGene_gn=LIMS2;refGene_tx=XM_023259204.1;refGene_gn=LIMS2;refGene_tx
=XM_023259205.1;refGene_gn=LIMS2;refGene_tx=XM_023259206.1
C1 113579571 113579722
refGene_gn=LIMS2;refGene_tx=XM_023259204.1;refGene_gn=LIMS2;refGene_tx
=XM_023259205.1;refGene_gn=LIMS2;refGene_tx=XM_023259206.1
C1 113581187 113581280
refGene_gn=LIMS2;refGene_tx=XM_023259204.1;refGene_gn=LIMS2;refGene_tx
=XM_023259205.1;refGene_gn=LIMS2;refGene_tx=XM_023259206.1
C1 113581810 113581859
refGene_gn=LIMS2;refGene_tx=XM_023259204.1;refGene_gn=LIMS2;refGene_tx
=XM_023259205.1;refGene_gn=LIMS2;refGene_tx=XM_023259206.1
C1 113582010 113582086
refGene_gn=LIMS2;refGene_tx=XM_023259204.1;refGene_gn=LIMS2;refGene_tx
=XM_023259205.1;refGene_gn=LIMS2;refGene_tx=XM_023259206.1
C1 113582704 113582852
refGene_gn=LIMS2;refGene_tx=XM_023259204.1;refGene_gn=LIMS2;refGene_tx
=XM_023259205.1;refGene_gn=LIMS2;refGene_tx=XM_023259206.1
C1 113584785 113584881
refGene_gn=MYO7B;refGene_tx=XM_019837914.2
C1 113585285 113585405
refGene_gn=MYO7B;refGene_tx=XM_019837914.2
C1 113585624 113585708
refGene_gn=MYO7B;refGene_tx=XM_019837914.2
C1 113585927 113586044
refGene_gn=MYO7B;refGene_tx=XM_019837914.2
C1 113586351 113586537
refGene_gn=MYO7B;refGene_tx=XM_019837914.2
C1 113587013 113587120
refGene_gn=MYO7B;refGene_tx=XM_019837914.2
C1 113587695 113587783
refGene_gn=MYO7B;refGene_tx=XM_019837914.2
C1 113588092 113588206
refGene_gn=MYO7B;refGene_tx=XM_019837914.2
C1 113589157 113589263
refGene_gn=MYO7B;refGene_tx=XM_019837914.2
C1 113590189 113590345
refGene_gn=MYO7B;refGene_tx=XM_019837914.2
C1 113590844 113590998
refGene_gn=MYO7B;refGene_tx=XM_019837914.2
C1 113591191 113591349
refGene_gn=MYO7B;refGene_tx=XM_019837914.2
C1 113591623 113591754

refGene_gn=MYO7B;refGene_tx=XM_019837914.2
C1	113592597	113592779
refGene_gn=MYO7B;refGene_tx=XM_019837914.2
C1	113593687	113593854
refGene_gn=MYO7B;refGene_tx=XM_019837914.2
C1	113594862	113594989
refGene_gn=MYO7B;refGene_tx=XM_019837914.2
C1	113595084	113595202
refGene_gn=MYO7B;refGene_tx=XM_019837914.2
C1	113598612	113598780
refGene_gn=MYO7B;refGene_tx=XM_019837914.2
C1	113600072	113600300
refGene_gn=MYO7B;refGene_tx=XM_019837914.2
C1	113600903	113601077
refGene_gn=MYO7B;refGene_tx=XM_019837914.2
C1	113602115	113602235
refGene_gn=MYO7B;refGene_tx=XM_019837914.2
C1	113603528	113603655
refGene_gn=MYO7B;refGene_tx=XM_019837914.2
C1	113607880	113608008
refGene_gn=MYO7B;refGene_tx=XM_019837914.2
C1	113608218	113608386
refGene_gn=MYO7B;refGene_tx=XM_019837914.2
C1	113608517	113608727
refGene_gn=MYO7B;refGene_tx=XM_019837914.2
C1	113609389	113609560
refGene_gn=MYO7B;refGene_tx=XM_019837914.2
C1	113610869	113611088
refGene_gn=MYO7B;refGene_tx=XM_019837914.2
C1	113612607	113612692
refGene_gn=MYO7B;refGene_tx=XM_019837914.2
C1	113619948	113620043
refGene_gn=MYO7B;refGene_tx=XM_019837914.2
C1	113621813	113621912
refGene_gn=MYO7B;refGene_tx=XM_019837914.2
C1	113622568	113622721
refGene_gn=MYO7B;refGene_tx=XM_019837914.2
C1	113624549	113624687
refGene_gn=MYO7B;refGene_tx=XM_019837914.2
C1	113626362	113626526
refGene_gn=MYO7B;refGene_tx=XM_019837914.2
C1	113628609	113628745
refGene_gn=MYO7B;refGene_tx=XM_019837914.2
C1	113629433	113629644
refGene_gn=MYO7B;refGene_tx=XM_019837914.2
C1	113631625	113631768
refGene_gn=MYO7B;refGene_tx=XM_019837914.2
C1	113632072	113632192
refGene_gn=MYO7B;refGene_tx=XM_019837914.2
C1	113633319	113633396

refGene_gn=MYO7B;refGene_tx=XM_019837914.2
C1 113635526 113635680
refGene_gn=MYO7B;refGene_tx=XM_019837914.2
C1 113637139 113637253
refGene_gn=MYO7B;refGene_tx=XM_019837914.2
C1 113637937 113638080
refGene_gn=MYO7B;refGene_tx=XM_019837914.2
C1 113641509 113641631
refGene_gn=MYO7B;refGene_tx=XM_019837914.2
C1 113645093 113645278
refGene_gn=MYO7B;refGene_tx=XM_019837914.2
C1 113646683 113646836
refGene_gn=MYO7B;refGene_tx=XM_019837914.2
C1 113649706 113649820
refGene_gn=MYO7B;refGene_tx=XM_019837914.2
C1 113660948 113660966
refGene_gn=MYO7B;refGene_tx=XM_019837914.2
C1 113688541 113688575
refGene_gn=IWS1;refGene_tx=XM_006935195.4;refGene_gn=IWS1;refGene_tx=XM_006935196.4;refGene_gn=IWS1;refGene_tx=XM_006935198.3;refGene_gn=IWS1;refGene_tx=XM_006935199.4;refGene_gn=IWS1;refGene_tx=XM_019837917.2
C1 113690611 113690730
refGene_gn=IWS1;refGene_tx=XM_006935195.4;refGene_gn=IWS1;refGene_tx=XM_006935196.4;refGene_gn=IWS1;refGene_tx=XM_006935198.3;refGene_gn=IWS1;refGene_tx=XM_006935199.4;refGene_gn=IWS1;refGene_tx=XM_019837917.2
C1 113705148 113706376
refGene_gn=IWS1;refGene_tx=XM_006935195.4;refGene_gn=IWS1;refGene_tx=XM_006935196.4;refGene_gn=IWS1;refGene_tx=XM_006935198.3;refGene_gn=IWS1;refGene_tx=XM_006935199.4;refGene_gn=IWS1;refGene_tx=XM_019837917.2
C1 113707785 113707978
refGene_gn=IWS1;refGene_tx=XM_006935195.4;refGene_gn=IWS1;refGene_tx=XM_006935196.4;refGene_gn=IWS1;refGene_tx=XM_006935198.3;refGene_gn=IWS1;refGene_tx=XM_006935199.4;refGene_gn=IWS1;refGene_tx=XM_019837917.2
C1 113708518 113708576
refGene_gn=IWS1;refGene_tx=XM_006935195.4;refGene_gn=IWS1;refGene_tx=XM_006935196.4;refGene_gn=IWS1;refGene_tx=XM_006935198.3;refGene_gn=IWS1;refGene_tx=XM_006935199.4;refGene_gn=IWS1;refGene_tx=XM_019837917.2
C1 113714148 113714246
refGene_gn=IWS1;refGene_tx=XM_006935195.4;refGene_gn=IWS1;refGene_tx=XM_006935196.4;refGene_gn=IWS1;refGene_tx=XM_006935198.3;refGene_gn=IWS1;refGene_tx=XM_006935199.4;refGene_gn=IWS1;refGene_tx=XM_019837917.2
C1 113714843 113714994
refGene_gn=IWS1;refGene_tx=XM_006935195.4;refGene_gn=IWS1;refGene_tx=XM_006935196.4;refGene_gn=IWS1;refGene_tx=XM_006935198.3;refGene_gn=IWS1;refGene_tx=XM_006935199.4;refGene_gn=IWS1;refGene_tx=XM_019837917.2
C1 113715546 113715629
refGene_gn=IWS1;refGene_tx=XM_006935195.4;refGene_gn=IWS1;refGene_tx=XM_006935196.4;refGene_gn=IWS1;refGene_tx=XM_006935198.3;refGene_gn=IWS1;refGene_tx=XM_006935199.4;refGene_gn=IWS1;refGene_tx=XM_019837917.2
C1 113716930 113717060

refGene_gn=IWS1;refGene_tx=XM_006935195.4;refGene_gn=IWS1;refGene_tx=XM_006935196.4;refGene_gn=IWS1;refGene_tx=XM_006935198.3;refGene_gn=IWS1;refGene_tx=XM_006935199.4;refGene_gn=IWS1;refGene_tx=XM_019837917.2
C1 113717925 113718043
refGene_gn=IWS1;refGene_tx=XM_006935195.4;refGene_gn=IWS1;refGene_tx=XM_006935196.4;refGene_gn=IWS1;refGene_tx=XM_006935198.3;refGene_gn=IWS1;refGene_tx=XM_006935199.4
C1 113719692 113719804
refGene_gn=IWS1;refGene_tx=XM_006935195.4;refGene_gn=IWS1;refGene_tx=XM_006935196.4;refGene_gn=IWS1;refGene_tx=XM_006935198.3;refGene_gn=IWS1;refGene_tx=XM_006935199.4;refGene_gn=IWS1;refGene_tx=XM_019837917.2
C1 113720358 113720415
refGene_gn=IWS1;refGene_tx=XM_006935195.4;refGene_gn=IWS1;refGene_tx=XM_006935196.4;refGene_gn=IWS1;refGene_tx=XM_006935198.3;refGene_gn=IWS1;refGene_tx=XM_006935199.4
C1 113723416 113723528
refGene_gn=IWS1;refGene_tx=XM_006935195.4;refGene_gn=IWS1;refGene_tx=XM_006935196.4;refGene_gn=IWS1;refGene_tx=XM_006935198.3;refGene_gn=IWS1;refGene_tx=XM_006935199.4
C1 113724006 113724135
refGene_gn=IWS1;refGene_tx=XM_006935198.3
C1 113728126 113728258
refGene_gn=IWS1;refGene_tx=XM_006935195.4;refGene_gn=IWS1;refGene_tx=XM_006935196.4;refGene_gn=IWS1;refGene_tx=XM_006935199.4
C1 113787616 113788194
refGene_gn=PROC;refGene_tx=XM_023259207.1;refGene_gn=PROC;refGene_tx=XM_023259208.1;refGene_gn=PROC;refGene_tx=XM_023259209.1;refGene_gn=PROC;refGene_tx=XM_023259210.1
C1 113789345 113789463
refGene_gn=PROC;refGene_tx=XM_023259207.1;refGene_gn=PROC;refGene_tx=XM_023259208.1;refGene_gn=PROC;refGene_tx=XM_023259209.1;refGene_gn=PROC;refGene_tx=XM_023259210.1
C1 113790399 113790539
refGene_gn=PROC;refGene_tx=XM_023259207.1;refGene_gn=PROC;refGene_tx=XM_023259208.1;refGene_gn=PROC;refGene_tx=XM_023259209.1;refGene_gn=PROC;refGene_tx=XM_023259210.1
C1 113793563 113793698
refGene_gn=PROC;refGene_tx=XM_023259207.1;refGene_gn=PROC;refGene_tx=XM_023259208.1;refGene_gn=PROC;refGene_tx=XM_023259209.1;refGene_gn=PROC;refGene_tx=XM_023259210.1
C1 113793878 113794016
refGene_gn=PROC;refGene_tx=XM_023259207.1;refGene_gn=PROC;refGene_tx=XM_023259208.1;refGene_gn=PROC;refGene_tx=XM_023259209.1;refGene_gn=PROC;refGene_tx=XM_023259210.1
C1 113794131 113794156
refGene_gn=PROC;refGene_tx=XM_023259207.1;refGene_gn=PROC;refGene_tx=XM_023259208.1;refGene_gn=PROC;refGene_tx=XM_023259209.1;refGene_gn=PROC;refGene_tx=XM_023259210.1
C1 113795401 113795568
refGene_gn=PROC;refGene_tx=XM_023259207.1;refGene_gn=PROC;refGene_tx=X

M_023259208.1;refGene_gn=PROC;refGene_tx=XM_023259209.1;refGene_gn=PRO
C;refGene_tx=XM_023259210.1
C1 113796116 113796239
refGene_gn=PROC;refGene_tx=XM_023259207.1;refGene_gn=PROC;refGene_tx=X
M_023259208.1
C1 113796893 113796980
refGene_gn=PROC;refGene_tx=XM_023259207.1;refGene_gn=PROC;refGene_tx=X
M_023259208.1;refGene_gn=PROC;refGene_tx=XM_023259209.1;refGene_gn=PRO
C;refGene_tx=XM_023259210.1
C1 113798142 113798188
refGene_gn=PROC;refGene_tx=XM_023259207.1;refGene_gn=PROC;refGene_tx=X
M_023259209.1
C1 113884152 113884159
refGene_gn=MAP3K2;refGene_tx=XM_023259211.1
C1 113887783 113887902
refGene_gn=MAP3K2;refGene_tx=XM_023259211.1
C1 113888531 113888572
refGene_gn=MAP3K2;refGene_tx=XM_023259211.1
C1 113889816 113889916
refGene_gn=MAP3K2;refGene_tx=XM_023259211.1
C1 113896337 113896451
refGene_gn=MAP3K2;refGene_tx=XM_023259211.1
C1 113896832 113896920
refGene_gn=MAP3K2;refGene_tx=XM_023259211.1
C1 113901302 113901433
refGene_gn=MAP3K2;refGene_tx=XM_023259211.1
C1 113902550 113902630
refGene_gn=MAP3K2;refGene_tx=XM_023259211.1
C1 113904106 113904174
refGene_gn=MAP3K2;refGene_tx=XM_023259211.1
C1 113904346 113904439
refGene_gn=MAP3K2;refGene_tx=XM_023259211.1
C1 113906056 113906263
refGene_gn=MAP3K2;refGene_tx=XM_023259211.1
C1 113907878 113908027
refGene_gn=MAP3K2;refGene_tx=XM_023259211.1
C1 113908384 113908516
refGene_gn=MAP3K2;refGene_tx=XM_023259211.1
C1 113912922 113913052
refGene_gn=MAP3K2;refGene_tx=XM_023259211.1
C1 113918586 113918764
refGene_gn=MAP3K2;refGene_tx=XM_023259211.1
C1 113920463 113920689
refGene_gn=MAP3K2;refGene_tx=XM_023259211.1
C1 113937633 113937661
refGene_gn=ERCC3;refGene_tx=XM_003990686.4
C1 113938024 113938230
refGene_gn=ERCC3;refGene_tx=XM_003990686.4
C1 113938883 113939120
refGene_gn=ERCC3;refGene_tx=XM_003990686.4

C1 113940953 113941003
refGene_gn=ERCC3;refGene_tx=XM_003990686.4
C1 113941436 113941572
refGene_gn=ERCC3;refGene_tx=XM_003990686.4
C1 113941977 113942142
refGene_gn=ERCC3;refGene_tx=XM_003990686.4
C1 113942625 113942830
refGene_gn=ERCC3;refGene_tx=XM_003990686.4
C1 113944955 113945270
refGene_gn=ERCC3;refGene_tx=XM_003990686.4
C1 113950436 113950621
refGene_gn=ERCC3;refGene_tx=XM_003990686.4
C1 113953924 113954127
refGene_gn=ERCC3;refGene_tx=XM_003990686.4
C1 113954816 113954913
refGene_gn=ERCC3;refGene_tx=XM_003990686.4
C1 113956023 113956141
refGene_gn=ERCC3;refGene_tx=XM_003990686.4
C1 113960387 113960506
refGene_gn=ERCC3;refGene_tx=XM_003990686.4
C1 113962662 113962815
refGene_gn=ERCC3;refGene_tx=XM_003990686.4
C1 113964625 113964757
refGene_gn=ERCC3;refGene_tx=XM_003990686.4
C1 113994764 113994842
refGene_gn=LOC101096908;refGene_tx=XM_023259212.1;refGene_gn=LOC101096908;refGene_tx=XM_023259213.1;refGene_gn=LOC101096908;refGene_tx=XM_023259214.1
C1 114001627 114001822
refGene_gn=LOC101096908;refGene_tx=XM_023259212.1;refGene_gn=LOC101096908;refGene_tx=XM_023259213.1;refGene_gn=LOC101096908;refGene_tx=XM_023259214.1;refGene_gn=LOC101096908;refGene_tx=XM_023259215.1
C1 114002996 114003196
refGene_gn=LOC101096908;refGene_tx=XM_023259212.1;refGene_gn=LOC101096908;refGene_tx=XM_023259213.1;refGene_gn=LOC101096908;refGene_tx=XM_023259214.1;refGene_gn=LOC101096908;refGene_tx=XM_023259215.1
C1 114005112 114005322
refGene_gn=LOC101096908;refGene_tx=XM_023259212.1;refGene_gn=LOC101096908;refGene_tx=XM_023259213.1;refGene_gn=LOC101096908;refGene_tx=XM_023259214.1;refGene_gn=LOC101096908;refGene_tx=XM_023259215.1
C1 114006366 114006530
refGene_gn=LOC101096908;refGene_tx=XM_023259212.1;refGene_gn=LOC101096908;refGene_tx=XM_023259213.1;refGene_gn=LOC101096908;refGene_tx=XM_023259214.1;refGene_gn=LOC101096908;refGene_tx=XM_023259215.1
C1 114010452 114010619
refGene_gn=LOC101096908;refGene_tx=XM_023259212.1;refGene_gn=LOC101096908;refGene_tx=XM_023259213.1;refGene_gn=LOC101096908;refGene_tx=XM_023259214.1;refGene_gn=LOC101096908;refGene_tx=XM_023259215.1
C1 114011797 114011876
refGene_gn=LOC101096908;refGene_tx=XM_023259212.1;refGene_gn=LOC101096

908;refGene_tx=XM_023259213.1;refGene_gn=LOC101096908;refGene_tx=XM_023259214.1;refGene_gn=LOC101096908;refGene_tx=XM_023259215.1
C1 114012312 114012516
refGene_gn=LOC101096908;refGene_tx=XM_023259212.1;refGene_gn=LOC101096908;refGene_tx=XM_023259213.1;refGene_gn=LOC101096908;refGene_tx=XM_023259214.1;refGene_gn=LOC101096908;refGene_tx=XM_023259215.1
C1 114015928 114016045
refGene_gn=LOC101096908;refGene_tx=XM_023259213.1
C1 114030096 114030219
refGene_gn=LOC101096908;refGene_tx=XM_023259212.1;refGene_gn=LOC101096908;refGene_tx=XM_023259214.1;refGene_gn=LOC101096908;refGene_tx=XM_023259215.1
C1 114092774 114092858
refGene_gn=BIN1;refGene_tx=XM_023259217.1;refGene_gn=BIN1;refGene_tx=XM_023259219.1;refGene_gn=BIN1;refGene_tx=XM_023259223.1;refGene_gn=BIN1;refGene_tx=XM_023259224.1;refGene_gn=BIN1;refGene_tx=XM_023259227.1;refGene_gn=BIN1;refGene_tx=XM_023259228.1;refGene_gn=BIN1;refGene_tx=XM_023259229.1;refGene_gn=BIN1;refGene_tx=XM_023259230.1;refGene_gn=BIN1;refGene_tx=XM_023259232.1;refGene_gn=BIN1;refGene_tx=XM_023259233.1;refGene_gn=BIN1;refGene_tx=XM_023259234.1;refGene_gn=BIN1;refGene_tx=XM_023259235.1;refGene_gn=BIN1;refGene_tx=XM_023259236.1;refGene_gn=BIN1;refGene_tx=XM_023259237.1;refGene_gn=BIN1;refGene_tx=XM_023259238.1
C1 114101420 114101516
refGene_gn=BIN1;refGene_tx=XM_023259216.1;refGene_gn=BIN1;refGene_tx=XM_023259218.1;refGene_gn=BIN1;refGene_tx=XM_023259220.1;refGene_gn=BIN1;refGene_tx=XM_023259221.1;refGene_gn=BIN1;refGene_tx=XM_023259222.1;refGene_gn=BIN1;refGene_tx=XM_023259225.1;refGene_gn=BIN1;refGene_tx=XM_023259226.1;refGene_gn=BIN1;refGene_tx=XM_023259231.1
C1 114113987 114113999
refGene_gn=BIN1;refGene_tx=XM_019837921.2;refGene_gn=BIN1;refGene_tx=XM_019837935.2
C1 114119502 114119583
refGene_gn=BIN1;refGene_tx=XM_019837921.2;refGene_gn=BIN1;refGene_tx=XM_019837935.2;refGene_gn=BIN1;refGene_tx=XM_023259216.1;refGene_gn=BIN1;refGene_tx=XM_023259217.1;refGene_gn=BIN1;refGene_tx=XM_023259218.1;refGene_gn=BIN1;refGene_tx=XM_023259219.1;refGene_gn=BIN1;refGene_tx=XM_023259220.1;refGene_gn=BIN1;refGene_tx=XM_023259221.1;refGene_gn=BIN1;refGene_tx=XM_023259222.1;refGene_gn=BIN1;refGene_tx=XM_023259223.1;refGene_gn=BIN1;refGene_tx=XM_023259224.1;refGene_gn=BIN1;refGene_tx=XM_023259225.1;refGene_gn=BIN1;refGene_tx=XM_023259226.1;refGene_gn=BIN1;refGene_tx=XM_023259227.1;refGene_gn=BIN1;refGene_tx=XM_023259228.1;refGene_gn=BIN1;refGene_tx=XM_023259229.1;refGene_gn=BIN1;refGene_tx=XM_023259230.1;refGene_gn=BIN1;refGene_tx=XM_023259231.1;refGene_gn=BIN1;refGene_tx=XM_023259232.1;refGene_gn=BIN1;refGene_tx=XM_023259233.1;refGene_gn=BIN1;refGene_tx=XM_023259234.1;refGene_gn=BIN1;refGene_tx=XM_023259235.1;refGene_gn=BIN1;refGene_tx=XM_023259236.1;refGene_gn=BIN1;refGene_tx=XM_023259237.1;refGene_gn=BIN1;refGene_tx=XM_023259238.1
C1 114125527 114125582
refGene_gn=BIN1;refGene_tx=XM_019837921.2;refGene_gn=BIN1;refGene_tx=XM_019837935.2;refGene_gn=BIN1;refGene_tx=XM_023259216.1;refGene_gn=BIN

1;refGene_tx=XM_023259217.1;refGene_gn=BIN1;refGene_tx=XM_023259218.1;
refGene_gn=BIN1;refGene_tx=XM_023259219.1;refGene_gn=BIN1;refGene_tx=X
M_023259220.1;refGene_gn=BIN1;refGene_tx=XM_023259221.1;refGene_gn=BIN
1;refGene_tx=XM_023259222.1;refGene_gn=BIN1;refGene_tx=XM_023259223.1;
refGene_gn=BIN1;refGene_tx=XM_023259224.1;refGene_gn=BIN1;refGene_tx=X
M_023259225.1;refGene_gn=BIN1;refGene_tx=XM_023259226.1;refGene_gn=BIN
1;refGene_tx=XM_023259227.1;refGene_gn=BIN1;refGene_tx=XM_023259228.1;
refGene_gn=BIN1;refGene_tx=XM_023259229.1;refGene_gn=BIN1;refGene_tx=X
M_023259230.1;refGene_gn=BIN1;refGene_tx=XM_023259231.1;refGene_gn=BIN
1;refGene_tx=XM_023259232.1;refGene_gn=BIN1;refGene_tx=XM_023259233.1;
refGene_gn=BIN1;refGene_tx=XM_023259234.1;refGene_gn=BIN1;refGene_tx=X
M_023259235.1;refGene_gn=BIN1;refGene_tx=XM_023259236.1;refGene_gn=BIN
1;refGene_tx=XM_023259237.1;refGene_gn=BIN1;refGene_tx=XM_023259238.1
C1 114125815 114125910
refGene_gn=BIN1;refGene_tx=XM_019837921.2;refGene_gn=BIN1;refGene_tx=X
M_019837935.2;refGene_gn=BIN1;refGene_tx=XM_023259216.1;refGene_gn=BIN
1;refGene_tx=XM_023259217.1;refGene_gn=BIN1;refGene_tx=XM_023259218.1;
refGene_gn=BIN1;refGene_tx=XM_023259219.1;refGene_gn=BIN1;refGene_tx=X
M_023259220.1;refGene_gn=BIN1;refGene_tx=XM_023259221.1;refGene_gn=BIN
1;refGene_tx=XM_023259222.1;refGene_gn=BIN1;refGene_tx=XM_023259223.1;
refGene_gn=BIN1;refGene_tx=XM_023259224.1;refGene_gn=BIN1;refGene_tx=X
M_023259225.1;refGene_gn=BIN1;refGene_tx=XM_023259226.1;refGene_gn=BIN
1;refGene_tx=XM_023259227.1;refGene_gn=BIN1;refGene_tx=XM_023259228.1;
refGene_gn=BIN1;refGene_tx=XM_023259229.1;refGene_gn=BIN1;refGene_tx=X
M_023259230.1;refGene_gn=BIN1;refGene_tx=XM_023259231.1;refGene_gn=BIN
1;refGene_tx=XM_023259232.1;refGene_gn=BIN1;refGene_tx=XM_023259233.1;
refGene_gn=BIN1;refGene_tx=XM_023259234.1;refGene_gn=BIN1;refGene_tx=X
M_023259235.1;refGene_gn=BIN1;refGene_tx=XM_023259236.1;refGene_gn=BIN
1;refGene_tx=XM_023259237.1;refGene_gn=BIN1;refGene_tx=XM_023259238.1
C1 114126279 114126375
refGene_gn=BIN1;refGene_tx=XM_019837921.2;refGene_gn=BIN1;refGene_tx=X
M_019837935.2;refGene_gn=BIN1;refGene_tx=XM_023259216.1;refGene_gn=BIN
1;refGene_tx=XM_023259217.1;refGene_gn=BIN1;refGene_tx=XM_023259218.1;
refGene_gn=BIN1;refGene_tx=XM_023259219.1;refGene_gn=BIN1;refGene_tx=X
M_023259220.1;refGene_gn=BIN1;refGene_tx=XM_023259221.1;refGene_gn=BIN
1;refGene_tx=XM_023259222.1;refGene_gn=BIN1;refGene_tx=XM_023259223.1;
refGene_gn=BIN1;refGene_tx=XM_023259224.1;refGene_gn=BIN1;refGene_tx=X
M_023259225.1;refGene_gn=BIN1;refGene_tx=XM_023259226.1;refGene_gn=BIN
1;refGene_tx=XM_023259227.1;refGene_gn=BIN1;refGene_tx=XM_023259228.1;
refGene_gn=BIN1;refGene_tx=XM_023259229.1;refGene_gn=BIN1;refGene_tx=X
M_023259230.1;refGene_gn=BIN1;refGene_tx=XM_023259231.1;refGene_gn=BIN
1;refGene_tx=XM_023259232.1;refGene_gn=BIN1;refGene_tx=XM_023259233.1;
refGene_gn=BIN1;refGene_tx=XM_023259234.1;refGene_gn=BIN1;refGene_tx=X
M_023259235.1;refGene_gn=BIN1;refGene_tx=XM_023259236.1;refGene_gn=BIN
1;refGene_tx=XM_023259237.1;refGene_gn=BIN1;refGene_tx=XM_023259238.1
C1 114127465 114127573
refGene_gn=BIN1;refGene_tx=XM_019837921.2;refGene_gn=BIN1;refGene_tx=X
M_019837935.2;refGene_gn=BIN1;refGene_tx=XM_023259216.1;refGene_gn=BIN
1;refGene_tx=XM_023259217.1;refGene_gn=BIN1;refGene_tx=XM_023259218.1;
refGene_gn=BIN1;refGene_tx=XM_023259219.1;refGene_gn=BIN1;refGene_tx=X

M_023259220.1;refGene_gn=BIN1;refGene_tx=XM_023259221.1;refGene_gn=BIN1;refGene_tx=XM_023259222.1;refGene_gn=BIN1;refGene_tx=XM_023259223.1;refGene_gn=BIN1;refGene_tx=XM_023259224.1;refGene_gn=BIN1;refGene_tx=XM_023259225.1;refGene_gn=BIN1;refGene_tx=XM_023259226.1;refGene_gn=BIN1;refGene_tx=XM_023259227.1;refGene_gn=BIN1;refGene_tx=XM_023259228.1;refGene_gn=BIN1;refGene_tx=XM_023259229.1;refGene_gn=BIN1;refGene_tx=XM_023259230.1;refGene_gn=BIN1;refGene_tx=XM_023259231.1;refGene_gn=BIN1;refGene_tx=XM_023259232.1;refGene_gn=BIN1;refGene_tx=XM_023259233.1;refGene_gn=BIN1;refGene_tx=XM_023259234.1;refGene_gn=BIN1;refGene_tx=XM_023259235.1;refGene_gn=BIN1;refGene_tx=XM_023259236.1;refGene_gn=BIN1;refGene_tx=XM_023259237.1;refGene_gn=BIN1;refGene_tx=XM_023259238.1

C1 114128433 114128526

refGene_gn=BIN1;refGene_tx=XM_019837921.2;refGene_gn=BIN1;refGene_tx=XM_023259216.1;refGene_gn=BIN1;refGene_tx=XM_023259217.1;refGene_gn=BIN1;refGene_tx=XM_023259218.1;refGene_gn=BIN1;refGene_tx=XM_023259219.1;refGene_gn=BIN1;refGene_tx=XM_023259221.1;refGene_gn=BIN1;refGene_tx=XM_023259222.1;refGene_gn=BIN1;refGene_tx=XM_023259224.1;refGene_gn=BIN1;refGene_tx=XM_023259225.1;refGene_gn=BIN1;refGene_tx=XM_023259226.1;refGene_gn=BIN1;refGene_tx=XM_023259228.1;refGene_gn=BIN1;refGene_tx=XM_023259229.1;refGene_gn=BIN1;refGene_tx=XM_023259231.1;refGene_gn=BIN1;refGene_tx=XM_023259234.1

C1 114132762 114132848

refGene_gn=BIN1;refGene_tx=XM_019837921.2;refGene_gn=BIN1;refGene_tx=XM_019837935.2;refGene_gn=BIN1;refGene_tx=XM_023259216.1;refGene_gn=BIN1;refGene_tx=XM_023259217.1;refGene_gn=BIN1;refGene_tx=XM_023259218.1;refGene_gn=BIN1;refGene_tx=XM_023259219.1;refGene_gn=BIN1;refGene_tx=XM_023259220.1;refGene_gn=BIN1;refGene_tx=XM_023259221.1;refGene_gn=BIN1;refGene_tx=XM_023259222.1;refGene_gn=BIN1;refGene_tx=XM_023259223.1;refGene_gn=BIN1;refGene_tx=XM_023259224.1;refGene_gn=BIN1;refGene_tx=XM_023259225.1;refGene_gn=BIN1;refGene_tx=XM_023259226.1;refGene_gn=BIN1;refGene_tx=XM_023259227.1;refGene_gn=BIN1;refGene_tx=XM_023259228.1;refGene_gn=BIN1;refGene_tx=XM_023259229.1;refGene_gn=BIN1;refGene_tx=XM_023259230.1;refGene_gn=BIN1;refGene_tx=XM_023259231.1;refGene_gn=BIN1;refGene_tx=XM_023259232.1;refGene_gn=BIN1;refGene_tx=XM_023259233.1;refGene_gn=BIN1;refGene_tx=XM_023259234.1;refGene_gn=BIN1;refGene_tx=XM_023259235.1;refGene_gn=BIN1;refGene_tx=XM_023259236.1;refGene_gn=BIN1;refGene_tx=XM_023259237.1;refGene_gn=BIN1;refGene_tx=XM_023259238.1

C1 114133048 114133124

refGene_gn=BIN1;refGene_tx=XM_019837921.2;refGene_gn=BIN1;refGene_tx=XM_019837935.2;refGene_gn=BIN1;refGene_tx=XM_023259216.1;refGene_gn=BIN1;refGene_tx=XM_023259217.1;refGene_gn=BIN1;refGene_tx=XM_023259218.1;refGene_gn=BIN1;refGene_tx=XM_023259219.1;refGene_gn=BIN1;refGene_tx=XM_023259220.1;refGene_gn=BIN1;refGene_tx=XM_023259221.1;refGene_gn=BIN1;refGene_tx=XM_023259222.1;refGene_gn=BIN1;refGene_tx=XM_023259223.1;refGene_gn=BIN1;refGene_tx=XM_023259224.1;refGene_gn=BIN1;refGene_tx=XM_023259225.1;refGene_gn=BIN1;refGene_tx=XM_023259226.1;refGene_gn=BIN1;refGene_tx=XM_023259227.1;refGene_gn=BIN1;refGene_tx=XM_023259228.1;refGene_gn=BIN1;refGene_tx=XM_023259229.1;refGene_gn=BIN1;refGene_tx=XM_023259230.1;refGene_gn=BIN1;refGene_tx=XM_023259231.1;refGene_gn=BIN1;refGene_tx=XM_023259232.1;refGene_gn=BIN1;refGene_tx=XM_023259233.1;

refGene_gn=BIN1;refGene_tx=XM_023259234.1;refGene_gn=BIN1;refGene_tx=XM_023259235.1;refGene_gn=BIN1;refGene_tx=XM_023259236.1;refGene_gn=BIN1;refGene_tx=XM_023259237.1;refGene_gn=BIN1;refGene_tx=XM_023259238.1
C1 114134456 114134539
refGene_gn=BIN1;refGene_tx=XM_019837921.2;refGene_gn=BIN1;refGene_tx=XM_019837935.2;refGene_gn=BIN1;refGene_tx=XM_023259216.1;refGene_gn=BIN1;refGene_tx=XM_023259217.1;refGene_gn=BIN1;refGene_tx=XM_023259218.1;refGene_gn=BIN1;refGene_tx=XM_023259219.1;refGene_gn=BIN1;refGene_tx=XM_023259220.1;refGene_gn=BIN1;refGene_tx=XM_023259221.1;refGene_gn=BIN1;refGene_tx=XM_023259222.1;refGene_gn=BIN1;refGene_tx=XM_023259223.1;refGene_gn=BIN1;refGene_tx=XM_023259224.1;refGene_gn=BIN1;refGene_tx=XM_023259225.1;refGene_gn=BIN1;refGene_tx=XM_023259226.1;refGene_gn=BIN1;refGene_tx=XM_023259227.1;refGene_gn=BIN1;refGene_tx=XM_023259228.1;refGene_gn=BIN1;refGene_tx=XM_023259229.1;refGene_gn=BIN1;refGene_tx=XM_023259230.1;refGene_gn=BIN1;refGene_tx=XM_023259231.1;refGene_gn=BIN1;refGene_tx=XM_023259232.1;refGene_gn=BIN1;refGene_tx=XM_023259233.1;refGene_gn=BIN1;refGene_tx=XM_023259234.1;refGene_gn=BIN1;refGene_tx=XM_023259235.1;refGene_gn=BIN1;refGene_tx=XM_023259236.1;refGene_gn=BIN1;refGene_tx=XM_023259237.1;refGene_gn=BIN1;refGene_tx=XM_023259238.1
C1 114136046 114136091
refGene_gn=BIN1;refGene_tx=XM_019837921.2;refGene_gn=BIN1;refGene_tx=XM_023259216.1;refGene_gn=BIN1;refGene_tx=XM_023259217.1;refGene_gn=BIN1;refGene_tx=XM_023259220.1;refGene_gn=BIN1;refGene_tx=XM_023259221.1;refGene_gn=BIN1;refGene_tx=XM_023259222.1;refGene_gn=BIN1;refGene_tx=XM_023259225.1;refGene_gn=BIN1;refGene_tx=XM_023259226.1;refGene_gn=BIN1;refGene_tx=XM_023259230.1;refGene_gn=BIN1;refGene_tx=XM_023259231.1;refGene_gn=BIN1;refGene_tx=XM_023259235.1;refGene_gn=BIN1;refGene_tx=XM_023259237.1
C1 114137639 114137784
refGene_gn=BIN1;refGene_tx=XM_019837921.2;refGene_gn=BIN1;refGene_tx=XM_019837935.2;refGene_gn=BIN1;refGene_tx=XM_023259216.1;refGene_gn=BIN1;refGene_tx=XM_023259217.1;refGene_gn=BIN1;refGene_tx=XM_023259218.1;refGene_gn=BIN1;refGene_tx=XM_023259219.1;refGene_gn=BIN1;refGene_tx=XM_023259220.1;refGene_gn=BIN1;refGene_tx=XM_023259221.1;refGene_gn=BIN1;refGene_tx=XM_023259222.1;refGene_gn=BIN1;refGene_tx=XM_023259223.1;refGene_gn=BIN1;refGene_tx=XM_023259224.1;refGene_gn=BIN1;refGene_tx=XM_023259225.1;refGene_gn=BIN1;refGene_tx=XM_023259226.1;refGene_gn=BIN1;refGene_tx=XM_023259227.1;refGene_gn=BIN1;refGene_tx=XM_023259228.1;refGene_gn=BIN1;refGene_tx=XM_023259229.1;refGene_gn=BIN1;refGene_tx=XM_023259230.1;refGene_gn=BIN1;refGene_tx=XM_023259231.1;refGene_gn=BIN1;refGene_tx=XM_023259232.1;refGene_gn=BIN1;refGene_tx=XM_023259233.1;refGene_gn=BIN1;refGene_tx=XM_023259234.1;refGene_gn=BIN1;refGene_tx=XM_023259235.1;refGene_gn=BIN1;refGene_tx=XM_023259236.1;refGene_gn=BIN1;refGene_tx=XM_023259237.1;refGene_gn=BIN1;refGene_tx=XM_023259238.1
C1 114139116 114139245
refGene_gn=BIN1;refGene_tx=XM_019837921.2;refGene_gn=BIN1;refGene_tx=XM_023259216.1;refGene_gn=BIN1;refGene_tx=XM_023259217.1;refGene_gn=BIN1;refGene_tx=XM_023259218.1;refGene_gn=BIN1;refGene_tx=XM_023259219.1;refGene_gn=BIN1;refGene_tx=XM_023259220.1;refGene_gn=BIN1;refGene_tx=XM_023259222.1;refGene_gn=BIN1;refGene_tx=XM_023259223.1;refGene_gn=BIN

1;refGene_tx=XM_023259225.1;refGene_gn=BIN1;refGene_tx=XM_023259227.1;
refGene_gn=BIN1;refGene_tx=XM_023259228.1;refGene_gn=BIN1;refGene_tx=X
M_023259230.1;refGene_gn=BIN1;refGene_tx=XM_023259232.1
C1 114142076 114142184
refGene_gn=BIN1;refGene_tx=XM_019837921.2;refGene_gn=BIN1;refGene_tx=X
M_023259216.1;refGene_gn=BIN1;refGene_tx=XM_023259217.1;refGene_gn=BIN
1;refGene_tx=XM_023259218.1;refGene_gn=BIN1;refGene_tx=XM_023259219.1;
refGene_gn=BIN1;refGene_tx=XM_023259220.1;refGene_gn=BIN1;refGene_tx=X
M_023259221.1;refGene_gn=BIN1;refGene_tx=XM_023259223.1;refGene_gn=BIN
1;refGene_tx=XM_023259224.1
C1 114142686 114142710
refGene_gn=BIN1;refGene_tx=XM_019837921.2;refGene_gn=BIN1;refGene_tx=X
M_023259216.1;refGene_gn=BIN1;refGene_tx=XM_023259217.1;refGene_gn=BIN
1;refGene_tx=XM_023259218.1;refGene_gn=BIN1;refGene_tx=XM_023259219.1;
refGene_gn=BIN1;refGene_tx=XM_023259220.1;refGene_gn=BIN1;refGene_tx=X
M_023259221.1;refGene_gn=BIN1;refGene_tx=XM_023259223.1;refGene_gn=BIN
1;refGene_tx=XM_023259224.1
C1 114143674 114143782
refGene_gn=BIN1;refGene_tx=XM_019837921.2;refGene_gn=BIN1;refGene_tx=X
M_023259216.1;refGene_gn=BIN1;refGene_tx=XM_023259217.1;refGene_gn=BIN
1;refGene_tx=XM_023259218.1;refGene_gn=BIN1;refGene_tx=XM_023259219.1;
refGene_gn=BIN1;refGene_tx=XM_023259220.1;refGene_gn=BIN1;refGene_tx=X
M_023259221.1;refGene_gn=BIN1;refGene_tx=XM_023259222.1;refGene_gn=BIN
1;refGene_tx=XM_023259223.1;refGene_gn=BIN1;refGene_tx=XM_023259224.1;
refGene_gn=BIN1;refGene_tx=XM_023259226.1;refGene_gn=BIN1;refGene_tx=X
M_023259227.1;refGene_gn=BIN1;refGene_tx=XM_023259229.1;refGene_gn=BIN
1;refGene_tx=XM_023259233.1
C1 114144705 114144795
refGene_gn=BIN1;refGene_tx=XM_019837921.2;refGene_gn=BIN1;refGene_tx=X
M_023259216.1;refGene_gn=BIN1;refGene_tx=XM_023259217.1;refGene_gn=BIN
1;refGene_tx=XM_023259218.1;refGene_gn=BIN1;refGene_tx=XM_023259219.1;
refGene_gn=BIN1;refGene_tx=XM_023259220.1;refGene_gn=BIN1;refGene_tx=X
M_023259221.1;refGene_gn=BIN1;refGene_tx=XM_023259222.1;refGene_gn=BIN
1;refGene_tx=XM_023259223.1;refGene_gn=BIN1;refGene_tx=XM_023259224.1;
refGene_gn=BIN1;refGene_tx=XM_023259225.1;refGene_gn=BIN1;refGene_tx=X
M_023259226.1;refGene_gn=BIN1;refGene_tx=XM_023259227.1;refGene_gn=BIN
1;refGene_tx=XM_023259228.1;refGene_gn=BIN1;refGene_tx=XM_023259229.1;
refGene_gn=BIN1;refGene_tx=XM_023259230.1;refGene_gn=BIN1;refGene_tx=X
M_023259231.1;refGene_gn=BIN1;refGene_tx=XM_023259232.1;refGene_gn=BIN
1;refGene_tx=XM_023259233.1;refGene_gn=BIN1;refGene_tx=XM_023259234.1;
refGene_gn=BIN1;refGene_tx=XM_023259235.1;refGene_gn=BIN1;refGene_tx=X
M_023259236.1
C1 114145048 114145159
refGene_gn=BIN1;refGene_tx=XM_019837921.2;refGene_gn=BIN1;refGene_tx=X
M_019837935.2;refGene_gn=BIN1;refGene_tx=XM_023259216.1;refGene_gn=BIN
1;refGene_tx=XM_023259217.1;refGene_gn=BIN1;refGene_tx=XM_023259218.1;
refGene_gn=BIN1;refGene_tx=XM_023259219.1;refGene_gn=BIN1;refGene_tx=X
M_023259220.1;refGene_gn=BIN1;refGene_tx=XM_023259221.1;refGene_gn=BIN
1;refGene_tx=XM_023259222.1;refGene_gn=BIN1;refGene_tx=XM_023259223.1;
refGene_gn=BIN1;refGene_tx=XM_023259224.1;refGene_gn=BIN1;refGene_tx=X

M_023259225.1;refGene_gn=BIN1;refGene_tx=XM_023259226.1;refGene_gn=BIN1;refGene_tx=XM_023259227.1;refGene_gn=BIN1;refGene_tx=XM_023259228.1;refGene_gn=BIN1;refGene_tx=XM_023259229.1;refGene_gn=BIN1;refGene_tx=XM_023259230.1;refGene_gn=BIN1;refGene_tx=XM_023259231.1;refGene_gn=BIN1;refGene_tx=XM_023259232.1;refGene_gn=BIN1;refGene_tx=XM_023259233.1;refGene_gn=BIN1;refGene_tx=XM_023259234.1;refGene_gn=BIN1;refGene_tx=XM_023259235.1;refGene_gn=BIN1;refGene_tx=XM_023259236.1;refGene_gn=BIN1;refGene_tx=XM_023259237.1;refGene_gn=BIN1;refGene_tx=XM_023259238.1
C1 114145461 114145563
refGene_gn=BIN1;refGene_tx=XM_019837921.2;refGene_gn=BIN1;refGene_tx=XM_019837935.2;refGene_gn=BIN1;refGene_tx=XM_023259216.1;refGene_gn=BIN1;refGene_tx=XM_023259217.1;refGene_gn=BIN1;refGene_tx=XM_023259218.1;refGene_gn=BIN1;refGene_tx=XM_023259219.1;refGene_gn=BIN1;refGene_tx=XM_023259220.1;refGene_gn=BIN1;refGene_tx=XM_023259221.1;refGene_gn=BIN1;refGene_tx=XM_023259222.1;refGene_gn=BIN1;refGene_tx=XM_023259223.1;refGene_gn=BIN1;refGene_tx=XM_023259224.1;refGene_gn=BIN1;refGene_tx=XM_023259225.1;refGene_gn=BIN1;refGene_tx=XM_023259226.1;refGene_gn=BIN1;refGene_tx=XM_023259227.1;refGene_gn=BIN1;refGene_tx=XM_023259228.1;refGene_gn=BIN1;refGene_tx=XM_023259229.1;refGene_gn=BIN1;refGene_tx=XM_023259230.1;refGene_gn=BIN1;refGene_tx=XM_023259231.1;refGene_gn=BIN1;refGene_tx=XM_023259232.1;refGene_gn=BIN1;refGene_tx=XM_023259233.1;refGene_gn=BIN1;refGene_tx=XM_023259234.1;refGene_gn=BIN1;refGene_tx=XM_023259235.1;refGene_gn=BIN1;refGene_tx=XM_023259236.1;refGene_gn=BIN1;refGene_tx=XM_023259237.1;refGene_gn=BIN1;refGene_tx=XM_023259238.1
C1 114147114 114147222
refGene_gn=BIN1;refGene_tx=XM_019837921.2;refGene_gn=BIN1;refGene_tx=XM_019837935.2;refGene_gn=BIN1;refGene_tx=XM_023259216.1;refGene_gn=BIN1;refGene_tx=XM_023259217.1;refGene_gn=BIN1;refGene_tx=XM_023259218.1;refGene_gn=BIN1;refGene_tx=XM_023259219.1;refGene_gn=BIN1;refGene_tx=XM_023259220.1;refGene_gn=BIN1;refGene_tx=XM_023259221.1;refGene_gn=BIN1;refGene_tx=XM_023259222.1;refGene_gn=BIN1;refGene_tx=XM_023259223.1;refGene_gn=BIN1;refGene_tx=XM_023259224.1;refGene_gn=BIN1;refGene_tx=XM_023259225.1;refGene_gn=BIN1;refGene_tx=XM_023259226.1;refGene_gn=BIN1;refGene_tx=XM_023259227.1;refGene_gn=BIN1;refGene_tx=XM_023259228.1;refGene_gn=BIN1;refGene_tx=XM_023259229.1;refGene_gn=BIN1;refGene_tx=XM_023259230.1;refGene_gn=BIN1;refGene_tx=XM_023259231.1;refGene_gn=BIN1;refGene_tx=XM_023259232.1;refGene_gn=BIN1;refGene_tx=XM_023259233.1;refGene_gn=BIN1;refGene_tx=XM_023259234.1;refGene_gn=BIN1;refGene_tx=XM_023259235.1;refGene_gn=BIN1;refGene_tx=XM_023259236.1;refGene_gn=BIN1;refGene_tx=XM_023259237.1;refGene_gn=BIN1;refGene_tx=XM_023259238.1
C1 114294057 114294093
refGene_gn=TEX51;refGene_tx=XM_006935221.3;refGene_gn=TEX51;refGene_tx=XM_019837937.2
C1 114294496 114294547
refGene_gn=TEX51;refGene_tx=XM_006935221.3;refGene_gn=TEX51;refGene_tx=XM_019837937.2;refGene_gn=TEX51;refGene_tx=XM_019837939.2;refGene_gn=TEX51;refGene_tx=XM_019837940.2;refGene_gn=TEX51;refGene_tx=XM_023259239.1
C1 114294631 114294707
refGene_gn=TEX51;refGene_tx=XM_006935221.3;refGene_gn=TEX51;refGene_tx

=XM_019837937.2;refGene_gn=TEX51;refGene_tx=XM_019837939.2;refGene_gn=TEX51;refGene_tx=XM_019837940.2;refGene_gn=TEX51;refGene_tx=XM_023259239.1
C1 114295643 114295927
refGene_gn=TEX51;refGene_tx=XM_006935221.3;refGene_gn=TEX51;refGene_tx=XM_019837936.2;refGene_gn=TEX51;refGene_tx=XM_019837937.2;refGene_gn=TEX51;refGene_tx=XM_019837938.2;refGene_gn=TEX51;refGene_tx=XM_019837939.2;refGene_gn=TEX51;refGene_tx=XM_019837940.2;refGene_gn=TEX51;refGene_tx=XM_019837942.2;refGene_gn=TEX51;refGene_tx=XM_023259239.1
C1 114296014 114296032
refGene_gn=TEX51;refGene_tx=XM_006935221.3;refGene_gn=TEX51;refGene_tx=XM_019837936.2;refGene_gn=TEX51;refGene_tx=XM_019837937.2;refGene_gn=TEX51;refGene_tx=XM_019837938.2;refGene_gn=TEX51;refGene_tx=XM_019837939.2;refGene_gn=TEX51;refGene_tx=XM_019837940.2;refGene_gn=TEX51;refGene_tx=XM_019837942.2;refGene_gn=TEX51;refGene_tx=XM_023259239.1
C1 114296308 114296398
refGene_gn=TEX51;refGene_tx=XM_006935221.3;refGene_gn=TEX51;refGene_tx=XM_019837936.2;refGene_gn=TEX51;refGene_tx=XM_019837937.2;refGene_gn=TEX51;refGene_tx=XM_019837938.2;refGene_gn=TEX51;refGene_tx=XM_019837939.2;refGene_gn=TEX51;refGene_tx=XM_019837940.2;refGene_gn=TEX51;refGene_tx=XM_019837942.2;refGene_gn=TEX51;refGene_tx=XM_023259239.1
C1 114296636 114296711
refGene_gn=TEX51;refGene_tx=XM_006935221.3;refGene_gn=TEX51;refGene_tx=XM_019837936.2;refGene_gn=TEX51;refGene_tx=XM_019837937.2;refGene_gn=TEX51;refGene_tx=XM_019837939.2;refGene_gn=TEX51;refGene_tx=XM_019837940.2;refGene_gn=TEX51;refGene_tx=XM_019837942.2;refGene_gn=TEX51;refGene_tx=XM_023259239.1
C1 114296800 114297050
refGene_gn=TEX51;refGene_tx=XM_006935221.3;refGene_gn=TEX51;refGene_tx=XM_019837936.2;refGene_gn=TEX51;refGene_tx=XM_019837937.2;refGene_gn=TEX51;refGene_tx=XM_019837938.2;refGene_gn=TEX51;refGene_tx=XM_019837939.2;refGene_gn=TEX51;refGene_tx=XM_019837940.2;refGene_gn=TEX51;refGene_tx=XM_019837942.2;refGene_gn=TEX51;refGene_tx=XM_023259239.1
C1 114515869 114516069
refGene_gn=GYPC;refGene_tx=XM_003990688.5
C1 114517024 114517084
refGene_gn=GYPC;refGene_tx=XM_003990688.5
C1 114557975 114558012
refGene_gn=GYPC;refGene_tx=XM_003990688.5
C1 116158718 116158720
refGene_gn=CNTNAP5;refGene_tx=XM_011285200.3
C1 116162930 116163127
refGene_gn=CNTNAP5;refGene_tx=XM_011285199.3;refGene_gn=CNTNAP5;refGene_tx=XM_011285200.3
C1 116165694 116165766
refGene_gn=CNTNAP5;refGene_tx=XM_011285199.3;refGene_gn=CNTNAP5;refGene_tx=XM_011285200.3
C1 116167872 116168091
refGene_gn=CNTNAP5;refGene_tx=XM_011285199.3;refGene_gn=CNTNAP5;refGene_tx=XM_011285200.3

C1 116203279 116203367
refGene_gn=CNTNAP5;refGene_tx=XM_011285199.3;refGene_gn=CNTNAP5;refGene_tx=XM_011285200.3
C1 116210639 116210770
refGene_gn=CNTNAP5;refGene_tx=XM_011285199.3;refGene_gn=CNTNAP5;refGene_tx=XM_011285200.3
C1 116254231 116254323
refGene_gn=CNTNAP5;refGene_tx=XM_019837943.2
C1 116257613 116257838
refGene_gn=CNTNAP5;refGene_tx=XM_011285199.3;refGene_gn=CNTNAP5;refGene_tx=XM_011285200.3;refGene_gn=CNTNAP5;refGene_tx=XM_019837943.2
C1 116265808 116266048
refGene_gn=CNTNAP5;refGene_tx=XM_011285199.3;refGene_gn=CNTNAP5;refGene_tx=XM_011285200.3;refGene_gn=CNTNAP5;refGene_tx=XM_019837943.2
C1 116285754 116285973
refGene_gn=CNTNAP5;refGene_tx=XM_011285199.3;refGene_gn=CNTNAP5;refGene_tx=XM_011285200.3;refGene_gn=CNTNAP5;refGene_tx=XM_019837943.2
C1 116296017 116296188
refGene_gn=CNTNAP5;refGene_tx=XM_011285199.3;refGene_gn=CNTNAP5;refGene_tx=XM_011285200.3;refGene_gn=CNTNAP5;refGene_tx=XM_019837943.2
C1 116296367 116296495
refGene_gn=CNTNAP5;refGene_tx=XM_011285199.3;refGene_gn=CNTNAP5;refGene_tx=XM_011285200.3;refGene_gn=CNTNAP5;refGene_tx=XM_019837943.2
C1 116312133 116312290
refGene_gn=CNTNAP5;refGene_tx=XM_011285199.3;refGene_gn=CNTNAP5;refGene_tx=XM_011285200.3;refGene_gn=CNTNAP5;refGene_tx=XM_019837943.2
C1 116403621 116403822
refGene_gn=CNTNAP5;refGene_tx=XM_011285199.3;refGene_gn=CNTNAP5;refGene_tx=XM_011285200.3;refGene_gn=CNTNAP5;refGene_tx=XM_019837943.2
C1 116429198 116429318
refGene_gn=CNTNAP5;refGene_tx=XM_011285199.3;refGene_gn=CNTNAP5;refGene_tx=XM_011285200.3;refGene_gn=CNTNAP5;refGene_tx=XM_019837943.2
C1 116468294 116468401
refGene_gn=CNTNAP5;refGene_tx=XM_011285199.3;refGene_gn=CNTNAP5;refGene_tx=XM_011285200.3;refGene_gn=CNTNAP5;refGene_tx=XM_019837943.2
C1 116501541 116501713
refGene_gn=CNTNAP5;refGene_tx=XM_011285199.3;refGene_gn=CNTNAP5;refGene_tx=XM_011285200.3;refGene_gn=CNTNAP5;refGene_tx=XM_019837943.2
C1 116504538 116504688
refGene_gn=CNTNAP5;refGene_tx=XM_011285199.3;refGene_gn=CNTNAP5;refGene_tx=XM_011285200.3;refGene_gn=CNTNAP5;refGene_tx=XM_019837943.2
C1 116523414 116523679
refGene_gn=CNTNAP5;refGene_tx=XM_011285199.3;refGene_gn=CNTNAP5;refGene_tx=XM_011285200.3;refGene_gn=CNTNAP5;refGene_tx=XM_019837943.2
C1 116553182 116553326
refGene_gn=CNTNAP5;refGene_tx=XM_011285199.3;refGene_gn=CNTNAP5;refGene_tx=XM_011285200.3;refGene_gn=CNTNAP5;refGene_tx=XM_019837943.2
C1 116572640 116572825
refGene_gn=CNTNAP5;refGene_tx=XM_011285199.3;refGene_gn=CNTNAP5;refGene_tx=XM_011285200.3;refGene_gn=CNTNAP5;refGene_tx=XM_019837943.2

C1 116587477 116587681
refGene_gn=CNTNAP5;refGene_tx=XM_011285199.3;refGene_gn=CNTNAP5;refGene_tx=XM_011285200.3;refGene_gn=CNTNAP5;refGene_tx=XM_019837943.2
C1 116610479 116610627
refGene_gn=CNTNAP5;refGene_tx=XM_011285199.3;refGene_gn=CNTNAP5;refGene_tx=XM_011285200.3;refGene_gn=CNTNAP5;refGene_tx=XM_019837943.2
C1 116759611 116759805
refGene_gn=CNTNAP5;refGene_tx=XM_011285199.3;refGene_gn=CNTNAP5;refGene_tx=XM_011285200.3;refGene_gn=CNTNAP5;refGene_tx=XM_019837943.2
C1 116777724 116777829
refGene_gn=CNTNAP5;refGene_tx=XM_011285199.3;refGene_gn=CNTNAP5;refGene_tx=XM_011285200.3;refGene_gn=CNTNAP5;refGene_tx=XM_019837943.2
C1 116959323 116959405
refGene_gn=CNTNAP5;refGene_tx=XM_011285199.3;refGene_gn=CNTNAP5;refGene_tx=XM_011285200.3;refGene_gn=CNTNAP5;refGene_tx=XM_019837943.2
C1 117969607 117971981
refGene_gn=LOC109503035;refGene_tx=XM_023259240.1
C1 117972443 117973525
refGene_gn=LOC109503035;refGene_tx=XM_023259240.1
C1 119091004 119091238
refGene_gn=TSN;refGene_tx=XM_003990692.5;refGene_gn=TSN;refGene_tx=XM_003990693.5
C1 119091891 119091971
refGene_gn=TSN;refGene_tx=XM_003990692.5
C1 119093322 119093438
refGene_gn=TSN;refGene_tx=XM_003990692.5;refGene_gn=TSN;refGene_tx=XM_003990693.5
C1 119095118 119095215
refGene_gn=TSN;refGene_tx=XM_003990692.5;refGene_gn=TSN;refGene_tx=XM_003990693.5
C1 119096832 119096926
refGene_gn=TSN;refGene_tx=XM_003990692.5;refGene_gn=TSN;refGene_tx=XM_003990693.5
C1 119098082 119098361
refGene_gn=NIFK;refGene_tx=XM_023257851.1;refGene_gn=TSN;refGene_tx=XM_003990692.5;refGene_gn=TSN;refGene_tx=XM_003990693.5
C1 119106211 119106347
refGene_gn=NIFK;refGene_tx=XM_023257851.1
C1 119107716 119107764
refGene_gn=NIFK;refGene_tx=XM_023257851.1
C1 119108735 119108873
refGene_gn=NIFK;refGene_tx=XM_023257851.1
C1 119115274 119115383
refGene_gn=NIFK;refGene_tx=XM_023257851.1
C1 119117045 119117257
refGene_gn=NIFK;refGene_tx=XM_023257851.1
C1 119118518 119118584
refGene_gn=NIFK;refGene_tx=XM_023257851.1
C1 119118708 119118780
refGene_gn=NIFK;refGene_tx=XM_023257851.1

C1 119119639 119119828
refGene_gn=NIFK;refGene_tx=XM_023257851.1
C1 119222091 119222286
refGene_gn=CLASP1;refGene_tx=XM_006935226.4;refGene_gn=CLASP1;refGene_tx=XM_006935227.4;refGene_gn=CLASP1;refGene_tx=XM_006935228.4;refGene_gn=CLASP1;refGene_tx=XM_006935229.4;refGene_gn=CLASP1;refGene_tx=XM_019837945.2;refGene_gn=CLASP1;refGene_tx=XM_019837946.2;refGene_gn=CLASP1;refGene_tx=XM_019837947.2;refGene_gn=CLASP1;refGene_tx=XM_019837948.2;refGene_gn=CLASP1;refGene_tx=XM_019837949.2;refGene_gn=CLASP1;refGene_tx=XM_019837951.2;refGene_gn=CLASP1;refGene_tx=XM_019837952.2;refGene_gn=CLASP1;refGene_tx=XM_019837953.2;refGene_gn=CLASP1;refGene_tx=XM_019837954.2;refGene_gn=CLASP1;refGene_tx=XM_019837955.2;refGene_gn=CLASP1;refGene_tx=XM_019837956.2;refGene_gn=CLASP1;refGene_tx=XM_019837957.2;refGene_gn=CLASP1;refGene_tx=XM_019837958.2;refGene_gn=CLASP1;refGene_tx=XM_019837959.2
C1 119279596 119279675
refGene_gn=CLASP1;refGene_tx=XM_006935226.4;refGene_gn=CLASP1;refGene_tx=XM_006935227.4;refGene_gn=CLASP1;refGene_tx=XM_006935228.4;refGene_gn=CLASP1;refGene_tx=XM_006935229.4;refGene_gn=CLASP1;refGene_tx=XM_019837945.2;refGene_gn=CLASP1;refGene_tx=XM_019837946.2;refGene_gn=CLASP1;refGene_tx=XM_019837947.2;refGene_gn=CLASP1;refGene_tx=XM_019837948.2;refGene_gn=CLASP1;refGene_tx=XM_019837949.2;refGene_gn=CLASP1;refGene_tx=XM_019837951.2;refGene_gn=CLASP1;refGene_tx=XM_019837952.2;refGene_gn=CLASP1;refGene_tx=XM_019837953.2;refGene_gn=CLASP1;refGene_tx=XM_019837954.2;refGene_gn=CLASP1;refGene_tx=XM_019837955.2;refGene_gn=CLASP1;refGene_tx=XM_019837956.2;refGene_gn=CLASP1;refGene_tx=XM_019837957.2;refGene_gn=CLASP1;refGene_tx=XM_019837958.2;refGene_gn=CLASP1;refGene_tx=XM_019837959.2
C1 119281449 119281553
refGene_gn=CLASP1;refGene_tx=XM_006935226.4;refGene_gn=CLASP1;refGene_tx=XM_006935227.4;refGene_gn=CLASP1;refGene_tx=XM_006935228.4;refGene_gn=CLASP1;refGene_tx=XM_006935229.4;refGene_gn=CLASP1;refGene_tx=XM_019837945.2;refGene_gn=CLASP1;refGene_tx=XM_019837946.2;refGene_gn=CLASP1;refGene_tx=XM_019837947.2;refGene_gn=CLASP1;refGene_tx=XM_019837948.2;refGene_gn=CLASP1;refGene_tx=XM_019837949.2;refGene_gn=CLASP1;refGene_tx=XM_019837951.2;refGene_gn=CLASP1;refGene_tx=XM_019837952.2;refGene_gn=CLASP1;refGene_tx=XM_019837953.2;refGene_gn=CLASP1;refGene_tx=XM_019837954.2;refGene_gn=CLASP1;refGene_tx=XM_019837955.2;refGene_gn=CLASP1;refGene_tx=XM_019837956.2;refGene_gn=CLASP1;refGene_tx=XM_019837957.2;refGene_gn=CLASP1;refGene_tx=XM_019837958.2;refGene_gn=CLASP1;refGene_tx=XM_019837959.2
C1 119282417 119282509
refGene_gn=CLASP1;refGene_tx=XM_006935226.4;refGene_gn=CLASP1;refGene_tx=XM_006935227.4;refGene_gn=CLASP1;refGene_tx=XM_006935228.4;refGene_gn=CLASP1;refGene_tx=XM_006935229.4;refGene_gn=CLASP1;refGene_tx=XM_019837945.2;refGene_gn=CLASP1;refGene_tx=XM_019837946.2;refGene_gn=CLASP1;refGene_tx=XM_019837947.2;refGene_gn=CLASP1;refGene_tx=XM_019837948.2;refGene_gn=CLASP1;refGene_tx=XM_019837949.2;refGene_gn=CLASP1;refGene_tx=XM_019837951.2;refGene_gn=CLASP1;refGene_tx=XM_019837952.2;refGene_gn=CLASP1;refGene_tx=XM_019837953.2;refGene_gn=CLASP1;refGene_tx=XM_

019837954.2;refGene_gn=CLASP1;refGene_tx=XM_019837955.2;refGene_gn=CLASP1;refGene_tx=XM_019837956.2;refGene_gn=CLASP1;refGene_tx=XM_019837957.2;refGene_gn=CLASP1;refGene_tx=XM_019837958.2;refGene_gn=CLASP1;refGene_tx=XM_019837959.2
C1 119284640 119284716
refGene_gn=CLASP1;refGene_tx=XM_006935226.4;refGene_gn=CLASP1;refGene_tx=XM_006935227.4;refGene_gn=CLASP1;refGene_tx=XM_006935228.4;refGene_gn=CLASP1;refGene_tx=XM_006935229.4;refGene_gn=CLASP1;refGene_tx=XM_019837945.2;refGene_gn=CLASP1;refGene_tx=XM_019837946.2;refGene_gn=CLASP1;refGene_tx=XM_019837947.2;refGene_gn=CLASP1;refGene_tx=XM_019837948.2;refGene_gn=CLASP1;refGene_tx=XM_019837949.2;refGene_gn=CLASP1;refGene_tx=XM_019837951.2;refGene_gn=CLASP1;refGene_tx=XM_019837952.2;refGene_gn=CLASP1;refGene_tx=XM_019837953.2;refGene_gn=CLASP1;refGene_tx=XM_019837954.2;refGene_gn=CLASP1;refGene_tx=XM_019837955.2;refGene_gn=CLASP1;refGene_tx=XM_019837956.2;refGene_gn=CLASP1;refGene_tx=XM_019837957.2;refGene_gn=CLASP1;refGene_tx=XM_019837958.2;refGene_gn=CLASP1;refGene_tx=XM_019837959.2
C1 119289981 119290079
refGene_gn=CLASP1;refGene_tx=XM_006935226.4;refGene_gn=CLASP1;refGene_tx=XM_006935227.4;refGene_gn=CLASP1;refGene_tx=XM_006935228.4;refGene_gn=CLASP1;refGene_tx=XM_006935229.4;refGene_gn=CLASP1;refGene_tx=XM_019837945.2;refGene_gn=CLASP1;refGene_tx=XM_019837946.2;refGene_gn=CLASP1;refGene_tx=XM_019837947.2;refGene_gn=CLASP1;refGene_tx=XM_019837948.2;refGene_gn=CLASP1;refGene_tx=XM_019837949.2;refGene_gn=CLASP1;refGene_tx=XM_019837951.2;refGene_gn=CLASP1;refGene_tx=XM_019837952.2;refGene_gn=CLASP1;refGene_tx=XM_019837953.2;refGene_gn=CLASP1;refGene_tx=XM_019837954.2;refGene_gn=CLASP1;refGene_tx=XM_019837955.2;refGene_gn=CLASP1;refGene_tx=XM_019837956.2;refGene_gn=CLASP1;refGene_tx=XM_019837957.2;refGene_gn=CLASP1;refGene_tx=XM_019837958.2;refGene_gn=CLASP1;refGene_tx=XM_019837959.2
C1 119302947 119303015
refGene_gn=CLASP1;refGene_tx=XM_006935226.4;refGene_gn=CLASP1;refGene_tx=XM_006935227.4;refGene_gn=CLASP1;refGene_tx=XM_006935228.4;refGene_gn=CLASP1;refGene_tx=XM_006935229.4;refGene_gn=CLASP1;refGene_tx=XM_019837945.2;refGene_gn=CLASP1;refGene_tx=XM_019837946.2;refGene_gn=CLASP1;refGene_tx=XM_019837947.2;refGene_gn=CLASP1;refGene_tx=XM_019837948.2;refGene_gn=CLASP1;refGene_tx=XM_019837949.2;refGene_gn=CLASP1;refGene_tx=XM_019837951.2;refGene_gn=CLASP1;refGene_tx=XM_019837952.2;refGene_gn=CLASP1;refGene_tx=XM_019837953.2;refGene_gn=CLASP1;refGene_tx=XM_019837954.2;refGene_gn=CLASP1;refGene_tx=XM_019837955.2;refGene_gn=CLASP1;refGene_tx=XM_019837956.2;refGene_gn=CLASP1;refGene_tx=XM_019837957.2;refGene_gn=CLASP1;refGene_tx=XM_019837958.2;refGene_gn=CLASP1;refGene_tx=XM_019837959.2
C1 119334358 119334511
refGene_gn=CLASP1;refGene_tx=XM_006935226.4;refGene_gn=CLASP1;refGene_tx=XM_006935227.4;refGene_gn=CLASP1;refGene_tx=XM_006935228.4;refGene_gn=CLASP1;refGene_tx=XM_006935229.4;refGene_gn=CLASP1;refGene_tx=XM_019837945.2;refGene_gn=CLASP1;refGene_tx=XM_019837946.2;refGene_gn=CLASP1;refGene_tx=XM_019837947.2;refGene_gn=CLASP1;refGene_tx=XM_019837948.2;refGene_gn=CLASP1;refGene_tx=XM_019837949.2;refGene_gn=CLASP1;refGen

e_tx=XM_019837951.2;refGene_gn=CLASP1;refGene_tx=XM_019837952.2;refGene_gn=CLASP1;refGene_tx=XM_019837953.2;refGene_gn=CLASP1;refGene_tx=XM_019837954.2;refGene_gn=CLASP1;refGene_tx=XM_019837955.2;refGene_gn=CLASP1;refGene_tx=XM_019837956.2;refGene_gn=CLASP1;refGene_tx=XM_019837957.2;refGene_gn=CLASP1;refGene_tx=XM_019837958.2;refGene_gn=CLASP1;refGene_tx=XM_019837959.2
C1 119337438 119337512
refGene_gn=CLASP1;refGene_tx=XM_006935226.4;refGene_gn=CLASP1;refGene_tx=XM_006935227.4;refGene_gn=CLASP1;refGene_tx=XM_006935228.4;refGene_gn=CLASP1;refGene_tx=XM_006935229.4;refGene_gn=CLASP1;refGene_tx=XM_019837945.2;refGene_gn=CLASP1;refGene_tx=XM_019837946.2;refGene_gn=CLASP1;refGene_tx=XM_019837947.2;refGene_gn=CLASP1;refGene_tx=XM_019837948.2;refGene_gn=CLASP1;refGene_tx=XM_019837949.2;refGene_gn=CLASP1;refGene_tx=XM_019837951.2;refGene_gn=CLASP1;refGene_tx=XM_019837952.2;refGene_gn=CLASP1;refGene_tx=XM_019837953.2;refGene_gn=CLASP1;refGene_tx=XM_019837954.2;refGene_gn=CLASP1;refGene_tx=XM_019837955.2;refGene_gn=CLASP1;refGene_tx=XM_019837956.2;refGene_gn=CLASP1;refGene_tx=XM_019837957.2;refGene_gn=CLASP1;refGene_tx=XM_019837958.2;refGene_gn=CLASP1;refGene_tx=XM_019837959.2
C1 119338618 119338711
refGene_gn=CLASP1;refGene_tx=XM_006935226.4;refGene_gn=CLASP1;refGene_tx=XM_006935227.4;refGene_gn=CLASP1;refGene_tx=XM_006935228.4;refGene_gn=CLASP1;refGene_tx=XM_006935229.4;refGene_gn=CLASP1;refGene_tx=XM_019837945.2;refGene_gn=CLASP1;refGene_tx=XM_019837946.2;refGene_gn=CLASP1;refGene_tx=XM_019837947.2;refGene_gn=CLASP1;refGene_tx=XM_019837948.2;refGene_gn=CLASP1;refGene_tx=XM_019837949.2;refGene_gn=CLASP1;refGene_tx=XM_019837951.2;refGene_gn=CLASP1;refGene_tx=XM_019837952.2;refGene_gn=CLASP1;refGene_tx=XM_019837953.2;refGene_gn=CLASP1;refGene_tx=XM_019837954.2;refGene_gn=CLASP1;refGene_tx=XM_019837955.2;refGene_gn=CLASP1;refGene_tx=XM_019837956.2;refGene_gn=CLASP1;refGene_tx=XM_019837957.2;refGene_gn=CLASP1;refGene_tx=XM_019837958.2;refGene_gn=CLASP1;refGene_tx=XM_019837959.2
C1 119339619 119339765
refGene_gn=CLASP1;refGene_tx=XM_006935226.4;refGene_gn=CLASP1;refGene_tx=XM_006935227.4;refGene_gn=CLASP1;refGene_tx=XM_006935228.4;refGene_gn=CLASP1;refGene_tx=XM_006935229.4;refGene_gn=CLASP1;refGene_tx=XM_019837945.2;refGene_gn=CLASP1;refGene_tx=XM_019837946.2;refGene_gn=CLASP1;refGene_tx=XM_019837947.2;refGene_gn=CLASP1;refGene_tx=XM_019837948.2;refGene_gn=CLASP1;refGene_tx=XM_019837949.2;refGene_gn=CLASP1;refGene_tx=XM_019837951.2;refGene_gn=CLASP1;refGene_tx=XM_019837952.2;refGene_gn=CLASP1;refGene_tx=XM_019837953.2;refGene_gn=CLASP1;refGene_tx=XM_019837954.2;refGene_gn=CLASP1;refGene_tx=XM_019837955.2;refGene_gn=CLASP1;refGene_tx=XM_019837956.2;refGene_gn=CLASP1;refGene_tx=XM_019837957.2;refGene_gn=CLASP1;refGene_tx=XM_019837958.2;refGene_gn=CLASP1;refGene_tx=XM_019837959.2
C1 119340952 119341088
refGene_gn=CLASP1;refGene_tx=XM_006935226.4;refGene_gn=CLASP1;refGene_tx=XM_006935227.4;refGene_gn=CLASP1;refGene_tx=XM_006935228.4;refGene_gn=CLASP1;refGene_tx=XM_006935229.4;refGene_gn=CLASP1;refGene_tx=XM_019837945.2;refGene_gn=CLASP1;refGene_tx=XM_019837946.2;refGene_gn=CLASP

1;refGene_tx=XM_019837947.2;refGene_gn=CLASP1;refGene_tx=XM_019837948.
2;refGene_gn=CLASP1;refGene_tx=XM_019837949.2;refGene_gn=CLASP1;refGen
e_tx=XM_019837951.2;refGene_gn=CLASP1;refGene_tx=XM_019837952.2;refGen
e_gn=CLASP1;refGene_tx=XM_019837953.2;refGene_gn=CLASP1;refGene_tx=XM_
019837954.2;refGene_gn=CLASP1;refGene_tx=XM_019837955.2;refGene_gn=CLA
SP1;refGene_tx=XM_019837956.2;refGene_gn=CLASP1;refGene_tx=XM_01983795
7.2;refGene_gn=CLASP1;refGene_tx=XM_019837958.2;refGene_gn=CLASP1;refG
ene_tx=XM_019837959.2
C1 119342107 119342178
refGene_gn=CLASP1;refGene_tx=XM_006935226.4;refGene_gn=CLASP1;refGene_
tx=XM_006935227.4;refGene_gn=CLASP1;refGene_tx=XM_006935228.4;refGene_
gn=CLASP1;refGene_tx=XM_006935229.4;refGene_gn=CLASP1;refGene_tx=XM_01
9837945.2;refGene_gn=CLASP1;refGene_tx=XM_019837946.2;refGene_gn=CLASP
1;refGene_tx=XM_019837947.2;refGene_gn=CLASP1;refGene_tx=XM_019837948.
2;refGene_gn=CLASP1;refGene_tx=XM_019837949.2;refGene_gn=CLASP1;refGen
e_tx=XM_019837951.2;refGene_gn=CLASP1;refGene_tx=XM_019837952.2;refGen
e_gn=CLASP1;refGene_tx=XM_019837953.2;refGene_gn=CLASP1;refGene_tx=XM_
019837954.2;refGene_gn=CLASP1;refGene_tx=XM_019837955.2;refGene_gn=CLA
SP1;refGene_tx=XM_019837956.2;refGene_gn=CLASP1;refGene_tx=XM_01983795
7.2;refGene_gn=CLASP1;refGene_tx=XM_019837958.2;refGene_gn=CLASP1;refG
ene_tx=XM_019837959.2
C1 119345967 119346027
refGene_gn=CLASP1;refGene_tx=XM_006935226.4;refGene_gn=CLASP1;refGene_
tx=XM_006935227.4;refGene_gn=CLASP1;refGene_tx=XM_006935228.4;refGene_
gn=CLASP1;refGene_tx=XM_006935229.4;refGene_gn=CLASP1;refGene_tx=XM_01
9837945.2;refGene_gn=CLASP1;refGene_tx=XM_019837946.2;refGene_gn=CLASP
1;refGene_tx=XM_019837947.2;refGene_gn=CLASP1;refGene_tx=XM_019837948.
2;refGene_gn=CLASP1;refGene_tx=XM_019837949.2;refGene_gn=CLASP1;refGen
e_tx=XM_019837951.2;refGene_gn=CLASP1;refGene_tx=XM_019837952.2;refGen
e_gn=CLASP1;refGene_tx=XM_019837953.2;refGene_gn=CLASP1;refGene_tx=XM_
019837954.2;refGene_gn=CLASP1;refGene_tx=XM_019837955.2;refGene_gn=CLA
SP1;refGene_tx=XM_019837956.2;refGene_gn=CLASP1;refGene_tx=XM_01983795
7.2;refGene_gn=CLASP1;refGene_tx=XM_019837958.2;refGene_gn=CLASP1;refG
ene_tx=XM_019837959.2
C1 119352643 119352721
refGene_gn=CLASP1;refGene_tx=XM_006935226.4;refGene_gn=CLASP1;refGene_
tx=XM_006935227.4;refGene_gn=CLASP1;refGene_tx=XM_006935228.4;refGene_
gn=CLASP1;refGene_tx=XM_006935229.4;refGene_gn=CLASP1;refGene_tx=XM_01
9837945.2;refGene_gn=CLASP1;refGene_tx=XM_019837946.2;refGene_gn=CLASP
1;refGene_tx=XM_019837947.2;refGene_gn=CLASP1;refGene_tx=XM_019837948.
2;refGene_gn=CLASP1;refGene_tx=XM_019837949.2;refGene_gn=CLASP1;refGen
e_tx=XM_019837951.2;refGene_gn=CLASP1;refGene_tx=XM_019837952.2;refGen
e_gn=CLASP1;refGene_tx=XM_019837953.2;refGene_gn=CLASP1;refGene_tx=XM_
019837954.2;refGene_gn=CLASP1;refGene_tx=XM_019837955.2;refGene_gn=CLA
SP1;refGene_tx=XM_019837956.2;refGene_gn=CLASP1;refGene_tx=XM_01983795
7.2;refGene_gn=CLASP1;refGene_tx=XM_019837958.2;refGene_gn=CLASP1;refG
ene_tx=XM_019837959.2
C1 119354904 119355072
refGene_gn=CLASP1;refGene_tx=XM_006935226.4;refGene_gn=CLASP1;refGene_
tx=XM_006935227.4;refGene_gn=CLASP1;refGene_tx=XM_006935228.4;refGene_

gn=CLASP1;refGene_tx=XM_006935229.4;refGene_gn=CLASP1;refGene_tx=XM_01
9837945.2;refGene_gn=CLASP1;refGene_tx=XM_019837946.2;refGene_gn=CLASP
1;refGene_tx=XM_019837947.2;refGene_gn=CLASP1;refGene_tx=XM_019837948.
2;refGene_gn=CLASP1;refGene_tx=XM_019837949.2;refGene_gn=CLASP1;refGen
e_tx=XM_019837951.2;refGene_gn=CLASP1;refGene_tx=XM_019837952.2;refGen
e_gn=CLASP1;refGene_tx=XM_019837953.2;refGene_gn=CLASP1;refGene_tx=XM_
019837954.2;refGene_gn=CLASP1;refGene_tx=XM_019837955.2;refGene_gn=CLA
SP1;refGene_tx=XM_019837956.2;refGene_gn=CLASP1;refGene_tx=XM_01983795
7.2;refGene_gn=CLASP1;refGene_tx=XM_019837958.2;refGene_gn=CLASP1;refG
ene_tx=XM_019837959.2
C1	119355672	119355722
refGene_gn=CLASP1;refGene_tx=XM_006935226.4;refGene_gn=CLASP1;refGene_
tx=XM_006935227.4;refGene_gn=CLASP1;refGene_tx=XM_006935228.4;refGene_
gn=CLASP1;refGene_tx=XM_006935229.4;refGene_gn=CLASP1;refGene_tx=XM_01
9837945.2;refGene_gn=CLASP1;refGene_tx=XM_019837946.2;refGene_gn=CLASP
1;refGene_tx=XM_019837947.2;refGene_gn=CLASP1;refGene_tx=XM_019837948.
2;refGene_gn=CLASP1;refGene_tx=XM_019837949.2;refGene_gn=CLASP1;refGen
e_tx=XM_019837951.2;refGene_gn=CLASP1;refGene_tx=XM_019837952.2;refGen
e_gn=CLASP1;refGene_tx=XM_019837953.2;refGene_gn=CLASP1;refGene_tx=XM_
019837954.2;refGene_gn=CLASP1;refGene_tx=XM_019837955.2;refGene_gn=CLA
SP1;refGene_tx=XM_019837956.2;refGene_gn=CLASP1;refGene_tx=XM_01983795
7.2;refGene_gn=CLASP1;refGene_tx=XM_019837958.2;refGene_gn=CLASP1;refG
ene_tx=XM_019837959.2
C1	119356462	119356633
refGene_gn=CLASP1;refGene_tx=XM_006935226.4;refGene_gn=CLASP1;refGene_
tx=XM_006935227.4;refGene_gn=CLASP1;refGene_tx=XM_006935228.4;refGene_
gn=CLASP1;refGene_tx=XM_006935229.4;refGene_gn=CLASP1;refGene_tx=XM_01
9837945.2;refGene_gn=CLASP1;refGene_tx=XM_019837946.2;refGene_gn=CLASP
1;refGene_tx=XM_019837947.2;refGene_gn=CLASP1;refGene_tx=XM_019837948.
2;refGene_gn=CLASP1;refGene_tx=XM_019837949.2;refGene_gn=CLASP1;refGen
e_tx=XM_019837951.2;refGene_gn=CLASP1;refGene_tx=XM_019837952.2;refGen
e_gn=CLASP1;refGene_tx=XM_019837953.2;refGene_gn=CLASP1;refGene_tx=XM_
019837954.2;refGene_gn=CLASP1;refGene_tx=XM_019837955.2;refGene_gn=CLA
SP1;refGene_tx=XM_019837956.2;refGene_gn=CLASP1;refGene_tx=XM_01983795
7.2;refGene_gn=CLASP1;refGene_tx=XM_019837958.2;refGene_gn=CLASP1;refG
ene_tx=XM_019837959.2
C1	119358406	119358454
refGene_gn=CLASP1;refGene_tx=XM_006935227.4;refGene_gn=CLASP1;refGene_
tx=XM_019837945.2;refGene_gn=CLASP1;refGene_tx=XM_019837946.2;refGene_
gn=CLASP1;refGene_tx=XM_019837948.2;refGene_gn=CLASP1;refGene_tx=XM_01
9837951.2;refGene_gn=CLASP1;refGene_tx=XM_019837952.2;refGene_gn=CLASP
1;refGene_tx=XM_019837953.2;refGene_gn=CLASP1;refGene_tx=XM_019837954.
2;refGene_gn=CLASP1;refGene_tx=XM_019837955.2;refGene_gn=CLASP1;refGen
e_tx=XM_019837956.2;refGene_gn=CLASP1;refGene_tx=XM_019837959.2
C1	119358998	119359022
refGene_gn=CLASP1;refGene_tx=XM_019837945.2;refGene_gn=CLASP1;refGene_
tx=XM_019837947.2;refGene_gn=CLASP1;refGene_tx=XM_019837948.2;refGene_
gn=CLASP1;refGene_tx=XM_019837951.2;refGene_gn=CLASP1;refGene_tx=XM_01
9837952.2;refGene_gn=CLASP1;refGene_tx=XM_019837953.2;refGene_gn=CLASP
1;refGene_tx=XM_019837954.2;refGene_gn=CLASP1;refGene_tx=XM_019837955.

2;refGene_gn=CLASP1;refGene_tx=XM_019837958.2;refGene_gn=CLASP1;refGene_tx=XM_019837959.2
C1 119371680 119371785
refGene_gn=CLASP1;refGene_tx=XM_006935226.4;refGene_gn=CLASP1;refGene_tx=XM_006935227.4;refGene_gn=CLASP1;refGene_tx=XM_006935228.4;refGene_gn=CLASP1;refGene_tx=XM_006935229.4;refGene_gn=CLASP1;refGene_tx=XM_019837945.2;refGene_gn=CLASP1;refGene_tx=XM_019837946.2;refGene_gn=CLASP1;refGene_tx=XM_019837947.2;refGene_gn=CLASP1;refGene_tx=XM_019837948.2;refGene_gn=CLASP1;refGene_tx=XM_019837949.2;refGene_gn=CLASP1;refGene_tx=XM_019837951.2;refGene_gn=CLASP1;refGene_tx=XM_019837952.2;refGene_gn=CLASP1;refGene_tx=XM_019837953.2;refGene_gn=CLASP1;refGene_tx=XM_019837954.2;refGene_gn=CLASP1;refGene_tx=XM_019837955.2;refGene_gn=CLASP1;refGene_tx=XM_019837956.2;refGene_gn=CLASP1;refGene_tx=XM_019837957.2;refGene_gn=CLASP1;refGene_tx=XM_019837958.2;refGene_gn=CLASP1;refGene_tx=XM_019837959.2
C1 119374064 119374091
refGene_gn=CLASP1;refGene_tx=XM_006935226.4;refGene_gn=CLASP1;refGene_tx=XM_019837945.2;refGene_gn=CLASP1;refGene_tx=XM_019837946.2;refGene_gn=CLASP1;refGene_tx=XM_019837947.2;refGene_gn=CLASP1;refGene_tx=XM_019837948.2;refGene_gn=CLASP1;refGene_tx=XM_019837949.2;refGene_gn=CLASP1;refGene_tx=XM_019837951.2;refGene_gn=CLASP1;refGene_tx=XM_019837952.2;refGene_gn=CLASP1;refGene_tx=XM_019837953.2;refGene_gn=CLASP1;refGene_tx=XM_019837954.2;refGene_gn=CLASP1;refGene_tx=XM_019837955.2;refGene_gn=CLASP1;refGene_tx=XM_019837957.2;refGene_gn=CLASP1;refGene_tx=XM_019837958.2
C1 119375408 119375576
refGene_gn=CLASP1;refGene_tx=XM_006935226.4;refGene_gn=CLASP1;refGene_tx=XM_006935227.4;refGene_gn=CLASP1;refGene_tx=XM_006935228.4;refGene_gn=CLASP1;refGene_tx=XM_006935229.4;refGene_gn=CLASP1;refGene_tx=XM_019837945.2;refGene_gn=CLASP1;refGene_tx=XM_019837946.2;refGene_gn=CLASP1;refGene_tx=XM_019837947.2;refGene_gn=CLASP1;refGene_tx=XM_019837948.2;refGene_gn=CLASP1;refGene_tx=XM_019837949.2;refGene_gn=CLASP1;refGene_tx=XM_019837951.2;refGene_gn=CLASP1;refGene_tx=XM_019837952.2;refGene_gn=CLASP1;refGene_tx=XM_019837953.2;refGene_gn=CLASP1;refGene_tx=XM_019837954.2;refGene_gn=CLASP1;refGene_tx=XM_019837955.2;refGene_gn=CLASP1;refGene_tx=XM_019837956.2;refGene_gn=CLASP1;refGene_tx=XM_019837957.2;refGene_gn=CLASP1;refGene_tx=XM_019837958.2;refGene_gn=CLASP1;refGene_tx=XM_019837959.2
C1 119381593 119381701
refGene_gn=CLASP1;refGene_tx=XM_019837945.2;refGene_gn=CLASP1;refGene_tx=XM_019837946.2;refGene_gn=CLASP1;refGene_tx=XM_019837947.2;refGene_gn=CLASP1;refGene_tx=XM_019837948.2;refGene_gn=CLASP1;refGene_tx=XM_019837949.2;refGene_gn=CLASP1;refGene_tx=XM_019837951.2;refGene_gn=CLASP1;refGene_tx=XM_019837952.2;refGene_gn=CLASP1;refGene_tx=XM_019837954.2;refGene_gn=CLASP1;refGene_tx=XM_019837955.2;refGene_gn=CLASP1;refGene_tx=XM_019837957.2;refGene_gn=CLASP1;refGene_tx=XM_019837959.2
C1 119386400 119386463
refGene_gn=CLASP1;refGene_tx=XM_019837945.2;refGene_gn=CLASP1;refGene_tx=XM_019837946.2;refGene_gn=CLASP1;refGene_tx=XM_019837947.2;refGene_gn=CLASP1;refGene_tx=XM_019837949.2;refGene_gn=CLASP1;refGene_tx=XM_01

9837951.2;refGene_gn=CLASP1;refGene_tx=XM_019837952.2;refGene_gn=CLASP1;refGene_tx=XM_019837954.2;refGene_gn=CLASP1;refGene_tx=XM_019837955.2;refGene_gn=CLASP1;refGene_tx=XM_019837959.2
C1 119389384 119389488
refGene_gn=CLASP1;refGene_tx=XM_006935226.4;refGene_gn=CLASP1;refGene_tx=XM_006935227.4;refGene_gn=CLASP1;refGene_tx=XM_006935228.4;refGene_gn=CLASP1;refGene_tx=XM_006935229.4;refGene_gn=CLASP1;refGene_tx=XM_019837945.2;refGene_gn=CLASP1;refGene_tx=XM_019837946.2;refGene_gn=CLASP1;refGene_tx=XM_019837947.2;refGene_gn=CLASP1;refGene_tx=XM_019837948.2;refGene_gn=CLASP1;refGene_tx=XM_019837949.2;refGene_gn=CLASP1;refGene_tx=XM_019837951.2;refGene_gn=CLASP1;refGene_tx=XM_019837952.2;refGene_gn=CLASP1;refGene_tx=XM_019837953.2;refGene_gn=CLASP1;refGene_tx=XM_019837954.2;refGene_gn=CLASP1;refGene_tx=XM_019837955.2;refGene_gn=CLASP1;refGene_tx=XM_019837956.2;refGene_gn=CLASP1;refGene_tx=XM_019837957.2;refGene_gn=CLASP1;refGene_tx=XM_019837958.2;refGene_gn=CLASP1;refGene_tx=XM_019837959.2
C1 119391413 119391437
refGene_gn=CLASP1;refGene_tx=XM_006935226.4;refGene_gn=CLASP1;refGene_tx=XM_006935228.4;refGene_gn=CLASP1;refGene_tx=XM_019837945.2;refGene_gn=CLASP1;refGene_tx=XM_019837946.2;refGene_gn=CLASP1;refGene_tx=XM_019837947.2;refGene_gn=CLASP1;refGene_tx=XM_019837948.2;refGene_gn=CLASP1;refGene_tx=XM_019837949.2;refGene_gn=CLASP1;refGene_tx=XM_019837951.2;refGene_gn=CLASP1;refGene_tx=XM_019837952.2;refGene_gn=CLASP1;refGene_tx=XM_019837953.2;refGene_gn=CLASP1;refGene_tx=XM_019837954.2;refGene_gn=CLASP1;refGene_tx=XM_019837956.2;refGene_gn=CLASP1;refGene_tx=XM_019837958.2
C1 119392762 119393007
refGene_gn=CLASP1;refGene_tx=XM_006935226.4;refGene_gn=CLASP1;refGene_tx=XM_006935227.4;refGene_gn=CLASP1;refGene_tx=XM_006935228.4;refGene_gn=CLASP1;refGene_tx=XM_006935229.4;refGene_gn=CLASP1;refGene_tx=XM_019837945.2;refGene_gn=CLASP1;refGene_tx=XM_019837946.2;refGene_gn=CLASP1;refGene_tx=XM_019837947.2;refGene_gn=CLASP1;refGene_tx=XM_019837948.2;refGene_gn=CLASP1;refGene_tx=XM_019837949.2;refGene_gn=CLASP1;refGene_tx=XM_019837951.2;refGene_gn=CLASP1;refGene_tx=XM_019837952.2;refGene_gn=CLASP1;refGene_tx=XM_019837953.2;refGene_gn=CLASP1;refGene_tx=XM_019837954.2;refGene_gn=CLASP1;refGene_tx=XM_019837955.2;refGene_gn=CLASP1;refGene_tx=XM_019837956.2;refGene_gn=CLASP1;refGene_tx=XM_019837957.2;refGene_gn=CLASP1;refGene_tx=XM_019837958.2;refGene_gn=CLASP1;refGene_tx=XM_019837959.2
C1 119395594 119395658
refGene_gn=CLASP1;refGene_tx=XM_006935226.4;refGene_gn=CLASP1;refGene_tx=XM_006935227.4;refGene_gn=CLASP1;refGene_tx=XM_006935228.4;refGene_gn=CLASP1;refGene_tx=XM_006935229.4;refGene_gn=CLASP1;refGene_tx=XM_019837945.2;refGene_gn=CLASP1;refGene_tx=XM_019837946.2;refGene_gn=CLASP1;refGene_tx=XM_019837947.2;refGene_gn=CLASP1;refGene_tx=XM_019837948.2;refGene_gn=CLASP1;refGene_tx=XM_019837949.2;refGene_gn=CLASP1;refGene_tx=XM_019837951.2;refGene_gn=CLASP1;refGene_tx=XM_019837952.2;refGene_gn=CLASP1;refGene_tx=XM_019837953.2;refGene_gn=CLASP1;refGene_tx=XM_019837954.2;refGene_gn=CLASP1;refGene_tx=XM_019837955.2;refGene_gn=CLASP1;refGene_tx=XM_019837956.2;refGene_gn=CLASP1;refGene_tx=XM_01983795

7.2;refGene_gn=CLASP1;refGene_tx=XM_019837958.2;refGene_gn=CLASP1;refGene_tx=XM_019837959.2
C1 119397036 119397117
refGene_gn=CLASP1;refGene_tx=XM_019837945.2;refGene_gn=CLASP1;refGene_tx=XM_019837946.2;refGene_gn=CLASP1;refGene_tx=XM_019837947.2;refGene_gn=CLASP1;refGene_tx=XM_019837948.2;refGene_gn=CLASP1;refGene_tx=XM_019837949.2;refGene_gn=CLASP1;refGene_tx=XM_019837953.2;refGene_gn=CLASP1;refGene_tx=XM_019837956.2
C1 119397858 119397861
refGene_gn=CLASP1;refGene_tx=XM_019837945.2;refGene_gn=CLASP1;refGene_tx=XM_019837946.2;refGene_gn=CLASP1;refGene_tx=XM_019837947.2;refGene_gn=CLASP1;refGene_tx=XM_019837948.2;refGene_gn=CLASP1;refGene_tx=XM_019837949.2;refGene_gn=CLASP1;refGene_tx=XM_019837953.2;refGene_gn=CLASP1;refGene_tx=XM_019837956.2
C1 119397984 119398148
refGene_gn=CLASP1;refGene_tx=XM_006935226.4;refGene_gn=CLASP1;refGene_tx=XM_006935227.4;refGene_gn=CLASP1;refGene_tx=XM_006935228.4;refGene_gn=CLASP1;refGene_tx=XM_006935229.4;refGene_gn=CLASP1;refGene_tx=XM_019837945.2;refGene_gn=CLASP1;refGene_tx=XM_019837946.2;refGene_gn=CLASP1;refGene_tx=XM_019837947.2;refGene_gn=CLASP1;refGene_tx=XM_019837948.2;refGene_gn=CLASP1;refGene_tx=XM_019837949.2;refGene_gn=CLASP1;refGene_tx=XM_019837951.2;refGene_gn=CLASP1;refGene_tx=XM_019837952.2;refGene_gn=CLASP1;refGene_tx=XM_019837953.2;refGene_gn=CLASP1;refGene_tx=XM_019837954.2;refGene_gn=CLASP1;refGene_tx=XM_019837955.2;refGene_gn=CLASP1;refGene_tx=XM_019837956.2;refGene_gn=CLASP1;refGene_tx=XM_019837957.2;refGene_gn=CLASP1;refGene_tx=XM_019837958.2;refGene_gn=CLASP1;refGene_tx=XM_019837959.2
C1 119404536 119404615
refGene_gn=CLASP1;refGene_tx=XM_006935226.4;refGene_gn=CLASP1;refGene_tx=XM_006935227.4;refGene_gn=CLASP1;refGene_tx=XM_006935228.4;refGene_gn=CLASP1;refGene_tx=XM_006935229.4;refGene_gn=CLASP1;refGene_tx=XM_019837945.2;refGene_gn=CLASP1;refGene_tx=XM_019837946.2;refGene_gn=CLASP1;refGene_tx=XM_019837947.2;refGene_gn=CLASP1;refGene_tx=XM_019837948.2;refGene_gn=CLASP1;refGene_tx=XM_019837949.2;refGene_gn=CLASP1;refGene_tx=XM_019837951.2;refGene_gn=CLASP1;refGene_tx=XM_019837952.2;refGene_gn=CLASP1;refGene_tx=XM_019837953.2;refGene_gn=CLASP1;refGene_tx=XM_019837954.2;refGene_gn=CLASP1;refGene_tx=XM_019837955.2;refGene_gn=CLASP1;refGene_tx=XM_019837956.2;refGene_gn=CLASP1;refGene_tx=XM_019837957.2;refGene_gn=CLASP1;refGene_tx=XM_019837958.2;refGene_gn=CLASP1;refGene_tx=XM_019837959.2
C1 119405634 119405778
refGene_gn=CLASP1;refGene_tx=XM_006935226.4;refGene_gn=CLASP1;refGene_tx=XM_006935227.4;refGene_gn=CLASP1;refGene_tx=XM_006935228.4;refGene_gn=CLASP1;refGene_tx=XM_006935229.4;refGene_gn=CLASP1;refGene_tx=XM_019837945.2;refGene_gn=CLASP1;refGene_tx=XM_019837946.2;refGene_gn=CLASP1;refGene_tx=XM_019837947.2;refGene_gn=CLASP1;refGene_tx=XM_019837948.2;refGene_gn=CLASP1;refGene_tx=XM_019837949.2;refGene_gn=CLASP1;refGene_tx=XM_019837951.2;refGene_gn=CLASP1;refGene_tx=XM_019837952.2;refGene_gn=CLASP1;refGene_tx=XM_019837953.2;refGene_gn=CLASP1;refGene_tx=XM_019837954.2;refGene_gn=CLASP1;refGene_tx=XM_019837955.2;refGene_gn=CLA

SP1;refGene_tx=XM_019837956.2;refGene_gn=CLASP1;refGene_tx=XM_01983795
7.2;refGene_gn=CLASP1;refGene_tx=XM_019837958.2;refGene_gn=CLASP1;refG
ene_tx=XM_019837959.2
C1 119415778 119415922
refGene_gn=CLASP1;refGene_tx=XM_006935226.4;refGene_gn=CLASP1;refGene_
tx=XM_006935227.4;refGene_gn=CLASP1;refGene_tx=XM_006935228.4;refGene_
gn=CLASP1;refGene_tx=XM_006935229.4;refGene_gn=CLASP1;refGene_tx=XM_01
9837945.2;refGene_gn=CLASP1;refGene_tx=XM_019837946.2;refGene_gn=CLASP
1;refGene_tx=XM_019837947.2;refGene_gn=CLASP1;refGene_tx=XM_019837948.
2;refGene_gn=CLASP1;refGene_tx=XM_019837949.2;refGene_gn=CLASP1;refGen
e_tx=XM_019837951.2;refGene_gn=CLASP1;refGene_tx=XM_019837952.2;refGen
e_gn=CLASP1;refGene_tx=XM_019837953.2;refGene_gn=CLASP1;refGene_tx=XM_
019837954.2;refGene_gn=CLASP1;refGene_tx=XM_019837955.2;refGene_gn=CLA
SP1;refGene_tx=XM_019837956.2;refGene_gn=CLASP1;refGene_tx=XM_01983795
7.2;refGene_gn=CLASP1;refGene_tx=XM_019837958.2;refGene_gn=CLASP1;refG
ene_tx=XM_019837959.2
C1 119416405 119416512
refGene_gn=CLASP1;refGene_tx=XM_006935226.4;refGene_gn=CLASP1;refGene_
tx=XM_006935227.4;refGene_gn=CLASP1;refGene_tx=XM_006935228.4;refGene_
gn=CLASP1;refGene_tx=XM_006935229.4;refGene_gn=CLASP1;refGene_tx=XM_01
9837945.2;refGene_gn=CLASP1;refGene_tx=XM_019837946.2;refGene_gn=CLASP
1;refGene_tx=XM_019837947.2;refGene_gn=CLASP1;refGene_tx=XM_019837948.
2;refGene_gn=CLASP1;refGene_tx=XM_019837949.2;refGene_gn=CLASP1;refGen
e_tx=XM_019837951.2;refGene_gn=CLASP1;refGene_tx=XM_019837952.2;refGen
e_gn=CLASP1;refGene_tx=XM_019837953.2;refGene_gn=CLASP1;refGene_tx=XM_
019837954.2;refGene_gn=CLASP1;refGene_tx=XM_019837955.2;refGene_gn=CLA
SP1;refGene_tx=XM_019837956.2;refGene_gn=CLASP1;refGene_tx=XM_01983795
7.2;refGene_gn=CLASP1;refGene_tx=XM_019837958.2;refGene_gn=CLASP1;refG
ene_tx=XM_019837959.2
C1 119420515 119420632
refGene_gn=CLASP1;refGene_tx=XM_019837945.2;refGene_gn=CLASP1;refGene_
tx=XM_019837946.2;refGene_gn=CLASP1;refGene_tx=XM_019837947.2;refGene_
gn=CLASP1;refGene_tx=XM_019837948.2;refGene_gn=CLASP1;refGene_tx=XM_01
9837949.2;refGene_gn=CLASP1;refGene_tx=XM_019837951.2;refGene_gn=CLASP
1;refGene_tx=XM_019837952.2;refGene_gn=CLASP1;refGene_tx=XM_019837953.
2;refGene_gn=CLASP1;refGene_tx=XM_019837956.2;refGene_gn=CLASP1;refGen
e_tx=XM_019837957.2
C1 119425059 119425210
refGene_gn=CLASP1;refGene_tx=XM_006935226.4;refGene_gn=CLASP1;refGene_
tx=XM_006935227.4;refGene_gn=CLASP1;refGene_tx=XM_006935228.4;refGene_
gn=CLASP1;refGene_tx=XM_006935229.4;refGene_gn=CLASP1;refGene_tx=XM_01
9837945.2;refGene_gn=CLASP1;refGene_tx=XM_019837946.2;refGene_gn=CLASP
1;refGene_tx=XM_019837947.2;refGene_gn=CLASP1;refGene_tx=XM_019837948.
2;refGene_gn=CLASP1;refGene_tx=XM_019837949.2;refGene_gn=CLASP1;refGen
e_tx=XM_019837951.2;refGene_gn=CLASP1;refGene_tx=XM_019837952.2;refGen
e_gn=CLASP1;refGene_tx=XM_019837953.2;refGene_gn=CLASP1;refGene_tx=XM_
019837954.2;refGene_gn=CLASP1;refGene_tx=XM_019837955.2;refGene_gn=CLA
SP1;refGene_tx=XM_019837956.2;refGene_gn=CLASP1;refGene_tx=XM_01983795
7.2;refGene_gn=CLASP1;refGene_tx=XM_019837958.2;refGene_gn=CLASP1;refG
ene_tx=XM_019837959.2

C1	119429210	119429457
refGene_gn=CLASP1;refGene_tx=XM_006935226.4;refGene_gn=CLASP1;refGene_tx=XM_006935227.4;refGene_gn=CLASP1;refGene_tx=XM_006935228.4;refGene_gn=CLASP1;refGene_tx=XM_006935229.4;refGene_gn=CLASP1;refGene_tx=XM_019837945.2;refGene_gn=CLASP1;refGene_tx=XM_019837946.2;refGene_gn=CLASP1;refGene_tx=XM_019837947.2;refGene_gn=CLASP1;refGene_tx=XM_019837948.2;refGene_gn=CLASP1;refGene_tx=XM_019837949.2;refGene_gn=CLASP1;refGene_tx=XM_019837951.2;refGene_gn=CLASP1;refGene_tx=XM_019837952.2;refGene_gn=CLASP1;refGene_tx=XM_019837953.2;refGene_gn=CLASP1;refGene_tx=XM_019837954.2;refGene_gn=CLASP1;refGene_tx=XM_019837955.2;refGene_gn=CLASP1;refGene_tx=XM_019837956.2;refGene_gn=CLASP1;refGene_tx=XM_019837957.2;refGene_gn=CLASP1;refGene_tx=XM_019837958.2;refGene_gn=CLASP1;refGene_tx=XM_019837959.2
C1	119431904	119432095
refGene_gn=CLASP1;refGene_tx=XM_006935226.4;refGene_gn=CLASP1;refGene_tx=XM_006935227.4;refGene_gn=CLASP1;refGene_tx=XM_006935228.4;refGene_gn=CLASP1;refGene_tx=XM_006935229.4;refGene_gn=CLASP1;refGene_tx=XM_019837945.2;refGene_gn=CLASP1;refGene_tx=XM_019837946.2;refGene_gn=CLASP1;refGene_tx=XM_019837947.2;refGene_gn=CLASP1;refGene_tx=XM_019837948.2;refGene_gn=CLASP1;refGene_tx=XM_019837949.2;refGene_gn=CLASP1;refGene_tx=XM_019837951.2;refGene_gn=CLASP1;refGene_tx=XM_019837952.2;refGene_gn=CLASP1;refGene_tx=XM_019837953.2;refGene_gn=CLASP1;refGene_tx=XM_019837954.2;refGene_gn=CLASP1;refGene_tx=XM_019837955.2;refGene_gn=CLASP1;refGene_tx=XM_019837956.2;refGene_gn=CLASP1;refGene_tx=XM_019837957.2;refGene_gn=CLASP1;refGene_tx=XM_019837958.2;refGene_gn=CLASP1;refGene_tx=XM_019837959.2
C1	119433858	119433987
refGene_gn=CLASP1;refGene_tx=XM_006935226.4;refGene_gn=CLASP1;refGene_tx=XM_006935227.4;refGene_gn=CLASP1;refGene_tx=XM_006935228.4;refGene_gn=CLASP1;refGene_tx=XM_006935229.4;refGene_gn=CLASP1;refGene_tx=XM_019837945.2;refGene_gn=CLASP1;refGene_tx=XM_019837946.2;refGene_gn=CLASP1;refGene_tx=XM_019837947.2;refGene_gn=CLASP1;refGene_tx=XM_019837948.2;refGene_gn=CLASP1;refGene_tx=XM_019837949.2;refGene_gn=CLASP1;refGene_tx=XM_019837951.2;refGene_gn=CLASP1;refGene_tx=XM_019837952.2;refGene_gn=CLASP1;refGene_tx=XM_019837953.2;refGene_gn=CLASP1;refGene_tx=XM_019837954.2;refGene_gn=CLASP1;refGene_tx=XM_019837955.2;refGene_gn=CLASP1;refGene_tx=XM_019837956.2;refGene_gn=CLASP1;refGene_tx=XM_019837957.2;refGene_gn=CLASP1;refGene_tx=XM_019837958.2;refGene_gn=CLASP1;refGene_tx=XM_019837959.2
C1	119449214	119449421
refGene_gn=CLASP1;refGene_tx=XM_006935226.4;refGene_gn=CLASP1;refGene_tx=XM_006935227.4;refGene_gn=CLASP1;refGene_tx=XM_006935228.4;refGene_gn=CLASP1;refGene_tx=XM_006935229.4;refGene_gn=CLASP1;refGene_tx=XM_019837945.2;refGene_gn=CLASP1;refGene_tx=XM_019837946.2;refGene_gn=CLASP1;refGene_tx=XM_019837947.2;refGene_gn=CLASP1;refGene_tx=XM_019837948.2;refGene_gn=CLASP1;refGene_tx=XM_019837949.2;refGene_gn=CLASP1;refGene_tx=XM_019837951.2;refGene_gn=CLASP1;refGene_tx=XM_019837952.2;refGene_gn=CLASP1;refGene_tx=XM_019837953.2;refGene_gn=CLASP1;refGene_tx=XM_019837954.2;refGene_gn=CLASP1;refGene_tx=XM_019837955.2;refGene_gn=CLASP1;refGene_tx=XM_019837956.2;refGene_gn=CLASP1;refGene_tx=XM_01983795

7.2;refGene_gn=CLASP1;refGene_tx=XM_019837958.2;refGene_gn=CLASP1;refGene_tx=XM_019837959.2
C1 119450907 119451024
refGene_gn=CLASP1;refGene_tx=XM_006935226.4;refGene_gn=CLASP1;refGene_tx=XM_006935227.4;refGene_gn=CLASP1;refGene_tx=XM_006935228.4;refGene_gn=CLASP1;refGene_tx=XM_006935229.4;refGene_gn=CLASP1;refGene_tx=XM_019837945.2;refGene_gn=CLASP1;refGene_tx=XM_019837946.2;refGene_gn=CLASP1;refGene_tx=XM_019837947.2;refGene_gn=CLASP1;refGene_tx=XM_019837948.2;refGene_gn=CLASP1;refGene_tx=XM_019837949.2;refGene_gn=CLASP1;refGene_tx=XM_019837951.2;refGene_gn=CLASP1;refGene_tx=XM_019837952.2;refGene_gn=CLASP1;refGene_tx=XM_019837953.2;refGene_gn=CLASP1;refGene_tx=XM_019837954.2;refGene_gn=CLASP1;refGene_tx=XM_019837955.2;refGene_gn=CLASP1;refGene_tx=XM_019837956.2;refGene_gn=CLASP1;refGene_tx=XM_019837957.2;refGene_gn=CLASP1;refGene_tx=XM_019837958.2;refGene_gn=CLASP1;refGene_tx=XM_019837959.2
C1 119457961 119458048
refGene_gn=CLASP1;refGene_tx=XM_006935226.4;refGene_gn=CLASP1;refGene_tx=XM_006935227.4;refGene_gn=CLASP1;refGene_tx=XM_006935228.4;refGene_gn=CLASP1;refGene_tx=XM_006935229.4;refGene_gn=CLASP1;refGene_tx=XM_019837945.2;refGene_gn=CLASP1;refGene_tx=XM_019837946.2;refGene_gn=CLASP1;refGene_tx=XM_019837947.2;refGene_gn=CLASP1;refGene_tx=XM_019837948.2;refGene_gn=CLASP1;refGene_tx=XM_019837949.2;refGene_gn=CLASP1;refGene_tx=XM_019837951.2;refGene_gn=CLASP1;refGene_tx=XM_019837952.2;refGene_gn=CLASP1;refGene_tx=XM_019837953.2;refGene_gn=CLASP1;refGene_tx=XM_019837954.2;refGene_gn=CLASP1;refGene_tx=XM_019837955.2;refGene_gn=CLASP1;refGene_tx=XM_019837956.2;refGene_gn=CLASP1;refGene_tx=XM_019837957.2;refGene_gn=CLASP1;refGene_tx=XM_019837958.2;refGene_gn=CLASP1;refGene_tx=XM_019837959.2
C1 119495775 119495837
refGene_gn=TFCP2L1;refGene_tx=XM_023259241.1;refGene_gn=TFCP2L1;refGene_tx=XM_023259243.1
C1 119496444 119496474
refGene_gn=TFCP2L1;refGene_tx=XM_023259241.1
C1 119499262 119499414
refGene_gn=TFCP2L1;refGene_tx=XM_023259241.1;refGene_gn=TFCP2L1;refGene_tx=XM_023259243.1
C1 119528116 119528193
refGene_gn=TFCP2L1;refGene_tx=XM_023259241.1;refGene_gn=TFCP2L1;refGene_tx=XM_023259243.1
C1 119528725 119528831
refGene_gn=TFCP2L1;refGene_tx=XM_023259241.1;refGene_gn=TFCP2L1;refGene_tx=XM_023259243.1
C1 119529519 119529626
refGene_gn=TFCP2L1;refGene_tx=XM_023259241.1;refGene_gn=TFCP2L1;refGene_tx=XM_023259243.1
C1 119530846 119530999
refGene_gn=TFCP2L1;refGene_tx=XM_023259241.1;refGene_gn=TFCP2L1;refGene_tx=XM_023259243.1
C1 119534158 119534269
refGene_gn=TFCP2L1;refGene_tx=XM_023259241.1;refGene_gn=TFCP2L1;refGen

e_tx=XM_023259243.1
C1 119536708 119536800
refGene_gn=TFCP2L1;refGene_tx=XM_023259241.1;refGene_gn=TFCP2L1;refGen
e_tx=XM_023259243.1
C1 119539130 119539179
refGene_gn=TFCP2L1;refGene_tx=XM_023259241.1;refGene_gn=TFCP2L1;refGen
e_tx=XM_023259243.1
C1 119539266 119539360
refGene_gn=TFCP2L1;refGene_tx=XM_023259241.1;refGene_gn=TFCP2L1;refGen
e_tx=XM_023259243.1
C1 119541967 119542058
refGene_gn=TFCP2L1;refGene_tx=XM_023259241.1;refGene_gn=TFCP2L1;refGen
e_tx=XM_023259243.1
C1 119543035 119543139
refGene_gn=TFCP2L1;refGene_tx=XM_023259241.1;refGene_gn=TFCP2L1;refGen
e_tx=XM_023259243.1
C1 119545552 119545695
refGene_gn=TFCP2L1;refGene_tx=XM_023259241.1;refGene_gn=TFCP2L1;refGen
e_tx=XM_023259243.1
C1 119549866 119549918
refGene_gn=TFCP2L1;refGene_tx=XM_023259241.1;refGene_gn=TFCP2L1;refGen
e_tx=XM_023259243.1
C1 119550709 119550756
refGene_gn=TFCP2L1;refGene_tx=XM_023259241.1;refGene_gn=TFCP2L1;refGen
e_tx=XM_023259243.1
C1 119768063 119770483
refGene_gn=GLI2;refGene_tx=XM_023259244.1
C1 119771752 119772089
refGene_gn=GLI2;refGene_tx=XM_023259244.1
C1 119773652 119773934
refGene_gn=GLI2;refGene_tx=XM_023259244.1
C1 119775195 119775360
refGene_gn=GLI2;refGene_tx=XM_023259244.1
C1 119781041 119781191
refGene_gn=GLI2;refGene_tx=XM_023259244.1
C1 119784742 119784877
refGene_gn=GLI2;refGene_tx=XM_023259244.1
C1 119788835 119788958
refGene_gn=GLI2;refGene_tx=XM_023259244.1
C1 119790133 119790347
refGene_gn=GLI2;refGene_tx=XM_023259244.1
C1 119791916 119792118
refGene_gn=GLI2;refGene_tx=XM_023259244.1
C1 119802445 119802631
refGene_gn=GLI2;refGene_tx=XM_023259244.1
C1 119806203 119806406
refGene_gn=GLI2;refGene_tx=XM_023259244.1
C1 119830604 119830710
refGene_gn=GLI2;refGene_tx=XM_023259244.1
C1 119955676 119955824

refGene_gn=GLI2;refGene_tx=XM_023259244.1
C1	120403432	120404208
refGene_gn=INHBB;refGene_tx=XM_003990713.5
C1	120406725	120407173
refGene_gn=INHBB;refGene_tx=XM_003990713.5
C1	120458156	120458276
refGene_gn=RALB;refGene_tx=XM_003990699.4
C1	120464346	120464524
refGene_gn=RALB;refGene_tx=XM_003990699.4
C1	120468589	120468798
refGene_gn=RALB;refGene_tx=XM_003990699.4
C1	120475925	120476039
refGene_gn=RALB;refGene_tx=XM_003990699.4
C1	120525900	120526953
refGene_gn=TMEM185B;refGene_tx=XM_003990700.4
C1	120554314	120554381
refGene_gn=EPB41L5;refGene_tx=XM_019837962.2;refGene_gn=EPB41L5;refGene_tx=XM_019837963.2;refGene_gn=EPB41L5;refGene_tx=XM_023259246.1;refGene_gn=EPB41L5;refGene_tx=XM_023259247.1
C1	120559199	120559221
refGene_gn=EPB41L5;refGene_tx=XM_019837965.2
C1	120559552	120559683
refGene_gn=EPB41L5;refGene_tx=XM_019837962.2;refGene_gn=EPB41L5;refGene_tx=XM_019837963.2;refGene_gn=EPB41L5;refGene_tx=XM_019837965.2;refGene_gn=EPB41L5;refGene_tx=XM_023259246.1;refGene_gn=EPB41L5;refGene_tx=XM_023259247.1
C1	120560050	120560092
refGene_gn=EPB41L5;refGene_tx=XM_019837962.2;refGene_gn=EPB41L5;refGene_tx=XM_019837963.2;refGene_gn=EPB41L5;refGene_tx=XM_019837965.2;refGene_gn=EPB41L5;refGene_tx=XM_023259246.1;refGene_gn=EPB41L5;refGene_tx=XM_023259247.1
C1	120566301	120566376
refGene_gn=EPB41L5;refGene_tx=XM_019837962.2;refGene_gn=EPB41L5;refGene_tx=XM_019837963.2;refGene_gn=EPB41L5;refGene_tx=XM_019837965.2;refGene_gn=EPB41L5;refGene_tx=XM_023259246.1;refGene_gn=EPB41L5;refGene_tx=XM_023259247.1
C1	120568122	120568213
refGene_gn=EPB41L5;refGene_tx=XM_019837962.2;refGene_gn=EPB41L5;refGene_tx=XM_019837963.2;refGene_gn=EPB41L5;refGene_tx=XM_019837965.2;refGene_gn=EPB41L5;refGene_tx=XM_023259246.1;refGene_gn=EPB41L5;refGene_tx=XM_023259247.1
C1	120572744	120572809
refGene_gn=EPB41L5;refGene_tx=XM_019837962.2;refGene_gn=EPB41L5;refGene_tx=XM_019837963.2;refGene_gn=EPB41L5;refGene_tx=XM_019837965.2;refGene_gn=EPB41L5;refGene_tx=XM_023259246.1;refGene_gn=EPB41L5;refGene_tx=XM_023259247.1
C1	120576285	120576414
refGene_gn=EPB41L5;refGene_tx=XM_019837962.2;refGene_gn=EPB41L5;refGene_tx=XM_019837963.2;refGene_gn=EPB41L5;refGene_tx=XM_019837965.2;refGene_gn=EPB41L5;refGene_tx=XM_023259246.1;refGene_gn=EPB41L5;refGene_tx=

XM_023259247.1
C1	120584897	120584995
refGene_gn=EPB41L5;refGene_tx=XM_019837962.2;refGene_gn=EPB41L5;refGene_tx=XM_019837963.2;refGene_gn=EPB41L5;refGene_tx=XM_019837965.2;refGene_gn=EPB41L5;refGene_tx=XM_023259246.1;refGene_gn=EPB41L5;refGene_tx=XM_023259247.1
C1	120588200	120588364
refGene_gn=EPB41L5;refGene_tx=XM_019837962.2;refGene_gn=EPB41L5;refGene_tx=XM_019837963.2;refGene_gn=EPB41L5;refGene_tx=XM_019837965.2;refGene_gn=EPB41L5;refGene_tx=XM_023259246.1;refGene_gn=EPB41L5;refGene_tx=XM_023259247.1
C1	120617504	120617685
refGene_gn=EPB41L5;refGene_tx=XM_019837968.2
C1	120621187	120621303
refGene_gn=EPB41L5;refGene_tx=XM_019837962.2;refGene_gn=EPB41L5;refGene_tx=XM_019837963.2;refGene_gn=EPB41L5;refGene_tx=XM_019837965.2;refGene_gn=EPB41L5;refGene_tx=XM_019837968.2;refGene_gn=EPB41L5;refGene_tx=XM_023259246.1;refGene_gn=EPB41L5;refGene_tx=XM_023259247.1
C1	120621748	120621791
refGene_gn=EPB41L5;refGene_tx=XM_019837962.2;refGene_gn=EPB41L5;refGene_tx=XM_019837963.2;refGene_gn=EPB41L5;refGene_tx=XM_019837965.2;refGene_gn=EPB41L5;refGene_tx=XM_019837968.2;refGene_gn=EPB41L5;refGene_tx=XM_023259246.1;refGene_gn=EPB41L5;refGene_tx=XM_023259247.1
C1	120629352	120629380
refGene_gn=EPB41L5;refGene_tx=XM_019837962.2;refGene_gn=EPB41L5;refGene_tx=XM_019837963.2;refGene_gn=EPB41L5;refGene_tx=XM_019837965.2;refGene_gn=EPB41L5;refGene_tx=XM_019837968.2;refGene_gn=EPB41L5;refGene_tx=XM_023259246.1;refGene_gn=EPB41L5;refGene_tx=XM_023259247.1
C1	120631417	120631524
refGene_gn=EPB41L5;refGene_tx=XM_019837962.2;refGene_gn=EPB41L5;refGene_tx=XM_019837963.2;refGene_gn=EPB41L5;refGene_tx=XM_019837965.2;refGene_gn=EPB41L5;refGene_tx=XM_019837968.2;refGene_gn=EPB41L5;refGene_tx=XM_023259246.1;refGene_gn=EPB41L5;refGene_tx=XM_023259247.1
C1	120632687	120632857
refGene_gn=EPB41L5;refGene_tx=XM_019837962.2;refGene_gn=EPB41L5;refGene_tx=XM_019837963.2;refGene_gn=EPB41L5;refGene_tx=XM_019837965.2;refGene_gn=EPB41L5;refGene_tx=XM_019837968.2;refGene_gn=EPB41L5;refGene_tx=XM_023259246.1;refGene_gn=EPB41L5;refGene_tx=XM_023259247.1
C1	120638402	120638472
refGene_gn=EPB41L5;refGene_tx=XM_019837962.2;refGene_gn=EPB41L5;refGene_tx=XM_019837963.2;refGene_gn=EPB41L5;refGene_tx=XM_019837965.2;refGene_gn=EPB41L5;refGene_tx=XM_019837968.2;refGene_gn=EPB41L5;refGene_tx=XM_023259246.1;refGene_gn=EPB41L5;refGene_tx=XM_023259247.1
C1	120641375	120641464
refGene_gn=EPB41L5;refGene_tx=XM_019837962.2;refGene_gn=EPB41L5;refGene_tx=XM_019837963.2;refGene_gn=EPB41L5;refGene_tx=XM_019837965.2;refGene_gn=EPB41L5;refGene_tx=XM_019837968.2;refGene_gn=EPB41L5;refGene_tx=XM_023259246.1;refGene_gn=EPB41L5;refGene_tx=XM_023259247.1
C1	120642671	120642759
refGene_gn=EPB41L5;refGene_tx=XM_019837962.2;refGene_gn=EPB41L5;refGen

e_tx=XM_019837963.2;refGene_gn=EPB41L5;refGene_tx=XM_019837965.2;refGene_gn=EPB41L5;refGene_tx=XM_019837968.2;refGene_gn=EPB41L5;refGene_tx=XM_023259246.1;refGene_gn=EPB41L5;refGene_tx=XM_023259247.1
C1 120642886 120643007
refGene_gn=EPB41L5;refGene_tx=XM_019837962.2;refGene_gn=EPB41L5;refGene_tx=XM_019837963.2;refGene_gn=EPB41L5;refGene_tx=XM_019837965.2;refGene_gn=EPB41L5;refGene_tx=XM_019837968.2;refGene_gn=EPB41L5;refGene_tx=XM_023259246.1;refGene_gn=EPB41L5;refGene_tx=XM_023259247.1
C1 120645445 120645498
refGene_gn=EPB41L5;refGene_tx=XM_019837962.2;refGene_gn=EPB41L5;refGene_tx=XM_019837963.2;refGene_gn=EPB41L5;refGene_tx=XM_019837965.2;refGene_gn=EPB41L5;refGene_tx=XM_019837968.2;refGene_gn=EPB41L5;refGene_tx=XM_023259246.1;refGene_gn=EPB41L5;refGene_tx=XM_023259247.1
C1 120645676 120645721
refGene_gn=EPB41L5;refGene_tx=XM_019837962.2;refGene_gn=EPB41L5;refGene_tx=XM_019837963.2;refGene_gn=EPB41L5;refGene_tx=XM_019837965.2;refGene_gn=EPB41L5;refGene_tx=XM_019837968.2;refGene_gn=EPB41L5;refGene_tx=XM_023259246.1;refGene_gn=EPB41L5;refGene_tx=XM_023259247.1
C1 120646936 120647015
refGene_gn=EPB41L5;refGene_tx=XM_019837962.2;refGene_gn=EPB41L5;refGene_tx=XM_019837963.2;refGene_gn=EPB41L5;refGene_tx=XM_019837965.2;refGene_gn=EPB41L5;refGene_tx=XM_019837968.2;refGene_gn=EPB41L5;refGene_tx=XM_023259246.1;refGene_gn=EPB41L5;refGene_tx=XM_023259247.1
C1 120648008 120648051
refGene_gn=EPB41L5;refGene_tx=XM_019837962.2;refGene_gn=EPB41L5;refGene_tx=XM_019837963.2;refGene_gn=EPB41L5;refGene_tx=XM_019837965.2;refGene_gn=EPB41L5;refGene_tx=XM_019837968.2;refGene_gn=EPB41L5;refGene_tx=XM_023259246.1;refGene_gn=EPB41L5;refGene_tx=XM_023259247.1
C1 120666809 120666914
refGene_gn=EPB41L5;refGene_tx=XM_019837962.2;refGene_gn=EPB41L5;refGene_tx=XM_019837963.2;refGene_gn=EPB41L5;refGene_tx=XM_019837965.2;refGene_gn=EPB41L5;refGene_tx=XM_019837968.2;refGene_gn=EPB41L5;refGene_tx=XM_023259246.1;refGene_gn=EPB41L5;refGene_tx=XM_023259247.1
C1 120681802 120681990
refGene_gn=EPB41L5;refGene_tx=XM_019837962.2;refGene_gn=EPB41L5;refGene_tx=XM_019837963.2;refGene_gn=EPB41L5;refGene_tx=XM_019837965.2;refGene_gn=EPB41L5;refGene_tx=XM_019837968.2;refGene_gn=EPB41L5;refGene_tx=XM_023259246.1;refGene_gn=EPB41L5;refGene_tx=XM_023259247.1
C1 120690065 120690132
refGene_gn=EPB41L5;refGene_tx=XM_019837962.2;refGene_gn=EPB41L5;refGene_tx=XM_019837963.2;refGene_gn=EPB41L5;refGene_tx=XM_019837965.2;refGene_gn=EPB41L5;refGene_tx=XM_019837968.2
C1 120715020 120715107
refGene_gn=PTPN4;refGene_tx=XM_011285247.3
C1 120722547 120722683
refGene_gn=PTPN4;refGene_tx=XM_011285247.3
C1 120724232 120724381
refGene_gn=PTPN4;refGene_tx=XM_011285247.3
C1 120726837 120726966
refGene_gn=PTPN4;refGene_tx=XM_011285247.3

C1 120728981 120729128
refGene_gn=PTPN4;refGene_tx=XM_011285247.3
C1 120731806 120731868
refGene_gn=PTPN4;refGene_tx=XM_011285247.3
C1 120731953 120732044
refGene_gn=PTPN4;refGene_tx=XM_011285247.3
C1 120732802 120732969
refGene_gn=PTPN4;refGene_tx=XM_011285247.3
C1 120743530 120743687
refGene_gn=PTPN4;refGene_tx=XM_011285247.3
C1 120749535 120749592
refGene_gn=PTPN4;refGene_tx=XM_011285247.3
C1 120749685 120749769
refGene_gn=PTPN4;refGene_tx=XM_011285247.3
C1 120753608 120753768
refGene_gn=PTPN4;refGene_tx=XM_011285247.3
C1 120765829 120765988
refGene_gn=PTPN4;refGene_tx=XM_011285247.3
C1 120769312 120769438
refGene_gn=PTPN4;refGene_tx=XM_011285247.3
C1 120775886 120775955
refGene_gn=PTPN4;refGene_tx=XM_011285247.3
C1 120782156 120782170
refGene_gn=PTPN4;refGene_tx=XM_011285247.3
C1 120782172 120782331
refGene_gn=PTPN4;refGene_tx=XM_011285247.3
C1 120786016 120786080
refGene_gn=PTPN4;refGene_tx=XM_011285247.3
C1 120796712 120796801
refGene_gn=PTPN4;refGene_tx=XM_011285247.3
C1 120812596 120812684
refGene_gn=PTPN4;refGene_tx=XM_011285247.3
C1 120821514 120821635
refGene_gn=PTPN4;refGene_tx=XM_011285247.3
C1 120821999 120822052
refGene_gn=PTPN4;refGene_tx=XM_011285247.3
C1 120822302 120822347
refGene_gn=PTPN4;refGene_tx=XM_011285247.3
C1 120826400 120826479
refGene_gn=PTPN4;refGene_tx=XM_011285247.3
C1 120826577 120826620
refGene_gn=PTPN4;refGene_tx=XM_011285247.3
C1 120841293 120841401
refGene_gn=PTPN4;refGene_tx=XM_011285247.3
C1 120887497 120887635
refGene_gn=PTPN4;refGene_tx=XM_011285247.3
C1 121025700 121026636
refGene_gn=TMEM177;refGene_tx=XM_006935234.2;refGene_gn=TMEM177;refGen
e_tx=XM_006935235.2;refGene_gn=TMEM177;refGene_tx=XM_006935236.2
C1 121050761 121050870

refGene_gn=CFAP221;refGene_tx=XM_023257852.1
C1 121057054 121057150
refGene_gn=CFAP221;refGene_tx=XM_023257852.1
C1 121060504 121060597
refGene_gn=CFAP221;refGene_tx=XM_023257852.1
C1 121065904 121065996
refGene_gn=CFAP221;refGene_tx=XM_023257852.1
C1 121067568 121067727
refGene_gn=CFAP221;refGene_tx=XM_023257852.1
C1 121080654 121080789
refGene_gn=CFAP221;refGene_tx=XM_023257852.1
C1 121081093 121081201
refGene_gn=CFAP221;refGene_tx=XM_023257852.1
C1 121081831 121081912
refGene_gn=CFAP221;refGene_tx=XM_023257852.1
C1 121082166 121082177
refGene_gn=CFAP221;refGene_tx=XM_023257852.1
C1 121087652 121087758
refGene_gn=CFAP221;refGene_tx=XM_023257852.1
C1 121089984 121090083
refGene_gn=CFAP221;refGene_tx=XM_023257852.1
C1 121092331 121092421
refGene_gn=CFAP221;refGene_tx=XM_023257852.1
C1 121097011 121097099
refGene_gn=CFAP221;refGene_tx=XM_023257852.1
C1 121100392 121100501
refGene_gn=CFAP221;refGene_tx=XM_023257852.1
C1 121101182 121101294
refGene_gn=CFAP221;refGene_tx=XM_023257852.1
C1 121101378 121101499
refGene_gn=CFAP221;refGene_tx=XM_023257852.1
C1 121106700 121106860
refGene_gn=CFAP221;refGene_tx=XM_023257852.1
C1 121122922 121123026
refGene_gn=CFAP221;refGene_tx=XM_023257852.1
C1 121142844 121142945
refGene_gn=CFAP221;refGene_tx=XM_023257852.1
C1 121145081 121145180
refGene_gn=CFAP221;refGene_tx=XM_023257852.1
C1 121145328 121145415
refGene_gn=CFAP221;refGene_tx=XM_023257852.1
C1 121152676 121152777
refGene_gn=CFAP221;refGene_tx=XM_023257852.1
C1 121156382 121156521
refGene_gn=CFAP221;refGene_tx=XM_023257852.1
C1 121174988 121175048
refGene_gn=SCTR;refGene_tx=XM_019839111.2;refGene_gn=SCTR;refGene_tx=XM_023259249.1
C1 121198803 121198924
refGene_gn=SCTR;refGene_tx=XM_019839111.2;refGene_gn=SCTR;refGene_tx=X

M_023259249.1
C1	121216938	121217046
refGene_gn=SCTR;refGene_tx=XM_019839111.2;refGene_gn=SCTR;refGene_tx=XM_023259249.1
C1	121223486	121223590
refGene_gn=SCTR;refGene_tx=XM_019839111.2;refGene_gn=SCTR;refGene_tx=XM_023259249.1
C1	121230638	121230736
refGene_gn=SCTR;refGene_tx=XM_019839111.2;refGene_gn=SCTR;refGene_tx=XM_023259249.1
C1	121231857	121231990
refGene_gn=SCTR;refGene_tx=XM_019839111.2;refGene_gn=SCTR;refGene_tx=XM_023259249.1
C1	121234112	121234266
refGene_gn=SCTR;refGene_tx=XM_019839111.2;refGene_gn=SCTR;refGene_tx=XM_023259249.1
C1	121242898	121242959
refGene_gn=SCTR;refGene_tx=XM_019839111.2;refGene_gn=SCTR;refGene_tx=XM_023259249.1
C1	121243275	121243345
refGene_gn=SCTR;refGene_tx=XM_019839111.2;refGene_gn=SCTR;refGene_tx=XM_023259249.1
C1	121247528	121247620
refGene_gn=SCTR;refGene_tx=XM_019839111.2;refGene_gn=SCTR;refGene_tx=XM_023259249.1
C1	121249052	121249179
refGene_gn=SCTR;refGene_tx=XM_019839111.2;refGene_gn=SCTR;refGene_tx=XM_023259249.1
C1	121254270	121254312
refGene_gn=SCTR;refGene_tx=XM_019839111.2;refGene_gn=SCTR;refGene_tx=XM_023259249.1
C1	121255384	121255555
refGene_gn=SCTR;refGene_tx=XM_019839111.2;refGene_gn=SCTR;refGene_tx=XM_023259249.1
C1	121258217	121258790
refGene_gn=TMEM37;refGene_tx=XM_003990716.5
C1	121264826	121264847
refGene_gn=TMEM37;refGene_tx=XM_003990716.5
C1	121318261	121318309
refGene_gn=CC1H2orf76;refGene_tx=XM_023259253.1
C1	121321104	121321178
refGene_gn=DBI;refGene_tx=XM_003990705.5
C1	121322580	121322643
refGene_gn=DBI;refGene_tx=XM_003990705.5
C1	121325063	121325181
refGene_gn=DBI;refGene_tx=XM_003990705.5
C1	121326217	121326489
refGene_gn=CC1H2orf76;refGene_tx=XM_023259250.1;refGene_gn=CC1H2orf76;refGene_tx=XM_023259251.1;refGene_gn=CC1H2orf76;refGene_tx=XM_023259252.1;refGene_gn=DBI;refGene_tx=XM_003990705.5

C1 121341286 121341490
refGene_gn=CC1H2orf76;refGene_tx=XM_023259250.1;refGene_gn=CC1H2orf76;
refGene_tx=XM_023259251.1
C1 121352848 121352993
refGene_gn=CC1H2orf76;refGene_tx=XM_023259253.1;refGene_gn=CC1H2orf76;
refGene_tx=XM_023259254.1;refGene_gn=CC1H2orf76;refGene_tx=XM_02325925
5.1;refGene_gn=CC1H2orf76;refGene_tx=XM_023259256.1;refGene_gn=CC1H2or
f76;refGene_tx=XM_023259257.1;refGene_gn=CC1H2orf76;refGene_tx=XM_0232
59258.1;refGene_gn=CC1H2orf76;refGene_tx=XM_023259259.1;refGene_gn=CC1
H2orf76;refGene_tx=XM_023259261.1
C1 121368686 121368737
refGene_gn=CC1H2orf76;refGene_tx=XM_023259250.1;refGene_gn=CC1H2orf76;
refGene_tx=XM_023259251.1;refGene_gn=CC1H2orf76;refGene_tx=XM_02325925
2.1;refGene_gn=CC1H2orf76;refGene_tx=XM_023259253.1;refGene_gn=CC1H2or
f76;refGene_tx=XM_023259254.1;refGene_gn=CC1H2orf76;refGene_tx=XM_0232
59255.1;refGene_gn=CC1H2orf76;refGene_tx=XM_023259256.1;refGene_gn=CC1
H2orf76;refGene_tx=XM_023259257.1;refGene_gn=CC1H2orf76;refGene_tx=XM_
023259258.1;refGene_gn=CC1H2orf76;refGene_tx=XM_023259259.1;refGene_gn
=CC1H2orf76;refGene_tx=XM_023259261.1
C1 121373571 121373609
refGene_gn=CC1H2orf76;refGene_tx=XM_023259250.1;refGene_gn=CC1H2orf76;
refGene_tx=XM_023259251.1;refGene_gn=CC1H2orf76;refGene_tx=XM_02325925
2.1;refGene_gn=CC1H2orf76;refGene_tx=XM_023259253.1;refGene_gn=CC1H2or
f76;refGene_tx=XM_023259254.1;refGene_gn=CC1H2orf76;refGene_tx=XM_0232
59255.1;refGene_gn=CC1H2orf76;refGene_tx=XM_023259256.1;refGene_gn=CC1
H2orf76;refGene_tx=XM_023259257.1;refGene_gn=CC1H2orf76;refGene_tx=XM_
023259258.1;refGene_gn=CC1H2orf76;refGene_tx=XM_023259259.1;refGene_gn
=CC1H2orf76;refGene_tx=XM_023259261.1
C1 121384002 121384084
refGene_gn=CC1H2orf76;refGene_tx=XM_023259250.1;refGene_gn=CC1H2orf76;
refGene_tx=XM_023259251.1;refGene_gn=CC1H2orf76;refGene_tx=XM_02325925
2.1;refGene_gn=CC1H2orf76;refGene_tx=XM_023259253.1;refGene_gn=CC1H2or
f76;refGene_tx=XM_023259254.1;refGene_gn=CC1H2orf76;refGene_tx=XM_0232
59255.1;refGene_gn=CC1H2orf76;refGene_tx=XM_023259256.1;refGene_gn=CC1
H2orf76;refGene_tx=XM_023259257.1;refGene_gn=CC1H2orf76;refGene_tx=XM_
023259258.1;refGene_gn=CC1H2orf76;refGene_tx=XM_023259259.1;refGene_gn
=CC1H2orf76;refGene_tx=XM_023259261.1
C1 121384883 121384954
refGene_gn=CC1H2orf76;refGene_tx=XM_023259251.1
C1 121392460 121392537
refGene_gn=CC1H2orf76;refGene_tx=XM_023259250.1;refGene_gn=CC1H2orf76;
refGene_tx=XM_023259252.1;refGene_gn=CC1H2orf76;refGene_tx=XM_02325925
3.1;refGene_gn=CC1H2orf76;refGene_tx=XM_023259254.1;refGene_gn=CC1H2or
f76;refGene_tx=XM_023259255.1;refGene_gn=CC1H2orf76;refGene_tx=XM_0232
59256.1;refGene_gn=CC1H2orf76;refGene_tx=XM_023259257.1;refGene_gn=CC1
H2orf76;refGene_tx=XM_023259258.1;refGene_gn=CC1H2orf76;refGene_tx=XM_
023259259.1;refGene_gn=CC1H2orf76;refGene_tx=XM_023259261.1
C1 121439622 121439904
refGene_gn=STEAP3;refGene_tx=XM_003990708.4;refGene_gn=STEAP3;refGene_
tx=XM_019837996.2;refGene_gn=STEAP3;refGene_tx=XM_019837997.2

C1 121452894 121453059
refGene_gn=STEAP3;refGene_tx=XM_003990708.4;refGene_gn=STEAP3;refGene_tx=XM_019837996.2;refGene_gn=STEAP3;refGene_tx=XM_019837997.2
C1 121460247 121460775
refGene_gn=STEAP3;refGene_tx=XM_003990708.4;refGene_gn=STEAP3;refGene_tx=XM_019837996.2;refGene_gn=STEAP3;refGene_tx=XM_019837997.2
C1 121462434 121462934
refGene_gn=STEAP3;refGene_tx=XM_003990708.4;refGene_gn=STEAP3;refGene_tx=XM_019837996.2;refGene_gn=STEAP3;refGene_tx=XM_019837997.2
C1 121481709 121481731
refGene_gn=STEAP3;refGene_tx=XM_019837997.2
C1 121493962 121494056
refGene_gn=STEAP3;refGene_tx=XM_019837996.2
C1 121575201 121575793
refGene_gn=C1QL2;refGene_tx=XM_023259262.1
C1 121575795 121575884
refGene_gn=C1QL2;refGene_tx=XM_023259262.1
C1 121576574 121576754
refGene_gn=C1QL2;refGene_tx=XM_023259262.1
C1 121726485 121726616
refGene_gn=MARCO;refGene_tx=XM_019837998.2
C1 121728773 121728950
refGene_gn=MARCO;refGene_tx=XM_019837998.2
C1 121729450 121729495
refGene_gn=MARCO;refGene_tx=XM_019837998.2
C1 121730084 121730183
refGene_gn=MARCO;refGene_tx=XM_019837998.2
C1 121731412 121731457
refGene_gn=MARCO;refGene_tx=XM_019837998.2
C1 121739885 121739948
refGene_gn=MARCO;refGene_tx=XM_019837998.2
C1 121740040 121740139
refGene_gn=MARCO;refGene_tx=XM_019837998.2
C1 121740623 121740659
refGene_gn=MARCO;refGene_tx=XM_019837998.2
C1 121740755 121740854
refGene_gn=MARCO;refGene_tx=XM_019837998.2
C1 121744272 121744380
refGene_gn=MARCO;refGene_tx=XM_019837998.2
C1 121744672 121744717
refGene_gn=MARCO;refGene_tx=XM_019837998.2
C1 121745041 121745086
refGene_gn=MARCO;refGene_tx=XM_019837998.2
C1 121750346 121750382
refGene_gn=MARCO;refGene_tx=XM_019837998.2
C1 121751708 121751930
refGene_gn=MARCO;refGene_tx=XM_019837998.2
C1 121753046 121753148
refGene_gn=MARCO;refGene_tx=XM_019837998.2
C1 121771127 121771236

refGene_gn=MARCO;refGene_tx=XM_019837998.2
C1 121857449 121858323
refGene_gn=EN1;refGene_tx=XM_006935242.2
C1 121861385 121861702
refGene_gn=EN1;refGene_tx=XM_006935242.2
C1 122550326 122550368
refGene_gn=INSIG2;refGene_tx=XM_003990710.5
C1 122551342 122551442
refGene_gn=INSIG2;refGene_tx=XM_003990710.5
C1 122551607 122551774
refGene_gn=INSIG2;refGene_tx=XM_003990710.5
C1 122555217 122555342
refGene_gn=INSIG2;refGene_tx=XM_003990710.5
C1 122562213 122562457
refGene_gn=INSIG2;refGene_tx=XM_003990710.5
C1 122635944 122635986
refGene_gn=CCDC93;refGene_tx=XM_023259263.1;refGene_gn=CCDC93;refGene_tx=XM_023259264.1;refGene_gn=CCDC93;refGene_tx=XM_023259265.1;refGene_gn=CCDC93;refGene_tx=XM_023259266.1
C1 122640174 122640288
refGene_gn=CCDC93;refGene_tx=XM_023259263.1;refGene_gn=CCDC93;refGene_tx=XM_023259264.1;refGene_gn=CCDC93;refGene_tx=XM_023259265.1;refGene_gn=CCDC93;refGene_tx=XM_023259266.1
C1 122642259 122642354
refGene_gn=CCDC93;refGene_tx=XM_023259263.1;refGene_gn=CCDC93;refGene_tx=XM_023259264.1;refGene_gn=CCDC93;refGene_tx=XM_023259265.1;refGene_gn=CCDC93;refGene_tx=XM_023259266.1
C1 122650509 122650621
refGene_gn=CCDC93;refGene_tx=XM_023259263.1;refGene_gn=CCDC93;refGene_tx=XM_023259264.1;refGene_gn=CCDC93;refGene_tx=XM_023259265.1;refGene_gn=CCDC93;refGene_tx=XM_023259266.1
C1 122655015 122655114
refGene_gn=CCDC93;refGene_tx=XM_023259263.1;refGene_gn=CCDC93;refGene_tx=XM_023259264.1;refGene_gn=CCDC93;refGene_tx=XM_023259265.1;refGene_gn=CCDC93;refGene_tx=XM_023259266.1
C1 122655926 122655983
refGene_gn=CCDC93;refGene_tx=XM_023259263.1;refGene_gn=CCDC93;refGene_tx=XM_023259264.1;refGene_gn=CCDC93;refGene_tx=XM_023259265.1;refGene_gn=CCDC93;refGene_tx=XM_023259266.1
C1 122666155 122666256
refGene_gn=CCDC93;refGene_tx=XM_023259263.1;refGene_gn=CCDC93;refGene_tx=XM_023259264.1;refGene_gn=CCDC93;refGene_tx=XM_023259265.1;refGene_gn=CCDC93;refGene_tx=XM_023259266.1
C1 122666864 122666882
refGene_gn=CCDC93;refGene_tx=XM_023259263.1;refGene_gn=CCDC93;refGene_tx=XM_023259264.1
C1 122673766 122673803
refGene_gn=CCDC93;refGene_tx=XM_023259263.1;refGene_gn=CCDC93;refGene_tx=XM_023259264.1;refGene_gn=CCDC93;refGene_tx=XM_023259265.1;refGene_gn=CCDC93;refGene_tx=XM_023259266.1

C1 122676890 122676983
refGene_gn=CCDC93;refGene_tx=XM_023259263.1;refGene_gn=CCDC93;refGene_tx=XM_023259264.1;refGene_gn=CCDC93;refGene_tx=XM_023259265.1;refGene_gn=CCDC93;refGene_tx=XM_023259266.1
C1 122677290 122677341
refGene_gn=CCDC93;refGene_tx=XM_023259263.1;refGene_gn=CCDC93;refGene_tx=XM_023259264.1;refGene_gn=CCDC93;refGene_tx=XM_023259265.1;refGene_gn=CCDC93;refGene_tx=XM_023259266.1
C1 122678217 122678304
refGene_gn=CCDC93;refGene_tx=XM_023259263.1;refGene_gn=CCDC93;refGene_tx=XM_023259264.1;refGene_gn=CCDC93;refGene_tx=XM_023259265.1;refGene_gn=CCDC93;refGene_tx=XM_023259266.1
C1 122693949 122694066
refGene_gn=CCDC93;refGene_tx=XM_023259263.1;refGene_gn=CCDC93;refGene_tx=XM_023259264.1;refGene_gn=CCDC93;refGene_tx=XM_023259265.1;refGene_gn=CCDC93;refGene_tx=XM_023259266.1
C1 122699989 122700052
refGene_gn=CCDC93;refGene_tx=XM_023259263.1;refGene_gn=CCDC93;refGene_tx=XM_023259264.1;refGene_gn=CCDC93;refGene_tx=XM_023259265.1;refGene_gn=CCDC93;refGene_tx=XM_023259266.1
C1 122703557 122703631
refGene_gn=CCDC93;refGene_tx=XM_023259263.1;refGene_gn=CCDC93;refGene_tx=XM_023259264.1;refGene_gn=CCDC93;refGene_tx=XM_023259265.1;refGene_gn=CCDC93;refGene_tx=XM_023259266.1
C1 122705092 122705174
refGene_gn=CCDC93;refGene_tx=XM_023259263.1;refGene_gn=CCDC93;refGene_tx=XM_023259264.1;refGene_gn=CCDC93;refGene_tx=XM_023259265.1;refGene_gn=CCDC93;refGene_tx=XM_023259266.1
C1 122705629 122705701
refGene_gn=CCDC93;refGene_tx=XM_023259263.1;refGene_gn=CCDC93;refGene_tx=XM_023259264.1;refGene_gn=CCDC93;refGene_tx=XM_023259265.1;refGene_gn=CCDC93;refGene_tx=XM_023259266.1
C1 122707229 122707283
refGene_gn=CCDC93;refGene_tx=XM_023259263.1;refGene_gn=CCDC93;refGene_tx=XM_023259264.1;refGene_gn=CCDC93;refGene_tx=XM_023259265.1;refGene_gn=CCDC93;refGene_tx=XM_023259266.1
C1 122708662 122708725
refGene_gn=CCDC93;refGene_tx=XM_023259263.1;refGene_gn=CCDC93;refGene_tx=XM_023259264.1;refGene_gn=CCDC93;refGene_tx=XM_023259265.1;refGene_gn=CCDC93;refGene_tx=XM_023259266.1
C1 122711417 122711526
refGene_gn=CCDC93;refGene_tx=XM_023259263.1;refGene_gn=CCDC93;refGene_tx=XM_023259264.1;refGene_gn=CCDC93;refGene_tx=XM_023259265.1;refGene_gn=CCDC93;refGene_tx=XM_023259266.1
C1 122715199 122715282
refGene_gn=CCDC93;refGene_tx=XM_023259263.1;refGene_gn=CCDC93;refGene_tx=XM_023259264.1;refGene_gn=CCDC93;refGene_tx=XM_023259265.1;refGene_gn=CCDC93;refGene_tx=XM_023259266.1
C1 122717940 122717978
refGene_gn=CCDC93;refGene_tx=XM_023259263.1;refGene_gn=CCDC93;refGene_

tx=XM_023259264.1;refGene_gn=CCDC93;refGene_tx=XM_023259265.1;refGene_gn=CCDC93;refGene_tx=XM_023259266.1
C1 122719391 122719476
refGene_gn=CCDC93;refGene_tx=XM_023259263.1;refGene_gn=CCDC93;refGene_tx=XM_023259264.1;refGene_gn=CCDC93;refGene_tx=XM_023259265.1;refGene_gn=CCDC93;refGene_tx=XM_023259266.1
C1 122723663 122723777
refGene_gn=CCDC93;refGene_tx=XM_023259263.1;refGene_gn=CCDC93;refGene_tx=XM_023259264.1;refGene_gn=CCDC93;refGene_tx=XM_023259265.1;refGene_gn=CCDC93;refGene_tx=XM_023259266.1
C1 122728503 122728557
refGene_gn=CCDC93;refGene_tx=XM_023259263.1;refGene_gn=CCDC93;refGene_tx=XM_023259264.1;refGene_gn=CCDC93;refGene_tx=XM_023259265.1;refGene_gn=CCDC93;refGene_tx=XM_023259266.1
C1 122738846 122739179
refGene_gn=LOC101084129;refGene_tx=XM_003990711.3
C1 122754101 122754878
refGene_gn=LOC101084129;refGene_tx=XM_003990711.3
C1 122781766 122781909
refGene_gn=DDX18;refGene_tx=XM_011285241.3
C1 122782959 122783137
refGene_gn=DDX18;refGene_tx=XM_011285241.3
C1 122783507 122783564
refGene_gn=DDX18;refGene_tx=XM_011285241.3
C1 122785067 122785181
refGene_gn=DDX18;refGene_tx=XM_011285241.3
C1 122786264 122786417
refGene_gn=DDX18;refGene_tx=XM_011285241.3
C1 122786723 122786885
refGene_gn=DDX18;refGene_tx=XM_011285241.3
C1 122788471 122788611
refGene_gn=DDX18;refGene_tx=XM_011285241.3
C1 122791478 122791593
refGene_gn=DDX18;refGene_tx=XM_011285241.3
C1 122791688 122791888
refGene_gn=DDX18;refGene_tx=XM_011285241.3
C1 122792007 122792108
refGene_gn=DDX18;refGene_tx=XM_011285241.3
C1 122792464 122792600
refGene_gn=DDX18;refGene_tx=XM_011285241.3
C1 122794241 122794382
refGene_gn=DDX18;refGene_tx=XM_011285241.3
C1 122797386 122797686
refGene_gn=DDX18;refGene_tx=XM_011285241.3
C1 122799210 122799295
refGene_gn=DDX18;refGene_tx=XM_011285241.3
C1 124205029 124205164
refGene_gn=DPP10;refGene_tx=XM_019837999.2;refGene_gn=DPP10;refGene_tx=XM_019838002.2;refGene_gn=DPP10;refGene_tx=XM_019838003.2
C1 124206886 124206960

refGene_gn=DPP10;refGene_tx=XM_019837999.2;refGene_gn=DPP10;refGene_tx=XM_019838002.2;refGene_gn=DPP10;refGene_tx=XM_019838003.2
C1 124212527 124212600
refGene_gn=DPP10;refGene_tx=XM_019837999.2;refGene_gn=DPP10;refGene_tx=XM_019838002.2;refGene_gn=DPP10;refGene_tx=XM_019838003.2
C1 124212706 124212765
refGene_gn=DPP10;refGene_tx=XM_019837999.2;refGene_gn=DPP10;refGene_tx=XM_019838002.2;refGene_gn=DPP10;refGene_tx=XM_019838003.2
C1 124213017 124213117
refGene_gn=DPP10;refGene_tx=XM_019837999.2;refGene_gn=DPP10;refGene_tx=XM_019838002.2;refGene_gn=DPP10;refGene_tx=XM_019838003.2
C1 124222465 124222520
refGene_gn=DPP10;refGene_tx=XM_019837999.2;refGene_gn=DPP10;refGene_tx=XM_019838002.2;refGene_gn=DPP10;refGene_tx=XM_019838003.2
C1 124223176 124223371
refGene_gn=DPP10;refGene_tx=XM_019837999.2;refGene_gn=DPP10;refGene_tx=XM_019838002.2;refGene_gn=DPP10;refGene_tx=XM_019838003.2
C1 124248982 124249052
refGene_gn=DPP10;refGene_tx=XM_019837999.2;refGene_gn=DPP10;refGene_tx=XM_019838002.2;refGene_gn=DPP10;refGene_tx=XM_019838003.2
C1 124249157 124249256
refGene_gn=DPP10;refGene_tx=XM_019837999.2;refGene_gn=DPP10;refGene_tx=XM_019838002.2;refGene_gn=DPP10;refGene_tx=XM_019838003.2
C1 124266432 124266480
refGene_gn=DPP10;refGene_tx=XM_019837999.2;refGene_gn=DPP10;refGene_tx=XM_019838002.2;refGene_gn=DPP10;refGene_tx=XM_019838003.2
C1 124267784 124267906
refGene_gn=DPP10;refGene_tx=XM_019837999.2;refGene_gn=DPP10;refGene_tx=XM_019838002.2;refGene_gn=DPP10;refGene_tx=XM_019838003.2
C1 124270948 124270996
refGene_gn=DPP10;refGene_tx=XM_019837999.2;refGene_gn=DPP10;refGene_tx=XM_019838002.2;refGene_gn=DPP10;refGene_tx=XM_019838003.2
C1 124271385 124271477
refGene_gn=DPP10;refGene_tx=XM_019837999.2;refGene_gn=DPP10;refGene_tx=XM_019838002.2;refGene_gn=DPP10;refGene_tx=XM_019838003.2
C1 124291309 124291417
refGene_gn=DPP10;refGene_tx=XM_019837999.2;refGene_gn=DPP10;refGene_tx=XM_019838002.2;refGene_gn=DPP10;refGene_tx=XM_019838003.2
C1 124296056 124296095
refGene_gn=DPP10;refGene_tx=XM_019837999.2;refGene_gn=DPP10;refGene_tx=XM_019838002.2;refGene_gn=DPP10;refGene_tx=XM_019838003.2
C1 124306344 124306468
refGene_gn=DPP10;refGene_tx=XM_019837999.2;refGene_gn=DPP10;refGene_tx=XM_019838002.2;refGene_gn=DPP10;refGene_tx=XM_019838003.2
C1 124314649 124314747
refGene_gn=DPP10;refGene_tx=XM_019837999.2;refGene_gn=DPP10;refGene_tx=XM_019838002.2;refGene_gn=DPP10;refGene_tx=XM_019838003.2
C1 124321879 124322034
refGene_gn=DPP10;refGene_tx=XM_019837999.2;refGene_gn=DPP10;refGene_tx=XM_019838002.2;refGene_gn=DPP10;refGene_tx=XM_019838003.2

C1 124332626 124332747
refGene_gn=DPP10;refGene_tx=XM_019837999.2;refGene_gn=DPP10;refGene_tx=XM_019838002.2;refGene_gn=DPP10;refGene_tx=XM_019838003.2
C1 124368730 124368812
refGene_gn=DPP10;refGene_tx=XM_019837999.2;refGene_gn=DPP10;refGene_tx=XM_019838002.2;refGene_gn=DPP10;refGene_tx=XM_019838003.2
C1 124368910 124368963
refGene_gn=DPP10;refGene_tx=XM_019837999.2;refGene_gn=DPP10;refGene_tx=XM_019838002.2;refGene_gn=DPP10;refGene_tx=XM_019838003.2
C1 124509630 124509705
refGene_gn=DPP10;refGene_tx=XM_019837999.2;refGene_gn=DPP10;refGene_tx=XM_019838002.2;refGene_gn=DPP10;refGene_tx=XM_019838003.2
C1 124536626 124536721
refGene_gn=DPP10;refGene_tx=XM_019837999.2;refGene_gn=DPP10;refGene_tx=XM_019838002.2;refGene_gn=DPP10;refGene_tx=XM_019838003.2
C1 124676095 124676191
refGene_gn=DPP10;refGene_tx=XM_019837999.2;refGene_gn=DPP10;refGene_tx=XM_019838002.2;refGene_gn=DPP10;refGene_tx=XM_019838003.2
C1 124706060 124706175
refGene_gn=DPP10;refGene_tx=XM_019837999.2;refGene_gn=DPP10;refGene_tx=XM_019838002.2;refGene_gn=DPP10;refGene_tx=XM_019838003.2
C1 124853209 124853248
refGene_gn=DPP10;refGene_tx=XM_019838003.2
C1 124853421 124853493
refGene_gn=DPP10;refGene_tx=XM_019838002.2
C1 125570371 125570431
refGene_gn=DPP10;refGene_tx=XM_019837999.2
C1 125996118 125996214
refGene_gn=ACTR3;refGene_tx=XM_023259267.1
C1 125997813 125997897
refGene_gn=ACTR3;refGene_tx=XM_023259267.1
C1 126000980 126001106
refGene_gn=ACTR3;refGene_tx=XM_023259267.1
C1 126001313 126001406
refGene_gn=ACTR3;refGene_tx=XM_023259267.1
C1 126007885 126008032
refGene_gn=ACTR3;refGene_tx=XM_023259267.1
C1 126008033 126008060
refGene_gn=ACTR3;refGene_tx=XM_023259267.1
C1 126010056 126010200
refGene_gn=ACTR3;refGene_tx=XM_023259267.1
C1 126015291 126015399
refGene_gn=ACTR3;refGene_tx=XM_023259267.1
C1 126017157 126017253
refGene_gn=ACTR3;refGene_tx=XM_023259267.1
C1 126021475 126021586
refGene_gn=ACTR3;refGene_tx=XM_023259267.1
C1 126028513 126028638
refGene_gn=ACTR3;refGene_tx=XM_023259267.1
C1 126032631 126032687

refGene_gn=ACTR3;refGene_tx=XM_023259267.1
C1 126062743 126062787
refGene_gn=ACTR3;refGene_tx=XM_023259267.1
C1 126169568 126169608
refGene_gn=SLC35F5;refGene_tx=XM_011285239.3
C1 126170481 126170572
refGene_gn=SLC35F5;refGene_tx=XM_011285239.3
C1 126170718 126170860
refGene_gn=SLC35F5;refGene_tx=XM_011285239.3
C1 126174511 126174655
refGene_gn=SLC35F5;refGene_tx=XM_011285239.3
C1 126177746 126177809
refGene_gn=SLC35F5;refGene_tx=XM_011285239.3
C1 126179879 126179961
refGene_gn=SLC35F5;refGene_tx=XM_011285239.3
C1 126180745 126180936
refGene_gn=SLC35F5;refGene_tx=XM_011285239.3
C1 126186160 126186242
refGene_gn=SLC35F5;refGene_tx=XM_011285239.3
C1 126186843 126186931
refGene_gn=SLC35F5;refGene_tx=XM_011285239.3
C1 126188652 126188717
refGene_gn=SLC35F5;refGene_tx=XM_011285239.3
C1 126190647 126190752
refGene_gn=SLC35F5;refGene_tx=XM_011285239.3
C1 126193905 126194065
refGene_gn=SLC35F5;refGene_tx=XM_011285239.3
C1 126198718 126198809
refGene_gn=SLC35F5;refGene_tx=XM_011285239.3
C1 126203266 126203421
refGene_gn=SLC35F5;refGene_tx=XM_011285239.3
C1 126203934 126204010
refGene_gn=SLC35F5;refGene_tx=XM_011285239.3
C1 126304089 126304945
refGene_gn=GPR39;refGene_tx=XM_003990722.5
C1 126502409 126502915
refGene_gn=GPR39;refGene_tx=XM_003990722.5
C1 126503340 126503576
refGene_gn=LYPD1;refGene_tx=XM_011285238.3
C1 126537809 126537947
refGene_gn=LYPD1;refGene_tx=XM_011285238.3
C1 126539269 126539321
refGene_gn=LYPD1;refGene_tx=XM_011285238.3
C1 126542631 126542648
refGene_gn=NCKAP5;refGene_tx=XM_023259268.1
C1 126594670 126594803
refGene_gn=NCKAP5;refGene_tx=XM_023259268.1
C1 126598025 126598162
refGene_gn=NCKAP5;refGene_tx=XM_023259268.1
C1 126600938 126601253

refGene_gn=NCKAP5;refGene_tx=XM_023259268.1
C1 126638752 126638828
refGene_gn=NCKAP5;refGene_tx=XM_023259268.1
C1 126646105 126646283
refGene_gn=NCKAP5;refGene_tx=XM_023259268.1
C1 126647039 126648483
refGene_gn=NCKAP5;refGene_tx=XM_023259268.1
C1 126648485 126650811
refGene_gn=NCKAP5;refGene_tx=XM_023259268.1
C1 126655498 126655687
refGene_gn=NCKAP5;refGene_tx=XM_023259268.1
C1 126667288 126667390
refGene_gn=NCKAP5;refGene_tx=XM_023259268.1
C1 126730680 126730800
refGene_gn=NCKAP5;refGene_tx=XM_023259268.1
C1 126739508 126739547
refGene_gn=NCKAP5;refGene_tx=XM_023259268.1
C1 126749685 126749754
refGene_gn=NCKAP5;refGene_tx=XM_023259268.1
C1 126829113 126829263
refGene_gn=NCKAP5;refGene_tx=XM_023259268.1
C1 126857892 126857980
refGene_gn=NCKAP5;refGene_tx=XM_023259268.1
C1 126982521 126982655
refGene_gn=NCKAP5;refGene_tx=XM_023259268.1
C1 127068735 127068799
refGene_gn=NCKAP5;refGene_tx=XM_023259268.1
C1 127160742 127160816
refGene_gn=NCKAP5;refGene_tx=XM_023259268.1
C1 127369716 127369785
refGene_gn=NCKAP5;refGene_tx=XM_023259268.1
C1 127450175 127450257
refGene_gn=LOC111561638;refGene_tx=XM_023257853.1
C1 127644365 127644429
refGene_gn=LOC111561639;refGene_tx=XM_023257855.1
C1 127645663 127647282
refGene_gn=LOC111561639;refGene_tx=XM_023257855.1
C1 127699903 127699937
refGene_gn=LOC105260851;refGene_tx=XM_023259269.1
C1 127703992 127704014
refGene_gn=LOC105260851;refGene_tx=XM_023259269.1
C1 127705790 127705866
refGene_gn=LOC105260851;refGene_tx=XM_023259269.1
C1 127705956 127706067
refGene_gn=LOC105260851;refGene_tx=XM_023259269.1
C1 127706659 127706692
refGene_gn=LOC105260851;refGene_tx=XM_023259269.1
C1 128315899 128316140
refGene_gn=MGAT5;refGene_tx=XM_019838021.2;refGene_gn=MGAT5;refGene_tx=XM_023259270.1;refGene_gn=MGAT5;refGene_tx=XM_023259271.1;refGene_gn=

MGAT5;refGene_tx=XM_023259272.1
C1	128334492	128334657
refGene_gn=MGAT5;refGene_tx=XM_019838021.2;refGene_gn=MGAT5;refGene_tx=XM_023259270.1;refGene_gn=MGAT5;refGene_tx=XM_023259271.1;refGene_gn=MGAT5;refGene_tx=XM_023259272.1
C1	128382127	128382244
refGene_gn=MGAT5;refGene_tx=XM_019838021.2;refGene_gn=MGAT5;refGene_tx=XM_023259270.1;refGene_gn=MGAT5;refGene_tx=XM_023259271.1
C1	128388678	128388755
refGene_gn=MGAT5;refGene_tx=XM_019838021.2;refGene_gn=MGAT5;refGene_tx=XM_023259270.1;refGene_gn=MGAT5;refGene_tx=XM_023259271.1;refGene_gn=MGAT5;refGene_tx=XM_023259272.1
C1	128389810	128389900
refGene_gn=MGAT5;refGene_tx=XM_019838021.2;refGene_gn=MGAT5;refGene_tx=XM_023259270.1;refGene_gn=MGAT5;refGene_tx=XM_023259271.1;refGene_gn=MGAT5;refGene_tx=XM_023259272.1
C1	128405826	128405898
refGene_gn=MGAT5;refGene_tx=XM_019838021.2;refGene_gn=MGAT5;refGene_tx=XM_023259270.1;refGene_gn=MGAT5;refGene_tx=XM_023259271.1;refGene_gn=MGAT5;refGene_tx=XM_023259272.1
C1	128408293	128408455
refGene_gn=MGAT5;refGene_tx=XM_019838021.2;refGene_gn=MGAT5;refGene_tx=XM_023259270.1;refGene_gn=MGAT5;refGene_tx=XM_023259271.1;refGene_gn=MGAT5;refGene_tx=XM_023259272.1
C1	128412066	128412236
refGene_gn=MGAT5;refGene_tx=XM_019838021.2;refGene_gn=MGAT5;refGene_tx=XM_023259270.1;refGene_gn=MGAT5;refGene_tx=XM_023259271.1;refGene_gn=MGAT5;refGene_tx=XM_023259272.1
C1	128415523	128415658
refGene_gn=MGAT5;refGene_tx=XM_019838021.2;refGene_gn=MGAT5;refGene_tx=XM_023259270.1;refGene_gn=MGAT5;refGene_tx=XM_023259271.1;refGene_gn=MGAT5;refGene_tx=XM_023259272.1
C1	128422040	128422174
refGene_gn=MGAT5;refGene_tx=XM_019838021.2;refGene_gn=MGAT5;refGene_tx=XM_023259270.1;refGene_gn=MGAT5;refGene_tx=XM_023259271.1;refGene_gn=MGAT5;refGene_tx=XM_023259272.1
C1	128438046	128438180
refGene_gn=MGAT5;refGene_tx=XM_019838021.2;refGene_gn=MGAT5;refGene_tx=XM_023259270.1;refGene_gn=MGAT5;refGene_tx=XM_023259271.1;refGene_gn=MGAT5;refGene_tx=XM_023259272.1
C1	128483086	128483236
refGene_gn=MGAT5;refGene_tx=XM_019838021.2;refGene_gn=MGAT5;refGene_tx=XM_023259270.1;refGene_gn=MGAT5;refGene_tx=XM_023259271.1;refGene_gn=MGAT5;refGene_tx=XM_023259272.1
C1	128492962	128493109
refGene_gn=MGAT5;refGene_tx=XM_019838021.2;refGene_gn=MGAT5;refGene_tx=XM_023259270.1;refGene_gn=MGAT5;refGene_tx=XM_023259271.1;refGene_gn=MGAT5;refGene_tx=XM_023259272.1
C1	128500710	128500827
refGene_gn=MGAT5;refGene_tx=XM_019838021.2;refGene_gn=MGAT5;refGene_tx

=XM_023259270.1;refGene_gn=MGAT5;refGene_tx=XM_023259271.1;refGene_gn=MGAT5;refGene_tx=XM_023259272.1
C1 128508915 128508990
refGene_gn=MGAT5;refGene_tx=XM_019838021.2;refGene_gn=MGAT5;refGene_tx=XM_023259270.1;refGene_gn=MGAT5;refGene_tx=XM_023259271.1;refGene_gn=MGAT5;refGene_tx=XM_023259272.1
C1 128521791 128521949
refGene_gn=MGAT5;refGene_tx=XM_019838021.2;refGene_gn=MGAT5;refGene_tx=XM_023259270.1;refGene_gn=MGAT5;refGene_tx=XM_023259271.1;refGene_gn=MGAT5;refGene_tx=XM_023259272.1
C1 128527556 128527755
refGene_gn=MGAT5;refGene_tx=XM_019838021.2;refGene_gn=MGAT5;refGene_tx=XM_023259270.1;refGene_gn=MGAT5;refGene_tx=XM_023259271.1;refGene_gn=MGAT5;refGene_tx=XM_023259272.1
C1 128535567 128535628
refGene_gn=TMEM163;refGene_tx=XM_023259273.1
C1 128536909 128537051
refGene_gn=TMEM163;refGene_tx=XM_023259273.1
C1 128545336 128545448
refGene_gn=TMEM163;refGene_tx=XM_023259273.1
C1 128583070 128583167
refGene_gn=TMEM163;refGene_tx=XM_023259273.1
C1 128636137 128636229
refGene_gn=TMEM163;refGene_tx=XM_023259273.1
C1 128637605 128637649
refGene_gn=TMEM163;refGene_tx=XM_023259273.1
C1 128767881 128768001
refGene_gn=TMEM163;refGene_tx=XM_023259273.1
C1 128773518 128773534
refGene_gn=TMEM163;refGene_tx=XM_023259273.1
C1 128773535 128773721
refGene_gn=TMEM163;refGene_tx=XM_023259273.1
C1 128877801 128877858
refGene_gn=ACMSD;refGene_tx=XM_023259275.1;refGene_gn=ACMSD;refGene_tx=XM_023259276.1
C1 128883949 128883994
refGene_gn=ACMSD;refGene_tx=XM_023259275.1;refGene_gn=ACMSD;refGene_tx=XM_023259276.1
C1 128892757 128892854
refGene_gn=ACMSD;refGene_tx=XM_023259275.1;refGene_gn=ACMSD;refGene_tx=XM_023259276.1;refGene_gn=ACMSD;refGene_tx=XM_023259277.1
C1 128898105 128898155
refGene_gn=ACMSD;refGene_tx=XM_023259275.1;refGene_gn=ACMSD;refGene_tx=XM_023259276.1
C1 128899584 128899821
refGene_gn=ACMSD;refGene_tx=XM_023259275.1;refGene_gn=ACMSD;refGene_tx=XM_023259276.1;refGene_gn=ACMSD;refGene_tx=XM_023259277.1
C1 128904119 128904213
refGene_gn=ACMSD;refGene_tx=XM_023259275.1;refGene_gn=ACMSD;refGene_tx=XM_023259276.1;refGene_gn=ACMSD;refGene_tx=XM_023259277.1

C1 128906788 128906884
refGene_gn=ACMSD;refGene_tx=XM_023259275.1;refGene_gn=ACMSD;refGene_tx=XM_023259276.1;refGene_gn=ACMSD;refGene_tx=XM_023259277.1
C1 128907915 128908088
refGene_gn=ACMSD;refGene_tx=XM_023259275.1;refGene_gn=ACMSD;refGene_tx=XM_023259276.1;refGene_gn=ACMSD;refGene_tx=XM_023259277.1
C1 128927423 128927522
refGene_gn=ACMSD;refGene_tx=XM_023259275.1;refGene_gn=ACMSD;refGene_tx=XM_023259276.1;refGene_gn=ACMSD;refGene_tx=XM_023259277.1
C1 128931781 128931844
refGene_gn=ACMSD;refGene_tx=XM_023259275.1;refGene_gn=ACMSD;refGene_tx=XM_023259277.1
C1 128934529 128934571
refGene_gn=ACMSD;refGene_tx=XM_023259276.1
C1 128941627 128941785
refGene_gn=CCNT2;refGene_tx=XM_003990727.5;refGene_gn=CCNT2;refGene_tx=XM_011285232.3;refGene_gn=CCNT2;refGene_tx=XM_019838023.2;refGene_gn=CCNT2;refGene_tx=XM_019838024.2
C1 128942589 128942671
refGene_gn=CCNT2;refGene_tx=XM_003990727.5;refGene_gn=CCNT2;refGene_tx=XM_011285232.3;refGene_gn=CCNT2;refGene_tx=XM_019838023.2;refGene_gn=CCNT2;refGene_tx=XM_019838024.2
C1 128944601 128944628
refGene_gn=CCNT2;refGene_tx=XM_019838023.2;refGene_gn=CCNT2;refGene_tx=XM_019838024.2
C1 128963321 128963450
refGene_gn=CCNT2;refGene_tx=XM_003990727.5;refGene_gn=CCNT2;refGene_tx=XM_011285232.3;refGene_gn=CCNT2;refGene_tx=XM_019838023.2;refGene_gn=CCNT2;refGene_tx=XM_019838024.2
C1 128964841 128964902
refGene_gn=CCNT2;refGene_tx=XM_003990727.5;refGene_gn=CCNT2;refGene_tx=XM_011285232.3;refGene_gn=CCNT2;refGene_tx=XM_019838023.2;refGene_gn=CCNT2;refGene_tx=XM_019838024.2;refGene_gn=CCNT2;refGene_tx=XM_019838026.2;refGene_gn=CCNT2;refGene_tx=XM_019838028.2;refGene_gn=CCNT2;refGene_tx=XM_023259274.1
C1 128967179 128967242
refGene_gn=CCNT2;refGene_tx=XM_003990727.5;refGene_gn=CCNT2;refGene_tx=XM_011285232.3;refGene_gn=CCNT2;refGene_tx=XM_019838023.2;refGene_gn=CCNT2;refGene_tx=XM_019838024.2;refGene_gn=CCNT2;refGene_tx=XM_019838026.2;refGene_gn=CCNT2;refGene_tx=XM_019838028.2;refGene_gn=CCNT2;refGene_tx=XM_023259274.1
C1 128970237 128970298
refGene_gn=CCNT2;refGene_tx=XM_019838031.2
C1 128970420 128970466
refGene_gn=CCNT2;refGene_tx=XM_003990727.5;refGene_gn=CCNT2;refGene_tx=XM_011285232.3;refGene_gn=CCNT2;refGene_tx=XM_019838023.2;refGene_gn=CCNT2;refGene_tx=XM_019838024.2;refGene_gn=CCNT2;refGene_tx=XM_019838026.2;refGene_gn=CCNT2;refGene_tx=XM_019838028.2;refGene_gn=CCNT2;refGene_tx=XM_019838031.2;refGene_gn=CCNT2;refGene_tx=XM_023259274.1
C1 128972074 128972238

refGene_gn=CCNT2;refGene_tx=XM_003990727.5;refGene_gn=CCNT2;refGene_tx=XM_011285232.3;refGene_gn=CCNT2;refGene_tx=XM_019838023.2;refGene_gn=CCNT2;refGene_tx=XM_019838024.2;refGene_gn=CCNT2;refGene_tx=XM_019838026.2;refGene_gn=CCNT2;refGene_tx=XM_019838028.2;refGene_gn=CCNT2;refGene_tx=XM_019838031.2;refGene_gn=CCNT2;refGene_tx=XM_023259274.1
C1 128976201 128976272
refGene_gn=CCNT2;refGene_tx=XM_003990727.5;refGene_gn=CCNT2;refGene_tx=XM_011285232.3;refGene_gn=CCNT2;refGene_tx=XM_019838023.2;refGene_gn=CCNT2;refGene_tx=XM_019838024.2;refGene_gn=CCNT2;refGene_tx=XM_019838026.2;refGene_gn=CCNT2;refGene_tx=XM_019838028.2;refGene_gn=CCNT2;refGene_tx=XM_019838031.2;refGene_gn=CCNT2;refGene_tx=XM_023259274.1
C1 128976794 128978222
refGene_gn=CCNT2;refGene_tx=XM_003990727.5;refGene_gn=CCNT2;refGene_tx=XM_011285232.3;refGene_gn=CCNT2;refGene_tx=XM_019838023.2;refGene_gn=CCNT2;refGene_tx=XM_019838024.2;refGene_gn=CCNT2;refGene_tx=XM_019838026.2;refGene_gn=CCNT2;refGene_tx=XM_019838028.2;refGene_gn=CCNT2;refGene_tx=XM_019838031.2;refGene_gn=CCNT2;refGene_tx=XM_023259274.1
C1 128986724 128986791
refGene_gn=MAP3K19;refGene_tx=XM_023259278.1
C1 128991652 128992350
refGene_gn=MAP3K19;refGene_tx=XM_023259278.1
C1 128993430 128993580
refGene_gn=MAP3K19;refGene_tx=XM_023259278.1
C1 128995805 128996747
refGene_gn=MAP3K19;refGene_tx=XM_023259278.1
C1 128996748 128998254
refGene_gn=MAP3K19;refGene_tx=XM_023259278.1
C1 129000250 129000294
refGene_gn=MAP3K19;refGene_tx=XM_023259278.1
C1 129004158 129004409
refGene_gn=MAP3K19;refGene_tx=XM_023259278.1
C1 129005407 129005486
refGene_gn=MAP3K19;refGene_tx=XM_023259278.1
C1 129013545 129013642
refGene_gn=MAP3K19;refGene_tx=XM_023259278.1
C1 129023000 129023116
refGene_gn=MAP3K19;refGene_tx=XM_023259278.1
C1 129025157 129025273
refGene_gn=MAP3K19;refGene_tx=XM_023259278.1
C1 129027664 129027819
refGene_gn=MAP3K19;refGene_tx=XM_023259278.1
C1 129030290 129030479
refGene_gn=MAP3K19;refGene_tx=XM_023259278.1
C1 129042232 129042406
refGene_gn=MAP3K19;refGene_tx=XM_023259278.1
C1 129059592 129059610
refGene_gn=RAB3GAP1;refGene_tx=XM_023259279.1;refGene_gn=RAB3GAP1;refGene_tx=XM_023259280.1
C1 129059710 129059766
refGene_gn=RAB3GAP1;refGene_tx=XM_023259279.1;refGene_gn=RAB3GAP1;refG

ene_tx=XM_023259280.1;refGene_gn=RAB3GAP1;refGene_tx=XM_023259281.1
C1 129064206 129064282
refGene_gn=RAB3GAP1;refGene_tx=XM_023259279.1;refGene_gn=RAB3GAP1;refGene_tx=XM_023259280.1
C1 129081671 129081804
refGene_gn=RAB3GAP1;refGene_tx=XM_023259279.1;refGene_gn=RAB3GAP1;refGene_tx=XM_023259280.1;refGene_gn=RAB3GAP1;refGene_tx=XM_023259281.1
C1 129085748 129085827
refGene_gn=RAB3GAP1;refGene_tx=XM_023259279.1;refGene_gn=RAB3GAP1;refGene_tx=XM_023259280.1;refGene_gn=RAB3GAP1;refGene_tx=XM_023259281.1;refGene_gn=RAB3GAP1;refGene_tx=XM_023259282.1;refGene_gn=RAB3GAP1;refGene_tx=XM_023259283.1
C1 129110109 129110229
refGene_gn=RAB3GAP1;refGene_tx=XM_023259279.1;refGene_gn=RAB3GAP1;refGene_tx=XM_023259280.1;refGene_gn=RAB3GAP1;refGene_tx=XM_023259281.1;refGene_gn=RAB3GAP1;refGene_tx=XM_023259282.1;refGene_gn=RAB3GAP1;refGene_tx=XM_023259283.1
C1 129111727 129111893
refGene_gn=RAB3GAP1;refGene_tx=XM_023259279.1;refGene_gn=RAB3GAP1;refGene_tx=XM_023259280.1;refGene_gn=RAB3GAP1;refGene_tx=XM_023259281.1;refGene_gn=RAB3GAP1;refGene_tx=XM_023259282.1;refGene_gn=RAB3GAP1;refGene_tx=XM_023259283.1;refGene_gn=RAB3GAP1;refGene_tx=XM_023259284.1
C1 129116475 129116575
refGene_gn=RAB3GAP1;refGene_tx=XM_023259279.1;refGene_gn=RAB3GAP1;refGene_tx=XM_023259280.1;refGene_gn=RAB3GAP1;refGene_tx=XM_023259281.1;refGene_gn=RAB3GAP1;refGene_tx=XM_023259282.1;refGene_gn=RAB3GAP1;refGene_tx=XM_023259283.1;refGene_gn=RAB3GAP1;refGene_tx=XM_023259284.1
C1 129119627 129119709
refGene_gn=RAB3GAP1;refGene_tx=XM_023259279.1;refGene_gn=RAB3GAP1;refGene_tx=XM_023259280.1;refGene_gn=RAB3GAP1;refGene_tx=XM_023259281.1;refGene_gn=RAB3GAP1;refGene_tx=XM_023259282.1;refGene_gn=RAB3GAP1;refGene_tx=XM_023259283.1;refGene_gn=RAB3GAP1;refGene_tx=XM_023259284.1
C1 129120593 129120884
refGene_gn=LOC101091220;refGene_tx=XM_019838039.2
C1 129126048 129126117
refGene_gn=RAB3GAP1;refGene_tx=XM_023259279.1;refGene_gn=RAB3GAP1;refGene_tx=XM_023259280.1;refGene_gn=RAB3GAP1;refGene_tx=XM_023259281.1;refGene_gn=RAB3GAP1;refGene_tx=XM_023259282.1;refGene_gn=RAB3GAP1;refGene_tx=XM_023259283.1;refGene_gn=RAB3GAP1;refGene_tx=XM_023259284.1
C1 129127637 129127711
refGene_gn=RAB3GAP1;refGene_tx=XM_023259279.1;refGene_gn=RAB3GAP1;refGene_tx=XM_023259280.1;refGene_gn=RAB3GAP1;refGene_tx=XM_023259281.1;refGene_gn=RAB3GAP1;refGene_tx=XM_023259282.1;refGene_gn=RAB3GAP1;refGene_tx=XM_023259283.1;refGene_gn=RAB3GAP1;refGene_tx=XM_023259284.1
C1 129130366 129130459
refGene_gn=RAB3GAP1;refGene_tx=XM_023259279.1;refGene_gn=RAB3GAP1;refGene_tx=XM_023259280.1;refGene_gn=RAB3GAP1;refGene_tx=XM_023259281.1;refGene_gn=RAB3GAP1;refGene_tx=XM_023259282.1;refGene_gn=RAB3GAP1;refGene_tx=XM_023259283.1;refGene_gn=RAB3GAP1;refGene_tx=XM_023259284.1
C1 129130939 129131109

refGene_gn=RAB3GAP1;refGene_tx=XM_023259279.1;refGene_gn=RAB3GAP1;refGene_tx=XM_023259280.1;refGene_gn=RAB3GAP1;refGene_tx=XM_023259281.1;refGene_gn=RAB3GAP1;refGene_tx=XM_023259282.1;refGene_gn=RAB3GAP1;refGene_tx=XM_023259283.1;refGene_gn=RAB3GAP1;refGene_tx=XM_023259284.1
C1 129132882 129132972
refGene_gn=RAB3GAP1;refGene_tx=XM_023259279.1;refGene_gn=RAB3GAP1;refGene_tx=XM_023259280.1;refGene_gn=RAB3GAP1;refGene_tx=XM_023259281.1;refGene_gn=RAB3GAP1;refGene_tx=XM_023259282.1;refGene_gn=RAB3GAP1;refGene_tx=XM_023259283.1;refGene_gn=RAB3GAP1;refGene_tx=XM_023259284.1
C1 129134146 129134319
refGene_gn=RAB3GAP1;refGene_tx=XM_023259279.1;refGene_gn=RAB3GAP1;refGene_tx=XM_023259280.1;refGene_gn=RAB3GAP1;refGene_tx=XM_023259281.1;refGene_gn=RAB3GAP1;refGene_tx=XM_023259282.1;refGene_gn=RAB3GAP1;refGene_tx=XM_023259283.1;refGene_gn=RAB3GAP1;refGene_tx=XM_023259284.1
C1 129135505 129135560
refGene_gn=RAB3GAP1;refGene_tx=XM_023259279.1;refGene_gn=RAB3GAP1;refGene_tx=XM_023259280.1;refGene_gn=RAB3GAP1;refGene_tx=XM_023259281.1;refGene_gn=RAB3GAP1;refGene_tx=XM_023259282.1;refGene_gn=RAB3GAP1;refGene_tx=XM_023259283.1;refGene_gn=RAB3GAP1;refGene_tx=XM_023259284.1
C1 129135805 129136165
refGene_gn=RAB3GAP1;refGene_tx=XM_023259279.1;refGene_gn=RAB3GAP1;refGene_tx=XM_023259280.1;refGene_gn=RAB3GAP1;refGene_tx=XM_023259281.1;refGene_gn=RAB3GAP1;refGene_tx=XM_023259282.1;refGene_gn=RAB3GAP1;refGene_tx=XM_023259283.1;refGene_gn=RAB3GAP1;refGene_tx=XM_023259284.1
C1 129145587 129145725
refGene_gn=RAB3GAP1;refGene_tx=XM_023259279.1;refGene_gn=RAB3GAP1;refGene_tx=XM_023259280.1;refGene_gn=RAB3GAP1;refGene_tx=XM_023259281.1;refGene_gn=RAB3GAP1;refGene_tx=XM_023259282.1;refGene_gn=RAB3GAP1;refGene_tx=XM_023259283.1;refGene_gn=RAB3GAP1;refGene_tx=XM_023259284.1
C1 129149408 129149636
refGene_gn=RAB3GAP1;refGene_tx=XM_023259279.1;refGene_gn=RAB3GAP1;refGene_tx=XM_023259280.1;refGene_gn=RAB3GAP1;refGene_tx=XM_023259281.1;refGene_gn=RAB3GAP1;refGene_tx=XM_023259282.1;refGene_gn=RAB3GAP1;refGene_tx=XM_023259283.1;refGene_gn=RAB3GAP1;refGene_tx=XM_023259284.1
C1 129163489 129163586
refGene_gn=RAB3GAP1;refGene_tx=XM_023259279.1;refGene_gn=RAB3GAP1;refGene_tx=XM_023259280.1;refGene_gn=RAB3GAP1;refGene_tx=XM_023259281.1;refGene_gn=RAB3GAP1;refGene_tx=XM_023259282.1;refGene_gn=RAB3GAP1;refGene_tx=XM_023259283.1;refGene_gn=RAB3GAP1;refGene_tx=XM_023259284.1
C1 129163679 129163783
refGene_gn=RAB3GAP1;refGene_tx=XM_023259279.1;refGene_gn=RAB3GAP1;refGene_tx=XM_023259280.1;refGene_gn=RAB3GAP1;refGene_tx=XM_023259281.1;refGene_gn=RAB3GAP1;refGene_tx=XM_023259282.1;refGene_gn=RAB3GAP1;refGene_tx=XM_023259283.1;refGene_gn=RAB3GAP1;refGene_tx=XM_023259284.1
C1 129163915 129164031
refGene_gn=RAB3GAP1;refGene_tx=XM_023259279.1;refGene_gn=RAB3GAP1;refGene_tx=XM_023259280.1;refGene_gn=RAB3GAP1;refGene_tx=XM_023259281.1;refGene_gn=RAB3GAP1;refGene_tx=XM_023259282.1;refGene_gn=RAB3GAP1;refGene_tx=XM_023259283.1;refGene_gn=RAB3GAP1;refGene_tx=XM_023259284.1
C1 129165443 129165546

refGene_gn=RAB3GAP1;refGene_tx=XM_023259279.1;refGene_gn=RAB3GAP1;refGene_tx=XM_023259280.1;refGene_gn=RAB3GAP1;refGene_tx=XM_023259281.1;refGene_gn=RAB3GAP1;refGene_tx=XM_023259282.1;refGene_gn=RAB3GAP1;refGene_tx=XM_023259283.1;refGene_gn=RAB3GAP1;refGene_tx=XM_023259284.1
C1 129168667 129168688
refGene_gn=RAB3GAP1;refGene_tx=XM_023259279.1;refGene_gn=RAB3GAP1;refGene_tx=XM_023259281.1;refGene_gn=RAB3GAP1;refGene_tx=XM_023259282.1;refGene_gn=RAB3GAP1;refGene_tx=XM_023259283.1;refGene_gn=RAB3GAP1;refGene_tx=XM_023259284.1
C1 129169548 129169785
refGene_gn=RAB3GAP1;refGene_tx=XM_023259279.1;refGene_gn=RAB3GAP1;refGene_tx=XM_023259280.1;refGene_gn=RAB3GAP1;refGene_tx=XM_023259281.1;refGene_gn=RAB3GAP1;refGene_tx=XM_023259282.1;refGene_gn=RAB3GAP1;refGene_tx=XM_023259283.1;refGene_gn=RAB3GAP1;refGene_tx=XM_023259284.1
C1 129207460 129207559
refGene_gn=ZRANB3;refGene_tx=XM_023259285.1;refGene_gn=ZRANB3;refGene_tx=XM_023259286.1;refGene_gn=ZRANB3;refGene_tx=XM_023259287.1;refGene_gn=ZRANB3;refGene_tx=XM_023259288.1
C1 129209320 129209452
refGene_gn=ZRANB3;refGene_tx=XM_023259285.1;refGene_gn=ZRANB3;refGene_tx=XM_023259286.1;refGene_gn=ZRANB3;refGene_tx=XM_023259287.1;refGene_gn=ZRANB3;refGene_tx=XM_023259288.1
C1 129212786 129213189
refGene_gn=ZRANB3;refGene_tx=XM_023259285.1;refGene_gn=ZRANB3;refGene_tx=XM_023259286.1;refGene_gn=ZRANB3;refGene_tx=XM_023259287.1;refGene_gn=ZRANB3;refGene_tx=XM_023259288.1
C1 129214229 129214340
refGene_gn=ZRANB3;refGene_tx=XM_023259285.1;refGene_gn=ZRANB3;refGene_tx=XM_023259286.1;refGene_gn=ZRANB3;refGene_tx=XM_023259287.1;refGene_gn=ZRANB3;refGene_tx=XM_023259288.1
C1 129222521 129222664
refGene_gn=ZRANB3;refGene_tx=XM_023259285.1;refGene_gn=ZRANB3;refGene_tx=XM_023259286.1;refGene_gn=ZRANB3;refGene_tx=XM_023259287.1;refGene_gn=ZRANB3;refGene_tx=XM_023259288.1
C1 129223845 129223947
refGene_gn=ZRANB3;refGene_tx=XM_023259285.1;refGene_gn=ZRANB3;refGene_tx=XM_023259286.1;refGene_gn=ZRANB3;refGene_tx=XM_023259287.1;refGene_gn=ZRANB3;refGene_tx=XM_023259288.1
C1 129227125 129227217
refGene_gn=ZRANB3;refGene_tx=XM_023259285.1;refGene_gn=ZRANB3;refGene_tx=XM_023259286.1;refGene_gn=ZRANB3;refGene_tx=XM_023259287.1;refGene_gn=ZRANB3;refGene_tx=XM_023259288.1
C1 129231131 129231329
refGene_gn=ZRANB3;refGene_tx=XM_023259285.1;refGene_gn=ZRANB3;refGene_tx=XM_023259286.1;refGene_gn=ZRANB3;refGene_tx=XM_023259288.1
C1 129234304 129234692
refGene_gn=ZRANB3;refGene_tx=XM_023259285.1;refGene_gn=ZRANB3;refGene_tx=XM_023259286.1;refGene_gn=ZRANB3;refGene_tx=XM_023259287.1;refGene_gn=ZRANB3;refGene_tx=XM_023259288.1
C1 129263107 129263260

refGene_gn=ZRANB3;refGene_tx=XM_023259285.1;refGene_gn=ZRANB3;refGene_tx=XM_023259286.1;refGene_gn=ZRANB3;refGene_tx=XM_023259287.1;refGene_gn=ZRANB3;refGene_tx=XM_023259288.1
C1 129264045 129264225
refGene_gn=ZRANB3;refGene_tx=XM_023259285.1;refGene_gn=ZRANB3;refGene_tx=XM_023259286.1;refGene_gn=ZRANB3;refGene_tx=XM_023259287.1;refGene_gn=ZRANB3;refGene_tx=XM_023259288.1
C1 129264996 129264999
refGene_gn=ZRANB3;refGene_tx=XM_023259288.1
C1 129266812 129266932
refGene_gn=ZRANB3;refGene_tx=XM_023259285.1;refGene_gn=ZRANB3;refGene_tx=XM_023259286.1;refGene_gn=ZRANB3;refGene_tx=XM_023259287.1
C1 129268469 129268589
refGene_gn=ZRANB3;refGene_tx=XM_023259285.1;refGene_gn=ZRANB3;refGene_tx=XM_023259286.1;refGene_gn=ZRANB3;refGene_tx=XM_023259287.1
C1 129289542 129289659
refGene_gn=ZRANB3;refGene_tx=XM_023259285.1;refGene_gn=ZRANB3;refGene_tx=XM_023259286.1;refGene_gn=ZRANB3;refGene_tx=XM_023259287.1
C1 129291469 129291641
refGene_gn=ZRANB3;refGene_tx=XM_023259285.1;refGene_gn=ZRANB3;refGene_tx=XM_023259286.1;refGene_gn=ZRANB3;refGene_tx=XM_023259287.1
C1 129319802 129319888
refGene_gn=ZRANB3;refGene_tx=XM_023259285.1;refGene_gn=ZRANB3;refGene_tx=XM_023259286.1;refGene_gn=ZRANB3;refGene_tx=XM_023259287.1
C1 129323987 129324219
refGene_gn=ZRANB3;refGene_tx=XM_023259285.1;refGene_gn=ZRANB3;refGene_tx=XM_023259286.1;refGene_gn=ZRANB3;refGene_tx=XM_023259287.1
C1 129332512 129332691
refGene_gn=ZRANB3;refGene_tx=XM_023259285.1;refGene_gn=ZRANB3;refGene_tx=XM_023259286.1;refGene_gn=ZRANB3;refGene_tx=XM_023259287.1
C1 129382598 129382617
refGene_gn=ZRANB3;refGene_tx=XM_023259285.1;refGene_gn=ZRANB3;refGene_tx=XM_023259286.1;refGene_gn=ZRANB3;refGene_tx=XM_023259287.1
C1 129488871 129489035
refGene_gn=ZRANB3;refGene_tx=XM_023259285.1;refGene_gn=ZRANB3;refGene_tx=XM_023259286.1;refGene_gn=ZRANB3;refGene_tx=XM_023259287.1
C1 129590782 129590953
refGene_gn=R3HDM1;refGene_tx=XM_011285215.3;refGene_gn=R3HDM1;refGene_tx=XM_011285216.3;refGene_gn=R3HDM1;refGene_tx=XM_011285217.3;refGene_gn=R3HDM1;refGene_tx=XM_019838041.2;refGene_gn=R3HDM1;refGene_tx=XM_019838042.2;refGene_gn=R3HDM1;refGene_tx=XM_019838043.2;refGene_gn=R3HDM1;refGene_tx=XM_019838044.2;refGene_gn=R3HDM1;refGene_tx=XM_019838045.2;refGene_gn=R3HDM1;refGene_tx=XM_019838046.2;refGene_gn=R3HDM1;refGene_tx=XM_019838048.2;refGene_gn=R3HDM1;refGene_tx=XM_019838049.2;refGene_gn=R3HDM1;refGene_tx=XM_019838050.2;refGene_gn=R3HDM1;refGene_tx=XM_019838051.2
C1 129602443 129602485
refGene_gn=R3HDM1;refGene_tx=XM_011285215.3;refGene_gn=R3HDM1;refGene_tx=XM_011285216.3;refGene_gn=R3HDM1;refGene_tx=XM_011285217.3;refGene_gn=R3HDM1;refGene_tx=XM_019838041.2;refGene_gn=R3HDM1;refGene_tx=XM_01

9838042.2;refGene_gn=R3HDM1;refGene_tx=XM_019838043.2;refGene_gn=R3HDM1;refGene_tx=XM_019838044.2;refGene_gn=R3HDM1;refGene_tx=XM_019838045.2;refGene_gn=R3HDM1;refGene_tx=XM_019838048.2;refGene_gn=R3HDM1;refGene_tx=XM_019838049.2;refGene_gn=R3HDM1;refGene_tx=XM_019838050.2
C1 129602948 129603038
refGene_gn=R3HDM1;refGene_tx=XM_011285215.3;refGene_gn=R3HDM1;refGene_tx=XM_011285216.3;refGene_gn=R3HDM1;refGene_tx=XM_011285217.3;refGene_gn=R3HDM1;refGene_tx=XM_019838041.2;refGene_gn=R3HDM1;refGene_tx=XM_019838042.2;refGene_gn=R3HDM1;refGene_tx=XM_019838043.2;refGene_gn=R3HDM1;refGene_tx=XM_019838044.2;refGene_gn=R3HDM1;refGene_tx=XM_019838045.2;refGene_gn=R3HDM1;refGene_tx=XM_019838048.2;refGene_gn=R3HDM1;refGene_tx=XM_019838049.2;refGene_gn=R3HDM1;refGene_tx=XM_019838050.2
C1 129607168 129607283
refGene_gn=R3HDM1;refGene_tx=XM_011285215.3;refGene_gn=R3HDM1;refGene_tx=XM_011285216.3;refGene_gn=R3HDM1;refGene_tx=XM_011285217.3;refGene_gn=R3HDM1;refGene_tx=XM_019838041.2;refGene_gn=R3HDM1;refGene_tx=XM_019838042.2;refGene_gn=R3HDM1;refGene_tx=XM_019838043.2;refGene_gn=R3HDM1;refGene_tx=XM_019838044.2;refGene_gn=R3HDM1;refGene_tx=XM_019838045.2;refGene_gn=R3HDM1;refGene_tx=XM_019838046.2;refGene_gn=R3HDM1;refGene_tx=XM_019838048.2;refGene_gn=R3HDM1;refGene_tx=XM_019838049.2;refGene_gn=R3HDM1;refGene_tx=XM_019838050.2;refGene_gn=R3HDM1;refGene_tx=XM_019838051.2
C1 129608344 129608423
refGene_gn=R3HDM1;refGene_tx=XM_011285215.3;refGene_gn=R3HDM1;refGene_tx=XM_011285216.3;refGene_gn=R3HDM1;refGene_tx=XM_011285217.3;refGene_gn=R3HDM1;refGene_tx=XM_019838041.2;refGene_gn=R3HDM1;refGene_tx=XM_019838042.2;refGene_gn=R3HDM1;refGene_tx=XM_019838043.2;refGene_gn=R3HDM1;refGene_tx=XM_019838044.2;refGene_gn=R3HDM1;refGene_tx=XM_019838045.2;refGene_gn=R3HDM1;refGene_tx=XM_019838046.2;refGene_gn=R3HDM1;refGene_tx=XM_019838048.2;refGene_gn=R3HDM1;refGene_tx=XM_019838049.2;refGene_gn=R3HDM1;refGene_tx=XM_019838050.2;refGene_gn=R3HDM1;refGene_tx=XM_019838051.2
C1 129615903 129615963
refGene_gn=R3HDM1;refGene_tx=XM_011285215.3;refGene_gn=R3HDM1;refGene_tx=XM_011285216.3;refGene_gn=R3HDM1;refGene_tx=XM_011285217.3;refGene_gn=R3HDM1;refGene_tx=XM_019838041.2;refGene_gn=R3HDM1;refGene_tx=XM_019838042.2;refGene_gn=R3HDM1;refGene_tx=XM_019838043.2;refGene_gn=R3HDM1;refGene_tx=XM_019838044.2;refGene_gn=R3HDM1;refGene_tx=XM_019838045.2;refGene_gn=R3HDM1;refGene_tx=XM_019838046.2;refGene_gn=R3HDM1;refGene_tx=XM_019838048.2;refGene_gn=R3HDM1;refGene_tx=XM_019838049.2;refGene_gn=R3HDM1;refGene_tx=XM_019838050.2;refGene_gn=R3HDM1;refGene_tx=XM_019838051.2
C1 129616046 129616187
refGene_gn=R3HDM1;refGene_tx=XM_011285215.3;refGene_gn=R3HDM1;refGene_tx=XM_011285216.3;refGene_gn=R3HDM1;refGene_tx=XM_011285217.3;refGene_gn=R3HDM1;refGene_tx=XM_019838041.2;refGene_gn=R3HDM1;refGene_tx=XM_019838042.2;refGene_gn=R3HDM1;refGene_tx=XM_019838043.2;refGene_gn=R3HDM1;refGene_tx=XM_019838044.2;refGene_gn=R3HDM1;refGene_tx=XM_019838045.2;refGene_gn=R3HDM1;refGene_tx=XM_019838046.2;refGene_gn=R3HDM1;refGene_tx=XM_019838048.2;refGene_gn=R3HDM1;refGene_tx=XM_019838049.2;refGen

e_gn=R3HDM1;refGene_tx=XM_019838050.2;refGene_gn=R3HDM1;refGene_tx=XM_019838051.2
C1 129620955 129621014
refGene_gn=R3HDM1;refGene_tx=XM_019838052.2
C1 129622810 129622919
refGene_gn=R3HDM1;refGene_tx=XM_011285215.3;refGene_gn=R3HDM1;refGene_tx=XM_011285216.3;refGene_gn=R3HDM1;refGene_tx=XM_011285217.3;refGene_gn=R3HDM1;refGene_tx=XM_019838041.2;refGene_gn=R3HDM1;refGene_tx=XM_019838042.2;refGene_gn=R3HDM1;refGene_tx=XM_019838043.2;refGene_gn=R3HDM1;refGene_tx=XM_019838044.2;refGene_gn=R3HDM1;refGene_tx=XM_019838045.2;refGene_gn=R3HDM1;refGene_tx=XM_019838046.2;refGene_gn=R3HDM1;refGene_tx=XM_019838048.2;refGene_gn=R3HDM1;refGene_tx=XM_019838049.2;refGene_gn=R3HDM1;refGene_tx=XM_019838050.2;refGene_gn=R3HDM1;refGene_tx=XM_019838051.2;refGene_gn=R3HDM1;refGene_tx=XM_019838052.2
C1 129623010 129623106
refGene_gn=R3HDM1;refGene_tx=XM_011285215.3;refGene_gn=R3HDM1;refGene_tx=XM_011285216.3;refGene_gn=R3HDM1;refGene_tx=XM_011285217.3;refGene_gn=R3HDM1;refGene_tx=XM_019838041.2;refGene_gn=R3HDM1;refGene_tx=XM_019838042.2;refGene_gn=R3HDM1;refGene_tx=XM_019838043.2;refGene_gn=R3HDM1;refGene_tx=XM_019838044.2;refGene_gn=R3HDM1;refGene_tx=XM_019838045.2;refGene_gn=R3HDM1;refGene_tx=XM_019838046.2;refGene_gn=R3HDM1;refGene_tx=XM_019838048.2;refGene_gn=R3HDM1;refGene_tx=XM_019838049.2;refGene_gn=R3HDM1;refGene_tx=XM_019838050.2;refGene_gn=R3HDM1;refGene_tx=XM_019838051.2;refGene_gn=R3HDM1;refGene_tx=XM_019838052.2
C1 129625254 129625292
refGene_gn=R3HDM1;refGene_tx=XM_011285215.3;refGene_gn=R3HDM1;refGene_tx=XM_011285216.3;refGene_gn=R3HDM1;refGene_tx=XM_011285217.3;refGene_gn=R3HDM1;refGene_tx=XM_019838041.2;refGene_gn=R3HDM1;refGene_tx=XM_019838042.2;refGene_gn=R3HDM1;refGene_tx=XM_019838043.2;refGene_gn=R3HDM1;refGene_tx=XM_019838044.2;refGene_gn=R3HDM1;refGene_tx=XM_019838045.2;refGene_gn=R3HDM1;refGene_tx=XM_019838046.2;refGene_gn=R3HDM1;refGene_tx=XM_019838048.2;refGene_gn=R3HDM1;refGene_tx=XM_019838049.2;refGene_gn=R3HDM1;refGene_tx=XM_019838050.2;refGene_gn=R3HDM1;refGene_tx=XM_019838051.2;refGene_gn=R3HDM1;refGene_tx=XM_019838052.2
C1 129625374 129625425
refGene_gn=R3HDM1;refGene_tx=XM_011285215.3;refGene_gn=R3HDM1;refGene_tx=XM_011285216.3;refGene_gn=R3HDM1;refGene_tx=XM_011285217.3;refGene_gn=R3HDM1;refGene_tx=XM_019838041.2;refGene_gn=R3HDM1;refGene_tx=XM_019838042.2;refGene_gn=R3HDM1;refGene_tx=XM_019838044.2;refGene_gn=R3HDM1;refGene_tx=XM_019838045.2;refGene_gn=R3HDM1;refGene_tx=XM_019838046.2;refGene_gn=R3HDM1;refGene_tx=XM_019838048.2;refGene_gn=R3HDM1;refGene_tx=XM_019838049.2;refGene_gn=R3HDM1;refGene_tx=XM_019838050.2;refGene_gn=R3HDM1;refGene_tx=XM_019838051.2;refGene_gn=R3HDM1;refGene_tx=XM_019838052.2
C1 129625533 129625760
refGene_gn=R3HDM1;refGene_tx=XM_011285215.3;refGene_gn=R3HDM1;refGene_tx=XM_011285216.3;refGene_gn=R3HDM1;refGene_tx=XM_011285217.3;refGene_gn=R3HDM1;refGene_tx=XM_019838041.2;refGene_gn=R3HDM1;refGene_tx=XM_019838042.2;refGene_gn=R3HDM1;refGene_tx=XM_019838043.2;refGene_gn=R3HDM1;refGene_tx=XM_019838044.2;refGene_gn=R3HDM1;refGene_tx=XM_019838045.

2;refGene_gn=R3HDM1;refGene_tx=XM_019838046.2;refGene_gn=R3HDM1;refGene_tx=XM_019838048.2;refGene_gn=R3HDM1;refGene_tx=XM_019838049.2;refGene_gn=R3HDM1;refGene_tx=XM_019838050.2;refGene_gn=R3HDM1;refGene_tx=XM_019838051.2;refGene_gn=R3HDM1;refGene_tx=XM_019838052.2
C1 129627828 129628083
refGene_gn=R3HDM1;refGene_tx=XM_011285215.3;refGene_gn=R3HDM1;refGene_tx=XM_011285216.3;refGene_gn=R3HDM1;refGene_tx=XM_011285217.3;refGene_gn=R3HDM1;refGene_tx=XM_019838041.2;refGene_gn=R3HDM1;refGene_tx=XM_019838042.2;refGene_gn=R3HDM1;refGene_tx=XM_019838043.2;refGene_gn=R3HDM1;refGene_tx=XM_019838044.2;refGene_gn=R3HDM1;refGene_tx=XM_019838045.2;refGene_gn=R3HDM1;refGene_tx=XM_019838046.2;refGene_gn=R3HDM1;refGene_tx=XM_019838048.2;refGene_gn=R3HDM1;refGene_tx=XM_019838052.2
C1 129631042 129631194
refGene_gn=R3HDM1;refGene_tx=XM_011285215.3;refGene_gn=R3HDM1;refGene_tx=XM_011285216.3;refGene_gn=R3HDM1;refGene_tx=XM_011285217.3;refGene_gn=R3HDM1;refGene_tx=XM_019838041.2;refGene_gn=R3HDM1;refGene_tx=XM_019838042.2;refGene_gn=R3HDM1;refGene_tx=XM_019838043.2;refGene_gn=R3HDM1;refGene_tx=XM_019838044.2;refGene_gn=R3HDM1;refGene_tx=XM_019838045.2;refGene_gn=R3HDM1;refGene_tx=XM_019838046.2;refGene_gn=R3HDM1;refGene_tx=XM_019838048.2;refGene_gn=R3HDM1;refGene_tx=XM_019838049.2;refGene_gn=R3HDM1;refGene_tx=XM_019838050.2;refGene_gn=R3HDM1;refGene_tx=XM_019838051.2;refGene_gn=R3HDM1;refGene_tx=XM_019838052.2
C1 129635019 129635121
refGene_gn=R3HDM1;refGene_tx=XM_011285215.3;refGene_gn=R3HDM1;refGene_tx=XM_011285216.3;refGene_gn=R3HDM1;refGene_tx=XM_011285217.3;refGene_gn=R3HDM1;refGene_tx=XM_019838041.2;refGene_gn=R3HDM1;refGene_tx=XM_019838042.2;refGene_gn=R3HDM1;refGene_tx=XM_019838043.2;refGene_gn=R3HDM1;refGene_tx=XM_019838046.2;refGene_gn=R3HDM1;refGene_tx=XM_019838048.2;refGene_gn=R3HDM1;refGene_tx=XM_019838049.2;refGene_gn=R3HDM1;refGene_tx=XM_019838050.2;refGene_gn=R3HDM1;refGene_tx=XM_019838052.2
C1 129637364 129637655
refGene_gn=R3HDM1;refGene_tx=XM_011285215.3;refGene_gn=R3HDM1;refGene_tx=XM_011285216.3;refGene_gn=R3HDM1;refGene_tx=XM_011285217.3;refGene_gn=R3HDM1;refGene_tx=XM_019838041.2;refGene_gn=R3HDM1;refGene_tx=XM_019838042.2;refGene_gn=R3HDM1;refGene_tx=XM_019838043.2;refGene_gn=R3HDM1;refGene_tx=XM_019838044.2;refGene_gn=R3HDM1;refGene_tx=XM_019838045.2;refGene_gn=R3HDM1;refGene_tx=XM_019838046.2;refGene_gn=R3HDM1;refGene_tx=XM_019838048.2;refGene_gn=R3HDM1;refGene_tx=XM_019838049.2;refGene_gn=R3HDM1;refGene_tx=XM_019838050.2;refGene_gn=R3HDM1;refGene_tx=XM_019838051.2;refGene_gn=R3HDM1;refGene_tx=XM_019838052.2
C1 129644878 129645002
refGene_gn=R3HDM1;refGene_tx=XM_011285215.3;refGene_gn=R3HDM1;refGene_tx=XM_011285216.3;refGene_gn=R3HDM1;refGene_tx=XM_011285217.3;refGene_gn=R3HDM1;refGene_tx=XM_019838041.2;refGene_gn=R3HDM1;refGene_tx=XM_019838042.2;refGene_gn=R3HDM1;refGene_tx=XM_019838043.2;refGene_gn=R3HDM1;refGene_tx=XM_019838044.2;refGene_gn=R3HDM1;refGene_tx=XM_019838045.2;refGene_gn=R3HDM1;refGene_tx=XM_019838046.2;refGene_gn=R3HDM1;refGene_tx=XM_019838048.2;refGene_gn=R3HDM1;refGene_tx=XM_019838049.2;refGene_gn=R3HDM1;refGene_tx=XM_019838050.2;refGene_gn=R3HDM1;refGene_tx=XM_019838051.2;refGene_gn=R3HDM1;refGene_tx=XM_019838052.2

C1 129665457 129665612
refGene_gn=R3HDM1;refGene_tx=XM_011285215.3;refGene_gn=R3HDM1;refGene_tx=XM_011285216.3;refGene_gn=R3HDM1;refGene_tx=XM_011285217.3;refGene_gn=R3HDM1;refGene_tx=XM_019838041.2;refGene_gn=R3HDM1;refGene_tx=XM_019838042.2;refGene_gn=R3HDM1;refGene_tx=XM_019838043.2;refGene_gn=R3HDM1;refGene_tx=XM_019838044.2;refGene_gn=R3HDM1;refGene_tx=XM_019838045.2;refGene_gn=R3HDM1;refGene_tx=XM_019838046.2;refGene_gn=R3HDM1;refGene_tx=XM_019838048.2;refGene_gn=R3HDM1;refGene_tx=XM_019838049.2;refGene_gn=R3HDM1;refGene_tx=XM_019838050.2;refGene_gn=R3HDM1;refGene_tx=XM_019838051.2;refGene_gn=R3HDM1;refGene_tx=XM_019838052.2
C1 129670147 129670299
refGene_gn=R3HDM1;refGene_tx=XM_011285215.3;refGene_gn=R3HDM1;refGene_tx=XM_011285216.3;refGene_gn=R3HDM1;refGene_tx=XM_011285217.3;refGene_gn=R3HDM1;refGene_tx=XM_019838041.2;refGene_gn=R3HDM1;refGene_tx=XM_019838042.2;refGene_gn=R3HDM1;refGene_tx=XM_019838043.2;refGene_gn=R3HDM1;refGene_tx=XM_019838044.2;refGene_gn=R3HDM1;refGene_tx=XM_019838045.2;refGene_gn=R3HDM1;refGene_tx=XM_019838046.2;refGene_gn=R3HDM1;refGene_tx=XM_019838048.2;refGene_gn=R3HDM1;refGene_tx=XM_019838049.2;refGene_gn=R3HDM1;refGene_tx=XM_019838050.2;refGene_gn=R3HDM1;refGene_tx=XM_019838051.2;refGene_gn=R3HDM1;refGene_tx=XM_019838052.2
C1 129700418 129700522
refGene_gn=R3HDM1;refGene_tx=XM_011285215.3;refGene_gn=R3HDM1;refGene_tx=XM_011285216.3;refGene_gn=R3HDM1;refGene_tx=XM_011285217.3;refGene_gn=R3HDM1;refGene_tx=XM_019838041.2;refGene_gn=R3HDM1;refGene_tx=XM_019838042.2;refGene_gn=R3HDM1;refGene_tx=XM_019838043.2;refGene_gn=R3HDM1;refGene_tx=XM_019838044.2;refGene_gn=R3HDM1;refGene_tx=XM_019838045.2;refGene_gn=R3HDM1;refGene_tx=XM_019838046.2;refGene_gn=R3HDM1;refGene_tx=XM_019838048.2;refGene_gn=R3HDM1;refGene_tx=XM_019838049.2;refGene_gn=R3HDM1;refGene_tx=XM_019838050.2;refGene_gn=R3HDM1;refGene_tx=XM_019838051.2;refGene_gn=R3HDM1;refGene_tx=XM_019838052.2
C1 129701904 129702077
refGene_gn=R3HDM1;refGene_tx=XM_011285215.3;refGene_gn=R3HDM1;refGene_tx=XM_011285216.3;refGene_gn=R3HDM1;refGene_tx=XM_011285217.3;refGene_gn=R3HDM1;refGene_tx=XM_019838041.2;refGene_gn=R3HDM1;refGene_tx=XM_019838042.2;refGene_gn=R3HDM1;refGene_tx=XM_019838043.2;refGene_gn=R3HDM1;refGene_tx=XM_019838044.2;refGene_gn=R3HDM1;refGene_tx=XM_019838045.2;refGene_gn=R3HDM1;refGene_tx=XM_019838046.2;refGene_gn=R3HDM1;refGene_tx=XM_019838048.2;refGene_gn=R3HDM1;refGene_tx=XM_019838049.2;refGene_gn=R3HDM1;refGene_tx=XM_019838050.2;refGene_gn=R3HDM1;refGene_tx=XM_019838051.2;refGene_gn=R3HDM1;refGene_tx=XM_019838052.2
C1 129710676 129710821
refGene_gn=R3HDM1;refGene_tx=XM_011285215.3;refGene_gn=R3HDM1;refGene_tx=XM_011285216.3;refGene_gn=R3HDM1;refGene_tx=XM_011285217.3;refGene_gn=R3HDM1;refGene_tx=XM_019838041.2;refGene_gn=R3HDM1;refGene_tx=XM_019838042.2;refGene_gn=R3HDM1;refGene_tx=XM_019838043.2;refGene_gn=R3HDM1;refGene_tx=XM_019838044.2;refGene_gn=R3HDM1;refGene_tx=XM_019838045.2;refGene_gn=R3HDM1;refGene_tx=XM_019838046.2;refGene_gn=R3HDM1;refGene_tx=XM_019838048.2;refGene_gn=R3HDM1;refGene_tx=XM_019838049.2;refGene_gn=R3HDM1;refGene_tx=XM_019838050.2;refGene_gn=R3HDM1;refGene_tx=XM_019838051.2;refGene_gn=R3HDM1;refGene_tx=XM_019838052.2

C1 129714416 129714499
refGene_gn=R3HDM1;refGene_tx=XM_011285215.3;refGene_gn=R3HDM1;refGene_tx=XM_011285216.3;refGene_gn=R3HDM1;refGene_tx=XM_011285217.3;refGene_gn=R3HDM1;refGene_tx=XM_019838041.2;refGene_gn=R3HDM1;refGene_tx=XM_019838042.2;refGene_gn=R3HDM1;refGene_tx=XM_019838043.2;refGene_gn=R3HDM1;refGene_tx=XM_019838044.2;refGene_gn=R3HDM1;refGene_tx=XM_019838045.2;refGene_gn=R3HDM1;refGene_tx=XM_019838046.2;refGene_gn=R3HDM1;refGene_tx=XM_019838048.2;refGene_gn=R3HDM1;refGene_tx=XM_019838049.2;refGene_gn=R3HDM1;refGene_tx=XM_019838050.2;refGene_gn=R3HDM1;refGene_tx=XM_019838051.2;refGene_gn=R3HDM1;refGene_tx=XM_019838052.2
C1 129714957 129715042
refGene_gn=R3HDM1;refGene_tx=XM_011285215.3;refGene_gn=R3HDM1;refGene_tx=XM_011285216.3;refGene_gn=R3HDM1;refGene_tx=XM_011285217.3;refGene_gn=R3HDM1;refGene_tx=XM_019838041.2;refGene_gn=R3HDM1;refGene_tx=XM_019838042.2;refGene_gn=R3HDM1;refGene_tx=XM_019838043.2;refGene_gn=R3HDM1;refGene_tx=XM_019838044.2;refGene_gn=R3HDM1;refGene_tx=XM_019838045.2;refGene_gn=R3HDM1;refGene_tx=XM_019838046.2;refGene_gn=R3HDM1;refGene_tx=XM_019838048.2;refGene_gn=R3HDM1;refGene_tx=XM_019838049.2;refGene_gn=R3HDM1;refGene_tx=XM_019838050.2;refGene_gn=R3HDM1;refGene_tx=XM_019838051.2;refGene_gn=R3HDM1;refGene_tx=XM_019838052.2
C1 129715756 129716118
refGene_gn=R3HDM1;refGene_tx=XM_011285215.3;refGene_gn=R3HDM1;refGene_tx=XM_011285216.3;refGene_gn=R3HDM1;refGene_tx=XM_011285217.3;refGene_gn=R3HDM1;refGene_tx=XM_019838041.2;refGene_gn=R3HDM1;refGene_tx=XM_019838042.2;refGene_gn=R3HDM1;refGene_tx=XM_019838043.2;refGene_gn=R3HDM1;refGene_tx=XM_019838044.2;refGene_gn=R3HDM1;refGene_tx=XM_019838045.2;refGene_gn=R3HDM1;refGene_tx=XM_019838046.2;refGene_gn=R3HDM1;refGene_tx=XM_019838048.2;refGene_gn=R3HDM1;refGene_tx=XM_019838049.2;refGene_gn=R3HDM1;refGene_tx=XM_019838050.2;refGene_gn=R3HDM1;refGene_tx=XM_019838051.2;refGene_gn=R3HDM1;refGene_tx=XM_019838052.2
C1 129730937 129731019
refGene_gn=UBXN4;refGene_tx=XM_011285214.2
C1 129740733 129740836
refGene_gn=UBXN4;refGene_tx=XM_011285214.2
C1 129744077 129744106
refGene_gn=UBXN4;refGene_tx=XM_011285214.2
C1 129744608 129744727
refGene_gn=UBXN4;refGene_tx=XM_011285214.2
C1 129745326 129745501
refGene_gn=UBXN4;refGene_tx=XM_011285214.2
C1 129753839 129753933
refGene_gn=UBXN4;refGene_tx=XM_011285214.2
C1 129760158 129760213
refGene_gn=UBXN4;refGene_tx=XM_011285214.2
C1 129760980 129761145
refGene_gn=UBXN4;refGene_tx=XM_011285214.2
C1 129763185 129763313
refGene_gn=UBXN4;refGene_tx=XM_011285214.2
C1 129767722 129767825
refGene_gn=UBXN4;refGene_tx=XM_011285214.2

C1	129770752	129770884
refGene_gn=UBXN4;refGene_tx=XM_011285214.2
C1	129772201	129772401
refGene_gn=UBXN4;refGene_tx=XM_011285214.2
C1	129774494	129774633
refGene_gn=UBXN4;refGene_tx=XM_011285214.2
C1	129780194	129780415
refGene_gn=LCT;refGene_tx=XM_003990733.3
C1	129781022	129781250
refGene_gn=LCT;refGene_tx=XM_003990733.3
C1	129782058	129782282
refGene_gn=LCT;refGene_tx=XM_003990733.3
C1	129783877	129784012
refGene_gn=LCT;refGene_tx=XM_003990733.3
C1	129786626	129786736
refGene_gn=LCT;refGene_tx=XM_003990733.3
C1	129788804	129789007
refGene_gn=LCT;refGene_tx=XM_003990733.3
C1	129790308	129790507
refGene_gn=LCT;refGene_tx=XM_003990733.3
C1	129792913	129793204
refGene_gn=LCT;refGene_tx=XM_003990733.3
C1	129797141	129797410
refGene_gn=LCT;refGene_tx=XM_003990733.3
C1	129798082	129799633
refGene_gn=LCT;refGene_tx=XM_003990733.3
C1	129803555	129804201
refGene_gn=LCT;refGene_tx=XM_003990733.3
C1	129808151	129808881
refGene_gn=LCT;refGene_tx=XM_003990733.3
C1	129813171	129813247
refGene_gn=LCT;refGene_tx=XM_003990733.3
C1	129814515	129814618
refGene_gn=LCT;refGene_tx=XM_003990733.3
C1	129819037	129819121
refGene_gn=LCT;refGene_tx=XM_003990733.3
C1	129822366	129822446
refGene_gn=LCT;refGene_tx=XM_003990733.3
C1	129830141	129830781
refGene_gn=LCT;refGene_tx=XM_003990733.3
C1	129837038	129837155
refGene_gn=MCM6;refGene_tx=XM_003990734.4
C1	129841665	129841805
refGene_gn=MCM6;refGene_tx=XM_003990734.4
C1	129843629	129843785
refGene_gn=MCM6;refGene_tx=XM_003990734.4
C1	129844901	129845037
refGene_gn=MCM6;refGene_tx=XM_003990734.4
C1	129846710	129846872
refGene_gn=MCM6;refGene_tx=XM_003990734.4

C1 129848543 129848672
refGene_gn=MCM6;refGene_tx=XM_003990734.4
C1 129850836 129850992
refGene_gn=MCM6;refGene_tx=XM_003990734.4
C1 129851907 129852015
refGene_gn=MCM6;refGene_tx=XM_003990734.4
C1 129853343 129853485
refGene_gn=MCM6;refGene_tx=XM_003990734.4
C1 129858208 129858350
refGene_gn=MCM6;refGene_tx=XM_003990734.4
C1 129861465 129861616
refGene_gn=MCM6;refGene_tx=XM_003990734.4
C1 129862525 129862671
refGene_gn=MCM6;refGene_tx=XM_003990734.4
C1 129862953 129863119
refGene_gn=MCM6;refGene_tx=XM_003990734.4
C1 129867323 129867573
refGene_gn=MCM6;refGene_tx=XM_003990734.4
C1 129868947 129869058
refGene_gn=MCM6;refGene_tx=XM_003990734.4
C1 129873309 129873456
refGene_gn=MCM6;refGene_tx=XM_003990734.4
C1 129876914 129877021
refGene_gn=MCM6;refGene_tx=XM_003990734.4
C1 129901739 129901831
refGene_gn=DARS;refGene_tx=XM_011285210.3;refGene_gn=DARS;refGene_tx=XM_011285212.3;refGene_gn=DARS;refGene_tx=XM_019838057.2
C1 129905115 129905187
refGene_gn=DARS;refGene_tx=XM_011285210.3;refGene_gn=DARS;refGene_tx=XM_011285212.3;refGene_gn=DARS;refGene_tx=XM_019838057.2
C1 129905389 129905501
refGene_gn=DARS;refGene_tx=XM_011285210.3;refGene_gn=DARS;refGene_tx=XM_011285212.3;refGene_gn=DARS;refGene_tx=XM_019838057.2
C1 129906582 129906663
refGene_gn=DARS;refGene_tx=XM_011285210.3;refGene_gn=DARS;refGene_tx=XM_011285212.3;refGene_gn=DARS;refGene_tx=XM_019838057.2
C1 129907978 129908021
refGene_gn=DARS;refGene_tx=XM_011285210.3;refGene_gn=DARS;refGene_tx=XM_011285212.3;refGene_gn=DARS;refGene_tx=XM_019838057.2
C1 129909522 129909669
refGene_gn=DARS;refGene_tx=XM_011285210.3;refGene_gn=DARS;refGene_tx=XM_011285212.3;refGene_gn=DARS;refGene_tx=XM_019838057.2
C1 129912969 129913117
refGene_gn=DARS;refGene_tx=XM_011285210.3;refGene_gn=DARS;refGene_tx=XM_011285212.3;refGene_gn=DARS;refGene_tx=XM_019838057.2
C1 129915629 129915764
refGene_gn=DARS;refGene_tx=XM_011285210.3;refGene_gn=DARS;refGene_tx=XM_011285212.3;refGene_gn=DARS;refGene_tx=XM_019838057.2
C1 129916621 129916733
refGene_gn=DARS;refGene_tx=XM_011285210.3;refGene_gn=DARS;refGene_tx=X

M_011285212.3;refGene_gn=DARS;refGene_tx=XM_019838057.2
C1	129918192	129918252
refGene_gn=DARS;refGene_tx=XM_011285210.3;refGene_gn=DARS;refGene_tx=XM_011285212.3;refGene_gn=DARS;refGene_tx=XM_019838057.2
C1	129919234	129919315
refGene_gn=DARS;refGene_tx=XM_011285210.3;refGene_gn=DARS;refGene_tx=XM_011285212.3;refGene_gn=DARS;refGene_tx=XM_019838057.2
C1	129922536	129922639
refGene_gn=DARS;refGene_tx=XM_011285210.3;refGene_gn=DARS;refGene_tx=XM_011285212.3;refGene_gn=DARS;refGene_tx=XM_019838057.2
C1	129949524	129949627
refGene_gn=DARS;refGene_tx=XM_011285210.3;refGene_gn=DARS;refGene_tx=XM_011285212.3;refGene_gn=DARS;refGene_tx=XM_019838057.2
C1	129960668	129960761
refGene_gn=DARS;refGene_tx=XM_011285210.3;refGene_gn=DARS;refGene_tx=XM_011285212.3;refGene_gn=DARS;refGene_tx=XM_019838057.2
C1	129964013	129964071
refGene_gn=DARS;refGene_tx=XM_011285210.3;refGene_gn=DARS;refGene_tx=XM_011285212.3;refGene_gn=DARS;refGene_tx=XM_019838057.2
C1	129965826	129965883
refGene_gn=DARS;refGene_tx=XM_019838057.2
C1	129966189	129966408
refGene_gn=DARS;refGene_tx=XM_011285210.3;refGene_gn=DARS;refGene_tx=XM_011285212.3
C1	130089335	130090382
refGene_gn=CXCR4;refGene_tx=NM_001009826.1
C1	130092512	130092527
refGene_gn=CXCR4;refGene_tx=NM_001009826.1
C1	130808310	130808449
refGene_gn=THSD7B;refGene_tx=XM_023259291.1;refGene_gn=THSD7B;refGene_tx=XM_023259292.1
C1	130970358	130971169
refGene_gn=THSD7B;refGene_tx=XM_023259291.1;refGene_gn=THSD7B;refGene_tx=XM_023259292.1
C1	131008227	131008473
refGene_gn=THSD7B;refGene_tx=XM_023259291.1;refGene_gn=THSD7B;refGene_tx=XM_023259292.1
C1	131022083	131022253
refGene_gn=THSD7B;refGene_tx=XM_023259291.1;refGene_gn=THSD7B;refGene_tx=XM_023259292.1
C1	131073620	131073776
refGene_gn=THSD7B;refGene_tx=XM_023259291.1;refGene_gn=THSD7B;refGene_tx=XM_023259292.1
C1	131084054	131084252
refGene_gn=THSD7B;refGene_tx=XM_023259291.1;refGene_gn=THSD7B;refGene_tx=XM_023259292.1
C1	131161058	131161250
refGene_gn=THSD7B;refGene_tx=XM_023259291.1;refGene_gn=THSD7B;refGene_tx=XM_023259292.1
C1	131163274	131163509

refGene_gn=THSD7B;refGene_tx=XM_023259291.1;refGene_gn=THSD7B;refGene_tx=XM_023259292.1
C1	131171718	131171834
refGene_gn=THSD7B;refGene_tx=XM_023259291.1;refGene_gn=THSD7B;refGene_tx=XM_023259292.1
C1	131199908	131200038
refGene_gn=THSD7B;refGene_tx=XM_023259291.1;refGene_gn=THSD7B;refGene_tx=XM_023259292.1
C1	131204287	131204391
refGene_gn=THSD7B;refGene_tx=XM_023259291.1;refGene_gn=THSD7B;refGene_tx=XM_023259292.1
C1	131304075	131304270
refGene_gn=THSD7B;refGene_tx=XM_023259291.1;refGene_gn=THSD7B;refGene_tx=XM_023259292.1
C1	131310085	131310352
refGene_gn=THSD7B;refGene_tx=XM_023259291.1;refGene_gn=THSD7B;refGene_tx=XM_023259292.1
C1	131360578	131360757
refGene_gn=THSD7B;refGene_tx=XM_023259291.1;refGene_gn=THSD7B;refGene_tx=XM_023259292.1
C1	131450229	131450363
refGene_gn=THSD7B;refGene_tx=XM_023259291.1;refGene_gn=THSD7B;refGene_tx=XM_023259292.1
C1	131457303	131457454
refGene_gn=THSD7B;refGene_tx=XM_023259291.1;refGene_gn=THSD7B;refGene_tx=XM_023259292.1
C1	131509480	131509622
refGene_gn=THSD7B;refGene_tx=XM_023259291.1;refGene_gn=THSD7B;refGene_tx=XM_023259292.1
C1	131511634	131511750
refGene_gn=THSD7B;refGene_tx=XM_023259291.1;refGene_gn=THSD7B;refGene_tx=XM_023259292.1
C1	131513893	131514011
refGene_gn=THSD7B;refGene_tx=XM_023259291.1;refGene_gn=THSD7B;refGene_tx=XM_023259292.1
C1	131529631	131529777
refGene_gn=THSD7B;refGene_tx=XM_023259291.1;refGene_gn=THSD7B;refGene_tx=XM_023259292.1
C1	131534872	131535032
refGene_gn=THSD7B;refGene_tx=XM_023259291.1;refGene_gn=THSD7B;refGene_tx=XM_023259292.1
C1	131536037	131536211
refGene_gn=THSD7B;refGene_tx=XM_023259291.1;refGene_gn=THSD7B;refGene_tx=XM_023259292.1
C1	131536311	131536407
refGene_gn=THSD7B;refGene_tx=XM_023259291.1;refGene_gn=THSD7B;refGene_tx=XM_023259292.1
C1	131539060	131539143
refGene_gn=THSD7B;refGene_tx=XM_023259291.1;refGene_gn=THSD7B;refGene_tx=XM_023259292.1

C1 131541190 131541383
refGene_gn=THSD7B;refGene_tx=XM_023259291.1;refGene_gn=THSD7B;refGene_tx=XM_023259292.1
C1 131544668 131544756
refGene_gn=THSD7B;refGene_tx=XM_023259291.1;refGene_gn=THSD7B;refGene_tx=XM_023259292.1
C1 131552459 131552541
refGene_gn=THSD7B;refGene_tx=XM_023259291.1;refGene_gn=THSD7B;refGene_tx=XM_023259292.1
C1 131894645 131894782
refGene_gn=HNMT;refGene_tx=XM_003990738.4;refGene_gn=HNMT;refGene_tx=XM_006935262.3
C1 131908384 131908437
refGene_gn=HNMT;refGene_tx=XM_003990738.4;refGene_gn=HNMT;refGene_tx=XM_006935262.3
C1 131929096 131929204
refGene_gn=HNMT;refGene_tx=XM_003990738.4;refGene_gn=HNMT;refGene_tx=XM_006935262.3
C1 131930246 131930377
refGene_gn=HNMT;refGene_tx=XM_003990738.4;refGene_gn=HNMT;refGene_tx=XM_006935262.3
C1 131932820 131932914
refGene_gn=HNMT;refGene_tx=XM_003990738.4;refGene_gn=HNMT;refGene_tx=XM_006935262.3
C1 131940505 131940861
refGene_gn=HNMT;refGene_tx=XM_003990738.4;refGene_gn=HNMT;refGene_tx=XM_006935262.3
C1 132470288 132470366
refGene_gn=SPOPL;refGene_tx=XM_019838058.2;refGene_gn=SPOPL;refGene_tx=XM_019838059.2;refGene_gn=SPOPL;refGene_tx=XM_023259293.1;refGene_gn=SPOPL;refGene_tx=XM_023259294.1;refGene_gn=SPOPL;refGene_tx=XM_023259296.1
C1 132470557 132470679
refGene_gn=SPOPL;refGene_tx=XM_019838058.2;refGene_gn=SPOPL;refGene_tx=XM_019838059.2;refGene_gn=SPOPL;refGene_tx=XM_023259293.1;refGene_gn=SPOPL;refGene_tx=XM_023259294.1;refGene_gn=SPOPL;refGene_tx=XM_023259296.1
C1 132471170 132471322
refGene_gn=SPOPL;refGene_tx=XM_019838058.2;refGene_gn=SPOPL;refGene_tx=XM_019838059.2;refGene_gn=SPOPL;refGene_tx=XM_023259293.1;refGene_gn=SPOPL;refGene_tx=XM_023259294.1;refGene_gn=SPOPL;refGene_tx=XM_023259296.1
C1 132473358 132473486
refGene_gn=SPOPL;refGene_tx=XM_019838058.2;refGene_gn=SPOPL;refGene_tx=XM_019838059.2;refGene_gn=SPOPL;refGene_tx=XM_023259293.1;refGene_gn=SPOPL;refGene_tx=XM_023259294.1;refGene_gn=SPOPL;refGene_tx=XM_023259296.1
C1 132478898 132479076
refGene_gn=SPOPL;refGene_tx=XM_019838058.2;refGene_gn=SPOPL;refGene_tx=XM_019838059.2;refGene_gn=SPOPL;refGene_tx=XM_023259293.1;refGene_gn=

SPOPL;refGene_tx=XM_023259294.1;refGene_gn=SPOPL;refGene_tx=XM_0232592
96.1
C1 132479156 132479212
refGene_gn=SPOPL;refGene_tx=XM_019838058.2;refGene_gn=SPOPL;refGene_tx
=XM_019838059.2;refGene_gn=SPOPL;refGene_tx=XM_023259293.1;refGene_gn=
SPOPL;refGene_tx=XM_023259294.1;refGene_gn=SPOPL;refGene_tx=XM_0232592
96.1
C1 132480579 132480702
refGene_gn=SPOPL;refGene_tx=XM_019838058.2;refGene_gn=SPOPL;refGene_tx
=XM_019838059.2;refGene_gn=SPOPL;refGene_tx=XM_023259293.1;refGene_gn=
SPOPL;refGene_tx=XM_023259294.1;refGene_gn=SPOPL;refGene_tx=XM_0232592
96.1
C1 132490699 132490842
refGene_gn=SPOPL;refGene_tx=XM_019838058.2;refGene_gn=SPOPL;refGene_tx
=XM_019838059.2;refGene_gn=SPOPL;refGene_tx=XM_023259293.1;refGene_gn=
SPOPL;refGene_tx=XM_023259294.1;refGene_gn=SPOPL;refGene_tx=XM_0232592
96.1
C1 132490931 132490985
refGene_gn=SPOPL;refGene_tx=XM_019838058.2;refGene_gn=SPOPL;refGene_tx
=XM_019838059.2;refGene_gn=SPOPL;refGene_tx=XM_023259293.1;refGene_gn=
SPOPL;refGene_tx=XM_023259294.1;refGene_gn=SPOPL;refGene_tx=XM_0232592
96.1
C1 132494449 132494594
refGene_gn=SPOPL;refGene_tx=XM_019838058.2;refGene_gn=SPOPL;refGene_tx
=XM_019838059.2;refGene_gn=SPOPL;refGene_tx=XM_023259293.1;refGene_gn=
SPOPL;refGene_tx=XM_023259294.1;refGene_gn=SPOPL;refGene_tx=XM_0232592
96.1
C1 132595799 132596543
refGene_gn=NXPH2;refGene_tx=XM_023259297.1
C1 132707811 132707862
refGene_gn=NXPH2;refGene_tx=XM_023259297.1
C1 133437311 133437408
refGene_gn=LOC101095597;refGene_tx=XM_023257856.1
C1 133437639 133438046
refGene_gn=LOC101095597;refGene_tx=XM_023257856.1
C1 133463415 133463565
refGene_gn=LOC101095597;refGene_tx=XM_023257856.1
C1 133465432 133465600
refGene_gn=LOC101095597;refGene_tx=XM_023257856.1
C1 133971240 133971381
refGene_gn=LRP1B;refGene_tx=XM_023259298.1;refGene_gn=LRP1B;refGene_tx
=XM_023259299.1;refGene_gn=LRP1B;refGene_tx=XM_023259300.1
C1 133972837 133972936
refGene_gn=LRP1B;refGene_tx=XM_023259298.1;refGene_gn=LRP1B;refGene_tx
=XM_023259300.1
C1 133975787 133975932
refGene_gn=LRP1B;refGene_tx=XM_023259298.1;refGene_gn=LRP1B;refGene_tx
=XM_023259299.1;refGene_gn=LRP1B;refGene_tx=XM_023259300.1
C1 133977060 133977151
refGene_gn=LRP1B;refGene_tx=XM_023259298.1;refGene_gn=LRP1B;refGene_tx

=XM_023259299.1;refGene_gn=LRP1B;refGene_tx=XM_023259300.1
C1 133984289 133984366
refGene_gn=LRP1B;refGene_tx=XM_023259298.1;refGene_gn=LRP1B;refGene_tx=XM_023259299.1;refGene_gn=LRP1B;refGene_tx=XM_023259300.1
C1 134006909 134007023
refGene_gn=LRP1B;refGene_tx=XM_023259298.1;refGene_gn=LRP1B;refGene_tx=XM_023259299.1
C1 134007886 134007991
refGene_gn=LRP1B;refGene_tx=XM_023259298.1;refGene_gn=LRP1B;refGene_tx=XM_023259299.1;refGene_gn=LRP1B;refGene_tx=XM_023259300.1
C1 134012903 134013078
refGene_gn=LRP1B;refGene_tx=XM_023259298.1;refGene_gn=LRP1B;refGene_tx=XM_023259299.1;refGene_gn=LRP1B;refGene_tx=XM_023259300.1
C1 134041514 134041676
refGene_gn=LRP1B;refGene_tx=XM_023259298.1;refGene_gn=LRP1B;refGene_tx=XM_023259299.1;refGene_gn=LRP1B;refGene_tx=XM_023259300.1
C1 134055762 134055927
refGene_gn=LRP1B;refGene_tx=XM_023259298.1;refGene_gn=LRP1B;refGene_tx=XM_023259299.1;refGene_gn=LRP1B;refGene_tx=XM_023259300.1
C1 134060908 134061034
refGene_gn=LRP1B;refGene_tx=XM_023259298.1;refGene_gn=LRP1B;refGene_tx=XM_023259299.1;refGene_gn=LRP1B;refGene_tx=XM_023259300.1
C1 134062674 134062848
refGene_gn=LRP1B;refGene_tx=XM_023259298.1;refGene_gn=LRP1B;refGene_tx=XM_023259299.1;refGene_gn=LRP1B;refGene_tx=XM_023259300.1
C1 134064740 134064857
refGene_gn=LRP1B;refGene_tx=XM_023259298.1;refGene_gn=LRP1B;refGene_tx=XM_023259299.1;refGene_gn=LRP1B;refGene_tx=XM_023259300.1
C1 134074915 134075022
refGene_gn=LRP1B;refGene_tx=XM_023259298.1;refGene_gn=LRP1B;refGene_tx=XM_023259299.1;refGene_gn=LRP1B;refGene_tx=XM_023259300.1
C1 134075741 134075965
refGene_gn=LRP1B;refGene_tx=XM_023259298.1;refGene_gn=LRP1B;refGene_tx=XM_023259299.1;refGene_gn=LRP1B;refGene_tx=XM_023259300.1
C1 134098475 134098717
refGene_gn=LRP1B;refGene_tx=XM_023259298.1;refGene_gn=LRP1B;refGene_tx=XM_023259299.1;refGene_gn=LRP1B;refGene_tx=XM_023259300.1;refGene_gn=LRP1B;refGene_tx=XM_023259301.1
C1 134100225 134100345
refGene_gn=LRP1B;refGene_tx=XM_023259298.1;refGene_gn=LRP1B;refGene_tx=XM_023259299.1;refGene_gn=LRP1B;refGene_tx=XM_023259300.1;refGene_gn=LRP1B;refGene_tx=XM_023259301.1
C1 134103840 134103975
refGene_gn=LRP1B;refGene_tx=XM_023259298.1;refGene_gn=LRP1B;refGene_tx=XM_023259299.1;refGene_gn=LRP1B;refGene_tx=XM_023259300.1;refGene_gn=LRP1B;refGene_tx=XM_023259301.1
C1 134105618 134105756
refGene_gn=LRP1B;refGene_tx=XM_023259298.1;refGene_gn=LRP1B;refGene_tx=XM_023259299.1;refGene_gn=LRP1B;refGene_tx=XM_023259300.1;refGene_gn=LRP1B;refGene_tx=XM_023259301.1

C1 134106723 134106849
refGene_gn=LRP1B;refGene_tx=XM_023259298.1;refGene_gn=LRP1B;refGene_tx=XM_023259299.1;refGene_gn=LRP1B;refGene_tx=XM_023259300.1;refGene_gn=LRP1B;refGene_tx=XM_023259301.1
C1 134112004 134112127
refGene_gn=LRP1B;refGene_tx=XM_023259298.1;refGene_gn=LRP1B;refGene_tx=XM_023259299.1;refGene_gn=LRP1B;refGene_tx=XM_023259300.1;refGene_gn=LRP1B;refGene_tx=XM_023259301.1
C1 134119163 134119296
refGene_gn=LRP1B;refGene_tx=XM_023259298.1;refGene_gn=LRP1B;refGene_tx=XM_023259299.1;refGene_gn=LRP1B;refGene_tx=XM_023259300.1;refGene_gn=LRP1B;refGene_tx=XM_023259301.1
C1 134119634 134119741
refGene_gn=LRP1B;refGene_tx=XM_023259298.1;refGene_gn=LRP1B;refGene_tx=XM_023259299.1;refGene_gn=LRP1B;refGene_tx=XM_023259300.1;refGene_gn=LRP1B;refGene_tx=XM_023259301.1
C1 134121346 134121476
refGene_gn=LRP1B;refGene_tx=XM_023259298.1;refGene_gn=LRP1B;refGene_tx=XM_023259299.1;refGene_gn=LRP1B;refGene_tx=XM_023259300.1;refGene_gn=LRP1B;refGene_tx=XM_023259301.1
C1 134127499 134127606
refGene_gn=LRP1B;refGene_tx=XM_023259298.1;refGene_gn=LRP1B;refGene_tx=XM_023259299.1;refGene_gn=LRP1B;refGene_tx=XM_023259300.1;refGene_gn=LRP1B;refGene_tx=XM_023259301.1
C1 134134825 134134942
refGene_gn=LRP1B;refGene_tx=XM_023259298.1;refGene_gn=LRP1B;refGene_tx=XM_023259299.1;refGene_gn=LRP1B;refGene_tx=XM_023259300.1;refGene_gn=LRP1B;refGene_tx=XM_023259301.1
C1 134180742 134180862
refGene_gn=LRP1B;refGene_tx=XM_023259298.1;refGene_gn=LRP1B;refGene_tx=XM_023259299.1;refGene_gn=LRP1B;refGene_tx=XM_023259300.1;refGene_gn=LRP1B;refGene_tx=XM_023259301.1
C1 134182264 134182384
refGene_gn=LRP1B;refGene_tx=XM_023259298.1;refGene_gn=LRP1B;refGene_tx=XM_023259299.1;refGene_gn=LRP1B;refGene_tx=XM_023259300.1;refGene_gn=LRP1B;refGene_tx=XM_023259301.1
C1 134182502 134182619
refGene_gn=LRP1B;refGene_tx=XM_023259298.1;refGene_gn=LRP1B;refGene_tx=XM_023259299.1;refGene_gn=LRP1B;refGene_tx=XM_023259300.1;refGene_gn=LRP1B;refGene_tx=XM_023259301.1
C1 134186085 134186179
refGene_gn=LRP1B;refGene_tx=XM_023259298.1;refGene_gn=LRP1B;refGene_tx=XM_023259299.1;refGene_gn=LRP1B;refGene_tx=XM_023259300.1;refGene_gn=LRP1B;refGene_tx=XM_023259301.1
C1 134192529 134192678
refGene_gn=LRP1B;refGene_tx=XM_023259298.1;refGene_gn=LRP1B;refGene_tx=XM_023259299.1;refGene_gn=LRP1B;refGene_tx=XM_023259300.1;refGene_gn=LRP1B;refGene_tx=XM_023259301.1
C1 134193516 134193705
refGene_gn=LRP1B;refGene_tx=XM_023259298.1;refGene_gn=LRP1B;refGene_tx

=XM_023259299.1;refGene_gn=LRP1B;refGene_tx=XM_023259300.1;refGene_gn=LRP1B;refGene_tx=XM_023259301.1
C1 134210102 134210302
refGene_gn=LRP1B;refGene_tx=XM_023259298.1;refGene_gn=LRP1B;refGene_tx=XM_023259299.1;refGene_gn=LRP1B;refGene_tx=XM_023259300.1;refGene_gn=LRP1B;refGene_tx=XM_023259301.1
C1 134219302 134219484
refGene_gn=LRP1B;refGene_tx=XM_023259298.1;refGene_gn=LRP1B;refGene_tx=XM_023259299.1;refGene_gn=LRP1B;refGene_tx=XM_023259300.1;refGene_gn=LRP1B;refGene_tx=XM_023259301.1
C1 134221734 134221857
refGene_gn=LRP1B;refGene_tx=XM_023259298.1;refGene_gn=LRP1B;refGene_tx=XM_023259299.1;refGene_gn=LRP1B;refGene_tx=XM_023259300.1;refGene_gn=LRP1B;refGene_tx=XM_023259301.1
C1 134227874 134227960
refGene_gn=LRP1B;refGene_tx=XM_023259298.1;refGene_gn=LRP1B;refGene_tx=XM_023259299.1;refGene_gn=LRP1B;refGene_tx=XM_023259300.1;refGene_gn=LRP1B;refGene_tx=XM_023259301.1
C1 134232835 134233019
refGene_gn=LRP1B;refGene_tx=XM_023259298.1;refGene_gn=LRP1B;refGene_tx=XM_023259299.1;refGene_gn=LRP1B;refGene_tx=XM_023259300.1;refGene_gn=LRP1B;refGene_tx=XM_023259301.1
C1 134240340 134240528
refGene_gn=LRP1B;refGene_tx=XM_023259298.1;refGene_gn=LRP1B;refGene_tx=XM_023259299.1;refGene_gn=LRP1B;refGene_tx=XM_023259300.1;refGene_gn=LRP1B;refGene_tx=XM_023259301.1
C1 134241898 134242039
refGene_gn=LRP1B;refGene_tx=XM_023259298.1;refGene_gn=LRP1B;refGene_tx=XM_023259299.1;refGene_gn=LRP1B;refGene_tx=XM_023259300.1;refGene_gn=LRP1B;refGene_tx=XM_023259301.1
C1 134245604 134245727
refGene_gn=LRP1B;refGene_tx=XM_023259298.1;refGene_gn=LRP1B;refGene_tx=XM_023259299.1;refGene_gn=LRP1B;refGene_tx=XM_023259300.1;refGene_gn=LRP1B;refGene_tx=XM_023259301.1
C1 134249858 134249987
refGene_gn=LRP1B;refGene_tx=XM_023259298.1;refGene_gn=LRP1B;refGene_tx=XM_023259299.1;refGene_gn=LRP1B;refGene_tx=XM_023259300.1;refGene_gn=LRP1B;refGene_tx=XM_023259301.1
C1 134254409 134254529
refGene_gn=LRP1B;refGene_tx=XM_023259298.1;refGene_gn=LRP1B;refGene_tx=XM_023259299.1;refGene_gn=LRP1B;refGene_tx=XM_023259300.1;refGene_gn=LRP1B;refGene_tx=XM_023259301.1
C1 134256296 134256419
refGene_gn=LRP1B;refGene_tx=XM_023259298.1;refGene_gn=LRP1B;refGene_tx=XM_023259299.1;refGene_gn=LRP1B;refGene_tx=XM_023259300.1;refGene_gn=LRP1B;refGene_tx=XM_023259301.1
C1 134259915 134260065
refGene_gn=LRP1B;refGene_tx=XM_023259298.1;refGene_gn=LRP1B;refGene_tx=XM_023259299.1;refGene_gn=LRP1B;refGene_tx=XM_023259300.1;refGene_gn=LRP1B;refGene_tx=XM_023259301.1

C1 134260308 134260422
refGene_gn=LRP1B;refGene_tx=XM_023259298.1;refGene_gn=LRP1B;refGene_tx=XM_023259299.1;refGene_gn=LRP1B;refGene_tx=XM_023259300.1;refGene_gn=LRP1B;refGene_tx=XM_023259301.1
C1 134269504 134269624
refGene_gn=LRP1B;refGene_tx=XM_023259298.1;refGene_gn=LRP1B;refGene_tx=XM_023259299.1;refGene_gn=LRP1B;refGene_tx=XM_023259300.1;refGene_gn=LRP1B;refGene_tx=XM_023259301.1
C1 134272271 134272400
refGene_gn=LRP1B;refGene_tx=XM_023259298.1;refGene_gn=LRP1B;refGene_tx=XM_023259299.1;refGene_gn=LRP1B;refGene_tx=XM_023259300.1;refGene_gn=LRP1B;refGene_tx=XM_023259301.1
C1 134276586 134276712
refGene_gn=LRP1B;refGene_tx=XM_023259298.1;refGene_gn=LRP1B;refGene_tx=XM_023259299.1;refGene_gn=LRP1B;refGene_tx=XM_023259300.1;refGene_gn=LRP1B;refGene_tx=XM_023259301.1
C1 134277342 134277535
refGene_gn=LRP1B;refGene_tx=XM_023259298.1;refGene_gn=LRP1B;refGene_tx=XM_023259299.1;refGene_gn=LRP1B;refGene_tx=XM_023259300.1;refGene_gn=LRP1B;refGene_tx=XM_023259301.1
C1 134326127 134326332
refGene_gn=LRP1B;refGene_tx=XM_023259298.1;refGene_gn=LRP1B;refGene_tx=XM_023259299.1;refGene_gn=LRP1B;refGene_tx=XM_023259300.1;refGene_gn=LRP1B;refGene_tx=XM_023259301.1
C1 134329518 134329708
refGene_gn=LRP1B;refGene_tx=XM_023259298.1;refGene_gn=LRP1B;refGene_tx=XM_023259299.1;refGene_gn=LRP1B;refGene_tx=XM_023259300.1;refGene_gn=LRP1B;refGene_tx=XM_023259301.1
C1 134431170 134431542
refGene_gn=LRP1B;refGene_tx=XM_023259298.1;refGene_gn=LRP1B;refGene_tx=XM_023259299.1;refGene_gn=LRP1B;refGene_tx=XM_023259300.1;refGene_gn=LRP1B;refGene_tx=XM_023259301.1
C1 134433092 134433217
refGene_gn=LRP1B;refGene_tx=XM_023259298.1;refGene_gn=LRP1B;refGene_tx=XM_023259299.1;refGene_gn=LRP1B;refGene_tx=XM_023259300.1;refGene_gn=LRP1B;refGene_tx=XM_023259301.1
C1 134433513 134433665
refGene_gn=LRP1B;refGene_tx=XM_023259298.1;refGene_gn=LRP1B;refGene_tx=XM_023259299.1;refGene_gn=LRP1B;refGene_tx=XM_023259300.1;refGene_gn=LRP1B;refGene_tx=XM_023259301.1
C1 134433801 134433928
refGene_gn=LRP1B;refGene_tx=XM_023259298.1;refGene_gn=LRP1B;refGene_tx=XM_023259299.1;refGene_gn=LRP1B;refGene_tx=XM_023259300.1;refGene_gn=LRP1B;refGene_tx=XM_023259301.1
C1 134450257 134450387
refGene_gn=LRP1B;refGene_tx=XM_023259298.1;refGene_gn=LRP1B;refGene_tx=XM_023259299.1;refGene_gn=LRP1B;refGene_tx=XM_023259300.1;refGene_gn=LRP1B;refGene_tx=XM_023259301.1
C1 134450966 134451101
refGene_gn=LRP1B;refGene_tx=XM_023259298.1;refGene_gn=LRP1B;refGene_tx

=XM_023259299.1;refGene_gn=LRP1B;refGene_tx=XM_023259300.1;refGene_gn=LRP1B;refGene_tx=XM_023259301.1
C1 134493081 134493213
refGene_gn=LRP1B;refGene_tx=XM_023259298.1;refGene_gn=LRP1B;refGene_tx=XM_023259299.1;refGene_gn=LRP1B;refGene_tx=XM_023259300.1;refGene_gn=LRP1B;refGene_tx=XM_023259301.1
C1 134494679 134494805
refGene_gn=LRP1B;refGene_tx=XM_023259298.1;refGene_gn=LRP1B;refGene_tx=XM_023259299.1;refGene_gn=LRP1B;refGene_tx=XM_023259300.1;refGene_gn=LRP1B;refGene_tx=XM_023259301.1
C1 134499202 134499343
refGene_gn=LRP1B;refGene_tx=XM_023259298.1;refGene_gn=LRP1B;refGene_tx=XM_023259299.1;refGene_gn=LRP1B;refGene_tx=XM_023259300.1;refGene_gn=LRP1B;refGene_tx=XM_023259301.1
C1 134533193 134533343
refGene_gn=LRP1B;refGene_tx=XM_023259298.1;refGene_gn=LRP1B;refGene_tx=XM_023259299.1;refGene_gn=LRP1B;refGene_tx=XM_023259300.1;refGene_gn=LRP1B;refGene_tx=XM_023259301.1
C1 134552763 134552858
refGene_gn=LRP1B;refGene_tx=XM_023259298.1;refGene_gn=LRP1B;refGene_tx=XM_023259299.1;refGene_gn=LRP1B;refGene_tx=XM_023259300.1;refGene_gn=LRP1B;refGene_tx=XM_023259301.1
C1 134553659 134553834
refGene_gn=LRP1B;refGene_tx=XM_023259298.1;refGene_gn=LRP1B;refGene_tx=XM_023259299.1;refGene_gn=LRP1B;refGene_tx=XM_023259300.1;refGene_gn=LRP1B;refGene_tx=XM_023259301.1
C1 134566490 134566718
refGene_gn=LRP1B;refGene_tx=XM_023259298.1;refGene_gn=LRP1B;refGene_tx=XM_023259299.1;refGene_gn=LRP1B;refGene_tx=XM_023259300.1;refGene_gn=LRP1B;refGene_tx=XM_023259301.1
C1 134568061 134568193
refGene_gn=LRP1B;refGene_tx=XM_023259298.1;refGene_gn=LRP1B;refGene_tx=XM_023259299.1;refGene_gn=LRP1B;refGene_tx=XM_023259300.1;refGene_gn=LRP1B;refGene_tx=XM_023259301.1
C1 134593934 134594179
refGene_gn=LRP1B;refGene_tx=XM_023259298.1;refGene_gn=LRP1B;refGene_tx=XM_023259299.1;refGene_gn=LRP1B;refGene_tx=XM_023259300.1;refGene_gn=LRP1B;refGene_tx=XM_023259301.1
C1 134594511 134594676
refGene_gn=LRP1B;refGene_tx=XM_023259298.1;refGene_gn=LRP1B;refGene_tx=XM_023259299.1;refGene_gn=LRP1B;refGene_tx=XM_023259300.1;refGene_gn=LRP1B;refGene_tx=XM_023259301.1
C1 134604199 134604404
refGene_gn=LRP1B;refGene_tx=XM_023259298.1;refGene_gn=LRP1B;refGene_tx=XM_023259299.1;refGene_gn=LRP1B;refGene_tx=XM_023259300.1;refGene_gn=LRP1B;refGene_tx=XM_023259301.1
C1 134605890 134606088
refGene_gn=LRP1B;refGene_tx=XM_023259298.1;refGene_gn=LRP1B;refGene_tx=XM_023259299.1;refGene_gn=LRP1B;refGene_tx=XM_023259300.1;refGene_gn=LRP1B;refGene_tx=XM_023259301.1

C1	134620795	134621041
refGene_gn=LRP1B;refGene_tx=XM_023259298.1;refGene_gn=LRP1B;refGene_tx=XM_023259299.1;refGene_gn=LRP1B;refGene_tx=XM_023259300.1;refGene_gn=LRP1B;refGene_tx=XM_023259301.1
C1	134626796	134626997
refGene_gn=LRP1B;refGene_tx=XM_023259298.1;refGene_gn=LRP1B;refGene_tx=XM_023259299.1;refGene_gn=LRP1B;refGene_tx=XM_023259300.1;refGene_gn=LRP1B;refGene_tx=XM_023259301.1
C1	134645227	134645410
refGene_gn=LRP1B;refGene_tx=XM_023259298.1;refGene_gn=LRP1B;refGene_tx=XM_023259299.1;refGene_gn=LRP1B;refGene_tx=XM_023259300.1;refGene_gn=LRP1B;refGene_tx=XM_023259301.1
C1	134662231	134662399
refGene_gn=LRP1B;refGene_tx=XM_023259298.1;refGene_gn=LRP1B;refGene_tx=XM_023259299.1;refGene_gn=LRP1B;refGene_tx=XM_023259300.1;refGene_gn=LRP1B;refGene_tx=XM_023259301.1
C1	134663518	134663599
refGene_gn=LRP1B;refGene_tx=XM_023259298.1;refGene_gn=LRP1B;refGene_tx=XM_023259299.1;refGene_gn=LRP1B;refGene_tx=XM_023259300.1;refGene_gn=LRP1B;refGene_tx=XM_023259301.1
C1	134687935	134688052
refGene_gn=LRP1B;refGene_tx=XM_023259298.1;refGene_gn=LRP1B;refGene_tx=XM_023259299.1;refGene_gn=LRP1B;refGene_tx=XM_023259300.1;refGene_gn=LRP1B;refGene_tx=XM_023259301.1
C1	134695559	134695685
refGene_gn=LRP1B;refGene_tx=XM_023259298.1;refGene_gn=LRP1B;refGene_tx=XM_023259299.1;refGene_gn=LRP1B;refGene_tx=XM_023259300.1;refGene_gn=LRP1B;refGene_tx=XM_023259301.1
C1	134700184	134700325
refGene_gn=LRP1B;refGene_tx=XM_023259298.1;refGene_gn=LRP1B;refGene_tx=XM_023259299.1;refGene_gn=LRP1B;refGene_tx=XM_023259300.1;refGene_gn=LRP1B;refGene_tx=XM_023259301.1
C1	134711756	134711879
refGene_gn=LRP1B;refGene_tx=XM_023259298.1;refGene_gn=LRP1B;refGene_tx=XM_023259299.1;refGene_gn=LRP1B;refGene_tx=XM_023259300.1;refGene_gn=LRP1B;refGene_tx=XM_023259301.1
C1	134721263	134721453
refGene_gn=LRP1B;refGene_tx=XM_023259298.1;refGene_gn=LRP1B;refGene_tx=XM_023259299.1;refGene_gn=LRP1B;refGene_tx=XM_023259300.1;refGene_gn=LRP1B;refGene_tx=XM_023259301.1
C1	134723365	134723585
refGene_gn=LRP1B;refGene_tx=XM_023259298.1;refGene_gn=LRP1B;refGene_tx=XM_023259299.1;refGene_gn=LRP1B;refGene_tx=XM_023259300.1;refGene_gn=LRP1B;refGene_tx=XM_023259301.1
C1	134729490	134729671
refGene_gn=LRP1B;refGene_tx=XM_023259298.1;refGene_gn=LRP1B;refGene_tx=XM_023259299.1;refGene_gn=LRP1B;refGene_tx=XM_023259300.1;refGene_gn=LRP1B;refGene_tx=XM_023259301.1
C1	134771543	134771780
refGene_gn=LRP1B;refGene_tx=XM_023259298.1;refGene_gn=LRP1B;refGene_tx

=XM_023259299.1;refGene_gn=LRP1B;refGene_tx=XM_023259300.1;refGene_gn=LRP1B;refGene_tx=XM_023259301.1
C1 134776666 134776810
refGene_gn=LRP1B;refGene_tx=XM_023259298.1;refGene_gn=LRP1B;refGene_tx=XM_023259299.1;refGene_gn=LRP1B;refGene_tx=XM_023259300.1;refGene_gn=LRP1B;refGene_tx=XM_023259301.1
C1 134779328 134779500
refGene_gn=LRP1B;refGene_tx=XM_023259298.1;refGene_gn=LRP1B;refGene_tx=XM_023259299.1;refGene_gn=LRP1B;refGene_tx=XM_023259300.1;refGene_gn=LRP1B;refGene_tx=XM_023259301.1
C1 134783011 134783234
refGene_gn=LRP1B;refGene_tx=XM_023259298.1;refGene_gn=LRP1B;refGene_tx=XM_023259299.1;refGene_gn=LRP1B;refGene_tx=XM_023259300.1;refGene_gn=LRP1B;refGene_tx=XM_023259301.1
C1 134927526 134927689
refGene_gn=LRP1B;refGene_tx=XM_023259298.1;refGene_gn=LRP1B;refGene_tx=XM_023259299.1;refGene_gn=LRP1B;refGene_tx=XM_023259300.1;refGene_gn=LRP1B;refGene_tx=XM_023259301.1
C1 134962524 134962782
refGene_gn=LRP1B;refGene_tx=XM_023259298.1;refGene_gn=LRP1B;refGene_tx=XM_023259299.1;refGene_gn=LRP1B;refGene_tx=XM_023259300.1;refGene_gn=LRP1B;refGene_tx=XM_023259301.1
C1 134980337 134980466
refGene_gn=LRP1B;refGene_tx=XM_023259298.1;refGene_gn=LRP1B;refGene_tx=XM_023259299.1;refGene_gn=LRP1B;refGene_tx=XM_023259300.1;refGene_gn=LRP1B;refGene_tx=XM_023259301.1
C1 134984502 134984622
refGene_gn=LRP1B;refGene_tx=XM_023259298.1;refGene_gn=LRP1B;refGene_tx=XM_023259299.1;refGene_gn=LRP1B;refGene_tx=XM_023259300.1;refGene_gn=LRP1B;refGene_tx=XM_023259301.1
C1 135190359 135190497
refGene_gn=LRP1B;refGene_tx=XM_023259298.1;refGene_gn=LRP1B;refGene_tx=XM_023259299.1;refGene_gn=LRP1B;refGene_tx=XM_023259300.1;refGene_gn=LRP1B;refGene_tx=XM_023259301.1
C1 135522950 135523073
refGene_gn=LRP1B;refGene_tx=XM_023259298.1;refGene_gn=LRP1B;refGene_tx=XM_023259299.1;refGene_gn=LRP1B;refGene_tx=XM_023259300.1;refGene_gn=LRP1B;refGene_tx=XM_023259301.1
C1 135875440 135875621
refGene_gn=LRP1B;refGene_tx=XM_023259298.1;refGene_gn=LRP1B;refGene_tx=XM_023259299.1;refGene_gn=LRP1B;refGene_tx=XM_023259300.1;refGene_gn=LRP1B;refGene_tx=XM_023259301.1
C1 136479701 136479870
refGene_gn=KYNU;refGene_tx=XM_011285295.3
C1 136493427 136493548
refGene_gn=KYNU;refGene_tx=XM_011285295.3
C1 136503046 136503129
refGene_gn=KYNU;refGene_tx=XM_011285295.3
C1 136511325 136511387
refGene_gn=KYNU;refGene_tx=XM_011285295.3

C1 136512942 136513014
refGene_gn=KYNU;refGene_tx=XM_011285295.3
C1 136514271 136514346
refGene_gn=KYNU;refGene_tx=XM_011285295.3
C1 136516894 136517041
refGene_gn=KYNU;refGene_tx=XM_011285295.3
C1 136536244 136536343
refGene_gn=KYNU;refGene_tx=XM_011285295.3
C1 136537125 136537199
refGene_gn=KYNU;refGene_tx=XM_011285295.3
C1 136572071 136572124
refGene_gn=KYNU;refGene_tx=XM_011285295.3
C1 136574034 136574120
refGene_gn=KYNU;refGene_tx=XM_011285295.3
C1 136580216 136580447
refGene_gn=KYNU;refGene_tx=XM_011285295.3
C1 136582538 136582676
refGene_gn=KYNU;refGene_tx=XM_011285295.3
C1 136666878 136667043
refGene_gn=ARHGAP15;refGene_tx=XM_003990745.5;refGene_gn=ARHGAP15;refGene_tx=XM_019838066.2;refGene_gn=ARHGAP15;refGene_tx=XM_019838067.2
C1 136713325 136713394
refGene_gn=ARHGAP15;refGene_tx=XM_003990745.5;refGene_gn=ARHGAP15;refGene_tx=XM_019838066.2;refGene_gn=ARHGAP15;refGene_tx=XM_019838067.2
C1 136727908 136727970
refGene_gn=ARHGAP15;refGene_tx=XM_003990745.5;refGene_gn=ARHGAP15;refGene_tx=XM_019838066.2;refGene_gn=ARHGAP15;refGene_tx=XM_019838067.2
C1 136740553 136740641
refGene_gn=ARHGAP15;refGene_tx=XM_003990745.5;refGene_gn=ARHGAP15;refGene_tx=XM_019838066.2;refGene_gn=ARHGAP15;refGene_tx=XM_019838067.2
C1 136760009 136760099
refGene_gn=ARHGAP15;refGene_tx=XM_003990745.5;refGene_gn=ARHGAP15;refGene_tx=XM_019838066.2;refGene_gn=ARHGAP15;refGene_tx=XM_019838067.2
C1 136945735 136945834
refGene_gn=ARHGAP15;refGene_tx=XM_003990745.5;refGene_gn=ARHGAP15;refGene_tx=XM_019838066.2;refGene_gn=ARHGAP15;refGene_tx=XM_019838067.2
C1 136947100 136947230
refGene_gn=ARHGAP15;refGene_tx=XM_003990745.5;refGene_gn=ARHGAP15;refGene_tx=XM_019838066.2;refGene_gn=ARHGAP15;refGene_tx=XM_019838067.2
C1 136995387 136995510
refGene_gn=ARHGAP15;refGene_tx=XM_003990745.5;refGene_gn=ARHGAP15;refGene_tx=XM_019838066.2;refGene_gn=ARHGAP15;refGene_tx=XM_019838067.2
C1 137025936 137026035
refGene_gn=ARHGAP15;refGene_tx=XM_003990745.5;refGene_gn=ARHGAP15;refGene_tx=XM_019838066.2;refGene_gn=ARHGAP15;refGene_tx=XM_019838067.2
C1 137056292 137056370
refGene_gn=ARHGAP15;refGene_tx=XM_003990745.5;refGene_gn=ARHGAP15;refGene_tx=XM_019838066.2;refGene_gn=ARHGAP15;refGene_tx=XM_019838067.2
C1 137122057 137122192
refGene_gn=ARHGAP15;refGene_tx=XM_003990745.5;refGene_gn=ARHGAP15;refG

ene_tx=XM_019838066.2;refGene_gn=ARHGAP15;refGene_tx=XM_019838067.2
C1	137193043	137193149
refGene_gn=ARHGAP15;refGene_tx=XM_003990745.5;refGene_gn=ARHGAP15;refGene_tx=XM_019838066.2;refGene_gn=ARHGAP15;refGene_tx=XM_019838067.2
C1	137208099	137208146
refGene_gn=ARHGAP15;refGene_tx=XM_019838066.2;refGene_gn=ARHGAP15;refGene_tx=XM_019838067.2
C1	137255433	137255617
refGene_gn=ARHGAP15;refGene_tx=XM_003990745.5
C1	137428743	137428827
refGene_gn=GTDC1;refGene_tx=XM_006935266.4;refGene_gn=GTDC1;refGene_tx=XM_011285294.3;refGene_gn=GTDC1;refGene_tx=XM_019838068.2;refGene_gn=GTDC1;refGene_tx=XM_019838069.2;refGene_gn=GTDC1;refGene_tx=XM_023259302.1;refGene_gn=GTDC1;refGene_tx=XM_023259303.1;refGene_gn=GTDC1;refGene_tx=XM_023259304.1;refGene_gn=GTDC1;refGene_tx=XM_023259305.1;refGene_gn=GTDC1;refGene_tx=XM_023259308.1
C1	137434055	137434147
refGene_gn=GTDC1;refGene_tx=XM_006935266.4;refGene_gn=GTDC1;refGene_tx=XM_011285294.3;refGene_gn=GTDC1;refGene_tx=XM_019838068.2;refGene_gn=GTDC1;refGene_tx=XM_019838069.2;refGene_gn=GTDC1;refGene_tx=XM_023259302.1;refGene_gn=GTDC1;refGene_tx=XM_023259303.1;refGene_gn=GTDC1;refGene_tx=XM_023259304.1;refGene_gn=GTDC1;refGene_tx=XM_023259305.1;refGene_gn=GTDC1;refGene_tx=XM_023259306.1;refGene_gn=GTDC1;refGene_tx=XM_023259307.1;refGene_gn=GTDC1;refGene_tx=XM_023259308.1
C1	137434849	137434920
refGene_gn=GTDC1;refGene_tx=XM_006935266.4;refGene_gn=GTDC1;refGene_tx=XM_011285294.3;refGene_gn=GTDC1;refGene_tx=XM_019838068.2;refGene_gn=GTDC1;refGene_tx=XM_019838069.2;refGene_gn=GTDC1;refGene_tx=XM_023259302.1;refGene_gn=GTDC1;refGene_tx=XM_023259303.1;refGene_gn=GTDC1;refGene_tx=XM_023259304.1;refGene_gn=GTDC1;refGene_tx=XM_023259305.1;refGene_gn=GTDC1;refGene_tx=XM_023259306.1;refGene_gn=GTDC1;refGene_tx=XM_023259307.1;refGene_gn=GTDC1;refGene_tx=XM_023259308.1
C1	137439358	137439503
refGene_gn=GTDC1;refGene_tx=XM_006935266.4;refGene_gn=GTDC1;refGene_tx=XM_011285294.3;refGene_gn=GTDC1;refGene_tx=XM_019838068.2;refGene_gn=GTDC1;refGene_tx=XM_019838069.2;refGene_gn=GTDC1;refGene_tx=XM_023259302.1;refGene_gn=GTDC1;refGene_tx=XM_023259303.1;refGene_gn=GTDC1;refGene_tx=XM_023259304.1;refGene_gn=GTDC1;refGene_tx=XM_023259305.1;refGene_gn=GTDC1;refGene_tx=XM_023259308.1
C1	137454580	137454690
refGene_gn=GTDC1;refGene_tx=XM_006935266.4;refGene_gn=GTDC1;refGene_tx=XM_011285294.3;refGene_gn=GTDC1;refGene_tx=XM_019838068.2;refGene_gn=GTDC1;refGene_tx=XM_019838069.2;refGene_gn=GTDC1;refGene_tx=XM_023259302.1;refGene_gn=GTDC1;refGene_tx=XM_023259303.1;refGene_gn=GTDC1;refGene_tx=XM_023259304.1;refGene_gn=GTDC1;refGene_tx=XM_023259305.1;refGene_gn=GTDC1;refGene_tx=XM_023259306.1;refGene_gn=GTDC1;refGene_tx=XM_023259307.1;refGene_gn=GTDC1;refGene_tx=XM_023259308.1
C1	137490475	137490820
refGene_gn=GTDC1;refGene_tx=XM_006935266.4;refGene_gn=GTDC1;refGene_tx=XM_011285294.3;refGene_gn=GTDC1;refGene_tx=XM_019838068.2;refGene_gn=

GTDC1;refGene_tx=XM_019838069.2;refGene_gn=GTDC1;refGene_tx=XM_0232593
02.1;refGene_gn=GTDC1;refGene_tx=XM_023259303.1;refGene_gn=GTDC1;refGe
ne_tx=XM_023259304.1;refGene_gn=GTDC1;refGene_tx=XM_023259305.1;refGen
e_gn=GTDC1;refGene_tx=XM_023259306.1;refGene_gn=GTDC1;refGene_tx=XM_02
3259307.1
C1 137494949 137495111
refGene_gn=GTDC1;refGene_tx=XM_006935266.4;refGene_gn=GTDC1;refGene_tx
=XM_011285294.3;refGene_gn=GTDC1;refGene_tx=XM_019838068.2;refGene_gn=
GTDC1;refGene_tx=XM_019838069.2;refGene_gn=GTDC1;refGene_tx=XM_0232593
02.1;refGene_gn=GTDC1;refGene_tx=XM_023259303.1;refGene_gn=GTDC1;refGe
ne_tx=XM_023259304.1;refGene_gn=GTDC1;refGene_tx=XM_023259306.1
C1 137624335 137624509
refGene_gn=GTDC1;refGene_tx=XM_006935266.4;refGene_gn=GTDC1;refGene_tx
=XM_011285294.3;refGene_gn=GTDC1;refGene_tx=XM_019838068.2;refGene_gn=
GTDC1;refGene_tx=XM_019838069.2;refGene_gn=GTDC1;refGene_tx=XM_0232593
02.1;refGene_gn=GTDC1;refGene_tx=XM_023259303.1;refGene_gn=GTDC1;refGe
ne_tx=XM_023259304.1;refGene_gn=GTDC1;refGene_tx=XM_023259305.1;refGen
e_gn=GTDC1;refGene_tx=XM_023259306.1;refGene_gn=GTDC1;refGene_tx=XM_02
3259307.1;refGene_gn=GTDC1;refGene_tx=XM_023259308.1
C1 137628542 137628710
refGene_gn=GTDC1;refGene_tx=XM_006935266.4;refGene_gn=GTDC1;refGene_tx
=XM_011285294.3;refGene_gn=GTDC1;refGene_tx=XM_019838068.2;refGene_gn=
GTDC1;refGene_tx=XM_019838069.2;refGene_gn=GTDC1;refGene_tx=XM_0232593
02.1;refGene_gn=GTDC1;refGene_tx=XM_023259303.1;refGene_gn=GTDC1;refGe
ne_tx=XM_023259304.1;refGene_gn=GTDC1;refGene_tx=XM_023259305.1;refGen
e_gn=GTDC1;refGene_tx=XM_023259306.1;refGene_gn=GTDC1;refGene_tx=XM_02
3259307.1;refGene_gn=GTDC1;refGene_tx=XM_023259308.1
C1 137681327 137681413
refGene_gn=GTDC1;refGene_tx=XM_011285294.3
C1 137695062 137695241
refGene_gn=GTDC1;refGene_tx=XM_006935266.4;refGene_gn=GTDC1;refGene_tx
=XM_019838068.2;refGene_gn=GTDC1;refGene_tx=XM_019838069.2;refGene_gn=
GTDC1;refGene_tx=XM_023259302.1;refGene_gn=GTDC1;refGene_tx=XM_0232593
03.1;refGene_gn=GTDC1;refGene_tx=XM_023259304.1;refGene_gn=GTDC1;refGe
ne_tx=XM_023259305.1;refGene_gn=GTDC1;refGene_tx=XM_023259306.1;refGen
e_gn=GTDC1;refGene_tx=XM_023259307.1;refGene_gn=GTDC1;refGene_tx=XM_02
3259308.1
C1 137884901 137885479
refGene_gn=ZEB2;refGene_tx=XM_023259309.1;refGene_gn=ZEB2;refGene_tx=X
M_023259310.1;refGene_gn=ZEB2;refGene_tx=XM_023259311.1
C1 137891268 137891449
refGene_gn=ZEB2;refGene_tx=XM_023259309.1;refGene_gn=ZEB2;refGene_tx=X
M_023259310.1
C1 137892860 137894830
refGene_gn=ZEB2;refGene_tx=XM_023259309.1;refGene_gn=ZEB2;refGene_tx=X
M_023259310.1;refGene_gn=ZEB2;refGene_tx=XM_023259311.1
C1 137895761 137895870
refGene_gn=ZEB2;refGene_tx=XM_023259309.1;refGene_gn=ZEB2;refGene_tx=X
M_023259310.1;refGene_gn=ZEB2;refGene_tx=XM_023259311.1
C1 137898496 137898711

refGene_gn=ZEB2;refGene_tx=XM_023259309.1;refGene_gn=ZEB2;refGene_tx=XM_023259310.1;refGene_gn=ZEB2;refGene_tx=XM_023259311.1
C1 137899432 137899621
refGene_gn=ZEB2;refGene_tx=XM_023259309.1;refGene_gn=ZEB2;refGene_tx=XM_023259310.1;refGene_gn=ZEB2;refGene_tx=XM_023259311.1
C1 137920025 137920097
refGene_gn=ZEB2;refGene_tx=XM_023259309.1;refGene_gn=ZEB2;refGene_tx=XM_023259310.1;refGene_gn=ZEB2;refGene_tx=XM_023259311.1
C1 137925240 137925498
refGene_gn=ZEB2;refGene_tx=XM_023259309.1;refGene_gn=ZEB2;refGene_tx=XM_023259310.1;refGene_gn=ZEB2;refGene_tx=XM_023259311.1
C1 138010975 138011048
refGene_gn=ZEB2;refGene_tx=XM_023259309.1;refGene_gn=ZEB2;refGene_tx=XM_023259310.1;refGene_gn=ZEB2;refGene_tx=XM_023259311.1
C1 138126302 138126515
refGene_gn=LOC101097506;refGene_tx=XM_006935269.1
C1 141282475 141283356
refGene_gn=LOC111561641;refGene_tx=XM_023257857.1
C1 141283397 141283401
refGene_gn=LOC111561641;refGene_tx=XM_023257857.1
C1 141283490 141283511
refGene_gn=LOC111561641;refGene_tx=XM_023257857.1
C1 141283707 141283762
refGene_gn=ACVR2A;refGene_tx=XM_003990748.5;refGene_gn=ACVR2A;refGene_tx=XM_019838079.2
C1 141331457 141331665
refGene_gn=ACVR2A;refGene_tx=XM_003990748.5;refGene_gn=ACVR2A;refGene_tx=XM_019838079.2
C1 141334185 141334295
refGene_gn=ACVR2A;refGene_tx=XM_003990748.5;refGene_gn=ACVR2A;refGene_tx=XM_019838079.2
C1 141334474 141334629
refGene_gn=ACVR2A;refGene_tx=XM_003990748.5;refGene_gn=ACVR2A;refGene_tx=XM_019838079.2
C1 141346282 141346306
refGene_gn=ACVR2A;refGene_tx=XM_019838079.2
C1 141350608 141350752
refGene_gn=ACVR2A;refGene_tx=XM_003990748.5;refGene_gn=ACVR2A;refGene_tx=XM_019838079.2
C1 141353041 141353185
refGene_gn=ACVR2A;refGene_tx=XM_003990748.5;refGene_gn=ACVR2A;refGene_tx=XM_019838079.2
C1 141354185 141354331
refGene_gn=ACVR2A;refGene_tx=XM_003990748.5;refGene_gn=ACVR2A;refGene_tx=XM_019838079.2
C1 141356275 141356390
refGene_gn=ACVR2A;refGene_tx=XM_003990748.5;refGene_gn=ACVR2A;refGene_tx=XM_019838079.2
C1 141358864 141359003
refGene_gn=ACVR2A;refGene_tx=XM_003990748.5;refGene_gn=ACVR2A;refGene_

tx=XM_019838079.2
C1	141362371	141362502
refGene_gn=ACVR2A;refGene_tx=XM_003990748.5;refGene_gn=ACVR2A;refGene_tx=XM_019838079.2
C1	141363399	141363594
refGene_gn=ACVR2A;refGene_tx=XM_003990748.5;refGene_gn=ACVR2A;refGene_tx=XM_019838079.2
C1	141370309	141370498
refGene_gn=ORC4;refGene_tx=XM_003990749.5;refGene_gn=ORC4;refGene_tx=XM_006935270.3;refGene_gn=ORC4;refGene_tx=XM_006935271.4;refGene_gn=ORC4;refGene_tx=XM_006935272.3;refGene_gn=ORC4;refGene_tx=XM_006935273.4;refGene_gn=ORC4;refGene_tx=XM_011285292.3;refGene_gn=ORC4;refGene_tx=XM_019838083.2;refGene_gn=ORC4;refGene_tx=XM_019838084.1;refGene_gn=ORC4;refGene_tx=XM_019838085.1;refGene_gn=ORC4;refGene_tx=XM_023259312.1;refGene_gn=ORC4;refGene_tx=XM_023259313.1
C1	141372780	141372848
refGene_gn=ORC4;refGene_tx=XM_003990749.5;refGene_gn=ORC4;refGene_tx=XM_006935270.3;refGene_gn=ORC4;refGene_tx=XM_006935271.4;refGene_gn=ORC4;refGene_tx=XM_006935272.3;refGene_gn=ORC4;refGene_tx=XM_006935273.4;refGene_gn=ORC4;refGene_tx=XM_011285292.3;refGene_gn=ORC4;refGene_tx=XM_019838083.2;refGene_gn=ORC4;refGene_tx=XM_019838084.1;refGene_gn=ORC4;refGene_tx=XM_019838085.1;refGene_gn=ORC4;refGene_tx=XM_023259312.1;refGene_gn=ORC4;refGene_tx=XM_023259313.1
C1	141372933	141373029
refGene_gn=ORC4;refGene_tx=XM_003990749.5;refGene_gn=ORC4;refGene_tx=XM_006935270.3;refGene_gn=ORC4;refGene_tx=XM_006935271.4;refGene_gn=ORC4;refGene_tx=XM_006935272.3;refGene_gn=ORC4;refGene_tx=XM_006935273.4;refGene_gn=ORC4;refGene_tx=XM_011285292.3;refGene_gn=ORC4;refGene_tx=XM_019838083.2;refGene_gn=ORC4;refGene_tx=XM_019838084.1;refGene_gn=ORC4;refGene_tx=XM_019838085.1;refGene_gn=ORC4;refGene_tx=XM_023259312.1;refGene_gn=ORC4;refGene_tx=XM_023259313.1
C1	141373738	141373847
refGene_gn=ORC4;refGene_tx=XM_003990749.5;refGene_gn=ORC4;refGene_tx=XM_006935270.3;refGene_gn=ORC4;refGene_tx=XM_006935271.4;refGene_gn=ORC4;refGene_tx=XM_006935272.3;refGene_gn=ORC4;refGene_tx=XM_006935273.4;refGene_gn=ORC4;refGene_tx=XM_011285292.3;refGene_gn=ORC4;refGene_tx=XM_019838083.2;refGene_gn=ORC4;refGene_tx=XM_019838084.1;refGene_gn=ORC4;refGene_tx=XM_019838085.1;refGene_gn=ORC4;refGene_tx=XM_023259312.1;refGene_gn=ORC4;refGene_tx=XM_023259313.1
C1	141378973	141379060
refGene_gn=ORC4;refGene_tx=XM_003990749.5;refGene_gn=ORC4;refGene_tx=XM_006935270.3;refGene_gn=ORC4;refGene_tx=XM_006935271.4;refGene_gn=ORC4;refGene_tx=XM_006935272.3;refGene_gn=ORC4;refGene_tx=XM_006935273.4;refGene_gn=ORC4;refGene_tx=XM_011285292.3;refGene_gn=ORC4;refGene_tx=XM_019838083.2;refGene_gn=ORC4;refGene_tx=XM_019838084.1;refGene_gn=ORC4;refGene_tx=XM_019838085.1;refGene_gn=ORC4;refGene_tx=XM_023259312.1;refGene_gn=ORC4;refGene_tx=XM_023259313.1
C1	141384314	141384488
refGene_gn=ORC4;refGene_tx=XM_003990749.5;refGene_gn=ORC4;refGene_tx=XM_006935270.3;refGene_gn=ORC4;refGene_tx=XM_006935271.4;refGene_gn=ORC

4;refGene_tx=XM_006935272.3;refGene_gn=ORC4;refGene_tx=XM_006935273.4;
refGene_gn=ORC4;refGene_tx=XM_011285292.3;refGene_gn=ORC4;refGene_tx=X
M_019838083.2;refGene_gn=ORC4;refGene_tx=XM_019838084.1;refGene_gn=ORC
4;refGene_tx=XM_019838085.1;refGene_gn=ORC4;refGene_tx=XM_023259312.1;
refGene_gn=ORC4;refGene_tx=XM_023259313.1
C1 141387130 141387282
refGene_gn=ORC4;refGene_tx=XM_003990749.5;refGene_gn=ORC4;refGene_tx=X
M_006935270.3;refGene_gn=ORC4;refGene_tx=XM_006935271.4;refGene_gn=ORC
4;refGene_tx=XM_006935272.3;refGene_gn=ORC4;refGene_tx=XM_006935273.4;
refGene_gn=ORC4;refGene_tx=XM_011285292.3;refGene_gn=ORC4;refGene_tx=X
M_019838083.2;refGene_gn=ORC4;refGene_tx=XM_019838084.1;refGene_gn=ORC
4;refGene_tx=XM_019838085.1;refGene_gn=ORC4;refGene_tx=XM_023259312.1;
refGene_gn=ORC4;refGene_tx=XM_023259313.1
C1 141388973 141389022
refGene_gn=ORC4;refGene_tx=XM_003990749.5;refGene_gn=ORC4;refGene_tx=X
M_006935270.3;refGene_gn=ORC4;refGene_tx=XM_006935271.4;refGene_gn=ORC
4;refGene_tx=XM_006935272.3;refGene_gn=ORC4;refGene_tx=XM_006935273.4;
refGene_gn=ORC4;refGene_tx=XM_011285292.3;refGene_gn=ORC4;refGene_tx=X
M_019838083.2;refGene_gn=ORC4;refGene_tx=XM_019838084.1;refGene_gn=ORC
4;refGene_tx=XM_019838085.1;refGene_gn=ORC4;refGene_tx=XM_023259312.1;
refGene_gn=ORC4;refGene_tx=XM_023259313.1
C1 141391783 141391869
refGene_gn=ORC4;refGene_tx=XM_003990749.5;refGene_gn=ORC4;refGene_tx=X
M_006935270.3;refGene_gn=ORC4;refGene_tx=XM_006935271.4;refGene_gn=ORC
4;refGene_tx=XM_006935272.3;refGene_gn=ORC4;refGene_tx=XM_006935273.4;
refGene_gn=ORC4;refGene_tx=XM_011285292.3;refGene_gn=ORC4;refGene_tx=X
M_019838083.2;refGene_gn=ORC4;refGene_tx=XM_019838084.1;refGene_gn=ORC
4;refGene_tx=XM_019838085.1;refGene_gn=ORC4;refGene_tx=XM_023259312.1;
refGene_gn=ORC4;refGene_tx=XM_023259313.1
C1 141392348 141392424
refGene_gn=ORC4;refGene_tx=XM_003990749.5;refGene_gn=ORC4;refGene_tx=X
M_006935270.3;refGene_gn=ORC4;refGene_tx=XM_006935271.4;refGene_gn=ORC
4;refGene_tx=XM_006935272.3;refGene_gn=ORC4;refGene_tx=XM_006935273.4;
refGene_gn=ORC4;refGene_tx=XM_011285292.3;refGene_gn=ORC4;refGene_tx=X
M_019838083.2;refGene_gn=ORC4;refGene_tx=XM_019838084.1;refGene_gn=ORC
4;refGene_tx=XM_019838085.1;refGene_gn=ORC4;refGene_tx=XM_023259312.1;
refGene_gn=ORC4;refGene_tx=XM_023259313.1
C1 141410995 141411086
refGene_gn=ORC4;refGene_tx=XM_003990749.5;refGene_gn=ORC4;refGene_tx=X
M_006935270.3;refGene_gn=ORC4;refGene_tx=XM_006935271.4;refGene_gn=ORC
4;refGene_tx=XM_006935272.3;refGene_gn=ORC4;refGene_tx=XM_011285292.3;
refGene_gn=ORC4;refGene_tx=XM_019838083.2;refGene_gn=ORC4;refGene_tx=X
M_019838084.1;refGene_gn=ORC4;refGene_tx=XM_023259312.1;refGene_gn=ORC
4;refGene_tx=XM_023259313.1
C1 141411668 141411745
refGene_gn=ORC4;refGene_tx=XM_003990749.5;refGene_gn=ORC4;refGene_tx=X
M_006935270.3;refGene_gn=ORC4;refGene_tx=XM_006935271.4;refGene_gn=ORC
4;refGene_tx=XM_019838083.2;refGene_gn=ORC4;refGene_tx=XM_019838084.1;
refGene_gn=ORC4;refGene_tx=XM_023259312.1;refGene_gn=ORC4;refGene_tx=X
M_023259313.1

C1	141414545	141414602
refGene_gn=ORC4;refGene_tx=XM_003990749.5;refGene_gn=ORC4;refGene_tx=XM_006935270.3;refGene_gn=ORC4;refGene_tx=XM_006935271.4;refGene_gn=ORC4;refGene_tx=XM_006935272.3;refGene_gn=ORC4;refGene_tx=XM_011285292.3;refGene_gn=ORC4;refGene_tx=XM_019838083.2;refGene_gn=ORC4;refGene_tx=XM_019838084.1;refGene_gn=ORC4;refGene_tx=XM_023259312.1;refGene_gn=ORC4;refGene_tx=XM_023259313.1
C1	141839748	141839861
refGene_gn=MBD5;refGene_tx=XM_019838080.2;refGene_gn=MBD5;refGene_tx=XM_019838081.2;refGene_gn=MBD5;refGene_tx=XM_019838082.2
C1	141848992	141849095
refGene_gn=MBD5;refGene_tx=XM_019838080.2;refGene_gn=MBD5;refGene_tx=XM_019838081.2;refGene_gn=MBD5;refGene_tx=XM_019838082.2
C1	141850219	141850400
refGene_gn=MBD5;refGene_tx=XM_019838080.2;refGene_gn=MBD5;refGene_tx=XM_019838081.2;refGene_gn=MBD5;refGene_tx=XM_019838082.2
C1	141855229	141857365
refGene_gn=MBD5;refGene_tx=XM_019838080.2;refGene_gn=MBD5;refGene_tx=XM_019838081.2;refGene_gn=MBD5;refGene_tx=XM_019838082.2
C1	141869562	141870588
refGene_gn=MBD5;refGene_tx=XM_019838080.2;refGene_gn=MBD5;refGene_tx=XM_019838081.2;refGene_gn=MBD5;refGene_tx=XM_019838082.2
C1	141872469	141872681
refGene_gn=MBD5;refGene_tx=XM_019838080.2;refGene_gn=MBD5;refGene_tx=XM_019838081.2;refGene_gn=MBD5;refGene_tx=XM_019838082.2
C1	141876693	141877902
refGene_gn=MBD5;refGene_tx=XM_019838080.2;refGene_gn=MBD5;refGene_tx=XM_019838081.2;refGene_gn=MBD5;refGene_tx=XM_019838082.2
C1	141888653	141888727
refGene_gn=MBD5;refGene_tx=XM_019838080.2;refGene_gn=MBD5;refGene_tx=XM_019838081.2;refGene_gn=MBD5;refGene_tx=XM_019838082.2
C1	141898005	141898081
refGene_gn=MBD5;refGene_tx=XM_019838080.2;refGene_gn=MBD5;refGene_tx=XM_019838081.2;refGene_gn=MBD5;refGene_tx=XM_019838082.2
C1	141900964	141901036
refGene_gn=MBD5;refGene_tx=XM_019838080.2;refGene_gn=MBD5;refGene_tx=XM_019838081.2;refGene_gn=MBD5;refGene_tx=XM_019838082.2
C1	142019056	142019209
refGene_gn=EPC2;refGene_tx=XM_003990752.5;refGene_gn=EPC2;refGene_tx=XM_006935275.4;refGene_gn=EPC2;refGene_tx=XM_006935276.4
C1	142067277	142067437
refGene_gn=EPC2;refGene_tx=XM_003990752.5;refGene_gn=EPC2;refGene_tx=XM_006935275.4;refGene_gn=EPC2;refGene_tx=XM_006935276.4
C1	142099546	142099692
refGene_gn=EPC2;refGene_tx=XM_003990752.5;refGene_gn=EPC2;refGene_tx=XM_006935275.4;refGene_gn=EPC2;refGene_tx=XM_006935276.4
C1	142105871	142106078
refGene_gn=EPC2;refGene_tx=XM_003990752.5;refGene_gn=EPC2;refGene_tx=XM_006935275.4;refGene_gn=EPC2;refGene_tx=XM_006935276.4
C1	142112129	142112278

refGene_gn=EPC2;refGene_tx=XM_003990752.5;refGene_gn=EPC2;refGene_tx=XM_006935275.4;refGene_gn=EPC2;refGene_tx=XM_006935276.4
C1 142113053 142113186
refGene_gn=EPC2;refGene_tx=XM_003990752.5;refGene_gn=EPC2;refGene_tx=XM_006935275.4;refGene_gn=EPC2;refGene_tx=XM_006935276.4
C1 142115368 142115560
refGene_gn=EPC2;refGene_tx=XM_003990752.5;refGene_gn=EPC2;refGene_tx=XM_006935275.4
C1 142119810 142119900
refGene_gn=EPC2;refGene_tx=XM_003990752.5;refGene_gn=EPC2;refGene_tx=XM_006935275.4;refGene_gn=EPC2;refGene_tx=XM_006935276.4
C1 142121154 142121300
refGene_gn=EPC2;refGene_tx=XM_003990752.5;refGene_gn=EPC2;refGene_tx=XM_006935275.4;refGene_gn=EPC2;refGene_tx=XM_006935276.4
C1 142121391 142121735
refGene_gn=EPC2;refGene_tx=XM_003990752.5;refGene_gn=EPC2;refGene_tx=XM_006935275.4;refGene_gn=EPC2;refGene_tx=XM_006935276.4
C1 142133614 142133751
refGene_gn=EPC2;refGene_tx=XM_003990752.5;refGene_gn=EPC2;refGene_tx=XM_006935275.4;refGene_gn=EPC2;refGene_tx=XM_006935276.4
C1 142135154 142135314
refGene_gn=EPC2;refGene_tx=XM_003990752.5;refGene_gn=EPC2;refGene_tx=XM_006935275.4;refGene_gn=EPC2;refGene_tx=XM_006935276.4
C1 142136476 142136810
refGene_gn=EPC2;refGene_tx=XM_003990752.5;refGene_gn=EPC2;refGene_tx=XM_006935275.4;refGene_gn=EPC2;refGene_tx=XM_006935276.4
C1 142138054 142138127
refGene_gn=EPC2;refGene_tx=XM_003990752.5;refGene_gn=EPC2;refGene_tx=XM_006935275.4;refGene_gn=EPC2;refGene_tx=XM_006935276.4
C1 142198020 142198146
refGene_gn=KIF5C;refGene_tx=XM_011285287.3
C1 142250546 142250637
refGene_gn=KIF5C;refGene_tx=XM_011285287.3
C1 142258139 142258213
refGene_gn=KIF5C;refGene_tx=XM_011285287.3
C1 142264981 142265086
refGene_gn=KIF5C;refGene_tx=XM_011285287.3
C1 142269454 142269503
refGene_gn=KIF5C;refGene_tx=XM_011285287.3
C1 142269777 142269833
refGene_gn=KIF5C;refGene_tx=XM_011285287.3
C1 142270528 142270616
refGene_gn=KIF5C;refGene_tx=XM_011285287.3
C1 142276330 142276455
refGene_gn=KIF5C;refGene_tx=XM_011285287.3
C1 142279332 142279437
refGene_gn=KIF5C;refGene_tx=XM_011285287.3
C1 142279807 142279956
refGene_gn=KIF5C;refGene_tx=XM_011285287.3
C1 142290953 142291102

refGene_gn=KIF5C;refGene_tx=XM_011285287.3
C1 142301053 142301229
refGene_gn=KIF5C;refGene_tx=XM_011285287.3
C1 142305997 142306066
refGene_gn=KIF5C;refGene_tx=XM_011285287.3
C1 142308214 142308421
refGene_gn=KIF5C;refGene_tx=XM_011285287.3
C1 142310720 142310867
refGene_gn=KIF5C;refGene_tx=XM_011285287.3
C1 142317307 142317496
refGene_gn=KIF5C;refGene_tx=XM_011285287.3
C1 142320758 142320876
refGene_gn=KIF5C;refGene_tx=XM_011285287.3
C1 142323085 142323162
refGene_gn=KIF5C;refGene_tx=XM_011285287.3
C1 142324495 142324605
refGene_gn=KIF5C;refGene_tx=XM_011285287.3
C1 142326910 142327012
refGene_gn=KIF5C;refGene_tx=XM_011285287.3
C1 142327236 142327297
refGene_gn=KIF5C;refGene_tx=XM_011285287.3
C1 142332672 142332744
refGene_gn=KIF5C;refGene_tx=XM_011285287.3
C1 142335339 142335444
refGene_gn=KIF5C;refGene_tx=XM_011285287.3
C1 142337389 142337606
refGene_gn=KIF5C;refGene_tx=XM_011285287.3
C1 142338805 142338912
refGene_gn=KIF5C;refGene_tx=XM_011285287.3
C1 142546894 142547024
refGene_gn=LYPD6B;refGene_tx=XM_003990774.3;refGene_gn=LYPD6B;refGene_tx=XM_019838086.2;refGene_gn=LYPD6B;refGene_tx=XM_019838087.2;refGene_gn=LYPD6B;refGene_tx=XM_019838088.2;refGene_gn=LYPD6B;refGene_tx=XM_019838089.2;refGene_gn=LYPD6B;refGene_tx=XM_023259314.1
C1 142549561 142549660
refGene_gn=LYPD6B;refGene_tx=XM_003990774.3;refGene_gn=LYPD6B;refGene_tx=XM_019838086.2;refGene_gn=LYPD6B;refGene_tx=XM_019838087.2;refGene_gn=LYPD6B;refGene_tx=XM_019838088.2;refGene_gn=LYPD6B;refGene_tx=XM_019838089.2;refGene_gn=LYPD6B;refGene_tx=XM_023259314.1
C1 142553299 142553430
refGene_gn=LYPD6B;refGene_tx=XM_003990774.3;refGene_gn=LYPD6B;refGene_tx=XM_019838086.2;refGene_gn=LYPD6B;refGene_tx=XM_019838087.2;refGene_gn=LYPD6B;refGene_tx=XM_019838088.2;refGene_gn=LYPD6B;refGene_tx=XM_019838089.2;refGene_gn=LYPD6B;refGene_tx=XM_023259314.1
C1 142555186 142555351
refGene_gn=LYPD6B;refGene_tx=XM_003990774.3;refGene_gn=LYPD6B;refGene_tx=XM_019838086.2;refGene_gn=LYPD6B;refGene_tx=XM_019838087.2;refGene_gn=LYPD6B;refGene_tx=XM_019838088.2;refGene_gn=LYPD6B;refGene_tx=XM_019838089.2;refGene_gn=LYPD6B;refGene_tx=XM_023259314.1
C1 142753389 142753507

refGene_gn=LYPD6;refGene_tx=XM_003990754.5;refGene_gn=LYPD6;refGene_tx=XM_006935277.4;refGene_gn=LYPD6;refGene_tx=XM_019838090.2;refGene_gn=LYPD6;refGene_tx=XM_019838091.1;refGene_gn=LYPD6;refGene_tx=XM_019838092.2
C1 142761058 142761157
refGene_gn=LYPD6;refGene_tx=XM_003990754.5;refGene_gn=LYPD6;refGene_tx=XM_006935277.4;refGene_gn=LYPD6;refGene_tx=XM_019838090.2;refGene_gn=LYPD6;refGene_tx=XM_019838091.1;refGene_gn=LYPD6;refGene_tx=XM_019838092.2
C1 142776680 142776836
refGene_gn=LYPD6;refGene_tx=XM_003990754.5;refGene_gn=LYPD6;refGene_tx=XM_006935277.4;refGene_gn=LYPD6;refGene_tx=XM_019838090.2;refGene_gn=LYPD6;refGene_tx=XM_019838091.1;refGene_gn=LYPD6;refGene_tx=XM_019838092.2
C1 142778747 142778915
refGene_gn=LYPD6;refGene_tx=XM_003990754.5;refGene_gn=LYPD6;refGene_tx=XM_006935277.4;refGene_gn=LYPD6;refGene_tx=XM_019838090.2;refGene_gn=LYPD6;refGene_tx=XM_019838091.1;refGene_gn=LYPD6;refGene_tx=XM_019838092.2
C1 142871624 142871819
refGene_gn=MMADHC;refGene_tx=XM_006935278.4;refGene_gn=MMADHC;refGene_tx=XM_006935279.4;refGene_gn=MMADHC;refGene_tx=XM_011285286.3;refGene_gn=MMADHC;refGene_tx=XM_019838093.2;refGene_gn=MMADHC;refGene_tx=XM_019838094.2
C1 142872567 142872654
refGene_gn=MMADHC;refGene_tx=XM_006935278.4;refGene_gn=MMADHC;refGene_tx=XM_006935279.4;refGene_gn=MMADHC;refGene_tx=XM_011285286.3;refGene_gn=MMADHC;refGene_tx=XM_019838093.2;refGene_gn=MMADHC;refGene_tx=XM_019838094.2
C1 142876456 142876587
refGene_gn=MMADHC;refGene_tx=XM_006935278.4;refGene_gn=MMADHC;refGene_tx=XM_006935279.4;refGene_gn=MMADHC;refGene_tx=XM_011285286.3;refGene_gn=MMADHC;refGene_tx=XM_019838093.2;refGene_gn=MMADHC;refGene_tx=XM_019838094.2
C1 142877393 142877499
refGene_gn=MMADHC;refGene_tx=XM_006935278.4;refGene_gn=MMADHC;refGene_tx=XM_006935279.4;refGene_gn=MMADHC;refGene_tx=XM_011285286.3;refGene_gn=MMADHC;refGene_tx=XM_019838093.2;refGene_gn=MMADHC;refGene_tx=XM_019838094.2
C1 142879825 142880043
refGene_gn=MMADHC;refGene_tx=XM_006935278.4;refGene_gn=MMADHC;refGene_tx=XM_006935279.4;refGene_gn=MMADHC;refGene_tx=XM_011285286.3;refGene_gn=MMADHC;refGene_tx=XM_019838093.2;refGene_gn=MMADHC;refGene_tx=XM_019838094.2
C1 142881693 142881729
refGene_gn=MMADHC;refGene_tx=XM_006935278.4;refGene_gn=MMADHC;refGene_tx=XM_006935279.4;refGene_gn=MMADHC;refGene_tx=XM_019838093.2;refGene_gn=MMADHC;refGene_tx=XM_019838094.2
C1 142903720 142903865
refGene_gn=MMADHC;refGene_tx=XM_006935278.4;refGene_gn=MMADHC;refGene_

tx=XM_006935279.4;refGene_gn=MMADHC;refGene_tx=XM_011285286.3;refGene_gn=MMADHC;refGene_tx=XM_019838093.2
C1 142908673 142908682
refGene_gn=MMADHC;refGene_tx=XM_006935278.4;refGene_gn=MMADHC;refGene_tx=XM_006935279.4;refGene_gn=MMADHC;refGene_tx=XM_011285286.3;refGene_gn=MMADHC;refGene_tx=XM_019838093.2
C1 143659522 143659774
refGene_gn=RND3;refGene_tx=XM_003990756.5
C1 143661215 143661350
refGene_gn=RND3;refGene_tx=XM_003990756.5
C1 143664524 143664634
refGene_gn=RND3;refGene_tx=XM_003990756.5
C1 143676639 143676727
refGene_gn=RND3;refGene_tx=XM_003990756.5
C1 143677212 143677362
refGene_gn=RND3;refGene_tx=XM_003990756.5
C1 144346199 144346949
refGene_gn=RBM43;refGene_tx=XM_011285285.3;refGene_gn=RBM43;refGene_tx=XM_019838098.2
C1 144347731 144347820
refGene_gn=RBM43;refGene_tx=XM_011285285.3;refGene_gn=RBM43;refGene_tx=XM_019838098.2
C1 144350502 144350719
refGene_gn=RBM43;refGene_tx=XM_011285285.3;refGene_gn=RBM43;refGene_tx=XM_019838098.2
C1 144356674 144356677
refGene_gn=RBM43;refGene_tx=XM_011285285.3
C1 144362808 144362994
refGene_gn=NMI;refGene_tx=XM_023259315.1
C1 144363743 144363850
refGene_gn=NMI;refGene_tx=XM_023259315.1
C1 144366446 144366633
refGene_gn=NMI;refGene_tx=XM_023259315.1
C1 144366720 144366827
refGene_gn=NMI;refGene_tx=XM_023259315.1
C1 144370669 144370834
refGene_gn=NMI;refGene_tx=XM_023259315.1
C1 144373502 144373596
refGene_gn=NMI;refGene_tx=XM_023259315.1
C1 144374310 144374394
refGene_gn=NMI;refGene_tx=XM_023259315.1
C1 144427226 144427320
refGene_gn=TNFAIP6;refGene_tx=XM_003990758.4
C1 144431999 144432137
refGene_gn=TNFAIP6;refGene_tx=XM_003990758.4
C1 144433832 144433994
refGene_gn=TNFAIP6;refGene_tx=XM_003990758.4
C1 144437754 144437983
refGene_gn=TNFAIP6;refGene_tx=XM_003990758.4
C1 144440196 144440237

refGene_gn=TNFAIP6;refGene_tx=XM_003990758.4
C1 144443728 144443898
refGene_gn=TNFAIP6;refGene_tx=XM_003990758.4
C1 144474049 144474153
refGene_gn=RIF1;refGene_tx=XM_019838099.2;refGene_gn=RIF1;refGene_tx=XM_023259316.1;refGene_gn=RIF1;refGene_tx=XM_023259317.1;refGene_gn=RIF1;refGene_tx=XM_023259318.1;refGene_gn=RIF1;refGene_tx=XM_023259319.1
C1 144474928 144475007
refGene_gn=RIF1;refGene_tx=XM_019838099.2;refGene_gn=RIF1;refGene_tx=XM_023259316.1;refGene_gn=RIF1;refGene_tx=XM_023259317.1;refGene_gn=RIF1;refGene_tx=XM_023259318.1;refGene_gn=RIF1;refGene_tx=XM_023259319.1
C1 144477777 144477874
refGene_gn=RIF1;refGene_tx=XM_019838099.2;refGene_gn=RIF1;refGene_tx=XM_023259316.1;refGene_gn=RIF1;refGene_tx=XM_023259317.1;refGene_gn=RIF1;refGene_tx=XM_023259318.1;refGene_gn=RIF1;refGene_tx=XM_023259319.1
C1 144478966 144479094
refGene_gn=RIF1;refGene_tx=XM_019838099.2;refGene_gn=RIF1;refGene_tx=XM_023259316.1;refGene_gn=RIF1;refGene_tx=XM_023259317.1;refGene_gn=RIF1;refGene_tx=XM_023259318.1;refGene_gn=RIF1;refGene_tx=XM_023259319.1
C1 144479187 144479282
refGene_gn=RIF1;refGene_tx=XM_019838099.2;refGene_gn=RIF1;refGene_tx=XM_023259316.1;refGene_gn=RIF1;refGene_tx=XM_023259317.1;refGene_gn=RIF1;refGene_tx=XM_023259318.1;refGene_gn=RIF1;refGene_tx=XM_023259319.1
C1 144480385 144480575
refGene_gn=RIF1;refGene_tx=XM_019838099.2;refGene_gn=RIF1;refGene_tx=XM_023259316.1;refGene_gn=RIF1;refGene_tx=XM_023259317.1;refGene_gn=RIF1;refGene_tx=XM_023259318.1;refGene_gn=RIF1;refGene_tx=XM_023259319.1
C1 144482358 144482451
refGene_gn=RIF1;refGene_tx=XM_019838099.2;refGene_gn=RIF1;refGene_tx=XM_023259316.1;refGene_gn=RIF1;refGene_tx=XM_023259317.1;refGene_gn=RIF1;refGene_tx=XM_023259318.1;refGene_gn=RIF1;refGene_tx=XM_023259319.1
C1 144487836 144487975
refGene_gn=RIF1;refGene_tx=XM_019838099.2;refGene_gn=RIF1;refGene_tx=XM_023259316.1;refGene_gn=RIF1;refGene_tx=XM_023259317.1;refGene_gn=RIF1;refGene_tx=XM_023259318.1;refGene_gn=RIF1;refGene_tx=XM_023259319.1
C1 144490447 144490599
refGene_gn=RIF1;refGene_tx=XM_019838099.2;refGene_gn=RIF1;refGene_tx=XM_023259316.1;refGene_gn=RIF1;refGene_tx=XM_023259317.1;refGene_gn=RIF1;refGene_tx=XM_023259318.1;refGene_gn=RIF1;refGene_tx=XM_023259319.1
C1 144491978 144492096
refGene_gn=RIF1;refGene_tx=XM_019838099.2;refGene_gn=RIF1;refGene_tx=XM_023259316.1;refGene_gn=RIF1;refGene_tx=XM_023259317.1;refGene_gn=RIF1;refGene_tx=XM_023259318.1;refGene_gn=RIF1;refGene_tx=XM_023259319.1
C1 144492994 144493168
refGene_gn=RIF1;refGene_tx=XM_019838099.2;refGene_gn=RIF1;refGene_tx=XM_023259316.1;refGene_gn=RIF1;refGene_tx=XM_023259317.1;refGene_gn=RIF1;refGene_tx=XM_023259318.1;refGene_gn=RIF1;refGene_tx=XM_023259319.1
C1 144493382 144493493
refGene_gn=RIF1;refGene_tx=XM_019838099.2;refGene_gn=RIF1;refGene_tx=XM_023259316.1;refGene_gn=RIF1;refGene_tx=XM_023259317.1;refGene_gn=RIF

1;refGene_tx=XM_023259318.1;refGene_gn=RIF1;refGene_tx=XM_023259319.1
C1 144494160 144494223
refGene_gn=RIF1;refGene_tx=XM_019838099.2;refGene_gn=RIF1;refGene_tx=XM_023259316.1;refGene_gn=RIF1;refGene_tx=XM_023259317.1;refGene_gn=RIF1;refGene_tx=XM_023259318.1;refGene_gn=RIF1;refGene_tx=XM_023259319.1
C1 144494488 144494589
refGene_gn=RIF1;refGene_tx=XM_019838099.2;refGene_gn=RIF1;refGene_tx=XM_023259316.1;refGene_gn=RIF1;refGene_tx=XM_023259317.1;refGene_gn=RIF1;refGene_tx=XM_023259318.1;refGene_gn=RIF1;refGene_tx=XM_023259319.1
C1 144502209 144502296
refGene_gn=RIF1;refGene_tx=XM_019838099.2;refGene_gn=RIF1;refGene_tx=XM_023259316.1;refGene_gn=RIF1;refGene_tx=XM_023259317.1;refGene_gn=RIF1;refGene_tx=XM_023259318.1;refGene_gn=RIF1;refGene_tx=XM_023259319.1
C1 144502931 144503005
refGene_gn=RIF1;refGene_tx=XM_019838099.2;refGene_gn=RIF1;refGene_tx=XM_023259316.1;refGene_gn=RIF1;refGene_tx=XM_023259317.1;refGene_gn=RIF1;refGene_tx=XM_023259318.1;refGene_gn=RIF1;refGene_tx=XM_023259319.1
C1 144503453 144503634
refGene_gn=RIF1;refGene_tx=XM_019838099.2;refGene_gn=RIF1;refGene_tx=XM_023259316.1;refGene_gn=RIF1;refGene_tx=XM_023259317.1;refGene_gn=RIF1;refGene_tx=XM_023259318.1;refGene_gn=RIF1;refGene_tx=XM_023259319.1
C1 144504944 144505052
refGene_gn=RIF1;refGene_tx=XM_019838099.2;refGene_gn=RIF1;refGene_tx=XM_023259316.1;refGene_gn=RIF1;refGene_tx=XM_023259317.1;refGene_gn=RIF1;refGene_tx=XM_023259318.1;refGene_gn=RIF1;refGene_tx=XM_023259319.1
C1 144505409 144505559
refGene_gn=RIF1;refGene_tx=XM_019838099.2;refGene_gn=RIF1;refGene_tx=XM_023259316.1;refGene_gn=RIF1;refGene_tx=XM_023259317.1;refGene_gn=RIF1;refGene_tx=XM_023259318.1;refGene_gn=RIF1;refGene_tx=XM_023259319.1
C1 144522458 144522558
refGene_gn=RIF1;refGene_tx=XM_019838099.2;refGene_gn=RIF1;refGene_tx=XM_023259316.1;refGene_gn=RIF1;refGene_tx=XM_023259317.1;refGene_gn=RIF1;refGene_tx=XM_023259318.1;refGene_gn=RIF1;refGene_tx=XM_023259319.1
C1 144524427 144524692
refGene_gn=RIF1;refGene_tx=XM_019838099.2;refGene_gn=RIF1;refGene_tx=XM_023259316.1;refGene_gn=RIF1;refGene_tx=XM_023259317.1;refGene_gn=RIF1;refGene_tx=XM_023259318.1;refGene_gn=RIF1;refGene_tx=XM_023259319.1
C1 144526030 144526073
refGene_gn=RIF1;refGene_tx=XM_019838099.2;refGene_gn=RIF1;refGene_tx=XM_023259316.1;refGene_gn=RIF1;refGene_tx=XM_023259317.1;refGene_gn=RIF1;refGene_tx=XM_023259318.1;refGene_gn=RIF1;refGene_tx=XM_023259319.1
C1 144526613 144526816
refGene_gn=RIF1;refGene_tx=XM_019838099.2;refGene_gn=RIF1;refGene_tx=XM_023259316.1;refGene_gn=RIF1;refGene_tx=XM_023259317.1;refGene_gn=RIF1;refGene_tx=XM_023259318.1;refGene_gn=RIF1;refGene_tx=XM_023259319.1
C1 144527366 144527466
refGene_gn=RIF1;refGene_tx=XM_019838099.2;refGene_gn=RIF1;refGene_tx=XM_023259316.1;refGene_gn=RIF1;refGene_tx=XM_023259317.1;refGene_gn=RIF1;refGene_tx=XM_023259318.1;refGene_gn=RIF1;refGene_tx=XM_023259319.1
C1 144530545 144530665

refGene_gn=RIF1;refGene_tx=XM_019838099.2;refGene_gn=RIF1;refGene_tx=XM_023259316.1;refGene_gn=RIF1;refGene_tx=XM_023259317.1;refGene_gn=RIF1;refGene_tx=XM_023259318.1;refGene_gn=RIF1;refGene_tx=XM_023259319.1
C1 144531631 144531783
refGene_gn=RIF1;refGene_tx=XM_019838099.2;refGene_gn=RIF1;refGene_tx=XM_023259316.1;refGene_gn=RIF1;refGene_tx=XM_023259317.1;refGene_gn=RIF1;refGene_tx=XM_023259318.1;refGene_gn=RIF1;refGene_tx=XM_023259319.1
C1 144532159 144532240
refGene_gn=RIF1;refGene_tx=XM_019838099.2;refGene_gn=RIF1;refGene_tx=XM_023259316.1;refGene_gn=RIF1;refGene_tx=XM_023259317.1;refGene_gn=RIF1;refGene_tx=XM_023259318.1;refGene_gn=RIF1;refGene_tx=XM_023259319.1
C1 144532532 144532587
refGene_gn=RIF1;refGene_tx=XM_019838099.2;refGene_gn=RIF1;refGene_tx=XM_023259316.1;refGene_gn=RIF1;refGene_tx=XM_023259317.1;refGene_gn=RIF1;refGene_tx=XM_023259318.1;refGene_gn=RIF1;refGene_tx=XM_023259319.1
C1 144532944 144536205
refGene_gn=RIF1;refGene_tx=XM_019838099.2;refGene_gn=RIF1;refGene_tx=XM_023259316.1;refGene_gn=RIF1;refGene_tx=XM_023259317.1;refGene_gn=RIF1;refGene_tx=XM_023259318.1;refGene_gn=RIF1;refGene_tx=XM_023259319.1
C1 144538152 144538296
refGene_gn=RIF1;refGene_tx=XM_019838099.2;refGene_gn=RIF1;refGene_tx=XM_023259316.1;refGene_gn=RIF1;refGene_tx=XM_023259317.1;refGene_gn=RIF1;refGene_tx=XM_023259318.1;refGene_gn=RIF1;refGene_tx=XM_023259319.1
C1 144538609 144538687
refGene_gn=RIF1;refGene_tx=XM_019838099.2;refGene_gn=RIF1;refGene_tx=XM_023259316.1;refGene_gn=RIF1;refGene_tx=XM_023259317.1;refGene_gn=RIF1;refGene_tx=XM_023259318.1
C1 144538775 144538891
refGene_gn=RIF1;refGene_tx=XM_019838099.2;refGene_gn=RIF1;refGene_tx=XM_023259316.1;refGene_gn=RIF1;refGene_tx=XM_023259317.1;refGene_gn=RIF1;refGene_tx=XM_023259318.1;refGene_gn=RIF1;refGene_tx=XM_023259319.1
C1 144539986 144540140
refGene_gn=RIF1;refGene_tx=XM_019838099.2;refGene_gn=RIF1;refGene_tx=XM_023259316.1;refGene_gn=RIF1;refGene_tx=XM_023259317.1;refGene_gn=RIF1;refGene_tx=XM_023259318.1;refGene_gn=RIF1;refGene_tx=XM_023259319.1
C1 144541734 144541843
refGene_gn=RIF1;refGene_tx=XM_019838099.2;refGene_gn=RIF1;refGene_tx=XM_023259316.1;refGene_gn=RIF1;refGene_tx=XM_023259317.1;refGene_gn=RIF1;refGene_tx=XM_023259318.1;refGene_gn=RIF1;refGene_tx=XM_023259319.1
C1 144542313 144542528
refGene_gn=RIF1;refGene_tx=XM_019838099.2;refGene_gn=RIF1;refGene_tx=XM_023259316.1;refGene_gn=RIF1;refGene_tx=XM_023259317.1;refGene_gn=RIF1;refGene_tx=XM_023259318.1;refGene_gn=RIF1;refGene_tx=XM_023259319.1
C1 144823902 144823951
refGene_gn=ARL5A;refGene_tx=XM_003990761.5
C1 144826869 144827021
refGene_gn=ARL5A;refGene_tx=XM_003990761.5
C1 144832281 144832365
refGene_gn=ARL5A;refGene_tx=XM_003990761.5
C1 144834220 144834368

refGene_gn=ARL5A;refGene_tx=XM_003990761.5
C1	144835062	144835123
refGene_gn=ARL5A;refGene_tx=XM_003990761.5
C1	144846378	144846424
refGene_gn=ARL5A;refGene_tx=XM_003990761.5
C1	144857423	144857684
refGene_gn=CACNB4;refGene_tx=XM_003990762.5;refGene_gn=CACNB4;refGene_tx=XM_019838110.2;refGene_gn=CACNB4;refGene_tx=XM_019838111.2;refGene_gn=CACNB4;refGene_tx=XM_019838112.2;refGene_gn=CACNB4;refGene_tx=XM_019838113.2;refGene_gn=CACNB4;refGene_tx=XM_019838114.2
C1	144859944	144860130
refGene_gn=CACNB4;refGene_tx=XM_003990762.5;refGene_gn=CACNB4;refGene_tx=XM_019838110.2;refGene_gn=CACNB4;refGene_tx=XM_019838111.2;refGene_gn=CACNB4;refGene_tx=XM_019838112.2;refGene_gn=CACNB4;refGene_tx=XM_019838113.2;refGene_gn=CACNB4;refGene_tx=XM_019838114.2
C1	144873240	144873336
refGene_gn=CACNB4;refGene_tx=XM_003990762.5;refGene_gn=CACNB4;refGene_tx=XM_019838110.2;refGene_gn=CACNB4;refGene_tx=XM_019838111.2;refGene_gn=CACNB4;refGene_tx=XM_019838112.2;refGene_gn=CACNB4;refGene_tx=XM_019838113.2;refGene_gn=CACNB4;refGene_tx=XM_019838114.2
C1	144875671	144875823
refGene_gn=CACNB4;refGene_tx=XM_003990762.5;refGene_gn=CACNB4;refGene_tx=XM_019838110.2;refGene_gn=CACNB4;refGene_tx=XM_019838111.2;refGene_gn=CACNB4;refGene_tx=XM_019838112.2;refGene_gn=CACNB4;refGene_tx=XM_019838113.2;refGene_gn=CACNB4;refGene_tx=XM_019838114.2
C1	144881423	144881533
refGene_gn=CACNB4;refGene_tx=XM_003990762.5;refGene_gn=CACNB4;refGene_tx=XM_019838110.2;refGene_gn=CACNB4;refGene_tx=XM_019838111.2;refGene_gn=CACNB4;refGene_tx=XM_019838112.2;refGene_gn=CACNB4;refGene_tx=XM_019838113.2;refGene_gn=CACNB4;refGene_tx=XM_019838114.2
C1	144890652	144890711
refGene_gn=CACNB4;refGene_tx=XM_003990762.5;refGene_gn=CACNB4;refGene_tx=XM_019838110.2;refGene_gn=CACNB4;refGene_tx=XM_019838111.2;refGene_gn=CACNB4;refGene_tx=XM_019838112.2;refGene_gn=CACNB4;refGene_tx=XM_019838113.2;refGene_gn=CACNB4;refGene_tx=XM_019838114.2
C1	144892025	144892106
refGene_gn=CACNB4;refGene_tx=XM_003990762.5;refGene_gn=CACNB4;refGene_tx=XM_019838110.2;refGene_gn=CACNB4;refGene_tx=XM_019838111.2;refGene_gn=CACNB4;refGene_tx=XM_019838112.2;refGene_gn=CACNB4;refGene_tx=XM_019838113.2;refGene_gn=CACNB4;refGene_tx=XM_019838114.2
C1	144892328	144892348
refGene_gn=CACNB4;refGene_tx=XM_003990762.5;refGene_gn=CACNB4;refGene_tx=XM_019838110.2;refGene_gn=CACNB4;refGene_tx=XM_019838111.2;refGene_gn=CACNB4;refGene_tx=XM_019838112.2;refGene_gn=CACNB4;refGene_tx=XM_019838113.2
C1	144893885	144893962
refGene_gn=CACNB4;refGene_tx=XM_003990762.5;refGene_gn=CACNB4;refGene_tx=XM_019838110.2;refGene_gn=CACNB4;refGene_tx=XM_019838111.2;refGene_gn=CACNB4;refGene_tx=XM_019838112.2;refGene_gn=CACNB4;refGene_tx=XM_019838113.2

C1	144896362	144896493
refGene_gn=CACNB4;refGene_tx=XM_003990762.5;refGene_gn=CACNB4;refGene_tx=XM_019838110.2;refGene_gn=CACNB4;refGene_tx=XM_019838111.2;refGene_gn=CACNB4;refGene_tx=XM_019838112.2;refGene_gn=CACNB4;refGene_tx=XM_019838113.2
C1	144905834	144905957
refGene_gn=CACNB4;refGene_tx=XM_003990762.5;refGene_gn=CACNB4;refGene_tx=XM_019838110.2;refGene_gn=CACNB4;refGene_tx=XM_019838111.2;refGene_gn=CACNB4;refGene_tx=XM_019838112.2;refGene_gn=CACNB4;refGene_tx=XM_019838113.2
C1	144907794	144907914
refGene_gn=CACNB4;refGene_tx=XM_003990762.5;refGene_gn=CACNB4;refGene_tx=XM_019838110.2;refGene_gn=CACNB4;refGene_tx=XM_019838111.2;refGene_gn=CACNB4;refGene_tx=XM_019838112.2;refGene_gn=CACNB4;refGene_tx=XM_019838113.2
C1	144992551	144992557
refGene_gn=CACNB4;refGene_tx=XM_019838111.2;refGene_gn=CACNB4;refGene_tx=XM_019838112.2;refGene_gn=CACNB4;refGene_tx=XM_019838113.2
C1	144994649	144994694
refGene_gn=CACNB4;refGene_tx=XM_019838110.2
C1	145106538	145106622
refGene_gn=CACNB4;refGene_tx=XM_003990762.5
C1	145106848	145106857
refGene_gn=CACNB4;refGene_tx=XM_003990762.5
C1	145123610	145123839
refGene_gn=STAM2;refGene_tx=XM_003990764.4;refGene_gn=STAM2;refGene_tx=XM_006935282.3;refGene_gn=STAM2;refGene_tx=XM_006935283.3;refGene_gn=STAM2;refGene_tx=XM_019838115.2
C1	145125262	145125432
refGene_gn=STAM2;refGene_tx=XM_003990764.4;refGene_gn=STAM2;refGene_tx=XM_006935282.3;refGene_gn=STAM2;refGene_tx=XM_006935283.3;refGene_gn=STAM2;refGene_tx=XM_019838115.2
C1	145128061	145128215
refGene_gn=STAM2;refGene_tx=XM_003990764.4;refGene_gn=STAM2;refGene_tx=XM_006935282.3;refGene_gn=STAM2;refGene_tx=XM_006935283.3;refGene_gn=STAM2;refGene_tx=XM_019838115.2
C1	145132124	145132179
refGene_gn=STAM2;refGene_tx=XM_003990764.4;refGene_gn=STAM2;refGene_tx=XM_006935282.3;refGene_gn=STAM2;refGene_tx=XM_006935283.3;refGene_gn=STAM2;refGene_tx=XM_019838115.2
C1	145133190	145133278
refGene_gn=STAM2;refGene_tx=XM_003990764.4;refGene_gn=STAM2;refGene_tx=XM_006935282.3;refGene_gn=STAM2;refGene_tx=XM_006935283.3;refGene_gn=STAM2;refGene_tx=XM_019838115.2
C1	145133418	145133501
refGene_gn=STAM2;refGene_tx=XM_003990764.4;refGene_gn=STAM2;refGene_tx=XM_006935282.3;refGene_gn=STAM2;refGene_tx=XM_006935283.3;refGene_gn=STAM2;refGene_tx=XM_019838115.2
C1	145136073	145136168
refGene_gn=STAM2;refGene_tx=XM_003990764.4;refGene_gn=STAM2;refGene_tx

=XM_006935282.3;refGene_gn=STAM2;refGene_tx=XM_006935283.3;refGene_gn=STAM2;refGene_tx=XM_019838115.2
C1	145145350	145145537
refGene_gn=STAM2;refGene_tx=XM_003990764.4;refGene_gn=STAM2;refGene_tx=XM_006935282.3;refGene_gn=STAM2;refGene_tx=XM_006935283.3;refGene_gn=STAM2;refGene_tx=XM_019838115.2
C1	145146130	145146200
refGene_gn=STAM2;refGene_tx=XM_003990764.4;refGene_gn=STAM2;refGene_tx=XM_006935282.3;refGene_gn=STAM2;refGene_tx=XM_006935283.3;refGene_gn=STAM2;refGene_tx=XM_019838115.2
C1	145148353	145148500
refGene_gn=STAM2;refGene_tx=XM_003990764.4;refGene_gn=STAM2;refGene_tx=XM_006935282.3;refGene_gn=STAM2;refGene_tx=XM_006935283.3;refGene_gn=STAM2;refGene_tx=XM_019838115.2
C1	145149429	145149528
refGene_gn=STAM2;refGene_tx=XM_003990764.4;refGene_gn=STAM2;refGene_tx=XM_006935282.3;refGene_gn=STAM2;refGene_tx=XM_006935283.3;refGene_gn=STAM2;refGene_tx=XM_019838115.2
C1	145149646	145149722
refGene_gn=STAM2;refGene_tx=XM_003990764.4;refGene_gn=STAM2;refGene_tx=XM_006935282.3;refGene_gn=STAM2;refGene_tx=XM_006935283.3;refGene_gn=STAM2;refGene_tx=XM_019838115.2
C1	145151087	145151172
refGene_gn=STAM2;refGene_tx=XM_003990764.4;refGene_gn=STAM2;refGene_tx=XM_006935282.3;refGene_gn=STAM2;refGene_tx=XM_006935283.3;refGene_gn=STAM2;refGene_tx=XM_019838115.2
C1	145172468	145172508
refGene_gn=STAM2;refGene_tx=XM_003990764.4;refGene_gn=STAM2;refGene_tx=XM_006935282.3
C1	145323984	145324101
refGene_gn=FMNL2;refGene_tx=XM_003990765.5;refGene_gn=FMNL2;refGene_tx=XM_023259320.1;refGene_gn=FMNL2;refGene_tx=XM_023259321.1;refGene_gn=FMNL2;refGene_tx=XM_023259322.1;refGene_gn=FMNL2;refGene_tx=XM_023259323.1;refGene_gn=FMNL2;refGene_tx=XM_023259324.1;refGene_gn=FMNL2;refGene_tx=XM_023259325.1;refGene_gn=FMNL2;refGene_tx=XM_023259326.1;refGene_gn=FMNL2;refGene_tx=XM_023259327.1
C1	145505893	145505977
refGene_gn=FMNL2;refGene_tx=XM_003990765.5;refGene_gn=FMNL2;refGene_tx=XM_023259320.1;refGene_gn=FMNL2;refGene_tx=XM_023259321.1;refGene_gn=FMNL2;refGene_tx=XM_023259322.1;refGene_gn=FMNL2;refGene_tx=XM_023259323.1;refGene_gn=FMNL2;refGene_tx=XM_023259324.1;refGene_gn=FMNL2;refGene_tx=XM_023259325.1;refGene_gn=FMNL2;refGene_tx=XM_023259326.1;refGene_gn=FMNL2;refGene_tx=XM_023259327.1
C1	145530380	145530461
refGene_gn=FMNL2;refGene_tx=XM_003990765.5;refGene_gn=FMNL2;refGene_tx=XM_023259320.1;refGene_gn=FMNL2;refGene_tx=XM_023259321.1;refGene_gn=FMNL2;refGene_tx=XM_023259322.1;refGene_gn=FMNL2;refGene_tx=XM_023259323.1;refGene_gn=FMNL2;refGene_tx=XM_023259324.1;refGene_gn=FMNL2;refGene_tx=XM_023259325.1;refGene_gn=FMNL2;refGene_tx=XM_023259326.1;refGene_gn=FMNL2;refGene_tx=XM_023259327.1

C1 145535748 145535825
refGene_gn=FMNL2;refGene_tx=XM_003990765.5;refGene_gn=FMNL2;refGene_tx=XM_023259320.1;refGene_gn=FMNL2;refGene_tx=XM_023259321.1;refGene_gn=FMNL2;refGene_tx=XM_023259322.1;refGene_gn=FMNL2;refGene_tx=XM_023259323.1;refGene_gn=FMNL2;refGene_tx=XM_023259324.1;refGene_gn=FMNL2;refGene_tx=XM_023259325.1;refGene_gn=FMNL2;refGene_tx=XM_023259326.1;refGene_gn=FMNL2;refGene_tx=XM_023259327.1
C1 145545590 145545674
refGene_gn=FMNL2;refGene_tx=XM_003990765.5;refGene_gn=FMNL2;refGene_tx=XM_023259320.1;refGene_gn=FMNL2;refGene_tx=XM_023259321.1;refGene_gn=FMNL2;refGene_tx=XM_023259322.1;refGene_gn=FMNL2;refGene_tx=XM_023259323.1;refGene_gn=FMNL2;refGene_tx=XM_023259324.1;refGene_gn=FMNL2;refGene_tx=XM_023259325.1;refGene_gn=FMNL2;refGene_tx=XM_023259326.1;refGene_gn=FMNL2;refGene_tx=XM_023259327.1
C1 145548698 145548851
refGene_gn=FMNL2;refGene_tx=XM_003990765.5;refGene_gn=FMNL2;refGene_tx=XM_023259320.1;refGene_gn=FMNL2;refGene_tx=XM_023259321.1;refGene_gn=FMNL2;refGene_tx=XM_023259322.1;refGene_gn=FMNL2;refGene_tx=XM_023259323.1;refGene_gn=FMNL2;refGene_tx=XM_023259324.1;refGene_gn=FMNL2;refGene_tx=XM_023259325.1;refGene_gn=FMNL2;refGene_tx=XM_023259326.1;refGene_gn=FMNL2;refGene_tx=XM_023259327.1
C1 145552671 145552811
refGene_gn=FMNL2;refGene_tx=XM_023259328.1
C1 145570741 145570850
refGene_gn=FMNL2;refGene_tx=XM_003990765.5;refGene_gn=FMNL2;refGene_tx=XM_023259320.1;refGene_gn=FMNL2;refGene_tx=XM_023259321.1;refGene_gn=FMNL2;refGene_tx=XM_023259322.1;refGene_gn=FMNL2;refGene_tx=XM_023259323.1;refGene_gn=FMNL2;refGene_tx=XM_023259324.1;refGene_gn=FMNL2;refGene_tx=XM_023259325.1;refGene_gn=FMNL2;refGene_tx=XM_023259326.1;refGene_gn=FMNL2;refGene_tx=XM_023259327.1;refGene_gn=FMNL2;refGene_tx=XM_023259328.1
C1 145574992 145575069
refGene_gn=FMNL2;refGene_tx=XM_003990765.5;refGene_gn=FMNL2;refGene_tx=XM_023259320.1;refGene_gn=FMNL2;refGene_tx=XM_023259321.1;refGene_gn=FMNL2;refGene_tx=XM_023259322.1;refGene_gn=FMNL2;refGene_tx=XM_023259323.1;refGene_gn=FMNL2;refGene_tx=XM_023259324.1;refGene_gn=FMNL2;refGene_tx=XM_023259325.1;refGene_gn=FMNL2;refGene_tx=XM_023259326.1;refGene_gn=FMNL2;refGene_tx=XM_023259327.1;refGene_gn=FMNL2;refGene_tx=XM_023259328.1
C1 145576789 145576883
refGene_gn=FMNL2;refGene_tx=XM_003990765.5;refGene_gn=FMNL2;refGene_tx=XM_023259320.1;refGene_gn=FMNL2;refGene_tx=XM_023259321.1;refGene_gn=FMNL2;refGene_tx=XM_023259322.1;refGene_gn=FMNL2;refGene_tx=XM_023259323.1;refGene_gn=FMNL2;refGene_tx=XM_023259324.1;refGene_gn=FMNL2;refGene_tx=XM_023259325.1;refGene_gn=FMNL2;refGene_tx=XM_023259326.1;refGene_gn=FMNL2;refGene_tx=XM_023259327.1;refGene_gn=FMNL2;refGene_tx=XM_023259328.1
C1 145600063 145600138
refGene_gn=FMNL2;refGene_tx=XM_003990765.5;refGene_gn=FMNL2;refGene_tx=XM_023259320.1;refGene_gn=FMNL2;refGene_tx=XM_023259321.1;refGene_gn=

FMNL2;refGene_tx=XM_023259322.1;refGene_gn=FMNL2;refGene_tx=XM_023259323.1;refGene_gn=FMNL2;refGene_tx=XM_023259324.1;refGene_gn=FMNL2;refGene_tx=XM_023259325.1;refGene_gn=FMNL2;refGene_tx=XM_023259326.1;refGene_gn=FMNL2;refGene_tx=XM_023259327.1;refGene_gn=FMNL2;refGene_tx=XM_023259328.1
C1 145605122 145605233
refGene_gn=FMNL2;refGene_tx=XM_003990765.5;refGene_gn=FMNL2;refGene_tx=XM_023259320.1;refGene_gn=FMNL2;refGene_tx=XM_023259321.1;refGene_gn=FMNL2;refGene_tx=XM_023259322.1;refGene_gn=FMNL2;refGene_tx=XM_023259323.1;refGene_gn=FMNL2;refGene_tx=XM_023259324.1;refGene_gn=FMNL2;refGene_tx=XM_023259325.1;refGene_gn=FMNL2;refGene_tx=XM_023259326.1;refGene_gn=FMNL2;refGene_tx=XM_023259327.1;refGene_gn=FMNL2;refGene_tx=XM_023259328.1
C1 145607887 145608037
refGene_gn=FMNL2;refGene_tx=XM_003990765.5;refGene_gn=FMNL2;refGene_tx=XM_023259320.1;refGene_gn=FMNL2;refGene_tx=XM_023259321.1;refGene_gn=FMNL2;refGene_tx=XM_023259322.1;refGene_gn=FMNL2;refGene_tx=XM_023259323.1;refGene_gn=FMNL2;refGene_tx=XM_023259324.1;refGene_gn=FMNL2;refGene_tx=XM_023259325.1;refGene_gn=FMNL2;refGene_tx=XM_023259326.1;refGene_gn=FMNL2;refGene_tx=XM_023259327.1;refGene_gn=FMNL2;refGene_tx=XM_023259328.1
C1 145610519 145610621
refGene_gn=FMNL2;refGene_tx=XM_003990765.5;refGene_gn=FMNL2;refGene_tx=XM_023259320.1;refGene_gn=FMNL2;refGene_tx=XM_023259321.1;refGene_gn=FMNL2;refGene_tx=XM_023259322.1;refGene_gn=FMNL2;refGene_tx=XM_023259323.1;refGene_gn=FMNL2;refGene_tx=XM_023259324.1;refGene_gn=FMNL2;refGene_tx=XM_023259325.1;refGene_gn=FMNL2;refGene_tx=XM_023259326.1;refGene_gn=FMNL2;refGene_tx=XM_023259327.1;refGene_gn=FMNL2;refGene_tx=XM_023259328.1
C1 145612551 145612864
refGene_gn=FMNL2;refGene_tx=XM_003990765.5;refGene_gn=FMNL2;refGene_tx=XM_023259320.1;refGene_gn=FMNL2;refGene_tx=XM_023259321.1;refGene_gn=FMNL2;refGene_tx=XM_023259322.1;refGene_gn=FMNL2;refGene_tx=XM_023259323.1;refGene_gn=FMNL2;refGene_tx=XM_023259324.1;refGene_gn=FMNL2;refGene_tx=XM_023259325.1;refGene_gn=FMNL2;refGene_tx=XM_023259326.1;refGene_gn=FMNL2;refGene_tx=XM_023259327.1;refGene_gn=FMNL2;refGene_tx=XM_023259328.1
C1 145613204 145613417
refGene_gn=FMNL2;refGene_tx=XM_003990765.5;refGene_gn=FMNL2;refGene_tx=XM_023259320.1;refGene_gn=FMNL2;refGene_tx=XM_023259321.1;refGene_gn=FMNL2;refGene_tx=XM_023259322.1;refGene_gn=FMNL2;refGene_tx=XM_023259323.1;refGene_gn=FMNL2;refGene_tx=XM_023259324.1;refGene_gn=FMNL2;refGene_tx=XM_023259325.1;refGene_gn=FMNL2;refGene_tx=XM_023259326.1;refGene_gn=FMNL2;refGene_tx=XM_023259327.1;refGene_gn=FMNL2;refGene_tx=XM_023259328.1
C1 145616033 145616054
refGene_gn=FMNL2;refGene_tx=XM_023259320.1;refGene_gn=FMNL2;refGene_tx=XM_023259321.1;refGene_gn=FMNL2;refGene_tx=XM_023259322.1;refGene_gn=FMNL2;refGene_tx=XM_023259323.1;refGene_gn=FMNL2;refGene_tx=XM_023259324.1;refGene_gn=FMNL2;refGene_tx=XM_023259328.1

C1	145618217	145618342
refGene_gn=FMNL2;refGene_tx=XM_003990765.5;refGene_gn=FMNL2;refGene_tx=XM_023259320.1;refGene_gn=FMNL2;refGene_tx=XM_023259321.1;refGene_gn=FMNL2;refGene_tx=XM_023259322.1;refGene_gn=FMNL2;refGene_tx=XM_023259323.1;refGene_gn=FMNL2;refGene_tx=XM_023259324.1;refGene_gn=FMNL2;refGene_tx=XM_023259325.1;refGene_gn=FMNL2;refGene_tx=XM_023259326.1;refGene_gn=FMNL2;refGene_tx=XM_023259327.1;refGene_gn=FMNL2;refGene_tx=XM_023259328.1
C1	145618975	145619178
refGene_gn=FMNL2;refGene_tx=XM_003990765.5;refGene_gn=FMNL2;refGene_tx=XM_023259320.1;refGene_gn=FMNL2;refGene_tx=XM_023259321.1;refGene_gn=FMNL2;refGene_tx=XM_023259322.1;refGene_gn=FMNL2;refGene_tx=XM_023259323.1;refGene_gn=FMNL2;refGene_tx=XM_023259324.1;refGene_gn=FMNL2;refGene_tx=XM_023259325.1;refGene_gn=FMNL2;refGene_tx=XM_023259326.1;refGene_gn=FMNL2;refGene_tx=XM_023259327.1;refGene_gn=FMNL2;refGene_tx=XM_023259328.1
C1	145620969	145621204
refGene_gn=FMNL2;refGene_tx=XM_003990765.5;refGene_gn=FMNL2;refGene_tx=XM_023259320.1;refGene_gn=FMNL2;refGene_tx=XM_023259321.1;refGene_gn=FMNL2;refGene_tx=XM_023259322.1;refGene_gn=FMNL2;refGene_tx=XM_023259323.1;refGene_gn=FMNL2;refGene_tx=XM_023259324.1;refGene_gn=FMNL2;refGene_tx=XM_023259325.1;refGene_gn=FMNL2;refGene_tx=XM_023259326.1;refGene_gn=FMNL2;refGene_tx=XM_023259327.1;refGene_gn=FMNL2;refGene_tx=XM_023259328.1
C1	145622144	145622213
refGene_gn=FMNL2;refGene_tx=XM_003990765.5;refGene_gn=FMNL2;refGene_tx=XM_023259320.1;refGene_gn=FMNL2;refGene_tx=XM_023259321.1;refGene_gn=FMNL2;refGene_tx=XM_023259322.1;refGene_gn=FMNL2;refGene_tx=XM_023259323.1;refGene_gn=FMNL2;refGene_tx=XM_023259324.1;refGene_gn=FMNL2;refGene_tx=XM_023259325.1;refGene_gn=FMNL2;refGene_tx=XM_023259326.1;refGene_gn=FMNL2;refGene_tx=XM_023259327.1;refGene_gn=FMNL2;refGene_tx=XM_023259328.1
C1	145622357	145622438
refGene_gn=FMNL2;refGene_tx=XM_003990765.5;refGene_gn=FMNL2;refGene_tx=XM_023259320.1;refGene_gn=FMNL2;refGene_tx=XM_023259321.1;refGene_gn=FMNL2;refGene_tx=XM_023259322.1;refGene_gn=FMNL2;refGene_tx=XM_023259323.1;refGene_gn=FMNL2;refGene_tx=XM_023259324.1;refGene_gn=FMNL2;refGene_tx=XM_023259325.1;refGene_gn=FMNL2;refGene_tx=XM_023259326.1;refGene_gn=FMNL2;refGene_tx=XM_023259327.1;refGene_gn=FMNL2;refGene_tx=XM_023259328.1
C1	145624410	145624540
refGene_gn=FMNL2;refGene_tx=XM_003990765.5;refGene_gn=FMNL2;refGene_tx=XM_023259320.1;refGene_gn=FMNL2;refGene_tx=XM_023259321.1;refGene_gn=FMNL2;refGene_tx=XM_023259322.1;refGene_gn=FMNL2;refGene_tx=XM_023259323.1;refGene_gn=FMNL2;refGene_tx=XM_023259324.1;refGene_gn=FMNL2;refGene_tx=XM_023259325.1;refGene_gn=FMNL2;refGene_tx=XM_023259326.1;refGene_gn=FMNL2;refGene_tx=XM_023259327.1;refGene_gn=FMNL2;refGene_tx=XM_023259328.1
C1	145626852	145627016
refGene_gn=FMNL2;refGene_tx=XM_003990765.5;refGene_gn=FMNL2;refGene_tx

=XM_023259320.1;refGene_gn=FMNL2;refGene_tx=XM_023259321.1;refGene_gn=FMNL2;refGene_tx=XM_023259322.1;refGene_gn=FMNL2;refGene_tx=XM_023259323.1;refGene_gn=FMNL2;refGene_tx=XM_023259324.1;refGene_gn=FMNL2;refGene_tx=XM_023259325.1;refGene_gn=FMNL2;refGene_tx=XM_023259326.1;refGene_gn=FMNL2;refGene_tx=XM_023259327.1;refGene_gn=FMNL2;refGene_tx=XM_023259328.1
C1 145627929 145628031
refGene_gn=FMNL2;refGene_tx=XM_003990765.5;refGene_gn=FMNL2;refGene_tx=XM_023259320.1;refGene_gn=FMNL2;refGene_tx=XM_023259321.1;refGene_gn=FMNL2;refGene_tx=XM_023259322.1;refGene_gn=FMNL2;refGene_tx=XM_023259323.1;refGene_gn=FMNL2;refGene_tx=XM_023259324.1;refGene_gn=FMNL2;refGene_tx=XM_023259325.1;refGene_gn=FMNL2;refGene_tx=XM_023259326.1;refGene_gn=FMNL2;refGene_tx=XM_023259327.1;refGene_gn=FMNL2;refGene_tx=XM_023259328.1
C1 145630010 145630109
refGene_gn=FMNL2;refGene_tx=XM_003990765.5;refGene_gn=FMNL2;refGene_tx=XM_023259320.1;refGene_gn=FMNL2;refGene_tx=XM_023259321.1;refGene_gn=FMNL2;refGene_tx=XM_023259322.1;refGene_gn=FMNL2;refGene_tx=XM_023259323.1;refGene_gn=FMNL2;refGene_tx=XM_023259324.1;refGene_gn=FMNL2;refGene_tx=XM_023259325.1;refGene_gn=FMNL2;refGene_tx=XM_023259326.1;refGene_gn=FMNL2;refGene_tx=XM_023259327.1;refGene_gn=FMNL2;refGene_tx=XM_023259328.1
C1 145630831 145630955
refGene_gn=FMNL2;refGene_tx=XM_003990765.5;refGene_gn=FMNL2;refGene_tx=XM_023259320.1;refGene_gn=FMNL2;refGene_tx=XM_023259321.1;refGene_gn=FMNL2;refGene_tx=XM_023259322.1;refGene_gn=FMNL2;refGene_tx=XM_023259323.1;refGene_gn=FMNL2;refGene_tx=XM_023259324.1;refGene_gn=FMNL2;refGene_tx=XM_023259325.1;refGene_gn=FMNL2;refGene_tx=XM_023259326.1;refGene_gn=FMNL2;refGene_tx=XM_023259327.1;refGene_gn=FMNL2;refGene_tx=XM_023259328.1
C1 145632908 145633000
refGene_gn=FMNL2;refGene_tx=XM_023259322.1;refGene_gn=FMNL2;refGene_tx=XM_023259323.1;refGene_gn=FMNL2;refGene_tx=XM_023259326.1
C1 145634594 145634688
refGene_gn=FMNL2;refGene_tx=XM_023259321.1;refGene_gn=FMNL2;refGene_tx=XM_023259324.1;refGene_gn=FMNL2;refGene_tx=XM_023259325.1;refGene_gn=FMNL2;refGene_tx=XM_023259327.1
C1 145636506 145636616
refGene_gn=FMNL2;refGene_tx=XM_003990765.5;refGene_gn=FMNL2;refGene_tx=XM_023259320.1;refGene_gn=FMNL2;refGene_tx=XM_023259321.1;refGene_gn=FMNL2;refGene_tx=XM_023259324.1;refGene_gn=FMNL2;refGene_tx=XM_023259325.1;refGene_gn=FMNL2;refGene_tx=XM_023259327.1;refGene_gn=FMNL2;refGene_tx=XM_023259328.1
C1 145645206 145645299
refGene_gn=PRPF40A;refGene_tx=XM_011285260.3;refGene_gn=PRPF40A;refGene_tx=XM_011285262.3;refGene_gn=PRPF40A;refGene_tx=XM_011285264.3;refGene_gn=PRPF40A;refGene_tx=XM_019838116.2;refGene_gn=PRPF40A;refGene_tx=XM_023259329.1;refGene_gn=PRPF40A;refGene_tx=XM_023259330.1;refGene_gn=PRPF40A;refGene_tx=XM_023259331.1
C1 145646663 145646795

refGene_gn=PRPF40A;refGene_tx=XM_011285260.3;refGene_gn=PRPF40A;refGene_tx=XM_011285262.3;refGene_gn=PRPF40A;refGene_tx=XM_011285264.3;refGene_gn=PRPF40A;refGene_tx=XM_019838116.2;refGene_gn=PRPF40A;refGene_tx=XM_023259329.1;refGene_gn=PRPF40A;refGene_tx=XM_023259330.1;refGene_gn=PRPF40A;refGene_tx=XM_023259331.1
C1 145647625 145647687
refGene_gn=PRPF40A;refGene_tx=XM_011285260.3;refGene_gn=PRPF40A;refGene_tx=XM_011285262.3;refGene_gn=PRPF40A;refGene_tx=XM_011285264.3;refGene_gn=PRPF40A;refGene_tx=XM_019838116.2;refGene_gn=PRPF40A;refGene_tx=XM_023259329.1;refGene_gn=PRPF40A;refGene_tx=XM_023259330.1;refGene_gn=PRPF40A;refGene_tx=XM_023259331.1
C1 145647779 145647876
refGene_gn=PRPF40A;refGene_tx=XM_011285260.3;refGene_gn=PRPF40A;refGene_tx=XM_011285262.3;refGene_gn=PRPF40A;refGene_tx=XM_011285264.3;refGene_gn=PRPF40A;refGene_tx=XM_019838116.2;refGene_gn=PRPF40A;refGene_tx=XM_023259329.1;refGene_gn=PRPF40A;refGene_tx=XM_023259330.1;refGene_gn=PRPF40A;refGene_tx=XM_023259331.1
C1 145647964 145648051
refGene_gn=PRPF40A;refGene_tx=XM_011285260.3;refGene_gn=PRPF40A;refGene_tx=XM_011285262.3;refGene_gn=PRPF40A;refGene_tx=XM_011285264.3;refGene_gn=PRPF40A;refGene_tx=XM_019838116.2;refGene_gn=PRPF40A;refGene_tx=XM_023259329.1;refGene_gn=PRPF40A;refGene_tx=XM_023259330.1;refGene_gn=PRPF40A;refGene_tx=XM_023259331.1
C1 145650191 145650287
refGene_gn=PRPF40A;refGene_tx=XM_011285260.3;refGene_gn=PRPF40A;refGene_tx=XM_011285262.3;refGene_gn=PRPF40A;refGene_tx=XM_011285264.3;refGene_gn=PRPF40A;refGene_tx=XM_019838116.2;refGene_gn=PRPF40A;refGene_tx=XM_023259329.1;refGene_gn=PRPF40A;refGene_tx=XM_023259330.1;refGene_gn=PRPF40A;refGene_tx=XM_023259331.1
C1 145650622 145650760
refGene_gn=PRPF40A;refGene_tx=XM_011285260.3;refGene_gn=PRPF40A;refGene_tx=XM_011285262.3;refGene_gn=PRPF40A;refGene_tx=XM_011285264.3;refGene_gn=PRPF40A;refGene_tx=XM_019838116.2;refGene_gn=PRPF40A;refGene_tx=XM_023259329.1;refGene_gn=PRPF40A;refGene_tx=XM_023259330.1;refGene_gn=PRPF40A;refGene_tx=XM_023259331.1
C1 145651080 145651197
refGene_gn=PRPF40A;refGene_tx=XM_011285260.3;refGene_gn=PRPF40A;refGene_tx=XM_011285262.3;refGene_gn=PRPF40A;refGene_tx=XM_011285264.3;refGene_gn=PRPF40A;refGene_tx=XM_019838116.2;refGene_gn=PRPF40A;refGene_tx=XM_023259329.1;refGene_gn=PRPF40A;refGene_tx=XM_023259330.1;refGene_gn=PRPF40A;refGene_tx=XM_023259331.1
C1 145651309 145651413
refGene_gn=PRPF40A;refGene_tx=XM_011285260.3;refGene_gn=PRPF40A;refGene_tx=XM_011285262.3;refGene_gn=PRPF40A;refGene_tx=XM_011285264.3;refGene_gn=PRPF40A;refGene_tx=XM_019838116.2;refGene_gn=PRPF40A;refGene_tx=XM_023259329.1;refGene_gn=PRPF40A;refGene_tx=XM_023259330.1;refGene_gn=PRPF40A;refGene_tx=XM_023259331.1
C1 145651573 145651688
refGene_gn=PRPF40A;refGene_tx=XM_011285260.3;refGene_gn=PRPF40A;refGene_tx=XM_011285262.3;refGene_gn=PRPF40A;refGene_tx=XM_011285264.3;refGe

ne_gn=PRPF40A;refGene_tx=XM_019838116.2;refGene_gn=PRPF40A;refGene_tx=XM_023259329.1;refGene_gn=PRPF40A;refGene_tx=XM_023259330.1;refGene_gn=PRPF40A;refGene_tx=XM_023259331.1
C1 145655610 145655744
refGene_gn=PRPF40A;refGene_tx=XM_011285260.3;refGene_gn=PRPF40A;refGene_tx=XM_011285262.3;refGene_gn=PRPF40A;refGene_tx=XM_011285264.3;refGene_gn=PRPF40A;refGene_tx=XM_019838116.2;refGene_gn=PRPF40A;refGene_tx=XM_023259329.1;refGene_gn=PRPF40A;refGene_tx=XM_023259330.1;refGene_gn=PRPF40A;refGene_tx=XM_023259331.1
C1 145656493 145656644
refGene_gn=PRPF40A;refGene_tx=XM_011285260.3;refGene_gn=PRPF40A;refGene_tx=XM_011285262.3;refGene_gn=PRPF40A;refGene_tx=XM_011285264.3;refGene_gn=PRPF40A;refGene_tx=XM_019838116.2;refGene_gn=PRPF40A;refGene_tx=XM_023259329.1;refGene_gn=PRPF40A;refGene_tx=XM_023259330.1;refGene_gn=PRPF40A;refGene_tx=XM_023259331.1
C1 145657703 145657812
refGene_gn=PRPF40A;refGene_tx=XM_011285260.3;refGene_gn=PRPF40A;refGene_tx=XM_011285262.3;refGene_gn=PRPF40A;refGene_tx=XM_011285264.3;refGene_gn=PRPF40A;refGene_tx=XM_019838116.2;refGene_gn=PRPF40A;refGene_tx=XM_023259329.1;refGene_gn=PRPF40A;refGene_tx=XM_023259330.1;refGene_gn=PRPF40A;refGene_tx=XM_023259331.1
C1 145658549 145658714
refGene_gn=PRPF40A;refGene_tx=XM_011285260.3;refGene_gn=PRPF40A;refGene_tx=XM_011285262.3;refGene_gn=PRPF40A;refGene_tx=XM_011285264.3;refGene_gn=PRPF40A;refGene_tx=XM_019838116.2;refGene_gn=PRPF40A;refGene_tx=XM_023259329.1;refGene_gn=PRPF40A;refGene_tx=XM_023259330.1;refGene_gn=PRPF40A;refGene_tx=XM_023259331.1
C1 145659064 145659129
refGene_gn=PRPF40A;refGene_tx=XM_011285260.3;refGene_gn=PRPF40A;refGene_tx=XM_011285262.3;refGene_gn=PRPF40A;refGene_tx=XM_011285264.3;refGene_gn=PRPF40A;refGene_tx=XM_019838116.2;refGene_gn=PRPF40A;refGene_tx=XM_023259329.1;refGene_gn=PRPF40A;refGene_tx=XM_023259330.1;refGene_gn=PRPF40A;refGene_tx=XM_023259331.1
C1 145659541 145659650
refGene_gn=PRPF40A;refGene_tx=XM_011285260.3;refGene_gn=PRPF40A;refGene_tx=XM_011285262.3;refGene_gn=PRPF40A;refGene_tx=XM_011285264.3;refGene_gn=PRPF40A;refGene_tx=XM_019838116.2;refGene_gn=PRPF40A;refGene_tx=XM_023259329.1;refGene_gn=PRPF40A;refGene_tx=XM_023259330.1;refGene_gn=PRPF40A;refGene_tx=XM_023259331.1
C1 145662364 145662739
refGene_gn=PRPF40A;refGene_tx=XM_011285260.3;refGene_gn=PRPF40A;refGene_tx=XM_011285262.3;refGene_gn=PRPF40A;refGene_tx=XM_011285264.3;refGene_gn=PRPF40A;refGene_tx=XM_019838116.2;refGene_gn=PRPF40A;refGene_tx=XM_023259329.1;refGene_gn=PRPF40A;refGene_tx=XM_023259330.1;refGene_gn=PRPF40A;refGene_tx=XM_023259331.1
C1 145663414 145663439
refGene_gn=PRPF40A;refGene_tx=XM_011285260.3;refGene_gn=PRPF40A;refGene_tx=XM_011285262.3;refGene_gn=PRPF40A;refGene_tx=XM_011285264.3;refGene_gn=PRPF40A;refGene_tx=XM_019838116.2;refGene_gn=PRPF40A;refGene_tx=XM_023259329.1;refGene_gn=PRPF40A;refGene_tx=XM_023259330.1;refGene_gn

=PRPF40A;refGene_tx=XM_023259331.1
C1 145665284 145665338
refGene_gn=PRPF40A;refGene_tx=XM_011285260.3;refGene_gn=PRPF40A;refGene_tx=XM_011285264.3;refGene_gn=PRPF40A;refGene_tx=XM_019838116.2;refGene_gn=PRPF40A;refGene_tx=XM_023259329.1;refGene_gn=PRPF40A;refGene_tx=XM_023259331.1
C1 145665572 145665693
refGene_gn=PRPF40A;refGene_tx=XM_011285260.3;refGene_gn=PRPF40A;refGene_tx=XM_011285262.3;refGene_gn=PRPF40A;refGene_tx=XM_011285264.3;refGene_gn=PRPF40A;refGene_tx=XM_019838116.2;refGene_gn=PRPF40A;refGene_tx=XM_023259329.1;refGene_gn=PRPF40A;refGene_tx=XM_023259330.1;refGene_gn=PRPF40A;refGene_tx=XM_023259331.1
C1 145666994 145667105
refGene_gn=PRPF40A;refGene_tx=XM_011285260.3;refGene_gn=PRPF40A;refGene_tx=XM_011285262.3;refGene_gn=PRPF40A;refGene_tx=XM_011285264.3;refGene_gn=PRPF40A;refGene_tx=XM_019838116.2;refGene_gn=PRPF40A;refGene_tx=XM_023259329.1;refGene_gn=PRPF40A;refGene_tx=XM_023259330.1;refGene_gn=PRPF40A;refGene_tx=XM_023259331.1
C1 145680677 145680694
refGene_gn=PRPF40A;refGene_tx=XM_011285260.3;refGene_gn=PRPF40A;refGene_tx=XM_011285262.3;refGene_gn=PRPF40A;refGene_tx=XM_011285264.3;refGene_gn=PRPF40A;refGene_tx=XM_019838116.2;refGene_gn=PRPF40A;refGene_tx=XM_023259329.1;refGene_gn=PRPF40A;refGene_tx=XM_023259330.1;refGene_gn=PRPF40A;refGene_tx=XM_023259331.1
C1 145682785 145682816
refGene_gn=PRPF40A;refGene_tx=XM_011285260.3;refGene_gn=PRPF40A;refGene_tx=XM_011285262.3;refGene_gn=PRPF40A;refGene_tx=XM_011285264.3;refGene_gn=PRPF40A;refGene_tx=XM_019838116.2;refGene_gn=PRPF40A;refGene_tx=XM_023259329.1;refGene_gn=PRPF40A;refGene_tx=XM_023259330.1;refGene_gn=PRPF40A;refGene_tx=XM_023259331.1
C1 145684212 145684293
refGene_gn=PRPF40A;refGene_tx=XM_011285260.3;refGene_gn=PRPF40A;refGene_tx=XM_011285262.3;refGene_gn=PRPF40A;refGene_tx=XM_011285264.3;refGene_gn=PRPF40A;refGene_tx=XM_019838116.2;refGene_gn=PRPF40A;refGene_tx=XM_023259329.1;refGene_gn=PRPF40A;refGene_tx=XM_023259330.1;refGene_gn=PRPF40A;refGene_tx=XM_023259331.1
C1 145694774 145694822
refGene_gn=PRPF40A;refGene_tx=XM_011285264.3
C1 145696235 145696367
refGene_gn=PRPF40A;refGene_tx=XM_011285260.3;refGene_gn=PRPF40A;refGene_tx=XM_011285262.3;refGene_gn=PRPF40A;refGene_tx=XM_019838116.2;refGene_gn=PRPF40A;refGene_tx=XM_023259329.1;refGene_gn=PRPF40A;refGene_tx=XM_023259330.1;refGene_gn=PRPF40A;refGene_tx=XM_023259331.1
C1 145697528 145697930
refGene_gn=PRPF40A;refGene_tx=XM_011285260.3;refGene_gn=PRPF40A;refGene_tx=XM_011285262.3;refGene_gn=PRPF40A;refGene_tx=XM_019838116.2;refGene_gn=PRPF40A;refGene_tx=XM_023259329.1;refGene_gn=PRPF40A;refGene_tx=XM_023259330.1;refGene_gn=PRPF40A;refGene_tx=XM_023259331.1
C1 145698949 145699349
refGene_gn=ARL6IP6;refGene_tx=XM_003990770.5;refGene_gn=ARL6IP6;refGen

e_tx=XM_006935285.4;refGene_gn=ARL6IP6;refGene_tx=XM_019838117.2;refGene_gn=ARL6IP6;refGene_tx=XM_019838118.2
C1 145700898 145700949
refGene_gn=ARL6IP6;refGene_tx=XM_003990770.5;refGene_gn=ARL6IP6;refGene_tx=XM_006935285.4;refGene_gn=ARL6IP6;refGene_tx=XM_019838118.2
C1 145707270 145707434
refGene_gn=ARL6IP6;refGene_tx=XM_019838118.2
C1 145711746 145711879
refGene_gn=ARL6IP6;refGene_tx=XM_003990770.5;refGene_gn=ARL6IP6;refGene_tx=XM_006935285.4;refGene_gn=ARL6IP6;refGene_tx=XM_019838117.2
C1 145735534 145735609
refGene_gn=ARL6IP6;refGene_tx=XM_006935285.4;refGene_gn=ARL6IP6;refGene_tx=XM_019838117.2
C1 145737363 145737457
refGene_gn=ARL6IP6;refGene_tx=XM_003990770.5;refGene_gn=ARL6IP6;refGene_tx=XM_006935285.4;refGene_gn=ARL6IP6;refGene_tx=XM_019838117.2
C1 146333961 146334291
refGene_gn=RPRM;refGene_tx=XM_011285259.3
C1 146683123 146683265
refGene_gn=GALNT13;refGene_tx=XM_011285258.3;refGene_gn=GALNT13;refGene_tx=XM_019838119.2;refGene_gn=GALNT13;refGene_tx=XM_019838120.2;refGene_gn=GALNT13;refGene_tx=XM_023259332.1
C1 146837477 146837646
refGene_gn=GALNT13;refGene_tx=XM_011285258.3;refGene_gn=GALNT13;refGene_tx=XM_019838119.2;refGene_gn=GALNT13;refGene_tx=XM_019838120.2;refGene_gn=GALNT13;refGene_tx=XM_023259332.1
C1 146938379 146938546
refGene_gn=GALNT13;refGene_tx=XM_011285258.3;refGene_gn=GALNT13;refGene_tx=XM_019838119.2;refGene_gn=GALNT13;refGene_tx=XM_019838120.2;refGene_gn=GALNT13;refGene_tx=XM_023259332.1
C1 146939154 146939362
refGene_gn=GALNT13;refGene_tx=XM_011285258.3;refGene_gn=GALNT13;refGene_tx=XM_019838119.2;refGene_gn=GALNT13;refGene_tx=XM_019838120.2;refGene_gn=GALNT13;refGene_tx=XM_023259332.1
C1 146942447 146942618
refGene_gn=GALNT13;refGene_tx=XM_011285258.3;refGene_gn=GALNT13;refGene_tx=XM_019838119.2;refGene_gn=GALNT13;refGene_tx=XM_019838120.2;refGene_gn=GALNT13;refGene_tx=XM_023259332.1
C1 146956821 146956939
refGene_gn=GALNT13;refGene_tx=XM_011285258.3;refGene_gn=GALNT13;refGene_tx=XM_019838119.2;refGene_gn=GALNT13;refGene_tx=XM_019838120.2;refGene_gn=GALNT13;refGene_tx=XM_023259332.1
C1 146987333 146987514
refGene_gn=GALNT13;refGene_tx=XM_011285258.3;refGene_gn=GALNT13;refGene_tx=XM_019838119.2;refGene_gn=GALNT13;refGene_tx=XM_019838120.2;refGene_gn=GALNT13;refGene_tx=XM_023259332.1
C1 147096363 147096503
refGene_gn=GALNT13;refGene_tx=XM_011285258.3;refGene_gn=GALNT13;refGene_tx=XM_019838119.2;refGene_gn=GALNT13;refGene_tx=XM_019838120.2;refGene_gn=GALNT13;refGene_tx=XM_023259332.1

C1 147109052 147109151
refGene_gn=GALNT13;refGene_tx=XM_011285258.3;refGene_gn=GALNT13;refGen
e_tx=XM_019838119.2;refGene_gn=GALNT13;refGene_tx=XM_019838120.2;refGe
ne_gn=GALNT13;refGene_tx=XM_023259332.1
C1 147147596 147147731
refGene_gn=GALNT13;refGene_tx=XM_011285258.3;refGene_gn=GALNT13;refGen
e_tx=XM_019838119.2;refGene_gn=GALNT13;refGene_tx=XM_019838120.2;refGe
ne_gn=GALNT13;refGene_tx=XM_023259332.1
C1 147151982 147152012
refGene_gn=GALNT13;refGene_tx=XM_019838119.2;refGene_gn=GALNT13;refGen
e_tx=XM_023259332.1
C1 147155640 147155826
refGene_gn=GALNT13;refGene_tx=XM_019838119.2;refGene_gn=GALNT13;refGen
e_tx=XM_019838120.2;refGene_gn=GALNT13;refGene_tx=XM_023259332.1
C1 147159725 147159866
refGene_gn=GALNT13;refGene_tx=XM_011285258.3
C1 147372808 147373510
refGene_gn=KCNJ3;refGene_tx=XM_003990778.5;refGene_gn=KCNJ3;refGene_tx
=XM_019838121.2
C1 147383952 147384169
refGene_gn=KCNJ3;refGene_tx=XM_003990778.5
C1 147524528 147525115
refGene_gn=KCNJ3;refGene_tx=XM_003990778.5;refGene_gn=KCNJ3;refGene_tx
=XM_019838121.2
C1 148712927 148713184
refGene_gn=NR4A2;refGene_tx=XM_006935294.4;refGene_gn=NR4A2;refGene_tx
=XM_019838122.2;refGene_gn=NR4A2;refGene_tx=XM_019838123.2;refGene_gn=
NR4A2;refGene_tx=XM_019838124.2
C1 148713330 148713509
refGene_gn=NR4A2;refGene_tx=XM_006935294.4;refGene_gn=NR4A2;refGene_tx
=XM_019838122.2;refGene_gn=NR4A2;refGene_tx=XM_019838123.2;refGene_gn=
NR4A2;refGene_tx=XM_019838124.2
C1 148713906 148714109
refGene_gn=NR4A2;refGene_tx=XM_006935294.4;refGene_gn=NR4A2;refGene_tx
=XM_019838122.2;refGene_gn=NR4A2;refGene_tx=XM_019838123.2;refGene_gn=
NR4A2;refGene_tx=XM_019838124.2
C1 148715002 148715166
refGene_gn=NR4A2;refGene_tx=XM_006935294.4;refGene_gn=NR4A2;refGene_tx
=XM_019838122.2;refGene_gn=NR4A2;refGene_tx=XM_019838123.2;refGene_gn=
NR4A2;refGene_tx=XM_019838124.2
C1 148715582 148715712
refGene_gn=NR4A2;refGene_tx=XM_006935294.4;refGene_gn=NR4A2;refGene_tx
=XM_019838122.2;refGene_gn=NR4A2;refGene_tx=XM_019838123.2;refGene_gn=
NR4A2;refGene_tx=XM_019838124.2
C1 148716509 148717373
refGene_gn=NR4A2;refGene_tx=XM_006935294.4;refGene_gn=NR4A2;refGene_tx
=XM_019838122.2;refGene_gn=NR4A2;refGene_tx=XM_019838123.2;refGene_gn=
NR4A2;refGene_tx=XM_019838124.2
C1 148849986 148850088
refGene_gn=GPD2;refGene_tx=XM_003990784.5;refGene_gn=GPD2;refGene_tx=X

M_011285364.3;refGene_gn=GPD2;refGene_tx=XM_011285365.3
C1 148871612 148871784
refGene_gn=GPD2;refGene_tx=XM_003990784.5;refGene_gn=GPD2;refGene_tx=XM_011285364.3;refGene_gn=GPD2;refGene_tx=XM_011285365.3
C1 148884625 148884750
refGene_gn=GPD2;refGene_tx=XM_003990784.5;refGene_gn=GPD2;refGene_tx=XM_011285364.3;refGene_gn=GPD2;refGene_tx=XM_011285365.3
C1 148886055 148886153
refGene_gn=GPD2;refGene_tx=XM_003990784.5;refGene_gn=GPD2;refGene_tx=XM_011285364.3;refGene_gn=GPD2;refGene_tx=XM_011285365.3
C1 148887233 148887397
refGene_gn=GPD2;refGene_tx=XM_003990784.5;refGene_gn=GPD2;refGene_tx=XM_011285364.3;refGene_gn=GPD2;refGene_tx=XM_011285365.3
C1 148918269 148918434
refGene_gn=GPD2;refGene_tx=XM_003990784.5;refGene_gn=GPD2;refGene_tx=XM_011285364.3;refGene_gn=GPD2;refGene_tx=XM_011285365.3
C1 148919263 148919408
refGene_gn=GPD2;refGene_tx=XM_003990784.5;refGene_gn=GPD2;refGene_tx=XM_011285364.3;refGene_gn=GPD2;refGene_tx=XM_011285365.3
C1 148926349 148926543
refGene_gn=GPD2;refGene_tx=XM_003990784.5;refGene_gn=GPD2;refGene_tx=XM_011285364.3;refGene_gn=GPD2;refGene_tx=XM_011285365.3
C1 148939374 148939509
refGene_gn=GPD2;refGene_tx=XM_003990784.5;refGene_gn=GPD2;refGene_tx=XM_011285364.3;refGene_gn=GPD2;refGene_tx=XM_011285365.3
C1 148939793 148939969
refGene_gn=GPD2;refGene_tx=XM_003990784.5;refGene_gn=GPD2;refGene_tx=XM_011285364.3;refGene_gn=GPD2;refGene_tx=XM_011285365.3
C1 148940490 148940622
refGene_gn=GPD2;refGene_tx=XM_003990784.5;refGene_gn=GPD2;refGene_tx=XM_011285364.3;refGene_gn=GPD2;refGene_tx=XM_011285365.3
C1 148941545 148941704
refGene_gn=GPD2;refGene_tx=XM_003990784.5;refGene_gn=GPD2;refGene_tx=XM_011285364.3;refGene_gn=GPD2;refGene_tx=XM_011285365.3
C1 148948749 148948862
refGene_gn=GPD2;refGene_tx=XM_003990784.5;refGene_gn=GPD2;refGene_tx=XM_011285364.3;refGene_gn=GPD2;refGene_tx=XM_011285365.3
C1 148948945 148949024
refGene_gn=GPD2;refGene_tx=XM_003990784.5;refGene_gn=GPD2;refGene_tx=XM_011285364.3;refGene_gn=GPD2;refGene_tx=XM_011285365.3
C1 148949214 148949313
refGene_gn=GPD2;refGene_tx=XM_003990784.5;refGene_gn=GPD2;refGene_tx=XM_011285364.3;refGene_gn=GPD2;refGene_tx=XM_011285365.3
C1 148952326 148952452
refGene_gn=GPD2;refGene_tx=XM_003990784.5;refGene_gn=GPD2;refGene_tx=XM_011285364.3;refGene_gn=GPD2;refGene_tx=XM_011285365.3
C1 149525426 149526877
refGene_gn=GALNT5;refGene_tx=XM_003990785.4
C1 149540976 149541143
refGene_gn=GALNT5;refGene_tx=XM_003990785.4

C1	149542654	149542774
refGene_gn=GALNT5;refGene_tx=XM_003990785.4
C1	149550789	149550925
refGene_gn=GALNT5;refGene_tx=XM_003990785.4
C1	149551510	149551630
refGene_gn=GALNT5;refGene_tx=XM_003990785.4
C1	149553845	149553963
refGene_gn=GALNT5;refGene_tx=XM_003990785.4
C1	149554958	149555282
refGene_gn=GALNT5;refGene_tx=XM_003990785.4
C1	149560187	149560268
refGene_gn=GALNT5;refGene_tx=XM_003990785.4
C1	149563395	149563557
refGene_gn=GALNT5;refGene_tx=XM_003990785.4
C1	149566762	149566903
refGene_gn=GALNT5;refGene_tx=XM_003990785.4
C1	149575854	149576372
refGene_gn=ERMN;refGene_tx=XM_006935296.3
C1	149579379	149579472
refGene_gn=ERMN;refGene_tx=XM_006935296.3
C1	149580147	149580388
refGene_gn=ERMN;refGene_tx=XM_006935296.3
C1	149641059	149641517
refGene_gn=CYTIP;refGene_tx=XM_011285360.3;refGene_gn=CYTIP;refGene_tx
=XM_011285361.2;refGene_gn=CYTIP;refGene_tx=XM_011285362.2;refGene_gn=
CYTIP;refGene_tx=XM_011285363.2;refGene_gn=CYTIP;refGene_tx=XM_0198381
28.1
C1	149644345	149644412
refGene_gn=CYTIP;refGene_tx=XM_011285360.3;refGene_gn=CYTIP;refGene_tx
=XM_011285361.2;refGene_gn=CYTIP;refGene_tx=XM_011285362.2;refGene_gn=
CYTIP;refGene_tx=XM_011285363.2;refGene_gn=CYTIP;refGene_tx=XM_0198381
28.1
C1	149656280	149656350
refGene_gn=CYTIP;refGene_tx=XM_011285360.3;refGene_gn=CYTIP;refGene_tx
=XM_011285361.2;refGene_gn=CYTIP;refGene_tx=XM_011285362.2;refGene_gn=
CYTIP;refGene_tx=XM_011285363.2;refGene_gn=CYTIP;refGene_tx=XM_0198381
28.1
C1	149659183	149659277
refGene_gn=CYTIP;refGene_tx=XM_011285360.3;refGene_gn=CYTIP;refGene_tx
=XM_011285361.2;refGene_gn=CYTIP;refGene_tx=XM_011285362.2;refGene_gn=
CYTIP;refGene_tx=XM_011285363.2;refGene_gn=CYTIP;refGene_tx=XM_0198381
28.1
C1	149659490	149659593
refGene_gn=CYTIP;refGene_tx=XM_011285360.3;refGene_gn=CYTIP;refGene_tx
=XM_011285361.2;refGene_gn=CYTIP;refGene_tx=XM_011285362.2;refGene_gn=
CYTIP;refGene_tx=XM_011285363.2;refGene_gn=CYTIP;refGene_tx=XM_0198381
28.1
C1	149663566	149663621
refGene_gn=CYTIP;refGene_tx=XM_011285360.3;refGene_gn=CYTIP;refGene_tx
=XM_011285361.2;refGene_gn=CYTIP;refGene_tx=XM_011285362.2;refGene_gn=

CYTIP;refGene_tx=XM_011285363.2;refGene_gn=CYTIP;refGene_tx=XM_0198381
28.1
C1 149663859 149663909
refGene_gn=CYTIP;refGene_tx=XM_011285360.3;refGene_gn=CYTIP;refGene_tx
=XM_011285361.2;refGene_gn=CYTIP;refGene_tx=XM_011285362.2;refGene_gn=
CYTIP;refGene_tx=XM_011285363.2;refGene_gn=CYTIP;refGene_tx=XM_0198381
28.1
C1 149671131 149671305
refGene_gn=CYTIP;refGene_tx=XM_011285360.3;refGene_gn=CYTIP;refGene_tx
=XM_011285361.2;refGene_gn=CYTIP;refGene_tx=XM_011285362.2;refGene_gn=
CYTIP;refGene_tx=XM_019838128.1
C1 149712696 149712759
refGene_gn=CYTIP;refGene_tx=XM_011285363.2
C1 149747601 149747727
refGene_gn=ACVR1C;refGene_tx=XM_023259333.1
C1 149753075 149753206
refGene_gn=ACVR1C;refGene_tx=XM_023259333.1
C1 149755818 149755943
refGene_gn=ACVR1C;refGene_tx=XM_023259333.1
C1 149757710 149757867
refGene_gn=ACVR1C;refGene_tx=XM_023259333.1
C1 149759233 149759401
refGene_gn=ACVR1C;refGene_tx=XM_023259333.1
C1 149762245 149762476
refGene_gn=ACVR1C;refGene_tx=XM_023259333.1
C1 149767114 149767354
refGene_gn=ACVR1C;refGene_tx=XM_023259333.1
C1 149788257 149788488
refGene_gn=ACVR1C;refGene_tx=XM_023259333.1
C1 149823865 149823938
refGene_gn=ACVR1C;refGene_tx=XM_023259333.1
C1 149972112 149972247
refGene_gn=ACVR1;refGene_tx=XM_019838129.2;refGene_gn=ACVR1;refGene_tx
=XM_019838130.2;refGene_gn=ACVR1;refGene_tx=XM_019838131.2;refGene_gn=
ACVR1;refGene_tx=XM_023259334.1;refGene_gn=ACVR1;refGene_tx=XM_0232593
35.1
C1 149972955 149973086
refGene_gn=ACVR1;refGene_tx=XM_019838129.2;refGene_gn=ACVR1;refGene_tx
=XM_019838130.2;refGene_gn=ACVR1;refGene_tx=XM_019838131.2;refGene_gn=
ACVR1;refGene_tx=XM_023259334.1;refGene_gn=ACVR1;refGene_tx=XM_0232593
35.1
C1 149999406 149999604
refGene_gn=ACVR1;refGene_tx=XM_019838129.2;refGene_gn=ACVR1;refGene_tx
=XM_019838130.2;refGene_gn=ACVR1;refGene_tx=XM_019838131.2;refGene_gn=
ACVR1;refGene_tx=XM_023259334.1;refGene_gn=ACVR1;refGene_tx=XM_0232593
35.1
C1 150003554 150003830
refGene_gn=ACVR1;refGene_tx=XM_019838129.2;refGene_gn=ACVR1;refGene_tx
=XM_019838130.2;refGene_gn=ACVR1;refGene_tx=XM_019838131.2;refGene_gn=
ACVR1;refGene_tx=XM_023259334.1;refGene_gn=ACVR1;refGene_tx=XM_0232593

35.1
C1	150010510	150010657
refGene_gn=ACVR1;refGene_tx=XM_019838129.2;refGene_gn=ACVR1;refGene_tx=XM_019838130.2;refGene_gn=ACVR1;refGene_tx=XM_019838131.2;refGene_gn=ACVR1;refGene_tx=XM_023259334.1;refGene_gn=ACVR1;refGene_tx=XM_023259335.1
C1	150014327	150014427
refGene_gn=ACVR1;refGene_tx=XM_019838129.2;refGene_gn=ACVR1;refGene_tx=XM_019838130.2;refGene_gn=ACVR1;refGene_tx=XM_019838131.2;refGene_gn=ACVR1;refGene_tx=XM_023259334.1;refGene_gn=ACVR1;refGene_tx=XM_023259335.1
C1	150017916	150018128
refGene_gn=ACVR1;refGene_tx=XM_019838129.2;refGene_gn=ACVR1;refGene_tx=XM_019838130.2;refGene_gn=ACVR1;refGene_tx=XM_019838131.2;refGene_gn=ACVR1;refGene_tx=XM_023259334.1;refGene_gn=ACVR1;refGene_tx=XM_023259335.1
C1	150020578	150020842
refGene_gn=ACVR1;refGene_tx=XM_019838129.2;refGene_gn=ACVR1;refGene_tx=XM_019838130.2;refGene_gn=ACVR1;refGene_tx=XM_019838131.2;refGene_gn=ACVR1;refGene_tx=XM_023259334.1;refGene_gn=ACVR1;refGene_tx=XM_023259335.1
C1	150037678	150037745
refGene_gn=ACVR1;refGene_tx=XM_019838129.2;refGene_gn=ACVR1;refGene_tx=XM_019838130.2;refGene_gn=ACVR1;refGene_tx=XM_019838131.2;refGene_gn=ACVR1;refGene_tx=XM_023259334.1;refGene_gn=ACVR1;refGene_tx=XM_023259335.1
C1	150341213	150341360
refGene_gn=CCDC148;refGene_tx=XM_019838135.2
C1	150345196	150345337
refGene_gn=CCDC148;refGene_tx=XM_019838135.2
C1	150347526	150347644
refGene_gn=CCDC148;refGene_tx=XM_019838135.2
C1	150409366	150409485
refGene_gn=CCDC148;refGene_tx=XM_019838135.2
C1	150437248	150437389
refGene_gn=CCDC148;refGene_tx=XM_019838135.2
C1	150483747	150483954
refGene_gn=CCDC148;refGene_tx=XM_019838135.2
C1	150486230	150486369
refGene_gn=CCDC148;refGene_tx=XM_019838135.2
C1	150503703	150503885
refGene_gn=CCDC148;refGene_tx=XM_019838135.2
C1	150503963	150504059
refGene_gn=CCDC148;refGene_tx=XM_019838135.2
C1	150506076	150506228
refGene_gn=CCDC148;refGene_tx=XM_019838135.2
C1	150506698	150506781
refGene_gn=CCDC148;refGene_tx=XM_019838135.2
C1	150516043	150516147
refGene_gn=CCDC148;refGene_tx=XM_019838135.2

C1	150524114	150524236
refGene_gn=CCDC148;refGene_tx=XM_019838135.2
C1	150525423	150525484
refGene_gn=CCDC148;refGene_tx=XM_019838135.2
C1	150672018	150672150
refGene_gn=PKP4;refGene_tx=XM_023259337.1;refGene_gn=PKP4;refGene_tx=XM_023259338.1;refGene_gn=PKP4;refGene_tx=XM_023259339.1;refGene_gn=PKP4;refGene_tx=XM_023259340.1;refGene_gn=PKP4;refGene_tx=XM_023259341.1;refGene_gn=PKP4;refGene_tx=XM_023259342.1;refGene_gn=PKP4;refGene_tx=XM_023259343.1;refGene_gn=PKP4;refGene_tx=XM_023259344.1;refGene_gn=PKP4;refGene_tx=XM_023259345.1;refGene_gn=PKP4;refGene_tx=XM_023259346.1
C1	150728206	150728319
refGene_gn=PKP4;refGene_tx=XM_023259337.1;refGene_gn=PKP4;refGene_tx=XM_023259338.1;refGene_gn=PKP4;refGene_tx=XM_023259339.1;refGene_gn=PKP4;refGene_tx=XM_023259340.1;refGene_gn=PKP4;refGene_tx=XM_023259341.1;refGene_gn=PKP4;refGene_tx=XM_023259342.1;refGene_gn=PKP4;refGene_tx=XM_023259343.1;refGene_gn=PKP4;refGene_tx=XM_023259344.1;refGene_gn=PKP4;refGene_tx=XM_023259345.1;refGene_gn=PKP4;refGene_tx=XM_023259346.1
C1	150755829	150755864
refGene_gn=PKP4;refGene_tx=XM_023259337.1;refGene_gn=PKP4;refGene_tx=XM_023259338.1;refGene_gn=PKP4;refGene_tx=XM_023259339.1;refGene_gn=PKP4;refGene_tx=XM_023259340.1;refGene_gn=PKP4;refGene_tx=XM_023259341.1;refGene_gn=PKP4;refGene_tx=XM_023259342.1;refGene_gn=PKP4;refGene_tx=XM_023259343.1;refGene_gn=PKP4;refGene_tx=XM_023259344.1;refGene_gn=PKP4;refGene_tx=XM_023259345.1;refGene_gn=PKP4;refGene_tx=XM_023259346.1
C1	150773374	150773506
refGene_gn=PKP4;refGene_tx=XM_023259337.1;refGene_gn=PKP4;refGene_tx=XM_023259338.1;refGene_gn=PKP4;refGene_tx=XM_023259339.1;refGene_gn=PKP4;refGene_tx=XM_023259340.1;refGene_gn=PKP4;refGene_tx=XM_023259341.1;refGene_gn=PKP4;refGene_tx=XM_023259342.1;refGene_gn=PKP4;refGene_tx=XM_023259343.1;refGene_gn=PKP4;refGene_tx=XM_023259344.1;refGene_gn=PKP4;refGene_tx=XM_023259345.1;refGene_gn=PKP4;refGene_tx=XM_023259346.1
C1	150773617	150773808
refGene_gn=PKP4;refGene_tx=XM_023259337.1;refGene_gn=PKP4;refGene_tx=XM_023259338.1;refGene_gn=PKP4;refGene_tx=XM_023259339.1;refGene_gn=PKP4;refGene_tx=XM_023259340.1;refGene_gn=PKP4;refGene_tx=XM_023259341.1;refGene_gn=PKP4;refGene_tx=XM_023259342.1;refGene_gn=PKP4;refGene_tx=XM_023259343.1;refGene_gn=PKP4;refGene_tx=XM_023259344.1;refGene_gn=PKP4;refGene_tx=XM_023259345.1;refGene_gn=PKP4;refGene_tx=XM_023259346.1;refGene_gn=PKP4;refGene_tx=XM_023259347.1
C1	150775982	150776535
refGene_gn=PKP4;refGene_tx=XM_023259337.1;refGene_gn=PKP4;refGene_tx=XM_023259338.1;refGene_gn=PKP4;refGene_tx=XM_023259339.1;refGene_gn=PKP4;refGene_tx=XM_023259340.1;refGene_gn=PKP4;refGene_tx=XM_023259341.1;refGene_gn=PKP4;refGene_tx=XM_023259342.1;refGene_gn=PKP4;refGene_tx=XM_023259343.1;refGene_gn=PKP4;refGene_tx=XM_023259344.1;refGene_gn=PKP4;refGene_tx=XM_023259345.1;refGene_gn=PKP4;refGene_tx=XM_023259346.1;refGene_gn=PKP4;refGene_tx=XM_023259347.1
C1	150781627	150781816
refGene_gn=PKP4;refGene_tx=XM_023259337.1;refGene_gn=PKP4;refGene_tx=X

M_023259338.1;refGene_gn=PKP4;refGene_tx=XM_023259339.1;refGene_gn=PKP4;refGene_tx=XM_023259340.1;refGene_gn=PKP4;refGene_tx=XM_023259341.1;refGene_gn=PKP4;refGene_tx=XM_023259342.1;refGene_gn=PKP4;refGene_tx=XM_023259343.1;refGene_gn=PKP4;refGene_tx=XM_023259344.1;refGene_gn=PKP4;refGene_tx=XM_023259345.1;refGene_gn=PKP4;refGene_tx=XM_023259346.1;refGene_gn=PKP4;refGene_tx=XM_023259347.1
C1	150783765	150783985
refGene_gn=PKP4;refGene_tx=XM_023259337.1;refGene_gn=PKP4;refGene_tx=XM_023259338.1;refGene_gn=PKP4;refGene_tx=XM_023259339.1;refGene_gn=PKP4;refGene_tx=XM_023259340.1;refGene_gn=PKP4;refGene_tx=XM_023259341.1;refGene_gn=PKP4;refGene_tx=XM_023259342.1;refGene_gn=PKP4;refGene_tx=XM_023259343.1;refGene_gn=PKP4;refGene_tx=XM_023259344.1;refGene_gn=PKP4;refGene_tx=XM_023259345.1;refGene_gn=PKP4;refGene_tx=XM_023259346.1;refGene_gn=PKP4;refGene_tx=XM_023259347.1
C1	150791311	150791444
refGene_gn=PKP4;refGene_tx=XM_023259337.1;refGene_gn=PKP4;refGene_tx=XM_023259338.1;refGene_gn=PKP4;refGene_tx=XM_023259339.1;refGene_gn=PKP4;refGene_tx=XM_023259340.1;refGene_gn=PKP4;refGene_tx=XM_023259341.1;refGene_gn=PKP4;refGene_tx=XM_023259342.1;refGene_gn=PKP4;refGene_tx=XM_023259343.1;refGene_gn=PKP4;refGene_tx=XM_023259344.1;refGene_gn=PKP4;refGene_tx=XM_023259345.1;refGene_gn=PKP4;refGene_tx=XM_023259346.1;refGene_gn=PKP4;refGene_tx=XM_023259347.1
C1	150792446	150792660
refGene_gn=PKP4;refGene_tx=XM_023259337.1;refGene_gn=PKP4;refGene_tx=XM_023259338.1;refGene_gn=PKP4;refGene_tx=XM_023259339.1;refGene_gn=PKP4;refGene_tx=XM_023259340.1;refGene_gn=PKP4;refGene_tx=XM_023259341.1;refGene_gn=PKP4;refGene_tx=XM_023259342.1;refGene_gn=PKP4;refGene_tx=XM_023259343.1;refGene_gn=PKP4;refGene_tx=XM_023259344.1;refGene_gn=PKP4;refGene_tx=XM_023259345.1;refGene_gn=PKP4;refGene_tx=XM_023259346.1;refGene_gn=PKP4;refGene_tx=XM_023259347.1
C1	150808323	150808507
refGene_gn=PKP4;refGene_tx=XM_023259337.1;refGene_gn=PKP4;refGene_tx=XM_023259338.1;refGene_gn=PKP4;refGene_tx=XM_023259339.1;refGene_gn=PKP4;refGene_tx=XM_023259340.1;refGene_gn=PKP4;refGene_tx=XM_023259341.1;refGene_gn=PKP4;refGene_tx=XM_023259342.1;refGene_gn=PKP4;refGene_tx=XM_023259343.1;refGene_gn=PKP4;refGene_tx=XM_023259344.1;refGene_gn=PKP4;refGene_tx=XM_023259345.1;refGene_gn=PKP4;refGene_tx=XM_023259346.1;refGene_gn=PKP4;refGene_tx=XM_023259347.1
C1	150811606	150811724
refGene_gn=PKP4;refGene_tx=XM_023259337.1;refGene_gn=PKP4;refGene_tx=XM_023259338.1;refGene_gn=PKP4;refGene_tx=XM_023259339.1;refGene_gn=PKP4;refGene_tx=XM_023259340.1;refGene_gn=PKP4;refGene_tx=XM_023259341.1;refGene_gn=PKP4;refGene_tx=XM_023259342.1;refGene_gn=PKP4;refGene_tx=XM_023259343.1;refGene_gn=PKP4;refGene_tx=XM_023259344.1;refGene_gn=PKP4;refGene_tx=XM_023259345.1;refGene_gn=PKP4;refGene_tx=XM_023259346.1;refGene_gn=PKP4;refGene_tx=XM_023259347.1
C1	150813188	150813380
refGene_gn=PKP4;refGene_tx=XM_023259337.1;refGene_gn=PKP4;refGene_tx=XM_023259338.1;refGene_gn=PKP4;refGene_tx=XM_023259339.1;refGene_gn=PKP4;refGene_tx=XM_023259340.1;refGene_gn=PKP4;refGene_tx=XM_023259341.1;

refGene_gn=PKP4;refGene_tx=XM_023259342.1;refGene_gn=PKP4;refGene_tx=XM_023259343.1;refGene_gn=PKP4;refGene_tx=XM_023259344.1;refGene_gn=PKP4;refGene_tx=XM_023259345.1;refGene_gn=PKP4;refGene_tx=XM_023259346.1;refGene_gn=PKP4;refGene_tx=XM_023259347.1
C1 150813572 150813746
refGene_gn=PKP4;refGene_tx=XM_023259337.1;refGene_gn=PKP4;refGene_tx=XM_023259338.1;refGene_gn=PKP4;refGene_tx=XM_023259339.1;refGene_gn=PKP4;refGene_tx=XM_023259340.1;refGene_gn=PKP4;refGene_tx=XM_023259341.1;refGene_gn=PKP4;refGene_tx=XM_023259342.1;refGene_gn=PKP4;refGene_tx=XM_023259343.1;refGene_gn=PKP4;refGene_tx=XM_023259344.1;refGene_gn=PKP4;refGene_tx=XM_023259345.1;refGene_gn=PKP4;refGene_tx=XM_023259346.1;refGene_gn=PKP4;refGene_tx=XM_023259347.1
C1 150816766 150816917
refGene_gn=PKP4;refGene_tx=XM_023259337.1;refGene_gn=PKP4;refGene_tx=XM_023259338.1;refGene_gn=PKP4;refGene_tx=XM_023259339.1;refGene_gn=PKP4;refGene_tx=XM_023259340.1;refGene_gn=PKP4;refGene_tx=XM_023259341.1;refGene_gn=PKP4;refGene_tx=XM_023259342.1;refGene_gn=PKP4;refGene_tx=XM_023259343.1;refGene_gn=PKP4;refGene_tx=XM_023259344.1;refGene_gn=PKP4;refGene_tx=XM_023259345.1;refGene_gn=PKP4;refGene_tx=XM_023259346.1;refGene_gn=PKP4;refGene_tx=XM_023259347.1
C1 150820278 150820474
refGene_gn=PKP4;refGene_tx=XM_023259337.1;refGene_gn=PKP4;refGene_tx=XM_023259338.1;refGene_gn=PKP4;refGene_tx=XM_023259339.1;refGene_gn=PKP4;refGene_tx=XM_023259340.1;refGene_gn=PKP4;refGene_tx=XM_023259341.1;refGene_gn=PKP4;refGene_tx=XM_023259342.1;refGene_gn=PKP4;refGene_tx=XM_023259343.1;refGene_gn=PKP4;refGene_tx=XM_023259344.1;refGene_gn=PKP4;refGene_tx=XM_023259345.1;refGene_gn=PKP4;refGene_tx=XM_023259346.1;refGene_gn=PKP4;refGene_tx=XM_023259347.1
C1 150824497 150824582
refGene_gn=PKP4;refGene_tx=XM_023259337.1;refGene_gn=PKP4;refGene_tx=XM_023259338.1;refGene_gn=PKP4;refGene_tx=XM_023259339.1;refGene_gn=PKP4;refGene_tx=XM_023259340.1;refGene_gn=PKP4;refGene_tx=XM_023259341.1;refGene_gn=PKP4;refGene_tx=XM_023259342.1;refGene_gn=PKP4;refGene_tx=XM_023259343.1;refGene_gn=PKP4;refGene_tx=XM_023259344.1;refGene_gn=PKP4;refGene_tx=XM_023259345.1;refGene_gn=PKP4;refGene_tx=XM_023259346.1;refGene_gn=PKP4;refGene_tx=XM_023259347.1
C1 150824706 150824824
refGene_gn=PKP4;refGene_tx=XM_023259337.1;refGene_gn=PKP4;refGene_tx=XM_023259338.1;refGene_gn=PKP4;refGene_tx=XM_023259339.1;refGene_gn=PKP4;refGene_tx=XM_023259340.1;refGene_gn=PKP4;refGene_tx=XM_023259341.1;refGene_gn=PKP4;refGene_tx=XM_023259342.1;refGene_gn=PKP4;refGene_tx=XM_023259343.1;refGene_gn=PKP4;refGene_tx=XM_023259344.1;refGene_gn=PKP4;refGene_tx=XM_023259345.1;refGene_gn=PKP4;refGene_tx=XM_023259346.1;refGene_gn=PKP4;refGene_tx=XM_023259347.1
C1 150827390 150827519
refGene_gn=PKP4;refGene_tx=XM_023259337.1;refGene_gn=PKP4;refGene_tx=XM_023259338.1;refGene_gn=PKP4;refGene_tx=XM_023259339.1;refGene_gn=PKP4;refGene_tx=XM_023259340.1;refGene_gn=PKP4;refGene_tx=XM_023259341.1;refGene_gn=PKP4;refGene_tx=XM_023259342.1;refGene_gn=PKP4;refGene_tx=XM_023259344.1;refGene_gn=PKP4;refGene_tx=XM_023259347.1

C1 150829217 150829291
refGene_gn=PKP4;refGene_tx=XM_023259337.1;refGene_gn=PKP4;refGene_tx=XM_023259338.1;refGene_gn=PKP4;refGene_tx=XM_023259339.1;refGene_gn=PKP4;refGene_tx=XM_023259340.1;refGene_gn=PKP4;refGene_tx=XM_023259341.1;refGene_gn=PKP4;refGene_tx=XM_023259342.1;refGene_gn=PKP4;refGene_tx=XM_023259343.1;refGene_gn=PKP4;refGene_tx=XM_023259344.1;refGene_gn=PKP4;refGene_tx=XM_023259345.1;refGene_gn=PKP4;refGene_tx=XM_023259347.1
C1 150830498 150830537
refGene_gn=PKP4;refGene_tx=XM_023259344.1;refGene_gn=PKP4;refGene_tx=XM_023259345.1
C1 150831044 150831293
refGene_gn=PKP4;refGene_tx=XM_023259337.1;refGene_gn=PKP4;refGene_tx=XM_023259338.1;refGene_gn=PKP4;refGene_tx=XM_023259339.1;refGene_gn=PKP4;refGene_tx=XM_023259340.1;refGene_gn=PKP4;refGene_tx=XM_023259341.1;refGene_gn=PKP4;refGene_tx=XM_023259342.1;refGene_gn=PKP4;refGene_tx=XM_023259343.1;refGene_gn=PKP4;refGene_tx=XM_023259346.1;refGene_gn=PKP4;refGene_tx=XM_023259347.1
C1 150938425 150938483
refGene_gn=DAPL1;refGene_tx=XM_023259348.1
C1 150948648 150948736
refGene_gn=DAPL1;refGene_tx=XM_023259348.1
C1 150951704 150951765
refGene_gn=DAPL1;refGene_tx=XM_023259348.1
C1 150956727 150956757
refGene_gn=DAPL1;refGene_tx=XM_023259348.1
C1 150960465 150960582
refGene_gn=DAPL1;refGene_tx=XM_023259348.1
C1 151136085 151136146
refGene_gn=TANC1;refGene_tx=XM_003990793.4;refGene_gn=TANC1;refGene_tx=XM_023259370.1;refGene_gn=TANC1;refGene_tx=XM_023259371.1;refGene_gn=TANC1;refGene_tx=XM_023259372.1;refGene_gn=TANC1;refGene_tx=XM_023259373.1;refGene_gn=TANC1;refGene_tx=XM_023259374.1;refGene_gn=TANC1;refGene_tx=XM_023259375.1;refGene_gn=TANC1;refGene_tx=XM_023259376.1;refGene_gn=TANC1;refGene_tx=XM_023259377.1;refGene_gn=TANC1;refGene_tx=XM_023259378.1
C1 151166659 151166860
refGene_gn=TANC1;refGene_tx=XM_003990793.4;refGene_gn=TANC1;refGene_tx=XM_023259370.1;refGene_gn=TANC1;refGene_tx=XM_023259371.1;refGene_gn=TANC1;refGene_tx=XM_023259372.1;refGene_gn=TANC1;refGene_tx=XM_023259373.1;refGene_gn=TANC1;refGene_tx=XM_023259374.1;refGene_gn=TANC1;refGene_tx=XM_023259375.1;refGene_gn=TANC1;refGene_tx=XM_023259376.1;refGene_gn=TANC1;refGene_tx=XM_023259377.1;refGene_gn=TANC1;refGene_tx=XM_023259378.1
C1 151208504 151208609
refGene_gn=TANC1;refGene_tx=XM_003990793.4;refGene_gn=TANC1;refGene_tx=XM_023259370.1;refGene_gn=TANC1;refGene_tx=XM_023259371.1;refGene_gn=TANC1;refGene_tx=XM_023259372.1;refGene_gn=TANC1;refGene_tx=XM_023259373.1;refGene_gn=TANC1;refGene_tx=XM_023259375.1;refGene_gn=TANC1;refGene_tx=XM_023259376.1;refGene_gn=TANC1;refGene_tx=XM_023259378.1
C1 151219857 151219988

refGene_gn=TANC1;refGene_tx=XM_003990793.4;refGene_gn=TANC1;refGene_tx
=XM_019838143.1;refGene_gn=TANC1;refGene_tx=XM_023259370.1;refGene_gn=
TANC1;refGene_tx=XM_023259371.1;refGene_gn=TANC1;refGene_tx=XM_0232593
72.1;refGene_gn=TANC1;refGene_tx=XM_023259373.1;refGene_gn=TANC1;refGe
ne_tx=XM_023259374.1;refGene_gn=TANC1;refGene_tx=XM_023259375.1;refGen
e_gn=TANC1;refGene_tx=XM_023259377.1;refGene_gn=TANC1;refGene_tx=XM_02
3259378.1
C1 151221033 151221220
refGene_gn=TANC1;refGene_tx=XM_003990793.4;refGene_gn=TANC1;refGene_tx
=XM_019838143.1;refGene_gn=TANC1;refGene_tx=XM_023259370.1;refGene_gn=
TANC1;refGene_tx=XM_023259371.1;refGene_gn=TANC1;refGene_tx=XM_0232593
72.1;refGene_gn=TANC1;refGene_tx=XM_023259373.1;refGene_gn=TANC1;refGe
ne_tx=XM_023259374.1;refGene_gn=TANC1;refGene_tx=XM_023259375.1;refGen
e_gn=TANC1;refGene_tx=XM_023259377.1;refGene_gn=TANC1;refGene_tx=XM_02
3259378.1
C1 151228326 151228498
refGene_gn=TANC1;refGene_tx=XM_019838144.1
C1 151234390 151234654
refGene_gn=TANC1;refGene_tx=XM_003990793.4;refGene_gn=TANC1;refGene_tx
=XM_019838143.1;refGene_gn=TANC1;refGene_tx=XM_019838144.1;refGene_gn=
TANC1;refGene_tx=XM_023259370.1;refGene_gn=TANC1;refGene_tx=XM_0232593
71.1;refGene_gn=TANC1;refGene_tx=XM_023259372.1;refGene_gn=TANC1;refGe
ne_tx=XM_023259373.1;refGene_gn=TANC1;refGene_tx=XM_023259374.1;refGen
e_gn=TANC1;refGene_tx=XM_023259376.1;refGene_gn=TANC1;refGene_tx=XM_02
3259378.1
C1 151240133 151240256
refGene_gn=TANC1;refGene_tx=XM_003990793.4;refGene_gn=TANC1;refGene_tx
=XM_019838143.1;refGene_gn=TANC1;refGene_tx=XM_019838144.1;refGene_gn=
TANC1;refGene_tx=XM_023259370.1;refGene_gn=TANC1;refGene_tx=XM_0232593
71.1;refGene_gn=TANC1;refGene_tx=XM_023259372.1;refGene_gn=TANC1;refGe
ne_tx=XM_023259373.1;refGene_gn=TANC1;refGene_tx=XM_023259374.1;refGen
e_gn=TANC1;refGene_tx=XM_023259375.1;refGene_gn=TANC1;refGene_tx=XM_02
3259376.1;refGene_gn=TANC1;refGene_tx=XM_023259377.1;refGene_gn=TANC1;
refGene_tx=XM_023259378.1
C1 151241042 151241324
refGene_gn=TANC1;refGene_tx=XM_003990793.4;refGene_gn=TANC1;refGene_tx
=XM_019838143.1;refGene_gn=TANC1;refGene_tx=XM_019838144.1;refGene_gn=
TANC1;refGene_tx=XM_023259370.1;refGene_gn=TANC1;refGene_tx=XM_0232593
71.1;refGene_gn=TANC1;refGene_tx=XM_023259372.1;refGene_gn=TANC1;refGe
ne_tx=XM_023259373.1;refGene_gn=TANC1;refGene_tx=XM_023259374.1;refGen
e_gn=TANC1;refGene_tx=XM_023259375.1;refGene_gn=TANC1;refGene_tx=XM_02
3259376.1;refGene_gn=TANC1;refGene_tx=XM_023259377.1;refGene_gn=TANC1;
refGene_tx=XM_023259378.1
C1 151242204 151242362
refGene_gn=TANC1;refGene_tx=XM_003990793.4;refGene_gn=TANC1;refGene_tx
=XM_019838143.1;refGene_gn=TANC1;refGene_tx=XM_019838144.1;refGene_gn=
TANC1;refGene_tx=XM_023259370.1;refGene_gn=TANC1;refGene_tx=XM_0232593
71.1;refGene_gn=TANC1;refGene_tx=XM_023259372.1;refGene_gn=TANC1;refGe
ne_tx=XM_023259373.1;refGene_gn=TANC1;refGene_tx=XM_023259374.1;refGen
e_gn=TANC1;refGene_tx=XM_023259375.1;refGene_gn=TANC1;refGene_tx=XM_02

3259376.1;refGene_gn=TANC1;refGene_tx=XM_023259377.1;refGene_gn=TANC1;
refGene_tx=XM_023259378.1
C1 151245552 151245784
refGene_gn=TANC1;refGene_tx=XM_003990793.4;refGene_gn=TANC1;refGene_tx
=XM_019838143.1;refGene_gn=TANC1;refGene_tx=XM_019838144.1;refGene_gn=
TANC1;refGene_tx=XM_023259370.1;refGene_gn=TANC1;refGene_tx=XM_0232593
71.1;refGene_gn=TANC1;refGene_tx=XM_023259372.1;refGene_gn=TANC1;refGe
ne_tx=XM_023259373.1;refGene_gn=TANC1;refGene_tx=XM_023259374.1;refGen
e_gn=TANC1;refGene_tx=XM_023259375.1;refGene_gn=TANC1;refGene_tx=XM_02
3259376.1;refGene_gn=TANC1;refGene_tx=XM_023259377.1
C1 151246923 151247090
refGene_gn=TANC1;refGene_tx=XM_003990793.4;refGene_gn=TANC1;refGene_tx
=XM_019838143.1;refGene_gn=TANC1;refGene_tx=XM_019838144.1;refGene_gn=
TANC1;refGene_tx=XM_023259370.1;refGene_gn=TANC1;refGene_tx=XM_0232593
71.1;refGene_gn=TANC1;refGene_tx=XM_023259372.1;refGene_gn=TANC1;refGe
ne_tx=XM_023259373.1;refGene_gn=TANC1;refGene_tx=XM_023259374.1;refGen
e_gn=TANC1;refGene_tx=XM_023259375.1;refGene_gn=TANC1;refGene_tx=XM_02
3259376.1;refGene_gn=TANC1;refGene_tx=XM_023259377.1
C1 151249083 151249691
refGene_gn=TANC1;refGene_tx=XM_003990793.4;refGene_gn=TANC1;refGene_tx
=XM_019838143.1;refGene_gn=TANC1;refGene_tx=XM_019838144.1;refGene_gn=
TANC1;refGene_tx=XM_023259370.1;refGene_gn=TANC1;refGene_tx=XM_0232593
71.1;refGene_gn=TANC1;refGene_tx=XM_023259372.1;refGene_gn=TANC1;refGe
ne_tx=XM_023259373.1;refGene_gn=TANC1;refGene_tx=XM_023259374.1;refGen
e_gn=TANC1;refGene_tx=XM_023259375.1;refGene_gn=TANC1;refGene_tx=XM_02
3259376.1;refGene_gn=TANC1;refGene_tx=XM_023259377.1;refGene_gn=TANC1;
refGene_tx=XM_023259378.1
C1 151255962 151256071
refGene_gn=TANC1;refGene_tx=XM_003990793.4;refGene_gn=TANC1;refGene_tx
=XM_019838143.1;refGene_gn=TANC1;refGene_tx=XM_019838144.1;refGene_gn=
TANC1;refGene_tx=XM_023259370.1;refGene_gn=TANC1;refGene_tx=XM_0232593
71.1;refGene_gn=TANC1;refGene_tx=XM_023259372.1;refGene_gn=TANC1;refGe
ne_tx=XM_023259373.1;refGene_gn=TANC1;refGene_tx=XM_023259374.1;refGen
e_gn=TANC1;refGene_tx=XM_023259375.1;refGene_gn=TANC1;refGene_tx=XM_02
3259376.1;refGene_gn=TANC1;refGene_tx=XM_023259377.1;refGene_gn=TANC1;
refGene_tx=XM_023259378.1
C1 151257104 151257227
refGene_gn=TANC1;refGene_tx=XM_003990793.4;refGene_gn=TANC1;refGene_tx
=XM_019838143.1;refGene_gn=TANC1;refGene_tx=XM_019838144.1;refGene_gn=
TANC1;refGene_tx=XM_023259370.1;refGene_gn=TANC1;refGene_tx=XM_0232593
71.1;refGene_gn=TANC1;refGene_tx=XM_023259372.1;refGene_gn=TANC1;refGe
ne_tx=XM_023259373.1;refGene_gn=TANC1;refGene_tx=XM_023259374.1;refGen
e_gn=TANC1;refGene_tx=XM_023259375.1;refGene_gn=TANC1;refGene_tx=XM_02
3259376.1;refGene_gn=TANC1;refGene_tx=XM_023259377.1;refGene_gn=TANC1;
refGene_tx=XM_023259378.1
C1 151263635 151263872
refGene_gn=TANC1;refGene_tx=XM_003990793.4;refGene_gn=TANC1;refGene_tx
=XM_019838143.1;refGene_gn=TANC1;refGene_tx=XM_019838144.1;refGene_gn=
TANC1;refGene_tx=XM_023259370.1;refGene_gn=TANC1;refGene_tx=XM_0232593
71.1;refGene_gn=TANC1;refGene_tx=XM_023259372.1;refGene_gn=TANC1;refGe

ne_tx=XM_023259373.1;refGene_gn=TANC1;refGene_tx=XM_023259374.1;refGene_gn=TANC1;refGene_tx=XM_023259375.1;refGene_gn=TANC1;refGene_tx=XM_023259376.1;refGene_gn=TANC1;refGene_tx=XM_023259377.1;refGene_gn=TANC1;refGene_tx=XM_023259378.1
C1 151265747 151265933
refGene_gn=TANC1;refGene_tx=XM_003990793.4;refGene_gn=TANC1;refGene_tx=XM_019838143.1;refGene_gn=TANC1;refGene_tx=XM_019838144.1;refGene_gn=TANC1;refGene_tx=XM_023259370.1;refGene_gn=TANC1;refGene_tx=XM_023259371.1;refGene_gn=TANC1;refGene_tx=XM_023259372.1;refGene_gn=TANC1;refGene_tx=XM_023259373.1;refGene_gn=TANC1;refGene_tx=XM_023259374.1;refGene_gn=TANC1;refGene_tx=XM_023259375.1;refGene_gn=TANC1;refGene_tx=XM_023259376.1;refGene_gn=TANC1;refGene_tx=XM_023259377.1;refGene_gn=TANC1;refGene_tx=XM_023259378.1
C1 151269175 151269254
refGene_gn=TANC1;refGene_tx=XM_003990793.4;refGene_gn=TANC1;refGene_tx=XM_019838143.1;refGene_gn=TANC1;refGene_tx=XM_019838144.1;refGene_gn=TANC1;refGene_tx=XM_023259370.1;refGene_gn=TANC1;refGene_tx=XM_023259371.1;refGene_gn=TANC1;refGene_tx=XM_023259372.1;refGene_gn=TANC1;refGene_tx=XM_023259373.1;refGene_gn=TANC1;refGene_tx=XM_023259374.1;refGene_gn=TANC1;refGene_tx=XM_023259375.1;refGene_gn=TANC1;refGene_tx=XM_023259376.1;refGene_gn=TANC1;refGene_tx=XM_023259377.1;refGene_gn=TANC1;refGene_tx=XM_023259378.1
C1 151285170 151285304
refGene_gn=TANC1;refGene_tx=XM_003990793.4;refGene_gn=TANC1;refGene_tx=XM_019838143.1;refGene_gn=TANC1;refGene_tx=XM_019838144.1;refGene_gn=TANC1;refGene_tx=XM_023259370.1;refGene_gn=TANC1;refGene_tx=XM_023259371.1;refGene_gn=TANC1;refGene_tx=XM_023259372.1;refGene_gn=TANC1;refGene_tx=XM_023259373.1;refGene_gn=TANC1;refGene_tx=XM_023259374.1;refGene_gn=TANC1;refGene_tx=XM_023259375.1;refGene_gn=TANC1;refGene_tx=XM_023259376.1;refGene_gn=TANC1;refGene_tx=XM_023259377.1;refGene_gn=TANC1;refGene_tx=XM_023259378.1
C1 151286837 151286961
refGene_gn=TANC1;refGene_tx=XM_003990793.4;refGene_gn=TANC1;refGene_tx=XM_019838143.1;refGene_gn=TANC1;refGene_tx=XM_019838144.1;refGene_gn=TANC1;refGene_tx=XM_023259370.1;refGene_gn=TANC1;refGene_tx=XM_023259371.1;refGene_gn=TANC1;refGene_tx=XM_023259372.1;refGene_gn=TANC1;refGene_tx=XM_023259373.1;refGene_gn=TANC1;refGene_tx=XM_023259374.1;refGene_gn=TANC1;refGene_tx=XM_023259375.1;refGene_gn=TANC1;refGene_tx=XM_023259376.1;refGene_gn=TANC1;refGene_tx=XM_023259377.1;refGene_gn=TANC1;refGene_tx=XM_023259378.1
C1 151287284 151287460
refGene_gn=TANC1;refGene_tx=XM_003990793.4;refGene_gn=TANC1;refGene_tx=XM_019838143.1;refGene_gn=TANC1;refGene_tx=XM_019838144.1;refGene_gn=TANC1;refGene_tx=XM_023259370.1;refGene_gn=TANC1;refGene_tx=XM_023259371.1;refGene_gn=TANC1;refGene_tx=XM_023259372.1;refGene_gn=TANC1;refGene_tx=XM_023259373.1;refGene_gn=TANC1;refGene_tx=XM_023259374.1;refGene_gn=TANC1;refGene_tx=XM_023259375.1;refGene_gn=TANC1;refGene_tx=XM_023259376.1;refGene_gn=TANC1;refGene_tx=XM_023259377.1;refGene_gn=TANC1;refGene_tx=XM_023259378.1
C1 151291225 151291358

refGene_gn=TANC1;refGene_tx=XM_003990793.4;refGene_gn=TANC1;refGene_tx=XM_019838143.1;refGene_gn=TANC1;refGene_tx=XM_019838144.1;refGene_gn=TANC1;refGene_tx=XM_023259370.1;refGene_gn=TANC1;refGene_tx=XM_023259371.1;refGene_gn=TANC1;refGene_tx=XM_023259372.1;refGene_gn=TANC1;refGene_tx=XM_023259373.1;refGene_gn=TANC1;refGene_tx=XM_023259374.1;refGene_gn=TANC1;refGene_tx=XM_023259375.1;refGene_gn=TANC1;refGene_tx=XM_023259376.1;refGene_gn=TANC1;refGene_tx=XM_023259377.1;refGene_gn=TANC1;refGene_tx=XM_023259378.1
C1 151292782 151292874
refGene_gn=TANC1;refGene_tx=XM_003990793.4;refGene_gn=TANC1;refGene_tx=XM_019838143.1;refGene_gn=TANC1;refGene_tx=XM_019838144.1;refGene_gn=TANC1;refGene_tx=XM_023259370.1;refGene_gn=TANC1;refGene_tx=XM_023259371.1;refGene_gn=TANC1;refGene_tx=XM_023259372.1;refGene_gn=TANC1;refGene_tx=XM_023259373.1;refGene_gn=TANC1;refGene_tx=XM_023259374.1;refGene_gn=TANC1;refGene_tx=XM_023259375.1;refGene_gn=TANC1;refGene_tx=XM_023259376.1;refGene_gn=TANC1;refGene_tx=XM_023259377.1;refGene_gn=TANC1;refGene_tx=XM_023259378.1
C1 151294590 151294737
refGene_gn=TANC1;refGene_tx=XM_003990793.4;refGene_gn=TANC1;refGene_tx=XM_019838143.1;refGene_gn=TANC1;refGene_tx=XM_019838144.1;refGene_gn=TANC1;refGene_tx=XM_023259370.1;refGene_gn=TANC1;refGene_tx=XM_023259371.1;refGene_gn=TANC1;refGene_tx=XM_023259372.1;refGene_gn=TANC1;refGene_tx=XM_023259373.1;refGene_gn=TANC1;refGene_tx=XM_023259374.1;refGene_gn=TANC1;refGene_tx=XM_023259375.1;refGene_gn=TANC1;refGene_tx=XM_023259376.1;refGene_gn=TANC1;refGene_tx=XM_023259377.1;refGene_gn=TANC1;refGene_tx=XM_023259378.1
C1 151295629 151295730
refGene_gn=TANC1;refGene_tx=XM_003990793.4;refGene_gn=TANC1;refGene_tx=XM_019838143.1;refGene_gn=TANC1;refGene_tx=XM_019838144.1;refGene_gn=TANC1;refGene_tx=XM_023259370.1;refGene_gn=TANC1;refGene_tx=XM_023259371.1;refGene_gn=TANC1;refGene_tx=XM_023259372.1;refGene_gn=TANC1;refGene_tx=XM_023259373.1;refGene_gn=TANC1;refGene_tx=XM_023259374.1;refGene_gn=TANC1;refGene_tx=XM_023259375.1;refGene_gn=TANC1;refGene_tx=XM_023259376.1;refGene_gn=TANC1;refGene_tx=XM_023259377.1;refGene_gn=TANC1;refGene_tx=XM_023259378.1
C1 151296515 151297950
refGene_gn=TANC1;refGene_tx=XM_003990793.4;refGene_gn=TANC1;refGene_tx=XM_019838143.1;refGene_gn=TANC1;refGene_tx=XM_019838144.1;refGene_gn=TANC1;refGene_tx=XM_023259370.1;refGene_gn=TANC1;refGene_tx=XM_023259371.1;refGene_gn=TANC1;refGene_tx=XM_023259372.1;refGene_gn=TANC1;refGene_tx=XM_023259373.1;refGene_gn=TANC1;refGene_tx=XM_023259374.1;refGene_gn=TANC1;refGene_tx=XM_023259375.1;refGene_gn=TANC1;refGene_tx=XM_023259376.1;refGene_gn=TANC1;refGene_tx=XM_023259377.1;refGene_gn=TANC1;refGene_tx=XM_023259378.1
C1 151301974 151302132
refGene_gn=BAZ2B;refGene_tx=XM_011285357.2;refGene_gn=BAZ2B;refGene_tx=XM_011285358.2;refGene_gn=BAZ2B;refGene_tx=XM_019838146.1;refGene_gn=BAZ2B;refGene_tx=XM_019838147.2;refGene_gn=BAZ2B;refGene_tx=XM_019838148.1;refGene_gn=BAZ2B;refGene_tx=XM_023259349.1;refGene_gn=BAZ2B;refGene_tx=XM_023259350.1;refGene_gn=BAZ2B;refGene_tx=XM_023259351.1;refGen

e_gn=BAZ2B;refGene_tx=XM_023259352.1;refGene_gn=BAZ2B;refGene_tx=XM_02
3259353.1;refGene_gn=BAZ2B;refGene_tx=XM_023259354.1;refGene_gn=BAZ2B;
refGene_tx=XM_023259355.1;refGene_gn=BAZ2B;refGene_tx=XM_023259356.1;r
efGene_gn=BAZ2B;refGene_tx=XM_023259357.1;refGene_gn=BAZ2B;refGene_tx=
XM_023259358.1;refGene_gn=BAZ2B;refGene_tx=XM_023259359.1;refGene_gn=B
AZ2B;refGene_tx=XM_023259361.1;refGene_gn=BAZ2B;refGene_tx=XM_02325936
2.1
C1 151309168 151309182
refGene_gn=BAZ2B;refGene_tx=XM_023259360.1
C1 151321050 151321191
refGene_gn=BAZ2B;refGene_tx=XM_011285357.2;refGene_gn=BAZ2B;refGene_tx
=XM_011285358.2;refGene_gn=BAZ2B;refGene_tx=XM_019838146.1;refGene_gn=
BAZ2B;refGene_tx=XM_019838147.2;refGene_gn=BAZ2B;refGene_tx=XM_0198381
48.1;refGene_gn=BAZ2B;refGene_tx=XM_023259349.1;refGene_gn=BAZ2B;refGe
ne_tx=XM_023259350.1;refGene_gn=BAZ2B;refGene_tx=XM_023259351.1;refGen
e_gn=BAZ2B;refGene_tx=XM_023259352.1;refGene_gn=BAZ2B;refGene_tx=XM_02
3259353.1;refGene_gn=BAZ2B;refGene_tx=XM_023259354.1;refGene_gn=BAZ2B;
refGene_tx=XM_023259355.1;refGene_gn=BAZ2B;refGene_tx=XM_023259356.1;r
efGene_gn=BAZ2B;refGene_tx=XM_023259357.1;refGene_gn=BAZ2B;refGene_tx=
XM_023259358.1;refGene_gn=BAZ2B;refGene_tx=XM_023259359.1;refGene_gn=B
AZ2B;refGene_tx=XM_023259360.1;refGene_gn=BAZ2B;refGene_tx=XM_02325936
1.1;refGene_gn=BAZ2B;refGene_tx=XM_023259362.1
C1 151330020 151330200
refGene_gn=BAZ2B;refGene_tx=XM_011285357.2;refGene_gn=BAZ2B;refGene_tx
=XM_011285358.2;refGene_gn=BAZ2B;refGene_tx=XM_019838146.1;refGene_gn=
BAZ2B;refGene_tx=XM_019838147.2;refGene_gn=BAZ2B;refGene_tx=XM_0198381
48.1;refGene_gn=BAZ2B;refGene_tx=XM_023259349.1;refGene_gn=BAZ2B;refGe
ne_tx=XM_023259350.1;refGene_gn=BAZ2B;refGene_tx=XM_023259351.1;refGen
e_gn=BAZ2B;refGene_tx=XM_023259352.1;refGene_gn=BAZ2B;refGene_tx=XM_02
3259353.1;refGene_gn=BAZ2B;refGene_tx=XM_023259354.1;refGene_gn=BAZ2B;
refGene_tx=XM_023259355.1;refGene_gn=BAZ2B;refGene_tx=XM_023259356.1;r
efGene_gn=BAZ2B;refGene_tx=XM_023259357.1;refGene_gn=BAZ2B;refGene_tx=
XM_023259358.1;refGene_gn=BAZ2B;refGene_tx=XM_023259359.1;refGene_gn=B
AZ2B;refGene_tx=XM_023259360.1;refGene_gn=BAZ2B;refGene_tx=XM_02325936
1.1;refGene_gn=BAZ2B;refGene_tx=XM_023259362.1
C1 151331115 151331222
refGene_gn=BAZ2B;refGene_tx=XM_011285357.2;refGene_gn=BAZ2B;refGene_tx
=XM_011285358.2;refGene_gn=BAZ2B;refGene_tx=XM_019838146.1;refGene_gn=
BAZ2B;refGene_tx=XM_019838147.2;refGene_gn=BAZ2B;refGene_tx=XM_0198381
48.1;refGene_gn=BAZ2B;refGene_tx=XM_023259349.1;refGene_gn=BAZ2B;refGe
ne_tx=XM_023259350.1;refGene_gn=BAZ2B;refGene_tx=XM_023259351.1;refGen
e_gn=BAZ2B;refGene_tx=XM_023259352.1;refGene_gn=BAZ2B;refGene_tx=XM_02
3259353.1;refGene_gn=BAZ2B;refGene_tx=XM_023259354.1;refGene_gn=BAZ2B;
refGene_tx=XM_023259355.1;refGene_gn=BAZ2B;refGene_tx=XM_023259356.1;r
efGene_gn=BAZ2B;refGene_tx=XM_023259357.1;refGene_gn=BAZ2B;refGene_tx=
XM_023259358.1;refGene_gn=BAZ2B;refGene_tx=XM_023259359.1;refGene_gn=B
AZ2B;refGene_tx=XM_023259360.1;refGene_gn=BAZ2B;refGene_tx=XM_02325936
1.1;refGene_gn=BAZ2B;refGene_tx=XM_023259362.1
C1 151331302 151331343
refGene_gn=BAZ2B;refGene_tx=XM_011285357.2;refGene_gn=BAZ2B;refGene_tx

=XM_011285358.2;refGene_gn=BAZ2B;refGene_tx=XM_019838146.1;refGene_gn=
BAZ2B;refGene_tx=XM_019838147.2;refGene_gn=BAZ2B;refGene_tx=XM_0198381
48.1;refGene_gn=BAZ2B;refGene_tx=XM_023259349.1;refGene_gn=BAZ2B;refGe
ne_tx=XM_023259350.1;refGene_gn=BAZ2B;refGene_tx=XM_023259351.1;refGen
e_gn=BAZ2B;refGene_tx=XM_023259352.1;refGene_gn=BAZ2B;refGene_tx=XM_02
3259353.1;refGene_gn=BAZ2B;refGene_tx=XM_023259354.1;refGene_gn=BAZ2B;
refGene_tx=XM_023259355.1;refGene_gn=BAZ2B;refGene_tx=XM_023259356.1;r
efGene_gn=BAZ2B;refGene_tx=XM_023259357.1;refGene_gn=BAZ2B;refGene_tx=
XM_023259358.1;refGene_gn=BAZ2B;refGene_tx=XM_023259359.1;refGene_gn=B
AZ2B;refGene_tx=XM_023259360.1;refGene_gn=BAZ2B;refGene_tx=XM_02325936
1.1;refGene_gn=BAZ2B;refGene_tx=XM_023259362.1
C1 151332715 151332749
refGene_gn=BAZ2B;refGene_tx=XM_011285357.2;refGene_gn=BAZ2B;refGene_tx
=XM_011285358.2;refGene_gn=BAZ2B;refGene_tx=XM_019838146.1;refGene_gn=
BAZ2B;refGene_tx=XM_019838147.2;refGene_gn=BAZ2B;refGene_tx=XM_0198381
48.1;refGene_gn=BAZ2B;refGene_tx=XM_023259349.1;refGene_gn=BAZ2B;refGe
ne_tx=XM_023259350.1;refGene_gn=BAZ2B;refGene_tx=XM_023259351.1;refGen
e_gn=BAZ2B;refGene_tx=XM_023259352.1;refGene_gn=BAZ2B;refGene_tx=XM_02
3259353.1;refGene_gn=BAZ2B;refGene_tx=XM_023259354.1;refGene_gn=BAZ2B;
refGene_tx=XM_023259355.1;refGene_gn=BAZ2B;refGene_tx=XM_023259356.1;r
efGene_gn=BAZ2B;refGene_tx=XM_023259357.1;refGene_gn=BAZ2B;refGene_tx=
XM_023259358.1;refGene_gn=BAZ2B;refGene_tx=XM_023259359.1;refGene_gn=B
AZ2B;refGene_tx=XM_023259360.1;refGene_gn=BAZ2B;refGene_tx=XM_02325936
1.1;refGene_gn=BAZ2B;refGene_tx=XM_023259362.1
C1 151343172 151343266
refGene_gn=BAZ2B;refGene_tx=XM_011285357.2;refGene_gn=BAZ2B;refGene_tx
=XM_011285358.2;refGene_gn=BAZ2B;refGene_tx=XM_019838146.1;refGene_gn=
BAZ2B;refGene_tx=XM_019838147.2;refGene_gn=BAZ2B;refGene_tx=XM_0232593
49.1;refGene_gn=BAZ2B;refGene_tx=XM_023259350.1;refGene_gn=BAZ2B;refGe
ne_tx=XM_023259351.1;refGene_gn=BAZ2B;refGene_tx=XM_023259352.1;refGen
e_gn=BAZ2B;refGene_tx=XM_023259353.1;refGene_gn=BAZ2B;refGene_tx=XM_02
3259354.1;refGene_gn=BAZ2B;refGene_tx=XM_023259355.1;refGene_gn=BAZ2B;
refGene_tx=XM_023259356.1;refGene_gn=BAZ2B;refGene_tx=XM_023259357.1;r
efGene_gn=BAZ2B;refGene_tx=XM_023259358.1;refGene_gn=BAZ2B;refGene_tx=
XM_023259359.1;refGene_gn=BAZ2B;refGene_tx=XM_023259360.1;refGene_gn=B
AZ2B;refGene_tx=XM_023259361.1;refGene_gn=BAZ2B;refGene_tx=XM_02325936
2.1
C1 151345740 151345833
refGene_gn=BAZ2B;refGene_tx=XM_011285357.2;refGene_gn=BAZ2B;refGene_tx
=XM_011285358.2;refGene_gn=BAZ2B;refGene_tx=XM_019838146.1;refGene_gn=
BAZ2B;refGene_tx=XM_019838147.2;refGene_gn=BAZ2B;refGene_tx=XM_0198381
48.1;refGene_gn=BAZ2B;refGene_tx=XM_023259349.1;refGene_gn=BAZ2B;refGe
ne_tx=XM_023259350.1;refGene_gn=BAZ2B;refGene_tx=XM_023259351.1;refGen
e_gn=BAZ2B;refGene_tx=XM_023259352.1;refGene_gn=BAZ2B;refGene_tx=XM_02
3259353.1;refGene_gn=BAZ2B;refGene_tx=XM_023259354.1;refGene_gn=BAZ2B;
refGene_tx=XM_023259355.1;refGene_gn=BAZ2B;refGene_tx=XM_023259356.1;r
efGene_gn=BAZ2B;refGene_tx=XM_023259357.1;refGene_gn=BAZ2B;refGene_tx=
XM_023259358.1;refGene_gn=BAZ2B;refGene_tx=XM_023259359.1;refGene_gn=B
AZ2B;refGene_tx=XM_023259360.1;refGene_gn=BAZ2B;refGene_tx=XM_02325936
1.1;refGene_gn=BAZ2B;refGene_tx=XM_023259362.1

C1 151348476 151348661
refGene_gn=BAZ2B;refGene_tx=XM_011285357.2;refGene_gn=BAZ2B;refGene_tx=XM_011285358.2;refGene_gn=BAZ2B;refGene_tx=XM_019838146.1;refGene_gn=BAZ2B;refGene_tx=XM_019838147.2;refGene_gn=BAZ2B;refGene_tx=XM_023259349.1;refGene_gn=BAZ2B;refGene_tx=XM_023259350.1;refGene_gn=BAZ2B;refGene_tx=XM_023259351.1;refGene_gn=BAZ2B;refGene_tx=XM_023259352.1;refGene_gn=BAZ2B;refGene_tx=XM_023259353.1;refGene_gn=BAZ2B;refGene_tx=XM_023259354.1;refGene_gn=BAZ2B;refGene_tx=XM_023259355.1;refGene_gn=BAZ2B;refGene_tx=XM_023259356.1;refGene_gn=BAZ2B;refGene_tx=XM_023259357.1;refGene_gn=BAZ2B;refGene_tx=XM_023259358.1;refGene_gn=BAZ2B;refGene_tx=XM_023259359.1;refGene_gn=BAZ2B;refGene_tx=XM_023259360.1;refGene_gn=BAZ2B;refGene_tx=XM_023259361.1;refGene_gn=BAZ2B;refGene_tx=XM_023259362.1
C1 151350541 151350963
refGene_gn=BAZ2B;refGene_tx=XM_011285357.2;refGene_gn=BAZ2B;refGene_tx=XM_011285358.2;refGene_gn=BAZ2B;refGene_tx=XM_019838146.1;refGene_gn=BAZ2B;refGene_tx=XM_019838147.2;refGene_gn=BAZ2B;refGene_tx=XM_023259349.1;refGene_gn=BAZ2B;refGene_tx=XM_023259350.1;refGene_gn=BAZ2B;refGene_tx=XM_023259351.1;refGene_gn=BAZ2B;refGene_tx=XM_023259352.1;refGene_gn=BAZ2B;refGene_tx=XM_023259353.1;refGene_gn=BAZ2B;refGene_tx=XM_023259354.1;refGene_gn=BAZ2B;refGene_tx=XM_023259355.1;refGene_gn=BAZ2B;refGene_tx=XM_023259356.1;refGene_gn=BAZ2B;refGene_tx=XM_023259357.1;refGene_gn=BAZ2B;refGene_tx=XM_023259358.1;refGene_gn=BAZ2B;refGene_tx=XM_023259359.1;refGene_gn=BAZ2B;refGene_tx=XM_023259360.1;refGene_gn=BAZ2B;refGene_tx=XM_023259361.1;refGene_gn=BAZ2B;refGene_tx=XM_023259362.1
C1 151358135 151358150
refGene_gn=BAZ2B;refGene_tx=XM_023259363.1
C1 151366839 151366993
refGene_gn=BAZ2B;refGene_tx=XM_023259349.1;refGene_gn=BAZ2B;refGene_tx=XM_023259350.1;refGene_gn=BAZ2B;refGene_tx=XM_023259351.1;refGene_gn=BAZ2B;refGene_tx=XM_023259352.1;refGene_gn=BAZ2B;refGene_tx=XM_023259353.1;refGene_gn=BAZ2B;refGene_tx=XM_023259354.1;refGene_gn=BAZ2B;refGene_tx=XM_023259355.1;refGene_gn=BAZ2B;refGene_tx=XM_023259356.1;refGene_gn=BAZ2B;refGene_tx=XM_023259357.1;refGene_gn=BAZ2B;refGene_tx=XM_023259358.1;refGene_gn=BAZ2B;refGene_tx=XM_023259359.1;refGene_gn=BAZ2B;refGene_tx=XM_023259360.1;refGene_gn=BAZ2B;refGene_tx=XM_023259361.1;refGene_gn=BAZ2B;refGene_tx=XM_023259362.1;refGene_gn=BAZ2B;refGene_tx=XM_023259363.1;refGene_gn=BAZ2B;refGene_tx=XM_023259364.1;refGene_gn=BAZ2B;refGene_tx=XM_023259365.1;refGene_gn=BAZ2B;refGene_tx=XM_023259366.1;refGene_gn=BAZ2B;refGene_tx=XM_023259367.1;refGene_gn=BAZ2B;refGene_tx=XM_023259368.1;refGene_gn=BAZ2B;refGene_tx=XM_023259369.1
C1 151370468 151370612
refGene_gn=BAZ2B;refGene_tx=XM_023259349.1;refGene_gn=BAZ2B;refGene_tx=XM_023259350.1;refGene_gn=BAZ2B;refGene_tx=XM_023259351.1;refGene_gn=BAZ2B;refGene_tx=XM_023259352.1;refGene_gn=BAZ2B;refGene_tx=XM_023259353.1;refGene_gn=BAZ2B;refGene_tx=XM_023259354.1;refGene_gn=BAZ2B;refGene_tx=XM_023259355.1;refGene_gn=BAZ2B;refGene_tx=XM_023259356.1;refGene_gn=BAZ2B;refGene_tx=XM_023259357.1;refGene_gn=BAZ2B;refGene_tx=XM_023259358.1;refGene_gn=BAZ2B;refGene_tx=XM_023259359.1;refGene_gn=BAZ2B;

refGene_tx=XM_023259360.1;refGene_gn=BAZ2B;refGene_tx=XM_023259361.1;refGene_gn=BAZ2B;refGene_tx=XM_023259362.1;refGene_gn=BAZ2B;refGene_tx=XM_023259363.1;refGene_gn=BAZ2B;refGene_tx=XM_023259364.1;refGene_gn=BAZ2B;refGene_tx=XM_023259365.1;refGene_gn=BAZ2B;refGene_tx=XM_023259366.1;refGene_gn=BAZ2B;refGene_tx=XM_023259367.1;refGene_gn=BAZ2B;refGene_tx=XM_023259368.1;refGene_gn=BAZ2B;refGene_tx=XM_023259369.1
C1 151370864 151371130
refGene_gn=BAZ2B;refGene_tx=XM_023259349.1;refGene_gn=BAZ2B;refGene_tx=XM_023259350.1;refGene_gn=BAZ2B;refGene_tx=XM_023259351.1;refGene_gn=BAZ2B;refGene_tx=XM_023259352.1;refGene_gn=BAZ2B;refGene_tx=XM_023259353.1;refGene_gn=BAZ2B;refGene_tx=XM_023259354.1;refGene_gn=BAZ2B;refGene_tx=XM_023259355.1;refGene_gn=BAZ2B;refGene_tx=XM_023259356.1;refGene_gn=BAZ2B;refGene_tx=XM_023259357.1;refGene_gn=BAZ2B;refGene_tx=XM_023259358.1;refGene_gn=BAZ2B;refGene_tx=XM_023259359.1;refGene_gn=BAZ2B;refGene_tx=XM_023259360.1;refGene_gn=BAZ2B;refGene_tx=XM_023259361.1;refGene_gn=BAZ2B;refGene_tx=XM_023259362.1;refGene_gn=BAZ2B;refGene_tx=XM_023259363.1;refGene_gn=BAZ2B;refGene_tx=XM_023259364.1;refGene_gn=BAZ2B;refGene_tx=XM_023259365.1;refGene_gn=BAZ2B;refGene_tx=XM_023259366.1;refGene_gn=BAZ2B;refGene_tx=XM_023259367.1;refGene_gn=BAZ2B;refGene_tx=XM_023259368.1;refGene_gn=BAZ2B;refGene_tx=XM_023259369.1
C1 151376445 151376592
refGene_gn=BAZ2B;refGene_tx=XM_023259349.1;refGene_gn=BAZ2B;refGene_tx=XM_023259350.1;refGene_gn=BAZ2B;refGene_tx=XM_023259351.1;refGene_gn=BAZ2B;refGene_tx=XM_023259352.1;refGene_gn=BAZ2B;refGene_tx=XM_023259353.1;refGene_gn=BAZ2B;refGene_tx=XM_023259354.1;refGene_gn=BAZ2B;refGene_tx=XM_023259355.1;refGene_gn=BAZ2B;refGene_tx=XM_023259356.1;refGene_gn=BAZ2B;refGene_tx=XM_023259357.1;refGene_gn=BAZ2B;refGene_tx=XM_023259358.1;refGene_gn=BAZ2B;refGene_tx=XM_023259359.1;refGene_gn=BAZ2B;refGene_tx=XM_023259360.1;refGene_gn=BAZ2B;refGene_tx=XM_023259361.1;refGene_gn=BAZ2B;refGene_tx=XM_023259362.1;refGene_gn=BAZ2B;refGene_tx=XM_023259363.1;refGene_gn=BAZ2B;refGene_tx=XM_023259364.1;refGene_gn=BAZ2B;refGene_tx=XM_023259365.1;refGene_gn=BAZ2B;refGene_tx=XM_023259366.1;refGene_gn=BAZ2B;refGene_tx=XM_023259367.1;refGene_gn=BAZ2B;refGene_tx=XM_023259368.1;refGene_gn=BAZ2B;refGene_tx=XM_023259369.1
C1 151380243 151380379
refGene_gn=BAZ2B;refGene_tx=XM_023259349.1;refGene_gn=BAZ2B;refGene_tx=XM_023259350.1;refGene_gn=BAZ2B;refGene_tx=XM_023259351.1;refGene_gn=BAZ2B;refGene_tx=XM_023259352.1;refGene_gn=BAZ2B;refGene_tx=XM_023259353.1;refGene_gn=BAZ2B;refGene_tx=XM_023259354.1;refGene_gn=BAZ2B;refGene_tx=XM_023259355.1;refGene_gn=BAZ2B;refGene_tx=XM_023259356.1;refGene_gn=BAZ2B;refGene_tx=XM_023259357.1;refGene_gn=BAZ2B;refGene_tx=XM_023259358.1;refGene_gn=BAZ2B;refGene_tx=XM_023259359.1;refGene_gn=BAZ2B;refGene_tx=XM_023259360.1;refGene_gn=BAZ2B;refGene_tx=XM_023259361.1;refGene_gn=BAZ2B;refGene_tx=XM_023259362.1;refGene_gn=BAZ2B;refGene_tx=XM_023259363.1;refGene_gn=BAZ2B;refGene_tx=XM_023259364.1;refGene_gn=BAZ2B;refGene_tx=XM_023259365.1;refGene_gn=BAZ2B;refGene_tx=XM_023259366.1;refGene_gn=BAZ2B;refGene_tx=XM_023259367.1;refGene_gn=BAZ2B;refGene_tx=XM_023259368.1;refGene_gn=BAZ2B;refGene_tx=XM_023259369.1
C1 151380766 151381077
refGene_gn=BAZ2B;refGene_tx=XM_023259349.1;refGene_gn=BAZ2B;refGene_tx

=XM_023259350.1;refGene_gn=BAZ2B;refGene_tx=XM_023259351.1;refGene_gn=BAZ2B;refGene_tx=XM_023259352.1;refGene_gn=BAZ2B;refGene_tx=XM_023259353.1;refGene_gn=BAZ2B;refGene_tx=XM_023259354.1;refGene_gn=BAZ2B;refGene_tx=XM_023259355.1;refGene_gn=BAZ2B;refGene_tx=XM_023259356.1;refGene_gn=BAZ2B;refGene_tx=XM_023259357.1;refGene_gn=BAZ2B;refGene_tx=XM_023259358.1;refGene_gn=BAZ2B;refGene_tx=XM_023259359.1;refGene_gn=BAZ2B;refGene_tx=XM_023259360.1;refGene_gn=BAZ2B;refGene_tx=XM_023259361.1;refGene_gn=BAZ2B;refGene_tx=XM_023259362.1;refGene_gn=BAZ2B;refGene_tx=XM_023259363.1;refGene_gn=BAZ2B;refGene_tx=XM_023259364.1;refGene_gn=BAZ2B;refGene_tx=XM_023259365.1;refGene_gn=BAZ2B;refGene_tx=XM_023259366.1;refGene_gn=BAZ2B;refGene_tx=XM_023259367.1;refGene_gn=BAZ2B;refGene_tx=XM_023259368.1;refGene_gn=BAZ2B;refGene_tx=XM_023259369.1
C1 151384942 151385103
refGene_gn=BAZ2B;refGene_tx=XM_023259349.1;refGene_gn=BAZ2B;refGene_tx=XM_023259350.1;refGene_gn=BAZ2B;refGene_tx=XM_023259351.1;refGene_gn=BAZ2B;refGene_tx=XM_023259352.1;refGene_gn=BAZ2B;refGene_tx=XM_023259353.1;refGene_gn=BAZ2B;refGene_tx=XM_023259354.1;refGene_gn=BAZ2B;refGene_tx=XM_023259355.1;refGene_gn=BAZ2B;refGene_tx=XM_023259356.1;refGene_gn=BAZ2B;refGene_tx=XM_023259357.1;refGene_gn=BAZ2B;refGene_tx=XM_023259358.1;refGene_gn=BAZ2B;refGene_tx=XM_023259359.1;refGene_gn=BAZ2B;refGene_tx=XM_023259360.1;refGene_gn=BAZ2B;refGene_tx=XM_023259361.1;refGene_gn=BAZ2B;refGene_tx=XM_023259362.1;refGene_gn=BAZ2B;refGene_tx=XM_023259363.1;refGene_gn=BAZ2B;refGene_tx=XM_023259364.1;refGene_gn=BAZ2B;refGene_tx=XM_023259365.1;refGene_gn=BAZ2B;refGene_tx=XM_023259366.1;refGene_gn=BAZ2B;refGene_tx=XM_023259367.1;refGene_gn=BAZ2B;refGene_tx=XM_023259368.1;refGene_gn=BAZ2B;refGene_tx=XM_023259369.1
C1 151385479 151385635
refGene_gn=BAZ2B;refGene_tx=XM_023259349.1;refGene_gn=BAZ2B;refGene_tx=XM_023259350.1;refGene_gn=BAZ2B;refGene_tx=XM_023259351.1;refGene_gn=BAZ2B;refGene_tx=XM_023259352.1;refGene_gn=BAZ2B;refGene_tx=XM_023259353.1;refGene_gn=BAZ2B;refGene_tx=XM_023259354.1;refGene_gn=BAZ2B;refGene_tx=XM_023259355.1;refGene_gn=BAZ2B;refGene_tx=XM_023259356.1;refGene_gn=BAZ2B;refGene_tx=XM_023259357.1;refGene_gn=BAZ2B;refGene_tx=XM_023259358.1;refGene_gn=BAZ2B;refGene_tx=XM_023259359.1;refGene_gn=BAZ2B;refGene_tx=XM_023259360.1;refGene_gn=BAZ2B;refGene_tx=XM_023259361.1;refGene_gn=BAZ2B;refGene_tx=XM_023259362.1;refGene_gn=BAZ2B;refGene_tx=XM_023259363.1;refGene_gn=BAZ2B;refGene_tx=XM_023259364.1;refGene_gn=BAZ2B;refGene_tx=XM_023259365.1;refGene_gn=BAZ2B;refGene_tx=XM_023259366.1;refGene_gn=BAZ2B;refGene_tx=XM_023259367.1;refGene_gn=BAZ2B;refGene_tx=XM_023259368.1;refGene_gn=BAZ2B;refGene_tx=XM_023259369.1
C1 151385811 151386085
refGene_gn=BAZ2B;refGene_tx=XM_023259349.1;refGene_gn=BAZ2B;refGene_tx=XM_023259350.1;refGene_gn=BAZ2B;refGene_tx=XM_023259351.1;refGene_gn=BAZ2B;refGene_tx=XM_023259352.1;refGene_gn=BAZ2B;refGene_tx=XM_023259353.1;refGene_gn=BAZ2B;refGene_tx=XM_023259354.1;refGene_gn=BAZ2B;refGene_tx=XM_023259355.1;refGene_gn=BAZ2B;refGene_tx=XM_023259356.1;refGene_gn=BAZ2B;refGene_tx=XM_023259357.1;refGene_gn=BAZ2B;refGene_tx=XM_023259358.1;refGene_gn=BAZ2B;refGene_tx=XM_023259359.1;refGene_gn=BAZ2B;refGene_tx=XM_023259360.1;refGene_gn=BAZ2B;refGene_tx=XM_023259361.1;refGene_gn=BAZ2B;refGene_tx=XM_023259362.1;refGene_gn=BAZ2B;refGene_tx=

XM_023259363.1;refGene_gn=BAZ2B;refGene_tx=XM_023259364.1;refGene_gn=BAZ2B;refGene_tx=XM_023259365.1;refGene_gn=BAZ2B;refGene_tx=XM_023259366.1;refGene_gn=BAZ2B;refGene_tx=XM_023259367.1;refGene_gn=BAZ2B;refGene_tx=XM_023259368.1;refGene_gn=BAZ2B;refGene_tx=XM_023259369.1
C1 151386362 151387018
refGene_gn=BAZ2B;refGene_tx=XM_023259349.1;refGene_gn=BAZ2B;refGene_tx=XM_023259350.1;refGene_gn=BAZ2B;refGene_tx=XM_023259351.1;refGene_gn=BAZ2B;refGene_tx=XM_023259352.1;refGene_gn=BAZ2B;refGene_tx=XM_023259353.1;refGene_gn=BAZ2B;refGene_tx=XM_023259354.1;refGene_gn=BAZ2B;refGene_tx=XM_023259355.1;refGene_gn=BAZ2B;refGene_tx=XM_023259356.1;refGene_gn=BAZ2B;refGene_tx=XM_023259357.1;refGene_gn=BAZ2B;refGene_tx=XM_023259358.1;refGene_gn=BAZ2B;refGene_tx=XM_023259359.1;refGene_gn=BAZ2B;refGene_tx=XM_023259360.1;refGene_gn=BAZ2B;refGene_tx=XM_023259361.1;refGene_gn=BAZ2B;refGene_tx=XM_023259362.1;refGene_gn=BAZ2B;refGene_tx=XM_023259363.1;refGene_gn=BAZ2B;refGene_tx=XM_023259364.1;refGene_gn=BAZ2B;refGene_tx=XM_023259365.1;refGene_gn=BAZ2B;refGene_tx=XM_023259366.1;refGene_gn=BAZ2B;refGene_tx=XM_023259367.1;refGene_gn=BAZ2B;refGene_tx=XM_023259368.1;refGene_gn=BAZ2B;refGene_tx=XM_023259369.1
C1 151405144 151405289
refGene_gn=BAZ2B;refGene_tx=XM_023259349.1;refGene_gn=BAZ2B;refGene_tx=XM_023259350.1;refGene_gn=BAZ2B;refGene_tx=XM_023259351.1;refGene_gn=BAZ2B;refGene_tx=XM_023259352.1;refGene_gn=BAZ2B;refGene_tx=XM_023259353.1;refGene_gn=BAZ2B;refGene_tx=XM_023259354.1;refGene_gn=BAZ2B;refGene_tx=XM_023259355.1;refGene_gn=BAZ2B;refGene_tx=XM_023259356.1;refGene_gn=BAZ2B;refGene_tx=XM_023259357.1;refGene_gn=BAZ2B;refGene_tx=XM_023259358.1;refGene_gn=BAZ2B;refGene_tx=XM_023259359.1;refGene_gn=BAZ2B;refGene_tx=XM_023259360.1;refGene_gn=BAZ2B;refGene_tx=XM_023259361.1;refGene_gn=BAZ2B;refGene_tx=XM_023259362.1;refGene_gn=BAZ2B;refGene_tx=XM_023259363.1;refGene_gn=BAZ2B;refGene_tx=XM_023259364.1;refGene_gn=BAZ2B;refGene_tx=XM_023259365.1;refGene_gn=BAZ2B;refGene_tx=XM_023259366.1;refGene_gn=BAZ2B;refGene_tx=XM_023259367.1;refGene_gn=BAZ2B;refGene_tx=XM_023259368.1;refGene_gn=BAZ2B;refGene_tx=XM_023259369.1
C1 151406291 151406354
refGene_gn=BAZ2B;refGene_tx=XM_023259349.1;refGene_gn=BAZ2B;refGene_tx=XM_023259350.1;refGene_gn=BAZ2B;refGene_tx=XM_023259351.1;refGene_gn=BAZ2B;refGene_tx=XM_023259352.1;refGene_gn=BAZ2B;refGene_tx=XM_023259353.1;refGene_gn=BAZ2B;refGene_tx=XM_023259354.1;refGene_gn=BAZ2B;refGene_tx=XM_023259355.1;refGene_gn=BAZ2B;refGene_tx=XM_023259356.1;refGene_gn=BAZ2B;refGene_tx=XM_023259357.1;refGene_gn=BAZ2B;refGene_tx=XM_023259358.1;refGene_gn=BAZ2B;refGene_tx=XM_023259359.1;refGene_gn=BAZ2B;refGene_tx=XM_023259360.1;refGene_gn=BAZ2B;refGene_tx=XM_023259361.1;refGene_gn=BAZ2B;refGene_tx=XM_023259362.1;refGene_gn=BAZ2B;refGene_tx=XM_023259363.1;refGene_gn=BAZ2B;refGene_tx=XM_023259364.1;refGene_gn=BAZ2B;refGene_tx=XM_023259365.1;refGene_gn=BAZ2B;refGene_tx=XM_023259366.1;refGene_gn=BAZ2B;refGene_tx=XM_023259367.1;refGene_gn=BAZ2B;refGene_tx=XM_023259368.1;refGene_gn=BAZ2B;refGene_tx=XM_023259369.1
C1 151413217 151413461
refGene_gn=BAZ2B;refGene_tx=XM_023259349.1;refGene_gn=BAZ2B;refGene_tx=XM_023259350.1;refGene_gn=BAZ2B;refGene_tx=XM_023259351.1;refGene_gn=BAZ2B;refGene_tx=XM_023259352.1;refGene_gn=BAZ2B;refGene_tx=XM_0232593

53.1;refGene_gn=BAZ2B;refGene_tx=XM_023259354.1;refGene_gn=BAZ2B;refGene_tx=XM_023259355.1;refGene_gn=BAZ2B;refGene_tx=XM_023259356.1;refGene_gn=BAZ2B;refGene_tx=XM_023259357.1;refGene_gn=BAZ2B;refGene_tx=XM_023259358.1;refGene_gn=BAZ2B;refGene_tx=XM_023259359.1;refGene_gn=BAZ2B;refGene_tx=XM_023259360.1;refGene_gn=BAZ2B;refGene_tx=XM_023259361.1;refGene_gn=BAZ2B;refGene_tx=XM_023259362.1;refGene_gn=BAZ2B;refGene_tx=XM_023259363.1;refGene_gn=BAZ2B;refGene_tx=XM_023259364.1;refGene_gn=BAZ2B;refGene_tx=XM_023259365.1;refGene_gn=BAZ2B;refGene_tx=XM_023259366.1;refGene_gn=BAZ2B;refGene_tx=XM_023259367.1;refGene_gn=BAZ2B;refGene_tx=XM_023259368.1;refGene_gn=BAZ2B;refGene_tx=XM_023259369.1
C1 151414517 151414592
refGene_gn=BAZ2B;refGene_tx=XM_023259349.1;refGene_gn=BAZ2B;refGene_tx=XM_023259350.1;refGene_gn=BAZ2B;refGene_tx=XM_023259351.1;refGene_gn=BAZ2B;refGene_tx=XM_023259352.1;refGene_gn=BAZ2B;refGene_tx=XM_023259353.1;refGene_gn=BAZ2B;refGene_tx=XM_023259354.1;refGene_gn=BAZ2B;refGene_tx=XM_023259355.1;refGene_gn=BAZ2B;refGene_tx=XM_023259356.1;refGene_gn=BAZ2B;refGene_tx=XM_023259357.1;refGene_gn=BAZ2B;refGene_tx=XM_023259358.1;refGene_gn=BAZ2B;refGene_tx=XM_023259359.1;refGene_gn=BAZ2B;refGene_tx=XM_023259360.1;refGene_gn=BAZ2B;refGene_tx=XM_023259361.1;refGene_gn=BAZ2B;refGene_tx=XM_023259362.1;refGene_gn=BAZ2B;refGene_tx=XM_023259363.1;refGene_gn=BAZ2B;refGene_tx=XM_023259364.1;refGene_gn=BAZ2B;refGene_tx=XM_023259365.1;refGene_gn=BAZ2B;refGene_tx=XM_023259366.1;refGene_gn=BAZ2B;refGene_tx=XM_023259367.1;refGene_gn=BAZ2B;refGene_tx=XM_023259368.1;refGene_gn=BAZ2B;refGene_tx=XM_023259369.1
C1 151415522 151415737
refGene_gn=BAZ2B;refGene_tx=XM_023259349.1;refGene_gn=BAZ2B;refGene_tx=XM_023259350.1;refGene_gn=BAZ2B;refGene_tx=XM_023259351.1;refGene_gn=BAZ2B;refGene_tx=XM_023259352.1;refGene_gn=BAZ2B;refGene_tx=XM_023259353.1;refGene_gn=BAZ2B;refGene_tx=XM_023259354.1;refGene_gn=BAZ2B;refGene_tx=XM_023259355.1;refGene_gn=BAZ2B;refGene_tx=XM_023259356.1;refGene_gn=BAZ2B;refGene_tx=XM_023259357.1;refGene_gn=BAZ2B;refGene_tx=XM_023259358.1;refGene_gn=BAZ2B;refGene_tx=XM_023259359.1;refGene_gn=BAZ2B;refGene_tx=XM_023259360.1;refGene_gn=BAZ2B;refGene_tx=XM_023259361.1;refGene_gn=BAZ2B;refGene_tx=XM_023259362.1;refGene_gn=BAZ2B;refGene_tx=XM_023259363.1;refGene_gn=BAZ2B;refGene_tx=XM_023259364.1;refGene_gn=BAZ2B;refGene_tx=XM_023259365.1;refGene_gn=BAZ2B;refGene_tx=XM_023259366.1;refGene_gn=BAZ2B;refGene_tx=XM_023259367.1;refGene_gn=BAZ2B;refGene_tx=XM_023259368.1;refGene_gn=BAZ2B;refGene_tx=XM_023259369.1
C1 151416476 151416731
refGene_gn=BAZ2B;refGene_tx=XM_023259349.1;refGene_gn=BAZ2B;refGene_tx=XM_023259350.1;refGene_gn=BAZ2B;refGene_tx=XM_023259351.1;refGene_gn=BAZ2B;refGene_tx=XM_023259352.1;refGene_gn=BAZ2B;refGene_tx=XM_023259353.1;refGene_gn=BAZ2B;refGene_tx=XM_023259354.1;refGene_gn=BAZ2B;refGene_tx=XM_023259355.1;refGene_gn=BAZ2B;refGene_tx=XM_023259356.1;refGene_gn=BAZ2B;refGene_tx=XM_023259357.1;refGene_gn=BAZ2B;refGene_tx=XM_023259358.1;refGene_gn=BAZ2B;refGene_tx=XM_023259359.1;refGene_gn=BAZ2B;refGene_tx=XM_023259360.1;refGene_gn=BAZ2B;refGene_tx=XM_023259361.1;refGene_gn=BAZ2B;refGene_tx=XM_023259362.1;refGene_gn=BAZ2B;refGene_tx=XM_023259363.1;refGene_gn=BAZ2B;refGene_tx=XM_023259364.1;refGene_gn=BAZ2B;refGene_tx=XM_023259365.1;refGene_gn=BAZ2B;refGene_tx=XM_0232593 6

6.1;refGene_gn=BAZ2B;refGene_tx=XM_023259367.1;refGene_gn=BAZ2B;refGene_tx=XM_023259368.1;refGene_gn=BAZ2B;refGene_tx=XM_023259369.1
C1 151419120 151419261
refGene_gn=BAZ2B;refGene_tx=XM_023259349.1;refGene_gn=BAZ2B;refGene_tx=XM_023259350.1;refGene_gn=BAZ2B;refGene_tx=XM_023259351.1;refGene_gn=BAZ2B;refGene_tx=XM_023259352.1;refGene_gn=BAZ2B;refGene_tx=XM_023259353.1;refGene_gn=BAZ2B;refGene_tx=XM_023259354.1;refGene_gn=BAZ2B;refGene_tx=XM_023259355.1;refGene_gn=BAZ2B;refGene_tx=XM_023259356.1;refGene_gn=BAZ2B;refGene_tx=XM_023259357.1;refGene_gn=BAZ2B;refGene_tx=XM_023259358.1;refGene_gn=BAZ2B;refGene_tx=XM_023259359.1;refGene_gn=BAZ2B;refGene_tx=XM_023259360.1;refGene_gn=BAZ2B;refGene_tx=XM_023259361.1;refGene_gn=BAZ2B;refGene_tx=XM_023259362.1;refGene_gn=BAZ2B;refGene_tx=XM_023259363.1;refGene_gn=BAZ2B;refGene_tx=XM_023259364.1;refGene_gn=BAZ2B;refGene_tx=XM_023259365.1;refGene_gn=BAZ2B;refGene_tx=XM_023259366.1;refGene_gn=BAZ2B;refGene_tx=XM_023259367.1;refGene_gn=BAZ2B;refGene_tx=XM_023259368.1;refGene_gn=BAZ2B;refGene_tx=XM_023259369.1
C1 151429303 151429369
refGene_gn=BAZ2B;refGene_tx=XM_023259349.1;refGene_gn=BAZ2B;refGene_tx=XM_023259350.1;refGene_gn=BAZ2B;refGene_tx=XM_023259351.1;refGene_gn=BAZ2B;refGene_tx=XM_023259352.1;refGene_gn=BAZ2B;refGene_tx=XM_023259353.1;refGene_gn=BAZ2B;refGene_tx=XM_023259354.1;refGene_gn=BAZ2B;refGene_tx=XM_023259355.1;refGene_gn=BAZ2B;refGene_tx=XM_023259356.1;refGene_gn=BAZ2B;refGene_tx=XM_023259357.1;refGene_gn=BAZ2B;refGene_tx=XM_023259358.1;refGene_gn=BAZ2B;refGene_tx=XM_023259359.1;refGene_gn=BAZ2B;refGene_tx=XM_023259360.1;refGene_gn=BAZ2B;refGene_tx=XM_023259361.1;refGene_gn=BAZ2B;refGene_tx=XM_023259362.1;refGene_gn=BAZ2B;refGene_tx=XM_023259363.1;refGene_gn=BAZ2B;refGene_tx=XM_023259364.1;refGene_gn=BAZ2B;refGene_tx=XM_023259365.1;refGene_gn=BAZ2B;refGene_tx=XM_023259366.1;refGene_gn=BAZ2B;refGene_tx=XM_023259367.1;refGene_gn=BAZ2B;refGene_tx=XM_023259368.1;refGene_gn=BAZ2B;refGene_tx=XM_023259369.1
C1 151430607 151430640
refGene_gn=BAZ2B;refGene_tx=XM_023259349.1;refGene_gn=BAZ2B;refGene_tx=XM_023259350.1;refGene_gn=BAZ2B;refGene_tx=XM_023259351.1;refGene_gn=BAZ2B;refGene_tx=XM_023259352.1;refGene_gn=BAZ2B;refGene_tx=XM_023259353.1;refGene_gn=BAZ2B;refGene_tx=XM_023259354.1;refGene_gn=BAZ2B;refGene_tx=XM_023259355.1;refGene_gn=BAZ2B;refGene_tx=XM_023259356.1;refGene_gn=BAZ2B;refGene_tx=XM_023259358.1;refGene_gn=BAZ2B;refGene_tx=XM_023259359.1;refGene_gn=BAZ2B;refGene_tx=XM_023259360.1;refGene_gn=BAZ2B;refGene_tx=XM_023259361.1;refGene_gn=BAZ2B;refGene_tx=XM_023259362.1;refGene_gn=BAZ2B;refGene_tx=XM_023259363.1;refGene_gn=BAZ2B;refGene_tx=XM_023259364.1;refGene_gn=BAZ2B;refGene_tx=XM_023259367.1;refGene_gn=BAZ2B;refGene_tx=XM_023259368.1
C1 151430882 151430927
refGene_gn=BAZ2B;refGene_tx=XM_023259349.1;refGene_gn=BAZ2B;refGene_tx=XM_023259350.1;refGene_gn=BAZ2B;refGene_tx=XM_023259351.1;refGene_gn=BAZ2B;refGene_tx=XM_023259352.1;refGene_gn=BAZ2B;refGene_tx=XM_023259353.1;refGene_gn=BAZ2B;refGene_tx=XM_023259354.1;refGene_gn=BAZ2B;refGene_tx=XM_023259355.1;refGene_gn=BAZ2B;refGene_tx=XM_023259356.1;refGene_gn=BAZ2B;refGene_tx=XM_023259357.1;refGene_gn=BAZ2B;refGene_tx=XM_023259358.1;refGene_gn=BAZ2B;refGene_tx=XM_023259359.1;refGene_gn=BAZ2B;

refGene_tx=XM_023259360.1;refGene_gn=BAZ2B;refGene_tx=XM_023259361.1;refGene_gn=BAZ2B;refGene_tx=XM_023259362.1;refGene_gn=BAZ2B;refGene_tx=XM_023259363.1;refGene_gn=BAZ2B;refGene_tx=XM_023259364.1;refGene_gn=BAZ2B;refGene_tx=XM_023259365.1;refGene_gn=BAZ2B;refGene_tx=XM_023259366.1;refGene_gn=BAZ2B;refGene_tx=XM_023259367.1;refGene_gn=BAZ2B;refGene_tx=XM_023259368.1;refGene_gn=BAZ2B;refGene_tx=XM_023259369.1
C1 151432312 151432378
refGene_gn=BAZ2B;refGene_tx=XM_023259349.1;refGene_gn=BAZ2B;refGene_tx=XM_023259350.1;refGene_gn=BAZ2B;refGene_tx=XM_023259351.1;refGene_gn=BAZ2B;refGene_tx=XM_023259352.1;refGene_gn=BAZ2B;refGene_tx=XM_023259353.1;refGene_gn=BAZ2B;refGene_tx=XM_023259354.1;refGene_gn=BAZ2B;refGene_tx=XM_023259355.1;refGene_gn=BAZ2B;refGene_tx=XM_023259356.1;refGene_gn=BAZ2B;refGene_tx=XM_023259357.1;refGene_gn=BAZ2B;refGene_tx=XM_023259358.1;refGene_gn=BAZ2B;refGene_tx=XM_023259359.1;refGene_gn=BAZ2B;refGene_tx=XM_023259360.1;refGene_gn=BAZ2B;refGene_tx=XM_023259361.1;refGene_gn=BAZ2B;refGene_tx=XM_023259362.1;refGene_gn=BAZ2B;refGene_tx=XM_023259363.1;refGene_gn=BAZ2B;refGene_tx=XM_023259364.1;refGene_gn=BAZ2B;refGene_tx=XM_023259365.1;refGene_gn=BAZ2B;refGene_tx=XM_023259366.1;refGene_gn=BAZ2B;refGene_tx=XM_023259367.1;refGene_gn=BAZ2B;refGene_tx=XM_023259368.1;refGene_gn=BAZ2B;refGene_tx=XM_023259369.1
C1 151434270 151434336
refGene_gn=BAZ2B;refGene_tx=XM_023259349.1;refGene_gn=BAZ2B;refGene_tx=XM_023259350.1;refGene_gn=BAZ2B;refGene_tx=XM_023259351.1;refGene_gn=BAZ2B;refGene_tx=XM_023259352.1;refGene_gn=BAZ2B;refGene_tx=XM_023259353.1;refGene_gn=BAZ2B;refGene_tx=XM_023259354.1;refGene_gn=BAZ2B;refGene_tx=XM_023259355.1;refGene_gn=BAZ2B;refGene_tx=XM_023259356.1;refGene_gn=BAZ2B;refGene_tx=XM_023259357.1;refGene_gn=BAZ2B;refGene_tx=XM_023259358.1;refGene_gn=BAZ2B;refGene_tx=XM_023259359.1;refGene_gn=BAZ2B;refGene_tx=XM_023259360.1;refGene_gn=BAZ2B;refGene_tx=XM_023259361.1;refGene_gn=BAZ2B;refGene_tx=XM_023259362.1;refGene_gn=BAZ2B;refGene_tx=XM_023259363.1;refGene_gn=BAZ2B;refGene_tx=XM_023259364.1;refGene_gn=BAZ2B;refGene_tx=XM_023259365.1;refGene_gn=BAZ2B;refGene_tx=XM_023259366.1;refGene_gn=BAZ2B;refGene_tx=XM_023259367.1;refGene_gn=BAZ2B;refGene_tx=XM_023259368.1;refGene_gn=BAZ2B;refGene_tx=XM_023259369.1
C1 151438749 151438811
refGene_gn=BAZ2B;refGene_tx=XM_023259349.1;refGene_gn=BAZ2B;refGene_tx=XM_023259350.1;refGene_gn=BAZ2B;refGene_tx=XM_023259351.1;refGene_gn=BAZ2B;refGene_tx=XM_023259352.1;refGene_gn=BAZ2B;refGene_tx=XM_023259353.1;refGene_gn=BAZ2B;refGene_tx=XM_023259354.1;refGene_gn=BAZ2B;refGene_tx=XM_023259355.1;refGene_gn=BAZ2B;refGene_tx=XM_023259356.1;refGene_gn=BAZ2B;refGene_tx=XM_023259357.1;refGene_gn=BAZ2B;refGene_tx=XM_023259358.1;refGene_gn=BAZ2B;refGene_tx=XM_023259359.1;refGene_gn=BAZ2B;refGene_tx=XM_023259360.1;refGene_gn=BAZ2B;refGene_tx=XM_023259361.1;refGene_gn=BAZ2B;refGene_tx=XM_023259362.1;refGene_gn=BAZ2B;refGene_tx=XM_023259363.1;refGene_gn=BAZ2B;refGene_tx=XM_023259364.1;refGene_gn=BAZ2B;refGene_tx=XM_023259365.1;refGene_gn=BAZ2B;refGene_tx=XM_023259366.1;refGene_gn=BAZ2B;refGene_tx=XM_023259367.1;refGene_gn=BAZ2B;refGene_tx=XM_023259368.1;refGene_gn=BAZ2B;refGene_tx=XM_023259369.1
C1 151438926 151439019
refGene_gn=BAZ2B;refGene_tx=XM_023259349.1;refGene_gn=BAZ2B;refGene_tx

=XM_023259350.1;refGene_gn=BAZ2B;refGene_tx=XM_023259351.1;refGene_gn=BAZ2B;refGene_tx=XM_023259352.1;refGene_gn=BAZ2B;refGene_tx=XM_023259353.1;refGene_gn=BAZ2B;refGene_tx=XM_023259354.1;refGene_gn=BAZ2B;refGene_tx=XM_023259355.1;refGene_gn=BAZ2B;refGene_tx=XM_023259356.1;refGene_gn=BAZ2B;refGene_tx=XM_023259357.1;refGene_gn=BAZ2B;refGene_tx=XM_023259358.1;refGene_gn=BAZ2B;refGene_tx=XM_023259359.1;refGene_gn=BAZ2B;refGene_tx=XM_023259360.1;refGene_gn=BAZ2B;refGene_tx=XM_023259361.1;refGene_gn=BAZ2B;refGene_tx=XM_023259362.1;refGene_gn=BAZ2B;refGene_tx=XM_023259363.1;refGene_gn=BAZ2B;refGene_tx=XM_023259364.1;refGene_gn=BAZ2B;refGene_tx=XM_023259365.1;refGene_gn=BAZ2B;refGene_tx=XM_023259366.1;refGene_gn=BAZ2B;refGene_tx=XM_023259367.1;refGene_gn=BAZ2B;refGene_tx=XM_023259368.1;refGene_gn=BAZ2B;refGene_tx=XM_023259369.1
C1 151443960 151444171
refGene_gn=BAZ2B;refGene_tx=XM_023259349.1;refGene_gn=BAZ2B;refGene_tx=XM_023259350.1;refGene_gn=BAZ2B;refGene_tx=XM_023259351.1;refGene_gn=BAZ2B;refGene_tx=XM_023259352.1;refGene_gn=BAZ2B;refGene_tx=XM_023259353.1;refGene_gn=BAZ2B;refGene_tx=XM_023259354.1;refGene_gn=BAZ2B;refGene_tx=XM_023259355.1;refGene_gn=BAZ2B;refGene_tx=XM_023259356.1;refGene_gn=BAZ2B;refGene_tx=XM_023259357.1;refGene_gn=BAZ2B;refGene_tx=XM_023259358.1;refGene_gn=BAZ2B;refGene_tx=XM_023259359.1;refGene_gn=BAZ2B;refGene_tx=XM_023259360.1;refGene_gn=BAZ2B;refGene_tx=XM_023259361.1;refGene_gn=BAZ2B;refGene_tx=XM_023259362.1;refGene_gn=BAZ2B;refGene_tx=XM_023259363.1;refGene_gn=BAZ2B;refGene_tx=XM_023259364.1;refGene_gn=BAZ2B;refGene_tx=XM_023259365.1;refGene_gn=BAZ2B;refGene_tx=XM_023259366.1;refGene_gn=BAZ2B;refGene_tx=XM_023259367.1;refGene_gn=BAZ2B;refGene_tx=XM_023259368.1;refGene_gn=BAZ2B;refGene_tx=XM_023259369.1
C1 151455761 151455863
refGene_gn=BAZ2B;refGene_tx=XM_023259349.1;refGene_gn=BAZ2B;refGene_tx=XM_023259350.1;refGene_gn=BAZ2B;refGene_tx=XM_023259351.1;refGene_gn=BAZ2B;refGene_tx=XM_023259352.1;refGene_gn=BAZ2B;refGene_tx=XM_023259353.1;refGene_gn=BAZ2B;refGene_tx=XM_023259354.1;refGene_gn=BAZ2B;refGene_tx=XM_023259355.1;refGene_gn=BAZ2B;refGene_tx=XM_023259356.1;refGene_gn=BAZ2B;refGene_tx=XM_023259357.1;refGene_gn=BAZ2B;refGene_tx=XM_023259359.1;refGene_gn=BAZ2B;refGene_tx=XM_023259360.1;refGene_gn=BAZ2B;refGene_tx=XM_023259361.1;refGene_gn=BAZ2B;refGene_tx=XM_023259362.1;refGene_gn=BAZ2B;refGene_tx=XM_023259363.1;refGene_gn=BAZ2B;refGene_tx=XM_023259366.1;refGene_gn=BAZ2B;refGene_tx=XM_023259367.1
C1 151456113 151456222
refGene_gn=BAZ2B;refGene_tx=XM_023259349.1;refGene_gn=BAZ2B;refGene_tx=XM_023259350.1;refGene_gn=BAZ2B;refGene_tx=XM_023259351.1;refGene_gn=BAZ2B;refGene_tx=XM_023259352.1;refGene_gn=BAZ2B;refGene_tx=XM_023259353.1;refGene_gn=BAZ2B;refGene_tx=XM_023259354.1;refGene_gn=BAZ2B;refGene_tx=XM_023259355.1;refGene_gn=BAZ2B;refGene_tx=XM_023259356.1;refGene_gn=BAZ2B;refGene_tx=XM_023259357.1;refGene_gn=BAZ2B;refGene_tx=XM_023259358.1;refGene_gn=BAZ2B;refGene_tx=XM_023259359.1;refGene_gn=BAZ2B;refGene_tx=XM_023259360.1;refGene_gn=BAZ2B;refGene_tx=XM_023259361.1;refGene_gn=BAZ2B;refGene_tx=XM_023259362.1;refGene_gn=BAZ2B;refGene_tx=XM_023259363.1;refGene_gn=BAZ2B;refGene_tx=XM_023259364.1;refGene_gn=BAZ2B;refGene_tx=XM_023259365.1;refGene_gn=BAZ2B;refGene_tx=XM_023259366.1;refGene_gn=BAZ2B;refGene_tx=XM_023259367.1;refGene_gn=BAZ2B;refGen

e_tx=XM_023259368.1;refGene_gn=BAZ2B;refGene_tx=XM_023259369.1
C1 151457008 151457069
refGene_gn=BAZ2B;refGene_tx=XM_023259349.1;refGene_gn=BAZ2B;refGene_tx=XM_023259350.1;refGene_gn=BAZ2B;refGene_tx=XM_023259351.1;refGene_gn=BAZ2B;refGene_tx=XM_023259352.1;refGene_gn=BAZ2B;refGene_tx=XM_023259353.1;refGene_gn=BAZ2B;refGene_tx=XM_023259354.1;refGene_gn=BAZ2B;refGene_tx=XM_023259355.1;refGene_gn=BAZ2B;refGene_tx=XM_023259356.1;refGene_gn=BAZ2B;refGene_tx=XM_023259357.1;refGene_gn=BAZ2B;refGene_tx=XM_023259358.1;refGene_gn=BAZ2B;refGene_tx=XM_023259359.1;refGene_gn=BAZ2B;refGene_tx=XM_023259360.1;refGene_gn=BAZ2B;refGene_tx=XM_023259361.1;refGene_gn=BAZ2B;refGene_tx=XM_023259362.1;refGene_gn=BAZ2B;refGene_tx=XM_023259363.1;refGene_gn=BAZ2B;refGene_tx=XM_023259364.1;refGene_gn=BAZ2B;refGene_tx=XM_023259365.1;refGene_gn=BAZ2B;refGene_tx=XM_023259366.1;refGene_gn=BAZ2B;refGene_tx=XM_023259367.1;refGene_gn=BAZ2B;refGene_tx=XM_023259368.1;refGene_gn=BAZ2B;refGene_tx=XM_023259369.1
C1 151458256 151458550
refGene_gn=BAZ2B;refGene_tx=XM_023259349.1;refGene_gn=BAZ2B;refGene_tx=XM_023259350.1;refGene_gn=BAZ2B;refGene_tx=XM_023259351.1;refGene_gn=BAZ2B;refGene_tx=XM_023259352.1;refGene_gn=BAZ2B;refGene_tx=XM_023259353.1;refGene_gn=BAZ2B;refGene_tx=XM_023259354.1;refGene_gn=BAZ2B;refGene_tx=XM_023259355.1;refGene_gn=BAZ2B;refGene_tx=XM_023259356.1;refGene_gn=BAZ2B;refGene_tx=XM_023259357.1;refGene_gn=BAZ2B;refGene_tx=XM_023259358.1;refGene_gn=BAZ2B;refGene_tx=XM_023259359.1;refGene_gn=BAZ2B;refGene_tx=XM_023259360.1;refGene_gn=BAZ2B;refGene_tx=XM_023259361.1;refGene_gn=BAZ2B;refGene_tx=XM_023259362.1;refGene_gn=BAZ2B;refGene_tx=XM_023259363.1;refGene_gn=BAZ2B;refGene_tx=XM_023259364.1;refGene_gn=BAZ2B;refGene_tx=XM_023259365.1;refGene_gn=BAZ2B;refGene_tx=XM_023259366.1;refGene_gn=BAZ2B;refGene_tx=XM_023259367.1;refGene_gn=BAZ2B;refGene_tx=XM_023259368.1;refGene_gn=BAZ2B;refGene_tx=XM_023259369.1
C1 151460159 151460766
refGene_gn=BAZ2B;refGene_tx=XM_023259349.1;refGene_gn=BAZ2B;refGene_tx=XM_023259350.1;refGene_gn=BAZ2B;refGene_tx=XM_023259351.1;refGene_gn=BAZ2B;refGene_tx=XM_023259352.1;refGene_gn=BAZ2B;refGene_tx=XM_023259353.1;refGene_gn=BAZ2B;refGene_tx=XM_023259354.1;refGene_gn=BAZ2B;refGene_tx=XM_023259355.1;refGene_gn=BAZ2B;refGene_tx=XM_023259356.1;refGene_gn=BAZ2B;refGene_tx=XM_023259357.1;refGene_gn=BAZ2B;refGene_tx=XM_023259358.1;refGene_gn=BAZ2B;refGene_tx=XM_023259359.1;refGene_gn=BAZ2B;refGene_tx=XM_023259360.1;refGene_gn=BAZ2B;refGene_tx=XM_023259361.1;refGene_gn=BAZ2B;refGene_tx=XM_023259362.1;refGene_gn=BAZ2B;refGene_tx=XM_023259363.1;refGene_gn=BAZ2B;refGene_tx=XM_023259364.1;refGene_gn=BAZ2B;refGene_tx=XM_023259365.1;refGene_gn=BAZ2B;refGene_tx=XM_023259366.1;refGene_gn=BAZ2B;refGene_tx=XM_023259367.1;refGene_gn=BAZ2B;refGene_tx=XM_023259368.1;refGene_gn=BAZ2B;refGene_tx=XM_023259369.1
C1 151465641 151466040
refGene_gn=BAZ2B;refGene_tx=XM_023259349.1;refGene_gn=BAZ2B;refGene_tx=XM_023259350.1;refGene_gn=BAZ2B;refGene_tx=XM_023259351.1;refGene_gn=BAZ2B;refGene_tx=XM_023259352.1;refGene_gn=BAZ2B;refGene_tx=XM_023259353.1;refGene_gn=BAZ2B;refGene_tx=XM_023259354.1;refGene_gn=BAZ2B;refGene_tx=XM_023259355.1;refGene_gn=BAZ2B;refGene_tx=XM_023259356.1;refGene_gn=BAZ2B;refGene_tx=XM_023259357.1;refGene_gn=BAZ2B;refGene_tx=XM_02

3259358.1;refGene_gn=BAZ2B;refGene_tx=XM_023259359.1;refGene_gn=BAZ2B;
refGene_tx=XM_023259360.1;refGene_gn=BAZ2B;refGene_tx=XM_023259361.1;r
efGene_gn=BAZ2B;refGene_tx=XM_023259362.1;refGene_gn=BAZ2B;refGene_tx=
XM_023259363.1;refGene_gn=BAZ2B;refGene_tx=XM_023259364.1;refGene_gn=B
AZ2B;refGene_tx=XM_023259365.1;refGene_gn=BAZ2B;refGene_tx=XM_02325936
6.1;refGene_gn=BAZ2B;refGene_tx=XM_023259367.1;refGene_gn=BAZ2B;refGen
e_tx=XM_023259368.1;refGene_gn=BAZ2B;refGene_tx=XM_023259369.1
C1 151466576 151466780
refGene_gn=BAZ2B;refGene_tx=XM_023259349.1;refGene_gn=BAZ2B;refGene_tx
=XM_023259350.1;refGene_gn=BAZ2B;refGene_tx=XM_023259351.1;refGene_gn=
BAZ2B;refGene_tx=XM_023259352.1;refGene_gn=BAZ2B;refGene_tx=XM_0232593
53.1;refGene_gn=BAZ2B;refGene_tx=XM_023259354.1;refGene_gn=BAZ2B;refGe
ne_tx=XM_023259355.1;refGene_gn=BAZ2B;refGene_tx=XM_023259356.1;refGen
e_gn=BAZ2B;refGene_tx=XM_023259357.1;refGene_gn=BAZ2B;refGene_tx=XM_02
3259358.1;refGene_gn=BAZ2B;refGene_tx=XM_023259359.1;refGene_gn=BAZ2B;
refGene_tx=XM_023259360.1;refGene_gn=BAZ2B;refGene_tx=XM_023259361.1;r
efGene_gn=BAZ2B;refGene_tx=XM_023259362.1;refGene_gn=BAZ2B;refGene_tx=
XM_023259363.1;refGene_gn=BAZ2B;refGene_tx=XM_023259364.1;refGene_gn=B
AZ2B;refGene_tx=XM_023259365.1;refGene_gn=BAZ2B;refGene_tx=XM_02325936
6.1;refGene_gn=BAZ2B;refGene_tx=XM_023259367.1;refGene_gn=BAZ2B;refGen
e_tx=XM_023259368.1;refGene_gn=BAZ2B;refGene_tx=XM_023259369.1
C1 151479843 151480037
refGene_gn=BAZ2B;refGene_tx=XM_023259349.1;refGene_gn=BAZ2B;refGene_tx
=XM_023259350.1;refGene_gn=BAZ2B;refGene_tx=XM_023259351.1;refGene_gn=
BAZ2B;refGene_tx=XM_023259352.1;refGene_gn=BAZ2B;refGene_tx=XM_0232593
53.1;refGene_gn=BAZ2B;refGene_tx=XM_023259354.1;refGene_gn=BAZ2B;refGe
ne_tx=XM_023259355.1;refGene_gn=BAZ2B;refGene_tx=XM_023259356.1;refGen
e_gn=BAZ2B;refGene_tx=XM_023259357.1;refGene_gn=BAZ2B;refGene_tx=XM_02
3259358.1;refGene_gn=BAZ2B;refGene_tx=XM_023259359.1;refGene_gn=BAZ2B;
refGene_tx=XM_023259360.1;refGene_gn=BAZ2B;refGene_tx=XM_023259361.1;r
efGene_gn=BAZ2B;refGene_tx=XM_023259362.1;refGene_gn=BAZ2B;refGene_tx=
XM_023259363.1;refGene_gn=BAZ2B;refGene_tx=XM_023259364.1;refGene_gn=B
AZ2B;refGene_tx=XM_023259365.1;refGene_gn=BAZ2B;refGene_tx=XM_02325936
6.1;refGene_gn=BAZ2B;refGene_tx=XM_023259367.1;refGene_gn=BAZ2B;refGen
e_tx=XM_023259368.1;refGene_gn=BAZ2B;refGene_tx=XM_023259369.1
C1 151481373 151481541
refGene_gn=BAZ2B;refGene_tx=XM_023259349.1;refGene_gn=BAZ2B;refGene_tx
=XM_023259350.1;refGene_gn=BAZ2B;refGene_tx=XM_023259351.1;refGene_gn=
BAZ2B;refGene_tx=XM_023259352.1;refGene_gn=BAZ2B;refGene_tx=XM_0232593
53.1;refGene_gn=BAZ2B;refGene_tx=XM_023259354.1;refGene_gn=BAZ2B;refGe
ne_tx=XM_023259355.1;refGene_gn=BAZ2B;refGene_tx=XM_023259356.1;refGen
e_gn=BAZ2B;refGene_tx=XM_023259357.1;refGene_gn=BAZ2B;refGene_tx=XM_02
3259358.1;refGene_gn=BAZ2B;refGene_tx=XM_023259359.1;refGene_gn=BAZ2B;
refGene_tx=XM_023259360.1;refGene_gn=BAZ2B;refGene_tx=XM_023259361.1;r
efGene_gn=BAZ2B;refGene_tx=XM_023259362.1;refGene_gn=BAZ2B;refGene_tx=
XM_023259363.1;refGene_gn=BAZ2B;refGene_tx=XM_023259364.1;refGene_gn=B
AZ2B;refGene_tx=XM_023259365.1;refGene_gn=BAZ2B;refGene_tx=XM_02325936
6.1;refGene_gn=BAZ2B;refGene_tx=XM_023259367.1;refGene_gn=BAZ2B;refGen
e_tx=XM_023259368.1;refGene_gn=BAZ2B;refGene_tx=XM_023259369.1
C1 151485876 151486065

refGene_gn=BAZ2B;refGene_tx=XM_023259349.1;refGene_gn=BAZ2B;refGene_tx=XM_023259350.1;refGene_gn=BAZ2B;refGene_tx=XM_023259351.1;refGene_gn=BAZ2B;refGene_tx=XM_023259352.1;refGene_gn=BAZ2B;refGene_tx=XM_023259353.1;refGene_gn=BAZ2B;refGene_tx=XM_023259354.1;refGene_gn=BAZ2B;refGene_tx=XM_023259355.1;refGene_gn=BAZ2B;refGene_tx=XM_023259356.1;refGene_gn=BAZ2B;refGene_tx=XM_023259357.1;refGene_gn=BAZ2B;refGene_tx=XM_023259358.1;refGene_gn=BAZ2B;refGene_tx=XM_023259359.1;refGene_gn=BAZ2B;refGene_tx=XM_023259360.1;refGene_gn=BAZ2B;refGene_tx=XM_023259363.1;refGene_gn=BAZ2B;refGene_tx=XM_023259364.1;refGene_gn=BAZ2B;refGene_tx=XM_023259365.1;refGene_gn=BAZ2B;refGene_tx=XM_023259366.1;refGene_gn=BAZ2B;refGene_tx=XM_023259368.1;refGene_gn=BAZ2B;refGene_tx=XM_023259369.1
C1 151507403 151507545
refGene_gn=BAZ2B;refGene_tx=XM_023259349.1;refGene_gn=BAZ2B;refGene_tx=XM_023259350.1;refGene_gn=BAZ2B;refGene_tx=XM_023259351.1;refGene_gn=BAZ2B;refGene_tx=XM_023259352.1;refGene_gn=BAZ2B;refGene_tx=XM_023259353.1;refGene_gn=BAZ2B;refGene_tx=XM_023259354.1;refGene_gn=BAZ2B;refGene_tx=XM_023259355.1;refGene_gn=BAZ2B;refGene_tx=XM_023259356.1;refGene_gn=BAZ2B;refGene_tx=XM_023259357.1;refGene_gn=BAZ2B;refGene_tx=XM_023259358.1;refGene_gn=BAZ2B;refGene_tx=XM_023259359.1;refGene_gn=BAZ2B;refGene_tx=XM_023259360.1;refGene_gn=BAZ2B;refGene_tx=XM_023259361.1;refGene_gn=BAZ2B;refGene_tx=XM_023259362.1;refGene_gn=BAZ2B;refGene_tx=XM_023259363.1;refGene_gn=BAZ2B;refGene_tx=XM_023259364.1;refGene_gn=BAZ2B;refGene_tx=XM_023259365.1;refGene_gn=BAZ2B;refGene_tx=XM_023259366.1;refGene_gn=BAZ2B;refGene_tx=XM_023259367.1;refGene_gn=BAZ2B;refGene_tx=XM_023259368.1;refGene_gn=BAZ2B;refGene_tx=XM_023259369.1
C1 151823924 151824077
refGene_gn=MARCH7;refGene_tx=XM_003990797.5
C1 151840463 151840653
refGene_gn=MARCH7;refGene_tx=XM_003990797.5
C1 151842281 151842449
refGene_gn=MARCH7;refGene_tx=XM_003990797.5
C1 151845253 151845367
refGene_gn=MARCH7;refGene_tx=XM_003990797.5
C1 151845369 151846360
refGene_gn=MARCH7;refGene_tx=XM_003990797.5
C1 151848903 151849073
refGene_gn=MARCH7;refGene_tx=XM_003990797.5
C1 151853314 151853424
refGene_gn=MARCH7;refGene_tx=XM_003990797.5
C1 151856611 151856725
refGene_gn=MARCH7;refGene_tx=XM_003990797.5
C1 151858288 151858337
refGene_gn=MARCH7;refGene_tx=XM_003990797.5
C1 151860874 151860933
refGene_gn=MARCH7;refGene_tx=XM_003990797.5
C1 151865316 151865519
refGene_gn=LY75;refGene_tx=XM_003990798.4
C1 151869688 151869715
refGene_gn=LY75;refGene_tx=XM_003990798.4

C1 151871022 151871196
refGene_gn=LY75;refGene_tx=XM_003990798.4
C1 151872066 151872183
refGene_gn=LY75;refGene_tx=XM_003990798.4
C1 151878520 151878631
refGene_gn=LY75;refGene_tx=XM_003990798.4
C1 151898914 151899093
refGene_gn=LY75;refGene_tx=XM_011285355.3
C1 151900769 151900937
refGene_gn=LY75;refGene_tx=XM_003990798.4;refGene_gn=LY75;refGene_tx=X
M_011285355.3
C1 151902624 151902747
refGene_gn=LY75;refGene_tx=XM_003990798.4;refGene_gn=LY75;refGene_tx=X
M_011285355.3
C1 151905104 151905254
refGene_gn=LY75;refGene_tx=XM_003990798.4;refGene_gn=LY75;refGene_tx=X
M_011285355.3
C1 151908558 151908727
refGene_gn=LY75;refGene_tx=XM_003990798.4;refGene_gn=LY75;refGene_tx=X
M_011285355.3
C1 151910084 151910311
refGene_gn=LY75;refGene_tx=XM_003990798.4;refGene_gn=LY75;refGene_tx=X
M_011285355.3
C1 151912589 151912784
refGene_gn=LY75;refGene_tx=XM_003990798.4;refGene_gn=LY75;refGene_tx=X
M_011285355.3
C1 151917238 151917355
refGene_gn=LY75;refGene_tx=XM_003990798.4;refGene_gn=LY75;refGene_tx=X
M_011285355.3
C1 151918691 151918859
refGene_gn=LY75;refGene_tx=XM_003990798.4;refGene_gn=LY75;refGene_tx=X
M_011285355.3
C1 151920527 151920693
refGene_gn=LY75;refGene_tx=XM_003990798.4;refGene_gn=LY75;refGene_tx=X
M_011285355.3
C1 151924569 151924796
refGene_gn=LY75;refGene_tx=XM_003990798.4;refGene_gn=LY75;refGene_tx=X
M_011285355.3
C1 151926016 151926146
refGene_gn=LY75;refGene_tx=XM_003990798.4;refGene_gn=LY75;refGene_tx=X
M_011285355.3
C1 151933312 151933473
refGene_gn=LY75;refGene_tx=XM_003990798.4;refGene_gn=LY75;refGene_tx=X
M_011285355.3
C1 151933903 151934009
refGene_gn=LY75;refGene_tx=XM_003990798.4;refGene_gn=LY75;refGene_tx=X
M_011285355.3
C1 151935951 151936091
refGene_gn=LY75;refGene_tx=XM_003990798.4;refGene_gn=LY75;refGene_tx=X
M_011285355.3

C1	151936672	151936752
refGene_gn=LY75;refGene_tx=XM_003990798.4;refGene_gn=LY75;refGene_tx=XM_011285355.3
C1	151937257	151937325
refGene_gn=LY75;refGene_tx=XM_003990798.4;refGene_gn=LY75;refGene_tx=XM_011285355.3
C1	151938203	151938379
refGene_gn=LY75;refGene_tx=XM_003990798.4;refGene_gn=LY75;refGene_tx=XM_011285355.3
C1	151938759	151938795
refGene_gn=LY75;refGene_tx=XM_003990798.4;refGene_gn=LY75;refGene_tx=XM_011285355.3
C1	151943963	151944078
refGene_gn=LY75;refGene_tx=XM_003990798.4;refGene_gn=LY75;refGene_tx=XM_011285355.3
C1	151946393	151946462
refGene_gn=LY75;refGene_tx=XM_003990798.4;refGene_gn=LY75;refGene_tx=XM_011285355.3
C1	151947655	151947737
refGene_gn=LY75;refGene_tx=XM_003990798.4;refGene_gn=LY75;refGene_tx=XM_011285355.3
C1	151962971	151963114
refGene_gn=LY75;refGene_tx=XM_003990798.4;refGene_gn=LY75;refGene_tx=XM_011285355.3
C1	151965169	151965369
refGene_gn=LY75;refGene_tx=XM_003990798.4;refGene_gn=LY75;refGene_tx=XM_011285355.3
C1	151967859	151968029
refGene_gn=LY75;refGene_tx=XM_003990798.4;refGene_gn=LY75;refGene_tx=XM_011285355.3
C1	151968174	151968263
refGene_gn=LY75;refGene_tx=XM_003990798.4;refGene_gn=LY75;refGene_tx=XM_011285355.3
C1	151968827	151968938
refGene_gn=LY75;refGene_tx=XM_003990798.4;refGene_gn=LY75;refGene_tx=XM_011285355.3
C1	151971096	151971254
refGene_gn=LY75;refGene_tx=XM_003990798.4;refGene_gn=LY75;refGene_tx=XM_011285355.3
C1	151972573	151972765
refGene_gn=LY75;refGene_tx=XM_003990798.4;refGene_gn=LY75;refGene_tx=XM_011285355.3
C1	151974981	151975125
refGene_gn=LY75;refGene_tx=XM_003990798.4;refGene_gn=LY75;refGene_tx=XM_011285355.3
C1	151976518	151976629
refGene_gn=LY75;refGene_tx=XM_003990798.4;refGene_gn=LY75;refGene_tx=XM_011285355.3
C1	151984477	151984642
refGene_gn=LY75;refGene_tx=XM_003990798.4;refGene_gn=LY75;refGene_tx=X

M_011285355.3
C1 151988169 151988340
refGene_gn=LY75;refGene_tx=XM_003990798.4;refGene_gn=LY75;refGene_tx=XM_011285355.3
C1 151994085 151994457
refGene_gn=LY75;refGene_tx=XM_003990798.4;refGene_gn=LY75;refGene_tx=XM_011285355.3
C1 151999890 151999984
refGene_gn=LY75;refGene_tx=XM_003990798.4;refGene_gn=LY75;refGene_tx=XM_011285355.3
C1 152036592 152036807
refGene_gn=PLA2R1;refGene_tx=XM_006935305.4
C1 152036924 152036957
refGene_gn=PLA2R1;refGene_tx=XM_006935305.4
C1 152039796 152039973
refGene_gn=PLA2R1;refGene_tx=XM_006935305.4
C1 152041562 152041679
refGene_gn=PLA2R1;refGene_tx=XM_006935305.4
C1 152042178 152042319
refGene_gn=PLA2R1;refGene_tx=XM_006935305.4
C1 152042602 152042616
refGene_gn=PLA2R1;refGene_tx=XM_019838149.2
C1 152044630 152044799
refGene_gn=PLA2R1;refGene_tx=XM_006935305.4;refGene_gn=PLA2R1;refGene_tx=XM_019838149.2
C1 152046355 152046594
refGene_gn=PLA2R1;refGene_tx=XM_006935305.4;refGene_gn=PLA2R1;refGene_tx=XM_019838149.2
C1 152050770 152050918
refGene_gn=PLA2R1;refGene_tx=XM_006935305.4;refGene_gn=PLA2R1;refGene_tx=XM_019838149.2
C1 152051276 152051404
refGene_gn=PLA2R1;refGene_tx=XM_006935305.4;refGene_gn=PLA2R1;refGene_tx=XM_019838149.2
C1 152052540 152052658
refGene_gn=PLA2R1;refGene_tx=XM_006935305.4;refGene_gn=PLA2R1;refGene_tx=XM_019838149.2
C1 152061152 152061292
refGene_gn=PLA2R1;refGene_tx=XM_006935305.4;refGene_gn=PLA2R1;refGene_tx=XM_019838149.2
C1 152062879 152062983
refGene_gn=PLA2R1;refGene_tx=XM_006935305.4;refGene_gn=PLA2R1;refGene_tx=XM_019838149.2
C1 152063254 152063319
refGene_gn=PLA2R1;refGene_tx=XM_006935305.4;refGene_gn=PLA2R1;refGene_tx=XM_019838149.2
C1 152068419 152068577
refGene_gn=PLA2R1;refGene_tx=XM_006935305.4;refGene_gn=PLA2R1;refGene_tx=XM_019838149.2
C1 152069051 152069087

refGene_gn=PLA2R1;refGene_tx=XM_006935305.4;refGene_gn=PLA2R1;refGene_tx=XM_019838149.2
C1 152069920 152070053
refGene_gn=PLA2R1;refGene_tx=XM_006935305.4;refGene_gn=PLA2R1;refGene_tx=XM_019838149.2
C1 152072692 152072777
refGene_gn=PLA2R1;refGene_tx=XM_006935305.4;refGene_gn=PLA2R1;refGene_tx=XM_019838149.2
C1 152076076 152076222
refGene_gn=PLA2R1;refGene_tx=XM_006935305.4;refGene_gn=PLA2R1;refGene_tx=XM_019838149.2
C1 152077930 152078133
refGene_gn=PLA2R1;refGene_tx=XM_006935305.4;refGene_gn=PLA2R1;refGene_tx=XM_019838149.2
C1 152090752 152090925
refGene_gn=PLA2R1;refGene_tx=XM_006935305.4;refGene_gn=PLA2R1;refGene_tx=XM_019838149.2
C1 152099796 152099909
refGene_gn=PLA2R1;refGene_tx=XM_006935305.4;refGene_gn=PLA2R1;refGene_tx=XM_019838149.2
C1 152104588 152104687
refGene_gn=PLA2R1;refGene_tx=XM_006935305.4;refGene_gn=PLA2R1;refGene_tx=XM_019838149.2
C1 152109005 152109163
refGene_gn=PLA2R1;refGene_tx=XM_006935305.4;refGene_gn=PLA2R1;refGene_tx=XM_019838149.2
C1 152110333 152110528
refGene_gn=PLA2R1;refGene_tx=XM_006935305.4;refGene_gn=PLA2R1;refGene_tx=XM_019838149.2
C1 152112791 152112935
refGene_gn=PLA2R1;refGene_tx=XM_006935305.4;refGene_gn=PLA2R1;refGene_tx=XM_019838149.2
C1 152113462 152113576
refGene_gn=PLA2R1;refGene_tx=XM_006935305.4;refGene_gn=PLA2R1;refGene_tx=XM_019838149.2
C1 152118337 152118511
refGene_gn=PLA2R1;refGene_tx=XM_006935305.4;refGene_gn=PLA2R1;refGene_tx=XM_019838149.2
C1 152133239 152133413
refGene_gn=PLA2R1;refGene_tx=XM_006935305.4;refGene_gn=PLA2R1;refGene_tx=XM_019838149.2
C1 152136043 152136427
refGene_gn=PLA2R1;refGene_tx=XM_006935305.4;refGene_gn=PLA2R1;refGene_tx=XM_019838149.2
C1 152153571 152153680
refGene_gn=PLA2R1;refGene_tx=XM_006935305.4;refGene_gn=PLA2R1;refGene_tx=XM_019838149.2
C1 152197043 152197056
refGene_gn=ITGB6;refGene_tx=XM_023259379.1;refGene_gn=ITGB6;refGene_tx=XM_023259380.1;refGene_gn=ITGB6;refGene_tx=XM_023259382.1;refGene_gn=

ITGB6;refGene_tx=XM_023259383.1
C1 152199830 152200057
refGene_gn=ITGB6;refGene_tx=XM_023259379.1;refGene_gn=ITGB6;refGene_tx=XM_023259380.1;refGene_gn=ITGB6;refGene_tx=XM_023259381.1;refGene_gn=ITGB6;refGene_tx=XM_023259382.1;refGene_gn=ITGB6;refGene_tx=XM_023259383.1
C1 152206271 152206689
refGene_gn=ITGB6;refGene_tx=XM_023259379.1;refGene_gn=ITGB6;refGene_tx=XM_023259380.1;refGene_gn=ITGB6;refGene_tx=XM_023259381.1;refGene_gn=ITGB6;refGene_tx=XM_023259382.1;refGene_gn=ITGB6;refGene_tx=XM_023259383.1
C1 152206910 152207045
refGene_gn=ITGB6;refGene_tx=XM_023259379.1;refGene_gn=ITGB6;refGene_tx=XM_023259380.1;refGene_gn=ITGB6;refGene_tx=XM_023259381.1;refGene_gn=ITGB6;refGene_tx=XM_023259382.1;refGene_gn=ITGB6;refGene_tx=XM_023259383.1
C1 152213691 152213781
refGene_gn=ITGB6;refGene_tx=XM_023259379.1;refGene_gn=ITGB6;refGene_tx=XM_023259380.1;refGene_gn=ITGB6;refGene_tx=XM_023259381.1;refGene_gn=ITGB6;refGene_tx=XM_023259382.1;refGene_gn=ITGB6;refGene_tx=XM_023259383.1
C1 152231331 152231427
refGene_gn=ITGB6;refGene_tx=XM_023259379.1;refGene_gn=ITGB6;refGene_tx=XM_023259380.1;refGene_gn=ITGB6;refGene_tx=XM_023259381.1;refGene_gn=ITGB6;refGene_tx=XM_023259382.1;refGene_gn=ITGB6;refGene_tx=XM_023259383.1
C1 152234395 152234557
refGene_gn=ITGB6;refGene_tx=XM_023259379.1;refGene_gn=ITGB6;refGene_tx=XM_023259380.1;refGene_gn=ITGB6;refGene_tx=XM_023259381.1;refGene_gn=ITGB6;refGene_tx=XM_023259382.1;refGene_gn=ITGB6;refGene_tx=XM_023259383.1
C1 152235781 152235947
refGene_gn=ITGB6;refGene_tx=XM_023259379.1;refGene_gn=ITGB6;refGene_tx=XM_023259380.1;refGene_gn=ITGB6;refGene_tx=XM_023259381.1;refGene_gn=ITGB6;refGene_tx=XM_023259382.1;refGene_gn=ITGB6;refGene_tx=XM_023259383.1
C1 152247986 152248233
refGene_gn=ITGB6;refGene_tx=XM_023259379.1;refGene_gn=ITGB6;refGene_tx=XM_023259380.1;refGene_gn=ITGB6;refGene_tx=XM_023259381.1;refGene_gn=ITGB6;refGene_tx=XM_023259382.1;refGene_gn=ITGB6;refGene_tx=XM_023259383.1
C1 152249092 152249297
refGene_gn=ITGB6;refGene_tx=XM_023259379.1;refGene_gn=ITGB6;refGene_tx=XM_023259380.1;refGene_gn=ITGB6;refGene_tx=XM_023259381.1;refGene_gn=ITGB6;refGene_tx=XM_023259382.1;refGene_gn=ITGB6;refGene_tx=XM_023259383.1
C1 152251853 152251933
refGene_gn=ITGB6;refGene_tx=XM_023259379.1;refGene_gn=ITGB6;refGene_tx=XM_023259380.1;refGene_gn=ITGB6;refGene_tx=XM_023259381.1;refGene_gn=ITGB6;refGene_tx=XM_0232593

83.1
C1	152252670	152252746
refGene_gn=ITGB6;refGene_tx=XM_023259379.1;refGene_gn=ITGB6;refGene_tx=XM_023259380.1;refGene_gn=ITGB6;refGene_tx=XM_023259381.1;refGene_gn=ITGB6;refGene_tx=XM_023259382.1;refGene_gn=ITGB6;refGene_tx=XM_023259383.1
C1	152252854	152252899
refGene_gn=ITGB6;refGene_tx=XM_023259379.1;refGene_gn=ITGB6;refGene_tx=XM_023259380.1;refGene_gn=ITGB6;refGene_tx=XM_023259381.1
C1	152330963	152331041
refGene_gn=RBMS1;refGene_tx=XM_019838158.2;refGene_gn=RBMS1;refGene_tx=XM_019838159.2;refGene_gn=RBMS1;refGene_tx=XM_023259384.1;refGene_gn=RBMS1;refGene_tx=XM_023259385.1;refGene_gn=RBMS1;refGene_tx=XM_023259386.1;refGene_gn=RBMS1;refGene_tx=XM_023259387.1;refGene_gn=RBMS1;refGene_tx=XM_023259388.1;refGene_gn=RBMS1;refGene_tx=XM_023259389.1;refGene_gn=RBMS1;refGene_tx=XM_023259390.1
C1	152332795	152332876
refGene_gn=RBMS1;refGene_tx=XM_019838158.2;refGene_gn=RBMS1;refGene_tx=XM_019838159.2;refGene_gn=RBMS1;refGene_tx=XM_023259384.1;refGene_gn=RBMS1;refGene_tx=XM_023259385.1;refGene_gn=RBMS1;refGene_tx=XM_023259386.1;refGene_gn=RBMS1;refGene_tx=XM_023259387.1;refGene_gn=RBMS1;refGene_tx=XM_023259388.1;refGene_gn=RBMS1;refGene_tx=XM_023259389.1;refGene_gn=RBMS1;refGene_tx=XM_023259390.1
C1	152333995	152334106
refGene_gn=RBMS1;refGene_tx=XM_019838158.2;refGene_gn=RBMS1;refGene_tx=XM_019838159.2;refGene_gn=RBMS1;refGene_tx=XM_023259384.1;refGene_gn=RBMS1;refGene_tx=XM_023259385.1;refGene_gn=RBMS1;refGene_tx=XM_023259386.1;refGene_gn=RBMS1;refGene_tx=XM_023259387.1;refGene_gn=RBMS1;refGene_tx=XM_023259388.1;refGene_gn=RBMS1;refGene_tx=XM_023259389.1;refGene_gn=RBMS1;refGene_tx=XM_023259390.1
C1	152337001	152337052
refGene_gn=RBMS1;refGene_tx=XM_019838158.2;refGene_gn=RBMS1;refGene_tx=XM_019838159.2;refGene_gn=RBMS1;refGene_tx=XM_023259384.1;refGene_gn=RBMS1;refGene_tx=XM_023259385.1;refGene_gn=RBMS1;refGene_tx=XM_023259386.1;refGene_gn=RBMS1;refGene_tx=XM_023259387.1;refGene_gn=RBMS1;refGene_tx=XM_023259388.1;refGene_gn=RBMS1;refGene_tx=XM_023259389.1;refGene_gn=RBMS1;refGene_tx=XM_023259390.1
C1	152337935	152337983
refGene_gn=RBMS1;refGene_tx=XM_019838158.2;refGene_gn=RBMS1;refGene_tx=XM_019838159.2;refGene_gn=RBMS1;refGene_tx=XM_023259384.1;refGene_gn=RBMS1;refGene_tx=XM_023259385.1;refGene_gn=RBMS1;refGene_tx=XM_023259386.1;refGene_gn=RBMS1;refGene_tx=XM_023259387.1;refGene_gn=RBMS1;refGene_tx=XM_023259388.1
C1	152340472	152340566
refGene_gn=RBMS1;refGene_tx=XM_019838158.2;refGene_gn=RBMS1;refGene_tx=XM_019838159.2;refGene_gn=RBMS1;refGene_tx=XM_023259384.1;refGene_gn=RBMS1;refGene_tx=XM_023259385.1;refGene_gn=RBMS1;refGene_tx=XM_023259386.1;refGene_gn=RBMS1;refGene_tx=XM_023259387.1;refGene_gn=RBMS1;refGene_tx=XM_023259388.1;refGene_gn=RBMS1;refGene_tx=XM_023259389.1;refGene_gn=RBMS1;refGene_tx=XM_023259390.1

C1 152340692 152340742
refGene_gn=RBMS1;refGene_tx=XM_019838158.2;refGene_gn=RBMS1;refGene_tx=XM_019838159.2;refGene_gn=RBMS1;refGene_tx=XM_023259384.1;refGene_gn=RBMS1;refGene_tx=XM_023259385.1;refGene_gn=RBMS1;refGene_tx=XM_023259386.1;refGene_gn=RBMS1;refGene_tx=XM_023259387.1;refGene_gn=RBMS1;refGene_tx=XM_023259388.1;refGene_gn=RBMS1;refGene_tx=XM_023259389.1;refGene_gn=RBMS1;refGene_tx=XM_023259390.1
C1 152341860 152341976
refGene_gn=RBMS1;refGene_tx=XM_019838158.2;refGene_gn=RBMS1;refGene_tx=XM_019838159.2;refGene_gn=RBMS1;refGene_tx=XM_023259384.1;refGene_gn=RBMS1;refGene_tx=XM_023259385.1;refGene_gn=RBMS1;refGene_tx=XM_023259386.1;refGene_gn=RBMS1;refGene_tx=XM_023259387.1;refGene_gn=RBMS1;refGene_tx=XM_023259388.1;refGene_gn=RBMS1;refGene_tx=XM_023259389.1;refGene_gn=RBMS1;refGene_tx=XM_023259390.1
C1 152355347 152355427
refGene_gn=RBMS1;refGene_tx=XM_019838158.2;refGene_gn=RBMS1;refGene_tx=XM_019838159.2;refGene_gn=RBMS1;refGene_tx=XM_023259384.1;refGene_gn=RBMS1;refGene_tx=XM_023259385.1;refGene_gn=RBMS1;refGene_tx=XM_023259386.1;refGene_gn=RBMS1;refGene_tx=XM_023259387.1;refGene_gn=RBMS1;refGene_tx=XM_023259388.1;refGene_gn=RBMS1;refGene_tx=XM_023259389.1;refGene_gn=RBMS1;refGene_tx=XM_023259390.1
C1 152357628 152357786
refGene_gn=RBMS1;refGene_tx=XM_019838158.2;refGene_gn=RBMS1;refGene_tx=XM_019838159.2;refGene_gn=RBMS1;refGene_tx=XM_023259384.1;refGene_gn=RBMS1;refGene_tx=XM_023259385.1;refGene_gn=RBMS1;refGene_tx=XM_023259386.1;refGene_gn=RBMS1;refGene_tx=XM_023259387.1;refGene_gn=RBMS1;refGene_tx=XM_023259388.1;refGene_gn=RBMS1;refGene_tx=XM_023259389.1;refGene_gn=RBMS1;refGene_tx=XM_023259390.1
C1 152366637 152366729
refGene_gn=RBMS1;refGene_tx=XM_019838158.2;refGene_gn=RBMS1;refGene_tx=XM_019838159.2;refGene_gn=RBMS1;refGene_tx=XM_023259384.1;refGene_gn=RBMS1;refGene_tx=XM_023259385.1;refGene_gn=RBMS1;refGene_tx=XM_023259386.1;refGene_gn=RBMS1;refGene_tx=XM_023259387.1;refGene_gn=RBMS1;refGene_tx=XM_023259388.1;refGene_gn=RBMS1;refGene_tx=XM_023259389.1;refGene_gn=RBMS1;refGene_tx=XM_023259390.1
C1 152371613 152371672
refGene_gn=RBMS1;refGene_tx=XM_019838158.2;refGene_gn=RBMS1;refGene_tx=XM_019838159.2;refGene_gn=RBMS1;refGene_tx=XM_023259384.1;refGene_gn=RBMS1;refGene_tx=XM_023259385.1;refGene_gn=RBMS1;refGene_tx=XM_023259386.1;refGene_gn=RBMS1;refGene_tx=XM_023259387.1;refGene_gn=RBMS1;refGene_tx=XM_023259388.1;refGene_gn=RBMS1;refGene_tx=XM_023259389.1;refGene_gn=RBMS1;refGene_tx=XM_023259390.1
C1 152420188 152420364
refGene_gn=RBMS1;refGene_tx=XM_019838158.2;refGene_gn=RBMS1;refGene_tx=XM_019838159.2;refGene_gn=RBMS1;refGene_tx=XM_023259384.1;refGene_gn=RBMS1;refGene_tx=XM_023259385.1;refGene_gn=RBMS1;refGene_tx=XM_023259386.1;refGene_gn=RBMS1;refGene_tx=XM_023259387.1;refGene_gn=RBMS1;refGene_tx=XM_023259388.1;refGene_gn=RBMS1;refGene_tx=XM_023259389.1;refGene_gn=RBMS1;refGene_tx=XM_023259390.1
C1 152442669 152442696

refGene_gn=RBMS1;refGene_tx=XM_019838158.2
C1 152459331 152459400
refGene_gn=RBMS1;refGene_tx=XM_023259386.1;refGene_gn=RBMS1;refGene_tx=XM_023259388.1
C1 152547177 152547252
refGene_gn=RBMS1;refGene_tx=XM_023259384.1;refGene_gn=RBMS1;refGene_tx=XM_023259385.1;refGene_gn=RBMS1;refGene_tx=XM_023259387.1;refGene_gn=RBMS1;refGene_tx=XM_023259389.1;refGene_gn=RBMS1;refGene_tx=XM_023259390.1
C1 153114241 153114249
refGene_gn=TANK;refGene_tx=XM_003990801.5
C1 153133452 153133600
refGene_gn=TANK;refGene_tx=XM_003990801.5;refGene_gn=TANK;refGene_tx=XM_006935314.3;refGene_gn=TANK;refGene_tx=XM_006935315.4
C1 153158845 153158954
refGene_gn=TANK;refGene_tx=XM_003990801.5;refGene_gn=TANK;refGene_tx=XM_006935314.3;refGene_gn=TANK;refGene_tx=XM_006935315.4
C1 153160533 153160652
refGene_gn=TANK;refGene_tx=XM_003990801.5;refGene_gn=TANK;refGene_tx=XM_006935314.3;refGene_gn=TANK;refGene_tx=XM_006935315.4
C1 153182082 153182159
refGene_gn=TANK;refGene_tx=XM_003990801.5;refGene_gn=TANK;refGene_tx=XM_006935314.3;refGene_gn=TANK;refGene_tx=XM_006935315.4
C1 153182599 153182715
refGene_gn=TANK;refGene_tx=XM_003990801.5;refGene_gn=TANK;refGene_tx=XM_006935314.3;refGene_gn=TANK;refGene_tx=XM_006935315.4
C1 153188254 153188835
refGene_gn=TANK;refGene_tx=XM_003990801.5;refGene_gn=TANK;refGene_tx=XM_006935314.3;refGene_gn=TANK;refGene_tx=XM_006935315.4
C1 153192281 153192461
refGene_gn=TANK;refGene_tx=XM_003990801.5;refGene_gn=TANK;refGene_tx=XM_006935314.3;refGene_gn=TANK;refGene_tx=XM_006935315.4
C1 153264076 153264124
refGene_gn=PSMD14;refGene_tx=XM_006935316.3
C1 153307862 153307934
refGene_gn=PSMD14;refGene_tx=XM_006935316.3
C1 153308156 153308276
refGene_gn=PSMD14;refGene_tx=XM_006935316.3
C1 153311037 153311108
refGene_gn=PSMD14;refGene_tx=XM_006935316.3
C1 153312097 153312248
refGene_gn=PSMD14;refGene_tx=XM_006935316.3
C1 153326193 153326301
refGene_gn=PSMD14;refGene_tx=XM_006935316.3
C1 153332870 153332945
refGene_gn=PSMD14;refGene_tx=XM_006935316.3
C1 153337433 153337559
refGene_gn=PSMD14;refGene_tx=XM_006935316.3
C1 153348715 153348778
refGene_gn=PSMD14;refGene_tx=XM_006935316.3

C1 153351259 153351358
refGene_gn=PSMD14;refGene_tx=XM_006935316.3
C1 153356608 153357300
refGene_gn=TBR1;refGene_tx=XM_023259391.1
C1 153357891 153358046
refGene_gn=TBR1;refGene_tx=XM_023259391.1
C1 153358422 153358544
refGene_gn=TBR1;refGene_tx=XM_023259391.1
C1 153359115 153359274
refGene_gn=TBR1;refGene_tx=XM_023259391.1
C1 153360456 153360518
refGene_gn=TBR1;refGene_tx=XM_023259391.1
C1 153363761 153364620
refGene_gn=TBR1;refGene_tx=XM_023259391.1
C1 153484052 153484100
refGene_gn=SLC4A10;refGene_tx=XM_019838163.2
C1 153564947 153565028
refGene_gn=SLC4A10;refGene_tx=XM_011285348.2;refGene_gn=SLC4A10;refGene_tx=XM_019838161.2;refGene_gn=SLC4A10;refGene_tx=XM_019838162.2;refGene_gn=SLC4A10;refGene_tx=XM_019838165.2;refGene_gn=SLC4A10;refGene_tx=XM_019838166.2;refGene_gn=SLC4A10;refGene_tx=XM_019838167.2
C1 153590427 153590509
refGene_gn=SLC4A10;refGene_tx=XM_011285348.2;refGene_gn=SLC4A10;refGene_tx=XM_019838161.2;refGene_gn=SLC4A10;refGene_tx=XM_019838162.2;refGene_gn=SLC4A10;refGene_tx=XM_019838163.2;refGene_gn=SLC4A10;refGene_tx=XM_019838165.2;refGene_gn=SLC4A10;refGene_tx=XM_019838166.2;refGene_gn=SLC4A10;refGene_tx=XM_019838167.2
C1 153623277 153623424
refGene_gn=SLC4A10;refGene_tx=XM_011285348.2;refGene_gn=SLC4A10;refGene_tx=XM_019838161.2;refGene_gn=SLC4A10;refGene_tx=XM_019838162.2;refGene_gn=SLC4A10;refGene_tx=XM_019838163.2;refGene_gn=SLC4A10;refGene_tx=XM_019838165.2;refGene_gn=SLC4A10;refGene_tx=XM_019838166.2;refGene_gn=SLC4A10;refGene_tx=XM_019838167.2
C1 153645538 153645677
refGene_gn=SLC4A10;refGene_tx=XM_011285348.2;refGene_gn=SLC4A10;refGene_tx=XM_019838161.2;refGene_gn=SLC4A10;refGene_tx=XM_019838162.2;refGene_gn=SLC4A10;refGene_tx=XM_019838163.2;refGene_gn=SLC4A10;refGene_tx=XM_019838165.2;refGene_gn=SLC4A10;refGene_tx=XM_019838166.2;refGene_gn=SLC4A10;refGene_tx=XM_019838167.2
C1 153652594 153652755
refGene_gn=SLC4A10;refGene_tx=XM_011285348.2;refGene_gn=SLC4A10;refGene_tx=XM_019838161.2;refGene_gn=SLC4A10;refGene_tx=XM_019838162.2;refGene_gn=SLC4A10;refGene_tx=XM_019838163.2;refGene_gn=SLC4A10;refGene_tx=XM_019838165.2;refGene_gn=SLC4A10;refGene_tx=XM_019838166.2;refGene_gn=SLC4A10;refGene_tx=XM_019838167.2
C1 153667880 153668069
refGene_gn=SLC4A10;refGene_tx=XM_011285348.2;refGene_gn=SLC4A10;refGene_tx=XM_019838161.2;refGene_gn=SLC4A10;refGene_tx=XM_019838162.2;refGene_gn=SLC4A10;refGene_tx=XM_019838163.2;refGene_gn=SLC4A10;refGene_tx=XM_019838165.2;refGene_gn=SLC4A10;refGene_tx=XM_019838166.2;refGene_gn

=SLC4A10;refGene_tx=XM_019838167.2
C1	153677588	153677680
refGene_gn=SLC4A10;refGene_tx=XM_011285348.2;refGene_gn=SLC4A10;refGene_tx=XM_019838161.2;refGene_gn=SLC4A10;refGene_tx=XM_019838162.2;refGene_gn=SLC4A10;refGene_tx=XM_019838163.2;refGene_gn=SLC4A10;refGene_tx=XM_019838165.2;refGene_gn=SLC4A10;refGene_tx=XM_019838166.2;refGene_gn=SLC4A10;refGene_tx=XM_019838167.2
C1	153678634	153678724
refGene_gn=SLC4A10;refGene_tx=XM_019838161.2;refGene_gn=SLC4A10;refGene_tx=XM_019838162.2;refGene_gn=SLC4A10;refGene_tx=XM_019838163.2;refGene_gn=SLC4A10;refGene_tx=XM_019838165.2;refGene_gn=SLC4A10;refGene_tx=XM_019838166.2
C1	153683687	153683845
refGene_gn=SLC4A10;refGene_tx=XM_011285348.2;refGene_gn=SLC4A10;refGene_tx=XM_019838161.2;refGene_gn=SLC4A10;refGene_tx=XM_019838162.2;refGene_gn=SLC4A10;refGene_tx=XM_019838163.2;refGene_gn=SLC4A10;refGene_tx=XM_019838165.2;refGene_gn=SLC4A10;refGene_tx=XM_019838166.2;refGene_gn=SLC4A10;refGene_tx=XM_019838167.2
C1	153686925	153687013
refGene_gn=SLC4A10;refGene_tx=XM_011285348.2;refGene_gn=SLC4A10;refGene_tx=XM_019838161.2;refGene_gn=SLC4A10;refGene_tx=XM_019838162.2;refGene_gn=SLC4A10;refGene_tx=XM_019838163.2;refGene_gn=SLC4A10;refGene_tx=XM_019838165.2;refGene_gn=SLC4A10;refGene_tx=XM_019838166.2;refGene_gn=SLC4A10;refGene_tx=XM_019838167.2
C1	153695361	153695508
refGene_gn=SLC4A10;refGene_tx=XM_011285348.2;refGene_gn=SLC4A10;refGene_tx=XM_019838161.2;refGene_gn=SLC4A10;refGene_tx=XM_019838162.2;refGene_gn=SLC4A10;refGene_tx=XM_019838163.2;refGene_gn=SLC4A10;refGene_tx=XM_019838165.2;refGene_gn=SLC4A10;refGene_tx=XM_019838166.2;refGene_gn=SLC4A10;refGene_tx=XM_019838167.2
C1	153703564	153703665
refGene_gn=SLC4A10;refGene_tx=XM_011285348.2;refGene_gn=SLC4A10;refGene_tx=XM_019838161.2;refGene_gn=SLC4A10;refGene_tx=XM_019838162.2;refGene_gn=SLC4A10;refGene_tx=XM_019838163.2;refGene_gn=SLC4A10;refGene_tx=XM_019838165.2;refGene_gn=SLC4A10;refGene_tx=XM_019838166.2;refGene_gn=SLC4A10;refGene_tx=XM_019838167.2
C1	153706744	153706919
refGene_gn=SLC4A10;refGene_tx=XM_011285348.2;refGene_gn=SLC4A10;refGene_tx=XM_019838161.2;refGene_gn=SLC4A10;refGene_tx=XM_019838162.2;refGene_gn=SLC4A10;refGene_tx=XM_019838163.2;refGene_gn=SLC4A10;refGene_tx=XM_019838165.2;refGene_gn=SLC4A10;refGene_tx=XM_019838166.2;refGene_gn=SLC4A10;refGene_tx=XM_019838167.2
C1	153707468	153707602
refGene_gn=SLC4A10;refGene_tx=XM_011285348.2;refGene_gn=SLC4A10;refGene_tx=XM_019838161.2;refGene_gn=SLC4A10;refGene_tx=XM_019838162.2;refGene_gn=SLC4A10;refGene_tx=XM_019838163.2;refGene_gn=SLC4A10;refGene_tx=XM_019838165.2;refGene_gn=SLC4A10;refGene_tx=XM_019838166.2;refGene_gn=SLC4A10;refGene_tx=XM_019838167.2
C1	153708001	153708247
refGene_gn=SLC4A10;refGene_tx=XM_011285348.2;refGene_gn=SLC4A10;refGen

e_tx=XM_019838161.2;refGene_gn=SLC4A10;refGene_tx=XM_019838162.2;refGene_gn=SLC4A10;refGene_tx=XM_019838163.2;refGene_gn=SLC4A10;refGene_tx=XM_019838165.2;refGene_gn=SLC4A10;refGene_tx=XM_019838166.2;refGene_gn=SLC4A10;refGene_tx=XM_019838167.2
C1 153735438 153735544
refGene_gn=SLC4A10;refGene_tx=XM_011285348.2;refGene_gn=SLC4A10;refGene_tx=XM_019838161.2;refGene_gn=SLC4A10;refGene_tx=XM_019838162.2;refGene_gn=SLC4A10;refGene_tx=XM_019838163.2;refGene_gn=SLC4A10;refGene_tx=XM_019838165.2;refGene_gn=SLC4A10;refGene_tx=XM_019838166.2;refGene_gn=SLC4A10;refGene_tx=XM_019838167.2
C1 153741253 153741415
refGene_gn=SLC4A10;refGene_tx=XM_011285348.2;refGene_gn=SLC4A10;refGene_tx=XM_019838161.2;refGene_gn=SLC4A10;refGene_tx=XM_019838162.2;refGene_gn=SLC4A10;refGene_tx=XM_019838163.2;refGene_gn=SLC4A10;refGene_tx=XM_019838165.2;refGene_gn=SLC4A10;refGene_tx=XM_019838166.2;refGene_gn=SLC4A10;refGene_tx=XM_019838167.2
C1 153743130 153743244
refGene_gn=SLC4A10;refGene_tx=XM_011285348.2;refGene_gn=SLC4A10;refGene_tx=XM_019838161.2;refGene_gn=SLC4A10;refGene_tx=XM_019838162.2;refGene_gn=SLC4A10;refGene_tx=XM_019838163.2;refGene_gn=SLC4A10;refGene_tx=XM_019838165.2;refGene_gn=SLC4A10;refGene_tx=XM_019838166.2;refGene_gn=SLC4A10;refGene_tx=XM_019838167.2
C1 153744283 153744445
refGene_gn=SLC4A10;refGene_tx=XM_011285348.2;refGene_gn=SLC4A10;refGene_tx=XM_019838161.2;refGene_gn=SLC4A10;refGene_tx=XM_019838162.2;refGene_gn=SLC4A10;refGene_tx=XM_019838163.2;refGene_gn=SLC4A10;refGene_tx=XM_019838165.2;refGene_gn=SLC4A10;refGene_tx=XM_019838166.2;refGene_gn=SLC4A10;refGene_tx=XM_019838167.2
C1 153752808 153753060
refGene_gn=SLC4A10;refGene_tx=XM_011285348.2;refGene_gn=SLC4A10;refGene_tx=XM_019838161.2;refGene_gn=SLC4A10;refGene_tx=XM_019838162.2;refGene_gn=SLC4A10;refGene_tx=XM_019838163.2;refGene_gn=SLC4A10;refGene_tx=XM_019838165.2;refGene_gn=SLC4A10;refGene_tx=XM_019838166.2;refGene_gn=SLC4A10;refGene_tx=XM_019838167.2
C1 153755514 153755583
refGene_gn=SLC4A10;refGene_tx=XM_011285348.2;refGene_gn=SLC4A10;refGene_tx=XM_019838161.2;refGene_gn=SLC4A10;refGene_tx=XM_019838162.2;refGene_gn=SLC4A10;refGene_tx=XM_019838163.2;refGene_gn=SLC4A10;refGene_tx=XM_019838165.2;refGene_gn=SLC4A10;refGene_tx=XM_019838166.2;refGene_gn=SLC4A10;refGene_tx=XM_019838167.2
C1 153760841 153761015
refGene_gn=SLC4A10;refGene_tx=XM_011285348.2;refGene_gn=SLC4A10;refGene_tx=XM_019838161.2;refGene_gn=SLC4A10;refGene_tx=XM_019838162.2;refGene_gn=SLC4A10;refGene_tx=XM_019838163.2;refGene_gn=SLC4A10;refGene_tx=XM_019838165.2;refGene_gn=SLC4A10;refGene_tx=XM_019838166.2;refGene_gn=SLC4A10;refGene_tx=XM_019838167.2
C1 153761765 153761888
refGene_gn=SLC4A10;refGene_tx=XM_011285348.2;refGene_gn=SLC4A10;refGene_tx=XM_019838161.2;refGene_gn=SLC4A10;refGene_tx=XM_019838162.2;refGene_gn=SLC4A10;refGene_tx=XM_019838163.2;refGene_gn=SLC4A10;refGene_tx=

XM_019838165.2;refGene_gn=SLC4A10;refGene_tx=XM_019838166.2;refGene_gn=SLC4A10;refGene_tx=XM_019838167.2
C1 153771151 153771219
refGene_gn=SLC4A10;refGene_tx=XM_011285348.2;refGene_gn=SLC4A10;refGene_tx=XM_019838161.2;refGene_gn=SLC4A10;refGene_tx=XM_019838162.2;refGene_gn=SLC4A10;refGene_tx=XM_019838163.2;refGene_gn=SLC4A10;refGene_tx=XM_019838165.2;refGene_gn=SLC4A10;refGene_tx=XM_019838166.2;refGene_gn=SLC4A10;refGene_tx=XM_019838167.2
C1 153774377 153774494
refGene_gn=SLC4A10;refGene_tx=XM_011285348.2;refGene_gn=SLC4A10;refGene_tx=XM_019838161.2;refGene_gn=SLC4A10;refGene_tx=XM_019838162.2;refGene_gn=SLC4A10;refGene_tx=XM_019838163.2;refGene_gn=SLC4A10;refGene_tx=XM_019838165.2;refGene_gn=SLC4A10;refGene_tx=XM_019838166.2;refGene_gn=SLC4A10;refGene_tx=XM_019838167.2
C1 153775329 153775342
refGene_gn=SLC4A10;refGene_tx=XM_019838165.2
C1 153775351 153775358
refGene_gn=SLC4A10;refGene_tx=XM_019838166.2
C1 153780506 153780573
refGene_gn=SLC4A10;refGene_tx=XM_011285348.2;refGene_gn=SLC4A10;refGene_tx=XM_019838161.2;refGene_gn=SLC4A10;refGene_tx=XM_019838162.2;refGene_gn=SLC4A10;refGene_tx=XM_019838163.2;refGene_gn=SLC4A10;refGene_tx=XM_019838167.2
C1 153788901 153789003
refGene_gn=DPP4;refGene_tx=NM_001009838.1;refGene_gn=DPP4;refGene_tx=XM_006935319.4;refGene_gn=DPP4;refGene_tx=XM_019835951.1
C1 153790522 153790596
refGene_gn=DPP4;refGene_tx=NM_001009838.1;refGene_gn=DPP4;refGene_tx=XM_006935319.4;refGene_gn=DPP4;refGene_tx=XM_019835951.1
C1 153790858 153790931
refGene_gn=DPP4;refGene_tx=NM_001009838.1;refGene_gn=DPP4;refGene_tx=XM_006935319.4;refGene_gn=DPP4;refGene_tx=XM_019835951.1
C1 153799963 153800028
refGene_gn=DPP4;refGene_tx=NM_001009838.1;refGene_gn=DPP4;refGene_tx=XM_006935319.4;refGene_gn=DPP4;refGene_tx=XM_019835951.1
C1 153802405 153802505
refGene_gn=DPP4;refGene_tx=NM_001009838.1;refGene_gn=DPP4;refGene_tx=XM_006935319.4;refGene_gn=DPP4;refGene_tx=XM_019835951.1
C1 153803094 153803149
refGene_gn=DPP4;refGene_tx=NM_001009838.1;refGene_gn=DPP4;refGene_tx=XM_006935319.4;refGene_gn=DPP4;refGene_tx=XM_019835951.1
C1 153806109 153806304
refGene_gn=DPP4;refGene_tx=NM_001009838.1;refGene_gn=DPP4;refGene_tx=XM_006935319.4;refGene_gn=DPP4;refGene_tx=XM_019835951.1
C1 153808626 153808696
refGene_gn=DPP4;refGene_tx=NM_001009838.1;refGene_gn=DPP4;refGene_tx=XM_006935319.4;refGene_gn=DPP4;refGene_tx=XM_019835951.1
C1 153811110 153811209
refGene_gn=DPP4;refGene_tx=NM_001009838.1;refGene_gn=DPP4;refGene_tx=XM_006935319.4;refGene_gn=DPP4;refGene_tx=XM_019835951.1

C1	153811702	153811750
refGene_gn=DPP4;refGene_tx=NM_001009838.1;refGene_gn=DPP4;refGene_tx=XM_019835951.1
C1	153813538	153813660
refGene_gn=DPP4;refGene_tx=NM_001009838.1;refGene_gn=DPP4;refGene_tx=XM_006935319.4;refGene_gn=DPP4;refGene_tx=XM_019835951.1
C1	153814050	153814104
refGene_gn=DPP4;refGene_tx=NM_001009838.1;refGene_gn=DPP4;refGene_tx=XM_006935319.4;refGene_gn=DPP4;refGene_tx=XM_019835951.1
C1	153815040	153815108
refGene_gn=DPP4;refGene_tx=NM_001009838.1;refGene_gn=DPP4;refGene_tx=XM_006935319.4;refGene_gn=DPP4;refGene_tx=XM_019835951.1
C1	153815377	153815485
refGene_gn=DPP4;refGene_tx=NM_001009838.1;refGene_gn=DPP4;refGene_tx=XM_006935319.4;refGene_gn=DPP4;refGene_tx=XM_019835951.1
C1	153817898	153817943
refGene_gn=DPP4;refGene_tx=NM_001009838.1;refGene_gn=DPP4;refGene_tx=XM_006935319.4;refGene_gn=DPP4;refGene_tx=XM_019835951.1
C1	153819993	153820129
refGene_gn=DPP4;refGene_tx=NM_001009838.1;refGene_gn=DPP4;refGene_tx=XM_006935319.4;refGene_gn=DPP4;refGene_tx=XM_019835951.1
C1	153827929	153828042
refGene_gn=DPP4;refGene_tx=NM_001009838.1;refGene_gn=DPP4;refGene_tx=XM_006935319.4;refGene_gn=DPP4;refGene_tx=XM_019835951.1
C1	153829726	153829887
refGene_gn=DPP4;refGene_tx=NM_001009838.1;refGene_gn=DPP4;refGene_tx=XM_006935319.4;refGene_gn=DPP4;refGene_tx=XM_019835951.1
C1	153834099	153834220
refGene_gn=DPP4;refGene_tx=NM_001009838.1;refGene_gn=DPP4;refGene_tx=XM_006935319.4;refGene_gn=DPP4;refGene_tx=XM_019835951.1
C1	153835038	153835111
refGene_gn=DPP4;refGene_tx=NM_001009838.1;refGene_gn=DPP4;refGene_tx=XM_006935319.4;refGene_gn=DPP4;refGene_tx=XM_019835951.1
C1	153835236	153835289
refGene_gn=DPP4;refGene_tx=NM_001009838.1;refGene_gn=DPP4;refGene_tx=XM_006935319.4;refGene_gn=DPP4;refGene_tx=XM_019835951.1
C1	153842342	153842423
refGene_gn=DPP4;refGene_tx=NM_001009838.1;refGene_gn=DPP4;refGene_tx=XM_006935319.4;refGene_gn=DPP4;refGene_tx=XM_019835951.1
C1	153843773	153843865
refGene_gn=DPP4;refGene_tx=NM_001009838.1;refGene_gn=DPP4;refGene_tx=XM_006935319.4;refGene_gn=DPP4;refGene_tx=XM_019835951.1
C1	153844271	153844367
refGene_gn=DPP4;refGene_tx=NM_001009838.1;refGene_gn=DPP4;refGene_tx=XM_006935319.4;refGene_gn=DPP4;refGene_tx=XM_019835951.1
C1	153865518	153865606
refGene_gn=DPP4;refGene_tx=NM_001009838.1;refGene_gn=DPP4;refGene_tx=XM_006935319.4
C1	153866132	153866138
refGene_gn=DPP4;refGene_tx=NM_001009838.1;refGene_gn=DPP4;refGene_tx=X

M_006935319.4
C1 153896435 153896487
refGene_gn=DPP4;refGene_tx=XM_019835951.1
C1 153936051 153936058
refGene_gn=GCG;refGene_tx=XM_019838170.2
C1 153936730 153936878
refGene_gn=GCG;refGene_tx=XM_006935320.3;refGene_gn=GCG;refGene_tx=XM_019838170.2
C1 153938238 153938376
refGene_gn=GCG;refGene_tx=XM_006935320.3;refGene_gn=GCG;refGene_tx=XM_019838170.2
C1 153940260 153940422
refGene_gn=GCG;refGene_tx=XM_006935320.3;refGene_gn=GCG;refGene_tx=XM_019838170.2
C1 153942183 153942275
refGene_gn=GCG;refGene_tx=XM_006935320.3;refGene_gn=GCG;refGene_tx=XM_019838170.2
C1 153969302 153969404
refGene_gn=FAP;refGene_tx=XM_011285347.2;refGene_gn=FAP;refGene_tx=XM_019838169.1
C1 153970845 153970919
refGene_gn=FAP;refGene_tx=XM_011285347.2;refGene_gn=FAP;refGene_tx=XM_019838169.1
C1 153971181 153971254
refGene_gn=FAP;refGene_tx=XM_011285347.2;refGene_gn=FAP;refGene_tx=XM_019838169.1
C1 153971757 153971822
refGene_gn=FAP;refGene_tx=XM_011285347.2;refGene_gn=FAP;refGene_tx=XM_019838169.1
C1 153972904 153973004
refGene_gn=FAP;refGene_tx=XM_011285347.2;refGene_gn=FAP;refGene_tx=XM_019838169.1
C1 153981077 153981132
refGene_gn=FAP;refGene_tx=XM_011285347.2;refGene_gn=FAP;refGene_tx=XM_019838169.1
C1 153986205 153986400
refGene_gn=FAP;refGene_tx=XM_011285347.2;refGene_gn=FAP;refGene_tx=XM_019838169.1
C1 153987145 153987215
refGene_gn=FAP;refGene_tx=XM_011285347.2;refGene_gn=FAP;refGene_tx=XM_019838169.1
C1 153987674 153987773
refGene_gn=FAP;refGene_tx=XM_011285347.2;refGene_gn=FAP;refGene_tx=XM_019838169.1
C1 153992737 153992785
refGene_gn=FAP;refGene_tx=XM_011285347.2;refGene_gn=FAP;refGene_tx=XM_019838169.1
C1 153996725 153996850
refGene_gn=FAP;refGene_tx=XM_011285347.2;refGene_gn=FAP;refGene_tx=XM_019838169.1

C1 153998572 153998626
refGene_gn=FAP;refGene_tx=XM_011285347.2;refGene_gn=FAP;refGene_tx=XM_019838169.1
C1 154001401 154001472
refGene_gn=FAP;refGene_tx=XM_011285347.2;refGene_gn=FAP;refGene_tx=XM_019838169.1
C1 154001570 154001675
refGene_gn=FAP;refGene_tx=XM_011285347.2;refGene_gn=FAP;refGene_tx=XM_019838169.1
C1 154007533 154007578
refGene_gn=FAP;refGene_tx=XM_011285347.2;refGene_gn=FAP;refGene_tx=XM_019838169.1
C1 154011509 154011645
refGene_gn=FAP;refGene_tx=XM_011285347.2;refGene_gn=FAP;refGene_tx=XM_019838169.1
C1 154013797 154013901
refGene_gn=FAP;refGene_tx=XM_011285347.2;refGene_gn=FAP;refGene_tx=XM_019838169.1
C1 154015939 154016094
refGene_gn=FAP;refGene_tx=XM_011285347.2;refGene_gn=FAP;refGene_tx=XM_019838169.1
C1 154017039 154017160
refGene_gn=FAP;refGene_tx=XM_011285347.2;refGene_gn=FAP;refGene_tx=XM_019838169.1
C1 154017833 154017906
refGene_gn=FAP;refGene_tx=XM_011285347.2;refGene_gn=FAP;refGene_tx=XM_019838169.1
C1 154021590 154021643
refGene_gn=FAP;refGene_tx=XM_011285347.2;refGene_gn=FAP;refGene_tx=XM_019838169.1
C1 154023106 154023181
refGene_gn=FAP;refGene_tx=XM_011285347.2;refGene_gn=FAP;refGene_tx=XM_019838169.1
C1 154024105 154024200
refGene_gn=FAP;refGene_tx=XM_011285347.2
C1 154025156 154025255
refGene_gn=FAP;refGene_tx=XM_011285347.2;refGene_gn=FAP;refGene_tx=XM_019838169.1
C1 154042273 154042358
refGene_gn=FAP;refGene_tx=XM_011285347.2
C1 154042687 154042693
refGene_gn=FAP;refGene_tx=XM_011285347.2
C1 154070924 154071104
refGene_gn=IFIH1;refGene_tx=XM_006935321.4;refGene_gn=IFIH1;refGene_tx=XM_011285346.3;refGene_gn=IFIH1;refGene_tx=XM_019838171.2
C1 154071202 154071296
refGene_gn=IFIH1;refGene_tx=XM_006935321.4;refGene_gn=IFIH1;refGene_tx=XM_011285346.3;refGene_gn=IFIH1;refGene_tx=XM_019838171.2
C1 154071809 154072000
refGene_gn=IFIH1;refGene_tx=XM_006935321.4;refGene_gn=IFIH1;refGene_tx

=XM_011285346.3;refGene_gn=IFIH1;refGene_tx=XM_019838171.2
C1 154075474 154075636
refGene_gn=IFIH1;refGene_tx=XM_006935321.4;refGene_gn=IFIH1;refGene_tx=XM_011285346.3;refGene_gn=IFIH1;refGene_tx=XM_019838171.2
C1 154077310 154077460
refGene_gn=IFIH1;refGene_tx=XM_006935321.4;refGene_gn=IFIH1;refGene_tx=XM_011285346.3
C1 154080053 154080313
refGene_gn=IFIH1;refGene_tx=XM_006935321.4;refGene_gn=IFIH1;refGene_tx=XM_011285346.3;refGene_gn=IFIH1;refGene_tx=XM_019838171.2
C1 154080902 154081181
refGene_gn=IFIH1;refGene_tx=XM_006935321.4;refGene_gn=IFIH1;refGene_tx=XM_011285346.3;refGene_gn=IFIH1;refGene_tx=XM_019838171.2
C1 154081687 154081811
refGene_gn=IFIH1;refGene_tx=XM_006935321.4;refGene_gn=IFIH1;refGene_tx=XM_011285346.3;refGene_gn=IFIH1;refGene_tx=XM_019838171.2
C1 154084012 154084129
refGene_gn=IFIH1;refGene_tx=XM_006935321.4;refGene_gn=IFIH1;refGene_tx=XM_019838171.2
C1 154085656 154085874
refGene_gn=IFIH1;refGene_tx=XM_006935321.4;refGene_gn=IFIH1;refGene_tx=XM_011285346.3;refGene_gn=IFIH1;refGene_tx=XM_019838171.2
C1 154086769 154086980
refGene_gn=IFIH1;refGene_tx=XM_006935321.4;refGene_gn=IFIH1;refGene_tx=XM_011285346.3;refGene_gn=IFIH1;refGene_tx=XM_019838171.2
C1 154093022 154093243
refGene_gn=IFIH1;refGene_tx=XM_006935321.4;refGene_gn=IFIH1;refGene_tx=XM_011285346.3;refGene_gn=IFIH1;refGene_tx=XM_019838171.2
C1 154098445 154098550
refGene_gn=IFIH1;refGene_tx=XM_006935321.4;refGene_gn=IFIH1;refGene_tx=XM_011285346.3;refGene_gn=IFIH1;refGene_tx=XM_019838171.2
C1 154112296 154112443
refGene_gn=IFIH1;refGene_tx=XM_006935321.4;refGene_gn=IFIH1;refGene_tx=XM_011285346.3;refGene_gn=IFIH1;refGene_tx=XM_019838171.2
C1 154116173 154116342
refGene_gn=IFIH1;refGene_tx=XM_006935321.4;refGene_gn=IFIH1;refGene_tx=XM_011285346.3;refGene_gn=IFIH1;refGene_tx=XM_019838171.2
C1 154125213 154125666
refGene_gn=IFIH1;refGene_tx=XM_006935321.4;refGene_gn=IFIH1;refGene_tx=XM_011285346.3;refGene_gn=IFIH1;refGene_tx=XM_019838171.2
C1 154141038 154141158
refGene_gn=GCA;refGene_tx=XM_019838173.2
C1 154147770 154147797
refGene_gn=GCA;refGene_tx=XM_003990808.5
C1 154149864 154150029
refGene_gn=GCA;refGene_tx=XM_003990808.5;refGene_gn=GCA;refGene_tx=XM_019838173.2
C1 154153829 154153899
refGene_gn=GCA;refGene_tx=XM_003990808.5;refGene_gn=GCA;refGene_tx=XM_019838173.2

C1 154160410 154160454
refGene_gn=GCA;refGene_tx=XM_003990808.5;refGene_gn=GCA;refGene_tx=XM_019838173.2
C1 154160739 154160887
refGene_gn=GCA;refGene_tx=XM_003990808.5;refGene_gn=GCA;refGene_tx=XM_019838173.2
C1 154165297 154165411
refGene_gn=GCA;refGene_tx=XM_003990808.5;refGene_gn=GCA;refGene_tx=XM_019838173.2
C1 154171255 154171314
refGene_gn=GCA;refGene_tx=XM_003990808.5;refGene_gn=GCA;refGene_tx=XM_019838173.2
C1 154171949 154171976
refGene_gn=GCA;refGene_tx=XM_003990808.5;refGene_gn=GCA;refGene_tx=XM_019838173.2
C1 154183434 154183701
refGene_gn=KCNH7;refGene_tx=XM_023259392.1;refGene_gn=KCNH7;refGene_tx=XM_023259393.1;refGene_gn=KCNH7;refGene_tx=XM_023259394.1;refGene_gn=KCNH7;refGene_tx=XM_023259395.1;refGene_gn=KCNH7;refGene_tx=XM_023259396.1;refGene_gn=KCNH7;refGene_tx=XM_023259397.1
C1 154185784 154185977
refGene_gn=KCNH7;refGene_tx=XM_023259392.1;refGene_gn=KCNH7;refGene_tx=XM_023259393.1;refGene_gn=KCNH7;refGene_tx=XM_023259394.1;refGene_gn=KCNH7;refGene_tx=XM_023259395.1;refGene_gn=KCNH7;refGene_tx=XM_023259396.1;refGene_gn=KCNH7;refGene_tx=XM_023259397.1
C1 154193819 154193988
refGene_gn=KCNH7;refGene_tx=XM_023259392.1;refGene_gn=KCNH7;refGene_tx=XM_023259393.1;refGene_gn=KCNH7;refGene_tx=XM_023259394.1;refGene_gn=KCNH7;refGene_tx=XM_023259395.1;refGene_gn=KCNH7;refGene_tx=XM_023259396.1;refGene_gn=KCNH7;refGene_tx=XM_023259397.1
C1 154199730 154199994
refGene_gn=KCNH7;refGene_tx=XM_023259392.1;refGene_gn=KCNH7;refGene_tx=XM_023259393.1;refGene_gn=KCNH7;refGene_tx=XM_023259394.1;refGene_gn=KCNH7;refGene_tx=XM_023259395.1;refGene_gn=KCNH7;refGene_tx=XM_023259396.1
C1 154208797 154208897
refGene_gn=KCNH7;refGene_tx=XM_023259392.1;refGene_gn=KCNH7;refGene_tx=XM_023259393.1;refGene_gn=KCNH7;refGene_tx=XM_023259394.1;refGene_gn=KCNH7;refGene_tx=XM_023259395.1;refGene_gn=KCNH7;refGene_tx=XM_023259396.1;refGene_gn=KCNH7;refGene_tx=XM_023259397.1
C1 154211440 154211646
refGene_gn=KCNH7;refGene_tx=XM_023259392.1;refGene_gn=KCNH7;refGene_tx=XM_023259393.1;refGene_gn=KCNH7;refGene_tx=XM_023259394.1;refGene_gn=KCNH7;refGene_tx=XM_023259395.1;refGene_gn=KCNH7;refGene_tx=XM_023259396.1;refGene_gn=KCNH7;refGene_tx=XM_023259397.1
C1 154214582 154214835
refGene_gn=KCNH7;refGene_tx=XM_023259392.1;refGene_gn=KCNH7;refGene_tx=XM_023259393.1;refGene_gn=KCNH7;refGene_tx=XM_023259394.1;refGene_gn=KCNH7;refGene_tx=XM_023259395.1;refGene_gn=KCNH7;refGene_tx=XM_023259396.1;refGene_gn=KCNH7;refGene_tx=XM_023259397.1

C1 154233280 154233480
refGene_gn=KCNH7;refGene_tx=XM_023259392.1;refGene_gn=KCNH7;refGene_tx
=XM_023259393.1;refGene_gn=KCNH7;refGene_tx=XM_023259394.1;refGene_gn=
KCNH7;refGene_tx=XM_023259395.1;refGene_gn=KCNH7;refGene_tx=XM_0232593
96.1;refGene_gn=KCNH7;refGene_tx=XM_023259397.1
C1 154240515 154240915
refGene_gn=KCNH7;refGene_tx=XM_023259392.1;refGene_gn=KCNH7;refGene_tx
=XM_023259393.1;refGene_gn=KCNH7;refGene_tx=XM_023259394.1;refGene_gn=
KCNH7;refGene_tx=XM_023259395.1;refGene_gn=KCNH7;refGene_tx=XM_0232593
96.1;refGene_gn=KCNH7;refGene_tx=XM_023259397.1
C1 154254789 154255215
refGene_gn=KCNH7;refGene_tx=XM_023259392.1;refGene_gn=KCNH7;refGene_tx
=XM_023259393.1;refGene_gn=KCNH7;refGene_tx=XM_023259394.1;refGene_gn=
KCNH7;refGene_tx=XM_023259395.1;refGene_gn=KCNH7;refGene_tx=XM_0232593
96.1;refGene_gn=KCNH7;refGene_tx=XM_023259397.1
C1 154301833 154302048
refGene_gn=KCNH7;refGene_tx=XM_023259392.1;refGene_gn=KCNH7;refGene_tx
=XM_023259393.1;refGene_gn=KCNH7;refGene_tx=XM_023259394.1;refGene_gn=
KCNH7;refGene_tx=XM_023259395.1;refGene_gn=KCNH7;refGene_tx=XM_0232593
96.1;refGene_gn=KCNH7;refGene_tx=XM_023259397.1
C1 154309651 154309675
refGene_gn=KCNH7;refGene_tx=XM_023259392.1;refGene_gn=KCNH7;refGene_tx
=XM_023259393.1;refGene_gn=KCNH7;refGene_tx=XM_023259394.1;refGene_gn=
KCNH7;refGene_tx=XM_023259397.1
C1 154310099 154310120
refGene_gn=KCNH7;refGene_tx=XM_023259392.1;refGene_gn=KCNH7;refGene_tx
=XM_023259393.1;refGene_gn=KCNH7;refGene_tx=XM_023259395.1;refGene_gn=
KCNH7;refGene_tx=XM_023259397.1
C1 154314864 154315293
refGene_gn=KCNH7;refGene_tx=XM_023259392.1;refGene_gn=KCNH7;refGene_tx
=XM_023259393.1;refGene_gn=KCNH7;refGene_tx=XM_023259394.1;refGene_gn=
KCNH7;refGene_tx=XM_023259395.1;refGene_gn=KCNH7;refGene_tx=XM_0232593
96.1;refGene_gn=KCNH7;refGene_tx=XM_023259397.1
C1 154335089 154335245
refGene_gn=KCNH7;refGene_tx=XM_023259392.1;refGene_gn=KCNH7;refGene_tx
=XM_023259393.1;refGene_gn=KCNH7;refGene_tx=XM_023259394.1;refGene_gn=
KCNH7;refGene_tx=XM_023259395.1;refGene_gn=KCNH7;refGene_tx=XM_0232593
96.1;refGene_gn=KCNH7;refGene_tx=XM_023259397.1
C1 154660993 154661224
refGene_gn=KCNH7;refGene_tx=XM_023259392.1;refGene_gn=KCNH7;refGene_tx
=XM_023259393.1;refGene_gn=KCNH7;refGene_tx=XM_023259394.1;refGene_gn=
KCNH7;refGene_tx=XM_023259395.1;refGene_gn=KCNH7;refGene_tx=XM_0232593
96.1;refGene_gn=KCNH7;refGene_tx=XM_023259397.1
C1 154662893 154662969
refGene_gn=KCNH7;refGene_tx=XM_023259392.1;refGene_gn=KCNH7;refGene_tx
=XM_023259393.1;refGene_gn=KCNH7;refGene_tx=XM_023259394.1;refGene_gn=
KCNH7;refGene_tx=XM_023259395.1;refGene_gn=KCNH7;refGene_tx=XM_0232593
96.1;refGene_gn=KCNH7;refGene_tx=XM_023259397.1
C1 155352776 155355031
refGene_gn=FIGN;refGene_tx=XM_003990813.5;refGene_gn=FIGN;refGene_tx=X

M_006935322.4;refGene_gn=FIGN;refGene_tx=XM_019838179.2
C1 155480783 155480808
refGene_gn=FIGN;refGene_tx=XM_003990813.5
C1 156186600 156186747
refGene_gn=GRB14;refGene_tx=XM_006935323.3;refGene_gn=GRB14;refGene_tx=XM_023259398.1;refGene_gn=GRB14;refGene_tx=XM_023259399.1
C1 156188097 156188191
refGene_gn=GRB14;refGene_tx=XM_006935323.3;refGene_gn=GRB14;refGene_tx=XM_023259398.1;refGene_gn=GRB14;refGene_tx=XM_023259399.1
C1 156190379 156190467
refGene_gn=GRB14;refGene_tx=XM_006935323.3;refGene_gn=GRB14;refGene_tx=XM_023259398.1;refGene_gn=GRB14;refGene_tx=XM_023259399.1
C1 156190624 156190697
refGene_gn=GRB14;refGene_tx=XM_006935323.3;refGene_gn=GRB14;refGene_tx=XM_023259398.1;refGene_gn=GRB14;refGene_tx=XM_023259399.1
C1 156190788 156190905
refGene_gn=GRB14;refGene_tx=XM_006935323.3;refGene_gn=GRB14;refGene_tx=XM_023259398.1;refGene_gn=GRB14;refGene_tx=XM_023259399.1
C1 156197020 156197101
refGene_gn=GRB14;refGene_tx=XM_006935323.3;refGene_gn=GRB14;refGene_tx=XM_023259398.1;refGene_gn=GRB14;refGene_tx=XM_023259399.1
C1 156203223 156203319
refGene_gn=GRB14;refGene_tx=XM_006935323.3;refGene_gn=GRB14;refGene_tx=XM_023259398.1
C1 156203510 156203621
refGene_gn=GRB14;refGene_tx=XM_006935323.3;refGene_gn=GRB14;refGene_tx=XM_023259398.1;refGene_gn=GRB14;refGene_tx=XM_023259399.1
C1 156208786 156208924
refGene_gn=GRB14;refGene_tx=XM_006935323.3;refGene_gn=GRB14;refGene_tx=XM_023259398.1;refGene_gn=GRB14;refGene_tx=XM_023259399.1
C1 156211828 156211903
refGene_gn=GRB14;refGene_tx=XM_006935323.3;refGene_gn=GRB14;refGene_tx=XM_023259398.1;refGene_gn=GRB14;refGene_tx=XM_023259399.1
C1 156213824 156213946
refGene_gn=GRB14;refGene_tx=XM_006935323.3;refGene_gn=GRB14;refGene_tx=XM_023259398.1;refGene_gn=GRB14;refGene_tx=XM_023259399.1
C1 156230311 156230468
refGene_gn=GRB14;refGene_tx=XM_006935323.3;refGene_gn=GRB14;refGene_tx=XM_023259398.1;refGene_gn=GRB14;refGene_tx=XM_023259399.1
C1 156249624 156249687
refGene_gn=GRB14;refGene_tx=XM_006935323.3
C1 156302691 156302824
refGene_gn=GRB14;refGene_tx=XM_023259398.1;refGene_gn=GRB14;refGene_tx=XM_023259399.1
C1 156304112 156304303
refGene_gn=GRB14;refGene_tx=XM_023259398.1;refGene_gn=GRB14;refGene_tx=XM_023259399.1
C1 156363598 156363685
refGene_gn=COBLL1;refGene_tx=XM_019838187.2;refGene_gn=COBLL1;refGene_tx=XM_019838188.1;refGene_gn=COBLL1;refGene_tx=XM_019838190.2;refGene_

gn=COBLL1;refGene_tx=XM_023259400.1;refGene_gn=COBLL1;refGene_tx=XM_02
3259401.1;refGene_gn=COBLL1;refGene_tx=XM_023259402.1;refGene_gn=COBLL
1;refGene_tx=XM_023259403.1;refGene_gn=COBLL1;refGene_tx=XM_023259404.
1;refGene_gn=COBLL1;refGene_tx=XM_023259405.1;refGene_gn=COBLL1;refGen
e_tx=XM_023259406.1;refGene_gn=COBLL1;refGene_tx=XM_023259407.1
C1 156369556 156369733
refGene_gn=COBLL1;refGene_tx=XM_019838187.2;refGene_gn=COBLL1;refGene_
tx=XM_019838188.1;refGene_gn=COBLL1;refGene_tx=XM_019838190.2;refGene_
gn=COBLL1;refGene_tx=XM_023259400.1;refGene_gn=COBLL1;refGene_tx=XM_02
3259401.1;refGene_gn=COBLL1;refGene_tx=XM_023259402.1;refGene_gn=COBLL
1;refGene_tx=XM_023259403.1;refGene_gn=COBLL1;refGene_tx=XM_023259404.
1;refGene_gn=COBLL1;refGene_tx=XM_023259405.1;refGene_gn=COBLL1;refGen
e_tx=XM_023259407.1
C1 156371862 156373433
refGene_gn=COBLL1;refGene_tx=XM_019838187.2;refGene_gn=COBLL1;refGene_
tx=XM_019838188.1;refGene_gn=COBLL1;refGene_tx=XM_019838190.2;refGene_
gn=COBLL1;refGene_tx=XM_023259400.1;refGene_gn=COBLL1;refGene_tx=XM_02
3259401.1;refGene_gn=COBLL1;refGene_tx=XM_023259402.1;refGene_gn=COBLL
1;refGene_tx=XM_023259403.1;refGene_gn=COBLL1;refGene_tx=XM_023259404.
1;refGene_gn=COBLL1;refGene_tx=XM_023259405.1;refGene_gn=COBLL1;refGen
e_tx=XM_023259406.1;refGene_gn=COBLL1;refGene_tx=XM_023259407.1
C1 156378446 156378541
refGene_gn=COBLL1;refGene_tx=XM_019838187.2;refGene_gn=COBLL1;refGene_
tx=XM_019838188.1;refGene_gn=COBLL1;refGene_tx=XM_019838190.2;refGene_
gn=COBLL1;refGene_tx=XM_023259400.1;refGene_gn=COBLL1;refGene_tx=XM_02
3259401.1;refGene_gn=COBLL1;refGene_tx=XM_023259402.1;refGene_gn=COBLL
1;refGene_tx=XM_023259403.1;refGene_gn=COBLL1;refGene_tx=XM_023259404.
1;refGene_gn=COBLL1;refGene_tx=XM_023259405.1;refGene_gn=COBLL1;refGen
e_tx=XM_023259406.1;refGene_gn=COBLL1;refGene_tx=XM_023259407.1
C1 156379760 156379995
refGene_gn=COBLL1;refGene_tx=XM_019838187.2;refGene_gn=COBLL1;refGene_
tx=XM_019838188.1;refGene_gn=COBLL1;refGene_tx=XM_019838190.2;refGene_
gn=COBLL1;refGene_tx=XM_023259400.1;refGene_gn=COBLL1;refGene_tx=XM_02
3259401.1;refGene_gn=COBLL1;refGene_tx=XM_023259402.1;refGene_gn=COBLL
1;refGene_tx=XM_023259403.1;refGene_gn=COBLL1;refGene_tx=XM_023259404.
1;refGene_gn=COBLL1;refGene_tx=XM_023259405.1;refGene_gn=COBLL1;refGen
e_tx=XM_023259406.1;refGene_gn=COBLL1;refGene_tx=XM_023259407.1
C1 156384802 156384940
refGene_gn=COBLL1;refGene_tx=XM_019838187.2;refGene_gn=COBLL1;refGene_
tx=XM_019838188.1;refGene_gn=COBLL1;refGene_tx=XM_019838190.2;refGene_
gn=COBLL1;refGene_tx=XM_023259400.1;refGene_gn=COBLL1;refGene_tx=XM_02
3259401.1;refGene_gn=COBLL1;refGene_tx=XM_023259402.1;refGene_gn=COBLL
1;refGene_tx=XM_023259403.1;refGene_gn=COBLL1;refGene_tx=XM_023259406.
1;refGene_gn=COBLL1;refGene_tx=XM_023259407.1
C1 156386119 156386194
refGene_gn=COBLL1;refGene_tx=XM_019838187.2;refGene_gn=COBLL1;refGene_
tx=XM_019838188.1;refGene_gn=COBLL1;refGene_tx=XM_019838190.2;refGene_
gn=COBLL1;refGene_tx=XM_023259400.1;refGene_gn=COBLL1;refGene_tx=XM_02
3259401.1;refGene_gn=COBLL1;refGene_tx=XM_023259402.1;refGene_gn=COBLL
1;refGene_tx=XM_023259403.1;refGene_gn=COBLL1;refGene_tx=XM_023259404.

1;refGene_gn=COBLL1;refGene_tx=XM_023259405.1;refGene_gn=COBLL1;refGene_tx=XM_023259406.1;refGene_gn=COBLL1;refGene_tx=XM_023259407.1
C1 156392666 156392820
refGene_gn=COBLL1;refGene_tx=XM_019838187.2;refGene_gn=COBLL1;refGene_tx=XM_019838188.1;refGene_gn=COBLL1;refGene_tx=XM_019838190.2;refGene_gn=COBLL1;refGene_tx=XM_023259400.1;refGene_gn=COBLL1;refGene_tx=XM_023259401.1;refGene_gn=COBLL1;refGene_tx=XM_023259402.1;refGene_gn=COBLL1;refGene_tx=XM_023259403.1;refGene_gn=COBLL1;refGene_tx=XM_023259404.1;refGene_gn=COBLL1;refGene_tx=XM_023259405.1;refGene_gn=COBLL1;refGene_tx=XM_023259406.1;refGene_gn=COBLL1;refGene_tx=XM_023259407.1
C1 156412337 156412574
refGene_gn=COBLL1;refGene_tx=XM_019838187.2;refGene_gn=COBLL1;refGene_tx=XM_019838188.1;refGene_gn=COBLL1;refGene_tx=XM_019838190.2;refGene_gn=COBLL1;refGene_tx=XM_023259400.1;refGene_gn=COBLL1;refGene_tx=XM_023259401.1;refGene_gn=COBLL1;refGene_tx=XM_023259402.1;refGene_gn=COBLL1;refGene_tx=XM_023259403.1;refGene_gn=COBLL1;refGene_tx=XM_023259404.1;refGene_gn=COBLL1;refGene_tx=XM_023259405.1;refGene_gn=COBLL1;refGene_tx=XM_023259406.1;refGene_gn=COBLL1;refGene_tx=XM_023259407.1
C1 156412686 156412784
refGene_gn=COBLL1;refGene_tx=XM_019838187.2;refGene_gn=COBLL1;refGene_tx=XM_019838188.1;refGene_gn=COBLL1;refGene_tx=XM_019838190.2;refGene_gn=COBLL1;refGene_tx=XM_023259400.1;refGene_gn=COBLL1;refGene_tx=XM_023259401.1;refGene_gn=COBLL1;refGene_tx=XM_023259402.1;refGene_gn=COBLL1;refGene_tx=XM_023259403.1;refGene_gn=COBLL1;refGene_tx=XM_023259404.1;refGene_gn=COBLL1;refGene_tx=XM_023259405.1;refGene_gn=COBLL1;refGene_tx=XM_023259406.1;refGene_gn=COBLL1;refGene_tx=XM_023259407.1
C1 156417217 156417256
refGene_gn=COBLL1;refGene_tx=XM_019838187.2;refGene_gn=COBLL1;refGene_tx=XM_019838188.1;refGene_gn=COBLL1;refGene_tx=XM_019838190.2;refGene_gn=COBLL1;refGene_tx=XM_023259400.1;refGene_gn=COBLL1;refGene_tx=XM_023259401.1;refGene_gn=COBLL1;refGene_tx=XM_023259402.1;refGene_gn=COBLL1;refGene_tx=XM_023259404.1;refGene_gn=COBLL1;refGene_tx=XM_023259406.1
C1 156418047 156418276
refGene_gn=COBLL1;refGene_tx=XM_019838187.2;refGene_gn=COBLL1;refGene_tx=XM_019838188.1;refGene_gn=COBLL1;refGene_tx=XM_019838190.2;refGene_gn=COBLL1;refGene_tx=XM_023259400.1;refGene_gn=COBLL1;refGene_tx=XM_023259401.1;refGene_gn=COBLL1;refGene_tx=XM_023259402.1;refGene_gn=COBLL1;refGene_tx=XM_023259403.1;refGene_gn=COBLL1;refGene_tx=XM_023259404.1;refGene_gn=COBLL1;refGene_tx=XM_023259405.1;refGene_gn=COBLL1;refGene_tx=XM_023259406.1;refGene_gn=COBLL1;refGene_tx=XM_023259407.1
C1 156420108 156420310
refGene_gn=COBLL1;refGene_tx=XM_019838187.2;refGene_gn=COBLL1;refGene_tx=XM_019838188.1;refGene_gn=COBLL1;refGene_tx=XM_019838190.2;refGene_gn=COBLL1;refGene_tx=XM_023259400.1;refGene_gn=COBLL1;refGene_tx=XM_023259401.1;refGene_gn=COBLL1;refGene_tx=XM_023259402.1;refGene_gn=COBLL1;refGene_tx=XM_023259403.1;refGene_gn=COBLL1;refGene_tx=XM_023259404.1;refGene_gn=COBLL1;refGene_tx=XM_023259405.1;refGene_gn=COBLL1;refGene_tx=XM_023259406.1;refGene_gn=COBLL1;refGene_tx=XM_023259407.1
C1 156434475 156434664

refGene_gn=COBLL1;refGene_tx=XM_019838187.2;refGene_gn=COBLL1;refGene_tx=XM_019838190.2;refGene_gn=COBLL1;refGene_tx=XM_023259400.1;refGene_gn=COBLL1;refGene_tx=XM_023259401.1;refGene_gn=COBLL1;refGene_tx=XM_023259402.1;refGene_gn=COBLL1;refGene_tx=XM_023259403.1;refGene_gn=COBLL1;refGene_tx=XM_023259404.1;refGene_gn=COBLL1;refGene_tx=XM_023259405.1;refGene_gn=COBLL1;refGene_tx=XM_023259406.1;refGene_gn=COBLL1;refGene_tx=XM_023259407.1
C1 156544687 156544805
refGene_gn=COBLL1;refGene_tx=XM_023259400.1;refGene_gn=COBLL1;refGene_tx=XM_023259401.1;refGene_gn=COBLL1;refGene_tx=XM_023259402.1;refGene_gn=COBLL1;refGene_tx=XM_023259403.1;refGene_gn=COBLL1;refGene_tx=XM_023259404.1;refGene_gn=COBLL1;refGene_tx=XM_023259405.1;refGene_gn=COBLL1;refGene_tx=XM_023259406.1;refGene_gn=COBLL1;refGene_tx=XM_023259407.1
C1 156545495 156545584
refGene_gn=COBLL1;refGene_tx=XM_019838190.2;refGene_gn=COBLL1;refGene_tx=XM_023259400.1;refGene_gn=COBLL1;refGene_tx=XM_023259401.1;refGene_gn=COBLL1;refGene_tx=XM_023259402.1;refGene_gn=COBLL1;refGene_tx=XM_023259403.1;refGene_gn=COBLL1;refGene_tx=XM_023259404.1;refGene_gn=COBLL1;refGene_tx=XM_023259405.1;refGene_gn=COBLL1;refGene_tx=XM_023259406.1;refGene_gn=COBLL1;refGene_tx=XM_023259407.1
C1 156613263 156613557
refGene_gn=SLC38A11;refGene_tx=XM_006935325.4;refGene_gn=SLC38A11;refGene_tx=XM_019838195.1
C1 156623538 156623670
refGene_gn=SLC38A11;refGene_tx=XM_006935325.4
C1 156626541 156626654
refGene_gn=SLC38A11;refGene_tx=XM_006935325.4;refGene_gn=SLC38A11;refGene_tx=XM_019838195.1
C1 156629906 156630068
refGene_gn=SLC38A11;refGene_tx=XM_006935325.4;refGene_gn=SLC38A11;refGene_tx=XM_019838195.1
C1 156630748 156630819
refGene_gn=SLC38A11;refGene_tx=XM_006935325.4;refGene_gn=SLC38A11;refGene_tx=XM_019838195.1
C1 156650237 156650317
refGene_gn=SLC38A11;refGene_tx=XM_006935325.4;refGene_gn=SLC38A11;refGene_tx=XM_019838195.1
C1 156652508 156652615
refGene_gn=SLC38A11;refGene_tx=XM_006935325.4;refGene_gn=SLC38A11;refGene_tx=XM_019838195.1
C1 156657524 156657590
refGene_gn=SLC38A11;refGene_tx=XM_006935325.4;refGene_gn=SLC38A11;refGene_tx=XM_019838195.1
C1 156659004 156659139
refGene_gn=SLC38A11;refGene_tx=XM_006935325.4;refGene_gn=SLC38A11;refGene_tx=XM_019838195.1
C1 156666114 156666189
refGene_gn=SLC38A11;refGene_tx=XM_006935325.4;refGene_gn=SLC38A11;refGene_tx=XM_019838195.1

C1 156668137 156668255
refGene_gn=SLC38A11;refGene_tx=XM_006935325.4;refGene_gn=SLC38A11;refGene_tx=XM_019838195.1
C1 156668744 156668783
refGene_gn=SLC38A11;refGene_tx=XM_006935325.4;refGene_gn=SLC38A11;refGene_tx=XM_019838195.1
C1 156719179 156719404
refGene_gn=LOC101101656;refGene_tx=XM_019838196.2
C1 156789720 156790916
refGene_gn=SCN3A;refGene_tx=XM_011285345.3;refGene_gn=SCN3A;refGene_tx=XM_019838191.2;refGene_gn=SCN3A;refGene_tx=XM_019838192.2;refGene_gn=SCN3A;refGene_tx=XM_019838194.2
C1 156791431 156791702
refGene_gn=SCN3A;refGene_tx=XM_011285345.3;refGene_gn=SCN3A;refGene_tx=XM_019838191.2;refGene_gn=SCN3A;refGene_tx=XM_019838192.2;refGene_gn=SCN3A;refGene_tx=XM_019838194.2
C1 156793488 156793593
refGene_gn=SCN3A;refGene_tx=XM_011285345.3;refGene_gn=SCN3A;refGene_tx=XM_019838191.2;refGene_gn=SCN3A;refGene_tx=XM_019838192.2;refGene_gn=SCN3A;refGene_tx=XM_019838194.2
C1 156794825 156794963
refGene_gn=SCN3A;refGene_tx=XM_011285345.3;refGene_gn=SCN3A;refGene_tx=XM_019838191.2;refGene_gn=SCN3A;refGene_tx=XM_019838192.2;refGene_gn=SCN3A;refGene_tx=XM_019838194.2
C1 156795771 156795825
refGene_gn=SCN3A;refGene_tx=XM_011285345.3;refGene_gn=SCN3A;refGene_tx=XM_019838191.2;refGene_gn=SCN3A;refGene_tx=XM_019838192.2;refGene_gn=SCN3A;refGene_tx=XM_019838194.2
C1 156797224 156797497
refGene_gn=SCN3A;refGene_tx=XM_011285345.3;refGene_gn=SCN3A;refGene_tx=XM_019838191.2;refGene_gn=SCN3A;refGene_tx=XM_019838192.2;refGene_gn=SCN3A;refGene_tx=XM_019838194.2
C1 156800000 156800123
refGene_gn=SCN3A;refGene_tx=XM_011285345.3;refGene_gn=SCN3A;refGene_tx=XM_019838191.2;refGene_gn=SCN3A;refGene_tx=XM_019838192.2;refGene_gn=SCN3A;refGene_tx=XM_019838194.2
C1 156806602 156806776
refGene_gn=SCN3A;refGene_tx=XM_011285345.3;refGene_gn=SCN3A;refGene_tx=XM_019838191.2;refGene_gn=SCN3A;refGene_tx=XM_019838192.2;refGene_gn=SCN3A;refGene_tx=XM_019838194.2
C1 156807520 156807675
refGene_gn=SCN3A;refGene_tx=XM_011285345.3;refGene_gn=SCN3A;refGene_tx=XM_019838191.2;refGene_gn=SCN3A;refGene_tx=XM_019838192.2;refGene_gn=SCN3A;refGene_tx=XM_019838194.2
C1 156809057 156809178
refGene_gn=SCN3A;refGene_tx=XM_011285345.3;refGene_gn=SCN3A;refGene_tx=XM_019838191.2;refGene_gn=SCN3A;refGene_tx=XM_019838192.2;refGene_gn=SCN3A;refGene_tx=XM_019838194.2
C1 156819842 156820313
refGene_gn=SCN3A;refGene_tx=XM_011285345.3;refGene_gn=SCN3A;refGene_tx

=XM_019838191.2;refGene_gn=SCN3A;refGene_tx=XM_019838192.2;refGene_gn=SCN3A;refGene_tx=XM_019838194.2
C1 156821981 156822338
refGene_gn=SCN3A;refGene_tx=XM_011285345.3;refGene_gn=SCN3A;refGene_tx=XM_019838191.2;refGene_gn=SCN3A;refGene_tx=XM_019838192.2;refGene_gn=SCN3A;refGene_tx=XM_019838194.2
C1 156823518 156823692
refGene_gn=SCN3A;refGene_tx=XM_011285345.3;refGene_gn=SCN3A;refGene_tx=XM_019838191.2;refGene_gn=SCN3A;refGene_tx=XM_019838192.2;refGene_gn=SCN3A;refGene_tx=XM_019838194.2
C1 156831306 156831545
refGene_gn=SCN3A;refGene_tx=XM_011285345.3;refGene_gn=SCN3A;refGene_tx=XM_019838191.2;refGene_gn=SCN3A;refGene_tx=XM_019838192.2;refGene_gn=SCN3A;refGene_tx=XM_019838194.2
C1 156833050 156833183
refGene_gn=SCN3A;refGene_tx=XM_011285345.3;refGene_gn=SCN3A;refGene_tx=XM_019838191.2;refGene_gn=SCN3A;refGene_tx=XM_019838192.2;refGene_gn=SCN3A;refGene_tx=XM_019838194.2
C1 156834195 156834543
refGene_gn=SCN3A;refGene_tx=XM_011285345.3;refGene_gn=SCN3A;refGene_tx=XM_019838191.2;refGene_gn=SCN3A;refGene_tx=XM_019838192.2;refGene_gn=SCN3A;refGene_tx=XM_019838194.2
C1 156840436 156840727
refGene_gn=SCN3A;refGene_tx=XM_011285345.3;refGene_gn=SCN3A;refGene_tx=XM_019838191.2;refGene_gn=SCN3A;refGene_tx=XM_019838192.2;refGene_gn=SCN3A;refGene_tx=XM_019838194.2
C1 156847689 156847896
refGene_gn=SCN3A;refGene_tx=XM_011285345.3;refGene_gn=SCN3A;refGene_tx=XM_019838191.2;refGene_gn=SCN3A;refGene_tx=XM_019838192.2;refGene_gn=SCN3A;refGene_tx=XM_019838194.2
C1 156848930 156849072
refGene_gn=SCN3A;refGene_tx=XM_011285345.3;refGene_gn=SCN3A;refGene_tx=XM_019838191.2;refGene_gn=SCN3A;refGene_tx=XM_019838192.2;refGene_gn=SCN3A;refGene_tx=XM_019838194.2
C1 156856546 156856610
refGene_gn=SCN3A;refGene_tx=XM_011285345.3;refGene_gn=SCN3A;refGene_tx=XM_019838191.2;refGene_gn=SCN3A;refGene_tx=XM_019838192.2;refGene_gn=SCN3A;refGene_tx=XM_019838194.2
C1 156856817 156857090
refGene_gn=SCN3A;refGene_tx=XM_011285345.3;refGene_gn=SCN3A;refGene_tx=XM_019838191.2;refGene_gn=SCN3A;refGene_tx=XM_019838192.2;refGene_gn=SCN3A;refGene_tx=XM_019838194.2
C1 156858149 156858241
refGene_gn=SCN3A;refGene_tx=XM_011285345.3;refGene_gn=SCN3A;refGene_tx=XM_019838191.2;refGene_gn=SCN3A;refGene_tx=XM_019838192.2;refGene_gn=SCN3A;refGene_tx=XM_019838194.2
C1 156858888 156859017
refGene_gn=SCN3A;refGene_tx=XM_011285345.3;refGene_gn=SCN3A;refGene_tx=XM_019838191.2;refGene_gn=SCN3A;refGene_tx=XM_019838192.2;refGene_gn=SCN3A;refGene_tx=XM_019838194.2

C1 156863151 156863241
refGene_gn=SCN3A;refGene_tx=XM_011285345.3;refGene_gn=SCN3A;refGene_tx=XM_019838191.2;refGene_gn=SCN3A;refGene_tx=XM_019838192.2;refGene_gn=SCN3A;refGene_tx=XM_019838194.2
C1 156865836 156865955
refGene_gn=SCN3A;refGene_tx=XM_011285345.3;refGene_gn=SCN3A;refGene_tx=XM_019838191.2;refGene_gn=SCN3A;refGene_tx=XM_019838192.2;refGene_gn=SCN3A;refGene_tx=XM_019838194.2
C1 156871461 156871725
refGene_gn=SCN3A;refGene_tx=XM_011285345.3;refGene_gn=SCN3A;refGene_tx=XM_019838191.2;refGene_gn=SCN3A;refGene_tx=XM_019838192.2;refGene_gn=SCN3A;refGene_tx=XM_019838194.2
C1 156947748 156947774
refGene_gn=LOC101080472;refGene_tx=XM_023259412.1
C1 156998937 156999204
refGene_gn=LOC101080472;refGene_tx=XM_023259408.1;refGene_gn=LOC101080472;refGene_tx=XM_023259409.1;refGene_gn=LOC101080472;refGene_tx=XM_023259410.1;refGene_gn=LOC101080472;refGene_tx=XM_023259411.1;refGene_gn=LOC101080472;refGene_tx=XM_023259412.1
C1 157000106 157000225
refGene_gn=LOC101080472;refGene_tx=XM_023259408.1;refGene_gn=LOC101080472;refGene_tx=XM_023259409.1;refGene_gn=LOC101080472;refGene_tx=XM_023259410.1;refGene_gn=LOC101080472;refGene_tx=XM_023259411.1;refGene_gn=LOC101080472;refGene_tx=XM_023259412.1
C1 157011896 157011986
refGene_gn=LOC101080472;refGene_tx=XM_023259408.1;refGene_gn=LOC101080472;refGene_tx=XM_023259409.1;refGene_gn=LOC101080472;refGene_tx=XM_023259410.1;refGene_gn=LOC101080472;refGene_tx=XM_023259411.1;refGene_gn=LOC101080472;refGene_tx=XM_023259412.1
C1 157012751 157012880
refGene_gn=LOC101080472;refGene_tx=XM_023259408.1;refGene_gn=LOC101080472;refGene_tx=XM_023259409.1;refGene_gn=LOC101080472;refGene_tx=XM_023259410.1;refGene_gn=LOC101080472;refGene_tx=XM_023259411.1;refGene_gn=LOC101080472;refGene_tx=XM_023259412.1
C1 157013263 157013355
refGene_gn=LOC101080472;refGene_tx=XM_023259411.1
C1 157013449 157013541
refGene_gn=LOC101080472;refGene_tx=XM_023259408.1;refGene_gn=LOC101080472;refGene_tx=XM_023259409.1;refGene_gn=LOC101080472;refGene_tx=XM_023259410.1;refGene_gn=LOC101080472;refGene_tx=XM_023259412.1
C1 157014419 157014692
refGene_gn=LOC101080472;refGene_tx=XM_023259408.1;refGene_gn=LOC101080472;refGene_tx=XM_023259409.1;refGene_gn=LOC101080472;refGene_tx=XM_023259410.1;refGene_gn=LOC101080472;refGene_tx=XM_023259411.1;refGene_gn=LOC101080472;refGene_tx=XM_023259412.1;refGene_gn=LOC101080472;refGene_tx=XM_023259413.1
C1 157017582 157017646
refGene_gn=LOC101080472;refGene_tx=XM_023259408.1;refGene_gn=LOC101080472;refGene_tx=XM_023259409.1;refGene_gn=LOC101080472;refGene_tx=XM_023259410.1;refGene_gn=LOC101080472;refGene_tx=XM_023259411.1;refGene_gn

=LOC101080472;refGene_tx=XM_023259412.1;refGene_gn=LOC101080472;refGene_tx=XM_023259413.1
C1 157019664 157019806
refGene_gn=LOC101080472;refGene_tx=XM_023259408.1;refGene_gn=LOC101080472;refGene_tx=XM_023259409.1;refGene_gn=LOC101080472;refGene_tx=XM_023259410.1;refGene_gn=LOC101080472;refGene_tx=XM_023259411.1;refGene_gn=LOC101080472;refGene_tx=XM_023259412.1;refGene_gn=LOC101080472;refGene_tx=XM_023259413.1
C1 157019922 157020129
refGene_gn=LOC101080472;refGene_tx=XM_023259408.1;refGene_gn=LOC101080472;refGene_tx=XM_023259409.1;refGene_gn=LOC101080472;refGene_tx=XM_023259410.1;refGene_gn=LOC101080472;refGene_tx=XM_023259411.1;refGene_gn=LOC101080472;refGene_tx=XM_023259412.1;refGene_gn=LOC101080472;refGene_tx=XM_023259413.1
C1 157021501 157021789
refGene_gn=LOC101080472;refGene_tx=XM_023259408.1;refGene_gn=LOC101080472;refGene_tx=XM_023259409.1;refGene_gn=LOC101080472;refGene_tx=XM_023259410.1;refGene_gn=LOC101080472;refGene_tx=XM_023259411.1;refGene_gn=LOC101080472;refGene_tx=XM_023259412.1;refGene_gn=LOC101080472;refGene_tx=XM_023259413.1
C1 157027401 157027749
refGene_gn=LOC101080472;refGene_tx=XM_023259408.1;refGene_gn=LOC101080472;refGene_tx=XM_023259409.1;refGene_gn=LOC101080472;refGene_tx=XM_023259410.1;refGene_gn=LOC101080472;refGene_tx=XM_023259411.1;refGene_gn=LOC101080472;refGene_tx=XM_023259412.1;refGene_gn=LOC101080472;refGene_tx=XM_023259413.1
C1 157031724 157031857
refGene_gn=LOC101080472;refGene_tx=XM_023259408.1;refGene_gn=LOC101080472;refGene_tx=XM_023259409.1;refGene_gn=LOC101080472;refGene_tx=XM_023259410.1;refGene_gn=LOC101080472;refGene_tx=XM_023259411.1;refGene_gn=LOC101080472;refGene_tx=XM_023259412.1;refGene_gn=LOC101080472;refGene_tx=XM_023259413.1
C1 157036159 157036398
refGene_gn=LOC101080472;refGene_tx=XM_023259408.1;refGene_gn=LOC101080472;refGene_tx=XM_023259409.1;refGene_gn=LOC101080472;refGene_tx=XM_023259410.1;refGene_gn=LOC101080472;refGene_tx=XM_023259411.1;refGene_gn=LOC101080472;refGene_tx=XM_023259412.1;refGene_gn=LOC101080472;refGene_tx=XM_023259413.1;refGene_gn=LOC101080472;refGene_tx=XM_023259414.1
C1 157042868 157043042
refGene_gn=LOC101080472;refGene_tx=XM_023259408.1;refGene_gn=LOC101080472;refGene_tx=XM_023259409.1;refGene_gn=LOC101080472;refGene_tx=XM_023259410.1;refGene_gn=LOC101080472;refGene_tx=XM_023259411.1;refGene_gn=LOC101080472;refGene_tx=XM_023259412.1;refGene_gn=LOC101080472;refGene_tx=XM_023259413.1;refGene_gn=LOC101080472;refGene_tx=XM_023259414.1
C1 157045710 157046067
refGene_gn=LOC101080472;refGene_tx=XM_023259408.1;refGene_gn=LOC101080472;refGene_tx=XM_023259409.1;refGene_gn=LOC101080472;refGene_tx=XM_023259410.1;refGene_gn=LOC101080472;refGene_tx=XM_023259411.1;refGene_gn=LOC101080472;refGene_tx=XM_023259412.1;refGene_gn=LOC101080472;refGene_tx=XM_023259413.1;refGene_gn=LOC101080472;refGene_tx=XM_023259414.1

C1	157050005	157050485
refGene_gn=LOC101080472;refGene_tx=XM_023259408.1;refGene_gn=LOC101080472;refGene_tx=XM_023259409.1;refGene_gn=LOC101080472;refGene_tx=XM_023259410.1;refGene_gn=LOC101080472;refGene_tx=XM_023259411.1;refGene_gn=LOC101080472;refGene_tx=XM_023259412.1;refGene_gn=LOC101080472;refGene_tx=XM_023259413.1;refGene_gn=LOC101080472;refGene_tx=XM_023259414.1
C1	157060320	157060441
refGene_gn=LOC101080472;refGene_tx=XM_023259408.1;refGene_gn=LOC101080472;refGene_tx=XM_023259409.1;refGene_gn=LOC101080472;refGene_tx=XM_023259410.1;refGene_gn=LOC101080472;refGene_tx=XM_023259411.1;refGene_gn=LOC101080472;refGene_tx=XM_023259412.1;refGene_gn=LOC101080472;refGene_tx=XM_023259413.1;refGene_gn=LOC101080472;refGene_tx=XM_023259414.1
C1	157064396	157064551
refGene_gn=LOC101080472;refGene_tx=XM_023259408.1;refGene_gn=LOC101080472;refGene_tx=XM_023259409.1;refGene_gn=LOC101080472;refGene_tx=XM_023259410.1;refGene_gn=LOC101080472;refGene_tx=XM_023259411.1;refGene_gn=LOC101080472;refGene_tx=XM_023259412.1;refGene_gn=LOC101080472;refGene_tx=XM_023259413.1;refGene_gn=LOC101080472;refGene_tx=XM_023259414.1
C1	157066138	157066312
refGene_gn=LOC101080472;refGene_tx=XM_023259408.1;refGene_gn=LOC101080472;refGene_tx=XM_023259409.1;refGene_gn=LOC101080472;refGene_tx=XM_023259410.1;refGene_gn=LOC101080472;refGene_tx=XM_023259411.1;refGene_gn=LOC101080472;refGene_tx=XM_023259412.1;refGene_gn=LOC101080472;refGene_tx=XM_023259413.1;refGene_gn=LOC101080472;refGene_tx=XM_023259414.1
C1	157070773	157070896
refGene_gn=LOC101080472;refGene_tx=XM_023259408.1;refGene_gn=LOC101080472;refGene_tx=XM_023259409.1;refGene_gn=LOC101080472;refGene_tx=XM_023259410.1;refGene_gn=LOC101080472;refGene_tx=XM_023259411.1;refGene_gn=LOC101080472;refGene_tx=XM_023259412.1;refGene_gn=LOC101080472;refGene_tx=XM_023259413.1;refGene_gn=LOC101080472;refGene_tx=XM_023259414.1
C1	157073182	157073464
refGene_gn=LOC101080472;refGene_tx=XM_023259408.1;refGene_gn=LOC101080472;refGene_tx=XM_023259409.1;refGene_gn=LOC101080472;refGene_tx=XM_023259410.1;refGene_gn=LOC101080472;refGene_tx=XM_023259411.1;refGene_gn=LOC101080472;refGene_tx=XM_023259412.1;refGene_gn=LOC101080472;refGene_tx=XM_023259413.1;refGene_gn=LOC101080472;refGene_tx=XM_023259414.1
C1	157074730	157074784
refGene_gn=LOC101080472;refGene_tx=XM_023259408.1;refGene_gn=LOC101080472;refGene_tx=XM_023259409.1;refGene_gn=LOC101080472;refGene_tx=XM_023259410.1;refGene_gn=LOC101080472;refGene_tx=XM_023259411.1;refGene_gn=LOC101080472;refGene_tx=XM_023259412.1;refGene_gn=LOC101080472;refGene_tx=XM_023259413.1;refGene_gn=LOC101080472;refGene_tx=XM_023259414.1
C1	157077814	157077952
refGene_gn=LOC101080472;refGene_tx=XM_023259408.1;refGene_gn=LOC101080472;refGene_tx=XM_023259409.1;refGene_gn=LOC101080472;refGene_tx=XM_023259410.1;refGene_gn=LOC101080472;refGene_tx=XM_023259411.1;refGene_gn=LOC101080472;refGene_tx=XM_023259412.1;refGene_gn=LOC101080472;refGene_tx=XM_023259413.1;refGene_gn=LOC101080472;refGene_tx=XM_023259414.1
C1	157078319	157078424
refGene_gn=LOC101080472;refGene_tx=XM_023259408.1;refGene_gn=LOC101080

472;refGene_tx=XM_023259409.1;refGene_gn=LOC101080472;refGene_tx=XM_02
3259410.1;refGene_gn=LOC101080472;refGene_tx=XM_023259411.1;refGene_gn
=LOC101080472;refGene_tx=XM_023259412.1;refGene_gn=LOC101080472;refGen
e_tx=XM_023259413.1;refGene_gn=LOC101080472;refGene_tx=XM_023259414.1
C1 157081749 157082020
refGene_gn=LOC101080472;refGene_tx=XM_023259408.1;refGene_gn=LOC101080
472;refGene_tx=XM_023259409.1;refGene_gn=LOC101080472;refGene_tx=XM_02
3259410.1;refGene_gn=LOC101080472;refGene_tx=XM_023259411.1;refGene_gn
=LOC101080472;refGene_tx=XM_023259412.1;refGene_gn=LOC101080472;refGen
e_tx=XM_023259413.1;refGene_gn=LOC101080472;refGene_tx=XM_023259414.1
C1 157083628 157084824
refGene_gn=LOC101080472;refGene_tx=XM_023259408.1;refGene_gn=LOC101080
472;refGene_tx=XM_023259409.1;refGene_gn=LOC101080472;refGene_tx=XM_02
3259410.1;refGene_gn=LOC101080472;refGene_tx=XM_023259411.1;refGene_gn
=LOC101080472;refGene_tx=XM_023259412.1;refGene_gn=LOC101080472;refGen
e_tx=XM_023259413.1;refGene_gn=LOC101080472;refGene_tx=XM_023259414.1
C1 157210392 157210405
refGene_gn=CSRNP3;refGene_tx=XM_023259415.1
C1 157290233 157290404
refGene_gn=CSRNP3;refGene_tx=XM_006935332.4;refGene_gn=CSRNP3;refGene_
tx=XM_019838201.2;refGene_gn=CSRNP3;refGene_tx=XM_023259415.1
C1 157348025 157348285
refGene_gn=CSRNP3;refGene_tx=XM_006935332.4;refGene_gn=CSRNP3;refGene_
tx=XM_019838201.2;refGene_gn=CSRNP3;refGene_tx=XM_023259415.1
C1 157362975 157363272
refGene_gn=CSRNP3;refGene_tx=XM_006935332.4;refGene_gn=CSRNP3;refGene_
tx=XM_019838201.2;refGene_gn=CSRNP3;refGene_tx=XM_023259415.1
C1 157365652 157366711
refGene_gn=CSRNP3;refGene_tx=XM_006935332.4;refGene_gn=CSRNP3;refGene_
tx=XM_019838201.2;refGene_gn=CSRNP3;refGene_tx=XM_023259415.1
C1 157412755 157412878
refGene_gn=GALNT3;refGene_tx=XM_019838203.2;refGene_gn=GALNT3;refGene_
tx=XM_023259416.1;refGene_gn=GALNT3;refGene_tx=XM_023259417.1;refGene_
gn=GALNT3;refGene_tx=XM_023259418.1;refGene_gn=GALNT3;refGene_tx=XM_02
3259419.1
C1 157413984 157414137
refGene_gn=GALNT3;refGene_tx=XM_019838203.2;refGene_gn=GALNT3;refGene_
tx=XM_023259416.1;refGene_gn=GALNT3;refGene_tx=XM_023259417.1;refGene_
gn=GALNT3;refGene_tx=XM_023259418.1;refGene_gn=GALNT3;refGene_tx=XM_02
3259419.1
C1 157419706 157419808
refGene_gn=GALNT3;refGene_tx=XM_019838203.2;refGene_gn=GALNT3;refGene_
tx=XM_023259416.1;refGene_gn=GALNT3;refGene_tx=XM_023259417.1;refGene_
gn=GALNT3;refGene_tx=XM_023259418.1;refGene_gn=GALNT3;refGene_tx=XM_02
3259419.1
C1 157420019 157420151
refGene_gn=GALNT3;refGene_tx=XM_019838203.2;refGene_gn=GALNT3;refGene_
tx=XM_023259416.1;refGene_gn=GALNT3;refGene_tx=XM_023259417.1;refGene_
gn=GALNT3;refGene_tx=XM_023259418.1;refGene_gn=GALNT3;refGene_tx=XM_02
3259419.1

C1	157422378	157422579
refGene_gn=GALNT3;refGene_tx=XM_019838203.2;refGene_gn=GALNT3;refGene_tx=XM_023259416.1;refGene_gn=GALNT3;refGene_tx=XM_023259417.1;refGene_gn=GALNT3;refGene_tx=XM_023259418.1;refGene_gn=GALNT3;refGene_tx=XM_023259419.1
C1	157424094	157424212
refGene_gn=GALNT3;refGene_tx=XM_019838203.2;refGene_gn=GALNT3;refGene_tx=XM_023259416.1;refGene_gn=GALNT3;refGene_tx=XM_023259417.1;refGene_gn=GALNT3;refGene_tx=XM_023259418.1;refGene_gn=GALNT3;refGene_tx=XM_023259419.1
C1	157424697	157424932
refGene_gn=GALNT3;refGene_tx=XM_019838203.2;refGene_gn=GALNT3;refGene_tx=XM_023259416.1;refGene_gn=GALNT3;refGene_tx=XM_023259417.1;refGene_gn=GALNT3;refGene_tx=XM_023259418.1;refGene_gn=GALNT3;refGene_tx=XM_023259419.1
C1	157426824	157426974
refGene_gn=GALNT3;refGene_tx=XM_019838203.2;refGene_gn=GALNT3;refGene_tx=XM_023259416.1;refGene_gn=GALNT3;refGene_tx=XM_023259417.1;refGene_gn=GALNT3;refGene_tx=XM_023259418.1;refGene_gn=GALNT3;refGene_tx=XM_023259419.1
C1	157431309	157431482
refGene_gn=GALNT3;refGene_tx=XM_019838203.2;refGene_gn=GALNT3;refGene_tx=XM_023259416.1;refGene_gn=GALNT3;refGene_tx=XM_023259417.1;refGene_gn=GALNT3;refGene_tx=XM_023259418.1;refGene_gn=GALNT3;refGene_tx=XM_023259419.1
C1	157435572	157436087
refGene_gn=GALNT3;refGene_tx=XM_019838203.2;refGene_gn=GALNT3;refGene_tx=XM_023259416.1;refGene_gn=GALNT3;refGene_tx=XM_023259417.1;refGene_gn=GALNT3;refGene_tx=XM_023259418.1;refGene_gn=GALNT3;refGene_tx=XM_023259419.1
C1	157527090	157527168
refGene_gn=TTC21B;refGene_tx=XM_003990824.4
C1	157528573	157528641
refGene_gn=TTC21B;refGene_tx=XM_003990824.4
C1	157532578	157532699
refGene_gn=TTC21B;refGene_tx=XM_003990824.4
C1	157538227	157538452
refGene_gn=TTC21B;refGene_tx=XM_003990824.4
C1	157542486	157542682
refGene_gn=TTC21B;refGene_tx=XM_003990824.4
C1	157544558	157544720
refGene_gn=TTC21B;refGene_tx=XM_003990824.4
C1	157545183	157545334
refGene_gn=TTC21B;refGene_tx=XM_003990824.4
C1	157553092	157553174
refGene_gn=TTC21B;refGene_tx=XM_003990824.4
C1	157554403	157554514
refGene_gn=TTC21B;refGene_tx=XM_003990824.4
C1	157556154	157556343
refGene_gn=TTC21B;refGene_tx=XM_003990824.4

C1 157563237 157563344
refGene_gn=TTC21B;refGene_tx=XM_003990824.4
C1 157569025 157569164
refGene_gn=TTC21B;refGene_tx=XM_003990824.4
C1 157569854 157569965
refGene_gn=TTC21B;refGene_tx=XM_003990824.4
C1 157570661 157570734
refGene_gn=TTC21B;refGene_tx=XM_003990824.4
C1 157572045 157572284
refGene_gn=TTC21B;refGene_tx=XM_003990824.4
C1 157573811 157574036
refGene_gn=TTC21B;refGene_tx=XM_003990824.4
C1 157575790 157575948
refGene_gn=TTC21B;refGene_tx=XM_003990824.4
C1 157579462 157579592
refGene_gn=TTC21B;refGene_tx=XM_003990824.4
C1 157584214 157584415
refGene_gn=TTC21B;refGene_tx=XM_003990824.4
C1 157584966 157585064
refGene_gn=TTC21B;refGene_tx=XM_003990824.4
C1 157586122 157586315
refGene_gn=TTC21B;refGene_tx=XM_003990824.4
C1 157589182 157589281
refGene_gn=TTC21B;refGene_tx=XM_003990824.4
C1 157590621 157590706
refGene_gn=TTC21B;refGene_tx=XM_003990824.4
C1 157595107 157595265
refGene_gn=TTC21B;refGene_tx=XM_003990824.4
C1 157598794 157598917
refGene_gn=TTC21B;refGene_tx=XM_003990824.4
C1 157601206 157601373
refGene_gn=TTC21B;refGene_tx=XM_003990824.4
C1 157604872 157604983
refGene_gn=TTC21B;refGene_tx=XM_003990824.4
C1 157605071 157605201
refGene_gn=TTC21B;refGene_tx=XM_003990824.4
C1 157609200 157609221
refGene_gn=TTC21B;refGene_tx=XM_003990824.4
C1 157629021 157630199
refGene_gn=SCN1A;refGene_tx=XM_003990825.5;refGene_gn=SCN1A;refGene_tx
=XM_019838204.2;refGene_gn=SCN1A;refGene_tx=XM_019838205.2;refGene_gn=
SCN1A;refGene_tx=XM_019838206.2;refGene_gn=SCN1A;refGene_tx=XM_0198382
07.2
C1 157632298 157632569
refGene_gn=SCN1A;refGene_tx=XM_003990825.5;refGene_gn=SCN1A;refGene_tx
=XM_019838204.2;refGene_gn=SCN1A;refGene_tx=XM_019838205.2;refGene_gn=
SCN1A;refGene_tx=XM_019838206.2;refGene_gn=SCN1A;refGene_tx=XM_0198382
07.2
C1 157634163 157634268
refGene_gn=SCN1A;refGene_tx=XM_003990825.5;refGene_gn=SCN1A;refGene_tx

=XM_019838204.2;refGene_gn=SCN1A;refGene_tx=XM_019838205.2;refGene_gn=SCN1A;refGene_tx=XM_019838206.2;refGene_gn=SCN1A;refGene_tx=XM_019838207.2
C1 157636186 157636324
refGene_gn=SCN1A;refGene_tx=XM_003990825.5;refGene_gn=SCN1A;refGene_tx=XM_019838204.2;refGene_gn=SCN1A;refGene_tx=XM_019838205.2;refGene_gn=SCN1A;refGene_tx=XM_019838206.2;refGene_gn=SCN1A;refGene_tx=XM_019838207.2
C1 157637899 157637953
refGene_gn=SCN1A;refGene_tx=XM_003990825.5;refGene_gn=SCN1A;refGene_tx=XM_019838204.2;refGene_gn=SCN1A;refGene_tx=XM_019838205.2;refGene_gn=SCN1A;refGene_tx=XM_019838206.2;refGene_gn=SCN1A;refGene_tx=XM_019838207.2
C1 157640425 157640707
refGene_gn=SCN1A;refGene_tx=XM_003990825.5;refGene_gn=SCN1A;refGene_tx=XM_019838204.2;refGene_gn=SCN1A;refGene_tx=XM_019838205.2;refGene_gn=SCN1A;refGene_tx=XM_019838206.2;refGene_gn=SCN1A;refGene_tx=XM_019838207.2
C1 157647785 157647908
refGene_gn=SCN1A;refGene_tx=XM_003990825.5;refGene_gn=SCN1A;refGene_tx=XM_019838204.2;refGene_gn=SCN1A;refGene_tx=XM_019838205.2;refGene_gn=SCN1A;refGene_tx=XM_019838206.2;refGene_gn=SCN1A;refGene_tx=XM_019838207.2
C1 157650747 157650921
refGene_gn=SCN1A;refGene_tx=XM_003990825.5;refGene_gn=SCN1A;refGene_tx=XM_019838204.2;refGene_gn=SCN1A;refGene_tx=XM_019838205.2;refGene_gn=SCN1A;refGene_tx=XM_019838206.2;refGene_gn=SCN1A;refGene_tx=XM_019838207.2
C1 157652650 157652805
refGene_gn=SCN1A;refGene_tx=XM_003990825.5;refGene_gn=SCN1A;refGene_tx=XM_019838204.2;refGene_gn=SCN1A;refGene_tx=XM_019838205.2;refGene_gn=SCN1A;refGene_tx=XM_019838206.2;refGene_gn=SCN1A;refGene_tx=XM_019838207.2
C1 157655118 157655239
refGene_gn=SCN1A;refGene_tx=XM_003990825.5;refGene_gn=SCN1A;refGene_tx=XM_019838204.2;refGene_gn=SCN1A;refGene_tx=XM_019838205.2;refGene_gn=SCN1A;refGene_tx=XM_019838206.2;refGene_gn=SCN1A;refGene_tx=XM_019838207.2
C1 157674885 157675368
refGene_gn=SCN1A;refGene_tx=XM_003990825.5;refGene_gn=SCN1A;refGene_tx=XM_019838204.2;refGene_gn=SCN1A;refGene_tx=XM_019838205.2;refGene_gn=SCN1A;refGene_tx=XM_019838206.2;refGene_gn=SCN1A;refGene_tx=XM_019838207.2
C1 157676749 157677106
refGene_gn=SCN1A;refGene_tx=XM_003990825.5;refGene_gn=SCN1A;refGene_tx=XM_019838204.2;refGene_gn=SCN1A;refGene_tx=XM_019838205.2;refGene_gn=SCN1A;refGene_tx=XM_019838206.2;refGene_gn=SCN1A;refGene_tx=XM_019838207.2
C1 157678099 157678273
refGene_gn=SCN1A;refGene_tx=XM_003990825.5;refGene_gn=SCN1A;refGene_tx

=XM_019838204.2;refGene_gn=SCN1A;refGene_tx=XM_019838205.2;refGene_gn=SCN1A;refGene_tx=XM_019838206.2;refGene_gn=SCN1A;refGene_tx=XM_019838207.2
C1 157679882 157680121
refGene_gn=SCN1A;refGene_tx=XM_003990825.5;refGene_gn=SCN1A;refGene_tx=XM_019838204.2;refGene_gn=SCN1A;refGene_tx=XM_019838205.2;refGene_gn=SCN1A;refGene_tx=XM_019838206.2;refGene_gn=SCN1A;refGene_tx=XM_019838207.2
C1 157680931 157681061
refGene_gn=SCN1A;refGene_tx=XM_003990825.5;refGene_gn=SCN1A;refGene_tx=XM_019838204.2;refGene_gn=SCN1A;refGene_tx=XM_019838205.2;refGene_gn=SCN1A;refGene_tx=XM_019838206.2;refGene_gn=SCN1A;refGene_tx=XM_019838207.2
C1 157682310 157682691
refGene_gn=SCN1A;refGene_tx=XM_003990825.5;refGene_gn=SCN1A;refGene_tx=XM_019838204.2;refGene_gn=SCN1A;refGene_tx=XM_019838205.2;refGene_gn=SCN1A;refGene_tx=XM_019838206.2;refGene_gn=SCN1A;refGene_tx=XM_019838207.2
C1 157683626 157683911
refGene_gn=SCN1A;refGene_tx=XM_003990825.5;refGene_gn=SCN1A;refGene_tx=XM_019838204.2;refGene_gn=SCN1A;refGene_tx=XM_019838205.2;refGene_gn=SCN1A;refGene_tx=XM_019838206.2;refGene_gn=SCN1A;refGene_tx=XM_019838207.2
C1 157685385 157685592
refGene_gn=SCN1A;refGene_tx=XM_003990825.5;refGene_gn=SCN1A;refGene_tx=XM_019838204.2;refGene_gn=SCN1A;refGene_tx=XM_019838205.2;refGene_gn=SCN1A;refGene_tx=XM_019838206.2;refGene_gn=SCN1A;refGene_tx=XM_019838207.2
C1 157686243 157686385
refGene_gn=SCN1A;refGene_tx=XM_003990825.5;refGene_gn=SCN1A;refGene_tx=XM_019838204.2;refGene_gn=SCN1A;refGene_tx=XM_019838205.2;refGene_gn=SCN1A;refGene_tx=XM_019838206.2;refGene_gn=SCN1A;refGene_tx=XM_019838207.2
C1 157686885 157686949
refGene_gn=SCN1A;refGene_tx=XM_003990825.5;refGene_gn=SCN1A;refGene_tx=XM_019838204.2;refGene_gn=SCN1A;refGene_tx=XM_019838205.2;refGene_gn=SCN1A;refGene_tx=XM_019838206.2;refGene_gn=SCN1A;refGene_tx=XM_019838207.2
C1 157690234 157690504
refGene_gn=SCN1A;refGene_tx=XM_003990825.5;refGene_gn=SCN1A;refGene_tx=XM_019838204.2;refGene_gn=SCN1A;refGene_tx=XM_019838205.2;refGene_gn=SCN1A;refGene_tx=XM_019838206.2;refGene_gn=SCN1A;refGene_tx=XM_019838207.2
C1 157691377 157691469
refGene_gn=SCN1A;refGene_tx=XM_003990825.5;refGene_gn=SCN1A;refGene_tx=XM_019838204.2;refGene_gn=SCN1A;refGene_tx=XM_019838205.2;refGene_gn=SCN1A;refGene_tx=XM_019838206.2;refGene_gn=SCN1A;refGene_tx=XM_019838207.2
C1 157693172 157693301
refGene_gn=SCN1A;refGene_tx=XM_003990825.5;refGene_gn=SCN1A;refGene_tx

=XM_019838204.2;refGene_gn=SCN1A;refGene_tx=XM_019838205.2;refGene_gn=SCN1A;refGene_tx=XM_019838206.2;refGene_gn=SCN1A;refGene_tx=XM_019838207.2
C1 157694906 157694996
refGene_gn=SCN1A;refGene_tx=XM_003990825.5;refGene_gn=SCN1A;refGene_tx=XM_019838204.2;refGene_gn=SCN1A;refGene_tx=XM_019838205.2;refGene_gn=SCN1A;refGene_tx=XM_019838206.2;refGene_gn=SCN1A;refGene_tx=XM_019838207.2
C1 157697125 157697244
refGene_gn=SCN1A;refGene_tx=XM_003990825.5;refGene_gn=SCN1A;refGene_tx=XM_019838204.2;refGene_gn=SCN1A;refGene_tx=XM_019838205.2;refGene_gn=SCN1A;refGene_tx=XM_019838206.2;refGene_gn=SCN1A;refGene_tx=XM_019838207.2
C1 157709426 157709690
refGene_gn=SCN1A;refGene_tx=XM_003990825.5;refGene_gn=SCN1A;refGene_tx=XM_019838204.2;refGene_gn=SCN1A;refGene_tx=XM_019838205.2;refGene_gn=SCN1A;refGene_tx=XM_019838206.2;refGene_gn=SCN1A;refGene_tx=XM_019838207.2
C1 157823200 157824393
refGene_gn=SCN9A;refGene_tx=XM_019838208.2
C1 157827245 157827516
refGene_gn=SCN9A;refGene_tx=XM_019838208.2
C1 157829240 157829345
refGene_gn=SCN9A;refGene_tx=XM_019838208.2
C1 157835410 157835548
refGene_gn=SCN9A;refGene_tx=XM_019838208.2
C1 157836481 157836535
refGene_gn=SCN9A;refGene_tx=XM_019838208.2
C1 157836977 157837259
refGene_gn=SCN9A;refGene_tx=XM_019838208.2
C1 157841166 157841289
refGene_gn=SCN9A;refGene_tx=XM_019838208.2
C1 157844837 157845011
refGene_gn=SCN9A;refGene_tx=XM_019838208.2
C1 157848892 157849047
refGene_gn=SCN9A;refGene_tx=XM_019838208.2
C1 157858419 157858540
refGene_gn=SCN9A;refGene_tx=XM_019838208.2
C1 157881421 157881898
refGene_gn=SCN9A;refGene_tx=XM_019838208.2
C1 157886225 157886582
refGene_gn=SCN9A;refGene_tx=XM_019838208.2
C1 157887993 157888167
refGene_gn=SCN9A;refGene_tx=XM_019838208.2
C1 157890243 157890482
refGene_gn=SCN9A;refGene_tx=XM_019838208.2
C1 157891828 157891958
refGene_gn=SCN9A;refGene_tx=XM_019838208.2
C1 157894882 157895254
refGene_gn=SCN9A;refGene_tx=XM_019838208.2

C1 157897497 157897785
refGene_gn=SCN9A;refGene_tx=XM_019838208.2
C1 157899614 157899821
refGene_gn=SCN9A;refGene_tx=XM_019838208.2
C1 157902571 157902713
refGene_gn=SCN9A;refGene_tx=XM_019838208.2
C1 157904182 157904246
refGene_gn=SCN9A;refGene_tx=XM_019838208.2
C1 157912040 157912250
refGene_gn=SCN9A;refGene_tx=XM_019838208.2
C1 157913230 157913322
refGene_gn=SCN9A;refGene_tx=XM_019838208.2
C1 157914804 157914933
refGene_gn=SCN9A;refGene_tx=XM_019838208.2
C1 157915483 157915573
refGene_gn=SCN9A;refGene_tx=XM_019838208.2
C1 157915922 157916041
refGene_gn=SCN9A;refGene_tx=XM_019838208.2
C1 157923419 157923677
refGene_gn=SCN9A;refGene_tx=XM_019838208.2
C1 158024033 158025100
refGene_gn=SCN7A;refGene_tx=XM_019838209.2;refGene_gn=SCN7A;refGene_tx
=XM_023259420.1;refGene_gn=SCN7A;refGene_tx=XM_023259422.1;refGene_gn=
SCN7A;refGene_tx=XM_023259423.1
C1 158027695 158027966
refGene_gn=SCN7A;refGene_tx=XM_019838209.2;refGene_gn=SCN7A;refGene_tx
=XM_023259420.1;refGene_gn=SCN7A;refGene_tx=XM_023259422.1;refGene_gn=
SCN7A;refGene_tx=XM_023259423.1
C1 158028543 158028648
refGene_gn=SCN7A;refGene_tx=XM_019838209.2;refGene_gn=SCN7A;refGene_tx
=XM_023259420.1;refGene_gn=SCN7A;refGene_tx=XM_023259422.1;refGene_gn=
SCN7A;refGene_tx=XM_023259423.1
C1 158030185 158030323
refGene_gn=SCN7A;refGene_tx=XM_019838209.2;refGene_gn=SCN7A;refGene_tx
=XM_023259420.1;refGene_gn=SCN7A;refGene_tx=XM_023259422.1;refGene_gn=
SCN7A;refGene_tx=XM_023259423.1
C1 158031108 158031162
refGene_gn=SCN7A;refGene_tx=XM_019838209.2;refGene_gn=SCN7A;refGene_tx
=XM_023259420.1;refGene_gn=SCN7A;refGene_tx=XM_023259422.1;refGene_gn=
SCN7A;refGene_tx=XM_023259423.1
C1 158031979 158032258
refGene_gn=SCN7A;refGene_tx=XM_019838209.2;refGene_gn=SCN7A;refGene_tx
=XM_023259420.1;refGene_gn=SCN7A;refGene_tx=XM_023259422.1;refGene_gn=
SCN7A;refGene_tx=XM_023259423.1
C1 158036155 158036263
refGene_gn=SCN7A;refGene_tx=XM_019838209.2;refGene_gn=SCN7A;refGene_tx
=XM_023259420.1;refGene_gn=SCN7A;refGene_tx=XM_023259422.1;refGene_gn=
SCN7A;refGene_tx=XM_023259423.1
C1 158038263 158038437
refGene_gn=SCN7A;refGene_tx=XM_019838209.2;refGene_gn=SCN7A;refGene_tx

=XM_023259420.1;refGene_gn=SCN7A;refGene_tx=XM_023259422.1;refGene_gn=SCN7A;refGene_tx=XM_023259423.1
C1 158041298 158041453 refGene_gn=SCN7A;refGene_tx=XM_019838209.2;refGene_gn=SCN7A;refGene_tx=XM_023259420.1;refGene_gn=SCN7A;refGene_tx=XM_023259422.1;refGene_gn=SCN7A;refGene_tx=XM_023259423.1
C1 158043036 158043139 refGene_gn=SCN7A;refGene_tx=XM_019838209.2;refGene_gn=SCN7A;refGene_tx=XM_023259420.1;refGene_gn=SCN7A;refGene_tx=XM_023259422.1;refGene_gn=SCN7A;refGene_tx=XM_023259423.1
C1 158046231 158046669 refGene_gn=SCN7A;refGene_tx=XM_019838209.2;refGene_gn=SCN7A;refGene_tx=XM_023259420.1;refGene_gn=SCN7A;refGene_tx=XM_023259422.1;refGene_gn=SCN7A;refGene_tx=XM_023259423.1
C1 158051692 158052049 refGene_gn=SCN7A;refGene_tx=XM_019838209.2;refGene_gn=SCN7A;refGene_tx=XM_023259420.1;refGene_gn=SCN7A;refGene_tx=XM_023259422.1;refGene_gn=SCN7A;refGene_tx=XM_023259423.1
C1 158053749 158053923 refGene_gn=SCN7A;refGene_tx=XM_019838209.2;refGene_gn=SCN7A;refGene_tx=XM_023259420.1;refGene_gn=SCN7A;refGene_tx=XM_023259422.1;refGene_gn=SCN7A;refGene_tx=XM_023259423.1
C1 158055227 158055466 refGene_gn=SCN7A;refGene_tx=XM_019838209.2;refGene_gn=SCN7A;refGene_tx=XM_023259420.1;refGene_gn=SCN7A;refGene_tx=XM_023259422.1;refGene_gn=SCN7A;refGene_tx=XM_023259423.1
C1 158056142 158056245 refGene_gn=SCN7A;refGene_tx=XM_019838209.2;refGene_gn=SCN7A;refGene_tx=XM_023259420.1;refGene_gn=SCN7A;refGene_tx=XM_023259422.1;refGene_gn=SCN7A;refGene_tx=XM_023259423.1
C1 158062893 158063100 refGene_gn=SCN7A;refGene_tx=XM_019838209.2;refGene_gn=SCN7A;refGene_tx=XM_023259420.1;refGene_gn=SCN7A;refGene_tx=XM_023259422.1;refGene_gn=SCN7A;refGene_tx=XM_023259423.1
C1 158066885 158067027 refGene_gn=SCN7A;refGene_tx=XM_019838209.2;refGene_gn=SCN7A;refGene_tx=XM_023259420.1;refGene_gn=SCN7A;refGene_tx=XM_023259422.1;refGene_gn=SCN7A;refGene_tx=XM_023259423.1
C1 158068754 158068824 refGene_gn=SCN7A;refGene_tx=XM_019838209.2;refGene_gn=SCN7A;refGene_tx=XM_023259420.1;refGene_gn=SCN7A;refGene_tx=XM_023259422.1;refGene_gn=SCN7A;refGene_tx=XM_023259423.1
C1 158071572 158071779 refGene_gn=SCN7A;refGene_tx=XM_019838209.2;refGene_gn=SCN7A;refGene_tx=XM_023259420.1;refGene_gn=SCN7A;refGene_tx=XM_023259422.1;refGene_gn=SCN7A;refGene_tx=XM_023259423.1
C1 158076044 158076136 refGene_gn=SCN7A;refGene_tx=XM_019838209.2;refGene_gn=SCN7A;refGene_tx=XM_023259420.1;refGene_gn=SCN7A;refGene_tx=XM_023259422.1;refGene_gn=SCN7A;refGene_tx=XM_023259423.1

C1 158077547 158077676
refGene_gn=SCN7A;refGene_tx=XM_019838209.2;refGene_gn=SCN7A;refGene_tx
=XM_023259420.1;refGene_gn=SCN7A;refGene_tx=XM_023259422.1;refGene_gn=
SCN7A;refGene_tx=XM_023259423.1
C1 158078495 158078585
refGene_gn=SCN7A;refGene_tx=XM_019838209.2;refGene_gn=SCN7A;refGene_tx
=XM_023259420.1;refGene_gn=SCN7A;refGene_tx=XM_023259422.1;refGene_gn=
SCN7A;refGene_tx=XM_023259423.1
C1 158078939 158079052
refGene_gn=SCN7A;refGene_tx=XM_019838209.2;refGene_gn=SCN7A;refGene_tx
=XM_023259420.1;refGene_gn=SCN7A;refGene_tx=XM_023259422.1;refGene_gn=
SCN7A;refGene_tx=XM_023259423.1
C1 158081966 158082205
refGene_gn=SCN7A;refGene_tx=XM_019838209.2;refGene_gn=SCN7A;refGene_tx
=XM_023259420.1;refGene_gn=SCN7A;refGene_tx=XM_023259422.1;refGene_gn=
SCN7A;refGene_tx=XM_023259423.1
C1 158101293 158101303
refGene_gn=SCN7A;refGene_tx=XM_019838209.2
C1 158511204 158511345
refGene_gn=LOC111561645;refGene_tx=XM_023257860.1
C1 158514665 158515085
refGene_gn=LOC111561645;refGene_tx=XM_023257860.1
C1 158515381 158515459
refGene_gn=LOC111561645;refGene_tx=XM_023257860.1
C1 158780084 158780141
refGene_gn=XIRP2;refGene_tx=XM_011285335.2
C1 158784802 158784940
refGene_gn=XIRP2;refGene_tx=XM_011285335.2;refGene_gn=XIRP2;refGene_tx
=XM_019838210.1
C1 158806867 158806978
refGene_gn=XIRP2;refGene_tx=XM_011285335.2;refGene_gn=XIRP2;refGene_tx
=XM_019838210.1
C1 158807627 158807700
refGene_gn=XIRP2;refGene_tx=XM_011285335.2;refGene_gn=XIRP2;refGene_tx
=XM_019838210.1
C1 158808626 158808757
refGene_gn=XIRP2;refGene_tx=XM_011285335.2;refGene_gn=XIRP2;refGene_tx
=XM_019838210.1
C1 158809404 158818711
refGene_gn=XIRP2;refGene_tx=XM_011285335.2;refGene_gn=XIRP2;refGene_tx
=XM_019838210.1
C1 158820617 158820712
refGene_gn=XIRP2;refGene_tx=XM_011285335.2;refGene_gn=XIRP2;refGene_tx
=XM_019838210.1
C1 159373163 159374144
refGene_gn=B3GALT1;refGene_tx=XM_019838212.2;refGene_gn=B3GALT1;refGen
e_tx=XM_023259424.1;refGene_gn=B3GALT1;refGene_tx=XM_023259425.1;refGe
ne_gn=B3GALT1;refGene_tx=XM_023259426.1
C1 159499293 159499368
refGene_gn=STK39;refGene_tx=XM_023259427.1

C1 159513452 159513517
refGene_gn=STK39;refGene_tx=XM_023259427.1
C1 159563410 159563479
refGene_gn=STK39;refGene_tx=XM_023259427.1
C1 159567762 159567815
refGene_gn=STK39;refGene_tx=XM_023259427.1
C1 159626473 159626541
refGene_gn=STK39;refGene_tx=XM_023259427.1
C1 159628329 159628392
refGene_gn=STK39;refGene_tx=XM_023259427.1
C1 159637207 159637237
refGene_gn=STK39;refGene_tx=XM_023259427.1
C1 159637334 159637457
refGene_gn=STK39;refGene_tx=XM_023259427.1
C1 159689517 159689583
refGene_gn=STK39;refGene_tx=XM_023259427.1
C1 159689685 159689734
refGene_gn=STK39;refGene_tx=XM_023259427.1
C1 159701497 159701631
refGene_gn=STK39;refGene_tx=XM_023259427.1
C1 159703959 159704061
refGene_gn=STK39;refGene_tx=XM_023259427.1
C1 159704368 159704449
refGene_gn=STK39;refGene_tx=XM_023259427.1
C1 159704450 159704479
refGene_gn=STK39;refGene_tx=XM_023259427.1
C1 159729272 159729328
refGene_gn=STK39;refGene_tx=XM_023259427.1
C1 159730743 159730885
refGene_gn=STK39;refGene_tx=XM_023259427.1
C1 159734319 159734428
refGene_gn=STK39;refGene_tx=XM_023259427.1
C1 159751580 159751693
refGene_gn=STK39;refGene_tx=XM_023259427.1
C1 159810835 159811040
refGene_gn=STK39;refGene_tx=XM_023259427.1
C1 159983086 159983256
refGene_gn=CERS6;refGene_tx=XM_023259428.1;refGene_gn=CERS6;refGene_tx
=XM_023259429.1
C1 160077065 160077171
refGene_gn=CERS6;refGene_tx=XM_023259428.1;refGene_gn=CERS6;refGene_tx
=XM_023259429.1
C1 160089191 160089322
refGene_gn=CERS6;refGene_tx=XM_023259428.1;refGene_gn=CERS6;refGene_tx
=XM_023259429.1
C1 160161765 160161823
refGene_gn=CERS6;refGene_tx=XM_023259428.1;refGene_gn=CERS6;refGene_tx
=XM_023259429.1
C1 160222463 160222514
refGene_gn=CERS6;refGene_tx=XM_023259428.1;refGene_gn=CERS6;refGene_tx

=XM_023259429.1
C1 160226364 160226457
refGene_gn=CERS6;refGene_tx=XM_023259428.1;refGene_gn=CERS6;refGene_tx
=XM_023259429.1
C1 160243500 160243629
refGene_gn=CERS6;refGene_tx=XM_023259428.1;refGene_gn=CERS6;refGene_tx
=XM_023259429.1
C1 160246328 160246435
refGene_gn=CERS6;refGene_tx=XM_023259428.1;refGene_gn=CERS6;refGene_tx
=XM_023259429.1
C1 160292465 160292622
refGene_gn=CERS6;refGene_tx=XM_023259428.1;refGene_gn=CERS6;refGene_tx
=XM_023259429.1
C1 160293164 160293188
refGene_gn=CERS6;refGene_tx=XM_023259428.1
C1 160296824 160296977
refGene_gn=CERS6;refGene_tx=XM_023259428.1;refGene_gn=CERS6;refGene_tx
=XM_023259429.1
C1 160323885 160323912
refGene_gn=NOSTRIN;refGene_tx=XM_003990832.3;refGene_gn=NOSTRIN;refGen
e_tx=XM_006935337.3;refGene_gn=NOSTRIN;refGene_tx=XM_006935338.3;refGe
ne_gn=NOSTRIN;refGene_tx=XM_019838219.2;refGene_gn=NOSTRIN;refGene_tx=
XM_019838220.2
C1 160332607 160332693
refGene_gn=NOSTRIN;refGene_tx=XM_003990832.3;refGene_gn=NOSTRIN;refGen
e_tx=XM_006935337.3;refGene_gn=NOSTRIN;refGene_tx=XM_006935338.3;refGe
ne_gn=NOSTRIN;refGene_tx=XM_019838219.2;refGene_gn=NOSTRIN;refGene_tx=
XM_019838220.2
C1 160345698 160345782
refGene_gn=NOSTRIN;refGene_tx=XM_003990832.3;refGene_gn=NOSTRIN;refGen
e_tx=XM_006935337.3;refGene_gn=NOSTRIN;refGene_tx=XM_006935338.3;refGe
ne_gn=NOSTRIN;refGene_tx=XM_019838219.2;refGene_gn=NOSTRIN;refGene_tx=
XM_019838220.2
C1 160346547 160346664
refGene_gn=NOSTRIN;refGene_tx=XM_006935337.3;refGene_gn=NOSTRIN;refGen
e_tx=XM_006935338.3;refGene_gn=NOSTRIN;refGene_tx=XM_019838219.2;refGe
ne_gn=NOSTRIN;refGene_tx=XM_019838220.2
C1 160349214 160349277
refGene_gn=NOSTRIN;refGene_tx=XM_003990832.3;refGene_gn=NOSTRIN;refGen
e_tx=XM_006935337.3;refGene_gn=NOSTRIN;refGene_tx=XM_006935338.3;refGe
ne_gn=NOSTRIN;refGene_tx=XM_019838219.2;refGene_gn=NOSTRIN;refGene_tx=
XM_019838220.2
C1 160349476 160349558
refGene_gn=NOSTRIN;refGene_tx=XM_003990832.3;refGene_gn=NOSTRIN;refGen
e_tx=XM_006935337.3;refGene_gn=NOSTRIN;refGene_tx=XM_006935338.3;refGe
ne_gn=NOSTRIN;refGene_tx=XM_019838219.2;refGene_gn=NOSTRIN;refGene_tx=
XM_019838220.2
C1 160351988 160352051
refGene_gn=NOSTRIN;refGene_tx=XM_003990832.3;refGene_gn=NOSTRIN;refGen
e_tx=XM_006935337.3;refGene_gn=NOSTRIN;refGene_tx=XM_006935338.3;refGe

ne_gn=NOSTRIN;refGene_tx=XM_019838219.2;refGene_gn=NOSTRIN;refGene_tx=XM_019838220.2
C1 160354601 160354700
refGene_gn=NOSTRIN;refGene_tx=XM_003990832.3;refGene_gn=NOSTRIN;refGene_tx=XM_006935337.3;refGene_gn=NOSTRIN;refGene_tx=XM_006935338.3;refGene_gn=NOSTRIN;refGene_tx=XM_019838219.2;refGene_gn=NOSTRIN;refGene_tx=XM_019838220.2
C1 160360536 160360662
refGene_gn=NOSTRIN;refGene_tx=XM_003990832.3;refGene_gn=NOSTRIN;refGene_tx=XM_006935337.3;refGene_gn=NOSTRIN;refGene_tx=XM_006935338.3;refGene_gn=NOSTRIN;refGene_tx=XM_019838219.2;refGene_gn=NOSTRIN;refGene_tx=XM_019838220.2
C1 160368556 160368655
refGene_gn=NOSTRIN;refGene_tx=XM_003990832.3;refGene_gn=NOSTRIN;refGene_tx=XM_006935337.3;refGene_gn=NOSTRIN;refGene_tx=XM_006935338.3;refGene_gn=NOSTRIN;refGene_tx=XM_019838219.2;refGene_gn=NOSTRIN;refGene_tx=XM_019838220.2
C1 160368750 160368876
refGene_gn=NOSTRIN;refGene_tx=XM_003990832.3;refGene_gn=NOSTRIN;refGene_tx=XM_006935337.3;refGene_gn=NOSTRIN;refGene_tx=XM_006935338.3;refGene_gn=NOSTRIN;refGene_tx=XM_019838219.2;refGene_gn=NOSTRIN;refGene_tx=XM_019838220.2
C1 160372505 160372614
refGene_gn=NOSTRIN;refGene_tx=XM_003990832.3;refGene_gn=NOSTRIN;refGene_tx=XM_006935337.3;refGene_gn=NOSTRIN;refGene_tx=XM_006935338.3;refGene_gn=NOSTRIN;refGene_tx=XM_019838219.2;refGene_gn=NOSTRIN;refGene_tx=XM_019838220.2
C1 160373472 160373561
refGene_gn=NOSTRIN;refGene_tx=XM_003990832.3;refGene_gn=NOSTRIN;refGene_tx=XM_006935337.3;refGene_gn=NOSTRIN;refGene_tx=XM_006935338.3;refGene_gn=NOSTRIN;refGene_tx=XM_019838219.2;refGene_gn=NOSTRIN;refGene_tx=XM_019838220.2
C1 160376640 160376766
refGene_gn=NOSTRIN;refGene_tx=XM_003990832.3;refGene_gn=NOSTRIN;refGene_tx=XM_006935337.3;refGene_gn=NOSTRIN;refGene_tx=XM_006935338.3;refGene_gn=NOSTRIN;refGene_tx=XM_019838219.2;refGene_gn=NOSTRIN;refGene_tx=XM_019838220.2
C1 160377692 160377792
refGene_gn=NOSTRIN;refGene_tx=XM_003990832.3;refGene_gn=NOSTRIN;refGene_tx=XM_006935337.3;refGene_gn=NOSTRIN;refGene_tx=XM_006935338.3;refGene_gn=NOSTRIN;refGene_tx=XM_019838219.2;refGene_gn=NOSTRIN;refGene_tx=XM_019838220.2
C1 160378869 160378959
refGene_gn=NOSTRIN;refGene_tx=XM_003990832.3;refGene_gn=NOSTRIN;refGene_tx=XM_006935337.3;refGene_gn=NOSTRIN;refGene_tx=XM_006935338.3;refGene_gn=NOSTRIN;refGene_tx=XM_019838219.2;refGene_gn=NOSTRIN;refGene_tx=XM_019838220.2
C1 160379901 160379930
refGene_gn=SPC25;refGene_tx=XM_019838222.2
C1 160380213 160380275

```
refGene_gn=NOSTRIN;refGene_tx=XM_019838219.2
C1	160381810	160381947
refGene_gn=NOSTRIN;refGene_tx=XM_003990832.3;refGene_gn=NOSTRIN;refGene_tx=XM_006935337.3;refGene_gn=NOSTRIN;refGene_tx=XM_006935338.3
C1	160394702	160394827
refGene_gn=SPC25;refGene_tx=XM_003990834.4
C1	160396826	160396925
refGene_gn=SPC25;refGene_tx=XM_003990834.4;refGene_gn=SPC25;refGene_tx=XM_019838222.2
C1	160402342	160402447
refGene_gn=SPC25;refGene_tx=XM_003990834.4;refGene_gn=SPC25;refGene_tx=XM_019838222.2
C1	160402884	160403031
refGene_gn=SPC25;refGene_tx=XM_003990834.4;refGene_gn=SPC25;refGene_tx=XM_019838222.2
C1	160407628	160407694
refGene_gn=SPC25;refGene_tx=XM_003990834.4;refGene_gn=SPC25;refGene_tx=XM_019838222.2
C1	160407799	160407938
refGene_gn=SPC25;refGene_tx=XM_003990834.4;refGene_gn=SPC25;refGene_tx=XM_019838222.2
C1	160426492	160426710
refGene_gn=G6PC2;refGene_tx=XM_003990835.3
C1	160427263	160427373
refGene_gn=G6PC2;refGene_tx=XM_003990835.3
C1	160429266	160429378
refGene_gn=G6PC2;refGene_tx=XM_003990835.3
C1	160431633	160431749
refGene_gn=G6PC2;refGene_tx=XM_003990835.3
C1	160432534	160433046
refGene_gn=G6PC2;refGene_tx=XM_003990835.3
C1	160448774	160448975
refGene_gn=ABCB11;refGene_tx=NM_001287551.1;refGene_gn=ABCB11;refGene_tx=XM_019835934.1
C1	160449806	160449953
refGene_gn=ABCB11;refGene_tx=NM_001287551.1;refGene_gn=ABCB11;refGene_tx=XM_019835934.1
C1	160451924	160452131
refGene_gn=ABCB11;refGene_tx=NM_001287551.1;refGene_gn=ABCB11;refGene_tx=XM_019835934.1
C1	160455740	160455938
refGene_gn=ABCB11;refGene_tx=NM_001287551.1;refGene_gn=ABCB11;refGene_tx=XM_019835934.1
C1	160457562	160457719
refGene_gn=ABCB11;refGene_tx=NM_001287551.1;refGene_gn=ABCB11;refGene_tx=XM_019835934.1
C1	160460210	160460452
refGene_gn=ABCB11;refGene_tx=NM_001287551.1;refGene_gn=ABCB11;refGene_tx=XM_019835934.1
C1	160461729	160461933
```

refGene_gn=ABCB11;refGene_tx=NM_001287551.1;refGene_gn=ABCB11;refGene_tx=XM_019835934.1
C1 160466586 160466748
refGene_gn=ABCB11;refGene_tx=NM_001287551.1;refGene_gn=ABCB11;refGene_tx=XM_019835934.1
C1 160466835 160466940
refGene_gn=ABCB11;refGene_tx=NM_001287551.1;refGene_gn=ABCB11;refGene_tx=XM_019835934.1
C1 160472649 160472664
refGene_gn=ABCB11;refGene_tx=XM_019835935.1
C1 160472821 160472986
refGene_gn=ABCB11;refGene_tx=NM_001287551.1;refGene_gn=ABCB11;refGene_tx=XM_019835934.1;refGene_gn=ABCB11;refGene_tx=XM_019835935.1
C1 160480362 160480465
refGene_gn=ABCB11;refGene_tx=NM_001287551.1;refGene_gn=ABCB11;refGene_tx=XM_019835934.1;refGene_gn=ABCB11;refGene_tx=XM_019835935.1
C1 160484893 160484969
refGene_gn=ABCB11;refGene_tx=NM_001287551.1;refGene_gn=ABCB11;refGene_tx=XM_019835934.1;refGene_gn=ABCB11;refGene_tx=XM_019835935.1
C1 160485976 160486178
refGene_gn=ABCB11;refGene_tx=NM_001287551.1;refGene_gn=ABCB11;refGene_tx=XM_019835934.1;refGene_gn=ABCB11;refGene_tx=XM_019835935.1
C1 160486678 160486849
refGene_gn=ABCB11;refGene_tx=NM_001287551.1;refGene_gn=ABCB11;refGene_tx=XM_019835934.1;refGene_gn=ABCB11;refGene_tx=XM_019835935.1
C1 160488468 160488672
refGene_gn=ABCB11;refGene_tx=NM_001287551.1;refGene_gn=ABCB11;refGene_tx=XM_019835934.1;refGene_gn=ABCB11;refGene_tx=XM_019835935.1
C1 160490404 160490530
refGene_gn=ABCB11;refGene_tx=NM_001287551.1;refGene_gn=ABCB11;refGene_tx=XM_019835934.1;refGene_gn=ABCB11;refGene_tx=XM_019835935.1
C1 160492545 160492656
refGene_gn=ABCB11;refGene_tx=NM_001287551.1;refGene_gn=ABCB11;refGene_tx=XM_019835934.1;refGene_gn=ABCB11;refGene_tx=XM_019835935.1
C1 160495726 160495840
refGene_gn=ABCB11;refGene_tx=NM_001287551.1;refGene_gn=ABCB11;refGene_tx=XM_019835934.1;refGene_gn=ABCB11;refGene_tx=XM_019835935.1
C1 160502710 160502885
refGene_gn=ABCB11;refGene_tx=NM_001287551.1;refGene_gn=ABCB11;refGene_tx=XM_019835934.1;refGene_gn=ABCB11;refGene_tx=XM_019835935.1
C1 160504735 160504860
refGene_gn=ABCB11;refGene_tx=NM_001287551.1;refGene_gn=ABCB11;refGene_tx=XM_019835934.1;refGene_gn=ABCB11;refGene_tx=XM_019835935.1
C1 160507669 160507841
refGene_gn=ABCB11;refGene_tx=NM_001287551.1;refGene_gn=ABCB11;refGene_tx=XM_019835934.1;refGene_gn=ABCB11;refGene_tx=XM_019835935.1
C1 160509466 160509600
refGene_gn=ABCB11;refGene_tx=NM_001287551.1;refGene_gn=ABCB11;refGene_tx=XM_019835934.1;refGene_gn=ABCB11;refGene_tx=XM_019835935.1
C1 160511842 160511930

refGene_gn=ABCB11;refGene_tx=NM_001287551.1;refGene_gn=ABCB11;refGene_tx=XM_019835934.1;refGene_gn=ABCB11;refGene_tx=XM_019835935.1
C1 160524588 160524827
refGene_gn=ABCB11;refGene_tx=NM_001287551.1;refGene_gn=ABCB11;refGene_tx=XM_019835934.1;refGene_gn=ABCB11;refGene_tx=XM_019835935.1
C1 160525587 160525639
refGene_gn=ABCB11;refGene_tx=NM_001287551.1;refGene_gn=ABCB11;refGene_tx=XM_019835935.1
C1 160528158 160528180
refGene_gn=ABCB11;refGene_tx=NM_001287551.1;refGene_gn=ABCB11;refGene_tx=XM_019835935.1
C1 160529701 160529777
refGene_gn=ABCB11;refGene_tx=NM_001287551.1;refGene_gn=ABCB11;refGene_tx=XM_019835935.1
C1 160569883 160570196
refGene_gn=DHRS9;refGene_tx=XM_003990837.4
C1 160571606 160571865
refGene_gn=DHRS9;refGene_tx=XM_003990837.4
C1 160575216 160575380
refGene_gn=DHRS9;refGene_tx=XM_003990837.4
C1 160578080 160578304
refGene_gn=DHRS9;refGene_tx=XM_003990837.4
C1 160615477 160615639
refGene_gn=LRP2;refGene_tx=XM_019839119.2;refGene_gn=LRP2;refGene_tx=XM_023259430.1;refGene_gn=LRP2;refGene_tx=XM_023259431.1
C1 160615828 160615900
refGene_gn=LRP2;refGene_tx=XM_019839119.2;refGene_gn=LRP2;refGene_tx=XM_023259430.1;refGene_gn=LRP2;refGene_tx=XM_023259431.1
C1 160619468 160619576
refGene_gn=LRP2;refGene_tx=XM_019839119.2;refGene_gn=LRP2;refGene_tx=XM_023259430.1;refGene_gn=LRP2;refGene_tx=XM_023259431.1
C1 160621432 160621537
refGene_gn=LRP2;refGene_tx=XM_019839119.2;refGene_gn=LRP2;refGene_tx=XM_023259430.1;refGene_gn=LRP2;refGene_tx=XM_023259431.1
C1 160622527 160622657
refGene_gn=LRP2;refGene_tx=XM_019839119.2;refGene_gn=LRP2;refGene_tx=XM_023259430.1;refGene_gn=LRP2;refGene_tx=XM_023259431.1
C1 160623188 160623306
refGene_gn=LRP2;refGene_tx=XM_019839119.2;refGene_gn=LRP2;refGene_tx=XM_023259430.1;refGene_gn=LRP2;refGene_tx=XM_023259431.1
C1 160623683 160623751
refGene_gn=LRP2;refGene_tx=XM_019839119.2;refGene_gn=LRP2;refGene_tx=XM_023259430.1;refGene_gn=LRP2;refGene_tx=XM_023259431.1
C1 160624552 160624643
refGene_gn=LRP2;refGene_tx=XM_019839119.2;refGene_gn=LRP2;refGene_tx=XM_023259430.1;refGene_gn=LRP2;refGene_tx=XM_023259431.1
C1 160627151 160627271
refGene_gn=LRP2;refGene_tx=XM_019839119.2;refGene_gn=LRP2;refGene_tx=XM_023259430.1;refGene_gn=LRP2;refGene_tx=XM_023259431.1
C1 160629717 160629894

refGene_gn=LRP2;refGene_tx=XM_019839119.2;refGene_gn=LRP2;refGene_tx=XM_023259430.1;refGene_gn=LRP2;refGene_tx=XM_023259431.1
C1 160630695 160630916
refGene_gn=LRP2;refGene_tx=XM_019839119.2;refGene_gn=LRP2;refGene_tx=XM_023259430.1;refGene_gn=LRP2;refGene_tx=XM_023259431.1
C1 160632370 160632499
refGene_gn=LRP2;refGene_tx=XM_019839119.2;refGene_gn=LRP2;refGene_tx=XM_023259430.1;refGene_gn=LRP2;refGene_tx=XM_023259431.1
C1 160634154 160634320
refGene_gn=LRP2;refGene_tx=XM_019839119.2;refGene_gn=LRP2;refGene_tx=XM_023259430.1;refGene_gn=LRP2;refGene_tx=XM_023259431.1
C1 160635426 160635570
refGene_gn=LRP2;refGene_tx=XM_019839119.2;refGene_gn=LRP2;refGene_tx=XM_023259430.1;refGene_gn=LRP2;refGene_tx=XM_023259431.1
C1 160637267 160637399
refGene_gn=LRP2;refGene_tx=XM_019839119.2;refGene_gn=LRP2;refGene_tx=XM_023259430.1;refGene_gn=LRP2;refGene_tx=XM_023259431.1
C1 160638474 160638606
refGene_gn=LRP2;refGene_tx=XM_019839119.2;refGene_gn=LRP2;refGene_tx=XM_023259430.1;refGene_gn=LRP2;refGene_tx=XM_023259431.1
C1 160642833 160642962
refGene_gn=LRP2;refGene_tx=XM_019839119.2;refGene_gn=LRP2;refGene_tx=XM_023259430.1;refGene_gn=LRP2;refGene_tx=XM_023259431.1
C1 160646190 160646313
refGene_gn=LRP2;refGene_tx=XM_019839119.2;refGene_gn=LRP2;refGene_tx=XM_023259430.1;refGene_gn=LRP2;refGene_tx=XM_023259431.1
C1 160648261 160648399
refGene_gn=LRP2;refGene_tx=XM_019839119.2;refGene_gn=LRP2;refGene_tx=XM_023259430.1;refGene_gn=LRP2;refGene_tx=XM_023259431.1
C1 160649272 160649389
refGene_gn=LRP2;refGene_tx=XM_019839119.2;refGene_gn=LRP2;refGene_tx=XM_023259430.1;refGene_gn=LRP2;refGene_tx=XM_023259431.1
C1 160650095 160650212
refGene_gn=LRP2;refGene_tx=XM_019839119.2;refGene_gn=LRP2;refGene_tx=XM_023259430.1;refGene_gn=LRP2;refGene_tx=XM_023259431.1
C1 160651449 160651569
refGene_gn=LRP2;refGene_tx=XM_019839119.2;refGene_gn=LRP2;refGene_tx=XM_023259430.1;refGene_gn=LRP2;refGene_tx=XM_023259431.1
C1 160652567 160652690
refGene_gn=LRP2;refGene_tx=XM_019839119.2;refGene_gn=LRP2;refGene_tx=XM_023259430.1;refGene_gn=LRP2;refGene_tx=XM_023259431.1
C1 160653317 160653563
refGene_gn=LRP2;refGene_tx=XM_019839119.2;refGene_gn=LRP2;refGene_tx=XM_023259430.1;refGene_gn=LRP2;refGene_tx=XM_023259431.1
C1 160654482 160654679
refGene_gn=LRP2;refGene_tx=XM_019839119.2;refGene_gn=LRP2;refGene_tx=XM_023259430.1;refGene_gn=LRP2;refGene_tx=XM_023259431.1
C1 160655645 160655823
refGene_gn=LRP2;refGene_tx=XM_019839119.2;refGene_gn=LRP2;refGene_tx=XM_023259430.1;refGene_gn=LRP2;refGene_tx=XM_023259431.1

C1 160657710 160657934
refGene_gn=LRP2;refGene_tx=XM_019839119.2;refGene_gn=LRP2;refGene_tx=XM_023259430.1;refGene_gn=LRP2;refGene_tx=XM_023259431.1
C1 160659575 160659746
refGene_gn=LRP2;refGene_tx=XM_019839119.2;refGene_gn=LRP2;refGene_tx=XM_023259430.1;refGene_gn=LRP2;refGene_tx=XM_023259431.1
C1 160660263 160660416
refGene_gn=LRP2;refGene_tx=XM_019839119.2;refGene_gn=LRP2;refGene_tx=XM_023259430.1;refGene_gn=LRP2;refGene_tx=XM_023259431.1
C1 160661766 160662283
refGene_gn=LRP2;refGene_tx=XM_019839119.2;refGene_gn=LRP2;refGene_tx=XM_023259430.1;refGene_gn=LRP2;refGene_tx=XM_023259431.1
C1 160663358 160663654
refGene_gn=LRP2;refGene_tx=XM_019839119.2;refGene_gn=LRP2;refGene_tx=XM_023259430.1;refGene_gn=LRP2;refGene_tx=XM_023259431.1
C1 160667118 160667320
refGene_gn=LRP2;refGene_tx=XM_019839119.2;refGene_gn=LRP2;refGene_tx=XM_023259430.1;refGene_gn=LRP2;refGene_tx=XM_023259431.1
C1 160669352 160669484
refGene_gn=LRP2;refGene_tx=XM_019839119.2;refGene_gn=LRP2;refGene_tx=XM_023259430.1;refGene_gn=LRP2;refGene_tx=XM_023259431.1
C1 160672411 160672531
refGene_gn=LRP2;refGene_tx=XM_019839119.2;refGene_gn=LRP2;refGene_tx=XM_023259430.1;refGene_gn=LRP2;refGene_tx=XM_023259431.1
C1 160673877 160674003
refGene_gn=LRP2;refGene_tx=XM_019839119.2;refGene_gn=LRP2;refGene_tx=XM_023259430.1;refGene_gn=LRP2;refGene_tx=XM_023259431.1
C1 160676055 160676298
refGene_gn=LRP2;refGene_tx=XM_019839119.2;refGene_gn=LRP2;refGene_tx=XM_023259430.1;refGene_gn=LRP2;refGene_tx=XM_023259431.1
C1 160677402 160677606
refGene_gn=LRP2;refGene_tx=XM_019839119.2;refGene_gn=LRP2;refGene_tx=XM_023259430.1;refGene_gn=LRP2;refGene_tx=XM_023259431.1
C1 160678543 160678830
refGene_gn=LRP2;refGene_tx=XM_019839119.2;refGene_gn=LRP2;refGene_tx=XM_023259430.1;refGene_gn=LRP2;refGene_tx=XM_023259431.1
C1 160679992 160680151
refGene_gn=LRP2;refGene_tx=XM_019839119.2;refGene_gn=LRP2;refGene_tx=XM_023259430.1;refGene_gn=LRP2;refGene_tx=XM_023259431.1
C1 160680533 160680699
refGene_gn=LRP2;refGene_tx=XM_019839119.2;refGene_gn=LRP2;refGene_tx=XM_023259430.1;refGene_gn=LRP2;refGene_tx=XM_023259431.1
C1 160680831 160681752
refGene_gn=LRP2;refGene_tx=XM_019839119.2;refGene_gn=LRP2;refGene_tx=XM_023259430.1;refGene_gn=LRP2;refGene_tx=XM_023259431.1
C1 160684721 160684910
refGene_gn=LRP2;refGene_tx=XM_019839119.2;refGene_gn=LRP2;refGene_tx=XM_023259430.1;refGene_gn=LRP2;refGene_tx=XM_023259431.1
C1 160686489 160686723
refGene_gn=LRP2;refGene_tx=XM_019839119.2;refGene_gn=LRP2;refGene_tx=X

M_023259430.1;refGene_gn=LRP2;refGene_tx=XM_023259431.1
C1 160688041 160688255
refGene_gn=LRP2;refGene_tx=XM_019839119.2;refGene_gn=LRP2;refGene_tx=XM_023259430.1;refGene_gn=LRP2;refGene_tx=XM_023259431.1
C1 160690944 160691122
refGene_gn=LRP2;refGene_tx=XM_019839119.2;refGene_gn=LRP2;refGene_tx=XM_023259430.1;refGene_gn=LRP2;refGene_tx=XM_023259431.1
C1 160692914 160693024
refGene_gn=LRP2;refGene_tx=XM_019839119.2;refGene_gn=LRP2;refGene_tx=XM_023259430.1;refGene_gn=LRP2;refGene_tx=XM_023259431.1
C1 160697009 160697153
refGene_gn=LRP2;refGene_tx=XM_019839119.2;refGene_gn=LRP2;refGene_tx=XM_023259430.1;refGene_gn=LRP2;refGene_tx=XM_023259431.1
C1 160698100 160698267
refGene_gn=LRP2;refGene_tx=XM_019839119.2;refGene_gn=LRP2;refGene_tx=XM_023259430.1;refGene_gn=LRP2;refGene_tx=XM_023259431.1
C1 160704075 160704204
refGene_gn=LRP2;refGene_tx=XM_019839119.2;refGene_gn=LRP2;refGene_tx=XM_023259430.1;refGene_gn=LRP2;refGene_tx=XM_023259431.1
C1 160706025 160706203
refGene_gn=LRP2;refGene_tx=XM_019839119.2;refGene_gn=LRP2;refGene_tx=XM_023259430.1;refGene_gn=LRP2;refGene_tx=XM_023259431.1
C1 160707232 160707461
refGene_gn=LRP2;refGene_tx=XM_019839119.2;refGene_gn=LRP2;refGene_tx=XM_023259430.1;refGene_gn=LRP2;refGene_tx=XM_023259431.1
C1 160708462 160708647
refGene_gn=LRP2;refGene_tx=XM_019839119.2;refGene_gn=LRP2;refGene_tx=XM_023259430.1;refGene_gn=LRP2;refGene_tx=XM_023259431.1
C1 160709478 160709690
refGene_gn=LRP2;refGene_tx=XM_019839119.2;refGene_gn=LRP2;refGene_tx=XM_023259430.1;refGene_gn=LRP2;refGene_tx=XM_023259431.1
C1 160710685 160710934
refGene_gn=LRP2;refGene_tx=XM_019839119.2;refGene_gn=LRP2;refGene_tx=XM_023259430.1;refGene_gn=LRP2;refGene_tx=XM_023259431.1
C1 160711662 160711677
refGene_gn=LRP2;refGene_tx=XM_019839119.2;refGene_gn=LRP2;refGene_tx=XM_023259430.1
C1 160712131 160712509
refGene_gn=LRP2;refGene_tx=XM_019839119.2;refGene_gn=LRP2;refGene_tx=XM_023259430.1;refGene_gn=LRP2;refGene_tx=XM_023259431.1
C1 160713978 160714095
refGene_gn=LRP2;refGene_tx=XM_019839119.2;refGene_gn=LRP2;refGene_tx=XM_023259430.1;refGene_gn=LRP2;refGene_tx=XM_023259431.1
C1 160714196 160714316
refGene_gn=LRP2;refGene_tx=XM_019839119.2;refGene_gn=LRP2;refGene_tx=XM_023259430.1;refGene_gn=LRP2;refGene_tx=XM_023259431.1
C1 160715550 160715790
refGene_gn=LRP2;refGene_tx=XM_019839119.2;refGene_gn=LRP2;refGene_tx=XM_023259430.1;refGene_gn=LRP2;refGene_tx=XM_023259431.1
C1 160716942 160717224

refGene_gn=LRP2;refGene_tx=XM_019839119.2;refGene_gn=LRP2;refGene_tx=XM_023259430.1;refGene_gn=LRP2;refGene_tx=XM_023259431.1
C1 160717600 160717738
refGene_gn=LRP2;refGene_tx=XM_019839119.2;refGene_gn=LRP2;refGene_tx=XM_023259430.1;refGene_gn=LRP2;refGene_tx=XM_023259431.1
C1 160720303 160720434
refGene_gn=LRP2;refGene_tx=XM_019839119.2;refGene_gn=LRP2;refGene_tx=XM_023259430.1;refGene_gn=LRP2;refGene_tx=XM_023259431.1
C1 160721332 160721458
refGene_gn=LRP2;refGene_tx=XM_019839119.2;refGene_gn=LRP2;refGene_tx=XM_023259430.1;refGene_gn=LRP2;refGene_tx=XM_023259431.1
C1 160723212 160723405
refGene_gn=LRP2;refGene_tx=XM_019839119.2;refGene_gn=LRP2;refGene_tx=XM_023259430.1;refGene_gn=LRP2;refGene_tx=XM_023259431.1
C1 160729309 160729513
refGene_gn=LRP2;refGene_tx=XM_019839119.2;refGene_gn=LRP2;refGene_tx=XM_023259430.1;refGene_gn=LRP2;refGene_tx=XM_023259431.1
C1 160731615 160731756
refGene_gn=LRP2;refGene_tx=XM_019839119.2;refGene_gn=LRP2;refGene_tx=XM_023259430.1;refGene_gn=LRP2;refGene_tx=XM_023259431.1
C1 160733444 160733647
refGene_gn=LRP2;refGene_tx=XM_019839119.2;refGene_gn=LRP2;refGene_tx=XM_023259430.1;refGene_gn=LRP2;refGene_tx=XM_023259431.1
C1 160736482 160736689
refGene_gn=LRP2;refGene_tx=XM_019839119.2;refGene_gn=LRP2;refGene_tx=XM_023259430.1;refGene_gn=LRP2;refGene_tx=XM_023259431.1
C1 160737749 160737967
refGene_gn=LRP2;refGene_tx=XM_019839119.2;refGene_gn=LRP2;refGene_tx=XM_023259430.1;refGene_gn=LRP2;refGene_tx=XM_023259431.1
C1 160738975 160739145
refGene_gn=LRP2;refGene_tx=XM_019839119.2;refGene_gn=LRP2;refGene_tx=XM_023259430.1;refGene_gn=LRP2;refGene_tx=XM_023259431.1
C1 160740797 160740926
refGene_gn=LRP2;refGene_tx=XM_019839119.2;refGene_gn=LRP2;refGene_tx=XM_023259430.1;refGene_gn=LRP2;refGene_tx=XM_023259431.1
C1 160746932 160747052
refGene_gn=LRP2;refGene_tx=XM_019839119.2;refGene_gn=LRP2;refGene_tx=XM_023259430.1;refGene_gn=LRP2;refGene_tx=XM_023259431.1
C1 160748632 160748785
refGene_gn=LRP2;refGene_tx=XM_019839119.2;refGene_gn=LRP2;refGene_tx=XM_023259430.1;refGene_gn=LRP2;refGene_tx=XM_023259431.1
C1 160749811 160749928
refGene_gn=LRP2;refGene_tx=XM_019839119.2;refGene_gn=LRP2;refGene_tx=XM_023259430.1;refGene_gn=LRP2;refGene_tx=XM_023259431.1
C1 160751618 160751732
refGene_gn=LRP2;refGene_tx=XM_019839119.2;refGene_gn=LRP2;refGene_tx=XM_023259430.1;refGene_gn=LRP2;refGene_tx=XM_023259431.1
C1 160752064 160752175
refGene_gn=LRP2;refGene_tx=XM_019839119.2;refGene_gn=LRP2;refGene_tx=XM_023259430.1;refGene_gn=LRP2;refGene_tx=XM_023259431.1

C1	160764455	160764572
refGene_gn=LRP2;refGene_tx=XM_019839119.2;refGene_gn=LRP2;refGene_tx=XM_023259430.1;refGene_gn=LRP2;refGene_tx=XM_023259431.1
C1	160775111	160775234
refGene_gn=LRP2;refGene_tx=XM_019839119.2;refGene_gn=LRP2;refGene_tx=XM_023259430.1;refGene_gn=LRP2;refGene_tx=XM_023259431.1
C1	160777039	160777147
refGene_gn=LRP2;refGene_tx=XM_019839119.2;refGene_gn=LRP2;refGene_tx=XM_023259430.1;refGene_gn=LRP2;refGene_tx=XM_023259431.1
C1	160814072	160814151
refGene_gn=LRP2;refGene_tx=XM_019839119.2;refGene_gn=LRP2;refGene_tx=XM_023259430.1;refGene_gn=LRP2;refGene_tx=XM_023259431.1
C1	160868376	160868589
refGene_gn=LOC101086493;refGene_tx=XM_019839187.1
C1	160910051	160910110
refGene_gn=BBS5;refGene_tx=XM_011285332.3;refGene_gn=BBS5;refGene_tx=XM_011285333.3;refGene_gn=BBS5;refGene_tx=XM_019838225.2
C1	160912193	160912276
refGene_gn=BBS5;refGene_tx=XM_011285332.3;refGene_gn=BBS5;refGene_tx=XM_011285333.3;refGene_gn=BBS5;refGene_tx=XM_019838224.2;refGene_gn=BBS5;refGene_tx=XM_019838225.2
C1	160915717	160915783
refGene_gn=BBS5;refGene_tx=XM_011285332.3;refGene_gn=BBS5;refGene_tx=XM_019838224.2;refGene_gn=BBS5;refGene_tx=XM_019838225.2
C1	160916417	160916467
refGene_gn=BBS5;refGene_tx=XM_011285332.3;refGene_gn=BBS5;refGene_tx=XM_011285333.3;refGene_gn=BBS5;refGene_tx=XM_019838224.2;refGene_gn=BBS5;refGene_tx=XM_019838225.2
C1	160916598	160916726
refGene_gn=BBS5;refGene_tx=XM_011285332.3;refGene_gn=BBS5;refGene_tx=XM_011285333.3;refGene_gn=BBS5;refGene_tx=XM_019838224.2;refGene_gn=BBS5;refGene_tx=XM_019838225.2
C1	160920792	160920928
refGene_gn=BBS5;refGene_tx=XM_011285332.3;refGene_gn=BBS5;refGene_tx=XM_011285333.3;refGene_gn=BBS5;refGene_tx=XM_019838224.2;refGene_gn=BBS5;refGene_tx=XM_019838225.2
C1	160923595	160923691
refGene_gn=BBS5;refGene_tx=XM_011285332.3;refGene_gn=BBS5;refGene_tx=XM_011285333.3;refGene_gn=BBS5;refGene_tx=XM_019838224.2;refGene_gn=BBS5;refGene_tx=XM_019838225.2
C1	160925847	160925910
refGene_gn=BBS5;refGene_tx=XM_011285332.3;refGene_gn=BBS5;refGene_tx=XM_011285333.3;refGene_gn=BBS5;refGene_tx=XM_019838224.2;refGene_gn=BBS5;refGene_tx=XM_019838225.2
C1	160927677	160927812
refGene_gn=BBS5;refGene_tx=XM_011285332.3;refGene_gn=BBS5;refGene_tx=XM_011285333.3;refGene_gn=BBS5;refGene_tx=XM_019838224.2
C1	160931728	160931812
refGene_gn=BBS5;refGene_tx=XM_011285332.3;refGene_gn=BBS5;refGene_tx=XM_011285333.3;refGene_gn=BBS5;refGene_tx=XM_019838224.2;refGene_gn=BBS

5;refGene_tx=XM_019838225.2
C1	160932755	160932779
refGene_gn=BBS5;refGene_tx=XM_011285332.3;refGene_gn=BBS5;refGene_tx=XM_011285333.3;refGene_gn=BBS5;refGene_tx=XM_019838224.2;refGene_gn=BBS5;refGene_tx=XM_019838225.2
C1	160932934	160933036
refGene_gn=BBS5;refGene_tx=XM_011285332.3;refGene_gn=BBS5;refGene_tx=XM_011285333.3;refGene_gn=BBS5;refGene_tx=XM_019838224.2;refGene_gn=BBS5;refGene_tx=XM_019838225.2
C1	160935925	160937035
refGene_gn=KLHL41;refGene_tx=XM_004001577.5
C1	160941279	160941437
refGene_gn=KLHL41;refGene_tx=XM_004001577.5
C1	160941581	160941689
refGene_gn=KLHL41;refGene_tx=XM_004001577.5
C1	160944767	160944953
refGene_gn=KLHL41;refGene_tx=XM_004001577.5
C1	160946574	160946721
refGene_gn=KLHL41;refGene_tx=XM_004001577.5
C1	160951532	160951644
refGene_gn=KLHL41;refGene_tx=XM_004001577.5
C1	160955872	160955968
refGene_gn=FASTKD1;refGene_tx=XM_011285330.3;refGene_gn=FASTKD1;refGene_tx=XM_011285331.3;refGene_gn=FASTKD1;refGene_tx=XM_019838228.2
C1	160957188	160957303
refGene_gn=FASTKD1;refGene_tx=XM_011285330.3;refGene_gn=FASTKD1;refGene_tx=XM_011285331.3;refGene_gn=FASTKD1;refGene_tx=XM_019838228.2
C1	160957947	160958086
refGene_gn=FASTKD1;refGene_tx=XM_011285330.3;refGene_gn=FASTKD1;refGene_tx=XM_011285331.3;refGene_gn=FASTKD1;refGene_tx=XM_019838228.2
C1	160966079	160966193
refGene_gn=FASTKD1;refGene_tx=XM_011285330.3;refGene_gn=FASTKD1;refGene_tx=XM_011285331.3;refGene_gn=FASTKD1;refGene_tx=XM_019838227.2;refGene_gn=FASTKD1;refGene_tx=XM_019838228.2
C1	160966555	160966687
refGene_gn=FASTKD1;refGene_tx=XM_011285330.3;refGene_gn=FASTKD1;refGene_tx=XM_011285331.3;refGene_gn=FASTKD1;refGene_tx=XM_019838227.2;refGene_gn=FASTKD1;refGene_tx=XM_019838228.2
C1	160967478	160967607
refGene_gn=FASTKD1;refGene_tx=XM_011285330.3;refGene_gn=FASTKD1;refGene_tx=XM_011285331.3;refGene_gn=FASTKD1;refGene_tx=XM_019838227.2;refGene_gn=FASTKD1;refGene_tx=XM_019838228.2
C1	160969980	160970095
refGene_gn=FASTKD1;refGene_tx=XM_011285330.3;refGene_gn=FASTKD1;refGene_tx=XM_011285331.3;refGene_gn=FASTKD1;refGene_tx=XM_019838227.2;refGene_gn=FASTKD1;refGene_tx=XM_019838228.2
C1	160971140	160971627
refGene_gn=FASTKD1;refGene_tx=XM_011285330.3;refGene_gn=FASTKD1;refGene_tx=XM_011285331.3;refGene_gn=FASTKD1;refGene_tx=XM_019838227.2;refGene_gn=FASTKD1;refGene_tx=XM_019838228.2

C1 160976082 160976214
refGene_gn=FASTKD1;refGene_tx=XM_011285330.3;refGene_gn=FASTKD1;refGene_tx=XM_011285331.3;refGene_gn=FASTKD1;refGene_tx=XM_019838227.2;refGene_gn=FASTKD1;refGene_tx=XM_019838228.2
C1 160977323 160977434
refGene_gn=FASTKD1;refGene_tx=XM_011285330.3;refGene_gn=FASTKD1;refGene_tx=XM_011285331.3;refGene_gn=FASTKD1;refGene_tx=XM_019838227.2;refGene_gn=FASTKD1;refGene_tx=XM_019838228.2
C1 160979183 160979582
refGene_gn=FASTKD1;refGene_tx=XM_011285330.3;refGene_gn=FASTKD1;refGene_tx=XM_011285331.3;refGene_gn=FASTKD1;refGene_tx=XM_019838227.2;refGene_gn=FASTKD1;refGene_tx=XM_019838228.2
C1 160982743 160982869
refGene_gn=FASTKD1;refGene_tx=XM_011285330.3;refGene_gn=FASTKD1;refGene_tx=XM_011285331.3;refGene_gn=FASTKD1;refGene_tx=XM_019838227.2
C1 160998751 160998820
refGene_gn=FASTKD1;refGene_tx=XM_011285330.3;refGene_gn=FASTKD1;refGene_tx=XM_011285331.3;refGene_gn=FASTKD1;refGene_tx=XM_019838227.2;refGene_gn=FASTKD1;refGene_tx=XM_019838228.2
C1 161001454 161001789
refGene_gn=FASTKD1;refGene_tx=XM_011285330.3;refGene_gn=FASTKD1;refGene_tx=XM_011285331.3;refGene_gn=FASTKD1;refGene_tx=XM_019838227.2;refGene_gn=FASTKD1;refGene_tx=XM_019838228.2
C1 161027166 161027227
refGene_gn=PPIG;refGene_tx=XM_006935342.4
C1 161027318 161027393
refGene_gn=PPIG;refGene_tx=XM_006935342.4
C1 161027992 161028100
refGene_gn=PPIG;refGene_tx=XM_006935342.4
C1 161028712 161028757
refGene_gn=PPIG;refGene_tx=XM_006935342.4
C1 161030038 161030126
refGene_gn=PPIG;refGene_tx=XM_006935342.4
C1 161035323 161035353
refGene_gn=PPIG;refGene_tx=XM_006935342.4
C1 161035438 161035578
refGene_gn=PPIG;refGene_tx=XM_006935342.4
C1 161046149 161046363
refGene_gn=PPIG;refGene_tx=XM_006935342.4
C1 161047380 161047507
refGene_gn=PPIG;refGene_tx=XM_006935342.4
C1 161047508 161047549
refGene_gn=PPIG;refGene_tx=XM_006935342.4
C1 161048973 161049061
refGene_gn=PPIG;refGene_tx=XM_006935342.4
C1 161051530 161051667
refGene_gn=PPIG;refGene_tx=XM_006935342.4
C1 161051855 161052966
refGene_gn=PPIG;refGene_tx=XM_006935342.4
C1 161059554 161059893

refGene_gn=CCDC173;refGene_tx=XM_011285329.3
C1 161065368 161065533
refGene_gn=CCDC173;refGene_tx=XM_011285329.3
C1 161066780 161066973
refGene_gn=CCDC173;refGene_tx=XM_011285329.3
C1 161069858 161069998
refGene_gn=CCDC173;refGene_tx=XM_011285329.3
C1 161073257 161073414
refGene_gn=CCDC173;refGene_tx=XM_011285329.3
C1 161081043 161081237
refGene_gn=CCDC173;refGene_tx=XM_011285329.3
C1 161081346 161081520
refGene_gn=CCDC173;refGene_tx=XM_011285329.3
C1 161085857 161086084
refGene_gn=CCDC173;refGene_tx=XM_011285329.3
C1 161091715 161091776
refGene_gn=CCDC173;refGene_tx=XM_011285329.3
C1 161097904 161098630
refGene_gn=PHOSPHO2;refGene_tx=XM_019838231.2;refGene_gn=PHOSPHO2;refGene_tx=XM_019838232.2;refGene_gn=PHOSPHO2;refGene_tx=XM_019838233.2;refGene_gn=PHOSPHO2;refGene_tx=XM_019838234.2;refGene_gn=PHOSPHO2;refGene_tx=XM_019838235.2;refGene_gn=PHOSPHO2;refGene_tx=XM_019838236.2
C1 161114088 161115301
refGene_gn=KLHL23;refGene_tx=XM_006935345.3;refGene_gn=KLHL23;refGene_tx=XM_006935346.4
C1 161119018 161119171
refGene_gn=KLHL23;refGene_tx=XM_006935345.3;refGene_gn=KLHL23;refGene_tx=XM_006935346.4
C1 161130263 161130574
refGene_gn=KLHL23;refGene_tx=XM_006935345.3;refGene_gn=KLHL23;refGene_tx=XM_006935346.4
C1 161156678 161156744
refGene_gn=SSB;refGene_tx=XM_011285323.3
C1 161158717 161158821
refGene_gn=SSB;refGene_tx=XM_011285323.3
C1 161158909 161159084
refGene_gn=SSB;refGene_tx=XM_011285323.3
C1 161159749 161159857
refGene_gn=SSB;refGene_tx=XM_011285323.3
C1 161159947 161160048
refGene_gn=SSB;refGene_tx=XM_011285323.3
C1 161161276 161161348
refGene_gn=SSB;refGene_tx=XM_011285323.3
C1 161161662 161161705
refGene_gn=SSB;refGene_tx=XM_011285323.3
C1 161162094 161162235
refGene_gn=SSB;refGene_tx=XM_011285323.3
C1 161162844 161163031
refGene_gn=SSB;refGene_tx=XM_011285323.3
C1 161163147 161163279

refGene_gn=SSB;refGene_tx=XM_011285323.3
C1 161163593 161163685
refGene_gn=SSB;refGene_tx=XM_011285323.3
C1 161163750 161163789
refGene_gn=METTL5;refGene_tx=XM_006935349.4;refGene_gn=METTL5;refGene_tx=XM_023259435.1;refGene_gn=METTL5;refGene_tx=XM_023259436.1;refGene_gn=METTL5;refGene_tx=XM_023259437.1;refGene_gn=METTL5;refGene_tx=XM_023259438.1;refGene_gn=METTL5;refGene_tx=XM_023259439.1
C1 161164182 161164250
refGene_gn=METTL5;refGene_tx=XM_006935349.4;refGene_gn=METTL5;refGene_tx=XM_023259435.1;refGene_gn=METTL5;refGene_tx=XM_023259436.1;refGene_gn=METTL5;refGene_tx=XM_023259437.1;refGene_gn=METTL5;refGene_tx=XM_023259438.1
C1 161166126 161166178
refGene_gn=METTL5;refGene_tx=XM_023259435.1;refGene_gn=METTL5;refGene_tx=XM_023259436.1;refGene_gn=METTL5;refGene_tx=XM_023259437.1;refGene_gn=METTL5;refGene_tx=XM_023259438.1;refGene_gn=METTL5;refGene_tx=XM_023259439.1
C1 161166966 161167049
refGene_gn=METTL5;refGene_tx=XM_006935349.4;refGene_gn=METTL5;refGene_tx=XM_023259435.1;refGene_gn=METTL5;refGene_tx=XM_023259436.1;refGene_gn=METTL5;refGene_tx=XM_023259437.1;refGene_gn=METTL5;refGene_tx=XM_023259438.1;refGene_gn=METTL5;refGene_tx=XM_023259439.1
C1 161168275 161168457
refGene_gn=METTL5;refGene_tx=XM_006935349.4;refGene_gn=METTL5;refGene_tx=XM_023259435.1;refGene_gn=METTL5;refGene_tx=XM_023259436.1;refGene_gn=METTL5;refGene_tx=XM_023259437.1;refGene_gn=METTL5;refGene_tx=XM_023259438.1;refGene_gn=METTL5;refGene_tx=XM_023259439.1
C1 161168828 161168943
refGene_gn=METTL5;refGene_tx=XM_006935349.4;refGene_gn=METTL5;refGene_tx=XM_023259435.1;refGene_gn=METTL5;refGene_tx=XM_023259436.1;refGene_gn=METTL5;refGene_tx=XM_023259437.1;refGene_gn=METTL5;refGene_tx=XM_023259438.1;refGene_gn=METTL5;refGene_tx=XM_023259439.1
C1 161170808 161170917
refGene_gn=METTL5;refGene_tx=XM_006935349.4;refGene_gn=METTL5;refGene_tx=XM_023259435.1;refGene_gn=METTL5;refGene_tx=XM_023259436.1;refGene_gn=METTL5;refGene_tx=XM_023259437.1;refGene_gn=METTL5;refGene_tx=XM_023259438.1;refGene_gn=METTL5;refGene_tx=XM_023259439.1
C1 161195729 161196274
refGene_gn=UBR3;refGene_tx=XM_023259440.1;refGene_gn=UBR3;refGene_tx=XM_023259442.1;refGene_gn=UBR3;refGene_tx=XM_023259443.1;refGene_gn=UBR3;refGene_tx=XM_023259444.1;refGene_gn=UBR3;refGene_tx=XM_023259445.1
C1 161235047 161235187
refGene_gn=UBR3;refGene_tx=XM_023259440.1;refGene_gn=UBR3;refGene_tx=XM_023259442.1;refGene_gn=UBR3;refGene_tx=XM_023259443.1;refGene_gn=UBR3;refGene_tx=XM_023259444.1;refGene_gn=UBR3;refGene_tx=XM_023259445.1
C1 161238172 161238331
refGene_gn=UBR3;refGene_tx=XM_023259440.1;refGene_gn=UBR3;refGene_tx=XM_023259442.1;refGene_gn=UBR3;refGene_tx=XM_023259443.1;refGene_gn=UBR3;refGene_tx=XM_023259444.1;refGene_gn=UBR3;refGene_tx=XM_023259445.1

C1	161244374	161244518
refGene_gn=UBR3;refGene_tx=XM_023259440.1;refGene_gn=UBR3;refGene_tx=XM_023259442.1;refGene_gn=UBR3;refGene_tx=XM_023259443.1;refGene_gn=UBR3;refGene_tx=XM_023259444.1;refGene_gn=UBR3;refGene_tx=XM_023259445.1
C1	161245058	161245108
refGene_gn=UBR3;refGene_tx=XM_023259440.1;refGene_gn=UBR3;refGene_tx=XM_023259442.1;refGene_gn=UBR3;refGene_tx=XM_023259443.1;refGene_gn=UBR3;refGene_tx=XM_023259444.1;refGene_gn=UBR3;refGene_tx=XM_023259445.1
C1	161275074	161275141
refGene_gn=UBR3;refGene_tx=XM_023259440.1;refGene_gn=UBR3;refGene_tx=XM_023259442.1;refGene_gn=UBR3;refGene_tx=XM_023259443.1;refGene_gn=UBR3;refGene_tx=XM_023259444.1;refGene_gn=UBR3;refGene_tx=XM_023259445.1
C1	161281983	161282114
refGene_gn=UBR3;refGene_tx=XM_023259440.1;refGene_gn=UBR3;refGene_tx=XM_023259442.1;refGene_gn=UBR3;refGene_tx=XM_023259443.1;refGene_gn=UBR3;refGene_tx=XM_023259444.1;refGene_gn=UBR3;refGene_tx=XM_023259445.1
C1	161283643	161283872
refGene_gn=UBR3;refGene_tx=XM_023259440.1;refGene_gn=UBR3;refGene_tx=XM_023259442.1;refGene_gn=UBR3;refGene_tx=XM_023259443.1;refGene_gn=UBR3;refGene_tx=XM_023259444.1;refGene_gn=UBR3;refGene_tx=XM_023259445.1
C1	161288367	161288547
refGene_gn=UBR3;refGene_tx=XM_023259440.1;refGene_gn=UBR3;refGene_tx=XM_023259442.1;refGene_gn=UBR3;refGene_tx=XM_023259443.1;refGene_gn=UBR3;refGene_tx=XM_023259444.1;refGene_gn=UBR3;refGene_tx=XM_023259445.1
C1	161289121	161289255
refGene_gn=UBR3;refGene_tx=XM_023259440.1;refGene_gn=UBR3;refGene_tx=XM_023259442.1;refGene_gn=UBR3;refGene_tx=XM_023259443.1;refGene_gn=UBR3;refGene_tx=XM_023259444.1;refGene_gn=UBR3;refGene_tx=XM_023259445.1
C1	161295896	161295983
refGene_gn=UBR3;refGene_tx=XM_023259440.1;refGene_gn=UBR3;refGene_tx=XM_023259442.1;refGene_gn=UBR3;refGene_tx=XM_023259443.1;refGene_gn=UBR3;refGene_tx=XM_023259444.1;refGene_gn=UBR3;refGene_tx=XM_023259445.1
C1	161299013	161299079
refGene_gn=UBR3;refGene_tx=XM_023259440.1;refGene_gn=UBR3;refGene_tx=XM_023259442.1;refGene_gn=UBR3;refGene_tx=XM_023259443.1;refGene_gn=UBR3;refGene_tx=XM_023259444.1;refGene_gn=UBR3;refGene_tx=XM_023259445.1
C1	161299172	161299262
refGene_gn=UBR3;refGene_tx=XM_023259440.1;refGene_gn=UBR3;refGene_tx=XM_023259442.1;refGene_gn=UBR3;refGene_tx=XM_023259443.1;refGene_gn=UBR3;refGene_tx=XM_023259444.1;refGene_gn=UBR3;refGene_tx=XM_023259445.1
C1	161300597	161300726
refGene_gn=UBR3;refGene_tx=XM_023259440.1;refGene_gn=UBR3;refGene_tx=XM_023259442.1;refGene_gn=UBR3;refGene_tx=XM_023259443.1;refGene_gn=UBR3;refGene_tx=XM_023259444.1;refGene_gn=UBR3;refGene_tx=XM_023259445.1
C1	161301392	161301445
refGene_gn=UBR3;refGene_tx=XM_023259440.1;refGene_gn=UBR3;refGene_tx=XM_023259442.1;refGene_gn=UBR3;refGene_tx=XM_023259443.1;refGene_gn=UBR3;refGene_tx=XM_023259444.1;refGene_gn=UBR3;refGene_tx=XM_023259445.1
C1	161301547	161301681
refGene_gn=UBR3;refGene_tx=XM_023259440.1;refGene_gn=UBR3;refGene_tx=X

M_023259442.1;refGene_gn=UBR3;refGene_tx=XM_023259443.1;refGene_gn=UBR3;refGene_tx=XM_023259444.1;refGene_gn=UBR3;refGene_tx=XM_023259445.1
C1 161302044 161302130
refGene_gn=UBR3;refGene_tx=XM_023259440.1;refGene_gn=UBR3;refGene_tx=XM_023259442.1;refGene_gn=UBR3;refGene_tx=XM_023259443.1;refGene_gn=UBR3;refGene_tx=XM_023259444.1;refGene_gn=UBR3;refGene_tx=XM_023259445.1
C1 161303662 161303804
refGene_gn=UBR3;refGene_tx=XM_023259440.1;refGene_gn=UBR3;refGene_tx=XM_023259442.1;refGene_gn=UBR3;refGene_tx=XM_023259443.1;refGene_gn=UBR3;refGene_tx=XM_023259444.1;refGene_gn=UBR3;refGene_tx=XM_023259445.1
C1 161306250 161306347
refGene_gn=UBR3;refGene_tx=XM_023259440.1;refGene_gn=UBR3;refGene_tx=XM_023259442.1;refGene_gn=UBR3;refGene_tx=XM_023259443.1;refGene_gn=UBR3;refGene_tx=XM_023259444.1;refGene_gn=UBR3;refGene_tx=XM_023259445.1
C1 161320057 161320199
refGene_gn=UBR3;refGene_tx=XM_023259440.1;refGene_gn=UBR3;refGene_tx=XM_023259442.1;refGene_gn=UBR3;refGene_tx=XM_023259443.1;refGene_gn=UBR3;refGene_tx=XM_023259444.1;refGene_gn=UBR3;refGene_tx=XM_023259445.1
C1 161323732 161323837
refGene_gn=UBR3;refGene_tx=XM_023259440.1;refGene_gn=UBR3;refGene_tx=XM_023259442.1;refGene_gn=UBR3;refGene_tx=XM_023259443.1;refGene_gn=UBR3;refGene_tx=XM_023259444.1;refGene_gn=UBR3;refGene_tx=XM_023259445.1
C1 161325512 161325686
refGene_gn=UBR3;refGene_tx=XM_023259440.1;refGene_gn=UBR3;refGene_tx=XM_023259442.1;refGene_gn=UBR3;refGene_tx=XM_023259443.1;refGene_gn=UBR3;refGene_tx=XM_023259444.1;refGene_gn=UBR3;refGene_tx=XM_023259445.1
C1 161325894 161325927
refGene_gn=UBR3;refGene_tx=XM_023259440.1;refGene_gn=UBR3;refGene_tx=XM_023259442.1;refGene_gn=UBR3;refGene_tx=XM_023259445.1
C1 161327555 161328016
refGene_gn=UBR3;refGene_tx=XM_023259440.1;refGene_gn=UBR3;refGene_tx=XM_023259442.1;refGene_gn=UBR3;refGene_tx=XM_023259443.1;refGene_gn=UBR3;refGene_tx=XM_023259444.1;refGene_gn=UBR3;refGene_tx=XM_023259445.1
C1 161334409 161334498
refGene_gn=UBR3;refGene_tx=XM_023259440.1;refGene_gn=UBR3;refGene_tx=XM_023259442.1;refGene_gn=UBR3;refGene_tx=XM_023259443.1;refGene_gn=UBR3;refGene_tx=XM_023259444.1;refGene_gn=UBR3;refGene_tx=XM_023259445.1
C1 161347412 161347562
refGene_gn=UBR3;refGene_tx=XM_023259440.1;refGene_gn=UBR3;refGene_tx=XM_023259442.1;refGene_gn=UBR3;refGene_tx=XM_023259443.1;refGene_gn=UBR3;refGene_tx=XM_023259444.1;refGene_gn=UBR3;refGene_tx=XM_023259445.1
C1 161355825 161355959
refGene_gn=UBR3;refGene_tx=XM_023259440.1;refGene_gn=UBR3;refGene_tx=XM_023259442.1;refGene_gn=UBR3;refGene_tx=XM_023259443.1;refGene_gn=UBR3;refGene_tx=XM_023259444.1;refGene_gn=UBR3;refGene_tx=XM_023259445.1
C1 161362053 161362164
refGene_gn=UBR3;refGene_tx=XM_023259440.1;refGene_gn=UBR3;refGene_tx=XM_023259442.1;refGene_gn=UBR3;refGene_tx=XM_023259443.1;refGene_gn=UBR3;refGene_tx=XM_023259444.1;refGene_gn=UBR3;refGene_tx=XM_023259445.1
C1 161366984 161367197

refGene_gn=UBR3;refGene_tx=XM_023259440.1;refGene_gn=UBR3;refGene_tx=XM_023259442.1;refGene_gn=UBR3;refGene_tx=XM_023259443.1;refGene_gn=UBR3;refGene_tx=XM_023259444.1;refGene_gn=UBR3;refGene_tx=XM_023259445.1
C1 161368657 161368794
refGene_gn=UBR3;refGene_tx=XM_023259440.1;refGene_gn=UBR3;refGene_tx=XM_023259442.1;refGene_gn=UBR3;refGene_tx=XM_023259443.1;refGene_gn=UBR3;refGene_tx=XM_023259444.1;refGene_gn=UBR3;refGene_tx=XM_023259445.1
C1 161374873 161374959
refGene_gn=UBR3;refGene_tx=XM_023259440.1;refGene_gn=UBR3;refGene_tx=XM_023259442.1;refGene_gn=UBR3;refGene_tx=XM_023259443.1;refGene_gn=UBR3;refGene_tx=XM_023259444.1;refGene_gn=UBR3;refGene_tx=XM_023259445.1
C1 161377584 161377688
refGene_gn=UBR3;refGene_tx=XM_023259445.1
C1 161386691 161386797
refGene_gn=UBR3;refGene_tx=XM_023259440.1;refGene_gn=UBR3;refGene_tx=XM_023259442.1;refGene_gn=UBR3;refGene_tx=XM_023259443.1;refGene_gn=UBR3;refGene_tx=XM_023259444.1;refGene_gn=UBR3;refGene_tx=XM_023259445.1
C1 161399190 161399294
refGene_gn=UBR3;refGene_tx=XM_023259440.1;refGene_gn=UBR3;refGene_tx=XM_023259442.1;refGene_gn=UBR3;refGene_tx=XM_023259443.1;refGene_gn=UBR3;refGene_tx=XM_023259444.1
C1 161411016 161411141
refGene_gn=UBR3;refGene_tx=XM_023259440.1;refGene_gn=UBR3;refGene_tx=XM_023259442.1;refGene_gn=UBR3;refGene_tx=XM_023259443.1;refGene_gn=UBR3;refGene_tx=XM_023259444.1
C1 161417013 161417121
refGene_gn=UBR3;refGene_tx=XM_023259440.1;refGene_gn=UBR3;refGene_tx=XM_023259442.1;refGene_gn=UBR3;refGene_tx=XM_023259443.1;refGene_gn=UBR3;refGene_tx=XM_023259444.1
C1 161417241 161417367
refGene_gn=UBR3;refGene_tx=XM_023259440.1;refGene_gn=UBR3;refGene_tx=XM_023259442.1;refGene_gn=UBR3;refGene_tx=XM_023259443.1;refGene_gn=UBR3;refGene_tx=XM_023259444.1
C1 161432759 161432939
refGene_gn=UBR3;refGene_tx=XM_023259440.1;refGene_gn=UBR3;refGene_tx=XM_023259442.1;refGene_gn=UBR3;refGene_tx=XM_023259443.1;refGene_gn=UBR3;refGene_tx=XM_023259444.1
C1 161437902 161438112
refGene_gn=UBR3;refGene_tx=XM_023259440.1;refGene_gn=UBR3;refGene_tx=XM_023259442.1;refGene_gn=UBR3;refGene_tx=XM_023259443.1;refGene_gn=UBR3;refGene_tx=XM_023259444.1
C1 161438871 161439011
refGene_gn=UBR3;refGene_tx=XM_023259440.1;refGene_gn=UBR3;refGene_tx=XM_023259442.1;refGene_gn=UBR3;refGene_tx=XM_023259443.1;refGene_gn=UBR3;refGene_tx=XM_023259444.1
C1 161439315 161439433
refGene_gn=UBR3;refGene_tx=XM_023259440.1;refGene_gn=UBR3;refGene_tx=XM_023259442.1;refGene_gn=UBR3;refGene_tx=XM_023259443.1;refGene_gn=UBR3;refGene_tx=XM_023259444.1
C1 161535669 161535746

refGene_gn=MYO3B;refGene_tx=XM_011285321.3
C1	161554553	161554737
refGene_gn=MYO3B;refGene_tx=XM_011285321.3
C1	161555458	161555593
refGene_gn=MYO3B;refGene_tx=XM_011285321.3
C1	161566113	161566218
refGene_gn=MYO3B;refGene_tx=XM_011285321.3
C1	161566462	161566562
refGene_gn=MYO3B;refGene_tx=XM_011285321.3
C1	161569527	161569604
refGene_gn=MYO3B;refGene_tx=XM_011285321.3
C1	161588987	161589133
refGene_gn=MYO3B;refGene_tx=XM_011285321.3
C1	161673390	161673456
refGene_gn=MYO3B;refGene_tx=XM_011285321.3
C1	161703860	161704016
refGene_gn=MYO3B;refGene_tx=XM_011285321.3
C1	161715715	161715812
refGene_gn=MYO3B;refGene_tx=XM_011285321.3
C1	161716574	161716691
refGene_gn=MYO3B;refGene_tx=XM_011285321.3
C1	161717206	161717311
refGene_gn=MYO3B;refGene_tx=XM_011285321.3
C1	161719918	161720002
refGene_gn=MYO3B;refGene_tx=XM_011285321.3
C1	161720802	161721005
refGene_gn=MYO3B;refGene_tx=XM_011285321.3
C1	161725228	161725327
refGene_gn=MYO3B;refGene_tx=XM_011285321.3
C1	161725700	161725815
refGene_gn=MYO3B;refGene_tx=XM_011285321.3
C1	161732241	161732368
refGene_gn=MYO3B;refGene_tx=XM_011285321.3
C1	161733140	161733351
refGene_gn=MYO3B;refGene_tx=XM_011285321.3
C1	161734193	161734341
refGene_gn=MYO3B;refGene_tx=XM_011285321.3
C1	161735603	161735757
refGene_gn=MYO3B;refGene_tx=XM_011285321.3
C1	161736941	161737030
refGene_gn=MYO3B;refGene_tx=XM_011285321.3
C1	161738972	161739102
refGene_gn=MYO3B;refGene_tx=XM_011285321.3
C1	161772532	161772612
refGene_gn=MYO3B;refGene_tx=XM_011285321.3
C1	161779478	161779556
refGene_gn=MYO3B;refGene_tx=XM_011285321.3
C1	161781676	161781882
refGene_gn=MYO3B;refGene_tx=XM_011285321.3
C1	161804976	161805088

refGene_gn=MYO3B;refGene_tx=XM_011285321.3
C1 161806273 161806436
refGene_gn=MYO3B;refGene_tx=XM_011285321.3
C1 161807819 161807900
refGene_gn=MYO3B;refGene_tx=XM_011285321.3
C1 161822186 161822288
refGene_gn=MYO3B;refGene_tx=XM_011285321.3
C1 161826332 161826435
refGene_gn=MYO3B;refGene_tx=XM_011285321.3
C1 161853995 161854056
refGene_gn=MYO3B;refGene_tx=XM_011285321.3
C1 161854982 161855082
refGene_gn=MYO3B;refGene_tx=XM_011285321.3
C1 161941954 161942061
refGene_gn=MYO3B;refGene_tx=XM_011285321.3
C1 161942435 161942482
refGene_gn=MYO3B;refGene_tx=XM_011285321.3
C1 161943294 161943433
refGene_gn=MYO3B;refGene_tx=XM_011285321.3
C1 161994480 161994531
refGene_gn=SP5;refGene_tx=XM_023259446.1
C1 161995240 161996389
refGene_gn=SP5;refGene_tx=XM_023259446.1;refGene_gn=SP5;refGene_tx=XM_023259447.1
C1 162030284 162030850
refGene_gn=ERICH2;refGene_tx=XM_019838242.2;refGene_gn=ERICH2;refGene_tx=XM_019838243.2;refGene_gn=ERICH2;refGene_tx=XM_023259448.1;refGene_gn=ERICH2;refGene_tx=XM_023259449.1;refGene_gn=ERICH2;refGene_tx=XM_023259450.1;refGene_gn=ERICH2;refGene_tx=XM_023259451.1
C1 162030964 162031010
refGene_gn=ERICH2;refGene_tx=XM_019838242.2;refGene_gn=ERICH2;refGene_tx=XM_019838243.2;refGene_gn=ERICH2;refGene_tx=XM_023259448.1;refGene_gn=ERICH2;refGene_tx=XM_023259449.1;refGene_gn=ERICH2;refGene_tx=XM_023259450.1;refGene_gn=ERICH2;refGene_tx=XM_023259451.1
C1 162038385 162038434
refGene_gn=ERICH2;refGene_tx=XM_023259448.1;refGene_gn=ERICH2;refGene_tx=XM_023259449.1
C1 162038667 162038704
refGene_gn=ERICH2;refGene_tx=XM_006935356.4;refGene_gn=ERICH2;refGene_tx=XM_011285317.3;refGene_gn=ERICH2;refGene_tx=XM_011285318.3;refGene_gn=ERICH2;refGene_tx=XM_023259448.1;refGene_gn=ERICH2;refGene_tx=XM_023259449.1;refGene_gn=ERICH2;refGene_tx=XM_023259450.1
C1 162041098 162041194
refGene_gn=ERICH2;refGene_tx=XM_011285317.3;refGene_gn=ERICH2;refGene_tx=XM_019838242.2;refGene_gn=ERICH2;refGene_tx=XM_019838243.2;refGene_gn=ERICH2;refGene_tx=XM_023259448.1;refGene_gn=ERICH2;refGene_tx=XM_023259449.1;refGene_gn=ERICH2;refGene_tx=XM_023259451.1
C1 162043268 162043417
refGene_gn=ERICH2;refGene_tx=XM_011285317.3;refGene_gn=ERICH2;refGene_tx=XM_019838242.2;refGene_gn=ERICH2;refGene_tx=XM_023259448.1

C1	162043934	162044019
refGene_gn=ERICH2;refGene_tx=XM_006935354.3
C1	162046721	162046909
refGene_gn=ERICH2;refGene_tx=XM_006935354.3;refGene_gn=ERICH2;refGene_tx=XM_006935356.4;refGene_gn=ERICH2;refGene_tx=XM_011285317.3;refGene_gn=ERICH2;refGene_tx=XM_011285318.3;refGene_gn=ERICH2;refGene_tx=XM_019838242.2;refGene_gn=ERICH2;refGene_tx=XM_019838243.2;refGene_gn=ERICH2;refGene_tx=XM_023259448.1;refGene_gn=ERICH2;refGene_tx=XM_023259449.1
C1	162058996	162059054
refGene_gn=ERICH2;refGene_tx=XM_006935354.3;refGene_gn=ERICH2;refGene_tx=XM_006935356.4;refGene_gn=ERICH2;refGene_tx=XM_011285317.3;refGene_gn=ERICH2;refGene_tx=XM_011285318.3;refGene_gn=ERICH2;refGene_tx=XM_019838242.2;refGene_gn=ERICH2;refGene_tx=XM_023259448.1
C1	162066315	162066387
refGene_gn=ERICH2;refGene_tx=XM_006935354.3;refGene_gn=ERICH2;refGene_tx=XM_006935356.4;refGene_gn=ERICH2;refGene_tx=XM_011285317.3;refGene_gn=ERICH2;refGene_tx=XM_011285318.3;refGene_gn=ERICH2;refGene_tx=XM_019838242.2;refGene_gn=ERICH2;refGene_tx=XM_023259448.1
C1	162067032	162067154
refGene_gn=ERICH2;refGene_tx=XM_006935354.3;refGene_gn=ERICH2;refGene_tx=XM_006935356.4;refGene_gn=ERICH2;refGene_tx=XM_011285317.3;refGene_gn=ERICH2;refGene_tx=XM_011285318.3;refGene_gn=ERICH2;refGene_tx=XM_019838242.2;refGene_gn=ERICH2;refGene_tx=XM_023259448.1
C1	162085404	162085486
refGene_gn=GAD1;refGene_tx=NM_001009225.1;refGene_gn=GAD1;refGene_tx=XM_019835946.2;refGene_gn=GAD1;refGene_tx=XM_019835947.2
C1	162088429	162088541
refGene_gn=GAD1;refGene_tx=XM_019835945.2
C1	162088901	162088964
refGene_gn=GAD1;refGene_tx=NM_001009225.1;refGene_gn=GAD1;refGene_tx=XM_019835945.2;refGene_gn=GAD1;refGene_tx=XM_019835946.2;refGene_gn=GAD1;refGene_tx=XM_019835947.2
C1	162095016	162095175
refGene_gn=GAD1;refGene_tx=NM_001009225.1;refGene_gn=GAD1;refGene_tx=XM_019835945.2;refGene_gn=GAD1;refGene_tx=XM_019835946.2;refGene_gn=GAD1;refGene_tx=XM_019835947.2
C1	162096537	162096780
refGene_gn=GAD1;refGene_tx=NM_001009225.1;refGene_gn=GAD1;refGene_tx=XM_019835945.2;refGene_gn=GAD1;refGene_tx=XM_019835946.2;refGene_gn=GAD1;refGene_tx=XM_019835947.2
C1	162102927	162103018
refGene_gn=GAD1;refGene_tx=NM_001009225.1;refGene_gn=GAD1;refGene_tx=XM_019835945.2;refGene_gn=GAD1;refGene_tx=XM_019835946.2;refGene_gn=GAD1;refGene_tx=XM_019835947.2
C1	162109672	162109785
refGene_gn=GAD1;refGene_tx=NM_001009225.1;refGene_gn=GAD1;refGene_tx=XM_019835945.2;refGene_gn=GAD1;refGene_tx=XM_019835946.2;refGene_gn=GAD1;refGene_tx=XM_019835947.2
C1	162110932	162111048

refGene_gn=GAD1;refGene_tx=NM_001009225.1;refGene_gn=GAD1;refGene_tx=XM_019835945.2;refGene_gn=GAD1;refGene_tx=XM_019835946.2;refGene_gn=GAD1;refGene_tx=XM_019835947.2
C1 162111131 162111211
refGene_gn=GAD1;refGene_tx=NM_001009225.1;refGene_gn=GAD1;refGene_tx=XM_019835945.2;refGene_gn=GAD1;refGene_tx=XM_019835946.2;refGene_gn=GAD1;refGene_tx=XM_019835947.2
C1 162111434 162111489
refGene_gn=GAD1;refGene_tx=NM_001009225.1;refGene_gn=GAD1;refGene_tx=XM_019835945.2;refGene_gn=GAD1;refGene_tx=XM_019835946.2;refGene_gn=GAD1;refGene_tx=XM_019835947.2
C1 162113002 162113119
refGene_gn=GAD1;refGene_tx=NM_001009225.1;refGene_gn=GAD1;refGene_tx=XM_019835945.2;refGene_gn=GAD1;refGene_tx=XM_019835946.2;refGene_gn=GAD1;refGene_tx=XM_019835947.2
C1 162114287 162114352
refGene_gn=GAD1;refGene_tx=NM_001009225.1;refGene_gn=GAD1;refGene_tx=XM_019835945.2;refGene_gn=GAD1;refGene_tx=XM_019835946.2;refGene_gn=GAD1;refGene_tx=XM_019835947.2
C1 162118440 162118519
refGene_gn=GAD1;refGene_tx=NM_001009225.1;refGene_gn=GAD1;refGene_tx=XM_019835945.2;refGene_gn=GAD1;refGene_tx=XM_019835946.2;refGene_gn=GAD1;refGene_tx=XM_019835947.2
C1 162119648 162119798
refGene_gn=GAD1;refGene_tx=NM_001009225.1;refGene_gn=GAD1;refGene_tx=XM_019835945.2;refGene_gn=GAD1;refGene_tx=XM_019835946.2;refGene_gn=GAD1;refGene_tx=XM_019835947.2
C1 162122176 162122284
refGene_gn=GAD1;refGene_tx=NM_001009225.1;refGene_gn=GAD1;refGene_tx=XM_019835945.2;refGene_gn=GAD1;refGene_tx=XM_019835946.2;refGene_gn=GAD1;refGene_tx=XM_019835947.2
C1 162123820 162123910
refGene_gn=GAD1;refGene_tx=NM_001009225.1;refGene_gn=GAD1;refGene_tx=XM_019835945.2;refGene_gn=GAD1;refGene_tx=XM_019835946.2;refGene_gn=GAD1;refGene_tx=XM_019835947.2
C1 162124682 162124856
refGene_gn=GAD1;refGene_tx=NM_001009225.1;refGene_gn=GAD1;refGene_tx=XM_019835945.2;refGene_gn=GAD1;refGene_tx=XM_019835946.2;refGene_gn=GAD1;refGene_tx=XM_019835947.2
C1 162176038 162176101
refGene_gn=GORASP2;refGene_tx=XM_003990844.5
C1 162195764 162195845
refGene_gn=GORASP2;refGene_tx=XM_003990844.5
C1 162198152 162198356
refGene_gn=GORASP2;refGene_tx=XM_003990844.5
C1 162199274 162199361
refGene_gn=GORASP2;refGene_tx=XM_003990844.5
C1 162200077 162200208
refGene_gn=GORASP2;refGene_tx=XM_003990844.5
C1 162203238 162203371

refGene_gn=GORASP2;refGene_tx=XM_003990844.5
C1 162205931 162206055
refGene_gn=GORASP2;refGene_tx=XM_003990844.5
C1 162209938 162210025
refGene_gn=GORASP2;refGene_tx=XM_003990844.5
C1 162211120 162211231
refGene_gn=GORASP2;refGene_tx=XM_003990844.5
C1 162212834 162213175
refGene_gn=GORASP2;refGene_tx=XM_003990844.5
C1 162263255 162263432
refGene_gn=TLK1;refGene_tx=XM_011285316.3
C1 162269130 162269238
refGene_gn=TLK1;refGene_tx=XM_011285316.3
C1 162270034 162270146
refGene_gn=TLK1;refGene_tx=XM_011285316.3
C1 162272985 162273121
refGene_gn=TLK1;refGene_tx=XM_011285316.3
C1 162273399 162273569
refGene_gn=TLK1;refGene_tx=XM_011285316.3
C1 162273713 162273803
refGene_gn=TLK1;refGene_tx=XM_011285316.3
C1 162273887 162273979
refGene_gn=TLK1;refGene_tx=XM_011285316.3
C1 162278964 162279046
refGene_gn=TLK1;refGene_tx=XM_011285316.3
C1 162281615 162281713
refGene_gn=TLK1;refGene_tx=XM_011285316.3
C1 162293960 162294027
refGene_gn=TLK1;refGene_tx=XM_011285316.3
C1 162308962 162309151
refGene_gn=TLK1;refGene_tx=XM_011285316.3
C1 162313395 162313532
refGene_gn=TLK1;refGene_tx=XM_011285316.3
C1 162313633 162313744
refGene_gn=TLK1;refGene_tx=XM_011285316.3
C1 162317540 162317573
refGene_gn=TLK1;refGene_tx=XM_011285316.3
C1 162317574 162317634
refGene_gn=TLK1;refGene_tx=XM_011285316.3
C1 162318991 162319081
refGene_gn=TLK1;refGene_tx=XM_011285316.3
C1 162320390 162320486
refGene_gn=TLK1;refGene_tx=XM_011285316.3
C1 162322153 162322200
refGene_gn=TLK1;refGene_tx=XM_011285316.3
C1 162326772 162326848
refGene_gn=TLK1;refGene_tx=XM_011285316.3
C1 162344044 162344116
refGene_gn=TLK1;refGene_tx=XM_011285316.3
C1 162366595 162366714

refGene_gn=TLK1;refGene_tx=XM_011285316.3
C1 162409477 162409616
refGene_gn=TLK1;refGene_tx=XM_011285316.3
C1 162545553 162545576
refGene_gn=METTL8;refGene_tx=XM_019838253.2
C1 162555725 162555916
refGene_gn=METTL8;refGene_tx=XM_019838252.2;refGene_gn=METTL8;refGene_tx=XM_023259454.1;refGene_gn=METTL8;refGene_tx=XM_023259455.1;refGene_gn=METTL8;refGene_tx=XM_023259456.1
C1 162556881 162556947
refGene_gn=METTL8;refGene_tx=XM_019838252.2;refGene_gn=METTL8;refGene_tx=XM_019838253.2;refGene_gn=METTL8;refGene_tx=XM_023259454.1;refGene_gn=METTL8;refGene_tx=XM_023259455.1;refGene_gn=METTL8;refGene_tx=XM_023259456.1
C1 162557068 162557175
refGene_gn=METTL8;refGene_tx=XM_019838252.2;refGene_gn=METTL8;refGene_tx=XM_019838253.2;refGene_gn=METTL8;refGene_tx=XM_023259454.1;refGene_gn=METTL8;refGene_tx=XM_023259455.1;refGene_gn=METTL8;refGene_tx=XM_023259456.1
C1 162559041 162559162
refGene_gn=METTL8;refGene_tx=XM_019838254.2
C1 162561491 162561631
refGene_gn=METTL8;refGene_tx=XM_019838252.2;refGene_gn=METTL8;refGene_tx=XM_019838253.2;refGene_gn=METTL8;refGene_tx=XM_019838254.2;refGene_gn=METTL8;refGene_tx=XM_023259454.1;refGene_gn=METTL8;refGene_tx=XM_023259455.1;refGene_gn=METTL8;refGene_tx=XM_023259456.1
C1 162563275 162563339
refGene_gn=METTL8;refGene_tx=XM_019838252.2;refGene_gn=METTL8;refGene_tx=XM_019838253.2;refGene_gn=METTL8;refGene_tx=XM_019838254.2;refGene_gn=METTL8;refGene_tx=XM_023259454.1;refGene_gn=METTL8;refGene_tx=XM_023259455.1;refGene_gn=METTL8;refGene_tx=XM_023259456.1
C1 162568861 162568911
refGene_gn=METTL8;refGene_tx=XM_019838252.2;refGene_gn=METTL8;refGene_tx=XM_019838253.2;refGene_gn=METTL8;refGene_tx=XM_019838254.2;refGene_gn=METTL8;refGene_tx=XM_023259454.1;refGene_gn=METTL8;refGene_tx=XM_023259455.1;refGene_gn=METTL8;refGene_tx=XM_023259456.1
C1 162570694 162571065
refGene_gn=METTL8;refGene_tx=XM_019838252.2;refGene_gn=METTL8;refGene_tx=XM_019838253.2;refGene_gn=METTL8;refGene_tx=XM_019838254.2;refGene_gn=METTL8;refGene_tx=XM_023259454.1;refGene_gn=METTL8;refGene_tx=XM_023259455.1;refGene_gn=METTL8;refGene_tx=XM_023259456.1
C1 162585078 162585170
refGene_gn=METTL8;refGene_tx=XM_019838252.2;refGene_gn=METTL8;refGene_tx=XM_019838253.2;refGene_gn=METTL8;refGene_tx=XM_019838254.2;refGene_gn=METTL8;refGene_tx=XM_023259454.1;refGene_gn=METTL8;refGene_tx=XM_023259455.1;refGene_gn=METTL8;refGene_tx=XM_023259456.1
C1 162600703 162600858
refGene_gn=METTL8;refGene_tx=XM_019838252.2;refGene_gn=METTL8;refGene_tx=XM_019838253.2;refGene_gn=METTL8;refGene_tx=XM_019838254.2;refGene_gn=METTL8;refGene_tx=XM_023259454.1;refGene_gn=METTL8;refGene_tx=XM_02

3259455.1;refGene_gn=METTL8;refGene_tx=XM_023259456.1
C1 162623953 162624001
refGene_gn=METTL8;refGene_tx=XM_019838252.2;refGene_gn=METTL8;refGene_tx=XM_019838253.2;refGene_gn=METTL8;refGene_tx=XM_019838254.2
C1 162643654 162643780
refGene_gn=DCAF17;refGene_tx=XM_023259453.1
C1 162644184 162644288
refGene_gn=DCAF17;refGene_tx=XM_023259453.1
C1 162646730 162646821
refGene_gn=DCAF17;refGene_tx=XM_023259453.1
C1 162650927 162651064
refGene_gn=DCAF17;refGene_tx=XM_023259453.1
C1 162652410 162652489
refGene_gn=DCAF17;refGene_tx=XM_023259453.1
C1 162653308 162653398
refGene_gn=DCAF17;refGene_tx=XM_023259453.1
C1 162655841 162655946
refGene_gn=DCAF17;refGene_tx=XM_023259453.1
C1 162656245 162656351
refGene_gn=DCAF17;refGene_tx=XM_023259453.1
C1 162664405 162664548
refGene_gn=DCAF17;refGene_tx=XM_023259453.1
C1 162667877 162667987
refGene_gn=DCAF17;refGene_tx=XM_023259453.1
C1 162671748 162671810
refGene_gn=DCAF17;refGene_tx=XM_023259453.1
C1 162671812 162671838
refGene_gn=DCAF17;refGene_tx=XM_023259453.1
C1 162672473 162672557
refGene_gn=DCAF17;refGene_tx=XM_023259453.1
C1 162674591 162674747
refGene_gn=DCAF17;refGene_tx=XM_023259453.1
C1 162675522 162675669
refGene_gn=DCAF17;refGene_tx=XM_023259453.1
C1 162703033 162703226
refGene_gn=LOC101089456;refGene_tx=XM_003990848.5
C1 162719760 162719969
refGene_gn=LOC101089456;refGene_tx=XM_003990848.5
C1 162735132 162735287
refGene_gn=LOC101089456;refGene_tx=XM_003990848.5
C1 162736053 162736351
refGene_gn=LOC101089456;refGene_tx=XM_003990848.5
C1 162847415 162847523
refGene_gn=DYNC1I2;refGene_tx=XM_006935360.4;refGene_gn=DYNC1I2;refGene_tx=XM_019838256.2;refGene_gn=DYNC1I2;refGene_tx=XM_019838257.2;refGene_gn=DYNC1I2;refGene_tx=XM_019838258.2;refGene_gn=DYNC1I2;refGene_tx=XM_019838259.2;refGene_gn=DYNC1I2;refGene_tx=XM_019838260.2
C1 162849483 162849601
refGene_gn=DYNC1I2;refGene_tx=XM_006935360.4;refGene_gn=DYNC1I2;refGene_tx=XM_019838256.2;refGene_gn=DYNC1I2;refGene_tx=XM_019838257.2;refGe

ne_gn=DYNC1I2;refGene_tx=XM_019838258.2;refGene_gn=DYNC1I2;refGene_tx=XM_019838259.2;refGene_gn=DYNC1I2;refGene_tx=XM_019838260.2
C1 162861441 162861586
refGene_gn=DYNC1I2;refGene_tx=XM_006935360.4;refGene_gn=DYNC1I2;refGene_tx=XM_019838256.2;refGene_gn=DYNC1I2;refGene_tx=XM_019838257.2;refGene_gn=DYNC1I2;refGene_tx=XM_019838258.2;refGene_gn=DYNC1I2;refGene_tx=XM_019838259.2;refGene_gn=DYNC1I2;refGene_tx=XM_019838260.2
C1 162868256 162868316
refGene_gn=DYNC1I2;refGene_tx=XM_019838256.2;refGene_gn=DYNC1I2;refGene_tx=XM_019838257.2;refGene_gn=DYNC1I2;refGene_tx=XM_019838258.2;refGene_gn=DYNC1I2;refGene_tx=XM_019838259.2
C1 162870109 162870225
refGene_gn=DYNC1I2;refGene_tx=XM_006935360.4;refGene_gn=DYNC1I2;refGene_tx=XM_019838256.2;refGene_gn=DYNC1I2;refGene_tx=XM_019838257.2;refGene_gn=DYNC1I2;refGene_tx=XM_019838258.2;refGene_gn=DYNC1I2;refGene_tx=XM_019838259.2;refGene_gn=DYNC1I2;refGene_tx=XM_019838260.2
C1 162881535 162881631
refGene_gn=DYNC1I2;refGene_tx=XM_006935360.4;refGene_gn=DYNC1I2;refGene_tx=XM_019838256.2;refGene_gn=DYNC1I2;refGene_tx=XM_019838257.2;refGene_gn=DYNC1I2;refGene_tx=XM_019838258.2;refGene_gn=DYNC1I2;refGene_tx=XM_019838259.2;refGene_gn=DYNC1I2;refGene_tx=XM_019838260.2
C1 162881845 162882008
refGene_gn=DYNC1I2;refGene_tx=XM_006935360.4;refGene_gn=DYNC1I2;refGene_tx=XM_019838256.2;refGene_gn=DYNC1I2;refGene_tx=XM_019838257.2;refGene_gn=DYNC1I2;refGene_tx=XM_019838258.2;refGene_gn=DYNC1I2;refGene_tx=XM_019838259.2;refGene_gn=DYNC1I2;refGene_tx=XM_019838260.2
C1 162882132 162882232
refGene_gn=DYNC1I2;refGene_tx=XM_006935360.4;refGene_gn=DYNC1I2;refGene_tx=XM_019838256.2;refGene_gn=DYNC1I2;refGene_tx=XM_019838257.2;refGene_gn=DYNC1I2;refGene_tx=XM_019838258.2;refGene_gn=DYNC1I2;refGene_tx=XM_019838259.2;refGene_gn=DYNC1I2;refGene_tx=XM_019838260.2
C1 162882760 162882886
refGene_gn=DYNC1I2;refGene_tx=XM_006935360.4;refGene_gn=DYNC1I2;refGene_tx=XM_019838256.2;refGene_gn=DYNC1I2;refGene_tx=XM_019838257.2;refGene_gn=DYNC1I2;refGene_tx=XM_019838258.2;refGene_gn=DYNC1I2;refGene_tx=XM_019838259.2;refGene_gn=DYNC1I2;refGene_tx=XM_019838260.2
C1 162883772 162883919
refGene_gn=DYNC1I2;refGene_tx=XM_006935360.4;refGene_gn=DYNC1I2;refGene_tx=XM_019838256.2;refGene_gn=DYNC1I2;refGene_tx=XM_019838257.2;refGene_gn=DYNC1I2;refGene_tx=XM_019838258.2;refGene_gn=DYNC1I2;refGene_tx=XM_019838259.2;refGene_gn=DYNC1I2;refGene_tx=XM_019838260.2
C1 162884299 162884413
refGene_gn=DYNC1I2;refGene_tx=XM_006935360.4;refGene_gn=DYNC1I2;refGene_tx=XM_019838256.2;refGene_gn=DYNC1I2;refGene_tx=XM_019838257.2;refGene_gn=DYNC1I2;refGene_tx=XM_019838258.2;refGene_gn=DYNC1I2;refGene_tx=XM_019838259.2;refGene_gn=DYNC1I2;refGene_tx=XM_019838260.2
C1 162884691 162884825
refGene_gn=DYNC1I2;refGene_tx=XM_006935360.4;refGene_gn=DYNC1I2;refGene_tx=XM_019838256.2;refGene_gn=DYNC1I2;refGene_tx=XM_019838257.2;refGene_gn=DYNC1I2;refGene_tx=XM_019838258.2;refGene_gn=DYNC1I2;refGene_tx=

XM_019838259.2;refGene_gn=DYNC1I2;refGene_tx=XM_019838260.2
C1 162885920 162886065
refGene_gn=DYNC1I2;refGene_tx=XM_006935360.4;refGene_gn=DYNC1I2;refGene_tx=XM_019838256.2;refGene_gn=DYNC1I2;refGene_tx=XM_019838257.2;refGene_gn=DYNC1I2;refGene_tx=XM_019838258.2;refGene_gn=DYNC1I2;refGene_tx=XM_019838259.2;refGene_gn=DYNC1I2;refGene_tx=XM_019838260.2
C1 162900643 162900784
refGene_gn=DYNC1I2;refGene_tx=XM_006935360.4;refGene_gn=DYNC1I2;refGene_tx=XM_019838256.2;refGene_gn=DYNC1I2;refGene_tx=XM_019838257.2;refGene_gn=DYNC1I2;refGene_tx=XM_019838258.2;refGene_gn=DYNC1I2;refGene_tx=XM_019838259.2;refGene_gn=DYNC1I2;refGene_tx=XM_019838260.2
C1 162902622 162902748
refGene_gn=DYNC1I2;refGene_tx=XM_006935360.4;refGene_gn=DYNC1I2;refGene_tx=XM_019838256.2;refGene_gn=DYNC1I2;refGene_tx=XM_019838257.2;refGene_gn=DYNC1I2;refGene_tx=XM_019838258.2;refGene_gn=DYNC1I2;refGene_tx=XM_019838259.2;refGene_gn=DYNC1I2;refGene_tx=XM_019838260.2
C1 162904730 162904844
refGene_gn=DYNC1I2;refGene_tx=XM_006935360.4;refGene_gn=DYNC1I2;refGene_tx=XM_019838256.2;refGene_gn=DYNC1I2;refGene_tx=XM_019838257.2;refGene_gn=DYNC1I2;refGene_tx=XM_019838258.2;refGene_gn=DYNC1I2;refGene_tx=XM_019838259.2;refGene_gn=DYNC1I2;refGene_tx=XM_019838260.2
C1 162933852 162934054
refGene_gn=SLC25A12;refGene_tx=XM_011285309.3;refGene_gn=SLC25A12;refGene_tx=XM_011285310.3;refGene_gn=SLC25A12;refGene_tx=XM_019838261.2;refGene_gn=SLC25A12;refGene_tx=XM_019838262.2
C1 162935705 162935796
refGene_gn=SLC25A12;refGene_tx=XM_011285309.3;refGene_gn=SLC25A12;refGene_tx=XM_011285310.3;refGene_gn=SLC25A12;refGene_tx=XM_019838261.2;refGene_gn=SLC25A12;refGene_tx=XM_019838262.2
C1 162935909 162936068
refGene_gn=SLC25A12;refGene_tx=XM_011285309.3;refGene_gn=SLC25A12;refGene_tx=XM_011285310.3;refGene_gn=SLC25A12;refGene_tx=XM_019838261.2;refGene_gn=SLC25A12;refGene_tx=XM_019838262.2
C1 162938640 162938779
refGene_gn=SLC25A12;refGene_tx=XM_011285309.3;refGene_gn=SLC25A12;refGene_tx=XM_011285310.3;refGene_gn=SLC25A12;refGene_tx=XM_019838261.2;refGene_gn=SLC25A12;refGene_tx=XM_019838262.2
C1 162940877 162941018
refGene_gn=SLC25A12;refGene_tx=XM_011285309.3;refGene_gn=SLC25A12;refGene_tx=XM_011285310.3;refGene_gn=SLC25A12;refGene_tx=XM_019838261.2;refGene_gn=SLC25A12;refGene_tx=XM_019838262.2
C1 162964361 162964442
refGene_gn=SLC25A12;refGene_tx=XM_011285309.3;refGene_gn=SLC25A12;refGene_tx=XM_011285310.3;refGene_gn=SLC25A12;refGene_tx=XM_019838261.2;refGene_gn=SLC25A12;refGene_tx=XM_019838262.2
C1 162964927 162964980
refGene_gn=SLC25A12;refGene_tx=XM_011285309.3;refGene_gn=SLC25A12;refGene_tx=XM_011285310.3;refGene_gn=SLC25A12;refGene_tx=XM_019838261.2;refGene_gn=SLC25A12;refGene_tx=XM_019838262.2
C1 162968055 162968214

refGene_gn=SLC25A12;refGene_tx=XM_011285309.3;refGene_gn=SLC25A12;refGene_tx=XM_011285310.3;refGene_gn=SLC25A12;refGene_tx=XM_019838261.2;refGene_gn=SLC25A12;refGene_tx=XM_019838262.2
C1 162969451 162969533
refGene_gn=SLC25A12;refGene_tx=XM_011285309.3;refGene_gn=SLC25A12;refGene_tx=XM_011285310.3;refGene_gn=SLC25A12;refGene_tx=XM_019838261.2;refGene_gn=SLC25A12;refGene_tx=XM_019838262.2
C1 162976389 162976474
refGene_gn=SLC25A12;refGene_tx=XM_011285309.3;refGene_gn=SLC25A12;refGene_tx=XM_011285310.3;refGene_gn=SLC25A12;refGene_tx=XM_019838261.2;refGene_gn=SLC25A12;refGene_tx=XM_019838262.2
C1 162979505 162979599
refGene_gn=SLC25A12;refGene_tx=XM_011285309.3;refGene_gn=SLC25A12;refGene_tx=XM_011285310.3;refGene_gn=SLC25A12;refGene_tx=XM_019838261.2;refGene_gn=SLC25A12;refGene_tx=XM_019838262.2
C1 162980252 162980391
refGene_gn=SLC25A12;refGene_tx=XM_011285309.3;refGene_gn=SLC25A12;refGene_tx=XM_011285310.3;refGene_gn=SLC25A12;refGene_tx=XM_019838261.2;refGene_gn=SLC25A12;refGene_tx=XM_019838262.2
C1 162982564 162982711
refGene_gn=SLC25A12;refGene_tx=XM_011285309.3;refGene_gn=SLC25A12;refGene_tx=XM_011285310.3;refGene_gn=SLC25A12;refGene_tx=XM_019838261.2;refGene_gn=SLC25A12;refGene_tx=XM_019838262.2
C1 162996590 162996730
refGene_gn=SLC25A12;refGene_tx=XM_011285309.3;refGene_gn=SLC25A12;refGene_tx=XM_011285310.3;refGene_gn=SLC25A12;refGene_tx=XM_019838261.2;refGene_gn=SLC25A12;refGene_tx=XM_019838262.2
C1 163000950 163001002
refGene_gn=SLC25A12;refGene_tx=XM_011285310.3
C1 163005640 163005756
refGene_gn=SLC25A12;refGene_tx=XM_011285309.3;refGene_gn=SLC25A12;refGene_tx=XM_019838261.2;refGene_gn=SLC25A12;refGene_tx=XM_019838262.2
C1 163016289 163016432
refGene_gn=SLC25A12;refGene_tx=XM_011285309.3;refGene_gn=SLC25A12;refGene_tx=XM_019838261.2;refGene_gn=SLC25A12;refGene_tx=XM_019838262.2
C1 163036247 163036301
refGene_gn=SLC25A12;refGene_tx=XM_011285309.3;refGene_gn=SLC25A12;refGene_tx=XM_019838262.2
C1 163037264 163037276
refGene_gn=SLC25A12;refGene_tx=XM_011285309.3
C1 163062511 163062626
refGene_gn=HAT1;refGene_tx=XM_003990857.3
C1 163064819 163064924
refGene_gn=HAT1;refGene_tx=XM_003990857.3;refGene_gn=HAT1;refGene_tx=XM_023259457.1
C1 163079781 163079857
refGene_gn=HAT1;refGene_tx=XM_003990857.3;refGene_gn=HAT1;refGene_tx=XM_023259457.1
C1 163083916 163084037
refGene_gn=HAT1;refGene_tx=XM_003990857.3;refGene_gn=HAT1;refGene_tx=X

M_023259457.1
C1	163092938	163093118
refGene_gn=HAT1;refGene_tx=XM_003990857.3;refGene_gn=HAT1;refGene_tx=X
M_023259457.1
C1	163093392	163093514
refGene_gn=HAT1;refGene_tx=XM_003990857.3;refGene_gn=HAT1;refGene_tx=X
M_023259457.1
C1	163094014	163094119
refGene_gn=HAT1;refGene_tx=XM_003990857.3;refGene_gn=HAT1;refGene_tx=X
M_023259457.1
C1	163094443	163094550
refGene_gn=HAT1;refGene_tx=XM_003990857.3;refGene_gn=HAT1;refGene_tx=X
M_023259457.1
C1	163106729	163106881
refGene_gn=HAT1;refGene_tx=XM_003990857.3;refGene_gn=HAT1;refGene_tx=X
M_023259457.1
C1	163109391	163109508
refGene_gn=HAT1;refGene_tx=XM_003990857.3;refGene_gn=HAT1;refGene_tx=X
M_023259457.1
C1	163120484	163120652
refGene_gn=HAT1;refGene_tx=XM_003990857.3;refGene_gn=HAT1;refGene_tx=X
M_023259457.1
C1	163141872	163141890
refGene_gn=SLC25A12;refGene_tx=XM_019838262.2
C1	163142171	163142211
refGene_gn=METAP1D;refGene_tx=XM_003990859.4;refGene_gn=METAP1D;refGen
e_tx=XM_019838263.2
C1	163212626	163212784
refGene_gn=METAP1D;refGene_tx=XM_003990859.4;refGene_gn=METAP1D;refGen
e_tx=XM_019838263.2
C1	163214800	163214950
refGene_gn=METAP1D;refGene_tx=XM_003990859.4;refGene_gn=METAP1D;refGen
e_tx=XM_019838263.2
C1	163216371	163216520
refGene_gn=METAP1D;refGene_tx=XM_003990859.4;refGene_gn=METAP1D;refGen
e_tx=XM_019838263.2
C1	163217099	163217142
refGene_gn=METAP1D;refGene_tx=XM_003990859.4;refGene_gn=METAP1D;refGen
e_tx=XM_019838263.2
C1	163220730	163220894
refGene_gn=METAP1D;refGene_tx=XM_003990859.4;refGene_gn=METAP1D;refGen
e_tx=XM_019838263.2
C1	163223506	163223618
refGene_gn=METAP1D;refGene_tx=XM_019838263.2
C1	163227573	163227671
refGene_gn=METAP1D;refGene_tx=XM_003990859.4
C1	163228983	163229031
refGene_gn=METAP1D;refGene_tx=XM_003990859.4
C1	163230003	163230082
refGene_gn=METAP1D;refGene_tx=XM_003990859.4

C1	163230236	163230315
refGene_gn=METAP1D;refGene_tx=XM_003990859.4
C1	163235595	163235908
refGene_gn=DLX1;refGene_tx=XM_003990860.5;refGene_gn=DLX1;refGene_tx=XM_023259458.1
C1	163236596	163236796
refGene_gn=DLX1;refGene_tx=XM_003990860.5
C1	163237997	163238252
refGene_gn=DLX1;refGene_tx=XM_003990860.5;refGene_gn=DLX1;refGene_tx=XM_023259458.1
C1	163250530	163250947
refGene_gn=DLX2;refGene_tx=XM_003990862.4
C1	163251442	163251627
refGene_gn=DLX2;refGene_tx=XM_003990862.4
C1	163252118	163252512
refGene_gn=DLX2;refGene_tx=XM_003990862.4
C1	163411022	163411349
refGene_gn=LOC111561647;refGene_tx=XM_023257861.1
C1	163414258	163414401
refGene_gn=LOC111561647;refGene_tx=XM_023257861.1
C1	163432200	163432278
refGene_gn=LOC111561647;refGene_tx=XM_023257861.1
C1	163437150	163437217
refGene_gn=LOC111561647;refGene_tx=XM_023257861.1
C1	163536477	163536659
refGene_gn=ITGA6;refGene_tx=XM_023259459.1
C1	163575788	163575879
refGene_gn=ITGA6;refGene_tx=XM_023259459.1
C1	163575880	163575914
refGene_gn=ITGA6;refGene_tx=XM_023259459.1
C1	163577344	163577424
refGene_gn=ITGA6;refGene_tx=XM_023259459.1
C1	163578853	163579109
refGene_gn=ITGA6;refGene_tx=XM_023259459.1
C1	163581236	163581368
refGene_gn=ITGA6;refGene_tx=XM_023259459.1
C1	163584474	163584685
refGene_gn=ITGA6;refGene_tx=XM_023259459.1
C1	163585420	163585614
refGene_gn=ITGA6;refGene_tx=XM_023259459.1
C1	163585760	163585849
refGene_gn=ITGA6;refGene_tx=XM_023259459.1
C1	163586546	163586665
refGene_gn=ITGA6;refGene_tx=XM_023259459.1
C1	163589519	163589618
refGene_gn=ITGA6;refGene_tx=XM_023259459.1
C1	163590075	163590137
refGene_gn=ITGA6;refGene_tx=XM_023259459.1
C1	163594170	163594331
refGene_gn=ITGA6;refGene_tx=XM_023259459.1

C1 163594720 163594864
refGene_gn=ITGA6;refGene_tx=XM_023259459.1
C1 163596470 163596586
refGene_gn=ITGA6;refGene_tx=XM_023259459.1
C1 163596713 163596903
refGene_gn=ITGA6;refGene_tx=XM_023259459.1
C1 163596992 163597076
refGene_gn=ITGA6;refGene_tx=XM_023259459.1
C1 163597173 163597253
refGene_gn=ITGA6;refGene_tx=XM_023259459.1
C1 163597402 163597480
refGene_gn=ITGA6;refGene_tx=XM_023259459.1
C1 163597566 163597669
refGene_gn=ITGA6;refGene_tx=XM_023259459.1
C1 163598964 163599138
refGene_gn=ITGA6;refGene_tx=XM_023259459.1
C1 163600914 163601013
refGene_gn=ITGA6;refGene_tx=XM_023259459.1
C1 163601113 163601224
refGene_gn=ITGA6;refGene_tx=XM_023259459.1
C1 163601323 163601422
refGene_gn=ITGA6;refGene_tx=XM_023259459.1
C1 163607694 163607820
refGene_gn=ITGA6;refGene_tx=XM_023259459.1
C1 163613273 163613435
refGene_gn=ITGA6;refGene_tx=XM_023259459.1
C1 163671448 163671650
refGene_gn=PDK1;refGene_tx=XM_003990864.5;refGene_gn=PDK1;refGene_tx=XM_006935366.4;refGene_gn=PDK1;refGene_tx=XM_019838267.2
C1 163673506 163673648
refGene_gn=PDK1;refGene_tx=XM_003990864.5;refGene_gn=PDK1;refGene_tx=XM_006935366.4;refGene_gn=PDK1;refGene_tx=XM_019838267.2
C1 163676253 163676325
refGene_gn=PDK1;refGene_tx=XM_003990864.5;refGene_gn=PDK1;refGene_tx=XM_006935366.4;refGene_gn=PDK1;refGene_tx=XM_019838267.2
C1 163678467 163678652
refGene_gn=PDK1;refGene_tx=XM_003990864.5;refGene_gn=PDK1;refGene_tx=XM_006935366.4;refGene_gn=PDK1;refGene_tx=XM_019838267.2
C1 163678932 163679028
refGene_gn=PDK1;refGene_tx=XM_003990864.5;refGene_gn=PDK1;refGene_tx=XM_006935366.4;refGene_gn=PDK1;refGene_tx=XM_019838267.2
C1 163680244 163680322
refGene_gn=PDK1;refGene_tx=XM_003990864.5;refGene_gn=PDK1;refGene_tx=XM_006935366.4;refGene_gn=PDK1;refGene_tx=XM_019838267.2
C1 163682123 163682200
refGene_gn=PDK1;refGene_tx=XM_003990864.5;refGene_gn=PDK1;refGene_tx=XM_006935366.4;refGene_gn=PDK1;refGene_tx=XM_019838267.2
C1 163683979 163684111
refGene_gn=PDK1;refGene_tx=XM_003990864.5;refGene_gn=PDK1;refGene_tx=XM_006935366.4;refGene_gn=PDK1;refGene_tx=XM_019838267.2

C1	163695492	163695603
refGene_gn=PDK1;refGene_tx=XM_003990864.5;refGene_gn=PDK1;refGene_tx=XM_006935366.4
C1	163699371	163699485
refGene_gn=PDK1;refGene_tx=XM_003990864.5;refGene_gn=PDK1;refGene_tx=XM_006935366.4;refGene_gn=PDK1;refGene_tx=XM_019838267.2
C1	163702214	163702355
refGene_gn=PDK1;refGene_tx=XM_003990864.5;refGene_gn=PDK1;refGene_tx=XM_006935366.4;refGene_gn=PDK1;refGene_tx=XM_019838267.2
C1	163783983	163784048
refGene_gn=RAPGEF4;refGene_tx=XM_023259461.1;refGene_gn=RAPGEF4;refGene_tx=XM_023259462.1
C1	163832113	163832256
refGene_gn=RAPGEF4;refGene_tx=XM_023259461.1;refGene_gn=RAPGEF4;refGene_tx=XM_023259462.1
C1	163835646	163835735
refGene_gn=RAPGEF4;refGene_tx=XM_023259461.1;refGene_gn=RAPGEF4;refGene_tx=XM_023259462.1
C1	163851351	163851498
refGene_gn=RAPGEF4;refGene_tx=XM_019838268.2;refGene_gn=RAPGEF4;refGene_tx=XM_023259461.1;refGene_gn=RAPGEF4;refGene_tx=XM_023259462.1
C1	163858232	163858244
refGene_gn=RAPGEF4;refGene_tx=XM_019838270.2
C1	163886029	163886047
refGene_gn=RAPGEF4;refGene_tx=XM_019838269.2
C1	163947213	163947286
refGene_gn=RAPGEF4;refGene_tx=XM_019838268.2;refGene_gn=RAPGEF4;refGene_tx=XM_019838269.2;refGene_gn=RAPGEF4;refGene_tx=XM_019838270.2;refGene_gn=RAPGEF4;refGene_tx=XM_019838271.2;refGene_gn=RAPGEF4;refGene_tx=XM_019838272.2;refGene_gn=RAPGEF4;refGene_tx=XM_019838273.2;refGene_gn=RAPGEF4;refGene_tx=XM_023259461.1;refGene_gn=RAPGEF4;refGene_tx=XM_023259462.1
C1	163952090	163952110
refGene_gn=RAPGEF4;refGene_tx=XM_019838269.2;refGene_gn=RAPGEF4;refGene_tx=XM_019838270.2;refGene_gn=RAPGEF4;refGene_tx=XM_019838271.2;refGene_gn=RAPGEF4;refGene_tx=XM_019838272.2;refGene_gn=RAPGEF4;refGene_tx=XM_019838273.2;refGene_gn=RAPGEF4;refGene_tx=XM_023259461.1;refGene_gn=RAPGEF4;refGene_tx=XM_023259462.1
C1	163985084	163985138
refGene_gn=RAPGEF4;refGene_tx=XM_019838268.2;refGene_gn=RAPGEF4;refGene_tx=XM_019838269.2;refGene_gn=RAPGEF4;refGene_tx=XM_019838270.2;refGene_gn=RAPGEF4;refGene_tx=XM_019838271.2;refGene_gn=RAPGEF4;refGene_tx=XM_019838272.2;refGene_gn=RAPGEF4;refGene_tx=XM_019838273.2;refGene_gn=RAPGEF4;refGene_tx=XM_023259461.1
C1	163985445	163985552
refGene_gn=RAPGEF4;refGene_tx=XM_019838268.2;refGene_gn=RAPGEF4;refGene_tx=XM_019838269.2;refGene_gn=RAPGEF4;refGene_tx=XM_019838270.2;refGene_gn=RAPGEF4;refGene_tx=XM_019838271.2;refGene_gn=RAPGEF4;refGene_tx=XM_019838272.2;refGene_gn=RAPGEF4;refGene_tx=XM_019838273.2;refGene_gn=RAPGEF4;refGene_tx=XM_023259461.1;refGene_gn=RAPGEF4;refGene_tx=XM_02

3259462.1
C1 163989529 163989651
refGene_gn=RAPGEF4;refGene_tx=XM_019838268.2;refGene_gn=RAPGEF4;refGene_tx=XM_019838269.2;refGene_gn=RAPGEF4;refGene_tx=XM_019838270.2;refGene_gn=RAPGEF4;refGene_tx=XM_019838271.2;refGene_gn=RAPGEF4;refGene_tx=XM_019838272.2;refGene_gn=RAPGEF4;refGene_tx=XM_019838273.2;refGene_gn=RAPGEF4;refGene_tx=XM_023259461.1;refGene_gn=RAPGEF4;refGene_tx=XM_023259462.1
C1 163991320 163991504
refGene_gn=RAPGEF4;refGene_tx=XM_019838268.2;refGene_gn=RAPGEF4;refGene_tx=XM_019838269.2;refGene_gn=RAPGEF4;refGene_tx=XM_019838270.2;refGene_gn=RAPGEF4;refGene_tx=XM_019838271.2;refGene_gn=RAPGEF4;refGene_tx=XM_019838272.2;refGene_gn=RAPGEF4;refGene_tx=XM_019838273.2;refGene_gn=RAPGEF4;refGene_tx=XM_023259461.1;refGene_gn=RAPGEF4;refGene_tx=XM_023259462.1
C1 164007844 164007929
refGene_gn=RAPGEF4;refGene_tx=XM_019838268.2;refGene_gn=RAPGEF4;refGene_tx=XM_019838269.2;refGene_gn=RAPGEF4;refGene_tx=XM_019838270.2;refGene_gn=RAPGEF4;refGene_tx=XM_019838271.2;refGene_gn=RAPGEF4;refGene_tx=XM_019838272.2;refGene_gn=RAPGEF4;refGene_tx=XM_019838273.2;refGene_gn=RAPGEF4;refGene_tx=XM_023259461.1;refGene_gn=RAPGEF4;refGene_tx=XM_023259462.1
C1 164009913 164009974
refGene_gn=RAPGEF4;refGene_tx=XM_019838268.2;refGene_gn=RAPGEF4;refGene_tx=XM_019838269.2;refGene_gn=RAPGEF4;refGene_tx=XM_019838270.2;refGene_gn=RAPGEF4;refGene_tx=XM_019838271.2;refGene_gn=RAPGEF4;refGene_tx=XM_019838272.2;refGene_gn=RAPGEF4;refGene_tx=XM_019838273.2;refGene_gn=RAPGEF4;refGene_tx=XM_023259461.1;refGene_gn=RAPGEF4;refGene_tx=XM_023259462.1
C1 164012259 164012336
refGene_gn=RAPGEF4;refGene_tx=XM_019838268.2;refGene_gn=RAPGEF4;refGene_tx=XM_019838269.2;refGene_gn=RAPGEF4;refGene_tx=XM_019838270.2;refGene_gn=RAPGEF4;refGene_tx=XM_019838271.2;refGene_gn=RAPGEF4;refGene_tx=XM_019838272.2;refGene_gn=RAPGEF4;refGene_tx=XM_019838273.2;refGene_gn=RAPGEF4;refGene_tx=XM_023259461.1;refGene_gn=RAPGEF4;refGene_tx=XM_023259462.1
C1 164012773 164012920
refGene_gn=RAPGEF4;refGene_tx=XM_019838268.2;refGene_gn=RAPGEF4;refGene_tx=XM_019838269.2;refGene_gn=RAPGEF4;refGene_tx=XM_019838270.2;refGene_gn=RAPGEF4;refGene_tx=XM_019838271.2;refGene_gn=RAPGEF4;refGene_tx=XM_019838272.2;refGene_gn=RAPGEF4;refGene_tx=XM_019838273.2;refGene_gn=RAPGEF4;refGene_tx=XM_023259461.1;refGene_gn=RAPGEF4;refGene_tx=XM_023259462.1
C1 164014798 164014914
refGene_gn=RAPGEF4;refGene_tx=XM_019838268.2;refGene_gn=RAPGEF4;refGene_tx=XM_019838269.2;refGene_gn=RAPGEF4;refGene_tx=XM_019838270.2;refGene_gn=RAPGEF4;refGene_tx=XM_019838271.2;refGene_gn=RAPGEF4;refGene_tx=XM_019838272.2;refGene_gn=RAPGEF4;refGene_tx=XM_019838273.2;refGene_gn=RAPGEF4;refGene_tx=XM_023259461.1;refGene_gn=RAPGEF4;refGene_tx=XM_023259462.1

C1 164020849 164020938
refGene_gn=RAPGEF4;refGene_tx=XM_019838268.2;refGene_gn=RAPGEF4;refGene_tx=XM_019838269.2;refGene_gn=RAPGEF4;refGene_tx=XM_019838270.2;refGene_gn=RAPGEF4;refGene_tx=XM_019838271.2;refGene_gn=RAPGEF4;refGene_tx=XM_019838272.2;refGene_gn=RAPGEF4;refGene_tx=XM_019838273.2;refGene_gn=RAPGEF4;refGene_tx=XM_023259461.1;refGene_gn=RAPGEF4;refGene_tx=XM_023259462.1
C1 164025587 164025666
refGene_gn=RAPGEF4;refGene_tx=XM_019838268.2;refGene_gn=RAPGEF4;refGene_tx=XM_019838269.2;refGene_gn=RAPGEF4;refGene_tx=XM_019838270.2;refGene_gn=RAPGEF4;refGene_tx=XM_019838271.2;refGene_gn=RAPGEF4;refGene_tx=XM_019838272.2;refGene_gn=RAPGEF4;refGene_tx=XM_019838273.2;refGene_gn=RAPGEF4;refGene_tx=XM_023259461.1;refGene_gn=RAPGEF4;refGene_tx=XM_023259462.1
C1 164039495 164039646
refGene_gn=RAPGEF4;refGene_tx=XM_019838268.2;refGene_gn=RAPGEF4;refGene_tx=XM_019838269.2;refGene_gn=RAPGEF4;refGene_tx=XM_019838270.2;refGene_gn=RAPGEF4;refGene_tx=XM_019838271.2;refGene_gn=RAPGEF4;refGene_tx=XM_019838272.2;refGene_gn=RAPGEF4;refGene_tx=XM_019838273.2;refGene_gn=RAPGEF4;refGene_tx=XM_023259461.1;refGene_gn=RAPGEF4;refGene_tx=XM_023259462.1
C1 164041359 164041448
refGene_gn=RAPGEF4;refGene_tx=XM_019838268.2;refGene_gn=RAPGEF4;refGene_tx=XM_019838269.2;refGene_gn=RAPGEF4;refGene_tx=XM_019838270.2;refGene_gn=RAPGEF4;refGene_tx=XM_019838271.2;refGene_gn=RAPGEF4;refGene_tx=XM_019838272.2;refGene_gn=RAPGEF4;refGene_tx=XM_019838273.2;refGene_gn=RAPGEF4;refGene_tx=XM_023259461.1;refGene_gn=RAPGEF4;refGene_tx=XM_023259462.1
C1 164042412 164042443
refGene_gn=RAPGEF4;refGene_tx=XM_019838268.2;refGene_gn=RAPGEF4;refGene_tx=XM_019838269.2;refGene_gn=RAPGEF4;refGene_tx=XM_019838270.2;refGene_gn=RAPGEF4;refGene_tx=XM_019838271.2;refGene_gn=RAPGEF4;refGene_tx=XM_019838272.2;refGene_gn=RAPGEF4;refGene_tx=XM_019838273.2;refGene_gn=RAPGEF4;refGene_tx=XM_023259461.1;refGene_gn=RAPGEF4;refGene_tx=XM_023259462.1
C1 164042648 164042727
refGene_gn=RAPGEF4;refGene_tx=XM_019838268.2;refGene_gn=RAPGEF4;refGene_tx=XM_019838269.2;refGene_gn=RAPGEF4;refGene_tx=XM_019838270.2;refGene_gn=RAPGEF4;refGene_tx=XM_019838271.2;refGene_gn=RAPGEF4;refGene_tx=XM_019838272.2;refGene_gn=RAPGEF4;refGene_tx=XM_019838273.2;refGene_gn=RAPGEF4;refGene_tx=XM_023259461.1;refGene_gn=RAPGEF4;refGene_tx=XM_023259462.1
C1 164044133 164044280
refGene_gn=RAPGEF4;refGene_tx=XM_019838268.2;refGene_gn=RAPGEF4;refGene_tx=XM_019838269.2;refGene_gn=RAPGEF4;refGene_tx=XM_019838270.2;refGene_gn=RAPGEF4;refGene_tx=XM_019838271.2;refGene_gn=RAPGEF4;refGene_tx=XM_019838272.2;refGene_gn=RAPGEF4;refGene_tx=XM_019838273.2;refGene_gn=RAPGEF4;refGene_tx=XM_023259461.1;refGene_gn=RAPGEF4;refGene_tx=XM_023259462.1
C1 164046021 164046119

refGene_gn=RAPGEF4;refGene_tx=XM_019838268.2;refGene_gn=RAPGEF4;refGene_tx=XM_019838269.2;refGene_gn=RAPGEF4;refGene_tx=XM_019838270.2;refGene_gn=RAPGEF4;refGene_tx=XM_019838271.2;refGene_gn=RAPGEF4;refGene_tx=XM_019838272.2;refGene_gn=RAPGEF4;refGene_tx=XM_019838273.2;refGene_gn=RAPGEF4;refGene_tx=XM_023259461.1;refGene_gn=RAPGEF4;refGene_tx=XM_023259462.1
C1 164051680 164051806
refGene_gn=RAPGEF4;refGene_tx=XM_019838268.2;refGene_gn=RAPGEF4;refGene_tx=XM_019838269.2;refGene_gn=RAPGEF4;refGene_tx=XM_019838270.2;refGene_gn=RAPGEF4;refGene_tx=XM_019838271.2;refGene_gn=RAPGEF4;refGene_tx=XM_019838272.2;refGene_gn=RAPGEF4;refGene_tx=XM_019838273.2;refGene_gn=RAPGEF4;refGene_tx=XM_023259461.1;refGene_gn=RAPGEF4;refGene_tx=XM_023259462.1
C1 164052227 164052406
refGene_gn=RAPGEF4;refGene_tx=XM_019838268.2;refGene_gn=RAPGEF4;refGene_tx=XM_019838269.2;refGene_gn=RAPGEF4;refGene_tx=XM_019838270.2;refGene_gn=RAPGEF4;refGene_tx=XM_019838271.2;refGene_gn=RAPGEF4;refGene_tx=XM_019838272.2;refGene_gn=RAPGEF4;refGene_tx=XM_019838273.2;refGene_gn=RAPGEF4;refGene_tx=XM_023259461.1;refGene_gn=RAPGEF4;refGene_tx=XM_023259462.1
C1 164055289 164055380
refGene_gn=RAPGEF4;refGene_tx=XM_019838268.2;refGene_gn=RAPGEF4;refGene_tx=XM_019838269.2;refGene_gn=RAPGEF4;refGene_tx=XM_019838270.2;refGene_gn=RAPGEF4;refGene_tx=XM_019838271.2;refGene_gn=RAPGEF4;refGene_tx=XM_019838272.2;refGene_gn=RAPGEF4;refGene_tx=XM_019838273.2;refGene_gn=RAPGEF4;refGene_tx=XM_023259461.1;refGene_gn=RAPGEF4;refGene_tx=XM_023259462.1
C1 164059202 164059253
refGene_gn=RAPGEF4;refGene_tx=XM_019838268.2;refGene_gn=RAPGEF4;refGene_tx=XM_019838269.2;refGene_gn=RAPGEF4;refGene_tx=XM_019838270.2;refGene_gn=RAPGEF4;refGene_tx=XM_019838271.2;refGene_gn=RAPGEF4;refGene_tx=XM_019838272.2;refGene_gn=RAPGEF4;refGene_tx=XM_019838273.2;refGene_gn=RAPGEF4;refGene_tx=XM_023259461.1;refGene_gn=RAPGEF4;refGene_tx=XM_023259462.1
C1 164061234 164061322
refGene_gn=RAPGEF4;refGene_tx=XM_019838268.2;refGene_gn=RAPGEF4;refGene_tx=XM_019838269.2;refGene_gn=RAPGEF4;refGene_tx=XM_019838270.2;refGene_gn=RAPGEF4;refGene_tx=XM_019838271.2;refGene_gn=RAPGEF4;refGene_tx=XM_019838272.2;refGene_gn=RAPGEF4;refGene_tx=XM_019838273.2;refGene_gn=RAPGEF4;refGene_tx=XM_023259461.1;refGene_gn=RAPGEF4;refGene_tx=XM_023259462.1
C1 164061730 164061795
refGene_gn=RAPGEF4;refGene_tx=XM_019838268.2;refGene_gn=RAPGEF4;refGene_tx=XM_019838269.2;refGene_gn=RAPGEF4;refGene_tx=XM_019838270.2;refGene_gn=RAPGEF4;refGene_tx=XM_019838271.2;refGene_gn=RAPGEF4;refGene_tx=XM_019838272.2;refGene_gn=RAPGEF4;refGene_tx=XM_019838273.2;refGene_gn=RAPGEF4;refGene_tx=XM_023259461.1;refGene_gn=RAPGEF4;refGene_tx=XM_023259462.1
C1 164071672 164071727
refGene_gn=RAPGEF4;refGene_tx=XM_019838268.2;refGene_gn=RAPGEF4;refGen

e_tx=XM_019838269.2;refGene_gn=RAPGEF4;refGene_tx=XM_019838270.2;refGene_gn=RAPGEF4;refGene_tx=XM_019838271.2;refGene_gn=RAPGEF4;refGene_tx=XM_019838272.2;refGene_gn=RAPGEF4;refGene_tx=XM_019838273.2;refGene_gn=RAPGEF4;refGene_tx=XM_023259461.1;refGene_gn=RAPGEF4;refGene_tx=XM_023259462.1
C1 164074851 164074979
refGene_gn=RAPGEF4;refGene_tx=XM_019838268.2;refGene_gn=RAPGEF4;refGene_tx=XM_019838269.2;refGene_gn=RAPGEF4;refGene_tx=XM_019838270.2;refGene_gn=RAPGEF4;refGene_tx=XM_019838271.2;refGene_gn=RAPGEF4;refGene_tx=XM_019838272.2;refGene_gn=RAPGEF4;refGene_tx=XM_019838273.2;refGene_gn=RAPGEF4;refGene_tx=XM_023259461.1;refGene_gn=RAPGEF4;refGene_tx=XM_023259462.1
C1 164108572 164108731
refGene_gn=MAP3K20;refGene_tx=XM_003990872.5;refGene_gn=MAP3K20;refGene_tx=XM_006935367.4;refGene_gn=MAP3K20;refGene_tx=XM_023259463.1;refGene_gn=MAP3K20;refGene_tx=XM_023259464.1;refGene_gn=MAP3K20;refGene_tx=XM_023259465.1;refGene_gn=MAP3K20;refGene_tx=XM_023259466.1
C1 164185945 164186033
refGene_gn=MAP3K20;refGene_tx=XM_003990872.5;refGene_gn=MAP3K20;refGene_tx=XM_006935367.4;refGene_gn=MAP3K20;refGene_tx=XM_023259463.1;refGene_gn=MAP3K20;refGene_tx=XM_023259464.1;refGene_gn=MAP3K20;refGene_tx=XM_023259465.1;refGene_gn=MAP3K20;refGene_tx=XM_023259466.1
C1 164200260 164200362
refGene_gn=MAP3K20;refGene_tx=XM_003990872.5;refGene_gn=MAP3K20;refGene_tx=XM_006935367.4;refGene_gn=MAP3K20;refGene_tx=XM_023259463.1;refGene_gn=MAP3K20;refGene_tx=XM_023259464.1;refGene_gn=MAP3K20;refGene_tx=XM_023259465.1;refGene_gn=MAP3K20;refGene_tx=XM_023259466.1
C1 164204771 164204837
refGene_gn=MAP3K20;refGene_tx=XM_003990872.5;refGene_gn=MAP3K20;refGene_tx=XM_006935367.4;refGene_gn=MAP3K20;refGene_tx=XM_023259463.1;refGene_gn=MAP3K20;refGene_tx=XM_023259464.1;refGene_gn=MAP3K20;refGene_tx=XM_023259465.1;refGene_gn=MAP3K20;refGene_tx=XM_023259466.1
C1 164208385 164208414
refGene_gn=MAP3K20;refGene_tx=XM_003990872.5;refGene_gn=MAP3K20;refGene_tx=XM_006935367.4;refGene_gn=MAP3K20;refGene_tx=XM_023259463.1;refGene_gn=MAP3K20;refGene_tx=XM_023259464.1;refGene_gn=MAP3K20;refGene_tx=XM_023259465.1;refGene_gn=MAP3K20;refGene_tx=XM_023259466.1
C1 164208536 164208674
refGene_gn=MAP3K20;refGene_tx=XM_003990872.5;refGene_gn=MAP3K20;refGene_tx=XM_006935367.4;refGene_gn=MAP3K20;refGene_tx=XM_023259463.1;refGene_gn=MAP3K20;refGene_tx=XM_023259464.1;refGene_gn=MAP3K20;refGene_tx=XM_023259465.1;refGene_gn=MAP3K20;refGene_tx=XM_023259466.1
C1 164215805 164215892
refGene_gn=MAP3K20;refGene_tx=XM_003990872.5;refGene_gn=MAP3K20;refGene_tx=XM_006935367.4;refGene_gn=MAP3K20;refGene_tx=XM_023259463.1;refGene_gn=MAP3K20;refGene_tx=XM_023259464.1;refGene_gn=MAP3K20;refGene_tx=XM_023259465.1;refGene_gn=MAP3K20;refGene_tx=XM_023259466.1
C1 164221736 164221811
refGene_gn=MAP3K20;refGene_tx=XM_003990872.5;refGene_gn=MAP3K20;refGene_tx=XM_006935367.4;refGene_gn=MAP3K20;refGene_tx=XM_023259463.1;refGe

ne_gn=MAP3K20;refGene_tx=XM_023259464.1;refGene_gn=MAP3K20;refGene_tx=XM_023259465.1;refGene_gn=MAP3K20;refGene_tx=XM_023259466.1
C1 164226596 164226703
refGene_gn=MAP3K20;refGene_tx=XM_003990872.5;refGene_gn=MAP3K20;refGene_tx=XM_006935367.4;refGene_gn=MAP3K20;refGene_tx=XM_023259463.1;refGene_gn=MAP3K20;refGene_tx=XM_023259464.1;refGene_gn=MAP3K20;refGene_tx=XM_023259465.1;refGene_gn=MAP3K20;refGene_tx=XM_023259466.1
C1 164235065 164235201
refGene_gn=MAP3K20;refGene_tx=XM_003990872.5;refGene_gn=MAP3K20;refGene_tx=XM_006935367.4;refGene_gn=MAP3K20;refGene_tx=XM_023259463.1;refGene_gn=MAP3K20;refGene_tx=XM_023259464.1;refGene_gn=MAP3K20;refGene_tx=XM_023259465.1;refGene_gn=MAP3K20;refGene_tx=XM_023259466.1
C1 164239044 164239434
refGene_gn=MAP3K20;refGene_tx=XM_023259466.1
C1 164247905 164247950
refGene_gn=MAP3K20;refGene_tx=XM_003990872.5;refGene_gn=MAP3K20;refGene_tx=XM_006935367.4;refGene_gn=MAP3K20;refGene_tx=XM_023259463.1;refGene_gn=MAP3K20;refGene_tx=XM_023259464.1;refGene_gn=MAP3K20;refGene_tx=XM_023259465.1
C1 164249768 164249799
refGene_gn=MAP3K20;refGene_tx=XM_003990872.5;refGene_gn=MAP3K20;refGene_tx=XM_006935367.4;refGene_gn=MAP3K20;refGene_tx=XM_023259463.1;refGene_gn=MAP3K20;refGene_tx=XM_023259464.1;refGene_gn=MAP3K20;refGene_tx=XM_023259465.1
C1 164249896 164250036
refGene_gn=MAP3K20;refGene_tx=XM_003990872.5;refGene_gn=MAP3K20;refGene_tx=XM_006935367.4;refGene_gn=MAP3K20;refGene_tx=XM_023259463.1;refGene_gn=MAP3K20;refGene_tx=XM_023259464.1;refGene_gn=MAP3K20;refGene_tx=XM_023259465.1
C1 164254546 164254609
refGene_gn=MAP3K20;refGene_tx=XM_003990872.5;refGene_gn=MAP3K20;refGene_tx=XM_006935367.4;refGene_gn=MAP3K20;refGene_tx=XM_023259463.1;refGene_gn=MAP3K20;refGene_tx=XM_023259464.1;refGene_gn=MAP3K20;refGene_tx=XM_023259465.1
C1 164255586 164255679
refGene_gn=MAP3K20;refGene_tx=XM_003990872.5;refGene_gn=MAP3K20;refGene_tx=XM_006935367.4;refGene_gn=MAP3K20;refGene_tx=XM_023259463.1;refGene_gn=MAP3K20;refGene_tx=XM_023259464.1;refGene_gn=MAP3K20;refGene_tx=XM_023259465.1
C1 164259397 164259439
refGene_gn=MAP3K20;refGene_tx=XM_023259465.1
C1 164271510 164271627
refGene_gn=MAP3K20;refGene_tx=XM_003990872.5;refGene_gn=MAP3K20;refGene_tx=XM_006935367.4;refGene_gn=MAP3K20;refGene_tx=XM_023259463.1;refGene_gn=MAP3K20;refGene_tx=XM_023259464.1
C1 164274120 164274195
refGene_gn=MAP3K20;refGene_tx=XM_003990872.5;refGene_gn=MAP3K20;refGene_tx=XM_006935367.4;refGene_gn=MAP3K20;refGene_tx=XM_023259463.1;refGene_gn=MAP3K20;refGene_tx=XM_023259464.1
C1 164277045 164277196

refGene_gn=MAP3K20;refGene_tx=XM_003990872.5;refGene_gn=MAP3K20;refGene_tx=XM_006935367.4;refGene_gn=MAP3K20;refGene_tx=XM_023259463.1
C1 164280934 164281635
refGene_gn=MAP3K20;refGene_tx=XM_003990872.5;refGene_gn=MAP3K20;refGene_tx=XM_006935367.4;refGene_gn=MAP3K20;refGene_tx=XM_023259463.1;refGene_gn=MAP3K20;refGene_tx=XM_023259464.1
C1 164340919 164340940
refGene_gn=CDCA7;refGene_tx=XM_003990873.5;refGene_gn=CDCA7;refGene_tx=XM_003990875.5;refGene_gn=CDCA7;refGene_tx=XM_023259467.1
C1 164343811 164343937
refGene_gn=CDCA7;refGene_tx=XM_003990873.5;refGene_gn=CDCA7;refGene_tx=XM_003990875.5;refGene_gn=CDCA7;refGene_tx=XM_023259467.1
C1 164344355 164344592
refGene_gn=CDCA7;refGene_tx=XM_003990875.5;refGene_gn=CDCA7;refGene_tx=XM_023259467.1
C1 164347256 164347493
refGene_gn=CDCA7;refGene_tx=XM_003990873.5;refGene_gn=CDCA7;refGene_tx=XM_003990875.5;refGene_gn=CDCA7;refGene_tx=XM_023259467.1
C1 164347784 164347862
refGene_gn=CDCA7;refGene_tx=XM_003990873.5;refGene_gn=CDCA7;refGene_tx=XM_003990875.5;refGene_gn=CDCA7;refGene_tx=XM_023259467.1
C1 164349470 164349671
refGene_gn=CDCA7;refGene_tx=XM_003990873.5;refGene_gn=CDCA7;refGene_tx=XM_003990875.5;refGene_gn=CDCA7;refGene_tx=XM_023259467.1
C1 164350071 164350209
refGene_gn=CDCA7;refGene_tx=XM_003990873.5;refGene_gn=CDCA7;refGene_tx=XM_003990875.5;refGene_gn=CDCA7;refGene_tx=XM_023259467.1
C1 164350844 164350994
refGene_gn=CDCA7;refGene_tx=XM_003990873.5;refGene_gn=CDCA7;refGene_tx=XM_003990875.5;refGene_gn=CDCA7;refGene_tx=XM_023259467.1
C1 164351454 164351591
refGene_gn=CDCA7;refGene_tx=XM_003990873.5;refGene_gn=CDCA7;refGene_tx=XM_003990875.5;refGene_gn=CDCA7;refGene_tx=XM_023259467.1
C1 164352175 164352206
refGene_gn=CDCA7;refGene_tx=XM_003990873.5;refGene_gn=CDCA7;refGene_tx=XM_003990875.5;refGene_gn=CDCA7;refGene_tx=XM_023259467.1
C1 164819929 164820246
refGene_gn=SP3;refGene_tx=XM_023259468.1
C1 164822971 164823168
refGene_gn=SP3;refGene_tx=XM_023259468.1
C1 164828537 164828730
refGene_gn=SP3;refGene_tx=XM_023259468.1
C1 164867319 164868679
refGene_gn=SP3;refGene_tx=XM_023259468.1
C1 164876616 164876739
refGene_gn=SP3;refGene_tx=XM_023259468.1
C1 164877269 164877392
refGene_gn=SP3;refGene_tx=XM_023259468.1
C1 164984704 164984806
refGene_gn=OLA1;refGene_tx=XM_023259469.1

C1 164988112 164988235
refGene_gn=OLA1;refGene_tx=XM_023259469.1
C1 164990289 164990386
refGene_gn=OLA1;refGene_tx=XM_023259469.1
C1 164991074 164991215
refGene_gn=OLA1;refGene_tx=XM_023259469.1
C1 165029200 165029298
refGene_gn=OLA1;refGene_tx=XM_023259469.1
C1 165029622 165029703
refGene_gn=OLA1;refGene_tx=XM_023259469.1
C1 165052229 165052405
refGene_gn=OLA1;refGene_tx=XM_023259469.1
C1 165138546 165138674
refGene_gn=OLA1;refGene_tx=XM_023259469.1
C1 165142380 165142524
refGene_gn=OLA1;refGene_tx=XM_023259469.1
C1 165156167 165156268
refGene_gn=OLA1;refGene_tx=XM_023259469.1
C1 165228116 165228137
refGene_gn=SP9;refGene_tx=XM_023259470.1
C1 165229159 165230596
refGene_gn=SP9;refGene_tx=XM_023259470.1
C1 165241099 165241838
refGene_gn=CIR1;refGene_tx=XM_003990878.5
C1 165243187 165243273
refGene_gn=CIR1;refGene_tx=XM_003990878.5
C1 165244377 165244484
refGene_gn=CIR1;refGene_tx=XM_003990878.5
C1 165274889 165275007
refGene_gn=CIR1;refGene_tx=XM_003990878.5
C1 165276156 165276205
refGene_gn=CIR1;refGene_tx=XM_003990878.5
C1 165277013 165277078
refGene_gn=CIR1;refGene_tx=XM_003990878.5
C1 165277167 165277178
refGene_gn=CIR1;refGene_tx=XM_003990878.5
C1 165277767 165277841
refGene_gn=CIR1;refGene_tx=XM_003990878.5
C1 165283489 165283584
refGene_gn=CIR1;refGene_tx=XM_003990878.5
C1 165289874 165289946
refGene_gn=CIR1;refGene_tx=XM_003990878.5
C1 165292201 165292360
refGene_gn=SCRN3;refGene_tx=XM_023259471.1;refGene_gn=SCRN3;refGene_tx=XM_023259472.1;refGene_gn=SCRN3;refGene_tx=XM_023259473.1
C1 165293846 165294028
refGene_gn=SCRN3;refGene_tx=XM_023259471.1;refGene_gn=SCRN3;refGene_tx=XM_023259472.1;refGene_gn=SCRN3;refGene_tx=XM_023259473.1;refGene_gn=SCRN3;refGene_tx=XM_023259474.1
C1 165296001 165296201

refGene_gn=SCRN3;refGene_tx=XM_023259471.1;refGene_gn=SCRN3;refGene_tx=XM_023259472.1;refGene_gn=SCRN3;refGene_tx=XM_023259473.1;refGene_gn=SCRN3;refGene_tx=XM_023259474.1
C1 165298186 165298399
refGene_gn=SCRN3;refGene_tx=XM_023259471.1;refGene_gn=SCRN3;refGene_tx=XM_023259472.1;refGene_gn=SCRN3;refGene_tx=XM_023259473.1;refGene_gn=SCRN3;refGene_tx=XM_023259474.1
C1 165312822 165312861
refGene_gn=SCRN3;refGene_tx=XM_023259471.1;refGene_gn=SCRN3;refGene_tx=XM_023259472.1;refGene_gn=SCRN3;refGene_tx=XM_023259474.1
C1 165318937 165319100
refGene_gn=SCRN3;refGene_tx=XM_023259471.1;refGene_gn=SCRN3;refGene_tx=XM_023259472.1;refGene_gn=SCRN3;refGene_tx=XM_023259473.1;refGene_gn=SCRN3;refGene_tx=XM_023259474.1
C1 165319946 165320127
refGene_gn=SCRN3;refGene_tx=XM_023259471.1;refGene_gn=SCRN3;refGene_tx=XM_023259472.1;refGene_gn=SCRN3;refGene_tx=XM_023259473.1;refGene_gn=SCRN3;refGene_tx=XM_023259474.1
C1 165324215 165324386
refGene_gn=SCRN3;refGene_tx=XM_023259471.1;refGene_gn=SCRN3;refGene_tx=XM_023259472.1;refGene_gn=SCRN3;refGene_tx=XM_023259473.1;refGene_gn=SCRN3;refGene_tx=XM_023259474.1
C1 165333616 165333908
refGene_gn=GPR155;refGene_tx=XM_003990880.5;refGene_gn=GPR155;refGene_tx=XM_019838276.2
C1 165336838 165336976
refGene_gn=GPR155;refGene_tx=XM_003990880.5;refGene_gn=GPR155;refGene_tx=XM_019838276.2
C1 165338823 165338888
refGene_gn=GPR155;refGene_tx=XM_003990880.5;refGene_gn=GPR155;refGene_tx=XM_019838276.2
C1 165340927 165341023
refGene_gn=GPR155;refGene_tx=XM_003990880.5;refGene_gn=GPR155;refGene_tx=XM_019838276.2
C1 165342906 165343043
refGene_gn=GPR155;refGene_tx=XM_003990880.5;refGene_gn=GPR155;refGene_tx=XM_019838276.2
C1 165347162 165347267
refGene_gn=GPR155;refGene_tx=XM_003990880.5;refGene_gn=GPR155;refGene_tx=XM_019838276.2
C1 165351761 165351972
refGene_gn=GPR155;refGene_tx=XM_003990880.5;refGene_gn=GPR155;refGene_tx=XM_019838276.2
C1 165353223 165353314
refGene_gn=GPR155;refGene_tx=XM_003990880.5;refGene_gn=GPR155;refGene_tx=XM_019838276.2
C1 165353402 165353487
refGene_gn=GPR155;refGene_tx=XM_003990880.5;refGene_gn=GPR155;refGene_tx=XM_019838276.2
C1 165358018 165358136

refGene_gn=GPR155;refGene_tx=XM_003990880.5;refGene_gn=GPR155;refGene_tx=XM_019838276.2
C1 165358756 165358840
refGene_gn=GPR155;refGene_tx=XM_003990880.5;refGene_gn=GPR155;refGene_tx=XM_019838276.2
C1 165360798 165360954
refGene_gn=GPR155;refGene_tx=XM_003990880.5;refGene_gn=GPR155;refGene_tx=XM_019838276.2
C1 165361835 165362001
refGene_gn=GPR155;refGene_tx=XM_003990880.5;refGene_gn=GPR155;refGene_tx=XM_019838276.2
C1 165365268 165365668
refGene_gn=GPR155;refGene_tx=XM_003990880.5;refGene_gn=GPR155;refGene_tx=XM_019838276.2
C1 165373893 165374359
refGene_gn=GPR155;refGene_tx=XM_003990880.5;refGene_gn=GPR155;refGene_tx=XM_019838276.2
C1 165433986 165434042
refGene_gn=WIPF1;refGene_tx=XM_023259475.1
C1 165440056 165440170
refGene_gn=WIPF1;refGene_tx=XM_023259475.1
C1 165440879 165441092
refGene_gn=WIPF1;refGene_tx=XM_023259475.1
C1 165446289 165446911
refGene_gn=WIPF1;refGene_tx=XM_023259475.1
C1 165446912 165447064
refGene_gn=WIPF1;refGene_tx=XM_023259475.1
C1 165450179 165450341
refGene_gn=WIPF1;refGene_tx=XM_023259475.1
C1 165456068 165456198
refGene_gn=WIPF1;refGene_tx=XM_023259475.1
C1 165460184 165460235
refGene_gn=WIPF1;refGene_tx=XM_023259475.1
C1 165587627 165587759
refGene_gn=CHRNA1;refGene_tx=XM_003990883.5
C1 165588053 165588293
refGene_gn=CHRNA1;refGene_tx=XM_003990883.5
C1 165589443 165589667
refGene_gn=CHRNA1;refGene_tx=XM_003990883.5
C1 165592984 165593222
refGene_gn=CHRNA1;refGene_tx=XM_003990883.5
C1 165593717 165593913
refGene_gn=CHRNA1;refGene_tx=XM_003990883.5
C1 165595977 165596087
refGene_gn=CHRNA1;refGene_tx=XM_003990883.5
C1 165597150 165597195
refGene_gn=CHRNA1;refGene_tx=XM_003990883.5
C1 165597297 165597443
refGene_gn=CHRNA1;refGene_tx=XM_003990883.5
C1 165602940 165602983

refGene_gn=CHRNA1;refGene_tx=XM_003990883.5
C1 165650833 165651005
refGene_gn=CHN1;refGene_tx=NM_001193650.1
C1 165652388 165652494
refGene_gn=CHN1;refGene_tx=NM_001193650.1
C1 165659512 165659650
refGene_gn=CHN1;refGene_tx=NM_001193650.1
C1 165662515 165662593
refGene_gn=CHN1;refGene_tx=NM_001193650.1
C1 165663351 165663525
refGene_gn=CHN1;refGene_tx=NM_001193650.1
C1 165674151 165674236
refGene_gn=CHN1;refGene_tx=NM_001193650.1
C1 165698865 165699117
refGene_gn=CHN1;refGene_tx=NM_001193650.1
C1 165937813 165938040
refGene_gn=ATF2;refGene_tx=XM_023259476.1
C1 165942822 165942928
refGene_gn=ATF2;refGene_tx=XM_023259476.1
C1 165951135 165951342
refGene_gn=ATF2;refGene_tx=XM_023259476.1
C1 165955551 165955701
refGene_gn=ATF2;refGene_tx=XM_023259476.1
C1 165969786 165969873
refGene_gn=ATF2;refGene_tx=XM_023259476.1
C1 165970602 165970717
refGene_gn=ATF2;refGene_tx=XM_023259476.1
C1 165971423 165971602
refGene_gn=ATF2;refGene_tx=XM_023259476.1
C1 165976085 165976214
refGene_gn=ATF2;refGene_tx=XM_023259476.1
C1 165976357 165976445
refGene_gn=ATF2;refGene_tx=XM_023259476.1
C1 165976446 165976474
refGene_gn=ATF2;refGene_tx=XM_023259476.1
C1 165979529 165979626
refGene_gn=ATF2;refGene_tx=XM_023259476.1
C1 165985413 165985483
refGene_gn=ATF2;refGene_tx=XM_023259476.1
C1 165987339 165987371
refGene_gn=ATF2;refGene_tx=XM_023259476.1
C1 166034220 166034335
refGene_gn=ATP5MC3;refGene_tx=XM_003990884.5
C1 166035024 166035218
refGene_gn=ATP5MC3;refGene_tx=XM_003990884.5
C1 166035748 166035826
refGene_gn=ATP5MC3;refGene_tx=XM_003990884.5
C1 166037011 166037050
refGene_gn=ATP5MC3;refGene_tx=XM_003990884.5
C1 166696584 166696820

refGene_gn=LNPK;refGene_tx=XM_003990886.4
C1 166704742 166704913
refGene_gn=LNPK;refGene_tx=XM_003990886.4
C1 166706060 166706131
refGene_gn=LNPK;refGene_tx=XM_003990886.4
C1 166707654 166707719
refGene_gn=LNPK;refGene_tx=XM_003990886.4
C1 166707720 166707761
refGene_gn=LNPK;refGene_tx=XM_003990886.4
C1 166718454 166718667
refGene_gn=LNPK;refGene_tx=XM_003990886.4
C1 166733362 166733414
refGene_gn=LNPK;refGene_tx=XM_003990886.4
C1 166733496 166733580
refGene_gn=LNPK;refGene_tx=XM_003990886.4
C1 166740806 166740847
refGene_gn=LNPK;refGene_tx=XM_003990886.4
C1 166749044 166749103
refGene_gn=LNPK;refGene_tx=XM_003990886.4
C1 166760987 166761175
refGene_gn=LNPK;refGene_tx=XM_003990886.4
C1 166762201 166762243
refGene_gn=LNPK;refGene_tx=XM_003990886.4
C1 166763797 166763824
refGene_gn=LNPK;refGene_tx=XM_003990886.4
C1 166849206 166849929
refGene_gn=EVX2;refGene_tx=XM_019839122.2
C1 166851267 166851566
refGene_gn=EVX2;refGene_tx=XM_019839122.2
C1 166852494 166852921
refGene_gn=EVX2;refGene_tx=XM_019839122.2
C1 166861915 166862690
refGene_gn=HOXD13;refGene_tx=XM_003990941.5
C1 166863495 166863746
refGene_gn=HOXD13;refGene_tx=XM_003990941.5
C1 166868881 166869085
refGene_gn=HOXD12;refGene_tx=XM_011285399.3
C1 166869086 166869456
refGene_gn=HOXD12;refGene_tx=XM_011285399.3
C1 166869606 166869845
refGene_gn=HOXD12;refGene_tx=XM_011285399.3
C1 166876460 166877241
refGene_gn=HOXD11;refGene_tx=XM_023259477.1
C1 166877993 166878229
refGene_gn=HOXD11;refGene_tx=XM_023259477.1
C1 166885841 166886796
refGene_gn=HOXD10;refGene_tx=XM_003990888.4
C1 166888142 166888420
refGene_gn=HOXD10;refGene_tx=XM_003990888.4
C1 166892001 166892797

refGene_gn=HOXD9;refGene_tx=XM_019839123.2
C1 166893142 166893384
refGene_gn=HOXD9;refGene_tx=XM_019839123.2
C1 166899642 166900240
refGene_gn=HOXD8;refGene_tx=XM_019838286.2;refGene_gn=HOXD8;refGene_tx=XM_023259478.1;refGene_gn=HOXD8;refGene_tx=XM_023259479.1
C1 166900631 166900927
refGene_gn=HOXD8;refGene_tx=XM_019838286.2;refGene_gn=HOXD8;refGene_tx=XM_023259478.1;refGene_gn=HOXD8;refGene_tx=XM_023259479.1
C1 166921130 166921566
refGene_gn=HOXD4;refGene_tx=XM_023259481.1
C1 166922107 166922448
refGene_gn=HOXD4;refGene_tx=XM_023259481.1
C1 166937900 166938441
refGene_gn=HOXD3;refGene_tx=XM_023259480.1
C1 166940310 166941068
refGene_gn=HOXD3;refGene_tx=XM_023259480.1
C1 166955982 166956628
refGene_gn=HOXD1;refGene_tx=XM_003990947.5
C1 166957001 166957336
refGene_gn=HOXD1;refGene_tx=XM_003990947.5
C1 167024571 167024611
refGene_gn=MTX2;refGene_tx=XM_023259483.1
C1 167033760 167033797
refGene_gn=MTX2;refGene_tx=XM_023259482.1
C1 167046458 167046506
refGene_gn=MTX2;refGene_tx=XM_023259482.1;refGene_gn=MTX2;refGene_tx=XM_023259483.1
C1 167047499 167047546
refGene_gn=MTX2;refGene_tx=XM_023259482.1;refGene_gn=MTX2;refGene_tx=XM_023259483.1
C1 167069268 167069341
refGene_gn=MTX2;refGene_tx=XM_023259482.1;refGene_gn=MTX2;refGene_tx=XM_023259483.1
C1 167073825 167073902
refGene_gn=MTX2;refGene_tx=XM_023259482.1;refGene_gn=MTX2;refGene_tx=XM_023259483.1
C1 167075166 167075259
refGene_gn=MTX2;refGene_tx=XM_023259482.1;refGene_gn=MTX2;refGene_tx=XM_023259483.1
C1 167075766 167075805
refGene_gn=MTX2;refGene_tx=XM_023259482.1;refGene_gn=MTX2;refGene_tx=XM_023259483.1
C1 167076190 167076316
refGene_gn=MTX2;refGene_tx=XM_023259482.1;refGene_gn=MTX2;refGene_tx=XM_023259483.1
C1 167077397 167077474
refGene_gn=MTX2;refGene_tx=XM_023259482.1;refGene_gn=MTX2;refGene_tx=XM_023259483.1
C1 167088804 167088988

refGene_gn=MTX2;refGene_tx=XM_023259482.1;refGene_gn=MTX2;refGene_tx=XM_023259483.1
C1 167387450 167387454
refGene_gn=LOC109502751;refGene_tx=XM_023257862.1
C1 167389395 167389454
refGene_gn=LOC109502751;refGene_tx=XM_023257862.1
C1 167393145 167393970
refGene_gn=LOC109502751;refGene_tx=XM_023257862.1
C1 167398428 167398554
refGene_gn=LOC109502751;refGene_tx=XM_023257862.1
C1 167946085 167946157
refGene_gn=HNRNPA3;refGene_tx=XM_011285395.2;refGene_gn=HNRNPA3;refGene_tx=XM_011285396.2;refGene_gn=HNRNPA3;refGene_tx=XM_011285397.2;refGene_gn=HNRNPA3;refGene_tx=XM_019838288.1
C1 167948290 167948413
refGene_gn=HNRNPA3;refGene_tx=XM_011285395.2;refGene_gn=HNRNPA3;refGene_tx=XM_011285396.2;refGene_gn=HNRNPA3;refGene_tx=XM_011285397.2;refGene_gn=HNRNPA3;refGene_tx=XM_019838288.1
C1 167948502 167948649
refGene_gn=HNRNPA3;refGene_tx=XM_011285395.2;refGene_gn=HNRNPA3;refGene_tx=XM_011285396.2;refGene_gn=HNRNPA3;refGene_tx=XM_011285397.2;refGene_gn=HNRNPA3;refGene_tx=XM_019838288.1;refGene_gn=HNRNPA3;refGene_tx=XM_019838289.2
C1 167948731 167948942
refGene_gn=HNRNPA3;refGene_tx=XM_011285395.2;refGene_gn=HNRNPA3;refGene_tx=XM_011285396.2;refGene_gn=HNRNPA3;refGene_tx=XM_011285397.2;refGene_gn=HNRNPA3;refGene_tx=XM_019838288.1;refGene_gn=HNRNPA3;refGene_tx=XM_019838289.2
C1 167949268 167949358
refGene_gn=HNRNPA3;refGene_tx=XM_011285395.2;refGene_gn=HNRNPA3;refGene_tx=XM_011285396.2;refGene_gn=HNRNPA3;refGene_tx=XM_011285397.2;refGene_gn=HNRNPA3;refGene_tx=XM_019838288.1;refGene_gn=HNRNPA3;refGene_tx=XM_019838289.2
C1 167949441 167949534
refGene_gn=HNRNPA3;refGene_tx=XM_011285395.2;refGene_gn=HNRNPA3;refGene_tx=XM_011285396.2;refGene_gn=HNRNPA3;refGene_tx=XM_011285397.2;refGene_gn=HNRNPA3;refGene_tx=XM_019838288.1;refGene_gn=HNRNPA3;refGene_tx=XM_019838289.2
C1 167949615 167949696
refGene_gn=HNRNPA3;refGene_tx=XM_011285395.2;refGene_gn=HNRNPA3;refGene_tx=XM_011285396.2;refGene_gn=HNRNPA3;refGene_tx=XM_011285397.2;refGene_gn=HNRNPA3;refGene_tx=XM_019838288.1;refGene_gn=HNRNPA3;refGene_tx=XM_019838289.2
C1 167950193 167950337
refGene_gn=HNRNPA3;refGene_tx=XM_011285395.2;refGene_gn=HNRNPA3;refGene_tx=XM_011285396.2;refGene_gn=HNRNPA3;refGene_tx=XM_011285397.2;refGene_gn=HNRNPA3;refGene_tx=XM_019838288.1;refGene_gn=HNRNPA3;refGene_tx=XM_019838289.2
C1 167951453 167951576
refGene_gn=HNRNPA3;refGene_tx=XM_011285395.2;refGene_gn=HNRNPA3;refGen

e_tx=XM_011285396.2;refGene_gn=HNRNPA3;refGene_tx=XM_011285397.2;refGene_gn=HNRNPA3;refGene_tx=XM_019838288.1;refGene_gn=HNRNPA3;refGene_tx=XM_019838289.2
C1 167951671 167951724
refGene_gn=HNRNPA3;refGene_tx=XM_011285395.2;refGene_gn=HNRNPA3;refGene_tx=XM_011285396.2;refGene_gn=HNRNPA3;refGene_tx=XM_011285397.2;refGene_gn=HNRNPA3;refGene_tx=XM_019838288.1;refGene_gn=HNRNPA3;refGene_tx=XM_019838289.2
C1 167961917 167963144
refGene_gn=NFE2L2;refGene_tx=XM_003990892.5;refGene_gn=NFE2L2;refGene_tx=XM_003990893.5
C1 167963843 167964035
refGene_gn=NFE2L2;refGene_tx=XM_003990892.5;refGene_gn=NFE2L2;refGene_tx=XM_003990893.5
C1 167964903 167964993
refGene_gn=NFE2L2;refGene_tx=XM_003990892.5;refGene_gn=NFE2L2;refGene_tx=XM_003990893.5
C1 167965532 167965799
refGene_gn=NFE2L2;refGene_tx=XM_003990892.5;refGene_gn=NFE2L2;refGene_tx=XM_003990893.5
C1 167996723 167996768
refGene_gn=NFE2L2;refGene_tx=XM_003990893.5
C1 168111981 168112241
refGene_gn=AGPS;refGene_tx=XM_023259484.1
C1 168146926 168147016
refGene_gn=AGPS;refGene_tx=XM_023259484.1
C1 168155628 168155719
refGene_gn=AGPS;refGene_tx=XM_023259484.1
C1 168158532 168158653
refGene_gn=AGPS;refGene_tx=XM_023259484.1
C1 168158744 168158819
refGene_gn=AGPS;refGene_tx=XM_023259484.1
C1 168162011 168162083
refGene_gn=AGPS;refGene_tx=XM_023259484.1
C1 168163431 168163511
refGene_gn=AGPS;refGene_tx=XM_023259484.1
C1 168168257 168168338
refGene_gn=AGPS;refGene_tx=XM_023259484.1
C1 168182436 168182562
refGene_gn=AGPS;refGene_tx=XM_023259484.1
C1 168185290 168185399
refGene_gn=AGPS;refGene_tx=XM_023259484.1
C1 168196247 168196375
refGene_gn=AGPS;refGene_tx=XM_023259484.1
C1 168205988 168206040
refGene_gn=AGPS;refGene_tx=XM_023259484.1
C1 168211000 168211077
refGene_gn=AGPS;refGene_tx=XM_023259484.1
C1 168213670 168213783
refGene_gn=AGPS;refGene_tx=XM_023259484.1

C1 168221178 168221248
refGene_gn=AGPS;refGene_tx=XM_023259484.1
C1 168223374 168223436
refGene_gn=AGPS;refGene_tx=XM_023259484.1
C1 168228270 168228360
refGene_gn=AGPS;refGene_tx=XM_023259484.1
C1 168234198 168234298
refGene_gn=AGPS;refGene_tx=XM_023259484.1
C1 168236488 168236546
refGene_gn=AGPS;refGene_tx=XM_023259484.1
C1 168249801 168249923
refGene_gn=AGPS;refGene_tx=XM_023259484.1
C1 168261158 168263156
refGene_gn=LOC101099574;refGene_tx=XM_003990896.4
C1 168317787 168319785
refGene_gn=LOC101099822;refGene_tx=XM_003990897.5
C1 168330920 168331076
refGene_gn=PDE11A;refGene_tx=XM_003990898.5
C1 168360773 168360857
refGene_gn=PDE11A;refGene_tx=XM_003990898.5
C1 168366478 168366553
refGene_gn=PDE11A;refGene_tx=XM_003990898.5
C1 168372588 168372652
refGene_gn=PDE11A;refGene_tx=XM_003990898.5
C1 168377872 168377950
refGene_gn=PDE11A;refGene_tx=XM_003990898.5
C1 168393201 168393302
refGene_gn=PDE11A;refGene_tx=XM_003990898.5
C1 168396914 168397005
refGene_gn=PDE11A;refGene_tx=XM_003990898.5
C1 168404599 168404709
refGene_gn=PDE11A;refGene_tx=XM_003990898.5
C1 168419063 168419171
refGene_gn=PDE11A;refGene_tx=XM_003990898.5
C1 168419413 168419560
refGene_gn=PDE11A;refGene_tx=XM_003990898.5
C1 168465085 168465136
refGene_gn=PDE11A;refGene_tx=XM_003990898.5
C1 168491800 168491893
refGene_gn=PDE11A;refGene_tx=XM_003990898.5
C1 168492635 168492703
refGene_gn=PDE11A;refGene_tx=XM_003990898.5
C1 168497293 168497384
refGene_gn=PDE11A;refGene_tx=XM_003990898.5
C1 168513433 168513566
refGene_gn=PDE11A;refGene_tx=XM_003990898.5
C1 168538770 168538835
refGene_gn=PDE11A;refGene_tx=XM_003990898.5
C1 168556391 168556532
refGene_gn=PDE11A;refGene_tx=XM_003990898.5

C1       168562348        168562438
refGene_gn=PDE11A;refGene_tx=XM_003990898.5
C1       168649839        168649998
refGene_gn=PDE11A;refGene_tx=XM_003990898.5
C1       168701235        168702147
refGene_gn=PDE11A;refGene_tx=XM_003990898.5
C1       168719322        168719470
refGene_gn=LOC101094614;refGene_tx=XM_003990950.1
C1       168737436        168737615
refGene_gn=LOC101094614;refGene_tx=XM_003990950.1
C1       168745888        168745960
refGene_gn=LOC101094614;refGene_tx=XM_003990950.1
C1       168750676        168750982
refGene_gn=RBM45;refGene_tx=XM_003990900.5;refGene_gn=RBM45;refGene_tx
=XM_006935378.4;refGene_gn=RBM45;refGene_tx=XM_006935381.4;refGene_gn=
RBM45;refGene_tx=XM_019838293.2;refGene_gn=RBM45;refGene_tx=XM_0198382
94.2;refGene_gn=RBM45;refGene_tx=XM_019838295.2;refGene_gn=RBM45;refGe
ne_tx=XM_019838296.2
C1       168755299        168755422
refGene_gn=RBM45;refGene_tx=XM_003990900.5;refGene_gn=RBM45;refGene_tx
=XM_006935378.4;refGene_gn=RBM45;refGene_tx=XM_006935381.4;refGene_gn=
RBM45;refGene_tx=XM_019838293.2;refGene_gn=RBM45;refGene_tx=XM_0198382
94.2;refGene_gn=RBM45;refGene_tx=XM_019838295.2;refGene_gn=RBM45;refGe
ne_tx=XM_019838296.2
C1       168757773        168757900
refGene_gn=RBM45;refGene_tx=XM_003990900.5;refGene_gn=RBM45;refGene_tx
=XM_006935378.4;refGene_gn=RBM45;refGene_tx=XM_006935381.4;refGene_gn=
RBM45;refGene_tx=XM_019838293.2;refGene_gn=RBM45;refGene_tx=XM_0198382
94.2;refGene_gn=RBM45;refGene_tx=XM_019838295.2;refGene_gn=RBM45;refGe
ne_tx=XM_019838296.2
C1       168759999        168760122
refGene_gn=RBM45;refGene_tx=XM_003990900.5;refGene_gn=RBM45;refGene_tx
=XM_006935378.4;refGene_gn=RBM45;refGene_tx=XM_006935381.4;refGene_gn=
RBM45;refGene_tx=XM_019838293.2;refGene_gn=RBM45;refGene_tx=XM_0198382
94.2;refGene_gn=RBM45;refGene_tx=XM_019838295.2;refGene_gn=RBM45;refGe
ne_tx=XM_019838296.2
C1       168760890        168761070
refGene_gn=RBM45;refGene_tx=XM_003990900.5;refGene_gn=RBM45;refGene_tx
=XM_006935378.4;refGene_gn=RBM45;refGene_tx=XM_006935381.4;refGene_gn=
RBM45;refGene_tx=XM_019838293.2;refGene_gn=RBM45;refGene_tx=XM_0198382
94.2;refGene_gn=RBM45;refGene_tx=XM_019838295.2;refGene_gn=RBM45;refGe
ne_tx=XM_019838296.2
C1       168763657        168763787
refGene_gn=RBM45;refGene_tx=XM_003990900.5;refGene_gn=RBM45;refGene_tx
=XM_006935378.4;refGene_gn=RBM45;refGene_tx=XM_006935381.4;refGene_gn=
RBM45;refGene_tx=XM_019838293.2;refGene_gn=RBM45;refGene_tx=XM_0198382
94.2;refGene_gn=RBM45;refGene_tx=XM_019838295.2;refGene_gn=RBM45;refGe
ne_tx=XM_019838296.2
C1       168763953        168764038
refGene_gn=RBM45;refGene_tx=XM_003990900.5;refGene_gn=RBM45;refGene_tx

=XM_006935378.4;refGene_gn=RBM45;refGene_tx=XM_006935381.4;refGene_gn=
RBM45;refGene_tx=XM_019838293.2;refGene_gn=RBM45;refGene_tx=XM_0198382
94.2;refGene_gn=RBM45;refGene_tx=XM_019838295.2;refGene_gn=RBM45;refGe
ne_tx=XM_019838296.2
C1 168764247 168764411
refGene_gn=RBM45;refGene_tx=XM_003990900.5;refGene_gn=RBM45;refGene_tx
=XM_006935378.4;refGene_gn=RBM45;refGene_tx=XM_006935381.4;refGene_gn=
RBM45;refGene_tx=XM_019838293.2;refGene_gn=RBM45;refGene_tx=XM_0198382
94.2;refGene_gn=RBM45;refGene_tx=XM_019838295.2;refGene_gn=RBM45;refGe
ne_tx=XM_019838296.2
C1 168765886 168766079
refGene_gn=RBM45;refGene_tx=XM_003990900.5;refGene_gn=RBM45;refGene_tx
=XM_006935378.4;refGene_gn=RBM45;refGene_tx=XM_006935381.4;refGene_gn=
RBM45;refGene_tx=XM_019838293.2;refGene_gn=RBM45;refGene_tx=XM_0198382
94.2;refGene_gn=RBM45;refGene_tx=XM_019838295.2;refGene_gn=RBM45;refGe
ne_tx=XM_019838296.2
C1 168944302 168944404
refGene_gn=OSBPL6;refGene_tx=XM_019838297.2;refGene_gn=OSBPL6;refGene_
tx=XM_023259486.1;refGene_gn=OSBPL6;refGene_tx=XM_023259487.1;refGene_
gn=OSBPL6;refGene_tx=XM_023259488.1;refGene_gn=OSBPL6;refGene_tx=XM_02
3259489.1;refGene_gn=OSBPL6;refGene_tx=XM_023259490.1;refGene_gn=OSBPL
6;refGene_tx=XM_023259491.1;refGene_gn=OSBPL6;refGene_tx=XM_023259492.
1;refGene_gn=OSBPL6;refGene_tx=XM_023259493.1;refGene_gn=OSBPL6;refGen
e_tx=XM_023259494.1;refGene_gn=OSBPL6;refGene_tx=XM_023259495.1;refGen
e_gn=OSBPL6;refGene_tx=XM_023259496.1;refGene_gn=OSBPL6;refGene_tx=XM_
023259497.1;refGene_gn=OSBPL6;refGene_tx=XM_023259498.1;refGene_gn=OSB
PL6;refGene_tx=XM_023259499.1;refGene_gn=OSBPL6;refGene_tx=XM_02325950
0.1;refGene_gn=OSBPL6;refGene_tx=XM_023259501.1;refGene_gn=OSBPL6;refG
ene_tx=XM_023259502.1
C1 168964823 168964916
refGene_gn=OSBPL6;refGene_tx=XM_019838297.2;refGene_gn=OSBPL6;refGene_
tx=XM_023259486.1;refGene_gn=OSBPL6;refGene_tx=XM_023259487.1;refGene_
gn=OSBPL6;refGene_tx=XM_023259488.1;refGene_gn=OSBPL6;refGene_tx=XM_02
3259489.1;refGene_gn=OSBPL6;refGene_tx=XM_023259490.1;refGene_gn=OSBPL
6;refGene_tx=XM_023259491.1;refGene_gn=OSBPL6;refGene_tx=XM_023259492.
1;refGene_gn=OSBPL6;refGene_tx=XM_023259493.1;refGene_gn=OSBPL6;refGen
e_tx=XM_023259494.1;refGene_gn=OSBPL6;refGene_tx=XM_023259495.1;refGen
e_gn=OSBPL6;refGene_tx=XM_023259496.1;refGene_gn=OSBPL6;refGene_tx=XM_
023259497.1;refGene_gn=OSBPL6;refGene_tx=XM_023259498.1;refGene_gn=OSB
PL6;refGene_tx=XM_023259499.1;refGene_gn=OSBPL6;refGene_tx=XM_02325950
0.1;refGene_gn=OSBPL6;refGene_tx=XM_023259501.1;refGene_gn=OSBPL6;refG
ene_tx=XM_023259502.1
C1 168973397 168973520
refGene_gn=OSBPL6;refGene_tx=XM_019838297.2;refGene_gn=OSBPL6;refGene_
tx=XM_023259486.1;refGene_gn=OSBPL6;refGene_tx=XM_023259487.1;refGene_
gn=OSBPL6;refGene_tx=XM_023259488.1;refGene_gn=OSBPL6;refGene_tx=XM_02
3259489.1;refGene_gn=OSBPL6;refGene_tx=XM_023259490.1;refGene_gn=OSBPL
6;refGene_tx=XM_023259491.1;refGene_gn=OSBPL6;refGene_tx=XM_023259492.
1;refGene_gn=OSBPL6;refGene_tx=XM_023259493.1;refGene_gn=OSBPL6;refGen
e_tx=XM_023259494.1;refGene_gn=OSBPL6;refGene_tx=XM_023259495.1;refGen

e_gn=OSBPL6;refGene_tx=XM_023259496.1;refGene_gn=OSBPL6;refGene_tx=XM_023259497.1;refGene_gn=OSBPL6;refGene_tx=XM_023259498.1;refGene_gn=OSBPL6;refGene_tx=XM_023259499.1;refGene_gn=OSBPL6;refGene_tx=XM_023259500.1;refGene_gn=OSBPL6;refGene_tx=XM_023259501.1;refGene_gn=OSBPL6;refGene_tx=XM_023259502.1
C1 168976081 168976135
refGene_gn=OSBPL6;refGene_tx=XM_019838297.2;refGene_gn=OSBPL6;refGene_tx=XM_023259486.1;refGene_gn=OSBPL6;refGene_tx=XM_023259487.1;refGene_gn=OSBPL6;refGene_tx=XM_023259488.1;refGene_gn=OSBPL6;refGene_tx=XM_023259489.1;refGene_gn=OSBPL6;refGene_tx=XM_023259490.1;refGene_gn=OSBPL6;refGene_tx=XM_023259491.1;refGene_gn=OSBPL6;refGene_tx=XM_023259492.1;refGene_gn=OSBPL6;refGene_tx=XM_023259493.1;refGene_gn=OSBPL6;refGene_tx=XM_023259494.1;refGene_gn=OSBPL6;refGene_tx=XM_023259495.1;refGene_gn=OSBPL6;refGene_tx=XM_023259496.1;refGene_gn=OSBPL6;refGene_tx=XM_023259497.1;refGene_gn=OSBPL6;refGene_tx=XM_023259498.1;refGene_gn=OSBPL6;refGene_tx=XM_023259499.1;refGene_gn=OSBPL6;refGene_tx=XM_023259500.1;refGene_gn=OSBPL6;refGene_tx=XM_023259501.1;refGene_gn=OSBPL6;refGene_tx=XM_023259502.1
C1 168977430 168977544
refGene_gn=OSBPL6;refGene_tx=XM_019838297.2;refGene_gn=OSBPL6;refGene_tx=XM_023259486.1;refGene_gn=OSBPL6;refGene_tx=XM_023259487.1;refGene_gn=OSBPL6;refGene_tx=XM_023259488.1;refGene_gn=OSBPL6;refGene_tx=XM_023259489.1;refGene_gn=OSBPL6;refGene_tx=XM_023259490.1;refGene_gn=OSBPL6;refGene_tx=XM_023259491.1;refGene_gn=OSBPL6;refGene_tx=XM_023259492.1;refGene_gn=OSBPL6;refGene_tx=XM_023259493.1;refGene_gn=OSBPL6;refGene_tx=XM_023259494.1;refGene_gn=OSBPL6;refGene_tx=XM_023259495.1;refGene_gn=OSBPL6;refGene_tx=XM_023259496.1;refGene_gn=OSBPL6;refGene_tx=XM_023259497.1;refGene_gn=OSBPL6;refGene_tx=XM_023259498.1;refGene_gn=OSBPL6;refGene_tx=XM_023259499.1;refGene_gn=OSBPL6;refGene_tx=XM_023259500.1;refGene_gn=OSBPL6;refGene_tx=XM_023259501.1;refGene_gn=OSBPL6;refGene_tx=XM_023259502.1
C1 168977660 168977831
refGene_gn=OSBPL6;refGene_tx=XM_019838297.2;refGene_gn=OSBPL6;refGene_tx=XM_023259486.1;refGene_gn=OSBPL6;refGene_tx=XM_023259487.1;refGene_gn=OSBPL6;refGene_tx=XM_023259488.1;refGene_gn=OSBPL6;refGene_tx=XM_023259489.1;refGene_gn=OSBPL6;refGene_tx=XM_023259490.1;refGene_gn=OSBPL6;refGene_tx=XM_023259491.1;refGene_gn=OSBPL6;refGene_tx=XM_023259492.1;refGene_gn=OSBPL6;refGene_tx=XM_023259493.1;refGene_gn=OSBPL6;refGene_tx=XM_023259494.1;refGene_gn=OSBPL6;refGene_tx=XM_023259495.1;refGene_gn=OSBPL6;refGene_tx=XM_023259496.1;refGene_gn=OSBPL6;refGene_tx=XM_023259497.1;refGene_gn=OSBPL6;refGene_tx=XM_023259498.1;refGene_gn=OSBPL6;refGene_tx=XM_023259499.1;refGene_gn=OSBPL6;refGene_tx=XM_023259500.1;refGene_gn=OSBPL6;refGene_tx=XM_023259501.1;refGene_gn=OSBPL6;refGene_tx=XM_023259502.1
C1 168980883 168981016
refGene_gn=OSBPL6;refGene_tx=XM_019838297.2;refGene_gn=OSBPL6;refGene_tx=XM_023259486.1;refGene_gn=OSBPL6;refGene_tx=XM_023259487.1;refGene_gn=OSBPL6;refGene_tx=XM_023259488.1;refGene_gn=OSBPL6;refGene_tx=XM_023259489.1;refGene_gn=OSBPL6;refGene_tx=XM_023259490.1;refGene_gn=OSBPL6;refGene_tx=XM_023259491.1;refGene_gn=OSBPL6;refGene_tx=XM_023259492.

1;refGene_gn=OSBPL6;refGene_tx=XM_023259493.1;refGene_gn=OSBPL6;refGene_tx=XM_023259494.1;refGene_gn=OSBPL6;refGene_tx=XM_023259495.1;refGene_gn=OSBPL6;refGene_tx=XM_023259496.1;refGene_gn=OSBPL6;refGene_tx=XM_023259497.1;refGene_gn=OSBPL6;refGene_tx=XM_023259498.1;refGene_gn=OSBPL6;refGene_tx=XM_023259499.1;refGene_gn=OSBPL6;refGene_tx=XM_023259500.1;refGene_gn=OSBPL6;refGene_tx=XM_023259501.1;refGene_gn=OSBPL6;refGene_tx=XM_023259502.1
C1 168988144 168988248
refGene_gn=OSBPL6;refGene_tx=XM_019838297.2;refGene_gn=OSBPL6;refGene_tx=XM_023259486.1;refGene_gn=OSBPL6;refGene_tx=XM_023259487.1;refGene_gn=OSBPL6;refGene_tx=XM_023259488.1;refGene_gn=OSBPL6;refGene_tx=XM_023259489.1;refGene_gn=OSBPL6;refGene_tx=XM_023259490.1;refGene_gn=OSBPL6;refGene_tx=XM_023259491.1;refGene_gn=OSBPL6;refGene_tx=XM_023259492.1;refGene_gn=OSBPL6;refGene_tx=XM_023259493.1;refGene_gn=OSBPL6;refGene_tx=XM_023259494.1;refGene_gn=OSBPL6;refGene_tx=XM_023259495.1;refGene_gn=OSBPL6;refGene_tx=XM_023259496.1;refGene_gn=OSBPL6;refGene_tx=XM_023259497.1;refGene_gn=OSBPL6;refGene_tx=XM_023259498.1;refGene_gn=OSBPL6;refGene_tx=XM_023259499.1;refGene_gn=OSBPL6;refGene_tx=XM_023259500.1;refGene_gn=OSBPL6;refGene_tx=XM_023259501.1;refGene_gn=OSBPL6;refGene_tx=XM_023259502.1
C1 168988823 168988916
refGene_gn=OSBPL6;refGene_tx=XM_019838297.2;refGene_gn=OSBPL6;refGene_tx=XM_023259486.1;refGene_gn=OSBPL6;refGene_tx=XM_023259487.1;refGene_gn=OSBPL6;refGene_tx=XM_023259488.1;refGene_gn=OSBPL6;refGene_tx=XM_023259489.1;refGene_gn=OSBPL6;refGene_tx=XM_023259490.1;refGene_gn=OSBPL6;refGene_tx=XM_023259491.1;refGene_gn=OSBPL6;refGene_tx=XM_023259492.1;refGene_gn=OSBPL6;refGene_tx=XM_023259493.1;refGene_gn=OSBPL6;refGene_tx=XM_023259494.1;refGene_gn=OSBPL6;refGene_tx=XM_023259495.1;refGene_gn=OSBPL6;refGene_tx=XM_023259497.1;refGene_gn=OSBPL6;refGene_tx=XM_023259498.1;refGene_gn=OSBPL6;refGene_tx=XM_023259500.1;refGene_gn=OSBPL6;refGene_tx=XM_023259502.1
C1 168993872 168993947
refGene_gn=OSBPL6;refGene_tx=XM_019838297.2;refGene_gn=OSBPL6;refGene_tx=XM_023259486.1;refGene_gn=OSBPL6;refGene_tx=XM_023259487.1;refGene_gn=OSBPL6;refGene_tx=XM_023259488.1;refGene_gn=OSBPL6;refGene_tx=XM_023259489.1;refGene_gn=OSBPL6;refGene_tx=XM_023259490.1;refGene_gn=OSBPL6;refGene_tx=XM_023259491.1;refGene_gn=OSBPL6;refGene_tx=XM_023259492.1;refGene_gn=OSBPL6;refGene_tx=XM_023259493.1;refGene_gn=OSBPL6;refGene_tx=XM_023259494.1;refGene_gn=OSBPL6;refGene_tx=XM_023259496.1;refGene_gn=OSBPL6;refGene_tx=XM_023259498.1;refGene_gn=OSBPL6;refGene_tx=XM_023259500.1
C1 168999221 168999387
refGene_gn=OSBPL6;refGene_tx=XM_019838297.2;refGene_gn=OSBPL6;refGene_tx=XM_023259486.1;refGene_gn=OSBPL6;refGene_tx=XM_023259487.1;refGene_gn=OSBPL6;refGene_tx=XM_023259488.1;refGene_gn=OSBPL6;refGene_tx=XM_023259489.1;refGene_gn=OSBPL6;refGene_tx=XM_023259490.1;refGene_gn=OSBPL6;refGene_tx=XM_023259491.1;refGene_gn=OSBPL6;refGene_tx=XM_023259492.1;refGene_gn=OSBPL6;refGene_tx=XM_023259493.1;refGene_gn=OSBPL6;refGene_tx=XM_023259494.1;refGene_gn=OSBPL6;refGene_tx=XM_023259495.1;refGene_gn=OSBPL6;refGene_tx=XM_023259496.1;refGene_gn=OSBPL6;refGene_tx=XM_

023259497.1;refGene_gn=OSBPL6;refGene_tx=XM_023259498.1;refGene_gn=OSBPL6;refGene_tx=XM_023259499.1;refGene_gn=OSBPL6;refGene_tx=XM_023259500.1;refGene_gn=OSBPL6;refGene_tx=XM_023259501.1;refGene_gn=OSBPL6;refGene_tx=XM_023259502.1
C1 169002901 169003035
refGene_gn=OSBPL6;refGene_tx=XM_019838297.2;refGene_gn=OSBPL6;refGene_tx=XM_023259486.1;refGene_gn=OSBPL6;refGene_tx=XM_023259487.1;refGene_gn=OSBPL6;refGene_tx=XM_023259488.1;refGene_gn=OSBPL6;refGene_tx=XM_023259489.1;refGene_gn=OSBPL6;refGene_tx=XM_023259490.1;refGene_gn=OSBPL6;refGene_tx=XM_023259491.1;refGene_gn=OSBPL6;refGene_tx=XM_023259492.1;refGene_gn=OSBPL6;refGene_tx=XM_023259493.1;refGene_gn=OSBPL6;refGene_tx=XM_023259494.1;refGene_gn=OSBPL6;refGene_tx=XM_023259495.1;refGene_gn=OSBPL6;refGene_tx=XM_023259496.1;refGene_gn=OSBPL6;refGene_tx=XM_023259497.1;refGene_gn=OSBPL6;refGene_tx=XM_023259498.1;refGene_gn=OSBPL6;refGene_tx=XM_023259499.1;refGene_gn=OSBPL6;refGene_tx=XM_023259500.1;refGene_gn=OSBPL6;refGene_tx=XM_023259501.1;refGene_gn=OSBPL6;refGene_tx=XM_023259502.1
C1 169003884 169003908
refGene_gn=OSBPL6;refGene_tx=XM_019838297.2;refGene_gn=OSBPL6;refGene_tx=XM_023259486.1;refGene_gn=OSBPL6;refGene_tx=XM_023259487.1;refGene_gn=OSBPL6;refGene_tx=XM_023259488.1;refGene_gn=OSBPL6;refGene_tx=XM_023259489.1;refGene_gn=OSBPL6;refGene_tx=XM_023259490.1;refGene_gn=OSBPL6;refGene_tx=XM_023259491.1;refGene_gn=OSBPL6;refGene_tx=XM_023259493.1;refGene_gn=OSBPL6;refGene_tx=XM_023259494.1;refGene_gn=OSBPL6;refGene_tx=XM_023259495.1;refGene_gn=OSBPL6;refGene_tx=XM_023259496.1;refGene_gn=OSBPL6;refGene_tx=XM_023259499.1
C1 169012540 169012648
refGene_gn=OSBPL6;refGene_tx=XM_019838297.2;refGene_gn=OSBPL6;refGene_tx=XM_023259486.1;refGene_gn=OSBPL6;refGene_tx=XM_023259487.1;refGene_gn=OSBPL6;refGene_tx=XM_023259488.1;refGene_gn=OSBPL6;refGene_tx=XM_023259489.1;refGene_gn=OSBPL6;refGene_tx=XM_023259490.1;refGene_gn=OSBPL6;refGene_tx=XM_023259491.1;refGene_gn=OSBPL6;refGene_tx=XM_023259492.1;refGene_gn=OSBPL6;refGene_tx=XM_023259493.1;refGene_gn=OSBPL6;refGene_tx=XM_023259494.1;refGene_gn=OSBPL6;refGene_tx=XM_023259495.1;refGene_gn=OSBPL6;refGene_tx=XM_023259496.1;refGene_gn=OSBPL6;refGene_tx=XM_023259497.1;refGene_gn=OSBPL6;refGene_tx=XM_023259499.1;refGene_gn=OSBPL6;refGene_tx=XM_023259501.1
C1 169014601 169014739
refGene_gn=OSBPL6;refGene_tx=XM_019838297.2;refGene_gn=OSBPL6;refGene_tx=XM_023259486.1;refGene_gn=OSBPL6;refGene_tx=XM_023259487.1;refGene_gn=OSBPL6;refGene_tx=XM_023259488.1;refGene_gn=OSBPL6;refGene_tx=XM_023259489.1;refGene_gn=OSBPL6;refGene_tx=XM_023259490.1;refGene_gn=OSBPL6;refGene_tx=XM_023259491.1;refGene_gn=OSBPL6;refGene_tx=XM_023259492.1;refGene_gn=OSBPL6;refGene_tx=XM_023259493.1;refGene_gn=OSBPL6;refGene_tx=XM_023259494.1;refGene_gn=OSBPL6;refGene_tx=XM_023259495.1;refGene_gn=OSBPL6;refGene_tx=XM_023259496.1;refGene_gn=OSBPL6;refGene_tx=XM_023259497.1;refGene_gn=OSBPL6;refGene_tx=XM_023259498.1;refGene_gn=OSBPL6;refGene_tx=XM_023259499.1;refGene_gn=OSBPL6;refGene_tx=XM_023259500.1;refGene_gn=OSBPL6;refGene_tx=XM_023259501.1;refGene_gn=OSBPL6;refGene_tx=XM_023259502.1

C1	169024216	169024304
refGene_gn=OSBPL6;refGene_tx=XM_019838297.2;refGene_gn=OSBPL6;refGene_tx=XM_023259486.1;refGene_gn=OSBPL6;refGene_tx=XM_023259487.1;refGene_gn=OSBPL6;refGene_tx=XM_023259488.1;refGene_gn=OSBPL6;refGene_tx=XM_023259489.1;refGene_gn=OSBPL6;refGene_tx=XM_023259490.1;refGene_gn=OSBPL6;refGene_tx=XM_023259491.1;refGene_gn=OSBPL6;refGene_tx=XM_023259492.1;refGene_gn=OSBPL6;refGene_tx=XM_023259493.1;refGene_gn=OSBPL6;refGene_tx=XM_023259494.1;refGene_gn=OSBPL6;refGene_tx=XM_023259495.1;refGene_gn=OSBPL6;refGene_tx=XM_023259496.1;refGene_gn=OSBPL6;refGene_tx=XM_023259497.1;refGene_gn=OSBPL6;refGene_tx=XM_023259498.1;refGene_gn=OSBPL6;refGene_tx=XM_023259499.1;refGene_gn=OSBPL6;refGene_tx=XM_023259500.1;refGene_gn=OSBPL6;refGene_tx=XM_023259501.1;refGene_gn=OSBPL6;refGene_tx=XM_023259502.1
C1	169024807	169025061
refGene_gn=OSBPL6;refGene_tx=XM_019838297.2;refGene_gn=OSBPL6;refGene_tx=XM_023259486.1;refGene_gn=OSBPL6;refGene_tx=XM_023259487.1;refGene_gn=OSBPL6;refGene_tx=XM_023259488.1;refGene_gn=OSBPL6;refGene_tx=XM_023259489.1;refGene_gn=OSBPL6;refGene_tx=XM_023259490.1;refGene_gn=OSBPL6;refGene_tx=XM_023259491.1;refGene_gn=OSBPL6;refGene_tx=XM_023259492.1;refGene_gn=OSBPL6;refGene_tx=XM_023259493.1;refGene_gn=OSBPL6;refGene_tx=XM_023259494.1;refGene_gn=OSBPL6;refGene_tx=XM_023259495.1;refGene_gn=OSBPL6;refGene_tx=XM_023259496.1;refGene_gn=OSBPL6;refGene_tx=XM_023259497.1;refGene_gn=OSBPL6;refGene_tx=XM_023259498.1;refGene_gn=OSBPL6;refGene_tx=XM_023259499.1;refGene_gn=OSBPL6;refGene_tx=XM_023259500.1;refGene_gn=OSBPL6;refGene_tx=XM_023259501.1;refGene_gn=OSBPL6;refGene_tx=XM_023259502.1
C1	169025808	169025946
refGene_gn=OSBPL6;refGene_tx=XM_019838297.2;refGene_gn=OSBPL6;refGene_tx=XM_023259486.1;refGene_gn=OSBPL6;refGene_tx=XM_023259487.1;refGene_gn=OSBPL6;refGene_tx=XM_023259488.1;refGene_gn=OSBPL6;refGene_tx=XM_023259489.1;refGene_gn=OSBPL6;refGene_tx=XM_023259490.1;refGene_gn=OSBPL6;refGene_tx=XM_023259491.1;refGene_gn=OSBPL6;refGene_tx=XM_023259492.1;refGene_gn=OSBPL6;refGene_tx=XM_023259493.1;refGene_gn=OSBPL6;refGene_tx=XM_023259494.1;refGene_gn=OSBPL6;refGene_tx=XM_023259495.1;refGene_gn=OSBPL6;refGene_tx=XM_023259496.1;refGene_gn=OSBPL6;refGene_tx=XM_023259497.1;refGene_gn=OSBPL6;refGene_tx=XM_023259498.1;refGene_gn=OSBPL6;refGene_tx=XM_023259499.1;refGene_gn=OSBPL6;refGene_tx=XM_023259500.1;refGene_gn=OSBPL6;refGene_tx=XM_023259501.1;refGene_gn=OSBPL6;refGene_tx=XM_023259502.1
C1	169026310	169026374
refGene_gn=OSBPL6;refGene_tx=XM_019838297.2;refGene_gn=OSBPL6;refGene_tx=XM_023259486.1;refGene_gn=OSBPL6;refGene_tx=XM_023259487.1;refGene_gn=OSBPL6;refGene_tx=XM_023259488.1;refGene_gn=OSBPL6;refGene_tx=XM_023259489.1;refGene_gn=OSBPL6;refGene_tx=XM_023259490.1;refGene_gn=OSBPL6;refGene_tx=XM_023259491.1;refGene_gn=OSBPL6;refGene_tx=XM_023259492.1;refGene_gn=OSBPL6;refGene_tx=XM_023259493.1;refGene_gn=OSBPL6;refGene_tx=XM_023259494.1;refGene_gn=OSBPL6;refGene_tx=XM_023259495.1;refGene_gn=OSBPL6;refGene_tx=XM_023259496.1;refGene_gn=OSBPL6;refGene_tx=XM_023259497.1;refGene_gn=OSBPL6;refGene_tx=XM_023259498.1;refGene_gn=OSBPL6;refGene_tx=XM_023259499.1;refGene_gn=OSBPL6;refGene_tx=XM_02325950

0.1;refGene_gn=OSBPL6;refGene_tx=XM_023259501.1;refGene_gn=OSBPL6;refGene_tx=XM_023259502.1
C1 169028403 169028482
refGene_gn=OSBPL6;refGene_tx=XM_019838297.2;refGene_gn=OSBPL6;refGene_tx=XM_023259486.1;refGene_gn=OSBPL6;refGene_tx=XM_023259487.1;refGene_gn=OSBPL6;refGene_tx=XM_023259488.1;refGene_gn=OSBPL6;refGene_tx=XM_023259489.1;refGene_gn=OSBPL6;refGene_tx=XM_023259490.1;refGene_gn=OSBPL6;refGene_tx=XM_023259491.1;refGene_gn=OSBPL6;refGene_tx=XM_023259492.1;refGene_gn=OSBPL6;refGene_tx=XM_023259493.1;refGene_gn=OSBPL6;refGene_tx=XM_023259494.1;refGene_gn=OSBPL6;refGene_tx=XM_023259495.1;refGene_gn=OSBPL6;refGene_tx=XM_023259496.1;refGene_gn=OSBPL6;refGene_tx=XM_023259497.1;refGene_gn=OSBPL6;refGene_tx=XM_023259498.1;refGene_gn=OSBPL6;refGene_tx=XM_023259499.1;refGene_gn=OSBPL6;refGene_tx=XM_023259500.1;refGene_gn=OSBPL6;refGene_tx=XM_023259501.1;refGene_gn=OSBPL6;refGene_tx=XM_023259502.1
C1 169030328 169030473
refGene_gn=OSBPL6;refGene_tx=XM_019838297.2;refGene_gn=OSBPL6;refGene_tx=XM_023259486.1;refGene_gn=OSBPL6;refGene_tx=XM_023259487.1;refGene_gn=OSBPL6;refGene_tx=XM_023259488.1;refGene_gn=OSBPL6;refGene_tx=XM_023259489.1;refGene_gn=OSBPL6;refGene_tx=XM_023259490.1;refGene_gn=OSBPL6;refGene_tx=XM_023259491.1;refGene_gn=OSBPL6;refGene_tx=XM_023259492.1;refGene_gn=OSBPL6;refGene_tx=XM_023259493.1;refGene_gn=OSBPL6;refGene_tx=XM_023259494.1;refGene_gn=OSBPL6;refGene_tx=XM_023259495.1;refGene_gn=OSBPL6;refGene_tx=XM_023259496.1;refGene_gn=OSBPL6;refGene_tx=XM_023259497.1;refGene_gn=OSBPL6;refGene_tx=XM_023259498.1;refGene_gn=OSBPL6;refGene_tx=XM_023259499.1;refGene_gn=OSBPL6;refGene_tx=XM_023259500.1;refGene_gn=OSBPL6;refGene_tx=XM_023259501.1;refGene_gn=OSBPL6;refGene_tx=XM_023259502.1
C1 169033194 169033339
refGene_gn=OSBPL6;refGene_tx=XM_019838297.2;refGene_gn=OSBPL6;refGene_tx=XM_023259486.1;refGene_gn=OSBPL6;refGene_tx=XM_023259487.1;refGene_gn=OSBPL6;refGene_tx=XM_023259488.1;refGene_gn=OSBPL6;refGene_tx=XM_023259489.1;refGene_gn=OSBPL6;refGene_tx=XM_023259490.1;refGene_gn=OSBPL6;refGene_tx=XM_023259491.1;refGene_gn=OSBPL6;refGene_tx=XM_023259492.1;refGene_gn=OSBPL6;refGene_tx=XM_023259493.1;refGene_gn=OSBPL6;refGene_tx=XM_023259494.1;refGene_gn=OSBPL6;refGene_tx=XM_023259495.1;refGene_gn=OSBPL6;refGene_tx=XM_023259496.1;refGene_gn=OSBPL6;refGene_tx=XM_023259497.1;refGene_gn=OSBPL6;refGene_tx=XM_023259498.1;refGene_gn=OSBPL6;refGene_tx=XM_023259499.1;refGene_gn=OSBPL6;refGene_tx=XM_023259500.1;refGene_gn=OSBPL6;refGene_tx=XM_023259501.1;refGene_gn=OSBPL6;refGene_tx=XM_023259502.1
C1 169034744 169034871
refGene_gn=OSBPL6;refGene_tx=XM_019838297.2;refGene_gn=OSBPL6;refGene_tx=XM_023259486.1;refGene_gn=OSBPL6;refGene_tx=XM_023259487.1;refGene_gn=OSBPL6;refGene_tx=XM_023259488.1;refGene_gn=OSBPL6;refGene_tx=XM_023259489.1;refGene_gn=OSBPL6;refGene_tx=XM_023259490.1;refGene_gn=OSBPL6;refGene_tx=XM_023259491.1;refGene_gn=OSBPL6;refGene_tx=XM_023259492.1;refGene_gn=OSBPL6;refGene_tx=XM_023259493.1;refGene_gn=OSBPL6;refGene_tx=XM_023259494.1;refGene_gn=OSBPL6;refGene_tx=XM_023259495.1;refGene_gn=OSBPL6;refGene_tx=XM_023259496.1;refGene_gn=OSBPL6;refGene_tx=XM_

023259497.1;refGene_gn=OSBPL6;refGene_tx=XM_023259498.1;refGene_gn=OSBPL6;refGene_tx=XM_023259499.1;refGene_gn=OSBPL6;refGene_tx=XM_023259500.1;refGene_gn=OSBPL6;refGene_tx=XM_023259501.1;refGene_gn=OSBPL6;refGene_tx=XM_023259502.1
C1 169036070 169036193
refGene_gn=OSBPL6;refGene_tx=XM_019838297.2;refGene_gn=OSBPL6;refGene_tx=XM_023259486.1;refGene_gn=OSBPL6;refGene_tx=XM_023259487.1;refGene_gn=OSBPL6;refGene_tx=XM_023259488.1;refGene_gn=OSBPL6;refGene_tx=XM_023259489.1;refGene_gn=OSBPL6;refGene_tx=XM_023259490.1;refGene_gn=OSBPL6;refGene_tx=XM_023259491.1;refGene_gn=OSBPL6;refGene_tx=XM_023259492.1;refGene_gn=OSBPL6;refGene_tx=XM_023259493.1;refGene_gn=OSBPL6;refGene_tx=XM_023259494.1;refGene_gn=OSBPL6;refGene_tx=XM_023259495.1;refGene_gn=OSBPL6;refGene_tx=XM_023259496.1;refGene_gn=OSBPL6;refGene_tx=XM_023259497.1;refGene_gn=OSBPL6;refGene_tx=XM_023259498.1;refGene_gn=OSBPL6;refGene_tx=XM_023259499.1;refGene_gn=OSBPL6;refGene_tx=XM_023259500.1;refGene_gn=OSBPL6;refGene_tx=XM_023259501.1;refGene_gn=OSBPL6;refGene_tx=XM_023259502.1
C1 169037241 169037350
refGene_gn=OSBPL6;refGene_tx=XM_019838297.2;refGene_gn=OSBPL6;refGene_tx=XM_023259486.1;refGene_gn=OSBPL6;refGene_tx=XM_023259487.1;refGene_gn=OSBPL6;refGene_tx=XM_023259488.1;refGene_gn=OSBPL6;refGene_tx=XM_023259489.1;refGene_gn=OSBPL6;refGene_tx=XM_023259490.1;refGene_gn=OSBPL6;refGene_tx=XM_023259491.1;refGene_gn=OSBPL6;refGene_tx=XM_023259492.1;refGene_gn=OSBPL6;refGene_tx=XM_023259493.1;refGene_gn=OSBPL6;refGene_tx=XM_023259494.1;refGene_gn=OSBPL6;refGene_tx=XM_023259495.1;refGene_gn=OSBPL6;refGene_tx=XM_023259496.1;refGene_gn=OSBPL6;refGene_tx=XM_023259497.1;refGene_gn=OSBPL6;refGene_tx=XM_023259498.1;refGene_gn=OSBPL6;refGene_tx=XM_023259499.1;refGene_gn=OSBPL6;refGene_tx=XM_023259500.1;refGene_gn=OSBPL6;refGene_tx=XM_023259501.1;refGene_gn=OSBPL6;refGene_tx=XM_023259502.1
C1 169076790 169076948
refGene_gn=PRKRA;refGene_tx=XM_019838312.2;refGene_gn=PRKRA;refGene_tx=XM_019838313.2;refGene_gn=PRKRA;refGene_tx=XM_023259504.1;refGene_gn=PRKRA;refGene_tx=XM_023259505.1;refGene_gn=PRKRA;refGene_tx=XM_023259506.1
C1 169081510 169081685
refGene_gn=PRKRA;refGene_tx=XM_019838312.2;refGene_gn=PRKRA;refGene_tx=XM_019838313.2;refGene_gn=PRKRA;refGene_tx=XM_023259504.1;refGene_gn=PRKRA;refGene_tx=XM_023259505.1;refGene_gn=PRKRA;refGene_tx=XM_023259506.1
C1 169090982 169091077
refGene_gn=PRKRA;refGene_tx=XM_019838312.2;refGene_gn=PRKRA;refGene_tx=XM_019838313.2;refGene_gn=PRKRA;refGene_tx=XM_023259504.1;refGene_gn=PRKRA;refGene_tx=XM_023259505.1;refGene_gn=PRKRA;refGene_tx=XM_023259506.1
C1 169092570 169092688
refGene_gn=PRKRA;refGene_tx=XM_019838312.2;refGene_gn=PRKRA;refGene_tx=XM_019838313.2;refGene_gn=PRKRA;refGene_tx=XM_023259504.1;refGene_gn=PRKRA;refGene_tx=XM_023259505.1;refGene_gn=PRKRA;refGene_tx=XM_023259506.1

C1 169093746 169093825
refGene_gn=PRKRA;refGene_tx=XM_019838312.2;refGene_gn=PRKRA;refGene_tx=XM_019838313.2;refGene_gn=PRKRA;refGene_tx=XM_023259504.1;refGene_gn=PRKRA;refGene_tx=XM_023259505.1;refGene_gn=PRKRA;refGene_tx=XM_023259506.1
C1 169095872 169096003
refGene_gn=PRKRA;refGene_tx=XM_019838312.2;refGene_gn=PRKRA;refGene_tx=XM_023259504.1;refGene_gn=PRKRA;refGene_tx=XM_023259505.1
C1 169099327 169099497
refGene_gn=PRKRA;refGene_tx=XM_023259504.1
C1 169100049 169100114
refGene_gn=PRKRA;refGene_tx=XM_023259504.1
C1 169102293 169102504
refGene_gn=PJVK;refGene_tx=XM_023259503.1
C1 169103350 169103546
refGene_gn=PJVK;refGene_tx=XM_023259503.1
C1 169104078 169104220
refGene_gn=PJVK;refGene_tx=XM_023259503.1
C1 169106136 169106254
refGene_gn=PJVK;refGene_tx=XM_023259503.1
C1 169107662 169107761
refGene_gn=PJVK;refGene_tx=XM_023259503.1
C1 169107976 169108269
refGene_gn=PJVK;refGene_tx=XM_023259503.1
C1 169111381 169111543
refGene_gn=FKBP7;refGene_tx=XM_011285393.3;refGene_gn=FKBP7;refGene_tx=XM_023259508.1
C1 169115207 169115341
refGene_gn=FKBP7;refGene_tx=XM_011285393.3;refGene_gn=FKBP7;refGene_tx=XM_023259508.1
C1 169119867 169120019
refGene_gn=FKBP7;refGene_tx=XM_011285393.3;refGene_gn=FKBP7;refGene_tx=XM_023259508.1
C1 169121339 169121548
refGene_gn=FKBP7;refGene_tx=XM_011285393.3;refGene_gn=FKBP7;refGene_tx=XM_023259508.1
C1 169123856 169123896
refGene_gn=PLEKHA3;refGene_tx=XM_003990953.5
C1 169127961 169128078
refGene_gn=PLEKHA3;refGene_tx=XM_003990953.5;refGene_gn=PLEKHA3;refGene_tx=XM_023259507.1
C1 169136557 169136713
refGene_gn=PLEKHA3;refGene_tx=XM_003990953.5;refGene_gn=PLEKHA3;refGene_tx=XM_023259507.1
C1 169138866 169139003
refGene_gn=PLEKHA3;refGene_tx=XM_003990953.5;refGene_gn=PLEKHA3;refGene_tx=XM_023259507.1
C1 169140521 169140686
refGene_gn=PLEKHA3;refGene_tx=XM_003990953.5;refGene_gn=PLEKHA3;refGene_tx=XM_023259507.1

C1	169144044	169144088
refGene_gn=PLEKHA3;refGene_tx=XM_003990953.5;refGene_gn=PLEKHA3;refGene_tx=XM_023259507.1
C1	169146070	169146186
refGene_gn=PLEKHA3;refGene_tx=XM_003990953.5;refGene_gn=PLEKHA3;refGene_tx=XM_023259507.1
C1	169149101	169149229
refGene_gn=PLEKHA3;refGene_tx=XM_003990953.5;refGene_gn=PLEKHA3;refGene_tx=XM_023259507.1
C1	169169341	169169640
refGene_gn=TTN;refGene_tx=XM_023259509.1
C1	169169772	169170075
refGene_gn=TTN;refGene_tx=XM_023259509.1
C1	169170610	169170764
refGene_gn=TTN;refGene_tx=XM_023259509.1
C1	169170867	169171559
refGene_gn=TTN;refGene_tx=XM_023259509.1
C1	169172681	169172838
refGene_gn=TTN;refGene_tx=XM_023259509.1
C1	169172964	169178579
refGene_gn=TTN;refGene_tx=XM_023259509.1
C1	169178715	169179309
refGene_gn=TTN;refGene_tx=XM_023259509.1
C1	169179680	169179986
refGene_gn=TTN;refGene_tx=XM_023259509.1
C1	169180107	169180683
refGene_gn=TTN;refGene_tx=XM_023259509.1
C1	169181423	169181723
refGene_gn=TTN;refGene_tx=XM_023259509.1
C1	169181836	169182142
refGene_gn=TTN;refGene_tx=XM_023259509.1
C1	169182261	169182846
refGene_gn=TTN;refGene_tx=XM_023259509.1
C1	169182949	169183252
refGene_gn=TTN;refGene_tx=XM_023259509.1
C1	169183798	169184101
refGene_gn=TTN;refGene_tx=XM_023259509.1
C1	169184578	169184878
refGene_gn=TTN;refGene_tx=XM_023259509.1
C1	169184980	169185268
refGene_gn=TTN;refGene_tx=XM_023259509.1
C1	169185388	169185982
refGene_gn=TTN;refGene_tx=XM_023259509.1
C1	169186194	169186476
refGene_gn=TTN;refGene_tx=XM_023259509.1
C1	169186561	169186867
refGene_gn=TTN;refGene_tx=XM_023259509.1
C1	169188012	169188318
refGene_gn=TTN;refGene_tx=XM_023259509.1
C1	169188445	169188742

refGene_gn=TTN;refGene_tx=XM_023259509.1
C1 169188838 169189129
refGene_gn=TTN;refGene_tx=XM_023259509.1
C1 169189246 169189552
refGene_gn=TTN;refGene_tx=XM_023259509.1
C1 169189653 169189956
refGene_gn=TTN;refGene_tx=XM_023259509.1
C1 169190052 169192119
refGene_gn=TTN;refGene_tx=XM_023259509.1
C1 169192202 169192502
refGene_gn=TTN;refGene_tx=XM_023259509.1
C1 169192615 169192903
refGene_gn=TTN;refGene_tx=XM_023259509.1
C1 169193611 169193905
refGene_gn=TTN;refGene_tx=XM_023259509.1
C1 169194222 169195989
refGene_gn=TTN;refGene_tx=XM_023259509.1
C1 169196099 169196405
refGene_gn=TTN;refGene_tx=XM_023259509.1
C1 169196502 169196805
refGene_gn=TTN;refGene_tx=XM_023259509.1
C1 169197047 169197347
refGene_gn=TTN;refGene_tx=XM_023259509.1
C1 169197458 169197746
refGene_gn=TTN;refGene_tx=XM_023259509.1
C1 169199413 169199710
refGene_gn=TTN;refGene_tx=XM_023259509.1
C1 169199827 169200130
refGene_gn=TTN;refGene_tx=XM_023259509.1
C1 169200223 169200811
refGene_gn=TTN;refGene_tx=XM_023259509.1
C1 169200908 169201205
refGene_gn=TTN;refGene_tx=XM_023259509.1
C1 169201784 169218890
refGene_gn=TTN;refGene_tx=XM_023259509.1
C1 169219002 169219305
refGene_gn=TTN;refGene_tx=XM_023259509.1
C1 169219396 169219984
refGene_gn=TTN;refGene_tx=XM_023259509.1
C1 169220073 169220370
refGene_gn=TTN;refGene_tx=XM_023259509.1
C1 169220472 169220670
refGene_gn=TTN;refGene_tx=XM_023259509.1
C1 169221130 169221235
refGene_gn=TTN;refGene_tx=XM_023259509.1
C1 169221325 169221913
refGene_gn=TTN;refGene_tx=XM_023259509.1
C1 169222073 169222361
refGene_gn=TTN;refGene_tx=XM_023259509.1
C1 169222451 169222742

refGene_gn=TTN;refGene_tx=XM_023259509.1
C1	169222834	169223122
refGene_gn=TTN;refGene_tx=XM_023259509.1
C1	169224189	169224495
refGene_gn=TTN;refGene_tx=XM_023259509.1
C1	169224593	169224896
refGene_gn=TTN;refGene_tx=XM_023259509.1
C1	169224985	169225282
refGene_gn=TTN;refGene_tx=XM_023259509.1
C1	169225667	169225955
refGene_gn=TTN;refGene_tx=XM_023259509.1
C1	169226651	169226951
refGene_gn=TTN;refGene_tx=XM_023259509.1
C1	169227122	169227425
refGene_gn=TTN;refGene_tx=XM_023259509.1
C1	169227514	169227814
refGene_gn=TTN;refGene_tx=XM_023259509.1
C1	169227920	169228196
refGene_gn=TTN;refGene_tx=XM_023259509.1
C1	169228450	169228753
refGene_gn=TTN;refGene_tx=XM_023259509.1
C1	169229045	169229345
refGene_gn=TTN;refGene_tx=XM_023259509.1
C1	169229433	169229718
refGene_gn=TTN;refGene_tx=XM_023259509.1
C1	169229818	169230139
refGene_gn=TTN;refGene_tx=XM_023259509.1
C1	169230678	169233645
refGene_gn=TTN;refGene_tx=XM_023259509.1
C1	169233741	169234035
refGene_gn=TTN;refGene_tx=XM_023259509.1
C1	169234122	169234422
refGene_gn=TTN;refGene_tx=XM_023259509.1
C1	169234541	169234823
refGene_gn=TTN;refGene_tx=XM_023259509.1
C1	169234933	169235242
refGene_gn=TTN;refGene_tx=XM_023259509.1
C1	169235333	169235636
refGene_gn=TTN;refGene_tx=XM_023259509.1
C1	169235726	169236026
refGene_gn=TTN;refGene_tx=XM_023259509.1
C1	169236121	169236403
refGene_gn=TTN;refGene_tx=XM_023259509.1
C1	169236504	169236807
refGene_gn=TTN;refGene_tx=XM_023259509.1
C1	169237346	169237649
refGene_gn=TTN;refGene_tx=XM_023259509.1
C1	169239275	169239557
refGene_gn=TTN;refGene_tx=XM_023259509.1
C1	169239646	169239797

refGene_gn=TTN;refGene_tx=XM_023259509.1
C1 169240237 169240386
refGene_gn=TTN;refGene_tx=XM_023259509.1
C1 169240471 169240786
refGene_gn=TTN;refGene_tx=XM_023259509.1
C1 169240873 169241173
refGene_gn=TTN;refGene_tx=XM_023259509.1
C1 169241283 169241580
refGene_gn=TTN;refGene_tx=XM_023259509.1
C1 169242517 169242835
refGene_gn=TTN;refGene_tx=XM_023259509.1
C1 169242927 169243227
refGene_gn=TTN;refGene_tx=XM_023259509.1
C1 169243316 169243446
refGene_gn=TTN;refGene_tx=XM_023259509.1
C1 169243539 169243572
refGene_gn=TTN;refGene_tx=XM_023259509.1
C1 169243654 169243803
refGene_gn=TTN;refGene_tx=XM_023259509.1
C1 169243930 169244239
refGene_gn=TTN;refGene_tx=XM_023259509.1
C1 169245747 169246177
refGene_gn=TTN;refGene_tx=XM_023259509.1
C1 169246577 169246768
refGene_gn=TTN;refGene_tx=XM_023259509.1
C1 169246856 169247165
refGene_gn=TTN;refGene_tx=XM_023259509.1
C1 169247277 169247577
refGene_gn=TTN;refGene_tx=XM_023259509.1
C1 169248785 169249079
refGene_gn=TTN;refGene_tx=XM_023259509.1
C1 169249170 169249455
refGene_gn=TTN;refGene_tx=XM_023259509.1
C1 169249565 169249862
refGene_gn=TTN;refGene_tx=XM_023259509.1
C1 169250307 169250607
refGene_gn=TTN;refGene_tx=XM_023259509.1
C1 169250738 169251041
refGene_gn=TTN;refGene_tx=XM_023259509.1
C1 169251361 169251724
refGene_gn=TTN;refGene_tx=XM_023259509.1
C1 169251836 169252139
refGene_gn=TTN;refGene_tx=XM_023259509.1
C1 169252252 169252552
refGene_gn=TTN;refGene_tx=XM_023259509.1
C1 169253150 169253429
refGene_gn=TTN;refGene_tx=XM_023259509.1
C1 169253525 169253831
refGene_gn=TTN;refGene_tx=XM_023259509.1
C1 169253914 169254111

refGene_gn=TTN;refGene_tx=XM_023259509.1
C1 169254211 169254317
refGene_gn=TTN;refGene_tx=XM_023259509.1
C1 169254406 169254706
refGene_gn=TTN;refGene_tx=XM_023259509.1
C1 169254901 169255201
refGene_gn=TTN;refGene_tx=XM_023259509.1
C1 169255296 169255412
refGene_gn=TTN;refGene_tx=XM_023259509.1
C1 169255870 169256057
refGene_gn=TTN;refGene_tx=XM_023259509.1
C1 169256174 169256471
refGene_gn=TTN;refGene_tx=XM_023259509.1
C1 169256581 169256869
refGene_gn=TTN;refGene_tx=XM_023259509.1
C1 169256963 169257085
refGene_gn=TTN;refGene_tx=XM_023259509.1
C1 169257475 169257653
refGene_gn=TTN;refGene_tx=XM_023259509.1
C1 169257807 169257955
refGene_gn=TTN;refGene_tx=XM_023259509.1
C1 169258639 169258791
refGene_gn=TTN;refGene_tx=XM_023259509.1
C1 169258891 169259176
refGene_gn=TTN;refGene_tx=XM_023259509.1
C1 169259271 169259386
refGene_gn=TTN;refGene_tx=XM_023259509.1
C1 169259508 169259696
refGene_gn=TTN;refGene_tx=XM_023259509.1
C1 169259957 169260260
refGene_gn=TTN;refGene_tx=XM_023259509.1
C1 169260355 169260658
refGene_gn=TTN;refGene_tx=XM_023259509.1
C1 169260748 169261018
refGene_gn=TTN;refGene_tx=XM_023259509.1
C1 169261788 169262055
refGene_gn=TTN;refGene_tx=XM_023259509.1
C1 169262156 169262281
refGene_gn=TTN;refGene_tx=XM_023259509.1
C1 169262382 169262791
refGene_gn=TTN;refGene_tx=XM_023259509.1
C1 169262882 169263161
refGene_gn=TTN;refGene_tx=XM_023259509.1
C1 169263251 169263518
refGene_gn=TTN;refGene_tx=XM_023259509.1
C1 169263618 169263885
refGene_gn=TTN;refGene_tx=XM_023259509.1
C1 169263971 169264140
refGene_gn=TTN;refGene_tx=XM_023259509.1
C1 169264771 169264869

refGene_gn=TTN;refGene_tx=XM_023259509.1
C1 169266599 169266866
refGene_gn=TTN;refGene_tx=XM_023259509.1
C1 169267391 169267515
refGene_gn=TTN;refGene_tx=XM_023259509.1
C1 169271739 169271882
refGene_gn=TTN;refGene_tx=XM_023259509.1
C1 169272673 169272800
refGene_gn=TTN;refGene_tx=XM_023259509.1
C1 169273218 169273358
refGene_gn=TTN;refGene_tx=XM_023259509.1
C1 169273449 169273716
refGene_gn=TTN;refGene_tx=XM_023259509.1
C1 169274546 169274813
refGene_gn=TTN;refGene_tx=XM_023259509.1
C1 169274922 169275189
refGene_gn=TTN;refGene_tx=XM_023259509.1
C1 169275312 169275439
refGene_gn=TTN;refGene_tx=XM_023259509.1
C1 169275602 169275742
refGene_gn=TTN;refGene_tx=XM_023259509.1
C1 169275830 169276094
refGene_gn=TTN;refGene_tx=XM_023259509.1
C1 169276232 169276499
refGene_gn=TTN;refGene_tx=XM_023259509.1
C1 169277656 169277920
refGene_gn=TTN;refGene_tx=XM_023259509.1
C1 169278015 169278142
refGene_gn=TTN;refGene_tx=XM_023259509.1
C1 169278659 169278799
refGene_gn=TTN;refGene_tx=XM_023259509.1
C1 169279000 169279276
refGene_gn=TTN;refGene_tx=XM_023259509.1
C1 169279521 169279800
refGene_gn=TTN;refGene_tx=XM_023259509.1
C1 169279978 169280380
refGene_gn=TTN;refGene_tx=XM_023259509.1
C1 169281106 169281157
refGene_gn=TTN;refGene_tx=XM_023259509.1
C1 169283185 169283275
refGene_gn=TTN;refGene_tx=XM_023259509.1
C1 169284073 169284136
refGene_gn=TTN;refGene_tx=XM_023259509.1
C1 169284574 169284664
refGene_gn=TTN;refGene_tx=XM_023259509.1
C1 169285281 169285353
refGene_gn=TTN;refGene_tx=XM_023259509.1
C1 169286266 169286347
refGene_gn=TTN;refGene_tx=XM_023259509.1
C1 169288639 169288708

refGene_gn=TTN;refGene_tx=XM_023259509.1
C1	169289979	169290093
refGene_gn=TTN;refGene_tx=XM_023259509.1
C1	169293715	169293793
refGene_gn=TTN;refGene_tx=XM_023259509.1
C1	169294041	169294185
refGene_gn=TTN;refGene_tx=XM_023259509.1
C1	169294644	169294728
refGene_gn=TTN;refGene_tx=XM_023259509.1
C1	169295125	169295203
refGene_gn=TTN;refGene_tx=XM_023259509.1
C1	169295328	169295412
refGene_gn=TTN;refGene_tx=XM_023259509.1
C1	169295542	169295626
refGene_gn=TTN;refGene_tx=XM_023259509.1
C1	169296730	169296814
refGene_gn=TTN;refGene_tx=XM_023259509.1
C1	169297001	169297085
refGene_gn=TTN;refGene_tx=XM_023259509.1
C1	169297169	169297253
refGene_gn=TTN;refGene_tx=XM_023259509.1
C1	169297602	169297689
refGene_gn=TTN;refGene_tx=XM_023259509.1
C1	169297801	169297888
refGene_gn=TTN;refGene_tx=XM_023259509.1
C1	169298000	169298087
refGene_gn=TTN;refGene_tx=XM_023259509.1
C1	169298189	169298276
refGene_gn=TTN;refGene_tx=XM_023259509.1
C1	169298389	169298476
refGene_gn=TTN;refGene_tx=XM_023259509.1
C1	169298587	169298671
refGene_gn=TTN;refGene_tx=XM_023259509.1
C1	169298783	169298867
refGene_gn=TTN;refGene_tx=XM_023259509.1
C1	169298980	169299064
refGene_gn=TTN;refGene_tx=XM_023259509.1
C1	169299180	169299264
refGene_gn=TTN;refGene_tx=XM_023259509.1
C1	169299379	169299463
refGene_gn=TTN;refGene_tx=XM_023259509.1
C1	169299622	169299709
refGene_gn=TTN;refGene_tx=XM_023259509.1
C1	169299917	169300001
refGene_gn=TTN;refGene_tx=XM_023259509.1
C1	169300086	169300170
refGene_gn=TTN;refGene_tx=XM_023259509.1
C1	169300284	169300371
refGene_gn=TTN;refGene_tx=XM_023259509.1
C1	169300483	169300567

refGene_gn=TTN;refGene_tx=XM_023259509.1
C1 169306650 169306728
refGene_gn=TTN;refGene_tx=XM_023259509.1
C1 169307523 169307604
refGene_gn=TTN;refGene_tx=XM_023259509.1
C1 169308484 169308778
refGene_gn=TTN;refGene_tx=XM_023259509.1
C1 169308933 169309023
refGene_gn=TTN;refGene_tx=XM_023259509.1
C1 169309300 169309369
refGene_gn=TTN;refGene_tx=XM_023259509.1
C1 169310339 169310411
refGene_gn=TTN;refGene_tx=XM_023259509.1
C1 169310989 169311082
refGene_gn=TTN;refGene_tx=XM_023259509.1
C1 169311724 169311799
refGene_gn=TTN;refGene_tx=XM_023259509.1
C1 169312326 169312410
refGene_gn=TTN;refGene_tx=XM_023259509.1
C1 169312572 169312647
refGene_gn=TTN;refGene_tx=XM_023259509.1
C1 169314170 169314254
refGene_gn=TTN;refGene_tx=XM_023259509.1
C1 169314576 169314879
refGene_gn=TTN;refGene_tx=XM_023259509.1
C1 169315086 169315170
refGene_gn=TTN;refGene_tx=XM_023259509.1
C1 169315359 169315443
refGene_gn=TTN;refGene_tx=XM_023259509.1
C1 169315693 169315777
refGene_gn=TTN;refGene_tx=XM_023259509.1
C1 169316248 169316326
refGene_gn=TTN;refGene_tx=XM_023259509.1
C1 169316498 169316582
refGene_gn=TTN;refGene_tx=XM_023259509.1
C1 169316794 169316953
refGene_gn=TTN;refGene_tx=XM_023259509.1
C1 169317141 169317219
refGene_gn=TTN;refGene_tx=XM_023259509.1
C1 169317967 169318060
refGene_gn=TTN;refGene_tx=XM_023259509.1
C1 169318264 169318339
refGene_gn=TTN;refGene_tx=XM_023259509.1
C1 169318544 169318622
refGene_gn=TTN;refGene_tx=XM_023259509.1
C1 169319937 169320153
refGene_gn=TTN;refGene_tx=XM_023259509.1
C1 169320442 169320523
refGene_gn=TTN;refGene_tx=XM_023259509.1
C1 169321487 169321571

refGene_gn=TTN;refGene_tx=XM_023259509.1
C1 169321796 169321880
refGene_gn=TTN;refGene_tx=XM_023259509.1
C1 169322171 169322255
refGene_gn=TTN;refGene_tx=XM_023259509.1
C1 169322415 169322499
refGene_gn=TTN;refGene_tx=XM_023259509.1
C1 169322714 169322792
refGene_gn=TTN;refGene_tx=XM_023259509.1
C1 169323205 169323286
refGene_gn=TTN;refGene_tx=XM_023259509.1
C1 169325345 169325453
refGene_gn=TTN;refGene_tx=XM_023259509.1
C1 169325879 169325981
refGene_gn=TTN;refGene_tx=XM_023259509.1
C1 169326229 169326313
refGene_gn=TTN;refGene_tx=XM_023259509.1
C1 169326482 169326563
refGene_gn=TTN;refGene_tx=XM_023259509.1
C1 169326707 169326788
refGene_gn=TTN;refGene_tx=XM_023259509.1
C1 169327015 169327099
refGene_gn=TTN;refGene_tx=XM_023259509.1
C1 169329108 169329192
refGene_gn=TTN;refGene_tx=XM_023259509.1
C1 169329586 169329670
refGene_gn=TTN;refGene_tx=XM_023259509.1
C1 169330667 169330692
refGene_gn=TTN;refGene_tx=XM_023259509.1
C1 169330693 169330749
refGene_gn=TTN;refGene_tx=XM_023259509.1
C1 169330985 169331072
refGene_gn=TTN;refGene_tx=XM_023259509.1
C1 169331672 169331750
refGene_gn=TTN;refGene_tx=XM_023259509.1
C1 169331911 169331989
refGene_gn=TTN;refGene_tx=XM_023259509.1
C1 169332394 169332454
refGene_gn=TTN;refGene_tx=XM_023259509.1
C1 169332890 169333295
refGene_gn=TTN;refGene_tx=XM_023259509.1
C1 169334413 169334461
refGene_gn=TTN;refGene_tx=XM_023259509.1
C1 169335444 169335513
refGene_gn=TTN;refGene_tx=XM_023259509.1
C1 169337176 169337260
refGene_gn=TTN;refGene_tx=XM_023259509.1
C1 169337552 169337612
refGene_gn=TTN;refGene_tx=XM_023259509.1
C1 169338210 169338288

refGene_gn=TTN;refGene_tx=XM_023259509.1
C1 169338517 169338727
refGene_gn=TTN;refGene_tx=XM_023259509.1
C1 169340319 169340580
refGene_gn=TTN;refGene_tx=XM_023259509.1
C1 169340688 169340956
refGene_gn=TTN;refGene_tx=XM_023259509.1
C1 169341052 169341142
refGene_gn=TTN;refGene_tx=XM_023259509.1
C1 169341586 169341770
refGene_gn=TTN;refGene_tx=XM_023259509.1
C1 169342305 169342591
refGene_gn=TTN;refGene_tx=XM_023259509.1
C1 169342712 169342805
refGene_gn=TTN;refGene_tx=XM_023259509.1
C1 169343402 169343690
refGene_gn=TTN;refGene_tx=XM_023259509.1
C1 169345916 169346207
refGene_gn=TTN;refGene_tx=XM_023259509.1
C1 169346678 169346966
refGene_gn=TTN;refGene_tx=XM_023259509.1
C1 169347093 169347381
refGene_gn=TTN;refGene_tx=XM_023259509.1
C1 169347957 169348236
refGene_gn=TTN;refGene_tx=XM_023259509.1
C1 169348320 169348599
refGene_gn=TTN;refGene_tx=XM_023259509.1
C1 169348711 169348990
refGene_gn=TTN;refGene_tx=XM_023259509.1
C1 169349332 169349620
refGene_gn=TTN;refGene_tx=XM_023259509.1
C1 169350110 169350389
refGene_gn=TTN;refGene_tx=XM_023259509.1
C1 169350505 169350787
refGene_gn=TTN;refGene_tx=XM_023259509.1
C1 169352248 169352527
refGene_gn=TTN;refGene_tx=XM_023259509.1
C1 169353133 169353415
refGene_gn=TTN;refGene_tx=XM_023259509.1
C1 169354943 169355231
refGene_gn=TTN;refGene_tx=XM_023259509.1
C1 169355367 169355655
refGene_gn=TTN;refGene_tx=XM_023259509.1
C1 169355795 169356074
refGene_gn=TTN;refGene_tx=XM_023259509.1
C1 169356177 169356456
refGene_gn=TTN;refGene_tx=XM_023259509.1
C1 169356550 169356829
refGene_gn=TTN;refGene_tx=XM_023259509.1
C1 169357020 169357308

refGene_gn=TTN;refGene_tx=XM_023259509.1
C1	169357408	169357687
refGene_gn=TTN;refGene_tx=XM_023259509.1
C1	169357833	169358115
refGene_gn=TTN;refGene_tx=XM_023259509.1
C1	169358233	169358512
refGene_gn=TTN;refGene_tx=XM_023259509.1
C1	169358777	169359059
refGene_gn=TTN;refGene_tx=XM_023259509.1
C1	169359832	169360120
refGene_gn=TTN;refGene_tx=XM_023259509.1
C1	169360246	169360534
refGene_gn=TTN;refGene_tx=XM_023259509.1
C1	169360647	169360926
refGene_gn=TTN;refGene_tx=XM_023259509.1
C1	169361036	169361315
refGene_gn=TTN;refGene_tx=XM_023259509.1
C1	169361407	169361686
refGene_gn=TTN;refGene_tx=XM_023259509.1
C1	169361835	169362123
refGene_gn=TTN;refGene_tx=XM_023259509.1
C1	169362239	169362518
refGene_gn=TTN;refGene_tx=XM_023259509.1
C1	169363344	169363626
refGene_gn=TTN;refGene_tx=XM_023259509.1
C1	169363748	169364027
refGene_gn=TTN;refGene_tx=XM_023259509.1
C1	169365270	169365552
refGene_gn=TTN;refGene_tx=XM_023259509.1
C1	169365761	169366040
refGene_gn=TTN;refGene_tx=XM_023259509.1
C1	169366278	169366566
refGene_gn=TTN;refGene_tx=XM_023259509.1
C1	169366677	169366956
refGene_gn=TTN;refGene_tx=XM_023259509.1
C1	169367069	169367348
refGene_gn=TTN;refGene_tx=XM_023259509.1
C1	169367470	169367749
refGene_gn=TTN;refGene_tx=XM_023259509.1
C1	169367849	169368131
refGene_gn=TTN;refGene_tx=XM_023259509.1
C1	169368260	169368539
refGene_gn=TTN;refGene_tx=XM_023259509.1
C1	169368664	169368952
refGene_gn=TTN;refGene_tx=XM_023259509.1
C1	169369067	169369346
refGene_gn=TTN;refGene_tx=XM_023259509.1
C1	169369447	169369726
refGene_gn=TTN;refGene_tx=XM_023259509.1
C1	169369843	169370122

refGene_gn=TTN;refGene_tx=XM_023259509.1
C1	169370214	169370496
refGene_gn=TTN;refGene_tx=XM_023259509.1
C1	169370588	169370867
refGene_gn=TTN;refGene_tx=XM_023259509.1
C1	169370981	169371269
refGene_gn=TTN;refGene_tx=XM_023259509.1
C1	169371387	169371666
refGene_gn=TTN;refGene_tx=XM_023259509.1
C1	169371763	169372042
refGene_gn=TTN;refGene_tx=XM_023259509.1
C1	169372493	169372772
refGene_gn=TTN;refGene_tx=XM_023259509.1
C1	169372868	169373150
refGene_gn=TTN;refGene_tx=XM_023259509.1
C1	169373719	169374283
refGene_gn=TTN;refGene_tx=XM_023259509.1
C1	169376310	169376589
refGene_gn=TTN;refGene_tx=XM_023259509.1
C1	169377367	169380139
refGene_gn=TTN;refGene_tx=XM_023259509.1
C1	169383801	169390116
refGene_gn=LOC109502758;refGene_tx=XM_019838314.2
C1	169391299	169391356
refGene_gn=TTN;refGene_tx=XM_023259509.1
C1	169396805	169396994
refGene_gn=TTN;refGene_tx=XM_023259509.1
C1	169402420	169402546
refGene_gn=TTN;refGene_tx=XM_023259509.1
C1	169402777	169403062
refGene_gn=TTN;refGene_tx=XM_023259509.1
C1	169404819	169405051
refGene_gn=TTN;refGene_tx=XM_023259509.1
C1	169406158	169406324
refGene_gn=TTN;refGene_tx=XM_023259509.1
C1	169406413	169406555
refGene_gn=TTN;refGene_tx=XM_023259509.1
C1	169407064	169407325
refGene_gn=TTN;refGene_tx=XM_023259509.1
C1	169407737	169407998
refGene_gn=TTN;refGene_tx=XM_023259509.1
C1	169408129	169408390
refGene_gn=TTN;refGene_tx=XM_023259509.1
C1	169408483	169408747
refGene_gn=TTN;refGene_tx=XM_023259509.1
C1	169409286	169409547
refGene_gn=TTN;refGene_tx=XM_023259509.1
C1	169411167	169411428
refGene_gn=TTN;refGene_tx=XM_023259509.1
C1	169411519	169411783

```
refGene_gn=TTN;refGene_tx=XM_023259509.1
C1	169411947	169412220
refGene_gn=TTN;refGene_tx=XM_023259509.1
C1	169412317	169412584
refGene_gn=TTN;refGene_tx=XM_023259509.1
C1	169413090	169413372
refGene_gn=TTN;refGene_tx=XM_023259509.1
C1	169413516	169415210
refGene_gn=TTN;refGene_tx=XM_023259509.1
C1	169415311	169415480
refGene_gn=TTN;refGene_tx=XM_023259509.1
C1	169415574	169415739
refGene_gn=TTN;refGene_tx=XM_023259509.1
C1	169415862	169416134
refGene_gn=TTN;refGene_tx=XM_023259509.1
C1	169417009	169417254
refGene_gn=TTN;refGene_tx=XM_023259509.1
C1	169417357	169417591
refGene_gn=TTN;refGene_tx=XM_023259509.1
C1	169418356	169418562
refGene_gn=TTN;refGene_tx=XM_023259509.1
C1	169419821	169419964
refGene_gn=TTN;refGene_tx=XM_023259509.1
C1	169420902	169421118
refGene_gn=TTN;refGene_tx=XM_023259509.1
C1	169421214	169421278
refGene_gn=TTN;refGene_tx=XM_023259509.1
C1	169421494	169421753
refGene_gn=TTN;refGene_tx=XM_023259509.1
C1	169422459	169422525
refGene_gn=TTN;refGene_tx=XM_023259509.1
C1	169422785	169423067
refGene_gn=TTN;refGene_tx=XM_023259509.1
C1	169424516	169424639
refGene_gn=TTN;refGene_tx=XM_023259509.1
C1	169424743	169425037
refGene_gn=TTN;refGene_tx=XM_023259509.1
C1	169428362	169428500
refGene_gn=TTN;refGene_tx=XM_023259509.1
C1	169428989	169429127
refGene_gn=TTN;refGene_tx=XM_023259509.1
C1	169429745	169429883
refGene_gn=TTN;refGene_tx=XM_023259509.1
C1	169431264	169431396
refGene_gn=TTN;refGene_tx=XM_023259509.1
C1	169432445	169432586
refGene_gn=TTN;refGene_tx=XM_023259509.1
C1	169433260	169433413
refGene_gn=TTN;refGene_tx=XM_023259509.1
C1	169433989	169434317
```

refGene_gn=TTN;refGene_tx=XM_023259509.1
C1 169438803 169439048
refGene_gn=TTN;refGene_tx=XM_023259509.1
C1 169439143 169439229
refGene_gn=TTN;refGene_tx=XM_023259509.1
C1 169439701 169439989
refGene_gn=TTN;refGene_tx=XM_023259509.1
C1 169441441 169441645
refGene_gn=TTN;refGene_tx=XM_023259509.1
C1 169443828 169443919
refGene_gn=TTN;refGene_tx=XM_023259509.1
C1 169477463 169477731
refGene_gn=CCDC141;refGene_tx=XM_019838317.2;refGene_gn=CCDC141;refGene_tx=XM_019838318.2;refGene_gn=CCDC141;refGene_tx=XM_019838319.2;refGene_gn=CCDC141;refGene_tx=XM_019838320.2;refGene_gn=CCDC141;refGene_tx=XM_023259510.1
C1 169479879 169480761
refGene_gn=CCDC141;refGene_tx=XM_019838317.2;refGene_gn=CCDC141;refGene_tx=XM_019838318.2;refGene_gn=CCDC141;refGene_tx=XM_019838319.2;refGene_gn=CCDC141;refGene_tx=XM_019838320.2;refGene_gn=CCDC141;refGene_tx=XM_019838321.2;refGene_gn=CCDC141;refGene_tx=XM_023259510.1
C1 169494270 169494387
refGene_gn=CCDC141;refGene_tx=XM_019838317.2;refGene_gn=CCDC141;refGene_tx=XM_019838318.2;refGene_gn=CCDC141;refGene_tx=XM_019838319.2;refGene_gn=CCDC141;refGene_tx=XM_019838320.2;refGene_gn=CCDC141;refGene_tx=XM_019838321.2;refGene_gn=CCDC141;refGene_tx=XM_023259510.1
C1 169498001 169498114
refGene_gn=CCDC141;refGene_tx=XM_019838317.2;refGene_gn=CCDC141;refGene_tx=XM_019838318.2;refGene_gn=CCDC141;refGene_tx=XM_019838319.2;refGene_gn=CCDC141;refGene_tx=XM_019838320.2;refGene_gn=CCDC141;refGene_tx=XM_019838321.2;refGene_gn=CCDC141;refGene_tx=XM_023259510.1
C1 169500707 169500891
refGene_gn=CCDC141;refGene_tx=XM_019838317.2;refGene_gn=CCDC141;refGene_tx=XM_019838318.2;refGene_gn=CCDC141;refGene_tx=XM_019838319.2;refGene_gn=CCDC141;refGene_tx=XM_019838320.2;refGene_gn=CCDC141;refGene_tx=XM_019838321.2;refGene_gn=CCDC141;refGene_tx=XM_023259510.1
C1 169502269 169502464
refGene_gn=CCDC141;refGene_tx=XM_019838317.2;refGene_gn=CCDC141;refGene_tx=XM_019838318.2;refGene_gn=CCDC141;refGene_tx=XM_019838319.2;refGene_gn=CCDC141;refGene_tx=XM_019838320.2;refGene_gn=CCDC141;refGene_tx=XM_019838321.2;refGene_gn=CCDC141;refGene_tx=XM_023259510.1
C1 169503264 169503405
refGene_gn=CCDC141;refGene_tx=XM_019838317.2;refGene_gn=CCDC141;refGene_tx=XM_019838318.2;refGene_gn=CCDC141;refGene_tx=XM_019838319.2;refGene_gn=CCDC141;refGene_tx=XM_019838320.2;refGene_gn=CCDC141;refGene_tx=XM_019838321.2;refGene_gn=CCDC141;refGene_tx=XM_023259510.1
C1 169513294 169513444
refGene_gn=CCDC141;refGene_tx=XM_019838317.2;refGene_gn=CCDC141;refGene_tx=XM_019838318.2;refGene_gn=CCDC141;refGene_tx=XM_019838319.2;refGene_gn=CCDC141;refGene_tx=XM_019838320.2;refGene_gn=CCDC141;refGene_tx=

XM_019838321.2;refGene_gn=CCDC141;refGene_tx=XM_023259510.1
C1 169515154 169515334
refGene_gn=CCDC141;refGene_tx=XM_019838317.2;refGene_gn=CCDC141;refGene_tx=XM_019838318.2;refGene_gn=CCDC141;refGene_tx=XM_019838319.2;refGene_gn=CCDC141;refGene_tx=XM_019838320.2;refGene_gn=CCDC141;refGene_tx=XM_019838321.2;refGene_gn=CCDC141;refGene_tx=XM_023259510.1
C1 169516862 169517051
refGene_gn=CCDC141;refGene_tx=XM_019838317.2;refGene_gn=CCDC141;refGene_tx=XM_019838318.2;refGene_gn=CCDC141;refGene_tx=XM_019838319.2;refGene_gn=CCDC141;refGene_tx=XM_019838320.2;refGene_gn=CCDC141;refGene_tx=XM_019838321.2;refGene_gn=CCDC141;refGene_tx=XM_023259510.1
C1 169519336 169519462
refGene_gn=CCDC141;refGene_tx=XM_019838317.2;refGene_gn=CCDC141;refGene_tx=XM_019838319.2;refGene_gn=CCDC141;refGene_tx=XM_019838320.2;refGene_gn=CCDC141;refGene_tx=XM_019838321.2;refGene_gn=CCDC141;refGene_tx=XM_023259510.1
C1 169520060 169520240
refGene_gn=CCDC141;refGene_tx=XM_019838317.2;refGene_gn=CCDC141;refGene_tx=XM_019838318.2;refGene_gn=CCDC141;refGene_tx=XM_019838319.2;refGene_gn=CCDC141;refGene_tx=XM_019838320.2;refGene_gn=CCDC141;refGene_tx=XM_019838321.2;refGene_gn=CCDC141;refGene_tx=XM_023259510.1
C1 169523427 169523445
refGene_gn=CCDC141;refGene_tx=XM_019838322.2
C1 169525545 169525725
refGene_gn=CCDC141;refGene_tx=XM_019838317.2;refGene_gn=CCDC141;refGene_tx=XM_019838318.2;refGene_gn=CCDC141;refGene_tx=XM_019838320.2;refGene_gn=CCDC141;refGene_tx=XM_019838321.2;refGene_gn=CCDC141;refGene_tx=XM_019838322.2;refGene_gn=CCDC141;refGene_tx=XM_023259510.1
C1 169531202 169531394
refGene_gn=CCDC141;refGene_tx=XM_019838317.2;refGene_gn=CCDC141;refGene_tx=XM_019838318.2;refGene_gn=CCDC141;refGene_tx=XM_019838319.2;refGene_gn=CCDC141;refGene_tx=XM_019838320.2;refGene_gn=CCDC141;refGene_tx=XM_019838321.2;refGene_gn=CCDC141;refGene_tx=XM_019838322.2;refGene_gn=CCDC141;refGene_tx=XM_023259510.1
C1 169533080 169533200
refGene_gn=CCDC141;refGene_tx=XM_019838317.2;refGene_gn=CCDC141;refGene_tx=XM_019838318.2;refGene_gn=CCDC141;refGene_tx=XM_019838319.2;refGene_gn=CCDC141;refGene_tx=XM_019838320.2;refGene_gn=CCDC141;refGene_tx=XM_019838321.2;refGene_gn=CCDC141;refGene_tx=XM_019838322.2;refGene_gn=CCDC141;refGene_tx=XM_023259510.1
C1 169534946 169535088
refGene_gn=CCDC141;refGene_tx=XM_019838317.2;refGene_gn=CCDC141;refGene_tx=XM_019838318.2;refGene_gn=CCDC141;refGene_tx=XM_019838319.2;refGene_gn=CCDC141;refGene_tx=XM_019838320.2;refGene_gn=CCDC141;refGene_tx=XM_019838321.2;refGene_gn=CCDC141;refGene_tx=XM_019838322.2;refGene_gn=CCDC141;refGene_tx=XM_023259510.1
C1 169549722 169549895
refGene_gn=CCDC141;refGene_tx=XM_019838317.2;refGene_gn=CCDC141;refGene_tx=XM_019838318.2;refGene_gn=CCDC141;refGene_tx=XM_019838319.2;refGene_gn=CCDC141;refGene_tx=XM_019838320.2;refGene_gn=CCDC141;refGene_tx=

XM_019838321.2;refGene_gn=CCDC141;refGene_tx=XM_019838322.2;refGene_gn=CCDC141;refGene_tx=XM_023259510.1
C1 169565334 169565529
refGene_gn=CCDC141;refGene_tx=XM_019838317.2;refGene_gn=CCDC141;refGene_tx=XM_019838318.2;refGene_gn=CCDC141;refGene_tx=XM_019838319.2;refGene_gn=CCDC141;refGene_tx=XM_019838321.2;refGene_gn=CCDC141;refGene_tx=XM_019838322.2;refGene_gn=CCDC141;refGene_tx=XM_023259510.1
C1 169597499 169597616
refGene_gn=CCDC141;refGene_tx=XM_019838317.2;refGene_gn=CCDC141;refGene_tx=XM_019838318.2;refGene_gn=CCDC141;refGene_tx=XM_019838319.2;refGene_gn=CCDC141;refGene_tx=XM_019838320.2;refGene_gn=CCDC141;refGene_tx=XM_019838321.2;refGene_gn=CCDC141;refGene_tx=XM_019838322.2;refGene_gn=CCDC141;refGene_tx=XM_023259510.1
C1 169614904 169615158
refGene_gn=CCDC141;refGene_tx=XM_019838317.2;refGene_gn=CCDC141;refGene_tx=XM_019838318.2;refGene_gn=CCDC141;refGene_tx=XM_019838319.2;refGene_gn=CCDC141;refGene_tx=XM_019838320.2;refGene_gn=CCDC141;refGene_tx=XM_019838321.2;refGene_gn=CCDC141;refGene_tx=XM_019838322.2;refGene_gn=CCDC141;refGene_tx=XM_023259510.1
C1 169619243 169619352
refGene_gn=CCDC141;refGene_tx=XM_019838317.2;refGene_gn=CCDC141;refGene_tx=XM_019838318.2;refGene_gn=CCDC141;refGene_tx=XM_019838319.2;refGene_gn=CCDC141;refGene_tx=XM_019838320.2;refGene_gn=CCDC141;refGene_tx=XM_019838321.2;refGene_gn=CCDC141;refGene_tx=XM_019838322.2;refGene_gn=CCDC141;refGene_tx=XM_023259510.1
C1 169623168 169623360
refGene_gn=CCDC141;refGene_tx=XM_019838317.2;refGene_gn=CCDC141;refGene_tx=XM_019838318.2;refGene_gn=CCDC141;refGene_tx=XM_019838319.2;refGene_gn=CCDC141;refGene_tx=XM_019838320.2;refGene_gn=CCDC141;refGene_tx=XM_019838321.2;refGene_gn=CCDC141;refGene_tx=XM_019838322.2;refGene_gn=CCDC141;refGene_tx=XM_023259510.1
C1 169674117 169674240
refGene_gn=CCDC141;refGene_tx=XM_019838317.2;refGene_gn=CCDC141;refGene_tx=XM_019838318.2;refGene_gn=CCDC141;refGene_tx=XM_019838319.2;refGene_gn=CCDC141;refGene_tx=XM_019838320.2;refGene_gn=CCDC141;refGene_tx=XM_019838321.2;refGene_gn=CCDC141;refGene_tx=XM_019838322.2;refGene_gn=CCDC141;refGene_tx=XM_023259510.1
C1 169676865 169676967
refGene_gn=CCDC141;refGene_tx=XM_019838317.2;refGene_gn=CCDC141;refGene_tx=XM_019838318.2;refGene_gn=CCDC141;refGene_tx=XM_019838319.2;refGene_gn=CCDC141;refGene_tx=XM_019838320.2;refGene_gn=CCDC141;refGene_tx=XM_019838321.2;refGene_gn=CCDC141;refGene_tx=XM_019838322.2;refGene_gn=CCDC141;refGene_tx=XM_023259510.1
C1 169707940 169708070
refGene_gn=SESTD1;refGene_tx=XM_011285390.3;refGene_gn=SESTD1;refGene_tx=XM_019838323.2;refGene_gn=SESTD1;refGene_tx=XM_023259511.1;refGene_gn=SESTD1;refGene_tx=XM_023259512.1
C1 169710605 169710727
refGene_gn=SESTD1;refGene_tx=XM_011285390.3;refGene_gn=SESTD1;refGene_tx=XM_019838323.2;refGene_gn=SESTD1;refGene_tx=XM_023259511.1;refGene_

gn=SESTD1;refGene_tx=XM_023259512.1
C1 169714459 169714651
refGene_gn=SESTD1;refGene_tx=XM_011285390.3;refGene_gn=SESTD1;refGene_tx=XM_019838323.2;refGene_gn=SESTD1;refGene_tx=XM_023259511.1;refGene_gn=SESTD1;refGene_tx=XM_023259512.1
C1 169715517 169715688
refGene_gn=SESTD1;refGene_tx=XM_011285390.3;refGene_gn=SESTD1;refGene_tx=XM_019838323.2;refGene_gn=SESTD1;refGene_tx=XM_023259511.1;refGene_gn=SESTD1;refGene_tx=XM_023259512.1;refGene_gn=SESTD1;refGene_tx=XM_023259513.1
C1 169716783 169716865
refGene_gn=SESTD1;refGene_tx=XM_011285390.3;refGene_gn=SESTD1;refGene_tx=XM_019838323.2;refGene_gn=SESTD1;refGene_tx=XM_023259511.1;refGene_gn=SESTD1;refGene_tx=XM_023259512.1;refGene_gn=SESTD1;refGene_tx=XM_023259513.1
C1 169721148 169721308
refGene_gn=SESTD1;refGene_tx=XM_011285390.3;refGene_gn=SESTD1;refGene_tx=XM_019838323.2;refGene_gn=SESTD1;refGene_tx=XM_023259511.1;refGene_gn=SESTD1;refGene_tx=XM_023259512.1;refGene_gn=SESTD1;refGene_tx=XM_023259513.1
C1 169722740 169722855
refGene_gn=SESTD1;refGene_tx=XM_011285390.3;refGene_gn=SESTD1;refGene_tx=XM_019838323.2;refGene_gn=SESTD1;refGene_tx=XM_023259511.1;refGene_gn=SESTD1;refGene_tx=XM_023259512.1;refGene_gn=SESTD1;refGene_tx=XM_023259513.1
C1 169723381 169723576
refGene_gn=SESTD1;refGene_tx=XM_011285390.3;refGene_gn=SESTD1;refGene_tx=XM_019838323.2;refGene_gn=SESTD1;refGene_tx=XM_023259511.1;refGene_gn=SESTD1;refGene_tx=XM_023259512.1;refGene_gn=SESTD1;refGene_tx=XM_023259513.1
C1 169737261 169737384
refGene_gn=SESTD1;refGene_tx=XM_011285390.3;refGene_gn=SESTD1;refGene_tx=XM_019838323.2;refGene_gn=SESTD1;refGene_tx=XM_023259511.1;refGene_gn=SESTD1;refGene_tx=XM_023259512.1;refGene_gn=SESTD1;refGene_tx=XM_023259513.1
C1 169751496 169751708
refGene_gn=SESTD1;refGene_tx=XM_011285390.3;refGene_gn=SESTD1;refGene_tx=XM_019838323.2;refGene_gn=SESTD1;refGene_tx=XM_023259511.1;refGene_gn=SESTD1;refGene_tx=XM_023259512.1;refGene_gn=SESTD1;refGene_tx=XM_023259513.1
C1 169754390 169754446
refGene_gn=SESTD1;refGene_tx=XM_011285390.3;refGene_gn=SESTD1;refGene_tx=XM_019838323.2;refGene_gn=SESTD1;refGene_tx=XM_023259511.1;refGene_gn=SESTD1;refGene_tx=XM_023259512.1;refGene_gn=SESTD1;refGene_tx=XM_023259513.1
C1 169757489 169757587
refGene_gn=SESTD1;refGene_tx=XM_011285390.3;refGene_gn=SESTD1;refGene_tx=XM_019838323.2;refGene_gn=SESTD1;refGene_tx=XM_023259511.1;refGene_gn=SESTD1;refGene_tx=XM_023259512.1;refGene_gn=SESTD1;refGene_tx=XM_023259513.1

C1 169759216 169759330
refGene_gn=SESTD1;refGene_tx=XM_011285390.3;refGene_gn=SESTD1;refGene_tx=XM_019838323.2;refGene_gn=SESTD1;refGene_tx=XM_023259511.1;refGene_gn=SESTD1;refGene_tx=XM_023259512.1;refGene_gn=SESTD1;refGene_tx=XM_023259513.1
C1 169776430 169776544
refGene_gn=SESTD1;refGene_tx=XM_011285390.3;refGene_gn=SESTD1;refGene_tx=XM_019838323.2;refGene_gn=SESTD1;refGene_tx=XM_023259511.1;refGene_gn=SESTD1;refGene_tx=XM_023259512.1;refGene_gn=SESTD1;refGene_tx=XM_023259513.1
C1 169781202 169781293
refGene_gn=SESTD1;refGene_tx=XM_011285390.3;refGene_gn=SESTD1;refGene_tx=XM_019838323.2;refGene_gn=SESTD1;refGene_tx=XM_023259511.1;refGene_gn=SESTD1;refGene_tx=XM_023259512.1;refGene_gn=SESTD1;refGene_tx=XM_023259513.1
C1 169804240 169804349
refGene_gn=SESTD1;refGene_tx=XM_011285390.3;refGene_gn=SESTD1;refGene_tx=XM_023259511.1;refGene_gn=SESTD1;refGene_tx=XM_023259512.1;refGene_gn=SESTD1;refGene_tx=XM_023259513.1
C1 169811490 169811545
refGene_gn=SESTD1;refGene_tx=XM_011285390.3;refGene_gn=SESTD1;refGene_tx=XM_023259511.1;refGene_gn=SESTD1;refGene_tx=XM_023259512.1;refGene_gn=SESTD1;refGene_tx=XM_023259513.1
C1 170004254 170004473
refGene_gn=ZNF385B;refGene_tx=XM_023259514.1;refGene_gn=ZNF385B;refGene_tx=XM_023259515.1;refGene_gn=ZNF385B;refGene_tx=XM_023259516.1;refGene_gn=ZNF385B;refGene_tx=XM_023259517.1;refGene_gn=ZNF385B;refGene_tx=XM_023259518.1;refGene_gn=ZNF385B;refGene_tx=XM_023259519.1;refGene_gn=ZNF385B;refGene_tx=XM_023259520.1;refGene_gn=ZNF385B;refGene_tx=XM_023259521.1;refGene_gn=ZNF385B;refGene_tx=XM_023259522.1
C1 170005944 170006046
refGene_gn=ZNF385B;refGene_tx=XM_023259514.1;refGene_gn=ZNF385B;refGene_tx=XM_023259515.1;refGene_gn=ZNF385B;refGene_tx=XM_023259516.1;refGene_gn=ZNF385B;refGene_tx=XM_023259517.1;refGene_gn=ZNF385B;refGene_tx=XM_023259518.1;refGene_gn=ZNF385B;refGene_tx=XM_023259519.1;refGene_gn=ZNF385B;refGene_tx=XM_023259520.1;refGene_gn=ZNF385B;refGene_tx=XM_023259521.1;refGene_gn=ZNF385B;refGene_tx=XM_023259522.1
C1 170006852 170007031
refGene_gn=ZNF385B;refGene_tx=XM_023259514.1;refGene_gn=ZNF385B;refGene_tx=XM_023259515.1;refGene_gn=ZNF385B;refGene_tx=XM_023259516.1;refGene_gn=ZNF385B;refGene_tx=XM_023259517.1;refGene_gn=ZNF385B;refGene_tx=XM_023259518.1;refGene_gn=ZNF385B;refGene_tx=XM_023259519.1;refGene_gn=ZNF385B;refGene_tx=XM_023259520.1;refGene_gn=ZNF385B;refGene_tx=XM_023259521.1;refGene_gn=ZNF385B;refGene_tx=XM_023259522.1
C1 170008366 170008618
refGene_gn=ZNF385B;refGene_tx=XM_023259514.1;refGene_gn=ZNF385B;refGene_tx=XM_023259515.1;refGene_gn=ZNF385B;refGene_tx=XM_023259516.1;refGene_gn=ZNF385B;refGene_tx=XM_023259517.1;refGene_gn=ZNF385B;refGene_tx=XM_023259518.1;refGene_gn=ZNF385B;refGene_tx=XM_023259519.1;refGene_gn=ZNF385B;refGene_tx=XM_023259520.1;refGene_gn=ZNF385B;refGene_tx=XM_02

3259521.1;refGene_gn=ZNF385B;refGene_tx=XM_023259522.1
C1 170034554 170034717
refGene_gn=ZNF385B;refGene_tx=XM_023259514.1;refGene_gn=ZNF385B;refGene_tx=XM_023259515.1;refGene_gn=ZNF385B;refGene_tx=XM_023259516.1;refGene_gn=ZNF385B;refGene_tx=XM_023259517.1;refGene_gn=ZNF385B;refGene_tx=XM_023259518.1;refGene_gn=ZNF385B;refGene_tx=XM_023259519.1;refGene_gn=ZNF385B;refGene_tx=XM_023259520.1;refGene_gn=ZNF385B;refGene_tx=XM_023259521.1;refGene_gn=ZNF385B;refGene_tx=XM_023259522.1
C1 170064237 170064348
refGene_gn=ZNF385B;refGene_tx=XM_023259514.1;refGene_gn=ZNF385B;refGene_tx=XM_023259515.1;refGene_gn=ZNF385B;refGene_tx=XM_023259517.1;refGene_gn=ZNF385B;refGene_tx=XM_023259518.1;refGene_gn=ZNF385B;refGene_tx=XM_023259519.1;refGene_gn=ZNF385B;refGene_tx=XM_023259520.1;refGene_gn=ZNF385B;refGene_tx=XM_023259521.1;refGene_gn=ZNF385B;refGene_tx=XM_023259522.1
C1 170068454 170068493
refGene_gn=ZNF385B;refGene_tx=XM_023259514.1;refGene_gn=ZNF385B;refGene_tx=XM_023259517.1;refGene_gn=ZNF385B;refGene_tx=XM_023259518.1;refGene_gn=ZNF385B;refGene_tx=XM_023259519.1;refGene_gn=ZNF385B;refGene_tx=XM_023259520.1;refGene_gn=ZNF385B;refGene_tx=XM_023259521.1;refGene_gn=ZNF385B;refGene_tx=XM_023259522.1
C1 170090060 170090206
refGene_gn=ZNF385B;refGene_tx=XM_023259514.1;refGene_gn=ZNF385B;refGene_tx=XM_023259515.1;refGene_gn=ZNF385B;refGene_tx=XM_023259516.1;refGene_gn=ZNF385B;refGene_tx=XM_023259517.1;refGene_gn=ZNF385B;refGene_tx=XM_023259518.1;refGene_gn=ZNF385B;refGene_tx=XM_023259519.1;refGene_gn=ZNF385B;refGene_tx=XM_023259520.1;refGene_gn=ZNF385B;refGene_tx=XM_023259521.1;refGene_gn=ZNF385B;refGene_tx=XM_023259522.1
C1 170201451 170201461
refGene_gn=ZNF385B;refGene_tx=XM_023259517.1
C1 170296105 170296130
refGene_gn=ZNF385B;refGene_tx=XM_023259522.1
C1 170319208 170319524
refGene_gn=ZNF385B;refGene_tx=XM_023259514.1;refGene_gn=ZNF385B;refGene_tx=XM_023259515.1;refGene_gn=ZNF385B;refGene_tx=XM_023259516.1
C1 170485312 170485899
refGene_gn=CWC22;refGene_tx=XM_003990908.2;refGene_gn=CWC22;refGene_tx=XM_006935392.3
C1 170490115 170490336
refGene_gn=CWC22;refGene_tx=XM_003990908.2;refGene_gn=CWC22;refGene_tx=XM_006935392.3
C1 170490432 170490534
refGene_gn=CWC22;refGene_tx=XM_003990908.2;refGene_gn=CWC22;refGene_tx=XM_006935392.3
C1 170492055 170492183
refGene_gn=CWC22;refGene_tx=XM_003990908.2;refGene_gn=CWC22;refGene_tx=XM_006935392.3
C1 170493861 170494014
refGene_gn=CWC22;refGene_tx=XM_003990908.2;refGene_gn=CWC22;refGene_tx=XM_006935392.3

C1 170494684 170494762
refGene_gn=CWC22;refGene_tx=XM_003990908.2;refGene_gn=CWC22;refGene_tx=XM_006935392.3
C1 170498545 170498606
refGene_gn=CWC22;refGene_tx=XM_003990908.2;refGene_gn=CWC22;refGene_tx=XM_006935392.3
C1 170502340 170502422
refGene_gn=CWC22;refGene_tx=XM_003990908.2;refGene_gn=CWC22;refGene_tx=XM_006935392.3
C1 170503805 170503910
refGene_gn=CWC22;refGene_tx=XM_003990908.2;refGene_gn=CWC22;refGene_tx=XM_006935392.3
C1 170508316 170508379
refGene_gn=CWC22;refGene_tx=XM_003990908.2;refGene_gn=CWC22;refGene_tx=XM_006935392.3
C1 170508471 170508678
refGene_gn=CWC22;refGene_tx=XM_003990908.2;refGene_gn=CWC22;refGene_tx=XM_006935392.3
C1 170508771 170508907
refGene_gn=CWC22;refGene_tx=XM_003990908.2;refGene_gn=CWC22;refGene_tx=XM_006935392.3
C1 170510703 170510757
refGene_gn=CWC22;refGene_tx=XM_003990908.2;refGene_gn=CWC22;refGene_tx=XM_006935392.3
C1 170511136 170511305
refGene_gn=CWC22;refGene_tx=XM_003990908.2;refGene_gn=CWC22;refGene_tx=XM_006935392.3
C1 170512822 170512951
refGene_gn=CWC22;refGene_tx=XM_003990908.2;refGene_gn=CWC22;refGene_tx=XM_006935392.3
C1 170516503 170516749
refGene_gn=CWC22;refGene_tx=XM_003990908.2;refGene_gn=CWC22;refGene_tx=XM_006935392.3
C1 170519668 170519779
refGene_gn=CWC22;refGene_tx=XM_003990908.2;refGene_gn=CWC22;refGene_tx=XM_006935392.3
C1 170521709 170521777
refGene_gn=CWC22;refGene_tx=XM_003990908.2;refGene_gn=CWC22;refGene_tx=XM_006935392.3
C1 170526354 170526381
refGene_gn=CWC22;refGene_tx=XM_003990908.2;refGene_gn=CWC22;refGene_tx=XM_006935392.3
C1 171396257 171396451
refGene_gn=UBE2E3;refGene_tx=XM_006935393.4;refGene_gn=UBE2E3;refGene_tx=XM_019838326.2;refGene_gn=UBE2E3;refGene_tx=XM_023259523.1;refGene_gn=UBE2E3;refGene_tx=XM_023259524.1
C1 171398257 171398308
refGene_gn=UBE2E3;refGene_tx=XM_006935393.4;refGene_gn=UBE2E3;refGene_tx=XM_019838326.2;refGene_gn=UBE2E3;refGene_tx=XM_023259523.1;refGene_gn=UBE2E3;refGene_tx=XM_023259524.1

C1	171483302	171483435
refGene_gn=UBE2E3;refGene_tx=XM_006935393.4;refGene_gn=UBE2E3;refGene_tx=XM_019838326.2;refGene_gn=UBE2E3;refGene_tx=XM_023259523.1;refGene_gn=UBE2E3;refGene_tx=XM_023259524.1
C1	171486787	171486935
refGene_gn=UBE2E3;refGene_tx=XM_006935393.4;refGene_gn=UBE2E3;refGene_tx=XM_019838326.2;refGene_gn=UBE2E3;refGene_tx=XM_023259523.1;refGene_gn=UBE2E3;refGene_tx=XM_023259524.1
C1	171488927	171489025
refGene_gn=UBE2E3;refGene_tx=XM_006935393.4;refGene_gn=UBE2E3;refGene_tx=XM_019838326.2;refGene_gn=UBE2E3;refGene_tx=XM_023259523.1;refGene_gn=UBE2E3;refGene_tx=XM_023259524.1
C1	171880003	171880209
refGene_gn=ITGA4;refGene_tx=XM_003990911.5
C1	171880568	171880690
refGene_gn=ITGA4;refGene_tx=XM_003990911.5
C1	171900122	171900229
refGene_gn=ITGA4;refGene_tx=XM_003990911.5
C1	171900325	171900361
refGene_gn=ITGA4;refGene_tx=XM_003990911.5
C1	171900362	171900456
refGene_gn=ITGA4;refGene_tx=XM_003990911.5
C1	171902227	171902295
refGene_gn=ITGA4;refGene_tx=XM_003990911.5
C1	171903544	171903674
refGene_gn=ITGA4;refGene_tx=XM_003990911.5
C1	171905232	171905318
refGene_gn=ITGA4;refGene_tx=XM_003990911.5
C1	171906239	171906302
refGene_gn=ITGA4;refGene_tx=XM_003990911.5
C1	171906391	171906529
refGene_gn=ITGA4;refGene_tx=XM_003990911.5
C1	171910391	171910503
refGene_gn=ITGA4;refGene_tx=XM_003990911.5
C1	171916688	171916783
refGene_gn=ITGA4;refGene_tx=XM_003990911.5
C1	171918095	171918186
refGene_gn=ITGA4;refGene_tx=XM_003990911.5
C1	171918759	171918805
refGene_gn=ITGA4;refGene_tx=XM_003990911.5
C1	171919175	171919330
refGene_gn=ITGA4;refGene_tx=XM_003990911.5
C1	171921837	171921992
refGene_gn=ITGA4;refGene_tx=XM_003990911.5
C1	171945651	171945801
refGene_gn=ITGA4;refGene_tx=XM_003990911.5
C1	171947915	171947992
refGene_gn=ITGA4;refGene_tx=XM_003990911.5
C1	171959449	171959600
refGene_gn=ITGA4;refGene_tx=XM_003990911.5

C1 171960948 171961044
refGene_gn=ITGA4;refGene_tx=XM_003990911.5
C1 171961733 171961813
refGene_gn=ITGA4;refGene_tx=XM_003990911.5
C1 171962173 171962263
refGene_gn=ITGA4;refGene_tx=XM_003990911.5
C1 171964573 171964664
refGene_gn=ITGA4;refGene_tx=XM_003990911.5
C1 171967218 171967326
refGene_gn=ITGA4;refGene_tx=XM_003990911.5
C1 171968345 171968471
refGene_gn=ITGA4;refGene_tx=XM_003990911.5
C1 171969074 171969194
refGene_gn=ITGA4;refGene_tx=XM_003990911.5
C1 171971008 171971107
refGene_gn=ITGA4;refGene_tx=XM_003990911.5
C1 171971491 171971611
refGene_gn=ITGA4;refGene_tx=XM_003990911.5
C1 171972223 171972322
refGene_gn=ITGA4;refGene_tx=XM_003990911.5
C1 171974770 171974852
refGene_gn=CERKL;refGene_tx=XM_023259525.1;refGene_gn=CERKL;refGene_tx=XM_023259526.1;refGene_gn=CERKL;refGene_tx=XM_023259528.1
C1 171975492 171975668
refGene_gn=CERKL;refGene_tx=XM_023259525.1;refGene_gn=CERKL;refGene_tx=XM_023259526.1
C1 171977114 171977243
refGene_gn=CERKL;refGene_tx=XM_023259527.1
C1 171980774 171980871
refGene_gn=CERKL;refGene_tx=XM_023259525.1;refGene_gn=CERKL;refGene_tx=XM_023259526.1;refGene_gn=CERKL;refGene_tx=XM_023259527.1;refGene_gn=CERKL;refGene_tx=XM_023259528.1
C1 171983523 171983632
refGene_gn=CERKL;refGene_tx=XM_023259525.1;refGene_gn=CERKL;refGene_tx=XM_023259526.1;refGene_gn=CERKL;refGene_tx=XM_023259527.1;refGene_gn=CERKL;refGene_tx=XM_023259528.1
C1 171983727 171983753
refGene_gn=CERKL;refGene_tx=XM_023259525.1;refGene_gn=CERKL;refGene_tx=XM_023259526.1;refGene_gn=CERKL;refGene_tx=XM_023259527.1;refGene_gn=CERKL;refGene_tx=XM_023259528.1
C1 171984343 171984403
refGene_gn=CERKL;refGene_tx=XM_023259525.1;refGene_gn=CERKL;refGene_tx=XM_023259526.1;refGene_gn=CERKL;refGene_tx=XM_023259527.1;refGene_gn=CERKL;refGene_tx=XM_023259528.1
C1 171984483 171984661
refGene_gn=CERKL;refGene_tx=XM_023259525.1;refGene_gn=CERKL;refGene_tx=XM_023259526.1;refGene_gn=CERKL;refGene_tx=XM_023259527.1;refGene_gn=CERKL;refGene_tx=XM_023259528.1
C1 171985638 171985713
refGene_gn=CERKL;refGene_tx=XM_023259525.1;refGene_gn=CERKL;refGene_tx

=XM_023259527.1;refGene_gn=CERKL;refGene_tx=XM_023259528.1
C1 171990816 171990959
refGene_gn=CERKL;refGene_tx=XM_023259525.1;refGene_gn=CERKL;refGene_tx=XM_023259526.1;refGene_gn=CERKL;refGene_tx=XM_023259527.1;refGene_gn=CERKL;refGene_tx=XM_023259528.1
C1 171997953 171998017
refGene_gn=CERKL;refGene_tx=XM_023259525.1;refGene_gn=CERKL;refGene_tx=XM_023259526.1;refGene_gn=CERKL;refGene_tx=XM_023259527.1;refGene_gn=CERKL;refGene_tx=XM_023259528.1
C1 172005485 172005617
refGene_gn=CERKL;refGene_tx=XM_023259525.1;refGene_gn=CERKL;refGene_tx=XM_023259526.1;refGene_gn=CERKL;refGene_tx=XM_023259527.1;refGene_gn=CERKL;refGene_tx=XM_023259528.1
C1 172041306 172041549
refGene_gn=CERKL;refGene_tx=XM_023259525.1;refGene_gn=CERKL;refGene_tx=XM_023259526.1;refGene_gn=CERKL;refGene_tx=XM_023259527.1;refGene_gn=CERKL;refGene_tx=XM_023259528.1
C1 172080843 172081081
refGene_gn=CERKL;refGene_tx=XM_023259525.1;refGene_gn=CERKL;refGene_tx=XM_023259526.1;refGene_gn=CERKL;refGene_tx=XM_023259527.1;refGene_gn=CERKL;refGene_tx=XM_023259528.1
C1 172096262 172097333
refGene_gn=NEUROD1;refGene_tx=XM_003990912.4
C1 172290242 172290453
refGene_gn=SSFA2;refGene_tx=XM_011285388.2;refGene_gn=SSFA2;refGene_tx=XM_011285389.2
C1 172290811 172290857
refGene_gn=SSFA2;refGene_tx=XM_011285388.2;refGene_gn=SSFA2;refGene_tx=XM_011285389.2
C1 172293834 172293884
refGene_gn=SSFA2;refGene_tx=XM_011285388.2;refGene_gn=SSFA2;refGene_tx=XM_011285389.2
C1 172294736 172294793
refGene_gn=SSFA2;refGene_tx=XM_011285388.2;refGene_gn=SSFA2;refGene_tx=XM_011285389.2
C1 172297095 172297135
refGene_gn=SSFA2;refGene_tx=XM_011285388.2;refGene_gn=SSFA2;refGene_tx=XM_011285389.2
C1 172297225 172297324
refGene_gn=SSFA2;refGene_tx=XM_011285388.2;refGene_gn=SSFA2;refGene_tx=XM_011285389.2
C1 172298506 172298715
refGene_gn=SSFA2;refGene_tx=XM_011285388.2;refGene_gn=SSFA2;refGene_tx=XM_011285389.2
C1 172299522 172300223
refGene_gn=SSFA2;refGene_tx=XM_011285388.2;refGene_gn=SSFA2;refGene_tx=XM_011285389.2
C1 172308486 172308559
refGene_gn=SSFA2;refGene_tx=XM_011285388.2;refGene_gn=SSFA2;refGene_tx=XM_011285389.2

C1 172312665 172312754
refGene_gn=SSFA2;refGene_tx=XM_011285388.2;refGene_gn=SSFA2;refGene_tx=XM_011285389.2
C1 172314376 172315585
refGene_gn=SSFA2;refGene_tx=XM_011285388.2;refGene_gn=SSFA2;refGene_tx=XM_011285389.2
C1 172317630 172317708
refGene_gn=SSFA2;refGene_tx=XM_011285388.2;refGene_gn=SSFA2;refGene_tx=XM_011285389.2
C1 172317787 172317915
refGene_gn=SSFA2;refGene_tx=XM_011285388.2;refGene_gn=SSFA2;refGene_tx=XM_011285389.2
C1 172318306 172318457
refGene_gn=SSFA2;refGene_tx=XM_011285388.2;refGene_gn=SSFA2;refGene_tx=XM_011285389.2
C1 172319537 172319603
refGene_gn=SSFA2;refGene_tx=XM_011285388.2
C1 172320529 172320994
refGene_gn=SSFA2;refGene_tx=XM_011285388.2;refGene_gn=SSFA2;refGene_tx=XM_011285389.2
C1 172328192 172328297
refGene_gn=SSFA2;refGene_tx=XM_011285388.2;refGene_gn=SSFA2;refGene_tx=XM_011285389.2
C1 172387130 172387211
refGene_gn=PPP1R1C;refGene_tx=XM_003990915.5;refGene_gn=PPP1R1C;refGene_tx=XM_006935395.4;refGene_gn=PPP1R1C;refGene_tx=XM_019838341.2
C1 172388868 172388929
refGene_gn=PPP1R1C;refGene_tx=XM_003990915.5;refGene_gn=PPP1R1C;refGene_tx=XM_006935395.4;refGene_gn=PPP1R1C;refGene_tx=XM_019838341.2
C1 172443911 172443949
refGene_gn=PPP1R1C;refGene_tx=XM_003990915.5;refGene_gn=PPP1R1C;refGene_tx=XM_006935395.4;refGene_gn=PPP1R1C;refGene_tx=XM_019838341.2
C1 172446207 172446279
refGene_gn=PPP1R1C;refGene_tx=XM_003990915.5;refGene_gn=PPP1R1C;refGene_tx=XM_006935395.4;refGene_gn=PPP1R1C;refGene_tx=XM_019838341.2
C1 172505052 172505141
refGene_gn=PPP1R1C;refGene_tx=XM_003990915.5;refGene_gn=PPP1R1C;refGene_tx=XM_006935395.4
C1 172510647 172510712
refGene_gn=PPP1R1C;refGene_tx=XM_019838341.2
C1 172526309 172526383
refGene_gn=PDE1A;refGene_tx=XM_003990917.5;refGene_gn=PDE1A;refGene_tx=XM_011285381.3;refGene_gn=PDE1A;refGene_tx=XM_011285382.3;refGene_gn=PDE1A;refGene_tx=XM_011285383.3;refGene_gn=PDE1A;refGene_tx=XM_011285384.3;refGene_gn=PDE1A;refGene_tx=XM_011285385.3;refGene_gn=PDE1A;refGene_tx=XM_011285386.3;refGene_gn=PDE1A;refGene_tx=XM_011285387.3;refGene_gn=PDE1A;refGene_tx=XM_019838342.2
C1 172551892 172552053
refGene_gn=PDE1A;refGene_tx=XM_003990917.5;refGene_gn=PDE1A;refGene_tx=XM_011285381.3;refGene_gn=PDE1A;refGene_tx=XM_011285382.3;refGene_gn=

PDE1A;refGene_tx=XM_011285383.3;refGene_gn=PDE1A;refGene_tx=XM_0112853
84.3;refGene_gn=PDE1A;refGene_tx=XM_011285385.3;refGene_gn=PDE1A;refGe
ne_tx=XM_011285386.3;refGene_gn=PDE1A;refGene_tx=XM_011285387.3;refGen
e_gn=PDE1A;refGene_tx=XM_019838342.2
C1 172553129 172553250
refGene_gn=PDE1A;refGene_tx=XM_003990917.5;refGene_gn=PDE1A;refGene_tx
=XM_011285381.3;refGene_gn=PDE1A;refGene_tx=XM_011285382.3;refGene_gn=
PDE1A;refGene_tx=XM_011285383.3;refGene_gn=PDE1A;refGene_tx=XM_0112853
84.3;refGene_gn=PDE1A;refGene_tx=XM_011285385.3;refGene_gn=PDE1A;refGe
ne_tx=XM_011285386.3;refGene_gn=PDE1A;refGene_tx=XM_011285387.3;refGen
e_gn=PDE1A;refGene_tx=XM_019838342.2
C1 172555254 172555336
refGene_gn=PDE1A;refGene_tx=XM_003990917.5;refGene_gn=PDE1A;refGene_tx
=XM_011285381.3;refGene_gn=PDE1A;refGene_tx=XM_011285382.3;refGene_gn=
PDE1A;refGene_tx=XM_011285383.3;refGene_gn=PDE1A;refGene_tx=XM_0112853
84.3;refGene_gn=PDE1A;refGene_tx=XM_011285385.3;refGene_gn=PDE1A;refGe
ne_tx=XM_011285386.3;refGene_gn=PDE1A;refGene_tx=XM_011285387.3;refGen
e_gn=PDE1A;refGene_tx=XM_019838342.2
C1 172566034 172566155
refGene_gn=PDE1A;refGene_tx=XM_003990917.5;refGene_gn=PDE1A;refGene_tx
=XM_011285381.3;refGene_gn=PDE1A;refGene_tx=XM_011285382.3;refGene_gn=
PDE1A;refGene_tx=XM_011285383.3;refGene_gn=PDE1A;refGene_tx=XM_0112853
84.3;refGene_gn=PDE1A;refGene_tx=XM_011285385.3;refGene_gn=PDE1A;refGe
ne_tx=XM_011285386.3;refGene_gn=PDE1A;refGene_tx=XM_011285387.3;refGen
e_gn=PDE1A;refGene_tx=XM_019838342.2
C1 172566808 172566910
refGene_gn=PDE1A;refGene_tx=XM_003990917.5;refGene_gn=PDE1A;refGene_tx
=XM_011285381.3;refGene_gn=PDE1A;refGene_tx=XM_011285382.3;refGene_gn=
PDE1A;refGene_tx=XM_011285383.3;refGene_gn=PDE1A;refGene_tx=XM_0112853
84.3;refGene_gn=PDE1A;refGene_tx=XM_011285385.3;refGene_gn=PDE1A;refGe
ne_tx=XM_011285386.3;refGene_gn=PDE1A;refGene_tx=XM_011285387.3;refGen
e_gn=PDE1A;refGene_tx=XM_019838342.2
C1 172570418 172570544
refGene_gn=PDE1A;refGene_tx=XM_003990917.5;refGene_gn=PDE1A;refGene_tx
=XM_011285381.3;refGene_gn=PDE1A;refGene_tx=XM_011285382.3;refGene_gn=
PDE1A;refGene_tx=XM_011285383.3;refGene_gn=PDE1A;refGene_tx=XM_0112853
84.3;refGene_gn=PDE1A;refGene_tx=XM_011285385.3;refGene_gn=PDE1A;refGe
ne_tx=XM_011285386.3;refGene_gn=PDE1A;refGene_tx=XM_011285387.3;refGen
e_gn=PDE1A;refGene_tx=XM_019838342.2
C1 172577739 172577840
refGene_gn=PDE1A;refGene_tx=XM_003990917.5;refGene_gn=PDE1A;refGene_tx
=XM_011285381.3;refGene_gn=PDE1A;refGene_tx=XM_011285382.3;refGene_gn=
PDE1A;refGene_tx=XM_011285383.3;refGene_gn=PDE1A;refGene_tx=XM_0112853
84.3;refGene_gn=PDE1A;refGene_tx=XM_011285385.3;refGene_gn=PDE1A;refGe
ne_tx=XM_011285386.3;refGene_gn=PDE1A;refGene_tx=XM_011285387.3;refGen
e_gn=PDE1A;refGene_tx=XM_019838342.2
C1 172585656 172585797
refGene_gn=PDE1A;refGene_tx=XM_003990917.5;refGene_gn=PDE1A;refGene_tx
=XM_011285381.3;refGene_gn=PDE1A;refGene_tx=XM_011285382.3;refGene_gn=
PDE1A;refGene_tx=XM_011285383.3;refGene_gn=PDE1A;refGene_tx=XM_0112853

84.3;refGene_gn=PDE1A;refGene_tx=XM_011285385.3;refGene_gn=PDE1A;refGene_tx=XM_011285386.3;refGene_gn=PDE1A;refGene_tx=XM_011285387.3;refGene_gn=PDE1A;refGene_tx=XM_019838342.2
C1 172586660 172586777
refGene_gn=PDE1A;refGene_tx=XM_003990917.5;refGene_gn=PDE1A;refGene_tx=XM_011285381.3;refGene_gn=PDE1A;refGene_tx=XM_011285382.3;refGene_gn=PDE1A;refGene_tx=XM_011285383.3;refGene_gn=PDE1A;refGene_tx=XM_011285384.3;refGene_gn=PDE1A;refGene_tx=XM_011285385.3;refGene_gn=PDE1A;refGene_tx=XM_011285386.3;refGene_gn=PDE1A;refGene_tx=XM_011285387.3;refGene_gn=PDE1A;refGene_tx=XM_019838342.2
C1 172589863 172589930
refGene_gn=PDE1A;refGene_tx=XM_003990917.5;refGene_gn=PDE1A;refGene_tx=XM_011285381.3;refGene_gn=PDE1A;refGene_tx=XM_011285382.3;refGene_gn=PDE1A;refGene_tx=XM_011285383.3;refGene_gn=PDE1A;refGene_tx=XM_011285384.3;refGene_gn=PDE1A;refGene_tx=XM_011285385.3;refGene_gn=PDE1A;refGene_tx=XM_011285386.3;refGene_gn=PDE1A;refGene_tx=XM_011285387.3;refGene_gn=PDE1A;refGene_tx=XM_019838342.2
C1 172595358 172595541
refGene_gn=PDE1A;refGene_tx=XM_003990917.5;refGene_gn=PDE1A;refGene_tx=XM_011285381.3;refGene_gn=PDE1A;refGene_tx=XM_011285382.3;refGene_gn=PDE1A;refGene_tx=XM_011285383.3;refGene_gn=PDE1A;refGene_tx=XM_011285384.3;refGene_gn=PDE1A;refGene_tx=XM_011285385.3;refGene_gn=PDE1A;refGene_tx=XM_011285386.3;refGene_gn=PDE1A;refGene_tx=XM_011285387.3;refGene_gn=PDE1A;refGene_tx=XM_019838342.2
C1 172597066 172597179
refGene_gn=PDE1A;refGene_tx=XM_011285387.3
C1 172618197 172618311
refGene_gn=PDE1A;refGene_tx=XM_003990917.5;refGene_gn=PDE1A;refGene_tx=XM_011285381.3;refGene_gn=PDE1A;refGene_tx=XM_011285382.3;refGene_gn=PDE1A;refGene_tx=XM_011285383.3;refGene_gn=PDE1A;refGene_tx=XM_011285384.3;refGene_gn=PDE1A;refGene_tx=XM_011285385.3;refGene_gn=PDE1A;refGene_tx=XM_011285386.3;refGene_gn=PDE1A;refGene_tx=XM_019838342.2
C1 172710097 172710150
refGene_gn=PDE1A;refGene_tx=XM_019838342.2
C1 172760769 172760822
refGene_gn=PDE1A;refGene_tx=XM_003990917.5;refGene_gn=PDE1A;refGene_tx=XM_011285386.3
C1 172848664 172848765
refGene_gn=PDE1A;refGene_tx=XM_011285381.3;refGene_gn=PDE1A;refGene_tx=XM_011285382.3;refGene_gn=PDE1A;refGene_tx=XM_011285383.3;refGene_gn=PDE1A;refGene_tx=XM_011285384.3;refGene_gn=PDE1A;refGene_tx=XM_011285385.3
C1 173052946 173052999
refGene_gn=DNAJC10;refGene_tx=XM_006935397.3
C1 173053000 173053148
refGene_gn=DNAJC10;refGene_tx=XM_006935397.3
C1 173054589 173054755
refGene_gn=DNAJC10;refGene_tx=XM_006935397.3
C1 173056346 173056397
refGene_gn=DNAJC10;refGene_tx=XM_006935397.3

C1	173069359	173069442
refGene_gn=DNAJC10;refGene_tx=XM_006935397.3
C1	173069744	173069876
refGene_gn=DNAJC10;refGene_tx=XM_006935397.3
C1	173070502	173070596
refGene_gn=DNAJC10;refGene_tx=XM_006935397.3
C1	173071631	173071709
refGene_gn=DNAJC10;refGene_tx=XM_006935397.3
C1	173074986	173075030
refGene_gn=DNAJC10;refGene_tx=XM_006935397.3
C1	173075680	173075818
refGene_gn=DNAJC10;refGene_tx=XM_006935397.3
C1	173079397	173079487
refGene_gn=DNAJC10;refGene_tx=XM_006935397.3
C1	173080240	173080354
refGene_gn=DNAJC10;refGene_tx=XM_006935397.3
C1	173082789	173082904
refGene_gn=DNAJC10;refGene_tx=XM_006935397.3
C1	173087752	173087880
refGene_gn=DNAJC10;refGene_tx=XM_006935397.3
C1	173088195	173088312
refGene_gn=DNAJC10;refGene_tx=XM_006935397.3
C1	173091366	173091468
refGene_gn=DNAJC10;refGene_tx=XM_006935397.3
C1	173092497	173092653
refGene_gn=DNAJC10;refGene_tx=XM_006935397.3
C1	173093604	173093738
refGene_gn=DNAJC10;refGene_tx=XM_006935397.3
C1	173094818	173094872
refGene_gn=DNAJC10;refGene_tx=XM_006935397.3
C1	173095134	173095282
refGene_gn=DNAJC10;refGene_tx=XM_006935397.3
C1	173098565	173098685
refGene_gn=DNAJC10;refGene_tx=XM_006935397.3
C1	173100500	173100605
refGene_gn=DNAJC10;refGene_tx=XM_006935397.3
C1	173101927	173101939
refGene_gn=DNAJC10;refGene_tx=XM_006935397.3
C1	173188685	173188802
refGene_gn=FRZB;refGene_tx=XM_003990920.5
C1	173192239	173192303
refGene_gn=FRZB;refGene_tx=XM_003990920.5
C1	173192691	173192896
refGene_gn=FRZB;refGene_tx=XM_003990920.5
C1	173197161	173197227
refGene_gn=FRZB;refGene_tx=XM_003990920.5
C1	173211721	173211766
refGene_gn=FRZB;refGene_tx=XM_003990920.5
C1	173219019	173219497
refGene_gn=FRZB;refGene_tx=XM_003990920.5

C1	173276319	173276436
refGene_gn=NCKAP1;refGene_tx=XM_023259529.1;refGene_gn=NCKAP1;refGene_tx=XM_023259530.1;refGene_gn=NCKAP1;refGene_tx=XM_023259531.1;refGene_gn=NCKAP1;refGene_tx=XM_023259532.1;refGene_gn=NCKAP1;refGene_tx=XM_023259533.1
C1	173277248	173277338
refGene_gn=NCKAP1;refGene_tx=XM_023259529.1;refGene_gn=NCKAP1;refGene_tx=XM_023259530.1;refGene_gn=NCKAP1;refGene_tx=XM_023259531.1;refGene_gn=NCKAP1;refGene_tx=XM_023259532.1;refGene_gn=NCKAP1;refGene_tx=XM_023259533.1
C1	173277674	173277784
refGene_gn=NCKAP1;refGene_tx=XM_023259529.1;refGene_gn=NCKAP1;refGene_tx=XM_023259530.1;refGene_gn=NCKAP1;refGene_tx=XM_023259531.1;refGene_gn=NCKAP1;refGene_tx=XM_023259532.1;refGene_gn=NCKAP1;refGene_tx=XM_023259533.1
C1	173278389	173278506
refGene_gn=NCKAP1;refGene_tx=XM_023259529.1;refGene_gn=NCKAP1;refGene_tx=XM_023259530.1;refGene_gn=NCKAP1;refGene_tx=XM_023259531.1;refGene_gn=NCKAP1;refGene_tx=XM_023259532.1;refGene_gn=NCKAP1;refGene_tx=XM_023259533.1
C1	173281170	173281264
refGene_gn=NCKAP1;refGene_tx=XM_023259529.1;refGene_gn=NCKAP1;refGene_tx=XM_023259530.1;refGene_gn=NCKAP1;refGene_tx=XM_023259531.1;refGene_gn=NCKAP1;refGene_tx=XM_023259532.1;refGene_gn=NCKAP1;refGene_tx=XM_023259533.1
C1	173285458	173285539
refGene_gn=NCKAP1;refGene_tx=XM_023259529.1;refGene_gn=NCKAP1;refGene_tx=XM_023259530.1;refGene_gn=NCKAP1;refGene_tx=XM_023259531.1;refGene_gn=NCKAP1;refGene_tx=XM_023259532.1;refGene_gn=NCKAP1;refGene_tx=XM_023259533.1
C1	173285960	173286043
refGene_gn=NCKAP1;refGene_tx=XM_023259529.1;refGene_gn=NCKAP1;refGene_tx=XM_023259530.1;refGene_gn=NCKAP1;refGene_tx=XM_023259531.1;refGene_gn=NCKAP1;refGene_tx=XM_023259532.1;refGene_gn=NCKAP1;refGene_tx=XM_023259533.1
C1	173293542	173293636
refGene_gn=NCKAP1;refGene_tx=XM_023259529.1;refGene_gn=NCKAP1;refGene_tx=XM_023259530.1;refGene_gn=NCKAP1;refGene_tx=XM_023259531.1;refGene_gn=NCKAP1;refGene_tx=XM_023259532.1;refGene_gn=NCKAP1;refGene_tx=XM_023259533.1
C1	173306328	173306426
refGene_gn=NCKAP1;refGene_tx=XM_023259529.1;refGene_gn=NCKAP1;refGene_tx=XM_023259530.1;refGene_gn=NCKAP1;refGene_tx=XM_023259531.1;refGene_gn=NCKAP1;refGene_tx=XM_023259532.1;refGene_gn=NCKAP1;refGene_tx=XM_023259533.1
C1	173306702	173306833
refGene_gn=NCKAP1;refGene_tx=XM_023259529.1;refGene_gn=NCKAP1;refGene_tx=XM_023259530.1;refGene_gn=NCKAP1;refGene_tx=XM_023259531.1;refGene_gn=NCKAP1;refGene_tx=XM_023259532.1;refGene_gn=NCKAP1;refGene_tx=XM_023259533.1

C1 173307261 173307480
refGene_gn=NCKAP1;refGene_tx=XM_023259529.1;refGene_gn=NCKAP1;refGene_tx=XM_023259530.1;refGene_gn=NCKAP1;refGene_tx=XM_023259531.1;refGene_gn=NCKAP1;refGene_tx=XM_023259532.1;refGene_gn=NCKAP1;refGene_tx=XM_023259533.1
C1 173310000 173310132
refGene_gn=NCKAP1;refGene_tx=XM_023259529.1;refGene_gn=NCKAP1;refGene_tx=XM_023259530.1;refGene_gn=NCKAP1;refGene_tx=XM_023259531.1;refGene_gn=NCKAP1;refGene_tx=XM_023259532.1;refGene_gn=NCKAP1;refGene_tx=XM_023259533.1
C1 173310965 173311105
refGene_gn=NCKAP1;refGene_tx=XM_023259529.1;refGene_gn=NCKAP1;refGene_tx=XM_023259530.1;refGene_gn=NCKAP1;refGene_tx=XM_023259531.1;refGene_gn=NCKAP1;refGene_tx=XM_023259532.1;refGene_gn=NCKAP1;refGene_tx=XM_023259533.1
C1 173311762 173311882
refGene_gn=NCKAP1;refGene_tx=XM_023259529.1;refGene_gn=NCKAP1;refGene_tx=XM_023259530.1;refGene_gn=NCKAP1;refGene_tx=XM_023259531.1;refGene_gn=NCKAP1;refGene_tx=XM_023259532.1;refGene_gn=NCKAP1;refGene_tx=XM_023259533.1
C1 173315152 173315285
refGene_gn=NCKAP1;refGene_tx=XM_023259529.1;refGene_gn=NCKAP1;refGene_tx=XM_023259530.1;refGene_gn=NCKAP1;refGene_tx=XM_023259531.1;refGene_gn=NCKAP1;refGene_tx=XM_023259532.1;refGene_gn=NCKAP1;refGene_tx=XM_023259533.1
C1 173317181 173317327
refGene_gn=NCKAP1;refGene_tx=XM_023259529.1;refGene_gn=NCKAP1;refGene_tx=XM_023259530.1;refGene_gn=NCKAP1;refGene_tx=XM_023259531.1;refGene_gn=NCKAP1;refGene_tx=XM_023259532.1;refGene_gn=NCKAP1;refGene_tx=XM_023259533.1
C1 173325652 173325711
refGene_gn=NCKAP1;refGene_tx=XM_023259529.1;refGene_gn=NCKAP1;refGene_tx=XM_023259530.1;refGene_gn=NCKAP1;refGene_tx=XM_023259531.1;refGene_gn=NCKAP1;refGene_tx=XM_023259532.1;refGene_gn=NCKAP1;refGene_tx=XM_023259533.1
C1 173327240 173327322
refGene_gn=NCKAP1;refGene_tx=XM_023259529.1;refGene_gn=NCKAP1;refGene_tx=XM_023259530.1;refGene_gn=NCKAP1;refGene_tx=XM_023259531.1;refGene_gn=NCKAP1;refGene_tx=XM_023259532.1;refGene_gn=NCKAP1;refGene_tx=XM_023259533.1
C1 173329883 173330016
refGene_gn=NCKAP1;refGene_tx=XM_023259529.1;refGene_gn=NCKAP1;refGene_tx=XM_023259530.1;refGene_gn=NCKAP1;refGene_tx=XM_023259531.1;refGene_gn=NCKAP1;refGene_tx=XM_023259532.1
C1 173330734 173330841
refGene_gn=NCKAP1;refGene_tx=XM_023259529.1;refGene_gn=NCKAP1;refGene_tx=XM_023259530.1;refGene_gn=NCKAP1;refGene_tx=XM_023259531.1;refGene_gn=NCKAP1;refGene_tx=XM_023259532.1
C1 173331490 173331587
refGene_gn=NCKAP1;refGene_tx=XM_023259529.1;refGene_gn=NCKAP1;refGene_

tx=XM_023259530.1;refGene_gn=NCKAP1;refGene_tx=XM_023259531.1;refGene_gn=NCKAP1;refGene_tx=XM_023259532.1;refGene_gn=NCKAP1;refGene_tx=XM_023259533.1
C1 173333654 173333711
refGene_gn=NCKAP1;refGene_tx=XM_023259529.1;refGene_gn=NCKAP1;refGene_tx=XM_023259530.1;refGene_gn=NCKAP1;refGene_tx=XM_023259531.1;refGene_gn=NCKAP1;refGene_tx=XM_023259532.1;refGene_gn=NCKAP1;refGene_tx=XM_023259533.1
C1 173336726 173336883
refGene_gn=NCKAP1;refGene_tx=XM_023259529.1;refGene_gn=NCKAP1;refGene_tx=XM_023259530.1;refGene_gn=NCKAP1;refGene_tx=XM_023259531.1;refGene_gn=NCKAP1;refGene_tx=XM_023259532.1;refGene_gn=NCKAP1;refGene_tx=XM_023259533.1
C1 173340788 173340837
refGene_gn=NCKAP1;refGene_tx=XM_023259529.1;refGene_gn=NCKAP1;refGene_tx=XM_023259530.1;refGene_gn=NCKAP1;refGene_tx=XM_023259531.1;refGene_gn=NCKAP1;refGene_tx=XM_023259532.1;refGene_gn=NCKAP1;refGene_tx=XM_023259533.1
C1 173341965 173342103
refGene_gn=NCKAP1;refGene_tx=XM_023259529.1;refGene_gn=NCKAP1;refGene_tx=XM_023259530.1;refGene_gn=NCKAP1;refGene_tx=XM_023259531.1;refGene_gn=NCKAP1;refGene_tx=XM_023259532.1;refGene_gn=NCKAP1;refGene_tx=XM_023259533.1
C1 173344531 173344622
refGene_gn=NCKAP1;refGene_tx=XM_023259529.1;refGene_gn=NCKAP1;refGene_tx=XM_023259530.1;refGene_gn=NCKAP1;refGene_tx=XM_023259531.1;refGene_gn=NCKAP1;refGene_tx=XM_023259532.1;refGene_gn=NCKAP1;refGene_tx=XM_023259533.1
C1 173344705 173344848
refGene_gn=NCKAP1;refGene_tx=XM_023259529.1;refGene_gn=NCKAP1;refGene_tx=XM_023259530.1;refGene_gn=NCKAP1;refGene_tx=XM_023259531.1;refGene_gn=NCKAP1;refGene_tx=XM_023259532.1;refGene_gn=NCKAP1;refGene_tx=XM_023259533.1
C1 173345832 173345889
refGene_gn=NCKAP1;refGene_tx=XM_023259529.1;refGene_gn=NCKAP1;refGene_tx=XM_023259530.1;refGene_gn=NCKAP1;refGene_tx=XM_023259531.1;refGene_gn=NCKAP1;refGene_tx=XM_023259532.1;refGene_gn=NCKAP1;refGene_tx=XM_023259533.1
C1 173346087 173346180
refGene_gn=NCKAP1;refGene_tx=XM_023259529.1;refGene_gn=NCKAP1;refGene_tx=XM_023259530.1;refGene_gn=NCKAP1;refGene_tx=XM_023259531.1;refGene_gn=NCKAP1;refGene_tx=XM_023259532.1;refGene_gn=NCKAP1;refGene_tx=XM_023259533.1
C1 173366185 173366296
refGene_gn=NCKAP1;refGene_tx=XM_023259529.1;refGene_gn=NCKAP1;refGene_tx=XM_023259530.1;refGene_gn=NCKAP1;refGene_tx=XM_023259531.1;refGene_gn=NCKAP1;refGene_tx=XM_023259532.1;refGene_gn=NCKAP1;refGene_tx=XM_023259533.1
C1 173367351 173367369
refGene_gn=NCKAP1;refGene_tx=XM_023259529.1;refGene_gn=NCKAP1;refGene_

tx=XM_023259530.1;refGene_gn=NCKAP1;refGene_tx=XM_023259533.1
C1	173383811	173383919
refGene_gn=NCKAP1;refGene_tx=XM_023259529.1;refGene_gn=NCKAP1;refGene_tx=XM_023259530.1;refGene_gn=NCKAP1;refGene_tx=XM_023259531.1;refGene_gn=NCKAP1;refGene_tx=XM_023259532.1;refGene_gn=NCKAP1;refGene_tx=XM_023259533.1
C1	173414490	173414716
refGene_gn=DUSP19;refGene_tx=XM_003990922.4
C1	173420395	173420442
refGene_gn=DUSP19;refGene_tx=XM_003990922.4
C1	173428397	173428550
refGene_gn=DUSP19;refGene_tx=XM_003990922.4
C1	173431782	173432019
refGene_gn=DUSP19;refGene_tx=XM_003990922.4
C1	173459998	173460038
refGene_gn=NUP35;refGene_tx=XM_003990924.3;refGene_gn=NUP35;refGene_tx=XM_019838348.1
C1	173463505	173463676
refGene_gn=NUP35;refGene_tx=XM_003990924.3;refGene_gn=NUP35;refGene_tx=XM_011285380.3;refGene_gn=NUP35;refGene_tx=XM_019838348.1
C1	173467214	173467342
refGene_gn=NUP35;refGene_tx=XM_003990924.3;refGene_gn=NUP35;refGene_tx=XM_011285380.3;refGene_gn=NUP35;refGene_tx=XM_019838348.1
C1	173470189	173470256
refGene_gn=NUP35;refGene_tx=XM_003990924.3;refGene_gn=NUP35;refGene_tx=XM_011285380.3;refGene_gn=NUP35;refGene_tx=XM_019838348.1
C1	173488542	173488684
refGene_gn=NUP35;refGene_tx=XM_003990924.3;refGene_gn=NUP35;refGene_tx=XM_011285380.3;refGene_gn=NUP35;refGene_tx=XM_019838348.1
C1	173491214	173491284
refGene_gn=NUP35;refGene_tx=XM_003990924.3;refGene_gn=NUP35;refGene_tx=XM_011285380.3;refGene_gn=NUP35;refGene_tx=XM_019838348.1
C1	173492081	173492210
refGene_gn=NUP35;refGene_tx=XM_003990924.3;refGene_gn=NUP35;refGene_tx=XM_011285380.3
C1	173493268	173493433
refGene_gn=NUP35;refGene_tx=XM_003990924.3;refGene_gn=NUP35;refGene_tx=XM_011285380.3;refGene_gn=NUP35;refGene_tx=XM_019838348.1
C1	173494485	173494563
refGene_gn=NUP35;refGene_tx=XM_003990924.3;refGene_gn=NUP35;refGene_tx=XM_011285380.3;refGene_gn=NUP35;refGene_tx=XM_019838348.1
C1	174716810	174716921
refGene_gn=ZNF804A;refGene_tx=XM_023259534.1
C1	174941402	174941546
refGene_gn=ZNF804A;refGene_tx=XM_023259534.1
C1	174999059	174999190
refGene_gn=ZNF804A;refGene_tx=XM_023259534.1
C1	175001210	175004466
refGene_gn=ZNF804A;refGene_tx=XM_023259534.1
C1	175793743	175793842

refGene_gn=FSIP2;refGene_tx=XM_023259535.1
C1 175794186 175794312
refGene_gn=FSIP2;refGene_tx=XM_023259535.1
C1 175798786 175798948
refGene_gn=FSIP2;refGene_tx=XM_023259535.1
C1 175799672 175799762
refGene_gn=FSIP2;refGene_tx=XM_023259535.1
C1 175800829 175800969
refGene_gn=FSIP2;refGene_tx=XM_023259535.1
C1 175801611 175801685
refGene_gn=FSIP2;refGene_tx=XM_023259535.1
C1 175801687 175801746
refGene_gn=FSIP2;refGene_tx=XM_023259535.1
C1 175802246 175802357
refGene_gn=FSIP2;refGene_tx=XM_023259535.1
C1 175805613 175805740
refGene_gn=FSIP2;refGene_tx=XM_023259535.1
C1 175808652 175808742
refGene_gn=FSIP2;refGene_tx=XM_023259535.1
C1 175818299 175818412
refGene_gn=FSIP2;refGene_tx=XM_023259535.1
C1 175819192 175819235
refGene_gn=FSIP2;refGene_tx=XM_023259535.1
C1 175820530 175820637
refGene_gn=FSIP2;refGene_tx=XM_023259535.1
C1 175822894 175822958
refGene_gn=FSIP2;refGene_tx=XM_023259535.1
C1 175828935 175828993
refGene_gn=FSIP2;refGene_tx=XM_023259535.1
C1 175832865 175832902
refGene_gn=FSIP2;refGene_tx=XM_023259535.1
C1 175835161 175844111
refGene_gn=FSIP2;refGene_tx=XM_023259535.1
C1 175846176 175853252
refGene_gn=FSIP2;refGene_tx=XM_023259535.1
C1 175853253 175855689
refGene_gn=FSIP2;refGene_tx=XM_023259535.1
C1 175858595 175859090
refGene_gn=FSIP2;refGene_tx=XM_023259535.1
C1 175860663 175860764
refGene_gn=FSIP2;refGene_tx=XM_023259535.1
C1 175880033 175880080
refGene_gn=FSIP2;refGene_tx=XM_023259535.1
C1 175883706 175883750
refGene_gn=FSIP2;refGene_tx=XM_023259535.1
C1 175887189 175887259
refGene_gn=FSIP2;refGene_tx=XM_023259535.1
C1 175890033 175890170
refGene_gn=FSIP2;refGene_tx=XM_023259535.1
C1 176397264 176397339

refGene_gn=ZC3H15;refGene_tx=XM_003990925.5
C1 176405067 176405169
refGene_gn=ZC3H15;refGene_tx=XM_003990925.5
C1 176409530 176409642
refGene_gn=ZC3H15;refGene_tx=XM_003990925.5
C1 176410688 176410841
refGene_gn=ZC3H15;refGene_tx=XM_003990925.5
C1 176411283 176411375
refGene_gn=ZC3H15;refGene_tx=XM_003990925.5
C1 176412522 176412705
refGene_gn=ZC3H15;refGene_tx=XM_003990925.5
C1 176414315 176414462
refGene_gn=ZC3H15;refGene_tx=XM_003990925.5
C1 176414626 176414728
refGene_gn=ZC3H15;refGene_tx=XM_003990925.5
C1 176415416 176415540
refGene_gn=ZC3H15;refGene_tx=XM_003990925.5
C1 176416984 176417175
refGene_gn=ZC3H15;refGene_tx=XM_003990925.5
C1 176487620 176487805
refGene_gn=ITGAV;refGene_tx=XM_023259536.1
C1 176499397 176499525
refGene_gn=ITGAV;refGene_tx=XM_023259536.1
C1 176526238 176526330
refGene_gn=ITGAV;refGene_tx=XM_023259536.1
C1 176528724 176528839
refGene_gn=ITGAV;refGene_tx=XM_023259536.1
C1 176533742 176533804
refGene_gn=ITGAV;refGene_tx=XM_023259536.1
C1 176536246 176536292
refGene_gn=ITGAV;refGene_tx=XM_023259536.1
C1 176539280 176539406
refGene_gn=ITGAV;refGene_tx=XM_023259536.1
C1 176541421 176541466
refGene_gn=ITGAV;refGene_tx=XM_023259536.1
C1 176542252 176542296
refGene_gn=ITGAV;refGene_tx=XM_023259536.1
C1 176542384 176542441
refGene_gn=ITGAV;refGene_tx=XM_023259536.1
C1 176544882 176544935
refGene_gn=ITGAV;refGene_tx=XM_023259536.1
C1 176545289 176545492
refGene_gn=ITGAV;refGene_tx=XM_023259536.1
C1 176550546 176550738
refGene_gn=ITGAV;refGene_tx=XM_023259536.1
C1 176552976 176553022
refGene_gn=ITGAV;refGene_tx=XM_023259536.1
C1 176554381 176554489
refGene_gn=ITGAV;refGene_tx=XM_023259536.1
C1 176556412 176556471

refGene_gn=ITGAV;refGene_tx=XM_023259536.1
C1 176557978 176558133
refGene_gn=ITGAV;refGene_tx=XM_023259536.1
C1 176559871 176560009
refGene_gn=ITGAV;refGene_tx=XM_023259536.1
C1 176563876 176563944
refGene_gn=ITGAV;refGene_tx=XM_023259536.1
C1 176565781 176565929
refGene_gn=ITGAV;refGene_tx=XM_023259536.1
C1 176566448 176566541
refGene_gn=ITGAV;refGene_tx=XM_023259536.1
C1 176568869 176568949
refGene_gn=ITGAV;refGene_tx=XM_023259536.1
C1 176569327 176569408
refGene_gn=ITGAV;refGene_tx=XM_023259536.1
C1 176570166 176570272
refGene_gn=ITGAV;refGene_tx=XM_023259536.1
C1 176570908 176571067
refGene_gn=ITGAV;refGene_tx=XM_023259536.1
C1 176571762 176571801
refGene_gn=ITGAV;refGene_tx=XM_023259536.1
C1 176571802 176571874
refGene_gn=ITGAV;refGene_tx=XM_023259536.1
C1 176576345 176576459
refGene_gn=ITGAV;refGene_tx=XM_023259536.1
C1 176576560 176576668
refGene_gn=ITGAV;refGene_tx=XM_023259536.1
C1 176577264 176577387
refGene_gn=ITGAV;refGene_tx=XM_023259536.1
C1 176578106 176578202
refGene_gn=ITGAV;refGene_tx=XM_023259536.1
C1 176605653 176605864
refGene_gn=FAM171B;refGene_tx=XM_023259537.1
C1 176647191 176647425
refGene_gn=FAM171B;refGene_tx=XM_023259537.1
C1 176650335 176650428
refGene_gn=FAM171B;refGene_tx=XM_023259537.1
C1 176653540 176653699
refGene_gn=FAM171B;refGene_tx=XM_023259537.1
C1 176657520 176657691
refGene_gn=FAM171B;refGene_tx=XM_023259537.1
C1 176661250 176661367
refGene_gn=FAM171B;refGene_tx=XM_023259537.1
C1 176666765 176666889
refGene_gn=FAM171B;refGene_tx=XM_023259537.1
C1 176667158 176668503
refGene_gn=FAM171B;refGene_tx=XM_023259537.1
C1 176728268 176729075
refGene_gn=ZSWIM2;refGene_tx=XM_003990927.4;refGene_gn=ZSWIM2;refGene_
tx=XM_006935399.3;refGene_gn=ZSWIM2;refGene_tx=XM_011285417.3;refGene_

gn=ZSWIM2;refGene_tx=XM_023259538.1
C1 176730008 176730162
refGene_gn=ZSWIM2;refGene_tx=XM_003990927.4;refGene_gn=ZSWIM2;refGene_tx=XM_006935399.3;refGene_gn=ZSWIM2;refGene_tx=XM_011285417.3;refGene_gn=ZSWIM2;refGene_tx=XM_023259538.1
C1 176736524 176736637
refGene_gn=ZSWIM2;refGene_tx=XM_003990927.4;refGene_gn=ZSWIM2;refGene_tx=XM_006935399.3;refGene_gn=ZSWIM2;refGene_tx=XM_011285417.3;refGene_gn=ZSWIM2;refGene_tx=XM_023259538.1
C1 176737329 176737414
refGene_gn=ZSWIM2;refGene_tx=XM_003990927.4;refGene_gn=ZSWIM2;refGene_tx=XM_006935399.3;refGene_gn=ZSWIM2;refGene_tx=XM_011285417.3;refGene_gn=ZSWIM2;refGene_tx=XM_023259538.1
C1 176740882 176741134
refGene_gn=ZSWIM2;refGene_tx=XM_003990927.4;refGene_gn=ZSWIM2;refGene_tx=XM_011285417.3;refGene_gn=ZSWIM2;refGene_tx=XM_023259538.1
C1 176751120 176751331
refGene_gn=ZSWIM2;refGene_tx=XM_003990927.4;refGene_gn=ZSWIM2;refGene_tx=XM_006935399.3;refGene_gn=ZSWIM2;refGene_tx=XM_011285417.3;refGene_gn=ZSWIM2;refGene_tx=XM_023259538.1
C1 176756819 176756860
refGene_gn=ZSWIM2;refGene_tx=XM_003990927.4;refGene_gn=ZSWIM2;refGene_tx=XM_006935399.3;refGene_gn=ZSWIM2;refGene_tx=XM_011285417.3;refGene_gn=ZSWIM2;refGene_tx=XM_023259538.1
C1 176758914 176758991
refGene_gn=ZSWIM2;refGene_tx=XM_003990927.4;refGene_gn=ZSWIM2;refGene_tx=XM_006935399.3;refGene_gn=ZSWIM2;refGene_tx=XM_011285417.3;refGene_gn=ZSWIM2;refGene_tx=XM_023259538.1
C1 176759896 176759914
refGene_gn=ZSWIM2;refGene_tx=XM_011285417.3
C1 176760168 176760333
refGene_gn=ZSWIM2;refGene_tx=XM_003990927.4;refGene_gn=ZSWIM2;refGene_tx=XM_006935399.3
C1 176764049 176764094
refGene_gn=ZSWIM2;refGene_tx=XM_023259538.1
C1 177181014 177181230
refGene_gn=CALCRL;refGene_tx=XM_011285418.2;refGene_gn=CALCRL;refGene_tx=XM_011285419.2
C1 177186954 177186996
refGene_gn=CALCRL;refGene_tx=XM_011285418.2;refGene_gn=CALCRL;refGene_tx=XM_011285419.2
C1 177187150 177187369
refGene_gn=CALCRL;refGene_tx=XM_011285418.2;refGene_gn=CALCRL;refGene_tx=XM_011285419.2
C1 177192085 177192152
refGene_gn=CALCRL;refGene_tx=XM_011285418.2;refGene_gn=CALCRL;refGene_tx=XM_011285419.2
C1 177192235 177192296
refGene_gn=CALCRL;refGene_tx=XM_011285418.2;refGene_gn=CALCRL;refGene_tx=XM_011285419.2

C1	177194294	177194448
refGene_gn=CALCRL;refGene_tx=XM_011285418.2;refGene_gn=CALCRL;refGene_tx=XM_011285419.2
C1	177197765	177197892
refGene_gn=CALCRL;refGene_tx=XM_011285418.2;refGene_gn=CALCRL;refGene_tx=XM_011285419.2
C1	177214961	177215053
refGene_gn=CALCRL;refGene_tx=XM_011285418.2;refGene_gn=CALCRL;refGene_tx=XM_011285419.2
C1	177216452	177216565
refGene_gn=CALCRL;refGene_tx=XM_011285418.2;refGene_gn=CALCRL;refGene_tx=XM_011285419.2
C1	177216658	177216769
refGene_gn=CALCRL;refGene_tx=XM_011285418.2;refGene_gn=CALCRL;refGene_tx=XM_011285419.2
C1	177219516	177219652
refGene_gn=CALCRL;refGene_tx=XM_011285418.2;refGene_gn=CALCRL;refGene_tx=XM_011285419.2
C1	177222372	177222426
refGene_gn=CALCRL;refGene_tx=XM_011285418.2;refGene_gn=CALCRL;refGene_tx=XM_011285419.2
C1	177301625	177301732
refGene_gn=TFPI;refGene_tx=XM_006935416.3;refGene_gn=TFPI;refGene_tx=XM_019838357.2;refGene_gn=TFPI;refGene_tx=XM_023259539.1
C1	177302428	177302608
refGene_gn=TFPI;refGene_tx=XM_006935416.3;refGene_gn=TFPI;refGene_tx=XM_019838357.2;refGene_gn=TFPI;refGene_tx=XM_023259539.1
C1	177319610	177319796
refGene_gn=TFPI;refGene_tx=XM_006935416.3;refGene_gn=TFPI;refGene_tx=XM_019838357.2;refGene_gn=TFPI;refGene_tx=XM_023259539.1
C1	177320655	177320832
refGene_gn=TFPI;refGene_tx=XM_006935416.3;refGene_gn=TFPI;refGene_tx=XM_019838357.2;refGene_gn=TFPI;refGene_tx=XM_023259539.1
C1	177324940	177324979
refGene_gn=TFPI;refGene_tx=XM_006935416.3;refGene_gn=TFPI;refGene_tx=XM_019838357.2;refGene_gn=TFPI;refGene_tx=XM_023259539.1
C1	177332963	177333161
refGene_gn=TFPI;refGene_tx=XM_006935416.3;refGene_gn=TFPI;refGene_tx=XM_019838357.2;refGene_gn=TFPI;refGene_tx=XM_023259539.1
C1	177340099	177340208
refGene_gn=TFPI;refGene_tx=XM_006935416.3;refGene_gn=TFPI;refGene_tx=XM_019838357.2;refGene_gn=TFPI;refGene_tx=XM_023259539.1
C1	178140312	178140340
refGene_gn=GULP1;refGene_tx=XM_011285420.3;refGene_gn=GULP1;refGene_tx=XM_011285421.3;refGene_gn=GULP1;refGene_tx=XM_019838359.2;refGene_gn=GULP1;refGene_tx=XM_019838360.2;refGene_gn=GULP1;refGene_tx=XM_019838361.2;refGene_gn=GULP1;refGene_tx=XM_019838363.2;refGene_gn=GULP1;refGene_tx=XM_019838364.2;refGene_gn=GULP1;refGene_tx=XM_023259540.1;refGene_gn=GULP1;refGene_tx=XM_023259541.1
C1	178146756	178146818

refGene_gn=GULP1;refGene_tx=XM_011285420.3;refGene_gn=GULP1;refGene_tx=XM_011285421.3;refGene_gn=GULP1;refGene_tx=XM_019838359.2;refGene_gn=GULP1;refGene_tx=XM_019838360.2;refGene_gn=GULP1;refGene_tx=XM_019838361.2;refGene_gn=GULP1;refGene_tx=XM_019838363.2;refGene_gn=GULP1;refGene_tx=XM_019838364.2;refGene_gn=GULP1;refGene_tx=XM_023259540.1;refGene_gn=GULP1;refGene_tx=XM_023259541.1
C1 178185523 178185595
refGene_gn=GULP1;refGene_tx=XM_011285420.3;refGene_gn=GULP1;refGene_tx=XM_011285421.3;refGene_gn=GULP1;refGene_tx=XM_019838359.2;refGene_gn=GULP1;refGene_tx=XM_019838360.2;refGene_gn=GULP1;refGene_tx=XM_019838361.2;refGene_gn=GULP1;refGene_tx=XM_019838363.2;refGene_gn=GULP1;refGene_tx=XM_019838364.2;refGene_gn=GULP1;refGene_tx=XM_023259540.1;refGene_gn=GULP1;refGene_tx=XM_023259541.1
C1 178191644 178191743
refGene_gn=GULP1;refGene_tx=XM_011285420.3;refGene_gn=GULP1;refGene_tx=XM_011285421.3;refGene_gn=GULP1;refGene_tx=XM_019838359.2;refGene_gn=GULP1;refGene_tx=XM_019838360.2;refGene_gn=GULP1;refGene_tx=XM_019838361.2;refGene_gn=GULP1;refGene_tx=XM_019838363.2;refGene_gn=GULP1;refGene_tx=XM_019838364.2;refGene_gn=GULP1;refGene_tx=XM_023259540.1;refGene_gn=GULP1;refGene_tx=XM_023259541.1
C1 178199361 178199499
refGene_gn=GULP1;refGene_tx=XM_011285420.3;refGene_gn=GULP1;refGene_tx=XM_011285421.3;refGene_gn=GULP1;refGene_tx=XM_019838359.2;refGene_gn=GULP1;refGene_tx=XM_019838360.2;refGene_gn=GULP1;refGene_tx=XM_019838361.2;refGene_gn=GULP1;refGene_tx=XM_019838363.2;refGene_gn=GULP1;refGene_tx=XM_019838364.2;refGene_gn=GULP1;refGene_tx=XM_023259540.1;refGene_gn=GULP1;refGene_tx=XM_023259541.1
C1 178214807 178214924
refGene_gn=GULP1;refGene_tx=XM_011285420.3;refGene_gn=GULP1;refGene_tx=XM_011285421.3;refGene_gn=GULP1;refGene_tx=XM_019838359.2;refGene_gn=GULP1;refGene_tx=XM_019838360.2;refGene_gn=GULP1;refGene_tx=XM_019838361.2;refGene_gn=GULP1;refGene_tx=XM_019838363.2;refGene_gn=GULP1;refGene_tx=XM_019838364.2;refGene_gn=GULP1;refGene_tx=XM_023259540.1;refGene_gn=GULP1;refGene_tx=XM_023259541.1
C1 178215120 178215201
refGene_gn=GULP1;refGene_tx=XM_011285420.3;refGene_gn=GULP1;refGene_tx=XM_019838359.2;refGene_gn=GULP1;refGene_tx=XM_019838360.2;refGene_gn=GULP1;refGene_tx=XM_019838361.2;refGene_gn=GULP1;refGene_tx=XM_019838363.2;refGene_gn=GULP1;refGene_tx=XM_023259540.1;refGene_gn=GULP1;refGene_tx=XM_023259541.1
C1 178215600 178215693
refGene_gn=GULP1;refGene_tx=XM_011285420.3;refGene_gn=GULP1;refGene_tx=XM_011285421.3;refGene_gn=GULP1;refGene_tx=XM_019838359.2;refGene_gn=GULP1;refGene_tx=XM_019838360.2;refGene_gn=GULP1;refGene_tx=XM_019838361.2;refGene_gn=GULP1;refGene_tx=XM_019838363.2;refGene_gn=GULP1;refGene_tx=XM_019838364.2;refGene_gn=GULP1;refGene_tx=XM_023259540.1;refGene_gn=GULP1;refGene_tx=XM_023259541.1
C1 178227707 178227935
refGene_gn=GULP1;refGene_tx=XM_019838359.2;refGene_gn=GULP1;refGene_tx=XM_019838360.2;refGene_gn=GULP1;refGene_tx=XM_019838361.2;refGene_gn=

GULP1;refGene_tx=XM_019838363.2;refGene_gn=GULP1;refGene_tx=XM_0198383
64.2;refGene_gn=GULP1;refGene_tx=XM_023259540.1;refGene_gn=GULP1;refGe
ne_tx=XM_023259541.1
C1	178229564	178229703
refGene_gn=GULP1;refGene_tx=XM_011285420.3;refGene_gn=GULP1;refGene_tx
=XM_011285421.3;refGene_gn=GULP1;refGene_tx=XM_019838359.2;refGene_gn=
GULP1;refGene_tx=XM_019838360.2;refGene_gn=GULP1;refGene_tx=XM_0198383
61.2;refGene_gn=GULP1;refGene_tx=XM_019838363.2;refGene_gn=GULP1;refGe
ne_tx=XM_019838364.2;refGene_gn=GULP1;refGene_tx=XM_023259540.1;refGen
e_gn=GULP1;refGene_tx=XM_023259541.1
C1	178232666	178232761
refGene_gn=GULP1;refGene_tx=XM_011285420.3;refGene_gn=GULP1;refGene_tx
=XM_011285421.3;refGene_gn=GULP1;refGene_tx=XM_019838359.2;refGene_gn=
GULP1;refGene_tx=XM_019838360.2;refGene_gn=GULP1;refGene_tx=XM_0198383
61.2;refGene_gn=GULP1;refGene_tx=XM_019838363.2;refGene_gn=GULP1;refGe
ne_tx=XM_019838364.2;refGene_gn=GULP1;refGene_tx=XM_023259540.1;refGen
e_gn=GULP1;refGene_tx=XM_023259541.1
C1	178234588	178234648
refGene_gn=GULP1;refGene_tx=XM_019838363.2
C1	178239754	178239826
refGene_gn=GULP1;refGene_tx=XM_011285420.3;refGene_gn=GULP1;refGene_tx
=XM_011285421.3;refGene_gn=GULP1;refGene_tx=XM_019838359.2;refGene_gn=
GULP1;refGene_tx=XM_019838360.2;refGene_gn=GULP1;refGene_tx=XM_0198383
61.2;refGene_gn=GULP1;refGene_tx=XM_019838364.2;refGene_gn=GULP1;refGe
ne_tx=XM_023259540.1;refGene_gn=GULP1;refGene_tx=XM_023259541.1
C1	178525408	178525487
refGene_gn=COL3A1;refGene_tx=XM_003990961.3
C1	178535809	178536012
refGene_gn=COL3A1;refGene_tx=XM_003990961.3
C1	178536496	178536547
refGene_gn=COL3A1;refGene_tx=XM_003990961.3
C1	178536985	178537099
refGene_gn=COL3A1;refGene_tx=XM_003990961.3
C1	178538410	178538491
refGene_gn=COL3A1;refGene_tx=XM_003990961.3
C1	178540508	178540562
refGene_gn=COL3A1;refGene_tx=XM_003990961.3
C1	178541221	178541275
refGene_gn=COL3A1;refGene_tx=XM_003990961.3
C1	178542059	178542113
refGene_gn=COL3A1;refGene_tx=XM_003990961.3
C1	178542758	178542812
refGene_gn=COL3A1;refGene_tx=XM_003990961.3
C1	178542979	178543033
refGene_gn=COL3A1;refGene_tx=XM_003990961.3
C1	178543726	178543780
refGene_gn=COL3A1;refGene_tx=XM_003990961.3
C1	178544202	178544247
refGene_gn=COL3A1;refGene_tx=XM_003990961.3
C1	178544385	178544439

refGene_gn=COL3A1;refGene_tx=XM_003990961.3
C1 178544901 178544946
refGene_gn=COL3A1;refGene_tx=XM_003990961.3
C1 178545516 178545570
refGene_gn=COL3A1;refGene_tx=XM_003990961.3
C1 178545971 178546070
refGene_gn=COL3A1;refGene_tx=XM_003990961.3
C1 178546616 178546661
refGene_gn=COL3A1;refGene_tx=XM_003990961.3
C1 178546815 178546914
refGene_gn=COL3A1;refGene_tx=XM_003990961.3
C1 178547114 178547168
refGene_gn=COL3A1;refGene_tx=XM_003990961.3
C1 178547257 178547365
refGene_gn=COL3A1;refGene_tx=XM_003990961.3
C1 178547579 178547633
refGene_gn=COL3A1;refGene_tx=XM_003990961.3
C1 178548235 178548334
refGene_gn=COL3A1;refGene_tx=XM_003990961.3
C1 178548646 178548700
refGene_gn=COL3A1;refGene_tx=XM_003990961.3
C1 178548858 178548957
refGene_gn=COL3A1;refGene_tx=XM_003990961.3
C1 178549335 178549389
refGene_gn=COL3A1;refGene_tx=XM_003990961.3
C1 178549497 178549551
refGene_gn=COL3A1;refGene_tx=XM_003990961.3
C1 178549882 178549936
refGene_gn=COL3A1;refGene_tx=XM_003990961.3
C1 178550711 178550765
refGene_gn=COL3A1;refGene_tx=XM_003990961.3
C1 178551093 178551138
refGene_gn=COL3A1;refGene_tx=XM_003990961.3
C1 178551726 178551825
refGene_gn=COL3A1;refGene_tx=XM_003990961.3
C1 178551912 178552020
refGene_gn=COL3A1;refGene_tx=XM_003990961.3
C1 178552279 178552333
refGene_gn=COL3A1;refGene_tx=XM_003990961.3
C1 178554050 178554104
refGene_gn=COL3A1;refGene_tx=XM_003990961.3
C1 178554190 178554244
refGene_gn=COL3A1;refGene_tx=XM_003990961.3
C1 178554940 178554994
refGene_gn=COL3A1;refGene_tx=XM_003990961.3
C1 178555815 178555923
refGene_gn=COL3A1;refGene_tx=XM_003990961.3
C1 178556359 178556413
refGene_gn=COL3A1;refGene_tx=XM_003990961.3
C1 178556638 178556692

refGene_gn=COL3A1;refGene_tx=XM_003990961.3
C1 178556867 178557029
refGene_gn=COL3A1;refGene_tx=XM_003990961.3
C1 178557143 178557251
refGene_gn=COL3A1;refGene_tx=XM_003990961.3
C1 178558175 178558283
refGene_gn=COL3A1;refGene_tx=XM_003990961.3
C1 178559436 178559490
refGene_gn=COL3A1;refGene_tx=XM_003990961.3
C1 178559585 178559693
refGene_gn=COL3A1;refGene_tx=XM_003990961.3
C1 178560127 178560181
refGene_gn=COL3A1;refGene_tx=XM_003990961.3
C1 178560558 178560666
refGene_gn=COL3A1;refGene_tx=XM_003990961.3
C1 178560965 178561019
refGene_gn=COL3A1;refGene_tx=XM_003990961.3
C1 178561115 178561223
refGene_gn=COL3A1;refGene_tx=XM_003990961.3
C1 178562085 178562383
refGene_gn=COL3A1;refGene_tx=XM_003990961.3
C1 178563452 178563640
refGene_gn=COL3A1;refGene_tx=XM_003990961.3
C1 178564133 178564376
refGene_gn=COL3A1;refGene_tx=XM_003990961.3
C1 178565129 178565276
refGene_gn=COL3A1;refGene_tx=XM_003990961.3
C1 178582903 178583050
refGene_gn=COL5A2;refGene_tx=XM_003990962.4
C1 178583809 178584049
refGene_gn=COL5A2;refGene_tx=XM_003990962.4
C1 178586330 178586518
refGene_gn=COL5A2;refGene_tx=XM_003990962.4
C1 178587240 178587532
refGene_gn=COL5A2;refGene_tx=XM_003990962.4
C1 178589667 178589775
refGene_gn=COL5A2;refGene_tx=XM_003990962.4
C1 178590788 178590842
refGene_gn=COL5A2;refGene_tx=XM_003990962.4
C1 178591210 178591318
refGene_gn=COL5A2;refGene_tx=XM_003990962.4
C1 178593391 178593445
refGene_gn=COL5A2;refGene_tx=XM_003990962.4
C1 178593941 178594049
refGene_gn=COL5A2;refGene_tx=XM_003990962.4
C1 178595940 178595994
refGene_gn=COL5A2;refGene_tx=XM_003990962.4
C1 178597465 178597573
refGene_gn=COL5A2;refGene_tx=XM_003990962.4
C1 178599097 178599205

refGene_gn=COL5A2;refGene_tx=XM_003990962.4
C1 178600307 178600469
refGene_gn=COL5A2;refGene_tx=XM_003990962.4
C1 178601197 178601251
refGene_gn=COL5A2;refGene_tx=XM_003990962.4
C1 178601784 178601838
refGene_gn=COL5A2;refGene_tx=XM_003990962.4
C1 178601955 178602063
refGene_gn=COL5A2;refGene_tx=XM_003990962.4
C1 178602471 178602525
refGene_gn=COL5A2;refGene_tx=XM_003990962.4
C1 178602944 178602998
refGene_gn=COL5A2;refGene_tx=XM_003990962.4
C1 178603216 178603270
refGene_gn=COL5A2;refGene_tx=XM_003990962.4
C1 178605472 178605526
refGene_gn=COL5A2;refGene_tx=XM_003990962.4
C1 178605862 178605970
refGene_gn=COL5A2;refGene_tx=XM_003990962.4
C1 178606977 178607076
refGene_gn=COL5A2;refGene_tx=XM_003990962.4
C1 178607864 178607909
refGene_gn=COL5A2;refGene_tx=XM_003990962.4
C1 178610212 178610266
refGene_gn=COL5A2;refGene_tx=XM_003990962.4
C1 178611022 178611076
refGene_gn=COL5A2;refGene_tx=XM_003990962.4
C1 178612468 178612522
refGene_gn=COL5A2;refGene_tx=XM_003990962.4
C1 178612634 178612688
refGene_gn=COL5A2;refGene_tx=XM_003990962.4
C1 178612796 178612895
refGene_gn=COL5A2;refGene_tx=XM_003990962.4
C1 178613613 178613667
refGene_gn=COL5A2;refGene_tx=XM_003990962.4
C1 178615318 178615417
refGene_gn=COL5A2;refGene_tx=XM_003990962.4
C1 178615978 178616032
refGene_gn=COL5A2;refGene_tx=XM_003990962.4
C1 178617109 178617217
refGene_gn=COL5A2;refGene_tx=XM_003990962.4
C1 178617496 178617550
refGene_gn=COL5A2;refGene_tx=XM_003990962.4
C1 178618765 178618864
refGene_gn=COL5A2;refGene_tx=XM_003990962.4
C1 178618981 178619026
refGene_gn=COL5A2;refGene_tx=XM_003990962.4
C1 178619514 178619613
refGene_gn=COL5A2;refGene_tx=XM_003990962.4
C1 178622089 178622143

refGene_gn=COL5A2;refGene_tx=XM_003990962.4
C1 178623242 178623287
refGene_gn=COL5A2;refGene_tx=XM_003990962.4
C1 178626206 178626260
refGene_gn=COL5A2;refGene_tx=XM_003990962.4
C1 178626682 178626727
refGene_gn=COL5A2;refGene_tx=XM_003990962.4
C1 178627895 178627949
refGene_gn=COL5A2;refGene_tx=XM_003990962.4
C1 178631093 178631147
refGene_gn=COL5A2;refGene_tx=XM_003990962.4
C1 178634039 178634093
refGene_gn=COL5A2;refGene_tx=XM_003990962.4
C1 178635167 178635221
refGene_gn=COL5A2;refGene_tx=XM_003990962.4
C1 178635863 178635917
refGene_gn=COL5A2;refGene_tx=XM_003990962.4
C1 178636866 178636911
refGene_gn=COL5A2;refGene_tx=XM_003990962.4
C1 178639295 178639373
refGene_gn=COL5A2;refGene_tx=XM_003990962.4
C1 178640938 178641049
refGene_gn=COL5A2;refGene_tx=XM_003990962.4
C1 178646204 178646258
refGene_gn=COL5A2;refGene_tx=XM_003990962.4
C1 178647226 178647259
refGene_gn=COL5A2;refGene_tx=XM_003990962.4
C1 178648552 178648585
refGene_gn=COL5A2;refGene_tx=XM_003990962.4
C1 178652894 178652908
refGene_gn=COL5A2;refGene_tx=XM_003990962.4
C1 178655901 178656126
refGene_gn=COL5A2;refGene_tx=XM_003990962.4
C1 178729239 178729336
refGene_gn=COL5A2;refGene_tx=XM_003990962.4
C1 178971309 178971395
refGene_gn=WDR75;refGene_tx=XM_003990963.5
C1 178976974 178977104
refGene_gn=WDR75;refGene_tx=XM_003990963.5
C1 178977468 178977480
refGene_gn=WDR75;refGene_tx=XM_006935420.3
C1 178979869 178979935
refGene_gn=WDR75;refGene_tx=XM_003990963.5;refGene_gn=WDR75;refGene_tx
=XM_006935420.3
C1 178980807 178980898
refGene_gn=WDR75;refGene_tx=XM_003990963.5;refGene_gn=WDR75;refGene_tx
=XM_006935420.3
C1 178984069 178984194
refGene_gn=WDR75;refGene_tx=XM_003990963.5;refGene_gn=WDR75;refGene_tx
=XM_006935420.3

C1 178985762 178985833
refGene_gn=WDR75;refGene_tx=XM_003990963.5;refGene_gn=WDR75;refGene_tx=XM_006935420.3
C1 178986824 178986944
refGene_gn=WDR75;refGene_tx=XM_003990963.5;refGene_gn=WDR75;refGene_tx=XM_006935420.3
C1 178987452 178987541
refGene_gn=WDR75;refGene_tx=XM_003990963.5;refGene_gn=WDR75;refGene_tx=XM_006935420.3
C1 178989454 178989613
refGene_gn=WDR75;refGene_tx=XM_003990963.5;refGene_gn=WDR75;refGene_tx=XM_006935420.3
C1 178990420 178990480
refGene_gn=WDR75;refGene_tx=XM_003990963.5;refGene_gn=WDR75;refGene_tx=XM_006935420.3
C1 178990565 178990681
refGene_gn=WDR75;refGene_tx=XM_003990963.5;refGene_gn=WDR75;refGene_tx=XM_006935420.3
C1 178991721 178991897
refGene_gn=WDR75;refGene_tx=XM_003990963.5;refGene_gn=WDR75;refGene_tx=XM_006935420.3
C1 178993044 178993202
refGene_gn=WDR75;refGene_tx=XM_003990963.5;refGene_gn=WDR75;refGene_tx=XM_006935420.3
C1 178993588 178993769
refGene_gn=WDR75;refGene_tx=XM_003990963.5;refGene_gn=WDR75;refGene_tx=XM_006935420.3
C1 178994598 178994693
refGene_gn=WDR75;refGene_tx=XM_003990963.5;refGene_gn=WDR75;refGene_tx=XM_006935420.3
C1 178995506 178995602
refGene_gn=WDR75;refGene_tx=XM_003990963.5;refGene_gn=WDR75;refGene_tx=XM_006935420.3
C1 178996144 178996314
refGene_gn=WDR75;refGene_tx=XM_003990963.5;refGene_gn=WDR75;refGene_tx=XM_006935420.3
C1 178996848 178996908
refGene_gn=WDR75;refGene_tx=XM_003990963.5;refGene_gn=WDR75;refGene_tx=XM_006935420.3
C1 178999787 178999934
refGene_gn=WDR75;refGene_tx=XM_003990963.5;refGene_gn=WDR75;refGene_tx=XM_006935420.3
C1 179000255 179000347
refGene_gn=WDR75;refGene_tx=XM_003990963.5;refGene_gn=WDR75;refGene_tx=XM_006935420.3
C1 179001487 179001692
refGene_gn=WDR75;refGene_tx=XM_003990963.5;refGene_gn=WDR75;refGene_tx=XM_006935420.3
C1 179106412 179106726
refGene_gn=SLC40A1;refGene_tx=XM_003990965.5

C1 179108669 179109326
refGene_gn=SLC40A1;refGene_tx=XM_003990965.5
C1 179110238 179110484
refGene_gn=SLC40A1;refGene_tx=XM_003990965.5
C1 179117440 179117567
refGene_gn=SLC40A1;refGene_tx=XM_003990965.5
C1 179118596 179118712
refGene_gn=SLC40A1;refGene_tx=XM_003990965.5
C1 179120909 179121069
refGene_gn=SLC40A1;refGene_tx=XM_003990965.5
C1 179126905 179126973
refGene_gn=SLC40A1;refGene_tx=XM_003990965.5
C1 179127466 179127509
refGene_gn=SLC40A1;refGene_tx=XM_003990965.5
C1 179236472 179236562
refGene_gn=LOC101101577;refGene_tx=XM_023259543.1
C1 179238713 179238767
refGene_gn=LOC101101577;refGene_tx=XM_023259543.1
C1 179239975 179240051
refGene_gn=LOC101101577;refGene_tx=XM_023259543.1
C1 179240865 179240936
refGene_gn=LOC101101577;refGene_tx=XM_023259543.1
C1 179240976 179242425
refGene_gn=ASNSD1;refGene_tx=XM_023259542.1
C1 179242426 179242477
refGene_gn=ASNSD1;refGene_tx=XM_023259542.1
C1 179242754 179242936
refGene_gn=ASNSD1;refGene_tx=XM_023259542.1
C1 179245130 179245416
refGene_gn=ASNSD1;refGene_tx=XM_023259542.1
C1 179250899 179251500
refGene_gn=ANKAR;refGene_tx=XM_019838366.2
C1 179260833 179261271
refGene_gn=ANKAR;refGene_tx=XM_019838366.2;refGene_gn=ANKAR;refGene_tx
=XM_023259544.1
C1 179262887 179263051
refGene_gn=ANKAR;refGene_tx=XM_019838366.2;refGene_gn=ANKAR;refGene_tx
=XM_023259544.1
C1 179263971 179264075
refGene_gn=ANKAR;refGene_tx=XM_019838366.2;refGene_gn=ANKAR;refGene_tx
=XM_023259544.1
C1 179267962 179268143
refGene_gn=ANKAR;refGene_tx=XM_019838366.2;refGene_gn=ANKAR;refGene_tx
=XM_023259544.1
C1 179269309 179269529
refGene_gn=ANKAR;refGene_tx=XM_019838366.2;refGene_gn=ANKAR;refGene_tx
=XM_023259544.1
C1 179271293 179271329
refGene_gn=ANKAR;refGene_tx=XM_019838366.2;refGene_gn=ANKAR;refGene_tx
=XM_023259544.1

C1 179278295 179278497
refGene_gn=ANKAR;refGene_tx=XM_019838366.2;refGene_gn=ANKAR;refGene_tx
=XM_023259544.1
C1 179281213 179281422
refGene_gn=ANKAR;refGene_tx=XM_019838366.2;refGene_gn=ANKAR;refGene_tx
=XM_023259544.1
C1 179283572 179283677
refGene_gn=ANKAR;refGene_tx=XM_019838366.2;refGene_gn=ANKAR;refGene_tx
=XM_023259544.1
C1 179285825 179286067
refGene_gn=ANKAR;refGene_tx=XM_019838366.2;refGene_gn=ANKAR;refGene_tx
=XM_023259544.1
C1 179290428 179290597
refGene_gn=ANKAR;refGene_tx=XM_019838366.2;refGene_gn=ANKAR;refGene_tx
=XM_023259544.1
C1 179296497 179296739
refGene_gn=ANKAR;refGene_tx=XM_019838366.2;refGene_gn=ANKAR;refGene_tx
=XM_023259544.1
C1 179296908 179297062
refGene_gn=ANKAR;refGene_tx=XM_019838366.2;refGene_gn=ANKAR;refGene_tx
=XM_023259544.1
C1 179297146 179297308
refGene_gn=ANKAR;refGene_tx=XM_019838366.2;refGene_gn=ANKAR;refGene_tx
=XM_023259544.1
C1 179302671 179302778
refGene_gn=ANKAR;refGene_tx=XM_019838366.2;refGene_gn=ANKAR;refGene_tx
=XM_023259544.1
C1 179304428 179304551
refGene_gn=ANKAR;refGene_tx=XM_019838366.2;refGene_gn=ANKAR;refGene_tx
=XM_023259544.1
C1 179311340 179311499
refGene_gn=ANKAR;refGene_tx=XM_019838366.2;refGene_gn=ANKAR;refGene_tx
=XM_023259544.1
C1 179312181 179312299
refGene_gn=ANKAR;refGene_tx=XM_019838366.2;refGene_gn=ANKAR;refGene_tx
=XM_023259544.1
C1 179314942 179315052
refGene_gn=ANKAR;refGene_tx=XM_019838366.2;refGene_gn=ANKAR;refGene_tx
=XM_023259544.1
C1 179316561 179316758
refGene_gn=ANKAR;refGene_tx=XM_019838366.2;refGene_gn=ANKAR;refGene_tx
=XM_023259544.1
C1 179319329 179319376
refGene_gn=ANKAR;refGene_tx=XM_019838366.2;refGene_gn=ANKAR;refGene_tx
=XM_023259544.1
C1 179320656 179320904
refGene_gn=ANKAR;refGene_tx=XM_019838366.2;refGene_gn=ANKAR;refGene_tx
=XM_023259544.1
C1 179323374 179323453
refGene_gn=LOC101080649;refGene_tx=XM_019838369.2

C1 179324876 179324948
refGene_gn=LOC101080649;refGene_tx=XM_019838369.2
C1 179325071 179325202
refGene_gn=LOC101080649;refGene_tx=XM_019838369.2
C1 179326011 179326160
refGene_gn=LOC101080649;refGene_tx=XM_019838369.2
C1 179326236 179326441
refGene_gn=LOC101080649;refGene_tx=XM_019838369.2
C1 179327219 179327607
refGene_gn=LOC101080649;refGene_tx=XM_019838369.2
C1 179331830 179332071
refGene_gn=LOC101080649;refGene_tx=XM_019838369.2
C1 179333211 179333366
refGene_gn=LOC101080649;refGene_tx=XM_019838369.2
C1 179342163 179342299
refGene_gn=LOC101080649;refGene_tx=XM_003990969.5
C1 179345860 179346012
refGene_gn=LOC101080649;refGene_tx=XM_003990969.5;refGene_gn=LOC101080649;refGene_tx=XM_019838369.2
C1 179350767 179350941
refGene_gn=LOC101080649;refGene_tx=XM_003990969.5;refGene_gn=LOC101080649;refGene_tx=XM_019838369.2
C1 179361004 179361136
refGene_gn=PMS1;refGene_tx=XM_006935421.3;refGene_gn=PMS1;refGene_tx=XM_011285422.3;refGene_gn=PMS1;refGene_tx=XM_011285424.2;refGene_gn=PMS1;refGene_tx=XM_011285427.2
C1 179371176 179371359
refGene_gn=PMS1;refGene_tx=XM_006935421.3;refGene_gn=PMS1;refGene_tx=XM_011285422.3;refGene_gn=PMS1;refGene_tx=XM_011285424.2;refGene_gn=PMS1;refGene_tx=XM_011285427.2
C1 179381385 179381488
refGene_gn=PMS1;refGene_tx=XM_006935421.3;refGene_gn=PMS1;refGene_tx=XM_011285422.3;refGene_gn=PMS1;refGene_tx=XM_011285424.2;refGene_gn=PMS1;refGene_tx=XM_011285427.2
C1 179391586 179391750
refGene_gn=PMS1;refGene_tx=XM_006935421.3;refGene_gn=PMS1;refGene_tx=XM_011285422.3;refGene_gn=PMS1;refGene_tx=XM_011285424.2;refGene_gn=PMS1;refGene_tx=XM_011285427.2
C1 179420067 179420184
refGene_gn=PMS1;refGene_tx=XM_006935421.3;refGene_gn=PMS1;refGene_tx=XM_011285422.3;refGene_gn=PMS1;refGene_tx=XM_011285424.2;refGene_gn=PMS1;refGene_tx=XM_011285427.2
C1 179430662 179430785
refGene_gn=PMS1;refGene_tx=XM_006935421.3;refGene_gn=PMS1;refGene_tx=XM_011285422.3;refGene_gn=PMS1;refGene_tx=XM_011285424.2;refGene_gn=PMS1;refGene_tx=XM_011285427.2
C1 179431321 179431465
refGene_gn=PMS1;refGene_tx=XM_006935421.3;refGene_gn=PMS1;refGene_tx=XM_011285422.3;refGene_gn=PMS1;refGene_tx=XM_011285424.2;refGene_gn=PMS1;refGene_tx=XM_011285427.2

C1	179431623	179432495
refGene_gn=PMS1;refGene_tx=XM_006935421.3;refGene_gn=PMS1;refGene_tx=XM_011285422.3;refGene_gn=PMS1;refGene_tx=XM_011285424.2;refGene_gn=PMS1;refGene_tx=XM_011285427.2
C1	179440731	179441214
refGene_gn=PMS1;refGene_tx=XM_006935421.3;refGene_gn=PMS1;refGene_tx=XM_011285422.3;refGene_gn=PMS1;refGene_tx=XM_011285424.2
C1	179444941	179445072
refGene_gn=PMS1;refGene_tx=XM_006935421.3;refGene_gn=PMS1;refGene_tx=XM_011285422.3;refGene_gn=PMS1;refGene_tx=XM_011285424.2;refGene_gn=PMS1;refGene_tx=XM_011285427.2
C1	179449449	179449610
refGene_gn=PMS1;refGene_tx=XM_006935421.3;refGene_gn=PMS1;refGene_tx=XM_011285422.3;refGene_gn=PMS1;refGene_tx=XM_011285424.2;refGene_gn=PMS1;refGene_tx=XM_011285427.2
C1	179452993	179453158
refGene_gn=PMS1;refGene_tx=XM_006935421.3;refGene_gn=PMS1;refGene_tx=XM_011285422.3;refGene_gn=PMS1;refGene_tx=XM_011285424.2;refGene_gn=PMS1;refGene_tx=XM_011285427.2
C1	179583073	179583454
refGene_gn=MSTN;refGene_tx=XM_003990972.4
C1	179586848	179587222
refGene_gn=MSTN;refGene_tx=XM_003990972.4
C1	179588993	179589366
refGene_gn=MSTN;refGene_tx=XM_003990972.4
C1	179715299	179715587
refGene_gn=CC1H2orf88;refGene_tx=XM_006935430.2;refGene_gn=CC1H2orf88;refGene_tx=XM_006935431.4;refGene_gn=CC1H2orf88;refGene_tx=XM_006935432.3;refGene_gn=CC1H2orf88;refGene_tx=XM_011285429.3;refGene_gn=CC1H2orf88;refGene_tx=XM_019838375.2;refGene_gn=CC1H2orf88;refGene_tx=XM_019838376.2;refGene_gn=CC1H2orf88;refGene_tx=XM_023259551.1
C1	179720418	179720534
refGene_gn=HIBCH;refGene_tx=XM_003990974.4;refGene_gn=HIBCH;refGene_tx=XM_019838371.2;refGene_gn=HIBCH;refGene_tx=XM_019838372.2;refGene_gn=HIBCH;refGene_tx=XM_019838373.2
C1	179723843	179723877
refGene_gn=HIBCH;refGene_tx=XM_003990974.4;refGene_gn=HIBCH;refGene_tx=XM_019838371.2;refGene_gn=HIBCH;refGene_tx=XM_019838372.2;refGene_gn=HIBCH;refGene_tx=XM_019838373.2
C1	179728810	179728930
refGene_gn=HIBCH;refGene_tx=XM_003990974.4;refGene_gn=HIBCH;refGene_tx=XM_019838371.2;refGene_gn=HIBCH;refGene_tx=XM_019838372.2;refGene_gn=HIBCH;refGene_tx=XM_019838373.2
C1	179743217	179743299
refGene_gn=HIBCH;refGene_tx=XM_003990974.4;refGene_gn=HIBCH;refGene_tx=XM_019838371.2;refGene_gn=HIBCH;refGene_tx=XM_019838372.2;refGene_gn=HIBCH;refGene_tx=XM_019838373.2
C1	179743988	179744047
refGene_gn=HIBCH;refGene_tx=XM_003990974.4;refGene_gn=HIBCH;refGene_tx=XM_019838371.2;refGene_gn=HIBCH;refGene_tx=XM_019838372.2;refGene_gn=

HIBCH;refGene_tx=XM_019838373.2
C1 179745003 179745099
refGene_gn=HIBCH;refGene_tx=XM_019838371.2;refGene_gn=HIBCH;refGene_tx=XM_019838372.2;refGene_gn=HIBCH;refGene_tx=XM_019838373.2
C1 179748877 179748964
refGene_gn=HIBCH;refGene_tx=XM_003990974.4;refGene_gn=HIBCH;refGene_tx=XM_019838371.2;refGene_gn=HIBCH;refGene_tx=XM_019838372.2;refGene_gn=HIBCH;refGene_tx=XM_019838373.2
C1 179751044 179751190
refGene_gn=HIBCH;refGene_tx=XM_003990974.4;refGene_gn=HIBCH;refGene_tx=XM_019838371.2;refGene_gn=HIBCH;refGene_tx=XM_019838372.2;refGene_gn=HIBCH;refGene_tx=XM_019838373.2
C1 179759102 179759181
refGene_gn=HIBCH;refGene_tx=XM_003990974.4;refGene_gn=HIBCH;refGene_tx=XM_019838371.2;refGene_gn=HIBCH;refGene_tx=XM_019838372.2;refGene_gn=HIBCH;refGene_tx=XM_019838373.2
C1 179773528 179773581
refGene_gn=HIBCH;refGene_tx=XM_003990974.4;refGene_gn=HIBCH;refGene_tx=XM_019838371.2;refGene_gn=HIBCH;refGene_tx=XM_019838372.2;refGene_gn=HIBCH;refGene_tx=XM_019838373.2
C1 179779122 179779203
refGene_gn=HIBCH;refGene_tx=XM_003990974.4;refGene_gn=HIBCH;refGene_tx=XM_019838371.2;refGene_gn=HIBCH;refGene_tx=XM_019838372.2;refGene_gn=HIBCH;refGene_tx=XM_019838373.2
C1 179782122 179782207
refGene_gn=HIBCH;refGene_tx=XM_003990974.4;refGene_gn=HIBCH;refGene_tx=XM_019838371.2;refGene_gn=HIBCH;refGene_tx=XM_019838372.2;refGene_gn=HIBCH;refGene_tx=XM_019838373.2
C1 179783922 179784063
refGene_gn=HIBCH;refGene_tx=XM_003990974.4;refGene_gn=HIBCH;refGene_tx=XM_019838371.2;refGene_gn=HIBCH;refGene_tx=XM_019838372.2;refGene_gn=HIBCH;refGene_tx=XM_019838373.2
C1 179804058 179804076
refGene_gn=HIBCH;refGene_tx=XM_019838371.2
C1 179805457 179805500
refGene_gn=HIBCH;refGene_tx=XM_003990974.4
C1 179814428 179814463
refGene_gn=HIBCH;refGene_tx=XM_003990974.4
C1 179847773 179847977
refGene_gn=INPP1;refGene_tx=XM_023259545.1;refGene_gn=INPP1;refGene_tx=XM_023259546.1;refGene_gn=INPP1;refGene_tx=XM_023259547.1;refGene_gn=INPP1;refGene_tx=XM_023259548.1
C1 179850370 179850431
refGene_gn=INPP1;refGene_tx=XM_023259545.1;refGene_gn=INPP1;refGene_tx=XM_023259546.1;refGene_gn=INPP1;refGene_tx=XM_023259547.1;refGene_gn=INPP1;refGene_tx=XM_023259548.1;refGene_gn=INPP1;refGene_tx=XM_023259549.1
C1 179853472 179853689
refGene_gn=INPP1;refGene_tx=XM_023259545.1;refGene_gn=INPP1;refGene_tx=XM_023259546.1;refGene_gn=INPP1;refGene_tx=XM_023259547.1;refGene_gn=

INPP1;refGene_tx=XM_023259548.1;refGene_gn=INPP1;refGene_tx=XM_0232595
49.1
C1 179855953 179856128
refGene_gn=INPP1;refGene_tx=XM_023259545.1;refGene_gn=INPP1;refGene_tx
=XM_023259546.1;refGene_gn=INPP1;refGene_tx=XM_023259547.1;refGene_gn=
INPP1;refGene_tx=XM_023259548.1;refGene_gn=INPP1;refGene_tx=XM_0232595
49.1
C1 179857616 179858184
refGene_gn=INPP1;refGene_tx=XM_023259545.1;refGene_gn=INPP1;refGene_tx
=XM_023259546.1;refGene_gn=INPP1;refGene_tx=XM_023259547.1;refGene_gn=
INPP1;refGene_tx=XM_023259548.1;refGene_gn=INPP1;refGene_tx=XM_0232595
49.1
C1 179925737 179927284
refGene_gn=MFSD6;refGene_tx=XM_003990976.4;refGene_gn=MFSD6;refGene_tx
=XM_023259552.1;refGene_gn=MFSD6;refGene_tx=XM_023259553.1
C1 179951120 179951218
refGene_gn=MFSD6;refGene_tx=XM_003990976.4;refGene_gn=MFSD6;refGene_tx
=XM_023259552.1;refGene_gn=MFSD6;refGene_tx=XM_023259553.1
C1 179969735 179969897
refGene_gn=MFSD6;refGene_tx=XM_003990976.4;refGene_gn=MFSD6;refGene_tx
=XM_023259552.1;refGene_gn=MFSD6;refGene_tx=XM_023259553.1
C1 179970804 179970903
refGene_gn=MFSD6;refGene_tx=XM_003990976.4;refGene_gn=MFSD6;refGene_tx
=XM_023259552.1;refGene_gn=MFSD6;refGene_tx=XM_023259553.1
C1 179972033 179972314
refGene_gn=MFSD6;refGene_tx=XM_003990976.4;refGene_gn=MFSD6;refGene_tx
=XM_023259552.1;refGene_gn=MFSD6;refGene_tx=XM_023259553.1
C1 179974292 179974317
refGene_gn=MFSD6;refGene_tx=XM_023259553.1
C1 179974472 179974673
refGene_gn=MFSD6;refGene_tx=XM_003990976.4;refGene_gn=MFSD6;refGene_tx
=XM_023259552.1;refGene_gn=MFSD6;refGene_tx=XM_023259553.1
C1 179983663 179983787
refGene_gn=NEMP2;refGene_tx=XM_023259554.1
C1 179984483 179984660
refGene_gn=NEMP2;refGene_tx=XM_023259554.1;refGene_gn=NEMP2;refGene_tx
=XM_023259555.1
C1 179988256 179988482
refGene_gn=NEMP2;refGene_tx=XM_023259554.1;refGene_gn=NEMP2;refGene_tx
=XM_023259555.1
C1 179990525 179990640
refGene_gn=NEMP2;refGene_tx=XM_023259554.1;refGene_gn=NEMP2;refGene_tx
=XM_023259555.1
C1 179991728 179991822
refGene_gn=NEMP2;refGene_tx=XM_023259554.1;refGene_gn=NEMP2;refGene_tx
=XM_023259555.1
C1 179993508 179993581
refGene_gn=NEMP2;refGene_tx=XM_023259554.1;refGene_gn=NEMP2;refGene_tx
=XM_023259555.1
C1 179993715 179993947

refGene_gn=NEMP2;refGene_tx=XM_023259554.1;refGene_gn=NEMP2;refGene_tx=XM_023259555.1
C1 179998965 179999081
refGene_gn=NEMP2;refGene_tx=XM_023259554.1;refGene_gn=NEMP2;refGene_tx=XM_023259555.1
C1 180008699 180008808
refGene_gn=NEMP2;refGene_tx=XM_023259554.1;refGene_gn=NEMP2;refGene_tx=XM_023259555.1
C1 180114908 180115727
refGene_gn=NAB1;refGene_tx=XM_019838383.2;refGene_gn=NAB1;refGene_tx=XM_023259556.1;refGene_gn=NAB1;refGene_tx=XM_023259557.1
C1 180126047 180126181
refGene_gn=NAB1;refGene_tx=XM_019838383.2;refGene_gn=NAB1;refGene_tx=XM_023259556.1;refGene_gn=NAB1;refGene_tx=XM_023259557.1
C1 180128655 180128707
refGene_gn=NAB1;refGene_tx=XM_019838383.2;refGene_gn=NAB1;refGene_tx=XM_023259556.1;refGene_gn=NAB1;refGene_tx=XM_023259557.1
C1 180139973 180140063
refGene_gn=NAB1;refGene_tx=XM_019838383.2;refGene_gn=NAB1;refGene_tx=XM_023259556.1;refGene_gn=NAB1;refGene_tx=XM_023259557.1
C1 180141712 180141875
refGene_gn=NAB1;refGene_tx=XM_019838383.2;refGene_gn=NAB1;refGene_tx=XM_023259556.1;refGene_gn=NAB1;refGene_tx=XM_023259557.1
C1 180143837 180143951
refGene_gn=NAB1;refGene_tx=XM_019838383.2;refGene_gn=NAB1;refGene_tx=XM_023259556.1;refGene_gn=NAB1;refGene_tx=XM_023259557.1
C1 180145656 180145745
refGene_gn=NAB1;refGene_tx=XM_019838383.2;refGene_gn=NAB1;refGene_tx=XM_023259556.1;refGene_gn=NAB1;refGene_tx=XM_023259557.1
C1 180206575 180206725
refGene_gn=LOC101084304;refGene_tx=XM_019838384.2;refGene_gn=LOC101084304;refGene_tx=XM_023259561.1;refGene_gn=LOC101084304;refGene_tx=XM_023259562.1;refGene_gn=LOC101084304;refGene_tx=XM_023259563.1;refGene_gn=LOC101084304;refGene_tx=XM_023259564.1;refGene_gn=LOC101084304;refGene_tx=XM_023259565.1;refGene_gn=LOC101084304;refGene_tx=XM_023259566.1;refGene_gn=LOC101084304;refGene_tx=XM_023259567.1;refGene_gn=LOC101084304;refGene_tx=XM_023259568.1;refGene_gn=LOC101084304;refGene_tx=XM_023259569.1;refGene_gn=LOC101084304;refGene_tx=XM_023259570.1
C1 180207094 180207243
refGene_gn=LOC101084304;refGene_tx=XM_019838384.2;refGene_gn=LOC101084304;refGene_tx=XM_023259561.1;refGene_gn=LOC101084304;refGene_tx=XM_023259562.1;refGene_gn=LOC101084304;refGene_tx=XM_023259563.1;refGene_gn=LOC101084304;refGene_tx=XM_023259564.1;refGene_gn=LOC101084304;refGene_tx=XM_023259565.1;refGene_gn=LOC101084304;refGene_tx=XM_023259566.1;refGene_gn=LOC101084304;refGene_tx=XM_023259567.1;refGene_gn=LOC101084304;refGene_tx=XM_023259568.1;refGene_gn=LOC101084304;refGene_tx=XM_023259569.1;refGene_gn=LOC101084304;refGene_tx=XM_023259570.1
C1 180207842 180208096
refGene_gn=LOC101084304;refGene_tx=XM_019838384.2;refGene_gn=LOC101084304;refGene_tx=XM_023259561.1;refGene_gn=LOC101084304;refGene_tx=XM_02

3259562.1;refGene_gn=LOC101084304;refGene_tx=XM_023259563.1;refGene_gn
=LOC101084304;refGene_tx=XM_023259564.1;refGene_gn=LOC101084304;refGen
e_tx=XM_023259565.1;refGene_gn=LOC101084304;refGene_tx=XM_023259566.1;
refGene_gn=LOC101084304;refGene_tx=XM_023259567.1;refGene_gn=LOC101084
304;refGene_tx=XM_023259568.1;refGene_gn=LOC101084304;refGene_tx=XM_02
3259569.1;refGene_gn=LOC101084304;refGene_tx=XM_023259570.1
C1 180208534 180208715
refGene_gn=LOC101084304;refGene_tx=XM_019838384.2;refGene_gn=LOC101084
304;refGene_tx=XM_023259561.1;refGene_gn=LOC101084304;refGene_tx=XM_02
3259562.1;refGene_gn=LOC101084304;refGene_tx=XM_023259563.1;refGene_gn
=LOC101084304;refGene_tx=XM_023259564.1;refGene_gn=LOC101084304;refGen
e_tx=XM_023259565.1;refGene_gn=LOC101084304;refGene_tx=XM_023259566.1;
refGene_gn=LOC101084304;refGene_tx=XM_023259567.1;refGene_gn=LOC101084
304;refGene_tx=XM_023259568.1;refGene_gn=LOC101084304;refGene_tx=XM_02
3259569.1;refGene_gn=LOC101084304;refGene_tx=XM_023259570.1
C1 180209252 180209310
refGene_gn=LOC101084304;refGene_tx=XM_023259564.1;refGene_gn=LOC101084
304;refGene_tx=XM_023259565.1
C1 180223478 180223532
refGene_gn=LOC101084304;refGene_tx=XM_019838384.2;refGene_gn=LOC101084
304;refGene_tx=XM_023259561.1;refGene_gn=LOC101084304;refGene_tx=XM_02
3259562.1;refGene_gn=LOC101084304;refGene_tx=XM_023259563.1
C1 180223617 180223633
refGene_gn=LOC101084304;refGene_tx=XM_019838384.2;refGene_gn=LOC101084
304;refGene_tx=XM_023259561.1;refGene_gn=LOC101084304;refGene_tx=XM_02
3259562.1;refGene_gn=LOC101084304;refGene_tx=XM_023259563.1
C1 180343785 180344180
refGene_gn=GLS;refGene_tx=XM_023259558.1;refGene_gn=GLS;refGene_tx=XM_
023259559.1;refGene_gn=GLS;refGene_tx=XM_023259560.1
C1 180358572 180358669
refGene_gn=GLS;refGene_tx=XM_023259558.1;refGene_gn=GLS;refGene_tx=XM_
023259559.1;refGene_gn=GLS;refGene_tx=XM_023259560.1
C1 180359030 180359152
refGene_gn=GLS;refGene_tx=XM_023259558.1;refGene_gn=GLS;refGene_tx=XM_
023259559.1;refGene_gn=GLS;refGene_tx=XM_023259560.1
C1 180363589 180363719
refGene_gn=GLS;refGene_tx=XM_019838386.1;refGene_gn=GLS;refGene_tx=XM_
023259558.1;refGene_gn=GLS;refGene_tx=XM_023259559.1;refGene_gn=GLS;re
fGene_tx=XM_023259560.1
C1 180364420 180364500
refGene_gn=GLS;refGene_tx=XM_019838386.1;refGene_gn=GLS;refGene_tx=XM_
023259558.1;refGene_gn=GLS;refGene_tx=XM_023259559.1;refGene_gn=GLS;re
fGene_tx=XM_023259560.1
C1 180367793 180367957
refGene_gn=GLS;refGene_tx=XM_019838386.1;refGene_gn=GLS;refGene_tx=XM_
023259558.1;refGene_gn=GLS;refGene_tx=XM_023259559.1;refGene_gn=GLS;re
fGene_tx=XM_023259560.1
C1 180377180 180377239
refGene_gn=GLS;refGene_tx=XM_019838386.1;refGene_gn=GLS;refGene_tx=XM_
023259558.1;refGene_gn=GLS;refGene_tx=XM_023259559.1;refGene_gn=GLS;re

fGene_tx=XM_023259560.1
C1 180385442 180385475
refGene_gn=GLS;refGene_tx=XM_019838386.1;refGene_gn=GLS;refGene_tx=XM_023259558.1;refGene_gn=GLS;refGene_tx=XM_023259559.1;refGene_gn=GLS;refGene_tx=XM_023259560.1
C1 180385569 180385628
refGene_gn=GLS;refGene_tx=XM_019838386.1;refGene_gn=GLS;refGene_tx=XM_023259558.1;refGene_gn=GLS;refGene_tx=XM_023259559.1;refGene_gn=GLS;refGene_tx=XM_023259560.1
C1 180388771 180388838
refGene_gn=GLS;refGene_tx=XM_019838386.1;refGene_gn=GLS;refGene_tx=XM_023259558.1;refGene_gn=GLS;refGene_tx=XM_023259559.1;refGene_gn=GLS;refGene_tx=XM_023259560.1
C1 180389407 180389458
refGene_gn=GLS;refGene_tx=XM_019838386.1;refGene_gn=GLS;refGene_tx=XM_023259558.1;refGene_gn=GLS;refGene_tx=XM_023259559.1;refGene_gn=GLS;refGene_tx=XM_023259560.1
C1 180392108 180392285
refGene_gn=GLS;refGene_tx=XM_019838386.1;refGene_gn=GLS;refGene_tx=XM_023259558.1;refGene_gn=GLS;refGene_tx=XM_023259559.1;refGene_gn=GLS;refGene_tx=XM_023259560.1
C1 180394275 180394407
refGene_gn=GLS;refGene_tx=XM_019838386.1;refGene_gn=GLS;refGene_tx=XM_023259558.1;refGene_gn=GLS;refGene_tx=XM_023259559.1;refGene_gn=GLS;refGene_tx=XM_023259560.1
C1 180397735 180397828
refGene_gn=GLS;refGene_tx=XM_019838386.1;refGene_gn=GLS;refGene_tx=XM_023259558.1;refGene_gn=GLS;refGene_tx=XM_023259559.1;refGene_gn=GLS;refGene_tx=XM_023259560.1
C1 180398887 180398917
refGene_gn=GLS;refGene_tx=XM_023259560.1
C1 180398946 180399093
refGene_gn=GLS;refGene_tx=XM_023259559.1
C1 180423594 180423656
refGene_gn=GLS;refGene_tx=XM_019838386.1;refGene_gn=GLS;refGene_tx=XM_023259558.1
C1 180424714 180424791
refGene_gn=GLS;refGene_tx=XM_019838386.1;refGene_gn=GLS;refGene_tx=XM_023259558.1
C1 180424869 180424933
refGene_gn=GLS;refGene_tx=XM_019838386.1;refGene_gn=GLS;refGene_tx=XM_023259558.1
C1 180427902 180428059
refGene_gn=GLS;refGene_tx=XM_019838386.1;refGene_gn=GLS;refGene_tx=XM_023259558.1
C1 180435844 180435859
refGene_gn=STAT1;refGene_tx=XM_006935443.3
C1 180439421 180439524
refGene_gn=STAT1;refGene_tx=XM_006935443.3
C1 180440418 180440494

refGene_gn=STAT1;refGene_tx=XM_006935443.3
C1 180441548 180441734
refGene_gn=STAT1;refGene_tx=XM_006935443.3
C1 180443541 180443687
refGene_gn=STAT1;refGene_tx=XM_006935443.3
C1 180444489 180444584
refGene_gn=STAT1;refGene_tx=XM_006935443.3
C1 180445296 180445346
refGene_gn=STAT1;refGene_tx=XM_006935443.3
C1 180446758 180446894
refGene_gn=STAT1;refGene_tx=XM_006935443.3
C1 180448241 180448340
refGene_gn=STAT1;refGene_tx=XM_006935443.3
C1 180448757 180448841
refGene_gn=STAT1;refGene_tx=XM_006935443.3
C1 180450071 180450113
refGene_gn=STAT1;refGene_tx=XM_006935443.3
C1 180451218 180451312
refGene_gn=STAT1;refGene_tx=XM_006935443.3
C1 180451402 180451432
refGene_gn=STAT1;refGene_tx=XM_006935443.3
C1 180453736 180453796
refGene_gn=STAT1;refGene_tx=XM_006935443.3
C1 180456262 180456355
refGene_gn=STAT1;refGene_tx=XM_006935443.3
C1 180457992 180458151
refGene_gn=STAT1;refGene_tx=XM_006935443.3
C1 180460358 180460510
refGene_gn=STAT1;refGene_tx=XM_006935443.3
C1 180460920 180461012
refGene_gn=STAT1;refGene_tx=XM_006935443.3
C1 180462104 180462183
refGene_gn=STAT1;refGene_tx=XM_006935443.3
C1 180463193 180463283
refGene_gn=STAT1;refGene_tx=XM_006935443.3
C1 180469274 180469373
refGene_gn=STAT1;refGene_tx=XM_006935443.3
C1 180470646 180470791
refGene_gn=STAT1;refGene_tx=XM_006935443.3
C1 180472244 180472372
refGene_gn=STAT1;refGene_tx=XM_006935443.3
C1 180491058 180491085
refGene_gn=STAT4;refGene_tx=XM_023259577.1;refGene_gn=STAT4;refGene_tx=XM_023259578.1
C1 180492676 180492878
refGene_gn=STAT4;refGene_tx=XM_019838390.2;refGene_gn=STAT4;refGene_tx=XM_023259571.1;refGene_gn=STAT4;refGene_tx=XM_023259572.1;refGene_gn=STAT4;refGene_tx=XM_023259573.1;refGene_gn=STAT4;refGene_tx=XM_023259574.1;refGene_gn=STAT4;refGene_tx=XM_023259575.1;refGene_gn=STAT4;refGene_tx=XM_023259576.1;refGene_gn=STAT4;refGene_tx=XM_023259577.1

C1 180493229 180493296
refGene_gn=STAT4;refGene_tx=XM_019838390.2;refGene_gn=STAT4;refGene_tx=XM_023259571.1;refGene_gn=STAT4;refGene_tx=XM_023259572.1;refGene_gn=STAT4;refGene_tx=XM_023259573.1;refGene_gn=STAT4;refGene_tx=XM_023259574.1;refGene_gn=STAT4;refGene_tx=XM_023259575.1;refGene_gn=STAT4;refGene_tx=XM_023259576.1;refGene_gn=STAT4;refGene_tx=XM_023259577.1;refGene_gn=STAT4;refGene_tx=XM_023259578.1
C1 180494447 180494639
refGene_gn=STAT4;refGene_tx=XM_019838390.2;refGene_gn=STAT4;refGene_tx=XM_023259571.1;refGene_gn=STAT4;refGene_tx=XM_023259572.1;refGene_gn=STAT4;refGene_tx=XM_023259573.1;refGene_gn=STAT4;refGene_tx=XM_023259574.1;refGene_gn=STAT4;refGene_tx=XM_023259575.1;refGene_gn=STAT4;refGene_tx=XM_023259576.1;refGene_gn=STAT4;refGene_tx=XM_023259577.1;refGene_gn=STAT4;refGene_tx=XM_023259578.1
C1 180495197 180495334
refGene_gn=STAT4;refGene_tx=XM_019838390.2;refGene_gn=STAT4;refGene_tx=XM_023259571.1;refGene_gn=STAT4;refGene_tx=XM_023259572.1;refGene_gn=STAT4;refGene_tx=XM_023259573.1;refGene_gn=STAT4;refGene_tx=XM_023259574.1;refGene_gn=STAT4;refGene_tx=XM_023259575.1;refGene_gn=STAT4;refGene_tx=XM_023259576.1;refGene_gn=STAT4;refGene_tx=XM_023259577.1;refGene_gn=STAT4;refGene_tx=XM_023259578.1
C1 180495626 180495721
refGene_gn=STAT4;refGene_tx=XM_019838390.2;refGene_gn=STAT4;refGene_tx=XM_023259571.1;refGene_gn=STAT4;refGene_tx=XM_023259572.1;refGene_gn=STAT4;refGene_tx=XM_023259573.1;refGene_gn=STAT4;refGene_tx=XM_023259574.1;refGene_gn=STAT4;refGene_tx=XM_023259575.1;refGene_gn=STAT4;refGene_tx=XM_023259576.1;refGene_gn=STAT4;refGene_tx=XM_023259577.1;refGene_gn=STAT4;refGene_tx=XM_023259578.1
C1 180495983 180496033
refGene_gn=STAT4;refGene_tx=XM_019838390.2;refGene_gn=STAT4;refGene_tx=XM_023259571.1;refGene_gn=STAT4;refGene_tx=XM_023259572.1;refGene_gn=STAT4;refGene_tx=XM_023259573.1;refGene_gn=STAT4;refGene_tx=XM_023259574.1;refGene_gn=STAT4;refGene_tx=XM_023259575.1;refGene_gn=STAT4;refGene_tx=XM_023259576.1;refGene_gn=STAT4;refGene_tx=XM_023259577.1;refGene_gn=STAT4;refGene_tx=XM_023259578.1
C1 180497831 180497967
refGene_gn=STAT4;refGene_tx=XM_019838390.2;refGene_gn=STAT4;refGene_tx=XM_023259571.1;refGene_gn=STAT4;refGene_tx=XM_023259572.1;refGene_gn=STAT4;refGene_tx=XM_023259573.1;refGene_gn=STAT4;refGene_tx=XM_023259574.1;refGene_gn=STAT4;refGene_tx=XM_023259575.1;refGene_gn=STAT4;refGene_tx=XM_023259576.1;refGene_gn=STAT4;refGene_tx=XM_023259577.1;refGene_gn=STAT4;refGene_tx=XM_023259578.1
C1 180500645 180500744
refGene_gn=STAT4;refGene_tx=XM_019838390.2;refGene_gn=STAT4;refGene_tx=XM_023259571.1;refGene_gn=STAT4;refGene_tx=XM_023259572.1;refGene_gn=STAT4;refGene_tx=XM_023259573.1;refGene_gn=STAT4;refGene_tx=XM_023259574.1;refGene_gn=STAT4;refGene_tx=XM_023259575.1;refGene_gn=STAT4;refGene_tx=XM_023259576.1;refGene_gn=STAT4;refGene_tx=XM_023259577.1;refGene_gn=STAT4;refGene_tx=XM_023259578.1
C1 180502231 180502315

refGene_gn=STAT4;refGene_tx=XM_019838390.2;refGene_gn=STAT4;refGene_tx
=XM_023259571.1;refGene_gn=STAT4;refGene_tx=XM_023259572.1;refGene_gn=
STAT4;refGene_tx=XM_023259573.1;refGene_gn=STAT4;refGene_tx=XM_0232595
74.1;refGene_gn=STAT4;refGene_tx=XM_023259575.1;refGene_gn=STAT4;refGe
ne_tx=XM_023259576.1;refGene_gn=STAT4;refGene_tx=XM_023259577.1;refGen
e_gn=STAT4;refGene_tx=XM_023259578.1
C1 180512429 180512474
refGene_gn=STAT4;refGene_tx=XM_019838390.2;refGene_gn=STAT4;refGene_tx
=XM_023259571.1;refGene_gn=STAT4;refGene_tx=XM_023259572.1;refGene_gn=
STAT4;refGene_tx=XM_023259573.1;refGene_gn=STAT4;refGene_tx=XM_0232595
74.1;refGene_gn=STAT4;refGene_tx=XM_023259575.1;refGene_gn=STAT4;refGe
ne_tx=XM_023259576.1;refGene_gn=STAT4;refGene_tx=XM_023259577.1;refGen
e_gn=STAT4;refGene_tx=XM_023259578.1
C1 180514476 180514570
refGene_gn=STAT4;refGene_tx=XM_019838390.2;refGene_gn=STAT4;refGene_tx
=XM_023259571.1;refGene_gn=STAT4;refGene_tx=XM_023259572.1;refGene_gn=
STAT4;refGene_tx=XM_023259573.1;refGene_gn=STAT4;refGene_tx=XM_0232595
74.1;refGene_gn=STAT4;refGene_tx=XM_023259575.1;refGene_gn=STAT4;refGe
ne_tx=XM_023259576.1;refGene_gn=STAT4;refGene_tx=XM_023259577.1;refGen
e_gn=STAT4;refGene_tx=XM_023259578.1
C1 180514657 180514675
refGene_gn=STAT4;refGene_tx=XM_019838390.2;refGene_gn=STAT4;refGene_tx
=XM_023259571.1;refGene_gn=STAT4;refGene_tx=XM_023259572.1;refGene_gn=
STAT4;refGene_tx=XM_023259573.1;refGene_gn=STAT4;refGene_tx=XM_0232595
74.1;refGene_gn=STAT4;refGene_tx=XM_023259575.1;refGene_gn=STAT4;refGe
ne_tx=XM_023259576.1;refGene_gn=STAT4;refGene_tx=XM_023259577.1;refGen
e_gn=STAT4;refGene_tx=XM_023259578.1
C1 180514984 180515044
refGene_gn=STAT4;refGene_tx=XM_019838390.2;refGene_gn=STAT4;refGene_tx
=XM_023259571.1;refGene_gn=STAT4;refGene_tx=XM_023259572.1;refGene_gn=
STAT4;refGene_tx=XM_023259573.1;refGene_gn=STAT4;refGene_tx=XM_0232595
74.1;refGene_gn=STAT4;refGene_tx=XM_023259575.1;refGene_gn=STAT4;refGe
ne_tx=XM_023259576.1;refGene_gn=STAT4;refGene_tx=XM_023259577.1;refGen
e_gn=STAT4;refGene_tx=XM_023259578.1
C1 180517963 180518056
refGene_gn=STAT4;refGene_tx=XM_019838390.2;refGene_gn=STAT4;refGene_tx
=XM_023259571.1;refGene_gn=STAT4;refGene_tx=XM_023259572.1;refGene_gn=
STAT4;refGene_tx=XM_023259573.1;refGene_gn=STAT4;refGene_tx=XM_0232595
74.1;refGene_gn=STAT4;refGene_tx=XM_023259575.1;refGene_gn=STAT4;refGe
ne_tx=XM_023259576.1;refGene_gn=STAT4;refGene_tx=XM_023259577.1;refGen
e_gn=STAT4;refGene_tx=XM_023259578.1
C1 180518771 180518930
refGene_gn=STAT4;refGene_tx=XM_019838390.2;refGene_gn=STAT4;refGene_tx
=XM_023259571.1;refGene_gn=STAT4;refGene_tx=XM_023259572.1;refGene_gn=
STAT4;refGene_tx=XM_023259573.1;refGene_gn=STAT4;refGene_tx=XM_0232595
74.1;refGene_gn=STAT4;refGene_tx=XM_023259575.1;refGene_gn=STAT4;refGe
ne_tx=XM_023259576.1;refGene_gn=STAT4;refGene_tx=XM_023259577.1;refGen
e_gn=STAT4;refGene_tx=XM_023259578.1
C1 180520724 180520876
refGene_gn=STAT4;refGene_tx=XM_019838390.2;refGene_gn=STAT4;refGene_tx

=XM_023259571.1;refGene_gn=STAT4;refGene_tx=XM_023259572.1;refGene_gn=STAT4;refGene_tx=XM_023259573.1;refGene_gn=STAT4;refGene_tx=XM_023259574.1;refGene_gn=STAT4;refGene_tx=XM_023259575.1;refGene_gn=STAT4;refGene_tx=XM_023259576.1;refGene_gn=STAT4;refGene_tx=XM_023259577.1;refGene_gn=STAT4;refGene_tx=XM_023259578.1
C1 180522342 180522428
refGene_gn=STAT4;refGene_tx=XM_019838390.2;refGene_gn=STAT4;refGene_tx=XM_023259571.1;refGene_gn=STAT4;refGene_tx=XM_023259572.1;refGene_gn=STAT4;refGene_tx=XM_023259573.1;refGene_gn=STAT4;refGene_tx=XM_023259574.1;refGene_gn=STAT4;refGene_tx=XM_023259575.1;refGene_gn=STAT4;refGene_tx=XM_023259576.1;refGene_gn=STAT4;refGene_tx=XM_023259577.1;refGene_gn=STAT4;refGene_tx=XM_023259578.1
C1 180525967 180526046
refGene_gn=STAT4;refGene_tx=XM_019838390.2;refGene_gn=STAT4;refGene_tx=XM_023259571.1;refGene_gn=STAT4;refGene_tx=XM_023259572.1;refGene_gn=STAT4;refGene_tx=XM_023259573.1;refGene_gn=STAT4;refGene_tx=XM_023259574.1;refGene_gn=STAT4;refGene_tx=XM_023259575.1;refGene_gn=STAT4;refGene_tx=XM_023259576.1;refGene_gn=STAT4;refGene_tx=XM_023259577.1;refGene_gn=STAT4;refGene_tx=XM_023259578.1
C1 180526919 180527012
refGene_gn=STAT4;refGene_tx=XM_023259571.1;refGene_gn=STAT4;refGene_tx=XM_023259572.1;refGene_gn=STAT4;refGene_tx=XM_023259573.1;refGene_gn=STAT4;refGene_tx=XM_023259574.1;refGene_gn=STAT4;refGene_tx=XM_023259575.1;refGene_gn=STAT4;refGene_tx=XM_023259576.1;refGene_gn=STAT4;refGene_tx=XM_023259577.1;refGene_gn=STAT4;refGene_tx=XM_023259578.1
C1 180529397 180529496
refGene_gn=STAT4;refGene_tx=XM_023259571.1;refGene_gn=STAT4;refGene_tx=XM_023259572.1;refGene_gn=STAT4;refGene_tx=XM_023259573.1;refGene_gn=STAT4;refGene_tx=XM_023259574.1;refGene_gn=STAT4;refGene_tx=XM_023259575.1;refGene_gn=STAT4;refGene_tx=XM_023259576.1;refGene_gn=STAT4;refGene_tx=XM_023259577.1;refGene_gn=STAT4;refGene_tx=XM_023259578.1
C1 180579036 180579181
refGene_gn=STAT4;refGene_tx=XM_023259571.1;refGene_gn=STAT4;refGene_tx=XM_023259572.1;refGene_gn=STAT4;refGene_tx=XM_023259573.1;refGene_gn=STAT4;refGene_tx=XM_023259574.1;refGene_gn=STAT4;refGene_tx=XM_023259575.1;refGene_gn=STAT4;refGene_tx=XM_023259576.1;refGene_gn=STAT4;refGene_tx=XM_023259577.1;refGene_gn=STAT4;refGene_tx=XM_023259578.1
C1 180580480 180580609
refGene_gn=STAT4;refGene_tx=XM_023259571.1;refGene_gn=STAT4;refGene_tx=XM_023259572.1;refGene_gn=STAT4;refGene_tx=XM_023259573.1;refGene_gn=STAT4;refGene_tx=XM_023259574.1;refGene_gn=STAT4;refGene_tx=XM_023259575.1;refGene_gn=STAT4;refGene_tx=XM_023259576.1;refGene_gn=STAT4;refGene_tx=XM_023259577.1;refGene_gn=STAT4;refGene_tx=XM_023259578.1
C1 180606385 180606414
refGene_gn=STAT4;refGene_tx=XM_023259571.1;refGene_gn=STAT4;refGene_tx=XM_023259572.1;refGene_gn=STAT4;refGene_tx=XM_023259577.1;refGene_gn=STAT4;refGene_tx=XM_023259578.1
C1 180708105 180708240
refGene_gn=MYO1B;refGene_tx=XM_011285438.2;refGene_gn=MYO1B;refGene_tx=XM_011285439.2;refGene_gn=MYO1B;refGene_tx=XM_011285440.2;refGene_gn=

MYO1B;refGene_tx=XM_019838392.2;refGene_gn=MYO1B;refGene_tx=XM_0198383
93.1;refGene_gn=MYO1B;refGene_tx=XM_019838394.1;refGene_gn=MYO1B;refGe
ne_tx=XM_019838395.2
C1 180728879 180728995
refGene_gn=MYO1B;refGene_tx=XM_011285438.2;refGene_gn=MYO1B;refGene_tx
=XM_011285439.2;refGene_gn=MYO1B;refGene_tx=XM_011285440.2;refGene_gn=
MYO1B;refGene_tx=XM_019838392.2;refGene_gn=MYO1B;refGene_tx=XM_0198383
93.1;refGene_gn=MYO1B;refGene_tx=XM_019838394.1;refGene_gn=MYO1B;refGe
ne_tx=XM_019838395.2
C1 180761260 180761355
refGene_gn=MYO1B;refGene_tx=XM_011285438.2;refGene_gn=MYO1B;refGene_tx
=XM_011285439.2;refGene_gn=MYO1B;refGene_tx=XM_011285440.2;refGene_gn=
MYO1B;refGene_tx=XM_019838392.2;refGene_gn=MYO1B;refGene_tx=XM_0198383
93.1;refGene_gn=MYO1B;refGene_tx=XM_019838394.1;refGene_gn=MYO1B;refGe
ne_tx=XM_019838395.2
C1 180777950 180778055
refGene_gn=MYO1B;refGene_tx=XM_011285438.2;refGene_gn=MYO1B;refGene_tx
=XM_011285439.2;refGene_gn=MYO1B;refGene_tx=XM_011285440.2;refGene_gn=
MYO1B;refGene_tx=XM_019838392.2;refGene_gn=MYO1B;refGene_tx=XM_0198383
93.1;refGene_gn=MYO1B;refGene_tx=XM_019838394.1;refGene_gn=MYO1B;refGe
ne_tx=XM_019838395.2
C1 180782221 180782268
refGene_gn=MYO1B;refGene_tx=XM_011285438.2;refGene_gn=MYO1B;refGene_tx
=XM_011285439.2;refGene_gn=MYO1B;refGene_tx=XM_011285440.2;refGene_gn=
MYO1B;refGene_tx=XM_019838392.2;refGene_gn=MYO1B;refGene_tx=XM_0198383
93.1;refGene_gn=MYO1B;refGene_tx=XM_019838394.1;refGene_gn=MYO1B;refGe
ne_tx=XM_019838395.2
C1 180788208 180788272
refGene_gn=MYO1B;refGene_tx=XM_011285438.2;refGene_gn=MYO1B;refGene_tx
=XM_011285439.2;refGene_gn=MYO1B;refGene_tx=XM_011285440.2;refGene_gn=
MYO1B;refGene_tx=XM_019838392.2;refGene_gn=MYO1B;refGene_tx=XM_0198383
93.1;refGene_gn=MYO1B;refGene_tx=XM_019838394.1;refGene_gn=MYO1B;refGe
ne_tx=XM_019838395.2
C1 180799887 180799986
refGene_gn=MYO1B;refGene_tx=XM_011285438.2;refGene_gn=MYO1B;refGene_tx
=XM_011285439.2;refGene_gn=MYO1B;refGene_tx=XM_011285440.2;refGene_gn=
MYO1B;refGene_tx=XM_019838392.2;refGene_gn=MYO1B;refGene_tx=XM_0198383
93.1;refGene_gn=MYO1B;refGene_tx=XM_019838394.1;refGene_gn=MYO1B;refGe
ne_tx=XM_019838395.2
C1 180801613 180801717
refGene_gn=MYO1B;refGene_tx=XM_011285438.2;refGene_gn=MYO1B;refGene_tx
=XM_011285439.2;refGene_gn=MYO1B;refGene_tx=XM_011285440.2;refGene_gn=
MYO1B;refGene_tx=XM_019838392.2;refGene_gn=MYO1B;refGene_tx=XM_0198383
93.1;refGene_gn=MYO1B;refGene_tx=XM_019838394.1;refGene_gn=MYO1B;refGe
ne_tx=XM_019838395.2
C1 180803720 180803868
refGene_gn=MYO1B;refGene_tx=XM_011285438.2;refGene_gn=MYO1B;refGene_tx
=XM_011285439.2;refGene_gn=MYO1B;refGene_tx=XM_011285440.2;refGene_gn=
MYO1B;refGene_tx=XM_019838392.2;refGene_gn=MYO1B;refGene_tx=XM_0198383
93.1;refGene_gn=MYO1B;refGene_tx=XM_019838394.1;refGene_gn=MYO1B;refGe

ne_tx=XM_019838395.2
C1	180804148	180804267
refGene_gn=MYO1B;refGene_tx=XM_011285438.2;refGene_gn=MYO1B;refGene_tx=XM_011285439.2;refGene_gn=MYO1B;refGene_tx=XM_011285440.2;refGene_gn=MYO1B;refGene_tx=XM_019838392.2;refGene_gn=MYO1B;refGene_tx=XM_019838393.1;refGene_gn=MYO1B;refGene_tx=XM_019838394.1;refGene_gn=MYO1B;refGene_tx=XM_019838395.2
C1	180810130	180810217
refGene_gn=MYO1B;refGene_tx=XM_011285438.2;refGene_gn=MYO1B;refGene_tx=XM_011285439.2;refGene_gn=MYO1B;refGene_tx=XM_011285440.2;refGene_gn=MYO1B;refGene_tx=XM_019838392.2;refGene_gn=MYO1B;refGene_tx=XM_019838393.1;refGene_gn=MYO1B;refGene_tx=XM_019838394.1;refGene_gn=MYO1B;refGene_tx=XM_019838395.2
C1	180810844	180810910
refGene_gn=MYO1B;refGene_tx=XM_011285438.2;refGene_gn=MYO1B;refGene_tx=XM_011285439.2;refGene_gn=MYO1B;refGene_tx=XM_011285440.2;refGene_gn=MYO1B;refGene_tx=XM_019838392.2;refGene_gn=MYO1B;refGene_tx=XM_019838393.1;refGene_gn=MYO1B;refGene_tx=XM_019838394.1;refGene_gn=MYO1B;refGene_tx=XM_019838395.2
C1	180818491	180818497
refGene_gn=MYO1B;refGene_tx=XM_019838392.2;refGene_gn=MYO1B;refGene_tx=XM_019838393.1;refGene_gn=MYO1B;refGene_tx=XM_019838394.1;refGene_gn=MYO1B;refGene_tx=XM_019838395.2
C1	180820657	180820762
refGene_gn=MYO1B;refGene_tx=XM_011285438.2;refGene_gn=MYO1B;refGene_tx=XM_011285439.2;refGene_gn=MYO1B;refGene_tx=XM_011285440.2;refGene_gn=MYO1B;refGene_tx=XM_019838392.2;refGene_gn=MYO1B;refGene_tx=XM_019838393.1;refGene_gn=MYO1B;refGene_tx=XM_019838394.1;refGene_gn=MYO1B;refGene_tx=XM_019838395.2
C1	180823861	180823924
refGene_gn=MYO1B;refGene_tx=XM_011285438.2;refGene_gn=MYO1B;refGene_tx=XM_011285439.2;refGene_gn=MYO1B;refGene_tx=XM_011285440.2;refGene_gn=MYO1B;refGene_tx=XM_019838392.2;refGene_gn=MYO1B;refGene_tx=XM_019838393.1;refGene_gn=MYO1B;refGene_tx=XM_019838394.1;refGene_gn=MYO1B;refGene_tx=XM_019838395.2
C1	180827548	180827749
refGene_gn=MYO1B;refGene_tx=XM_011285438.2;refGene_gn=MYO1B;refGene_tx=XM_011285439.2;refGene_gn=MYO1B;refGene_tx=XM_011285440.2;refGene_gn=MYO1B;refGene_tx=XM_019838392.2;refGene_gn=MYO1B;refGene_tx=XM_019838393.1;refGene_gn=MYO1B;refGene_tx=XM_019838394.1;refGene_gn=MYO1B;refGene_tx=XM_019838395.2
C1	180828922	180829149
refGene_gn=MYO1B;refGene_tx=XM_011285438.2;refGene_gn=MYO1B;refGene_tx=XM_011285439.2;refGene_gn=MYO1B;refGene_tx=XM_011285440.2;refGene_gn=MYO1B;refGene_tx=XM_019838392.2;refGene_gn=MYO1B;refGene_tx=XM_019838393.1;refGene_gn=MYO1B;refGene_tx=XM_019838394.1;refGene_gn=MYO1B;refGene_tx=XM_019838395.2
C1	180832078	180832279
refGene_gn=MYO1B;refGene_tx=XM_011285438.2;refGene_gn=MYO1B;refGene_tx=XM_011285439.2;refGene_gn=MYO1B;refGene_tx=XM_011285440.2;refGene_gn=

MYO1B;refGene_tx=XM_019838392.2;refGene_gn=MYO1B;refGene_tx=XM_0198383
93.1;refGene_gn=MYO1B;refGene_tx=XM_019838394.1;refGene_gn=MYO1B;refGe
ne_tx=XM_019838395.2
C1	180833910	180834004
refGene_gn=MYO1B;refGene_tx=XM_011285438.2;refGene_gn=MYO1B;refGene_tx
=XM_011285439.2;refGene_gn=MYO1B;refGene_tx=XM_011285440.2;refGene_gn=
MYO1B;refGene_tx=XM_019838392.2;refGene_gn=MYO1B;refGene_tx=XM_0198383
93.1;refGene_gn=MYO1B;refGene_tx=XM_019838394.1;refGene_gn=MYO1B;refGe
ne_tx=XM_019838395.2
C1	180834863	180835013
refGene_gn=MYO1B;refGene_tx=XM_011285438.2;refGene_gn=MYO1B;refGene_tx
=XM_011285439.2;refGene_gn=MYO1B;refGene_tx=XM_011285440.2;refGene_gn=
MYO1B;refGene_tx=XM_019838392.2;refGene_gn=MYO1B;refGene_tx=XM_0198383
93.1;refGene_gn=MYO1B;refGene_tx=XM_019838394.1;refGene_gn=MYO1B;refGe
ne_tx=XM_019838395.2
C1	180838177	180838246
refGene_gn=MYO1B;refGene_tx=XM_011285438.2;refGene_gn=MYO1B;refGene_tx
=XM_011285439.2;refGene_gn=MYO1B;refGene_tx=XM_011285440.2;refGene_gn=
MYO1B;refGene_tx=XM_019838392.2;refGene_gn=MYO1B;refGene_tx=XM_0198383
93.1;refGene_gn=MYO1B;refGene_tx=XM_019838394.1;refGene_gn=MYO1B;refGe
ne_tx=XM_019838395.2
C1	180839981	180840068
refGene_gn=MYO1B;refGene_tx=XM_011285438.2;refGene_gn=MYO1B;refGene_tx
=XM_011285439.2;refGene_gn=MYO1B;refGene_tx=XM_011285440.2;refGene_gn=
MYO1B;refGene_tx=XM_019838392.2;refGene_gn=MYO1B;refGene_tx=XM_0198383
93.1;refGene_gn=MYO1B;refGene_tx=XM_019838394.1;refGene_gn=MYO1B;refGe
ne_tx=XM_019838395.2
C1	180840314	180840401
refGene_gn=MYO1B;refGene_tx=XM_011285438.2;refGene_gn=MYO1B;refGene_tx
=XM_019838392.2;refGene_gn=MYO1B;refGene_tx=XM_019838395.2
C1	180842198	180842285
refGene_gn=MYO1B;refGene_tx=XM_011285438.2;refGene_gn=MYO1B;refGene_tx
=XM_011285439.2;refGene_gn=MYO1B;refGene_tx=XM_019838392.2;refGene_gn=
MYO1B;refGene_tx=XM_019838393.1;refGene_gn=MYO1B;refGene_tx=XM_0198383
95.2
C1	180845713	180845788
refGene_gn=MYO1B;refGene_tx=XM_011285438.2;refGene_gn=MYO1B;refGene_tx
=XM_011285439.2;refGene_gn=MYO1B;refGene_tx=XM_011285440.2;refGene_gn=
MYO1B;refGene_tx=XM_019838392.2;refGene_gn=MYO1B;refGene_tx=XM_0198383
93.1;refGene_gn=MYO1B;refGene_tx=XM_019838394.1;refGene_gn=MYO1B;refGe
ne_tx=XM_019838395.2
C1	180847111	180847246
refGene_gn=MYO1B;refGene_tx=XM_011285438.2;refGene_gn=MYO1B;refGene_tx
=XM_011285439.2;refGene_gn=MYO1B;refGene_tx=XM_011285440.2;refGene_gn=
MYO1B;refGene_tx=XM_019838392.2;refGene_gn=MYO1B;refGene_tx=XM_0198383
93.1;refGene_gn=MYO1B;refGene_tx=XM_019838394.1;refGene_gn=MYO1B;refGe
ne_tx=XM_019838395.2
C1	180850309	180850416
refGene_gn=MYO1B;refGene_tx=XM_011285438.2;refGene_gn=MYO1B;refGene_tx
=XM_011285439.2;refGene_gn=MYO1B;refGene_tx=XM_011285440.2;refGene_gn=

MYO1B;refGene_tx=XM_019838392.2;refGene_gn=MYO1B;refGene_tx=XM_0198383
93.1;refGene_gn=MYO1B;refGene_tx=XM_019838394.1;refGene_gn=MYO1B;refGe
ne_tx=XM_019838395.2
C1	180853399	180853532
refGene_gn=MYO1B;refGene_tx=XM_011285438.2;refGene_gn=MYO1B;refGene_tx
=XM_011285439.2;refGene_gn=MYO1B;refGene_tx=XM_011285440.2;refGene_gn=
MYO1B;refGene_tx=XM_019838392.2;refGene_gn=MYO1B;refGene_tx=XM_0198383
93.1;refGene_gn=MYO1B;refGene_tx=XM_019838394.1;refGene_gn=MYO1B;refGe
ne_tx=XM_019838395.2
C1	180853900	180854053
refGene_gn=MYO1B;refGene_tx=XM_011285438.2;refGene_gn=MYO1B;refGene_tx
=XM_011285439.2;refGene_gn=MYO1B;refGene_tx=XM_011285440.2;refGene_gn=
MYO1B;refGene_tx=XM_019838392.2;refGene_gn=MYO1B;refGene_tx=XM_0198383
93.1;refGene_gn=MYO1B;refGene_tx=XM_019838394.1;refGene_gn=MYO1B;refGe
ne_tx=XM_019838395.2
C1	180856314	180856442
refGene_gn=MYO1B;refGene_tx=XM_011285438.2;refGene_gn=MYO1B;refGene_tx
=XM_011285439.2;refGene_gn=MYO1B;refGene_tx=XM_011285440.2;refGene_gn=
MYO1B;refGene_tx=XM_019838392.2;refGene_gn=MYO1B;refGene_tx=XM_0198383
93.1;refGene_gn=MYO1B;refGene_tx=XM_019838394.1;refGene_gn=MYO1B;refGe
ne_tx=XM_019838395.2
C1	180862443	180862567
refGene_gn=MYO1B;refGene_tx=XM_011285438.2;refGene_gn=MYO1B;refGene_tx
=XM_011285439.2;refGene_gn=MYO1B;refGene_tx=XM_011285440.2;refGene_gn=
MYO1B;refGene_tx=XM_019838392.2;refGene_gn=MYO1B;refGene_tx=XM_0198383
93.1;refGene_gn=MYO1B;refGene_tx=XM_019838394.1;refGene_gn=MYO1B;refGe
ne_tx=XM_019838395.2
C1	181101573	181101664
refGene_gn=NABP1;refGene_tx=XM_003990982.4
C1	181101964	181102103
refGene_gn=NABP1;refGene_tx=XM_003990982.4
C1	181104718	181104790
refGene_gn=NABP1;refGene_tx=XM_003990982.4;refGene_gn=NABP1;refGene_tx
=XM_019838396.2;refGene_gn=NABP1;refGene_tx=XM_019838397.2
C1	181106461	181106537
refGene_gn=NABP1;refGene_tx=XM_003990982.4;refGene_gn=NABP1;refGene_tx
=XM_019838396.2;refGene_gn=NABP1;refGene_tx=XM_019838397.2
C1	181106990	181107057
refGene_gn=NABP1;refGene_tx=XM_003990982.4;refGene_gn=NABP1;refGene_tx
=XM_019838396.2;refGene_gn=NABP1;refGene_tx=XM_019838397.2
C1	181108293	181108463
refGene_gn=NABP1;refGene_tx=XM_003990982.4;refGene_gn=NABP1;refGene_tx
=XM_019838396.2;refGene_gn=NABP1;refGene_tx=XM_019838397.2
C1	181255173	181255968
refGene_gn=CAVIN2;refGene_tx=XM_003990983.5
C1	181266484	181266967
refGene_gn=CAVIN2;refGene_tx=XM_003990983.5
C1	181369757	181369854
refGene_gn=TMEFF2;refGene_tx=XM_003990984.4;refGene_gn=TMEFF2;refGene_
tx=XM_006935444.3

C1	181373219	181373378
refGene_gn=TMEFF2;refGene_tx=XM_003990984.4;refGene_gn=TMEFF2;refGene_tx=XM_006935444.3
C1	181375205	181375329
refGene_gn=TMEFF2;refGene_tx=XM_003990984.4;refGene_gn=TMEFF2;refGene_tx=XM_006935444.3
C1	181412466	181412526
refGene_gn=TMEFF2;refGene_tx=XM_006935444.3
C1	181417420	181417480
refGene_gn=TMEFF2;refGene_tx=XM_003990984.4;refGene_gn=TMEFF2;refGene_tx=XM_006935444.3
C1	181418213	181418362
refGene_gn=TMEFF2;refGene_tx=XM_003990984.4;refGene_gn=TMEFF2;refGene_tx=XM_006935444.3
C1	181468261	181468358
refGene_gn=TMEFF2;refGene_tx=XM_003990984.4;refGene_gn=TMEFF2;refGene_tx=XM_006935444.3
C1	181584746	181584773
refGene_gn=TMEFF2;refGene_tx=XM_003990984.4;refGene_gn=TMEFF2;refGene_tx=XM_006935444.3
C1	181589270	181589400
refGene_gn=TMEFF2;refGene_tx=XM_003990984.4;refGene_gn=TMEFF2;refGene_tx=XM_006935444.3
C1	181597942	181598052
refGene_gn=TMEFF2;refGene_tx=XM_003990984.4;refGene_gn=TMEFF2;refGene_tx=XM_006935444.3
C1	181600424	181600596
refGene_gn=TMEFF2;refGene_tx=XM_003990984.4;refGene_gn=TMEFF2;refGene_tx=XM_006935444.3
C1	181743116	181743389
refGene_gn=LOC101085738;refGene_tx=XM_003990986.4
C1	182974950	182975054
refGene_gn=LOC109503051;refGene_tx=XM_019839134.1
C1	182979795	182980677
refGene_gn=LOC109503051;refGene_tx=XM_019839134.1
C1	184738127	184739141
refGene_gn=SLC39A10;refGene_tx=XM_003990987.4;refGene_gn=SLC39A10;refGene_tx=XM_011285442.2;refGene_gn=SLC39A10;refGene_tx=XM_019838398.1;refGene_gn=SLC39A10;refGene_tx=XM_019838399.1
C1	184741050	184741258
refGene_gn=SLC39A10;refGene_tx=XM_003990987.4;refGene_gn=SLC39A10;refGene_tx=XM_011285442.2;refGene_gn=SLC39A10;refGene_tx=XM_019838398.1;refGene_gn=SLC39A10;refGene_tx=XM_019838399.1
C1	184753734	184753904
refGene_gn=SLC39A10;refGene_tx=XM_003990987.4;refGene_gn=SLC39A10;refGene_tx=XM_011285442.2;refGene_gn=SLC39A10;refGene_tx=XM_019838398.1;refGene_gn=SLC39A10;refGene_tx=XM_019838399.1
C1	184756148	184756337
refGene_gn=SLC39A10;refGene_tx=XM_003990987.4;refGene_gn=SLC39A10;refGene_tx=XM_011285442.2;refGene_gn=SLC39A10;refGene_tx=XM_019838398.1;re

fGene_gn=SLC39A10;refGene_tx=XM_019838399.1
C1 184759674 184759795
refGene_gn=SLC39A10;refGene_tx=XM_003990987.4;refGene_gn=SLC39A10;refGene_tx=XM_011285442.2;refGene_gn=SLC39A10;refGene_tx=XM_019838398.1;refGene_gn=SLC39A10;refGene_tx=XM_019838399.1
C1 184760970 184761339
refGene_gn=SLC39A10;refGene_tx=XM_003990987.4;refGene_gn=SLC39A10;refGene_tx=XM_011285442.2;refGene_gn=SLC39A10;refGene_tx=XM_019838398.1;refGene_gn=SLC39A10;refGene_tx=XM_019838399.1
C1 184762678 184762759
refGene_gn=SLC39A10;refGene_tx=XM_003990987.4;refGene_gn=SLC39A10;refGene_tx=XM_011285442.2;refGene_gn=SLC39A10;refGene_tx=XM_019838398.1;refGene_gn=SLC39A10;refGene_tx=XM_019838399.1
C1 184772555 184772746
refGene_gn=SLC39A10;refGene_tx=XM_003990987.4;refGene_gn=SLC39A10;refGene_tx=XM_011285442.2;refGene_gn=SLC39A10;refGene_tx=XM_019838398.1;refGene_gn=SLC39A10;refGene_tx=XM_019838399.1
C1 184778204 184778363
refGene_gn=SLC39A10;refGene_tx=XM_003990987.4;refGene_gn=SLC39A10;refGene_tx=XM_011285442.2;refGene_gn=SLC39A10;refGene_tx=XM_019838398.1;refGene_gn=SLC39A10;refGene_tx=XM_019838399.1
C1 184781202 184781409
refGene_gn=DNAH7;refGene_tx=XM_011285443.3
C1 184786006 184786110
refGene_gn=DNAH7;refGene_tx=XM_011285443.3
C1 184791128 184791306
refGene_gn=DNAH7;refGene_tx=XM_011285443.3
C1 184792699 184792852
refGene_gn=DNAH7;refGene_tx=XM_011285443.3
C1 184796086 184796317
refGene_gn=DNAH7;refGene_tx=XM_011285443.3
C1 184799793 184799931
refGene_gn=DNAH7;refGene_tx=XM_011285443.3
C1 184801386 184801572
refGene_gn=DNAH7;refGene_tx=XM_011285443.3
C1 184814766 184814928
refGene_gn=DNAH7;refGene_tx=XM_011285443.3
C1 184819055 184819256
refGene_gn=DNAH7;refGene_tx=XM_011285443.3
C1 184828459 184828621
refGene_gn=DNAH7;refGene_tx=XM_011285443.3
C1 184831503 184831680
refGene_gn=DNAH7;refGene_tx=XM_011285443.3
C1 184836867 184836960
refGene_gn=DNAH7;refGene_tx=XM_011285443.3
C1 184839743 184839938
refGene_gn=DNAH7;refGene_tx=XM_011285443.3
C1 184841183 184841310
refGene_gn=DNAH7;refGene_tx=XM_011285443.3
C1 184849310 184849646

refGene_gn=DNAH7;refGene_tx=XM_011285443.3
C1	184850344	184850478
refGene_gn=DNAH7;refGene_tx=XM_011285443.3
C1	184854436	184854627
refGene_gn=DNAH7;refGene_tx=XM_011285443.3
C1	184864722	184864877
refGene_gn=DNAH7;refGene_tx=XM_011285443.3
C1	184882800	184882964
refGene_gn=DNAH7;refGene_tx=XM_011285443.3
C1	184887662	184887848
refGene_gn=DNAH7;refGene_tx=XM_011285443.3
C1	184890954	184891135
refGene_gn=DNAH7;refGene_tx=XM_011285443.3
C1	184892783	184893130
refGene_gn=DNAH7;refGene_tx=XM_011285443.3
C1	184895801	184896132
refGene_gn=DNAH7;refGene_tx=XM_011285443.3
C1	184900620	184900850
refGene_gn=DNAH7;refGene_tx=XM_011285443.3
C1	184902868	184903741
refGene_gn=DNAH7;refGene_tx=XM_011285443.3
C1	184908324	184908544
refGene_gn=DNAH7;refGene_tx=XM_011285443.3
C1	184910017	184910144
refGene_gn=DNAH7;refGene_tx=XM_011285443.3
C1	184913396	184913565
refGene_gn=DNAH7;refGene_tx=XM_011285443.3
C1	184914272	184914428
refGene_gn=DNAH7;refGene_tx=XM_011285443.3
C1	184916487	184916685
refGene_gn=DNAH7;refGene_tx=XM_011285443.3
C1	184919741	184919966
refGene_gn=DNAH7;refGene_tx=XM_011285443.3
C1	184921748	184921880
refGene_gn=DNAH7;refGene_tx=XM_011285443.3
C1	184923947	184924124
refGene_gn=DNAH7;refGene_tx=XM_011285443.3
C1	184924457	184924640
refGene_gn=DNAH7;refGene_tx=XM_011285443.3
C1	184927109	184927259
refGene_gn=DNAH7;refGene_tx=XM_011285443.3
C1	184929897	184930146
refGene_gn=DNAH7;refGene_tx=XM_011285443.3
C1	184935093	184935192
refGene_gn=DNAH7;refGene_tx=XM_011285443.3
C1	184937080	184937293
refGene_gn=DNAH7;refGene_tx=XM_011285443.3
C1	184942282	184942410
refGene_gn=DNAH7;refGene_tx=XM_011285443.3
C1	184942527	184942630

refGene_gn=DNAH7;refGene_tx=XM_011285443.3
C1 184944919 184945088
refGene_gn=DNAH7;refGene_tx=XM_011285443.3
C1 184949274 184949384
refGene_gn=DNAH7;refGene_tx=XM_011285443.3
C1 184954376 184954589
refGene_gn=DNAH7;refGene_tx=XM_011285443.3
C1 184959531 184959672
refGene_gn=DNAH7;refGene_tx=XM_011285443.3
C1 184962804 184963003
refGene_gn=DNAH7;refGene_tx=XM_011285443.3
C1 184967222 184967416
refGene_gn=DNAH7;refGene_tx=XM_011285443.3
C1 184975845 184976032
refGene_gn=DNAH7;refGene_tx=XM_011285443.3
C1 184978442 184979128
refGene_gn=DNAH7;refGene_tx=XM_011285443.3
C1 184980597 184980744
refGene_gn=DNAH7;refGene_tx=XM_011285443.3
C1 184983543 184983569
refGene_gn=DNAH7;refGene_tx=XM_011285443.3
C1 184983571 184983767
refGene_gn=DNAH7;refGene_tx=XM_011285443.3
C1 184985482 184985561
refGene_gn=DNAH7;refGene_tx=XM_011285443.3
C1 184988522 184988650
refGene_gn=DNAH7;refGene_tx=XM_011285443.3
C1 184989430 184989703
refGene_gn=DNAH7;refGene_tx=XM_011285443.3
C1 184993886 184994066
refGene_gn=DNAH7;refGene_tx=XM_011285443.3
C1 184994667 184994851
refGene_gn=DNAH7;refGene_tx=XM_011285443.3
C1 184999368 184999488
refGene_gn=DNAH7;refGene_tx=XM_011285443.3
C1 185005962 185006088
refGene_gn=DNAH7;refGene_tx=XM_011285443.3
C1 185007581 185007657
refGene_gn=DNAH7;refGene_tx=XM_011285443.3
C1 185009733 185009914
refGene_gn=DNAH7;refGene_tx=XM_011285443.3
C1 185010999 185011029
refGene_gn=DNAH7;refGene_tx=XM_011285443.3
C1 185011030 185011088
refGene_gn=DNAH7;refGene_tx=XM_011285443.3
C1 185019544 185019692
refGene_gn=DNAH7;refGene_tx=XM_011285443.3
C1 185020424 185020533
refGene_gn=DNAH7;refGene_tx=XM_011285443.3
C1 185023853 185023916

refGene_gn=DNAH7;refGene_tx=XM_011285443.3
C1 185030683 185030746
refGene_gn=DNAH7;refGene_tx=XM_011285443.3
C1 185043227 185043242
refGene_gn=DNAH7;refGene_tx=XM_011285443.3
C1 185089286 185089569
refGene_gn=STK17B;refGene_tx=XM_003990988.4;refGene_gn=STK17B;refGene_tx=XM_006935445.3
C1 185091065 185091245
refGene_gn=STK17B;refGene_tx=XM_003990988.4;refGene_gn=STK17B;refGene_tx=XM_006935445.3
C1 185092431 185092480
refGene_gn=STK17B;refGene_tx=XM_003990988.4;refGene_gn=STK17B;refGene_tx=XM_006935445.3
C1 185094129 185094256
refGene_gn=STK17B;refGene_tx=XM_003990988.4;refGene_gn=STK17B;refGene_tx=XM_006935445.3
C1 185096047 185096192
refGene_gn=STK17B;refGene_tx=XM_003990988.4;refGene_gn=STK17B;refGene_tx=XM_006935445.3
C1 185101231 185101444
refGene_gn=STK17B;refGene_tx=XM_003990988.4;refGene_gn=STK17B;refGene_tx=XM_006935445.3
C1 185107824 185107946
refGene_gn=STK17B;refGene_tx=XM_003990988.4;refGene_gn=STK17B;refGene_tx=XM_006935445.3
C1 185144996 185145069
refGene_gn=HECW2;refGene_tx=XM_003990989.4
C1 185161031 185161144
refGene_gn=HECW2;refGene_tx=XM_003990989.4
C1 185162124 185162210
refGene_gn=HECW2;refGene_tx=XM_003990989.4
C1 185165049 185165164
refGene_gn=HECW2;refGene_tx=XM_003990989.4
C1 185165811 185165958
refGene_gn=HECW2;refGene_tx=XM_003990989.4
C1 185166971 185167101
refGene_gn=HECW2;refGene_tx=XM_003990989.4
C1 185170405 185170504
refGene_gn=HECW2;refGene_tx=XM_003990989.4
C1 185172277 185172430
refGene_gn=HECW2;refGene_tx=XM_003990989.4
C1 185182520 185182634
refGene_gn=HECW2;refGene_tx=XM_003990989.4
C1 185184094 185184215
refGene_gn=HECW2;refGene_tx=XM_003990989.4
C1 185195261 185195371
refGene_gn=HECW2;refGene_tx=XM_003990989.4
C1 185200642 185200726
refGene_gn=HECW2;refGene_tx=XM_003990989.4

C1 185209410 185209507
refGene_gn=HECW2;refGene_tx=XM_003990989.4
C1 185213599 185213702
refGene_gn=HECW2;refGene_tx=XM_003990989.4
C1 185218166 185218301
refGene_gn=HECW2;refGene_tx=XM_003990989.4
C1 185240513 185240699
refGene_gn=HECW2;refGene_tx=XM_003990989.4
C1 185253340 185253465
refGene_gn=HECW2;refGene_tx=XM_003990989.4
C1 185254193 185254297
refGene_gn=HECW2;refGene_tx=XM_003990989.4
C1 185254978 185255129
refGene_gn=HECW2;refGene_tx=XM_003990989.4
C1 185265150 185265246
refGene_gn=HECW2;refGene_tx=XM_003990989.4
C1 185266269 185267619
refGene_gn=HECW2;refGene_tx=XM_003990989.4
C1 185268306 185268407
refGene_gn=HECW2;refGene_tx=XM_003990989.4
C1 185273064 185273207
refGene_gn=HECW2;refGene_tx=XM_003990989.4
C1 185275820 185275990
refGene_gn=HECW2;refGene_tx=XM_003990989.4
C1 185280413 185280489
refGene_gn=HECW2;refGene_tx=XM_003990989.4
C1 185286304 185286399
refGene_gn=HECW2;refGene_tx=XM_003990989.4
C1 185296573 185296681
refGene_gn=HECW2;refGene_tx=XM_003990989.4
C1 185375501 185375793
refGene_gn=HECW2;refGene_tx=XM_003990989.4
C1 185552533 185552569
refGene_gn=CCDC150;refGene_tx=XM_019838401.1;refGene_gn=CCDC150;refGene_tx=XM_019838402.1;refGene_gn=CCDC150;refGene_tx=XM_019838403.1;refGene_gn=CCDC150;refGene_tx=XM_019838406.2
C1 185552640 185552652
refGene_gn=CCDC150;refGene_tx=XM_019838404.1
C1 185560123 185560287
refGene_gn=CCDC150;refGene_tx=XM_019838401.1;refGene_gn=CCDC150;refGene_tx=XM_019838402.1;refGene_gn=CCDC150;refGene_tx=XM_019838403.1;refGene_gn=CCDC150;refGene_tx=XM_019838404.1;refGene_gn=CCDC150;refGene_tx=XM_019838406.2
C1 185561511 185561732
refGene_gn=CCDC150;refGene_tx=XM_019838401.1;refGene_gn=CCDC150;refGene_tx=XM_019838402.1;refGene_gn=CCDC150;refGene_tx=XM_019838403.1;refGene_gn=CCDC150;refGene_tx=XM_019838404.1;refGene_gn=CCDC150;refGene_tx=XM_019838406.2;refGene_gn=CCDC150;refGene_tx=XM_019838407.1;refGene_gn=CCDC150;refGene_tx=XM_023259580.1
C1 185562602 185562781

refGene_gn=CCDC150;refGene_tx=XM_019838401.1;refGene_gn=CCDC150;refGene_tx=XM_019838402.1;refGene_gn=CCDC150;refGene_tx=XM_019838403.1;refGene_gn=CCDC150;refGene_tx=XM_019838404.1;refGene_gn=CCDC150;refGene_tx=XM_019838406.2;refGene_gn=CCDC150;refGene_tx=XM_019838407.1;refGene_gn=CCDC150;refGene_tx=XM_023259580.1
C1 185564614 185564683
refGene_gn=CCDC150;refGene_tx=XM_019838401.1;refGene_gn=CCDC150;refGene_tx=XM_019838402.1;refGene_gn=CCDC150;refGene_tx=XM_019838403.1;refGene_gn=CCDC150;refGene_tx=XM_019838404.1;refGene_gn=CCDC150;refGene_tx=XM_019838406.2;refGene_gn=CCDC150;refGene_tx=XM_019838407.1;refGene_gn=CCDC150;refGene_tx=XM_019838408.1;refGene_gn=CCDC150;refGene_tx=XM_023259580.1
C1 185573283 185573400
refGene_gn=CCDC150;refGene_tx=XM_019838401.1;refGene_gn=CCDC150;refGene_tx=XM_019838402.1;refGene_gn=CCDC150;refGene_tx=XM_019838403.1;refGene_gn=CCDC150;refGene_tx=XM_019838404.1;refGene_gn=CCDC150;refGene_tx=XM_019838406.2;refGene_gn=CCDC150;refGene_tx=XM_019838407.1;refGene_gn=CCDC150;refGene_tx=XM_019838408.1;refGene_gn=CCDC150;refGene_tx=XM_023259580.1
C1 185574033 185574163
refGene_gn=CCDC150;refGene_tx=XM_019838401.1;refGene_gn=CCDC150;refGene_tx=XM_019838402.1;refGene_gn=CCDC150;refGene_tx=XM_019838403.1;refGene_gn=CCDC150;refGene_tx=XM_019838404.1;refGene_gn=CCDC150;refGene_tx=XM_019838406.2;refGene_gn=CCDC150;refGene_tx=XM_019838407.1;refGene_gn=CCDC150;refGene_tx=XM_019838408.1;refGene_gn=CCDC150;refGene_tx=XM_023259580.1
C1 185577023 185577067
refGene_gn=CCDC150;refGene_tx=XM_019838401.1;refGene_gn=CCDC150;refGene_tx=XM_019838402.1;refGene_gn=CCDC150;refGene_tx=XM_019838403.1;refGene_gn=CCDC150;refGene_tx=XM_019838404.1;refGene_gn=CCDC150;refGene_tx=XM_019838406.2;refGene_gn=CCDC150;refGene_tx=XM_019838407.1;refGene_gn=CCDC150;refGene_tx=XM_019838408.1;refGene_gn=CCDC150;refGene_tx=XM_023259580.1
C1 185578550 185578643
refGene_gn=CCDC150;refGene_tx=XM_019838401.1;refGene_gn=CCDC150;refGene_tx=XM_019838402.1;refGene_gn=CCDC150;refGene_tx=XM_019838403.1;refGene_gn=CCDC150;refGene_tx=XM_019838404.1;refGene_gn=CCDC150;refGene_tx=XM_019838406.2;refGene_gn=CCDC150;refGene_tx=XM_019838407.1;refGene_gn=CCDC150;refGene_tx=XM_019838408.1;refGene_gn=CCDC150;refGene_tx=XM_023259580.1
C1 185582201 185582309
refGene_gn=CCDC150;refGene_tx=XM_019838401.1;refGene_gn=CCDC150;refGene_tx=XM_019838402.1;refGene_gn=CCDC150;refGene_tx=XM_019838403.1;refGene_gn=CCDC150;refGene_tx=XM_019838404.1;refGene_gn=CCDC150;refGene_tx=XM_019838406.2;refGene_gn=CCDC150;refGene_tx=XM_019838407.1;refGene_gn=CCDC150;refGene_tx=XM_019838408.1;refGene_gn=CCDC150;refGene_tx=XM_023259580.1
C1 185587159 185587284
refGene_gn=CCDC150;refGene_tx=XM_019838401.1;refGene_gn=CCDC150;refGene_tx=XM_019838402.1;refGene_gn=CCDC150;refGene_tx=XM_019838403.1;refGe

ne_gn=CCDC150;refGene_tx=XM_019838404.1;refGene_gn=CCDC150;refGene_tx=XM_019838406.2;refGene_gn=CCDC150;refGene_tx=XM_019838407.1;refGene_gn=CCDC150;refGene_tx=XM_019838408.1;refGene_gn=CCDC150;refGene_tx=XM_023259580.1
C1 185587526 185587704
refGene_gn=CCDC150;refGene_tx=XM_019838401.1;refGene_gn=CCDC150;refGene_tx=XM_019838402.1;refGene_gn=CCDC150;refGene_tx=XM_019838403.1;refGene_gn=CCDC150;refGene_tx=XM_019838404.1;refGene_gn=CCDC150;refGene_tx=XM_019838406.2;refGene_gn=CCDC150;refGene_tx=XM_019838407.1;refGene_gn=CCDC150;refGene_tx=XM_019838408.1;refGene_gn=CCDC150;refGene_tx=XM_023259580.1
C1 185588256 185588325
refGene_gn=CCDC150;refGene_tx=XM_019838401.1;refGene_gn=CCDC150;refGene_tx=XM_019838402.1;refGene_gn=CCDC150;refGene_tx=XM_019838403.1;refGene_gn=CCDC150;refGene_tx=XM_019838404.1;refGene_gn=CCDC150;refGene_tx=XM_019838406.2;refGene_gn=CCDC150;refGene_tx=XM_019838407.1;refGene_gn=CCDC150;refGene_tx=XM_019838408.1;refGene_gn=CCDC150;refGene_tx=XM_023259580.1
C1 185602242 185602356
refGene_gn=CCDC150;refGene_tx=XM_019838401.1;refGene_gn=CCDC150;refGene_tx=XM_019838402.1;refGene_gn=CCDC150;refGene_tx=XM_019838403.1;refGene_gn=CCDC150;refGene_tx=XM_019838404.1;refGene_gn=CCDC150;refGene_tx=XM_019838406.2;refGene_gn=CCDC150;refGene_tx=XM_019838407.1;refGene_gn=CCDC150;refGene_tx=XM_019838408.1;refGene_gn=CCDC150;refGene_tx=XM_023259580.1
C1 185608683 185608767
refGene_gn=CCDC150;refGene_tx=XM_019838401.1;refGene_gn=CCDC150;refGene_tx=XM_019838402.1;refGene_gn=CCDC150;refGene_tx=XM_019838403.1;refGene_gn=CCDC150;refGene_tx=XM_019838404.1;refGene_gn=CCDC150;refGene_tx=XM_019838406.2;refGene_gn=CCDC150;refGene_tx=XM_019838407.1;refGene_gn=CCDC150;refGene_tx=XM_019838408.1;refGene_gn=CCDC150;refGene_tx=XM_023259580.1
C1 185621939 185622047
refGene_gn=CCDC150;refGene_tx=XM_019838401.1;refGene_gn=CCDC150;refGene_tx=XM_019838402.1;refGene_gn=CCDC150;refGene_tx=XM_019838403.1;refGene_gn=CCDC150;refGene_tx=XM_019838404.1;refGene_gn=CCDC150;refGene_tx=XM_019838406.2;refGene_gn=CCDC150;refGene_tx=XM_019838407.1;refGene_gn=CCDC150;refGene_tx=XM_019838408.1;refGene_gn=CCDC150;refGene_tx=XM_023259580.1
C1 185623059 185623383
refGene_gn=CCDC150;refGene_tx=XM_023259582.1
C1 185628557 185628686
refGene_gn=CCDC150;refGene_tx=XM_019838401.1;refGene_gn=CCDC150;refGene_tx=XM_019838402.1;refGene_gn=CCDC150;refGene_tx=XM_019838403.1;refGene_gn=CCDC150;refGene_tx=XM_019838404.1;refGene_gn=CCDC150;refGene_tx=XM_019838406.2;refGene_gn=CCDC150;refGene_tx=XM_019838407.1;refGene_gn=CCDC150;refGene_tx=XM_019838408.1;refGene_gn=CCDC150;refGene_tx=XM_023259580.1;refGene_gn=CCDC150;refGene_tx=XM_023259582.1
C1 185630080 185630250
refGene_gn=CCDC150;refGene_tx=XM_019838401.1;refGene_gn=CCDC150;refGen

e_tx=XM_019838402.1;refGene_gn=CCDC150;refGene_tx=XM_019838403.1;refGene_gn=CCDC150;refGene_tx=XM_019838404.1;refGene_gn=CCDC150;refGene_tx=XM_019838406.2;refGene_gn=CCDC150;refGene_tx=XM_019838407.1;refGene_gn=CCDC150;refGene_tx=XM_019838408.1;refGene_gn=CCDC150;refGene_tx=XM_023259580.1;refGene_gn=CCDC150;refGene_tx=XM_023259582.1
C1 185632536 185632630
refGene_gn=CCDC150;refGene_tx=XM_019838401.1;refGene_gn=CCDC150;refGene_tx=XM_019838402.1;refGene_gn=CCDC150;refGene_tx=XM_019838403.1;refGene_gn=CCDC150;refGene_tx=XM_019838404.1;refGene_gn=CCDC150;refGene_tx=XM_019838406.2;refGene_gn=CCDC150;refGene_tx=XM_019838407.1;refGene_gn=CCDC150;refGene_tx=XM_019838408.1;refGene_gn=CCDC150;refGene_tx=XM_023259580.1;refGene_gn=CCDC150;refGene_tx=XM_023259582.1
C1 185633426 185633596
refGene_gn=CCDC150;refGene_tx=XM_019838401.1;refGene_gn=CCDC150;refGene_tx=XM_019838402.1;refGene_gn=CCDC150;refGene_tx=XM_019838403.1;refGene_gn=CCDC150;refGene_tx=XM_019838404.1;refGene_gn=CCDC150;refGene_tx=XM_019838406.2;refGene_gn=CCDC150;refGene_tx=XM_019838407.1;refGene_gn=CCDC150;refGene_tx=XM_019838408.1;refGene_gn=CCDC150;refGene_tx=XM_023259580.1;refGene_gn=CCDC150;refGene_tx=XM_023259582.1
C1 185634756 185634883
refGene_gn=CCDC150;refGene_tx=XM_019838401.1;refGene_gn=CCDC150;refGene_tx=XM_019838402.1;refGene_gn=CCDC150;refGene_tx=XM_019838403.1;refGene_gn=CCDC150;refGene_tx=XM_019838404.1;refGene_gn=CCDC150;refGene_tx=XM_019838406.2;refGene_gn=CCDC150;refGene_tx=XM_019838407.1;refGene_gn=CCDC150;refGene_tx=XM_019838408.1;refGene_gn=CCDC150;refGene_tx=XM_023259580.1;refGene_gn=CCDC150;refGene_tx=XM_023259582.1
C1 185643694 185643889
refGene_gn=CCDC150;refGene_tx=XM_019838401.1;refGene_gn=CCDC150;refGene_tx=XM_019838402.1;refGene_gn=CCDC150;refGene_tx=XM_019838403.1;refGene_gn=CCDC150;refGene_tx=XM_019838404.1;refGene_gn=CCDC150;refGene_tx=XM_019838406.2;refGene_gn=CCDC150;refGene_tx=XM_019838407.1;refGene_gn=CCDC150;refGene_tx=XM_019838408.1;refGene_gn=CCDC150;refGene_tx=XM_023259580.1;refGene_gn=CCDC150;refGene_tx=XM_023259582.1
C1 185644290 185644359
refGene_gn=CCDC150;refGene_tx=XM_019838401.1;refGene_gn=CCDC150;refGene_tx=XM_019838402.1;refGene_gn=CCDC150;refGene_tx=XM_019838403.1;refGene_gn=CCDC150;refGene_tx=XM_019838404.1;refGene_gn=CCDC150;refGene_tx=XM_019838406.2;refGene_gn=CCDC150;refGene_tx=XM_019838407.1;refGene_gn=CCDC150;refGene_tx=XM_019838408.1;refGene_gn=CCDC150;refGene_tx=XM_023259580.1;refGene_gn=CCDC150;refGene_tx=XM_023259582.1
C1 185644453 185644615
refGene_gn=CCDC150;refGene_tx=XM_019838401.1;refGene_gn=CCDC150;refGene_tx=XM_019838402.1;refGene_gn=CCDC150;refGene_tx=XM_019838403.1;refGene_gn=CCDC150;refGene_tx=XM_019838404.1;refGene_gn=CCDC150;refGene_tx=XM_019838406.2;refGene_gn=CCDC150;refGene_tx=XM_019838407.1;refGene_gn=CCDC150;refGene_tx=XM_019838408.1;refGene_gn=CCDC150;refGene_tx=XM_023259580.1;refGene_gn=CCDC150;refGene_tx=XM_023259582.1
C1 185645292 185645379
refGene_gn=CCDC150;refGene_tx=XM_019838401.1;refGene_gn=CCDC150;refGene_tx=XM_019838402.1;refGene_gn=CCDC150;refGene_tx=XM_019838403.1;refGe

ne_gn=CCDC150;refGene_tx=XM_019838404.1;refGene_gn=CCDC150;refGene_tx=XM_019838406.2;refGene_gn=CCDC150;refGene_tx=XM_019838407.1;refGene_gn=CCDC150;refGene_tx=XM_019838408.1;refGene_gn=CCDC150;refGene_tx=XM_023259580.1;refGene_gn=CCDC150;refGene_tx=XM_023259582.1
C1 185645766 185645838
refGene_gn=CCDC150;refGene_tx=XM_019838406.2
C1 185647515 185647635
refGene_gn=CCDC150;refGene_tx=XM_019838401.1;refGene_gn=CCDC150;refGene_tx=XM_019838402.1;refGene_gn=CCDC150;refGene_tx=XM_019838403.1;refGene_gn=CCDC150;refGene_tx=XM_019838404.1;refGene_gn=CCDC150;refGene_tx=XM_019838407.1;refGene_gn=CCDC150;refGene_tx=XM_019838408.1;refGene_gn=CCDC150;refGene_tx=XM_023259580.1;refGene_gn=CCDC150;refGene_tx=XM_023259582.1
C1 185648020 185648163
refGene_gn=CCDC150;refGene_tx=XM_019838401.1;refGene_gn=CCDC150;refGene_tx=XM_019838402.1;refGene_gn=CCDC150;refGene_tx=XM_019838403.1;refGene_gn=CCDC150;refGene_tx=XM_019838404.1;refGene_gn=CCDC150;refGene_tx=XM_019838407.1;refGene_gn=CCDC150;refGene_tx=XM_019838408.1;refGene_gn=CCDC150;refGene_tx=XM_023259580.1;refGene_gn=CCDC150;refGene_tx=XM_023259582.1
C1 185671581 185671704
refGene_gn=GTF3C3;refGene_tx=XM_003990991.4
C1 185673482 185673635
refGene_gn=GTF3C3;refGene_tx=XM_003990991.4
C1 185677049 185677174
refGene_gn=GTF3C3;refGene_tx=XM_003990991.4
C1 185679214 185679405
refGene_gn=GTF3C3;refGene_tx=XM_003990991.4
C1 185680013 185680251
refGene_gn=GTF3C3;refGene_tx=XM_003990991.4
C1 185682440 185682576
refGene_gn=GTF3C3;refGene_tx=XM_003990991.4
C1 185683651 185683753
refGene_gn=GTF3C3;refGene_tx=XM_003990991.4
C1 185684308 185684511
refGene_gn=GTF3C3;refGene_tx=XM_003990991.4
C1 185686804 185686976
refGene_gn=GTF3C3;refGene_tx=XM_003990991.4
C1 185688304 185688408
refGene_gn=GTF3C3;refGene_tx=XM_003990991.4
C1 185692380 185692422
refGene_gn=GTF3C3;refGene_tx=XM_003990991.4
C1 185692424 185692452
refGene_gn=GTF3C3;refGene_tx=XM_003990991.4
C1 185692999 185693147
refGene_gn=GTF3C3;refGene_tx=XM_003990991.4
C1 185695705 185695871
refGene_gn=GTF3C3;refGene_tx=XM_003990991.4
C1 185696400 185696592
refGene_gn=GTF3C3;refGene_tx=XM_003990991.4

C1	185704860	185704984
refGene_gn=GTF3C3;refGene_tx=XM_003990991.4
C1	185705860	185706051
refGene_gn=GTF3C3;refGene_tx=XM_003990991.4
C1	185708440	185708552
refGene_gn=GTF3C3;refGene_tx=XM_003990991.4
C1	185709418	185709520
refGene_gn=GTF3C3;refGene_tx=XM_003990991.4
C1	185740632	185740675
refGene_gn=PGAP1;refGene_tx=XM_019838410.2
C1	185760073	185760212
refGene_gn=PGAP1;refGene_tx=XM_011285444.2;refGene_gn=PGAP1;refGene_tx=XM_019838409.2
C1	185761449	185761554
refGene_gn=PGAP1;refGene_tx=XM_011285444.2;refGene_gn=PGAP1;refGene_tx=XM_019838409.2;refGene_gn=PGAP1;refGene_tx=XM_019838410.2
C1	185763504	185763692
refGene_gn=PGAP1;refGene_tx=XM_011285444.2;refGene_gn=PGAP1;refGene_tx=XM_019838409.2;refGene_gn=PGAP1;refGene_tx=XM_019838410.2
C1	185764165	185764216
refGene_gn=PGAP1;refGene_tx=XM_011285444.2;refGene_gn=PGAP1;refGene_tx=XM_019838409.2;refGene_gn=PGAP1;refGene_tx=XM_019838410.2
C1	185765446	185765582
refGene_gn=PGAP1;refGene_tx=XM_011285444.2;refGene_gn=PGAP1;refGene_tx=XM_019838409.2;refGene_gn=PGAP1;refGene_tx=XM_019838410.2
C1	185766656	185766854
refGene_gn=PGAP1;refGene_tx=XM_011285444.2;refGene_gn=PGAP1;refGene_tx=XM_019838409.2;refGene_gn=PGAP1;refGene_tx=XM_019838410.2
C1	185767665	185767756
refGene_gn=PGAP1;refGene_tx=XM_011285444.2;refGene_gn=PGAP1;refGene_tx=XM_019838409.2;refGene_gn=PGAP1;refGene_tx=XM_019838410.2
C1	185769567	185769661
refGene_gn=PGAP1;refGene_tx=XM_011285444.2;refGene_gn=PGAP1;refGene_tx=XM_019838409.2;refGene_gn=PGAP1;refGene_tx=XM_019838410.2
C1	185775091	185775130
refGene_gn=PGAP1;refGene_tx=XM_011285444.2;refGene_gn=PGAP1;refGene_tx=XM_019838409.2;refGene_gn=PGAP1;refGene_tx=XM_019838410.2
C1	185776827	185776936
refGene_gn=PGAP1;refGene_tx=XM_011285444.2;refGene_gn=PGAP1;refGene_tx=XM_019838409.2;refGene_gn=PGAP1;refGene_tx=XM_019838410.2
C1	185778878	185778945
refGene_gn=PGAP1;refGene_tx=XM_011285444.2;refGene_gn=PGAP1;refGene_tx=XM_019838409.2;refGene_gn=PGAP1;refGene_tx=XM_019838410.2
C1	185779431	185779483
refGene_gn=PGAP1;refGene_tx=XM_011285444.2;refGene_gn=PGAP1;refGene_tx=XM_019838409.2;refGene_gn=PGAP1;refGene_tx=XM_019838410.2
C1	185779575	185779649
refGene_gn=PGAP1;refGene_tx=XM_011285444.2;refGene_gn=PGAP1;refGene_tx=XM_019838409.2;refGene_gn=PGAP1;refGene_tx=XM_019838410.2
C1	185782762	185782838

refGene_gn=PGAP1;refGene_tx=XM_011285444.2;refGene_gn=PGAP1;refGene_tx=XM_019838409.2;refGene_gn=PGAP1;refGene_tx=XM_019838410.2
C1 185786981 185787059
refGene_gn=PGAP1;refGene_tx=XM_011285444.2;refGene_gn=PGAP1;refGene_tx=XM_019838410.2
C1 185791682 185791734
refGene_gn=PGAP1;refGene_tx=XM_011285444.2;refGene_gn=PGAP1;refGene_tx=XM_019838409.2;refGene_gn=PGAP1;refGene_tx=XM_019838410.2
C1 185792123 185792170
refGene_gn=PGAP1;refGene_tx=XM_011285444.2;refGene_gn=PGAP1;refGene_tx=XM_019838409.2;refGene_gn=PGAP1;refGene_tx=XM_019838410.2
C1 185796061 185796145
refGene_gn=PGAP1;refGene_tx=XM_011285444.2;refGene_gn=PGAP1;refGene_tx=XM_019838409.2;refGene_gn=PGAP1;refGene_tx=XM_019838410.2
C1 185797772 185797828
refGene_gn=PGAP1;refGene_tx=XM_011285444.2;refGene_gn=PGAP1;refGene_tx=XM_019838409.2;refGene_gn=PGAP1;refGene_tx=XM_019838410.2
C1 185798505 185798611
refGene_gn=PGAP1;refGene_tx=XM_011285444.2;refGene_gn=PGAP1;refGene_tx=XM_019838409.2;refGene_gn=PGAP1;refGene_tx=XM_019838410.2
C1 185802101 185802168
refGene_gn=PGAP1;refGene_tx=XM_011285444.2;refGene_gn=PGAP1;refGene_tx=XM_019838409.2;refGene_gn=PGAP1;refGene_tx=XM_019838410.2
C1 185802967 185803020
refGene_gn=PGAP1;refGene_tx=XM_011285444.2;refGene_gn=PGAP1;refGene_tx=XM_019838409.2;refGene_gn=PGAP1;refGene_tx=XM_019838410.2
C1 185806446 185806604
refGene_gn=PGAP1;refGene_tx=XM_011285444.2;refGene_gn=PGAP1;refGene_tx=XM_019838409.2;refGene_gn=PGAP1;refGene_tx=XM_019838410.2
C1 185810086 185810258
refGene_gn=PGAP1;refGene_tx=XM_011285444.2;refGene_gn=PGAP1;refGene_tx=XM_019838409.2;refGene_gn=PGAP1;refGene_tx=XM_019838410.2
C1 185830331 185830507
refGene_gn=PGAP1;refGene_tx=XM_011285444.2;refGene_gn=PGAP1;refGene_tx=XM_019838409.2;refGene_gn=PGAP1;refGene_tx=XM_019838410.2
C1 185831915 185832069
refGene_gn=PGAP1;refGene_tx=XM_011285444.2;refGene_gn=PGAP1;refGene_tx=XM_019838409.2;refGene_gn=PGAP1;refGene_tx=XM_019838410.2
C1 185838492 185838639
refGene_gn=PGAP1;refGene_tx=XM_011285444.2;refGene_gn=PGAP1;refGene_tx=XM_019838409.2;refGene_gn=PGAP1;refGene_tx=XM_019838410.2
C1 185875097 185875279
refGene_gn=ANKRD44;refGene_tx=XM_023259585.1
C1 185878585 185878590
refGene_gn=ANKRD44;refGene_tx=XM_023259586.1
C1 185894554 185894802
refGene_gn=ANKRD44;refGene_tx=XM_023259583.1;refGene_gn=ANKRD44;refGene_tx=XM_023259584.1
C1 185896293 185896329
refGene_gn=ANKRD44;refGene_tx=XM_023259584.1

C1 185897114 185897206
refGene_gn=ANKRD44;refGene_tx=XM_023259583.1;refGene_gn=ANKRD44;refGene_tx=XM_023259584.1;refGene_gn=ANKRD44;refGene_tx=XM_023259585.1;refGene_gn=ANKRD44;refGene_tx=XM_023259586.1
C1 185898199 185898282
refGene_gn=ANKRD44;refGene_tx=XM_023259583.1;refGene_gn=ANKRD44;refGene_tx=XM_023259584.1;refGene_gn=ANKRD44;refGene_tx=XM_023259585.1;refGene_gn=ANKRD44;refGene_tx=XM_023259586.1
C1 185900402 185900485
refGene_gn=ANKRD44;refGene_tx=XM_023259583.1;refGene_gn=ANKRD44;refGene_tx=XM_023259584.1;refGene_gn=ANKRD44;refGene_tx=XM_023259585.1;refGene_gn=ANKRD44;refGene_tx=XM_023259586.1
C1 185900938 185901084
refGene_gn=ANKRD44;refGene_tx=XM_023259583.1;refGene_gn=ANKRD44;refGene_tx=XM_023259584.1;refGene_gn=ANKRD44;refGene_tx=XM_023259585.1;refGene_gn=ANKRD44;refGene_tx=XM_023259586.1
C1 185902592 185902676
refGene_gn=ANKRD44;refGene_tx=XM_023259583.1;refGene_gn=ANKRD44;refGene_tx=XM_023259584.1;refGene_gn=ANKRD44;refGene_tx=XM_023259585.1;refGene_gn=ANKRD44;refGene_tx=XM_023259586.1
C1 185910057 185910145
refGene_gn=ANKRD44;refGene_tx=XM_023259583.1;refGene_gn=ANKRD44;refGene_tx=XM_023259584.1;refGene_gn=ANKRD44;refGene_tx=XM_023259585.1;refGene_gn=ANKRD44;refGene_tx=XM_023259586.1
C1 185913766 185913983
refGene_gn=ANKRD44;refGene_tx=XM_023259583.1;refGene_gn=ANKRD44;refGene_tx=XM_023259584.1;refGene_gn=ANKRD44;refGene_tx=XM_023259585.1;refGene_gn=ANKRD44;refGene_tx=XM_023259586.1
C1 185915639 185915757
refGene_gn=ANKRD44;refGene_tx=XM_023259583.1;refGene_gn=ANKRD44;refGene_tx=XM_023259584.1;refGene_gn=ANKRD44;refGene_tx=XM_023259585.1;refGene_gn=ANKRD44;refGene_tx=XM_023259586.1
C1 185916799 185916887
refGene_gn=ANKRD44;refGene_tx=XM_023259583.1;refGene_gn=ANKRD44;refGene_tx=XM_023259584.1;refGene_gn=ANKRD44;refGene_tx=XM_023259585.1;refGene_gn=ANKRD44;refGene_tx=XM_023259586.1
C1 185921427 185921629
refGene_gn=ANKRD44;refGene_tx=XM_023259583.1;refGene_gn=ANKRD44;refGene_tx=XM_023259584.1;refGene_gn=ANKRD44;refGene_tx=XM_023259585.1;refGene_gn=ANKRD44;refGene_tx=XM_023259586.1
C1 185931973 185932045
refGene_gn=ANKRD44;refGene_tx=XM_023259583.1;refGene_gn=ANKRD44;refGene_tx=XM_023259584.1;refGene_gn=ANKRD44;refGene_tx=XM_023259585.1;refGene_gn=ANKRD44;refGene_tx=XM_023259586.1
C1 185986913 185987025
refGene_gn=ANKRD44;refGene_tx=XM_023259583.1;refGene_gn=ANKRD44;refGene_tx=XM_023259584.1;refGene_gn=ANKRD44;refGene_tx=XM_023259585.1;refGene_gn=ANKRD44;refGene_tx=XM_023259586.1
C1 185990856 185990940
refGene_gn=ANKRD44;refGene_tx=XM_023259583.1;refGene_gn=ANKRD44;refGen

e_tx=XM_023259584.1;refGene_gn=ANKRD44;refGene_tx=XM_023259585.1;refGene_gn=ANKRD44;refGene_tx=XM_023259586.1
C1	185992623	185992764	refGene_gn=ANKRD44;refGene_tx=XM_023259583.1;refGene_gn=ANKRD44;refGene_tx=XM_023259584.1;refGene_gn=ANKRD44;refGene_tx=XM_023259585.1;refGene_gn=ANKRD44;refGene_tx=XM_023259586.1
C1	185994298	185994367	refGene_gn=ANKRD44;refGene_tx=XM_023259583.1;refGene_gn=ANKRD44;refGene_tx=XM_023259584.1;refGene_gn=ANKRD44;refGene_tx=XM_023259585.1;refGene_gn=ANKRD44;refGene_tx=XM_023259586.1
C1	185996366	185996484	refGene_gn=ANKRD44;refGene_tx=XM_023259583.1;refGene_gn=ANKRD44;refGene_tx=XM_023259584.1;refGene_gn=ANKRD44;refGene_tx=XM_023259585.1;refGene_gn=ANKRD44;refGene_tx=XM_023259586.1
C1	185997575	185997658	refGene_gn=ANKRD44;refGene_tx=XM_023259583.1;refGene_gn=ANKRD44;refGene_tx=XM_023259584.1;refGene_gn=ANKRD44;refGene_tx=XM_023259585.1;refGene_gn=ANKRD44;refGene_tx=XM_023259586.1
C1	186007083	186007198	refGene_gn=ANKRD44;refGene_tx=XM_023259583.1;refGene_gn=ANKRD44;refGene_tx=XM_023259584.1;refGene_gn=ANKRD44;refGene_tx=XM_023259585.1;refGene_gn=ANKRD44;refGene_tx=XM_023259586.1
C1	186014291	186014370	refGene_gn=ANKRD44;refGene_tx=XM_023259583.1;refGene_gn=ANKRD44;refGene_tx=XM_023259584.1;refGene_gn=ANKRD44;refGene_tx=XM_023259585.1;refGene_gn=ANKRD44;refGene_tx=XM_023259586.1
C1	186023109	186023322	refGene_gn=ANKRD44;refGene_tx=XM_023259583.1;refGene_gn=ANKRD44;refGene_tx=XM_023259584.1;refGene_gn=ANKRD44;refGene_tx=XM_023259585.1;refGene_gn=ANKRD44;refGene_tx=XM_023259586.1
C1	186024395	186024538	refGene_gn=ANKRD44;refGene_tx=XM_023259583.1;refGene_gn=ANKRD44;refGene_tx=XM_023259584.1;refGene_gn=ANKRD44;refGene_tx=XM_023259585.1;refGene_gn=ANKRD44;refGene_tx=XM_023259586.1
C1	186027181	186027269	refGene_gn=ANKRD44;refGene_tx=XM_023259583.1;refGene_gn=ANKRD44;refGene_tx=XM_023259584.1;refGene_gn=ANKRD44;refGene_tx=XM_023259585.1;refGene_gn=ANKRD44;refGene_tx=XM_023259586.1
C1	186027591	186027792	refGene_gn=ANKRD44;refGene_tx=XM_023259583.1;refGene_gn=ANKRD44;refGene_tx=XM_023259584.1;refGene_gn=ANKRD44;refGene_tx=XM_023259585.1;refGene_gn=ANKRD44;refGene_tx=XM_023259586.1
C1	186041293	186041364	refGene_gn=ANKRD44;refGene_tx=XM_023259583.1;refGene_gn=ANKRD44;refGene_tx=XM_023259584.1;refGene_gn=ANKRD44;refGene_tx=XM_023259585.1;refGene_gn=ANKRD44;refGene_tx=XM_023259586.1
C1	186053643	186053722	refGene_gn=ANKRD44;refGene_tx=XM_023259583.1;refGene_gn=ANKRD44;refGene_tx=XM_023259584.1;refGene_gn=ANKRD44;refGene_tx=XM_023259585.1;refGene_gn=ANKRD44;refGene_tx=XM_023259586.1

C1 186099097 186099133
refGene_gn=ANKRD44;refGene_tx=XM_023259583.1;refGene_gn=ANKRD44;refGene_tx=XM_023259584.1;refGene_gn=ANKRD44;refGene_tx=XM_023259585.1;refGene_gn=ANKRD44;refGene_tx=XM_023259586.1
C1 186282296 186282455
refGene_gn=SF3B1;refGene_tx=XM_003990992.4;refGene_gn=SF3B1;refGene_tx=XM_011285447.3
C1 186282745 186282962
refGene_gn=SF3B1;refGene_tx=XM_003990992.4;refGene_gn=SF3B1;refGene_tx=XM_011285447.3
C1 186285574 186285847
refGene_gn=SF3B1;refGene_tx=XM_003990992.4;refGene_gn=SF3B1;refGene_tx=XM_011285447.3
C1 186286957 186287089
refGene_gn=SF3B1;refGene_tx=XM_003990992.4;refGene_gn=SF3B1;refGene_tx=XM_011285447.3
C1 186287379 186287500
refGene_gn=SF3B1;refGene_tx=XM_003990992.4;refGene_gn=SF3B1;refGene_tx=XM_011285447.3;refGene_gn=SF3B1;refGene_tx=XM_011285448.2
C1 186287895 186287965
refGene_gn=SF3B1;refGene_tx=XM_011285448.2
C1 186288986 186289098
refGene_gn=SF3B1;refGene_tx=XM_003990992.4;refGene_gn=SF3B1;refGene_tx=XM_011285447.3;refGene_gn=SF3B1;refGene_tx=XM_011285448.2
C1 186289192 186289375
refGene_gn=SF3B1;refGene_tx=XM_003990992.4;refGene_gn=SF3B1;refGene_tx=XM_011285447.3;refGene_gn=SF3B1;refGene_tx=XM_011285448.2
C1 186289867 186290089
refGene_gn=SF3B1;refGene_tx=XM_003990992.4;refGene_gn=SF3B1;refGene_tx=XM_011285447.3;refGene_gn=SF3B1;refGene_tx=XM_011285448.2
C1 186290826 186290952
refGene_gn=SF3B1;refGene_tx=XM_003990992.4;refGene_gn=SF3B1;refGene_tx=XM_011285447.3;refGene_gn=SF3B1;refGene_tx=XM_011285448.2
C1 186291090 186291237
refGene_gn=SF3B1;refGene_tx=XM_003990992.4;refGene_gn=SF3B1;refGene_tx=XM_011285447.3;refGene_gn=SF3B1;refGene_tx=XM_011285448.2
C1 186291350 186291496
refGene_gn=SF3B1;refGene_tx=XM_003990992.4;refGene_gn=SF3B1;refGene_tx=XM_011285447.3;refGene_gn=SF3B1;refGene_tx=XM_011285448.2
C1 186292022 186292293
refGene_gn=SF3B1;refGene_tx=XM_003990992.4;refGene_gn=SF3B1;refGene_tx=XM_011285447.3;refGene_gn=SF3B1;refGene_tx=XM_011285448.2
C1 186292416 186292503
refGene_gn=SF3B1;refGene_tx=XM_003990992.4;refGene_gn=SF3B1;refGene_tx=XM_011285447.3;refGene_gn=SF3B1;refGene_tx=XM_011285448.2
C1 186293462 186293642
refGene_gn=SF3B1;refGene_tx=XM_003990992.4;refGene_gn=SF3B1;refGene_tx=XM_011285447.3;refGene_gn=SF3B1;refGene_tx=XM_011285448.2
C1 186294192 186294294
refGene_gn=SF3B1;refGene_tx=XM_003990992.4;refGene_gn=SF3B1;refGene_tx

=XM_011285447.3;refGene_gn=SF3B1;refGene_tx=XM_011285448.2
C1 186294395 186294593
refGene_gn=SF3B1;refGene_tx=XM_003990992.4;refGene_gn=SF3B1;refGene_tx
=XM_011285447.3;refGene_gn=SF3B1;refGene_tx=XM_011285448.2
C1 186296213 186296335
refGene_gn=SF3B1;refGene_tx=XM_003990992.4;refGene_gn=SF3B1;refGene_tx
=XM_011285447.3;refGene_gn=SF3B1;refGene_tx=XM_011285448.2
C1 186296575 186296788
refGene_gn=SF3B1;refGene_tx=XM_003990992.4;refGene_gn=SF3B1;refGene_tx
=XM_011285447.3;refGene_gn=SF3B1;refGene_tx=XM_011285448.2
C1 186298324 186298562
refGene_gn=SF3B1;refGene_tx=XM_003990992.4;refGene_gn=SF3B1;refGene_tx
=XM_011285447.3;refGene_gn=SF3B1;refGene_tx=XM_011285448.2
C1 186302157 186302328
refGene_gn=SF3B1;refGene_tx=XM_003990992.4;refGene_gn=SF3B1;refGene_tx
=XM_011285447.3;refGene_gn=SF3B1;refGene_tx=XM_011285448.2
C1 186303697 186303777
refGene_gn=SF3B1;refGene_tx=XM_003990992.4;refGene_gn=SF3B1;refGene_tx
=XM_011285447.3;refGene_gn=SF3B1;refGene_tx=XM_011285448.2
C1 186304120 186304140
refGene_gn=SF3B1;refGene_tx=XM_006935449.4
C1 186305618 186305733
refGene_gn=SF3B1;refGene_tx=XM_003990992.4;refGene_gn=SF3B1;refGene_tx
=XM_006935449.4;refGene_gn=SF3B1;refGene_tx=XM_011285448.2
C1 186306125 186306230
refGene_gn=SF3B1;refGene_tx=XM_003990992.4;refGene_gn=SF3B1;refGene_tx
=XM_006935449.4;refGene_gn=SF3B1;refGene_tx=XM_011285448.2
C1 186307906 186308073
refGene_gn=SF3B1;refGene_tx=XM_003990992.4;refGene_gn=SF3B1;refGene_tx
=XM_006935449.4;refGene_gn=SF3B1;refGene_tx=XM_011285448.2
C1 186319830 186319858
refGene_gn=SF3B1;refGene_tx=XM_003990992.4;refGene_gn=SF3B1;refGene_tx
=XM_006935449.4;refGene_gn=SF3B1;refGene_tx=XM_011285448.2
C1 186339980 186340303
refGene_gn=COQ10B;refGene_tx=XM_011285449.2
C1 186346216 186346369
refGene_gn=COQ10B;refGene_tx=XM_011285449.2;refGene_gn=COQ10B;refGene_
tx=XM_019838415.1
C1 186352859 186353052
refGene_gn=COQ10B;refGene_tx=XM_011285449.2;refGene_gn=COQ10B;refGene_
tx=XM_019838415.1
C1 186358408 186358510
refGene_gn=COQ10B;refGene_tx=XM_011285449.2;refGene_gn=COQ10B;refGene_
tx=XM_019838415.1
C1 186360785 186360953
refGene_gn=COQ10B;refGene_tx=XM_011285449.2;refGene_gn=COQ10B;refGene_
tx=XM_019838415.1
C1 186369730 186369883
refGene_gn=HSPD1;refGene_tx=XM_003990995.4;refGene_gn=HSPD1;refGene_tx
=XM_006935450.3;refGene_gn=HSPD1;refGene_tx=XM_019838416.2

C1 186369985 186370164
refGene_gn=HSPD1;refGene_tx=XM_003990995.4;refGene_gn=HSPD1;refGene_tx=XM_006935450.3;refGene_gn=HSPD1;refGene_tx=XM_019838416.2
C1 186370449 186370624
refGene_gn=HSPD1;refGene_tx=XM_003990995.4;refGene_gn=HSPD1;refGene_tx=XM_006935450.3;refGene_gn=HSPD1;refGene_tx=XM_019838416.2
C1 186371265 186371511
refGene_gn=HSPD1;refGene_tx=XM_003990995.4;refGene_gn=HSPD1;refGene_tx=XM_006935450.3;refGene_gn=HSPD1;refGene_tx=XM_019838416.2
C1 186372290 186372390
refGene_gn=HSPD1;refGene_tx=XM_003990995.4;refGene_gn=HSPD1;refGene_tx=XM_006935450.3;refGene_gn=HSPD1;refGene_tx=XM_019838416.2
C1 186374192 186374361
refGene_gn=HSPD1;refGene_tx=XM_003990995.4;refGene_gn=HSPD1;refGene_tx=XM_006935450.3;refGene_gn=HSPD1;refGene_tx=XM_019838416.2
C1 186374626 186374720
refGene_gn=HSPD1;refGene_tx=XM_003990995.4;refGene_gn=HSPD1;refGene_tx=XM_006935450.3;refGene_gn=HSPD1;refGene_tx=XM_019838416.2
C1 186375218 186375314
refGene_gn=HSPD1;refGene_tx=XM_003990995.4;refGene_gn=HSPD1;refGene_tx=XM_006935450.3;refGene_gn=HSPD1;refGene_tx=XM_019838416.2
C1 186375830 186375913
refGene_gn=HSPD1;refGene_tx=XM_003990995.4;refGene_gn=HSPD1;refGene_tx=XM_006935450.3;refGene_gn=HSPD1;refGene_tx=XM_019838416.2
C1 186377717 186377970
refGene_gn=HSPD1;refGene_tx=XM_003990995.4;refGene_gn=HSPD1;refGene_tx=XM_006935450.3;refGene_gn=HSPD1;refGene_tx=XM_019838416.2
C1 186379106 186379280
refGene_gn=HSPD1;refGene_tx=XM_003990995.4;refGene_gn=HSPD1;refGene_tx=XM_006935450.3;refGene_gn=HSPD1;refGene_tx=XM_019838416.2
C1 186380246 186380258
refGene_gn=HSPE1;refGene_tx=XM_019839136.1
C1 186380480 186380876
refGene_gn=HSPE1;refGene_tx=XM_019839136.1
C1 186381482 186381647
refGene_gn=HSPE1;refGene_tx=XM_019839136.1
C1 186382712 186382802
refGene_gn=HSPE1;refGene_tx=XM_019839136.1
C1 186382879 186382930
refGene_gn=HSPE1;refGene_tx=XM_019839136.1
C1 186395634 186395975
refGene_gn=MOB4;refGene_tx=XM_019839137.1
C1 186396180 186396263
refGene_gn=MOB4;refGene_tx=XM_019839137.1
C1 186396335 186396511
refGene_gn=MOB4;refGene_tx=XM_019839137.1
C1 186399790 186399853
refGene_gn=MOB4;refGene_tx=XM_019839137.1
C1 186411181 186411282
refGene_gn=MOB4;refGene_tx=XM_019839137.1

C1 186415577 186415620
refGene_gn=MOB4;refGene_tx=XM_019839137.1
C1 186415808 186415937
refGene_gn=MOB4;refGene_tx=XM_019839137.1
C1 186444595 186444868
refGene_gn=RFTN2;refGene_tx=XM_019838417.2;refGene_gn=RFTN2;refGene_tx
=XM_019838418.2
C1 186456867 186456946
refGene_gn=RFTN2;refGene_tx=XM_019838417.2
C1 186471265 186471369
refGene_gn=RFTN2;refGene_tx=XM_019838417.2;refGene_gn=RFTN2;refGene_tx
=XM_019838418.2
C1 186472628 186472750
refGene_gn=RFTN2;refGene_tx=XM_019838417.2;refGene_gn=RFTN2;refGene_tx
=XM_019838418.2
C1 186482311 186482515
refGene_gn=RFTN2;refGene_tx=XM_019838417.2;refGene_gn=RFTN2;refGene_tx
=XM_019838418.2
C1 186485338 186485618
refGene_gn=RFTN2;refGene_tx=XM_019838417.2;refGene_gn=RFTN2;refGene_tx
=XM_019838418.2
C1 186489218 186489333
refGene_gn=RFTN2;refGene_tx=XM_019838417.2;refGene_gn=RFTN2;refGene_tx
=XM_019838418.2
C1 186490813 186490997
refGene_gn=RFTN2;refGene_tx=XM_019838417.2;refGene_gn=RFTN2;refGene_tx
=XM_019838418.2
C1 186513985 186514124
refGene_gn=RFTN2;refGene_tx=XM_019838417.2;refGene_gn=RFTN2;refGene_tx
=XM_019838418.2
C1 186522715 186524497
refGene_gn=MARS2;refGene_tx=XM_003990998.5
C1 186545760 186545784
refGene_gn=BOLL;refGene_tx=XM_011285451.2;refGene_gn=BOLL;refGene_tx=X
M_011285452.2
C1 186562046 186562145
refGene_gn=BOLL;refGene_tx=XM_011285451.2;refGene_gn=BOLL;refGene_tx=X
M_011285452.2
C1 186564074 186564203
refGene_gn=BOLL;refGene_tx=XM_011285451.2;refGene_gn=BOLL;refGene_tx=X
M_011285452.2
C1 186565222 186565270
refGene_gn=BOLL;refGene_tx=XM_011285451.2;refGene_gn=BOLL;refGene_tx=X
M_011285452.2
C1 186567430 186567502
refGene_gn=BOLL;refGene_tx=XM_011285451.2;refGene_gn=BOLL;refGene_tx=X
M_011285452.2
C1 186580289 186580417
refGene_gn=BOLL;refGene_tx=XM_011285451.2;refGene_gn=BOLL;refGene_tx=X
M_011285452.2

C1 186583430 186583506
refGene_gn=BOLL;refGene_tx=XM_011285451.2;refGene_gn=BOLL;refGene_tx=XM_011285452.2
C1 186585689 186585744
refGene_gn=BOLL;refGene_tx=XM_011285451.2;refGene_gn=BOLL;refGene_tx=XM_011285452.2
C1 186587606 186587698
refGene_gn=BOLL;refGene_tx=XM_011285451.2;refGene_gn=BOLL;refGene_tx=XM_011285452.2
C1 186590198 186590342
refGene_gn=BOLL;refGene_tx=XM_011285451.2;refGene_gn=BOLL;refGene_tx=XM_011285452.2
C1 186592932 186592947
refGene_gn=BOLL;refGene_tx=XM_011285452.2
C1 186593935 186593956
refGene_gn=BOLL;refGene_tx=XM_011285451.2
C1 186870025 186872446
refGene_gn=PLCL1;refGene_tx=XM_023259587.1
C1 186874785 186874989
refGene_gn=PLCL1;refGene_tx=XM_023259587.1
C1 186893896 186893972
refGene_gn=PLCL1;refGene_tx=XM_023259587.1
C1 186896054 186896230
refGene_gn=PLCL1;refGene_tx=XM_023259587.1
C1 187244603 187244642
refGene_gn=SPATS2L;refGene_tx=XM_019838432.1;refGene_gn=SPATS2L;refGene_tx=XM_023259588.1;refGene_gn=SPATS2L;refGene_tx=XM_023259589.1;refGene_gn=SPATS2L;refGene_tx=XM_023259590.1
C1 187267889 187267998
refGene_gn=SPATS2L;refGene_tx=XM_019838432.1;refGene_gn=SPATS2L;refGene_tx=XM_023259588.1;refGene_gn=SPATS2L;refGene_tx=XM_023259589.1;refGene_gn=SPATS2L;refGene_tx=XM_023259590.1
C1 187271429 187271479
refGene_gn=SPATS2L;refGene_tx=XM_019838432.1;refGene_gn=SPATS2L;refGene_tx=XM_023259588.1;refGene_gn=SPATS2L;refGene_tx=XM_023259589.1;refGene_gn=SPATS2L;refGene_tx=XM_023259590.1
C1 187274093 187274340
refGene_gn=SPATS2L;refGene_tx=XM_019838432.1;refGene_gn=SPATS2L;refGene_tx=XM_023259588.1;refGene_gn=SPATS2L;refGene_tx=XM_023259589.1;refGene_gn=SPATS2L;refGene_tx=XM_023259590.1
C1 187290590 187290797
refGene_gn=SPATS2L;refGene_tx=XM_023259588.1;refGene_gn=SPATS2L;refGene_tx=XM_023259589.1;refGene_gn=SPATS2L;refGene_tx=XM_023259590.1
C1 187292145 187292281
refGene_gn=SPATS2L;refGene_tx=XM_019838432.1;refGene_gn=SPATS2L;refGene_tx=XM_023259588.1;refGene_gn=SPATS2L;refGene_tx=XM_023259589.1;refGene_gn=SPATS2L;refGene_tx=XM_023259590.1
C1 187312454 187312513
refGene_gn=SPATS2L;refGene_tx=XM_019838432.1;refGene_gn=SPATS2L;refGene_tx=XM_023259588.1;refGene_gn=SPATS2L;refGene_tx=XM_023259589.1;refGe

ne_gn=SPATS2L;refGene_tx=XM_023259590.1
C1	187318981	187319091
refGene_gn=SPATS2L;refGene_tx=XM_019838432.1;refGene_gn=SPATS2L;refGene_tx=XM_023259588.1;refGene_gn=SPATS2L;refGene_tx=XM_023259589.1;refGene_gn=SPATS2L;refGene_tx=XM_023259590.1
C1	187321660	187321763
refGene_gn=SPATS2L;refGene_tx=XM_019838432.1;refGene_gn=SPATS2L;refGene_tx=XM_023259588.1;refGene_gn=SPATS2L;refGene_tx=XM_023259589.1;refGene_gn=SPATS2L;refGene_tx=XM_023259590.1
C1	187324622	187324843
refGene_gn=SPATS2L;refGene_tx=XM_019838432.1;refGene_gn=SPATS2L;refGene_tx=XM_023259588.1;refGene_gn=SPATS2L;refGene_tx=XM_023259589.1;refGene_gn=SPATS2L;refGene_tx=XM_023259590.1
C1	187327428	187327824
refGene_gn=SPATS2L;refGene_tx=XM_019838432.1;refGene_gn=SPATS2L;refGene_tx=XM_023259588.1;refGene_gn=SPATS2L;refGene_tx=XM_023259589.1;refGene_gn=SPATS2L;refGene_tx=XM_023259590.1
C1	187339033	187339514
refGene_gn=KCTD18;refGene_tx=XM_006935458.4;refGene_gn=KCTD18;refGene_tx=XM_019838433.2;refGene_gn=KCTD18;refGene_tx=XM_019838435.2
C1	187346280	187346383
refGene_gn=KCTD18;refGene_tx=XM_006935458.4;refGene_gn=KCTD18;refGene_tx=XM_019838433.2;refGene_gn=KCTD18;refGene_tx=XM_019838435.2
C1	187351333	187351428
refGene_gn=KCTD18;refGene_tx=XM_006935458.4;refGene_gn=KCTD18;refGene_tx=XM_019838433.2;refGene_gn=KCTD18;refGene_tx=XM_019838435.2
C1	187352494	187352688
refGene_gn=KCTD18;refGene_tx=XM_006935458.4;refGene_gn=KCTD18;refGene_tx=XM_019838433.2
C1	187354962	187355174
refGene_gn=KCTD18;refGene_tx=XM_006935458.4;refGene_gn=KCTD18;refGene_tx=XM_019838433.2;refGene_gn=KCTD18;refGene_tx=XM_019838435.2
C1	187356715	187356884
refGene_gn=KCTD18;refGene_tx=XM_006935458.4;refGene_gn=KCTD18;refGene_tx=XM_019838433.2;refGene_gn=KCTD18;refGene_tx=XM_019838435.2
C1	187390535	187390665
refGene_gn=SGO2;refGene_tx=XM_011285458.3;refGene_gn=SGO2;refGene_tx=XM_019838427.2;refGene_gn=SGO2;refGene_tx=XM_019838428.2;refGene_gn=SGO2;refGene_tx=XM_019838429.2;refGene_gn=SGO2;refGene_tx=XM_019838430.2
C1	187392225	187392401
refGene_gn=SGO2;refGene_tx=XM_011285458.3;refGene_gn=SGO2;refGene_tx=XM_019838427.2;refGene_gn=SGO2;refGene_tx=XM_019838428.2;refGene_gn=SGO2;refGene_tx=XM_019838429.2;refGene_gn=SGO2;refGene_tx=XM_019838430.2
C1	187393205	187393283
refGene_gn=SGO2;refGene_tx=XM_011285458.3;refGene_gn=SGO2;refGene_tx=XM_019838427.2;refGene_gn=SGO2;refGene_tx=XM_019838428.2;refGene_gn=SGO2;refGene_tx=XM_019838429.2;refGene_gn=SGO2;refGene_tx=XM_019838430.2
C1	187401029	187401115
refGene_gn=SGO2;refGene_tx=XM_011285458.3;refGene_gn=SGO2;refGene_tx=XM_019838427.2;refGene_gn=SGO2;refGene_tx=XM_019838428.2;refGene_gn=SGO

2;refGene_tx=XM_019838429.2;refGene_gn=SGO2;refGene_tx=XM_019838430.2
C1 187413842 187413970
refGene_gn=SGO2;refGene_tx=XM_011285458.3;refGene_gn=SGO2;refGene_tx=XM_019838427.2;refGene_gn=SGO2;refGene_tx=XM_019838428.2;refGene_gn=SGO2;refGene_tx=XM_019838429.2;refGene_gn=SGO2;refGene_tx=XM_019838430.2
C1 187415141 187418330
refGene_gn=SGO2;refGene_tx=XM_011285458.3;refGene_gn=SGO2;refGene_tx=XM_019838427.2;refGene_gn=SGO2;refGene_tx=XM_019838428.2;refGene_gn=SGO2;refGene_tx=XM_019838429.2;refGene_gn=SGO2;refGene_tx=XM_019838430.2
C1 187419312 187419463
refGene_gn=SGO2;refGene_tx=XM_011285458.3;refGene_gn=SGO2;refGene_tx=XM_019838427.2;refGene_gn=SGO2;refGene_tx=XM_019838428.2;refGene_gn=SGO2;refGene_tx=XM_019838429.2;refGene_gn=SGO2;refGene_tx=XM_019838430.2
C1 187429832 187429848
refGene_gn=SGO2;refGene_tx=XM_011285458.3;refGene_gn=SGO2;refGene_tx=XM_019838427.2;refGene_gn=SGO2;refGene_tx=XM_019838428.2;refGene_gn=SGO2;refGene_tx=XM_019838429.2;refGene_gn=SGO2;refGene_tx=XM_019838430.2
C1 187683671 187683725
refGene_gn=AOX1;refGene_tx=NM_001290238.1
C1 187685698 187685756
refGene_gn=AOX1;refGene_tx=NM_001290238.1
C1 187687247 187687344
refGene_gn=AOX1;refGene_tx=NM_001290238.1
C1 187688546 187688655
refGene_gn=AOX1;refGene_tx=NM_001290238.1
C1 187690471 187690598
refGene_gn=AOX1;refGene_tx=NM_001290238.1
C1 187693834 187693896
refGene_gn=AOX1;refGene_tx=NM_001290238.1
C1 187696278 187696368
refGene_gn=AOX1;refGene_tx=NM_001290238.1
C1 187698710 187698791
refGene_gn=AOX1;refGene_tx=NM_001290238.1
C1 187701155 187701300
refGene_gn=AOX1;refGene_tx=NM_001290238.1
C1 187705834 187705927
refGene_gn=AOX1;refGene_tx=NM_001290238.1
C1 187709854 187710006
refGene_gn=AOX1;refGene_tx=NM_001290238.1
C1 187711992 187712086
refGene_gn=AOX1;refGene_tx=NM_001290238.1
C1 187714092 187714202
refGene_gn=AOX1;refGene_tx=NM_001290238.1
C1 187715792 187715977
refGene_gn=AOX1;refGene_tx=NM_001290238.1
C1 187716647 187716810
refGene_gn=AOX1;refGene_tx=NM_001290238.1
C1 187718353 187718446
refGene_gn=AOX1;refGene_tx=NM_001290238.1
C1 187719587 187719757

refGene_gn=AOX1;refGene_tx=NM_001290238.1
C1 187720365 187720480
refGene_gn=AOX1;refGene_tx=NM_001290238.1
C1 187722042 187722165
refGene_gn=AOX1;refGene_tx=NM_001290238.1
C1 187723920 187724017
refGene_gn=AOX1;refGene_tx=NM_001290238.1
C1 187724492 187724614
refGene_gn=AOX1;refGene_tx=NM_001290238.1
C1 187725514 187725648
refGene_gn=AOX1;refGene_tx=NM_001290238.1
C1 187727871 187727931
refGene_gn=AOX1;refGene_tx=NM_001290238.1
C1 187727932 187727960
refGene_gn=AOX1;refGene_tx=NM_001290238.1
C1 187730846 187730933
refGene_gn=AOX1;refGene_tx=NM_001290238.1
C1 187731182 187731374
refGene_gn=AOX1;refGene_tx=NM_001290238.1
C1 187734214 187734442
refGene_gn=AOX1;refGene_tx=NM_001290238.1
C1 187738532 187738628
refGene_gn=AOX1;refGene_tx=NM_001290238.1
C1 187742835 187742964
refGene_gn=AOX1;refGene_tx=NM_001290238.1
C1 187744684 187744759
refGene_gn=AOX1;refGene_tx=NM_001290238.1
C1 187745511 187745564
refGene_gn=AOX1;refGene_tx=NM_001290238.1
C1 187747483 187747598
refGene_gn=AOX1;refGene_tx=NM_001290238.1
C1 187751185 187751251
refGene_gn=AOX1;refGene_tx=NM_001290238.1
C1 187751362 187751500
refGene_gn=AOX1;refGene_tx=NM_001290238.1
C1 187754637 187754805
refGene_gn=AOX1;refGene_tx=NM_001290238.1
C1 187768729 187768783
refGene_gn=AOX1;refGene_tx=NM_001290238.1
C1 187788969 187789055
refGene_gn=BZW1;refGene_tx=XM_019838438.2;refGene_gn=BZW1;refGene_tx=X
M_023259592.1
C1 187789761 187789835
refGene_gn=BZW1;refGene_tx=XM_019838438.2;refGene_gn=BZW1;refGene_tx=X
M_023259592.1;refGene_gn=BZW1;refGene_tx=XM_023259593.1;refGene_gn=BZW
1;refGene_tx=XM_023259594.1;refGene_gn=BZW1;refGene_tx=XM_023259595.1;
refGene_gn=BZW1;refGene_tx=XM_023259596.1
C1 187791600 187791783
refGene_gn=BZW1;refGene_tx=XM_019838438.2;refGene_gn=BZW1;refGene_tx=X
M_019838439.2;refGene_gn=BZW1;refGene_tx=XM_023259592.1;refGene_gn=BZW

1;refGene_tx=XM_023259593.1;refGene_gn=BZW1;refGene_tx=XM_023259594.1;refGene_gn=BZW1;refGene_tx=XM_023259595.1;refGene_gn=BZW1;refGene_tx=XM_023259596.1
C1 187793753 187793848
refGene_gn=BZW1;refGene_tx=XM_019838438.2;refGene_gn=BZW1;refGene_tx=XM_023259592.1;refGene_gn=BZW1;refGene_tx=XM_023259593.1;refGene_gn=BZW1;refGene_tx=XM_023259594.1;refGene_gn=BZW1;refGene_tx=XM_023259595.1;refGene_gn=BZW1;refGene_tx=XM_023259596.1
C1 187794826 187794892
refGene_gn=BZW1;refGene_tx=XM_019838438.2;refGene_gn=BZW1;refGene_tx=XM_019838439.2;refGene_gn=BZW1;refGene_tx=XM_023259592.1;refGene_gn=BZW1;refGene_tx=XM_023259593.1;refGene_gn=BZW1;refGene_tx=XM_023259594.1;refGene_gn=BZW1;refGene_tx=XM_023259595.1;refGene_gn=BZW1;refGene_tx=XM_023259596.1
C1 187795516 187795652
refGene_gn=BZW1;refGene_tx=XM_019838438.2;refGene_gn=BZW1;refGene_tx=XM_019838439.2;refGene_gn=BZW1;refGene_tx=XM_023259592.1;refGene_gn=BZW1;refGene_tx=XM_023259593.1;refGene_gn=BZW1;refGene_tx=XM_023259594.1;refGene_gn=BZW1;refGene_tx=XM_023259595.1;refGene_gn=BZW1;refGene_tx=XM_023259596.1
C1 187796322 187796432
refGene_gn=BZW1;refGene_tx=XM_019838438.2;refGene_gn=BZW1;refGene_tx=XM_019838439.2;refGene_gn=BZW1;refGene_tx=XM_023259592.1;refGene_gn=BZW1;refGene_tx=XM_023259593.1;refGene_gn=BZW1;refGene_tx=XM_023259594.1;refGene_gn=BZW1;refGene_tx=XM_023259595.1;refGene_gn=BZW1;refGene_tx=XM_023259596.1
C1 187796576 187796747
refGene_gn=BZW1;refGene_tx=XM_019838438.2;refGene_gn=BZW1;refGene_tx=XM_019838439.2;refGene_gn=BZW1;refGene_tx=XM_023259592.1;refGene_gn=BZW1;refGene_tx=XM_023259593.1;refGene_gn=BZW1;refGene_tx=XM_023259594.1;refGene_gn=BZW1;refGene_tx=XM_023259595.1;refGene_gn=BZW1;refGene_tx=XM_023259596.1
C1 187797225 187797372
refGene_gn=BZW1;refGene_tx=XM_019838438.2;refGene_gn=BZW1;refGene_tx=XM_019838439.2;refGene_gn=BZW1;refGene_tx=XM_023259592.1;refGene_gn=BZW1;refGene_tx=XM_023259593.1;refGene_gn=BZW1;refGene_tx=XM_023259594.1;refGene_gn=BZW1;refGene_tx=XM_023259595.1;refGene_gn=BZW1;refGene_tx=XM_023259596.1
C1 187797730 187797869
refGene_gn=BZW1;refGene_tx=XM_019838438.2;refGene_gn=BZW1;refGene_tx=XM_019838439.2;refGene_gn=BZW1;refGene_tx=XM_023259592.1;refGene_gn=BZW1;refGene_tx=XM_023259593.1;refGene_gn=BZW1;refGene_tx=XM_023259594.1;refGene_gn=BZW1;refGene_tx=XM_023259595.1;refGene_gn=BZW1;refGene_tx=XM_023259596.1
C1 187799269 187799392
refGene_gn=BZW1;refGene_tx=XM_019838438.2;refGene_gn=BZW1;refGene_tx=XM_019838439.2;refGene_gn=BZW1;refGene_tx=XM_023259592.1;refGene_gn=BZW1;refGene_tx=XM_023259593.1;refGene_gn=BZW1;refGene_tx=XM_023259594.1;refGene_gn=BZW1;refGene_tx=XM_023259595.1;refGene_gn=BZW1;refGene_tx=XM_023259596.1

C1	187799811	187799843
refGene_gn=BZW1;refGene_tx=XM_019838438.2;refGene_gn=BZW1;refGene_tx=XM_019838439.2;refGene_gn=BZW1;refGene_tx=XM_023259592.1;refGene_gn=BZW1;refGene_tx=XM_023259593.1;refGene_gn=BZW1;refGene_tx=XM_023259594.1;refGene_gn=BZW1;refGene_tx=XM_023259595.1;refGene_gn=BZW1;refGene_tx=XM_023259596.1
C1	187804099	187804179
refGene_gn=BZW1;refGene_tx=XM_019838438.2
C1	187806283	187806380
refGene_gn=BZW1;refGene_tx=XM_023259592.1;refGene_gn=BZW1;refGene_tx=XM_023259593.1;refGene_gn=BZW1;refGene_tx=XM_023259594.1;refGene_gn=BZW1;refGene_tx=XM_023259595.1;refGene_gn=BZW1;refGene_tx=XM_023259596.1
C1	187806481	187806497
refGene_gn=BZW1;refGene_tx=XM_023259592.1;refGene_gn=BZW1;refGene_tx=XM_023259593.1;refGene_gn=BZW1;refGene_tx=XM_023259594.1;refGene_gn=BZW1;refGene_tx=XM_023259595.1;refGene_gn=BZW1;refGene_tx=XM_023259596.1
C1	187826472	187826616
refGene_gn=CLK1;refGene_tx=XM_003991010.5;refGene_gn=CLK1;refGene_tx=XM_019838441.2;refGene_gn=CLK1;refGene_tx=XM_019838442.1
C1	187826747	187826838
refGene_gn=CLK1;refGene_tx=XM_003991010.5;refGene_gn=CLK1;refGene_tx=XM_019838441.2;refGene_gn=CLK1;refGene_tx=XM_019838442.1
C1	187827073	187827153
refGene_gn=CLK1;refGene_tx=XM_003991010.5;refGene_gn=CLK1;refGene_tx=XM_019838441.2;refGene_gn=CLK1;refGene_tx=XM_019838442.1
C1	187827634	187827717
refGene_gn=CLK1;refGene_tx=XM_003991010.5;refGene_gn=CLK1;refGene_tx=XM_019838441.2;refGene_gn=CLK1;refGene_tx=XM_019838442.1
C1	187827860	187827990
refGene_gn=CLK1;refGene_tx=XM_003991010.5;refGene_gn=CLK1;refGene_tx=XM_019838441.2;refGene_gn=CLK1;refGene_tx=XM_019838442.1
C1	187828086	187828181
refGene_gn=CLK1;refGene_tx=XM_003991010.5;refGene_gn=CLK1;refGene_tx=XM_019838441.2;refGene_gn=CLK1;refGene_tx=XM_019838442.1
C1	187828707	187828874
refGene_gn=CLK1;refGene_tx=XM_003991010.5;refGene_gn=CLK1;refGene_tx=XM_019838441.2;refGene_gn=CLK1;refGene_tx=XM_019838442.1
C1	187828957	187829074
refGene_gn=CLK1;refGene_tx=XM_003991010.5;refGene_gn=CLK1;refGene_tx=XM_019838441.2
C1	187830176	187830243
refGene_gn=CLK1;refGene_tx=XM_003991010.5;refGene_gn=CLK1;refGene_tx=XM_019838441.2
C1	187830641	187830732
refGene_gn=CLK1;refGene_tx=XM_003991010.5;refGene_gn=CLK1;refGene_tx=XM_019838441.2
C1	187831712	187831941
refGene_gn=CLK1;refGene_tx=XM_003991010.5;refGene_gn=CLK1;refGene_tx=XM_019838441.2
C1	187832234	187832395

refGene_gn=CLK1;refGene_tx=XM_003991010.5;refGene_gn=CLK1;refGene_tx=XM_019838441.2
C1 187839646 187839773
refGene_gn=PPIL3;refGene_tx=XM_006935466.4
C1 187843080 187843199
refGene_gn=PPIL3;refGene_tx=XM_006935466.4
C1 187846388 187846456
refGene_gn=PPIL3;refGene_tx=XM_006935466.4
C1 187847433 187847527
refGene_gn=PPIL3;refGene_tx=XM_006935466.4
C1 187849001 187849076
refGene_gn=PPIL3;refGene_tx=XM_006935466.4
C1 187850335 187850338
refGene_gn=PPIL3;refGene_tx=XM_006935466.4
C1 187853940 187854373
refGene_gn=NIF3L1;refGene_tx=XM_023259597.1;refGene_gn=NIF3L1;refGene_tx=XM_023259598.1
C1 187855192 187855355
refGene_gn=NIF3L1;refGene_tx=XM_023259597.1;refGene_gn=NIF3L1;refGene_tx=XM_023259598.1
C1 187858723 187858850
refGene_gn=NIF3L1;refGene_tx=XM_023259597.1;refGene_gn=NIF3L1;refGene_tx=XM_023259598.1
C1 187860749 187860888
refGene_gn=NIF3L1;refGene_tx=XM_023259597.1
C1 187870566 187870650
refGene_gn=NIF3L1;refGene_tx=XM_023259597.1;refGene_gn=NIF3L1;refGene_tx=XM_023259598.1
C1 187874522 187874707
refGene_gn=NIF3L1;refGene_tx=XM_023259597.1
C1 187880007 187880094
refGene_gn=ORC2;refGene_tx=XM_011285460.3;refGene_gn=ORC2;refGene_tx=XM_011285461.3
C1 187882098 187882217
refGene_gn=ORC2;refGene_tx=XM_011285460.3;refGene_gn=ORC2;refGene_tx=XM_011285461.3
C1 187883334 187883396
refGene_gn=ORC2;refGene_tx=XM_011285460.3;refGene_gn=ORC2;refGene_tx=XM_011285461.3
C1 187886898 187887070
refGene_gn=ORC2;refGene_tx=XM_011285460.3;refGene_gn=ORC2;refGene_tx=XM_011285461.3
C1 187887688 187887835
refGene_gn=ORC2;refGene_tx=XM_011285460.3;refGene_gn=ORC2;refGene_tx=XM_011285461.3
C1 187891023 187891120
refGene_gn=ORC2;refGene_tx=XM_011285460.3;refGene_gn=ORC2;refGene_tx=XM_011285461.3
C1 187891897 187892030
refGene_gn=ORC2;refGene_tx=XM_011285460.3;refGene_gn=ORC2;refGene_tx=X

M_011285461.3
C1	187894294	187894404
refGene_gn=ORC2;refGene_tx=XM_011285460.3;refGene_gn=ORC2;refGene_tx=X
M_011285461.3
C1	187900593	187900692
refGene_gn=ORC2;refGene_tx=XM_011285460.3;refGene_gn=ORC2;refGene_tx=X
M_011285461.3
C1	187901946	187902140
refGene_gn=ORC2;refGene_tx=XM_011285460.3;refGene_gn=ORC2;refGene_tx=X
M_011285461.3
C1	187904297	187904358
refGene_gn=ORC2;refGene_tx=XM_011285460.3;refGene_gn=ORC2;refGene_tx=X
M_011285461.3
C1	187907676	187907708
refGene_gn=ORC2;refGene_tx=XM_011285460.3;refGene_gn=ORC2;refGene_tx=X
M_011285461.3
C1	187908383	187908476
refGene_gn=ORC2;refGene_tx=XM_011285460.3;refGene_gn=ORC2;refGene_tx=X
M_011285461.3
C1	187912100	187912190
refGene_gn=ORC2;refGene_tx=XM_011285460.3;refGene_gn=ORC2;refGene_tx=X
M_011285461.3
C1	187920734	187920878
refGene_gn=ORC2;refGene_tx=XM_011285460.3;refGene_gn=ORC2;refGene_tx=X
M_011285461.3
C1	187921342	187921439
refGene_gn=ORC2;refGene_tx=XM_011285460.3;refGene_gn=ORC2;refGene_tx=X
M_011285461.3
C1	187939847	187940449
refGene_gn=FAM126B;refGene_tx=XM_006935468.4;refGene_gn=FAM126B;refGen
e_tx=XM_006935469.4;refGene_gn=FAM126B;refGene_tx=XM_019838443.2;refGe
ne_gn=FAM126B;refGene_tx=XM_019838445.2;refGene_gn=FAM126B;refGene_tx=
XM_019838446.2;refGene_gn=FAM126B;refGene_tx=XM_019838447.2;refGene_gn
=FAM126B;refGene_tx=XM_019838448.2;refGene_gn=FAM126B;refGene_tx=XM_01
9838449.2
C1	187944574	187944742
refGene_gn=FAM126B;refGene_tx=XM_006935469.4;refGene_gn=FAM126B;refGen
e_tx=XM_019838443.2;refGene_gn=FAM126B;refGene_tx=XM_019838445.2;refGe
ne_gn=FAM126B;refGene_tx=XM_019838446.2;refGene_gn=FAM126B;refGene_tx=
XM_019838447.2;refGene_gn=FAM126B;refGene_tx=XM_019838448.2;refGene_gn
=FAM126B;refGene_tx=XM_019838449.2
C1	187945495	187945655
refGene_gn=FAM126B;refGene_tx=XM_006935468.4;refGene_gn=FAM126B;refGen
e_tx=XM_006935469.4;refGene_gn=FAM126B;refGene_tx=XM_019838443.2;refGe
ne_gn=FAM126B;refGene_tx=XM_019838445.2;refGene_gn=FAM126B;refGene_tx=
XM_019838446.2;refGene_gn=FAM126B;refGene_tx=XM_019838447.2;refGene_gn
=FAM126B;refGene_tx=XM_019838448.2;refGene_gn=FAM126B;refGene_tx=XM_01
9838449.2
C1	187949844	187949932
refGene_gn=FAM126B;refGene_tx=XM_006935468.4;refGene_gn=FAM126B;refGen

e_tx=XM_006935469.4;refGene_gn=FAM126B;refGene_tx=XM_019838443.2;refGene_gn=FAM126B;refGene_tx=XM_019838445.2;refGene_gn=FAM126B;refGene_tx=XM_019838446.2;refGene_gn=FAM126B;refGene_tx=XM_019838447.2;refGene_gn=FAM126B;refGene_tx=XM_019838448.2;refGene_gn=FAM126B;refGene_tx=XM_019838449.2
C1 187953429 187953546
refGene_gn=FAM126B;refGene_tx=XM_006935468.4;refGene_gn=FAM126B;refGene_tx=XM_006935469.4;refGene_gn=FAM126B;refGene_tx=XM_019838443.2;refGene_gn=FAM126B;refGene_tx=XM_019838445.2;refGene_gn=FAM126B;refGene_tx=XM_019838446.2;refGene_gn=FAM126B;refGene_tx=XM_019838447.2;refGene_gn=FAM126B;refGene_tx=XM_019838448.2;refGene_gn=FAM126B;refGene_tx=XM_019838449.2
C1 187957801 187957897
refGene_gn=FAM126B;refGene_tx=XM_006935468.4;refGene_gn=FAM126B;refGene_tx=XM_006935469.4;refGene_gn=FAM126B;refGene_tx=XM_019838443.2;refGene_gn=FAM126B;refGene_tx=XM_019838445.2;refGene_gn=FAM126B;refGene_tx=XM_019838446.2;refGene_gn=FAM126B;refGene_tx=XM_019838447.2;refGene_gn=FAM126B;refGene_tx=XM_019838448.2;refGene_gn=FAM126B;refGene_tx=XM_019838449.2
C1 187968683 187968799
refGene_gn=FAM126B;refGene_tx=XM_006935468.4;refGene_gn=FAM126B;refGene_tx=XM_006935469.4;refGene_gn=FAM126B;refGene_tx=XM_019838443.2;refGene_gn=FAM126B;refGene_tx=XM_019838445.2;refGene_gn=FAM126B;refGene_tx=XM_019838446.2;refGene_gn=FAM126B;refGene_tx=XM_019838447.2;refGene_gn=FAM126B;refGene_tx=XM_019838448.2;refGene_gn=FAM126B;refGene_tx=XM_019838449.2
C1 187970951 187971032
refGene_gn=FAM126B;refGene_tx=XM_006935468.4;refGene_gn=FAM126B;refGene_tx=XM_006935469.4;refGene_gn=FAM126B;refGene_tx=XM_019838443.2;refGene_gn=FAM126B;refGene_tx=XM_019838445.2;refGene_gn=FAM126B;refGene_tx=XM_019838446.2;refGene_gn=FAM126B;refGene_tx=XM_019838447.2;refGene_gn=FAM126B;refGene_tx=XM_019838448.2;refGene_gn=FAM126B;refGene_tx=XM_019838449.2
C1 187974200 187974380
refGene_gn=FAM126B;refGene_tx=XM_006935468.4;refGene_gn=FAM126B;refGene_tx=XM_006935469.4;refGene_gn=FAM126B;refGene_tx=XM_019838443.2;refGene_gn=FAM126B;refGene_tx=XM_019838445.2;refGene_gn=FAM126B;refGene_tx=XM_019838446.2;refGene_gn=FAM126B;refGene_tx=XM_019838447.2;refGene_gn=FAM126B;refGene_tx=XM_019838448.2;refGene_gn=FAM126B;refGene_tx=XM_019838449.2
C1 187978823 187978925
refGene_gn=FAM126B;refGene_tx=XM_006935468.4;refGene_gn=FAM126B;refGene_tx=XM_019838443.2;refGene_gn=FAM126B;refGene_tx=XM_019838445.2;refGene_gn=FAM126B;refGene_tx=XM_019838446.2;refGene_gn=FAM126B;refGene_tx=XM_019838447.2;refGene_gn=FAM126B;refGene_tx=XM_019838448.2;refGene_gn=FAM126B;refGene_tx=XM_019838449.2
C1 187980073 187980124
refGene_gn=FAM126B;refGene_tx=XM_006935468.4;refGene_gn=FAM126B;refGene_tx=XM_019838443.2;refGene_gn=FAM126B;refGene_tx=XM_019838445.2;refGene_gn=FAM126B;refGene_tx=XM_019838446.2;refGene_gn=FAM126B;refGene_tx=

XM_019838447.2;refGene_gn=FAM126B;refGene_tx=XM_019838448.2;refGene_gn=FAM126B;refGene_tx=XM_019838449.2
C1 188026636 188026649
refGene_gn=NDUFB3;refGene_tx=XM_006935471.4
C1 188033140 188033282
refGene_gn=NDUFB3;refGene_tx=XM_003991014.4;refGene_gn=NDUFB3;refGene_tx=XM_006935471.4
C1 188035414 188035571
refGene_gn=NDUFB3;refGene_tx=XM_003991014.4;refGene_gn=NDUFB3;refGene_tx=XM_006935471.4
C1 188088204 188088566
refGene_gn=CFLAR;refGene_tx=XM_011285463.3;refGene_gn=CFLAR;refGene_tx=XM_011285464.3;refGene_gn=CFLAR;refGene_tx=XM_011285465.3;refGene_gn=CFLAR;refGene_tx=XM_019838451.2;refGene_gn=CFLAR;refGene_tx=XM_019838452.2;refGene_gn=CFLAR;refGene_tx=XM_019838453.2;refGene_gn=CFLAR;refGene_tx=XM_019838454.2;refGene_gn=CFLAR;refGene_tx=XM_019838455.2;refGene_gn=CFLAR;refGene_tx=XM_023259600.1
C1 188090825 188090931
refGene_gn=CFLAR;refGene_tx=XM_011285463.3;refGene_gn=CFLAR;refGene_tx=XM_011285464.3;refGene_gn=CFLAR;refGene_tx=XM_011285465.3;refGene_gn=CFLAR;refGene_tx=XM_019838451.2;refGene_gn=CFLAR;refGene_tx=XM_019838452.2;refGene_gn=CFLAR;refGene_tx=XM_019838453.2;refGene_gn=CFLAR;refGene_tx=XM_019838454.2;refGene_gn=CFLAR;refGene_tx=XM_019838455.2;refGene_gn=CFLAR;refGene_tx=XM_019838456.2;refGene_gn=CFLAR;refGene_tx=XM_023259600.1
C1 188092597 188092733
refGene_gn=CFLAR;refGene_tx=XM_011285463.3;refGene_gn=CFLAR;refGene_tx=XM_011285464.3;refGene_gn=CFLAR;refGene_tx=XM_011285465.3;refGene_gn=CFLAR;refGene_tx=XM_019838451.2;refGene_gn=CFLAR;refGene_tx=XM_019838452.2;refGene_gn=CFLAR;refGene_tx=XM_019838453.2;refGene_gn=CFLAR;refGene_tx=XM_019838454.2;refGene_gn=CFLAR;refGene_tx=XM_019838455.2;refGene_gn=CFLAR;refGene_tx=XM_019838456.2;refGene_gn=CFLAR;refGene_tx=XM_023259600.1
C1 188093581 188093703
refGene_gn=CFLAR;refGene_tx=XM_011285463.3;refGene_gn=CFLAR;refGene_tx=XM_011285464.3;refGene_gn=CFLAR;refGene_tx=XM_011285465.3;refGene_gn=CFLAR;refGene_tx=XM_019838451.2;refGene_gn=CFLAR;refGene_tx=XM_019838452.2;refGene_gn=CFLAR;refGene_tx=XM_019838453.2;refGene_gn=CFLAR;refGene_tx=XM_019838454.2;refGene_gn=CFLAR;refGene_tx=XM_019838456.2;refGene_gn=CFLAR;refGene_tx=XM_023259600.1
C1 188098394 188098452
refGene_gn=CFLAR;refGene_tx=XM_011285463.3;refGene_gn=CFLAR;refGene_tx=XM_011285464.3;refGene_gn=CFLAR;refGene_tx=XM_011285465.3;refGene_gn=CFLAR;refGene_tx=XM_019838451.2;refGene_gn=CFLAR;refGene_tx=XM_019838452.2;refGene_gn=CFLAR;refGene_tx=XM_019838453.2;refGene_gn=CFLAR;refGene_tx=XM_019838454.2;refGene_gn=CFLAR;refGene_tx=XM_019838456.2
C1 188101670 188101720
refGene_gn=CFLAR;refGene_tx=XM_011285463.3;refGene_gn=CFLAR;refGene_tx=XM_011285464.3;refGene_gn=CFLAR;refGene_tx=XM_011285465.3;refGene_gn=CFLAR;refGene_tx=XM_019838451.2;refGene_gn=CFLAR;refGene_tx=XM_0198384

52.2;refGene_gn=CFLAR;refGene_tx=XM_019838453.2;refGene_gn=CFLAR;refGene_tx=XM_019838454.2;refGene_gn=CFLAR;refGene_tx=XM_019838455.2;refGene_gn=CFLAR;refGene_tx=XM_019838456.2
C1 188102454 188102533
refGene_gn=CFLAR;refGene_tx=XM_011285463.3;refGene_gn=CFLAR;refGene_tx=XM_011285464.3;refGene_gn=CFLAR;refGene_tx=XM_011285465.3;refGene_gn=CFLAR;refGene_tx=XM_011285466.3;refGene_gn=CFLAR;refGene_tx=XM_019838451.2;refGene_gn=CFLAR;refGene_tx=XM_019838452.2;refGene_gn=CFLAR;refGene_tx=XM_019838453.2;refGene_gn=CFLAR;refGene_tx=XM_019838454.2;refGene_gn=CFLAR;refGene_tx=XM_019838455.2;refGene_gn=CFLAR;refGene_tx=XM_019838456.2;refGene_gn=CFLAR;refGene_tx=XM_019838457.2
C1 188116468 188116979
refGene_gn=CFLAR;refGene_tx=XM_011285463.3;refGene_gn=CFLAR;refGene_tx=XM_011285464.3;refGene_gn=CFLAR;refGene_tx=XM_011285465.3;refGene_gn=CFLAR;refGene_tx=XM_011285466.3;refGene_gn=CFLAR;refGene_tx=XM_019838451.2;refGene_gn=CFLAR;refGene_tx=XM_019838452.2;refGene_gn=CFLAR;refGene_tx=XM_019838453.2;refGene_gn=CFLAR;refGene_tx=XM_019838454.2;refGene_gn=CFLAR;refGene_tx=XM_019838455.2;refGene_gn=CFLAR;refGene_tx=XM_019838456.2;refGene_gn=CFLAR;refGene_tx=XM_019838457.2
C1 188118753 188118892
refGene_gn=CFLAR;refGene_tx=XM_011285463.3;refGene_gn=CFLAR;refGene_tx=XM_011285464.3;refGene_gn=CFLAR;refGene_tx=XM_011285465.3;refGene_gn=CFLAR;refGene_tx=XM_011285466.3;refGene_gn=CFLAR;refGene_tx=XM_019838451.2;refGene_gn=CFLAR;refGene_tx=XM_019838452.2;refGene_gn=CFLAR;refGene_tx=XM_019838453.2;refGene_gn=CFLAR;refGene_tx=XM_019838454.2;refGene_gn=CFLAR;refGene_tx=XM_019838455.2;refGene_gn=CFLAR;refGene_tx=XM_019838456.2;refGene_gn=CFLAR;refGene_tx=XM_019838457.2
C1 188149487 188149834
refGene_gn=CASP10;refGene_tx=XM_019838459.2;refGene_gn=CASP10;refGene_tx=XM_019838460.2
C1 188151632 188151726
refGene_gn=CASP10;refGene_tx=XM_019838459.2;refGene_gn=CASP10;refGene_tx=XM_019838460.2
C1 188157268 188157404
refGene_gn=CASP10;refGene_tx=XM_019838459.2;refGene_gn=CASP10;refGene_tx=XM_019838460.2
C1 188160925 188161035
refGene_gn=CASP10;refGene_tx=XM_019838459.2;refGene_gn=CASP10;refGene_tx=XM_019838460.2
C1 188170047 188170156
refGene_gn=CASP10;refGene_tx=XM_019838459.2;refGene_gn=CASP10;refGene_tx=XM_019838460.2
C1 188171554 188172053
refGene_gn=CASP10;refGene_tx=XM_019838459.2;refGene_gn=CASP10;refGene_tx=XM_019838460.2
C1 188175178 188175332
refGene_gn=CASP10;refGene_tx=XM_019838459.2;refGene_gn=CASP10;refGene_tx=XM_019838460.2
C1 188235013 188235318
refGene_gn=CASP8;refGene_tx=XM_006935473.4;refGene_gn=CASP8;refGene_tx

=XM_019838462.1;refGene_gn=CASP8;refGene_tx=XM_019838463.1;refGene_gn=CASP8;refGene_tx=XM_019838464.1;refGene_gn=CASP8;refGene_tx=XM_019838465.2
C1 188239781 188239887
refGene_gn=CASP8;refGene_tx=XM_006935473.4;refGene_gn=CASP8;refGene_tx=XM_019838462.1;refGene_gn=CASP8;refGene_tx=XM_019838463.1;refGene_gn=CASP8;refGene_tx=XM_019838464.1;refGene_gn=CASP8;refGene_tx=XM_019838465.2
C1 188240629 188240768
refGene_gn=CASP8;refGene_tx=XM_006935473.4;refGene_gn=CASP8;refGene_tx=XM_019838462.1;refGene_gn=CASP8;refGene_tx=XM_019838463.1;refGene_gn=CASP8;refGene_tx=XM_019838464.1;refGene_gn=CASP8;refGene_tx=XM_019838465.2
C1 188240876 188240921
refGene_gn=CASP8;refGene_tx=XM_006935473.4;refGene_gn=CASP8;refGene_tx=XM_019838462.1;refGene_gn=CASP8;refGene_tx=XM_019838463.1;refGene_gn=CASP8;refGene_tx=XM_019838464.1;refGene_gn=CASP8;refGene_tx=XM_019838465.2
C1 188241317 188241382
refGene_gn=CASP8;refGene_tx=XM_006935473.4;refGene_gn=CASP8;refGene_tx=XM_019838462.1;refGene_gn=CASP8;refGene_tx=XM_019838463.1;refGene_gn=CASP8;refGene_tx=XM_019838464.1;refGene_gn=CASP8;refGene_tx=XM_019838465.2
C1 188243115 188243257
refGene_gn=CASP8;refGene_tx=XM_006935473.4;refGene_gn=CASP8;refGene_tx=XM_019838462.1;refGene_gn=CASP8;refGene_tx=XM_019838463.1;refGene_gn=CASP8;refGene_tx=XM_019838464.1;refGene_gn=CASP8;refGene_tx=XM_019838465.2
C1 188243458 188243475
refGene_gn=CASP8;refGene_tx=XM_019838465.2
C1 188246774 188247276
refGene_gn=CASP8;refGene_tx=XM_006935473.4;refGene_gn=CASP8;refGene_tx=XM_019838462.1;refGene_gn=CASP8;refGene_tx=XM_019838463.1;refGene_gn=CASP8;refGene_tx=XM_019838464.1
C1 188248411 188248547
refGene_gn=CASP8;refGene_tx=XM_006935473.4;refGene_gn=CASP8;refGene_tx=XM_019838462.1;refGene_gn=CASP8;refGene_tx=XM_019838463.1;refGene_gn=CASP8;refGene_tx=XM_019838464.1
C1 188252117 188252238
refGene_gn=ALS2CR12;refGene_tx=XM_003991061.5;refGene_gn=ALS2CR12;refGene_tx=XM_019838468.2;refGene_gn=ALS2CR12;refGene_tx=XM_019838469.2;refGene_gn=ALS2CR12;refGene_tx=XM_019838470.2
C1 188254206 188254316
refGene_gn=ALS2CR12;refGene_tx=XM_003991061.5;refGene_gn=ALS2CR12;refGene_tx=XM_019838468.2;refGene_gn=ALS2CR12;refGene_tx=XM_019838469.2;refGene_gn=ALS2CR12;refGene_tx=XM_019838470.2
C1 188254447 188254537
refGene_gn=ALS2CR12;refGene_tx=XM_003991061.5;refGene_gn=ALS2CR12;refGene_tx=XM_019838468.2;refGene_gn=ALS2CR12;refGene_tx=XM_019838469.2;refGene_gn=ALS2CR12;refGene_tx=XM_019838470.2

C1 188257698 188257761
refGene_gn=ALS2CR12;refGene_tx=XM_003991061.5;refGene_gn=ALS2CR12;refGene_tx=XM_019838468.2;refGene_gn=ALS2CR12;refGene_tx=XM_019838469.2;refGene_gn=ALS2CR12;refGene_tx=XM_019838470.2
C1 188263111 188263215
refGene_gn=ALS2CR12;refGene_tx=XM_003991061.5;refGene_gn=ALS2CR12;refGene_tx=XM_019838468.2;refGene_gn=ALS2CR12;refGene_tx=XM_019838469.2;refGene_gn=ALS2CR12;refGene_tx=XM_019838470.2
C1 188264977 188265077
refGene_gn=ALS2CR12;refGene_tx=XM_003991061.5;refGene_gn=ALS2CR12;refGene_tx=XM_019838468.2;refGene_gn=ALS2CR12;refGene_tx=XM_019838469.2;refGene_gn=ALS2CR12;refGene_tx=XM_019838470.2
C1 188266957 188267010
refGene_gn=ALS2CR12;refGene_tx=XM_003991061.5;refGene_gn=ALS2CR12;refGene_tx=XM_019838468.2;refGene_gn=ALS2CR12;refGene_tx=XM_019838469.2;refGene_gn=ALS2CR12;refGene_tx=XM_019838470.2
C1 188268313 188268411
refGene_gn=ALS2CR12;refGene_tx=XM_003991061.5;refGene_gn=ALS2CR12;refGene_tx=XM_019838468.2;refGene_gn=ALS2CR12;refGene_tx=XM_019838469.2;refGene_gn=ALS2CR12;refGene_tx=XM_019838470.2
C1 188269926 188269988
refGene_gn=ALS2CR12;refGene_tx=XM_003991061.5;refGene_gn=ALS2CR12;refGene_tx=XM_019838468.2;refGene_gn=ALS2CR12;refGene_tx=XM_019838469.2;refGene_gn=ALS2CR12;refGene_tx=XM_019838470.2
C1 188271674 188271768
refGene_gn=ALS2CR12;refGene_tx=XM_003991061.5;refGene_gn=ALS2CR12;refGene_tx=XM_019838468.2;refGene_gn=ALS2CR12;refGene_tx=XM_019838469.2
C1 188273002 188273136
refGene_gn=ALS2CR12;refGene_tx=XM_003991061.5;refGene_gn=ALS2CR12;refGene_tx=XM_019838468.2;refGene_gn=ALS2CR12;refGene_tx=XM_019838469.2
C1 188274137 188274164
refGene_gn=ALS2CR12;refGene_tx=XM_019838469.2
C1 188274644 188274693
refGene_gn=ALS2CR12;refGene_tx=XM_003991061.5;refGene_gn=ALS2CR12;refGene_tx=XM_019838468.2;refGene_gn=ALS2CR12;refGene_tx=XM_019838470.2
C1 188286190 188286265
refGene_gn=ALS2CR12;refGene_tx=XM_003991061.5;refGene_gn=ALS2CR12;refGene_tx=XM_019838468.2;refGene_gn=ALS2CR12;refGene_tx=XM_019838470.2
C1 188287041 188287154
refGene_gn=ALS2CR12;refGene_tx=XM_003991061.5;refGene_gn=ALS2CR12;refGene_tx=XM_019838468.2;refGene_gn=ALS2CR12;refGene_tx=XM_019838470.2
C1 188311635 188312314
refGene_gn=TRAK2;refGene_tx=XM_011285469.3;refGene_gn=TRAK2;refGene_tx=XM_019838466.2
C1 188315833 188315939
refGene_gn=TRAK2;refGene_tx=XM_011285469.3;refGene_gn=TRAK2;refGene_tx=XM_019838466.2
C1 188317827 188318076
refGene_gn=TRAK2;refGene_tx=XM_011285469.3;refGene_gn=TRAK2;refGene_tx=XM_019838466.2

C1	188319435	188319734
refGene_gn=TRAK2;refGene_tx=XM_011285469.3;refGene_gn=TRAK2;refGene_tx=XM_019838466.2
C1	188321662	188321866
refGene_gn=TRAK2;refGene_tx=XM_011285469.3;refGene_gn=TRAK2;refGene_tx=XM_019838466.2
C1	188322410	188322490
refGene_gn=TRAK2;refGene_tx=XM_011285469.3;refGene_gn=TRAK2;refGene_tx=XM_019838466.2
C1	188324999	188325137
refGene_gn=TRAK2;refGene_tx=XM_011285469.3;refGene_gn=TRAK2;refGene_tx=XM_019838466.2
C1	188326193	188326268
refGene_gn=TRAK2;refGene_tx=XM_011285469.3;refGene_gn=TRAK2;refGene_tx=XM_019838466.2
C1	188326944	188327075
refGene_gn=TRAK2;refGene_tx=XM_011285469.3;refGene_gn=TRAK2;refGene_tx=XM_019838466.2
C1	188328776	188328855
refGene_gn=TRAK2;refGene_tx=XM_011285469.3;refGene_gn=TRAK2;refGene_tx=XM_019838466.2
C1	188329737	188329947
refGene_gn=TRAK2;refGene_tx=XM_011285469.3;refGene_gn=TRAK2;refGene_tx=XM_019838466.2
C1	188331209	188331326
refGene_gn=TRAK2;refGene_tx=XM_011285469.3;refGene_gn=TRAK2;refGene_tx=XM_019838466.2
C1	188333294	188333371
refGene_gn=TRAK2;refGene_tx=XM_011285469.3;refGene_gn=TRAK2;refGene_tx=XM_019838466.2
C1	188338796	188338991
refGene_gn=TRAK2;refGene_tx=XM_011285469.3;refGene_gn=TRAK2;refGene_tx=XM_019838466.2
C1	188343472	188343563
refGene_gn=TRAK2;refGene_tx=XM_011285469.3;refGene_gn=TRAK2;refGene_tx=XM_019838466.2
C1	188375948	188375960
refGene_gn=STRADB;refGene_tx=XM_003991018.5;refGene_gn=STRADB;refGene_tx=XM_011285471.3
C1	188380258	188380339
refGene_gn=STRADB;refGene_tx=XM_003991018.5;refGene_gn=STRADB;refGene_tx=XM_011285471.3
C1	188396287	188396387
refGene_gn=STRADB;refGene_tx=XM_003991018.5;refGene_gn=STRADB;refGene_tx=XM_011285471.3
C1	188400668	188400790
refGene_gn=STRADB;refGene_tx=XM_003991018.5;refGene_gn=STRADB;refGene_tx=XM_011285471.3
C1	188403112	188403221
refGene_gn=STRADB;refGene_tx=XM_003991018.5;refGene_gn=STRADB;refGene_

tx=XM_011285471.3
C1	188404248	188404372
refGene_gn=STRADB;refGene_tx=XM_003991018.5;refGene_gn=STRADB;refGene_tx=XM_011285471.3
C1	188405558	188405730
refGene_gn=STRADB;refGene_tx=XM_003991018.5;refGene_gn=STRADB;refGene_tx=XM_011285471.3
C1	188406025	188406130
refGene_gn=STRADB;refGene_tx=XM_003991018.5;refGene_gn=STRADB;refGene_tx=XM_011285471.3
C1	188406281	188406526
refGene_gn=STRADB;refGene_tx=XM_003991018.5;refGene_gn=STRADB;refGene_tx=XM_011285471.3
C1	188407139	188407182
refGene_gn=STRADB;refGene_tx=XM_003991018.5;refGene_gn=STRADB;refGene_tx=XM_011285471.3
C1	188407649	188407784
refGene_gn=STRADB;refGene_tx=XM_003991018.5;refGene_gn=STRADB;refGene_tx=XM_011285471.3
C1	188429576	188429924
refGene_gn=C2CD6;refGene_tx=XM_019838477.2;refGene_gn=C2CD6;refGene_tx=XM_023259601.1;refGene_gn=C2CD6;refGene_tx=XM_023259602.1;refGene_gn=C2CD6;refGene_tx=XM_023259603.1;refGene_gn=C2CD6;refGene_tx=XM_023259604.1;refGene_gn=C2CD6;refGene_tx=XM_023259605.1;refGene_gn=C2CD6;refGene_tx=XM_023259606.1;refGene_gn=C2CD6;refGene_tx=XM_023259608.1
C1	188432769	188436273
refGene_gn=C2CD6;refGene_tx=XM_019838477.2;refGene_gn=C2CD6;refGene_tx=XM_023259601.1;refGene_gn=C2CD6;refGene_tx=XM_023259602.1;refGene_gn=C2CD6;refGene_tx=XM_023259603.1;refGene_gn=C2CD6;refGene_tx=XM_023259604.1;refGene_gn=C2CD6;refGene_tx=XM_023259605.1;refGene_gn=C2CD6;refGene_tx=XM_023259606.1;refGene_gn=C2CD6;refGene_tx=XM_023259608.1
C1	188437390	188437465
refGene_gn=C2CD6;refGene_tx=XM_019838477.2;refGene_gn=C2CD6;refGene_tx=XM_023259601.1;refGene_gn=C2CD6;refGene_tx=XM_023259602.1;refGene_gn=C2CD6;refGene_tx=XM_023259603.1;refGene_gn=C2CD6;refGene_tx=XM_023259604.1;refGene_gn=C2CD6;refGene_tx=XM_023259605.1;refGene_gn=C2CD6;refGene_tx=XM_023259606.1;refGene_gn=C2CD6;refGene_tx=XM_023259608.1
C1	188461468	188461527
refGene_gn=C2CD6;refGene_tx=XM_019838477.2;refGene_gn=C2CD6;refGene_tx=XM_023259601.1;refGene_gn=C2CD6;refGene_tx=XM_023259602.1;refGene_gn=C2CD6;refGene_tx=XM_023259603.1;refGene_gn=C2CD6;refGene_tx=XM_023259604.1;refGene_gn=C2CD6;refGene_tx=XM_023259605.1;refGene_gn=C2CD6;refGene_tx=XM_023259606.1;refGene_gn=C2CD6;refGene_tx=XM_023259608.1
C1	188479015	188479073
refGene_gn=C2CD6;refGene_tx=XM_019838477.2;refGene_gn=C2CD6;refGene_tx=XM_023259601.1;refGene_gn=C2CD6;refGene_tx=XM_023259602.1;refGene_gn=C2CD6;refGene_tx=XM_023259603.1;refGene_gn=C2CD6;refGene_tx=XM_023259604.1;refGene_gn=C2CD6;refGene_tx=XM_023259605.1;refGene_gn=C2CD6;refGene_tx=XM_023259606.1;refGene_gn=C2CD6;refGene_tx=XM_023259608.1
C1	188509671	188509980

refGene_gn=C2CD6;refGene_tx=XM_023259601.1;refGene_gn=C2CD6;refGene_tx=XM_023259602.1;refGene_gn=C2CD6;refGene_tx=XM_023259603.1;refGene_gn=C2CD6;refGene_tx=XM_023259604.1;refGene_gn=C2CD6;refGene_tx=XM_023259605.1;refGene_gn=C2CD6;refGene_tx=XM_023259606.1;refGene_gn=C2CD6;refGene_tx=XM_023259608.1
C1 188517062 188517106
refGene_gn=C2CD6;refGene_tx=XM_019838477.2;refGene_gn=C2CD6;refGene_tx=XM_023259601.1;refGene_gn=C2CD6;refGene_tx=XM_023259602.1;refGene_gn=C2CD6;refGene_tx=XM_023259603.1;refGene_gn=C2CD6;refGene_tx=XM_023259604.1;refGene_gn=C2CD6;refGene_tx=XM_023259605.1;refGene_gn=C2CD6;refGene_tx=XM_023259606.1;refGene_gn=C2CD6;refGene_tx=XM_023259608.1
C1 188519384 188519451
refGene_gn=C2CD6;refGene_tx=XM_019838477.2;refGene_gn=C2CD6;refGene_tx=XM_023259601.1;refGene_gn=C2CD6;refGene_tx=XM_023259602.1;refGene_gn=C2CD6;refGene_tx=XM_023259603.1;refGene_gn=C2CD6;refGene_tx=XM_023259604.1;refGene_gn=C2CD6;refGene_tx=XM_023259605.1;refGene_gn=C2CD6;refGene_tx=XM_023259606.1;refGene_gn=C2CD6;refGene_tx=XM_023259608.1
C1 188521691 188521797
refGene_gn=C2CD6;refGene_tx=XM_019838477.2;refGene_gn=C2CD6;refGene_tx=XM_023259601.1;refGene_gn=C2CD6;refGene_tx=XM_023259602.1;refGene_gn=C2CD6;refGene_tx=XM_023259603.1;refGene_gn=C2CD6;refGene_tx=XM_023259604.1;refGene_gn=C2CD6;refGene_tx=XM_023259605.1;refGene_gn=C2CD6;refGene_tx=XM_023259606.1;refGene_gn=C2CD6;refGene_tx=XM_023259608.1
C1 188532026 188532173
refGene_gn=C2CD6;refGene_tx=XM_019838477.2;refGene_gn=C2CD6;refGene_tx=XM_023259601.1;refGene_gn=C2CD6;refGene_tx=XM_023259602.1;refGene_gn=C2CD6;refGene_tx=XM_023259603.1;refGene_gn=C2CD6;refGene_tx=XM_023259604.1;refGene_gn=C2CD6;refGene_tx=XM_023259605.1;refGene_gn=C2CD6;refGene_tx=XM_023259606.1;refGene_gn=C2CD6;refGene_tx=XM_023259608.1
C1 188536020 188536100
refGene_gn=C2CD6;refGene_tx=XM_019838477.2;refGene_gn=C2CD6;refGene_tx=XM_023259601.1;refGene_gn=C2CD6;refGene_tx=XM_023259602.1;refGene_gn=C2CD6;refGene_tx=XM_023259603.1;refGene_gn=C2CD6;refGene_tx=XM_023259604.1;refGene_gn=C2CD6;refGene_tx=XM_023259605.1;refGene_gn=C2CD6;refGene_tx=XM_023259606.1;refGene_gn=C2CD6;refGene_tx=XM_023259608.1
C1 188537721 188537790
refGene_gn=C2CD6;refGene_tx=XM_019838477.2;refGene_gn=C2CD6;refGene_tx=XM_023259601.1;refGene_gn=C2CD6;refGene_tx=XM_023259602.1;refGene_gn=C2CD6;refGene_tx=XM_023259603.1;refGene_gn=C2CD6;refGene_tx=XM_023259604.1;refGene_gn=C2CD6;refGene_tx=XM_023259605.1;refGene_gn=C2CD6;refGene_tx=XM_023259606.1;refGene_gn=C2CD6;refGene_tx=XM_023259608.1
C1 188538703 188538748
refGene_gn=C2CD6;refGene_tx=XM_019838477.2;refGene_gn=C2CD6;refGene_tx=XM_023259601.1;refGene_gn=C2CD6;refGene_tx=XM_023259603.1;refGene_gn=C2CD6;refGene_tx=XM_023259604.1;refGene_gn=C2CD6;refGene_tx=XM_023259605.1;refGene_gn=C2CD6;refGene_tx=XM_023259606.1;refGene_gn=C2CD6;refGene_tx=XM_023259608.1
C1 188545756 188545879
refGene_gn=C2CD6;refGene_tx=XM_019838477.2;refGene_gn=C2CD6;refGene_tx=XM_023259601.1;refGene_gn=C2CD6;refGene_tx=XM_023259602.1;refGene_gn=

C2CD6;refGene_tx=XM_023259603.1;refGene_gn=C2CD6;refGene_tx=XM_0232596
04.1;refGene_gn=C2CD6;refGene_tx=XM_023259605.1;refGene_gn=C2CD6;refGe
ne_tx=XM_023259606.1;refGene_gn=C2CD6;refGene_tx=XM_023259608.1
C1 188553511 188553657
refGene_gn=C2CD6;refGene_tx=XM_019838477.2;refGene_gn=C2CD6;refGene_tx
=XM_023259601.1;refGene_gn=C2CD6;refGene_tx=XM_023259602.1;refGene_gn=
C2CD6;refGene_tx=XM_023259604.1;refGene_gn=C2CD6;refGene_tx=XM_0232596
08.1
C1 188554947 188555019
refGene_gn=C2CD6;refGene_tx=XM_023259606.1
C1 188555234 188555286
refGene_gn=C2CD6;refGene_tx=XM_019838477.2;refGene_gn=C2CD6;refGene_tx
=XM_023259601.1;refGene_gn=C2CD6;refGene_tx=XM_023259602.1;refGene_gn=
C2CD6;refGene_tx=XM_023259604.1;refGene_gn=C2CD6;refGene_tx=XM_0232596
08.1
C1 188558217 188558304
refGene_gn=C2CD6;refGene_tx=XM_019838477.2;refGene_gn=C2CD6;refGene_tx
=XM_023259601.1;refGene_gn=C2CD6;refGene_tx=XM_023259602.1;refGene_gn=
C2CD6;refGene_tx=XM_023259603.1;refGene_gn=C2CD6;refGene_tx=XM_0232596
04.1;refGene_gn=C2CD6;refGene_tx=XM_023259605.1;refGene_gn=C2CD6;refGe
ne_tx=XM_023259608.1
C1 188601321 188601534
refGene_gn=C2CD6;refGene_tx=XM_019838477.2;refGene_gn=C2CD6;refGene_tx
=XM_023259601.1;refGene_gn=C2CD6;refGene_tx=XM_023259602.1;refGene_gn=
C2CD6;refGene_tx=XM_023259603.1;refGene_gn=C2CD6;refGene_tx=XM_0232596
08.1
C1 188608532 188608600
refGene_gn=MPP4;refGene_tx=XM_011285473.3;refGene_gn=MPP4;refGene_tx=X
M_011285474.3;refGene_gn=MPP4;refGene_tx=XM_011285475.3;refGene_gn=MPP
4;refGene_tx=XM_019838481.2;refGene_gn=MPP4;refGene_tx=XM_019838482.2
C1 188610573 188610695
refGene_gn=MPP4;refGene_tx=XM_011285473.3;refGene_gn=MPP4;refGene_tx=X
M_011285474.3;refGene_gn=MPP4;refGene_tx=XM_011285475.3;refGene_gn=MPP
4;refGene_tx=XM_019838481.2;refGene_gn=MPP4;refGene_tx=XM_019838482.2
C1 188611516 188611610
refGene_gn=MPP4;refGene_tx=XM_011285473.3;refGene_gn=MPP4;refGene_tx=X
M_011285474.3;refGene_gn=MPP4;refGene_tx=XM_011285475.3;refGene_gn=MPP
4;refGene_tx=XM_019838481.2;refGene_gn=MPP4;refGene_tx=XM_019838482.2
C1 188612264 188612338
refGene_gn=MPP4;refGene_tx=XM_011285473.3;refGene_gn=MPP4;refGene_tx=X
M_011285474.3;refGene_gn=MPP4;refGene_tx=XM_011285475.3;refGene_gn=MPP
4;refGene_tx=XM_019838481.2;refGene_gn=MPP4;refGene_tx=XM_019838482.2
C1 188613739 188613931
refGene_gn=MPP4;refGene_tx=XM_011285473.3;refGene_gn=MPP4;refGene_tx=X
M_011285474.3;refGene_gn=MPP4;refGene_tx=XM_011285475.3;refGene_gn=MPP
4;refGene_tx=XM_019838481.2;refGene_gn=MPP4;refGene_tx=XM_019838482.2
C1 188614227 188614351
refGene_gn=MPP4;refGene_tx=XM_011285473.3;refGene_gn=MPP4;refGene_tx=X
M_011285474.3;refGene_gn=MPP4;refGene_tx=XM_011285475.3;refGene_gn=MPP
4;refGene_tx=XM_019838481.2;refGene_gn=MPP4;refGene_tx=XM_019838482.2

C1 188616480 188616638
refGene_gn=MPP4;refGene_tx=XM_011285473.3;refGene_gn=MPP4;refGene_tx=XM_011285474.3;refGene_gn=MPP4;refGene_tx=XM_011285475.3;refGene_gn=MPP4;refGene_tx=XM_019838481.2;refGene_gn=MPP4;refGene_tx=XM_019838482.2
C1 188617378 188617499
refGene_gn=MPP4;refGene_tx=XM_011285473.3;refGene_gn=MPP4;refGene_tx=XM_011285474.3;refGene_gn=MPP4;refGene_tx=XM_011285475.3;refGene_gn=MPP4;refGene_tx=XM_019838481.2;refGene_gn=MPP4;refGene_tx=XM_019838482.2
C1 188624824 188624965
refGene_gn=MPP4;refGene_tx=XM_011285473.3;refGene_gn=MPP4;refGene_tx=XM_011285474.3;refGene_gn=MPP4;refGene_tx=XM_011285475.3;refGene_gn=MPP4;refGene_tx=XM_019838481.2;refGene_gn=MPP4;refGene_tx=XM_019838482.2
C1 188625921 188625978
refGene_gn=MPP4;refGene_tx=XM_011285473.3;refGene_gn=MPP4;refGene_tx=XM_011285474.3;refGene_gn=MPP4;refGene_tx=XM_011285475.3;refGene_gn=MPP4;refGene_tx=XM_019838481.2;refGene_gn=MPP4;refGene_tx=XM_019838482.2
C1 188627151 188627162
refGene_gn=MPP4;refGene_tx=XM_011285473.3;refGene_gn=MPP4;refGene_tx=XM_011285474.3
C1 188627777 188627809
refGene_gn=MPP4;refGene_tx=XM_011285473.3;refGene_gn=MPP4;refGene_tx=XM_011285474.3;refGene_gn=MPP4;refGene_tx=XM_011285475.3;refGene_gn=MPP4;refGene_tx=XM_019838481.2;refGene_gn=MPP4;refGene_tx=XM_019838482.2
C1 188629289 188629307
refGene_gn=MPP4;refGene_tx=XM_011285474.3
C1 188630066 188630105
refGene_gn=MPP4;refGene_tx=XM_011285473.3
C1 188632350 188632542
refGene_gn=MPP4;refGene_tx=XM_011285475.3;refGene_gn=MPP4;refGene_tx=XM_019838481.2;refGene_gn=MPP4;refGene_tx=XM_019838482.2;refGene_gn=MPP4;refGene_tx=XM_023259609.1
C1 188634926 188635061
refGene_gn=MPP4;refGene_tx=XM_011285475.3;refGene_gn=MPP4;refGene_tx=XM_019838481.2;refGene_gn=MPP4;refGene_tx=XM_019838482.2;refGene_gn=MPP4;refGene_tx=XM_023259609.1
C1 188636014 188636123
refGene_gn=MPP4;refGene_tx=XM_011285475.3;refGene_gn=MPP4;refGene_tx=XM_019838481.2;refGene_gn=MPP4;refGene_tx=XM_019838482.2;refGene_gn=MPP4;refGene_tx=XM_023259609.1
C1 188636520 188636614
refGene_gn=MPP4;refGene_tx=XM_011285475.3;refGene_gn=MPP4;refGene_tx=XM_019838481.2;refGene_gn=MPP4;refGene_tx=XM_019838482.2;refGene_gn=MPP4;refGene_tx=XM_023259609.1
C1 188639982 188640063
refGene_gn=MPP4;refGene_tx=XM_011285475.3;refGene_gn=MPP4;refGene_tx=XM_019838481.2;refGene_gn=MPP4;refGene_tx=XM_019838482.2;refGene_gn=MPP4;refGene_tx=XM_023259609.1
C1 188641708 188641879
refGene_gn=MPP4;refGene_tx=XM_011285475.3;refGene_gn=MPP4;refGene_tx=XM_019838481.2;refGene_gn=MPP4;refGene_tx=XM_019838482.2;refGene_gn=MPP

4;refGene_tx=XM_023259609.1
C1 188643959 188644001
refGene_gn=MPP4;refGene_tx=XM_011285475.3;refGene_gn=MPP4;refGene_tx=XM_019838481.2;refGene_gn=MPP4;refGene_tx=XM_019838482.2;refGene_gn=MPP4;refGene_tx=XM_023259609.1
C1 188645130 188645145
refGene_gn=MPP4;refGene_tx=XM_011285475.3;refGene_gn=MPP4;refGene_tx=XM_019838481.2;refGene_gn=MPP4;refGene_tx=XM_019838482.2;refGene_gn=MPP4;refGene_tx=XM_023259609.1
C1 188647269 188647290
refGene_gn=MPP4;refGene_tx=XM_011285475.3;refGene_gn=MPP4;refGene_tx=XM_019838481.2;refGene_gn=MPP4;refGene_tx=XM_019838482.2;refGene_gn=MPP4;refGene_tx=XM_023259609.1
C1 188649290 188649329
refGene_gn=MPP4;refGene_tx=XM_011285475.3;refGene_gn=MPP4;refGene_tx=XM_019838481.2;refGene_gn=MPP4;refGene_tx=XM_019838482.2;refGene_gn=MPP4;refGene_tx=XM_023259609.1
C1 188651879 188651897
refGene_gn=MPP4;refGene_tx=XM_011285475.3;refGene_gn=MPP4;refGene_tx=XM_019838481.2;refGene_gn=MPP4;refGene_tx=XM_019838482.2;refGene_gn=MPP4;refGene_tx=XM_023259609.1
C1 188654311 188654376
refGene_gn=MPP4;refGene_tx=XM_011285475.3;refGene_gn=MPP4;refGene_tx=XM_019838481.2;refGene_gn=MPP4;refGene_tx=XM_019838482.2;refGene_gn=MPP4;refGene_tx=XM_023259609.1
C1 188656580 188656777
refGene_gn=MPP4;refGene_tx=XM_011285475.3;refGene_gn=MPP4;refGene_tx=XM_019838481.2;refGene_gn=MPP4;refGene_tx=XM_019838482.2;refGene_gn=MPP4;refGene_tx=XM_023259609.1
C1 188657465 188657537
refGene_gn=MPP4;refGene_tx=XM_011285475.3;refGene_gn=MPP4;refGene_tx=XM_019838481.2;refGene_gn=MPP4;refGene_tx=XM_019838482.2;refGene_gn=MPP4;refGene_tx=XM_023259609.1
C1 188658986 188659072
refGene_gn=MPP4;refGene_tx=XM_011285475.3;refGene_gn=MPP4;refGene_tx=XM_019838481.2;refGene_gn=MPP4;refGene_tx=XM_019838482.2;refGene_gn=MPP4;refGene_tx=XM_023259609.1
C1 188660183 188660265
refGene_gn=MPP4;refGene_tx=XM_011285475.3;refGene_gn=MPP4;refGene_tx=XM_019838481.2;refGene_gn=MPP4;refGene_tx=XM_019838482.2;refGene_gn=MPP4;refGene_tx=XM_023259609.1
C1 188660973 188661105
refGene_gn=MPP4;refGene_tx=XM_011285475.3;refGene_gn=MPP4;refGene_tx=XM_019838481.2;refGene_gn=MPP4;refGene_tx=XM_019838482.2;refGene_gn=MPP4;refGene_tx=XM_023259609.1
C1 188662258 188662339
refGene_gn=MPP4;refGene_tx=XM_011285475.3;refGene_gn=MPP4;refGene_tx=XM_019838481.2;refGene_gn=MPP4;refGene_tx=XM_019838482.2;refGene_gn=MPP4;refGene_tx=XM_023259609.1
C1 188664434 188664512

refGene_gn=MPP4;refGene_tx=XM_011285475.3;refGene_gn=MPP4;refGene_tx=XM_019838481.2;refGene_gn=MPP4;refGene_tx=XM_019838482.2;refGene_gn=MPP4;refGene_tx=XM_023259609.1
C1 188668093 188668215
refGene_gn=MPP4;refGene_tx=XM_011285475.3;refGene_gn=MPP4;refGene_tx=XM_019838481.2;refGene_gn=MPP4;refGene_tx=XM_019838482.2;refGene_gn=MPP4;refGene_tx=XM_023259609.1
C1 188669305 188669339
refGene_gn=MPP4;refGene_tx=XM_011285475.3;refGene_gn=MPP4;refGene_tx=XM_019838481.2;refGene_gn=MPP4;refGene_tx=XM_019838482.2;refGene_gn=MPP4;refGene_tx=XM_023259609.1
C1 188678204 188678243
refGene_gn=ALS2;refGene_tx=XM_011285476.3;refGene_gn=ALS2;refGene_tx=XM_019838483.2;refGene_gn=ALS2;refGene_tx=XM_019838484.2;refGene_gn=ALS2;refGene_tx=XM_019838485.2
C1 188680177 188680274
refGene_gn=ALS2;refGene_tx=XM_011285476.3;refGene_gn=ALS2;refGene_tx=XM_019838483.2;refGene_gn=ALS2;refGene_tx=XM_019838484.2;refGene_gn=ALS2;refGene_tx=XM_019838485.2
C1 188680486 188680636
refGene_gn=ALS2;refGene_tx=XM_011285476.3;refGene_gn=ALS2;refGene_tx=XM_019838483.2;refGene_gn=ALS2;refGene_tx=XM_019838484.2;refGene_gn=ALS2;refGene_tx=XM_019838485.2
C1 188681176 188681238
refGene_gn=ALS2;refGene_tx=XM_011285476.3;refGene_gn=ALS2;refGene_tx=XM_019838483.2;refGene_gn=ALS2;refGene_tx=XM_019838484.2;refGene_gn=ALS2;refGene_tx=XM_019838485.2
C1 188681441 188681487
refGene_gn=ALS2;refGene_tx=XM_011285476.3;refGene_gn=ALS2;refGene_tx=XM_019838483.2;refGene_gn=ALS2;refGene_tx=XM_019838484.2;refGene_gn=ALS2;refGene_tx=XM_019838485.2
C1 188682826 188683003
refGene_gn=ALS2;refGene_tx=XM_011285476.3;refGene_gn=ALS2;refGene_tx=XM_019838483.2;refGene_gn=ALS2;refGene_tx=XM_019838484.2;refGene_gn=ALS2;refGene_tx=XM_019838485.2
C1 188683506 188683629
refGene_gn=ALS2;refGene_tx=XM_011285476.3;refGene_gn=ALS2;refGene_tx=XM_019838483.2;refGene_gn=ALS2;refGene_tx=XM_019838484.2;refGene_gn=ALS2;refGene_tx=XM_019838485.2
C1 188685407 188685565
refGene_gn=ALS2;refGene_tx=XM_011285476.3;refGene_gn=ALS2;refGene_tx=XM_019838483.2;refGene_gn=ALS2;refGene_tx=XM_019838484.2;refGene_gn=ALS2;refGene_tx=XM_019838485.2
C1 188686314 188686432
refGene_gn=ALS2;refGene_tx=XM_011285476.3;refGene_gn=ALS2;refGene_tx=XM_019838483.2;refGene_gn=ALS2;refGene_tx=XM_019838484.2;refGene_gn=ALS2;refGene_tx=XM_019838485.2
C1 188690110 188690278
refGene_gn=ALS2;refGene_tx=XM_011285476.3;refGene_gn=ALS2;refGene_tx=XM_019838483.2;refGene_gn=ALS2;refGene_tx=XM_019838484.2;refGene_gn=ALS

2;refGene_tx=XM_019838485.2
C1	188691159	188691293
refGene_gn=ALS2;refGene_tx=XM_011285476.3;refGene_gn=ALS2;refGene_tx=XM_019838483.2;refGene_gn=ALS2;refGene_tx=XM_019838484.2;refGene_gn=ALS2;refGene_tx=XM_019838485.2
C1	188696229	188696307
refGene_gn=ALS2;refGene_tx=XM_011285476.3;refGene_gn=ALS2;refGene_tx=XM_019838483.2;refGene_gn=ALS2;refGene_tx=XM_019838484.2;refGene_gn=ALS2;refGene_tx=XM_019838485.2
C1	188696514	188696626
refGene_gn=ALS2;refGene_tx=XM_011285476.3;refGene_gn=ALS2;refGene_tx=XM_019838483.2;refGene_gn=ALS2;refGene_tx=XM_019838484.2;refGene_gn=ALS2;refGene_tx=XM_019838485.2
C1	188697863	188698028
refGene_gn=ALS2;refGene_tx=XM_011285476.3;refGene_gn=ALS2;refGene_tx=XM_019838483.2;refGene_gn=ALS2;refGene_tx=XM_019838484.2;refGene_gn=ALS2;refGene_tx=XM_019838485.2
C1	188698604	188698703
refGene_gn=ALS2;refGene_tx=XM_011285476.3;refGene_gn=ALS2;refGene_tx=XM_019838483.2;refGene_gn=ALS2;refGene_tx=XM_019838484.2;refGene_gn=ALS2;refGene_tx=XM_019838485.2
C1	188699756	188699822
refGene_gn=ALS2;refGene_tx=XM_011285476.3;refGene_gn=ALS2;refGene_tx=XM_019838483.2;refGene_gn=ALS2;refGene_tx=XM_019838484.2;refGene_gn=ALS2;refGene_tx=XM_019838485.2
C1	188699940	188700143
refGene_gn=ALS2;refGene_tx=XM_011285476.3;refGene_gn=ALS2;refGene_tx=XM_019838483.2;refGene_gn=ALS2;refGene_tx=XM_019838484.2;refGene_gn=ALS2;refGene_tx=XM_019838485.2
C1	188700528	188700595
refGene_gn=ALS2;refGene_tx=XM_011285476.3;refGene_gn=ALS2;refGene_tx=XM_019838483.2;refGene_gn=ALS2;refGene_tx=XM_019838484.2;refGene_gn=ALS2;refGene_tx=XM_019838485.2
C1	188700938	188701009
refGene_gn=ALS2;refGene_tx=XM_011285476.3;refGene_gn=ALS2;refGene_tx=XM_019838483.2;refGene_gn=ALS2;refGene_tx=XM_019838484.2;refGene_gn=ALS2;refGene_tx=XM_019838485.2
C1	188701374	188701503
refGene_gn=ALS2;refGene_tx=XM_011285476.3;refGene_gn=ALS2;refGene_tx=XM_019838483.2;refGene_gn=ALS2;refGene_tx=XM_019838484.2;refGene_gn=ALS2;refGene_tx=XM_019838485.2
C1	188702147	188702279
refGene_gn=ALS2;refGene_tx=XM_011285476.3;refGene_gn=ALS2;refGene_tx=XM_019838483.2;refGene_gn=ALS2;refGene_tx=XM_019838484.2;refGene_gn=ALS2;refGene_tx=XM_019838485.2
C1	188705843	188706006
refGene_gn=ALS2;refGene_tx=XM_011285476.3;refGene_gn=ALS2;refGene_tx=XM_019838483.2;refGene_gn=ALS2;refGene_tx=XM_019838484.2;refGene_gn=ALS2;refGene_tx=XM_019838485.2
C1	188707159	188707264

refGene_gn=ALS2;refGene_tx=XM_019838483.2;refGene_gn=ALS2;refGene_tx=XM_019838484.2;refGene_gn=ALS2;refGene_tx=XM_019838485.2
C1 188710924 188710990
refGene_gn=ALS2;refGene_tx=XM_011285476.3;refGene_gn=ALS2;refGene_tx=XM_019838483.2;refGene_gn=ALS2;refGene_tx=XM_019838484.2;refGene_gn=ALS2;refGene_tx=XM_019838485.2
C1 188714079 188714348
refGene_gn=ALS2;refGene_tx=XM_011285476.3;refGene_gn=ALS2;refGene_tx=XM_019838483.2;refGene_gn=ALS2;refGene_tx=XM_019838484.2;refGene_gn=ALS2;refGene_tx=XM_019838485.2;refGene_gn=ALS2;refGene_tx=XM_019838486.2
C1 188716572 188716744
refGene_gn=ALS2;refGene_tx=XM_011285476.3;refGene_gn=ALS2;refGene_tx=XM_019838483.2;refGene_gn=ALS2;refGene_tx=XM_019838484.2;refGene_gn=ALS2;refGene_tx=XM_019838485.2;refGene_gn=ALS2;refGene_tx=XM_019838486.2;refGene_gn=ALS2;refGene_tx=XM_019838487.2
C1 188718864 188719047
refGene_gn=ALS2;refGene_tx=XM_011285476.3;refGene_gn=ALS2;refGene_tx=XM_019838483.2;refGene_gn=ALS2;refGene_tx=XM_019838484.2;refGene_gn=ALS2;refGene_tx=XM_019838485.2;refGene_gn=ALS2;refGene_tx=XM_019838486.2;refGene_gn=ALS2;refGene_tx=XM_019838487.2
C1 188722041 188722119
refGene_gn=ALS2;refGene_tx=XM_011285476.3;refGene_gn=ALS2;refGene_tx=XM_019838483.2;refGene_gn=ALS2;refGene_tx=XM_019838484.2;refGene_gn=ALS2;refGene_tx=XM_019838485.2;refGene_gn=ALS2;refGene_tx=XM_019838486.2;refGene_gn=ALS2;refGene_tx=XM_019838487.2
C1 188724911 188725008
refGene_gn=ALS2;refGene_tx=XM_011285476.3;refGene_gn=ALS2;refGene_tx=XM_019838483.2;refGene_gn=ALS2;refGene_tx=XM_019838484.2;refGene_gn=ALS2;refGene_tx=XM_019838485.2;refGene_gn=ALS2;refGene_tx=XM_019838486.2;refGene_gn=ALS2;refGene_tx=XM_019838487.2
C1 188726345 188726514
refGene_gn=ALS2;refGene_tx=XM_011285476.3;refGene_gn=ALS2;refGene_tx=XM_019838483.2;refGene_gn=ALS2;refGene_tx=XM_019838484.2;refGene_gn=ALS2;refGene_tx=XM_019838485.2;refGene_gn=ALS2;refGene_tx=XM_019838486.2;refGene_gn=ALS2;refGene_tx=XM_019838487.2
C1 188728002 188728360
refGene_gn=ALS2;refGene_tx=XM_011285476.3;refGene_gn=ALS2;refGene_tx=XM_019838483.2;refGene_gn=ALS2;refGene_tx=XM_019838484.2;refGene_gn=ALS2;refGene_tx=XM_019838485.2;refGene_gn=ALS2;refGene_tx=XM_019838486.2;refGene_gn=ALS2;refGene_tx=XM_019838487.2
C1 188731440 188732357
refGene_gn=ALS2;refGene_tx=XM_011285476.3;refGene_gn=ALS2;refGene_tx=XM_019838483.2;refGene_gn=ALS2;refGene_tx=XM_019838484.2;refGene_gn=ALS2;refGene_tx=XM_019838485.2;refGene_gn=ALS2;refGene_tx=XM_019838486.2;refGene_gn=ALS2;refGene_tx=XM_019838487.2
C1 188735519 188735674
refGene_gn=ALS2;refGene_tx=XM_011285476.3;refGene_gn=ALS2;refGene_tx=XM_019838483.2;refGene_gn=ALS2;refGene_tx=XM_019838484.2;refGene_gn=ALS2;refGene_tx=XM_019838485.2;refGene_gn=ALS2;refGene_tx=XM_019838486.2;refGene_gn=ALS2;refGene_tx=XM_019838487.2

C1 188736881 188736901
refGene_gn=ALS2;refGene_tx=XM_011285476.3;refGene_gn=ALS2;refGene_tx=XM_019838483.2;refGene_gn=ALS2;refGene_tx=XM_019838484.2;refGene_gn=ALS2;refGene_tx=XM_019838485.2;refGene_gn=ALS2;refGene_tx=XM_019838486.2;refGene_gn=ALS2;refGene_tx=XM_019838487.2
C1 188770048 188770171
refGene_gn=CDK15;refGene_tx=XM_006935476.3;refGene_gn=CDK15;refGene_tx=XM_019838488.2;refGene_gn=CDK15;refGene_tx=XM_019838489.1
C1 188770879 188771029
refGene_gn=CDK15;refGene_tx=XM_006935476.3;refGene_gn=CDK15;refGene_tx=XM_019838488.2;refGene_gn=CDK15;refGene_tx=XM_019838489.1
C1 188771226 188771324
refGene_gn=CDK15;refGene_tx=XM_006935476.3;refGene_gn=CDK15;refGene_tx=XM_019838488.2;refGene_gn=CDK15;refGene_tx=XM_019838489.1
C1 188778440 188778520
refGene_gn=CDK15;refGene_tx=XM_006935476.3;refGene_gn=CDK15;refGene_tx=XM_019838488.2;refGene_gn=CDK15;refGene_tx=XM_019838489.1
C1 188788543 188788638
refGene_gn=CDK15;refGene_tx=XM_006935476.3;refGene_gn=CDK15;refGene_tx=XM_019838488.2;refGene_gn=CDK15;refGene_tx=XM_019838489.1
C1 188789382 188789445
refGene_gn=CDK15;refGene_tx=XM_006935476.3;refGene_gn=CDK15;refGene_tx=XM_019838488.2;refGene_gn=CDK15;refGene_tx=XM_019838489.1
C1 188795103 188795227
refGene_gn=CDK15;refGene_tx=XM_006935476.3;refGene_gn=CDK15;refGene_tx=XM_019838488.2;refGene_gn=CDK15;refGene_tx=XM_019838489.1
C1 188796868 188796989
refGene_gn=CDK15;refGene_tx=XM_006935476.3;refGene_gn=CDK15;refGene_tx=XM_019838488.2;refGene_gn=CDK15;refGene_tx=XM_019838489.1
C1 188808573 188808667
refGene_gn=CDK15;refGene_tx=XM_006935476.3;refGene_gn=CDK15;refGene_tx=XM_019838488.2;refGene_gn=CDK15;refGene_tx=XM_019838489.1
C1 188813422 188813486
refGene_gn=CDK15;refGene_tx=XM_006935476.3;refGene_gn=CDK15;refGene_tx=XM_019838488.2;refGene_gn=CDK15;refGene_tx=XM_019838489.1
C1 188828804 188828853
refGene_gn=CDK15;refGene_tx=XM_006935476.3;refGene_gn=CDK15;refGene_tx=XM_019838488.2;refGene_gn=CDK15;refGene_tx=XM_019838489.1
C1 188835200 188835340
refGene_gn=CDK15;refGene_tx=XM_006935476.3;refGene_gn=CDK15;refGene_tx=XM_019838488.2;refGene_gn=CDK15;refGene_tx=XM_019838489.1
C1 188839962 188840066
refGene_gn=CDK15;refGene_tx=XM_019838489.1
C1 188846862 188846972
refGene_gn=CDK15;refGene_tx=XM_006935476.3;refGene_gn=CDK15;refGene_tx=XM_019838488.2
C1 188981896 188983615
refGene_gn=FZD7;refGene_tx=XM_019839139.2
C1 188993117 188993199
refGene_gn=FZD7;refGene_tx=XM_019839139.2

C1 189005074 189005082
refGene_gn=FZD7;refGene_tx=XM_019839139.2
C1 189016842 189016926
refGene_gn=KIAA2012;refGene_tx=XM_019838490.1;refGene_gn=KIAA2012;refGene_tx=XM_019838491.1;refGene_gn=KIAA2012;refGene_tx=XM_019838492.1;refGene_gn=KIAA2012;refGene_tx=XM_019838493.1
C1 189018333 189018618
refGene_gn=KIAA2012;refGene_tx=XM_019838490.1;refGene_gn=KIAA2012;refGene_tx=XM_019838491.1;refGene_gn=KIAA2012;refGene_tx=XM_019838492.1;refGene_gn=KIAA2012;refGene_tx=XM_019838493.1
C1 189034507 189034667
refGene_gn=KIAA2012;refGene_tx=XM_019838490.1;refGene_gn=KIAA2012;refGene_tx=XM_019838491.1;refGene_gn=KIAA2012;refGene_tx=XM_019838492.1
C1 189036624 189036780
refGene_gn=KIAA2012;refGene_tx=XM_019838490.1;refGene_gn=KIAA2012;refGene_tx=XM_019838491.1;refGene_gn=KIAA2012;refGene_tx=XM_019838492.1
C1 189040625 189040765
refGene_gn=KIAA2012;refGene_tx=XM_019838490.1;refGene_gn=KIAA2012;refGene_tx=XM_019838491.1;refGene_gn=KIAA2012;refGene_tx=XM_019838492.1
C1 189041865 189042049
refGene_gn=KIAA2012;refGene_tx=XM_019838490.1;refGene_gn=KIAA2012;refGene_tx=XM_019838491.1;refGene_gn=KIAA2012;refGene_tx=XM_019838492.1;refGene_gn=KIAA2012;refGene_tx=XM_019838493.1
C1 189042669 189042812
refGene_gn=KIAA2012;refGene_tx=XM_019838490.1;refGene_gn=KIAA2012;refGene_tx=XM_019838491.1;refGene_gn=KIAA2012;refGene_tx=XM_019838492.1;refGene_gn=KIAA2012;refGene_tx=XM_019838493.1
C1 189045608 189045777
refGene_gn=KIAA2012;refGene_tx=XM_019838490.1;refGene_gn=KIAA2012;refGene_tx=XM_019838491.1;refGene_gn=KIAA2012;refGene_tx=XM_019838492.1;refGene_gn=KIAA2012;refGene_tx=XM_019838493.1
C1 189046796 189046946
refGene_gn=KIAA2012;refGene_tx=XM_019838490.1;refGene_gn=KIAA2012;refGene_tx=XM_019838491.1;refGene_gn=KIAA2012;refGene_tx=XM_019838492.1;refGene_gn=KIAA2012;refGene_tx=XM_019838493.1
C1 189049981 189050152
refGene_gn=KIAA2012;refGene_tx=XM_019838490.1;refGene_gn=KIAA2012;refGene_tx=XM_019838491.1;refGene_gn=KIAA2012;refGene_tx=XM_019838492.1;refGene_gn=KIAA2012;refGene_tx=XM_019838493.1
C1 189052772 189052883
refGene_gn=KIAA2012;refGene_tx=XM_019838490.1;refGene_gn=KIAA2012;refGene_tx=XM_019838491.1;refGene_gn=KIAA2012;refGene_tx=XM_019838492.1;refGene_gn=KIAA2012;refGene_tx=XM_019838493.1
C1 189064569 189064638
refGene_gn=KIAA2012;refGene_tx=XM_019838490.1;refGene_gn=KIAA2012;refGene_tx=XM_019838491.1;refGene_gn=KIAA2012;refGene_tx=XM_019838493.1
C1 189073289 189073366
refGene_gn=KIAA2012;refGene_tx=XM_019838490.1;refGene_gn=KIAA2012;refGene_tx=XM_019838491.1;refGene_gn=KIAA2012;refGene_tx=XM_019838492.1;refGene_gn=KIAA2012;refGene_tx=XM_019838493.1

C1	189087819	189087957
refGene_gn=KIAA2012;refGene_tx=XM_019838490.1;refGene_gn=KIAA2012;refGene_tx=XM_019838491.1;refGene_gn=KIAA2012;refGene_tx=XM_019838492.1;refGene_gn=KIAA2012;refGene_tx=XM_019838493.1
C1	189096785	189096858
refGene_gn=KIAA2012;refGene_tx=XM_019838490.1;refGene_gn=KIAA2012;refGene_tx=XM_019838491.1;refGene_gn=KIAA2012;refGene_tx=XM_019838492.1;refGene_gn=KIAA2012;refGene_tx=XM_019838493.1
C1	189111918	189112009
refGene_gn=KIAA2012;refGene_tx=XM_019838490.1;refGene_gn=KIAA2012;refGene_tx=XM_019838491.1;refGene_gn=KIAA2012;refGene_tx=XM_019838492.1;refGene_gn=KIAA2012;refGene_tx=XM_019838493.1
C1	189114932	189115089
refGene_gn=KIAA2012;refGene_tx=XM_019838490.1;refGene_gn=KIAA2012;refGene_tx=XM_019838491.1;refGene_gn=KIAA2012;refGene_tx=XM_019838492.1;refGene_gn=KIAA2012;refGene_tx=XM_019838493.1
C1	189117092	189117207
refGene_gn=KIAA2012;refGene_tx=XM_019838490.1;refGene_gn=KIAA2012;refGene_tx=XM_019838491.1;refGene_gn=KIAA2012;refGene_tx=XM_019838492.1;refGene_gn=KIAA2012;refGene_tx=XM_019838493.1
C1	189118628	189118966
refGene_gn=KIAA2012;refGene_tx=XM_019838490.1;refGene_gn=KIAA2012;refGene_tx=XM_019838491.1;refGene_gn=KIAA2012;refGene_tx=XM_019838492.1;refGene_gn=KIAA2012;refGene_tx=XM_019838493.1
C1	189120900	189121100
refGene_gn=KIAA2012;refGene_tx=XM_019838490.1;refGene_gn=KIAA2012;refGene_tx=XM_019838491.1;refGene_gn=KIAA2012;refGene_tx=XM_019838492.1;refGene_gn=KIAA2012;refGene_tx=XM_019838493.1
C1	189121717	189121890
refGene_gn=KIAA2012;refGene_tx=XM_019838490.1;refGene_gn=KIAA2012;refGene_tx=XM_019838491.1;refGene_gn=KIAA2012;refGene_tx=XM_019838492.1;refGene_gn=KIAA2012;refGene_tx=XM_019838493.1
C1	189122694	189122914
refGene_gn=KIAA2012;refGene_tx=XM_019838490.1;refGene_gn=KIAA2012;refGene_tx=XM_019838491.1;refGene_gn=KIAA2012;refGene_tx=XM_019838492.1;refGene_gn=KIAA2012;refGene_tx=XM_019838493.1
C1	189124971	189125110
refGene_gn=KIAA2012;refGene_tx=XM_019838490.1;refGene_gn=KIAA2012;refGene_tx=XM_019838491.1;refGene_gn=KIAA2012;refGene_tx=XM_019838492.1;refGene_gn=KIAA2012;refGene_tx=XM_019838493.1
C1	189128498	189128567
refGene_gn=SUMO1;refGene_tx=XM_023257866.1
C1	189135661	189135733
refGene_gn=SUMO1;refGene_tx=XM_023257866.1
C1	189138275	189138353
refGene_gn=SUMO1;refGene_tx=XM_023257866.1
C1	189142262	189142337
refGene_gn=SUMO1;refGene_tx=XM_023257866.1
C1	189156980	189157492
refGene_gn=SUMO1;refGene_tx=XM_023257866.1

C1 189159961 189159992
refGene_gn=SUMO1;refGene_tx=XM_023257866.1
C1 189181357 189181402
refGene_gn=NOP58;refGene_tx=XM_023257867.1
C1 189189596 189189673
refGene_gn=NOP58;refGene_tx=XM_023257867.1
C1 189192248 189192301
refGene_gn=NOP58;refGene_tx=XM_023257867.1
C1 189197399 189197522
refGene_gn=NOP58;refGene_tx=XM_023257867.1
C1 189198642 189198653
refGene_gn=NOP58;refGene_tx=XM_023257867.1
C1 189199966 189200103
refGene_gn=NOP58;refGene_tx=XM_023257867.1
C1 189204492 189204557
refGene_gn=NOP58;refGene_tx=XM_023257867.1
C1 189205518 189205653
refGene_gn=NOP58;refGene_tx=XM_023257867.1
C1 189206759 189206905
refGene_gn=NOP58;refGene_tx=XM_023257867.1
C1 189207831 189207958
refGene_gn=NOP58;refGene_tx=XM_023257867.1
C1 189209340 189209505
refGene_gn=NOP58;refGene_tx=XM_023257867.1
C1 189211025 189211171
refGene_gn=NOP58;refGene_tx=XM_023257867.1
C1 189211363 189211425
refGene_gn=NOP58;refGene_tx=XM_023257867.1
C1 189213131 189213265
refGene_gn=NOP58;refGene_tx=XM_023257867.1
C1 189216122 189216295
refGene_gn=NOP58;refGene_tx=XM_023257867.1
C1 189216723 189216774
refGene_gn=NOP58;refGene_tx=XM_023257867.1
C1 189272447 189272523
refGene_gn=BMPR2;refGene_tx=XM_003991026.5
C1 189384797 189384968
refGene_gn=BMPR2;refGene_tx=XM_003991026.5
C1 189386100 189386271
refGene_gn=BMPR2;refGene_tx=XM_003991026.5
C1 189428411 189428522
refGene_gn=BMPR2;refGene_tx=XM_003991026.5
C1 189429557 189429649
refGene_gn=BMPR2;refGene_tx=XM_003991026.5
C1 189431492 189431723
refGene_gn=BMPR2;refGene_tx=XM_003991026.5
C1 189432476 189432591
refGene_gn=BMPR2;refGene_tx=XM_003991026.5
C1 189436861 189437022
refGene_gn=BMPR2;refGene_tx=XM_003991026.5

C1 189438270 189438418
refGene_gn=BMPR2;refGene_tx=XM_003991026.5
C1 189444806 189444943
refGene_gn=BMPR2;refGene_tx=XM_003991026.5
C1 189460101 189460274
refGene_gn=BMPR2;refGene_tx=XM_003991026.5
C1 189462484 189463764
refGene_gn=BMPR2;refGene_tx=XM_003991026.5
C1 189466112 189466363
refGene_gn=BMPR2;refGene_tx=XM_003991026.5
C1 189528615 189529207
refGene_gn=FAM117B;refGene_tx=XM_023259611.1
C1 189584537 189584689
refGene_gn=FAM117B;refGene_tx=XM_023259611.1
C1 189598398 189598491
refGene_gn=FAM117B;refGene_tx=XM_023259611.1
C1 189599460 189599574
refGene_gn=FAM117B;refGene_tx=XM_023259611.1
C1 189610959 189611103
refGene_gn=FAM117B;refGene_tx=XM_023259611.1
C1 189612417 189612643
refGene_gn=FAM117B;refGene_tx=XM_023259611.1
C1 189614710 189614831
refGene_gn=FAM117B;refGene_tx=XM_023259611.1
C1 189620836 189621155
refGene_gn=FAM117B;refGene_tx=XM_023259611.1
C1 189640907 189641023
refGene_gn=ICA1L;refGene_tx=XM_006935481.4;refGene_gn=ICA1L;refGene_tx=XM_019838496.1;refGene_gn=ICA1L;refGene_tx=XM_019838499.2;refGene_gn=ICA1L;refGene_tx=XM_019838500.2;refGene_gn=ICA1L;refGene_tx=XM_023259613.1
C1 189657670 189657760
refGene_gn=ICA1L;refGene_tx=XM_006935481.4;refGene_gn=ICA1L;refGene_tx=XM_019838496.1;refGene_gn=ICA1L;refGene_tx=XM_019838499.2;refGene_gn=ICA1L;refGene_tx=XM_019838500.2;refGene_gn=ICA1L;refGene_tx=XM_023259613.1
C1 189667877 189668132
refGene_gn=ICA1L;refGene_tx=XM_006935481.4;refGene_gn=ICA1L;refGene_tx=XM_019838496.1;refGene_gn=ICA1L;refGene_tx=XM_019838499.2;refGene_gn=ICA1L;refGene_tx=XM_019838500.2
C1 189688971 189689015
refGene_gn=ICA1L;refGene_tx=XM_006935481.4;refGene_gn=ICA1L;refGene_tx=XM_019838496.1;refGene_gn=ICA1L;refGene_tx=XM_019838499.2;refGene_gn=ICA1L;refGene_tx=XM_019838500.2;refGene_gn=ICA1L;refGene_tx=XM_023259613.1
C1 189691228 189691314
refGene_gn=ICA1L;refGene_tx=XM_006935481.4;refGene_gn=ICA1L;refGene_tx=XM_019838496.1;refGene_gn=ICA1L;refGene_tx=XM_019838499.2;refGene_gn=ICA1L;refGene_tx=XM_019838500.2;refGene_gn=ICA1L;refGene_tx=XM_023259613.1

C1	189692581	189692680
refGene_gn=ICA1L;refGene_tx=XM_006935481.4;refGene_gn=ICA1L;refGene_tx=XM_019838496.1;refGene_gn=ICA1L;refGene_tx=XM_019838499.2;refGene_gn=ICA1L;refGene_tx=XM_019838500.2;refGene_gn=ICA1L;refGene_tx=XM_023259613.1
C1	189694052	189694178
refGene_gn=ICA1L;refGene_tx=XM_006935481.4;refGene_gn=ICA1L;refGene_tx=XM_019838496.1;refGene_gn=ICA1L;refGene_tx=XM_019838499.2;refGene_gn=ICA1L;refGene_tx=XM_019838500.2;refGene_gn=ICA1L;refGene_tx=XM_023259613.1
C1	189696570	189696769
refGene_gn=ICA1L;refGene_tx=XM_006935481.4;refGene_gn=ICA1L;refGene_tx=XM_019838496.1;refGene_gn=ICA1L;refGene_tx=XM_019838499.2;refGene_gn=ICA1L;refGene_tx=XM_019838500.2;refGene_gn=ICA1L;refGene_tx=XM_023259613.1
C1	189697320	189697444
refGene_gn=ICA1L;refGene_tx=XM_006935481.4;refGene_gn=ICA1L;refGene_tx=XM_019838496.1;refGene_gn=ICA1L;refGene_tx=XM_019838499.2;refGene_gn=ICA1L;refGene_tx=XM_019838500.2;refGene_gn=ICA1L;refGene_tx=XM_023259613.1
C1	189698724	189698797
refGene_gn=ICA1L;refGene_tx=XM_006935481.4;refGene_gn=ICA1L;refGene_tx=XM_019838496.1;refGene_gn=ICA1L;refGene_tx=XM_019838499.2;refGene_gn=ICA1L;refGene_tx=XM_019838500.2;refGene_gn=ICA1L;refGene_tx=XM_023259613.1
C1	189701731	189701914
refGene_gn=ICA1L;refGene_tx=XM_006935481.4;refGene_gn=ICA1L;refGene_tx=XM_019838496.1;refGene_gn=ICA1L;refGene_tx=XM_019838499.2;refGene_gn=ICA1L;refGene_tx=XM_019838500.2;refGene_gn=ICA1L;refGene_tx=XM_023259613.1
C1	189717214	189717240
refGene_gn=ICA1L;refGene_tx=XM_006935481.4;refGene_gn=ICA1L;refGene_tx=XM_019838496.1;refGene_gn=ICA1L;refGene_tx=XM_019838499.2;refGene_gn=ICA1L;refGene_tx=XM_019838500.2;refGene_gn=ICA1L;refGene_tx=XM_023259613.1
C1	189733130	189733208
refGene_gn=WDR12;refGene_tx=XM_003991029.4;refGene_gn=WDR12;refGene_tx=XM_006935484.3;refGene_gn=WDR12;refGene_tx=XM_011285480.2
C1	189734187	189734260
refGene_gn=WDR12;refGene_tx=XM_003991029.4;refGene_gn=WDR12;refGene_tx=XM_006935484.3;refGene_gn=WDR12;refGene_tx=XM_011285480.2
C1	189736329	189736462
refGene_gn=WDR12;refGene_tx=XM_003991029.4;refGene_gn=WDR12;refGene_tx=XM_006935484.3;refGene_gn=WDR12;refGene_tx=XM_011285480.2
C1	189736626	189736732
refGene_gn=WDR12;refGene_tx=XM_003991029.4;refGene_gn=WDR12;refGene_tx=XM_006935484.3;refGene_gn=WDR12;refGene_tx=XM_011285480.2
C1	189736831	189736972
refGene_gn=WDR12;refGene_tx=XM_003991029.4;refGene_gn=WDR12;refGene_tx=XM_006935484.3;refGene_gn=WDR12;refGene_tx=XM_011285480.2

C1	189742753	189742839
refGene_gn=WDR12;refGene_tx=XM_003991029.4;refGene_gn=WDR12;refGene_tx
=XM_006935484.3;refGene_gn=WDR12;refGene_tx=XM_011285480.2
C1	189745013	189745059
refGene_gn=WDR12;refGene_tx=XM_003991029.4;refGene_gn=WDR12;refGene_tx
=XM_006935484.3;refGene_gn=WDR12;refGene_tx=XM_011285480.2
C1	189746782	189746937
refGene_gn=WDR12;refGene_tx=XM_003991029.4;refGene_gn=WDR12;refGene_tx
=XM_006935484.3;refGene_gn=WDR12;refGene_tx=XM_011285480.2
C1	189747374	189747490
refGene_gn=WDR12;refGene_tx=XM_003991029.4;refGene_gn=WDR12;refGene_tx
=XM_006935484.3;refGene_gn=WDR12;refGene_tx=XM_011285480.2
C1	189748589	189748696
refGene_gn=WDR12;refGene_tx=XM_003991029.4;refGene_gn=WDR12;refGene_tx
=XM_006935484.3;refGene_gn=WDR12;refGene_tx=XM_011285480.2
C1	189749796	189749891
refGene_gn=WDR12;refGene_tx=XM_003991029.4;refGene_gn=WDR12;refGene_tx
=XM_006935484.3;refGene_gn=WDR12;refGene_tx=XM_011285480.2
C1	189753183	189753278
refGene_gn=WDR12;refGene_tx=XM_003991029.4;refGene_gn=WDR12;refGene_tx
=XM_006935484.3;refGene_gn=WDR12;refGene_tx=XM_011285480.2
C1	189756622	189756663
refGene_gn=WDR12;refGene_tx=XM_003991029.4;refGene_gn=WDR12;refGene_tx
=XM_006935484.3;refGene_gn=WDR12;refGene_tx=XM_011285480.2
C1	189775092	189775170
refGene_gn=CARF;refGene_tx=XM_023259614.1;refGene_gn=CARF;refGene_tx=X
M_023259615.1;refGene_gn=CARF;refGene_tx=XM_023259616.1;refGene_gn=CAR
F;refGene_tx=XM_023259617.1;refGene_gn=CARF;refGene_tx=XM_023259618.1;
refGene_gn=CARF;refGene_tx=XM_023259619.1;refGene_gn=CARF;refGene_tx=X
M_023259620.1;refGene_gn=CARF;refGene_tx=XM_023259621.1;refGene_gn=CAR
F;refGene_tx=XM_023259623.1;refGene_gn=CARF;refGene_tx=XM_023259624.1;
refGene_gn=CARF;refGene_tx=XM_023259625.1
C1	189776414	189776642
refGene_gn=CARF;refGene_tx=XM_023259614.1;refGene_gn=CARF;refGene_tx=X
M_023259615.1;refGene_gn=CARF;refGene_tx=XM_023259616.1;refGene_gn=CAR
F;refGene_tx=XM_023259617.1;refGene_gn=CARF;refGene_tx=XM_023259618.1;
refGene_gn=CARF;refGene_tx=XM_023259619.1;refGene_gn=CARF;refGene_tx=X
M_023259620.1;refGene_gn=CARF;refGene_tx=XM_023259621.1;refGene_gn=CAR
F;refGene_tx=XM_023259622.1;refGene_gn=CARF;refGene_tx=XM_023259623.1;
refGene_gn=CARF;refGene_tx=XM_023259624.1;refGene_gn=CARF;refGene_tx=X
M_023259625.1;refGene_gn=CARF;refGene_tx=XM_023259626.1;refGene_gn=CAR
F;refGene_tx=XM_023259627.1;refGene_gn=CARF;refGene_tx=XM_023259628.1
C1	189780448	189780569
refGene_gn=CARF;refGene_tx=XM_023259614.1;refGene_gn=CARF;refGene_tx=X
M_023259615.1;refGene_gn=CARF;refGene_tx=XM_023259616.1;refGene_gn=CAR
F;refGene_tx=XM_023259617.1;refGene_gn=CARF;refGene_tx=XM_023259618.1;
refGene_gn=CARF;refGene_tx=XM_023259619.1;refGene_gn=CARF;refGene_tx=X
M_023259620.1;refGene_gn=CARF;refGene_tx=XM_023259621.1;refGene_gn=CAR
F;refGene_tx=XM_023259622.1;refGene_gn=CARF;refGene_tx=XM_023259623.1;
refGene_gn=CARF;refGene_tx=XM_023259624.1;refGene_gn=CARF;refGene_tx=X

M_023259625.1;refGene_gn=CARF;refGene_tx=XM_023259626.1;refGene_gn=CARF;refGene_tx=XM_023259627.1;refGene_gn=CARF;refGene_tx=XM_023259628.1;refGene_gn=CARF;refGene_tx=XM_023259629.1
C1 189780802 189780809
refGene_gn=CARF;refGene_tx=XM_023259630.1
C1 189781566 189781696
refGene_gn=CARF;refGene_tx=XM_023259614.1;refGene_gn=CARF;refGene_tx=XM_023259615.1;refGene_gn=CARF;refGene_tx=XM_023259616.1;refGene_gn=CARF;refGene_tx=XM_023259617.1;refGene_gn=CARF;refGene_tx=XM_023259618.1;refGene_gn=CARF;refGene_tx=XM_023259619.1;refGene_gn=CARF;refGene_tx=XM_023259620.1;refGene_gn=CARF;refGene_tx=XM_023259621.1;refGene_gn=CARF;refGene_tx=XM_023259622.1;refGene_gn=CARF;refGene_tx=XM_023259623.1;refGene_gn=CARF;refGene_tx=XM_023259624.1;refGene_gn=CARF;refGene_tx=XM_023259625.1;refGene_gn=CARF;refGene_tx=XM_023259626.1;refGene_gn=CARF;refGene_tx=XM_023259627.1;refGene_gn=CARF;refGene_tx=XM_023259628.1;refGene_gn=CARF;refGene_tx=XM_023259629.1;refGene_gn=CARF;refGene_tx=XM_023259630.1
C1 189782963 189783048
refGene_gn=CARF;refGene_tx=XM_023259614.1;refGene_gn=CARF;refGene_tx=XM_023259615.1;refGene_gn=CARF;refGene_tx=XM_023259616.1;refGene_gn=CARF;refGene_tx=XM_023259617.1;refGene_gn=CARF;refGene_tx=XM_023259618.1;refGene_gn=CARF;refGene_tx=XM_023259619.1;refGene_gn=CARF;refGene_tx=XM_023259620.1;refGene_gn=CARF;refGene_tx=XM_023259621.1;refGene_gn=CARF;refGene_tx=XM_023259622.1;refGene_gn=CARF;refGene_tx=XM_023259623.1;refGene_gn=CARF;refGene_tx=XM_023259624.1;refGene_gn=CARF;refGene_tx=XM_023259625.1;refGene_gn=CARF;refGene_tx=XM_023259626.1;refGene_gn=CARF;refGene_tx=XM_023259627.1;refGene_gn=CARF;refGene_tx=XM_023259628.1;refGene_gn=CARF;refGene_tx=XM_023259629.1;refGene_gn=CARF;refGene_tx=XM_023259630.1
C1 189787229 189787419
refGene_gn=CARF;refGene_tx=XM_023259614.1;refGene_gn=CARF;refGene_tx=XM_023259615.1;refGene_gn=CARF;refGene_tx=XM_023259616.1;refGene_gn=CARF;refGene_tx=XM_023259617.1;refGene_gn=CARF;refGene_tx=XM_023259618.1;refGene_gn=CARF;refGene_tx=XM_023259619.1;refGene_gn=CARF;refGene_tx=XM_023259620.1;refGene_gn=CARF;refGene_tx=XM_023259621.1;refGene_gn=CARF;refGene_tx=XM_023259622.1;refGene_gn=CARF;refGene_tx=XM_023259623.1;refGene_gn=CARF;refGene_tx=XM_023259624.1;refGene_gn=CARF;refGene_tx=XM_023259625.1;refGene_gn=CARF;refGene_tx=XM_023259626.1;refGene_gn=CARF;refGene_tx=XM_023259627.1;refGene_gn=CARF;refGene_tx=XM_023259628.1;refGene_gn=CARF;refGene_tx=XM_023259629.1;refGene_gn=CARF;refGene_tx=XM_023259630.1
C1 189792249 189792370
refGene_gn=CARF;refGene_tx=XM_023259614.1;refGene_gn=CARF;refGene_tx=XM_023259615.1;refGene_gn=CARF;refGene_tx=XM_023259616.1;refGene_gn=CARF;refGene_tx=XM_023259617.1;refGene_gn=CARF;refGene_tx=XM_023259618.1;refGene_gn=CARF;refGene_tx=XM_023259619.1;refGene_gn=CARF;refGene_tx=XM_023259620.1;refGene_gn=CARF;refGene_tx=XM_023259621.1;refGene_gn=CARF;refGene_tx=XM_023259622.1;refGene_gn=CARF;refGene_tx=XM_023259623.1;refGene_gn=CARF;refGene_tx=XM_023259624.1;refGene_gn=CARF;refGene_tx=XM_023259625.1;refGene_gn=CARF;refGene_tx=XM_023259626.1;refGene_gn=CAR

F;refGene_tx=XM_023259627.1;refGene_gn=CARF;refGene_tx=XM_023259628.1;
refGene_gn=CARF;refGene_tx=XM_023259629.1;refGene_gn=CARF;refGene_tx=X
M_023259630.1
C1 189796541 189796685
refGene_gn=CARF;refGene_tx=XM_023259614.1;refGene_gn=CARF;refGene_tx=X
M_023259615.1;refGene_gn=CARF;refGene_tx=XM_023259616.1;refGene_gn=CAR
F;refGene_tx=XM_023259617.1;refGene_gn=CARF;refGene_tx=XM_023259618.1;
refGene_gn=CARF;refGene_tx=XM_023259619.1;refGene_gn=CARF;refGene_tx=X
M_023259620.1;refGene_gn=CARF;refGene_tx=XM_023259621.1;refGene_gn=CAR
F;refGene_tx=XM_023259622.1;refGene_gn=CARF;refGene_tx=XM_023259623.1;
refGene_gn=CARF;refGene_tx=XM_023259624.1;refGene_gn=CARF;refGene_tx=X
M_023259625.1;refGene_gn=CARF;refGene_tx=XM_023259626.1;refGene_gn=CAR
F;refGene_tx=XM_023259627.1;refGene_gn=CARF;refGene_tx=XM_023259628.1;
refGene_gn=CARF;refGene_tx=XM_023259629.1;refGene_gn=CARF;refGene_tx=X
M_023259630.1
C1 189799672 189799906
refGene_gn=CARF;refGene_tx=XM_023259614.1;refGene_gn=CARF;refGene_tx=X
M_023259615.1;refGene_gn=CARF;refGene_tx=XM_023259616.1;refGene_gn=CAR
F;refGene_tx=XM_023259617.1;refGene_gn=CARF;refGene_tx=XM_023259618.1;
refGene_gn=CARF;refGene_tx=XM_023259619.1;refGene_gn=CARF;refGene_tx=X
M_023259620.1;refGene_gn=CARF;refGene_tx=XM_023259621.1;refGene_gn=CAR
F;refGene_tx=XM_023259622.1;refGene_gn=CARF;refGene_tx=XM_023259623.1;
refGene_gn=CARF;refGene_tx=XM_023259624.1;refGene_gn=CARF;refGene_tx=X
M_023259625.1;refGene_gn=CARF;refGene_tx=XM_023259626.1;refGene_gn=CAR
F;refGene_tx=XM_023259627.1;refGene_gn=CARF;refGene_tx=XM_023259628.1;
refGene_gn=CARF;refGene_tx=XM_023259629.1;refGene_gn=CARF;refGene_tx=X
M_023259630.1
C1 189802315 189802478
refGene_gn=CARF;refGene_tx=XM_023259614.1;refGene_gn=CARF;refGene_tx=X
M_023259615.1;refGene_gn=CARF;refGene_tx=XM_023259616.1;refGene_gn=CAR
F;refGene_tx=XM_023259617.1;refGene_gn=CARF;refGene_tx=XM_023259618.1;
refGene_gn=CARF;refGene_tx=XM_023259619.1;refGene_gn=CARF;refGene_tx=X
M_023259620.1;refGene_gn=CARF;refGene_tx=XM_023259621.1;refGene_gn=CAR
F;refGene_tx=XM_023259622.1;refGene_gn=CARF;refGene_tx=XM_023259623.1;
refGene_gn=CARF;refGene_tx=XM_023259624.1;refGene_gn=CARF;refGene_tx=X
M_023259625.1;refGene_gn=CARF;refGene_tx=XM_023259626.1;refGene_gn=CAR
F;refGene_tx=XM_023259627.1;refGene_gn=CARF;refGene_tx=XM_023259628.1;
refGene_gn=CARF;refGene_tx=XM_023259629.1;refGene_gn=CARF;refGene_tx=X
M_023259630.1
C1 189803456 189803520
refGene_gn=CARF;refGene_tx=XM_023259614.1;refGene_gn=CARF;refGene_tx=X
M_023259615.1;refGene_gn=CARF;refGene_tx=XM_023259616.1;refGene_gn=CAR
F;refGene_tx=XM_023259617.1;refGene_gn=CARF;refGene_tx=XM_023259618.1;
refGene_gn=CARF;refGene_tx=XM_023259619.1;refGene_gn=CARF;refGene_tx=X
M_023259620.1;refGene_gn=CARF;refGene_tx=XM_023259621.1;refGene_gn=CAR
F;refGene_tx=XM_023259622.1;refGene_gn=CARF;refGene_tx=XM_023259623.1;
refGene_gn=CARF;refGene_tx=XM_023259624.1;refGene_gn=CARF;refGene_tx=X
M_023259625.1;refGene_gn=CARF;refGene_tx=XM_023259626.1;refGene_gn=CAR
F;refGene_tx=XM_023259627.1;refGene_gn=CARF;refGene_tx=XM_023259628.1;
refGene_gn=CARF;refGene_tx=XM_023259629.1;refGene_gn=CARF;refGene_tx=X

M_023259630.1
C1	189808449	189808580
refGene_gn=CARF;refGene_tx=XM_023259614.1;refGene_gn=CARF;refGene_tx=XM_023259615.1;refGene_gn=CARF;refGene_tx=XM_023259616.1;refGene_gn=CARF;refGene_tx=XM_023259617.1;refGene_gn=CARF;refGene_tx=XM_023259618.1;refGene_gn=CARF;refGene_tx=XM_023259619.1;refGene_gn=CARF;refGene_tx=XM_023259620.1;refGene_gn=CARF;refGene_tx=XM_023259621.1;refGene_gn=CARF;refGene_tx=XM_023259622.1;refGene_gn=CARF;refGene_tx=XM_023259623.1;refGene_gn=CARF;refGene_tx=XM_023259624.1;refGene_gn=CARF;refGene_tx=XM_023259625.1;refGene_gn=CARF;refGene_tx=XM_023259626.1;refGene_gn=CARF;refGene_tx=XM_023259627.1;refGene_gn=CARF;refGene_tx=XM_023259628.1;refGene_gn=CARF;refGene_tx=XM_023259629.1;refGene_gn=CARF;refGene_tx=XM_023259630.1
C1	189808955	189809328
refGene_gn=CARF;refGene_tx=XM_023259614.1;refGene_gn=CARF;refGene_tx=XM_023259615.1;refGene_gn=CARF;refGene_tx=XM_023259616.1;refGene_gn=CARF;refGene_tx=XM_023259617.1;refGene_gn=CARF;refGene_tx=XM_023259618.1;refGene_gn=CARF;refGene_tx=XM_023259619.1;refGene_gn=CARF;refGene_tx=XM_023259620.1;refGene_gn=CARF;refGene_tx=XM_023259621.1;refGene_gn=CARF;refGene_tx=XM_023259622.1;refGene_gn=CARF;refGene_tx=XM_023259623.1;refGene_gn=CARF;refGene_tx=XM_023259624.1;refGene_gn=CARF;refGene_tx=XM_023259625.1;refGene_gn=CARF;refGene_tx=XM_023259626.1;refGene_gn=CARF;refGene_tx=XM_023259627.1;refGene_gn=CARF;refGene_tx=XM_023259628.1;refGene_gn=CARF;refGene_tx=XM_023259629.1;refGene_gn=CARF;refGene_tx=XM_023259630.1
C1	189812281	189812377
refGene_gn=CARF;refGene_tx=XM_023259614.1;refGene_gn=CARF;refGene_tx=XM_023259615.1;refGene_gn=CARF;refGene_tx=XM_023259616.1;refGene_gn=CARF;refGene_tx=XM_023259617.1;refGene_gn=CARF;refGene_tx=XM_023259622.1;refGene_gn=CARF;refGene_tx=XM_023259623.1;refGene_gn=CARF;refGene_tx=XM_023259626.1;refGene_gn=CARF;refGene_tx=XM_023259627.1;refGene_gn=CARF;refGene_tx=XM_023259628.1;refGene_gn=CARF;refGene_tx=XM_023259629.1;refGene_gn=CARF;refGene_tx=XM_023259630.1
C1	189814575	189814721
refGene_gn=CARF;refGene_tx=XM_023259614.1;refGene_gn=CARF;refGene_tx=XM_023259615.1;refGene_gn=CARF;refGene_tx=XM_023259616.1;refGene_gn=CARF;refGene_tx=XM_023259617.1;refGene_gn=CARF;refGene_tx=XM_023259622.1;refGene_gn=CARF;refGene_tx=XM_023259623.1;refGene_gn=CARF;refGene_tx=XM_023259624.1;refGene_gn=CARF;refGene_tx=XM_023259625.1;refGene_gn=CARF;refGene_tx=XM_023259626.1;refGene_gn=CARF;refGene_tx=XM_023259627.1;refGene_gn=CARF;refGene_tx=XM_023259628.1;refGene_gn=CARF;refGene_tx=XM_023259629.1;refGene_gn=CARF;refGene_tx=XM_023259630.1
C1	189825377	189825517
refGene_gn=CARF;refGene_tx=XM_023259618.1;refGene_gn=CARF;refGene_tx=XM_023259619.1;refGene_gn=CARF;refGene_tx=XM_023259620.1;refGene_gn=CARF;refGene_tx=XM_023259621.1;refGene_gn=CARF;refGene_tx=XM_023259623.1;refGene_gn=CARF;refGene_tx=XM_023259625.1
C1	189829724	189829748
refGene_gn=CARF;refGene_tx=XM_023259621.1
C1	189840470	189840521

refGene_gn=CARF;refGene_tx=XM_023259618.1
C1 189840734 189840782
refGene_gn=CARF;refGene_tx=XM_023259619.1
C1 189858970 189858997
refGene_gn=CARF;refGene_tx=XM_023259620.1
C1 189859205 189859256
refGene_gn=NBEAL1;refGene_tx=XM_019838502.2
C1 189884335 189884427
refGene_gn=NBEAL1;refGene_tx=XM_019838502.2
C1 189889083 189889245
refGene_gn=NBEAL1;refGene_tx=XM_019838502.2
C1 189890683 189890765
refGene_gn=NBEAL1;refGene_tx=XM_019838502.2
C1 189891320 189891448
refGene_gn=NBEAL1;refGene_tx=XM_019838502.2;refGene_gn=NBEAL1;refGene_tx=XM_019838503.1
C1 189897995 189898078
refGene_gn=NBEAL1;refGene_tx=XM_019838502.2;refGene_gn=NBEAL1;refGene_tx=XM_019838503.1
C1 189903520 189903606
refGene_gn=NBEAL1;refGene_tx=XM_019838502.2
C1 189906935 189907242
refGene_gn=NBEAL1;refGene_tx=XM_019838502.2;refGene_gn=NBEAL1;refGene_tx=XM_019838503.1
C1 189908933 189909040
refGene_gn=NBEAL1;refGene_tx=XM_019838502.2;refGene_gn=NBEAL1;refGene_tx=XM_019838503.1
C1 189915944 189916031
refGene_gn=NBEAL1;refGene_tx=XM_019838503.1
C1 189918315 189918399
refGene_gn=NBEAL1;refGene_tx=XM_019838502.2;refGene_gn=NBEAL1;refGene_tx=XM_019838503.1
C1 189921262 189921361
refGene_gn=NBEAL1;refGene_tx=XM_019838502.2;refGene_gn=NBEAL1;refGene_tx=XM_019838503.1
C1 189921448 189922029
refGene_gn=NBEAL1;refGene_tx=XM_019838502.2;refGene_gn=NBEAL1;refGene_tx=XM_019838503.1
C1 189922755 189922888
refGene_gn=NBEAL1;refGene_tx=XM_019838502.2;refGene_gn=NBEAL1;refGene_tx=XM_019838503.1
C1 189923511 189923631
refGene_gn=NBEAL1;refGene_tx=XM_019838502.2;refGene_gn=NBEAL1;refGene_tx=XM_019838503.1
C1 189925012 189925316
refGene_gn=NBEAL1;refGene_tx=XM_019838502.2;refGene_gn=NBEAL1;refGene_tx=XM_019838503.1
C1 189936202 189936288
refGene_gn=NBEAL1;refGene_tx=XM_019838502.2;refGene_gn=NBEAL1;refGene_tx=XM_019838503.1

C1 189942610 189942700
refGene_gn=NBEAL1;refGene_tx=XM_019838502.2;refGene_gn=NBEAL1;refGene_
tx=XM_019838503.1
C1 189945677 189945846
refGene_gn=NBEAL1;refGene_tx=XM_019838502.2;refGene_gn=NBEAL1;refGene_
tx=XM_019838503.1
C1 189946304 189946438
refGene_gn=NBEAL1;refGene_tx=XM_019838502.2;refGene_gn=NBEAL1;refGene_
tx=XM_019838503.1
C1 189946901 189947061
refGene_gn=NBEAL1;refGene_tx=XM_019838502.2;refGene_gn=NBEAL1;refGene_
tx=XM_019838503.1
C1 189947170 189947273
refGene_gn=NBEAL1;refGene_tx=XM_019838502.2;refGene_gn=NBEAL1;refGene_
tx=XM_019838503.1
C1 189948024 189948181
refGene_gn=NBEAL1;refGene_tx=XM_019838502.2;refGene_gn=NBEAL1;refGene_
tx=XM_019838503.1
C1 189951385 189951544
refGene_gn=NBEAL1;refGene_tx=XM_019838502.2;refGene_gn=NBEAL1;refGene_
tx=XM_019838503.1
C1 189953028 189953188
refGene_gn=NBEAL1;refGene_tx=XM_019838502.2;refGene_gn=NBEAL1;refGene_
tx=XM_019838503.1
C1 189956031 189956120
refGene_gn=NBEAL1;refGene_tx=XM_019838502.2;refGene_gn=NBEAL1;refGene_
tx=XM_019838503.1
C1 189958827 189959403
refGene_gn=NBEAL1;refGene_tx=XM_019838502.2;refGene_gn=NBEAL1;refGene_
tx=XM_019838503.1
C1 189959750 189959926
refGene_gn=NBEAL1;refGene_tx=XM_019838502.2;refGene_gn=NBEAL1;refGene_
tx=XM_019838503.1
C1 189960660 189960814
refGene_gn=NBEAL1;refGene_tx=XM_019838502.2;refGene_gn=NBEAL1;refGene_
tx=XM_019838503.1
C1 189961106 189961235
refGene_gn=NBEAL1;refGene_tx=XM_019838502.2;refGene_gn=NBEAL1;refGene_
tx=XM_019838503.1
C1 189966982 189967288
refGene_gn=NBEAL1;refGene_tx=XM_019838502.2;refGene_gn=NBEAL1;refGene_
tx=XM_019838503.1
C1 189967379 189967529
refGene_gn=NBEAL1;refGene_tx=XM_019838502.2;refGene_gn=NBEAL1;refGene_
tx=XM_019838503.1
C1 189971007 189971165
refGene_gn=NBEAL1;refGene_tx=XM_019838502.2;refGene_gn=NBEAL1;refGene_
tx=XM_019838503.1
C1 189973791 189973916
refGene_gn=NBEAL1;refGene_tx=XM_019838502.2;refGene_gn=NBEAL1;refGene_

tx=XM_019838503.1
C1	189979577	189979704
refGene_gn=NBEAL1;refGene_tx=XM_019838502.2;refGene_gn=NBEAL1;refGene_
tx=XM_019838503.1
C1	189985070	189985219
refGene_gn=NBEAL1;refGene_tx=XM_019838502.2;refGene_gn=NBEAL1;refGene_
tx=XM_019838503.1
C1	189985807	189985941
refGene_gn=NBEAL1;refGene_tx=XM_019838502.2;refGene_gn=NBEAL1;refGene_
tx=XM_019838503.1
C1	189988221	189988323
refGene_gn=NBEAL1;refGene_tx=XM_019838502.2;refGene_gn=NBEAL1;refGene_
tx=XM_019838503.1
C1	189990380	189990476
refGene_gn=NBEAL1;refGene_tx=XM_019838502.2;refGene_gn=NBEAL1;refGene_
tx=XM_019838503.1
C1	189990873	189990998
refGene_gn=NBEAL1;refGene_tx=XM_019838502.2;refGene_gn=NBEAL1;refGene_
tx=XM_019838503.1
C1	189993905	189994046
refGene_gn=NBEAL1;refGene_tx=XM_019838502.2;refGene_gn=NBEAL1;refGene_
tx=XM_019838503.1
C1	189997391	189997522
refGene_gn=NBEAL1;refGene_tx=XM_019838502.2;refGene_gn=NBEAL1;refGene_
tx=XM_019838503.1
C1	189999988	190000098
refGene_gn=NBEAL1;refGene_tx=XM_019838502.2;refGene_gn=NBEAL1;refGene_
tx=XM_019838503.1
C1	190004071	190004189
refGene_gn=NBEAL1;refGene_tx=XM_019838502.2;refGene_gn=NBEAL1;refGene_
tx=XM_019838503.1
C1	190009185	190009283
refGene_gn=NBEAL1;refGene_tx=XM_019838502.2;refGene_gn=NBEAL1;refGene_
tx=XM_019838503.1
C1	190011624	190011741
refGene_gn=NBEAL1;refGene_tx=XM_019838502.2;refGene_gn=NBEAL1;refGene_
tx=XM_019838503.1
C1	190016002	190016092
refGene_gn=NBEAL1;refGene_tx=XM_019838502.2;refGene_gn=NBEAL1;refGene_
tx=XM_019838503.1
C1	190023142	190023252
refGene_gn=NBEAL1;refGene_tx=XM_019838502.2;refGene_gn=NBEAL1;refGene_
tx=XM_019838503.1
C1	190024328	190024501
refGene_gn=NBEAL1;refGene_tx=XM_019838502.2;refGene_gn=NBEAL1;refGene_
tx=XM_019838503.1
C1	190025500	190025595
refGene_gn=NBEAL1;refGene_tx=XM_019838502.2;refGene_gn=NBEAL1;refGene_
tx=XM_019838503.1
C1	190028767	190028884

refGene_gn=NBEAL1;refGene_tx=XM_019838502.2;refGene_gn=NBEAL1;refGene_tx=XM_019838503.1
C1 190029322 190029484
refGene_gn=NBEAL1;refGene_tx=XM_019838502.2;refGene_gn=NBEAL1;refGene_tx=XM_019838503.1
C1 190030622 190030771
refGene_gn=NBEAL1;refGene_tx=XM_019838502.2;refGene_gn=NBEAL1;refGene_tx=XM_019838503.1
C1 190032649 190032923
refGene_gn=NBEAL1;refGene_tx=XM_019838502.2;refGene_gn=NBEAL1;refGene_tx=XM_019838503.1
C1 190035233 190035335
refGene_gn=NBEAL1;refGene_tx=XM_019838502.2
C1 190059726 190059798
refGene_gn=LOC101088261;refGene_tx=XM_011285483.3
C1 190060405 190060417
refGene_gn=LOC101088261;refGene_tx=XM_023259631.1
C1 190067109 190067159
refGene_gn=LOC101088261;refGene_tx=XM_011285483.3;refGene_gn=LOC101088261;refGene_tx=XM_023259631.1
C1 190067964 190068131
refGene_gn=LOC101088261;refGene_tx=XM_011285483.3;refGene_gn=LOC101088261;refGene_tx=XM_023259631.1
C1 190070652 190070795
refGene_gn=LOC101088261;refGene_tx=XM_011285483.3;refGene_gn=LOC101088261;refGene_tx=XM_023259631.1
C1 190099207 190099375
refGene_gn=LOC101088261;refGene_tx=XM_011285483.3;refGene_gn=LOC101088261;refGene_tx=XM_023259631.1
C1 190103167 190103246
refGene_gn=LOC101088261;refGene_tx=XM_011285483.3;refGene_gn=LOC101088261;refGene_tx=XM_023259631.1
C1 190109550 190109666
refGene_gn=LOC101088261;refGene_tx=XM_011285483.3;refGene_gn=LOC101088261;refGene_tx=XM_023259631.1
C1 190110715 190110770
refGene_gn=LOC101088261;refGene_tx=XM_011285483.3;refGene_gn=LOC101088261;refGene_tx=XM_023259631.1
C1 190113690 190113811
refGene_gn=LOC101088261;refGene_tx=XM_011285483.3;refGene_gn=LOC101088261;refGene_tx=XM_023259631.1
C1 190118831 190118943
refGene_gn=LOC101088261;refGene_tx=XM_011285483.3;refGene_gn=LOC101088261;refGene_tx=XM_023259631.1
C1 190120630 190120695
refGene_gn=LOC101088261;refGene_tx=XM_011285483.3;refGene_gn=LOC101088261;refGene_tx=XM_023259631.1
C1 190138098 190138188
refGene_gn=LOC101088261;refGene_tx=XM_011285483.3;refGene_gn=LOC101088261;refGene_tx=XM_023259631.1

C1	190138351	190138502
refGene_gn=LOC101088261;refGene_tx=XM_011285483.3;refGene_gn=LOC101088261;refGene_tx=XM_023259631.1
C1	190165446	190165563
refGene_gn=ABI2;refGene_tx=XM_011285484.3;refGene_gn=ABI2;refGene_tx=XM_011285485.3;refGene_gn=ABI2;refGene_tx=XM_011285486.2;refGene_gn=ABI2;refGene_tx=XM_011285487.2;refGene_gn=ABI2;refGene_tx=XM_011285488.2;refGene_gn=ABI2;refGene_tx=XM_011285489.3;refGene_gn=ABI2;refGene_tx=XM_011285490.2;refGene_gn=ABI2;refGene_tx=XM_011285491.2;refGene_gn=ABI2;refGene_tx=XM_011285492.3;refGene_gn=ABI2;refGene_tx=XM_011285493.2;refGene_gn=ABI2;refGene_tx=XM_019838505.1;refGene_gn=ABI2;refGene_tx=XM_019838506.1;refGene_gn=ABI2;refGene_tx=XM_019838509.2;refGene_gn=ABI2;refGene_tx=XM_019838510.2
C1	190165572	190165602
refGene_gn=ABI2;refGene_tx=XM_019838508.2
C1	190221450	190221501
refGene_gn=ABI2;refGene_tx=XM_019838507.1
C1	190223930	190224098
refGene_gn=ABI2;refGene_tx=XM_011285484.3;refGene_gn=ABI2;refGene_tx=XM_011285485.3;refGene_gn=ABI2;refGene_tx=XM_011285486.2;refGene_gn=ABI2;refGene_tx=XM_011285487.2;refGene_gn=ABI2;refGene_tx=XM_011285488.2;refGene_gn=ABI2;refGene_tx=XM_011285489.3;refGene_gn=ABI2;refGene_tx=XM_011285490.2;refGene_gn=ABI2;refGene_tx=XM_011285491.2;refGene_gn=ABI2;refGene_tx=XM_011285492.3;refGene_gn=ABI2;refGene_tx=XM_011285493.2;refGene_gn=ABI2;refGene_tx=XM_019838505.1;refGene_gn=ABI2;refGene_tx=XM_019838506.1;refGene_gn=ABI2;refGene_tx=XM_019838507.1;refGene_gn=ABI2;refGene_tx=XM_019838508.2;refGene_gn=ABI2;refGene_tx=XM_019838509.2
C1	190235864	190236041
refGene_gn=ABI2;refGene_tx=XM_011285484.3;refGene_gn=ABI2;refGene_tx=XM_011285485.3;refGene_gn=ABI2;refGene_tx=XM_011285486.2;refGene_gn=ABI2;refGene_tx=XM_011285487.2;refGene_gn=ABI2;refGene_tx=XM_011285488.2;refGene_gn=ABI2;refGene_tx=XM_011285489.3;refGene_gn=ABI2;refGene_tx=XM_011285490.2;refGene_gn=ABI2;refGene_tx=XM_011285491.2;refGene_gn=ABI2;refGene_tx=XM_011285492.3;refGene_gn=ABI2;refGene_tx=XM_011285493.2;refGene_gn=ABI2;refGene_tx=XM_019838505.1;refGene_gn=ABI2;refGene_tx=XM_019838506.1;refGene_gn=ABI2;refGene_tx=XM_019838507.1;refGene_gn=ABI2;refGene_tx=XM_019838508.2;refGene_gn=ABI2;refGene_tx=XM_019838509.2;refGene_gn=ABI2;refGene_tx=XM_019838510.2
C1	190237502	190237520
refGene_gn=ABI2;refGene_tx=XM_011285484.3;refGene_gn=ABI2;refGene_tx=XM_011285485.3;refGene_gn=ABI2;refGene_tx=XM_011285487.2;refGene_gn=ABI2;refGene_tx=XM_011285489.3;refGene_gn=ABI2;refGene_tx=XM_011285490.2;refGene_gn=ABI2;refGene_tx=XM_019838505.1;refGene_gn=ABI2;refGene_tx=XM_019838506.1;refGene_gn=ABI2;refGene_tx=XM_019838507.1;refGene_gn=ABI2;refGene_tx=XM_019838508.2;refGene_gn=ABI2;refGene_tx=XM_019838510.2
C1	190244226	190244324
refGene_gn=ABI2;refGene_tx=XM_011285484.3;refGene_gn=ABI2;refGene_tx=XM_011285485.3;refGene_gn=ABI2;refGene_tx=XM_011285486.2;refGene_gn=ABI2;refGene_tx=XM_011285487.2;refGene_gn=ABI2;refGene_tx=XM_011285488.2;refGene_gn=ABI2;refGene_tx=XM_011285489.3;refGene_gn=ABI2;refGene_tx=X

M_011285490.2;refGene_gn=ABI2;refGene_tx=XM_011285491.2;refGene_gn=ABI2;refGene_tx=XM_011285492.3;refGene_gn=ABI2;refGene_tx=XM_011285493.2;refGene_gn=ABI2;refGene_tx=XM_019838505.1;refGene_gn=ABI2;refGene_tx=XM_019838506.1;refGene_gn=ABI2;refGene_tx=XM_019838507.1;refGene_gn=ABI2;refGene_tx=XM_019838508.2;refGene_gn=ABI2;refGene_tx=XM_019838509.2;refGene_gn=ABI2;refGene_tx=XM_019838510.2
C1 190248793 190248940
refGene_gn=ABI2;refGene_tx=XM_011285484.3;refGene_gn=ABI2;refGene_tx=XM_011285485.3;refGene_gn=ABI2;refGene_tx=XM_011285486.2;refGene_gn=ABI2;refGene_tx=XM_011285487.2;refGene_gn=ABI2;refGene_tx=XM_011285488.2;refGene_gn=ABI2;refGene_tx=XM_011285490.2;refGene_gn=ABI2;refGene_tx=XM_011285491.2;refGene_gn=ABI2;refGene_tx=XM_011285493.2;refGene_gn=ABI2;refGene_tx=XM_019838505.1;refGene_gn=ABI2;refGene_tx=XM_019838506.1;refGene_gn=ABI2;refGene_tx=XM_019838507.1;refGene_gn=ABI2;refGene_tx=XM_019838508.2;refGene_gn=ABI2;refGene_tx=XM_019838510.2
C1 190249698 190249823
refGene_gn=ABI2;refGene_tx=XM_011285484.3;refGene_gn=ABI2;refGene_tx=XM_011285485.3;refGene_gn=ABI2;refGene_tx=XM_011285486.2;refGene_gn=ABI2;refGene_tx=XM_011285487.2;refGene_gn=ABI2;refGene_tx=XM_011285488.2;refGene_gn=ABI2;refGene_tx=XM_011285489.3;refGene_gn=ABI2;refGene_tx=XM_011285490.2;refGene_gn=ABI2;refGene_tx=XM_011285491.2;refGene_gn=ABI2;refGene_tx=XM_011285492.3;refGene_gn=ABI2;refGene_tx=XM_011285493.2;refGene_gn=ABI2;refGene_tx=XM_019838505.1;refGene_gn=ABI2;refGene_tx=XM_019838506.1;refGene_gn=ABI2;refGene_tx=XM_019838507.1;refGene_gn=ABI2;refGene_tx=XM_019838508.2;refGene_gn=ABI2;refGene_tx=XM_019838509.2;refGene_gn=ABI2;refGene_tx=XM_019838510.2
C1 190251297 190251480
refGene_gn=ABI2;refGene_tx=XM_011285484.3;refGene_gn=ABI2;refGene_tx=XM_011285485.3;refGene_gn=ABI2;refGene_tx=XM_011285486.2;refGene_gn=ABI2;refGene_tx=XM_011285487.2;refGene_gn=ABI2;refGene_tx=XM_011285488.2;refGene_gn=ABI2;refGene_tx=XM_011285489.3;refGene_gn=ABI2;refGene_tx=XM_011285492.3;refGene_gn=ABI2;refGene_tx=XM_019838505.1;refGene_gn=ABI2;refGene_tx=XM_019838506.1;refGene_gn=ABI2;refGene_tx=XM_019838507.1;refGene_gn=ABI2;refGene_tx=XM_019838508.2;refGene_gn=ABI2;refGene_tx=XM_019838509.2;refGene_gn=ABI2;refGene_tx=XM_019838510.2
C1 190257248 190257407
refGene_gn=ABI2;refGene_tx=XM_011285484.3;refGene_gn=ABI2;refGene_tx=XM_011285485.3;refGene_gn=ABI2;refGene_tx=XM_011285486.2;refGene_gn=ABI2;refGene_tx=XM_011285487.2;refGene_gn=ABI2;refGene_tx=XM_011285488.2;refGene_gn=ABI2;refGene_tx=XM_011285489.3;refGene_gn=ABI2;refGene_tx=XM_011285490.2;refGene_gn=ABI2;refGene_tx=XM_011285491.2;refGene_gn=ABI2;refGene_tx=XM_011285492.3;refGene_gn=ABI2;refGene_tx=XM_011285493.2;refGene_gn=ABI2;refGene_tx=XM_019838505.1;refGene_gn=ABI2;refGene_tx=XM_019838506.1;refGene_gn=ABI2;refGene_tx=XM_019838507.1;refGene_gn=ABI2;refGene_tx=XM_019838508.2;refGene_gn=ABI2;refGene_tx=XM_019838509.2;refGene_gn=ABI2;refGene_tx=XM_019838510.2
C1 190271264 190271351
refGene_gn=ABI2;refGene_tx=XM_011285484.3;refGene_gn=ABI2;refGene_tx=XM_011285485.3;refGene_gn=ABI2;refGene_tx=XM_011285486.2;refGene_gn=ABI2;refGene_tx=XM_011285489.3;refGene_gn=ABI2;refGene_tx=XM_011285490.2;

refGene_gn=ABI2;refGene_tx=XM_011285491.2;refGene_gn=ABI2;refGene_tx=XM_019838505.1;refGene_gn=ABI2;refGene_tx=XM_019838506.1;refGene_gn=ABI2;refGene_tx=XM_019838507.1;refGene_gn=ABI2;refGene_tx=XM_019838508.2;refGene_gn=ABI2;refGene_tx=XM_019838509.2;refGene_gn=ABI2;refGene_tx=XM_019838510.2
C1 190275421 190275595
refGene_gn=ABI2;refGene_tx=XM_011285484.3;refGene_gn=ABI2;refGene_tx=XM_011285485.3;refGene_gn=ABI2;refGene_tx=XM_011285486.2;refGene_gn=ABI2;refGene_tx=XM_011285487.2;refGene_gn=ABI2;refGene_tx=XM_011285488.2;refGene_gn=ABI2;refGene_tx=XM_011285489.3;refGene_gn=ABI2;refGene_tx=XM_011285490.2;refGene_gn=ABI2;refGene_tx=XM_011285491.2;refGene_gn=ABI2;refGene_tx=XM_011285492.3;refGene_gn=ABI2;refGene_tx=XM_011285493.2;refGene_gn=ABI2;refGene_tx=XM_019838505.1;refGene_gn=ABI2;refGene_tx=XM_019838506.1;refGene_gn=ABI2;refGene_tx=XM_019838507.1;refGene_gn=ABI2;refGene_tx=XM_019838508.2;refGene_gn=ABI2;refGene_tx=XM_019838509.2;refGene_gn=ABI2;refGene_tx=XM_019838510.2
C1 190284333 190284509
refGene_gn=ABI2;refGene_tx=XM_011285484.3;refGene_gn=ABI2;refGene_tx=XM_011285485.3;refGene_gn=ABI2;refGene_tx=XM_011285486.2;refGene_gn=ABI2;refGene_tx=XM_011285487.2;refGene_gn=ABI2;refGene_tx=XM_011285488.2;refGene_gn=ABI2;refGene_tx=XM_011285489.3;refGene_gn=ABI2;refGene_tx=XM_011285490.2;refGene_gn=ABI2;refGene_tx=XM_011285491.2;refGene_gn=ABI2;refGene_tx=XM_011285492.3;refGene_gn=ABI2;refGene_tx=XM_011285493.2;refGene_gn=ABI2;refGene_tx=XM_019838505.1;refGene_gn=ABI2;refGene_tx=XM_019838506.1;refGene_gn=ABI2;refGene_tx=XM_019838507.1;refGene_gn=ABI2;refGene_tx=XM_019838508.2;refGene_gn=ABI2;refGene_tx=XM_019838509.2;refGene_gn=ABI2;refGene_tx=XM_019838510.2
C1 190284825 190284899
refGene_gn=ABI2;refGene_tx=XM_019838506.1
C1 190294917 190296576
refGene_gn=RAPH1;refGene_tx=XM_023259632.1
C1 190296578 190296995
refGene_gn=RAPH1;refGene_tx=XM_023259632.1
C1 190299746 190299889
refGene_gn=RAPH1;refGene_tx=XM_023259632.1
C1 190304268 190304389
refGene_gn=RAPH1;refGene_tx=XM_023259632.1
C1 190305433 190305532
refGene_gn=RAPH1;refGene_tx=XM_023259632.1
C1 190314847 190314958
refGene_gn=RAPH1;refGene_tx=XM_023259632.1
C1 190315822 190315966
refGene_gn=RAPH1;refGene_tx=XM_023259632.1
C1 190317943 190318009
refGene_gn=RAPH1;refGene_tx=XM_023259632.1
C1 190319194 190319316
refGene_gn=RAPH1;refGene_tx=XM_023259632.1
C1 190321687 190321847
refGene_gn=RAPH1;refGene_tx=XM_023259632.1
C1 190322248 190322326

refGene_gn=RAPH1;refGene_tx=XM_023259632.1
C1 190350037 190350543
refGene_gn=RAPH1;refGene_tx=XM_023259632.1
C1 190351352 190351458
refGene_gn=RAPH1;refGene_tx=XM_023259632.1
C1 190355513 190355633
refGene_gn=RAPH1;refGene_tx=XM_023259632.1
C1 190521176 190521231
refGene_gn=CD28;refGene_tx=NM_001009844.1
C1 190542271 190542628
refGene_gn=CD28;refGene_tx=NM_001009844.1
C1 190542857 190542982
refGene_gn=CD28;refGene_tx=NM_001009844.1
C1 190548340 190548469
refGene_gn=CD28;refGene_tx=NM_001009844.1
C1 190643354 190643366
refGene_gn=CTLA4;refGene_tx=XM_011285413.2
C1 190645855 190645976
refGene_gn=CTLA4;refGene_tx=XM_011285413.2;refGene_gn=CTLA4;refGene_tx=XM_011285414.2;refGene_gn=CTLA4;refGene_tx=XM_019835948.2;refGene_gn=CTLA4;refGene_tx=XM_023257815.1
C1 190654992 190655126
refGene_gn=CTLA4;refGene_tx=NM_001009236.1;refGene_gn=CTLA4;refGene_tx=XM_019835948.2;refGene_gn=CTLA4;refGene_tx=XM_023257815.1
C1 190657761 190658109
refGene_gn=CTLA4;refGene_tx=NM_001009236.1;refGene_gn=CTLA4;refGene_tx=XM_011285413.2;refGene_gn=CTLA4;refGene_tx=XM_011285414.2;refGene_gn=CTLA4;refGene_tx=XM_019835948.2;refGene_gn=CTLA4;refGene_tx=XM_023257815.1
C1 190658549 190658659
refGene_gn=CTLA4;refGene_tx=NM_001009236.1;refGene_gn=CTLA4;refGene_tx=XM_011285413.2;refGene_gn=CTLA4;refGene_tx=XM_011285414.2;refGene_gn=CTLA4;refGene_tx=XM_019835948.2;refGene_gn=CTLA4;refGene_tx=XM_023257815.1
C1 190659956 190660061
refGene_gn=CTLA4;refGene_tx=NM_001009236.1;refGene_gn=CTLA4;refGene_tx=XM_011285413.2;refGene_gn=CTLA4;refGene_tx=XM_011285414.2;refGene_gn=CTLA4;refGene_tx=XM_019835948.2;refGene_gn=CTLA4;refGene_tx=XM_023257815.1
C1 190722858 190722916
refGene_gn=ICOS;refGene_tx=XM_006935485.3
C1 190738784 190739123
refGene_gn=ICOS;refGene_tx=XM_006935485.3
C1 190740193 190740300
refGene_gn=ICOS;refGene_tx=XM_006935485.3
C1 190741250 190741335
refGene_gn=ICOS;refGene_tx=XM_006935485.3
C1 190743547 190743588
refGene_gn=ICOS;refGene_tx=XM_006935485.3
C1 191284943 191285063

refGene_gn=PARD3B;refGene_tx=XM_011285494.2;refGene_gn=PARD3B;refGene_tx=XM_019838513.1;refGene_gn=PARD3B;refGene_tx=XM_019838515.1
C1 191429529 191429631
refGene_gn=PARD3B;refGene_tx=XM_011285494.2;refGene_gn=PARD3B;refGene_tx=XM_019838513.1;refGene_gn=PARD3B;refGene_tx=XM_019838515.1
C1 191686932 191687104
refGene_gn=PARD3B;refGene_tx=XM_011285494.2;refGene_gn=PARD3B;refGene_tx=XM_019838513.1;refGene_gn=PARD3B;refGene_tx=XM_019838515.1
C1 191758202 191758312
refGene_gn=PARD3B;refGene_tx=XM_011285494.2;refGene_gn=PARD3B;refGene_tx=XM_019838513.1;refGene_gn=PARD3B;refGene_tx=XM_019838515.1
C1 191825455 191825544
refGene_gn=PARD3B;refGene_tx=XM_011285494.2;refGene_gn=PARD3B;refGene_tx=XM_019838513.1;refGene_gn=PARD3B;refGene_tx=XM_019838515.1
C1 191834242 191834329
refGene_gn=PARD3B;refGene_tx=XM_011285494.2;refGene_gn=PARD3B;refGene_tx=XM_019838513.1;refGene_gn=PARD3B;refGene_tx=XM_019838515.1;refGene_gn=PARD3B;refGene_tx=XM_019838516.2
C1 191839834 191839960
refGene_gn=PARD3B;refGene_tx=XM_011285494.2;refGene_gn=PARD3B;refGene_tx=XM_019838513.1;refGene_gn=PARD3B;refGene_tx=XM_019838515.1;refGene_gn=PARD3B;refGene_tx=XM_019838516.2
C1 191842385 191842738
refGene_gn=PARD3B;refGene_tx=XM_011285494.2;refGene_gn=PARD3B;refGene_tx=XM_019838513.1;refGene_gn=PARD3B;refGene_tx=XM_019838515.1;refGene_gn=PARD3B;refGene_tx=XM_019838516.2
C1 191845077 191845217
refGene_gn=PARD3B;refGene_tx=XM_011285494.2;refGene_gn=PARD3B;refGene_tx=XM_019838513.1;refGene_gn=PARD3B;refGene_tx=XM_019838515.1;refGene_gn=PARD3B;refGene_tx=XM_019838516.2
C1 191846519 191846648
refGene_gn=PARD3B;refGene_tx=XM_011285494.2;refGene_gn=PARD3B;refGene_tx=XM_019838513.1;refGene_gn=PARD3B;refGene_tx=XM_019838515.1;refGene_gn=PARD3B;refGene_tx=XM_019838516.2
C1 191877487 191877673
refGene_gn=PARD3B;refGene_tx=XM_011285494.2;refGene_gn=PARD3B;refGene_tx=XM_019838513.1;refGene_gn=PARD3B;refGene_tx=XM_019838515.1;refGene_gn=PARD3B;refGene_tx=XM_019838516.2
C1 191890835 191891006
refGene_gn=PARD3B;refGene_tx=XM_011285494.2;refGene_gn=PARD3B;refGene_tx=XM_019838513.1;refGene_gn=PARD3B;refGene_tx=XM_019838515.1;refGene_gn=PARD3B;refGene_tx=XM_019838516.2
C1 191895232 191895365
refGene_gn=PARD3B;refGene_tx=XM_011285494.2;refGene_gn=PARD3B;refGene_tx=XM_019838513.1;refGene_gn=PARD3B;refGene_tx=XM_019838515.1;refGene_gn=PARD3B;refGene_tx=XM_019838516.2
C1 191903375 191903475
refGene_gn=PARD3B;refGene_tx=XM_011285494.2;refGene_gn=PARD3B;refGene_tx=XM_019838513.1;refGene_gn=PARD3B;refGene_tx=XM_019838515.1;refGene_gn=PARD3B;refGene_tx=XM_019838516.2

C1	191909868	191909984
refGene_gn=PARD3B;refGene_tx=XM_011285494.2;refGene_gn=PARD3B;refGene_tx=XM_019838513.1;refGene_gn=PARD3B;refGene_tx=XM_019838515.1;refGene_gn=PARD3B;refGene_tx=XM_019838516.2
C1	191944742	191944787
refGene_gn=PARD3B;refGene_tx=XM_011285494.2;refGene_gn=PARD3B;refGene_tx=XM_019838513.1;refGene_gn=PARD3B;refGene_tx=XM_019838515.1;refGene_gn=PARD3B;refGene_tx=XM_019838516.2
C1	191998247	191998454
refGene_gn=PARD3B;refGene_tx=XM_011285494.2;refGene_gn=PARD3B;refGene_tx=XM_019838513.1;refGene_gn=PARD3B;refGene_tx=XM_019838515.1;refGene_gn=PARD3B;refGene_tx=XM_019838516.2
C1	191999192	191999430
refGene_gn=PARD3B;refGene_tx=XM_011285494.2;refGene_gn=PARD3B;refGene_tx=XM_019838513.1;refGene_gn=PARD3B;refGene_tx=XM_019838515.1;refGene_gn=PARD3B;refGene_tx=XM_019838516.2
C1	192094106	192094217
refGene_gn=PARD3B;refGene_tx=XM_011285494.2;refGene_gn=PARD3B;refGene_tx=XM_019838513.1;refGene_gn=PARD3B;refGene_tx=XM_019838515.1;refGene_gn=PARD3B;refGene_tx=XM_019838516.2
C1	192133298	192133601
refGene_gn=PARD3B;refGene_tx=XM_011285494.2;refGene_gn=PARD3B;refGene_tx=XM_019838513.1;refGene_gn=PARD3B;refGene_tx=XM_019838515.1;refGene_gn=PARD3B;refGene_tx=XM_019838516.2
C1	192189298	192189431
refGene_gn=PARD3B;refGene_tx=XM_011285494.2;refGene_gn=PARD3B;refGene_tx=XM_019838513.1;refGene_gn=PARD3B;refGene_tx=XM_019838515.1;refGene_gn=PARD3B;refGene_tx=XM_019838516.2
C1	192190804	192190927
refGene_gn=PARD3B;refGene_tx=XM_019838513.1
C1	192215196	192215244
refGene_gn=PARD3B;refGene_tx=XM_019838515.1
C1	192237722	192237802
refGene_gn=PARD3B;refGene_tx=XM_011285494.2;refGene_gn=PARD3B;refGene_tx=XM_019838516.2
C1	192290109	192290467
refGene_gn=PARD3B;refGene_tx=XM_011285494.2;refGene_gn=PARD3B;refGene_tx=XM_019838516.2
C1	192346583	192346656
refGene_gn=NRP2;refGene_tx=XM_003991039.5;refGene_gn=NRP2;refGene_tx=XM_006935486.4;refGene_gn=NRP2;refGene_tx=XM_019838518.2
C1	192361619	192361797
refGene_gn=NRP2;refGene_tx=XM_003991039.5;refGene_gn=NRP2;refGene_tx=XM_006935486.4;refGene_gn=NRP2;refGene_tx=XM_019838518.2
C1	192380975	192381157
refGene_gn=NRP2;refGene_tx=XM_003991039.5;refGene_gn=NRP2;refGene_tx=XM_006935486.4;refGene_gn=NRP2;refGene_tx=XM_019838518.2
C1	192387547	192387778
refGene_gn=NRP2;refGene_tx=XM_003991039.5;refGene_gn=NRP2;refGene_tx=XM_006935486.4;refGene_gn=NRP2;refGene_tx=XM_019838518.2

C1 192389186 192389342
refGene_gn=NRP2;refGene_tx=XM_003991039.5;refGene_gn=NRP2;refGene_tx=XM_006935486.4;refGene_gn=NRP2;refGene_tx=XM_019838518.2
C1 192391099 192391269
refGene_gn=NRP2;refGene_tx=XM_003991039.5;refGene_gn=NRP2;refGene_tx=XM_006935486.4;refGene_gn=NRP2;refGene_tx=XM_019838518.2
C1 192393156 192393312
refGene_gn=NRP2;refGene_tx=XM_003991039.5;refGene_gn=NRP2;refGene_tx=XM_006935486.4;refGene_gn=NRP2;refGene_tx=XM_019838518.2
C1 192406051 192406196
refGene_gn=NRP2;refGene_tx=XM_003991039.5;refGene_gn=NRP2;refGene_tx=XM_006935486.4;refGene_gn=NRP2;refGene_tx=XM_019838518.2
C1 192407326 192407676
refGene_gn=NRP2;refGene_tx=XM_003991039.5;refGene_gn=NRP2;refGene_tx=XM_006935486.4;refGene_gn=NRP2;refGene_tx=XM_019838518.2
C1 192409525 192409670
refGene_gn=NRP2;refGene_tx=XM_003991039.5;refGene_gn=NRP2;refGene_tx=XM_006935486.4;refGene_gn=NRP2;refGene_tx=XM_019838518.2
C1 192414203 192414320
refGene_gn=NRP2;refGene_tx=XM_003991039.5;refGene_gn=NRP2;refGene_tx=XM_006935486.4;refGene_gn=NRP2;refGene_tx=XM_019838518.2
C1 192417192 192417333
refGene_gn=NRP2;refGene_tx=XM_003991039.5;refGene_gn=NRP2;refGene_tx=XM_006935486.4;refGene_gn=NRP2;refGene_tx=XM_019838518.2
C1 192428692 192428955
refGene_gn=NRP2;refGene_tx=XM_003991039.5;refGene_gn=NRP2;refGene_tx=XM_006935486.4;refGene_gn=NRP2;refGene_tx=XM_019838518.2
C1 192430519 192430616
refGene_gn=NRP2;refGene_tx=XM_003991039.5;refGene_gn=NRP2;refGene_tx=XM_006935486.4;refGene_gn=NRP2;refGene_tx=XM_019838518.2
C1 192431793 192431829
refGene_gn=NRP2;refGene_tx=XM_003991039.5;refGene_gn=NRP2;refGene_tx=XM_006935486.4;refGene_gn=NRP2;refGene_tx=XM_019838518.2
C1 192441085 192441366
refGene_gn=NRP2;refGene_tx=XM_003991039.5;refGene_gn=NRP2;refGene_tx=XM_019838518.2
C1 192456861 192456909
refGene_gn=NRP2;refGene_tx=XM_006935486.4
C1 192459510 192460013
refGene_gn=NRP2;refGene_tx=XM_006935486.4
C1 192646257 192647423
refGene_gn=INO80D;refGene_tx=XM_019838519.2
C1 192649026 192649184
refGene_gn=INO80D;refGene_tx=XM_019838519.2
C1 192649874 192650092
refGene_gn=INO80D;refGene_tx=XM_019838519.2
C1 192655735 192655869
refGene_gn=INO80D;refGene_tx=XM_019838519.2
C1 192657711 192657821
refGene_gn=INO80D;refGene_tx=XM_019838519.2

C1 192663511 192663736
refGene_gn=INO80D;refGene_tx=XM_019838519.2
C1 192674767 192674876
refGene_gn=INO80D;refGene_tx=XM_019838519.2
C1 192682764 192683516
refGene_gn=INO80D;refGene_tx=XM_019838519.2
C1 192689360 192689578
refGene_gn=INO80D;refGene_tx=XM_019838519.2
C1 192739687 192739779
refGene_gn=NDUFS1;refGene_tx=XM_003991043.4;refGene_gn=NDUFS1;refGene_tx=XM_006935488.4;refGene_gn=NDUFS1;refGene_tx=XM_006935489.4
C1 192742019 192742092
refGene_gn=NDUFS1;refGene_tx=XM_003991043.4;refGene_gn=NDUFS1;refGene_tx=XM_006935488.4;refGene_gn=NDUFS1;refGene_tx=XM_006935489.4
C1 192742193 192742328
refGene_gn=NDUFS1;refGene_tx=XM_003991043.4;refGene_gn=NDUFS1;refGene_tx=XM_006935488.4;refGene_gn=NDUFS1;refGene_tx=XM_006935489.4
C1 192742886 192743062
refGene_gn=NDUFS1;refGene_tx=XM_003991043.4;refGene_gn=NDUFS1;refGene_tx=XM_006935488.4;refGene_gn=NDUFS1;refGene_tx=XM_006935489.4
C1 192745039 192745194
refGene_gn=NDUFS1;refGene_tx=XM_003991043.4;refGene_gn=NDUFS1;refGene_tx=XM_006935488.4;refGene_gn=NDUFS1;refGene_tx=XM_006935489.4
C1 192750217 192750378
refGene_gn=NDUFS1;refGene_tx=XM_003991043.4;refGene_gn=NDUFS1;refGene_tx=XM_006935488.4;refGene_gn=NDUFS1;refGene_tx=XM_006935489.4
C1 192752860 192752990
refGene_gn=NDUFS1;refGene_tx=XM_003991043.4;refGene_gn=NDUFS1;refGene_tx=XM_006935488.4;refGene_gn=NDUFS1;refGene_tx=XM_006935489.4
C1 192753930 192754059
refGene_gn=NDUFS1;refGene_tx=XM_003991043.4;refGene_gn=NDUFS1;refGene_tx=XM_006935488.4;refGene_gn=NDUFS1;refGene_tx=XM_006935489.4
C1 192754764 192754910
refGene_gn=NDUFS1;refGene_tx=XM_003991043.4;refGene_gn=NDUFS1;refGene_tx=XM_006935488.4;refGene_gn=NDUFS1;refGene_tx=XM_006935489.4
C1 192755905 192756020
refGene_gn=NDUFS1;refGene_tx=XM_003991043.4;refGene_gn=NDUFS1;refGene_tx=XM_006935488.4;refGene_gn=NDUFS1;refGene_tx=XM_006935489.4
C1 192756699 192756834
refGene_gn=NDUFS1;refGene_tx=XM_003991043.4;refGene_gn=NDUFS1;refGene_tx=XM_006935488.4;refGene_gn=NDUFS1;refGene_tx=XM_006935489.4
C1 192758630 192758816
refGene_gn=NDUFS1;refGene_tx=XM_003991043.4;refGene_gn=NDUFS1;refGene_tx=XM_006935488.4;refGene_gn=NDUFS1;refGene_tx=XM_006935489.4
C1 192759261 192759392
refGene_gn=NDUFS1;refGene_tx=XM_003991043.4;refGene_gn=NDUFS1;refGene_tx=XM_006935488.4;refGene_gn=NDUFS1;refGene_tx=XM_006935489.4
C1 192759483 192759565
refGene_gn=NDUFS1;refGene_tx=XM_003991043.4;refGene_gn=NDUFS1;refGene_tx=XM_006935488.4;refGene_gn=NDUFS1;refGene_tx=XM_006935489.4

C1 192760626 192760703
refGene_gn=NDUFS1;refGene_tx=XM_003991043.4;refGene_gn=NDUFS1;refGene_tx=XM_006935488.4;refGene_gn=NDUFS1;refGene_tx=XM_006935489.4
C1 192761404 192761512
refGene_gn=NDUFS1;refGene_tx=XM_003991043.4;refGene_gn=NDUFS1;refGene_tx=XM_006935488.4;refGene_gn=NDUFS1;refGene_tx=XM_006935489.4
C1 192763215 192763307
refGene_gn=NDUFS1;refGene_tx=XM_003991043.4;refGene_gn=NDUFS1;refGene_tx=XM_006935488.4;refGene_gn=NDUFS1;refGene_tx=XM_006935489.4
C1 192763829 192763890
refGene_gn=NDUFS1;refGene_tx=XM_003991043.4;refGene_gn=NDUFS1;refGene_tx=XM_006935488.4;refGene_gn=NDUFS1;refGene_tx=XM_006935489.4
C1 192769064 192769144
refGene_gn=EEF1B2;refGene_tx=XM_003991046.4
C1 192769592 192769715
refGene_gn=EEF1B2;refGene_tx=XM_003991046.4
C1 192770356 192770483
refGene_gn=EEF1B2;refGene_tx=XM_003991046.4
C1 192770738 192770805
refGene_gn=EEF1B2;refGene_tx=XM_003991046.4
C1 192771209 192771335
refGene_gn=EEF1B2;refGene_tx=XM_003991046.4
C1 192771442 192771597
refGene_gn=EEF1B2;refGene_tx=XM_003991046.4
C1 192775485 192775492
refGene_gn=GPR1;refGene_tx=XM_011285498.3;refGene_gn=GPR1;refGene_tx=XM_011285501.3;refGene_gn=GPR1;refGene_tx=XM_019838523.2;refGene_gn=GPR1;refGene_tx=XM_019838524.2;refGene_gn=GPR1;refGene_tx=XM_019838525.2;refGene_gn=GPR1;refGene_tx=XM_019838529.2;refGene_gn=GPR1;refGene_tx=XM_019838530.2;refGene_gn=GPR1;refGene_tx=XM_019838531.2;refGene_gn=GPR1;refGene_tx=XM_019838532.1;refGene_gn=GPR1;refGene_tx=XM_019838533.1
C1 192776403 192776523
refGene_gn=GPR1;refGene_tx=XM_011285498.3;refGene_gn=GPR1;refGene_tx=XM_011285501.3;refGene_gn=GPR1;refGene_tx=XM_019838523.2;refGene_gn=GPR1;refGene_tx=XM_019838524.2;refGene_gn=GPR1;refGene_tx=XM_019838525.2;refGene_gn=GPR1;refGene_tx=XM_019838529.2;refGene_gn=GPR1;refGene_tx=XM_019838530.2;refGene_gn=GPR1;refGene_tx=XM_019838531.2;refGene_gn=GPR1;refGene_tx=XM_019838532.1;refGene_gn=GPR1;refGene_tx=XM_019838533.1
C1 192780851 192781915
refGene_gn=GPR1;refGene_tx=XM_011285498.3;refGene_gn=GPR1;refGene_tx=XM_011285501.3;refGene_gn=GPR1;refGene_tx=XM_019838523.2;refGene_gn=GPR1;refGene_tx=XM_019838524.2;refGene_gn=GPR1;refGene_tx=XM_019838525.2;refGene_gn=GPR1;refGene_tx=XM_019838529.2;refGene_gn=GPR1;refGene_tx=XM_019838530.2;refGene_gn=GPR1;refGene_tx=XM_019838531.2;refGene_gn=GPR1;refGene_tx=XM_019838532.1;refGene_gn=GPR1;refGene_tx=XM_019838533.1
C1 192913662 192913722
refGene_gn=ZDBF2;refGene_tx=XM_023259635.1
C1 192918961 192919089
refGene_gn=ZDBF2;refGene_tx=XM_023259635.1
C1 192926085 192930379

refGene_gn=ZDBF2;refGene_tx=XM_023259635.1
C1 192931381 192934438
refGene_gn=ZDBF2;refGene_tx=XM_023259635.1
C1 193064341 193064555
refGene_gn=ADAM23;refGene_tx=XM_011285539.3;refGene_gn=ADAM23;refGene_tx=XM_023259636.1;refGene_gn=ADAM23;refGene_tx=XM_023259637.1;refGene_gn=ADAM23;refGene_tx=XM_023259638.1
C1 193065717 193065935
refGene_gn=ADAM23;refGene_tx=XM_011285539.3;refGene_gn=ADAM23;refGene_tx=XM_023259636.1;refGene_gn=ADAM23;refGene_tx=XM_023259637.1;refGene_gn=ADAM23;refGene_tx=XM_023259638.1
C1 193114526 193114603
refGene_gn=ADAM23;refGene_tx=XM_011285539.3;refGene_gn=ADAM23;refGene_tx=XM_023259636.1;refGene_gn=ADAM23;refGene_tx=XM_023259637.1;refGene_gn=ADAM23;refGene_tx=XM_023259638.1
C1 193158884 193158948
refGene_gn=ADAM23;refGene_tx=XM_011285539.3;refGene_gn=ADAM23;refGene_tx=XM_023259636.1;refGene_gn=ADAM23;refGene_tx=XM_023259637.1;refGene_gn=ADAM23;refGene_tx=XM_023259638.1
C1 193168921 193169004
refGene_gn=ADAM23;refGene_tx=XM_011285539.3;refGene_gn=ADAM23;refGene_tx=XM_023259636.1;refGene_gn=ADAM23;refGene_tx=XM_023259637.1;refGene_gn=ADAM23;refGene_tx=XM_023259638.1
C1 193170206 193170270
refGene_gn=ADAM23;refGene_tx=XM_011285539.3;refGene_gn=ADAM23;refGene_tx=XM_023259636.1;refGene_gn=ADAM23;refGene_tx=XM_023259637.1;refGene_gn=ADAM23;refGene_tx=XM_023259638.1
C1 193175379 193175452
refGene_gn=ADAM23;refGene_tx=XM_011285539.3;refGene_gn=ADAM23;refGene_tx=XM_023259636.1;refGene_gn=ADAM23;refGene_tx=XM_023259637.1;refGene_gn=ADAM23;refGene_tx=XM_023259638.1
C1 193178926 193179003
refGene_gn=ADAM23;refGene_tx=XM_011285539.3;refGene_gn=ADAM23;refGene_tx=XM_023259636.1;refGene_gn=ADAM23;refGene_tx=XM_023259637.1;refGene_gn=ADAM23;refGene_tx=XM_023259638.1
C1 193180940 193181006
refGene_gn=ADAM23;refGene_tx=XM_011285539.3;refGene_gn=ADAM23;refGene_tx=XM_023259636.1;refGene_gn=ADAM23;refGene_tx=XM_023259637.1;refGene_gn=ADAM23;refGene_tx=XM_023259638.1
C1 193187867 193187939
refGene_gn=ADAM23;refGene_tx=XM_011285539.3;refGene_gn=ADAM23;refGene_tx=XM_023259636.1;refGene_gn=ADAM23;refGene_tx=XM_023259637.1;refGene_gn=ADAM23;refGene_tx=XM_023259638.1
C1 193190051 193190215
refGene_gn=ADAM23;refGene_tx=XM_011285539.3;refGene_gn=ADAM23;refGene_tx=XM_023259636.1;refGene_gn=ADAM23;refGene_tx=XM_023259637.1;refGene_gn=ADAM23;refGene_tx=XM_023259638.1
C1 193191341 193191426
refGene_gn=ADAM23;refGene_tx=XM_011285539.3;refGene_gn=ADAM23;refGene_tx=XM_023259636.1;refGene_gn=ADAM23;refGene_tx=XM_023259637.1;refGene_

gn=ADAM23;refGene_tx=XM_023259638.1
C1 193192419 193192510
refGene_gn=ADAM23;refGene_tx=XM_011285539.3;refGene_gn=ADAM23;refGene_tx=XM_023259636.1;refGene_gn=ADAM23;refGene_tx=XM_023259637.1;refGene_gn=ADAM23;refGene_tx=XM_023259638.1
C1 193195058 193195107
refGene_gn=ADAM23;refGene_tx=XM_011285539.3;refGene_gn=ADAM23;refGene_tx=XM_023259636.1;refGene_gn=ADAM23;refGene_tx=XM_023259637.1;refGene_gn=ADAM23;refGene_tx=XM_023259638.1
C1 193197424 193197524
refGene_gn=ADAM23;refGene_tx=XM_011285539.3;refGene_gn=ADAM23;refGene_tx=XM_023259636.1;refGene_gn=ADAM23;refGene_tx=XM_023259637.1;refGene_gn=ADAM23;refGene_tx=XM_023259638.1
C1 193201535 193201607
refGene_gn=ADAM23;refGene_tx=XM_011285539.3;refGene_gn=ADAM23;refGene_tx=XM_023259636.1;refGene_gn=ADAM23;refGene_tx=XM_023259637.1;refGene_gn=ADAM23;refGene_tx=XM_023259638.1
C1 193202333 193202423
refGene_gn=ADAM23;refGene_tx=XM_011285539.3;refGene_gn=ADAM23;refGene_tx=XM_023259636.1;refGene_gn=ADAM23;refGene_tx=XM_023259637.1;refGene_gn=ADAM23;refGene_tx=XM_023259638.1
C1 193203658 193203739
refGene_gn=ADAM23;refGene_tx=XM_011285539.3;refGene_gn=ADAM23;refGene_tx=XM_023259636.1;refGene_gn=ADAM23;refGene_tx=XM_023259637.1;refGene_gn=ADAM23;refGene_tx=XM_023259638.1
C1 193208993 193209044
refGene_gn=ADAM23;refGene_tx=XM_011285539.3;refGene_gn=ADAM23;refGene_tx=XM_023259636.1;refGene_gn=ADAM23;refGene_tx=XM_023259637.1;refGene_gn=ADAM23;refGene_tx=XM_023259638.1
C1 193209747 193209811
refGene_gn=ADAM23;refGene_tx=XM_011285539.3;refGene_gn=ADAM23;refGene_tx=XM_023259636.1;refGene_gn=ADAM23;refGene_tx=XM_023259637.1;refGene_gn=ADAM23;refGene_tx=XM_023259638.1
C1 193211572 193211678
refGene_gn=ADAM23;refGene_tx=XM_011285539.3;refGene_gn=ADAM23;refGene_tx=XM_023259636.1;refGene_gn=ADAM23;refGene_tx=XM_023259637.1;refGene_gn=ADAM23;refGene_tx=XM_023259638.1
C1 193214391 193214511
refGene_gn=ADAM23;refGene_tx=XM_011285539.3;refGene_gn=ADAM23;refGene_tx=XM_023259636.1;refGene_gn=ADAM23;refGene_tx=XM_023259637.1;refGene_gn=ADAM23;refGene_tx=XM_023259638.1
C1 193215761 193215930
refGene_gn=ADAM23;refGene_tx=XM_011285539.3;refGene_gn=ADAM23;refGene_tx=XM_023259636.1;refGene_gn=ADAM23;refGene_tx=XM_023259637.1;refGene_gn=ADAM23;refGene_tx=XM_023259638.1
C1 193218220 193218332
refGene_gn=ADAM23;refGene_tx=XM_011285539.3;refGene_gn=ADAM23;refGene_tx=XM_023259636.1;refGene_gn=ADAM23;refGene_tx=XM_023259637.1;refGene_gn=ADAM23;refGene_tx=XM_023259638.1
C1 193223639 193223650

refGene_gn=ADAM23;refGene_tx=XM_023259638.1
C1 193230910 193231001
refGene_gn=ADAM23;refGene_tx=XM_011285539.3
C1 193238079 193238219
refGene_gn=ADAM23;refGene_tx=XM_011285539.3;refGene_gn=ADAM23;refGene_tx=XM_023259636.1
C1 193251840 193251944
refGene_gn=ADAM23;refGene_tx=XM_023259637.1
C1 193263434 193263846
refGene_gn=FAM237A;refGene_tx=XM_023259639.1
C1 193267780 193267920
refGene_gn=FAM237A;refGene_tx=XM_023259639.1
C1 193272590 193272694
refGene_gn=DYTN;refGene_tx=XM_023260056.1
C1 193278034 193278539
refGene_gn=DYTN;refGene_tx=XM_023260056.1
C1 193281189 193281349
refGene_gn=DYTN;refGene_tx=XM_023260056.1
C1 193304734 193304883
refGene_gn=DYTN;refGene_tx=XM_023260056.1
C1 193306802 193306914
refGene_gn=DYTN;refGene_tx=XM_023260056.1
C1 193311475 193311639
refGene_gn=DYTN;refGene_tx=XM_023260056.1
C1 193312134 193312206
refGene_gn=DYTN;refGene_tx=XM_023260056.1
C1 193317011 193317112
refGene_gn=DYTN;refGene_tx=XM_023260056.1
C1 193317628 193317714
refGene_gn=DYTN;refGene_tx=XM_023260056.1
C1 193319494 193319549
refGene_gn=DYTN;refGene_tx=XM_023260056.1
C1 193319551 193319695
refGene_gn=DYTN;refGene_tx=XM_023260056.1
C1 193323324 193323399
refGene_gn=DYTN;refGene_tx=XM_023260056.1
C1 193327092 193327111
refGene_gn=DYTN;refGene_tx=XM_023260056.1
C1 193340262 193340294
refGene_gn=MDH1B;refGene_tx=XM_006935504.3;refGene_gn=MDH1B;refGene_tx=XM_019838538.1;refGene_gn=MDH1B;refGene_tx=XM_023259640.1
C1 193344119 193344200
refGene_gn=MDH1B;refGene_tx=XM_006935504.3;refGene_gn=MDH1B;refGene_tx=XM_019838538.1;refGene_gn=MDH1B;refGene_tx=XM_023259640.1
C1 193346758 193346810
refGene_gn=MDH1B;refGene_tx=XM_006935504.3;refGene_gn=MDH1B;refGene_tx=XM_019838538.1;refGene_gn=MDH1B;refGene_tx=XM_023259640.1
C1 193347349 193347489
refGene_gn=MDH1B;refGene_tx=XM_006935504.3;refGene_gn=MDH1B;refGene_tx=XM_019838538.1;refGene_gn=MDH1B;refGene_tx=XM_023259640.1

C1 193350147 193350311
refGene_gn=MDH1B;refGene_tx=XM_006935504.3;refGene_gn=MDH1B;refGene_tx=XM_019838538.1;refGene_gn=MDH1B;refGene_tx=XM_023259640.1
C1 193351631 193351773
refGene_gn=MDH1B;refGene_tx=XM_006935504.3;refGene_gn=MDH1B;refGene_tx=XM_019838538.1;refGene_gn=MDH1B;refGene_tx=XM_023259640.1
C1 193355055 193355552
refGene_gn=MDH1B;refGene_tx=XM_006935504.3;refGene_gn=MDH1B;refGene_tx=XM_019838538.1;refGene_gn=MDH1B;refGene_tx=XM_023259640.1
C1 193357120 193357263
refGene_gn=MDH1B;refGene_tx=XM_006935504.3;refGene_gn=MDH1B;refGene_tx=XM_019838538.1;refGene_gn=MDH1B;refGene_tx=XM_023259640.1
C1 193357458 193357593
refGene_gn=MDH1B;refGene_tx=XM_006935504.3;refGene_gn=MDH1B;refGene_tx=XM_023259640.1
C1 193360846 193360959
refGene_gn=MDH1B;refGene_tx=XM_006935504.3;refGene_gn=MDH1B;refGene_tx=XM_023259640.1
C1 193364428 193364680
refGene_gn=MDH1B;refGene_tx=XM_023259640.1
C1 193364779 193364801
refGene_gn=MDH1B;refGene_tx=XM_006935504.3;refGene_gn=MDH1B;refGene_tx=XM_023259640.1
C1 193366058 193366781
refGene_gn=FASTKD2;refGene_tx=XM_011285507.3
C1 193370187 193370291
refGene_gn=FASTKD2;refGene_tx=XM_011285507.3
C1 193370982 193371091
refGene_gn=FASTKD2;refGene_tx=XM_011285507.3
C1 193371381 193371505
refGene_gn=FASTKD2;refGene_tx=XM_011285507.3
C1 193371777 193371917
refGene_gn=FASTKD2;refGene_tx=XM_011285507.3
C1 193373050 193373223
refGene_gn=FASTKD2;refGene_tx=XM_011285507.3
C1 193377179 193377343
refGene_gn=FASTKD2;refGene_tx=XM_011285507.3
C1 193378570 193378780
refGene_gn=FASTKD2;refGene_tx=XM_011285507.3
C1 193379179 193379267
refGene_gn=FASTKD2;refGene_tx=XM_011285507.3
C1 193381210 193381325
refGene_gn=FASTKD2;refGene_tx=XM_011285507.3
C1 193382686 193382806
refGene_gn=FASTKD2;refGene_tx=XM_011285507.3
C1 193610339 193610391
refGene_gn=KLF7;refGene_tx=XM_003991071.5;refGene_gn=KLF7;refGene_tx=XM_006935507.4
C1 193617534 193617658
refGene_gn=KLF7;refGene_tx=XM_003991071.5;refGene_gn=KLF7;refGene_tx=X

M_006935507.4
C1	193652151	193652782
refGene_gn=KLF7;refGene_tx=XM_003991071.5;refGene_gn=KLF7;refGene_tx=X
M_006935507.4
C1	193695838	193695940
refGene_gn=KLF7;refGene_tx=XM_003991071.5;refGene_gn=KLF7;refGene_tx=X
M_006935507.4
C1	194032646	194032760
refGene_gn=CREB1;refGene_tx=XM_023259641.1;refGene_gn=CREB1;refGene_tx
=XM_023259642.1;refGene_gn=CREB1;refGene_tx=XM_023259643.1;refGene_gn=
CREB1;refGene_tx=XM_023259644.1
C1	194038433	194038580
refGene_gn=CREB1;refGene_tx=XM_023259641.1;refGene_gn=CREB1;refGene_tx
=XM_023259642.1;refGene_gn=CREB1;refGene_tx=XM_023259643.1;refGene_gn=
CREB1;refGene_tx=XM_023259644.1
C1	194039326	194039368
refGene_gn=CREB1;refGene_tx=XM_023259641.1;refGene_gn=CREB1;refGene_tx
=XM_023259642.1;refGene_gn=CREB1;refGene_tx=XM_023259643.1
C1	194045359	194045460
refGene_gn=CREB1;refGene_tx=XM_023259641.1;refGene_gn=CREB1;refGene_tx
=XM_023259642.1;refGene_gn=CREB1;refGene_tx=XM_023259643.1;refGene_gn=
CREB1;refGene_tx=XM_023259644.1
C1	194049319	194049462
refGene_gn=CREB1;refGene_tx=XM_023259641.1;refGene_gn=CREB1;refGene_tx
=XM_023259642.1;refGene_gn=CREB1;refGene_tx=XM_023259643.1;refGene_gn=
CREB1;refGene_tx=XM_023259644.1
C1	194054370	194054553
refGene_gn=CREB1;refGene_tx=XM_023259641.1;refGene_gn=CREB1;refGene_tx
=XM_023259642.1;refGene_gn=CREB1;refGene_tx=XM_023259643.1;refGene_gn=
CREB1;refGene_tx=XM_023259644.1
C1	194055670	194055710
refGene_gn=CREB1;refGene_tx=XM_019838539.2
C1	194056542	194056693
refGene_gn=CREB1;refGene_tx=XM_019838539.2;refGene_gn=CREB1;refGene_tx
=XM_023259641.1;refGene_gn=CREB1;refGene_tx=XM_023259642.1;refGene_gn=
CREB1;refGene_tx=XM_023259643.1;refGene_gn=CREB1;refGene_tx=XM_0232596
44.1
C1	194069869	194070014
refGene_gn=CREB1;refGene_tx=XM_019838539.2;refGene_gn=CREB1;refGene_tx
=XM_023259641.1;refGene_gn=CREB1;refGene_tx=XM_023259642.1;refGene_gn=
CREB1;refGene_tx=XM_023259643.1;refGene_gn=CREB1;refGene_tx=XM_0232596
44.1
C1	194081506	194081754
refGene_gn=METTL21A;refGene_tx=XM_006935509.4
C1	194082475	194082873
refGene_gn=METTL21A;refGene_tx=XM_003991074.5;refGene_gn=METTL21A;refG
ene_tx=XM_019838540.2;refGene_gn=METTL21A;refGene_tx=XM_019838542.2;re
fGene_gn=METTL21A;refGene_tx=XM_023259645.1
C1	194091971	194092130
refGene_gn=METTL21A;refGene_tx=XM_003991074.5;refGene_gn=METTL21A;refG

ene_tx=XM_006935509.4;refGene_gn=METTL21A;refGene_tx=XM_019838540.2;refGene_gn=METTL21A;refGene_tx=XM_019838542.2;refGene_gn=METTL21A;refGene_tx=XM_019838545.2;refGene_gn=METTL21A;refGene_tx=XM_023259645.1
C1 194093804 194093951
refGene_gn=METTL21A;refGene_tx=XM_003991074.5;refGene_gn=METTL21A;refGene_tx=XM_006935509.4;refGene_gn=METTL21A;refGene_tx=XM_019838540.2;refGene_gn=METTL21A;refGene_tx=XM_019838542.2;refGene_gn=METTL21A;refGene_tx=XM_019838545.2;refGene_gn=METTL21A;refGene_tx=XM_023259645.1
C1 194159435 194159661
refGene_gn=CCNYL1;refGene_tx=XM_003991075.5;refGene_gn=CCNYL1;refGene_tx=XM_003991076.5
C1 194171093 194171168
refGene_gn=CCNYL1;refGene_tx=XM_003991075.5;refGene_gn=CCNYL1;refGene_tx=XM_003991076.5
C1 194172840 194172875
refGene_gn=CCNYL1;refGene_tx=XM_003991075.5;refGene_gn=CCNYL1;refGene_tx=XM_003991076.5
C1 194178357 194178458
refGene_gn=CCNYL1;refGene_tx=XM_003991075.5
C1 194181790 194181826
refGene_gn=CCNYL1;refGene_tx=XM_003991075.5;refGene_gn=CCNYL1;refGene_tx=XM_003991076.5
C1 194184067 194184119
refGene_gn=CCNYL1;refGene_tx=XM_003991075.5
C1 194184967 194185087
refGene_gn=CCNYL1;refGene_tx=XM_003991075.5;refGene_gn=CCNYL1;refGene_tx=XM_003991076.5
C1 194188784 194188951
refGene_gn=CCNYL1;refGene_tx=XM_003991075.5;refGene_gn=CCNYL1;refGene_tx=XM_003991076.5
C1 194192197 194192360
refGene_gn=CCNYL1;refGene_tx=XM_003991075.5;refGene_gn=CCNYL1;refGene_tx=XM_003991076.5
C1 194193549 194193666
refGene_gn=CCNYL1;refGene_tx=XM_003991075.5;refGene_gn=CCNYL1;refGene_tx=XM_003991076.5
C1 194205404 194207177
refGene_gn=FZD5;refGene_tx=XM_023259646.1
C1 194263696 194263874
refGene_gn=PLEKHM3;refGene_tx=XM_003991077.5;refGene_gn=PLEKHM3;refGene_tx=XM_006935510.4;refGene_gn=PLEKHM3;refGene_tx=XM_019838547.2;refGene_gn=PLEKHM3;refGene_tx=XM_019838548.2;refGene_gn=PLEKHM3;refGene_tx=XM_019838552.2
C1 194284517 194284587
refGene_gn=PLEKHM3;refGene_tx=XM_019838549.2
C1 194290104 194290262
refGene_gn=PLEKHM3;refGene_tx=XM_003991077.5;refGene_gn=PLEKHM3;refGene_tx=XM_006935510.4;refGene_gn=PLEKHM3;refGene_tx=XM_019838547.2;refGene_gn=PLEKHM3;refGene_tx=XM_019838548.2;refGene_gn=PLEKHM3;refGene_tx=XM_019838549.2

C1	194317296	194317452
refGene_gn=PLEKHM3;refGene_tx=XM_019838550.2
C1	194336996	194337060
refGene_gn=PLEKHM3;refGene_tx=XM_003991077.5;refGene_gn=PLEKHM3;refGene_tx=XM_006935510.4;refGene_gn=PLEKHM3;refGene_tx=XM_019838547.2;refGene_gn=PLEKHM3;refGene_tx=XM_019838548.2;refGene_gn=PLEKHM3;refGene_tx=XM_019838549.2;refGene_gn=PLEKHM3;refGene_tx=XM_019838550.2;refGene_gn=PLEKHM3;refGene_tx=XM_019838552.2
C1	194368973	194368986
refGene_gn=PLEKHM3;refGene_tx=XM_006935511.4
C1	194369131	194369325
refGene_gn=PLEKHM3;refGene_tx=XM_003991077.5;refGene_gn=PLEKHM3;refGene_tx=XM_006935510.4;refGene_gn=PLEKHM3;refGene_tx=XM_006935511.4;refGene_gn=PLEKHM3;refGene_tx=XM_019838547.2;refGene_gn=PLEKHM3;refGene_tx=XM_019838548.2;refGene_gn=PLEKHM3;refGene_tx=XM_019838549.2;refGene_gn=PLEKHM3;refGene_tx=XM_019838550.2;refGene_gn=PLEKHM3;refGene_tx=XM_019838552.2
C1	194380927	194381073
refGene_gn=PLEKHM3;refGene_tx=XM_003991077.5;refGene_gn=PLEKHM3;refGene_tx=XM_006935510.4;refGene_gn=PLEKHM3;refGene_tx=XM_006935511.4;refGene_gn=PLEKHM3;refGene_tx=XM_019838547.2;refGene_gn=PLEKHM3;refGene_tx=XM_019838548.2;refGene_gn=PLEKHM3;refGene_tx=XM_019838549.2;refGene_gn=PLEKHM3;refGene_tx=XM_019838550.2;refGene_gn=PLEKHM3;refGene_tx=XM_019838552.2
C1	194409884	194410820
refGene_gn=PLEKHM3;refGene_tx=XM_003991077.5;refGene_gn=PLEKHM3;refGene_tx=XM_006935510.4;refGene_gn=PLEKHM3;refGene_tx=XM_006935511.4;refGene_gn=PLEKHM3;refGene_tx=XM_019838547.2;refGene_gn=PLEKHM3;refGene_tx=XM_019838548.2;refGene_gn=PLEKHM3;refGene_tx=XM_019838549.2;refGene_gn=PLEKHM3;refGene_tx=XM_019838550.2;refGene_gn=PLEKHM3;refGene_tx=XM_019838552.2
C1	194434230	194434831
refGene_gn=PLEKHM3;refGene_tx=XM_003991077.5;refGene_gn=PLEKHM3;refGene_tx=XM_006935510.4;refGene_gn=PLEKHM3;refGene_tx=XM_006935511.4;refGene_gn=PLEKHM3;refGene_tx=XM_019838547.2;refGene_gn=PLEKHM3;refGene_tx=XM_019838548.2;refGene_gn=PLEKHM3;refGene_tx=XM_019838549.2;refGene_gn=PLEKHM3;refGene_tx=XM_019838550.2;refGene_gn=PLEKHM3;refGene_tx=XM_019838552.2
C1	194510097	194510370
refGene_gn=LOC101092631;refGene_tx=XM_003991078.3
C1	194512235	194512478
refGene_gn=LOC101092631;refGene_tx=XM_003991078.3
C1	194512569	194512578
refGene_gn=LOC101092631;refGene_tx=XM_003991078.3
C1	194518787	194519060
refGene_gn=LOC101092875;refGene_tx=XM_003991079.3
C1	194520156	194520399
refGene_gn=LOC101092875;refGene_tx=XM_003991079.3
C1	194520490	194520499
refGene_gn=LOC101092875;refGene_tx=XM_003991079.3

C1 194525211 194525484
refGene_gn=LOC101093129;refGene_tx=XM_003991080.3
C1 194526451 194526694
refGene_gn=LOC101093129;refGene_tx=XM_003991080.3
C1 194526804 194526813
refGene_gn=LOC101093129;refGene_tx=XM_003991080.3
C1 194534372 194534648
refGene_gn=LOC101093383;refGene_tx=XM_003991081.4
C1 194536314 194536557
refGene_gn=LOC101093383;refGene_tx=XM_003991081.4
C1 194536652 194536661
refGene_gn=LOC101093383;refGene_tx=XM_003991081.4
C1 194546616 194546889
refGene_gn=LOC101093630;refGene_tx=XM_003991082.4
C1 194548092 194548335
refGene_gn=LOC101093630;refGene_tx=XM_003991082.4
C1 194548439 194548448
refGene_gn=LOC101093630;refGene_tx=XM_003991082.4
C1 194550510 194550587
refGene_gn=CC1H2orf80;refGene_tx=XM_006935514.4;refGene_gn=CC1H2orf80;
refGene_tx=XM_019838554.2;refGene_gn=CC1H2orf80;refGene_tx=XM_01983855
5.2;refGene_gn=CC1H2orf80;refGene_tx=XM_019838557.2
C1 194553786 194553843
refGene_gn=CC1H2orf80;refGene_tx=XM_019838554.2;refGene_gn=CC1H2orf80;
refGene_tx=XM_019838555.2;refGene_gn=CC1H2orf80;refGene_tx=XM_01983855
7.2;refGene_gn=CC1H2orf80;refGene_tx=XM_019838558.2
C1 194554735 194554769
refGene_gn=CC1H2orf80;refGene_tx=XM_019838554.2;refGene_gn=CC1H2orf80;
refGene_tx=XM_019838555.2;refGene_gn=CC1H2orf80;refGene_tx=XM_01983855
7.2
C1 194556072 194556165
refGene_gn=CC1H2orf80;refGene_tx=XM_023259647.1
C1 194556697 194556816
refGene_gn=CC1H2orf80;refGene_tx=XM_006935514.4;refGene_gn=CC1H2orf80;
refGene_tx=XM_019838554.2;refGene_gn=CC1H2orf80;refGene_tx=XM_01983855
5.2;refGene_gn=CC1H2orf80;refGene_tx=XM_019838557.2;refGene_gn=CC1H2or
f80;refGene_tx=XM_019838558.2;refGene_gn=CC1H2orf80;refGene_tx=XM_0232
59647.1
C1 194557597 194557685
refGene_gn=CC1H2orf80;refGene_tx=XM_006935514.4;refGene_gn=CC1H2orf80;
refGene_tx=XM_019838554.2;refGene_gn=CC1H2orf80;refGene_tx=XM_01983855
5.2;refGene_gn=CC1H2orf80;refGene_tx=XM_019838557.2;refGene_gn=CC1H2or
f80;refGene_tx=XM_019838558.2;refGene_gn=CC1H2orf80;refGene_tx=XM_0232
59647.1
C1 194568658 194568730
refGene_gn=CC1H2orf80;refGene_tx=XM_006935514.4;refGene_gn=CC1H2orf80;
refGene_tx=XM_019838554.2;refGene_gn=CC1H2orf80;refGene_tx=XM_01983855
5.2;refGene_gn=CC1H2orf80;refGene_tx=XM_019838558.2;refGene_gn=CC1H2or
f80;refGene_tx=XM_023259647.1
C1 194569405 194569493

refGene_gn=CC1H2orf80;refGene_tx=XM_006935514.4;refGene_gn=CC1H2orf80;
refGene_tx=XM_019838554.2;refGene_gn=CC1H2orf80;refGene_tx=XM_01983855
5.2;refGene_gn=CC1H2orf80;refGene_tx=XM_019838557.2;refGene_gn=CC1H2or
f80;refGene_tx=XM_019838558.2;refGene_gn=CC1H2orf80;refGene_tx=XM_0232
59647.1
C1 194571087 194571170
refGene_gn=CC1H2orf80;refGene_tx=XM_006935514.4;refGene_gn=CC1H2orf80;
refGene_tx=XM_019838554.2;refGene_gn=CC1H2orf80;refGene_tx=XM_01983855
5.2;refGene_gn=CC1H2orf80;refGene_tx=XM_019838557.2;refGene_gn=CC1H2or
f80;refGene_tx=XM_019838558.2;refGene_gn=CC1H2orf80;refGene_tx=XM_0232
59647.1
C1 194573219 194573301
refGene_gn=CC1H2orf80;refGene_tx=XM_006935514.4;refGene_gn=CC1H2orf80;
refGene_tx=XM_019838554.2;refGene_gn=CC1H2orf80;refGene_tx=XM_01983855
5.2;refGene_gn=CC1H2orf80;refGene_tx=XM_019838557.2;refGene_gn=CC1H2or
f80;refGene_tx=XM_019838558.2;refGene_gn=CC1H2orf80;refGene_tx=XM_0232
59647.1
C1 194575053 194575094
refGene_gn=CC1H2orf80;refGene_tx=XM_006935514.4;refGene_gn=CC1H2orf80;
refGene_tx=XM_019838554.2;refGene_gn=CC1H2orf80;refGene_tx=XM_01983855
5.2;refGene_gn=CC1H2orf80;refGene_tx=XM_019838557.2;refGene_gn=CC1H2or
f80;refGene_tx=XM_019838558.2;refGene_gn=CC1H2orf80;refGene_tx=XM_0232
59647.1
C1 194609636 194609727
refGene_gn=IDH1;refGene_tx=XM_006935515.4;refGene_gn=IDH1;refGene_tx=X
M_006935516.4;refGene_gn=IDH1;refGene_tx=XM_006935517.3
C1 194610920 194611083
refGene_gn=IDH1;refGene_tx=XM_006935515.4;refGene_gn=IDH1;refGene_tx=X
M_006935516.4;refGene_gn=IDH1;refGene_tx=XM_006935517.3
C1 194612187 194612328
refGene_gn=IDH1;refGene_tx=XM_006935515.4;refGene_gn=IDH1;refGene_tx=X
M_006935516.4;refGene_gn=IDH1;refGene_tx=XM_006935517.3
C1 194614496 194614648
refGene_gn=IDH1;refGene_tx=XM_006935515.4;refGene_gn=IDH1;refGene_tx=X
M_006935516.4;refGene_gn=IDH1;refGene_tx=XM_006935517.3
C1 194615668 194615846
refGene_gn=IDH1;refGene_tx=XM_006935515.4;refGene_gn=IDH1;refGene_tx=X
M_006935516.4;refGene_gn=IDH1;refGene_tx=XM_006935517.3
C1 194617369 194617475
refGene_gn=IDH1;refGene_tx=XM_006935515.4;refGene_gn=IDH1;refGene_tx=X
M_006935516.4;refGene_gn=IDH1;refGene_tx=XM_006935517.3
C1 194620036 194620328
refGene_gn=IDH1;refGene_tx=XM_006935515.4;refGene_gn=IDH1;refGene_tx=X
M_006935516.4;refGene_gn=IDH1;refGene_tx=XM_006935517.3
C1 194622955 194623077
refGene_gn=IDH1;refGene_tx=XM_006935515.4;refGene_gn=IDH1;refGene_tx=X
M_006935516.4;refGene_gn=IDH1;refGene_tx=XM_006935517.3
C1 194637179 194637351
refGene_gn=PIKFYVE;refGene_tx=XM_019838560.1;refGene_gn=PIKFYVE;refGen
e_tx=XM_019838561.1;refGene_gn=PIKFYVE;refGene_tx=XM_019838562.1;refGe

ne_gn=PIKFYVE;refGene_tx=XM_019838563.2;refGene_gn=PIKFYVE;refGene_tx=
XM_019838564.2;refGene_gn=PIKFYVE;refGene_tx=XM_019838565.1;refGene_gn
=PIKFYVE;refGene_tx=XM_019838566.1;refGene_gn=PIKFYVE;refGene_tx=XM_01
9838569.1
C1 194639636 194639789
refGene_gn=PIKFYVE;refGene_tx=XM_019838560.1;refGene_gn=PIKFYVE;refGen
e_tx=XM_019838561.1;refGene_gn=PIKFYVE;refGene_tx=XM_019838562.1;refGe
ne_gn=PIKFYVE;refGene_tx=XM_019838563.2;refGene_gn=PIKFYVE;refGene_tx=
XM_019838564.2;refGene_gn=PIKFYVE;refGene_tx=XM_019838565.1;refGene_gn
=PIKFYVE;refGene_tx=XM_019838566.1;refGene_gn=PIKFYVE;refGene_tx=XM_01
9838569.1
C1 194640055 194640091
refGene_gn=PIKFYVE;refGene_tx=XM_019838560.1;refGene_gn=PIKFYVE;refGen
e_tx=XM_019838561.1;refGene_gn=PIKFYVE;refGene_tx=XM_019838562.1;refGe
ne_gn=PIKFYVE;refGene_tx=XM_019838563.2;refGene_gn=PIKFYVE;refGene_tx=
XM_019838564.2;refGene_gn=PIKFYVE;refGene_tx=XM_019838565.1;refGene_gn
=PIKFYVE;refGene_tx=XM_019838569.1
C1 194643234 194643353
refGene_gn=PIKFYVE;refGene_tx=XM_019838560.1;refGene_gn=PIKFYVE;refGen
e_tx=XM_019838561.1;refGene_gn=PIKFYVE;refGene_tx=XM_019838562.1;refGe
ne_gn=PIKFYVE;refGene_tx=XM_019838563.2;refGene_gn=PIKFYVE;refGene_tx=
XM_019838564.2;refGene_gn=PIKFYVE;refGene_tx=XM_019838565.1;refGene_gn
=PIKFYVE;refGene_tx=XM_019838566.1;refGene_gn=PIKFYVE;refGene_tx=XM_01
9838567.1;refGene_gn=PIKFYVE;refGene_tx=XM_019838569.1
C1 194644060 194644232
refGene_gn=PIKFYVE;refGene_tx=XM_019838560.1;refGene_gn=PIKFYVE;refGen
e_tx=XM_019838561.1;refGene_gn=PIKFYVE;refGene_tx=XM_019838562.1;refGe
ne_gn=PIKFYVE;refGene_tx=XM_019838563.2;refGene_gn=PIKFYVE;refGene_tx=
XM_019838564.2;refGene_gn=PIKFYVE;refGene_tx=XM_019838565.1;refGene_gn
=PIKFYVE;refGene_tx=XM_019838566.1;refGene_gn=PIKFYVE;refGene_tx=XM_01
9838567.1;refGene_gn=PIKFYVE;refGene_tx=XM_019838569.1
C1 194649169 194649377
refGene_gn=PIKFYVE;refGene_tx=XM_019838560.1;refGene_gn=PIKFYVE;refGen
e_tx=XM_019838561.1;refGene_gn=PIKFYVE;refGene_tx=XM_019838562.1;refGe
ne_gn=PIKFYVE;refGene_tx=XM_019838563.2;refGene_gn=PIKFYVE;refGene_tx=
XM_019838564.2;refGene_gn=PIKFYVE;refGene_tx=XM_019838565.1;refGene_gn
=PIKFYVE;refGene_tx=XM_019838566.1;refGene_gn=PIKFYVE;refGene_tx=XM_01
9838567.1;refGene_gn=PIKFYVE;refGene_tx=XM_019838569.1
C1 194651554 194651644
refGene_gn=PIKFYVE;refGene_tx=XM_019838560.1;refGene_gn=PIKFYVE;refGen
e_tx=XM_019838561.1;refGene_gn=PIKFYVE;refGene_tx=XM_019838562.1;refGe
ne_gn=PIKFYVE;refGene_tx=XM_019838563.2;refGene_gn=PIKFYVE;refGene_tx=
XM_019838564.2;refGene_gn=PIKFYVE;refGene_tx=XM_019838565.1;refGene_gn
=PIKFYVE;refGene_tx=XM_019838566.1;refGene_gn=PIKFYVE;refGene_tx=XM_01
9838567.1;refGene_gn=PIKFYVE;refGene_tx=XM_019838569.1
C1 194654130 194654269
refGene_gn=PIKFYVE;refGene_tx=XM_019838560.1;refGene_gn=PIKFYVE;refGen
e_tx=XM_019838561.1;refGene_gn=PIKFYVE;refGene_tx=XM_019838562.1;refGe
ne_gn=PIKFYVE;refGene_tx=XM_019838563.2;refGene_gn=PIKFYVE;refGene_tx=
XM_019838564.2;refGene_gn=PIKFYVE;refGene_tx=XM_019838565.1;refGene_gn

=PIKFYVE;refGene_tx=XM_019838566.1;refGene_gn=PIKFYVE;refGene_tx=XM_019838567.1;refGene_gn=PIKFYVE;refGene_tx=XM_019838569.1
C1 194655560 194655718
refGene_gn=PIKFYVE;refGene_tx=XM_019838560.1;refGene_gn=PIKFYVE;refGene_tx=XM_019838561.1;refGene_gn=PIKFYVE;refGene_tx=XM_019838562.1;refGene_gn=PIKFYVE;refGene_tx=XM_019838563.2;refGene_gn=PIKFYVE;refGene_tx=XM_019838564.2;refGene_gn=PIKFYVE;refGene_tx=XM_019838565.1;refGene_gn=PIKFYVE;refGene_tx=XM_019838566.1;refGene_gn=PIKFYVE;refGene_tx=XM_019838567.1;refGene_gn=PIKFYVE;refGene_tx=XM_019838569.1
C1 194656918 194657030
refGene_gn=PIKFYVE;refGene_tx=XM_019838560.1;refGene_gn=PIKFYVE;refGene_tx=XM_019838561.1;refGene_gn=PIKFYVE;refGene_tx=XM_019838562.1;refGene_gn=PIKFYVE;refGene_tx=XM_019838563.2;refGene_gn=PIKFYVE;refGene_tx=XM_019838564.2;refGene_gn=PIKFYVE;refGene_tx=XM_019838565.1;refGene_gn=PIKFYVE;refGene_tx=XM_019838566.1;refGene_gn=PIKFYVE;refGene_tx=XM_019838567.1;refGene_gn=PIKFYVE;refGene_tx=XM_019838569.1
C1 194657961 194658109
refGene_gn=PIKFYVE;refGene_tx=XM_019838560.1;refGene_gn=PIKFYVE;refGene_tx=XM_019838561.1;refGene_gn=PIKFYVE;refGene_tx=XM_019838562.1;refGene_gn=PIKFYVE;refGene_tx=XM_019838563.2;refGene_gn=PIKFYVE;refGene_tx=XM_019838564.2;refGene_gn=PIKFYVE;refGene_tx=XM_019838565.1;refGene_gn=PIKFYVE;refGene_tx=XM_019838566.1;refGene_gn=PIKFYVE;refGene_tx=XM_019838567.1;refGene_gn=PIKFYVE;refGene_tx=XM_019838569.1
C1 194658650 194658829
refGene_gn=PIKFYVE;refGene_tx=XM_019838560.1;refGene_gn=PIKFYVE;refGene_tx=XM_019838561.1;refGene_gn=PIKFYVE;refGene_tx=XM_019838562.1;refGene_gn=PIKFYVE;refGene_tx=XM_019838563.2;refGene_gn=PIKFYVE;refGene_tx=XM_019838564.2;refGene_gn=PIKFYVE;refGene_tx=XM_019838565.1;refGene_gn=PIKFYVE;refGene_tx=XM_019838566.1;refGene_gn=PIKFYVE;refGene_tx=XM_019838567.1;refGene_gn=PIKFYVE;refGene_tx=XM_019838569.1
C1 194666756 194666816
refGene_gn=PIKFYVE;refGene_tx=XM_019838560.1;refGene_gn=PIKFYVE;refGene_tx=XM_019838561.1;refGene_gn=PIKFYVE;refGene_tx=XM_019838562.1;refGene_gn=PIKFYVE;refGene_tx=XM_019838563.2;refGene_gn=PIKFYVE;refGene_tx=XM_019838564.2;refGene_gn=PIKFYVE;refGene_tx=XM_019838565.1;refGene_gn=PIKFYVE;refGene_tx=XM_019838566.1;refGene_gn=PIKFYVE;refGene_tx=XM_019838567.1
C1 194668441 194668571
refGene_gn=PIKFYVE;refGene_tx=XM_019838560.1;refGene_gn=PIKFYVE;refGene_tx=XM_019838561.1;refGene_gn=PIKFYVE;refGene_tx=XM_019838562.1;refGene_gn=PIKFYVE;refGene_tx=XM_019838563.2;refGene_gn=PIKFYVE;refGene_tx=XM_019838564.2;refGene_gn=PIKFYVE;refGene_tx=XM_019838565.1;refGene_gn=PIKFYVE;refGene_tx=XM_019838566.1;refGene_gn=PIKFYVE;refGene_tx=XM_019838567.1
C1 194668987 194669168
refGene_gn=PIKFYVE;refGene_tx=XM_019838560.1;refGene_gn=PIKFYVE;refGene_tx=XM_019838561.1;refGene_gn=PIKFYVE;refGene_tx=XM_019838562.1;refGene_gn=PIKFYVE;refGene_tx=XM_019838563.2;refGene_gn=PIKFYVE;refGene_tx=XM_019838564.2;refGene_gn=PIKFYVE;refGene_tx=XM_019838565.1;refGene_gn=PIKFYVE;refGene_tx=XM_019838566.1;refGene_gn=PIKFYVE;refGene_tx=XM_01

9838567.1
C1	194669816	194669891
refGene_gn=PIKFYVE;refGene_tx=XM_019838560.1;refGene_gn=PIKFYVE;refGene_tx=XM_019838561.1;refGene_gn=PIKFYVE;refGene_tx=XM_019838562.1;refGene_gn=PIKFYVE;refGene_tx=XM_019838563.2;refGene_gn=PIKFYVE;refGene_tx=XM_019838564.2;refGene_gn=PIKFYVE;refGene_tx=XM_019838565.1;refGene_gn=PIKFYVE;refGene_tx=XM_019838566.1;refGene_gn=PIKFYVE;refGene_tx=XM_019838567.1
C1	194671846	194671954
refGene_gn=PIKFYVE;refGene_tx=XM_019838560.1;refGene_gn=PIKFYVE;refGene_tx=XM_019838561.1;refGene_gn=PIKFYVE;refGene_tx=XM_019838562.1;refGene_gn=PIKFYVE;refGene_tx=XM_019838563.2;refGene_gn=PIKFYVE;refGene_tx=XM_019838564.2;refGene_gn=PIKFYVE;refGene_tx=XM_019838565.1;refGene_gn=PIKFYVE;refGene_tx=XM_019838566.1;refGene_gn=PIKFYVE;refGene_tx=XM_019838567.1
C1	194676388	194676529
refGene_gn=PIKFYVE;refGene_tx=XM_019838560.1;refGene_gn=PIKFYVE;refGene_tx=XM_019838561.1;refGene_gn=PIKFYVE;refGene_tx=XM_019838562.1;refGene_gn=PIKFYVE;refGene_tx=XM_019838563.2;refGene_gn=PIKFYVE;refGene_tx=XM_019838564.2;refGene_gn=PIKFYVE;refGene_tx=XM_019838565.1;refGene_gn=PIKFYVE;refGene_tx=XM_019838566.1;refGene_gn=PIKFYVE;refGene_tx=XM_019838567.1
C1	194677148	194677275
refGene_gn=PIKFYVE;refGene_tx=XM_019838560.1;refGene_gn=PIKFYVE;refGene_tx=XM_019838561.1;refGene_gn=PIKFYVE;refGene_tx=XM_019838562.1;refGene_gn=PIKFYVE;refGene_tx=XM_019838563.2;refGene_gn=PIKFYVE;refGene_tx=XM_019838564.2;refGene_gn=PIKFYVE;refGene_tx=XM_019838565.1;refGene_gn=PIKFYVE;refGene_tx=XM_019838566.1;refGene_gn=PIKFYVE;refGene_tx=XM_019838567.1
C1	194677557	194678717
refGene_gn=PIKFYVE;refGene_tx=XM_019838560.1;refGene_gn=PIKFYVE;refGene_tx=XM_019838561.1;refGene_gn=PIKFYVE;refGene_tx=XM_019838562.1;refGene_gn=PIKFYVE;refGene_tx=XM_019838563.2;refGene_gn=PIKFYVE;refGene_tx=XM_019838564.2;refGene_gn=PIKFYVE;refGene_tx=XM_019838565.1;refGene_gn=PIKFYVE;refGene_tx=XM_019838566.1;refGene_gn=PIKFYVE;refGene_tx=XM_019838567.1
C1	194682032	194682133
refGene_gn=PIKFYVE;refGene_tx=XM_019838560.1;refGene_gn=PIKFYVE;refGene_tx=XM_019838561.1;refGene_gn=PIKFYVE;refGene_tx=XM_019838562.1;refGene_gn=PIKFYVE;refGene_tx=XM_019838563.2;refGene_gn=PIKFYVE;refGene_tx=XM_019838564.2;refGene_gn=PIKFYVE;refGene_tx=XM_019838565.1;refGene_gn=PIKFYVE;refGene_tx=XM_019838566.1;refGene_gn=PIKFYVE;refGene_tx=XM_019838567.1
C1	194683406	194683478
refGene_gn=PIKFYVE;refGene_tx=XM_019838560.1;refGene_gn=PIKFYVE;refGene_tx=XM_019838561.1;refGene_gn=PIKFYVE;refGene_tx=XM_019838562.1;refGene_gn=PIKFYVE;refGene_tx=XM_019838563.2;refGene_gn=PIKFYVE;refGene_tx=XM_019838564.2;refGene_gn=PIKFYVE;refGene_tx=XM_019838565.1;refGene_gn=PIKFYVE;refGene_tx=XM_019838566.1;refGene_gn=PIKFYVE;refGene_tx=XM_019838567.1

C1 194684148 194684320
refGene_gn=PIKFYVE;refGene_tx=XM_019838560.1;refGene_gn=PIKFYVE;refGene_tx=XM_019838561.1;refGene_gn=PIKFYVE;refGene_tx=XM_019838562.1;refGene_gn=PIKFYVE;refGene_tx=XM_019838563.2;refGene_gn=PIKFYVE;refGene_tx=XM_019838564.2;refGene_gn=PIKFYVE;refGene_tx=XM_019838565.1;refGene_gn=PIKFYVE;refGene_tx=XM_019838566.1;refGene_gn=PIKFYVE;refGene_tx=XM_019838567.1
C1 194686313 194686492
refGene_gn=PIKFYVE;refGene_tx=XM_019838560.1;refGene_gn=PIKFYVE;refGene_tx=XM_019838561.1;refGene_gn=PIKFYVE;refGene_tx=XM_019838562.1;refGene_gn=PIKFYVE;refGene_tx=XM_019838563.2;refGene_gn=PIKFYVE;refGene_tx=XM_019838564.2;refGene_gn=PIKFYVE;refGene_tx=XM_019838565.1;refGene_gn=PIKFYVE;refGene_tx=XM_019838566.1;refGene_gn=PIKFYVE;refGene_tx=XM_019838567.1
C1 194688099 194688213
refGene_gn=PIKFYVE;refGene_tx=XM_019838560.1;refGene_gn=PIKFYVE;refGene_tx=XM_019838561.1;refGene_gn=PIKFYVE;refGene_tx=XM_019838562.1;refGene_gn=PIKFYVE;refGene_tx=XM_019838563.2;refGene_gn=PIKFYVE;refGene_tx=XM_019838564.2;refGene_gn=PIKFYVE;refGene_tx=XM_019838565.1;refGene_gn=PIKFYVE;refGene_tx=XM_019838566.1;refGene_gn=PIKFYVE;refGene_tx=XM_019838567.1
C1 194688924 194689033
refGene_gn=PIKFYVE;refGene_tx=XM_019838560.1;refGene_gn=PIKFYVE;refGene_tx=XM_019838561.1;refGene_gn=PIKFYVE;refGene_tx=XM_019838562.1;refGene_gn=PIKFYVE;refGene_tx=XM_019838563.2;refGene_gn=PIKFYVE;refGene_tx=XM_019838564.2;refGene_gn=PIKFYVE;refGene_tx=XM_019838565.1;refGene_gn=PIKFYVE;refGene_tx=XM_019838566.1;refGene_gn=PIKFYVE;refGene_tx=XM_019838567.1
C1 194689143 194689298
refGene_gn=PIKFYVE;refGene_tx=XM_019838560.1;refGene_gn=PIKFYVE;refGene_tx=XM_019838561.1;refGene_gn=PIKFYVE;refGene_tx=XM_019838562.1;refGene_gn=PIKFYVE;refGene_tx=XM_019838563.2;refGene_gn=PIKFYVE;refGene_tx=XM_019838564.2;refGene_gn=PIKFYVE;refGene_tx=XM_019838565.1;refGene_gn=PIKFYVE;refGene_tx=XM_019838566.1;refGene_gn=PIKFYVE;refGene_tx=XM_019838567.1
C1 194690861 194690952
refGene_gn=PIKFYVE;refGene_tx=XM_019838560.1;refGene_gn=PIKFYVE;refGene_tx=XM_019838561.1;refGene_gn=PIKFYVE;refGene_tx=XM_019838562.1;refGene_gn=PIKFYVE;refGene_tx=XM_019838563.2;refGene_gn=PIKFYVE;refGene_tx=XM_019838564.2;refGene_gn=PIKFYVE;refGene_tx=XM_019838565.1;refGene_gn=PIKFYVE;refGene_tx=XM_019838566.1;refGene_gn=PIKFYVE;refGene_tx=XM_019838567.1
C1 194693915 194693976
refGene_gn=PIKFYVE;refGene_tx=XM_019838560.1;refGene_gn=PIKFYVE;refGene_tx=XM_019838561.1;refGene_gn=PIKFYVE;refGene_tx=XM_019838562.1;refGene_gn=PIKFYVE;refGene_tx=XM_019838563.2;refGene_gn=PIKFYVE;refGene_tx=XM_019838564.2;refGene_gn=PIKFYVE;refGene_tx=XM_019838565.1;refGene_gn=PIKFYVE;refGene_tx=XM_019838566.1;refGene_gn=PIKFYVE;refGene_tx=XM_019838567.1
C1 194695056 194695194

refGene_gn=PIKFYVE;refGene_tx=XM_019838560.1;refGene_gn=PIKFYVE;refGene_tx=XM_019838561.1;refGene_gn=PIKFYVE;refGene_tx=XM_019838562.1;refGene_gn=PIKFYVE;refGene_tx=XM_019838563.2;refGene_gn=PIKFYVE;refGene_tx=XM_019838564.2;refGene_gn=PIKFYVE;refGene_tx=XM_019838565.1;refGene_gn=PIKFYVE;refGene_tx=XM_019838566.1;refGene_gn=PIKFYVE;refGene_tx=XM_019838567.1
C1 194696248 194696369
refGene_gn=PIKFYVE;refGene_tx=XM_019838560.1;refGene_gn=PIKFYVE;refGene_tx=XM_019838561.1;refGene_gn=PIKFYVE;refGene_tx=XM_019838562.1;refGene_gn=PIKFYVE;refGene_tx=XM_019838563.2;refGene_gn=PIKFYVE;refGene_tx=XM_019838564.2;refGene_gn=PIKFYVE;refGene_tx=XM_019838565.1;refGene_gn=PIKFYVE;refGene_tx=XM_019838566.1;refGene_gn=PIKFYVE;refGene_tx=XM_019838567.1
C1 194698557 194698653
refGene_gn=PIKFYVE;refGene_tx=XM_019838560.1;refGene_gn=PIKFYVE;refGene_tx=XM_019838561.1;refGene_gn=PIKFYVE;refGene_tx=XM_019838562.1;refGene_gn=PIKFYVE;refGene_tx=XM_019838563.2;refGene_gn=PIKFYVE;refGene_tx=XM_019838564.2;refGene_gn=PIKFYVE;refGene_tx=XM_019838565.1;refGene_gn=PIKFYVE;refGene_tx=XM_019838566.1;refGene_gn=PIKFYVE;refGene_tx=XM_019838567.1
C1 194700356 194700440
refGene_gn=PIKFYVE;refGene_tx=XM_019838560.1;refGene_gn=PIKFYVE;refGene_tx=XM_019838561.1;refGene_gn=PIKFYVE;refGene_tx=XM_019838562.1;refGene_gn=PIKFYVE;refGene_tx=XM_019838563.2;refGene_gn=PIKFYVE;refGene_tx=XM_019838564.2;refGene_gn=PIKFYVE;refGene_tx=XM_019838565.1;refGene_gn=PIKFYVE;refGene_tx=XM_019838566.1;refGene_gn=PIKFYVE;refGene_tx=XM_019838567.1
C1 194701514 194701612
refGene_gn=PIKFYVE;refGene_tx=XM_019838560.1;refGene_gn=PIKFYVE;refGene_tx=XM_019838561.1;refGene_gn=PIKFYVE;refGene_tx=XM_019838562.1;refGene_gn=PIKFYVE;refGene_tx=XM_019838563.2;refGene_gn=PIKFYVE;refGene_tx=XM_019838564.2;refGene_gn=PIKFYVE;refGene_tx=XM_019838565.1;refGene_gn=PIKFYVE;refGene_tx=XM_019838566.1;refGene_gn=PIKFYVE;refGene_tx=XM_019838567.1
C1 194703347 194703512
refGene_gn=PIKFYVE;refGene_tx=XM_019838560.1;refGene_gn=PIKFYVE;refGene_tx=XM_019838561.1;refGene_gn=PIKFYVE;refGene_tx=XM_019838562.1;refGene_gn=PIKFYVE;refGene_tx=XM_019838563.2;refGene_gn=PIKFYVE;refGene_tx=XM_019838564.2;refGene_gn=PIKFYVE;refGene_tx=XM_019838565.1;refGene_gn=PIKFYVE;refGene_tx=XM_019838566.1;refGene_gn=PIKFYVE;refGene_tx=XM_019838567.1
C1 194705942 194706002
refGene_gn=PIKFYVE;refGene_tx=XM_019838560.1;refGene_gn=PIKFYVE;refGene_tx=XM_019838561.1;refGene_gn=PIKFYVE;refGene_tx=XM_019838562.1;refGene_gn=PIKFYVE;refGene_tx=XM_019838563.2;refGene_gn=PIKFYVE;refGene_tx=XM_019838564.2;refGene_gn=PIKFYVE;refGene_tx=XM_019838565.1;refGene_gn=PIKFYVE;refGene_tx=XM_019838566.1;refGene_gn=PIKFYVE;refGene_tx=XM_019838567.1
C1 194706663 194706840
refGene_gn=PIKFYVE;refGene_tx=XM_019838560.1;refGene_gn=PIKFYVE;refGen

e_tx=XM_019838561.1;refGene_gn=PIKFYVE;refGene_tx=XM_019838562.1;refGe
ne_gn=PIKFYVE;refGene_tx=XM_019838563.2;refGene_gn=PIKFYVE;refGene_tx=
XM_019838564.2;refGene_gn=PIKFYVE;refGene_tx=XM_019838565.1;refGene_gn
=PIKFYVE;refGene_tx=XM_019838566.1;refGene_gn=PIKFYVE;refGene_tx=XM_01
9838567.1
C1 194707838 194707942
refGene_gn=PIKFYVE;refGene_tx=XM_019838560.1;refGene_gn=PIKFYVE;refGen
e_tx=XM_019838561.1;refGene_gn=PIKFYVE;refGene_tx=XM_019838562.1;refGe
ne_gn=PIKFYVE;refGene_tx=XM_019838563.2;refGene_gn=PIKFYVE;refGene_tx=
XM_019838564.2;refGene_gn=PIKFYVE;refGene_tx=XM_019838565.1;refGene_gn
=PIKFYVE;refGene_tx=XM_019838566.1;refGene_gn=PIKFYVE;refGene_tx=XM_01
9838567.1
C1 194708715 194708844
refGene_gn=PIKFYVE;refGene_tx=XM_019838560.1;refGene_gn=PIKFYVE;refGen
e_tx=XM_019838561.1;refGene_gn=PIKFYVE;refGene_tx=XM_019838562.1;refGe
ne_gn=PIKFYVE;refGene_tx=XM_019838563.2;refGene_gn=PIKFYVE;refGene_tx=
XM_019838564.2;refGene_gn=PIKFYVE;refGene_tx=XM_019838565.1;refGene_gn
=PIKFYVE;refGene_tx=XM_019838566.1;refGene_gn=PIKFYVE;refGene_tx=XM_01
9838567.1
C1 194710168 194710430
refGene_gn=PIKFYVE;refGene_tx=XM_019838560.1;refGene_gn=PIKFYVE;refGen
e_tx=XM_019838561.1;refGene_gn=PIKFYVE;refGene_tx=XM_019838562.1;refGe
ne_gn=PIKFYVE;refGene_tx=XM_019838563.2;refGene_gn=PIKFYVE;refGene_tx=
XM_019838564.2;refGene_gn=PIKFYVE;refGene_tx=XM_019838565.1;refGene_gn
=PIKFYVE;refGene_tx=XM_019838566.1;refGene_gn=PIKFYVE;refGene_tx=XM_01
9838567.1
C1 194710835 194710910
refGene_gn=PIKFYVE;refGene_tx=XM_019838560.1;refGene_gn=PIKFYVE;refGen
e_tx=XM_019838561.1;refGene_gn=PIKFYVE;refGene_tx=XM_019838562.1;refGe
ne_gn=PIKFYVE;refGene_tx=XM_019838563.2;refGene_gn=PIKFYVE;refGene_tx=
XM_019838564.2;refGene_gn=PIKFYVE;refGene_tx=XM_019838565.1;refGene_gn
=PIKFYVE;refGene_tx=XM_019838566.1;refGene_gn=PIKFYVE;refGene_tx=XM_01
9838567.1
C1 194712020 194712136
refGene_gn=PIKFYVE;refGene_tx=XM_019838560.1;refGene_gn=PIKFYVE;refGen
e_tx=XM_019838561.1;refGene_gn=PIKFYVE;refGene_tx=XM_019838562.1;refGe
ne_gn=PIKFYVE;refGene_tx=XM_019838563.2;refGene_gn=PIKFYVE;refGene_tx=
XM_019838564.2;refGene_gn=PIKFYVE;refGene_tx=XM_019838565.1;refGene_gn
=PIKFYVE;refGene_tx=XM_019838566.1;refGene_gn=PIKFYVE;refGene_tx=XM_01
9838567.1
C1 194729280 194729328
refGene_gn=PTH2R;refGene_tx=XM_003991086.5
C1 194760833 194760908
refGene_gn=PTH2R;refGene_tx=XM_023259649.1
C1 194786686 194786789
refGene_gn=PTH2R;refGene_tx=XM_003991086.5;refGene_gn=PTH2R;refGene_tx
=XM_023259649.1
C1 194795232 194795343
refGene_gn=PTH2R;refGene_tx=XM_003991086.5;refGene_gn=PTH2R;refGene_tx
=XM_023259649.1

C1 194795451 194795573
refGene_gn=PTH2R;refGene_tx=XM_003991086.5;refGene_gn=PTH2R;refGene_tx=XM_023259649.1
C1 194803931 194804029
refGene_gn=PTH2R;refGene_tx=XM_003991086.5;refGene_gn=PTH2R;refGene_tx=XM_023259649.1
C1 194807902 194808092
refGene_gn=PTH2R;refGene_tx=XM_003991086.5;refGene_gn=PTH2R;refGene_tx=XM_023259649.1
C1 194809278 194809432
refGene_gn=PTH2R;refGene_tx=XM_003991086.5;refGene_gn=PTH2R;refGene_tx=XM_023259649.1
C1 194818015 194818076
refGene_gn=PTH2R;refGene_tx=XM_003991086.5;refGene_gn=PTH2R;refGene_tx=XM_023259649.1
C1 194826121 194826188
refGene_gn=PTH2R;refGene_tx=XM_003991086.5;refGene_gn=PTH2R;refGene_tx=XM_023259649.1
C1 194835283 194835378
refGene_gn=PTH2R;refGene_tx=XM_003991086.5;refGene_gn=PTH2R;refGene_tx=XM_023259649.1
C1 194843319 194843458
refGene_gn=PTH2R;refGene_tx=XM_003991086.5;refGene_gn=PTH2R;refGene_tx=XM_023259649.1
C1 194844939 194844981
refGene_gn=PTH2R;refGene_tx=XM_003991086.5;refGene_gn=PTH2R;refGene_tx=XM_023259649.1
C1 194846033 194846429
refGene_gn=PTH2R;refGene_tx=XM_003991086.5;refGene_gn=PTH2R;refGene_tx=XM_023259649.1
C1 195271863 195271872
refGene_gn=LOC101094779;refGene_tx=XM_003991087.3
C1 195271963 195272206
refGene_gn=LOC101094779;refGene_tx=XM_003991087.3
C1 195274063 195274336
refGene_gn=LOC101094779;refGene_tx=XM_003991087.3
C1 195677312 195677574
refGene_gn=MAP2;refGene_tx=XM_011285517.3;refGene_gn=MAP2;refGene_tx=XM_019838574.2;refGene_gn=MAP2;refGene_tx=XM_019838575.2;refGene_gn=MAP2;refGene_tx=XM_019838576.2;refGene_gn=MAP2;refGene_tx=XM_019838577.2;refGene_gn=MAP2;refGene_tx=XM_019838578.2;refGene_gn=MAP2;refGene_tx=XM_019838579.1;refGene_gn=MAP2;refGene_tx=XM_019838580.1;refGene_gn=MAP2;refGene_tx=XM_019838581.1;refGene_gn=MAP2;refGene_tx=XM_019838582.2;refGene_gn=MAP2;refGene_tx=XM_019838583.2;refGene_gn=MAP2;refGene_tx=XM_019838584.1;refGene_gn=MAP2;refGene_tx=XM_019838585.2;refGene_gn=MAP2;refGene_tx=XM_019838586.2;refGene_gn=MAP2;refGene_tx=XM_019838587.2;refGene_gn=MAP2;refGene_tx=XM_019838588.2;refGene_gn=MAP2;refGene_tx=XM_019838589.2;refGene_gn=MAP2;refGene_tx=XM_019838590.2;refGene_gn=MAP2;refGene_tx=XM_019838591.2;refGene_gn=MAP2;refGene_tx=XM_019838592.2;refGene_gn=MAP2;refGene_tx=XM_019838594.2;refGene_gn=MAP2;refGene_tx=X

M_019838595.2;refGene_gn=MAP2;refGene_tx=XM_019838596.2;refGene_gn=MAP2;refGene_tx=XM_019838597.2;refGene_gn=MAP2;refGene_tx=XM_019838598.2;refGene_gn=MAP2;refGene_tx=XM_019838599.2;refGene_gn=MAP2;refGene_tx=XM_019838600.2;refGene_gn=MAP2;refGene_tx=XM_019838601.2;refGene_gn=MAP2;refGene_tx=XM_023259651.1
C1 195701017 195701131
refGene_gn=MAP2;refGene_tx=XM_011285517.3;refGene_gn=MAP2;refGene_tx=XM_019838574.2;refGene_gn=MAP2;refGene_tx=XM_019838575.2;refGene_gn=MAP2;refGene_tx=XM_019838576.2;refGene_gn=MAP2;refGene_tx=XM_019838577.2;refGene_gn=MAP2;refGene_tx=XM_019838578.2;refGene_gn=MAP2;refGene_tx=XM_019838579.1;refGene_gn=MAP2;refGene_tx=XM_019838580.1;refGene_gn=MAP2;refGene_tx=XM_019838581.1;refGene_gn=MAP2;refGene_tx=XM_019838582.2;refGene_gn=MAP2;refGene_tx=XM_019838583.2;refGene_gn=MAP2;refGene_tx=XM_019838584.1;refGene_gn=MAP2;refGene_tx=XM_019838585.2;refGene_gn=MAP2;refGene_tx=XM_019838586.2;refGene_gn=MAP2;refGene_tx=XM_019838587.2;refGene_gn=MAP2;refGene_tx=XM_019838588.2;refGene_gn=MAP2;refGene_tx=XM_019838589.2;refGene_gn=MAP2;refGene_tx=XM_019838590.2;refGene_gn=MAP2;refGene_tx=XM_019838591.2;refGene_gn=MAP2;refGene_tx=XM_019838592.2;refGene_gn=MAP2;refGene_tx=XM_019838594.2;refGene_gn=MAP2;refGene_tx=XM_019838595.2;refGene_gn=MAP2;refGene_tx=XM_019838596.2;refGene_gn=MAP2;refGene_tx=XM_019838597.2;refGene_gn=MAP2;refGene_tx=XM_019838598.2;refGene_gn=MAP2;refGene_tx=XM_019838599.2;refGene_gn=MAP2;refGene_tx=XM_019838600.2;refGene_gn=MAP2;refGene_tx=XM_019838601.2;refGene_gn=MAP2;refGene_tx=XM_023259651.1
C1 195703382 195703460
refGene_gn=MAP2;refGene_tx=XM_011285517.3;refGene_gn=MAP2;refGene_tx=XM_019838574.2;refGene_gn=MAP2;refGene_tx=XM_019838575.2;refGene_gn=MAP2;refGene_tx=XM_019838576.2;refGene_gn=MAP2;refGene_tx=XM_019838577.2;refGene_gn=MAP2;refGene_tx=XM_019838578.2;refGene_gn=MAP2;refGene_tx=XM_019838579.1;refGene_gn=MAP2;refGene_tx=XM_019838580.1;refGene_gn=MAP2;refGene_tx=XM_019838581.1;refGene_gn=MAP2;refGene_tx=XM_019838582.2;refGene_gn=MAP2;refGene_tx=XM_019838583.2;refGene_gn=MAP2;refGene_tx=XM_019838584.1;refGene_gn=MAP2;refGene_tx=XM_019838585.2;refGene_gn=MAP2;refGene_tx=XM_019838586.2;refGene_gn=MAP2;refGene_tx=XM_019838587.2;refGene_gn=MAP2;refGene_tx=XM_019838588.2;refGene_gn=MAP2;refGene_tx=XM_019838589.2;refGene_gn=MAP2;refGene_tx=XM_019838590.2;refGene_gn=MAP2;refGene_tx=XM_019838591.2;refGene_gn=MAP2;refGene_tx=XM_019838592.2;refGene_gn=MAP2;refGene_tx=XM_019838594.2;refGene_gn=MAP2;refGene_tx=XM_019838595.2;refGene_gn=MAP2;refGene_tx=XM_019838596.2;refGene_gn=MAP2;refGene_tx=XM_019838597.2;refGene_gn=MAP2;refGene_tx=XM_019838598.2;refGene_gn=MAP2;refGene_tx=XM_019838599.2;refGene_gn=MAP2;refGene_tx=XM_019838600.2;refGene_gn=MAP2;refGene_tx=XM_019838601.2;refGene_gn=MAP2;refGene_tx=XM_023259651.1
C1 195712990 195713236
refGene_gn=MAP2;refGene_tx=XM_019838574.2;refGene_gn=MAP2;refGene_tx=XM_019838575.2;refGene_gn=MAP2;refGene_tx=XM_019838576.2;refGene_gn=MAP2;refGene_tx=XM_019838577.2;refGene_gn=MAP2;refGene_tx=XM_019838578.2;refGene_gn=MAP2;refGene_tx=XM_019838579.1;refGene_gn=MAP2;refGene_tx=XM_019838580.1;refGene_gn=MAP2;refGene_tx=XM_019838581.1;refGene_gn=MAP2;refGene_tx=XM_019838582.2;refGene_gn=MAP2;refGene_tx=XM_019838583.2;

refGene_gn=MAP2;refGene_tx=XM_019838584.1;refGene_gn=MAP2;refGene_tx=X
M_019838585.2;refGene_gn=MAP2;refGene_tx=XM_019838586.2;refGene_gn=MAP
2;refGene_tx=XM_019838587.2;refGene_gn=MAP2;refGene_tx=XM_019838588.2;
refGene_gn=MAP2;refGene_tx=XM_019838589.2;refGene_gn=MAP2;refGene_tx=X
M_019838590.2;refGene_gn=MAP2;refGene_tx=XM_019838591.2;refGene_gn=MAP
2;refGene_tx=XM_019838592.2;refGene_gn=MAP2;refGene_tx=XM_019838593.2
C1 195715024 195718783
refGene_gn=MAP2;refGene_tx=XM_011285517.3;refGene_gn=MAP2;refGene_tx=X
M_019838574.2;refGene_gn=MAP2;refGene_tx=XM_019838575.2;refGene_gn=MAP
2;refGene_tx=XM_019838576.2;refGene_gn=MAP2;refGene_tx=XM_019838577.2;
refGene_gn=MAP2;refGene_tx=XM_019838578.2;refGene_gn=MAP2;refGene_tx=X
M_019838579.1;refGene_gn=MAP2;refGene_tx=XM_019838580.1;refGene_gn=MAP
2;refGene_tx=XM_019838581.1;refGene_gn=MAP2;refGene_tx=XM_019838582.2;
refGene_gn=MAP2;refGene_tx=XM_019838583.2;refGene_gn=MAP2;refGene_tx=X
M_019838584.1;refGene_gn=MAP2;refGene_tx=XM_019838585.2;refGene_gn=MAP
2;refGene_tx=XM_019838586.2;refGene_gn=MAP2;refGene_tx=XM_019838587.2;
refGene_gn=MAP2;refGene_tx=XM_019838588.2;refGene_gn=MAP2;refGene_tx=X
M_019838589.2;refGene_gn=MAP2;refGene_tx=XM_019838590.2;refGene_gn=MAP
2;refGene_tx=XM_019838591.2;refGene_gn=MAP2;refGene_tx=XM_019838592.2;
refGene_gn=MAP2;refGene_tx=XM_019838593.2;refGene_gn=MAP2;refGene_tx=X
M_019838594.2;refGene_gn=MAP2;refGene_tx=XM_019838595.2;refGene_gn=MAP
2;refGene_tx=XM_019838596.2;refGene_gn=MAP2;refGene_tx=XM_019838597.2;
refGene_gn=MAP2;refGene_tx=XM_019838598.2;refGene_gn=MAP2;refGene_tx=X
M_023259651.1
C1 195718981 195719188
refGene_gn=MAP2;refGene_tx=XM_011285517.3;refGene_gn=MAP2;refGene_tx=X
M_019838574.2;refGene_gn=MAP2;refGene_tx=XM_019838575.2;refGene_gn=MAP
2;refGene_tx=XM_019838576.2;refGene_gn=MAP2;refGene_tx=XM_019838577.2;
refGene_gn=MAP2;refGene_tx=XM_019838578.2;refGene_gn=MAP2;refGene_tx=X
M_019838579.1;refGene_gn=MAP2;refGene_tx=XM_019838580.1;refGene_gn=MAP
2;refGene_tx=XM_019838581.1;refGene_gn=MAP2;refGene_tx=XM_019838582.2;
refGene_gn=MAP2;refGene_tx=XM_019838583.2;refGene_gn=MAP2;refGene_tx=X
M_019838584.1;refGene_gn=MAP2;refGene_tx=XM_019838585.2;refGene_gn=MAP
2;refGene_tx=XM_019838586.2;refGene_gn=MAP2;refGene_tx=XM_019838587.2;
refGene_gn=MAP2;refGene_tx=XM_019838588.2;refGene_gn=MAP2;refGene_tx=X
M_019838589.2;refGene_gn=MAP2;refGene_tx=XM_019838590.2;refGene_gn=MAP
2;refGene_tx=XM_019838591.2;refGene_gn=MAP2;refGene_tx=XM_019838592.2;
refGene_gn=MAP2;refGene_tx=XM_019838593.2;refGene_gn=MAP2;refGene_tx=X
M_019838594.2;refGene_gn=MAP2;refGene_tx=XM_019838595.2;refGene_gn=MAP
2;refGene_tx=XM_019838596.2;refGene_gn=MAP2;refGene_tx=XM_019838597.2;
refGene_gn=MAP2;refGene_tx=XM_019838598.2;refGene_gn=MAP2;refGene_tx=X
M_023259651.1
C1 195719362 195719497
refGene_gn=MAP2;refGene_tx=XM_011285517.3;refGene_gn=MAP2;refGene_tx=X
M_019838574.2;refGene_gn=MAP2;refGene_tx=XM_019838575.2;refGene_gn=MAP
2;refGene_tx=XM_019838576.2;refGene_gn=MAP2;refGene_tx=XM_019838577.2;
refGene_gn=MAP2;refGene_tx=XM_019838578.2;refGene_gn=MAP2;refGene_tx=X
M_019838579.1;refGene_gn=MAP2;refGene_tx=XM_019838580.1;refGene_gn=MAP
2;refGene_tx=XM_019838581.1;refGene_gn=MAP2;refGene_tx=XM_019838582.2;
refGene_gn=MAP2;refGene_tx=XM_019838583.2;refGene_gn=MAP2;refGene_tx=X

M_019838584.1;refGene_gn=MAP2;refGene_tx=XM_019838585.2;refGene_gn=MAP2;refGene_tx=XM_019838586.2;refGene_gn=MAP2;refGene_tx=XM_019838587.2;refGene_gn=MAP2;refGene_tx=XM_019838588.2;refGene_gn=MAP2;refGene_tx=XM_019838589.2;refGene_gn=MAP2;refGene_tx=XM_019838590.2;refGene_gn=MAP2;refGene_tx=XM_019838591.2;refGene_gn=MAP2;refGene_tx=XM_019838592.2;refGene_gn=MAP2;refGene_tx=XM_019838593.2;refGene_gn=MAP2;refGene_tx=XM_019838594.2;refGene_gn=MAP2;refGene_tx=XM_019838595.2;refGene_gn=MAP2;refGene_tx=XM_019838596.2;refGene_gn=MAP2;refGene_tx=XM_019838597.2;refGene_gn=MAP2;refGene_tx=XM_019838598.2;refGene_gn=MAP2;refGene_tx=XM_023259651.1

C1 195722348 195722410
refGene_gn=MAP2;refGene_tx=XM_011285517.3;refGene_gn=MAP2;refGene_tx=XM_019838574.2;refGene_gn=MAP2;refGene_tx=XM_019838575.2;refGene_gn=MAP2;refGene_tx=XM_019838576.2;refGene_gn=MAP2;refGene_tx=XM_019838577.2;refGene_gn=MAP2;refGene_tx=XM_019838578.2;refGene_gn=MAP2;refGene_tx=XM_019838579.1;refGene_gn=MAP2;refGene_tx=XM_019838580.1;refGene_gn=MAP2;refGene_tx=XM_019838581.1;refGene_gn=MAP2;refGene_tx=XM_019838582.2;refGene_gn=MAP2;refGene_tx=XM_019838583.2;refGene_gn=MAP2;refGene_tx=XM_019838584.1;refGene_gn=MAP2;refGene_tx=XM_019838585.2;refGene_gn=MAP2;refGene_tx=XM_019838586.2;refGene_gn=MAP2;refGene_tx=XM_019838587.2;refGene_gn=MAP2;refGene_tx=XM_019838588.2;refGene_gn=MAP2;refGene_tx=XM_019838589.2;refGene_gn=MAP2;refGene_tx=XM_019838590.2;refGene_gn=MAP2;refGene_tx=XM_019838591.2;refGene_gn=MAP2;refGene_tx=XM_019838592.2;refGene_gn=MAP2;refGene_tx=XM_019838593.2;refGene_gn=MAP2;refGene_tx=XM_019838594.2;refGene_gn=MAP2;refGene_tx=XM_019838595.2;refGene_gn=MAP2;refGene_tx=XM_019838596.2;refGene_gn=MAP2;refGene_tx=XM_019838597.2;refGene_gn=MAP2;refGene_tx=XM_019838598.2;refGene_gn=MAP2;refGene_tx=XM_019838599.2;refGene_gn=MAP2;refGene_tx=XM_019838600.2;refGene_gn=MAP2;refGene_tx=XM_019838601.2;refGene_gn=MAP2;refGene_tx=XM_023259651.1

C1 195726305 195726476
refGene_gn=MAP2;refGene_tx=XM_019838574.2;refGene_gn=MAP2;refGene_tx=XM_019838575.2;refGene_gn=MAP2;refGene_tx=XM_019838576.2;refGene_gn=MAP2;refGene_tx=XM_019838577.2;refGene_gn=MAP2;refGene_tx=XM_019838578.2;refGene_gn=MAP2;refGene_tx=XM_019838579.1;refGene_gn=MAP2;refGene_tx=XM_019838580.1;refGene_gn=MAP2;refGene_tx=XM_019838581.1;refGene_gn=MAP2;refGene_tx=XM_019838582.2;refGene_gn=MAP2;refGene_tx=XM_019838583.2;refGene_gn=MAP2;refGene_tx=XM_019838584.1;refGene_gn=MAP2;refGene_tx=XM_019838585.2;refGene_gn=MAP2;refGene_tx=XM_019838586.2;refGene_gn=MAP2;refGene_tx=XM_019838587.2;refGene_gn=MAP2;refGene_tx=XM_019838588.2;refGene_gn=MAP2;refGene_tx=XM_019838589.2;refGene_gn=MAP2;refGene_tx=XM_019838590.2;refGene_gn=MAP2;refGene_tx=XM_019838593.2;refGene_gn=MAP2;refGene_tx=XM_019838594.2;refGene_gn=MAP2;refGene_tx=XM_019838595.2;refGene_gn=MAP2;refGene_tx=XM_019838596.2;refGene_gn=MAP2;refGene_tx=XM_019838597.2;refGene_gn=MAP2;refGene_tx=XM_019838599.2

C1 195727452 195727600
refGene_gn=MAP2;refGene_tx=XM_011285517.3;refGene_gn=MAP2;refGene_tx=XM_019838574.2;refGene_gn=MAP2;refGene_tx=XM_019838575.2;refGene_gn=MAP2;refGene_tx=XM_019838576.2;refGene_gn=MAP2;refGene_tx=XM_019838577.2;refGene_gn=MAP2;refGene_tx=XM_019838578.2;refGene_gn=MAP2;refGene_tx=XM_019838579.1;refGene_gn=MAP2;refGene_tx=XM_019838580.1;refGene_gn=MAP

2;refGene_tx=XM_019838581.1;refGene_gn=MAP2;refGene_tx=XM_019838582.2;
refGene_gn=MAP2;refGene_tx=XM_019838583.2;refGene_gn=MAP2;refGene_tx=X
M_019838584.1;refGene_gn=MAP2;refGene_tx=XM_019838585.2;refGene_gn=MAP
2;refGene_tx=XM_019838586.2;refGene_gn=MAP2;refGene_tx=XM_019838587.2;
refGene_gn=MAP2;refGene_tx=XM_019838588.2;refGene_gn=MAP2;refGene_tx=X
M_019838589.2;refGene_gn=MAP2;refGene_tx=XM_019838590.2;refGene_gn=MAP
2;refGene_tx=XM_019838591.2;refGene_gn=MAP2;refGene_tx=XM_019838592.2;
refGene_gn=MAP2;refGene_tx=XM_019838593.2;refGene_gn=MAP2;refGene_tx=X
M_019838594.2;refGene_gn=MAP2;refGene_tx=XM_019838595.2;refGene_gn=MAP
2;refGene_tx=XM_019838596.2;refGene_gn=MAP2;refGene_tx=XM_019838597.2;
refGene_gn=MAP2;refGene_tx=XM_019838598.2;refGene_gn=MAP2;refGene_tx=X
M_019838599.2;refGene_gn=MAP2;refGene_tx=XM_019838600.2;refGene_gn=MAP
2;refGene_tx=XM_019838601.2;refGene_gn=MAP2;refGene_tx=XM_023259651.1
C1 195732548 195732889
refGene_gn=MAP2;refGene_tx=XM_011285517.3;refGene_gn=MAP2;refGene_tx=X
M_019838574.2;refGene_gn=MAP2;refGene_tx=XM_019838575.2;refGene_gn=MAP
2;refGene_tx=XM_019838576.2;refGene_gn=MAP2;refGene_tx=XM_019838577.2;
refGene_gn=MAP2;refGene_tx=XM_019838578.2;refGene_gn=MAP2;refGene_tx=X
M_019838579.1;refGene_gn=MAP2;refGene_tx=XM_019838580.1;refGene_gn=MAP
2;refGene_tx=XM_019838581.1;refGene_gn=MAP2;refGene_tx=XM_019838582.2;
refGene_gn=MAP2;refGene_tx=XM_019838583.2;refGene_gn=MAP2;refGene_tx=X
M_019838584.1;refGene_gn=MAP2;refGene_tx=XM_019838585.2;refGene_gn=MAP
2;refGene_tx=XM_019838586.2;refGene_gn=MAP2;refGene_tx=XM_019838587.2;
refGene_gn=MAP2;refGene_tx=XM_019838588.2;refGene_gn=MAP2;refGene_tx=X
M_019838589.2;refGene_gn=MAP2;refGene_tx=XM_019838590.2;refGene_gn=MAP
2;refGene_tx=XM_019838591.2;refGene_gn=MAP2;refGene_tx=XM_019838592.2;
refGene_gn=MAP2;refGene_tx=XM_019838593.2;refGene_gn=MAP2;refGene_tx=X
M_019838594.2;refGene_gn=MAP2;refGene_tx=XM_019838595.2;refGene_gn=MAP
2;refGene_tx=XM_019838596.2;refGene_gn=MAP2;refGene_tx=XM_019838597.2;
refGene_gn=MAP2;refGene_tx=XM_019838598.2;refGene_gn=MAP2;refGene_tx=X
M_019838599.2;refGene_gn=MAP2;refGene_tx=XM_019838600.2;refGene_gn=MAP
2;refGene_tx=XM_019838601.2;refGene_gn=MAP2;refGene_tx=XM_023259651.1
C1 195744053 195744146
refGene_gn=MAP2;refGene_tx=XM_019838574.2;refGene_gn=MAP2;refGene_tx=X
M_019838575.2;refGene_gn=MAP2;refGene_tx=XM_019838576.2;refGene_gn=MAP
2;refGene_tx=XM_019838577.2;refGene_gn=MAP2;refGene_tx=XM_019838578.2;
refGene_gn=MAP2;refGene_tx=XM_019838579.1;refGene_gn=MAP2;refGene_tx=X
M_019838580.1;refGene_gn=MAP2;refGene_tx=XM_019838581.1;refGene_gn=MAP
2;refGene_tx=XM_019838582.2;refGene_gn=MAP2;refGene_tx=XM_019838583.2;
refGene_gn=MAP2;refGene_tx=XM_019838584.1;refGene_gn=MAP2;refGene_tx=X
M_019838585.2;refGene_gn=MAP2;refGene_tx=XM_019838586.2;refGene_gn=MAP
2;refGene_tx=XM_019838587.2;refGene_gn=MAP2;refGene_tx=XM_019838588.2;
refGene_gn=MAP2;refGene_tx=XM_019838589.2;refGene_gn=MAP2;refGene_tx=X
M_019838591.2;refGene_gn=MAP2;refGene_tx=XM_019838592.2;refGene_gn=MAP
2;refGene_tx=XM_019838593.2;refGene_gn=MAP2;refGene_tx=XM_019838594.2;
refGene_gn=MAP2;refGene_tx=XM_019838595.2;refGene_gn=MAP2;refGene_tx=X
M_019838596.2;refGene_gn=MAP2;refGene_tx=XM_019838597.2;refGene_gn=MAP
2;refGene_tx=XM_019838599.2;refGene_gn=MAP2;refGene_tx=XM_019838600.2;
refGene_gn=MAP2;refGene_tx=XM_023259651.1
C1 195746113 195746195

refGene_gn=MAP2;refGene_tx=XM_011285517.3;refGene_gn=MAP2;refGene_tx=X
M_019838574.2;refGene_gn=MAP2;refGene_tx=XM_019838575.2;refGene_gn=MAP
2;refGene_tx=XM_019838576.2;refGene_gn=MAP2;refGene_tx=XM_019838577.2;
refGene_gn=MAP2;refGene_tx=XM_019838578.2;refGene_gn=MAP2;refGene_tx=X
M_019838579.1;refGene_gn=MAP2;refGene_tx=XM_019838580.1;refGene_gn=MAP
2;refGene_tx=XM_019838581.1;refGene_gn=MAP2;refGene_tx=XM_019838582.2;
refGene_gn=MAP2;refGene_tx=XM_019838583.2;refGene_gn=MAP2;refGene_tx=X
M_019838584.1;refGene_gn=MAP2;refGene_tx=XM_019838585.2;refGene_gn=MAP
2;refGene_tx=XM_019838586.2;refGene_gn=MAP2;refGene_tx=XM_019838587.2;
refGene_gn=MAP2;refGene_tx=XM_019838588.2;refGene_gn=MAP2;refGene_tx=X
M_019838589.2;refGene_gn=MAP2;refGene_tx=XM_019838590.2;refGene_gn=MAP
2;refGene_tx=XM_019838591.2;refGene_gn=MAP2;refGene_tx=XM_019838592.2;
refGene_gn=MAP2;refGene_tx=XM_019838593.2;refGene_gn=MAP2;refGene_tx=X
M_019838594.2;refGene_gn=MAP2;refGene_tx=XM_019838595.2;refGene_gn=MAP
2;refGene_tx=XM_019838596.2;refGene_gn=MAP2;refGene_tx=XM_019838597.2;
refGene_gn=MAP2;refGene_tx=XM_019838598.2;refGene_gn=MAP2;refGene_tx=X
M_019838599.2;refGene_gn=MAP2;refGene_tx=XM_019838600.2;refGene_gn=MAP
2;refGene_tx=XM_019838601.2;refGene_gn=MAP2;refGene_tx=XM_023259651.1
C1 195750328 195750441
refGene_gn=MAP2;refGene_tx=XM_011285517.3;refGene_gn=MAP2;refGene_tx=X
M_019838574.2;refGene_gn=MAP2;refGene_tx=XM_019838575.2;refGene_gn=MAP
2;refGene_tx=XM_019838576.2;refGene_gn=MAP2;refGene_tx=XM_019838577.2;
refGene_gn=MAP2;refGene_tx=XM_019838578.2;refGene_gn=MAP2;refGene_tx=X
M_019838579.1;refGene_gn=MAP2;refGene_tx=XM_019838580.1;refGene_gn=MAP
2;refGene_tx=XM_019838581.1;refGene_gn=MAP2;refGene_tx=XM_019838582.2;
refGene_gn=MAP2;refGene_tx=XM_019838583.2;refGene_gn=MAP2;refGene_tx=X
M_019838584.1;refGene_gn=MAP2;refGene_tx=XM_019838585.2;refGene_gn=MAP
2;refGene_tx=XM_019838586.2;refGene_gn=MAP2;refGene_tx=XM_019838587.2;
refGene_gn=MAP2;refGene_tx=XM_019838588.2;refGene_gn=MAP2;refGene_tx=X
M_019838589.2;refGene_gn=MAP2;refGene_tx=XM_019838590.2;refGene_gn=MAP
2;refGene_tx=XM_019838591.2;refGene_gn=MAP2;refGene_tx=XM_019838592.2;
refGene_gn=MAP2;refGene_tx=XM_019838593.2;refGene_gn=MAP2;refGene_tx=X
M_019838594.2;refGene_gn=MAP2;refGene_tx=XM_019838595.2;refGene_gn=MAP
2;refGene_tx=XM_019838596.2;refGene_gn=MAP2;refGene_tx=XM_019838597.2;
refGene_gn=MAP2;refGene_tx=XM_019838598.2;refGene_gn=MAP2;refGene_tx=X
M_019838599.2;refGene_gn=MAP2;refGene_tx=XM_019838600.2;refGene_gn=MAP
2;refGene_tx=XM_019838601.2;refGene_gn=MAP2;refGene_tx=XM_023259651.1
C1 195750668 195750884
refGene_gn=MAP2;refGene_tx=XM_011285517.3;refGene_gn=MAP2;refGene_tx=X
M_019838574.2;refGene_gn=MAP2;refGene_tx=XM_019838575.2;refGene_gn=MAP
2;refGene_tx=XM_019838576.2;refGene_gn=MAP2;refGene_tx=XM_019838577.2;
refGene_gn=MAP2;refGene_tx=XM_019838578.2;refGene_gn=MAP2;refGene_tx=X
M_019838579.1;refGene_gn=MAP2;refGene_tx=XM_019838580.1;refGene_gn=MAP
2;refGene_tx=XM_019838581.1;refGene_gn=MAP2;refGene_tx=XM_019838582.2;
refGene_gn=MAP2;refGene_tx=XM_019838583.2;refGene_gn=MAP2;refGene_tx=X
M_019838584.1;refGene_gn=MAP2;refGene_tx=XM_019838585.2;refGene_gn=MAP
2;refGene_tx=XM_019838586.2;refGene_gn=MAP2;refGene_tx=XM_019838587.2;
refGene_gn=MAP2;refGene_tx=XM_019838588.2;refGene_gn=MAP2;refGene_tx=X
M_019838589.2;refGene_gn=MAP2;refGene_tx=XM_019838590.2;refGene_gn=MAP
2;refGene_tx=XM_019838591.2;refGene_gn=MAP2;refGene_tx=XM_019838592.2;

refGene_gn=MAP2;refGene_tx=XM_019838593.2;refGene_gn=MAP2;refGene_tx=XM_019838594.2;refGene_gn=MAP2;refGene_tx=XM_019838595.2;refGene_gn=MAP2;refGene_tx=XM_019838596.2;refGene_gn=MAP2;refGene_tx=XM_019838597.2;refGene_gn=MAP2;refGene_tx=XM_019838598.2;refGene_gn=MAP2;refGene_tx=XM_019838599.2;refGene_gn=MAP2;refGene_tx=XM_019838600.2;refGene_gn=MAP2;refGene_tx=XM_019838601.2;refGene_gn=MAP2;refGene_tx=XM_023259651.1
C1 195786649 195786741
refGene_gn=UNC80;refGene_tx=XM_011285519.3;refGene_gn=UNC80;refGene_tx=XM_011285520.3;refGene_gn=UNC80;refGene_tx=XM_023259652.1
C1 195787706 195787755
refGene_gn=UNC80;refGene_tx=XM_011285519.3;refGene_gn=UNC80;refGene_tx=XM_011285520.3;refGene_gn=UNC80;refGene_tx=XM_023259652.1
C1 195790616 195790773
refGene_gn=UNC80;refGene_tx=XM_011285519.3;refGene_gn=UNC80;refGene_tx=XM_011285520.3;refGene_gn=UNC80;refGene_tx=XM_023259652.1
C1 195791900 195792202
refGene_gn=UNC80;refGene_tx=XM_011285519.3;refGene_gn=UNC80;refGene_tx=XM_011285520.3;refGene_gn=UNC80;refGene_tx=XM_023259652.1
C1 195801478 195801602
refGene_gn=UNC80;refGene_tx=XM_011285519.3;refGene_gn=UNC80;refGene_tx=XM_011285520.3;refGene_gn=UNC80;refGene_tx=XM_023259652.1
C1 195805459 195805533
refGene_gn=UNC80;refGene_tx=XM_011285519.3;refGene_gn=UNC80;refGene_tx=XM_011285520.3;refGene_gn=UNC80;refGene_tx=XM_023259652.1
C1 195809431 195809571
refGene_gn=UNC80;refGene_tx=XM_011285519.3;refGene_gn=UNC80;refGene_tx=XM_011285520.3;refGene_gn=UNC80;refGene_tx=XM_023259652.1
C1 195830354 195830616
refGene_gn=UNC80;refGene_tx=XM_011285519.3;refGene_gn=UNC80;refGene_tx=XM_011285520.3;refGene_gn=UNC80;refGene_tx=XM_023259652.1
C1 195831745 195831880
refGene_gn=UNC80;refGene_tx=XM_011285519.3;refGene_gn=UNC80;refGene_tx=XM_011285520.3;refGene_gn=UNC80;refGene_tx=XM_023259652.1
C1 195834052 195834269
refGene_gn=UNC80;refGene_tx=XM_011285519.3;refGene_gn=UNC80;refGene_tx=XM_011285520.3;refGene_gn=UNC80;refGene_tx=XM_023259652.1
C1 195834902 195835043
refGene_gn=UNC80;refGene_tx=XM_011285519.3;refGene_gn=UNC80;refGene_tx=XM_011285520.3;refGene_gn=UNC80;refGene_tx=XM_023259652.1
C1 195836074 195836343
refGene_gn=UNC80;refGene_tx=XM_011285519.3;refGene_gn=UNC80;refGene_tx=XM_011285520.3;refGene_gn=UNC80;refGene_tx=XM_023259652.1
C1 195837427 195837805
refGene_gn=UNC80;refGene_tx=XM_011285519.3;refGene_gn=UNC80;refGene_tx=XM_011285520.3;refGene_gn=UNC80;refGene_tx=XM_023259652.1
C1 195843282 195843429
refGene_gn=UNC80;refGene_tx=XM_011285519.3;refGene_gn=UNC80;refGene_tx=XM_011285520.3;refGene_gn=UNC80;refGene_tx=XM_023259652.1
C1 195847305 195847453
refGene_gn=UNC80;refGene_tx=XM_011285519.3;refGene_gn=UNC80;refGene_tx

=XM_011285520.3;refGene_gn=UNC80;refGene_tx=XM_023259652.1
C1 195849513 195849662
refGene_gn=UNC80;refGene_tx=XM_011285519.3;refGene_gn=UNC80;refGene_tx
=XM_011285520.3;refGene_gn=UNC80;refGene_tx=XM_023259652.1
C1 195851814 195851981
refGene_gn=UNC80;refGene_tx=XM_011285519.3;refGene_gn=UNC80;refGene_tx
=XM_011285520.3;refGene_gn=UNC80;refGene_tx=XM_023259652.1
C1 195852733 195852832
refGene_gn=UNC80;refGene_tx=XM_011285519.3;refGene_gn=UNC80;refGene_tx
=XM_011285520.3;refGene_gn=UNC80;refGene_tx=XM_023259652.1
C1 195856094 195856309
refGene_gn=UNC80;refGene_tx=XM_011285519.3;refGene_gn=UNC80;refGene_tx
=XM_011285520.3;refGene_gn=UNC80;refGene_tx=XM_023259652.1
C1 195857699 195857806
refGene_gn=UNC80;refGene_tx=XM_011285519.3;refGene_gn=UNC80;refGene_tx
=XM_011285520.3;refGene_gn=UNC80;refGene_tx=XM_023259652.1
C1 195859644 195859741
refGene_gn=UNC80;refGene_tx=XM_011285519.3;refGene_gn=UNC80;refGene_tx
=XM_011285520.3;refGene_gn=UNC80;refGene_tx=XM_023259652.1
C1 195865632 195865805
refGene_gn=UNC80;refGene_tx=XM_011285519.3;refGene_gn=UNC80;refGene_tx
=XM_011285520.3;refGene_gn=UNC80;refGene_tx=XM_023259652.1
C1 195883542 195883755
refGene_gn=UNC80;refGene_tx=XM_011285519.3;refGene_gn=UNC80;refGene_tx
=XM_011285520.3;refGene_gn=UNC80;refGene_tx=XM_023259652.1
C1 195887064 195887200
refGene_gn=UNC80;refGene_tx=XM_011285519.3;refGene_gn=UNC80;refGene_tx
=XM_011285520.3;refGene_gn=UNC80;refGene_tx=XM_023259652.1
C1 195890280 195890414
refGene_gn=UNC80;refGene_tx=XM_011285519.3;refGene_gn=UNC80;refGene_tx
=XM_011285520.3;refGene_gn=UNC80;refGene_tx=XM_023259652.1
C1 195896938 195897104
refGene_gn=UNC80;refGene_tx=XM_011285519.3;refGene_gn=UNC80;refGene_tx
=XM_011285520.3;refGene_gn=UNC80;refGene_tx=XM_023259652.1
C1 195902177 195902381
refGene_gn=UNC80;refGene_tx=XM_023259652.1
C1 195904338 195904439
refGene_gn=UNC80;refGene_tx=XM_011285519.3;refGene_gn=UNC80;refGene_tx
=XM_011285520.3;refGene_gn=UNC80;refGene_tx=XM_023259652.1
C1 195909250 195909451
refGene_gn=UNC80;refGene_tx=XM_011285519.3;refGene_gn=UNC80;refGene_tx
=XM_011285520.3;refGene_gn=UNC80;refGene_tx=XM_023259652.1
C1 195917446 195917554
refGene_gn=UNC80;refGene_tx=XM_011285519.3;refGene_gn=UNC80;refGene_tx
=XM_011285520.3;refGene_gn=UNC80;refGene_tx=XM_023259652.1
C1 195918760 195918899
refGene_gn=UNC80;refGene_tx=XM_011285519.3;refGene_gn=UNC80;refGene_tx
=XM_011285520.3;refGene_gn=UNC80;refGene_tx=XM_023259652.1
C1 195927988 195928170
refGene_gn=UNC80;refGene_tx=XM_011285519.3;refGene_gn=UNC80;refGene_tx

=XM_011285520.3;refGene_gn=UNC80;refGene_tx=XM_023259652.1
C1 195928678 195928810
refGene_gn=UNC80;refGene_tx=XM_011285519.3;refGene_gn=UNC80;refGene_tx
=XM_011285520.3;refGene_gn=UNC80;refGene_tx=XM_023259652.1
C1 195930812 195930999
refGene_gn=UNC80;refGene_tx=XM_011285519.3;refGene_gn=UNC80;refGene_tx
=XM_011285520.3;refGene_gn=UNC80;refGene_tx=XM_023259652.1
C1 195931617 195931749
refGene_gn=UNC80;refGene_tx=XM_011285519.3;refGene_gn=UNC80;refGene_tx
=XM_011285520.3;refGene_gn=UNC80;refGene_tx=XM_023259652.1
C1 195933442 195933586
refGene_gn=UNC80;refGene_tx=XM_011285519.3;refGene_gn=UNC80;refGene_tx
=XM_011285520.3;refGene_gn=UNC80;refGene_tx=XM_023259652.1
C1 195936627 195936728
refGene_gn=UNC80;refGene_tx=XM_011285519.3;refGene_gn=UNC80;refGene_tx
=XM_011285520.3;refGene_gn=UNC80;refGene_tx=XM_023259652.1
C1 195937753 195937840
refGene_gn=UNC80;refGene_tx=XM_011285519.3;refGene_gn=UNC80;refGene_tx
=XM_011285520.3;refGene_gn=UNC80;refGene_tx=XM_023259652.1
C1 195941685 195941869
refGene_gn=UNC80;refGene_tx=XM_011285519.3;refGene_gn=UNC80;refGene_tx
=XM_011285520.3;refGene_gn=UNC80;refGene_tx=XM_023259652.1
C1 195943811 195943906
refGene_gn=UNC80;refGene_tx=XM_011285519.3;refGene_gn=UNC80;refGene_tx
=XM_011285520.3;refGene_gn=UNC80;refGene_tx=XM_023259652.1
C1 195945556 195945646
refGene_gn=UNC80;refGene_tx=XM_011285519.3;refGene_gn=UNC80;refGene_tx
=XM_011285520.3;refGene_gn=UNC80;refGene_tx=XM_023259652.1
C1 195946369 195946471
refGene_gn=UNC80;refGene_tx=XM_011285519.3;refGene_gn=UNC80;refGene_tx
=XM_011285520.3;refGene_gn=UNC80;refGene_tx=XM_023259652.1
C1 195955172 195955353
refGene_gn=UNC80;refGene_tx=XM_011285519.3;refGene_gn=UNC80;refGene_tx
=XM_011285520.3;refGene_gn=UNC80;refGene_tx=XM_023259652.1
C1 195956680 195956949
refGene_gn=UNC80;refGene_tx=XM_011285519.3;refGene_gn=UNC80;refGene_tx
=XM_011285520.3;refGene_gn=UNC80;refGene_tx=XM_023259652.1
C1 195958378 195958513
refGene_gn=UNC80;refGene_tx=XM_011285519.3;refGene_gn=UNC80;refGene_tx
=XM_011285520.3;refGene_gn=UNC80;refGene_tx=XM_023259652.1
C1 195960034 195960173
refGene_gn=UNC80;refGene_tx=XM_011285519.3;refGene_gn=UNC80;refGene_tx
=XM_011285520.3;refGene_gn=UNC80;refGene_tx=XM_023259652.1
C1 195960817 195960914
refGene_gn=UNC80;refGene_tx=XM_011285519.3;refGene_gn=UNC80;refGene_tx
=XM_011285520.3;refGene_gn=UNC80;refGene_tx=XM_023259652.1
C1 195964811 195964982
refGene_gn=UNC80;refGene_tx=XM_011285519.3;refGene_gn=UNC80;refGene_tx
=XM_011285520.3;refGene_gn=UNC80;refGene_tx=XM_023259652.1
C1 195968561 195968654

refGene_gn=UNC80;refGene_tx=XM_011285519.3;refGene_gn=UNC80;refGene_tx=XM_011285520.3;refGene_gn=UNC80;refGene_tx=XM_023259652.1
C1 195971194 195971230
refGene_gn=UNC80;refGene_tx=XM_011285519.3;refGene_gn=UNC80;refGene_tx=XM_011285520.3;refGene_gn=UNC80;refGene_tx=XM_023259652.1
C1 195971591 195971810
refGene_gn=UNC80;refGene_tx=XM_011285519.3;refGene_gn=UNC80;refGene_tx=XM_011285520.3;refGene_gn=UNC80;refGene_tx=XM_023259652.1
C1 195983352 195983553
refGene_gn=UNC80;refGene_tx=XM_011285519.3;refGene_gn=UNC80;refGene_tx=XM_011285520.3;refGene_gn=UNC80;refGene_tx=XM_023259652.1
C1 195985714 195985838
refGene_gn=UNC80;refGene_tx=XM_011285519.3;refGene_gn=UNC80;refGene_tx=XM_011285520.3;refGene_gn=UNC80;refGene_tx=XM_023259652.1
C1 195986802 195986928
refGene_gn=UNC80;refGene_tx=XM_011285519.3;refGene_gn=UNC80;refGene_tx=XM_011285520.3;refGene_gn=UNC80;refGene_tx=XM_023259652.1
C1 195988165 195988289
refGene_gn=UNC80;refGene_tx=XM_011285519.3;refGene_gn=UNC80;refGene_tx=XM_011285520.3;refGene_gn=UNC80;refGene_tx=XM_023259652.1
C1 195989026 195989233
refGene_gn=UNC80;refGene_tx=XM_011285519.3;refGene_gn=UNC80;refGene_tx=XM_011285520.3;refGene_gn=UNC80;refGene_tx=XM_023259652.1
C1 195992207 195992392
refGene_gn=UNC80;refGene_tx=XM_011285519.3;refGene_gn=UNC80;refGene_tx=XM_011285520.3;refGene_gn=UNC80;refGene_tx=XM_023259652.1
C1 195993238 195993404
refGene_gn=UNC80;refGene_tx=XM_011285519.3;refGene_gn=UNC80;refGene_tx=XM_011285520.3;refGene_gn=UNC80;refGene_tx=XM_023259652.1
C1 195994901 195995081
refGene_gn=UNC80;refGene_tx=XM_011285519.3;refGene_gn=UNC80;refGene_tx=XM_011285520.3;refGene_gn=UNC80;refGene_tx=XM_023259652.1
C1 195997872 195998011
refGene_gn=UNC80;refGene_tx=XM_011285519.3;refGene_gn=UNC80;refGene_tx=XM_011285520.3;refGene_gn=UNC80;refGene_tx=XM_023259652.1
C1 196000693 196000750
refGene_gn=UNC80;refGene_tx=XM_011285519.3;refGene_gn=UNC80;refGene_tx=XM_023259652.1
C1 196007833 196007915
refGene_gn=UNC80;refGene_tx=XM_011285519.3;refGene_gn=UNC80;refGene_tx=XM_011285520.3;refGene_gn=UNC80;refGene_tx=XM_023259652.1
C1 196008802 196008914
refGene_gn=UNC80;refGene_tx=XM_011285519.3;refGene_gn=UNC80;refGene_tx=XM_011285520.3;refGene_gn=UNC80;refGene_tx=XM_023259652.1
C1 196009522 196009722
refGene_gn=UNC80;refGene_tx=XM_011285519.3;refGene_gn=UNC80;refGene_tx=XM_011285520.3;refGene_gn=UNC80;refGene_tx=XM_023259652.1
C1 196010580 196010847
refGene_gn=UNC80;refGene_tx=XM_011285519.3;refGene_gn=UNC80;refGene_tx=XM_011285520.3;refGene_gn=UNC80;refGene_tx=XM_023259652.1

C1 196015801 196015923
refGene_gn=RPE;refGene_tx=XM_003991092.4;refGene_gn=RPE;refGene_tx=XM_006935518.3;refGene_gn=RPE;refGene_tx=XM_006935519.3;refGene_gn=RPE;refGene_tx=XM_006935520.3;refGene_gn=RPE;refGene_tx=XM_019838606.1;refGene_gn=RPE;refGene_tx=XM_023259660.1
C1 196016582 196016614
refGene_gn=RPE;refGene_tx=XM_006935518.3;refGene_gn=RPE;refGene_tx=XM_006935519.3;refGene_gn=RPE;refGene_tx=XM_019838606.1;refGene_gn=RPE;refGene_tx=XM_023259660.1
C1 196022676 196022756
refGene_gn=RPE;refGene_tx=XM_003991092.4;refGene_gn=RPE;refGene_tx=XM_006935518.3;refGene_gn=RPE;refGene_tx=XM_006935520.3;refGene_gn=RPE;refGene_tx=XM_019838606.1
C1 196027161 196027301
refGene_gn=RPE;refGene_tx=XM_003991092.4;refGene_gn=RPE;refGene_tx=XM_006935518.3;refGene_gn=RPE;refGene_tx=XM_006935519.3;refGene_gn=RPE;refGene_tx=XM_006935520.3;refGene_gn=RPE;refGene_tx=XM_006935521.3;refGene_gn=RPE;refGene_tx=XM_006935522.3;refGene_gn=RPE;refGene_tx=XM_019838606.1;refGene_gn=RPE;refGene_tx=XM_023259660.1;refGene_gn=RPE;refGene_tx=XM_023259661.1
C1 196027909 196028044
refGene_gn=RPE;refGene_tx=XM_003991092.4;refGene_gn=RPE;refGene_tx=XM_006935518.3;refGene_gn=RPE;refGene_tx=XM_006935519.3;refGene_gn=RPE;refGene_tx=XM_006935520.3;refGene_gn=RPE;refGene_tx=XM_006935521.3;refGene_gn=RPE;refGene_tx=XM_006935522.3;refGene_gn=RPE;refGene_tx=XM_019838606.1;refGene_gn=RPE;refGene_tx=XM_023259660.1;refGene_gn=RPE;refGene_tx=XM_023259661.1
C1 196028740 196028827
refGene_gn=RPE;refGene_tx=XM_003991092.4;refGene_gn=RPE;refGene_tx=XM_006935518.3;refGene_gn=RPE;refGene_tx=XM_006935519.3;refGene_gn=RPE;refGene_tx=XM_006935520.3;refGene_gn=RPE;refGene_tx=XM_006935521.3;refGene_gn=RPE;refGene_tx=XM_006935522.3;refGene_gn=RPE;refGene_tx=XM_019838606.1;refGene_gn=RPE;refGene_tx=XM_023259660.1;refGene_gn=RPE;refGene_tx=XM_023259661.1
C1 196031031 196031154
refGene_gn=RPE;refGene_tx=XM_003991092.4;refGene_gn=RPE;refGene_tx=XM_006935518.3;refGene_gn=RPE;refGene_tx=XM_006935519.3;refGene_gn=RPE;refGene_tx=XM_006935520.3;refGene_gn=RPE;refGene_tx=XM_006935521.3;refGene_gn=RPE;refGene_tx=XM_006935522.3;refGene_gn=RPE;refGene_tx=XM_019838606.1;refGene_gn=RPE;refGene_tx=XM_023259660.1;refGene_gn=RPE;refGene_tx=XM_023259661.1
C1 196034203 196034434
refGene_gn=KANSL1L;refGene_tx=XM_019838602.2;refGene_gn=KANSL1L;refGene_tx=XM_023259653.1;refGene_gn=KANSL1L;refGene_tx=XM_023259654.1;refGene_gn=KANSL1L;refGene_tx=XM_023259655.1;refGene_gn=KANSL1L;refGene_tx=XM_023259656.1;refGene_gn=KANSL1L;refGene_tx=XM_023259657.1;refGene_gn=KANSL1L;refGene_tx=XM_023259658.1;refGene_gn=KANSL1L;refGene_tx=XM_023259659.1
C1 196035295 196035464
refGene_gn=KANSL1L;refGene_tx=XM_019838602.2;refGene_gn=KANSL1L;refGen

e_tx=XM_023259653.1;refGene_gn=KANSL1L;refGene_tx=XM_023259654.1;refGene_gn=KANSL1L;refGene_tx=XM_023259655.1;refGene_gn=KANSL1L;refGene_tx=XM_023259656.1;refGene_gn=KANSL1L;refGene_tx=XM_023259657.1;refGene_gn=KANSL1L;refGene_tx=XM_023259658.1;refGene_gn=KANSL1L;refGene_tx=XM_023259659.1
C1 196037468 196037581
refGene_gn=KANSL1L;refGene_tx=XM_019838602.2;refGene_gn=KANSL1L;refGene_tx=XM_023259653.1;refGene_gn=KANSL1L;refGene_tx=XM_023259654.1;refGene_gn=KANSL1L;refGene_tx=XM_023259655.1;refGene_gn=KANSL1L;refGene_tx=XM_023259656.1;refGene_gn=KANSL1L;refGene_tx=XM_023259657.1;refGene_gn=KANSL1L;refGene_tx=XM_023259658.1;refGene_gn=KANSL1L;refGene_tx=XM_023259659.1
C1 196038748 196038800
refGene_gn=KANSL1L;refGene_tx=XM_019838602.2;refGene_gn=KANSL1L;refGene_tx=XM_023259653.1;refGene_gn=KANSL1L;refGene_tx=XM_023259654.1;refGene_gn=KANSL1L;refGene_tx=XM_023259655.1;refGene_gn=KANSL1L;refGene_tx=XM_023259656.1;refGene_gn=KANSL1L;refGene_tx=XM_023259657.1;refGene_gn=KANSL1L;refGene_tx=XM_023259658.1;refGene_gn=KANSL1L;refGene_tx=XM_023259659.1
C1 196040260 196040385
refGene_gn=KANSL1L;refGene_tx=XM_019838602.2;refGene_gn=KANSL1L;refGene_tx=XM_023259653.1;refGene_gn=KANSL1L;refGene_tx=XM_023259654.1;refGene_gn=KANSL1L;refGene_tx=XM_023259655.1;refGene_gn=KANSL1L;refGene_tx=XM_023259656.1;refGene_gn=KANSL1L;refGene_tx=XM_023259657.1;refGene_gn=KANSL1L;refGene_tx=XM_023259658.1;refGene_gn=KANSL1L;refGene_tx=XM_023259659.1
C1 196041371 196041490
refGene_gn=KANSL1L;refGene_tx=XM_019838602.2;refGene_gn=KANSL1L;refGene_tx=XM_023259653.1;refGene_gn=KANSL1L;refGene_tx=XM_023259654.1;refGene_gn=KANSL1L;refGene_tx=XM_023259655.1;refGene_gn=KANSL1L;refGene_tx=XM_023259656.1;refGene_gn=KANSL1L;refGene_tx=XM_023259657.1;refGene_gn=KANSL1L;refGene_tx=XM_023259658.1;refGene_gn=KANSL1L;refGene_tx=XM_023259659.1
C1 196043779 196043908
refGene_gn=KANSL1L;refGene_tx=XM_019838602.2;refGene_gn=KANSL1L;refGene_tx=XM_023259653.1;refGene_gn=KANSL1L;refGene_tx=XM_023259654.1;refGene_gn=KANSL1L;refGene_tx=XM_023259655.1;refGene_gn=KANSL1L;refGene_tx=XM_023259656.1;refGene_gn=KANSL1L;refGene_tx=XM_023259658.1
C1 196051685 196051793
refGene_gn=KANSL1L;refGene_tx=XM_019838602.2;refGene_gn=KANSL1L;refGene_tx=XM_023259653.1;refGene_gn=KANSL1L;refGene_tx=XM_023259654.1;refGene_gn=KANSL1L;refGene_tx=XM_023259655.1;refGene_gn=KANSL1L;refGene_tx=XM_023259656.1;refGene_gn=KANSL1L;refGene_tx=XM_023259657.1;refGene_gn=KANSL1L;refGene_tx=XM_023259658.1;refGene_gn=KANSL1L;refGene_tx=XM_023259659.1
C1 196054438 196054604
refGene_gn=KANSL1L;refGene_tx=XM_019838602.2;refGene_gn=KANSL1L;refGene_tx=XM_023259653.1;refGene_gn=KANSL1L;refGene_tx=XM_023259654.1;refGene_gn=KANSL1L;refGene_tx=XM_023259655.1;refGene_gn=KANSL1L;refGene_tx=XM_023259656.1;refGene_gn=KANSL1L;refGene_tx=XM_023259657.1;refGene_gn

=KANSL1L;refGene_tx=XM_023259658.1;refGene_gn=KANSL1L;refGene_tx=XM_023259659.1
C1 196056499 196056646
refGene_gn=KANSL1L;refGene_tx=XM_019838602.2;refGene_gn=KANSL1L;refGene_tx=XM_023259653.1;refGene_gn=KANSL1L;refGene_tx=XM_023259654.1;refGene_gn=KANSL1L;refGene_tx=XM_023259655.1;refGene_gn=KANSL1L;refGene_tx=XM_023259656.1;refGene_gn=KANSL1L;refGene_tx=XM_023259657.1
C1 196093750 196093955
refGene_gn=KANSL1L;refGene_tx=XM_019838602.2;refGene_gn=KANSL1L;refGene_tx=XM_023259653.1;refGene_gn=KANSL1L;refGene_tx=XM_023259654.1;refGene_gn=KANSL1L;refGene_tx=XM_023259655.1;refGene_gn=KANSL1L;refGene_tx=XM_023259656.1;refGene_gn=KANSL1L;refGene_tx=XM_023259657.1;refGene_gn=KANSL1L;refGene_tx=XM_023259658.1;refGene_gn=KANSL1L;refGene_tx=XM_023259659.1
C1 196112331 196112453
refGene_gn=KANSL1L;refGene_tx=XM_019838602.2;refGene_gn=KANSL1L;refGene_tx=XM_023259653.1;refGene_gn=KANSL1L;refGene_tx=XM_023259654.1;refGene_gn=KANSL1L;refGene_tx=XM_023259655.1;refGene_gn=KANSL1L;refGene_tx=XM_023259656.1;refGene_gn=KANSL1L;refGene_tx=XM_023259657.1;refGene_gn=KANSL1L;refGene_tx=XM_023259658.1;refGene_gn=KANSL1L;refGene_tx=XM_023259659.1
C1 196116217 196116415
refGene_gn=KANSL1L;refGene_tx=XM_019838602.2;refGene_gn=KANSL1L;refGene_tx=XM_023259653.1;refGene_gn=KANSL1L;refGene_tx=XM_023259654.1;refGene_gn=KANSL1L;refGene_tx=XM_023259655.1;refGene_gn=KANSL1L;refGene_tx=XM_023259656.1;refGene_gn=KANSL1L;refGene_tx=XM_023259657.1;refGene_gn=KANSL1L;refGene_tx=XM_023259658.1;refGene_gn=KANSL1L;refGene_tx=XM_023259659.1
C1 196125037 196125179
refGene_gn=KANSL1L;refGene_tx=XM_019838602.2;refGene_gn=KANSL1L;refGene_tx=XM_023259653.1;refGene_gn=KANSL1L;refGene_tx=XM_023259654.1;refGene_gn=KANSL1L;refGene_tx=XM_023259655.1;refGene_gn=KANSL1L;refGene_tx=XM_023259656.1;refGene_gn=KANSL1L;refGene_tx=XM_023259657.1;refGene_gn=KANSL1L;refGene_tx=XM_023259658.1;refGene_gn=KANSL1L;refGene_tx=XM_023259659.1
C1 196145383 196146471
refGene_gn=KANSL1L;refGene_tx=XM_019838602.2;refGene_gn=KANSL1L;refGene_tx=XM_023259653.1;refGene_gn=KANSL1L;refGene_tx=XM_023259654.1;refGene_gn=KANSL1L;refGene_tx=XM_023259655.1;refGene_gn=KANSL1L;refGene_tx=XM_023259656.1;refGene_gn=KANSL1L;refGene_tx=XM_023259657.1;refGene_gn=KANSL1L;refGene_tx=XM_023259658.1;refGene_gn=KANSL1L;refGene_tx=XM_023259659.1
C1 196182528 196182622
refGene_gn=ACADL;refGene_tx=XM_003991101.4;refGene_gn=ACADL;refGene_tx=XM_006935526.3
C1 196188300 196188387
refGene_gn=ACADL;refGene_tx=XM_003991101.4;refGene_gn=ACADL;refGene_tx=XM_006935526.3
C1 196192242 196192370
refGene_gn=ACADL;refGene_tx=XM_003991101.4;refGene_gn=ACADL;refGene_tx

=XM_006935526.3
C1	196203362	196203476
refGene_gn=ACADL;refGene_tx=XM_003991101.4;refGene_gn=ACADL;refGene_tx=XM_006935526.3
C1	196204578	196204680
refGene_gn=ACADL;refGene_tx=XM_003991101.4;refGene_gn=ACADL;refGene_tx=XM_006935526.3
C1	196205836	196206001
refGene_gn=ACADL;refGene_tx=XM_003991101.4;refGene_gn=ACADL;refGene_tx=XM_006935526.3
C1	196210953	196211020
refGene_gn=ACADL;refGene_tx=XM_003991101.4;refGene_gn=ACADL;refGene_tx=XM_006935526.3
C1	196218236	196218401
refGene_gn=ACADL;refGene_tx=XM_003991101.4
C1	196220161	196220299
refGene_gn=ACADL;refGene_tx=XM_003991101.4;refGene_gn=ACADL;refGene_tx=XM_006935526.3
C1	196222470	196222626
refGene_gn=ACADL;refGene_tx=XM_003991101.4;refGene_gn=ACADL;refGene_tx=XM_006935526.3
C1	196227937	196228014
refGene_gn=ACADL;refGene_tx=XM_003991101.4;refGene_gn=ACADL;refGene_tx=XM_006935526.3
C1	196294915	196294944
refGene_gn=MYL1;refGene_tx=XM_003991102.3;refGene_gn=MYL1;refGene_tx=XM_003991103.4
C1	196297161	196297239
refGene_gn=MYL1;refGene_tx=XM_003991102.3;refGene_gn=MYL1;refGene_tx=XM_003991103.4
C1	196297692	196297866
refGene_gn=MYL1;refGene_tx=XM_003991102.3;refGene_gn=MYL1;refGene_tx=XM_003991103.4
C1	196299994	196300138
refGene_gn=MYL1;refGene_tx=XM_003991102.3;refGene_gn=MYL1;refGene_tx=XM_003991103.4
C1	196303963	196303991
refGene_gn=MYL1;refGene_tx=XM_003991103.4
C1	196304244	196304269
refGene_gn=MYL1;refGene_tx=XM_003991102.3
C1	196305027	196305030
refGene_gn=MYL1;refGene_tx=XM_003991102.3
C1	196316013	196316139
refGene_gn=MYL1;refGene_tx=XM_003991103.4
C1	196371617	196371694
refGene_gn=LANCL1;refGene_tx=XM_003991104.5;refGene_gn=LANCL1;refGene_tx=XM_006935527.4;refGene_gn=LANCL1;refGene_tx=XM_006935528.4;refGene_gn=LANCL1;refGene_tx=XM_019838608.2
C1	196372315	196372388
refGene_gn=LANCL1;refGene_tx=XM_003991104.5;refGene_gn=LANCL1;refGene_

tx=XM_006935527.4;refGene_gn=LANCL1;refGene_tx=XM_006935528.4;refGene_gn=LANCL1;refGene_tx=XM_019838608.2
C1	196372827	196373004
refGene_gn=LANCL1;refGene_tx=XM_003991104.5;refGene_gn=LANCL1;refGene_tx=XM_006935527.4;refGene_gn=LANCL1;refGene_tx=XM_006935528.4;refGene_gn=LANCL1;refGene_tx=XM_019838608.2
C1	196374790	196374973
refGene_gn=LANCL1;refGene_tx=XM_003991104.5;refGene_gn=LANCL1;refGene_tx=XM_006935527.4;refGene_gn=LANCL1;refGene_tx=XM_006935528.4;refGene_gn=LANCL1;refGene_tx=XM_019838608.2
C1	196377182	196377329
refGene_gn=LANCL1;refGene_tx=XM_003991104.5;refGene_gn=LANCL1;refGene_tx=XM_006935527.4;refGene_gn=LANCL1;refGene_tx=XM_006935528.4;refGene_gn=LANCL1;refGene_tx=XM_019838608.2
C1	196377882	196378018
refGene_gn=LANCL1;refGene_tx=XM_003991104.5;refGene_gn=LANCL1;refGene_tx=XM_006935527.4;refGene_gn=LANCL1;refGene_tx=XM_006935528.4;refGene_gn=LANCL1;refGene_tx=XM_019838608.2
C1	196386213	196386421
refGene_gn=LANCL1;refGene_tx=XM_003991104.5;refGene_gn=LANCL1;refGene_tx=XM_006935527.4;refGene_gn=LANCL1;refGene_tx=XM_006935528.4;refGene_gn=LANCL1;refGene_tx=XM_019838608.2
C1	196401558	196401676
refGene_gn=LANCL1;refGene_tx=XM_003991104.5;refGene_gn=LANCL1;refGene_tx=XM_006935527.4;refGene_gn=LANCL1;refGene_tx=XM_006935528.4;refGene_gn=LANCL1;refGene_tx=XM_019838608.2
C1	196405239	196405320
refGene_gn=LANCL1;refGene_tx=XM_003991104.5;refGene_gn=LANCL1;refGene_tx=XM_006935527.4;refGene_gn=LANCL1;refGene_tx=XM_006935528.4;refGene_gn=LANCL1;refGene_tx=XM_019838608.2
C1	196470808	196470934
refGene_gn=CPS1;refGene_tx=XM_019838609.2
C1	196492808	196492918
refGene_gn=CPS1;refGene_tx=XM_019838609.2
C1	196496644	196496789
refGene_gn=CPS1;refGene_tx=XM_019838609.2
C1	196498314	196498404
refGene_gn=CPS1;refGene_tx=XM_019838609.2
C1	196500339	196500396
refGene_gn=CPS1;refGene_tx=XM_019838609.2
C1	196503762	196503855
refGene_gn=CPS1;refGene_tx=XM_019838609.2
C1	196506664	196506754
refGene_gn=CPS1;refGene_tx=XM_019838609.2
C1	196508470	196508599
refGene_gn=CPS1;refGene_tx=XM_019838609.2
C1	196509605	196509712
refGene_gn=CPS1;refGene_tx=XM_019838609.2
C1	196510563	196510702
refGene_gn=CPS1;refGene_tx=XM_019838609.2

C1	196511331	196511409
refGene_gn=CPS1;refGene_tx=XM_019838609.2
C1	196513252	196513351
refGene_gn=CPS1;refGene_tx=XM_019838609.2
C1	196515008	196515104
refGene_gn=CPS1;refGene_tx=XM_019838609.2
C1	196518099	196518289
refGene_gn=CPS1;refGene_tx=XM_019838609.2
C1	196519288	196519446
refGene_gn=CPS1;refGene_tx=XM_019838609.2
C1	196520964	196521093
refGene_gn=CPS1;refGene_tx=XM_019838609.2
C1	196523648	196523793
refGene_gn=CPS1;refGene_tx=XM_019838609.2
C1	196525261	196525472
refGene_gn=CPS1;refGene_tx=XM_019838609.2
C1	196527119	196527318
refGene_gn=CPS1;refGene_tx=XM_019838609.2
C1	196531301	196531478
refGene_gn=CPS1;refGene_tx=XM_019838609.2
C1	196534923	196535042
refGene_gn=CPS1;refGene_tx=XM_019838609.2
C1	196555837	196555979
refGene_gn=CPS1;refGene_tx=XM_019838609.2
C1	196557296	196557362
refGene_gn=CPS1;refGene_tx=XM_019838609.2
C1	196557813	196557877
refGene_gn=CPS1;refGene_tx=XM_019838609.2
C1	196559797	196559979
refGene_gn=CPS1;refGene_tx=XM_019838609.2
C1	196566646	196566841
refGene_gn=CPS1;refGene_tx=XM_019838609.2
C1	196567263	196567331
refGene_gn=CPS1;refGene_tx=XM_019838609.2
C1	196568997	196569073
refGene_gn=CPS1;refGene_tx=XM_019838609.2
C1	196574292	196574370
refGene_gn=CPS1;refGene_tx=XM_019838609.2
C1	196577187	196577295
refGene_gn=CPS1;refGene_tx=XM_019838609.2
C1	196579179	196579269
refGene_gn=CPS1;refGene_tx=XM_019838609.2
C1	196580822	196580993
refGene_gn=CPS1;refGene_tx=XM_019838609.2
C1	196584303	196584378
refGene_gn=CPS1;refGene_tx=XM_019838609.2
C1	196587053	196587152
refGene_gn=CPS1;refGene_tx=XM_019838609.2
C1	196594886	196594946
refGene_gn=CPS1;refGene_tx=XM_019838609.2

C1 196595691 196595804
refGene_gn=CPS1;refGene_tx=XM_019838609.2
C1 196597285 196597415
refGene_gn=CPS1;refGene_tx=XM_019838609.2
C1 196598141 196598240
refGene_gn=CPS1;refGene_tx=XM_019838609.2
C1 197231161 197231607
refGene_gn=ERBB4;refGene_tx=XM_003991109.5;refGene_gn=ERBB4;refGene_tx=XM_019838610.2;refGene_gn=ERBB4;refGene_tx=XM_019838611.2;refGene_gn=ERBB4;refGene_tx=XM_019838612.2;refGene_gn=ERBB4;refGene_tx=XM_019838613.2;refGene_gn=ERBB4;refGene_tx=XM_019838614.2
C1 197234387 197234685
refGene_gn=ERBB4;refGene_tx=XM_003991109.5;refGene_gn=ERBB4;refGene_tx=XM_019838610.2;refGene_gn=ERBB4;refGene_tx=XM_019838611.2;refGene_gn=ERBB4;refGene_tx=XM_019838612.2;refGene_gn=ERBB4;refGene_tx=XM_019838613.2;refGene_gn=ERBB4;refGene_tx=XM_019838614.2
C1 197235471 197235519
refGene_gn=ERBB4;refGene_tx=XM_003991109.5;refGene_gn=ERBB4;refGene_tx=XM_019838610.2;refGene_gn=ERBB4;refGene_tx=XM_019838612.2
C1 197266073 197266244
refGene_gn=ERBB4;refGene_tx=XM_003991109.5;refGene_gn=ERBB4;refGene_tx=XM_019838610.2;refGene_gn=ERBB4;refGene_tx=XM_019838611.2;refGene_gn=ERBB4;refGene_tx=XM_019838612.2;refGene_gn=ERBB4;refGene_tx=XM_019838613.2;refGene_gn=ERBB4;refGene_tx=XM_019838614.2
C1 197267803 197267901
refGene_gn=ERBB4;refGene_tx=XM_003991109.5;refGene_gn=ERBB4;refGene_tx=XM_019838610.2;refGene_gn=ERBB4;refGene_tx=XM_019838611.2;refGene_gn=ERBB4;refGene_tx=XM_019838612.2;refGene_gn=ERBB4;refGene_tx=XM_019838613.2;refGene_gn=ERBB4;refGene_tx=XM_019838614.2
C1 197270063 197270210
refGene_gn=ERBB4;refGene_tx=XM_003991109.5;refGene_gn=ERBB4;refGene_tx=XM_019838610.2;refGene_gn=ERBB4;refGene_tx=XM_019838611.2;refGene_gn=ERBB4;refGene_tx=XM_019838612.2;refGene_gn=ERBB4;refGene_tx=XM_019838613.2;refGene_gn=ERBB4;refGene_tx=XM_019838614.2
C1 197274023 197274099
refGene_gn=ERBB4;refGene_tx=XM_003991109.5;refGene_gn=ERBB4;refGene_tx=XM_019838610.2;refGene_gn=ERBB4;refGene_tx=XM_019838611.2;refGene_gn=ERBB4;refGene_tx=XM_019838612.2;refGene_gn=ERBB4;refGene_tx=XM_019838613.2;refGene_gn=ERBB4;refGene_tx=XM_019838614.2
C1 197277074 197277230
refGene_gn=ERBB4;refGene_tx=XM_003991109.5;refGene_gn=ERBB4;refGene_tx=XM_019838610.2;refGene_gn=ERBB4;refGene_tx=XM_019838611.2;refGene_gn=ERBB4;refGene_tx=XM_019838612.2;refGene_gn=ERBB4;refGene_tx=XM_019838613.2;refGene_gn=ERBB4;refGene_tx=XM_019838614.2
C1 197397464 197397650
refGene_gn=ERBB4;refGene_tx=XM_003991109.5;refGene_gn=ERBB4;refGene_tx=XM_019838610.2;refGene_gn=ERBB4;refGene_tx=XM_019838611.2;refGene_gn=ERBB4;refGene_tx=XM_019838612.2;refGene_gn=ERBB4;refGene_tx=XM_019838613.2;refGene_gn=ERBB4;refGene_tx=XM_019838614.2
C1 197436665 197436764

refGene_gn=ERBB4;refGene_tx=XM_003991109.5;refGene_gn=ERBB4;refGene_tx=XM_019838610.2;refGene_gn=ERBB4;refGene_tx=XM_019838611.2;refGene_gn=ERBB4;refGene_tx=XM_019838612.2;refGene_gn=ERBB4;refGene_tx=XM_019838613.2;refGene_gn=ERBB4;refGene_tx=XM_019838614.2
C1 197440630 197440753
refGene_gn=ERBB4;refGene_tx=XM_003991109.5;refGene_gn=ERBB4;refGene_tx=XM_019838610.2;refGene_gn=ERBB4;refGene_tx=XM_019838611.2;refGene_gn=ERBB4;refGene_tx=XM_019838612.2;refGene_gn=ERBB4;refGene_tx=XM_019838613.2;refGene_gn=ERBB4;refGene_tx=XM_019838614.2
C1 197444982 197445115
refGene_gn=ERBB4;refGene_tx=XM_003991109.5;refGene_gn=ERBB4;refGene_tx=XM_019838610.2;refGene_gn=ERBB4;refGene_tx=XM_019838611.2;refGene_gn=ERBB4;refGene_tx=XM_019838612.2;refGene_gn=ERBB4;refGene_tx=XM_019838613.2;refGene_gn=ERBB4;refGene_tx=XM_019838614.2
C1 197481432 197481507
refGene_gn=ERBB4;refGene_tx=XM_003991109.5;refGene_gn=ERBB4;refGene_tx=XM_019838610.2;refGene_gn=ERBB4;refGene_tx=XM_019838611.2;refGene_gn=ERBB4;refGene_tx=XM_019838613.2
C1 197481628 197481673
refGene_gn=ERBB4;refGene_tx=XM_019838610.2;refGene_gn=ERBB4;refGene_tx=XM_019838611.2;refGene_gn=ERBB4;refGene_tx=XM_019838612.2;refGene_gn=ERBB4;refGene_tx=XM_019838614.2
C1 197488463 197488618
refGene_gn=ERBB4;refGene_tx=XM_003991109.5;refGene_gn=ERBB4;refGene_tx=XM_019838610.2;refGene_gn=ERBB4;refGene_tx=XM_019838611.2;refGene_gn=ERBB4;refGene_tx=XM_019838612.2;refGene_gn=ERBB4;refGene_tx=XM_019838613.2;refGene_gn=ERBB4;refGene_tx=XM_019838614.2
C1 197494579 197494673
refGene_gn=ERBB4;refGene_tx=XM_003991109.5;refGene_gn=ERBB4;refGene_tx=XM_019838610.2;refGene_gn=ERBB4;refGene_tx=XM_019838611.2;refGene_gn=ERBB4;refGene_tx=XM_019838612.2;refGene_gn=ERBB4;refGene_tx=XM_019838613.2;refGene_gn=ERBB4;refGene_tx=XM_019838614.2
C1 197499578 197499711
refGene_gn=ERBB4;refGene_tx=XM_003991109.5;refGene_gn=ERBB4;refGene_tx=XM_019838610.2;refGene_gn=ERBB4;refGene_tx=XM_019838611.2;refGene_gn=ERBB4;refGene_tx=XM_019838612.2;refGene_gn=ERBB4;refGene_tx=XM_019838613.2;refGene_gn=ERBB4;refGene_tx=XM_019838614.2
C1 197520856 197521056
refGene_gn=ERBB4;refGene_tx=XM_003991109.5;refGene_gn=ERBB4;refGene_tx=XM_019838610.2;refGene_gn=ERBB4;refGene_tx=XM_019838611.2;refGene_gn=ERBB4;refGene_tx=XM_019838612.2;refGene_gn=ERBB4;refGene_tx=XM_019838613.2;refGene_gn=ERBB4;refGene_tx=XM_019838614.2
C1 197524242 197524333
refGene_gn=ERBB4;refGene_tx=XM_003991109.5;refGene_gn=ERBB4;refGene_tx=XM_019838610.2;refGene_gn=ERBB4;refGene_tx=XM_019838611.2;refGene_gn=ERBB4;refGene_tx=XM_019838612.2;refGene_gn=ERBB4;refGene_tx=XM_019838613.2;refGene_gn=ERBB4;refGene_tx=XM_019838614.2
C1 197525245 197525319
refGene_gn=ERBB4;refGene_tx=XM_003991109.5;refGene_gn=ERBB4;refGene_tx=XM_019838610.2;refGene_gn=ERBB4;refGene_tx=XM_019838611.2;refGene_gn=

ERBB4;refGene_tx=XM_019838612.2;refGene_gn=ERBB4;refGene_tx=XM_0198386
13.2;refGene_gn=ERBB4;refGene_tx=XM_019838614.2
C1 197532822 197532949
refGene_gn=ERBB4;refGene_tx=XM_003991109.5;refGene_gn=ERBB4;refGene_tx
=XM_019838610.2;refGene_gn=ERBB4;refGene_tx=XM_019838611.2;refGene_gn=
ERBB4;refGene_tx=XM_019838612.2;refGene_gn=ERBB4;refGene_tx=XM_0198386
13.2;refGene_gn=ERBB4;refGene_tx=XM_019838614.2
C1 197534357 197534471
refGene_gn=ERBB4;refGene_tx=XM_003991109.5;refGene_gn=ERBB4;refGene_tx
=XM_019838610.2;refGene_gn=ERBB4;refGene_tx=XM_019838611.2;refGene_gn=
ERBB4;refGene_tx=XM_019838612.2;refGene_gn=ERBB4;refGene_tx=XM_0198386
13.2;refGene_gn=ERBB4;refGene_tx=XM_019838614.2
C1 197538003 197538145
refGene_gn=ERBB4;refGene_tx=XM_003991109.5;refGene_gn=ERBB4;refGene_tx
=XM_019838610.2;refGene_gn=ERBB4;refGene_tx=XM_019838611.2;refGene_gn=
ERBB4;refGene_tx=XM_019838612.2;refGene_gn=ERBB4;refGene_tx=XM_0198386
13.2;refGene_gn=ERBB4;refGene_tx=XM_019838614.2
C1 197540872 197540991
refGene_gn=ERBB4;refGene_tx=XM_003991109.5;refGene_gn=ERBB4;refGene_tx
=XM_019838610.2;refGene_gn=ERBB4;refGene_tx=XM_019838611.2;refGene_gn=
ERBB4;refGene_tx=XM_019838612.2;refGene_gn=ERBB4;refGene_tx=XM_0198386
13.2;refGene_gn=ERBB4;refGene_tx=XM_019838614.2
C1 197576297 197576363
refGene_gn=ERBB4;refGene_tx=XM_003991109.5;refGene_gn=ERBB4;refGene_tx
=XM_019838610.2;refGene_gn=ERBB4;refGene_tx=XM_019838611.2;refGene_gn=
ERBB4;refGene_tx=XM_019838612.2;refGene_gn=ERBB4;refGene_tx=XM_0198386
13.2;refGene_gn=ERBB4;refGene_tx=XM_019838614.2
C1 197605709 197605844
refGene_gn=ERBB4;refGene_tx=XM_003991109.5;refGene_gn=ERBB4;refGene_tx
=XM_019838610.2;refGene_gn=ERBB4;refGene_tx=XM_019838611.2;refGene_gn=
ERBB4;refGene_tx=XM_019838612.2;refGene_gn=ERBB4;refGene_tx=XM_0198386
13.2;refGene_gn=ERBB4;refGene_tx=XM_019838614.2
C1 197757818 197758005
refGene_gn=ERBB4;refGene_tx=XM_003991109.5;refGene_gn=ERBB4;refGene_tx
=XM_019838610.2;refGene_gn=ERBB4;refGene_tx=XM_019838611.2;refGene_gn=
ERBB4;refGene_tx=XM_019838612.2;refGene_gn=ERBB4;refGene_tx=XM_0198386
13.2;refGene_gn=ERBB4;refGene_tx=XM_019838614.2
C1 197945785 197945937
refGene_gn=ERBB4;refGene_tx=XM_003991109.5;refGene_gn=ERBB4;refGene_tx
=XM_019838610.2;refGene_gn=ERBB4;refGene_tx=XM_019838611.2;refGene_gn=
ERBB4;refGene_tx=XM_019838612.2;refGene_gn=ERBB4;refGene_tx=XM_0198386
13.2;refGene_gn=ERBB4;refGene_tx=XM_019838614.2
C1 198363789 198363871
refGene_gn=ERBB4;refGene_tx=XM_003991109.5;refGene_gn=ERBB4;refGene_tx
=XM_019838610.2;refGene_gn=ERBB4;refGene_tx=XM_019838611.2;refGene_gn=
ERBB4;refGene_tx=XM_019838612.2;refGene_gn=ERBB4;refGene_tx=XM_0198386
13.2;refGene_gn=ERBB4;refGene_tx=XM_019838614.2
C1 198773601 198774326
refGene_gn=IKZF2;refGene_tx=XM_006935530.4;refGene_gn=IKZF2;refGene_tx
=XM_019838616.2;refGene_gn=IKZF2;refGene_tx=XM_019838617.2;refGene_gn=

```
IKZF2;refGene_tx=XM_019838618.2;refGene_gn=IKZF2;refGene_tx=XM_0198386
19.2;refGene_gn=IKZF2;refGene_tx=XM_019838620.2;refGene_gn=IKZF2;refGe
ne_tx=XM_019838621.2;refGene_gn=IKZF2;refGene_tx=XM_019838622.2;refGen
e_gn=IKZF2;refGene_tx=XM_019838623.2;refGene_gn=IKZF2;refGene_tx=XM_01
9838624.2;refGene_gn=IKZF2;refGene_tx=XM_019838625.2;refGene_gn=IKZF2;
refGene_tx=XM_019838626.2;refGene_gn=IKZF2;refGene_tx=XM_019838627.2
C1	198780212	198780356
refGene_gn=IKZF2;refGene_tx=XM_006935530.4;refGene_gn=IKZF2;refGene_tx
=XM_019838616.2;refGene_gn=IKZF2;refGene_tx=XM_019838617.2;refGene_gn=
IKZF2;refGene_tx=XM_019838618.2;refGene_gn=IKZF2;refGene_tx=XM_0198386
19.2;refGene_gn=IKZF2;refGene_tx=XM_019838620.2;refGene_gn=IKZF2;refGe
ne_tx=XM_019838621.2;refGene_gn=IKZF2;refGene_tx=XM_019838622.2;refGen
e_gn=IKZF2;refGene_tx=XM_019838623.2;refGene_gn=IKZF2;refGene_tx=XM_01
9838624.2;refGene_gn=IKZF2;refGene_tx=XM_019838625.2;refGene_gn=IKZF2;
refGene_tx=XM_019838626.2;refGene_gn=IKZF2;refGene_tx=XM_019838627.2
C1	198788270	198788408
refGene_gn=IKZF2;refGene_tx=XM_006935530.4;refGene_gn=IKZF2;refGene_tx
=XM_019838616.2;refGene_gn=IKZF2;refGene_tx=XM_019838617.2;refGene_gn=
IKZF2;refGene_tx=XM_019838618.2;refGene_gn=IKZF2;refGene_tx=XM_0198386
19.2;refGene_gn=IKZF2;refGene_tx=XM_019838620.2;refGene_gn=IKZF2;refGe
ne_tx=XM_019838621.2;refGene_gn=IKZF2;refGene_tx=XM_019838622.2;refGen
e_gn=IKZF2;refGene_tx=XM_019838623.2;refGene_gn=IKZF2;refGene_tx=XM_01
9838624.2;refGene_gn=IKZF2;refGene_tx=XM_019838625.2;refGene_gn=IKZF2;
refGene_tx=XM_019838626.2;refGene_gn=IKZF2;refGene_tx=XM_019838627.2
C1	198818361	198818529
refGene_gn=IKZF2;refGene_tx=XM_006935530.4;refGene_gn=IKZF2;refGene_tx
=XM_019838616.2;refGene_gn=IKZF2;refGene_tx=XM_019838617.2;refGene_gn=
IKZF2;refGene_tx=XM_019838618.2;refGene_gn=IKZF2;refGene_tx=XM_0198386
19.2;refGene_gn=IKZF2;refGene_tx=XM_019838620.2;refGene_gn=IKZF2;refGe
ne_tx=XM_019838621.2;refGene_gn=IKZF2;refGene_tx=XM_019838622.2;refGen
e_gn=IKZF2;refGene_tx=XM_019838623.2;refGene_gn=IKZF2;refGene_tx=XM_01
9838624.2;refGene_gn=IKZF2;refGene_tx=XM_019838625.2;refGene_gn=IKZF2;
refGene_tx=XM_019838626.2;refGene_gn=IKZF2;refGene_tx=XM_019838627.2
C1	198825262	198825529
refGene_gn=IKZF2;refGene_tx=XM_006935530.4;refGene_gn=IKZF2;refGene_tx
=XM_019838616.2;refGene_gn=IKZF2;refGene_tx=XM_019838617.2;refGene_gn=
IKZF2;refGene_tx=XM_019838618.2;refGene_gn=IKZF2;refGene_tx=XM_0198386
19.2;refGene_gn=IKZF2;refGene_tx=XM_019838620.2;refGene_gn=IKZF2;refGe
ne_tx=XM_019838621.2;refGene_gn=IKZF2;refGene_tx=XM_019838622.2;refGen
e_gn=IKZF2;refGene_tx=XM_019838623.2;refGene_gn=IKZF2;refGene_tx=XM_01
9838624.2;refGene_gn=IKZF2;refGene_tx=XM_019838625.2;refGene_gn=IKZF2;
refGene_tx=XM_019838626.2;refGene_gn=IKZF2;refGene_tx=XM_019838627.2
C1	198924721	198924788
refGene_gn=IKZF2;refGene_tx=XM_019838627.2
C1	198924955	198925063
refGene_gn=IKZF2;refGene_tx=XM_006935530.4;refGene_gn=IKZF2;refGene_tx
=XM_019838616.2;refGene_gn=IKZF2;refGene_tx=XM_019838617.2;refGene_gn=
IKZF2;refGene_tx=XM_019838618.2;refGene_gn=IKZF2;refGene_tx=XM_0198386
19.2;refGene_gn=IKZF2;refGene_tx=XM_019838620.2;refGene_gn=IKZF2;refGe
ne_tx=XM_019838621.2;refGene_gn=IKZF2;refGene_tx=XM_019838622.2;refGen
```

e_gn=IKZF2;refGene_tx=XM_019838623.2;refGene_gn=IKZF2;refGene_tx=XM_019838624.2;refGene_gn=IKZF2;refGene_tx=XM_019838625.2;refGene_gn=IKZF2;refGene_tx=XM_019838626.2
C1 198925860 198925909
refGene_gn=IKZF2;refGene_tx=XM_006935530.4;refGene_gn=IKZF2;refGene_tx=XM_019838616.2;refGene_gn=IKZF2;refGene_tx=XM_019838617.2;refGene_gn=IKZF2;refGene_tx=XM_019838618.2;refGene_gn=IKZF2;refGene_tx=XM_019838619.2;refGene_gn=IKZF2;refGene_tx=XM_019838620.2;refGene_gn=IKZF2;refGene_tx=XM_019838621.2;refGene_gn=IKZF2;refGene_tx=XM_019838622.2;refGene_gn=IKZF2;refGene_tx=XM_019838623.2;refGene_gn=IKZF2;refGene_tx=XM_019838624.2;refGene_gn=IKZF2;refGene_tx=XM_019838625.2;refGene_gn=IKZF2;refGene_tx=XM_019838626.2
C1 198927444 198927447
refGene_gn=IKZF2;refGene_tx=XM_019838616.2;refGene_gn=IKZF2;refGene_tx=XM_019838617.2;refGene_gn=IKZF2;refGene_tx=XM_019838618.2;refGene_gn=IKZF2;refGene_tx=XM_019838625.2;refGene_gn=IKZF2;refGene_tx=XM_019838626.2
C1 199063151 199063278
refGene_gn=SPAG16;refGene_tx=XM_019838628.2;refGene_gn=SPAG16;refGene_tx=XM_019838629.2
C1 199073411 199073458
refGene_gn=SPAG16;refGene_tx=XM_019838628.2;refGene_gn=SPAG16;refGene_tx=XM_019838629.2
C1 199074600 199074696
refGene_gn=SPAG16;refGene_tx=XM_019838628.2;refGene_gn=SPAG16;refGene_tx=XM_019838629.2
C1 199077886 199078005
refGene_gn=SPAG16;refGene_tx=XM_019838628.2;refGene_gn=SPAG16;refGene_tx=XM_019838629.2
C1 199084255 199084393
refGene_gn=SPAG16;refGene_tx=XM_019838628.2
C1 199103742 199103850
refGene_gn=SPAG16;refGene_tx=XM_019838628.2;refGene_gn=SPAG16;refGene_tx=XM_019838629.2
C1 199111475 199111593
refGene_gn=SPAG16;refGene_tx=XM_019838628.2;refGene_gn=SPAG16;refGene_tx=XM_019838629.2
C1 199118951 199119021
refGene_gn=SPAG16;refGene_tx=XM_019838628.2;refGene_gn=SPAG16;refGene_tx=XM_019838629.2
C1 199124435 199124545
refGene_gn=SPAG16;refGene_tx=XM_019838628.2;refGene_gn=SPAG16;refGene_tx=XM_019838629.2
C1 199180843 199180889
refGene_gn=LOC101101657;refGene_tx=XM_011285525.3
C1 199180891 199181448
refGene_gn=LOC101101657;refGene_tx=XM_011285525.3
C1 199222135 199222263
refGene_gn=SPAG16;refGene_tx=XM_019838628.2;refGene_gn=SPAG16;refGene_tx=XM_019838629.2

C1 199548468 199548612
refGene_gn=SPAG16;refGene_tx=XM_019838628.2;refGene_gn=SPAG16;refGene_tx=XM_019838629.2
C1 199596812 199596998
refGene_gn=SPAG16;refGene_tx=XM_019838628.2;refGene_gn=SPAG16;refGene_tx=XM_019838629.2
C1 199685699 199685826
refGene_gn=SPAG16;refGene_tx=XM_019838628.2;refGene_gn=SPAG16;refGene_tx=XM_019838629.2
C1 199771059 199771125
refGene_gn=SPAG16;refGene_tx=XM_019838628.2;refGene_gn=SPAG16;refGene_tx=XM_019838629.2
C1 199808392 199808519
refGene_gn=SPAG16;refGene_tx=XM_019838628.2;refGene_gn=SPAG16;refGene_tx=XM_019838629.2
C1 200067945 200068121
refGene_gn=SPAG16;refGene_tx=XM_019838628.2;refGene_gn=SPAG16;refGene_tx=XM_019838629.2
C1 200071966 200072356
refGene_gn=VWC2L;refGene_tx=XM_006935531.3
C1 200093968 200094098
refGene_gn=VWC2L;refGene_tx=XM_006935531.3
C1 200232794 200232943
refGene_gn=VWC2L;refGene_tx=XM_006935531.3
C1 200369763 200370096
refGene_gn=BARD1;refGene_tx=XM_006935532.4;refGene_gn=BARD1;refGene_tx=XM_019838636.2;refGene_gn=BARD1;refGene_tx=XM_019838637.2;refGene_gn=BARD1;refGene_tx=XM_023259666.1
C1 200371174 200371272
refGene_gn=BARD1;refGene_tx=XM_006935532.4;refGene_gn=BARD1;refGene_tx=XM_019838636.2;refGene_gn=BARD1;refGene_tx=XM_019838637.2;refGene_gn=BARD1;refGene_tx=XM_023259666.1
C1 200381576 200381669
refGene_gn=BARD1;refGene_tx=XM_006935532.4;refGene_gn=BARD1;refGene_tx=XM_019838636.2;refGene_gn=BARD1;refGene_tx=XM_019838637.2;refGene_gn=BARD1;refGene_tx=XM_023259666.1
C1 200382203 200382336
refGene_gn=BARD1;refGene_tx=XM_006935532.4;refGene_gn=BARD1;refGene_tx=XM_019838636.2;refGene_gn=BARD1;refGene_tx=XM_019838637.2;refGene_gn=BARD1;refGene_tx=XM_023259666.1
C1 200389836 200389945
refGene_gn=BARD1;refGene_tx=XM_006935532.4;refGene_gn=BARD1;refGene_tx=XM_019838636.2;refGene_gn=BARD1;refGene_tx=XM_019838637.2;refGene_gn=BARD1;refGene_tx=XM_023259666.1
C1 200411151 200411324
refGene_gn=BARD1;refGene_tx=XM_006935532.4;refGene_gn=BARD1;refGene_tx=XM_019838636.2;refGene_gn=BARD1;refGene_tx=XM_019838637.2;refGene_gn=BARD1;refGene_tx=XM_023259666.1
C1 200413168 200413249
refGene_gn=BARD1;refGene_tx=XM_006935532.4;refGene_gn=BARD1;refGene_tx

=XM_019838636.2;refGene_gn=BARD1;refGene_tx=XM_019838637.2;refGene_gn=BARD1;refGene_tx=XM_023259666.1
C1 200420970 200421923
refGene_gn=BARD1;refGene_tx=XM_006935532.4;refGene_gn=BARD1;refGene_tx=XM_019838636.2;refGene_gn=BARD1;refGene_tx=XM_019838637.2;refGene_gn=BARD1;refGene_tx=XM_023259666.1
C1 200435357 200435506
refGene_gn=BARD1;refGene_tx=XM_006935532.4;refGene_gn=BARD1;refGene_tx=XM_019838636.2
C1 200439991 200440048
refGene_gn=BARD1;refGene_tx=XM_006935532.4
C1 200451528 200451671
refGene_gn=BARD1;refGene_tx=XM_006935532.4
C1 200567003 200567111
refGene_gn=ABCA12;refGene_tx=XM_019838638.1;refGene_gn=ABCA12;refGene_tx=XM_019838639.1;refGene_gn=ABCA12;refGene_tx=XM_019838640.1
C1 200568858 200568996
refGene_gn=ABCA12;refGene_tx=XM_019838638.1;refGene_gn=ABCA12;refGene_tx=XM_019838639.1;refGene_gn=ABCA12;refGene_tx=XM_019838640.1
C1 200571728 200571834
refGene_gn=ABCA12;refGene_tx=XM_019838638.1;refGene_gn=ABCA12;refGene_tx=XM_019838639.1;refGene_gn=ABCA12;refGene_tx=XM_019838640.1
C1 200572906 200572999
refGene_gn=ABCA12;refGene_tx=XM_019838638.1;refGene_gn=ABCA12;refGene_tx=XM_019838639.1;refGene_gn=ABCA12;refGene_tx=XM_019838640.1
C1 200574423 200574527
refGene_gn=ABCA12;refGene_tx=XM_019838638.1;refGene_gn=ABCA12;refGene_tx=XM_019838639.1;refGene_gn=ABCA12;refGene_tx=XM_019838640.1
C1 200576425 200576560
refGene_gn=ABCA12;refGene_tx=XM_019838638.1;refGene_gn=ABCA12;refGene_tx=XM_019838639.1;refGene_gn=ABCA12;refGene_tx=XM_019838640.1
C1 200577750 200577892
refGene_gn=ABCA12;refGene_tx=XM_019838638.1;refGene_gn=ABCA12;refGene_tx=XM_019838639.1;refGene_gn=ABCA12;refGene_tx=XM_019838640.1
C1 200578172 200578282
refGene_gn=ABCA12;refGene_tx=XM_019838638.1;refGene_gn=ABCA12;refGene_tx=XM_019838639.1;refGene_gn=ABCA12;refGene_tx=XM_019838640.1
C1 200579657 200579862
refGene_gn=ABCA12;refGene_tx=XM_019838638.1;refGene_gn=ABCA12;refGene_tx=XM_019838639.1;refGene_gn=ABCA12;refGene_tx=XM_019838640.1
C1 200581675 200581929
refGene_gn=ABCA12;refGene_tx=XM_019838638.1;refGene_gn=ABCA12;refGene_tx=XM_019838639.1;refGene_gn=ABCA12;refGene_tx=XM_019838640.1
C1 200583063 200583223
refGene_gn=ABCA12;refGene_tx=XM_019838638.1;refGene_gn=ABCA12;refGene_tx=XM_019838639.1;refGene_gn=ABCA12;refGene_tx=XM_019838640.1
C1 200584987 200585103
refGene_gn=ABCA12;refGene_tx=XM_019838638.1;refGene_gn=ABCA12;refGene_tx=XM_019838639.1;refGene_gn=ABCA12;refGene_tx=XM_019838640.1
C1 200587017 200587195

refGene_gn=ABCA12;refGene_tx=XM_019838638.1;refGene_gn=ABCA12;refGene_tx=XM_019838639.1;refGene_gn=ABCA12;refGene_tx=XM_019838640.1
C1	200587783	200587838	refGene_gn=ABCA12;refGene_tx=XM_019838638.1;refGene_gn=ABCA12;refGene_tx=XM_019838639.1;refGene_gn=ABCA12;refGene_tx=XM_019838640.1
C1	200591914	200592020	refGene_gn=ABCA12;refGene_tx=XM_019838638.1;refGene_gn=ABCA12;refGene_tx=XM_019838639.1;refGene_gn=ABCA12;refGene_tx=XM_019838640.1
C1	200594756	200594844	refGene_gn=ABCA12;refGene_tx=XM_019838638.1;refGene_gn=ABCA12;refGene_tx=XM_019838639.1;refGene_gn=ABCA12;refGene_tx=XM_019838640.1
C1	200596863	200596991	refGene_gn=ABCA12;refGene_tx=XM_019838638.1;refGene_gn=ABCA12;refGene_tx=XM_019838639.1;refGene_gn=ABCA12;refGene_tx=XM_019838640.1
C1	200599863	200599957	refGene_gn=ABCA12;refGene_tx=XM_019838638.1;refGene_gn=ABCA12;refGene_tx=XM_019838639.1;refGene_gn=ABCA12;refGene_tx=XM_019838640.1
C1	200600754	200600841	refGene_gn=ABCA12;refGene_tx=XM_019838638.1;refGene_gn=ABCA12;refGene_tx=XM_019838639.1;refGene_gn=ABCA12;refGene_tx=XM_019838640.1
C1	200601487	200601740	refGene_gn=ABCA12;refGene_tx=XM_019838638.1;refGene_gn=ABCA12;refGene_tx=XM_019838639.1;refGene_gn=ABCA12;refGene_tx=XM_019838640.1
C1	200604536	200604687	refGene_gn=ABCA12;refGene_tx=XM_019838638.1;refGene_gn=ABCA12;refGene_tx=XM_019838639.1;refGene_gn=ABCA12;refGene_tx=XM_019838640.1
C1	200605194	200605431	refGene_gn=ABCA12;refGene_tx=XM_019838638.1;refGene_gn=ABCA12;refGene_tx=XM_019838639.1;refGene_gn=ABCA12;refGene_tx=XM_019838640.1
C1	200607013	200607174	refGene_gn=ABCA12;refGene_tx=XM_019838638.1;refGene_gn=ABCA12;refGene_tx=XM_019838639.1;refGene_gn=ABCA12;refGene_tx=XM_019838640.1
C1	200607959	200608156	refGene_gn=ABCA12;refGene_tx=XM_019838638.1;refGene_gn=ABCA12;refGene_tx=XM_019838639.1;refGene_gn=ABCA12;refGene_tx=XM_019838640.1
C1	200609612	200609831	refGene_gn=ABCA12;refGene_tx=XM_019838638.1;refGene_gn=ABCA12;refGene_tx=XM_019838639.1;refGene_gn=ABCA12;refGene_tx=XM_019838640.1
C1	200611924	200612111	refGene_gn=ABCA12;refGene_tx=XM_019838638.1;refGene_gn=ABCA12;refGene_tx=XM_019838639.1;refGene_gn=ABCA12;refGene_tx=XM_019838640.1
C1	200613015	200613171	refGene_gn=ABCA12;refGene_tx=XM_019838638.1;refGene_gn=ABCA12;refGene_tx=XM_019838639.1;refGene_gn=ABCA12;refGene_tx=XM_019838640.1
C1	200614068	200614203	refGene_gn=ABCA12;refGene_tx=XM_019838638.1;refGene_gn=ABCA12;refGene_tx=XM_019838639.1;refGene_gn=ABCA12;refGene_tx=XM_019838640.1
C1	200614291	200614361	refGene_gn=ABCA12;refGene_tx=XM_019838638.1;refGene_gn=ABCA12;refGene_tx=XM_019838639.1;refGene_gn=ABCA12;refGene_tx=XM_019838640.1

C1	200615481	200615811
refGene_gn=ABCA12;refGene_tx=XM_019838638.1;refGene_gn=ABCA12;refGene_tx=XM_019838639.1;refGene_gn=ABCA12;refGene_tx=XM_019838640.1
C1	200622442	200622557
refGene_gn=ABCA12;refGene_tx=XM_019838638.1;refGene_gn=ABCA12;refGene_tx=XM_019838639.1;refGene_gn=ABCA12;refGene_tx=XM_019838640.1
C1	200625395	200625711
refGene_gn=ABCA12;refGene_tx=XM_019838638.1;refGene_gn=ABCA12;refGene_tx=XM_019838639.1;refGene_gn=ABCA12;refGene_tx=XM_019838640.1
C1	200626287	200626467
refGene_gn=ABCA12;refGene_tx=XM_019838638.1;refGene_gn=ABCA12;refGene_tx=XM_019838639.1;refGene_gn=ABCA12;refGene_tx=XM_019838640.1
C1	200628610	200628701
refGene_gn=ABCA12;refGene_tx=XM_019838638.1;refGene_gn=ABCA12;refGene_tx=XM_019838639.1;refGene_gn=ABCA12;refGene_tx=XM_019838640.1
C1	200631894	200632014
refGene_gn=ABCA12;refGene_tx=XM_019838638.1;refGene_gn=ABCA12;refGene_tx=XM_019838639.1;refGene_gn=ABCA12;refGene_tx=XM_019838640.1
C1	200634049	200634189
refGene_gn=ABCA12;refGene_tx=XM_019838638.1;refGene_gn=ABCA12;refGene_tx=XM_019838639.1;refGene_gn=ABCA12;refGene_tx=XM_019838640.1
C1	200635141	200635352
refGene_gn=ABCA12;refGene_tx=XM_019838638.1;refGene_gn=ABCA12;refGene_tx=XM_019838639.1;refGene_gn=ABCA12;refGene_tx=XM_019838640.1
C1	200635676	200635841
refGene_gn=ABCA12;refGene_tx=XM_019838638.1;refGene_gn=ABCA12;refGene_tx=XM_019838639.1;refGene_gn=ABCA12;refGene_tx=XM_019838640.1
C1	200639146	200639320
refGene_gn=ABCA12;refGene_tx=XM_019838638.1;refGene_gn=ABCA12;refGene_tx=XM_019838639.1;refGene_gn=ABCA12;refGene_tx=XM_019838640.1
C1	200641159	200641284
refGene_gn=ABCA12;refGene_tx=XM_019838638.1;refGene_gn=ABCA12;refGene_tx=XM_019838639.1;refGene_gn=ABCA12;refGene_tx=XM_019838640.1
C1	200642505	200642618
refGene_gn=ABCA12;refGene_tx=XM_019838638.1;refGene_gn=ABCA12;refGene_tx=XM_019838639.1;refGene_gn=ABCA12;refGene_tx=XM_019838640.1
C1	200642720	200642977
refGene_gn=ABCA12;refGene_tx=XM_019838638.1;refGene_gn=ABCA12;refGene_tx=XM_019838639.1;refGene_gn=ABCA12;refGene_tx=XM_019838640.1
C1	200649243	200649350
refGene_gn=ABCA12;refGene_tx=XM_019838638.1;refGene_gn=ABCA12;refGene_tx=XM_019838639.1;refGene_gn=ABCA12;refGene_tx=XM_019838640.1
C1	200650450	200650569
refGene_gn=ABCA12;refGene_tx=XM_019838638.1;refGene_gn=ABCA12;refGene_tx=XM_019838639.1;refGene_gn=ABCA12;refGene_tx=XM_019838640.1
C1	200654216	200654292
refGene_gn=ABCA12;refGene_tx=XM_019838638.1;refGene_gn=ABCA12;refGene_tx=XM_019838639.1;refGene_gn=ABCA12;refGene_tx=XM_019838640.1
C1	200658710	200658823
refGene_gn=ABCA12;refGene_tx=XM_019838638.1;refGene_gn=ABCA12;refGene_

tx=XM_019838639.1;refGene_gn=ABCA12;refGene_tx=XM_019838640.1
C1	200664662	200664841
refGene_gn=ABCA12;refGene_tx=XM_019838638.1;refGene_gn=ABCA12;refGene_tx=XM_019838639.1;refGene_gn=ABCA12;refGene_tx=XM_019838640.1
C1	200666495	200666681
refGene_gn=ABCA12;refGene_tx=XM_019838638.1;refGene_gn=ABCA12;refGene_tx=XM_019838639.1;refGene_gn=ABCA12;refGene_tx=XM_019838640.1
C1	200669804	200669902
refGene_gn=ABCA12;refGene_tx=XM_019838638.1;refGene_gn=ABCA12;refGene_tx=XM_019838639.1;refGene_gn=ABCA12;refGene_tx=XM_019838640.1
C1	200673044	200673136
refGene_gn=ABCA12;refGene_tx=XM_019838638.1;refGene_gn=ABCA12;refGene_tx=XM_019838639.1;refGene_gn=ABCA12;refGene_tx=XM_019838640.1
C1	200685845	200685999
refGene_gn=ABCA12;refGene_tx=XM_019838638.1;refGene_gn=ABCA12;refGene_tx=XM_019838639.1;refGene_gn=ABCA12;refGene_tx=XM_019838640.1
C1	200727257	200727351
refGene_gn=ABCA12;refGene_tx=XM_019838638.1;refGene_gn=ABCA12;refGene_tx=XM_019838639.1;refGene_gn=ABCA12;refGene_tx=XM_019838640.1
C1	200740979	200741048
refGene_gn=ABCA12;refGene_tx=XM_019838638.1;refGene_gn=ABCA12;refGene_tx=XM_019838639.1
C1	200784782	200784835
refGene_gn=ABCA12;refGene_tx=XM_019838640.1
C1	200842528	200842532
refGene_gn=ABCA12;refGene_tx=XM_019838640.1
C1	200853705	200853724
refGene_gn=ATIC;refGene_tx=XM_003991117.3
C1	200854027	200854154
refGene_gn=ATIC;refGene_tx=XM_003991117.3
C1	200857415	200857492
refGene_gn=ATIC;refGene_tx=XM_003991117.3
C1	200858398	200858465
refGene_gn=ATIC;refGene_tx=XM_003991117.3
C1	200861904	200861993
refGene_gn=ATIC;refGene_tx=XM_003991117.3
C1	200862787	200862939
refGene_gn=ATIC;refGene_tx=XM_003991117.3
C1	200863266	200863423
refGene_gn=ATIC;refGene_tx=XM_003991117.3
C1	200867981	200868107
refGene_gn=ATIC;refGene_tx=XM_003991117.3
C1	200869070	200869178
refGene_gn=ATIC;refGene_tx=XM_003991117.3
C1	200870275	200870361
refGene_gn=ATIC;refGene_tx=XM_003991117.3
C1	200871736	200871826
refGene_gn=ATIC;refGene_tx=XM_003991117.3
C1	200873078	200873207
refGene_gn=ATIC;refGene_tx=XM_003991117.3

C1 200877966 200878059
refGene_gn=ATIC;refGene_tx=XM_003991117.3
C1 200879263 200879446
refGene_gn=ATIC;refGene_tx=XM_003991117.3
C1 200881184 200881340
refGene_gn=ATIC;refGene_tx=XM_003991117.3
C1 200881633 200881753
refGene_gn=ATIC;refGene_tx=XM_003991117.3
C1 200894239 200894311
refGene_gn=FN1;refGene_tx=XM_003991118.4;refGene_gn=FN1;refGene_tx=XM_003991119.5;refGene_gn=FN1;refGene_tx=XM_003991120.4;refGene_gn=FN1;refGene_tx=XM_003991121.4;refGene_gn=FN1;refGene_tx=XM_006935534.3;refGene_gn=FN1;refGene_tx=XM_006935535.3;refGene_gn=FN1;refGene_tx=XM_006935536.3;refGene_gn=FN1;refGene_tx=XM_006935537.3;refGene_gn=FN1;refGene_tx=XM_006935538.3;refGene_gn=FN1;refGene_tx=XM_006935539.3;refGene_gn=FN1;refGene_tx=XM_019838641.1
C1 200894897 200895005
refGene_gn=FN1;refGene_tx=XM_003991118.4;refGene_gn=FN1;refGene_tx=XM_003991119.5;refGene_gn=FN1;refGene_tx=XM_003991120.4;refGene_gn=FN1;refGene_tx=XM_003991121.4;refGene_gn=FN1;refGene_tx=XM_006935534.3;refGene_gn=FN1;refGene_tx=XM_006935535.3;refGene_gn=FN1;refGene_tx=XM_006935536.3;refGene_gn=FN1;refGene_tx=XM_006935537.3;refGene_gn=FN1;refGene_tx=XM_006935538.3;refGene_gn=FN1;refGene_tx=XM_006935539.3;refGene_gn=FN1;refGene_tx=XM_019838641.1
C1 200898107 200898214
refGene_gn=FN1;refGene_tx=XM_003991118.4;refGene_gn=FN1;refGene_tx=XM_003991119.5;refGene_gn=FN1;refGene_tx=XM_003991120.4;refGene_gn=FN1;refGene_tx=XM_003991121.4;refGene_gn=FN1;refGene_tx=XM_006935534.3;refGene_gn=FN1;refGene_tx=XM_006935535.3;refGene_gn=FN1;refGene_tx=XM_006935536.3;refGene_gn=FN1;refGene_tx=XM_006935537.3;refGene_gn=FN1;refGene_tx=XM_006935538.3;refGene_gn=FN1;refGene_tx=XM_006935539.3;refGene_gn=FN1;refGene_tx=XM_019838641.1
C1 200898716 200898842
refGene_gn=FN1;refGene_tx=XM_003991118.4;refGene_gn=FN1;refGene_tx=XM_003991119.5;refGene_gn=FN1;refGene_tx=XM_003991120.4;refGene_gn=FN1;refGene_tx=XM_003991121.4;refGene_gn=FN1;refGene_tx=XM_006935534.3;refGene_gn=FN1;refGene_tx=XM_006935535.3;refGene_gn=FN1;refGene_tx=XM_006935536.3;refGene_gn=FN1;refGene_tx=XM_006935537.3;refGene_gn=FN1;refGene_tx=XM_006935538.3;refGene_gn=FN1;refGene_tx=XM_006935539.3;refGene_gn=FN1;refGene_tx=XM_019838641.1
C1 200900707 200900872
refGene_gn=FN1;refGene_tx=XM_003991118.4;refGene_gn=FN1;refGene_tx=XM_003991119.5;refGene_gn=FN1;refGene_tx=XM_003991120.4;refGene_gn=FN1;refGene_tx=XM_003991121.4;refGene_gn=FN1;refGene_tx=XM_006935534.3;refGene_gn=FN1;refGene_tx=XM_006935535.3;refGene_gn=FN1;refGene_tx=XM_006935536.3;refGene_gn=FN1;refGene_tx=XM_006935537.3;refGene_gn=FN1;refGene_tx=XM_006935538.3;refGene_gn=FN1;refGene_tx=XM_006935539.3
C1 200903319 200903458
refGene_gn=FN1;refGene_tx=XM_003991118.4;refGene_gn=FN1;refGene_tx=XM_003991119.5;refGene_gn=FN1;refGene_tx=XM_003991120.4;refGene_gn=FN1;re

fGene_tx=XM_003991121.4;refGene_gn=FN1;refGene_tx=XM_006935534.3;refGene_gn=FN1;refGene_tx=XM_006935535.3;refGene_gn=FN1;refGene_tx=XM_006935536.3;refGene_gn=FN1;refGene_tx=XM_006935537.3;refGene_gn=FN1;refGene_tx=XM_006935538.3;refGene_gn=FN1;refGene_tx=XM_006935539.3
C1 200904584 200905048
refGene_gn=FN1;refGene_tx=XM_003991118.4;refGene_gn=FN1;refGene_tx=XM_003991119.5;refGene_gn=FN1;refGene_tx=XM_003991120.4;refGene_gn=FN1;refGene_tx=XM_003991121.4;refGene_gn=FN1;refGene_tx=XM_006935534.3;refGene_gn=FN1;refGene_tx=XM_006935535.3;refGene_gn=FN1;refGene_tx=XM_006935536.3;refGene_gn=FN1;refGene_tx=XM_006935537.3;refGene_gn=FN1;refGene_tx=XM_006935538.3;refGene_gn=FN1;refGene_tx=XM_006935539.3
C1 200905988 200906078
refGene_gn=FN1;refGene_tx=XM_003991118.4;refGene_gn=FN1;refGene_tx=XM_003991119.5;refGene_gn=FN1;refGene_tx=XM_003991120.4;refGene_gn=FN1;refGene_tx=XM_003991121.4;refGene_gn=FN1;refGene_tx=XM_006935534.3;refGene_gn=FN1;refGene_tx=XM_006935535.3;refGene_gn=FN1;refGene_tx=XM_006935536.3;refGene_gn=FN1;refGene_tx=XM_006935537.3;refGene_gn=FN1;refGene_tx=XM_006935538.3;refGene_gn=FN1;refGene_tx=XM_006935539.3;refGene_gn=FN1;refGene_tx=XM_019838641.1
C1 200907221 200907401
refGene_gn=FN1;refGene_tx=XM_003991118.4;refGene_gn=FN1;refGene_tx=XM_003991119.5;refGene_gn=FN1;refGene_tx=XM_003991120.4;refGene_gn=FN1;refGene_tx=XM_003991121.4;refGene_gn=FN1;refGene_tx=XM_006935534.3;refGene_gn=FN1;refGene_tx=XM_006935535.3;refGene_gn=FN1;refGene_tx=XM_006935536.3;refGene_gn=FN1;refGene_tx=XM_006935537.3;refGene_gn=FN1;refGene_tx=XM_006935538.3;refGene_gn=FN1;refGene_tx=XM_006935539.3;refGene_gn=FN1;refGene_tx=XM_019838641.1
C1 200907654 200907744
refGene_gn=FN1;refGene_tx=XM_003991118.4;refGene_gn=FN1;refGene_tx=XM_003991119.5;refGene_gn=FN1;refGene_tx=XM_003991120.4;refGene_gn=FN1;refGene_tx=XM_003991121.4;refGene_gn=FN1;refGene_tx=XM_006935534.3;refGene_gn=FN1;refGene_tx=XM_006935535.3;refGene_gn=FN1;refGene_tx=XM_006935536.3;refGene_gn=FN1;refGene_tx=XM_006935537.3;refGene_gn=FN1;refGene_tx=XM_006935538.3;refGene_gn=FN1;refGene_tx=XM_006935539.3;refGene_gn=FN1;refGene_tx=XM_019838641.1
C1 200908495 200908672
refGene_gn=FN1;refGene_tx=XM_003991118.4;refGene_gn=FN1;refGene_tx=XM_003991119.5;refGene_gn=FN1;refGene_tx=XM_003991120.4;refGene_gn=FN1;refGene_tx=XM_003991121.4;refGene_gn=FN1;refGene_tx=XM_006935534.3;refGene_gn=FN1;refGene_tx=XM_006935535.3;refGene_gn=FN1;refGene_tx=XM_006935536.3;refGene_gn=FN1;refGene_tx=XM_006935537.3;refGene_gn=FN1;refGene_tx=XM_006935538.3;refGene_gn=FN1;refGene_tx=XM_006935539.3;refGene_gn=FN1;refGene_tx=XM_019838641.1
C1 200909560 200909648
refGene_gn=FN1;refGene_tx=XM_003991118.4;refGene_gn=FN1;refGene_tx=XM_003991119.5;refGene_gn=FN1;refGene_tx=XM_003991120.4;refGene_gn=FN1;refGene_tx=XM_003991121.4;refGene_gn=FN1;refGene_tx=XM_006935534.3;refGene_gn=FN1;refGene_tx=XM_006935535.3;refGene_gn=FN1;refGene_tx=XM_006935536.3;refGene_gn=FN1;refGene_tx=XM_006935537.3;refGene_gn=FN1;refGene_tx=XM_006935538.3;refGene_gn=FN1;refGene_tx=XM_006935539.3;refGene_gn

=FN1;refGene_tx=XM_019838641.1
C1	200910891	200911079
refGene_gn=FN1;refGene_tx=XM_003991118.4;refGene_gn=FN1;refGene_tx=XM_003991119.5;refGene_gn=FN1;refGene_tx=XM_003991120.4;refGene_gn=FN1;refGene_tx=XM_003991121.4;refGene_gn=FN1;refGene_tx=XM_006935534.3;refGene_gn=FN1;refGene_tx=XM_006935535.3;refGene_gn=FN1;refGene_tx=XM_006935536.3;refGene_gn=FN1;refGene_tx=XM_006935537.3;refGene_gn=FN1;refGene_tx=XM_006935538.3;refGene_gn=FN1;refGene_tx=XM_006935539.3;refGene_gn=FN1;refGene_tx=XM_019838641.1
C1	200912309	200912579
refGene_gn=FN1;refGene_tx=XM_003991118.4;refGene_gn=FN1;refGene_tx=XM_006935534.3;refGene_gn=FN1;refGene_tx=XM_006935536.3;refGene_gn=FN1;refGene_tx=XM_006935537.3;refGene_gn=FN1;refGene_tx=XM_006935538.3;refGene_gn=FN1;refGene_tx=XM_019838641.1
C1	200913431	200913545
refGene_gn=FN1;refGene_tx=XM_003991118.4;refGene_gn=FN1;refGene_tx=XM_003991119.5;refGene_gn=FN1;refGene_tx=XM_003991120.4;refGene_gn=FN1;refGene_tx=XM_003991121.4;refGene_gn=FN1;refGene_tx=XM_006935534.3;refGene_gn=FN1;refGene_tx=XM_006935535.3;refGene_gn=FN1;refGene_tx=XM_006935536.3;refGene_gn=FN1;refGene_tx=XM_006935537.3;refGene_gn=FN1;refGene_tx=XM_006935538.3;refGene_gn=FN1;refGene_tx=XM_006935539.3;refGene_gn=FN1;refGene_tx=XM_019838641.1
C1	200914219	200914375
refGene_gn=FN1;refGene_tx=XM_003991118.4;refGene_gn=FN1;refGene_tx=XM_003991119.5;refGene_gn=FN1;refGene_tx=XM_003991120.4;refGene_gn=FN1;refGene_tx=XM_003991121.4;refGene_gn=FN1;refGene_tx=XM_006935534.3;refGene_gn=FN1;refGene_tx=XM_006935535.3;refGene_gn=FN1;refGene_tx=XM_006935536.3;refGene_gn=FN1;refGene_tx=XM_006935537.3;refGene_gn=FN1;refGene_tx=XM_006935538.3;refGene_gn=FN1;refGene_tx=XM_006935539.3;refGene_gn=FN1;refGene_tx=XM_019838641.1
C1	200915135	200915300
refGene_gn=FN1;refGene_tx=XM_003991118.4;refGene_gn=FN1;refGene_tx=XM_003991119.5;refGene_gn=FN1;refGene_tx=XM_003991120.4;refGene_gn=FN1;refGene_tx=XM_003991121.4;refGene_gn=FN1;refGene_tx=XM_006935534.3;refGene_gn=FN1;refGene_tx=XM_006935535.3;refGene_gn=FN1;refGene_tx=XM_006935536.3;refGene_gn=FN1;refGene_tx=XM_006935537.3;refGene_gn=FN1;refGene_tx=XM_006935538.3;refGene_gn=FN1;refGene_tx=XM_006935539.3;refGene_gn=FN1;refGene_tx=XM_019838641.1
C1	200916188	200916305
refGene_gn=FN1;refGene_tx=XM_003991118.4;refGene_gn=FN1;refGene_tx=XM_003991119.5;refGene_gn=FN1;refGene_tx=XM_003991120.4;refGene_gn=FN1;refGene_tx=XM_003991121.4;refGene_gn=FN1;refGene_tx=XM_006935534.3;refGene_gn=FN1;refGene_tx=XM_006935535.3;refGene_gn=FN1;refGene_tx=XM_006935536.3;refGene_gn=FN1;refGene_tx=XM_006935537.3;refGene_gn=FN1;refGene_tx=XM_006935538.3;refGene_gn=FN1;refGene_tx=XM_006935539.3;refGene_gn=FN1;refGene_tx=XM_019838641.1
C1	200917300	200917570
refGene_gn=FN1;refGene_tx=XM_003991118.4;refGene_gn=FN1;refGene_tx=XM_003991119.5;refGene_gn=FN1;refGene_tx=XM_003991120.4;refGene_gn=FN1;refGene_tx=XM_003991121.4;refGene_gn=FN1;refGene_tx=XM_006935534.3;refGe

ne_gn=FN1;refGene_tx=XM_006935535.3;refGene_gn=FN1;refGene_tx=XM_00693
5536.3;refGene_gn=FN1;refGene_tx=XM_006935537.3;refGene_gn=FN1;refGene
_tx=XM_006935538.3;refGene_gn=FN1;refGene_tx=XM_006935539.3;refGene_gn
=FN1;refGene_tx=XM_019838641.1
C1	200918857	200918947
refGene_gn=FN1;refGene_tx=XM_003991118.4;refGene_gn=FN1;refGene_tx=XM_
003991119.5;refGene_gn=FN1;refGene_tx=XM_003991120.4;refGene_gn=FN1;re
fGene_tx=XM_003991121.4;refGene_gn=FN1;refGene_tx=XM_006935534.3;refGe
ne_gn=FN1;refGene_tx=XM_006935535.3;refGene_gn=FN1;refGene_tx=XM_00693
5536.3;refGene_gn=FN1;refGene_tx=XM_006935537.3;refGene_gn=FN1;refGene
_tx=XM_006935538.3;refGene_gn=FN1;refGene_tx=XM_006935539.3;refGene_gn
=FN1;refGene_tx=XM_019838641.1
C1	200921075	200921258
refGene_gn=FN1;refGene_tx=XM_003991118.4;refGene_gn=FN1;refGene_tx=XM_
003991119.5;refGene_gn=FN1;refGene_tx=XM_003991120.4;refGene_gn=FN1;re
fGene_tx=XM_003991121.4;refGene_gn=FN1;refGene_tx=XM_006935534.3;refGe
ne_gn=FN1;refGene_tx=XM_006935535.3;refGene_gn=FN1;refGene_tx=XM_00693
5536.3;refGene_gn=FN1;refGene_tx=XM_006935537.3;refGene_gn=FN1;refGene
_tx=XM_006935538.3;refGene_gn=FN1;refGene_tx=XM_006935539.3;refGene_gn
=FN1;refGene_tx=XM_019838641.1
C1	200922380	200922653
refGene_gn=FN1;refGene_tx=XM_003991118.4;refGene_gn=FN1;refGene_tx=XM_
003991121.4;refGene_gn=FN1;refGene_tx=XM_006935534.3;refGene_gn=FN1;re
fGene_tx=XM_006935535.3;refGene_gn=FN1;refGene_tx=XM_006935538.3;refGe
ne_gn=FN1;refGene_tx=XM_019838641.1
C1	200923955	200924147
refGene_gn=FN1;refGene_tx=XM_003991118.4;refGene_gn=FN1;refGene_tx=XM_
003991119.5;refGene_gn=FN1;refGene_tx=XM_003991120.4;refGene_gn=FN1;re
fGene_tx=XM_003991121.4;refGene_gn=FN1;refGene_tx=XM_006935534.3;refGe
ne_gn=FN1;refGene_tx=XM_006935535.3;refGene_gn=FN1;refGene_tx=XM_00693
5536.3;refGene_gn=FN1;refGene_tx=XM_006935537.3;refGene_gn=FN1;refGene
_tx=XM_006935538.3;refGene_gn=FN1;refGene_tx=XM_006935539.3;refGene_gn
=FN1;refGene_tx=XM_019838641.1
C1	200925397	200925484
refGene_gn=FN1;refGene_tx=XM_003991118.4;refGene_gn=FN1;refGene_tx=XM_
003991119.5;refGene_gn=FN1;refGene_tx=XM_003991120.4;refGene_gn=FN1;re
fGene_tx=XM_003991121.4;refGene_gn=FN1;refGene_tx=XM_006935534.3;refGe
ne_gn=FN1;refGene_tx=XM_006935535.3;refGene_gn=FN1;refGene_tx=XM_00693
5536.3;refGene_gn=FN1;refGene_tx=XM_006935537.3;refGene_gn=FN1;refGene
_tx=XM_006935538.3;refGene_gn=FN1;refGene_tx=XM_006935539.3;refGene_gn
=FN1;refGene_tx=XM_019838641.1
C1	200926026	200926195
refGene_gn=FN1;refGene_tx=XM_003991118.4;refGene_gn=FN1;refGene_tx=XM_
003991119.5;refGene_gn=FN1;refGene_tx=XM_003991120.4;refGene_gn=FN1;re
fGene_tx=XM_003991121.4;refGene_gn=FN1;refGene_tx=XM_006935534.3;refGe
ne_gn=FN1;refGene_tx=XM_006935535.3;refGene_gn=FN1;refGene_tx=XM_00693
5536.3;refGene_gn=FN1;refGene_tx=XM_006935537.3;refGene_gn=FN1;refGene
_tx=XM_006935538.3;refGene_gn=FN1;refGene_tx=XM_006935539.3;refGene_gn
=FN1;refGene_tx=XM_019838641.1
C1	200927892	200927987

refGene_gn=FN1;refGene_tx=XM_003991118.4;refGene_gn=FN1;refGene_tx=XM_
003991119.5;refGene_gn=FN1;refGene_tx=XM_003991120.4;refGene_gn=FN1;re
fGene_tx=XM_003991121.4;refGene_gn=FN1;refGene_tx=XM_006935534.3;refGe
ne_gn=FN1;refGene_tx=XM_006935535.3;refGene_gn=FN1;refGene_tx=XM_00693
5536.3;refGene_gn=FN1;refGene_tx=XM_006935537.3;refGene_gn=FN1;refGene
_tx=XM_006935538.3;refGene_gn=FN1;refGene_tx=XM_006935539.3;refGene_gn
=FN1;refGene_tx=XM_019838641.1
C1 200933060 200933327
refGene_gn=FN1;refGene_tx=XM_003991118.4;refGene_gn=FN1;refGene_tx=XM_
003991119.5;refGene_gn=FN1;refGene_tx=XM_003991120.4;refGene_gn=FN1;re
fGene_tx=XM_003991121.4;refGene_gn=FN1;refGene_tx=XM_006935534.3;refGe
ne_gn=FN1;refGene_tx=XM_006935535.3;refGene_gn=FN1;refGene_tx=XM_00693
5536.3;refGene_gn=FN1;refGene_tx=XM_006935537.3;refGene_gn=FN1;refGene
_tx=XM_006935538.3;refGene_gn=FN1;refGene_tx=XM_006935539.3;refGene_gn
=FN1;refGene_tx=XM_019838641.1
C1 200934646 200934919
refGene_gn=FN1;refGene_tx=XM_003991118.4;refGene_gn=FN1;refGene_tx=XM_
003991119.5;refGene_gn=FN1;refGene_tx=XM_003991120.4;refGene_gn=FN1;re
fGene_tx=XM_003991121.4;refGene_gn=FN1;refGene_tx=XM_006935534.3;refGe
ne_gn=FN1;refGene_tx=XM_006935535.3;refGene_gn=FN1;refGene_tx=XM_00693
5536.3;refGene_gn=FN1;refGene_tx=XM_006935537.3;refGene_gn=FN1;refGene
_tx=XM_006935538.3;refGene_gn=FN1;refGene_tx=XM_006935539.3;refGene_gn
=FN1;refGene_tx=XM_019838641.1
C1 200935707 200935902
refGene_gn=FN1;refGene_tx=XM_003991118.4;refGene_gn=FN1;refGene_tx=XM_
003991119.5;refGene_gn=FN1;refGene_tx=XM_003991120.4;refGene_gn=FN1;re
fGene_tx=XM_003991121.4;refGene_gn=FN1;refGene_tx=XM_006935534.3;refGe
ne_gn=FN1;refGene_tx=XM_006935535.3;refGene_gn=FN1;refGene_tx=XM_00693
5536.3;refGene_gn=FN1;refGene_tx=XM_006935537.3;refGene_gn=FN1;refGene
_tx=XM_006935538.3;refGene_gn=FN1;refGene_tx=XM_006935539.3;refGene_gn
=FN1;refGene_tx=XM_019838641.1
C1 200936919 200937009
refGene_gn=FN1;refGene_tx=XM_003991118.4;refGene_gn=FN1;refGene_tx=XM_
003991119.5;refGene_gn=FN1;refGene_tx=XM_003991120.4;refGene_gn=FN1;re
fGene_tx=XM_003991121.4;refGene_gn=FN1;refGene_tx=XM_006935534.3;refGe
ne_gn=FN1;refGene_tx=XM_006935535.3;refGene_gn=FN1;refGene_tx=XM_00693
5536.3;refGene_gn=FN1;refGene_tx=XM_006935537.3;refGene_gn=FN1;refGene
_tx=XM_006935538.3;refGene_gn=FN1;refGene_tx=XM_006935539.3;refGene_gn
=FN1;refGene_tx=XM_019838641.1
C1 200937116 200937245
refGene_gn=FN1;refGene_tx=XM_003991118.4;refGene_gn=FN1;refGene_tx=XM_
003991119.5;refGene_gn=FN1;refGene_tx=XM_003991120.4;refGene_gn=FN1;re
fGene_tx=XM_003991121.4;refGene_gn=FN1;refGene_tx=XM_006935534.3;refGe
ne_gn=FN1;refGene_tx=XM_006935535.3;refGene_gn=FN1;refGene_tx=XM_00693
5536.3;refGene_gn=FN1;refGene_tx=XM_006935537.3;refGene_gn=FN1;refGene
_tx=XM_006935538.3;refGene_gn=FN1;refGene_tx=XM_006935539.3;refGene_gn
=FN1;refGene_tx=XM_019838641.1
C1 200938268 200938445
refGene_gn=FN1;refGene_tx=XM_003991118.4;refGene_gn=FN1;refGene_tx=XM_
003991119.5;refGene_gn=FN1;refGene_tx=XM_003991120.4;refGene_gn=FN1;re

fGene_tx=XM_003991121.4;refGene_gn=FN1;refGene_tx=XM_006935534.3;refGene_gn=FN1;refGene_tx=XM_006935535.3;refGene_gn=FN1;refGene_tx=XM_006935536.3;refGene_gn=FN1;refGene_tx=XM_006935537.3;refGene_gn=FN1;refGene_tx=XM_006935538.3;refGene_gn=FN1;refGene_tx=XM_006935539.3;refGene_gn=FN1;refGene_tx=XM_019838641.1
C1 200938643 200938824
refGene_gn=FN1;refGene_tx=XM_003991118.4;refGene_gn=FN1;refGene_tx=XM_003991119.5;refGene_gn=FN1;refGene_tx=XM_003991120.4;refGene_gn=FN1;refGene_tx=XM_003991121.4;refGene_gn=FN1;refGene_tx=XM_006935534.3;refGene_gn=FN1;refGene_tx=XM_006935535.3;refGene_gn=FN1;refGene_tx=XM_006935536.3;refGene_gn=FN1;refGene_tx=XM_006935537.3;refGene_gn=FN1;refGene_tx=XM_006935538.3;refGene_gn=FN1;refGene_tx=XM_006935539.3;refGene_gn=FN1;refGene_tx=XM_019838641.1
C1 200941472 200941594
refGene_gn=FN1;refGene_tx=XM_003991118.4;refGene_gn=FN1;refGene_tx=XM_003991119.5;refGene_gn=FN1;refGene_tx=XM_003991120.4;refGene_gn=FN1;refGene_tx=XM_003991121.4;refGene_gn=FN1;refGene_tx=XM_006935534.3;refGene_gn=FN1;refGene_tx=XM_006935535.3;refGene_gn=FN1;refGene_tx=XM_006935536.3;refGene_gn=FN1;refGene_tx=XM_006935537.3;refGene_gn=FN1;refGene_tx=XM_006935538.3;refGene_gn=FN1;refGene_tx=XM_006935539.3;refGene_gn=FN1;refGene_tx=XM_019838641.1
C1 200946338 200946482
refGene_gn=FN1;refGene_tx=XM_003991118.4;refGene_gn=FN1;refGene_tx=XM_003991119.5;refGene_gn=FN1;refGene_tx=XM_003991120.4;refGene_gn=FN1;refGene_tx=XM_003991121.4;refGene_gn=FN1;refGene_tx=XM_006935534.3;refGene_gn=FN1;refGene_tx=XM_006935535.3;refGene_gn=FN1;refGene_tx=XM_006935536.3;refGene_gn=FN1;refGene_tx=XM_006935537.3;refGene_gn=FN1;refGene_tx=XM_006935538.3;refGene_gn=FN1;refGene_tx=XM_006935539.3;refGene_gn=FN1;refGene_tx=XM_019838641.1
C1 200948325 200948454
refGene_gn=FN1;refGene_tx=XM_003991118.4;refGene_gn=FN1;refGene_tx=XM_003991119.5;refGene_gn=FN1;refGene_tx=XM_003991120.4;refGene_gn=FN1;refGene_tx=XM_003991121.4;refGene_gn=FN1;refGene_tx=XM_006935534.3;refGene_gn=FN1;refGene_tx=XM_006935535.3;refGene_gn=FN1;refGene_tx=XM_006935536.3;refGene_gn=FN1;refGene_tx=XM_006935537.3;refGene_gn=FN1;refGene_tx=XM_006935538.3;refGene_gn=FN1;refGene_tx=XM_006935539.3;refGene_gn=FN1;refGene_tx=XM_019838641.1
C1 200949728 200949881
refGene_gn=FN1;refGene_tx=XM_003991118.4;refGene_gn=FN1;refGene_tx=XM_003991119.5;refGene_gn=FN1;refGene_tx=XM_003991120.4;refGene_gn=FN1;refGene_tx=XM_003991121.4;refGene_gn=FN1;refGene_tx=XM_006935534.3;refGene_gn=FN1;refGene_tx=XM_006935535.3;refGene_gn=FN1;refGene_tx=XM_006935536.3;refGene_gn=FN1;refGene_tx=XM_006935537.3;refGene_gn=FN1;refGene_tx=XM_006935538.3;refGene_gn=FN1;refGene_tx=XM_006935539.3;refGene_gn=FN1;refGene_tx=XM_019838641.1
C1 200950938 200951115
refGene_gn=FN1;refGene_tx=XM_003991118.4;refGene_gn=FN1;refGene_tx=XM_003991119.5;refGene_gn=FN1;refGene_tx=XM_003991120.4;refGene_gn=FN1;refGene_tx=XM_003991121.4;refGene_gn=FN1;refGene_tx=XM_006935534.3;refGene_gn=FN1;refGene_tx=XM_006935535.3;refGene_gn=FN1;refGene_tx=XM_00693

5536.3;refGene_gn=FN1;refGene_tx=XM_006935537.3;refGene_gn=FN1;refGene_tx=XM_006935538.3;refGene_gn=FN1;refGene_tx=XM_006935539.3;refGene_gn=FN1;refGene_tx=XM_019838641.1
C1 200951774 200951954
refGene_gn=FN1;refGene_tx=XM_003991118.4;refGene_gn=FN1;refGene_tx=XM_003991119.5;refGene_gn=FN1;refGene_tx=XM_003991120.4;refGene_gn=FN1;refGene_tx=XM_003991121.4;refGene_gn=FN1;refGene_tx=XM_006935534.3;refGene_gn=FN1;refGene_tx=XM_006935535.3;refGene_gn=FN1;refGene_tx=XM_006935536.3;refGene_gn=FN1;refGene_tx=XM_006935537.3;refGene_gn=FN1;refGene_tx=XM_006935538.3;refGene_gn=FN1;refGene_tx=XM_006935539.3;refGene_gn=FN1;refGene_tx=XM_019838641.1
C1 200952782 200952974
refGene_gn=FN1;refGene_tx=XM_003991118.4;refGene_gn=FN1;refGene_tx=XM_003991119.5;refGene_gn=FN1;refGene_tx=XM_003991120.4;refGene_gn=FN1;refGene_tx=XM_003991121.4;refGene_gn=FN1;refGene_tx=XM_006935534.3;refGene_gn=FN1;refGene_tx=XM_006935535.3;refGene_gn=FN1;refGene_tx=XM_006935536.3;refGene_gn=FN1;refGene_tx=XM_006935537.3;refGene_gn=FN1;refGene_tx=XM_006935538.3;refGene_gn=FN1;refGene_tx=XM_006935539.3;refGene_gn=FN1;refGene_tx=XM_019838641.1
C1 200955472 200955631
refGene_gn=FN1;refGene_tx=XM_003991118.4;refGene_gn=FN1;refGene_tx=XM_003991119.5;refGene_gn=FN1;refGene_tx=XM_003991120.4;refGene_gn=FN1;refGene_tx=XM_003991121.4;refGene_gn=FN1;refGene_tx=XM_006935534.3;refGene_gn=FN1;refGene_tx=XM_006935535.3;refGene_gn=FN1;refGene_tx=XM_006935536.3;refGene_gn=FN1;refGene_tx=XM_006935537.3;refGene_gn=FN1;refGene_tx=XM_006935538.3;refGene_gn=FN1;refGene_tx=XM_006935539.3;refGene_gn=FN1;refGene_tx=XM_019838641.1
C1 200957841 200957979
refGene_gn=FN1;refGene_tx=XM_003991118.4;refGene_gn=FN1;refGene_tx=XM_003991119.5;refGene_gn=FN1;refGene_tx=XM_003991120.4;refGene_gn=FN1;refGene_tx=XM_003991121.4;refGene_gn=FN1;refGene_tx=XM_006935534.3;refGene_gn=FN1;refGene_tx=XM_006935535.3;refGene_gn=FN1;refGene_tx=XM_006935536.3;refGene_gn=FN1;refGene_tx=XM_006935537.3;refGene_gn=FN1;refGene_tx=XM_006935538.3;refGene_gn=FN1;refGene_tx=XM_006935539.3;refGene_gn=FN1;refGene_tx=XM_019838641.1
C1 200958917 200959049
refGene_gn=FN1;refGene_tx=XM_003991118.4;refGene_gn=FN1;refGene_tx=XM_003991119.5;refGene_gn=FN1;refGene_tx=XM_003991120.4;refGene_gn=FN1;refGene_tx=XM_003991121.4;refGene_gn=FN1;refGene_tx=XM_006935534.3;refGene_gn=FN1;refGene_tx=XM_006935535.3;refGene_gn=FN1;refGene_tx=XM_006935536.3;refGene_gn=FN1;refGene_tx=XM_006935537.3;refGene_gn=FN1;refGene_tx=XM_006935538.3;refGene_gn=FN1;refGene_tx=XM_006935539.3;refGene_gn=FN1;refGene_tx=XM_019838641.1
C1 200960640 200960778
refGene_gn=FN1;refGene_tx=XM_003991118.4;refGene_gn=FN1;refGene_tx=XM_003991119.5;refGene_gn=FN1;refGene_tx=XM_003991120.4;refGene_gn=FN1;refGene_tx=XM_003991121.4;refGene_gn=FN1;refGene_tx=XM_006935534.3;refGene_gn=FN1;refGene_tx=XM_006935535.3;refGene_gn=FN1;refGene_tx=XM_006935536.3;refGene_gn=FN1;refGene_tx=XM_006935537.3;refGene_gn=FN1;refGene_tx=XM_006935538.3;refGene_gn=FN1;refGene_tx=XM_006935539.3;refGene_gn

=FN1;refGene_tx=XM_019838641.1
C1	200961707	200961836
refGene_gn=FN1;refGene_tx=XM_003991118.4;refGene_gn=FN1;refGene_tx=XM_003991119.5;refGene_gn=FN1;refGene_tx=XM_003991120.4;refGene_gn=FN1;refGene_tx=XM_003991121.4;refGene_gn=FN1;refGene_tx=XM_006935534.3;refGene_gn=FN1;refGene_tx=XM_006935535.3;refGene_gn=FN1;refGene_tx=XM_006935536.3;refGene_gn=FN1;refGene_tx=XM_006935537.3;refGene_gn=FN1;refGene_tx=XM_006935538.3;refGene_gn=FN1;refGene_tx=XM_006935539.3;refGene_gn=FN1;refGene_tx=XM_019838641.1
C1	200962828	200962976
refGene_gn=FN1;refGene_tx=XM_003991118.4;refGene_gn=FN1;refGene_tx=XM_003991119.5;refGene_gn=FN1;refGene_tx=XM_003991120.4;refGene_gn=FN1;refGene_tx=XM_003991121.4;refGene_gn=FN1;refGene_tx=XM_006935534.3;refGene_gn=FN1;refGene_tx=XM_006935535.3;refGene_gn=FN1;refGene_tx=XM_006935536.3;refGene_gn=FN1;refGene_tx=XM_006935537.3;refGene_gn=FN1;refGene_tx=XM_006935538.3;refGene_gn=FN1;refGene_tx=XM_006935539.3;refGene_gn=FN1;refGene_tx=XM_019838641.1
C1	201411898	201412033
refGene_gn=MREG;refGene_tx=XM_023259667.1
C1	201413326	201413490
refGene_gn=MREG;refGene_tx=XM_023259667.1
C1	201415054	201415145
refGene_gn=MREG;refGene_tx=XM_023259667.1
C1	201451835	201451995
refGene_gn=MREG;refGene_tx=XM_023259667.1
C1	201469174	201469269
refGene_gn=MREG;refGene_tx=XM_023259667.1
C1	201489239	201489325
refGene_gn=PECR;refGene_tx=XM_011285577.3;refGene_gn=PECR;refGene_tx=XM_011285578.3
C1	201492457	201492569
refGene_gn=PECR;refGene_tx=XM_011285577.3
C1	201499976	201500087
refGene_gn=PECR;refGene_tx=XM_011285577.3;refGene_gn=PECR;refGene_tx=XM_011285578.3
C1	201502159	201502256
refGene_gn=PECR;refGene_tx=XM_011285577.3;refGene_gn=PECR;refGene_tx=XM_011285578.3
C1	201506791	201506873
refGene_gn=PECR;refGene_tx=XM_011285577.3;refGene_gn=PECR;refGene_tx=XM_011285578.3
C1	201512535	201512701
refGene_gn=PECR;refGene_tx=XM_011285577.3;refGene_gn=PECR;refGene_tx=XM_011285578.3
C1	201513409	201513543
refGene_gn=PECR;refGene_tx=XM_011285577.3;refGene_gn=PECR;refGene_tx=XM_011285578.3
C1	201517429	201517553
refGene_gn=PECR;refGene_tx=XM_011285577.3;refGene_gn=PECR;refGene_tx=XM_011285578.3

C1	201535208	201535479
refGene_gn=TMEM169;refGene_tx=XM_003991122.5;refGene_gn=TMEM169;refGene_tx=XM_019838646.2;refGene_gn=TMEM169;refGene_tx=XM_019838648.2
C1	201537842	201538465
refGene_gn=TMEM169;refGene_tx=XM_003991122.5;refGene_gn=TMEM169;refGene_tx=XM_019838646.2;refGene_gn=TMEM169;refGene_tx=XM_019838648.2
C1	201543880	201543901
refGene_gn=XRCC5;refGene_tx=XM_006935540.4;refGene_gn=XRCC5;refGene_tx=XM_006935541.3;refGene_gn=XRCC5;refGene_tx=XM_019838644.1;refGene_gn=XRCC5;refGene_tx=XM_023259668.1
C1	201547983	201548097
refGene_gn=XRCC5;refGene_tx=XM_006935540.4;refGene_gn=XRCC5;refGene_tx=XM_006935541.3;refGene_gn=XRCC5;refGene_tx=XM_019838644.1;refGene_gn=XRCC5;refGene_tx=XM_023259668.1
C1	201551775	201551959
refGene_gn=XRCC5;refGene_tx=XM_006935540.4;refGene_gn=XRCC5;refGene_tx=XM_006935541.3;refGene_gn=XRCC5;refGene_tx=XM_019838644.1;refGene_gn=XRCC5;refGene_tx=XM_023259668.1
C1	201552873	201552922
refGene_gn=XRCC5;refGene_tx=XM_006935540.4;refGene_gn=XRCC5;refGene_tx=XM_006935541.3;refGene_gn=XRCC5;refGene_tx=XM_019838644.1;refGene_gn=XRCC5;refGene_tx=XM_023259668.1
C1	201554201	201554324
refGene_gn=XRCC5;refGene_tx=XM_006935540.4;refGene_gn=XRCC5;refGene_tx=XM_006935541.3;refGene_gn=XRCC5;refGene_tx=XM_019838644.1;refGene_gn=XRCC5;refGene_tx=XM_023259668.1
C1	201557317	201557509
refGene_gn=XRCC5;refGene_tx=XM_006935540.4;refGene_gn=XRCC5;refGene_tx=XM_006935541.3;refGene_gn=XRCC5;refGene_tx=XM_019838644.1;refGene_gn=XRCC5;refGene_tx=XM_023259668.1
C1	201559603	201559718
refGene_gn=XRCC5;refGene_tx=XM_006935540.4;refGene_gn=XRCC5;refGene_tx=XM_006935541.3;refGene_gn=XRCC5;refGene_tx=XM_019838644.1;refGene_gn=XRCC5;refGene_tx=XM_023259668.1
C1	201561194	201561333
refGene_gn=XRCC5;refGene_tx=XM_006935540.4;refGene_gn=XRCC5;refGene_tx=XM_006935541.3;refGene_gn=XRCC5;refGene_tx=XM_019838644.1;refGene_gn=XRCC5;refGene_tx=XM_023259668.1
C1	201564326	201564439
refGene_gn=XRCC5;refGene_tx=XM_006935540.4;refGene_gn=XRCC5;refGene_tx=XM_006935541.3;refGene_gn=XRCC5;refGene_tx=XM_019838644.1;refGene_gn=XRCC5;refGene_tx=XM_023259668.1
C1	201569011	201569074
refGene_gn=XRCC5;refGene_tx=XM_006935540.4;refGene_gn=XRCC5;refGene_tx=XM_006935541.3;refGene_gn=XRCC5;refGene_tx=XM_019838644.1;refGene_gn=XRCC5;refGene_tx=XM_023259668.1
C1	201574699	201574837
refGene_gn=XRCC5;refGene_tx=XM_006935540.4;refGene_gn=XRCC5;refGene_tx=XM_006935541.3;refGene_gn=XRCC5;refGene_tx=XM_019838644.1;refGene_gn=XRCC5;refGene_tx=XM_023259668.1

C1	201575879	201575970
refGene_gn=XRCC5;refGene_tx=XM_006935540.4;refGene_gn=XRCC5;refGene_tx=XM_006935541.3;refGene_gn=XRCC5;refGene_tx=XM_019838644.1;refGene_gn=XRCC5;refGene_tx=XM_023259668.1
C1	201578306	201578440
refGene_gn=XRCC5;refGene_tx=XM_006935540.4;refGene_gn=XRCC5;refGene_tx=XM_006935541.3;refGene_gn=XRCC5;refGene_tx=XM_019838644.1;refGene_gn=XRCC5;refGene_tx=XM_023259668.1
C1	201584335	201584532
refGene_gn=XRCC5;refGene_tx=XM_006935540.4;refGene_gn=XRCC5;refGene_tx=XM_006935541.3;refGene_gn=XRCC5;refGene_tx=XM_019838644.1;refGene_gn=XRCC5;refGene_tx=XM_023259668.1
C1	201593067	201593161
refGene_gn=XRCC5;refGene_tx=XM_006935540.4;refGene_gn=XRCC5;refGene_tx=XM_006935541.3;refGene_gn=XRCC5;refGene_tx=XM_019838644.1;refGene_gn=XRCC5;refGene_tx=XM_023259668.1
C1	201594859	201594929
refGene_gn=XRCC5;refGene_tx=XM_006935540.4;refGene_gn=XRCC5;refGene_tx=XM_006935541.3;refGene_gn=XRCC5;refGene_tx=XM_019838644.1;refGene_gn=XRCC5;refGene_tx=XM_023259668.1
C1	201619563	201619673
refGene_gn=XRCC5;refGene_tx=XM_006935540.4;refGene_gn=XRCC5;refGene_tx=XM_006935541.3;refGene_gn=XRCC5;refGene_tx=XM_019838644.1;refGene_gn=XRCC5;refGene_tx=XM_023259668.1
C1	201621604	201621701
refGene_gn=XRCC5;refGene_tx=XM_006935540.4;refGene_gn=XRCC5;refGene_tx=XM_006935541.3;refGene_gn=XRCC5;refGene_tx=XM_019838644.1;refGene_gn=XRCC5;refGene_tx=XM_023259668.1
C1	201623725	201623793
refGene_gn=XRCC5;refGene_tx=XM_006935540.4;refGene_gn=XRCC5;refGene_tx=XM_006935541.3;refGene_gn=XRCC5;refGene_tx=XM_019838644.1;refGene_gn=XRCC5;refGene_tx=XM_023259668.1
C1	201630628	201630661
refGene_gn=XRCC5;refGene_tx=XM_023259668.1
C1	201633287	201633362
refGene_gn=XRCC5;refGene_tx=XM_006935540.4
C1	201633386	201633476
refGene_gn=XRCC5;refGene_tx=XM_006935541.3
C1	201634126	201634141
refGene_gn=XRCC5;refGene_tx=XM_006935540.4
C1	201667168	201667536
refGene_gn=MARCH4;refGene_tx=XM_003991123.2;refGene_gn=MARCH4;refGene_tx=XM_019838645.1
C1	201673892	201673895
refGene_gn=XRCC5;refGene_tx=XM_019838644.1
C1	201686134	201686327
refGene_gn=MARCH4;refGene_tx=XM_003991123.2;refGene_gn=MARCH4;refGene_tx=XM_019838645.1
C1	201692003	201692159
refGene_gn=MARCH4;refGene_tx=XM_003991123.2

C1 201774164 201774680
refGene_gn=MARCH4;refGene_tx=XM_003991123.2;refGene_gn=MARCH4;refGene_tx=XM_019838645.1
C1 201811206 201812023
refGene_gn=SMARCAL1;refGene_tx=XM_011285571.2;refGene_gn=SMARCAL1;refGene_tx=XM_011285572.2;refGene_gn=SMARCAL1;refGene_tx=XM_011285573.3;refGene_gn=SMARCAL1;refGene_tx=XM_011285574.3;refGene_gn=SMARCAL1;refGene_tx=XM_011285575.3;refGene_gn=SMARCAL1;refGene_tx=XM_023259669.1
C1 201812783 201812834
refGene_gn=SMARCAL1;refGene_tx=XM_011285571.2;refGene_gn=SMARCAL1;refGene_tx=XM_011285572.2;refGene_gn=SMARCAL1;refGene_tx=XM_011285573.3;refGene_gn=SMARCAL1;refGene_tx=XM_011285574.3;refGene_gn=SMARCAL1;refGene_tx=XM_011285575.3;refGene_gn=SMARCAL1;refGene_tx=XM_023259669.1
C1 201816824 201817058
refGene_gn=SMARCAL1;refGene_tx=XM_011285571.2;refGene_gn=SMARCAL1;refGene_tx=XM_011285572.2;refGene_gn=SMARCAL1;refGene_tx=XM_011285573.3;refGene_gn=SMARCAL1;refGene_tx=XM_011285574.3;refGene_gn=SMARCAL1;refGene_tx=XM_011285575.3;refGene_gn=SMARCAL1;refGene_tx=XM_023259669.1
C1 201819191 201819242
refGene_gn=SMARCAL1;refGene_tx=XM_011285571.2;refGene_gn=SMARCAL1;refGene_tx=XM_011285572.2;refGene_gn=SMARCAL1;refGene_tx=XM_011285573.3;refGene_gn=SMARCAL1;refGene_tx=XM_011285574.3;refGene_gn=SMARCAL1;refGene_tx=XM_011285575.3;refGene_gn=SMARCAL1;refGene_tx=XM_023259669.1
C1 201824087 201824274
refGene_gn=SMARCAL1;refGene_tx=XM_011285571.2;refGene_gn=SMARCAL1;refGene_tx=XM_011285572.2;refGene_gn=SMARCAL1;refGene_tx=XM_011285573.3;refGene_gn=SMARCAL1;refGene_tx=XM_011285574.3;refGene_gn=SMARCAL1;refGene_tx=XM_011285575.3;refGene_gn=SMARCAL1;refGene_tx=XM_023259669.1
C1 201827256 201827407
refGene_gn=SMARCAL1;refGene_tx=XM_011285571.2;refGene_gn=SMARCAL1;refGene_tx=XM_011285572.2;refGene_gn=SMARCAL1;refGene_tx=XM_011285573.3;refGene_gn=SMARCAL1;refGene_tx=XM_011285574.3;refGene_gn=SMARCAL1;refGene_tx=XM_011285575.3;refGene_gn=SMARCAL1;refGene_tx=XM_023259669.1
C1 201829001 201829160
refGene_gn=SMARCAL1;refGene_tx=XM_011285571.2;refGene_gn=SMARCAL1;refGene_tx=XM_011285572.2;refGene_gn=SMARCAL1;refGene_tx=XM_011285573.3;refGene_gn=SMARCAL1;refGene_tx=XM_011285574.3;refGene_gn=SMARCAL1;refGene_tx=XM_011285575.3;refGene_gn=SMARCAL1;refGene_tx=XM_023259669.1
C1 201832254 201832320
refGene_gn=SMARCAL1;refGene_tx=XM_011285571.2;refGene_gn=SMARCAL1;refGene_tx=XM_011285572.2;refGene_gn=SMARCAL1;refGene_tx=XM_011285573.3;refGene_gn=SMARCAL1;refGene_tx=XM_011285574.3;refGene_gn=SMARCAL1;refGene_tx=XM_011285575.3;refGene_gn=SMARCAL1;refGene_tx=XM_023259669.1
C1 201839913 201840054
refGene_gn=SMARCAL1;refGene_tx=XM_011285571.2;refGene_gn=SMARCAL1;refGene_tx=XM_011285572.2;refGene_gn=SMARCAL1;refGene_tx=XM_011285573.3;refGene_gn=SMARCAL1;refGene_tx=XM_011285574.3;refGene_gn=SMARCAL1;refGene_tx=XM_011285575.3;refGene_gn=SMARCAL1;refGene_tx=XM_023259669.1
C1 201844425 201844644
refGene_gn=SMARCAL1;refGene_tx=XM_011285571.2;refGene_gn=SMARCAL1;refG

ene_tx=XM_011285572.2;refGene_gn=SMARCAL1;refGene_tx=XM_011285573.3;refGene_gn=SMARCAL1;refGene_tx=XM_011285574.3;refGene_gn=SMARCAL1;refGene_tx=XM_011285575.3;refGene_gn=SMARCAL1;refGene_tx=XM_023259669.1
C1 201850958 201851029
refGene_gn=SMARCAL1;refGene_tx=XM_011285571.2;refGene_gn=SMARCAL1;refGene_tx=XM_011285572.2;refGene_gn=SMARCAL1;refGene_tx=XM_011285573.3;refGene_gn=SMARCAL1;refGene_tx=XM_011285574.3;refGene_gn=SMARCAL1;refGene_tx=XM_011285575.3;refGene_gn=SMARCAL1;refGene_tx=XM_023259669.1
C1 201854358 201854461
refGene_gn=SMARCAL1;refGene_tx=XM_011285571.2;refGene_gn=SMARCAL1;refGene_tx=XM_011285572.2;refGene_gn=SMARCAL1;refGene_tx=XM_011285573.3;refGene_gn=SMARCAL1;refGene_tx=XM_011285574.3;refGene_gn=SMARCAL1;refGene_tx=XM_011285575.3;refGene_gn=SMARCAL1;refGene_tx=XM_023259669.1
C1 201857713 201857896
refGene_gn=SMARCAL1;refGene_tx=XM_011285571.2;refGene_gn=SMARCAL1;refGene_tx=XM_011285572.2;refGene_gn=SMARCAL1;refGene_tx=XM_011285573.3;refGene_gn=SMARCAL1;refGene_tx=XM_011285574.3;refGene_gn=SMARCAL1;refGene_tx=XM_011285575.3;refGene_gn=SMARCAL1;refGene_tx=XM_023259669.1
C1 201858986 201859087
refGene_gn=SMARCAL1;refGene_tx=XM_011285571.2;refGene_gn=SMARCAL1;refGene_tx=XM_011285572.2;refGene_gn=SMARCAL1;refGene_tx=XM_011285573.3;refGene_gn=SMARCAL1;refGene_tx=XM_011285574.3;refGene_gn=SMARCAL1;refGene_tx=XM_011285575.3;refGene_gn=SMARCAL1;refGene_tx=XM_023259669.1
C1 201860250 201860347
refGene_gn=SMARCAL1;refGene_tx=XM_011285571.2;refGene_gn=SMARCAL1;refGene_tx=XM_011285572.2;refGene_gn=SMARCAL1;refGene_tx=XM_011285573.3;refGene_gn=SMARCAL1;refGene_tx=XM_011285574.3;refGene_gn=SMARCAL1;refGene_tx=XM_011285575.3;refGene_gn=SMARCAL1;refGene_tx=XM_023259669.1
C1 201863223 201863442
refGene_gn=SMARCAL1;refGene_tx=XM_011285571.2;refGene_gn=SMARCAL1;refGene_tx=XM_011285572.2;refGene_gn=SMARCAL1;refGene_tx=XM_011285573.3;refGene_gn=SMARCAL1;refGene_tx=XM_011285574.3;refGene_gn=SMARCAL1;refGene_tx=XM_011285575.3;refGene_gn=SMARCAL1;refGene_tx=XM_023259669.1
C1 201873274 201873277
refGene_gn=RPL37A;refGene_tx=XM_004001367.5
C1 201873655 201873784
refGene_gn=RPL37A;refGene_tx=XM_004001367.5
C1 201874319 201874402
refGene_gn=RPL37A;refGene_tx=XM_004001367.5
C1 201875949 201876013
refGene_gn=RPL37A;refGene_tx=XM_004001367.5
C1 201986124 201986581
refGene_gn=IGFBP2;refGene_tx=XM_011285583.3
C1 202007974 202008204
refGene_gn=IGFBP2;refGene_tx=XM_011285583.3
C1 202008948 202009089
refGene_gn=IGFBP2;refGene_tx=XM_011285583.3
C1 202010721 202010886
refGene_gn=IGFBP2;refGene_tx=XM_011285583.3
C1 202022429 202022561

refGene_gn=IGFBP5;refGene_tx=XM_003991128.5
C1 202023688 202023808
refGene_gn=IGFBP5;refGene_tx=XM_003991128.5
C1 202024403 202024633
refGene_gn=IGFBP5;refGene_tx=XM_003991128.5
C1 202038278 202038612
refGene_gn=IGFBP5;refGene_tx=XM_003991128.5
C1 202191000 202191029
refGene_gn=TNP1;refGene_tx=XM_003991129.5
C1 202191236 202191375
refGene_gn=TNP1;refGene_tx=XM_003991129.5
C1 203027540 203027685
refGene_gn=TNS1;refGene_tx=XM_006935547.3;refGene_gn=TNS1;refGene_tx=XM_023259670.1;refGene_gn=TNS1;refGene_tx=XM_023259671.1;refGene_gn=TNS1;refGene_tx=XM_023259672.1;refGene_gn=TNS1;refGene_tx=XM_023259673.1;refGene_gn=TNS1;refGene_tx=XM_023259674.1;refGene_gn=TNS1;refGene_tx=XM_023259675.1;refGene_gn=TNS1;refGene_tx=XM_023259676.1
C1 203030768 203030801
refGene_gn=TNS1;refGene_tx=XM_006935547.3;refGene_gn=TNS1;refGene_tx=XM_023259670.1;refGene_gn=TNS1;refGene_tx=XM_023259671.1;refGene_gn=TNS1;refGene_tx=XM_023259672.1;refGene_gn=TNS1;refGene_tx=XM_023259673.1;refGene_gn=TNS1;refGene_tx=XM_023259674.1;refGene_gn=TNS1;refGene_tx=XM_023259675.1;refGene_gn=TNS1;refGene_tx=XM_023259676.1
C1 203031268 203031337
refGene_gn=TNS1;refGene_tx=XM_006935547.3;refGene_gn=TNS1;refGene_tx=XM_023259670.1;refGene_gn=TNS1;refGene_tx=XM_023259671.1;refGene_gn=TNS1;refGene_tx=XM_023259672.1;refGene_gn=TNS1;refGene_tx=XM_023259673.1;refGene_gn=TNS1;refGene_tx=XM_023259674.1;refGene_gn=TNS1;refGene_tx=XM_023259675.1;refGene_gn=TNS1;refGene_tx=XM_023259676.1
C1 203031966 203032135
refGene_gn=TNS1;refGene_tx=XM_006935547.3;refGene_gn=TNS1;refGene_tx=XM_023259670.1;refGene_gn=TNS1;refGene_tx=XM_023259671.1;refGene_gn=TNS1;refGene_tx=XM_023259672.1;refGene_gn=TNS1;refGene_tx=XM_023259673.1;refGene_gn=TNS1;refGene_tx=XM_023259674.1;refGene_gn=TNS1;refGene_tx=XM_023259675.1;refGene_gn=TNS1;refGene_tx=XM_023259676.1
C1 203032361 203032433
refGene_gn=TNS1;refGene_tx=XM_006935547.3;refGene_gn=TNS1;refGene_tx=XM_023259670.1;refGene_gn=TNS1;refGene_tx=XM_023259671.1;refGene_gn=TNS1;refGene_tx=XM_023259672.1;refGene_gn=TNS1;refGene_tx=XM_023259673.1;refGene_gn=TNS1;refGene_tx=XM_023259674.1;refGene_gn=TNS1;refGene_tx=XM_023259675.1;refGene_gn=TNS1;refGene_tx=XM_023259676.1
C1 203034226 203034304
refGene_gn=TNS1;refGene_tx=XM_006935547.3;refGene_gn=TNS1;refGene_tx=XM_023259670.1;refGene_gn=TNS1;refGene_tx=XM_023259671.1;refGene_gn=TNS1;refGene_tx=XM_023259672.1;refGene_gn=TNS1;refGene_tx=XM_023259673.1;refGene_gn=TNS1;refGene_tx=XM_023259674.1;refGene_gn=TNS1;refGene_tx=XM_023259675.1;refGene_gn=TNS1;refGene_tx=XM_023259676.1
C1 203035012 203035105
refGene_gn=TNS1;refGene_tx=XM_006935547.3;refGene_gn=TNS1;refGene_tx=XM_023259670.1;refGene_gn=TNS1;refGene_tx=XM_023259671.1;refGene_gn=TNS

1;refGene_tx=XM_023259672.1;refGene_gn=TNS1;refGene_tx=XM_023259673.1;
refGene_gn=TNS1;refGene_tx=XM_023259674.1;refGene_gn=TNS1;refGene_tx=X
M_023259675.1;refGene_gn=TNS1;refGene_tx=XM_023259676.1
C1 203035458 203035590
refGene_gn=TNS1;refGene_tx=XM_006935547.3;refGene_gn=TNS1;refGene_tx=X
M_023259670.1;refGene_gn=TNS1;refGene_tx=XM_023259671.1;refGene_gn=TNS
1;refGene_tx=XM_023259672.1;refGene_gn=TNS1;refGene_tx=XM_023259673.1;
refGene_gn=TNS1;refGene_tx=XM_023259674.1;refGene_gn=TNS1;refGene_tx=X
M_023259675.1;refGene_gn=TNS1;refGene_tx=XM_023259676.1
C1 203036551 203036638
refGene_gn=TNS1;refGene_tx=XM_006935547.3;refGene_gn=TNS1;refGene_tx=X
M_023259670.1;refGene_gn=TNS1;refGene_tx=XM_023259671.1;refGene_gn=TNS
1;refGene_tx=XM_023259672.1;refGene_gn=TNS1;refGene_tx=XM_023259673.1;
refGene_gn=TNS1;refGene_tx=XM_023259674.1;refGene_gn=TNS1;refGene_tx=X
M_023259675.1;refGene_gn=TNS1;refGene_tx=XM_023259676.1
C1 203039480 203040547
refGene_gn=TNS1;refGene_tx=XM_006935547.3;refGene_gn=TNS1;refGene_tx=X
M_023259670.1;refGene_gn=TNS1;refGene_tx=XM_023259671.1;refGene_gn=TNS
1;refGene_tx=XM_023259672.1;refGene_gn=TNS1;refGene_tx=XM_023259673.1;
refGene_gn=TNS1;refGene_tx=XM_023259674.1;refGene_gn=TNS1;refGene_tx=X
M_023259675.1;refGene_gn=TNS1;refGene_tx=XM_023259676.1
C1 203043417 203043616
refGene_gn=TNS1;refGene_tx=XM_006935547.3;refGene_gn=TNS1;refGene_tx=X
M_023259670.1;refGene_gn=TNS1;refGene_tx=XM_023259671.1;refGene_gn=TNS
1;refGene_tx=XM_023259672.1;refGene_gn=TNS1;refGene_tx=XM_023259673.1;
refGene_gn=TNS1;refGene_tx=XM_023259674.1;refGene_gn=TNS1;refGene_tx=X
M_023259675.1;refGene_gn=TNS1;refGene_tx=XM_023259676.1
C1 203048420 203048459
refGene_gn=TNS1;refGene_tx=XM_006935547.3;refGene_gn=TNS1;refGene_tx=X
M_023259674.1;refGene_gn=TNS1;refGene_tx=XM_023259675.1;refGene_gn=TNS
1;refGene_tx=XM_023259676.1
C1 203048910 203048934
refGene_gn=TNS1;refGene_tx=XM_023259670.1;refGene_gn=TNS1;refGene_tx=X
M_023259672.1
C1 203049533 203049626
refGene_gn=TNS1;refGene_tx=XM_006935547.3;refGene_gn=TNS1;refGene_tx=X
M_023259670.1;refGene_gn=TNS1;refGene_tx=XM_023259671.1;refGene_gn=TNS
1;refGene_tx=XM_023259672.1;refGene_gn=TNS1;refGene_tx=XM_023259673.1;
refGene_gn=TNS1;refGene_tx=XM_023259674.1;refGene_gn=TNS1;refGene_tx=X
M_023259675.1;refGene_gn=TNS1;refGene_tx=XM_023259676.1
C1 203053340 203053416
refGene_gn=TNS1;refGene_tx=XM_006935547.3;refGene_gn=TNS1;refGene_tx=X
M_023259670.1;refGene_gn=TNS1;refGene_tx=XM_023259671.1;refGene_gn=TNS
1;refGene_tx=XM_023259672.1;refGene_gn=TNS1;refGene_tx=XM_023259673.1;
refGene_gn=TNS1;refGene_tx=XM_023259674.1;refGene_gn=TNS1;refGene_tx=X
M_023259675.1;refGene_gn=TNS1;refGene_tx=XM_023259676.1
C1 203054305 203054502
refGene_gn=TNS1;refGene_tx=XM_006935547.3;refGene_gn=TNS1;refGene_tx=X
M_023259670.1;refGene_gn=TNS1;refGene_tx=XM_023259671.1;refGene_gn=TNS
1;refGene_tx=XM_023259672.1;refGene_gn=TNS1;refGene_tx=XM_023259673.1;

refGene_gn=TNS1;refGene_tx=XM_023259674.1;refGene_gn=TNS1;refGene_tx=XM_023259675.1;refGene_gn=TNS1;refGene_tx=XM_023259676.1
C1 203058969 203059053
refGene_gn=TNS1;refGene_tx=XM_006935547.3;refGene_gn=TNS1;refGene_tx=XM_023259670.1;refGene_gn=TNS1;refGene_tx=XM_023259671.1;refGene_gn=TNS1;refGene_tx=XM_023259674.1;refGene_gn=TNS1;refGene_tx=XM_023259675.1
C1 203064909 203066505
refGene_gn=TNS1;refGene_tx=XM_006935547.3;refGene_gn=TNS1;refGene_tx=XM_023259670.1;refGene_gn=TNS1;refGene_tx=XM_023259671.1;refGene_gn=TNS1;refGene_tx=XM_023259672.1;refGene_gn=TNS1;refGene_tx=XM_023259673.1;refGene_gn=TNS1;refGene_tx=XM_023259674.1;refGene_gn=TNS1;refGene_tx=XM_023259675.1;refGene_gn=TNS1;refGene_tx=XM_023259676.1
C1 203098187 203098304
refGene_gn=TNS1;refGene_tx=XM_006935547.3;refGene_gn=TNS1;refGene_tx=XM_023259670.1;refGene_gn=TNS1;refGene_tx=XM_023259671.1;refGene_gn=TNS1;refGene_tx=XM_023259672.1;refGene_gn=TNS1;refGene_tx=XM_023259673.1;refGene_gn=TNS1;refGene_tx=XM_023259674.1;refGene_gn=TNS1;refGene_tx=XM_023259675.1;refGene_gn=TNS1;refGene_tx=XM_023259676.1
C1 203099438 203099504
refGene_gn=TNS1;refGene_tx=XM_006935547.3;refGene_gn=TNS1;refGene_tx=XM_023259670.1;refGene_gn=TNS1;refGene_tx=XM_023259671.1;refGene_gn=TNS1;refGene_tx=XM_023259672.1;refGene_gn=TNS1;refGene_tx=XM_023259673.1;refGene_gn=TNS1;refGene_tx=XM_023259674.1;refGene_gn=TNS1;refGene_tx=XM_023259675.1;refGene_gn=TNS1;refGene_tx=XM_023259676.1
C1 203101239 203101369
refGene_gn=TNS1;refGene_tx=XM_006935547.3;refGene_gn=TNS1;refGene_tx=XM_023259670.1;refGene_gn=TNS1;refGene_tx=XM_023259671.1;refGene_gn=TNS1;refGene_tx=XM_023259672.1;refGene_gn=TNS1;refGene_tx=XM_023259673.1;refGene_gn=TNS1;refGene_tx=XM_023259674.1;refGene_gn=TNS1;refGene_tx=XM_023259675.1;refGene_gn=TNS1;refGene_tx=XM_023259676.1
C1 203101965 203102041
refGene_gn=TNS1;refGene_tx=XM_006935547.3;refGene_gn=TNS1;refGene_tx=XM_023259670.1;refGene_gn=TNS1;refGene_tx=XM_023259671.1;refGene_gn=TNS1;refGene_tx=XM_023259672.1;refGene_gn=TNS1;refGene_tx=XM_023259673.1;refGene_gn=TNS1;refGene_tx=XM_023259674.1;refGene_gn=TNS1;refGene_tx=XM_023259675.1;refGene_gn=TNS1;refGene_tx=XM_023259676.1
C1 203102278 203102339
refGene_gn=TNS1;refGene_tx=XM_006935547.3;refGene_gn=TNS1;refGene_tx=XM_023259670.1;refGene_gn=TNS1;refGene_tx=XM_023259671.1;refGene_gn=TNS1;refGene_tx=XM_023259672.1;refGene_gn=TNS1;refGene_tx=XM_023259673.1;refGene_gn=TNS1;refGene_tx=XM_023259674.1;refGene_gn=TNS1;refGene_tx=XM_023259675.1;refGene_gn=TNS1;refGene_tx=XM_023259676.1
C1 203102740 203102853
refGene_gn=TNS1;refGene_tx=XM_006935547.3;refGene_gn=TNS1;refGene_tx=XM_023259670.1;refGene_gn=TNS1;refGene_tx=XM_023259671.1;refGene_gn=TNS1;refGene_tx=XM_023259672.1;refGene_gn=TNS1;refGene_tx=XM_023259673.1;refGene_gn=TNS1;refGene_tx=XM_023259674.1;refGene_gn=TNS1;refGene_tx=XM_023259675.1;refGene_gn=TNS1;refGene_tx=XM_023259676.1
C1 203106435 203106519
refGene_gn=TNS1;refGene_tx=XM_006935547.3;refGene_gn=TNS1;refGene_tx=X

M_023259670.1;refGene_gn=TNS1;refGene_tx=XM_023259671.1;refGene_gn=TNS1;refGene_tx=XM_023259672.1;refGene_gn=TNS1;refGene_tx=XM_023259673.1;refGene_gn=TNS1;refGene_tx=XM_023259674.1;refGene_gn=TNS1;refGene_tx=XM_023259675.1;refGene_gn=TNS1;refGene_tx=XM_023259676.1
C1 203107474 203107539
refGene_gn=TNS1;refGene_tx=XM_006935547.3;refGene_gn=TNS1;refGene_tx=XM_023259670.1;refGene_gn=TNS1;refGene_tx=XM_023259671.1;refGene_gn=TNS1;refGene_tx=XM_023259672.1;refGene_gn=TNS1;refGene_tx=XM_023259673.1;refGene_gn=TNS1;refGene_tx=XM_023259674.1;refGene_gn=TNS1;refGene_tx=XM_023259675.1;refGene_gn=TNS1;refGene_tx=XM_023259676.1
C1 203107930 203108053
refGene_gn=TNS1;refGene_tx=XM_006935547.3;refGene_gn=TNS1;refGene_tx=XM_023259670.1;refGene_gn=TNS1;refGene_tx=XM_023259671.1;refGene_gn=TNS1;refGene_tx=XM_023259672.1;refGene_gn=TNS1;refGene_tx=XM_023259673.1;refGene_gn=TNS1;refGene_tx=XM_023259674.1;refGene_gn=TNS1;refGene_tx=XM_023259675.1;refGene_gn=TNS1;refGene_tx=XM_023259676.1
C1 203108964 203109015
refGene_gn=TNS1;refGene_tx=XM_006935547.3;refGene_gn=TNS1;refGene_tx=XM_023259670.1;refGene_gn=TNS1;refGene_tx=XM_023259671.1;refGene_gn=TNS1;refGene_tx=XM_023259672.1;refGene_gn=TNS1;refGene_tx=XM_023259673.1;refGene_gn=TNS1;refGene_tx=XM_023259674.1;refGene_gn=TNS1;refGene_tx=XM_023259675.1;refGene_gn=TNS1;refGene_tx=XM_023259676.1
C1 203111470 203111642
refGene_gn=TNS1;refGene_tx=XM_006935547.3;refGene_gn=TNS1;refGene_tx=XM_023259670.1;refGene_gn=TNS1;refGene_tx=XM_023259671.1;refGene_gn=TNS1;refGene_tx=XM_023259672.1;refGene_gn=TNS1;refGene_tx=XM_023259673.1;refGene_gn=TNS1;refGene_tx=XM_023259674.1;refGene_gn=TNS1;refGene_tx=XM_023259675.1;refGene_gn=TNS1;refGene_tx=XM_023259676.1
C1 203113826 203113876
refGene_gn=TNS1;refGene_tx=XM_006935547.3;refGene_gn=TNS1;refGene_tx=XM_023259674.1
C1 203118843 203118879
refGene_gn=TNS1;refGene_tx=XM_006935547.3
C1 203119935 203119986
refGene_gn=TNS1;refGene_tx=XM_023259674.1
C1 203121000 203121042
refGene_gn=TNS1;refGene_tx=XM_023259674.1
C1 203134418 203134460
refGene_gn=TNS1;refGene_tx=XM_023259674.1
C1 203193541 203193579
refGene_gn=TNS1;refGene_tx=XM_023259674.1
C1 203204847 203204962
refGene_gn=TNS1;refGene_tx=XM_023259674.1
C1 203217094 203217127
refGene_gn=TNS1;refGene_tx=XM_023259674.1
C1 203252244 203252284
refGene_gn=RUFY4;refGene_tx=XM_003991145.4;refGene_gn=RUFY4;refGene_tx=XM_019838654.2;refGene_gn=RUFY4;refGene_tx=XM_019838655.1;refGene_gn=RUFY4;refGene_tx=XM_019838656.1;refGene_gn=RUFY4;refGene_tx=XM_023259677.1;refGene_gn=RUFY4;refGene_tx=XM_023259678.1

C1 203252413 203252520
refGene_gn=RUFY4;refGene_tx=XM_003991145.4;refGene_gn=RUFY4;refGene_tx=XM_019838654.2;refGene_gn=RUFY4;refGene_tx=XM_019838655.1;refGene_gn=RUFY4;refGene_tx=XM_019838656.1;refGene_gn=RUFY4;refGene_tx=XM_023259677.1;refGene_gn=RUFY4;refGene_tx=XM_023259678.1
C1 203253986 203254112
refGene_gn=RUFY4;refGene_tx=XM_003991145.4;refGene_gn=RUFY4;refGene_tx=XM_019838654.2;refGene_gn=RUFY4;refGene_tx=XM_019838655.1;refGene_gn=RUFY4;refGene_tx=XM_019838656.1;refGene_gn=RUFY4;refGene_tx=XM_023259677.1;refGene_gn=RUFY4;refGene_tx=XM_023259678.1
C1 203254424 203254531
refGene_gn=RUFY4;refGene_tx=XM_003991145.4;refGene_gn=RUFY4;refGene_tx=XM_019838654.2;refGene_gn=RUFY4;refGene_tx=XM_019838655.1;refGene_gn=RUFY4;refGene_tx=XM_019838656.1;refGene_gn=RUFY4;refGene_tx=XM_023259677.1;refGene_gn=RUFY4;refGene_tx=XM_023259678.1
C1 203254875 203255019
refGene_gn=RUFY4;refGene_tx=XM_003991145.4;refGene_gn=RUFY4;refGene_tx=XM_019838654.2;refGene_gn=RUFY4;refGene_tx=XM_019838655.1;refGene_gn=RUFY4;refGene_tx=XM_019838656.1;refGene_gn=RUFY4;refGene_tx=XM_023259677.1;refGene_gn=RUFY4;refGene_tx=XM_023259678.1
C1 203255484 203255548
refGene_gn=RUFY4;refGene_tx=XM_003991145.4;refGene_gn=RUFY4;refGene_tx=XM_019838654.2;refGene_gn=RUFY4;refGene_tx=XM_019838655.1;refGene_gn=RUFY4;refGene_tx=XM_019838656.1;refGene_gn=RUFY4;refGene_tx=XM_023259677.1;refGene_gn=RUFY4;refGene_tx=XM_023259678.1
C1 203256087 203256228
refGene_gn=RUFY4;refGene_tx=XM_019838654.2;refGene_gn=RUFY4;refGene_tx=XM_019838655.1;refGene_gn=RUFY4;refGene_tx=XM_019838656.1;refGene_gn=RUFY4;refGene_tx=XM_023259677.1;refGene_gn=RUFY4;refGene_tx=XM_023259678.1
C1 203256632 203257259
refGene_gn=RUFY4;refGene_tx=XM_003991145.4;refGene_gn=RUFY4;refGene_tx=XM_019838654.2;refGene_gn=RUFY4;refGene_tx=XM_019838655.1;refGene_gn=RUFY4;refGene_tx=XM_019838656.1;refGene_gn=RUFY4;refGene_tx=XM_023259677.1;refGene_gn=RUFY4;refGene_tx=XM_023259678.1
C1 203258017 203258124
refGene_gn=RUFY4;refGene_tx=XM_003991145.4;refGene_gn=RUFY4;refGene_tx=XM_019838654.2;refGene_gn=RUFY4;refGene_tx=XM_019838655.1;refGene_gn=RUFY4;refGene_tx=XM_019838656.1;refGene_gn=RUFY4;refGene_tx=XM_023259677.1;refGene_gn=RUFY4;refGene_tx=XM_023259678.1
C1 203263399 203263514
refGene_gn=RUFY4;refGene_tx=XM_019838656.1
C1 203263836 203263983
refGene_gn=RUFY4;refGene_tx=XM_003991145.4;refGene_gn=RUFY4;refGene_tx=XM_019838654.2;refGene_gn=RUFY4;refGene_tx=XM_019838655.1;refGene_gn=RUFY4;refGene_tx=XM_023259677.1;refGene_gn=RUFY4;refGene_tx=XM_023259678.1
C1 203264117 203264175
refGene_gn=RUFY4;refGene_tx=XM_019838655.1
C1 203265584 203265783

refGene_gn=RUFY4;refGene_tx=XM_019838654.2;refGene_gn=RUFY4;refGene_tx=XM_023259678.1
C1	203266417	203266437
refGene_gn=RUFY4;refGene_tx=XM_019838654.2
C1	203268337	203268448
refGene_gn=RUFY4;refGene_tx=XM_003991145.4;refGene_gn=RUFY4;refGene_tx=XM_023259677.1
C1	203268972	203269078
refGene_gn=RUFY4;refGene_tx=XM_003991145.4;refGene_gn=RUFY4;refGene_tx=XM_023259677.1
C1	203282350	203283421
refGene_gn=CXCR2;refGene_tx=XM_019838658.2;refGene_gn=CXCR2;refGene_tx=XM_019838659.2
C1	203308394	203309468
refGene_gn=LOC101085650;refGene_tx=XM_011285563.3
C1	203348466	203348540
refGene_gn=ARPC2;refGene_tx=XM_011285562.3
C1	203356484	203356519
refGene_gn=ARPC2;refGene_tx=XM_011285562.3
C1	203358276	203358389
refGene_gn=ARPC2;refGene_tx=XM_011285562.3
C1	203362247	203362293
refGene_gn=ARPC2;refGene_tx=XM_011285562.3
C1	203365269	203365456
refGene_gn=ARPC2;refGene_tx=XM_011285562.3
C1	203366347	203366441
refGene_gn=ARPC2;refGene_tx=XM_011285562.3
C1	203371858	203371985
refGene_gn=ARPC2;refGene_tx=XM_011285562.3
C1	203373467	203373568
refGene_gn=ARPC2;refGene_tx=XM_011285562.3
C1	203374021	203374122
refGene_gn=ARPC2;refGene_tx=XM_011285562.3
C1	203376689	203376714
refGene_gn=ARPC2;refGene_tx=XM_011285562.3
C1	203383072	203384065
refGene_gn=GPBAR1;refGene_tx=XM_023259679.1
C1	203384852	203384928
refGene_gn=AAMP;refGene_tx=XM_003991135.5;refGene_gn=AAMP;refGene_tx=XM_006935549.4
C1	203385424	203385579
refGene_gn=AAMP;refGene_tx=XM_003991135.5;refGene_gn=AAMP;refGene_tx=XM_006935549.4
C1	203385709	203385800
refGene_gn=AAMP;refGene_tx=XM_003991135.5;refGene_gn=AAMP;refGene_tx=XM_006935549.4
C1	203385952	203386056
refGene_gn=AAMP;refGene_tx=XM_003991135.5;refGene_gn=AAMP;refGene_tx=XM_006935549.4
C1	203386204	203386320

refGene_gn=AAMP;refGene_tx=XM_003991135.5;refGene_gn=AAMP;refGene_tx=XM_006935549.4
C1 203386655 203386739
refGene_gn=AAMP;refGene_tx=XM_003991135.5;refGene_gn=AAMP;refGene_tx=XM_006935549.4
C1 203387010 203387155
refGene_gn=AAMP;refGene_tx=XM_003991135.5;refGene_gn=AAMP;refGene_tx=XM_006935549.4
C1 203387408 203387548
refGene_gn=AAMP;refGene_tx=XM_003991135.5;refGene_gn=AAMP;refGene_tx=XM_006935549.4
C1 203388308 203388428
refGene_gn=AAMP;refGene_tx=XM_003991135.5;refGene_gn=AAMP;refGene_tx=XM_006935549.4
C1 203389092 203389251
refGene_gn=AAMP;refGene_tx=XM_003991135.5;refGene_gn=AAMP;refGene_tx=XM_006935549.4
C1 203389709 203389830
refGene_gn=AAMP;refGene_tx=XM_003991135.5;refGene_gn=AAMP;refGene_tx=XM_006935549.4
C1 203390277 203390344
refGene_gn=PNKD;refGene_tx=XM_006935550.4;refGene_gn=PNKD;refGene_tx=XM_019838661.2;refGene_gn=PNKD;refGene_tx=XM_019838662.2;refGene_gn=PNKD;refGene_tx=XM_019838663.2
C1 203391083 203391252
refGene_gn=PNKD;refGene_tx=XM_006935550.4;refGene_gn=PNKD;refGene_tx=XM_019838661.2;refGene_gn=PNKD;refGene_tx=XM_019838662.2;refGene_gn=PNKD;refGene_tx=XM_019838663.2;refGene_gn=PNKD;refGene_tx=XM_019838664.2
C1 203392613 203392806
refGene_gn=PNKD;refGene_tx=XM_006935550.4
C1 203395056 203395203
refGene_gn=TMBIM1;refGene_tx=XM_006935551.2;refGene_gn=TMBIM1;refGene_tx=XM_019838665.1
C1 203395613 203395667
refGene_gn=TMBIM1;refGene_tx=XM_006935551.2;refGene_gn=TMBIM1;refGene_tx=XM_019838665.1
C1 203396672 203396768
refGene_gn=TMBIM1;refGene_tx=XM_006935551.2;refGene_gn=TMBIM1;refGene_tx=XM_019838665.1
C1 203397089 203397177
refGene_gn=TMBIM1;refGene_tx=XM_006935551.2;refGene_gn=TMBIM1;refGene_tx=XM_019838665.1
C1 203397362 203397400
refGene_gn=TMBIM1;refGene_tx=XM_006935551.2;refGene_gn=TMBIM1;refGene_tx=XM_019838665.1
C1 203397662 203397702
refGene_gn=TMBIM1;refGene_tx=XM_006935551.2;refGene_gn=TMBIM1;refGene_tx=XM_019838665.1
C1 203398219 203398270
refGene_gn=TMBIM1;refGene_tx=XM_006935551.2;refGene_gn=TMBIM1;refGene_

tx=XM_019838665.1
C1 203398797 203398851
refGene_gn=TMBIM1;refGene_tx=XM_006935551.2;refGene_gn=TMBIM1;refGene_
tx=XM_019838665.1
C1 203399066 203399131
refGene_gn=TMBIM1;refGene_tx=XM_006935551.2;refGene_gn=TMBIM1;refGene_
tx=XM_019838665.1
C1 203399951 203400052
refGene_gn=TMBIM1;refGene_tx=XM_006935551.2;refGene_gn=TMBIM1;refGene_
tx=XM_019838665.1
C1 203401303 203401505
refGene_gn=TMBIM1;refGene_tx=XM_006935551.2;refGene_gn=TMBIM1;refGene_
tx=XM_019838665.1
C1 203430971 203431132
refGene_gn=PNKD;refGene_tx=XM_023259680.1;refGene_gn=PNKD;refGene_tx=X
M_023259681.1
C1 203444599 203444715
refGene_gn=PNKD;refGene_tx=XM_019838661.2;refGene_gn=PNKD;refGene_tx=X
M_019838662.2;refGene_gn=PNKD;refGene_tx=XM_019838663.2;refGene_gn=PNK
D;refGene_tx=XM_019838664.2;refGene_gn=PNKD;refGene_tx=XM_023259680.1;
refGene_gn=PNKD;refGene_tx=XM_023259681.1
C1 203444834 203444965
refGene_gn=PNKD;refGene_tx=XM_019838661.2;refGene_gn=PNKD;refGene_tx=X
M_019838662.2;refGene_gn=PNKD;refGene_tx=XM_019838663.2;refGene_gn=PNK
D;refGene_tx=XM_019838664.2;refGene_gn=PNKD;refGene_tx=XM_023259680.1;
refGene_gn=PNKD;refGene_tx=XM_023259681.1
C1 203445556 203445615
refGene_gn=PNKD;refGene_tx=XM_019838661.2;refGene_gn=PNKD;refGene_tx=X
M_019838662.2;refGene_gn=PNKD;refGene_tx=XM_019838663.2;refGene_gn=PNK
D;refGene_tx=XM_019838664.2;refGene_gn=PNKD;refGene_tx=XM_023259680.1;
refGene_gn=PNKD;refGene_tx=XM_023259681.1
C1 203446338 203446431
refGene_gn=PNKD;refGene_tx=XM_019838661.2;refGene_gn=PNKD;refGene_tx=X
M_019838662.2;refGene_gn=PNKD;refGene_tx=XM_019838663.2;refGene_gn=PNK
D;refGene_tx=XM_019838664.2;refGene_gn=PNKD;refGene_tx=XM_023259680.1;
refGene_gn=PNKD;refGene_tx=XM_023259681.1
C1 203446773 203446937
refGene_gn=PNKD;refGene_tx=XM_019838661.2;refGene_gn=PNKD;refGene_tx=X
M_019838662.2;refGene_gn=PNKD;refGene_tx=XM_019838663.2;refGene_gn=PNK
D;refGene_tx=XM_019838664.2;refGene_gn=PNKD;refGene_tx=XM_023259680.1;
refGene_gn=PNKD;refGene_tx=XM_023259681.1
C1 203447665 203447752
refGene_gn=PNKD;refGene_tx=XM_019838661.2;refGene_gn=PNKD;refGene_tx=X
M_019838662.2;refGene_gn=PNKD;refGene_tx=XM_019838663.2;refGene_gn=PNK
D;refGene_tx=XM_019838664.2;refGene_gn=PNKD;refGene_tx=XM_023259680.1;
refGene_gn=PNKD;refGene_tx=XM_023259681.1
C1 203448689 203448805
refGene_gn=PNKD;refGene_tx=XM_019838661.2;refGene_gn=PNKD;refGene_tx=X
M_019838662.2;refGene_gn=PNKD;refGene_tx=XM_019838663.2;refGene_gn=PNK
D;refGene_tx=XM_019838664.2;refGene_gn=PNKD;refGene_tx=XM_023259680.1;

refGene_gn=PNKD;refGene_tx=XM_023259681.1
C1 203449042 203449216
refGene_gn=PNKD;refGene_tx=XM_019838661.2;refGene_gn=PNKD;refGene_tx=XM_019838662.2;refGene_gn=PNKD;refGene_tx=XM_019838663.2;refGene_gn=PNKD;refGene_tx=XM_019838664.2;refGene_gn=PNKD;refGene_tx=XM_023259680.1;refGene_gn=PNKD;refGene_tx=XM_023259681.1
C1 203461245 203461270
refGene_gn=CATIP;refGene_tx=XM_003991147.4;refGene_gn=CATIP;refGene_tx=XM_006935552.3;refGene_gn=CATIP;refGene_tx=XM_019838666.1
C1 203461422 203461515
refGene_gn=CATIP;refGene_tx=XM_003991147.4;refGene_gn=CATIP;refGene_tx=XM_006935552.3;refGene_gn=CATIP;refGene_tx=XM_019838666.1
C1 203461861 203462062
refGene_gn=CATIP;refGene_tx=XM_003991147.4;refGene_gn=CATIP;refGene_tx=XM_019838666.1
C1 203462366 203462422
refGene_gn=CATIP;refGene_tx=XM_003991147.4;refGene_gn=CATIP;refGene_tx=XM_006935552.3;refGene_gn=CATIP;refGene_tx=XM_019838666.1
C1 203463551 203463638
refGene_gn=CATIP;refGene_tx=XM_003991147.4;refGene_gn=CATIP;refGene_tx=XM_006935552.3
C1 203463826 203463994
refGene_gn=CATIP;refGene_tx=XM_003991147.4;refGene_gn=CATIP;refGene_tx=XM_006935552.3;refGene_gn=CATIP;refGene_tx=XM_019838666.1
C1 203465286 203465411
refGene_gn=CATIP;refGene_tx=XM_003991147.4;refGene_gn=CATIP;refGene_tx=XM_006935552.3;refGene_gn=CATIP;refGene_tx=XM_019838666.1
C1 203466169 203466246
refGene_gn=CATIP;refGene_tx=XM_003991147.4;refGene_gn=CATIP;refGene_tx=XM_006935552.3;refGene_gn=CATIP;refGene_tx=XM_019838666.1
C1 203466560 203466649
refGene_gn=CATIP;refGene_tx=XM_003991147.4;refGene_gn=CATIP;refGene_tx=XM_006935552.3;refGene_gn=CATIP;refGene_tx=XM_019838666.1
C1 203466831 203467062
refGene_gn=CATIP;refGene_tx=XM_003991147.4;refGene_gn=CATIP;refGene_tx=XM_006935552.3;refGene_gn=CATIP;refGene_tx=XM_019838666.1
C1 203477573 203477580
refGene_gn=SLC11A1;refGene_tx=XM_006935553.3
C1 203478140 203478274
refGene_gn=SLC11A1;refGene_tx=XM_006935553.3
C1 203479121 203479244
refGene_gn=SLC11A1;refGene_tx=XM_006935553.3
C1 203479586 203479706
refGene_gn=SLC11A1;refGene_tx=XM_006935553.3
C1 203480469 203480576
refGene_gn=SLC11A1;refGene_tx=XM_006935553.3
C1 203480957 203481028
refGene_gn=SLC11A1;refGene_tx=XM_006935553.3
C1 203481366 203481434
refGene_gn=SLC11A1;refGene_tx=XM_006935553.3

C1 203481568 203481724
refGene_gn=SLC11A1;refGene_tx=XM_006935553.3
C1 203482646 203482805
refGene_gn=SLC11A1;refGene_tx=XM_006935553.3
C1 203483858 203483948
refGene_gn=SLC11A1;refGene_tx=XM_006935553.3
C1 203484071 203484191
refGene_gn=SLC11A1;refGene_tx=XM_006935553.3
C1 203484620 203484770
refGene_gn=SLC11A1;refGene_tx=XM_006935553.3
C1 203485257 203485331
refGene_gn=SLC11A1;refGene_tx=XM_006935553.3
C1 203485753 203485907
refGene_gn=SLC11A1;refGene_tx=XM_006935553.3
C1 203486056 203486167
refGene_gn=SLC11A1;refGene_tx=XM_006935553.3
C1 203490953 203491020
refGene_gn=CTDSP1;refGene_tx=XM_011285560.2;refGene_gn=CTDSP1;refGene_tx=XM_011285561.2;refGene_gn=CTDSP1;refGene_tx=XM_019838673.2
C1 203492447 203492596
refGene_gn=CTDSP1;refGene_tx=XM_011285560.2;refGene_gn=CTDSP1;refGene_tx=XM_011285561.2;refGene_gn=CTDSP1;refGene_tx=XM_019838673.2
C1 203493040 203493145
refGene_gn=CTDSP1;refGene_tx=XM_011285560.2;refGene_gn=CTDSP1;refGene_tx=XM_011285561.2;refGene_gn=CTDSP1;refGene_tx=XM_019838673.2
C1 203493254 203493311
refGene_gn=CTDSP1;refGene_tx=XM_011285560.2;refGene_gn=CTDSP1;refGene_tx=XM_011285561.2
C1 203493936 203494029
refGene_gn=CTDSP1;refGene_tx=XM_011285560.2;refGene_gn=CTDSP1;refGene_tx=XM_011285561.2;refGene_gn=CTDSP1;refGene_tx=XM_019838673.2
C1 203494159 203494345
refGene_gn=CTDSP1;refGene_tx=XM_011285560.2;refGene_gn=CTDSP1;refGene_tx=XM_011285561.2;refGene_gn=CTDSP1;refGene_tx=XM_019838673.2
C1 203495114 203495243
refGene_gn=CTDSP1;refGene_tx=XM_011285560.2;refGene_gn=CTDSP1;refGene_tx=XM_011285561.2;refGene_gn=CTDSP1;refGene_tx=XM_019838673.2
C1 203510263 203510338
refGene_gn=VIL1;refGene_tx=XM_006935554.3;refGene_gn=VIL1;refGene_tx=XM_019838672.2
C1 203510665 203510740
refGene_gn=VIL1;refGene_tx=XM_006935554.3;refGene_gn=VIL1;refGene_tx=XM_019838672.2
C1 203511560 203511757
refGene_gn=VIL1;refGene_tx=XM_006935554.3;refGene_gn=VIL1;refGene_tx=XM_019838672.2
C1 203513223 203513332
refGene_gn=VIL1;refGene_tx=XM_006935554.3;refGene_gn=VIL1;refGene_tx=XM_019838672.2
C1 203513500 203513611

refGene_gn=VIL1;refGene_tx=XM_006935554.3;refGene_gn=VIL1;refGene_tx=XM_019838672.2
C1 203515181 203515384
refGene_gn=VIL1;refGene_tx=XM_006935554.3;refGene_gn=VIL1;refGene_tx=XM_019838672.2
C1 203515494 203515573
refGene_gn=VIL1;refGene_tx=XM_006935554.3;refGene_gn=VIL1;refGene_tx=XM_019838672.2
C1 203515766 203515865
refGene_gn=VIL1;refGene_tx=XM_006935554.3;refGene_gn=VIL1;refGene_tx=XM_019838672.2
C1 203516512 203516666
refGene_gn=VIL1;refGene_tx=XM_006935554.3;refGene_gn=VIL1;refGene_tx=XM_019838672.2
C1 203517192 203517293
refGene_gn=VIL1;refGene_tx=XM_006935554.3;refGene_gn=VIL1;refGene_tx=XM_019838672.2
C1 203517387 203517525
refGene_gn=VIL1;refGene_tx=XM_006935554.3;refGene_gn=VIL1;refGene_tx=XM_019838672.2
C1 203518125 203518284
refGene_gn=VIL1;refGene_tx=XM_006935554.3;refGene_gn=VIL1;refGene_tx=XM_019838672.2
C1 203519315 203519495
refGene_gn=VIL1;refGene_tx=XM_006935554.3;refGene_gn=VIL1;refGene_tx=XM_019838672.2
C1 203519978 203520124
refGene_gn=VIL1;refGene_tx=XM_006935554.3;refGene_gn=VIL1;refGene_tx=XM_019838672.2
C1 203520825 203520970
refGene_gn=VIL1;refGene_tx=XM_006935554.3;refGene_gn=VIL1;refGene_tx=XM_019838672.2
C1 203521419 203521608
refGene_gn=VIL1;refGene_tx=XM_006935554.3;refGene_gn=VIL1;refGene_tx=XM_019838672.2
C1 203523704 203523773
refGene_gn=VIL1;refGene_tx=XM_006935554.3;refGene_gn=VIL1;refGene_tx=XM_019838672.2
C1 203524868 203525009
refGene_gn=VIL1;refGene_tx=XM_006935554.3;refGene_gn=VIL1;refGene_tx=XM_019838672.2
C1 203529600 203529714
refGene_gn=VIL1;refGene_tx=XM_006935554.3;refGene_gn=VIL1;refGene_tx=XM_019838672.2
C1 203535163 203535251
refGene_gn=USP37;refGene_tx=XM_003991142.5;refGene_gn=USP37;refGene_tx=XM_019838668.2;refGene_gn=USP37;refGene_tx=XM_019838669.2;refGene_gn=USP37;refGene_tx=XM_019838670.2;refGene_gn=USP37;refGene_tx=XM_019838671.2
C1 203535768 203535907

refGene_gn=USP37;refGene_tx=XM_003991142.5;refGene_gn=USP37;refGene_tx=XM_019838668.2;refGene_gn=USP37;refGene_tx=XM_019838669.2;refGene_gn=USP37;refGene_tx=XM_019838670.2;refGene_gn=USP37;refGene_tx=XM_019838671.2
C1 203536423 203536493
refGene_gn=USP37;refGene_tx=XM_003991142.5;refGene_gn=USP37;refGene_tx=XM_019838668.2;refGene_gn=USP37;refGene_tx=XM_019838669.2;refGene_gn=USP37;refGene_tx=XM_019838670.2;refGene_gn=USP37;refGene_tx=XM_019838671.2
C1 203538395 203538511
refGene_gn=USP37;refGene_tx=XM_003991142.5;refGene_gn=USP37;refGene_tx=XM_019838668.2;refGene_gn=USP37;refGene_tx=XM_019838669.2;refGene_gn=USP37;refGene_tx=XM_019838670.2;refGene_gn=USP37;refGene_tx=XM_019838671.2
C1 203543061 203543122
refGene_gn=USP37;refGene_tx=XM_003991142.5;refGene_gn=USP37;refGene_tx=XM_019838668.2;refGene_gn=USP37;refGene_tx=XM_019838669.2;refGene_gn=USP37;refGene_tx=XM_019838670.2;refGene_gn=USP37;refGene_tx=XM_019838671.2
C1 203545726 203545893
refGene_gn=USP37;refGene_tx=XM_003991142.5;refGene_gn=USP37;refGene_tx=XM_019838668.2;refGene_gn=USP37;refGene_tx=XM_019838669.2;refGene_gn=USP37;refGene_tx=XM_019838670.2;refGene_gn=USP37;refGene_tx=XM_019838671.2
C1 203558627 203558883
refGene_gn=USP37;refGene_tx=XM_003991142.5;refGene_gn=USP37;refGene_tx=XM_019838668.2;refGene_gn=USP37;refGene_tx=XM_019838669.2;refGene_gn=USP37;refGene_tx=XM_019838670.2;refGene_gn=USP37;refGene_tx=XM_019838671.2
C1 203561565 203561707
refGene_gn=USP37;refGene_tx=XM_003991142.5;refGene_gn=USP37;refGene_tx=XM_019838668.2;refGene_gn=USP37;refGene_tx=XM_019838669.2;refGene_gn=USP37;refGene_tx=XM_019838670.2;refGene_gn=USP37;refGene_tx=XM_019838671.2
C1 203564365 203564431
refGene_gn=USP37;refGene_tx=XM_003991142.5;refGene_gn=USP37;refGene_tx=XM_019838668.2;refGene_gn=USP37;refGene_tx=XM_019838669.2;refGene_gn=USP37;refGene_tx=XM_019838670.2
C1 203566093 203566258
refGene_gn=USP37;refGene_tx=XM_003991142.5;refGene_gn=USP37;refGene_tx=XM_019838668.2;refGene_gn=USP37;refGene_tx=XM_019838669.2;refGene_gn=USP37;refGene_tx=XM_019838670.2;refGene_gn=USP37;refGene_tx=XM_019838671.2
C1 203568770 203568850
refGene_gn=USP37;refGene_tx=XM_003991142.5;refGene_gn=USP37;refGene_tx=XM_019838668.2;refGene_gn=USP37;refGene_tx=XM_019838669.2;refGene_gn=USP37;refGene_tx=XM_019838670.2;refGene_gn=USP37;refGene_tx=XM_019838671.2
C1 203570147 203570265
refGene_gn=USP37;refGene_tx=XM_003991142.5;refGene_gn=USP37;refGene_tx

=XM_019838668.2;refGene_gn=USP37;refGene_tx=XM_019838669.2;refGene_gn=USP37;refGene_tx=XM_019838670.2;refGene_gn=USP37;refGene_tx=XM_019838671.2
C1 203575694 203575885
refGene_gn=USP37;refGene_tx=XM_003991142.5;refGene_gn=USP37;refGene_tx=XM_019838668.2;refGene_gn=USP37;refGene_tx=XM_019838669.2;refGene_gn=USP37;refGene_tx=XM_019838670.2;refGene_gn=USP37;refGene_tx=XM_019838671.2
C1 203576372 203576496
refGene_gn=USP37;refGene_tx=XM_003991142.5;refGene_gn=USP37;refGene_tx=XM_019838668.2;refGene_gn=USP37;refGene_tx=XM_019838669.2;refGene_gn=USP37;refGene_tx=XM_019838670.2;refGene_gn=USP37;refGene_tx=XM_019838671.2
C1 203576887 203577019
refGene_gn=USP37;refGene_tx=XM_003991142.5;refGene_gn=USP37;refGene_tx=XM_019838668.2;refGene_gn=USP37;refGene_tx=XM_019838669.2;refGene_gn=USP37;refGene_tx=XM_019838670.2;refGene_gn=USP37;refGene_tx=XM_019838671.2
C1 203591934 203592096
refGene_gn=USP37;refGene_tx=XM_003991142.5;refGene_gn=USP37;refGene_tx=XM_019838668.2;refGene_gn=USP37;refGene_tx=XM_019838669.2;refGene_gn=USP37;refGene_tx=XM_019838670.2;refGene_gn=USP37;refGene_tx=XM_019838671.2
C1 203599607 203599692
refGene_gn=USP37;refGene_tx=XM_003991142.5;refGene_gn=USP37;refGene_tx=XM_019838668.2;refGene_gn=USP37;refGene_tx=XM_019838669.2;refGene_gn=USP37;refGene_tx=XM_019838670.2;refGene_gn=USP37;refGene_tx=XM_019838671.2
C1 203601005 203601103
refGene_gn=USP37;refGene_tx=XM_003991142.5;refGene_gn=USP37;refGene_tx=XM_019838668.2;refGene_gn=USP37;refGene_tx=XM_019838669.2;refGene_gn=USP37;refGene_tx=XM_019838670.2;refGene_gn=USP37;refGene_tx=XM_019838671.2
C1 203608894 203608972
refGene_gn=USP37;refGene_tx=XM_003991142.5;refGene_gn=USP37;refGene_tx=XM_019838668.2;refGene_gn=USP37;refGene_tx=XM_019838669.2;refGene_gn=USP37;refGene_tx=XM_019838670.2;refGene_gn=USP37;refGene_tx=XM_019838671.2
C1 203609321 203609500
refGene_gn=USP37;refGene_tx=XM_003991142.5;refGene_gn=USP37;refGene_tx=XM_019838668.2;refGene_gn=USP37;refGene_tx=XM_019838669.2;refGene_gn=USP37;refGene_tx=XM_019838670.2;refGene_gn=USP37;refGene_tx=XM_019838671.2
C1 203612321 203612419
refGene_gn=USP37;refGene_tx=XM_003991142.5;refGene_gn=USP37;refGene_tx=XM_019838668.2;refGene_gn=USP37;refGene_tx=XM_019838669.2;refGene_gn=USP37;refGene_tx=XM_019838670.2;refGene_gn=USP37;refGene_tx=XM_019838671.2
C1 203614764 203614936
refGene_gn=USP37;refGene_tx=XM_003991142.5;refGene_gn=USP37;refGene_tx

=XM_019838668.2;refGene_gn=USP37;refGene_tx=XM_019838669.2;refGene_gn=USP37;refGene_tx=XM_019838670.2;refGene_gn=USP37;refGene_tx=XM_019838671.2
C1 203618579 203618735
refGene_gn=USP37;refGene_tx=XM_003991142.5;refGene_gn=USP37;refGene_tx=XM_019838668.2;refGene_gn=USP37;refGene_tx=XM_019838669.2;refGene_gn=USP37;refGene_tx=XM_019838670.2;refGene_gn=USP37;refGene_tx=XM_019838671.2
C1 203628773 203628992
refGene_gn=CNOT9;refGene_tx=XM_011285554.3;refGene_gn=CNOT9;refGene_tx=XM_011285555.3
C1 203629163 203629187
refGene_gn=CNOT9;refGene_tx=XM_011285556.2
C1 203640565 203640745
refGene_gn=CNOT9;refGene_tx=XM_011285554.3;refGene_gn=CNOT9;refGene_tx=XM_011285555.3;refGene_gn=CNOT9;refGene_tx=XM_011285556.2;refGene_gn=CNOT9;refGene_tx=XM_011285558.3
C1 203642799 203642915
refGene_gn=CNOT9;refGene_tx=XM_011285554.3;refGene_gn=CNOT9;refGene_tx=XM_011285555.3;refGene_gn=CNOT9;refGene_tx=XM_011285556.2;refGene_gn=CNOT9;refGene_tx=XM_011285558.3
C1 203645104 203645214
refGene_gn=CNOT9;refGene_tx=XM_011285554.3;refGene_gn=CNOT9;refGene_tx=XM_011285555.3;refGene_gn=CNOT9;refGene_tx=XM_011285556.2;refGene_gn=CNOT9;refGene_tx=XM_011285558.3
C1 203650561 203650671
refGene_gn=CNOT9;refGene_tx=XM_011285554.3;refGene_gn=CNOT9;refGene_tx=XM_011285555.3;refGene_gn=CNOT9;refGene_tx=XM_011285556.2;refGene_gn=CNOT9;refGene_tx=XM_011285558.3
C1 203653573 203653672
refGene_gn=CNOT9;refGene_tx=XM_011285554.3;refGene_gn=CNOT9;refGene_tx=XM_011285555.3;refGene_gn=CNOT9;refGene_tx=XM_011285556.2;refGene_gn=CNOT9;refGene_tx=XM_011285558.3
C1 203653868 203653960
refGene_gn=CNOT9;refGene_tx=XM_011285554.3;refGene_gn=CNOT9;refGene_tx=XM_011285555.3;refGene_gn=CNOT9;refGene_tx=XM_011285556.2;refGene_gn=CNOT9;refGene_tx=XM_011285558.3
C1 203655088 203655110
refGene_gn=CNOT9;refGene_tx=XM_011285555.3
C1 203656043 203656212
refGene_gn=CNOT9;refGene_tx=XM_011285554.3;refGene_gn=CNOT9;refGene_tx=XM_011285556.2;refGene_gn=CNOT9;refGene_tx=XM_011285558.3
C1 203669865 203669887
refGene_gn=PLCD4;refGene_tx=XM_006935566.4;refGene_gn=PLCD4;refGene_tx=XM_006935568.4;refGene_gn=PLCD4;refGene_tx=XM_006935569.4;refGene_gn=PLCD4;refGene_tx=XM_006935570.4;refGene_gn=PLCD4;refGene_tx=XM_019838682.2;refGene_gn=PLCD4;refGene_tx=XM_019838683.2
C1 203670029 203670188
refGene_gn=PLCD4;refGene_tx=XM_006935566.4;refGene_gn=PLCD4;refGene_tx=XM_006935568.4;refGene_gn=PLCD4;refGene_tx=XM_006935569.4;refGene_gn=

PLCD4;refGene_tx=XM_006935570.4;refGene_gn=PLCD4;refGene_tx=XM_0198386
82.2;refGene_gn=PLCD4;refGene_tx=XM_019838683.2
C1 203671476 203671705
refGene_gn=PLCD4;refGene_tx=XM_006935566.4;refGene_gn=PLCD4;refGene_tx
=XM_006935568.4;refGene_gn=PLCD4;refGene_tx=XM_006935569.4;refGene_gn=
PLCD4;refGene_tx=XM_006935570.4;refGene_gn=PLCD4;refGene_tx=XM_0198386
82.2;refGene_gn=PLCD4;refGene_tx=XM_019838683.2
C1 203672828 203672958
refGene_gn=PLCD4;refGene_tx=XM_006935566.4;refGene_gn=PLCD4;refGene_tx
=XM_006935568.4;refGene_gn=PLCD4;refGene_tx=XM_006935569.4;refGene_gn=
PLCD4;refGene_tx=XM_006935570.4;refGene_gn=PLCD4;refGene_tx=XM_0198386
82.2;refGene_gn=PLCD4;refGene_tx=XM_019838683.2
C1 203673778 203674010
refGene_gn=PLCD4;refGene_tx=XM_006935566.4;refGene_gn=PLCD4;refGene_tx
=XM_006935568.4;refGene_gn=PLCD4;refGene_tx=XM_006935569.4;refGene_gn=
PLCD4;refGene_tx=XM_006935570.4;refGene_gn=PLCD4;refGene_tx=XM_0198386
82.2;refGene_gn=PLCD4;refGene_tx=XM_019838683.2
C1 203678509 203678711
refGene_gn=PLCD4;refGene_tx=XM_006935566.4;refGene_gn=PLCD4;refGene_tx
=XM_006935568.4;refGene_gn=PLCD4;refGene_tx=XM_006935569.4;refGene_gn=
PLCD4;refGene_tx=XM_006935570.4;refGene_gn=PLCD4;refGene_tx=XM_0198386
82.2;refGene_gn=PLCD4;refGene_tx=XM_019838683.2
C1 203680228 203680373
refGene_gn=PLCD4;refGene_tx=XM_006935566.4;refGene_gn=PLCD4;refGene_tx
=XM_006935568.4;refGene_gn=PLCD4;refGene_tx=XM_006935569.4;refGene_gn=
PLCD4;refGene_tx=XM_006935570.4;refGene_gn=PLCD4;refGene_tx=XM_0198386
82.2;refGene_gn=PLCD4;refGene_tx=XM_019838683.2
C1 203681266 203681419
refGene_gn=PLCD4;refGene_tx=XM_006935566.4;refGene_gn=PLCD4;refGene_tx
=XM_006935568.4;refGene_gn=PLCD4;refGene_tx=XM_006935569.4;refGene_gn=
PLCD4;refGene_tx=XM_006935570.4;refGene_gn=PLCD4;refGene_tx=XM_0198386
82.2;refGene_gn=PLCD4;refGene_tx=XM_019838683.2
C1 203684436 203684631
refGene_gn=PLCD4;refGene_tx=XM_006935566.4;refGene_gn=PLCD4;refGene_tx
=XM_006935568.4;refGene_gn=PLCD4;refGene_tx=XM_006935569.4;refGene_gn=
PLCD4;refGene_tx=XM_006935570.4;refGene_gn=PLCD4;refGene_tx=XM_0198386
82.2;refGene_gn=PLCD4;refGene_tx=XM_019838683.2
C1 203686192 203686288
refGene_gn=PLCD4;refGene_tx=XM_006935566.4;refGene_gn=PLCD4;refGene_tx
=XM_006935568.4;refGene_gn=PLCD4;refGene_tx=XM_019838682.2;refGene_gn=
PLCD4;refGene_tx=XM_019838683.2
C1 203686429 203686586
refGene_gn=PLCD4;refGene_tx=XM_006935566.4;refGene_gn=PLCD4;refGene_tx
=XM_006935568.4;refGene_gn=PLCD4;refGene_tx=XM_006935569.4;refGene_gn=
PLCD4;refGene_tx=XM_006935570.4;refGene_gn=PLCD4;refGene_tx=XM_0198386
82.2;refGene_gn=PLCD4;refGene_tx=XM_019838683.2
C1 203686926 203687043
refGene_gn=PLCD4;refGene_tx=XM_006935566.4;refGene_gn=PLCD4;refGene_tx
=XM_006935568.4;refGene_gn=PLCD4;refGene_tx=XM_006935569.4;refGene_gn=
PLCD4;refGene_tx=XM_006935570.4;refGene_gn=PLCD4;refGene_tx=XM_0198386

82.2
C1	203687257	203687430
refGene_gn=PLCD4;refGene_tx=XM_006935566.4;refGene_gn=PLCD4;refGene_tx=XM_006935568.4;refGene_gn=PLCD4;refGene_tx=XM_006935569.4;refGene_gn=PLCD4;refGene_tx=XM_006935570.4;refGene_gn=PLCD4;refGene_tx=XM_019838682.2;refGene_gn=PLCD4;refGene_tx=XM_019838683.2
C1	203687969	203688105
refGene_gn=PLCD4;refGene_tx=XM_006935566.4;refGene_gn=PLCD4;refGene_tx=XM_006935568.4;refGene_gn=PLCD4;refGene_tx=XM_006935569.4;refGene_gn=PLCD4;refGene_tx=XM_006935570.4;refGene_gn=PLCD4;refGene_tx=XM_019838682.2;refGene_gn=PLCD4;refGene_tx=XM_019838683.2
C1	203688447	203688782
refGene_gn=PLCD4;refGene_tx=XM_006935566.4;refGene_gn=PLCD4;refGene_tx=XM_006935568.4;refGene_gn=PLCD4;refGene_tx=XM_006935569.4;refGene_gn=PLCD4;refGene_tx=XM_006935570.4;refGene_gn=PLCD4;refGene_tx=XM_019838682.2;refGene_gn=PLCD4;refGene_tx=XM_019838683.2
C1	203690495	203690965
refGene_gn=ZNF142;refGene_tx=XM_006935559.3;refGene_gn=ZNF142;refGene_tx=XM_019838674.1;refGene_gn=ZNF142;refGene_tx=XM_019838675.1;refGene_gn=ZNF142;refGene_tx=XM_019838676.1;refGene_gn=ZNF142;refGene_tx=XM_019838677.1
C1	203691891	203691997
refGene_gn=ZNF142;refGene_tx=XM_006935559.3;refGene_gn=ZNF142;refGene_tx=XM_019838674.1;refGene_gn=ZNF142;refGene_tx=XM_019838675.1;refGene_gn=ZNF142;refGene_tx=XM_019838676.1;refGene_gn=ZNF142;refGene_tx=XM_019838677.1
C1	203692541	203695458
refGene_gn=ZNF142;refGene_tx=XM_006935559.3;refGene_gn=ZNF142;refGene_tx=XM_019838674.1;refGene_gn=ZNF142;refGene_tx=XM_019838675.1;refGene_gn=ZNF142;refGene_tx=XM_019838676.1;refGene_gn=ZNF142;refGene_tx=XM_019838677.1
C1	203696204	203696382
refGene_gn=ZNF142;refGene_tx=XM_006935559.3;refGene_gn=ZNF142;refGene_tx=XM_019838674.1;refGene_gn=ZNF142;refGene_tx=XM_019838675.1;refGene_gn=ZNF142;refGene_tx=XM_019838676.1;refGene_gn=ZNF142;refGene_tx=XM_019838677.1
C1	203698273	203699098
refGene_gn=ZNF142;refGene_tx=XM_006935559.3;refGene_gn=ZNF142;refGene_tx=XM_019838674.1;refGene_gn=ZNF142;refGene_tx=XM_019838675.1;refGene_gn=ZNF142;refGene_tx=XM_019838676.1;refGene_gn=ZNF142;refGene_tx=XM_019838677.1
C1	203700007	203700202
refGene_gn=ZNF142;refGene_tx=XM_006935559.3;refGene_gn=ZNF142;refGene_tx=XM_019838674.1;refGene_gn=ZNF142;refGene_tx=XM_019838675.1;refGene_gn=ZNF142;refGene_tx=XM_019838677.1
C1	203701302	203701902
refGene_gn=ZNF142;refGene_tx=XM_006935559.3;refGene_gn=ZNF142;refGene_tx=XM_019838674.1;refGene_gn=ZNF142;refGene_tx=XM_019838675.1;refGene_gn=ZNF142;refGene_tx=XM_019838676.1
C1	203704735	203705015

refGene_gn=ZNF142;refGene_tx=XM_006935559.3;refGene_gn=ZNF142;refGene_tx=XM_019838674.1;refGene_gn=ZNF142;refGene_tx=XM_019838675.1;refGene_gn=ZNF142;refGene_tx=XM_019838676.1;refGene_gn=ZNF142;refGene_tx=XM_019838677.1
C1 203708651 203708971
refGene_gn=LOC101090709;refGene_tx=XM_003991151.4;refGene_gn=LOC101090709;refGene_tx=XM_003991152.5
C1 203709074 203709214
refGene_gn=LOC101090709;refGene_tx=XM_003991151.4;refGene_gn=LOC101090709;refGene_tx=XM_003991152.5
C1 203709415 203709610
refGene_gn=LOC101090709;refGene_tx=XM_003991151.4;refGene_gn=LOC101090709;refGene_tx=XM_003991152.5
C1 203709872 203709936
refGene_gn=LOC101090709;refGene_tx=XM_003991151.4;refGene_gn=LOC101090709;refGene_tx=XM_003991152.5
C1 203710174 203710344
refGene_gn=LOC101090709;refGene_tx=XM_003991151.4;refGene_gn=LOC101090709;refGene_tx=XM_003991152.5
C1 203710569 203710687
refGene_gn=LOC101090709;refGene_tx=XM_003991151.4;refGene_gn=LOC101090709;refGene_tx=XM_003991152.5
C1 203710833 203711086
refGene_gn=LOC101090709;refGene_tx=XM_003991151.4;refGene_gn=LOC101090709;refGene_tx=XM_003991152.5
C1 203711647 203712226
refGene_gn=RNF25;refGene_tx=XM_003991153.4;refGene_gn=RNF25;refGene_tx=XM_006935574.3
C1 203712425 203712560
refGene_gn=RNF25;refGene_tx=XM_003991153.4;refGene_gn=RNF25;refGene_tx=XM_006935574.3
C1 203713043 203713136
refGene_gn=RNF25;refGene_tx=XM_003991153.4;refGene_gn=RNF25;refGene_tx=XM_006935574.3
C1 203713490 203713628
refGene_gn=RNF25;refGene_tx=XM_003991153.4;refGene_gn=RNF25;refGene_tx=XM_006935574.3
C1 203713725 203713797
refGene_gn=RNF25;refGene_tx=XM_003991153.4;refGene_gn=RNF25;refGene_tx=XM_006935574.3
C1 203714946 203715016
refGene_gn=RNF25;refGene_tx=XM_003991153.4;refGene_gn=RNF25;refGene_tx=XM_006935574.3
C1 203715108 203715176
refGene_gn=RNF25;refGene_tx=XM_003991153.4;refGene_gn=RNF25;refGene_tx=XM_006935574.3
C1 203715268 203715371
refGene_gn=RNF25;refGene_tx=XM_003991153.4;refGene_gn=RNF25;refGene_tx=XM_006935574.3
C1 203715614 203715689

refGene_gn=RNF25;refGene_tx=XM_003991153.4;refGene_gn=RNF25;refGene_tx
=XM_006935574.3
C1 203717951 203717992
refGene_gn=RNF25;refGene_tx=XM_003991153.4;refGene_gn=RNF25;refGene_tx
=XM_006935574.3
C1 203718929 203719013
refGene_gn=STK36;refGene_tx=XM_003991154.5;refGene_gn=STK36;refGene_tx
=XM_019838678.2;refGene_gn=STK36;refGene_tx=XM_019838679.2;refGene_gn=
STK36;refGene_tx=XM_019838680.1
C1 203719704 203719845
refGene_gn=STK36;refGene_tx=XM_003991154.5;refGene_gn=STK36;refGene_tx
=XM_019838678.2;refGene_gn=STK36;refGene_tx=XM_019838679.2;refGene_gn=
STK36;refGene_tx=XM_019838680.1
C1 203719955 203720033
refGene_gn=STK36;refGene_tx=XM_003991154.5;refGene_gn=STK36;refGene_tx
=XM_019838678.2;refGene_gn=STK36;refGene_tx=XM_019838679.2;refGene_gn=
STK36;refGene_tx=XM_019838680.1
C1 203722087 203722218
refGene_gn=STK36;refGene_tx=XM_003991154.5;refGene_gn=STK36;refGene_tx
=XM_019838678.2;refGene_gn=STK36;refGene_tx=XM_019838679.2;refGene_gn=
STK36;refGene_tx=XM_019838680.1
C1 203722448 203722698
refGene_gn=STK36;refGene_tx=XM_003991154.5;refGene_gn=STK36;refGene_tx
=XM_019838678.2;refGene_gn=STK36;refGene_tx=XM_019838679.2;refGene_gn=
STK36;refGene_tx=XM_019838680.1
C1 203724351 203724445
refGene_gn=STK36;refGene_tx=XM_003991154.5;refGene_gn=STK36;refGene_tx
=XM_019838678.2;refGene_gn=STK36;refGene_tx=XM_019838679.2;refGene_gn=
STK36;refGene_tx=XM_019838680.1
C1 203724738 203724908
refGene_gn=STK36;refGene_tx=XM_003991154.5;refGene_gn=STK36;refGene_tx
=XM_019838678.2;refGene_gn=STK36;refGene_tx=XM_019838679.2;refGene_gn=
STK36;refGene_tx=XM_019838680.1
C1 203725308 203725496
refGene_gn=STK36;refGene_tx=XM_003991154.5;refGene_gn=STK36;refGene_tx
=XM_019838678.2;refGene_gn=STK36;refGene_tx=XM_019838679.2;refGene_gn=
STK36;refGene_tx=XM_019838680.1
C1 203726000 203726100
refGene_gn=STK36;refGene_tx=XM_003991154.5;refGene_gn=STK36;refGene_tx
=XM_019838678.2;refGene_gn=STK36;refGene_tx=XM_019838679.2;refGene_gn=
STK36;refGene_tx=XM_019838680.1
C1 203727765 203727909
refGene_gn=STK36;refGene_tx=XM_003991154.5;refGene_gn=STK36;refGene_tx
=XM_019838678.2;refGene_gn=STK36;refGene_tx=XM_019838679.2;refGene_gn=
STK36;refGene_tx=XM_019838680.1
C1 203732408 203732588
refGene_gn=STK36;refGene_tx=XM_003991154.5;refGene_gn=STK36;refGene_tx
=XM_019838678.2;refGene_gn=STK36;refGene_tx=XM_019838679.2;refGene_gn=
STK36;refGene_tx=XM_019838680.1
C1 203733414 203733614

refGene_gn=STK36;refGene_tx=XM_003991154.5;refGene_gn=STK36;refGene_tx
=XM_019838678.2;refGene_gn=STK36;refGene_tx=XM_019838679.2;refGene_gn=
STK36;refGene_tx=XM_019838680.1
C1 203734002 203734108
refGene_gn=STK36;refGene_tx=XM_003991154.5;refGene_gn=STK36;refGene_tx
=XM_019838678.2;refGene_gn=STK36;refGene_tx=XM_019838679.2;refGene_gn=
STK36;refGene_tx=XM_019838680.1
C1 203735612 203735763
refGene_gn=STK36;refGene_tx=XM_003991154.5;refGene_gn=STK36;refGene_tx
=XM_019838678.2;refGene_gn=STK36;refGene_tx=XM_019838679.2;refGene_gn=
STK36;refGene_tx=XM_019838680.1
C1 203735954 203736082
refGene_gn=STK36;refGene_tx=XM_003991154.5;refGene_gn=STK36;refGene_tx
=XM_019838678.2;refGene_gn=STK36;refGene_tx=XM_019838679.2;refGene_gn=
STK36;refGene_tx=XM_019838680.1
C1 203736617 203736722
refGene_gn=STK36;refGene_tx=XM_003991154.5;refGene_gn=STK36;refGene_tx
=XM_019838678.2;refGene_gn=STK36;refGene_tx=XM_019838679.2;refGene_gn=
STK36;refGene_tx=XM_019838680.1
C1 203737069 203737168
refGene_gn=STK36;refGene_tx=XM_003991154.5;refGene_gn=STK36;refGene_tx
=XM_019838678.2;refGene_gn=STK36;refGene_tx=XM_019838679.2;refGene_gn=
STK36;refGene_tx=XM_019838680.1
C1 203737254 203737343
refGene_gn=STK36;refGene_tx=XM_003991154.5;refGene_gn=STK36;refGene_tx
=XM_019838678.2;refGene_gn=STK36;refGene_tx=XM_019838679.2;refGene_gn=
STK36;refGene_tx=XM_019838680.1
C1 203737627 203737691
refGene_gn=STK36;refGene_tx=XM_003991154.5;refGene_gn=STK36;refGene_tx
=XM_019838678.2;refGene_gn=STK36;refGene_tx=XM_019838679.2;refGene_gn=
STK36;refGene_tx=XM_019838680.1
C1 203737877 203737988
refGene_gn=STK36;refGene_tx=XM_003991154.5;refGene_gn=STK36;refGene_tx
=XM_019838678.2;refGene_gn=STK36;refGene_tx=XM_019838679.2;refGene_gn=
STK36;refGene_tx=XM_019838680.1
C1 203739613 203739688
refGene_gn=STK36;refGene_tx=XM_003991154.5;refGene_gn=STK36;refGene_tx
=XM_019838678.2;refGene_gn=STK36;refGene_tx=XM_019838679.2;refGene_gn=
STK36;refGene_tx=XM_019838680.1
C1 203740133 203740296
refGene_gn=STK36;refGene_tx=XM_003991154.5;refGene_gn=STK36;refGene_tx
=XM_019838678.2;refGene_gn=STK36;refGene_tx=XM_019838679.2;refGene_gn=
STK36;refGene_tx=XM_019838680.1
C1 203740512 203740660
refGene_gn=STK36;refGene_tx=XM_003991154.5;refGene_gn=STK36;refGene_tx
=XM_019838678.2;refGene_gn=STK36;refGene_tx=XM_019838679.2;refGene_gn=
STK36;refGene_tx=XM_019838680.1
C1 203740889 203741037
refGene_gn=STK36;refGene_tx=XM_003991154.5;refGene_gn=STK36;refGene_tx
=XM_019838678.2;refGene_gn=STK36;refGene_tx=XM_019838679.2;refGene_gn=

STK36;refGene_tx=XM_019838680.1
C1	203741599	203742346
refGene_gn=STK36;refGene_tx=XM_003991154.5;refGene_gn=STK36;refGene_tx
=XM_019838678.2;refGene_gn=STK36;refGene_tx=XM_019838679.2;refGene_gn=
STK36;refGene_tx=XM_019838680.1
C1	203743406	203743550
refGene_gn=STK36;refGene_tx=XM_003991154.5;refGene_gn=STK36;refGene_tx
=XM_019838678.2;refGene_gn=STK36;refGene_tx=XM_019838679.2;refGene_gn=
STK36;refGene_tx=XM_019838680.1
C1	203767563	203768189
refGene_gn=TTLL4;refGene_tx=XM_023259683.1
C1	203768294	203769158
refGene_gn=TTLL4;refGene_tx=XM_023259683.1
C1	203770248	203770358
refGene_gn=TTLL4;refGene_tx=XM_023259683.1
C1	203770711	203770775
refGene_gn=TTLL4;refGene_tx=XM_023259683.1
C1	203775994	203776116
refGene_gn=TTLL4;refGene_tx=XM_023259683.1
C1	203776551	203776662
refGene_gn=TTLL4;refGene_tx=XM_023259683.1
C1	203777020	203777097
refGene_gn=TTLL4;refGene_tx=XM_023259683.1
C1	203777795	203777987
refGene_gn=TTLL4;refGene_tx=XM_023259683.1
C1	203778080	203778163
refGene_gn=TTLL4;refGene_tx=XM_023259683.1
C1	203778386	203778515
refGene_gn=TTLL4;refGene_tx=XM_023259683.1
C1	203780185	203780308
refGene_gn=TTLL4;refGene_tx=XM_023259683.1
C1	203780607	203780706
refGene_gn=TTLL4;refGene_tx=XM_023259683.1
C1	203781026	203781161
refGene_gn=TTLL4;refGene_tx=XM_023259683.1
C1	203781585	203781723
refGene_gn=TTLL4;refGene_tx=XM_023259683.1
C1	203783133	203783236
refGene_gn=TTLL4;refGene_tx=XM_023259683.1
C1	203783904	203784115
refGene_gn=TTLL4;refGene_tx=XM_023259683.1
C1	203784255	203784326
refGene_gn=TTLL4;refGene_tx=XM_023259683.1
C1	203784729	203784815
refGene_gn=TTLL4;refGene_tx=XM_023259683.1
C1	203785291	203785550
refGene_gn=TTLL4;refGene_tx=XM_023259683.1
C1	203815508	203815772
refGene_gn=LOC101091962;refGene_tx=XM_003991157.5
C1	203820177	203820222

refGene_gn=LOC101091962;refGene_tx=XM_006935575.4
C1	203840552	203840743
refGene_gn=LOC101091962;refGene_tx=XM_003991157.5;refGene_gn=LOC101091
962;refGene_tx=XM_006935575.4
C1	203843792	203843992
refGene_gn=LOC101091962;refGene_tx=XM_003991157.5;refGene_gn=LOC101091
962;refGene_tx=XM_006935575.4
C1	203844119	203844317
refGene_gn=LOC101091962;refGene_tx=XM_003991157.5;refGene_gn=LOC101091
962;refGene_tx=XM_006935575.4
C1	203844483	203844656
refGene_gn=LOC101091962;refGene_tx=XM_003991157.5;refGene_gn=LOC101091
962;refGene_tx=XM_006935575.4
C1	203845455	203845622
refGene_gn=LOC101091962;refGene_tx=XM_003991157.5;refGene_gn=LOC101091
962;refGene_tx=XM_006935575.4
C1	203845780	203845862
refGene_gn=LOC101091962;refGene_tx=XM_003991157.5;refGene_gn=LOC101091
962;refGene_tx=XM_006935575.4
C1	203845942	203846155
refGene_gn=LOC101091962;refGene_tx=XM_003991157.5;refGene_gn=LOC101091
962;refGene_tx=XM_006935575.4
C1	203846310	203846430
refGene_gn=LOC101091962;refGene_tx=XM_003991157.5;refGene_gn=LOC101091
962;refGene_tx=XM_006935575.4
C1	203852470	203852587
refGene_gn=PRKAG3;refGene_tx=XM_019838684.2;refGene_gn=PRKAG3;refGene_
tx=XM_019838685.2;refGene_gn=PRKAG3;refGene_tx=XM_019838686.2;refGene_
gn=PRKAG3;refGene_tx=XM_019838687.2
C1	203852958	203853105
refGene_gn=PRKAG3;refGene_tx=XM_019838684.2;refGene_gn=PRKAG3;refGene_
tx=XM_019838685.2;refGene_gn=PRKAG3;refGene_tx=XM_019838686.2;refGene_
gn=PRKAG3;refGene_tx=XM_019838687.2
C1	203853278	203853316
refGene_gn=PRKAG3;refGene_tx=XM_019838684.2;refGene_gn=PRKAG3;refGene_
tx=XM_019838685.2;refGene_gn=PRKAG3;refGene_tx=XM_019838686.2;refGene_
gn=PRKAG3;refGene_tx=XM_019838687.2
C1	203855075	203855241
refGene_gn=PRKAG3;refGene_tx=XM_019838684.2;refGene_gn=PRKAG3;refGene_
tx=XM_019838685.2;refGene_gn=PRKAG3;refGene_tx=XM_019838686.2;refGene_
gn=PRKAG3;refGene_tx=XM_019838687.2
C1	203855374	203855501
refGene_gn=PRKAG3;refGene_tx=XM_019838684.2;refGene_gn=PRKAG3;refGene_
tx=XM_019838685.2;refGene_gn=PRKAG3;refGene_tx=XM_019838686.2;refGene_
gn=PRKAG3;refGene_tx=XM_019838687.2
C1	203855685	203855740
refGene_gn=PRKAG3;refGene_tx=XM_019838684.2;refGene_gn=PRKAG3;refGene_
tx=XM_019838685.2;refGene_gn=PRKAG3;refGene_tx=XM_019838686.2;refGene_
gn=PRKAG3;refGene_tx=XM_019838687.2
C1	203855954	203856000

refGene_gn=PRKAG3;refGene_tx=XM_019838684.2;refGene_gn=PRKAG3;refGene_tx=XM_019838685.2;refGene_gn=PRKAG3;refGene_tx=XM_019838686.2;refGene_gn=PRKAG3;refGene_tx=XM_019838687.2
C1 203856119 203856178
refGene_gn=PRKAG3;refGene_tx=XM_019838684.2;refGene_gn=PRKAG3;refGene_tx=XM_019838685.2;refGene_gn=PRKAG3;refGene_tx=XM_019838686.2;refGene_gn=PRKAG3;refGene_tx=XM_019838687.2
C1 203856667 203856749
refGene_gn=PRKAG3;refGene_tx=XM_019838684.2;refGene_gn=PRKAG3;refGene_tx=XM_019838685.2;refGene_gn=PRKAG3;refGene_tx=XM_019838686.2;refGene_gn=PRKAG3;refGene_tx=XM_019838687.2
C1 203857766 203858170
refGene_gn=PRKAG3;refGene_tx=XM_019838684.2;refGene_gn=PRKAG3;refGene_tx=XM_019838685.2;refGene_gn=PRKAG3;refGene_tx=XM_019838686.2;refGene_gn=PRKAG3;refGene_tx=XM_019838687.2
C1 203858520 203858676
refGene_gn=PRKAG3;refGene_tx=XM_019838684.2;refGene_gn=PRKAG3;refGene_tx=XM_019838685.2;refGene_gn=PRKAG3;refGene_tx=XM_019838686.2;refGene_gn=PRKAG3;refGene_tx=XM_019838687.2
C1 203859145 203859185
refGene_gn=PRKAG3;refGene_tx=XM_019838684.2
C1 203859562 203859583
refGene_gn=PRKAG3;refGene_tx=XM_019838684.2
C1 203887966 203888046
refGene_gn=WNT6;refGene_tx=XM_023259684.1
C1 203897809 203898030
refGene_gn=WNT6;refGene_tx=XM_023259684.1
C1 203898261 203898596
refGene_gn=WNT6;refGene_tx=XM_023259684.1
C1 203900174 203900636
refGene_gn=WNT6;refGene_tx=XM_023259684.1
C1 203907770 203907880
refGene_gn=WNT10A;refGene_tx=XM_003991159.4
C1 203908921 203909184
refGene_gn=WNT10A;refGene_tx=XM_003991159.4
C1 203915633 203916013
refGene_gn=WNT10A;refGene_tx=XM_003991159.4
C1 203918422 203918920
refGene_gn=WNT10A;refGene_tx=XM_003991159.4
C1 203968654 203969767
refGene_gn=CDK5R2;refGene_tx=XM_011285580.3
C1 203988936 203989523
refGene_gn=FEV;refGene_tx=XM_011285579.2
C1 203991560 203991635
refGene_gn=FEV;refGene_tx=XM_011285579.2
C1 203992383 203992435
refGene_gn=FEV;refGene_tx=XM_011285579.2
C1 203997052 203997200
refGene_gn=CRYBA2;refGene_tx=XM_006935581.2
C1 203997605 203997748

refGene_gn=CRYBA2;refGene_tx=XM_006935581.2
C1 203998893 203999035
refGene_gn=CRYBA2;refGene_tx=XM_006935581.2
C1 203999805 203999966
refGene_gn=CRYBA2;refGene_tx=XM_006935581.2
C1 204010034 204010167
refGene_gn=CFAP65;refGene_tx=XM_011285549.2;refGene_gn=CFAP65;refGene_tx=XM_019838690.1;refGene_gn=CFAP65;refGene_tx=XM_019838691.2;refGene_gn=CFAP65;refGene_tx=XM_019838693.2;refGene_gn=CFAP65;refGene_tx=XM_019838694.2;refGene_gn=CFAP65;refGene_tx=XM_019838695.2
C1 204010271 204010409
refGene_gn=CFAP65;refGene_tx=XM_011285549.2;refGene_gn=CFAP65;refGene_tx=XM_019838690.1;refGene_gn=CFAP65;refGene_tx=XM_019838691.2;refGene_gn=CFAP65;refGene_tx=XM_019838693.2;refGene_gn=CFAP65;refGene_tx=XM_019838694.2;refGene_gn=CFAP65;refGene_tx=XM_019838695.2
C1 204011101 204011590
refGene_gn=CFAP65;refGene_tx=XM_011285549.2;refGene_gn=CFAP65;refGene_tx=XM_019838690.1;refGene_gn=CFAP65;refGene_tx=XM_019838691.2;refGene_gn=CFAP65;refGene_tx=XM_019838693.2;refGene_gn=CFAP65;refGene_tx=XM_019838694.2;refGene_gn=CFAP65;refGene_tx=XM_019838695.2
C1 204012095 204012224
refGene_gn=CFAP65;refGene_tx=XM_011285549.2;refGene_gn=CFAP65;refGene_tx=XM_019838690.1;refGene_gn=CFAP65;refGene_tx=XM_019838691.2;refGene_gn=CFAP65;refGene_tx=XM_019838693.2;refGene_gn=CFAP65;refGene_tx=XM_019838694.2;refGene_gn=CFAP65;refGene_tx=XM_019838695.2
C1 204012510 204012713
refGene_gn=CFAP65;refGene_tx=XM_011285549.2;refGene_gn=CFAP65;refGene_tx=XM_019838690.1;refGene_gn=CFAP65;refGene_tx=XM_019838691.2;refGene_gn=CFAP65;refGene_tx=XM_019838693.2;refGene_gn=CFAP65;refGene_tx=XM_019838694.2;refGene_gn=CFAP65;refGene_tx=XM_019838695.2
C1 204012918 204012963
refGene_gn=CFAP65;refGene_tx=XM_011285549.2;refGene_gn=CFAP65;refGene_tx=XM_019838690.1;refGene_gn=CFAP65;refGene_tx=XM_019838691.2;refGene_gn=CFAP65;refGene_tx=XM_019838693.2;refGene_gn=CFAP65;refGene_tx=XM_019838694.2;refGene_gn=CFAP65;refGene_tx=XM_019838695.2
C1 204014251 204014359
refGene_gn=CFAP65;refGene_tx=XM_011285549.2;refGene_gn=CFAP65;refGene_tx=XM_019838690.1;refGene_gn=CFAP65;refGene_tx=XM_019838691.2;refGene_gn=CFAP65;refGene_tx=XM_019838693.2;refGene_gn=CFAP65;refGene_tx=XM_019838694.2;refGene_gn=CFAP65;refGene_tx=XM_019838695.2
C1 204014537 204014651
refGene_gn=CFAP65;refGene_tx=XM_011285549.2;refGene_gn=CFAP65;refGene_tx=XM_019838690.1;refGene_gn=CFAP65;refGene_tx=XM_019838691.2;refGene_gn=CFAP65;refGene_tx=XM_019838693.2;refGene_gn=CFAP65;refGene_tx=XM_019838694.2;refGene_gn=CFAP65;refGene_tx=XM_019838695.2
C1 204014949 204015093
refGene_gn=CFAP65;refGene_tx=XM_011285549.2;refGene_gn=CFAP65;refGene_tx=XM_019838690.1;refGene_gn=CFAP65;refGene_tx=XM_019838691.2;refGene_gn=CFAP65;refGene_tx=XM_019838693.2;refGene_gn=CFAP65;refGene_tx=XM_019838694.2;refGene_gn=CFAP65;refGene_tx=XM_019838695.2

C1 204016189 204016348
refGene_gn=CFAP65;refGene_tx=XM_011285549.2;refGene_gn=CFAP65;refGene_tx=XM_019838690.1;refGene_gn=CFAP65;refGene_tx=XM_019838691.2;refGene_gn=CFAP65;refGene_tx=XM_019838693.2;refGene_gn=CFAP65;refGene_tx=XM_019838694.2;refGene_gn=CFAP65;refGene_tx=XM_019838695.2
C1 204016448 204016640
refGene_gn=CFAP65;refGene_tx=XM_011285549.2;refGene_gn=CFAP65;refGene_tx=XM_019838690.1;refGene_gn=CFAP65;refGene_tx=XM_019838691.2;refGene_gn=CFAP65;refGene_tx=XM_019838693.2;refGene_gn=CFAP65;refGene_tx=XM_019838694.2;refGene_gn=CFAP65;refGene_tx=XM_019838695.2
C1 204018723 204018834
refGene_gn=CFAP65;refGene_tx=XM_011285549.2;refGene_gn=CFAP65;refGene_tx=XM_019838690.1;refGene_gn=CFAP65;refGene_tx=XM_019838691.2;refGene_gn=CFAP65;refGene_tx=XM_019838693.2;refGene_gn=CFAP65;refGene_tx=XM_019838694.2
C1 204018983 204019050
refGene_gn=CFAP65;refGene_tx=XM_011285549.2;refGene_gn=CFAP65;refGene_tx=XM_019838690.1;refGene_gn=CFAP65;refGene_tx=XM_019838691.2;refGene_gn=CFAP65;refGene_tx=XM_019838693.2;refGene_gn=CFAP65;refGene_tx=XM_019838694.2;refGene_gn=CFAP65;refGene_tx=XM_019838695.2
C1 204019345 204019522
refGene_gn=CFAP65;refGene_tx=XM_011285549.2;refGene_gn=CFAP65;refGene_tx=XM_019838690.1;refGene_gn=CFAP65;refGene_tx=XM_019838691.2;refGene_gn=CFAP65;refGene_tx=XM_019838693.2;refGene_gn=CFAP65;refGene_tx=XM_019838694.2;refGene_gn=CFAP65;refGene_tx=XM_019838695.2;refGene_gn=CFAP65;refGene_tx=XM_019838696.2
C1 204023452 204023581
refGene_gn=CFAP65;refGene_tx=XM_011285549.2;refGene_gn=CFAP65;refGene_tx=XM_019838690.1;refGene_gn=CFAP65;refGene_tx=XM_019838691.2;refGene_gn=CFAP65;refGene_tx=XM_019838693.2;refGene_gn=CFAP65;refGene_tx=XM_019838694.2;refGene_gn=CFAP65;refGene_tx=XM_019838695.2;refGene_gn=CFAP65;refGene_tx=XM_019838696.2
C1 204023902 204024116
refGene_gn=CFAP65;refGene_tx=XM_011285549.2;refGene_gn=CFAP65;refGene_tx=XM_019838690.1;refGene_gn=CFAP65;refGene_tx=XM_019838691.2;refGene_gn=CFAP65;refGene_tx=XM_019838693.2;refGene_gn=CFAP65;refGene_tx=XM_019838694.2;refGene_gn=CFAP65;refGene_tx=XM_019838695.2;refGene_gn=CFAP65;refGene_tx=XM_019838696.2
C1 204025647 204025776
refGene_gn=CFAP65;refGene_tx=XM_011285549.2;refGene_gn=CFAP65;refGene_tx=XM_019838690.1;refGene_gn=CFAP65;refGene_tx=XM_019838691.2;refGene_gn=CFAP65;refGene_tx=XM_019838693.2;refGene_gn=CFAP65;refGene_tx=XM_019838694.2;refGene_gn=CFAP65;refGene_tx=XM_019838695.2;refGene_gn=CFAP65;refGene_tx=XM_019838696.2
C1 204026444 204026595
refGene_gn=CFAP65;refGene_tx=XM_011285549.2;refGene_gn=CFAP65;refGene_tx=XM_019838690.1;refGene_gn=CFAP65;refGene_tx=XM_019838691.2;refGene_gn=CFAP65;refGene_tx=XM_019838693.2;refGene_gn=CFAP65;refGene_tx=XM_019838694.2;refGene_gn=CFAP65;refGene_tx=XM_019838695.2;refGene_gn=CFAP65;refGene_tx=XM_019838696.2

C1	204026806	204026965
refGene_gn=CFAP65;refGene_tx=XM_011285549.2;refGene_gn=CFAP65;refGene_tx=XM_019838690.1;refGene_gn=CFAP65;refGene_tx=XM_019838691.2;refGene_gn=CFAP65;refGene_tx=XM_019838693.2;refGene_gn=CFAP65;refGene_tx=XM_019838694.2;refGene_gn=CFAP65;refGene_tx=XM_019838695.2;refGene_gn=CFAP65;refGene_tx=XM_019838696.2
C1	204028237	204028462
refGene_gn=CFAP65;refGene_tx=XM_011285549.2;refGene_gn=CFAP65;refGene_tx=XM_019838690.1;refGene_gn=CFAP65;refGene_tx=XM_019838691.2;refGene_gn=CFAP65;refGene_tx=XM_019838693.2;refGene_gn=CFAP65;refGene_tx=XM_019838694.2;refGene_gn=CFAP65;refGene_tx=XM_019838695.2;refGene_gn=CFAP65;refGene_tx=XM_019838696.2
C1	204029039	204029285
refGene_gn=CFAP65;refGene_tx=XM_011285549.2;refGene_gn=CFAP65;refGene_tx=XM_019838690.1;refGene_gn=CFAP65;refGene_tx=XM_019838691.2;refGene_gn=CFAP65;refGene_tx=XM_019838693.2;refGene_gn=CFAP65;refGene_tx=XM_019838694.2;refGene_gn=CFAP65;refGene_tx=XM_019838695.2;refGene_gn=CFAP65;refGene_tx=XM_019838696.2
C1	204031428	204031566
refGene_gn=CFAP65;refGene_tx=XM_011285549.2;refGene_gn=CFAP65;refGene_tx=XM_019838690.1;refGene_gn=CFAP65;refGene_tx=XM_019838691.2;refGene_gn=CFAP65;refGene_tx=XM_019838693.2;refGene_gn=CFAP65;refGene_tx=XM_019838694.2;refGene_gn=CFAP65;refGene_tx=XM_019838695.2;refGene_gn=CFAP65;refGene_tx=XM_019838696.2
C1	204033012	204033372
refGene_gn=CFAP65;refGene_tx=XM_011285549.2;refGene_gn=CFAP65;refGene_tx=XM_019838690.1;refGene_gn=CFAP65;refGene_tx=XM_019838691.2;refGene_gn=CFAP65;refGene_tx=XM_019838693.2;refGene_gn=CFAP65;refGene_tx=XM_019838694.2;refGene_gn=CFAP65;refGene_tx=XM_019838695.2;refGene_gn=CFAP65;refGene_tx=XM_019838696.2
C1	204033608	204033809
refGene_gn=CFAP65;refGene_tx=XM_011285549.2;refGene_gn=CFAP65;refGene_tx=XM_019838690.1;refGene_gn=CFAP65;refGene_tx=XM_019838691.2;refGene_gn=CFAP65;refGene_tx=XM_019838693.2;refGene_gn=CFAP65;refGene_tx=XM_019838694.2;refGene_gn=CFAP65;refGene_tx=XM_019838695.2;refGene_gn=CFAP65;refGene_tx=XM_019838696.2
C1	204034268	204034534
refGene_gn=CFAP65;refGene_tx=XM_011285549.2;refGene_gn=CFAP65;refGene_tx=XM_019838690.1;refGene_gn=CFAP65;refGene_tx=XM_019838691.2;refGene_gn=CFAP65;refGene_tx=XM_019838693.2;refGene_gn=CFAP65;refGene_tx=XM_019838694.2;refGene_gn=CFAP65;refGene_tx=XM_019838695.2;refGene_gn=CFAP65;refGene_tx=XM_019838696.2
C1	204034912	204035135
refGene_gn=CFAP65;refGene_tx=XM_011285549.2;refGene_gn=CFAP65;refGene_tx=XM_019838690.1;refGene_gn=CFAP65;refGene_tx=XM_019838691.2;refGene_gn=CFAP65;refGene_tx=XM_019838693.2;refGene_gn=CFAP65;refGene_tx=XM_019838694.2;refGene_gn=CFAP65;refGene_tx=XM_019838695.2;refGene_gn=CFAP65;refGene_tx=XM_019838696.2
C1	204035833	204035979
refGene_gn=CFAP65;refGene_tx=XM_011285549.2;refGene_gn=CFAP65;refGene_

tx=XM_019838690.1;refGene_gn=CFAP65;refGene_tx=XM_019838691.2;refGene_gn=CFAP65;refGene_tx=XM_019838693.2;refGene_gn=CFAP65;refGene_tx=XM_019838694.2;refGene_gn=CFAP65;refGene_tx=XM_019838695.2;refGene_gn=CFAP65;refGene_tx=XM_019838696.2
C1 204036254 204036454
refGene_gn=CFAP65;refGene_tx=XM_011285549.2;refGene_gn=CFAP65;refGene_tx=XM_019838690.1;refGene_gn=CFAP65;refGene_tx=XM_019838691.2;refGene_gn=CFAP65;refGene_tx=XM_019838693.2;refGene_gn=CFAP65;refGene_tx=XM_019838694.2;refGene_gn=CFAP65;refGene_tx=XM_019838695.2;refGene_gn=CFAP65;refGene_tx=XM_019838696.2
C1 204036631 204036798
refGene_gn=CFAP65;refGene_tx=XM_011285549.2;refGene_gn=CFAP65;refGene_tx=XM_019838690.1;refGene_gn=CFAP65;refGene_tx=XM_019838691.2;refGene_gn=CFAP65;refGene_tx=XM_019838693.2;refGene_gn=CFAP65;refGene_tx=XM_019838694.2;refGene_gn=CFAP65;refGene_tx=XM_019838695.2;refGene_gn=CFAP65;refGene_tx=XM_019838696.2
C1 204037362 204037465
refGene_gn=CFAP65;refGene_tx=XM_011285549.2;refGene_gn=CFAP65;refGene_tx=XM_019838690.1;refGene_gn=CFAP65;refGene_tx=XM_019838691.2;refGene_gn=CFAP65;refGene_tx=XM_019838693.2;refGene_gn=CFAP65;refGene_tx=XM_019838694.2;refGene_gn=CFAP65;refGene_tx=XM_019838695.2;refGene_gn=CFAP65;refGene_tx=XM_019838696.2
C1 204040804 204040989
refGene_gn=CFAP65;refGene_tx=XM_011285549.2;refGene_gn=CFAP65;refGene_tx=XM_019838690.1;refGene_gn=CFAP65;refGene_tx=XM_019838691.2;refGene_gn=CFAP65;refGene_tx=XM_019838693.2;refGene_gn=CFAP65;refGene_tx=XM_019838694.2;refGene_gn=CFAP65;refGene_tx=XM_019838695.2;refGene_gn=CFAP65;refGene_tx=XM_019838696.2
C1 204042827 204042992
refGene_gn=CFAP65;refGene_tx=XM_011285549.2;refGene_gn=CFAP65;refGene_tx=XM_019838690.1;refGene_gn=CFAP65;refGene_tx=XM_019838691.2;refGene_gn=CFAP65;refGene_tx=XM_019838693.2;refGene_gn=CFAP65;refGene_tx=XM_019838694.2;refGene_gn=CFAP65;refGene_tx=XM_019838695.2;refGene_gn=CFAP65;refGene_tx=XM_019838696.2
C1 204058356 204059015
refGene_gn=IHH;refGene_tx=XM_019839142.2
C1 204060696 204060958
refGene_gn=IHH;refGene_tx=XM_019839142.2
C1 204063387 204063699
refGene_gn=IHH;refGene_tx=XM_019839142.2
C1 204078697 204078772
refGene_gn=NHEJ1;refGene_tx=XM_006935583.3;refGene_gn=NHEJ1;refGene_tx=XM_011285546.3;refGene_gn=NHEJ1;refGene_tx=XM_019838698.2;refGene_gn=NHEJ1;refGene_tx=XM_019838699.2
C1 204079188 204079307
refGene_gn=NHEJ1;refGene_tx=XM_006935583.3;refGene_gn=NHEJ1;refGene_tx=XM_011285546.3;refGene_gn=NHEJ1;refGene_tx=XM_019838698.2;refGene_gn=NHEJ1;refGene_tx=XM_019838699.2
C1 204079903 204080021
refGene_gn=NHEJ1;refGene_tx=XM_006935583.3;refGene_gn=NHEJ1;refGene_tx

=XM_011285546.3;refGene_gn=NHEJ1;refGene_tx=XM_019838698.2;refGene_gn=NHEJ1;refGene_tx=XM_019838699.2
C1 204148901 204148960
refGene_gn=NHEJ1;refGene_tx=XM_006935583.3;refGene_gn=NHEJ1;refGene_tx=XM_011285546.3;refGene_gn=NHEJ1;refGene_tx=XM_019838698.2;refGene_gn=NHEJ1;refGene_tx=XM_019838699.2
C1 204149535 204149674
refGene_gn=NHEJ1;refGene_tx=XM_006935583.3;refGene_gn=NHEJ1;refGene_tx=XM_011285546.3;refGene_gn=NHEJ1;refGene_tx=XM_019838698.2;refGene_gn=NHEJ1;refGene_tx=XM_019838699.2
C1 204156323 204156536
refGene_gn=NHEJ1;refGene_tx=XM_006935583.3;refGene_gn=NHEJ1;refGene_tx=XM_011285546.3;refGene_gn=NHEJ1;refGene_tx=XM_019838698.2;refGene_gn=NHEJ1;refGene_tx=XM_019838699.2
C1 204156948 204157125
refGene_gn=NHEJ1;refGene_tx=XM_006935583.3;refGene_gn=NHEJ1;refGene_tx=XM_011285546.3;refGene_gn=NHEJ1;refGene_tx=XM_019838698.2;refGene_gn=NHEJ1;refGene_tx=XM_019838699.2
C1 204159421 204159442
refGene_gn=NHEJ1;refGene_tx=XM_019838698.2;refGene_gn=NHEJ1;refGene_tx=XM_019838699.2
C1 204161424 204161714
refGene_gn=SLC23A3;refGene_tx=XM_003991163.4;refGene_gn=SLC23A3;refGene_tx=XM_019838700.2;refGene_gn=SLC23A3;refGene_tx=XM_019838701.1;refGene_gn=SLC23A3;refGene_tx=XM_019838702.1;refGene_gn=SLC23A3;refGene_tx=XM_019838704.1
C1 204161806 204161902
refGene_gn=SLC23A3;refGene_tx=XM_003991163.4;refGene_gn=SLC23A3;refGene_tx=XM_019838700.2;refGene_gn=SLC23A3;refGene_tx=XM_019838701.1;refGene_gn=SLC23A3;refGene_tx=XM_019838702.1
C1 204162898 204163063
refGene_gn=SLC23A3;refGene_tx=XM_003991163.4;refGene_gn=SLC23A3;refGene_tx=XM_019838700.2;refGene_gn=SLC23A3;refGene_tx=XM_019838702.1
C1 204163369 204163475
refGene_gn=SLC23A3;refGene_tx=XM_003991163.4;refGene_gn=SLC23A3;refGene_tx=XM_019838700.2;refGene_gn=SLC23A3;refGene_tx=XM_019838701.1;refGene_gn=SLC23A3;refGene_tx=XM_019838702.1;refGene_gn=SLC23A3;refGene_tx=XM_019838704.1
C1 204164046 204164300
refGene_gn=SLC23A3;refGene_tx=XM_019838700.2;refGene_gn=SLC23A3;refGene_tx=XM_019838701.1;refGene_gn=SLC23A3;refGene_tx=XM_019838702.1;refGene_gn=SLC23A3;refGene_tx=XM_019838704.1
C1 204165262 204165377
refGene_gn=SLC23A3;refGene_tx=XM_003991163.4;refGene_gn=SLC23A3;refGene_tx=XM_019838700.2;refGene_gn=SLC23A3;refGene_tx=XM_019838701.1;refGene_gn=SLC23A3;refGene_tx=XM_019838702.1;refGene_gn=SLC23A3;refGene_tx=XM_019838704.1
C1 204165515 204165639
refGene_gn=SLC23A3;refGene_tx=XM_003991163.4;refGene_gn=SLC23A3;refGene_tx=XM_019838700.2;refGene_gn=SLC23A3;refGene_tx=XM_019838701.1;refGe

ne_gn=SLC23A3;refGene_tx=XM_019838702.1;refGene_gn=SLC23A3;refGene_tx=XM_019838704.1
C1 204165953 204166139 refGene_gn=SLC23A3;refGene_tx=XM_003991163.4;refGene_gn=SLC23A3;refGene_tx=XM_019838700.2;refGene_gn=SLC23A3;refGene_tx=XM_019838701.1;refGene_gn=SLC23A3;refGene_tx=XM_019838702.1;refGene_gn=SLC23A3;refGene_tx=XM_019838704.1
C1 204166345 204166419 refGene_gn=SLC23A3;refGene_tx=XM_003991163.4;refGene_gn=SLC23A3;refGene_tx=XM_019838700.2;refGene_gn=SLC23A3;refGene_tx=XM_019838701.1;refGene_gn=SLC23A3;refGene_tx=XM_019838704.1
C1 204166630 204166728 refGene_gn=SLC23A3;refGene_tx=XM_003991163.4;refGene_gn=SLC23A3;refGene_tx=XM_019838700.2;refGene_gn=SLC23A3;refGene_tx=XM_019838701.1;refGene_gn=SLC23A3;refGene_tx=XM_019838702.1;refGene_gn=SLC23A3;refGene_tx=XM_019838704.1
C1 204166889 204167047 refGene_gn=SLC23A3;refGene_tx=XM_003991163.4;refGene_gn=SLC23A3;refGene_tx=XM_019838700.2;refGene_gn=SLC23A3;refGene_tx=XM_019838701.1;refGene_gn=SLC23A3;refGene_tx=XM_019838702.1;refGene_gn=SLC23A3;refGene_tx=XM_019838704.1
C1 204167208 204167388 refGene_gn=SLC23A3;refGene_tx=XM_003991163.4;refGene_gn=SLC23A3;refGene_tx=XM_019838700.2;refGene_gn=SLC23A3;refGene_tx=XM_019838701.1;refGene_gn=SLC23A3;refGene_tx=XM_019838702.1;refGene_gn=SLC23A3;refGene_tx=XM_019838704.1
C1 204170113 204170659 refGene_gn=CNPPD1;refGene_tx=XM_003991165.5;refGene_gn=CNPPD1;refGene_tx=XM_006935585.2;refGene_gn=CNPPD1;refGene_tx=XM_006935586.3
C1 204170884 204171002 refGene_gn=CNPPD1;refGene_tx=XM_003991165.5;refGene_gn=CNPPD1;refGene_tx=XM_006935585.2;refGene_gn=CNPPD1;refGene_tx=XM_006935586.3
C1 204171348 204171410 refGene_gn=CNPPD1;refGene_tx=XM_003991165.5;refGene_gn=CNPPD1;refGene_tx=XM_006935585.2;refGene_gn=CNPPD1;refGene_tx=XM_006935586.3
C1 204172010 204172139 refGene_gn=CNPPD1;refGene_tx=XM_003991165.5;refGene_gn=CNPPD1;refGene_tx=XM_006935585.2;refGene_gn=CNPPD1;refGene_tx=XM_006935586.3
C1 204172224 204172345 refGene_gn=CNPPD1;refGene_tx=XM_003991165.5;refGene_gn=CNPPD1;refGene_tx=XM_006935585.2;refGene_gn=CNPPD1;refGene_tx=XM_006935586.3
C1 204172511 204172593 refGene_gn=CNPPD1;refGene_tx=XM_003991165.5;refGene_gn=CNPPD1;refGene_tx=XM_006935585.2;refGene_gn=CNPPD1;refGene_tx=XM_006935586.3
C1 204173384 204173493 refGene_gn=CNPPD1;refGene_tx=XM_003991165.5;refGene_gn=CNPPD1;refGene_tx=XM_006935585.2;refGene_gn=CNPPD1;refGene_tx=XM_006935586.3
C1 204173927 204173996 refGene_gn=CNPPD1;refGene_tx=XM_003991165.5;refGene_gn=CNPPD1;refGene_tx=XM_006935585.2;refGene_gn=CNPPD1;refGene_tx=XM_006935586.3

C1	204175589	204175870
refGene_gn=RETREG2;refGene_tx=XM_003991166.5;refGene_gn=RETREG2;refGene_tx=XM_019838705.2
C1	204176153	204176260
refGene_gn=RETREG2;refGene_tx=XM_003991166.5;refGene_gn=RETREG2;refGene_tx=XM_019838705.2
C1	204177253	204177284
refGene_gn=RETREG2;refGene_tx=XM_003991166.5;refGene_gn=RETREG2;refGene_tx=XM_019838705.2
C1	204177635	204177771
refGene_gn=RETREG2;refGene_tx=XM_003991166.5;refGene_gn=RETREG2;refGene_tx=XM_019838705.2
C1	204178184	204178269
refGene_gn=RETREG2;refGene_tx=XM_003991166.5;refGene_gn=RETREG2;refGene_tx=XM_019838705.2
C1	204178596	204178740
refGene_gn=RETREG2;refGene_tx=XM_003991166.5;refGene_gn=RETREG2;refGene_tx=XM_019838705.2
C1	204178896	204178991
refGene_gn=RETREG2;refGene_tx=XM_003991166.5;refGene_gn=RETREG2;refGene_tx=XM_019838705.2
C1	204179150	204179286
refGene_gn=RETREG2;refGene_tx=XM_003991166.5;refGene_gn=RETREG2;refGene_tx=XM_019838705.2
C1	204179545	204180171
refGene_gn=RETREG2;refGene_tx=XM_003991166.5;refGene_gn=RETREG2;refGene_tx=XM_019838705.2
C1	204202188	204202243
refGene_gn=ZFAND2B;refGene_tx=XM_003991167.4;refGene_gn=ZFAND2B;refGene_tx=XM_019838714.1;refGene_gn=ZFAND2B;refGene_tx=XM_019838715.1
C1	204202518	204202613
refGene_gn=ZFAND2B;refGene_tx=XM_003991167.4;refGene_gn=ZFAND2B;refGene_tx=XM_019838714.1;refGene_gn=ZFAND2B;refGene_tx=XM_019838715.1
C1	204202849	204202981
refGene_gn=ZFAND2B;refGene_tx=XM_003991167.4;refGene_gn=ZFAND2B;refGene_tx=XM_019838714.1;refGene_gn=ZFAND2B;refGene_tx=XM_019838715.1
C1	204203114	204203266
refGene_gn=ZFAND2B;refGene_tx=XM_003991167.4;refGene_gn=ZFAND2B;refGene_tx=XM_019838714.1;refGene_gn=ZFAND2B;refGene_tx=XM_019838715.1
C1	204203520	204203610
refGene_gn=ZFAND2B;refGene_tx=XM_003991167.4;refGene_gn=ZFAND2B;refGene_tx=XM_019838714.1;refGene_gn=ZFAND2B;refGene_tx=XM_019838715.1
C1	204203664	204203746
refGene_gn=ZFAND2B;refGene_tx=XM_003991167.4;refGene_gn=ZFAND2B;refGene_tx=XM_019838714.1;refGene_gn=ZFAND2B;refGene_tx=XM_019838715.1
C1	204203831	204203899
refGene_gn=ZFAND2B;refGene_tx=XM_003991167.4;refGene_gn=ZFAND2B;refGene_tx=XM_019838714.1;refGene_gn=ZFAND2B;refGene_tx=XM_019838715.1
C1	204204237	204204310
refGene_gn=ZFAND2B;refGene_tx=XM_003991167.4;refGene_gn=ZFAND2B;refGen

e_tx=XM_019838714.1;refGene_gn=ZFAND2B;refGene_tx=XM_019838715.1
C1	204204534	204204579
refGene_gn=ZFAND2B;refGene_tx=XM_003991167.4;refGene_gn=ZFAND2B;refGene_tx=XM_019838714.1
C1	204204638	204204680
refGene_gn=ZFAND2B;refGene_tx=XM_019838715.1
C1	204205234	204205346
refGene_gn=ABCB6;refGene_tx=XM_003991168.5
C1	204205529	204205598
refGene_gn=ABCB6;refGene_tx=XM_003991168.5
C1	204205673	204205768
refGene_gn=ABCB6;refGene_tx=XM_003991168.5
C1	204206011	204206124
refGene_gn=ABCB6;refGene_tx=XM_003991168.5
C1	204206276	204206451
refGene_gn=ABCB6;refGene_tx=XM_003991168.5
C1	204206826	204206931
refGene_gn=ABCB6;refGene_tx=XM_003991168.5
C1	204207147	204207205
refGene_gn=ABCB6;refGene_tx=XM_003991168.5
C1	204207439	204207525
refGene_gn=ABCB6;refGene_tx=XM_003991168.5
C1	204207644	204207708
refGene_gn=ABCB6;refGene_tx=XM_003991168.5
C1	204207796	204207873
refGene_gn=ABCB6;refGene_tx=XM_003991168.5
C1	204208002	204208128
refGene_gn=ABCB6;refGene_tx=XM_003991168.5
C1	204208319	204208385
refGene_gn=ABCB6;refGene_tx=XM_003991168.5
C1	204208582	204208692
refGene_gn=ABCB6;refGene_tx=XM_003991168.5
C1	204209213	204209335
refGene_gn=ABCB6;refGene_tx=XM_003991168.5
C1	204210179	204210363
refGene_gn=ABCB6;refGene_tx=XM_003991168.5
C1	204210535	204210637
refGene_gn=ABCB6;refGene_tx=XM_003991168.5
C1	204210839	204211026
refGene_gn=ABCB6;refGene_tx=XM_003991168.5
C1	204211534	204211672
refGene_gn=ABCB6;refGene_tx=XM_003991168.5
C1	204211982	204212531
refGene_gn=ABCB6;refGene_tx=XM_003991168.5
C1	204214264	204214270
refGene_gn=ATG9A;refGene_tx=XM_003991169.5;refGene_gn=ATG9A;refGene_tx=XM_006935589.4;refGene_gn=ATG9A;refGene_tx=XM_006935590.4;refGene_gn=ATG9A;refGene_tx=XM_006935591.4;refGene_gn=ATG9A;refGene_tx=XM_006935592.3;refGene_gn=ATG9A;refGene_tx=XM_011285544.3;refGene_gn=ATG9A;refGene_tx=XM_019838706.2

C1 204214531 204214677
refGene_gn=ATG9A;refGene_tx=XM_003991169.5;refGene_gn=ATG9A;refGene_tx=XM_006935589.4;refGene_gn=ATG9A;refGene_tx=XM_006935590.4;refGene_gn=ATG9A;refGene_tx=XM_006935591.4;refGene_gn=ATG9A;refGene_tx=XM_006935592.3;refGene_gn=ATG9A;refGene_tx=XM_011285544.3;refGene_gn=ATG9A;refGene_tx=XM_019838706.2
C1 204214867 204215090
refGene_gn=ATG9A;refGene_tx=XM_003991169.5;refGene_gn=ATG9A;refGene_tx=XM_006935589.4;refGene_gn=ATG9A;refGene_tx=XM_006935590.4;refGene_gn=ATG9A;refGene_tx=XM_006935591.4;refGene_gn=ATG9A;refGene_tx=XM_006935592.3;refGene_gn=ATG9A;refGene_tx=XM_011285544.3;refGene_gn=ATG9A;refGene_tx=XM_019838706.2
C1 204215875 204215993
refGene_gn=ATG9A;refGene_tx=XM_003991169.5;refGene_gn=ATG9A;refGene_tx=XM_006935589.4;refGene_gn=ATG9A;refGene_tx=XM_006935590.4;refGene_gn=ATG9A;refGene_tx=XM_006935591.4;refGene_gn=ATG9A;refGene_tx=XM_006935592.3;refGene_gn=ATG9A;refGene_tx=XM_011285544.3;refGene_gn=ATG9A;refGene_tx=XM_019838706.2
C1 204216098 204216277
refGene_gn=ATG9A;refGene_tx=XM_003991169.5;refGene_gn=ATG9A;refGene_tx=XM_006935589.4;refGene_gn=ATG9A;refGene_tx=XM_006935590.4;refGene_gn=ATG9A;refGene_tx=XM_006935591.4;refGene_gn=ATG9A;refGene_tx=XM_006935592.3;refGene_gn=ATG9A;refGene_tx=XM_011285544.3;refGene_gn=ATG9A;refGene_tx=XM_019838706.2
C1 204216488 204216737
refGene_gn=ATG9A;refGene_tx=XM_003991169.5;refGene_gn=ATG9A;refGene_tx=XM_006935589.4;refGene_gn=ATG9A;refGene_tx=XM_006935590.4;refGene_gn=ATG9A;refGene_tx=XM_006935591.4;refGene_gn=ATG9A;refGene_tx=XM_006935592.3;refGene_gn=ATG9A;refGene_tx=XM_011285544.3;refGene_gn=ATG9A;refGene_tx=XM_019838706.2
C1 204217122 204217302
refGene_gn=ATG9A;refGene_tx=XM_003991169.5;refGene_gn=ATG9A;refGene_tx=XM_006935589.4;refGene_gn=ATG9A;refGene_tx=XM_006935590.4;refGene_gn=ATG9A;refGene_tx=XM_006935591.4;refGene_gn=ATG9A;refGene_tx=XM_006935592.3;refGene_gn=ATG9A;refGene_tx=XM_011285544.3;refGene_gn=ATG9A;refGene_tx=XM_019838706.2
C1 204217407 204217561
refGene_gn=ATG9A;refGene_tx=XM_003991169.5;refGene_gn=ATG9A;refGene_tx=XM_006935589.4;refGene_gn=ATG9A;refGene_tx=XM_006935590.4;refGene_gn=ATG9A;refGene_tx=XM_006935591.4;refGene_gn=ATG9A;refGene_tx=XM_006935592.3;refGene_gn=ATG9A;refGene_tx=XM_011285544.3;refGene_gn=ATG9A;refGene_tx=XM_019838706.2
C1 204217645 204218394
refGene_gn=ATG9A;refGene_tx=XM_003991169.5;refGene_gn=ATG9A;refGene_tx=XM_006935589.4;refGene_gn=ATG9A;refGene_tx=XM_006935590.4;refGene_gn=ATG9A;refGene_tx=XM_006935591.4;refGene_gn=ATG9A;refGene_tx=XM_006935592.3;refGene_gn=ATG9A;refGene_tx=XM_011285544.3;refGene_gn=ATG9A;refGene_tx=XM_019838706.2
C1 204218623 204218765
refGene_gn=ATG9A;refGene_tx=XM_003991169.5;refGene_gn=ATG9A;refGene_tx

=XM_006935589.4;refGene_gn=ATG9A;refGene_tx=XM_006935590.4;refGene_gn=ATG9A;refGene_tx=XM_006935591.4;refGene_gn=ATG9A;refGene_tx=XM_006935592.3;refGene_gn=ATG9A;refGene_tx=XM_011285544.3;refGene_gn=ATG9A;refGene_tx=XM_019838706.2
C1 204218934 204219096
refGene_gn=ATG9A;refGene_tx=XM_003991169.5;refGene_gn=ATG9A;refGene_tx=XM_006935589.4;refGene_gn=ATG9A;refGene_tx=XM_006935590.4;refGene_gn=ATG9A;refGene_tx=XM_006935591.4;refGene_gn=ATG9A;refGene_tx=XM_006935592.3;refGene_gn=ATG9A;refGene_tx=XM_011285544.3;refGene_gn=ATG9A;refGene_tx=XM_019838706.2
C1 204219900 204219965
refGene_gn=ATG9A;refGene_tx=XM_003991169.5;refGene_gn=ATG9A;refGene_tx=XM_006935589.4;refGene_gn=ATG9A;refGene_tx=XM_006935590.4;refGene_gn=ATG9A;refGene_tx=XM_006935591.4;refGene_gn=ATG9A;refGene_tx=XM_006935592.3;refGene_gn=ATG9A;refGene_tx=XM_011285544.3;refGene_gn=ATG9A;refGene_tx=XM_019838706.2
C1 204220516 204220560
refGene_gn=ATG9A;refGene_tx=XM_003991169.5;refGene_gn=ATG9A;refGene_tx=XM_006935589.4;refGene_gn=ATG9A;refGene_tx=XM_006935590.4;refGene_gn=ATG9A;refGene_tx=XM_006935591.4;refGene_gn=ATG9A;refGene_tx=XM_006935592.3;refGene_gn=ATG9A;refGene_tx=XM_011285544.3;refGene_gn=ATG9A;refGene_tx=XM_019838706.2
C1 204220681 204220784
refGene_gn=ATG9A;refGene_tx=XM_003991169.5;refGene_gn=ATG9A;refGene_tx=XM_006935589.4;refGene_gn=ATG9A;refGene_tx=XM_006935590.4;refGene_gn=ATG9A;refGene_tx=XM_006935591.4;refGene_gn=ATG9A;refGene_tx=XM_006935592.3;refGene_gn=ATG9A;refGene_tx=XM_011285544.3;refGene_gn=ATG9A;refGene_tx=XM_019838706.2
C1 204222623 204222750
refGene_gn=ANKZF1;refGene_tx=XM_019838709.2
C1 204223066 204223214
refGene_gn=ANKZF1;refGene_tx=XM_006935599.2;refGene_gn=ANKZF1;refGene_tx=XM_019838707.1;refGene_gn=ANKZF1;refGene_tx=XM_019838708.1
C1 204224176 204224286
refGene_gn=ANKZF1;refGene_tx=XM_006935599.2;refGene_gn=ANKZF1;refGene_tx=XM_019838707.1;refGene_gn=ANKZF1;refGene_tx=XM_019838708.1;refGene_gn=ANKZF1;refGene_tx=XM_019838709.2
C1 204224505 204224608
refGene_gn=ANKZF1;refGene_tx=XM_006935599.2;refGene_gn=ANKZF1;refGene_tx=XM_019838707.1;refGene_gn=ANKZF1;refGene_tx=XM_019838708.1;refGene_gn=ANKZF1;refGene_tx=XM_019838709.2
C1 204224724 204224918
refGene_gn=ANKZF1;refGene_tx=XM_006935599.2;refGene_gn=ANKZF1;refGene_tx=XM_019838707.1;refGene_gn=ANKZF1;refGene_tx=XM_019838708.1;refGene_gn=ANKZF1;refGene_tx=XM_019838709.2
C1 204225317 204225430
refGene_gn=ANKZF1;refGene_tx=XM_006935599.2;refGene_gn=ANKZF1;refGene_tx=XM_019838707.1;refGene_gn=ANKZF1;refGene_tx=XM_019838708.1;refGene_gn=ANKZF1;refGene_tx=XM_019838709.2
C1 204225524 204225672

refGene_gn=ANKZF1;refGene_tx=XM_006935599.2;refGene_gn=ANKZF1;refGene_tx=XM_019838707.1;refGene_gn=ANKZF1;refGene_tx=XM_019838708.1;refGene_gn=ANKZF1;refGene_tx=XM_019838709.2
C1 204225961 204226190
refGene_gn=ANKZF1;refGene_tx=XM_006935599.2;refGene_gn=ANKZF1;refGene_tx=XM_019838707.1;refGene_gn=ANKZF1;refGene_tx=XM_019838708.1;refGene_gn=ANKZF1;refGene_tx=XM_019838709.2
C1 204226351 204226498
refGene_gn=ANKZF1;refGene_tx=XM_006935599.2;refGene_gn=ANKZF1;refGene_tx=XM_019838707.1;refGene_gn=ANKZF1;refGene_tx=XM_019838708.1;refGene_gn=ANKZF1;refGene_tx=XM_019838709.2
C1 204227327 204227802
refGene_gn=ANKZF1;refGene_tx=XM_006935599.2;refGene_gn=ANKZF1;refGene_tx=XM_019838707.1;refGene_gn=ANKZF1;refGene_tx=XM_019838708.1;refGene_gn=ANKZF1;refGene_tx=XM_019838709.2
C1 204227970 204228082
refGene_gn=ANKZF1;refGene_tx=XM_006935599.2;refGene_gn=ANKZF1;refGene_tx=XM_019838707.1;refGene_gn=ANKZF1;refGene_tx=XM_019838708.1;refGene_gn=ANKZF1;refGene_tx=XM_019838709.2
C1 204228215 204228606
refGene_gn=ANKZF1;refGene_tx=XM_006935599.2;refGene_gn=ANKZF1;refGene_tx=XM_019838707.1;refGene_gn=ANKZF1;refGene_tx=XM_019838708.1;refGene_gn=ANKZF1;refGene_tx=XM_019838709.2
C1 204228819 204229001
refGene_gn=ANKZF1;refGene_tx=XM_006935599.2;refGene_gn=ANKZF1;refGene_tx=XM_019838707.1;refGene_gn=ANKZF1;refGene_tx=XM_019838708.1;refGene_gn=ANKZF1;refGene_tx=XM_019838709.2
C1 204229260 204229530
refGene_gn=GLB1L;refGene_tx=XM_003991175.5;refGene_gn=GLB1L;refGene_tx=XM_006935596.4;refGene_gn=GLB1L;refGene_tx=XM_006935597.4;refGene_gn=GLB1L;refGene_tx=XM_006935598.4;refGene_gn=GLB1L;refGene_tx=XM_019838710.2;refGene_gn=GLB1L;refGene_tx=XM_019838711.2;refGene_gn=GLB1L;refGene_tx=XM_019838712.2
C1 204229677 204229893
refGene_gn=GLB1L;refGene_tx=XM_003991175.5;refGene_gn=GLB1L;refGene_tx=XM_006935596.4;refGene_gn=GLB1L;refGene_tx=XM_006935597.4;refGene_gn=GLB1L;refGene_tx=XM_006935598.4;refGene_gn=GLB1L;refGene_tx=XM_019838710.2;refGene_gn=GLB1L;refGene_tx=XM_019838711.2
C1 204230127 204230259
refGene_gn=GLB1L;refGene_tx=XM_003991175.5;refGene_gn=GLB1L;refGene_tx=XM_006935596.4;refGene_gn=GLB1L;refGene_tx=XM_006935597.4;refGene_gn=GLB1L;refGene_tx=XM_006935598.4;refGene_gn=GLB1L;refGene_tx=XM_019838710.2;refGene_gn=GLB1L;refGene_tx=XM_019838711.2;refGene_gn=GLB1L;refGene_tx=XM_019838712.2
C1 204230538 204230652
refGene_gn=GLB1L;refGene_tx=XM_003991175.5;refGene_gn=GLB1L;refGene_tx=XM_006935596.4;refGene_gn=GLB1L;refGene_tx=XM_006935597.4;refGene_gn=GLB1L;refGene_tx=XM_006935598.4;refGene_gn=GLB1L;refGene_tx=XM_019838710.2;refGene_gn=GLB1L;refGene_tx=XM_019838711.2;refGene_gn=GLB1L;refGene_tx=XM_019838712.2

C1	204230794	204230884
refGene_gn=GLB1L;refGene_tx=XM_003991175.5;refGene_gn=GLB1L;refGene_tx=XM_006935596.4;refGene_gn=GLB1L;refGene_tx=XM_006935597.4;refGene_gn=GLB1L;refGene_tx=XM_006935598.4;refGene_gn=GLB1L;refGene_tx=XM_019838710.2;refGene_gn=GLB1L;refGene_tx=XM_019838711.2;refGene_gn=GLB1L;refGene_tx=XM_019838712.2
C1	204231006	204231081
refGene_gn=GLB1L;refGene_tx=XM_003991175.5;refGene_gn=GLB1L;refGene_tx=XM_006935596.4;refGene_gn=GLB1L;refGene_tx=XM_006935597.4;refGene_gn=GLB1L;refGene_tx=XM_006935598.4;refGene_gn=GLB1L;refGene_tx=XM_019838710.2;refGene_gn=GLB1L;refGene_tx=XM_019838712.2
C1	204231386	204231505
refGene_gn=GLB1L;refGene_tx=XM_003991175.5;refGene_gn=GLB1L;refGene_tx=XM_006935596.4;refGene_gn=GLB1L;refGene_tx=XM_006935597.4;refGene_gn=GLB1L;refGene_tx=XM_006935598.4;refGene_gn=GLB1L;refGene_tx=XM_019838710.2;refGene_gn=GLB1L;refGene_tx=XM_019838711.2;refGene_gn=GLB1L;refGene_tx=XM_019838712.2
C1	204231685	204231726
refGene_gn=GLB1L;refGene_tx=XM_003991175.5;refGene_gn=GLB1L;refGene_tx=XM_006935596.4;refGene_gn=GLB1L;refGene_tx=XM_006935597.4;refGene_gn=GLB1L;refGene_tx=XM_006935598.4;refGene_gn=GLB1L;refGene_tx=XM_019838710.2;refGene_gn=GLB1L;refGene_tx=XM_019838711.2;refGene_gn=GLB1L;refGene_tx=XM_019838712.2
C1	204231834	204231956
refGene_gn=GLB1L;refGene_tx=XM_003991175.5;refGene_gn=GLB1L;refGene_tx=XM_006935596.4;refGene_gn=GLB1L;refGene_tx=XM_006935597.4;refGene_gn=GLB1L;refGene_tx=XM_006935598.4;refGene_gn=GLB1L;refGene_tx=XM_019838710.2;refGene_gn=GLB1L;refGene_tx=XM_019838711.2;refGene_gn=GLB1L;refGene_tx=XM_019838712.2
C1	204232060	204232119
refGene_gn=GLB1L;refGene_tx=XM_003991175.5;refGene_gn=GLB1L;refGene_tx=XM_006935596.4;refGene_gn=GLB1L;refGene_tx=XM_006935597.4;refGene_gn=GLB1L;refGene_tx=XM_006935598.4;refGene_gn=GLB1L;refGene_tx=XM_019838710.2;refGene_gn=GLB1L;refGene_tx=XM_019838711.2;refGene_gn=GLB1L;refGene_tx=XM_019838712.2
C1	204232203	204232378
refGene_gn=GLB1L;refGene_tx=XM_003991175.5;refGene_gn=GLB1L;refGene_tx=XM_006935596.4;refGene_gn=GLB1L;refGene_tx=XM_006935597.4;refGene_gn=GLB1L;refGene_tx=XM_006935598.4;refGene_gn=GLB1L;refGene_tx=XM_019838710.2;refGene_gn=GLB1L;refGene_tx=XM_019838711.2;refGene_gn=GLB1L;refGene_tx=XM_019838712.2
C1	204232481	204232576
refGene_gn=GLB1L;refGene_tx=XM_003991175.5;refGene_gn=GLB1L;refGene_tx=XM_006935596.4;refGene_gn=GLB1L;refGene_tx=XM_006935597.4;refGene_gn=GLB1L;refGene_tx=XM_006935598.4;refGene_gn=GLB1L;refGene_tx=XM_019838710.2;refGene_gn=GLB1L;refGene_tx=XM_019838711.2;refGene_gn=GLB1L;refGene_tx=XM_019838712.2
C1	204236702	204236763
refGene_gn=GLB1L;refGene_tx=XM_003991175.5;refGene_gn=GLB1L;refGene_tx=XM_006935596.4;refGene_gn=GLB1L;refGene_tx=XM_006935597.4;refGene_gn=

GLB1L;refGene_tx=XM_006935598.4;refGene_gn=GLB1L;refGene_tx=XM_019838710.2;refGene_gn=GLB1L;refGene_tx=XM_019838711.2;refGene_gn=GLB1L;refGene_tx=XM_019838712.2
C1 204236907 204237058
refGene_gn=GLB1L;refGene_tx=XM_003991175.5;refGene_gn=GLB1L;refGene_tx=XM_006935596.4;refGene_gn=GLB1L;refGene_tx=XM_006935597.4;refGene_gn=GLB1L;refGene_tx=XM_006935598.4;refGene_gn=GLB1L;refGene_tx=XM_019838710.2;refGene_gn=GLB1L;refGene_tx=XM_019838711.2;refGene_gn=GLB1L;refGene_tx=XM_019838712.2
C1 204237302 204237469
refGene_gn=GLB1L;refGene_tx=XM_003991175.5;refGene_gn=GLB1L;refGene_tx=XM_006935596.4;refGene_gn=GLB1L;refGene_tx=XM_006935597.4;refGene_gn=GLB1L;refGene_tx=XM_006935598.4;refGene_gn=GLB1L;refGene_tx=XM_019838710.2;refGene_gn=GLB1L;refGene_tx=XM_019838711.2;refGene_gn=GLB1L;refGene_tx=XM_019838712.2
C1 204237695 204237764
refGene_gn=GLB1L;refGene_tx=XM_003991175.5;refGene_gn=GLB1L;refGene_tx=XM_006935596.4;refGene_gn=GLB1L;refGene_tx=XM_006935597.4;refGene_gn=GLB1L;refGene_tx=XM_019838710.2;refGene_gn=GLB1L;refGene_tx=XM_019838711.2;refGene_gn=GLB1L;refGene_tx=XM_019838712.2
C1 204239003 204239012
refGene_gn=GLB1L;refGene_tx=XM_006935598.4
C1 204239757 204239843
refGene_gn=STK16;refGene_tx=XM_003991173.2;refGene_gn=STK16;refGene_tx=XM_006935593.3;refGene_gn=STK16;refGene_tx=XM_006935594.4;refGene_gn=STK16;refGene_tx=XM_019838713.2
C1 204240437 204240657
refGene_gn=STK16;refGene_tx=XM_003991173.2;refGene_gn=STK16;refGene_tx=XM_006935593.3;refGene_gn=STK16;refGene_tx=XM_006935594.4;refGene_gn=STK16;refGene_tx=XM_019838713.2
C1 204240921 204241055
refGene_gn=STK16;refGene_tx=XM_003991173.2;refGene_gn=STK16;refGene_tx=XM_006935593.3;refGene_gn=STK16;refGene_tx=XM_006935594.4;refGene_gn=STK16;refGene_tx=XM_019838713.2
C1 204241225 204241346
refGene_gn=STK16;refGene_tx=XM_003991173.2;refGene_gn=STK16;refGene_tx=XM_006935593.3;refGene_gn=STK16;refGene_tx=XM_006935594.4;refGene_gn=STK16;refGene_tx=XM_019838713.2
C1 204241472 204241568
refGene_gn=STK16;refGene_tx=XM_003991173.2;refGene_gn=STK16;refGene_tx=XM_006935593.3;refGene_gn=STK16;refGene_tx=XM_006935594.4
C1 204241992 204242114
refGene_gn=STK16;refGene_tx=XM_003991173.2;refGene_gn=STK16;refGene_tx=XM_006935593.3;refGene_gn=STK16;refGene_tx=XM_006935594.4;refGene_gn=STK16;refGene_tx=XM_019838713.2
C1 204242229 204242368
refGene_gn=STK16;refGene_tx=XM_003991173.2;refGene_gn=STK16;refGene_tx=XM_006935593.3;refGene_gn=STK16;refGene_tx=XM_006935594.4;refGene_gn=STK16;refGene_tx=XM_019838713.2
C1 204244248 204245220

refGene_gn=TUBA4A;refGene_tx=XM_003991171.4
C1 204245456 204245605
refGene_gn=TUBA4A;refGene_tx=XM_003991171.4
C1 204245918 204246141
refGene_gn=TUBA4A;refGene_tx=XM_003991171.4
C1 204247946 204247949
refGene_gn=TUBA4A;refGene_tx=XM_003991171.4
C1 204270154 204270219
refGene_gn=DNAJB2;refGene_tx=XM_003991176.5
C1 204270871 204270981
refGene_gn=DNAJB2;refGene_tx=XM_003991176.5
C1 204272067 204272121
refGene_gn=DNAJB2;refGene_tx=XM_003991176.5
C1 204272293 204272416
refGene_gn=DNAJB2;refGene_tx=XM_003991176.5
C1 204273139 204273232
refGene_gn=DNAJB2;refGene_tx=XM_003991176.5
C1 204273467 204273570
refGene_gn=DNAJB2;refGene_tx=XM_003991176.5
C1 204273760 204273831
refGene_gn=DNAJB2;refGene_tx=XM_003991176.5
C1 204275123 204275330
refGene_gn=DNAJB2;refGene_tx=XM_003991176.5
C1 204276464 204276475
refGene_gn=DNAJB2;refGene_tx=XM_003991176.5
C1 204279723 204279795
refGene_gn=PTPRN;refGene_tx=XM_011285542.3;refGene_gn=PTPRN;refGene_tx
=XM_019838716.2
C1 204279955 204280029
refGene_gn=PTPRN;refGene_tx=XM_011285542.3;refGene_gn=PTPRN;refGene_tx
=XM_019838716.2
C1 204280223 204280288
refGene_gn=PTPRN;refGene_tx=XM_011285542.3;refGene_gn=PTPRN;refGene_tx
=XM_019838716.2
C1 204281421 204281475
refGene_gn=PTPRN;refGene_tx=XM_011285542.3;refGene_gn=PTPRN;refGene_tx
=XM_019838716.2
C1 204284265 204284432
refGene_gn=PTPRN;refGene_tx=XM_011285542.3;refGene_gn=PTPRN;refGene_tx
=XM_019838716.2
C1 204285641 204285761
refGene_gn=PTPRN;refGene_tx=XM_011285542.3;refGene_gn=PTPRN;refGene_tx
=XM_019838716.2
C1 204285863 204285941
refGene_gn=PTPRN;refGene_tx=XM_011285542.3;refGene_gn=PTPRN;refGene_tx
=XM_019838716.2
C1 204286179 204286253
refGene_gn=PTPRN;refGene_tx=XM_011285542.3;refGene_gn=PTPRN;refGene_tx
=XM_019838716.2
C1 204286413 204286561

refGene_gn=PTPRN;refGene_tx=XM_011285542.3;refGene_gn=PTPRN;refGene_tx=XM_019838716.2
C1 204286661 204286862
refGene_gn=PTPRN;refGene_tx=XM_011285542.3;refGene_gn=PTPRN;refGene_tx=XM_019838716.2
C1 204287299 204287518
refGene_gn=PTPRN;refGene_tx=XM_011285542.3;refGene_gn=PTPRN;refGene_tx=XM_019838716.2
C1 204288442 204288507
refGene_gn=PTPRN;refGene_tx=XM_011285542.3;refGene_gn=PTPRN;refGene_tx=XM_019838716.2
C1 204288684 204288764
refGene_gn=PTPRN;refGene_tx=XM_011285542.3;refGene_gn=PTPRN;refGene_tx=XM_019838716.2
C1 204289083 204289170
refGene_gn=PTPRN;refGene_tx=XM_011285542.3;refGene_gn=PTPRN;refGene_tx=XM_019838716.2
C1 204289391 204289666
refGene_gn=PTPRN;refGene_tx=XM_011285542.3;refGene_gn=PTPRN;refGene_tx=XM_019838716.2
C1 204290566 204290601
refGene_gn=PTPRN;refGene_tx=XM_011285542.3;refGene_gn=PTPRN;refGene_tx=XM_019838716.2
C1 204291206 204291341
refGene_gn=PTPRN;refGene_tx=XM_011285542.3;refGene_gn=PTPRN;refGene_tx=XM_019838716.2
C1 204291759 204292114
refGene_gn=PTPRN;refGene_tx=XM_011285542.3;refGene_gn=PTPRN;refGene_tx=XM_019838716.2
C1 204292199 204292461
refGene_gn=PTPRN;refGene_tx=XM_011285542.3;refGene_gn=PTPRN;refGene_tx=XM_019838716.2
C1 204293343 204293440
refGene_gn=PTPRN;refGene_tx=XM_011285542.3;refGene_gn=PTPRN;refGene_tx=XM_019838716.2
C1 204296292 204296406
refGene_gn=PTPRN;refGene_tx=XM_011285542.3;refGene_gn=PTPRN;refGene_tx=XM_019838716.2
C1 204296638 204296689
refGene_gn=PTPRN;refGene_tx=XM_011285542.3;refGene_gn=PTPRN;refGene_tx=XM_019838716.2
C1 204298129 204298244
refGene_gn=PTPRN;refGene_tx=XM_011285542.3;refGene_gn=PTPRN;refGene_tx=XM_019838716.2
C1 204308697 204308748
refGene_gn=RESP18;refGene_tx=XM_019839143.1
C1 204310635 204310723
refGene_gn=RESP18;refGene_tx=XM_019839143.1
C1 204311110 204311243
refGene_gn=RESP18;refGene_tx=XM_019839143.1

C1 204312255 204312360
refGene_gn=RESP18;refGene_tx=XM_019839143.1
C1 204314072 204314288
refGene_gn=RESP18;refGene_tx=XM_019839143.1
C1 204314718 204314740
refGene_gn=RESP18;refGene_tx=XM_019839143.1
C1 204328552 204328588
refGene_gn=LOC109502880;refGene_tx=XM_023259686.1
C1 204329826 204329930
refGene_gn=LOC109502880;refGene_tx=XM_023259686.1
C1 204330103 204330154
refGene_gn=LOC109502880;refGene_tx=XM_023259686.1
C1 204331034 204331152
refGene_gn=LOC109502880;refGene_tx=XM_023259686.1
C1 204344152 204344203
refGene_gn=DNPEP;refGene_tx=XM_006935601.4;refGene_gn=DNPEP;refGene_tx
=XM_006935602.3;refGene_gn=DNPEP;refGene_tx=XM_006935603.4;refGene_gn=
DNPEP;refGene_tx=XM_006935604.4;refGene_gn=DNPEP;refGene_tx=XM_0232596
87.1
C1 204344509 204344677
refGene_gn=DNPEP;refGene_tx=XM_006935601.4;refGene_gn=DNPEP;refGene_tx
=XM_006935602.3;refGene_gn=DNPEP;refGene_tx=XM_006935603.4;refGene_gn=
DNPEP;refGene_tx=XM_006935604.4;refGene_gn=DNPEP;refGene_tx=XM_0232596
87.1
C1 204348807 204348909
refGene_gn=DNPEP;refGene_tx=XM_006935601.4;refGene_gn=DNPEP;refGene_tx
=XM_006935602.3;refGene_gn=DNPEP;refGene_tx=XM_006935603.4;refGene_gn=
DNPEP;refGene_tx=XM_006935604.4;refGene_gn=DNPEP;refGene_tx=XM_0232596
87.1
C1 204349012 204349052
refGene_gn=DNPEP;refGene_tx=XM_006935601.4;refGene_gn=DNPEP;refGene_tx
=XM_006935602.3;refGene_gn=DNPEP;refGene_tx=XM_006935603.4;refGene_gn=
DNPEP;refGene_tx=XM_006935604.4;refGene_gn=DNPEP;refGene_tx=XM_0232596
87.1
C1 204349430 204349591
refGene_gn=DNPEP;refGene_tx=XM_006935601.4;refGene_gn=DNPEP;refGene_tx
=XM_006935602.3;refGene_gn=DNPEP;refGene_tx=XM_006935603.4;refGene_gn=
DNPEP;refGene_tx=XM_006935604.4;refGene_gn=DNPEP;refGene_tx=XM_0232596
87.1
C1 204350609 204350693
refGene_gn=DNPEP;refGene_tx=XM_006935601.4;refGene_gn=DNPEP;refGene_tx
=XM_006935602.3;refGene_gn=DNPEP;refGene_tx=XM_006935603.4;refGene_gn=
DNPEP;refGene_tx=XM_006935604.4;refGene_gn=DNPEP;refGene_tx=XM_0232596
87.1
C1 204351891 204351969
refGene_gn=DNPEP;refGene_tx=XM_006935601.4;refGene_gn=DNPEP;refGene_tx
=XM_006935602.3;refGene_gn=DNPEP;refGene_tx=XM_006935603.4;refGene_gn=
DNPEP;refGene_tx=XM_006935604.4;refGene_gn=DNPEP;refGene_tx=XM_0232596
87.1
C1 204352713 204352821

refGene_gn=DNPEP;refGene_tx=XM_006935601.4;refGene_gn=DNPEP;refGene_tx=XM_006935602.3;refGene_gn=DNPEP;refGene_tx=XM_006935603.4;refGene_gn=DNPEP;refGene_tx=XM_006935604.4;refGene_gn=DNPEP;refGene_tx=XM_023259687.1
C1 204352928 204353004
refGene_gn=DNPEP;refGene_tx=XM_006935601.4;refGene_gn=DNPEP;refGene_tx=XM_006935602.3;refGene_gn=DNPEP;refGene_tx=XM_006935603.4;refGene_gn=DNPEP;refGene_tx=XM_006935604.4;refGene_gn=DNPEP;refGene_tx=XM_023259687.1
C1 204353265 204353396
refGene_gn=DNPEP;refGene_tx=XM_006935601.4;refGene_gn=DNPEP;refGene_tx=XM_006935602.3;refGene_gn=DNPEP;refGene_tx=XM_006935603.4;refGene_gn=DNPEP;refGene_tx=XM_006935604.4;refGene_gn=DNPEP;refGene_tx=XM_023259687.1
C1 204353600 204353726
refGene_gn=DNPEP;refGene_tx=XM_006935601.4;refGene_gn=DNPEP;refGene_tx=XM_006935602.3;refGene_gn=DNPEP;refGene_tx=XM_006935603.4;refGene_gn=DNPEP;refGene_tx=XM_006935604.4;refGene_gn=DNPEP;refGene_tx=XM_023259687.1
C1 204353953 204354067
refGene_gn=DNPEP;refGene_tx=XM_006935601.4;refGene_gn=DNPEP;refGene_tx=XM_006935602.3;refGene_gn=DNPEP;refGene_tx=XM_006935603.4;refGene_gn=DNPEP;refGene_tx=XM_006935604.4;refGene_gn=DNPEP;refGene_tx=XM_023259687.1
C1 204354196 204354285
refGene_gn=DNPEP;refGene_tx=XM_006935601.4;refGene_gn=DNPEP;refGene_tx=XM_006935602.3;refGene_gn=DNPEP;refGene_tx=XM_006935603.4;refGene_gn=DNPEP;refGene_tx=XM_006935604.4;refGene_gn=DNPEP;refGene_tx=XM_023259687.1
C1 204354371 204354465
refGene_gn=DNPEP;refGene_tx=XM_006935601.4;refGene_gn=DNPEP;refGene_tx=XM_006935602.3;refGene_gn=DNPEP;refGene_tx=XM_006935603.4;refGene_gn=DNPEP;refGene_tx=XM_006935604.4;refGene_gn=DNPEP;refGene_tx=XM_023259687.1
C1 204355084 204355090
refGene_gn=DNPEP;refGene_tx=XM_006935602.3
C1 204355374 204355422
refGene_gn=DNPEP;refGene_tx=XM_006935601.4
C1 204363274 204363334
refGene_gn=DNPEP;refGene_tx=XM_006935601.4
C1 204379746 204380330
refGene_gn=DES;refGene_tx=XM_003991178.4
C1 204381354 204381415
refGene_gn=DES;refGene_tx=XM_003991178.4
C1 204381534 204381630
refGene_gn=DES;refGene_tx=XM_003991178.4
C1 204381782 204381944
refGene_gn=DES;refGene_tx=XM_003991178.4
C1 204382115 204382241
refGene_gn=DES;refGene_tx=XM_003991178.4

C1	204382654	204382875
refGene_gn=DES;refGene_tx=XM_003991178.4
C1	204384419	204384463
refGene_gn=DES;refGene_tx=XM_003991178.4
C1	204385903	204385986
refGene_gn=DES;refGene_tx=XM_003991178.4
C1	204386201	204386243
refGene_gn=DES;refGene_tx=XM_003991178.4
C1	204393574	204393953
refGene_gn=SPEG;refGene_tx=XM_011285666.3;refGene_gn=SPEG;refGene_tx=XM_023259689.1;refGene_gn=SPEG;refGene_tx=XM_023259691.1;refGene_gn=SPEG;refGene_tx=XM_023259694.1;refGene_gn=SPEG;refGene_tx=XM_023259701.1;refGene_gn=SPEG;refGene_tx=XM_023259702.1;refGene_gn=SPEG;refGene_tx=XM_023259703.1
C1	204400738	204400771
refGene_gn=SPEG;refGene_tx=XM_023259693.1
C1	204401825	204401868
refGene_gn=SPEG;refGene_tx=XM_023259693.1
C1	204402461	204402813
refGene_gn=SPEG;refGene_tx=XM_023259689.1;refGene_gn=SPEG;refGene_tx=XM_023259690.1;refGene_gn=SPEG;refGene_tx=XM_023259691.1;refGene_gn=SPEG;refGene_tx=XM_023259693.1;refGene_gn=SPEG;refGene_tx=XM_023259694.1;refGene_gn=SPEG;refGene_tx=XM_023259701.1;refGene_gn=SPEG;refGene_tx=XM_023259702.1;refGene_gn=SPEG;refGene_tx=XM_023259703.1
C1	204403402	204403492
refGene_gn=SPEG;refGene_tx=XM_011285666.3;refGene_gn=SPEG;refGene_tx=XM_023259689.1;refGene_gn=SPEG;refGene_tx=XM_023259690.1;refGene_gn=SPEG;refGene_tx=XM_023259691.1;refGene_gn=SPEG;refGene_tx=XM_023259693.1;refGene_gn=SPEG;refGene_tx=XM_023259694.1;refGene_gn=SPEG;refGene_tx=XM_023259701.1;refGene_gn=SPEG;refGene_tx=XM_023259702.1;refGene_gn=SPEG;refGene_tx=XM_023259703.1
C1	204403575	204403912
refGene_gn=SPEG;refGene_tx=XM_011285666.3;refGene_gn=SPEG;refGene_tx=XM_023259689.1;refGene_gn=SPEG;refGene_tx=XM_023259690.1;refGene_gn=SPEG;refGene_tx=XM_023259691.1;refGene_gn=SPEG;refGene_tx=XM_023259693.1;refGene_gn=SPEG;refGene_tx=XM_023259694.1;refGene_gn=SPEG;refGene_tx=XM_023259701.1;refGene_gn=SPEG;refGene_tx=XM_023259702.1;refGene_gn=SPEG;refGene_tx=XM_023259703.1
C1	204403919	204404455
refGene_gn=SPEG;refGene_tx=XM_023259692.1
C1	204406777	204408081
refGene_gn=SPEG;refGene_tx=XM_011285666.3;refGene_gn=SPEG;refGene_tx=XM_023259689.1;refGene_gn=SPEG;refGene_tx=XM_023259690.1;refGene_gn=SPEG;refGene_tx=XM_023259691.1;refGene_gn=SPEG;refGene_tx=XM_023259692.1;refGene_gn=SPEG;refGene_tx=XM_023259693.1;refGene_gn=SPEG;refGene_tx=XM_023259694.1;refGene_gn=SPEG;refGene_tx=XM_023259701.1;refGene_gn=SPEG;refGene_tx=XM_023259702.1;refGene_gn=SPEG;refGene_tx=XM_023259703.1
C1	204409925	204410069
refGene_gn=SPEG;refGene_tx=XM_011285666.3;refGene_gn=SPEG;refGene_tx=XM_023259689.1;refGene_gn=SPEG;refGene_tx=XM_023259690.1;refGene_gn=SPE

G;refGene_tx=XM_023259691.1;refGene_gn=SPEG;refGene_tx=XM_023259692.1;
refGene_gn=SPEG;refGene_tx=XM_023259693.1;refGene_gn=SPEG;refGene_tx=X
M_023259701.1;refGene_gn=SPEG;refGene_tx=XM_023259702.1;refGene_gn=SPE
G;refGene_tx=XM_023259703.1
C1 204410376 204410559
refGene_gn=SPEG;refGene_tx=XM_011285666.3;refGene_gn=SPEG;refGene_tx=X
M_023259689.1;refGene_gn=SPEG;refGene_tx=XM_023259690.1;refGene_gn=SPE
G;refGene_tx=XM_023259691.1;refGene_gn=SPEG;refGene_tx=XM_023259692.1;
refGene_gn=SPEG;refGene_tx=XM_023259693.1;refGene_gn=SPEG;refGene_tx=X
M_023259701.1;refGene_gn=SPEG;refGene_tx=XM_023259702.1;refGene_gn=SPE
G;refGene_tx=XM_023259703.1
C1 204419806 204419982
refGene_gn=SPEG;refGene_tx=XM_011285666.3;refGene_gn=SPEG;refGene_tx=X
M_023259689.1;refGene_gn=SPEG;refGene_tx=XM_023259690.1;refGene_gn=SPE
G;refGene_tx=XM_023259691.1;refGene_gn=SPEG;refGene_tx=XM_023259692.1;
refGene_gn=SPEG;refGene_tx=XM_023259693.1;refGene_gn=SPEG;refGene_tx=X
M_023259694.1;refGene_gn=SPEG;refGene_tx=XM_023259695.1;refGene_gn=SPE
G;refGene_tx=XM_023259697.1;refGene_gn=SPEG;refGene_tx=XM_023259698.1;
refGene_gn=SPEG;refGene_tx=XM_023259699.1;refGene_gn=SPEG;refGene_tx=X
M_023259700.1;refGene_gn=SPEG;refGene_tx=XM_023259701.1;refGene_gn=SPE
G;refGene_tx=XM_023259702.1;refGene_gn=SPEG;refGene_tx=XM_023259703.1
C1 204420219 204420308
refGene_gn=SPEG;refGene_tx=XM_011285666.3;refGene_gn=SPEG;refGene_tx=X
M_023259689.1;refGene_gn=SPEG;refGene_tx=XM_023259690.1;refGene_gn=SPE
G;refGene_tx=XM_023259691.1;refGene_gn=SPEG;refGene_tx=XM_023259692.1;
refGene_gn=SPEG;refGene_tx=XM_023259693.1;refGene_gn=SPEG;refGene_tx=X
M_023259694.1;refGene_gn=SPEG;refGene_tx=XM_023259695.1;refGene_gn=SPE
G;refGene_tx=XM_023259697.1;refGene_gn=SPEG;refGene_tx=XM_023259698.1;
refGene_gn=SPEG;refGene_tx=XM_023259699.1;refGene_gn=SPEG;refGene_tx=X
M_023259700.1;refGene_gn=SPEG;refGene_tx=XM_023259701.1;refGene_gn=SPE
G;refGene_tx=XM_023259702.1;refGene_gn=SPEG;refGene_tx=XM_023259703.1
C1 204421203 204421379
refGene_gn=SPEG;refGene_tx=XM_011285666.3;refGene_gn=SPEG;refGene_tx=X
M_023259689.1;refGene_gn=SPEG;refGene_tx=XM_023259690.1;refGene_gn=SPE
G;refGene_tx=XM_023259691.1;refGene_gn=SPEG;refGene_tx=XM_023259692.1;
refGene_gn=SPEG;refGene_tx=XM_023259693.1;refGene_gn=SPEG;refGene_tx=X
M_023259694.1;refGene_gn=SPEG;refGene_tx=XM_023259695.1;refGene_gn=SPE
G;refGene_tx=XM_023259697.1;refGene_gn=SPEG;refGene_tx=XM_023259698.1;
refGene_gn=SPEG;refGene_tx=XM_023259699.1;refGene_gn=SPEG;refGene_tx=X
M_023259700.1;refGene_gn=SPEG;refGene_tx=XM_023259701.1;refGene_gn=SPE
G;refGene_tx=XM_023259702.1;refGene_gn=SPEG;refGene_tx=XM_023259703.1
C1 204422742 204422750
refGene_gn=SPEG;refGene_tx=XM_023259703.1
C1 204423896 204424157
refGene_gn=SPEG;refGene_tx=XM_011285666.3;refGene_gn=SPEG;refGene_tx=X
M_023259689.1;refGene_gn=SPEG;refGene_tx=XM_023259690.1;refGene_gn=SPE
G;refGene_tx=XM_023259691.1;refGene_gn=SPEG;refGene_tx=XM_023259692.1;
refGene_gn=SPEG;refGene_tx=XM_023259693.1;refGene_gn=SPEG;refGene_tx=X
M_023259694.1;refGene_gn=SPEG;refGene_tx=XM_023259695.1;refGene_gn=SPE
G;refGene_tx=XM_023259697.1;refGene_gn=SPEG;refGene_tx=XM_023259698.1;

refGene_gn=SPEG;refGene_tx=XM_023259699.1;refGene_gn=SPEG;refGene_tx=XM_023259700.1;refGene_gn=SPEG;refGene_tx=XM_023259701.1;refGene_gn=SPEG;refGene_tx=XM_023259702.1
C1 204425185 204425344
refGene_gn=SPEG;refGene_tx=XM_011285666.3;refGene_gn=SPEG;refGene_tx=XM_023259689.1;refGene_gn=SPEG;refGene_tx=XM_023259690.1;refGene_gn=SPEG;refGene_tx=XM_023259691.1;refGene_gn=SPEG;refGene_tx=XM_023259692.1;refGene_gn=SPEG;refGene_tx=XM_023259693.1;refGene_gn=SPEG;refGene_tx=XM_023259694.1;refGene_gn=SPEG;refGene_tx=XM_023259695.1;refGene_gn=SPEG;refGene_tx=XM_023259697.1;refGene_gn=SPEG;refGene_tx=XM_023259698.1;refGene_gn=SPEG;refGene_tx=XM_023259699.1;refGene_gn=SPEG;refGene_tx=XM_023259700.1;refGene_gn=SPEG;refGene_tx=XM_023259701.1
C1 204425450 204425640
refGene_gn=SPEG;refGene_tx=XM_011285666.3;refGene_gn=SPEG;refGene_tx=XM_023259689.1;refGene_gn=SPEG;refGene_tx=XM_023259690.1;refGene_gn=SPEG;refGene_tx=XM_023259691.1;refGene_gn=SPEG;refGene_tx=XM_023259692.1;refGene_gn=SPEG;refGene_tx=XM_023259693.1;refGene_gn=SPEG;refGene_tx=XM_023259694.1;refGene_gn=SPEG;refGene_tx=XM_023259695.1;refGene_gn=SPEG;refGene_tx=XM_023259697.1;refGene_gn=SPEG;refGene_tx=XM_023259698.1;refGene_gn=SPEG;refGene_tx=XM_023259699.1;refGene_gn=SPEG;refGene_tx=XM_023259700.1;refGene_gn=SPEG;refGene_tx=XM_023259701.1
C1 204425743 204425967
refGene_gn=SPEG;refGene_tx=XM_011285666.3;refGene_gn=SPEG;refGene_tx=XM_023259689.1;refGene_gn=SPEG;refGene_tx=XM_023259690.1;refGene_gn=SPEG;refGene_tx=XM_023259691.1;refGene_gn=SPEG;refGene_tx=XM_023259692.1;refGene_gn=SPEG;refGene_tx=XM_023259693.1;refGene_gn=SPEG;refGene_tx=XM_023259694.1;refGene_gn=SPEG;refGene_tx=XM_023259695.1;refGene_gn=SPEG;refGene_tx=XM_023259697.1;refGene_gn=SPEG;refGene_tx=XM_023259698.1;refGene_gn=SPEG;refGene_tx=XM_023259699.1;refGene_gn=SPEG;refGene_tx=XM_023259700.1;refGene_gn=SPEG;refGene_tx=XM_023259701.1
C1 204428457 204428577
refGene_gn=SPEG;refGene_tx=XM_011285666.3;refGene_gn=SPEG;refGene_tx=XM_023259689.1;refGene_gn=SPEG;refGene_tx=XM_023259690.1;refGene_gn=SPEG;refGene_tx=XM_023259691.1;refGene_gn=SPEG;refGene_tx=XM_023259692.1;refGene_gn=SPEG;refGene_tx=XM_023259693.1;refGene_gn=SPEG;refGene_tx=XM_023259694.1;refGene_gn=SPEG;refGene_tx=XM_023259695.1;refGene_gn=SPEG;refGene_tx=XM_023259697.1;refGene_gn=SPEG;refGene_tx=XM_023259698.1;refGene_gn=SPEG;refGene_tx=XM_023259699.1;refGene_gn=SPEG;refGene_tx=XM_023259700.1;refGene_gn=SPEG;refGene_tx=XM_023259701.1
C1 204428796 204428901
refGene_gn=SPEG;refGene_tx=XM_011285666.3;refGene_gn=SPEG;refGene_tx=XM_023259689.1;refGene_gn=SPEG;refGene_tx=XM_023259690.1;refGene_gn=SPEG;refGene_tx=XM_023259691.1;refGene_gn=SPEG;refGene_tx=XM_023259692.1;refGene_gn=SPEG;refGene_tx=XM_023259693.1;refGene_gn=SPEG;refGene_tx=XM_023259694.1;refGene_gn=SPEG;refGene_tx=XM_023259695.1;refGene_gn=SPEG;refGene_tx=XM_023259697.1;refGene_gn=SPEG;refGene_tx=XM_023259698.1;refGene_gn=SPEG;refGene_tx=XM_023259699.1;refGene_gn=SPEG;refGene_tx=XM_023259700.1;refGene_gn=SPEG;refGene_tx=XM_023259701.1
C1 204429502 204429709
refGene_gn=SPEG;refGene_tx=XM_011285666.3;refGene_gn=SPEG;refGene_tx=X

M_023259689.1;refGene_gn=SPEG;refGene_tx=XM_023259690.1;refGene_gn=SPEG;refGene_tx=XM_023259691.1;refGene_gn=SPEG;refGene_tx=XM_023259692.1;refGene_gn=SPEG;refGene_tx=XM_023259693.1;refGene_gn=SPEG;refGene_tx=XM_023259694.1;refGene_gn=SPEG;refGene_tx=XM_023259695.1;refGene_gn=SPEG;refGene_tx=XM_023259697.1;refGene_gn=SPEG;refGene_tx=XM_023259698.1;refGene_gn=SPEG;refGene_tx=XM_023259699.1;refGene_gn=SPEG;refGene_tx=XM_023259700.1;refGene_gn=SPEG;refGene_tx=XM_023259701.1
C1 204430111 204430235
refGene_gn=SPEG;refGene_tx=XM_011285666.3;refGene_gn=SPEG;refGene_tx=XM_023259689.1;refGene_gn=SPEG;refGene_tx=XM_023259690.1;refGene_gn=SPEG;refGene_tx=XM_023259691.1;refGene_gn=SPEG;refGene_tx=XM_023259692.1;refGene_gn=SPEG;refGene_tx=XM_023259693.1;refGene_gn=SPEG;refGene_tx=XM_023259694.1;refGene_gn=SPEG;refGene_tx=XM_023259695.1;refGene_gn=SPEG;refGene_tx=XM_023259697.1;refGene_gn=SPEG;refGene_tx=XM_023259698.1;refGene_gn=SPEG;refGene_tx=XM_023259699.1;refGene_gn=SPEG;refGene_tx=XM_023259700.1;refGene_gn=SPEG;refGene_tx=XM_023259701.1
C1 204430352 204430528
refGene_gn=SPEG;refGene_tx=XM_011285666.3;refGene_gn=SPEG;refGene_tx=XM_023259689.1;refGene_gn=SPEG;refGene_tx=XM_023259690.1;refGene_gn=SPEG;refGene_tx=XM_023259691.1;refGene_gn=SPEG;refGene_tx=XM_023259692.1;refGene_gn=SPEG;refGene_tx=XM_023259693.1;refGene_gn=SPEG;refGene_tx=XM_023259694.1;refGene_gn=SPEG;refGene_tx=XM_023259695.1;refGene_gn=SPEG;refGene_tx=XM_023259697.1;refGene_gn=SPEG;refGene_tx=XM_023259698.1;refGene_gn=SPEG;refGene_tx=XM_023259699.1;refGene_gn=SPEG;refGene_tx=XM_023259700.1;refGene_gn=SPEG;refGene_tx=XM_023259701.1
C1 204432883 204432996
refGene_gn=SPEG;refGene_tx=XM_011285666.3;refGene_gn=SPEG;refGene_tx=XM_023259689.1;refGene_gn=SPEG;refGene_tx=XM_023259690.1;refGene_gn=SPEG;refGene_tx=XM_023259691.1;refGene_gn=SPEG;refGene_tx=XM_023259692.1;refGene_gn=SPEG;refGene_tx=XM_023259693.1;refGene_gn=SPEG;refGene_tx=XM_023259694.1;refGene_gn=SPEG;refGene_tx=XM_023259695.1;refGene_gn=SPEG;refGene_tx=XM_023259697.1;refGene_gn=SPEG;refGene_tx=XM_023259698.1;refGene_gn=SPEG;refGene_tx=XM_023259699.1;refGene_gn=SPEG;refGene_tx=XM_023259700.1;refGene_gn=SPEG;refGene_tx=XM_023259701.1
C1 204433440 204433609
refGene_gn=SPEG;refGene_tx=XM_011285666.3;refGene_gn=SPEG;refGene_tx=XM_023259689.1;refGene_gn=SPEG;refGene_tx=XM_023259690.1;refGene_gn=SPEG;refGene_tx=XM_023259691.1;refGene_gn=SPEG;refGene_tx=XM_023259692.1;refGene_gn=SPEG;refGene_tx=XM_023259693.1;refGene_gn=SPEG;refGene_tx=XM_023259694.1;refGene_gn=SPEG;refGene_tx=XM_023259695.1;refGene_gn=SPEG;refGene_tx=XM_023259697.1;refGene_gn=SPEG;refGene_tx=XM_023259698.1;refGene_gn=SPEG;refGene_tx=XM_023259699.1;refGene_gn=SPEG;refGene_tx=XM_023259700.1;refGene_gn=SPEG;refGene_tx=XM_023259701.1
C1 204433859 204433956
refGene_gn=SPEG;refGene_tx=XM_011285666.3;refGene_gn=SPEG;refGene_tx=XM_023259689.1;refGene_gn=SPEG;refGene_tx=XM_023259690.1;refGene_gn=SPEG;refGene_tx=XM_023259691.1;refGene_gn=SPEG;refGene_tx=XM_023259692.1;refGene_gn=SPEG;refGene_tx=XM_023259693.1;refGene_gn=SPEG;refGene_tx=XM_023259694.1;refGene_gn=SPEG;refGene_tx=XM_023259695.1;refGene_gn=SPEG;refGene_tx=XM_023259697.1;refGene_gn=SPEG;refGene_tx=XM_023259698.1;

refGene_gn=SPEG;refGene_tx=XM_023259699.1;refGene_gn=SPEG;refGene_tx=XM_023259700.1;refGene_gn=SPEG;refGene_tx=XM_023259701.1
C1 204434081 204434282
refGene_gn=SPEG;refGene_tx=XM_011285666.3;refGene_gn=SPEG;refGene_tx=XM_023259689.1;refGene_gn=SPEG;refGene_tx=XM_023259690.1;refGene_gn=SPEG;refGene_tx=XM_023259691.1;refGene_gn=SPEG;refGene_tx=XM_023259692.1;refGene_gn=SPEG;refGene_tx=XM_023259693.1;refGene_gn=SPEG;refGene_tx=XM_023259694.1;refGene_gn=SPEG;refGene_tx=XM_023259695.1;refGene_gn=SPEG;refGene_tx=XM_023259697.1;refGene_gn=SPEG;refGene_tx=XM_023259698.1;refGene_gn=SPEG;refGene_tx=XM_023259699.1;refGene_gn=SPEG;refGene_tx=XM_023259700.1;refGene_gn=SPEG;refGene_tx=XM_023259701.1
C1 204435572 204435630
refGene_gn=SPEG;refGene_tx=XM_011285666.3;refGene_gn=SPEG;refGene_tx=XM_023259689.1;refGene_gn=SPEG;refGene_tx=XM_023259690.1;refGene_gn=SPEG;refGene_tx=XM_023259691.1;refGene_gn=SPEG;refGene_tx=XM_023259692.1;refGene_gn=SPEG;refGene_tx=XM_023259693.1;refGene_gn=SPEG;refGene_tx=XM_023259694.1;refGene_gn=SPEG;refGene_tx=XM_023259695.1;refGene_gn=SPEG;refGene_tx=XM_023259697.1;refGene_gn=SPEG;refGene_tx=XM_023259698.1;refGene_gn=SPEG;refGene_tx=XM_023259699.1;refGene_gn=SPEG;refGene_tx=XM_023259700.1;refGene_gn=SPEG;refGene_tx=XM_023259701.1
C1 204436213 204436291
refGene_gn=SPEG;refGene_tx=XM_011285666.3;refGene_gn=SPEG;refGene_tx=XM_023259689.1;refGene_gn=SPEG;refGene_tx=XM_023259690.1;refGene_gn=SPEG;refGene_tx=XM_023259691.1;refGene_gn=SPEG;refGene_tx=XM_023259692.1;refGene_gn=SPEG;refGene_tx=XM_023259693.1;refGene_gn=SPEG;refGene_tx=XM_023259694.1;refGene_gn=SPEG;refGene_tx=XM_023259695.1;refGene_gn=SPEG;refGene_tx=XM_023259697.1;refGene_gn=SPEG;refGene_tx=XM_023259698.1;refGene_gn=SPEG;refGene_tx=XM_023259699.1;refGene_gn=SPEG;refGene_tx=XM_023259700.1;refGene_gn=SPEG;refGene_tx=XM_023259701.1
C1 204436397 204436576
refGene_gn=SPEG;refGene_tx=XM_011285666.3;refGene_gn=SPEG;refGene_tx=XM_023259689.1;refGene_gn=SPEG;refGene_tx=XM_023259690.1;refGene_gn=SPEG;refGene_tx=XM_023259691.1;refGene_gn=SPEG;refGene_tx=XM_023259692.1;refGene_gn=SPEG;refGene_tx=XM_023259693.1;refGene_gn=SPEG;refGene_tx=XM_023259694.1;refGene_gn=SPEG;refGene_tx=XM_023259695.1;refGene_gn=SPEG;refGene_tx=XM_023259697.1;refGene_gn=SPEG;refGene_tx=XM_023259698.1;refGene_gn=SPEG;refGene_tx=XM_023259699.1;refGene_gn=SPEG;refGene_tx=XM_023259700.1;refGene_gn=SPEG;refGene_tx=XM_023259701.1
C1 204437087 204437114
refGene_gn=SPEG;refGene_tx=XM_011285666.3;refGene_gn=SPEG;refGene_tx=XM_023259689.1;refGene_gn=SPEG;refGene_tx=XM_023259690.1;refGene_gn=SPEG;refGene_tx=XM_023259691.1;refGene_gn=SPEG;refGene_tx=XM_023259692.1;refGene_gn=SPEG;refGene_tx=XM_023259693.1;refGene_gn=SPEG;refGene_tx=XM_023259694.1;refGene_gn=SPEG;refGene_tx=XM_023259695.1;refGene_gn=SPEG;refGene_tx=XM_023259697.1;refGene_gn=SPEG;refGene_tx=XM_023259698.1;refGene_gn=SPEG;refGene_tx=XM_023259699.1;refGene_gn=SPEG;refGene_tx=XM_023259700.1
C1 204437777 204437930
refGene_gn=SPEG;refGene_tx=XM_011285666.3;refGene_gn=SPEG;refGene_tx=XM_023259689.1;refGene_gn=SPEG;refGene_tx=XM_023259690.1;refGene_gn=SPE

G;refGene_tx=XM_023259691.1;refGene_gn=SPEG;refGene_tx=XM_023259692.1;
refGene_gn=SPEG;refGene_tx=XM_023259693.1;refGene_gn=SPEG;refGene_tx=X
M_023259694.1;refGene_gn=SPEG;refGene_tx=XM_023259695.1;refGene_gn=SPE
G;refGene_tx=XM_023259697.1;refGene_gn=SPEG;refGene_tx=XM_023259698.1;
refGene_gn=SPEG;refGene_tx=XM_023259699.1;refGene_gn=SPEG;refGene_tx=X
M_023259700.1
C1 204438114 204438157
refGene_gn=SPEG;refGene_tx=XM_011285666.3;refGene_gn=SPEG;refGene_tx=X
M_023259689.1;refGene_gn=SPEG;refGene_tx=XM_023259690.1;refGene_gn=SPE
G;refGene_tx=XM_023259691.1;refGene_gn=SPEG;refGene_tx=XM_023259692.1;
refGene_gn=SPEG;refGene_tx=XM_023259693.1;refGene_gn=SPEG;refGene_tx=X
M_023259694.1;refGene_gn=SPEG;refGene_tx=XM_023259695.1;refGene_gn=SPE
G;refGene_tx=XM_023259697.1;refGene_gn=SPEG;refGene_tx=XM_023259698.1;
refGene_gn=SPEG;refGene_tx=XM_023259699.1;refGene_gn=SPEG;refGene_tx=X
M_023259700.1
C1 204439103 204439172
refGene_gn=SPEG;refGene_tx=XM_011285666.3;refGene_gn=SPEG;refGene_tx=X
M_023259689.1;refGene_gn=SPEG;refGene_tx=XM_023259690.1;refGene_gn=SPE
G;refGene_tx=XM_023259691.1;refGene_gn=SPEG;refGene_tx=XM_023259692.1;
refGene_gn=SPEG;refGene_tx=XM_023259693.1;refGene_gn=SPEG;refGene_tx=X
M_023259694.1;refGene_gn=SPEG;refGene_tx=XM_023259695.1;refGene_gn=SPE
G;refGene_tx=XM_023259697.1;refGene_gn=SPEG;refGene_tx=XM_023259698.1;
refGene_gn=SPEG;refGene_tx=XM_023259699.1;refGene_gn=SPEG;refGene_tx=X
M_023259700.1
C1 204439439 204441417
refGene_gn=SPEG;refGene_tx=XM_011285666.3;refGene_gn=SPEG;refGene_tx=X
M_023259689.1;refGene_gn=SPEG;refGene_tx=XM_023259690.1;refGene_gn=SPE
G;refGene_tx=XM_023259691.1;refGene_gn=SPEG;refGene_tx=XM_023259692.1;
refGene_gn=SPEG;refGene_tx=XM_023259693.1;refGene_gn=SPEG;refGene_tx=X
M_023259694.1;refGene_gn=SPEG;refGene_tx=XM_023259695.1;refGene_gn=SPE
G;refGene_tx=XM_023259697.1;refGene_gn=SPEG;refGene_tx=XM_023259698.1;
refGene_gn=SPEG;refGene_tx=XM_023259699.1;refGene_gn=SPEG;refGene_tx=X
M_023259700.1
C1 204441683 204441815
refGene_gn=SPEG;refGene_tx=XM_011285666.3;refGene_gn=SPEG;refGene_tx=X
M_023259689.1;refGene_gn=SPEG;refGene_tx=XM_023259690.1;refGene_gn=SPE
G;refGene_tx=XM_023259691.1;refGene_gn=SPEG;refGene_tx=XM_023259692.1;
refGene_gn=SPEG;refGene_tx=XM_023259693.1;refGene_gn=SPEG;refGene_tx=X
M_023259694.1;refGene_gn=SPEG;refGene_tx=XM_023259695.1;refGene_gn=SPE
G;refGene_tx=XM_023259697.1;refGene_gn=SPEG;refGene_tx=XM_023259698.1;
refGene_gn=SPEG;refGene_tx=XM_023259699.1;refGene_gn=SPEG;refGene_tx=X
M_023259700.1
C1 204444359 204444476
refGene_gn=SPEG;refGene_tx=XM_011285666.3;refGene_gn=SPEG;refGene_tx=X
M_023259689.1;refGene_gn=SPEG;refGene_tx=XM_023259690.1;refGene_gn=SPE
G;refGene_tx=XM_023259691.1;refGene_gn=SPEG;refGene_tx=XM_023259692.1;
refGene_gn=SPEG;refGene_tx=XM_023259693.1;refGene_gn=SPEG;refGene_tx=X
M_023259694.1;refGene_gn=SPEG;refGene_tx=XM_023259695.1;refGene_gn=SPE
G;refGene_tx=XM_023259697.1;refGene_gn=SPEG;refGene_tx=XM_023259698.1;
refGene_gn=SPEG;refGene_tx=XM_023259699.1;refGene_gn=SPEG;refGene_tx=X

M_023259700.1
C1	204444616	204444784
refGene_gn=SPEG;refGene_tx=XM_011285666.3;refGene_gn=SPEG;refGene_tx=XM_023259689.1;refGene_gn=SPEG;refGene_tx=XM_023259690.1;refGene_gn=SPEG;refGene_tx=XM_023259691.1;refGene_gn=SPEG;refGene_tx=XM_023259692.1;refGene_gn=SPEG;refGene_tx=XM_023259693.1;refGene_gn=SPEG;refGene_tx=XM_023259694.1;refGene_gn=SPEG;refGene_tx=XM_023259695.1;refGene_gn=SPEG;refGene_tx=XM_023259697.1;refGene_gn=SPEG;refGene_tx=XM_023259698.1;refGene_gn=SPEG;refGene_tx=XM_023259699.1;refGene_gn=SPEG;refGene_tx=XM_023259700.1
C1	204444896	204445019
refGene_gn=SPEG;refGene_tx=XM_011285666.3;refGene_gn=SPEG;refGene_tx=XM_023259689.1;refGene_gn=SPEG;refGene_tx=XM_023259690.1;refGene_gn=SPEG;refGene_tx=XM_023259691.1;refGene_gn=SPEG;refGene_tx=XM_023259692.1;refGene_gn=SPEG;refGene_tx=XM_023259693.1;refGene_gn=SPEG;refGene_tx=XM_023259694.1;refGene_gn=SPEG;refGene_tx=XM_023259695.1;refGene_gn=SPEG;refGene_tx=XM_023259697.1;refGene_gn=SPEG;refGene_tx=XM_023259698.1;refGene_gn=SPEG;refGene_tx=XM_023259699.1;refGene_gn=SPEG;refGene_tx=XM_023259700.1
C1	204445174	204445342
refGene_gn=SPEG;refGene_tx=XM_011285666.3;refGene_gn=SPEG;refGene_tx=XM_023259689.1;refGene_gn=SPEG;refGene_tx=XM_023259690.1;refGene_gn=SPEG;refGene_tx=XM_023259691.1;refGene_gn=SPEG;refGene_tx=XM_023259692.1;refGene_gn=SPEG;refGene_tx=XM_023259693.1;refGene_gn=SPEG;refGene_tx=XM_023259694.1;refGene_gn=SPEG;refGene_tx=XM_023259695.1;refGene_gn=SPEG;refGene_tx=XM_023259697.1;refGene_gn=SPEG;refGene_tx=XM_023259698.1;refGene_gn=SPEG;refGene_tx=XM_023259699.1;refGene_gn=SPEG;refGene_tx=XM_023259700.1
C1	204445429	204446045
refGene_gn=SPEG;refGene_tx=XM_011285666.3;refGene_gn=SPEG;refGene_tx=XM_023259689.1;refGene_gn=SPEG;refGene_tx=XM_023259690.1;refGene_gn=SPEG;refGene_tx=XM_023259691.1;refGene_gn=SPEG;refGene_tx=XM_023259692.1;refGene_gn=SPEG;refGene_tx=XM_023259693.1;refGene_gn=SPEG;refGene_tx=XM_023259694.1;refGene_gn=SPEG;refGene_tx=XM_023259695.1;refGene_gn=SPEG;refGene_tx=XM_023259697.1;refGene_gn=SPEG;refGene_tx=XM_023259698.1;refGene_gn=SPEG;refGene_tx=XM_023259699.1;refGene_gn=SPEG;refGene_tx=XM_023259700.1
C1	204446503	204446743
refGene_gn=SPEG;refGene_tx=XM_011285666.3;refGene_gn=SPEG;refGene_tx=XM_023259689.1;refGene_gn=SPEG;refGene_tx=XM_023259690.1;refGene_gn=SPEG;refGene_tx=XM_023259691.1;refGene_gn=SPEG;refGene_tx=XM_023259692.1;refGene_gn=SPEG;refGene_tx=XM_023259693.1;refGene_gn=SPEG;refGene_tx=XM_023259694.1;refGene_gn=SPEG;refGene_tx=XM_023259695.1;refGene_gn=SPEG;refGene_tx=XM_023259697.1;refGene_gn=SPEG;refGene_tx=XM_023259698.1;refGene_gn=SPEG;refGene_tx=XM_023259699.1;refGene_gn=SPEG;refGene_tx=XM_023259700.1
C1	204446824	204447048
refGene_gn=SPEG;refGene_tx=XM_011285666.3;refGene_gn=SPEG;refGene_tx=XM_023259689.1;refGene_gn=SPEG;refGene_tx=XM_023259690.1;refGene_gn=SPEG;refGene_tx=XM_023259691.1;refGene_gn=SPEG;refGene_tx=XM_023259692.1;

refGene_gn=SPEG;refGene_tx=XM_023259693.1;refGene_gn=SPEG;refGene_tx=X
M_023259694.1;refGene_gn=SPEG;refGene_tx=XM_023259695.1;refGene_gn=SPE
G;refGene_tx=XM_023259697.1;refGene_gn=SPEG;refGene_tx=XM_023259698.1;
refGene_gn=SPEG;refGene_tx=XM_023259699.1;refGene_gn=SPEG;refGene_tx=X
M_023259700.1
C1 204448116 204448192
refGene_gn=SPEG;refGene_tx=XM_011285666.3;refGene_gn=SPEG;refGene_tx=X
M_023259689.1;refGene_gn=SPEG;refGene_tx=XM_023259690.1;refGene_gn=SPE
G;refGene_tx=XM_023259691.1;refGene_gn=SPEG;refGene_tx=XM_023259692.1;
refGene_gn=SPEG;refGene_tx=XM_023259693.1;refGene_gn=SPEG;refGene_tx=X
M_023259694.1;refGene_gn=SPEG;refGene_tx=XM_023259695.1;refGene_gn=SPE
G;refGene_tx=XM_023259697.1;refGene_gn=SPEG;refGene_tx=XM_023259698.1;
refGene_gn=SPEG;refGene_tx=XM_023259699.1;refGene_gn=SPEG;refGene_tx=X
M_023259700.1
C1 204448394 204448544
refGene_gn=SPEG;refGene_tx=XM_011285666.3;refGene_gn=SPEG;refGene_tx=X
M_023259689.1;refGene_gn=SPEG;refGene_tx=XM_023259690.1;refGene_gn=SPE
G;refGene_tx=XM_023259691.1;refGene_gn=SPEG;refGene_tx=XM_023259692.1;
refGene_gn=SPEG;refGene_tx=XM_023259693.1;refGene_gn=SPEG;refGene_tx=X
M_023259694.1;refGene_gn=SPEG;refGene_tx=XM_023259695.1;refGene_gn=SPE
G;refGene_tx=XM_023259697.1;refGene_gn=SPEG;refGene_tx=XM_023259698.1;
refGene_gn=SPEG;refGene_tx=XM_023259699.1;refGene_gn=SPEG;refGene_tx=X
M_023259700.1
C1 204448868 204449061
refGene_gn=SPEG;refGene_tx=XM_011285666.3;refGene_gn=SPEG;refGene_tx=X
M_023259689.1;refGene_gn=SPEG;refGene_tx=XM_023259690.1;refGene_gn=SPE
G;refGene_tx=XM_023259691.1;refGene_gn=SPEG;refGene_tx=XM_023259692.1;
refGene_gn=SPEG;refGene_tx=XM_023259693.1;refGene_gn=SPEG;refGene_tx=X
M_023259694.1;refGene_gn=SPEG;refGene_tx=XM_023259695.1;refGene_gn=SPE
G;refGene_tx=XM_023259697.1;refGene_gn=SPEG;refGene_tx=XM_023259698.1;
refGene_gn=SPEG;refGene_tx=XM_023259699.1;refGene_gn=SPEG;refGene_tx=X
M_023259700.1
C1 204452398 204452526
refGene_gn=LOC101101404;refGene_tx=XM_023259706.1
C1 204452842 204453150
refGene_gn=LOC101101404;refGene_tx=XM_023259706.1
C1 204453706 204453848
refGene_gn=LOC101101404;refGene_tx=XM_023259706.1
C1 204454017 204454156
refGene_gn=LOC101101404;refGene_tx=XM_023259706.1
C1 204455839 204455879
refGene_gn=GMPPA;refGene_tx=XM_006935606.4;refGene_gn=GMPPA;refGene_tx
=XM_006935607.4;refGene_gn=GMPPA;refGene_tx=XM_006935611.4;refGene_gn=
GMPPA;refGene_tx=XM_023259704.1
C1 204455989 204456087
refGene_gn=GMPPA;refGene_tx=XM_006935606.4;refGene_gn=GMPPA;refGene_tx
=XM_006935607.4;refGene_gn=GMPPA;refGene_tx=XM_006935611.4;refGene_gn=
GMPPA;refGene_tx=XM_023259704.1
C1 204457011 204457115
refGene_gn=GMPPA;refGene_tx=XM_006935606.4;refGene_gn=GMPPA;refGene_tx

=XM_006935607.4;refGene_gn=GMPPA;refGene_tx=XM_006935611.4;refGene_gn=GMPPA;refGene_tx=XM_006935612.4;refGene_gn=GMPPA;refGene_tx=XM_023259704.1;refGene_gn=GMPPA;refGene_tx=XM_023259705.1
C1 204457370 204457557 refGene_gn=GMPPA;refGene_tx=XM_006935606.4;refGene_gn=GMPPA;refGene_tx=XM_006935607.4;refGene_gn=GMPPA;refGene_tx=XM_006935611.4;refGene_gn=GMPPA;refGene_tx=XM_006935612.4;refGene_gn=GMPPA;refGene_tx=XM_023259704.1;refGene_gn=GMPPA;refGene_tx=XM_023259705.1
C1 204457908 204457968 refGene_gn=GMPPA;refGene_tx=XM_006935606.4;refGene_gn=GMPPA;refGene_tx=XM_006935607.4;refGene_gn=GMPPA;refGene_tx=XM_006935611.4;refGene_gn=GMPPA;refGene_tx=XM_006935612.4;refGene_gn=GMPPA;refGene_tx=XM_023259704.1;refGene_gn=GMPPA;refGene_tx=XM_023259705.1
C1 204459269 204459400 refGene_gn=GMPPA;refGene_tx=XM_006935606.4;refGene_gn=GMPPA;refGene_tx=XM_006935607.4;refGene_gn=GMPPA;refGene_tx=XM_006935611.4;refGene_gn=GMPPA;refGene_tx=XM_006935612.4;refGene_gn=GMPPA;refGene_tx=XM_023259704.1;refGene_gn=GMPPA;refGene_tx=XM_023259705.1
C1 204460109 204460124 refGene_gn=GMPPA;refGene_tx=XM_006935606.4;refGene_gn=GMPPA;refGene_tx=XM_006935607.4;refGene_gn=GMPPA;refGene_tx=XM_006935612.4;refGene_gn=GMPPA;refGene_tx=XM_023259704.1;refGene_gn=GMPPA;refGene_tx=XM_023259705.1
C1 204460486 204460621 refGene_gn=GMPPA;refGene_tx=XM_006935606.4;refGene_gn=GMPPA;refGene_tx=XM_006935607.4;refGene_gn=GMPPA;refGene_tx=XM_006935611.4;refGene_gn=GMPPA;refGene_tx=XM_006935612.4;refGene_gn=GMPPA;refGene_tx=XM_023259704.1;refGene_gn=GMPPA;refGene_tx=XM_023259705.1
C1 204460727 204460825 refGene_gn=GMPPA;refGene_tx=XM_006935606.4;refGene_gn=GMPPA;refGene_tx=XM_006935607.4;refGene_gn=GMPPA;refGene_tx=XM_006935611.4;refGene_gn=GMPPA;refGene_tx=XM_006935612.4;refGene_gn=GMPPA;refGene_tx=XM_023259704.1;refGene_gn=GMPPA;refGene_tx=XM_023259705.1
C1 204460992 204461039 refGene_gn=GMPPA;refGene_tx=XM_006935606.4;refGene_gn=GMPPA;refGene_tx=XM_006935607.4;refGene_gn=GMPPA;refGene_tx=XM_006935611.4;refGene_gn=GMPPA;refGene_tx=XM_006935612.4;refGene_gn=GMPPA;refGene_tx=XM_023259704.1;refGene_gn=GMPPA;refGene_tx=XM_023259705.1
C1 204461233 204461326 refGene_gn=GMPPA;refGene_tx=XM_006935606.4;refGene_gn=GMPPA;refGene_tx=XM_006935607.4;refGene_gn=GMPPA;refGene_tx=XM_006935611.4;refGene_gn=GMPPA;refGene_tx=XM_006935612.4;refGene_gn=GMPPA;refGene_tx=XM_023259704.1;refGene_gn=GMPPA;refGene_tx=XM_023259705.1
C1 204461447 204461616 refGene_gn=GMPPA;refGene_tx=XM_006935606.4;refGene_gn=GMPPA;refGene_tx=XM_006935607.4;refGene_gn=GMPPA;refGene_tx=XM_006935611.4;refGene_gn=GMPPA;refGene_tx=XM_006935612.4;refGene_gn=GMPPA;refGene_tx=XM_023259704.1;refGene_gn=GMPPA;refGene_tx=XM_023259705.1
C1 204461926 204462027 refGene_gn=GMPPA;refGene_tx=XM_006935606.4;refGene_gn=GMPPA;refGene_tx

=XM_006935607.4;refGene_gn=GMPPA;refGene_tx=XM_006935611.4;refGene_gn=GMPPA;refGene_tx=XM_006935612.4;refGene_gn=GMPPA;refGene_tx=XM_023259704.1;refGene_gn=GMPPA;refGene_tx=XM_023259705.1
C1 204470023 204470605
refGene_gn=ASIC4;refGene_tx=XM_019838718.1;refGene_gn=ASIC4;refGene_tx=XM_019838719.2;refGene_gn=ASIC4;refGene_tx=XM_019838720.1;refGene_gn=ASIC4;refGene_tx=XM_023259714.1
C1 204486158 204486303
refGene_gn=ASIC4;refGene_tx=XM_019838718.1;refGene_gn=ASIC4;refGene_tx=XM_019838719.2;refGene_gn=ASIC4;refGene_tx=XM_019838720.1;refGene_gn=ASIC4;refGene_tx=XM_023259714.1
C1 204486398 204486526
refGene_gn=ASIC4;refGene_tx=XM_019838718.1;refGene_gn=ASIC4;refGene_tx=XM_019838719.2;refGene_gn=ASIC4;refGene_tx=XM_019838720.1;refGene_gn=ASIC4;refGene_tx=XM_023259714.1
C1 204486715 204486878
refGene_gn=ASIC4;refGene_tx=XM_019838718.1;refGene_gn=ASIC4;refGene_tx=XM_019838719.2;refGene_gn=ASIC4;refGene_tx=XM_019838720.1;refGene_gn=ASIC4;refGene_tx=XM_023259714.1
C1 204487411 204487468
refGene_gn=ASIC4;refGene_tx=XM_019838718.1;refGene_gn=ASIC4;refGene_tx=XM_019838719.2;refGene_gn=ASIC4;refGene_tx=XM_019838720.1;refGene_gn=ASIC4;refGene_tx=XM_023259714.1
C1 204489482 204489636
refGene_gn=ASIC4;refGene_tx=XM_019838718.1;refGene_gn=ASIC4;refGene_tx=XM_019838719.2;refGene_gn=ASIC4;refGene_tx=XM_019838720.1;refGene_gn=ASIC4;refGene_tx=XM_023259714.1
C1 204491007 204491099
refGene_gn=ASIC4;refGene_tx=XM_019838718.1;refGene_gn=ASIC4;refGene_tx=XM_019838719.2;refGene_gn=ASIC4;refGene_tx=XM_019838720.1;refGene_gn=ASIC4;refGene_tx=XM_023259714.1
C1 204491180 204491260
refGene_gn=ASIC4;refGene_tx=XM_019838718.1;refGene_gn=ASIC4;refGene_tx=XM_019838719.2;refGene_gn=ASIC4;refGene_tx=XM_019838720.1;refGene_gn=ASIC4;refGene_tx=XM_023259714.1
C1 204491586 204491691
refGene_gn=ASIC4;refGene_tx=XM_019838718.1;refGene_gn=ASIC4;refGene_tx=XM_019838719.2;refGene_gn=ASIC4;refGene_tx=XM_019838720.1;refGene_gn=ASIC4;refGene_tx=XM_023259714.1
C1 204491914 204492028
refGene_gn=ASIC4;refGene_tx=XM_019838718.1;refGene_gn=ASIC4;refGene_tx=XM_019838719.2;refGene_gn=ASIC4;refGene_tx=XM_019838720.1;refGene_gn=ASIC4;refGene_tx=XM_023259714.1
C1 204493351 204494611
refGene_gn=CHPF;refGene_tx=XM_006935614.4
C1 204494929 204495109
refGene_gn=CHPF;refGene_tx=XM_006935614.4
C1 204495623 204496197
refGene_gn=CHPF;refGene_tx=XM_006935614.4
C1 204497238 204497552

refGene_gn=CHPF;refGene_tx=XM_006935614.4
C1	204497796	204497840
refGene_gn=OBSL1;refGene_tx=XM_023259709.1
C1	204498716	204498882
refGene_gn=TMEM198;refGene_tx=XM_003991183.5;refGene_gn=TMEM198;refGene_tx=XM_019838722.2;refGene_gn=TMEM198;refGene_tx=XM_023259715.1
C1	204501402	204501978
refGene_gn=TMEM198;refGene_tx=XM_003991183.5;refGene_gn=TMEM198;refGene_tx=XM_019838722.2;refGene_gn=TMEM198;refGene_tx=XM_023259715.1
C1	204503060	204503263
refGene_gn=TMEM198;refGene_tx=XM_003991183.5;refGene_gn=TMEM198;refGene_tx=XM_019838722.2;refGene_gn=TMEM198;refGene_tx=XM_023259715.1
C1	204503636	204503774
refGene_gn=TMEM198;refGene_tx=XM_003991183.5;refGene_gn=TMEM198;refGene_tx=XM_019838722.2;refGene_gn=TMEM198;refGene_tx=XM_023259715.1
C1	204504690	204504807
refGene_gn=OBSL1;refGene_tx=XM_023259708.1;refGene_gn=OBSL1;refGene_tx=XM_023259710.1;refGene_gn=OBSL1;refGene_tx=XM_023259712.1
C1	204505504	204505774
refGene_gn=OBSL1;refGene_tx=XM_023259708.1;refGene_gn=OBSL1;refGene_tx=XM_023259709.1;refGene_gn=OBSL1;refGene_tx=XM_023259710.1;refGene_gn=OBSL1;refGene_tx=XM_023259711.1;refGene_gn=OBSL1;refGene_tx=XM_023259712.1
C1	204506077	204506182
refGene_gn=OBSL1;refGene_tx=XM_023259708.1;refGene_gn=OBSL1;refGene_tx=XM_023259709.1;refGene_gn=OBSL1;refGene_tx=XM_023259710.1;refGene_gn=OBSL1;refGene_tx=XM_023259711.1;refGene_gn=OBSL1;refGene_tx=XM_023259712.1
C1	204506496	204506658
refGene_gn=OBSL1;refGene_tx=XM_023259708.1;refGene_gn=OBSL1;refGene_tx=XM_023259709.1;refGene_gn=OBSL1;refGene_tx=XM_023259710.1;refGene_gn=OBSL1;refGene_tx=XM_023259711.1;refGene_gn=OBSL1;refGene_tx=XM_023259712.1
C1	204506830	204506987
refGene_gn=OBSL1;refGene_tx=XM_023259708.1;refGene_gn=OBSL1;refGene_tx=XM_023259709.1;refGene_gn=OBSL1;refGene_tx=XM_023259710.1;refGene_gn=OBSL1;refGene_tx=XM_023259711.1;refGene_gn=OBSL1;refGene_tx=XM_023259712.1
C1	204507536	204507649
refGene_gn=OBSL1;refGene_tx=XM_023259708.1;refGene_gn=OBSL1;refGene_tx=XM_023259709.1;refGene_gn=OBSL1;refGene_tx=XM_023259710.1;refGene_gn=OBSL1;refGene_tx=XM_023259711.1;refGene_gn=OBSL1;refGene_tx=XM_023259712.1
C1	204508517	204508784
refGene_gn=OBSL1;refGene_tx=XM_023259708.1;refGene_gn=OBSL1;refGene_tx=XM_023259709.1;refGene_gn=OBSL1;refGene_tx=XM_023259710.1;refGene_gn=OBSL1;refGene_tx=XM_023259711.1;refGene_gn=OBSL1;refGene_tx=XM_023259712.1
C1	204509782	204510055
refGene_gn=OBSL1;refGene_tx=XM_023259708.1;refGene_gn=OBSL1;refGene_tx

=XM_023259709.1;refGene_gn=OBSL1;refGene_tx=XM_023259710.1;refGene_gn=OBSL1;refGene_tx=XM_023259711.1;refGene_gn=OBSL1;refGene_tx=XM_023259712.1
C1 204510218 204510488
refGene_gn=OBSL1;refGene_tx=XM_023259708.1;refGene_gn=OBSL1;refGene_tx=XM_023259709.1;refGene_gn=OBSL1;refGene_tx=XM_023259710.1;refGene_gn=OBSL1;refGene_tx=XM_023259711.1;refGene_gn=OBSL1;refGene_tx=XM_023259712.1
C1 204511083 204511359
refGene_gn=OBSL1;refGene_tx=XM_023259708.1;refGene_gn=OBSL1;refGene_tx=XM_023259709.1;refGene_gn=OBSL1;refGene_tx=XM_023259710.1;refGene_gn=OBSL1;refGene_tx=XM_023259711.1
C1 204511532 204511820
refGene_gn=OBSL1;refGene_tx=XM_023259708.1;refGene_gn=OBSL1;refGene_tx=XM_023259709.1;refGene_gn=OBSL1;refGene_tx=XM_023259710.1;refGene_gn=OBSL1;refGene_tx=XM_023259711.1;refGene_gn=OBSL1;refGene_tx=XM_023259712.1
C1 204511894 204512170
refGene_gn=OBSL1;refGene_tx=XM_023259708.1;refGene_gn=OBSL1;refGene_tx=XM_023259709.1;refGene_gn=OBSL1;refGene_tx=XM_023259710.1;refGene_gn=OBSL1;refGene_tx=XM_023259711.1;refGene_gn=OBSL1;refGene_tx=XM_023259712.1
C1 204512920 204513193
refGene_gn=OBSL1;refGene_tx=XM_023259708.1;refGene_gn=OBSL1;refGene_tx=XM_023259709.1;refGene_gn=OBSL1;refGene_tx=XM_023259710.1;refGene_gn=OBSL1;refGene_tx=XM_023259711.1;refGene_gn=OBSL1;refGene_tx=XM_023259712.1
C1 204515532 204515657
refGene_gn=OBSL1;refGene_tx=XM_023259713.1
C1 204516067 204516340
refGene_gn=OBSL1;refGene_tx=XM_023259708.1;refGene_gn=OBSL1;refGene_tx=XM_023259709.1;refGene_gn=OBSL1;refGene_tx=XM_023259710.1;refGene_gn=OBSL1;refGene_tx=XM_023259711.1;refGene_gn=OBSL1;refGene_tx=XM_023259712.1;refGene_gn=OBSL1;refGene_tx=XM_023259713.1
C1 204517063 204517336
refGene_gn=OBSL1;refGene_tx=XM_023259708.1;refGene_gn=OBSL1;refGene_tx=XM_023259709.1;refGene_gn=OBSL1;refGene_tx=XM_023259710.1;refGene_gn=OBSL1;refGene_tx=XM_023259711.1;refGene_gn=OBSL1;refGene_tx=XM_023259712.1;refGene_gn=OBSL1;refGene_tx=XM_023259713.1
C1 204518502 204518775
refGene_gn=OBSL1;refGene_tx=XM_023259708.1;refGene_gn=OBSL1;refGene_tx=XM_023259709.1;refGene_gn=OBSL1;refGene_tx=XM_023259710.1;refGene_gn=OBSL1;refGene_tx=XM_023259711.1;refGene_gn=OBSL1;refGene_tx=XM_023259712.1;refGene_gn=OBSL1;refGene_tx=XM_023259713.1
C1 204519782 204520079
refGene_gn=OBSL1;refGene_tx=XM_023259708.1;refGene_gn=OBSL1;refGene_tx=XM_023259709.1;refGene_gn=OBSL1;refGene_tx=XM_023259710.1;refGene_gn=OBSL1;refGene_tx=XM_023259711.1;refGene_gn=OBSL1;refGene_tx=XM_023259712.1;refGene_gn=OBSL1;refGene_tx=XM_023259713.1
C1 204520256 204520559

refGene_gn=OBSL1;refGene_tx=XM_023259708.1;refGene_gn=OBSL1;refGene_tx=XM_023259709.1;refGene_gn=OBSL1;refGene_tx=XM_023259710.1;refGene_gn=OBSL1;refGene_tx=XM_023259711.1;refGene_gn=OBSL1;refGene_tx=XM_023259712.1;refGene_gn=OBSL1;refGene_tx=XM_023259713.1
C1 204520728 204520980
refGene_gn=OBSL1;refGene_tx=XM_023259708.1;refGene_gn=OBSL1;refGene_tx=XM_023259709.1;refGene_gn=OBSL1;refGene_tx=XM_023259710.1;refGene_gn=OBSL1;refGene_tx=XM_023259711.1;refGene_gn=OBSL1;refGene_tx=XM_023259712.1;refGene_gn=OBSL1;refGene_tx=XM_023259713.1
C1 204521064 204521334
refGene_gn=OBSL1;refGene_tx=XM_023259708.1;refGene_gn=OBSL1;refGene_tx=XM_023259709.1;refGene_gn=OBSL1;refGene_tx=XM_023259710.1;refGene_gn=OBSL1;refGene_tx=XM_023259711.1;refGene_gn=OBSL1;refGene_tx=XM_023259712.1;refGene_gn=OBSL1;refGene_tx=XM_023259713.1
C1 204523497 204524509
refGene_gn=OBSL1;refGene_tx=XM_023259708.1;refGene_gn=OBSL1;refGene_tx=XM_023259709.1;refGene_gn=OBSL1;refGene_tx=XM_023259710.1;refGene_gn=OBSL1;refGene_tx=XM_023259711.1;refGene_gn=OBSL1;refGene_tx=XM_023259712.1;refGene_gn=OBSL1;refGene_tx=XM_023259713.1
C1 204525662 204525936
refGene_gn=INHA;refGene_tx=XM_003991182.4
C1 204527891 204528724
refGene_gn=INHA;refGene_tx=XM_003991182.4
C1 204545612 204545738
refGene_gn=STK11IP;refGene_tx=XM_023259716.1;refGene_gn=STK11IP;refGene_tx=XM_023259717.1
C1 204546243 204546329
refGene_gn=STK11IP;refGene_tx=XM_011285660.3;refGene_gn=STK11IP;refGene_tx=XM_023259716.1;refGene_gn=STK11IP;refGene_tx=XM_023259717.1;refGene_gn=STK11IP;refGene_tx=XM_023259718.1
C1 204546526 204546603
refGene_gn=STK11IP;refGene_tx=XM_023259716.1;refGene_gn=STK11IP;refGene_tx=XM_023259717.1
C1 204548827 204549033
refGene_gn=STK11IP;refGene_tx=XM_011285660.3;refGene_gn=STK11IP;refGene_tx=XM_023259716.1;refGene_gn=STK11IP;refGene_tx=XM_023259717.1;refGene_gn=STK11IP;refGene_tx=XM_023259718.1
C1 204549231 204549306
refGene_gn=STK11IP;refGene_tx=XM_011285660.3;refGene_gn=STK11IP;refGene_tx=XM_023259716.1;refGene_gn=STK11IP;refGene_tx=XM_023259717.1;refGene_gn=STK11IP;refGene_tx=XM_023259718.1
C1 204549596 204549692
refGene_gn=STK11IP;refGene_tx=XM_011285660.3;refGene_gn=STK11IP;refGene_tx=XM_023259716.1;refGene_gn=STK11IP;refGene_tx=XM_023259717.1;refGene_gn=STK11IP;refGene_tx=XM_023259718.1
C1 204550074 204550182
refGene_gn=STK11IP;refGene_tx=XM_011285660.3;refGene_gn=STK11IP;refGene_tx=XM_023259716.1;refGene_gn=STK11IP;refGene_tx=XM_023259717.1;refGene_gn=STK11IP;refGene_tx=XM_023259718.1
C1 204550299 204550371

refGene_gn=STK11IP;refGene_tx=XM_011285660.3;refGene_gn=STK11IP;refGene_tx=XM_023259716.1;refGene_gn=STK11IP;refGene_tx=XM_023259717.1;refGene_gn=STK11IP;refGene_tx=XM_023259718.1
C1 204551453 204551580
refGene_gn=STK11IP;refGene_tx=XM_011285660.3;refGene_gn=STK11IP;refGene_tx=XM_023259716.1;refGene_gn=STK11IP;refGene_tx=XM_023259717.1;refGene_gn=STK11IP;refGene_tx=XM_023259718.1
C1 204551813 204551917
refGene_gn=STK11IP;refGene_tx=XM_011285660.3;refGene_gn=STK11IP;refGene_tx=XM_023259716.1;refGene_gn=STK11IP;refGene_tx=XM_023259717.1;refGene_gn=STK11IP;refGene_tx=XM_023259718.1
C1 204552051 204552147
refGene_gn=STK11IP;refGene_tx=XM_011285660.3;refGene_gn=STK11IP;refGene_tx=XM_023259716.1;refGene_gn=STK11IP;refGene_tx=XM_023259718.1
C1 204552380 204552422
refGene_gn=STK11IP;refGene_tx=XM_011285660.3;refGene_gn=STK11IP;refGene_tx=XM_023259716.1;refGene_gn=STK11IP;refGene_tx=XM_023259717.1;refGene_gn=STK11IP;refGene_tx=XM_023259718.1
C1 204552597 204552738
refGene_gn=STK11IP;refGene_tx=XM_011285660.3;refGene_gn=STK11IP;refGene_tx=XM_023259716.1;refGene_gn=STK11IP;refGene_tx=XM_023259717.1;refGene_gn=STK11IP;refGene_tx=XM_023259718.1
C1 204552936 204553021
refGene_gn=STK11IP;refGene_tx=XM_011285660.3;refGene_gn=STK11IP;refGene_tx=XM_023259716.1;refGene_gn=STK11IP;refGene_tx=XM_023259717.1;refGene_gn=STK11IP;refGene_tx=XM_023259718.1
C1 204553594 204553936
refGene_gn=STK11IP;refGene_tx=XM_011285660.3;refGene_gn=STK11IP;refGene_tx=XM_023259716.1;refGene_gn=STK11IP;refGene_tx=XM_023259717.1;refGene_gn=STK11IP;refGene_tx=XM_023259718.1
C1 204554341 204554547
refGene_gn=STK11IP;refGene_tx=XM_011285660.3;refGene_gn=STK11IP;refGene_tx=XM_023259716.1;refGene_gn=STK11IP;refGene_tx=XM_023259717.1;refGene_gn=STK11IP;refGene_tx=XM_023259718.1
C1 204555223 204555340
refGene_gn=STK11IP;refGene_tx=XM_011285660.3;refGene_gn=STK11IP;refGene_tx=XM_023259716.1;refGene_gn=STK11IP;refGene_tx=XM_023259717.1;refGene_gn=STK11IP;refGene_tx=XM_023259718.1
C1 204555441 204555619
refGene_gn=STK11IP;refGene_tx=XM_011285660.3;refGene_gn=STK11IP;refGene_tx=XM_023259716.1;refGene_gn=STK11IP;refGene_tx=XM_023259717.1;refGene_gn=STK11IP;refGene_tx=XM_023259718.1
C1 204557362 204557593
refGene_gn=STK11IP;refGene_tx=XM_011285660.3;refGene_gn=STK11IP;refGene_tx=XM_023259716.1;refGene_gn=STK11IP;refGene_tx=XM_023259717.1;refGene_gn=STK11IP;refGene_tx=XM_023259718.1
C1 204557722 204557826
refGene_gn=STK11IP;refGene_tx=XM_011285660.3;refGene_gn=STK11IP;refGene_tx=XM_023259716.1;refGene_gn=STK11IP;refGene_tx=XM_023259717.1;refGene_gn=STK11IP;refGene_tx=XM_023259718.1

C1 204558792 204558890
refGene_gn=STK11IP;refGene_tx=XM_011285660.3;refGene_gn=STK11IP;refGen
e_tx=XM_023259716.1;refGene_gn=STK11IP;refGene_tx=XM_023259717.1;refGe
ne_gn=STK11IP;refGene_tx=XM_023259718.1
C1 204558983 204559162
refGene_gn=STK11IP;refGene_tx=XM_011285660.3;refGene_gn=STK11IP;refGen
e_tx=XM_023259716.1;refGene_gn=STK11IP;refGene_tx=XM_023259717.1;refGe
ne_gn=STK11IP;refGene_tx=XM_023259718.1
C1 204559383 204559465
refGene_gn=STK11IP;refGene_tx=XM_011285660.3;refGene_gn=STK11IP;refGen
e_tx=XM_023259716.1;refGene_gn=STK11IP;refGene_tx=XM_023259717.1;refGe
ne_gn=STK11IP;refGene_tx=XM_023259718.1
C1 204559679 204559750
refGene_gn=STK11IP;refGene_tx=XM_011285660.3;refGene_gn=STK11IP;refGen
e_tx=XM_023259716.1;refGene_gn=STK11IP;refGene_tx=XM_023259717.1;refGe
ne_gn=STK11IP;refGene_tx=XM_023259718.1
C1 204560302 204560550
refGene_gn=STK11IP;refGene_tx=XM_011285660.3;refGene_gn=STK11IP;refGen
e_tx=XM_023259716.1;refGene_gn=STK11IP;refGene_tx=XM_023259717.1;refGe
ne_gn=STK11IP;refGene_tx=XM_023259718.1
C1 204561272 204561422
refGene_gn=STK11IP;refGene_tx=XM_011285660.3;refGene_gn=STK11IP;refGen
e_tx=XM_023259716.1;refGene_gn=STK11IP;refGene_tx=XM_023259717.1;refGe
ne_gn=STK11IP;refGene_tx=XM_023259718.1
C1 204575882 204575933
refGene_gn=SLC4A3;refGene_tx=XM_023259719.1;refGene_gn=SLC4A3;refGene_
tx=XM_023259720.1
C1 204576282 204576448
refGene_gn=SLC4A3;refGene_tx=XM_023259719.1;refGene_gn=SLC4A3;refGene_
tx=XM_023259720.1
C1 204577037 204577318
refGene_gn=SLC4A3;refGene_tx=XM_023259719.1;refGene_gn=SLC4A3;refGene_
tx=XM_023259720.1
C1 204577481 204577597
refGene_gn=SLC4A3;refGene_tx=XM_023259719.1;refGene_gn=SLC4A3;refGene_
tx=XM_023259720.1
C1 204577976 204578203
refGene_gn=SLC4A3;refGene_tx=XM_023259719.1;refGene_gn=SLC4A3;refGene_
tx=XM_023259720.1
C1 204579069 204579286
refGene_gn=SLC4A3;refGene_tx=XM_019838726.2
C1 204579766 204579915
refGene_gn=SLC4A3;refGene_tx=XM_019838726.2;refGene_gn=SLC4A3;refGene_
tx=XM_023259719.1;refGene_gn=SLC4A3;refGene_tx=XM_023259720.1
C1 204580056 204580237
refGene_gn=SLC4A3;refGene_tx=XM_019838726.2;refGene_gn=SLC4A3;refGene_
tx=XM_023259719.1;refGene_gn=SLC4A3;refGene_tx=XM_023259720.1
C1 204580706 204580842
refGene_gn=SLC4A3;refGene_tx=XM_019838726.2;refGene_gn=SLC4A3;refGene_
tx=XM_023259719.1;refGene_gn=SLC4A3;refGene_tx=XM_023259720.1

C1 204581556 204581740
refGene_gn=SLC4A3;refGene_tx=XM_019838726.2;refGene_gn=SLC4A3;refGene_tx=XM_023259719.1;refGene_gn=SLC4A3;refGene_tx=XM_023259720.1
C1 204582173 204582273
refGene_gn=SLC4A3;refGene_tx=XM_019838726.2;refGene_gn=SLC4A3;refGene_tx=XM_023259719.1;refGene_gn=SLC4A3;refGene_tx=XM_023259720.1
C1 204582737 204582922
refGene_gn=SLC4A3;refGene_tx=XM_019838726.2;refGene_gn=SLC4A3;refGene_tx=XM_023259719.1;refGene_gn=SLC4A3;refGene_tx=XM_023259720.1
C1 204583593 204583819
refGene_gn=SLC4A3;refGene_tx=XM_019838726.2;refGene_gn=SLC4A3;refGene_tx=XM_023259719.1;refGene_gn=SLC4A3;refGene_tx=XM_023259720.1
C1 204583996 204584215
refGene_gn=SLC4A3;refGene_tx=XM_019838726.2;refGene_gn=SLC4A3;refGene_tx=XM_023259719.1;refGene_gn=SLC4A3;refGene_tx=XM_023259720.1
C1 204584503 204584652
refGene_gn=SLC4A3;refGene_tx=XM_019838726.2;refGene_gn=SLC4A3;refGene_tx=XM_023259719.1;refGene_gn=SLC4A3;refGene_tx=XM_023259720.1
C1 204584862 204585057
refGene_gn=SLC4A3;refGene_tx=XM_019838726.2;refGene_gn=SLC4A3;refGene_tx=XM_023259719.1;refGene_gn=SLC4A3;refGene_tx=XM_023259720.1
C1 204585538 204585757
refGene_gn=SLC4A3;refGene_tx=XM_019838726.2;refGene_gn=SLC4A3;refGene_tx=XM_023259719.1;refGene_gn=SLC4A3;refGene_tx=XM_023259720.1
C1 204586105 204586195
refGene_gn=SLC4A3;refGene_tx=XM_019838726.2;refGene_gn=SLC4A3;refGene_tx=XM_023259719.1;refGene_gn=SLC4A3;refGene_tx=XM_023259720.1
C1 204586704 204586871
refGene_gn=SLC4A3;refGene_tx=XM_019838726.2;refGene_gn=SLC4A3;refGene_tx=XM_023259719.1;refGene_gn=SLC4A3;refGene_tx=XM_023259720.1
C1 204587361 204587615
refGene_gn=SLC4A3;refGene_tx=XM_019838726.2;refGene_gn=SLC4A3;refGene_tx=XM_023259719.1;refGene_gn=SLC4A3;refGene_tx=XM_023259720.1
C1 204587942 204588112
refGene_gn=SLC4A3;refGene_tx=XM_019838726.2;refGene_gn=SLC4A3;refGene_tx=XM_023259719.1;refGene_gn=SLC4A3;refGene_tx=XM_023259720.1
C1 204588319 204588493
refGene_gn=SLC4A3;refGene_tx=XM_019838726.2;refGene_gn=SLC4A3;refGene_tx=XM_023259719.1;refGene_gn=SLC4A3;refGene_tx=XM_023259720.1
C1 204589157 204589235
refGene_gn=SLC4A3;refGene_tx=XM_019838726.2;refGene_gn=SLC4A3;refGene_tx=XM_023259719.1;refGene_gn=SLC4A3;refGene_tx=XM_023259720.1
C1 206200430 206200437
refGene_gn=EPHA4;refGene_tx=XM_019838727.1
C1 206205960 206206120
refGene_gn=EPHA4;refGene_tx=XM_019838727.1;refGene_gn=EPHA4;refGene_tx=XM_019838728.1
C1 206209707 206209901
refGene_gn=EPHA4;refGene_tx=XM_019838727.1;refGene_gn=EPHA4;refGene_tx=XM_019838728.1

C1	206213833	206213983
refGene_gn=EPHA4;refGene_tx=XM_019838727.1;refGene_gn=EPHA4;refGene_tx=XM_019838728.1
C1	206216159	206216369
refGene_gn=EPHA4;refGene_tx=XM_019838727.1;refGene_gn=EPHA4;refGene_tx=XM_019838728.1
C1	206216825	206216887
refGene_gn=EPHA4;refGene_tx=XM_019838727.1;refGene_gn=EPHA4;refGene_tx=XM_019838728.1
C1	206221280	206221466
refGene_gn=EPHA4;refGene_tx=XM_019838727.1;refGene_gn=EPHA4;refGene_tx=XM_019838728.1
C1	206221727	206221841
refGene_gn=EPHA4;refGene_tx=XM_019838727.1;refGene_gn=EPHA4;refGene_tx=XM_019838728.1
C1	206223640	206223699
refGene_gn=EPHA4;refGene_tx=XM_019838727.1;refGene_gn=EPHA4;refGene_tx=XM_019838728.1
C1	206232746	206232858
refGene_gn=EPHA4;refGene_tx=XM_019838727.1;refGene_gn=EPHA4;refGene_tx=XM_019838728.1
C1	206234052	206234212
refGene_gn=EPHA4;refGene_tx=XM_019838727.1;refGene_gn=EPHA4;refGene_tx=XM_019838728.1
C1	206235239	206235364
refGene_gn=EPHA4;refGene_tx=XM_019838727.1;refGene_gn=EPHA4;refGene_tx=XM_019838728.1
C1	206262883	206263222
refGene_gn=EPHA4;refGene_tx=XM_019838727.1;refGene_gn=EPHA4;refGene_tx=XM_019838728.1
C1	206279756	206279912
refGene_gn=EPHA4;refGene_tx=XM_019838727.1;refGene_gn=EPHA4;refGene_tx=XM_019838728.1
C1	206341952	206342616
refGene_gn=EPHA4;refGene_tx=XM_019838727.1;refGene_gn=EPHA4;refGene_tx=XM_019838728.1
C1	206346921	206346989
refGene_gn=EPHA4;refGene_tx=XM_019838727.1;refGene_gn=EPHA4;refGene_tx=XM_019838728.1
C1	206350047	206350138
refGene_gn=EPHA4;refGene_tx=XM_019838727.1;refGene_gn=EPHA4;refGene_tx=XM_019838728.1
C1	206350759	206351557
refGene_gn=LOC109502886;refGene_tx=XM_019838729.2
C1	206352779	206352842
refGene_gn=LOC109502886;refGene_tx=XM_019838729.2
C1	206907557	206907592
refGene_gn=PAX3;refGene_tx=XM_011285657.3;refGene_gn=PAX3;refGene_tx=XM_019838731.2
C1	206908092	206908339

refGene_gn=PAX3;refGene_tx=XM_011285657.3;refGene_gn=PAX3;refGene_tx=XM_019838731.2
C1 206926824 206927039
refGene_gn=PAX3;refGene_tx=XM_011285657.3;refGene_gn=PAX3;refGene_tx=XM_019838731.2
C1 206928136 206928302
refGene_gn=PAX3;refGene_tx=XM_011285657.3;refGene_gn=PAX3;refGene_tx=XM_019838731.2
C1 206938370 206938576
refGene_gn=PAX3;refGene_tx=XM_011285657.3;refGene_gn=PAX3;refGene_tx=XM_019838731.2
C1 206998945 206999080
refGene_gn=PAX3;refGene_tx=XM_011285657.3;refGene_gn=PAX3;refGene_tx=XM_019838731.2
C1 207000231 207000361
refGene_gn=PAX3;refGene_tx=XM_011285657.3;refGene_gn=PAX3;refGene_tx=XM_019838731.2
C1 207001673 207001909
refGene_gn=PAX3;refGene_tx=XM_011285657.3;refGene_gn=PAX3;refGene_tx=XM_019838731.2
C1 207003195 207003280
refGene_gn=PAX3;refGene_tx=XM_011285657.3;refGene_gn=PAX3;refGene_tx=XM_019838731.2
C1 207100082 207100301
refGene_gn=SGPP2;refGene_tx=XM_023259721.1
C1 207147739 207147898
refGene_gn=SGPP2;refGene_tx=XM_023259721.1
C1 207183694 207183874
refGene_gn=SGPP2;refGene_tx=XM_019838733.2;refGene_gn=SGPP2;refGene_tx=XM_019838734.2;refGene_gn=SGPP2;refGene_tx=XM_023259721.1;refGene_gn=SGPP2;refGene_tx=XM_023259722.1;refGene_gn=SGPP2;refGene_tx=XM_023259723.1
C1 207186949 207187039
refGene_gn=SGPP2;refGene_tx=XM_019838733.2;refGene_gn=SGPP2;refGene_tx=XM_019838734.2;refGene_gn=SGPP2;refGene_tx=XM_023259721.1;refGene_gn=SGPP2;refGene_tx=XM_023259722.1;refGene_gn=SGPP2;refGene_tx=XM_023259723.1
C1 207210887 207211439
refGene_gn=SGPP2;refGene_tx=XM_019838733.2;refGene_gn=SGPP2;refGene_tx=XM_019838734.2;refGene_gn=SGPP2;refGene_tx=XM_023259721.1;refGene_gn=SGPP2;refGene_tx=XM_023259722.1;refGene_gn=SGPP2;refGene_tx=XM_023259723.1
C1 207222439 207222591
refGene_gn=FARSB;refGene_tx=XM_003991201.5
C1 207247231 207247387
refGene_gn=FARSB;refGene_tx=XM_003991201.5
C1 207259406 207259524
refGene_gn=FARSB;refGene_tx=XM_003991201.5
C1 207263183 207263276
refGene_gn=FARSB;refGene_tx=XM_003991201.5

C1 207266411 207266492
refGene_gn=FARSB;refGene_tx=XM_003991201.5
C1 207266979 207267187
refGene_gn=FARSB;refGene_tx=XM_003991201.5
C1 207267658 207267720
refGene_gn=FARSB;refGene_tx=XM_003991201.5
C1 207271127 207271179
refGene_gn=FARSB;refGene_tx=XM_003991201.5
C1 207272081 207272143
refGene_gn=FARSB;refGene_tx=XM_003991201.5
C1 207273607 207273678
refGene_gn=FARSB;refGene_tx=XM_003991201.5
C1 207275487 207275596
refGene_gn=FARSB;refGene_tx=XM_003991201.5
C1 207276853 207277004
refGene_gn=FARSB;refGene_tx=XM_003991201.5
C1 207282063 207282179
refGene_gn=FARSB;refGene_tx=XM_003991201.5
C1 207282813 207282883
refGene_gn=FARSB;refGene_tx=XM_003991201.5
C1 207284892 207285047
refGene_gn=FARSB;refGene_tx=XM_003991201.5
C1 207287964 207288020
refGene_gn=FARSB;refGene_tx=XM_003991201.5
C1 207294272 207294330
refGene_gn=FARSB;refGene_tx=XM_003991201.5
C1 207305125 207305219
refGene_gn=MOGAT1;refGene_tx=XM_023259724.1;refGene_gn=MOGAT1;refGene_
tx=XM_023259725.1
C1 207318822 207319001
refGene_gn=MOGAT1;refGene_tx=XM_023259724.1;refGene_gn=MOGAT1;refGene_
tx=XM_023259725.1;refGene_gn=MOGAT1;refGene_tx=XM_023259726.1
C1 207319339 207319544
refGene_gn=MOGAT1;refGene_tx=XM_023259724.1;refGene_gn=MOGAT1;refGene_
tx=XM_023259725.1;refGene_gn=MOGAT1;refGene_tx=XM_023259726.1
C1 207323900 207324075
refGene_gn=MOGAT1;refGene_tx=XM_023259724.1;refGene_gn=MOGAT1;refGene_
tx=XM_023259725.1;refGene_gn=MOGAT1;refGene_tx=XM_023259726.1
C1 207325381 207325581
refGene_gn=MOGAT1;refGene_tx=XM_023259724.1;refGene_gn=MOGAT1;refGene_
tx=XM_023259725.1;refGene_gn=MOGAT1;refGene_tx=XM_023259726.1
C1 207327421 207327426
refGene_gn=MOGAT1;refGene_tx=XM_023259725.1
C1 207339318 207339473
refGene_gn=MOGAT1;refGene_tx=XM_023259724.1;refGene_gn=MOGAT1;refGene_
tx=XM_023259726.1
C1 207497157 207497535
refGene_gn=ACSL3;refGene_tx=XM_019838737.2;refGene_gn=ACSL3;refGene_tx
=XM_023259727.1;refGene_gn=ACSL3;refGene_tx=XM_023259728.1;refGene_gn=
ACSL3;refGene_tx=XM_023259729.1;refGene_gn=ACSL3;refGene_tx=XM_0232597

30.1
C1 207505121 207505182
refGene_gn=ACSL3;refGene_tx=XM_019838737.2
C1 207506227 207506405
refGene_gn=ACSL3;refGene_tx=XM_019838737.2;refGene_gn=ACSL3;refGene_tx=XM_023259727.1;refGene_gn=ACSL3;refGene_tx=XM_023259728.1;refGene_gn=ACSL3;refGene_tx=XM_023259729.1;refGene_gn=ACSL3;refGene_tx=XM_023259730.1
C1 207507873 207507983
refGene_gn=ACSL3;refGene_tx=XM_019838737.2;refGene_gn=ACSL3;refGene_tx=XM_023259727.1;refGene_gn=ACSL3;refGene_tx=XM_023259728.1;refGene_gn=ACSL3;refGene_tx=XM_023259729.1;refGene_gn=ACSL3;refGene_tx=XM_023259730.1
C1 207509204 207509343
refGene_gn=ACSL3;refGene_tx=XM_019838737.2;refGene_gn=ACSL3;refGene_tx=XM_023259727.1;refGene_gn=ACSL3;refGene_tx=XM_023259728.1;refGene_gn=ACSL3;refGene_tx=XM_023259729.1;refGene_gn=ACSL3;refGene_tx=XM_023259730.1
C1 207511542 207511693
refGene_gn=ACSL3;refGene_tx=XM_019838737.2;refGene_gn=ACSL3;refGene_tx=XM_023259727.1;refGene_gn=ACSL3;refGene_tx=XM_023259728.1;refGene_gn=ACSL3;refGene_tx=XM_023259729.1;refGene_gn=ACSL3;refGene_tx=XM_023259730.1
C1 207512938 207513062
refGene_gn=ACSL3;refGene_tx=XM_019838737.2;refGene_gn=ACSL3;refGene_tx=XM_023259727.1;refGene_gn=ACSL3;refGene_tx=XM_023259728.1;refGene_gn=ACSL3;refGene_tx=XM_023259729.1;refGene_gn=ACSL3;refGene_tx=XM_023259730.1
C1 207513291 207513363
refGene_gn=ACSL3;refGene_tx=XM_019838737.2;refGene_gn=ACSL3;refGene_tx=XM_023259727.1;refGene_gn=ACSL3;refGene_tx=XM_023259728.1;refGene_gn=ACSL3;refGene_tx=XM_023259729.1;refGene_gn=ACSL3;refGene_tx=XM_023259730.1
C1 207515351 207515491
refGene_gn=ACSL3;refGene_tx=XM_019838737.2;refGene_gn=ACSL3;refGene_tx=XM_023259727.1;refGene_gn=ACSL3;refGene_tx=XM_023259728.1;refGene_gn=ACSL3;refGene_tx=XM_023259729.1;refGene_gn=ACSL3;refGene_tx=XM_023259730.1
C1 207518273 207518446
refGene_gn=ACSL3;refGene_tx=XM_019838737.2;refGene_gn=ACSL3;refGene_tx=XM_023259727.1;refGene_gn=ACSL3;refGene_tx=XM_023259728.1;refGene_gn=ACSL3;refGene_tx=XM_023259729.1;refGene_gn=ACSL3;refGene_tx=XM_023259730.1
C1 207520089 207520164
refGene_gn=ACSL3;refGene_tx=XM_019838737.2;refGene_gn=ACSL3;refGene_tx=XM_023259727.1;refGene_gn=ACSL3;refGene_tx=XM_023259728.1;refGene_gn=ACSL3;refGene_tx=XM_023259729.1;refGene_gn=ACSL3;refGene_tx=XM_023259730.1
C1 207521719 207521911
refGene_gn=ACSL3;refGene_tx=XM_019838737.2;refGene_gn=ACSL3;refGene_tx

=XM_023259727.1;refGene_gn=ACSL3;refGene_tx=XM_023259728.1;refGene_gn=ACSL3;refGene_tx=XM_023259729.1;refGene_gn=ACSL3;refGene_tx=XM_023259730.1
C1 207523214 207523329
refGene_gn=ACSL3;refGene_tx=XM_019838737.2;refGene_gn=ACSL3;refGene_tx=XM_023259727.1;refGene_gn=ACSL3;refGene_tx=XM_023259728.1;refGene_gn=ACSL3;refGene_tx=XM_023259729.1;refGene_gn=ACSL3;refGene_tx=XM_023259730.1
C1 207524929 207525087
refGene_gn=ACSL3;refGene_tx=XM_019838737.2;refGene_gn=ACSL3;refGene_tx=XM_023259727.1;refGene_gn=ACSL3;refGene_tx=XM_023259728.1;refGene_gn=ACSL3;refGene_tx=XM_023259729.1;refGene_gn=ACSL3;refGene_tx=XM_023259730.1
C1 207530261 207530419
refGene_gn=ACSL3;refGene_tx=XM_019838737.2;refGene_gn=ACSL3;refGene_tx=XM_023259727.1;refGene_gn=ACSL3;refGene_tx=XM_023259728.1;refGene_gn=ACSL3;refGene_tx=XM_023259729.1;refGene_gn=ACSL3;refGene_tx=XM_023259730.1
C1 207615011 207615515
refGene_gn=KCNE4;refGene_tx=XM_023259731.1
C1 208060476 208062348
refGene_gn=SCG2;refGene_tx=XM_006935628.3
C1 208065397 208065433
refGene_gn=SCG2;refGene_tx=XM_006935628.3
C1 208198146 208198182
refGene_gn=AP1S3;refGene_tx=XM_011285654.2;refGene_gn=AP1S3;refGene_tx=XM_023259732.1
C1 208203602 208203740
refGene_gn=AP1S3;refGene_tx=XM_011285654.2;refGene_gn=AP1S3;refGene_tx=XM_023259732.1
C1 208211282 208211391
refGene_gn=AP1S3;refGene_tx=XM_011285654.2;refGene_gn=AP1S3;refGene_tx=XM_023259732.1
C1 208213562 208213741
refGene_gn=AP1S3;refGene_tx=XM_011285654.2;refGene_gn=AP1S3;refGene_tx=XM_023259732.1
C1 208244377 208244461
refGene_gn=AP1S3;refGene_tx=XM_023259732.1
C1 208256854 208256857
refGene_gn=AP1S3;refGene_tx=XM_011285654.2
C1 208286565 208286625
refGene_gn=WDFY1;refGene_tx=XM_023259733.1;refGene_gn=WDFY1;refGene_tx=XM_023259734.1
C1 208287444 208287553
refGene_gn=WDFY1;refGene_tx=XM_023259733.1;refGene_gn=WDFY1;refGene_tx=XM_023259734.1
C1 208288617 208288748
refGene_gn=WDFY1;refGene_tx=XM_023259733.1;refGene_gn=WDFY1;refGene_tx=XM_023259734.1
C1 208290217 208290319

refGene_gn=WDFY1;refGene_tx=XM_023259733.1;refGene_gn=WDFY1;refGene_tx=XM_023259734.1
C1 208295309 208295415
refGene_gn=WDFY1;refGene_tx=XM_023259733.1;refGene_gn=WDFY1;refGene_tx=XM_023259734.1
C1 208296404 208296531
refGene_gn=WDFY1;refGene_tx=XM_023259733.1;refGene_gn=WDFY1;refGene_tx=XM_023259734.1
C1 208298296 208298409
refGene_gn=WDFY1;refGene_tx=XM_023259733.1;refGene_gn=WDFY1;refGene_tx=XM_023259734.1
C1 208300048 208300199
refGene_gn=WDFY1;refGene_tx=XM_023259733.1;refGene_gn=WDFY1;refGene_tx=XM_023259734.1
C1 208301830 208301885
refGene_gn=WDFY1;refGene_tx=XM_023259733.1;refGene_gn=WDFY1;refGene_tx=XM_023259734.1
C1 208306522 208306596
refGene_gn=WDFY1;refGene_tx=XM_023259733.1;refGene_gn=WDFY1;refGene_tx=XM_023259734.1
C1 208312856 208312924
refGene_gn=WDFY1;refGene_tx=XM_023259733.1;refGene_gn=WDFY1;refGene_tx=XM_023259734.1
C1 208328962 208329099
refGene_gn=WDFY1;refGene_tx=XM_023259733.1
C1 208334293 208334472
refGene_gn=MRPL44;refGene_tx=XM_003991213.5
C1 208336442 208336911
refGene_gn=MRPL44;refGene_tx=XM_003991213.5
C1 208342826 208343005
refGene_gn=MRPL44;refGene_tx=XM_003991213.5
C1 208346206 208346396
refGene_gn=MRPL44;refGene_tx=XM_003991213.5
C1 208355150 208355188
refGene_gn=SERPINE2;refGene_tx=XM_011285653.3;refGene_gn=SERPINE2;refGene_tx=XM_023259735.1
C1 208357393 208357477
refGene_gn=SERPINE2;refGene_tx=XM_011285653.3;refGene_gn=SERPINE2;refGene_tx=XM_023259735.1
C1 208360034 208360121
refGene_gn=SERPINE2;refGene_tx=XM_011285653.3;refGene_gn=SERPINE2;refGene_tx=XM_023259735.1
C1 208362000 208362101
refGene_gn=SERPINE2;refGene_tx=XM_011285653.3;refGene_gn=SERPINE2;refGene_tx=XM_023259735.1
C1 208363742 208363941
refGene_gn=SERPINE2;refGene_tx=XM_011285653.3;refGene_gn=SERPINE2;refGene_tx=XM_023259735.1
C1 208371330 208371528
refGene_gn=SERPINE2;refGene_tx=XM_011285653.3;refGene_gn=SERPINE2;refG

ene_tx=XM_023259735.1
C1 208377400 208377628
refGene_gn=SERPINE2;refGene_tx=XM_011285653.3;refGene_gn=SERPINE2;refGene_tx=XM_023259735.1
C1 208380555 208380814
refGene_gn=SERPINE2;refGene_tx=XM_011285653.3;refGene_gn=SERPINE2;refGene_tx=XM_023259735.1
C1 208694492 208695143
refGene_gn=FAM124B;refGene_tx=XM_003991214.5
C1 208712322 208713054
refGene_gn=FAM124B;refGene_tx=XM_003991214.5
C1 208781125 208781257
refGene_gn=CUL3;refGene_tx=XM_019838740.1;refGene_gn=CUL3;refGene_tx=XM_023259737.1;refGene_gn=CUL3;refGene_tx=XM_023259738.1
C1 208783884 208784030
refGene_gn=CUL3;refGene_tx=XM_019838740.1;refGene_gn=CUL3;refGene_tx=XM_023259737.1;refGene_gn=CUL3;refGene_tx=XM_023259738.1
C1 208788737 208788924
refGene_gn=CUL3;refGene_tx=XM_019838740.1;refGene_gn=CUL3;refGene_tx=XM_023259737.1;refGene_gn=CUL3;refGene_tx=XM_023259738.1
C1 208795617 208795752
refGene_gn=CUL3;refGene_tx=XM_019838740.1;refGene_gn=CUL3;refGene_tx=XM_023259737.1;refGene_gn=CUL3;refGene_tx=XM_023259738.1
C1 208797209 208797306
refGene_gn=CUL3;refGene_tx=XM_019838740.1;refGene_gn=CUL3;refGene_tx=XM_023259737.1;refGene_gn=CUL3;refGene_tx=XM_023259738.1
C1 208797761 208797886
refGene_gn=CUL3;refGene_tx=XM_019838740.1;refGene_gn=CUL3;refGene_tx=XM_023259737.1;refGene_gn=CUL3;refGene_tx=XM_023259738.1
C1 208801483 208801591
refGene_gn=CUL3;refGene_tx=XM_019838740.1;refGene_gn=CUL3;refGene_tx=XM_023259737.1;refGene_gn=CUL3;refGene_tx=XM_023259738.1
C1 208802225 208802396
refGene_gn=CUL3;refGene_tx=XM_019838740.1;refGene_gn=CUL3;refGene_tx=XM_023259737.1;refGene_gn=CUL3;refGene_tx=XM_023259738.1
C1 208803535 208803712
refGene_gn=CUL3;refGene_tx=XM_019838740.1;refGene_gn=CUL3;refGene_tx=XM_023259737.1;refGene_gn=CUL3;refGene_tx=XM_023259738.1
C1 208804692 208804838
refGene_gn=CUL3;refGene_tx=XM_019838740.1;refGene_gn=CUL3;refGene_tx=XM_023259737.1;refGene_gn=CUL3;refGene_tx=XM_023259738.1
C1 208810575 208810804
refGene_gn=CUL3;refGene_tx=XM_019838740.1;refGene_gn=CUL3;refGene_tx=XM_023259737.1;refGene_gn=CUL3;refGene_tx=XM_023259738.1
C1 208813010 208813125
refGene_gn=CUL3;refGene_tx=XM_019838740.1;refGene_gn=CUL3;refGene_tx=XM_023259737.1;refGene_gn=CUL3;refGene_tx=XM_023259738.1
C1 208814133 208814294
refGene_gn=CUL3;refGene_tx=XM_019838740.1;refGene_gn=CUL3;refGene_tx=XM_023259737.1;refGene_gn=CUL3;refGene_tx=XM_023259738.1

C1	208835591	208835705
refGene_gn=CUL3;refGene_tx=XM_019838740.1;refGene_gn=CUL3;refGene_tx=XM_023259737.1;refGene_gn=CUL3;refGene_tx=XM_023259738.1
C1	208854464	208854512
refGene_gn=CUL3;refGene_tx=XM_019838740.1
C1	208855796	208855994
refGene_gn=CUL3;refGene_tx=XM_023259737.1
C1	208877219	208877285
refGene_gn=CUL3;refGene_tx=XM_023259737.1;refGene_gn=CUL3;refGene_tx=XM_023259738.1
C1	208959079	208959210
refGene_gn=LOC111561649;refGene_tx=XM_023257868.1
C1	208964688	208964930
refGene_gn=LOC111561649;refGene_tx=XM_023257868.1
C1	208975982	208976035
refGene_gn=LOC111561649;refGene_tx=XM_023257868.1
C1	208999222	208999338
refGene_gn=LOC111561649;refGene_tx=XM_023257868.1
C1	209000809	209000908
refGene_gn=LOC111561649;refGene_tx=XM_023257868.1
C1	209002139	209002170
refGene_gn=LOC111561649;refGene_tx=XM_023257868.1
C1	209026073	209026190
refGene_gn=DOCK10;refGene_tx=XM_011285652.3;refGene_gn=DOCK10;refGene_tx=XM_019838741.2;refGene_gn=DOCK10;refGene_tx=XM_019838742.2;refGene_gn=DOCK10;refGene_tx=XM_019838743.2;refGene_gn=DOCK10;refGene_tx=XM_019838744.2;refGene_gn=DOCK10;refGene_tx=XM_019838745.2;refGene_gn=DOCK10;refGene_tx=XM_019838746.2;refGene_gn=DOCK10;refGene_tx=XM_019838748.2;refGene_gn=DOCK10;refGene_tx=XM_019838749.2;refGene_gn=DOCK10;refGene_tx=XM_019838750.2;refGene_gn=DOCK10;refGene_tx=XM_023259739.1;refGene_gn=DOCK10;refGene_tx=XM_023259740.1
C1	209029332	209029408
refGene_gn=DOCK10;refGene_tx=XM_019838741.2;refGene_gn=DOCK10;refGene_tx=XM_019838742.2;refGene_gn=DOCK10;refGene_tx=XM_019838743.2;refGene_gn=DOCK10;refGene_tx=XM_019838745.2;refGene_gn=DOCK10;refGene_tx=XM_019838746.2;refGene_gn=DOCK10;refGene_tx=XM_019838749.2;refGene_gn=DOCK10;refGene_tx=XM_019838750.2;refGene_gn=DOCK10;refGene_tx=XM_023259739.1
C1	209031211	209031350
refGene_gn=DOCK10;refGene_tx=XM_011285652.3;refGene_gn=DOCK10;refGene_tx=XM_019838741.2;refGene_gn=DOCK10;refGene_tx=XM_019838742.2;refGene_gn=DOCK10;refGene_tx=XM_019838743.2;refGene_gn=DOCK10;refGene_tx=XM_019838744.2;refGene_gn=DOCK10;refGene_tx=XM_019838745.2;refGene_gn=DOCK10;refGene_tx=XM_019838746.2;refGene_gn=DOCK10;refGene_tx=XM_019838748.2;refGene_gn=DOCK10;refGene_tx=XM_019838749.2;refGene_gn=DOCK10;refGene_tx=XM_019838750.2;refGene_gn=DOCK10;refGene_tx=XM_023259739.1;refGene_gn=DOCK10;refGene_tx=XM_023259740.1
C1	209031502	209031603
refGene_gn=DOCK10;refGene_tx=XM_011285652.3;refGene_gn=DOCK10;refGene_tx=XM_019838741.2;refGene_gn=DOCK10;refGene_tx=XM_019838742.2;refGene_

gn=DOCK10;refGene_tx=XM_019838743.2;refGene_gn=DOCK10;refGene_tx=XM_019838744.2;refGene_gn=DOCK10;refGene_tx=XM_019838745.2;refGene_gn=DOCK10;refGene_tx=XM_019838746.2;refGene_gn=DOCK10;refGene_tx=XM_019838748.2;refGene_gn=DOCK10;refGene_tx=XM_019838749.2;refGene_gn=DOCK10;refGene_tx=XM_019838750.2;refGene_gn=DOCK10;refGene_tx=XM_023259739.1;refGene_gn=DOCK10;refGene_tx=XM_023259740.1
C1 209033947 209034138
refGene_gn=DOCK10;refGene_tx=XM_011285652.3;refGene_gn=DOCK10;refGene_tx=XM_019838741.2;refGene_gn=DOCK10;refGene_tx=XM_019838742.2;refGene_gn=DOCK10;refGene_tx=XM_019838743.2;refGene_gn=DOCK10;refGene_tx=XM_019838744.2;refGene_gn=DOCK10;refGene_tx=XM_019838745.2;refGene_gn=DOCK10;refGene_tx=XM_019838746.2;refGene_gn=DOCK10;refGene_tx=XM_019838748.2;refGene_gn=DOCK10;refGene_tx=XM_019838749.2;refGene_gn=DOCK10;refGene_tx=XM_019838750.2;refGene_gn=DOCK10;refGene_tx=XM_023259739.1;refGene_gn=DOCK10;refGene_tx=XM_023259740.1
C1 209035564 209035775
refGene_gn=DOCK10;refGene_tx=XM_011285652.3;refGene_gn=DOCK10;refGene_tx=XM_019838741.2;refGene_gn=DOCK10;refGene_tx=XM_019838742.2;refGene_gn=DOCK10;refGene_tx=XM_019838743.2;refGene_gn=DOCK10;refGene_tx=XM_019838744.2;refGene_gn=DOCK10;refGene_tx=XM_019838745.2;refGene_gn=DOCK10;refGene_tx=XM_019838746.2;refGene_gn=DOCK10;refGene_tx=XM_019838748.2;refGene_gn=DOCK10;refGene_tx=XM_019838749.2;refGene_gn=DOCK10;refGene_tx=XM_019838750.2;refGene_gn=DOCK10;refGene_tx=XM_023259739.1;refGene_gn=DOCK10;refGene_tx=XM_023259740.1
C1 209038305 209038452
refGene_gn=DOCK10;refGene_tx=XM_011285652.3;refGene_gn=DOCK10;refGene_tx=XM_019838741.2;refGene_gn=DOCK10;refGene_tx=XM_019838742.2;refGene_gn=DOCK10;refGene_tx=XM_019838743.2;refGene_gn=DOCK10;refGene_tx=XM_019838744.2;refGene_gn=DOCK10;refGene_tx=XM_019838745.2;refGene_gn=DOCK10;refGene_tx=XM_019838746.2;refGene_gn=DOCK10;refGene_tx=XM_019838748.2;refGene_gn=DOCK10;refGene_tx=XM_019838749.2;refGene_gn=DOCK10;refGene_tx=XM_019838750.2;refGene_gn=DOCK10;refGene_tx=XM_023259739.1;refGene_gn=DOCK10;refGene_tx=XM_023259740.1
C1 209044164 209044170
refGene_gn=DOCK10;refGene_tx=XM_019838741.2;refGene_gn=DOCK10;refGene_tx=XM_019838742.2;refGene_gn=DOCK10;refGene_tx=XM_019838744.2;refGene_gn=DOCK10;refGene_tx=XM_019838745.2;refGene_gn=DOCK10;refGene_tx=XM_019838746.2;refGene_gn=DOCK10;refGene_tx=XM_019838749.2;refGene_gn=DOCK10;refGene_tx=XM_023259739.1
C1 209045749 209045863
refGene_gn=DOCK10;refGene_tx=XM_011285652.3;refGene_gn=DOCK10;refGene_tx=XM_019838741.2;refGene_gn=DOCK10;refGene_tx=XM_019838742.2;refGene_gn=DOCK10;refGene_tx=XM_019838743.2;refGene_gn=DOCK10;refGene_tx=XM_019838744.2;refGene_gn=DOCK10;refGene_tx=XM_019838745.2;refGene_gn=DOCK10;refGene_tx=XM_019838746.2;refGene_gn=DOCK10;refGene_tx=XM_019838748.2;refGene_gn=DOCK10;refGene_tx=XM_019838749.2;refGene_gn=DOCK10;refGene_tx=XM_019838750.2;refGene_gn=DOCK10;refGene_tx=XM_023259739.1;refGene_gn=DOCK10;refGene_tx=XM_023259740.1
C1 209046000 209046123
refGene_gn=DOCK10;refGene_tx=XM_011285652.3;refGene_gn=DOCK10;refGene_

tx=XM_019838741.2;refGene_gn=DOCK10;refGene_tx=XM_019838742.2;refGene_gn=DOCK10;refGene_tx=XM_019838743.2;refGene_gn=DOCK10;refGene_tx=XM_019838744.2;refGene_gn=DOCK10;refGene_tx=XM_019838745.2;refGene_gn=DOCK10;refGene_tx=XM_019838746.2;refGene_gn=DOCK10;refGene_tx=XM_019838748.2;refGene_gn=DOCK10;refGene_tx=XM_019838749.2;refGene_gn=DOCK10;refGene_tx=XM_019838750.2;refGene_gn=DOCK10;refGene_tx=XM_023259739.1;refGene_gn=DOCK10;refGene_tx=XM_023259740.1
C1 209047585 209047692
refGene_gn=DOCK10;refGene_tx=XM_011285652.3;refGene_gn=DOCK10;refGene_tx=XM_019838741.2;refGene_gn=DOCK10;refGene_tx=XM_019838742.2;refGene_gn=DOCK10;refGene_tx=XM_019838743.2;refGene_gn=DOCK10;refGene_tx=XM_019838744.2;refGene_gn=DOCK10;refGene_tx=XM_019838745.2;refGene_gn=DOCK10;refGene_tx=XM_019838746.2;refGene_gn=DOCK10;refGene_tx=XM_019838748.2;refGene_gn=DOCK10;refGene_tx=XM_019838749.2;refGene_gn=DOCK10;refGene_tx=XM_019838750.2;refGene_gn=DOCK10;refGene_tx=XM_023259739.1;refGene_gn=DOCK10;refGene_tx=XM_023259740.1
C1 209050762 209050861
refGene_gn=DOCK10;refGene_tx=XM_011285652.3;refGene_gn=DOCK10;refGene_tx=XM_019838741.2;refGene_gn=DOCK10;refGene_tx=XM_019838742.2;refGene_gn=DOCK10;refGene_tx=XM_019838743.2;refGene_gn=DOCK10;refGene_tx=XM_019838744.2;refGene_gn=DOCK10;refGene_tx=XM_019838745.2;refGene_gn=DOCK10;refGene_tx=XM_019838746.2;refGene_gn=DOCK10;refGene_tx=XM_019838748.2;refGene_gn=DOCK10;refGene_tx=XM_023259739.1;refGene_gn=DOCK10;refGene_tx=XM_023259740.1
C1 209051181 209051239
refGene_gn=DOCK10;refGene_tx=XM_011285652.3;refGene_gn=DOCK10;refGene_tx=XM_019838741.2;refGene_gn=DOCK10;refGene_tx=XM_019838742.2;refGene_gn=DOCK10;refGene_tx=XM_019838743.2;refGene_gn=DOCK10;refGene_tx=XM_019838744.2;refGene_gn=DOCK10;refGene_tx=XM_019838745.2;refGene_gn=DOCK10;refGene_tx=XM_019838746.2;refGene_gn=DOCK10;refGene_tx=XM_019838748.2;refGene_gn=DOCK10;refGene_tx=XM_019838749.2;refGene_gn=DOCK10;refGene_tx=XM_019838750.2;refGene_gn=DOCK10;refGene_tx=XM_023259739.1;refGene_gn=DOCK10;refGene_tx=XM_023259740.1
C1 209053149 209053365
refGene_gn=DOCK10;refGene_tx=XM_011285652.3;refGene_gn=DOCK10;refGene_tx=XM_019838741.2;refGene_gn=DOCK10;refGene_tx=XM_019838742.2;refGene_gn=DOCK10;refGene_tx=XM_019838743.2;refGene_gn=DOCK10;refGene_tx=XM_019838744.2;refGene_gn=DOCK10;refGene_tx=XM_019838745.2;refGene_gn=DOCK10;refGene_tx=XM_019838746.2;refGene_gn=DOCK10;refGene_tx=XM_019838748.2;refGene_gn=DOCK10;refGene_tx=XM_019838749.2;refGene_gn=DOCK10;refGene_tx=XM_019838750.2;refGene_gn=DOCK10;refGene_tx=XM_023259739.1;refGene_gn=DOCK10;refGene_tx=XM_023259740.1
C1 209055803 209055914
refGene_gn=DOCK10;refGene_tx=XM_011285652.3;refGene_gn=DOCK10;refGene_tx=XM_019838741.2;refGene_gn=DOCK10;refGene_tx=XM_019838742.2;refGene_gn=DOCK10;refGene_tx=XM_019838743.2;refGene_gn=DOCK10;refGene_tx=XM_019838744.2;refGene_gn=DOCK10;refGene_tx=XM_019838745.2;refGene_gn=DOCK10;refGene_tx=XM_019838746.2;refGene_gn=DOCK10;refGene_tx=XM_019838748.2;refGene_gn=DOCK10;refGene_tx=XM_019838749.2;refGene_gn=DOCK10;refGene_tx=XM_019838750.2;refGene_gn=DOCK10;refGene_tx=XM_023259739.1;refGen

e_gn=DOCK10;refGene_tx=XM_023259740.1
C1 209056444 209056627
refGene_gn=DOCK10;refGene_tx=XM_011285652.3;refGene_gn=DOCK10;refGene_tx=XM_019838741.2;refGene_gn=DOCK10;refGene_tx=XM_019838742.2;refGene_gn=DOCK10;refGene_tx=XM_019838743.2;refGene_gn=DOCK10;refGene_tx=XM_019838744.2;refGene_gn=DOCK10;refGene_tx=XM_019838745.2;refGene_gn=DOCK10;refGene_tx=XM_019838746.2;refGene_gn=DOCK10;refGene_tx=XM_019838748.2;refGene_gn=DOCK10;refGene_tx=XM_019838749.2;refGene_gn=DOCK10;refGene_tx=XM_019838750.2;refGene_gn=DOCK10;refGene_tx=XM_023259739.1;refGene_gn=DOCK10;refGene_tx=XM_023259740.1
C1 209057294 209057432
refGene_gn=DOCK10;refGene_tx=XM_011285652.3;refGene_gn=DOCK10;refGene_tx=XM_019838741.2;refGene_gn=DOCK10;refGene_tx=XM_019838742.2;refGene_gn=DOCK10;refGene_tx=XM_019838743.2;refGene_gn=DOCK10;refGene_tx=XM_019838744.2;refGene_gn=DOCK10;refGene_tx=XM_019838745.2;refGene_gn=DOCK10;refGene_tx=XM_019838746.2;refGene_gn=DOCK10;refGene_tx=XM_019838748.2;refGene_gn=DOCK10;refGene_tx=XM_019838749.2;refGene_gn=DOCK10;refGene_tx=XM_019838750.2;refGene_gn=DOCK10;refGene_tx=XM_023259739.1;refGene_gn=DOCK10;refGene_tx=XM_023259740.1
C1 209059232 209059345
refGene_gn=DOCK10;refGene_tx=XM_011285652.3;refGene_gn=DOCK10;refGene_tx=XM_019838741.2;refGene_gn=DOCK10;refGene_tx=XM_019838742.2;refGene_gn=DOCK10;refGene_tx=XM_019838743.2;refGene_gn=DOCK10;refGene_tx=XM_019838744.2;refGene_gn=DOCK10;refGene_tx=XM_019838745.2;refGene_gn=DOCK10;refGene_tx=XM_019838746.2;refGene_gn=DOCK10;refGene_tx=XM_019838748.2;refGene_gn=DOCK10;refGene_tx=XM_019838749.2;refGene_gn=DOCK10;refGene_tx=XM_019838750.2;refGene_gn=DOCK10;refGene_tx=XM_023259739.1;refGene_gn=DOCK10;refGene_tx=XM_023259740.1
C1 209060614 209060739
refGene_gn=DOCK10;refGene_tx=XM_011285652.3;refGene_gn=DOCK10;refGene_tx=XM_019838741.2;refGene_gn=DOCK10;refGene_tx=XM_019838742.2;refGene_gn=DOCK10;refGene_tx=XM_019838743.2;refGene_gn=DOCK10;refGene_tx=XM_019838744.2;refGene_gn=DOCK10;refGene_tx=XM_019838745.2;refGene_gn=DOCK10;refGene_tx=XM_019838746.2;refGene_gn=DOCK10;refGene_tx=XM_019838748.2;refGene_gn=DOCK10;refGene_tx=XM_019838749.2;refGene_gn=DOCK10;refGene_tx=XM_019838750.2;refGene_gn=DOCK10;refGene_tx=XM_023259739.1;refGene_gn=DOCK10;refGene_tx=XM_023259740.1
C1 209062797 209062899
refGene_gn=DOCK10;refGene_tx=XM_011285652.3;refGene_gn=DOCK10;refGene_tx=XM_019838741.2;refGene_gn=DOCK10;refGene_tx=XM_019838742.2;refGene_gn=DOCK10;refGene_tx=XM_019838743.2;refGene_gn=DOCK10;refGene_tx=XM_019838744.2;refGene_gn=DOCK10;refGene_tx=XM_019838745.2;refGene_gn=DOCK10;refGene_tx=XM_019838746.2;refGene_gn=DOCK10;refGene_tx=XM_019838748.2;refGene_gn=DOCK10;refGene_tx=XM_019838749.2;refGene_gn=DOCK10;refGene_tx=XM_019838750.2;refGene_gn=DOCK10;refGene_tx=XM_023259739.1;refGene_gn=DOCK10;refGene_tx=XM_023259740.1
C1 209063711 209063759
refGene_gn=DOCK10;refGene_tx=XM_011285652.3;refGene_gn=DOCK10;refGene_tx=XM_019838741.2;refGene_gn=DOCK10;refGene_tx=XM_019838742.2;refGene_gn=DOCK10;refGene_tx=XM_019838743.2;refGene_gn=DOCK10;refGene_tx=XM_01

9838744.2;refGene_gn=DOCK10;refGene_tx=XM_019838745.2;refGene_gn=DOCK1
0;refGene_tx=XM_019838746.2;refGene_gn=DOCK10;refGene_tx=XM_019838748.
2;refGene_gn=DOCK10;refGene_tx=XM_019838749.2;refGene_gn=DOCK10;refGen
e_tx=XM_019838750.2;refGene_gn=DOCK10;refGene_tx=XM_023259739.1;refGen
e_gn=DOCK10;refGene_tx=XM_023259740.1
C1 209063999 209064066
refGene_gn=DOCK10;refGene_tx=XM_011285652.3;refGene_gn=DOCK10;refGene_
tx=XM_019838741.2;refGene_gn=DOCK10;refGene_tx=XM_019838742.2;refGene_
gn=DOCK10;refGene_tx=XM_019838743.2;refGene_gn=DOCK10;refGene_tx=XM_01
9838744.2;refGene_gn=DOCK10;refGene_tx=XM_019838745.2;refGene_gn=DOCK1
0;refGene_tx=XM_019838746.2;refGene_gn=DOCK10;refGene_tx=XM_019838748.
2;refGene_gn=DOCK10;refGene_tx=XM_019838749.2;refGene_gn=DOCK10;refGen
e_tx=XM_019838750.2;refGene_gn=DOCK10;refGene_tx=XM_023259739.1;refGen
e_gn=DOCK10;refGene_tx=XM_023259740.1
C1 209064157 209064272
refGene_gn=DOCK10;refGene_tx=XM_011285652.3;refGene_gn=DOCK10;refGene_
tx=XM_019838741.2;refGene_gn=DOCK10;refGene_tx=XM_019838742.2;refGene_
gn=DOCK10;refGene_tx=XM_019838743.2;refGene_gn=DOCK10;refGene_tx=XM_01
9838744.2;refGene_gn=DOCK10;refGene_tx=XM_019838745.2;refGene_gn=DOCK1
0;refGene_tx=XM_019838746.2;refGene_gn=DOCK10;refGene_tx=XM_019838748.
2;refGene_gn=DOCK10;refGene_tx=XM_019838749.2;refGene_gn=DOCK10;refGen
e_tx=XM_019838750.2;refGene_gn=DOCK10;refGene_tx=XM_023259739.1;refGen
e_gn=DOCK10;refGene_tx=XM_023259740.1
C1 209064369 209064491
refGene_gn=DOCK10;refGene_tx=XM_011285652.3;refGene_gn=DOCK10;refGene_
tx=XM_019838741.2;refGene_gn=DOCK10;refGene_tx=XM_019838742.2;refGene_
gn=DOCK10;refGene_tx=XM_019838743.2;refGene_gn=DOCK10;refGene_tx=XM_01
9838744.2;refGene_gn=DOCK10;refGene_tx=XM_019838745.2;refGene_gn=DOCK1
0;refGene_tx=XM_019838746.2;refGene_gn=DOCK10;refGene_tx=XM_019838748.
2;refGene_gn=DOCK10;refGene_tx=XM_019838749.2;refGene_gn=DOCK10;refGen
e_tx=XM_019838750.2;refGene_gn=DOCK10;refGene_tx=XM_023259739.1;refGen
e_gn=DOCK10;refGene_tx=XM_023259740.1
C1 209065211 209065323
refGene_gn=DOCK10;refGene_tx=XM_011285652.3;refGene_gn=DOCK10;refGene_
tx=XM_019838741.2;refGene_gn=DOCK10;refGene_tx=XM_019838742.2;refGene_
gn=DOCK10;refGene_tx=XM_019838743.2;refGene_gn=DOCK10;refGene_tx=XM_01
9838744.2;refGene_gn=DOCK10;refGene_tx=XM_019838745.2;refGene_gn=DOCK1
0;refGene_tx=XM_019838746.2;refGene_gn=DOCK10;refGene_tx=XM_019838748.
2;refGene_gn=DOCK10;refGene_tx=XM_019838749.2;refGene_gn=DOCK10;refGen
e_tx=XM_019838750.2;refGene_gn=DOCK10;refGene_tx=XM_023259739.1;refGen
e_gn=DOCK10;refGene_tx=XM_023259740.1
C1 209066556 209066673
refGene_gn=DOCK10;refGene_tx=XM_011285652.3;refGene_gn=DOCK10;refGene_
tx=XM_019838741.2;refGene_gn=DOCK10;refGene_tx=XM_019838742.2;refGene_
gn=DOCK10;refGene_tx=XM_019838743.2;refGene_gn=DOCK10;refGene_tx=XM_01
9838744.2;refGene_gn=DOCK10;refGene_tx=XM_019838745.2;refGene_gn=DOCK1
0;refGene_tx=XM_019838746.2;refGene_gn=DOCK10;refGene_tx=XM_019838748.
2;refGene_gn=DOCK10;refGene_tx=XM_019838749.2;refGene_gn=DOCK10;refGen
e_tx=XM_019838750.2;refGene_gn=DOCK10;refGene_tx=XM_023259739.1;refGen
e_gn=DOCK10;refGene_tx=XM_023259740.1

C1	209066798	209066974
refGene_gn=DOCK10;refGene_tx=XM_011285652.3;refGene_gn=DOCK10;refGene_tx=XM_019838741.2;refGene_gn=DOCK10;refGene_tx=XM_019838742.2;refGene_gn=DOCK10;refGene_tx=XM_019838743.2;refGene_gn=DOCK10;refGene_tx=XM_019838744.2;refGene_gn=DOCK10;refGene_tx=XM_019838745.2;refGene_gn=DOCK10;refGene_tx=XM_019838746.2;refGene_gn=DOCK10;refGene_tx=XM_019838748.2;refGene_gn=DOCK10;refGene_tx=XM_019838749.2;refGene_gn=DOCK10;refGene_tx=XM_019838750.2;refGene_gn=DOCK10;refGene_tx=XM_023259739.1;refGene_gn=DOCK10;refGene_tx=XM_023259740.1
C1	209068967	209069012
refGene_gn=DOCK10;refGene_tx=XM_011285652.3;refGene_gn=DOCK10;refGene_tx=XM_019838741.2;refGene_gn=DOCK10;refGene_tx=XM_019838742.2;refGene_gn=DOCK10;refGene_tx=XM_019838743.2;refGene_gn=DOCK10;refGene_tx=XM_019838744.2;refGene_gn=DOCK10;refGene_tx=XM_019838745.2;refGene_gn=DOCK10;refGene_tx=XM_019838746.2;refGene_gn=DOCK10;refGene_tx=XM_019838748.2;refGene_gn=DOCK10;refGene_tx=XM_019838749.2;refGene_gn=DOCK10;refGene_tx=XM_019838750.2;refGene_gn=DOCK10;refGene_tx=XM_023259739.1;refGene_gn=DOCK10;refGene_tx=XM_023259740.1
C1	209070788	209070885
refGene_gn=DOCK10;refGene_tx=XM_011285652.3;refGene_gn=DOCK10;refGene_tx=XM_019838741.2;refGene_gn=DOCK10;refGene_tx=XM_019838742.2;refGene_gn=DOCK10;refGene_tx=XM_019838743.2;refGene_gn=DOCK10;refGene_tx=XM_019838744.2;refGene_gn=DOCK10;refGene_tx=XM_019838745.2;refGene_gn=DOCK10;refGene_tx=XM_019838746.2;refGene_gn=DOCK10;refGene_tx=XM_019838748.2;refGene_gn=DOCK10;refGene_tx=XM_019838749.2;refGene_gn=DOCK10;refGene_tx=XM_019838750.2;refGene_gn=DOCK10;refGene_tx=XM_023259739.1;refGene_gn=DOCK10;refGene_tx=XM_023259740.1
C1	209073292	209073376
refGene_gn=DOCK10;refGene_tx=XM_011285652.3;refGene_gn=DOCK10;refGene_tx=XM_019838741.2;refGene_gn=DOCK10;refGene_tx=XM_019838742.2;refGene_gn=DOCK10;refGene_tx=XM_019838743.2;refGene_gn=DOCK10;refGene_tx=XM_019838744.2;refGene_gn=DOCK10;refGene_tx=XM_019838745.2;refGene_gn=DOCK10;refGene_tx=XM_019838746.2;refGene_gn=DOCK10;refGene_tx=XM_019838748.2;refGene_gn=DOCK10;refGene_tx=XM_019838749.2;refGene_gn=DOCK10;refGene_tx=XM_019838750.2;refGene_gn=DOCK10;refGene_tx=XM_023259739.1;refGene_gn=DOCK10;refGene_tx=XM_023259740.1
C1	209080045	209080192
refGene_gn=DOCK10;refGene_tx=XM_011285652.3;refGene_gn=DOCK10;refGene_tx=XM_019838741.2;refGene_gn=DOCK10;refGene_tx=XM_019838742.2;refGene_gn=DOCK10;refGene_tx=XM_019838743.2;refGene_gn=DOCK10;refGene_tx=XM_019838744.2;refGene_gn=DOCK10;refGene_tx=XM_019838745.2;refGene_gn=DOCK10;refGene_tx=XM_019838746.2;refGene_gn=DOCK10;refGene_tx=XM_019838748.2;refGene_gn=DOCK10;refGene_tx=XM_019838749.2;refGene_gn=DOCK10;refGene_tx=XM_019838750.2;refGene_gn=DOCK10;refGene_tx=XM_023259739.1;refGene_gn=DOCK10;refGene_tx=XM_023259740.1
C1	209086204	209086276
refGene_gn=DOCK10;refGene_tx=XM_011285652.3;refGene_gn=DOCK10;refGene_tx=XM_019838741.2;refGene_gn=DOCK10;refGene_tx=XM_019838742.2;refGene_gn=DOCK10;refGene_tx=XM_019838743.2;refGene_gn=DOCK10;refGene_tx=XM_019838744.2;refGene_gn=DOCK10;refGene_tx=XM_019838745.2;refGene_gn=DOCK1

0;refGene_tx=XM_019838746.2;refGene_gn=DOCK10;refGene_tx=XM_019838748.2;refGene_gn=DOCK10;refGene_tx=XM_019838749.2;refGene_gn=DOCK10;refGene_tx=XM_019838750.2;refGene_gn=DOCK10;refGene_tx=XM_023259739.1;refGene_gn=DOCK10;refGene_tx=XM_023259740.1
C1 209089863 209089977
refGene_gn=DOCK10;refGene_tx=XM_011285652.3;refGene_gn=DOCK10;refGene_tx=XM_019838741.2;refGene_gn=DOCK10;refGene_tx=XM_019838742.2;refGene_gn=DOCK10;refGene_tx=XM_019838743.2;refGene_gn=DOCK10;refGene_tx=XM_019838744.2;refGene_gn=DOCK10;refGene_tx=XM_019838745.2;refGene_gn=DOCK10;refGene_tx=XM_019838746.2;refGene_gn=DOCK10;refGene_tx=XM_019838748.2;refGene_gn=DOCK10;refGene_tx=XM_019838749.2;refGene_gn=DOCK10;refGene_tx=XM_019838750.2;refGene_gn=DOCK10;refGene_tx=XM_023259739.1;refGene_gn=DOCK10;refGene_tx=XM_023259740.1
C1 209093365 209093435
refGene_gn=DOCK10;refGene_tx=XM_011285652.3;refGene_gn=DOCK10;refGene_tx=XM_019838741.2;refGene_gn=DOCK10;refGene_tx=XM_019838742.2;refGene_gn=DOCK10;refGene_tx=XM_019838743.2;refGene_gn=DOCK10;refGene_tx=XM_019838744.2;refGene_gn=DOCK10;refGene_tx=XM_019838745.2;refGene_gn=DOCK10;refGene_tx=XM_019838746.2;refGene_gn=DOCK10;refGene_tx=XM_019838748.2;refGene_gn=DOCK10;refGene_tx=XM_019838749.2;refGene_gn=DOCK10;refGene_tx=XM_019838750.2;refGene_gn=DOCK10;refGene_tx=XM_023259739.1;refGene_gn=DOCK10;refGene_tx=XM_023259740.1
C1 209095787 209095906
refGene_gn=DOCK10;refGene_tx=XM_011285652.3;refGene_gn=DOCK10;refGene_tx=XM_019838741.2;refGene_gn=DOCK10;refGene_tx=XM_019838742.2;refGene_gn=DOCK10;refGene_tx=XM_019838743.2;refGene_gn=DOCK10;refGene_tx=XM_019838744.2;refGene_gn=DOCK10;refGene_tx=XM_019838745.2;refGene_gn=DOCK10;refGene_tx=XM_019838746.2;refGene_gn=DOCK10;refGene_tx=XM_019838748.2;refGene_gn=DOCK10;refGene_tx=XM_019838749.2;refGene_gn=DOCK10;refGene_tx=XM_019838750.2;refGene_gn=DOCK10;refGene_tx=XM_023259739.1;refGene_gn=DOCK10;refGene_tx=XM_023259740.1
C1 209096344 209096437
refGene_gn=DOCK10;refGene_tx=XM_011285652.3;refGene_gn=DOCK10;refGene_tx=XM_019838741.2;refGene_gn=DOCK10;refGene_tx=XM_019838742.2;refGene_gn=DOCK10;refGene_tx=XM_019838743.2;refGene_gn=DOCK10;refGene_tx=XM_019838744.2;refGene_gn=DOCK10;refGene_tx=XM_019838745.2;refGene_gn=DOCK10;refGene_tx=XM_019838746.2;refGene_gn=DOCK10;refGene_tx=XM_019838748.2;refGene_gn=DOCK10;refGene_tx=XM_019838749.2;refGene_gn=DOCK10;refGene_tx=XM_019838750.2;refGene_gn=DOCK10;refGene_tx=XM_023259739.1;refGene_gn=DOCK10;refGene_tx=XM_023259740.1
C1 209099560 209099647
refGene_gn=DOCK10;refGene_tx=XM_011285652.3;refGene_gn=DOCK10;refGene_tx=XM_019838741.2;refGene_gn=DOCK10;refGene_tx=XM_019838742.2;refGene_gn=DOCK10;refGene_tx=XM_019838743.2;refGene_gn=DOCK10;refGene_tx=XM_019838744.2;refGene_gn=DOCK10;refGene_tx=XM_019838745.2;refGene_gn=DOCK10;refGene_tx=XM_019838746.2;refGene_gn=DOCK10;refGene_tx=XM_019838748.2;refGene_gn=DOCK10;refGene_tx=XM_019838749.2;refGene_gn=DOCK10;refGene_tx=XM_019838750.2;refGene_gn=DOCK10;refGene_tx=XM_023259739.1;refGene_gn=DOCK10;refGene_tx=XM_023259740.1
C1 209100022 209100144

refGene_gn=DOCK10;refGene_tx=XM_011285652.3;refGene_gn=DOCK10;refGene_tx=XM_019838741.2;refGene_gn=DOCK10;refGene_tx=XM_019838742.2;refGene_gn=DOCK10;refGene_tx=XM_019838743.2;refGene_gn=DOCK10;refGene_tx=XM_019838744.2;refGene_gn=DOCK10;refGene_tx=XM_019838745.2;refGene_gn=DOCK10;refGene_tx=XM_019838746.2;refGene_gn=DOCK10;refGene_tx=XM_019838748.2;refGene_gn=DOCK10;refGene_tx=XM_019838749.2;refGene_gn=DOCK10;refGene_tx=XM_019838750.2;refGene_gn=DOCK10;refGene_tx=XM_023259739.1;refGene_gn=DOCK10;refGene_tx=XM_023259740.1
C1 209100337 209100461
refGene_gn=DOCK10;refGene_tx=XM_011285652.3;refGene_gn=DOCK10;refGene_tx=XM_019838741.2;refGene_gn=DOCK10;refGene_tx=XM_019838742.2;refGene_gn=DOCK10;refGene_tx=XM_019838743.2;refGene_gn=DOCK10;refGene_tx=XM_019838744.2;refGene_gn=DOCK10;refGene_tx=XM_019838745.2;refGene_gn=DOCK10;refGene_tx=XM_019838746.2;refGene_gn=DOCK10;refGene_tx=XM_019838748.2;refGene_gn=DOCK10;refGene_tx=XM_019838749.2;refGene_gn=DOCK10;refGene_tx=XM_019838750.2;refGene_gn=DOCK10;refGene_tx=XM_023259739.1;refGene_gn=DOCK10;refGene_tx=XM_023259740.1
C1 209104559 209104652
refGene_gn=DOCK10;refGene_tx=XM_011285652.3;refGene_gn=DOCK10;refGene_tx=XM_019838741.2;refGene_gn=DOCK10;refGene_tx=XM_019838742.2;refGene_gn=DOCK10;refGene_tx=XM_019838743.2;refGene_gn=DOCK10;refGene_tx=XM_019838744.2;refGene_gn=DOCK10;refGene_tx=XM_019838745.2;refGene_gn=DOCK10;refGene_tx=XM_019838746.2;refGene_gn=DOCK10;refGene_tx=XM_019838748.2;refGene_gn=DOCK10;refGene_tx=XM_019838749.2;refGene_gn=DOCK10;refGene_tx=XM_019838750.2;refGene_gn=DOCK10;refGene_tx=XM_023259739.1;refGene_gn=DOCK10;refGene_tx=XM_023259740.1
C1 209106868 209106934
refGene_gn=DOCK10;refGene_tx=XM_011285652.3;refGene_gn=DOCK10;refGene_tx=XM_019838741.2;refGene_gn=DOCK10;refGene_tx=XM_019838742.2;refGene_gn=DOCK10;refGene_tx=XM_019838743.2;refGene_gn=DOCK10;refGene_tx=XM_019838744.2;refGene_gn=DOCK10;refGene_tx=XM_019838745.2;refGene_gn=DOCK10;refGene_tx=XM_019838746.2;refGene_gn=DOCK10;refGene_tx=XM_019838748.2;refGene_gn=DOCK10;refGene_tx=XM_019838749.2;refGene_gn=DOCK10;refGene_tx=XM_019838750.2;refGene_gn=DOCK10;refGene_tx=XM_023259739.1;refGene_gn=DOCK10;refGene_tx=XM_023259740.1
C1 209107681 209107869
refGene_gn=DOCK10;refGene_tx=XM_011285652.3;refGene_gn=DOCK10;refGene_tx=XM_019838741.2;refGene_gn=DOCK10;refGene_tx=XM_019838742.2;refGene_gn=DOCK10;refGene_tx=XM_019838743.2;refGene_gn=DOCK10;refGene_tx=XM_019838744.2;refGene_gn=DOCK10;refGene_tx=XM_019838745.2;refGene_gn=DOCK10;refGene_tx=XM_019838746.2;refGene_gn=DOCK10;refGene_tx=XM_019838748.2;refGene_gn=DOCK10;refGene_tx=XM_019838749.2;refGene_gn=DOCK10;refGene_tx=XM_019838750.2;refGene_gn=DOCK10;refGene_tx=XM_023259739.1;refGene_gn=DOCK10;refGene_tx=XM_023259740.1
C1 209109916 209109996
refGene_gn=DOCK10;refGene_tx=XM_011285652.3;refGene_gn=DOCK10;refGene_tx=XM_019838741.2;refGene_gn=DOCK10;refGene_tx=XM_019838742.2;refGene_gn=DOCK10;refGene_tx=XM_019838743.2;refGene_gn=DOCK10;refGene_tx=XM_019838744.2;refGene_gn=DOCK10;refGene_tx=XM_019838745.2;refGene_gn=DOCK10;refGene_tx=XM_019838746.2;refGene_gn=DOCK10;refGene_tx=XM_019838748.

2;refGene_gn=DOCK10;refGene_tx=XM_019838749.2;refGene_gn=DOCK10;refGene_tx=XM_019838750.2;refGene_gn=DOCK10;refGene_tx=XM_023259739.1;refGene_gn=DOCK10;refGene_tx=XM_023259740.1
C1 209112954 209113077
refGene_gn=DOCK10;refGene_tx=XM_011285652.3;refGene_gn=DOCK10;refGene_tx=XM_019838741.2;refGene_gn=DOCK10;refGene_tx=XM_019838742.2;refGene_gn=DOCK10;refGene_tx=XM_019838743.2;refGene_gn=DOCK10;refGene_tx=XM_019838744.2;refGene_gn=DOCK10;refGene_tx=XM_019838745.2;refGene_gn=DOCK10;refGene_tx=XM_019838746.2;refGene_gn=DOCK10;refGene_tx=XM_019838748.2;refGene_gn=DOCK10;refGene_tx=XM_019838749.2;refGene_gn=DOCK10;refGene_tx=XM_019838750.2;refGene_gn=DOCK10;refGene_tx=XM_023259739.1;refGene_gn=DOCK10;refGene_tx=XM_023259740.1
C1 209118605 209118688
refGene_gn=DOCK10;refGene_tx=XM_011285652.3;refGene_gn=DOCK10;refGene_tx=XM_019838741.2;refGene_gn=DOCK10;refGene_tx=XM_019838742.2;refGene_gn=DOCK10;refGene_tx=XM_019838743.2;refGene_gn=DOCK10;refGene_tx=XM_019838744.2;refGene_gn=DOCK10;refGene_tx=XM_019838745.2;refGene_gn=DOCK10;refGene_tx=XM_019838746.2;refGene_gn=DOCK10;refGene_tx=XM_019838748.2;refGene_gn=DOCK10;refGene_tx=XM_019838749.2;refGene_gn=DOCK10;refGene_tx=XM_019838750.2;refGene_gn=DOCK10;refGene_tx=XM_023259739.1;refGene_gn=DOCK10;refGene_tx=XM_023259740.1
C1 209119877 209120000
refGene_gn=DOCK10;refGene_tx=XM_011285652.3;refGene_gn=DOCK10;refGene_tx=XM_019838741.2;refGene_gn=DOCK10;refGene_tx=XM_019838742.2;refGene_gn=DOCK10;refGene_tx=XM_019838743.2;refGene_gn=DOCK10;refGene_tx=XM_019838744.2;refGene_gn=DOCK10;refGene_tx=XM_019838745.2;refGene_gn=DOCK10;refGene_tx=XM_019838746.2;refGene_gn=DOCK10;refGene_tx=XM_019838748.2;refGene_gn=DOCK10;refGene_tx=XM_019838749.2;refGene_gn=DOCK10;refGene_tx=XM_019838750.2;refGene_gn=DOCK10;refGene_tx=XM_023259739.1;refGene_gn=DOCK10;refGene_tx=XM_023259740.1
C1 209120187 209120409
refGene_gn=DOCK10;refGene_tx=XM_011285652.3;refGene_gn=DOCK10;refGene_tx=XM_019838741.2;refGene_gn=DOCK10;refGene_tx=XM_019838742.2;refGene_gn=DOCK10;refGene_tx=XM_019838743.2;refGene_gn=DOCK10;refGene_tx=XM_019838744.2;refGene_gn=DOCK10;refGene_tx=XM_019838745.2;refGene_gn=DOCK10;refGene_tx=XM_019838746.2;refGene_gn=DOCK10;refGene_tx=XM_019838748.2;refGene_gn=DOCK10;refGene_tx=XM_019838749.2;refGene_gn=DOCK10;refGene_tx=XM_019838750.2;refGene_gn=DOCK10;refGene_tx=XM_023259739.1;refGene_gn=DOCK10;refGene_tx=XM_023259740.1
C1 209127954 209128110
refGene_gn=DOCK10;refGene_tx=XM_011285652.3;refGene_gn=DOCK10;refGene_tx=XM_019838741.2;refGene_gn=DOCK10;refGene_tx=XM_019838742.2;refGene_gn=DOCK10;refGene_tx=XM_019838743.2;refGene_gn=DOCK10;refGene_tx=XM_019838744.2;refGene_gn=DOCK10;refGene_tx=XM_019838745.2;refGene_gn=DOCK10;refGene_tx=XM_019838746.2;refGene_gn=DOCK10;refGene_tx=XM_019838748.2;refGene_gn=DOCK10;refGene_tx=XM_019838749.2;refGene_gn=DOCK10;refGene_tx=XM_019838750.2;refGene_gn=DOCK10;refGene_tx=XM_023259739.1;refGene_gn=DOCK10;refGene_tx=XM_023259740.1
C1 209128231 209128315
refGene_gn=DOCK10;refGene_tx=XM_011285652.3;refGene_gn=DOCK10;refGene_

tx=XM_019838741.2;refGene_gn=DOCK10;refGene_tx=XM_019838742.2;refGene_gn=DOCK10;refGene_tx=XM_019838743.2;refGene_gn=DOCK10;refGene_tx=XM_019838744.2;refGene_gn=DOCK10;refGene_tx=XM_019838745.2;refGene_gn=DOCK10;refGene_tx=XM_019838746.2;refGene_gn=DOCK10;refGene_tx=XM_019838748.2;refGene_gn=DOCK10;refGene_tx=XM_019838749.2;refGene_gn=DOCK10;refGene_tx=XM_019838750.2;refGene_gn=DOCK10;refGene_tx=XM_023259739.1;refGene_gn=DOCK10;refGene_tx=XM_023259740.1
C1 209128590 209128676
refGene_gn=DOCK10;refGene_tx=XM_011285652.3;refGene_gn=DOCK10;refGene_tx=XM_019838741.2;refGene_gn=DOCK10;refGene_tx=XM_019838742.2;refGene_gn=DOCK10;refGene_tx=XM_019838743.2;refGene_gn=DOCK10;refGene_tx=XM_019838744.2;refGene_gn=DOCK10;refGene_tx=XM_019838745.2;refGene_gn=DOCK10;refGene_tx=XM_019838746.2;refGene_gn=DOCK10;refGene_tx=XM_019838748.2;refGene_gn=DOCK10;refGene_tx=XM_019838749.2;refGene_gn=DOCK10;refGene_tx=XM_019838750.2;refGene_gn=DOCK10;refGene_tx=XM_023259739.1;refGene_gn=DOCK10;refGene_tx=XM_023259740.1
C1 209129938 209130122
refGene_gn=DOCK10;refGene_tx=XM_011285652.3;refGene_gn=DOCK10;refGene_tx=XM_019838741.2;refGene_gn=DOCK10;refGene_tx=XM_019838742.2;refGene_gn=DOCK10;refGene_tx=XM_019838743.2;refGene_gn=DOCK10;refGene_tx=XM_019838744.2;refGene_gn=DOCK10;refGene_tx=XM_019838745.2;refGene_gn=DOCK10;refGene_tx=XM_019838746.2;refGene_gn=DOCK10;refGene_tx=XM_019838748.2;refGene_gn=DOCK10;refGene_tx=XM_019838749.2;refGene_gn=DOCK10;refGene_tx=XM_019838750.2;refGene_gn=DOCK10;refGene_tx=XM_023259739.1;refGene_gn=DOCK10;refGene_tx=XM_023259740.1
C1 209140522 209140657
refGene_gn=DOCK10;refGene_tx=XM_011285652.3;refGene_gn=DOCK10;refGene_tx=XM_019838741.2;refGene_gn=DOCK10;refGene_tx=XM_019838742.2;refGene_gn=DOCK10;refGene_tx=XM_019838743.2;refGene_gn=DOCK10;refGene_tx=XM_019838744.2;refGene_gn=DOCK10;refGene_tx=XM_019838745.2;refGene_gn=DOCK10;refGene_tx=XM_019838746.2;refGene_gn=DOCK10;refGene_tx=XM_019838748.2;refGene_gn=DOCK10;refGene_tx=XM_019838749.2;refGene_gn=DOCK10;refGene_tx=XM_019838750.2;refGene_gn=DOCK10;refGene_tx=XM_023259739.1;refGene_gn=DOCK10;refGene_tx=XM_023259740.1
C1 209140910 209141033
refGene_gn=DOCK10;refGene_tx=XM_011285652.3;refGene_gn=DOCK10;refGene_tx=XM_019838741.2;refGene_gn=DOCK10;refGene_tx=XM_019838742.2;refGene_gn=DOCK10;refGene_tx=XM_019838743.2;refGene_gn=DOCK10;refGene_tx=XM_019838744.2;refGene_gn=DOCK10;refGene_tx=XM_019838745.2;refGene_gn=DOCK10;refGene_tx=XM_019838746.2;refGene_gn=DOCK10;refGene_tx=XM_019838748.2;refGene_gn=DOCK10;refGene_tx=XM_019838749.2;refGene_gn=DOCK10;refGene_tx=XM_019838750.2;refGene_gn=DOCK10;refGene_tx=XM_023259739.1;refGene_gn=DOCK10;refGene_tx=XM_023259740.1
C1 209141315 209141388
refGene_gn=DOCK10;refGene_tx=XM_011285652.3;refGene_gn=DOCK10;refGene_tx=XM_019838741.2;refGene_gn=DOCK10;refGene_tx=XM_019838742.2;refGene_gn=DOCK10;refGene_tx=XM_019838743.2;refGene_gn=DOCK10;refGene_tx=XM_019838744.2;refGene_gn=DOCK10;refGene_tx=XM_019838745.2;refGene_gn=DOCK10;refGene_tx=XM_019838746.2;refGene_gn=DOCK10;refGene_tx=XM_019838748.2;refGene_gn=DOCK10;refGene_tx=XM_019838749.2;refGene_gn=DOCK10;refGen

e_tx=XM_019838750.2;refGene_gn=DOCK10;refGene_tx=XM_023259739.1;refGene_gn=DOCK10;refGene_tx=XM_023259740.1
C1 209149448 209149531
refGene_gn=DOCK10;refGene_tx=XM_011285652.3;refGene_gn=DOCK10;refGene_tx=XM_019838741.2;refGene_gn=DOCK10;refGene_tx=XM_019838742.2;refGene_gn=DOCK10;refGene_tx=XM_019838743.2;refGene_gn=DOCK10;refGene_tx=XM_019838744.2;refGene_gn=DOCK10;refGene_tx=XM_019838745.2;refGene_gn=DOCK10;refGene_tx=XM_019838746.2;refGene_gn=DOCK10;refGene_tx=XM_019838748.2;refGene_gn=DOCK10;refGene_tx=XM_019838749.2;refGene_gn=DOCK10;refGene_tx=XM_019838750.2;refGene_gn=DOCK10;refGene_tx=XM_023259739.1;refGene_gn=DOCK10;refGene_tx=XM_023259740.1
C1 209168119 209168209
refGene_gn=DOCK10;refGene_tx=XM_011285652.3;refGene_gn=DOCK10;refGene_tx=XM_019838741.2;refGene_gn=DOCK10;refGene_tx=XM_019838742.2;refGene_gn=DOCK10;refGene_tx=XM_019838743.2;refGene_gn=DOCK10;refGene_tx=XM_019838744.2;refGene_gn=DOCK10;refGene_tx=XM_019838745.2;refGene_gn=DOCK10;refGene_tx=XM_019838746.2;refGene_gn=DOCK10;refGene_tx=XM_019838748.2;refGene_gn=DOCK10;refGene_tx=XM_019838749.2;refGene_gn=DOCK10;refGene_tx=XM_019838750.2;refGene_gn=DOCK10;refGene_tx=XM_023259739.1;refGene_gn=DOCK10;refGene_tx=XM_023259740.1
C1 209182821 209182941
refGene_gn=DOCK10;refGene_tx=XM_011285652.3;refGene_gn=DOCK10;refGene_tx=XM_019838741.2;refGene_gn=DOCK10;refGene_tx=XM_019838742.2;refGene_gn=DOCK10;refGene_tx=XM_019838743.2;refGene_gn=DOCK10;refGene_tx=XM_019838744.2;refGene_gn=DOCK10;refGene_tx=XM_019838745.2;refGene_gn=DOCK10;refGene_tx=XM_019838746.2;refGene_gn=DOCK10;refGene_tx=XM_019838748.2;refGene_gn=DOCK10;refGene_tx=XM_019838749.2;refGene_gn=DOCK10;refGene_tx=XM_019838750.2;refGene_gn=DOCK10;refGene_tx=XM_023259739.1;refGene_gn=DOCK10;refGene_tx=XM_023259740.1
C1 209197910 209198135
refGene_gn=DOCK10;refGene_tx=XM_011285652.3;refGene_gn=DOCK10;refGene_tx=XM_019838741.2;refGene_gn=DOCK10;refGene_tx=XM_019838742.2;refGene_gn=DOCK10;refGene_tx=XM_019838743.2;refGene_gn=DOCK10;refGene_tx=XM_019838744.2;refGene_gn=DOCK10;refGene_tx=XM_019838745.2;refGene_gn=DOCK10;refGene_tx=XM_019838749.2;refGene_gn=DOCK10;refGene_tx=XM_019838750.2
C1 209235931 209236093
refGene_gn=DOCK10;refGene_tx=XM_019838746.2;refGene_gn=DOCK10;refGene_tx=XM_019838748.2
C1 209296789 209296912
refGene_gn=DOCK10;refGene_tx=XM_023259739.1;refGene_gn=DOCK10;refGene_tx=XM_023259740.1
C1 209602807 209603028
refGene_gn=NYAP2;refGene_tx=XM_003991216.4;refGene_gn=NYAP2;refGene_tx=XM_019838751.1;refGene_gn=NYAP2;refGene_tx=XM_019838752.2;refGene_gn=NYAP2;refGene_tx=XM_019838753.2;refGene_gn=NYAP2;refGene_tx=XM_019838754.2;refGene_gn=NYAP2;refGene_tx=XM_019838755.2;refGene_gn=NYAP2;refGene_tx=XM_019838756.2
C1 209705615 209705917
refGene_gn=NYAP2;refGene_tx=XM_003991216.4;refGene_gn=NYAP2;refGene_tx

=XM_019838751.1;refGene_gn=NYAP2;refGene_tx=XM_019838752.2;refGene_gn=NYAP2;refGene_tx=XM_019838753.2;refGene_gn=NYAP2;refGene_tx=XM_019838754.2;refGene_gn=NYAP2;refGene_tx=XM_019838755.2;refGene_gn=NYAP2;refGene_tx=XM_019838756.2;refGene_gn=NYAP2;refGene_tx=XM_019838759.1
C1 209752987 209754094
refGene_gn=NYAP2;refGene_tx=XM_003991216.4;refGene_gn=NYAP2;refGene_tx=XM_019838751.1;refGene_gn=NYAP2;refGene_tx=XM_019838752.2;refGene_gn=NYAP2;refGene_tx=XM_019838753.2;refGene_gn=NYAP2;refGene_tx=XM_019838754.2;refGene_gn=NYAP2;refGene_tx=XM_019838755.2;refGene_gn=NYAP2;refGene_tx=XM_019838756.2;refGene_gn=NYAP2;refGene_tx=XM_019838759.1
C1 209789889 209790096
refGene_gn=NYAP2;refGene_tx=XM_003991216.4;refGene_gn=NYAP2;refGene_tx=XM_019838751.1;refGene_gn=NYAP2;refGene_tx=XM_019838752.2;refGene_gn=NYAP2;refGene_tx=XM_019838753.2;refGene_gn=NYAP2;refGene_tx=XM_019838754.2;refGene_gn=NYAP2;refGene_tx=XM_019838756.2;refGene_gn=NYAP2;refGene_tx=XM_019838759.1
C1 209817009 209817143
refGene_gn=NYAP2;refGene_tx=XM_003991216.4;refGene_gn=NYAP2;refGene_tx=XM_019838751.1;refGene_gn=NYAP2;refGene_tx=XM_019838752.2;refGene_gn=NYAP2;refGene_tx=XM_019838753.2;refGene_gn=NYAP2;refGene_tx=XM_019838754.2;refGene_gn=NYAP2;refGene_tx=XM_019838755.2;refGene_gn=NYAP2;refGene_tx=XM_019838756.2;refGene_gn=NYAP2;refGene_tx=XM_019838759.1
C1 209817651 209817663
refGene_gn=NYAP2;refGene_tx=XM_019838756.2
C1 209852630 209852703
refGene_gn=NYAP2;refGene_tx=XM_019838753.2;refGene_gn=NYAP2;refGene_tx=XM_019838754.2
C1 209858360 209858588
refGene_gn=NYAP2;refGene_tx=XM_019838751.1;refGene_gn=NYAP2;refGene_tx=XM_019838755.2;refGene_gn=NYAP2;refGene_tx=XM_019838759.1
C1 209860829 209860889
refGene_gn=NYAP2;refGene_tx=XM_019838752.2
C1 210835204 210838909
refGene_gn=IRS1;refGene_tx=XM_003991219.3
C1 210899568 210900001
refGene_gn=RHBDD1;refGene_tx=XM_023259741.1;refGene_gn=RHBDD1;refGene_tx=XM_023259742.1;refGene_gn=RHBDD1;refGene_tx=XM_023259743.1;refGene_gn=RHBDD1;refGene_tx=XM_023259744.1;refGene_gn=RHBDD1;refGene_tx=XM_023259745.1;refGene_gn=RHBDD1;refGene_tx=XM_023259746.1;refGene_gn=RHBDD1;refGene_tx=XM_023259747.1;refGene_gn=RHBDD1;refGene_tx=XM_023259748.1
C1 210901074 210901207
refGene_gn=RHBDD1;refGene_tx=XM_023259741.1;refGene_gn=RHBDD1;refGene_tx=XM_023259742.1;refGene_gn=RHBDD1;refGene_tx=XM_023259743.1;refGene_gn=RHBDD1;refGene_tx=XM_023259744.1;refGene_gn=RHBDD1;refGene_tx=XM_023259745.1;refGene_gn=RHBDD1;refGene_tx=XM_023259746.1;refGene_gn=RHBDD1;refGene_tx=XM_023259747.1;refGene_gn=RHBDD1;refGene_tx=XM_023259748.1
C1 210938896 210938985
refGene_gn=RHBDD1;refGene_tx=XM_023259741.1;refGene_gn=RHBDD1;refGene_

tx=XM_023259742.1;refGene_gn=RHBDD1;refGene_tx=XM_023259743.1;refGene_gn=RHBDD1;refGene_tx=XM_023259744.1;refGene_gn=RHBDD1;refGene_tx=XM_023259745.1;refGene_gn=RHBDD1;refGene_tx=XM_023259746.1;refGene_gn=RHBDD1;refGene_tx=XM_023259747.1;refGene_gn=RHBDD1;refGene_tx=XM_023259748.1
C1 210942307 210942364
refGene_gn=RHBDD1;refGene_tx=XM_023259741.1;refGene_gn=RHBDD1;refGene_tx=XM_023259742.1;refGene_gn=RHBDD1;refGene_tx=XM_023259743.1;refGene_gn=RHBDD1;refGene_tx=XM_023259744.1;refGene_gn=RHBDD1;refGene_tx=XM_023259745.1;refGene_gn=RHBDD1;refGene_tx=XM_023259746.1;refGene_gn=RHBDD1;refGene_tx=XM_023259747.1;refGene_gn=RHBDD1;refGene_tx=XM_023259748.1
C1 210951910 210952054
refGene_gn=RHBDD1;refGene_tx=XM_023259741.1;refGene_gn=RHBDD1;refGene_tx=XM_023259742.1;refGene_gn=RHBDD1;refGene_tx=XM_023259743.1;refGene_gn=RHBDD1;refGene_tx=XM_023259744.1;refGene_gn=RHBDD1;refGene_tx=XM_023259745.1;refGene_gn=RHBDD1;refGene_tx=XM_023259746.1;refGene_gn=RHBDD1;refGene_tx=XM_023259747.1;refGene_gn=RHBDD1;refGene_tx=XM_023259748.1
C1 210954602 210954640
refGene_gn=RHBDD1;refGene_tx=XM_023259747.1
C1 210958248 210958355
refGene_gn=RHBDD1;refGene_tx=XM_023259741.1;refGene_gn=RHBDD1;refGene_tx=XM_023259742.1;refGene_gn=RHBDD1;refGene_tx=XM_023259743.1;refGene_gn=RHBDD1;refGene_tx=XM_023259744.1;refGene_gn=RHBDD1;refGene_tx=XM_023259745.1
C1 210959456 210959473
refGene_gn=RHBDD1;refGene_tx=XM_023259748.1
C1 211005458 211005553
refGene_gn=RHBDD1;refGene_tx=XM_023259746.1
C1 211017609 211017873
refGene_gn=COL4A4;refGene_tx=XM_023259749.1
C1 211018301 211018588
refGene_gn=COL4A4;refGene_tx=XM_023259749.1
C1 211020317 211020506
refGene_gn=COL4A4;refGene_tx=XM_023259749.1
C1 211022024 211022141
refGene_gn=COL4A4;refGene_tx=XM_023259749.1
C1 211027609 211027735
refGene_gn=COL4A4;refGene_tx=XM_023259749.1
C1 211031500 211031509
refGene_gn=COL4A4;refGene_tx=XM_023259749.1
C1 211032615 211032723
refGene_gn=COL4A4;refGene_tx=XM_023259749.1
C1 211036132 211036288
refGene_gn=COL4A4;refGene_tx=XM_023259749.1
C1 211037517 211037631
refGene_gn=COL4A4;refGene_tx=XM_023259749.1
C1 211037722 211037851
refGene_gn=COL4A4;refGene_tx=XM_023259749.1

C1 211039239 211039311
refGene_gn=COL4A4;refGene_tx=XM_023259749.1
C1 211045385 211045493
refGene_gn=COL4A4;refGene_tx=XM_023259749.1
C1 211046093 211046201
refGene_gn=COL4A4;refGene_tx=XM_023259749.1
C1 211050027 211050102
refGene_gn=COL4A4;refGene_tx=XM_023259749.1
C1 211054525 211054589
refGene_gn=COL4A4;refGene_tx=XM_023259749.1
C1 211055677 211055859
refGene_gn=COL4A4;refGene_tx=XM_023259749.1
C1 211056942 211057050
refGene_gn=COL4A4;refGene_tx=XM_023259749.1
C1 211059894 211060038
refGene_gn=COL4A4;refGene_tx=XM_023259749.1
C1 211060511 211060682
refGene_gn=COL4A4;refGene_tx=XM_023259749.1
C1 211061638 211061800
refGene_gn=COL4A4;refGene_tx=XM_023259749.1
C1 211064178 211064397
refGene_gn=COL4A4;refGene_tx=XM_023259749.1
C1 211064908 211065016
refGene_gn=COL4A4;refGene_tx=XM_023259749.1
C1 211066234 211066303
refGene_gn=COL4A4;refGene_tx=XM_023259749.1
C1 211067705 211067889
refGene_gn=COL4A4;refGene_tx=XM_023259749.1
C1 211070221 211070328
refGene_gn=COL4A4;refGene_tx=XM_023259749.1
C1 211071499 211071572
refGene_gn=COL4A4;refGene_tx=XM_023259749.1
C1 211076290 211076454
refGene_gn=COL4A4;refGene_tx=XM_023259749.1
C1 211077418 211077508
refGene_gn=COL4A4;refGene_tx=XM_023259749.1
C1 211080523 211080688
refGene_gn=COL4A4;refGene_tx=XM_023259749.1
C1 211083960 211083991
refGene_gn=COL4A4;refGene_tx=XM_023259749.1
C1 211083992 211084066
refGene_gn=COL4A4;refGene_tx=XM_023259749.1
C1 211084832 211084902
refGene_gn=COL4A4;refGene_tx=XM_023259749.1
C1 211085592 211085646
refGene_gn=COL4A4;refGene_tx=XM_023259749.1
C1 211085967 211086012
refGene_gn=COL4A4;refGene_tx=XM_023259749.1
C1 211086874 211086934
refGene_gn=COL4A4;refGene_tx=XM_023259749.1

C1 211087201 211087255
refGene_gn=COL4A4;refGene_tx=XM_023259749.1
C1 211088315 211088396
refGene_gn=COL4A4;refGene_tx=XM_023259749.1
C1 211091091 211091133
refGene_gn=COL4A4;refGene_tx=XM_023259749.1
C1 211091357 211091393
refGene_gn=COL4A4;refGene_tx=XM_023259749.1
C1 211091793 211091856
refGene_gn=COL4A4;refGene_tx=XM_023259749.1
C1 211093927 211093963
refGene_gn=COL4A4;refGene_tx=XM_023259749.1
C1 211096817 211096886
refGene_gn=COL4A4;refGene_tx=XM_023259749.1
C1 211100223 211100340
refGene_gn=COL4A4;refGene_tx=XM_023259749.1
C1 211100983 211101028
refGene_gn=COL4A4;refGene_tx=XM_023259749.1
C1 211101815 211101950
refGene_gn=COL4A4;refGene_tx=XM_023259749.1
C1 211122240 211122318
refGene_gn=COL4A4;refGene_tx=XM_023259749.1
C1 211126468 211126511
refGene_gn=COL4A4;refGene_tx=XM_023259749.1
C1 211129700 211129756
refGene_gn=COL4A4;refGene_tx=XM_023259749.1
C1 211148280 211148367
refGene_gn=COL4A3;refGene_tx=XM_019838774.2;refGene_gn=COL4A3;refGene_
tx=XM_019838775.2
C1 211213118 211213175
refGene_gn=COL4A3;refGene_tx=XM_019838774.2;refGene_gn=COL4A3;refGene_
tx=XM_019838775.2
C1 211215271 211215361
refGene_gn=COL4A3;refGene_tx=XM_019838774.2;refGene_gn=COL4A3;refGene_
tx=XM_019838775.2
C1 211218079 211218124
refGene_gn=COL4A3;refGene_tx=XM_019838774.2;refGene_gn=COL4A3;refGene_
tx=XM_019838775.2
C1 211218706 211218751
refGene_gn=COL4A3;refGene_tx=XM_019838774.2;refGene_gn=COL4A3;refGene_
tx=XM_019838775.2
C1 211220347 211220410
refGene_gn=COL4A3;refGene_tx=XM_019838774.2;refGene_gn=COL4A3;refGene_
tx=XM_019838775.2
C1 211221085 211221139
refGene_gn=COL4A3;refGene_tx=XM_019838774.2;refGene_gn=COL4A3;refGene_
tx=XM_019838775.2
C1 211221932 211221959
refGene_gn=COL4A3;refGene_tx=XM_019838774.2;refGene_gn=COL4A3;refGene_
tx=XM_019838775.2

C1	211222716	211222791
refGene_gn=COL4A3;refGene_tx=XM_019838774.2;refGene_gn=COL4A3;refGene_
tx=XM_019838775.2
C1	211224932	211224995
refGene_gn=COL4A3;refGene_tx=XM_019838774.2;refGene_gn=COL4A3;refGene_
tx=XM_019838775.2
C1	211225126	211225162
refGene_gn=COL4A3;refGene_tx=XM_019838774.2;refGene_gn=COL4A3;refGene_
tx=XM_019838775.2
C1	211225971	211226013
refGene_gn=COL4A3;refGene_tx=XM_019838774.2;refGene_gn=COL4A3;refGene_
tx=XM_019838775.2
C1	211226237	211226315
refGene_gn=COL4A3;refGene_tx=XM_019838774.2;refGene_gn=COL4A3;refGene_
tx=XM_019838775.2
C1	211228014	211228080
refGene_gn=COL4A3;refGene_tx=XM_019838774.2;refGene_gn=COL4A3;refGene_
tx=XM_019838775.2
C1	211229002	211229062
refGene_gn=COL4A3;refGene_tx=XM_019838774.2;refGene_gn=COL4A3;refGene_
tx=XM_019838775.2
C1	211230478	211230523
refGene_gn=COL4A3;refGene_tx=XM_019838774.2;refGene_gn=COL4A3;refGene_
tx=XM_019838775.2
C1	211230800	211230854
refGene_gn=COL4A3;refGene_tx=XM_019838774.2;refGene_gn=COL4A3;refGene_
tx=XM_019838775.2
C1	211231715	211231757
refGene_gn=COL4A3;refGene_tx=XM_019838774.2;refGene_gn=COL4A3;refGene_
tx=XM_019838775.2
C1	211234319	211234404
refGene_gn=COL4A3;refGene_tx=XM_019838774.2;refGene_gn=COL4A3;refGene_
tx=XM_019838775.2
C1	211235600	211235636
refGene_gn=COL4A3;refGene_tx=XM_019838774.2;refGene_gn=COL4A3;refGene_
tx=XM_019838775.2
C1	211237344	211237509
refGene_gn=COL4A3;refGene_tx=XM_019838774.2;refGene_gn=COL4A3;refGene_
tx=XM_019838775.2
C1	211241589	211241682
refGene_gn=COL4A3;refGene_tx=XM_019838774.2;refGene_gn=COL4A3;refGene_
tx=XM_019838775.2
C1	211242432	211242528
refGene_gn=COL4A3;refGene_tx=XM_019838774.2;refGene_gn=COL4A3;refGene_
tx=XM_019838775.2
C1	211245717	211245788
refGene_gn=COL4A3;refGene_tx=XM_019838774.2;refGene_gn=COL4A3;refGene_
tx=XM_019838775.2
C1	211246577	211246760
refGene_gn=COL4A3;refGene_tx=XM_019838774.2;refGene_gn=COL4A3;refGene_

tx=XM_019838775.2
C1	211248619	211248788
refGene_gn=COL4A3;refGene_tx=XM_019838774.2;refGene_gn=COL4A3;refGene_tx=XM_019838775.2
C1	211250732	211250825
refGene_gn=COL4A3;refGene_tx=XM_019838774.2;refGene_gn=COL4A3;refGene_tx=XM_019838775.2
C1	211251504	211251606
refGene_gn=COL4A3;refGene_tx=XM_019838774.2;refGene_gn=COL4A3;refGene_tx=XM_019838775.2
C1	211253712	211253810
refGene_gn=COL4A3;refGene_tx=XM_019838774.2;refGene_gn=COL4A3;refGene_tx=XM_019838775.2
C1	211254582	211254733
refGene_gn=COL4A3;refGene_tx=XM_019838774.2;refGene_gn=COL4A3;refGene_tx=XM_019838775.2
C1	211255072	211255186
refGene_gn=COL4A3;refGene_tx=XM_019838774.2;refGene_gn=COL4A3;refGene_tx=XM_019838775.2
C1	211256159	211256327
refGene_gn=COL4A3;refGene_tx=XM_019838774.2;refGene_gn=COL4A3;refGene_tx=XM_019838775.2
C1	211258208	211258298
refGene_gn=COL4A3;refGene_tx=XM_019838774.2;refGene_gn=COL4A3;refGene_tx=XM_019838775.2
C1	211258629	211258764
refGene_gn=COL4A3;refGene_tx=XM_019838774.2;refGene_gn=COL4A3;refGene_tx=XM_019838775.2
C1	211261208	211261307
refGene_gn=COL4A3;refGene_tx=XM_019838774.2;refGene_gn=COL4A3;refGene_tx=XM_019838775.2
C1	211262357	211262447
refGene_gn=COL4A3;refGene_tx=XM_019838774.2;refGene_gn=COL4A3;refGene_tx=XM_019838775.2
C1	211262746	211262886
refGene_gn=COL4A3;refGene_tx=XM_019838774.2;refGene_gn=COL4A3;refGene_tx=XM_019838775.2
C1	211263342	211263469
refGene_gn=COL4A3;refGene_tx=XM_019838774.2;refGene_gn=COL4A3;refGene_tx=XM_019838775.2
C1	211264764	211264845
refGene_gn=COL4A3;refGene_tx=XM_019838774.2;refGene_gn=COL4A3;refGene_tx=XM_019838775.2
C1	211265227	211265326
refGene_gn=COL4A3;refGene_tx=XM_019838774.2;refGene_gn=COL4A3;refGene_tx=XM_019838775.2
C1	211265560	211265608
refGene_gn=COL4A3;refGene_tx=XM_019838774.2;refGene_gn=COL4A3;refGene_tx=XM_019838775.2
C1	211268391	211268577

refGene_gn=COL4A3;refGene_tx=XM_019838774.2;refGene_gn=COL4A3;refGene_tx=XM_019838775.2
C1 211269742 211269873
refGene_gn=COL4A3;refGene_tx=XM_019838774.2;refGene_gn=COL4A3;refGene_tx=XM_019838775.2
C1 211272703 211272776
refGene_gn=COL4A3;refGene_tx=XM_019838774.2;refGene_gn=COL4A3;refGene_tx=XM_019838775.2
C1 211275083 211275155
refGene_gn=COL4A3;refGene_tx=XM_019838774.2;refGene_gn=COL4A3;refGene_tx=XM_019838775.2
C1 211275243 211275369
refGene_gn=COL4A3;refGene_tx=XM_019838774.2;refGene_gn=COL4A3;refGene_tx=XM_019838775.2
C1 211276157 211276256
refGene_gn=COL4A3;refGene_tx=XM_019838774.2;refGene_gn=COL4A3;refGene_tx=XM_019838775.2
C1 211279268 211279481
refGene_gn=COL4A3;refGene_tx=XM_019838774.2;refGene_gn=COL4A3;refGene_tx=XM_019838775.2
C1 211280114 211280292
refGene_gn=COL4A3;refGene_tx=XM_019838774.2;refGene_gn=COL4A3;refGene_tx=XM_019838775.2
C1 211280421 211280536
refGene_gn=COL4A3;refGene_tx=XM_019838774.2;refGene_gn=COL4A3;refGene_tx=XM_019838775.2
C1 211281548 211281721
refGene_gn=COL4A3;refGene_tx=XM_019838774.2;refGene_gn=COL4A3;refGene_tx=XM_019838775.2
C1 211282317 211282402
refGene_gn=COL4A3;refGene_tx=XM_019838774.2;refGene_gn=COL4A3;refGene_tx=XM_019838775.2
C1 211293996 211294034
refGene_gn=MFF;refGene_tx=XM_019838780.1;refGene_gn=MFF;refGene_tx=XM_019838782.1;refGene_gn=MFF;refGene_tx=XM_019838791.1
C1 211294881 211294931
refGene_gn=MFF;refGene_tx=XM_019838776.1;refGene_gn=MFF;refGene_tx=XM_019838779.2;refGene_gn=MFF;refGene_tx=XM_019838781.1;refGene_gn=MFF;refGene_tx=XM_019838783.1;refGene_gn=MFF;refGene_tx=XM_019838787.1;refGene_gn=MFF;refGene_tx=XM_019838788.1;refGene_gn=MFF;refGene_tx=XM_023259751.1;refGene_gn=MFF;refGene_tx=XM_023259752.1;refGene_gn=MFF;refGene_tx=XM_023259753.1
C1 211295020 211295241
refGene_gn=MFF;refGene_tx=XM_019838776.1;refGene_gn=MFF;refGene_tx=XM_019838779.2;refGene_gn=MFF;refGene_tx=XM_019838780.1;refGene_gn=MFF;refGene_tx=XM_019838781.1;refGene_gn=MFF;refGene_tx=XM_019838782.1;refGene_gn=MFF;refGene_tx=XM_019838783.1;refGene_gn=MFF;refGene_tx=XM_019838784.1;refGene_gn=MFF;refGene_tx=XM_019838785.1;refGene_gn=MFF;refGene_tx=XM_019838787.1;refGene_gn=MFF;refGene_tx=XM_019838788.1;refGene_gn=MFF;refGene_tx=XM_019838791.1;refGene_gn=MFF;refGene_tx=XM_023259751.

1;refGene_gn=MFF;refGene_tx=XM_023259752.1;refGene_gn=MFF;refGene_tx=XM_023259753.1
C1 211296076 211296246
refGene_gn=MFF;refGene_tx=XM_019838776.1;refGene_gn=MFF;refGene_tx=XM_019838779.2;refGene_gn=MFF;refGene_tx=XM_019838780.1;refGene_gn=MFF;refGene_tx=XM_019838781.1;refGene_gn=MFF;refGene_tx=XM_019838782.1;refGene_gn=MFF;refGene_tx=XM_019838783.1;refGene_gn=MFF;refGene_tx=XM_019838784.1;refGene_gn=MFF;refGene_tx=XM_019838785.1;refGene_gn=MFF;refGene_tx=XM_019838787.1;refGene_gn=MFF;refGene_tx=XM_019838788.1;refGene_gn=MFF;refGene_tx=XM_019838791.1;refGene_gn=MFF;refGene_tx=XM_023259751.1;refGene_gn=MFF;refGene_tx=XM_023259752.1;refGene_gn=MFF;refGene_tx=XM_023259753.1
C1 211304642 211304731
refGene_gn=MFF;refGene_tx=XM_019838776.1;refGene_gn=MFF;refGene_tx=XM_019838779.2;refGene_gn=MFF;refGene_tx=XM_019838780.1;refGene_gn=MFF;refGene_tx=XM_019838781.1;refGene_gn=MFF;refGene_tx=XM_019838782.1;refGene_gn=MFF;refGene_tx=XM_019838783.1;refGene_gn=MFF;refGene_tx=XM_019838784.1;refGene_gn=MFF;refGene_tx=XM_019838785.1;refGene_gn=MFF;refGene_tx=XM_019838787.1;refGene_gn=MFF;refGene_tx=XM_019838788.1;refGene_gn=MFF;refGene_tx=XM_019838791.1;refGene_gn=MFF;refGene_tx=XM_023259751.1;refGene_gn=MFF;refGene_tx=XM_023259752.1;refGene_gn=MFF;refGene_tx=XM_023259753.1
C1 211308042 211308117
refGene_gn=MFF;refGene_tx=XM_019838776.1;refGene_gn=MFF;refGene_tx=XM_019838779.2;refGene_gn=MFF;refGene_tx=XM_019838780.1;refGene_gn=MFF;refGene_tx=XM_019838781.1;refGene_gn=MFF;refGene_tx=XM_019838782.1;refGene_gn=MFF;refGene_tx=XM_019838784.1;refGene_gn=MFF;refGene_tx=XM_019838785.1;refGene_gn=MFF;refGene_tx=XM_019838787.1
C1 211312792 211312951
refGene_gn=MFF;refGene_tx=XM_019838776.1;refGene_gn=MFF;refGene_tx=XM_019838779.2;refGene_gn=MFF;refGene_tx=XM_019838780.1;refGene_gn=MFF;refGene_tx=XM_019838781.1;refGene_gn=MFF;refGene_tx=XM_019838782.1;refGene_gn=MFF;refGene_tx=XM_019838783.1;refGene_gn=MFF;refGene_tx=XM_019838784.1;refGene_gn=MFF;refGene_tx=XM_019838785.1;refGene_gn=MFF;refGene_tx=XM_023259751.1
C1 211316330 211316363
refGene_gn=MFF;refGene_tx=XM_019838776.1;refGene_gn=MFF;refGene_tx=XM_019838779.2;refGene_gn=MFF;refGene_tx=XM_019838780.1;refGene_gn=MFF;refGene_tx=XM_019838783.1;refGene_gn=MFF;refGene_tx=XM_019838784.1;refGene_gn=MFF;refGene_tx=XM_019838785.1;refGene_gn=MFF;refGene_tx=XM_019838787.1;refGene_gn=MFF;refGene_tx=XM_019838788.1
C1 211317894 211317954
refGene_gn=MFF;refGene_tx=XM_019838776.1;refGene_gn=MFF;refGene_tx=XM_019838779.2;refGene_gn=MFF;refGene_tx=XM_019838780.1;refGene_gn=MFF;refGene_tx=XM_019838781.1;refGene_gn=MFF;refGene_tx=XM_019838782.1;refGene_gn=MFF;refGene_tx=XM_019838783.1;refGene_gn=MFF;refGene_tx=XM_019838784.1;refGene_gn=MFF;refGene_tx=XM_019838785.1;refGene_gn=MFF;refGene_tx=XM_019838787.1;refGene_gn=MFF;refGene_tx=XM_019838788.1;refGene_gn=MFF;refGene_tx=XM_023259751.1;refGene_gn=MFF;refGene_tx=XM_023259752.1

C1	211320954	211321039
refGene_gn=MFF;refGene_tx=XM_019838776.1;refGene_gn=MFF;refGene_tx=XM_019838779.2;refGene_gn=MFF;refGene_tx=XM_019838780.1;refGene_gn=MFF;refGene_tx=XM_019838781.1;refGene_gn=MFF;refGene_tx=XM_019838782.1;refGene_gn=MFF;refGene_tx=XM_019838783.1;refGene_gn=MFF;refGene_tx=XM_019838784.1;refGene_gn=MFF;refGene_tx=XM_019838785.1;refGene_gn=MFF;refGene_tx=XM_019838787.1;refGene_gn=MFF;refGene_tx=XM_019838788.1;refGene_gn=MFF;refGene_tx=XM_019838791.1;refGene_gn=MFF;refGene_tx=XM_023259751.1;refGene_gn=MFF;refGene_tx=XM_023259752.1;refGene_gn=MFF;refGene_tx=XM_023259753.1
C1	211323756	211323888
refGene_gn=MFF;refGene_tx=XM_019838776.1;refGene_gn=MFF;refGene_tx=XM_019838779.2;refGene_gn=MFF;refGene_tx=XM_019838780.1;refGene_gn=MFF;refGene_tx=XM_019838781.1;refGene_gn=MFF;refGene_tx=XM_019838782.1;refGene_gn=MFF;refGene_tx=XM_019838783.1;refGene_gn=MFF;refGene_tx=XM_019838784.1;refGene_gn=MFF;refGene_tx=XM_019838785.1;refGene_gn=MFF;refGene_tx=XM_019838787.1;refGene_gn=MFF;refGene_tx=XM_019838788.1;refGene_gn=MFF;refGene_tx=XM_019838791.1;refGene_gn=MFF;refGene_tx=XM_023259751.1;refGene_gn=MFF;refGene_tx=XM_023259752.1;refGene_gn=MFF;refGene_tx=XM_023259753.1
C1	211330223	211330512
refGene_gn=TM4SF20;refGene_tx=XM_003991222.5
C1	211332685	211332837
refGene_gn=TM4SF20;refGene_tx=XM_003991222.5
C1	211336286	211336352
refGene_gn=TM4SF20;refGene_tx=XM_003991222.5
C1	211342246	211342429
refGene_gn=TM4SF20;refGene_tx=XM_003991222.5
C1	211423286	211423453
refGene_gn=AGFG1;refGene_tx=XM_023259755.1;refGene_gn=AGFG1;refGene_tx=XM_023259756.1;refGene_gn=AGFG1;refGene_tx=XM_023259757.1;refGene_gn=AGFG1;refGene_tx=XM_023259758.1;refGene_gn=AGFG1;refGene_tx=XM_023259760.1;refGene_gn=AGFG1;refGene_tx=XM_023259761.1;refGene_gn=AGFG1;refGene_tx=XM_023259762.1;refGene_gn=AGFG1;refGene_tx=XM_023259763.1;refGene_gn=AGFG1;refGene_tx=XM_023259764.1;refGene_gn=AGFG1;refGene_tx=XM_023259765.1
C1	211441226	211441237
refGene_gn=AGFG1;refGene_tx=XM_006935652.4
C1	211441471	211441565
refGene_gn=AGFG1;refGene_tx=XM_006935652.4;refGene_gn=AGFG1;refGene_tx=XM_023259755.1;refGene_gn=AGFG1;refGene_tx=XM_023259756.1;refGene_gn=AGFG1;refGene_tx=XM_023259757.1;refGene_gn=AGFG1;refGene_tx=XM_023259758.1;refGene_gn=AGFG1;refGene_tx=XM_023259760.1;refGene_gn=AGFG1;refGene_tx=XM_023259761.1;refGene_gn=AGFG1;refGene_tx=XM_023259762.1;refGene_gn=AGFG1;refGene_tx=XM_023259763.1;refGene_gn=AGFG1;refGene_tx=XM_023259764.1;refGene_gn=AGFG1;refGene_tx=XM_023259765.1
C1	211467118	211467234
refGene_gn=AGFG1;refGene_tx=XM_006935652.4;refGene_gn=AGFG1;refGene_tx=XM_023259755.1;refGene_gn=AGFG1;refGene_tx=XM_023259756.1;refGene_gn=AGFG1;refGene_tx=XM_023259757.1;refGene_gn=AGFG1;refGene_tx=XM_0232597

58.1;refGene_gn=AGFG1;refGene_tx=XM_023259760.1;refGene_gn=AGFG1;refGene_tx=XM_023259761.1;refGene_gn=AGFG1;refGene_tx=XM_023259762.1;refGene_gn=AGFG1;refGene_tx=XM_023259763.1;refGene_gn=AGFG1;refGene_tx=XM_023259764.1;refGene_gn=AGFG1;refGene_tx=XM_023259765.1
C1 211470485 211470648
refGene_gn=AGFG1;refGene_tx=XM_006935652.4;refGene_gn=AGFG1;refGene_tx=XM_023259755.1;refGene_gn=AGFG1;refGene_tx=XM_023259756.1;refGene_gn=AGFG1;refGene_tx=XM_023259757.1;refGene_gn=AGFG1;refGene_tx=XM_023259758.1;refGene_gn=AGFG1;refGene_tx=XM_023259760.1;refGene_gn=AGFG1;refGene_tx=XM_023259761.1;refGene_gn=AGFG1;refGene_tx=XM_023259762.1;refGene_gn=AGFG1;refGene_tx=XM_023259763.1;refGene_gn=AGFG1;refGene_tx=XM_023259764.1;refGene_gn=AGFG1;refGene_tx=XM_023259765.1
C1 211471649 211471803
refGene_gn=AGFG1;refGene_tx=XM_006935652.4;refGene_gn=AGFG1;refGene_tx=XM_023259755.1;refGene_gn=AGFG1;refGene_tx=XM_023259756.1;refGene_gn=AGFG1;refGene_tx=XM_023259757.1;refGene_gn=AGFG1;refGene_tx=XM_023259758.1;refGene_gn=AGFG1;refGene_tx=XM_023259760.1;refGene_gn=AGFG1;refGene_tx=XM_023259761.1;refGene_gn=AGFG1;refGene_tx=XM_023259762.1;refGene_gn=AGFG1;refGene_tx=XM_023259763.1;refGene_gn=AGFG1;refGene_tx=XM_023259764.1;refGene_gn=AGFG1;refGene_tx=XM_023259765.1
C1 211479307 211479427
refGene_gn=AGFG1;refGene_tx=XM_006935652.4;refGene_gn=AGFG1;refGene_tx=XM_023259755.1;refGene_gn=AGFG1;refGene_tx=XM_023259756.1;refGene_gn=AGFG1;refGene_tx=XM_023259757.1;refGene_gn=AGFG1;refGene_tx=XM_023259758.1;refGene_gn=AGFG1;refGene_tx=XM_023259760.1;refGene_gn=AGFG1;refGene_tx=XM_023259761.1;refGene_gn=AGFG1;refGene_tx=XM_023259762.1;refGene_gn=AGFG1;refGene_tx=XM_023259763.1
C1 211480104 211480176
refGene_gn=AGFG1;refGene_tx=XM_006935652.4;refGene_gn=AGFG1;refGene_tx=XM_023259755.1;refGene_gn=AGFG1;refGene_tx=XM_023259756.1;refGene_gn=AGFG1;refGene_tx=XM_023259757.1;refGene_gn=AGFG1;refGene_tx=XM_023259758.1;refGene_gn=AGFG1;refGene_tx=XM_023259760.1
C1 211481673 211481883
refGene_gn=AGFG1;refGene_tx=XM_006935652.4;refGene_gn=AGFG1;refGene_tx=XM_023259755.1;refGene_gn=AGFG1;refGene_tx=XM_023259756.1;refGene_gn=AGFG1;refGene_tx=XM_023259757.1;refGene_gn=AGFG1;refGene_tx=XM_023259758.1;refGene_gn=AGFG1;refGene_tx=XM_023259760.1;refGene_gn=AGFG1;refGene_tx=XM_023259761.1;refGene_gn=AGFG1;refGene_tx=XM_023259762.1;refGene_gn=AGFG1;refGene_tx=XM_023259763.1;refGene_gn=AGFG1;refGene_tx=XM_023259764.1;refGene_gn=AGFG1;refGene_tx=XM_023259765.1
C1 211483448 211483629
refGene_gn=AGFG1;refGene_tx=XM_006935652.4;refGene_gn=AGFG1;refGene_tx=XM_023259755.1;refGene_gn=AGFG1;refGene_tx=XM_023259756.1;refGene_gn=AGFG1;refGene_tx=XM_023259757.1;refGene_gn=AGFG1;refGene_tx=XM_023259758.1;refGene_gn=AGFG1;refGene_tx=XM_023259760.1;refGene_gn=AGFG1;refGene_tx=XM_023259761.1;refGene_gn=AGFG1;refGene_tx=XM_023259762.1;refGene_gn=AGFG1;refGene_tx=XM_023259763.1;refGene_gn=AGFG1;refGene_tx=XM_023259764.1;refGene_gn=AGFG1;refGene_tx=XM_023259765.1
C1 211486652 211486732
refGene_gn=AGFG1;refGene_tx=XM_006935652.4;refGene_gn=AGFG1;refGene_tx

=XM_023259755.1;refGene_gn=AGFG1;refGene_tx=XM_023259756.1;refGene_gn=AGFG1;refGene_tx=XM_023259757.1;refGene_gn=AGFG1;refGene_tx=XM_023259758.1;refGene_gn=AGFG1;refGene_tx=XM_023259760.1;refGene_gn=AGFG1;refGene_tx=XM_023259761.1;refGene_gn=AGFG1;refGene_tx=XM_023259762.1;refGene_gn=AGFG1;refGene_tx=XM_023259763.1;refGene_gn=AGFG1;refGene_tx=XM_023259764.1;refGene_gn=AGFG1;refGene_tx=XM_023259765.1
C1 211486941 211487034
refGene_gn=AGFG1;refGene_tx=XM_006935652.4;refGene_gn=AGFG1;refGene_tx=XM_023259755.1;refGene_gn=AGFG1;refGene_tx=XM_023259756.1;refGene_gn=AGFG1;refGene_tx=XM_023259757.1;refGene_gn=AGFG1;refGene_tx=XM_023259758.1;refGene_gn=AGFG1;refGene_tx=XM_023259760.1;refGene_gn=AGFG1;refGene_tx=XM_023259761.1;refGene_gn=AGFG1;refGene_tx=XM_023259762.1;refGene_gn=AGFG1;refGene_tx=XM_023259763.1;refGene_gn=AGFG1;refGene_tx=XM_023259764.1;refGene_gn=AGFG1;refGene_tx=XM_023259765.1
C1 211492025 211492073
refGene_gn=AGFG1;refGene_tx=XM_006935652.4;refGene_gn=AGFG1;refGene_tx=XM_023259755.1;refGene_gn=AGFG1;refGene_tx=XM_023259756.1;refGene_gn=AGFG1;refGene_tx=XM_023259760.1;refGene_gn=AGFG1;refGene_tx=XM_023259761.1;refGene_gn=AGFG1;refGene_tx=XM_023259764.1
C1 211493911 211494070
refGene_gn=AGFG1;refGene_tx=XM_006935652.4;refGene_gn=AGFG1;refGene_tx=XM_023259755.1;refGene_gn=AGFG1;refGene_tx=XM_023259756.1;refGene_gn=AGFG1;refGene_tx=XM_023259757.1;refGene_gn=AGFG1;refGene_tx=XM_023259758.1;refGene_gn=AGFG1;refGene_tx=XM_023259760.1;refGene_gn=AGFG1;refGene_tx=XM_023259761.1;refGene_gn=AGFG1;refGene_tx=XM_023259762.1;refGene_gn=AGFG1;refGene_tx=XM_023259763.1;refGene_gn=AGFG1;refGene_tx=XM_023259764.1;refGene_gn=AGFG1;refGene_tx=XM_023259765.1
C1 211495020 211495112
refGene_gn=AGFG1;refGene_tx=XM_006935652.4;refGene_gn=AGFG1;refGene_tx=XM_023259755.1;refGene_gn=AGFG1;refGene_tx=XM_023259756.1;refGene_gn=AGFG1;refGene_tx=XM_023259757.1;refGene_gn=AGFG1;refGene_tx=XM_023259758.1;refGene_gn=AGFG1;refGene_tx=XM_023259760.1;refGene_gn=AGFG1;refGene_tx=XM_023259761.1;refGene_gn=AGFG1;refGene_tx=XM_023259762.1;refGene_gn=AGFG1;refGene_tx=XM_023259763.1;refGene_gn=AGFG1;refGene_tx=XM_023259764.1;refGene_gn=AGFG1;refGene_tx=XM_023259765.1
C1 211495707 211495767
refGene_gn=AGFG1;refGene_tx=XM_006935652.4;refGene_gn=AGFG1;refGene_tx=XM_023259755.1;refGene_gn=AGFG1;refGene_tx=XM_023259756.1;refGene_gn=AGFG1;refGene_tx=XM_023259757.1;refGene_gn=AGFG1;refGene_tx=XM_023259758.1;refGene_gn=AGFG1;refGene_tx=XM_023259760.1;refGene_gn=AGFG1;refGene_tx=XM_023259761.1;refGene_gn=AGFG1;refGene_tx=XM_023259762.1;refGene_gn=AGFG1;refGene_tx=XM_023259763.1;refGene_gn=AGFG1;refGene_tx=XM_023259764.1;refGene_gn=AGFG1;refGene_tx=XM_023259765.1
C1 211528723 211529059
refGene_gn=LOC111561915;refGene_tx=XM_023259766.1
C1 211542454 211542814
refGene_gn=LOC109503062;refGene_tx=XM_019839145.2
C1 211559364 211559538
refGene_gn=LOC101091721;refGene_tx=XM_023259767.1;refGene_gn=LOC101091721;refGene_tx=XM_023259768.1

C1	211569978	211570120
refGene_gn=LOC101091721;refGene_tx=XM_023259767.1;refGene_gn=LOC101091
721;refGene_tx=XM_023259768.1
C1	211576650	211576843
refGene_gn=LOC101091721;refGene_tx=XM_023259767.1;refGene_gn=LOC101091
721;refGene_tx=XM_023259768.1
C1	211581020	211581837
refGene_gn=LOC101091721;refGene_tx=XM_023259767.1;refGene_gn=LOC101091
721;refGene_tx=XM_023259768.1
C1	211583827	211583977
refGene_gn=LOC101091721;refGene_tx=XM_023259767.1;refGene_gn=LOC101091
721;refGene_tx=XM_023259768.1
C1	211607362	211607539
refGene_gn=SLC19A3;refGene_tx=NM_001278820.1;refGene_gn=SLC19A3;refGen
e_tx=XM_006935634.4;refGene_gn=SLC19A3;refGene_tx=XM_006935635.4;refGe
ne_gn=SLC19A3;refGene_tx=XM_019835936.2;refGene_gn=SLC19A3;refGene_tx=
XM_019835937.2;refGene_gn=SLC19A3;refGene_tx=XM_019835938.2;refGene_gn
=SLC19A3;refGene_tx=XM_019835939.2;refGene_gn=SLC19A3;refGene_tx=XM_01
9835940.2;refGene_gn=SLC19A3;refGene_tx=XM_019835941.2;refGene_gn=SLC1
9A3;refGene_tx=XM_023257809.1;refGene_gn=SLC19A3;refGene_tx=XM_0232578
10.1
C1	211608387	211608529
refGene_gn=SLC19A3;refGene_tx=NM_001278820.1;refGene_gn=SLC19A3;refGen
e_tx=XM_006935634.4;refGene_gn=SLC19A3;refGene_tx=XM_006935635.4;refGe
ne_gn=SLC19A3;refGene_tx=XM_019835936.2;refGene_gn=SLC19A3;refGene_tx=
XM_019835937.2;refGene_gn=SLC19A3;refGene_tx=XM_019835938.2;refGene_gn
=SLC19A3;refGene_tx=XM_019835939.2;refGene_gn=SLC19A3;refGene_tx=XM_01
9835940.2;refGene_gn=SLC19A3;refGene_tx=XM_019835941.2;refGene_gn=SLC1
9A3;refGene_tx=XM_023257809.1;refGene_gn=SLC19A3;refGene_tx=XM_0232578
10.1
C1	211609951	211610144
refGene_gn=SLC19A3;refGene_tx=NM_001278820.1;refGene_gn=SLC19A3;refGen
e_tx=XM_006935634.4;refGene_gn=SLC19A3;refGene_tx=XM_006935635.4;refGe
ne_gn=SLC19A3;refGene_tx=XM_019835936.2;refGene_gn=SLC19A3;refGene_tx=
XM_019835937.2;refGene_gn=SLC19A3;refGene_tx=XM_019835938.2;refGene_gn
=SLC19A3;refGene_tx=XM_019835939.2;refGene_gn=SLC19A3;refGene_tx=XM_01
9835940.2;refGene_gn=SLC19A3;refGene_tx=XM_019835941.2;refGene_gn=SLC1
9A3;refGene_tx=XM_023257809.1;refGene_gn=SLC19A3;refGene_tx=XM_0232578
10.1
C1	211611969	211612795
refGene_gn=SLC19A3;refGene_tx=NM_001278820.1;refGene_gn=SLC19A3;refGen
e_tx=XM_006935634.4;refGene_gn=SLC19A3;refGene_tx=XM_006935635.4;refGe
ne_gn=SLC19A3;refGene_tx=XM_019835936.2;refGene_gn=SLC19A3;refGene_tx=
XM_019835937.2;refGene_gn=SLC19A3;refGene_tx=XM_019835938.2;refGene_gn
=SLC19A3;refGene_tx=XM_019835939.2;refGene_gn=SLC19A3;refGene_tx=XM_01
9835940.2;refGene_gn=SLC19A3;refGene_tx=XM_019835941.2;refGene_gn=SLC1
9A3;refGene_tx=XM_023257809.1;refGene_gn=SLC19A3;refGene_tx=XM_0232578
10.1
C1	211614545	211614695
refGene_gn=SLC19A3;refGene_tx=NM_001278820.1;refGene_gn=SLC19A3;refGen

e_tx=XM_006935634.4;refGene_gn=SLC19A3;refGene_tx=XM_006935635.4;refGene_gn=SLC19A3;refGene_tx=XM_019835936.2;refGene_gn=SLC19A3;refGene_tx=XM_019835937.2;refGene_gn=SLC19A3;refGene_tx=XM_019835938.2;refGene_gn=SLC19A3;refGene_tx=XM_019835939.2;refGene_gn=SLC19A3;refGene_tx=XM_019835940.2;refGene_gn=SLC19A3;refGene_tx=XM_019835941.2;refGene_gn=SLC19A3;refGene_tx=XM_023257809.1;refGene_gn=SLC19A3;refGene_tx=XM_023257810.1
C1 211647248 211647593
refGene_gn=LOC111561651;refGene_tx=XM_023257870.1
C1 211697425 211697504
refGene_gn=CCL20;refGene_tx=XM_003991225.3;refGene_gn=CCL20;refGene_tx=XM_011285647.3
C1 211698650 211698765
refGene_gn=CCL20;refGene_tx=XM_003991225.3;refGene_gn=CCL20;refGene_tx=XM_011285647.3
C1 211699458 211699536
refGene_gn=CCL20;refGene_tx=XM_003991225.3;refGene_gn=CCL20;refGene_tx=XM_011285647.3
C1 211700474 211700496
refGene_gn=CCL20;refGene_tx=XM_003991225.3;refGene_gn=CCL20;refGene_tx=XM_011285647.3
C1 211741774 211741895
refGene_gn=DAW1;refGene_tx=XM_011285645.3;refGene_gn=DAW1;refGene_tx=XM_011285646.3;refGene_gn=DAW1;refGene_tx=XM_019838796.2;refGene_gn=DAW1;refGene_tx=XM_019838798.2
C1 211753376 211753449
refGene_gn=DAW1;refGene_tx=XM_011285645.3;refGene_gn=DAW1;refGene_tx=XM_011285646.3;refGene_gn=DAW1;refGene_tx=XM_019838796.2;refGene_gn=DAW1;refGene_tx=XM_019838797.2;refGene_gn=DAW1;refGene_tx=XM_019838798.2
C1 211757092 211757237
refGene_gn=DAW1;refGene_tx=XM_011285645.3;refGene_gn=DAW1;refGene_tx=XM_011285646.3;refGene_gn=DAW1;refGene_tx=XM_019838796.2;refGene_gn=DAW1;refGene_tx=XM_019838797.2;refGene_gn=DAW1;refGene_tx=XM_019838798.2
C1 211759381 211759440
refGene_gn=DAW1;refGene_tx=XM_011285645.3;refGene_gn=DAW1;refGene_tx=XM_011285646.3;refGene_gn=DAW1;refGene_tx=XM_019838796.2;refGene_gn=DAW1;refGene_tx=XM_019838797.2;refGene_gn=DAW1;refGene_tx=XM_019838798.2
C1 211765911 211766034
refGene_gn=DAW1;refGene_tx=XM_011285645.3;refGene_gn=DAW1;refGene_tx=XM_011285646.3;refGene_gn=DAW1;refGene_tx=XM_019838796.2;refGene_gn=DAW1;refGene_tx=XM_019838797.2;refGene_gn=DAW1;refGene_tx=XM_019838798.2
C1 211769786 211769886
refGene_gn=DAW1;refGene_tx=XM_011285645.3;refGene_gn=DAW1;refGene_tx=XM_011285646.3;refGene_gn=DAW1;refGene_tx=XM_019838796.2;refGene_gn=DAW1;refGene_tx=XM_019838797.2;refGene_gn=DAW1;refGene_tx=XM_019838798.2
C1 211773705 211773813
refGene_gn=DAW1;refGene_tx=XM_011285645.3;refGene_gn=DAW1;refGene_tx=XM_011285646.3;refGene_gn=DAW1;refGene_tx=XM_019838796.2;refGene_gn=DAW1;refGene_tx=XM_019838797.2;refGene_gn=DAW1;refGene_tx=XM_019838798.2
C1 211776081 211776188

refGene_gn=DAW1;refGene_tx=XM_011285645.3;refGene_gn=DAW1;refGene_tx=XM_011285646.3;refGene_gn=DAW1;refGene_tx=XM_019838796.2;refGene_gn=DAW1;refGene_tx=XM_019838797.2;refGene_gn=DAW1;refGene_tx=XM_019838798.2
C1 211777290 211777393
refGene_gn=DAW1;refGene_tx=XM_011285645.3;refGene_gn=DAW1;refGene_tx=XM_011285646.3;refGene_gn=DAW1;refGene_tx=XM_019838796.2;refGene_gn=DAW1;refGene_tx=XM_019838797.2;refGene_gn=DAW1;refGene_tx=XM_019838798.2
C1 211778469 211778584
refGene_gn=DAW1;refGene_tx=XM_011285645.3;refGene_gn=DAW1;refGene_tx=XM_011285646.3;refGene_gn=DAW1;refGene_tx=XM_019838796.2;refGene_gn=DAW1;refGene_tx=XM_019838797.2;refGene_gn=DAW1;refGene_tx=XM_019838798.2
C1 211780612 211780713
refGene_gn=DAW1;refGene_tx=XM_019838798.2
C1 211785839 211785916
refGene_gn=DAW1;refGene_tx=XM_011285645.3;refGene_gn=DAW1;refGene_tx=XM_011285646.3;refGene_gn=DAW1;refGene_tx=XM_019838796.2;refGene_gn=DAW1;refGene_tx=XM_019838797.2
C1 211788160 211788323
refGene_gn=DAW1;refGene_tx=XM_011285645.3;refGene_gn=DAW1;refGene_tx=XM_011285646.3;refGene_gn=DAW1;refGene_tx=XM_019838796.2;refGene_gn=DAW1;refGene_tx=XM_019838797.2
C1 211788697 211788766
refGene_gn=DAW1;refGene_tx=XM_011285645.3;refGene_gn=DAW1;refGene_tx=XM_011285646.3;refGene_gn=DAW1;refGene_tx=XM_019838797.2
C1 211807607 211807636
refGene_gn=DAW1;refGene_tx=XM_019838796.2
C1 211828954 211829098
refGene_gn=SPHKAP;refGene_tx=XM_019838800.2;refGene_gn=SPHKAP;refGene_tx=XM_019838801.2
C1 211835871 211836056
refGene_gn=SPHKAP;refGene_tx=XM_019838800.2;refGene_gn=SPHKAP;refGene_tx=XM_019838801.2
C1 211836146 211836199
refGene_gn=SPHKAP;refGene_tx=XM_019838800.2;refGene_gn=SPHKAP;refGene_tx=XM_019838801.2
C1 211838377 211838464
refGene_gn=SPHKAP;refGene_tx=XM_019838800.2
C1 211839782 211839981
refGene_gn=SPHKAP;refGene_tx=XM_023259769.1
C1 211840316 211840493
refGene_gn=SPHKAP;refGene_tx=XM_019838800.2;refGene_gn=SPHKAP;refGene_tx=XM_019838801.2;refGene_gn=SPHKAP;refGene_tx=XM_023259769.1
C1 211854951 211858717
refGene_gn=SPHKAP;refGene_tx=XM_019838800.2;refGene_gn=SPHKAP;refGene_tx=XM_019838801.2;refGene_gn=SPHKAP;refGene_tx=XM_023259769.1
C1 211859794 211860050
refGene_gn=SPHKAP;refGene_tx=XM_019838800.2;refGene_gn=SPHKAP;refGene_tx=XM_019838801.2;refGene_gn=SPHKAP;refGene_tx=XM_023259769.1
C1 211866739 211866874
refGene_gn=SPHKAP;refGene_tx=XM_019838800.2;refGene_gn=SPHKAP;refGene_

tx=XM_019838801.2;refGene_gn=SPHKAP;refGene_tx=XM_023259769.1
C1	211868706	211868766
refGene_gn=SPHKAP;refGene_tx=XM_019838800.2;refGene_gn=SPHKAP;refGene_tx=XM_019838801.2;refGene_gn=SPHKAP;refGene_tx=XM_023259769.1
C1	211927571	211927679
refGene_gn=SPHKAP;refGene_tx=XM_019838800.2;refGene_gn=SPHKAP;refGene_tx=XM_019838801.2;refGene_gn=SPHKAP;refGene_tx=XM_023259769.1
C1	211950889	211950995
refGene_gn=SPHKAP;refGene_tx=XM_019838800.2;refGene_gn=SPHKAP;refGene_tx=XM_019838801.2;refGene_gn=SPHKAP;refGene_tx=XM_023259769.1
C1	211987776	211987808
refGene_gn=SPHKAP;refGene_tx=XM_019838800.2;refGene_gn=SPHKAP;refGene_tx=XM_019838801.2;refGene_gn=SPHKAP;refGene_tx=XM_023259769.1
C1	212615107	212615584
refGene_gn=PID1;refGene_tx=XM_023259770.1;refGene_gn=PID1;refGene_tx=XM_023259771.1
C1	212737158	212737305
refGene_gn=PID1;refGene_tx=XM_023259770.1
C1	212844402	212844432
refGene_gn=PID1;refGene_tx=XM_023259770.1;refGene_gn=PID1;refGene_tx=XM_023259771.1
C1	212903459	212903571
refGene_gn=DNER;refGene_tx=XM_023259772.1
C1	212910090	212910337
refGene_gn=DNER;refGene_tx=XM_023259772.1
C1	212920457	212920589
refGene_gn=DNER;refGene_tx=XM_023259772.1
C1	212930289	212930403
refGene_gn=DNER;refGene_tx=XM_023259772.1
C1	212942184	212942307
refGene_gn=DNER;refGene_tx=XM_023259772.1
C1	212969612	212969837
refGene_gn=DNER;refGene_tx=XM_023259772.1
C1	212997895	212998009
refGene_gn=DNER;refGene_tx=XM_023259772.1
C1	213021495	213021649
refGene_gn=DNER;refGene_tx=XM_023259772.1
C1	213052289	213052435
refGene_gn=DNER;refGene_tx=XM_023259772.1
C1	213079315	213079482
refGene_gn=DNER;refGene_tx=XM_023259772.1
C1	213082168	213082263
refGene_gn=DNER;refGene_tx=XM_023259772.1
C1	213084742	213085039
refGene_gn=DNER;refGene_tx=XM_023259772.1
C1	213223482	213223731
refGene_gn=DNER;refGene_tx=XM_023259772.1
C1	213261610	213261807
refGene_gn=TRIP12;refGene_tx=XM_011285634.3;refGene_gn=TRIP12;refGene_tx=XM_019838807.2;refGene_gn=TRIP12;refGene_tx=XM_019838808.2;refGene_

gn=TRIP12;refGene_tx=XM_019838809.2;refGene_gn=TRIP12;refGene_tx=XM_023259773.1;refGene_gn=TRIP12;refGene_tx=XM_023259774.1;refGene_gn=TRIP12;refGene_tx=XM_023259775.1;refGene_gn=TRIP12;refGene_tx=XM_023259776.1;refGene_gn=TRIP12;refGene_tx=XM_023259777.1;refGene_gn=TRIP12;refGene_tx=XM_023259778.1;refGene_gn=TRIP12;refGene_tx=XM_023259779.1;refGene_gn=TRIP12;refGene_tx=XM_023259780.1;refGene_gn=TRIP12;refGene_tx=XM_023259781.1;refGene_gn=TRIP12;refGene_tx=XM_023259782.1;refGene_gn=TRIP12;refGene_tx=XM_023259783.1;refGene_gn=TRIP12;refGene_tx=XM_023259784.1;refGene_gn=TRIP12;refGene_tx=XM_023259785.1;refGene_gn=TRIP12;refGene_tx=XM_023259786.1
C1 213262558 213262662
refGene_gn=TRIP12;refGene_tx=XM_011285634.3;refGene_gn=TRIP12;refGene_tx=XM_019838807.2;refGene_gn=TRIP12;refGene_tx=XM_019838808.2;refGene_gn=TRIP12;refGene_tx=XM_019838809.2;refGene_gn=TRIP12;refGene_tx=XM_023259773.1;refGene_gn=TRIP12;refGene_tx=XM_023259774.1;refGene_gn=TRIP12;refGene_tx=XM_023259775.1;refGene_gn=TRIP12;refGene_tx=XM_023259776.1;refGene_gn=TRIP12;refGene_tx=XM_023259777.1;refGene_gn=TRIP12;refGene_tx=XM_023259778.1;refGene_gn=TRIP12;refGene_tx=XM_023259779.1;refGene_gn=TRIP12;refGene_tx=XM_023259780.1;refGene_gn=TRIP12;refGene_tx=XM_023259781.1;refGene_gn=TRIP12;refGene_tx=XM_023259782.1;refGene_gn=TRIP12;refGene_tx=XM_023259783.1;refGene_gn=TRIP12;refGene_tx=XM_023259784.1;refGene_gn=TRIP12;refGene_tx=XM_023259785.1;refGene_gn=TRIP12;refGene_tx=XM_023259786.1
C1 213263191 213263286
refGene_gn=TRIP12;refGene_tx=XM_011285634.3;refGene_gn=TRIP12;refGene_tx=XM_019838807.2;refGene_gn=TRIP12;refGene_tx=XM_019838808.2;refGene_gn=TRIP12;refGene_tx=XM_019838809.2;refGene_gn=TRIP12;refGene_tx=XM_023259773.1;refGene_gn=TRIP12;refGene_tx=XM_023259774.1;refGene_gn=TRIP12;refGene_tx=XM_023259775.1;refGene_gn=TRIP12;refGene_tx=XM_023259776.1;refGene_gn=TRIP12;refGene_tx=XM_023259777.1;refGene_gn=TRIP12;refGene_tx=XM_023259778.1;refGene_gn=TRIP12;refGene_tx=XM_023259779.1;refGene_gn=TRIP12;refGene_tx=XM_023259780.1;refGene_gn=TRIP12;refGene_tx=XM_023259781.1;refGene_gn=TRIP12;refGene_tx=XM_023259782.1;refGene_gn=TRIP12;refGene_tx=XM_023259783.1;refGene_gn=TRIP12;refGene_tx=XM_023259784.1;refGene_gn=TRIP12;refGene_tx=XM_023259785.1;refGene_gn=TRIP12;refGene_tx=XM_023259786.1
C1 213265011 213265125
refGene_gn=TRIP12;refGene_tx=XM_011285634.3;refGene_gn=TRIP12;refGene_tx=XM_019838807.2;refGene_gn=TRIP12;refGene_tx=XM_019838808.2;refGene_gn=TRIP12;refGene_tx=XM_019838809.2;refGene_gn=TRIP12;refGene_tx=XM_023259773.1;refGene_gn=TRIP12;refGene_tx=XM_023259774.1;refGene_gn=TRIP12;refGene_tx=XM_023259775.1;refGene_gn=TRIP12;refGene_tx=XM_023259776.1;refGene_gn=TRIP12;refGene_tx=XM_023259777.1;refGene_gn=TRIP12;refGene_tx=XM_023259778.1;refGene_gn=TRIP12;refGene_tx=XM_023259779.1;refGene_gn=TRIP12;refGene_tx=XM_023259780.1;refGene_gn=TRIP12;refGene_tx=XM_023259781.1;refGene_gn=TRIP12;refGene_tx=XM_023259782.1;refGene_gn=TRIP12;refGene_tx=XM_023259783.1;refGene_gn=TRIP12;refGene_tx=XM_023259784.1;refGene_gn=TRIP12;refGene_tx=XM_023259785.1;refGene_gn=TRIP12;refGene_tx=XM_023259786.1
C1 213267179 213267344

refGene_gn=TRIP12;refGene_tx=XM_011285634.3;refGene_gn=TRIP12;refGene_tx=XM_019838807.2;refGene_gn=TRIP12;refGene_tx=XM_019838808.2;refGene_gn=TRIP12;refGene_tx=XM_019838809.2;refGene_gn=TRIP12;refGene_tx=XM_023259773.1;refGene_gn=TRIP12;refGene_tx=XM_023259774.1;refGene_gn=TRIP12;refGene_tx=XM_023259775.1;refGene_gn=TRIP12;refGene_tx=XM_023259776.1;refGene_gn=TRIP12;refGene_tx=XM_023259777.1;refGene_gn=TRIP12;refGene_tx=XM_023259778.1;refGene_gn=TRIP12;refGene_tx=XM_023259779.1;refGene_gn=TRIP12;refGene_tx=XM_023259780.1;refGene_gn=TRIP12;refGene_tx=XM_023259781.1;refGene_gn=TRIP12;refGene_tx=XM_023259782.1;refGene_gn=TRIP12;refGene_tx=XM_023259783.1;refGene_gn=TRIP12;refGene_tx=XM_023259784.1;refGene_gn=TRIP12;refGene_tx=XM_023259785.1;refGene_gn=TRIP12;refGene_tx=XM_023259786.1
C1 213270592 213270757
refGene_gn=TRIP12;refGene_tx=XM_011285634.3;refGene_gn=TRIP12;refGene_tx=XM_019838807.2;refGene_gn=TRIP12;refGene_tx=XM_019838808.2;refGene_gn=TRIP12;refGene_tx=XM_019838809.2;refGene_gn=TRIP12;refGene_tx=XM_023259773.1;refGene_gn=TRIP12;refGene_tx=XM_023259774.1;refGene_gn=TRIP12;refGene_tx=XM_023259775.1;refGene_gn=TRIP12;refGene_tx=XM_023259776.1;refGene_gn=TRIP12;refGene_tx=XM_023259777.1;refGene_gn=TRIP12;refGene_tx=XM_023259778.1;refGene_gn=TRIP12;refGene_tx=XM_023259779.1;refGene_gn=TRIP12;refGene_tx=XM_023259780.1;refGene_gn=TRIP12;refGene_tx=XM_023259781.1;refGene_gn=TRIP12;refGene_tx=XM_023259782.1;refGene_gn=TRIP12;refGene_tx=XM_023259783.1;refGene_gn=TRIP12;refGene_tx=XM_023259784.1;refGene_gn=TRIP12;refGene_tx=XM_023259785.1;refGene_gn=TRIP12;refGene_tx=XM_023259786.1
C1 213271490 213271645
refGene_gn=TRIP12;refGene_tx=XM_011285634.3;refGene_gn=TRIP12;refGene_tx=XM_019838807.2;refGene_gn=TRIP12;refGene_tx=XM_019838808.2;refGene_gn=TRIP12;refGene_tx=XM_019838809.2;refGene_gn=TRIP12;refGene_tx=XM_023259773.1;refGene_gn=TRIP12;refGene_tx=XM_023259774.1;refGene_gn=TRIP12;refGene_tx=XM_023259775.1;refGene_gn=TRIP12;refGene_tx=XM_023259776.1;refGene_gn=TRIP12;refGene_tx=XM_023259777.1;refGene_gn=TRIP12;refGene_tx=XM_023259778.1;refGene_gn=TRIP12;refGene_tx=XM_023259779.1;refGene_gn=TRIP12;refGene_tx=XM_023259780.1;refGene_gn=TRIP12;refGene_tx=XM_023259781.1;refGene_gn=TRIP12;refGene_tx=XM_023259782.1;refGene_gn=TRIP12;refGene_tx=XM_023259783.1;refGene_gn=TRIP12;refGene_tx=XM_023259784.1;refGene_gn=TRIP12;refGene_tx=XM_023259785.1;refGene_gn=TRIP12;refGene_tx=XM_023259786.1
C1 213271955 213272070
refGene_gn=TRIP12;refGene_tx=XM_011285634.3;refGene_gn=TRIP12;refGene_tx=XM_019838807.2;refGene_gn=TRIP12;refGene_tx=XM_019838808.2;refGene_gn=TRIP12;refGene_tx=XM_019838809.2;refGene_gn=TRIP12;refGene_tx=XM_023259773.1;refGene_gn=TRIP12;refGene_tx=XM_023259774.1;refGene_gn=TRIP12;refGene_tx=XM_023259775.1;refGene_gn=TRIP12;refGene_tx=XM_023259776.1;refGene_gn=TRIP12;refGene_tx=XM_023259777.1;refGene_gn=TRIP12;refGene_tx=XM_023259778.1;refGene_gn=TRIP12;refGene_tx=XM_023259779.1;refGene_gn=TRIP12;refGene_tx=XM_023259780.1;refGene_gn=TRIP12;refGene_tx=XM_023259781.1;refGene_gn=TRIP12;refGene_tx=XM_023259782.1;refGene_gn=TRIP12;refGene_tx=XM_023259783.1;refGene_gn=TRIP12;refGene_tx=XM_023259784.1;refGene_gn=TRIP12;refGene_tx=XM_023259785.1;refGene_gn=TRIP12;refG

ene_tx=XM_023259786.1
C1	213278682	213278781
refGene_gn=TRIP12;refGene_tx=XM_011285634.3;refGene_gn=TRIP12;refGene_tx=XM_019838807.2;refGene_gn=TRIP12;refGene_tx=XM_019838808.2;refGene_gn=TRIP12;refGene_tx=XM_019838809.2;refGene_gn=TRIP12;refGene_tx=XM_023259773.1;refGene_gn=TRIP12;refGene_tx=XM_023259774.1;refGene_gn=TRIP12;refGene_tx=XM_023259775.1;refGene_gn=TRIP12;refGene_tx=XM_023259776.1;refGene_gn=TRIP12;refGene_tx=XM_023259777.1;refGene_gn=TRIP12;refGene_tx=XM_023259778.1;refGene_gn=TRIP12;refGene_tx=XM_023259779.1;refGene_gn=TRIP12;refGene_tx=XM_023259780.1;refGene_gn=TRIP12;refGene_tx=XM_023259781.1;refGene_gn=TRIP12;refGene_tx=XM_023259782.1;refGene_gn=TRIP12;refGene_tx=XM_023259783.1;refGene_gn=TRIP12;refGene_tx=XM_023259784.1;refGene_gn=TRIP12;refGene_tx=XM_023259785.1;refGene_gn=TRIP12;refGene_tx=XM_023259786.1
C1	213280086	213280243
refGene_gn=TRIP12;refGene_tx=XM_011285634.3;refGene_gn=TRIP12;refGene_tx=XM_019838807.2;refGene_gn=TRIP12;refGene_tx=XM_019838808.2;refGene_gn=TRIP12;refGene_tx=XM_019838809.2;refGene_gn=TRIP12;refGene_tx=XM_023259773.1;refGene_gn=TRIP12;refGene_tx=XM_023259774.1;refGene_gn=TRIP12;refGene_tx=XM_023259775.1;refGene_gn=TRIP12;refGene_tx=XM_023259776.1;refGene_gn=TRIP12;refGene_tx=XM_023259777.1;refGene_gn=TRIP12;refGene_tx=XM_023259778.1;refGene_gn=TRIP12;refGene_tx=XM_023259779.1;refGene_gn=TRIP12;refGene_tx=XM_023259780.1;refGene_gn=TRIP12;refGene_tx=XM_023259781.1;refGene_gn=TRIP12;refGene_tx=XM_023259782.1;refGene_gn=TRIP12;refGene_tx=XM_023259783.1;refGene_gn=TRIP12;refGene_tx=XM_023259784.1;refGene_gn=TRIP12;refGene_tx=XM_023259785.1;refGene_gn=TRIP12;refGene_tx=XM_023259786.1
C1	213281330	213281473
refGene_gn=TRIP12;refGene_tx=XM_011285634.3;refGene_gn=TRIP12;refGene_tx=XM_019838807.2;refGene_gn=TRIP12;refGene_tx=XM_019838808.2;refGene_gn=TRIP12;refGene_tx=XM_019838809.2;refGene_gn=TRIP12;refGene_tx=XM_023259773.1;refGene_gn=TRIP12;refGene_tx=XM_023259774.1;refGene_gn=TRIP12;refGene_tx=XM_023259775.1;refGene_gn=TRIP12;refGene_tx=XM_023259776.1;refGene_gn=TRIP12;refGene_tx=XM_023259777.1;refGene_gn=TRIP12;refGene_tx=XM_023259778.1;refGene_gn=TRIP12;refGene_tx=XM_023259779.1;refGene_gn=TRIP12;refGene_tx=XM_023259780.1;refGene_gn=TRIP12;refGene_tx=XM_023259781.1;refGene_gn=TRIP12;refGene_tx=XM_023259782.1;refGene_gn=TRIP12;refGene_tx=XM_023259783.1;refGene_gn=TRIP12;refGene_tx=XM_023259784.1;refGene_gn=TRIP12;refGene_tx=XM_023259785.1;refGene_gn=TRIP12;refGene_tx=XM_023259786.1
C1	213282369	213282521
refGene_gn=TRIP12;refGene_tx=XM_011285634.3;refGene_gn=TRIP12;refGene_tx=XM_019838807.2;refGene_gn=TRIP12;refGene_tx=XM_019838808.2;refGene_gn=TRIP12;refGene_tx=XM_019838809.2;refGene_gn=TRIP12;refGene_tx=XM_023259773.1;refGene_gn=TRIP12;refGene_tx=XM_023259774.1;refGene_gn=TRIP12;refGene_tx=XM_023259775.1;refGene_gn=TRIP12;refGene_tx=XM_023259776.1;refGene_gn=TRIP12;refGene_tx=XM_023259777.1;refGene_gn=TRIP12;refGene_tx=XM_023259778.1;refGene_gn=TRIP12;refGene_tx=XM_023259779.1;refGene_gn=TRIP12;refGene_tx=XM_023259780.1;refGene_gn=TRIP12;refGene_tx=XM_023259781.1;refGene_gn=TRIP12;refGene_tx=XM_023259782.1;refGene_gn=TRI

P12;refGene_tx=XM_023259783.1;refGene_gn=TRIP12;refGene_tx=XM_02325978
4.1;refGene_gn=TRIP12;refGene_tx=XM_023259785.1;refGene_gn=TRIP12;refG
ene_tx=XM_023259786.1
C1 213283858 213283986
refGene_gn=TRIP12;refGene_tx=XM_011285634.3;refGene_gn=TRIP12;refGene_
tx=XM_019838807.2;refGene_gn=TRIP12;refGene_tx=XM_019838808.2;refGene_
gn=TRIP12;refGene_tx=XM_019838809.2;refGene_gn=TRIP12;refGene_tx=XM_02
3259773.1;refGene_gn=TRIP12;refGene_tx=XM_023259774.1;refGene_gn=TRIP1
2;refGene_tx=XM_023259775.1;refGene_gn=TRIP12;refGene_tx=XM_023259776.
1;refGene_gn=TRIP12;refGene_tx=XM_023259777.1;refGene_gn=TRIP12;refGen
e_tx=XM_023259778.1;refGene_gn=TRIP12;refGene_tx=XM_023259779.1;refGen
e_gn=TRIP12;refGene_tx=XM_023259780.1;refGene_gn=TRIP12;refGene_tx=XM_
023259781.1;refGene_gn=TRIP12;refGene_tx=XM_023259782.1;refGene_gn=TRI
P12;refGene_tx=XM_023259783.1;refGene_gn=TRIP12;refGene_tx=XM_02325978
4.1;refGene_gn=TRIP12;refGene_tx=XM_023259785.1;refGene_gn=TRIP12;refG
ene_tx=XM_023259786.1
C1 213284638 213284838
refGene_gn=TRIP12;refGene_tx=XM_011285634.3;refGene_gn=TRIP12;refGene_
tx=XM_019838807.2;refGene_gn=TRIP12;refGene_tx=XM_019838808.2;refGene_
gn=TRIP12;refGene_tx=XM_019838809.2;refGene_gn=TRIP12;refGene_tx=XM_02
3259773.1;refGene_gn=TRIP12;refGene_tx=XM_023259774.1;refGene_gn=TRIP1
2;refGene_tx=XM_023259775.1;refGene_gn=TRIP12;refGene_tx=XM_023259776.
1;refGene_gn=TRIP12;refGene_tx=XM_023259777.1;refGene_gn=TRIP12;refGen
e_tx=XM_023259778.1;refGene_gn=TRIP12;refGene_tx=XM_023259779.1;refGen
e_gn=TRIP12;refGene_tx=XM_023259780.1;refGene_gn=TRIP12;refGene_tx=XM_
023259781.1;refGene_gn=TRIP12;refGene_tx=XM_023259782.1;refGene_gn=TRI
P12;refGene_tx=XM_023259783.1;refGene_gn=TRIP12;refGene_tx=XM_02325978
4.1;refGene_gn=TRIP12;refGene_tx=XM_023259785.1;refGene_gn=TRIP12;refG
ene_tx=XM_023259786.1
C1 213284919 213284993
refGene_gn=TRIP12;refGene_tx=XM_011285634.3;refGene_gn=TRIP12;refGene_
tx=XM_019838807.2;refGene_gn=TRIP12;refGene_tx=XM_019838808.2;refGene_
gn=TRIP12;refGene_tx=XM_019838809.2;refGene_gn=TRIP12;refGene_tx=XM_02
3259773.1;refGene_gn=TRIP12;refGene_tx=XM_023259774.1;refGene_gn=TRIP1
2;refGene_tx=XM_023259775.1;refGene_gn=TRIP12;refGene_tx=XM_023259776.
1;refGene_gn=TRIP12;refGene_tx=XM_023259777.1;refGene_gn=TRIP12;refGen
e_tx=XM_023259778.1;refGene_gn=TRIP12;refGene_tx=XM_023259779.1;refGen
e_gn=TRIP12;refGene_tx=XM_023259780.1;refGene_gn=TRIP12;refGene_tx=XM_
023259781.1;refGene_gn=TRIP12;refGene_tx=XM_023259782.1;refGene_gn=TRI
P12;refGene_tx=XM_023259783.1;refGene_gn=TRIP12;refGene_tx=XM_02325978
4.1;refGene_gn=TRIP12;refGene_tx=XM_023259785.1;refGene_gn=TRIP12;refG
ene_tx=XM_023259786.1
C1 213285367 213285540
refGene_gn=TRIP12;refGene_tx=XM_011285634.3;refGene_gn=TRIP12;refGene_
tx=XM_019838807.2;refGene_gn=TRIP12;refGene_tx=XM_019838808.2;refGene_
gn=TRIP12;refGene_tx=XM_019838809.2;refGene_gn=TRIP12;refGene_tx=XM_02
3259773.1;refGene_gn=TRIP12;refGene_tx=XM_023259774.1;refGene_gn=TRIP1
2;refGene_tx=XM_023259775.1;refGene_gn=TRIP12;refGene_tx=XM_023259776.
1;refGene_gn=TRIP12;refGene_tx=XM_023259777.1;refGene_gn=TRIP12;refGen
e_tx=XM_023259778.1;refGene_gn=TRIP12;refGene_tx=XM_023259779.1;refGen

e_gn=TRIP12;refGene_tx=XM_023259780.1;refGene_gn=TRIP12;refGene_tx=XM_023259781.1;refGene_gn=TRIP12;refGene_tx=XM_023259782.1;refGene_gn=TRIP12;refGene_tx=XM_023259783.1;refGene_gn=TRIP12;refGene_tx=XM_023259784.1;refGene_gn=TRIP12;refGene_tx=XM_023259785.1;refGene_gn=TRIP12;refGene_tx=XM_023259786.1
C1 213287213 213287365
refGene_gn=TRIP12;refGene_tx=XM_011285634.3;refGene_gn=TRIP12;refGene_tx=XM_019838807.2;refGene_gn=TRIP12;refGene_tx=XM_019838808.2;refGene_gn=TRIP12;refGene_tx=XM_019838809.2;refGene_gn=TRIP12;refGene_tx=XM_023259773.1;refGene_gn=TRIP12;refGene_tx=XM_023259774.1;refGene_gn=TRIP12;refGene_tx=XM_023259775.1;refGene_gn=TRIP12;refGene_tx=XM_023259776.1;refGene_gn=TRIP12;refGene_tx=XM_023259777.1;refGene_gn=TRIP12;refGene_tx=XM_023259778.1;refGene_gn=TRIP12;refGene_tx=XM_023259779.1;refGene_gn=TRIP12;refGene_tx=XM_023259780.1;refGene_gn=TRIP12;refGene_tx=XM_023259781.1;refGene_gn=TRIP12;refGene_tx=XM_023259782.1;refGene_gn=TRIP12;refGene_tx=XM_023259783.1;refGene_gn=TRIP12;refGene_tx=XM_023259784.1;refGene_gn=TRIP12;refGene_tx=XM_023259785.1;refGene_gn=TRIP12;refGene_tx=XM_023259786.1
C1 213289097 213289289
refGene_gn=TRIP12;refGene_tx=XM_011285634.3;refGene_gn=TRIP12;refGene_tx=XM_019838807.2;refGene_gn=TRIP12;refGene_tx=XM_019838808.2;refGene_gn=TRIP12;refGene_tx=XM_019838809.2;refGene_gn=TRIP12;refGene_tx=XM_023259773.1;refGene_gn=TRIP12;refGene_tx=XM_023259774.1;refGene_gn=TRIP12;refGene_tx=XM_023259775.1;refGene_gn=TRIP12;refGene_tx=XM_023259776.1;refGene_gn=TRIP12;refGene_tx=XM_023259777.1;refGene_gn=TRIP12;refGene_tx=XM_023259778.1;refGene_gn=TRIP12;refGene_tx=XM_023259779.1;refGene_gn=TRIP12;refGene_tx=XM_023259780.1;refGene_gn=TRIP12;refGene_tx=XM_023259781.1;refGene_gn=TRIP12;refGene_tx=XM_023259782.1;refGene_gn=TRIP12;refGene_tx=XM_023259783.1;refGene_gn=TRIP12;refGene_tx=XM_023259784.1;refGene_gn=TRIP12;refGene_tx=XM_023259785.1;refGene_gn=TRIP12;refGene_tx=XM_023259786.1
C1 213290102 213290244
refGene_gn=TRIP12;refGene_tx=XM_011285634.3;refGene_gn=TRIP12;refGene_tx=XM_019838807.2;refGene_gn=TRIP12;refGene_tx=XM_019838808.2;refGene_gn=TRIP12;refGene_tx=XM_019838809.2;refGene_gn=TRIP12;refGene_tx=XM_023259773.1;refGene_gn=TRIP12;refGene_tx=XM_023259774.1;refGene_gn=TRIP12;refGene_tx=XM_023259775.1;refGene_gn=TRIP12;refGene_tx=XM_023259776.1;refGene_gn=TRIP12;refGene_tx=XM_023259777.1;refGene_gn=TRIP12;refGene_tx=XM_023259778.1;refGene_gn=TRIP12;refGene_tx=XM_023259779.1;refGene_gn=TRIP12;refGene_tx=XM_023259780.1;refGene_gn=TRIP12;refGene_tx=XM_023259781.1;refGene_gn=TRIP12;refGene_tx=XM_023259782.1;refGene_gn=TRIP12;refGene_tx=XM_023259783.1;refGene_gn=TRIP12;refGene_tx=XM_023259784.1;refGene_gn=TRIP12;refGene_tx=XM_023259785.1;refGene_gn=TRIP12;refGene_tx=XM_023259786.1
C1 213291415 213291590
refGene_gn=TRIP12;refGene_tx=XM_011285634.3;refGene_gn=TRIP12;refGene_tx=XM_019838807.2;refGene_gn=TRIP12;refGene_tx=XM_019838808.2;refGene_gn=TRIP12;refGene_tx=XM_019838809.2;refGene_gn=TRIP12;refGene_tx=XM_023259773.1;refGene_gn=TRIP12;refGene_tx=XM_023259774.1;refGene_gn=TRIP12;refGene_tx=XM_023259775.1;refGene_gn=TRIP12;refGene_tx=XM_023259776.

1;refGene_gn=TRIP12;refGene_tx=XM_023259777.1;refGene_gn=TRIP12;refGene_tx=XM_023259778.1;refGene_gn=TRIP12;refGene_tx=XM_023259779.1;refGene_gn=TRIP12;refGene_tx=XM_023259780.1;refGene_gn=TRIP12;refGene_tx=XM_023259781.1;refGene_gn=TRIP12;refGene_tx=XM_023259782.1;refGene_gn=TRIP12;refGene_tx=XM_023259783.1;refGene_gn=TRIP12;refGene_tx=XM_023259784.1;refGene_gn=TRIP12;refGene_tx=XM_023259785.1;refGene_gn=TRIP12;refGene_tx=XM_023259786.1
C1 213291820 213291921
refGene_gn=TRIP12;refGene_tx=XM_011285634.3;refGene_gn=TRIP12;refGene_tx=XM_019838807.2;refGene_gn=TRIP12;refGene_tx=XM_019838808.2;refGene_gn=TRIP12;refGene_tx=XM_019838809.2;refGene_gn=TRIP12;refGene_tx=XM_023259773.1;refGene_gn=TRIP12;refGene_tx=XM_023259774.1;refGene_gn=TRIP12;refGene_tx=XM_023259775.1;refGene_gn=TRIP12;refGene_tx=XM_023259776.1;refGene_gn=TRIP12;refGene_tx=XM_023259777.1;refGene_gn=TRIP12;refGene_tx=XM_023259778.1;refGene_gn=TRIP12;refGene_tx=XM_023259779.1;refGene_gn=TRIP12;refGene_tx=XM_023259780.1;refGene_gn=TRIP12;refGene_tx=XM_023259781.1;refGene_gn=TRIP12;refGene_tx=XM_023259782.1;refGene_gn=TRIP12;refGene_tx=XM_023259783.1;refGene_gn=TRIP12;refGene_tx=XM_023259784.1;refGene_gn=TRIP12;refGene_tx=XM_023259785.1;refGene_gn=TRIP12;refGene_tx=XM_023259786.1
C1 213294442 213294650
refGene_gn=TRIP12;refGene_tx=XM_011285634.3;refGene_gn=TRIP12;refGene_tx=XM_019838807.2;refGene_gn=TRIP12;refGene_tx=XM_019838808.2;refGene_gn=TRIP12;refGene_tx=XM_019838809.2;refGene_gn=TRIP12;refGene_tx=XM_023259773.1;refGene_gn=TRIP12;refGene_tx=XM_023259774.1;refGene_gn=TRIP12;refGene_tx=XM_023259775.1;refGene_gn=TRIP12;refGene_tx=XM_023259776.1;refGene_gn=TRIP12;refGene_tx=XM_023259777.1;refGene_gn=TRIP12;refGene_tx=XM_023259778.1;refGene_gn=TRIP12;refGene_tx=XM_023259779.1;refGene_gn=TRIP12;refGene_tx=XM_023259780.1;refGene_gn=TRIP12;refGene_tx=XM_023259781.1;refGene_gn=TRIP12;refGene_tx=XM_023259782.1;refGene_gn=TRIP12;refGene_tx=XM_023259783.1;refGene_gn=TRIP12;refGene_tx=XM_023259784.1;refGene_gn=TRIP12;refGene_tx=XM_023259785.1;refGene_gn=TRIP12;refGene_tx=XM_023259786.1
C1 213296966 213297085
refGene_gn=TRIP12;refGene_tx=XM_011285634.3;refGene_gn=TRIP12;refGene_tx=XM_019838807.2;refGene_gn=TRIP12;refGene_tx=XM_019838808.2;refGene_gn=TRIP12;refGene_tx=XM_019838809.2;refGene_gn=TRIP12;refGene_tx=XM_023259773.1;refGene_gn=TRIP12;refGene_tx=XM_023259774.1;refGene_gn=TRIP12;refGene_tx=XM_023259775.1;refGene_gn=TRIP12;refGene_tx=XM_023259776.1;refGene_gn=TRIP12;refGene_tx=XM_023259777.1;refGene_gn=TRIP12;refGene_tx=XM_023259778.1;refGene_gn=TRIP12;refGene_tx=XM_023259779.1;refGene_gn=TRIP12;refGene_tx=XM_023259780.1;refGene_gn=TRIP12;refGene_tx=XM_023259781.1;refGene_gn=TRIP12;refGene_tx=XM_023259782.1;refGene_gn=TRIP12;refGene_tx=XM_023259783.1;refGene_gn=TRIP12;refGene_tx=XM_023259784.1;refGene_gn=TRIP12;refGene_tx=XM_023259785.1;refGene_gn=TRIP12;refGene_tx=XM_023259786.1
C1 213297408 213297637
refGene_gn=TRIP12;refGene_tx=XM_011285634.3;refGene_gn=TRIP12;refGene_tx=XM_019838807.2;refGene_gn=TRIP12;refGene_tx=XM_019838808.2;refGene_gn=TRIP12;refGene_tx=XM_019838809.2;refGene_gn=TRIP12;refGene_tx=XM_02

3259773.1;refGene_gn=TRIP12;refGene_tx=XM_023259774.1;refGene_gn=TRIP12;refGene_tx=XM_023259775.1;refGene_gn=TRIP12;refGene_tx=XM_023259776.1;refGene_gn=TRIP12;refGene_tx=XM_023259777.1;refGene_gn=TRIP12;refGene_tx=XM_023259778.1;refGene_gn=TRIP12;refGene_tx=XM_023259779.1;refGene_gn=TRIP12;refGene_tx=XM_023259780.1;refGene_gn=TRIP12;refGene_tx=XM_023259781.1;refGene_gn=TRIP12;refGene_tx=XM_023259782.1;refGene_gn=TRIP12;refGene_tx=XM_023259783.1;refGene_gn=TRIP12;refGene_tx=XM_023259784.1;refGene_gn=TRIP12;refGene_tx=XM_023259785.1;refGene_gn=TRIP12;refGene_tx=XM_023259786.1
C1 213298740 213298894
refGene_gn=TRIP12;refGene_tx=XM_011285634.3;refGene_gn=TRIP12;refGene_tx=XM_019838807.2;refGene_gn=TRIP12;refGene_tx=XM_019838808.2;refGene_gn=TRIP12;refGene_tx=XM_019838809.2;refGene_gn=TRIP12;refGene_tx=XM_023259773.1;refGene_gn=TRIP12;refGene_tx=XM_023259774.1;refGene_gn=TRIP12;refGene_tx=XM_023259775.1;refGene_gn=TRIP12;refGene_tx=XM_023259776.1;refGene_gn=TRIP12;refGene_tx=XM_023259777.1;refGene_gn=TRIP12;refGene_tx=XM_023259778.1;refGene_gn=TRIP12;refGene_tx=XM_023259779.1;refGene_gn=TRIP12;refGene_tx=XM_023259780.1;refGene_gn=TRIP12;refGene_tx=XM_023259781.1;refGene_gn=TRIP12;refGene_tx=XM_023259782.1;refGene_gn=TRIP12;refGene_tx=XM_023259783.1;refGene_gn=TRIP12;refGene_tx=XM_023259784.1;refGene_gn=TRIP12;refGene_tx=XM_023259785.1;refGene_gn=TRIP12;refGene_tx=XM_023259786.1
C1 213300714 213300874
refGene_gn=TRIP12;refGene_tx=XM_011285634.3;refGene_gn=TRIP12;refGene_tx=XM_019838807.2;refGene_gn=TRIP12;refGene_tx=XM_019838808.2;refGene_gn=TRIP12;refGene_tx=XM_019838809.2;refGene_gn=TRIP12;refGene_tx=XM_023259773.1;refGene_gn=TRIP12;refGene_tx=XM_023259774.1;refGene_gn=TRIP12;refGene_tx=XM_023259775.1;refGene_gn=TRIP12;refGene_tx=XM_023259776.1;refGene_gn=TRIP12;refGene_tx=XM_023259777.1;refGene_gn=TRIP12;refGene_tx=XM_023259778.1;refGene_gn=TRIP12;refGene_tx=XM_023259779.1;refGene_gn=TRIP12;refGene_tx=XM_023259780.1;refGene_gn=TRIP12;refGene_tx=XM_023259781.1;refGene_gn=TRIP12;refGene_tx=XM_023259782.1;refGene_gn=TRIP12;refGene_tx=XM_023259783.1;refGene_gn=TRIP12;refGene_tx=XM_023259784.1;refGene_gn=TRIP12;refGene_tx=XM_023259785.1;refGene_gn=TRIP12;refGene_tx=XM_023259786.1
C1 213300988 213301106
refGene_gn=TRIP12;refGene_tx=XM_011285634.3;refGene_gn=TRIP12;refGene_tx=XM_019838807.2;refGene_gn=TRIP12;refGene_tx=XM_019838808.2;refGene_gn=TRIP12;refGene_tx=XM_019838809.2;refGene_gn=TRIP12;refGene_tx=XM_023259773.1;refGene_gn=TRIP12;refGene_tx=XM_023259774.1;refGene_gn=TRIP12;refGene_tx=XM_023259775.1;refGene_gn=TRIP12;refGene_tx=XM_023259776.1;refGene_gn=TRIP12;refGene_tx=XM_023259777.1;refGene_gn=TRIP12;refGene_tx=XM_023259778.1;refGene_gn=TRIP12;refGene_tx=XM_023259779.1;refGene_gn=TRIP12;refGene_tx=XM_023259780.1;refGene_gn=TRIP12;refGene_tx=XM_023259781.1;refGene_gn=TRIP12;refGene_tx=XM_023259782.1;refGene_gn=TRIP12;refGene_tx=XM_023259783.1;refGene_gn=TRIP12;refGene_tx=XM_023259784.1;refGene_gn=TRIP12;refGene_tx=XM_023259785.1;refGene_gn=TRIP12;refGene_tx=XM_023259786.1
C1 213304061 213304227
refGene_gn=TRIP12;refGene_tx=XM_011285634.3;refGene_gn=TRIP12;refGene_

tx=XM_019838807.2;refGene_gn=TRIP12;refGene_tx=XM_019838808.2;refGene_gn=TRIP12;refGene_tx=XM_019838809.2;refGene_gn=TRIP12;refGene_tx=XM_023259773.1;refGene_gn=TRIP12;refGene_tx=XM_023259774.1;refGene_gn=TRIP12;refGene_tx=XM_023259775.1;refGene_gn=TRIP12;refGene_tx=XM_023259776.1;refGene_gn=TRIP12;refGene_tx=XM_023259777.1;refGene_gn=TRIP12;refGene_tx=XM_023259778.1;refGene_gn=TRIP12;refGene_tx=XM_023259779.1;refGene_gn=TRIP12;refGene_tx=XM_023259780.1;refGene_gn=TRIP12;refGene_tx=XM_023259781.1;refGene_gn=TRIP12;refGene_tx=XM_023259782.1;refGene_gn=TRIP12;refGene_tx=XM_023259783.1;refGene_gn=TRIP12;refGene_tx=XM_023259784.1;refGene_gn=TRIP12;refGene_tx=XM_023259785.1;refGene_gn=TRIP12;refGene_tx=XM_023259786.1
C1 213304318 213304387
refGene_gn=TRIP12;refGene_tx=XM_011285634.3;refGene_gn=TRIP12;refGene_tx=XM_019838807.2;refGene_gn=TRIP12;refGene_tx=XM_019838808.2;refGene_gn=TRIP12;refGene_tx=XM_019838809.2;refGene_gn=TRIP12;refGene_tx=XM_023259773.1;refGene_gn=TRIP12;refGene_tx=XM_023259774.1;refGene_gn=TRIP12;refGene_tx=XM_023259775.1;refGene_gn=TRIP12;refGene_tx=XM_023259776.1;refGene_gn=TRIP12;refGene_tx=XM_023259777.1;refGene_gn=TRIP12;refGene_tx=XM_023259778.1;refGene_gn=TRIP12;refGene_tx=XM_023259779.1;refGene_gn=TRIP12;refGene_tx=XM_023259780.1;refGene_gn=TRIP12;refGene_tx=XM_023259781.1;refGene_gn=TRIP12;refGene_tx=XM_023259782.1;refGene_gn=TRIP12;refGene_tx=XM_023259783.1;refGene_gn=TRIP12;refGene_tx=XM_023259784.1;refGene_gn=TRIP12;refGene_tx=XM_023259785.1;refGene_gn=TRIP12;refGene_tx=XM_023259786.1
C1 213306260 213306422
refGene_gn=TRIP12;refGene_tx=XM_011285634.3;refGene_gn=TRIP12;refGene_tx=XM_019838807.2;refGene_gn=TRIP12;refGene_tx=XM_019838808.2;refGene_gn=TRIP12;refGene_tx=XM_019838809.2;refGene_gn=TRIP12;refGene_tx=XM_023259773.1;refGene_gn=TRIP12;refGene_tx=XM_023259774.1;refGene_gn=TRIP12;refGene_tx=XM_023259775.1;refGene_gn=TRIP12;refGene_tx=XM_023259776.1;refGene_gn=TRIP12;refGene_tx=XM_023259777.1;refGene_gn=TRIP12;refGene_tx=XM_023259778.1;refGene_gn=TRIP12;refGene_tx=XM_023259779.1;refGene_gn=TRIP12;refGene_tx=XM_023259780.1;refGene_gn=TRIP12;refGene_tx=XM_023259781.1;refGene_gn=TRIP12;refGene_tx=XM_023259782.1;refGene_gn=TRIP12;refGene_tx=XM_023259783.1;refGene_gn=TRIP12;refGene_tx=XM_023259784.1;refGene_gn=TRIP12;refGene_tx=XM_023259785.1;refGene_gn=TRIP12;refGene_tx=XM_023259786.1
C1 213306622 213306715
refGene_gn=TRIP12;refGene_tx=XM_011285634.3;refGene_gn=TRIP12;refGene_tx=XM_019838807.2;refGene_gn=TRIP12;refGene_tx=XM_019838808.2;refGene_gn=TRIP12;refGene_tx=XM_019838809.2;refGene_gn=TRIP12;refGene_tx=XM_023259773.1;refGene_gn=TRIP12;refGene_tx=XM_023259774.1;refGene_gn=TRIP12;refGene_tx=XM_023259775.1;refGene_gn=TRIP12;refGene_tx=XM_023259776.1;refGene_gn=TRIP12;refGene_tx=XM_023259777.1;refGene_gn=TRIP12;refGene_tx=XM_023259778.1;refGene_gn=TRIP12;refGene_tx=XM_023259779.1;refGene_gn=TRIP12;refGene_tx=XM_023259780.1;refGene_gn=TRIP12;refGene_tx=XM_023259781.1;refGene_gn=TRIP12;refGene_tx=XM_023259782.1;refGene_gn=TRIP12;refGene_tx=XM_023259783.1;refGene_gn=TRIP12;refGene_tx=XM_023259784.1;refGene_gn=TRIP12;refGene_tx=XM_023259785.1;refGene_gn=TRIP12;refGene_tx=XM_023259786.1

C1 213307846 213307942
refGene_gn=TRIP12;refGene_tx=XM_011285634.3;refGene_gn=TRIP12;refGene_tx=XM_019838807.2;refGene_gn=TRIP12;refGene_tx=XM_019838808.2;refGene_gn=TRIP12;refGene_tx=XM_019838809.2;refGene_gn=TRIP12;refGene_tx=XM_023259773.1;refGene_gn=TRIP12;refGene_tx=XM_023259774.1;refGene_gn=TRIP12;refGene_tx=XM_023259775.1;refGene_gn=TRIP12;refGene_tx=XM_023259776.1;refGene_gn=TRIP12;refGene_tx=XM_023259777.1;refGene_gn=TRIP12;refGene_tx=XM_023259778.1;refGene_gn=TRIP12;refGene_tx=XM_023259779.1;refGene_gn=TRIP12;refGene_tx=XM_023259780.1;refGene_gn=TRIP12;refGene_tx=XM_023259781.1;refGene_gn=TRIP12;refGene_tx=XM_023259782.1;refGene_gn=TRIP12;refGene_tx=XM_023259783.1;refGene_gn=TRIP12;refGene_tx=XM_023259784.1;refGene_gn=TRIP12;refGene_tx=XM_023259785.1;refGene_gn=TRIP12;refGene_tx=XM_023259786.1
C1 213308022 213308058
refGene_gn=TRIP12;refGene_tx=XM_011285634.3;refGene_gn=TRIP12;refGene_tx=XM_019838807.2;refGene_gn=TRIP12;refGene_tx=XM_019838808.2;refGene_gn=TRIP12;refGene_tx=XM_019838809.2;refGene_gn=TRIP12;refGene_tx=XM_023259773.1;refGene_gn=TRIP12;refGene_tx=XM_023259774.1;refGene_gn=TRIP12;refGene_tx=XM_023259775.1;refGene_gn=TRIP12;refGene_tx=XM_023259776.1;refGene_gn=TRIP12;refGene_tx=XM_023259777.1;refGene_gn=TRIP12;refGene_tx=XM_023259778.1;refGene_gn=TRIP12;refGene_tx=XM_023259779.1;refGene_gn=TRIP12;refGene_tx=XM_023259780.1;refGene_gn=TRIP12;refGene_tx=XM_023259781.1;refGene_gn=TRIP12;refGene_tx=XM_023259782.1;refGene_gn=TRIP12;refGene_tx=XM_023259783.1;refGene_gn=TRIP12;refGene_tx=XM_023259784.1;refGene_gn=TRIP12;refGene_tx=XM_023259785.1;refGene_gn=TRIP12;refGene_tx=XM_023259786.1
C1 213310612 213310761
refGene_gn=TRIP12;refGene_tx=XM_011285634.3;refGene_gn=TRIP12;refGene_tx=XM_019838807.2;refGene_gn=TRIP12;refGene_tx=XM_019838808.2;refGene_gn=TRIP12;refGene_tx=XM_019838809.2;refGene_gn=TRIP12;refGene_tx=XM_023259773.1;refGene_gn=TRIP12;refGene_tx=XM_023259774.1;refGene_gn=TRIP12;refGene_tx=XM_023259775.1;refGene_gn=TRIP12;refGene_tx=XM_023259776.1;refGene_gn=TRIP12;refGene_tx=XM_023259777.1;refGene_gn=TRIP12;refGene_tx=XM_023259778.1;refGene_gn=TRIP12;refGene_tx=XM_023259779.1;refGene_gn=TRIP12;refGene_tx=XM_023259780.1;refGene_gn=TRIP12;refGene_tx=XM_023259781.1;refGene_gn=TRIP12;refGene_tx=XM_023259782.1;refGene_gn=TRIP12;refGene_tx=XM_023259783.1;refGene_gn=TRIP12;refGene_tx=XM_023259784.1;refGene_gn=TRIP12;refGene_tx=XM_023259785.1;refGene_gn=TRIP12;refGene_tx=XM_023259786.1
C1 213319968 213320064
refGene_gn=TRIP12;refGene_tx=XM_011285634.3;refGene_gn=TRIP12;refGene_tx=XM_019838807.2;refGene_gn=TRIP12;refGene_tx=XM_019838808.2;refGene_gn=TRIP12;refGene_tx=XM_019838809.2;refGene_gn=TRIP12;refGene_tx=XM_023259773.1;refGene_gn=TRIP12;refGene_tx=XM_023259774.1;refGene_gn=TRIP12;refGene_tx=XM_023259775.1;refGene_gn=TRIP12;refGene_tx=XM_023259776.1;refGene_gn=TRIP12;refGene_tx=XM_023259777.1;refGene_gn=TRIP12;refGene_tx=XM_023259778.1;refGene_gn=TRIP12;refGene_tx=XM_023259779.1;refGene_gn=TRIP12;refGene_tx=XM_023259780.1;refGene_gn=TRIP12;refGene_tx=XM_023259781.1;refGene_gn=TRIP12;refGene_tx=XM_023259782.1;refGene_gn=TRIP12;refGene_tx=XM_023259783.1;refGene_gn=TRIP12;refGene_tx=XM_02325978

4.1;refGene_gn=TRIP12;refGene_tx=XM_023259785.1;refGene_gn=TRIP12;refGene_tx=XM_023259786.1
C1 213321155 213321239
refGene_gn=TRIP12;refGene_tx=XM_011285634.3;refGene_gn=TRIP12;refGene_tx=XM_019838807.2;refGene_gn=TRIP12;refGene_tx=XM_019838808.2;refGene_gn=TRIP12;refGene_tx=XM_019838809.2;refGene_gn=TRIP12;refGene_tx=XM_023259773.1;refGene_gn=TRIP12;refGene_tx=XM_023259774.1;refGene_gn=TRIP12;refGene_tx=XM_023259775.1;refGene_gn=TRIP12;refGene_tx=XM_023259776.1;refGene_gn=TRIP12;refGene_tx=XM_023259777.1;refGene_gn=TRIP12;refGene_tx=XM_023259778.1;refGene_gn=TRIP12;refGene_tx=XM_023259779.1;refGene_gn=TRIP12;refGene_tx=XM_023259780.1;refGene_gn=TRIP12;refGene_tx=XM_023259781.1;refGene_gn=TRIP12;refGene_tx=XM_023259782.1;refGene_gn=TRIP12;refGene_tx=XM_023259783.1;refGene_gn=TRIP12;refGene_tx=XM_023259784.1;refGene_gn=TRIP12;refGene_tx=XM_023259785.1;refGene_gn=TRIP12;refGene_tx=XM_023259786.1
C1 213327075 213327212
refGene_gn=TRIP12;refGene_tx=XM_011285634.3;refGene_gn=TRIP12;refGene_tx=XM_019838807.2;refGene_gn=TRIP12;refGene_tx=XM_019838808.2;refGene_gn=TRIP12;refGene_tx=XM_019838809.2;refGene_gn=TRIP12;refGene_tx=XM_023259773.1;refGene_gn=TRIP12;refGene_tx=XM_023259774.1;refGene_gn=TRIP12;refGene_tx=XM_023259775.1;refGene_gn=TRIP12;refGene_tx=XM_023259776.1;refGene_gn=TRIP12;refGene_tx=XM_023259777.1;refGene_gn=TRIP12;refGene_tx=XM_023259778.1;refGene_gn=TRIP12;refGene_tx=XM_023259779.1;refGene_gn=TRIP12;refGene_tx=XM_023259780.1;refGene_gn=TRIP12;refGene_tx=XM_023259781.1;refGene_gn=TRIP12;refGene_tx=XM_023259782.1;refGene_gn=TRIP12;refGene_tx=XM_023259783.1;refGene_gn=TRIP12;refGene_tx=XM_023259784.1;refGene_gn=TRIP12;refGene_tx=XM_023259785.1;refGene_gn=TRIP12;refGene_tx=XM_023259786.1
C1 213330456 213330562
refGene_gn=TRIP12;refGene_tx=XM_011285634.3;refGene_gn=TRIP12;refGene_tx=XM_019838807.2;refGene_gn=TRIP12;refGene_tx=XM_019838808.2;refGene_gn=TRIP12;refGene_tx=XM_019838809.2;refGene_gn=TRIP12;refGene_tx=XM_023259773.1;refGene_gn=TRIP12;refGene_tx=XM_023259774.1;refGene_gn=TRIP12;refGene_tx=XM_023259775.1;refGene_gn=TRIP12;refGene_tx=XM_023259776.1;refGene_gn=TRIP12;refGene_tx=XM_023259777.1;refGene_gn=TRIP12;refGene_tx=XM_023259778.1;refGene_gn=TRIP12;refGene_tx=XM_023259779.1;refGene_gn=TRIP12;refGene_tx=XM_023259780.1;refGene_gn=TRIP12;refGene_tx=XM_023259781.1;refGene_gn=TRIP12;refGene_tx=XM_023259782.1;refGene_gn=TRIP12;refGene_tx=XM_023259783.1;refGene_gn=TRIP12;refGene_tx=XM_023259784.1;refGene_gn=TRIP12;refGene_tx=XM_023259785.1;refGene_gn=TRIP12;refGene_tx=XM_023259786.1
C1 213344265 213345068
refGene_gn=TRIP12;refGene_tx=XM_011285634.3;refGene_gn=TRIP12;refGene_tx=XM_019838807.2;refGene_gn=TRIP12;refGene_tx=XM_019838808.2;refGene_gn=TRIP12;refGene_tx=XM_019838809.2;refGene_gn=TRIP12;refGene_tx=XM_023259773.1;refGene_gn=TRIP12;refGene_tx=XM_023259774.1;refGene_gn=TRIP12;refGene_tx=XM_023259775.1;refGene_gn=TRIP12;refGene_tx=XM_023259776.1;refGene_gn=TRIP12;refGene_tx=XM_023259777.1;refGene_gn=TRIP12;refGene_tx=XM_023259778.1;refGene_gn=TRIP12;refGene_tx=XM_023259779.1;refGene_gn=TRIP12;refGene_tx=XM_023259780.1;refGene_gn=TRIP12;refGene_tx=XM_

023259781.1;refGene_gn=TRIP12;refGene_tx=XM_023259782.1;refGene_gn=TRIP12;refGene_tx=XM_023259783.1;refGene_gn=TRIP12;refGene_tx=XM_023259784.1;refGene_gn=TRIP12;refGene_tx=XM_023259785.1;refGene_gn=TRIP12;refGene_tx=XM_023259786.1
C1 213345911 213346037
refGene_gn=TRIP12;refGene_tx=XM_011285634.3;refGene_gn=TRIP12;refGene_tx=XM_019838807.2;refGene_gn=TRIP12;refGene_tx=XM_019838808.2;refGene_gn=TRIP12;refGene_tx=XM_019838809.2;refGene_gn=TRIP12;refGene_tx=XM_023259773.1;refGene_gn=TRIP12;refGene_tx=XM_023259774.1;refGene_gn=TRIP12;refGene_tx=XM_023259775.1;refGene_gn=TRIP12;refGene_tx=XM_023259776.1;refGene_gn=TRIP12;refGene_tx=XM_023259777.1;refGene_gn=TRIP12;refGene_tx=XM_023259778.1;refGene_gn=TRIP12;refGene_tx=XM_023259779.1;refGene_gn=TRIP12;refGene_tx=XM_023259780.1;refGene_gn=TRIP12;refGene_tx=XM_023259781.1
C1 213361485 213361583
refGene_gn=TRIP12;refGene_tx=XM_011285634.3;refGene_gn=TRIP12;refGene_tx=XM_019838807.2;refGene_gn=TRIP12;refGene_tx=XM_019838808.2;refGene_gn=TRIP12;refGene_tx=XM_019838809.2;refGene_gn=TRIP12;refGene_tx=XM_023259773.1;refGene_gn=TRIP12;refGene_tx=XM_023259774.1;refGene_gn=TRIP12;refGene_tx=XM_023259775.1;refGene_gn=TRIP12;refGene_tx=XM_023259776.1;refGene_gn=TRIP12;refGene_tx=XM_023259777.1;refGene_gn=TRIP12;refGene_tx=XM_023259778.1;refGene_gn=TRIP12;refGene_tx=XM_023259779.1;refGene_gn=TRIP12;refGene_tx=XM_023259780.1;refGene_gn=TRIP12;refGene_tx=XM_023259781.1;refGene_gn=TRIP12;refGene_tx=XM_023259782.1;refGene_gn=TRIP12;refGene_tx=XM_023259783.1;refGene_gn=TRIP12;refGene_tx=XM_023259784.1;refGene_gn=TRIP12;refGene_tx=XM_023259785.1;refGene_gn=TRIP12;refGene_tx=XM_023259786.1
C1 213400575 213400671
refGene_gn=FBXO36;refGene_tx=XM_003991229.5;refGene_gn=FBXO36;refGene_tx=XM_019838812.2;refGene_gn=FBXO36;refGene_tx=XM_023259787.1
C1 213461968 213462077
refGene_gn=FBXO36;refGene_tx=XM_003991229.5;refGene_gn=FBXO36;refGene_tx=XM_019838812.2;refGene_gn=FBXO36;refGene_tx=XM_023259787.1
C1 213477968 213478141
refGene_gn=FBXO36;refGene_tx=XM_003991229.5;refGene_gn=FBXO36;refGene_tx=XM_019838812.2;refGene_gn=FBXO36;refGene_tx=XM_023259787.1
C1 213487557 213487684
refGene_gn=FBXO36;refGene_tx=XM_019838812.2
C1 213492589 213492778
refGene_gn=FBXO36;refGene_tx=XM_003991229.5;refGene_gn=FBXO36;refGene_tx=XM_019838812.2;refGene_gn=FBXO36;refGene_tx=XM_023259787.1
C1 213511039 213511191
refGene_gn=SLC16A14;refGene_tx=XM_023259789.1
C1 213518992 213519967
refGene_gn=SLC16A14;refGene_tx=XM_023259789.1
C1 213525233 213525377
refGene_gn=SLC16A14;refGene_tx=XM_023259789.1
C1 213532620 213532879
refGene_gn=SLC16A14;refGene_tx=XM_023259789.1
C1 213577830 213577850

refGene_gn=LOC101093204;refGene_tx=XM_023259790.1
C1	213585276	213586213
refGene_gn=LOC101093204;refGene_tx=XM_019838826.2;refGene_gn=LOC101093204;refGene_tx=XM_023259790.1;refGene_gn=LOC101093204;refGene_tx=XM_023259791.1
C1	213607065	213607176
refGene_gn=LOC101093454;refGene_tx=XM_011285632.3
C1	213608343	213608485
refGene_gn=LOC101093454;refGene_tx=XM_011285632.3
C1	213609062	213609133
refGene_gn=LOC101093454;refGene_tx=XM_011285632.3
C1	213609390	213609496
refGene_gn=LOC101093454;refGene_tx=XM_011285632.3
C1	213610198	213610225
refGene_gn=LOC101093454;refGene_tx=XM_011285632.3
C1	213610226	213610312
refGene_gn=LOC101093454;refGene_tx=XM_011285632.3
C1	213615532	213615577
refGene_gn=LOC101093454;refGene_tx=XM_011285632.3
C1	213616384	213616527
refGene_gn=LOC101093454;refGene_tx=XM_011285632.3
C1	213616967	213617066
refGene_gn=LOC101093454;refGene_tx=XM_011285632.3
C1	213621339	213621408
refGene_gn=LOC101093454;refGene_tx=XM_011285632.3
C1	213623900	213624050
refGene_gn=LOC101093454;refGene_tx=XM_011285632.3
C1	213626142	213626211
refGene_gn=LOC101093454;refGene_tx=XM_011285632.3
C1	213628313	213628391
refGene_gn=LOC101093454;refGene_tx=XM_011285632.3
C1	213630637	213630682
refGene_gn=LOC101093454;refGene_tx=XM_011285632.3
C1	213631027	213631294
refGene_gn=LOC101093454;refGene_tx=XM_011285632.3
C1	213633617	213633786
refGene_gn=LOC101093454;refGene_tx=XM_011285632.3
C1	213635410	213635557
refGene_gn=LOC101093454;refGene_tx=XM_011285632.3
C1	213654585	213654586
refGene_gn=LOC101093705;refGene_tx=XM_006935658.3;refGene_gn=LOC101093705;refGene_tx=XM_006935659.3;refGene_gn=LOC101093705;refGene_tx=XM_006935660.4;refGene_gn=LOC101093705;refGene_tx=XM_006935661.3;refGene_gn=LOC101093705;refGene_tx=XM_006935662.3;refGene_gn=LOC101093705;refGene_tx=XM_006935664.3;refGene_gn=LOC101093705;refGene_tx=XM_006935665.3;refGene_gn=LOC101093705;refGene_tx=XM_019838813.2;refGene_gn=LOC101093705;refGene_tx=XM_019838814.2;refGene_gn=LOC101093705;refGene_tx=XM_019838818.2;refGene_gn=LOC101093705;refGene_tx=XM_019838819.2;refGene_gn=LOC101093705;refGene_tx=XM_019838822.2;refGene_gn=LOC101093705;refGene_tx=XM_019838823.2;refGene_gn=LOC101093705;refGene_tx=XM_019838824.2

C1	213664055	213664139
refGene_gn=LOC101093705;refGene_tx=XM_006935658.3;refGene_gn=LOC101093
705;refGene_tx=XM_006935659.3;refGene_gn=LOC101093705;refGene_tx=XM_00
6935660.4;refGene_gn=LOC101093705;refGene_tx=XM_006935661.3;refGene_gn
=LOC101093705;refGene_tx=XM_006935662.3;refGene_gn=LOC101093705;refGen
e_tx=XM_006935663.3;refGene_gn=LOC101093705;refGene_tx=XM_006935664.3;
refGene_gn=LOC101093705;refGene_tx=XM_006935665.3;refGene_gn=LOC101093
705;refGene_tx=XM_019838813.2;refGene_gn=LOC101093705;refGene_tx=XM_01
9838814.2;refGene_gn=LOC101093705;refGene_tx=XM_019838816.2;refGene_gn
=LOC101093705;refGene_tx=XM_019838817.2;refGene_gn=LOC101093705;refGen
e_tx=XM_019838818.2;refGene_gn=LOC101093705;refGene_tx=XM_019838819.2;
refGene_gn=LOC101093705;refGene_tx=XM_019838822.2;refGene_gn=LOC101093
705;refGene_tx=XM_019838823.2;refGene_gn=LOC101093705;refGene_tx=XM_01
9838824.2
C1	213666544	213666676
refGene_gn=LOC101093705;refGene_tx=XM_006935658.3;refGene_gn=LOC101093
705;refGene_tx=XM_006935659.3;refGene_gn=LOC101093705;refGene_tx=XM_00
6935660.4;refGene_gn=LOC101093705;refGene_tx=XM_006935661.3;refGene_gn
=LOC101093705;refGene_tx=XM_006935662.3;refGene_gn=LOC101093705;refGen
e_tx=XM_006935663.3;refGene_gn=LOC101093705;refGene_tx=XM_006935664.3;
refGene_gn=LOC101093705;refGene_tx=XM_006935665.3;refGene_gn=LOC101093
705;refGene_tx=XM_006935666.3;refGene_gn=LOC101093705;refGene_tx=XM_01
9838813.2;refGene_gn=LOC101093705;refGene_tx=XM_019838814.2;refGene_gn
=LOC101093705;refGene_tx=XM_019838816.2;refGene_gn=LOC101093705;refGen
e_tx=XM_019838817.2;refGene_gn=LOC101093705;refGene_tx=XM_019838818.2;
refGene_gn=LOC101093705;refGene_tx=XM_019838819.2;refGene_gn=LOC101093
705;refGene_tx=XM_019838820.2;refGene_gn=LOC101093705;refGene_tx=XM_01
9838822.2;refGene_gn=LOC101093705;refGene_tx=XM_019838823.2;refGene_gn
=LOC101093705;refGene_tx=XM_019838824.2
C1	213668782	213668866
refGene_gn=LOC101093705;refGene_tx=XM_006935658.3;refGene_gn=LOC101093
705;refGene_tx=XM_006935659.3;refGene_gn=LOC101093705;refGene_tx=XM_00
6935660.4;refGene_gn=LOC101093705;refGene_tx=XM_006935661.3;refGene_gn
=LOC101093705;refGene_tx=XM_006935662.3;refGene_gn=LOC101093705;refGen
e_tx=XM_006935663.3;refGene_gn=LOC101093705;refGene_tx=XM_006935664.3;
refGene_gn=LOC101093705;refGene_tx=XM_006935665.3;refGene_gn=LOC101093
705;refGene_tx=XM_006935666.3;refGene_gn=LOC101093705;refGene_tx=XM_01
9838813.2;refGene_gn=LOC101093705;refGene_tx=XM_019838814.2;refGene_gn
=LOC101093705;refGene_tx=XM_019838816.2;refGene_gn=LOC101093705;refGen
e_tx=XM_019838817.2;refGene_gn=LOC101093705;refGene_tx=XM_019838818.2;
refGene_gn=LOC101093705;refGene_tx=XM_019838819.2;refGene_gn=LOC101093
705;refGene_tx=XM_019838820.2;refGene_gn=LOC101093705;refGene_tx=XM_01
9838822.2;refGene_gn=LOC101093705;refGene_tx=XM_019838823.2;refGene_gn
=LOC101093705;refGene_tx=XM_019838824.2
C1	213672983	213673067
refGene_gn=LOC101093705;refGene_tx=XM_006935658.3;refGene_gn=LOC101093
705;refGene_tx=XM_006935659.3;refGene_gn=LOC101093705;refGene_tx=XM_00
6935660.4;refGene_gn=LOC101093705;refGene_tx=XM_006935661.3;refGene_gn
=LOC101093705;refGene_tx=XM_006935662.3;refGene_gn=LOC101093705;refGen
e_tx=XM_006935663.3;refGene_gn=LOC101093705;refGene_tx=XM_006935664.3;

refGene_gn=LOC101093705;refGene_tx=XM_006935665.3;refGene_gn=LOC101093
705;refGene_tx=XM_006935666.3;refGene_gn=LOC101093705;refGene_tx=XM_01
9838813.2;refGene_gn=LOC101093705;refGene_tx=XM_019838814.2;refGene_gn
=LOC101093705;refGene_tx=XM_019838816.2;refGene_gn=LOC101093705;refGen
e_tx=XM_019838817.2;refGene_gn=LOC101093705;refGene_tx=XM_019838819.2;
refGene_gn=LOC101093705;refGene_tx=XM_019838820.2;refGene_gn=LOC101093
705;refGene_tx=XM_019838823.2;refGene_gn=LOC101093705;refGene_tx=XM_01
9838824.2
C1 213673833 213673917
refGene_gn=LOC101093705;refGene_tx=XM_006935658.3;refGene_gn=LOC101093
705;refGene_tx=XM_006935659.3;refGene_gn=LOC101093705;refGene_tx=XM_00
6935660.4;refGene_gn=LOC101093705;refGene_tx=XM_006935661.3;refGene_gn
=LOC101093705;refGene_tx=XM_006935662.3;refGene_gn=LOC101093705;refGen
e_tx=XM_006935663.3;refGene_gn=LOC101093705;refGene_tx=XM_006935664.3;
refGene_gn=LOC101093705;refGene_tx=XM_006935665.3;refGene_gn=LOC101093
705;refGene_tx=XM_006935666.3;refGene_gn=LOC101093705;refGene_tx=XM_01
9838813.2;refGene_gn=LOC101093705;refGene_tx=XM_019838814.2;refGene_gn
=LOC101093705;refGene_tx=XM_019838816.2;refGene_gn=LOC101093705;refGen
e_tx=XM_019838817.2;refGene_gn=LOC101093705;refGene_tx=XM_019838818.2;
refGene_gn=LOC101093705;refGene_tx=XM_019838819.2;refGene_gn=LOC101093
705;refGene_tx=XM_019838820.2;refGene_gn=LOC101093705;refGene_tx=XM_01
9838822.2;refGene_gn=LOC101093705;refGene_tx=XM_019838823.2;refGene_gn
=LOC101093705;refGene_tx=XM_019838824.2
C1 213676677 213676761
refGene_gn=LOC101093705;refGene_tx=XM_006935658.3;refGene_gn=LOC101093
705;refGene_tx=XM_006935659.3;refGene_gn=LOC101093705;refGene_tx=XM_00
6935660.4;refGene_gn=LOC101093705;refGene_tx=XM_006935661.3;refGene_gn
=LOC101093705;refGene_tx=XM_006935662.3;refGene_gn=LOC101093705;refGen
e_tx=XM_006935663.3;refGene_gn=LOC101093705;refGene_tx=XM_006935664.3;
refGene_gn=LOC101093705;refGene_tx=XM_006935666.3;refGene_gn=LOC101093
705;refGene_tx=XM_019838813.2;refGene_gn=LOC101093705;refGene_tx=XM_01
9838814.2;refGene_gn=LOC101093705;refGene_tx=XM_019838816.2;refGene_gn
=LOC101093705;refGene_tx=XM_019838817.2;refGene_gn=LOC101093705;refGen
e_tx=XM_019838818.2;refGene_gn=LOC101093705;refGene_tx=XM_019838819.2;
refGene_gn=LOC101093705;refGene_tx=XM_019838820.2;refGene_gn=LOC101093
705;refGene_tx=XM_019838823.2;refGene_gn=LOC101093705;refGene_tx=XM_01
9838824.2
C1 213678780 213678861
refGene_gn=LOC101093705;refGene_tx=XM_006935658.3;refGene_gn=LOC101093
705;refGene_tx=XM_006935659.3;refGene_gn=LOC101093705;refGene_tx=XM_00
6935660.4;refGene_gn=LOC101093705;refGene_tx=XM_006935661.3;refGene_gn
=LOC101093705;refGene_tx=XM_006935662.3;refGene_gn=LOC101093705;refGen
e_tx=XM_006935663.3;refGene_gn=LOC101093705;refGene_tx=XM_006935664.3;
refGene_gn=LOC101093705;refGene_tx=XM_006935665.3;refGene_gn=LOC101093
705;refGene_tx=XM_006935666.3;refGene_gn=LOC101093705;refGene_tx=XM_01
9838813.2;refGene_gn=LOC101093705;refGene_tx=XM_019838814.2;refGene_gn
=LOC101093705;refGene_tx=XM_019838816.2;refGene_gn=LOC101093705;refGen
e_tx=XM_019838817.2;refGene_gn=LOC101093705;refGene_tx=XM_019838818.2;
refGene_gn=LOC101093705;refGene_tx=XM_019838819.2;refGene_gn=LOC101093
705;refGene_tx=XM_019838820.2;refGene_gn=LOC101093705;refGene_tx=XM_01

9838822.2;refGene_gn=LOC101093705;refGene_tx=XM_019838823.2;refGene_gn=LOC101093705;refGene_tx=XM_019838824.2
C1 213680014 213680098
refGene_gn=LOC101093705;refGene_tx=XM_006935658.3;refGene_gn=LOC101093705;refGene_tx=XM_006935659.3;refGene_gn=LOC101093705;refGene_tx=XM_006935660.4;refGene_gn=LOC101093705;refGene_tx=XM_006935661.3;refGene_gn=LOC101093705;refGene_tx=XM_006935662.3;refGene_gn=LOC101093705;refGene_tx=XM_006935663.3;refGene_gn=LOC101093705;refGene_tx=XM_006935664.3;refGene_gn=LOC101093705;refGene_tx=XM_006935665.3;refGene_gn=LOC101093705;refGene_tx=XM_006935666.3;refGene_gn=LOC101093705;refGene_tx=XM_019838813.2;refGene_gn=LOC101093705;refGene_tx=XM_019838814.2;refGene_gn=LOC101093705;refGene_tx=XM_019838816.2;refGene_gn=LOC101093705;refGene_tx=XM_019838817.2;refGene_gn=LOC101093705;refGene_tx=XM_019838818.2;refGene_gn=LOC101093705;refGene_tx=XM_019838819.2;refGene_gn=LOC101093705;refGene_tx=XM_019838820.2;refGene_gn=LOC101093705;refGene_tx=XM_019838822.2;refGene_gn=LOC101093705;refGene_tx=XM_019838823.2;refGene_gn=LOC101093705;refGene_tx=XM_019838824.2
C1 213680497 213680578
refGene_gn=LOC101093705;refGene_tx=XM_006935658.3;refGene_gn=LOC101093705;refGene_tx=XM_006935659.3;refGene_gn=LOC101093705;refGene_tx=XM_006935660.4;refGene_gn=LOC101093705;refGene_tx=XM_006935661.3;refGene_gn=LOC101093705;refGene_tx=XM_006935662.3;refGene_gn=LOC101093705;refGene_tx=XM_006935663.3;refGene_gn=LOC101093705;refGene_tx=XM_006935664.3;refGene_gn=LOC101093705;refGene_tx=XM_006935665.3;refGene_gn=LOC101093705;refGene_tx=XM_006935666.3;refGene_gn=LOC101093705;refGene_tx=XM_019838813.2;refGene_gn=LOC101093705;refGene_tx=XM_019838814.2;refGene_gn=LOC101093705;refGene_tx=XM_019838816.2;refGene_gn=LOC101093705;refGene_tx=XM_019838817.2;refGene_gn=LOC101093705;refGene_tx=XM_019838818.2;refGene_gn=LOC101093705;refGene_tx=XM_019838819.2;refGene_gn=LOC101093705;refGene_tx=XM_019838820.2;refGene_gn=LOC101093705;refGene_tx=XM_019838822.2;refGene_gn=LOC101093705;refGene_tx=XM_019838823.2;refGene_gn=LOC101093705;refGene_tx=XM_019838824.2
C1 213683582 213683669
refGene_gn=LOC101093705;refGene_tx=XM_006935658.3;refGene_gn=LOC101093705;refGene_tx=XM_006935659.3;refGene_gn=LOC101093705;refGene_tx=XM_006935660.4;refGene_gn=LOC101093705;refGene_tx=XM_006935661.3;refGene_gn=LOC101093705;refGene_tx=XM_006935662.3;refGene_gn=LOC101093705;refGene_tx=XM_006935663.3;refGene_gn=LOC101093705;refGene_tx=XM_006935664.3;refGene_gn=LOC101093705;refGene_tx=XM_006935665.3;refGene_gn=LOC101093705;refGene_tx=XM_006935666.3;refGene_gn=LOC101093705;refGene_tx=XM_019838813.2;refGene_gn=LOC101093705;refGene_tx=XM_019838814.2;refGene_gn=LOC101093705;refGene_tx=XM_019838816.2;refGene_gn=LOC101093705;refGene_tx=XM_019838817.2;refGene_gn=LOC101093705;refGene_tx=XM_019838818.2;refGene_gn=LOC101093705;refGene_tx=XM_019838819.2;refGene_gn=LOC101093705;refGene_tx=XM_019838820.2;refGene_gn=LOC101093705;refGene_tx=XM_019838822.2;refGene_gn=LOC101093705;refGene_tx=XM_019838823.2;refGene_gn=LOC101093705;refGene_tx=XM_019838824.2
C1 213683886 213684003
refGene_gn=LOC101093705;refGene_tx=XM_006935658.3;refGene_gn=LOC101093705;refGene_tx=XM_006935659.3;refGene_gn=LOC101093705;refGene_tx=XM_00

6935660.4;refGene_gn=LOC101093705;refGene_tx=XM_006935661.3;refGene_gn=LOC101093705;refGene_tx=XM_006935662.3;refGene_gn=LOC101093705;refGene_tx=XM_006935663.3;refGene_gn=LOC101093705;refGene_tx=XM_006935664.3;refGene_gn=LOC101093705;refGene_tx=XM_006935665.3;refGene_gn=LOC101093705;refGene_tx=XM_006935666.3;refGene_gn=LOC101093705;refGene_tx=XM_019838813.2;refGene_gn=LOC101093705;refGene_tx=XM_019838814.2;refGene_gn=LOC101093705;refGene_tx=XM_019838816.2;refGene_gn=LOC101093705;refGene_tx=XM_019838817.2;refGene_gn=LOC101093705;refGene_tx=XM_019838818.2;refGene_gn=LOC101093705;refGene_tx=XM_019838820.2;refGene_gn=LOC101093705;refGene_tx=XM_019838822.2;refGene_gn=LOC101093705;refGene_tx=XM_019838823.2;refGene_gn=LOC101093705;refGene_tx=XM_019838824.2
C1 213684808 213684862
refGene_gn=LOC101093705;refGene_tx=XM_006935658.3;refGene_gn=LOC101093705;refGene_tx=XM_006935659.3;refGene_gn=LOC101093705;refGene_tx=XM_006935660.4;refGene_gn=LOC101093705;refGene_tx=XM_006935661.3;refGene_gn=LOC101093705;refGene_tx=XM_006935662.3;refGene_gn=LOC101093705;refGene_tx=XM_006935663.3;refGene_gn=LOC101093705;refGene_tx=XM_006935664.3;refGene_gn=LOC101093705;refGene_tx=XM_006935665.3;refGene_gn=LOC101093705;refGene_tx=XM_006935666.3;refGene_gn=LOC101093705;refGene_tx=XM_019838813.2;refGene_gn=LOC101093705;refGene_tx=XM_019838814.2;refGene_gn=LOC101093705;refGene_tx=XM_019838816.2;refGene_gn=LOC101093705;refGene_tx=XM_019838817.2;refGene_gn=LOC101093705;refGene_tx=XM_019838818.2;refGene_gn=LOC101093705;refGene_tx=XM_019838819.2;refGene_gn=LOC101093705;refGene_tx=XM_019838820.2;refGene_gn=LOC101093705;refGene_tx=XM_019838822.2;refGene_gn=LOC101093705;refGene_tx=XM_019838823.2;refGene_gn=LOC101093705;refGene_tx=XM_019838824.2
C1 213687590 213687656
refGene_gn=LOC101093705;refGene_tx=XM_006935658.3;refGene_gn=LOC101093705;refGene_tx=XM_006935659.3;refGene_gn=LOC101093705;refGene_tx=XM_006935660.4;refGene_gn=LOC101093705;refGene_tx=XM_006935661.3;refGene_gn=LOC101093705;refGene_tx=XM_006935662.3;refGene_gn=LOC101093705;refGene_tx=XM_006935663.3;refGene_gn=LOC101093705;refGene_tx=XM_006935664.3;refGene_gn=LOC101093705;refGene_tx=XM_006935665.3;refGene_gn=LOC101093705;refGene_tx=XM_006935666.3;refGene_gn=LOC101093705;refGene_tx=XM_019838813.2;refGene_gn=LOC101093705;refGene_tx=XM_019838814.2;refGene_gn=LOC101093705;refGene_tx=XM_019838816.2;refGene_gn=LOC101093705;refGene_tx=XM_019838817.2;refGene_gn=LOC101093705;refGene_tx=XM_019838818.2;refGene_gn=LOC101093705;refGene_tx=XM_019838819.2;refGene_gn=LOC101093705;refGene_tx=XM_019838820.2;refGene_gn=LOC101093705;refGene_tx=XM_019838822.2;refGene_gn=LOC101093705;refGene_tx=XM_019838823.2;refGene_gn=LOC101093705;refGene_tx=XM_019838824.2
C1 213689385 213689466
refGene_gn=LOC101093705;refGene_tx=XM_006935658.3;refGene_gn=LOC101093705;refGene_tx=XM_006935659.3;refGene_gn=LOC101093705;refGene_tx=XM_006935660.4;refGene_gn=LOC101093705;refGene_tx=XM_006935661.3;refGene_gn=LOC101093705;refGene_tx=XM_006935662.3;refGene_gn=LOC101093705;refGene_tx=XM_006935663.3;refGene_gn=LOC101093705;refGene_tx=XM_006935664.3;refGene_gn=LOC101093705;refGene_tx=XM_006935665.3;refGene_gn=LOC101093705;refGene_tx=XM_006935666.3;refGene_gn=LOC101093705;refGene_tx=XM_019838813.2;refGene_gn=LOC101093705;refGene_tx=XM_019838814.2;refGene_gn

=LOC101093705;refGene_tx=XM_019838816.2;refGene_gn=LOC101093705;refGene_tx=XM_019838817.2;refGene_gn=LOC101093705;refGene_tx=XM_019838818.2;refGene_gn=LOC101093705;refGene_tx=XM_019838819.2;refGene_gn=LOC101093705;refGene_tx=XM_019838820.2;refGene_gn=LOC101093705;refGene_tx=XM_019838822.2;refGene_gn=LOC101093705;refGene_tx=XM_019838823.2;refGene_gn=LOC101093705;refGene_tx=XM_019838824.2
C1 213692580 213692655
refGene_gn=LOC101093705;refGene_tx=XM_006935658.3;refGene_gn=LOC101093705;refGene_tx=XM_006935659.3;refGene_gn=LOC101093705;refGene_tx=XM_006935660.4;refGene_gn=LOC101093705;refGene_tx=XM_006935661.3;refGene_gn=LOC101093705;refGene_tx=XM_006935662.3;refGene_gn=LOC101093705;refGene_tx=XM_006935663.3;refGene_gn=LOC101093705;refGene_tx=XM_006935664.3;refGene_gn=LOC101093705;refGene_tx=XM_006935665.3;refGene_gn=LOC101093705;refGene_tx=XM_006935666.3;refGene_gn=LOC101093705;refGene_tx=XM_019838813.2;refGene_gn=LOC101093705;refGene_tx=XM_019838814.2;refGene_gn=LOC101093705;refGene_tx=XM_019838816.2;refGene_gn=LOC101093705;refGene_tx=XM_019838817.2;refGene_gn=LOC101093705;refGene_tx=XM_019838818.2;refGene_gn=LOC101093705;refGene_tx=XM_019838819.2;refGene_gn=LOC101093705;refGene_tx=XM_019838820.2;refGene_gn=LOC101093705;refGene_tx=XM_019838822.2;refGene_gn=LOC101093705;refGene_tx=XM_019838823.2;refGene_gn=LOC101093705;refGene_tx=XM_019838824.2
C1 213693878 213693983
refGene_gn=LOC101093705;refGene_tx=XM_006935658.3;refGene_gn=LOC101093705;refGene_tx=XM_006935659.3;refGene_gn=LOC101093705;refGene_tx=XM_006935660.4;refGene_gn=LOC101093705;refGene_tx=XM_006935661.3;refGene_gn=LOC101093705;refGene_tx=XM_006935662.3;refGene_gn=LOC101093705;refGene_tx=XM_006935663.3;refGene_gn=LOC101093705;refGene_tx=XM_006935664.3;refGene_gn=LOC101093705;refGene_tx=XM_006935665.3;refGene_gn=LOC101093705;refGene_tx=XM_006935666.3;refGene_gn=LOC101093705;refGene_tx=XM_019838813.2;refGene_gn=LOC101093705;refGene_tx=XM_019838814.2;refGene_gn=LOC101093705;refGene_tx=XM_019838816.2;refGene_gn=LOC101093705;refGene_tx=XM_019838817.2;refGene_gn=LOC101093705;refGene_tx=XM_019838818.2;refGene_gn=LOC101093705;refGene_tx=XM_019838819.2;refGene_gn=LOC101093705;refGene_tx=XM_019838820.2;refGene_gn=LOC101093705;refGene_tx=XM_019838822.2;refGene_gn=LOC101093705;refGene_tx=XM_019838823.2;refGene_gn=LOC101093705;refGene_tx=XM_019838824.2
C1 213696089 213696235
refGene_gn=LOC101093705;refGene_tx=XM_006935658.3;refGene_gn=LOC101093705;refGene_tx=XM_006935659.3;refGene_gn=LOC101093705;refGene_tx=XM_006935660.4;refGene_gn=LOC101093705;refGene_tx=XM_006935661.3;refGene_gn=LOC101093705;refGene_tx=XM_006935662.3;refGene_gn=LOC101093705;refGene_tx=XM_006935663.3;refGene_gn=LOC101093705;refGene_tx=XM_006935664.3;refGene_gn=LOC101093705;refGene_tx=XM_006935665.3;refGene_gn=LOC101093705;refGene_tx=XM_006935666.3;refGene_gn=LOC101093705;refGene_tx=XM_019838813.2;refGene_gn=LOC101093705;refGene_tx=XM_019838814.2;refGene_gn=LOC101093705;refGene_tx=XM_019838816.2;refGene_gn=LOC101093705;refGene_tx=XM_019838817.2;refGene_gn=LOC101093705;refGene_tx=XM_019838818.2;refGene_gn=LOC101093705;refGene_tx=XM_019838819.2;refGene_gn=LOC101093705;refGene_tx=XM_019838820.2;refGene_gn=LOC101093705;refGene_tx=XM_019838822.2;refGene_gn=LOC101093705;refGene_tx=XM_019838823.2;refGene_gn

=LOC101093705;refGene_tx=XM_019838824.2
C1 213697307 213697355
refGene_gn=LOC101093705;refGene_tx=XM_006935658.3;refGene_gn=LOC101093705;refGene_tx=XM_006935659.3;refGene_gn=LOC101093705;refGene_tx=XM_006935660.4;refGene_gn=LOC101093705;refGene_tx=XM_006935661.3;refGene_gn=LOC101093705;refGene_tx=XM_006935663.3;refGene_gn=LOC101093705;refGene_tx=XM_006935664.3;refGene_gn=LOC101093705;refGene_tx=XM_006935665.3;refGene_gn=LOC101093705;refGene_tx=XM_006935666.3;refGene_gn=LOC101093705;refGene_tx=XM_019838813.2;refGene_gn=LOC101093705;refGene_tx=XM_019838814.2;refGene_gn=LOC101093705;refGene_tx=XM_019838816.2;refGene_gn=LOC101093705;refGene_tx=XM_019838817.2;refGene_gn=LOC101093705;refGene_tx=XM_019838818.2;refGene_gn=LOC101093705;refGene_tx=XM_019838819.2;refGene_gn=LOC101093705;refGene_tx=XM_019838820.2;refGene_gn=LOC101093705;refGene_tx=XM_019838822.2;refGene_gn=LOC101093705;refGene_tx=XM_019838823.2;refGene_gn=LOC101093705;refGene_tx=XM_019838824.2
C1 213700161 213700203
refGene_gn=LOC101093705;refGene_tx=XM_006935658.3;refGene_gn=LOC101093705;refGene_tx=XM_006935659.3;refGene_gn=LOC101093705;refGene_tx=XM_006935660.4;refGene_gn=LOC101093705;refGene_tx=XM_006935661.3;refGene_gn=LOC101093705;refGene_tx=XM_006935662.3;refGene_gn=LOC101093705;refGene_tx=XM_006935663.3;refGene_gn=LOC101093705;refGene_tx=XM_006935664.3;refGene_gn=LOC101093705;refGene_tx=XM_006935665.3;refGene_gn=LOC101093705;refGene_tx=XM_006935666.3;refGene_gn=LOC101093705;refGene_tx=XM_019838813.2;refGene_gn=LOC101093705;refGene_tx=XM_019838814.2;refGene_gn=LOC101093705;refGene_tx=XM_019838816.2;refGene_gn=LOC101093705;refGene_tx=XM_019838817.2;refGene_gn=LOC101093705;refGene_tx=XM_019838818.2;refGene_gn=LOC101093705;refGene_tx=XM_019838819.2;refGene_gn=LOC101093705;refGene_tx=XM_019838820.2;refGene_gn=LOC101093705;refGene_tx=XM_019838822.2;refGene_gn=LOC101093705;refGene_tx=XM_019838823.2;refGene_gn=LOC101093705;refGene_tx=XM_019838824.2
C1 213702389 213702505
refGene_gn=LOC101093705;refGene_tx=XM_006935658.3;refGene_gn=LOC101093705;refGene_tx=XM_006935659.3;refGene_gn=LOC101093705;refGene_tx=XM_006935660.4;refGene_gn=LOC101093705;refGene_tx=XM_006935661.3;refGene_gn=LOC101093705;refGene_tx=XM_006935662.3;refGene_gn=LOC101093705;refGene_tx=XM_006935663.3;refGene_gn=LOC101093705;refGene_tx=XM_006935664.3;refGene_gn=LOC101093705;refGene_tx=XM_006935665.3;refGene_gn=LOC101093705;refGene_tx=XM_006935666.3;refGene_gn=LOC101093705;refGene_tx=XM_019838813.2;refGene_gn=LOC101093705;refGene_tx=XM_019838814.2;refGene_gn=LOC101093705;refGene_tx=XM_019838816.2;refGene_gn=LOC101093705;refGene_tx=XM_019838817.2;refGene_gn=LOC101093705;refGene_tx=XM_019838818.2;refGene_gn=LOC101093705;refGene_tx=XM_019838819.2;refGene_gn=LOC101093705;refGene_tx=XM_019838820.2;refGene_gn=LOC101093705;refGene_tx=XM_019838822.2;refGene_gn=LOC101093705;refGene_tx=XM_019838823.2;refGene_gn=LOC101093705;refGene_tx=XM_019838824.2
C1 213703207 213703313
refGene_gn=LOC101093705;refGene_tx=XM_006935658.3;refGene_gn=LOC101093705;refGene_tx=XM_006935659.3;refGene_gn=LOC101093705;refGene_tx=XM_006935660.4;refGene_gn=LOC101093705;refGene_tx=XM_006935661.3;refGene_gn=LOC101093705;refGene_tx=XM_006935662.3;refGene_gn=LOC101093705;refGen

e_tx=XM_006935663.3;refGene_gn=LOC101093705;refGene_tx=XM_006935664.3;refGene_gn=LOC101093705;refGene_tx=XM_006935665.3;refGene_gn=LOC101093705;refGene_tx=XM_006935666.3;refGene_gn=LOC101093705;refGene_tx=XM_019838813.2;refGene_gn=LOC101093705;refGene_tx=XM_019838814.2;refGene_gn=LOC101093705;refGene_tx=XM_019838816.2;refGene_gn=LOC101093705;refGene_tx=XM_019838817.2;refGene_gn=LOC101093705;refGene_tx=XM_019838818.2;refGene_gn=LOC101093705;refGene_tx=XM_019838819.2;refGene_gn=LOC101093705;refGene_tx=XM_019838820.2;refGene_gn=LOC101093705;refGene_tx=XM_019838822.2;refGene_gn=LOC101093705;refGene_tx=XM_019838823.2;refGene_gn=LOC101093705;refGene_tx=XM_019838824.2
C1 213703500 213703539
refGene_gn=LOC101093705;refGene_tx=XM_019838823.2
C1 213703617 213703695
refGene_gn=LOC101093705;refGene_tx=XM_006935658.3;refGene_gn=LOC101093705;refGene_tx=XM_006935659.3;refGene_gn=LOC101093705;refGene_tx=XM_006935660.4;refGene_gn=LOC101093705;refGene_tx=XM_006935661.3;refGene_gn=LOC101093705;refGene_tx=XM_006935662.3;refGene_gn=LOC101093705;refGene_tx=XM_006935663.3;refGene_gn=LOC101093705;refGene_tx=XM_006935665.3;refGene_gn=LOC101093705;refGene_tx=XM_006935666.3;refGene_gn=LOC101093705;refGene_tx=XM_019838813.2;refGene_gn=LOC101093705;refGene_tx=XM_019838814.2;refGene_gn=LOC101093705;refGene_tx=XM_019838816.2;refGene_gn=LOC101093705;refGene_tx=XM_019838817.2;refGene_gn=LOC101093705;refGene_tx=XM_019838818.2;refGene_gn=LOC101093705;refGene_tx=XM_019838819.2;refGene_gn=LOC101093705;refGene_tx=XM_019838820.2;refGene_gn=LOC101093705;refGene_tx=XM_019838822.2;refGene_gn=LOC101093705;refGene_tx=XM_019838824.2
C1 213703904 213704048
refGene_gn=LOC101093705;refGene_tx=XM_006935658.3;refGene_gn=LOC101093705;refGene_tx=XM_006935659.3;refGene_gn=LOC101093705;refGene_tx=XM_006935660.4;refGene_gn=LOC101093705;refGene_tx=XM_006935661.3;refGene_gn=LOC101093705;refGene_tx=XM_006935662.3;refGene_gn=LOC101093705;refGene_tx=XM_006935663.3;refGene_gn=LOC101093705;refGene_tx=XM_006935664.3;refGene_gn=LOC101093705;refGene_tx=XM_006935665.3;refGene_gn=LOC101093705;refGene_tx=XM_006935666.3;refGene_gn=LOC101093705;refGene_tx=XM_019838813.2;refGene_gn=LOC101093705;refGene_tx=XM_019838814.2;refGene_gn=LOC101093705;refGene_tx=XM_019838816.2;refGene_gn=LOC101093705;refGene_tx=XM_019838817.2;refGene_gn=LOC101093705;refGene_tx=XM_019838818.2;refGene_gn=LOC101093705;refGene_tx=XM_019838819.2;refGene_gn=LOC101093705;refGene_tx=XM_019838820.2;refGene_gn=LOC101093705;refGene_tx=XM_019838822.2;refGene_gn=LOC101093705;refGene_tx=XM_019838824.2
C1 213704824 213704932
refGene_gn=LOC101093705;refGene_tx=XM_006935658.3;refGene_gn=LOC101093705;refGene_tx=XM_006935659.3;refGene_gn=LOC101093705;refGene_tx=XM_006935660.4;refGene_gn=LOC101093705;refGene_tx=XM_006935661.3;refGene_gn=LOC101093705;refGene_tx=XM_006935662.3;refGene_gn=LOC101093705;refGene_tx=XM_006935663.3;refGene_gn=LOC101093705;refGene_tx=XM_006935664.3;refGene_gn=LOC101093705;refGene_tx=XM_006935665.3;refGene_gn=LOC101093705;refGene_tx=XM_006935666.3;refGene_gn=LOC101093705;refGene_tx=XM_019838813.2;refGene_gn=LOC101093705;refGene_tx=XM_019838814.2;refGene_gn=LOC101093705;refGene_tx=XM_019838816.2;refGene_gn=LOC101093705;refGen

e_tx=XM_019838817.2;refGene_gn=LOC101093705;refGene_tx=XM_019838818.2;refGene_gn=LOC101093705;refGene_tx=XM_019838819.2;refGene_gn=LOC101093705;refGene_tx=XM_019838820.2;refGene_gn=LOC101093705;refGene_tx=XM_019838822.2
C1 213716374 213716406
refGene_gn=SP100;refGene_tx=XM_006935668.3;refGene_gn=SP100;refGene_tx=XM_006935669.3;refGene_gn=SP100;refGene_tx=XM_006935670.3;refGene_gn=SP100;refGene_tx=XM_023259792.1
C1 213759113 213759276
refGene_gn=SP100;refGene_tx=XM_006935668.3;refGene_gn=SP100;refGene_tx=XM_006935669.3;refGene_gn=SP100;refGene_tx=XM_006935670.3;refGene_gn=SP100;refGene_tx=XM_023259792.1
C1 213760589 213760758
refGene_gn=SP100;refGene_tx=XM_006935668.3;refGene_gn=SP100;refGene_tx=XM_006935669.3;refGene_gn=SP100;refGene_tx=XM_006935670.3;refGene_gn=SP100;refGene_tx=XM_023259792.1
C1 213764178 213764262
refGene_gn=SP100;refGene_tx=XM_006935668.3;refGene_gn=SP100;refGene_tx=XM_006935669.3;refGene_gn=SP100;refGene_tx=XM_006935670.3;refGene_gn=SP100;refGene_tx=XM_023259792.1
C1 213766528 213766591
refGene_gn=SP100;refGene_tx=XM_006935668.3;refGene_gn=SP100;refGene_tx=XM_006935669.3;refGene_gn=SP100;refGene_tx=XM_006935670.3;refGene_gn=SP100;refGene_tx=XM_023259792.1
C1 213767224 213767374
refGene_gn=SP100;refGene_tx=XM_006935668.3;refGene_gn=SP100;refGene_tx=XM_006935669.3;refGene_gn=SP100;refGene_tx=XM_006935670.3;refGene_gn=SP100;refGene_tx=XM_023259792.1
C1 213767831 213767915
refGene_gn=SP100;refGene_tx=XM_006935668.3;refGene_gn=SP100;refGene_tx=XM_006935669.3;refGene_gn=SP100;refGene_tx=XM_006935670.3;refGene_gn=SP100;refGene_tx=XM_023259792.1
C1 213770727 213770874
refGene_gn=SP100;refGene_tx=XM_006935668.3;refGene_gn=SP100;refGene_tx=XM_006935669.3;refGene_gn=SP100;refGene_tx=XM_006935670.3;refGene_gn=SP100;refGene_tx=XM_023259792.1
C1 213771564 213771645
refGene_gn=SP100;refGene_tx=XM_006935668.3;refGene_gn=SP100;refGene_tx=XM_006935669.3;refGene_gn=SP100;refGene_tx=XM_006935670.3;refGene_gn=SP100;refGene_tx=XM_023259792.1
C1 213773175 213773259
refGene_gn=SP100;refGene_tx=XM_006935668.3;refGene_gn=SP100;refGene_tx=XM_006935669.3;refGene_gn=SP100;refGene_tx=XM_006935670.3;refGene_gn=SP100;refGene_tx=XM_023259792.1
C1 213774836 213774929
refGene_gn=SP100;refGene_tx=XM_006935668.3;refGene_gn=SP100;refGene_tx=XM_006935669.3;refGene_gn=SP100;refGene_tx=XM_023259792.1
C1 213776089 213776143
refGene_gn=SP100;refGene_tx=XM_006935668.3;refGene_gn=SP100;refGene_tx=XM_006935669.3;refGene_gn=SP100;refGene_tx=XM_006935670.3;refGene_gn=

SP100;refGene_tx=XM_023259792.1
C1 213776780 213776864
refGene_gn=SP100;refGene_tx=XM_006935668.3;refGene_gn=SP100;refGene_tx=XM_006935669.3;refGene_gn=SP100;refGene_tx=XM_006935670.3;refGene_gn=SP100;refGene_tx=XM_023259792.1
C1 213780469 213780529
refGene_gn=SP100;refGene_tx=XM_006935668.3;refGene_gn=SP100;refGene_tx=XM_006935669.3;refGene_gn=SP100;refGene_tx=XM_006935670.3;refGene_gn=SP100;refGene_tx=XM_023259792.1
C1 213781455 213781509
refGene_gn=SP100;refGene_tx=XM_006935668.3;refGene_gn=SP100;refGene_tx=XM_006935669.3;refGene_gn=SP100;refGene_tx=XM_006935670.3;refGene_gn=SP100;refGene_tx=XM_023259792.1
C1 213791389 213791440
refGene_gn=SP100;refGene_tx=XM_006935668.3;refGene_gn=SP100;refGene_tx=XM_006935670.3;refGene_gn=SP100;refGene_tx=XM_023259792.1
C1 213797523 213797592
refGene_gn=SP100;refGene_tx=XM_006935668.3;refGene_gn=SP100;refGene_tx=XM_006935669.3;refGene_gn=SP100;refGene_tx=XM_006935670.3;refGene_gn=SP100;refGene_tx=XM_023259792.1
C1 213800336 213800384
refGene_gn=SP100;refGene_tx=XM_006935668.3;refGene_gn=SP100;refGene_tx=XM_006935669.3;refGene_gn=SP100;refGene_tx=XM_006935670.3;refGene_gn=SP100;refGene_tx=XM_023259792.1
C1 213801614 213801719
refGene_gn=SP100;refGene_tx=XM_006935668.3;refGene_gn=SP100;refGene_tx=XM_006935669.3;refGene_gn=SP100;refGene_tx=XM_006935670.3;refGene_gn=SP100;refGene_tx=XM_023259792.1
C1 213803395 213803538
refGene_gn=SP100;refGene_tx=XM_006935668.3;refGene_gn=SP100;refGene_tx=XM_006935669.3;refGene_gn=SP100;refGene_tx=XM_006935670.3;refGene_gn=SP100;refGene_tx=XM_023259792.1
C1 213807333 213807375
refGene_gn=SP100;refGene_tx=XM_006935668.3;refGene_gn=SP100;refGene_tx=XM_006935669.3;refGene_gn=SP100;refGene_tx=XM_006935670.3
C1 213809314 213809347
refGene_gn=SP100;refGene_tx=XM_006935668.3;refGene_gn=SP100;refGene_tx=XM_006935669.3;refGene_gn=SP100;refGene_tx=XM_006935670.3;refGene_gn=SP100;refGene_tx=XM_023259792.1
C1 213813324 213813440
refGene_gn=SP100;refGene_tx=XM_006935668.3;refGene_gn=SP100;refGene_tx=XM_006935669.3;refGene_gn=SP100;refGene_tx=XM_006935670.3;refGene_gn=SP100;refGene_tx=XM_023259792.1
C1 213814875 213814996
refGene_gn=SP100;refGene_tx=XM_006935668.3;refGene_gn=SP100;refGene_tx=XM_006935669.3;refGene_gn=SP100;refGene_tx=XM_006935670.3;refGene_gn=SP100;refGene_tx=XM_023259792.1
C1 213815278 213815350
refGene_gn=SP100;refGene_tx=XM_006935668.3;refGene_gn=SP100;refGene_tx=XM_006935669.3;refGene_gn=SP100;refGene_tx=XM_006935670.3;refGene_gn=

SP100;refGene_tx=XM_023259792.1
C1	213815888	213816032
refGene_gn=SP100;refGene_tx=XM_006935668.3;refGene_gn=SP100;refGene_tx=XM_006935669.3;refGene_gn=SP100;refGene_tx=XM_006935670.3;refGene_gn=SP100;refGene_tx=XM_023259792.1
C1	213817194	213817320
refGene_gn=SP100;refGene_tx=XM_006935668.3;refGene_gn=SP100;refGene_tx=XM_006935669.3;refGene_gn=SP100;refGene_tx=XM_006935670.3;refGene_gn=SP100;refGene_tx=XM_023259792.1
C1	213941269	213941383
refGene_gn=CAB39;refGene_tx=XM_023259793.1
C1	213967467	213967632
refGene_gn=CAB39;refGene_tx=XM_023259793.1
C1	213969772	213969891
refGene_gn=CAB39;refGene_tx=XM_023259793.1
C1	213974805	213974974
refGene_gn=CAB39;refGene_tx=XM_023259793.1
C1	213982126	213982186
refGene_gn=CAB39;refGene_tx=XM_023259793.1
C1	213985032	213985098
refGene_gn=CAB39;refGene_tx=XM_023259793.1
C1	213989108	213989252
refGene_gn=CAB39;refGene_tx=XM_023259793.1
C1	213989857	213990046
refGene_gn=CAB39;refGene_tx=XM_023259793.1
C1	214021256	214021382
refGene_gn=ITM2C;refGene_tx=XM_003991232.4;refGene_gn=ITM2C;refGene_tx=XM_011285629.2
C1	214029037	214029178
refGene_gn=ITM2C;refGene_tx=XM_003991232.4;refGene_gn=ITM2C;refGene_tx=XM_011285629.2
C1	214031148	214031337
refGene_gn=ITM2C;refGene_tx=XM_003991232.4;refGene_gn=ITM2C;refGene_tx=XM_011285629.2
C1	214032316	214032427
refGene_gn=ITM2C;refGene_tx=XM_003991232.4;refGene_gn=ITM2C;refGene_tx=XM_011285629.2
C1	214032632	214032860
refGene_gn=ITM2C;refGene_tx=XM_011285629.2
C1	214032957	214033108
refGene_gn=ITM2C;refGene_tx=XM_003991232.4
C1	214033615	214033707
refGene_gn=ITM2C;refGene_tx=XM_003991232.4
C1	214061074	214062045
refGene_gn=GPR55;refGene_tx=XM_019838831.2;refGene_gn=GPR55;refGene_tx=XM_019838835.2;refGene_gn=GPR55;refGene_tx=XM_019838838.2;refGene_gn=GPR55;refGene_tx=XM_019838843.2
C1	214102612	214102631
refGene_gn=GPR55;refGene_tx=XM_019838843.2
C1	214132939	214133255

refGene_gn=SPATA3;refGene_tx=XM_003991246.4;refGene_gn=SPATA3;refGene_tx=XM_019838845.2
C1 214136603 214136851
refGene_gn=SPATA3;refGene_tx=XM_003991246.4;refGene_gn=SPATA3;refGene_tx=XM_019838845.2
C1 214138916 214139022
refGene_gn=SPATA3;refGene_tx=XM_003991246.4
C1 214169813 214170420
refGene_gn=CC1H2orf72;refGene_tx=XM_023259794.1
C1 214173200 214173314
refGene_gn=CC1H2orf72;refGene_tx=XM_023259794.1
C1 214178175 214178315
refGene_gn=CC1H2orf72;refGene_tx=XM_023259794.1
C1 214188796 214188812
refGene_gn=PSMD1;refGene_tx=XM_011285628.3
C1 214193636 214193680
refGene_gn=PSMD1;refGene_tx=XM_011285628.3
C1 214194312 214194386
refGene_gn=PSMD1;refGene_tx=XM_011285628.3
C1 214194564 214194734
refGene_gn=PSMD1;refGene_tx=XM_011285628.3
C1 214196666 214196872
refGene_gn=PSMD1;refGene_tx=XM_011285628.3
C1 214199370 214199514
refGene_gn=PSMD1;refGene_tx=XM_011285628.3
C1 214202736 214202963
refGene_gn=PSMD1;refGene_tx=XM_011285628.3
C1 214207704 214207765
refGene_gn=PSMD1;refGene_tx=XM_011285628.3
C1 214208671 214208800
refGene_gn=PSMD1;refGene_tx=XM_011285628.3
C1 214210304 214210393
refGene_gn=PSMD1;refGene_tx=XM_011285628.3
C1 214210829 214210908
refGene_gn=PSMD1;refGene_tx=XM_011285628.3
C1 214211405 214211579
refGene_gn=PSMD1;refGene_tx=XM_011285628.3
C1 214213181 214213293
refGene_gn=PSMD1;refGene_tx=XM_011285628.3
C1 214214241 214214438
refGene_gn=PSMD1;refGene_tx=XM_011285628.3
C1 214215394 214215490
refGene_gn=PSMD1;refGene_tx=XM_011285628.3
C1 214216537 214216602
refGene_gn=PSMD1;refGene_tx=XM_011285628.3
C1 214222698 214223591
refGene_gn=HTR2B;refGene_tx=XM_003991248.5
C1 214227855 214228056
refGene_gn=HTR2B;refGene_tx=XM_003991248.5
C1 214236903 214237321

refGene_gn=HTR2B;refGene_tx=XM_003991248.5
C1	214253305	214253420
refGene_gn=PSMD1;refGene_tx=XM_011285628.3
C1	214257960	214258077
refGene_gn=PSMD1;refGene_tx=XM_011285628.3
C1	214265692	214265795
refGene_gn=PSMD1;refGene_tx=XM_011285628.3
C1	214270970	214271140
refGene_gn=PSMD1;refGene_tx=XM_011285628.3
C1	214273119	214273212
refGene_gn=PSMD1;refGene_tx=XM_011285628.3
C1	214274769	214274856
refGene_gn=PSMD1;refGene_tx=XM_011285628.3
C1	214275626	214275773
refGene_gn=PSMD1;refGene_tx=XM_011285628.3
C1	214282214	214282361
refGene_gn=PSMD1;refGene_tx=XM_011285628.3
C1	214314368	214314419
refGene_gn=ARMC9;refGene_tx=XM_006935682.4;refGene_gn=ARMC9;refGene_tx=XM_006935683.4;refGene_gn=ARMC9;refGene_tx=XM_006935684.4;refGene_gn=ARMC9;refGene_tx=XM_019838847.2;refGene_gn=ARMC9;refGene_tx=XM_019838848.2;refGene_gn=ARMC9;refGene_tx=XM_019838849.2;refGene_gn=ARMC9;refGene_tx=XM_019838850.2;refGene_gn=ARMC9;refGene_tx=XM_019838852.2
C1	214316066	214316192
refGene_gn=ARMC9;refGene_tx=XM_006935682.4;refGene_gn=ARMC9;refGene_tx=XM_006935683.4;refGene_gn=ARMC9;refGene_tx=XM_006935684.4;refGene_gn=ARMC9;refGene_tx=XM_019838847.2;refGene_gn=ARMC9;refGene_tx=XM_019838848.2;refGene_gn=ARMC9;refGene_tx=XM_019838849.2;refGene_gn=ARMC9;refGene_tx=XM_019838850.2;refGene_gn=ARMC9;refGene_tx=XM_019838852.2
C1	214321547	214321718
refGene_gn=ARMC9;refGene_tx=XM_006935682.4;refGene_gn=ARMC9;refGene_tx=XM_006935683.4;refGene_gn=ARMC9;refGene_tx=XM_006935684.4;refGene_gn=ARMC9;refGene_tx=XM_019838847.2;refGene_gn=ARMC9;refGene_tx=XM_019838849.2;refGene_gn=ARMC9;refGene_tx=XM_019838852.2
C1	214323511	214323667
refGene_gn=ARMC9;refGene_tx=XM_006935682.4;refGene_gn=ARMC9;refGene_tx=XM_006935683.4;refGene_gn=ARMC9;refGene_tx=XM_006935684.4;refGene_gn=ARMC9;refGene_tx=XM_019838847.2;refGene_gn=ARMC9;refGene_tx=XM_019838848.2;refGene_gn=ARMC9;refGene_tx=XM_019838849.2;refGene_gn=ARMC9;refGene_tx=XM_019838852.2
C1	214329568	214329661
refGene_gn=ARMC9;refGene_tx=XM_006935682.4;refGene_gn=ARMC9;refGene_tx=XM_006935683.4;refGene_gn=ARMC9;refGene_tx=XM_006935684.4;refGene_gn=ARMC9;refGene_tx=XM_019838847.2;refGene_gn=ARMC9;refGene_tx=XM_019838848.2;refGene_gn=ARMC9;refGene_tx=XM_019838850.2;refGene_gn=ARMC9;refGene_tx=XM_019838852.2
C1	214331346	214331371
refGene_gn=ARMC9;refGene_tx=XM_006935682.4;refGene_gn=ARMC9;refGene_tx=XM_006935683.4;refGene_gn=ARMC9;refGene_tx=XM_006935684.4;refGene_gn=ARMC9;refGene_tx=XM_019838847.2;refGene_gn=ARMC9;refGene_tx=XM_0198388

48.2;refGene_gn=ARMC9;refGene_tx=XM_019838850.2;refGene_gn=ARMC9;refGene_tx=XM_019838852.2
C1 214338696 214338854
refGene_gn=ARMC9;refGene_tx=XM_006935682.4;refGene_gn=ARMC9;refGene_tx=XM_006935683.4;refGene_gn=ARMC9;refGene_tx=XM_006935684.4;refGene_gn=ARMC9;refGene_tx=XM_019838847.2;refGene_gn=ARMC9;refGene_tx=XM_019838848.2;refGene_gn=ARMC9;refGene_tx=XM_019838850.2;refGene_gn=ARMC9;refGene_tx=XM_019838852.2
C1 214345581 214345680
refGene_gn=ARMC9;refGene_tx=XM_006935682.4;refGene_gn=ARMC9;refGene_tx=XM_006935683.4;refGene_gn=ARMC9;refGene_tx=XM_006935684.4;refGene_gn=ARMC9;refGene_tx=XM_019838847.2;refGene_gn=ARMC9;refGene_tx=XM_019838848.2;refGene_gn=ARMC9;refGene_tx=XM_019838849.2;refGene_gn=ARMC9;refGene_tx=XM_019838850.2;refGene_gn=ARMC9;refGene_tx=XM_019838852.2
C1 214356574 214356609
refGene_gn=ARMC9;refGene_tx=XM_006935682.4;refGene_gn=ARMC9;refGene_tx=XM_006935683.4;refGene_gn=ARMC9;refGene_tx=XM_006935684.4;refGene_gn=ARMC9;refGene_tx=XM_019838848.2;refGene_gn=ARMC9;refGene_tx=XM_019838849.2;refGene_gn=ARMC9;refGene_tx=XM_019838850.2;refGene_gn=ARMC9;refGene_tx=XM_019838852.2
C1 214359919 214360034
refGene_gn=ARMC9;refGene_tx=XM_006935682.4;refGene_gn=ARMC9;refGene_tx=XM_006935683.4;refGene_gn=ARMC9;refGene_tx=XM_006935684.4;refGene_gn=ARMC9;refGene_tx=XM_019838848.2;refGene_gn=ARMC9;refGene_tx=XM_019838849.2;refGene_gn=ARMC9;refGene_tx=XM_019838850.2;refGene_gn=ARMC9;refGene_tx=XM_019838852.2
C1 214362837 214362930
refGene_gn=ARMC9;refGene_tx=XM_006935682.4;refGene_gn=ARMC9;refGene_tx=XM_006935683.4;refGene_gn=ARMC9;refGene_tx=XM_006935684.4;refGene_gn=ARMC9;refGene_tx=XM_019838847.2;refGene_gn=ARMC9;refGene_tx=XM_019838848.2;refGene_gn=ARMC9;refGene_tx=XM_019838849.2;refGene_gn=ARMC9;refGene_tx=XM_019838850.2;refGene_gn=ARMC9;refGene_tx=XM_019838852.2
C1 214366120 214366211
refGene_gn=ARMC9;refGene_tx=XM_006935682.4;refGene_gn=ARMC9;refGene_tx=XM_006935683.4;refGene_gn=ARMC9;refGene_tx=XM_006935684.4;refGene_gn=ARMC9;refGene_tx=XM_019838847.2;refGene_gn=ARMC9;refGene_tx=XM_019838848.2;refGene_gn=ARMC9;refGene_tx=XM_019838849.2;refGene_gn=ARMC9;refGene_tx=XM_019838850.2;refGene_gn=ARMC9;refGene_tx=XM_019838852.2
C1 214367465 214367589
refGene_gn=ARMC9;refGene_tx=XM_006935682.4;refGene_gn=ARMC9;refGene_tx=XM_006935683.4;refGene_gn=ARMC9;refGene_tx=XM_006935684.4;refGene_gn=ARMC9;refGene_tx=XM_019838847.2;refGene_gn=ARMC9;refGene_tx=XM_019838848.2;refGene_gn=ARMC9;refGene_tx=XM_019838849.2;refGene_gn=ARMC9;refGene_tx=XM_019838850.2;refGene_gn=ARMC9;refGene_tx=XM_019838852.2
C1 214370492 214370632
refGene_gn=ARMC9;refGene_tx=XM_006935682.4;refGene_gn=ARMC9;refGene_tx=XM_006935683.4;refGene_gn=ARMC9;refGene_tx=XM_006935684.4;refGene_gn=ARMC9;refGene_tx=XM_019838847.2;refGene_gn=ARMC9;refGene_tx=XM_019838848.2;refGene_gn=ARMC9;refGene_tx=XM_019838849.2;refGene_gn=ARMC9;refGene_tx=XM_019838850.2;refGene_gn=ARMC9;refGene_tx=XM_019838852.2

C1	214372024	214372101
refGene_gn=ARMC9;refGene_tx=XM_006935682.4;refGene_gn=ARMC9;refGene_tx=XM_006935683.4;refGene_gn=ARMC9;refGene_tx=XM_006935684.4;refGene_gn=ARMC9;refGene_tx=XM_019838847.2;refGene_gn=ARMC9;refGene_tx=XM_019838848.2;refGene_gn=ARMC9;refGene_tx=XM_019838849.2;refGene_gn=ARMC9;refGene_tx=XM_019838850.2;refGene_gn=ARMC9;refGene_tx=XM_019838852.2
C1	214374648	214374723
refGene_gn=ARMC9;refGene_tx=XM_006935682.4;refGene_gn=ARMC9;refGene_tx=XM_006935683.4;refGene_gn=ARMC9;refGene_tx=XM_006935684.4;refGene_gn=ARMC9;refGene_tx=XM_019838847.2;refGene_gn=ARMC9;refGene_tx=XM_019838848.2;refGene_gn=ARMC9;refGene_tx=XM_019838849.2;refGene_gn=ARMC9;refGene_tx=XM_019838850.2;refGene_gn=ARMC9;refGene_tx=XM_019838852.2
C1	214378512	214378603
refGene_gn=ARMC9;refGene_tx=XM_006935682.4;refGene_gn=ARMC9;refGene_tx=XM_006935683.4;refGene_gn=ARMC9;refGene_tx=XM_006935684.4;refGene_gn=ARMC9;refGene_tx=XM_019838847.2;refGene_gn=ARMC9;refGene_tx=XM_019838848.2;refGene_gn=ARMC9;refGene_tx=XM_019838849.2;refGene_gn=ARMC9;refGene_tx=XM_019838850.2;refGene_gn=ARMC9;refGene_tx=XM_019838852.2
C1	214382948	214383004
refGene_gn=ARMC9;refGene_tx=XM_006935682.4;refGene_gn=ARMC9;refGene_tx=XM_006935683.4;refGene_gn=ARMC9;refGene_tx=XM_006935684.4;refGene_gn=ARMC9;refGene_tx=XM_019838847.2;refGene_gn=ARMC9;refGene_tx=XM_019838848.2;refGene_gn=ARMC9;refGene_tx=XM_019838849.2;refGene_gn=ARMC9;refGene_tx=XM_019838850.2;refGene_gn=ARMC9;refGene_tx=XM_019838852.2
C1	214421873	214421978
refGene_gn=ARMC9;refGene_tx=XM_006935682.4;refGene_gn=ARMC9;refGene_tx=XM_006935683.4;refGene_gn=ARMC9;refGene_tx=XM_006935684.4;refGene_gn=ARMC9;refGene_tx=XM_019838847.2;refGene_gn=ARMC9;refGene_tx=XM_019838848.2;refGene_gn=ARMC9;refGene_tx=XM_019838849.2;refGene_gn=ARMC9;refGene_tx=XM_019838850.2;refGene_gn=ARMC9;refGene_tx=XM_019838852.2
C1	214429927	214430089
refGene_gn=ARMC9;refGene_tx=XM_019838852.2
C1	214433347	214433463
refGene_gn=ARMC9;refGene_tx=XM_006935682.4;refGene_gn=ARMC9;refGene_tx=XM_006935683.4;refGene_gn=ARMC9;refGene_tx=XM_006935684.4;refGene_gn=ARMC9;refGene_tx=XM_019838847.2;refGene_gn=ARMC9;refGene_tx=XM_019838848.2;refGene_gn=ARMC9;refGene_tx=XM_019838849.2;refGene_gn=ARMC9;refGene_tx=XM_019838850.2
C1	214437938	214438063
refGene_gn=ARMC9;refGene_tx=XM_006935682.4;refGene_gn=ARMC9;refGene_tx=XM_006935683.4;refGene_gn=ARMC9;refGene_tx=XM_006935684.4;refGene_gn=ARMC9;refGene_tx=XM_019838847.2;refGene_gn=ARMC9;refGene_tx=XM_019838848.2;refGene_gn=ARMC9;refGene_tx=XM_019838849.2;refGene_gn=ARMC9;refGene_tx=XM_019838850.2
C1	214442650	214442750
refGene_gn=ARMC9;refGene_tx=XM_006935682.4;refGene_gn=ARMC9;refGene_tx=XM_006935683.4;refGene_gn=ARMC9;refGene_tx=XM_006935684.4;refGene_gn=ARMC9;refGene_tx=XM_019838847.2;refGene_gn=ARMC9;refGene_tx=XM_019838848.2;refGene_gn=ARMC9;refGene_tx=XM_019838849.2;refGene_gn=ARMC9;refGene_tx=XM_019838850.2

C1	214451721	214451923
refGene_gn=ARMC9;refGene_tx=XM_006935682.4;refGene_gn=ARMC9;refGene_tx=XM_006935683.4;refGene_gn=ARMC9;refGene_tx=XM_006935684.4;refGene_gn=ARMC9;refGene_tx=XM_019838847.2;refGene_gn=ARMC9;refGene_tx=XM_019838848.2;refGene_gn=ARMC9;refGene_tx=XM_019838849.2;refGene_gn=ARMC9;refGene_tx=XM_019838850.2
C1	214453271	214453294
refGene_gn=ARMC9;refGene_tx=XM_006935684.4
C1	214477381	214477392
refGene_gn=B3GNT7;refGene_tx=XM_023259795.1
C1	214479484	214480679
refGene_gn=B3GNT7;refGene_tx=XM_023259795.1
C1	214511583	214511660
refGene_gn=NCL;refGene_tx=XM_003991250.5
C1	214511839	214512063
refGene_gn=NCL;refGene_tx=XM_003991250.5
C1	214512448	214512575
refGene_gn=NCL;refGene_tx=XM_003991250.5
C1	214513144	214513278
refGene_gn=NCL;refGene_tx=XM_003991250.5
C1	214513455	214513579
refGene_gn=NCL;refGene_tx=XM_003991250.5
C1	214514044	214514202
refGene_gn=NCL;refGene_tx=XM_003991250.5
C1	214514698	214514822
refGene_gn=NCL;refGene_tx=XM_003991250.5
C1	214515433	214515558
refGene_gn=NCL;refGene_tx=XM_003991250.5
C1	214516264	214516406
refGene_gn=NCL;refGene_tx=XM_003991250.5
C1	214516599	214516686
refGene_gn=NCL;refGene_tx=XM_003991250.5
C1	214516809	214517019
refGene_gn=NCL;refGene_tx=XM_003991250.5
C1	214517400	214517482
refGene_gn=NCL;refGene_tx=XM_003991250.5
C1	214517483	214517885
refGene_gn=NCL;refGene_tx=XM_003991250.5
C1	214519174	214519291
refGene_gn=NCL;refGene_tx=XM_003991250.5
C1	214520342	214520360
refGene_gn=NCL;refGene_tx=XM_003991250.5
C1	214538065	214538244
refGene_gn=LOC102900614;refGene_tx=XM_023259801.1
C1	214546593	214548378
refGene_gn=LOC102900614;refGene_tx=XM_019838855.2;refGene_gn=LOC102900614;refGene_tx=XM_019838856.2;refGene_gn=LOC102900614;refGene_tx=XM_019838857.2;refGene_gn=LOC102900614;refGene_tx=XM_019838859.2;refGene_gn=LOC102900614;refGene_tx=XM_019838861.2;refGene_gn=LOC102900614;refGene_tx=XM_019838862.2;refGene_gn=LOC102900614;refGene_tx=XM_023259797.1;

refGene_gn=LOC102900614;refGene_tx=XM_023259798.1;refGene_gn=LOC102900614;refGene_tx=XM_023259799.1;refGene_gn=LOC102900614;refGene_tx=XM_023259800.1
C1 214549293 214549451
refGene_gn=LOC102900614;refGene_tx=XM_019838855.2;refGene_gn=LOC102900614;refGene_tx=XM_019838856.2;refGene_gn=LOC102900614;refGene_tx=XM_023259797.1;refGene_gn=LOC102900614;refGene_tx=XM_023259798.1;refGene_gn=LOC102900614;refGene_tx=XM_023259799.1;refGene_gn=LOC102900614;refGene_tx=XM_023259800.1;refGene_gn=LOC102900614;refGene_tx=XM_023259801.1
C1 214557156 214557299
refGene_gn=LOC102900614;refGene_tx=XM_023259797.1;refGene_gn=LOC102900614;refGene_tx=XM_023259798.1;refGene_gn=LOC102900614;refGene_tx=XM_023259799.1;refGene_gn=LOC102900614;refGene_tx=XM_023259800.1;refGene_gn=LOC102900614;refGene_tx=XM_023259801.1
C1 214567899 214567940
refGene_gn=LOC102900614;refGene_tx=XM_019838855.2;refGene_gn=LOC102900614;refGene_tx=XM_019838856.2
C1 214577969 214578352
refGene_gn=NMUR1;refGene_tx=XM_011285674.3
C1 214580590 214581539
refGene_gn=NMUR1;refGene_tx=XM_011285674.3
C1 214625794 214626979
refGene_gn=TEX44;refGene_tx=XM_023259802.1
C1 214726072 214726117
refGene_gn=PTMA;refGene_tx=XM_003991251.4;refGene_gn=PTMA;refGene_tx=XM_006935686.3
C1 214728862 214728934
refGene_gn=PTMA;refGene_tx=XM_003991251.4;refGene_gn=PTMA;refGene_tx=XM_006935686.3
C1 214729200 214729297
refGene_gn=PTMA;refGene_tx=XM_003991251.4;refGene_gn=PTMA;refGene_tx=XM_006935686.3
C1 214729798 214729872
refGene_gn=PTMA;refGene_tx=XM_003991251.4;refGene_gn=PTMA;refGene_tx=XM_006935686.3
C1 214730129 214730177
refGene_gn=PTMA;refGene_tx=XM_003991251.4;refGene_gn=PTMA;refGene_tx=XM_006935686.3
C1 214745943 214746025
refGene_gn=PDE6D;refGene_tx=XM_019838864.2
C1 214747922 214748028
refGene_gn=PDE6D;refGene_tx=XM_019838864.2
C1 214748760 214748886
refGene_gn=PDE6D;refGene_tx=XM_019838864.2
C1 214749496 214749585
refGene_gn=PDE6D;refGene_tx=XM_019838864.2
C1 214798448 214798498
refGene_gn=PDE6D;refGene_tx=XM_019838864.2
C1 214799218 214799281
refGene_gn=COPS7B;refGene_tx=XM_003991255.4

C1 214805748 214805910
refGene_gn=COPS7B;refGene_tx=XM_003991253.4;refGene_gn=COPS7B;refGene_tx=XM_006935687.4
C1 214807259 214807274
refGene_gn=COPS7B;refGene_tx=XM_006935688.4
C1 214809662 214809738
refGene_gn=COPS7B;refGene_tx=XM_003991253.4;refGene_gn=COPS7B;refGene_tx=XM_003991255.4;refGene_gn=COPS7B;refGene_tx=XM_006935687.4;refGene_gn=COPS7B;refGene_tx=XM_006935688.4
C1 214811065 214811154
refGene_gn=COPS7B;refGene_tx=XM_003991253.4;refGene_gn=COPS7B;refGene_tx=XM_003991255.4;refGene_gn=COPS7B;refGene_tx=XM_006935687.4;refGene_gn=COPS7B;refGene_tx=XM_006935688.4
C1 214812813 214813016
refGene_gn=COPS7B;refGene_tx=XM_003991253.4;refGene_gn=COPS7B;refGene_tx=XM_003991255.4;refGene_gn=COPS7B;refGene_tx=XM_006935687.4;refGene_gn=COPS7B;refGene_tx=XM_006935688.4
C1 214815387 214815493
refGene_gn=COPS7B;refGene_tx=XM_003991253.4;refGene_gn=COPS7B;refGene_tx=XM_003991255.4;refGene_gn=COPS7B;refGene_tx=XM_006935687.4;refGene_gn=COPS7B;refGene_tx=XM_006935688.4
C1 214821491 214821650
refGene_gn=COPS7B;refGene_tx=XM_003991253.4;refGene_gn=COPS7B;refGene_tx=XM_003991255.4;refGene_gn=COPS7B;refGene_tx=XM_006935687.4;refGene_gn=COPS7B;refGene_tx=XM_006935688.4
C1 214881493 214881784
refGene_gn=NPPC;refGene_tx=XM_003991256.5
C1 214882244 214882334
refGene_gn=NPPC;refGene_tx=XM_003991256.5
C1 214983107 214983159
refGene_gn=DIS3L2;refGene_tx=XM_011285625.3;refGene_gn=DIS3L2;refGene_tx=XM_011285626.3;refGene_gn=DIS3L2;refGene_tx=XM_019838866.2;refGene_gn=DIS3L2;refGene_tx=XM_019838867.2
C1 214983702 214983860
refGene_gn=DIS3L2;refGene_tx=XM_011285625.3;refGene_gn=DIS3L2;refGene_tx=XM_011285626.3;refGene_gn=DIS3L2;refGene_tx=XM_019838866.2;refGene_gn=DIS3L2;refGene_tx=XM_019838867.2
C1 214992016 214992070
refGene_gn=DIS3L2;refGene_tx=XM_011285625.3;refGene_gn=DIS3L2;refGene_tx=XM_011285626.3;refGene_gn=DIS3L2;refGene_tx=XM_019838866.2;refGene_gn=DIS3L2;refGene_tx=XM_019838867.2
C1 214999038 214999140
refGene_gn=DIS3L2;refGene_tx=XM_011285625.3;refGene_gn=DIS3L2;refGene_tx=XM_011285626.3;refGene_gn=DIS3L2;refGene_tx=XM_019838866.2;refGene_gn=DIS3L2;refGene_tx=XM_019838867.2
C1 215055019 215055254
refGene_gn=DIS3L2;refGene_tx=XM_011285625.3;refGene_gn=DIS3L2;refGene_tx=XM_011285626.3;refGene_gn=DIS3L2;refGene_tx=XM_019838866.2;refGene_gn=DIS3L2;refGene_tx=XM_019838867.2
C1 215056183 215056188

refGene_gn=DIS3L2;refGene_tx=XM_011285627.3
C1	215057519	215057605
refGene_gn=DIS3L2;refGene_tx=XM_011285627.3
C1	215084244	215084345
refGene_gn=DIS3L2;refGene_tx=XM_011285625.3;refGene_gn=DIS3L2;refGene_tx=XM_011285626.3;refGene_gn=DIS3L2;refGene_tx=XM_011285627.3;refGene_gn=DIS3L2;refGene_tx=XM_019838866.2;refGene_gn=DIS3L2;refGene_tx=XM_019838867.2
C1	215089156	215089404
refGene_gn=DIS3L2;refGene_tx=XM_011285625.3;refGene_gn=DIS3L2;refGene_tx=XM_011285626.3;refGene_gn=DIS3L2;refGene_tx=XM_011285627.3;refGene_gn=DIS3L2;refGene_tx=XM_019838866.2;refGene_gn=DIS3L2;refGene_tx=XM_019838867.2
C1	215108794	215108968
refGene_gn=DIS3L2;refGene_tx=XM_011285625.3;refGene_gn=DIS3L2;refGene_tx=XM_011285626.3;refGene_gn=DIS3L2;refGene_tx=XM_011285627.3;refGene_gn=DIS3L2;refGene_tx=XM_019838866.2;refGene_gn=DIS3L2;refGene_tx=XM_019838867.2
C1	215141463	215141543
refGene_gn=DIS3L2;refGene_tx=XM_011285625.3;refGene_gn=DIS3L2;refGene_tx=XM_011285626.3;refGene_gn=DIS3L2;refGene_tx=XM_011285627.3;refGene_gn=DIS3L2;refGene_tx=XM_019838866.2;refGene_gn=DIS3L2;refGene_tx=XM_019838867.2
C1	215171016	215171129
refGene_gn=DIS3L2;refGene_tx=XM_011285625.3;refGene_gn=DIS3L2;refGene_tx=XM_011285626.3;refGene_gn=DIS3L2;refGene_tx=XM_011285627.3;refGene_gn=DIS3L2;refGene_tx=XM_019838866.2;refGene_gn=DIS3L2;refGene_tx=XM_019838867.2
C1	215182451	215182559
refGene_gn=DIS3L2;refGene_tx=XM_011285625.3;refGene_gn=DIS3L2;refGene_tx=XM_011285626.3;refGene_gn=DIS3L2;refGene_tx=XM_011285627.3;refGene_gn=DIS3L2;refGene_tx=XM_019838867.2
C1	215201266	215201500
refGene_gn=DIS3L2;refGene_tx=XM_011285625.3;refGene_gn=DIS3L2;refGene_tx=XM_011285626.3;refGene_gn=DIS3L2;refGene_tx=XM_011285627.3;refGene_gn=DIS3L2;refGene_tx=XM_019838866.2
C1	215236443	215236523
refGene_gn=DIS3L2;refGene_tx=XM_011285625.3;refGene_gn=DIS3L2;refGene_tx=XM_011285626.3;refGene_gn=DIS3L2;refGene_tx=XM_011285627.3;refGene_gn=DIS3L2;refGene_tx=XM_019838866.2;refGene_gn=DIS3L2;refGene_tx=XM_019838867.2
C1	215271100	215271284
refGene_gn=DIS3L2;refGene_tx=XM_011285625.3;refGene_gn=DIS3L2;refGene_tx=XM_011285626.3;refGene_gn=DIS3L2;refGene_tx=XM_011285627.3;refGene_gn=DIS3L2;refGene_tx=XM_019838866.2;refGene_gn=DIS3L2;refGene_tx=XM_019838867.2
C1	215271884	215271971
refGene_gn=DIS3L2;refGene_tx=XM_011285625.3;refGene_gn=DIS3L2;refGene_tx=XM_011285626.3;refGene_gn=DIS3L2;refGene_tx=XM_011285627.3;refGene_gn=DIS3L2;refGene_tx=XM_019838866.2;refGene_gn=DIS3L2;refGene_tx=XM_01

9838867.2
C1	215275210	215275358
refGene_gn=DIS3L2;refGene_tx=XM_011285625.3;refGene_gn=DIS3L2;refGene_tx=XM_011285626.3;refGene_gn=DIS3L2;refGene_tx=XM_011285627.3;refGene_gn=DIS3L2;refGene_tx=XM_019838866.2;refGene_gn=DIS3L2;refGene_tx=XM_019838867.2
C1	215275750	215275881
refGene_gn=DIS3L2;refGene_tx=XM_011285625.3;refGene_gn=DIS3L2;refGene_tx=XM_011285626.3;refGene_gn=DIS3L2;refGene_tx=XM_011285627.3;refGene_gn=DIS3L2;refGene_tx=XM_019838866.2;refGene_gn=DIS3L2;refGene_tx=XM_019838867.2
C1	215276052	215276157
refGene_gn=DIS3L2;refGene_tx=XM_011285625.3;refGene_gn=DIS3L2;refGene_tx=XM_011285626.3;refGene_gn=DIS3L2;refGene_tx=XM_011285627.3;refGene_gn=DIS3L2;refGene_tx=XM_019838866.2;refGene_gn=DIS3L2;refGene_tx=XM_019838867.2
C1	215277549	215277651
refGene_gn=DIS3L2;refGene_tx=XM_011285625.3;refGene_gn=DIS3L2;refGene_tx=XM_011285626.3;refGene_gn=DIS3L2;refGene_tx=XM_011285627.3;refGene_gn=DIS3L2;refGene_tx=XM_019838866.2;refGene_gn=DIS3L2;refGene_tx=XM_019838867.2
C1	215278334	215278463
refGene_gn=DIS3L2;refGene_tx=XM_011285625.3;refGene_gn=DIS3L2;refGene_tx=XM_011285626.3;refGene_gn=DIS3L2;refGene_tx=XM_011285627.3;refGene_gn=DIS3L2;refGene_tx=XM_019838866.2;refGene_gn=DIS3L2;refGene_tx=XM_019838867.2
C1	215291838	215291908
refGene_gn=ALPI;refGene_tx=XM_023259805.1
C1	215291990	215292107
refGene_gn=ALPI;refGene_tx=XM_023259805.1
C1	215292221	215292337
refGene_gn=ALPI;refGene_tx=XM_023259805.1
C1	215292533	215292708
refGene_gn=ALPI;refGene_tx=XM_023259805.1
C1	215292786	215292959
refGene_gn=ALPI;refGene_tx=XM_023259805.1
C1	215293219	215293354
refGene_gn=ALPI;refGene_tx=XM_023259805.1
C1	215293452	215293525
refGene_gn=ALPI;refGene_tx=XM_023259805.1
C1	215293651	215293786
refGene_gn=ALPI;refGene_tx=XM_023259805.1
C1	215293877	215294069
refGene_gn=ALPI;refGene_tx=XM_023259805.1
C1	215294290	215294410
refGene_gn=ALPI;refGene_tx=XM_023259805.1
C1	215294512	215294811
refGene_gn=ALPI;refGene_tx=XM_023259805.1
C1	215305185	215305285
refGene_gn=ECEL1;refGene_tx=XM_023259806.1;refGene_gn=ECEL1;refGene_tx

=XM_023259807.1;refGene_gn=ECEL1;refGene_tx=XM_023259808.1
C1 215305415 215305492
refGene_gn=ECEL1;refGene_tx=XM_023259806.1;refGene_gn=ECEL1;refGene_tx
=XM_023259807.1;refGene_gn=ECEL1;refGene_tx=XM_023259808.1;refGene_gn=
ECEL1;refGene_tx=XM_023259809.1
C1 215305713 215305829
refGene_gn=ECEL1;refGene_tx=XM_023259806.1;refGene_gn=ECEL1;refGene_tx
=XM_023259807.1;refGene_gn=ECEL1;refGene_tx=XM_023259808.1;refGene_gn=
ECEL1;refGene_tx=XM_023259809.1
C1 215306181 215306247
refGene_gn=ECEL1;refGene_tx=XM_023259806.1;refGene_gn=ECEL1;refGene_tx
=XM_023259807.1;refGene_gn=ECEL1;refGene_tx=XM_023259808.1;refGene_gn=
ECEL1;refGene_tx=XM_023259809.1
C1 215306538 215306663
refGene_gn=ECEL1;refGene_tx=XM_023259806.1;refGene_gn=ECEL1;refGene_tx
=XM_023259807.1;refGene_gn=ECEL1;refGene_tx=XM_023259808.1;refGene_gn=
ECEL1;refGene_tx=XM_023259809.1
C1 215306862 215306930
refGene_gn=ECEL1;refGene_tx=XM_023259806.1;refGene_gn=ECEL1;refGene_tx
=XM_023259807.1;refGene_gn=ECEL1;refGene_tx=XM_023259808.1;refGene_gn=
ECEL1;refGene_tx=XM_023259809.1
C1 215307502 215307554
refGene_gn=ECEL1;refGene_tx=XM_023259806.1;refGene_gn=ECEL1;refGene_tx
=XM_023259807.1;refGene_gn=ECEL1;refGene_tx=XM_023259808.1;refGene_gn=
ECEL1;refGene_tx=XM_023259809.1
C1 215307629 215307688
refGene_gn=ECEL1;refGene_tx=XM_023259806.1;refGene_gn=ECEL1;refGene_tx
=XM_023259807.1;refGene_gn=ECEL1;refGene_tx=XM_023259808.1;refGene_gn=
ECEL1;refGene_tx=XM_023259809.1
C1 215308012 215308116
refGene_gn=ECEL1;refGene_tx=XM_023259806.1;refGene_gn=ECEL1;refGene_tx
=XM_023259807.1;refGene_gn=ECEL1;refGene_tx=XM_023259808.1;refGene_gn=
ECEL1;refGene_tx=XM_023259809.1
C1 215308319 215308394
refGene_gn=ECEL1;refGene_tx=XM_023259806.1;refGene_gn=ECEL1;refGene_tx
=XM_023259807.1;refGene_gn=ECEL1;refGene_tx=XM_023259808.1;refGene_gn=
ECEL1;refGene_tx=XM_023259809.1
C1 215308612 215308711
refGene_gn=ECEL1;refGene_tx=XM_023259806.1;refGene_gn=ECEL1;refGene_tx
=XM_023259807.1;refGene_gn=ECEL1;refGene_tx=XM_023259808.1;refGene_gn=
ECEL1;refGene_tx=XM_023259809.1
C1 215309165 215309388
refGene_gn=ECEL1;refGene_tx=XM_023259806.1;refGene_gn=ECEL1;refGene_tx
=XM_023259807.1;refGene_gn=ECEL1;refGene_tx=XM_023259808.1;refGene_gn=
ECEL1;refGene_tx=XM_023259809.1
C1 215310045 215310170
refGene_gn=ECEL1;refGene_tx=XM_023259806.1;refGene_gn=ECEL1;refGene_tx
=XM_023259807.1;refGene_gn=ECEL1;refGene_tx=XM_023259808.1;refGene_gn=
ECEL1;refGene_tx=XM_023259809.1
C1 215310359 215310452

refGene_gn=ECEL1;refGene_tx=XM_023259806.1;refGene_gn=ECEL1;refGene_tx=XM_023259807.1;refGene_gn=ECEL1;refGene_tx=XM_023259808.1;refGene_gn=ECEL1;refGene_tx=XM_023259809.1
C1 215310570 215310681
refGene_gn=ECEL1;refGene_tx=XM_023259806.1;refGene_gn=ECEL1;refGene_tx=XM_023259807.1;refGene_gn=ECEL1;refGene_tx=XM_023259808.1;refGene_gn=ECEL1;refGene_tx=XM_023259809.1
C1 215310798 215310867
refGene_gn=ECEL1;refGene_tx=XM_023259806.1;refGene_gn=ECEL1;refGene_tx=XM_023259807.1;refGene_gn=ECEL1;refGene_tx=XM_023259808.1;refGene_gn=ECEL1;refGene_tx=XM_023259809.1
C1 215311476 215312268
refGene_gn=ECEL1;refGene_tx=XM_023259806.1;refGene_gn=ECEL1;refGene_tx=XM_023259807.1;refGene_gn=ECEL1;refGene_tx=XM_023259808.1;refGene_gn=ECEL1;refGene_tx=XM_023259809.1
C1 215315268 215315659
refGene_gn=LOC109502935;refGene_tx=XM_023259810.1
C1 215315914 215316399
refGene_gn=LOC109502935;refGene_tx=XM_023259810.1
C1 215344272 215344375
refGene_gn=PRSS56;refGene_tx=XM_019838872.2;refGene_gn=PRSS56;refGene_tx=XM_019838873.2;refGene_gn=PRSS56;refGene_tx=XM_019838874.2
C1 215344997 215345135
refGene_gn=PRSS56;refGene_tx=XM_019838872.2;refGene_gn=PRSS56;refGene_tx=XM_019838873.2;refGene_gn=PRSS56;refGene_tx=XM_019838874.2
C1 215345478 215345526
refGene_gn=PRSS56;refGene_tx=XM_019838872.2;refGene_gn=PRSS56;refGene_tx=XM_019838873.2;refGene_gn=PRSS56;refGene_tx=XM_019838874.2
C1 215345642 215345832
refGene_gn=PRSS56;refGene_tx=XM_019838872.2;refGene_gn=PRSS56;refGene_tx=XM_019838873.2;refGene_gn=PRSS56;refGene_tx=XM_019838874.2
C1 215346220 215346320
refGene_gn=PRSS56;refGene_tx=XM_019838872.2;refGene_gn=PRSS56;refGene_tx=XM_019838873.2;refGene_gn=PRSS56;refGene_tx=XM_019838874.2
C1 215346408 215346568
refGene_gn=PRSS56;refGene_tx=XM_019838872.2;refGene_gn=PRSS56;refGene_tx=XM_019838873.2;refGene_gn=PRSS56;refGene_tx=XM_019838874.2
C1 215346810 215346953
refGene_gn=PRSS56;refGene_tx=XM_019838872.2;refGene_gn=PRSS56;refGene_tx=XM_019838873.2;refGene_gn=PRSS56;refGene_tx=XM_019838874.2
C1 215347190 215347353
refGene_gn=PRSS56;refGene_tx=XM_019838872.2;refGene_gn=PRSS56;refGene_tx=XM_019838873.2;refGene_gn=PRSS56;refGene_tx=XM_019838874.2
C1 215347540 215347717
refGene_gn=PRSS56;refGene_tx=XM_019838872.2;refGene_gn=PRSS56;refGene_tx=XM_019838873.2;refGene_gn=PRSS56;refGene_tx=XM_019838874.2
C1 215347803 215347968
refGene_gn=PRSS56;refGene_tx=XM_019838872.2;refGene_gn=PRSS56;refGene_tx=XM_019838873.2;refGene_gn=PRSS56;refGene_tx=XM_019838874.2
C1 215348049 215348112

refGene_gn=PRSS56;refGene_tx=XM_019838872.2;refGene_gn=PRSS56;refGene_tx=XM_019838873.2;refGene_gn=PRSS56;refGene_tx=XM_019838874.2
C1 215348478 215348585
refGene_gn=PRSS56;refGene_tx=XM_019838872.2;refGene_gn=PRSS56;refGene_tx=XM_019838873.2;refGene_gn=PRSS56;refGene_tx=XM_019838874.2
C1 215348959 215349175
refGene_gn=PRSS56;refGene_tx=XM_019838872.2;refGene_gn=PRSS56;refGene_tx=XM_019838873.2;refGene_gn=PRSS56;refGene_tx=XM_019838874.2
C1 215349424 215349428
refGene_gn=PRSS56;refGene_tx=XM_019838873.2
C1 215349936 215349988
refGene_gn=CHRND;refGene_tx=XM_003991258.4
C1 215350252 215350398
refGene_gn=CHRND;refGene_tx=XM_003991258.4
C1 215350836 215350881
refGene_gn=CHRND;refGene_tx=XM_003991258.4
C1 215351381 215351491
refGene_gn=CHRND;refGene_tx=XM_003991258.4
C1 215351597 215351753
refGene_gn=CHRND;refGene_tx=XM_003991258.4
C1 215351957 215352067
refGene_gn=CHRND;refGene_tx=XM_003991258.4
C1 215352968 215353169
refGene_gn=CHRND;refGene_tx=XM_003991258.4
C1 215354337 215354449
refGene_gn=CHRND;refGene_tx=XM_003991258.4
C1 215354574 215354689
refGene_gn=CHRND;refGene_tx=XM_003991258.4
C1 215355969 215356174
refGene_gn=CHRND;refGene_tx=XM_003991258.4
C1 215356270 215356389
refGene_gn=CHRND;refGene_tx=XM_003991258.4
C1 215357068 215357251
refGene_gn=CHRND;refGene_tx=XM_003991258.4
C1 215362328 215362383
refGene_gn=CHRNG;refGene_tx=XM_003991260.4
C1 215362586 215362726
refGene_gn=CHRNG;refGene_tx=XM_003991260.4
C1 215362987 215363032
refGene_gn=CHRNG;refGene_tx=XM_003991260.4
C1 215363193 215363303
refGene_gn=CHRNG;refGene_tx=XM_003991260.4
C1 215363760 215363916
refGene_gn=CHRNG;refGene_tx=XM_003991260.4
C1 215364719 215364817
refGene_gn=CHRNG;refGene_tx=XM_003991260.4
C1 215365143 215365344
refGene_gn=CHRNG;refGene_tx=XM_003991260.4
C1 215365524 215365639
refGene_gn=CHRNG;refGene_tx=XM_003991260.4

C1 215365852 215365967
refGene_gn=CHRNG;refGene_tx=XM_003991260.4
C1 215366716 215366933
refGene_gn=CHRNG;refGene_tx=XM_003991260.4
C1 215367149 215367283
refGene_gn=CHRNG;refGene_tx=XM_003991260.4
C1 215367559 215367733
refGene_gn=CHRNG;refGene_tx=XM_003991260.4
C1 215369667 215369687
refGene_gn=EIF4E2;refGene_tx=XM_003991262.3;refGene_gn=EIF4E2;refGene_tx=XM_006935691.4;refGene_gn=EIF4E2;refGene_tx=XM_006935692.4;refGene_gn=EIF4E2;refGene_tx=XM_019838878.2;refGene_gn=EIF4E2;refGene_tx=XM_019838879.2
C1 215376689 215376804
refGene_gn=EIF4E2;refGene_tx=XM_003991262.3;refGene_gn=EIF4E2;refGene_tx=XM_006935691.4;refGene_gn=EIF4E2;refGene_tx=XM_006935692.4;refGene_gn=EIF4E2;refGene_tx=XM_019838878.2;refGene_gn=EIF4E2;refGene_tx=XM_019838879.2
C1 215378855 215378990
refGene_gn=EIF4E2;refGene_tx=XM_003991262.3;refGene_gn=EIF4E2;refGene_tx=XM_006935691.4;refGene_gn=EIF4E2;refGene_tx=XM_006935692.4;refGene_gn=EIF4E2;refGene_tx=XM_019838878.2;refGene_gn=EIF4E2;refGene_tx=XM_019838879.2
C1 215383058 215383163
refGene_gn=EIF4E2;refGene_tx=XM_003991262.3;refGene_gn=EIF4E2;refGene_tx=XM_006935691.4;refGene_gn=EIF4E2;refGene_tx=XM_006935692.4;refGene_gn=EIF4E2;refGene_tx=XM_019838878.2;refGene_gn=EIF4E2;refGene_tx=XM_019838879.2
C1 215384575 215384728
refGene_gn=EIF4E2;refGene_tx=XM_003991262.3;refGene_gn=EIF4E2;refGene_tx=XM_006935691.4;refGene_gn=EIF4E2;refGene_tx=XM_006935692.4;refGene_gn=EIF4E2;refGene_tx=XM_019838878.2;refGene_gn=EIF4E2;refGene_tx=XM_019838879.2
C1 215384825 215384962
refGene_gn=EIF4E2;refGene_tx=XM_003991262.3;refGene_gn=EIF4E2;refGene_tx=XM_006935691.4;refGene_gn=EIF4E2;refGene_tx=XM_006935692.4;refGene_gn=EIF4E2;refGene_tx=XM_019838878.2;refGene_gn=EIF4E2;refGene_tx=XM_019838879.2
C1 215386702 215386775
refGene_gn=EIF4E2;refGene_tx=XM_003991262.3
C1 215388873 215388949
refGene_gn=EIF4E2;refGene_tx=XM_019838878.2;refGene_gn=EIF4E2;refGene_tx=XM_019838879.2
C1 215391657 215391703
refGene_gn=EIF4E2;refGene_tx=XM_006935691.4
C1 215397633 215397673
refGene_gn=EIF4E2;refGene_tx=XM_006935692.4
C1 215445265 215445555
refGene_gn=EFHD1;refGene_tx=XM_023259812.1
C1 215467495 215467643

refGene_gn=EFHD1;refGene_tx=XM_023259812.1
C1 215476888 215477023
refGene_gn=EFHD1;refGene_tx=XM_023259812.1
C1 215487690 215487825
refGene_gn=EFHD1;refGene_tx=XM_023259812.1
C1 215500645 215500699
refGene_gn=LOC111561652;refGene_tx=XM_023257871.1
C1 215501074 215501701
refGene_gn=LOC111561652;refGene_tx=XM_023257871.1
C1 215532677 215532718
refGene_gn=GIGYF2;refGene_tx=XM_003991263.5;refGene_gn=GIGYF2;refGene_tx=XM_006935693.4;refGene_gn=GIGYF2;refGene_tx=XM_019838880.2;refGene_gn=GIGYF2;refGene_tx=XM_019838881.2
C1 215544118 215544248
refGene_gn=GIGYF2;refGene_tx=XM_003991263.5;refGene_gn=GIGYF2;refGene_tx=XM_006935693.4;refGene_gn=GIGYF2;refGene_tx=XM_019838880.2;refGene_gn=GIGYF2;refGene_tx=XM_019838881.2
C1 215545178 215545274
refGene_gn=GIGYF2;refGene_tx=XM_003991263.5;refGene_gn=GIGYF2;refGene_tx=XM_006935693.4;refGene_gn=GIGYF2;refGene_tx=XM_019838880.2;refGene_gn=GIGYF2;refGene_tx=XM_019838881.2
C1 215558508 215558620
refGene_gn=GIGYF2;refGene_tx=XM_003991263.5;refGene_gn=GIGYF2;refGene_tx=XM_006935693.4;refGene_gn=GIGYF2;refGene_tx=XM_019838880.2;refGene_gn=GIGYF2;refGene_tx=XM_019838881.2
C1 215563680 215563792
refGene_gn=GIGYF2;refGene_tx=XM_003991263.5;refGene_gn=GIGYF2;refGene_tx=XM_006935693.4;refGene_gn=GIGYF2;refGene_tx=XM_019838880.2;refGene_gn=GIGYF2;refGene_tx=XM_019838881.2
C1 215564652 215564693
refGene_gn=GIGYF2;refGene_tx=XM_003991263.5;refGene_gn=GIGYF2;refGene_tx=XM_006935693.4;refGene_gn=GIGYF2;refGene_tx=XM_019838880.2;refGene_gn=GIGYF2;refGene_tx=XM_019838881.2
C1 215571041 215571664
refGene_gn=KCNJ13;refGene_tx=XM_006935695.3;refGene_gn=KCNJ13;refGene_tx=XM_019838884.2;refGene_gn=KCNJ13;refGene_tx=XM_019838885.1
C1 215573451 215573911
refGene_gn=KCNJ13;refGene_tx=XM_006935695.3;refGene_gn=KCNJ13;refGene_tx=XM_019838884.2;refGene_gn=KCNJ13;refGene_tx=XM_019838885.1
C1 215589865 215590045
refGene_gn=GIGYF2;refGene_tx=XM_003991263.5;refGene_gn=GIGYF2;refGene_tx=XM_006935693.4;refGene_gn=GIGYF2;refGene_tx=XM_019838880.2;refGene_gn=GIGYF2;refGene_tx=XM_019838881.2
C1 215591736 215591794
refGene_gn=GIGYF2;refGene_tx=XM_006935694.4;refGene_gn=GIGYF2;refGene_tx=XM_019838882.2
C1 215594152 215594370
refGene_gn=GIGYF2;refGene_tx=XM_003991263.5;refGene_gn=GIGYF2;refGene_tx=XM_006935693.4;refGene_gn=GIGYF2;refGene_tx=XM_006935694.4;refGene_gn=GIGYF2;refGene_tx=XM_019838880.2;refGene_gn=GIGYF2;refGene_tx=XM_01

9838881.2;refGene_gn=GIGYF2;refGene_tx=XM_019838882.2;refGene_gn=GIGYF2;refGene_tx=XM_019838883.2;refGene_gn=GIGYF2;refGene_tx=XM_023259811.1
C1 215594462 215594625
refGene_gn=GIGYF2;refGene_tx=XM_003991263.5;refGene_gn=GIGYF2;refGene_tx=XM_006935693.4;refGene_gn=GIGYF2;refGene_tx=XM_006935694.4;refGene_gn=GIGYF2;refGene_tx=XM_019838880.2;refGene_gn=GIGYF2;refGene_tx=XM_019838881.2;refGene_gn=GIGYF2;refGene_tx=XM_019838882.2;refGene_gn=GIGYF2;refGene_tx=XM_019838883.2;refGene_gn=GIGYF2;refGene_tx=XM_023259811.1
C1 215594713 215594902
refGene_gn=GIGYF2;refGene_tx=XM_003991263.5;refGene_gn=GIGYF2;refGene_tx=XM_006935693.4;refGene_gn=GIGYF2;refGene_tx=XM_006935694.4;refGene_gn=GIGYF2;refGene_tx=XM_019838880.2;refGene_gn=GIGYF2;refGene_tx=XM_019838881.2;refGene_gn=GIGYF2;refGene_tx=XM_019838882.2;refGene_gn=GIGYF2;refGene_tx=XM_019838883.2;refGene_gn=GIGYF2;refGene_tx=XM_023259811.1
C1 215597433 215597630
refGene_gn=GIGYF2;refGene_tx=XM_003991263.5;refGene_gn=GIGYF2;refGene_tx=XM_006935693.4;refGene_gn=GIGYF2;refGene_tx=XM_006935694.4;refGene_gn=GIGYF2;refGene_tx=XM_019838880.2;refGene_gn=GIGYF2;refGene_tx=XM_019838881.2;refGene_gn=GIGYF2;refGene_tx=XM_019838882.2;refGene_gn=GIGYF2;refGene_tx=XM_019838883.2;refGene_gn=GIGYF2;refGene_tx=XM_023259811.1
C1 215598613 215598773
refGene_gn=GIGYF2;refGene_tx=XM_003991263.5;refGene_gn=GIGYF2;refGene_tx=XM_006935693.4;refGene_gn=GIGYF2;refGene_tx=XM_006935694.4;refGene_gn=GIGYF2;refGene_tx=XM_019838880.2;refGene_gn=GIGYF2;refGene_tx=XM_019838881.2;refGene_gn=GIGYF2;refGene_tx=XM_019838882.2;refGene_gn=GIGYF2;refGene_tx=XM_019838883.2;refGene_gn=GIGYF2;refGene_tx=XM_023259811.1
C1 215606357 215606524
refGene_gn=GIGYF2;refGene_tx=XM_003991263.5;refGene_gn=GIGYF2;refGene_tx=XM_006935693.4;refGene_gn=GIGYF2;refGene_tx=XM_006935694.4;refGene_gn=GIGYF2;refGene_tx=XM_019838880.2;refGene_gn=GIGYF2;refGene_tx=XM_019838881.2;refGene_gn=GIGYF2;refGene_tx=XM_019838882.2;refGene_gn=GIGYF2;refGene_tx=XM_019838883.2;refGene_gn=GIGYF2;refGene_tx=XM_023259811.1
C1 215610082 215610174
refGene_gn=GIGYF2;refGene_tx=XM_003991263.5;refGene_gn=GIGYF2;refGene_tx=XM_006935693.4;refGene_gn=GIGYF2;refGene_tx=XM_006935694.4;refGene_gn=GIGYF2;refGene_tx=XM_019838880.2;refGene_gn=GIGYF2;refGene_tx=XM_019838881.2;refGene_gn=GIGYF2;refGene_tx=XM_019838882.2;refGene_gn=GIGYF2;refGene_tx=XM_019838883.2;refGene_gn=GIGYF2;refGene_tx=XM_023259811.1
C1 215611506 215611614
refGene_gn=GIGYF2;refGene_tx=XM_003991263.5;refGene_gn=GIGYF2;refGene_tx=XM_006935693.4;refGene_gn=GIGYF2;refGene_tx=XM_006935694.4;refGene_gn=GIGYF2;refGene_tx=XM_019838880.2;refGene_gn=GIGYF2;refGene_tx=XM_019838881.2;refGene_gn=GIGYF2;refGene_tx=XM_019838882.2;refGene_gn=GIGYF

2;refGene_tx=XM_019838883.2;refGene_gn=GIGYF2;refGene_tx=XM_023259811.
1
C1 215612668 215612769
refGene_gn=GIGYF2;refGene_tx=XM_003991263.5;refGene_gn=GIGYF2;refGene_
tx=XM_006935693.4;refGene_gn=GIGYF2;refGene_tx=XM_006935694.4;refGene_
gn=GIGYF2;refGene_tx=XM_019838880.2;refGene_gn=GIGYF2;refGene_tx=XM_01
9838881.2;refGene_gn=GIGYF2;refGene_tx=XM_019838882.2;refGene_gn=GIGYF
2;refGene_tx=XM_019838883.2;refGene_gn=GIGYF2;refGene_tx=XM_023259811.
1
C1 215615609 215615710
refGene_gn=GIGYF2;refGene_tx=XM_003991263.5;refGene_gn=GIGYF2;refGene_
tx=XM_006935693.4;refGene_gn=GIGYF2;refGene_tx=XM_006935694.4;refGene_
gn=GIGYF2;refGene_tx=XM_019838880.2;refGene_gn=GIGYF2;refGene_tx=XM_01
9838881.2;refGene_gn=GIGYF2;refGene_tx=XM_019838882.2;refGene_gn=GIGYF
2;refGene_tx=XM_019838883.2;refGene_gn=GIGYF2;refGene_tx=XM_023259811.
1
C1 215618585 215618747
refGene_gn=GIGYF2;refGene_tx=XM_003991263.5;refGene_gn=GIGYF2;refGene_
tx=XM_006935693.4;refGene_gn=GIGYF2;refGene_tx=XM_006935694.4;refGene_
gn=GIGYF2;refGene_tx=XM_019838880.2;refGene_gn=GIGYF2;refGene_tx=XM_01
9838881.2;refGene_gn=GIGYF2;refGene_tx=XM_019838882.2;refGene_gn=GIGYF
2;refGene_tx=XM_019838883.2;refGene_gn=GIGYF2;refGene_tx=XM_023259811.
1
C1 215620026 215620185
refGene_gn=GIGYF2;refGene_tx=XM_003991263.5;refGene_gn=GIGYF2;refGene_
tx=XM_006935693.4;refGene_gn=GIGYF2;refGene_tx=XM_006935694.4;refGene_
gn=GIGYF2;refGene_tx=XM_019838880.2;refGene_gn=GIGYF2;refGene_tx=XM_01
9838881.2;refGene_gn=GIGYF2;refGene_tx=XM_019838882.2;refGene_gn=GIGYF
2;refGene_tx=XM_019838883.2;refGene_gn=GIGYF2;refGene_tx=XM_023259811.
1
C1 215623879 215624116
refGene_gn=GIGYF2;refGene_tx=XM_003991263.5;refGene_gn=GIGYF2;refGene_
tx=XM_006935693.4;refGene_gn=GIGYF2;refGene_tx=XM_006935694.4;refGene_
gn=GIGYF2;refGene_tx=XM_019838880.2;refGene_gn=GIGYF2;refGene_tx=XM_01
9838881.2;refGene_gn=GIGYF2;refGene_tx=XM_019838882.2;refGene_gn=GIGYF
2;refGene_tx=XM_019838883.2;refGene_gn=GIGYF2;refGene_tx=XM_023259811.
1
C1 215632022 215632145
refGene_gn=GIGYF2;refGene_tx=XM_003991263.5;refGene_gn=GIGYF2;refGene_
tx=XM_006935693.4;refGene_gn=GIGYF2;refGene_tx=XM_006935694.4;refGene_
gn=GIGYF2;refGene_tx=XM_019838880.2;refGene_gn=GIGYF2;refGene_tx=XM_01
9838881.2;refGene_gn=GIGYF2;refGene_tx=XM_019838882.2;refGene_gn=GIGYF
2;refGene_tx=XM_019838883.2;refGene_gn=GIGYF2;refGene_tx=XM_023259811.
1
C1 215635188 215635392
refGene_gn=GIGYF2;refGene_tx=XM_003991263.5;refGene_gn=GIGYF2;refGene_
tx=XM_006935693.4;refGene_gn=GIGYF2;refGene_tx=XM_006935694.4;refGene_
gn=GIGYF2;refGene_tx=XM_019838880.2;refGene_gn=GIGYF2;refGene_tx=XM_01
9838881.2;refGene_gn=GIGYF2;refGene_tx=XM_019838882.2;refGene_gn=GIGYF
2;refGene_tx=XM_019838883.2;refGene_gn=GIGYF2;refGene_tx=XM_023259811.

1
C1	215635522	215635728
refGene_gn=GIGYF2;refGene_tx=XM_003991263.5;refGene_gn=GIGYF2;refGene_tx=XM_006935693.4;refGene_gn=GIGYF2;refGene_tx=XM_006935694.4;refGene_gn=GIGYF2;refGene_tx=XM_019838880.2;refGene_gn=GIGYF2;refGene_tx=XM_019838881.2;refGene_gn=GIGYF2;refGene_tx=XM_019838882.2;refGene_gn=GIGYF2;refGene_tx=XM_019838883.2;refGene_gn=GIGYF2;refGene_tx=XM_023259811.1
C1	215637127	215637282
refGene_gn=GIGYF2;refGene_tx=XM_003991263.5;refGene_gn=GIGYF2;refGene_tx=XM_006935693.4;refGene_gn=GIGYF2;refGene_tx=XM_006935694.4;refGene_gn=GIGYF2;refGene_tx=XM_019838880.2;refGene_gn=GIGYF2;refGene_tx=XM_019838882.2;refGene_gn=GIGYF2;refGene_tx=XM_019838883.2;refGene_gn=GIGYF2;refGene_tx=XM_023259811.1
C1	215638843	215639061
refGene_gn=GIGYF2;refGene_tx=XM_003991263.5;refGene_gn=GIGYF2;refGene_tx=XM_006935693.4;refGene_gn=GIGYF2;refGene_tx=XM_006935694.4;refGene_gn=GIGYF2;refGene_tx=XM_019838880.2;refGene_gn=GIGYF2;refGene_tx=XM_019838881.2;refGene_gn=GIGYF2;refGene_tx=XM_019838882.2;refGene_gn=GIGYF2;refGene_tx=XM_019838883.2;refGene_gn=GIGYF2;refGene_tx=XM_023259811.1
C1	215641803	215641951
refGene_gn=GIGYF2;refGene_tx=XM_003991263.5;refGene_gn=GIGYF2;refGene_tx=XM_006935693.4;refGene_gn=GIGYF2;refGene_tx=XM_006935694.4;refGene_gn=GIGYF2;refGene_tx=XM_019838880.2;refGene_gn=GIGYF2;refGene_tx=XM_019838882.2;refGene_gn=GIGYF2;refGene_tx=XM_019838883.2;refGene_gn=GIGYF2;refGene_tx=XM_023259811.1
C1	215646525	215646593
refGene_gn=GIGYF2;refGene_tx=XM_003991263.5;refGene_gn=GIGYF2;refGene_tx=XM_006935693.4;refGene_gn=GIGYF2;refGene_tx=XM_006935694.4;refGene_gn=GIGYF2;refGene_tx=XM_019838880.2;refGene_gn=GIGYF2;refGene_tx=XM_019838882.2;refGene_gn=GIGYF2;refGene_tx=XM_019838883.2;refGene_gn=GIGYF2;refGene_tx=XM_023259811.1
C1	215662521	215662594
refGene_gn=SNORC;refGene_tx=XM_023259817.1
C1	215668159	215668342
refGene_gn=SNORC;refGene_tx=XM_023259817.1
C1	215668423	215668533
refGene_gn=SNORC;refGene_tx=XM_023259817.1
C1	215671237	215671428
refGene_gn=NGEF;refGene_tx=XM_023259813.1
C1	215671946	215672051
refGene_gn=NGEF;refGene_tx=XM_023259813.1
C1	215672778	215672858
refGene_gn=NGEF;refGene_tx=XM_023259813.1
C1	215673455	215673611
refGene_gn=NGEF;refGene_tx=XM_023259813.1
C1	215673864	215674028
refGene_gn=NGEF;refGene_tx=XM_023259813.1
C1	215675054	215675144

```
refGene_gn=NGEF;refGene_tx=XM_023259813.1
C1	215677129	215677204
refGene_gn=NGEF;refGene_tx=XM_023259813.1
C1	215679475	215679605
refGene_gn=NGEF;refGene_tx=XM_023259813.1
C1	215681021	215681174
refGene_gn=NGEF;refGene_tx=XM_023259813.1
C1	215682451	215682612
refGene_gn=NGEF;refGene_tx=XM_023259813.1
C1	215700324	215700629
refGene_gn=NGEF;refGene_tx=XM_023259813.1
C1	215706931	215707074
refGene_gn=NGEF;refGene_tx=XM_023259813.1
C1	215707772	215707879
refGene_gn=NGEF;refGene_tx=XM_023259813.1
C1	215794008	215794209
refGene_gn=NEU2;refGene_tx=XM_003991289.5;refGene_gn=NEU2;refGene_tx=X
M_019838892.2;refGene_gn=NEU2;refGene_tx=XM_019838893.2;refGene_gn=NEU
2;refGene_tx=XM_019838899.2;refGene_gn=NEU2;refGene_tx=XM_019838902.2;
refGene_gn=NEU2;refGene_tx=XM_023259814.1;refGene_gn=NEU2;refGene_tx=X
M_023259815.1;refGene_gn=NEU2;refGene_tx=XM_023259816.1
C1	215795529	215796540
refGene_gn=NEU2;refGene_tx=XM_003991289.5;refGene_gn=NEU2;refGene_tx=X
M_019838892.2;refGene_gn=NEU2;refGene_tx=XM_019838893.2;refGene_gn=NEU
2;refGene_tx=XM_019838899.2;refGene_gn=NEU2;refGene_tx=XM_019838902.2;
refGene_gn=NEU2;refGene_tx=XM_023259814.1;refGene_gn=NEU2;refGene_tx=X
M_023259815.1;refGene_gn=NEU2;refGene_tx=XM_023259816.1
C1	215812060	215812194
refGene_gn=INPP5D;refGene_tx=XM_023259818.1
C1	215829385	215829449
refGene_gn=INPP5D;refGene_tx=XM_023259818.1
C1	215864278	215864429
refGene_gn=INPP5D;refGene_tx=XM_023259818.1
C1	215866951	215867126
refGene_gn=INPP5D;refGene_tx=XM_023259818.1
C1	215871035	215871176
refGene_gn=INPP5D;refGene_tx=XM_023259818.1
C1	215881567	215881655
refGene_gn=INPP5D;refGene_tx=XM_023259818.1
C1	215885667	215885748
refGene_gn=INPP5D;refGene_tx=XM_023259818.1
C1	215885915	215885987
refGene_gn=INPP5D;refGene_tx=XM_023259818.1
C1	215886932	215887056
refGene_gn=INPP5D;refGene_tx=XM_023259818.1
C1	215894250	215894357
refGene_gn=INPP5D;refGene_tx=XM_023259818.1
C1	215896499	215896602
refGene_gn=INPP5D;refGene_tx=XM_023259818.1
C1	215898874	215899071
```

refGene_gn=INPP5D;refGene_tx=XM_023259818.1
C1 215899533 215899651
refGene_gn=INPP5D;refGene_tx=XM_023259818.1
C1 215904705 215904802
refGene_gn=INPP5D;refGene_tx=XM_023259818.1
C1 215905328 215905467
refGene_gn=INPP5D;refGene_tx=XM_023259818.1
C1 215907143 215907252
refGene_gn=INPP5D;refGene_tx=XM_023259818.1
C1 215907439 215907528
refGene_gn=INPP5D;refGene_tx=XM_023259818.1
C1 215911542 215911624
refGene_gn=INPP5D;refGene_tx=XM_023259818.1
C1 215915171 215915261
refGene_gn=INPP5D;refGene_tx=XM_023259818.1
C1 215917682 215917796
refGene_gn=INPP5D;refGene_tx=XM_023259818.1
C1 215919092 215919175
refGene_gn=INPP5D;refGene_tx=XM_023259818.1
C1 215920641 215920729
refGene_gn=INPP5D;refGene_tx=XM_023259818.1
C1 215923570 215923720
refGene_gn=INPP5D;refGene_tx=XM_023259818.1
C1 215925751 215925851
refGene_gn=INPP5D;refGene_tx=XM_023259818.1
C1 215928032 215928299
refGene_gn=INPP5D;refGene_tx=XM_023259818.1
C1 215932706 215933307
refGene_gn=INPP5D;refGene_tx=XM_023259818.1
C1 215934178 215934181
refGene_gn=INPP5D;refGene_tx=XM_023259818.1
C1 215965072 215965187
refGene_gn=ATG16L1;refGene_tx=XM_023259819.1;refGene_gn=ATG16L1;refGen
e_tx=XM_023259820.1;refGene_gn=ATG16L1;refGene_tx=XM_023259821.1
C1 215969112 215969206
refGene_gn=ATG16L1;refGene_tx=XM_023259819.1;refGene_gn=ATG16L1;refGen
e_tx=XM_023259820.1;refGene_gn=ATG16L1;refGene_tx=XM_023259821.1
C1 215973265 215973371
refGene_gn=ATG16L1;refGene_tx=XM_023259819.1;refGene_gn=ATG16L1;refGen
e_tx=XM_023259820.1;refGene_gn=ATG16L1;refGene_tx=XM_023259821.1
C1 215974101 215974175
refGene_gn=ATG16L1;refGene_tx=XM_023259819.1;refGene_gn=ATG16L1;refGen
e_tx=XM_023259820.1;refGene_gn=ATG16L1;refGene_tx=XM_023259821.1
C1 215975012 215975264
refGene_gn=ATG16L1;refGene_tx=XM_023259819.1;refGene_gn=ATG16L1;refGen
e_tx=XM_023259820.1;refGene_gn=ATG16L1;refGene_tx=XM_023259821.1
C1 215979446 215979512
refGene_gn=ATG16L1;refGene_tx=XM_023259819.1;refGene_gn=ATG16L1;refGen
e_tx=XM_023259820.1;refGene_gn=ATG16L1;refGene_tx=XM_023259821.1
C1 215981517 215981604

refGene_gn=ATG16L1;refGene_tx=XM_023259819.1;refGene_gn=ATG16L1;refGene_tx=XM_023259820.1;refGene_gn=ATG16L1;refGene_tx=XM_023259821.1
C1 215982261 215982318
refGene_gn=ATG16L1;refGene_tx=XM_023259819.1;refGene_gn=ATG16L1;refGene_tx=XM_023259820.1
C1 215982533 215982581
refGene_gn=ATG16L1;refGene_tx=XM_023259819.1
C1 215983233 215983336
refGene_gn=ATG16L1;refGene_tx=XM_023259819.1;refGene_gn=ATG16L1;refGene_tx=XM_023259820.1;refGene_gn=ATG16L1;refGene_tx=XM_023259821.1
C1 215986165 215986271
refGene_gn=ATG16L1;refGene_tx=XM_023259819.1;refGene_gn=ATG16L1;refGene_tx=XM_023259820.1;refGene_gn=ATG16L1;refGene_tx=XM_023259821.1
C1 215989652 215989723
refGene_gn=ATG16L1;refGene_tx=XM_023259819.1;refGene_gn=ATG16L1;refGene_tx=XM_023259820.1;refGene_gn=ATG16L1;refGene_tx=XM_023259821.1
C1 215990447 215990519
refGene_gn=ATG16L1;refGene_tx=XM_023259819.1;refGene_gn=ATG16L1;refGene_tx=XM_023259820.1;refGene_gn=ATG16L1;refGene_tx=XM_023259821.1
C1 215995631 215995752
refGene_gn=ATG16L1;refGene_tx=XM_023259819.1;refGene_gn=ATG16L1;refGene_tx=XM_023259820.1;refGene_gn=ATG16L1;refGene_tx=XM_023259821.1
C1 215996023 215996129
refGene_gn=ATG16L1;refGene_tx=XM_023259819.1;refGene_gn=ATG16L1;refGene_tx=XM_023259820.1;refGene_gn=ATG16L1;refGene_tx=XM_023259821.1
C1 215998111 215998261
refGene_gn=ATG16L1;refGene_tx=XM_023259819.1;refGene_gn=ATG16L1;refGene_tx=XM_023259820.1;refGene_gn=ATG16L1;refGene_tx=XM_023259821.1
C1 215998402 215998450
refGene_gn=ATG16L1;refGene_tx=XM_023259819.1;refGene_gn=ATG16L1;refGene_tx=XM_023259820.1;refGene_gn=ATG16L1;refGene_tx=XM_023259821.1
C1 215998804 215998906
refGene_gn=ATG16L1;refGene_tx=XM_023259819.1;refGene_gn=ATG16L1;refGene_tx=XM_023259820.1;refGene_gn=ATG16L1;refGene_tx=XM_023259821.1
C1 215999687 215999781
refGene_gn=ATG16L1;refGene_tx=XM_023259819.1;refGene_gn=ATG16L1;refGene_tx=XM_023259820.1;refGene_gn=ATG16L1;refGene_tx=XM_023259821.1
C1 216013322 216013397
refGene_gn=SAG;refGene_tx=XM_003991270.4
C1 216017414 216017475
refGene_gn=SAG;refGene_tx=XM_003991270.4
C1 216019832 216019877
refGene_gn=SAG;refGene_tx=XM_003991270.4
C1 216021311 216021396
refGene_gn=SAG;refGene_tx=XM_019838903.1
C1 216021517 216021711
refGene_gn=SAG;refGene_tx=XM_003991270.4;refGene_gn=SAG;refGene_tx=XM_019838903.1
C1 216023612 216023672
refGene_gn=SAG;refGene_tx=XM_003991270.4;refGene_gn=SAG;refGene_tx=XM_

019838903.1
C1	216025740	216025817
refGene_gn=SAG;refGene_tx=XM_003991270.4;refGene_gn=SAG;refGene_tx=XM_
019838903.1
C1	216026262	216026398
refGene_gn=SAG;refGene_tx=XM_003991270.4;refGene_gn=SAG;refGene_tx=XM_
019838903.1
C1	216027421	216027506
refGene_gn=SAG;refGene_tx=XM_003991270.4;refGene_gn=SAG;refGene_tx=XM_
019838903.1
C1	216029851	216029924
refGene_gn=SAG;refGene_tx=XM_003991270.4;refGene_gn=SAG;refGene_tx=XM_
019838903.1
C1	216032069	216032207
refGene_gn=SAG;refGene_tx=XM_003991270.4;refGene_gn=SAG;refGene_tx=XM_
019838903.1
C1	216035060	216035138
refGene_gn=SAG;refGene_tx=XM_003991270.4;refGene_gn=SAG;refGene_tx=XM_
019838903.1
C1	216036765	216036789
refGene_gn=SAG;refGene_tx=XM_003991270.4;refGene_gn=SAG;refGene_tx=XM_
019838903.1
C1	216038569	216038625
refGene_gn=SAG;refGene_tx=XM_003991270.4;refGene_gn=SAG;refGene_tx=XM_
019838903.1
C1	216041494	216041504
refGene_gn=SAG;refGene_tx=XM_003991270.4;refGene_gn=SAG;refGene_tx=XM_
019838903.1
C1	216041897	216042003
refGene_gn=SAG;refGene_tx=XM_003991270.4;refGene_gn=SAG;refGene_tx=XM_
019838903.1
C1	216048990	216049146
refGene_gn=DGKD;refGene_tx=XM_023259828.1
C1	216073276	216073300
refGene_gn=DGKD;refGene_tx=XM_006935700.4
C1	216074689	216074800
refGene_gn=DGKD;refGene_tx=XM_006935700.4;refGene_gn=DGKD;refGene_tx=X
M_019838907.2;refGene_gn=DGKD;refGene_tx=XM_023259828.1
C1	216076428	216076509
refGene_gn=DGKD;refGene_tx=XM_006935700.4;refGene_gn=DGKD;refGene_tx=X
M_019838907.2;refGene_gn=DGKD;refGene_tx=XM_023259828.1
C1	216120761	216120866
refGene_gn=DGKD;refGene_tx=XM_006935700.4;refGene_gn=DGKD;refGene_tx=X
M_006935702.4;refGene_gn=DGKD;refGene_tx=XM_019838907.2;refGene_gn=DGK
D;refGene_tx=XM_023259828.1;refGene_gn=DGKD;refGene_tx=XM_023259829.1
C1	216121253	216121386
refGene_gn=DGKD;refGene_tx=XM_006935700.4;refGene_gn=DGKD;refGene_tx=X
M_006935702.4;refGene_gn=DGKD;refGene_tx=XM_019838907.2;refGene_gn=DGK
D;refGene_tx=XM_023259828.1;refGene_gn=DGKD;refGene_tx=XM_023259829.1
C1	216122180	216122287

refGene_gn=DGKD;refGene_tx=XM_006935700.4;refGene_gn=DGKD;refGene_tx=XM_006935702.4;refGene_gn=DGKD;refGene_tx=XM_019838907.2;refGene_gn=DGKD;refGene_tx=XM_023259828.1;refGene_gn=DGKD;refGene_tx=XM_023259829.1
C1 216122604 216122730
refGene_gn=DGKD;refGene_tx=XM_006935700.4;refGene_gn=DGKD;refGene_tx=XM_006935702.4;refGene_gn=DGKD;refGene_tx=XM_019838907.2;refGene_gn=DGKD;refGene_tx=XM_023259828.1;refGene_gn=DGKD;refGene_tx=XM_023259829.1
C1 216122877 216122980
refGene_gn=DGKD;refGene_tx=XM_006935700.4;refGene_gn=DGKD;refGene_tx=XM_006935702.4;refGene_gn=DGKD;refGene_tx=XM_019838907.2;refGene_gn=DGKD;refGene_tx=XM_023259828.1;refGene_gn=DGKD;refGene_tx=XM_023259829.1
C1 216123600 216123763
refGene_gn=DGKD;refGene_tx=XM_006935700.4;refGene_gn=DGKD;refGene_tx=XM_006935702.4;refGene_gn=DGKD;refGene_tx=XM_019838907.2;refGene_gn=DGKD;refGene_tx=XM_023259828.1;refGene_gn=DGKD;refGene_tx=XM_023259829.1
C1 216126404 216126513
refGene_gn=DGKD;refGene_tx=XM_006935700.4;refGene_gn=DGKD;refGene_tx=XM_006935702.4;refGene_gn=DGKD;refGene_tx=XM_019838907.2;refGene_gn=DGKD;refGene_tx=XM_023259828.1;refGene_gn=DGKD;refGene_tx=XM_023259829.1
C1 216130044 216130184
refGene_gn=DGKD;refGene_tx=XM_006935700.4;refGene_gn=DGKD;refGene_tx=XM_006935702.4;refGene_gn=DGKD;refGene_tx=XM_019838907.2;refGene_gn=DGKD;refGene_tx=XM_023259828.1;refGene_gn=DGKD;refGene_tx=XM_023259829.1
C1 216131640 216131725
refGene_gn=DGKD;refGene_tx=XM_006935700.4;refGene_gn=DGKD;refGene_tx=XM_006935702.4;refGene_gn=DGKD;refGene_tx=XM_019838907.2;refGene_gn=DGKD;refGene_tx=XM_023259828.1;refGene_gn=DGKD;refGene_tx=XM_023259829.1
C1 216132858 216132953
refGene_gn=DGKD;refGene_tx=XM_006935700.4;refGene_gn=DGKD;refGene_tx=XM_006935702.4;refGene_gn=DGKD;refGene_tx=XM_019838907.2;refGene_gn=DGKD;refGene_tx=XM_023259828.1;refGene_gn=DGKD;refGene_tx=XM_023259829.1
C1 216133058 216133158
refGene_gn=DGKD;refGene_tx=XM_006935700.4;refGene_gn=DGKD;refGene_tx=XM_006935702.4;refGene_gn=DGKD;refGene_tx=XM_019838907.2;refGene_gn=DGKD;refGene_tx=XM_023259828.1;refGene_gn=DGKD;refGene_tx=XM_023259829.1
C1 216133234 216133508
refGene_gn=DGKD;refGene_tx=XM_006935700.4;refGene_gn=DGKD;refGene_tx=XM_006935702.4;refGene_gn=DGKD;refGene_tx=XM_019838907.2;refGene_gn=DGKD;refGene_tx=XM_023259828.1;refGene_gn=DGKD;refGene_tx=XM_023259829.1
C1 216134711 216134858
refGene_gn=DGKD;refGene_tx=XM_006935700.4;refGene_gn=DGKD;refGene_tx=XM_006935702.4;refGene_gn=DGKD;refGene_tx=XM_019838907.2;refGene_gn=DGKD;refGene_tx=XM_023259828.1;refGene_gn=DGKD;refGene_tx=XM_023259829.1
C1 216135717 216135846
refGene_gn=DGKD;refGene_tx=XM_006935700.4;refGene_gn=DGKD;refGene_tx=XM_006935702.4;refGene_gn=DGKD;refGene_tx=XM_019838907.2;refGene_gn=DGKD;refGene_tx=XM_023259828.1;refGene_gn=DGKD;refGene_tx=XM_023259829.1
C1 216137296 216137393
refGene_gn=DGKD;refGene_tx=XM_006935700.4;refGene_gn=DGKD;refGene_tx=XM_006935702.4;refGene_gn=DGKD;refGene_tx=XM_019838907.2;refGene_gn=DGK

D;refGene_tx=XM_023259828.1;refGene_gn=DGKD;refGene_tx=XM_023259829.1
C1 216140318 216140429
refGene_gn=DGKD;refGene_tx=XM_006935700.4;refGene_gn=DGKD;refGene_tx=XM_006935702.4;refGene_gn=DGKD;refGene_tx=XM_019838907.2;refGene_gn=DGKD;refGene_tx=XM_023259828.1;refGene_gn=DGKD;refGene_tx=XM_023259829.1
C1 216142364 216142461
refGene_gn=DGKD;refGene_tx=XM_006935700.4;refGene_gn=DGKD;refGene_tx=XM_006935702.4;refGene_gn=DGKD;refGene_tx=XM_019838907.2;refGene_gn=DGKD;refGene_tx=XM_023259828.1;refGene_gn=DGKD;refGene_tx=XM_023259829.1
C1 216142685 216142793
refGene_gn=DGKD;refGene_tx=XM_006935700.4;refGene_gn=DGKD;refGene_tx=XM_006935702.4;refGene_gn=DGKD;refGene_tx=XM_019838907.2;refGene_gn=DGKD;refGene_tx=XM_023259828.1;refGene_gn=DGKD;refGene_tx=XM_023259829.1
C1 216143891 216144005
refGene_gn=DGKD;refGene_tx=XM_006935700.4;refGene_gn=DGKD;refGene_tx=XM_006935702.4;refGene_gn=DGKD;refGene_tx=XM_019838907.2;refGene_gn=DGKD;refGene_tx=XM_023259828.1;refGene_gn=DGKD;refGene_tx=XM_023259829.1
C1 216145044 216145179
refGene_gn=DGKD;refGene_tx=XM_006935700.4;refGene_gn=DGKD;refGene_tx=XM_006935702.4;refGene_gn=DGKD;refGene_tx=XM_019838907.2;refGene_gn=DGKD;refGene_tx=XM_023259828.1;refGene_gn=DGKD;refGene_tx=XM_023259829.1
C1 216145503 216145655
refGene_gn=DGKD;refGene_tx=XM_006935700.4;refGene_gn=DGKD;refGene_tx=XM_006935702.4;refGene_gn=DGKD;refGene_tx=XM_019838907.2;refGene_gn=DGKD;refGene_tx=XM_023259828.1;refGene_gn=DGKD;refGene_tx=XM_023259829.1
C1 216147130 216147242
refGene_gn=DGKD;refGene_tx=XM_006935700.4;refGene_gn=DGKD;refGene_tx=XM_006935702.4;refGene_gn=DGKD;refGene_tx=XM_019838907.2;refGene_gn=DGKD;refGene_tx=XM_023259828.1;refGene_gn=DGKD;refGene_tx=XM_023259829.1
C1 216147326 216147419
refGene_gn=DGKD;refGene_tx=XM_006935700.4;refGene_gn=DGKD;refGene_tx=XM_006935702.4;refGene_gn=DGKD;refGene_tx=XM_019838907.2;refGene_gn=DGKD;refGene_tx=XM_023259828.1;refGene_gn=DGKD;refGene_tx=XM_023259829.1
C1 216148389 216148509
refGene_gn=DGKD;refGene_tx=XM_006935700.4;refGene_gn=DGKD;refGene_tx=XM_006935702.4;refGene_gn=DGKD;refGene_tx=XM_019838907.2;refGene_gn=DGKD;refGene_tx=XM_023259828.1;refGene_gn=DGKD;refGene_tx=XM_023259829.1
C1 216154952 216155064
refGene_gn=DGKD;refGene_tx=XM_006935700.4;refGene_gn=DGKD;refGene_tx=XM_006935702.4;refGene_gn=DGKD;refGene_tx=XM_019838907.2;refGene_gn=DGKD;refGene_tx=XM_023259828.1;refGene_gn=DGKD;refGene_tx=XM_023259829.1
C1 216156418 216156549
refGene_gn=DGKD;refGene_tx=XM_006935700.4;refGene_gn=DGKD;refGene_tx=XM_006935702.4;refGene_gn=DGKD;refGene_tx=XM_019838907.2;refGene_gn=DGKD;refGene_tx=XM_023259828.1;refGene_gn=DGKD;refGene_tx=XM_023259829.1
C1 216157844 216157934
refGene_gn=DGKD;refGene_tx=XM_006935700.4;refGene_gn=DGKD;refGene_tx=XM_006935702.4;refGene_gn=DGKD;refGene_tx=XM_019838907.2;refGene_gn=DGKD;refGene_tx=XM_023259828.1;refGene_gn=DGKD;refGene_tx=XM_023259829.1
C1 216165647 216165759

refGene_gn=USP40;refGene_tx=XM_023259822.1;refGene_gn=USP40;refGene_tx=XM_023259823.1;refGene_gn=USP40;refGene_tx=XM_023259824.1;refGene_gn=USP40;refGene_tx=XM_023259825.1;refGene_gn=USP40;refGene_tx=XM_023259826.1;refGene_gn=USP40;refGene_tx=XM_023259827.1
C1 216167284 216167379
refGene_gn=USP40;refGene_tx=XM_023259822.1;refGene_gn=USP40;refGene_tx=XM_023259823.1;refGene_gn=USP40;refGene_tx=XM_023259824.1;refGene_gn=USP40;refGene_tx=XM_023259825.1;refGene_gn=USP40;refGene_tx=XM_023259826.1;refGene_gn=USP40;refGene_tx=XM_023259827.1
C1 216170911 216171007
refGene_gn=USP40;refGene_tx=XM_023259822.1;refGene_gn=USP40;refGene_tx=XM_023259823.1;refGene_gn=USP40;refGene_tx=XM_023259824.1;refGene_gn=USP40;refGene_tx=XM_023259825.1;refGene_gn=USP40;refGene_tx=XM_023259826.1;refGene_gn=USP40;refGene_tx=XM_023259827.1
C1 216171147 216171358
refGene_gn=USP40;refGene_tx=XM_023259822.1;refGene_gn=USP40;refGene_tx=XM_023259823.1;refGene_gn=USP40;refGene_tx=XM_023259824.1;refGene_gn=USP40;refGene_tx=XM_023259825.1;refGene_gn=USP40;refGene_tx=XM_023259826.1;refGene_gn=USP40;refGene_tx=XM_023259827.1
C1 216173532 216173598
refGene_gn=USP40;refGene_tx=XM_023259822.1;refGene_gn=USP40;refGene_tx=XM_023259823.1;refGene_gn=USP40;refGene_tx=XM_023259824.1;refGene_gn=USP40;refGene_tx=XM_023259825.1;refGene_gn=USP40;refGene_tx=XM_023259826.1;refGene_gn=USP40;refGene_tx=XM_023259827.1
C1 216178121 216178237
refGene_gn=USP40;refGene_tx=XM_023259822.1;refGene_gn=USP40;refGene_tx=XM_023259823.1;refGene_gn=USP40;refGene_tx=XM_023259824.1;refGene_gn=USP40;refGene_tx=XM_023259825.1;refGene_gn=USP40;refGene_tx=XM_023259826.1;refGene_gn=USP40;refGene_tx=XM_023259827.1
C1 216179545 216179640
refGene_gn=USP40;refGene_tx=XM_023259822.1;refGene_gn=USP40;refGene_tx=XM_023259823.1;refGene_gn=USP40;refGene_tx=XM_023259824.1;refGene_gn=USP40;refGene_tx=XM_023259825.1;refGene_gn=USP40;refGene_tx=XM_023259826.1;refGene_gn=USP40;refGene_tx=XM_023259827.1
C1 216181648 216181775
refGene_gn=USP40;refGene_tx=XM_023259822.1;refGene_gn=USP40;refGene_tx=XM_023259823.1;refGene_gn=USP40;refGene_tx=XM_023259824.1;refGene_gn=USP40;refGene_tx=XM_023259825.1;refGene_gn=USP40;refGene_tx=XM_023259826.1;refGene_gn=USP40;refGene_tx=XM_023259827.1
C1 216184656 216184731
refGene_gn=USP40;refGene_tx=XM_023259822.1;refGene_gn=USP40;refGene_tx=XM_023259823.1;refGene_gn=USP40;refGene_tx=XM_023259824.1;refGene_gn=USP40;refGene_tx=XM_023259825.1;refGene_gn=USP40;refGene_tx=XM_023259826.1;refGene_gn=USP40;refGene_tx=XM_023259827.1
C1 216186483 216186548
refGene_gn=USP40;refGene_tx=XM_023259822.1;refGene_gn=USP40;refGene_tx=XM_023259823.1;refGene_gn=USP40;refGene_tx=XM_023259824.1;refGene_gn=USP40;refGene_tx=XM_023259825.1;refGene_gn=USP40;refGene_tx=XM_023259826.1;refGene_gn=USP40;refGene_tx=XM_023259827.1
C1 216188139 216188176

refGene_gn=USP40;refGene_tx=XM_023259822.1;refGene_gn=USP40;refGene_tx=XM_023259823.1;refGene_gn=USP40;refGene_tx=XM_023259824.1;refGene_gn=USP40;refGene_tx=XM_023259825.1;refGene_gn=USP40;refGene_tx=XM_023259826.1;refGene_gn=USP40;refGene_tx=XM_023259827.1
C1 216192863 216192950
refGene_gn=USP40;refGene_tx=XM_023259822.1;refGene_gn=USP40;refGene_tx=XM_023259823.1;refGene_gn=USP40;refGene_tx=XM_023259824.1;refGene_gn=USP40;refGene_tx=XM_023259825.1;refGene_gn=USP40;refGene_tx=XM_023259826.1;refGene_gn=USP40;refGene_tx=XM_023259827.1
C1 216194329 216194418
refGene_gn=USP40;refGene_tx=XM_023259822.1;refGene_gn=USP40;refGene_tx=XM_023259823.1;refGene_gn=USP40;refGene_tx=XM_023259824.1;refGene_gn=USP40;refGene_tx=XM_023259825.1;refGene_gn=USP40;refGene_tx=XM_023259826.1;refGene_gn=USP40;refGene_tx=XM_023259827.1
C1 216195137 216195191
refGene_gn=USP40;refGene_tx=XM_023259822.1;refGene_gn=USP40;refGene_tx=XM_023259823.1;refGene_gn=USP40;refGene_tx=XM_023259824.1;refGene_gn=USP40;refGene_tx=XM_023259825.1;refGene_gn=USP40;refGene_tx=XM_023259826.1;refGene_gn=USP40;refGene_tx=XM_023259827.1
C1 216201317 216201375
refGene_gn=USP40;refGene_tx=XM_023259822.1;refGene_gn=USP40;refGene_tx=XM_023259823.1;refGene_gn=USP40;refGene_tx=XM_023259824.1;refGene_gn=USP40;refGene_tx=XM_023259825.1;refGene_gn=USP40;refGene_tx=XM_023259826.1;refGene_gn=USP40;refGene_tx=XM_023259827.1
C1 216203046 216203170
refGene_gn=USP40;refGene_tx=XM_023259822.1;refGene_gn=USP40;refGene_tx=XM_023259823.1;refGene_gn=USP40;refGene_tx=XM_023259824.1;refGene_gn=USP40;refGene_tx=XM_023259825.1;refGene_gn=USP40;refGene_tx=XM_023259826.1;refGene_gn=USP40;refGene_tx=XM_023259827.1
C1 216204453 216204773
refGene_gn=USP40;refGene_tx=XM_023259822.1;refGene_gn=USP40;refGene_tx=XM_023259823.1;refGene_gn=USP40;refGene_tx=XM_023259824.1;refGene_gn=USP40;refGene_tx=XM_023259825.1;refGene_gn=USP40;refGene_tx=XM_023259826.1;refGene_gn=USP40;refGene_tx=XM_023259827.1
C1 216205726 216205797
refGene_gn=USP40;refGene_tx=XM_023259822.1;refGene_gn=USP40;refGene_tx=XM_023259823.1;refGene_gn=USP40;refGene_tx=XM_023259824.1;refGene_gn=USP40;refGene_tx=XM_023259825.1;refGene_gn=USP40;refGene_tx=XM_023259826.1;refGene_gn=USP40;refGene_tx=XM_023259827.1
C1 216206897 216206982
refGene_gn=USP40;refGene_tx=XM_023259822.1;refGene_gn=USP40;refGene_tx=XM_023259823.1;refGene_gn=USP40;refGene_tx=XM_023259824.1;refGene_gn=USP40;refGene_tx=XM_023259825.1;refGene_gn=USP40;refGene_tx=XM_023259826.1;refGene_gn=USP40;refGene_tx=XM_023259827.1
C1 216209464 216209636
refGene_gn=USP40;refGene_tx=XM_023259822.1;refGene_gn=USP40;refGene_tx=XM_023259823.1;refGene_gn=USP40;refGene_tx=XM_023259824.1;refGene_gn=USP40;refGene_tx=XM_023259825.1;refGene_gn=USP40;refGene_tx=XM_023259826.1;refGene_gn=USP40;refGene_tx=XM_023259827.1
C1 216211132 216211214

refGene_gn=USP40;refGene_tx=XM_023259822.1;refGene_gn=USP40;refGene_tx=XM_023259823.1;refGene_gn=USP40;refGene_tx=XM_023259824.1;refGene_gn=USP40;refGene_tx=XM_023259825.1;refGene_gn=USP40;refGene_tx=XM_023259826.1;refGene_gn=USP40;refGene_tx=XM_023259827.1
C1 216215251 216215552
refGene_gn=USP40;refGene_tx=XM_023259822.1;refGene_gn=USP40;refGene_tx=XM_023259823.1;refGene_gn=USP40;refGene_tx=XM_023259824.1;refGene_gn=USP40;refGene_tx=XM_023259825.1;refGene_gn=USP40;refGene_tx=XM_023259826.1;refGene_gn=USP40;refGene_tx=XM_023259827.1
C1 216230743 216230851
refGene_gn=USP40;refGene_tx=XM_023259822.1;refGene_gn=USP40;refGene_tx=XM_023259823.1;refGene_gn=USP40;refGene_tx=XM_023259824.1;refGene_gn=USP40;refGene_tx=XM_023259825.1;refGene_gn=USP40;refGene_tx=XM_023259826.1;refGene_gn=USP40;refGene_tx=XM_023259827.1
C1 216232288 216232372
refGene_gn=USP40;refGene_tx=XM_023259822.1;refGene_gn=USP40;refGene_tx=XM_023259823.1;refGene_gn=USP40;refGene_tx=XM_023259824.1;refGene_gn=USP40;refGene_tx=XM_023259825.1;refGene_gn=USP40;refGene_tx=XM_023259826.1;refGene_gn=USP40;refGene_tx=XM_023259827.1
C1 216239752 216239881
refGene_gn=USP40;refGene_tx=XM_023259822.1;refGene_gn=USP40;refGene_tx=XM_023259823.1;refGene_gn=USP40;refGene_tx=XM_023259824.1;refGene_gn=USP40;refGene_tx=XM_023259825.1;refGene_gn=USP40;refGene_tx=XM_023259826.1
C1 216241256 216241400
refGene_gn=USP40;refGene_tx=XM_023259822.1;refGene_gn=USP40;refGene_tx=XM_023259823.1;refGene_gn=USP40;refGene_tx=XM_023259824.1;refGene_gn=USP40;refGene_tx=XM_023259825.1;refGene_gn=USP40;refGene_tx=XM_023259826.1
C1 216243354 216243558
refGene_gn=USP40;refGene_tx=XM_023259822.1;refGene_gn=USP40;refGene_tx=XM_023259823.1;refGene_gn=USP40;refGene_tx=XM_023259824.1;refGene_gn=USP40;refGene_tx=XM_023259825.1;refGene_gn=USP40;refGene_tx=XM_023259826.1
C1 216245204 216245369
refGene_gn=USP40;refGene_tx=XM_023259822.1;refGene_gn=USP40;refGene_tx=XM_023259823.1;refGene_gn=USP40;refGene_tx=XM_023259824.1;refGene_gn=USP40;refGene_tx=XM_023259825.1;refGene_gn=USP40;refGene_tx=XM_023259826.1
C1 216248304 216248418
refGene_gn=USP40;refGene_tx=XM_023259822.1;refGene_gn=USP40;refGene_tx=XM_023259823.1;refGene_gn=USP40;refGene_tx=XM_023259824.1;refGene_gn=USP40;refGene_tx=XM_023259827.1
C1 216249825 216249893
refGene_gn=USP40;refGene_tx=XM_023259822.1;refGene_gn=USP40;refGene_tx=XM_023259823.1;refGene_gn=USP40;refGene_tx=XM_023259824.1;refGene_gn=USP40;refGene_tx=XM_023259827.1
C1 216252432 216252631
refGene_gn=USP40;refGene_tx=XM_023259822.1;refGene_gn=USP40;refGene_tx=XM_023259823.1;refGene_gn=USP40;refGene_tx=XM_023259824.1;refGene_gn=

USP40;refGene_tx=XM_023259827.1
C1 216306022 216306886
refGene_gn=UGT1A1;refGene_tx=XM_006935678.3
C1 216311957 216312680
refGene_gn=DNAJB3;refGene_tx=XM_011285622.3
C1 216317730 216318594
refGene_gn=UGT1A1;refGene_tx=NM_001009359.1
C1 216326733 216326865
refGene_gn=UGT1A1;refGene_tx=NM_001009359.1;refGene_gn=UGT1A1;refGene_tx=XM_006935678.3
C1 216327709 216327797
refGene_gn=UGT1A1;refGene_tx=NM_001009359.1;refGene_gn=UGT1A1;refGene_tx=XM_006935678.3
C1 216328085 216328305
refGene_gn=UGT1A1;refGene_tx=NM_001009359.1;refGene_gn=UGT1A1;refGene_tx=XM_006935678.3
C1 216330722 216331020
refGene_gn=UGT1A1;refGene_tx=NM_001009359.1;refGene_gn=UGT1A1;refGene_tx=XM_006935678.3
C1 216336020 216336170
refGene_gn=MROH2A;refGene_tx=XM_019838910.1;refGene_gn=MROH2A;refGene_tx=XM_019838911.1;refGene_gn=MROH2A;refGene_tx=XM_019838912.1;refGene_gn=MROH2A;refGene_tx=XM_019838913.2;refGene_gn=MROH2A;refGene_tx=XM_019838915.1
C1 216340644 216340776
refGene_gn=MROH2A;refGene_tx=XM_019838910.1;refGene_gn=MROH2A;refGene_tx=XM_019838911.1;refGene_gn=MROH2A;refGene_tx=XM_019838912.1;refGene_gn=MROH2A;refGene_tx=XM_019838913.2;refGene_gn=MROH2A;refGene_tx=XM_019838915.1;refGene_gn=MROH2A;refGene_tx=XM_019838918.1
C1 216341058 216341221
refGene_gn=MROH2A;refGene_tx=XM_019838910.1;refGene_gn=MROH2A;refGene_tx=XM_019838911.1;refGene_gn=MROH2A;refGene_tx=XM_019838912.1;refGene_gn=MROH2A;refGene_tx=XM_019838913.2;refGene_gn=MROH2A;refGene_tx=XM_019838915.1;refGene_gn=MROH2A;refGene_tx=XM_019838918.1
C1 216343323 216343422
refGene_gn=MROH2A;refGene_tx=XM_019838910.1;refGene_gn=MROH2A;refGene_tx=XM_019838911.1;refGene_gn=MROH2A;refGene_tx=XM_019838912.1;refGene_gn=MROH2A;refGene_tx=XM_019838913.2;refGene_gn=MROH2A;refGene_tx=XM_019838915.1;refGene_gn=MROH2A;refGene_tx=XM_019838918.1
C1 216344167 216344319
refGene_gn=MROH2A;refGene_tx=XM_019838910.1;refGene_gn=MROH2A;refGene_tx=XM_019838911.1;refGene_gn=MROH2A;refGene_tx=XM_019838912.1;refGene_gn=MROH2A;refGene_tx=XM_019838913.2;refGene_gn=MROH2A;refGene_tx=XM_019838915.1;refGene_gn=MROH2A;refGene_tx=XM_019838918.1
C1 216344791 216344935
refGene_gn=MROH2A;refGene_tx=XM_019838910.1;refGene_gn=MROH2A;refGene_tx=XM_019838911.1;refGene_gn=MROH2A;refGene_tx=XM_019838912.1;refGene_gn=MROH2A;refGene_tx=XM_019838913.2;refGene_gn=MROH2A;refGene_tx=XM_019838915.1;refGene_gn=MROH2A;refGene_tx=XM_019838918.1
C1 216346836 216346929

refGene_gn=MROH2A;refGene_tx=XM_019838910.1;refGene_gn=MROH2A;refGene_tx=XM_019838911.1;refGene_gn=MROH2A;refGene_tx=XM_019838912.1;refGene_gn=MROH2A;refGene_tx=XM_019838913.2;refGene_gn=MROH2A;refGene_tx=XM_019838915.1;refGene_gn=MROH2A;refGene_tx=XM_019838918.1
C1 216347163 216347242
refGene_gn=MROH2A;refGene_tx=XM_019838910.1;refGene_gn=MROH2A;refGene_tx=XM_019838911.1;refGene_gn=MROH2A;refGene_tx=XM_019838912.1;refGene_gn=MROH2A;refGene_tx=XM_019838913.2;refGene_gn=MROH2A;refGene_tx=XM_019838915.1;refGene_gn=MROH2A;refGene_tx=XM_019838918.1
C1 216347371 216347485
refGene_gn=MROH2A;refGene_tx=XM_019838910.1;refGene_gn=MROH2A;refGene_tx=XM_019838911.1;refGene_gn=MROH2A;refGene_tx=XM_019838912.1;refGene_gn=MROH2A;refGene_tx=XM_019838913.2;refGene_gn=MROH2A;refGene_tx=XM_019838915.1;refGene_gn=MROH2A;refGene_tx=XM_019838918.1
C1 216350172 216350249
refGene_gn=MROH2A;refGene_tx=XM_019838910.1;refGene_gn=MROH2A;refGene_tx=XM_019838911.1;refGene_gn=MROH2A;refGene_tx=XM_019838912.1;refGene_gn=MROH2A;refGene_tx=XM_019838913.2;refGene_gn=MROH2A;refGene_tx=XM_019838915.1;refGene_gn=MROH2A;refGene_tx=XM_019838918.1
C1 216351191 216351311
refGene_gn=MROH2A;refGene_tx=XM_019838910.1;refGene_gn=MROH2A;refGene_tx=XM_019838911.1;refGene_gn=MROH2A;refGene_tx=XM_019838912.1;refGene_gn=MROH2A;refGene_tx=XM_019838913.2;refGene_gn=MROH2A;refGene_tx=XM_019838915.1;refGene_gn=MROH2A;refGene_tx=XM_019838918.1
C1 216351613 216351724
refGene_gn=MROH2A;refGene_tx=XM_019838910.1;refGene_gn=MROH2A;refGene_tx=XM_019838911.1;refGene_gn=MROH2A;refGene_tx=XM_019838912.1;refGene_gn=MROH2A;refGene_tx=XM_019838913.2;refGene_gn=MROH2A;refGene_tx=XM_019838915.1;refGene_gn=MROH2A;refGene_tx=XM_019838918.1
C1 216352528 216352676
refGene_gn=MROH2A;refGene_tx=XM_019838910.1;refGene_gn=MROH2A;refGene_tx=XM_019838911.1;refGene_gn=MROH2A;refGene_tx=XM_019838912.1;refGene_gn=MROH2A;refGene_tx=XM_019838913.2;refGene_gn=MROH2A;refGene_tx=XM_019838915.1;refGene_gn=MROH2A;refGene_tx=XM_019838918.1
C1 216353441 216353482
refGene_gn=MROH2A;refGene_tx=XM_019838910.1;refGene_gn=MROH2A;refGene_tx=XM_019838911.1;refGene_gn=MROH2A;refGene_tx=XM_019838912.1;refGene_gn=MROH2A;refGene_tx=XM_019838913.2;refGene_gn=MROH2A;refGene_tx=XM_019838915.1;refGene_gn=MROH2A;refGene_tx=XM_019838918.1
C1 216353929 216354071
refGene_gn=MROH2A;refGene_tx=XM_019838910.1;refGene_gn=MROH2A;refGene_tx=XM_019838911.1;refGene_gn=MROH2A;refGene_tx=XM_019838912.1;refGene_gn=MROH2A;refGene_tx=XM_019838913.2;refGene_gn=MROH2A;refGene_tx=XM_019838915.1;refGene_gn=MROH2A;refGene_tx=XM_019838918.1
C1 216354256 216354309
refGene_gn=MROH2A;refGene_tx=XM_019838910.1;refGene_gn=MROH2A;refGene_tx=XM_019838911.1;refGene_gn=MROH2A;refGene_tx=XM_019838912.1;refGene_gn=MROH2A;refGene_tx=XM_019838913.2;refGene_gn=MROH2A;refGene_tx=XM_019838915.1;refGene_gn=MROH2A;refGene_tx=XM_019838918.1
C1 216354747 216354855

refGene_gn=MROH2A;refGene_tx=XM_019838910.1;refGene_gn=MROH2A;refGene_
tx=XM_019838911.1;refGene_gn=MROH2A;refGene_tx=XM_019838912.1;refGene_
gn=MROH2A;refGene_tx=XM_019838913.2;refGene_gn=MROH2A;refGene_tx=XM_01
9838915.1;refGene_gn=MROH2A;refGene_tx=XM_019838918.1
C1 216358171 216358291
refGene_gn=MROH2A;refGene_tx=XM_019838910.1;refGene_gn=MROH2A;refGene_
tx=XM_019838911.1;refGene_gn=MROH2A;refGene_tx=XM_019838912.1;refGene_
gn=MROH2A;refGene_tx=XM_019838913.2;refGene_gn=MROH2A;refGene_tx=XM_01
9838915.1;refGene_gn=MROH2A;refGene_tx=XM_019838918.1
C1 216358564 216358687
refGene_gn=MROH2A;refGene_tx=XM_019838910.1;refGene_gn=MROH2A;refGene_
tx=XM_019838911.1;refGene_gn=MROH2A;refGene_tx=XM_019838912.1;refGene_
gn=MROH2A;refGene_tx=XM_019838913.2;refGene_gn=MROH2A;refGene_tx=XM_01
9838915.1;refGene_gn=MROH2A;refGene_tx=XM_019838918.1
C1 216359707 216359860
refGene_gn=MROH2A;refGene_tx=XM_019838910.1;refGene_gn=MROH2A;refGene_
tx=XM_019838911.1;refGene_gn=MROH2A;refGene_tx=XM_019838912.1;refGene_
gn=MROH2A;refGene_tx=XM_019838913.2;refGene_gn=MROH2A;refGene_tx=XM_01
9838915.1;refGene_gn=MROH2A;refGene_tx=XM_019838918.1
C1 216361010 216361133
refGene_gn=MROH2A;refGene_tx=XM_019838910.1;refGene_gn=MROH2A;refGene_
tx=XM_019838911.1;refGene_gn=MROH2A;refGene_tx=XM_019838912.1;refGene_
gn=MROH2A;refGene_tx=XM_019838913.2;refGene_gn=MROH2A;refGene_tx=XM_01
9838915.1;refGene_gn=MROH2A;refGene_tx=XM_019838918.1
C1 216362067 216362147
refGene_gn=MROH2A;refGene_tx=XM_019838910.1;refGene_gn=MROH2A;refGene_
tx=XM_019838911.1;refGene_gn=MROH2A;refGene_tx=XM_019838912.1;refGene_
gn=MROH2A;refGene_tx=XM_019838913.2;refGene_gn=MROH2A;refGene_tx=XM_01
9838915.1;refGene_gn=MROH2A;refGene_tx=XM_019838918.1
C1 216363581 216363690
refGene_gn=MROH2A;refGene_tx=XM_019838910.1;refGene_gn=MROH2A;refGene_
tx=XM_019838911.1;refGene_gn=MROH2A;refGene_tx=XM_019838912.1;refGene_
gn=MROH2A;refGene_tx=XM_019838913.2;refGene_gn=MROH2A;refGene_tx=XM_01
9838915.1;refGene_gn=MROH2A;refGene_tx=XM_019838918.1
C1 216364494 216364590
refGene_gn=MROH2A;refGene_tx=XM_019838910.1;refGene_gn=MROH2A;refGene_
tx=XM_019838911.1;refGene_gn=MROH2A;refGene_tx=XM_019838912.1;refGene_
gn=MROH2A;refGene_tx=XM_019838913.2;refGene_gn=MROH2A;refGene_tx=XM_01
9838915.1;refGene_gn=MROH2A;refGene_tx=XM_019838918.1
C1 216367131 216367236
refGene_gn=MROH2A;refGene_tx=XM_019838910.1;refGene_gn=MROH2A;refGene_
tx=XM_019838911.1;refGene_gn=MROH2A;refGene_tx=XM_019838912.1;refGene_
gn=MROH2A;refGene_tx=XM_019838913.2;refGene_gn=MROH2A;refGene_tx=XM_01
9838915.1;refGene_gn=MROH2A;refGene_tx=XM_019838918.1
C1 216367929 216368053
refGene_gn=MROH2A;refGene_tx=XM_019838910.1;refGene_gn=MROH2A;refGene_
tx=XM_019838911.1;refGene_gn=MROH2A;refGene_tx=XM_019838912.1;refGene_
gn=MROH2A;refGene_tx=XM_019838913.2;refGene_gn=MROH2A;refGene_tx=XM_01
9838915.1;refGene_gn=MROH2A;refGene_tx=XM_019838918.1
C1 216368627 216368746

refGene_gn=MROH2A;refGene_tx=XM_019838910.1;refGene_gn=MROH2A;refGene_tx=XM_019838911.1;refGene_gn=MROH2A;refGene_tx=XM_019838912.1;refGene_gn=MROH2A;refGene_tx=XM_019838913.2;refGene_gn=MROH2A;refGene_tx=XM_019838915.1;refGene_gn=MROH2A;refGene_tx=XM_019838918.1
C1 216369595 216369748
refGene_gn=MROH2A;refGene_tx=XM_019838910.1;refGene_gn=MROH2A;refGene_tx=XM_019838911.1;refGene_gn=MROH2A;refGene_tx=XM_019838912.1;refGene_gn=MROH2A;refGene_tx=XM_019838913.2;refGene_gn=MROH2A;refGene_tx=XM_019838915.1;refGene_gn=MROH2A;refGene_tx=XM_019838918.1
C1 216370153 216370308
refGene_gn=MROH2A;refGene_tx=XM_019838910.1;refGene_gn=MROH2A;refGene_tx=XM_019838911.1;refGene_gn=MROH2A;refGene_tx=XM_019838912.1;refGene_gn=MROH2A;refGene_tx=XM_019838913.2;refGene_gn=MROH2A;refGene_tx=XM_019838915.1;refGene_gn=MROH2A;refGene_tx=XM_019838918.1
C1 216372271 216372431
refGene_gn=MROH2A;refGene_tx=XM_019838910.1;refGene_gn=MROH2A;refGene_tx=XM_019838911.1;refGene_gn=MROH2A;refGene_tx=XM_019838912.1;refGene_gn=MROH2A;refGene_tx=XM_019838913.2;refGene_gn=MROH2A;refGene_tx=XM_019838915.1;refGene_gn=MROH2A;refGene_tx=XM_019838918.1
C1 216372510 216372704
refGene_gn=MROH2A;refGene_tx=XM_019838910.1;refGene_gn=MROH2A;refGene_tx=XM_019838911.1;refGene_gn=MROH2A;refGene_tx=XM_019838912.1;refGene_gn=MROH2A;refGene_tx=XM_019838913.2;refGene_gn=MROH2A;refGene_tx=XM_019838915.1;refGene_gn=MROH2A;refGene_tx=XM_019838918.1
C1 216373050 216373188
refGene_gn=MROH2A;refGene_tx=XM_019838910.1;refGene_gn=MROH2A;refGene_tx=XM_019838911.1;refGene_gn=MROH2A;refGene_tx=XM_019838912.1;refGene_gn=MROH2A;refGene_tx=XM_019838913.2;refGene_gn=MROH2A;refGene_tx=XM_019838915.1;refGene_gn=MROH2A;refGene_tx=XM_019838918.1
C1 216373636 216373751
refGene_gn=MROH2A;refGene_tx=XM_019838910.1;refGene_gn=MROH2A;refGene_tx=XM_019838911.1;refGene_gn=MROH2A;refGene_tx=XM_019838912.1;refGene_gn=MROH2A;refGene_tx=XM_019838913.2;refGene_gn=MROH2A;refGene_tx=XM_019838915.1;refGene_gn=MROH2A;refGene_tx=XM_019838918.1
C1 216375058 216375205
refGene_gn=MROH2A;refGene_tx=XM_019838910.1;refGene_gn=MROH2A;refGene_tx=XM_019838911.1;refGene_gn=MROH2A;refGene_tx=XM_019838912.1;refGene_gn=MROH2A;refGene_tx=XM_019838913.2;refGene_gn=MROH2A;refGene_tx=XM_019838915.1;refGene_gn=MROH2A;refGene_tx=XM_019838918.1
C1 216375360 216375543
refGene_gn=MROH2A;refGene_tx=XM_019838910.1;refGene_gn=MROH2A;refGene_tx=XM_019838911.1;refGene_gn=MROH2A;refGene_tx=XM_019838912.1;refGene_gn=MROH2A;refGene_tx=XM_019838913.2;refGene_gn=MROH2A;refGene_tx=XM_019838915.1;refGene_gn=MROH2A;refGene_tx=XM_019838918.1
C1 216375998 216376154
refGene_gn=MROH2A;refGene_tx=XM_019838910.1;refGene_gn=MROH2A;refGene_tx=XM_019838911.1;refGene_gn=MROH2A;refGene_tx=XM_019838912.1;refGene_gn=MROH2A;refGene_tx=XM_019838913.2;refGene_gn=MROH2A;refGene_tx=XM_019838915.1;refGene_gn=MROH2A;refGene_tx=XM_019838918.1
C1 216377581 216377710

refGene_gn=MROH2A;refGene_tx=XM_019838910.1;refGene_gn=MROH2A;refGene_tx=XM_019838911.1;refGene_gn=MROH2A;refGene_tx=XM_019838912.1;refGene_gn=MROH2A;refGene_tx=XM_019838913.2;refGene_gn=MROH2A;refGene_tx=XM_019838915.1;refGene_gn=MROH2A;refGene_tx=XM_019838918.1
C1 216378356 216378459
refGene_gn=MROH2A;refGene_tx=XM_019838910.1;refGene_gn=MROH2A;refGene_tx=XM_019838911.1;refGene_gn=MROH2A;refGene_tx=XM_019838912.1;refGene_gn=MROH2A;refGene_tx=XM_019838913.2;refGene_gn=MROH2A;refGene_tx=XM_019838918.1
C1 216378717 216378791
refGene_gn=MROH2A;refGene_tx=XM_019838910.1;refGene_gn=MROH2A;refGene_tx=XM_019838911.1;refGene_gn=MROH2A;refGene_tx=XM_019838912.1;refGene_gn=MROH2A;refGene_tx=XM_019838913.2;refGene_gn=MROH2A;refGene_tx=XM_019838915.1;refGene_gn=MROH2A;refGene_tx=XM_019838918.1
C1 216379366 216379488
refGene_gn=MROH2A;refGene_tx=XM_019838910.1;refGene_gn=MROH2A;refGene_tx=XM_019838911.1;refGene_gn=MROH2A;refGene_tx=XM_019838912.1;refGene_gn=MROH2A;refGene_tx=XM_019838918.1
C1 216383799 216383839
refGene_gn=HJURP;refGene_tx=XM_023257872.1
C1 216386430 216388000
refGene_gn=HJURP;refGene_tx=XM_023257872.1
C1 216390159 216390238
refGene_gn=HJURP;refGene_tx=XM_023257872.1
C1 216391045 216391138
refGene_gn=HJURP;refGene_tx=XM_023257872.1
C1 216392463 216392525
refGene_gn=HJURP;refGene_tx=XM_023257872.1
C1 216394955 216395073
refGene_gn=HJURP;refGene_tx=XM_023257872.1
C1 216396340 216396396
refGene_gn=HJURP;refGene_tx=XM_023257872.1
C1 216399435 216399502
refGene_gn=HJURP;refGene_tx=XM_023257872.1
C1 216399844 216400048
refGene_gn=HJURP;refGene_tx=XM_023257872.1
C1 216400151 216400289
refGene_gn=HJURP;refGene_tx=XM_023257872.1
C1 216408967 216409075
refGene_gn=HJURP;refGene_tx=XM_023257872.1
C1 216409846 216409870
refGene_gn=HJURP;refGene_tx=XM_023257872.1
C1 216446413 216446530
refGene_gn=TRPM8;refGene_tx=XM_006935704.4
C1 216451392 216451466
refGene_gn=TRPM8;refGene_tx=XM_006935704.4
C1 216457083 216457240
refGene_gn=TRPM8;refGene_tx=XM_006935704.4
C1 216458208 216458386
refGene_gn=TRPM8;refGene_tx=XM_006935704.4

C1 216461781 216461954
refGene_gn=TRPM8;refGene_tx=XM_006935704.4
C1 216468250 216468425
refGene_gn=TRPM8;refGene_tx=XM_006935704.4
C1 216469698 216469766
refGene_gn=TRPM8;refGene_tx=XM_006935704.4
C1 216472026 216472224
refGene_gn=TRPM8;refGene_tx=XM_006935704.4
C1 216477132 216477235
refGene_gn=TRPM8;refGene_tx=XM_006935704.4
C1 216478328 216478447
refGene_gn=TRPM8;refGene_tx=XM_006935704.4
C1 216482529 216482820
refGene_gn=TRPM8;refGene_tx=XM_006935704.4
C1 216484831 216484927
refGene_gn=TRPM8;refGene_tx=XM_006935704.4
C1 216485744 216485874
refGene_gn=TRPM8;refGene_tx=XM_006935704.4
C1 216487657 216487803
refGene_gn=TRPM8;refGene_tx=XM_006935704.4
C1 216490528 216490641
refGene_gn=TRPM8;refGene_tx=XM_006935704.4
C1 216491046 216491263
refGene_gn=TRPM8;refGene_tx=XM_006935704.4
C1 216497248 216497340
refGene_gn=TRPM8;refGene_tx=XM_006935704.4
C1 216498570 216498712
refGene_gn=TRPM8;refGene_tx=XM_006935704.4
C1 216499771 216499943
refGene_gn=TRPM8;refGene_tx=XM_006935704.4
C1 216501073 216501251
refGene_gn=TRPM8;refGene_tx=XM_006935704.4
C1 216511939 216512130
refGene_gn=TRPM8;refGene_tx=XM_006935704.4
C1 216516972 216517072
refGene_gn=TRPM8;refGene_tx=XM_006935704.4
C1 216518164 216518198
refGene_gn=TRPM8;refGene_tx=XM_006935704.4
C1 216522626 216522677
refGene_gn=TRPM8;refGene_tx=XM_006935704.4
C1 216550935 216550993
refGene_gn=SPP2;refGene_tx=XM_003991272.5
C1 216551122 216551247
refGene_gn=SPP2;refGene_tx=XM_003991272.5
C1 216557532 216557655
refGene_gn=SPP2;refGene_tx=XM_003991272.5
C1 216558840 216558948
refGene_gn=SPP2;refGene_tx=XM_003991272.5
C1 216563205 216563260
refGene_gn=SPP2;refGene_tx=XM_003991272.5

C1 216564432 216564480
refGene_gn=SPP2;refGene_tx=XM_003991272.5
C1 216566665 216566751
refGene_gn=SPP2;refGene_tx=XM_003991272.5
C1 216895673 216896252
refGene_gn=ARL4C;refGene_tx=XM_011285620.3
C1 217263573 217264009
refGene_gn=LOC109502951;refGene_tx=XM_019838924.2
C1 217265078 217265704
refGene_gn=LOC109502951;refGene_tx=XM_019838924.2
C1 217319886 217320004
refGene_gn=SH3BP4;refGene_tx=XM_023259832.1;refGene_gn=SH3BP4;refGene_tx=XM_023259833.1;refGene_gn=SH3BP4;refGene_tx=XM_023259834.1
C1 217324291 217326648
refGene_gn=SH3BP4;refGene_tx=XM_023259832.1;refGene_gn=SH3BP4;refGene_tx=XM_023259833.1;refGene_gn=SH3BP4;refGene_tx=XM_023259834.1
C1 217329112 217329235
refGene_gn=SH3BP4;refGene_tx=XM_023259834.1
C1 217335482 217335671
refGene_gn=SH3BP4;refGene_tx=XM_023259832.1;refGene_gn=SH3BP4;refGene_tx=XM_023259833.1
C1 217336472 217336697
refGene_gn=SH3BP4;refGene_tx=XM_023259832.1;refGene_gn=SH3BP4;refGene_tx=XM_023259833.1
C1 217665916 217666079
refGene_gn=AGAP1;refGene_tx=XM_023259842.1;refGene_gn=AGAP1;refGene_tx=XM_023259844.1;refGene_gn=AGAP1;refGene_tx=XM_023259845.1;refGene_gn=AGAP1;refGene_tx=XM_023259846.1
C1 217818588 217819606
refGene_gn=AGAP1;refGene_tx=XM_023259835.1;refGene_gn=AGAP1;refGene_tx=XM_023259836.1;refGene_gn=AGAP1;refGene_tx=XM_023259837.1;refGene_gn=AGAP1;refGene_tx=XM_023259838.1;refGene_gn=AGAP1;refGene_tx=XM_023259839.1;refGene_gn=AGAP1;refGene_tx=XM_023259840.1;refGene_gn=AGAP1;refGene_tx=XM_023259841.1
C1 217856563 217856622
refGene_gn=AGAP1;refGene_tx=XM_023259835.1;refGene_gn=AGAP1;refGene_tx=XM_023259836.1;refGene_gn=AGAP1;refGene_tx=XM_023259837.1;refGene_gn=AGAP1;refGene_tx=XM_023259838.1;refGene_gn=AGAP1;refGene_tx=XM_023259839.1;refGene_gn=AGAP1;refGene_tx=XM_023259840.1;refGene_gn=AGAP1;refGene_tx=XM_023259841.1;refGene_gn=AGAP1;refGene_tx=XM_023259842.1;refGene_gn=AGAP1;refGene_tx=XM_023259844.1;refGene_gn=AGAP1;refGene_tx=XM_023259845.1;refGene_gn=AGAP1;refGene_tx=XM_023259846.1
C1 217862618 217862706
refGene_gn=AGAP1;refGene_tx=XM_023259835.1;refGene_gn=AGAP1;refGene_tx=XM_023259836.1;refGene_gn=AGAP1;refGene_tx=XM_023259837.1;refGene_gn=AGAP1;refGene_tx=XM_023259838.1;refGene_gn=AGAP1;refGene_tx=XM_023259839.1;refGene_gn=AGAP1;refGene_tx=XM_023259840.1;refGene_gn=AGAP1;refGene_tx=XM_023259841.1;refGene_gn=AGAP1;refGene_tx=XM_023259842.1;refGene_gn=AGAP1;refGene_tx=XM_023259844.1;refGene_gn=AGAP1;refGene_tx=XM_023259845.1;refGene_gn=AGAP1;refGene_tx=XM_023259846.1

C1 217863408 217863420
refGene_gn=AGAP1;refGene_tx=XM_023259835.1;refGene_gn=AGAP1;refGene_tx=XM_023259837.1;refGene_gn=AGAP1;refGene_tx=XM_023259838.1;refGene_gn=AGAP1;refGene_tx=XM_023259839.1;refGene_gn=AGAP1;refGene_tx=XM_023259840.1;refGene_gn=AGAP1;refGene_tx=XM_023259841.1;refGene_gn=AGAP1;refGene_tx=XM_023259842.1
C1 217884870 217884956
refGene_gn=AGAP1;refGene_tx=XM_023259835.1;refGene_gn=AGAP1;refGene_tx=XM_023259836.1;refGene_gn=AGAP1;refGene_tx=XM_023259837.1;refGene_gn=AGAP1;refGene_tx=XM_023259838.1;refGene_gn=AGAP1;refGene_tx=XM_023259839.1;refGene_gn=AGAP1;refGene_tx=XM_023259840.1;refGene_gn=AGAP1;refGene_tx=XM_023259841.1;refGene_gn=AGAP1;refGene_tx=XM_023259842.1;refGene_gn=AGAP1;refGene_tx=XM_023259844.1;refGene_gn=AGAP1;refGene_tx=XM_023259845.1;refGene_gn=AGAP1;refGene_tx=XM_023259846.1
C1 217886631 217886773
refGene_gn=AGAP1;refGene_tx=XM_023259835.1;refGene_gn=AGAP1;refGene_tx=XM_023259836.1;refGene_gn=AGAP1;refGene_tx=XM_023259837.1;refGene_gn=AGAP1;refGene_tx=XM_023259838.1;refGene_gn=AGAP1;refGene_tx=XM_023259839.1;refGene_gn=AGAP1;refGene_tx=XM_023259840.1;refGene_gn=AGAP1;refGene_tx=XM_023259841.1;refGene_gn=AGAP1;refGene_tx=XM_023259842.1;refGene_gn=AGAP1;refGene_tx=XM_023259844.1;refGene_gn=AGAP1;refGene_tx=XM_023259845.1;refGene_gn=AGAP1;refGene_tx=XM_023259846.1
C1 217891226 217891361
refGene_gn=AGAP1;refGene_tx=XM_023259835.1;refGene_gn=AGAP1;refGene_tx=XM_023259836.1;refGene_gn=AGAP1;refGene_tx=XM_023259837.1;refGene_gn=AGAP1;refGene_tx=XM_023259838.1;refGene_gn=AGAP1;refGene_tx=XM_023259839.1;refGene_gn=AGAP1;refGene_tx=XM_023259840.1;refGene_gn=AGAP1;refGene_tx=XM_023259841.1;refGene_gn=AGAP1;refGene_tx=XM_023259842.1;refGene_gn=AGAP1;refGene_tx=XM_023259844.1;refGene_gn=AGAP1;refGene_tx=XM_023259845.1;refGene_gn=AGAP1;refGene_tx=XM_023259846.1
C1 217937206 217937334
refGene_gn=AGAP1;refGene_tx=XM_023259835.1;refGene_gn=AGAP1;refGene_tx=XM_023259836.1;refGene_gn=AGAP1;refGene_tx=XM_023259837.1;refGene_gn=AGAP1;refGene_tx=XM_023259838.1;refGene_gn=AGAP1;refGene_tx=XM_023259839.1;refGene_gn=AGAP1;refGene_tx=XM_023259840.1;refGene_gn=AGAP1;refGene_tx=XM_023259841.1;refGene_gn=AGAP1;refGene_tx=XM_023259842.1;refGene_gn=AGAP1;refGene_tx=XM_023259844.1;refGene_gn=AGAP1;refGene_tx=XM_023259845.1;refGene_gn=AGAP1;refGene_tx=XM_023259846.1
C1 217938548 217938704
refGene_gn=AGAP1;refGene_tx=XM_023259835.1;refGene_gn=AGAP1;refGene_tx=XM_023259836.1;refGene_gn=AGAP1;refGene_tx=XM_023259837.1;refGene_gn=AGAP1;refGene_tx=XM_023259838.1;refGene_gn=AGAP1;refGene_tx=XM_023259839.1;refGene_gn=AGAP1;refGene_tx=XM_023259840.1;refGene_gn=AGAP1;refGene_tx=XM_023259841.1;refGene_gn=AGAP1;refGene_tx=XM_023259842.1;refGene_gn=AGAP1;refGene_tx=XM_023259844.1;refGene_gn=AGAP1;refGene_tx=XM_023259845.1;refGene_gn=AGAP1;refGene_tx=XM_023259846.1
C1 217945638 217945731
refGene_gn=AGAP1;refGene_tx=XM_023259835.1;refGene_gn=AGAP1;refGene_tx=XM_023259836.1;refGene_gn=AGAP1;refGene_tx=XM_023259837.1;refGene_gn=AGAP1;refGene_tx=XM_023259838.1;refGene_gn=AGAP1;refGene_tx=XM_0232598

39.1;refGene_gn=AGAP1;refGene_tx=XM_023259840.1;refGene_gn=AGAP1;refGene_tx=XM_023259841.1;refGene_gn=AGAP1;refGene_tx=XM_023259842.1;refGene_gn=AGAP1;refGene_tx=XM_023259844.1;refGene_gn=AGAP1;refGene_tx=XM_023259845.1;refGene_gn=AGAP1;refGene_tx=XM_023259846.1
C1 218009807 218009912
refGene_gn=AGAP1;refGene_tx=XM_023259835.1;refGene_gn=AGAP1;refGene_tx=XM_023259836.1;refGene_gn=AGAP1;refGene_tx=XM_023259837.1;refGene_gn=AGAP1;refGene_tx=XM_023259838.1;refGene_gn=AGAP1;refGene_tx=XM_023259839.1;refGene_gn=AGAP1;refGene_tx=XM_023259840.1;refGene_gn=AGAP1;refGene_tx=XM_023259841.1;refGene_gn=AGAP1;refGene_tx=XM_023259842.1;refGene_gn=AGAP1;refGene_tx=XM_023259844.1;refGene_gn=AGAP1;refGene_tx=XM_023259845.1;refGene_gn=AGAP1;refGene_tx=XM_023259846.1
C1 218031773 218031942
refGene_gn=AGAP1;refGene_tx=XM_023259835.1;refGene_gn=AGAP1;refGene_tx=XM_023259836.1;refGene_gn=AGAP1;refGene_tx=XM_023259837.1;refGene_gn=AGAP1;refGene_tx=XM_023259838.1;refGene_gn=AGAP1;refGene_tx=XM_023259839.1;refGene_gn=AGAP1;refGene_tx=XM_023259840.1;refGene_gn=AGAP1;refGene_tx=XM_023259841.1;refGene_gn=AGAP1;refGene_tx=XM_023259842.1;refGene_gn=AGAP1;refGene_tx=XM_023259844.1;refGene_gn=AGAP1;refGene_tx=XM_023259845.1;refGene_gn=AGAP1;refGene_tx=XM_023259846.1
C1 218052687 218052846
refGene_gn=AGAP1;refGene_tx=XM_023259835.1;refGene_gn=AGAP1;refGene_tx=XM_023259836.1;refGene_gn=AGAP1;refGene_tx=XM_023259837.1;refGene_gn=AGAP1;refGene_tx=XM_023259838.1;refGene_gn=AGAP1;refGene_tx=XM_023259840.1;refGene_gn=AGAP1;refGene_tx=XM_023259842.1;refGene_gn=AGAP1;refGene_tx=XM_023259844.1;refGene_gn=AGAP1;refGene_tx=XM_023259845.1
C1 218084682 218084844
refGene_gn=AGAP1;refGene_tx=XM_023259835.1;refGene_gn=AGAP1;refGene_tx=XM_023259836.1;refGene_gn=AGAP1;refGene_tx=XM_023259837.1;refGene_gn=AGAP1;refGene_tx=XM_023259838.1;refGene_gn=AGAP1;refGene_tx=XM_023259839.1;refGene_gn=AGAP1;refGene_tx=XM_023259840.1;refGene_gn=AGAP1;refGene_tx=XM_023259842.1;refGene_gn=AGAP1;refGene_tx=XM_023259844.1;refGene_gn=AGAP1;refGene_tx=XM_023259845.1;refGene_gn=AGAP1;refGene_tx=XM_023259846.1
C1 218120879 218120957
refGene_gn=AGAP1;refGene_tx=XM_023259835.1;refGene_gn=AGAP1;refGene_tx=XM_023259836.1;refGene_gn=AGAP1;refGene_tx=XM_023259837.1;refGene_gn=AGAP1;refGene_tx=XM_023259839.1;refGene_gn=AGAP1;refGene_tx=XM_023259840.1;refGene_gn=AGAP1;refGene_tx=XM_023259842.1;refGene_gn=AGAP1;refGene_tx=XM_023259844.1
C1 218139800 218139955
refGene_gn=AGAP1;refGene_tx=XM_023259835.1;refGene_gn=AGAP1;refGene_tx=XM_023259836.1;refGene_gn=AGAP1;refGene_tx=XM_023259837.1;refGene_gn=AGAP1;refGene_tx=XM_023259838.1;refGene_gn=AGAP1;refGene_tx=XM_023259839.1;refGene_gn=AGAP1;refGene_tx=XM_023259840.1;refGene_gn=AGAP1;refGene_tx=XM_023259841.1;refGene_gn=AGAP1;refGene_tx=XM_023259842.1;refGene_gn=AGAP1;refGene_tx=XM_023259844.1;refGene_gn=AGAP1;refGene_tx=XM_023259845.1;refGene_gn=AGAP1;refGene_tx=XM_023259846.1
C1 218143848 218143939
refGene_gn=AGAP1;refGene_tx=XM_023259835.1;refGene_gn=AGAP1;refGene_tx

=XM_023259836.1;refGene_gn=AGAP1;refGene_tx=XM_023259837.1;refGene_gn=
AGAP1;refGene_tx=XM_023259838.1;refGene_gn=AGAP1;refGene_tx=XM_0232598
39.1;refGene_gn=AGAP1;refGene_tx=XM_023259840.1;refGene_gn=AGAP1;refGe
ne_tx=XM_023259841.1;refGene_gn=AGAP1;refGene_tx=XM_023259842.1;refGen
e_gn=AGAP1;refGene_tx=XM_023259844.1;refGene_gn=AGAP1;refGene_tx=XM_02
3259845.1;refGene_gn=AGAP1;refGene_tx=XM_023259846.1
C1 218153233 218153456
refGene_gn=AGAP1;refGene_tx=XM_023259835.1;refGene_gn=AGAP1;refGene_tx
=XM_023259836.1;refGene_gn=AGAP1;refGene_tx=XM_023259837.1;refGene_gn=
AGAP1;refGene_tx=XM_023259838.1;refGene_gn=AGAP1;refGene_tx=XM_0232598
39.1;refGene_gn=AGAP1;refGene_tx=XM_023259840.1;refGene_gn=AGAP1;refGe
ne_tx=XM_023259841.1;refGene_gn=AGAP1;refGene_tx=XM_023259842.1;refGen
e_gn=AGAP1;refGene_tx=XM_023259844.1;refGene_gn=AGAP1;refGene_tx=XM_02
3259845.1;refGene_gn=AGAP1;refGene_tx=XM_023259846.1
C1 218158294 218158391
refGene_gn=AGAP1;refGene_tx=XM_023259840.1
C1 218212394 218212650
refGene_gn=AGAP1;refGene_tx=XM_023259835.1;refGene_gn=AGAP1;refGene_tx
=XM_023259836.1;refGene_gn=AGAP1;refGene_tx=XM_023259837.1;refGene_gn=
AGAP1;refGene_tx=XM_023259838.1;refGene_gn=AGAP1;refGene_tx=XM_0232598
39.1;refGene_gn=AGAP1;refGene_tx=XM_023259841.1;refGene_gn=AGAP1;refGe
ne_tx=XM_023259842.1;refGene_gn=AGAP1;refGene_tx=XM_023259844.1;refGen
e_gn=AGAP1;refGene_tx=XM_023259845.1;refGene_gn=AGAP1;refGene_tx=XM_02
3259846.1
C1 218213968 218214172
refGene_gn=AGAP1;refGene_tx=XM_023259835.1;refGene_gn=AGAP1;refGene_tx
=XM_023259836.1;refGene_gn=AGAP1;refGene_tx=XM_023259837.1;refGene_gn=
AGAP1;refGene_tx=XM_023259838.1;refGene_gn=AGAP1;refGene_tx=XM_0232598
39.1;refGene_gn=AGAP1;refGene_tx=XM_023259841.1;refGene_gn=AGAP1;refGe
ne_tx=XM_023259842.1;refGene_gn=AGAP1;refGene_tx=XM_023259844.1;refGen
e_gn=AGAP1;refGene_tx=XM_023259845.1;refGene_gn=AGAP1;refGene_tx=XM_02
3259846.1
C1 218246169 218246693
refGene_gn=GBX2;refGene_tx=XM_023259847.1;refGene_gn=GBX2;refGene_tx=X
M_023259848.1
C1 218247675 218248265
refGene_gn=GBX2;refGene_tx=XM_023259847.1;refGene_gn=GBX2;refGene_tx=X
M_023259848.1
C1 218266305 218266491
refGene_gn=ASB18;refGene_tx=XM_023259849.1
C1 218266986 218267100
refGene_gn=ASB18;refGene_tx=XM_023259849.1
C1 218284377 218284882
refGene_gn=ASB18;refGene_tx=XM_023259849.1
C1 218303591 218303859
refGene_gn=ASB18;refGene_tx=XM_023259849.1
C1 218307291 218307532
refGene_gn=ASB18;refGene_tx=XM_023259849.1
C1 218369317 218369440
refGene_gn=IQCA1;refGene_tx=XM_006935734.4

C1 218374860 218375099
refGene_gn=IQCA1;refGene_tx=XM_006935734.4
C1 218381688 218381866
refGene_gn=IQCA1;refGene_tx=XM_006935734.4
C1 218391847 218391934
refGene_gn=IQCA1;refGene_tx=XM_006935734.4
C1 218405515 218405715
refGene_gn=IQCA1;refGene_tx=XM_006935734.4
C1 218409777 218409885
refGene_gn=IQCA1;refGene_tx=XM_006935734.4
C1 218417412 218417495
refGene_gn=IQCA1;refGene_tx=XM_006935734.4
C1 218420695 218420754
refGene_gn=IQCA1;refGene_tx=XM_006935734.4
C1 218429175 218429277
refGene_gn=IQCA1;refGene_tx=XM_006935734.4
C1 218429436 218429543
refGene_gn=IQCA1;refGene_tx=XM_006935734.4
C1 218435685 218435761
refGene_gn=IQCA1;refGene_tx=XM_006935734.4
C1 218445082 218445208
refGene_gn=IQCA1;refGene_tx=XM_006935734.4
C1 218457739 218457824
refGene_gn=IQCA1;refGene_tx=XM_006935734.4
C1 218489926 218490132
refGene_gn=IQCA1;refGene_tx=XM_006935734.4
C1 218505972 218506044
refGene_gn=IQCA1;refGene_tx=XM_006935734.4
C1 218508841 218509033
refGene_gn=IQCA1;refGene_tx=XM_006935734.4
C1 218512189 218512309
refGene_gn=IQCA1;refGene_tx=XM_006935734.4
C1 218515443 218515768
refGene_gn=IQCA1;refGene_tx=XM_006935734.4
C1 218524945 218524956
refGene_gn=IQCA1;refGene_tx=XM_006935734.4
C1 218579537 218579565
refGene_gn=ACKR3;refGene_tx=XM_006935707.4
C1 218584722 218584828
refGene_gn=ACKR3;refGene_tx=XM_011285618.2
C1 218588051 218589166
refGene_gn=ACKR3;refGene_tx=XM_003991275.3;refGene_gn=ACKR3;refGene_tx
=XM_006935707.4;refGene_gn=ACKR3;refGene_tx=XM_006935708.2;refGene_gn=
ACKR3;refGene_tx=XM_006935709.3;refGene_gn=ACKR3;refGene_tx=XM_0112856
18.2
C1 219044753 219044831
refGene_gn=COPS8;refGene_tx=XM_023259851.1;refGene_gn=COPS8;refGene_tx
=XM_023259853.1
C1 219045893 219045964
refGene_gn=COPS8;refGene_tx=XM_023259851.1;refGene_gn=COPS8;refGene_tx

=XM_023259853.1;refGene_gn=COPS8;refGene_tx=XM_023259854.1
C1	219047284	219047333
refGene_gn=COPS8;refGene_tx=XM_023259851.1;refGene_gn=COPS8;refGene_tx=XM_023259853.1;refGene_gn=COPS8;refGene_tx=XM_023259854.1
C1	219048543	219048676
refGene_gn=COPS8;refGene_tx=XM_023259851.1;refGene_gn=COPS8;refGene_tx=XM_023259853.1;refGene_gn=COPS8;refGene_tx=XM_023259854.1
C1	219053193	219053301
refGene_gn=COPS8;refGene_tx=XM_023259851.1;refGene_gn=COPS8;refGene_tx=XM_023259853.1;refGene_gn=COPS8;refGene_tx=XM_023259854.1
C1	219055283	219055346
refGene_gn=COPS8;refGene_tx=XM_023259851.1;refGene_gn=COPS8;refGene_tx=XM_023259854.1
C1	219056536	219056584
refGene_gn=COPS8;refGene_tx=XM_023259851.1;refGene_gn=COPS8;refGene_tx=XM_023259853.1;refGene_gn=COPS8;refGene_tx=XM_023259854.1
C1	219057503	219057583
refGene_gn=COPS8;refGene_tx=XM_023259851.1;refGene_gn=COPS8;refGene_tx=XM_023259853.1;refGene_gn=COPS8;refGene_tx=XM_023259854.1
C1	219239079	219239120
refGene_gn=COL6A3;refGene_tx=XM_019838939.2;refGene_gn=COL6A3;refGene_tx=XM_023259855.1;refGene_gn=COL6A3;refGene_tx=XM_023259856.1;refGene_gn=COL6A3;refGene_tx=XM_023259857.1;refGene_gn=COL6A3;refGene_tx=XM_023259858.1;refGene_gn=COL6A3;refGene_tx=XM_023259859.1;refGene_gn=COL6A3;refGene_tx=XM_023259860.1
C1	219239778	219239943
refGene_gn=COL6A3;refGene_tx=XM_019838939.2;refGene_gn=COL6A3;refGene_tx=XM_023259855.1;refGene_gn=COL6A3;refGene_tx=XM_023259856.1;refGene_gn=COL6A3;refGene_tx=XM_023259857.1;refGene_gn=COL6A3;refGene_tx=XM_023259858.1;refGene_gn=COL6A3;refGene_tx=XM_023259859.1;refGene_gn=COL6A3;refGene_tx=XM_023259860.1
C1	219247087	219247189
refGene_gn=COL6A3;refGene_tx=XM_019838939.2;refGene_gn=COL6A3;refGene_tx=XM_023259855.1;refGene_gn=COL6A3;refGene_tx=XM_023259856.1;refGene_gn=COL6A3;refGene_tx=XM_023259857.1;refGene_gn=COL6A3;refGene_tx=XM_023259858.1;refGene_gn=COL6A3;refGene_tx=XM_023259859.1;refGene_gn=COL6A3;refGene_tx=XM_023259860.1
C1	219248208	219248472
refGene_gn=COL6A3;refGene_tx=XM_019838939.2;refGene_gn=COL6A3;refGene_tx=XM_023259855.1;refGene_gn=COL6A3;refGene_tx=XM_023259856.1;refGene_gn=COL6A3;refGene_tx=XM_023259857.1;refGene_gn=COL6A3;refGene_tx=XM_023259858.1;refGene_gn=COL6A3;refGene_tx=XM_023259859.1;refGene_gn=COL6A3;refGene_tx=XM_023259860.1
C1	219249762	219250160
refGene_gn=COL6A3;refGene_tx=XM_019838939.2;refGene_gn=COL6A3;refGene_tx=XM_023259855.1;refGene_gn=COL6A3;refGene_tx=XM_023259856.1;refGene_gn=COL6A3;refGene_tx=XM_023259857.1;refGene_gn=COL6A3;refGene_tx=XM_023259858.1;refGene_gn=COL6A3;refGene_tx=XM_023259859.1;refGene_gn=COL6A3;refGene_tx=XM_023259860.1
C1	219252999	219253102

refGene_gn=COL6A3;refGene_tx=XM_019838939.2;refGene_gn=COL6A3;refGene_tx=XM_023259855.1;refGene_gn=COL6A3;refGene_tx=XM_023259856.1;refGene_gn=COL6A3;refGene_tx=XM_023259857.1;refGene_gn=COL6A3;refGene_tx=XM_023259858.1;refGene_gn=COL6A3;refGene_tx=XM_023259859.1;refGene_gn=COL6A3;refGene_tx=XM_023259860.1
C1 219254418 219255117
refGene_gn=COL6A3;refGene_tx=XM_019838939.2;refGene_gn=COL6A3;refGene_tx=XM_023259855.1;refGene_gn=COL6A3;refGene_tx=XM_023259856.1;refGene_gn=COL6A3;refGene_tx=XM_023259857.1;refGene_gn=COL6A3;refGene_tx=XM_023259858.1;refGene_gn=COL6A3;refGene_tx=XM_023259859.1;refGene_gn=COL6A3;refGene_tx=XM_023259860.1
C1 219255886 219255983
refGene_gn=COL6A3;refGene_tx=XM_019838939.2;refGene_gn=COL6A3;refGene_tx=XM_023259855.1;refGene_gn=COL6A3;refGene_tx=XM_023259856.1;refGene_gn=COL6A3;refGene_tx=XM_023259857.1;refGene_gn=COL6A3;refGene_tx=XM_023259858.1;refGene_gn=COL6A3;refGene_tx=XM_023259859.1;refGene_gn=COL6A3;refGene_tx=XM_023259860.1
C1 219257470 219257964
refGene_gn=COL6A3;refGene_tx=XM_019838939.2;refGene_gn=COL6A3;refGene_tx=XM_023259855.1;refGene_gn=COL6A3;refGene_tx=XM_023259856.1;refGene_gn=COL6A3;refGene_tx=XM_023259857.1;refGene_gn=COL6A3;refGene_tx=XM_023259858.1;refGene_gn=COL6A3;refGene_tx=XM_023259859.1;refGene_gn=COL6A3;refGene_tx=XM_023259860.1
C1 219258060 219258072
refGene_gn=COL6A3;refGene_tx=XM_019838939.2;refGene_gn=COL6A3;refGene_tx=XM_023259855.1;refGene_gn=COL6A3;refGene_tx=XM_023259856.1;refGene_gn=COL6A3;refGene_tx=XM_023259857.1;refGene_gn=COL6A3;refGene_tx=XM_023259858.1;refGene_gn=COL6A3;refGene_tx=XM_023259859.1;refGene_gn=COL6A3;refGene_tx=XM_023259860.1
C1 219258173 219258210
refGene_gn=COL6A3;refGene_tx=XM_019838939.2;refGene_gn=COL6A3;refGene_tx=XM_023259855.1;refGene_gn=COL6A3;refGene_tx=XM_023259856.1;refGene_gn=COL6A3;refGene_tx=XM_023259857.1;refGene_gn=COL6A3;refGene_tx=XM_023259858.1;refGene_gn=COL6A3;refGene_tx=XM_023259859.1;refGene_gn=COL6A3;refGene_tx=XM_023259860.1
C1 219258294 219258327
refGene_gn=COL6A3;refGene_tx=XM_019838939.2;refGene_gn=COL6A3;refGene_tx=XM_023259855.1;refGene_gn=COL6A3;refGene_tx=XM_023259856.1;refGene_gn=COL6A3;refGene_tx=XM_023259857.1;refGene_gn=COL6A3;refGene_tx=XM_023259858.1;refGene_gn=COL6A3;refGene_tx=XM_023259859.1;refGene_gn=COL6A3;refGene_tx=XM_023259860.1
C1 219259603 219259666
refGene_gn=COL6A3;refGene_tx=XM_019838939.2;refGene_gn=COL6A3;refGene_tx=XM_023259855.1;refGene_gn=COL6A3;refGene_tx=XM_023259856.1;refGene_gn=COL6A3;refGene_tx=XM_023259857.1;refGene_gn=COL6A3;refGene_tx=XM_023259858.1;refGene_gn=COL6A3;refGene_tx=XM_023259859.1;refGene_gn=COL6A3;refGene_tx=XM_023259860.1
C1 219260524 219260587
refGene_gn=COL6A3;refGene_tx=XM_019838939.2;refGene_gn=COL6A3;refGene_tx=XM_023259855.1;refGene_gn=COL6A3;refGene_tx=XM_023259856.1;refGene_

gn=COL6A3;refGene_tx=XM_023259857.1;refGene_gn=COL6A3;refGene_tx=XM_023259858.1;refGene_gn=COL6A3;refGene_tx=XM_023259859.1;refGene_gn=COL6A3;refGene_tx=XM_023259860.1
C1 219261034 219261070
refGene_gn=COL6A3;refGene_tx=XM_019838939.2;refGene_gn=COL6A3;refGene_tx=XM_023259855.1;refGene_gn=COL6A3;refGene_tx=XM_023259856.1;refGene_gn=COL6A3;refGene_tx=XM_023259857.1;refGene_gn=COL6A3;refGene_tx=XM_023259858.1;refGene_gn=COL6A3;refGene_tx=XM_023259859.1;refGene_gn=COL6A3;refGene_tx=XM_023259860.1
C1 219261270 219261321
refGene_gn=COL6A3;refGene_tx=XM_019838939.2;refGene_gn=COL6A3;refGene_tx=XM_023259855.1;refGene_gn=COL6A3;refGene_tx=XM_023259856.1;refGene_gn=COL6A3;refGene_tx=XM_023259857.1;refGene_gn=COL6A3;refGene_tx=XM_023259858.1;refGene_gn=COL6A3;refGene_tx=XM_023259859.1;refGene_gn=COL6A3;refGene_tx=XM_023259860.1
C1 219262793 219262856
refGene_gn=COL6A3;refGene_tx=XM_019838939.2;refGene_gn=COL6A3;refGene_tx=XM_023259855.1;refGene_gn=COL6A3;refGene_tx=XM_023259856.1;refGene_gn=COL6A3;refGene_tx=XM_023259857.1;refGene_gn=COL6A3;refGene_tx=XM_023259858.1;refGene_gn=COL6A3;refGene_tx=XM_023259859.1;refGene_gn=COL6A3;refGene_tx=XM_023259860.1
C1 219263582 219263645
refGene_gn=COL6A3;refGene_tx=XM_019838939.2;refGene_gn=COL6A3;refGene_tx=XM_023259855.1;refGene_gn=COL6A3;refGene_tx=XM_023259856.1;refGene_gn=COL6A3;refGene_tx=XM_023259857.1;refGene_gn=COL6A3;refGene_tx=XM_023259858.1;refGene_gn=COL6A3;refGene_tx=XM_023259859.1;refGene_gn=COL6A3;refGene_tx=XM_023259860.1
C1 219264919 219264982
refGene_gn=COL6A3;refGene_tx=XM_019838939.2;refGene_gn=COL6A3;refGene_tx=XM_023259855.1;refGene_gn=COL6A3;refGene_tx=XM_023259856.1;refGene_gn=COL6A3;refGene_tx=XM_023259857.1;refGene_gn=COL6A3;refGene_tx=XM_023259858.1;refGene_gn=COL6A3;refGene_tx=XM_023259859.1;refGene_gn=COL6A3;refGene_tx=XM_023259860.1
C1 219265806 219265869
refGene_gn=COL6A3;refGene_tx=XM_019838939.2;refGene_gn=COL6A3;refGene_tx=XM_023259855.1;refGene_gn=COL6A3;refGene_tx=XM_023259856.1;refGene_gn=COL6A3;refGene_tx=XM_023259857.1;refGene_gn=COL6A3;refGene_tx=XM_023259858.1;refGene_gn=COL6A3;refGene_tx=XM_023259859.1;refGene_gn=COL6A3;refGene_tx=XM_023259860.1
C1 219266471 219266507
refGene_gn=COL6A3;refGene_tx=XM_019838939.2;refGene_gn=COL6A3;refGene_tx=XM_023259855.1;refGene_gn=COL6A3;refGene_tx=XM_023259856.1;refGene_gn=COL6A3;refGene_tx=XM_023259857.1;refGene_gn=COL6A3;refGene_tx=XM_023259858.1;refGene_gn=COL6A3;refGene_tx=XM_023259859.1;refGene_gn=COL6A3;refGene_tx=XM_023259860.1
C1 219268667 219268721
refGene_gn=COL6A3;refGene_tx=XM_019838939.2;refGene_gn=COL6A3;refGene_tx=XM_023259855.1;refGene_gn=COL6A3;refGene_tx=XM_023259856.1;refGene_gn=COL6A3;refGene_tx=XM_023259857.1;refGene_gn=COL6A3;refGene_tx=XM_023259858.1;refGene_gn=COL6A3;refGene_tx=XM_023259859.1;refGene_gn=COL6A

3;refGene_tx=XM_023259860.1
C1 219269079 219269145
refGene_gn=COL6A3;refGene_tx=XM_019838939.2;refGene_gn=COL6A3;refGene_tx=XM_023259855.1;refGene_gn=COL6A3;refGene_tx=XM_023259856.1;refGene_gn=COL6A3;refGene_tx=XM_023259857.1;refGene_gn=COL6A3;refGene_tx=XM_023259858.1;refGene_gn=COL6A3;refGene_tx=XM_023259859.1;refGene_gn=COL6A3;refGene_tx=XM_023259860.1
C1 219269916 219269979
refGene_gn=COL6A3;refGene_tx=XM_019838939.2;refGene_gn=COL6A3;refGene_tx=XM_023259855.1;refGene_gn=COL6A3;refGene_tx=XM_023259856.1;refGene_gn=COL6A3;refGene_tx=XM_023259857.1;refGene_gn=COL6A3;refGene_tx=XM_023259858.1;refGene_gn=COL6A3;refGene_tx=XM_023259859.1;refGene_gn=COL6A3;refGene_tx=XM_023259860.1
C1 219270435 219270489
refGene_gn=COL6A3;refGene_tx=XM_019838939.2;refGene_gn=COL6A3;refGene_tx=XM_023259855.1;refGene_gn=COL6A3;refGene_tx=XM_023259856.1;refGene_gn=COL6A3;refGene_tx=XM_023259857.1;refGene_gn=COL6A3;refGene_tx=XM_023259858.1;refGene_gn=COL6A3;refGene_tx=XM_023259859.1;refGene_gn=COL6A3;refGene_tx=XM_023259860.1
C1 219270608 219270653
refGene_gn=COL6A3;refGene_tx=XM_019838939.2;refGene_gn=COL6A3;refGene_tx=XM_023259855.1;refGene_gn=COL6A3;refGene_tx=XM_023259856.1;refGene_gn=COL6A3;refGene_tx=XM_023259857.1;refGene_gn=COL6A3;refGene_tx=XM_023259858.1;refGene_gn=COL6A3;refGene_tx=XM_023259859.1;refGene_gn=COL6A3;refGene_tx=XM_023259860.1
C1 219271000 219271027
refGene_gn=COL6A3;refGene_tx=XM_019838939.2;refGene_gn=COL6A3;refGene_tx=XM_023259855.1;refGene_gn=COL6A3;refGene_tx=XM_023259856.1;refGene_gn=COL6A3;refGene_tx=XM_023259857.1;refGene_gn=COL6A3;refGene_tx=XM_023259858.1;refGene_gn=COL6A3;refGene_tx=XM_023259859.1;refGene_gn=COL6A3;refGene_tx=XM_023259860.1
C1 219271991 219272063
refGene_gn=COL6A3;refGene_tx=XM_019838939.2;refGene_gn=COL6A3;refGene_tx=XM_023259855.1;refGene_gn=COL6A3;refGene_tx=XM_023259856.1;refGene_gn=COL6A3;refGene_tx=XM_023259857.1;refGene_gn=COL6A3;refGene_tx=XM_023259858.1;refGene_gn=COL6A3;refGene_tx=XM_023259859.1;refGene_gn=COL6A3;refGene_tx=XM_023259860.1
C1 219273013 219273067
refGene_gn=COL6A3;refGene_tx=XM_019838939.2;refGene_gn=COL6A3;refGene_tx=XM_023259855.1;refGene_gn=COL6A3;refGene_tx=XM_023259856.1;refGene_gn=COL6A3;refGene_tx=XM_023259857.1;refGene_gn=COL6A3;refGene_tx=XM_023259858.1;refGene_gn=COL6A3;refGene_tx=XM_023259859.1;refGene_gn=COL6A3;refGene_tx=XM_023259860.1
C1 219273709 219273802
refGene_gn=COL6A3;refGene_tx=XM_019838939.2;refGene_gn=COL6A3;refGene_tx=XM_023259855.1;refGene_gn=COL6A3;refGene_tx=XM_023259856.1;refGene_gn=COL6A3;refGene_tx=XM_023259857.1;refGene_gn=COL6A3;refGene_tx=XM_023259858.1;refGene_gn=COL6A3;refGene_tx=XM_023259859.1;refGene_gn=COL6A3;refGene_tx=XM_023259860.1
C1 219275233 219275379

refGene_gn=COL6A3;refGene_tx=XM_019838939.2;refGene_gn=COL6A3;refGene_tx=XM_023259855.1;refGene_gn=COL6A3;refGene_tx=XM_023259856.1;refGene_gn=COL6A3;refGene_tx=XM_023259857.1;refGene_gn=COL6A3;refGene_tx=XM_023259858.1;refGene_gn=COL6A3;refGene_tx=XM_023259859.1;refGene_gn=COL6A3;refGene_tx=XM_023259860.1
C1 219276485 219276564
refGene_gn=COL6A3;refGene_tx=XM_019838939.2;refGene_gn=COL6A3;refGene_tx=XM_023259855.1;refGene_gn=COL6A3;refGene_tx=XM_023259856.1;refGene_gn=COL6A3;refGene_tx=XM_023259857.1;refGene_gn=COL6A3;refGene_tx=XM_023259858.1;refGene_gn=COL6A3;refGene_tx=XM_023259859.1;refGene_gn=COL6A3;refGene_tx=XM_023259860.1
C1 219277693 219278031
refGene_gn=COL6A3;refGene_tx=XM_019838939.2;refGene_gn=COL6A3;refGene_tx=XM_023259855.1;refGene_gn=COL6A3;refGene_tx=XM_023259856.1;refGene_gn=COL6A3;refGene_tx=XM_023259857.1;refGene_gn=COL6A3;refGene_tx=XM_023259858.1;refGene_gn=COL6A3;refGene_tx=XM_023259859.1;refGene_gn=COL6A3;refGene_tx=XM_023259860.1
C1 219278835 219279435
refGene_gn=COL6A3;refGene_tx=XM_019838939.2;refGene_gn=COL6A3;refGene_tx=XM_023259855.1;refGene_gn=COL6A3;refGene_tx=XM_023259856.1;refGene_gn=COL6A3;refGene_tx=XM_023259857.1;refGene_gn=COL6A3;refGene_tx=XM_023259858.1;refGene_gn=COL6A3;refGene_tx=XM_023259859.1;refGene_gn=COL6A3;refGene_tx=XM_023259860.1
C1 219280322 219280937
refGene_gn=COL6A3;refGene_tx=XM_019838939.2;refGene_gn=COL6A3;refGene_tx=XM_023259855.1;refGene_gn=COL6A3;refGene_tx=XM_023259856.1;refGene_gn=COL6A3;refGene_tx=XM_023259857.1;refGene_gn=COL6A3;refGene_tx=XM_023259858.1;refGene_gn=COL6A3;refGene_tx=XM_023259859.1;refGene_gn=COL6A3;refGene_tx=XM_023259860.1
C1 219282820 219283426
refGene_gn=COL6A3;refGene_tx=XM_019838939.2;refGene_gn=COL6A3;refGene_tx=XM_023259855.1;refGene_gn=COL6A3;refGene_tx=XM_023259856.1;refGene_gn=COL6A3;refGene_tx=XM_023259857.1;refGene_gn=COL6A3;refGene_tx=XM_023259858.1;refGene_gn=COL6A3;refGene_tx=XM_023259859.1;refGene_gn=COL6A3;refGene_tx=XM_023259860.1
C1 219285463 219286072
refGene_gn=COL6A3;refGene_tx=XM_019838939.2;refGene_gn=COL6A3;refGene_tx=XM_023259855.1;refGene_gn=COL6A3;refGene_tx=XM_023259856.1;refGene_gn=COL6A3;refGene_tx=XM_023259857.1;refGene_gn=COL6A3;refGene_tx=XM_023259858.1;refGene_gn=COL6A3;refGene_tx=XM_023259859.1;refGene_gn=COL6A3;refGene_tx=XM_023259860.1
C1 219287666 219288239
refGene_gn=COL6A3;refGene_tx=XM_019838939.2;refGene_gn=COL6A3;refGene_tx=XM_023259855.1;refGene_gn=COL6A3;refGene_tx=XM_023259856.1;refGene_gn=COL6A3;refGene_tx=XM_023259857.1;refGene_gn=COL6A3;refGene_tx=XM_023259858.1;refGene_gn=COL6A3;refGene_tx=XM_023259859.1;refGene_gn=COL6A3;refGene_tx=XM_023259860.1
C1 219289567 219290167
refGene_gn=COL6A3;refGene_tx=XM_019838939.2;refGene_gn=COL6A3;refGene_tx=XM_023259856.1;refGene_gn=COL6A3;refGene_tx=XM_023259857.1;refGene_

gn=COL6A3;refGene_tx=XM_023259858.1
C1 219291849 219292434
refGene_gn=COL6A3;refGene_tx=XM_019838939.2;refGene_gn=COL6A3;refGene_tx=XM_023259855.1;refGene_gn=COL6A3;refGene_tx=XM_023259856.1;refGene_gn=COL6A3;refGene_tx=XM_023259857.1;refGene_gn=COL6A3;refGene_tx=XM_023259859.1
C1 219296217 219296820
refGene_gn=COL6A3;refGene_tx=XM_019838939.2;refGene_gn=COL6A3;refGene_tx=XM_023259855.1;refGene_gn=COL6A3;refGene_tx=XM_023259856.1
C1 219302493 219303111
refGene_gn=COL6A3;refGene_tx=XM_019838939.2;refGene_gn=COL6A3;refGene_tx=XM_023259855.1
C1 219304621 219304712
refGene_gn=COL6A3;refGene_tx=XM_019838939.2;refGene_gn=COL6A3;refGene_tx=XM_023259855.1;refGene_gn=COL6A3;refGene_tx=XM_023259856.1;refGene_gn=COL6A3;refGene_tx=XM_023259857.1;refGene_gn=COL6A3;refGene_tx=XM_023259858.1;refGene_gn=COL6A3;refGene_tx=XM_023259859.1;refGene_gn=COL6A3;refGene_tx=XM_023259860.1
C1 219396737 219396847
refGene_gn=MLPH;refGene_tx=NM_001079655.1;refGene_gn=MLPH;refGene_tx=XM_023257818.1;refGene_gn=MLPH;refGene_tx=XM_023257819.1
C1 219409737 219409959
refGene_gn=MLPH;refGene_tx=NM_001079655.1;refGene_gn=MLPH;refGene_tx=XM_023257818.1;refGene_gn=MLPH;refGene_tx=XM_023257819.1
C1 219410086 219410199
refGene_gn=MLPH;refGene_tx=NM_001079655.1;refGene_gn=MLPH;refGene_tx=XM_023257818.1;refGene_gn=MLPH;refGene_tx=XM_023257819.1
C1 219412737 219412776
refGene_gn=MLPH;refGene_tx=XM_023257818.1
C1 219414464 219414568
refGene_gn=MLPH;refGene_tx=NM_001079655.1;refGene_gn=MLPH;refGene_tx=XM_023257818.1;refGene_gn=MLPH;refGene_tx=XM_023257819.1
C1 219415712 219415829
refGene_gn=MLPH;refGene_tx=NM_001079655.1;refGene_gn=MLPH;refGene_tx=XM_023257818.1;refGene_gn=MLPH;refGene_tx=XM_023257819.1
C1 219416983 219417185
refGene_gn=MLPH;refGene_tx=NM_001079655.1;refGene_gn=MLPH;refGene_tx=XM_023257818.1;refGene_gn=MLPH;refGene_tx=XM_023257819.1
C1 219419255 219419398
refGene_gn=MLPH;refGene_tx=NM_001079655.1;refGene_gn=MLPH;refGene_tx=XM_023257818.1;refGene_gn=MLPH;refGene_tx=XM_023257819.1
C1 219424120 219424306
refGene_gn=MLPH;refGene_tx=NM_001079655.1;refGene_gn=MLPH;refGene_tx=XM_023257818.1;refGene_gn=MLPH;refGene_tx=XM_023257819.1
C1 219424581 219424737
refGene_gn=MLPH;refGene_tx=NM_001079655.1;refGene_gn=MLPH;refGene_tx=XM_023257818.1;refGene_gn=MLPH;refGene_tx=XM_023257819.1
C1 219426158 219426251
refGene_gn=MLPH;refGene_tx=NM_001079655.1;refGene_gn=MLPH;refGene_tx=XM_023257818.1;refGene_gn=MLPH;refGene_tx=XM_023257819.1

C1 219426785 219426863
refGene_gn=MLPH;refGene_tx=NM_001079655.1;refGene_gn=MLPH;refGene_tx=X
M_023257818.1;refGene_gn=MLPH;refGene_tx=XM_023257819.1
C1 219428400 219428458
refGene_gn=MLPH;refGene_tx=NM_001079655.1;refGene_gn=MLPH;refGene_tx=X
M_023257818.1;refGene_gn=MLPH;refGene_tx=XM_023257819.1
C1 219430567 219430668
refGene_gn=MLPH;refGene_tx=NM_001079655.1;refGene_gn=MLPH;refGene_tx=X
M_023257818.1;refGene_gn=MLPH;refGene_tx=XM_023257819.1
C1 219433995 219434022
refGene_gn=MLPH;refGene_tx=NM_001079655.1;refGene_gn=MLPH;refGene_tx=X
M_023257818.1;refGene_gn=MLPH;refGene_tx=XM_023257819.1
C1 219445158 219445258
refGene_gn=PRLH;refGene_tx=XM_006935710.3
C1 219445666 219445797
refGene_gn=PRLH;refGene_tx=XM_006935710.3
C1 219452670 219452783
refGene_gn=RAB17;refGene_tx=XM_006935712.4;refGene_gn=RAB17;refGene_tx
=XM_019838941.2;refGene_gn=RAB17;refGene_tx=XM_019838942.2;refGene_gn=
RAB17;refGene_tx=XM_019838943.2;refGene_gn=RAB17;refGene_tx=XM_0232598
63.1
C1 219453025 219453119
refGene_gn=RAB17;refGene_tx=XM_006935712.4;refGene_gn=RAB17;refGene_tx
=XM_019838941.2;refGene_gn=RAB17;refGene_tx=XM_019838942.2;refGene_gn=
RAB17;refGene_tx=XM_019838943.2;refGene_gn=RAB17;refGene_tx=XM_0232598
63.1
C1 219454762 219454888
refGene_gn=RAB17;refGene_tx=XM_006935712.4;refGene_gn=RAB17;refGene_tx
=XM_019838941.2;refGene_gn=RAB17;refGene_tx=XM_019838942.2;refGene_gn=
RAB17;refGene_tx=XM_019838943.2;refGene_gn=RAB17;refGene_tx=XM_0232598
63.1
C1 219455988 219456140
refGene_gn=RAB17;refGene_tx=XM_006935712.4;refGene_gn=RAB17;refGene_tx
=XM_019838941.2;refGene_gn=RAB17;refGene_tx=XM_019838942.2;refGene_gn=
RAB17;refGene_tx=XM_019838943.2;refGene_gn=RAB17;refGene_tx=XM_0232598
63.1
C1 219462010 219462167
refGene_gn=RAB17;refGene_tx=XM_006935712.4;refGene_gn=RAB17;refGene_tx
=XM_019838941.2;refGene_gn=RAB17;refGene_tx=XM_019838942.2;refGene_gn=
RAB17;refGene_tx=XM_019838943.2;refGene_gn=RAB17;refGene_tx=XM_0232598
63.1
C1 219493783 219493879
refGene_gn=LRRFIP1;refGene_tx=XM_023259864.1;refGene_gn=LRRFIP1;refGen
e_tx=XM_023259865.1;refGene_gn=LRRFIP1;refGene_tx=XM_023259866.1;refGe
ne_gn=LRRFIP1;refGene_tx=XM_023259867.1;refGene_gn=LRRFIP1;refGene_tx=
XM_023259869.1;refGene_gn=LRRFIP1;refGene_tx=XM_023259870.1;refGene_gn
=LRRFIP1;refGene_tx=XM_023259871.1;refGene_gn=LRRFIP1;refGene_tx=XM_02
3259872.1;refGene_gn=LRRFIP1;refGene_tx=XM_023259873.1;refGene_gn=LRRF
IP1;refGene_tx=XM_023259874.1;refGene_gn=LRRFIP1;refGene_tx=XM_0232598
75.1;refGene_gn=LRRFIP1;refGene_tx=XM_023259876.1;refGene_gn=LRRFIP1;r

efGene_tx=XM_023259878.1;refGene_gn=LRRFIP1;refGene_tx=XM_023259879.1;
refGene_gn=LRRFIP1;refGene_tx=XM_023259880.1;refGene_gn=LRRFIP1;refGen
e_tx=XM_023259881.1;refGene_gn=LRRFIP1;refGene_tx=XM_023259882.1;refGe
ne_gn=LRRFIP1;refGene_tx=XM_023259883.1;refGene_gn=LRRFIP1;refGene_tx=
XM_023259884.1;refGene_gn=LRRFIP1;refGene_tx=XM_023259885.1;refGene_gn
=LRRFIP1;refGene_tx=XM_023259886.1;refGene_gn=LRRFIP1;refGene_tx=XM_02
3259888.1;refGene_gn=LRRFIP1;refGene_tx=XM_023259889.1;refGene_gn=LRRF
IP1;refGene_tx=XM_023259890.1;refGene_gn=LRRFIP1;refGene_tx=XM_0232598
91.1;refGene_gn=LRRFIP1;refGene_tx=XM_023259892.1;refGene_gn=LRRFIP1;r
efGene_tx=XM_023259893.1;refGene_gn=LRRFIP1;refGene_tx=XM_023259894.1;
refGene_gn=LRRFIP1;refGene_tx=XM_023259895.1;refGene_gn=LRRFIP1;refGen
e_tx=XM_023259896.1;refGene_gn=LRRFIP1;refGene_tx=XM_023259897.1;refGe
ne_gn=LRRFIP1;refGene_tx=XM_023259898.1;refGene_gn=LRRFIP1;refGene_tx=
XM_023259899.1;refGene_gn=LRRFIP1;refGene_tx=XM_023259900.1;refGene_gn
=LRRFIP1;refGene_tx=XM_023259901.1;refGene_gn=LRRFIP1;refGene_tx=XM_02
3259902.1;refGene_gn=LRRFIP1;refGene_tx=XM_023259903.1;refGene_gn=LRRF
IP1;refGene_tx=XM_023259904.1;refGene_gn=LRRFIP1;refGene_tx=XM_0232599
06.1;refGene_gn=LRRFIP1;refGene_tx=XM_023259907.1;refGene_gn=LRRFIP1;r
efGene_tx=XM_023259908.1;refGene_gn=LRRFIP1;refGene_tx=XM_023259909.1;
refGene_gn=LRRFIP1;refGene_tx=XM_023259910.1
C1 219551162 219551228
refGene_gn=LRRFIP1;refGene_tx=XM_006935720.2;refGene_gn=LRRFIP1;refGen
e_tx=XM_011285602.2;refGene_gn=LRRFIP1;refGene_tx=XM_011285607.2;refGe
ne_gn=LRRFIP1;refGene_tx=XM_019838947.2;refGene_gn=LRRFIP1;refGene_tx=
XM_019838952.2;refGene_gn=LRRFIP1;refGene_tx=XM_019838960.2;refGene_gn
=LRRFIP1;refGene_tx=XM_023259905.1
C1 219566721 219566808
refGene_gn=LRRFIP1;refGene_tx=XM_006935720.2;refGene_gn=LRRFIP1;refGen
e_tx=XM_011285602.2;refGene_gn=LRRFIP1;refGene_tx=XM_011285607.2;refGe
ne_gn=LRRFIP1;refGene_tx=XM_019838947.2;refGene_gn=LRRFIP1;refGene_tx=
XM_019838952.2;refGene_gn=LRRFIP1;refGene_tx=XM_019838960.2;refGene_gn
=LRRFIP1;refGene_tx=XM_023259864.1;refGene_gn=LRRFIP1;refGene_tx=XM_02
3259865.1;refGene_gn=LRRFIP1;refGene_tx=XM_023259866.1;refGene_gn=LRRF
IP1;refGene_tx=XM_023259867.1;refGene_gn=LRRFIP1;refGene_tx=XM_0232598
69.1;refGene_gn=LRRFIP1;refGene_tx=XM_023259870.1;refGene_gn=LRRFIP1;r
efGene_tx=XM_023259871.1;refGene_gn=LRRFIP1;refGene_tx=XM_023259872.1;
refGene_gn=LRRFIP1;refGene_tx=XM_023259873.1;refGene_gn=LRRFIP1;refGen
e_tx=XM_023259874.1;refGene_gn=LRRFIP1;refGene_tx=XM_023259875.1;refGe
ne_gn=LRRFIP1;refGene_tx=XM_023259876.1;refGene_gn=LRRFIP1;refGene_tx=
XM_023259878.1;refGene_gn=LRRFIP1;refGene_tx=XM_023259879.1;refGene_gn
=LRRFIP1;refGene_tx=XM_023259880.1;refGene_gn=LRRFIP1;refGene_tx=XM_02
3259881.1;refGene_gn=LRRFIP1;refGene_tx=XM_023259882.1;refGene_gn=LRRF
IP1;refGene_tx=XM_023259883.1;refGene_gn=LRRFIP1;refGene_tx=XM_0232598
84.1;refGene_gn=LRRFIP1;refGene_tx=XM_023259885.1;refGene_gn=LRRFIP1;r
efGene_tx=XM_023259886.1;refGene_gn=LRRFIP1;refGene_tx=XM_023259888.1;
refGene_gn=LRRFIP1;refGene_tx=XM_023259889.1;refGene_gn=LRRFIP1;refGen
e_tx=XM_023259890.1;refGene_gn=LRRFIP1;refGene_tx=XM_023259891.1;refGe
ne_gn=LRRFIP1;refGene_tx=XM_023259892.1;refGene_gn=LRRFIP1;refGene_tx=
XM_023259893.1;refGene_gn=LRRFIP1;refGene_tx=XM_023259894.1;refGene_gn
=LRRFIP1;refGene_tx=XM_023259895.1;refGene_gn=LRRFIP1;refGene_tx=XM_02

3259896.1;refGene_gn=LRRFIP1;refGene_tx=XM_023259897.1;refGene_gn=LRRFIP1;refGene_tx=XM_023259898.1;refGene_gn=LRRFIP1;refGene_tx=XM_023259899.1;refGene_gn=LRRFIP1;refGene_tx=XM_023259900.1;refGene_gn=LRRFIP1;refGene_tx=XM_023259901.1;refGene_gn=LRRFIP1;refGene_tx=XM_023259902.1;refGene_gn=LRRFIP1;refGene_tx=XM_023259903.1;refGene_gn=LRRFIP1;refGene_tx=XM_023259904.1;refGene_gn=LRRFIP1;refGene_tx=XM_023259905.1;refGene_gn=LRRFIP1;refGene_tx=XM_023259906.1;refGene_gn=LRRFIP1;refGene_tx=XM_023259907.1;refGene_gn=LRRFIP1;refGene_tx=XM_023259908.1;refGene_gn=LRRFIP1;refGene_tx=XM_023259909.1;refGene_gn=LRRFIP1;refGene_tx=XM_023259910.1
C1 219574441 219574459
refGene_gn=LRRFIP1;refGene_tx=XM_011285602.2;refGene_gn=LRRFIP1;refGene_tx=XM_023259864.1;refGene_gn=LRRFIP1;refGene_tx=XM_023259865.1;refGene_gn=LRRFIP1;refGene_tx=XM_023259866.1;refGene_gn=LRRFIP1;refGene_tx=XM_023259869.1;refGene_gn=LRRFIP1;refGene_tx=XM_023259870.1;refGene_gn=LRRFIP1;refGene_tx=XM_023259871.1;refGene_gn=LRRFIP1;refGene_tx=XM_023259872.1;refGene_gn=LRRFIP1;refGene_tx=XM_023259874.1;refGene_gn=LRRFIP1;refGene_tx=XM_023259875.1;refGene_gn=LRRFIP1;refGene_tx=XM_023259876.1;refGene_gn=LRRFIP1;refGene_tx=XM_023259881.1;refGene_gn=LRRFIP1;refGene_tx=XM_023259882.1;refGene_gn=LRRFIP1;refGene_tx=XM_023259884.1;refGene_gn=LRRFIP1;refGene_tx=XM_023259886.1;refGene_gn=LRRFIP1;refGene_tx=XM_023259888.1;refGene_gn=LRRFIP1;refGene_tx=XM_023259899.1;refGene_gn=LRRFIP1;refGene_tx=XM_023259900.1;refGene_gn=LRRFIP1;refGene_tx=XM_023259901.1;refGene_gn=LRRFIP1;refGene_tx=XM_023259902.1;refGene_gn=LRRFIP1;refGene_tx=XM_023259906.1;refGene_gn=LRRFIP1;refGene_tx=XM_023259907.1
C1 219577543 219577591
refGene_gn=LRRFIP1;refGene_tx=XM_011285602.2;refGene_gn=LRRFIP1;refGene_tx=XM_023259864.1;refGene_gn=LRRFIP1;refGene_tx=XM_023259865.1;refGene_gn=LRRFIP1;refGene_tx=XM_023259866.1;refGene_gn=LRRFIP1;refGene_tx=XM_023259869.1;refGene_gn=LRRFIP1;refGene_tx=XM_023259870.1;refGene_gn=LRRFIP1;refGene_tx=XM_023259871.1;refGene_gn=LRRFIP1;refGene_tx=XM_023259872.1;refGene_gn=LRRFIP1;refGene_tx=XM_023259874.1;refGene_gn=LRRFIP1;refGene_tx=XM_023259875.1;refGene_gn=LRRFIP1;refGene_tx=XM_023259876.1;refGene_gn=LRRFIP1;refGene_tx=XM_023259881.1;refGene_gn=LRRFIP1;refGene_tx=XM_023259882.1;refGene_gn=LRRFIP1;refGene_tx=XM_023259884.1;refGene_gn=LRRFIP1;refGene_tx=XM_023259886.1;refGene_gn=LRRFIP1;refGene_tx=XM_023259888.1;refGene_gn=LRRFIP1;refGene_tx=XM_023259899.1;refGene_gn=LRRFIP1;refGene_tx=XM_023259900.1;refGene_gn=LRRFIP1;refGene_tx=XM_023259901.1;refGene_gn=LRRFIP1;refGene_tx=XM_023259902.1;refGene_gn=LRRFIP1;refGene_tx=XM_023259906.1;refGene_gn=LRRFIP1;refGene_tx=XM_023259907.1
C1 219579109 219579154
refGene_gn=LRRFIP1;refGene_tx=XM_011285602.2;refGene_gn=LRRFIP1;refGene_tx=XM_019838947.2;refGene_gn=LRRFIP1;refGene_tx=XM_023259864.1;refGene_gn=LRRFIP1;refGene_tx=XM_023259865.1;refGene_gn=LRRFIP1;refGene_tx=XM_023259866.1;refGene_gn=LRRFIP1;refGene_tx=XM_023259867.1;refGene_gn=LRRFIP1;refGene_tx=XM_023259869.1;refGene_gn=LRRFIP1;refGene_tx=XM_023259870.1;refGene_gn=LRRFIP1;refGene_tx=XM_023259871.1;refGene_gn=LRRFIP1;refGene_tx=XM_023259872.1;refGene_gn=LRRFIP1;refGene_tx=XM_0232598

74.1;refGene_gn=LRRFIP1;refGene_tx=XM_023259875.1;refGene_gn=LRRFIP1;refGene_tx=XM_023259876.1;refGene_gn=LRRFIP1;refGene_tx=XM_023259881.1;refGene_gn=LRRFIP1;refGene_tx=XM_023259882.1;refGene_gn=LRRFIP1;refGene_tx=XM_023259883.1;refGene_gn=LRRFIP1;refGene_tx=XM_023259884.1;refGene_gn=LRRFIP1;refGene_tx=XM_023259886.1;refGene_gn=LRRFIP1;refGene_tx=XM_023259888.1;refGene_gn=LRRFIP1;refGene_tx=XM_023259890.1;refGene_gn=LRRFIP1;refGene_tx=XM_023259896.1;refGene_gn=LRRFIP1;refGene_tx=XM_023259897.1;refGene_gn=LRRFIP1;refGene_tx=XM_023259899.1;refGene_gn=LRRFIP1;refGene_tx=XM_023259900.1;refGene_gn=LRRFIP1;refGene_tx=XM_023259901.1;refGene_gn=LRRFIP1;refGene_tx=XM_023259902.1;refGene_gn=LRRFIP1;refGene_tx=XM_023259906.1;refGene_gn=LRRFIP1;refGene_tx=XM_023259907.1;refGene_gn=LRRFIP1;refGene_tx=XM_023259910.1
C1	219579685	219579766
refGene_gn=LRRFIP1;refGene_tx=XM_011285602.2;refGene_gn=LRRFIP1;refGene_tx=XM_011285607.2;refGene_gn=LRRFIP1;refGene_tx=XM_019838947.2;refGene_gn=LRRFIP1;refGene_tx=XM_023259864.1;refGene_gn=LRRFIP1;refGene_tx=XM_023259865.1;refGene_gn=LRRFIP1;refGene_tx=XM_023259866.1;refGene_gn=LRRFIP1;refGene_tx=XM_023259867.1;refGene_gn=LRRFIP1;refGene_tx=XM_023259869.1;refGene_gn=LRRFIP1;refGene_tx=XM_023259870.1;refGene_gn=LRRFIP1;refGene_tx=XM_023259871.1;refGene_gn=LRRFIP1;refGene_tx=XM_023259872.1;refGene_gn=LRRFIP1;refGene_tx=XM_023259873.1;refGene_gn=LRRFIP1;refGene_tx=XM_023259874.1;refGene_gn=LRRFIP1;refGene_tx=XM_023259875.1;refGene_gn=LRRFIP1;refGene_tx=XM_023259876.1;refGene_gn=LRRFIP1;refGene_tx=XM_023259879.1;refGene_gn=LRRFIP1;refGene_tx=XM_023259881.1;refGene_gn=LRRFIP1;refGene_tx=XM_023259882.1;refGene_gn=LRRFIP1;refGene_tx=XM_023259883.1;refGene_gn=LRRFIP1;refGene_tx=XM_023259884.1;refGene_gn=LRRFIP1;refGene_tx=XM_023259885.1;refGene_gn=LRRFIP1;refGene_tx=XM_023259886.1;refGene_gn=LRRFIP1;refGene_tx=XM_023259888.1;refGene_gn=LRRFIP1;refGene_tx=XM_023259889.1;refGene_gn=LRRFIP1;refGene_tx=XM_023259890.1;refGene_gn=LRRFIP1;refGene_tx=XM_023259891.1;refGene_gn=LRRFIP1;refGene_tx=XM_023259893.1;refGene_gn=LRRFIP1;refGene_tx=XM_023259896.1;refGene_gn=LRRFIP1;refGene_tx=XM_023259897.1;refGene_gn=LRRFIP1;refGene_tx=XM_023259899.1;refGene_gn=LRRFIP1;refGene_tx=XM_023259900.1;refGene_gn=LRRFIP1;refGene_tx=XM_023259901.1;refGene_gn=LRRFIP1;refGene_tx=XM_023259902.1;refGene_gn=LRRFIP1;refGene_tx=XM_023259906.1;refGene_gn=LRRFIP1;refGene_tx=XM_023259907.1;refGene_gn=LRRFIP1;refGene_tx=XM_023259910.1
C1	219582532	219582571
refGene_gn=LRRFIP1;refGene_tx=XM_011285602.2;refGene_gn=LRRFIP1;refGene_tx=XM_011285607.2;refGene_gn=LRRFIP1;refGene_tx=XM_019838947.2;refGene_gn=LRRFIP1;refGene_tx=XM_023259864.1;refGene_gn=LRRFIP1;refGene_tx=XM_023259865.1;refGene_gn=LRRFIP1;refGene_tx=XM_023259866.1;refGene_gn=LRRFIP1;refGene_tx=XM_023259867.1;refGene_gn=LRRFIP1;refGene_tx=XM_023259870.1;refGene_gn=LRRFIP1;refGene_tx=XM_023259871.1;refGene_gn=LRRFIP1;refGene_tx=XM_023259873.1;refGene_gn=LRRFIP1;refGene_tx=XM_023259874.1;refGene_gn=LRRFIP1;refGene_tx=XM_023259876.1;refGene_gn=LRRFIP1;refGene_tx=XM_023259899.1;refGene_gn=LRRFIP1;refGene_tx=XM_023259900.1;refGene_gn=LRRFIP1;refGene_tx=XM_023259901.1;refGene_gn=LRRFIP1;refGene_tx=XM_023259902.1;refGene_gn=LRRFIP1;refGene_tx=XM_023259906.1;refGene_gn=LRRFIP1;refGene_tx=XM_023259910.1

C1	219587817	219587877
refGene_gn=LRRFIP1;refGene_tx=XM_011285602.2;refGene_gn=LRRFIP1;refGene_tx=XM_011285607.2;refGene_gn=LRRFIP1;refGene_tx=XM_019838947.2;refGene_gn=LRRFIP1;refGene_tx=XM_023259864.1;refGene_gn=LRRFIP1;refGene_tx=XM_023259865.1;refGene_gn=LRRFIP1;refGene_tx=XM_023259866.1;refGene_gn=LRRFIP1;refGene_tx=XM_023259867.1;refGene_gn=LRRFIP1;refGene_tx=XM_023259869.1;refGene_gn=LRRFIP1;refGene_tx=XM_023259870.1;refGene_gn=LRRFIP1;refGene_tx=XM_023259871.1;refGene_gn=LRRFIP1;refGene_tx=XM_023259873.1;refGene_gn=LRRFIP1;refGene_tx=XM_023259874.1;refGene_gn=LRRFIP1;refGene_tx=XM_023259876.1;refGene_gn=LRRFIP1;refGene_tx=XM_023259878.1;refGene_gn=LRRFIP1;refGene_tx=XM_023259899.1;refGene_gn=LRRFIP1;refGene_tx=XM_023259900.1;refGene_gn=LRRFIP1;refGene_tx=XM_023259901.1;refGene_gn=LRRFIP1;refGene_tx=XM_023259902.1;refGene_gn=LRRFIP1;refGene_tx=XM_023259906.1;refGene_gn=LRRFIP1;refGene_tx=XM_023259907.1;refGene_gn=LRRFIP1;refGene_tx=XM_023259910.1
C1	219590301	219590370
refGene_gn=LRRFIP1;refGene_tx=XM_011285602.2;refGene_gn=LRRFIP1;refGene_tx=XM_011285607.2;refGene_gn=LRRFIP1;refGene_tx=XM_019838947.2;refGene_gn=LRRFIP1;refGene_tx=XM_019838952.2;refGene_gn=LRRFIP1;refGene_tx=XM_023259864.1;refGene_gn=LRRFIP1;refGene_tx=XM_023259865.1;refGene_gn=LRRFIP1;refGene_tx=XM_023259866.1;refGene_gn=LRRFIP1;refGene_tx=XM_023259867.1;refGene_gn=LRRFIP1;refGene_tx=XM_023259869.1;refGene_gn=LRRFIP1;refGene_tx=XM_023259870.1;refGene_gn=LRRFIP1;refGene_tx=XM_023259871.1;refGene_gn=LRRFIP1;refGene_tx=XM_023259872.1;refGene_gn=LRRFIP1;refGene_tx=XM_023259873.1;refGene_gn=LRRFIP1;refGene_tx=XM_023259874.1;refGene_gn=LRRFIP1;refGene_tx=XM_023259875.1;refGene_gn=LRRFIP1;refGene_tx=XM_023259876.1;refGene_gn=LRRFIP1;refGene_tx=XM_023259878.1;refGene_gn=LRRFIP1;refGene_tx=XM_023259879.1;refGene_gn=LRRFIP1;refGene_tx=XM_023259880.1;refGene_gn=LRRFIP1;refGene_tx=XM_023259899.1;refGene_gn=LRRFIP1;refGene_tx=XM_023259900.1;refGene_gn=LRRFIP1;refGene_tx=XM_023259901.1;refGene_gn=LRRFIP1;refGene_tx=XM_023259902.1;refGene_gn=LRRFIP1;refGene_tx=XM_023259906.1;refGene_gn=LRRFIP1;refGene_tx=XM_023259907.1;refGene_gn=LRRFIP1;refGene_tx=XM_023259910.1
C1	219592894	219592966
refGene_gn=LRRFIP1;refGene_tx=XM_011285602.2;refGene_gn=LRRFIP1;refGene_tx=XM_011285607.2;refGene_gn=LRRFIP1;refGene_tx=XM_019838947.2;refGene_gn=LRRFIP1;refGene_tx=XM_019838952.2;refGene_gn=LRRFIP1;refGene_tx=XM_023259864.1;refGene_gn=LRRFIP1;refGene_tx=XM_023259865.1;refGene_gn=LRRFIP1;refGene_tx=XM_023259866.1;refGene_gn=LRRFIP1;refGene_tx=XM_023259867.1;refGene_gn=LRRFIP1;refGene_tx=XM_023259869.1;refGene_gn=LRRFIP1;refGene_tx=XM_023259870.1;refGene_gn=LRRFIP1;refGene_tx=XM_023259871.1;refGene_gn=LRRFIP1;refGene_tx=XM_023259872.1;refGene_gn=LRRFIP1;refGene_tx=XM_023259873.1;refGene_gn=LRRFIP1;refGene_tx=XM_023259874.1;refGene_gn=LRRFIP1;refGene_tx=XM_023259875.1;refGene_gn=LRRFIP1;refGene_tx=XM_023259876.1;refGene_gn=LRRFIP1;refGene_tx=XM_023259878.1;refGene_gn=LRRFIP1;refGene_tx=XM_023259879.1;refGene_gn=LRRFIP1;refGene_tx=XM_023259880.1;refGene_gn=LRRFIP1;refGene_tx=XM_023259899.1;refGene_gn=LRRFIP1;refGene_tx=XM_023259900.1;refGene_gn=LRRFIP1;refGene_tx=XM_023259901.1;refGene_gn=LRRFIP1;refGene_tx=XM_023259902.1;refGene_gn=LRRFIP1;refGene_tx=XM_023259906.1;refGene_gn=LRRFIP1;refGene_tx=XM_0232599

07.1;refGene_gn=LRRFIP1;refGene_tx=XM_023259910.1
C1 219594306 219594351
refGene_gn=LRRFIP1;refGene_tx=XM_011285602.2;refGene_gn=LRRFIP1;refGen
e_tx=XM_011285607.2;refGene_gn=LRRFIP1;refGene_tx=XM_019838947.2;refGe
ne_gn=LRRFIP1;refGene_tx=XM_019838952.2;refGene_gn=LRRFIP1;refGene_tx=
XM_023259864.1;refGene_gn=LRRFIP1;refGene_tx=XM_023259865.1;refGene_gn
=LRRFIP1;refGene_tx=XM_023259867.1;refGene_gn=LRRFIP1;refGene_tx=XM_02
3259869.1;refGene_gn=LRRFIP1;refGene_tx=XM_023259870.1;refGene_gn=LRRF
IP1;refGene_tx=XM_023259871.1;refGene_gn=LRRFIP1;refGene_tx=XM_0232598
72.1;refGene_gn=LRRFIP1;refGene_tx=XM_023259873.1;refGene_gn=LRRFIP1;r
efGene_tx=XM_023259875.1;refGene_gn=LRRFIP1;refGene_tx=XM_023259876.1;
refGene_gn=LRRFIP1;refGene_tx=XM_023259878.1;refGene_gn=LRRFIP1;refGen
e_tx=XM_023259879.1;refGene_gn=LRRFIP1;refGene_tx=XM_023259880.1;refGe
ne_gn=LRRFIP1;refGene_tx=XM_023259899.1;refGene_gn=LRRFIP1;refGene_tx=
XM_023259902.1;refGene_gn=LRRFIP1;refGene_tx=XM_023259906.1;refGene_gn
=LRRFIP1;refGene_tx=XM_023259910.1
C1 219595910 219595976
refGene_gn=LRRFIP1;refGene_tx=XM_011285602.2;refGene_gn=LRRFIP1;refGen
e_tx=XM_011285607.2;refGene_gn=LRRFIP1;refGene_tx=XM_019838947.2;refGe
ne_gn=LRRFIP1;refGene_tx=XM_019838952.2;refGene_gn=LRRFIP1;refGene_tx=
XM_023259864.1;refGene_gn=LRRFIP1;refGene_tx=XM_023259865.1;refGene_gn
=LRRFIP1;refGene_tx=XM_023259866.1;refGene_gn=LRRFIP1;refGene_tx=XM_02
3259867.1;refGene_gn=LRRFIP1;refGene_tx=XM_023259869.1;refGene_gn=LRRF
IP1;refGene_tx=XM_023259870.1;refGene_gn=LRRFIP1;refGene_tx=XM_0232598
71.1;refGene_gn=LRRFIP1;refGene_tx=XM_023259872.1;refGene_gn=LRRFIP1;r
efGene_tx=XM_023259873.1;refGene_gn=LRRFIP1;refGene_tx=XM_023259874.1;
refGene_gn=LRRFIP1;refGene_tx=XM_023259875.1;refGene_gn=LRRFIP1;refGen
e_tx=XM_023259876.1;refGene_gn=LRRFIP1;refGene_tx=XM_023259878.1;refGe
ne_gn=LRRFIP1;refGene_tx=XM_023259879.1;refGene_gn=LRRFIP1;refGene_tx=
XM_023259880.1;refGene_gn=LRRFIP1;refGene_tx=XM_023259881.1;refGene_gn
=LRRFIP1;refGene_tx=XM_023259883.1;refGene_gn=LRRFIP1;refGene_tx=XM_02
3259884.1;refGene_gn=LRRFIP1;refGene_tx=XM_023259885.1;refGene_gn=LRRF
IP1;refGene_tx=XM_023259899.1;refGene_gn=LRRFIP1;refGene_tx=XM_0232599
00.1;refGene_gn=LRRFIP1;refGene_tx=XM_023259901.1;refGene_gn=LRRFIP1;r
efGene_tx=XM_023259902.1;refGene_gn=LRRFIP1;refGene_tx=XM_023259906.1;
refGene_gn=LRRFIP1;refGene_tx=XM_023259907.1;refGene_gn=LRRFIP1;refGen
e_tx=XM_023259910.1
C1 219600063 219600141
refGene_gn=LRRFIP1;refGene_tx=XM_011285602.2;refGene_gn=LRRFIP1;refGen
e_tx=XM_011285607.2;refGene_gn=LRRFIP1;refGene_tx=XM_019838947.2;refGe
ne_gn=LRRFIP1;refGene_tx=XM_019838952.2;refGene_gn=LRRFIP1;refGene_tx=
XM_019838960.2;refGene_gn=LRRFIP1;refGene_tx=XM_023259864.1;refGene_gn
=LRRFIP1;refGene_tx=XM_023259865.1;refGene_gn=LRRFIP1;refGene_tx=XM_02
3259866.1;refGene_gn=LRRFIP1;refGene_tx=XM_023259867.1;refGene_gn=LRRF
IP1;refGene_tx=XM_023259869.1;refGene_gn=LRRFIP1;refGene_tx=XM_0232598
70.1;refGene_gn=LRRFIP1;refGene_tx=XM_023259872.1;refGene_gn=LRRFIP1;r
efGene_tx=XM_023259873.1;refGene_gn=LRRFIP1;refGene_tx=XM_023259875.1;
refGene_gn=LRRFIP1;refGene_tx=XM_023259876.1;refGene_gn=LRRFIP1;refGen
e_tx=XM_023259878.1;refGene_gn=LRRFIP1;refGene_tx=XM_023259879.1;refGe
ne_gn=LRRFIP1;refGene_tx=XM_023259880.1;refGene_gn=LRRFIP1;refGene_tx=

XM_023259881.1;refGene_gn=LRRFIP1;refGene_tx=XM_023259882.1;refGene_gn
=LRRFIP1;refGene_tx=XM_023259883.1;refGene_gn=LRRFIP1;refGene_tx=XM_02
3259885.1;refGene_gn=LRRFIP1;refGene_tx=XM_023259889.1;refGene_gn=LRRF
IP1;refGene_tx=XM_023259892.1;refGene_gn=LRRFIP1;refGene_tx=XM_0232598
99.1;refGene_gn=LRRFIP1;refGene_tx=XM_023259900.1;refGene_gn=LRRFIP1;r
efGene_tx=XM_023259901.1;refGene_gn=LRRFIP1;refGene_tx=XM_023259902.1;
refGene_gn=LRRFIP1;refGene_tx=XM_023259906.1;refGene_gn=LRRFIP1;refGen
e_tx=XM_023259907.1;refGene_gn=LRRFIP1;refGene_tx=XM_023259910.1
C1 219607964 219608000
refGene_gn=LRRFIP1;refGene_tx=XM_006935720.2;refGene_gn=LRRFIP1;refGen
e_tx=XM_011285602.2;refGene_gn=LRRFIP1;refGene_tx=XM_011285607.2;refGe
ne_gn=LRRFIP1;refGene_tx=XM_019838947.2;refGene_gn=LRRFIP1;refGene_tx=
XM_019838952.2;refGene_gn=LRRFIP1;refGene_tx=XM_019838960.2;refGene_gn
=LRRFIP1;refGene_tx=XM_023259864.1;refGene_gn=LRRFIP1;refGene_tx=XM_02
3259865.1;refGene_gn=LRRFIP1;refGene_tx=XM_023259866.1;refGene_gn=LRRF
IP1;refGene_tx=XM_023259867.1;refGene_gn=LRRFIP1;refGene_tx=XM_0232598
69.1;refGene_gn=LRRFIP1;refGene_tx=XM_023259870.1;refGene_gn=LRRFIP1;r
efGene_tx=XM_023259871.1;refGene_gn=LRRFIP1;refGene_tx=XM_023259872.1;
refGene_gn=LRRFIP1;refGene_tx=XM_023259873.1;refGene_gn=LRRFIP1;refGen
e_tx=XM_023259874.1;refGene_gn=LRRFIP1;refGene_tx=XM_023259875.1;refGe
ne_gn=LRRFIP1;refGene_tx=XM_023259876.1;refGene_gn=LRRFIP1;refGene_tx=
XM_023259878.1;refGene_gn=LRRFIP1;refGene_tx=XM_023259879.1;refGene_gn
=LRRFIP1;refGene_tx=XM_023259880.1;refGene_gn=LRRFIP1;refGene_tx=XM_02
3259881.1;refGene_gn=LRRFIP1;refGene_tx=XM_023259882.1;refGene_gn=LRRF
IP1;refGene_tx=XM_023259883.1;refGene_gn=LRRFIP1;refGene_tx=XM_0232598
84.1;refGene_gn=LRRFIP1;refGene_tx=XM_023259885.1;refGene_gn=LRRFIP1;r
efGene_tx=XM_023259886.1;refGene_gn=LRRFIP1;refGene_tx=XM_023259888.1;
refGene_gn=LRRFIP1;refGene_tx=XM_023259889.1;refGene_gn=LRRFIP1;refGen
e_tx=XM_023259890.1;refGene_gn=LRRFIP1;refGene_tx=XM_023259891.1;refGe
ne_gn=LRRFIP1;refGene_tx=XM_023259892.1;refGene_gn=LRRFIP1;refGene_tx=
XM_023259893.1;refGene_gn=LRRFIP1;refGene_tx=XM_023259894.1;refGene_gn
=LRRFIP1;refGene_tx=XM_023259895.1;refGene_gn=LRRFIP1;refGene_tx=XM_02
3259896.1;refGene_gn=LRRFIP1;refGene_tx=XM_023259897.1;refGene_gn=LRRF
IP1;refGene_tx=XM_023259898.1;refGene_gn=LRRFIP1;refGene_tx=XM_0232598
99.1;refGene_gn=LRRFIP1;refGene_tx=XM_023259900.1;refGene_gn=LRRFIP1;r
efGene_tx=XM_023259901.1;refGene_gn=LRRFIP1;refGene_tx=XM_023259902.1;
refGene_gn=LRRFIP1;refGene_tx=XM_023259903.1;refGene_gn=LRRFIP1;refGen
e_tx=XM_023259904.1;refGene_gn=LRRFIP1;refGene_tx=XM_023259905.1;refGe
ne_gn=LRRFIP1;refGene_tx=XM_023259906.1;refGene_gn=LRRFIP1;refGene_tx=
XM_023259907.1;refGene_gn=LRRFIP1;refGene_tx=XM_023259908.1;refGene_gn
=LRRFIP1;refGene_tx=XM_023259909.1;refGene_gn=LRRFIP1;refGene_tx=XM_02
3259910.1
C1 219608418 219608544
refGene_gn=LRRFIP1;refGene_tx=XM_006935720.2;refGene_gn=LRRFIP1;refGen
e_tx=XM_011285602.2;refGene_gn=LRRFIP1;refGene_tx=XM_011285607.2;refGe
ne_gn=LRRFIP1;refGene_tx=XM_019838947.2;refGene_gn=LRRFIP1;refGene_tx=
XM_019838952.2;refGene_gn=LRRFIP1;refGene_tx=XM_019838960.2;refGene_gn
=LRRFIP1;refGene_tx=XM_023259864.1;refGene_gn=LRRFIP1;refGene_tx=XM_02
3259865.1;refGene_gn=LRRFIP1;refGene_tx=XM_023259866.1;refGene_gn=LRRF
IP1;refGene_tx=XM_023259867.1;refGene_gn=LRRFIP1;refGene_tx=XM_0232598

69.1;refGene_gn=LRRFIP1;refGene_tx=XM_023259870.1;refGene_gn=LRRFIP1;refGene_tx=XM_023259871.1;refGene_gn=LRRFIP1;refGene_tx=XM_023259872.1;refGene_gn=LRRFIP1;refGene_tx=XM_023259873.1;refGene_gn=LRRFIP1;refGene_tx=XM_023259874.1;refGene_gn=LRRFIP1;refGene_tx=XM_023259875.1;refGene_gn=LRRFIP1;refGene_tx=XM_023259876.1;refGene_gn=LRRFIP1;refGene_tx=XM_023259878.1;refGene_gn=LRRFIP1;refGene_tx=XM_023259879.1;refGene_gn=LRRFIP1;refGene_tx=XM_023259880.1;refGene_gn=LRRFIP1;refGene_tx=XM_023259881.1;refGene_gn=LRRFIP1;refGene_tx=XM_023259882.1;refGene_gn=LRRFIP1;refGene_tx=XM_023259883.1;refGene_gn=LRRFIP1;refGene_tx=XM_023259884.1;refGene_gn=LRRFIP1;refGene_tx=XM_023259885.1;refGene_gn=LRRFIP1;refGene_tx=XM_023259886.1;refGene_gn=LRRFIP1;refGene_tx=XM_023259888.1;refGene_gn=LRRFIP1;refGene_tx=XM_023259889.1;refGene_gn=LRRFIP1;refGene_tx=XM_023259890.1;refGene_gn=LRRFIP1;refGene_tx=XM_023259891.1;refGene_gn=LRRFIP1;refGene_tx=XM_023259892.1;refGene_gn=LRRFIP1;refGene_tx=XM_023259893.1;refGene_gn=LRRFIP1;refGene_tx=XM_023259894.1;refGene_gn=LRRFIP1;refGene_tx=XM_023259895.1;refGene_gn=LRRFIP1;refGene_tx=XM_023259896.1;refGene_gn=LRRFIP1;refGene_tx=XM_023259897.1;refGene_gn=LRRFIP1;refGene_tx=XM_023259898.1;refGene_gn=LRRFIP1;refGene_tx=XM_023259899.1;refGene_gn=LRRFIP1;refGene_tx=XM_023259900.1;refGene_gn=LRRFIP1;refGene_tx=XM_023259901.1;refGene_gn=LRRFIP1;refGene_tx=XM_023259902.1;refGene_gn=LRRFIP1;refGene_tx=XM_023259903.1;refGene_gn=LRRFIP1;refGene_tx=XM_023259904.1;refGene_gn=LRRFIP1;refGene_tx=XM_023259905.1;refGene_gn=LRRFIP1;refGene_tx=XM_023259906.1;refGene_gn=LRRFIP1;refGene_tx=XM_023259907.1;refGene_gn=LRRFIP1;refGene_tx=XM_023259908.1;refGene_gn=LRRFIP1;refGene_tx=XM_023259909.1;refGene_gn=LRRFIP1;refGene_tx=XM_023259910.1

C1 219609971 219610043

refGene_gn=LRRFIP1;refGene_tx=XM_006935720.2;refGene_gn=LRRFIP1;refGene_tx=XM_011285602.2;refGene_gn=LRRFIP1;refGene_tx=XM_011285607.2;refGene_gn=LRRFIP1;refGene_tx=XM_019838947.2;refGene_gn=LRRFIP1;refGene_tx=XM_019838952.2;refGene_gn=LRRFIP1;refGene_tx=XM_019838960.2;refGene_gn=LRRFIP1;refGene_tx=XM_023259864.1;refGene_gn=LRRFIP1;refGene_tx=XM_023259865.1;refGene_gn=LRRFIP1;refGene_tx=XM_023259866.1;refGene_gn=LRRFIP1;refGene_tx=XM_023259867.1;refGene_gn=LRRFIP1;refGene_tx=XM_023259869.1;refGene_gn=LRRFIP1;refGene_tx=XM_023259871.1;refGene_gn=LRRFIP1;refGene_tx=XM_023259872.1;refGene_gn=LRRFIP1;refGene_tx=XM_023259873.1;refGene_gn=LRRFIP1;refGene_tx=XM_023259874.1;refGene_gn=LRRFIP1;refGene_tx=XM_023259875.1;refGene_gn=LRRFIP1;refGene_tx=XM_023259876.1;refGene_gn=LRRFIP1;refGene_tx=XM_023259878.1;refGene_gn=LRRFIP1;refGene_tx=XM_023259879.1;refGene_gn=LRRFIP1;refGene_tx=XM_023259880.1;refGene_gn=LRRFIP1;refGene_tx=XM_023259881.1;refGene_gn=LRRFIP1;refGene_tx=XM_023259882.1;refGene_gn=LRRFIP1;refGene_tx=XM_023259883.1;refGene_gn=LRRFIP1;refGene_tx=XM_023259884.1;refGene_gn=LRRFIP1;refGene_tx=XM_023259885.1;refGene_gn=LRRFIP1;refGene_tx=XM_023259886.1;refGene_gn=LRRFIP1;refGene_tx=XM_023259888.1;refGene_gn=LRRFIP1;refGene_tx=XM_023259889.1;refGene_gn=LRRFIP1;refGene_tx=XM_023259890.1;refGene_gn=LRRFIP1;refGene_tx=XM_023259891.1;refGene_gn=LRRFIP1;refGene_tx=XM_023259892.1;refGene_gn=LRRFIP1;refGene_tx=XM_023259893.1;refGene_gn=LRRFIP1;refGene_tx=XM_023259894.1;refGene_gn=LRRFIP1;refGene_tx=XM_023259896.1;refGene_gn=LRRFIP1;refGene_tx=XM_023259899.1;refGene_gn=LRRFIP1;refGene_tx=XM_02

3259900.1;refGene_gn=LRRFIP1;refGene_tx=XM_023259901.1;refGene_gn=LRRFIP1;refGene_tx=XM_023259903.1;refGene_gn=LRRFIP1;refGene_tx=XM_023259906.1;refGene_gn=LRRFIP1;refGene_tx=XM_023259907.1;refGene_gn=LRRFIP1;refGene_tx=XM_023259908.1
C1 219611775 219611946
refGene_gn=LRRFIP1;refGene_tx=XM_006935720.2;refGene_gn=LRRFIP1;refGene_tx=XM_011285602.2;refGene_gn=LRRFIP1;refGene_tx=XM_011285607.2;refGene_gn=LRRFIP1;refGene_tx=XM_019838947.2;refGene_gn=LRRFIP1;refGene_tx=XM_019838952.2;refGene_gn=LRRFIP1;refGene_tx=XM_019838960.2;refGene_gn=LRRFIP1;refGene_tx=XM_023259864.1;refGene_gn=LRRFIP1;refGene_tx=XM_023259865.1;refGene_gn=LRRFIP1;refGene_tx=XM_023259866.1;refGene_gn=LRRFIP1;refGene_tx=XM_023259867.1;refGene_gn=LRRFIP1;refGene_tx=XM_023259869.1;refGene_gn=LRRFIP1;refGene_tx=XM_023259870.1;refGene_gn=LRRFIP1;refGene_tx=XM_023259871.1;refGene_gn=LRRFIP1;refGene_tx=XM_023259872.1;refGene_gn=LRRFIP1;refGene_tx=XM_023259873.1;refGene_gn=LRRFIP1;refGene_tx=XM_023259874.1;refGene_gn=LRRFIP1;refGene_tx=XM_023259875.1;refGene_gn=LRRFIP1;refGene_tx=XM_023259876.1;refGene_gn=LRRFIP1;refGene_tx=XM_023259878.1;refGene_gn=LRRFIP1;refGene_tx=XM_023259879.1;refGene_gn=LRRFIP1;refGene_tx=XM_023259880.1;refGene_gn=LRRFIP1;refGene_tx=XM_023259881.1;refGene_gn=LRRFIP1;refGene_tx=XM_023259882.1;refGene_gn=LRRFIP1;refGene_tx=XM_023259883.1;refGene_gn=LRRFIP1;refGene_tx=XM_023259884.1;refGene_gn=LRRFIP1;refGene_tx=XM_023259885.1;refGene_gn=LRRFIP1;refGene_tx=XM_023259886.1;refGene_gn=LRRFIP1;refGene_tx=XM_023259888.1;refGene_gn=LRRFIP1;refGene_tx=XM_023259889.1;refGene_gn=LRRFIP1;refGene_tx=XM_023259890.1;refGene_gn=LRRFIP1;refGene_tx=XM_023259891.1;refGene_gn=LRRFIP1;refGene_tx=XM_023259892.1;refGene_gn=LRRFIP1;refGene_tx=XM_023259893.1;refGene_gn=LRRFIP1;refGene_tx=XM_023259894.1;refGene_gn=LRRFIP1;refGene_tx=XM_023259895.1;refGene_gn=LRRFIP1;refGene_tx=XM_023259896.1;refGene_gn=LRRFIP1;refGene_tx=XM_023259897.1;refGene_gn=LRRFIP1;refGene_tx=XM_023259898.1;refGene_gn=LRRFIP1;refGene_tx=XM_023259899.1;refGene_gn=LRRFIP1;refGene_tx=XM_023259900.1;refGene_gn=LRRFIP1;refGene_tx=XM_023259901.1;refGene_gn=LRRFIP1;refGene_tx=XM_023259902.1;refGene_gn=LRRFIP1;refGene_tx=XM_023259903.1;refGene_gn=LRRFIP1;refGene_tx=XM_023259904.1;refGene_gn=LRRFIP1;refGene_tx=XM_023259905.1;refGene_gn=LRRFIP1;refGene_tx=XM_023259906.1;refGene_gn=LRRFIP1;refGene_tx=XM_023259907.1;refGene_gn=LRRFIP1;refGene_tx=XM_023259908.1;refGene_gn=LRRFIP1;refGene_tx=XM_023259909.1;refGene_gn=LRRFIP1;refGene_tx=XM_023259910.1
C1 219614458 219614551
refGene_gn=LRRFIP1;refGene_tx=XM_006935720.2;refGene_gn=LRRFIP1;refGene_tx=XM_011285602.2;refGene_gn=LRRFIP1;refGene_tx=XM_011285607.2;refGene_gn=LRRFIP1;refGene_tx=XM_019838947.2;refGene_gn=LRRFIP1;refGene_tx=XM_019838952.2;refGene_gn=LRRFIP1;refGene_tx=XM_019838960.2;refGene_gn=LRRFIP1;refGene_tx=XM_023259864.1;refGene_gn=LRRFIP1;refGene_tx=XM_023259865.1;refGene_gn=LRRFIP1;refGene_tx=XM_023259866.1;refGene_gn=LRRFIP1;refGene_tx=XM_023259867.1;refGene_gn=LRRFIP1;refGene_tx=XM_023259869.1;refGene_gn=LRRFIP1;refGene_tx=XM_023259870.1;refGene_gn=LRRFIP1;refGene_tx=XM_023259871.1;refGene_gn=LRRFIP1;refGene_tx=XM_023259872.1;refGene_gn=LRRFIP1;refGene_tx=XM_023259873.1;refGene_gn=LRRFIP1;refGene_tx=XM_023259874.1;refGene_gn=LRRFIP1;refGene_tx=XM_023259875.1;refGe

ne_gn=LRRFIP1;refGene_tx=XM_023259876.1;refGene_gn=LRRFIP1;refGene_tx=XM_023259878.1;refGene_gn=LRRFIP1;refGene_tx=XM_023259879.1;refGene_gn=LRRFIP1;refGene_tx=XM_023259880.1;refGene_gn=LRRFIP1;refGene_tx=XM_023259881.1;refGene_gn=LRRFIP1;refGene_tx=XM_023259882.1;refGene_gn=LRRFIP1;refGene_tx=XM_023259883.1;refGene_gn=LRRFIP1;refGene_tx=XM_023259884.1;refGene_gn=LRRFIP1;refGene_tx=XM_023259885.1;refGene_gn=LRRFIP1;refGene_tx=XM_023259886.1;refGene_gn=LRRFIP1;refGene_tx=XM_023259888.1;refGene_gn=LRRFIP1;refGene_tx=XM_023259889.1;refGene_gn=LRRFIP1;refGene_tx=XM_023259890.1;refGene_gn=LRRFIP1;refGene_tx=XM_023259891.1;refGene_gn=LRRFIP1;refGene_tx=XM_023259892.1;refGene_gn=LRRFIP1;refGene_tx=XM_023259893.1;refGene_gn=LRRFIP1;refGene_tx=XM_023259894.1;refGene_gn=LRRFIP1;refGene_tx=XM_023259895.1;refGene_gn=LRRFIP1;refGene_tx=XM_023259896.1;refGene_gn=LRRFIP1;refGene_tx=XM_023259897.1;refGene_gn=LRRFIP1;refGene_tx=XM_023259898.1;refGene_gn=LRRFIP1;refGene_tx=XM_023259899.1;refGene_gn=LRRFIP1;refGene_tx=XM_023259900.1;refGene_gn=LRRFIP1;refGene_tx=XM_023259901.1;refGene_gn=LRRFIP1;refGene_tx=XM_023259902.1;refGene_gn=LRRFIP1;refGene_tx=XM_023259903.1;refGene_gn=LRRFIP1;refGene_tx=XM_023259904.1;refGene_gn=LRRFIP1;refGene_tx=XM_023259905.1;refGene_gn=LRRFIP1;refGene_tx=XM_023259906.1;refGene_gn=LRRFIP1;refGene_tx=XM_023259907.1;refGene_gn=LRRFIP1;refGene_tx=XM_023259908.1;refGene_gn=LRRFIP1;refGene_tx=XM_023259909.1;refGene_gn=LRRFIP1;refGene_tx=XM_023259910.1
C1 219615691 219615784
refGene_gn=LRRFIP1;refGene_tx=XM_006935720.2;refGene_gn=LRRFIP1;refGene_tx=XM_011285602.2;refGene_gn=LRRFIP1;refGene_tx=XM_011285607.2;refGene_gn=LRRFIP1;refGene_tx=XM_019838947.2;refGene_gn=LRRFIP1;refGene_tx=XM_019838952.2;refGene_gn=LRRFIP1;refGene_tx=XM_019838960.2;refGene_gn=LRRFIP1;refGene_tx=XM_023259864.1;refGene_gn=LRRFIP1;refGene_tx=XM_023259865.1;refGene_gn=LRRFIP1;refGene_tx=XM_023259866.1;refGene_gn=LRRFIP1;refGene_tx=XM_023259867.1;refGene_gn=LRRFIP1;refGene_tx=XM_023259869.1;refGene_gn=LRRFIP1;refGene_tx=XM_023259870.1;refGene_gn=LRRFIP1;refGene_tx=XM_023259871.1;refGene_gn=LRRFIP1;refGene_tx=XM_023259872.1;refGene_gn=LRRFIP1;refGene_tx=XM_023259873.1;refGene_gn=LRRFIP1;refGene_tx=XM_023259874.1;refGene_gn=LRRFIP1;refGene_tx=XM_023259875.1;refGene_gn=LRRFIP1;refGene_tx=XM_023259878.1;refGene_gn=LRRFIP1;refGene_tx=XM_023259879.1;refGene_gn=LRRFIP1;refGene_tx=XM_023259880.1;refGene_gn=LRRFIP1;refGene_tx=XM_023259881.1;refGene_gn=LRRFIP1;refGene_tx=XM_023259882.1;refGene_gn=LRRFIP1;refGene_tx=XM_023259883.1;refGene_gn=LRRFIP1;refGene_tx=XM_023259884.1;refGene_gn=LRRFIP1;refGene_tx=XM_023259885.1;refGene_gn=LRRFIP1;refGene_tx=XM_023259886.1;refGene_gn=LRRFIP1;refGene_tx=XM_023259888.1;refGene_gn=LRRFIP1;refGene_tx=XM_023259889.1;refGene_gn=LRRFIP1;refGene_tx=XM_023259890.1;refGene_gn=LRRFIP1;refGene_tx=XM_023259891.1;refGene_gn=LRRFIP1;refGene_tx=XM_023259892.1;refGene_gn=LRRFIP1;refGene_tx=XM_023259893.1;refGene_gn=LRRFIP1;refGene_tx=XM_023259894.1;refGene_gn=LRRFIP1;refGene_tx=XM_023259895.1;refGene_gn=LRRFIP1;refGene_tx=XM_023259899.1;refGene_gn=LRRFIP1;refGene_tx=XM_023259900.1;refGene_gn=LRRFIP1;refGene_tx=XM_023259901.1;refGene_gn=LRRFIP1;refGene_tx=XM_023259906.1;refGene_gn=LRRFIP1;refGene_tx=XM_023259907.1;refGene_gn=LRRFIP1;refGene_tx=XM_023259908.1;refGene_gn=LRRFIP1;refGene_tx=XM_023259909.1;refGene_gn=LRRFIP1;refGene_tx=XM_023259910.1

C1 219616790 219616883
refGene_gn=LRRFIP1;refGene_tx=XM_006935720.2;refGene_gn=LRRFIP1;refGene_tx=XM_011285602.2;refGene_gn=LRRFIP1;refGene_tx=XM_011285607.2;refGene_gn=LRRFIP1;refGene_tx=XM_019838947.2;refGene_gn=LRRFIP1;refGene_tx=XM_019838952.2;refGene_gn=LRRFIP1;refGene_tx=XM_019838960.2;refGene_gn=LRRFIP1;refGene_tx=XM_023259864.1;refGene_gn=LRRFIP1;refGene_tx=XM_023259865.1;refGene_gn=LRRFIP1;refGene_tx=XM_023259866.1;refGene_gn=LRRFIP1;refGene_tx=XM_023259867.1;refGene_gn=LRRFIP1;refGene_tx=XM_023259869.1;refGene_gn=LRRFIP1;refGene_tx=XM_023259870.1;refGene_gn=LRRFIP1;refGene_tx=XM_023259871.1;refGene_gn=LRRFIP1;refGene_tx=XM_023259872.1;refGene_gn=LRRFIP1;refGene_tx=XM_023259873.1;refGene_gn=LRRFIP1;refGene_tx=XM_023259874.1;refGene_gn=LRRFIP1;refGene_tx=XM_023259875.1;refGene_gn=LRRFIP1;refGene_tx=XM_023259878.1;refGene_gn=LRRFIP1;refGene_tx=XM_023259879.1;refGene_gn=LRRFIP1;refGene_tx=XM_023259880.1;refGene_gn=LRRFIP1;refGene_tx=XM_023259881.1;refGene_gn=LRRFIP1;refGene_tx=XM_023259882.1;refGene_gn=LRRFIP1;refGene_tx=XM_023259883.1;refGene_gn=LRRFIP1;refGene_tx=XM_023259884.1;refGene_gn=LRRFIP1;refGene_tx=XM_023259885.1;refGene_gn=LRRFIP1;refGene_tx=XM_023259886.1;refGene_gn=LRRFIP1;refGene_tx=XM_023259888.1;refGene_gn=LRRFIP1;refGene_tx=XM_023259889.1;refGene_gn=LRRFIP1;refGene_tx=XM_023259890.1;refGene_gn=LRRFIP1;refGene_tx=XM_023259891.1;refGene_gn=LRRFIP1;refGene_tx=XM_023259892.1;refGene_gn=LRRFIP1;refGene_tx=XM_023259893.1;refGene_gn=LRRFIP1;refGene_tx=XM_023259894.1;refGene_gn=LRRFIP1;refGene_tx=XM_023259895.1;refGene_gn=LRRFIP1;refGene_tx=XM_023259899.1;refGene_gn=LRRFIP1;refGene_tx=XM_023259900.1;refGene_gn=LRRFIP1;refGene_tx=XM_023259901.1;refGene_gn=LRRFIP1;refGene_tx=XM_023259906.1;refGene_gn=LRRFIP1;refGene_tx=XM_023259907.1;refGene_gn=LRRFIP1;refGene_tx=XM_023259908.1;refGene_gn=LRRFIP1;refGene_tx=XM_023259909.1;refGene_gn=LRRFIP1;refGene_tx=XM_023259910.1
C1 219618683 219618825
refGene_gn=LRRFIP1;refGene_tx=XM_006935720.2;refGene_gn=LRRFIP1;refGene_tx=XM_011285602.2;refGene_gn=LRRFIP1;refGene_tx=XM_011285607.2;refGene_gn=LRRFIP1;refGene_tx=XM_019838947.2;refGene_gn=LRRFIP1;refGene_tx=XM_019838952.2;refGene_gn=LRRFIP1;refGene_tx=XM_019838960.2;refGene_gn=LRRFIP1;refGene_tx=XM_023259864.1;refGene_gn=LRRFIP1;refGene_tx=XM_023259865.1;refGene_gn=LRRFIP1;refGene_tx=XM_023259866.1;refGene_gn=LRRFIP1;refGene_tx=XM_023259867.1;refGene_gn=LRRFIP1;refGene_tx=XM_023259869.1;refGene_gn=LRRFIP1;refGene_tx=XM_023259870.1;refGene_gn=LRRFIP1;refGene_tx=XM_023259871.1;refGene_gn=LRRFIP1;refGene_tx=XM_023259872.1;refGene_gn=LRRFIP1;refGene_tx=XM_023259873.1;refGene_gn=LRRFIP1;refGene_tx=XM_023259874.1;refGene_gn=LRRFIP1;refGene_tx=XM_023259875.1;refGene_gn=LRRFIP1;refGene_tx=XM_023259876.1;refGene_gn=LRRFIP1;refGene_tx=XM_023259878.1;refGene_gn=LRRFIP1;refGene_tx=XM_023259879.1;refGene_gn=LRRFIP1;refGene_tx=XM_023259880.1;refGene_gn=LRRFIP1;refGene_tx=XM_023259881.1;refGene_gn=LRRFIP1;refGene_tx=XM_023259882.1;refGene_gn=LRRFIP1;refGene_tx=XM_023259883.1;refGene_gn=LRRFIP1;refGene_tx=XM_023259884.1;refGene_gn=LRRFIP1;refGene_tx=XM_023259885.1;refGene_gn=LRRFIP1;refGene_tx=XM_023259886.1;refGene_gn=LRRFIP1;refGene_tx=XM_023259888.1;refGene_gn=LRRFIP1;refGene_tx=XM_023259889.1;refGene_gn=LRRFIP1;refGene_tx=XM_023259890.1;refGene_gn=LRRFIP1;refGene_tx=XM_023259891.1;refGene_gn=LRRFIP1;refGene_tx=XM_023259892.1;refGene_gn=LRRFIP1;refGene_tx=

XM_023259893.1;refGene_gn=LRRFIP1;refGene_tx=XM_023259894.1;refGene_gn
=LRRFIP1;refGene_tx=XM_023259895.1;refGene_gn=LRRFIP1;refGene_tx=XM_02
3259896.1;refGene_gn=LRRFIP1;refGene_tx=XM_023259897.1;refGene_gn=LRRF
IP1;refGene_tx=XM_023259898.1;refGene_gn=LRRFIP1;refGene_tx=XM_0232598
99.1;refGene_gn=LRRFIP1;refGene_tx=XM_023259900.1;refGene_gn=LRRFIP1;r
efGene_tx=XM_023259901.1;refGene_gn=LRRFIP1;refGene_tx=XM_023259902.1;
refGene_gn=LRRFIP1;refGene_tx=XM_023259903.1;refGene_gn=LRRFIP1;refGen
e_tx=XM_023259904.1;refGene_gn=LRRFIP1;refGene_tx=XM_023259905.1;refGe
ne_gn=LRRFIP1;refGene_tx=XM_023259906.1;refGene_gn=LRRFIP1;refGene_tx=
XM_023259907.1;refGene_gn=LRRFIP1;refGene_tx=XM_023259908.1;refGene_gn
=LRRFIP1;refGene_tx=XM_023259909.1;refGene_gn=LRRFIP1;refGene_tx=XM_02
3259910.1
C1 219621240 219622928
refGene_gn=LRRFIP1;refGene_tx=XM_006935720.2;refGene_gn=LRRFIP1;refGen
e_tx=XM_011285602.2;refGene_gn=LRRFIP1;refGene_tx=XM_011285607.2;refGe
ne_gn=LRRFIP1;refGene_tx=XM_019838947.2;refGene_gn=LRRFIP1;refGene_tx=
XM_019838952.2;refGene_gn=LRRFIP1;refGene_tx=XM_019838960.2;refGene_gn
=LRRFIP1;refGene_tx=XM_023259864.1;refGene_gn=LRRFIP1;refGene_tx=XM_02
3259865.1;refGene_gn=LRRFIP1;refGene_tx=XM_023259866.1;refGene_gn=LRRF
IP1;refGene_tx=XM_023259867.1;refGene_gn=LRRFIP1;refGene_tx=XM_0232598
69.1;refGene_gn=LRRFIP1;refGene_tx=XM_023259870.1;refGene_gn=LRRFIP1;r
efGene_tx=XM_023259871.1;refGene_gn=LRRFIP1;refGene_tx=XM_023259872.1;
refGene_gn=LRRFIP1;refGene_tx=XM_023259873.1;refGene_gn=LRRFIP1;refGen
e_tx=XM_023259874.1;refGene_gn=LRRFIP1;refGene_tx=XM_023259875.1;refGe
ne_gn=LRRFIP1;refGene_tx=XM_023259876.1;refGene_gn=LRRFIP1;refGene_tx=
XM_023259878.1;refGene_gn=LRRFIP1;refGene_tx=XM_023259879.1;refGene_gn
=LRRFIP1;refGene_tx=XM_023259880.1;refGene_gn=LRRFIP1;refGene_tx=XM_02
3259881.1;refGene_gn=LRRFIP1;refGene_tx=XM_023259882.1;refGene_gn=LRRF
IP1;refGene_tx=XM_023259883.1;refGene_gn=LRRFIP1;refGene_tx=XM_0232598
84.1;refGene_gn=LRRFIP1;refGene_tx=XM_023259885.1;refGene_gn=LRRFIP1;r
efGene_tx=XM_023259886.1;refGene_gn=LRRFIP1;refGene_tx=XM_023259888.1;
refGene_gn=LRRFIP1;refGene_tx=XM_023259889.1;refGene_gn=LRRFIP1;refGen
e_tx=XM_023259890.1;refGene_gn=LRRFIP1;refGene_tx=XM_023259891.1;refGe
ne_gn=LRRFIP1;refGene_tx=XM_023259892.1;refGene_gn=LRRFIP1;refGene_tx=
XM_023259893.1;refGene_gn=LRRFIP1;refGene_tx=XM_023259894.1;refGene_gn
=LRRFIP1;refGene_tx=XM_023259895.1;refGene_gn=LRRFIP1;refGene_tx=XM_02
3259896.1;refGene_gn=LRRFIP1;refGene_tx=XM_023259897.1;refGene_gn=LRRF
IP1;refGene_tx=XM_023259898.1
C1 219628900 219628950
refGene_gn=LRRFIP1;refGene_tx=XM_023259899.1;refGene_gn=LRRFIP1;refGen
e_tx=XM_023259900.1;refGene_gn=LRRFIP1;refGene_tx=XM_023259901.1;refGe
ne_gn=LRRFIP1;refGene_tx=XM_023259902.1;refGene_gn=LRRFIP1;refGene_tx=
XM_023259903.1;refGene_gn=LRRFIP1;refGene_tx=XM_023259904.1;refGene_gn
=LRRFIP1;refGene_tx=XM_023259905.1;refGene_gn=LRRFIP1;refGene_tx=XM_02
3259906.1;refGene_gn=LRRFIP1;refGene_tx=XM_023259907.1;refGene_gn=LRRF
IP1;refGene_tx=XM_023259908.1;refGene_gn=LRRFIP1;refGene_tx=XM_0232599
09.1;refGene_gn=LRRFIP1;refGene_tx=XM_023259910.1
C1 219630515 219630633
refGene_gn=LRRFIP1;refGene_tx=XM_023259899.1;refGene_gn=LRRFIP1;refGen
e_tx=XM_023259900.1;refGene_gn=LRRFIP1;refGene_tx=XM_023259901.1;refGe

ne_gn=LRRFIP1;refGene_tx=XM_023259902.1;refGene_gn=LRRFIP1;refGene_tx=XM_023259903.1;refGene_gn=LRRFIP1;refGene_tx=XM_023259904.1;refGene_gn=LRRFIP1;refGene_tx=XM_023259905.1;refGene_gn=LRRFIP1;refGene_tx=XM_023259906.1;refGene_gn=LRRFIP1;refGene_tx=XM_023259907.1;refGene_gn=LRRFIP1;refGene_tx=XM_023259908.1;refGene_gn=LRRFIP1;refGene_tx=XM_023259909.1;refGene_gn=LRRFIP1;refGene_tx=XM_023259910.1
C1 219631499 219631579
refGene_gn=LRRFIP1;refGene_tx=XM_023259899.1;refGene_gn=LRRFIP1;refGene_tx=XM_023259900.1;refGene_gn=LRRFIP1;refGene_tx=XM_023259901.1;refGene_gn=LRRFIP1;refGene_tx=XM_023259902.1;refGene_gn=LRRFIP1;refGene_tx=XM_023259903.1;refGene_gn=LRRFIP1;refGene_tx=XM_023259904.1;refGene_gn=LRRFIP1;refGene_tx=XM_023259905.1;refGene_gn=LRRFIP1;refGene_tx=XM_023259906.1;refGene_gn=LRRFIP1;refGene_tx=XM_023259907.1;refGene_gn=LRRFIP1;refGene_tx=XM_023259908.1;refGene_gn=LRRFIP1;refGene_tx=XM_023259909.1;refGene_gn=LRRFIP1;refGene_tx=XM_023259910.1
C1 219632665 219632770
refGene_gn=LRRFIP1;refGene_tx=XM_023259899.1;refGene_gn=LRRFIP1;refGene_tx=XM_023259900.1;refGene_gn=LRRFIP1;refGene_tx=XM_023259901.1;refGene_gn=LRRFIP1;refGene_tx=XM_023259902.1;refGene_gn=LRRFIP1;refGene_tx=XM_023259903.1;refGene_gn=LRRFIP1;refGene_tx=XM_023259904.1;refGene_gn=LRRFIP1;refGene_tx=XM_023259905.1;refGene_gn=LRRFIP1;refGene_tx=XM_023259906.1;refGene_gn=LRRFIP1;refGene_tx=XM_023259907.1;refGene_gn=LRRFIP1;refGene_tx=XM_023259908.1;refGene_gn=LRRFIP1;refGene_tx=XM_023259909.1;refGene_gn=LRRFIP1;refGene_tx=XM_023259910.1
C1 219634979 219635090
refGene_gn=LRRFIP1;refGene_tx=XM_023259899.1;refGene_gn=LRRFIP1;refGene_tx=XM_023259900.1;refGene_gn=LRRFIP1;refGene_tx=XM_023259901.1;refGene_gn=LRRFIP1;refGene_tx=XM_023259902.1;refGene_gn=LRRFIP1;refGene_tx=XM_023259903.1;refGene_gn=LRRFIP1;refGene_tx=XM_023259904.1;refGene_gn=LRRFIP1;refGene_tx=XM_023259905.1;refGene_gn=LRRFIP1;refGene_tx=XM_023259906.1;refGene_gn=LRRFIP1;refGene_tx=XM_023259907.1;refGene_gn=LRRFIP1;refGene_tx=XM_023259908.1;refGene_gn=LRRFIP1;refGene_tx=XM_023259909.1;refGene_gn=LRRFIP1;refGene_tx=XM_023259910.1
C1 219655534 219655607
refGene_gn=RBM44;refGene_tx=XM_011285600.3;refGene_gn=RBM44;refGene_tx=XM_023259911.1;refGene_gn=RBM44;refGene_tx=XM_023259912.1
C1 219658288 219659901
refGene_gn=RBM44;refGene_tx=XM_011285600.3;refGene_gn=RBM44;refGene_tx=XM_023259911.1;refGene_gn=RBM44;refGene_tx=XM_023259912.1
C1 219660188 219660247
refGene_gn=RBM44;refGene_tx=XM_011285600.3;refGene_gn=RBM44;refGene_tx=XM_023259911.1;refGene_gn=RBM44;refGene_tx=XM_023259912.1
C1 219661398 219661575
refGene_gn=RBM44;refGene_tx=XM_011285600.3;refGene_gn=RBM44;refGene_tx=XM_023259911.1;refGene_gn=RBM44;refGene_tx=XM_023259912.1
C1 219662086 219662270
refGene_gn=RBM44;refGene_tx=XM_011285600.3;refGene_gn=RBM44;refGene_tx=XM_023259911.1
C1 219662364 219662387
refGene_gn=RBM44;refGene_tx=XM_011285600.3;refGene_gn=RBM44;refGene_tx

=XM_023259911.1
C1	219662517	219662602
refGene_gn=RBM44;refGene_tx=XM_011285600.3;refGene_gn=RBM44;refGene_tx=XM_023259911.1;refGene_gn=RBM44;refGene_tx=XM_023259912.1
C1	219666620	219666735
refGene_gn=RBM44;refGene_tx=XM_011285600.3;refGene_gn=RBM44;refGene_tx=XM_023259911.1;refGene_gn=RBM44;refGene_tx=XM_023259912.1
C1	219667712	219667841
refGene_gn=RBM44;refGene_tx=XM_011285600.3;refGene_gn=RBM44;refGene_tx=XM_023259911.1;refGene_gn=RBM44;refGene_tx=XM_023259912.1
C1	219670022	219670102
refGene_gn=RBM44;refGene_tx=XM_011285600.3;refGene_gn=RBM44;refGene_tx=XM_023259911.1;refGene_gn=RBM44;refGene_tx=XM_023259912.1
C1	219670204	219670275
refGene_gn=RBM44;refGene_tx=XM_011285600.3;refGene_gn=RBM44;refGene_tx=XM_023259911.1;refGene_gn=RBM44;refGene_tx=XM_023259912.1
C1	219672230	219672477
refGene_gn=RBM44;refGene_tx=XM_011285600.3;refGene_gn=RBM44;refGene_tx=XM_023259911.1;refGene_gn=RBM44;refGene_tx=XM_023259912.1
C1	219678595	219678741
refGene_gn=RBM44;refGene_tx=XM_011285600.3;refGene_gn=RBM44;refGene_tx=XM_023259911.1;refGene_gn=RBM44;refGene_tx=XM_023259912.1
C1	219678879	219679003
refGene_gn=RBM44;refGene_tx=XM_011285600.3;refGene_gn=RBM44;refGene_tx=XM_023259911.1;refGene_gn=RBM44;refGene_tx=XM_023259912.1
C1	219699562	219699599
refGene_gn=RAMP1;refGene_tx=XM_023259914.1;refGene_gn=RAMP1;refGene_tx=XM_023259915.1
C1	219717173	219717312
refGene_gn=RAMP1;refGene_tx=XM_023259914.1;refGene_gn=RAMP1;refGene_tx=XM_023259915.1
C1	219720457	219720731
refGene_gn=RAMP1;refGene_tx=XM_023259914.1
C1	219746665	219746921
refGene_gn=RAMP1;refGene_tx=XM_023259915.1
C1	219789211	219789329
refGene_gn=UBE2F;refGene_tx=XM_023259916.1;refGene_gn=UBE2F;refGene_tx=XM_023259917.1;refGene_gn=UBE2F;refGene_tx=XM_023259918.1;refGene_gn=UBE2F;refGene_tx=XM_023259919.1;refGene_gn=UBE2F;refGene_tx=XM_023259920.1;refGene_gn=UBE2F;refGene_tx=XM_023259921.1
C1	219799172	219799202
refGene_gn=UBE2F;refGene_tx=XM_023259916.1;refGene_gn=UBE2F;refGene_tx=XM_023259917.1;refGene_gn=UBE2F;refGene_tx=XM_023259918.1;refGene_gn=UBE2F;refGene_tx=XM_023259921.1
C1	219803754	219803820
refGene_gn=UBE2F;refGene_tx=XM_023259916.1;refGene_gn=UBE2F;refGene_tx=XM_023259917.1;refGene_gn=UBE2F;refGene_tx=XM_023259918.1;refGene_gn=UBE2F;refGene_tx=XM_023259919.1;refGene_gn=UBE2F;refGene_tx=XM_023259921.1
C1	219818653	219818721

refGene_gn=UBE2F;refGene_tx=XM_023259916.1;refGene_gn=UBE2F;refGene_tx=XM_023259917.1;refGene_gn=UBE2F;refGene_tx=XM_023259918.1;refGene_gn=UBE2F;refGene_tx=XM_023259919.1;refGene_gn=UBE2F;refGene_tx=XM_023259920.1
C1 219820130 219820170
refGene_gn=UBE2F;refGene_tx=XM_023259918.1
C1 219824860 219824931
refGene_gn=UBE2F;refGene_tx=XM_023259916.1;refGene_gn=UBE2F;refGene_tx=XM_023259917.1;refGene_gn=UBE2F;refGene_tx=XM_023259918.1;refGene_gn=UBE2F;refGene_tx=XM_023259919.1;refGene_gn=UBE2F;refGene_tx=XM_023259920.1;refGene_gn=UBE2F;refGene_tx=XM_023259921.1
C1 219828824 219828882
refGene_gn=UBE2F;refGene_tx=XM_023259916.1;refGene_gn=UBE2F;refGene_tx=XM_023259917.1;refGene_gn=UBE2F;refGene_tx=XM_023259918.1;refGene_gn=UBE2F;refGene_tx=XM_023259919.1;refGene_gn=UBE2F;refGene_tx=XM_023259920.1;refGene_gn=UBE2F;refGene_tx=XM_023259921.1
C1 219829407 219829511
refGene_gn=UBE2F;refGene_tx=XM_023259918.1;refGene_gn=UBE2F;refGene_tx=XM_023259921.1
C1 219831016 219831049
refGene_gn=UBE2F;refGene_tx=XM_023259916.1;refGene_gn=UBE2F;refGene_tx=XM_023259917.1;refGene_gn=UBE2F;refGene_tx=XM_023259919.1;refGene_gn=UBE2F;refGene_tx=XM_023259920.1
C1 219833592 219833655
refGene_gn=UBE2F;refGene_tx=XM_023259916.1;refGene_gn=UBE2F;refGene_tx=XM_023259917.1;refGene_gn=UBE2F;refGene_tx=XM_023259919.1;refGene_gn=UBE2F;refGene_tx=XM_023259920.1
C1 219837096 219837147
refGene_gn=UBE2F;refGene_tx=XM_023259916.1;refGene_gn=UBE2F;refGene_tx=XM_023259917.1;refGene_gn=UBE2F;refGene_tx=XM_023259919.1;refGene_gn=UBE2F;refGene_tx=XM_023259920.1
C1 219849911 219850000
refGene_gn=SCLY;refGene_tx=XM_003991297.5;refGene_gn=SCLY;refGene_tx=XM_006935738.4
C1 219852554 219852667
refGene_gn=SCLY;refGene_tx=XM_003991297.5;refGene_gn=SCLY;refGene_tx=XM_006935738.4
C1 219858460 219858561
refGene_gn=SCLY;refGene_tx=XM_003991297.5;refGene_gn=SCLY;refGene_tx=XM_006935737.2
C1 219861536 219861690
refGene_gn=SCLY;refGene_tx=XM_003991297.5;refGene_gn=SCLY;refGene_tx=XM_006935737.2
C1 219867242 219867370
refGene_gn=SCLY;refGene_tx=XM_003991297.5;refGene_gn=SCLY;refGene_tx=XM_006935737.2;refGene_gn=SCLY;refGene_tx=XM_006935738.4
C1 219867574 219867739
refGene_gn=SCLY;refGene_tx=XM_003991297.5;refGene_gn=SCLY;refGene_tx=XM_006935737.2;refGene_gn=SCLY;refGene_tx=XM_006935738.4
C1 219868694 219868801

refGene_gn=SCLY;refGene_tx=XM_003991297.5;refGene_gn=SCLY;refGene_tx=XM_006935737.2;refGene_gn=SCLY;refGene_tx=XM_006935738.4
C1 219875532 219875569
refGene_gn=SCLY;refGene_tx=XM_003991297.5;refGene_gn=SCLY;refGene_tx=XM_006935737.2;refGene_gn=SCLY;refGene_tx=XM_006935738.4
C1 219877938 219878022
refGene_gn=SCLY;refGene_tx=XM_003991297.5;refGene_gn=SCLY;refGene_tx=XM_006935737.2;refGene_gn=SCLY;refGene_tx=XM_006935738.4
C1 219878545 219878648
refGene_gn=SCLY;refGene_tx=XM_003991297.5;refGene_gn=SCLY;refGene_tx=XM_006935737.2;refGene_gn=SCLY;refGene_tx=XM_006935738.4
C1 219880134 219880210
refGene_gn=SCLY;refGene_tx=XM_003991297.5;refGene_gn=SCLY;refGene_tx=XM_006935737.2;refGene_gn=SCLY;refGene_tx=XM_006935738.4
C1 219881836 219882008
refGene_gn=SCLY;refGene_tx=XM_003991297.5;refGene_gn=SCLY;refGene_tx=XM_006935737.2;refGene_gn=SCLY;refGene_tx=XM_006935738.4
C1 219884041 219884335
refGene_gn=LOC111561653;refGene_tx=XM_023257873.1
C1 219884953 219885061
refGene_gn=LOC111561653;refGene_tx=XM_023257873.1
C1 219885563 219885754
refGene_gn=LOC111561653;refGene_tx=XM_023257873.1
C1 219887324 219887511
refGene_gn=LOC111561653;refGene_tx=XM_023257873.1
C1 219888786 219888945
refGene_gn=LOC111561653;refGene_tx=XM_023257873.1
C1 219889891 219889966
refGene_gn=LOC101092386;refGene_tx=XM_023259923.1
C1 219899291 219899423
refGene_gn=LOC101092386;refGene_tx=XM_023259923.1
C1 219905930 219906258
refGene_gn=LOC101092386;refGene_tx=XM_023259922.1;refGene_gn=LOC101092386;refGene_tx=XM_023259923.1
C1 219907711 219907824
refGene_gn=LOC101092386;refGene_tx=XM_023259922.1;refGene_gn=LOC101092386;refGene_tx=XM_023259923.1
C1 219908588 219908783
refGene_gn=LOC101092386;refGene_tx=XM_023259922.1;refGene_gn=LOC101092386;refGene_tx=XM_023259923.1
C1 219910346 219911948
refGene_gn=LOC101092386;refGene_tx=XM_023259922.1;refGene_gn=LOC101092386;refGene_tx=XM_023259923.1
C1 219920439 219921213
refGene_gn=KLHL30;refGene_tx=XM_011285598.2
C1 219922487 219922620
refGene_gn=KLHL30;refGene_tx=XM_011285598.2
C1 219924335 219924422
refGene_gn=KLHL30;refGene_tx=XM_011285598.2
C1 219925159 219925315

refGene_gn=KLHL30;refGene_tx=XM_011285598.2
C1	219925982	219926171
refGene_gn=KLHL30;refGene_tx=XM_011285598.2
C1	219927082	219927228
refGene_gn=KLHL30;refGene_tx=XM_011285598.2
C1	219928600	219928852
refGene_gn=KLHL30;refGene_tx=XM_011285598.2
C1	219937412	219937598
refGene_gn=ERFE;refGene_tx=XM_023259924.1
C1	219939757	219939880
refGene_gn=ERFE;refGene_tx=XM_023259924.1
C1	219940459	219940562
refGene_gn=ERFE;refGene_tx=XM_023259924.1
C1	219941542	219941787
refGene_gn=ERFE;refGene_tx=XM_023259924.1
C1	219941863	219941972
refGene_gn=ERFE;refGene_tx=XM_023259924.1
C1	219942064	219942155
refGene_gn=ERFE;refGene_tx=XM_023259924.1
C1	219943011	219943090
refGene_gn=ERFE;refGene_tx=XM_023259924.1
C1	219943940	219944039
refGene_gn=ERFE;refGene_tx=XM_023259924.1
C1	219946638	219946779
refGene_gn=ILKAP;refGene_tx=XM_006935741.3;refGene_gn=ILKAP;refGene_tx
=XM_019838971.2;refGene_gn=ILKAP;refGene_tx=XM_023259925.1;refGene_gn=
ILKAP;refGene_tx=XM_023259926.1;refGene_gn=ILKAP;refGene_tx=XM_0232599
27.1
C1	219947001	219947083
refGene_gn=ILKAP;refGene_tx=XM_006935741.3;refGene_gn=ILKAP;refGene_tx
=XM_019838971.2;refGene_gn=ILKAP;refGene_tx=XM_023259925.1;refGene_gn=
ILKAP;refGene_tx=XM_023259926.1;refGene_gn=ILKAP;refGene_tx=XM_0232599
27.1
C1	219948580	219948700
refGene_gn=ILKAP;refGene_tx=XM_006935741.3;refGene_gn=ILKAP;refGene_tx
=XM_019838971.2;refGene_gn=ILKAP;refGene_tx=XM_023259925.1;refGene_gn=
ILKAP;refGene_tx=XM_023259926.1;refGene_gn=ILKAP;refGene_tx=XM_0232599
27.1
C1	219951721	219951843
refGene_gn=ILKAP;refGene_tx=XM_006935741.3;refGene_gn=ILKAP;refGene_tx
=XM_019838971.2;refGene_gn=ILKAP;refGene_tx=XM_023259925.1;refGene_gn=
ILKAP;refGene_tx=XM_023259926.1;refGene_gn=ILKAP;refGene_tx=XM_0232599
27.1
C1	219953182	219953270
refGene_gn=ILKAP;refGene_tx=XM_006935741.3;refGene_gn=ILKAP;refGene_tx
=XM_019838971.2;refGene_gn=ILKAP;refGene_tx=XM_023259925.1;refGene_gn=
ILKAP;refGene_tx=XM_023259926.1;refGene_gn=ILKAP;refGene_tx=XM_0232599
27.1
C1	219953462	219953556
refGene_gn=ILKAP;refGene_tx=XM_006935741.3;refGene_gn=ILKAP;refGene_tx

=XM_019838971.2;refGene_gn=ILKAP;refGene_tx=XM_023259925.1;refGene_gn=ILKAP;refGene_tx=XM_023259926.1;refGene_gn=ILKAP;refGene_tx=XM_023259927.1
C1 219953935 219954042
refGene_gn=ILKAP;refGene_tx=XM_006935741.3;refGene_gn=ILKAP;refGene_tx=XM_019838971.2;refGene_gn=ILKAP;refGene_tx=XM_023259925.1;refGene_gn=ILKAP;refGene_tx=XM_023259926.1;refGene_gn=ILKAP;refGene_tx=XM_023259927.1
C1 219955836 219955963
refGene_gn=ILKAP;refGene_tx=XM_006935741.3;refGene_gn=ILKAP;refGene_tx=XM_019838971.2;refGene_gn=ILKAP;refGene_tx=XM_023259925.1;refGene_gn=ILKAP;refGene_tx=XM_023259926.1;refGene_gn=ILKAP;refGene_tx=XM_023259927.1
C1 219957476 219957593
refGene_gn=ILKAP;refGene_tx=XM_006935741.3;refGene_gn=ILKAP;refGene_tx=XM_019838971.2;refGene_gn=ILKAP;refGene_tx=XM_023259925.1;refGene_gn=ILKAP;refGene_tx=XM_023259927.1
C1 219960106 219960163
refGene_gn=ILKAP;refGene_tx=XM_023259925.1;refGene_gn=ILKAP;refGene_tx=XM_023259926.1
C1 219960634 219960700
refGene_gn=ILKAP;refGene_tx=XM_023259925.1;refGene_gn=ILKAP;refGene_tx=XM_023259926.1
C1 219968188 219968243
refGene_gn=ILKAP;refGene_tx=XM_023259925.1;refGene_gn=ILKAP;refGene_tx=XM_023259926.1
C1 219989949 219990374
refGene_gn=HES6;refGene_tx=XM_023259928.1;refGene_gn=HES6;refGene_tx=XM_023259929.1
C1 219990593 219990675
refGene_gn=HES6;refGene_tx=XM_023259928.1;refGene_gn=HES6;refGene_tx=XM_023259929.1
C1 219990793 219990880
refGene_gn=HES6;refGene_tx=XM_023259928.1;refGene_gn=HES6;refGene_tx=XM_023259929.1
C1 219990954 219991035
refGene_gn=HES6;refGene_tx=XM_023259928.1;refGene_gn=HES6;refGene_tx=XM_023259929.1
C1 219997530 219997680
refGene_gn=PER2;refGene_tx=XM_023259930.1;refGene_gn=PER2;refGene_tx=XM_023259931.1;refGene_gn=PER2;refGene_tx=XM_023259932.1;refGene_gn=PER2;refGene_tx=XM_023259933.1
C1 219999722 219999873
refGene_gn=PER2;refGene_tx=XM_023259930.1;refGene_gn=PER2;refGene_tx=XM_023259931.1;refGene_gn=PER2;refGene_tx=XM_023259932.1;refGene_gn=PER2;refGene_tx=XM_023259933.1
C1 220001233 220001426
refGene_gn=PER2;refGene_tx=XM_023259930.1;refGene_gn=PER2;refGene_tx=XM_023259931.1;refGene_gn=PER2;refGene_tx=XM_023259932.1;refGene_gn=PER2;refGene_tx=XM_023259933.1

C1	220002327	220002505
refGene_gn=PER2;refGene_tx=XM_023259930.1;refGene_gn=PER2;refGene_tx=XM_023259931.1;refGene_gn=PER2;refGene_tx=XM_023259932.1;refGene_gn=PER2;refGene_tx=XM_023259933.1
C1	220003791	220004603
refGene_gn=PER2;refGene_tx=XM_023259930.1;refGene_gn=PER2;refGene_tx=XM_023259931.1;refGene_gn=PER2;refGene_tx=XM_023259932.1;refGene_gn=PER2;refGene_tx=XM_023259933.1
C1	220006490	220006730
refGene_gn=PER2;refGene_tx=XM_023259930.1;refGene_gn=PER2;refGene_tx=XM_023259931.1;refGene_gn=PER2;refGene_tx=XM_023259932.1;refGene_gn=PER2;refGene_tx=XM_023259933.1
C1	220007853	220008018
refGene_gn=PER2;refGene_tx=XM_023259930.1;refGene_gn=PER2;refGene_tx=XM_023259931.1;refGene_gn=PER2;refGene_tx=XM_023259932.1;refGene_gn=PER2;refGene_tx=XM_023259933.1
C1	220008814	220008939
refGene_gn=PER2;refGene_tx=XM_023259930.1;refGene_gn=PER2;refGene_tx=XM_023259931.1;refGene_gn=PER2;refGene_tx=XM_023259932.1;refGene_gn=PER2;refGene_tx=XM_023259933.1
C1	220009027	220009187
refGene_gn=PER2;refGene_tx=XM_023259930.1;refGene_gn=PER2;refGene_tx=XM_023259931.1;refGene_gn=PER2;refGene_tx=XM_023259932.1;refGene_gn=PER2;refGene_tx=XM_023259933.1
C1	220010704	220010792
refGene_gn=PER2;refGene_tx=XM_023259930.1;refGene_gn=PER2;refGene_tx=XM_023259931.1;refGene_gn=PER2;refGene_tx=XM_023259932.1;refGene_gn=PER2;refGene_tx=XM_023259933.1
C1	220011212	220011338
refGene_gn=PER2;refGene_tx=XM_023259930.1;refGene_gn=PER2;refGene_tx=XM_023259931.1;refGene_gn=PER2;refGene_tx=XM_023259932.1;refGene_gn=PER2;refGene_tx=XM_023259933.1
C1	220012115	220012224
refGene_gn=PER2;refGene_tx=XM_023259930.1;refGene_gn=PER2;refGene_tx=XM_023259931.1;refGene_gn=PER2;refGene_tx=XM_023259932.1;refGene_gn=PER2;refGene_tx=XM_023259933.1
C1	220012515	220012669
refGene_gn=PER2;refGene_tx=XM_023259930.1;refGene_gn=PER2;refGene_tx=XM_023259931.1;refGene_gn=PER2;refGene_tx=XM_023259932.1;refGene_gn=PER2;refGene_tx=XM_023259933.1
C1	220013211	220013318
refGene_gn=PER2;refGene_tx=XM_023259930.1;refGene_gn=PER2;refGene_tx=XM_023259931.1;refGene_gn=PER2;refGene_tx=XM_023259932.1;refGene_gn=PER2;refGene_tx=XM_023259933.1
C1	220015639	220015718
refGene_gn=PER2;refGene_tx=XM_023259930.1;refGene_gn=PER2;refGene_tx=XM_023259931.1;refGene_gn=PER2;refGene_tx=XM_023259932.1;refGene_gn=PER2;refGene_tx=XM_023259933.1
C1	220017338	220017481
refGene_gn=PER2;refGene_tx=XM_023259930.1;refGene_gn=PER2;refGene_tx=X

M_023259931.1;refGene_gn=PER2;refGene_tx=XM_023259932.1;refGene_gn=PER2;refGene_tx=XM_023259933.1
C1 220018625 220018677
refGene_gn=PER2;refGene_tx=XM_023259930.1;refGene_gn=PER2;refGene_tx=XM_023259931.1;refGene_gn=PER2;refGene_tx=XM_023259932.1;refGene_gn=PER2;refGene_tx=XM_023259933.1
C1 220020606 220020808
refGene_gn=PER2;refGene_tx=XM_023259930.1;refGene_gn=PER2;refGene_tx=XM_023259931.1;refGene_gn=PER2;refGene_tx=XM_023259932.1;refGene_gn=PER2;refGene_tx=XM_023259933.1
C1 220021622 220021744
refGene_gn=PER2;refGene_tx=XM_023259930.1;refGene_gn=PER2;refGene_tx=XM_023259931.1;refGene_gn=PER2;refGene_tx=XM_023259932.1;refGene_gn=PER2;refGene_tx=XM_023259933.1
C1 220024538 220024693
refGene_gn=PER2;refGene_tx=XM_023259930.1;refGene_gn=PER2;refGene_tx=XM_023259931.1;refGene_gn=PER2;refGene_tx=XM_023259932.1;refGene_gn=PER2;refGene_tx=XM_023259933.1
C1 220025500 220025563
refGene_gn=PER2;refGene_tx=XM_023259930.1;refGene_gn=PER2;refGene_tx=XM_023259931.1;refGene_gn=PER2;refGene_tx=XM_023259932.1;refGene_gn=PER2;refGene_tx=XM_023259933.1
C1 220026050 220026277
refGene_gn=PER2;refGene_tx=XM_023259930.1;refGene_gn=PER2;refGene_tx=XM_023259931.1;refGene_gn=PER2;refGene_tx=XM_023259932.1;refGene_gn=PER2;refGene_tx=XM_023259933.1
C1 220033538 220033583
refGene_gn=PER2;refGene_tx=XM_023259930.1;refGene_gn=PER2;refGene_tx=XM_023259931.1
C1 220056679 220056802
refGene_gn=TRAF3IP1;refGene_tx=XM_011285589.3;refGene_gn=TRAF3IP1;refGene_tx=XM_011285590.3;refGene_gn=TRAF3IP1;refGene_tx=XM_011285591.3;refGene_gn=TRAF3IP1;refGene_tx=XM_011285592.3;refGene_gn=TRAF3IP1;refGene_tx=XM_019838973.2;refGene_gn=TRAF3IP1;refGene_tx=XM_019838974.2;refGene_gn=TRAF3IP1;refGene_tx=XM_019838975.2
C1 220059562 220059631
refGene_gn=TRAF3IP1;refGene_tx=XM_011285589.3;refGene_gn=TRAF3IP1;refGene_tx=XM_011285590.3;refGene_gn=TRAF3IP1;refGene_tx=XM_011285591.3;refGene_gn=TRAF3IP1;refGene_tx=XM_011285592.3;refGene_gn=TRAF3IP1;refGene_tx=XM_019838973.2;refGene_gn=TRAF3IP1;refGene_tx=XM_019838974.2;refGene_gn=TRAF3IP1;refGene_tx=XM_019838975.2
C1 220060003 220060165
refGene_gn=TRAF3IP1;refGene_tx=XM_011285589.3;refGene_gn=TRAF3IP1;refGene_tx=XM_011285590.3;refGene_gn=TRAF3IP1;refGene_tx=XM_011285591.3;refGene_gn=TRAF3IP1;refGene_tx=XM_011285592.3;refGene_gn=TRAF3IP1;refGene_tx=XM_019838973.2;refGene_gn=TRAF3IP1;refGene_tx=XM_019838974.2;refGene_gn=TRAF3IP1;refGene_tx=XM_019838975.2
C1 220061907 220062051
refGene_gn=TRAF3IP1;refGene_tx=XM_011285589.3;refGene_gn=TRAF3IP1;refGene_tx=XM_011285590.3;refGene_gn=TRAF3IP1;refGene_tx=XM_011285591.3;re

fGene_gn=TRAF3IP1;refGene_tx=XM_011285592.3;refGene_gn=TRAF3IP1;refGene_tx=XM_019838973.2;refGene_gn=TRAF3IP1;refGene_tx=XM_019838974.2;refGene_gn=TRAF3IP1;refGene_tx=XM_019838975.2
C1 220062153 220062576
refGene_gn=TRAF3IP1;refGene_tx=XM_011285589.3;refGene_gn=TRAF3IP1;refGene_tx=XM_011285590.3;refGene_gn=TRAF3IP1;refGene_tx=XM_011285591.3;refGene_gn=TRAF3IP1;refGene_tx=XM_011285592.3;refGene_gn=TRAF3IP1;refGene_tx=XM_019838973.2;refGene_gn=TRAF3IP1;refGene_tx=XM_019838974.2;refGene_gn=TRAF3IP1;refGene_tx=XM_019838975.2
C1 220065006 220065078
refGene_gn=TRAF3IP1;refGene_tx=XM_011285589.3;refGene_gn=TRAF3IP1;refGene_tx=XM_011285590.3;refGene_gn=TRAF3IP1;refGene_tx=XM_011285591.3;refGene_gn=TRAF3IP1;refGene_tx=XM_011285592.3;refGene_gn=TRAF3IP1;refGene_tx=XM_019838973.2;refGene_gn=TRAF3IP1;refGene_tx=XM_019838974.2;refGene_gn=TRAF3IP1;refGene_tx=XM_019838975.2
C1 220065979 220066055
refGene_gn=TRAF3IP1;refGene_tx=XM_011285589.3;refGene_gn=TRAF3IP1;refGene_tx=XM_011285590.3;refGene_gn=TRAF3IP1;refGene_tx=XM_011285591.3;refGene_gn=TRAF3IP1;refGene_tx=XM_011285592.3;refGene_gn=TRAF3IP1;refGene_tx=XM_019838973.2;refGene_gn=TRAF3IP1;refGene_tx=XM_019838974.2;refGene_gn=TRAF3IP1;refGene_tx=XM_019838975.2
C1 220068393 220068495
refGene_gn=TRAF3IP1;refGene_tx=XM_011285589.3;refGene_gn=TRAF3IP1;refGene_tx=XM_019838973.2;refGene_gn=TRAF3IP1;refGene_tx=XM_019838975.2
C1 220069944 220070040
refGene_gn=TRAF3IP1;refGene_tx=XM_011285589.3;refGene_gn=TRAF3IP1;refGene_tx=XM_019838973.2;refGene_gn=TRAF3IP1;refGene_tx=XM_019838974.2;refGene_gn=TRAF3IP1;refGene_tx=XM_019838975.2
C1 220073234 220073336
refGene_gn=TRAF3IP1;refGene_tx=XM_011285589.3;refGene_gn=TRAF3IP1;refGene_tx=XM_011285590.3;refGene_gn=TRAF3IP1;refGene_tx=XM_019838973.2;refGene_gn=TRAF3IP1;refGene_tx=XM_019838974.2;refGene_gn=TRAF3IP1;refGene_tx=XM_019838975.2
C1 220075509 220075530
refGene_gn=TRAF3IP1;refGene_tx=XM_011285589.3;refGene_gn=TRAF3IP1;refGene_tx=XM_011285590.3;refGene_gn=TRAF3IP1;refGene_tx=XM_011285591.3;refGene_gn=TRAF3IP1;refGene_tx=XM_011285592.3;refGene_gn=TRAF3IP1;refGene_tx=XM_019838973.2;refGene_gn=TRAF3IP1;refGene_tx=XM_019838974.2;refGene_gn=TRAF3IP1;refGene_tx=XM_019838975.2
C1 220076393 220076457
refGene_gn=TRAF3IP1;refGene_tx=XM_011285589.3;refGene_gn=TRAF3IP1;refGene_tx=XM_011285590.3;refGene_gn=TRAF3IP1;refGene_tx=XM_011285591.3;refGene_gn=TRAF3IP1;refGene_tx=XM_011285592.3;refGene_gn=TRAF3IP1;refGene_tx=XM_019838973.2;refGene_gn=TRAF3IP1;refGene_tx=XM_019838974.2;refGene_gn=TRAF3IP1;refGene_tx=XM_019838975.2
C1 220076864 220076948
refGene_gn=TRAF3IP1;refGene_tx=XM_011285589.3;refGene_gn=TRAF3IP1;refGene_tx=XM_011285590.3;refGene_gn=TRAF3IP1;refGene_tx=XM_011285591.3;refGene_gn=TRAF3IP1;refGene_tx=XM_011285592.3;refGene_gn=TRAF3IP1;refGene_tx=XM_019838973.2;refGene_gn=TRAF3IP1;refGene_tx=XM_019838974.2;refG

ene_gn=TRAF3IP1;refGene_tx=XM_019838975.2
C1	220078041	220078165
refGene_gn=TRAF3IP1;refGene_tx=XM_011285589.3;refGene_gn=TRAF3IP1;refGene_tx=XM_011285590.3;refGene_gn=TRAF3IP1;refGene_tx=XM_011285591.3;refGene_gn=TRAF3IP1;refGene_tx=XM_011285592.3;refGene_gn=TRAF3IP1;refGene_tx=XM_019838973.2;refGene_gn=TRAF3IP1;refGene_tx=XM_019838974.2;refGene_gn=TRAF3IP1;refGene_tx=XM_019838975.2
C1	220078364	220078401
refGene_gn=TRAF3IP1;refGene_tx=XM_011285589.3;refGene_gn=TRAF3IP1;refGene_tx=XM_011285590.3;refGene_gn=TRAF3IP1;refGene_tx=XM_011285591.3;refGene_gn=TRAF3IP1;refGene_tx=XM_011285592.3;refGene_gn=TRAF3IP1;refGene_tx=XM_019838973.2;refGene_gn=TRAF3IP1;refGene_tx=XM_019838974.2;refGene_gn=TRAF3IP1;refGene_tx=XM_019838975.2
C1	220083068	220083145
refGene_gn=TRAF3IP1;refGene_tx=XM_011285589.3;refGene_gn=TRAF3IP1;refGene_tx=XM_011285590.3;refGene_gn=TRAF3IP1;refGene_tx=XM_011285591.3;refGene_gn=TRAF3IP1;refGene_tx=XM_011285592.3;refGene_gn=TRAF3IP1;refGene_tx=XM_019838973.2;refGene_gn=TRAF3IP1;refGene_tx=XM_019838974.2;refGene_gn=TRAF3IP1;refGene_tx=XM_019838975.2
C1	220110783	220110873
refGene_gn=TRAF3IP1;refGene_tx=XM_019838975.2
C1	220114026	220114250
refGene_gn=TRAF3IP1;refGene_tx=XM_011285589.3;refGene_gn=TRAF3IP1;refGene_tx=XM_011285590.3;refGene_gn=TRAF3IP1;refGene_tx=XM_011285591.3;refGene_gn=TRAF3IP1;refGene_tx=XM_011285592.3;refGene_gn=TRAF3IP1;refGene_tx=XM_019838973.2;refGene_gn=TRAF3IP1;refGene_tx=XM_019838974.2
C1	220115749	220115915
refGene_gn=TRAF3IP1;refGene_tx=XM_011285589.3;refGene_gn=TRAF3IP1;refGene_tx=XM_011285590.3;refGene_gn=TRAF3IP1;refGene_tx=XM_011285591.3;refGene_gn=TRAF3IP1;refGene_tx=XM_011285592.3;refGene_gn=TRAF3IP1;refGene_tx=XM_019838973.2;refGene_gn=TRAF3IP1;refGene_tx=XM_019838974.2
C1	220138022	220138266
refGene_gn=ASB1;refGene_tx=XM_023259934.1
C1	220138403	220138452
refGene_gn=ASB1;refGene_tx=XM_003991302.5
C1	220138777	220138907
refGene_gn=ASB1;refGene_tx=XM_019838976.2
C1	220142579	220142721
refGene_gn=ASB1;refGene_tx=XM_003991302.5;refGene_gn=ASB1;refGene_tx=XM_019838976.2;refGene_gn=ASB1;refGene_tx=XM_023259934.1
C1	220144955	220145258
refGene_gn=ASB1;refGene_tx=XM_003991302.5;refGene_gn=ASB1;refGene_tx=XM_019838976.2;refGene_gn=ASB1;refGene_tx=XM_023259934.1
C1	220152915	220153301
refGene_gn=ASB1;refGene_tx=XM_003991302.5;refGene_gn=ASB1;refGene_tx=XM_019838976.2;refGene_gn=ASB1;refGene_tx=XM_023259934.1
C1	220154567	220154695
refGene_gn=ASB1;refGene_tx=XM_003991302.5;refGene_gn=ASB1;refGene_tx=XM_019838976.2;refGene_gn=ASB1;refGene_tx=XM_023259934.1
C1	220474483	220474966

refGene_gn=TWIST2;refGene_tx=XM_023259935.1;refGene_gn=TWIST2;refGene_tx=XM_023259936.1
C1 220619752 220619799
refGene_gn=LOC109502971;refGene_tx=XM_023257874.1
C1 220622338 220622433
refGene_gn=LOC109502971;refGene_tx=XM_023257874.1
C1 220622795 220622933
refGene_gn=LOC109502971;refGene_tx=XM_023257874.1
C1 220623849 220624036
refGene_gn=LOC109502971;refGene_tx=XM_023257874.1
C1 220630532 220630704
refGene_gn=LOC109502971;refGene_tx=XM_023257874.1
C1 220635906 220635946
refGene_gn=HDAC4;refGene_tx=XM_019838979.2;refGene_gn=HDAC4;refGene_tx=XM_023259937.1;refGene_gn=HDAC4;refGene_tx=XM_023259938.1;refGene_gn=HDAC4;refGene_tx=XM_023259939.1;refGene_gn=HDAC4;refGene_tx=XM_023259940.1
C1 220636356 220636498
refGene_gn=HDAC4;refGene_tx=XM_019838979.2;refGene_gn=HDAC4;refGene_tx=XM_023259937.1;refGene_gn=HDAC4;refGene_tx=XM_023259938.1;refGene_gn=HDAC4;refGene_tx=XM_023259939.1;refGene_gn=HDAC4;refGene_tx=XM_023259940.1
C1 220636915 220637000
refGene_gn=HDAC4;refGene_tx=XM_019838979.2;refGene_gn=HDAC4;refGene_tx=XM_023259937.1;refGene_gn=HDAC4;refGene_tx=XM_023259938.1;refGene_gn=HDAC4;refGene_tx=XM_023259939.1;refGene_gn=HDAC4;refGene_tx=XM_023259940.1
C1 220649958 220650092
refGene_gn=HDAC4;refGene_tx=XM_019838979.2;refGene_gn=HDAC4;refGene_tx=XM_023259937.1;refGene_gn=HDAC4;refGene_tx=XM_023259938.1;refGene_gn=HDAC4;refGene_tx=XM_023259939.1;refGene_gn=HDAC4;refGene_tx=XM_023259940.1
C1 220651572 220651691
refGene_gn=HDAC4;refGene_tx=XM_019838979.2;refGene_gn=HDAC4;refGene_tx=XM_023259937.1;refGene_gn=HDAC4;refGene_tx=XM_023259938.1;refGene_gn=HDAC4;refGene_tx=XM_023259939.1;refGene_gn=HDAC4;refGene_tx=XM_023259940.1
C1 220660372 220660470
refGene_gn=HDAC4;refGene_tx=XM_019838979.2;refGene_gn=HDAC4;refGene_tx=XM_023259937.1;refGene_gn=HDAC4;refGene_tx=XM_023259938.1;refGene_gn=HDAC4;refGene_tx=XM_023259939.1;refGene_gn=HDAC4;refGene_tx=XM_023259940.1
C1 220661250 220661370
refGene_gn=HDAC4;refGene_tx=XM_019838979.2;refGene_gn=HDAC4;refGene_tx=XM_023259937.1;refGene_gn=HDAC4;refGene_tx=XM_023259938.1;refGene_gn=HDAC4;refGene_tx=XM_023259939.1;refGene_gn=HDAC4;refGene_tx=XM_023259940.1
C1 220663599 220663687
refGene_gn=HDAC4;refGene_tx=XM_019838979.2;refGene_gn=HDAC4;refGene_tx=XM_023259937.1;refGene_gn=HDAC4;refGene_tx=XM_023259938.1;refGene_gn=

HDAC4;refGene_tx=XM_023259939.1;refGene_gn=HDAC4;refGene_tx=XM_0232599
40.1
C1 220665429 220665485
refGene_gn=HDAC4;refGene_tx=XM_019838979.2;refGene_gn=HDAC4;refGene_tx
=XM_023259937.1;refGene_gn=HDAC4;refGene_tx=XM_023259938.1;refGene_gn=
HDAC4;refGene_tx=XM_023259939.1;refGene_gn=HDAC4;refGene_tx=XM_0232599
40.1
C1 220667359 220667467
refGene_gn=HDAC4;refGene_tx=XM_019838979.2;refGene_gn=HDAC4;refGene_tx
=XM_023259937.1;refGene_gn=HDAC4;refGene_tx=XM_023259938.1;refGene_gn=
HDAC4;refGene_tx=XM_023259939.1;refGene_gn=HDAC4;refGene_tx=XM_0232599
40.1
C1 220671131 220671178
refGene_gn=HDAC4;refGene_tx=XM_019838979.2;refGene_gn=HDAC4;refGene_tx
=XM_023259937.1;refGene_gn=HDAC4;refGene_tx=XM_023259938.1;refGene_gn=
HDAC4;refGene_tx=XM_023259939.1;refGene_gn=HDAC4;refGene_tx=XM_0232599
40.1
C1 220677947 220678068
refGene_gn=HDAC4;refGene_tx=XM_019838979.2;refGene_gn=HDAC4;refGene_tx
=XM_023259937.1;refGene_gn=HDAC4;refGene_tx=XM_023259938.1;refGene_gn=
HDAC4;refGene_tx=XM_023259939.1;refGene_gn=HDAC4;refGene_tx=XM_0232599
40.1
C1 220680443 220680577
refGene_gn=HDAC4;refGene_tx=XM_019838979.2;refGene_gn=HDAC4;refGene_tx
=XM_023259937.1;refGene_gn=HDAC4;refGene_tx=XM_023259938.1;refGene_gn=
HDAC4;refGene_tx=XM_023259939.1;refGene_gn=HDAC4;refGene_tx=XM_0232599
40.1
C1 220683617 220683804
refGene_gn=HDAC4;refGene_tx=XM_019838979.2;refGene_gn=HDAC4;refGene_tx
=XM_023259937.1;refGene_gn=HDAC4;refGene_tx=XM_023259938.1;refGene_gn=
HDAC4;refGene_tx=XM_023259939.1;refGene_gn=HDAC4;refGene_tx=XM_0232599
40.1
C1 220685615 220685873
refGene_gn=HDAC4;refGene_tx=XM_019838979.2;refGene_gn=HDAC4;refGene_tx
=XM_023259937.1;refGene_gn=HDAC4;refGene_tx=XM_023259938.1;refGene_gn=
HDAC4;refGene_tx=XM_023259939.1;refGene_gn=HDAC4;refGene_tx=XM_0232599
40.1
C1 220688816 220689037
refGene_gn=HDAC4;refGene_tx=XM_019838979.2;refGene_gn=HDAC4;refGene_tx
=XM_023259937.1;refGene_gn=HDAC4;refGene_tx=XM_023259938.1;refGene_gn=
HDAC4;refGene_tx=XM_023259939.1;refGene_gn=HDAC4;refGene_tx=XM_0232599
40.1
C1 220695279 220695478
refGene_gn=HDAC4;refGene_tx=XM_019838979.2;refGene_gn=HDAC4;refGene_tx
=XM_023259937.1;refGene_gn=HDAC4;refGene_tx=XM_023259938.1;refGene_gn=
HDAC4;refGene_tx=XM_023259939.1;refGene_gn=HDAC4;refGene_tx=XM_0232599
40.1
C1 220695569 220695686
refGene_gn=HDAC4;refGene_tx=XM_019838979.2;refGene_gn=HDAC4;refGene_tx
=XM_023259937.1;refGene_gn=HDAC4;refGene_tx=XM_023259938.1;refGene_gn=

HDAC4;refGene_tx=XM_023259939.1;refGene_gn=HDAC4;refGene_tx=XM_0232599
40.1
C1	220700402	220700515
refGene_gn=HDAC4;refGene_tx=XM_019838979.2;refGene_gn=HDAC4;refGene_tx
=XM_023259937.1;refGene_gn=HDAC4;refGene_tx=XM_023259938.1;refGene_gn=
HDAC4;refGene_tx=XM_023259939.1;refGene_gn=HDAC4;refGene_tx=XM_0232599
40.1
C1	220703729	220703861
refGene_gn=HDAC4;refGene_tx=XM_019838979.2;refGene_gn=HDAC4;refGene_tx
=XM_023259937.1;refGene_gn=HDAC4;refGene_tx=XM_023259938.1;refGene_gn=
HDAC4;refGene_tx=XM_023259939.1;refGene_gn=HDAC4;refGene_tx=XM_0232599
40.1
C1	220713224	220713346
refGene_gn=HDAC4;refGene_tx=XM_019838979.2;refGene_gn=HDAC4;refGene_tx
=XM_023259937.1;refGene_gn=HDAC4;refGene_tx=XM_023259938.1;refGene_gn=
HDAC4;refGene_tx=XM_023259939.1;refGene_gn=HDAC4;refGene_tx=XM_0232599
40.1
C1	220717313	220717434
refGene_gn=HDAC4;refGene_tx=XM_019838979.2;refGene_gn=HDAC4;refGene_tx
=XM_023259937.1;refGene_gn=HDAC4;refGene_tx=XM_023259938.1;refGene_gn=
HDAC4;refGene_tx=XM_023259939.1;refGene_gn=HDAC4;refGene_tx=XM_0232599
40.1
C1	220726724	220726875
refGene_gn=HDAC4;refGene_tx=XM_019838979.2;refGene_gn=HDAC4;refGene_tx
=XM_023259937.1;refGene_gn=HDAC4;refGene_tx=XM_023259938.1;refGene_gn=
HDAC4;refGene_tx=XM_023259939.1;refGene_gn=HDAC4;refGene_tx=XM_0232599
40.1
C1	220745897	220746142
refGene_gn=HDAC4;refGene_tx=XM_023259937.1;refGene_gn=HDAC4;refGene_tx
=XM_023259938.1;refGene_gn=HDAC4;refGene_tx=XM_023259939.1;refGene_gn=
HDAC4;refGene_tx=XM_023259940.1
C1	220783600	220783672
refGene_gn=HDAC4;refGene_tx=XM_023259937.1;refGene_gn=HDAC4;refGene_tx
=XM_023259938.1;refGene_gn=HDAC4;refGene_tx=XM_023259939.1;refGene_gn=
HDAC4;refGene_tx=XM_023259940.1
C1	220886922	220886944
refGene_gn=HDAC4;refGene_tx=XM_023259937.1;refGene_gn=HDAC4;refGene_tx
=XM_023259938.1;refGene_gn=HDAC4;refGene_tx=XM_023259939.1;refGene_gn=
HDAC4;refGene_tx=XM_023259940.1
C1	221307115	221307184
refGene_gn=NDUFA10;refGene_tx=XM_023259943.1
C1	221313573	221313680
refGene_gn=NDUFA10;refGene_tx=XM_023259941.1
C1	221316065	221316072
refGene_gn=NDUFA10;refGene_tx=XM_023259941.1
C1	221329406	221329415
refGene_gn=NDUFA10;refGene_tx=XM_023259941.1
C1	221331928	221332037
refGene_gn=NDUFA10;refGene_tx=XM_023259941.1;refGene_gn=NDUFA10;refGen
e_tx=XM_023259943.1

C1 221340335 221340421
refGene_gn=NDUFA10;refGene_tx=XM_023259941.1;refGene_gn=NDUFA10;refGene_tx=XM_023259943.1
C1 221341987 221342042
refGene_gn=NDUFA10;refGene_tx=XM_023259941.1;refGene_gn=NDUFA10;refGene_tx=XM_023259943.1
C1 221347384 221347464
refGene_gn=NDUFA10;refGene_tx=XM_023259941.1;refGene_gn=NDUFA10;refGene_tx=XM_023259943.1
C1 221348705 221348827
refGene_gn=NDUFA10;refGene_tx=XM_023259941.1;refGene_gn=NDUFA10;refGene_tx=XM_023259943.1
C1 221351560 221351647
refGene_gn=NDUFA10;refGene_tx=XM_023259941.1;refGene_gn=NDUFA10;refGene_tx=XM_023259943.1
C1 221354012 221354228
refGene_gn=NDUFA10;refGene_tx=XM_023259941.1;refGene_gn=NDUFA10;refGene_tx=XM_023259943.1
C1 221354956 221355125
refGene_gn=NDUFA10;refGene_tx=XM_023259941.1;refGene_gn=NDUFA10;refGene_tx=XM_023259943.1
C1 221358522 221358597
refGene_gn=NDUFA10;refGene_tx=XM_023259941.1;refGene_gn=NDUFA10;refGene_tx=XM_023259943.1
C1 221364829 221365750
refGene_gn=LOC101089458;refGene_tx=XM_019839197.2
C1 221430580 221430833
refGene_gn=LOC101093881;refGene_tx=XM_011285684.3
C1 221430835 221431142
refGene_gn=LOC101093881;refGene_tx=XM_011285684.3
C1 221431143 221431483
refGene_gn=LOC101093881;refGene_tx=XM_011285684.3
C1 221455932 221455970
refGene_gn=COPS9;refGene_tx=XM_023259944.1
C1 221458365 221458438
refGene_gn=COPS9;refGene_tx=XM_023259944.1
C1 221460362 221460425
refGene_gn=COPS9;refGene_tx=XM_023259944.1
C1 221463144 221463329
refGene_gn=OTOS;refGene_tx=XM_003991304.2;refGene_gn=OTOS;refGene_tx=XM_019839199.1;refGene_gn=OTOS;refGene_tx=XM_019839200.2;refGene_gn=OTOS;refGene_tx=XM_019839201.1
C1 221464009 221464036
refGene_gn=OTOS;refGene_tx=XM_003991304.2;refGene_gn=OTOS;refGene_tx=XM_019839200.2;refGene_gn=OTOS;refGene_tx=XM_019839201.1
C1 221464149 221464348
refGene_gn=OTOS;refGene_tx=XM_003991304.2;refGene_gn=OTOS;refGene_tx=XM_019839199.1;refGene_gn=OTOS;refGene_tx=XM_019839200.2;refGene_gn=OTOS;refGene_tx=XM_019839201.1
C1 221626715 221627132

refGene_gn=LOC109502978;refGene_tx=XM_023257875.1
C1 221627502 221627876
refGene_gn=LOC109502978;refGene_tx=XM_023257875.1
C1 221628091 221628158
refGene_gn=LOC109502978;refGene_tx=XM_023257875.1
C1 221628834 221628888
refGene_gn=LOC109502978;refGene_tx=XM_023257875.1
C1 221663423 221663589
refGene_gn=GPC1;refGene_tx=XM_023259964.1
C1 221681453 221681679
refGene_gn=GPC1;refGene_tx=XM_023259963.1
C1 221684719 221684878
refGene_gn=GPC1;refGene_tx=XM_023259963.1;refGene_gn=GPC1;refGene_tx=XM_023259964.1
C1 221686496 221686888
refGene_gn=GPC1;refGene_tx=XM_023259963.1;refGene_gn=GPC1;refGene_tx=XM_023259964.1
C1 221687700 221687866
refGene_gn=GPC1;refGene_tx=XM_023259963.1;refGene_gn=GPC1;refGene_tx=XM_023259964.1
C1 221688560 221688691
refGene_gn=GPC1;refGene_tx=XM_023259963.1;refGene_gn=GPC1;refGene_tx=XM_023259964.1
C1 221688777 221688897
refGene_gn=GPC1;refGene_tx=XM_023259963.1;refGene_gn=GPC1;refGene_tx=XM_023259964.1
C1 221688990 221689124
refGene_gn=GPC1;refGene_tx=XM_023259963.1;refGene_gn=GPC1;refGene_tx=XM_023259964.1
C1 221689465 221689641
refGene_gn=GPC1;refGene_tx=XM_023259963.1;refGene_gn=GPC1;refGene_tx=XM_023259964.1
C1 221690028 221690258
refGene_gn=GPC1;refGene_tx=XM_023259963.1;refGene_gn=GPC1;refGene_tx=XM_023259964.1
C1 221692889 221693032
refGene_gn=ANKMY1;refGene_tx=XM_023259945.1;refGene_gn=ANKMY1;refGene_tx=XM_023259946.1;refGene_gn=ANKMY1;refGene_tx=XM_023259947.1;refGene_gn=ANKMY1;refGene_tx=XM_023259948.1;refGene_gn=ANKMY1;refGene_tx=XM_023259950.1;refGene_gn=ANKMY1;refGene_tx=XM_023259951.1;refGene_gn=ANKMY1;refGene_tx=XM_023259952.1;refGene_gn=ANKMY1;refGene_tx=XM_023259953.1;refGene_gn=ANKMY1;refGene_tx=XM_023259954.1;refGene_gn=ANKMY1;refGene_tx=XM_023259955.1;refGene_gn=ANKMY1;refGene_tx=XM_023259957.1;refGene_gn=ANKMY1;refGene_tx=XM_023259958.1;refGene_gn=ANKMY1;refGene_tx=XM_023259959.1;refGene_gn=ANKMY1;refGene_tx=XM_023259960.1;refGene_gn=ANKMY1;refGene_tx=XM_023259961.1
C1 221701506 221701562
refGene_gn=ANKMY1;refGene_tx=XM_023259949.1
C1 221702633 221702794
refGene_gn=ANKMY1;refGene_tx=XM_023259945.1;refGene_gn=ANKMY1;refGene_

tx=XM_023259946.1;refGene_gn=ANKMY1;refGene_tx=XM_023259947.1;refGene_
gn=ANKMY1;refGene_tx=XM_023259948.1;refGene_gn=ANKMY1;refGene_tx=XM_02
3259949.1;refGene_gn=ANKMY1;refGene_tx=XM_023259950.1;refGene_gn=ANKMY
1;refGene_tx=XM_023259951.1;refGene_gn=ANKMY1;refGene_tx=XM_023259953.
1;refGene_gn=ANKMY1;refGene_tx=XM_023259954.1;refGene_gn=ANKMY1;refGen
e_tx=XM_023259955.1;refGene_gn=ANKMY1;refGene_tx=XM_023259957.1;refGen
e_gn=ANKMY1;refGene_tx=XM_023259958.1;refGene_gn=ANKMY1;refGene_tx=XM_
023259959.1;refGene_gn=ANKMY1;refGene_tx=XM_023259961.1
C1 221703534 221703613
refGene_gn=ANKMY1;refGene_tx=XM_023259945.1;refGene_gn=ANKMY1;refGene_
tx=XM_023259946.1;refGene_gn=ANKMY1;refGene_tx=XM_023259947.1;refGene_
gn=ANKMY1;refGene_tx=XM_023259948.1;refGene_gn=ANKMY1;refGene_tx=XM_02
3259949.1;refGene_gn=ANKMY1;refGene_tx=XM_023259950.1;refGene_gn=ANKMY
1;refGene_tx=XM_023259951.1;refGene_gn=ANKMY1;refGene_tx=XM_023259953.
1;refGene_gn=ANKMY1;refGene_tx=XM_023259954.1;refGene_gn=ANKMY1;refGen
e_tx=XM_023259955.1;refGene_gn=ANKMY1;refGene_tx=XM_023259957.1;refGen
e_gn=ANKMY1;refGene_tx=XM_023259958.1;refGene_gn=ANKMY1;refGene_tx=XM_
023259959.1;refGene_gn=ANKMY1;refGene_tx=XM_023259961.1
C1 221706956 221707122
refGene_gn=ANKMY1;refGene_tx=XM_023259945.1;refGene_gn=ANKMY1;refGene_
tx=XM_023259946.1;refGene_gn=ANKMY1;refGene_tx=XM_023259947.1;refGene_
gn=ANKMY1;refGene_tx=XM_023259948.1;refGene_gn=ANKMY1;refGene_tx=XM_02
3259949.1;refGene_gn=ANKMY1;refGene_tx=XM_023259950.1;refGene_gn=ANKMY
1;refGene_tx=XM_023259951.1;refGene_gn=ANKMY1;refGene_tx=XM_023259952.
1;refGene_gn=ANKMY1;refGene_tx=XM_023259953.1;refGene_gn=ANKMY1;refGen
e_tx=XM_023259954.1;refGene_gn=ANKMY1;refGene_tx=XM_023259955.1;refGen
e_gn=ANKMY1;refGene_tx=XM_023259957.1;refGene_gn=ANKMY1;refGene_tx=XM_
023259958.1;refGene_gn=ANKMY1;refGene_tx=XM_023259959.1;refGene_gn=ANK
MY1;refGene_tx=XM_023259961.1
C1 221707494 221707608
refGene_gn=ANKMY1;refGene_tx=XM_023259945.1;refGene_gn=ANKMY1;refGene_
tx=XM_023259946.1;refGene_gn=ANKMY1;refGene_tx=XM_023259947.1;refGene_
gn=ANKMY1;refGene_tx=XM_023259948.1;refGene_gn=ANKMY1;refGene_tx=XM_02
3259949.1;refGene_gn=ANKMY1;refGene_tx=XM_023259950.1;refGene_gn=ANKMY
1;refGene_tx=XM_023259951.1;refGene_gn=ANKMY1;refGene_tx=XM_023259952.
1;refGene_gn=ANKMY1;refGene_tx=XM_023259953.1;refGene_gn=ANKMY1;refGen
e_tx=XM_023259954.1;refGene_gn=ANKMY1;refGene_tx=XM_023259955.1;refGen
e_gn=ANKMY1;refGene_tx=XM_023259957.1;refGene_gn=ANKMY1;refGene_tx=XM_
023259958.1;refGene_gn=ANKMY1;refGene_tx=XM_023259959.1;refGene_gn=ANK
MY1;refGene_tx=XM_023259961.1
C1 221710126 221710381
refGene_gn=ANKMY1;refGene_tx=XM_023259956.1
C1 221710848 221710980
refGene_gn=ANKMY1;refGene_tx=XM_023259945.1;refGene_gn=ANKMY1;refGene_
tx=XM_023259946.1;refGene_gn=ANKMY1;refGene_tx=XM_023259947.1;refGene_
gn=ANKMY1;refGene_tx=XM_023259948.1;refGene_gn=ANKMY1;refGene_tx=XM_02
3259949.1;refGene_gn=ANKMY1;refGene_tx=XM_023259950.1;refGene_gn=ANKMY
1;refGene_tx=XM_023259951.1;refGene_gn=ANKMY1;refGene_tx=XM_023259952.
1;refGene_gn=ANKMY1;refGene_tx=XM_023259955.1;refGene_gn=ANKMY1;refGen
e_tx=XM_023259956.1;refGene_gn=ANKMY1;refGene_tx=XM_023259957.1;refGen

e_gn=ANKMY1;refGene_tx=XM_023259958.1;refGene_gn=ANKMY1;refGene_tx=XM_023259959.1;refGene_gn=ANKMY1;refGene_tx=XM_023259960.1
C1 221712357 221712465
refGene_gn=ANKMY1;refGene_tx=XM_023259945.1;refGene_gn=ANKMY1;refGene_tx=XM_023259946.1;refGene_gn=ANKMY1;refGene_tx=XM_023259947.1;refGene_gn=ANKMY1;refGene_tx=XM_023259948.1;refGene_gn=ANKMY1;refGene_tx=XM_023259949.1;refGene_gn=ANKMY1;refGene_tx=XM_023259950.1;refGene_gn=ANKMY1;refGene_tx=XM_023259951.1;refGene_gn=ANKMY1;refGene_tx=XM_023259952.1;refGene_gn=ANKMY1;refGene_tx=XM_023259955.1;refGene_gn=ANKMY1;refGene_tx=XM_023259956.1;refGene_gn=ANKMY1;refGene_tx=XM_023259957.1;refGene_gn=ANKMY1;refGene_tx=XM_023259958.1;refGene_gn=ANKMY1;refGene_tx=XM_023259959.1;refGene_gn=ANKMY1;refGene_tx=XM_023259960.1
C1 221714024 221714165
refGene_gn=ANKMY1;refGene_tx=XM_023259945.1;refGene_gn=ANKMY1;refGene_tx=XM_023259946.1;refGene_gn=ANKMY1;refGene_tx=XM_023259947.1;refGene_gn=ANKMY1;refGene_tx=XM_023259948.1;refGene_gn=ANKMY1;refGene_tx=XM_023259949.1;refGene_gn=ANKMY1;refGene_tx=XM_023259950.1;refGene_gn=ANKMY1;refGene_tx=XM_023259951.1;refGene_gn=ANKMY1;refGene_tx=XM_023259952.1;refGene_gn=ANKMY1;refGene_tx=XM_023259953.1;refGene_gn=ANKMY1;refGene_tx=XM_023259954.1;refGene_gn=ANKMY1;refGene_tx=XM_023259955.1;refGene_gn=ANKMY1;refGene_tx=XM_023259956.1;refGene_gn=ANKMY1;refGene_tx=XM_023259957.1;refGene_gn=ANKMY1;refGene_tx=XM_023259958.1;refGene_gn=ANKMY1;refGene_tx=XM_023259959.1;refGene_gn=ANKMY1;refGene_tx=XM_023259960.1
C1 221714553 221714694
refGene_gn=ANKMY1;refGene_tx=XM_023259945.1;refGene_gn=ANKMY1;refGene_tx=XM_023259946.1;refGene_gn=ANKMY1;refGene_tx=XM_023259947.1;refGene_gn=ANKMY1;refGene_tx=XM_023259948.1;refGene_gn=ANKMY1;refGene_tx=XM_023259949.1;refGene_gn=ANKMY1;refGene_tx=XM_023259950.1;refGene_gn=ANKMY1;refGene_tx=XM_023259951.1;refGene_gn=ANKMY1;refGene_tx=XM_023259952.1;refGene_gn=ANKMY1;refGene_tx=XM_023259953.1;refGene_gn=ANKMY1;refGene_tx=XM_023259954.1;refGene_gn=ANKMY1;refGene_tx=XM_023259955.1;refGene_gn=ANKMY1;refGene_tx=XM_023259956.1;refGene_gn=ANKMY1;refGene_tx=XM_023259957.1;refGene_gn=ANKMY1;refGene_tx=XM_023259958.1;refGene_gn=ANKMY1;refGene_tx=XM_023259959.1;refGene_gn=ANKMY1;refGene_tx=XM_023259960.1
C1 221714829 221715474
refGene_gn=ANKMY1;refGene_tx=XM_023259945.1;refGene_gn=ANKMY1;refGene_tx=XM_023259946.1;refGene_gn=ANKMY1;refGene_tx=XM_023259947.1;refGene_gn=ANKMY1;refGene_tx=XM_023259949.1;refGene_gn=ANKMY1;refGene_tx=XM_023259950.1;refGene_gn=ANKMY1;refGene_tx=XM_023259951.1;refGene_gn=ANKMY1;refGene_tx=XM_023259952.1;refGene_gn=ANKMY1;refGene_tx=XM_023259953.1;refGene_gn=ANKMY1;refGene_tx=XM_023259955.1;refGene_gn=ANKMY1;refGene_tx=XM_023259956.1;refGene_gn=ANKMY1;refGene_tx=XM_023259957.1;refGene_gn=ANKMY1;refGene_tx=XM_023259958.1;refGene_gn=ANKMY1;refGene_tx=XM_023259959.1;refGene_gn=ANKMY1;refGene_tx=XM_023259960.1;refGene_gn=ANKMY1;refGene_tx=XM_023259962.1
C1 221715749 221715921
refGene_gn=ANKMY1;refGene_tx=XM_023259945.1;refGene_gn=ANKMY1;refGene_tx=XM_023259946.1;refGene_gn=ANKMY1;refGene_tx=XM_023259947.1;refGene_

gn=ANKMY1;refGene_tx=XM_023259948.1;refGene_gn=ANKMY1;refGene_tx=XM_023259949.1;refGene_gn=ANKMY1;refGene_tx=XM_023259950.1;refGene_gn=ANKMY1;refGene_tx=XM_023259951.1;refGene_gn=ANKMY1;refGene_tx=XM_023259952.1;refGene_gn=ANKMY1;refGene_tx=XM_023259953.1;refGene_gn=ANKMY1;refGene_tx=XM_023259954.1;refGene_gn=ANKMY1;refGene_tx=XM_023259955.1;refGene_gn=ANKMY1;refGene_tx=XM_023259956.1;refGene_gn=ANKMY1;refGene_tx=XM_023259957.1;refGene_gn=ANKMY1;refGene_tx=XM_023259958.1;refGene_gn=ANKMY1;refGene_tx=XM_023259959.1;refGene_gn=ANKMY1;refGene_tx=XM_023259960.1;refGene_gn=ANKMY1;refGene_tx=XM_023259961.1;refGene_gn=ANKMY1;refGene_tx=XM_023259962.1
C1 221717051 221717515
refGene_gn=ANKMY1;refGene_tx=XM_023259945.1;refGene_gn=ANKMY1;refGene_tx=XM_023259946.1;refGene_gn=ANKMY1;refGene_tx=XM_023259947.1;refGene_gn=ANKMY1;refGene_tx=XM_023259948.1;refGene_gn=ANKMY1;refGene_tx=XM_023259949.1;refGene_gn=ANKMY1;refGene_tx=XM_023259950.1;refGene_gn=ANKMY1;refGene_tx=XM_023259951.1;refGene_gn=ANKMY1;refGene_tx=XM_023259952.1;refGene_gn=ANKMY1;refGene_tx=XM_023259953.1;refGene_gn=ANKMY1;refGene_tx=XM_023259954.1;refGene_gn=ANKMY1;refGene_tx=XM_023259955.1;refGene_gn=ANKMY1;refGene_tx=XM_023259956.1;refGene_gn=ANKMY1;refGene_tx=XM_023259957.1;refGene_gn=ANKMY1;refGene_tx=XM_023259958.1;refGene_gn=ANKMY1;refGene_tx=XM_023259959.1;refGene_gn=ANKMY1;refGene_tx=XM_023259960.1;refGene_gn=ANKMY1;refGene_tx=XM_023259961.1;refGene_gn=ANKMY1;refGene_tx=XM_023259962.1
C1 221720125 221720389
refGene_gn=ANKMY1;refGene_tx=XM_023259945.1;refGene_gn=ANKMY1;refGene_tx=XM_023259946.1;refGene_gn=ANKMY1;refGene_tx=XM_023259947.1;refGene_gn=ANKMY1;refGene_tx=XM_023259948.1;refGene_gn=ANKMY1;refGene_tx=XM_023259949.1;refGene_gn=ANKMY1;refGene_tx=XM_023259950.1;refGene_gn=ANKMY1;refGene_tx=XM_023259951.1;refGene_gn=ANKMY1;refGene_tx=XM_023259952.1;refGene_gn=ANKMY1;refGene_tx=XM_023259953.1;refGene_gn=ANKMY1;refGene_tx=XM_023259954.1;refGene_gn=ANKMY1;refGene_tx=XM_023259955.1;refGene_gn=ANKMY1;refGene_tx=XM_023259956.1;refGene_gn=ANKMY1;refGene_tx=XM_023259957.1;refGene_gn=ANKMY1;refGene_tx=XM_023259958.1;refGene_gn=ANKMY1;refGene_tx=XM_023259959.1;refGene_gn=ANKMY1;refGene_tx=XM_023259960.1;refGene_gn=ANKMY1;refGene_tx=XM_023259961.1;refGene_gn=ANKMY1;refGene_tx=XM_023259962.1
C1 221720495 221720712
refGene_gn=ANKMY1;refGene_tx=XM_023259945.1;refGene_gn=ANKMY1;refGene_tx=XM_023259946.1;refGene_gn=ANKMY1;refGene_tx=XM_023259947.1;refGene_gn=ANKMY1;refGene_tx=XM_023259948.1;refGene_gn=ANKMY1;refGene_tx=XM_023259949.1;refGene_gn=ANKMY1;refGene_tx=XM_023259950.1;refGene_gn=ANKMY1;refGene_tx=XM_023259951.1;refGene_gn=ANKMY1;refGene_tx=XM_023259952.1;refGene_gn=ANKMY1;refGene_tx=XM_023259953.1;refGene_gn=ANKMY1;refGene_tx=XM_023259954.1;refGene_gn=ANKMY1;refGene_tx=XM_023259955.1;refGene_gn=ANKMY1;refGene_tx=XM_023259956.1;refGene_gn=ANKMY1;refGene_tx=XM_023259957.1;refGene_gn=ANKMY1;refGene_tx=XM_023259958.1;refGene_gn=ANKMY1;refGene_tx=XM_023259959.1;refGene_gn=ANKMY1;refGene_tx=XM_023259960.1;refGene_gn=ANKMY1;refGene_tx=XM_023259961.1;refGene_gn=ANKMY1;refGene_tx=XM_023259962.1
C1 221722461 221722931

refGene_gn=ANKMY1;refGene_tx=XM_023259945.1;refGene_gn=ANKMY1;refGene_tx=XM_023259946.1;refGene_gn=ANKMY1;refGene_tx=XM_023259947.1;refGene_gn=ANKMY1;refGene_tx=XM_023259948.1;refGene_gn=ANKMY1;refGene_tx=XM_023259949.1;refGene_gn=ANKMY1;refGene_tx=XM_023259950.1;refGene_gn=ANKMY1;refGene_tx=XM_023259951.1;refGene_gn=ANKMY1;refGene_tx=XM_023259952.1;refGene_gn=ANKMY1;refGene_tx=XM_023259953.1;refGene_gn=ANKMY1;refGene_tx=XM_023259954.1;refGene_gn=ANKMY1;refGene_tx=XM_023259955.1;refGene_gn=ANKMY1;refGene_tx=XM_023259956.1;refGene_gn=ANKMY1;refGene_tx=XM_023259957.1;refGene_gn=ANKMY1;refGene_tx=XM_023259958.1;refGene_gn=ANKMY1;refGene_tx=XM_023259959.1;refGene_gn=ANKMY1;refGene_tx=XM_023259960.1;refGene_gn=ANKMY1;refGene_tx=XM_023259961.1;refGene_gn=ANKMY1;refGene_tx=XM_023259962.1
C1 221728527 221728671
refGene_gn=ANKMY1;refGene_tx=XM_023259945.1;refGene_gn=ANKMY1;refGene_tx=XM_023259946.1;refGene_gn=ANKMY1;refGene_tx=XM_023259947.1;refGene_gn=ANKMY1;refGene_tx=XM_023259948.1;refGene_gn=ANKMY1;refGene_tx=XM_023259949.1;refGene_gn=ANKMY1;refGene_tx=XM_023259951.1;refGene_gn=ANKMY1;refGene_tx=XM_023259952.1;refGene_gn=ANKMY1;refGene_tx=XM_023259953.1;refGene_gn=ANKMY1;refGene_tx=XM_023259954.1;refGene_gn=ANKMY1;refGene_tx=XM_023259955.1;refGene_gn=ANKMY1;refGene_tx=XM_023259956.1;refGene_gn=ANKMY1;refGene_tx=XM_023259960.1;refGene_gn=ANKMY1;refGene_tx=XM_023259961.1;refGene_gn=ANKMY1;refGene_tx=XM_023259962.1
C1 221731297 221731484
refGene_gn=ANKMY1;refGene_tx=XM_023259945.1;refGene_gn=ANKMY1;refGene_tx=XM_023259946.1;refGene_gn=ANKMY1;refGene_tx=XM_023259947.1;refGene_gn=ANKMY1;refGene_tx=XM_023259948.1;refGene_gn=ANKMY1;refGene_tx=XM_023259949.1;refGene_gn=ANKMY1;refGene_tx=XM_023259950.1;refGene_gn=ANKMY1;refGene_tx=XM_023259951.1;refGene_gn=ANKMY1;refGene_tx=XM_023259952.1;refGene_gn=ANKMY1;refGene_tx=XM_023259953.1;refGene_gn=ANKMY1;refGene_tx=XM_023259954.1;refGene_gn=ANKMY1;refGene_tx=XM_023259955.1;refGene_gn=ANKMY1;refGene_tx=XM_023259956.1;refGene_gn=ANKMY1;refGene_tx=XM_023259960.1;refGene_gn=ANKMY1;refGene_tx=XM_023259961.1;refGene_gn=ANKMY1;refGene_tx=XM_023259962.1
C1 221734300 221734437
refGene_gn=ANKMY1;refGene_tx=XM_023259945.1;refGene_gn=ANKMY1;refGene_tx=XM_023259946.1;refGene_gn=ANKMY1;refGene_tx=XM_023259947.1;refGene_gn=ANKMY1;refGene_tx=XM_023259948.1;refGene_gn=ANKMY1;refGene_tx=XM_023259949.1;refGene_gn=ANKMY1;refGene_tx=XM_023259950.1;refGene_gn=ANKMY1;refGene_tx=XM_023259951.1;refGene_gn=ANKMY1;refGene_tx=XM_023259952.1;refGene_gn=ANKMY1;refGene_tx=XM_023259953.1;refGene_gn=ANKMY1;refGene_tx=XM_023259954.1;refGene_gn=ANKMY1;refGene_tx=XM_023259956.1;refGene_gn=ANKMY1;refGene_tx=XM_023259960.1;refGene_gn=ANKMY1;refGene_tx=XM_023259961.1;refGene_gn=ANKMY1;refGene_tx=XM_023259962.1
C1 221737694 221738096
refGene_gn=DUSP28;refGene_tx=XM_023259965.1
C1 221738531 221738687
refGene_gn=DUSP28;refGene_tx=XM_023259965.1
C1 221739819 221740069
refGene_gn=LOC111561654;refGene_tx=XM_023257876.1
C1 221742889 221743224

refGene_gn=LOC111561654;refGene_tx=XM_023257876.1
C1 221743235 221743769
refGene_gn=RNPEPL1;refGene_tx=XM_019839157.2
C1 221747133 221747274
refGene_gn=RNPEPL1;refGene_tx=XM_019839157.2
C1 221747603 221747755
refGene_gn=RNPEPL1;refGene_tx=XM_019839157.2
C1 221748248 221748365
refGene_gn=RNPEPL1;refGene_tx=XM_019839157.2
C1 221748546 221748782
refGene_gn=RNPEPL1;refGene_tx=XM_019839157.2
C1 221749053 221749167
refGene_gn=RNPEPL1;refGene_tx=XM_019839157.2
C1 221749728 221749841
refGene_gn=RNPEPL1;refGene_tx=XM_019839157.2
C1 221750118 221750227
refGene_gn=RNPEPL1;refGene_tx=XM_019839157.2
C1 221751182 221751413
refGene_gn=RNPEPL1;refGene_tx=XM_019839157.2
C1 221751487 221751630
refGene_gn=RNPEPL1;refGene_tx=XM_019839157.2
C1 221752090 221752378
refGene_gn=RNPEPL1;refGene_tx=XM_019839157.2
C1 221759821 221760085
refGene_gn=CAPN10;refGene_tx=XM_003991305.4;refGene_gn=CAPN10;refGene_tx=XM_019838984.2
C1 221761530 221761583
refGene_gn=CAPN10;refGene_tx=XM_019838984.2
C1 221762315 221762447
refGene_gn=CAPN10;refGene_tx=XM_003991305.4;refGene_gn=CAPN10;refGene_tx=XM_019838984.2
C1 221763595 221763792
refGene_gn=CAPN10;refGene_tx=XM_003991305.4;refGene_gn=CAPN10;refGene_tx=XM_019838984.2
C1 221764582 221764800
refGene_gn=CAPN10;refGene_tx=XM_003991305.4;refGene_gn=CAPN10;refGene_tx=XM_019838984.2
C1 221766373 221766515
refGene_gn=CAPN10;refGene_tx=XM_003991305.4;refGene_gn=CAPN10;refGene_tx=XM_019838984.2
C1 221767216 221767383
refGene_gn=CAPN10;refGene_tx=XM_003991305.4;refGene_gn=CAPN10;refGene_tx=XM_019838984.2
C1 221767751 221768032
refGene_gn=CAPN10;refGene_tx=XM_003991305.4;refGene_gn=CAPN10;refGene_tx=XM_019838984.2
C1 221768484 221768687
refGene_gn=CAPN10;refGene_tx=XM_003991305.4;refGene_gn=CAPN10;refGene_tx=XM_019838984.2
C1 221768833 221769089

refGene_gn=CAPN10;refGene_tx=XM_003991305.4;refGene_gn=CAPN10;refGene_tx=XM_019838984.2
C1 221770384 221770584
refGene_gn=CAPN10;refGene_tx=XM_003991305.4;refGene_gn=CAPN10;refGene_tx=XM_019838984.2
C1 221770801 221770847
refGene_gn=CAPN10;refGene_tx=XM_003991305.4;refGene_gn=CAPN10;refGene_tx=XM_019838984.2
C1 221771117 221771147
refGene_gn=CAPN10;refGene_tx=XM_003991305.4;refGene_gn=CAPN10;refGene_tx=XM_019838984.2
C1 221786079 221786334
refGene_gn=GPR35;refGene_tx=XM_023259967.1;refGene_gn=GPR35;refGene_tx=XM_023259968.1
C1 221788456 221788569
refGene_gn=GPR35;refGene_tx=XM_023259969.1
C1 221790534 221791467
refGene_gn=GPR35;refGene_tx=XM_023259967.1;refGene_gn=GPR35;refGene_tx=XM_023259968.1;refGene_gn=GPR35;refGene_tx=XM_023259969.1;refGene_gn=GPR35;refGene_tx=XM_023259970.1
C1 221795082 221795374
refGene_gn=LOC111561928;refGene_tx=XM_023259971.1
C1 221796246 221797394
refGene_gn=LOC111561928;refGene_tx=XM_023259971.1
C1 221818959 221819165
refGene_gn=LOC101095121;refGene_tx=XM_019839202.1;refGene_gn=LOC101095121;refGene_tx=XM_019839203.1
C1 221820457 221820574
refGene_gn=LOC101095121;refGene_tx=XM_019839202.1
C1 221822136 221822743
refGene_gn=LOC101095121;refGene_tx=XM_019839202.1;refGene_gn=LOC101095121;refGene_tx=XM_019839203.1
C1 221829843 221829886
refGene_gn=KIF1A;refGene_tx=XM_023259972.1;refGene_gn=KIF1A;refGene_tx=XM_023259973.1;refGene_gn=KIF1A;refGene_tx=XM_023259974.1;refGene_gn=KIF1A;refGene_tx=XM_023259975.1;refGene_gn=KIF1A;refGene_tx=XM_023259976.1;refGene_gn=KIF1A;refGene_tx=XM_023259977.1;refGene_gn=KIF1A;refGene_tx=XM_023259978.1;refGene_gn=KIF1A;refGene_tx=XM_023259979.1;refGene_gn=KIF1A;refGene_tx=XM_023259980.1;refGene_gn=KIF1A;refGene_tx=XM_023259981.1;refGene_gn=KIF1A;refGene_tx=XM_023259982.1;refGene_gn=KIF1A;refGene_tx=XM_023259983.1;refGene_gn=KIF1A;refGene_tx=XM_023259984.1;refGene_gn=KIF1A;refGene_tx=XM_023259985.1;refGene_gn=KIF1A;refGene_tx=XM_023259986.1;refGene_gn=KIF1A;refGene_tx=XM_023259987.1;refGene_gn=KIF1A;refGene_tx=XM_023259988.1
C1 221830238 221830357
refGene_gn=KIF1A;refGene_tx=XM_023259972.1;refGene_gn=KIF1A;refGene_tx=XM_023259973.1;refGene_gn=KIF1A;refGene_tx=XM_023259974.1;refGene_gn=KIF1A;refGene_tx=XM_023259975.1;refGene_gn=KIF1A;refGene_tx=XM_023259976.1;refGene_gn=KIF1A;refGene_tx=XM_023259977.1;refGene_gn=KIF1A;refGene_tx=XM_023259978.1;refGene_gn=KIF1A;refGene_tx=XM_023259979.1;refGen

e_gn=KIF1A;refGene_tx=XM_023259980.1;refGene_gn=KIF1A;refGene_tx=XM_02
3259981.1;refGene_gn=KIF1A;refGene_tx=XM_023259982.1;refGene_gn=KIF1A;
refGene_tx=XM_023259983.1;refGene_gn=KIF1A;refGene_tx=XM_023259984.1;r
efGene_gn=KIF1A;refGene_tx=XM_023259985.1;refGene_gn=KIF1A;refGene_tx=
XM_023259986.1;refGene_gn=KIF1A;refGene_tx=XM_023259987.1;refGene_gn=K
IF1A;refGene_tx=XM_023259988.1
C1 221831198 221831391
refGene_gn=KIF1A;refGene_tx=XM_023259972.1;refGene_gn=KIF1A;refGene_tx
=XM_023259973.1;refGene_gn=KIF1A;refGene_tx=XM_023259974.1;refGene_gn=
KIF1A;refGene_tx=XM_023259975.1;refGene_gn=KIF1A;refGene_tx=XM_0232599
76.1;refGene_gn=KIF1A;refGene_tx=XM_023259977.1;refGene_gn=KIF1A;refGe
ne_tx=XM_023259978.1;refGene_gn=KIF1A;refGene_tx=XM_023259979.1;refGen
e_gn=KIF1A;refGene_tx=XM_023259980.1;refGene_gn=KIF1A;refGene_tx=XM_02
3259981.1;refGene_gn=KIF1A;refGene_tx=XM_023259982.1;refGene_gn=KIF1A;
refGene_tx=XM_023259983.1;refGene_gn=KIF1A;refGene_tx=XM_023259984.1;r
efGene_gn=KIF1A;refGene_tx=XM_023259985.1;refGene_gn=KIF1A;refGene_tx=
XM_023259986.1;refGene_gn=KIF1A;refGene_tx=XM_023259987.1;refGene_gn=K
IF1A;refGene_tx=XM_023259988.1
C1 221831950 221832103
refGene_gn=KIF1A;refGene_tx=XM_023259972.1;refGene_gn=KIF1A;refGene_tx
=XM_023259973.1;refGene_gn=KIF1A;refGene_tx=XM_023259974.1;refGene_gn=
KIF1A;refGene_tx=XM_023259975.1;refGene_gn=KIF1A;refGene_tx=XM_0232599
76.1;refGene_gn=KIF1A;refGene_tx=XM_023259977.1;refGene_gn=KIF1A;refGe
ne_tx=XM_023259978.1;refGene_gn=KIF1A;refGene_tx=XM_023259979.1;refGen
e_gn=KIF1A;refGene_tx=XM_023259980.1;refGene_gn=KIF1A;refGene_tx=XM_02
3259981.1;refGene_gn=KIF1A;refGene_tx=XM_023259982.1;refGene_gn=KIF1A;
refGene_tx=XM_023259983.1;refGene_gn=KIF1A;refGene_tx=XM_023259984.1;r
efGene_gn=KIF1A;refGene_tx=XM_023259985.1;refGene_gn=KIF1A;refGene_tx=
XM_023259986.1;refGene_gn=KIF1A;refGene_tx=XM_023259987.1;refGene_gn=K
IF1A;refGene_tx=XM_023259988.1
C1 221832882 221833007
refGene_gn=KIF1A;refGene_tx=XM_023259972.1;refGene_gn=KIF1A;refGene_tx
=XM_023259973.1;refGene_gn=KIF1A;refGene_tx=XM_023259974.1;refGene_gn=
KIF1A;refGene_tx=XM_023259975.1;refGene_gn=KIF1A;refGene_tx=XM_0232599
76.1;refGene_gn=KIF1A;refGene_tx=XM_023259977.1;refGene_gn=KIF1A;refGe
ne_tx=XM_023259978.1;refGene_gn=KIF1A;refGene_tx=XM_023259979.1;refGen
e_gn=KIF1A;refGene_tx=XM_023259980.1;refGene_gn=KIF1A;refGene_tx=XM_02
3259981.1;refGene_gn=KIF1A;refGene_tx=XM_023259982.1;refGene_gn=KIF1A;
refGene_tx=XM_023259983.1;refGene_gn=KIF1A;refGene_tx=XM_023259984.1;r
efGene_gn=KIF1A;refGene_tx=XM_023259985.1;refGene_gn=KIF1A;refGene_tx=
XM_023259986.1;refGene_gn=KIF1A;refGene_tx=XM_023259987.1;refGene_gn=K
IF1A;refGene_tx=XM_023259988.1
C1 221833551 221833629
refGene_gn=KIF1A;refGene_tx=XM_023259972.1;refGene_gn=KIF1A;refGene_tx
=XM_023259973.1;refGene_gn=KIF1A;refGene_tx=XM_023259974.1;refGene_gn=
KIF1A;refGene_tx=XM_023259975.1;refGene_gn=KIF1A;refGene_tx=XM_0232599
76.1;refGene_gn=KIF1A;refGene_tx=XM_023259977.1;refGene_gn=KIF1A;refGe
ne_tx=XM_023259978.1;refGene_gn=KIF1A;refGene_tx=XM_023259979.1;refGen
e_gn=KIF1A;refGene_tx=XM_023259980.1;refGene_gn=KIF1A;refGene_tx=XM_02
3259981.1;refGene_gn=KIF1A;refGene_tx=XM_023259982.1;refGene_gn=KIF1A;

refGene_tx=XM_023259983.1;refGene_gn=KIF1A;refGene_tx=XM_023259984.1;r
efGene_gn=KIF1A;refGene_tx=XM_023259985.1;refGene_gn=KIF1A;refGene_tx=
XM_023259986.1;refGene_gn=KIF1A;refGene_tx=XM_023259987.1;refGene_gn=K
IF1A;refGene_tx=XM_023259988.1
C1 221834092 221834293
refGene_gn=KIF1A;refGene_tx=XM_023259972.1;refGene_gn=KIF1A;refGene_tx
=XM_023259973.1;refGene_gn=KIF1A;refGene_tx=XM_023259974.1;refGene_gn=
KIF1A;refGene_tx=XM_023259975.1;refGene_gn=KIF1A;refGene_tx=XM_0232599
76.1;refGene_gn=KIF1A;refGene_tx=XM_023259977.1;refGene_gn=KIF1A;refGe
ne_tx=XM_023259978.1;refGene_gn=KIF1A;refGene_tx=XM_023259979.1;refGen
e_gn=KIF1A;refGene_tx=XM_023259980.1;refGene_gn=KIF1A;refGene_tx=XM_02
3259981.1;refGene_gn=KIF1A;refGene_tx=XM_023259982.1;refGene_gn=KIF1A;
refGene_tx=XM_023259983.1;refGene_gn=KIF1A;refGene_tx=XM_023259984.1;r
efGene_gn=KIF1A;refGene_tx=XM_023259985.1;refGene_gn=KIF1A;refGene_tx=
XM_023259986.1;refGene_gn=KIF1A;refGene_tx=XM_023259987.1;refGene_gn=K
IF1A;refGene_tx=XM_023259988.1
C1 221834561 221834707
refGene_gn=KIF1A;refGene_tx=XM_023259972.1;refGene_gn=KIF1A;refGene_tx
=XM_023259973.1;refGene_gn=KIF1A;refGene_tx=XM_023259974.1;refGene_gn=
KIF1A;refGene_tx=XM_023259975.1;refGene_gn=KIF1A;refGene_tx=XM_0232599
76.1;refGene_gn=KIF1A;refGene_tx=XM_023259977.1;refGene_gn=KIF1A;refGe
ne_tx=XM_023259978.1;refGene_gn=KIF1A;refGene_tx=XM_023259979.1;refGen
e_gn=KIF1A;refGene_tx=XM_023259980.1;refGene_gn=KIF1A;refGene_tx=XM_02
3259981.1;refGene_gn=KIF1A;refGene_tx=XM_023259982.1;refGene_gn=KIF1A;
refGene_tx=XM_023259983.1;refGene_gn=KIF1A;refGene_tx=XM_023259984.1;r
efGene_gn=KIF1A;refGene_tx=XM_023259985.1;refGene_gn=KIF1A;refGene_tx=
XM_023259986.1;refGene_gn=KIF1A;refGene_tx=XM_023259987.1;refGene_gn=K
IF1A;refGene_tx=XM_023259988.1
C1 221835026 221835088
refGene_gn=KIF1A;refGene_tx=XM_023259972.1;refGene_gn=KIF1A;refGene_tx
=XM_023259973.1;refGene_gn=KIF1A;refGene_tx=XM_023259974.1;refGene_gn=
KIF1A;refGene_tx=XM_023259975.1;refGene_gn=KIF1A;refGene_tx=XM_0232599
76.1;refGene_gn=KIF1A;refGene_tx=XM_023259977.1;refGene_gn=KIF1A;refGe
ne_tx=XM_023259978.1;refGene_gn=KIF1A;refGene_tx=XM_023259979.1;refGen
e_gn=KIF1A;refGene_tx=XM_023259980.1;refGene_gn=KIF1A;refGene_tx=XM_02
3259981.1;refGene_gn=KIF1A;refGene_tx=XM_023259982.1;refGene_gn=KIF1A;
refGene_tx=XM_023259983.1;refGene_gn=KIF1A;refGene_tx=XM_023259984.1;r
efGene_gn=KIF1A;refGene_tx=XM_023259985.1;refGene_gn=KIF1A;refGene_tx=
XM_023259986.1;refGene_gn=KIF1A;refGene_tx=XM_023259987.1;refGene_gn=K
IF1A;refGene_tx=XM_023259988.1
C1 221836068 221836202
refGene_gn=KIF1A;refGene_tx=XM_023259972.1;refGene_gn=KIF1A;refGene_tx
=XM_023259973.1;refGene_gn=KIF1A;refGene_tx=XM_023259974.1;refGene_gn=
KIF1A;refGene_tx=XM_023259975.1;refGene_gn=KIF1A;refGene_tx=XM_0232599
76.1;refGene_gn=KIF1A;refGene_tx=XM_023259977.1;refGene_gn=KIF1A;refGe
ne_tx=XM_023259978.1;refGene_gn=KIF1A;refGene_tx=XM_023259979.1;refGen
e_gn=KIF1A;refGene_tx=XM_023259980.1;refGene_gn=KIF1A;refGene_tx=XM_02
3259981.1;refGene_gn=KIF1A;refGene_tx=XM_023259982.1;refGene_gn=KIF1A;
refGene_tx=XM_023259983.1;refGene_gn=KIF1A;refGene_tx=XM_023259984.1;r
efGene_gn=KIF1A;refGene_tx=XM_023259985.1;refGene_gn=KIF1A;refGene_tx=

XM_023259986.1;refGene_gn=KIF1A;refGene_tx=XM_023259987.1;refGene_gn=K
IF1A;refGene_tx=XM_023259988.1
C1 221837170 221837285
refGene_gn=KIF1A;refGene_tx=XM_023259972.1;refGene_gn=KIF1A;refGene_tx
=XM_023259973.1;refGene_gn=KIF1A;refGene_tx=XM_023259974.1;refGene_gn=
KIF1A;refGene_tx=XM_023259975.1;refGene_gn=KIF1A;refGene_tx=XM_0232599
76.1;refGene_gn=KIF1A;refGene_tx=XM_023259977.1;refGene_gn=KIF1A;refGe
ne_tx=XM_023259978.1;refGene_gn=KIF1A;refGene_tx=XM_023259979.1;refGen
e_gn=KIF1A;refGene_tx=XM_023259980.1;refGene_gn=KIF1A;refGene_tx=XM_02
3259981.1;refGene_gn=KIF1A;refGene_tx=XM_023259982.1;refGene_gn=KIF1A;
refGene_tx=XM_023259983.1;refGene_gn=KIF1A;refGene_tx=XM_023259984.1;r
efGene_gn=KIF1A;refGene_tx=XM_023259985.1;refGene_gn=KIF1A;refGene_tx=
XM_023259986.1;refGene_gn=KIF1A;refGene_tx=XM_023259987.1;refGene_gn=K
IF1A;refGene_tx=XM_023259988.1
C1 221838516 221838540
refGene_gn=KIF1A;refGene_tx=XM_023259972.1;refGene_gn=KIF1A;refGene_tx
=XM_023259973.1;refGene_gn=KIF1A;refGene_tx=XM_023259975.1;refGene_gn=
KIF1A;refGene_tx=XM_023259976.1;refGene_gn=KIF1A;refGene_tx=XM_0232599
78.1;refGene_gn=KIF1A;refGene_tx=XM_023259979.1;refGene_gn=KIF1A;refGe
ne_tx=XM_023259981.1;refGene_gn=KIF1A;refGene_tx=XM_023259982.1;refGen
e_gn=KIF1A;refGene_tx=XM_023259986.1
C1 221845206 221845233
refGene_gn=KIF1A;refGene_tx=XM_023259972.1;refGene_gn=KIF1A;refGene_tx
=XM_023259973.1;refGene_gn=KIF1A;refGene_tx=XM_023259974.1;refGene_gn=
KIF1A;refGene_tx=XM_023259975.1;refGene_gn=KIF1A;refGene_tx=XM_0232599
78.1;refGene_gn=KIF1A;refGene_tx=XM_023259979.1;refGene_gn=KIF1A;refGe
ne_tx=XM_023259981.1;refGene_gn=KIF1A;refGene_tx=XM_023259982.1;refGen
e_gn=KIF1A;refGene_tx=XM_023259984.1
C1 221847627 221847733
refGene_gn=KIF1A;refGene_tx=XM_023259972.1;refGene_gn=KIF1A;refGene_tx
=XM_023259973.1;refGene_gn=KIF1A;refGene_tx=XM_023259974.1;refGene_gn=
KIF1A;refGene_tx=XM_023259975.1;refGene_gn=KIF1A;refGene_tx=XM_0232599
76.1;refGene_gn=KIF1A;refGene_tx=XM_023259977.1;refGene_gn=KIF1A;refGe
ne_tx=XM_023259978.1;refGene_gn=KIF1A;refGene_tx=XM_023259979.1;refGen
e_gn=KIF1A;refGene_tx=XM_023259980.1;refGene_gn=KIF1A;refGene_tx=XM_02
3259981.1;refGene_gn=KIF1A;refGene_tx=XM_023259982.1;refGene_gn=KIF1A;
refGene_tx=XM_023259983.1;refGene_gn=KIF1A;refGene_tx=XM_023259984.1;r
efGene_gn=KIF1A;refGene_tx=XM_023259985.1;refGene_gn=KIF1A;refGene_tx=
XM_023259986.1;refGene_gn=KIF1A;refGene_tx=XM_023259987.1;refGene_gn=K
IF1A;refGene_tx=XM_023259988.1
C1 221850047 221850132
refGene_gn=KIF1A;refGene_tx=XM_023259972.1;refGene_gn=KIF1A;refGene_tx
=XM_023259973.1;refGene_gn=KIF1A;refGene_tx=XM_023259974.1;refGene_gn=
KIF1A;refGene_tx=XM_023259975.1;refGene_gn=KIF1A;refGene_tx=XM_0232599
76.1;refGene_gn=KIF1A;refGene_tx=XM_023259977.1;refGene_gn=KIF1A;refGe
ne_tx=XM_023259978.1;refGene_gn=KIF1A;refGene_tx=XM_023259979.1;refGen
e_gn=KIF1A;refGene_tx=XM_023259980.1;refGene_gn=KIF1A;refGene_tx=XM_02
3259981.1;refGene_gn=KIF1A;refGene_tx=XM_023259982.1;refGene_gn=KIF1A;
refGene_tx=XM_023259983.1;refGene_gn=KIF1A;refGene_tx=XM_023259984.1;r
efGene_gn=KIF1A;refGene_tx=XM_023259985.1;refGene_gn=KIF1A;refGene_tx=

XM_023259986.1;refGene_gn=KIF1A;refGene_tx=XM_023259987.1;refGene_gn=KIF1A;refGene_tx=XM_023259988.1
C1	221850231	221850298
refGene_gn=KIF1A;refGene_tx=XM_023259972.1;refGene_gn=KIF1A;refGene_tx=XM_023259973.1;refGene_gn=KIF1A;refGene_tx=XM_023259974.1;refGene_gn=KIF1A;refGene_tx=XM_023259975.1;refGene_gn=KIF1A;refGene_tx=XM_023259976.1;refGene_gn=KIF1A;refGene_tx=XM_023259977.1;refGene_gn=KIF1A;refGene_tx=XM_023259978.1;refGene_gn=KIF1A;refGene_tx=XM_023259979.1;refGene_gn=KIF1A;refGene_tx=XM_023259980.1;refGene_gn=KIF1A;refGene_tx=XM_023259981.1;refGene_gn=KIF1A;refGene_tx=XM_023259982.1;refGene_gn=KIF1A;refGene_tx=XM_023259983.1;refGene_gn=KIF1A;refGene_tx=XM_023259984.1;refGene_gn=KIF1A;refGene_tx=XM_023259985.1;refGene_gn=KIF1A;refGene_tx=XM_023259986.1;refGene_gn=KIF1A;refGene_tx=XM_023259987.1;refGene_gn=KIF1A;refGene_tx=XM_023259988.1
C1	221850490	221850599
refGene_gn=KIF1A;refGene_tx=XM_023259972.1;refGene_gn=KIF1A;refGene_tx=XM_023259973.1;refGene_gn=KIF1A;refGene_tx=XM_023259974.1;refGene_gn=KIF1A;refGene_tx=XM_023259975.1;refGene_gn=KIF1A;refGene_tx=XM_023259976.1;refGene_gn=KIF1A;refGene_tx=XM_023259977.1;refGene_gn=KIF1A;refGene_tx=XM_023259978.1;refGene_gn=KIF1A;refGene_tx=XM_023259979.1;refGene_gn=KIF1A;refGene_tx=XM_023259980.1;refGene_gn=KIF1A;refGene_tx=XM_023259981.1;refGene_gn=KIF1A;refGene_tx=XM_023259982.1;refGene_gn=KIF1A;refGene_tx=XM_023259983.1;refGene_gn=KIF1A;refGene_tx=XM_023259984.1;refGene_gn=KIF1A;refGene_tx=XM_023259985.1;refGene_gn=KIF1A;refGene_tx=XM_023259986.1;refGene_gn=KIF1A;refGene_tx=XM_023259987.1;refGene_gn=KIF1A;refGene_tx=XM_023259988.1
C1	221850758	221850814
refGene_gn=KIF1A;refGene_tx=XM_023259972.1;refGene_gn=KIF1A;refGene_tx=XM_023259973.1;refGene_gn=KIF1A;refGene_tx=XM_023259974.1;refGene_gn=KIF1A;refGene_tx=XM_023259975.1;refGene_gn=KIF1A;refGene_tx=XM_023259976.1;refGene_gn=KIF1A;refGene_tx=XM_023259977.1;refGene_gn=KIF1A;refGene_tx=XM_023259978.1;refGene_gn=KIF1A;refGene_tx=XM_023259979.1;refGene_gn=KIF1A;refGene_tx=XM_023259980.1;refGene_gn=KIF1A;refGene_tx=XM_023259981.1;refGene_gn=KIF1A;refGene_tx=XM_023259982.1;refGene_gn=KIF1A;refGene_tx=XM_023259983.1;refGene_gn=KIF1A;refGene_tx=XM_023259984.1;refGene_gn=KIF1A;refGene_tx=XM_023259985.1;refGene_gn=KIF1A;refGene_tx=XM_023259986.1;refGene_gn=KIF1A;refGene_tx=XM_023259987.1;refGene_gn=KIF1A;refGene_tx=XM_023259988.1
C1	221851845	221851964
refGene_gn=KIF1A;refGene_tx=XM_023259972.1;refGene_gn=KIF1A;refGene_tx=XM_023259973.1;refGene_gn=KIF1A;refGene_tx=XM_023259974.1;refGene_gn=KIF1A;refGene_tx=XM_023259975.1;refGene_gn=KIF1A;refGene_tx=XM_023259976.1;refGene_gn=KIF1A;refGene_tx=XM_023259977.1;refGene_gn=KIF1A;refGene_tx=XM_023259978.1;refGene_gn=KIF1A;refGene_tx=XM_023259979.1;refGene_gn=KIF1A;refGene_tx=XM_023259980.1;refGene_gn=KIF1A;refGene_tx=XM_023259981.1;refGene_gn=KIF1A;refGene_tx=XM_023259982.1;refGene_gn=KIF1A;refGene_tx=XM_023259983.1;refGene_gn=KIF1A;refGene_tx=XM_023259984.1;refGene_gn=KIF1A;refGene_tx=XM_023259985.1;refGene_gn=KIF1A;refGene_tx=XM_023259986.1;refGene_gn=KIF1A;refGene_tx=XM_023259987.1;refGene_gn=KIF1A;refGene_tx=XM_023259988.1

C1 221853349 221853440
refGene_gn=KIF1A;refGene_tx=XM_023259972.1;refGene_gn=KIF1A;refGene_tx=XM_023259973.1;refGene_gn=KIF1A;refGene_tx=XM_023259974.1;refGene_gn=KIF1A;refGene_tx=XM_023259975.1;refGene_gn=KIF1A;refGene_tx=XM_023259976.1;refGene_gn=KIF1A;refGene_tx=XM_023259977.1;refGene_gn=KIF1A;refGene_tx=XM_023259978.1;refGene_gn=KIF1A;refGene_tx=XM_023259979.1;refGene_gn=KIF1A;refGene_tx=XM_023259980.1;refGene_gn=KIF1A;refGene_tx=XM_023259981.1;refGene_gn=KIF1A;refGene_tx=XM_023259982.1;refGene_gn=KIF1A;refGene_tx=XM_023259983.1;refGene_gn=KIF1A;refGene_tx=XM_023259984.1;refGene_gn=KIF1A;refGene_tx=XM_023259985.1;refGene_gn=KIF1A;refGene_tx=XM_023259986.1;refGene_gn=KIF1A;refGene_tx=XM_023259987.1;refGene_gn=KIF1A;refGene_tx=XM_023259988.1
C1 221853572 221853744
refGene_gn=KIF1A;refGene_tx=XM_023259972.1;refGene_gn=KIF1A;refGene_tx=XM_023259973.1;refGene_gn=KIF1A;refGene_tx=XM_023259974.1;refGene_gn=KIF1A;refGene_tx=XM_023259975.1;refGene_gn=KIF1A;refGene_tx=XM_023259976.1;refGene_gn=KIF1A;refGene_tx=XM_023259977.1;refGene_gn=KIF1A;refGene_tx=XM_023259978.1;refGene_gn=KIF1A;refGene_tx=XM_023259979.1;refGene_gn=KIF1A;refGene_tx=XM_023259980.1;refGene_gn=KIF1A;refGene_tx=XM_023259981.1;refGene_gn=KIF1A;refGene_tx=XM_023259982.1;refGene_gn=KIF1A;refGene_tx=XM_023259983.1;refGene_gn=KIF1A;refGene_tx=XM_023259984.1;refGene_gn=KIF1A;refGene_tx=XM_023259985.1;refGene_gn=KIF1A;refGene_tx=XM_023259986.1;refGene_gn=KIF1A;refGene_tx=XM_023259987.1;refGene_gn=KIF1A;refGene_tx=XM_023259988.1
C1 221853920 221854059
refGene_gn=KIF1A;refGene_tx=XM_023259972.1;refGene_gn=KIF1A;refGene_tx=XM_023259973.1;refGene_gn=KIF1A;refGene_tx=XM_023259974.1;refGene_gn=KIF1A;refGene_tx=XM_023259975.1;refGene_gn=KIF1A;refGene_tx=XM_023259976.1;refGene_gn=KIF1A;refGene_tx=XM_023259977.1;refGene_gn=KIF1A;refGene_tx=XM_023259978.1;refGene_gn=KIF1A;refGene_tx=XM_023259979.1;refGene_gn=KIF1A;refGene_tx=XM_023259980.1;refGene_gn=KIF1A;refGene_tx=XM_023259981.1;refGene_gn=KIF1A;refGene_tx=XM_023259982.1;refGene_gn=KIF1A;refGene_tx=XM_023259983.1;refGene_gn=KIF1A;refGene_tx=XM_023259984.1;refGene_gn=KIF1A;refGene_tx=XM_023259985.1;refGene_gn=KIF1A;refGene_tx=XM_023259986.1;refGene_gn=KIF1A;refGene_tx=XM_023259987.1;refGene_gn=KIF1A;refGene_tx=XM_023259988.1
C1 221854873 221854959
refGene_gn=KIF1A;refGene_tx=XM_023259972.1;refGene_gn=KIF1A;refGene_tx=XM_023259973.1;refGene_gn=KIF1A;refGene_tx=XM_023259974.1;refGene_gn=KIF1A;refGene_tx=XM_023259975.1;refGene_gn=KIF1A;refGene_tx=XM_023259976.1;refGene_gn=KIF1A;refGene_tx=XM_023259977.1;refGene_gn=KIF1A;refGene_tx=XM_023259978.1;refGene_gn=KIF1A;refGene_tx=XM_023259979.1;refGene_gn=KIF1A;refGene_tx=XM_023259980.1;refGene_gn=KIF1A;refGene_tx=XM_023259981.1;refGene_gn=KIF1A;refGene_tx=XM_023259982.1;refGene_gn=KIF1A;refGene_tx=XM_023259983.1;refGene_gn=KIF1A;refGene_tx=XM_023259984.1;refGene_gn=KIF1A;refGene_tx=XM_023259985.1;refGene_gn=KIF1A;refGene_tx=XM_023259986.1;refGene_gn=KIF1A;refGene_tx=XM_023259987.1;refGene_gn=KIF1A;refGene_tx=XM_023259988.1
C1 221857063 221857182
refGene_gn=KIF1A;refGene_tx=XM_023259972.1;refGene_gn=KIF1A;refGene_tx

=XM_023259973.1;refGene_gn=KIF1A;refGene_tx=XM_023259974.1;refGene_gn=KIF1A;refGene_tx=XM_023259975.1;refGene_gn=KIF1A;refGene_tx=XM_023259976.1;refGene_gn=KIF1A;refGene_tx=XM_023259977.1;refGene_gn=KIF1A;refGene_tx=XM_023259978.1;refGene_gn=KIF1A;refGene_tx=XM_023259979.1;refGene_gn=KIF1A;refGene_tx=XM_023259980.1;refGene_gn=KIF1A;refGene_tx=XM_023259981.1;refGene_gn=KIF1A;refGene_tx=XM_023259982.1;refGene_gn=KIF1A;refGene_tx=XM_023259983.1;refGene_gn=KIF1A;refGene_tx=XM_023259984.1;refGene_gn=KIF1A;refGene_tx=XM_023259985.1;refGene_gn=KIF1A;refGene_tx=XM_023259986.1;refGene_gn=KIF1A;refGene_tx=XM_023259987.1;refGene_gn=KIF1A;refGene_tx=XM_023259988.1

C1 221863903 221864176

refGene_gn=KIF1A;refGene_tx=XM_023259972.1;refGene_gn=KIF1A;refGene_tx=XM_023259973.1;refGene_gn=KIF1A;refGene_tx=XM_023259974.1;refGene_gn=KIF1A;refGene_tx=XM_023259975.1;refGene_gn=KIF1A;refGene_tx=XM_023259976.1;refGene_gn=KIF1A;refGene_tx=XM_023259977.1;refGene_gn=KIF1A;refGene_tx=XM_023259978.1;refGene_gn=KIF1A;refGene_tx=XM_023259979.1;refGene_gn=KIF1A;refGene_tx=XM_023259980.1;refGene_gn=KIF1A;refGene_tx=XM_023259981.1;refGene_gn=KIF1A;refGene_tx=XM_023259988.1

C1 221864819 221864957

refGene_gn=KIF1A;refGene_tx=XM_023259972.1;refGene_gn=KIF1A;refGene_tx=XM_023259973.1;refGene_gn=KIF1A;refGene_tx=XM_023259974.1;refGene_gn=KIF1A;refGene_tx=XM_023259975.1;refGene_gn=KIF1A;refGene_tx=XM_023259976.1;refGene_gn=KIF1A;refGene_tx=XM_023259977.1;refGene_gn=KIF1A;refGene_tx=XM_023259978.1;refGene_gn=KIF1A;refGene_tx=XM_023259979.1;refGene_gn=KIF1A;refGene_tx=XM_023259980.1;refGene_gn=KIF1A;refGene_tx=XM_023259981.1;refGene_gn=KIF1A;refGene_tx=XM_023259982.1;refGene_gn=KIF1A;refGene_tx=XM_023259983.1;refGene_gn=KIF1A;refGene_tx=XM_023259984.1;refGene_gn=KIF1A;refGene_tx=XM_023259985.1;refGene_gn=KIF1A;refGene_tx=XM_023259986.1;refGene_gn=KIF1A;refGene_tx=XM_023259987.1;refGene_gn=KIF1A;refGene_tx=XM_023259988.1

C1 221866600 221866779

refGene_gn=KIF1A;refGene_tx=XM_023259972.1;refGene_gn=KIF1A;refGene_tx=XM_023259973.1;refGene_gn=KIF1A;refGene_tx=XM_023259974.1;refGene_gn=KIF1A;refGene_tx=XM_023259975.1;refGene_gn=KIF1A;refGene_tx=XM_023259976.1;refGene_gn=KIF1A;refGene_tx=XM_023259977.1;refGene_gn=KIF1A;refGene_tx=XM_023259978.1;refGene_gn=KIF1A;refGene_tx=XM_023259979.1;refGene_gn=KIF1A;refGene_tx=XM_023259980.1;refGene_gn=KIF1A;refGene_tx=XM_023259981.1;refGene_gn=KIF1A;refGene_tx=XM_023259982.1;refGene_gn=KIF1A;refGene_tx=XM_023259983.1;refGene_gn=KIF1A;refGene_tx=XM_023259984.1;refGene_gn=KIF1A;refGene_tx=XM_023259985.1;refGene_gn=KIF1A;refGene_tx=XM_023259986.1;refGene_gn=KIF1A;refGene_tx=XM_023259987.1;refGene_gn=KIF1A;refGene_tx=XM_023259988.1

C1 221867028 221867177

refGene_gn=KIF1A;refGene_tx=XM_023259972.1;refGene_gn=KIF1A;refGene_tx=XM_023259973.1;refGene_gn=KIF1A;refGene_tx=XM_023259974.1;refGene_gn=KIF1A;refGene_tx=XM_023259975.1;refGene_gn=KIF1A;refGene_tx=XM_023259976.1;refGene_gn=KIF1A;refGene_tx=XM_023259977.1;refGene_gn=KIF1A;refGene_tx=XM_023259978.1;refGene_gn=KIF1A;refGene_tx=XM_023259979.1;refGene_gn=KIF1A;refGene_tx=XM_023259980.1;refGene_gn=KIF1A;refGene_tx=XM_023259981.1;refGene_gn=KIF1A;refGene_tx=XM_023259982.1;refGene_gn=KIF1A;

refGene_tx=XM_023259983.1;refGene_gn=KIF1A;refGene_tx=XM_023259984.1;r
efGene_gn=KIF1A;refGene_tx=XM_023259985.1;refGene_gn=KIF1A;refGene_tx=
XM_023259986.1;refGene_gn=KIF1A;refGene_tx=XM_023259987.1;refGene_gn=K
IF1A;refGene_tx=XM_023259988.1
C1 221868146 221868240
refGene_gn=KIF1A;refGene_tx=XM_023259972.1;refGene_gn=KIF1A;refGene_tx
=XM_023259973.1;refGene_gn=KIF1A;refGene_tx=XM_023259974.1;refGene_gn=
KIF1A;refGene_tx=XM_023259975.1;refGene_gn=KIF1A;refGene_tx=XM_0232599
76.1;refGene_gn=KIF1A;refGene_tx=XM_023259977.1;refGene_gn=KIF1A;refGe
ne_tx=XM_023259978.1;refGene_gn=KIF1A;refGene_tx=XM_023259979.1;refGen
e_gn=KIF1A;refGene_tx=XM_023259980.1;refGene_gn=KIF1A;refGene_tx=XM_02
3259981.1;refGene_gn=KIF1A;refGene_tx=XM_023259982.1;refGene_gn=KIF1A;
refGene_tx=XM_023259983.1;refGene_gn=KIF1A;refGene_tx=XM_023259984.1;r
efGene_gn=KIF1A;refGene_tx=XM_023259985.1;refGene_gn=KIF1A;refGene_tx=
XM_023259986.1;refGene_gn=KIF1A;refGene_tx=XM_023259987.1;refGene_gn=K
IF1A;refGene_tx=XM_023259988.1
C1 221868388 221868461
refGene_gn=KIF1A;refGene_tx=XM_023259972.1;refGene_gn=KIF1A;refGene_tx
=XM_023259973.1;refGene_gn=KIF1A;refGene_tx=XM_023259974.1;refGene_gn=
KIF1A;refGene_tx=XM_023259975.1;refGene_gn=KIF1A;refGene_tx=XM_0232599
76.1;refGene_gn=KIF1A;refGene_tx=XM_023259977.1;refGene_gn=KIF1A;refGe
ne_tx=XM_023259978.1;refGene_gn=KIF1A;refGene_tx=XM_023259979.1;refGen
e_gn=KIF1A;refGene_tx=XM_023259980.1;refGene_gn=KIF1A;refGene_tx=XM_02
3259981.1;refGene_gn=KIF1A;refGene_tx=XM_023259982.1;refGene_gn=KIF1A;
refGene_tx=XM_023259983.1;refGene_gn=KIF1A;refGene_tx=XM_023259984.1;r
efGene_gn=KIF1A;refGene_tx=XM_023259985.1;refGene_gn=KIF1A;refGene_tx=
XM_023259986.1;refGene_gn=KIF1A;refGene_tx=XM_023259987.1;refGene_gn=K
IF1A;refGene_tx=XM_023259988.1
C1 221868584 221868765
refGene_gn=KIF1A;refGene_tx=XM_023259972.1;refGene_gn=KIF1A;refGene_tx
=XM_023259973.1;refGene_gn=KIF1A;refGene_tx=XM_023259974.1;refGene_gn=
KIF1A;refGene_tx=XM_023259975.1;refGene_gn=KIF1A;refGene_tx=XM_0232599
76.1;refGene_gn=KIF1A;refGene_tx=XM_023259977.1;refGene_gn=KIF1A;refGe
ne_tx=XM_023259978.1;refGene_gn=KIF1A;refGene_tx=XM_023259979.1;refGen
e_gn=KIF1A;refGene_tx=XM_023259980.1;refGene_gn=KIF1A;refGene_tx=XM_02
3259981.1;refGene_gn=KIF1A;refGene_tx=XM_023259982.1;refGene_gn=KIF1A;
refGene_tx=XM_023259983.1;refGene_gn=KIF1A;refGene_tx=XM_023259984.1;r
efGene_gn=KIF1A;refGene_tx=XM_023259985.1;refGene_gn=KIF1A;refGene_tx=
XM_023259986.1;refGene_gn=KIF1A;refGene_tx=XM_023259987.1;refGene_gn=K
IF1A;refGene_tx=XM_023259988.1
C1 221869653 221869737
refGene_gn=KIF1A;refGene_tx=XM_023259972.1;refGene_gn=KIF1A;refGene_tx
=XM_023259973.1;refGene_gn=KIF1A;refGene_tx=XM_023259974.1;refGene_gn=
KIF1A;refGene_tx=XM_023259975.1;refGene_gn=KIF1A;refGene_tx=XM_0232599
76.1;refGene_gn=KIF1A;refGene_tx=XM_023259977.1;refGene_gn=KIF1A;refGe
ne_tx=XM_023259978.1;refGene_gn=KIF1A;refGene_tx=XM_023259979.1;refGen
e_gn=KIF1A;refGene_tx=XM_023259980.1;refGene_gn=KIF1A;refGene_tx=XM_02
3259981.1;refGene_gn=KIF1A;refGene_tx=XM_023259982.1;refGene_gn=KIF1A;
refGene_tx=XM_023259983.1;refGene_gn=KIF1A;refGene_tx=XM_023259984.1;r
efGene_gn=KIF1A;refGene_tx=XM_023259985.1;refGene_gn=KIF1A;refGene_tx=

XM_023259986.1;refGene_gn=KIF1A;refGene_tx=XM_023259987.1;refGene_gn=K
IF1A;refGene_tx=XM_023259988.1
C1	221870472	221870579
refGene_gn=KIF1A;refGene_tx=XM_023259972.1;refGene_gn=KIF1A;refGene_tx
=XM_023259973.1;refGene_gn=KIF1A;refGene_tx=XM_023259974.1;refGene_gn=
KIF1A;refGene_tx=XM_023259975.1;refGene_gn=KIF1A;refGene_tx=XM_0232599
76.1;refGene_gn=KIF1A;refGene_tx=XM_023259977.1;refGene_gn=KIF1A;refGe
ne_tx=XM_023259978.1;refGene_gn=KIF1A;refGene_tx=XM_023259979.1;refGen
e_gn=KIF1A;refGene_tx=XM_023259980.1;refGene_gn=KIF1A;refGene_tx=XM_02
3259981.1;refGene_gn=KIF1A;refGene_tx=XM_023259982.1;refGene_gn=KIF1A;
refGene_tx=XM_023259983.1;refGene_gn=KIF1A;refGene_tx=XM_023259984.1;r
efGene_gn=KIF1A;refGene_tx=XM_023259985.1;refGene_gn=KIF1A;refGene_tx=
XM_023259986.1;refGene_gn=KIF1A;refGene_tx=XM_023259987.1;refGene_gn=K
IF1A;refGene_tx=XM_023259988.1
C1	221870824	221870904
refGene_gn=KIF1A;refGene_tx=XM_023259972.1;refGene_gn=KIF1A;refGene_tx
=XM_023259973.1;refGene_gn=KIF1A;refGene_tx=XM_023259974.1;refGene_gn=
KIF1A;refGene_tx=XM_023259975.1;refGene_gn=KIF1A;refGene_tx=XM_0232599
76.1;refGene_gn=KIF1A;refGene_tx=XM_023259977.1;refGene_gn=KIF1A;refGe
ne_tx=XM_023259978.1;refGene_gn=KIF1A;refGene_tx=XM_023259979.1;refGen
e_gn=KIF1A;refGene_tx=XM_023259980.1;refGene_gn=KIF1A;refGene_tx=XM_02
3259981.1;refGene_gn=KIF1A;refGene_tx=XM_023259982.1;refGene_gn=KIF1A;
refGene_tx=XM_023259983.1;refGene_gn=KIF1A;refGene_tx=XM_023259984.1;r
efGene_gn=KIF1A;refGene_tx=XM_023259985.1;refGene_gn=KIF1A;refGene_tx=
XM_023259986.1;refGene_gn=KIF1A;refGene_tx=XM_023259987.1;refGene_gn=K
IF1A;refGene_tx=XM_023259988.1
C1	221872839	221872915
refGene_gn=KIF1A;refGene_tx=XM_023259972.1;refGene_gn=KIF1A;refGene_tx
=XM_023259973.1;refGene_gn=KIF1A;refGene_tx=XM_023259974.1;refGene_gn=
KIF1A;refGene_tx=XM_023259975.1;refGene_gn=KIF1A;refGene_tx=XM_0232599
76.1;refGene_gn=KIF1A;refGene_tx=XM_023259977.1;refGene_gn=KIF1A;refGe
ne_tx=XM_023259978.1;refGene_gn=KIF1A;refGene_tx=XM_023259979.1;refGen
e_gn=KIF1A;refGene_tx=XM_023259980.1;refGene_gn=KIF1A;refGene_tx=XM_02
3259981.1;refGene_gn=KIF1A;refGene_tx=XM_023259982.1;refGene_gn=KIF1A;
refGene_tx=XM_023259983.1;refGene_gn=KIF1A;refGene_tx=XM_023259984.1;r
efGene_gn=KIF1A;refGene_tx=XM_023259985.1;refGene_gn=KIF1A;refGene_tx=
XM_023259986.1;refGene_gn=KIF1A;refGene_tx=XM_023259987.1;refGene_gn=K
IF1A;refGene_tx=XM_023259988.1
C1	221873210	221873290
refGene_gn=KIF1A;refGene_tx=XM_023259972.1;refGene_gn=KIF1A;refGene_tx
=XM_023259973.1;refGene_gn=KIF1A;refGene_tx=XM_023259974.1;refGene_gn=
KIF1A;refGene_tx=XM_023259975.1;refGene_gn=KIF1A;refGene_tx=XM_0232599
76.1;refGene_gn=KIF1A;refGene_tx=XM_023259977.1;refGene_gn=KIF1A;refGe
ne_tx=XM_023259978.1;refGene_gn=KIF1A;refGene_tx=XM_023259979.1;refGen
e_gn=KIF1A;refGene_tx=XM_023259980.1;refGene_gn=KIF1A;refGene_tx=XM_02
3259981.1;refGene_gn=KIF1A;refGene_tx=XM_023259982.1;refGene_gn=KIF1A;
refGene_tx=XM_023259983.1;refGene_gn=KIF1A;refGene_tx=XM_023259984.1;r
efGene_gn=KIF1A;refGene_tx=XM_023259985.1;refGene_gn=KIF1A;refGene_tx=
XM_023259986.1;refGene_gn=KIF1A;refGene_tx=XM_023259987.1;refGene_gn=K
IF1A;refGene_tx=XM_023259988.1

C1	221875655	221875867
refGene_gn=KIF1A;refGene_tx=XM_023259972.1;refGene_gn=KIF1A;refGene_tx=XM_023259973.1;refGene_gn=KIF1A;refGene_tx=XM_023259974.1;refGene_gn=KIF1A;refGene_tx=XM_023259975.1;refGene_gn=KIF1A;refGene_tx=XM_023259976.1;refGene_gn=KIF1A;refGene_tx=XM_023259977.1;refGene_gn=KIF1A;refGene_tx=XM_023259978.1;refGene_gn=KIF1A;refGene_tx=XM_023259979.1;refGene_gn=KIF1A;refGene_tx=XM_023259980.1;refGene_gn=KIF1A;refGene_tx=XM_023259981.1;refGene_gn=KIF1A;refGene_tx=XM_023259982.1;refGene_gn=KIF1A;refGene_tx=XM_023259983.1;refGene_gn=KIF1A;refGene_tx=XM_023259984.1;refGene_gn=KIF1A;refGene_tx=XM_023259985.1;refGene_gn=KIF1A;refGene_tx=XM_023259986.1;refGene_gn=KIF1A;refGene_tx=XM_023259987.1;refGene_gn=KIF1A;refGene_tx=XM_023259988.1
C1	221877200	221877227
refGene_gn=KIF1A;refGene_tx=XM_023259972.1;refGene_gn=KIF1A;refGene_tx=XM_023259973.1;refGene_gn=KIF1A;refGene_tx=XM_023259974.1;refGene_gn=KIF1A;refGene_tx=XM_023259975.1;refGene_gn=KIF1A;refGene_tx=XM_023259976.1;refGene_gn=KIF1A;refGene_tx=XM_023259977.1;refGene_gn=KIF1A;refGene_tx=XM_023259978.1;refGene_gn=KIF1A;refGene_tx=XM_023259980.1;refGene_gn=KIF1A;refGene_tx=XM_023259982.1;refGene_gn=KIF1A;refGene_tx=XM_023259983.1;refGene_gn=KIF1A;refGene_tx=XM_023259984.1;refGene_gn=KIF1A;refGene_tx=XM_023259985.1
C1	221877672	221877815
refGene_gn=KIF1A;refGene_tx=XM_023259972.1;refGene_gn=KIF1A;refGene_tx=XM_023259973.1;refGene_gn=KIF1A;refGene_tx=XM_023259974.1;refGene_gn=KIF1A;refGene_tx=XM_023259975.1;refGene_gn=KIF1A;refGene_tx=XM_023259976.1;refGene_gn=KIF1A;refGene_tx=XM_023259977.1;refGene_gn=KIF1A;refGene_tx=XM_023259978.1;refGene_gn=KIF1A;refGene_tx=XM_023259979.1;refGene_gn=KIF1A;refGene_tx=XM_023259980.1;refGene_gn=KIF1A;refGene_tx=XM_023259981.1;refGene_gn=KIF1A;refGene_tx=XM_023259982.1;refGene_gn=KIF1A;refGene_tx=XM_023259983.1;refGene_gn=KIF1A;refGene_tx=XM_023259984.1;refGene_gn=KIF1A;refGene_tx=XM_023259985.1;refGene_gn=KIF1A;refGene_tx=XM_023259986.1;refGene_gn=KIF1A;refGene_tx=XM_023259987.1;refGene_gn=KIF1A;refGene_tx=XM_023259988.1
C1	221878182	221878261
refGene_gn=KIF1A;refGene_tx=XM_023259972.1;refGene_gn=KIF1A;refGene_tx=XM_023259973.1;refGene_gn=KIF1A;refGene_tx=XM_023259974.1;refGene_gn=KIF1A;refGene_tx=XM_023259975.1;refGene_gn=KIF1A;refGene_tx=XM_023259976.1;refGene_gn=KIF1A;refGene_tx=XM_023259977.1;refGene_gn=KIF1A;refGene_tx=XM_023259978.1;refGene_gn=KIF1A;refGene_tx=XM_023259979.1;refGene_gn=KIF1A;refGene_tx=XM_023259980.1;refGene_gn=KIF1A;refGene_tx=XM_023259981.1;refGene_gn=KIF1A;refGene_tx=XM_023259982.1;refGene_gn=KIF1A;refGene_tx=XM_023259983.1;refGene_gn=KIF1A;refGene_tx=XM_023259984.1;refGene_gn=KIF1A;refGene_tx=XM_023259985.1;refGene_gn=KIF1A;refGene_tx=XM_023259986.1;refGene_gn=KIF1A;refGene_tx=XM_023259987.1;refGene_gn=KIF1A;refGene_tx=XM_023259988.1
C1	221879763	221879839
refGene_gn=KIF1A;refGene_tx=XM_023259972.1;refGene_gn=KIF1A;refGene_tx=XM_023259973.1;refGene_gn=KIF1A;refGene_tx=XM_023259974.1;refGene_gn=KIF1A;refGene_tx=XM_023259975.1;refGene_gn=KIF1A;refGene_tx=XM_023259976.1;refGene_gn=KIF1A;refGene_tx=XM_023259977.1;refGene_gn=KIF1A;refGe

ne_tx=XM_023259978.1;refGene_gn=KIF1A;refGene_tx=XM_023259979.1;refGen
e_gn=KIF1A;refGene_tx=XM_023259980.1;refGene_gn=KIF1A;refGene_tx=XM_02
3259981.1;refGene_gn=KIF1A;refGene_tx=XM_023259982.1;refGene_gn=KIF1A;
refGene_tx=XM_023259983.1;refGene_gn=KIF1A;refGene_tx=XM_023259984.1;r
efGene_gn=KIF1A;refGene_tx=XM_023259985.1;refGene_gn=KIF1A;refGene_tx=
XM_023259986.1;refGene_gn=KIF1A;refGene_tx=XM_023259987.1;refGene_gn=K
IF1A;refGene_tx=XM_023259988.1
C1 221890686 221890704
refGene_gn=KIF1A;refGene_tx=XM_023259972.1;refGene_gn=KIF1A;refGene_tx
=XM_023259973.1;refGene_gn=KIF1A;refGene_tx=XM_023259974.1;refGene_gn=
KIF1A;refGene_tx=XM_023259975.1;refGene_gn=KIF1A;refGene_tx=XM_0232599
76.1;refGene_gn=KIF1A;refGene_tx=XM_023259977.1;refGene_gn=KIF1A;refGe
ne_tx=XM_023259978.1;refGene_gn=KIF1A;refGene_tx=XM_023259979.1;refGen
e_gn=KIF1A;refGene_tx=XM_023259980.1;refGene_gn=KIF1A;refGene_tx=XM_02
3259981.1;refGene_gn=KIF1A;refGene_tx=XM_023259982.1;refGene_gn=KIF1A;
refGene_tx=XM_023259983.1;refGene_gn=KIF1A;refGene_tx=XM_023259984.1;r
efGene_gn=KIF1A;refGene_tx=XM_023259985.1;refGene_gn=KIF1A;refGene_tx=
XM_023259986.1;refGene_gn=KIF1A;refGene_tx=XM_023259987.1;refGene_gn=K
IF1A;refGene_tx=XM_023259988.1
C1 221891226 221891292
refGene_gn=KIF1A;refGene_tx=XM_023259972.1;refGene_gn=KIF1A;refGene_tx
=XM_023259973.1;refGene_gn=KIF1A;refGene_tx=XM_023259974.1;refGene_gn=
KIF1A;refGene_tx=XM_023259975.1;refGene_gn=KIF1A;refGene_tx=XM_0232599
76.1;refGene_gn=KIF1A;refGene_tx=XM_023259977.1;refGene_gn=KIF1A;refGe
ne_tx=XM_023259978.1;refGene_gn=KIF1A;refGene_tx=XM_023259979.1;refGen
e_gn=KIF1A;refGene_tx=XM_023259980.1;refGene_gn=KIF1A;refGene_tx=XM_02
3259981.1;refGene_gn=KIF1A;refGene_tx=XM_023259982.1;refGene_gn=KIF1A;
refGene_tx=XM_023259983.1;refGene_gn=KIF1A;refGene_tx=XM_023259984.1;r
efGene_gn=KIF1A;refGene_tx=XM_023259985.1;refGene_gn=KIF1A;refGene_tx=
XM_023259986.1;refGene_gn=KIF1A;refGene_tx=XM_023259987.1;refGene_gn=K
IF1A;refGene_tx=XM_023259988.1
C1 221891994 221892072
refGene_gn=KIF1A;refGene_tx=XM_023259972.1;refGene_gn=KIF1A;refGene_tx
=XM_023259973.1;refGene_gn=KIF1A;refGene_tx=XM_023259974.1;refGene_gn=
KIF1A;refGene_tx=XM_023259975.1;refGene_gn=KIF1A;refGene_tx=XM_0232599
76.1;refGene_gn=KIF1A;refGene_tx=XM_023259977.1;refGene_gn=KIF1A;refGe
ne_tx=XM_023259978.1;refGene_gn=KIF1A;refGene_tx=XM_023259979.1;refGen
e_gn=KIF1A;refGene_tx=XM_023259980.1;refGene_gn=KIF1A;refGene_tx=XM_02
3259981.1;refGene_gn=KIF1A;refGene_tx=XM_023259982.1;refGene_gn=KIF1A;
refGene_tx=XM_023259983.1;refGene_gn=KIF1A;refGene_tx=XM_023259984.1;r
efGene_gn=KIF1A;refGene_tx=XM_023259985.1;refGene_gn=KIF1A;refGene_tx=
XM_023259986.1;refGene_gn=KIF1A;refGene_tx=XM_023259987.1;refGene_gn=K
IF1A;refGene_tx=XM_023259988.1
C1 221893318 221893430
refGene_gn=KIF1A;refGene_tx=XM_023259972.1;refGene_gn=KIF1A;refGene_tx
=XM_023259973.1;refGene_gn=KIF1A;refGene_tx=XM_023259974.1;refGene_gn=
KIF1A;refGene_tx=XM_023259975.1;refGene_gn=KIF1A;refGene_tx=XM_0232599
76.1;refGene_gn=KIF1A;refGene_tx=XM_023259977.1;refGene_gn=KIF1A;refGe
ne_tx=XM_023259978.1;refGene_gn=KIF1A;refGene_tx=XM_023259979.1;refGen
e_gn=KIF1A;refGene_tx=XM_023259980.1;refGene_gn=KIF1A;refGene_tx=XM_02

3259981.1;refGene_gn=KIF1A;refGene_tx=XM_023259982.1;refGene_gn=KIF1A;
refGene_tx=XM_023259983.1;refGene_gn=KIF1A;refGene_tx=XM_023259984.1;r
efGene_gn=KIF1A;refGene_tx=XM_023259985.1;refGene_gn=KIF1A;refGene_tx=
XM_023259986.1;refGene_gn=KIF1A;refGene_tx=XM_023259987.1;refGene_gn=K
IF1A;refGene_tx=XM_023259988.1
C1 221894473 221894652
refGene_gn=KIF1A;refGene_tx=XM_023259972.1;refGene_gn=KIF1A;refGene_tx
=XM_023259973.1;refGene_gn=KIF1A;refGene_tx=XM_023259974.1;refGene_gn=
KIF1A;refGene_tx=XM_023259975.1;refGene_gn=KIF1A;refGene_tx=XM_0232599
76.1;refGene_gn=KIF1A;refGene_tx=XM_023259977.1;refGene_gn=KIF1A;refGe
ne_tx=XM_023259978.1;refGene_gn=KIF1A;refGene_tx=XM_023259979.1;refGen
e_gn=KIF1A;refGene_tx=XM_023259980.1;refGene_gn=KIF1A;refGene_tx=XM_02
3259981.1;refGene_gn=KIF1A;refGene_tx=XM_023259982.1;refGene_gn=KIF1A;
refGene_tx=XM_023259983.1;refGene_gn=KIF1A;refGene_tx=XM_023259984.1;r
efGene_gn=KIF1A;refGene_tx=XM_023259985.1;refGene_gn=KIF1A;refGene_tx=
XM_023259986.1;refGene_gn=KIF1A;refGene_tx=XM_023259987.1;refGene_gn=K
IF1A;refGene_tx=XM_023259988.1
C1 221895131 221895197
refGene_gn=KIF1A;refGene_tx=XM_023259972.1;refGene_gn=KIF1A;refGene_tx
=XM_023259973.1;refGene_gn=KIF1A;refGene_tx=XM_023259974.1;refGene_gn=
KIF1A;refGene_tx=XM_023259975.1;refGene_gn=KIF1A;refGene_tx=XM_0232599
76.1;refGene_gn=KIF1A;refGene_tx=XM_023259977.1;refGene_gn=KIF1A;refGe
ne_tx=XM_023259978.1;refGene_gn=KIF1A;refGene_tx=XM_023259979.1;refGen
e_gn=KIF1A;refGene_tx=XM_023259980.1;refGene_gn=KIF1A;refGene_tx=XM_02
3259981.1;refGene_gn=KIF1A;refGene_tx=XM_023259982.1;refGene_gn=KIF1A;
refGene_tx=XM_023259983.1;refGene_gn=KIF1A;refGene_tx=XM_023259984.1;r
efGene_gn=KIF1A;refGene_tx=XM_023259985.1;refGene_gn=KIF1A;refGene_tx=
XM_023259986.1;refGene_gn=KIF1A;refGene_tx=XM_023259987.1;refGene_gn=K
IF1A;refGene_tx=XM_023259988.1
C1 221895981 221896161
refGene_gn=KIF1A;refGene_tx=XM_023259972.1;refGene_gn=KIF1A;refGene_tx
=XM_023259973.1;refGene_gn=KIF1A;refGene_tx=XM_023259974.1;refGene_gn=
KIF1A;refGene_tx=XM_023259975.1;refGene_gn=KIF1A;refGene_tx=XM_0232599
76.1;refGene_gn=KIF1A;refGene_tx=XM_023259977.1;refGene_gn=KIF1A;refGe
ne_tx=XM_023259978.1;refGene_gn=KIF1A;refGene_tx=XM_023259979.1;refGen
e_gn=KIF1A;refGene_tx=XM_023259980.1;refGene_gn=KIF1A;refGene_tx=XM_02
3259981.1;refGene_gn=KIF1A;refGene_tx=XM_023259982.1;refGene_gn=KIF1A;
refGene_tx=XM_023259983.1;refGene_gn=KIF1A;refGene_tx=XM_023259984.1;r
efGene_gn=KIF1A;refGene_tx=XM_023259985.1;refGene_gn=KIF1A;refGene_tx=
XM_023259986.1;refGene_gn=KIF1A;refGene_tx=XM_023259987.1;refGene_gn=K
IF1A;refGene_tx=XM_023259988.1
C1 221896805 221896882
refGene_gn=KIF1A;refGene_tx=XM_023259972.1;refGene_gn=KIF1A;refGene_tx
=XM_023259973.1;refGene_gn=KIF1A;refGene_tx=XM_023259974.1;refGene_gn=
KIF1A;refGene_tx=XM_023259975.1;refGene_gn=KIF1A;refGene_tx=XM_0232599
76.1;refGene_gn=KIF1A;refGene_tx=XM_023259977.1;refGene_gn=KIF1A;refGe
ne_tx=XM_023259978.1;refGene_gn=KIF1A;refGene_tx=XM_023259979.1;refGen
e_gn=KIF1A;refGene_tx=XM_023259980.1;refGene_gn=KIF1A;refGene_tx=XM_02
3259981.1;refGene_gn=KIF1A;refGene_tx=XM_023259982.1;refGene_gn=KIF1A;
refGene_tx=XM_023259983.1;refGene_gn=KIF1A;refGene_tx=XM_023259984.1;r

efGene_gn=KIF1A;refGene_tx=XM_023259985.1;refGene_gn=KIF1A;refGene_tx=XM_023259986.1;refGene_gn=KIF1A;refGene_tx=XM_023259987.1;refGene_gn=KIF1A;refGene_tx=XM_023259988.1
C1 221904679 221904785
refGene_gn=KIF1A;refGene_tx=XM_023259972.1;refGene_gn=KIF1A;refGene_tx=XM_023259973.1;refGene_gn=KIF1A;refGene_tx=XM_023259974.1;refGene_gn=KIF1A;refGene_tx=XM_023259975.1;refGene_gn=KIF1A;refGene_tx=XM_023259976.1;refGene_gn=KIF1A;refGene_tx=XM_023259977.1;refGene_gn=KIF1A;refGene_tx=XM_023259978.1;refGene_gn=KIF1A;refGene_tx=XM_023259979.1;refGene_gn=KIF1A;refGene_tx=XM_023259980.1;refGene_gn=KIF1A;refGene_tx=XM_023259981.1;refGene_gn=KIF1A;refGene_tx=XM_023259982.1;refGene_gn=KIF1A;refGene_tx=XM_023259983.1;refGene_gn=KIF1A;refGene_tx=XM_023259984.1;refGene_gn=KIF1A;refGene_tx=XM_023259985.1;refGene_gn=KIF1A;refGene_tx=XM_023259986.1;refGene_gn=KIF1A;refGene_tx=XM_023259987.1;refGene_gn=KIF1A;refGene_tx=XM_023259988.1
C1 221954318 221954549
refGene_gn=AGXT;refGene_tx=NM_001042566.1
C1 221954648 221954841
refGene_gn=AGXT;refGene_tx=NM_001042566.1
C1 221956039 221956104
refGene_gn=AGXT;refGene_tx=NM_001042566.1
C1 221956743 221956844
refGene_gn=AGXT;refGene_tx=NM_001042566.1
C1 221958081 221958152
refGene_gn=AGXT;refGene_tx=NM_001042566.1
C1 221959623 221959708
refGene_gn=AGXT;refGene_tx=NM_001042566.1
C1 221960630 221960726
refGene_gn=AGXT;refGene_tx=NM_001042566.1
C1 221961374 221961444
refGene_gn=AGXT;refGene_tx=NM_001042566.1
C1 221962628 221962724
refGene_gn=AGXT;refGene_tx=NM_001042566.1
C1 221964348 221964477
refGene_gn=AGXT;refGene_tx=NM_001042566.1
C1 221965054 221965162
refGene_gn=AGXT;refGene_tx=NM_001042566.1
C1 221971534 221971627
refGene_gn=CC1H2orf54;refGene_tx=XM_003991327.4;refGene_gn=CC1H2orf54;refGene_tx=XM_019838990.2
C1 221972261 221972328
refGene_gn=CC1H2orf54;refGene_tx=XM_019838990.2
C1 221972528 221972885
refGene_gn=CC1H2orf54;refGene_tx=XM_003991327.4;refGene_gn=CC1H2orf54;refGene_tx=XM_019838990.2
C1 221975507 221975661
refGene_gn=CC1H2orf54;refGene_tx=XM_003991327.4;refGene_gn=CC1H2orf54;refGene_tx=XM_019838990.2
C1 221976716 221976942
refGene_gn=CC1H2orf54;refGene_tx=XM_003991327.4;refGene_gn=CC1H2orf54;

refGene_tx=XM_019838990.2
C1 221979293 221979777
refGene_gn=CC1H2orf54;refGene_tx=XM_003991327.4;refGene_gn=CC1H2orf54;
refGene_tx=XM_019838990.2
C1 222001875 222002182
refGene_gn=CROCC2;refGene_tx=XM_023257877.1
C1 222003182 222003331
refGene_gn=CROCC2;refGene_tx=XM_023257877.1
C1 222005455 222005562
refGene_gn=CROCC2;refGene_tx=XM_023257877.1
C1 222006902 222007127
refGene_gn=CROCC2;refGene_tx=XM_023257877.1
C1 222007813 222007971
refGene_gn=CROCC2;refGene_tx=XM_023257877.1
C1 222010946 222011142
refGene_gn=CROCC2;refGene_tx=XM_023257877.1
C1 222011684 222011882
refGene_gn=CROCC2;refGene_tx=XM_023257877.1
C1 222013512 222013609
refGene_gn=CROCC2;refGene_tx=XM_023257877.1
C1 222013688 222013778
refGene_gn=CROCC2;refGene_tx=XM_023257877.1
C1 222013936 222014143
refGene_gn=CROCC2;refGene_tx=XM_023257877.1
C1 222014461 222014673
refGene_gn=CROCC2;refGene_tx=XM_023257877.1
C1 222015846 222016029
refGene_gn=CROCC2;refGene_tx=XM_023257877.1
C1 222016826 222016971
refGene_gn=CROCC2;refGene_tx=XM_023257877.1
C1 222017386 222017533
refGene_gn=CROCC2;refGene_tx=XM_023257877.1
C1 222017730 222017931
refGene_gn=CROCC2;refGene_tx=XM_023257877.1
C1 222025586 222025780
refGene_gn=CROCC2;refGene_tx=XM_023257877.1
C1 222029196 222029324
refGene_gn=CROCC2;refGene_tx=XM_023257877.1
C1 222029836 222030006
refGene_gn=CROCC2;refGene_tx=XM_023257877.1
C1 222030601 222030778
refGene_gn=CROCC2;refGene_tx=XM_023257877.1
C1 222033135 222033249
refGene_gn=CROCC2;refGene_tx=XM_023257877.1
C1 222036734 222036878
refGene_gn=CROCC2;refGene_tx=XM_023257877.1
C1 222040311 222040529
refGene_gn=CROCC2;refGene_tx=XM_023257877.1
C1 222041168 222041328
refGene_gn=CROCC2;refGene_tx=XM_023257877.1

C1	222043816	222044174
refGene_gn=CROCC2;refGene_tx=XM_023257877.1
C1	222044411	222044578
refGene_gn=CROCC2;refGene_tx=XM_023257877.1
C1	222044970	222045091
refGene_gn=CROCC2;refGene_tx=XM_023257877.1
C1	222045275	222045372
refGene_gn=CROCC2;refGene_tx=XM_023257877.1
C1	222045460	222045875
refGene_gn=CROCC2;refGene_tx=XM_023257877.1
C1	222053919	222054066
refGene_gn=CROCC2;refGene_tx=XM_023257877.1
C1	222058167	222058530
refGene_gn=CROCC2;refGene_tx=XM_023257877.1
C1	222059006	222059231
refGene_gn=CROCC2;refGene_tx=XM_023257877.1
C1	222060138	222060225
refGene_gn=CROCC2;refGene_tx=XM_023257877.1
C1	222066881	222067094
refGene_gn=SNED1;refGene_tx=XM_023259989.1;refGene_gn=SNED1;refGene_tx=XM_023259990.1;refGene_gn=SNED1;refGene_tx=XM_023259991.1;refGene_gn=SNED1;refGene_tx=XM_023259992.1;refGene_gn=SNED1;refGene_tx=XM_023259993.1;refGene_gn=SNED1;refGene_tx=XM_023259994.1;refGene_gn=SNED1;refGene_tx=XM_023259995.1;refGene_gn=SNED1;refGene_tx=XM_023259996.1
C1	222090464	222090752
refGene_gn=SNED1;refGene_tx=XM_023259989.1;refGene_gn=SNED1;refGene_tx=XM_023259990.1;refGene_gn=SNED1;refGene_tx=XM_023259991.1;refGene_gn=SNED1;refGene_tx=XM_023259992.1;refGene_gn=SNED1;refGene_tx=XM_023259993.1;refGene_gn=SNED1;refGene_tx=XM_023259995.1;refGene_gn=SNED1;refGene_tx=XM_023259996.1
C1	222091385	222091583
refGene_gn=MTERF4;refGene_tx=XM_023260001.1
C1	222093605	222093746
refGene_gn=SNED1;refGene_tx=XM_023259989.1;refGene_gn=SNED1;refGene_tx=XM_023259990.1;refGene_gn=SNED1;refGene_tx=XM_023259991.1;refGene_gn=SNED1;refGene_tx=XM_023259992.1;refGene_gn=SNED1;refGene_tx=XM_023259993.1;refGene_gn=SNED1;refGene_tx=XM_023259994.1;refGene_gn=SNED1;refGene_tx=XM_023259995.1;refGene_gn=SNED1;refGene_tx=XM_023259996.1
C1	222094364	222094527
refGene_gn=SNED1;refGene_tx=XM_023259989.1;refGene_gn=SNED1;refGene_tx=XM_023259990.1;refGene_gn=SNED1;refGene_tx=XM_023259991.1;refGene_gn=SNED1;refGene_tx=XM_023259992.1;refGene_gn=SNED1;refGene_tx=XM_023259993.1;refGene_gn=SNED1;refGene_tx=XM_023259994.1;refGene_gn=SNED1;refGene_tx=XM_023259995.1;refGene_gn=SNED1;refGene_tx=XM_023259996.1
C1	222097382	222097508
refGene_gn=SNED1;refGene_tx=XM_023259989.1;refGene_gn=SNED1;refGene_tx=XM_023259990.1;refGene_gn=SNED1;refGene_tx=XM_023259991.1;refGene_gn=SNED1;refGene_tx=XM_023259992.1;refGene_gn=SNED1;refGene_tx=XM_023259993.1;refGene_gn=SNED1;refGene_tx=XM_023259994.1;refGene_gn=SNED1;refGene_tx=XM_023259995.1;refGene_gn=SNED1;refGene_tx=XM_023259996.1

C1	222097747	222097861
refGene_gn=SNED1;refGene_tx=XM_023259989.1;refGene_gn=SNED1;refGene_tx=XM_023259990.1;refGene_gn=SNED1;refGene_tx=XM_023259991.1;refGene_gn=SNED1;refGene_tx=XM_023259992.1;refGene_gn=SNED1;refGene_tx=XM_023259993.1;refGene_gn=SNED1;refGene_tx=XM_023259994.1;refGene_gn=SNED1;refGene_tx=XM_023259995.1;refGene_gn=SNED1;refGene_tx=XM_023259996.1
C1	222099685	222099799
refGene_gn=SNED1;refGene_tx=XM_023259989.1;refGene_gn=SNED1;refGene_tx=XM_023259990.1;refGene_gn=SNED1;refGene_tx=XM_023259991.1;refGene_gn=SNED1;refGene_tx=XM_023259992.1;refGene_gn=SNED1;refGene_tx=XM_023259993.1;refGene_gn=SNED1;refGene_tx=XM_023259994.1;refGene_gn=SNED1;refGene_tx=XM_023259995.1;refGene_gn=SNED1;refGene_tx=XM_023259996.1
C1	222099904	222100018
refGene_gn=SNED1;refGene_tx=XM_023259989.1;refGene_gn=SNED1;refGene_tx=XM_023259990.1;refGene_gn=SNED1;refGene_tx=XM_023259991.1;refGene_gn=SNED1;refGene_tx=XM_023259992.1;refGene_gn=SNED1;refGene_tx=XM_023259993.1;refGene_gn=SNED1;refGene_tx=XM_023259994.1;refGene_gn=SNED1;refGene_tx=XM_023259995.1;refGene_gn=SNED1;refGene_tx=XM_023259996.1
C1	222108796	222108922
refGene_gn=SNED1;refGene_tx=XM_023259989.1;refGene_gn=SNED1;refGene_tx=XM_023259990.1;refGene_gn=SNED1;refGene_tx=XM_023259991.1;refGene_gn=SNED1;refGene_tx=XM_023259992.1;refGene_gn=SNED1;refGene_tx=XM_023259993.1;refGene_gn=SNED1;refGene_tx=XM_023259994.1;refGene_gn=SNED1;refGene_tx=XM_023259995.1;refGene_gn=SNED1;refGene_tx=XM_023259996.1
C1	222109977	222110082
refGene_gn=SNED1;refGene_tx=XM_023259989.1;refGene_gn=SNED1;refGene_tx=XM_023259990.1;refGene_gn=SNED1;refGene_tx=XM_023259991.1;refGene_gn=SNED1;refGene_tx=XM_023259992.1;refGene_gn=SNED1;refGene_tx=XM_023259993.1;refGene_gn=SNED1;refGene_tx=XM_023259994.1;refGene_gn=SNED1;refGene_tx=XM_023259995.1;refGene_gn=SNED1;refGene_tx=XM_023259996.1
C1	222110311	222110425
refGene_gn=SNED1;refGene_tx=XM_023259989.1;refGene_gn=SNED1;refGene_tx=XM_023259990.1;refGene_gn=SNED1;refGene_tx=XM_023259991.1;refGene_gn=SNED1;refGene_tx=XM_023259992.1;refGene_gn=SNED1;refGene_tx=XM_023259993.1;refGene_gn=SNED1;refGene_tx=XM_023259994.1;refGene_gn=SNED1;refGene_tx=XM_023259995.1;refGene_gn=SNED1;refGene_tx=XM_023259996.1
C1	222111003	222111120
refGene_gn=SNED1;refGene_tx=XM_023259989.1;refGene_gn=SNED1;refGene_tx=XM_023259990.1;refGene_gn=SNED1;refGene_tx=XM_023259991.1;refGene_gn=SNED1;refGene_tx=XM_023259992.1;refGene_gn=SNED1;refGene_tx=XM_023259993.1;refGene_gn=SNED1;refGene_tx=XM_023259994.1;refGene_gn=SNED1;refGene_tx=XM_023259995.1;refGene_gn=SNED1;refGene_tx=XM_023259996.1
C1	222116552	222116669
refGene_gn=SNED1;refGene_tx=XM_023259989.1;refGene_gn=SNED1;refGene_tx=XM_023259990.1;refGene_gn=SNED1;refGene_tx=XM_023259992.1;refGene_gn=SNED1;refGene_tx=XM_023259993.1;refGene_gn=SNED1;refGene_tx=XM_023259994.1;refGene_gn=SNED1;refGene_tx=XM_023259995.1;refGene_gn=SNED1;refGene_tx=XM_023259996.1
C1	222116820	222116937
refGene_gn=SNED1;refGene_tx=XM_023259989.1;refGene_gn=SNED1;refGene_tx

=XM_023259990.1;refGene_gn=SNED1;refGene_tx=XM_023259991.1;refGene_gn=SNED1;refGene_tx=XM_023259992.1;refGene_gn=SNED1;refGene_tx=XM_023259993.1;refGene_gn=SNED1;refGene_tx=XM_023259994.1;refGene_gn=SNED1;refGene_tx=XM_023259995.1;refGene_gn=SNED1;refGene_tx=XM_023259996.1
C1 222117096 222117210
refGene_gn=SNED1;refGene_tx=XM_023259989.1;refGene_gn=SNED1;refGene_tx=XM_023259990.1;refGene_gn=SNED1;refGene_tx=XM_023259991.1;refGene_gn=SNED1;refGene_tx=XM_023259992.1;refGene_gn=SNED1;refGene_tx=XM_023259993.1;refGene_gn=SNED1;refGene_tx=XM_023259994.1;refGene_gn=SNED1;refGene_tx=XM_023259995.1;refGene_gn=SNED1;refGene_tx=XM_023259996.1
C1 222117512 222117686
refGene_gn=SNED1;refGene_tx=XM_023259989.1;refGene_gn=SNED1;refGene_tx=XM_023259990.1;refGene_gn=SNED1;refGene_tx=XM_023259991.1;refGene_gn=SNED1;refGene_tx=XM_023259993.1;refGene_gn=SNED1;refGene_tx=XM_023259994.1;refGene_gn=SNED1;refGene_tx=XM_023259995.1;refGene_gn=SNED1;refGene_tx=XM_023259996.1
C1 222126978 222127092
refGene_gn=SNED1;refGene_tx=XM_023259989.1;refGene_gn=SNED1;refGene_tx=XM_023259990.1;refGene_gn=SNED1;refGene_tx=XM_023259991.1;refGene_gn=SNED1;refGene_tx=XM_023259992.1;refGene_gn=SNED1;refGene_tx=XM_023259993.1;refGene_gn=SNED1;refGene_tx=XM_023259994.1;refGene_gn=SNED1;refGene_tx=XM_023259995.1;refGene_gn=SNED1;refGene_tx=XM_023259996.1
C1 222127409 222127523
refGene_gn=SNED1;refGene_tx=XM_023259989.1;refGene_gn=SNED1;refGene_tx=XM_023259990.1;refGene_gn=SNED1;refGene_tx=XM_023259991.1;refGene_gn=SNED1;refGene_tx=XM_023259992.1;refGene_gn=SNED1;refGene_tx=XM_023259993.1;refGene_gn=SNED1;refGene_tx=XM_023259994.1;refGene_gn=SNED1;refGene_tx=XM_023259995.1;refGene_gn=SNED1;refGene_tx=XM_023259996.1
C1 222128503 222128617
refGene_gn=SNED1;refGene_tx=XM_023259989.1;refGene_gn=SNED1;refGene_tx=XM_023259990.1;refGene_gn=SNED1;refGene_tx=XM_023259991.1;refGene_gn=SNED1;refGene_tx=XM_023259992.1;refGene_gn=SNED1;refGene_tx=XM_023259993.1;refGene_gn=SNED1;refGene_tx=XM_023259994.1;refGene_gn=SNED1;refGene_tx=XM_023259995.1;refGene_gn=SNED1;refGene_tx=XM_023259996.1
C1 222129033 222129330
refGene_gn=SNED1;refGene_tx=XM_023259989.1;refGene_gn=SNED1;refGene_tx=XM_023259990.1;refGene_gn=SNED1;refGene_tx=XM_023259991.1;refGene_gn=SNED1;refGene_tx=XM_023259992.1;refGene_gn=SNED1;refGene_tx=XM_023259993.1;refGene_gn=SNED1;refGene_tx=XM_023259994.1;refGene_gn=SNED1;refGene_tx=XM_023259995.1;refGene_gn=SNED1;refGene_tx=XM_023259996.1
C1 222130810 222130994
refGene_gn=SNED1;refGene_tx=XM_023259989.1;refGene_gn=SNED1;refGene_tx=XM_023259990.1;refGene_gn=SNED1;refGene_tx=XM_023259991.1;refGene_gn=SNED1;refGene_tx=XM_023259992.1;refGene_gn=SNED1;refGene_tx=XM_023259993.1;refGene_gn=SNED1;refGene_tx=XM_023259994.1;refGene_gn=SNED1;refGene_tx=XM_023259995.1;refGene_gn=SNED1;refGene_tx=XM_023259996.1
C1 222132723 222132836
refGene_gn=SNED1;refGene_tx=XM_023259989.1;refGene_gn=SNED1;refGene_tx=XM_023259990.1;refGene_gn=SNED1;refGene_tx=XM_023259991.1;refGene_gn=SNED1;refGene_tx=XM_023259992.1;refGene_gn=SNED1;refGene_tx=XM_0232599

93.1;refGene_gn=SNED1;refGene_tx=XM_023259994.1;refGene_gn=SNED1;refGene_tx=XM_023259995.1;refGene_gn=SNED1;refGene_tx=XM_023259996.1
C1 222133672 222133954
refGene_gn=SNED1;refGene_tx=XM_023259989.1;refGene_gn=SNED1;refGene_tx=XM_023259990.1;refGene_gn=SNED1;refGene_tx=XM_023259991.1;refGene_gn=SNED1;refGene_tx=XM_023259992.1;refGene_gn=SNED1;refGene_tx=XM_023259993.1;refGene_gn=SNED1;refGene_tx=XM_023259994.1;refGene_gn=SNED1;refGene_tx=XM_023259995.1;refGene_gn=SNED1;refGene_tx=XM_023259996.1
C1 222135110 222135252
refGene_gn=SNED1;refGene_tx=XM_023259989.1;refGene_gn=SNED1;refGene_tx=XM_023259990.1;refGene_gn=SNED1;refGene_tx=XM_023259991.1;refGene_gn=SNED1;refGene_tx=XM_023259992.1;refGene_gn=SNED1;refGene_tx=XM_023259993.1;refGene_gn=SNED1;refGene_tx=XM_023259994.1;refGene_gn=SNED1;refGene_tx=XM_023259995.1;refGene_gn=SNED1;refGene_tx=XM_023259996.1
C1 222135335 222135418
refGene_gn=SNED1;refGene_tx=XM_023259989.1;refGene_gn=SNED1;refGene_tx=XM_023259990.1;refGene_gn=SNED1;refGene_tx=XM_023259991.1;refGene_gn=SNED1;refGene_tx=XM_023259992.1;refGene_gn=SNED1;refGene_tx=XM_023259993.1;refGene_gn=SNED1;refGene_tx=XM_023259994.1;refGene_gn=SNED1;refGene_tx=XM_023259995.1;refGene_gn=SNED1;refGene_tx=XM_023259996.1
C1 222136886 222137000
refGene_gn=SNED1;refGene_tx=XM_023259989.1;refGene_gn=SNED1;refGene_tx=XM_023259990.1;refGene_gn=SNED1;refGene_tx=XM_023259991.1;refGene_gn=SNED1;refGene_tx=XM_023259992.1;refGene_gn=SNED1;refGene_tx=XM_023259993.1;refGene_gn=SNED1;refGene_tx=XM_023259994.1;refGene_gn=SNED1;refGene_tx=XM_023259995.1;refGene_gn=SNED1;refGene_tx=XM_023259996.1
C1 222139375 222139407
refGene_gn=SNED1;refGene_tx=XM_023259995.1
C1 222139780 222139911
refGene_gn=SNED1;refGene_tx=XM_023259993.1
C1 222142459 222142479
refGene_gn=SNED1;refGene_tx=XM_023259996.1
C1 222144614 222144731
refGene_gn=SNED1;refGene_tx=XM_023259989.1;refGene_gn=SNED1;refGene_tx=XM_023259990.1;refGene_gn=SNED1;refGene_tx=XM_023259991.1;refGene_gn=SNED1;refGene_tx=XM_023259992.1;refGene_gn=SNED1;refGene_tx=XM_023259994.1
C1 222145239 222145327
refGene_gn=SNED1;refGene_tx=XM_023259989.1;refGene_gn=SNED1;refGene_tx=XM_023259990.1;refGene_gn=SNED1;refGene_tx=XM_023259991.1;refGene_gn=SNED1;refGene_tx=XM_023259992.1;refGene_gn=SNED1;refGene_tx=XM_023259994.1
C1 222149698 222149782
refGene_gn=SNED1;refGene_tx=XM_023259989.1;refGene_gn=SNED1;refGene_tx=XM_023259990.1;refGene_gn=SNED1;refGene_tx=XM_023259991.1;refGene_gn=SNED1;refGene_tx=XM_023259992.1;refGene_gn=SNED1;refGene_tx=XM_023259994.1
C1 222150625 222150665
refGene_gn=SNED1;refGene_tx=XM_023259989.1;refGene_gn=SNED1;refGene_tx=XM_023259990.1;refGene_gn=SNED1;refGene_tx=XM_023259991.1;refGene_gn=

SNED1;refGene_tx=XM_023259992.1;refGene_gn=SNED1;refGene_tx=XM_0232599
94.1
C1	222156348	222156702
refGene_gn=MTERF4;refGene_tx=XM_023260000.1
C1	222157392	222157577
refGene_gn=MTERF4;refGene_tx=XM_023260000.1;refGene_gn=MTERF4;refGene_
tx=XM_023260001.1
C1	222159826	222160322
refGene_gn=MTERF4;refGene_tx=XM_023260000.1;refGene_gn=MTERF4;refGene_
tx=XM_023260001.1
C1	222161373	222161394
refGene_gn=MTERF4;refGene_tx=XM_023260000.1;refGene_gn=MTERF4;refGene_
tx=XM_023260001.1
C1	222163779	222163886
refGene_gn=PASK;refGene_tx=XM_006935745.3;refGene_gn=PASK;refGene_tx=X
M_006935746.3;refGene_gn=PASK;refGene_tx=XM_006935747.3;refGene_gn=PAS
K;refGene_tx=XM_019838992.2
C1	222164483	222164630
refGene_gn=PASK;refGene_tx=XM_006935745.3;refGene_gn=PASK;refGene_tx=X
M_006935746.3;refGene_gn=PASK;refGene_tx=XM_006935747.3;refGene_gn=PAS
K;refGene_tx=XM_019838992.2
C1	222166244	222166378
refGene_gn=PASK;refGene_tx=XM_006935745.3;refGene_gn=PASK;refGene_tx=X
M_006935746.3;refGene_gn=PASK;refGene_tx=XM_006935747.3;refGene_gn=PAS
K;refGene_tx=XM_019838992.2
C1	222168393	222168593
refGene_gn=PASK;refGene_tx=XM_006935745.3;refGene_gn=PASK;refGene_tx=X
M_006935746.3;refGene_gn=PASK;refGene_tx=XM_006935747.3;refGene_gn=PAS
K;refGene_tx=XM_019838992.2
C1	222171311	222171446
refGene_gn=PASK;refGene_tx=XM_006935745.3;refGene_gn=PASK;refGene_tx=X
M_006935746.3;refGene_gn=PASK;refGene_tx=XM_006935747.3;refGene_gn=PAS
K;refGene_tx=XM_019838992.2
C1	222171561	222171687
refGene_gn=PASK;refGene_tx=XM_006935745.3;refGene_gn=PASK;refGene_tx=X
M_006935746.3;refGene_gn=PASK;refGene_tx=XM_006935747.3;refGene_gn=PAS
K;refGene_tx=XM_019838992.2
C1	222175274	222175442
refGene_gn=PASK;refGene_tx=XM_006935745.3;refGene_gn=PASK;refGene_tx=X
M_006935746.3;refGene_gn=PASK;refGene_tx=XM_006935747.3
C1	222175815	222175997
refGene_gn=PASK;refGene_tx=XM_006935745.3;refGene_gn=PASK;refGene_tx=X
M_006935746.3;refGene_gn=PASK;refGene_tx=XM_006935747.3;refGene_gn=PAS
K;refGene_tx=XM_019838992.2
C1	222176844	222178586
refGene_gn=PASK;refGene_tx=XM_006935745.3;refGene_gn=PASK;refGene_tx=X
M_006935746.3;refGene_gn=PASK;refGene_tx=XM_006935747.3;refGene_gn=PAS
K;refGene_tx=XM_019838992.2
C1	222182702	222182859
refGene_gn=PASK;refGene_tx=XM_006935745.3;refGene_gn=PASK;refGene_tx=X

M_006935746.3;refGene_gn=PASK;refGene_tx=XM_006935747.3;refGene_gn=PASK;refGene_tx=XM_019838992.2
C1 222184969 222185162
refGene_gn=PASK;refGene_tx=XM_006935745.3;refGene_gn=PASK;refGene_tx=XM_006935746.3;refGene_gn=PASK;refGene_tx=XM_006935747.3;refGene_gn=PASK;refGene_tx=XM_019838992.2
C1 222186932 222187193
refGene_gn=PASK;refGene_tx=XM_006935745.3;refGene_gn=PASK;refGene_tx=XM_006935746.3;refGene_gn=PASK;refGene_tx=XM_006935747.3;refGene_gn=PASK;refGene_tx=XM_019838992.2
C1 222188968 222189103
refGene_gn=PASK;refGene_tx=XM_006935745.3;refGene_gn=PASK;refGene_tx=XM_006935746.3;refGene_gn=PASK;refGene_tx=XM_006935747.3;refGene_gn=PASK;refGene_tx=XM_019838992.2
C1 222189562 222189703
refGene_gn=PASK;refGene_tx=XM_006935745.3;refGene_gn=PASK;refGene_tx=XM_006935746.3;refGene_gn=PASK;refGene_tx=XM_006935747.3;refGene_gn=PASK;refGene_tx=XM_019838992.2
C1 222190357 222190528
refGene_gn=PASK;refGene_tx=XM_006935745.3;refGene_gn=PASK;refGene_tx=XM_006935746.3;refGene_gn=PASK;refGene_tx=XM_006935747.3;refGene_gn=PASK;refGene_tx=XM_019838992.2
C1 222191079 222191312
refGene_gn=PASK;refGene_tx=XM_006935745.3;refGene_gn=PASK;refGene_tx=XM_006935746.3;refGene_gn=PASK;refGene_tx=XM_006935747.3;refGene_gn=PASK;refGene_tx=XM_019838992.2
C1 222192874 222193232
refGene_gn=PASK;refGene_tx=XM_006935745.3;refGene_gn=PASK;refGene_tx=XM_006935746.3;refGene_gn=PASK;refGene_tx=XM_006935747.3;refGene_gn=PASK;refGene_tx=XM_019838992.2
C1 222204382 222204434
refGene_gn=PPP1R7;refGene_tx=XM_003991306.3;refGene_gn=PPP1R7;refGene_tx=XM_006935748.2;refGene_gn=PPP1R7;refGene_tx=XM_023259999.1
C1 222207247 222207388
refGene_gn=PPP1R7;refGene_tx=XM_003991306.3;refGene_gn=PPP1R7;refGene_tx=XM_006935748.2;refGene_gn=PPP1R7;refGene_tx=XM_023259999.1
C1 222211488 222211544
refGene_gn=PPP1R7;refGene_tx=XM_003991306.3;refGene_gn=PPP1R7;refGene_tx=XM_006935748.2;refGene_gn=PPP1R7;refGene_tx=XM_023259999.1
C1 222212141 222212207
refGene_gn=PPP1R7;refGene_tx=XM_003991306.3;refGene_gn=PPP1R7;refGene_tx=XM_006935748.2;refGene_gn=PPP1R7;refGene_tx=XM_023259999.1
C1 222212745 222212876
refGene_gn=PPP1R7;refGene_tx=XM_003991306.3;refGene_gn=PPP1R7;refGene_tx=XM_006935748.2;refGene_gn=PPP1R7;refGene_tx=XM_023259999.1
C1 222213234 222213397
refGene_gn=PPP1R7;refGene_tx=XM_003991306.3;refGene_gn=PPP1R7;refGene_tx=XM_006935748.2;refGene_gn=PPP1R7;refGene_tx=XM_023259999.1
C1 222215217 222215334
refGene_gn=PPP1R7;refGene_tx=XM_003991306.3;refGene_gn=PPP1R7;refGene_

tx=XM_006935748.2;refGene_gn=PPP1R7;refGene_tx=XM_023259999.1
C1 222216173 222216278
refGene_gn=PPP1R7;refGene_tx=XM_003991306.3;refGene_gn=PPP1R7;refGene_
tx=XM_006935748.2;refGene_gn=PPP1R7;refGene_tx=XM_023259999.1
C1 222219462 222219549
refGene_gn=PPP1R7;refGene_tx=XM_003991306.3;refGene_gn=PPP1R7;refGene_
tx=XM_006935748.2;refGene_gn=PPP1R7;refGene_tx=XM_023259999.1
C1 222220298 222220310
refGene_gn=PPP1R7;refGene_tx=XM_023259999.1
C1 222232947 222233124
refGene_gn=PPP1R7;refGene_tx=XM_003991306.3;refGene_gn=PPP1R7;refGene_
tx=XM_006935748.2
C1 222235720 222235811
refGene_gn=ANO7;refGene_tx=XM_023257878.1
C1 222238816 222239207
refGene_gn=ANO7;refGene_tx=XM_023257878.1
C1 222240347 222240459
refGene_gn=ANO7;refGene_tx=XM_023257878.1
C1 222241742 222241800
refGene_gn=ANO7;refGene_tx=XM_023257878.1
C1 222246156 222246296
refGene_gn=ANO7;refGene_tx=XM_023257878.1
C1 222248203 222248311
refGene_gn=ANO7;refGene_tx=XM_023257878.1
C1 222248879 222249016
refGene_gn=ANO7;refGene_tx=XM_023257878.1
C1 222249760 222249818
refGene_gn=ANO7;refGene_tx=XM_023257878.1
C1 222250533 222250644
refGene_gn=ANO7;refGene_tx=XM_023257878.1
C1 222251557 222251723
refGene_gn=ANO7;refGene_tx=XM_023257878.1
C1 222253048 222253139
refGene_gn=ANO7;refGene_tx=XM_023257878.1
C1 222254042 222254139
refGene_gn=ANO7;refGene_tx=XM_023257878.1
C1 222255498 222255642
refGene_gn=ANO7;refGene_tx=XM_023257878.1
C1 222255713 222255851
refGene_gn=ANO7;refGene_tx=XM_023257878.1
C1 222256499 222256598
refGene_gn=ANO7;refGene_tx=XM_023257878.1
C1 222256689 222256792
refGene_gn=ANO7;refGene_tx=XM_023257878.1
C1 222257673 222257785
refGene_gn=ANO7;refGene_tx=XM_023257878.1
C1 222258024 222258079
refGene_gn=ANO7;refGene_tx=XM_023257878.1
C1 222259673 222259771
refGene_gn=ANO7;refGene_tx=XM_023257878.1

C1 222260831 222260977
refGene_gn=ANO7;refGene_tx=XM_023257878.1
C1 222262390 222262596
refGene_gn=ANO7;refGene_tx=XM_023257878.1
C1 222262675 222262818
refGene_gn=ANO7;refGene_tx=XM_023257878.1
C1 222266121 222266212
refGene_gn=ANO7;refGene_tx=XM_023257878.1
C1 222266678 222266798
refGene_gn=ANO7;refGene_tx=XM_023257878.1
C1 222266929 222267046
refGene_gn=ANO7;refGene_tx=XM_023257878.1
C1 222269847 222269934
refGene_gn=HDLBP;refGene_tx=XM_006935755.3;refGene_gn=HDLBP;refGene_tx
=XM_023259997.1;refGene_gn=HDLBP;refGene_tx=XM_023259998.1
C1 222270049 222270178
refGene_gn=HDLBP;refGene_tx=XM_006935755.3;refGene_gn=HDLBP;refGene_tx
=XM_023259997.1;refGene_gn=HDLBP;refGene_tx=XM_023259998.1
C1 222270391 222270508
refGene_gn=HDLBP;refGene_tx=XM_006935755.3;refGene_gn=HDLBP;refGene_tx
=XM_023259997.1;refGene_gn=HDLBP;refGene_tx=XM_023259998.1
C1 222270981 222271167
refGene_gn=HDLBP;refGene_tx=XM_006935755.3;refGene_gn=HDLBP;refGene_tx
=XM_023259997.1;refGene_gn=HDLBP;refGene_tx=XM_023259998.1
C1 222273124 222273268
refGene_gn=HDLBP;refGene_tx=XM_006935755.3;refGene_gn=HDLBP;refGene_tx
=XM_023259997.1;refGene_gn=HDLBP;refGene_tx=XM_023259998.1
C1 222274218 222274353
refGene_gn=HDLBP;refGene_tx=XM_006935755.3;refGene_gn=HDLBP;refGene_tx
=XM_023259997.1;refGene_gn=HDLBP;refGene_tx=XM_023259998.1
C1 222274529 222274634
refGene_gn=HDLBP;refGene_tx=XM_006935755.3;refGene_gn=HDLBP;refGene_tx
=XM_023259997.1;refGene_gn=HDLBP;refGene_tx=XM_023259998.1
C1 222275470 222275625
refGene_gn=HDLBP;refGene_tx=XM_006935755.3;refGene_gn=HDLBP;refGene_tx
=XM_023259997.1;refGene_gn=HDLBP;refGene_tx=XM_023259998.1
C1 222277090 222277229
refGene_gn=HDLBP;refGene_tx=XM_006935755.3;refGene_gn=HDLBP;refGene_tx
=XM_023259997.1;refGene_gn=HDLBP;refGene_tx=XM_023259998.1
C1 222278013 222278232
refGene_gn=HDLBP;refGene_tx=XM_006935755.3;refGene_gn=HDLBP;refGene_tx
=XM_023259997.1;refGene_gn=HDLBP;refGene_tx=XM_023259998.1
C1 222278323 222278545
refGene_gn=HDLBP;refGene_tx=XM_006935755.3;refGene_gn=HDLBP;refGene_tx
=XM_023259997.1;refGene_gn=HDLBP;refGene_tx=XM_023259998.1
C1 222281061 222281280
refGene_gn=HDLBP;refGene_tx=XM_006935755.3;refGene_gn=HDLBP;refGene_tx
=XM_023259997.1;refGene_gn=HDLBP;refGene_tx=XM_023259998.1
C1 222284564 222284696
refGene_gn=HDLBP;refGene_tx=XM_006935755.3;refGene_gn=HDLBP;refGene_tx

=XM_023259997.1;refGene_gn=HDLBP;refGene_tx=XM_023259998.1
C1 222284861 222284948
refGene_gn=HDLBP;refGene_tx=XM_006935755.3;refGene_gn=HDLBP;refGene_tx=XM_023259997.1;refGene_gn=HDLBP;refGene_tx=XM_023259998.1
C1 222285926 222286040
refGene_gn=HDLBP;refGene_tx=XM_006935755.3;refGene_gn=HDLBP;refGene_tx=XM_023259997.1;refGene_gn=HDLBP;refGene_tx=XM_023259998.1
C1 222286152 222286257
refGene_gn=HDLBP;refGene_tx=XM_006935755.3;refGene_gn=HDLBP;refGene_tx=XM_023259997.1;refGene_gn=HDLBP;refGene_tx=XM_023259998.1
C1 222287668 222287808
refGene_gn=HDLBP;refGene_tx=XM_006935755.3;refGene_gn=HDLBP;refGene_tx=XM_023259997.1;refGene_gn=HDLBP;refGene_tx=XM_023259998.1
C1 222289434 222289513
refGene_gn=HDLBP;refGene_tx=XM_006935755.3;refGene_gn=HDLBP;refGene_tx=XM_023259997.1;refGene_gn=HDLBP;refGene_tx=XM_023259998.1
C1 222289884 222289989
refGene_gn=HDLBP;refGene_tx=XM_006935755.3;refGene_gn=HDLBP;refGene_tx=XM_023259997.1;refGene_gn=HDLBP;refGene_tx=XM_023259998.1
C1 222291286 222291394
refGene_gn=HDLBP;refGene_tx=XM_006935755.3;refGene_gn=HDLBP;refGene_tx=XM_023259997.1;refGene_gn=HDLBP;refGene_tx=XM_023259998.1
C1 222291526 222291733
refGene_gn=HDLBP;refGene_tx=XM_006935755.3;refGene_gn=HDLBP;refGene_tx=XM_023259997.1;refGene_gn=HDLBP;refGene_tx=XM_023259998.1
C1 222292256 222292472
refGene_gn=HDLBP;refGene_tx=XM_006935755.3;refGene_gn=HDLBP;refGene_tx=XM_023259997.1;refGene_gn=HDLBP;refGene_tx=XM_023259998.1
C1 222292556 222292763
refGene_gn=HDLBP;refGene_tx=XM_006935755.3;refGene_gn=HDLBP;refGene_tx=XM_023259997.1;refGene_gn=HDLBP;refGene_tx=XM_023259998.1
C1 222299663 222299879
refGene_gn=HDLBP;refGene_tx=XM_006935755.3;refGene_gn=HDLBP;refGene_tx=XM_023259997.1;refGene_gn=HDLBP;refGene_tx=XM_023259998.1
C1 222301097 222301255
refGene_gn=HDLBP;refGene_tx=XM_006935755.3;refGene_gn=HDLBP;refGene_tx=XM_023259997.1;refGene_gn=HDLBP;refGene_tx=XM_023259998.1
C1 222302637 222302713
refGene_gn=HDLBP;refGene_tx=XM_006935755.3;refGene_gn=HDLBP;refGene_tx=XM_023259997.1;refGene_gn=HDLBP;refGene_tx=XM_023259998.1
C1 222341945 222341951
refGene_gn=SEPT2;refGene_tx=XM_019838994.1
C1 222349555 222349585
refGene_gn=SEPT2;refGene_tx=XM_006935750.3;refGene_gn=SEPT2;refGene_tx=XM_019838993.2;refGene_gn=SEPT2;refGene_tx=XM_019838994.1;refGene_gn=SEPT2;refGene_tx=XM_019838995.1
C1 222351042 222351163
refGene_gn=SEPT2;refGene_tx=XM_006935750.3;refGene_gn=SEPT2;refGene_tx=XM_019838993.2;refGene_gn=SEPT2;refGene_tx=XM_019838994.1;refGene_gn=SEPT2;refGene_tx=XM_019838995.1;refGene_gn=SEPT2;refGene_tx=XM_0198389

96.2
C1 222360683 222360806
refGene_gn=SEPT2;refGene_tx=XM_006935750.3;refGene_gn=SEPT2;refGene_tx=XM_019838993.2;refGene_gn=SEPT2;refGene_tx=XM_019838994.1;refGene_gn=SEPT2;refGene_tx=XM_019838995.1;refGene_gn=SEPT2;refGene_tx=XM_019838996.2
C1 222361397 222361521
refGene_gn=SEPT2;refGene_tx=XM_006935750.3;refGene_gn=SEPT2;refGene_tx=XM_019838993.2;refGene_gn=SEPT2;refGene_tx=XM_019838994.1;refGene_gn=SEPT2;refGene_tx=XM_019838995.1;refGene_gn=SEPT2;refGene_tx=XM_019838996.2
C1 222362811 222362946
refGene_gn=SEPT2;refGene_tx=XM_006935750.3;refGene_gn=SEPT2;refGene_tx=XM_019838993.2;refGene_gn=SEPT2;refGene_tx=XM_019838994.1;refGene_gn=SEPT2;refGene_tx=XM_019838995.1;refGene_gn=SEPT2;refGene_tx=XM_019838996.2
C1 222363125 222363243
refGene_gn=SEPT2;refGene_tx=XM_006935750.3;refGene_gn=SEPT2;refGene_tx=XM_019838993.2;refGene_gn=SEPT2;refGene_tx=XM_019838994.1;refGene_gn=SEPT2;refGene_tx=XM_019838995.1;refGene_gn=SEPT2;refGene_tx=XM_019838996.2
C1 222367020 222367122
refGene_gn=SEPT2;refGene_tx=XM_006935750.3;refGene_gn=SEPT2;refGene_tx=XM_019838993.2;refGene_gn=SEPT2;refGene_tx=XM_019838994.1;refGene_gn=SEPT2;refGene_tx=XM_019838995.1;refGene_gn=SEPT2;refGene_tx=XM_019838996.2
C1 222367495 222367641
refGene_gn=SEPT2;refGene_tx=XM_006935750.3;refGene_gn=SEPT2;refGene_tx=XM_019838993.2;refGene_gn=SEPT2;refGene_tx=XM_019838994.1;refGene_gn=SEPT2;refGene_tx=XM_019838995.1;refGene_gn=SEPT2;refGene_tx=XM_019838996.2
C1 222368996 222369080
refGene_gn=SEPT2;refGene_tx=XM_006935750.3;refGene_gn=SEPT2;refGene_tx=XM_019838993.2;refGene_gn=SEPT2;refGene_tx=XM_019838994.1;refGene_gn=SEPT2;refGene_tx=XM_019838995.1;refGene_gn=SEPT2;refGene_tx=XM_019838996.2
C1 222370614 222370672
refGene_gn=SEPT2;refGene_tx=XM_006935750.3;refGene_gn=SEPT2;refGene_tx=XM_019838993.2;refGene_gn=SEPT2;refGene_tx=XM_019838994.1;refGene_gn=SEPT2;refGene_tx=XM_019838995.1;refGene_gn=SEPT2;refGene_tx=XM_019838996.2
C1 222372014 222372116
refGene_gn=SEPT2;refGene_tx=XM_006935750.3;refGene_gn=SEPT2;refGene_tx=XM_019838993.2;refGene_gn=SEPT2;refGene_tx=XM_019838994.1;refGene_gn=SEPT2;refGene_tx=XM_019838995.1;refGene_gn=SEPT2;refGene_tx=XM_019838996.2
C1 222393148 222393352
refGene_gn=FARP2;refGene_tx=XM_023260002.1;refGene_gn=FARP2;refGene_tx=XM_023260003.1;refGene_gn=FARP2;refGene_tx=XM_023260004.1;refGene_gn=FARP2;refGene_tx=XM_023260005.1;refGene_gn=FARP2;refGene_tx=XM_0232600

06.1;refGene_gn=FARP2;refGene_tx=XM_023260008.1;refGene_gn=FARP2;refGene_tx=XM_023260009.1;refGene_gn=FARP2;refGene_tx=XM_023260010.1
C1 222426343 222426448
refGene_gn=FARP2;refGene_tx=XM_023260002.1;refGene_gn=FARP2;refGene_tx=XM_023260003.1;refGene_gn=FARP2;refGene_tx=XM_023260004.1;refGene_gn=FARP2;refGene_tx=XM_023260005.1;refGene_gn=FARP2;refGene_tx=XM_023260006.1;refGene_gn=FARP2;refGene_tx=XM_023260008.1;refGene_gn=FARP2;refGene_tx=XM_023260009.1;refGene_gn=FARP2;refGene_tx=XM_023260010.1
C1 222427490 222427533
refGene_gn=FARP2;refGene_tx=XM_023260002.1;refGene_gn=FARP2;refGene_tx=XM_023260003.1;refGene_gn=FARP2;refGene_tx=XM_023260004.1;refGene_gn=FARP2;refGene_tx=XM_023260005.1;refGene_gn=FARP2;refGene_tx=XM_023260006.1;refGene_gn=FARP2;refGene_tx=XM_023260008.1;refGene_gn=FARP2;refGene_tx=XM_023260009.1;refGene_gn=FARP2;refGene_tx=XM_023260010.1
C1 222428899 222428978
refGene_gn=FARP2;refGene_tx=XM_023260002.1;refGene_gn=FARP2;refGene_tx=XM_023260003.1;refGene_gn=FARP2;refGene_tx=XM_023260004.1;refGene_gn=FARP2;refGene_tx=XM_023260005.1;refGene_gn=FARP2;refGene_tx=XM_023260006.1;refGene_gn=FARP2;refGene_tx=XM_023260008.1;refGene_gn=FARP2;refGene_tx=XM_023260009.1;refGene_gn=FARP2;refGene_tx=XM_023260010.1
C1 222431021 222431119
refGene_gn=FARP2;refGene_tx=XM_023260002.1;refGene_gn=FARP2;refGene_tx=XM_023260003.1;refGene_gn=FARP2;refGene_tx=XM_023260004.1;refGene_gn=FARP2;refGene_tx=XM_023260005.1;refGene_gn=FARP2;refGene_tx=XM_023260006.1;refGene_gn=FARP2;refGene_tx=XM_023260008.1;refGene_gn=FARP2;refGene_tx=XM_023260009.1;refGene_gn=FARP2;refGene_tx=XM_023260010.1
C1 222432774 222432889
refGene_gn=FARP2;refGene_tx=XM_023260002.1;refGene_gn=FARP2;refGene_tx=XM_023260003.1;refGene_gn=FARP2;refGene_tx=XM_023260004.1;refGene_gn=FARP2;refGene_tx=XM_023260005.1;refGene_gn=FARP2;refGene_tx=XM_023260006.1;refGene_gn=FARP2;refGene_tx=XM_023260008.1;refGene_gn=FARP2;refGene_tx=XM_023260009.1;refGene_gn=FARP2;refGene_tx=XM_023260010.1
C1 222437546 222437694
refGene_gn=FARP2;refGene_tx=XM_023260002.1;refGene_gn=FARP2;refGene_tx=XM_023260003.1;refGene_gn=FARP2;refGene_tx=XM_023260004.1;refGene_gn=FARP2;refGene_tx=XM_023260005.1;refGene_gn=FARP2;refGene_tx=XM_023260006.1;refGene_gn=FARP2;refGene_tx=XM_023260008.1;refGene_gn=FARP2;refGene_tx=XM_023260009.1;refGene_gn=FARP2;refGene_tx=XM_023260010.1
C1 222442398 222442494
refGene_gn=FARP2;refGene_tx=XM_023260002.1;refGene_gn=FARP2;refGene_tx=XM_023260003.1;refGene_gn=FARP2;refGene_tx=XM_023260004.1;refGene_gn=FARP2;refGene_tx=XM_023260005.1;refGene_gn=FARP2;refGene_tx=XM_023260006.1;refGene_gn=FARP2;refGene_tx=XM_023260008.1;refGene_gn=FARP2;refGene_tx=XM_023260009.1;refGene_gn=FARP2;refGene_tx=XM_023260010.1
C1 222444677 222444841
refGene_gn=FARP2;refGene_tx=XM_023260002.1;refGene_gn=FARP2;refGene_tx=XM_023260003.1;refGene_gn=FARP2;refGene_tx=XM_023260004.1;refGene_gn=FARP2;refGene_tx=XM_023260005.1;refGene_gn=FARP2;refGene_tx=XM_023260006.1;refGene_gn=FARP2;refGene_tx=XM_023260008.1;refGene_gn=FARP2;refGene_tx=XM_023260009.1;refGene_gn=FARP2;refGene_tx=XM_023260010.1

C1	222445173	222445242
refGene_gn=FARP2;refGene_tx=XM_023260002.1;refGene_gn=FARP2;refGene_tx=XM_023260003.1;refGene_gn=FARP2;refGene_tx=XM_023260004.1;refGene_gn=FARP2;refGene_tx=XM_023260005.1;refGene_gn=FARP2;refGene_tx=XM_023260006.1;refGene_gn=FARP2;refGene_tx=XM_023260008.1;refGene_gn=FARP2;refGene_tx=XM_023260009.1;refGene_gn=FARP2;refGene_tx=XM_023260010.1
C1	222445754	222445812
refGene_gn=FARP2;refGene_tx=XM_023260002.1;refGene_gn=FARP2;refGene_tx=XM_023260003.1;refGene_gn=FARP2;refGene_tx=XM_023260004.1;refGene_gn=FARP2;refGene_tx=XM_023260005.1;refGene_gn=FARP2;refGene_tx=XM_023260006.1;refGene_gn=FARP2;refGene_tx=XM_023260008.1;refGene_gn=FARP2;refGene_tx=XM_023260009.1;refGene_gn=FARP2;refGene_tx=XM_023260010.1
C1	222448527	222448780
refGene_gn=FARP2;refGene_tx=XM_023260002.1;refGene_gn=FARP2;refGene_tx=XM_023260003.1;refGene_gn=FARP2;refGene_tx=XM_023260004.1;refGene_gn=FARP2;refGene_tx=XM_023260005.1;refGene_gn=FARP2;refGene_tx=XM_023260006.1;refGene_gn=FARP2;refGene_tx=XM_023260008.1;refGene_gn=FARP2;refGene_tx=XM_023260009.1;refGene_gn=FARP2;refGene_tx=XM_023260010.1
C1	222459721	222459900
refGene_gn=FARP2;refGene_tx=XM_023260002.1;refGene_gn=FARP2;refGene_tx=XM_023260003.1;refGene_gn=FARP2;refGene_tx=XM_023260004.1;refGene_gn=FARP2;refGene_tx=XM_023260005.1;refGene_gn=FARP2;refGene_tx=XM_023260006.1;refGene_gn=FARP2;refGene_tx=XM_023260008.1;refGene_gn=FARP2;refGene_tx=XM_023260009.1;refGene_gn=FARP2;refGene_tx=XM_023260010.1
C1	222464231	222464321
refGene_gn=FARP2;refGene_tx=XM_023260002.1;refGene_gn=FARP2;refGene_tx=XM_023260003.1;refGene_gn=FARP2;refGene_tx=XM_023260004.1;refGene_gn=FARP2;refGene_tx=XM_023260005.1;refGene_gn=FARP2;refGene_tx=XM_023260006.1;refGene_gn=FARP2;refGene_tx=XM_023260008.1;refGene_gn=FARP2;refGene_tx=XM_023260009.1;refGene_gn=FARP2;refGene_tx=XM_023260010.1
C1	222464688	222464822
refGene_gn=FARP2;refGene_tx=XM_023260002.1;refGene_gn=FARP2;refGene_tx=XM_023260003.1;refGene_gn=FARP2;refGene_tx=XM_023260004.1;refGene_gn=FARP2;refGene_tx=XM_023260005.1;refGene_gn=FARP2;refGene_tx=XM_023260006.1;refGene_gn=FARP2;refGene_tx=XM_023260008.1;refGene_gn=FARP2;refGene_tx=XM_023260009.1;refGene_gn=FARP2;refGene_tx=XM_023260010.1
C1	222465215	222465297
refGene_gn=FARP2;refGene_tx=XM_023260002.1;refGene_gn=FARP2;refGene_tx=XM_023260003.1;refGene_gn=FARP2;refGene_tx=XM_023260004.1;refGene_gn=FARP2;refGene_tx=XM_023260005.1;refGene_gn=FARP2;refGene_tx=XM_023260006.1;refGene_gn=FARP2;refGene_tx=XM_023260008.1;refGene_gn=FARP2;refGene_tx=XM_023260009.1;refGene_gn=FARP2;refGene_tx=XM_023260010.1
C1	222469358	222469596
refGene_gn=FARP2;refGene_tx=XM_023260002.1;refGene_gn=FARP2;refGene_tx=XM_023260003.1;refGene_gn=FARP2;refGene_tx=XM_023260004.1;refGene_gn=FARP2;refGene_tx=XM_023260005.1;refGene_gn=FARP2;refGene_tx=XM_023260006.1;refGene_gn=FARP2;refGene_tx=XM_023260008.1;refGene_gn=FARP2;refGene_tx=XM_023260009.1;refGene_gn=FARP2;refGene_tx=XM_023260010.1
C1	222472198	222472329
refGene_gn=FARP2;refGene_tx=XM_023260002.1;refGene_gn=FARP2;refGene_tx

=XM_023260003.1;refGene_gn=FARP2;refGene_tx=XM_023260004.1;refGene_gn=FARP2;refGene_tx=XM_023260005.1;refGene_gn=FARP2;refGene_tx=XM_023260006.1;refGene_gn=FARP2;refGene_tx=XM_023260008.1;refGene_gn=FARP2;refGene_tx=XM_023260009.1;refGene_gn=FARP2;refGene_tx=XM_023260010.1
C1 222477669 222477738
refGene_gn=FARP2;refGene_tx=XM_023260002.1;refGene_gn=FARP2;refGene_tx=XM_023260003.1;refGene_gn=FARP2;refGene_tx=XM_023260004.1;refGene_gn=FARP2;refGene_tx=XM_023260005.1;refGene_gn=FARP2;refGene_tx=XM_023260006.1;refGene_gn=FARP2;refGene_tx=XM_023260008.1;refGene_gn=FARP2;refGene_tx=XM_023260009.1;refGene_gn=FARP2;refGene_tx=XM_023260010.1
C1 222478327 222478417
refGene_gn=FARP2;refGene_tx=XM_023260002.1;refGene_gn=FARP2;refGene_tx=XM_023260003.1;refGene_gn=FARP2;refGene_tx=XM_023260004.1;refGene_gn=FARP2;refGene_tx=XM_023260005.1;refGene_gn=FARP2;refGene_tx=XM_023260006.1;refGene_gn=FARP2;refGene_tx=XM_023260008.1;refGene_gn=FARP2;refGene_tx=XM_023260009.1;refGene_gn=FARP2;refGene_tx=XM_023260010.1
C1 222481741 222482052
refGene_gn=FARP2;refGene_tx=XM_023260002.1;refGene_gn=FARP2;refGene_tx=XM_023260003.1;refGene_gn=FARP2;refGene_tx=XM_023260004.1;refGene_gn=FARP2;refGene_tx=XM_023260005.1;refGene_gn=FARP2;refGene_tx=XM_023260006.1;refGene_gn=FARP2;refGene_tx=XM_023260008.1;refGene_gn=FARP2;refGene_tx=XM_023260009.1;refGene_gn=FARP2;refGene_tx=XM_023260010.1
C1 222482793 222483424
refGene_gn=FARP2;refGene_tx=XM_023260002.1;refGene_gn=FARP2;refGene_tx=XM_023260003.1;refGene_gn=FARP2;refGene_tx=XM_023260004.1;refGene_gn=FARP2;refGene_tx=XM_023260005.1;refGene_gn=FARP2;refGene_tx=XM_023260006.1;refGene_gn=FARP2;refGene_tx=XM_023260008.1;refGene_gn=FARP2;refGene_tx=XM_023260009.1;refGene_gn=FARP2;refGene_tx=XM_023260010.1
C1 222484113 222484143
refGene_gn=FARP2;refGene_tx=XM_023260009.1
C1 222484336 222484444
refGene_gn=FARP2;refGene_tx=XM_023260002.1;refGene_gn=FARP2;refGene_tx=XM_023260003.1;refGene_gn=FARP2;refGene_tx=XM_023260004.1;refGene_gn=FARP2;refGene_tx=XM_023260005.1;refGene_gn=FARP2;refGene_tx=XM_023260006.1;refGene_gn=FARP2;refGene_tx=XM_023260008.1;refGene_gn=FARP2;refGene_tx=XM_023260010.1
C1 222484654 222484806
refGene_gn=FARP2;refGene_tx=XM_023260002.1;refGene_gn=FARP2;refGene_tx=XM_023260003.1;refGene_gn=FARP2;refGene_tx=XM_023260004.1;refGene_gn=FARP2;refGene_tx=XM_023260005.1;refGene_gn=FARP2;refGene_tx=XM_023260006.1;refGene_gn=FARP2;refGene_tx=XM_023260008.1;refGene_gn=FARP2;refGene_tx=XM_023260010.1
C1 222485046 222485116
refGene_gn=FARP2;refGene_tx=XM_023260002.1;refGene_gn=FARP2;refGene_tx=XM_023260003.1;refGene_gn=FARP2;refGene_tx=XM_023260004.1;refGene_gn=FARP2;refGene_tx=XM_023260005.1;refGene_gn=FARP2;refGene_tx=XM_023260006.1;refGene_gn=FARP2;refGene_tx=XM_023260008.1;refGene_gn=FARP2;refGene_tx=XM_023260010.1
C1 222485404 222485473
refGene_gn=FARP2;refGene_tx=XM_023260002.1;refGene_gn=FARP2;refGene_tx

=XM_023260003.1;refGene_gn=FARP2;refGene_tx=XM_023260005.1;refGene_gn=FARP2;refGene_tx=XM_023260006.1;refGene_gn=FARP2;refGene_tx=XM_023260008.1;refGene_gn=FARP2;refGene_tx=XM_023260010.1
C1 222486194 222486234
refGene_gn=STK25;refGene_tx=XM_023260011.1
C1 222486462 222486599
refGene_gn=STK25;refGene_tx=XM_023260011.1
C1 222488023 222488095
refGene_gn=STK25;refGene_tx=XM_023260011.1
C1 222488252 222488287
refGene_gn=STK25;refGene_tx=XM_023260011.1
C1 222488289 222488333
refGene_gn=STK25;refGene_tx=XM_023260011.1
C1 222488334 222488367
refGene_gn=STK25;refGene_tx=XM_023260011.1
C1 222488599 222488745
refGene_gn=STK25;refGene_tx=XM_023260011.1
C1 222489133 222489319
refGene_gn=STK25;refGene_tx=XM_023260011.1
C1 222489402 222489560
refGene_gn=STK25;refGene_tx=XM_023260011.1
C1 222490281 222490390
refGene_gn=STK25;refGene_tx=XM_023260011.1
C1 222490846 222490903
refGene_gn=STK25;refGene_tx=XM_023260011.1
C1 222491583 222491814
refGene_gn=STK25;refGene_tx=XM_023260011.1
C1 222497008 222497038
refGene_gn=STK25;refGene_tx=XM_023260011.1
C1 222515832 222516055
refGene_gn=BOK;refGene_tx=XM_003991311.5;refGene_gn=BOK;refGene_tx=XM_006935762.4
C1 222518683 222518812
refGene_gn=BOK;refGene_tx=XM_003991311.5;refGene_gn=BOK;refGene_tx=XM_006935762.4
C1 222523633 222523756
refGene_gn=BOK;refGene_tx=XM_006935762.4
C1 222526301 222526465
refGene_gn=BOK;refGene_tx=XM_003991311.5;refGene_gn=BOK;refGene_tx=XM_006935762.4
C1 222527464 222527590
refGene_gn=BOK;refGene_tx=XM_003991311.5;refGene_gn=BOK;refGene_tx=XM_006935762.4
C1 222536275 222536395
refGene_gn=THAP4;refGene_tx=XM_003991312.5;refGene_gn=THAP4;refGene_tx=XM_006935765.4;refGene_gn=THAP4;refGene_tx=XM_023260012.1
C1 222551015 222551119
refGene_gn=THAP4;refGene_tx=XM_003991312.5;refGene_gn=THAP4;refGene_tx=XM_006935765.4;refGene_gn=THAP4;refGene_tx=XM_023260012.1
C1 222551899 222552009

refGene_gn=THAP4;refGene_tx=XM_003991312.5;refGene_gn=THAP4;refGene_tx
=XM_006935765.4;refGene_gn=THAP4;refGene_tx=XM_023260012.1
C1	222552958	222553118
refGene_gn=THAP4;refGene_tx=XM_003991312.5;refGene_gn=THAP4;refGene_tx
=XM_006935765.4;refGene_gn=THAP4;refGene_tx=XM_023260012.1
C1	222560197	222560201
refGene_gn=THAP4;refGene_tx=XM_003991312.5
C1	222574174	222575340
refGene_gn=THAP4;refGene_tx=XM_006935765.4;refGene_gn=THAP4;refGene_tx
=XM_023260012.1
C1	222576725	222576826
refGene_gn=THAP4;refGene_tx=XM_006935765.4
C1	222577382	222577459
refGene_gn=THAP4;refGene_tx=XM_023260012.1
C1	222577601	222578124
refGene_gn=ATG4B;refGene_tx=XM_023260014.1;refGene_gn=ATG4B;refGene_tx
=XM_023260015.1
C1	222578705	222578712
refGene_gn=ATG4B;refGene_tx=XM_023260016.1
C1	222586786	222586888
refGene_gn=ATG4B;refGene_tx=XM_023260014.1;refGene_gn=ATG4B;refGene_tx
=XM_023260016.1
C1	222587032	222587104
refGene_gn=ATG4B;refGene_tx=XM_023260014.1;refGene_gn=ATG4B;refGene_tx
=XM_023260016.1
C1	222589282	222589381
refGene_gn=ATG4B;refGene_tx=XM_023260014.1;refGene_gn=ATG4B;refGene_tx
=XM_023260015.1;refGene_gn=ATG4B;refGene_tx=XM_023260016.1
C1	222589769	222589871
refGene_gn=ATG4B;refGene_tx=XM_023260014.1;refGene_gn=ATG4B;refGene_tx
=XM_023260015.1;refGene_gn=ATG4B;refGene_tx=XM_023260016.1
C1	222590349	222590422
refGene_gn=ATG4B;refGene_tx=XM_023260014.1;refGene_gn=ATG4B;refGene_tx
=XM_023260015.1;refGene_gn=ATG4B;refGene_tx=XM_023260016.1
C1	222593121	222593201
refGene_gn=ATG4B;refGene_tx=XM_023260014.1;refGene_gn=ATG4B;refGene_tx
=XM_023260015.1;refGene_gn=ATG4B;refGene_tx=XM_023260016.1
C1	222599174	222599368
refGene_gn=ATG4B;refGene_tx=XM_023260014.1;refGene_gn=ATG4B;refGene_tx
=XM_023260015.1;refGene_gn=ATG4B;refGene_tx=XM_023260016.1
C1	222599888	222599967
refGene_gn=ATG4B;refGene_tx=XM_023260014.1;refGene_gn=ATG4B;refGene_tx
=XM_023260015.1;refGene_gn=ATG4B;refGene_tx=XM_023260016.1
C1	222600253	222600399
refGene_gn=ATG4B;refGene_tx=XM_023260014.1;refGene_gn=ATG4B;refGene_tx
=XM_023260015.1;refGene_gn=ATG4B;refGene_tx=XM_023260016.1
C1	222601598	222601655
refGene_gn=ATG4B;refGene_tx=XM_023260014.1;refGene_gn=ATG4B;refGene_tx
=XM_023260015.1;refGene_gn=ATG4B;refGene_tx=XM_023260016.1
C1	222602058	222602152

refGene_gn=ATG4B;refGene_tx=XM_023260014.1;refGene_gn=ATG4B;refGene_tx=XM_023260015.1;refGene_gn=ATG4B;refGene_tx=XM_023260016.1
C1	222603036	222603110
refGene_gn=ATG4B;refGene_tx=XM_023260014.1;refGene_gn=ATG4B;refGene_tx=XM_023260015.1;refGene_gn=ATG4B;refGene_tx=XM_023260016.1
C1	222607342	222607453
refGene_gn=DTYMK;refGene_tx=XM_023260017.1
C1	222611687	222611885
refGene_gn=DTYMK;refGene_tx=XM_023260017.1
C1	222612637	222612728
refGene_gn=DTYMK;refGene_tx=XM_023260017.1
C1	222615200	222615309
refGene_gn=DTYMK;refGene_tx=XM_023260017.1
C1	222616075	222616205
refGene_gn=DTYMK;refGene_tx=XM_023260017.1
C1	222624000	222624037
refGene_gn=ING5;refGene_tx=XM_023260023.1;refGene_gn=ING5;refGene_tx=XM_023260024.1
C1	222626331	222626403
refGene_gn=ING5;refGene_tx=XM_023260023.1;refGene_gn=ING5;refGene_tx=XM_023260025.1
C1	222629398	222629565
refGene_gn=ING5;refGene_tx=XM_023260023.1;refGene_gn=ING5;refGene_tx=XM_023260024.1;refGene_gn=ING5;refGene_tx=XM_023260025.1
C1	222630188	222630300
refGene_gn=ING5;refGene_tx=XM_023260023.1;refGene_gn=ING5;refGene_tx=XM_023260024.1;refGene_gn=ING5;refGene_tx=XM_023260025.1
C1	222630467	222630561
refGene_gn=ING5;refGene_tx=XM_023260023.1;refGene_gn=ING5;refGene_tx=XM_023260024.1;refGene_gn=ING5;refGene_tx=XM_023260025.1
C1	222643075	222643211
refGene_gn=ING5;refGene_tx=XM_023260023.1;refGene_gn=ING5;refGene_tx=XM_023260024.1;refGene_gn=ING5;refGene_tx=XM_023260025.1
C1	222643330	222643392
refGene_gn=ING5;refGene_tx=XM_023260023.1;refGene_gn=ING5;refGene_tx=XM_023260024.1;refGene_gn=ING5;refGene_tx=XM_023260025.1
C1	222645073	222645116
refGene_gn=ING5;refGene_tx=XM_023260023.1;refGene_gn=ING5;refGene_tx=XM_023260024.1;refGene_gn=ING5;refGene_tx=XM_023260025.1
C1	222652390	222652742
refGene_gn=D2HGDH;refGene_tx=XM_019839001.2;refGene_gn=D2HGDH;refGene_tx=XM_019839002.1;refGene_gn=D2HGDH;refGene_tx=XM_023260018.1;refGene_gn=D2HGDH;refGene_tx=XM_023260019.1;refGene_gn=D2HGDH;refGene_tx=XM_023260020.1;refGene_gn=D2HGDH;refGene_tx=XM_023260021.1;refGene_gn=D2HGDH;refGene_tx=XM_023260022.1
C1	222659739	222659797
refGene_gn=D2HGDH;refGene_tx=XM_019839001.2;refGene_gn=D2HGDH;refGene_tx=XM_019839002.1;refGene_gn=D2HGDH;refGene_tx=XM_023260018.1;refGene_gn=D2HGDH;refGene_tx=XM_023260019.1;refGene_gn=D2HGDH;refGene_tx=XM_023260020.1;refGene_gn=D2HGDH;refGene_tx=XM_023260021.1;refGene_gn=D2HGD

H;refGene_tx=XM_023260022.1
C1 222661249 222661389
refGene_gn=D2HGDH;refGene_tx=XM_019839001.2;refGene_gn=D2HGDH;refGene_tx=XM_019839002.1;refGene_gn=D2HGDH;refGene_tx=XM_023260018.1;refGene_gn=D2HGDH;refGene_tx=XM_023260019.1;refGene_gn=D2HGDH;refGene_tx=XM_023260020.1;refGene_gn=D2HGDH;refGene_tx=XM_023260021.1;refGene_gn=D2HGDH;refGene_tx=XM_023260022.1
C1 222662048 222662242
refGene_gn=D2HGDH;refGene_tx=XM_019839001.2;refGene_gn=D2HGDH;refGene_tx=XM_019839002.1;refGene_gn=D2HGDH;refGene_tx=XM_023260018.1;refGene_gn=D2HGDH;refGene_tx=XM_023260019.1;refGene_gn=D2HGDH;refGene_tx=XM_023260020.1;refGene_gn=D2HGDH;refGene_tx=XM_023260021.1;refGene_gn=D2HGDH;refGene_tx=XM_023260022.1
C1 222662706 222662875
refGene_gn=D2HGDH;refGene_tx=XM_019839001.2;refGene_gn=D2HGDH;refGene_tx=XM_019839002.1;refGene_gn=D2HGDH;refGene_tx=XM_023260018.1;refGene_gn=D2HGDH;refGene_tx=XM_023260019.1;refGene_gn=D2HGDH;refGene_tx=XM_023260020.1;refGene_gn=D2HGDH;refGene_tx=XM_023260021.1;refGene_gn=D2HGDH;refGene_tx=XM_023260022.1
C1 222664302 222664446
refGene_gn=D2HGDH;refGene_tx=XM_019839001.2;refGene_gn=D2HGDH;refGene_tx=XM_019839002.1;refGene_gn=D2HGDH;refGene_tx=XM_023260018.1;refGene_gn=D2HGDH;refGene_tx=XM_023260019.1;refGene_gn=D2HGDH;refGene_tx=XM_023260020.1;refGene_gn=D2HGDH;refGene_tx=XM_023260021.1;refGene_gn=D2HGDH;refGene_tx=XM_023260022.1
C1 222664753 222664840
refGene_gn=D2HGDH;refGene_tx=XM_019839001.2;refGene_gn=D2HGDH;refGene_tx=XM_019839002.1;refGene_gn=D2HGDH;refGene_tx=XM_023260018.1;refGene_gn=D2HGDH;refGene_tx=XM_023260020.1;refGene_gn=D2HGDH;refGene_tx=XM_023260021.1;refGene_gn=D2HGDH;refGene_tx=XM_023260022.1
C1 222666697 222666840
refGene_gn=D2HGDH;refGene_tx=XM_019839001.2;refGene_gn=D2HGDH;refGene_tx=XM_019839002.1;refGene_gn=D2HGDH;refGene_tx=XM_023260018.1;refGene_gn=D2HGDH;refGene_tx=XM_023260019.1;refGene_gn=D2HGDH;refGene_tx=XM_023260020.1;refGene_gn=D2HGDH;refGene_tx=XM_023260021.1;refGene_gn=D2HGDH;refGene_tx=XM_023260022.1
C1 222667572 222667614
refGene_gn=D2HGDH;refGene_tx=XM_023260021.1
C1 222669556 222669722
refGene_gn=D2HGDH;refGene_tx=XM_019839001.2;refGene_gn=D2HGDH;refGene_tx=XM_019839002.1;refGene_gn=D2HGDH;refGene_tx=XM_023260018.1;refGene_gn=D2HGDH;refGene_tx=XM_023260019.1;refGene_gn=D2HGDH;refGene_tx=XM_023260020.1
C1 222677244 222677504
refGene_gn=D2HGDH;refGene_tx=XM_019839001.2;refGene_gn=D2HGDH;refGene_tx=XM_019839002.1;refGene_gn=D2HGDH;refGene_tx=XM_023260018.1;refGene_gn=D2HGDH;refGene_tx=XM_023260019.1;refGene_gn=D2HGDH;refGene_tx=XM_023260020.1;refGene_gn=D2HGDH;refGene_tx=XM_023260022.1
C1 222685874 222685903
refGene_gn=GAL3ST2;refGene_tx=XM_023260026.1

C1	222696575	222696659
refGene_gn=GAL3ST2;refGene_tx=XM_023260026.1
C1	222697754	222698019
refGene_gn=GAL3ST2;refGene_tx=XM_023260026.1
C1	222698863	222699682
refGene_gn=GAL3ST2;refGene_tx=XM_023260026.1
C1	222703761	222704031
refGene_gn=NEU4;refGene_tx=XM_023260027.1
C1	222706370	222706466
refGene_gn=NEU4;refGene_tx=XM_023260027.1
C1	222707014	222707218
refGene_gn=NEU4;refGene_tx=XM_023260027.1;refGene_gn=NEU4;refGene_tx=XM_023260028.1;refGene_gn=NEU4;refGene_tx=XM_023260029.1
C1	222707318	222707574
refGene_gn=NEU4;refGene_tx=XM_023260027.1;refGene_gn=NEU4;refGene_tx=XM_023260028.1;refGene_gn=NEU4;refGene_tx=XM_023260029.1
C1	222708555	222709541
refGene_gn=NEU4;refGene_tx=XM_023260027.1;refGene_gn=NEU4;refGene_tx=XM_023260028.1;refGene_gn=NEU4;refGene_tx=XM_023260029.1
C1	222716207	222716411
refGene_gn=PDCD1;refGene_tx=NM_001145510.1
C1	222716658	222716693
refGene_gn=PDCD1;refGene_tx=NM_001145510.1
C1	222718386	222718826
refGene_gn=PDCD1;refGene_tx=NM_001145510.1;refGene_gn=PDCD1;refGene_tx=XM_023257806.1
C1	222720850	222721210
refGene_gn=PDCD1;refGene_tx=NM_001145510.1;refGene_gn=PDCD1;refGene_tx=XM_023257806.1
C1	222728311	222728387
refGene_gn=PDCD1;refGene_tx=NM_001145510.1;refGene_gn=PDCD1;refGene_tx=XM_023257806.1
C1	222736483	222736635
refGene_gn=RTP5;refGene_tx=XM_023260030.1
C1	222739189	222739877
refGene_gn=RTP5;refGene_tx=XM_023260030.1
C1	222739878	222740052
refGene_gn=RTP5;refGene_tx=XM_023260030.1
C1	222740053	222740677
refGene_gn=RTP5;refGene_tx=XM_023260030.1
C2	57405	57409
refGene_gn=LOC111562359;refGene_tx=XM_023260860.1
C2	68194	68919
refGene_gn=LOC101101073;refGene_tx=XM_019839246.2;refGene_gn=LOC101101073;refGene_tx=XM_023238575.1;refGene_gn=LOC111562359;refGene_tx=XM_023260860.1
C2	79181	79369
refGene_gn=LOC101101073;refGene_tx=XM_019839246.2;refGene_gn=LOC101101073;refGene_tx=XM_023238575.1
C2	82699	82821

refGene_gn=LOC101101073;refGene_tx=XM_019839246.2;refGene_gn=LOC101101073;refGene_tx=XM_023238575.1
C2 110514 110604
refGene_gn=LOC101101073;refGene_tx=XM_019839246.2;refGene_gn=LOC101101073;refGene_tx=XM_023238575.1
C2 115434 115561
refGene_gn=LOC101101073;refGene_tx=XM_019839246.2;refGene_gn=LOC101101073;refGene_tx=XM_023238575.1
C2 118409 118552
refGene_gn=LOC101101073;refGene_tx=XM_019839246.2;refGene_gn=LOC101101073;refGene_tx=XM_023238575.1
C2 119180 119383
refGene_gn=LOC101101073;refGene_tx=XM_019839246.2;refGene_gn=LOC101101073;refGene_tx=XM_023238575.1
C2 120279 120389
refGene_gn=LOC101101073;refGene_tx=XM_019839246.2;refGene_gn=LOC101101073;refGene_tx=XM_023238575.1
C2 120546 120688
refGene_gn=LOC101101073;refGene_tx=XM_019839246.2;refGene_gn=LOC101101073;refGene_tx=XM_023238575.1
C2 130311 132163
refGene_gn=LOC101101073;refGene_tx=XM_019839246.2;refGene_gn=LOC101101073;refGene_tx=XM_023238575.1
C2 133092 133173
refGene_gn=LOC101101073;refGene_tx=XM_023238575.1
C2 167859 167892 refGene_gn=PRMT2;refGene_tx=XM_006935787.4
C2 168688 168860 refGene_gn=PRMT2;refGene_tx=XM_006935787.4
C2 170225 170362 refGene_gn=PRMT2;refGene_tx=XM_006935787.4
C2 171205 171335 refGene_gn=PRMT2;refGene_tx=XM_006935787.4
C2 174284 174460 refGene_gn=PRMT2;refGene_tx=XM_006935787.4
C2 189124 189289 refGene_gn=PRMT2;refGene_tx=XM_006935787.4
C2 190292 190358 refGene_gn=PRMT2;refGene_tx=XM_006935787.4
C2 190359 190455 refGene_gn=PRMT2;refGene_tx=XM_006935787.4
C2 194905 195088 refGene_gn=PRMT2;refGene_tx=XM_006935787.4
C2 195762 195870 refGene_gn=PRMT2;refGene_tx=XM_006935787.4
C2 202163 202202 refGene_gn=PRMT2;refGene_tx=XM_006935787.4
C2 235488 235626
refGene_gn=S100B;refGene_tx=XM_023238576.1;refGene_gn=S100B;refGene_tx=XM_023238577.1
C2 237879 238020
refGene_gn=S100B;refGene_tx=XM_023238576.1;refGene_gn=S100B;refGene_tx=XM_023238577.1
C2 246306 246559
refGene_gn=DIP2A;refGene_tx=XM_023238578.1;refGene_gn=DIP2A;refGene_tx=XM_023238579.1;refGene_gn=DIP2A;refGene_tx=XM_023238580.1;refGene_gn=DIP2A;refGene_tx=XM_023238581.1;refGene_gn=DIP2A;refGene_tx=XM_023238582.1;refGene_gn=DIP2A;refGene_tx=XM_023238583.1;refGene_gn=DIP2A;refGene_tx=XM_023238584.1;refGene_gn=DIP2A;refGene_tx=XM_023238585.1
C2 247072 247196
refGene_gn=DIP2A;refGene_tx=XM_023238578.1;refGene_gn=DIP2A;refGene_tx

=XM_023238579.1;refGene_gn=DIP2A;refGene_tx=XM_023238580.1;refGene_gn=
DIP2A;refGene_tx=XM_023238581.1;refGene_gn=DIP2A;refGene_tx=XM_0232385
82.1;refGene_gn=DIP2A;refGene_tx=XM_023238583.1;refGene_gn=DIP2A;refGe
ne_tx=XM_023238584.1;refGene_gn=DIP2A;refGene_tx=XM_023238585.1
C2 247861 248036
refGene_gn=DIP2A;refGene_tx=XM_023238578.1;refGene_gn=DIP2A;refGene_tx
=XM_023238579.1;refGene_gn=DIP2A;refGene_tx=XM_023238580.1;refGene_gn=
DIP2A;refGene_tx=XM_023238581.1;refGene_gn=DIP2A;refGene_tx=XM_0232385
82.1;refGene_gn=DIP2A;refGene_tx=XM_023238583.1;refGene_gn=DIP2A;refGe
ne_tx=XM_023238584.1;refGene_gn=DIP2A;refGene_tx=XM_023238585.1
C2 249234 249309
refGene_gn=DIP2A;refGene_tx=XM_023238578.1;refGene_gn=DIP2A;refGene_tx
=XM_023238579.1;refGene_gn=DIP2A;refGene_tx=XM_023238580.1;refGene_gn=
DIP2A;refGene_tx=XM_023238581.1;refGene_gn=DIP2A;refGene_tx=XM_0232385
82.1;refGene_gn=DIP2A;refGene_tx=XM_023238583.1;refGene_gn=DIP2A;refGe
ne_tx=XM_023238584.1;refGene_gn=DIP2A;refGene_tx=XM_023238585.1
C2 251149 251207
refGene_gn=DIP2A;refGene_tx=XM_023238578.1;refGene_gn=DIP2A;refGene_tx
=XM_023238579.1;refGene_gn=DIP2A;refGene_tx=XM_023238580.1;refGene_gn=
DIP2A;refGene_tx=XM_023238581.1;refGene_gn=DIP2A;refGene_tx=XM_0232385
82.1;refGene_gn=DIP2A;refGene_tx=XM_023238583.1;refGene_gn=DIP2A;refGe
ne_tx=XM_023238584.1;refGene_gn=DIP2A;refGene_tx=XM_023238585.1
C2 251882 251944
refGene_gn=DIP2A;refGene_tx=XM_023238578.1;refGene_gn=DIP2A;refGene_tx
=XM_023238579.1;refGene_gn=DIP2A;refGene_tx=XM_023238580.1;refGene_gn=
DIP2A;refGene_tx=XM_023238581.1;refGene_gn=DIP2A;refGene_tx=XM_0232385
82.1;refGene_gn=DIP2A;refGene_tx=XM_023238583.1;refGene_gn=DIP2A;refGe
ne_tx=XM_023238584.1;refGene_gn=DIP2A;refGene_tx=XM_023238585.1
C2 252791 252821
refGene_gn=DIP2A;refGene_tx=XM_023238578.1;refGene_gn=DIP2A;refGene_tx
=XM_023238579.1;refGene_gn=DIP2A;refGene_tx=XM_023238582.1;refGene_gn=
DIP2A;refGene_tx=XM_023238583.1;refGene_gn=DIP2A;refGene_tx=XM_0232385
84.1;refGene_gn=DIP2A;refGene_tx=XM_023238585.1
C2 253932 254103
refGene_gn=DIP2A;refGene_tx=XM_023238578.1;refGene_gn=DIP2A;refGene_tx
=XM_023238579.1;refGene_gn=DIP2A;refGene_tx=XM_023238580.1;refGene_gn=
DIP2A;refGene_tx=XM_023238581.1;refGene_gn=DIP2A;refGene_tx=XM_0232385
82.1;refGene_gn=DIP2A;refGene_tx=XM_023238583.1;refGene_gn=DIP2A;refGe
ne_tx=XM_023238584.1;refGene_gn=DIP2A;refGene_tx=XM_023238585.1
C2 254659 254828
refGene_gn=DIP2A;refGene_tx=XM_023238578.1;refGene_gn=DIP2A;refGene_tx
=XM_023238579.1;refGene_gn=DIP2A;refGene_tx=XM_023238580.1;refGene_gn=
DIP2A;refGene_tx=XM_023238581.1;refGene_gn=DIP2A;refGene_tx=XM_0232385
82.1;refGene_gn=DIP2A;refGene_tx=XM_023238583.1;refGene_gn=DIP2A;refGe
ne_tx=XM_023238584.1;refGene_gn=DIP2A;refGene_tx=XM_023238585.1
C2 255515 255646
refGene_gn=DIP2A;refGene_tx=XM_023238578.1;refGene_gn=DIP2A;refGene_tx
=XM_023238579.1;refGene_gn=DIP2A;refGene_tx=XM_023238580.1;refGene_gn=
DIP2A;refGene_tx=XM_023238581.1;refGene_gn=DIP2A;refGene_tx=XM_0232385
82.1;refGene_gn=DIP2A;refGene_tx=XM_023238583.1;refGene_gn=DIP2A;refGe

ne_tx=XM_023238584.1;refGene_gn=DIP2A;refGene_tx=XM_023238585.1
C2 256122 256232
refGene_gn=DIP2A;refGene_tx=XM_023238578.1;refGene_gn=DIP2A;refGene_tx=XM_023238579.1;refGene_gn=DIP2A;refGene_tx=XM_023238580.1;refGene_gn=DIP2A;refGene_tx=XM_023238581.1;refGene_gn=DIP2A;refGene_tx=XM_023238582.1;refGene_gn=DIP2A;refGene_tx=XM_023238583.1;refGene_gn=DIP2A;refGene_tx=XM_023238584.1;refGene_gn=DIP2A;refGene_tx=XM_023238585.1
C2 257657 257769
refGene_gn=DIP2A;refGene_tx=XM_023238578.1;refGene_gn=DIP2A;refGene_tx=XM_023238579.1;refGene_gn=DIP2A;refGene_tx=XM_023238580.1;refGene_gn=DIP2A;refGene_tx=XM_023238581.1;refGene_gn=DIP2A;refGene_tx=XM_023238582.1;refGene_gn=DIP2A;refGene_tx=XM_023238583.1;refGene_gn=DIP2A;refGene_tx=XM_023238584.1;refGene_gn=DIP2A;refGene_tx=XM_023238585.1
C2 257899 258021
refGene_gn=DIP2A;refGene_tx=XM_023238578.1;refGene_gn=DIP2A;refGene_tx=XM_023238579.1;refGene_gn=DIP2A;refGene_tx=XM_023238580.1;refGene_gn=DIP2A;refGene_tx=XM_023238581.1;refGene_gn=DIP2A;refGene_tx=XM_023238582.1;refGene_gn=DIP2A;refGene_tx=XM_023238583.1;refGene_gn=DIP2A;refGene_tx=XM_023238584.1;refGene_gn=DIP2A;refGene_tx=XM_023238585.1
C2 263624 263748
refGene_gn=DIP2A;refGene_tx=XM_023238578.1;refGene_gn=DIP2A;refGene_tx=XM_023238579.1;refGene_gn=DIP2A;refGene_tx=XM_023238580.1;refGene_gn=DIP2A;refGene_tx=XM_023238581.1;refGene_gn=DIP2A;refGene_tx=XM_023238582.1;refGene_gn=DIP2A;refGene_tx=XM_023238583.1;refGene_gn=DIP2A;refGene_tx=XM_023238584.1;refGene_gn=DIP2A;refGene_tx=XM_023238585.1
C2 266005 266086
refGene_gn=DIP2A;refGene_tx=XM_023238578.1;refGene_gn=DIP2A;refGene_tx=XM_023238579.1;refGene_gn=DIP2A;refGene_tx=XM_023238580.1;refGene_gn=DIP2A;refGene_tx=XM_023238581.1;refGene_gn=DIP2A;refGene_tx=XM_023238582.1;refGene_gn=DIP2A;refGene_tx=XM_023238583.1;refGene_gn=DIP2A;refGene_tx=XM_023238584.1;refGene_gn=DIP2A;refGene_tx=XM_023238585.1
C2 266169 266279
refGene_gn=DIP2A;refGene_tx=XM_023238578.1;refGene_gn=DIP2A;refGene_tx=XM_023238579.1;refGene_gn=DIP2A;refGene_tx=XM_023238580.1;refGene_gn=DIP2A;refGene_tx=XM_023238581.1;refGene_gn=DIP2A;refGene_tx=XM_023238582.1;refGene_gn=DIP2A;refGene_tx=XM_023238583.1;refGene_gn=DIP2A;refGene_tx=XM_023238584.1;refGene_gn=DIP2A;refGene_tx=XM_023238585.1
C2 267179 267381
refGene_gn=DIP2A;refGene_tx=XM_023238578.1;refGene_gn=DIP2A;refGene_tx=XM_023238579.1;refGene_gn=DIP2A;refGene_tx=XM_023238580.1;refGene_gn=DIP2A;refGene_tx=XM_023238581.1;refGene_gn=DIP2A;refGene_tx=XM_023238582.1;refGene_gn=DIP2A;refGene_tx=XM_023238583.1;refGene_gn=DIP2A;refGene_tx=XM_023238584.1;refGene_gn=DIP2A;refGene_tx=XM_023238585.1
C2 274184 274299
refGene_gn=DIP2A;refGene_tx=XM_023238578.1;refGene_gn=DIP2A;refGene_tx=XM_023238579.1;refGene_gn=DIP2A;refGene_tx=XM_023238580.1;refGene_gn=DIP2A;refGene_tx=XM_023238581.1;refGene_gn=DIP2A;refGene_tx=XM_023238582.1;refGene_gn=DIP2A;refGene_tx=XM_023238583.1;refGene_gn=DIP2A;refGene_tx=XM_023238584.1;refGene_gn=DIP2A;refGene_tx=XM_023238585.1
C2 276219 276351

refGene_gn=DIP2A;refGene_tx=XM_023238578.1;refGene_gn=DIP2A;refGene_tx=XM_023238579.1;refGene_gn=DIP2A;refGene_tx=XM_023238580.1;refGene_gn=DIP2A;refGene_tx=XM_023238581.1;refGene_gn=DIP2A;refGene_tx=XM_023238582.1;refGene_gn=DIP2A;refGene_tx=XM_023238583.1;refGene_gn=DIP2A;refGene_tx=XM_023238584.1;refGene_gn=DIP2A;refGene_tx=XM_023238585.1;refGene_gn=DIP2A;refGene_tx=XM_023238586.1
C2 276768 276849
refGene_gn=DIP2A;refGene_tx=XM_023238578.1;refGene_gn=DIP2A;refGene_tx=XM_023238579.1;refGene_gn=DIP2A;refGene_tx=XM_023238580.1;refGene_gn=DIP2A;refGene_tx=XM_023238581.1;refGene_gn=DIP2A;refGene_tx=XM_023238582.1;refGene_gn=DIP2A;refGene_tx=XM_023238583.1;refGene_gn=DIP2A;refGene_tx=XM_023238584.1;refGene_gn=DIP2A;refGene_tx=XM_023238585.1;refGene_gn=DIP2A;refGene_tx=XM_023238586.1
C2 277252 277389
refGene_gn=DIP2A;refGene_tx=XM_023238578.1;refGene_gn=DIP2A;refGene_tx=XM_023238579.1;refGene_gn=DIP2A;refGene_tx=XM_023238580.1;refGene_gn=DIP2A;refGene_tx=XM_023238581.1;refGene_gn=DIP2A;refGene_tx=XM_023238582.1;refGene_gn=DIP2A;refGene_tx=XM_023238583.1;refGene_gn=DIP2A;refGene_tx=XM_023238584.1;refGene_gn=DIP2A;refGene_tx=XM_023238585.1;refGene_gn=DIP2A;refGene_tx=XM_023238586.1
C2 279303 279443
refGene_gn=DIP2A;refGene_tx=XM_023238578.1;refGene_gn=DIP2A;refGene_tx=XM_023238579.1;refGene_gn=DIP2A;refGene_tx=XM_023238580.1;refGene_gn=DIP2A;refGene_tx=XM_023238581.1;refGene_gn=DIP2A;refGene_tx=XM_023238582.1;refGene_gn=DIP2A;refGene_tx=XM_023238583.1;refGene_gn=DIP2A;refGene_tx=XM_023238584.1;refGene_gn=DIP2A;refGene_tx=XM_023238585.1;refGene_gn=DIP2A;refGene_tx=XM_023238586.1
C2 280842 280957
refGene_gn=DIP2A;refGene_tx=XM_023238578.1;refGene_gn=DIP2A;refGene_tx=XM_023238579.1;refGene_gn=DIP2A;refGene_tx=XM_023238580.1;refGene_gn=DIP2A;refGene_tx=XM_023238581.1;refGene_gn=DIP2A;refGene_tx=XM_023238582.1;refGene_gn=DIP2A;refGene_tx=XM_023238583.1;refGene_gn=DIP2A;refGene_tx=XM_023238584.1;refGene_gn=DIP2A;refGene_tx=XM_023238585.1;refGene_gn=DIP2A;refGene_tx=XM_023238586.1
C2 282416 282536
refGene_gn=DIP2A;refGene_tx=XM_023238578.1;refGene_gn=DIP2A;refGene_tx=XM_023238579.1;refGene_gn=DIP2A;refGene_tx=XM_023238580.1;refGene_gn=DIP2A;refGene_tx=XM_023238581.1;refGene_gn=DIP2A;refGene_tx=XM_023238582.1;refGene_gn=DIP2A;refGene_tx=XM_023238583.1;refGene_gn=DIP2A;refGene_tx=XM_023238584.1;refGene_gn=DIP2A;refGene_tx=XM_023238585.1;refGene_gn=DIP2A;refGene_tx=XM_023238586.1
C2 283903 283997
refGene_gn=DIP2A;refGene_tx=XM_023238578.1;refGene_gn=DIP2A;refGene_tx=XM_023238579.1;refGene_gn=DIP2A;refGene_tx=XM_023238580.1;refGene_gn=DIP2A;refGene_tx=XM_023238581.1;refGene_gn=DIP2A;refGene_tx=XM_023238582.1;refGene_gn=DIP2A;refGene_tx=XM_023238583.1;refGene_gn=DIP2A;refGene_tx=XM_023238584.1;refGene_gn=DIP2A;refGene_tx=XM_023238585.1;refGene_gn=DIP2A;refGene_tx=XM_023238586.1
C2 284231 284296
refGene_gn=DIP2A;refGene_tx=XM_023238578.1;refGene_gn=DIP2A;refGene_tx

=XM_023238579.1;refGene_gn=DIP2A;refGene_tx=XM_023238580.1;refGene_gn=DIP2A;refGene_tx=XM_023238581.1;refGene_gn=DIP2A;refGene_tx=XM_023238582.1;refGene_gn=DIP2A;refGene_tx=XM_023238583.1;refGene_gn=DIP2A;refGene_tx=XM_023238584.1;refGene_gn=DIP2A;refGene_tx=XM_023238585.1;refGene_gn=DIP2A;refGene_tx=XM_023238586.1
C2 286003 286106
refGene_gn=DIP2A;refGene_tx=XM_023238578.1;refGene_gn=DIP2A;refGene_tx=XM_023238579.1;refGene_gn=DIP2A;refGene_tx=XM_023238580.1;refGene_gn=DIP2A;refGene_tx=XM_023238581.1;refGene_gn=DIP2A;refGene_tx=XM_023238582.1;refGene_gn=DIP2A;refGene_tx=XM_023238583.1;refGene_gn=DIP2A;refGene_tx=XM_023238584.1;refGene_gn=DIP2A;refGene_tx=XM_023238585.1;refGene_gn=DIP2A;refGene_tx=XM_023238586.1
C2 286546 286656
refGene_gn=DIP2A;refGene_tx=XM_023238578.1;refGene_gn=DIP2A;refGene_tx=XM_023238579.1;refGene_gn=DIP2A;refGene_tx=XM_023238580.1;refGene_gn=DIP2A;refGene_tx=XM_023238581.1;refGene_gn=DIP2A;refGene_tx=XM_023238582.1;refGene_gn=DIP2A;refGene_tx=XM_023238583.1;refGene_gn=DIP2A;refGene_tx=XM_023238584.1;refGene_gn=DIP2A;refGene_tx=XM_023238585.1;refGene_gn=DIP2A;refGene_tx=XM_023238586.1
C2 286997 287121
refGene_gn=DIP2A;refGene_tx=XM_023238578.1;refGene_gn=DIP2A;refGene_tx=XM_023238579.1;refGene_gn=DIP2A;refGene_tx=XM_023238580.1;refGene_gn=DIP2A;refGene_tx=XM_023238581.1;refGene_gn=DIP2A;refGene_tx=XM_023238582.1;refGene_gn=DIP2A;refGene_tx=XM_023238583.1;refGene_gn=DIP2A;refGene_tx=XM_023238584.1;refGene_gn=DIP2A;refGene_tx=XM_023238585.1;refGene_gn=DIP2A;refGene_tx=XM_023238586.1
C2 288378 288489
refGene_gn=DIP2A;refGene_tx=XM_023238578.1;refGene_gn=DIP2A;refGene_tx=XM_023238579.1;refGene_gn=DIP2A;refGene_tx=XM_023238580.1;refGene_gn=DIP2A;refGene_tx=XM_023238581.1;refGene_gn=DIP2A;refGene_tx=XM_023238582.1;refGene_gn=DIP2A;refGene_tx=XM_023238583.1;refGene_gn=DIP2A;refGene_tx=XM_023238584.1;refGene_gn=DIP2A;refGene_tx=XM_023238585.1;refGene_gn=DIP2A;refGene_tx=XM_023238586.1
C2 292116 292208
refGene_gn=DIP2A;refGene_tx=XM_023238578.1;refGene_gn=DIP2A;refGene_tx=XM_023238579.1;refGene_gn=DIP2A;refGene_tx=XM_023238580.1;refGene_gn=DIP2A;refGene_tx=XM_023238581.1;refGene_gn=DIP2A;refGene_tx=XM_023238582.1;refGene_gn=DIP2A;refGene_tx=XM_023238583.1;refGene_gn=DIP2A;refGene_tx=XM_023238584.1;refGene_gn=DIP2A;refGene_tx=XM_023238585.1;refGene_gn=DIP2A;refGene_tx=XM_023238586.1
C2 299594 299792
refGene_gn=DIP2A;refGene_tx=XM_023238578.1;refGene_gn=DIP2A;refGene_tx=XM_023238579.1;refGene_gn=DIP2A;refGene_tx=XM_023238580.1;refGene_gn=DIP2A;refGene_tx=XM_023238581.1;refGene_gn=DIP2A;refGene_tx=XM_023238582.1;refGene_gn=DIP2A;refGene_tx=XM_023238583.1;refGene_gn=DIP2A;refGene_tx=XM_023238584.1;refGene_gn=DIP2A;refGene_tx=XM_023238585.1;refGene_gn=DIP2A;refGene_tx=XM_023238586.1
C2 301121 301241
refGene_gn=DIP2A;refGene_tx=XM_023238578.1;refGene_gn=DIP2A;refGene_tx=XM_023238579.1;refGene_gn=DIP2A;refGene_tx=XM_023238580.1;refGene_gn=

DIP2A;refGene_tx=XM_023238581.1;refGene_gn=DIP2A;refGene_tx=XM_0232385
82.1;refGene_gn=DIP2A;refGene_tx=XM_023238583.1;refGene_gn=DIP2A;refGe
ne_tx=XM_023238584.1;refGene_gn=DIP2A;refGene_tx=XM_023238585.1;refGen
e_gn=DIP2A;refGene_tx=XM_023238586.1
C2 311338 311470
refGene_gn=DIP2A;refGene_tx=XM_023238578.1;refGene_gn=DIP2A;refGene_tx
=XM_023238579.1;refGene_gn=DIP2A;refGene_tx=XM_023238580.1;refGene_gn=
DIP2A;refGene_tx=XM_023238581.1;refGene_gn=DIP2A;refGene_tx=XM_0232385
83.1;refGene_gn=DIP2A;refGene_tx=XM_023238584.1;refGene_gn=DIP2A;refGe
ne_tx=XM_023238585.1
C2 315050 315281
refGene_gn=DIP2A;refGene_tx=XM_023238578.1;refGene_gn=DIP2A;refGene_tx
=XM_023238579.1;refGene_gn=DIP2A;refGene_tx=XM_023238580.1;refGene_gn=
DIP2A;refGene_tx=XM_023238581.1;refGene_gn=DIP2A;refGene_tx=XM_0232385
82.1;refGene_gn=DIP2A;refGene_tx=XM_023238584.1;refGene_gn=DIP2A;refGe
ne_tx=XM_023238585.1;refGene_gn=DIP2A;refGene_tx=XM_023238586.1
C2 316607 316727
refGene_gn=DIP2A;refGene_tx=XM_023238578.1;refGene_gn=DIP2A;refGene_tx
=XM_023238579.1;refGene_gn=DIP2A;refGene_tx=XM_023238580.1;refGene_gn=
DIP2A;refGene_tx=XM_023238581.1;refGene_gn=DIP2A;refGene_tx=XM_0232385
82.1;refGene_gn=DIP2A;refGene_tx=XM_023238583.1;refGene_gn=DIP2A;refGe
ne_tx=XM_023238584.1;refGene_gn=DIP2A;refGene_tx=XM_023238585.1;refGen
e_gn=DIP2A;refGene_tx=XM_023238586.1
C2 324222 324342
refGene_gn=DIP2A;refGene_tx=XM_023238578.1;refGene_gn=DIP2A;refGene_tx
=XM_023238579.1;refGene_gn=DIP2A;refGene_tx=XM_023238580.1;refGene_gn=
DIP2A;refGene_tx=XM_023238581.1;refGene_gn=DIP2A;refGene_tx=XM_0232385
82.1;refGene_gn=DIP2A;refGene_tx=XM_023238583.1;refGene_gn=DIP2A;refGe
ne_tx=XM_023238586.1
C2 328340 328412
refGene_gn=DIP2A;refGene_tx=XM_023238578.1;refGene_gn=DIP2A;refGene_tx
=XM_023238579.1;refGene_gn=DIP2A;refGene_tx=XM_023238580.1;refGene_gn=
DIP2A;refGene_tx=XM_023238581.1;refGene_gn=DIP2A;refGene_tx=XM_0232385
82.1;refGene_gn=DIP2A;refGene_tx=XM_023238583.1;refGene_gn=DIP2A;refGe
ne_tx=XM_023238586.1
C2 364458 364549
refGene_gn=DIP2A;refGene_tx=XM_023238578.1;refGene_gn=DIP2A;refGene_tx
=XM_023238579.1;refGene_gn=DIP2A;refGene_tx=XM_023238580.1;refGene_gn=
DIP2A;refGene_tx=XM_023238581.1;refGene_gn=DIP2A;refGene_tx=XM_0232385
82.1;refGene_gn=DIP2A;refGene_tx=XM_023238583.1;refGene_gn=DIP2A;refGe
ne_tx=XM_023238586.1
C2 377439 377483 refGene_gn=PCNT;refGene_tx=XM_023238587.1
C2 377927 378055 refGene_gn=PCNT;refGene_tx=XM_023238587.1
C2 379822 379943 refGene_gn=PCNT;refGene_tx=XM_023238587.1
C2 381448 381525 refGene_gn=PCNT;refGene_tx=XM_023238587.1
C2 383091 383324 refGene_gn=PCNT;refGene_tx=XM_023238587.1
C2 384524 384635 refGene_gn=PCNT;refGene_tx=XM_023238587.1
C2 390719 390893 refGene_gn=PCNT;refGene_tx=XM_023238587.1
C2 392135 392238 refGene_gn=PCNT;refGene_tx=XM_023238587.1
C2 393020 393235 refGene_gn=PCNT;refGene_tx=XM_023238587.1

C2	396589	397210	refGene_gn=PCNT;refGene_tx=XM_023238587.1
C2	397879	398030	refGene_gn=PCNT;refGene_tx=XM_023238587.1
C2	398192	398415	refGene_gn=PCNT;refGene_tx=XM_023238587.1
C2	399582	399766	refGene_gn=PCNT;refGene_tx=XM_023238587.1
C2	400368	400542	refGene_gn=PCNT;refGene_tx=XM_023238587.1
C2	401172	401367	refGene_gn=PCNT;refGene_tx=XM_023238587.1
C2	405165	405320	refGene_gn=PCNT;refGene_tx=XM_023238587.1
C2	408142	408245	refGene_gn=PCNT;refGene_tx=XM_023238587.1
C2	409070	409784	refGene_gn=PCNT;refGene_tx=XM_023238587.1
C2	417980	418172	refGene_gn=PCNT;refGene_tx=XM_023238587.1
C2	419693	420518	refGene_gn=PCNT;refGene_tx=XM_023238587.1
C2	422268	422361	refGene_gn=PCNT;refGene_tx=XM_023238587.1
C2	423271	423424	refGene_gn=PCNT;refGene_tx=XM_023238587.1
C2	423687	423846	refGene_gn=PCNT;refGene_tx=XM_023238587.1
C2	425956	426093	refGene_gn=PCNT;refGene_tx=XM_023238587.1
C2	426095	426162	refGene_gn=PCNT;refGene_tx=XM_023238587.1
C2	426899	426914	refGene_gn=PCNT;refGene_tx=XM_023238587.1
C2	427018	427141	refGene_gn=PCNT;refGene_tx=XM_023238587.1
C2	427715	427945	refGene_gn=PCNT;refGene_tx=XM_023238587.1
C2	433407	433620	refGene_gn=PCNT;refGene_tx=XM_023238587.1
C2	434248	434411	refGene_gn=PCNT;refGene_tx=XM_023238587.1
C2	436075	436308	refGene_gn=PCNT;refGene_tx=XM_023238587.1
C2	436523	436684	refGene_gn=PCNT;refGene_tx=XM_023238587.1
C2	439273	439425	refGene_gn=PCNT;refGene_tx=XM_023238587.1
C2	441665	441812	refGene_gn=PCNT;refGene_tx=XM_023238587.1
C2	446533	446671	refGene_gn=PCNT;refGene_tx=XM_023238587.1
C2	459162	459727	refGene_gn=PCNT;refGene_tx=XM_023238587.1
C2	461348	461559	refGene_gn=PCNT;refGene_tx=XM_023238587.1
C2	461561	461817	refGene_gn=PCNT;refGene_tx=XM_023238587.1
C2	468254	468472	refGene_gn=PCNT;refGene_tx=XM_023238587.1
C2	469549	469682	refGene_gn=PCNT;refGene_tx=XM_023238587.1
C2	473302	473372	refGene_gn=PCNT;refGene_tx=XM_023238587.1
C2	474005	474276	refGene_gn=PCNT;refGene_tx=XM_023238587.1
C2	475823	475935	refGene_gn=PCNT;refGene_tx=XM_023238587.1
C2	477088	477225	refGene_gn=PCNT;refGene_tx=XM_023238587.1
C2	478156	478331	refGene_gn=PCNT;refGene_tx=XM_023238587.1
C2	479486	479542	refGene_gn=PCNT;refGene_tx=XM_023238587.1
C2	481696	481952	refGene_gn=PCNT;refGene_tx=XM_023238587.1
C2	483264	483375	refGene_gn=PCNT;refGene_tx=XM_023238587.1
C2	483684	483762	refGene_gn=PCNT;refGene_tx=XM_023238587.1
C2	489184	489280	refGene_gn=PCNT;refGene_tx=XM_023238587.1
C2	507246	507450	refGene_gn=PCNT;refGene_tx=XM_023238587.1
C2	510256	510313	refGene_gn=PCNT;refGene_tx=XM_023238587.1
C2	512616	512636
refGene_gn=CC2H21orf58;refGene_tx=XM_023260862.1
C2	515072	515139
refGene_gn=CC2H21orf58;refGene_tx=XM_023260862.1
C2	516204	516265
refGene_gn=CC2H21orf58;refGene_tx=XM_023260862.1
C2	517413	517487

refGene_gn=CC2H21orf58;refGene_tx=XM_023260862.1
C2	518285	518453
refGene_gn=CC2H21orf58;refGene_tx=XM_023260862.1
C2	522016	522128
refGene_gn=CC2H21orf58;refGene_tx=XM_023260862.1
C2	528151	528381
refGene_gn=CC2H21orf58;refGene_tx=XM_023260862.1
C2	533913	534000	refGene_gn=YBEY;refGene_tx=XM_023238589.1
C2	535153	535222
refGene_gn=YBEY;refGene_tx=XM_023238589.1;refGene_gn=YBEY;refGene_tx=XM_023238590.1;refGene_gn=YBEY;refGene_tx=XM_023238594.1
C2	535676	535759
refGene_gn=YBEY;refGene_tx=XM_023238590.1;refGene_gn=YBEY;refGene_tx=XM_023238591.1;refGene_gn=YBEY;refGene_tx=XM_023238592.1
C2	536236	536365
refGene_gn=YBEY;refGene_tx=XM_023238589.1;refGene_gn=YBEY;refGene_tx=XM_023238590.1;refGene_gn=YBEY;refGene_tx=XM_023238591.1;refGene_gn=YBEY;refGene_tx=XM_023238592.1;refGene_gn=YBEY;refGene_tx=XM_023238593.1;refGene_gn=YBEY;refGene_tx=XM_023238594.1
C2	536576	536652	refGene_gn=YBEY;refGene_tx=XM_023238593.1
C2	538730	538981
refGene_gn=YBEY;refGene_tx=XM_023238589.1;refGene_gn=YBEY;refGene_tx=XM_023238590.1;refGene_gn=YBEY;refGene_tx=XM_023238591.1;refGene_gn=YBEY;refGene_tx=XM_023238592.1;refGene_gn=YBEY;refGene_tx=XM_023238593.1;refGene_gn=YBEY;refGene_tx=XM_023238594.1
C2	539704	540206
refGene_gn=YBEY;refGene_tx=XM_023238590.1;refGene_gn=YBEY;refGene_tx=XM_023238591.1;refGene_gn=YBEY;refGene_tx=XM_023238592.1;refGene_gn=YBEY;refGene_tx=XM_023238593.1;refGene_gn=YBEY;refGene_tx=XM_023238594.1
C2	540775	541988
refGene_gn=MCM3AP;refGene_tx=XM_003991346.5;refGene_gn=MCM3AP;refGene_tx=XM_011285716.3;refGene_gn=MCM3AP;refGene_tx=XM_011285717.3;refGene_gn=MCM3AP;refGene_tx=XM_019839256.2;refGene_gn=MCM3AP;refGene_tx=XM_023238588.1
C2	542213	542437
refGene_gn=MCM3AP;refGene_tx=XM_003991346.5;refGene_gn=MCM3AP;refGene_tx=XM_011285716.3;refGene_gn=MCM3AP;refGene_tx=XM_011285717.3;refGene_gn=MCM3AP;refGene_tx=XM_019839256.2;refGene_gn=MCM3AP;refGene_tx=XM_023238588.1
C2	544652	544731
refGene_gn=MCM3AP;refGene_tx=XM_003991346.5;refGene_gn=MCM3AP;refGene_tx=XM_011285716.3;refGene_gn=MCM3AP;refGene_tx=XM_011285717.3;refGene_gn=MCM3AP;refGene_tx=XM_019839256.2;refGene_gn=MCM3AP;refGene_tx=XM_023238588.1
C2	544963	545108
refGene_gn=MCM3AP;refGene_tx=XM_003991346.5;refGene_gn=MCM3AP;refGene_tx=XM_011285716.3;refGene_gn=MCM3AP;refGene_tx=XM_011285717.3;refGene_gn=MCM3AP;refGene_tx=XM_019839256.2;refGene_gn=MCM3AP;refGene_tx=XM_023238588.1
C2	546218	546409

refGene_gn=MCM3AP;refGene_tx=XM_003991346.5;refGene_gn=MCM3AP;refGene_tx=XM_011285716.3;refGene_gn=MCM3AP;refGene_tx=XM_011285717.3;refGene_gn=MCM3AP;refGene_tx=XM_019839256.2;refGene_gn=MCM3AP;refGene_tx=XM_023238588.1
C2 550311 550451
refGene_gn=MCM3AP;refGene_tx=XM_003991346.5;refGene_gn=MCM3AP;refGene_tx=XM_011285716.3;refGene_gn=MCM3AP;refGene_tx=XM_011285717.3;refGene_gn=MCM3AP;refGene_tx=XM_019839256.2;refGene_gn=MCM3AP;refGene_tx=XM_023238588.1
C2 552635 552833
refGene_gn=MCM3AP;refGene_tx=XM_003991346.5;refGene_gn=MCM3AP;refGene_tx=XM_011285716.3;refGene_gn=MCM3AP;refGene_tx=XM_011285717.3;refGene_gn=MCM3AP;refGene_tx=XM_019839256.2;refGene_gn=MCM3AP;refGene_tx=XM_023238588.1
C2 553371 553640
refGene_gn=MCM3AP;refGene_tx=XM_003991346.5;refGene_gn=MCM3AP;refGene_tx=XM_011285716.3;refGene_gn=MCM3AP;refGene_tx=XM_011285717.3;refGene_gn=MCM3AP;refGene_tx=XM_019839256.2;refGene_gn=MCM3AP;refGene_tx=XM_023238588.1
C2 554773 554936
refGene_gn=MCM3AP;refGene_tx=XM_003991346.5;refGene_gn=MCM3AP;refGene_tx=XM_011285716.3;refGene_gn=MCM3AP;refGene_tx=XM_011285717.3;refGene_gn=MCM3AP;refGene_tx=XM_019839256.2;refGene_gn=MCM3AP;refGene_tx=XM_023238588.1
C2 557263 557424
refGene_gn=MCM3AP;refGene_tx=XM_003991346.5;refGene_gn=MCM3AP;refGene_tx=XM_011285716.3;refGene_gn=MCM3AP;refGene_tx=XM_011285717.3;refGene_gn=MCM3AP;refGene_tx=XM_019839256.2;refGene_gn=MCM3AP;refGene_tx=XM_023238588.1
C2 558327 558566
refGene_gn=MCM3AP;refGene_tx=XM_003991346.5;refGene_gn=MCM3AP;refGene_tx=XM_011285716.3;refGene_gn=MCM3AP;refGene_tx=XM_011285717.3;refGene_gn=MCM3AP;refGene_tx=XM_019839256.2;refGene_gn=MCM3AP;refGene_tx=XM_023238588.1
C2 558866 559093
refGene_gn=MCM3AP;refGene_tx=XM_003991346.5;refGene_gn=MCM3AP;refGene_tx=XM_011285716.3;refGene_gn=MCM3AP;refGene_tx=XM_011285717.3;refGene_gn=MCM3AP;refGene_tx=XM_019839256.2;refGene_gn=MCM3AP;refGene_tx=XM_023238588.1
C2 560214 560315
refGene_gn=MCM3AP;refGene_tx=XM_003991346.5;refGene_gn=MCM3AP;refGene_tx=XM_011285716.3;refGene_gn=MCM3AP;refGene_tx=XM_011285717.3;refGene_gn=MCM3AP;refGene_tx=XM_019839256.2;refGene_gn=MCM3AP;refGene_tx=XM_023238588.1
C2 567085 567217
refGene_gn=MCM3AP;refGene_tx=XM_003991346.5;refGene_gn=MCM3AP;refGene_tx=XM_011285716.3;refGene_gn=MCM3AP;refGene_tx=XM_011285717.3;refGene_gn=MCM3AP;refGene_tx=XM_019839256.2;refGene_gn=MCM3AP;refGene_tx=XM_023238588.1
C2 567623 567737

refGene_gn=MCM3AP;refGene_tx=XM_003991346.5;refGene_gn=MCM3AP;refGene_tx=XM_011285716.3;refGene_gn=MCM3AP;refGene_tx=XM_011285717.3;refGene_gn=MCM3AP;refGene_tx=XM_019839256.2;refGene_gn=MCM3AP;refGene_tx=XM_023238588.1
C2 574123 574276
refGene_gn=MCM3AP;refGene_tx=XM_003991346.5;refGene_gn=MCM3AP;refGene_tx=XM_011285716.3;refGene_gn=MCM3AP;refGene_tx=XM_011285717.3;refGene_gn=MCM3AP;refGene_tx=XM_019839256.2;refGene_gn=MCM3AP;refGene_tx=XM_023238588.1
C2 575816 576014
refGene_gn=MCM3AP;refGene_tx=XM_003991346.5;refGene_gn=MCM3AP;refGene_tx=XM_011285716.3;refGene_gn=MCM3AP;refGene_tx=XM_011285717.3;refGene_gn=MCM3AP;refGene_tx=XM_019839256.2;refGene_gn=MCM3AP;refGene_tx=XM_023238588.1
C2 576591 576660
refGene_gn=MCM3AP;refGene_tx=XM_003991346.5;refGene_gn=MCM3AP;refGene_tx=XM_011285716.3;refGene_gn=MCM3AP;refGene_tx=XM_011285717.3;refGene_gn=MCM3AP;refGene_tx=XM_019839256.2;refGene_gn=MCM3AP;refGene_tx=XM_023238588.1
C2 576741 576876
refGene_gn=MCM3AP;refGene_tx=XM_003991346.5;refGene_gn=MCM3AP;refGene_tx=XM_011285716.3;refGene_gn=MCM3AP;refGene_tx=XM_011285717.3;refGene_gn=MCM3AP;refGene_tx=XM_019839256.2;refGene_gn=MCM3AP;refGene_tx=XM_023238588.1
C2 580184 580338
refGene_gn=MCM3AP;refGene_tx=XM_003991346.5;refGene_gn=MCM3AP;refGene_tx=XM_011285716.3;refGene_gn=MCM3AP;refGene_tx=XM_011285717.3;refGene_gn=MCM3AP;refGene_tx=XM_019839256.2;refGene_gn=MCM3AP;refGene_tx=XM_023238588.1
C2 586435 586706
refGene_gn=MCM3AP;refGene_tx=XM_003991346.5;refGene_gn=MCM3AP;refGene_tx=XM_011285716.3;refGene_gn=MCM3AP;refGene_tx=XM_011285717.3;refGene_gn=MCM3AP;refGene_tx=XM_019839256.2;refGene_gn=MCM3AP;refGene_tx=XM_023238588.1
C2 586918 587016
refGene_gn=MCM3AP;refGene_tx=XM_003991346.5;refGene_gn=MCM3AP;refGene_tx=XM_011285716.3;refGene_gn=MCM3AP;refGene_tx=XM_011285717.3;refGene_gn=MCM3AP;refGene_tx=XM_019839256.2;refGene_gn=MCM3AP;refGene_tx=XM_023238588.1
C2 587676 588067
refGene_gn=MCM3AP;refGene_tx=XM_003991346.5;refGene_gn=MCM3AP;refGene_tx=XM_011285716.3;refGene_gn=MCM3AP;refGene_tx=XM_011285717.3;refGene_gn=MCM3AP;refGene_tx=XM_019839256.2;refGene_gn=MCM3AP;refGene_tx=XM_023238588.1
C2 589689 589947
refGene_gn=MCM3AP;refGene_tx=XM_003991346.5;refGene_gn=MCM3AP;refGene_tx=XM_011285716.3;refGene_gn=MCM3AP;refGene_tx=XM_011285717.3;refGene_gn=MCM3AP;refGene_tx=XM_019839256.2;refGene_gn=MCM3AP;refGene_tx=XM_023238588.1
C2 590433 590563

refGene_gn=MCM3AP;refGene_tx=XM_003991346.5;refGene_gn=MCM3AP;refGene_tx=XM_011285716.3;refGene_gn=MCM3AP;refGene_tx=XM_011285717.3;refGene_gn=MCM3AP;refGene_tx=XM_019839256.2;refGene_gn=MCM3AP;refGene_tx=XM_023238588.1
C2 592391 592621
refGene_gn=MCM3AP;refGene_tx=XM_003991346.5;refGene_gn=MCM3AP;refGene_tx=XM_011285716.3;refGene_gn=MCM3AP;refGene_tx=XM_011285717.3;refGene_gn=MCM3AP;refGene_tx=XM_019839256.2;refGene_gn=MCM3AP;refGene_tx=XM_023238588.1
C2 595006 595157
refGene_gn=MCM3AP;refGene_tx=XM_003991346.5;refGene_gn=MCM3AP;refGene_tx=XM_011285716.3;refGene_gn=MCM3AP;refGene_tx=XM_011285717.3;refGene_gn=MCM3AP;refGene_tx=XM_023238588.1
C2 596257 596436
refGene_gn=MCM3AP;refGene_tx=XM_003991346.5;refGene_gn=MCM3AP;refGene_tx=XM_011285716.3;refGene_gn=MCM3AP;refGene_tx=XM_011285717.3;refGene_gn=MCM3AP;refGene_tx=XM_023238588.1
C2 603514 603528
refGene_gn=LSS;refGene_tx=XM_003991353.5;refGene_gn=LSS;refGene_tx=XM_006935796.3;refGene_gn=LSS;refGene_tx=XM_019839257.2
C2 603830 603999
refGene_gn=LSS;refGene_tx=XM_003991353.5;refGene_gn=LSS;refGene_tx=XM_006935796.3;refGene_gn=LSS;refGene_tx=XM_019839257.2
C2 604652 604791
refGene_gn=LSS;refGene_tx=XM_003991353.5;refGene_gn=LSS;refGene_tx=XM_006935796.3;refGene_gn=LSS;refGene_tx=XM_019839257.2
C2 606764 606786 refGene_gn=LSS;refGene_tx=XM_006935797.4
C2 607641 607750
refGene_gn=LSS;refGene_tx=XM_003991353.5;refGene_gn=LSS;refGene_tx=XM_006935796.3;refGene_gn=LSS;refGene_tx=XM_006935797.4;refGene_gn=LSS;refGene_tx=XM_019839257.2
C2 608792 608914
refGene_gn=LSS;refGene_tx=XM_003991353.5;refGene_gn=LSS;refGene_tx=XM_006935796.3;refGene_gn=LSS;refGene_tx=XM_006935797.4;refGene_gn=LSS;refGene_tx=XM_019839257.2;refGene_gn=LSS;refGene_tx=XM_019839258.2
C2 610287 610384
refGene_gn=LSS;refGene_tx=XM_003991353.5;refGene_gn=LSS;refGene_tx=XM_006935796.3;refGene_gn=LSS;refGene_tx=XM_006935797.4;refGene_gn=LSS;refGene_tx=XM_019839257.2;refGene_gn=LSS;refGene_tx=XM_019839258.2
C2 612486 612622
refGene_gn=LSS;refGene_tx=XM_003991353.5;refGene_gn=LSS;refGene_tx=XM_006935796.3;refGene_gn=LSS;refGene_tx=XM_006935797.4;refGene_gn=LSS;refGene_tx=XM_019839257.2;refGene_gn=LSS;refGene_tx=XM_019839258.2
C2 612920 613029
refGene_gn=LSS;refGene_tx=XM_003991353.5;refGene_gn=LSS;refGene_tx=XM_006935796.3;refGene_gn=LSS;refGene_tx=XM_006935797.4;refGene_gn=LSS;refGene_tx=XM_019839257.2;refGene_gn=LSS;refGene_tx=XM_019839258.2
C2 613593 613712
refGene_gn=LSS;refGene_tx=XM_003991353.5;refGene_gn=LSS;refGene_tx=XM_006935796.3;refGene_gn=LSS;refGene_tx=XM_006935797.4;refGene_gn=LSS;re

fGene_tx=XM_019839257.2;refGene_gn=LSS;refGene_tx=XM_019839258.2
C2	615494	615592
refGene_gn=LSS;refGene_tx=XM_003991353.5;refGene_gn=LSS;refGene_tx=XM_006935796.3;refGene_gn=LSS;refGene_tx=XM_006935797.4;refGene_gn=LSS;refGene_tx=XM_019839257.2;refGene_gn=LSS;refGene_tx=XM_019839258.2
C2	616060	616088
refGene_gn=LSS;refGene_tx=XM_003991353.5;refGene_gn=LSS;refGene_tx=XM_006935796.3;refGene_gn=LSS;refGene_tx=XM_006935797.4;refGene_gn=LSS;refGene_tx=XM_019839257.2;refGene_gn=LSS;refGene_tx=XM_019839258.2
C2	618079	618136
refGene_gn=LSS;refGene_tx=XM_003991353.5;refGene_gn=LSS;refGene_tx=XM_006935796.3;refGene_gn=LSS;refGene_tx=XM_006935797.4;refGene_gn=LSS;refGene_tx=XM_019839257.2;refGene_gn=LSS;refGene_tx=XM_019839258.2
C2	618816	618888
refGene_gn=LSS;refGene_tx=XM_003991353.5;refGene_gn=LSS;refGene_tx=XM_006935796.3;refGene_gn=LSS;refGene_tx=XM_006935797.4;refGene_gn=LSS;refGene_tx=XM_019839257.2;refGene_gn=LSS;refGene_tx=XM_019839258.2
C2	619216	619267
refGene_gn=LSS;refGene_tx=XM_003991353.5;refGene_gn=LSS;refGene_tx=XM_006935796.3;refGene_gn=LSS;refGene_tx=XM_006935797.4;refGene_gn=LSS;refGene_tx=XM_019839257.2;refGene_gn=LSS;refGene_tx=XM_019839258.2
C2	619879	620029
refGene_gn=LSS;refGene_tx=XM_003991353.5;refGene_gn=LSS;refGene_tx=XM_006935796.3;refGene_gn=LSS;refGene_tx=XM_006935797.4;refGene_gn=LSS;refGene_tx=XM_019839257.2;refGene_gn=LSS;refGene_tx=XM_019839258.2
C2	620645	620742
refGene_gn=LSS;refGene_tx=XM_003991353.5;refGene_gn=LSS;refGene_tx=XM_006935796.3;refGene_gn=LSS;refGene_tx=XM_006935797.4;refGene_gn=LSS;refGene_tx=XM_019839257.2;refGene_gn=LSS;refGene_tx=XM_019839258.2
C2	621598	621704
refGene_gn=LSS;refGene_tx=XM_003991353.5;refGene_gn=LSS;refGene_tx=XM_006935796.3;refGene_gn=LSS;refGene_tx=XM_006935797.4;refGene_gn=LSS;refGene_tx=XM_019839257.2;refGene_gn=LSS;refGene_tx=XM_019839258.2
C2	633417	633483
refGene_gn=LSS;refGene_tx=XM_003991353.5;refGene_gn=LSS;refGene_tx=XM_006935796.3;refGene_gn=LSS;refGene_tx=XM_006935797.4;refGene_gn=LSS;refGene_tx=XM_019839257.2;refGene_gn=LSS;refGene_tx=XM_019839258.2
C2	634011	634092
refGene_gn=LSS;refGene_tx=XM_003991353.5;refGene_gn=LSS;refGene_tx=XM_006935796.3;refGene_gn=LSS;refGene_tx=XM_006935797.4;refGene_gn=LSS;refGene_tx=XM_019839257.2;refGene_gn=LSS;refGene_tx=XM_019839258.2
C2	634629	634800
refGene_gn=LSS;refGene_tx=XM_003991353.5;refGene_gn=LSS;refGene_tx=XM_006935796.3;refGene_gn=LSS;refGene_tx=XM_006935797.4;refGene_gn=LSS;refGene_tx=XM_019839257.2;refGene_gn=LSS;refGene_tx=XM_019839258.2
C2	635782	635861
refGene_gn=LSS;refGene_tx=XM_003991353.5;refGene_gn=LSS;refGene_tx=XM_006935796.3;refGene_gn=LSS;refGene_tx=XM_006935797.4;refGene_gn=LSS;refGene_tx=XM_019839257.2;refGene_gn=LSS;refGene_tx=XM_019839258.2
C2	636424	636556

refGene_gn=LSS;refGene_tx=XM_003991353.5;refGene_gn=LSS;refGene_tx=XM_006935796.3;refGene_gn=LSS;refGene_tx=XM_006935797.4;refGene_gn=LSS;refGene_tx=XM_019839257.2;refGene_gn=LSS;refGene_tx=XM_019839258.2
C2 645276 645478
refGene_gn=SPATC1L;refGene_tx=XM_003991347.5;refGene_gn=SPATC1L;refGene_tx=XM_006935798.4
C2 649972 650323
refGene_gn=SPATC1L;refGene_tx=XM_003991347.5;refGene_gn=SPATC1L;refGene_tx=XM_006935798.4
C2 658497 658649
refGene_gn=SPATC1L;refGene_tx=XM_003991347.5;refGene_gn=SPATC1L;refGene_tx=XM_006935798.4
C2 658779 659106
refGene_gn=SPATC1L;refGene_tx=XM_003991347.5;refGene_gn=SPATC1L;refGene_tx=XM_006935798.4
C2 661821 661875
refGene_gn=FTCD;refGene_tx=XM_011285715.3;refGene_gn=FTCD;refGene_tx=XM_019839259.2;refGene_gn=FTCD;refGene_tx=XM_023238597.1
C2 662632 662816
refGene_gn=FTCD;refGene_tx=XM_011285715.3;refGene_gn=FTCD;refGene_tx=XM_019839259.2;refGene_gn=FTCD;refGene_tx=XM_023238597.1
C2 663636 663765
refGene_gn=FTCD;refGene_tx=XM_011285715.3;refGene_gn=FTCD;refGene_tx=XM_019839259.2;refGene_gn=FTCD;refGene_tx=XM_023238597.1
C2 665882 665971
refGene_gn=FTCD;refGene_tx=XM_011285715.3;refGene_gn=FTCD;refGene_tx=XM_019839259.2;refGene_gn=FTCD;refGene_tx=XM_023238597.1
C2 666090 666270
refGene_gn=FTCD;refGene_tx=XM_011285715.3;refGene_gn=FTCD;refGene_tx=XM_019839259.2;refGene_gn=FTCD;refGene_tx=XM_023238597.1
C2 667449 667587
refGene_gn=FTCD;refGene_tx=XM_011285715.3;refGene_gn=FTCD;refGene_tx=XM_019839259.2;refGene_gn=FTCD;refGene_tx=XM_023238597.1
C2 667842 667974
refGene_gn=FTCD;refGene_tx=XM_011285715.3;refGene_gn=FTCD;refGene_tx=XM_019839259.2;refGene_gn=FTCD;refGene_tx=XM_023238597.1
C2 673746 673808
refGene_gn=FTCD;refGene_tx=XM_011285715.3;refGene_gn=FTCD;refGene_tx=XM_019839259.2;refGene_gn=FTCD;refGene_tx=XM_023238597.1
C2 674261 674391
refGene_gn=FTCD;refGene_tx=XM_011285715.3;refGene_gn=FTCD;refGene_tx=XM_019839259.2;refGene_gn=FTCD;refGene_tx=XM_023238597.1
C2 674484 674646
refGene_gn=FTCD;refGene_tx=XM_011285715.3;refGene_gn=FTCD;refGene_tx=XM_019839259.2;refGene_gn=FTCD;refGene_tx=XM_023238597.1
C2 678260 678304
refGene_gn=FTCD;refGene_tx=XM_011285715.3;refGene_gn=FTCD;refGene_tx=XM_019839259.2;refGene_gn=FTCD;refGene_tx=XM_023238597.1
C2 678507 678646
refGene_gn=FTCD;refGene_tx=XM_011285715.3;refGene_gn=FTCD;refGene_tx=X

M_023238597.1
C2	679854	679950
refGene_gn=FTCD;refGene_tx=XM_011285715.3;refGene_gn=FTCD;refGene_tx=XM_019839259.2;refGene_gn=FTCD;refGene_tx=XM_023238597.1
C2	680919	681365
refGene_gn=FTCD;refGene_tx=XM_011285715.3;refGene_gn=FTCD;refGene_tx=XM_023238597.1
C2	684863	685462	refGene_gn=COL6A2;refGene_tx=XM_023238595.1
C2	687517	687813	refGene_gn=COL6A2;refGene_tx=XM_023238596.1
C2	690794	690833
refGene_gn=COL6A2;refGene_tx=XM_023238595.1;refGene_gn=COL6A2;refGene_tx=XM_023238596.1
C2	691002	691455
refGene_gn=COL6A2;refGene_tx=XM_023238595.1;refGene_gn=COL6A2;refGene_tx=XM_023238596.1
C2	691681	691834
refGene_gn=COL6A2;refGene_tx=XM_023238595.1;refGene_gn=COL6A2;refGene_tx=XM_023238596.1
C2	692054	692100
refGene_gn=COL6A2;refGene_tx=XM_023238595.1;refGene_gn=COL6A2;refGene_tx=XM_023238596.1
C2	692238	692274
refGene_gn=COL6A2;refGene_tx=XM_023238595.1;refGene_gn=COL6A2;refGene_tx=XM_023238596.1
C2	692458	692521
refGene_gn=COL6A2;refGene_tx=XM_023238595.1;refGene_gn=COL6A2;refGene_tx=XM_023238596.1
C2	692772	692835
refGene_gn=COL6A2;refGene_tx=XM_023238595.1;refGene_gn=COL6A2;refGene_tx=XM_023238596.1
C2	693149	693185
refGene_gn=COL6A2;refGene_tx=XM_023238595.1;refGene_gn=COL6A2;refGene_tx=XM_023238596.1
C2	693492	693543
refGene_gn=COL6A2;refGene_tx=XM_023238595.1;refGene_gn=COL6A2;refGene_tx=XM_023238596.1
C2	693830	693893
refGene_gn=COL6A2;refGene_tx=XM_023238595.1;refGene_gn=COL6A2;refGene_tx=XM_023238596.1
C2	694277	694340
refGene_gn=COL6A2;refGene_tx=XM_023238595.1;refGene_gn=COL6A2;refGene_tx=XM_023238596.1
C2	694531	694594
refGene_gn=COL6A2;refGene_tx=XM_023238595.1;refGene_gn=COL6A2;refGene_tx=XM_023238596.1
C2	695184	695247
refGene_gn=COL6A2;refGene_tx=XM_023238595.1;refGene_gn=COL6A2;refGene_tx=XM_023238596.1
C2	695948	696038
refGene_gn=COL6A2;refGene_tx=XM_023238595.1;refGene_gn=COL6A2;refGene_

tx=XM_023238596.1
C2	696252	696315
refGene_gn=COL6A2;refGene_tx=XM_023238595.1;refGene_gn=COL6A2;refGene_tx=XM_023238596.1
C2	697125	697188
refGene_gn=COL6A2;refGene_tx=XM_023238595.1;refGene_gn=COL6A2;refGene_tx=XM_023238596.1
C2	697579	697633
refGene_gn=COL6A2;refGene_tx=XM_023238595.1;refGene_gn=COL6A2;refGene_tx=XM_023238596.1
C2	698141	698186
refGene_gn=COL6A2;refGene_tx=XM_023238595.1;refGene_gn=COL6A2;refGene_tx=XM_023238596.1
C2	698270	698297
refGene_gn=COL6A2;refGene_tx=XM_023238595.1;refGene_gn=COL6A2;refGene_tx=XM_023238596.1
C2	698514	698541
refGene_gn=COL6A2;refGene_tx=XM_023238595.1;refGene_gn=COL6A2;refGene_tx=XM_023238596.1
C2	698778	698823
refGene_gn=COL6A2;refGene_tx=XM_023238595.1;refGene_gn=COL6A2;refGene_tx=XM_023238596.1
C2	698905	698959
refGene_gn=COL6A2;refGene_tx=XM_023238595.1;refGene_gn=COL6A2;refGene_tx=XM_023238596.1
C2	699730	699796
refGene_gn=COL6A2;refGene_tx=XM_023238595.1;refGene_gn=COL6A2;refGene_tx=XM_023238596.1
C2	700498	700519
refGene_gn=COL6A2;refGene_tx=XM_023238595.1;refGene_gn=COL6A2;refGene_tx=XM_023238596.1
C2	700779	701378
refGene_gn=COL6A2;refGene_tx=XM_023238595.1;refGene_gn=COL6A2;refGene_tx=XM_023238596.1
C2	701514	701641
refGene_gn=COL6A2;refGene_tx=XM_023238595.1;refGene_gn=COL6A2;refGene_tx=XM_023238596.1
C2	834575	835198	refGene_gn=COL6A1;refGene_tx=XM_011285711.3
C2	835327	835357	refGene_gn=COL6A1;refGene_tx=XM_011285711.3
C2	836187	836368	refGene_gn=COL6A1;refGene_tx=XM_011285711.3
C2	836470	836654	refGene_gn=COL6A1;refGene_tx=XM_011285711.3
C2	836773	836883	refGene_gn=COL6A1;refGene_tx=XM_011285711.3
C2	837102	837236	refGene_gn=COL6A1;refGene_tx=XM_011285711.3
C2	837506	837515	refGene_gn=COL6A1;refGene_tx=XM_011285711.3
C2	837873	837910	refGene_gn=COL6A1;refGene_tx=XM_011285711.3
C2	838180	838216	refGene_gn=COL6A1;refGene_tx=XM_011285711.3
C2	838779	838845	refGene_gn=COL6A1;refGene_tx=XM_011285711.3
C2	839041	839104	refGene_gn=COL6A1;refGene_tx=XM_011285711.3
C2	839689	839725	refGene_gn=COL6A1;refGene_tx=XM_011285711.3
C2	839888	839939	refGene_gn=COL6A1;refGene_tx=XM_011285711.3

C2 840315 840378 refGene_gn=COL6A1;refGene_tx=XM_011285711.3
C2 841080 841143 refGene_gn=COL6A1;refGene_tx=XM_011285711.3
C2 845228 845291 refGene_gn=COL6A1;refGene_tx=XM_011285711.3
C2 846827 846890 refGene_gn=COL6A1;refGene_tx=XM_011285711.3
C2 847090 847126 refGene_gn=COL6A1;refGene_tx=XM_011285711.3
C2 847218 847272 refGene_gn=COL6A1;refGene_tx=XM_011285711.3
C2 847462 847525 refGene_gn=COL6A1;refGene_tx=XM_011285711.3
C2 848315 848378 refGene_gn=COL6A1;refGene_tx=XM_011285711.3
C2 848571 848625 refGene_gn=COL6A1;refGene_tx=XM_011285711.3
C2 848812 848857 refGene_gn=COL6A1;refGene_tx=XM_011285711.3
C2 848945 848972 refGene_gn=COL6A1;refGene_tx=XM_011285711.3
C2 849161 849188 refGene_gn=COL6A1;refGene_tx=XM_011285711.3
C2 849320 849365 refGene_gn=COL6A1;refGene_tx=XM_011285711.3
C2 849869 849923 refGene_gn=COL6A1;refGene_tx=XM_011285711.3
C2 850722 850767 refGene_gn=COL6A1;refGene_tx=XM_011285711.3
C2 850844 850865 refGene_gn=COL6A1;refGene_tx=XM_011285711.3
C2 851061 851082 refGene_gn=COL6A1;refGene_tx=XM_011285711.3
C2 851180 851309 refGene_gn=COL6A1;refGene_tx=XM_011285711.3
C2 851472 851632 refGene_gn=COL6A1;refGene_tx=XM_011285711.3
C2 852973 853174 refGene_gn=COL6A1;refGene_tx=XM_011285711.3
C2 854700 854830 refGene_gn=COL6A1;refGene_tx=XM_011285711.3
C2 855370 855467 refGene_gn=COL6A1;refGene_tx=XM_011285711.3
C2 868274 868810
refGene_gn=LOC109496987;refGene_tx=XM_023238598.1
C2 869105 869355
refGene_gn=LOC109496987;refGene_tx=XM_023238598.1
C2 869767 869905
refGene_gn=LOC109496987;refGene_tx=XM_023238598.1
C2 897840 897877
refGene_gn=PCBP3;refGene_tx=XM_019839261.2;refGene_gn=PCBP3;refGene_tx=XM_023238599.1;refGene_gn=PCBP3;refGene_tx=XM_023238600.1;refGene_gn=PCBP3;refGene_tx=XM_023238601.1;refGene_gn=PCBP3;refGene_tx=XM_023238602.1;refGene_gn=PCBP3;refGene_tx=XM_023238603.1;refGene_gn=PCBP3;refGene_tx=XM_023238604.1;refGene_gn=PCBP3;refGene_tx=XM_023238606.1;refGene_gn=PCBP3;refGene_tx=XM_023238607.1;refGene_gn=PCBP3;refGene_tx=XM_023238608.1;refGene_gn=PCBP3;refGene_tx=XM_023238609.1;refGene_gn=PCBP3;refGene_tx=XM_023238610.1;refGene_gn=PCBP3;refGene_tx=XM_023238611.1;refGene_gn=PCBP3;refGene_tx=XM_023238612.1;refGene_gn=PCBP3;refGene_tx=XM_023238613.1
C2 898964 899134
refGene_gn=PCBP3;refGene_tx=XM_019839261.2;refGene_gn=PCBP3;refGene_tx=XM_023238599.1;refGene_gn=PCBP3;refGene_tx=XM_023238600.1;refGene_gn=PCBP3;refGene_tx=XM_023238601.1;refGene_gn=PCBP3;refGene_tx=XM_023238602.1;refGene_gn=PCBP3;refGene_tx=XM_023238603.1;refGene_gn=PCBP3;refGene_tx=XM_023238604.1;refGene_gn=PCBP3;refGene_tx=XM_023238606.1;refGene_gn=PCBP3;refGene_tx=XM_023238607.1;refGene_gn=PCBP3;refGene_tx=XM_023238608.1;refGene_gn=PCBP3;refGene_tx=XM_023238609.1;refGene_gn=PCBP3;refGene_tx=XM_023238610.1;refGene_gn=PCBP3;refGene_tx=XM_023238611.1;refGene_gn=PCBP3;refGene_tx=XM_023238612.1;refGene_gn=PCBP3;refGene_tx=XM_023238613.1

C2	902820	902873
refGene_gn=PCBP3;refGene_tx=XM_019839261.2;refGene_gn=PCBP3;refGene_tx=XM_023238599.1;refGene_gn=PCBP3;refGene_tx=XM_023238600.1;refGene_gn=PCBP3;refGene_tx=XM_023238601.1;refGene_gn=PCBP3;refGene_tx=XM_023238602.1;refGene_gn=PCBP3;refGene_tx=XM_023238603.1;refGene_gn=PCBP3;refGene_tx=XM_023238604.1;refGene_gn=PCBP3;refGene_tx=XM_023238606.1;refGene_gn=PCBP3;refGene_tx=XM_023238607.1;refGene_gn=PCBP3;refGene_tx=XM_023238608.1;refGene_gn=PCBP3;refGene_tx=XM_023238609.1;refGene_gn=PCBP3;refGene_tx=XM_023238610.1;refGene_gn=PCBP3;refGene_tx=XM_023238611.1;refGene_gn=PCBP3;refGene_tx=XM_023238612.1;refGene_gn=PCBP3;refGene_tx=XM_023238613.1
C2	906271	906331
refGene_gn=PCBP3;refGene_tx=XM_019839261.2;refGene_gn=PCBP3;refGene_tx=XM_023238599.1;refGene_gn=PCBP3;refGene_tx=XM_023238600.1;refGene_gn=PCBP3;refGene_tx=XM_023238601.1;refGene_gn=PCBP3;refGene_tx=XM_023238602.1;refGene_gn=PCBP3;refGene_tx=XM_023238606.1;refGene_gn=PCBP3;refGene_tx=XM_023238607.1;refGene_gn=PCBP3;refGene_tx=XM_023238608.1;refGene_gn=PCBP3;refGene_tx=XM_023238609.1;refGene_gn=PCBP3;refGene_tx=XM_023238612.1;refGene_gn=PCBP3;refGene_tx=XM_023238613.1
C2	907061	907140
refGene_gn=PCBP3;refGene_tx=XM_019839261.2;refGene_gn=PCBP3;refGene_tx=XM_023238599.1;refGene_gn=PCBP3;refGene_tx=XM_023238600.1;refGene_gn=PCBP3;refGene_tx=XM_023238601.1;refGene_gn=PCBP3;refGene_tx=XM_023238602.1;refGene_gn=PCBP3;refGene_tx=XM_023238603.1;refGene_gn=PCBP3;refGene_tx=XM_023238604.1;refGene_gn=PCBP3;refGene_tx=XM_023238606.1;refGene_gn=PCBP3;refGene_tx=XM_023238607.1;refGene_gn=PCBP3;refGene_tx=XM_023238608.1;refGene_gn=PCBP3;refGene_tx=XM_023238609.1;refGene_gn=PCBP3;refGene_tx=XM_023238610.1;refGene_gn=PCBP3;refGene_tx=XM_023238611.1;refGene_gn=PCBP3;refGene_tx=XM_023238612.1;refGene_gn=PCBP3;refGene_tx=XM_023238613.1
C2	916248	916290
refGene_gn=PCBP3;refGene_tx=XM_019839261.2;refGene_gn=PCBP3;refGene_tx=XM_023238599.1;refGene_gn=PCBP3;refGene_tx=XM_023238600.1;refGene_gn=PCBP3;refGene_tx=XM_023238601.1;refGene_gn=PCBP3;refGene_tx=XM_023238602.1;refGene_gn=PCBP3;refGene_tx=XM_023238603.1;refGene_gn=PCBP3;refGene_tx=XM_023238604.1;refGene_gn=PCBP3;refGene_tx=XM_023238606.1;refGene_gn=PCBP3;refGene_tx=XM_023238607.1;refGene_gn=PCBP3;refGene_tx=XM_023238608.1;refGene_gn=PCBP3;refGene_tx=XM_023238609.1;refGene_gn=PCBP3;refGene_tx=XM_023238610.1;refGene_gn=PCBP3;refGene_tx=XM_023238611.1;refGene_gn=PCBP3;refGene_tx=XM_023238612.1;refGene_gn=PCBP3;refGene_tx=XM_023238613.1
C2	919809	919953
refGene_gn=PCBP3;refGene_tx=XM_019839261.2;refGene_gn=PCBP3;refGene_tx=XM_023238599.1;refGene_gn=PCBP3;refGene_tx=XM_023238600.1;refGene_gn=PCBP3;refGene_tx=XM_023238601.1;refGene_gn=PCBP3;refGene_tx=XM_023238602.1;refGene_gn=PCBP3;refGene_tx=XM_023238603.1;refGene_gn=PCBP3;refGene_tx=XM_023238604.1;refGene_gn=PCBP3;refGene_tx=XM_023238606.1;refGene_gn=PCBP3;refGene_tx=XM_023238607.1;refGene_gn=PCBP3;refGene_tx=XM_023238608.1;refGene_gn=PCBP3;refGene_tx=XM_023238609.1;refGene_gn=PCBP3;refGene_tx=XM_023238610.1;refGene_gn=PCBP3;refGene_tx=XM_023238611.1

C2	923118	923247
refGene_gn=PCBP3;refGene_tx=XM_019839261.2;refGene_gn=PCBP3;refGene_tx=XM_023238599.1;refGene_gn=PCBP3;refGene_tx=XM_023238600.1;refGene_gn=PCBP3;refGene_tx=XM_023238601.1;refGene_gn=PCBP3;refGene_tx=XM_023238602.1;refGene_gn=PCBP3;refGene_tx=XM_023238603.1;refGene_gn=PCBP3;refGene_tx=XM_023238604.1;refGene_gn=PCBP3;refGene_tx=XM_023238606.1;refGene_gn=PCBP3;refGene_tx=XM_023238607.1;refGene_gn=PCBP3;refGene_tx=XM_023238608.1;refGene_gn=PCBP3;refGene_tx=XM_023238609.1;refGene_gn=PCBP3;refGene_tx=XM_023238610.1;refGene_gn=PCBP3;refGene_tx=XM_023238611.1;refGene_gn=PCBP3;refGene_tx=XM_023238612.1;refGene_gn=PCBP3;refGene_tx=XM_023238613.1
C2	924090	924222
refGene_gn=PCBP3;refGene_tx=XM_019839261.2;refGene_gn=PCBP3;refGene_tx=XM_023238599.1;refGene_gn=PCBP3;refGene_tx=XM_023238600.1;refGene_gn=PCBP3;refGene_tx=XM_023238601.1;refGene_gn=PCBP3;refGene_tx=XM_023238602.1;refGene_gn=PCBP3;refGene_tx=XM_023238603.1;refGene_gn=PCBP3;refGene_tx=XM_023238604.1;refGene_gn=PCBP3;refGene_tx=XM_023238606.1;refGene_gn=PCBP3;refGene_tx=XM_023238607.1;refGene_gn=PCBP3;refGene_tx=XM_023238608.1;refGene_gn=PCBP3;refGene_tx=XM_023238609.1;refGene_gn=PCBP3;refGene_tx=XM_023238610.1;refGene_gn=PCBP3;refGene_tx=XM_023238611.1;refGene_gn=PCBP3;refGene_tx=XM_023238612.1;refGene_gn=PCBP3;refGene_tx=XM_023238613.1
C2	925134	925251
refGene_gn=PCBP3;refGene_tx=XM_019839261.2;refGene_gn=PCBP3;refGene_tx=XM_023238599.1;refGene_gn=PCBP3;refGene_tx=XM_023238600.1;refGene_gn=PCBP3;refGene_tx=XM_023238601.1;refGene_gn=PCBP3;refGene_tx=XM_023238602.1;refGene_gn=PCBP3;refGene_tx=XM_023238603.1;refGene_gn=PCBP3;refGene_tx=XM_023238604.1;refGene_gn=PCBP3;refGene_tx=XM_023238606.1;refGene_gn=PCBP3;refGene_tx=XM_023238607.1;refGene_gn=PCBP3;refGene_tx=XM_023238608.1;refGene_gn=PCBP3;refGene_tx=XM_023238609.1;refGene_gn=PCBP3;refGene_tx=XM_023238610.1;refGene_gn=PCBP3;refGene_tx=XM_023238611.1;refGene_gn=PCBP3;refGene_tx=XM_023238612.1;refGene_gn=PCBP3;refGene_tx=XM_023238613.1
C2	925575	925608
refGene_gn=PCBP3;refGene_tx=XM_019839261.2;refGene_gn=PCBP3;refGene_tx=XM_023238599.1;refGene_gn=PCBP3;refGene_tx=XM_023238600.1;refGene_gn=PCBP3;refGene_tx=XM_023238601.1;refGene_gn=PCBP3;refGene_tx=XM_023238602.1;refGene_gn=PCBP3;refGene_tx=XM_023238603.1;refGene_gn=PCBP3;refGene_tx=XM_023238604.1;refGene_gn=PCBP3;refGene_tx=XM_023238606.1;refGene_gn=PCBP3;refGene_tx=XM_023238607.1;refGene_gn=PCBP3;refGene_tx=XM_023238608.1;refGene_gn=PCBP3;refGene_tx=XM_023238609.1;refGene_gn=PCBP3;refGene_tx=XM_023238610.1;refGene_gn=PCBP3;refGene_tx=XM_023238611.1;refGene_gn=PCBP3;refGene_tx=XM_023238612.1;refGene_gn=PCBP3;refGene_tx=XM_023238613.1
C2	926423	926447
refGene_gn=PCBP3;refGene_tx=XM_019839261.2;refGene_gn=PCBP3;refGene_tx=XM_023238599.1;refGene_gn=PCBP3;refGene_tx=XM_023238600.1;refGene_gn=PCBP3;refGene_tx=XM_023238601.1;refGene_gn=PCBP3;refGene_tx=XM_023238602.1;refGene_gn=PCBP3;refGene_tx=XM_023238603.1;refGene_gn=PCBP3;refGene_tx=XM_023238604.1;refGene_gn=PCBP3;refGene_tx=XM_023238606.1;refGen

e_gn=PCBP3;refGene_tx=XM_023238607.1;refGene_gn=PCBP3;refGene_tx=XM_02
3238608.1;refGene_gn=PCBP3;refGene_tx=XM_023238609.1;refGene_gn=PCBP3;
refGene_tx=XM_023238610.1;refGene_gn=PCBP3;refGene_tx=XM_023238611.1;r
efGene_gn=PCBP3;refGene_tx=XM_023238612.1;refGene_gn=PCBP3;refGene_tx=
XM_023238613.1
C2 927615 927770
refGene_gn=PCBP3;refGene_tx=XM_019839261.2;refGene_gn=PCBP3;refGene_tx
=XM_023238599.1;refGene_gn=PCBP3;refGene_tx=XM_023238600.1;refGene_gn=
PCBP3;refGene_tx=XM_023238601.1;refGene_gn=PCBP3;refGene_tx=XM_0232386
02.1;refGene_gn=PCBP3;refGene_tx=XM_023238603.1;refGene_gn=PCBP3;refGe
ne_tx=XM_023238604.1;refGene_gn=PCBP3;refGene_tx=XM_023238606.1;refGen
e_gn=PCBP3;refGene_tx=XM_023238607.1;refGene_gn=PCBP3;refGene_tx=XM_02
3238608.1;refGene_gn=PCBP3;refGene_tx=XM_023238609.1;refGene_gn=PCBP3;
refGene_tx=XM_023238610.1;refGene_gn=PCBP3;refGene_tx=XM_023238611.1;r
efGene_gn=PCBP3;refGene_tx=XM_023238612.1;refGene_gn=PCBP3;refGene_tx=
XM_023238613.1
C2 965512 965522
refGene_gn=PCBP3;refGene_tx=XM_019839261.2;refGene_gn=PCBP3;refGene_tx
=XM_023238600.1;refGene_gn=PCBP3;refGene_tx=XM_023238601.1;refGene_gn=
PCBP3;refGene_tx=XM_023238602.1
C2 1240241 1240262 refGene_gn=SLC19A1;refGene_tx=XM_023238624.1
C2 1246159 1246348
refGene_gn=SLC19A1;refGene_tx=XM_023238620.1;refGene_gn=SLC19A1;refGen
e_tx=XM_023238621.1;refGene_gn=SLC19A1;refGene_tx=XM_023238622.1;refGe
ne_gn=SLC19A1;refGene_tx=XM_023238623.1;refGene_gn=SLC19A1;refGene_tx=
XM_023238625.1;refGene_gn=SLC19A1;refGene_tx=XM_023238626.1
C2 1250341 1251113
refGene_gn=SLC19A1;refGene_tx=XM_023238620.1;refGene_gn=SLC19A1;refGen
e_tx=XM_023238621.1;refGene_gn=SLC19A1;refGene_tx=XM_023238622.1;refGe
ne_gn=SLC19A1;refGene_tx=XM_023238623.1;refGene_gn=SLC19A1;refGene_tx=
XM_023238624.1;refGene_gn=SLC19A1;refGene_tx=XM_023238625.1;refGene_gn
=SLC19A1;refGene_tx=XM_023238626.1
C2 1251190 1251383
refGene_gn=SLC19A1;refGene_tx=XM_023238620.1;refGene_gn=SLC19A1;refGen
e_tx=XM_023238621.1;refGene_gn=SLC19A1;refGene_tx=XM_023238622.1;refGe
ne_gn=SLC19A1;refGene_tx=XM_023238623.1;refGene_gn=SLC19A1;refGene_tx=
XM_023238624.1;refGene_gn=SLC19A1;refGene_tx=XM_023238625.1;refGene_gn
=SLC19A1;refGene_tx=XM_023238626.1
C2 1253813 1253956
refGene_gn=SLC19A1;refGene_tx=XM_023238620.1;refGene_gn=SLC19A1;refGen
e_tx=XM_023238621.1;refGene_gn=SLC19A1;refGene_tx=XM_023238622.1;refGe
ne_gn=SLC19A1;refGene_tx=XM_023238623.1;refGene_gn=SLC19A1;refGene_tx=
XM_023238624.1;refGene_gn=SLC19A1;refGene_tx=XM_023238625.1;refGene_gn
=SLC19A1;refGene_tx=XM_023238626.1
C2 1261620 1262069
refGene_gn=SLC19A1;refGene_tx=XM_023238620.1;refGene_gn=SLC19A1;refGen
e_tx=XM_023238621.1;refGene_gn=SLC19A1;refGene_tx=XM_023238622.1;refGe
ne_gn=SLC19A1;refGene_tx=XM_023238623.1;refGene_gn=SLC19A1;refGene_tx=
XM_023238624.1;refGene_gn=SLC19A1;refGene_tx=XM_023238625.1;refGene_gn
=SLC19A1;refGene_tx=XM_023238626.1

C2 1264998 1265212
refGene_gn=COL18A1;refGene_tx=XM_023238614.1;refGene_gn=COL18A1;refGen
e_tx=XM_023238615.1;refGene_gn=COL18A1;refGene_tx=XM_023238616.1;refGe
ne_gn=COL18A1;refGene_tx=XM_023238617.1;refGene_gn=COL18A1;refGene_tx=
XM_023238618.1;refGene_gn=COL18A1;refGene_tx=XM_023238619.1
C2 1265395 1265514
refGene_gn=COL18A1;refGene_tx=XM_023238614.1;refGene_gn=COL18A1;refGen
e_tx=XM_023238615.1;refGene_gn=COL18A1;refGene_tx=XM_023238616.1;refGe
ne_gn=COL18A1;refGene_tx=XM_023238617.1;refGene_gn=COL18A1;refGene_tx=
XM_023238618.1;refGene_gn=COL18A1;refGene_tx=XM_023238619.1
C2 1265867 1266065
refGene_gn=COL18A1;refGene_tx=XM_023238614.1;refGene_gn=COL18A1;refGen
e_tx=XM_023238615.1;refGene_gn=COL18A1;refGene_tx=XM_023238616.1;refGe
ne_gn=COL18A1;refGene_tx=XM_023238617.1;refGene_gn=COL18A1;refGene_tx=
XM_023238618.1;refGene_gn=COL18A1;refGene_tx=XM_023238619.1
C2 1266359 1266611
refGene_gn=COL18A1;refGene_tx=XM_023238614.1;refGene_gn=COL18A1;refGen
e_tx=XM_023238615.1;refGene_gn=COL18A1;refGene_tx=XM_023238616.1;refGe
ne_gn=COL18A1;refGene_tx=XM_023238617.1;refGene_gn=COL18A1;refGene_tx=
XM_023238618.1;refGene_gn=COL18A1;refGene_tx=XM_023238619.1
C2 1267472 1267505
refGene_gn=COL18A1;refGene_tx=XM_023238614.1;refGene_gn=COL18A1;refGen
e_tx=XM_023238615.1;refGene_gn=COL18A1;refGene_tx=XM_023238616.1;refGe
ne_gn=COL18A1;refGene_tx=XM_023238617.1;refGene_gn=COL18A1;refGene_tx=
XM_023238618.1;refGene_gn=COL18A1;refGene_tx=XM_023238619.1
C2 1267874 1268006
refGene_gn=COL18A1;refGene_tx=XM_023238614.1;refGene_gn=COL18A1;refGen
e_tx=XM_023238615.1;refGene_gn=COL18A1;refGene_tx=XM_023238616.1;refGe
ne_gn=COL18A1;refGene_tx=XM_023238617.1;refGene_gn=COL18A1;refGene_tx=
XM_023238618.1;refGene_gn=COL18A1;refGene_tx=XM_023238619.1
C2 1268477 1268551
refGene_gn=COL18A1;refGene_tx=XM_023238614.1;refGene_gn=COL18A1;refGen
e_tx=XM_023238615.1;refGene_gn=COL18A1;refGene_tx=XM_023238616.1;refGe
ne_gn=COL18A1;refGene_tx=XM_023238617.1;refGene_gn=COL18A1;refGene_tx=
XM_023238618.1;refGene_gn=COL18A1;refGene_tx=XM_023238619.1
C2 1268635 1268789
refGene_gn=COL18A1;refGene_tx=XM_023238614.1;refGene_gn=COL18A1;refGen
e_tx=XM_023238615.1;refGene_gn=COL18A1;refGene_tx=XM_023238616.1;refGe
ne_gn=COL18A1;refGene_tx=XM_023238617.1;refGene_gn=COL18A1;refGene_tx=
XM_023238618.1;refGene_gn=COL18A1;refGene_tx=XM_023238619.1
C2 1269256 1269379
refGene_gn=COL18A1;refGene_tx=XM_023238614.1;refGene_gn=COL18A1;refGen
e_tx=XM_023238615.1;refGene_gn=COL18A1;refGene_tx=XM_023238616.1;refGe
ne_gn=COL18A1;refGene_tx=XM_023238617.1;refGene_gn=COL18A1;refGene_tx=
XM_023238618.1;refGene_gn=COL18A1;refGene_tx=XM_023238619.1
C2 1269605 1269649
refGene_gn=COL18A1;refGene_tx=XM_023238614.1;refGene_gn=COL18A1;refGen
e_tx=XM_023238615.1;refGene_gn=COL18A1;refGene_tx=XM_023238616.1;refGe
ne_gn=COL18A1;refGene_tx=XM_023238617.1;refGene_gn=COL18A1;refGene_tx=
XM_023238618.1;refGene_gn=COL18A1;refGene_tx=XM_023238619.1

C2	1273210	1273273
refGene_gn=COL18A1;refGene_tx=XM_023238614.1;refGene_gn=COL18A1;refGene_tx=XM_023238615.1;refGene_gn=COL18A1;refGene_tx=XM_023238616.1;refGene_gn=COL18A1;refGene_tx=XM_023238617.1;refGene_gn=COL18A1;refGene_tx=XM_023238618.1;refGene_gn=COL18A1;refGene_tx=XM_023238619.1
C2	1273803	1273846
refGene_gn=COL18A1;refGene_tx=XM_023238614.1;refGene_gn=COL18A1;refGene_tx=XM_023238615.1;refGene_gn=COL18A1;refGene_tx=XM_023238616.1;refGene_gn=COL18A1;refGene_tx=XM_023238617.1;refGene_gn=COL18A1;refGene_tx=XM_023238618.1;refGene_gn=COL18A1;refGene_tx=XM_023238619.1
C2	1274292	1274361
refGene_gn=COL18A1;refGene_tx=XM_023238614.1;refGene_gn=COL18A1;refGene_tx=XM_023238615.1;refGene_gn=COL18A1;refGene_tx=XM_023238616.1;refGene_gn=COL18A1;refGene_tx=XM_023238617.1;refGene_gn=COL18A1;refGene_tx=XM_023238618.1;refGene_gn=COL18A1;refGene_tx=XM_023238619.1
C2	1275710	1275785
refGene_gn=COL18A1;refGene_tx=XM_023238614.1;refGene_gn=COL18A1;refGene_tx=XM_023238615.1;refGene_gn=COL18A1;refGene_tx=XM_023238616.1;refGene_gn=COL18A1;refGene_tx=XM_023238617.1;refGene_gn=COL18A1;refGene_tx=XM_023238618.1;refGene_gn=COL18A1;refGene_tx=XM_023238619.1
C2	1276047	1276101
refGene_gn=COL18A1;refGene_tx=XM_023238614.1;refGene_gn=COL18A1;refGene_tx=XM_023238615.1;refGene_gn=COL18A1;refGene_tx=XM_023238616.1;refGene_gn=COL18A1;refGene_tx=XM_023238617.1;refGene_gn=COL18A1;refGene_tx=XM_023238618.1;refGene_gn=COL18A1;refGene_tx=XM_023238619.1
C2	1276306	1276333
refGene_gn=COL18A1;refGene_tx=XM_023238614.1;refGene_gn=COL18A1;refGene_tx=XM_023238615.1;refGene_gn=COL18A1;refGene_tx=XM_023238616.1;refGene_gn=COL18A1;refGene_tx=XM_023238617.1;refGene_gn=COL18A1;refGene_tx=XM_023238618.1;refGene_gn=COL18A1;refGene_tx=XM_023238619.1
C2	1277022	1277097
refGene_gn=COL18A1;refGene_tx=XM_023238614.1;refGene_gn=COL18A1;refGene_tx=XM_023238615.1;refGene_gn=COL18A1;refGene_tx=XM_023238616.1;refGene_gn=COL18A1;refGene_tx=XM_023238617.1;refGene_gn=COL18A1;refGene_tx=XM_023238618.1;refGene_gn=COL18A1;refGene_tx=XM_023238619.1
C2	1277380	1277443
refGene_gn=COL18A1;refGene_tx=XM_023238614.1;refGene_gn=COL18A1;refGene_tx=XM_023238615.1;refGene_gn=COL18A1;refGene_tx=XM_023238616.1;refGene_gn=COL18A1;refGene_tx=XM_023238617.1;refGene_gn=COL18A1;refGene_tx=XM_023238618.1;refGene_gn=COL18A1;refGene_tx=XM_023238619.1
C2	1277665	1277692
refGene_gn=COL18A1;refGene_tx=XM_023238614.1;refGene_gn=COL18A1;refGene_tx=XM_023238615.1;refGene_gn=COL18A1;refGene_tx=XM_023238616.1;refGene_gn=COL18A1;refGene_tx=XM_023238617.1;refGene_gn=COL18A1;refGene_tx=XM_023238618.1;refGene_gn=COL18A1;refGene_tx=XM_023238619.1
C2	1277772	1277802
refGene_gn=COL18A1;refGene_tx=XM_023238614.1;refGene_gn=COL18A1;refGene_tx=XM_023238615.1;refGene_gn=COL18A1;refGene_tx=XM_023238616.1;refGene_gn=COL18A1;refGene_tx=XM_023238617.1;refGene_gn=COL18A1;refGene_tx=XM_023238618.1;refGene_gn=COL18A1;refGene_tx=XM_023238619.1

C2 1278363 1278453
refGene_gn=COL18A1;refGene_tx=XM_023238614.1;refGene_gn=COL18A1;refGene_tx=XM_023238615.1;refGene_gn=COL18A1;refGene_tx=XM_023238616.1;refGene_gn=COL18A1;refGene_tx=XM_023238617.1;refGene_gn=COL18A1;refGene_tx=XM_023238618.1;refGene_gn=COL18A1;refGene_tx=XM_023238619.1
C2 1278708 1278744
refGene_gn=COL18A1;refGene_tx=XM_023238614.1;refGene_gn=COL18A1;refGene_tx=XM_023238615.1;refGene_gn=COL18A1;refGene_tx=XM_023238616.1;refGene_gn=COL18A1;refGene_tx=XM_023238617.1;refGene_gn=COL18A1;refGene_tx=XM_023238618.1;refGene_gn=COL18A1;refGene_tx=XM_023238619.1
C2 1279046 1279118
refGene_gn=COL18A1;refGene_tx=XM_023238614.1;refGene_gn=COL18A1;refGene_tx=XM_023238615.1;refGene_gn=COL18A1;refGene_tx=XM_023238616.1;refGene_gn=COL18A1;refGene_tx=XM_023238617.1;refGene_gn=COL18A1;refGene_tx=XM_023238618.1;refGene_gn=COL18A1;refGene_tx=XM_023238619.1
C2 1279639 1279675
refGene_gn=COL18A1;refGene_tx=XM_023238614.1;refGene_gn=COL18A1;refGene_tx=XM_023238615.1;refGene_gn=COL18A1;refGene_tx=XM_023238616.1;refGene_gn=COL18A1;refGene_tx=XM_023238617.1;refGene_gn=COL18A1;refGene_tx=XM_023238618.1;refGene_gn=COL18A1;refGene_tx=XM_023238619.1
C2 1280431 1280458
refGene_gn=COL18A1;refGene_tx=XM_023238614.1;refGene_gn=COL18A1;refGene_tx=XM_023238615.1;refGene_gn=COL18A1;refGene_tx=XM_023238616.1;refGene_gn=COL18A1;refGene_tx=XM_023238617.1;refGene_gn=COL18A1;refGene_tx=XM_023238618.1;refGene_gn=COL18A1;refGene_tx=XM_023238619.1
C2 1281602 1281665
refGene_gn=COL18A1;refGene_tx=XM_023238614.1;refGene_gn=COL18A1;refGene_tx=XM_023238615.1;refGene_gn=COL18A1;refGene_tx=XM_023238616.1;refGene_gn=COL18A1;refGene_tx=XM_023238617.1;refGene_gn=COL18A1;refGene_tx=XM_023238618.1;refGene_gn=COL18A1;refGene_tx=XM_023238619.1
C2 1282180 1282312
refGene_gn=COL18A1;refGene_tx=XM_023238614.1;refGene_gn=COL18A1;refGene_tx=XM_023238615.1;refGene_gn=COL18A1;refGene_tx=XM_023238616.1;refGene_gn=COL18A1;refGene_tx=XM_023238617.1;refGene_gn=COL18A1;refGene_tx=XM_023238618.1;refGene_gn=COL18A1;refGene_tx=XM_023238619.1
C2 1284377 1284404
refGene_gn=COL18A1;refGene_tx=XM_023238614.1;refGene_gn=COL18A1;refGene_tx=XM_023238615.1;refGene_gn=COL18A1;refGene_tx=XM_023238616.1;refGene_gn=COL18A1;refGene_tx=XM_023238617.1;refGene_gn=COL18A1;refGene_tx=XM_023238618.1;refGene_gn=COL18A1;refGene_tx=XM_023238619.1
C2 1285243 1285306
refGene_gn=COL18A1;refGene_tx=XM_023238614.1;refGene_gn=COL18A1;refGene_tx=XM_023238615.1;refGene_gn=COL18A1;refGene_tx=XM_023238616.1;refGene_gn=COL18A1;refGene_tx=XM_023238617.1;refGene_gn=COL18A1;refGene_tx=XM_023238618.1;refGene_gn=COL18A1;refGene_tx=XM_023238619.1
C2 1286073 1286232
refGene_gn=COL18A1;refGene_tx=XM_023238614.1;refGene_gn=COL18A1;refGene_tx=XM_023238615.1;refGene_gn=COL18A1;refGene_tx=XM_023238616.1;refGene_gn=COL18A1;refGene_tx=XM_023238617.1;refGene_gn=COL18A1;refGene_tx=XM_023238618.1;refGene_gn=COL18A1;refGene_tx=XM_023238619.1

C2 1286364 1286418
refGene_gn=COL18A1;refGene_tx=XM_023238614.1;refGene_gn=COL18A1;refGene_tx=XM_023238615.1;refGene_gn=COL18A1;refGene_tx=XM_023238616.1;refGene_gn=COL18A1;refGene_tx=XM_023238617.1;refGene_gn=COL18A1;refGene_tx=XM_023238618.1;refGene_gn=COL18A1;refGene_tx=XM_023238619.1
C2 1286817 1286904
refGene_gn=COL18A1;refGene_tx=XM_023238614.1;refGene_gn=COL18A1;refGene_tx=XM_023238615.1;refGene_gn=COL18A1;refGene_tx=XM_023238616.1;refGene_gn=COL18A1;refGene_tx=XM_023238617.1;refGene_gn=COL18A1;refGene_tx=XM_023238618.1;refGene_gn=COL18A1;refGene_tx=XM_023238619.1
C2 1287011 1287074
refGene_gn=COL18A1;refGene_tx=XM_023238614.1;refGene_gn=COL18A1;refGene_tx=XM_023238615.1;refGene_gn=COL18A1;refGene_tx=XM_023238616.1;refGene_gn=COL18A1;refGene_tx=XM_023238617.1;refGene_gn=COL18A1;refGene_tx=XM_023238618.1;refGene_gn=COL18A1;refGene_tx=XM_023238619.1
C2 1288234 1288261
refGene_gn=COL18A1;refGene_tx=XM_023238614.1;refGene_gn=COL18A1;refGene_tx=XM_023238615.1;refGene_gn=COL18A1;refGene_tx=XM_023238616.1;refGene_gn=COL18A1;refGene_tx=XM_023238617.1;refGene_gn=COL18A1;refGene_tx=XM_023238618.1;refGene_gn=COL18A1;refGene_tx=XM_023238619.1
C2 1288790 1288910
refGene_gn=COL18A1;refGene_tx=XM_023238614.1;refGene_gn=COL18A1;refGene_tx=XM_023238615.1;refGene_gn=COL18A1;refGene_tx=XM_023238616.1;refGene_gn=COL18A1;refGene_tx=XM_023238617.1;refGene_gn=COL18A1;refGene_tx=XM_023238618.1;refGene_gn=COL18A1;refGene_tx=XM_023238619.1
C2 1289159 1289230
refGene_gn=COL18A1;refGene_tx=XM_023238614.1;refGene_gn=COL18A1;refGene_tx=XM_023238615.1;refGene_gn=COL18A1;refGene_tx=XM_023238616.1;refGene_gn=COL18A1;refGene_tx=XM_023238617.1;refGene_gn=COL18A1;refGene_tx=XM_023238618.1;refGene_gn=COL18A1;refGene_tx=XM_023238619.1
C2 1293363 1293493
refGene_gn=COL18A1;refGene_tx=XM_023238614.1;refGene_gn=COL18A1;refGene_tx=XM_023238615.1;refGene_gn=COL18A1;refGene_tx=XM_023238616.1;refGene_gn=COL18A1;refGene_tx=XM_023238617.1;refGene_gn=COL18A1;refGene_tx=XM_023238618.1;refGene_gn=COL18A1;refGene_tx=XM_023238619.1
C2 1295277 1295337
refGene_gn=COL18A1;refGene_tx=XM_023238614.1;refGene_gn=COL18A1;refGene_tx=XM_023238615.1;refGene_gn=COL18A1;refGene_tx=XM_023238616.1;refGene_gn=COL18A1;refGene_tx=XM_023238617.1;refGene_gn=COL18A1;refGene_tx=XM_023238618.1;refGene_gn=COL18A1;refGene_tx=XM_023238619.1
C2 1296902 1296989
refGene_gn=COL18A1;refGene_tx=XM_023238614.1;refGene_gn=COL18A1;refGene_tx=XM_023238615.1;refGene_gn=COL18A1;refGene_tx=XM_023238616.1;refGene_gn=COL18A1;refGene_tx=XM_023238617.1;refGene_gn=COL18A1;refGene_tx=XM_023238618.1;refGene_gn=COL18A1;refGene_tx=XM_023238619.1
C2 1303764 1304309
refGene_gn=COL18A1;refGene_tx=XM_023238614.1;refGene_gn=COL18A1;refGene_tx=XM_023238615.1;refGene_gn=COL18A1;refGene_tx=XM_023238616.1;refGene_gn=COL18A1;refGene_tx=XM_023238617.1;refGene_gn=COL18A1;refGene_tx=XM_023238618.1;refGene_gn=COL18A1;refGene_tx=XM_023238619.1

C2 1314478 1315886
refGene_gn=COL18A1;refGene_tx=XM_023238614.1;refGene_gn=COL18A1;refGene_tx=XM_023238615.1;refGene_gn=COL18A1;refGene_tx=XM_023238616.1;refGene_gn=COL18A1;refGene_tx=XM_023238618.1
C2 1355939 1356034
refGene_gn=COL18A1;refGene_tx=XM_023238617.1;refGene_gn=COL18A1;refGene_tx=XM_023238619.1
C2 1356124 1356135
refGene_gn=COL18A1;refGene_tx=XM_023238617.1;refGene_gn=COL18A1;refGene_tx=XM_023238619.1
C2 1450971 1451222
refGene_gn=POFUT2;refGene_tx=XM_023238627.1;refGene_gn=POFUT2;refGene_tx=XM_023238628.1
C2 1452578 1452829
refGene_gn=POFUT2;refGene_tx=XM_023238627.1;refGene_gn=POFUT2;refGene_tx=XM_023238628.1
C2 1454993 1455138
refGene_gn=POFUT2;refGene_tx=XM_023238627.1;refGene_gn=POFUT2;refGene_tx=XM_023238628.1
C2 1455721 1455832
refGene_gn=POFUT2;refGene_tx=XM_023238627.1;refGene_gn=POFUT2;refGene_tx=XM_023238628.1
C2 1457494 1457561
refGene_gn=POFUT2;refGene_tx=XM_023238627.1;refGene_gn=POFUT2;refGene_tx=XM_023238628.1
C2 1458225 1458351
refGene_gn=POFUT2;refGene_tx=XM_023238627.1;refGene_gn=POFUT2;refGene_tx=XM_023238628.1
C2 1459025 1459206
refGene_gn=POFUT2;refGene_tx=XM_023238627.1;refGene_gn=POFUT2;refGene_tx=XM_023238628.1
C2 1459403 1459527
refGene_gn=POFUT2;refGene_tx=XM_023238627.1;refGene_gn=POFUT2;refGene_tx=XM_023238628.1
C2 1460230 1460449
refGene_gn=POFUT2;refGene_tx=XM_023238627.1;refGene_gn=POFUT2;refGene_tx=XM_023238628.1
C2 1487301 1487481
refGene_gn=ADARB1;refGene_tx=XM_019839278.2;refGene_gn=ADARB1;refGene_tx=XM_019839279.2;refGene_gn=ADARB1;refGene_tx=XM_019839281.2;refGene_gn=ADARB1;refGene_tx=XM_019839283.2;refGene_gn=ADARB1;refGene_tx=XM_019839288.1;refGene_gn=ADARB1;refGene_tx=XM_023238629.1;refGene_gn=ADARB1;refGene_tx=XM_023238630.1;refGene_gn=ADARB1;refGene_tx=XM_023238631.1;refGene_gn=ADARB1;refGene_tx=XM_023238632.1;refGene_gn=ADARB1;refGene_tx=XM_023238633.1
C2 1488382 1488561
refGene_gn=ADARB1;refGene_tx=XM_019839278.2;refGene_gn=ADARB1;refGene_tx=XM_019839279.2;refGene_gn=ADARB1;refGene_tx=XM_019839281.2;refGene_gn=ADARB1;refGene_tx=XM_019839283.2;refGene_gn=ADARB1;refGene_tx=XM_019839288.1;refGene_gn=ADARB1;refGene_tx=XM_023238629.1;refGene_gn=ADARB

1;refGene_tx=XM_023238630.1;refGene_gn=ADARB1;refGene_tx=XM_023238631.
1;refGene_gn=ADARB1;refGene_tx=XM_023238632.1;refGene_gn=ADARB1;refGen
e_tx=XM_023238633.1
C2 1502954 1503136
refGene_gn=ADARB1;refGene_tx=XM_019839278.2;refGene_gn=ADARB1;refGene_
tx=XM_019839279.2;refGene_gn=ADARB1;refGene_tx=XM_019839281.2;refGene_
gn=ADARB1;refGene_tx=XM_019839283.2;refGene_gn=ADARB1;refGene_tx=XM_01
9839288.1;refGene_gn=ADARB1;refGene_tx=XM_023238629.1;refGene_gn=ADARB
1;refGene_tx=XM_023238630.1;refGene_gn=ADARB1;refGene_tx=XM_023238631.
1;refGene_gn=ADARB1;refGene_tx=XM_023238632.1;refGene_gn=ADARB1;refGen
e_tx=XM_023238633.1
C2 1516030 1516262
refGene_gn=ADARB1;refGene_tx=XM_019839278.2;refGene_gn=ADARB1;refGene_
tx=XM_019839279.2;refGene_gn=ADARB1;refGene_tx=XM_019839281.2;refGene_
gn=ADARB1;refGene_tx=XM_019839283.2;refGene_gn=ADARB1;refGene_tx=XM_01
9839288.1;refGene_gn=ADARB1;refGene_tx=XM_023238629.1;refGene_gn=ADARB
1;refGene_tx=XM_023238630.1;refGene_gn=ADARB1;refGene_tx=XM_023238631.
1;refGene_gn=ADARB1;refGene_tx=XM_023238632.1;refGene_gn=ADARB1;refGen
e_tx=XM_023238633.1
C2 1517303 1517452
refGene_gn=ADARB1;refGene_tx=XM_019839278.2;refGene_gn=ADARB1;refGene_
tx=XM_019839279.2;refGene_gn=ADARB1;refGene_tx=XM_019839281.2;refGene_
gn=ADARB1;refGene_tx=XM_019839283.2;refGene_gn=ADARB1;refGene_tx=XM_01
9839288.1;refGene_gn=ADARB1;refGene_tx=XM_023238629.1;refGene_gn=ADARB
1;refGene_tx=XM_023238630.1;refGene_gn=ADARB1;refGene_tx=XM_023238631.
1;refGene_gn=ADARB1;refGene_tx=XM_023238632.1;refGene_gn=ADARB1;refGen
e_tx=XM_023238633.1
C2 1518081 1518250
refGene_gn=ADARB1;refGene_tx=XM_019839278.2;refGene_gn=ADARB1;refGene_
tx=XM_019839279.2;refGene_gn=ADARB1;refGene_tx=XM_019839281.2;refGene_
gn=ADARB1;refGene_tx=XM_019839283.2;refGene_gn=ADARB1;refGene_tx=XM_01
9839288.1;refGene_gn=ADARB1;refGene_tx=XM_023238629.1;refGene_gn=ADARB
1;refGene_tx=XM_023238630.1;refGene_gn=ADARB1;refGene_tx=XM_023238631.
1;refGene_gn=ADARB1;refGene_tx=XM_023238632.1;refGene_gn=ADARB1;refGen
e_tx=XM_023238633.1
C2 1520289 1520404
refGene_gn=ADARB1;refGene_tx=XM_019839278.2;refGene_gn=ADARB1;refGene_
tx=XM_019839279.2;refGene_gn=ADARB1;refGene_tx=XM_019839281.2;refGene_
gn=ADARB1;refGene_tx=XM_019839283.2;refGene_gn=ADARB1;refGene_tx=XM_01
9839288.1;refGene_gn=ADARB1;refGene_tx=XM_023238629.1;refGene_gn=ADARB
1;refGene_tx=XM_023238630.1;refGene_gn=ADARB1;refGene_tx=XM_023238631.
1;refGene_gn=ADARB1;refGene_tx=XM_023238632.1;refGene_gn=ADARB1;refGen
e_tx=XM_023238633.1
C2 1524011 1524993
refGene_gn=ADARB1;refGene_tx=XM_019839278.2;refGene_gn=ADARB1;refGene_
tx=XM_019839279.2;refGene_gn=ADARB1;refGene_tx=XM_019839281.2;refGene_
gn=ADARB1;refGene_tx=XM_019839283.2;refGene_gn=ADARB1;refGene_tx=XM_01
9839288.1;refGene_gn=ADARB1;refGene_tx=XM_023238629.1;refGene_gn=ADARB
1;refGene_tx=XM_023238630.1;refGene_gn=ADARB1;refGene_tx=XM_023238631.
1;refGene_gn=ADARB1;refGene_tx=XM_023238632.1;refGene_gn=ADARB1;refGen

e_tx=XM_023238633.1
C2 1526094 1526171 refGene_gn=ADARB1;refGene_tx=XM_019839278.2
C2 1528441 1528516
refGene_gn=ADARB1;refGene_tx=XM_019839278.2;refGene_gn=ADARB1;refGene_tx=XM_019839279.2;refGene_gn=ADARB1;refGene_tx=XM_019839281.2;refGene_gn=ADARB1;refGene_tx=XM_019839283.2;refGene_gn=ADARB1;refGene_tx=XM_019839288.1;refGene_gn=ADARB1;refGene_tx=XM_023238629.1;refGene_gn=ADARB1;refGene_tx=XM_023238630.1;refGene_gn=ADARB1;refGene_tx=XM_023238631.1;refGene_gn=ADARB1;refGene_tx=XM_023238632.1;refGene_gn=ADARB1;refGene_tx=XM_023238633.1
C2 1546824 1546924
refGene_gn=ADARB1;refGene_tx=XM_019839278.2;refGene_gn=ADARB1;refGene_tx=XM_019839279.2;refGene_gn=ADARB1;refGene_tx=XM_019839281.2;refGene_gn=ADARB1;refGene_tx=XM_019839283.2;refGene_gn=ADARB1;refGene_tx=XM_019839288.1;refGene_gn=ADARB1;refGene_tx=XM_023238629.1;refGene_gn=ADARB1;refGene_tx=XM_023238630.1
C2 1711186 1711310 refGene_gn=FAM207A;refGene_tx=XM_006935857.4
C2 1712456 1712528 refGene_gn=FAM207A;refGene_tx=XM_006935857.4
C2 1715378 1715529 refGene_gn=FAM207A;refGene_tx=XM_006935857.4
C2 1721672 1721741 refGene_gn=FAM207A;refGene_tx=XM_006935857.4
C2 1738283 1738392 refGene_gn=FAM207A;refGene_tx=XM_006935857.4
C2 1744173 1744338 refGene_gn=FAM207A;refGene_tx=XM_006935857.4
C2 1772055 1772113 refGene_gn=ITGB2;refGene_tx=XM_011285804.3
C2 1772404 1772493 refGene_gn=ITGB2;refGene_tx=XM_011285804.3
C2 1775604 1775785 refGene_gn=ITGB2;refGene_tx=XM_011285804.3
C2 1776751 1776922 refGene_gn=ITGB2;refGene_tx=XM_011285804.3
C2 1777492 1777734 refGene_gn=ITGB2;refGene_tx=XM_011285804.3
C2 1778859 1779015 refGene_gn=ITGB2;refGene_tx=XM_011285804.3
C2 1779423 1779519 refGene_gn=ITGB2;refGene_tx=XM_011285804.3
C2 1781700 1781790 refGene_gn=ITGB2;refGene_tx=XM_011285804.3
C2 1782594 1782735 refGene_gn=ITGB2;refGene_tx=XM_011285804.3
C2 1784273 1784461 refGene_gn=ITGB2;refGene_tx=XM_011285804.3
C2 1784552 1784797 refGene_gn=ITGB2;refGene_tx=XM_011285804.3
C2 1785082 1785302 refGene_gn=ITGB2;refGene_tx=XM_011285804.3
C2 1785493 1785699 refGene_gn=ITGB2;refGene_tx=XM_011285804.3
C2 1787673 1787840 refGene_gn=ITGB2;refGene_tx=XM_011285804.3
C2 1787992 1788055 refGene_gn=ITGB2;refGene_tx=XM_011285804.3
C2 1797167 1797282 refGene_gn=PTTG1IP;refGene_tx=XM_023238634.1
C2 1804622 1804689 refGene_gn=PTTG1IP;refGene_tx=XM_019839290.2
C2 1804933 1804986
refGene_gn=PTTG1IP;refGene_tx=XM_019839290.2;refGene_gn=PTTG1IP;refGene_tx=XM_023238634.1
C2 1807325 1807434
refGene_gn=PTTG1IP;refGene_tx=XM_019839290.2;refGene_gn=PTTG1IP;refGene_tx=XM_023238634.1
C2 1811160 1811335
refGene_gn=PTTG1IP;refGene_tx=XM_019839290.2;refGene_gn=PTTG1IP;refGene_tx=XM_023238634.1
C2 1811874 1811921
refGene_gn=PTTG1IP;refGene_tx=XM_019839290.2;refGene_gn=PTTG1IP;refGen

e_tx=XM_023238634.1
C2	1814703	1814750
refGene_gn=PTTG1IP;refGene_tx=XM_019839290.2;refGene_gn=PTTG1IP;refGen
e_tx=XM_023238634.1
C2	1830819	1830840	refGene_gn=SUMO3;refGene_tx=XM_023238639.1
C2	1835624	1835753	refGene_gn=SUMO3;refGene_tx=XM_023238639.1
C2	1839318	1839390	refGene_gn=SUMO3;refGene_tx=XM_023238639.1
C2	1841319	1841397	refGene_gn=SUMO3;refGene_tx=XM_023238639.1
C2	1844810	1844853	refGene_gn=UBE2G2;refGene_tx=XM_023238636.1
C2	1861622	1861658	refGene_gn=UBE2G2;refGene_tx=XM_023238636.1
C2	1861752	1861798
refGene_gn=UBE2G2;refGene_tx=XM_019839291.2;refGene_gn=UBE2G2;refGene_
tx=XM_023238636.1;refGene_gn=UBE2G2;refGene_tx=XM_023238637.1
C2	1876574	1876693
refGene_gn=UBE2G2;refGene_tx=XM_019839291.2;refGene_gn=UBE2G2;refGene_
tx=XM_023238636.1;refGene_gn=UBE2G2;refGene_tx=XM_023238637.1;refGene_
gn=UBE2G2;refGene_tx=XM_023238638.1
C2	1880205	1880346
refGene_gn=UBE2G2;refGene_tx=XM_019839291.2;refGene_gn=UBE2G2;refGene_
tx=XM_023238636.1;refGene_gn=UBE2G2;refGene_tx=XM_023238637.1;refGene_
gn=UBE2G2;refGene_tx=XM_023238638.1
C2	1882770	1882883
refGene_gn=UBE2G2;refGene_tx=XM_019839291.2;refGene_gn=UBE2G2;refGene_
tx=XM_023238636.1;refGene_gn=UBE2G2;refGene_tx=XM_023238637.1;refGene_
gn=UBE2G2;refGene_tx=XM_023238638.1
C2	1925666	1925751
refGene_gn=TSPEAR;refGene_tx=XM_023238640.1;refGene_gn=TSPEAR;refGene_
tx=XM_023238641.1
C2	1931044	1932082
refGene_gn=LOC105260904;refGene_tx=XM_019839298.2
C2	1936197	1936869
refGene_gn=LOC101088263;refGene_tx=XM_019839300.1
C2	1949201	1949489
refGene_gn=LOC101088509;refGene_tx=XM_003991381.5
C2	1962631	1963057
refGene_gn=LOC105260893;refGene_tx=XM_011285802.3
C2	1965198	1965549
refGene_gn=LOC105260892;refGene_tx=XM_011285801.2
C2	1969210	1969656
refGene_gn=LOC111556338;refGene_tx=XM_023238643.1
C2	1969658	1970247
refGene_gn=LOC111556338;refGene_tx=XM_023238643.1
C2	1978252	1978440
refGene_gn=LOC105260903;refGene_tx=XM_023260828.1
C2	1989247	1989342
refGene_gn=LOC109496909;refGene_tx=XM_023260857.1
C2	1989446	1989637
refGene_gn=LOC109496909;refGene_tx=XM_023260857.1
C2	1989791	1989853
refGene_gn=LOC109496909;refGene_tx=XM_023260857.1

C2 1990914 1990976
refGene_gn=LOC109496909;refGene_tx=XM_023260857.1
C2 1992282 1993513
refGene_gn=LOC109496909;refGene_tx=XM_023260857.1
C2 2006940 2008011
refGene_gn=LOC101089539;refGene_tx=XM_019839299.2
C2 2012261 2013068
refGene_gn=LOC109503265;refGene_tx=XM_023238646.1
C2 2031003 2031224
refGene_gn=TSPEAR;refGene_tx=XM_023238640.1;refGene_gn=TSPEAR;refGene_tx=XM_023238641.1
C2 2036861 2037395
refGene_gn=LOC111556339;refGene_tx=XM_023238644.1
C2 2044099 2044277
refGene_gn=LOC111562356;refGene_tx=XM_023260858.1
C2 2046672 2046735
refGene_gn=LOC111562356;refGene_tx=XM_023260858.1
C2 2053910 2054795
refGene_gn=LOC111556340;refGene_tx=XM_023238645.1
C2 2057014 2057253
refGene_gn=TSPEAR;refGene_tx=XM_023238640.1;refGene_gn=TSPEAR;refGene_tx=XM_023238641.1
C2 2059141 2059232
refGene_gn=TSPEAR;refGene_tx=XM_023238640.1;refGene_gn=TSPEAR;refGene_tx=XM_023238641.1
C2 2061957 2062114
refGene_gn=TSPEAR;refGene_tx=XM_023238640.1;refGene_gn=TSPEAR;refGene_tx=XM_023238641.1
C2 2063475 2063607
refGene_gn=TSPEAR;refGene_tx=XM_023238640.1;refGene_gn=TSPEAR;refGene_tx=XM_023238641.1
C2 2064300 2064527
refGene_gn=TSPEAR;refGene_tx=XM_023238640.1;refGene_gn=TSPEAR;refGene_tx=XM_023238641.1
C2 2065963 2066150
refGene_gn=TSPEAR;refGene_tx=XM_023238640.1;refGene_gn=TSPEAR;refGene_tx=XM_023238641.1
C2 2067309 2067539
refGene_gn=TSPEAR;refGene_tx=XM_023238640.1;refGene_gn=TSPEAR;refGene_tx=XM_023238641.1
C2 2079299 2079487 refGene_gn=TSPEAR;refGene_tx=XM_023238640.1
C2 2081721 2081823
refGene_gn=TSPEAR;refGene_tx=XM_023238640.1;refGene_gn=TSPEAR;refGene_tx=XM_023238641.1
C2 2084592 2084746
refGene_gn=TSPEAR;refGene_tx=XM_023238640.1;refGene_gn=TSPEAR;refGene_tx=XM_023238641.1
C2 2097953 2098727 refGene_gn=LRRC3;refGene_tx=XM_023238648.1
C2 2111487 2111613
refGene_gn=TRPM2;refGene_tx=XM_011285792.3;refGene_gn=TRPM2;refGene_tx

=XM_019839301.2
C2 2112193 2112310
refGene_gn=TRPM2;refGene_tx=XM_011285792.3;refGene_gn=TRPM2;refGene_tx=XM_019839301.2
C2 2112523 2112568 refGene_gn=TRPM2;refGene_tx=XM_019839302.2
C2 2113727 2113829
refGene_gn=TRPM2;refGene_tx=XM_011285792.3;refGene_gn=TRPM2;refGene_tx=XM_019839301.2;refGene_gn=TRPM2;refGene_tx=XM_019839302.2;refGene_gn=TRPM2;refGene_tx=XM_019839303.2;refGene_gn=TRPM2;refGene_tx=XM_019839304.2;refGene_gn=TRPM2;refGene_tx=XM_019839305.2
C2 2115394 2115500
refGene_gn=TRPM2;refGene_tx=XM_011285792.3;refGene_gn=TRPM2;refGene_tx=XM_019839301.2;refGene_gn=TRPM2;refGene_tx=XM_019839302.2;refGene_gn=TRPM2;refGene_tx=XM_019839303.2;refGene_gn=TRPM2;refGene_tx=XM_019839304.2;refGene_gn=TRPM2;refGene_tx=XM_019839305.2
C2 2116077 2116164
refGene_gn=TRPM2;refGene_tx=XM_011285792.3;refGene_gn=TRPM2;refGene_tx=XM_019839301.2;refGene_gn=TRPM2;refGene_tx=XM_019839302.2;refGene_gn=TRPM2;refGene_tx=XM_019839303.2;refGene_gn=TRPM2;refGene_tx=XM_019839304.2;refGene_gn=TRPM2;refGene_tx=XM_019839305.2
C2 2116447 2116549
refGene_gn=TRPM2;refGene_tx=XM_011285792.3;refGene_gn=TRPM2;refGene_tx=XM_019839301.2;refGene_gn=TRPM2;refGene_tx=XM_019839302.2;refGene_gn=TRPM2;refGene_tx=XM_019839303.2;refGene_gn=TRPM2;refGene_tx=XM_019839304.2;refGene_gn=TRPM2;refGene_tx=XM_019839305.2;refGene_gn=TRPM2;refGene_tx=XM_019839306.2
C2 2116799 2116876
refGene_gn=TRPM2;refGene_tx=XM_011285792.3;refGene_gn=TRPM2;refGene_tx=XM_019839301.2;refGene_gn=TRPM2;refGene_tx=XM_019839302.2;refGene_gn=TRPM2;refGene_tx=XM_019839306.2
C2 2117571 2117729
refGene_gn=TRPM2;refGene_tx=XM_011285792.3;refGene_gn=TRPM2;refGene_tx=XM_019839301.2;refGene_gn=TRPM2;refGene_tx=XM_019839302.2;refGene_gn=TRPM2;refGene_tx=XM_019839303.2;refGene_gn=TRPM2;refGene_tx=XM_019839304.2;refGene_gn=TRPM2;refGene_tx=XM_019839305.2;refGene_gn=TRPM2;refGene_tx=XM_019839306.2
C2 2117773 2117781 refGene_gn=TRPM2;refGene_tx=XM_011285799.3
C2 2118620 2118708
refGene_gn=TRPM2;refGene_tx=XM_011285792.3;refGene_gn=TRPM2;refGene_tx=XM_011285799.3;refGene_gn=TRPM2;refGene_tx=XM_019839301.2;refGene_gn=TRPM2;refGene_tx=XM_019839302.2;refGene_gn=TRPM2;refGene_tx=XM_019839303.2;refGene_gn=TRPM2;refGene_tx=XM_019839304.2;refGene_gn=TRPM2;refGene_tx=XM_019839305.2;refGene_gn=TRPM2;refGene_tx=XM_019839306.2
C2 2120529 2120617
refGene_gn=TRPM2;refGene_tx=XM_011285792.3;refGene_gn=TRPM2;refGene_tx=XM_011285799.3;refGene_gn=TRPM2;refGene_tx=XM_019839301.2;refGene_gn=TRPM2;refGene_tx=XM_019839302.2;refGene_gn=TRPM2;refGene_tx=XM_019839303.2;refGene_gn=TRPM2;refGene_tx=XM_019839304.2;refGene_gn=TRPM2;refGene_tx=XM_019839305.2;refGene_gn=TRPM2;refGene_tx=XM_019839306.2
C2 2122201 2122334

refGene_gn=TRPM2;refGene_tx=XM_011285792.3;refGene_gn=TRPM2;refGene_tx
=XM_011285799.3;refGene_gn=TRPM2;refGene_tx=XM_019839301.2;refGene_gn=
TRPM2;refGene_tx=XM_019839302.2;refGene_gn=TRPM2;refGene_tx=XM_0198393
03.2;refGene_gn=TRPM2;refGene_tx=XM_019839304.2;refGene_gn=TRPM2;refGe
ne_tx=XM_019839305.2;refGene_gn=TRPM2;refGene_tx=XM_019839306.2
C2 2122568 2122750
refGene_gn=TRPM2;refGene_tx=XM_011285792.3;refGene_gn=TRPM2;refGene_tx
=XM_011285799.3;refGene_gn=TRPM2;refGene_tx=XM_019839301.2;refGene_gn=
TRPM2;refGene_tx=XM_019839302.2;refGene_gn=TRPM2;refGene_tx=XM_0198393
03.2;refGene_gn=TRPM2;refGene_tx=XM_019839304.2;refGene_gn=TRPM2;refGe
ne_tx=XM_019839305.2;refGene_gn=TRPM2;refGene_tx=XM_019839306.2
C2 2123137 2123321
refGene_gn=TRPM2;refGene_tx=XM_011285792.3;refGene_gn=TRPM2;refGene_tx
=XM_011285799.3;refGene_gn=TRPM2;refGene_tx=XM_019839301.2;refGene_gn=
TRPM2;refGene_tx=XM_019839302.2;refGene_gn=TRPM2;refGene_tx=XM_0198393
03.2;refGene_gn=TRPM2;refGene_tx=XM_019839304.2;refGene_gn=TRPM2;refGe
ne_tx=XM_019839305.2;refGene_gn=TRPM2;refGene_tx=XM_019839306.2
C2 2128211 2128383
refGene_gn=TRPM2;refGene_tx=XM_011285792.3;refGene_gn=TRPM2;refGene_tx
=XM_011285799.3;refGene_gn=TRPM2;refGene_tx=XM_019839301.2;refGene_gn=
TRPM2;refGene_tx=XM_019839302.2;refGene_gn=TRPM2;refGene_tx=XM_0198393
03.2;refGene_gn=TRPM2;refGene_tx=XM_019839304.2;refGene_gn=TRPM2;refGe
ne_tx=XM_019839305.2;refGene_gn=TRPM2;refGene_tx=XM_019839306.2
C2 2128793 2128926
refGene_gn=TRPM2;refGene_tx=XM_011285792.3;refGene_gn=TRPM2;refGene_tx
=XM_011285799.3;refGene_gn=TRPM2;refGene_tx=XM_019839301.2;refGene_gn=
TRPM2;refGene_tx=XM_019839302.2;refGene_gn=TRPM2;refGene_tx=XM_0198393
03.2;refGene_gn=TRPM2;refGene_tx=XM_019839304.2;refGene_gn=TRPM2;refGe
ne_tx=XM_019839305.2;refGene_gn=TRPM2;refGene_tx=XM_019839306.2
C2 2129435 2129554
refGene_gn=TRPM2;refGene_tx=XM_011285792.3;refGene_gn=TRPM2;refGene_tx
=XM_011285799.3;refGene_gn=TRPM2;refGene_tx=XM_019839301.2;refGene_gn=
TRPM2;refGene_tx=XM_019839302.2;refGene_gn=TRPM2;refGene_tx=XM_0198393
03.2;refGene_gn=TRPM2;refGene_tx=XM_019839304.2;refGene_gn=TRPM2;refGe
ne_tx=XM_019839305.2;refGene_gn=TRPM2;refGene_tx=XM_019839306.2
C2 2130246 2130460
refGene_gn=TRPM2;refGene_tx=XM_011285792.3;refGene_gn=TRPM2;refGene_tx
=XM_011285799.3;refGene_gn=TRPM2;refGene_tx=XM_019839301.2;refGene_gn=
TRPM2;refGene_tx=XM_019839302.2;refGene_gn=TRPM2;refGene_tx=XM_0198393
03.2;refGene_gn=TRPM2;refGene_tx=XM_019839304.2;refGene_gn=TRPM2;refGe
ne_tx=XM_019839305.2;refGene_gn=TRPM2;refGene_tx=XM_019839306.2
C2 2131427 2131540
refGene_gn=TRPM2;refGene_tx=XM_011285792.3;refGene_gn=TRPM2;refGene_tx
=XM_011285799.3;refGene_gn=TRPM2;refGene_tx=XM_019839301.2;refGene_gn=
TRPM2;refGene_tx=XM_019839302.2;refGene_gn=TRPM2;refGene_tx=XM_0198393
03.2;refGene_gn=TRPM2;refGene_tx=XM_019839304.2;refGene_gn=TRPM2;refGe
ne_tx=XM_019839305.2;refGene_gn=TRPM2;refGene_tx=XM_019839306.2
C2 2132015 2132161
refGene_gn=TRPM2;refGene_tx=XM_011285792.3;refGene_gn=TRPM2;refGene_tx
=XM_011285799.3;refGene_gn=TRPM2;refGene_tx=XM_019839301.2;refGene_gn=

TRPM2;refGene_tx=XM_019839302.2;refGene_gn=TRPM2;refGene_tx=XM_019839303.2;refGene_gn=TRPM2;refGene_tx=XM_019839304.2;refGene_gn=TRPM2;refGene_tx=XM_019839305.2;refGene_gn=TRPM2;refGene_tx=XM_019839306.2
C2 2132348 2132478
refGene_gn=TRPM2;refGene_tx=XM_011285792.3;refGene_gn=TRPM2;refGene_tx=XM_011285799.3;refGene_gn=TRPM2;refGene_tx=XM_019839301.2;refGene_gn=TRPM2;refGene_tx=XM_019839302.2;refGene_gn=TRPM2;refGene_tx=XM_019839303.2;refGene_gn=TRPM2;refGene_tx=XM_019839304.2;refGene_gn=TRPM2;refGene_tx=XM_019839305.2;refGene_gn=TRPM2;refGene_tx=XM_019839306.2
C2 2133812 2133950
refGene_gn=TRPM2;refGene_tx=XM_011285792.3;refGene_gn=TRPM2;refGene_tx=XM_011285799.3;refGene_gn=TRPM2;refGene_tx=XM_019839301.2;refGene_gn=TRPM2;refGene_tx=XM_019839302.2;refGene_gn=TRPM2;refGene_tx=XM_019839303.2;refGene_gn=TRPM2;refGene_tx=XM_019839304.2;refGene_gn=TRPM2;refGene_tx=XM_019839305.2;refGene_gn=TRPM2;refGene_tx=XM_019839306.2
C2 2136243 2136597
refGene_gn=TRPM2;refGene_tx=XM_011285792.3;refGene_gn=TRPM2;refGene_tx=XM_011285799.3;refGene_gn=TRPM2;refGene_tx=XM_019839301.2;refGene_gn=TRPM2;refGene_tx=XM_019839302.2;refGene_gn=TRPM2;refGene_tx=XM_019839303.2;refGene_gn=TRPM2;refGene_tx=XM_019839304.2;refGene_gn=TRPM2;refGene_tx=XM_019839305.2;refGene_gn=TRPM2;refGene_tx=XM_019839306.2
C2 2136777 2136899
refGene_gn=TRPM2;refGene_tx=XM_011285792.3;refGene_gn=TRPM2;refGene_tx=XM_011285799.3;refGene_gn=TRPM2;refGene_tx=XM_019839301.2;refGene_gn=TRPM2;refGene_tx=XM_019839302.2;refGene_gn=TRPM2;refGene_tx=XM_019839303.2;refGene_gn=TRPM2;refGene_tx=XM_019839304.2;refGene_gn=TRPM2;refGene_tx=XM_019839305.2;refGene_gn=TRPM2;refGene_tx=XM_019839306.2
C2 2140201 2140304
refGene_gn=TRPM2;refGene_tx=XM_011285792.3;refGene_gn=TRPM2;refGene_tx=XM_011285799.3;refGene_gn=TRPM2;refGene_tx=XM_019839301.2;refGene_gn=TRPM2;refGene_tx=XM_019839302.2;refGene_gn=TRPM2;refGene_tx=XM_019839303.2;refGene_gn=TRPM2;refGene_tx=XM_019839304.2;refGene_gn=TRPM2;refGene_tx=XM_019839305.2;refGene_gn=TRPM2;refGene_tx=XM_019839306.2
C2 2142505 2142706
refGene_gn=TRPM2;refGene_tx=XM_011285792.3;refGene_gn=TRPM2;refGene_tx=XM_011285799.3;refGene_gn=TRPM2;refGene_tx=XM_019839301.2;refGene_gn=TRPM2;refGene_tx=XM_019839302.2;refGene_gn=TRPM2;refGene_tx=XM_019839303.2;refGene_gn=TRPM2;refGene_tx=XM_019839304.2;refGene_gn=TRPM2;refGene_tx=XM_019839305.2;refGene_gn=TRPM2;refGene_tx=XM_019839306.2
C2 2143767 2143829
refGene_gn=TRPM2;refGene_tx=XM_011285792.3;refGene_gn=TRPM2;refGene_tx=XM_011285799.3;refGene_gn=TRPM2;refGene_tx=XM_019839301.2;refGene_gn=TRPM2;refGene_tx=XM_019839302.2;refGene_gn=TRPM2;refGene_tx=XM_019839303.2;refGene_gn=TRPM2;refGene_tx=XM_019839304.2;refGene_gn=TRPM2;refGene_tx=XM_019839305.2;refGene_gn=TRPM2;refGene_tx=XM_019839306.2
C2 2144748 2144929
refGene_gn=TRPM2;refGene_tx=XM_011285792.3;refGene_gn=TRPM2;refGene_tx=XM_011285799.3;refGene_gn=TRPM2;refGene_tx=XM_019839301.2;refGene_gn=TRPM2;refGene_tx=XM_019839302.2;refGene_gn=TRPM2;refGene_tx=XM_019839303.2;refGene_gn=TRPM2;refGene_tx=XM_019839304.2;refGene_gn=TRPM2;refGe

ne_tx=XM_019839305.2;refGene_gn=TRPM2;refGene_tx=XM_019839306.2
C2 2146227 2146394
refGene_gn=TRPM2;refGene_tx=XM_011285792.3;refGene_gn=TRPM2;refGene_tx=XM_011285799.3;refGene_gn=TRPM2;refGene_tx=XM_019839301.2;refGene_gn=TRPM2;refGene_tx=XM_019839302.2;refGene_gn=TRPM2;refGene_tx=XM_019839303.2;refGene_gn=TRPM2;refGene_tx=XM_019839304.2;refGene_gn=TRPM2;refGene_tx=XM_019839305.2;refGene_gn=TRPM2;refGene_tx=XM_019839306.2
C2 2148775 2148956
refGene_gn=TRPM2;refGene_tx=XM_011285792.3;refGene_gn=TRPM2;refGene_tx=XM_011285799.3;refGene_gn=TRPM2;refGene_tx=XM_019839301.2;refGene_gn=TRPM2;refGene_tx=XM_019839302.2;refGene_gn=TRPM2;refGene_tx=XM_019839303.2;refGene_gn=TRPM2;refGene_tx=XM_019839304.2;refGene_gn=TRPM2;refGene_tx=XM_019839305.2;refGene_gn=TRPM2;refGene_tx=XM_019839306.2
C2 2149471 2149640
refGene_gn=TRPM2;refGene_tx=XM_011285792.3;refGene_gn=TRPM2;refGene_tx=XM_011285799.3;refGene_gn=TRPM2;refGene_tx=XM_019839301.2;refGene_gn=TRPM2;refGene_tx=XM_019839302.2;refGene_gn=TRPM2;refGene_tx=XM_019839303.2;refGene_gn=TRPM2;refGene_tx=XM_019839304.2;refGene_gn=TRPM2;refGene_tx=XM_019839305.2;refGene_gn=TRPM2;refGene_tx=XM_019839306.2
C2 2158197 2158286
refGene_gn=TRPM2;refGene_tx=XM_011285792.3;refGene_gn=TRPM2;refGene_tx=XM_011285799.3;refGene_gn=TRPM2;refGene_tx=XM_019839301.2;refGene_gn=TRPM2;refGene_tx=XM_019839302.2;refGene_gn=TRPM2;refGene_tx=XM_019839303.2;refGene_gn=TRPM2;refGene_tx=XM_019839304.2;refGene_gn=TRPM2;refGene_tx=XM_019839305.2;refGene_gn=TRPM2;refGene_tx=XM_019839306.2
C2 2159031 2159196
refGene_gn=TRPM2;refGene_tx=XM_011285792.3;refGene_gn=TRPM2;refGene_tx=XM_011285799.3;refGene_gn=TRPM2;refGene_tx=XM_019839301.2;refGene_gn=TRPM2;refGene_tx=XM_019839302.2;refGene_gn=TRPM2;refGene_tx=XM_019839303.2;refGene_gn=TRPM2;refGene_tx=XM_019839304.2;refGene_gn=TRPM2;refGene_tx=XM_019839305.2;refGene_gn=TRPM2;refGene_tx=XM_019839306.2
C2 2170059 2170136
refGene_gn=CC2H21orf2;refGene_tx=XM_011285788.3;refGene_gn=CC2H21orf2;refGene_tx=XM_011285789.3
C2 2171422 2171441
refGene_gn=CC2H21orf2;refGene_tx=XM_011285788.3;refGene_gn=CC2H21orf2;refGene_tx=XM_011285789.3
C2 2172723 2172770
refGene_gn=CC2H21orf2;refGene_tx=XM_006935815.4;refGene_gn=CC2H21orf2;refGene_tx=XM_011285788.3;refGene_gn=CC2H21orf2;refGene_tx=XM_011285789.3;refGene_gn=CC2H21orf2;refGene_tx=XM_011285791.3;refGene_gn=CC2H21orf2;refGene_tx=XM_019839307.2
C2 2174922 2175152
refGene_gn=CC2H21orf2;refGene_tx=XM_006935815.4;refGene_gn=CC2H21orf2;refGene_tx=XM_011285788.3;refGene_gn=CC2H21orf2;refGene_tx=XM_011285789.3;refGene_gn=CC2H21orf2;refGene_tx=XM_011285791.3;refGene_gn=CC2H21orf2;refGene_tx=XM_019839307.2
C2 2176238 2176410
refGene_gn=CC2H21orf2;refGene_tx=XM_006935815.4;refGene_gn=CC2H21orf2;refGene_tx=XM_011285788.3;refGene_gn=CC2H21orf2;refGene_tx=XM_01128578

9.3;refGene_gn=CC2H21orf2;refGene_tx=XM_011285791.3;refGene_gn=CC2H21orf2;refGene_tx=XM_019839307.2
C2 2177297 2177391
refGene_gn=CC2H21orf2;refGene_tx=XM_006935815.4;refGene_gn=CC2H21orf2;refGene_tx=XM_011285788.3;refGene_gn=CC2H21orf2;refGene_tx=XM_011285789.3;refGene_gn=CC2H21orf2;refGene_tx=XM_011285791.3;refGene_gn=CC2H21orf2;refGene_tx=XM_019839307.2
C2 2179262 2179394
refGene_gn=CC2H21orf2;refGene_tx=XM_006935815.4;refGene_gn=CC2H21orf2;refGene_tx=XM_011285788.3;refGene_gn=CC2H21orf2;refGene_tx=XM_011285789.3;refGene_gn=CC2H21orf2;refGene_tx=XM_011285791.3;refGene_gn=CC2H21orf2;refGene_tx=XM_019839307.2
C2 2183091 2183239
refGene_gn=PFKL;refGene_tx=XM_019839309.2;refGene_gn=PFKL;refGene_tx=XM_023238651.1
C2 2183954 2184060
refGene_gn=PFKL;refGene_tx=XM_019839309.2;refGene_gn=PFKL;refGene_tx=XM_023238651.1
C2 2184148 2184248
refGene_gn=PFKL;refGene_tx=XM_019839309.2;refGene_gn=PFKL;refGene_tx=XM_023238651.1
C2 2184825 2184937
refGene_gn=PFKL;refGene_tx=XM_019839309.2;refGene_gn=PFKL;refGene_tx=XM_023238651.1
C2 2185074 2185136
refGene_gn=PFKL;refGene_tx=XM_019839309.2;refGene_gn=PFKL;refGene_tx=XM_023238651.1
C2 2185550 2185715
refGene_gn=PFKL;refGene_tx=XM_019839309.2;refGene_gn=PFKL;refGene_tx=XM_023238651.1
C2 2187162 2187315
refGene_gn=PFKL;refGene_tx=XM_019839309.2;refGene_gn=PFKL;refGene_tx=XM_023238651.1
C2 2188932 2189020
refGene_gn=PFKL;refGene_tx=XM_019839309.2;refGene_gn=PFKL;refGene_tx=XM_023238651.1
C2 2189457 2189528
refGene_gn=PFKL;refGene_tx=XM_019839309.2;refGene_gn=PFKL;refGene_tx=XM_023238651.1
C2 2189773 2189920
refGene_gn=PFKL;refGene_tx=XM_019839309.2;refGene_gn=PFKL;refGene_tx=XM_023238651.1
C2 2191113 2191177
refGene_gn=PFKL;refGene_tx=XM_019839309.2;refGene_gn=PFKL;refGene_tx=XM_023238651.1
C2 2191725 2191790
refGene_gn=PFKL;refGene_tx=XM_019839309.2;refGene_gn=PFKL;refGene_tx=XM_023238651.1
C2 2192190 2192316
refGene_gn=PFKL;refGene_tx=XM_019839309.2;refGene_gn=PFKL;refGene_tx=X

M_023238651.1
C2	2193513	2193606
refGene_gn=PFKL;refGene_tx=XM_019839309.2;refGene_gn=PFKL;refGene_tx=X
M_023238651.1
C2	2194546	2194642
refGene_gn=PFKL;refGene_tx=XM_019839309.2;refGene_gn=PFKL;refGene_tx=X
M_023238651.1
C2	2196643	2196752
refGene_gn=PFKL;refGene_tx=XM_019839309.2;refGene_gn=PFKL;refGene_tx=X
M_023238651.1
C2	2196972	2197017
refGene_gn=PFKL;refGene_tx=XM_019839309.2;refGene_gn=PFKL;refGene_tx=X
M_023238651.1
C2	2197467	2197633
refGene_gn=PFKL;refGene_tx=XM_019839309.2;refGene_gn=PFKL;refGene_tx=X
M_023238651.1
C2	2198341	2198531
refGene_gn=PFKL;refGene_tx=XM_019839309.2;refGene_gn=PFKL;refGene_tx=X
M_023238651.1
C2	2201006	2201084
refGene_gn=PFKL;refGene_tx=XM_019839309.2;refGene_gn=PFKL;refGene_tx=X
M_023238651.1
C2	2204013	2204087
refGene_gn=PFKL;refGene_tx=XM_019839309.2;refGene_gn=PFKL;refGene_tx=X
M_023238651.1
C2	2209355	2209440	refGene_gn=PFKL;refGene_tx=XM_023238651.1
C2	2219761	2219842	refGene_gn=AIRE;refGene_tx=XM_023260864.1
C2	2220860	2220929	refGene_gn=AIRE;refGene_tx=XM_023260864.1
C2	2222601	2222704	refGene_gn=AIRE;refGene_tx=XM_023260864.1
C2	2222788	2222907	refGene_gn=AIRE;refGene_tx=XM_023260864.1
C2	2223376	2223559	refGene_gn=AIRE;refGene_tx=XM_023260864.1
C2	2224086	2224186	refGene_gn=AIRE;refGene_tx=XM_023260864.1
C2	2225191	2225307	refGene_gn=AIRE;refGene_tx=XM_023260864.1
C2	2226235	2226316	refGene_gn=AIRE;refGene_tx=XM_023260864.1
C2	2226423	2226566	refGene_gn=AIRE;refGene_tx=XM_023260864.1
C2	2227499	2227616	refGene_gn=AIRE;refGene_tx=XM_023260864.1
C2	2228317	2228392	refGene_gn=AIRE;refGene_tx=XM_023260864.1
C2	2228670	2228826	refGene_gn=AIRE;refGene_tx=XM_023260864.1
C2	2229046	2229221	refGene_gn=AIRE;refGene_tx=XM_023260864.1
C2	2229758	2230004	refGene_gn=AIRE;refGene_tx=XM_023260864.1
C2	2247035	2247114	refGene_gn=DNMT3L;refGene_tx=XM_023260865.1
C2	2247853	2247906	refGene_gn=DNMT3L;refGene_tx=XM_023260865.1
C2	2248135	2248358	refGene_gn=DNMT3L;refGene_tx=XM_023260865.1
C2	2248535	2248580	refGene_gn=DNMT3L;refGene_tx=XM_023260865.1
C2	2249127	2249207	refGene_gn=DNMT3L;refGene_tx=XM_023260865.1
C2	2249390	2249503	refGene_gn=DNMT3L;refGene_tx=XM_023260865.1
C2	2250133	2250305	refGene_gn=DNMT3L;refGene_tx=XM_023260865.1
C2	2252629	2252717	refGene_gn=DNMT3L;refGene_tx=XM_023260865.1
C2	2253820	2253909	refGene_gn=DNMT3L;refGene_tx=XM_023260865.1
C2	2256389	2256465	refGene_gn=DNMT3L;refGene_tx=XM_023260865.1

C2 2256817 2256957 refGene_gn=DNMT3L;refGene_tx=XM_023260865.1
C2 2259212 2259304 refGene_gn=DNMT3L;refGene_tx=XM_023260865.1
C2 2261149 2261315 refGene_gn=DNMT3L;refGene_tx=XM_023260865.1
C2 2266413 2266427
refGene_gn=ICOSLG;refGene_tx=XM_023238652.1;refGene_gn=ICOSLG;refGene_tx=XM_023238653.1
C2 2268421 2268462
refGene_gn=ICOSLG;refGene_tx=XM_023238652.1;refGene_gn=ICOSLG;refGene_tx=XM_023238653.1
C2 2269335 2269698
refGene_gn=ICOSLG;refGene_tx=XM_023238652.1;refGene_gn=ICOSLG;refGene_tx=XM_023238653.1
C2 2270311 2270644
refGene_gn=ICOSLG;refGene_tx=XM_023238652.1;refGene_gn=ICOSLG;refGene_tx=XM_023238653.1
C2 2272727 2272893
refGene_gn=ICOSLG;refGene_tx=XM_023238652.1;refGene_gn=ICOSLG;refGene_tx=XM_023238653.1
C2 2273749 2273786
refGene_gn=ICOSLG;refGene_tx=XM_023238652.1;refGene_gn=ICOSLG;refGene_tx=XM_023238653.1
C2 2274687 2274738
refGene_gn=ICOSLG;refGene_tx=XM_023238652.1;refGene_gn=ICOSLG;refGene_tx=XM_023238653.1
C2 2335008 2335137
refGene_gn=CC2H21orf33;refGene_tx=XM_023238654.1
C2 2336354 2336511
refGene_gn=CC2H21orf33;refGene_tx=XM_023238654.1
C2 2338388 2338481
refGene_gn=CC2H21orf33;refGene_tx=XM_023238654.1
C2 2341160 2341280
refGene_gn=CC2H21orf33;refGene_tx=XM_023238654.1
C2 2342200 2342314
refGene_gn=CC2H21orf33;refGene_tx=XM_023238654.1
C2 2344022 2344075
refGene_gn=CC2H21orf33;refGene_tx=XM_023238654.1
C2 2344452 2344581
refGene_gn=CC2H21orf33;refGene_tx=XM_023238654.1
C2 2347539 2347714
refGene_gn=PWP2;refGene_tx=XM_011285785.3;refGene_gn=PWP2;refGene_tx=XM_011285786.3;refGene_gn=PWP2;refGene_tx=XM_023238655.1
C2 2349163 2349238
refGene_gn=PWP2;refGene_tx=XM_011285785.3;refGene_gn=PWP2;refGene_tx=XM_011285786.3;refGene_gn=PWP2;refGene_tx=XM_023238655.1
C2 2349440 2349591
refGene_gn=PWP2;refGene_tx=XM_011285785.3;refGene_gn=PWP2;refGene_tx=XM_011285786.3;refGene_gn=PWP2;refGene_tx=XM_023238655.1
C2 2349717 2349936
refGene_gn=PWP2;refGene_tx=XM_011285785.3;refGene_gn=PWP2;refGene_tx=XM_011285786.3;refGene_gn=PWP2;refGene_tx=XM_023238655.1

C2 2350685 2350751
refGene_gn=PWP2;refGene_tx=XM_011285785.3;refGene_gn=PWP2;refGene_tx=XM_011285786.3;refGene_gn=PWP2;refGene_tx=XM_023238655.1
C2 2352516 2352625
refGene_gn=PWP2;refGene_tx=XM_011285785.3;refGene_gn=PWP2;refGene_tx=XM_011285786.3;refGene_gn=PWP2;refGene_tx=XM_023238655.1
C2 2353049 2353197
refGene_gn=PWP2;refGene_tx=XM_011285785.3;refGene_gn=PWP2;refGene_tx=XM_011285786.3;refGene_gn=PWP2;refGene_tx=XM_023238655.1
C2 2354167 2354348
refGene_gn=PWP2;refGene_tx=XM_011285785.3;refGene_gn=PWP2;refGene_tx=XM_011285786.3;refGene_gn=PWP2;refGene_tx=XM_023238655.1
C2 2355151 2355302
refGene_gn=PWP2;refGene_tx=XM_011285785.3;refGene_gn=PWP2;refGene_tx=XM_011285786.3;refGene_gn=PWP2;refGene_tx=XM_023238655.1
C2 2355519 2355667
refGene_gn=PWP2;refGene_tx=XM_011285785.3;refGene_gn=PWP2;refGene_tx=XM_011285786.3;refGene_gn=PWP2;refGene_tx=XM_023238655.1
C2 2355817 2355966
refGene_gn=PWP2;refGene_tx=XM_011285785.3;refGene_gn=PWP2;refGene_tx=XM_011285786.3;refGene_gn=PWP2;refGene_tx=XM_023238655.1
C2 2356417 2356554
refGene_gn=PWP2;refGene_tx=XM_011285785.3;refGene_gn=PWP2;refGene_tx=XM_011285786.3;refGene_gn=PWP2;refGene_tx=XM_023238655.1
C2 2356975 2357046
refGene_gn=PWP2;refGene_tx=XM_011285785.3;refGene_gn=PWP2;refGene_tx=XM_011285786.3;refGene_gn=PWP2;refGene_tx=XM_023238655.1
C2 2357224 2357368
refGene_gn=PWP2;refGene_tx=XM_011285785.3;refGene_gn=PWP2;refGene_tx=XM_011285786.3;refGene_gn=PWP2;refGene_tx=XM_023238655.1
C2 2357646 2357867
refGene_gn=PWP2;refGene_tx=XM_011285785.3;refGene_gn=PWP2;refGene_tx=XM_011285786.3;refGene_gn=PWP2;refGene_tx=XM_023238655.1
C2 2358127 2358263
refGene_gn=PWP2;refGene_tx=XM_011285785.3;refGene_gn=PWP2;refGene_tx=XM_011285786.3;refGene_gn=PWP2;refGene_tx=XM_023238655.1
C2 2359118 2359262
refGene_gn=PWP2;refGene_tx=XM_011285785.3;refGene_gn=PWP2;refGene_tx=XM_011285786.3;refGene_gn=PWP2;refGene_tx=XM_023238655.1
C2 2360073 2360173
refGene_gn=PWP2;refGene_tx=XM_011285785.3;refGene_gn=PWP2;refGene_tx=XM_011285786.3;refGene_gn=PWP2;refGene_tx=XM_023238655.1
C2 2360383 2360478
refGene_gn=PWP2;refGene_tx=XM_011285785.3;refGene_gn=PWP2;refGene_tx=XM_011285786.3;refGene_gn=PWP2;refGene_tx=XM_023238655.1
C2 2361512 2361646
refGene_gn=PWP2;refGene_tx=XM_011285785.3;refGene_gn=PWP2;refGene_tx=XM_011285786.3;refGene_gn=PWP2;refGene_tx=XM_023238655.1
C2 2362410 2362788 refGene_gn=PWP2;refGene_tx=XM_011285785.3
C2 2363111 2363129 refGene_gn=PWP2;refGene_tx=XM_011285786.3

C2 2367427 2367666
refGene_gn=TRAPPC10;refGene_tx=XM_023238657.1;refGene_gn=TRAPPC10;refGene_tx=XM_023238658.1
C2 2368007 2368202
refGene_gn=TRAPPC10;refGene_tx=XM_023238657.1;refGene_gn=TRAPPC10;refGene_tx=XM_023238658.1
C2 2371154 2371332
refGene_gn=TRAPPC10;refGene_tx=XM_023238657.1;refGene_gn=TRAPPC10;refGene_tx=XM_023238658.1
C2 2374381 2374552
refGene_gn=TRAPPC10;refGene_tx=XM_023238657.1;refGene_gn=TRAPPC10;refGene_tx=XM_023238658.1
C2 2375424 2375551
refGene_gn=TRAPPC10;refGene_tx=XM_023238657.1;refGene_gn=TRAPPC10;refGene_tx=XM_023238658.1
C2 2376882 2376983
refGene_gn=TRAPPC10;refGene_tx=XM_023238657.1;refGene_gn=TRAPPC10;refGene_tx=XM_023238658.1
C2 2384194 2384424
refGene_gn=TRAPPC10;refGene_tx=XM_023238657.1;refGene_gn=TRAPPC10;refGene_tx=XM_023238658.1
C2 2385232 2385391
refGene_gn=TRAPPC10;refGene_tx=XM_023238657.1;refGene_gn=TRAPPC10;refGene_tx=XM_023238658.1
C2 2386421 2386563
refGene_gn=TRAPPC10;refGene_tx=XM_023238657.1;refGene_gn=TRAPPC10;refGene_tx=XM_023238658.1
C2 2387554 2388069
refGene_gn=TRAPPC10;refGene_tx=XM_023238657.1;refGene_gn=TRAPPC10;refGene_tx=XM_023238658.1
C2 2392029 2392142
refGene_gn=TRAPPC10;refGene_tx=XM_023238657.1;refGene_gn=TRAPPC10;refGene_tx=XM_023238658.1
C2 2392460 2392601
refGene_gn=TRAPPC10;refGene_tx=XM_023238657.1;refGene_gn=TRAPPC10;refGene_tx=XM_023238658.1
C2 2394476 2394568
refGene_gn=TRAPPC10;refGene_tx=XM_023238657.1;refGene_gn=TRAPPC10;refGene_tx=XM_023238658.1
C2 2395344 2395421
refGene_gn=TRAPPC10;refGene_tx=XM_023238657.1;refGene_gn=TRAPPC10;refGene_tx=XM_023238658.1
C2 2396703 2396818
refGene_gn=TRAPPC10;refGene_tx=XM_023238657.1;refGene_gn=TRAPPC10;refGene_tx=XM_023238658.1
C2 2397417 2397564
refGene_gn=TRAPPC10;refGene_tx=XM_023238657.1;refGene_gn=TRAPPC10;refGene_tx=XM_023238658.1
C2 2409826 2410074
refGene_gn=TRAPPC10;refGene_tx=XM_023238657.1;refGene_gn=TRAPPC10;refG

ene_tx=XM_023238658.1
C2 2412602 2412714
refGene_gn=TRAPPC10;refGene_tx=XM_023238657.1;refGene_gn=TRAPPC10;refGene_tx=XM_023238658.1
C2 2414305 2414501
refGene_gn=TRAPPC10;refGene_tx=XM_023238657.1;refGene_gn=TRAPPC10;refGene_tx=XM_023238658.1
C2 2418651 2418848
refGene_gn=TRAPPC10;refGene_tx=XM_023238657.1;refGene_gn=TRAPPC10;refGene_tx=XM_023238658.1
C2 2429842 2429978
refGene_gn=TRAPPC10;refGene_tx=XM_023238657.1;refGene_gn=TRAPPC10;refGene_tx=XM_023238658.1
C2 2434444 2434526
refGene_gn=TRAPPC10;refGene_tx=XM_023238657.1;refGene_gn=TRAPPC10;refGene_tx=XM_023238658.1
C2 2453738 2453805
refGene_gn=TRAPPC10;refGene_tx=XM_023238657.1;refGene_gn=TRAPPC10;refGene_tx=XM_023238658.1
C2 2481626 2481715
refGene_gn=AGPAT3;refGene_tx=XM_003991359.5;refGene_gn=AGPAT3;refGene_tx=XM_006935818.4;refGene_gn=AGPAT3;refGene_tx=XM_019839314.2;refGene_gn=AGPAT3;refGene_tx=XM_023238659.1
C2 2482317 2482516
refGene_gn=AGPAT3;refGene_tx=XM_003991359.5;refGene_gn=AGPAT3;refGene_tx=XM_006935818.4;refGene_gn=AGPAT3;refGene_tx=XM_019839314.2;refGene_gn=AGPAT3;refGene_tx=XM_023238659.1
C2 2487058 2487134
refGene_gn=AGPAT3;refGene_tx=XM_003991359.5;refGene_gn=AGPAT3;refGene_tx=XM_006935818.4;refGene_gn=AGPAT3;refGene_tx=XM_019839314.2;refGene_gn=AGPAT3;refGene_tx=XM_023238659.1
C2 2488866 2489008 refGene_gn=AGPAT3;refGene_tx=XM_023238660.1
C2 2489628 2489731
refGene_gn=AGPAT3;refGene_tx=XM_003991359.5;refGene_gn=AGPAT3;refGene_tx=XM_006935818.4;refGene_gn=AGPAT3;refGene_tx=XM_019839314.2;refGene_gn=AGPAT3;refGene_tx=XM_023238659.1;refGene_gn=AGPAT3;refGene_tx=XM_023238660.1
C2 2490307 2490461
refGene_gn=AGPAT3;refGene_tx=XM_003991359.5;refGene_gn=AGPAT3;refGene_tx=XM_006935818.4;refGene_gn=AGPAT3;refGene_tx=XM_019839314.2;refGene_gn=AGPAT3;refGene_tx=XM_023238659.1;refGene_gn=AGPAT3;refGene_tx=XM_023238660.1
C2 2491552 2491714
refGene_gn=AGPAT3;refGene_tx=XM_003991359.5;refGene_gn=AGPAT3;refGene_tx=XM_006935818.4;refGene_gn=AGPAT3;refGene_tx=XM_019839314.2;refGene_gn=AGPAT3;refGene_tx=XM_023238659.1;refGene_gn=AGPAT3;refGene_tx=XM_023238660.1
C2 2492469 2492639
refGene_gn=AGPAT3;refGene_tx=XM_003991359.5;refGene_gn=AGPAT3;refGene_tx=XM_006935818.4;refGene_gn=AGPAT3;refGene_tx=XM_019839314.2;refGene_

gn=AGPAT3;refGene_tx=XM_023238659.1;refGene_gn=AGPAT3;refGene_tx=XM_023238660.1
C2 2500003 2500181
refGene_gn=AGPAT3;refGene_tx=XM_003991359.5;refGene_gn=AGPAT3;refGene_tx=XM_006935818.4;refGene_gn=AGPAT3;refGene_tx=XM_019839314.2;refGene_gn=AGPAT3;refGene_tx=XM_023238659.1;refGene_gn=AGPAT3;refGene_tx=XM_023238660.1
C2 2627027 2627266
refGene_gn=RRP1;refGene_tx=XM_023238662.1;refGene_gn=RRP1;refGene_tx=XM_023238663.1
C2 2627808 2627922
refGene_gn=RRP1;refGene_tx=XM_023238662.1;refGene_gn=RRP1;refGene_tx=XM_023238663.1
C2 2628825 2628845
refGene_gn=RRP1;refGene_tx=XM_023238662.1;refGene_gn=RRP1;refGene_tx=XM_023238663.1
C2 2629029 2629127
refGene_gn=RRP1;refGene_tx=XM_023238662.1;refGene_gn=RRP1;refGene_tx=XM_023238663.1
C2 2629735 2629815
refGene_gn=RRP1;refGene_tx=XM_023238662.1;refGene_gn=RRP1;refGene_tx=XM_023238663.1
C2 2632452 2632619
refGene_gn=RRP1;refGene_tx=XM_023238662.1;refGene_gn=RRP1;refGene_tx=XM_023238663.1
C2 2632803 2632868
refGene_gn=RRP1;refGene_tx=XM_023238662.1;refGene_gn=RRP1;refGene_tx=XM_023238663.1
C2 2632956 2633086
refGene_gn=RRP1;refGene_tx=XM_023238662.1;refGene_gn=RRP1;refGene_tx=XM_023238663.1
C2 2635339 2635401
refGene_gn=RRP1;refGene_tx=XM_023238662.1;refGene_gn=RRP1;refGene_tx=XM_023238663.1
C2 2637119 2637205
refGene_gn=RRP1;refGene_tx=XM_023238662.1;refGene_gn=RRP1;refGene_tx=XM_023238663.1
C2 2638681 2638739
refGene_gn=RRP1;refGene_tx=XM_023238662.1;refGene_gn=RRP1;refGene_tx=XM_023238663.1
C2 2639683 2639766
refGene_gn=RRP1;refGene_tx=XM_023238662.1;refGene_gn=RRP1;refGene_tx=XM_023238663.1
C2 2640873 2641006
refGene_gn=RRP1;refGene_tx=XM_023238662.1;refGene_gn=RRP1;refGene_tx=XM_023238663.1
C2 2652055 2652121 refGene_gn=CSTB;refGene_tx=XM_023238664.1
C2 2654492 2654594 refGene_gn=CSTB;refGene_tx=XM_023238664.1
C2 2654750 2654879 refGene_gn=CSTB;refGene_tx=XM_023238664.1
C2 2664474 2664587 refGene_gn=PDXK;refGene_tx=XM_023238665.1

C2 2664871 2664938 refGene_gn=PDXK;refGene_tx=XM_023238665.1
C2 2666488 2666625 refGene_gn=PDXK;refGene_tx=XM_023238665.1
C2 2667567 2667679 refGene_gn=PDXK;refGene_tx=XM_023238665.1
C2 2670282 2670328 refGene_gn=PDXK;refGene_tx=XM_023238665.1
C2 2670905 2670991 refGene_gn=PDXK;refGene_tx=XM_023238665.1
C2 2671845 2671892 refGene_gn=PDXK;refGene_tx=XM_023238665.1
C2 2672900 2672984 refGene_gn=PDXK;refGene_tx=XM_023238665.1
C2 2674312 2674417 refGene_gn=PDXK;refGene_tx=XM_023238665.1
C2 2678864 2678919 refGene_gn=PDXK;refGene_tx=XM_023238665.1
C2 2692058 2692145 refGene_gn=PDXK;refGene_tx=XM_023238665.1
C2 2714437 2714631
refGene_gn=RRP1B;refGene_tx=XM_023238666.1;refGene_gn=RRP1B;refGene_tx=XM_023238667.1;refGene_gn=RRP1B;refGene_tx=XM_023238668.1;refGene_gn=RRP1B;refGene_tx=XM_023238669.1
C2 2715404 2715468
refGene_gn=RRP1B;refGene_tx=XM_023238666.1;refGene_gn=RRP1B;refGene_tx=XM_023238667.1;refGene_gn=RRP1B;refGene_tx=XM_023238668.1;refGene_gn=RRP1B;refGene_tx=XM_023238669.1
C2 2716485 2716641
refGene_gn=RRP1B;refGene_tx=XM_023238666.1;refGene_gn=RRP1B;refGene_tx=XM_023238667.1;refGene_gn=RRP1B;refGene_tx=XM_023238668.1;refGene_gn=RRP1B;refGene_tx=XM_023238669.1
C2 2718507 2719172
refGene_gn=RRP1B;refGene_tx=XM_023238666.1;refGene_gn=RRP1B;refGene_tx=XM_023238667.1;refGene_gn=RRP1B;refGene_tx=XM_023238668.1;refGene_gn=RRP1B;refGene_tx=XM_023238669.1
C2 2719684 2719816
refGene_gn=RRP1B;refGene_tx=XM_023238666.1;refGene_gn=RRP1B;refGene_tx=XM_023238667.1;refGene_gn=RRP1B;refGene_tx=XM_023238668.1;refGene_gn=RRP1B;refGene_tx=XM_023238669.1
C2 2720484 2720504
refGene_gn=RRP1B;refGene_tx=XM_023238666.1;refGene_gn=RRP1B;refGene_tx=XM_023238667.1;refGene_gn=RRP1B;refGene_tx=XM_023238668.1;refGene_gn=RRP1B;refGene_tx=XM_023238669.1
C2 2721407 2721505
refGene_gn=RRP1B;refGene_tx=XM_023238666.1;refGene_gn=RRP1B;refGene_tx=XM_023238667.1;refGene_gn=RRP1B;refGene_tx=XM_023238668.1;refGene_gn=RRP1B;refGene_tx=XM_023238669.1
C2 2722142 2722234
refGene_gn=RRP1B;refGene_tx=XM_023238666.1;refGene_gn=RRP1B;refGene_tx=XM_023238667.1;refGene_gn=RRP1B;refGene_tx=XM_023238668.1;refGene_gn=RRP1B;refGene_tx=XM_023238669.1
C2 2723931 2724113
refGene_gn=RRP1B;refGene_tx=XM_023238666.1;refGene_gn=RRP1B;refGene_tx=XM_023238667.1;refGene_gn=RRP1B;refGene_tx=XM_023238668.1;refGene_gn=RRP1B;refGene_tx=XM_023238669.1
C2 2724527 2724592
refGene_gn=RRP1B;refGene_tx=XM_023238666.1;refGene_gn=RRP1B;refGene_tx=XM_023238667.1;refGene_gn=RRP1B;refGene_tx=XM_023238668.1;refGene_gn=RRP1B;refGene_tx=XM_023238669.1

C2	2725452	2725582
refGene_gn=RRP1B;refGene_tx=XM_023238666.1;refGene_gn=RRP1B;refGene_tx=XM_023238667.1;refGene_gn=RRP1B;refGene_tx=XM_023238669.1
C2	2725931	2725993
refGene_gn=RRP1B;refGene_tx=XM_023238666.1;refGene_gn=RRP1B;refGene_tx=XM_023238667.1;refGene_gn=RRP1B;refGene_tx=XM_023238669.1
C2	2726552	2726638
refGene_gn=RRP1B;refGene_tx=XM_023238666.1;refGene_gn=RRP1B;refGene_tx=XM_023238667.1;refGene_gn=RRP1B;refGene_tx=XM_023238668.1;refGene_gn=RRP1B;refGene_tx=XM_023238669.1
C2	2727917	2727975
refGene_gn=RRP1B;refGene_tx=XM_023238666.1;refGene_gn=RRP1B;refGene_tx=XM_023238667.1;refGene_gn=RRP1B;refGene_tx=XM_023238668.1
C2	2729543	2729626
refGene_gn=RRP1B;refGene_tx=XM_023238666.1;refGene_gn=RRP1B;refGene_tx=XM_023238667.1;refGene_gn=RRP1B;refGene_tx=XM_023238668.1
C2	2736581	2736711
refGene_gn=RRP1B;refGene_tx=XM_023238666.1;refGene_gn=RRP1B;refGene_tx=XM_023238667.1;refGene_gn=RRP1B;refGene_tx=XM_023238668.1
C2	2738165	2738186	refGene_gn=HSF2BP;refGene_tx=XM_019839315.2
C2	2738319	2738355
refGene_gn=HSF2BP;refGene_tx=XM_003991360.4;refGene_gn=HSF2BP;refGene_tx=XM_011285784.3;refGene_gn=HSF2BP;refGene_tx=XM_019839316.2;refGene_gn=HSF2BP;refGene_tx=XM_019839318.2
C2	2739129	2739280
refGene_gn=HSF2BP;refGene_tx=XM_003991360.4;refGene_gn=HSF2BP;refGene_tx=XM_011285784.3;refGene_gn=HSF2BP;refGene_tx=XM_019839315.2;refGene_gn=HSF2BP;refGene_tx=XM_019839316.2;refGene_gn=HSF2BP;refGene_tx=XM_019839318.2
C2	2739636	2739697	refGene_gn=HSF2BP;refGene_tx=XM_006935821.4
C2	2751719	2751823
refGene_gn=HSF2BP;refGene_tx=XM_003991360.4;refGene_gn=HSF2BP;refGene_tx=XM_006935821.4;refGene_gn=HSF2BP;refGene_tx=XM_011285784.3;refGene_gn=HSF2BP;refGene_tx=XM_019839315.2;refGene_gn=HSF2BP;refGene_tx=XM_019839316.2;refGene_gn=HSF2BP;refGene_tx=XM_019839318.2
C2	2764268	2764418
refGene_gn=HSF2BP;refGene_tx=XM_003991360.4;refGene_gn=HSF2BP;refGene_tx=XM_006935821.4;refGene_gn=HSF2BP;refGene_tx=XM_011285784.3;refGene_gn=HSF2BP;refGene_tx=XM_019839315.2;refGene_gn=HSF2BP;refGene_tx=XM_019839316.2;refGene_gn=HSF2BP;refGene_tx=XM_019839318.2
C2	2766472	2766605
refGene_gn=HSF2BP;refGene_tx=XM_003991360.4;refGene_gn=HSF2BP;refGene_tx=XM_006935821.4;refGene_gn=HSF2BP;refGene_tx=XM_011285784.3;refGene_gn=HSF2BP;refGene_tx=XM_019839315.2;refGene_gn=HSF2BP;refGene_tx=XM_019839316.2;refGene_gn=HSF2BP;refGene_tx=XM_019839318.2
C2	2769357	2769475
refGene_gn=HSF2BP;refGene_tx=XM_003991360.4;refGene_gn=HSF2BP;refGene_tx=XM_006935821.4;refGene_gn=HSF2BP;refGene_tx=XM_011285784.3;refGene_gn=HSF2BP;refGene_tx=XM_019839315.2;refGene_gn=HSF2BP;refGene_tx=XM_019839316.2;refGene_gn=HSF2BP;refGene_tx=XM_019839318.2

C2	2789682	2789786
refGene_gn=HSF2BP;refGene_tx=XM_003991360.4;refGene_gn=HSF2BP;refGene_tx=XM_006935821.4;refGene_gn=HSF2BP;refGene_tx=XM_011285784.3;refGene_gn=HSF2BP;refGene_tx=XM_019839315.2;refGene_gn=HSF2BP;refGene_tx=XM_019839316.2;refGene_gn=HSF2BP;refGene_tx=XM_019839318.2
C2	2790017	2790160	refGene_gn=HSF2BP;refGene_tx=XM_019839316.2
C2	2791694	2791750	refGene_gn=HSF2BP;refGene_tx=XM_019839318.2
C2	2832761	2832970
refGene_gn=HSF2BP;refGene_tx=XM_003991360.4;refGene_gn=HSF2BP;refGene_tx=XM_006935821.4;refGene_gn=HSF2BP;refGene_tx=XM_011285784.3;refGene_gn=HSF2BP;refGene_tx=XM_019839315.2
C2	2928061	2928217	refGene_gn=SIK1;refGene_tx=XM_023238671.1
C2	2928644	2928660	refGene_gn=SIK1;refGene_tx=XM_023238671.1
C2	2928662	2928760	refGene_gn=SIK1;refGene_tx=XM_023238671.1
C2	2931171	2931235	refGene_gn=SIK1;refGene_tx=XM_023238671.1
C2	2931412	2931574	refGene_gn=SIK1;refGene_tx=XM_023238671.1
C2	2931736	2931861	refGene_gn=SIK1;refGene_tx=XM_023238671.1
C2	2931962	2932086	refGene_gn=SIK1;refGene_tx=XM_023238671.1
C2	2932570	2932794	refGene_gn=SIK1;refGene_tx=XM_023238671.1
C2	2933087	2933237	refGene_gn=SIK1;refGene_tx=XM_023238671.1
C2	2933708	2933831	refGene_gn=SIK1;refGene_tx=XM_023238671.1
C2	2933948	2934165	refGene_gn=SIK1;refGene_tx=XM_023238671.1
C2	2934764	2935043	refGene_gn=SIK1;refGene_tx=XM_023238671.1
C2	2935601	2935836	refGene_gn=SIK1;refGene_tx=XM_023238671.1
C2	2936242	2936603	refGene_gn=SIK1;refGene_tx=XM_023238671.1
C2	3002794	3002829
refGene_gn=LOC109491668;refGene_tx=XM_019811138.2
C2	3003017	3003618
refGene_gn=LOC109491668;refGene_tx=XM_019811138.2
C2	3093402	3093612	refGene_gn=CRYAA;refGene_tx=XM_003991362.3
C2	3094677	3094800	refGene_gn=CRYAA;refGene_tx=XM_003991362.3
C2	3095809	3095998	refGene_gn=CRYAA;refGene_tx=XM_003991362.3
C2	3133997	3134041
refGene_gn=U2AF1;refGene_tx=XM_023238672.1;refGene_gn=U2AF1;refGene_tx=XM_023238673.1;refGene_gn=U2AF1;refGene_tx=XM_023238674.1
C2	3137700	3137788
refGene_gn=U2AF1;refGene_tx=XM_023238672.1;refGene_gn=U2AF1;refGene_tx=XM_023238673.1;refGene_gn=U2AF1;refGene_tx=XM_023238674.1
C2	3140641	3140708	refGene_gn=U2AF1;refGene_tx=XM_023238674.1
C2	3141521	3141609
refGene_gn=U2AF1;refGene_tx=XM_023238672.1;refGene_gn=U2AF1;refGene_tx=XM_023238673.1
C2	3144972	3145022
refGene_gn=U2AF1;refGene_tx=XM_019839319.2;refGene_gn=U2AF1;refGene_tx=XM_019839320.2;refGene_gn=U2AF1;refGene_tx=XM_019839321.2;refGene_gn=U2AF1;refGene_tx=XM_019839323.2;refGene_gn=U2AF1;refGene_tx=XM_019839325.2;refGene_gn=U2AF1;refGene_tx=XM_023238672.1;refGene_gn=U2AF1;refGene_tx=XM_023238673.1;refGene_gn=U2AF1;refGene_tx=XM_023238674.1;refGene_gn=U2AF1;refGene_tx=XM_023238675.1;refGene_gn=U2AF1;refGene_tx=XM_023238676.1

C2 3145111 3145210
refGene_gn=U2AF1;refGene_tx=XM_019839319.2;refGene_gn=U2AF1;refGene_tx=XM_019839320.2;refGene_gn=U2AF1;refGene_tx=XM_019839321.2;refGene_gn=U2AF1;refGene_tx=XM_019839323.2;refGene_gn=U2AF1;refGene_tx=XM_019839325.2;refGene_gn=U2AF1;refGene_tx=XM_023238672.1;refGene_gn=U2AF1;refGene_tx=XM_023238673.1;refGene_gn=U2AF1;refGene_tx=XM_023238674.1;refGene_gn=U2AF1;refGene_tx=XM_023238675.1;refGene_gn=U2AF1;refGene_tx=XM_023238676.1
C2 3145707 3145841
refGene_gn=U2AF1;refGene_tx=XM_019839319.2;refGene_gn=U2AF1;refGene_tx=XM_019839320.2;refGene_gn=U2AF1;refGene_tx=XM_019839321.2;refGene_gn=U2AF1;refGene_tx=XM_019839323.2;refGene_gn=U2AF1;refGene_tx=XM_019839325.2;refGene_gn=U2AF1;refGene_tx=XM_023238672.1;refGene_gn=U2AF1;refGene_tx=XM_023238673.1;refGene_gn=U2AF1;refGene_tx=XM_023238674.1;refGene_gn=U2AF1;refGene_tx=XM_023238675.1;refGene_gn=U2AF1;refGene_tx=XM_023238676.1
C2 3145936 3146029
refGene_gn=U2AF1;refGene_tx=XM_019839319.2;refGene_gn=U2AF1;refGene_tx=XM_019839320.2;refGene_gn=U2AF1;refGene_tx=XM_019839321.2;refGene_gn=U2AF1;refGene_tx=XM_019839323.2;refGene_gn=U2AF1;refGene_tx=XM_019839325.2;refGene_gn=U2AF1;refGene_tx=XM_023238672.1;refGene_gn=U2AF1;refGene_tx=XM_023238673.1;refGene_gn=U2AF1;refGene_tx=XM_023238674.1;refGene_gn=U2AF1;refGene_tx=XM_023238675.1;refGene_gn=U2AF1;refGene_tx=XM_023238676.1
C2 3146826 3146974
refGene_gn=U2AF1;refGene_tx=XM_019839319.2;refGene_gn=U2AF1;refGene_tx=XM_019839320.2;refGene_gn=U2AF1;refGene_tx=XM_019839321.2;refGene_gn=U2AF1;refGene_tx=XM_019839323.2;refGene_gn=U2AF1;refGene_tx=XM_019839325.2;refGene_gn=U2AF1;refGene_tx=XM_023238672.1;refGene_gn=U2AF1;refGene_tx=XM_023238673.1;refGene_gn=U2AF1;refGene_tx=XM_023238674.1;refGene_gn=U2AF1;refGene_tx=XM_023238675.1;refGene_gn=U2AF1;refGene_tx=XM_023238676.1
C2 3163874 3164040
refGene_gn=CBS;refGene_tx=XM_011285781.2;refGene_gn=CBS;refGene_tx=XM_023238677.1;refGene_gn=CBS;refGene_tx=XM_023238678.1
C2 3166545 3166762
refGene_gn=CBS;refGene_tx=XM_011285781.2;refGene_gn=CBS;refGene_tx=XM_019839326.2;refGene_gn=CBS;refGene_tx=XM_023238677.1;refGene_gn=CBS;refGene_tx=XM_023238678.1;refGene_gn=CBS;refGene_tx=XM_023238679.1
C2 3168761 3168868
refGene_gn=CBS;refGene_tx=XM_011285781.2;refGene_gn=CBS;refGene_tx=XM_019839326.2;refGene_gn=CBS;refGene_tx=XM_023238677.1;refGene_gn=CBS;refGene_tx=XM_023238678.1;refGene_gn=CBS;refGene_tx=XM_023238679.1
C2 3169804 3169939
refGene_gn=CBS;refGene_tx=XM_011285781.2;refGene_gn=CBS;refGene_tx=XM_019839326.2;refGene_gn=CBS;refGene_tx=XM_023238677.1;refGene_gn=CBS;refGene_tx=XM_023238678.1;refGene_gn=CBS;refGene_tx=XM_023238679.1
C2 3170288 3170368
refGene_gn=CBS;refGene_tx=XM_011285781.2;refGene_gn=CBS;refGene_tx=XM_019839326.2;refGene_gn=CBS;refGene_tx=XM_023238677.1;refGene_gn=CBS;re

fGene_tx=XM_023238678.1;refGene_gn=CBS;refGene_tx=XM_023238679.1
C2 3170450 3170585
refGene_gn=CBS;refGene_tx=XM_011285781.2;refGene_gn=CBS;refGene_tx=XM_019839326.2;refGene_gn=CBS;refGene_tx=XM_023238677.1;refGene_gn=CBS;refGene_tx=XM_023238678.1;refGene_gn=CBS;refGene_tx=XM_023238679.1
C2 3171119 3171189
refGene_gn=CBS;refGene_tx=XM_011285781.2;refGene_gn=CBS;refGene_tx=XM_019839326.2;refGene_gn=CBS;refGene_tx=XM_023238677.1;refGene_gn=CBS;refGene_tx=XM_023238678.1;refGene_gn=CBS;refGene_tx=XM_023238679.1
C2 3172411 3172503
refGene_gn=CBS;refGene_tx=XM_011285781.2;refGene_gn=CBS;refGene_tx=XM_019839326.2;refGene_gn=CBS;refGene_tx=XM_023238677.1;refGene_gn=CBS;refGene_tx=XM_023238678.1;refGene_gn=CBS;refGene_tx=XM_023238679.1
C2 3173268 3173394
refGene_gn=CBS;refGene_tx=XM_011285781.2;refGene_gn=CBS;refGene_tx=XM_019839326.2;refGene_gn=CBS;refGene_tx=XM_023238677.1;refGene_gn=CBS;refGene_tx=XM_023238678.1;refGene_gn=CBS;refGene_tx=XM_023238679.1
C2 3174013 3174098
refGene_gn=CBS;refGene_tx=XM_011285781.2;refGene_gn=CBS;refGene_tx=XM_019839326.2;refGene_gn=CBS;refGene_tx=XM_023238677.1;refGene_gn=CBS;refGene_tx=XM_023238678.1;refGene_gn=CBS;refGene_tx=XM_023238679.1
C2 3175673 3175779
refGene_gn=CBS;refGene_tx=XM_011285781.2;refGene_gn=CBS;refGene_tx=XM_019839326.2;refGene_gn=CBS;refGene_tx=XM_023238677.1;refGene_gn=CBS;refGene_tx=XM_023238678.1;refGene_gn=CBS;refGene_tx=XM_023238679.1
C2 3177299 3177377
refGene_gn=CBS;refGene_tx=XM_011285781.2;refGene_gn=CBS;refGene_tx=XM_019839326.2;refGene_gn=CBS;refGene_tx=XM_023238677.1;refGene_gn=CBS;refGene_tx=XM_023238678.1;refGene_gn=CBS;refGene_tx=XM_023238679.1
C2 3177540 3177675
refGene_gn=CBS;refGene_tx=XM_011285781.2;refGene_gn=CBS;refGene_tx=XM_019839326.2;refGene_gn=CBS;refGene_tx=XM_023238677.1;refGene_gn=CBS;refGene_tx=XM_023238678.1;refGene_gn=CBS;refGene_tx=XM_023238679.1
C2 3178246 3178355
refGene_gn=CBS;refGene_tx=XM_011285781.2;refGene_gn=CBS;refGene_tx=XM_019839326.2;refGene_gn=CBS;refGene_tx=XM_023238677.1;refGene_gn=CBS;refGene_tx=XM_023238678.1;refGene_gn=CBS;refGene_tx=XM_023238679.1
C2 3182149 3182234
refGene_gn=CBS;refGene_tx=XM_011285781.2;refGene_gn=CBS;refGene_tx=XM_019839326.2;refGene_gn=CBS;refGene_tx=XM_023238677.1;refGene_gn=CBS;refGene_tx=XM_023238678.1;refGene_gn=CBS;refGene_tx=XM_023238679.1
C2 3182609 3182651
refGene_gn=CBS;refGene_tx=XM_019839326.2;refGene_gn=CBS;refGene_tx=XM_023238677.1;refGene_gn=CBS;refGene_tx=XM_023238678.1;refGene_gn=CBS;refGene_tx=XM_023238679.1
C2 3183956 3184114
refGene_gn=CBS;refGene_tx=XM_011285781.2;refGene_gn=CBS;refGene_tx=XM_019839326.2;refGene_gn=CBS;refGene_tx=XM_023238677.1;refGene_gn=CBS;refGene_tx=XM_023238678.1;refGene_gn=CBS;refGene_tx=XM_023238679.1
C2 3198373 3198585

refGene_gn=PKNOX1;refGene_tx=XM_006935828.4;refGene_gn=PKNOX1;refGene_tx=XM_023238680.1;refGene_gn=PKNOX1;refGene_tx=XM_023238681.1;refGene_gn=PKNOX1;refGene_tx=XM_023238682.1
C2 3200761 3200934 refGene_gn=PKNOX1;refGene_tx=XM_006935828.4;refGene_gn=PKNOX1;refGene_tx=XM_023238680.1;refGene_gn=PKNOX1;refGene_tx=XM_023238681.1;refGene_gn=PKNOX1;refGene_tx=XM_023238682.1
C2 3203382 3203459 refGene_gn=PKNOX1;refGene_tx=XM_006935828.4;refGene_gn=PKNOX1;refGene_tx=XM_023238680.1;refGene_gn=PKNOX1;refGene_tx=XM_023238681.1;refGene_gn=PKNOX1;refGene_tx=XM_023238682.1
C2 3207198 3207327 refGene_gn=PKNOX1;refGene_tx=XM_006935828.4;refGene_gn=PKNOX1;refGene_tx=XM_023238680.1;refGene_gn=PKNOX1;refGene_tx=XM_023238681.1;refGene_gn=PKNOX1;refGene_tx=XM_023238682.1
C2 3213649 3213747 refGene_gn=PKNOX1;refGene_tx=XM_006935828.4;refGene_gn=PKNOX1;refGene_tx=XM_023238680.1;refGene_gn=PKNOX1;refGene_tx=XM_023238681.1;refGene_gn=PKNOX1;refGene_tx=XM_023238682.1
C2 3214674 3214774 refGene_gn=PKNOX1;refGene_tx=XM_006935828.4;refGene_gn=PKNOX1;refGene_tx=XM_023238680.1;refGene_gn=PKNOX1;refGene_tx=XM_023238681.1;refGene_gn=PKNOX1;refGene_tx=XM_023238682.1
C2 3219020 3219191 refGene_gn=PKNOX1;refGene_tx=XM_006935828.4;refGene_gn=PKNOX1;refGene_tx=XM_023238680.1;refGene_gn=PKNOX1;refGene_tx=XM_023238681.1;refGene_gn=PKNOX1;refGene_tx=XM_023238682.1
C2 3220950 3221122 refGene_gn=PKNOX1;refGene_tx=XM_006935828.4;refGene_gn=PKNOX1;refGene_tx=XM_023238680.1;refGene_gn=PKNOX1;refGene_tx=XM_023238681.1;refGene_gn=PKNOX1;refGene_tx=XM_023238682.1
C2 3223932 3224060 refGene_gn=PKNOX1;refGene_tx=XM_006935828.4;refGene_gn=PKNOX1;refGene_tx=XM_023238680.1;refGene_gn=PKNOX1;refGene_tx=XM_023238681.1;refGene_gn=PKNOX1;refGene_tx=XM_023238682.1
C2 3227237 3227288 refGene_gn=PKNOX1;refGene_tx=XM_006935828.4;refGene_gn=PKNOX1;refGene_tx=XM_023238680.1;refGene_gn=PKNOX1;refGene_tx=XM_023238681.1;refGene_gn=PKNOX1;refGene_tx=XM_023238682.1
C2 3283786 3283950 refGene_gn=NDUFV3;refGene_tx=XM_011285780.3
C2 3285966 3287052 refGene_gn=NDUFV3;refGene_tx=XM_011285780.3
C2 3293244 3293365 refGene_gn=NDUFV3;refGene_tx=XM_011285780.3
C2 3294796 3294844 refGene_gn=NDUFV3;refGene_tx=XM_011285780.3
C2 3307252 3307489 refGene_gn=LOC105260869;refGene_tx=XM_023238691.1
C2 3313533 3313757 refGene_gn=WDR4;refGene_tx=XM_023238683.1;refGene_gn=WDR4;refGene_tx=XM_023238684.1
C2 3315768 3315834 refGene_gn=WDR4;refGene_tx=XM_023238683.1;refGene_gn=WDR4;refGene_tx=X

M_023238684.1
C2	3317131	3317151
refGene_gn=WDR4;refGene_tx=XM_019839329.2;refGene_gn=WDR4;refGene_tx=XM_019839330.2;refGene_gn=WDR4;refGene_tx=XM_023238685.1;refGene_gn=WDR4;refGene_tx=XM_023238686.1
C2	3317182	3317326
refGene_gn=WDR4;refGene_tx=XM_023238683.1;refGene_gn=WDR4;refGene_tx=XM_023238684.1
C2	3321952	3322109
refGene_gn=WDR4;refGene_tx=XM_006935831.3;refGene_gn=WDR4;refGene_tx=XM_006935833.4;refGene_gn=WDR4;refGene_tx=XM_019839329.2;refGene_gn=WDR4;refGene_tx=XM_019839330.2;refGene_gn=WDR4;refGene_tx=XM_019839331.1;refGene_gn=WDR4;refGene_tx=XM_019839332.2;refGene_gn=WDR4;refGene_tx=XM_019839333.2;refGene_gn=WDR4;refGene_tx=XM_023238683.1;refGene_gn=WDR4;refGene_tx=XM_023238684.1;refGene_gn=WDR4;refGene_tx=XM_023238685.1;refGene_gn=WDR4;refGene_tx=XM_023238686.1;refGene_gn=WDR4;refGene_tx=XM_023238687.1;refGene_gn=WDR4;refGene_tx=XM_023238688.1;refGene_gn=WDR4;refGene_tx=XM_023238689.1;refGene_gn=WDR4;refGene_tx=XM_023238690.1
C2	3322762	3322875
refGene_gn=WDR4;refGene_tx=XM_006935831.3;refGene_gn=WDR4;refGene_tx=XM_006935833.4;refGene_gn=WDR4;refGene_tx=XM_019839329.2;refGene_gn=WDR4;refGene_tx=XM_019839330.2;refGene_gn=WDR4;refGene_tx=XM_019839331.1;refGene_gn=WDR4;refGene_tx=XM_019839332.2;refGene_gn=WDR4;refGene_tx=XM_019839333.2;refGene_gn=WDR4;refGene_tx=XM_023238683.1;refGene_gn=WDR4;refGene_tx=XM_023238684.1;refGene_gn=WDR4;refGene_tx=XM_023238685.1;refGene_gn=WDR4;refGene_tx=XM_023238686.1;refGene_gn=WDR4;refGene_tx=XM_023238687.1;refGene_gn=WDR4;refGene_tx=XM_023238688.1;refGene_gn=WDR4;refGene_tx=XM_023238689.1;refGene_gn=WDR4;refGene_tx=XM_023238690.1
C2	3325881	3325942
refGene_gn=WDR4;refGene_tx=XM_006935831.3;refGene_gn=WDR4;refGene_tx=XM_006935833.4;refGene_gn=WDR4;refGene_tx=XM_019839329.2;refGene_gn=WDR4;refGene_tx=XM_019839330.2;refGene_gn=WDR4;refGene_tx=XM_019839331.1;refGene_gn=WDR4;refGene_tx=XM_019839332.2;refGene_gn=WDR4;refGene_tx=XM_019839333.2;refGene_gn=WDR4;refGene_tx=XM_023238683.1;refGene_gn=WDR4;refGene_tx=XM_023238684.1;refGene_gn=WDR4;refGene_tx=XM_023238685.1;refGene_gn=WDR4;refGene_tx=XM_023238686.1;refGene_gn=WDR4;refGene_tx=XM_023238687.1;refGene_gn=WDR4;refGene_tx=XM_023238688.1;refGene_gn=WDR4;refGene_tx=XM_023238689.1;refGene_gn=WDR4;refGene_tx=XM_023238690.1
C2	3329621	3329721
refGene_gn=WDR4;refGene_tx=XM_006935831.3;refGene_gn=WDR4;refGene_tx=XM_006935833.4;refGene_gn=WDR4;refGene_tx=XM_019839329.2;refGene_gn=WDR4;refGene_tx=XM_019839330.2;refGene_gn=WDR4;refGene_tx=XM_019839331.1;refGene_gn=WDR4;refGene_tx=XM_019839332.2;refGene_gn=WDR4;refGene_tx=XM_019839333.2;refGene_gn=WDR4;refGene_tx=XM_023238683.1;refGene_gn=WDR4;refGene_tx=XM_023238684.1;refGene_gn=WDR4;refGene_tx=XM_023238685.1;refGene_gn=WDR4;refGene_tx=XM_023238686.1;refGene_gn=WDR4;refGene_tx=XM_023238687.1;refGene_gn=WDR4;refGene_tx=XM_023238688.1;refGene_gn=WDR4;refGene_tx=XM_023238689.1;refGene_gn=WDR4;refGene_tx=XM_023238690.1
C2	3329799	3329961	refGene_gn=WDR4;refGene_tx=XM_023238684.1
C2	3331403	3331468

refGene_gn=WDR4;refGene_tx=XM_006935831.3;refGene_gn=WDR4;refGene_tx=XM_006935833.4;refGene_gn=WDR4;refGene_tx=XM_019839329.2;refGene_gn=WDR4;refGene_tx=XM_019839330.2;refGene_gn=WDR4;refGene_tx=XM_019839331.1;refGene_gn=WDR4;refGene_tx=XM_019839332.2;refGene_gn=WDR4;refGene_tx=XM_019839333.2;refGene_gn=WDR4;refGene_tx=XM_023238683.1;refGene_gn=WDR4;refGene_tx=XM_023238684.1;refGene_gn=WDR4;refGene_tx=XM_023238685.1;refGene_gn=WDR4;refGene_tx=XM_023238686.1;refGene_gn=WDR4;refGene_tx=XM_023238687.1;refGene_gn=WDR4;refGene_tx=XM_023238688.1;refGene_gn=WDR4;refGene_tx=XM_023238689.1;refGene_gn=WDR4;refGene_tx=XM_023238690.1
C2 3332207 3332391
refGene_gn=WDR4;refGene_tx=XM_006935831.3;refGene_gn=WDR4;refGene_tx=XM_006935833.4;refGene_gn=WDR4;refGene_tx=XM_019839329.2;refGene_gn=WDR4;refGene_tx=XM_019839330.2;refGene_gn=WDR4;refGene_tx=XM_019839331.1;refGene_gn=WDR4;refGene_tx=XM_019839332.2;refGene_gn=WDR4;refGene_tx=XM_019839333.2;refGene_gn=WDR4;refGene_tx=XM_023238683.1;refGene_gn=WDR4;refGene_tx=XM_023238685.1;refGene_gn=WDR4;refGene_tx=XM_023238686.1;refGene_gn=WDR4;refGene_tx=XM_023238687.1;refGene_gn=WDR4;refGene_tx=XM_023238688.1;refGene_gn=WDR4;refGene_tx=XM_023238689.1;refGene_gn=WDR4;refGene_tx=XM_023238690.1
C2 3333522 3333592
refGene_gn=WDR4;refGene_tx=XM_006935831.3;refGene_gn=WDR4;refGene_tx=XM_006935833.4;refGene_gn=WDR4;refGene_tx=XM_019839329.2;refGene_gn=WDR4;refGene_tx=XM_019839330.2;refGene_gn=WDR4;refGene_tx=XM_019839331.1;refGene_gn=WDR4;refGene_tx=XM_019839332.2;refGene_gn=WDR4;refGene_tx=XM_019839333.2;refGene_gn=WDR4;refGene_tx=XM_023238683.1;refGene_gn=WDR4;refGene_tx=XM_023238685.1;refGene_gn=WDR4;refGene_tx=XM_023238686.1;refGene_gn=WDR4;refGene_tx=XM_023238687.1;refGene_gn=WDR4;refGene_tx=XM_023238688.1;refGene_gn=WDR4;refGene_tx=XM_023238689.1;refGene_gn=WDR4;refGene_tx=XM_023238690.1
C2 3335537 3335719
refGene_gn=WDR4;refGene_tx=XM_006935831.3;refGene_gn=WDR4;refGene_tx=XM_006935833.4;refGene_gn=WDR4;refGene_tx=XM_019839329.2;refGene_gn=WDR4;refGene_tx=XM_019839330.2;refGene_gn=WDR4;refGene_tx=XM_019839331.1;refGene_gn=WDR4;refGene_tx=XM_019839332.2;refGene_gn=WDR4;refGene_tx=XM_019839333.2;refGene_gn=WDR4;refGene_tx=XM_023238683.1;refGene_gn=WDR4;refGene_tx=XM_023238685.1;refGene_gn=WDR4;refGene_tx=XM_023238686.1;refGene_gn=WDR4;refGene_tx=XM_023238687.1;refGene_gn=WDR4;refGene_tx=XM_023238688.1;refGene_gn=WDR4;refGene_tx=XM_023238689.1;refGene_gn=WDR4;refGene_tx=XM_023238690.1
C2 3383218 3383323 refGene_gn=PDE9A;refGene_tx=XM_023238692.1
C2 3384773 3384869 refGene_gn=PDE9A;refGene_tx=XM_023238692.1
C2 3386013 3386115 refGene_gn=PDE9A;refGene_tx=XM_023238692.1
C2 3386116 3386143 refGene_gn=PDE9A;refGene_tx=XM_023238692.1
C2 3386809 3386914 refGene_gn=PDE9A;refGene_tx=XM_023238692.1
C2 3388795 3388909 refGene_gn=PDE9A;refGene_tx=XM_023238692.1
C2 3391112 3391269 refGene_gn=PDE9A;refGene_tx=XM_023238692.1
C2 3392392 3392475 refGene_gn=PDE9A;refGene_tx=XM_023238692.1
C2 3392794 3392899 refGene_gn=PDE9A;refGene_tx=XM_023238692.1
C2 3393885 3393972 refGene_gn=PDE9A;refGene_tx=XM_023238692.1
C2 3396571 3396646 refGene_gn=PDE9A;refGene_tx=XM_023238692.1

C2	3398434	3398516	refGene_gn=PDE9A;refGene_tx=XM_023238692.1
C2	3404344	3404429	refGene_gn=PDE9A;refGene_tx=XM_023238692.1
C2	3407931	3408002	refGene_gn=PDE9A;refGene_tx=XM_023238692.1
C2	3409285	3409340	refGene_gn=PDE9A;refGene_tx=XM_023238692.1
C2	3437186	3437230	refGene_gn=PDE9A;refGene_tx=XM_023238692.1
C2	3441708	3441786	refGene_gn=PDE9A;refGene_tx=XM_023238692.1
C2	3443408	3443479	refGene_gn=PDE9A;refGene_tx=XM_023238692.1
C2	3471799	3471868	refGene_gn=PDE9A;refGene_tx=XM_023238692.1
C2	3535051	3535067
refGene_gn=SLC37A1;refGene_tx=XM_003991402.5;refGene_gn=SLC37A1;refGene_tx=XM_023238693.1;refGene_gn=SLC37A1;refGene_tx=XM_023238694.1;refGene_gn=SLC37A1;refGene_tx=XM_023238695.1;refGene_gn=SLC37A1;refGene_tx=XM_023238696.1;refGene_gn=SLC37A1;refGene_tx=XM_023238697.1;refGene_gn=SLC37A1;refGene_tx=XM_023238698.1;refGene_gn=SLC37A1;refGene_tx=XM_023238699.1
C2	3535585	3535650
refGene_gn=SLC37A1;refGene_tx=XM_003991402.5;refGene_gn=SLC37A1;refGene_tx=XM_023238693.1;refGene_gn=SLC37A1;refGene_tx=XM_023238694.1;refGene_gn=SLC37A1;refGene_tx=XM_023238695.1;refGene_gn=SLC37A1;refGene_tx=XM_023238696.1;refGene_gn=SLC37A1;refGene_tx=XM_023238697.1;refGene_gn=SLC37A1;refGene_tx=XM_023238698.1;refGene_gn=SLC37A1;refGene_tx=XM_023238699.1
C2	3535754	3535823	refGene_gn=SLC37A1;refGene_tx=XM_019839340.2
C2	3539358	3539456
refGene_gn=SLC37A1;refGene_tx=XM_003991402.5;refGene_gn=SLC37A1;refGene_tx=XM_019839340.2;refGene_gn=SLC37A1;refGene_tx=XM_023238693.1;refGene_gn=SLC37A1;refGene_tx=XM_023238694.1;refGene_gn=SLC37A1;refGene_tx=XM_023238695.1;refGene_gn=SLC37A1;refGene_tx=XM_023238696.1;refGene_gn=SLC37A1;refGene_tx=XM_023238697.1;refGene_gn=SLC37A1;refGene_tx=XM_023238698.1;refGene_gn=SLC37A1;refGene_tx=XM_023238699.1
C2	3547342	3547421
refGene_gn=SLC37A1;refGene_tx=XM_003991402.5;refGene_gn=SLC37A1;refGene_tx=XM_019839340.2;refGene_gn=SLC37A1;refGene_tx=XM_023238693.1;refGene_gn=SLC37A1;refGene_tx=XM_023238694.1;refGene_gn=SLC37A1;refGene_tx=XM_023238695.1;refGene_gn=SLC37A1;refGene_tx=XM_023238696.1;refGene_gn=SLC37A1;refGene_tx=XM_023238697.1;refGene_gn=SLC37A1;refGene_tx=XM_023238698.1;refGene_gn=SLC37A1;refGene_tx=XM_023238699.1
C2	3550022	3550096
refGene_gn=SLC37A1;refGene_tx=XM_003991402.5;refGene_gn=SLC37A1;refGene_tx=XM_019839340.2;refGene_gn=SLC37A1;refGene_tx=XM_023238693.1;refGene_gn=SLC37A1;refGene_tx=XM_023238694.1;refGene_gn=SLC37A1;refGene_tx=XM_023238695.1;refGene_gn=SLC37A1;refGene_tx=XM_023238696.1;refGene_gn=SLC37A1;refGene_tx=XM_023238697.1;refGene_gn=SLC37A1;refGene_tx=XM_023238698.1;refGene_gn=SLC37A1;refGene_tx=XM_023238699.1
C2	3551750	3551799
refGene_gn=SLC37A1;refGene_tx=XM_003991402.5;refGene_gn=SLC37A1;refGene_tx=XM_019839340.2;refGene_gn=SLC37A1;refGene_tx=XM_023238693.1;refGene_gn=SLC37A1;refGene_tx=XM_023238694.1;refGene_gn=SLC37A1;refGene_tx=XM_023238695.1;refGene_gn=SLC37A1;refGene_tx=XM_023238696.1;refGene_gn=SLC37A1;refGene_tx=XM_023238697.1;refGene_gn=SLC37A1;refGene_tx=XM_02

3238698.1;refGene_gn=SLC37A1;refGene_tx=XM_023238699.1
C2	3552755	3552841
refGene_gn=SLC37A1;refGene_tx=XM_003991402.5;refGene_gn=SLC37A1;refGene_tx=XM_019839340.2;refGene_gn=SLC37A1;refGene_tx=XM_023238693.1;refGene_gn=SLC37A1;refGene_tx=XM_023238694.1;refGene_gn=SLC37A1;refGene_tx=XM_023238695.1;refGene_gn=SLC37A1;refGene_tx=XM_023238696.1;refGene_gn=SLC37A1;refGene_tx=XM_023238697.1;refGene_gn=SLC37A1;refGene_tx=XM_023238698.1;refGene_gn=SLC37A1;refGene_tx=XM_023238699.1
C2	3553160	3553223
refGene_gn=SLC37A1;refGene_tx=XM_003991402.5;refGene_gn=SLC37A1;refGene_tx=XM_019839340.2;refGene_gn=SLC37A1;refGene_tx=XM_023238693.1;refGene_gn=SLC37A1;refGene_tx=XM_023238694.1;refGene_gn=SLC37A1;refGene_tx=XM_023238695.1;refGene_gn=SLC37A1;refGene_tx=XM_023238696.1;refGene_gn=SLC37A1;refGene_tx=XM_023238697.1;refGene_gn=SLC37A1;refGene_tx=XM_023238698.1;refGene_gn=SLC37A1;refGene_tx=XM_023238699.1
C2	3555507	3555598
refGene_gn=SLC37A1;refGene_tx=XM_003991402.5;refGene_gn=SLC37A1;refGene_tx=XM_019839340.2;refGene_gn=SLC37A1;refGene_tx=XM_023238693.1;refGene_gn=SLC37A1;refGene_tx=XM_023238694.1;refGene_gn=SLC37A1;refGene_tx=XM_023238695.1;refGene_gn=SLC37A1;refGene_tx=XM_023238696.1;refGene_gn=SLC37A1;refGene_tx=XM_023238697.1;refGene_gn=SLC37A1;refGene_tx=XM_023238698.1;refGene_gn=SLC37A1;refGene_tx=XM_023238699.1
C2	3557947	3558079
refGene_gn=SLC37A1;refGene_tx=XM_003991402.5;refGene_gn=SLC37A1;refGene_tx=XM_019839340.2;refGene_gn=SLC37A1;refGene_tx=XM_023238693.1;refGene_gn=SLC37A1;refGene_tx=XM_023238694.1;refGene_gn=SLC37A1;refGene_tx=XM_023238695.1;refGene_gn=SLC37A1;refGene_tx=XM_023238696.1;refGene_gn=SLC37A1;refGene_tx=XM_023238697.1;refGene_gn=SLC37A1;refGene_tx=XM_023238698.1;refGene_gn=SLC37A1;refGene_tx=XM_023238699.1
C2	3564687	3564768
refGene_gn=SLC37A1;refGene_tx=XM_003991402.5;refGene_gn=SLC37A1;refGene_tx=XM_019839340.2;refGene_gn=SLC37A1;refGene_tx=XM_023238693.1;refGene_gn=SLC37A1;refGene_tx=XM_023238694.1;refGene_gn=SLC37A1;refGene_tx=XM_023238695.1;refGene_gn=SLC37A1;refGene_tx=XM_023238696.1;refGene_gn=SLC37A1;refGene_tx=XM_023238697.1;refGene_gn=SLC37A1;refGene_tx=XM_023238698.1;refGene_gn=SLC37A1;refGene_tx=XM_023238699.1
C2	3566467	3566505
refGene_gn=SLC37A1;refGene_tx=XM_003991402.5;refGene_gn=SLC37A1;refGene_tx=XM_019839340.2;refGene_gn=SLC37A1;refGene_tx=XM_023238693.1;refGene_gn=SLC37A1;refGene_tx=XM_023238694.1;refGene_gn=SLC37A1;refGene_tx=XM_023238695.1;refGene_gn=SLC37A1;refGene_tx=XM_023238696.1;refGene_gn=SLC37A1;refGene_tx=XM_023238697.1;refGene_gn=SLC37A1;refGene_tx=XM_023238698.1;refGene_gn=SLC37A1;refGene_tx=XM_023238699.1
C2	3571394	3571561
refGene_gn=SLC37A1;refGene_tx=XM_003991402.5;refGene_gn=SLC37A1;refGene_tx=XM_019839340.2;refGene_gn=SLC37A1;refGene_tx=XM_023238693.1;refGene_gn=SLC37A1;refGene_tx=XM_023238694.1;refGene_gn=SLC37A1;refGene_tx=XM_023238695.1;refGene_gn=SLC37A1;refGene_tx=XM_023238696.1;refGene_gn=SLC37A1;refGene_tx=XM_023238697.1;refGene_gn=SLC37A1;refGene_tx=XM_023238698.1;refGene_gn=SLC37A1;refGene_tx=XM_023238699.1

C2 3572609 3572686
refGene_gn=SLC37A1;refGene_tx=XM_003991402.5;refGene_gn=SLC37A1;refGene_tx=XM_019839340.2;refGene_gn=SLC37A1;refGene_tx=XM_023238693.1;refGene_gn=SLC37A1;refGene_tx=XM_023238694.1;refGene_gn=SLC37A1;refGene_tx=XM_023238695.1;refGene_gn=SLC37A1;refGene_tx=XM_023238696.1;refGene_gn=SLC37A1;refGene_tx=XM_023238697.1;refGene_gn=SLC37A1;refGene_tx=XM_023238698.1;refGene_gn=SLC37A1;refGene_tx=XM_023238699.1
C2 3573900 3574036
refGene_gn=SLC37A1;refGene_tx=XM_003991402.5;refGene_gn=SLC37A1;refGene_tx=XM_019839340.2;refGene_gn=SLC37A1;refGene_tx=XM_023238693.1;refGene_gn=SLC37A1;refGene_tx=XM_023238694.1;refGene_gn=SLC37A1;refGene_tx=XM_023238695.1;refGene_gn=SLC37A1;refGene_tx=XM_023238696.1;refGene_gn=SLC37A1;refGene_tx=XM_023238697.1;refGene_gn=SLC37A1;refGene_tx=XM_023238698.1;refGene_gn=SLC37A1;refGene_tx=XM_023238699.1
C2 3577313 3577392
refGene_gn=SLC37A1;refGene_tx=XM_003991402.5;refGene_gn=SLC37A1;refGene_tx=XM_019839340.2;refGene_gn=SLC37A1;refGene_tx=XM_023238693.1;refGene_gn=SLC37A1;refGene_tx=XM_023238694.1;refGene_gn=SLC37A1;refGene_tx=XM_023238695.1;refGene_gn=SLC37A1;refGene_tx=XM_023238696.1;refGene_gn=SLC37A1;refGene_tx=XM_023238697.1;refGene_gn=SLC37A1;refGene_tx=XM_023238698.1;refGene_gn=SLC37A1;refGene_tx=XM_023238699.1
C2 3578091 3578224
refGene_gn=SLC37A1;refGene_tx=XM_003991402.5;refGene_gn=SLC37A1;refGene_tx=XM_019839340.2;refGene_gn=SLC37A1;refGene_tx=XM_023238693.1;refGene_gn=SLC37A1;refGene_tx=XM_023238694.1;refGene_gn=SLC37A1;refGene_tx=XM_023238695.1;refGene_gn=SLC37A1;refGene_tx=XM_023238696.1;refGene_gn=SLC37A1;refGene_tx=XM_023238697.1;refGene_gn=SLC37A1;refGene_tx=XM_023238698.1;refGene_gn=SLC37A1;refGene_tx=XM_023238699.1
C2 3586294 3586376
refGene_gn=SLC37A1;refGene_tx=XM_003991402.5;refGene_gn=SLC37A1;refGene_tx=XM_019839340.2;refGene_gn=SLC37A1;refGene_tx=XM_023238693.1;refGene_gn=SLC37A1;refGene_tx=XM_023238694.1;refGene_gn=SLC37A1;refGene_tx=XM_023238695.1;refGene_gn=SLC37A1;refGene_tx=XM_023238696.1;refGene_gn=SLC37A1;refGene_tx=XM_023238697.1;refGene_gn=SLC37A1;refGene_tx=XM_023238698.1;refGene_gn=SLC37A1;refGene_tx=XM_023238699.1
C2 3595889 3595945
refGene_gn=SLC37A1;refGene_tx=XM_003991402.5;refGene_gn=SLC37A1;refGene_tx=XM_019839340.2;refGene_gn=SLC37A1;refGene_tx=XM_023238693.1;refGene_gn=SLC37A1;refGene_tx=XM_023238694.1;refGene_gn=SLC37A1;refGene_tx=XM_023238695.1;refGene_gn=SLC37A1;refGene_tx=XM_023238696.1;refGene_gn=SLC37A1;refGene_tx=XM_023238697.1;refGene_gn=SLC37A1;refGene_tx=XM_023238698.1;refGene_gn=SLC37A1;refGene_tx=XM_023238699.1
C2 3617939 3617993
refGene_gn=RSPH1;refGene_tx=XM_003991403.4;refGene_gn=RSPH1;refGene_tx=XM_023238700.1
C2 3620288 3620402
refGene_gn=RSPH1;refGene_tx=XM_003991403.4;refGene_gn=RSPH1;refGene_tx=XM_023238700.1
C2 3620504 3620610
refGene_gn=RSPH1;refGene_tx=XM_003991403.4;refGene_gn=RSPH1;refGene_tx

=XM_023238700.1
C2 3627169 3627260
refGene_gn=RSPH1;refGene_tx=XM_003991403.4;refGene_gn=RSPH1;refGene_tx=XM_023238700.1
C2 3627752 3627888
refGene_gn=RSPH1;refGene_tx=XM_003991403.4;refGene_gn=RSPH1;refGene_tx=XM_023238700.1
C2 3631410 3631482
refGene_gn=RSPH1;refGene_tx=XM_003991403.4;refGene_gn=RSPH1;refGene_tx=XM_023238700.1
C2 3633942 3634105
refGene_gn=RSPH1;refGene_tx=XM_003991403.4;refGene_gn=RSPH1;refGene_tx=XM_023238700.1
C2 3634935 3635091
refGene_gn=RSPH1;refGene_tx=XM_003991403.4;refGene_gn=RSPH1;refGene_tx=XM_023238700.1
C2 3635321 3635326 refGene_gn=RSPH1;refGene_tx=XM_023238700.1
C2 3636335 3636388 refGene_gn=RSPH1;refGene_tx=XM_003991403.4
C2 3653570 3653708
refGene_gn=UBASH3A;refGene_tx=XM_003991367.3;refGene_gn=UBASH3A;refGene_tx=XM_003991368.3;refGene_gn=UBASH3A;refGene_tx=XM_006935839.2
C2 3657923 3658033
refGene_gn=UBASH3A;refGene_tx=XM_003991367.3;refGene_gn=UBASH3A;refGene_tx=XM_006935839.2
C2 3659060 3659092 refGene_gn=UBASH3A;refGene_tx=XM_019839345.2
C2 3660574 3660681
refGene_gn=UBASH3A;refGene_tx=XM_003991367.3;refGene_gn=UBASH3A;refGene_tx=XM_006935839.2;refGene_gn=UBASH3A;refGene_tx=XM_019839345.2
C2 3661406 3661551
refGene_gn=UBASH3A;refGene_tx=XM_003991367.3;refGene_gn=UBASH3A;refGene_tx=XM_003991368.3;refGene_gn=UBASH3A;refGene_tx=XM_006935839.2;refGene_gn=UBASH3A;refGene_tx=XM_019839345.2
C2 3664017 3664110
refGene_gn=UBASH3A;refGene_tx=XM_003991367.3;refGene_gn=UBASH3A;refGene_tx=XM_003991368.3;refGene_gn=UBASH3A;refGene_tx=XM_006935839.2;refGene_gn=UBASH3A;refGene_tx=XM_019839345.2
C2 3665757 3665880
refGene_gn=UBASH3A;refGene_tx=XM_003991367.3;refGene_gn=UBASH3A;refGene_tx=XM_003991368.3;refGene_gn=UBASH3A;refGene_tx=XM_006935839.2;refGene_gn=UBASH3A;refGene_tx=XM_019839345.2
C2 3668765 3668865
refGene_gn=UBASH3A;refGene_tx=XM_003991367.3;refGene_gn=UBASH3A;refGene_tx=XM_003991368.3;refGene_gn=UBASH3A;refGene_tx=XM_006935839.2;refGene_gn=UBASH3A;refGene_tx=XM_019839345.2
C2 3673598 3673725
refGene_gn=UBASH3A;refGene_tx=XM_003991367.3;refGene_gn=UBASH3A;refGene_tx=XM_003991368.3;refGene_gn=UBASH3A;refGene_tx=XM_006935839.2;refGene_gn=UBASH3A;refGene_tx=XM_019839345.2
C2 3682045 3682254
refGene_gn=UBASH3A;refGene_tx=XM_003991367.3;refGene_gn=UBASH3A;refGen

e_tx=XM_003991368.3;refGene_gn=UBASH3A;refGene_tx=XM_006935839.2;refGene_gn=UBASH3A;refGene_tx=XM_019839345.2
C2 3683300 3683470
refGene_gn=UBASH3A;refGene_tx=XM_003991367.3;refGene_gn=UBASH3A;refGene_tx=XM_003991368.3;refGene_gn=UBASH3A;refGene_tx=XM_006935839.2;refGene_gn=UBASH3A;refGene_tx=XM_019839345.2
C2 3686890 3687089
refGene_gn=UBASH3A;refGene_tx=XM_003991367.3;refGene_gn=UBASH3A;refGene_tx=XM_003991368.3;refGene_gn=UBASH3A;refGene_tx=XM_006935839.2;refGene_gn=UBASH3A;refGene_tx=XM_019839345.2
C2 3689284 3689471
refGene_gn=UBASH3A;refGene_tx=XM_003991367.3;refGene_gn=UBASH3A;refGene_tx=XM_003991368.3;refGene_gn=UBASH3A;refGene_tx=XM_006935839.2;refGene_gn=UBASH3A;refGene_tx=XM_019839345.2
C2 3692196 3692250
refGene_gn=UBASH3A;refGene_tx=XM_003991367.3;refGene_gn=UBASH3A;refGene_tx=XM_003991368.3;refGene_gn=UBASH3A;refGene_tx=XM_006935839.2;refGene_gn=UBASH3A;refGene_tx=XM_019839345.2
C2 3694027 3694140
refGene_gn=UBASH3A;refGene_tx=XM_003991367.3;refGene_gn=UBASH3A;refGene_tx=XM_003991368.3;refGene_gn=UBASH3A;refGene_tx=XM_006935839.2;refGene_gn=UBASH3A;refGene_tx=XM_019839345.2
C2 3702915 3703009 refGene_gn=TMPRSS3;refGene_tx=XM_023238701.1
C2 3707055 3707166 refGene_gn=TMPRSS3;refGene_tx=XM_023238701.1
C2 3707998 3708115 refGene_gn=TMPRSS3;refGene_tx=XM_023238701.1
C2 3708486 3708610 refGene_gn=TMPRSS3;refGene_tx=XM_023238701.1
C2 3711496 3711622 refGene_gn=TMPRSS3;refGene_tx=XM_023238701.1
C2 3712828 3712872 refGene_gn=TMPRSS3;refGene_tx=XM_023238701.1
C2 3713551 3713717 refGene_gn=TMPRSS3;refGene_tx=XM_023238701.1
C2 3714467 3714637 refGene_gn=TMPRSS3;refGene_tx=XM_023238701.1
C2 3716475 3716571 refGene_gn=TMPRSS3;refGene_tx=XM_023238701.1
C2 3719196 3719339 refGene_gn=TMPRSS3;refGene_tx=XM_023238701.1
C2 3719946 3720099 refGene_gn=TMPRSS3;refGene_tx=XM_023238701.1
C2 3723027 3723072 refGene_gn=TMPRSS3;refGene_tx=XM_023238701.1
C2 3741215 3741291 refGene_gn=TFF1;refGene_tx=NM_001130389.1
C2 3742754 3742907 refGene_gn=TFF1;refGene_tx=NM_001130389.1
C2 3743330 3743347 refGene_gn=TFF1;refGene_tx=NM_001130389.1
C2 3751218 3751406
refGene_gn=LOC109503282;refGene_tx=XM_019839346.1
C2 3753501 3753615
refGene_gn=LOC109503282;refGene_tx=XM_019839346.1
C2 3753699 3753938
refGene_gn=LOC109503282;refGene_tx=XM_019839346.1
C2 3754809 3755024
refGene_gn=LOC109503282;refGene_tx=XM_019839346.1
C2 3755359 3755438 refGene_gn=TFF2;refGene_tx=NM_001130388.1
C2 3756129 3756279 refGene_gn=TFF2;refGene_tx=NM_001130388.1
C2 3757179 3757326 refGene_gn=TFF2;refGene_tx=NM_001130388.1
C2 3758177 3758191 refGene_gn=TFF2;refGene_tx=NM_001130388.1
C2 3784799 3784881 refGene_gn=TFF3;refGene_tx=NM_001130390.1

C2	3786458	3786605	refGene_gn=TFF3;refGene_tx=NM_001130390.1
C2	3787773	3787787	refGene_gn=TFF3;refGene_tx=NM_001130390.1
C2	3805970	3806199	refGene_gn=ABCG1;refGene_tx=XM_023238702.1
C2	3807162	3807281	refGene_gn=ABCG1;refGene_tx=XM_023238702.1
C2	3808352	3808511	refGene_gn=ABCG1;refGene_tx=XM_023238702.1
C2	3808625	3808726	refGene_gn=ABCG1;refGene_tx=XM_023238702.1
C2	3809536	3809660	refGene_gn=ABCG1;refGene_tx=XM_023238702.1
C2	3809661	3809706	refGene_gn=ABCG1;refGene_tx=XM_023238702.1
C2	3810571	3810673	refGene_gn=ABCG1;refGene_tx=XM_023238702.1
C2	3810846	3810995	refGene_gn=ABCG1;refGene_tx=XM_023238702.1
C2	3812818	3812933	refGene_gn=ABCG1;refGene_tx=XM_023238702.1
C2	3813899	3814023	refGene_gn=ABCG1;refGene_tx=XM_023238702.1
C2	3814303	3814449	refGene_gn=ABCG1;refGene_tx=XM_023238702.1
C2	3818754	3818805	refGene_gn=ABCG1;refGene_tx=XM_023238702.1
C2	3820689	3820822	refGene_gn=ABCG1;refGene_tx=XM_023238702.1
C2	3822505	3822623	refGene_gn=ABCG1;refGene_tx=XM_023238702.1
C2	3855007	3855251	refGene_gn=ABCG1;refGene_tx=XM_023238702.1
C2	3861875	3861917	refGene_gn=ABCG1;refGene_tx=XM_023238702.1
C2	3914623	3914805	refGene_gn=UMODL1;refGene_tx=XM_023238703.1
C2	3920347	3920432	refGene_gn=UMODL1;refGene_tx=XM_023238703.1
C2	3928194	3928354	refGene_gn=UMODL1;refGene_tx=XM_023238703.1
C2	3928556	3928793	refGene_gn=UMODL1;refGene_tx=XM_023238703.1
C2	3929550	3929696	refGene_gn=UMODL1;refGene_tx=XM_023238703.1
C2	3934713	3935051	refGene_gn=UMODL1;refGene_tx=XM_023238703.1
C2	3936116	3936254	refGene_gn=UMODL1;refGene_tx=XM_023238703.1
C2	3938790	3939004	refGene_gn=UMODL1;refGene_tx=XM_023238703.1
C2	3941640	3941753	refGene_gn=UMODL1;refGene_tx=XM_023238703.1
C2	3943389	3943647	refGene_gn=UMODL1;refGene_tx=XM_023238703.1
C2	3944156	3944729	refGene_gn=UMODL1;refGene_tx=XM_023238703.1
C2	3945244	3945382	refGene_gn=UMODL1;refGene_tx=XM_023238703.1
C2	3949807	3950027	refGene_gn=UMODL1;refGene_tx=XM_023238703.1
C2	3951513	3951626	refGene_gn=UMODL1;refGene_tx=XM_023238703.1
C2	3953353	3953608	refGene_gn=UMODL1;refGene_tx=XM_023238703.1
C2	3966843	3967030	refGene_gn=UMODL1;refGene_tx=XM_023238703.1
C2	3969083	3969205	refGene_gn=UMODL1;refGene_tx=XM_023238703.1
C2	3970057	3970219	refGene_gn=UMODL1;refGene_tx=XM_023238703.1
C2	3977167	3977410	refGene_gn=UMODL1;refGene_tx=XM_023238703.1
C2	3980924	3981000	refGene_gn=UMODL1;refGene_tx=XM_023238703.1
C2	4050590	4053773
refGene_gn=ZBTB21;refGene_tx=XM_003991370.5;refGene_gn=ZBTB21;refGene_tx=XM_006935841.4;refGene_gn=ZBTB21;refGene_tx=XM_006935843.4;refGene_gn=ZBTB21;refGene_tx=XM_006935844.4;refGene_gn=ZBTB21;refGene_tx=XM_019839353.2;refGene_gn=ZBTB21;refGene_tx=XM_019839354.2;refGene_gn=ZBTB21;refGene_tx=XM_019839355.2
C2	4085248	4085521
refGene_gn=C2CD2;refGene_tx=XM_023238704.1;refGene_gn=C2CD2;refGene_tx=XM_023238705.1;refGene_gn=C2CD2;refGene_tx=XM_023238706.1;refGene_gn=C2CD2;refGene_tx=XM_023238707.1
C2	4087719	4087746	refGene_gn=C2CD2;refGene_tx=XM_011285776.2
C2	4093278	4093386

refGene_gn=C2CD2;refGene_tx=XM_023238704.1;refGene_gn=C2CD2;refGene_tx=XM_023238705.1;refGene_gn=C2CD2;refGene_tx=XM_023238707.1
C2 4094473 4094572
refGene_gn=C2CD2;refGene_tx=XM_011285776.2;refGene_gn=C2CD2;refGene_tx=XM_023238704.1;refGene_gn=C2CD2;refGene_tx=XM_023238705.1;refGene_gn=C2CD2;refGene_tx=XM_023238706.1;refGene_gn=C2CD2;refGene_tx=XM_023238707.1
C2 4107178 4107298
refGene_gn=C2CD2;refGene_tx=XM_011285776.2;refGene_gn=C2CD2;refGene_tx=XM_023238704.1;refGene_gn=C2CD2;refGene_tx=XM_023238705.1;refGene_gn=C2CD2;refGene_tx=XM_023238706.1;refGene_gn=C2CD2;refGene_tx=XM_023238707.1
C2 4110977 4111088
refGene_gn=C2CD2;refGene_tx=XM_011285776.2;refGene_gn=C2CD2;refGene_tx=XM_023238704.1;refGene_gn=C2CD2;refGene_tx=XM_023238705.1;refGene_gn=C2CD2;refGene_tx=XM_023238706.1;refGene_gn=C2CD2;refGene_tx=XM_023238707.1
C2 4112867 4112990
refGene_gn=C2CD2;refGene_tx=XM_011285776.2;refGene_gn=C2CD2;refGene_tx=XM_023238704.1;refGene_gn=C2CD2;refGene_tx=XM_023238705.1;refGene_gn=C2CD2;refGene_tx=XM_023238706.1;refGene_gn=C2CD2;refGene_tx=XM_023238707.1
C2 4114407 4114531
refGene_gn=C2CD2;refGene_tx=XM_011285776.2;refGene_gn=C2CD2;refGene_tx=XM_023238704.1;refGene_gn=C2CD2;refGene_tx=XM_023238705.1;refGene_gn=C2CD2;refGene_tx=XM_023238706.1;refGene_gn=C2CD2;refGene_tx=XM_023238707.1
C2 4116224 4116330
refGene_gn=C2CD2;refGene_tx=XM_011285776.2;refGene_gn=C2CD2;refGene_tx=XM_023238704.1;refGene_gn=C2CD2;refGene_tx=XM_023238705.1;refGene_gn=C2CD2;refGene_tx=XM_023238706.1;refGene_gn=C2CD2;refGene_tx=XM_023238707.1
C2 4117961 4118026
refGene_gn=C2CD2;refGene_tx=XM_011285776.2;refGene_gn=C2CD2;refGene_tx=XM_023238704.1;refGene_gn=C2CD2;refGene_tx=XM_023238705.1;refGene_gn=C2CD2;refGene_tx=XM_023238706.1;refGene_gn=C2CD2;refGene_tx=XM_023238707.1
C2 4119835 4119960
refGene_gn=C2CD2;refGene_tx=XM_011285776.2;refGene_gn=C2CD2;refGene_tx=XM_023238704.1;refGene_gn=C2CD2;refGene_tx=XM_023238705.1;refGene_gn=C2CD2;refGene_tx=XM_023238706.1;refGene_gn=C2CD2;refGene_tx=XM_023238707.1
C2 4120451 4120626
refGene_gn=C2CD2;refGene_tx=XM_011285776.2;refGene_gn=C2CD2;refGene_tx=XM_023238704.1;refGene_gn=C2CD2;refGene_tx=XM_023238705.1;refGene_gn=C2CD2;refGene_tx=XM_023238706.1;refGene_gn=C2CD2;refGene_tx=XM_023238707.1
C2 4121575 4121689
refGene_gn=C2CD2;refGene_tx=XM_011285776.2;refGene_gn=C2CD2;refGene_tx=XM_023238704.1;refGene_gn=C2CD2;refGene_tx=XM_023238705.1;refGene_gn=

C2CD2;refGene_tx=XM_023238706.1;refGene_gn=C2CD2;refGene_tx=XM_0232387
07.1
C2	4123661	4123789
refGene_gn=C2CD2;refGene_tx=XM_011285776.2;refGene_gn=C2CD2;refGene_tx
=XM_023238704.1;refGene_gn=C2CD2;refGene_tx=XM_023238705.1;refGene_gn=
C2CD2;refGene_tx=XM_023238706.1;refGene_gn=C2CD2;refGene_tx=XM_0232387
07.1
C2	4125381	4125691
refGene_gn=C2CD2;refGene_tx=XM_011285776.2;refGene_gn=C2CD2;refGene_tx
=XM_023238704.1;refGene_gn=C2CD2;refGene_tx=XM_023238705.1;refGene_gn=
C2CD2;refGene_tx=XM_023238706.1;refGene_gn=C2CD2;refGene_tx=XM_0232387
07.1
C2	4127484	4127618	refGene_gn=C2CD2;refGene_tx=XM_023238707.1
C2	4133124	4133387
refGene_gn=C2CD2;refGene_tx=XM_011285776.2;refGene_gn=C2CD2;refGene_tx
=XM_023238704.1;refGene_gn=C2CD2;refGene_tx=XM_023238705.1;refGene_gn=
C2CD2;refGene_tx=XM_023238706.1
C2	4143024	4143126	refGene_gn=PRDM15;refGene_tx=XM_023260867.1
C2	4163851	4163897	refGene_gn=PRDM15;refGene_tx=XM_023260867.1
C2	4164456	4164550	refGene_gn=PRDM15;refGene_tx=XM_023260867.1
C2	4166438	4166592	refGene_gn=PRDM15;refGene_tx=XM_023260867.1
C2	4168820	4169073	refGene_gn=PRDM15;refGene_tx=XM_023260867.1
C2	4175887	4175989	refGene_gn=PRDM15;refGene_tx=XM_023260867.1
C2	4177291	4177516	refGene_gn=PRDM15;refGene_tx=XM_023260867.1
C2	4179145	4179275	refGene_gn=PRDM15;refGene_tx=XM_023260867.1
C2	4180845	4181027	refGene_gn=PRDM15;refGene_tx=XM_023260867.1
C2	4181186	4181281	refGene_gn=PRDM15;refGene_tx=XM_023260867.1
C2	4181728	4181816	refGene_gn=PRDM15;refGene_tx=XM_023260867.1
C2	4184668	4184836	refGene_gn=PRDM15;refGene_tx=XM_023260867.1
C2	4187584	4187679	refGene_gn=PRDM15;refGene_tx=XM_023260867.1
C2	4188500	4188632	refGene_gn=PRDM15;refGene_tx=XM_023260867.1
C2	4190050	4190185	refGene_gn=PRDM15;refGene_tx=XM_023260867.1
C2	4190774	4190938	refGene_gn=PRDM15;refGene_tx=XM_023260867.1
C2	4191805	4191885	refGene_gn=PRDM15;refGene_tx=XM_023260867.1
C2	4192244	4192364	refGene_gn=PRDM15;refGene_tx=XM_023260867.1
C2	4196709	4196841	refGene_gn=PRDM15;refGene_tx=XM_023260867.1
C2	4198308	4198392	refGene_gn=PRDM15;refGene_tx=XM_023260867.1
C2	4198718	4198894	refGene_gn=PRDM15;refGene_tx=XM_023260867.1
C2	4205515	4205596	refGene_gn=PRDM15;refGene_tx=XM_023260867.1
C2	4207178	4207887	refGene_gn=PRDM15;refGene_tx=XM_023260867.1
C2	4226847	4226885	refGene_gn=PRDM15;refGene_tx=XM_023260867.1
C2	4242278	4242460	refGene_gn=RIPK4;refGene_tx=XM_023238708.1
C2	4252528	4252820	refGene_gn=RIPK4;refGene_tx=XM_023238708.1
C2	4259185	4259334	refGene_gn=RIPK4;refGene_tx=XM_023238708.1
C2	4261064	4261114	refGene_gn=RIPK4;refGene_tx=XM_023238708.1
C2	4263357	4263516	refGene_gn=RIPK4;refGene_tx=XM_023238708.1
C2	4264294	4264398	refGene_gn=RIPK4;refGene_tx=XM_023238708.1
C2	4265821	4266083	refGene_gn=RIPK4;refGene_tx=XM_023238708.1
C2	4268034	4269194	refGene_gn=RIPK4;refGene_tx=XM_023238708.1
C2	4393822	4394140

refGene_gn=LOC101095514;refGene_tx=XM_003991409.5
C2 4396284 4396635
refGene_gn=LOC111562335;refGene_tx=XM_023260843.1
C2 4474237 4474252
refGene_gn=TMPRSS2;refGene_tx=XM_023238709.1;refGene_gn=TMPRSS2;refGene_tx=XM_023238710.1;refGene_gn=TMPRSS2;refGene_tx=XM_023238711.1
C2 4476449 4476672
refGene_gn=TMPRSS2;refGene_tx=XM_023238709.1;refGene_gn=TMPRSS2;refGene_tx=XM_023238710.1;refGene_gn=TMPRSS2;refGene_tx=XM_023238711.1
C2 4481457 4481544
refGene_gn=TMPRSS2;refGene_tx=XM_023238709.1;refGene_gn=TMPRSS2;refGene_tx=XM_023238710.1;refGene_gn=TMPRSS2;refGene_tx=XM_023238711.1
C2 4482213 4482333
refGene_gn=TMPRSS2;refGene_tx=XM_023238709.1;refGene_gn=TMPRSS2;refGene_tx=XM_023238710.1;refGene_gn=TMPRSS2;refGene_tx=XM_023238711.1
C2 4487569 4487696
refGene_gn=TMPRSS2;refGene_tx=XM_023238709.1;refGene_gn=TMPRSS2;refGene_tx=XM_023238710.1;refGene_gn=TMPRSS2;refGene_tx=XM_023238711.1
C2 4488946 4489057
refGene_gn=TMPRSS2;refGene_tx=XM_023238709.1;refGene_gn=TMPRSS2;refGene_tx=XM_023238710.1;refGene_gn=TMPRSS2;refGene_tx=XM_023238711.1
C2 4491455 4491499
refGene_gn=TMPRSS2;refGene_tx=XM_023238709.1;refGene_gn=TMPRSS2;refGene_tx=XM_023238710.1;refGene_gn=TMPRSS2;refGene_tx=XM_023238711.1
C2 4493366 4493538
refGene_gn=TMPRSS2;refGene_tx=XM_023238709.1;refGene_gn=TMPRSS2;refGene_tx=XM_023238710.1;refGene_gn=TMPRSS2;refGene_tx=XM_023238711.1
C2 4494917 4495093
refGene_gn=TMPRSS2;refGene_tx=XM_023238709.1;refGene_gn=TMPRSS2;refGene_tx=XM_023238710.1;refGene_gn=TMPRSS2;refGene_tx=XM_023238711.1
C2 4496204 4496300
refGene_gn=TMPRSS2;refGene_tx=XM_023238709.1;refGene_gn=TMPRSS2;refGene_tx=XM_023238710.1;refGene_gn=TMPRSS2;refGene_tx=XM_023238711.1
C2 4498354 4498497
refGene_gn=TMPRSS2;refGene_tx=XM_023238709.1;refGene_gn=TMPRSS2;refGene_tx=XM_023238710.1;refGene_gn=TMPRSS2;refGene_tx=XM_023238711.1
C2 4498948 4499101
refGene_gn=TMPRSS2;refGene_tx=XM_023238709.1;refGene_gn=TMPRSS2;refGene_tx=XM_023238710.1;refGene_gn=TMPRSS2;refGene_tx=XM_023238711.1
C2 4499551 4499563
refGene_gn=TMPRSS2;refGene_tx=XM_023238709.1;refGene_gn=TMPRSS2;refGene_tx=XM_023238710.1;refGene_gn=TMPRSS2;refGene_tx=XM_023238711.1
C2 4506672 4506903
refGene_gn=LOC101085996;refGene_tx=XM_023238713.1;refGene_gn=LOC101085996;refGene_tx=XM_023238714.1
C2 4510429 4510678
refGene_gn=LOC101085996;refGene_tx=XM_023238713.1;refGene_gn=LOC101085996;refGene_tx=XM_023238714.1
C2 4512084 4512161
refGene_gn=LOC101085996;refGene_tx=XM_023238713.1;refGene_gn=LOC101085

996;refGene_tx=XM_023238714.1
C2 4513341 4513500
refGene_gn=LOC101085996;refGene_tx=XM_023238713.1;refGene_gn=LOC101085
996;refGene_tx=XM_023238714.1
C2 4516529 4516671
refGene_gn=LOC101085996;refGene_tx=XM_023238713.1;refGene_gn=LOC101085
996;refGene_tx=XM_023238714.1
C2 4517037 4517160
refGene_gn=LOC101085996;refGene_tx=XM_023238713.1;refGene_gn=LOC101085
996;refGene_tx=XM_023238714.1
C2 4518681 4518760
refGene_gn=LOC101085996;refGene_tx=XM_023238713.1;refGene_gn=LOC101085
996;refGene_tx=XM_023238714.1
C2 4520169 4520368
refGene_gn=LOC101085996;refGene_tx=XM_023238713.1;refGene_gn=LOC101085
996;refGene_tx=XM_023238714.1
C2 4520944 4521083
refGene_gn=LOC101085996;refGene_tx=XM_023238713.1;refGene_gn=LOC101085
996;refGene_tx=XM_023238714.1
C2 4521700 4521855
refGene_gn=LOC101085996;refGene_tx=XM_023238713.1;refGene_gn=LOC101085
996;refGene_tx=XM_023238714.1
C2 4523699 4523837
refGene_gn=LOC101085996;refGene_tx=XM_023238713.1;refGene_gn=LOC101085
996;refGene_tx=XM_023238714.1
C2 4524902 4525095
refGene_gn=LOC101085996;refGene_tx=XM_023238713.1;refGene_gn=LOC101085
996;refGene_tx=XM_023238714.1
C2 4529331 4529430
refGene_gn=LOC101085996;refGene_tx=XM_023238713.1;refGene_gn=LOC101085
996;refGene_tx=XM_023238714.1
C2 4580102 4580192 refGene_gn=FAM3B;refGene_tx=XM_023238715.1
C2 4584956 4585089 refGene_gn=FAM3B;refGene_tx=XM_023238715.1
C2 4586465 4586553 refGene_gn=FAM3B;refGene_tx=XM_023238715.1
C2 4587581 4587632 refGene_gn=FAM3B;refGene_tx=XM_023238715.1
C2 4589525 4589584 refGene_gn=FAM3B;refGene_tx=XM_023238715.1
C2 4606993 4607117 refGene_gn=FAM3B;refGene_tx=XM_023238715.1
C2 4616767 4616911 refGene_gn=FAM3B;refGene_tx=XM_023238715.1
C2 4623640 4623659 refGene_gn=FAM3B;refGene_tx=XM_023238715.1
C2 4669763 4670017
refGene_gn=BACE2;refGene_tx=XM_023238716.1;refGene_gn=BACE2;refGene_tx
=XM_023238717.1
C2 4690996 4691165 refGene_gn=BACE2;refGene_tx=XM_023238716.1
C2 4696180 4696330
refGene_gn=BACE2;refGene_tx=XM_023238716.1;refGene_gn=BACE2;refGene_tx
=XM_023238717.1
C2 4701060 4701162
refGene_gn=BACE2;refGene_tx=XM_023238716.1;refGene_gn=BACE2;refGene_tx
=XM_023238717.1
C2 4702528 4702663

refGene_gn=BACE2;refGene_tx=XM_023238716.1;refGene_gn=BACE2;refGene_tx
=XM_023238717.1
C2 4704053 4704182
refGene_gn=BACE2;refGene_tx=XM_023238716.1;refGene_gn=BACE2;refGene_tx
=XM_023238717.1
C2 4708273 4708490
refGene_gn=BACE2;refGene_tx=XM_023238716.1;refGene_gn=BACE2;refGene_tx
=XM_023238717.1
C2 4719398 4719487
refGene_gn=BACE2;refGene_tx=XM_023238716.1;refGene_gn=BACE2;refGene_tx
=XM_023238717.1
C2 4759118 4759409
refGene_gn=BACE2;refGene_tx=XM_023238716.1;refGene_gn=BACE2;refGene_tx
=XM_023238717.1
C2 5069418 5069584 refGene_gn=DSCAM;refGene_tx=XM_019811073.2
C2 5203128 5203446 refGene_gn=DSCAM;refGene_tx=XM_019811073.2
C2 5220174 5220321 refGene_gn=DSCAM;refGene_tx=XM_019811073.2
C2 5524379 5524526 refGene_gn=DSCAM;refGene_tx=XM_019811073.2
C2 5533742 5534021 refGene_gn=DSCAM;refGene_tx=XM_019811073.2
C2 5537685 5537961 refGene_gn=DSCAM;refGene_tx=XM_019811073.2
C2 5547618 5547915 refGene_gn=DSCAM;refGene_tx=XM_019811073.2
C2 5548727 5549003 refGene_gn=DSCAM;refGene_tx=XM_019811073.2
C2 5573026 5573305 refGene_gn=DSCAM;refGene_tx=XM_019811073.2
C2 5588270 5588390 refGene_gn=DSCAM;refGene_tx=XM_019811073.2
C2 5606679 5606853 refGene_gn=DSCAM;refGene_tx=XM_019811073.2
C2 5695675 5695872 refGene_gn=DSCAM;refGene_tx=XM_019811073.2
C2 5696909 5697006 refGene_gn=DSCAM;refGene_tx=XM_019811073.2
C2 5698361 5698490 refGene_gn=DSCAM;refGene_tx=XM_019811073.2
C2 5707378 5707546 refGene_gn=DSCAM;refGene_tx=XM_019811073.2
C2 5718999 5719070 refGene_gn=DSCAM;refGene_tx=XM_019811073.2
C2 5736754 5736995 refGene_gn=DSCAM;refGene_tx=XM_019811073.2
C2 5738482 5738629 refGene_gn=DSCAM;refGene_tx=XM_019811073.2
C2 5744520 5744676 refGene_gn=DSCAM;refGene_tx=XM_019811073.2
C2 5755283 5755417 refGene_gn=DSCAM;refGene_tx=XM_019811073.2
C2 5773262 5773416 refGene_gn=DSCAM;refGene_tx=XM_019811073.2
C2 5777907 5778025 refGene_gn=DSCAM;refGene_tx=XM_019811073.2
C2 5779398 5779562 refGene_gn=DSCAM;refGene_tx=XM_019811073.2
C2 5781076 5781175 refGene_gn=DSCAM;refGene_tx=XM_019811073.2
C2 5785237 5785426 refGene_gn=DSCAM;refGene_tx=XM_019811073.2
C2 5786837 5787128 refGene_gn=DSCAM;refGene_tx=XM_019811073.2
C2 5790468 5790645 refGene_gn=DSCAM;refGene_tx=XM_019811073.2
C2 5801705 5801736 refGene_gn=DSCAM;refGene_tx=XM_019811073.2
C2 5808481 5808597 refGene_gn=DSCAM;refGene_tx=XM_019811073.2
C2 5811985 5812135 refGene_gn=DSCAM;refGene_tx=XM_019811073.2
C2 5820304 5820502 refGene_gn=DSCAM;refGene_tx=XM_019811073.2
C2 5821828 5822131 refGene_gn=DSCAM;refGene_tx=XM_019811073.2
C2 5834618 5834971 refGene_gn=DSCAM;refGene_tx=XM_019811073.2
C2 5921470 5921598 refGene_gn=PCP4;refGene_tx=XM_006935852.4
C2 5955348 5955400 refGene_gn=PCP4;refGene_tx=XM_006935852.4
C2 5981990 5981999 refGene_gn=PCP4;refGene_tx=XM_006935852.4

C2 6030852 6030948
refGene_gn=IGSF5;refGene_tx=XM_006935853.4;refGene_gn=IGSF5;refGene_tx=XM_019839362.2;refGene_gn=IGSF5;refGene_tx=XM_019839363.2
C2 6038328 6038402
refGene_gn=IGSF5;refGene_tx=XM_006935853.4;refGene_gn=IGSF5;refGene_tx=XM_019839362.2;refGene_gn=IGSF5;refGene_tx=XM_019839363.2
C2 6039842 6039943
refGene_gn=IGSF5;refGene_tx=XM_006935853.4;refGene_gn=IGSF5;refGene_tx=XM_019839362.2;refGene_gn=IGSF5;refGene_tx=XM_019839363.2
C2 6043713 6043738
refGene_gn=IGSF5;refGene_tx=XM_006935853.4;refGene_gn=IGSF5;refGene_tx=XM_019839362.2;refGene_gn=IGSF5;refGene_tx=XM_019839363.2
C2 6046963 6047104
refGene_gn=IGSF5;refGene_tx=XM_006935853.4;refGene_gn=IGSF5;refGene_tx=XM_019839362.2;refGene_gn=IGSF5;refGene_tx=XM_019839363.2
C2 6057369 6057669
refGene_gn=IGSF5;refGene_tx=XM_006935853.4;refGene_gn=IGSF5;refGene_tx=XM_019839362.2;refGene_gn=IGSF5;refGene_tx=XM_019839363.2
C2 6063042 6063360
refGene_gn=IGSF5;refGene_tx=XM_006935853.4;refGene_gn=IGSF5;refGene_tx=XM_019839362.2;refGene_gn=IGSF5;refGene_tx=XM_019839363.2
C2 6076773 6076831
refGene_gn=IGSF5;refGene_tx=XM_006935853.4;refGene_gn=IGSF5;refGene_tx=XM_019839362.2;refGene_gn=IGSF5;refGene_tx=XM_019839363.2
C2 6166667 6167603
refGene_gn=B3GALT5;refGene_tx=XM_023238719.1;refGene_gn=B3GALT5;refGene_tx=XM_023238720.1;refGene_gn=B3GALT5;refGene_tx=XM_023238721.1
C2 6261680 6261809 refGene_gn=SH3BGR;refGene_tx=XM_023260869.1
C2 6269399 6269498 refGene_gn=SH3BGR;refGene_tx=XM_023260869.1
C2 6270151 6270226 refGene_gn=SH3BGR;refGene_tx=XM_023260869.1
C2 6293849 6293930 refGene_gn=SH3BGR;refGene_tx=XM_023260869.1
C2 6302544 6302730 refGene_gn=SH3BGR;refGene_tx=XM_023260869.1
C2 6311393 6311438 refGene_gn=SH3BGR;refGene_tx=XM_023260869.1
C2 6326584 6326616 refGene_gn=WRB;refGene_tx=XM_023238727.1
C2 6336559 6336893
refGene_gn=LCA5L;refGene_tx=XM_019839369.2;refGene_gn=LCA5L;refGene_tx=XM_023238722.1;refGene_gn=LCA5L;refGene_tx=XM_023238723.1
C2 6337160 6337307
refGene_gn=LCA5L;refGene_tx=XM_019839369.2;refGene_gn=LCA5L;refGene_tx=XM_019839371.2;refGene_gn=LCA5L;refGene_tx=XM_023238722.1
C2 6340971 6341486
refGene_gn=LCA5L;refGene_tx=XM_019839369.2;refGene_gn=LCA5L;refGene_tx=XM_019839371.2;refGene_gn=LCA5L;refGene_tx=XM_019839373.2;refGene_gn=LCA5L;refGene_tx=XM_023238722.1;refGene_gn=LCA5L;refGene_tx=XM_023238723.1
C2 6343463 6343601
refGene_gn=LCA5L;refGene_tx=XM_019839369.2;refGene_gn=LCA5L;refGene_tx=XM_019839371.2;refGene_gn=LCA5L;refGene_tx=XM_019839373.2;refGene_gn=LCA5L;refGene_tx=XM_023238722.1;refGene_gn=LCA5L;refGene_tx=XM_023238723.1

C2 6357180 6357262
refGene_gn=LCA5L;refGene_tx=XM_019839369.2;refGene_gn=LCA5L;refGene_tx
=XM_019839371.2;refGene_gn=LCA5L;refGene_tx=XM_019839373.2;refGene_gn=
LCA5L;refGene_tx=XM_023238722.1;refGene_gn=LCA5L;refGene_tx=XM_0232387
23.1
C2 6359415 6359519
refGene_gn=LCA5L;refGene_tx=XM_019839369.2;refGene_gn=LCA5L;refGene_tx
=XM_019839371.2;refGene_gn=LCA5L;refGene_tx=XM_019839373.2;refGene_gn=
LCA5L;refGene_tx=XM_023238722.1;refGene_gn=LCA5L;refGene_tx=XM_0232387
23.1
C2 6359689 6359807
refGene_gn=LCA5L;refGene_tx=XM_019839369.2;refGene_gn=LCA5L;refGene_tx
=XM_019839371.2;refGene_gn=LCA5L;refGene_tx=XM_019839373.2;refGene_gn=
LCA5L;refGene_tx=XM_023238722.1;refGene_gn=LCA5L;refGene_tx=XM_0232387
23.1
C2 6365386 6366075
refGene_gn=LCA5L;refGene_tx=XM_019839369.2;refGene_gn=LCA5L;refGene_tx
=XM_019839371.2;refGene_gn=LCA5L;refGene_tx=XM_019839373.2;refGene_gn=
LCA5L;refGene_tx=XM_023238722.1;refGene_gn=LCA5L;refGene_tx=XM_0232387
23.1
C2 6370055 6370111 refGene_gn=WRB;refGene_tx=XM_023238725.1
C2 6371818 6371892
refGene_gn=WRB;refGene_tx=XM_023238724.1;refGene_gn=WRB;refGene_tx=XM_
023238728.1;refGene_gn=WRB;refGene_tx=XM_023238729.1;refGene_gn=WRB;re
fGene_tx=XM_023238730.1
C2 6372070 6372117 refGene_gn=WRB;refGene_tx=XM_023238726.1
C2 6373184 6373299
refGene_gn=WRB;refGene_tx=XM_023238724.1;refGene_gn=WRB;refGene_tx=XM_
023238725.1;refGene_gn=WRB;refGene_tx=XM_023238726.1;refGene_gn=WRB;re
fGene_tx=XM_023238727.1;refGene_gn=WRB;refGene_tx=XM_023238728.1;refGe
ne_gn=WRB;refGene_tx=XM_023238729.1;refGene_gn=WRB;refGene_tx=XM_02323
8730.1
C2 6375708 6375776
refGene_gn=WRB;refGene_tx=XM_023238724.1;refGene_gn=WRB;refGene_tx=XM_
023238725.1;refGene_gn=WRB;refGene_tx=XM_023238726.1;refGene_gn=WRB;re
fGene_tx=XM_023238727.1;refGene_gn=WRB;refGene_tx=XM_023238728.1;refGe
ne_gn=WRB;refGene_tx=XM_023238729.1;refGene_gn=WRB;refGene_tx=XM_02323
8730.1
C2 6377126 6377292
refGene_gn=WRB;refGene_tx=XM_023238724.1;refGene_gn=WRB;refGene_tx=XM_
023238725.1;refGene_gn=WRB;refGene_tx=XM_023238726.1;refGene_gn=WRB;re
fGene_tx=XM_023238727.1;refGene_gn=WRB;refGene_tx=XM_023238728.1;refGe
ne_gn=WRB;refGene_tx=XM_023238729.1;refGene_gn=WRB;refGene_tx=XM_02323
8730.1
C2 6381505 6381667
refGene_gn=WRB;refGene_tx=XM_023238725.1;refGene_gn=WRB;refGene_tx=XM_
023238726.1;refGene_gn=WRB;refGene_tx=XM_023238727.1
C2 6386608 6386710 refGene_gn=WRB;refGene_tx=XM_023238728.1
C2 6409709 6409973 refGene_gn=HMGN1;refGene_tx=XM_023238731.1
C2 6410325 6410358 refGene_gn=HMGN1;refGene_tx=XM_023238731.1

C2	6410444	6410474	refGene_gn=HMGN1;refGene_tx=XM_023238731.1
C2	6410557	6410608	refGene_gn=HMGN1;refGene_tx=XM_023238731.1
C2	6413965	6414094	refGene_gn=HMGN1;refGene_tx=XM_023238731.1
C2	6415676	6415724	refGene_gn=HMGN1;refGene_tx=XM_023238731.1
C2	6455046	6455095	refGene_gn=BRWD1;refGene_tx=XM_023238732.1
C2	6455205	6455264	refGene_gn=BRWD1;refGene_tx=XM_023238732.1
C2	6455411	6455441	refGene_gn=BRWD1;refGene_tx=XM_023238732.1
C2	6455624	6455684	refGene_gn=BRWD1;refGene_tx=XM_023238732.1
C2	6476976	6477127	refGene_gn=BRWD1;refGene_tx=XM_023238732.1
C2	6479183	6479282	refGene_gn=BRWD1;refGene_tx=XM_023238732.1
C2	6479645	6479806	refGene_gn=BRWD1;refGene_tx=XM_023238732.1
C2	6480906	6481128	refGene_gn=BRWD1;refGene_tx=XM_023238732.1
C2	6493817	6493918	refGene_gn=BRWD1;refGene_tx=XM_023238732.1
C2	6495311	6495382	refGene_gn=BRWD1;refGene_tx=XM_023238732.1
C2	6496916	6497017	refGene_gn=BRWD1;refGene_tx=XM_023238732.1
C2	6498009	6498050	refGene_gn=BRWD1;refGene_tx=XM_023238732.1
C2	6498896	6498995	refGene_gn=BRWD1;refGene_tx=XM_023238732.1
C2	6502574	6502725	refGene_gn=BRWD1;refGene_tx=XM_023238732.1
C2	6502980	6503115	refGene_gn=BRWD1;refGene_tx=XM_023238732.1
C2	6507689	6507818	refGene_gn=BRWD1;refGene_tx=XM_023238732.1
C2	6508258	6508484	refGene_gn=BRWD1;refGene_tx=XM_023238732.1
C2	6514628	6514814	refGene_gn=BRWD1;refGene_tx=XM_023238732.1
C2	6517030	6517214	refGene_gn=BRWD1;refGene_tx=XM_023238732.1
C2	6521112	6521206	refGene_gn=BRWD1;refGene_tx=XM_023238732.1
C2	6525418	6525550	refGene_gn=BRWD1;refGene_tx=XM_023238732.1
C2	6535323	6535418	refGene_gn=BRWD1;refGene_tx=XM_023238732.1
C2	6537645	6537835	refGene_gn=BRWD1;refGene_tx=XM_023238732.1
C2	6539236	6539362	refGene_gn=BRWD1;refGene_tx=XM_023238732.1
C2	6539456	6539564	refGene_gn=BRWD1;refGene_tx=XM_023238732.1
C2	6541766	6541891	refGene_gn=BRWD1;refGene_tx=XM_023238732.1
C2	6542523	6542606	refGene_gn=BRWD1;refGene_tx=XM_023238732.1
C2	6545928	6546040	refGene_gn=BRWD1;refGene_tx=XM_023238732.1
C2	6546636	6546698	refGene_gn=BRWD1;refGene_tx=XM_023238732.1
C2	6547429	6547585	refGene_gn=BRWD1;refGene_tx=XM_023238732.1
C2	6548032	6548153	refGene_gn=BRWD1;refGene_tx=XM_023238732.1
C2	6549355	6549481	refGene_gn=BRWD1;refGene_tx=XM_023238732.1
C2	6550652	6550725	refGene_gn=BRWD1;refGene_tx=XM_023238732.1
C2	6552263	6552305	refGene_gn=BRWD1;refGene_tx=XM_023238732.1
C2	6554722	6554869	refGene_gn=BRWD1;refGene_tx=XM_023238732.1
C2	6555494	6555647	refGene_gn=BRWD1;refGene_tx=XM_023238732.1
C2	6558958	6559125	refGene_gn=BRWD1;refGene_tx=XM_023238732.1
C2	6564745	6564972	refGene_gn=BRWD1;refGene_tx=XM_023238732.1
C2	6567225	6567390	refGene_gn=BRWD1;refGene_tx=XM_023238732.1
C2	6569200	6570112	refGene_gn=BRWD1;refGene_tx=XM_023238732.1
C2	6571099	6572238	refGene_gn=BRWD1;refGene_tx=XM_023238732.1
C2	6586497	6586631
refGene_gn=PSMG1;refGene_tx=XM_023238733.1;refGene_gn=PSMG1;refGene_tx
=XM_023238734.1;refGene_gn=PSMG1;refGene_tx=XM_023238735.1
C2	6588087	6588194
refGene_gn=PSMG1;refGene_tx=XM_023238733.1;refGene_gn=PSMG1;refGene_tx

=XM_023238734.1
C2 6589419 6589571
refGene_gn=PSMG1;refGene_tx=XM_023238733.1;refGene_gn=PSMG1;refGene_tx
=XM_023238734.1;refGene_gn=PSMG1;refGene_tx=XM_023238735.1
C2 6591340 6591403
refGene_gn=PSMG1;refGene_tx=XM_023238733.1;refGene_gn=PSMG1;refGene_tx
=XM_023238735.1
C2 6594403 6594602
refGene_gn=PSMG1;refGene_tx=XM_023238733.1;refGene_gn=PSMG1;refGene_tx
=XM_023238734.1;refGene_gn=PSMG1;refGene_tx=XM_023238735.1
C2 6595479 6595616
refGene_gn=PSMG1;refGene_tx=XM_023238733.1;refGene_gn=PSMG1;refGene_tx
=XM_023238734.1;refGene_gn=PSMG1;refGene_tx=XM_023238735.1
C2 6597139 6597214
refGene_gn=PSMG1;refGene_tx=XM_023238733.1;refGene_gn=PSMG1;refGene_tx
=XM_023238734.1;refGene_gn=PSMG1;refGene_tx=XM_023238735.1
C2 6883948 6884164 refGene_gn=ETS2;refGene_tx=XM_023238736.1
C2 6884905 6885024 refGene_gn=ETS2;refGene_tx=XM_023238736.1
C2 6887529 6887793 refGene_gn=ETS2;refGene_tx=XM_023238736.1
C2 6888907 6889129 refGene_gn=ETS2;refGene_tx=XM_023238736.1
C2 6890044 6890128 refGene_gn=ETS2;refGene_tx=XM_023238736.1
C2 6892193 6892394 refGene_gn=ETS2;refGene_tx=XM_023238736.1
C2 6892749 6892869 refGene_gn=ETS2;refGene_tx=XM_023238736.1
C2 6894194 6894306 refGene_gn=ETS2;refGene_tx=XM_023238736.1
C2 6897076 6897148 refGene_gn=ETS2;refGene_tx=XM_023238736.1
C2 7120175 7120214 refGene_gn=ERG;refGene_tx=XM_023238738.1
C2 7195698 7195716 refGene_gn=ERG;refGene_tx=XM_003991424.5
C2 7242328 7242454 refGene_gn=ERG;refGene_tx=XM_023238739.1
C2 7245055 7245273
refGene_gn=ERG;refGene_tx=XM_003991424.5;refGene_gn=ERG;refGene_tx=XM_
011285773.3;refGene_gn=ERG;refGene_tx=XM_011285774.3;refGene_gn=ERG;re
fGene_tx=XM_019839379.2;refGene_gn=ERG;refGene_tx=XM_023238738.1;refGe
ne_gn=ERG;refGene_tx=XM_023238739.1
C2 7265163 7265315
refGene_gn=ERG;refGene_tx=XM_003991424.5;refGene_gn=ERG;refGene_tx=XM_
011285773.3;refGene_gn=ERG;refGene_tx=XM_011285774.3;refGene_gn=ERG;re
fGene_tx=XM_019839379.2;refGene_gn=ERG;refGene_tx=XM_023238738.1;refGe
ne_gn=ERG;refGene_tx=XM_023238739.1
C2 7287196 7287400
refGene_gn=ERG;refGene_tx=XM_003991424.5;refGene_gn=ERG;refGene_tx=XM_
011285773.3;refGene_gn=ERG;refGene_tx=XM_011285774.3;refGene_gn=ERG;re
fGene_tx=XM_019839379.2;refGene_gn=ERG;refGene_tx=XM_023238738.1
C2 7288269 7288350
refGene_gn=ERG;refGene_tx=XM_003991424.5;refGene_gn=ERG;refGene_tx=XM_
011285773.3;refGene_gn=ERG;refGene_tx=XM_023238738.1
C2 7290246 7290318
refGene_gn=ERG;refGene_tx=XM_003991424.5;refGene_gn=ERG;refGene_tx=XM_
019839379.2;refGene_gn=ERG;refGene_tx=XM_023238738.1
C2 7298037 7298106
refGene_gn=ERG;refGene_tx=XM_003991424.5;refGene_gn=ERG;refGene_tx=XM_

011285773.3;refGene_gn=ERG;refGene_tx=XM_011285774.3;refGene_gn=ERG;refGene_tx=XM_019839379.2;refGene_gn=ERG;refGene_tx=XM_023238738.1;refGene_gn=ERG;refGene_tx=XM_023238739.1
C2 7298644 7298701
refGene_gn=ERG;refGene_tx=XM_003991424.5;refGene_gn=ERG;refGene_tx=XM_011285773.3;refGene_gn=ERG;refGene_tx=XM_011285774.3;refGene_gn=ERG;refGene_tx=XM_019839379.2;refGene_gn=ERG;refGene_tx=XM_023238738.1;refGene_gn=ERG;refGene_tx=XM_023238739.1
C2 7299629 7299677
refGene_gn=ERG;refGene_tx=XM_003991424.5;refGene_gn=ERG;refGene_tx=XM_011285773.3;refGene_gn=ERG;refGene_tx=XM_011285774.3;refGene_gn=ERG;refGene_tx=XM_019839379.2;refGene_gn=ERG;refGene_tx=XM_023238738.1;refGene_gn=ERG;refGene_tx=XM_023238739.1
C2 7307328 7307849
refGene_gn=ERG;refGene_tx=XM_003991424.5;refGene_gn=ERG;refGene_tx=XM_011285773.3;refGene_gn=ERG;refGene_tx=XM_011285774.3;refGene_gn=ERG;refGene_tx=XM_019839379.2;refGene_gn=ERG;refGene_tx=XM_023238738.1;refGene_gn=ERG;refGene_tx=XM_023238739.1
C2 7371868 7372996
refGene_gn=KCNJ15;refGene_tx=XM_019839382.2;refGene_gn=KCNJ15;refGene_tx=XM_019839383.2;refGene_gn=KCNJ15;refGene_tx=XM_019839384.2;refGene_gn=KCNJ15;refGene_tx=XM_023238740.1
C2 7793936 7793961 refGene_gn=KCNJ6;refGene_tx=XM_003991434.5
C2 7911499 7912420 refGene_gn=KCNJ6;refGene_tx=XM_003991434.5
C2 7983255 7983581 refGene_gn=KCNJ6;refGene_tx=XM_003991434.5
C2 8098584 8099205
refGene_gn=DYRK1A;refGene_tx=XM_003991427.4;refGene_gn=DYRK1A;refGene_tx=XM_023238742.1;refGene_gn=DYRK1A;refGene_tx=XM_023238743.1;refGene_gn=DYRK1A;refGene_tx=XM_023238745.1
C2 8104187 8104312
refGene_gn=DYRK1A;refGene_tx=XM_003991427.4;refGene_gn=DYRK1A;refGene_tx=XM_023238742.1;refGene_gn=DYRK1A;refGene_tx=XM_023238743.1;refGene_gn=DYRK1A;refGene_tx=XM_023238745.1
C2 8105034 8105341
refGene_gn=DYRK1A;refGene_tx=XM_003991427.4;refGene_gn=DYRK1A;refGene_tx=XM_023238742.1;refGene_gn=DYRK1A;refGene_tx=XM_023238743.1;refGene_gn=DYRK1A;refGene_tx=XM_023238745.1
C2 8113823 8113964
refGene_gn=DYRK1A;refGene_tx=XM_003991427.4;refGene_gn=DYRK1A;refGene_tx=XM_023238742.1;refGene_gn=DYRK1A;refGene_tx=XM_023238743.1;refGene_gn=DYRK1A;refGene_tx=XM_023238745.1
C2 8116015 8116162
refGene_gn=DYRK1A;refGene_tx=XM_003991427.4;refGene_gn=DYRK1A;refGene_tx=XM_023238742.1;refGene_gn=DYRK1A;refGene_tx=XM_023238743.1;refGene_gn=DYRK1A;refGene_tx=XM_023238745.1
C2 8118702 8118989
refGene_gn=DYRK1A;refGene_tx=XM_003991427.4;refGene_gn=DYRK1A;refGene_tx=XM_023238742.1;refGene_gn=DYRK1A;refGene_tx=XM_023238743.1;refGene_gn=DYRK1A;refGene_tx=XM_023238745.1
C2 8123107 8123255

refGene_gn=DYRK1A;refGene_tx=XM_003991427.4;refGene_gn=DYRK1A;refGene_tx=XM_023238742.1;refGene_gn=DYRK1A;refGene_tx=XM_023238743.1;refGene_gn=DYRK1A;refGene_tx=XM_023238745.1
C2 8127634 8127823
refGene_gn=DYRK1A;refGene_tx=XM_003991427.4;refGene_gn=DYRK1A;refGene_tx=XM_023238742.1;refGene_gn=DYRK1A;refGene_tx=XM_023238743.1;refGene_gn=DYRK1A;refGene_tx=XM_023238745.1
C2 8129827 8129947
refGene_gn=DYRK1A;refGene_tx=XM_003991427.4;refGene_gn=DYRK1A;refGene_tx=XM_023238742.1;refGene_gn=DYRK1A;refGene_tx=XM_023238743.1;refGene_gn=DYRK1A;refGene_tx=XM_023238745.1
C2 8136613 8136810
refGene_gn=DYRK1A;refGene_tx=XM_003991427.4;refGene_gn=DYRK1A;refGene_tx=XM_023238742.1;refGene_gn=DYRK1A;refGene_tx=XM_023238743.1;refGene_gn=DYRK1A;refGene_tx=XM_023238745.1
C2 8192106 8192116
refGene_gn=DYRK1A;refGene_tx=XM_003991427.4;refGene_gn=DYRK1A;refGene_tx=XM_023238742.1;refGene_gn=DYRK1A;refGene_tx=XM_023238743.1
C2 8329938 8329995
refGene_gn=DSCR3;refGene_tx=XM_011285771.3;refGene_gn=DSCR3;refGene_tx=XM_019839392.2
C2 8354994 8355138
refGene_gn=DSCR3;refGene_tx=XM_011285771.3;refGene_gn=DSCR3;refGene_tx=XM_019839392.2
C2 8356323 8356473
refGene_gn=DSCR3;refGene_tx=XM_011285771.3;refGene_gn=DSCR3;refGene_tx=XM_019839392.2
C2 8363844 8363925
refGene_gn=DSCR3;refGene_tx=XM_011285771.3;refGene_gn=DSCR3;refGene_tx=XM_019839392.2
C2 8367345 8367420
refGene_gn=DSCR3;refGene_tx=XM_011285771.3;refGene_gn=DSCR3;refGene_tx=XM_019839392.2
C2 8372212 8372363
refGene_gn=DSCR3;refGene_tx=XM_011285771.3;refGene_gn=DSCR3;refGene_tx=XM_019839392.2
C2 8372603 8372756
refGene_gn=DSCR3;refGene_tx=XM_011285771.3;refGene_gn=DSCR3;refGene_tx=XM_019839392.2
C2 8375338 8375421 refGene_gn=DSCR3;refGene_tx=XM_011285771.3
C2 8378924 8379013 refGene_gn=DSCR3;refGene_tx=XM_019839392.2
C2 8424218 8424332 refGene_gn=TTC3;refGene_tx=XM_023238746.1
C2 8425509 8425602 refGene_gn=TTC3;refGene_tx=XM_023238746.1
C2 8429124 8429259 refGene_gn=TTC3;refGene_tx=XM_023238746.1
C2 8429433 8429560 refGene_gn=TTC3;refGene_tx=XM_023238746.1
C2 8431328 8431690 refGene_gn=TTC3;refGene_tx=XM_023238746.1
C2 8434330 8434432 refGene_gn=TTC3;refGene_tx=XM_023238746.1
C2 8434863 8434954 refGene_gn=TTC3;refGene_tx=XM_023238746.1
C2 8437680 8437781 refGene_gn=TTC3;refGene_tx=XM_023238746.1
C2 8439192 8439289 refGene_gn=TTC3;refGene_tx=XM_023238746.1

C2 8440719 8440788 refGene_gn=TTC3;refGene_tx=XM_023238746.1
C2 8444848 8444988 refGene_gn=TTC3;refGene_tx=XM_023238746.1
C2 8455730 8455880 refGene_gn=TTC3;refGene_tx=XM_023238746.1
C2 8461471 8461537 refGene_gn=TTC3;refGene_tx=XM_023238746.1
C2 8462466 8463535 refGene_gn=TTC3;refGene_tx=XM_023238746.1
C2 8468748 8468913 refGene_gn=TTC3;refGene_tx=XM_023238746.1
C2 8470266 8470340 refGene_gn=TTC3;refGene_tx=XM_023238746.1
C2 8470874 8470931 refGene_gn=TTC3;refGene_tx=XM_023238746.1
C2 8472098 8472145 refGene_gn=TTC3;refGene_tx=XM_023238746.1
C2 8474998 8475176 refGene_gn=TTC3;refGene_tx=XM_023238746.1
C2 8475177 8475251 refGene_gn=TTC3;refGene_tx=XM_023238746.1
C2 8477569 8477896 refGene_gn=TTC3;refGene_tx=XM_023238746.1
C2 8479704 8479841 refGene_gn=TTC3;refGene_tx=XM_023238746.1
C2 8480859 8480924 refGene_gn=TTC3;refGene_tx=XM_023238746.1
C2 8481552 8481645 refGene_gn=TTC3;refGene_tx=XM_023238746.1
C2 8483238 8483340 refGene_gn=TTC3;refGene_tx=XM_023238746.1
C2 8484351 8484474 refGene_gn=TTC3;refGene_tx=XM_023238746.1
C2 8486885 8487006 refGene_gn=TTC3;refGene_tx=XM_023238746.1
C2 8490156 8490269 refGene_gn=TTC3;refGene_tx=XM_023238746.1
C2 8492907 8492988 refGene_gn=TTC3;refGene_tx=XM_023238746.1
C2 8493644 8493779 refGene_gn=TTC3;refGene_tx=XM_023238746.1
C2 8498801 8498886 refGene_gn=TTC3;refGene_tx=XM_023238746.1
C2 8501840 8501901 refGene_gn=TTC3;refGene_tx=XM_023238746.1
C2 8504038 8504102 refGene_gn=TTC3;refGene_tx=XM_023238746.1
C2 8505208 8505329 refGene_gn=TTC3;refGene_tx=XM_023238746.1
C2 8506386 8506432 refGene_gn=TTC3;refGene_tx=XM_023238746.1
C2 8507367 8507530 refGene_gn=TTC3;refGene_tx=XM_023238746.1
C2 8515747 8515802 refGene_gn=TTC3;refGene_tx=XM_023238746.1
C2 8526388 8526451 refGene_gn=TTC3;refGene_tx=XM_023238746.1
C2 8527585 8527680 refGene_gn=TTC3;refGene_tx=XM_023238746.1
C2 8529129 8529215 refGene_gn=TTC3;refGene_tx=XM_023238746.1
C2 8531567 8531688 refGene_gn=TTC3;refGene_tx=XM_023238746.1
C2 8534102 8534156 refGene_gn=TTC3;refGene_tx=XM_023238746.1
C2 8535397 8535485 refGene_gn=TTC3;refGene_tx=XM_023238746.1
C2 8535957 8536108 refGene_gn=TTC3;refGene_tx=XM_023238746.1
C2 8536948 8536991 refGene_gn=TTC3;refGene_tx=XM_023238746.1
C2 8538095 8538239 refGene_gn=TTC3;refGene_tx=XM_023238746.1
C2 8551780 8551862 refGene_gn=PIGP;refGene_tx=XM_023238747.1
C2 8558296 8558343 refGene_gn=PIGP;refGene_tx=XM_023238747.1
C2 8558344 8558370 refGene_gn=PIGP;refGene_tx=XM_023238747.1
C2 8558910 8559029 refGene_gn=PIGP;refGene_tx=XM_023238747.1
C2 8560089 8560220 refGene_gn=PIGP;refGene_tx=XM_023238747.1
C2 8597642 8597929 refGene_gn=RIPPLY3;refGene_tx=XM_023238748.1
C2 8598165 8598428
refGene_gn=RIPPLY3;refGene_tx=XM_023238748.1;refGene_gn=RIPPLY3;refGene_tx=XM_023238749.1
C2 8602190 8602258
refGene_gn=RIPPLY3;refGene_tx=XM_023238748.1;refGene_gn=RIPPLY3;refGene_tx=XM_023238749.1
C2 8608475 8608542

refGene_gn=RIPPLY3;refGene_tx=XM_023238748.1;refGene_gn=RIPPLY3;refGene_tx=XM_023238749.1
C2	8609624	8609728	refGene_gn=RIPPLY3;refGene_tx=XM_023238748.1;refGene_gn=RIPPLY3;refGene_tx=XM_023238749.1
C2	8644073	8644268	refGene_gn=HLCS;refGene_tx=XM_023238750.1;refGene_gn=HLCS;refGene_tx=XM_023238751.1
C2	8645342	8645417	refGene_gn=HLCS;refGene_tx=XM_023238750.1
C2	8648125	8648263	refGene_gn=HLCS;refGene_tx=XM_023238750.1;refGene_gn=HLCS;refGene_tx=XM_023238751.1
C2	8661623	8661786	refGene_gn=HLCS;refGene_tx=XM_019839391.2;refGene_gn=HLCS;refGene_tx=XM_023238750.1;refGene_gn=HLCS;refGene_tx=XM_023238751.1;refGene_gn=HLCS;refGene_tx=XM_023238752.1;refGene_gn=HLCS;refGene_tx=XM_023238753.1
C2	8663965	8664912	refGene_gn=HLCS;refGene_tx=XM_019839391.2;refGene_gn=HLCS;refGene_tx=XM_023238750.1;refGene_gn=HLCS;refGene_tx=XM_023238751.1;refGene_gn=HLCS;refGene_tx=XM_023238752.1;refGene_gn=HLCS;refGene_tx=XM_023238753.1
C2	8669955	8670138	refGene_gn=HLCS;refGene_tx=XM_019839391.2;refGene_gn=HLCS;refGene_tx=XM_023238750.1;refGene_gn=HLCS;refGene_tx=XM_023238751.1;refGene_gn=HLCS;refGene_tx=XM_023238752.1;refGene_gn=HLCS;refGene_tx=XM_023238753.1
C2	8704653	8704925	refGene_gn=HLCS;refGene_tx=XM_019839391.2;refGene_gn=HLCS;refGene_tx=XM_023238750.1;refGene_gn=HLCS;refGene_tx=XM_023238751.1;refGene_gn=HLCS;refGene_tx=XM_023238752.1;refGene_gn=HLCS;refGene_tx=XM_023238753.1
C2	8845989	8846057	refGene_gn=HLCS;refGene_tx=XM_019839391.2;refGene_gn=HLCS;refGene_tx=XM_023238750.1;refGene_gn=HLCS;refGene_tx=XM_023238751.1;refGene_gn=HLCS;refGene_tx=XM_023238752.1;refGene_gn=HLCS;refGene_tx=XM_023238753.1
C2	8847741	8847902	refGene_gn=HLCS;refGene_tx=XM_019839391.2;refGene_gn=HLCS;refGene_tx=XM_023238750.1;refGene_gn=HLCS;refGene_tx=XM_023238751.1;refGene_gn=HLCS;refGene_tx=XM_023238752.1;refGene_gn=HLCS;refGene_tx=XM_023238753.1
C2	8853779	8853894	refGene_gn=HLCS;refGene_tx=XM_019839391.2;refGene_gn=HLCS;refGene_tx=XM_023238750.1;refGene_gn=HLCS;refGene_tx=XM_023238751.1;refGene_gn=HLCS;refGene_tx=XM_023238752.1;refGene_gn=HLCS;refGene_tx=XM_023238753.1
C2	8856076	8856290	refGene_gn=HLCS;refGene_tx=XM_019839391.2;refGene_gn=HLCS;refGene_tx=XM_023238750.1;refGene_gn=HLCS;refGene_tx=XM_023238751.1;refGene_gn=HLCS;refGene_tx=XM_023238752.1;refGene_gn=HLCS;refGene_tx=XM_023238753.1
C2	8857755	8857927	refGene_gn=HLCS;refGene_tx=XM_019839391.2;refGene_gn=HLCS;refGene_tx=XM_023238750.1;refGene_gn=HLCS;refGene_tx=XM_023238751.1;refGene_gn=HLCS;refGene_tx=XM_023238752.1;refGene_gn=HLCS;refGene_tx=XM_023238753.1
C2	8864269	8864721	refGene_gn=SIM2;refGene_tx=XM_023238754.1
C2	8866931	8867340	refGene_gn=SIM2;refGene_tx=XM_023238754.1

C2 8867927 8868096 refGene_gn=SIM2;refGene_tx=XM_023238754.1
C2 8869537 8869685 refGene_gn=SIM2;refGene_tx=XM_023238754.1
C2 8883603 8883710 refGene_gn=SIM2;refGene_tx=XM_023238754.1
C2 8888853 8889053 refGene_gn=SIM2;refGene_tx=XM_023238754.1
C2 8894031 8894117 refGene_gn=SIM2;refGene_tx=XM_023238754.1
C2 8897630 8897739 refGene_gn=SIM2;refGene_tx=XM_023238754.1
C2 8904928 8905018 refGene_gn=SIM2;refGene_tx=XM_023238754.1
C2 8908130 8908213 refGene_gn=SIM2;refGene_tx=XM_023238754.1
C2 8917393 8917568 refGene_gn=SIM2;refGene_tx=XM_023238754.1
C2 9100842 9100882 refGene_gn=CLDN14;refGene_tx=XM_019839394.2
C2 9111852 9112610
refGene_gn=CLDN14;refGene_tx=XM_003991439.3;refGene_gn=CLDN14;refGene_tx=XM_019839394.2
C2 9171678 9171770
refGene_gn=CHAF1B;refGene_tx=XM_006935899.3;refGene_gn=CHAF1B;refGene_tx=XM_019839397.2
C2 9172324 9172419
refGene_gn=CHAF1B;refGene_tx=XM_006935899.3;refGene_gn=CHAF1B;refGene_tx=XM_019839397.2
C2 9173859 9174222 refGene_gn=CHAF1B;refGene_tx=XM_006935899.3
C2 9175391 9175554
refGene_gn=CHAF1B;refGene_tx=XM_006935899.3;refGene_gn=CHAF1B;refGene_tx=XM_019839397.2
C2 9177344 9177436
refGene_gn=CHAF1B;refGene_tx=XM_006935899.3;refGene_gn=CHAF1B;refGene_tx=XM_019839397.2
C2 9177528 9177598
refGene_gn=CHAF1B;refGene_tx=XM_006935899.3;refGene_gn=CHAF1B;refGene_tx=XM_019839397.2
C2 9181068 9181162
refGene_gn=CHAF1B;refGene_tx=XM_006935899.3;refGene_gn=CHAF1B;refGene_tx=XM_019839397.2
C2 9182568 9182653
refGene_gn=CHAF1B;refGene_tx=XM_006935899.3;refGene_gn=CHAF1B;refGene_tx=XM_019839397.2
C2 9184460 9184557
refGene_gn=CHAF1B;refGene_tx=XM_006935899.3;refGene_gn=CHAF1B;refGene_tx=XM_019839397.2
C2 9186996 9187100
refGene_gn=CHAF1B;refGene_tx=XM_006935899.3;refGene_gn=CHAF1B;refGene_tx=XM_019839397.2
C2 9187768 9187886
refGene_gn=CHAF1B;refGene_tx=XM_006935899.3;refGene_gn=CHAF1B;refGene_tx=XM_019839397.2
C2 9190012 9190145
refGene_gn=CHAF1B;refGene_tx=XM_006935899.3;refGene_gn=CHAF1B;refGene_tx=XM_019839397.2
C2 9190596 9190722
refGene_gn=CHAF1B;refGene_tx=XM_006935899.3;refGene_gn=CHAF1B;refGene_tx=XM_019839397.2

C2 9204956 9205110 refGene_gn=MORC3;refGene_tx=XM_023238756.1
C2 9208499 9208657 refGene_gn=MORC3;refGene_tx=XM_023238756.1
C2 9209871 9210760 refGene_gn=MORC3;refGene_tx=XM_023238756.1
C2 9213756 9213923 refGene_gn=MORC3;refGene_tx=XM_023238756.1
C2 9215132 9215178 refGene_gn=MORC3;refGene_tx=XM_023238756.1
C2 9216206 9216281 refGene_gn=MORC3;refGene_tx=XM_023238756.1
C2 9216387 9216510 refGene_gn=MORC3;refGene_tx=XM_023238756.1
C2 9218496 9218601 refGene_gn=MORC3;refGene_tx=XM_023238756.1
C2 9222925 9223023 refGene_gn=MORC3;refGene_tx=XM_023238756.1
C2 9225618 9225738 refGene_gn=MORC3;refGene_tx=XM_023238756.1
C2 9225946 9226075 refGene_gn=MORC3;refGene_tx=XM_023238756.1
C2 9228197 9228345 refGene_gn=MORC3;refGene_tx=XM_023238756.1
C2 9229856 9230004 refGene_gn=MORC3;refGene_tx=XM_023238756.1
C2 9230585 9230800 refGene_gn=MORC3;refGene_tx=XM_023238756.1
C2 9231933 9232066 refGene_gn=MORC3;refGene_tx=XM_023238756.1
C2 9233490 9233563 refGene_gn=MORC3;refGene_tx=XM_023238756.1
C2 9243532 9243571 refGene_gn=MORC3;refGene_tx=XM_023238756.1
C2 9256526 9256778 refGene_gn=DOPEY2;refGene_tx=XM_023238757.1;refGene_gn=DOPEY2;refGene_tx=XM_023238758.1
C2 9257298 9257428 refGene_gn=DOPEY2;refGene_tx=XM_023238757.1;refGene_gn=DOPEY2;refGene_tx=XM_023238758.1
C2 9258942 9259104 refGene_gn=DOPEY2;refGene_tx=XM_023238757.1;refGene_gn=DOPEY2;refGene_tx=XM_023238758.1
C2 9259765 9259821 refGene_gn=DOPEY2;refGene_tx=XM_023238757.1;refGene_gn=DOPEY2;refGene_tx=XM_023238758.1
C2 9262955 9263092 refGene_gn=DOPEY2;refGene_tx=XM_023238757.1;refGene_gn=DOPEY2;refGene_tx=XM_023238758.1
C2 9270588 9270717 refGene_gn=DOPEY2;refGene_tx=XM_023238757.1;refGene_gn=DOPEY2;refGene_tx=XM_023238758.1
C2 9271227 9271289 refGene_gn=DOPEY2;refGene_tx=XM_023238757.1;refGene_gn=DOPEY2;refGene_tx=XM_023238758.1
C2 9272274 9272421 refGene_gn=DOPEY2;refGene_tx=XM_023238757.1;refGene_gn=DOPEY2;refGene_tx=XM_023238758.1
C2 9272525 9272635 refGene_gn=DOPEY2;refGene_tx=XM_023238757.1;refGene_gn=DOPEY2;refGene_tx=XM_023238758.1
C2 9272851 9272937 refGene_gn=DOPEY2;refGene_tx=XM_023238757.1;refGene_gn=DOPEY2;refGene_tx=XM_023238758.1
C2 9275395 9275540 refGene_gn=DOPEY2;refGene_tx=XM_023238757.1;refGene_gn=DOPEY2;refGene_tx=XM_023238758.1

C2	9277591	9277658
refGene_gn=DOPEY2;refGene_tx=XM_023238757.1;refGene_gn=DOPEY2;refGene_
tx=XM_023238758.1
C2	9277754	9277859
refGene_gn=DOPEY2;refGene_tx=XM_023238757.1;refGene_gn=DOPEY2;refGene_
tx=XM_023238758.1
C2	9279162	9279218
refGene_gn=DOPEY2;refGene_tx=XM_023238757.1;refGene_gn=DOPEY2;refGene_
tx=XM_023238758.1
C2	9283753	9283888
refGene_gn=DOPEY2;refGene_tx=XM_023238757.1;refGene_gn=DOPEY2;refGene_
tx=XM_023238758.1
C2	9285879	9286002
refGene_gn=DOPEY2;refGene_tx=XM_023238757.1;refGene_gn=DOPEY2;refGene_
tx=XM_023238758.1
C2	9287395	9287584
refGene_gn=DOPEY2;refGene_tx=XM_023238757.1;refGene_gn=DOPEY2;refGene_
tx=XM_023238758.1
C2	9288224	9288336
refGene_gn=DOPEY2;refGene_tx=XM_023238757.1;refGene_gn=DOPEY2;refGene_
tx=XM_023238758.1
C2	9288821	9290451
refGene_gn=DOPEY2;refGene_tx=XM_023238757.1;refGene_gn=DOPEY2;refGene_
tx=XM_023238758.1
C2	9294192	9294383
refGene_gn=DOPEY2;refGene_tx=XM_023238757.1;refGene_gn=DOPEY2;refGene_
tx=XM_023238758.1
C2	9295352	9295453
refGene_gn=DOPEY2;refGene_tx=XM_023238757.1;refGene_gn=DOPEY2;refGene_
tx=XM_023238758.1
C2	9296130	9296283
refGene_gn=DOPEY2;refGene_tx=XM_023238757.1;refGene_gn=DOPEY2;refGene_
tx=XM_023238758.1
C2	9303899	9304171
refGene_gn=DOPEY2;refGene_tx=XM_023238757.1;refGene_gn=DOPEY2;refGene_
tx=XM_023238758.1
C2	9305872	9306557
refGene_gn=DOPEY2;refGene_tx=XM_023238757.1;refGene_gn=DOPEY2;refGene_
tx=XM_023238758.1
C2	9308617	9308809
refGene_gn=DOPEY2;refGene_tx=XM_023238757.1;refGene_gn=DOPEY2;refGene_
tx=XM_023238758.1
C2	9309850	9309953
refGene_gn=DOPEY2;refGene_tx=XM_023238757.1;refGene_gn=DOPEY2;refGene_
tx=XM_023238758.1
C2	9312464	9312584
refGene_gn=DOPEY2;refGene_tx=XM_023238757.1;refGene_gn=DOPEY2;refGene_
tx=XM_023238758.1
C2	9314609	9314730
refGene_gn=DOPEY2;refGene_tx=XM_023238757.1;refGene_gn=DOPEY2;refGene_

tx=XM_023238758.1
C2 9316762 9316877
refGene_gn=DOPEY2;refGene_tx=XM_023238757.1;refGene_gn=DOPEY2;refGene_tx=XM_023238758.1
C2 9317186 9317296
refGene_gn=DOPEY2;refGene_tx=XM_023238757.1;refGene_gn=DOPEY2;refGene_tx=XM_023238758.1
C2 9319170 9319294
refGene_gn=DOPEY2;refGene_tx=XM_023238757.1;refGene_gn=DOPEY2;refGene_tx=XM_023238758.1
C2 9319629 9319728
refGene_gn=DOPEY2;refGene_tx=XM_023238757.1;refGene_gn=DOPEY2;refGene_tx=XM_023238758.1
C2 9321707 9321897
refGene_gn=DOPEY2;refGene_tx=XM_023238757.1;refGene_gn=DOPEY2;refGene_tx=XM_023238758.1
C2 9327951 9328122
refGene_gn=DOPEY2;refGene_tx=XM_023238757.1;refGene_gn=DOPEY2;refGene_tx=XM_023238758.1
C2 9328745 9328927
refGene_gn=DOPEY2;refGene_tx=XM_023238757.1;refGene_gn=DOPEY2;refGene_tx=XM_023238758.1
C2 9354125 9354263
refGene_gn=DOPEY2;refGene_tx=XM_023238757.1;refGene_gn=DOPEY2;refGene_tx=XM_023238758.1
C2 9366442 9366879 refGene_gn=CBR3;refGene_tx=XM_003991442.5
C2 9372039 9372147 refGene_gn=CBR3;refGene_tx=XM_003991442.5
C2 9373929 9374218 refGene_gn=CBR3;refGene_tx=XM_003991442.5
C2 9409612 9410094
refGene_gn=LOC101083875;refGene_tx=XM_003991445.3
C2 9412165 9412273
refGene_gn=LOC101083875;refGene_tx=XM_003991445.3
C2 9412823 9413112
refGene_gn=LOC101083875;refGene_tx=XM_003991445.3
C2 9415884 9416366
refGene_gn=LOC102901316;refGene_tx=XM_011285830.3
C2 9422641 9422749
refGene_gn=LOC102901316;refGene_tx=XM_011285830.3
C2 9423316 9423605
refGene_gn=LOC102901316;refGene_tx=XM_011285830.3
C2 9462442 9462924
refGene_gn=LOC101084134;refGene_tx=XM_003991446.4
C2 9468751 9468859
refGene_gn=LOC101084134;refGene_tx=XM_003991446.4
C2 9469403 9469692
refGene_gn=LOC101084134;refGene_tx=XM_003991446.4
C2 9481549 9482031
refGene_gn=LOC105260891;refGene_tx=XM_023238763.1
C2 9484002 9484105
refGene_gn=LOC105260891;refGene_tx=XM_023238763.1

C2 9490936 9491118
refGene_gn=LOC101084388;refGene_tx=XM_011285768.3
C2 9491120 9491372
refGene_gn=LOC101084388;refGene_tx=XM_011285768.3
C2 9493201 9493309
refGene_gn=LOC101084388;refGene_tx=XM_011285768.3
C2 9493875 9493998
refGene_gn=LOC101084388;refGene_tx=XM_011285768.3
C2 9494000 9494163
refGene_gn=LOC101084388;refGene_tx=XM_011285768.3
C2 9499875 9500550
refGene_gn=LOC109503322;refGene_tx=XM_023238761.1
C2 9502476 9502549 refGene_gn=SETD4;refGene_tx=XM_023238760.1
C2 9506970 9507066 refGene_gn=SETD4;refGene_tx=XM_023238760.1
C2 9510669 9510707 refGene_gn=SETD4;refGene_tx=XM_023238760.1
C2 9513029 9513118 refGene_gn=SETD4;refGene_tx=XM_023238760.1
C2 9514663 9515093 refGene_gn=SETD4;refGene_tx=XM_023238760.1
C2 9517172 9517347 refGene_gn=SETD4;refGene_tx=XM_023238760.1
C2 9518998 9519080 refGene_gn=SETD4;refGene_tx=XM_023238760.1
C2 9519962 9520043 refGene_gn=SETD4;refGene_tx=XM_023238760.1
C2 9522503 9522627 refGene_gn=SETD4;refGene_tx=XM_023238760.1
C2 9523266 9523401 refGene_gn=SETD4;refGene_tx=XM_023238760.1
C2 9732846 9733527
refGene_gn=LOC109491637;refGene_tx=XM_023260844.1
C2 9767162 9767243
refGene_gn=LOC109491637;refGene_tx=XM_023260844.1
C2 10437893 10437951
refGene_gn=RUNX1;refGene_tx=XM_023238764.1;refGene_gn=RUNX1;refGene_tx
=XM_023238765.1;refGene_gn=RUNX1;refGene_tx=XM_023238766.1
C2 10594095 10594134
refGene_gn=RUNX1;refGene_tx=XM_023238764.1;refGene_gn=RUNX1;refGene_tx
=XM_023238765.1;refGene_gn=RUNX1;refGene_tx=XM_023238766.1
C2 10600151 10600405
refGene_gn=RUNX1;refGene_tx=XM_023238764.1;refGene_gn=RUNX1;refGene_tx
=XM_023238765.1;refGene_gn=RUNX1;refGene_tx=XM_023238766.1
C2 10606507 10606664
refGene_gn=RUNX1;refGene_tx=XM_023238764.1;refGene_gn=RUNX1;refGene_tx
=XM_023238765.1;refGene_gn=RUNX1;refGene_tx=XM_023238766.1
C2 10628629 10628734
refGene_gn=RUNX1;refGene_tx=XM_023238764.1;refGene_gn=RUNX1;refGene_tx
=XM_023238765.1;refGene_gn=RUNX1;refGene_tx=XM_023238766.1
C2 10652421 10652613
refGene_gn=RUNX1;refGene_tx=XM_023238764.1;refGene_gn=RUNX1;refGene_tx
=XM_023238765.1
C2 10685488 10685650
refGene_gn=RUNX1;refGene_tx=XM_023238764.1;refGene_gn=RUNX1;refGene_tx
=XM_023238765.1;refGene_gn=RUNX1;refGene_tx=XM_023238766.1
C2 10691261 10691737
refGene_gn=RUNX1;refGene_tx=XM_023238764.1;refGene_gn=RUNX1;refGene_tx
=XM_023238765.1;refGene_gn=RUNX1;refGene_tx=XM_023238766.1

C2 10761785 10761947 refGene_gn=CLIC6;refGene_tx=XM_023238767.1
C2 10769035 10769217 refGene_gn=CLIC6;refGene_tx=XM_023238767.1
C2 10769789 10769896 refGene_gn=CLIC6;refGene_tx=XM_023238767.1
C2 10770466 10770592 refGene_gn=CLIC6;refGene_tx=XM_023238767.1
C2 10771146 10771256 refGene_gn=CLIC6;refGene_tx=XM_023238767.1
C2 10804603 10805686 refGene_gn=CLIC6;refGene_tx=XM_023238767.1
C2 10858113 10858356 refGene_gn=RCAN1;refGene_tx=XM_023238770.1
C2 10947510 10947597 refGene_gn=RCAN1;refGene_tx=XM_023238771.1
C2 10949243 10949510 refGene_gn=RCAN1;refGene_tx=XM_023238769.1
C2 10950781 10950955
refGene_gn=RCAN1;refGene_tx=XM_023238769.1;refGene_gn=RCAN1;refGene_tx=XM_023238770.1;refGene_gn=RCAN1;refGene_tx=XM_023238771.1
C2 10955235 10955395
refGene_gn=RCAN1;refGene_tx=XM_023238769.1;refGene_gn=RCAN1;refGene_tx=XM_023238770.1;refGene_gn=RCAN1;refGene_tx=XM_023238771.1
C2 10958228 10958401
refGene_gn=RCAN1;refGene_tx=XM_023238769.1;refGene_gn=RCAN1;refGene_tx=XM_023238770.1;refGene_gn=RCAN1;refGene_tx=XM_023238771.1
C2 11009599 11009989
refGene_gn=KCNE1;refGene_tx=NM_001009206.1;refGene_gn=KCNE1;refGene_tx=XM_023260085.1
C2 11033728 11033939
refGene_gn=LOC109503329;refGene_tx=XM_019839403.1
C2 11034058 11034186
refGene_gn=LOC109503329;refGene_tx=XM_019839403.1
C2 11036366 11036471
refGene_gn=LOC109503329;refGene_tx=XM_019839403.1
C2 11064269 11065025
refGene_gn=CC2H21orf140;refGene_tx=XM_006935906.3
C2 11075913 11076069
refGene_gn=SMIM11A;refGene_tx=XM_003991450.4;refGene_gn=SMIM11A;refGene_tx=XM_006935905.4;refGene_gn=SMIM11A;refGene_tx=XM_011285764.3;refGene_gn=SMIM11A;refGene_tx=XM_011285765.3;refGene_gn=SMIM11A;refGene_tx=XM_011285766.3;refGene_gn=SMIM11A;refGene_tx=XM_019839410.2
C2 11081596 11081608
refGene_gn=SMIM11A;refGene_tx=XM_003991450.4;refGene_gn=SMIM11A;refGene_tx=XM_006935905.4;refGene_gn=SMIM11A;refGene_tx=XM_011285764.3;refGene_gn=SMIM11A;refGene_tx=XM_011285765.3;refGene_gn=SMIM11A;refGene_tx=XM_011285766.3;refGene_gn=SMIM11A;refGene_tx=XM_019839410.2
C2 11089741 11090125
refGene_gn=KCNE2;refGene_tx=XM_006935910.4;refGene_gn=KCNE2;refGene_tx=XM_019839406.2;refGene_gn=KCNE2;refGene_tx=XM_019839408.2;refGene_gn=KCNE2;refGene_tx=XM_023238773.1;refGene_gn=KCNE2;refGene_tx=XM_023238774.1;refGene_gn=KCNE2;refGene_tx=XM_023238775.1;refGene_gn=KCNE2;refGene_tx=XM_023238776.1
C2 11121490 11121610
refGene_gn=KCNE2;refGene_tx=XM_006935910.4;refGene_gn=KCNE2;refGene_tx=XM_019839406.2;refGene_gn=KCNE2;refGene_tx=XM_023238773.1;refGene_gn=KCNE2;refGene_tx=XM_023238774.1;refGene_gn=KCNE2;refGene_tx=XM_023238775.1;refGene_gn=KCNE2;refGene_tx=XM_023238776.1

C2	11173591	11173639	refGene_gn=KCNE2;refGene_tx=XM_019839406.2
C2	11294595	11294785	refGene_gn=MRPS6;refGene_tx=XM_023238777.1
C2	11309560	11309700	refGene_gn=MRPS6;refGene_tx=XM_023238777.1
C2	11335156	11337313
refGene_gn=SLC5A3;refGene_tx=XM_003991452.5;refGene_gn=SLC5A3;refGene_tx=XM_019839404.2;refGene_gn=SLC5A3;refGene_tx=XM_019839405.2;refGene_gn=SLC5A3;refGene_tx=XM_023238772.1
C2	11360888	11360933	refGene_gn=KCNE2;refGene_tx=XM_023238773.1
C2	11361337	11361379	refGene_gn=MRPS6;refGene_tx=XM_023238777.1
C2	11491836	11491872	refGene_gn=ATP5PO;refGene_tx=XM_003991455.3
C2	11493034	11493085	refGene_gn=ATP5PO;refGene_tx=XM_003991455.3
C2	11494024	11494135	refGene_gn=ATP5PO;refGene_tx=XM_003991455.3
C2	11496801	11496931	refGene_gn=ATP5PO;refGene_tx=XM_003991455.3
C2	11498544	11498657	refGene_gn=ATP5PO;refGene_tx=XM_003991455.3
C2	11501069	11501156	refGene_gn=ATP5PO;refGene_tx=XM_003991455.3
C2	11501452	11501566	refGene_gn=ATP5PO;refGene_tx=XM_003991455.3
C2	11515943	11516092
refGene_gn=ITSN1;refGene_tx=XM_019839414.1;refGene_gn=ITSN1;refGene_tx=XM_023238778.1;refGene_gn=ITSN1;refGene_tx=XM_023238779.1;refGene_gn=ITSN1;refGene_tx=XM_023238780.1;refGene_gn=ITSN1;refGene_tx=XM_023238781.1
C2	11518451	11518625
refGene_gn=ITSN1;refGene_tx=XM_019839414.1;refGene_gn=ITSN1;refGene_tx=XM_023238778.1;refGene_gn=ITSN1;refGene_tx=XM_023238779.1;refGene_gn=ITSN1;refGene_tx=XM_023238780.1;refGene_gn=ITSN1;refGene_tx=XM_023238781.1
C2	11519171	11519255
refGene_gn=ITSN1;refGene_tx=XM_019839414.1;refGene_gn=ITSN1;refGene_tx=XM_023238778.1;refGene_gn=ITSN1;refGene_tx=XM_023238779.1;refGene_gn=ITSN1;refGene_tx=XM_023238780.1;refGene_gn=ITSN1;refGene_tx=XM_023238781.1
C2	11519564	11519647
refGene_gn=ITSN1;refGene_tx=XM_019839414.1;refGene_gn=ITSN1;refGene_tx=XM_023238778.1;refGene_gn=ITSN1;refGene_tx=XM_023238779.1;refGene_gn=ITSN1;refGene_tx=XM_023238780.1;refGene_gn=ITSN1;refGene_tx=XM_023238781.1
C2	11520992	11521114
refGene_gn=ITSN1;refGene_tx=XM_019839414.1;refGene_gn=ITSN1;refGene_tx=XM_023238778.1;refGene_gn=ITSN1;refGene_tx=XM_023238779.1;refGene_gn=ITSN1;refGene_tx=XM_023238780.1;refGene_gn=ITSN1;refGene_tx=XM_023238781.1
C2	11522090	11522303
refGene_gn=ITSN1;refGene_tx=XM_019839414.1;refGene_gn=ITSN1;refGene_tx=XM_023238778.1;refGene_gn=ITSN1;refGene_tx=XM_023238779.1;refGene_gn=ITSN1;refGene_tx=XM_023238780.1;refGene_gn=ITSN1;refGene_tx=XM_023238781.1
C2	11526304	11526472
refGene_gn=ITSN1;refGene_tx=XM_019839414.1;refGene_gn=ITSN1;refGene_tx=XM_023238778.1;refGene_gn=ITSN1;refGene_tx=XM_023238779.1;refGene_gn=ITSN1;refGene_tx=XM_023238780.1;refGene_gn=ITSN1;refGene_tx=XM_0232387

81.1
C2	11533336	11533435
refGene_gn=ITSN1;refGene_tx=XM_019839414.1;refGene_gn=ITSN1;refGene_tx=XM_023238778.1;refGene_gn=ITSN1;refGene_tx=XM_023238779.1;refGene_gn=ITSN1;refGene_tx=XM_023238780.1;refGene_gn=ITSN1;refGene_tx=XM_023238781.1
C2	11535393	11535577
refGene_gn=ITSN1;refGene_tx=XM_019839414.1;refGene_gn=ITSN1;refGene_tx=XM_023238778.1;refGene_gn=ITSN1;refGene_tx=XM_023238779.1;refGene_gn=ITSN1;refGene_tx=XM_023238780.1;refGene_gn=ITSN1;refGene_tx=XM_023238781.1
C2	11541190	11541297
refGene_gn=ITSN1;refGene_tx=XM_019839414.1;refGene_gn=ITSN1;refGene_tx=XM_023238778.1;refGene_gn=ITSN1;refGene_tx=XM_023238779.1;refGene_gn=ITSN1;refGene_tx=XM_023238780.1;refGene_gn=ITSN1;refGene_tx=XM_023238781.1
C2	11543122	11543244
refGene_gn=ITSN1;refGene_tx=XM_019839414.1;refGene_gn=ITSN1;refGene_tx=XM_023238778.1;refGene_gn=ITSN1;refGene_tx=XM_023238779.1;refGene_gn=ITSN1;refGene_tx=XM_023238780.1;refGene_gn=ITSN1;refGene_tx=XM_023238781.1
C2	11552437	11552439	refGene_gn=ITSN1;refGene_tx=XM_023238783.1
C2	11562393	11562395	refGene_gn=ITSN1;refGene_tx=XM_023238782.1
C2	11562771	11562963
refGene_gn=ITSN1;refGene_tx=XM_019839414.1;refGene_gn=ITSN1;refGene_tx=XM_023238778.1;refGene_gn=ITSN1;refGene_tx=XM_023238779.1;refGene_gn=ITSN1;refGene_tx=XM_023238780.1;refGene_gn=ITSN1;refGene_tx=XM_023238781.1;refGene_gn=ITSN1;refGene_tx=XM_023238782.1;refGene_gn=ITSN1;refGene_tx=XM_023238783.1
C2	11564596	11564714
refGene_gn=ITSN1;refGene_tx=XM_019839414.1;refGene_gn=ITSN1;refGene_tx=XM_023238778.1;refGene_gn=ITSN1;refGene_tx=XM_023238779.1;refGene_gn=ITSN1;refGene_tx=XM_023238780.1;refGene_gn=ITSN1;refGene_tx=XM_023238781.1;refGene_gn=ITSN1;refGene_tx=XM_023238782.1;refGene_gn=ITSN1;refGene_tx=XM_023238783.1
C2	11567803	11567925
refGene_gn=ITSN1;refGene_tx=XM_019839414.1;refGene_gn=ITSN1;refGene_tx=XM_023238778.1;refGene_gn=ITSN1;refGene_tx=XM_023238779.1;refGene_gn=ITSN1;refGene_tx=XM_023238780.1;refGene_gn=ITSN1;refGene_tx=XM_023238781.1;refGene_gn=ITSN1;refGene_tx=XM_023238782.1;refGene_gn=ITSN1;refGene_tx=XM_023238783.1
C2	11570241	11570287
refGene_gn=ITSN1;refGene_tx=XM_019839414.1;refGene_gn=ITSN1;refGene_tx=XM_023238778.1;refGene_gn=ITSN1;refGene_tx=XM_023238779.1;refGene_gn=ITSN1;refGene_tx=XM_023238782.1;refGene_gn=ITSN1;refGene_tx=XM_023238783.1
C2	11574027	11574194
refGene_gn=ITSN1;refGene_tx=XM_019839414.1;refGene_gn=ITSN1;refGene_tx=XM_023238778.1;refGene_gn=ITSN1;refGene_tx=XM_023238779.1;refGene_gn=ITSN1;refGene_tx=XM_023238782.1;refGene_gn=ITSN1;refGene_tx=XM_0232387

83.1
C2	11578041 11578124
refGene_gn=ITSN1;refGene_tx=XM_019839414.1;refGene_gn=ITSN1;refGene_tx=XM_023238778.1;refGene_gn=ITSN1;refGene_tx=XM_023238779.1;refGene_gn=ITSN1;refGene_tx=XM_023238780.1;refGene_gn=ITSN1;refGene_tx=XM_023238781.1;refGene_gn=ITSN1;refGene_tx=XM_023238782.1;refGene_gn=ITSN1;refGene_tx=XM_023238783.1
C2	11578913 11579119
refGene_gn=ITSN1;refGene_tx=XM_019839414.1;refGene_gn=ITSN1;refGene_tx=XM_023238778.1;refGene_gn=ITSN1;refGene_tx=XM_023238779.1;refGene_gn=ITSN1;refGene_tx=XM_023238780.1;refGene_gn=ITSN1;refGene_tx=XM_023238781.1;refGene_gn=ITSN1;refGene_tx=XM_023238782.1;refGene_gn=ITSN1;refGene_tx=XM_023238783.1
C2	11582512 11582672
refGene_gn=ITSN1;refGene_tx=XM_019839414.1;refGene_gn=ITSN1;refGene_tx=XM_023238778.1;refGene_gn=ITSN1;refGene_tx=XM_023238779.1;refGene_gn=ITSN1;refGene_tx=XM_023238780.1;refGene_gn=ITSN1;refGene_tx=XM_023238781.1;refGene_gn=ITSN1;refGene_tx=XM_023238782.1;refGene_gn=ITSN1;refGene_tx=XM_023238783.1
C2	11584975 11585223
refGene_gn=ITSN1;refGene_tx=XM_019839414.1;refGene_gn=ITSN1;refGene_tx=XM_023238778.1;refGene_gn=ITSN1;refGene_tx=XM_023238779.1;refGene_gn=ITSN1;refGene_tx=XM_023238780.1;refGene_gn=ITSN1;refGene_tx=XM_023238781.1;refGene_gn=ITSN1;refGene_tx=XM_023238782.1;refGene_gn=ITSN1;refGene_tx=XM_023238783.1
C2	11592046 11592061
refGene_gn=ITSN1;refGene_tx=XM_019839414.1;refGene_gn=ITSN1;refGene_tx=XM_023238778.1;refGene_gn=ITSN1;refGene_tx=XM_023238780.1;refGene_gn=ITSN1;refGene_tx=XM_023238782.1;refGene_gn=ITSN1;refGene_tx=XM_023238783.1
C2	11593822 11593944
refGene_gn=ITSN1;refGene_tx=XM_019839414.1;refGene_gn=ITSN1;refGene_tx=XM_023238778.1;refGene_gn=ITSN1;refGene_tx=XM_023238779.1;refGene_gn=ITSN1;refGene_tx=XM_023238780.1;refGene_gn=ITSN1;refGene_tx=XM_023238781.1;refGene_gn=ITSN1;refGene_tx=XM_023238782.1;refGene_gn=ITSN1;refGene_tx=XM_023238783.1
C2	11596204 11596434
refGene_gn=ITSN1;refGene_tx=XM_019839414.1;refGene_gn=ITSN1;refGene_tx=XM_023238778.1;refGene_gn=ITSN1;refGene_tx=XM_023238779.1;refGene_gn=ITSN1;refGene_tx=XM_023238780.1;refGene_gn=ITSN1;refGene_tx=XM_023238781.1;refGene_gn=ITSN1;refGene_tx=XM_023238782.1;refGene_gn=ITSN1;refGene_tx=XM_023238783.1
C2	11600205 11600333
refGene_gn=ITSN1;refGene_tx=XM_019839414.1;refGene_gn=ITSN1;refGene_tx=XM_023238778.1;refGene_gn=ITSN1;refGene_tx=XM_023238779.1;refGene_gn=ITSN1;refGene_tx=XM_023238780.1;refGene_gn=ITSN1;refGene_tx=XM_023238781.1;refGene_gn=ITSN1;refGene_tx=XM_023238782.1;refGene_gn=ITSN1;refGene_tx=XM_023238783.1
C2	11612134 11612274
refGene_gn=ITSN1;refGene_tx=XM_019839414.1;refGene_gn=ITSN1;refGene_tx

=XM_023238778.1;refGene_gn=ITSN1;refGene_tx=XM_023238779.1;refGene_gn=ITSN1;refGene_tx=XM_023238780.1;refGene_gn=ITSN1;refGene_tx=XM_023238781.1;refGene_gn=ITSN1;refGene_tx=XM_023238782.1;refGene_gn=ITSN1;refGene_tx=XM_023238783.1
C2 11612412 11612500
refGene_gn=ITSN1;refGene_tx=XM_019839414.1;refGene_gn=ITSN1;refGene_tx=XM_023238778.1;refGene_gn=ITSN1;refGene_tx=XM_023238779.1;refGene_gn=ITSN1;refGene_tx=XM_023238780.1;refGene_gn=ITSN1;refGene_tx=XM_023238781.1;refGene_gn=ITSN1;refGene_tx=XM_023238782.1;refGene_gn=ITSN1;refGene_tx=XM_023238783.1
C2 11619244 11619385
refGene_gn=ITSN1;refGene_tx=XM_019839414.1;refGene_gn=ITSN1;refGene_tx=XM_023238778.1;refGene_gn=ITSN1;refGene_tx=XM_023238779.1;refGene_gn=ITSN1;refGene_tx=XM_023238780.1;refGene_gn=ITSN1;refGene_tx=XM_023238781.1;refGene_gn=ITSN1;refGene_tx=XM_023238782.1;refGene_gn=ITSN1;refGene_tx=XM_023238783.1
C2 11620007 11620157
refGene_gn=ITSN1;refGene_tx=XM_019839414.1;refGene_gn=ITSN1;refGene_tx=XM_023238778.1;refGene_gn=ITSN1;refGene_tx=XM_023238779.1;refGene_gn=ITSN1;refGene_tx=XM_023238780.1;refGene_gn=ITSN1;refGene_tx=XM_023238781.1;refGene_gn=ITSN1;refGene_tx=XM_023238782.1;refGene_gn=ITSN1;refGene_tx=XM_023238783.1
C2 11622265 11622528
refGene_gn=ITSN1;refGene_tx=XM_019839414.1;refGene_gn=ITSN1;refGene_tx=XM_023238778.1;refGene_gn=ITSN1;refGene_tx=XM_023238779.1;refGene_gn=ITSN1;refGene_tx=XM_023238780.1;refGene_gn=ITSN1;refGene_tx=XM_023238781.1;refGene_gn=ITSN1;refGene_tx=XM_023238782.1;refGene_gn=ITSN1;refGene_tx=XM_023238783.1
C2 11628913 11629029
refGene_gn=ITSN1;refGene_tx=XM_019839414.1;refGene_gn=ITSN1;refGene_tx=XM_023238778.1;refGene_gn=ITSN1;refGene_tx=XM_023238779.1;refGene_gn=ITSN1;refGene_tx=XM_023238780.1;refGene_gn=ITSN1;refGene_tx=XM_023238781.1;refGene_gn=ITSN1;refGene_tx=XM_023238782.1;refGene_gn=ITSN1;refGene_tx=XM_023238783.1
C2 11630648 11630786
refGene_gn=ITSN1;refGene_tx=XM_019839414.1;refGene_gn=ITSN1;refGene_tx=XM_023238778.1;refGene_gn=ITSN1;refGene_tx=XM_023238779.1;refGene_gn=ITSN1;refGene_tx=XM_023238780.1;refGene_gn=ITSN1;refGene_tx=XM_023238781.1;refGene_gn=ITSN1;refGene_tx=XM_023238782.1;refGene_gn=ITSN1;refGene_tx=XM_023238783.1
C2 11633453 11633517
refGene_gn=ITSN1;refGene_tx=XM_023238778.1;refGene_gn=ITSN1;refGene_tx=XM_023238779.1;refGene_gn=ITSN1;refGene_tx=XM_023238780.1;refGene_gn=ITSN1;refGene_tx=XM_023238781.1;refGene_gn=ITSN1;refGene_tx=XM_023238782.1;refGene_gn=ITSN1;refGene_tx=XM_023238783.1
C2 11640127 11640228
refGene_gn=ITSN1;refGene_tx=XM_023238778.1;refGene_gn=ITSN1;refGene_tx=XM_023238779.1;refGene_gn=ITSN1;refGene_tx=XM_023238780.1;refGene_gn=ITSN1;refGene_tx=XM_023238781.1;refGene_gn=ITSN1;refGene_tx=XM_023238782.1;refGene_gn=ITSN1;refGene_tx=XM_023238783.1

C2 11642762 11642859
refGene_gn=ITSN1;refGene_tx=XM_023238778.1;refGene_gn=ITSN1;refGene_tx=XM_023238779.1;refGene_gn=ITSN1;refGene_tx=XM_023238780.1;refGene_gn=ITSN1;refGene_tx=XM_023238781.1;refGene_gn=ITSN1;refGene_tx=XM_023238782.1;refGene_gn=ITSN1;refGene_tx=XM_023238783.1
C2 11644665 11644845
refGene_gn=ITSN1;refGene_tx=XM_023238778.1;refGene_gn=ITSN1;refGene_tx=XM_023238779.1;refGene_gn=ITSN1;refGene_tx=XM_023238780.1;refGene_gn=ITSN1;refGene_tx=XM_023238781.1;refGene_gn=ITSN1;refGene_tx=XM_023238782.1;refGene_gn=ITSN1;refGene_tx=XM_023238783.1
C2 11652310 11652471
refGene_gn=ITSN1;refGene_tx=XM_023238778.1;refGene_gn=ITSN1;refGene_tx=XM_023238779.1;refGene_gn=ITSN1;refGene_tx=XM_023238780.1;refGene_gn=ITSN1;refGene_tx=XM_023238781.1;refGene_gn=ITSN1;refGene_tx=XM_023238782.1;refGene_gn=ITSN1;refGene_tx=XM_023238783.1
C2 11663550 11663614
refGene_gn=ITSN1;refGene_tx=XM_023238778.1;refGene_gn=ITSN1;refGene_tx=XM_023238779.1;refGene_gn=ITSN1;refGene_tx=XM_023238780.1;refGene_gn=ITSN1;refGene_tx=XM_023238781.1;refGene_gn=ITSN1;refGene_tx=XM_023238782.1;refGene_gn=ITSN1;refGene_tx=XM_023238783.1
C2 11664606 11664699
refGene_gn=ITSN1;refGene_tx=XM_023238778.1;refGene_gn=ITSN1;refGene_tx=XM_023238779.1;refGene_gn=ITSN1;refGene_tx=XM_023238780.1;refGene_gn=ITSN1;refGene_tx=XM_023238781.1;refGene_gn=ITSN1;refGene_tx=XM_023238782.1;refGene_gn=ITSN1;refGene_tx=XM_023238783.1
C2 11667263 11667291
refGene_gn=ITSN1;refGene_tx=XM_023238778.1;refGene_gn=ITSN1;refGene_tx=XM_023238779.1;refGene_gn=ITSN1;refGene_tx=XM_023238780.1;refGene_gn=ITSN1;refGene_tx=XM_023238781.1;refGene_gn=ITSN1;refGene_tx=XM_023238782.1;refGene_gn=ITSN1;refGene_tx=XM_023238783.1
C2 11746633 11746699 refGene_gn=CRYZL1;refGene_tx=XM_003991454.5
C2 11748338 11748416 refGene_gn=CRYZL1;refGene_tx=XM_003991454.5
C2 11750975 11751048 refGene_gn=CRYZL1;refGene_tx=XM_003991454.5
C2 11754679 11754724 refGene_gn=CRYZL1;refGene_tx=XM_003991454.5
C2 11756445 11756471 refGene_gn=CRYZL1;refGene_tx=XM_003991454.5
C2 11756472 11756515 refGene_gn=CRYZL1;refGene_tx=XM_003991454.5
C2 11761159 11761293 refGene_gn=CRYZL1;refGene_tx=XM_003991454.5
C2 11762570 11762682 refGene_gn=CRYZL1;refGene_tx=XM_003991454.5
C2 11765884 11765983 refGene_gn=CRYZL1;refGene_tx=XM_003991454.5
C2 11767035 11767157 refGene_gn=CRYZL1;refGene_tx=XM_003991454.5
C2 11767937 11768043 refGene_gn=CRYZL1;refGene_tx=XM_003991454.5
C2 11770531 11770577 refGene_gn=CRYZL1;refGene_tx=XM_003991454.5
C2 11771106 11771206 refGene_gn=CRYZL1;refGene_tx=XM_003991454.5
C2 11772284 11772593
refGene_gn=DONSON;refGene_tx=XM_023238784.1;refGene_gn=DONSON;refGene_tx=XM_023238785.1;refGene_gn=DONSON;refGene_tx=XM_023238786.1
C2 11773320 11773401
refGene_gn=DONSON;refGene_tx=XM_023238784.1;refGene_gn=DONSON;refGene_tx=XM_023238785.1;refGene_gn=DONSON;refGene_tx=XM_023238786.1
C2 11774719 11774923

refGene_gn=DONSON;refGene_tx=XM_023238784.1;refGene_gn=DONSON;refGene_tx=XM_023238785.1;refGene_gn=DONSON;refGene_tx=XM_023238786.1
C2 11775874 11776053
refGene_gn=DONSON;refGene_tx=XM_023238784.1;refGene_gn=DONSON;refGene_tx=XM_023238785.1;refGene_gn=DONSON;refGene_tx=XM_023238786.1
C2 11776297 11776354
refGene_gn=DONSON;refGene_tx=XM_023238784.1;refGene_gn=DONSON;refGene_tx=XM_023238785.1
C2 11776771 11776950
refGene_gn=DONSON;refGene_tx=XM_023238784.1;refGene_gn=DONSON;refGene_tx=XM_023238785.1;refGene_gn=DONSON;refGene_tx=XM_023238786.1
C2 11778132 11778214
refGene_gn=DONSON;refGene_tx=XM_023238784.1;refGene_gn=DONSON;refGene_tx=XM_023238785.1;refGene_gn=DONSON;refGene_tx=XM_023238786.1
C2 11778323 11778428
refGene_gn=DONSON;refGene_tx=XM_023238784.1;refGene_gn=DONSON;refGene_tx=XM_023238785.1;refGene_gn=DONSON;refGene_tx=XM_023238786.1
C2 11779354 11779553
refGene_gn=DONSON;refGene_tx=XM_023238784.1;refGene_gn=DONSON;refGene_tx=XM_023238785.1;refGene_gn=DONSON;refGene_tx=XM_023238786.1
C2 11780627 11780840
refGene_gn=DONSON;refGene_tx=XM_023238784.1;refGene_gn=DONSON;refGene_tx=XM_023238785.1;refGene_gn=DONSON;refGene_tx=XM_023238786.1
C2 11781710 11781848
refGene_gn=DONSON;refGene_tx=XM_023238784.1;refGene_gn=DONSON;refGene_tx=XM_023238785.1;refGene_gn=DONSON;refGene_tx=XM_023238786.1
C2 11783620 11783832
refGene_gn=SON;refGene_tx=XM_023238787.1;refGene_gn=SON;refGene_tx=XM_023238789.1
C2 11784359 11784475
refGene_gn=SON;refGene_tx=XM_023238787.1;refGene_gn=SON;refGene_tx=XM_023238789.1
C2 11784584 11784656
refGene_gn=SON;refGene_tx=XM_023238787.1;refGene_gn=SON;refGene_tx=XM_023238789.1
C2 11787234 11787382
refGene_gn=SON;refGene_tx=XM_023238787.1;refGene_gn=SON;refGene_tx=XM_023238789.1
C2 11791546 11791663
refGene_gn=SON;refGene_tx=XM_023238787.1;refGene_gn=SON;refGene_tx=XM_023238789.1
C2 11792833 11792944
refGene_gn=SON;refGene_tx=XM_023238787.1;refGene_gn=SON;refGene_tx=XM_023238789.1
C2 11799619 11799874 refGene_gn=SON;refGene_tx=XM_023238788.1
C2 11799981 11800170
refGene_gn=SON;refGene_tx=XM_023238787.1;refGene_gn=SON;refGene_tx=XM_023238788.1;refGene_gn=SON;refGene_tx=XM_023238789.1
C2 11800347 11800494
refGene_gn=SON;refGene_tx=XM_023238787.1;refGene_gn=SON;refGene_tx=XM_

023238788.1;refGene_gn=SON;refGene_tx=XM_023238789.1
C2 11802406 11802567
refGene_gn=SON;refGene_tx=XM_023238787.1;refGene_gn=SON;refGene_tx=XM_023238788.1;refGene_gn=SON;refGene_tx=XM_023238789.1
C2 11804417 11808975
refGene_gn=SON;refGene_tx=XM_023238787.1;refGene_gn=SON;refGene_tx=XM_023238788.1;refGene_gn=SON;refGene_tx=XM_023238789.1
C2 11809095 11810729
refGene_gn=SON;refGene_tx=XM_023238787.1;refGene_gn=SON;refGene_tx=XM_023238788.1;refGene_gn=SON;refGene_tx=XM_023238789.1
C2 11812747 11812914
refGene_gn=SON;refGene_tx=XM_023238787.1;refGene_gn=SON;refGene_tx=XM_023238788.1;refGene_gn=SON;refGene_tx=XM_023238789.1
C2 11815953 11816030
refGene_gn=SON;refGene_tx=XM_023238787.1;refGene_gn=SON;refGene_tx=XM_023238788.1;refGene_gn=SON;refGene_tx=XM_023238789.1
C2 11819928 11820073
refGene_gn=GART;refGene_tx=XM_003991457.5;refGene_gn=GART;refGene_tx=XM_011285758.3;refGene_gn=GART;refGene_tx=XM_011285759.3
C2 11823146 11823242
refGene_gn=GART;refGene_tx=XM_003991457.5;refGene_gn=GART;refGene_tx=XM_011285758.3;refGene_gn=GART;refGene_tx=XM_011285759.3
C2 11823708 11823883
refGene_gn=GART;refGene_tx=XM_003991457.5;refGene_gn=GART;refGene_tx=XM_011285758.3;refGene_gn=GART;refGene_tx=XM_011285759.3
C2 11825120 11825232
refGene_gn=GART;refGene_tx=XM_003991457.5;refGene_gn=GART;refGene_tx=XM_011285758.3;refGene_gn=GART;refGene_tx=XM_011285759.3
C2 11826076 11826145
refGene_gn=GART;refGene_tx=XM_003991457.5;refGene_gn=GART;refGene_tx=XM_011285758.3;refGene_gn=GART;refGene_tx=XM_011285759.3
C2 11826624 11826750
refGene_gn=GART;refGene_tx=XM_003991457.5;refGene_gn=GART;refGene_tx=XM_011285758.3;refGene_gn=GART;refGene_tx=XM_011285759.3
C2 11829191 11829279
refGene_gn=GART;refGene_tx=XM_003991457.5;refGene_gn=GART;refGene_tx=XM_011285758.3;refGene_gn=GART;refGene_tx=XM_011285759.3
C2 11829503 11829589
refGene_gn=GART;refGene_tx=XM_003991457.5;refGene_gn=GART;refGene_tx=XM_011285758.3;refGene_gn=GART;refGene_tx=XM_011285759.3
C2 11830015 11830184
refGene_gn=GART;refGene_tx=XM_003991457.5;refGene_gn=GART;refGene_tx=XM_011285758.3;refGene_gn=GART;refGene_tx=XM_011285759.3
C2 11832470 11832702
refGene_gn=GART;refGene_tx=XM_003991457.5;refGene_gn=GART;refGene_tx=XM_011285758.3;refGene_gn=GART;refGene_tx=XM_011285759.3
C2 11834613 11834708
refGene_gn=GART;refGene_tx=XM_003991457.5;refGene_gn=GART;refGene_tx=XM_011285758.3;refGene_gn=GART;refGene_tx=XM_011285759.3
C2 11835455 11835565

refGene_gn=GART;refGene_tx=XM_003991457.5;refGene_gn=GART;refGene_tx=XM_011285758.3;refGene_gn=GART;refGene_tx=XM_011285759.3
C2 11835927 11836126
refGene_gn=GART;refGene_tx=XM_003991457.5;refGene_gn=GART;refGene_tx=XM_011285758.3;refGene_gn=GART;refGene_tx=XM_011285759.3
C2 11838039 11838291
refGene_gn=GART;refGene_tx=XM_003991457.5;refGene_gn=GART;refGene_tx=XM_011285758.3;refGene_gn=GART;refGene_tx=XM_011285759.3
C2 11838540 11838693
refGene_gn=GART;refGene_tx=XM_003991457.5;refGene_gn=GART;refGene_tx=XM_011285758.3;refGene_gn=GART;refGene_tx=XM_011285759.3
C2 11839542 11839749
refGene_gn=GART;refGene_tx=XM_003991457.5;refGene_gn=GART;refGene_tx=XM_011285758.3;refGene_gn=GART;refGene_tx=XM_011285759.3
C2 11840781 11840919
refGene_gn=GART;refGene_tx=XM_003991457.5;refGene_gn=GART;refGene_tx=XM_011285758.3;refGene_gn=GART;refGene_tx=XM_011285759.3
C2 11842614 11842745
refGene_gn=GART;refGene_tx=XM_003991457.5;refGene_gn=GART;refGene_tx=XM_011285758.3;refGene_gn=GART;refGene_tx=XM_011285759.3
C2 11843013 11843155
refGene_gn=GART;refGene_tx=XM_003991457.5;refGene_gn=GART;refGene_tx=XM_011285758.3;refGene_gn=GART;refGene_tx=XM_011285759.3
C2 11844160 11844276
refGene_gn=GART;refGene_tx=XM_003991457.5;refGene_gn=GART;refGene_tx=XM_011285758.3;refGene_gn=GART;refGene_tx=XM_011285759.3
C2 11844372 11844561
refGene_gn=GART;refGene_tx=XM_003991457.5;refGene_gn=GART;refGene_tx=XM_011285758.3;refGene_gn=GART;refGene_tx=XM_011285759.3
C2 11860841 11862002 refGene_gn=DNAJC28;refGene_tx=XM_006935914.4
C2 11898414 11898513
refGene_gn=TMEM50B;refGene_tx=XM_019839421.2;refGene_gn=TMEM50B;refGene_tx=XM_019839422.2;refGene_gn=TMEM50B;refGene_tx=XM_019839423.2;refGene_gn=TMEM50B;refGene_tx=XM_023238790.1;refGene_gn=TMEM50B;refGene_tx=XM_023238791.1;refGene_gn=TMEM50B;refGene_tx=XM_023238792.1;refGene_gn=TMEM50B;refGene_tx=XM_023238793.1
C2 11900214 11900327
refGene_gn=TMEM50B;refGene_tx=XM_019839421.2;refGene_gn=TMEM50B;refGene_tx=XM_019839422.2;refGene_gn=TMEM50B;refGene_tx=XM_019839423.2;refGene_gn=TMEM50B;refGene_tx=XM_023238790.1;refGene_gn=TMEM50B;refGene_tx=XM_023238791.1;refGene_gn=TMEM50B;refGene_tx=XM_023238792.1;refGene_gn=TMEM50B;refGene_tx=XM_023238793.1
C2 11901845 11901913
refGene_gn=TMEM50B;refGene_tx=XM_019839421.2;refGene_gn=TMEM50B;refGene_tx=XM_019839422.2;refGene_gn=TMEM50B;refGene_tx=XM_019839423.2;refGene_gn=TMEM50B;refGene_tx=XM_023238790.1;refGene_gn=TMEM50B;refGene_tx=XM_023238791.1;refGene_gn=TMEM50B;refGene_tx=XM_023238792.1;refGene_gn=TMEM50B;refGene_tx=XM_023238793.1
C2 11902522 11902615
refGene_gn=TMEM50B;refGene_tx=XM_019839421.2;refGene_gn=TMEM50B;refGen

e_tx=XM_019839422.2;refGene_gn=TMEM50B;refGene_tx=XM_019839423.2;refGene_gn=TMEM50B;refGene_tx=XM_023238790.1;refGene_gn=TMEM50B;refGene_tx=XM_023238791.1;refGene_gn=TMEM50B;refGene_tx=XM_023238792.1;refGene_gn=TMEM50B;refGene_tx=XM_023238793.1
C2	11905414	11905461	refGene_gn=TMEM50B;refGene_tx=XM_023238793.1
C2	11906359	11906417	refGene_gn=TMEM50B;refGene_tx=XM_023238792.1
C2	11911561	11911607	refGene_gn=TMEM50B;refGene_tx=XM_023238792.1
C2	11918180	11918362	refGene_gn=TMEM50B;refGene_tx=XM_019839421.2;refGene_gn=TMEM50B;refGene_tx=XM_019839422.2;refGene_gn=TMEM50B;refGene_tx=XM_019839423.2;refGene_gn=TMEM50B;refGene_tx=XM_023238790.1;refGene_gn=TMEM50B;refGene_tx=XM_023238791.1
C2	11922278	11922452	refGene_gn=IFNGR2;refGene_tx=XM_006935924.4;refGene_gn=IFNGR2;refGene_tx=XM_023238794.1
C2	11924917	11925075	refGene_gn=IFNGR2;refGene_tx=XM_006935924.4;refGene_gn=IFNGR2;refGene_tx=XM_023238794.1
C2	11925463	11925623	refGene_gn=IFNGR2;refGene_tx=XM_006935924.4;refGene_gn=IFNGR2;refGene_tx=XM_023238794.1
C2	11929660	11929806	refGene_gn=IFNGR2;refGene_tx=XM_006935924.4;refGene_gn=IFNGR2;refGene_tx=XM_023238794.1
C2	11934022	11934228	refGene_gn=IFNGR2;refGene_tx=XM_006935924.4;refGene_gn=IFNGR2;refGene_tx=XM_023238794.1
C2	11940848	11940981	refGene_gn=IFNGR2;refGene_tx=XM_006935924.4;refGene_gn=IFNGR2;refGene_tx=XM_023238794.1
C2	11946468	11946484	refGene_gn=IFNGR2;refGene_tx=XM_023238794.1
C2	11949228	11949310	refGene_gn=IFNGR2;refGene_tx=XM_006935924.4
C2	12006973	12007213	refGene_gn=IFNAR1;refGene_tx=XM_023238795.1
C2	12008341	12008487	refGene_gn=IFNAR1;refGene_tx=XM_023238795.1
C2	12010142	12010293	refGene_gn=IFNAR1;refGene_tx=XM_023238795.1
C2	12013021	12013179	refGene_gn=IFNAR1;refGene_tx=XM_023238795.1
C2	12013290	12013493	refGene_gn=IFNAR1;refGene_tx=XM_023238795.1
C2	12016963	12017078	refGene_gn=IFNAR1;refGene_tx=XM_023238795.1
C2	12018227	12018366	refGene_gn=IFNAR1;refGene_tx=XM_023238795.1
C2	12018475	12018633	refGene_gn=IFNAR1;refGene_tx=XM_023238795.1
C2	12020001	12020180	refGene_gn=IFNAR1;refGene_tx=XM_023238795.1
C2	12025229	12025353	refGene_gn=IFNAR1;refGene_tx=XM_023238795.1
C2	12029032	12029105	refGene_gn=IFNAR1;refGene_tx=XM_023238795.1
C2	12074556	12074871	refGene_gn=IL10RB;refGene_tx=XM_003991463.3
C2	12081184	12081339	refGene_gn=IL10RB;refGene_tx=XM_003991463.3
C2	12087044	12087192	refGene_gn=IL10RB;refGene_tx=XM_003991463.3
C2	12091571	12091738	refGene_gn=IL10RB;refGene_tx=XM_003991463.3
C2	12093820	12093978	refGene_gn=IL10RB;refGene_tx=XM_003991463.3
C2	12097517	12097641	refGene_gn=IL10RB;refGene_tx=XM_003991463.3
C2	12099266	12099315	refGene_gn=IL10RB;refGene_tx=XM_003991463.3

C2 12101709 12102414
refGene_gn=IFNAR2;refGene_tx=NM_001278859.1;refGene_gn=IFNAR2;refGene_tx=XM_019839220.2;refGene_gn=IFNAR2;refGene_tx=XM_023260081.1;refGene_gn=IFNAR2;refGene_tx=XM_023260082.1;refGene_gn=IFNAR2;refGene_tx=XM_023260083.1;refGene_gn=IFNAR2;refGene_tx=XM_023260084.1
C2 12106053 12106184
refGene_gn=IFNAR2;refGene_tx=NM_001278859.1;refGene_gn=IFNAR2;refGene_tx=XM_019839220.2;refGene_gn=IFNAR2;refGene_tx=XM_023260081.1;refGene_gn=IFNAR2;refGene_tx=XM_023260082.1;refGene_gn=IFNAR2;refGene_tx=XM_023260083.1
C2 12109502 12109671
refGene_gn=IFNAR2;refGene_tx=NM_001278859.1;refGene_gn=IFNAR2;refGene_tx=XM_019839220.2;refGene_gn=IFNAR2;refGene_tx=XM_023260081.1;refGene_gn=IFNAR2;refGene_tx=XM_023260082.1;refGene_gn=IFNAR2;refGene_tx=XM_023260083.1;refGene_gn=IFNAR2;refGene_tx=XM_023260084.1
C2 12114188 12114349
refGene_gn=IFNAR2;refGene_tx=NM_001278859.1;refGene_gn=IFNAR2;refGene_tx=XM_019839220.2;refGene_gn=IFNAR2;refGene_tx=XM_023260081.1;refGene_gn=IFNAR2;refGene_tx=XM_023260082.1;refGene_gn=IFNAR2;refGene_tx=XM_023260083.1;refGene_gn=IFNAR2;refGene_tx=XM_023260084.1
C2 12116771 12116944
refGene_gn=IFNAR2;refGene_tx=NM_001278859.1;refGene_gn=IFNAR2;refGene_tx=XM_019839220.2;refGene_gn=IFNAR2;refGene_tx=XM_023260081.1;refGene_gn=IFNAR2;refGene_tx=XM_023260082.1;refGene_gn=IFNAR2;refGene_tx=XM_023260083.1;refGene_gn=IFNAR2;refGene_tx=XM_023260084.1
C2 12118779 12118903
refGene_gn=IFNAR2;refGene_tx=NM_001278859.1;refGene_gn=IFNAR2;refGene_tx=XM_023260081.1;refGene_gn=IFNAR2;refGene_tx=XM_023260082.1;refGene_gn=IFNAR2;refGene_tx=XM_023260083.1;refGene_gn=IFNAR2;refGene_tx=XM_023260084.1
C2 12121016 12121063
refGene_gn=IFNAR2;refGene_tx=NM_001278859.1;refGene_gn=IFNAR2;refGene_tx=XM_019839220.2;refGene_gn=IFNAR2;refGene_tx=XM_023260081.1;refGene_gn=IFNAR2;refGene_tx=XM_023260082.1;refGene_gn=IFNAR2;refGene_tx=XM_023260083.1;refGene_gn=IFNAR2;refGene_tx=XM_023260084.1
C2 12121064 12121074 refGene_gn=IFNAR2;refGene_tx=NM_001278859.1
C2 12122145 12122200
refGene_gn=IFNAR2;refGene_tx=NM_001278859.1;refGene_gn=IFNAR2;refGene_tx=XM_023260081.1;refGene_gn=IFNAR2;refGene_tx=XM_023260082.1;refGene_gn=IFNAR2;refGene_tx=XM_023260083.1;refGene_gn=IFNAR2;refGene_tx=XM_023260084.1
C2 12286778 12287597 refGene_gn=OLIG1;refGene_tx=XM_023238796.1
C2 12330246 12330399 refGene_gn=OLIG2;refGene_tx=XM_023238797.1
C2 12330400 12330589 refGene_gn=OLIG2;refGene_tx=XM_023238797.1
C2 12330590 12331313 refGene_gn=OLIG2;refGene_tx=XM_023238797.1
C2 12332055 12332304 refGene_gn=OLIG2;refGene_tx=XM_023238797.1
C2 12550682 12551342
refGene_gn=CC2H21orf62;refGene_tx=XM_006935919.3;refGene_gn=CC2H21orf62;refGene_tx=XM_019839425.2
C2 12569639 12570129 refGene_gn=PAXBP1;refGene_tx=XM_023238798.1

C2 12571831 12571960 refGene_gn=PAXBP1;refGene_tx=XM_023238798.1
C2 12576196 12576373 refGene_gn=PAXBP1;refGene_tx=XM_023238798.1
C2 12577768 12577990 refGene_gn=PAXBP1;refGene_tx=XM_023238798.1
C2 12578535 12578639 refGene_gn=PAXBP1;refGene_tx=XM_023238798.1
C2 12579619 12579837 refGene_gn=PAXBP1;refGene_tx=XM_023238798.1
C2 12580349 12580539 refGene_gn=PAXBP1;refGene_tx=XM_023238798.1
C2 12584441 12584565 refGene_gn=PAXBP1;refGene_tx=XM_023238798.1
C2 12587844 12587944 refGene_gn=PAXBP1;refGene_tx=XM_023238798.1
C2 12588031 12588147 refGene_gn=PAXBP1;refGene_tx=XM_023238798.1
C2 12589961 12590161 refGene_gn=PAXBP1;refGene_tx=XM_023238798.1
C2 12591233 12591378 refGene_gn=PAXBP1;refGene_tx=XM_023238798.1
C2 12592070 12592192 refGene_gn=PAXBP1;refGene_tx=XM_023238798.1
C2 12592985 12593062 refGene_gn=PAXBP1;refGene_tx=XM_023238798.1
C2 12593548 12593615 refGene_gn=PAXBP1;refGene_tx=XM_023238798.1
C2 12598315 12598462 refGene_gn=PAXBP1;refGene_tx=XM_023238798.1
C2 12600458 12600613 refGene_gn=PAXBP1;refGene_tx=XM_023238798.1
C2 12602957 12603075 refGene_gn=PAXBP1;refGene_tx=XM_023238798.1
C2 12611233 12611357
refGene_gn=SYNJ1;refGene_tx=XM_023238799.1;refGene_gn=SYNJ1;refGene_tx=XM_023238800.1;refGene_gn=SYNJ1;refGene_tx=XM_023238801.1;refGene_gn=SYNJ1;refGene_tx=XM_023238802.1;refGene_gn=SYNJ1;refGene_tx=XM_023238803.1;refGene_gn=SYNJ1;refGene_tx=XM_023238804.1;refGene_gn=SYNJ1;refGene_tx=XM_023238805.1;refGene_gn=SYNJ1;refGene_tx=XM_023238806.1;refGene_gn=SYNJ1;refGene_tx=XM_023238807.1;refGene_gn=SYNJ1;refGene_tx=XM_023238808.1
C2 12635300 12635387
refGene_gn=SYNJ1;refGene_tx=XM_023238799.1;refGene_gn=SYNJ1;refGene_tx=XM_023238800.1;refGene_gn=SYNJ1;refGene_tx=XM_023238801.1;refGene_gn=SYNJ1;refGene_tx=XM_023238802.1;refGene_gn=SYNJ1;refGene_tx=XM_023238803.1;refGene_gn=SYNJ1;refGene_tx=XM_023238804.1;refGene_gn=SYNJ1;refGene_tx=XM_023238805.1;refGene_gn=SYNJ1;refGene_tx=XM_023238806.1;refGene_gn=SYNJ1;refGene_tx=XM_023238807.1;refGene_gn=SYNJ1;refGene_tx=XM_023238808.1
C2 12636989 12637257
refGene_gn=SYNJ1;refGene_tx=XM_023238799.1;refGene_gn=SYNJ1;refGene_tx=XM_023238800.1;refGene_gn=SYNJ1;refGene_tx=XM_023238801.1;refGene_gn=SYNJ1;refGene_tx=XM_023238802.1;refGene_gn=SYNJ1;refGene_tx=XM_023238803.1;refGene_gn=SYNJ1;refGene_tx=XM_023238804.1;refGene_gn=SYNJ1;refGene_tx=XM_023238805.1;refGene_gn=SYNJ1;refGene_tx=XM_023238806.1;refGene_gn=SYNJ1;refGene_tx=XM_023238807.1;refGene_gn=SYNJ1;refGene_tx=XM_023238808.1
C2 12640808 12641034
refGene_gn=SYNJ1;refGene_tx=XM_023238799.1;refGene_gn=SYNJ1;refGene_tx=XM_023238800.1;refGene_gn=SYNJ1;refGene_tx=XM_023238801.1;refGene_gn=SYNJ1;refGene_tx=XM_023238802.1;refGene_gn=SYNJ1;refGene_tx=XM_023238803.1;refGene_gn=SYNJ1;refGene_tx=XM_023238804.1;refGene_gn=SYNJ1;refGene_tx=XM_023238805.1;refGene_gn=SYNJ1;refGene_tx=XM_023238806.1;refGene_gn=SYNJ1;refGene_tx=XM_023238807.1;refGene_gn=SYNJ1;refGene_tx=XM_023238808.1
C2 12642120 12642204

refGene_gn=SYNJ1;refGene_tx=XM_023238799.1;refGene_gn=SYNJ1;refGene_tx=XM_023238800.1;refGene_gn=SYNJ1;refGene_tx=XM_023238801.1;refGene_gn=SYNJ1;refGene_tx=XM_023238802.1;refGene_gn=SYNJ1;refGene_tx=XM_023238803.1;refGene_gn=SYNJ1;refGene_tx=XM_023238804.1;refGene_gn=SYNJ1;refGene_tx=XM_023238805.1;refGene_gn=SYNJ1;refGene_tx=XM_023238806.1;refGene_gn=SYNJ1;refGene_tx=XM_023238807.1;refGene_gn=SYNJ1;refGene_tx=XM_023238808.1
C2 12648024 12648086
refGene_gn=SYNJ1;refGene_tx=XM_023238799.1;refGene_gn=SYNJ1;refGene_tx=XM_023238800.1;refGene_gn=SYNJ1;refGene_tx=XM_023238801.1;refGene_gn=SYNJ1;refGene_tx=XM_023238802.1;refGene_gn=SYNJ1;refGene_tx=XM_023238803.1;refGene_gn=SYNJ1;refGene_tx=XM_023238804.1;refGene_gn=SYNJ1;refGene_tx=XM_023238805.1;refGene_gn=SYNJ1;refGene_tx=XM_023238806.1;refGene_gn=SYNJ1;refGene_tx=XM_023238807.1;refGene_gn=SYNJ1;refGene_tx=XM_023238808.1
C2 12649109 12649206
refGene_gn=SYNJ1;refGene_tx=XM_023238799.1;refGene_gn=SYNJ1;refGene_tx=XM_023238800.1;refGene_gn=SYNJ1;refGene_tx=XM_023238801.1;refGene_gn=SYNJ1;refGene_tx=XM_023238802.1;refGene_gn=SYNJ1;refGene_tx=XM_023238803.1;refGene_gn=SYNJ1;refGene_tx=XM_023238804.1;refGene_gn=SYNJ1;refGene_tx=XM_023238805.1;refGene_gn=SYNJ1;refGene_tx=XM_023238806.1;refGene_gn=SYNJ1;refGene_tx=XM_023238807.1;refGene_gn=SYNJ1;refGene_tx=XM_023238808.1
C2 12650320 12650490
refGene_gn=SYNJ1;refGene_tx=XM_023238799.1;refGene_gn=SYNJ1;refGene_tx=XM_023238800.1;refGene_gn=SYNJ1;refGene_tx=XM_023238801.1;refGene_gn=SYNJ1;refGene_tx=XM_023238802.1;refGene_gn=SYNJ1;refGene_tx=XM_023238803.1;refGene_gn=SYNJ1;refGene_tx=XM_023238804.1;refGene_gn=SYNJ1;refGene_tx=XM_023238805.1;refGene_gn=SYNJ1;refGene_tx=XM_023238806.1;refGene_gn=SYNJ1;refGene_tx=XM_023238807.1;refGene_gn=SYNJ1;refGene_tx=XM_023238808.1
C2 12651544 12651626
refGene_gn=SYNJ1;refGene_tx=XM_023238799.1;refGene_gn=SYNJ1;refGene_tx=XM_023238800.1;refGene_gn=SYNJ1;refGene_tx=XM_023238801.1;refGene_gn=SYNJ1;refGene_tx=XM_023238802.1;refGene_gn=SYNJ1;refGene_tx=XM_023238803.1;refGene_gn=SYNJ1;refGene_tx=XM_023238804.1;refGene_gn=SYNJ1;refGene_tx=XM_023238805.1;refGene_gn=SYNJ1;refGene_tx=XM_023238806.1;refGene_gn=SYNJ1;refGene_tx=XM_023238807.1;refGene_gn=SYNJ1;refGene_tx=XM_023238808.1
C2 12654117 12654270
refGene_gn=SYNJ1;refGene_tx=XM_023238799.1;refGene_gn=SYNJ1;refGene_tx=XM_023238800.1;refGene_gn=SYNJ1;refGene_tx=XM_023238801.1;refGene_gn=SYNJ1;refGene_tx=XM_023238802.1;refGene_gn=SYNJ1;refGene_tx=XM_023238803.1;refGene_gn=SYNJ1;refGene_tx=XM_023238804.1;refGene_gn=SYNJ1;refGene_tx=XM_023238805.1;refGene_gn=SYNJ1;refGene_tx=XM_023238806.1;refGene_gn=SYNJ1;refGene_tx=XM_023238807.1;refGene_gn=SYNJ1;refGene_tx=XM_023238808.1
C2 12656789 12656955
refGene_gn=SYNJ1;refGene_tx=XM_023238799.1;refGene_gn=SYNJ1;refGene_tx=XM_023238800.1;refGene_gn=SYNJ1;refGene_tx=XM_023238801.1;refGene_gn=

SYNJ1;refGene_tx=XM_023238802.1;refGene_gn=SYNJ1;refGene_tx=XM_0232388
03.1;refGene_gn=SYNJ1;refGene_tx=XM_023238804.1;refGene_gn=SYNJ1;refGe
ne_tx=XM_023238805.1;refGene_gn=SYNJ1;refGene_tx=XM_023238806.1;refGen
e_gn=SYNJ1;refGene_tx=XM_023238807.1;refGene_gn=SYNJ1;refGene_tx=XM_02
3238808.1
C2 12659431 12659455
refGene_gn=SYNJ1;refGene_tx=XM_023238799.1;refGene_gn=SYNJ1;refGene_tx
=XM_023238801.1;refGene_gn=SYNJ1;refGene_tx=XM_023238802.1;refGene_gn=
SYNJ1;refGene_tx=XM_023238803.1;refGene_gn=SYNJ1;refGene_tx=XM_0232388
05.1;refGene_gn=SYNJ1;refGene_tx=XM_023238806.1;refGene_gn=SYNJ1;refGe
ne_tx=XM_023238807.1;refGene_gn=SYNJ1;refGene_tx=XM_023238808.1
C2 12662143 12662335
refGene_gn=SYNJ1;refGene_tx=XM_023238799.1;refGene_gn=SYNJ1;refGene_tx
=XM_023238800.1;refGene_gn=SYNJ1;refGene_tx=XM_023238801.1;refGene_gn=
SYNJ1;refGene_tx=XM_023238802.1;refGene_gn=SYNJ1;refGene_tx=XM_0232388
03.1;refGene_gn=SYNJ1;refGene_tx=XM_023238804.1;refGene_gn=SYNJ1;refGe
ne_tx=XM_023238805.1;refGene_gn=SYNJ1;refGene_tx=XM_023238806.1;refGen
e_gn=SYNJ1;refGene_tx=XM_023238807.1;refGene_gn=SYNJ1;refGene_tx=XM_02
3238808.1
C2 12664246 12664331
refGene_gn=SYNJ1;refGene_tx=XM_023238799.1;refGene_gn=SYNJ1;refGene_tx
=XM_023238800.1;refGene_gn=SYNJ1;refGene_tx=XM_023238801.1;refGene_gn=
SYNJ1;refGene_tx=XM_023238802.1;refGene_gn=SYNJ1;refGene_tx=XM_0232388
03.1;refGene_gn=SYNJ1;refGene_tx=XM_023238804.1;refGene_gn=SYNJ1;refGe
ne_tx=XM_023238805.1;refGene_gn=SYNJ1;refGene_tx=XM_023238806.1;refGen
e_gn=SYNJ1;refGene_tx=XM_023238807.1;refGene_gn=SYNJ1;refGene_tx=XM_02
3238808.1
C2 12668936 12669077
refGene_gn=SYNJ1;refGene_tx=XM_023238799.1;refGene_gn=SYNJ1;refGene_tx
=XM_023238800.1;refGene_gn=SYNJ1;refGene_tx=XM_023238801.1;refGene_gn=
SYNJ1;refGene_tx=XM_023238802.1;refGene_gn=SYNJ1;refGene_tx=XM_0232388
03.1;refGene_gn=SYNJ1;refGene_tx=XM_023238804.1;refGene_gn=SYNJ1;refGe
ne_tx=XM_023238805.1;refGene_gn=SYNJ1;refGene_tx=XM_023238806.1;refGen
e_gn=SYNJ1;refGene_tx=XM_023238807.1;refGene_gn=SYNJ1;refGene_tx=XM_02
3238808.1
C2 12669353 12669546
refGene_gn=SYNJ1;refGene_tx=XM_023238799.1;refGene_gn=SYNJ1;refGene_tx
=XM_023238800.1;refGene_gn=SYNJ1;refGene_tx=XM_023238801.1;refGene_gn=
SYNJ1;refGene_tx=XM_023238802.1;refGene_gn=SYNJ1;refGene_tx=XM_0232388
03.1;refGene_gn=SYNJ1;refGene_tx=XM_023238804.1;refGene_gn=SYNJ1;refGe
ne_tx=XM_023238805.1;refGene_gn=SYNJ1;refGene_tx=XM_023238806.1;refGen
e_gn=SYNJ1;refGene_tx=XM_023238807.1;refGene_gn=SYNJ1;refGene_tx=XM_02
3238808.1
C2 12669822 12669981
refGene_gn=SYNJ1;refGene_tx=XM_023238799.1;refGene_gn=SYNJ1;refGene_tx
=XM_023238800.1;refGene_gn=SYNJ1;refGene_tx=XM_023238801.1;refGene_gn=
SYNJ1;refGene_tx=XM_023238802.1;refGene_gn=SYNJ1;refGene_tx=XM_0232388
03.1;refGene_gn=SYNJ1;refGene_tx=XM_023238804.1;refGene_gn=SYNJ1;refGe
ne_tx=XM_023238805.1;refGene_gn=SYNJ1;refGene_tx=XM_023238806.1;refGen
e_gn=SYNJ1;refGene_tx=XM_023238807.1;refGene_gn=SYNJ1;refGene_tx=XM_02

3238808.1
C2 12671152 12671309
refGene_gn=SYNJ1;refGene_tx=XM_023238799.1;refGene_gn=SYNJ1;refGene_tx=XM_023238800.1;refGene_gn=SYNJ1;refGene_tx=XM_023238801.1;refGene_gn=SYNJ1;refGene_tx=XM_023238802.1;refGene_gn=SYNJ1;refGene_tx=XM_023238803.1;refGene_gn=SYNJ1;refGene_tx=XM_023238804.1;refGene_gn=SYNJ1;refGene_tx=XM_023238805.1;refGene_gn=SYNJ1;refGene_tx=XM_023238806.1;refGene_gn=SYNJ1;refGene_tx=XM_023238807.1;refGene_gn=SYNJ1;refGene_tx=XM_023238808.1
C2 12671924 12672042
refGene_gn=SYNJ1;refGene_tx=XM_023238799.1;refGene_gn=SYNJ1;refGene_tx=XM_023238800.1;refGene_gn=SYNJ1;refGene_tx=XM_023238801.1;refGene_gn=SYNJ1;refGene_tx=XM_023238802.1;refGene_gn=SYNJ1;refGene_tx=XM_023238803.1;refGene_gn=SYNJ1;refGene_tx=XM_023238804.1;refGene_gn=SYNJ1;refGene_tx=XM_023238805.1;refGene_gn=SYNJ1;refGene_tx=XM_023238806.1;refGene_gn=SYNJ1;refGene_tx=XM_023238807.1;refGene_gn=SYNJ1;refGene_tx=XM_023238808.1
C2 12672142 12672358
refGene_gn=SYNJ1;refGene_tx=XM_023238799.1;refGene_gn=SYNJ1;refGene_tx=XM_023238800.1;refGene_gn=SYNJ1;refGene_tx=XM_023238801.1;refGene_gn=SYNJ1;refGene_tx=XM_023238802.1;refGene_gn=SYNJ1;refGene_tx=XM_023238803.1;refGene_gn=SYNJ1;refGene_tx=XM_023238804.1;refGene_gn=SYNJ1;refGene_tx=XM_023238805.1;refGene_gn=SYNJ1;refGene_tx=XM_023238806.1;refGene_gn=SYNJ1;refGene_tx=XM_023238807.1;refGene_gn=SYNJ1;refGene_tx=XM_023238808.1
C2 12675510 12675589
refGene_gn=SYNJ1;refGene_tx=XM_023238799.1;refGene_gn=SYNJ1;refGene_tx=XM_023238800.1;refGene_gn=SYNJ1;refGene_tx=XM_023238801.1;refGene_gn=SYNJ1;refGene_tx=XM_023238802.1;refGene_gn=SYNJ1;refGene_tx=XM_023238803.1;refGene_gn=SYNJ1;refGene_tx=XM_023238804.1;refGene_gn=SYNJ1;refGene_tx=XM_023238805.1;refGene_gn=SYNJ1;refGene_tx=XM_023238806.1;refGene_gn=SYNJ1;refGene_tx=XM_023238807.1;refGene_gn=SYNJ1;refGene_tx=XM_023238808.1
C2 12679427 12679590
refGene_gn=SYNJ1;refGene_tx=XM_023238799.1;refGene_gn=SYNJ1;refGene_tx=XM_023238800.1;refGene_gn=SYNJ1;refGene_tx=XM_023238801.1;refGene_gn=SYNJ1;refGene_tx=XM_023238802.1;refGene_gn=SYNJ1;refGene_tx=XM_023238803.1;refGene_gn=SYNJ1;refGene_tx=XM_023238804.1;refGene_gn=SYNJ1;refGene_tx=XM_023238805.1;refGene_gn=SYNJ1;refGene_tx=XM_023238806.1;refGene_gn=SYNJ1;refGene_tx=XM_023238807.1;refGene_gn=SYNJ1;refGene_tx=XM_023238808.1
C2 12683141 12683351
refGene_gn=SYNJ1;refGene_tx=XM_023238799.1;refGene_gn=SYNJ1;refGene_tx=XM_023238800.1;refGene_gn=SYNJ1;refGene_tx=XM_023238801.1;refGene_gn=SYNJ1;refGene_tx=XM_023238802.1;refGene_gn=SYNJ1;refGene_tx=XM_023238803.1;refGene_gn=SYNJ1;refGene_tx=XM_023238804.1;refGene_gn=SYNJ1;refGene_tx=XM_023238805.1;refGene_gn=SYNJ1;refGene_tx=XM_023238806.1;refGene_gn=SYNJ1;refGene_tx=XM_023238807.1;refGene_gn=SYNJ1;refGene_tx=XM_023238808.1
C2 12684469 12684610

refGene_gn=SYNJ1;refGene_tx=XM_023238799.1;refGene_gn=SYNJ1;refGene_tx=XM_023238800.1;refGene_gn=SYNJ1;refGene_tx=XM_023238801.1;refGene_gn=SYNJ1;refGene_tx=XM_023238802.1;refGene_gn=SYNJ1;refGene_tx=XM_023238803.1;refGene_gn=SYNJ1;refGene_tx=XM_023238804.1;refGene_gn=SYNJ1;refGene_tx=XM_023238805.1;refGene_gn=SYNJ1;refGene_tx=XM_023238806.1;refGene_gn=SYNJ1;refGene_tx=XM_023238807.1;refGene_gn=SYNJ1;refGene_tx=XM_023238808.1
C2 12685181 12685220
refGene_gn=SYNJ1;refGene_tx=XM_023238799.1;refGene_gn=SYNJ1;refGene_tx=XM_023238800.1;refGene_gn=SYNJ1;refGene_tx=XM_023238802.1;refGene_gn=SYNJ1;refGene_tx=XM_023238805.1;refGene_gn=SYNJ1;refGene_tx=XM_023238806.1;refGene_gn=SYNJ1;refGene_tx=XM_023238807.1;refGene_gn=SYNJ1;refGene_tx=XM_023238808.1
C2 12686909 12686957
refGene_gn=SYNJ1;refGene_tx=XM_023238799.1;refGene_gn=SYNJ1;refGene_tx=XM_023238800.1;refGene_gn=SYNJ1;refGene_tx=XM_023238805.1;refGene_gn=SYNJ1;refGene_tx=XM_023238806.1;refGene_gn=SYNJ1;refGene_tx=XM_023238807.1
C2 12688236 12688275
refGene_gn=SYNJ1;refGene_tx=XM_023238799.1;refGene_gn=SYNJ1;refGene_tx=XM_023238800.1;refGene_gn=SYNJ1;refGene_tx=XM_023238801.1;refGene_gn=SYNJ1;refGene_tx=XM_023238805.1;refGene_gn=SYNJ1;refGene_tx=XM_023238806.1;refGene_gn=SYNJ1;refGene_tx=XM_023238807.1;refGene_gn=SYNJ1;refGene_tx=XM_023238808.1
C2 12688409 12688480
refGene_gn=SYNJ1;refGene_tx=XM_023238799.1;refGene_gn=SYNJ1;refGene_tx=XM_023238800.1;refGene_gn=SYNJ1;refGene_tx=XM_023238801.1;refGene_gn=SYNJ1;refGene_tx=XM_023238802.1;refGene_gn=SYNJ1;refGene_tx=XM_023238803.1;refGene_gn=SYNJ1;refGene_tx=XM_023238804.1;refGene_gn=SYNJ1;refGene_tx=XM_023238805.1;refGene_gn=SYNJ1;refGene_tx=XM_023238806.1;refGene_gn=SYNJ1;refGene_tx=XM_023238807.1;refGene_gn=SYNJ1;refGene_tx=XM_023238808.1
C2 12690447 12690556
refGene_gn=SYNJ1;refGene_tx=XM_023238799.1;refGene_gn=SYNJ1;refGene_tx=XM_023238800.1;refGene_gn=SYNJ1;refGene_tx=XM_023238801.1;refGene_gn=SYNJ1;refGene_tx=XM_023238802.1;refGene_gn=SYNJ1;refGene_tx=XM_023238803.1;refGene_gn=SYNJ1;refGene_tx=XM_023238804.1;refGene_gn=SYNJ1;refGene_tx=XM_023238805.1;refGene_gn=SYNJ1;refGene_tx=XM_023238806.1;refGene_gn=SYNJ1;refGene_tx=XM_023238807.1;refGene_gn=SYNJ1;refGene_tx=XM_023238808.1
C2 12690781 12691008
refGene_gn=SYNJ1;refGene_tx=XM_023238799.1;refGene_gn=SYNJ1;refGene_tx=XM_023238800.1;refGene_gn=SYNJ1;refGene_tx=XM_023238801.1;refGene_gn=SYNJ1;refGene_tx=XM_023238802.1;refGene_gn=SYNJ1;refGene_tx=XM_023238803.1;refGene_gn=SYNJ1;refGene_tx=XM_023238804.1;refGene_gn=SYNJ1;refGene_tx=XM_023238805.1;refGene_gn=SYNJ1;refGene_tx=XM_023238806.1;refGene_gn=SYNJ1;refGene_tx=XM_023238807.1;refGene_gn=SYNJ1;refGene_tx=XM_023238808.1
C2 12694682 12694703
refGene_gn=SYNJ1;refGene_tx=XM_023238806.1;refGene_gn=SYNJ1;refGene_tx

=XM_023238807.1;refGene_gn=SYNJ1;refGene_tx=XM_023238808.1
C2 12697630 12698422
refGene_gn=SYNJ1;refGene_tx=XM_023238799.1;refGene_gn=SYNJ1;refGene_tx=XM_023238800.1;refGene_gn=SYNJ1;refGene_tx=XM_023238801.1;refGene_gn=SYNJ1;refGene_tx=XM_023238802.1;refGene_gn=SYNJ1;refGene_tx=XM_023238803.1;refGene_gn=SYNJ1;refGene_tx=XM_023238804.1;refGene_gn=SYNJ1;refGene_tx=XM_023238805.1
C2 12705575 12705714
refGene_gn=CC2H21orf59;refGene_tx=XM_023238809.1
C2 12707472 12707502
refGene_gn=CC2H21orf59;refGene_tx=XM_023238809.1
C2 12707504 12707585
refGene_gn=CC2H21orf59;refGene_tx=XM_023238809.1
C2 12707587 12707638
refGene_gn=CC2H21orf59;refGene_tx=XM_023238809.1
C2 12709572 12709640
refGene_gn=CC2H21orf59;refGene_tx=XM_023238809.1
C2 12714532 12714571
refGene_gn=CC2H21orf59;refGene_tx=XM_023238809.1
C2 12714572 12714696
refGene_gn=CC2H21orf59;refGene_tx=XM_023238809.1
C2 12715548 12715676
refGene_gn=CC2H21orf59;refGene_tx=XM_023238809.1
C2 12716340 12716436
refGene_gn=CC2H21orf59;refGene_tx=XM_023238809.1
C2 12716739 12716850
refGene_gn=CC2H21orf59;refGene_tx=XM_023238809.1
C2 12752297 12752455 refGene_gn=EVA1C;refGene_tx=XM_023238810.1
C2 12752457 12752493 refGene_gn=EVA1C;refGene_tx=XM_023238810.1
C2 12752495 12752672 refGene_gn=EVA1C;refGene_tx=XM_023238810.1
C2 12760976 12761066 refGene_gn=EVA1C;refGene_tx=XM_023238810.1
C2 12762425 12762506 refGene_gn=EVA1C;refGene_tx=XM_023238810.1
C2 12769420 12769564 refGene_gn=EVA1C;refGene_tx=XM_023238810.1
C2 12781812 12781965 refGene_gn=EVA1C;refGene_tx=XM_023238810.1
C2 12790648 12790772 refGene_gn=EVA1C;refGene_tx=XM_023238810.1
C2 12793852 12794049 refGene_gn=EVA1C;refGene_tx=XM_023238810.1
C2 12817060 12817171 refGene_gn=EVA1C;refGene_tx=XM_023238810.1
C2 12830071 12830231 refGene_gn=EVA1C;refGene_tx=XM_023238810.1
C2 12846509 12846651
refGene_gn=URB1;refGene_tx=XM_011285748.3;refGene_gn=URB1;refGene_tx=XM_019839441.2
C2 12849601 12849741
refGene_gn=URB1;refGene_tx=XM_011285748.3;refGene_gn=URB1;refGene_tx=XM_019839441.2
C2 12850581 12850733
refGene_gn=URB1;refGene_tx=XM_011285748.3;refGene_gn=URB1;refGene_tx=XM_019839441.2
C2 12851697 12851830
refGene_gn=URB1;refGene_tx=XM_011285748.3;refGene_gn=URB1;refGene_tx=XM_019839441.2

C2 12856974 12857071
refGene_gn=URB1;refGene_tx=XM_011285748.3;refGene_gn=URB1;refGene_tx=XM_019839441.2
C2 12859310 12859396
refGene_gn=URB1;refGene_tx=XM_011285748.3;refGene_gn=URB1;refGene_tx=XM_019839441.2
C2 12860775 12860901
refGene_gn=URB1;refGene_tx=XM_011285748.3;refGene_gn=URB1;refGene_tx=XM_019839441.2
C2 12865581 12865706
refGene_gn=URB1;refGene_tx=XM_011285748.3;refGene_gn=URB1;refGene_tx=XM_019839441.2
C2 12869691 12869887
refGene_gn=URB1;refGene_tx=XM_011285748.3;refGene_gn=URB1;refGene_tx=XM_019839441.2
C2 12871139 12871277
refGene_gn=URB1;refGene_tx=XM_011285748.3;refGene_gn=URB1;refGene_tx=XM_019839441.2
C2 12873292 12873466
refGene_gn=URB1;refGene_tx=XM_011285748.3;refGene_gn=URB1;refGene_tx=XM_019839441.2
C2 12874082 12874212
refGene_gn=URB1;refGene_tx=XM_011285748.3;refGene_gn=URB1;refGene_tx=XM_019839441.2
C2 12874766 12874883
refGene_gn=URB1;refGene_tx=XM_011285748.3;refGene_gn=URB1;refGene_tx=XM_019839441.2
C2 12875972 12876085
refGene_gn=URB1;refGene_tx=XM_011285748.3;refGene_gn=URB1;refGene_tx=XM_019839441.2
C2 12879632 12879752
refGene_gn=URB1;refGene_tx=XM_011285748.3;refGene_gn=URB1;refGene_tx=XM_019839441.2
C2 12881181 12881298
refGene_gn=URB1;refGene_tx=XM_011285748.3;refGene_gn=URB1;refGene_tx=XM_019839441.2
C2 12881759 12881898
refGene_gn=URB1;refGene_tx=XM_011285748.3;refGene_gn=URB1;refGene_tx=XM_019839441.2
C2 12882678 12882849
refGene_gn=URB1;refGene_tx=XM_011285748.3;refGene_gn=URB1;refGene_tx=XM_019839441.2
C2 12883874 12884068
refGene_gn=URB1;refGene_tx=XM_011285748.3;refGene_gn=URB1;refGene_tx=XM_019839441.2
C2 12885895 12886114
refGene_gn=URB1;refGene_tx=XM_011285748.3;refGene_gn=URB1;refGene_tx=XM_019839441.2
C2 12886882 12887062
refGene_gn=URB1;refGene_tx=XM_011285748.3;refGene_gn=URB1;refGene_tx=X

M_019839441.2
C2	12888729	12889585
refGene_gn=URB1;refGene_tx=XM_011285748.3;refGene_gn=URB1;refGene_tx=XM_019839441.2
C2	12890287	12890489
refGene_gn=URB1;refGene_tx=XM_011285748.3;refGene_gn=URB1;refGene_tx=XM_019839441.2
C2	12891167	12891354
refGene_gn=URB1;refGene_tx=XM_011285748.3;refGene_gn=URB1;refGene_tx=XM_019839441.2
C2	12893919	12893978
refGene_gn=URB1;refGene_tx=XM_011285748.3;refGene_gn=URB1;refGene_tx=XM_019839441.2
C2	12896249	12896443
refGene_gn=URB1;refGene_tx=XM_011285748.3;refGene_gn=URB1;refGene_tx=XM_019839441.2
C2	12897453	12897564
refGene_gn=URB1;refGene_tx=XM_011285748.3;refGene_gn=URB1;refGene_tx=XM_019839441.2
C2	12897947	12898011
refGene_gn=URB1;refGene_tx=XM_011285748.3;refGene_gn=URB1;refGene_tx=XM_019839441.2
C2	12901502	12901674
refGene_gn=URB1;refGene_tx=XM_011285748.3;refGene_gn=URB1;refGene_tx=XM_019839441.2
C2	12902418	12902521
refGene_gn=URB1;refGene_tx=XM_011285748.3;refGene_gn=URB1;refGene_tx=XM_019839441.2
C2	12912192	12912353
refGene_gn=URB1;refGene_tx=XM_011285748.3;refGene_gn=URB1;refGene_tx=XM_019839441.2
C2	12912897	12913009
refGene_gn=URB1;refGene_tx=XM_011285748.3;refGene_gn=URB1;refGene_tx=XM_019839441.2
C2	12913935	12914042
refGene_gn=URB1;refGene_tx=XM_011285748.3;refGene_gn=URB1;refGene_tx=XM_019839441.2
C2	12914568	12914712
refGene_gn=URB1;refGene_tx=XM_011285748.3;refGene_gn=URB1;refGene_tx=XM_019839441.2
C2	12915817	12915927
refGene_gn=URB1;refGene_tx=XM_011285748.3;refGene_gn=URB1;refGene_tx=XM_019839441.2
C2	12916308	12916506
refGene_gn=URB1;refGene_tx=XM_011285748.3;refGene_gn=URB1;refGene_tx=XM_019839441.2
C2	12917680	12917922
refGene_gn=URB1;refGene_tx=XM_011285748.3;refGene_gn=URB1;refGene_tx=XM_019839441.2
C2	12918348	12918945

refGene_gn=URB1;refGene_tx=XM_011285748.3;refGene_gn=URB1;refGene_tx=XM_019839441.2
C2 12920010 12920192
refGene_gn=URB1;refGene_tx=XM_011285748.3;refGene_gn=URB1;refGene_tx=XM_019839441.2
C2 12921965 12922340
refGene_gn=MRAP;refGene_tx=XM_011285747.3;refGene_gn=MRAP;refGene_tx=XM_019839443.2
C2 12924672 12924772
refGene_gn=MRAP;refGene_tx=XM_011285747.3;refGene_gn=MRAP;refGene_tx=XM_019839443.2
C2 12931495 12931533 refGene_gn=MRAP;refGene_tx=XM_019839443.2
C2 12945464 12945570 refGene_gn=MRAP;refGene_tx=XM_011285747.3
C2 12963818 12964161
refGene_gn=MIS18A;refGene_tx=XM_019839446.2;refGene_gn=MIS18A;refGene_tx=XM_023238811.1;refGene_gn=MIS18A;refGene_tx=XM_023238812.1;refGene_gn=MIS18A;refGene_tx=XM_023238813.1
C2 12967129 12967196
refGene_gn=MIS18A;refGene_tx=XM_019839446.2;refGene_gn=MIS18A;refGene_tx=XM_023238811.1;refGene_gn=MIS18A;refGene_tx=XM_023238812.1;refGene_gn=MIS18A;refGene_tx=XM_023238813.1
C2 12971524 12971710
refGene_gn=MIS18A;refGene_tx=XM_019839446.2;refGene_gn=MIS18A;refGene_tx=XM_023238811.1;refGene_gn=MIS18A;refGene_tx=XM_023238812.1;refGene_gn=MIS18A;refGene_tx=XM_023238813.1
C2 12973909 12974014
refGene_gn=MIS18A;refGene_tx=XM_019839446.2;refGene_gn=MIS18A;refGene_tx=XM_023238811.1
C2 12974446 12974543
refGene_gn=MIS18A;refGene_tx=XM_019839446.2;refGene_gn=MIS18A;refGene_tx=XM_023238811.1;refGene_gn=MIS18A;refGene_tx=XM_023238812.1;refGene_gn=MIS18A;refGene_tx=XM_023238813.1
C2 12975325 12975406
refGene_gn=MIS18A;refGene_tx=XM_019839446.2;refGene_gn=MIS18A;refGene_tx=XM_023238811.1;refGene_gn=MIS18A;refGene_tx=XM_023238812.1;refGene_gn=MIS18A;refGene_tx=XM_023238813.1
C2 13174530 13175189 refGene_gn=HUNK;refGene_tx=XM_023238814.1
C2 13177951 13178132 refGene_gn=HUNK;refGene_tx=XM_023238814.1
C2 13182840 13182888 refGene_gn=HUNK;refGene_tx=XM_023238814.1
C2 13188024 13188108 refGene_gn=HUNK;refGene_tx=XM_023238814.1
C2 13192060 13192223 refGene_gn=HUNK;refGene_tx=XM_023238814.1
C2 13197648 13197784 refGene_gn=HUNK;refGene_tx=XM_023238814.1
C2 13204368 13204496 refGene_gn=HUNK;refGene_tx=XM_023238814.1
C2 13217519 13217655 refGene_gn=HUNK;refGene_tx=XM_023238814.1
C2 13223676 13223732 refGene_gn=HUNK;refGene_tx=XM_023238814.1
C2 13237158 13237451 refGene_gn=HUNK;refGene_tx=XM_023238814.1
C2 13283739 13284000 refGene_gn=HUNK;refGene_tx=XM_023238814.1
C2 13418237 13418267
refGene_gn=SCAF4;refGene_tx=XM_023238818.1;refGene_gn=SCAF4;refGene_tx=XM_023238819.1;refGene_gn=SCAF4;refGene_tx=XM_023238820.1

C2	13441243	13441327
refGene_gn=SCAF4;refGene_tx=XM_023238818.1;refGene_gn=SCAF4;refGene_tx
=XM_023238819.1;refGene_gn=SCAF4;refGene_tx=XM_023238820.1
C2	13442479	13442524
refGene_gn=SCAF4;refGene_tx=XM_023238818.1;refGene_gn=SCAF4;refGene_tx
=XM_023238819.1;refGene_gn=SCAF4;refGene_tx=XM_023238820.1
C2	13444023	13444185
refGene_gn=SCAF4;refGene_tx=XM_023238818.1;refGene_gn=SCAF4;refGene_tx
=XM_023238819.1;refGene_gn=SCAF4;refGene_tx=XM_023238820.1
C2	13445335	13445471
refGene_gn=SCAF4;refGene_tx=XM_023238818.1;refGene_gn=SCAF4;refGene_tx
=XM_023238819.1;refGene_gn=SCAF4;refGene_tx=XM_023238820.1
C2	13445778	13445918
refGene_gn=SCAF4;refGene_tx=XM_023238818.1;refGene_gn=SCAF4;refGene_tx
=XM_023238819.1;refGene_gn=SCAF4;refGene_tx=XM_023238820.1
C2	13446529	13446706
refGene_gn=SCAF4;refGene_tx=XM_023238818.1;refGene_gn=SCAF4;refGene_tx
=XM_023238819.1;refGene_gn=SCAF4;refGene_tx=XM_023238820.1
C2	13452229	13452426
refGene_gn=SCAF4;refGene_tx=XM_023238818.1;refGene_gn=SCAF4;refGene_tx
=XM_023238819.1;refGene_gn=SCAF4;refGene_tx=XM_023238820.1
C2	13453073	13453182
refGene_gn=SCAF4;refGene_tx=XM_023238818.1;refGene_gn=SCAF4;refGene_tx
=XM_023238819.1;refGene_gn=SCAF4;refGene_tx=XM_023238820.1
C2	13454792	13454957
refGene_gn=SCAF4;refGene_tx=XM_023238818.1;refGene_gn=SCAF4;refGene_tx
=XM_023238819.1;refGene_gn=SCAF4;refGene_tx=XM_023238820.1
C2	13455473	13455559
refGene_gn=SCAF4;refGene_tx=XM_023238818.1;refGene_gn=SCAF4;refGene_tx
=XM_023238819.1;refGene_gn=SCAF4;refGene_tx=XM_023238820.1
C2	13456258	13456449
refGene_gn=SCAF4;refGene_tx=XM_023238818.1;refGene_gn=SCAF4;refGene_tx
=XM_023238819.1;refGene_gn=SCAF4;refGene_tx=XM_023238820.1
C2	13457809	13457910
refGene_gn=SCAF4;refGene_tx=XM_023238818.1;refGene_gn=SCAF4;refGene_tx
=XM_023238819.1;refGene_gn=SCAF4;refGene_tx=XM_023238820.1
C2	13458354	13458468
refGene_gn=SCAF4;refGene_tx=XM_023238818.1;refGene_gn=SCAF4;refGene_tx
=XM_023238819.1;refGene_gn=SCAF4;refGene_tx=XM_023238820.1
C2	13459211	13459368
refGene_gn=SCAF4;refGene_tx=XM_023238818.1;refGene_gn=SCAF4;refGene_tx
=XM_023238819.1;refGene_gn=SCAF4;refGene_tx=XM_023238820.1
C2	13460802	13460960
refGene_gn=SCAF4;refGene_tx=XM_023238818.1;refGene_gn=SCAF4;refGene_tx
=XM_023238819.1;refGene_gn=SCAF4;refGene_tx=XM_023238820.1
C2	13462643	13462809
refGene_gn=SCAF4;refGene_tx=XM_023238818.1;refGene_gn=SCAF4;refGene_tx
=XM_023238819.1;refGene_gn=SCAF4;refGene_tx=XM_023238820.1
C2	13462896	13462983
refGene_gn=SCAF4;refGene_tx=XM_023238818.1;refGene_gn=SCAF4;refGene_tx

=XM_023238819.1;refGene_gn=SCAF4;refGene_tx=XM_023238820.1
C2 13463137 13463329
refGene_gn=SCAF4;refGene_tx=XM_023238818.1;refGene_gn=SCAF4;refGene_tx
=XM_023238819.1;refGene_gn=SCAF4;refGene_tx=XM_023238820.1
C2 13477142 13478074
refGene_gn=SCAF4;refGene_tx=XM_023238818.1;refGene_gn=SCAF4;refGene_tx
=XM_023238819.1;refGene_gn=SCAF4;refGene_tx=XM_023238820.1
C2 13480926 13481034 refGene_gn=SOD1;refGene_tx=XM_006935922.3
C2 13482068 13482186 refGene_gn=SOD1;refGene_tx=XM_006935922.3
C2 13483048 13483118 refGene_gn=SOD1;refGene_tx=XM_006935922.3
C2 13484545 13484639 refGene_gn=SOD1;refGene_tx=XM_006935922.3
C2 13489067 13489139 refGene_gn=SOD1;refGene_tx=XM_006935922.3
C2 13589457 13589576
refGene_gn=LOC105260899;refGene_tx=XM_023238821.1;refGene_gn=LOC105260
899;refGene_tx=XM_023238822.1
C2 13589876 13589916
refGene_gn=LOC105260899;refGene_tx=XM_023238821.1;refGene_gn=LOC105260
899;refGene_tx=XM_023238822.1
C2 13590342 13590664
refGene_gn=LOC105260899;refGene_tx=XM_023238821.1;refGene_gn=LOC105260
899;refGene_tx=XM_023238822.1
C2 13606211 13606291
refGene_gn=LOC105260899;refGene_tx=XM_023238821.1;refGene_gn=LOC105260
899;refGene_tx=XM_023238822.1
C2 13838501 13838555 refGene_gn=TIAM1;refGene_tx=XM_023238816.1
C2 13847501 13848464
refGene_gn=TIAM1;refGene_tx=XM_003991478.4;refGene_gn=TIAM1;refGene_tx
=XM_019839450.2;refGene_gn=TIAM1;refGene_tx=XM_019839453.2;refGene_gn=
TIAM1;refGene_tx=XM_019839454.2;refGene_gn=TIAM1;refGene_tx=XM_0232388
15.1
C2 13864809 13865257
refGene_gn=TIAM1;refGene_tx=XM_003991478.4;refGene_gn=TIAM1;refGene_tx
=XM_019839450.2;refGene_gn=TIAM1;refGene_tx=XM_019839453.2;refGene_gn=
TIAM1;refGene_tx=XM_019839454.2;refGene_gn=TIAM1;refGene_tx=XM_0232388
15.1;refGene_gn=TIAM1;refGene_tx=XM_023238816.1
C2 13870499 13870672
refGene_gn=TIAM1;refGene_tx=XM_003991478.4;refGene_gn=TIAM1;refGene_tx
=XM_019839450.2;refGene_gn=TIAM1;refGene_tx=XM_019839453.2;refGene_gn=
TIAM1;refGene_tx=XM_019839454.2;refGene_gn=TIAM1;refGene_tx=XM_0232388
15.1;refGene_gn=TIAM1;refGene_tx=XM_023238816.1
C2 13889101 13889326
refGene_gn=TIAM1;refGene_tx=XM_003991478.4;refGene_gn=TIAM1;refGene_tx
=XM_019839450.2;refGene_gn=TIAM1;refGene_tx=XM_019839453.2;refGene_gn=
TIAM1;refGene_tx=XM_019839454.2;refGene_gn=TIAM1;refGene_tx=XM_0232388
15.1;refGene_gn=TIAM1;refGene_tx=XM_023238816.1
C2 13891178 13891364
refGene_gn=TIAM1;refGene_tx=XM_003991478.4;refGene_gn=TIAM1;refGene_tx
=XM_019839450.2;refGene_gn=TIAM1;refGene_tx=XM_019839453.2;refGene_gn=
TIAM1;refGene_tx=XM_019839454.2;refGene_gn=TIAM1;refGene_tx=XM_0232388
15.1;refGene_gn=TIAM1;refGene_tx=XM_023238816.1

C2 13896853 13897000
refGene_gn=TIAM1;refGene_tx=XM_003991478.4;refGene_gn=TIAM1;refGene_tx=XM_019839450.2;refGene_gn=TIAM1;refGene_tx=XM_019839453.2;refGene_gn=TIAM1;refGene_tx=XM_019839454.2;refGene_gn=TIAM1;refGene_tx=XM_023238815.1;refGene_gn=TIAM1;refGene_tx=XM_023238816.1
C2 13899375 13899450
refGene_gn=TIAM1;refGene_tx=XM_003991478.4;refGene_gn=TIAM1;refGene_tx=XM_019839450.2;refGene_gn=TIAM1;refGene_tx=XM_019839453.2;refGene_gn=TIAM1;refGene_tx=XM_019839454.2;refGene_gn=TIAM1;refGene_tx=XM_023238816.1
C2 13902173 13902344
refGene_gn=TIAM1;refGene_tx=XM_003991478.4;refGene_gn=TIAM1;refGene_tx=XM_019839450.2;refGene_gn=TIAM1;refGene_tx=XM_019839453.2;refGene_gn=TIAM1;refGene_tx=XM_019839454.2;refGene_gn=TIAM1;refGene_tx=XM_023238815.1;refGene_gn=TIAM1;refGene_tx=XM_023238816.1
C2 13908984 13909089
refGene_gn=TIAM1;refGene_tx=XM_003991478.4;refGene_gn=TIAM1;refGene_tx=XM_019839450.2;refGene_gn=TIAM1;refGene_tx=XM_019839453.2;refGene_gn=TIAM1;refGene_tx=XM_019839454.2;refGene_gn=TIAM1;refGene_tx=XM_023238815.1;refGene_gn=TIAM1;refGene_tx=XM_023238816.1
C2 13916194 13916276
refGene_gn=TIAM1;refGene_tx=XM_003991478.4;refGene_gn=TIAM1;refGene_tx=XM_019839450.2;refGene_gn=TIAM1;refGene_tx=XM_019839453.2;refGene_gn=TIAM1;refGene_tx=XM_019839454.2;refGene_gn=TIAM1;refGene_tx=XM_023238815.1;refGene_gn=TIAM1;refGene_tx=XM_023238816.1
C2 13923273 13923360
refGene_gn=TIAM1;refGene_tx=XM_003991478.4;refGene_gn=TIAM1;refGene_tx=XM_019839450.2;refGene_gn=TIAM1;refGene_tx=XM_019839453.2;refGene_gn=TIAM1;refGene_tx=XM_019839454.2;refGene_gn=TIAM1;refGene_tx=XM_023238815.1;refGene_gn=TIAM1;refGene_tx=XM_023238816.1
C2 13928877 13929102
refGene_gn=TIAM1;refGene_tx=XM_003991478.4;refGene_gn=TIAM1;refGene_tx=XM_019839450.2;refGene_gn=TIAM1;refGene_tx=XM_019839453.2;refGene_gn=TIAM1;refGene_tx=XM_019839454.2;refGene_gn=TIAM1;refGene_tx=XM_023238815.1;refGene_gn=TIAM1;refGene_tx=XM_023238816.1
C2 13942653 13942757
refGene_gn=TIAM1;refGene_tx=XM_003991478.4;refGene_gn=TIAM1;refGene_tx=XM_019839450.2;refGene_gn=TIAM1;refGene_tx=XM_019839453.2;refGene_gn=TIAM1;refGene_tx=XM_019839454.2;refGene_gn=TIAM1;refGene_tx=XM_023238815.1;refGene_gn=TIAM1;refGene_tx=XM_023238816.1
C2 13947089 13947179 refGene_gn=TIAM1;refGene_tx=XM_023238817.1
C2 13952341 13952521
refGene_gn=TIAM1;refGene_tx=XM_003991478.4;refGene_gn=TIAM1;refGene_tx=XM_019839450.2;refGene_gn=TIAM1;refGene_tx=XM_019839453.2;refGene_gn=TIAM1;refGene_tx=XM_019839454.2;refGene_gn=TIAM1;refGene_tx=XM_023238815.1;refGene_gn=TIAM1;refGene_tx=XM_023238816.1;refGene_gn=TIAM1;refGene_tx=XM_023238817.1
C2 13953493 13953562
refGene_gn=TIAM1;refGene_tx=XM_003991478.4;refGene_gn=TIAM1;refGene_tx=XM_019839450.2;refGene_gn=TIAM1;refGene_tx=XM_019839453.2;refGene_gn=

TIAM1;refGene_tx=XM_019839454.2;refGene_gn=TIAM1;refGene_tx=XM_0232388
15.1;refGene_gn=TIAM1;refGene_tx=XM_023238816.1;refGene_gn=TIAM1;refGe
ne_tx=XM_023238817.1
C2 13953885 13954011
refGene_gn=TIAM1;refGene_tx=XM_003991478.4;refGene_gn=TIAM1;refGene_tx
=XM_019839450.2;refGene_gn=TIAM1;refGene_tx=XM_019839453.2;refGene_gn=
TIAM1;refGene_tx=XM_019839454.2;refGene_gn=TIAM1;refGene_tx=XM_0232388
15.1;refGene_gn=TIAM1;refGene_tx=XM_023238816.1;refGene_gn=TIAM1;refGe
ne_tx=XM_023238817.1
C2 13959845 13959954
refGene_gn=TIAM1;refGene_tx=XM_003991478.4;refGene_gn=TIAM1;refGene_tx
=XM_019839450.2;refGene_gn=TIAM1;refGene_tx=XM_019839453.2;refGene_gn=
TIAM1;refGene_tx=XM_019839454.2;refGene_gn=TIAM1;refGene_tx=XM_0232388
15.1;refGene_gn=TIAM1;refGene_tx=XM_023238816.1;refGene_gn=TIAM1;refGe
ne_tx=XM_023238817.1
C2 13964063 13964243
refGene_gn=TIAM1;refGene_tx=XM_003991478.4;refGene_gn=TIAM1;refGene_tx
=XM_019839450.2;refGene_gn=TIAM1;refGene_tx=XM_019839453.2;refGene_gn=
TIAM1;refGene_tx=XM_019839454.2;refGene_gn=TIAM1;refGene_tx=XM_0232388
15.1;refGene_gn=TIAM1;refGene_tx=XM_023238816.1;refGene_gn=TIAM1;refGe
ne_tx=XM_023238817.1
C2 13964340 13964459
refGene_gn=TIAM1;refGene_tx=XM_003991478.4;refGene_gn=TIAM1;refGene_tx
=XM_019839450.2;refGene_gn=TIAM1;refGene_tx=XM_019839453.2;refGene_gn=
TIAM1;refGene_tx=XM_019839454.2;refGene_gn=TIAM1;refGene_tx=XM_0232388
15.1;refGene_gn=TIAM1;refGene_tx=XM_023238816.1;refGene_gn=TIAM1;refGe
ne_tx=XM_023238817.1
C2 13970083 13970192
refGene_gn=TIAM1;refGene_tx=XM_003991478.4;refGene_gn=TIAM1;refGene_tx
=XM_019839450.2;refGene_gn=TIAM1;refGene_tx=XM_019839453.2;refGene_gn=
TIAM1;refGene_tx=XM_019839454.2;refGene_gn=TIAM1;refGene_tx=XM_0232388
15.1;refGene_gn=TIAM1;refGene_tx=XM_023238816.1;refGene_gn=TIAM1;refGe
ne_tx=XM_023238817.1
C2 13975466 13975525
refGene_gn=TIAM1;refGene_tx=XM_003991478.4;refGene_gn=TIAM1;refGene_tx
=XM_019839450.2;refGene_gn=TIAM1;refGene_tx=XM_019839453.2;refGene_gn=
TIAM1;refGene_tx=XM_019839454.2;refGene_gn=TIAM1;refGene_tx=XM_0232388
15.1;refGene_gn=TIAM1;refGene_tx=XM_023238816.1;refGene_gn=TIAM1;refGe
ne_tx=XM_023238817.1
C2 13976114 13976217
refGene_gn=TIAM1;refGene_tx=XM_003991478.4;refGene_gn=TIAM1;refGene_tx
=XM_019839450.2;refGene_gn=TIAM1;refGene_tx=XM_019839453.2;refGene_gn=
TIAM1;refGene_tx=XM_019839454.2;refGene_gn=TIAM1;refGene_tx=XM_0232388
15.1;refGene_gn=TIAM1;refGene_tx=XM_023238816.1;refGene_gn=TIAM1;refGe
ne_tx=XM_023238817.1
C2 13978289 13978377
refGene_gn=TIAM1;refGene_tx=XM_003991478.4;refGene_gn=TIAM1;refGene_tx
=XM_019839450.2;refGene_gn=TIAM1;refGene_tx=XM_019839453.2;refGene_gn=
TIAM1;refGene_tx=XM_019839454.2;refGene_gn=TIAM1;refGene_tx=XM_0232388
15.1;refGene_gn=TIAM1;refGene_tx=XM_023238816.1;refGene_gn=TIAM1;refGe

ne_tx=XM_023238817.1
C2 13980197 13980370
refGene_gn=TIAM1;refGene_tx=XM_003991478.4;refGene_gn=TIAM1;refGene_tx=XM_019839450.2;refGene_gn=TIAM1;refGene_tx=XM_019839453.2;refGene_gn=TIAM1;refGene_tx=XM_019839454.2;refGene_gn=TIAM1;refGene_tx=XM_023238815.1;refGene_gn=TIAM1;refGene_tx=XM_023238816.1;refGene_gn=TIAM1;refGene_tx=XM_023238817.1
C2 13983968 13984438
refGene_gn=TIAM1;refGene_tx=XM_003991478.4;refGene_gn=TIAM1;refGene_tx=XM_019839450.2;refGene_gn=TIAM1;refGene_tx=XM_019839453.2;refGene_gn=TIAM1;refGene_tx=XM_019839454.2;refGene_gn=TIAM1;refGene_tx=XM_023238815.1;refGene_gn=TIAM1;refGene_tx=XM_023238816.1;refGene_gn=TIAM1;refGene_tx=XM_023238817.1
C2 14175816 14176278
refGene_gn=LOC101094945;refGene_tx=XM_003991480.4
C2 14218612 14218876
refGene_gn=LOC101095189;refGene_tx=XM_006935932.2
C2 14236798 14236990
refGene_gn=LOC101095431;refGene_tx=XM_003991482.5
C2 14318646 14318671
refGene_gn=LOC111562339;refGene_tx=XM_023260845.1
C2 14319644 14319696
refGene_gn=LOC111562339;refGene_tx=XM_023260845.1
C2 14374725 14374823
refGene_gn=LOC111556324;refGene_tx=XM_023238567.1
C2 14375148 14375404
refGene_gn=LOC111556324;refGene_tx=XM_023238567.1
C2 14379840 14379914
refGene_gn=LOC111562340;refGene_tx=XM_023260846.1
C2 14380261 14380481
refGene_gn=LOC111562340;refGene_tx=XM_023260846.1
C2 14507866 14508118
refGene_gn=LOC109491684;refGene_tx=XM_019811144.2
C2 14522300 14522606
refGene_gn=LOC105260898;refGene_tx=XM_023238823.1
C2 14537645 14537692
refGene_gn=LOC105260890;refGene_tx=XM_023238568.1
C2 14542285 14542685
refGene_gn=LOC105260890;refGene_tx=XM_023238568.1
C2 14549451 14549598
refGene_gn=LOC111556359;refGene_tx=XM_023238824.1
C2 14554850 14555099
refGene_gn=LOC102899561;refGene_tx=XM_006935954.2
C2 14562223 14562472
refGene_gn=LOC111562341;refGene_tx=XM_023260847.1
C2 14572062 14572305
refGene_gn=LOC105260889;refGene_tx=XM_011285740.2
C2 14576052 14576199
refGene_gn=LOC111556335;refGene_tx=XM_023238574.1
C2 14584109 14584424

refGene_gn=LOC105260897;refGene_tx=XM_019839464.1
C2 14627255 14627405
refGene_gn=LOC111562342;refGene_tx=XM_023260848.1
C2 14652982 14653174
refGene_gn=LOC105260888;refGene_tx=XM_011285739.2
C2 14728669 14728861
refGene_gn=LOC111562343;refGene_tx=XM_023260849.1
C2 14733156 14733348
refGene_gn=LOC111562344;refGene_tx=XM_023260850.1
C2 14744209 14744401
refGene_gn=LOC105260887;refGene_tx=XM_011285738.3
C2 14775072 14775525
refGene_gn=LOC101080730;refGene_tx=XM_006935933.3
C2 14779020 14779524
refGene_gn=LOC101080990;refGene_tx=XM_006935934.2
C2 14786132 14786618
refGene_gn=LOC101081242;refGene_tx=XM_003991505.5
C2 14791488 14791920
refGene_gn=LOC101081486;refGene_tx=XM_023260851.1
C2 14808253 14808323
refGene_gn=LOC111562345;refGene_tx=XM_023260852.1
C2 14812625 14812741
refGene_gn=LOC111562345;refGene_tx=XM_023260852.1
C2 14817408 14817936
refGene_gn=LOC101082256;refGene_tx=XM_003991507.5
C2 14823548 14824034
refGene_gn=LOC101082509;refGene_tx=XM_003991508.5
C2 14844760 14845321
refGene_gn=LOC101082768;refGene_tx=XM_003991509.5
C2 14864356 14864887
refGene_gn=LOC101083025;refGene_tx=XM_023260854.1
C2 14872415 14873030
refGene_gn=LOC101083286;refGene_tx=XM_023260855.1
C2 14893636 14894188
refGene_gn=LOC101096407;refGene_tx=XM_003991483.3
C2 14930736 14931375
refGene_gn=LOC101083536;refGene_tx=XM_003991511.3
C2 14938283 14939039
refGene_gn=LOC101096663;refGene_tx=XM_003991484.3
C2 14999753 15000431 refGene_gn=CLDN8;refGene_tx=XM_019839465.2
C2 15049126 15049786 refGene_gn=CLDN17;refGene_tx=XM_003991512.2
C2 15263173 15263291
refGene_gn=GRIK1;refGene_tx=XM_019839466.2;refGene_gn=GRIK1;refGene_tx
=XM_023238825.1;refGene_gn=GRIK1;refGene_tx=XM_023238826.1;refGene_gn=
GRIK1;refGene_tx=XM_023238827.1;refGene_gn=GRIK1;refGene_tx=XM_0232388
28.1
C2 15494747 15494915
refGene_gn=GRIK1;refGene_tx=XM_019839466.2;refGene_gn=GRIK1;refGene_tx
=XM_023238825.1;refGene_gn=GRIK1;refGene_tx=XM_023238826.1;refGene_gn=
GRIK1;refGene_tx=XM_023238827.1;refGene_gn=GRIK1;refGene_tx=XM_0232388

28.1
C2 15498483 15498741
refGene_gn=GRIK1;refGene_tx=XM_019839466.2;refGene_gn=GRIK1;refGene_tx=XM_023238825.1;refGene_gn=GRIK1;refGene_tx=XM_023238826.1;refGene_gn=GRIK1;refGene_tx=XM_023238827.1;refGene_gn=GRIK1;refGene_tx=XM_023238828.1
C2 15511507 15511689
refGene_gn=GRIK1;refGene_tx=XM_019839466.2;refGene_gn=GRIK1;refGene_tx=XM_023238825.1;refGene_gn=GRIK1;refGene_tx=XM_023238826.1;refGene_gn=GRIK1;refGene_tx=XM_023238827.1;refGene_gn=GRIK1;refGene_tx=XM_023238828.1
C2 15528635 15528689
refGene_gn=GRIK1;refGene_tx=XM_019839466.2;refGene_gn=GRIK1;refGene_tx=XM_023238825.1;refGene_gn=GRIK1;refGene_tx=XM_023238826.1;refGene_gn=GRIK1;refGene_tx=XM_023238827.1;refGene_gn=GRIK1;refGene_tx=XM_023238828.1
C2 15531617 15531791
refGene_gn=GRIK1;refGene_tx=XM_019839466.2;refGene_gn=GRIK1;refGene_tx=XM_023238825.1;refGene_gn=GRIK1;refGene_tx=XM_023238826.1;refGene_gn=GRIK1;refGene_tx=XM_023238827.1;refGene_gn=GRIK1;refGene_tx=XM_023238828.1
C2 15540546 15540690
refGene_gn=GRIK1;refGene_tx=XM_019839466.2;refGene_gn=GRIK1;refGene_tx=XM_023238825.1;refGene_gn=GRIK1;refGene_tx=XM_023238826.1;refGene_gn=GRIK1;refGene_tx=XM_023238827.1;refGene_gn=GRIK1;refGene_tx=XM_023238828.1
C2 15579042 15579150
refGene_gn=GRIK1;refGene_tx=XM_019839466.2;refGene_gn=GRIK1;refGene_tx=XM_023238825.1;refGene_gn=GRIK1;refGene_tx=XM_023238826.1;refGene_gn=GRIK1;refGene_tx=XM_023238827.1;refGene_gn=GRIK1;refGene_tx=XM_023238828.1
C2 15581853 15581898
refGene_gn=GRIK1;refGene_tx=XM_019839466.2;refGene_gn=GRIK1;refGene_tx=XM_023238826.1;refGene_gn=GRIK1;refGene_tx=XM_023238827.1
C2 15587097 15587211
refGene_gn=GRIK1;refGene_tx=XM_019839466.2;refGene_gn=GRIK1;refGene_tx=XM_023238825.1;refGene_gn=GRIK1;refGene_tx=XM_023238826.1;refGene_gn=GRIK1;refGene_tx=XM_023238827.1;refGene_gn=GRIK1;refGene_tx=XM_023238828.1
C2 15588960 15589164
refGene_gn=GRIK1;refGene_tx=XM_019839466.2;refGene_gn=GRIK1;refGene_tx=XM_023238825.1;refGene_gn=GRIK1;refGene_tx=XM_023238826.1;refGene_gn=GRIK1;refGene_tx=XM_023238827.1;refGene_gn=GRIK1;refGene_tx=XM_023238828.1
C2 15589821 15590045
refGene_gn=GRIK1;refGene_tx=XM_019839466.2;refGene_gn=GRIK1;refGene_tx=XM_023238825.1;refGene_gn=GRIK1;refGene_tx=XM_023238826.1;refGene_gn=GRIK1;refGene_tx=XM_023238827.1;refGene_gn=GRIK1;refGene_tx=XM_023238828.1
C2 15595731 15595850

refGene_gn=GRIK1;refGene_tx=XM_019839466.2;refGene_gn=GRIK1;refGene_tx=XM_023238825.1;refGene_gn=GRIK1;refGene_tx=XM_023238826.1;refGene_gn=GRIK1;refGene_tx=XM_023238827.1;refGene_gn=GRIK1;refGene_tx=XM_023238828.1
C2 15599949 15600167 refGene_gn=GRIK1;refGene_tx=XM_019839466.2;refGene_gn=GRIK1;refGene_tx=XM_023238825.1;refGene_gn=GRIK1;refGene_tx=XM_023238826.1;refGene_gn=GRIK1;refGene_tx=XM_023238827.1;refGene_gn=GRIK1;refGene_tx=XM_023238828.1
C2 15612414 15612640 refGene_gn=GRIK1;refGene_tx=XM_019839466.2;refGene_gn=GRIK1;refGene_tx=XM_023238825.1;refGene_gn=GRIK1;refGene_tx=XM_023238826.1;refGene_gn=GRIK1;refGene_tx=XM_023238827.1;refGene_gn=GRIK1;refGene_tx=XM_023238828.1
C2 15618411 15618662 refGene_gn=GRIK1;refGene_tx=XM_019839466.2;refGene_gn=GRIK1;refGene_tx=XM_023238825.1;refGene_gn=GRIK1;refGene_tx=XM_023238826.1;refGene_gn=GRIK1;refGene_tx=XM_023238827.1;refGene_gn=GRIK1;refGene_tx=XM_023238828.1
C2 15620016 15620166 refGene_gn=GRIK1;refGene_tx=XM_023238827.1
C2 15636160 15636247 refGene_gn=GRIK1;refGene_tx=XM_019839466.2;refGene_gn=GRIK1;refGene_tx=XM_023238825.1
C2 15636645 15636801 refGene_gn=GRIK1;refGene_tx=XM_019839466.2;refGene_gn=GRIK1;refGene_tx=XM_023238825.1;refGene_gn=GRIK1;refGene_tx=XM_023238826.1;refGene_gn=GRIK1;refGene_tx=XM_023238828.1
C2 15820080 15820536 refGene_gn=BACH1;refGene_tx=XM_019839467.2;refGene_gn=BACH1;refGene_tx=XM_023238829.1
C2 15823994 15824045 refGene_gn=BACH1;refGene_tx=XM_023238831.1
C2 15824334 15824415 refGene_gn=BACH1;refGene_tx=XM_023238830.1
C2 15831364 15831571 refGene_gn=BACH1;refGene_tx=XM_019839467.2;refGene_gn=BACH1;refGene_tx=XM_023238829.1;refGene_gn=BACH1;refGene_tx=XM_023238830.1;refGene_gn=BACH1;refGene_tx=XM_023238831.1
C2 15833254 15834604 refGene_gn=BACH1;refGene_tx=XM_019839467.2;refGene_gn=BACH1;refGene_tx=XM_023238829.1;refGene_gn=BACH1;refGene_tx=XM_023238830.1;refGene_gn=BACH1;refGene_tx=XM_023238831.1
C2 15836123 15836357 refGene_gn=BACH1;refGene_tx=XM_019839467.2;refGene_gn=BACH1;refGene_tx=XM_023238829.1;refGene_gn=BACH1;refGene_tx=XM_023238830.1;refGene_gn=BACH1;refGene_tx=XM_023238831.1
C2 15977776 15977957 refGene_gn=MAP3K7CL;refGene_tx=XM_003991488.5;refGene_gn=MAP3K7CL;refGene_tx=XM_006935936.3;refGene_gn=MAP3K7CL;refGene_tx=XM_019839472.2
C2 15989745 15989861 refGene_gn=MAP3K7CL;refGene_tx=XM_003991488.5;refGene_gn=MAP3K7CL;refGene_tx=XM_006935936.3;refGene_gn=MAP3K7CL;refGene_tx=XM_019839472.2

C2	15999743	15999805
refGene_gn=MAP3K7CL;refGene_tx=XM_003991488.5;refGene_gn=MAP3K7CL;refGene_tx=XM_006935936.3
C2	16014072	16014181
refGene_gn=MAP3K7CL;refGene_tx=XM_003991488.5;refGene_gn=MAP3K7CL;refGene_tx=XM_006935936.3
C2	16015670	16015706	refGene_gn=MAP3K7CL;refGene_tx=XM_006935936.3
C2	16076750	16076810
refGene_gn=CCT8;refGene_tx=XM_003991489.5;refGene_gn=CCT8;refGene_tx=XM_006935938.3
C2	16079919	16080010
refGene_gn=CCT8;refGene_tx=XM_003991489.5;refGene_gn=CCT8;refGene_tx=XM_006935938.3
C2	16080847	16080927
refGene_gn=CCT8;refGene_tx=XM_003991489.5;refGene_gn=CCT8;refGene_tx=XM_006935938.3
C2	16083005	16083155
refGene_gn=CCT8;refGene_tx=XM_003991489.5;refGene_gn=CCT8;refGene_tx=XM_006935938.3
C2	16083613	16083794
refGene_gn=CCT8;refGene_tx=XM_003991489.5;refGene_gn=CCT8;refGene_tx=XM_006935938.3
C2	16083907	16083969
refGene_gn=CCT8;refGene_tx=XM_003991489.5;refGene_gn=CCT8;refGene_tx=XM_006935938.3
C2	16084848	16084986
refGene_gn=CCT8;refGene_tx=XM_003991489.5;refGene_gn=CCT8;refGene_tx=XM_006935938.3
C2	16086361	16086540
refGene_gn=CCT8;refGene_tx=XM_003991489.5;refGene_gn=CCT8;refGene_tx=XM_006935938.3
C2	16087254	16087321
refGene_gn=CCT8;refGene_tx=XM_003991489.5;refGene_gn=CCT8;refGene_tx=XM_006935938.3
C2	16087399	16087487
refGene_gn=CCT8;refGene_tx=XM_003991489.5;refGene_gn=CCT8;refGene_tx=XM_006935938.3
C2	16087568	16087684
refGene_gn=CCT8;refGene_tx=XM_003991489.5;refGene_gn=CCT8;refGene_tx=XM_006935938.3
C2	16088037	16088109
refGene_gn=CCT8;refGene_tx=XM_003991489.5;refGene_gn=CCT8;refGene_tx=XM_006935938.3
C2	16088187	16088352
refGene_gn=CCT8;refGene_tx=XM_003991489.5;refGene_gn=CCT8;refGene_tx=XM_006935938.3
C2	16088655	16088775
refGene_gn=CCT8;refGene_tx=XM_003991489.5;refGene_gn=CCT8;refGene_tx=XM_006935938.3
C2	16091316	16091409	refGene_gn=CCT8;refGene_tx=XM_006935938.3

C2	16091742	16091820	refGene_gn=CCT8;refGene_tx=XM_003991489.5
C2	16093812	16093937
refGene_gn=USP16;refGene_tx=XM_003991492.4;refGene_gn=USP16;refGene_tx=XM_011285733.2;refGene_gn=USP16;refGene_tx=XM_011285734.2;refGene_gn=USP16;refGene_tx=XM_011285735.2;refGene_gn=USP16;refGene_tx=XM_019839474.2;refGene_gn=USP16;refGene_tx=XM_019839475.1;refGene_gn=USP16;refGene_tx=XM_019839477.2;refGene_gn=USP16;refGene_tx=XM_023238832.1
C2	16094026	16094183
refGene_gn=USP16;refGene_tx=XM_003991492.4;refGene_gn=USP16;refGene_tx=XM_011285733.2;refGene_gn=USP16;refGene_tx=XM_011285734.2;refGene_gn=USP16;refGene_tx=XM_011285735.2;refGene_gn=USP16;refGene_tx=XM_019839474.2;refGene_gn=USP16;refGene_tx=XM_019839475.1;refGene_gn=USP16;refGene_tx=XM_019839477.2;refGene_gn=USP16;refGene_tx=XM_023238832.1
C2	16098403	16098490
refGene_gn=USP16;refGene_tx=XM_003991492.4;refGene_gn=USP16;refGene_tx=XM_011285733.2;refGene_gn=USP16;refGene_tx=XM_011285734.2;refGene_gn=USP16;refGene_tx=XM_011285735.2;refGene_gn=USP16;refGene_tx=XM_019839474.2;refGene_gn=USP16;refGene_tx=XM_019839475.1;refGene_gn=USP16;refGene_tx=XM_019839477.2;refGene_gn=USP16;refGene_tx=XM_023238832.1
C2	16099634	16099729
refGene_gn=USP16;refGene_tx=XM_003991492.4;refGene_gn=USP16;refGene_tx=XM_011285733.2;refGene_gn=USP16;refGene_tx=XM_011285734.2;refGene_gn=USP16;refGene_tx=XM_011285735.2;refGene_gn=USP16;refGene_tx=XM_019839474.2;refGene_gn=USP16;refGene_tx=XM_019839475.1;refGene_gn=USP16;refGene_tx=XM_019839477.2;refGene_gn=USP16;refGene_tx=XM_023238832.1
C2	16100813	16101477
refGene_gn=USP16;refGene_tx=XM_003991492.4;refGene_gn=USP16;refGene_tx=XM_011285733.2;refGene_gn=USP16;refGene_tx=XM_011285734.2;refGene_gn=USP16;refGene_tx=XM_011285735.2;refGene_gn=USP16;refGene_tx=XM_019839474.2;refGene_gn=USP16;refGene_tx=XM_019839475.1;refGene_gn=USP16;refGene_tx=XM_019839477.2;refGene_gn=USP16;refGene_tx=XM_023238832.1
C2	16103108	16103285
refGene_gn=USP16;refGene_tx=XM_003991492.4;refGene_gn=USP16;refGene_tx=XM_011285733.2;refGene_gn=USP16;refGene_tx=XM_011285734.2;refGene_gn=USP16;refGene_tx=XM_011285735.2;refGene_gn=USP16;refGene_tx=XM_019839474.2;refGene_gn=USP16;refGene_tx=XM_019839475.1;refGene_gn=USP16;refGene_tx=XM_019839477.2;refGene_gn=USP16;refGene_tx=XM_023238832.1
C2	16104190	16104247
refGene_gn=USP16;refGene_tx=XM_003991492.4;refGene_gn=USP16;refGene_tx=XM_011285733.2;refGene_gn=USP16;refGene_tx=XM_011285734.2;refGene_gn=USP16;refGene_tx=XM_011285735.2;refGene_gn=USP16;refGene_tx=XM_019839474.2;refGene_gn=USP16;refGene_tx=XM_019839477.2;refGene_gn=USP16;refGene_tx=XM_023238832.1
C2	16104611	16104703
refGene_gn=USP16;refGene_tx=XM_003991492.4;refGene_gn=USP16;refGene_tx=XM_011285733.2;refGene_gn=USP16;refGene_tx=XM_011285734.2;refGene_gn=USP16;refGene_tx=XM_011285735.2;refGene_gn=USP16;refGene_tx=XM_019839474.2;refGene_gn=USP16;refGene_tx=XM_019839475.1;refGene_gn=USP16;refGene_tx=XM_019839477.2;refGene_gn=USP16;refGene_tx=XM_023238832.1
C2	16106487	16106566

refGene_gn=USP16;refGene_tx=XM_003991492.4;refGene_gn=USP16;refGene_tx=XM_011285733.2;refGene_gn=USP16;refGene_tx=XM_011285734.2;refGene_gn=USP16;refGene_tx=XM_011285735.2;refGene_gn=USP16;refGene_tx=XM_019839474.2;refGene_gn=USP16;refGene_tx=XM_019839475.1;refGene_gn=USP16;refGene_tx=XM_019839477.2;refGene_gn=USP16;refGene_tx=XM_023238832.1
C2 16107641 16107729
refGene_gn=USP16;refGene_tx=XM_003991492.4;refGene_gn=USP16;refGene_tx=XM_011285733.2;refGene_gn=USP16;refGene_tx=XM_011285734.2;refGene_gn=USP16;refGene_tx=XM_011285735.2;refGene_gn=USP16;refGene_tx=XM_019839474.2;refGene_gn=USP16;refGene_tx=XM_019839475.1;refGene_gn=USP16;refGene_tx=XM_019839477.2;refGene_gn=USP16;refGene_tx=XM_023238832.1
C2 16108062 16108193
refGene_gn=USP16;refGene_tx=XM_003991492.4;refGene_gn=USP16;refGene_tx=XM_011285733.2;refGene_gn=USP16;refGene_tx=XM_011285734.2;refGene_gn=USP16;refGene_tx=XM_011285735.2;refGene_gn=USP16;refGene_tx=XM_019839474.2;refGene_gn=USP16;refGene_tx=XM_019839475.1;refGene_gn=USP16;refGene_tx=XM_019839477.2;refGene_gn=USP16;refGene_tx=XM_023238832.1
C2 16109454 16109550
refGene_gn=USP16;refGene_tx=XM_003991492.4;refGene_gn=USP16;refGene_tx=XM_011285733.2;refGene_gn=USP16;refGene_tx=XM_011285734.2;refGene_gn=USP16;refGene_tx=XM_011285735.2;refGene_gn=USP16;refGene_tx=XM_019839474.2;refGene_gn=USP16;refGene_tx=XM_019839475.1;refGene_gn=USP16;refGene_tx=XM_019839477.2;refGene_gn=USP16;refGene_tx=XM_023238832.1
C2 16110328 16110513
refGene_gn=USP16;refGene_tx=XM_003991492.4;refGene_gn=USP16;refGene_tx=XM_011285733.2;refGene_gn=USP16;refGene_tx=XM_011285734.2;refGene_gn=USP16;refGene_tx=XM_011285735.2;refGene_gn=USP16;refGene_tx=XM_019839474.2;refGene_gn=USP16;refGene_tx=XM_019839475.1;refGene_gn=USP16;refGene_tx=XM_019839477.2;refGene_gn=USP16;refGene_tx=XM_023238832.1
C2 16111435 16111539
refGene_gn=USP16;refGene_tx=XM_003991492.4;refGene_gn=USP16;refGene_tx=XM_011285733.2;refGene_gn=USP16;refGene_tx=XM_011285734.2;refGene_gn=USP16;refGene_tx=XM_011285735.2;refGene_gn=USP16;refGene_tx=XM_019839474.2;refGene_gn=USP16;refGene_tx=XM_019839475.1;refGene_gn=USP16;refGene_tx=XM_019839477.2;refGene_gn=USP16;refGene_tx=XM_023238832.1
C2 16112353 16112457
refGene_gn=USP16;refGene_tx=XM_003991492.4;refGene_gn=USP16;refGene_tx=XM_011285733.2;refGene_gn=USP16;refGene_tx=XM_011285734.2;refGene_gn=USP16;refGene_tx=XM_011285735.2;refGene_gn=USP16;refGene_tx=XM_019839474.2;refGene_gn=USP16;refGene_tx=XM_019839475.1;refGene_gn=USP16;refGene_tx=XM_019839477.2;refGene_gn=USP16;refGene_tx=XM_023238832.1
C2 16115354 16115533
refGene_gn=USP16;refGene_tx=XM_003991492.4;refGene_gn=USP16;refGene_tx=XM_011285733.2;refGene_gn=USP16;refGene_tx=XM_011285734.2;refGene_gn=USP16;refGene_tx=XM_011285735.2;refGene_gn=USP16;refGene_tx=XM_019839474.2;refGene_gn=USP16;refGene_tx=XM_019839475.1;refGene_gn=USP16;refGene_tx=XM_023238832.1
C2 16117474 16117535
refGene_gn=USP16;refGene_tx=XM_003991492.4;refGene_gn=USP16;refGene_tx=XM_011285733.2;refGene_gn=USP16;refGene_tx=XM_011285734.2;refGene_gn=

USP16;refGene_tx=XM_019839474.2;refGene_gn=USP16;refGene_tx=XM_0198394
75.1;refGene_gn=USP16;refGene_tx=XM_023238832.1
C2 16120788 16120807 refGene_gn=USP16;refGene_tx=XM_011285735.2
C2 16133671 16133878
refGene_gn=RWDD2B;refGene_tx=XM_003991493.3;refGene_gn=RWDD2B;refGene_
tx=XM_011285736.2
C2 16133960 16134028
refGene_gn=RWDD2B;refGene_tx=XM_003991493.3;refGene_gn=RWDD2B;refGene_
tx=XM_011285736.2
C2 16134152 16134515
refGene_gn=RWDD2B;refGene_tx=XM_003991493.3;refGene_gn=RWDD2B;refGene_
tx=XM_011285736.2
C2 16135122 16135357
refGene_gn=RWDD2B;refGene_tx=XM_003991493.3;refGene_gn=RWDD2B;refGene_
tx=XM_011285736.2
C2 16147040 16147074
refGene_gn=LTN1;refGene_tx=XM_011285732.2;refGene_gn=LTN1;refGene_tx=X
M_023238833.1
C2 16147155 16147268
refGene_gn=LTN1;refGene_tx=XM_011285732.2;refGene_gn=LTN1;refGene_tx=X
M_019839473.2;refGene_gn=LTN1;refGene_tx=XM_023238833.1
C2 16152105 16152309
refGene_gn=LTN1;refGene_tx=XM_011285732.2;refGene_gn=LTN1;refGene_tx=X
M_019839473.2;refGene_gn=LTN1;refGene_tx=XM_023238833.1
C2 16152815 16152914
refGene_gn=LTN1;refGene_tx=XM_011285732.2;refGene_gn=LTN1;refGene_tx=X
M_019839473.2;refGene_gn=LTN1;refGene_tx=XM_023238833.1
C2 16153421 16153652
refGene_gn=LTN1;refGene_tx=XM_011285732.2;refGene_gn=LTN1;refGene_tx=X
M_019839473.2;refGene_gn=LTN1;refGene_tx=XM_023238833.1
C2 16156028 16156081
refGene_gn=LTN1;refGene_tx=XM_011285732.2;refGene_gn=LTN1;refGene_tx=X
M_019839473.2;refGene_gn=LTN1;refGene_tx=XM_023238833.1
C2 16156794 16156975
refGene_gn=LTN1;refGene_tx=XM_011285732.2;refGene_gn=LTN1;refGene_tx=X
M_019839473.2;refGene_gn=LTN1;refGene_tx=XM_023238833.1
C2 16164290 16164464
refGene_gn=LTN1;refGene_tx=XM_011285732.2;refGene_gn=LTN1;refGene_tx=X
M_019839473.2;refGene_gn=LTN1;refGene_tx=XM_023238833.1
C2 16165033 16165224
refGene_gn=LTN1;refGene_tx=XM_011285732.2;refGene_gn=LTN1;refGene_tx=X
M_019839473.2;refGene_gn=LTN1;refGene_tx=XM_023238833.1
C2 16165969 16166105
refGene_gn=LTN1;refGene_tx=XM_011285732.2;refGene_gn=LTN1;refGene_tx=X
M_019839473.2;refGene_gn=LTN1;refGene_tx=XM_023238833.1
C2 16167171 16167981
refGene_gn=LTN1;refGene_tx=XM_011285732.2;refGene_gn=LTN1;refGene_tx=X
M_019839473.2;refGene_gn=LTN1;refGene_tx=XM_023238833.1
C2 16169974 16170016
refGene_gn=LTN1;refGene_tx=XM_011285732.2;refGene_gn=LTN1;refGene_tx=X

M_019839473.2;refGene_gn=LTN1;refGene_tx=XM_023238833.1
C2 16172534 16172724
refGene_gn=LTN1;refGene_tx=XM_011285732.2;refGene_gn=LTN1;refGene_tx=XM_019839473.2;refGene_gn=LTN1;refGene_tx=XM_023238833.1
C2 16173212 16173452
refGene_gn=LTN1;refGene_tx=XM_011285732.2;refGene_gn=LTN1;refGene_tx=XM_019839473.2;refGene_gn=LTN1;refGene_tx=XM_023238833.1
C2 16174261 16174415
refGene_gn=LTN1;refGene_tx=XM_011285732.2;refGene_gn=LTN1;refGene_tx=XM_019839473.2;refGene_gn=LTN1;refGene_tx=XM_023238833.1
C2 16175187 16175332
refGene_gn=LTN1;refGene_tx=XM_011285732.2;refGene_gn=LTN1;refGene_tx=XM_019839473.2;refGene_gn=LTN1;refGene_tx=XM_023238833.1
C2 16175827 16176014
refGene_gn=LTN1;refGene_tx=XM_011285732.2;refGene_gn=LTN1;refGene_tx=XM_019839473.2;refGene_gn=LTN1;refGene_tx=XM_023238833.1
C2 16177500 16177660
refGene_gn=LTN1;refGene_tx=XM_011285732.2;refGene_gn=LTN1;refGene_tx=XM_019839473.2;refGene_gn=LTN1;refGene_tx=XM_023238833.1
C2 16178527 16178632
refGene_gn=LTN1;refGene_tx=XM_011285732.2;refGene_gn=LTN1;refGene_tx=XM_019839473.2;refGene_gn=LTN1;refGene_tx=XM_023238833.1
C2 16181733 16181876
refGene_gn=LTN1;refGene_tx=XM_011285732.2;refGene_gn=LTN1;refGene_tx=XM_019839473.2;refGene_gn=LTN1;refGene_tx=XM_023238833.1
C2 16194013 16194149
refGene_gn=LTN1;refGene_tx=XM_011285732.2;refGene_gn=LTN1;refGene_tx=XM_019839473.2;refGene_gn=LTN1;refGene_tx=XM_023238833.1
C2 16194352 16194497
refGene_gn=LTN1;refGene_tx=XM_011285732.2;refGene_gn=LTN1;refGene_tx=XM_019839473.2;refGene_gn=LTN1;refGene_tx=XM_023238833.1
C2 16195366 16195580
refGene_gn=LTN1;refGene_tx=XM_011285732.2;refGene_gn=LTN1;refGene_tx=XM_019839473.2;refGene_gn=LTN1;refGene_tx=XM_023238833.1
C2 16196441 16196679
refGene_gn=LTN1;refGene_tx=XM_011285732.2;refGene_gn=LTN1;refGene_tx=XM_019839473.2;refGene_gn=LTN1;refGene_tx=XM_023238833.1
C2 16197014 16197089
refGene_gn=LTN1;refGene_tx=XM_011285732.2;refGene_gn=LTN1;refGene_tx=XM_019839473.2;refGene_gn=LTN1;refGene_tx=XM_023238833.1
C2 16199362 16199549
refGene_gn=LTN1;refGene_tx=XM_011285732.2;refGene_gn=LTN1;refGene_tx=XM_019839473.2;refGene_gn=LTN1;refGene_tx=XM_023238833.1
C2 16202668 16202840
refGene_gn=LTN1;refGene_tx=XM_011285732.2;refGene_gn=LTN1;refGene_tx=XM_019839473.2;refGene_gn=LTN1;refGene_tx=XM_023238833.1
C2 16203908 16204129
refGene_gn=LTN1;refGene_tx=XM_011285732.2;refGene_gn=LTN1;refGene_tx=XM_019839473.2
C2 16205505 16205700

refGene_gn=LTN1;refGene_tx=XM_011285732.2;refGene_gn=LTN1;refGene_tx=XM_019839473.2;refGene_gn=LTN1;refGene_tx=XM_023238833.1
C2 16206389 16206557
refGene_gn=LTN1;refGene_tx=XM_011285732.2;refGene_gn=LTN1;refGene_tx=XM_019839473.2
C2 16207050 16207113
refGene_gn=LTN1;refGene_tx=XM_011285732.2;refGene_gn=LTN1;refGene_tx=XM_019839473.2
C2 16246802 16246921
refGene_gn=N6AMT1;refGene_tx=XM_006935939.3;refGene_gn=N6AMT1;refGene_tx=XM_019839479.2;refGene_gn=N6AMT1;refGene_tx=XM_019839480.2
C2 16247961 16248103
refGene_gn=N6AMT1;refGene_tx=XM_006935939.3;refGene_gn=N6AMT1;refGene_tx=XM_019839479.2;refGene_gn=N6AMT1;refGene_tx=XM_019839480.2
C2 16250864 16250948
refGene_gn=N6AMT1;refGene_tx=XM_006935939.3;refGene_gn=N6AMT1;refGene_tx=XM_019839479.2;refGene_gn=N6AMT1;refGene_tx=XM_019839480.2
C2 16255411 16255502
refGene_gn=N6AMT1;refGene_tx=XM_006935939.3;refGene_gn=N6AMT1;refGene_tx=XM_019839479.2;refGene_gn=N6AMT1;refGene_tx=XM_019839480.2
C2 16257243 16257330
refGene_gn=N6AMT1;refGene_tx=XM_006935939.3;refGene_gn=N6AMT1;refGene_tx=XM_019839479.2;refGene_gn=N6AMT1;refGene_tx=XM_019839480.2
C2 16259285 16259419
refGene_gn=N6AMT1;refGene_tx=XM_006935939.3;refGene_gn=N6AMT1;refGene_tx=XM_019839479.2;refGene_gn=N6AMT1;refGene_tx=XM_019839480.2
C2 17942573 17943677
refGene_gn=ADAMTS5;refGene_tx=XM_023238834.1;refGene_gn=ADAMTS5;refGene_tx=XM_023238835.1
C2 17955510 17955643
refGene_gn=ADAMTS5;refGene_tx=XM_023238834.1;refGene_gn=ADAMTS5;refGene_tx=XM_023238835.1
C2 17967585 17967753
refGene_gn=ADAMTS5;refGene_tx=XM_023238834.1;refGene_gn=ADAMTS5;refGene_tx=XM_023238835.1
C2 17977144 17977428
refGene_gn=ADAMTS5;refGene_tx=XM_023238834.1;refGene_gn=ADAMTS5;refGene_tx=XM_023238835.1
C2 17978444 17978628
refGene_gn=ADAMTS5;refGene_tx=XM_023238834.1;refGene_gn=ADAMTS5;refGene_tx=XM_023238835.1
C2 17978999 17979175
refGene_gn=ADAMTS5;refGene_tx=XM_023238834.1;refGene_gn=ADAMTS5;refGene_tx=XM_023238835.1
C2 17981501 17981677
refGene_gn=ADAMTS5;refGene_tx=XM_023238834.1;refGene_gn=ADAMTS5;refGene_tx=XM_023238835.1
C2 17985826 17986394
refGene_gn=ADAMTS5;refGene_tx=XM_023238834.1;refGene_gn=ADAMTS5;refGene_tx=XM_023238835.1

C2 18071813 18072290 refGene_gn=ADAMTS1;refGene_tx=XM_019839482.2
C2 18072292 18072503 refGene_gn=ADAMTS1;refGene_tx=XM_019839482.2
C2 18074159 18074506 refGene_gn=ADAMTS1;refGene_tx=XM_019839482.2
C2 18074852 18074985 refGene_gn=ADAMTS1;refGene_tx=XM_019839482.2
C2 18075269 18075437 refGene_gn=ADAMTS1;refGene_tx=XM_019839482.2
C2 18076120 18076407 refGene_gn=ADAMTS1;refGene_tx=XM_019839482.2
C2 18076658 18076845 refGene_gn=ADAMTS1;refGene_tx=XM_019839482.2
C2 18076935 18077111 refGene_gn=ADAMTS1;refGene_tx=XM_019839482.2
C2 18078040 18078216 refGene_gn=ADAMTS1;refGene_tx=XM_019839482.2
C2 18078378 18079078 refGene_gn=ADAMTS1;refGene_tx=XM_019839482.2
C2 18299449 18299494 refGene_gn=CYYR1;refGene_tx=XM_003991494.5
C2 18299495 18299835 refGene_gn=CYYR1;refGene_tx=XM_003991494.5
C2 18305119 18305222 refGene_gn=CYYR1;refGene_tx=XM_003991494.5
C2 18390578 18390736 refGene_gn=CYYR1;refGene_tx=XM_003991494.5
C2 18403337 18403522 refGene_gn=CYYR1;refGene_tx=XM_003991494.5
C2 18421683 18421787
refGene_gn=LOC111562223;refGene_tx=XM_023260154.1
C2 18565598 18565636
refGene_gn=LOC111562223;refGene_tx=XM_023260154.1
C2 18656785 18656980
refGene_gn=LOC111562223;refGene_tx=XM_023260154.1
C2 18661686 18661870
refGene_gn=LOC111562223;refGene_tx=XM_023260154.1
C2 18661923 18662017
refGene_gn=LOC111562223;refGene_tx=XM_023260154.1
C2 18662109 18662166
refGene_gn=APP;refGene_tx=XM_023238836.1;refGene_gn=APP;refGene_tx=XM_023238837.1;refGene_gn=APP;refGene_tx=XM_023238838.1;refGene_gn=APP;refGene_tx=XM_023238839.1;refGene_gn=APP;refGene_tx=XM_023238840.1;refGene_gn=APP;refGene_tx=XM_023238841.1;refGene_gn=APP;refGene_tx=XM_023238842.1
C2 18724586 18724754
refGene_gn=APP;refGene_tx=XM_023238836.1;refGene_gn=APP;refGene_tx=XM_023238837.1;refGene_gn=APP;refGene_tx=XM_023238838.1;refGene_gn=APP;refGene_tx=XM_023238839.1;refGene_gn=APP;refGene_tx=XM_023238840.1;refGene_gn=APP;refGene_tx=XM_023238841.1;refGene_gn=APP;refGene_tx=XM_023238842.1
C2 18748858 18748988
refGene_gn=APP;refGene_tx=XM_023238836.1;refGene_gn=APP;refGene_tx=XM_023238837.1;refGene_gn=APP;refGene_tx=XM_023238838.1;refGene_gn=APP;refGene_tx=XM_023238839.1;refGene_gn=APP;refGene_tx=XM_023238840.1;refGene_gn=APP;refGene_tx=XM_023238841.1;refGene_gn=APP;refGene_tx=XM_023238842.1
C2 18780227 18780340
refGene_gn=APP;refGene_tx=XM_023238836.1;refGene_gn=APP;refGene_tx=XM_023238837.1;refGene_gn=APP;refGene_tx=XM_023238838.1;refGene_gn=APP;refGene_tx=XM_023238839.1;refGene_gn=APP;refGene_tx=XM_023238840.1;refGene_gn=APP;refGene_tx=XM_023238841.1;refGene_gn=APP;refGene_tx=XM_023238842.1
C2 18782714 18782908

refGene_gn=APP;refGene_tx=XM_023238836.1;refGene_gn=APP;refGene_tx=XM_023238837.1;refGene_gn=APP;refGene_tx=XM_023238838.1;refGene_gn=APP;refGene_tx=XM_023238839.1;refGene_gn=APP;refGene_tx=XM_023238840.1;refGene_gn=APP;refGene_tx=XM_023238841.1;refGene_gn=APP;refGene_tx=XM_023238842.1
C2 18819366 18819569
refGene_gn=APP;refGene_tx=XM_023238836.1;refGene_gn=APP;refGene_tx=XM_023238837.1;refGene_gn=APP;refGene_tx=XM_023238838.1;refGene_gn=APP;refGene_tx=XM_023238839.1;refGene_gn=APP;refGene_tx=XM_023238840.1;refGene_gn=APP;refGene_tx=XM_023238841.1;refGene_gn=APP;refGene_tx=XM_023238842.1
C2 18836771 18836939
refGene_gn=APP;refGene_tx=XM_023238836.1;refGene_gn=APP;refGene_tx=XM_023238837.1;refGene_gn=APP;refGene_tx=XM_023238838.1;refGene_gn=APP;refGene_tx=XM_023238839.1
C2 18839235 18839292
refGene_gn=APP;refGene_tx=XM_023238836.1;refGene_gn=APP;refGene_tx=XM_023238837.1;refGene_gn=APP;refGene_tx=XM_023238840.1
C2 18853368 18853502
refGene_gn=APP;refGene_tx=XM_023238836.1;refGene_gn=APP;refGene_tx=XM_023238837.1;refGene_gn=APP;refGene_tx=XM_023238838.1;refGene_gn=APP;refGene_tx=XM_023238839.1;refGene_gn=APP;refGene_tx=XM_023238840.1;refGene_gn=APP;refGene_tx=XM_023238841.1;refGene_gn=APP;refGene_tx=XM_023238842.1
C2 18859825 18859900
refGene_gn=APP;refGene_tx=XM_023238836.1;refGene_gn=APP;refGene_tx=XM_023238837.1;refGene_gn=APP;refGene_tx=XM_023238838.1;refGene_gn=APP;refGene_tx=XM_023238839.1;refGene_gn=APP;refGene_tx=XM_023238840.1;refGene_gn=APP;refGene_tx=XM_023238841.1;refGene_gn=APP;refGene_tx=XM_023238842.1
C2 18860567 18860726
refGene_gn=APP;refGene_tx=XM_023238836.1;refGene_gn=APP;refGene_tx=XM_023238837.1;refGene_gn=APP;refGene_tx=XM_023238838.1;refGene_gn=APP;refGene_tx=XM_023238839.1;refGene_gn=APP;refGene_tx=XM_023238840.1;refGene_gn=APP;refGene_tx=XM_023238841.1;refGene_gn=APP;refGene_tx=XM_023238842.1
C2 18877477 18877606
refGene_gn=APP;refGene_tx=XM_023238836.1;refGene_gn=APP;refGene_tx=XM_023238837.1;refGene_gn=APP;refGene_tx=XM_023238838.1;refGene_gn=APP;refGene_tx=XM_023238839.1;refGene_gn=APP;refGene_tx=XM_023238840.1;refGene_gn=APP;refGene_tx=XM_023238841.1;refGene_gn=APP;refGene_tx=XM_023238842.1
C2 18878489 18878589
refGene_gn=APP;refGene_tx=XM_023238836.1;refGene_gn=APP;refGene_tx=XM_023238837.1;refGene_gn=APP;refGene_tx=XM_023238838.1;refGene_gn=APP;refGene_tx=XM_023238839.1;refGene_gn=APP;refGene_tx=XM_023238840.1;refGene_gn=APP;refGene_tx=XM_023238841.1;refGene_gn=APP;refGene_tx=XM_023238842.1
C2 18912597 18912819
refGene_gn=APP;refGene_tx=XM_023238836.1;refGene_gn=APP;refGene_tx=XM_

023238837.1;refGene_gn=APP;refGene_tx=XM_023238838.1;refGene_gn=APP;refGene_tx=XM_023238839.1;refGene_gn=APP;refGene_tx=XM_023238840.1;refGene_gn=APP;refGene_tx=XM_023238841.1;refGene_gn=APP;refGene_tx=XM_023238842.1
C2 18921667 18921721
refGene_gn=APP;refGene_tx=XM_023238836.1;refGene_gn=APP;refGene_tx=XM_023238838.1;refGene_gn=APP;refGene_tx=XM_023238840.1;refGene_gn=APP;refGene_tx=XM_023238841.1
C2 18928313 18928414
refGene_gn=APP;refGene_tx=XM_023238836.1;refGene_gn=APP;refGene_tx=XM_023238837.1;refGene_gn=APP;refGene_tx=XM_023238838.1;refGene_gn=APP;refGene_tx=XM_023238839.1;refGene_gn=APP;refGene_tx=XM_023238840.1;refGene_gn=APP;refGene_tx=XM_023238841.1;refGene_gn=APP;refGene_tx=XM_023238842.1
C2 18932172 18932319
refGene_gn=APP;refGene_tx=XM_023238836.1;refGene_gn=APP;refGene_tx=XM_023238837.1;refGene_gn=APP;refGene_tx=XM_023238838.1;refGene_gn=APP;refGene_tx=XM_023238839.1;refGene_gn=APP;refGene_tx=XM_023238840.1;refGene_gn=APP;refGene_tx=XM_023238841.1;refGene_gn=APP;refGene_tx=XM_023238842.1
C2 18940693 18940795
refGene_gn=APP;refGene_tx=XM_023238836.1;refGene_gn=APP;refGene_tx=XM_023238837.1;refGene_gn=APP;refGene_tx=XM_023238838.1;refGene_gn=APP;refGene_tx=XM_023238839.1;refGene_gn=APP;refGene_tx=XM_023238840.1;refGene_gn=APP;refGene_tx=XM_023238841.1;refGene_gn=APP;refGene_tx=XM_023238842.1
C2 19052217 19052446
refGene_gn=GABPA;refGene_tx=XM_003991500.4;refGene_gn=GABPA;refGene_tx=XM_006935951.3
C2 19056488 19056681
refGene_gn=GABPA;refGene_tx=XM_003991500.4;refGene_gn=GABPA;refGene_tx=XM_006935951.3
C2 19056915 19057056
refGene_gn=GABPA;refGene_tx=XM_003991500.4;refGene_gn=GABPA;refGene_tx=XM_006935951.3
C2 19057998 19058052
refGene_gn=GABPA;refGene_tx=XM_003991500.4;refGene_gn=GABPA;refGene_tx=XM_006935951.3
C2 19062429 19062624
refGene_gn=GABPA;refGene_tx=XM_003991500.4;refGene_gn=GABPA;refGene_tx=XM_006935951.3
C2 19071605 19071851
refGene_gn=GABPA;refGene_tx=XM_003991500.4;refGene_gn=GABPA;refGene_tx=XM_006935951.3
C2 19075489 19075574
refGene_gn=GABPA;refGene_tx=XM_003991500.4;refGene_gn=GABPA;refGene_tx=XM_006935951.3
C2 19079565 19079710
refGene_gn=GABPA;refGene_tx=XM_003991500.4;refGene_gn=GABPA;refGene_tx=XM_006935951.3

C2	19082287	19082364
refGene_gn=GABPA;refGene_tx=XM_003991500.4;refGene_gn=GABPA;refGene_tx=XM_006935951.3
C2	19094210	19094374
refGene_gn=ATP5PF;refGene_tx=XM_006935949.3;refGene_gn=ATP5PF;refGene_tx=XM_006935950.3
C2	19099188	19099313
refGene_gn=ATP5PF;refGene_tx=XM_006935949.3;refGene_gn=ATP5PF;refGene_tx=XM_006935950.3
C2	19099865	19099903
refGene_gn=ATP5PF;refGene_tx=XM_006935949.3;refGene_gn=ATP5PF;refGene_tx=XM_006935950.3
C2	19106206	19106239	refGene_gn=JAM2;refGene_tx=XM_023238846.1
C2	19108372	19108415
refGene_gn=JAM2;refGene_tx=XM_023238845.1;refGene_gn=JAM2;refGene_tx=XM_023238846.1
C2	19110543	19110687	refGene_gn=JAM2;refGene_tx=XM_023238845.1
C2	19111036	19111052	refGene_gn=JAM2;refGene_tx=XM_023238846.1
C2	19112972	19113080
refGene_gn=JAM2;refGene_tx=XM_023238845.1;refGene_gn=JAM2;refGene_tx=XM_023238846.1
C2	19119173	19119273
refGene_gn=JAM2;refGene_tx=XM_023238845.1;refGene_gn=JAM2;refGene_tx=XM_023238846.1
C2	19121176	19121379
refGene_gn=JAM2;refGene_tx=XM_023238845.1;refGene_gn=JAM2;refGene_tx=XM_023238846.1
C2	19122993	19123146
refGene_gn=JAM2;refGene_tx=XM_023238845.1;refGene_gn=JAM2;refGene_tx=XM_023238846.1
C2	19125330	19125438
refGene_gn=JAM2;refGene_tx=XM_023238845.1;refGene_gn=JAM2;refGene_tx=XM_023238846.1
C2	19131078	19131144
refGene_gn=JAM2;refGene_tx=XM_023238845.1;refGene_gn=JAM2;refGene_tx=XM_023238846.1
C2	19165102	19165169
refGene_gn=JAM2;refGene_tx=XM_023238845.1;refGene_gn=JAM2;refGene_tx=XM_023238846.1
C2	19193452	19193516
refGene_gn=MRPL39;refGene_tx=XM_011285730.2;refGene_gn=MRPL39;refGene_tx=XM_019839485.1
C2	19194278	19194485
refGene_gn=MRPL39;refGene_tx=XM_011285730.2;refGene_gn=MRPL39;refGene_tx=XM_019839485.1;refGene_gn=MRPL39;refGene_tx=XM_019839486.1
C2	19196218	19196358
refGene_gn=MRPL39;refGene_tx=XM_011285730.2;refGene_gn=MRPL39;refGene_tx=XM_019839485.1;refGene_gn=MRPL39;refGene_tx=XM_019839486.1
C2	19198755	19198855
refGene_gn=MRPL39;refGene_tx=XM_011285730.2;refGene_gn=MRPL39;refGene_

tx=XM_019839485.1;refGene_gn=MRPL39;refGene_tx=XM_019839486.1
C2 19200269 19200337
refGene_gn=MRPL39;refGene_tx=XM_011285730.2;refGene_gn=MRPL39;refGene_
tx=XM_019839485.1;refGene_gn=MRPL39;refGene_tx=XM_019839486.1
C2 19203189 19203302
refGene_gn=MRPL39;refGene_tx=XM_011285730.2;refGene_gn=MRPL39;refGene_
tx=XM_019839485.1;refGene_gn=MRPL39;refGene_tx=XM_019839486.1
C2 19205815 19205881
refGene_gn=MRPL39;refGene_tx=XM_011285730.2;refGene_gn=MRPL39;refGene_
tx=XM_019839485.1;refGene_gn=MRPL39;refGene_tx=XM_019839486.1
C2 19206789 19206943
refGene_gn=MRPL39;refGene_tx=XM_011285730.2;refGene_gn=MRPL39;refGene_
tx=XM_019839485.1;refGene_gn=MRPL39;refGene_tx=XM_019839486.1
C2 19211452 19211500
refGene_gn=MRPL39;refGene_tx=XM_011285730.2;refGene_gn=MRPL39;refGene_
tx=XM_019839485.1;refGene_gn=MRPL39;refGene_tx=XM_019839486.1
C2 19213984 19214035
refGene_gn=MRPL39;refGene_tx=XM_011285730.2;refGene_gn=MRPL39;refGene_
tx=XM_019839485.1;refGene_gn=MRPL39;refGene_tx=XM_019839486.1
C2 22873109 22873221
refGene_gn=NCAM2;refGene_tx=XM_011285729.3;refGene_gn=NCAM2;refGene_tx
=XM_023238847.1
C2 22876044 22876164
refGene_gn=NCAM2;refGene_tx=XM_011285729.3;refGene_gn=NCAM2;refGene_tx
=XM_023238847.1
C2 22887291 22887496
refGene_gn=NCAM2;refGene_tx=XM_011285729.3;refGene_gn=NCAM2;refGene_tx
=XM_023238847.1
C2 22909106 22909213 refGene_gn=NCAM2;refGene_tx=XM_023238848.1
C2 22915212 22915393
refGene_gn=NCAM2;refGene_tx=XM_011285729.3;refGene_gn=NCAM2;refGene_tx
=XM_023238847.1;refGene_gn=NCAM2;refGene_tx=XM_023238848.1
C2 22925029 22925151
refGene_gn=NCAM2;refGene_tx=XM_011285729.3;refGene_gn=NCAM2;refGene_tx
=XM_023238847.1;refGene_gn=NCAM2;refGene_tx=XM_023238848.1
C2 22928784 22928904
refGene_gn=NCAM2;refGene_tx=XM_011285729.3;refGene_gn=NCAM2;refGene_tx
=XM_023238847.1;refGene_gn=NCAM2;refGene_tx=XM_023238848.1
C2 22956277 22956451
refGene_gn=NCAM2;refGene_tx=XM_011285729.3;refGene_gn=NCAM2;refGene_tx
=XM_023238847.1;refGene_gn=NCAM2;refGene_tx=XM_023238848.1
C2 22975870 22975967
refGene_gn=NCAM2;refGene_tx=XM_011285729.3;refGene_gn=NCAM2;refGene_tx
=XM_023238847.1;refGene_gn=NCAM2;refGene_tx=XM_023238848.1
C2 22980602 22980790
refGene_gn=NCAM2;refGene_tx=XM_011285729.3;refGene_gn=NCAM2;refGene_tx
=XM_023238847.1;refGene_gn=NCAM2;refGene_tx=XM_023238848.1
C2 23006807 23006958
refGene_gn=NCAM2;refGene_tx=XM_011285729.3;refGene_gn=NCAM2;refGene_tx
=XM_023238847.1;refGene_gn=NCAM2;refGene_tx=XM_023238848.1

C2 23037049 23037195
refGene_gn=NCAM2;refGene_tx=XM_011285729.3;refGene_gn=NCAM2;refGene_tx=XM_023238847.1;refGene_gn=NCAM2;refGene_tx=XM_023238848.1
C2 23039325 23039486
refGene_gn=NCAM2;refGene_tx=XM_011285729.3;refGene_gn=NCAM2;refGene_tx=XM_023238847.1;refGene_gn=NCAM2;refGene_tx=XM_023238848.1
C2 23052423 23052541
refGene_gn=NCAM2;refGene_tx=XM_011285729.3;refGene_gn=NCAM2;refGene_tx=XM_023238847.1;refGene_gn=NCAM2;refGene_tx=XM_023238848.1
C2 23082099 23082237
refGene_gn=NCAM2;refGene_tx=XM_011285729.3;refGene_gn=NCAM2;refGene_tx=XM_023238847.1;refGene_gn=NCAM2;refGene_tx=XM_023238848.1
C2 23089136 23089280
refGene_gn=NCAM2;refGene_tx=XM_011285729.3;refGene_gn=NCAM2;refGene_tx=XM_023238848.1
C2 23091910 23092117
refGene_gn=NCAM2;refGene_tx=XM_011285729.3;refGene_gn=NCAM2;refGene_tx=XM_023238847.1;refGene_gn=NCAM2;refGene_tx=XM_023238848.1
C2 23095809 23095884
refGene_gn=NCAM2;refGene_tx=XM_011285729.3;refGene_gn=NCAM2;refGene_tx=XM_023238847.1;refGene_gn=NCAM2;refGene_tx=XM_023238848.1
C2 23396091 23396146
refGene_gn=NCAM2;refGene_tx=XM_011285729.3;refGene_gn=NCAM2;refGene_tx=XM_023238847.1;refGene_gn=NCAM2;refGene_tx=XM_023238848.1
C2 25671840 25671985 refGene_gn=TMPRSS15;refGene_tx=XM_023260155.1
C2 25674516 25674647 refGene_gn=TMPRSS15;refGene_tx=XM_023260155.1
C2 25674955 25675023 refGene_gn=TMPRSS15;refGene_tx=XM_023260155.1
C2 25685880 25686032 refGene_gn=TMPRSS15;refGene_tx=XM_023260155.1
C2 25698222 25698354 refGene_gn=TMPRSS15;refGene_tx=XM_023260155.1
C2 25704566 25704675 refGene_gn=TMPRSS15;refGene_tx=XM_023260155.1
C2 25709886 25709993 refGene_gn=TMPRSS15;refGene_tx=XM_023260155.1
C2 25711074 25711215 refGene_gn=TMPRSS15;refGene_tx=XM_023260155.1
C2 25711831 25711981 refGene_gn=TMPRSS15;refGene_tx=XM_023260155.1
C2 25728462 25728568 refGene_gn=TMPRSS15;refGene_tx=XM_023260155.1
C2 25728863 25729014 refGene_gn=TMPRSS15;refGene_tx=XM_023260155.1
C2 25731428 25731564 refGene_gn=TMPRSS15;refGene_tx=XM_023260155.1
C2 25739028 25739118 refGene_gn=TMPRSS15;refGene_tx=XM_023260155.1
C2 25740560 25740686 refGene_gn=TMPRSS15;refGene_tx=XM_023260155.1
C2 25742648 25742789 refGene_gn=TMPRSS15;refGene_tx=XM_023260155.1
C2 25754897 25755008 refGene_gn=TMPRSS15;refGene_tx=XM_023260155.1
C2 25756536 25756669 refGene_gn=TMPRSS15;refGene_tx=XM_023260155.1
C2 25768189 25768285 refGene_gn=TMPRSS15;refGene_tx=XM_023260155.1
C2 25772516 25772566 refGene_gn=TMPRSS15;refGene_tx=XM_023260155.1
C2 25772744 25772919 refGene_gn=TMPRSS15;refGene_tx=XM_023260155.1
C2 25781584 25781766 refGene_gn=TMPRSS15;refGene_tx=XM_023260155.1
C2 25783015 25783111 refGene_gn=TMPRSS15;refGene_tx=XM_023260155.1
C2 25786650 25786790 refGene_gn=TMPRSS15;refGene_tx=XM_023260155.1
C2 25791864 25792020 refGene_gn=TMPRSS15;refGene_tx=XM_023260155.1
C2 25795541 25795626
refGene_gn=CHODL;refGene_tx=XM_011285862.3;refGene_gn=CHODL;refGene_tx

=XM_023238849.1
C2	25796944	25797023	refGene_gn=CHODL;refGene_tx=XM_023238849.1
C2	25799461	25799564
refGene_gn=CHODL;refGene_tx=XM_011285862.3;refGene_gn=CHODL;refGene_tx
=XM_023238849.1
C2	25801427	25801514
refGene_gn=CHODL;refGene_tx=XM_011285862.3;refGene_gn=CHODL;refGene_tx
=XM_023238849.1
C2	25805351	25805509
refGene_gn=CHODL;refGene_tx=XM_011285862.3;refGene_gn=CHODL;refGene_tx
=XM_023238849.1
C2	25806387	25806697
refGene_gn=CHODL;refGene_tx=XM_011285862.3;refGene_gn=CHODL;refGene_tx
=XM_023238849.1
C2	25818181	25818260
refGene_gn=CHODL;refGene_tx=XM_011285862.3;refGene_gn=CHODL;refGene_tx
=XM_023238849.1
C2	26199331	26199458
refGene_gn=CC2H21orf91;refGene_tx=XM_006935961.4;refGene_gn=CC2H21orf9
1;refGene_tx=XM_006935962.4;refGene_gn=CC2H21orf91;refGene_tx=XM_02323
8850.1;refGene_gn=CC2H21orf91;refGene_tx=XM_023238851.1
C2	26222938	26223475
refGene_gn=CC2H21orf91;refGene_tx=XM_006935961.4;refGene_gn=CC2H21orf9
1;refGene_tx=XM_006935962.4;refGene_gn=CC2H21orf91;refGene_tx=XM_02323
8850.1;refGene_gn=CC2H21orf91;refGene_tx=XM_023238851.1
C2	26224493	26224556
refGene_gn=CC2H21orf91;refGene_tx=XM_006935961.4;refGene_gn=CC2H21orf9
1;refGene_tx=XM_006935962.4;refGene_gn=CC2H21orf91;refGene_tx=XM_02323
8850.1;refGene_gn=CC2H21orf91;refGene_tx=XM_023238851.1
C2	26225733	26225900
refGene_gn=CC2H21orf91;refGene_tx=XM_006935961.4;refGene_gn=CC2H21orf9
1;refGene_tx=XM_006935962.4;refGene_gn=CC2H21orf91;refGene_tx=XM_02323
8850.1;refGene_gn=CC2H21orf91;refGene_tx=XM_023238851.1
C2	26355325	26355498	refGene_gn=BTG3;refGene_tx=XM_004001326.5
C2	26359313	26359451	refGene_gn=BTG3;refGene_tx=XM_004001326.5
C2	26364953	26365161	refGene_gn=BTG3;refGene_tx=XM_004001326.5
C2	26371689	26371929	refGene_gn=BTG3;refGene_tx=XM_004001326.5
C2	26372825	26372867	refGene_gn=CXADR;refGene_tx=XM_019839492.1
C2	26389946	26390211
refGene_gn=CXADR;refGene_tx=XM_011285859.2;refGene_gn=CXADR;refGene_tx
=XM_019839491.1;refGene_gn=CXADR;refGene_tx=XM_019839492.1
C2	26391728	26391867
refGene_gn=CXADR;refGene_tx=XM_011285859.2;refGene_gn=CXADR;refGene_tx
=XM_019839491.1;refGene_gn=CXADR;refGene_tx=XM_019839492.1
C2	26392399	26392522
refGene_gn=CXADR;refGene_tx=XM_011285859.2;refGene_gn=CXADR;refGene_tx
=XM_019839491.1;refGene_gn=CXADR;refGene_tx=XM_019839492.1
C2	26393892	26394048
refGene_gn=CXADR;refGene_tx=XM_011285859.2;refGene_gn=CXADR;refGene_tx
=XM_019839491.1;refGene_gn=CXADR;refGene_tx=XM_019839492.1

C2 26400397 26400602
refGene_gn=CXADR;refGene_tx=XM_011285859.2;refGene_gn=CXADR;refGene_tx
=XM_019839491.1;refGene_gn=CXADR;refGene_tx=XM_019839492.1
C2 26404672 26404839
refGene_gn=CXADR;refGene_tx=XM_011285859.2;refGene_gn=CXADR;refGene_tx
=XM_019839491.1;refGene_gn=CXADR;refGene_tx=XM_019839492.1
C2 26421221 26421297
refGene_gn=CXADR;refGene_tx=XM_019839491.1;refGene_gn=CXADR;refGene_tx
=XM_019839492.1
C2 26427014 26427057 refGene_gn=CXADR;refGene_tx=XM_011285859.2
C2 27794781 27794954
refGene_gn=USP25;refGene_tx=XM_019839497.2;refGene_gn=USP25;refGene_tx
=XM_023238852.1;refGene_gn=USP25;refGene_tx=XM_023238853.1;refGene_gn=
USP25;refGene_tx=XM_023238854.1;refGene_gn=USP25;refGene_tx=XM_0232388
55.1
C2 27795294 27795490
refGene_gn=USP25;refGene_tx=XM_019839497.2;refGene_gn=USP25;refGene_tx
=XM_023238852.1;refGene_gn=USP25;refGene_tx=XM_023238853.1;refGene_gn=
USP25;refGene_tx=XM_023238854.1;refGene_gn=USP25;refGene_tx=XM_0232388
55.1
C2 27799135 27799259
refGene_gn=USP25;refGene_tx=XM_019839497.2;refGene_gn=USP25;refGene_tx
=XM_023238852.1;refGene_gn=USP25;refGene_tx=XM_023238853.1;refGene_gn=
USP25;refGene_tx=XM_023238854.1;refGene_gn=USP25;refGene_tx=XM_0232388
55.1
C2 27804503 27804583
refGene_gn=USP25;refGene_tx=XM_019839497.2;refGene_gn=USP25;refGene_tx
=XM_023238852.1;refGene_gn=USP25;refGene_tx=XM_023238853.1;refGene_gn=
USP25;refGene_tx=XM_023238854.1;refGene_gn=USP25;refGene_tx=XM_0232388
55.1
C2 27807550 27807629
refGene_gn=USP25;refGene_tx=XM_019839497.2;refGene_gn=USP25;refGene_tx
=XM_023238852.1;refGene_gn=USP25;refGene_tx=XM_023238853.1;refGene_gn=
USP25;refGene_tx=XM_023238854.1;refGene_gn=USP25;refGene_tx=XM_0232388
55.1
C2 27810237 27810416
refGene_gn=USP25;refGene_tx=XM_019839497.2;refGene_gn=USP25;refGene_tx
=XM_023238852.1;refGene_gn=USP25;refGene_tx=XM_023238853.1;refGene_gn=
USP25;refGene_tx=XM_023238854.1;refGene_gn=USP25;refGene_tx=XM_0232388
55.1
C2 27821509 27821605
refGene_gn=USP25;refGene_tx=XM_019839497.2;refGene_gn=USP25;refGene_tx
=XM_023238852.1;refGene_gn=USP25;refGene_tx=XM_023238853.1
C2 27823573 27823687
refGene_gn=USP25;refGene_tx=XM_019839497.2;refGene_gn=USP25;refGene_tx
=XM_023238852.1;refGene_gn=USP25;refGene_tx=XM_023238853.1
C2 27829211 27829351
refGene_gn=USP25;refGene_tx=XM_019839497.2;refGene_gn=USP25;refGene_tx
=XM_023238852.1;refGene_gn=USP25;refGene_tx=XM_023238853.1;refGene_gn=
USP25;refGene_tx=XM_023238854.1;refGene_gn=USP25;refGene_tx=XM_0232388

55.1
C2 27838769 27838970
refGene_gn=USP25;refGene_tx=XM_019839497.2;refGene_gn=USP25;refGene_tx=XM_023238852.1;refGene_gn=USP25;refGene_tx=XM_023238853.1;refGene_gn=USP25;refGene_tx=XM_023238854.1;refGene_gn=USP25;refGene_tx=XM_023238855.1
C2 27840620 27840849
refGene_gn=USP25;refGene_tx=XM_019839497.2;refGene_gn=USP25;refGene_tx=XM_023238852.1;refGene_gn=USP25;refGene_tx=XM_023238853.1;refGene_gn=USP25;refGene_tx=XM_023238854.1;refGene_gn=USP25;refGene_tx=XM_023238855.1
C2 27841516 27841587
refGene_gn=USP25;refGene_tx=XM_019839497.2;refGene_gn=USP25;refGene_tx=XM_023238852.1;refGene_gn=USP25;refGene_tx=XM_023238853.1;refGene_gn=USP25;refGene_tx=XM_023238854.1;refGene_gn=USP25;refGene_tx=XM_023238855.1
C2 27845154 27845381
refGene_gn=USP25;refGene_tx=XM_019839497.2;refGene_gn=USP25;refGene_tx=XM_023238852.1;refGene_gn=USP25;refGene_tx=XM_023238853.1;refGene_gn=USP25;refGene_tx=XM_023238854.1;refGene_gn=USP25;refGene_tx=XM_023238855.1
C2 27845966 27846125
refGene_gn=USP25;refGene_tx=XM_019839497.2;refGene_gn=USP25;refGene_tx=XM_023238852.1;refGene_gn=USP25;refGene_tx=XM_023238853.1;refGene_gn=USP25;refGene_tx=XM_023238854.1;refGene_gn=USP25;refGene_tx=XM_023238855.1
C2 27847440 27847536
refGene_gn=USP25;refGene_tx=XM_019839497.2;refGene_gn=USP25;refGene_tx=XM_023238852.1;refGene_gn=USP25;refGene_tx=XM_023238853.1;refGene_gn=USP25;refGene_tx=XM_023238854.1;refGene_gn=USP25;refGene_tx=XM_023238855.1
C2 27847931 27848059
refGene_gn=USP25;refGene_tx=XM_019839497.2;refGene_gn=USP25;refGene_tx=XM_023238852.1;refGene_gn=USP25;refGene_tx=XM_023238853.1;refGene_gn=USP25;refGene_tx=XM_023238854.1;refGene_gn=USP25;refGene_tx=XM_023238855.1
C2 27854860 27855009
refGene_gn=USP25;refGene_tx=XM_019839497.2;refGene_gn=USP25;refGene_tx=XM_023238852.1;refGene_gn=USP25;refGene_tx=XM_023238853.1;refGene_gn=USP25;refGene_tx=XM_023238854.1;refGene_gn=USP25;refGene_tx=XM_023238855.1
C2 27859882 27859956
refGene_gn=USP25;refGene_tx=XM_019839497.2;refGene_gn=USP25;refGene_tx=XM_023238852.1;refGene_gn=USP25;refGene_tx=XM_023238853.1;refGene_gn=USP25;refGene_tx=XM_023238854.1;refGene_gn=USP25;refGene_tx=XM_023238855.1
C2 27866825 27866902
refGene_gn=USP25;refGene_tx=XM_019839497.2;refGene_gn=USP25;refGene_tx=XM_023238852.1;refGene_gn=USP25;refGene_tx=XM_023238853.1;refGene_gn=USP25;refGene_tx=XM_023238854.1;refGene_gn=USP25;refGene_tx=XM_0232388

55.1
C2 27870287 27870425
refGene_gn=USP25;refGene_tx=XM_019839497.2;refGene_gn=USP25;refGene_tx=XM_023238852.1;refGene_gn=USP25;refGene_tx=XM_023238853.1;refGene_gn=USP25;refGene_tx=XM_023238854.1;refGene_gn=USP25;refGene_tx=XM_023238855.1
C2 27880513 27880600
refGene_gn=USP25;refGene_tx=XM_019839497.2;refGene_gn=USP25;refGene_tx=XM_023238852.1;refGene_gn=USP25;refGene_tx=XM_023238853.1;refGene_gn=USP25;refGene_tx=XM_023238854.1;refGene_gn=USP25;refGene_tx=XM_023238855.1
C2 27892885 27893048
refGene_gn=USP25;refGene_tx=XM_019839497.2;refGene_gn=USP25;refGene_tx=XM_023238852.1;refGene_gn=USP25;refGene_tx=XM_023238853.1;refGene_gn=USP25;refGene_tx=XM_023238854.1;refGene_gn=USP25;refGene_tx=XM_023238855.1
C2 27903407 27903531
refGene_gn=USP25;refGene_tx=XM_019839497.2;refGene_gn=USP25;refGene_tx=XM_023238852.1;refGene_gn=USP25;refGene_tx=XM_023238853.1;refGene_gn=USP25;refGene_tx=XM_023238854.1;refGene_gn=USP25;refGene_tx=XM_023238855.1
C2 27911306 27911387
refGene_gn=USP25;refGene_tx=XM_019839497.2;refGene_gn=USP25;refGene_tx=XM_023238852.1;refGene_gn=USP25;refGene_tx=XM_023238854.1
C2 27915509 27915654
refGene_gn=USP25;refGene_tx=XM_023238852.1;refGene_gn=USP25;refGene_tx=XM_023238853.1;refGene_gn=USP25;refGene_tx=XM_023238854.1;refGene_gn=USP25;refGene_tx=XM_023238855.1
C2 27919371 27919449
refGene_gn=USP25;refGene_tx=XM_023238852.1;refGene_gn=USP25;refGene_tx=XM_023238853.1;refGene_gn=USP25;refGene_tx=XM_023238854.1;refGene_gn=USP25;refGene_tx=XM_023238855.1
C2 27946041 27946086
refGene_gn=USP25;refGene_tx=XM_023238852.1;refGene_gn=USP25;refGene_tx=XM_023238853.1;refGene_gn=USP25;refGene_tx=XM_023238854.1;refGene_gn=USP25;refGene_tx=XM_023238855.1
C2 28667501 28670969
refGene_gn=NRIP1;refGene_tx=XM_006935964.4;refGene_gn=NRIP1;refGene_tx=XM_019839501.2;refGene_gn=NRIP1;refGene_tx=XM_023238856.1
C2 28929477 28929501
refGene_gn=SAMSN1;refGene_tx=XM_019839502.2;refGene_gn=SAMSN1;refGene_tx=XM_019839505.2
C2 28935513 28935645
refGene_gn=SAMSN1;refGene_tx=XM_019839502.2;refGene_gn=SAMSN1;refGene_tx=XM_019839503.2;refGene_gn=SAMSN1;refGene_tx=XM_019839505.2
C2 28953839 28953877
refGene_gn=SAMSN1;refGene_tx=XM_019839502.2;refGene_gn=SAMSN1;refGene_tx=XM_019839505.2
C2 28956546 28956584
refGene_gn=SAMSN1;refGene_tx=XM_019839502.2;refGene_gn=SAMSN1;refGene_

tx=XM_019839503.2;refGene_gn=SAMSN1;refGene_tx=XM_019839504.2;refGene_gn=SAMSN1;refGene_tx=XM_019839505.2
C2 28958313 28958400 refGene_gn=SAMSN1;refGene_tx=XM_019839502.2;refGene_gn=SAMSN1;refGene_tx=XM_019839503.2;refGene_gn=SAMSN1;refGene_tx=XM_019839504.2;refGene_gn=SAMSN1;refGene_tx=XM_019839505.2
C2 28966110 28966190 refGene_gn=SAMSN1;refGene_tx=XM_019839502.2;refGene_gn=SAMSN1;refGene_tx=XM_019839503.2;refGene_gn=SAMSN1;refGene_tx=XM_019839504.2;refGene_gn=SAMSN1;refGene_tx=XM_019839505.2
C2 28984655 28984782 refGene_gn=SAMSN1;refGene_tx=XM_019839502.2;refGene_gn=SAMSN1;refGene_tx=XM_019839503.2;refGene_gn=SAMSN1;refGene_tx=XM_019839504.2
C2 28986211 28986564 refGene_gn=SAMSN1;refGene_tx=XM_019839502.2;refGene_gn=SAMSN1;refGene_tx=XM_019839503.2;refGene_gn=SAMSN1;refGene_tx=XM_019839504.2
C2 29019919 29019976 refGene_gn=SAMSN1;refGene_tx=XM_003991527.5
C2 29041959 29042031 refGene_gn=SAMSN1;refGene_tx=XM_003991527.5;refGene_gn=SAMSN1;refGene_tx=XM_019839502.2;refGene_gn=SAMSN1;refGene_tx=XM_019839503.2;refGene_gn=SAMSN1;refGene_tx=XM_019839504.2;refGene_gn=SAMSN1;refGene_tx=XM_019839505.2
C2 29045460 29045610 refGene_gn=SAMSN1;refGene_tx=XM_003991527.5;refGene_gn=SAMSN1;refGene_tx=XM_019839502.2;refGene_gn=SAMSN1;refGene_tx=XM_019839503.2;refGene_gn=SAMSN1;refGene_tx=XM_019839504.2;refGene_gn=SAMSN1;refGene_tx=XM_019839505.2
C2 29049860 29049990 refGene_gn=SAMSN1;refGene_tx=XM_003991527.5;refGene_gn=SAMSN1;refGene_tx=XM_019839502.2;refGene_gn=SAMSN1;refGene_tx=XM_019839503.2;refGene_gn=SAMSN1;refGene_tx=XM_019839504.2;refGene_gn=SAMSN1;refGene_tx=XM_019839505.2
C2 29052547 29052699 refGene_gn=SAMSN1;refGene_tx=XM_003991527.5;refGene_gn=SAMSN1;refGene_tx=XM_019839502.2;refGene_gn=SAMSN1;refGene_tx=XM_019839503.2;refGene_gn=SAMSN1;refGene_tx=XM_019839504.2;refGene_gn=SAMSN1;refGene_tx=XM_019839505.2
C2 29055818 29056025 refGene_gn=SAMSN1;refGene_tx=XM_003991527.5;refGene_gn=SAMSN1;refGene_tx=XM_019839502.2;refGene_gn=SAMSN1;refGene_tx=XM_019839503.2;refGene_gn=SAMSN1;refGene_tx=XM_019839504.2;refGene_gn=SAMSN1;refGene_tx=XM_019839505.2
C2 29057944 29058095 refGene_gn=SAMSN1;refGene_tx=XM_003991527.5;refGene_gn=SAMSN1;refGene_tx=XM_019839502.2;refGene_gn=SAMSN1;refGene_tx=XM_019839503.2;refGene_gn=SAMSN1;refGene_tx=XM_019839504.2;refGene_gn=SAMSN1;refGene_tx=XM_019839505.2
C2 29069132 29069335 refGene_gn=SAMSN1;refGene_tx=XM_003991527.5;refGene_gn=SAMSN1;refGene_tx=XM_019839502.2;refGene_gn=SAMSN1;refGene_tx=XM_019839503.2;refGene_

gn=SAMSN1;refGene_tx=XM_019839504.2;refGene_gn=SAMSN1;refGene_tx=XM_019839505.2
C2 29165020 29165045 refGene_gn=HSPA13;refGene_tx=XM_003991529.5
C2 29166652 29166993 refGene_gn=HSPA13;refGene_tx=XM_003991529.5
C2 29170678 29170892 refGene_gn=HSPA13;refGene_tx=XM_003991529.5
C2 29172974 29173142 refGene_gn=HSPA13;refGene_tx=XM_003991529.5
C2 29174682 29175350 refGene_gn=HSPA13;refGene_tx=XM_003991529.5
C2 29184255 29184382
refGene_gn=LOC101095601;refGene_tx=XM_011285865.3;refGene_gn=LOC101095601;refGene_tx=XM_019839508.2;refGene_gn=LOC101095601;refGene_tx=XM_019839509.2;refGene_gn=LOC101095601;refGene_tx=XM_019839510.2;refGene_gn=LOC101095601;refGene_tx=XM_019839511.2
C2 29184715 29184910
refGene_gn=LOC101095601;refGene_tx=XM_011285865.3;refGene_gn=LOC101095601;refGene_tx=XM_019839508.2;refGene_gn=LOC101095601;refGene_tx=XM_019839509.2;refGene_gn=LOC101095601;refGene_tx=XM_019839510.2
C2 29185868 29186035
refGene_gn=LOC101095601;refGene_tx=XM_011285865.3;refGene_gn=LOC101095601;refGene_tx=XM_019839508.2;refGene_gn=LOC101095601;refGene_tx=XM_019839509.2;refGene_gn=LOC101095601;refGene_tx=XM_019839510.2;refGene_gn=LOC101095601;refGene_tx=XM_019839511.2
C2 29187299 29187458
refGene_gn=LOC101095601;refGene_tx=XM_011285865.3;refGene_gn=LOC101095601;refGene_tx=XM_019839508.2;refGene_gn=LOC101095601;refGene_tx=XM_019839509.2;refGene_gn=LOC101095601;refGene_tx=XM_019839510.2;refGene_gn=LOC101095601;refGene_tx=XM_019839511.2
C2 29189939 29190083
refGene_gn=LOC101095601;refGene_tx=XM_011285865.3;refGene_gn=LOC101095601;refGene_tx=XM_019839508.2;refGene_gn=LOC101095601;refGene_tx=XM_019839509.2;refGene_gn=LOC101095601;refGene_tx=XM_019839510.2;refGene_gn=LOC101095601;refGene_tx=XM_019839511.2
C2 29193807 29193909
refGene_gn=LOC101095601;refGene_tx=XM_011285865.3;refGene_gn=LOC101095601;refGene_tx=XM_019839509.2;refGene_gn=LOC101095601;refGene_tx=XM_019839510.2;refGene_gn=LOC101095601;refGene_tx=XM_019839511.2
C2 29195226 29195353
refGene_gn=LOC101095601;refGene_tx=XM_011285865.3;refGene_gn=LOC101095601;refGene_tx=XM_019839508.2;refGene_gn=LOC101095601;refGene_tx=XM_019839509.2;refGene_gn=LOC101095601;refGene_tx=XM_019839510.2;refGene_gn=LOC101095601;refGene_tx=XM_019839511.2
C2 29196098 29196298
refGene_gn=LOC101095601;refGene_tx=XM_011285865.3;refGene_gn=LOC101095601;refGene_tx=XM_019839508.2;refGene_gn=LOC101095601;refGene_tx=XM_019839509.2;refGene_gn=LOC101095601;refGene_tx=XM_019839510.2;refGene_gn=LOC101095601;refGene_tx=XM_019839511.2
C2 29199306 29199462
refGene_gn=LOC101095601;refGene_tx=XM_011285865.3;refGene_gn=LOC101095601;refGene_tx=XM_019839508.2;refGene_gn=LOC101095601;refGene_tx=XM_019839509.2;refGene_gn=LOC101095601;refGene_tx=XM_019839510.2;refGene_gn=LOC101095601;refGene_tx=XM_019839511.2

C2 29201671 29201826
refGene_gn=LOC101095601;refGene_tx=XM_011285865.3;refGene_gn=LOC101095601;refGene_tx=XM_019839508.2;refGene_gn=LOC101095601;refGene_tx=XM_019839509.2;refGene_gn=LOC101095601;refGene_tx=XM_019839510.2;refGene_gn=LOC101095601;refGene_tx=XM_019839511.2
C2 29208897 29209117
refGene_gn=LOC101095601;refGene_tx=XM_011285865.3;refGene_gn=LOC101095601;refGene_tx=XM_019839508.2;refGene_gn=LOC101095601;refGene_tx=XM_019839509.2;refGene_gn=LOC101095601;refGene_tx=XM_019839510.2;refGene_gn=LOC101095601;refGene_tx=XM_019839511.2
C2 29209573 29209634
refGene_gn=LOC101095601;refGene_tx=XM_011285865.3;refGene_gn=LOC101095601;refGene_tx=XM_019839508.2;refGene_gn=LOC101095601;refGene_tx=XM_019839509.2;refGene_gn=LOC101095601;refGene_tx=XM_019839510.2;refGene_gn=LOC101095601;refGene_tx=XM_019839511.2
C2 29217829 29217905
refGene_gn=LOC101095601;refGene_tx=XM_011285865.3;refGene_gn=LOC101095601;refGene_tx=XM_019839508.2;refGene_gn=LOC101095601;refGene_tx=XM_019839509.2;refGene_gn=LOC101095601;refGene_tx=XM_019839510.2;refGene_gn=LOC101095601;refGene_tx=XM_019839511.2
C2 29222493 29222656
refGene_gn=LOC101095601;refGene_tx=XM_011285865.3;refGene_gn=LOC101095601;refGene_tx=XM_019839508.2;refGene_gn=LOC101095601;refGene_tx=XM_019839509.2;refGene_gn=LOC101095601;refGene_tx=XM_019839510.2;refGene_gn=LOC101095601;refGene_tx=XM_019839511.2
C2 29224864 29225032
refGene_gn=LOC101095601;refGene_tx=XM_011285865.3;refGene_gn=LOC101095601;refGene_tx=XM_019839508.2;refGene_gn=LOC101095601;refGene_tx=XM_019839509.2;refGene_gn=LOC101095601;refGene_tx=XM_019839510.2;refGene_gn=LOC101095601;refGene_tx=XM_019839511.2
C2 29228148 29228325
refGene_gn=LOC101095601;refGene_tx=XM_011285865.3;refGene_gn=LOC101095601;refGene_tx=XM_019839508.2;refGene_gn=LOC101095601;refGene_tx=XM_019839509.2;refGene_gn=LOC101095601;refGene_tx=XM_019839510.2;refGene_gn=LOC101095601;refGene_tx=XM_019839511.2
C2 29232341 29232468
refGene_gn=LOC101095601;refGene_tx=XM_011285865.3;refGene_gn=LOC101095601;refGene_tx=XM_019839508.2;refGene_gn=LOC101095601;refGene_tx=XM_019839509.2;refGene_gn=LOC101095601;refGene_tx=XM_019839510.2;refGene_gn=LOC101095601;refGene_tx=XM_019839511.2
C2 29233074 29233141
refGene_gn=LOC101095601;refGene_tx=XM_011285865.3;refGene_gn=LOC101095601;refGene_tx=XM_019839508.2;refGene_gn=LOC101095601;refGene_tx=XM_019839509.2;refGene_gn=LOC101095601;refGene_tx=XM_019839510.2;refGene_gn=LOC101095601;refGene_tx=XM_019839511.2
C2 29233234 29233319
refGene_gn=LOC101095601;refGene_tx=XM_011285865.3;refGene_gn=LOC101095601;refGene_tx=XM_019839508.2;refGene_gn=LOC101095601;refGene_tx=XM_019839509.2;refGene_gn=LOC101095601;refGene_tx=XM_019839510.2;refGene_gn=LOC101095601;refGene_tx=XM_019839511.2

C2	29235633	29235780
refGene_gn=LOC101095601;refGene_tx=XM_011285865.3;refGene_gn=LOC101095601;refGene_tx=XM_019839508.2;refGene_gn=LOC101095601;refGene_tx=XM_019839509.2;refGene_gn=LOC101095601;refGene_tx=XM_019839511.2
C2	29237315	29237453
refGene_gn=LOC101095601;refGene_tx=XM_011285865.3;refGene_gn=LOC101095601;refGene_tx=XM_019839508.2;refGene_gn=LOC101095601;refGene_tx=XM_019839509.2;refGene_gn=LOC101095601;refGene_tx=XM_019839510.2;refGene_gn=LOC101095601;refGene_tx=XM_019839511.2
C2	29240372	29240438
refGene_gn=LOC101095601;refGene_tx=XM_011285865.3;refGene_gn=LOC101095601;refGene_tx=XM_019839508.2;refGene_gn=LOC101095601;refGene_tx=XM_019839509.2;refGene_gn=LOC101095601;refGene_tx=XM_019839510.2;refGene_gn=LOC101095601;refGene_tx=XM_019839511.2
C2	29242956	29243211
refGene_gn=LOC101095601;refGene_tx=XM_011285865.3;refGene_gn=LOC101095601;refGene_tx=XM_019839508.2;refGene_gn=LOC101095601;refGene_tx=XM_019839509.2;refGene_gn=LOC101095601;refGene_tx=XM_019839510.2;refGene_gn=LOC101095601;refGene_tx=XM_019839511.2
C2	29252400	29252578
refGene_gn=LOC101095601;refGene_tx=XM_011285865.3;refGene_gn=LOC101095601;refGene_tx=XM_019839508.2;refGene_gn=LOC101095601;refGene_tx=XM_019839509.2;refGene_gn=LOC101095601;refGene_tx=XM_019839510.2;refGene_gn=LOC101095601;refGene_tx=XM_019839511.2
C2	29258991	29259134
refGene_gn=LOC101095601;refGene_tx=XM_011285865.3;refGene_gn=LOC101095601;refGene_tx=XM_019839508.2;refGene_gn=LOC101095601;refGene_tx=XM_019839509.2;refGene_gn=LOC101095601;refGene_tx=XM_019839510.2;refGene_gn=LOC101095601;refGene_tx=XM_019839511.2
C2	29260316	29260464
refGene_gn=LOC101095601;refGene_tx=XM_011285865.3;refGene_gn=LOC101095601;refGene_tx=XM_019839508.2;refGene_gn=LOC101095601;refGene_tx=XM_019839509.2;refGene_gn=LOC101095601;refGene_tx=XM_019839510.2;refGene_gn=LOC101095601;refGene_tx=XM_019839511.2
C2	29261449	29261460
refGene_gn=LOC101095601;refGene_tx=XM_019839509.2
C2	29267312	29267374
refGene_gn=LOC101095601;refGene_tx=XM_011285865.3;refGene_gn=LOC101095601;refGene_tx=XM_019839508.2;refGene_gn=LOC101095601;refGene_tx=XM_019839510.2;refGene_gn=LOC101095601;refGene_tx=XM_019839511.2
C2	29272539	29272611
refGene_gn=LOC101095601;refGene_tx=XM_011285865.3;refGene_gn=LOC101095601;refGene_tx=XM_019839508.2;refGene_gn=LOC101095601;refGene_tx=XM_019839510.2;refGene_gn=LOC101095601;refGene_tx=XM_019839511.2
C2	29315254	29315539	refGene_gn=RBM11;refGene_tx=XM_023238863.1
C2	29317286	29317383	refGene_gn=RBM11;refGene_tx=XM_023238863.1
C2	29328093	29328166	refGene_gn=RBM11;refGene_tx=XM_023238863.1
C2	29329501	29329664	refGene_gn=RBM11;refGene_tx=XM_023238863.1
C2	29333245	29333341	refGene_gn=RBM11;refGene_tx=XM_023238863.1
C2	29339280	29339389

refGene_gn=LIPI;refGene_tx=XM_023238857.1;refGene_gn=LIPI;refGene_tx=X
M_023238858.1;refGene_gn=LIPI;refGene_tx=XM_023238859.1;refGene_gn=LIP
I;refGene_tx=XM_023238860.1
C2 29359563 29359949
refGene_gn=LIPI;refGene_tx=XM_019839519.2;refGene_gn=LIPI;refGene_tx=X
M_023238857.1;refGene_gn=LIPI;refGene_tx=XM_023238858.1;refGene_gn=LIP
I;refGene_tx=XM_023238859.1;refGene_gn=LIPI;refGene_tx=XM_023238860.1;
refGene_gn=LIPI;refGene_tx=XM_023238861.1;refGene_gn=LIPI;refGene_tx=X
M_023238862.1
C2 29362402 29362511
refGene_gn=LIPI;refGene_tx=XM_019839519.2;refGene_gn=LIPI;refGene_tx=X
M_023238857.1;refGene_gn=LIPI;refGene_tx=XM_023238858.1;refGene_gn=LIP
I;refGene_tx=XM_023238859.1;refGene_gn=LIPI;refGene_tx=XM_023238860.1;
refGene_gn=LIPI;refGene_tx=XM_023238861.1;refGene_gn=LIPI;refGene_tx=X
M_023238862.1
C2 29364778 29364880
refGene_gn=LIPI;refGene_tx=XM_019839519.2;refGene_gn=LIPI;refGene_tx=X
M_023238857.1;refGene_gn=LIPI;refGene_tx=XM_023238858.1;refGene_gn=LIP
I;refGene_tx=XM_023238859.1;refGene_gn=LIPI;refGene_tx=XM_023238860.1;
refGene_gn=LIPI;refGene_tx=XM_023238861.1;refGene_gn=LIPI;refGene_tx=X
M_023238862.1
C2 29369773 29369863
refGene_gn=LIPI;refGene_tx=XM_019839519.2;refGene_gn=LIPI;refGene_tx=X
M_023238857.1;refGene_gn=LIPI;refGene_tx=XM_023238858.1;refGene_gn=LIP
I;refGene_tx=XM_023238860.1;refGene_gn=LIPI;refGene_tx=XM_023238861.1;
refGene_gn=LIPI;refGene_tx=XM_023238862.1
C2 29370840 29371008
refGene_gn=LIPI;refGene_tx=XM_019839519.2;refGene_gn=LIPI;refGene_tx=X
M_023238857.1;refGene_gn=LIPI;refGene_tx=XM_023238858.1;refGene_gn=LIP
I;refGene_tx=XM_023238859.1;refGene_gn=LIPI;refGene_tx=XM_023238860.1;
refGene_gn=LIPI;refGene_tx=XM_023238861.1;refGene_gn=LIPI;refGene_tx=X
M_023238862.1
C2 29372771 29372876
refGene_gn=LIPI;refGene_tx=XM_019839519.2;refGene_gn=LIPI;refGene_tx=X
M_023238857.1;refGene_gn=LIPI;refGene_tx=XM_023238858.1;refGene_gn=LIP
I;refGene_tx=XM_023238859.1;refGene_gn=LIPI;refGene_tx=XM_023238860.1;
refGene_gn=LIPI;refGene_tx=XM_023238861.1;refGene_gn=LIPI;refGene_tx=X
M_023238862.1
C2 29387208 29387320
refGene_gn=LIPI;refGene_tx=XM_019839519.2;refGene_gn=LIPI;refGene_tx=X
M_023238857.1;refGene_gn=LIPI;refGene_tx=XM_023238858.1;refGene_gn=LIP
I;refGene_tx=XM_023238859.1;refGene_gn=LIPI;refGene_tx=XM_023238860.1;
refGene_gn=LIPI;refGene_tx=XM_023238861.1;refGene_gn=LIPI;refGene_tx=X
M_023238862.1
C2 29393514 29393691
refGene_gn=LIPI;refGene_tx=XM_019839519.2;refGene_gn=LIPI;refGene_tx=X
M_023238857.1;refGene_gn=LIPI;refGene_tx=XM_023238858.1;refGene_gn=LIP
I;refGene_tx=XM_023238859.1;refGene_gn=LIPI;refGene_tx=XM_023238860.1;
refGene_gn=LIPI;refGene_tx=XM_023238861.1;refGene_gn=LIPI;refGene_tx=X
M_023238862.1

C2	29400981 29401012 refGene_gn=LIPI;refGene_tx=XM_023238858.1
C2	29407350 29407441
refGene_gn=LIPI;refGene_tx=XM_019839519.2;refGene_gn=LIPI;refGene_tx=XM_023238857.1;refGene_gn=LIPI;refGene_tx=XM_023238859.1;refGene_gn=LIPI;refGene_tx=XM_023238860.1;refGene_gn=LIPI;refGene_tx=XM_023238861.1;refGene_gn=LIPI;refGene_tx=XM_023238862.1
C2	29529379 29529488 refGene_gn=ROBO2;refGene_tx=XM_019839528.1
C2	29903492 29903622
refGene_gn=ROBO2;refGene_tx=XM_019839525.1;refGene_gn=ROBO2;refGene_tx=XM_019839526.1;refGene_gn=ROBO2;refGene_tx=XM_019839527.1;refGene_gn=ROBO2;refGene_tx=XM_019839531.1;refGene_gn=ROBO2;refGene_tx=XM_019839532.1;refGene_gn=ROBO2;refGene_tx=XM_019839533.1;refGene_gn=ROBO2;refGene_tx=XM_019839534.1;refGene_gn=ROBO2;refGene_tx=XM_019839536.1;refGene_gn=ROBO2;refGene_tx=XM_019839537.1;refGene_gn=ROBO2;refGene_tx=XM_019839538.1;refGene_gn=ROBO2;refGene_tx=XM_019839539.1;refGene_gn=ROBO2;refGene_tx=XM_019839540.1
C2	30600583 30600644 refGene_gn=ROBO2;refGene_tx=XM_019839529.2
C2	30666447 30666774
refGene_gn=ROBO2;refGene_tx=XM_019839525.1;refGene_gn=ROBO2;refGene_tx=XM_019839526.1;refGene_gn=ROBO2;refGene_tx=XM_019839527.1;refGene_gn=ROBO2;refGene_tx=XM_019839528.1;refGene_gn=ROBO2;refGene_tx=XM_019839529.2;refGene_gn=ROBO2;refGene_tx=XM_019839531.1;refGene_gn=ROBO2;refGene_tx=XM_019839532.1;refGene_gn=ROBO2;refGene_tx=XM_019839533.1;refGene_gn=ROBO2;refGene_tx=XM_019839534.1;refGene_gn=ROBO2;refGene_tx=XM_019839536.1;refGene_gn=ROBO2;refGene_tx=XM_019839537.1;refGene_gn=ROBO2;refGene_tx=XM_019839538.1;refGene_gn=ROBO2;refGene_tx=XM_019839539.1;refGene_gn=ROBO2;refGene_tx=XM_019839540.1
C2	31017326 31017484
refGene_gn=ROBO2;refGene_tx=XM_019839525.1;refGene_gn=ROBO2;refGene_tx=XM_019839526.1;refGene_gn=ROBO2;refGene_tx=XM_019839527.1;refGene_gn=ROBO2;refGene_tx=XM_019839528.1;refGene_gn=ROBO2;refGene_tx=XM_019839529.2;refGene_gn=ROBO2;refGene_tx=XM_019839531.1;refGene_gn=ROBO2;refGene_tx=XM_019839532.1;refGene_gn=ROBO2;refGene_tx=XM_019839533.1;refGene_gn=ROBO2;refGene_tx=XM_019839534.1;refGene_gn=ROBO2;refGene_tx=XM_019839536.1;refGene_gn=ROBO2;refGene_tx=XM_019839537.1;refGene_gn=ROBO2;refGene_tx=XM_019839538.1;refGene_gn=ROBO2;refGene_tx=XM_019839539.1;refGene_gn=ROBO2;refGene_tx=XM_019839540.1
C2	31020311 31020432
refGene_gn=ROBO2;refGene_tx=XM_019839525.1;refGene_gn=ROBO2;refGene_tx=XM_019839526.1;refGene_gn=ROBO2;refGene_tx=XM_019839527.1;refGene_gn=ROBO2;refGene_tx=XM_019839528.1;refGene_gn=ROBO2;refGene_tx=XM_019839529.2;refGene_gn=ROBO2;refGene_tx=XM_019839531.1;refGene_gn=ROBO2;refGene_tx=XM_019839532.1;refGene_gn=ROBO2;refGene_tx=XM_019839533.1;refGene_gn=ROBO2;refGene_tx=XM_019839534.1;refGene_gn=ROBO2;refGene_tx=XM_019839536.1;refGene_gn=ROBO2;refGene_tx=XM_019839537.1;refGene_gn=ROBO2;refGene_tx=XM_019839538.1;refGene_gn=ROBO2;refGene_tx=XM_019839539.1;refGene_gn=ROBO2;refGene_tx=XM_019839540.1
C2	31032673 31032812
refGene_gn=ROBO2;refGene_tx=XM_019839525.1;refGene_gn=ROBO2;refGene_tx=XM_019839526.1;refGene_gn=ROBO2;refGene_tx=XM_019839527.1;refGene_gn=

ROBO2;refGene_tx=XM_019839528.1;refGene_gn=ROBO2;refGene_tx=XM_0198395
29.2;refGene_gn=ROBO2;refGene_tx=XM_019839531.1;refGene_gn=ROBO2;refGe
ne_tx=XM_019839532.1;refGene_gn=ROBO2;refGene_tx=XM_019839533.1;refGen
e_gn=ROBO2;refGene_tx=XM_019839534.1;refGene_gn=ROBO2;refGene_tx=XM_01
9839536.1;refGene_gn=ROBO2;refGene_tx=XM_019839537.1;refGene_gn=ROBO2;
refGene_tx=XM_019839538.1;refGene_gn=ROBO2;refGene_tx=XM_019839539.1;r
efGene_gn=ROBO2;refGene_tx=XM_019839540.1
C2 31061837 31061965
refGene_gn=ROBO2;refGene_tx=XM_019839525.1;refGene_gn=ROBO2;refGene_tx
=XM_019839526.1;refGene_gn=ROBO2;refGene_tx=XM_019839527.1;refGene_gn=
ROBO2;refGene_tx=XM_019839528.1;refGene_gn=ROBO2;refGene_tx=XM_0198395
29.2;refGene_gn=ROBO2;refGene_tx=XM_019839531.1;refGene_gn=ROBO2;refGe
ne_tx=XM_019839532.1;refGene_gn=ROBO2;refGene_tx=XM_019839533.1;refGen
e_gn=ROBO2;refGene_tx=XM_019839534.1;refGene_gn=ROBO2;refGene_tx=XM_01
9839536.1;refGene_gn=ROBO2;refGene_tx=XM_019839537.1;refGene_gn=ROBO2;
refGene_tx=XM_019839538.1;refGene_gn=ROBO2;refGene_tx=XM_019839539.1;r
efGene_gn=ROBO2;refGene_tx=XM_019839540.1;refGene_gn=ROBO2;refGene_tx=
XM_019839541.2
C2 31066539 31066551
refGene_gn=ROBO2;refGene_tx=XM_019839525.1;refGene_gn=ROBO2;refGene_tx
=XM_019839526.1;refGene_gn=ROBO2;refGene_tx=XM_019839528.1;refGene_gn=
ROBO2;refGene_tx=XM_019839529.2;refGene_gn=ROBO2;refGene_tx=XM_0198395
31.1;refGene_gn=ROBO2;refGene_tx=XM_019839532.1;refGene_gn=ROBO2;refGe
ne_tx=XM_019839533.1;refGene_gn=ROBO2;refGene_tx=XM_019839534.1;refGen
e_gn=ROBO2;refGene_tx=XM_019839537.1;refGene_gn=ROBO2;refGene_tx=XM_01
9839538.1;refGene_gn=ROBO2;refGene_tx=XM_019839539.1;refGene_gn=ROBO2;
refGene_tx=XM_019839541.2
C2 31085686 31085811
refGene_gn=ROBO2;refGene_tx=XM_019839525.1;refGene_gn=ROBO2;refGene_tx
=XM_019839526.1;refGene_gn=ROBO2;refGene_tx=XM_019839527.1;refGene_gn=
ROBO2;refGene_tx=XM_019839528.1;refGene_gn=ROBO2;refGene_tx=XM_0198395
29.2;refGene_gn=ROBO2;refGene_tx=XM_019839531.1;refGene_gn=ROBO2;refGe
ne_tx=XM_019839532.1;refGene_gn=ROBO2;refGene_tx=XM_019839533.1;refGen
e_gn=ROBO2;refGene_tx=XM_019839534.1;refGene_gn=ROBO2;refGene_tx=XM_01
9839536.1;refGene_gn=ROBO2;refGene_tx=XM_019839537.1;refGene_gn=ROBO2;
refGene_tx=XM_019839538.1;refGene_gn=ROBO2;refGene_tx=XM_019839539.1;r
efGene_gn=ROBO2;refGene_tx=XM_019839540.1;refGene_gn=ROBO2;refGene_tx=
XM_019839541.2
C2 31090602 31090774
refGene_gn=ROBO2;refGene_tx=XM_019839525.1;refGene_gn=ROBO2;refGene_tx
=XM_019839526.1;refGene_gn=ROBO2;refGene_tx=XM_019839527.1;refGene_gn=
ROBO2;refGene_tx=XM_019839528.1;refGene_gn=ROBO2;refGene_tx=XM_0198395
29.2;refGene_gn=ROBO2;refGene_tx=XM_019839531.1;refGene_gn=ROBO2;refGe
ne_tx=XM_019839532.1;refGene_gn=ROBO2;refGene_tx=XM_019839533.1;refGen
e_gn=ROBO2;refGene_tx=XM_019839534.1;refGene_gn=ROBO2;refGene_tx=XM_01
9839536.1;refGene_gn=ROBO2;refGene_tx=XM_019839537.1;refGene_gn=ROBO2;
refGene_tx=XM_019839538.1;refGene_gn=ROBO2;refGene_tx=XM_019839539.1;r
efGene_gn=ROBO2;refGene_tx=XM_019839540.1;refGene_gn=ROBO2;refGene_tx=
XM_019839541.2
C2 31095468 31095674

refGene_gn=ROBO2;refGene_tx=XM_019839525.1;refGene_gn=ROBO2;refGene_tx=XM_019839526.1;refGene_gn=ROBO2;refGene_tx=XM_019839527.1;refGene_gn=ROBO2;refGene_tx=XM_019839528.1;refGene_gn=ROBO2;refGene_tx=XM_019839529.2;refGene_gn=ROBO2;refGene_tx=XM_019839531.1;refGene_gn=ROBO2;refGene_tx=XM_019839532.1;refGene_gn=ROBO2;refGene_tx=XM_019839533.1;refGene_gn=ROBO2;refGene_tx=XM_019839534.1;refGene_gn=ROBO2;refGene_tx=XM_019839536.1;refGene_gn=ROBO2;refGene_tx=XM_019839537.1;refGene_gn=ROBO2;refGene_tx=XM_019839538.1;refGene_gn=ROBO2;refGene_tx=XM_019839539.1;refGene_gn=ROBO2;refGene_tx=XM_019839540.1;refGene_gn=ROBO2;refGene_tx=XM_019839541.2
C2 31100010 31100092
refGene_gn=ROBO2;refGene_tx=XM_019839525.1;refGene_gn=ROBO2;refGene_tx=XM_019839526.1;refGene_gn=ROBO2;refGene_tx=XM_019839527.1;refGene_gn=ROBO2;refGene_tx=XM_019839528.1;refGene_gn=ROBO2;refGene_tx=XM_019839529.2;refGene_gn=ROBO2;refGene_tx=XM_019839531.1;refGene_gn=ROBO2;refGene_tx=XM_019839532.1;refGene_gn=ROBO2;refGene_tx=XM_019839533.1;refGene_gn=ROBO2;refGene_tx=XM_019839534.1;refGene_gn=ROBO2;refGene_tx=XM_019839536.1;refGene_gn=ROBO2;refGene_tx=XM_019839537.1;refGene_gn=ROBO2;refGene_tx=XM_019839538.1;refGene_gn=ROBO2;refGene_tx=XM_019839539.1;refGene_gn=ROBO2;refGene_tx=XM_019839540.1;refGene_gn=ROBO2;refGene_tx=XM_019839541.2
C2 31100545 31100708
refGene_gn=ROBO2;refGene_tx=XM_019839525.1;refGene_gn=ROBO2;refGene_tx=XM_019839526.1;refGene_gn=ROBO2;refGene_tx=XM_019839527.1;refGene_gn=ROBO2;refGene_tx=XM_019839528.1;refGene_gn=ROBO2;refGene_tx=XM_019839529.2;refGene_gn=ROBO2;refGene_tx=XM_019839531.1;refGene_gn=ROBO2;refGene_tx=XM_019839532.1;refGene_gn=ROBO2;refGene_tx=XM_019839533.1;refGene_gn=ROBO2;refGene_tx=XM_019839534.1;refGene_gn=ROBO2;refGene_tx=XM_019839536.1;refGene_gn=ROBO2;refGene_tx=XM_019839537.1;refGene_gn=ROBO2;refGene_tx=XM_019839538.1;refGene_gn=ROBO2;refGene_tx=XM_019839539.1;refGene_gn=ROBO2;refGene_tx=XM_019839540.1;refGene_gn=ROBO2;refGene_tx=XM_019839541.2
C2 31102336 31102503
refGene_gn=ROBO2;refGene_tx=XM_019839525.1;refGene_gn=ROBO2;refGene_tx=XM_019839526.1;refGene_gn=ROBO2;refGene_tx=XM_019839527.1;refGene_gn=ROBO2;refGene_tx=XM_019839528.1;refGene_gn=ROBO2;refGene_tx=XM_019839529.2;refGene_gn=ROBO2;refGene_tx=XM_019839531.1;refGene_gn=ROBO2;refGene_tx=XM_019839532.1;refGene_gn=ROBO2;refGene_tx=XM_019839533.1;refGene_gn=ROBO2;refGene_tx=XM_019839534.1;refGene_gn=ROBO2;refGene_tx=XM_019839536.1;refGene_gn=ROBO2;refGene_tx=XM_019839537.1;refGene_gn=ROBO2;refGene_tx=XM_019839538.1;refGene_gn=ROBO2;refGene_tx=XM_019839539.1;refGene_gn=ROBO2;refGene_tx=XM_019839540.1;refGene_gn=ROBO2;refGene_tx=XM_019839541.2
C2 31106774 31106896
refGene_gn=ROBO2;refGene_tx=XM_019839525.1;refGene_gn=ROBO2;refGene_tx=XM_019839526.1;refGene_gn=ROBO2;refGene_tx=XM_019839527.1;refGene_gn=ROBO2;refGene_tx=XM_019839528.1;refGene_gn=ROBO2;refGene_tx=XM_019839529.2;refGene_gn=ROBO2;refGene_tx=XM_019839531.1;refGene_gn=ROBO2;refGene_tx=XM_019839532.1;refGene_gn=ROBO2;refGene_tx=XM_019839533.1;refGene_gn=ROBO2;refGene_tx=XM_01

9839536.1;refGene_gn=ROBO2;refGene_tx=XM_019839537.1;refGene_gn=ROBO2;
refGene_tx=XM_019839538.1;refGene_gn=ROBO2;refGene_tx=XM_019839539.1;r
efGene_gn=ROBO2;refGene_tx=XM_019839540.1;refGene_gn=ROBO2;refGene_tx=
XM_019839541.2
C2 31114980 31115212
refGene_gn=ROBO2;refGene_tx=XM_019839525.1;refGene_gn=ROBO2;refGene_tx
=XM_019839526.1;refGene_gn=ROBO2;refGene_tx=XM_019839527.1;refGene_gn=
ROBO2;refGene_tx=XM_019839528.1;refGene_gn=ROBO2;refGene_tx=XM_0198395
29.2;refGene_gn=ROBO2;refGene_tx=XM_019839531.1;refGene_gn=ROBO2;refGe
ne_tx=XM_019839532.1;refGene_gn=ROBO2;refGene_tx=XM_019839533.1;refGen
e_gn=ROBO2;refGene_tx=XM_019839534.1;refGene_gn=ROBO2;refGene_tx=XM_01
9839536.1;refGene_gn=ROBO2;refGene_tx=XM_019839537.1;refGene_gn=ROBO2;
refGene_tx=XM_019839538.1;refGene_gn=ROBO2;refGene_tx=XM_019839539.1;r
efGene_gn=ROBO2;refGene_tx=XM_019839540.1;refGene_gn=ROBO2;refGene_tx=
XM_019839541.2
C2 31117931 31118056
refGene_gn=ROBO2;refGene_tx=XM_019839525.1;refGene_gn=ROBO2;refGene_tx
=XM_019839526.1;refGene_gn=ROBO2;refGene_tx=XM_019839527.1;refGene_gn=
ROBO2;refGene_tx=XM_019839528.1;refGene_gn=ROBO2;refGene_tx=XM_0198395
29.2;refGene_gn=ROBO2;refGene_tx=XM_019839531.1;refGene_gn=ROBO2;refGe
ne_tx=XM_019839532.1;refGene_gn=ROBO2;refGene_tx=XM_019839533.1;refGen
e_gn=ROBO2;refGene_tx=XM_019839534.1;refGene_gn=ROBO2;refGene_tx=XM_01
9839536.1;refGene_gn=ROBO2;refGene_tx=XM_019839537.1;refGene_gn=ROBO2;
refGene_tx=XM_019839538.1;refGene_gn=ROBO2;refGene_tx=XM_019839539.1;r
efGene_gn=ROBO2;refGene_tx=XM_019839540.1;refGene_gn=ROBO2;refGene_tx=
XM_019839541.2
C2 31120289 31120461
refGene_gn=ROBO2;refGene_tx=XM_019839525.1;refGene_gn=ROBO2;refGene_tx
=XM_019839526.1;refGene_gn=ROBO2;refGene_tx=XM_019839527.1;refGene_gn=
ROBO2;refGene_tx=XM_019839528.1;refGene_gn=ROBO2;refGene_tx=XM_0198395
29.2;refGene_gn=ROBO2;refGene_tx=XM_019839531.1;refGene_gn=ROBO2;refGe
ne_tx=XM_019839532.1;refGene_gn=ROBO2;refGene_tx=XM_019839533.1;refGen
e_gn=ROBO2;refGene_tx=XM_019839534.1;refGene_gn=ROBO2;refGene_tx=XM_01
9839536.1;refGene_gn=ROBO2;refGene_tx=XM_019839537.1;refGene_gn=ROBO2;
refGene_tx=XM_019839538.1;refGene_gn=ROBO2;refGene_tx=XM_019839539.1;r
efGene_gn=ROBO2;refGene_tx=XM_019839540.1;refGene_gn=ROBO2;refGene_tx=
XM_019839541.2
C2 31130934 31131117
refGene_gn=ROBO2;refGene_tx=XM_019839525.1;refGene_gn=ROBO2;refGene_tx
=XM_019839526.1;refGene_gn=ROBO2;refGene_tx=XM_019839527.1;refGene_gn=
ROBO2;refGene_tx=XM_019839528.1;refGene_gn=ROBO2;refGene_tx=XM_0198395
29.2;refGene_gn=ROBO2;refGene_tx=XM_019839531.1;refGene_gn=ROBO2;refGe
ne_tx=XM_019839532.1;refGene_gn=ROBO2;refGene_tx=XM_019839533.1;refGen
e_gn=ROBO2;refGene_tx=XM_019839534.1;refGene_gn=ROBO2;refGene_tx=XM_01
9839536.1;refGene_gn=ROBO2;refGene_tx=XM_019839537.1;refGene_gn=ROBO2;
refGene_tx=XM_019839538.1;refGene_gn=ROBO2;refGene_tx=XM_019839539.1;r
efGene_gn=ROBO2;refGene_tx=XM_019839540.1;refGene_gn=ROBO2;refGene_tx=
XM_019839541.2
C2 31141226 31141269
refGene_gn=ROBO2;refGene_tx=XM_019839525.1;refGene_gn=ROBO2;refGene_tx

=XM_019839526.1;refGene_gn=ROBO2;refGene_tx=XM_019839527.1;refGene_gn=ROBO2;refGene_tx=XM_019839528.1;refGene_gn=ROBO2;refGene_tx=XM_019839529.2;refGene_gn=ROBO2;refGene_tx=XM_019839531.1;refGene_gn=ROBO2;refGene_tx=XM_019839532.1;refGene_gn=ROBO2;refGene_tx=XM_019839534.1;refGene_gn=ROBO2;refGene_tx=XM_019839536.1;refGene_gn=ROBO2;refGene_tx=XM_019839537.1;refGene_gn=ROBO2;refGene_tx=XM_019839538.1;refGene_gn=ROBO2;refGene_tx=XM_019839539.1;refGene_gn=ROBO2;refGene_tx=XM_019839540.1;refGene_gn=ROBO2;refGene_tx=XM_019839541.2
C2 31142691 31142819
refGene_gn=ROBO2;refGene_tx=XM_019839525.1;refGene_gn=ROBO2;refGene_tx=XM_019839526.1;refGene_gn=ROBO2;refGene_tx=XM_019839527.1;refGene_gn=ROBO2;refGene_tx=XM_019839528.1;refGene_gn=ROBO2;refGene_tx=XM_019839529.2;refGene_gn=ROBO2;refGene_tx=XM_019839531.1;refGene_gn=ROBO2;refGene_tx=XM_019839532.1;refGene_gn=ROBO2;refGene_tx=XM_019839534.1;refGene_gn=ROBO2;refGene_tx=XM_019839536.1;refGene_gn=ROBO2;refGene_tx=XM_019839537.1;refGene_gn=ROBO2;refGene_tx=XM_019839538.1;refGene_gn=ROBO2;refGene_tx=XM_019839539.1;refGene_gn=ROBO2;refGene_tx=XM_019839540.1;refGene_gn=ROBO2;refGene_tx=XM_019839541.2
C2 31148984 31149266
refGene_gn=ROBO2;refGene_tx=XM_019839525.1;refGene_gn=ROBO2;refGene_tx=XM_019839526.1;refGene_gn=ROBO2;refGene_tx=XM_019839527.1;refGene_gn=ROBO2;refGene_tx=XM_019839528.1;refGene_gn=ROBO2;refGene_tx=XM_019839529.2;refGene_gn=ROBO2;refGene_tx=XM_019839531.1;refGene_gn=ROBO2;refGene_tx=XM_019839532.1;refGene_gn=ROBO2;refGene_tx=XM_019839533.1;refGene_gn=ROBO2;refGene_tx=XM_019839534.1;refGene_gn=ROBO2;refGene_tx=XM_019839536.1;refGene_gn=ROBO2;refGene_tx=XM_019839537.1;refGene_gn=ROBO2;refGene_tx=XM_019839538.1;refGene_gn=ROBO2;refGene_tx=XM_019839539.1;refGene_gn=ROBO2;refGene_tx=XM_019839540.1;refGene_gn=ROBO2;refGene_tx=XM_019839541.2
C2 31149665 31149791
refGene_gn=ROBO2;refGene_tx=XM_019839525.1;refGene_gn=ROBO2;refGene_tx=XM_019839526.1;refGene_gn=ROBO2;refGene_tx=XM_019839527.1;refGene_gn=ROBO2;refGene_tx=XM_019839528.1;refGene_gn=ROBO2;refGene_tx=XM_019839529.2;refGene_gn=ROBO2;refGene_tx=XM_019839533.1;refGene_gn=ROBO2;refGene_tx=XM_019839534.1;refGene_gn=ROBO2;refGene_tx=XM_019839536.1;refGene_gn=ROBO2;refGene_tx=XM_019839537.1;refGene_gn=ROBO2;refGene_tx=XM_019839541.2
C2 31155400 31155557
refGene_gn=ROBO2;refGene_tx=XM_019839525.1;refGene_gn=ROBO2;refGene_tx=XM_019839526.1;refGene_gn=ROBO2;refGene_tx=XM_019839527.1;refGene_gn=ROBO2;refGene_tx=XM_019839528.1;refGene_gn=ROBO2;refGene_tx=XM_019839529.2;refGene_gn=ROBO2;refGene_tx=XM_019839531.1;refGene_gn=ROBO2;refGene_tx=XM_019839532.1;refGene_gn=ROBO2;refGene_tx=XM_019839533.1;refGene_gn=ROBO2;refGene_tx=XM_019839534.1;refGene_gn=ROBO2;refGene_tx=XM_019839536.1;refGene_gn=ROBO2;refGene_tx=XM_019839537.1;refGene_gn=ROBO2;refGene_tx=XM_019839538.1;refGene_gn=ROBO2;refGene_tx=XM_019839539.1;refGene_gn=ROBO2;refGene_tx=XM_019839540.1;refGene_gn=ROBO2;refGene_tx=XM_019839541.2
C2 31166689 31166950
refGene_gn=ROBO2;refGene_tx=XM_019839525.1;refGene_gn=ROBO2;refGene_tx

=XM_019839526.1;refGene_gn=ROBO2;refGene_tx=XM_019839527.1;refGene_gn=ROBO2;refGene_tx=XM_019839528.1;refGene_gn=ROBO2;refGene_tx=XM_019839529.2;refGene_gn=ROBO2;refGene_tx=XM_019839531.1;refGene_gn=ROBO2;refGene_tx=XM_019839532.1;refGene_gn=ROBO2;refGene_tx=XM_019839533.1;refGene_gn=ROBO2;refGene_tx=XM_019839534.1;refGene_gn=ROBO2;refGene_tx=XM_019839536.1;refGene_gn=ROBO2;refGene_tx=XM_019839537.1;refGene_gn=ROBO2;refGene_tx=XM_019839538.1;refGene_gn=ROBO2;refGene_tx=XM_019839539.1;refGene_gn=ROBO2;refGene_tx=XM_019839540.1;refGene_gn=ROBO2;refGene_tx=XM_019839541.2
C2 31171594 31171800
refGene_gn=ROBO2;refGene_tx=XM_019839525.1;refGene_gn=ROBO2;refGene_tx=XM_019839526.1;refGene_gn=ROBO2;refGene_tx=XM_019839527.1;refGene_gn=ROBO2;refGene_tx=XM_019839528.1;refGene_gn=ROBO2;refGene_tx=XM_019839529.2;refGene_gn=ROBO2;refGene_tx=XM_019839531.1;refGene_gn=ROBO2;refGene_tx=XM_019839532.1;refGene_gn=ROBO2;refGene_tx=XM_019839533.1;refGene_gn=ROBO2;refGene_tx=XM_019839534.1;refGene_gn=ROBO2;refGene_tx=XM_019839536.1;refGene_gn=ROBO2;refGene_tx=XM_019839537.1;refGene_gn=ROBO2;refGene_tx=XM_019839538.1;refGene_gn=ROBO2;refGene_tx=XM_019839539.1;refGene_gn=ROBO2;refGene_tx=XM_019839540.1;refGene_gn=ROBO2;refGene_tx=XM_019839541.2
C2 31181506 31181689
refGene_gn=ROBO2;refGene_tx=XM_019839525.1;refGene_gn=ROBO2;refGene_tx=XM_019839526.1;refGene_gn=ROBO2;refGene_tx=XM_019839527.1;refGene_gn=ROBO2;refGene_tx=XM_019839528.1;refGene_gn=ROBO2;refGene_tx=XM_019839529.2;refGene_gn=ROBO2;refGene_tx=XM_019839531.1;refGene_gn=ROBO2;refGene_tx=XM_019839532.1;refGene_gn=ROBO2;refGene_tx=XM_019839533.1;refGene_gn=ROBO2;refGene_tx=XM_019839537.1;refGene_gn=ROBO2;refGene_tx=XM_019839538.1;refGene_gn=ROBO2;refGene_tx=XM_019839541.2
C2 31183906 31184080
refGene_gn=ROBO2;refGene_tx=XM_019839525.1;refGene_gn=ROBO2;refGene_tx=XM_019839526.1;refGene_gn=ROBO2;refGene_tx=XM_019839527.1;refGene_gn=ROBO2;refGene_tx=XM_019839528.1;refGene_gn=ROBO2;refGene_tx=XM_019839529.2;refGene_gn=ROBO2;refGene_tx=XM_019839531.1;refGene_gn=ROBO2;refGene_tx=XM_019839532.1;refGene_gn=ROBO2;refGene_tx=XM_019839533.1;refGene_gn=ROBO2;refGene_tx=XM_019839534.1;refGene_gn=ROBO2;refGene_tx=XM_019839536.1;refGene_gn=ROBO2;refGene_tx=XM_019839537.1;refGene_gn=ROBO2;refGene_tx=XM_019839538.1;refGene_gn=ROBO2;refGene_tx=XM_019839539.1;refGene_gn=ROBO2;refGene_tx=XM_019839540.1;refGene_gn=ROBO2;refGene_tx=XM_019839541.2
C2 31190806 31191079
refGene_gn=ROBO2;refGene_tx=XM_019839525.1;refGene_gn=ROBO2;refGene_tx=XM_019839526.1;refGene_gn=ROBO2;refGene_tx=XM_019839527.1;refGene_gn=ROBO2;refGene_tx=XM_019839528.1;refGene_gn=ROBO2;refGene_tx=XM_019839529.2;refGene_gn=ROBO2;refGene_tx=XM_019839531.1;refGene_gn=ROBO2;refGene_tx=XM_019839532.1;refGene_gn=ROBO2;refGene_tx=XM_019839533.1;refGene_gn=ROBO2;refGene_tx=XM_019839534.1;refGene_gn=ROBO2;refGene_tx=XM_019839536.1;refGene_gn=ROBO2;refGene_tx=XM_019839541.2
C2 31192819 31193020
refGene_gn=ROBO2;refGene_tx=XM_019839525.1;refGene_gn=ROBO2;refGene_tx=XM_019839526.1;refGene_gn=ROBO2;refGene_tx=XM_019839527.1;refGene_gn=

ROBO2;refGene_tx=XM_019839528.1;refGene_gn=ROBO2;refGene_tx=XM_0198395
29.2;refGene_gn=ROBO2;refGene_tx=XM_019839531.1;refGene_gn=ROBO2;refGe
ne_tx=XM_019839532.1;refGene_gn=ROBO2;refGene_tx=XM_019839533.1;refGen
e_gn=ROBO2;refGene_tx=XM_019839534.1;refGene_gn=ROBO2;refGene_tx=XM_01
9839536.1;refGene_gn=ROBO2;refGene_tx=XM_019839537.1;refGene_gn=ROBO2;
refGene_tx=XM_019839538.1;refGene_gn=ROBO2;refGene_tx=XM_019839539.1;r
efGene_gn=ROBO2;refGene_tx=XM_019839540.1;refGene_gn=ROBO2;refGene_tx=
XM_019839541.2
C2 31195273 31195278
refGene_gn=ROBO2;refGene_tx=XM_019839525.1;refGene_gn=ROBO2;refGene_tx
=XM_019839526.1;refGene_gn=ROBO2;refGene_tx=XM_019839527.1;refGene_gn=
ROBO2;refGene_tx=XM_019839528.1;refGene_gn=ROBO2;refGene_tx=XM_0198395
29.2;refGene_gn=ROBO2;refGene_tx=XM_019839531.1;refGene_gn=ROBO2;refGe
ne_tx=XM_019839532.1;refGene_gn=ROBO2;refGene_tx=XM_019839533.1;refGen
e_gn=ROBO2;refGene_tx=XM_019839534.1;refGene_gn=ROBO2;refGene_tx=XM_01
9839536.1;refGene_gn=ROBO2;refGene_tx=XM_019839537.1;refGene_gn=ROBO2;
refGene_tx=XM_019839538.1;refGene_gn=ROBO2;refGene_tx=XM_019839539.1;r
efGene_gn=ROBO2;refGene_tx=XM_019839540.1;refGene_gn=ROBO2;refGene_tx=
XM_019839541.2
C2 32073865 32073880 refGene_gn=ROBO1;refGene_tx=XM_019839543.2
C2 32075124 32075321 refGene_gn=ROBO1;refGene_tx=XM_019839543.2
C2 32080725 32080777 refGene_gn=ROBO1;refGene_tx=XM_019839543.2
C2 32080778 32081014 refGene_gn=ROBO1;refGene_tx=XM_019839543.2
C2 32081016 32081034 refGene_gn=ROBO1;refGene_tx=XM_019839543.2
C2 32088166 32088319 refGene_gn=ROBO1;refGene_tx=XM_019839543.2
C2 32091392 32091799 refGene_gn=ROBO1;refGene_tx=XM_019839543.2
C2 32101603 32101852 refGene_gn=ROBO1;refGene_tx=XM_019839543.2
C2 32104998 32105143 refGene_gn=ROBO1;refGene_tx=XM_019839543.2
C2 32107818 32107926 refGene_gn=ROBO1;refGene_tx=XM_019839543.2
C2 32109789 32110080 refGene_gn=ROBO1;refGene_tx=XM_019839543.2
C2 32110082 32110124 refGene_gn=ROBO1;refGene_tx=XM_019839543.2
C2 32113213 32113368 refGene_gn=ROBO1;refGene_tx=XM_019839543.2
C2 32119953 32119996 refGene_gn=ROBO1;refGene_tx=XM_019839543.2
C2 32121656 32121683 refGene_gn=ROBO1;refGene_tx=XM_019839543.2
C2 32126698 32126896 refGene_gn=ROBO1;refGene_tx=XM_019839543.2
C2 32131307 32131479 refGene_gn=ROBO1;refGene_tx=XM_019839543.2
C2 32134506 32134628 refGene_gn=ROBO1;refGene_tx=XM_019839543.2
C2 32135586 32135818 refGene_gn=ROBO1;refGene_tx=XM_019839543.2
C2 32136814 32136936 refGene_gn=ROBO1;refGene_tx=XM_019839543.2
C2 32142242 32142409 refGene_gn=ROBO1;refGene_tx=XM_019839543.2
C2 32142510 32142679 refGene_gn=ROBO1;refGene_tx=XM_019839543.2
C2 32142849 32142931 refGene_gn=ROBO1;refGene_tx=XM_019839543.2
C2 32144719 32144925 refGene_gn=ROBO1;refGene_tx=XM_019839543.2
C2 32151183 32151355 refGene_gn=ROBO1;refGene_tx=XM_019839543.2
C2 32154209 32154334 refGene_gn=ROBO1;refGene_tx=XM_019839543.2
C2 32181398 32181526 refGene_gn=ROBO1;refGene_tx=XM_019839543.2
C2 32183927 32184066 refGene_gn=ROBO1;refGene_tx=XM_019839543.2
C2 32184418 32184539 refGene_gn=ROBO1;refGene_tx=XM_019839543.2
C2 32213086 32213244 refGene_gn=ROBO1;refGene_tx=XM_019839543.2
C2 32403655 32403982 refGene_gn=ROBO1;refGene_tx=XM_019839543.2

C2 32581370 32581454 refGene_gn=ROBO1;refGene_tx=XM_019839543.2
C2 33045007 33045095 refGene_gn=ROBO1;refGene_tx=XM_019839543.2
C2 34697813 34697858 refGene_gn=GBE1;refGene_tx=XM_019839239.2
C2 34712956 34713013 refGene_gn=GBE1;refGene_tx=NM_001009872.1
C2 34720742 34720860
refGene_gn=GBE1;refGene_tx=NM_001009872.1;refGene_gn=GBE1;refGene_tx=XM_019839239.2
C2 34741234 34741365
refGene_gn=GBE1;refGene_tx=NM_001009872.1;refGene_gn=GBE1;refGene_tx=XM_019839239.2
C2 34742990 34743175
refGene_gn=GBE1;refGene_tx=NM_001009872.1;refGene_gn=GBE1;refGene_tx=XM_019839239.2
C2 34779385 34779557
refGene_gn=GBE1;refGene_tx=NM_001009872.1;refGene_gn=GBE1;refGene_tx=XM_019839239.2
C2 34783447 34783558
refGene_gn=GBE1;refGene_tx=NM_001009872.1;refGene_gn=GBE1;refGene_tx=XM_019839239.2
C2 34788694 34788793
refGene_gn=GBE1;refGene_tx=NM_001009872.1;refGene_gn=GBE1;refGene_tx=XM_019839239.2
C2 34793091 34793219
refGene_gn=GBE1;refGene_tx=NM_001009872.1;refGene_gn=GBE1;refGene_tx=XM_019839239.2
C2 34797530 34797646
refGene_gn=GBE1;refGene_tx=NM_001009872.1;refGene_gn=GBE1;refGene_tx=XM_019839239.2
C2 34844767 34844977
refGene_gn=GBE1;refGene_tx=NM_001009872.1;refGene_gn=GBE1;refGene_tx=XM_019839239.2
C2 34848425 34848516
refGene_gn=GBE1;refGene_tx=NM_001009872.1;refGene_gn=GBE1;refGene_tx=XM_019839239.2
C2 34850673 34850809
refGene_gn=GBE1;refGene_tx=NM_001009872.1;refGene_gn=GBE1;refGene_tx=XM_019839239.2
C2 34851611 34851737
refGene_gn=GBE1;refGene_tx=NM_001009872.1;refGene_gn=GBE1;refGene_tx=XM_019839239.2
C2 34894332 34894448
refGene_gn=GBE1;refGene_tx=NM_001009872.1;refGene_gn=GBE1;refGene_tx=XM_019839239.2
C2 34927474 34927644
refGene_gn=GBE1;refGene_tx=NM_001009872.1;refGene_gn=GBE1;refGene_tx=XM_019839239.2
C2 34980995 34981129
refGene_gn=GBE1;refGene_tx=NM_001009872.1;refGene_gn=GBE1;refGene_tx=XM_019839239.2
C2 37936378 37936439

refGene_gn=CADM2;refGene_tx=XM_003991537.5;refGene_gn=CADM2;refGene_tx
=XM_019839555.2;refGene_gn=CADM2;refGene_tx=XM_019839556.2
C2 38156185 38156730
refGene_gn=LOC109503392;refGene_tx=XM_019839557.1
C2 38156732 38156970
refGene_gn=LOC109503392;refGene_tx=XM_019839557.1
C2 38664319 38664346
refGene_gn=CADM2;refGene_tx=XM_019839555.2;refGene_gn=CADM2;refGene_tx
=XM_019839556.2
C2 38736536 38736686
refGene_gn=CADM2;refGene_tx=XM_003991537.5;refGene_gn=CADM2;refGene_tx
=XM_019839555.2;refGene_gn=CADM2;refGene_tx=XM_019839556.2
C2 38814053 38814206
refGene_gn=CADM2;refGene_tx=XM_003991537.5;refGene_gn=CADM2;refGene_tx
=XM_019839555.2;refGene_gn=CADM2;refGene_tx=XM_019839556.2
C2 38817270 38817408
refGene_gn=CADM2;refGene_tx=XM_003991537.5;refGene_gn=CADM2;refGene_tx
=XM_019839555.2;refGene_gn=CADM2;refGene_tx=XM_019839556.2
C2 38847413 38847584
refGene_gn=CADM2;refGene_tx=XM_003991537.5;refGene_gn=CADM2;refGene_tx
=XM_019839555.2;refGene_gn=CADM2;refGene_tx=XM_019839556.2
C2 38872367 38872458
refGene_gn=CADM2;refGene_tx=XM_003991537.5;refGene_gn=CADM2;refGene_tx
=XM_019839555.2;refGene_gn=CADM2;refGene_tx=XM_019839556.2
C2 38897952 38898131
refGene_gn=CADM2;refGene_tx=XM_003991537.5;refGene_gn=CADM2;refGene_tx
=XM_019839555.2;refGene_gn=CADM2;refGene_tx=XM_019839556.2
C2 38914892 38915012
refGene_gn=CADM2;refGene_tx=XM_003991537.5;refGene_gn=CADM2;refGene_tx
=XM_019839555.2
C2 38998255 38998381
refGene_gn=CADM2;refGene_tx=XM_003991537.5;refGene_gn=CADM2;refGene_tx
=XM_019839555.2;refGene_gn=CADM2;refGene_tx=XM_019839556.2
C2 38999169 38999288
refGene_gn=CADM2;refGene_tx=XM_003991537.5;refGene_gn=CADM2;refGene_tx
=XM_019839555.2;refGene_gn=CADM2;refGene_tx=XM_019839556.2
C2 39823987 39824031 refGene_gn=VGLL3;refGene_tx=XM_023238867.1
C2 39842879 39843039 refGene_gn=VGLL3;refGene_tx=XM_023238867.1
C2 39843040 39843271 refGene_gn=VGLL3;refGene_tx=XM_023238867.1
C2 39843272 39843418 refGene_gn=VGLL3;refGene_tx=XM_023238867.1
C2 39851965 39852251 refGene_gn=VGLL3;refGene_tx=XM_023238867.1
C2 39865813 39865939 refGene_gn=VGLL3;refGene_tx=XM_023238867.1
C2 40070684 40070718 refGene_gn=CHMP2B;refGene_tx=XM_003991539.5
C2 40084402 40084494 refGene_gn=CHMP2B;refGene_tx=XM_003991539.5
C2 40090421 40090616 refGene_gn=CHMP2B;refGene_tx=XM_003991539.5
C2 40094021 40094124 refGene_gn=CHMP2B;refGene_tx=XM_003991539.5
C2 40098644 40098751 refGene_gn=CHMP2B;refGene_tx=XM_003991539.5
C2 40098939 40099050 refGene_gn=CHMP2B;refGene_tx=XM_003991539.5
C2 40104860 40105071 refGene_gn=POU1F1;refGene_tx=XM_023238868.1
C2 40105986 40106047 refGene_gn=POU1F1;refGene_tx=XM_023238868.1

C2 40106976 40107141 refGene_gn=POU1F1;refGene_tx=XM_023238868.1
C2 40109743 40109968 refGene_gn=POU1F1;refGene_tx=XM_023238868.1
C2 40119764 40119836 refGene_gn=POU1F1;refGene_tx=XM_023238868.1
C2 40122204 40122346 refGene_gn=POU1F1;refGene_tx=XM_023238868.1
C2 40693804 40694905 refGene_gn=HTR1F;refGene_tx=XM_023238870.1
C2 40758589 40759093
refGene_gn=CGGBP1;refGene_tx=XM_006935969.4;refGene_gn=CGGBP1;refGene_tx=XM_011285989.3
C2 40762656 40762845 refGene_gn=ZNF654;refGene_tx=XM_023238871.1
C2 40791825 40791971 refGene_gn=ZNF654;refGene_tx=XM_023238871.1
C2 40818712 40818794 refGene_gn=ZNF654;refGene_tx=XM_023238871.1
C2 40833249 40833385 refGene_gn=ZNF654;refGene_tx=XM_023238871.1
C2 40836004 40836207 refGene_gn=ZNF654;refGene_tx=XM_023238871.1
C2 40836883 40837023 refGene_gn=ZNF654;refGene_tx=XM_023238871.1
C2 40847276 40847418 refGene_gn=ZNF654;refGene_tx=XM_023238871.1
C2 40851640 40853987 refGene_gn=ZNF654;refGene_tx=XM_023238871.1
C2 40854579 40854587 refGene_gn=ZNF654;refGene_tx=XM_023238871.1
C2 40863103 40863230
refGene_gn=CC2H3orf38;refGene_tx=XM_003991549.4
C2 40865976 40866215
refGene_gn=CC2H3orf38;refGene_tx=XM_003991549.4
C2 40868007 40868688
refGene_gn=CC2H3orf38;refGene_tx=XM_003991549.4
C2 41682203 41682291
refGene_gn=EPHA3;refGene_tx=XM_011285988.3;refGene_gn=EPHA3;refGene_tx=XM_023238873.1
C2 41701296 41701361
refGene_gn=EPHA3;refGene_tx=XM_011285988.3;refGene_gn=EPHA3;refGene_tx=XM_023238873.1
C2 41779993 41780654
refGene_gn=EPHA3;refGene_tx=XM_011285988.3;refGene_gn=EPHA3;refGene_tx=XM_023238873.1
C2 41907320 41907476
refGene_gn=EPHA3;refGene_tx=XM_011285988.3;refGene_gn=EPHA3;refGene_tx=XM_023238873.1
C2 41908174 41908510
refGene_gn=EPHA3;refGene_tx=XM_011285988.3;refGene_gn=EPHA3;refGene_tx=XM_023238873.1
C2 41959882 41960007
refGene_gn=EPHA3;refGene_tx=XM_011285988.3;refGene_gn=EPHA3;refGene_tx=XM_023238873.1
C2 41962913 41963076
refGene_gn=EPHA3;refGene_tx=XM_011285988.3;refGene_gn=EPHA3;refGene_tx=XM_023238873.1
C2 41970785 41970888
refGene_gn=EPHA3;refGene_tx=XM_011285988.3;refGene_gn=EPHA3;refGene_tx=XM_023238873.1
C2 41971592 41971660
refGene_gn=EPHA3;refGene_tx=XM_011285988.3;refGene_gn=EPHA3;refGene_tx=XM_023238873.1

C2 41976912 41977038
refGene_gn=EPHA3;refGene_tx=XM_011285988.3;refGene_gn=EPHA3;refGene_tx=XM_023238873.1
C2 41983419 41983605
refGene_gn=EPHA3;refGene_tx=XM_011285988.3;refGene_gn=EPHA3;refGene_tx=XM_023238873.1
C2 41993967 41994029
refGene_gn=EPHA3;refGene_tx=XM_011285988.3;refGene_gn=EPHA3;refGene_tx=XM_023238873.1
C2 41996434 41996644
refGene_gn=EPHA3;refGene_tx=XM_011285988.3;refGene_gn=EPHA3;refGene_tx=XM_023238873.1
C2 42012492 42012642
refGene_gn=EPHA3;refGene_tx=XM_011285988.3;refGene_gn=EPHA3;refGene_tx=XM_023238873.1
C2 42014229 42014423
refGene_gn=EPHA3;refGene_tx=XM_011285988.3;refGene_gn=EPHA3;refGene_tx=XM_023238873.1
C2 42030560 42030716
refGene_gn=EPHA3;refGene_tx=XM_011285988.3;refGene_gn=EPHA3;refGene_tx=XM_023238873.1
C2 42036854 42036960
refGene_gn=EPHA3;refGene_tx=XM_011285988.3;refGene_gn=EPHA3;refGene_tx=XM_023238873.1
C2 42636050 42636211
refGene_gn=PROS1;refGene_tx=XM_011285987.3;refGene_gn=PROS1;refGene_tx=XM_023238875.1
C2 42640062 42640288
refGene_gn=PROS1;refGene_tx=XM_011285987.3;refGene_gn=PROS1;refGene_tx=XM_023238875.1
C2 42642337 42642486
refGene_gn=PROS1;refGene_tx=XM_011285987.3;refGene_gn=PROS1;refGene_tx=XM_023238875.1
C2 42648080 42648249
refGene_gn=PROS1;refGene_tx=XM_011285987.3;refGene_gn=PROS1;refGene_tx=XM_023238875.1
C2 42649523 42649691
refGene_gn=PROS1;refGene_tx=XM_011285987.3;refGene_gn=PROS1;refGene_tx=XM_023238875.1
C2 42656185 42656375
refGene_gn=PROS1;refGene_tx=XM_011285987.3;refGene_gn=PROS1;refGene_tx=XM_023238875.1
C2 42659392 42659508
refGene_gn=PROS1;refGene_tx=XM_011285987.3;refGene_gn=PROS1;refGene_tx=XM_023238875.1
C2 42662147 42662269
refGene_gn=PROS1;refGene_tx=XM_011285987.3;refGene_gn=PROS1;refGene_tx=XM_023238875.1
C2 42664737 42664863
refGene_gn=PROS1;refGene_tx=XM_011285987.3;refGene_gn=PROS1;refGene_tx

=XM_023238875.1
C2	42668491	42668623
refGene_gn=PROS1;refGene_tx=XM_011285987.3;refGene_gn=PROS1;refGene_tx
=XM_023238875.1
C2	42668722	42668845
refGene_gn=PROS1;refGene_tx=XM_011285987.3;refGene_gn=PROS1;refGene_tx
=XM_023238875.1
C2	42671606	42671693
refGene_gn=PROS1;refGene_tx=XM_011285987.3;refGene_gn=PROS1;refGene_tx
=XM_023238875.1
C2	42679975	42680000
refGene_gn=PROS1;refGene_tx=XM_011285987.3;refGene_gn=PROS1;refGene_tx
=XM_023238875.1
C2	42682737	42682895
refGene_gn=PROS1;refGene_tx=XM_011285987.3;refGene_gn=PROS1;refGene_tx
=XM_023238875.1
C2	42688269	42688363	refGene_gn=PROS1;refGene_tx=XM_023238875.1
C2	42705092	42705168	refGene_gn=PROS1;refGene_tx=XM_011285987.3
C2	42710958	42711017
refGene_gn=ARL13B;refGene_tx=XM_003991550.4;refGene_gn=ARL13B;refGene_
tx=XM_019839570.2;refGene_gn=ARL13B;refGene_tx=XM_019839571.2
C2	42724590	42724661
refGene_gn=ARL13B;refGene_tx=XM_003991550.4;refGene_gn=ARL13B;refGene_
tx=XM_019839570.2;refGene_gn=ARL13B;refGene_tx=XM_019839571.2
C2	42731108	42731358
refGene_gn=ARL13B;refGene_tx=XM_003991550.4;refGene_gn=ARL13B;refGene_
tx=XM_019839570.2;refGene_gn=ARL13B;refGene_tx=XM_019839571.2;refGene_
gn=ARL13B;refGene_tx=XM_019839572.2
C2	42739891	42740776
refGene_gn=STX19;refGene_tx=XM_019839573.2;refGene_gn=STX19;refGene_tx
=XM_019839574.2
C2	42757235	42757341
refGene_gn=ARL13B;refGene_tx=XM_003991550.4;refGene_gn=ARL13B;refGene_
tx=XM_019839570.2;refGene_gn=ARL13B;refGene_tx=XM_019839571.2;refGene_
gn=ARL13B;refGene_tx=XM_019839572.2
C2	42758402	42758605
refGene_gn=ARL13B;refGene_tx=XM_003991550.4;refGene_gn=ARL13B;refGene_
tx=XM_019839570.2;refGene_gn=ARL13B;refGene_tx=XM_019839571.2;refGene_
gn=ARL13B;refGene_tx=XM_019839572.2
C2	42763871	42763986
refGene_gn=ARL13B;refGene_tx=XM_003991550.4;refGene_gn=ARL13B;refGene_
tx=XM_019839570.2;refGene_gn=ARL13B;refGene_tx=XM_019839571.2;refGene_
gn=ARL13B;refGene_tx=XM_019839572.2
C2	42767157	42767383
refGene_gn=ARL13B;refGene_tx=XM_003991550.4;refGene_gn=ARL13B;refGene_
tx=XM_019839570.2;refGene_gn=ARL13B;refGene_tx=XM_019839571.2;refGene_
gn=ARL13B;refGene_tx=XM_019839572.2
C2	42770425	42770542
refGene_gn=ARL13B;refGene_tx=XM_003991550.4;refGene_gn=ARL13B;refGene_
tx=XM_019839570.2;refGene_gn=ARL13B;refGene_tx=XM_019839571.2;refGene_

gn=ARL13B;refGene_tx=XM_019839572.2
C2	42771499	42771568
refGene_gn=ARL13B;refGene_tx=XM_003991550.4;refGene_gn=ARL13B;refGene_tx=XM_019839570.2;refGene_gn=ARL13B;refGene_tx=XM_019839572.2
C2	42775238	42775306	refGene_gn=ARL13B;refGene_tx=XM_019839570.2
C2	42780750	42780827
refGene_gn=ARL13B;refGene_tx=XM_003991550.4;refGene_gn=ARL13B;refGene_tx=XM_019839571.2;refGene_gn=ARL13B;refGene_tx=XM_019839572.2
C2	42787999	42788011	refGene_gn=NSUN3;refGene_tx=XM_003991552.4
C2	42788781	42788891	refGene_gn=NSUN3;refGene_tx=XM_003991552.4
C2	42800041	42800385	refGene_gn=NSUN3;refGene_tx=XM_003991552.4
C2	42815071	42815226	refGene_gn=NSUN3;refGene_tx=XM_003991552.4
C2	42815941	42816063	refGene_gn=NSUN3;refGene_tx=XM_003991552.4
C2	42846612	42846883	refGene_gn=NSUN3;refGene_tx=XM_003991552.4
C2	45123046	45123425
refGene_gn=EPHA6;refGene_tx=XM_019839578.2;refGene_gn=EPHA6;refGene_tx=XM_019839585.2;refGene_gn=EPHA6;refGene_tx=XM_023238877.1;refGene_gn=EPHA6;refGene_tx=XM_023238880.1
C2	45191620	45191685
refGene_gn=EPHA6;refGene_tx=XM_019839578.2;refGene_gn=EPHA6;refGene_tx=XM_019839585.2;refGene_gn=EPHA6;refGene_tx=XM_023238877.1;refGene_gn=EPHA6;refGene_tx=XM_023238880.1
C2	45250102	45250150	refGene_gn=EPHA6;refGene_tx=XM_023238878.1
C2	45295138	45295802
refGene_gn=EPHA6;refGene_tx=XM_019839578.2;refGene_gn=EPHA6;refGene_tx=XM_019839585.2;refGene_gn=EPHA6;refGene_tx=XM_023238877.1;refGene_gn=EPHA6;refGene_tx=XM_023238878.1;refGene_gn=EPHA6;refGene_tx=XM_023238880.1
C2	45490620	45490776
refGene_gn=EPHA6;refGene_tx=XM_019839578.2;refGene_gn=EPHA6;refGene_tx=XM_019839585.2;refGene_gn=EPHA6;refGene_tx=XM_023238877.1;refGene_gn=EPHA6;refGene_tx=XM_023238878.1;refGene_gn=EPHA6;refGene_tx=XM_023238879.1;refGene_gn=EPHA6;refGene_tx=XM_023238880.1
C2	45507691	45508027
refGene_gn=EPHA6;refGene_tx=XM_019839578.2;refGene_gn=EPHA6;refGene_tx=XM_019839585.2;refGene_gn=EPHA6;refGene_tx=XM_023238877.1;refGene_gn=EPHA6;refGene_tx=XM_023238878.1;refGene_gn=EPHA6;refGene_tx=XM_023238879.1;refGene_gn=EPHA6;refGene_tx=XM_023238880.1
C2	45657573	45657698
refGene_gn=EPHA6;refGene_tx=XM_019839578.2;refGene_gn=EPHA6;refGene_tx=XM_019839585.2;refGene_gn=EPHA6;refGene_tx=XM_023238877.1;refGene_gn=EPHA6;refGene_tx=XM_023238878.1;refGene_gn=EPHA6;refGene_tx=XM_023238879.1;refGene_gn=EPHA6;refGene_tx=XM_023238880.1
C2	45699516	45699564
refGene_gn=EPHA6;refGene_tx=XM_019839578.2;refGene_gn=EPHA6;refGene_tx=XM_023238877.1;refGene_gn=EPHA6;refGene_tx=XM_023238878.1;refGene_gn=EPHA6;refGene_tx=XM_023238879.1
C2	45699974	45700137
refGene_gn=EPHA6;refGene_tx=XM_019839578.2;refGene_gn=EPHA6;refGene_tx=XM_019839585.2;refGene_gn=EPHA6;refGene_tx=XM_023238877.1;refGene_gn=

EPHA6;refGene_tx=XM_023238878.1;refGene_gn=EPHA6;refGene_tx=XM_0232388
79.1;refGene_gn=EPHA6;refGene_tx=XM_023238880.1
C2	45733601	45733710
refGene_gn=EPHA6;refGene_tx=XM_019839578.2;refGene_gn=EPHA6;refGene_tx
=XM_019839585.2;refGene_gn=EPHA6;refGene_tx=XM_023238877.1;refGene_gn=
EPHA6;refGene_tx=XM_023238878.1;refGene_gn=EPHA6;refGene_tx=XM_0232388
79.1;refGene_gn=EPHA6;refGene_tx=XM_023238880.1
C2	45737429	45737503
refGene_gn=EPHA6;refGene_tx=XM_019839578.2;refGene_gn=EPHA6;refGene_tx
=XM_019839585.2;refGene_gn=EPHA6;refGene_tx=XM_023238877.1;refGene_gn=
EPHA6;refGene_tx=XM_023238878.1;refGene_gn=EPHA6;refGene_tx=XM_0232388
79.1;refGene_gn=EPHA6;refGene_tx=XM_023238880.1
C2	45740148	45740274
refGene_gn=EPHA6;refGene_tx=XM_019839578.2;refGene_gn=EPHA6;refGene_tx
=XM_019839585.2;refGene_gn=EPHA6;refGene_tx=XM_023238877.1;refGene_gn=
EPHA6;refGene_tx=XM_023238878.1;refGene_gn=EPHA6;refGene_tx=XM_0232388
79.1;refGene_gn=EPHA6;refGene_tx=XM_023238880.1
C2	45778573	45778759
refGene_gn=EPHA6;refGene_tx=XM_019839578.2;refGene_gn=EPHA6;refGene_tx
=XM_019839585.2;refGene_gn=EPHA6;refGene_tx=XM_023238877.1;refGene_gn=
EPHA6;refGene_tx=XM_023238878.1;refGene_gn=EPHA6;refGene_tx=XM_0232388
79.1;refGene_gn=EPHA6;refGene_tx=XM_023238880.1
C2	45825987	45826113
refGene_gn=EPHA6;refGene_tx=XM_019839578.2;refGene_gn=EPHA6;refGene_tx
=XM_019839585.2;refGene_gn=EPHA6;refGene_tx=XM_023238878.1;refGene_gn=
EPHA6;refGene_tx=XM_023238879.1
C2	45848832	45848894
refGene_gn=EPHA6;refGene_tx=XM_019839578.2;refGene_gn=EPHA6;refGene_tx
=XM_019839585.2;refGene_gn=EPHA6;refGene_tx=XM_023238877.1;refGene_gn=
EPHA6;refGene_tx=XM_023238878.1;refGene_gn=EPHA6;refGene_tx=XM_0232388
79.1;refGene_gn=EPHA6;refGene_tx=XM_023238880.1
C2	45875534	45875744
refGene_gn=EPHA6;refGene_tx=XM_019839578.2;refGene_gn=EPHA6;refGene_tx
=XM_019839585.2;refGene_gn=EPHA6;refGene_tx=XM_023238877.1;refGene_gn=
EPHA6;refGene_tx=XM_023238878.1;refGene_gn=EPHA6;refGene_tx=XM_0232388
79.1;refGene_gn=EPHA6;refGene_tx=XM_023238880.1
C2	45948756	45948906
refGene_gn=EPHA6;refGene_tx=XM_019839578.2;refGene_gn=EPHA6;refGene_tx
=XM_019839585.2;refGene_gn=EPHA6;refGene_tx=XM_023238877.1;refGene_gn=
EPHA6;refGene_tx=XM_023238878.1;refGene_gn=EPHA6;refGene_tx=XM_0232388
79.1;refGene_gn=EPHA6;refGene_tx=XM_023238880.1
C2	45964902	45965096
refGene_gn=EPHA6;refGene_tx=XM_019839578.2;refGene_gn=EPHA6;refGene_tx
=XM_019839585.2;refGene_gn=EPHA6;refGene_tx=XM_023238877.1;refGene_gn=
EPHA6;refGene_tx=XM_023238878.1;refGene_gn=EPHA6;refGene_tx=XM_0232388
79.1;refGene_gn=EPHA6;refGene_tx=XM_023238880.1
C2	45967419	45967491
refGene_gn=EPHA6;refGene_tx=XM_019839578.2;refGene_gn=EPHA6;refGene_tx
=XM_019839585.2;refGene_gn=EPHA6;refGene_tx=XM_023238877.1;refGene_gn=
EPHA6;refGene_tx=XM_023238878.1;refGene_gn=EPHA6;refGene_tx=XM_0232388

79.1;refGene_gn=EPHA6;refGene_tx=XM_023238880.1
C2 45977123 45977273
refGene_gn=EPHA6;refGene_tx=XM_019839578.2;refGene_gn=EPHA6;refGene_tx=XM_019839585.2;refGene_gn=EPHA6;refGene_tx=XM_023238877.1;refGene_gn=EPHA6;refGene_tx=XM_023238878.1;refGene_gn=EPHA6;refGene_tx=XM_023238879.1;refGene_gn=EPHA6;refGene_tx=XM_023238880.1
C2 45978225 45978340
refGene_gn=EPHA6;refGene_tx=XM_019839578.2;refGene_gn=EPHA6;refGene_tx=XM_019839585.2;refGene_gn=EPHA6;refGene_tx=XM_023238877.1;refGene_gn=EPHA6;refGene_tx=XM_023238878.1;refGene_gn=EPHA6;refGene_tx=XM_023238879.1;refGene_gn=EPHA6;refGene_tx=XM_023238880.1
C2 45996741 45996800 refGene_gn=ARL6;refGene_tx=XM_023238881.1
C2 45996801 45996862 refGene_gn=ARL6;refGene_tx=XM_023238881.1
C2 46004626 46004688 refGene_gn=ARL6;refGene_tx=XM_023238881.1
C2 46005125 46005194 refGene_gn=ARL6;refGene_tx=XM_023238881.1
C2 46009204 46009299 refGene_gn=ARL6;refGene_tx=XM_023238881.1
C2 46023121 46023251 refGene_gn=ARL6;refGene_tx=XM_023238881.1
C2 46026876 46026932 refGene_gn=ARL6;refGene_tx=XM_023238881.1
C2 46027063 46027076 refGene_gn=ARL6;refGene_tx=XM_023238881.1
C2 46032698 46032732 refGene_gn=ARL6;refGene_tx=XM_023238881.1
C2 46065322 46065471
refGene_gn=CRYBG3;refGene_tx=XM_023238882.1;refGene_gn=CRYBG3;refGene_tx=XM_023238884.1;refGene_gn=CRYBG3;refGene_tx=XM_023238885.1
C2 46083181 46083248
refGene_gn=CRYBG3;refGene_tx=XM_023238882.1;refGene_gn=CRYBG3;refGene_tx=XM_023238884.1;refGene_gn=CRYBG3;refGene_tx=XM_023238885.1
C2 46100257 46100688
refGene_gn=CRYBG3;refGene_tx=XM_023238882.1;refGene_gn=CRYBG3;refGene_tx=XM_023238884.1;refGene_gn=CRYBG3;refGene_tx=XM_023238885.1
C2 46110535 46116755
refGene_gn=CRYBG3;refGene_tx=XM_023238882.1;refGene_gn=CRYBG3;refGene_tx=XM_023238883.1;refGene_gn=CRYBG3;refGene_tx=XM_023238884.1;refGene_gn=CRYBG3;refGene_tx=XM_023238885.1
C2 46118118 46118163
refGene_gn=CRYBG3;refGene_tx=XM_023238882.1;refGene_gn=CRYBG3;refGene_tx=XM_023238883.1;refGene_gn=CRYBG3;refGene_tx=XM_023238884.1;refGene_gn=CRYBG3;refGene_tx=XM_023238885.1
C2 46118518 46118634
refGene_gn=CRYBG3;refGene_tx=XM_023238882.1;refGene_gn=CRYBG3;refGene_tx=XM_023238883.1;refGene_gn=CRYBG3;refGene_tx=XM_023238884.1;refGene_gn=CRYBG3;refGene_tx=XM_023238885.1
C2 46119599 46119747
refGene_gn=CRYBG3;refGene_tx=XM_023238882.1;refGene_gn=CRYBG3;refGene_tx=XM_023238883.1;refGene_gn=CRYBG3;refGene_tx=XM_023238884.1;refGene_gn=CRYBG3;refGene_tx=XM_023238885.1
C2 46125228 46125365
refGene_gn=CRYBG3;refGene_tx=XM_023238882.1;refGene_gn=CRYBG3;refGene_tx=XM_023238883.1;refGene_gn=CRYBG3;refGene_tx=XM_023238884.1;refGene_gn=CRYBG3;refGene_tx=XM_023238885.1
C2 46126915 46127030

refGene_gn=CRYBG3;refGene_tx=XM_023238882.1;refGene_gn=CRYBG3;refGene_tx=XM_023238883.1;refGene_gn=CRYBG3;refGene_tx=XM_023238884.1;refGene_gn=CRYBG3;refGene_tx=XM_023238885.1
C2 46128157 46128193
refGene_gn=CRYBG3;refGene_tx=XM_023238882.1;refGene_gn=CRYBG3;refGene_tx=XM_023238883.1;refGene_gn=CRYBG3;refGene_tx=XM_023238884.1;refGene_gn=CRYBG3;refGene_tx=XM_023238885.1
C2 46130287 46130421
refGene_gn=CRYBG3;refGene_tx=XM_023238882.1;refGene_gn=CRYBG3;refGene_tx=XM_023238883.1;refGene_gn=CRYBG3;refGene_tx=XM_023238884.1;refGene_gn=CRYBG3;refGene_tx=XM_023238885.1
C2 46133749 46133876
refGene_gn=CRYBG3;refGene_tx=XM_023238882.1;refGene_gn=CRYBG3;refGene_tx=XM_023238883.1;refGene_gn=CRYBG3;refGene_tx=XM_023238884.1;refGene_gn=CRYBG3;refGene_tx=XM_023238885.1
C2 46154254 46154397
refGene_gn=CRYBG3;refGene_tx=XM_023238882.1;refGene_gn=CRYBG3;refGene_tx=XM_023238883.1;refGene_gn=CRYBG3;refGene_tx=XM_023238884.1;refGene_gn=CRYBG3;refGene_tx=XM_023238885.1
C2 46154508 46154635
refGene_gn=CRYBG3;refGene_tx=XM_023238882.1;refGene_gn=CRYBG3;refGene_tx=XM_023238883.1;refGene_gn=CRYBG3;refGene_tx=XM_023238884.1;refGene_gn=CRYBG3;refGene_tx=XM_023238885.1
C2 46155527 46155560
refGene_gn=CRYBG3;refGene_tx=XM_023238882.1;refGene_gn=CRYBG3;refGene_tx=XM_023238883.1;refGene_gn=CRYBG3;refGene_tx=XM_023238884.1;refGene_gn=CRYBG3;refGene_tx=XM_023238885.1
C2 46161160 46161270
refGene_gn=CRYBG3;refGene_tx=XM_023238882.1;refGene_gn=CRYBG3;refGene_tx=XM_023238883.1;refGene_gn=CRYBG3;refGene_tx=XM_023238884.1;refGene_gn=CRYBG3;refGene_tx=XM_023238885.1
C2 46171490 46171617
refGene_gn=CRYBG3;refGene_tx=XM_023238882.1;refGene_gn=CRYBG3;refGene_tx=XM_023238883.1;refGene_gn=CRYBG3;refGene_tx=XM_023238884.1;refGene_gn=CRYBG3;refGene_tx=XM_023238885.1
C2 46184959 46185099
refGene_gn=CRYBG3;refGene_tx=XM_023238882.1;refGene_gn=CRYBG3;refGene_tx=XM_023238883.1;refGene_gn=CRYBG3;refGene_tx=XM_023238884.1;refGene_gn=CRYBG3;refGene_tx=XM_023238885.1
C2 46187914 46188038
refGene_gn=CRYBG3;refGene_tx=XM_023238882.1;refGene_gn=CRYBG3;refGene_tx=XM_023238883.1;refGene_gn=CRYBG3;refGene_tx=XM_023238884.1;refGene_gn=CRYBG3;refGene_tx=XM_023238885.1
C2 46191536 46191695
refGene_gn=CRYBG3;refGene_tx=XM_023238882.1;refGene_gn=CRYBG3;refGene_tx=XM_023238883.1;refGene_gn=CRYBG3;refGene_tx=XM_023238884.1;refGene_gn=CRYBG3;refGene_tx=XM_023238885.1
C2 46192819 46192979
refGene_gn=CRYBG3;refGene_tx=XM_023238882.1;refGene_gn=CRYBG3;refGene_tx=XM_023238883.1;refGene_gn=CRYBG3;refGene_tx=XM_023238884.1;refGene_

gn=CRYBG3;refGene_tx=XM_023238885.1
C2 46193736 46193825
refGene_gn=CRYBG3;refGene_tx=XM_023238882.1;refGene_gn=CRYBG3;refGene_tx=XM_023238883.1;refGene_gn=CRYBG3;refGene_tx=XM_023238884.1;refGene_gn=CRYBG3;refGene_tx=XM_023238885.1
C2 46195545 46195704
refGene_gn=RIOX2;refGene_tx=XM_003991555.5;refGene_gn=RIOX2;refGene_tx=XM_006935973.4;refGene_gn=RIOX2;refGene_tx=XM_006935974.4;refGene_gn=RIOX2;refGene_tx=XM_006935975.4;refGene_gn=RIOX2;refGene_tx=XM_006935976.4;refGene_gn=RIOX2;refGene_tx=XM_023238886.1
C2 46196131 46196218
refGene_gn=RIOX2;refGene_tx=XM_003991555.5;refGene_gn=RIOX2;refGene_tx=XM_006935973.4;refGene_gn=RIOX2;refGene_tx=XM_006935974.4;refGene_gn=RIOX2;refGene_tx=XM_006935975.4;refGene_gn=RIOX2;refGene_tx=XM_006935976.4;refGene_gn=RIOX2;refGene_tx=XM_023238886.1
C2 46197812 46197901
refGene_gn=RIOX2;refGene_tx=XM_003991555.5;refGene_gn=RIOX2;refGene_tx=XM_006935973.4;refGene_gn=RIOX2;refGene_tx=XM_006935974.4;refGene_gn=RIOX2;refGene_tx=XM_006935975.4;refGene_gn=RIOX2;refGene_tx=XM_006935976.4;refGene_gn=RIOX2;refGene_tx=XM_023238886.1
C2 46199867 46200036
refGene_gn=RIOX2;refGene_tx=XM_003991555.5;refGene_gn=RIOX2;refGene_tx=XM_006935973.4;refGene_gn=RIOX2;refGene_tx=XM_006935974.4;refGene_gn=RIOX2;refGene_tx=XM_006935975.4;refGene_gn=RIOX2;refGene_tx=XM_006935976.4;refGene_gn=RIOX2;refGene_tx=XM_023238886.1
C2 46200829 46200932
refGene_gn=RIOX2;refGene_tx=XM_003991555.5;refGene_gn=RIOX2;refGene_tx=XM_006935973.4;refGene_gn=RIOX2;refGene_tx=XM_006935974.4;refGene_gn=RIOX2;refGene_tx=XM_006935975.4;refGene_gn=RIOX2;refGene_tx=XM_006935976.4;refGene_gn=RIOX2;refGene_tx=XM_023238886.1
C2 46204133 46204237
refGene_gn=RIOX2;refGene_tx=XM_003991555.5;refGene_gn=RIOX2;refGene_tx=XM_006935973.4;refGene_gn=RIOX2;refGene_tx=XM_006935974.4;refGene_gn=RIOX2;refGene_tx=XM_006935975.4;refGene_gn=RIOX2;refGene_tx=XM_006935976.4;refGene_gn=RIOX2;refGene_tx=XM_023238886.1
C2 46209980 46210109
refGene_gn=RIOX2;refGene_tx=XM_003991555.5;refGene_gn=RIOX2;refGene_tx=XM_006935973.4;refGene_gn=RIOX2;refGene_tx=XM_006935974.4;refGene_gn=RIOX2;refGene_tx=XM_006935975.4;refGene_gn=RIOX2;refGene_tx=XM_006935976.4;refGene_gn=RIOX2;refGene_tx=XM_023238886.1
C2 46211173 46211293
refGene_gn=RIOX2;refGene_tx=XM_003991555.5;refGene_gn=RIOX2;refGene_tx=XM_006935973.4;refGene_gn=RIOX2;refGene_tx=XM_006935974.4;refGene_gn=RIOX2;refGene_tx=XM_006935975.4;refGene_gn=RIOX2;refGene_tx=XM_006935976.4;refGene_gn=RIOX2;refGene_tx=XM_023238886.1
C2 46217124 46217559
refGene_gn=RIOX2;refGene_tx=XM_003991555.5;refGene_gn=RIOX2;refGene_tx=XM_006935973.4;refGene_gn=RIOX2;refGene_tx=XM_006935974.4;refGene_gn=RIOX2;refGene_tx=XM_006935975.4;refGene_gn=RIOX2;refGene_tx=XM_006935976.4;refGene_gn=RIOX2;refGene_tx=XM_023238886.1

C2 46234606 46234909
refGene_gn=GABRR3;refGene_tx=XM_006935978.3;refGene_gn=GABRR3;refGene_tx=XM_019839590.2;refGene_gn=GABRR3;refGene_tx=XM_019839592.2
C2 46239953 46240150
refGene_gn=GABRR3;refGene_tx=XM_006935978.3;refGene_gn=GABRR3;refGene_tx=XM_019839590.2;refGene_gn=GABRR3;refGene_tx=XM_019839592.2
C2 46249204 46249357
refGene_gn=GABRR3;refGene_tx=XM_006935978.3;refGene_gn=GABRR3;refGene_tx=XM_019839590.2;refGene_gn=GABRR3;refGene_tx=XM_019839592.2
C2 46255470 46255611
refGene_gn=GABRR3;refGene_tx=XM_006935978.3;refGene_gn=GABRR3;refGene_tx=XM_019839590.2;refGene_gn=GABRR3;refGene_tx=XM_019839592.2
C2 46256618 46256701
refGene_gn=GABRR3;refGene_tx=XM_006935978.3;refGene_gn=GABRR3;refGene_tx=XM_019839590.2;refGene_gn=GABRR3;refGene_tx=XM_019839592.2
C2 46262165 46262389
refGene_gn=GABRR3;refGene_tx=XM_006935978.3;refGene_gn=GABRR3;refGene_tx=XM_019839590.2;refGene_gn=GABRR3;refGene_tx=XM_019839592.2
C2 46264857 46264925
refGene_gn=GABRR3;refGene_tx=XM_006935978.3;refGene_gn=GABRR3;refGene_tx=XM_019839590.2;refGene_gn=GABRR3;refGene_tx=XM_019839592.2
C2 46271825 46271944
refGene_gn=GABRR3;refGene_tx=XM_006935978.3;refGene_gn=GABRR3;refGene_tx=XM_019839590.2;refGene_gn=GABRR3;refGene_tx=XM_019839592.2
C2 46282242 46282367
refGene_gn=GABRR3;refGene_tx=XM_006935978.3;refGene_gn=GABRR3;refGene_tx=XM_019839590.2;refGene_gn=GABRR3;refGene_tx=XM_019839592.2
C2 46303020 46303944
refGene_gn=LOC101088100;refGene_tx=XM_011285985.1
C2 46317049 46317187
refGene_gn=LOC101088354;refGene_tx=XM_011286021.2
C2 46317188 46317324
refGene_gn=LOC101088354;refGene_tx=XM_011286021.2
C2 46317326 46318000
refGene_gn=LOC101088354;refGene_tx=XM_011286021.2
C2 46325974 46326898
refGene_gn=LOC101083026;refGene_tx=XM_003991557.1
C2 46364007 46364311
refGene_gn=LOC101088606;refGene_tx=XM_011286020.2
C2 46364313 46364533
refGene_gn=LOC101088606;refGene_tx=XM_011286020.2
C2 46364591 46364617
refGene_gn=LOC101088606;refGene_tx=XM_011286020.2
C2 46364619 46364942
refGene_gn=LOC101088606;refGene_tx=XM_011286020.2
C2 46364943 46364973
refGene_gn=LOC101088606;refGene_tx=XM_011286020.2
C2 46373157 46373241
refGene_gn=LOC101088869;refGene_tx=XM_011286019.2
C2 46373768 46373868

refGene_gn=LOC101088869;refGene_tx=XM_011286019.2
C2	46378855	46379796
refGene_gn=LOC101088869;refGene_tx=XM_011286019.2
C2	46401937	46402879
refGene_gn=LOC101089133;refGene_tx=XM_003991567.1
C2	46428170	46429100
refGene_gn=LOC101089384;refGene_tx=XM_011286018.2
C2	46447262	46448192
refGene_gn=LOC101089637;refGene_tx=XM_006935982.3
C2	46483048	46483978
refGene_gn=LOC101090130;refGene_tx=XM_006935979.4
C2	46547244	46548174
refGene_gn=LOC101083287;refGene_tx=XM_011285984.1
C2	46564779	46565709
refGene_gn=LOC101090380;refGene_tx=XM_019811147.1
C2	46600103	46601057
refGene_gn=LOC101083537;refGene_tx=XM_003991559.3
C2	46620752	46620973	refGene_gn=CLDND1;refGene_tx=XM_023238887.1
C2	46621117	46621183	refGene_gn=CLDND1;refGene_tx=XM_023238887.1
C2	46621185	46621254	refGene_gn=CLDND1;refGene_tx=XM_023238887.1
C2	46623422	46623530	refGene_gn=CLDND1;refGene_tx=XM_023238887.1
C2	46625615	46625925	refGene_gn=CLDND1;refGene_tx=XM_023238887.1
C2	46626146	46626190	refGene_gn=CLDND1;refGene_tx=XM_023238887.1
C2	46627603	46627610	refGene_gn=CLDND1;refGene_tx=XM_023238887.1
C2	46635807	46636662	refGene_gn=GPR15;refGene_tx=XM_023238888.1
C2	46636727	46636781	refGene_gn=GPR15;refGene_tx=XM_023238888.1
C2	46686137	46686225	refGene_gn=CPOX;refGene_tx=XM_003991594.5
C2	46689051	46689156	refGene_gn=CPOX;refGene_tx=XM_003991594.5
C2	46693416	46693635	refGene_gn=CPOX;refGene_tx=XM_003991594.5
C2	46695893	46696035	refGene_gn=CPOX;refGene_tx=XM_003991594.5
C2	46697852	46697963	refGene_gn=CPOX;refGene_tx=XM_003991594.5
C2	46698224	46698368	refGene_gn=CPOX;refGene_tx=XM_003991594.5
C2	46700656	46701185	refGene_gn=CPOX;refGene_tx=XM_003991594.5
C2	46852709	46852798
refGene_gn=ST3GAL6;refGene_tx=XM_006935985.3;refGene_gn=ST3GAL6;refGene_tx=XM_006935986.4;refGene_gn=ST3GAL6;refGene_tx=XM_006935988.4;refGene_gn=ST3GAL6;refGene_tx=XM_023238893.1;refGene_gn=ST3GAL6;refGene_tx=XM_023238894.1;refGene_gn=ST3GAL6;refGene_tx=XM_023238895.1;refGene_gn=ST3GAL6;refGene_tx=XM_023238896.1
C2	46855807	46855885
refGene_gn=ST3GAL6;refGene_tx=XM_006935985.3;refGene_gn=ST3GAL6;refGene_tx=XM_006935986.4;refGene_gn=ST3GAL6;refGene_tx=XM_006935988.4;refGene_gn=ST3GAL6;refGene_tx=XM_023238893.1;refGene_gn=ST3GAL6;refGene_tx=XM_023238894.1;refGene_gn=ST3GAL6;refGene_tx=XM_023238895.1;refGene_gn=ST3GAL6;refGene_tx=XM_023238896.1
C2	46858270	46858374
refGene_gn=ST3GAL6;refGene_tx=XM_006935985.3;refGene_gn=ST3GAL6;refGene_tx=XM_006935986.4;refGene_gn=ST3GAL6;refGene_tx=XM_006935988.4;refGene_gn=ST3GAL6;refGene_tx=XM_023238893.1;refGene_gn=ST3GAL6;refGene_tx=XM_023238894.1;refGene_gn=ST3GAL6;refGene_tx=XM_023238895.1;refGene_gn

=ST3GAL6;refGene_tx=XM_023238896.1
C2 46859679 46859743
refGene_gn=ST3GAL6;refGene_tx=XM_006935985.3;refGene_gn=ST3GAL6;refGene_tx=XM_006935986.4;refGene_gn=ST3GAL6;refGene_tx=XM_006935988.4;refGene_gn=ST3GAL6;refGene_tx=XM_023238893.1;refGene_gn=ST3GAL6;refGene_tx=XM_023238894.1;refGene_gn=ST3GAL6;refGene_tx=XM_023238895.1;refGene_gn=ST3GAL6;refGene_tx=XM_023238896.1
C2 46869981 46870080
refGene_gn=ST3GAL6;refGene_tx=XM_006935985.3;refGene_gn=ST3GAL6;refGene_tx=XM_006935986.4;refGene_gn=ST3GAL6;refGene_tx=XM_006935988.4;refGene_gn=ST3GAL6;refGene_tx=XM_023238893.1;refGene_gn=ST3GAL6;refGene_tx=XM_023238894.1;refGene_gn=ST3GAL6;refGene_tx=XM_023238895.1
C2 46873422 46873609
refGene_gn=ST3GAL6;refGene_tx=XM_006935985.3;refGene_gn=ST3GAL6;refGene_tx=XM_006935986.4;refGene_gn=ST3GAL6;refGene_tx=XM_006935988.4;refGene_gn=ST3GAL6;refGene_tx=XM_023238893.1;refGene_gn=ST3GAL6;refGene_tx=XM_023238894.1;refGene_gn=ST3GAL6;refGene_tx=XM_023238895.1;refGene_gn=ST3GAL6;refGene_tx=XM_023238896.1
C2 46873722 46873860
refGene_gn=ST3GAL6;refGene_tx=XM_006935985.3;refGene_gn=ST3GAL6;refGene_tx=XM_006935986.4;refGene_gn=ST3GAL6;refGene_tx=XM_006935988.4;refGene_gn=ST3GAL6;refGene_tx=XM_023238893.1;refGene_gn=ST3GAL6;refGene_tx=XM_023238894.1;refGene_gn=ST3GAL6;refGene_tx=XM_023238895.1;refGene_gn=ST3GAL6;refGene_tx=XM_023238896.1
C2 46877254 46877407
refGene_gn=ST3GAL6;refGene_tx=XM_006935985.3;refGene_gn=ST3GAL6;refGene_tx=XM_006935986.4;refGene_gn=ST3GAL6;refGene_tx=XM_006935988.4;refGene_gn=ST3GAL6;refGene_tx=XM_023238893.1;refGene_gn=ST3GAL6;refGene_tx=XM_023238894.1;refGene_gn=ST3GAL6;refGene_tx=XM_023238895.1;refGene_gn=ST3GAL6;refGene_tx=XM_023238896.1
C2 46878898 46878985
refGene_gn=ST3GAL6;refGene_tx=XM_006935985.3;refGene_gn=ST3GAL6;refGene_tx=XM_006935986.4;refGene_gn=ST3GAL6;refGene_tx=XM_006935988.4;refGene_gn=ST3GAL6;refGene_tx=XM_023238893.1;refGene_gn=ST3GAL6;refGene_tx=XM_023238896.1
C2 46883246 46883314 refGene_gn=DCBLD2;refGene_tx=XM_023238890.1
C2 46883632 46884102
refGene_gn=DCBLD2;refGene_tx=XM_023238889.1;refGene_gn=DCBLD2;refGene_tx=XM_023238890.1;refGene_gn=DCBLD2;refGene_tx=XM_023238891.1;refGene_gn=DCBLD2;refGene_tx=XM_023238892.1
C2 46885183 46885321
refGene_gn=DCBLD2;refGene_tx=XM_023238889.1;refGene_gn=DCBLD2;refGene_tx=XM_023238890.1;refGene_gn=DCBLD2;refGene_tx=XM_023238891.1
C2 46888470 46888566
refGene_gn=DCBLD2;refGene_tx=XM_023238889.1;refGene_gn=DCBLD2;refGene_tx=XM_023238890.1;refGene_gn=DCBLD2;refGene_tx=XM_023238891.1;refGene_gn=DCBLD2;refGene_tx=XM_023238892.1;refGene_gn=ST3GAL6;refGene_tx=XM_023238894.1;refGene_gn=ST3GAL6;refGene_tx=XM_023238895.1
C2 46891870 46891964
refGene_gn=DCBLD2;refGene_tx=XM_023238889.1;refGene_gn=DCBLD2;refGene_

tx=XM_023238890.1;refGene_gn=DCBLD2;refGene_tx=XM_023238891.1;refGene_gn=DCBLD2;refGene_tx=XM_023238892.1
C2 46895110 46895236
refGene_gn=DCBLD2;refGene_tx=XM_023238889.1;refGene_gn=DCBLD2;refGene_tx=XM_023238890.1;refGene_gn=DCBLD2;refGene_tx=XM_023238891.1;refGene_gn=DCBLD2;refGene_tx=XM_023238892.1
C2 46895382 46895469
refGene_gn=DCBLD2;refGene_tx=XM_023238889.1;refGene_gn=DCBLD2;refGene_tx=XM_023238890.1;refGene_gn=DCBLD2;refGene_tx=XM_023238891.1;refGene_gn=DCBLD2;refGene_tx=XM_023238892.1
C2 46896258 46896409
refGene_gn=DCBLD2;refGene_tx=XM_023238889.1;refGene_gn=DCBLD2;refGene_tx=XM_023238890.1;refGene_gn=DCBLD2;refGene_tx=XM_023238891.1;refGene_gn=DCBLD2;refGene_tx=XM_023238892.1
C2 46901804 46901929
refGene_gn=DCBLD2;refGene_tx=XM_023238889.1;refGene_gn=DCBLD2;refGene_tx=XM_023238890.1;refGene_gn=DCBLD2;refGene_tx=XM_023238891.1;refGene_gn=DCBLD2;refGene_tx=XM_023238892.1
C2 46902900 46903116
refGene_gn=DCBLD2;refGene_tx=XM_023238889.1;refGene_gn=DCBLD2;refGene_tx=XM_023238890.1;refGene_gn=DCBLD2;refGene_tx=XM_023238891.1;refGene_gn=DCBLD2;refGene_tx=XM_023238892.1
C2 46903929 46903970
refGene_gn=DCBLD2;refGene_tx=XM_023238889.1;refGene_gn=DCBLD2;refGene_tx=XM_023238890.1;refGene_gn=DCBLD2;refGene_tx=XM_023238891.1;refGene_gn=DCBLD2;refGene_tx=XM_023238892.1
C2 46905701 46905835
refGene_gn=DCBLD2;refGene_tx=XM_023238889.1;refGene_gn=DCBLD2;refGene_tx=XM_023238890.1;refGene_gn=DCBLD2;refGene_tx=XM_023238891.1;refGene_gn=DCBLD2;refGene_tx=XM_023238892.1
C2 46906149 46906222
refGene_gn=DCBLD2;refGene_tx=XM_023238889.1;refGene_gn=DCBLD2;refGene_tx=XM_023238890.1;refGene_gn=DCBLD2;refGene_tx=XM_023238891.1;refGene_gn=DCBLD2;refGene_tx=XM_023238892.1
C2 46908410 46908462
refGene_gn=DCBLD2;refGene_tx=XM_023238889.1;refGene_gn=DCBLD2;refGene_tx=XM_023238890.1;refGene_gn=DCBLD2;refGene_tx=XM_023238891.1;refGene_gn=DCBLD2;refGene_tx=XM_023238892.1
C2 46921922 46922060
refGene_gn=DCBLD2;refGene_tx=XM_023238889.1;refGene_gn=DCBLD2;refGene_tx=XM_023238890.1;refGene_gn=DCBLD2;refGene_tx=XM_023238892.1
C2 46951422 46951650
refGene_gn=DCBLD2;refGene_tx=XM_023238889.1;refGene_gn=DCBLD2;refGene_tx=XM_023238890.1;refGene_gn=DCBLD2;refGene_tx=XM_023238891.1;refGene_gn=DCBLD2;refGene_tx=XM_023238892.1
C2 46969360 46969571
refGene_gn=DCBLD2;refGene_tx=XM_023238889.1;refGene_gn=DCBLD2;refGene_tx=XM_023238890.1;refGene_gn=DCBLD2;refGene_tx=XM_023238891.1;refGene_gn=DCBLD2;refGene_tx=XM_023238892.1
C2 47743614 47743942

refGene_gn=COL8A1;refGene_tx=XM_011285982.3;refGene_gn=COL8A1;refGene_tx=XM_011285983.3;refGene_gn=COL8A1;refGene_tx=XM_019839605.2
C2 47746707 47748614
refGene_gn=COL8A1;refGene_tx=XM_011285982.3;refGene_gn=COL8A1;refGene_tx=XM_011285983.3;refGene_gn=COL8A1;refGene_tx=XM_019839605.2
C2 47771040 47771104 refGene_gn=CMSS1;refGene_tx=XM_006935991.4
C2 47781330 47781522 refGene_gn=FILIP1L;refGene_tx=XM_003991577.5
C2 47799101 47800675 refGene_gn=FILIP1L;refGene_tx=XM_003991577.5
C2 47800677 47801876 refGene_gn=FILIP1L;refGene_tx=XM_003991577.5
C2 47866785 47866903 refGene_gn=CMSS1;refGene_tx=XM_019839608.2
C2 47882456 47882635 refGene_gn=FILIP1L;refGene_tx=XM_003991577.5
C2 47892545 47892719 refGene_gn=FILIP1L;refGene_tx=XM_003991577.5
C2 47893446 47893689 refGene_gn=FILIP1L;refGene_tx=XM_003991577.5
C2 48010389 48010425 refGene_gn=CMSS1;refGene_tx=XM_019839612.2
C2 48118633 48118722
refGene_gn=CMSS1;refGene_tx=XM_003991578.5;refGene_gn=CMSS1;refGene_tx=XM_006935991.4;refGene_gn=CMSS1;refGene_tx=XM_019839608.2;refGene_gn=CMSS1;refGene_tx=XM_019839612.2;refGene_gn=CMSS1;refGene_tx=XM_019839613.2
C2 48134420 48134492
refGene_gn=CMSS1;refGene_tx=XM_003991578.5;refGene_gn=CMSS1;refGene_tx=XM_006935991.4;refGene_gn=CMSS1;refGene_tx=XM_019839608.2;refGene_gn=CMSS1;refGene_tx=XM_019839612.2;refGene_gn=CMSS1;refGene_tx=XM_019839613.2
C2 48136280 48136410
refGene_gn=CMSS1;refGene_tx=XM_003991578.5;refGene_gn=CMSS1;refGene_tx=XM_006935991.4;refGene_gn=CMSS1;refGene_tx=XM_019839608.2;refGene_gn=CMSS1;refGene_tx=XM_019839612.2;refGene_gn=CMSS1;refGene_tx=XM_019839613.2
C2 48139841 48139901
refGene_gn=CMSS1;refGene_tx=XM_003991578.5;refGene_gn=CMSS1;refGene_tx=XM_006935991.4;refGene_gn=CMSS1;refGene_tx=XM_019839608.2;refGene_gn=CMSS1;refGene_tx=XM_019839612.2;refGene_gn=CMSS1;refGene_tx=XM_019839613.2
C2 48140984 48141087
refGene_gn=CMSS1;refGene_tx=XM_003991578.5;refGene_gn=CMSS1;refGene_tx=XM_006935991.4;refGene_gn=CMSS1;refGene_tx=XM_019839608.2;refGene_gn=CMSS1;refGene_tx=XM_019839612.2;refGene_gn=CMSS1;refGene_tx=XM_019839613.2
C2 48144232 48144293
refGene_gn=CMSS1;refGene_tx=XM_003991578.5;refGene_gn=CMSS1;refGene_tx=XM_006935991.4;refGene_gn=CMSS1;refGene_tx=XM_019839608.2;refGene_gn=CMSS1;refGene_tx=XM_019839612.2;refGene_gn=CMSS1;refGene_tx=XM_019839613.2
C2 48144727 48144815
refGene_gn=CMSS1;refGene_tx=XM_003991578.5;refGene_gn=CMSS1;refGene_tx=XM_006935991.4;refGene_gn=CMSS1;refGene_tx=XM_019839608.2;refGene_gn=CMSS1;refGene_tx=XM_019839612.2;refGene_gn=CMSS1;refGene_tx=XM_019839613.2
C2 48149992 48150081

refGene_gn=CMSS1;refGene_tx=XM_003991578.5;refGene_gn=CMSS1;refGene_tx=XM_006935991.4;refGene_gn=CMSS1;refGene_tx=XM_019839608.2;refGene_gn=CMSS1;refGene_tx=XM_019839612.2;refGene_gn=CMSS1;refGene_tx=XM_019839613.2
C2 48151473 48151557
refGene_gn=CMSS1;refGene_tx=XM_003991578.5;refGene_gn=CMSS1;refGene_tx=XM_006935991.4;refGene_gn=CMSS1;refGene_tx=XM_019839608.2;refGene_gn=CMSS1;refGene_tx=XM_019839612.2;refGene_gn=CMSS1;refGene_tx=XM_019839613.2
C2 48153194 48153196
refGene_gn=LOC101098325;refGene_tx=XM_006935995.4
C2 48154784 48154983
refGene_gn=LOC101098325;refGene_tx=XM_003991597.5;refGene_gn=LOC101098325;refGene_tx=XM_006935995.4
C2 48159004 48159115
refGene_gn=LOC101098325;refGene_tx=XM_003991597.5;refGene_gn=LOC101098325;refGene_tx=XM_006935995.4
C2 48167352 48167454
refGene_gn=LOC101098325;refGene_tx=XM_003991597.5;refGene_gn=LOC101098325;refGene_tx=XM_006935995.4
C2 48168449 48168537
refGene_gn=LOC101098325;refGene_tx=XM_003991597.5;refGene_gn=LOC101098325;refGene_tx=XM_006935995.4
C2 48168930 48169071
refGene_gn=LOC101098325;refGene_tx=XM_003991597.5;refGene_gn=LOC101098325;refGene_tx=XM_006935995.4
C2 48176240 48176450
refGene_gn=LOC101098325;refGene_tx=XM_003991597.5;refGene_gn=LOC101098325;refGene_tx=XM_006935995.4
C2 48179204 48179395
refGene_gn=LOC101098325;refGene_tx=XM_003991597.5;refGene_gn=LOC101098325;refGene_tx=XM_006935995.4
C2 48216557 48216610
refGene_gn=TBC1D23;refGene_tx=XM_003991580.5;refGene_gn=TBC1D23;refGene_tx=XM_019839614.2
C2 48238327 48238439
refGene_gn=TBC1D23;refGene_tx=XM_003991580.5;refGene_gn=TBC1D23;refGene_tx=XM_019839614.2
C2 48239807 48239913
refGene_gn=TBC1D23;refGene_tx=XM_003991580.5;refGene_gn=TBC1D23;refGene_tx=XM_019839614.2
C2 48240925 48241130
refGene_gn=TBC1D23;refGene_tx=XM_003991580.5;refGene_gn=TBC1D23;refGene_tx=XM_019839614.2
C2 48244987 48245111
refGene_gn=TBC1D23;refGene_tx=XM_003991580.5;refGene_gn=TBC1D23;refGene_tx=XM_019839614.2
C2 48248167 48248292
refGene_gn=TBC1D23;refGene_tx=XM_003991580.5;refGene_gn=TBC1D23;refGene_tx=XM_019839614.2

C2 48248385 48248432
refGene_gn=TBC1D23;refGene_tx=XM_003991580.5;refGene_gn=TBC1D23;refGene_tx=XM_019839614.2
C2 48249257 48249361
refGene_gn=TBC1D23;refGene_tx=XM_003991580.5;refGene_gn=TBC1D23;refGene_tx=XM_019839614.2
C2 48250968 48251091
refGene_gn=TBC1D23;refGene_tx=XM_003991580.5;refGene_gn=TBC1D23;refGene_tx=XM_019839614.2
C2 48252382 48252475
refGene_gn=TBC1D23;refGene_tx=XM_003991580.5;refGene_gn=TBC1D23;refGene_tx=XM_019839614.2
C2 48256065 48256236
refGene_gn=TBC1D23;refGene_tx=XM_003991580.5;refGene_gn=TBC1D23;refGene_tx=XM_019839614.2
C2 48258521 48258564
refGene_gn=TBC1D23;refGene_tx=XM_003991580.5;refGene_gn=TBC1D23;refGene_tx=XM_019839614.2
C2 48260261 48260371
refGene_gn=TBC1D23;refGene_tx=XM_003991580.5;refGene_gn=TBC1D23;refGene_tx=XM_019839614.2
C2 48263529 48263669
refGene_gn=TBC1D23;refGene_tx=XM_003991580.5;refGene_gn=TBC1D23;refGene_tx=XM_019839614.2
C2 48265055 48265100 refGene_gn=TBC1D23;refGene_tx=XM_003991580.5
C2 48268657 48268746
refGene_gn=TBC1D23;refGene_tx=XM_003991580.5;refGene_gn=TBC1D23;refGene_tx=XM_019839614.2
C2 48271077 48271213
refGene_gn=TBC1D23;refGene_tx=XM_003991580.5;refGene_gn=TBC1D23;refGene_tx=XM_019839614.2
C2 48272747 48272942
refGene_gn=TBC1D23;refGene_tx=XM_003991580.5;refGene_gn=TBC1D23;refGene_tx=XM_019839614.2
C2 48273764 48273846
refGene_gn=TBC1D23;refGene_tx=XM_003991580.5;refGene_gn=TBC1D23;refGene_tx=XM_019839614.2
C2 48279688 48279716 refGene_gn=NIT2;refGene_tx=XM_019839616.2
C2 48282990 48283109 refGene_gn=NIT2;refGene_tx=XM_019839616.2
C2 48284588 48284709 refGene_gn=NIT2;refGene_tx=XM_019839616.2
C2 48285771 48285860 refGene_gn=NIT2;refGene_tx=XM_019839616.2
C2 48291947 48292041 refGene_gn=NIT2;refGene_tx=XM_019839616.2
C2 48292557 48292632 refGene_gn=NIT2;refGene_tx=XM_019839616.2
C2 48293694 48293773 refGene_gn=NIT2;refGene_tx=XM_019839616.2
C2 48295498 48295597 refGene_gn=NIT2;refGene_tx=XM_019839616.2
C2 48297732 48297788 refGene_gn=NIT2;refGene_tx=XM_019839616.2
C2 48298136 48298228 refGene_gn=NIT2;refGene_tx=XM_019839616.2
C2 48322465 48322619 refGene_gn=TOMM70;refGene_tx=XM_003991582.4
C2 48323614 48323737 refGene_gn=TOMM70;refGene_tx=XM_003991582.4
C2 48324303 48324401 refGene_gn=TOMM70;refGene_tx=XM_003991582.4

C2 48327665 48327782 refGene_gn=TOMM70;refGene_tx=XM_003991582.4
C2 48328685 48328793 refGene_gn=TOMM70;refGene_tx=XM_003991582.4
C2 48329803 48329938 refGene_gn=TOMM70;refGene_tx=XM_003991582.4
C2 48331710 48331918 refGene_gn=TOMM70;refGene_tx=XM_003991582.4
C2 48333521 48333670 refGene_gn=TOMM70;refGene_tx=XM_003991582.4
C2 48336606 48336716 refGene_gn=TOMM70;refGene_tx=XM_003991582.4
C2 48338319 48338446 refGene_gn=TOMM70;refGene_tx=XM_003991582.4
C2 48340657 48340831 refGene_gn=TOMM70;refGene_tx=XM_003991582.4
C2 48353916 48354243 refGene_gn=TOMM70;refGene_tx=XM_003991582.4
C2 48375418 48375580
refGene_gn=LNP1;refGene_tx=XM_006935996.4;refGene_gn=LNP1;refGene_tx=XM_006936000.4;refGene_gn=LNP1;refGene_tx=XM_011285976.3;refGene_gn=LNP1;refGene_tx=XM_019839617.2;refGene_gn=LNP1;refGene_tx=XM_019839618.2;refGene_gn=LNP1;refGene_tx=XM_019839619.2;refGene_gn=LNP1;refGene_tx=XM_023260089.1;refGene_gn=LNP1;refGene_tx=XM_023260090.1;refGene_gn=LNP1;refGene_tx=XM_023260091.1
C2 48376328 48376386 refGene_gn=LNP1;refGene_tx=XM_019839619.2
C2 48385918 48386149
refGene_gn=LNP1;refGene_tx=XM_006935996.4;refGene_gn=LNP1;refGene_tx=XM_006936000.4;refGene_gn=LNP1;refGene_tx=XM_011285976.3;refGene_gn=LNP1;refGene_tx=XM_019839617.2;refGene_gn=LNP1;refGene_tx=XM_019839618.2;refGene_gn=LNP1;refGene_tx=XM_019839619.2;refGene_gn=LNP1;refGene_tx=XM_023260089.1;refGene_gn=LNP1;refGene_tx=XM_023260090.1;refGene_gn=LNP1;refGene_tx=XM_023260091.1
C2 48391076 48391226
refGene_gn=LNP1;refGene_tx=XM_006935996.4;refGene_gn=LNP1;refGene_tx=XM_006936000.4;refGene_gn=LNP1;refGene_tx=XM_011285976.3;refGene_gn=LNP1;refGene_tx=XM_019839617.2;refGene_gn=LNP1;refGene_tx=XM_019839618.2;refGene_gn=LNP1;refGene_tx=XM_019839619.2;refGene_gn=LNP1;refGene_tx=XM_023260089.1;refGene_gn=LNP1;refGene_tx=XM_023260090.1;refGene_gn=LNP1;refGene_tx=XM_023260091.1
C2 48478079 48478269
refGene_gn=TMEM45A;refGene_tx=XM_003991584.5;refGene_gn=TMEM45A;refGene_tx=XM_019839620.2;refGene_gn=TMEM45A;refGene_tx=XM_019839621.2
C2 48479828 48480041
refGene_gn=TMEM45A;refGene_tx=XM_003991584.5;refGene_gn=TMEM45A;refGene_tx=XM_019839620.2;refGene_gn=TMEM45A;refGene_tx=XM_019839621.2
C2 48481448 48481633
refGene_gn=TMEM45A;refGene_tx=XM_003991584.5;refGene_gn=TMEM45A;refGene_tx=XM_019839620.2;refGene_gn=TMEM45A;refGene_tx=XM_019839621.2
C2 48488951 48489097
refGene_gn=TMEM45A;refGene_tx=XM_003991584.5;refGene_gn=TMEM45A;refGene_tx=XM_019839620.2;refGene_gn=TMEM45A;refGene_tx=XM_019839621.2
C2 48493393 48493487
refGene_gn=TMEM45A;refGene_tx=XM_003991584.5;refGene_gn=TMEM45A;refGene_tx=XM_019839620.2;refGene_gn=TMEM45A;refGene_tx=XM_019839621.2
C2 48514970 48515085 refGene_gn=ADGRG7;refGene_tx=XM_023260092.1
C2 48527703 48527817 refGene_gn=ADGRG7;refGene_tx=XM_023260092.1
C2 48529509 48529614 refGene_gn=ADGRG7;refGene_tx=XM_023260092.1
C2 48532102 48532215 refGene_gn=ADGRG7;refGene_tx=XM_023260092.1

C2 48535078 48535228 refGene_gn=ADGRG7;refGene_tx=XM_023260092.1
C2 48535680 48535784 refGene_gn=ADGRG7;refGene_tx=XM_023260092.1
C2 48541601 48541726 refGene_gn=ADGRG7;refGene_tx=XM_023260092.1
C2 48541841 48541949 refGene_gn=ADGRG7;refGene_tx=XM_023260092.1
C2 48545176 48545337 refGene_gn=ADGRG7;refGene_tx=XM_023260092.1
C2 48545770 48545926 refGene_gn=ADGRG7;refGene_tx=XM_023260092.1
C2 48556967 48557080 refGene_gn=ADGRG7;refGene_tx=XM_023260092.1
C2 48563616 48563963 refGene_gn=ADGRG7;refGene_tx=XM_023260092.1
C2 48564654 48564751 refGene_gn=ADGRG7;refGene_tx=XM_023260092.1
C2 48572462 48572618 refGene_gn=ADGRG7;refGene_tx=XM_023260092.1
C2 48574673 48574830 refGene_gn=ADGRG7;refGene_tx=XM_023260092.1
C2 48585188 48585449 refGene_gn=ADGRG7;refGene_tx=XM_023260092.1
C2 48603754 48603938 refGene_gn=TFG;refGene_tx=XM_023260094.1
C2 48605080 48605264
refGene_gn=TFG;refGene_tx=XM_006936010.3;refGene_gn=TFG;refGene_tx=XM_006936011.3;refGene_gn=TFG;refGene_tx=XM_011285975.3;refGene_gn=TFG;refGene_tx=XM_019839622.2
C2 48608709 48608793
refGene_gn=TFG;refGene_tx=XM_006936010.3;refGene_gn=TFG;refGene_tx=XM_006936011.3;refGene_gn=TFG;refGene_tx=XM_011285975.3;refGene_gn=TFG;refGene_tx=XM_019839622.2;refGene_gn=TFG;refGene_tx=XM_023260094.1
C2 48615076 48615223
refGene_gn=TFG;refGene_tx=XM_006936010.3;refGene_gn=TFG;refGene_tx=XM_006936011.3;refGene_gn=TFG;refGene_tx=XM_011285975.3;refGene_gn=TFG;refGene_tx=XM_019839622.2;refGene_gn=TFG;refGene_tx=XM_023260094.1
C2 48618995 48619163
refGene_gn=TFG;refGene_tx=XM_006936010.3;refGene_gn=TFG;refGene_tx=XM_006936011.3;refGene_gn=TFG;refGene_tx=XM_011285975.3;refGene_gn=TFG;refGene_tx=XM_019839622.2;refGene_gn=TFG;refGene_tx=XM_023260094.1
C2 48623092 48623233
refGene_gn=TFG;refGene_tx=XM_006936010.3;refGene_gn=TFG;refGene_tx=XM_006936011.3;refGene_gn=TFG;refGene_tx=XM_011285975.3;refGene_gn=TFG;refGene_tx=XM_019839622.2;refGene_gn=TFG;refGene_tx=XM_023260094.1
C2 48628581 48628680
refGene_gn=TFG;refGene_tx=XM_006936010.3;refGene_gn=TFG;refGene_tx=XM_006936011.3;refGene_gn=TFG;refGene_tx=XM_011285975.3;refGene_gn=TFG;refGene_tx=XM_019839622.2;refGene_gn=TFG;refGene_tx=XM_023260094.1
C2 48632010 48632393
refGene_gn=TFG;refGene_tx=XM_006936010.3;refGene_gn=TFG;refGene_tx=XM_006936011.3;refGene_gn=TFG;refGene_tx=XM_011285975.3;refGene_gn=TFG;refGene_tx=XM_019839622.2;refGene_gn=TFG;refGene_tx=XM_023260094.1
C2 48634124 48634240 refGene_gn=ABI3BP;refGene_tx=XM_023260093.1
C2 48635462 48635585 refGene_gn=ABI3BP;refGene_tx=XM_023260093.1
C2 48636791 48636953 refGene_gn=ABI3BP;refGene_tx=XM_023260093.1
C2 48637745 48637775 refGene_gn=ABI3BP;refGene_tx=XM_023260093.1
C2 48638718 48638798 refGene_gn=ABI3BP;refGene_tx=XM_023260093.1
C2 48654413 48654522 refGene_gn=ABI3BP;refGene_tx=XM_023260093.1
C2 48659948 48660158 refGene_gn=ABI3BP;refGene_tx=XM_023260093.1
C2 48676649 48676718 refGene_gn=ABI3BP;refGene_tx=XM_023260093.1
C2 48677493 48677622 refGene_gn=ABI3BP;refGene_tx=XM_023260093.1

| | | | |
|---|---|---|---|
| C2 | 48680843 | 48680948 | refGene_gn=ABI3BP;refGene_tx=XM_023260093.1 |
| C2 | 48682615 | 48682693 | refGene_gn=ABI3BP;refGene_tx=XM_023260093.1 |
| C2 | 48689172 | 48689247 | refGene_gn=ABI3BP;refGene_tx=XM_023260093.1 |
| C2 | 48690936 | 48691002 | refGene_gn=ABI3BP;refGene_tx=XM_023260093.1 |
| C2 | 48701626 | 48701704 | refGene_gn=ABI3BP;refGene_tx=XM_023260093.1 |
| C2 | 48703826 | 48703907 | refGene_gn=ABI3BP;refGene_tx=XM_023260093.1 |
| C2 | 48704783 | 48704831 | refGene_gn=ABI3BP;refGene_tx=XM_023260093.1 |
| C2 | 48705428 | 48705488 | refGene_gn=ABI3BP;refGene_tx=XM_023260093.1 |
| C2 | 48711933 | 48712008 | refGene_gn=ABI3BP;refGene_tx=XM_023260093.1 |
| C2 | 48715815 | 48715890 | refGene_gn=ABI3BP;refGene_tx=XM_023260093.1 |
| C2 | 48717957 | 48718023 | refGene_gn=ABI3BP;refGene_tx=XM_023260093.1 |
| C2 | 48718774 | 48718822 | refGene_gn=ABI3BP;refGene_tx=XM_023260093.1 |
| C2 | 48719289 | 48719361 | refGene_gn=ABI3BP;refGene_tx=XM_023260093.1 |
| C2 | 48720032 | 48720089 | refGene_gn=ABI3BP;refGene_tx=XM_023260093.1 |
| C2 | 48720950 | 48721040 | refGene_gn=ABI3BP;refGene_tx=XM_023260093.1 |
| C2 | 48723462 | 48723522 | refGene_gn=ABI3BP;refGene_tx=XM_023260093.1 |
| C2 | 48724137 | 48724221 | refGene_gn=ABI3BP;refGene_tx=XM_023260093.1 |
| C2 | 48724861 | 48724921 | refGene_gn=ABI3BP;refGene_tx=XM_023260093.1 |
| C2 | 48725924 | 48725981 | refGene_gn=ABI3BP;refGene_tx=XM_023260093.1 |
| C2 | 48727322 | 48727406 | refGene_gn=ABI3BP;refGene_tx=XM_023260093.1 |
| C2 | 48728141 | 48728201 | refGene_gn=ABI3BP;refGene_tx=XM_023260093.1 |
| C2 | 48729322 | 48729406 | refGene_gn=ABI3BP;refGene_tx=XM_023260093.1 |
| C2 | 48731601 | 48731685 | refGene_gn=ABI3BP;refGene_tx=XM_023260093.1 |
| C2 | 48732488 | 48732548 | refGene_gn=ABI3BP;refGene_tx=XM_023260093.1 |
| C2 | 48735157 | 48735217 | refGene_gn=ABI3BP;refGene_tx=XM_023260093.1 |
| C2 | 48736456 | 48736540 | refGene_gn=ABI3BP;refGene_tx=XM_023260093.1 |
| C2 | 48737387 | 48737447 | refGene_gn=ABI3BP;refGene_tx=XM_023260093.1 |
| C2 | 48738602 | 48738683 | refGene_gn=ABI3BP;refGene_tx=XM_023260093.1 |
| C2 | 48741067 | 48741157 | refGene_gn=ABI3BP;refGene_tx=XM_023260093.1 |
| C2 | 48741951 | 48742011 | refGene_gn=ABI3BP;refGene_tx=XM_023260093.1 |
| C2 | 48743501 | 48743549 | refGene_gn=ABI3BP;refGene_tx=XM_023260093.1 |
| C2 | 48744238 | 48744313 | refGene_gn=ABI3BP;refGene_tx=XM_023260093.1 |
| C2 | 48744429 | 48744492 | refGene_gn=ABI3BP;refGene_tx=XM_023260093.1 |
| C2 | 48745564 | 48745612 | refGene_gn=ABI3BP;refGene_tx=XM_023260093.1 |
| C2 | 48746046 | 48746127 | refGene_gn=ABI3BP;refGene_tx=XM_023260093.1 |
| C2 | 48746750 | 48746789 | refGene_gn=ABI3BP;refGene_tx=XM_023260093.1 |
| C2 | 48747646 | 48747688 | refGene_gn=ABI3BP;refGene_tx=XM_023260093.1 |
| C2 | 48752487 | 48752562 | refGene_gn=ABI3BP;refGene_tx=XM_023260093.1 |
| C2 | 48753366 | 48753438 | refGene_gn=ABI3BP;refGene_tx=XM_023260093.1 |
| C2 | 48754319 | 48754394 | refGene_gn=ABI3BP;refGene_tx=XM_023260093.1 |
| C2 | 48755267 | 48755342 | refGene_gn=ABI3BP;refGene_tx=XM_023260093.1 |
| C2 | 48755894 | 48755969 | refGene_gn=ABI3BP;refGene_tx=XM_023260093.1 |
| C2 | 48757102 | 48757168 | refGene_gn=ABI3BP;refGene_tx=XM_023260093.1 |
| C2 | 48768207 | 48768282 | refGene_gn=ABI3BP;refGene_tx=XM_023260093.1 |
| C2 | 48768731 | 48768803 | refGene_gn=ABI3BP;refGene_tx=XM_023260093.1 |
| C2 | 48769652 | 48769727 | refGene_gn=ABI3BP;refGene_tx=XM_023260093.1 |
| C2 | 48770468 | 48770543 | refGene_gn=ABI3BP;refGene_tx=XM_023260093.1 |
| C2 | 48772454 | 48772532 | refGene_gn=ABI3BP;refGene_tx=XM_023260093.1 |
| C2 | 48777774 | 48777867 | refGene_gn=ABI3BP;refGene_tx=XM_023260093.1 |
| C2 | 48778638 | 48778710 | refGene_gn=ABI3BP;refGene_tx=XM_023260093.1 |

C2 48779711 48779760 refGene_gn=ABI3BP;refGene_tx=XM_023260093.1
C2 48790289 48790345 refGene_gn=ABI3BP;refGene_tx=XM_023260093.1
C2 48790903 48791085 refGene_gn=ABI3BP;refGene_tx=XM_023260093.1
C2 48804228 48804361 refGene_gn=ABI3BP;refGene_tx=XM_023260093.1
C2 48807743 48807812 refGene_gn=ABI3BP;refGene_tx=XM_023260093.1
C2 48833916 48834096 refGene_gn=ABI3BP;refGene_tx=XM_023260093.1
C2 48899142 48899344 refGene_gn=ABI3BP;refGene_tx=XM_023260093.1
C2 49122011 49122024 refGene_gn=IMPG2;refGene_tx=XM_023260158.1
C2 49124423 49124503 refGene_gn=IMPG2;refGene_tx=XM_023260158.1
C2 49125001 49125200 refGene_gn=IMPG2;refGene_tx=XM_023260158.1
C2 49126794 49126983 refGene_gn=IMPG2;refGene_tx=XM_023260158.1
C2 49128477 49128688 refGene_gn=IMPG2;refGene_tx=XM_023260158.1
C2 49133617 49133837 refGene_gn=IMPG2;refGene_tx=XM_023260158.1
C2 49134695 49135978 refGene_gn=IMPG2;refGene_tx=XM_023260158.1
C2 49136966 49137270 refGene_gn=IMPG2;refGene_tx=XM_023260158.1
C2 49155341 49155427 refGene_gn=IMPG2;refGene_tx=XM_023260158.1
C2 49159205 49159450 refGene_gn=IMPG2;refGene_tx=XM_023260158.1
C2 49167382 49167403 refGene_gn=IMPG2;refGene_tx=XM_023260158.1
C2 49168779 49168838 refGene_gn=IMPG2;refGene_tx=XM_023260158.1
C2 49178791 49179015 refGene_gn=IMPG2;refGene_tx=XM_023260158.1
C2 49179926 49179976 refGene_gn=IMPG2;refGene_tx=XM_023260158.1
C2 49187851 49187883 refGene_gn=IMPG2;refGene_tx=XM_023260158.1
C2 49191063 49191230 refGene_gn=IMPG2;refGene_tx=XM_023260158.1
C2 49204505 49204757 refGene_gn=IMPG2;refGene_tx=XM_023260158.1
C2 49205217 49205302 refGene_gn=IMPG2;refGene_tx=XM_023260158.1
C2 49209559 49209697
refGene_gn=SENP7;refGene_tx=XM_023260096.1;refGene_gn=SENP7;refGene_tx=XM_023260097.1;refGene_gn=SENP7;refGene_tx=XM_023260098.1;refGene_gn=SENP7;refGene_tx=XM_023260099.1;refGene_gn=SENP7;refGene_tx=XM_023260100.1;refGene_gn=SENP7;refGene_tx=XM_023260102.1;refGene_gn=SENP7;refGene_tx=XM_023260103.1
C2 49212665 49212816
refGene_gn=SENP7;refGene_tx=XM_023260096.1;refGene_gn=SENP7;refGene_tx=XM_023260097.1;refGene_gn=SENP7;refGene_tx=XM_023260098.1;refGene_gn=SENP7;refGene_tx=XM_023260099.1;refGene_gn=SENP7;refGene_tx=XM_023260100.1;refGene_gn=SENP7;refGene_tx=XM_023260102.1;refGene_gn=SENP7;refGene_tx=XM_023260103.1
C2 49213677 49213746
refGene_gn=SENP7;refGene_tx=XM_023260096.1;refGene_gn=SENP7;refGene_tx=XM_023260097.1;refGene_gn=SENP7;refGene_tx=XM_023260098.1;refGene_gn=SENP7;refGene_tx=XM_023260099.1;refGene_gn=SENP7;refGene_tx=XM_023260100.1;refGene_gn=SENP7;refGene_tx=XM_023260102.1;refGene_gn=SENP7;refGene_tx=XM_023260103.1
C2 49213840 49213890
refGene_gn=SENP7;refGene_tx=XM_023260096.1;refGene_gn=SENP7;refGene_tx=XM_023260097.1;refGene_gn=SENP7;refGene_tx=XM_023260098.1;refGene_gn=SENP7;refGene_tx=XM_023260099.1;refGene_gn=SENP7;refGene_tx=XM_023260100.1;refGene_gn=SENP7;refGene_tx=XM_023260102.1;refGene_gn=SENP7;refGene_tx=XM_023260103.1
C2 49215319 49215372

refGene_gn=SENP7;refGene_tx=XM_023260096.1;refGene_gn=SENP7;refGene_tx=XM_023260097.1;refGene_gn=SENP7;refGene_tx=XM_023260098.1;refGene_gn=SENP7;refGene_tx=XM_023260099.1;refGene_gn=SENP7;refGene_tx=XM_023260100.1;refGene_gn=SENP7;refGene_tx=XM_023260102.1;refGene_gn=SENP7;refGene_tx=XM_023260103.1
C2 49217220 49217345
refGene_gn=SENP7;refGene_tx=XM_023260096.1;refGene_gn=SENP7;refGene_tx=XM_023260097.1;refGene_gn=SENP7;refGene_tx=XM_023260098.1;refGene_gn=SENP7;refGene_tx=XM_023260099.1;refGene_gn=SENP7;refGene_tx=XM_023260100.1;refGene_gn=SENP7;refGene_tx=XM_023260102.1;refGene_gn=SENP7;refGene_tx=XM_023260103.1
C2 49217985 49218078
refGene_gn=SENP7;refGene_tx=XM_023260096.1;refGene_gn=SENP7;refGene_tx=XM_023260097.1;refGene_gn=SENP7;refGene_tx=XM_023260098.1;refGene_gn=SENP7;refGene_tx=XM_023260099.1;refGene_gn=SENP7;refGene_tx=XM_023260100.1;refGene_gn=SENP7;refGene_tx=XM_023260102.1;refGene_gn=SENP7;refGene_tx=XM_023260103.1
C2 49221701 49221824
refGene_gn=SENP7;refGene_tx=XM_023260096.1;refGene_gn=SENP7;refGene_tx=XM_023260097.1;refGene_gn=SENP7;refGene_tx=XM_023260098.1;refGene_gn=SENP7;refGene_tx=XM_023260099.1;refGene_gn=SENP7;refGene_tx=XM_023260100.1;refGene_gn=SENP7;refGene_tx=XM_023260102.1;refGene_gn=SENP7;refGene_tx=XM_023260103.1
C2 49226656 49226773
refGene_gn=SENP7;refGene_tx=XM_023260096.1;refGene_gn=SENP7;refGene_tx=XM_023260097.1;refGene_gn=SENP7;refGene_tx=XM_023260099.1;refGene_gn=SENP7;refGene_tx=XM_023260100.1;refGene_gn=SENP7;refGene_tx=XM_023260102.1;refGene_gn=SENP7;refGene_tx=XM_023260103.1
C2 49228371 49228505
refGene_gn=SENP7;refGene_tx=XM_023260096.1;refGene_gn=SENP7;refGene_tx=XM_023260097.1;refGene_gn=SENP7;refGene_tx=XM_023260098.1;refGene_gn=SENP7;refGene_tx=XM_023260099.1;refGene_gn=SENP7;refGene_tx=XM_023260100.1;refGene_gn=SENP7;refGene_tx=XM_023260102.1;refGene_gn=SENP7;refGene_tx=XM_023260103.1
C2 49230211 49230474
refGene_gn=SENP7;refGene_tx=XM_023260096.1;refGene_gn=SENP7;refGene_tx=XM_023260097.1;refGene_gn=SENP7;refGene_tx=XM_023260098.1;refGene_gn=SENP7;refGene_tx=XM_023260099.1;refGene_gn=SENP7;refGene_tx=XM_023260100.1;refGene_gn=SENP7;refGene_tx=XM_023260102.1;refGene_gn=SENP7;refGene_tx=XM_023260103.1
C2 49232068 49232248
refGene_gn=SENP7;refGene_tx=XM_023260096.1;refGene_gn=SENP7;refGene_tx=XM_023260097.1;refGene_gn=SENP7;refGene_tx=XM_023260098.1;refGene_gn=SENP7;refGene_tx=XM_023260099.1;refGene_gn=SENP7;refGene_tx=XM_023260100.1;refGene_gn=SENP7;refGene_tx=XM_023260102.1;refGene_gn=SENP7;refGene_tx=XM_023260103.1
C2 49246522 49246556
refGene_gn=SENP7;refGene_tx=XM_023260096.1;refGene_gn=SENP7;refGene_tx=XM_023260097.1;refGene_gn=SENP7;refGene_tx=XM_023260098.1;refGene_gn=SENP7;refGene_tx=XM_023260099.1;refGene_gn=SENP7;refGene_tx=XM_0232601

00.1;refGene_gn=SENP7;refGene_tx=XM_023260102.1;refGene_gn=SENP7;refGene_tx=XM_023260103.1
C2 49251745 49251892
refGene_gn=SENP7;refGene_tx=XM_023260096.1;refGene_gn=SENP7;refGene_tx=XM_023260097.1;refGene_gn=SENP7;refGene_tx=XM_023260098.1;refGene_gn=SENP7;refGene_tx=XM_023260099.1;refGene_gn=SENP7;refGene_tx=XM_023260100.1;refGene_gn=SENP7;refGene_tx=XM_023260102.1;refGene_gn=SENP7;refGene_tx=XM_023260103.1
C2 49254489 49254644
refGene_gn=SENP7;refGene_tx=XM_023260096.1;refGene_gn=SENP7;refGene_tx=XM_023260097.1;refGene_gn=SENP7;refGene_tx=XM_023260098.1;refGene_gn=SENP7;refGene_tx=XM_023260099.1;refGene_gn=SENP7;refGene_tx=XM_023260100.1;refGene_gn=SENP7;refGene_tx=XM_023260102.1;refGene_gn=SENP7;refGene_tx=XM_023260103.1
C2 49255698 49256038
refGene_gn=SENP7;refGene_tx=XM_023260096.1;refGene_gn=SENP7;refGene_tx=XM_023260097.1;refGene_gn=SENP7;refGene_tx=XM_023260098.1;refGene_gn=SENP7;refGene_tx=XM_023260099.1;refGene_gn=SENP7;refGene_tx=XM_023260100.1;refGene_gn=SENP7;refGene_tx=XM_023260102.1;refGene_gn=SENP7;refGene_tx=XM_023260103.1
C2 49257101 49257286
refGene_gn=SENP7;refGene_tx=XM_023260096.1;refGene_gn=SENP7;refGene_tx=XM_023260097.1;refGene_gn=SENP7;refGene_tx=XM_023260098.1;refGene_gn=SENP7;refGene_tx=XM_023260099.1;refGene_gn=SENP7;refGene_tx=XM_023260100.1;refGene_gn=SENP7;refGene_tx=XM_023260102.1;refGene_gn=SENP7;refGene_tx=XM_023260103.1
C2 49261014 49261141
refGene_gn=SENP7;refGene_tx=XM_011285974.3;refGene_gn=SENP7;refGene_tx=XM_023260096.1;refGene_gn=SENP7;refGene_tx=XM_023260097.1;refGene_gn=SENP7;refGene_tx=XM_023260098.1;refGene_gn=SENP7;refGene_tx=XM_023260099.1;refGene_gn=SENP7;refGene_tx=XM_023260100.1;refGene_gn=SENP7;refGene_tx=XM_023260102.1;refGene_gn=SENP7;refGene_tx=XM_023260103.1
C2 49284481 49284676
refGene_gn=SENP7;refGene_tx=XM_023260096.1;refGene_gn=SENP7;refGene_tx=XM_023260097.1;refGene_gn=SENP7;refGene_tx=XM_023260098.1;refGene_gn=SENP7;refGene_tx=XM_023260100.1
C2 49292750 49292948
refGene_gn=SENP7;refGene_tx=XM_011285974.3;refGene_gn=SENP7;refGene_tx=XM_023260096.1;refGene_gn=SENP7;refGene_tx=XM_023260097.1;refGene_gn=SENP7;refGene_tx=XM_023260098.1;refGene_gn=SENP7;refGene_tx=XM_023260099.1
C2 49332379 49332477
refGene_gn=SENP7;refGene_tx=XM_011285974.3;refGene_gn=SENP7;refGene_tx=XM_023260096.1;refGene_gn=SENP7;refGene_tx=XM_023260097.1;refGene_gn=SENP7;refGene_tx=XM_023260098.1;refGene_gn=SENP7;refGene_tx=XM_023260099.1;refGene_gn=SENP7;refGene_tx=XM_023260100.1;refGene_gn=SENP7;refGene_tx=XM_023260102.1
C2 49350472 49350571
refGene_gn=SENP7;refGene_tx=XM_011285974.3;refGene_gn=SENP7;refGene_tx=XM_023260096.1;refGene_gn=SENP7;refGene_tx=XM_023260097.1;refGene_gn=

SENP7;refGene_tx=XM_023260098.1;refGene_gn=SENP7;refGene_tx=XM_023260099.1;refGene_gn=SENP7;refGene_tx=XM_023260100.1;refGene_gn=SENP7;refGene_tx=XM_023260102.1;refGene_gn=SENP7;refGene_tx=XM_023260103.1
C2 49360766 49360819
refGene_gn=SENP7;refGene_tx=XM_011285974.3;refGene_gn=SENP7;refGene_tx=XM_023260096.1;refGene_gn=SENP7;refGene_tx=XM_023260098.1;refGene_gn=SENP7;refGene_tx=XM_023260099.1;refGene_gn=SENP7;refGene_tx=XM_023260100.1;refGene_gn=SENP7;refGene_tx=XM_023260102.1
C2 49365596 49365796
refGene_gn=LOC101099826;refGene_tx=XM_023260159.1
C2 49366273 49366553
refGene_gn=LOC101099826;refGene_tx=XM_023260159.1;refGene_gn=SENP7;refGene_tx=XM_011285974.3;refGene_gn=SENP7;refGene_tx=XM_023260096.1;refGene_gn=SENP7;refGene_tx=XM_023260098.1;refGene_gn=SENP7;refGene_tx=XM_023260099.1;refGene_gn=SENP7;refGene_tx=XM_023260100.1;refGene_gn=SENP7;refGene_tx=XM_023260102.1
C2 49377511 49378606
refGene_gn=LOC101099826;refGene_tx=XM_023260159.1
C2 49390999 49392271 refGene_gn=TRMT10C;refGene_tx=XM_003991587.4
C2 49401073 49401137 refGene_gn=PCNP;refGene_tx=XM_023260104.1
C2 49407246 49407461 refGene_gn=PCNP;refGene_tx=XM_023260104.1
C2 49410533 49410608 refGene_gn=PCNP;refGene_tx=XM_023260104.1
C2 49415692 49415748 refGene_gn=PCNP;refGene_tx=XM_023260104.1
C2 49416443 49416570 refGene_gn=PCNP;refGene_tx=XM_023260104.1
C2 49438136 49438654 refGene_gn=ZBTB11;refGene_tx=XM_003991589.5
C2 49439032 49439208 refGene_gn=ZBTB11;refGene_tx=XM_003991589.5
C2 49439319 49439478 refGene_gn=ZBTB11;refGene_tx=XM_003991589.5
C2 49441980 49442098 refGene_gn=ZBTB11;refGene_tx=XM_003991589.5
C2 49442940 49443085 refGene_gn=ZBTB11;refGene_tx=XM_003991589.5
C2 49444412 49444658 refGene_gn=ZBTB11;refGene_tx=XM_003991589.5
C2 49447769 49447949 refGene_gn=ZBTB11;refGene_tx=XM_003991589.5
C2 49448187 49449029 refGene_gn=ZBTB11;refGene_tx=XM_003991589.5
C2 49458940 49459172 refGene_gn=ZBTB11;refGene_tx=XM_003991589.5
C2 49460616 49460852 refGene_gn=ZBTB11;refGene_tx=XM_003991589.5
C2 49468809 49468903 refGene_gn=ZBTB11;refGene_tx=XM_003991589.5
C2 49468904 49469120 refGene_gn=ZBTB11;refGene_tx=XM_003991589.5
C2 49473577 49473658 refGene_gn=RPL24;refGene_tx=NM_001128841.1
C2 49474577 49474641 refGene_gn=RPL24;refGene_tx=NM_001128841.1
C2 49475301 49475438 refGene_gn=RPL24;refGene_tx=NM_001128841.1
C2 49477466 49477577 refGene_gn=RPL24;refGene_tx=NM_001128841.1
C2 49478107 49478183 refGene_gn=RPL24;refGene_tx=NM_001128841.1
C2 49478307 49478312 refGene_gn=RPL24;refGene_tx=NM_001128841.1
C2 49492839 49492882
refGene_gn=CEP97;refGene_tx=XM_006936013.4;refGene_gn=CEP97;refGene_tx=XM_023260106.1
C2 49494716 49494859
refGene_gn=CEP97;refGene_tx=XM_006936013.4;refGene_gn=CEP97;refGene_tx=XM_023260106.1
C2 49495513 49495672
refGene_gn=CEP97;refGene_tx=XM_006936013.4;refGene_gn=CEP97;refGene_tx

=XM_023260106.1
C2	49496986	49497088
refGene_gn=CEP97;refGene_tx=XM_006936013.4;refGene_gn=CEP97;refGene_tx=XM_023260106.1
C2	49498686	49498800
refGene_gn=CEP97;refGene_tx=XM_006936013.4;refGene_gn=CEP97;refGene_tx=XM_023260106.1
C2	49499359	49499526
refGene_gn=CEP97;refGene_tx=XM_006936013.4;refGene_gn=CEP97;refGene_tx=XM_023260106.1
C2	49507067	49507232
refGene_gn=CEP97;refGene_tx=XM_006936013.4;refGene_gn=CEP97;refGene_tx=XM_023260106.1
C2	49508968	49509102
refGene_gn=CEP97;refGene_tx=XM_006936013.4;refGene_gn=CEP97;refGene_tx=XM_023260106.1
C2	49510661	49511436
refGene_gn=CEP97;refGene_tx=XM_006936013.4;refGene_gn=CEP97;refGene_tx=XM_023260106.1
C2	49515721	49515797
refGene_gn=CEP97;refGene_tx=XM_006936013.4;refGene_gn=CEP97;refGene_tx=XM_023260106.1
C2	49517698	49518403
refGene_gn=CEP97;refGene_tx=XM_006936013.4;refGene_gn=CEP97;refGene_tx=XM_023260106.1
C2	49533558	49533603	refGene_gn=NXPE3;refGene_tx=XM_006936015.4
C2	49540349	49540442
refGene_gn=NXPE3;refGene_tx=XM_003991591.5;refGene_gn=NXPE3;refGene_tx=XM_006936014.4
C2	49554975	49555730
refGene_gn=NXPE3;refGene_tx=XM_003991591.5;refGene_gn=NXPE3;refGene_tx=XM_006936014.4;refGene_gn=NXPE3;refGene_tx=XM_006936015.4
C2	49562249	49562323
refGene_gn=NXPE3;refGene_tx=XM_003991591.5;refGene_gn=NXPE3;refGene_tx=XM_006936014.4;refGene_gn=NXPE3;refGene_tx=XM_006936015.4
C2	49577662	49577869
refGene_gn=NXPE3;refGene_tx=XM_003991591.5;refGene_gn=NXPE3;refGene_tx=XM_006936014.4;refGene_gn=NXPE3;refGene_tx=XM_006936015.4
C2	49580215	49580766
refGene_gn=NXPE3;refGene_tx=XM_003991591.5;refGene_gn=NXPE3;refGene_tx=XM_006936014.4;refGene_gn=NXPE3;refGene_tx=XM_006936015.4
C2	49602284	49603212
refGene_gn=NFKBIZ;refGene_tx=XM_023260107.1;refGene_gn=NFKBIZ;refGene_tx=XM_023260108.1
C2	49605542	49605682
refGene_gn=NFKBIZ;refGene_tx=XM_023260107.1;refGene_gn=NFKBIZ;refGene_tx=XM_023260108.1
C2	49606161	49606192
refGene_gn=NFKBIZ;refGene_tx=XM_023260107.1;refGene_gn=NFKBIZ;refGene_tx=XM_023260108.1

C2 49606311 49606415
refGene_gn=NFKBIZ;refGene_tx=XM_023260107.1;refGene_gn=NFKBIZ;refGene_tx=XM_023260108.1
C2 49606506 49607276
refGene_gn=NFKBIZ;refGene_tx=XM_023260107.1;refGene_gn=NFKBIZ;refGene_tx=XM_023260108.1
C2 49607968 49608074
refGene_gn=NFKBIZ;refGene_tx=XM_023260107.1;refGene_gn=NFKBIZ;refGene_tx=XM_023260108.1
C2 49608462 49608609
refGene_gn=NFKBIZ;refGene_tx=XM_023260107.1;refGene_gn=NFKBIZ;refGene_tx=XM_023260108.1
C2 49608796 49608860
refGene_gn=NFKBIZ;refGene_tx=XM_023260107.1;refGene_gn=NFKBIZ;refGene_tx=XM_023260108.1
C2 49609158 49609328
refGene_gn=NFKBIZ;refGene_tx=XM_023260107.1;refGene_gn=NFKBIZ;refGene_tx=XM_023260108.1
C2 49610185 49610296
refGene_gn=NFKBIZ;refGene_tx=XM_023260107.1;refGene_gn=NFKBIZ;refGene_tx=XM_023260108.1
C2 49610423 49610675
refGene_gn=NFKBIZ;refGene_tx=XM_023260107.1;refGene_gn=NFKBIZ;refGene_tx=XM_023260108.1
C2 49613137 49613191 refGene_gn=NFKBIZ;refGene_tx=XM_023260108.1
C2 50149982 50150088 refGene_gn=ZPLD1;refGene_tx=XM_023260109.1
C2 50164292 50164513 refGene_gn=ZPLD1;refGene_tx=XM_023260109.1
C2 50168136 50168318 refGene_gn=ZPLD1;refGene_tx=XM_023260109.1
C2 50169740 50169813 refGene_gn=ZPLD1;refGene_tx=XM_023260109.1
C2 50174630 50174728 refGene_gn=ZPLD1;refGene_tx=XM_023260109.1
C2 50176052 50176133 refGene_gn=ZPLD1;refGene_tx=XM_023260109.1
C2 50185032 50185204 refGene_gn=ZPLD1;refGene_tx=XM_023260109.1
C2 50186513 50186622 refGene_gn=ZPLD1;refGene_tx=XM_023260109.1
C2 50191976 50192006 refGene_gn=ZPLD1;refGene_tx=XM_023260109.1
C2 50192422 50192598 refGene_gn=ZPLD1;refGene_tx=XM_023260109.1
C2 52667664 52667737
refGene_gn=ALCAM;refGene_tx=XM_011285971.3;refGene_gn=ALCAM;refGene_tx=XM_011285972.3
C2 52823384 52823485
refGene_gn=ALCAM;refGene_tx=XM_011285971.3;refGene_gn=ALCAM;refGene_tx=XM_011285972.3
C2 52827194 52827414
refGene_gn=ALCAM;refGene_tx=XM_011285971.3;refGene_gn=ALCAM;refGene_tx=XM_011285972.3
C2 52834250 52834315
refGene_gn=ALCAM;refGene_tx=XM_011285971.3;refGene_gn=ALCAM;refGene_tx=XM_011285972.3
C2 52835745 52835833
refGene_gn=ALCAM;refGene_tx=XM_011285971.3;refGene_gn=ALCAM;refGene_tx=XM_011285972.3

C2 52836292 52836475
refGene_gn=ALCAM;refGene_tx=XM_011285971.3;refGene_gn=ALCAM;refGene_tx
=XM_011285972.3
C2 52845935 52846063
refGene_gn=ALCAM;refGene_tx=XM_011285971.3;refGene_gn=ALCAM;refGene_tx
=XM_011285972.3
C2 52847651 52847784
refGene_gn=ALCAM;refGene_tx=XM_011285971.3;refGene_gn=ALCAM;refGene_tx
=XM_011285972.3
C2 52851576 52851689
refGene_gn=ALCAM;refGene_tx=XM_011285971.3;refGene_gn=ALCAM;refGene_tx
=XM_011285972.3
C2 52853927 52854063
refGene_gn=ALCAM;refGene_tx=XM_011285971.3;refGene_gn=ALCAM;refGene_tx
=XM_011285972.3
C2 52854170 52854304
refGene_gn=ALCAM;refGene_tx=XM_011285971.3;refGene_gn=ALCAM;refGene_tx
=XM_011285972.3
C2 52856940 52857073
refGene_gn=ALCAM;refGene_tx=XM_011285971.3;refGene_gn=ALCAM;refGene_tx
=XM_011285972.3
C2 52858907 52858946 refGene_gn=ALCAM;refGene_tx=XM_011285971.3
C2 52859238 52859356
refGene_gn=ALCAM;refGene_tx=XM_011285971.3;refGene_gn=ALCAM;refGene_tx
=XM_011285972.3
C2 52879290 52879378
refGene_gn=ALCAM;refGene_tx=XM_011285971.3;refGene_gn=ALCAM;refGene_tx
=XM_011285972.3
C2 52958590 52958850
refGene_gn=CBLB;refGene_tx=XM_006936019.4;refGene_gn=CBLB;refGene_tx=X
M_006936020.4;refGene_gn=CBLB;refGene_tx=XM_006936021.4;refGene_gn=CBL
B;refGene_tx=XM_019839645.2
C2 52975182 52975302
refGene_gn=CBLB;refGene_tx=XM_006936019.4;refGene_gn=CBLB;refGene_tx=X
M_006936020.4;refGene_gn=CBLB;refGene_tx=XM_006936021.4
C2 52983195 52983336
refGene_gn=CBLB;refGene_tx=XM_006936019.4;refGene_gn=CBLB;refGene_tx=X
M_006936020.4;refGene_gn=CBLB;refGene_tx=XM_006936021.4;refGene_gn=CBL
B;refGene_tx=XM_019839645.2
C2 52985932 52986064
refGene_gn=CBLB;refGene_tx=XM_006936019.4;refGene_gn=CBLB;refGene_tx=X
M_006936020.4;refGene_gn=CBLB;refGene_tx=XM_019839645.2
C2 52986179 52986277
refGene_gn=CBLB;refGene_tx=XM_006936019.4;refGene_gn=CBLB;refGene_tx=X
M_006936020.4;refGene_gn=CBLB;refGene_tx=XM_006936021.4;refGene_gn=CBL
B;refGene_tx=XM_019839645.2
C2 52989547 52989694
refGene_gn=CBLB;refGene_tx=XM_006936019.4;refGene_gn=CBLB;refGene_tx=X
M_006936020.4;refGene_gn=CBLB;refGene_tx=XM_006936021.4;refGene_gn=CBL
B;refGene_tx=XM_019839645.2

C2 53002056 53002151
refGene_gn=CBLB;refGene_tx=XM_006936019.4;refGene_gn=CBLB;refGene_tx=XM_006936020.4;refGene_gn=CBLB;refGene_tx=XM_006936021.4;refGene_gn=CBLB;refGene_tx=XM_019839645.2
C2 53009771 53010140
refGene_gn=CBLB;refGene_tx=XM_006936019.4;refGene_gn=CBLB;refGene_tx=XM_006936020.4;refGene_gn=CBLB;refGene_tx=XM_006936021.4;refGene_gn=CBLB;refGene_tx=XM_019839645.2
C2 53011925 53012111
refGene_gn=CBLB;refGene_tx=XM_006936019.4;refGene_gn=CBLB;refGene_tx=XM_006936020.4;refGene_gn=CBLB;refGene_tx=XM_006936021.4;refGene_gn=CBLB;refGene_tx=XM_019839645.2
C2 53029325 53029529
refGene_gn=CBLB;refGene_tx=XM_006936019.4;refGene_gn=CBLB;refGene_tx=XM_006936020.4;refGene_gn=CBLB;refGene_tx=XM_006936021.4;refGene_gn=CBLB;refGene_tx=XM_019839645.2
C2 53040962 53041094
refGene_gn=CBLB;refGene_tx=XM_006936019.4;refGene_gn=CBLB;refGene_tx=XM_006936020.4;refGene_gn=CBLB;refGene_tx=XM_006936021.4;refGene_gn=CBLB;refGene_tx=XM_019839645.2
C2 53043857 53043945
refGene_gn=CBLB;refGene_tx=XM_006936019.4;refGene_gn=CBLB;refGene_tx=XM_006936020.4;refGene_gn=CBLB;refGene_tx=XM_006936021.4;refGene_gn=CBLB;refGene_tx=XM_019839645.2
C2 53047244 53047382
refGene_gn=CBLB;refGene_tx=XM_006936019.4;refGene_gn=CBLB;refGene_tx=XM_006936020.4;refGene_gn=CBLB;refGene_tx=XM_006936021.4;refGene_gn=CBLB;refGene_tx=XM_019839645.2
C2 53050478 53050600
refGene_gn=CBLB;refGene_tx=XM_006936019.4;refGene_gn=CBLB;refGene_tx=XM_006936020.4;refGene_gn=CBLB;refGene_tx=XM_006936021.4;refGene_gn=CBLB;refGene_tx=XM_019839645.2
C2 53058442 53058599
refGene_gn=CBLB;refGene_tx=XM_006936019.4;refGene_gn=CBLB;refGene_tx=XM_006936020.4;refGene_gn=CBLB;refGene_tx=XM_006936021.4;refGene_gn=CBLB;refGene_tx=XM_019839645.2
C2 53077605 53077752
refGene_gn=CBLB;refGene_tx=XM_006936019.4;refGene_gn=CBLB;refGene_tx=XM_006936020.4;refGene_gn=CBLB;refGene_tx=XM_006936021.4;refGene_gn=CBLB;refGene_tx=XM_019839645.2
C2 53154671 53154922
refGene_gn=CBLB;refGene_tx=XM_006936019.4;refGene_gn=CBLB;refGene_tx=XM_006936020.4;refGene_gn=CBLB;refGene_tx=XM_006936021.4;refGene_gn=CBLB;refGene_tx=XM_019839645.2
C2 53167795 53167963
refGene_gn=CBLB;refGene_tx=XM_006936019.4;refGene_gn=CBLB;refGene_tx=XM_006936020.4;refGene_gn=CBLB;refGene_tx=XM_006936021.4;refGene_gn=CBLB;refGene_tx=XM_019839645.2
C2 54462632 54462978 refGene_gn=CCDC54;refGene_tx=XM_019839652.2
C2 54462980 54463042 refGene_gn=CCDC54;refGene_tx=XM_019839652.2

C2	54463284 54463526 refGene_gn=CCDC54;refGene_tx=XM_019839652.2
C2	54463527 54463855 refGene_gn=CCDC54;refGene_tx=XM_019839652.2
C2	54792226 54792388
refGene_gn=BBX;refGene_tx=XM_003991606.4;refGene_gn=BBX;refGene_tx=XM_019839646.1;refGene_gn=BBX;refGene_tx=XM_019839647.1;refGene_gn=BBX;refGene_tx=XM_019839649.2;refGene_gn=BBX;refGene_tx=XM_019839650.1;refGene_gn=BBX;refGene_tx=XM_019839651.1
C2	54798082 54798322
refGene_gn=BBX;refGene_tx=XM_003991606.4;refGene_gn=BBX;refGene_tx=XM_019839646.1;refGene_gn=BBX;refGene_tx=XM_019839647.1;refGene_gn=BBX;refGene_tx=XM_019839649.2;refGene_gn=BBX;refGene_tx=XM_019839650.1;refGene_gn=BBX;refGene_tx=XM_019839651.1
C2	54810251 54810447
refGene_gn=BBX;refGene_tx=XM_003991606.4;refGene_gn=BBX;refGene_tx=XM_019839646.1;refGene_gn=BBX;refGene_tx=XM_019839647.1;refGene_gn=BBX;refGene_tx=XM_019839649.2;refGene_gn=BBX;refGene_tx=XM_019839650.1;refGene_gn=BBX;refGene_tx=XM_019839651.1
C2	54818340 54818408
refGene_gn=BBX;refGene_tx=XM_003991606.4;refGene_gn=BBX;refGene_tx=XM_019839646.1;refGene_gn=BBX;refGene_tx=XM_019839647.1;refGene_gn=BBX;refGene_tx=XM_019839649.2;refGene_gn=BBX;refGene_tx=XM_019839650.1;refGene_gn=BBX;refGene_tx=XM_019839651.1
C2	54831766 54831847
refGene_gn=BBX;refGene_tx=XM_003991606.4;refGene_gn=BBX;refGene_tx=XM_019839646.1;refGene_gn=BBX;refGene_tx=XM_019839647.1;refGene_gn=BBX;refGene_tx=XM_019839649.2;refGene_gn=BBX;refGene_tx=XM_019839650.1;refGene_gn=BBX;refGene_tx=XM_019839651.1
C2	54834935 54835010
refGene_gn=BBX;refGene_tx=XM_003991606.4;refGene_gn=BBX;refGene_tx=XM_019839646.1;refGene_gn=BBX;refGene_tx=XM_019839647.1;refGene_gn=BBX;refGene_tx=XM_019839649.2;refGene_gn=BBX;refGene_tx=XM_019839650.1;refGene_gn=BBX;refGene_tx=XM_019839651.1
C2	54843787 54843868
refGene_gn=BBX;refGene_tx=XM_003991606.4;refGene_gn=BBX;refGene_tx=XM_019839646.1;refGene_gn=BBX;refGene_tx=XM_019839647.1;refGene_gn=BBX;refGene_tx=XM_019839649.2;refGene_gn=BBX;refGene_tx=XM_019839650.1;refGene_gn=BBX;refGene_tx=XM_019839651.1
C2	54858420 54859429
refGene_gn=BBX;refGene_tx=XM_003991606.4;refGene_gn=BBX;refGene_tx=XM_019839646.1;refGene_gn=BBX;refGene_tx=XM_019839647.1;refGene_gn=BBX;refGene_tx=XM_019839649.2;refGene_gn=BBX;refGene_tx=XM_019839650.1;refGene_gn=BBX;refGene_tx=XM_019839651.1
C2	54860415 54860554
refGene_gn=BBX;refGene_tx=XM_003991606.4;refGene_gn=BBX;refGene_tx=XM_019839646.1;refGene_gn=BBX;refGene_tx=XM_019839647.1;refGene_gn=BBX;refGene_tx=XM_019839649.2;refGene_gn=BBX;refGene_tx=XM_019839650.1;refGene_gn=BBX;refGene_tx=XM_019839651.1
C2	54863884 54864033
refGene_gn=BBX;refGene_tx=XM_003991606.4;refGene_gn=BBX;refGene_tx=XM_019839646.1;refGene_gn=BBX;refGene_tx=XM_019839647.1;refGene_gn=BBX;re

fGene_tx=XM_019839649.2;refGene_gn=BBX;refGene_tx=XM_019839650.1;refGene_gn=BBX;refGene_tx=XM_019839651.1
C2 54874798 54874888
refGene_gn=BBX;refGene_tx=XM_003991606.4;refGene_gn=BBX;refGene_tx=XM_019839646.1;refGene_gn=BBX;refGene_tx=XM_019839647.1;refGene_gn=BBX;refGene_tx=XM_019839649.2;refGene_gn=BBX;refGene_tx=XM_019839650.1
C2 54876241 54876301
refGene_gn=BBX;refGene_tx=XM_003991606.4;refGene_gn=BBX;refGene_tx=XM_019839646.1;refGene_gn=BBX;refGene_tx=XM_019839647.1;refGene_gn=BBX;refGene_tx=XM_019839649.2;refGene_gn=BBX;refGene_tx=XM_019839650.1;refGene_gn=BBX;refGene_tx=XM_019839651.1
C2 54883516 54883714
refGene_gn=BBX;refGene_tx=XM_003991606.4;refGene_gn=BBX;refGene_tx=XM_019839646.1;refGene_gn=BBX;refGene_tx=XM_019839647.1;refGene_gn=BBX;refGene_tx=XM_019839649.2;refGene_gn=BBX;refGene_tx=XM_019839650.1;refGene_gn=BBX;refGene_tx=XM_019839651.1
C2 54886106 54886293
refGene_gn=BBX;refGene_tx=XM_003991606.4;refGene_gn=BBX;refGene_tx=XM_019839646.1;refGene_gn=BBX;refGene_tx=XM_019839647.1;refGene_gn=BBX;refGene_tx=XM_019839649.2;refGene_gn=BBX;refGene_tx=XM_019839650.1;refGene_gn=BBX;refGene_tx=XM_019839651.1
C2 54890538 54890626
refGene_gn=BBX;refGene_tx=XM_003991606.4;refGene_gn=BBX;refGene_tx=XM_019839646.1;refGene_gn=BBX;refGene_tx=XM_019839647.1;refGene_gn=BBX;refGene_tx=XM_019839649.2;refGene_gn=BBX;refGene_tx=XM_019839650.1;refGene_gn=BBX;refGene_tx=XM_019839651.1
C2 55122284 55122293
refGene_gn=CD47;refGene_tx=XM_023260110.1;refGene_gn=CD47;refGene_tx=XM_023260111.1;refGene_gn=CD47;refGene_tx=XM_023260112.1;refGene_gn=CD47;refGene_tx=XM_023260113.1;refGene_gn=CD47;refGene_tx=XM_023260114.1;refGene_gn=CD47;refGene_tx=XM_023260115.1
C2 55124174 55124207
refGene_gn=CD47;refGene_tx=XM_023260110.1;refGene_gn=CD47;refGene_tx=XM_023260113.1;refGene_gn=CD47;refGene_tx=XM_023260114.1;refGene_gn=CD47;refGene_tx=XM_023260115.1
C2 55124818 55124843
refGene_gn=CD47;refGene_tx=XM_023260110.1;refGene_gn=CD47;refGene_tx=XM_023260111.1;refGene_gn=CD47;refGene_tx=XM_023260113.1;refGene_gn=CD47;refGene_tx=XM_023260114.1;refGene_gn=CD47;refGene_tx=XM_023260115.1
C2 55126225 55126257
refGene_gn=CD47;refGene_tx=XM_023260110.1;refGene_gn=CD47;refGene_tx=XM_023260111.1;refGene_gn=CD47;refGene_tx=XM_023260112.1;refGene_gn=CD47;refGene_tx=XM_023260113.1;refGene_gn=CD47;refGene_tx=XM_023260114.1;refGene_gn=CD47;refGene_tx=XM_023260115.1
C2 55132166 55132259
refGene_gn=CD47;refGene_tx=XM_023260110.1;refGene_gn=CD47;refGene_tx=XM_023260111.1;refGene_gn=CD47;refGene_tx=XM_023260112.1;refGene_gn=CD47;refGene_tx=XM_023260113.1;refGene_gn=CD47;refGene_tx=XM_023260114.1;refGene_gn=CD47;refGene_tx=XM_023260115.1
C2 55133076 55133169

refGene_gn=CD47;refGene_tx=XM_023260110.1;refGene_gn=CD47;refGene_tx=XM_023260111.1;refGene_gn=CD47;refGene_tx=XM_023260112.1;refGene_gn=CD47;refGene_tx=XM_023260113.1;refGene_gn=CD47;refGene_tx=XM_023260114.1;refGene_gn=CD47;refGene_tx=XM_023260115.1
C2 55134202 55134295
refGene_gn=CD47;refGene_tx=XM_023260110.1;refGene_gn=CD47;refGene_tx=XM_023260111.1;refGene_gn=CD47;refGene_tx=XM_023260112.1;refGene_gn=CD47;refGene_tx=XM_023260113.1;refGene_gn=CD47;refGene_tx=XM_023260114.1;refGene_gn=CD47;refGene_tx=XM_023260115.1
C2 55135916 55136024
refGene_gn=CD47;refGene_tx=XM_023260110.1;refGene_gn=CD47;refGene_tx=XM_023260111.1;refGene_gn=CD47;refGene_tx=XM_023260112.1;refGene_gn=CD47;refGene_tx=XM_023260113.1;refGene_gn=CD47;refGene_tx=XM_023260114.1;refGene_gn=CD47;refGene_tx=XM_023260115.1
C2 55148860 55148950
refGene_gn=CD47;refGene_tx=XM_023260110.1;refGene_gn=CD47;refGene_tx=XM_023260111.1;refGene_gn=CD47;refGene_tx=XM_023260112.1;refGene_gn=CD47;refGene_tx=XM_023260114.1;refGene_gn=CD47;refGene_tx=XM_023260115.1
C2 55156490 55156838
refGene_gn=CD47;refGene_tx=XM_023260110.1;refGene_gn=CD47;refGene_tx=XM_023260111.1;refGene_gn=CD47;refGene_tx=XM_023260112.1;refGene_gn=CD47;refGene_tx=XM_023260113.1;refGene_gn=CD47;refGene_tx=XM_023260114.1;refGene_gn=CD47;refGene_tx=XM_023260115.1
C2 55168268 55168314
refGene_gn=CD47;refGene_tx=XM_023260110.1;refGene_gn=CD47;refGene_tx=XM_023260111.1;refGene_gn=CD47;refGene_tx=XM_023260112.1;refGene_gn=CD47;refGene_tx=XM_023260113.1
C2 55246004 55246183 refGene_gn=IFT57;refGene_tx=XM_003991608.4
C2 55247171 55247238 refGene_gn=IFT57;refGene_tx=XM_003991608.4
C2 55248905 55248968 refGene_gn=IFT57;refGene_tx=XM_003991608.4
C2 55250327 55250459 refGene_gn=IFT57;refGene_tx=XM_003991608.4
C2 55251385 55251457 refGene_gn=IFT57;refGene_tx=XM_003991608.4
C2 55271096 55271219 refGene_gn=IFT57;refGene_tx=XM_003991608.4
C2 55287613 55287682 refGene_gn=IFT57;refGene_tx=XM_003991608.4
C2 55292005 55292096 refGene_gn=IFT57;refGene_tx=XM_003991608.4
C2 55294488 55294607 refGene_gn=IFT57;refGene_tx=XM_003991608.4
C2 55295435 55295598 refGene_gn=IFT57;refGene_tx=XM_003991608.4
C2 55298073 55298285 refGene_gn=IFT57;refGene_tx=XM_003991608.4
C2 55421116 55421180 refGene_gn=HHLA2;refGene_tx=XM_023260116.1
C2 55422601 55422771 refGene_gn=HHLA2;refGene_tx=XM_023260116.1
C2 55422773 55422822 refGene_gn=HHLA2;refGene_tx=XM_023260116.1
C2 55422824 55422944 refGene_gn=HHLA2;refGene_tx=XM_023260116.1
C2 55423858 55424112 refGene_gn=HHLA2;refGene_tx=XM_023260116.1
C2 55434957 55434972 refGene_gn=MYH15;refGene_tx=XM_011285970.3
C2 55437022 55437157 refGene_gn=MYH15;refGene_tx=XM_011285970.3
C2 55438066 55438162 refGene_gn=MYH15;refGene_tx=XM_011285970.3
C2 55453115 55453220 refGene_gn=MYH15;refGene_tx=XM_011285970.3
C2 55456531 55456702 refGene_gn=MYH15;refGene_tx=XM_011285970.3
C2 55458793 55458919 refGene_gn=MYH15;refGene_tx=XM_011285970.3
C2 55462734 55462938 refGene_gn=MYH15;refGene_tx=XM_011285970.3

C2 55463140 55463333 refGene_gn=MYH15;refGene_tx=XM_011285970.3
C2 55473400 55473516 refGene_gn=MYH15;refGene_tx=XM_011285970.3
C2 55476401 55476526 refGene_gn=MYH15;refGene_tx=XM_011285970.3
C2 55479490 55479840 refGene_gn=MYH15;refGene_tx=XM_011285970.3
C2 55483277 55483474 refGene_gn=MYH15;refGene_tx=XM_011285970.3
C2 55486312 55486431 refGene_gn=MYH15;refGene_tx=XM_011285970.3
C2 55490927 55491054 refGene_gn=MYH15;refGene_tx=XM_011285970.3
C2 55499505 55499895 refGene_gn=MYH15;refGene_tx=XM_011285970.3
C2 55502155 55502246 refGene_gn=MYH15;refGene_tx=XM_011285970.3
C2 55505790 55505936 refGene_gn=MYH15;refGene_tx=XM_011285970.3
C2 55507635 55507812 refGene_gn=MYH15;refGene_tx=XM_011285970.3
C2 55508860 55509103 refGene_gn=MYH15;refGene_tx=XM_011285970.3
C2 55512252 55512508 refGene_gn=MYH15;refGene_tx=XM_011285970.3
C2 55515968 55516105 refGene_gn=MYH15;refGene_tx=XM_011285970.3
C2 55520170 55520294 refGene_gn=MYH15;refGene_tx=XM_011285970.3
C2 55521438 55521556 refGene_gn=MYH15;refGene_tx=XM_011285970.3
C2 55524382 55524470 refGene_gn=MYH15;refGene_tx=XM_011285970.3
C2 55525299 55525367 refGene_gn=MYH15;refGene_tx=XM_011285970.3
C2 55528405 55528538 refGene_gn=MYH15;refGene_tx=XM_011285970.3
C2 55529609 55529786 refGene_gn=MYH15;refGene_tx=XM_011285970.3
C2 55536981 55537152 refGene_gn=MYH15;refGene_tx=XM_011285970.3
C2 55537753 55537903 refGene_gn=MYH15;refGene_tx=XM_011285970.3
C2 55542636 55542755 refGene_gn=MYH15;refGene_tx=XM_011285970.3
C2 55552879 55553018 refGene_gn=MYH15;refGene_tx=XM_011285970.3
C2 55554278 55554382 refGene_gn=MYH15;refGene_tx=XM_011285970.3
C2 55563622 55563718 refGene_gn=MYH15;refGene_tx=XM_011285970.3
C2 55563808 55563872 refGene_gn=MYH15;refGene_tx=XM_011285970.3
C2 55566822 55566915 refGene_gn=MYH15;refGene_tx=XM_011285970.3
C2 55568667 55568761 refGene_gn=MYH15;refGene_tx=XM_011285970.3
C2 55570057 55570085 refGene_gn=MYH15;refGene_tx=XM_011285970.3
C2 55570724 55570881 refGene_gn=MYH15;refGene_tx=XM_011285970.3
C2 55572564 55572708 refGene_gn=MYH15;refGene_tx=XM_011285970.3
C2 55576458 55576565 refGene_gn=MYH15;refGene_tx=XM_011285970.3
C2 55580862 55580950 refGene_gn=MYH15;refGene_tx=XM_011285970.3
C2 55624112 55624283
refGene_gn=CIP2A;refGene_tx=XM_003991609.4;refGene_gn=CIP2A;refGene_tx
=XM_023260117.1
C2 55625280 55625420
refGene_gn=CIP2A;refGene_tx=XM_003991609.4;refGene_gn=CIP2A;refGene_tx
=XM_023260117.1
C2 55626153 55626236
refGene_gn=CIP2A;refGene_tx=XM_003991609.4;refGene_gn=CIP2A;refGene_tx
=XM_023260117.1
C2 55626480 55626594
refGene_gn=CIP2A;refGene_tx=XM_003991609.4;refGene_gn=CIP2A;refGene_tx
=XM_023260117.1
C2 55628989 55629186
refGene_gn=CIP2A;refGene_tx=XM_003991609.4;refGene_gn=CIP2A;refGene_tx
=XM_023260117.1
C2 55631492 55631603

refGene_gn=CIP2A;refGene_tx=XM_003991609.4;refGene_gn=CIP2A;refGene_tx
=XM_023260117.1
C2	55631699	55631774
refGene_gn=CIP2A;refGene_tx=XM_003991609.4;refGene_gn=CIP2A;refGene_tx
=XM_023260117.1
C2	55632485	55632681
refGene_gn=CIP2A;refGene_tx=XM_003991609.4;refGene_gn=CIP2A;refGene_tx
=XM_023260117.1
C2	55635766	55635885
refGene_gn=CIP2A;refGene_tx=XM_003991609.4;refGene_gn=CIP2A;refGene_tx
=XM_023260117.1
C2	55637387	55637487
refGene_gn=CIP2A;refGene_tx=XM_003991609.4;refGene_gn=CIP2A;refGene_tx
=XM_023260117.1
C2	55638591	55638733
refGene_gn=CIP2A;refGene_tx=XM_003991609.4;refGene_gn=CIP2A;refGene_tx
=XM_023260117.1
C2	55640240	55640400
refGene_gn=CIP2A;refGene_tx=XM_003991609.4;refGene_gn=CIP2A;refGene_tx
=XM_023260117.1
C2	55641673	55641892
refGene_gn=CIP2A;refGene_tx=XM_003991609.4;refGene_gn=CIP2A;refGene_tx
=XM_023260117.1
C2	55643492	55643568
refGene_gn=CIP2A;refGene_tx=XM_003991609.4;refGene_gn=CIP2A;refGene_tx
=XM_023260117.1
C2	55647061	55647207
refGene_gn=CIP2A;refGene_tx=XM_003991609.4;refGene_gn=CIP2A;refGene_tx
=XM_023260117.1
C2	55647353	55647476
refGene_gn=CIP2A;refGene_tx=XM_003991609.4;refGene_gn=CIP2A;refGene_tx
=XM_023260117.1
C2	55648358	55648455
refGene_gn=CIP2A;refGene_tx=XM_003991609.4;refGene_gn=CIP2A;refGene_tx
=XM_023260117.1
C2	55648961	55649056	refGene_gn=CIP2A;refGene_tx=XM_003991609.4
C2	55649908	55650021	refGene_gn=CIP2A;refGene_tx=XM_003991609.4
C2	55652201	55652349	refGene_gn=CIP2A;refGene_tx=XM_003991609.4
C2	55656638	55656740	refGene_gn=CIP2A;refGene_tx=XM_003991609.4
C2	55672907	55672939
refGene_gn=DZIP3;refGene_tx=XM_011285969.3;refGene_gn=DZIP3;refGene_tx
=XM_019839661.2;refGene_gn=DZIP3;refGene_tx=XM_019839662.2;refGene_gn=
DZIP3;refGene_tx=XM_019839663.2;refGene_gn=DZIP3;refGene_tx=XM_0198396
64.2;refGene_gn=DZIP3;refGene_tx=XM_019839665.2;refGene_gn=DZIP3;refGe
ne_tx=XM_019839666.2;refGene_gn=DZIP3;refGene_tx=XM_019839667.2
C2	55674856	55674929
refGene_gn=DZIP3;refGene_tx=XM_011285969.3;refGene_gn=DZIP3;refGene_tx
=XM_019839661.2;refGene_gn=DZIP3;refGene_tx=XM_019839662.2;refGene_gn=
DZIP3;refGene_tx=XM_019839663.2;refGene_gn=DZIP3;refGene_tx=XM_0198396
64.2;refGene_gn=DZIP3;refGene_tx=XM_019839665.2;refGene_gn=DZIP3;refGe

ne_tx=XM_019839666.2;refGene_gn=DZIP3;refGene_tx=XM_019839667.2
C2	55677216	55677372
refGene_gn=DZIP3;refGene_tx=XM_011285969.3;refGene_gn=DZIP3;refGene_tx=XM_019839661.2;refGene_gn=DZIP3;refGene_tx=XM_019839662.2;refGene_gn=DZIP3;refGene_tx=XM_019839663.2;refGene_gn=DZIP3;refGene_tx=XM_019839664.2;refGene_gn=DZIP3;refGene_tx=XM_019839665.2;refGene_gn=DZIP3;refGene_tx=XM_019839666.2;refGene_gn=DZIP3;refGene_tx=XM_019839667.2
C2	55682286	55682403
refGene_gn=DZIP3;refGene_tx=XM_011285969.3;refGene_gn=DZIP3;refGene_tx=XM_019839661.2;refGene_gn=DZIP3;refGene_tx=XM_019839662.2;refGene_gn=DZIP3;refGene_tx=XM_019839663.2;refGene_gn=DZIP3;refGene_tx=XM_019839664.2;refGene_gn=DZIP3;refGene_tx=XM_019839665.2;refGene_gn=DZIP3;refGene_tx=XM_019839666.2;refGene_gn=DZIP3;refGene_tx=XM_019839667.2
C2	55685441	55685522
refGene_gn=DZIP3;refGene_tx=XM_011285969.3;refGene_gn=DZIP3;refGene_tx=XM_019839661.2;refGene_gn=DZIP3;refGene_tx=XM_019839662.2;refGene_gn=DZIP3;refGene_tx=XM_019839663.2;refGene_gn=DZIP3;refGene_tx=XM_019839664.2;refGene_gn=DZIP3;refGene_tx=XM_019839665.2;refGene_gn=DZIP3;refGene_tx=XM_019839666.2;refGene_gn=DZIP3;refGene_tx=XM_019839667.2
C2	55686022	55686147
refGene_gn=DZIP3;refGene_tx=XM_011285969.3;refGene_gn=DZIP3;refGene_tx=XM_019839661.2;refGene_gn=DZIP3;refGene_tx=XM_019839662.2;refGene_gn=DZIP3;refGene_tx=XM_019839663.2;refGene_gn=DZIP3;refGene_tx=XM_019839664.2;refGene_gn=DZIP3;refGene_tx=XM_019839665.2;refGene_gn=DZIP3;refGene_tx=XM_019839666.2;refGene_gn=DZIP3;refGene_tx=XM_019839667.2
C2	55689098	55689213
refGene_gn=DZIP3;refGene_tx=XM_011285969.3;refGene_gn=DZIP3;refGene_tx=XM_019839661.2;refGene_gn=DZIP3;refGene_tx=XM_019839662.2;refGene_gn=DZIP3;refGene_tx=XM_019839663.2;refGene_gn=DZIP3;refGene_tx=XM_019839664.2;refGene_gn=DZIP3;refGene_tx=XM_019839665.2;refGene_gn=DZIP3;refGene_tx=XM_019839666.2;refGene_gn=DZIP3;refGene_tx=XM_019839667.2
C2	55694449	55694569
refGene_gn=DZIP3;refGene_tx=XM_011285969.3;refGene_gn=DZIP3;refGene_tx=XM_019839661.2;refGene_gn=DZIP3;refGene_tx=XM_019839662.2;refGene_gn=DZIP3;refGene_tx=XM_019839663.2;refGene_gn=DZIP3;refGene_tx=XM_019839664.2;refGene_gn=DZIP3;refGene_tx=XM_019839665.2;refGene_gn=DZIP3;refGene_tx=XM_019839666.2;refGene_gn=DZIP3;refGene_tx=XM_019839667.2
C2	55695955	55696057
refGene_gn=DZIP3;refGene_tx=XM_011285969.3;refGene_gn=DZIP3;refGene_tx=XM_019839661.2;refGene_gn=DZIP3;refGene_tx=XM_019839662.2;refGene_gn=DZIP3;refGene_tx=XM_019839663.2;refGene_gn=DZIP3;refGene_tx=XM_019839664.2;refGene_gn=DZIP3;refGene_tx=XM_019839665.2;refGene_gn=DZIP3;refGene_tx=XM_019839666.2;refGene_gn=DZIP3;refGene_tx=XM_019839667.2
C2	55703804	55703897
refGene_gn=DZIP3;refGene_tx=XM_011285969.3;refGene_gn=DZIP3;refGene_tx=XM_019839661.2;refGene_gn=DZIP3;refGene_tx=XM_019839662.2;refGene_gn=DZIP3;refGene_tx=XM_019839663.2;refGene_gn=DZIP3;refGene_tx=XM_019839664.2;refGene_gn=DZIP3;refGene_tx=XM_019839665.2;refGene_gn=DZIP3;refGene_tx=XM_019839666.2;refGene_gn=DZIP3;refGene_tx=XM_019839667.2
C2	55704946	55704999

refGene_gn=DZIP3;refGene_tx=XM_011285969.3;refGene_gn=DZIP3;refGene_tx=XM_019839661.2;refGene_gn=DZIP3;refGene_tx=XM_019839662.2;refGene_gn=DZIP3;refGene_tx=XM_019839663.2;refGene_gn=DZIP3;refGene_tx=XM_019839664.2;refGene_gn=DZIP3;refGene_tx=XM_019839665.2;refGene_gn=DZIP3;refGene_tx=XM_019839666.2;refGene_gn=DZIP3;refGene_tx=XM_019839667.2
C2 55713202 55713279
refGene_gn=DZIP3;refGene_tx=XM_011285969.3;refGene_gn=DZIP3;refGene_tx=XM_019839661.2;refGene_gn=DZIP3;refGene_tx=XM_019839662.2;refGene_gn=DZIP3;refGene_tx=XM_019839663.2;refGene_gn=DZIP3;refGene_tx=XM_019839664.2;refGene_gn=DZIP3;refGene_tx=XM_019839665.2;refGene_gn=DZIP3;refGene_tx=XM_019839666.2;refGene_gn=DZIP3;refGene_tx=XM_019839667.2
C2 55714936 55715554
refGene_gn=DZIP3;refGene_tx=XM_011285969.3;refGene_gn=DZIP3;refGene_tx=XM_019839661.2;refGene_gn=DZIP3;refGene_tx=XM_019839662.2;refGene_gn=DZIP3;refGene_tx=XM_019839663.2;refGene_gn=DZIP3;refGene_tx=XM_019839664.2;refGene_gn=DZIP3;refGene_tx=XM_019839665.2
C2 55717604 55717637
refGene_gn=DZIP3;refGene_tx=XM_011285969.3;refGene_gn=DZIP3;refGene_tx=XM_019839661.2;refGene_gn=DZIP3;refGene_tx=XM_019839662.2;refGene_gn=DZIP3;refGene_tx=XM_019839663.2;refGene_gn=DZIP3;refGene_tx=XM_019839664.2;refGene_gn=DZIP3;refGene_tx=XM_019839665.2;refGene_gn=DZIP3;refGene_tx=XM_019839666.2
C2 55718503 55718673
refGene_gn=DZIP3;refGene_tx=XM_011285969.3;refGene_gn=DZIP3;refGene_tx=XM_019839661.2;refGene_gn=DZIP3;refGene_tx=XM_019839662.2;refGene_gn=DZIP3;refGene_tx=XM_019839663.2;refGene_gn=DZIP3;refGene_tx=XM_019839664.2;refGene_gn=DZIP3;refGene_tx=XM_019839665.2;refGene_gn=DZIP3;refGene_tx=XM_019839666.2;refGene_gn=DZIP3;refGene_tx=XM_019839667.2
C2 55719485 55719530
refGene_gn=DZIP3;refGene_tx=XM_011285969.3;refGene_gn=DZIP3;refGene_tx=XM_019839661.2;refGene_gn=DZIP3;refGene_tx=XM_019839662.2;refGene_gn=DZIP3;refGene_tx=XM_019839663.2;refGene_gn=DZIP3;refGene_tx=XM_019839664.2;refGene_gn=DZIP3;refGene_tx=XM_019839665.2;refGene_gn=DZIP3;refGene_tx=XM_019839666.2;refGene_gn=DZIP3;refGene_tx=XM_019839667.2
C2 55724252 55724278
refGene_gn=DZIP3;refGene_tx=XM_011285969.3;refGene_gn=DZIP3;refGene_tx=XM_019839661.2;refGene_gn=DZIP3;refGene_tx=XM_019839662.2;refGene_gn=DZIP3;refGene_tx=XM_019839663.2;refGene_gn=DZIP3;refGene_tx=XM_019839664.2;refGene_gn=DZIP3;refGene_tx=XM_019839665.2;refGene_gn=DZIP3;refGene_tx=XM_019839666.2;refGene_gn=DZIP3;refGene_tx=XM_019839667.2
C2 55726510 55726676
refGene_gn=DZIP3;refGene_tx=XM_011285969.3;refGene_gn=DZIP3;refGene_tx=XM_019839661.2;refGene_gn=DZIP3;refGene_tx=XM_019839662.2;refGene_gn=DZIP3;refGene_tx=XM_019839663.2;refGene_gn=DZIP3;refGene_tx=XM_019839664.2;refGene_gn=DZIP3;refGene_tx=XM_019839665.2;refGene_gn=DZIP3;refGene_tx=XM_019839666.2;refGene_gn=DZIP3;refGene_tx=XM_019839667.2
C2 55728074 55728170
refGene_gn=DZIP3;refGene_tx=XM_011285969.3;refGene_gn=DZIP3;refGene_tx=XM_019839661.2;refGene_gn=DZIP3;refGene_tx=XM_019839662.2;refGene_gn=DZIP3;refGene_tx=XM_019839663.2;refGene_gn=DZIP3;refGene_tx=XM_0198396

64.2;refGene_gn=DZIP3;refGene_tx=XM_019839665.2;refGene_gn=DZIP3;refGene_tx=XM_019839666.2;refGene_gn=DZIP3;refGene_tx=XM_019839667.2
C2 55728333 55728461
refGene_gn=DZIP3;refGene_tx=XM_011285969.3;refGene_gn=DZIP3;refGene_tx=XM_019839661.2;refGene_gn=DZIP3;refGene_tx=XM_019839662.2;refGene_gn=DZIP3;refGene_tx=XM_019839663.2;refGene_gn=DZIP3;refGene_tx=XM_019839664.2;refGene_gn=DZIP3;refGene_tx=XM_019839665.2;refGene_gn=DZIP3;refGene_tx=XM_019839666.2;refGene_gn=DZIP3;refGene_tx=XM_019839667.2
C2 55740220 55740289
refGene_gn=DZIP3;refGene_tx=XM_011285969.3;refGene_gn=DZIP3;refGene_tx=XM_019839661.2;refGene_gn=DZIP3;refGene_tx=XM_019839662.2;refGene_gn=DZIP3;refGene_tx=XM_019839663.2;refGene_gn=DZIP3;refGene_tx=XM_019839664.2;refGene_gn=DZIP3;refGene_tx=XM_019839665.2;refGene_gn=DZIP3;refGene_tx=XM_019839666.2;refGene_gn=DZIP3;refGene_tx=XM_019839667.2
C2 55744065 55744162
refGene_gn=DZIP3;refGene_tx=XM_011285969.3;refGene_gn=DZIP3;refGene_tx=XM_019839661.2;refGene_gn=DZIP3;refGene_tx=XM_019839662.2;refGene_gn=DZIP3;refGene_tx=XM_019839663.2;refGene_gn=DZIP3;refGene_tx=XM_019839664.2;refGene_gn=DZIP3;refGene_tx=XM_019839665.2;refGene_gn=DZIP3;refGene_tx=XM_019839666.2;refGene_gn=DZIP3;refGene_tx=XM_019839667.2
C2 55745684 55745788
refGene_gn=DZIP3;refGene_tx=XM_011285969.3;refGene_gn=DZIP3;refGene_tx=XM_019839661.2;refGene_gn=DZIP3;refGene_tx=XM_019839662.2;refGene_gn=DZIP3;refGene_tx=XM_019839663.2;refGene_gn=DZIP3;refGene_tx=XM_019839664.2;refGene_gn=DZIP3;refGene_tx=XM_019839665.2;refGene_gn=DZIP3;refGene_tx=XM_019839666.2;refGene_gn=DZIP3;refGene_tx=XM_019839667.2
C2 55747332 55747420
refGene_gn=DZIP3;refGene_tx=XM_011285969.3;refGene_gn=DZIP3;refGene_tx=XM_019839661.2;refGene_gn=DZIP3;refGene_tx=XM_019839662.2;refGene_gn=DZIP3;refGene_tx=XM_019839663.2;refGene_gn=DZIP3;refGene_tx=XM_019839664.2;refGene_gn=DZIP3;refGene_tx=XM_019839665.2;refGene_gn=DZIP3;refGene_tx=XM_019839666.2;refGene_gn=DZIP3;refGene_tx=XM_019839667.2
C2 55749088 55749190
refGene_gn=DZIP3;refGene_tx=XM_011285969.3;refGene_gn=DZIP3;refGene_tx=XM_019839661.2;refGene_gn=DZIP3;refGene_tx=XM_019839662.2;refGene_gn=DZIP3;refGene_tx=XM_019839663.2;refGene_gn=DZIP3;refGene_tx=XM_019839664.2;refGene_gn=DZIP3;refGene_tx=XM_019839665.2;refGene_gn=DZIP3;refGene_tx=XM_019839666.2;refGene_gn=DZIP3;refGene_tx=XM_019839667.2
C2 55752809 55752935
refGene_gn=DZIP3;refGene_tx=XM_011285969.3;refGene_gn=DZIP3;refGene_tx=XM_019839661.2;refGene_gn=DZIP3;refGene_tx=XM_019839662.2;refGene_gn=DZIP3;refGene_tx=XM_019839663.2;refGene_gn=DZIP3;refGene_tx=XM_019839664.2;refGene_gn=DZIP3;refGene_tx=XM_019839665.2;refGene_gn=DZIP3;refGene_tx=XM_019839666.2;refGene_gn=DZIP3;refGene_tx=XM_019839667.2
C2 55755772 55755915
refGene_gn=DZIP3;refGene_tx=XM_011285969.3;refGene_gn=DZIP3;refGene_tx=XM_019839661.2;refGene_gn=DZIP3;refGene_tx=XM_019839662.2;refGene_gn=DZIP3;refGene_tx=XM_019839663.2;refGene_gn=DZIP3;refGene_tx=XM_019839664.2;refGene_gn=DZIP3;refGene_tx=XM_019839665.2;refGene_gn=DZIP3;refGene_tx=XM_019839666.2;refGene_gn=DZIP3;refGene_tx=XM_019839667.2

C2	55757206	55757327
refGene_gn=DZIP3;refGene_tx=XM_011285969.3;refGene_gn=DZIP3;refGene_tx=XM_019839661.2;refGene_gn=DZIP3;refGene_tx=XM_019839662.2;refGene_gn=DZIP3;refGene_tx=XM_019839663.2;refGene_gn=DZIP3;refGene_tx=XM_019839664.2;refGene_gn=DZIP3;refGene_tx=XM_019839665.2;refGene_gn=DZIP3;refGene_tx=XM_019839666.2;refGene_gn=DZIP3;refGene_tx=XM_019839667.2
C2	55757791	55757935
refGene_gn=DZIP3;refGene_tx=XM_011285969.3;refGene_gn=DZIP3;refGene_tx=XM_019839661.2;refGene_gn=DZIP3;refGene_tx=XM_019839662.2;refGene_gn=DZIP3;refGene_tx=XM_019839663.2;refGene_gn=DZIP3;refGene_tx=XM_019839664.2;refGene_gn=DZIP3;refGene_tx=XM_019839665.2;refGene_gn=DZIP3;refGene_tx=XM_019839666.2;refGene_gn=DZIP3;refGene_tx=XM_019839667.2
C2	55758023	55758125
refGene_gn=DZIP3;refGene_tx=XM_011285969.3;refGene_gn=DZIP3;refGene_tx=XM_019839661.2;refGene_gn=DZIP3;refGene_tx=XM_019839662.2;refGene_gn=DZIP3;refGene_tx=XM_019839663.2;refGene_gn=DZIP3;refGene_tx=XM_019839664.2;refGene_gn=DZIP3;refGene_tx=XM_019839665.2;refGene_gn=DZIP3;refGene_tx=XM_019839666.2;refGene_gn=DZIP3;refGene_tx=XM_019839667.2
C2	55759965	55760076
refGene_gn=DZIP3;refGene_tx=XM_011285969.3;refGene_gn=DZIP3;refGene_tx=XM_019839661.2;refGene_gn=DZIP3;refGene_tx=XM_019839662.2;refGene_gn=DZIP3;refGene_tx=XM_019839663.2;refGene_gn=DZIP3;refGene_tx=XM_019839664.2;refGene_gn=DZIP3;refGene_tx=XM_019839665.2;refGene_gn=DZIP3;refGene_tx=XM_019839666.2;refGene_gn=DZIP3;refGene_tx=XM_019839667.2
C2	55837091	55837230	refGene_gn=RETNLB;refGene_tx=XM_023260161.1
C2	55837818	55837874	refGene_gn=RETNLB;refGene_tx=XM_023260161.1
C2	55838355	55838438	refGene_gn=RETNLB;refGene_tx=XM_023260161.1
C2	55838620	55838747	refGene_gn=RETNLB;refGene_tx=XM_023260161.1
C2	55888075	55888082
refGene_gn=LOC101084050;refGene_tx=XM_006936023.4
C2	55901301	55901412
refGene_gn=LOC101084050;refGene_tx=XM_006936023.4
C2	55909730	55909767
refGene_gn=LOC101084050;refGene_tx=XM_006936023.4
C2	55916789	55916851
refGene_gn=LOC101084050;refGene_tx=XM_006936023.4
C2	55919830	55919919
refGene_gn=LOC101084050;refGene_tx=XM_006936023.4
C2	55925157	55925439
refGene_gn=LOC101084050;refGene_tx=XM_006936023.4
C2	55968949	55969119	refGene_gn=GUCA1C;refGene_tx=XM_003991614.4
C2	55977002	55977090	refGene_gn=GUCA1C;refGene_tx=XM_003991614.4
C2	55981496	55981646	refGene_gn=GUCA1C;refGene_tx=XM_003991614.4
C2	56000496	56000700	refGene_gn=GUCA1C;refGene_tx=XM_003991614.4
C2	56004413	56004551	refGene_gn=MORC1;refGene_tx=XM_019839671.2
C2	56007257	56007452	refGene_gn=MORC1;refGene_tx=XM_019839671.2
C2	56013991	56014045	refGene_gn=MORC1;refGene_tx=XM_019839671.2
C2	56015798	56015886	refGene_gn=MORC1;refGene_tx=XM_019839671.2
C2	56024504	56024651	refGene_gn=MORC1;refGene_tx=XM_019839671.2
C2	56028234	56028301	refGene_gn=MORC1;refGene_tx=XM_019839671.2

C2 56030395 56030465 refGene_gn=MORC1;refGene_tx=XM_019839671.2
C2 56045040 56045145 refGene_gn=MORC1;refGene_tx=XM_019839671.2
C2 56047770 56048016 refGene_gn=MORC1;refGene_tx=XM_019839671.2
C2 56049724 56049790 refGene_gn=MORC1;refGene_tx=XM_019839671.2
C2 56074065 56074204 refGene_gn=MORC1;refGene_tx=XM_019839671.2
C2 56079391 56079497 refGene_gn=MORC1;refGene_tx=XM_019839671.2
C2 56082653 56082782 refGene_gn=MORC1;refGene_tx=XM_019839671.2
C2 56102448 56102603 refGene_gn=MORC1;refGene_tx=XM_019839671.2
C2 56104896 56105040 refGene_gn=MORC1;refGene_tx=XM_019839671.2
C2 56108024 56108089 refGene_gn=MORC1;refGene_tx=XM_019839671.2
C2 56110229 56110300 refGene_gn=MORC1;refGene_tx=XM_019839671.2
C2 56111014 56111094 refGene_gn=MORC1;refGene_tx=XM_019839671.2
C2 56125870 56125996 refGene_gn=MORC1;refGene_tx=XM_019839671.2
C2 56143075 56143181 refGene_gn=MORC1;refGene_tx=XM_019839671.2
C2 56145336 56145496 refGene_gn=MORC1;refGene_tx=XM_019839671.2
C2 56149907 56150016 refGene_gn=MORC1;refGene_tx=XM_019839671.2
C2 56151228 56151319 refGene_gn=MORC1;refGene_tx=XM_019839671.2
C2 56155019 56155088 refGene_gn=MORC1;refGene_tx=XM_019839671.2
C2 56161527 56161562 refGene_gn=MORC1;refGene_tx=XM_019839671.2
C2 56164211 56164265 refGene_gn=MORC1;refGene_tx=XM_019839671.2
C2 56168141 56168206 refGene_gn=MORC1;refGene_tx=XM_019839671.2
C2 56306950 56307089 refGene_gn=DPPA2;refGene_tx=XM_023260162.1
C2 56331992 56332188 refGene_gn=DPPA2;refGene_tx=XM_023260162.1
C2 56333863 56334125 refGene_gn=DPPA2;refGene_tx=XM_023260162.1
C2 56335054 56335108 refGene_gn=DPPA2;refGene_tx=XM_023260162.1
C2 56335196 56335360 refGene_gn=DPPA2;refGene_tx=XM_023260162.1
C2 56337249 56337397 refGene_gn=DPPA2;refGene_tx=XM_023260162.1
C2 56339596 56339629 refGene_gn=DPPA2;refGene_tx=XM_023260162.1
C2 56359816 56360016 refGene_gn=DPPA4;refGene_tx=XM_023260163.1
C2 56362271 56362584 refGene_gn=DPPA4;refGene_tx=XM_023260163.1
C2 56362863 56362914 refGene_gn=DPPA4;refGene_tx=XM_023260163.1
C2 56363000 56363149 refGene_gn=DPPA4;refGene_tx=XM_023260163.1
C2 56364445 56364686 refGene_gn=DPPA4;refGene_tx=XM_023260163.1
C2 57779963 57780278
refGene_gn=LOC102900940;refGene_tx=XM_019839684.2
C2 57780507 57780744
refGene_gn=LOC102900940;refGene_tx=XM_019839684.2
C2 57907293 57907453
refGene_gn=NECTIN3;refGene_tx=XM_023260118.1;refGene_gn=NECTIN3;refGene_tx=XM_023260119.1
C2 57947672 57948014
refGene_gn=NECTIN3;refGene_tx=XM_023260118.1;refGene_gn=NECTIN3;refGene_tx=XM_023260119.1
C2 57955384 57955681
refGene_gn=NECTIN3;refGene_tx=XM_023260118.1;refGene_gn=NECTIN3;refGene_tx=XM_023260119.1
C2 57958318 57958436
refGene_gn=NECTIN3;refGene_tx=XM_023260118.1;refGene_gn=NECTIN3;refGene_tx=XM_023260119.1
C2 57962614 57962766

refGene_gn=NECTIN3;refGene_tx=XM_023260118.1;refGene_gn=NECTIN3;refGene_tx=XM_023260119.1
C2 57970559 57971140 refGene_gn=NECTIN3;refGene_tx=XM_023260118.1
C2 57981723 57981862 refGene_gn=NECTIN3;refGene_tx=XM_023260119.1
C2 57984225 57984310 refGene_gn=NECTIN3;refGene_tx=XM_023260119.1
C2 58033330 58033393 refGene_gn=NECTIN3;refGene_tx=XM_023260119.1
C2 58034155 58034335 refGene_gn=NECTIN3;refGene_tx=XM_023260119.1
C2 58301932 58301993
refGene_gn=CD96;refGene_tx=XM_019839674.2;refGene_gn=CD96;refGene_tx=XM_023260120.1;refGene_gn=CD96;refGene_tx=XM_023260121.1;refGene_gn=CD96;refGene_tx=XM_023260122.1;refGene_gn=CD96;refGene_tx=XM_023260123.1;refGene_gn=CD96;refGene_tx=XM_023260124.1;refGene_gn=CD96;refGene_tx=XM_023260125.1;refGene_gn=CD96;refGene_tx=XM_023260126.1
C2 58304663 58305017
refGene_gn=CD96;refGene_tx=XM_019839674.2;refGene_gn=CD96;refGene_tx=XM_023260120.1;refGene_gn=CD96;refGene_tx=XM_023260121.1;refGene_gn=CD96;refGene_tx=XM_023260122.1;refGene_gn=CD96;refGene_tx=XM_023260123.1;refGene_gn=CD96;refGene_tx=XM_023260124.1;refGene_gn=CD96;refGene_tx=XM_023260125.1;refGene_gn=CD96;refGene_tx=XM_023260126.1
C2 58322447 58322572
refGene_gn=CD96;refGene_tx=XM_019839674.2;refGene_gn=CD96;refGene_tx=XM_023260120.1;refGene_gn=CD96;refGene_tx=XM_023260121.1;refGene_gn=CD96;refGene_tx=XM_023260122.1;refGene_gn=CD96;refGene_tx=XM_023260123.1;refGene_gn=CD96;refGene_tx=XM_023260124.1;refGene_gn=CD96;refGene_tx=XM_023260125.1;refGene_gn=CD96;refGene_tx=XM_023260126.1
C2 58332753 58332964
refGene_gn=CD96;refGene_tx=XM_019839674.2;refGene_gn=CD96;refGene_tx=XM_023260120.1;refGene_gn=CD96;refGene_tx=XM_023260121.1;refGene_gn=CD96;refGene_tx=XM_023260122.1;refGene_gn=CD96;refGene_tx=XM_023260123.1;refGene_gn=CD96;refGene_tx=XM_023260124.1;refGene_gn=CD96;refGene_tx=XM_023260125.1;refGene_gn=CD96;refGene_tx=XM_023260126.1
C2 58336003 58336059
refGene_gn=CD96;refGene_tx=XM_019839674.2;refGene_gn=CD96;refGene_tx=XM_023260120.1;refGene_gn=CD96;refGene_tx=XM_023260121.1;refGene_gn=CD96;refGene_tx=XM_023260122.1;refGene_gn=CD96;refGene_tx=XM_023260123.1;refGene_gn=CD96;refGene_tx=XM_023260124.1;refGene_gn=CD96;refGene_tx=XM_023260125.1;refGene_gn=CD96;refGene_tx=XM_023260126.1
C2 58341711 58342365 refGene_gn=ZBED2;refGene_tx=XM_006936029.4
C2 58347262 58347353
refGene_gn=CD96;refGene_tx=XM_019839674.2;refGene_gn=CD96;refGene_tx=XM_023260120.1;refGene_gn=CD96;refGene_tx=XM_023260121.1;refGene_gn=CD96;refGene_tx=XM_023260122.1;refGene_gn=CD96;refGene_tx=XM_023260123.1;refGene_gn=CD96;refGene_tx=XM_023260124.1;refGene_gn=CD96;refGene_tx=XM_023260125.1;refGene_gn=CD96;refGene_tx=XM_023260126.1
C2 58350183 58350372
refGene_gn=CD96;refGene_tx=XM_019839674.2;refGene_gn=CD96;refGene_tx=XM_023260120.1;refGene_gn=CD96;refGene_tx=XM_023260121.1;refGene_gn=CD96;refGene_tx=XM_023260122.1;refGene_gn=CD96;refGene_tx=XM_023260123.1;refGene_gn=CD96;refGene_tx=XM_023260124.1;refGene_gn=CD96;refGene_tx=XM_023260125.1;refGene_gn=CD96;refGene_tx=XM_023260126.1

C2 58357830 58357923
refGene_gn=CD96;refGene_tx=XM_019839674.2;refGene_gn=CD96;refGene_tx=XM_023260120.1;refGene_gn=CD96;refGene_tx=XM_023260121.1;refGene_gn=CD96;refGene_tx=XM_023260122.1;refGene_gn=CD96;refGene_tx=XM_023260123.1;refGene_gn=CD96;refGene_tx=XM_023260124.1;refGene_gn=CD96;refGene_tx=XM_023260125.1;refGene_gn=CD96;refGene_tx=XM_023260126.1
C2 58381192 58381261
refGene_gn=CD96;refGene_tx=XM_019839674.2;refGene_gn=CD96;refGene_tx=XM_023260120.1;refGene_gn=CD96;refGene_tx=XM_023260121.1;refGene_gn=CD96;refGene_tx=XM_023260122.1;refGene_gn=CD96;refGene_tx=XM_023260123.1;refGene_gn=CD96;refGene_tx=XM_023260124.1;refGene_gn=CD96;refGene_tx=XM_023260125.1;refGene_gn=CD96;refGene_tx=XM_023260126.1
C2 58381776 58381857
refGene_gn=CD96;refGene_tx=XM_019839674.2;refGene_gn=CD96;refGene_tx=XM_023260120.1;refGene_gn=CD96;refGene_tx=XM_023260121.1;refGene_gn=CD96;refGene_tx=XM_023260122.1;refGene_gn=CD96;refGene_tx=XM_023260123.1;refGene_gn=CD96;refGene_tx=XM_023260124.1;refGene_gn=CD96;refGene_tx=XM_023260125.1;refGene_gn=CD96;refGene_tx=XM_023260126.1
C2 58387280 58387348 refGene_gn=CD96;refGene_tx=XM_023260126.1
C2 58390598 58390664
refGene_gn=CD96;refGene_tx=XM_019839674.2;refGene_gn=CD96;refGene_tx=XM_023260120.1;refGene_gn=CD96;refGene_tx=XM_023260121.1;refGene_gn=CD96;refGene_tx=XM_023260122.1;refGene_gn=CD96;refGene_tx=XM_023260123.1;refGene_gn=CD96;refGene_tx=XM_023260124.1
C2 58391465 58391552
refGene_gn=CD96;refGene_tx=XM_019839674.2;refGene_gn=CD96;refGene_tx=XM_023260120.1;refGene_gn=CD96;refGene_tx=XM_023260121.1;refGene_gn=CD96;refGene_tx=XM_023260122.1
C2 58395371 58395495
refGene_gn=CD96;refGene_tx=XM_019839674.2;refGene_gn=CD96;refGene_tx=XM_023260120.1;refGene_gn=CD96;refGene_tx=XM_023260121.1;refGene_gn=CD96;refGene_tx=XM_023260122.1;refGene_gn=CD96;refGene_tx=XM_023260123.1;refGene_gn=CD96;refGene_tx=XM_023260124.1;refGene_gn=CD96;refGene_tx=XM_023260125.1
C2 58397417 58397526
refGene_gn=CD96;refGene_tx=XM_019839674.2;refGene_gn=CD96;refGene_tx=XM_023260120.1;refGene_gn=CD96;refGene_tx=XM_023260121.1;refGene_gn=CD96;refGene_tx=XM_023260122.1;refGene_gn=CD96;refGene_tx=XM_023260123.1;refGene_gn=CD96;refGene_tx=XM_023260124.1;refGene_gn=CD96;refGene_tx=XM_023260125.1
C2 58418484 58418647
refGene_gn=PLCXD2;refGene_tx=XM_003991618.5;refGene_gn=PLCXD2;refGene_tx=XM_019839700.2;refGene_gn=PLCXD2;refGene_tx=XM_019839701.2
C2 58426736 58426830 refGene_gn=PLCXD2;refGene_tx=XM_019839702.1
C2 58449453 58449914
refGene_gn=PLCXD2;refGene_tx=XM_003991618.5;refGene_gn=PLCXD2;refGene_tx=XM_019839700.2;refGene_gn=PLCXD2;refGene_tx=XM_019839701.2;refGene_gn=PLCXD2;refGene_tx=XM_019839702.1
C2 58455357 58455599
refGene_gn=PLCXD2;refGene_tx=XM_003991618.5;refGene_gn=PLCXD2;refGene_

tx=XM_019839700.2;refGene_gn=PLCXD2;refGene_tx=XM_019839701.2;refGene_gn=PLCXD2;refGene_tx=XM_019839702.1
C2 58462507 58462667
refGene_gn=PLCXD2;refGene_tx=XM_003991618.5;refGene_gn=PLCXD2;refGene_tx=XM_019839700.2;refGene_gn=PLCXD2;refGene_tx=XM_019839701.2;refGene_gn=PLCXD2;refGene_tx=XM_019839702.1
C2 58614640 58614682 refGene_gn=PHLDB2;refGene_tx=XM_011285963.2
C2 58614683 58614764 refGene_gn=PHLDB2;refGene_tx=XM_011285963.2
C2 58614766 58615978 refGene_gn=PHLDB2;refGene_tx=XM_011285963.2
C2 58638157 58638520 refGene_gn=PHLDB2;refGene_tx=XM_011285963.2
C2 58643936 58644080 refGene_gn=PHLDB2;refGene_tx=XM_011285963.2
C2 58645104 58645242 refGene_gn=PHLDB2;refGene_tx=XM_011285963.2
C2 58657664 58657793 refGene_gn=PHLDB2;refGene_tx=XM_011285963.2
C2 58663548 58663704 refGene_gn=PHLDB2;refGene_tx=XM_011285963.2
C2 58664636 58664747 refGene_gn=PHLDB2;refGene_tx=XM_011285963.2
C2 58667903 58667993 refGene_gn=PHLDB2;refGene_tx=XM_011285963.2
C2 58671665 58671809 refGene_gn=PHLDB2;refGene_tx=XM_011285963.2
C2 58674660 58674801 refGene_gn=PHLDB2;refGene_tx=XM_011285963.2
C2 58675993 58676093 refGene_gn=PHLDB2;refGene_tx=XM_011285963.2
C2 58683567 58683775 refGene_gn=PHLDB2;refGene_tx=XM_011285963.2
C2 58694632 58694723 refGene_gn=PHLDB2;refGene_tx=XM_011285963.2
C2 58695630 58695777 refGene_gn=PHLDB2;refGene_tx=XM_011285963.2
C2 58697794 58698014 refGene_gn=PHLDB2;refGene_tx=XM_011285963.2
C2 58703024 58703110 refGene_gn=PHLDB2;refGene_tx=XM_011285963.2
C2 58703712 58703853 refGene_gn=PHLDB2;refGene_tx=XM_011285963.2
C2 58708347 58708696 refGene_gn=ABHD10;refGene_tx=XM_003991621.5
C2 58711592 58711776 refGene_gn=ABHD10;refGene_tx=XM_003991621.5
C2 58717490 58717602 refGene_gn=ABHD10;refGene_tx=XM_003991621.5
C2 58718121 58718259 refGene_gn=ABHD10;refGene_tx=XM_003991621.5
C2 58722836 58723181 refGene_gn=ABHD10;refGene_tx=XM_003991621.5
C2 58731243 58731423 refGene_gn=TAGLN3;refGene_tx=XM_011285960.2
C2 58732564 58732739 refGene_gn=TAGLN3;refGene_tx=XM_011285960.2
C2 58742526 58742629 refGene_gn=TAGLN3;refGene_tx=XM_011285960.2
C2 58744401 58744543 refGene_gn=TAGLN3;refGene_tx=XM_011285960.2
C2 58767011 58767104 refGene_gn=TMPRSS7;refGene_tx=XM_023260128.1
C2 58769704 58769772 refGene_gn=TMPRSS7;refGene_tx=XM_023260128.1
C2 58771154 58771345 refGene_gn=TMPRSS7;refGene_tx=XM_023260128.1
C2 58773213 58773442 refGene_gn=TMPRSS7;refGene_tx=XM_023260128.1
C2 58774742 58774873 refGene_gn=TMPRSS7;refGene_tx=XM_023260128.1
C2 58776802 58776915 refGene_gn=TMPRSS7;refGene_tx=XM_023260128.1
C2 58783960 58784067 refGene_gn=TMPRSS7;refGene_tx=XM_023260128.1
C2 58787531 58787668 refGene_gn=TMPRSS7;refGene_tx=XM_023260128.1
C2 58790477 58790585 refGene_gn=TMPRSS7;refGene_tx=XM_023260128.1
C2 58792971 58793082 refGene_gn=TMPRSS7;refGene_tx=XM_023260128.1
C2 58794082 58794199 refGene_gn=TMPRSS7;refGene_tx=XM_023260128.1
C2 58795347 58795519 refGene_gn=TMPRSS7;refGene_tx=XM_023260128.1
C2 58798256 58798525 refGene_gn=TMPRSS7;refGene_tx=XM_023260128.1
C2 58800293 58800430 refGene_gn=TMPRSS7;refGene_tx=XM_023260128.1
C2 58802687 58802858 refGene_gn=TMPRSS7;refGene_tx=XM_023260128.1
C2 58811399 58811537

refGene_gn=CC2H3orf52;refGene_tx=XM_006936064.3;refGene_gn=CC2H3orf52;
refGene_tx=XM_019839704.2;refGene_gn=CC2H3orf52;refGene_tx=XM_01983970
5.2
C2 58818084 58818214
refGene_gn=CC2H3orf52;refGene_tx=XM_006936064.3;refGene_gn=CC2H3orf52;
refGene_tx=XM_019839704.2;refGene_gn=CC2H3orf52;refGene_tx=XM_01983970
5.2
C2 58828836 58828964
refGene_gn=CC2H3orf52;refGene_tx=XM_006936064.3;refGene_gn=CC2H3orf52;
refGene_tx=XM_019839704.2;refGene_gn=CC2H3orf52;refGene_tx=XM_01983970
5.2
C2 58830621 58830840
refGene_gn=CC2H3orf52;refGene_tx=XM_019839705.2
C2 58835155 58835226
refGene_gn=CC2H3orf52;refGene_tx=XM_006936064.3;refGene_gn=CC2H3orf52;
refGene_tx=XM_019839704.2
C2 58838689 58838871
refGene_gn=CC2H3orf52;refGene_tx=XM_006936064.3;refGene_gn=CC2H3orf52;
refGene_tx=XM_019839704.2
C2 58843223 58843228
refGene_gn=CC2H3orf52;refGene_tx=XM_006936064.3;refGene_gn=CC2H3orf52;
refGene_tx=XM_019839704.2
C2 58846129 58846233
refGene_gn=GCSAM;refGene_tx=XM_006936038.3;refGene_gn=GCSAM;refGene_tx
=XM_011285959.3;refGene_gn=GCSAM;refGene_tx=XM_019839707.2;refGene_gn=
GCSAM;refGene_tx=XM_019839708.2
C2 58849923 58850161
refGene_gn=GCSAM;refGene_tx=XM_006936038.3;refGene_gn=GCSAM;refGene_tx
=XM_011285959.3;refGene_gn=GCSAM;refGene_tx=XM_019839707.2;refGene_gn=
GCSAM;refGene_tx=XM_019839708.2
C2 58851052 58851081
refGene_gn=GCSAM;refGene_tx=XM_006936038.3;refGene_gn=GCSAM;refGene_tx
=XM_011285959.3;refGene_gn=GCSAM;refGene_tx=XM_019839707.2;refGene_gn=
GCSAM;refGene_tx=XM_019839708.2
C2 58853290 58853337
refGene_gn=GCSAM;refGene_tx=XM_006936038.3;refGene_gn=GCSAM;refGene_tx
=XM_011285959.3;refGene_gn=GCSAM;refGene_tx=XM_019839707.2;refGene_gn=
GCSAM;refGene_tx=XM_019839708.2
C2 58854334 58854379
refGene_gn=GCSAM;refGene_tx=XM_006936038.3;refGene_gn=GCSAM;refGene_tx
=XM_011285959.3
C2 58861707 58861736
refGene_gn=GCSAM;refGene_tx=XM_006936038.3;refGene_gn=GCSAM;refGene_tx
=XM_011285959.3;refGene_gn=GCSAM;refGene_tx=XM_019839707.2;refGene_gn=
GCSAM;refGene_tx=XM_019839708.2
C2 58875377 58875443
refGene_gn=SLC9C1;refGene_tx=XM_011285957.3;refGene_gn=SLC9C1;refGene_
tx=XM_019839709.2;refGene_gn=SLC9C1;refGene_tx=XM_019839711.1
C2 58882554 58882601
refGene_gn=SLC9C1;refGene_tx=XM_011285957.3;refGene_gn=SLC9C1;refGene_

tx=XM_019839709.2;refGene_gn=SLC9C1;refGene_tx=XM_019839711.1
C2 58883452 58883579
refGene_gn=SLC9C1;refGene_tx=XM_011285957.3;refGene_gn=SLC9C1;refGene_
tx=XM_019839709.2;refGene_gn=SLC9C1;refGene_tx=XM_019839711.1
C2 58885226 58885412
refGene_gn=SLC9C1;refGene_tx=XM_011285957.3;refGene_gn=SLC9C1;refGene_
tx=XM_019839709.2;refGene_gn=SLC9C1;refGene_tx=XM_019839711.1
C2 58885545 58885677
refGene_gn=SLC9C1;refGene_tx=XM_011285957.3;refGene_gn=SLC9C1;refGene_
tx=XM_019839709.2;refGene_gn=SLC9C1;refGene_tx=XM_019839711.1
C2 58892632 58892806
refGene_gn=SLC9C1;refGene_tx=XM_011285957.3;refGene_gn=SLC9C1;refGene_
tx=XM_019839709.2;refGene_gn=SLC9C1;refGene_tx=XM_019839711.1
C2 58893330 58893429
refGene_gn=SLC9C1;refGene_tx=XM_011285957.3;refGene_gn=SLC9C1;refGene_
tx=XM_019839709.2;refGene_gn=SLC9C1;refGene_tx=XM_019839711.1
C2 58894788 58894914
refGene_gn=SLC9C1;refGene_tx=XM_011285957.3;refGene_gn=SLC9C1;refGene_
tx=XM_019839709.2;refGene_gn=SLC9C1;refGene_tx=XM_019839711.1
C2 58899444 58899593
refGene_gn=SLC9C1;refGene_tx=XM_011285957.3;refGene_gn=SLC9C1;refGene_
tx=XM_019839709.2;refGene_gn=SLC9C1;refGene_tx=XM_019839711.1
C2 58900843 58900895
refGene_gn=SLC9C1;refGene_tx=XM_011285957.3;refGene_gn=SLC9C1;refGene_
tx=XM_019839709.2;refGene_gn=SLC9C1;refGene_tx=XM_019839711.1
C2 58902635 58902785
refGene_gn=SLC9C1;refGene_tx=XM_011285957.3;refGene_gn=SLC9C1;refGene_
tx=XM_019839709.2;refGene_gn=SLC9C1;refGene_tx=XM_019839711.1
C2 58904648 58904834
refGene_gn=SLC9C1;refGene_tx=XM_011285957.3;refGene_gn=SLC9C1;refGene_
tx=XM_019839709.2;refGene_gn=SLC9C1;refGene_tx=XM_019839711.1
C2 58906308 58906504
refGene_gn=SLC9C1;refGene_tx=XM_011285957.3;refGene_gn=SLC9C1;refGene_
tx=XM_019839709.2;refGene_gn=SLC9C1;refGene_tx=XM_019839711.1
C2 58914018 58914138
refGene_gn=SLC9C1;refGene_tx=XM_011285957.3;refGene_gn=SLC9C1;refGene_
tx=XM_019839709.2;refGene_gn=SLC9C1;refGene_tx=XM_019839711.1
C2 58919701 58919799
refGene_gn=SLC9C1;refGene_tx=XM_011285957.3;refGene_gn=SLC9C1;refGene_
tx=XM_019839709.2;refGene_gn=SLC9C1;refGene_tx=XM_019839711.1
C2 58923281 58923407
refGene_gn=SLC9C1;refGene_tx=XM_011285957.3;refGene_gn=SLC9C1;refGene_
tx=XM_019839709.2;refGene_gn=SLC9C1;refGene_tx=XM_019839711.1
C2 58925512 58925679
refGene_gn=SLC9C1;refGene_tx=XM_011285957.3;refGene_gn=SLC9C1;refGene_
tx=XM_019839709.2;refGene_gn=SLC9C1;refGene_tx=XM_019839711.1
C2 58927837 58927919
refGene_gn=SLC9C1;refGene_tx=XM_011285957.3;refGene_gn=SLC9C1;refGene_
tx=XM_019839709.2;refGene_gn=SLC9C1;refGene_tx=XM_019839711.1
C2 58931493 58931668

refGene_gn=SLC9C1;refGene_tx=XM_011285957.3;refGene_gn=SLC9C1;refGene_tx=XM_019839709.2;refGene_gn=SLC9C1;refGene_tx=XM_019839711.1
C2 58932887 58933031
refGene_gn=SLC9C1;refGene_tx=XM_011285957.3;refGene_gn=SLC9C1;refGene_tx=XM_019839709.2;refGene_gn=SLC9C1;refGene_tx=XM_019839711.1
C2 58938626 58938726
refGene_gn=SLC9C1;refGene_tx=XM_011285957.3;refGene_gn=SLC9C1;refGene_tx=XM_019839709.2;refGene_gn=SLC9C1;refGene_tx=XM_019839711.1
C2 58943222 58943384
refGene_gn=SLC9C1;refGene_tx=XM_011285957.3;refGene_gn=SLC9C1;refGene_tx=XM_019839709.2;refGene_gn=SLC9C1;refGene_tx=XM_019839711.1
C2 58947800 58947926
refGene_gn=SLC9C1;refGene_tx=XM_011285957.3;refGene_gn=SLC9C1;refGene_tx=XM_019839709.2;refGene_gn=SLC9C1;refGene_tx=XM_019839711.1
C2 58950901 58951067
refGene_gn=SLC9C1;refGene_tx=XM_011285957.3;refGene_gn=SLC9C1;refGene_tx=XM_019839709.2;refGene_gn=SLC9C1;refGene_tx=XM_019839711.1
C2 58953380 58953509
refGene_gn=SLC9C1;refGene_tx=XM_011285957.3;refGene_gn=SLC9C1;refGene_tx=XM_019839709.2;refGene_gn=SLC9C1;refGene_tx=XM_019839711.1
C2 58956207 58956308
refGene_gn=SLC9C1;refGene_tx=XM_011285957.3;refGene_gn=SLC9C1;refGene_tx=XM_019839709.2
C2 58957429 58957580
refGene_gn=SLC9C1;refGene_tx=XM_011285957.3;refGene_gn=SLC9C1;refGene_tx=XM_019839709.2
C2 58984748 58984842
refGene_gn=LOC109491644;refGene_tx=XM_023260129.1
C2 58994094 58994433
refGene_gn=LOC109491644;refGene_tx=XM_023260129.1
C2 58995134 58995434
refGene_gn=LOC109491644;refGene_tx=XM_023260129.1
C2 59001939 59002042
refGene_gn=LOC109491644;refGene_tx=XM_023260129.1
C2 59003063 59003147
refGene_gn=LOC109491644;refGene_tx=XM_023260129.1
C2 59010224 59010249
refGene_gn=LOC109491644;refGene_tx=XM_023260129.1
C2 59011436 59011448
refGene_gn=LOC101096094;refGene_tx=XM_023260130.1
C2 59014645 59014730
refGene_gn=LOC101096094;refGene_tx=XM_023260130.1
C2 59026111 59026432
refGene_gn=LOC101096094;refGene_tx=XM_019839712.1;refGene_gn=LOC101096094;refGene_tx=XM_023260130.1;refGene_gn=LOC101096094;refGene_tx=XM_023260131.1
C2 59029061 59029334
refGene_gn=LOC101096094;refGene_tx=XM_019839712.1;refGene_gn=LOC101096094;refGene_tx=XM_023260130.1;refGene_gn=LOC101096094;refGene_tx=XM_023260131.1

C2 59031341 59031449
refGene_gn=LOC101096094;refGene_tx=XM_019839712.1;refGene_gn=LOC101096094;refGene_tx=XM_023260130.1;refGene_gn=LOC101096094;refGene_tx=XM_023260131.1
C2 59036414 59036422
refGene_gn=LOC101096094;refGene_tx=XM_019839712.1;refGene_gn=LOC101096094;refGene_tx=XM_023260130.1;refGene_gn=LOC101096094;refGene_tx=XM_023260131.1
C2 59096469 59096496 refGene_gn=BTLA;refGene_tx=XM_023260135.1
C2 59096762 59096956
refGene_gn=BTLA;refGene_tx=XM_023260133.1;refGene_gn=BTLA;refGene_tx=XM_023260135.1
C2 59105152 59105428
refGene_gn=BTLA;refGene_tx=XM_003991628.4;refGene_gn=BTLA;refGene_tx=XM_006936039.3;refGene_gn=BTLA;refGene_tx=XM_023260133.1;refGene_gn=BTLA;refGene_tx=XM_023260134.1;refGene_gn=BTLA;refGene_tx=XM_023260135.1
C2 59109100 59109147
refGene_gn=BTLA;refGene_tx=XM_003991628.4;refGene_gn=BTLA;refGene_tx=XM_006936039.3;refGene_gn=BTLA;refGene_tx=XM_023260133.1;refGene_gn=BTLA;refGene_tx=XM_023260134.1;refGene_gn=BTLA;refGene_tx=XM_023260135.1
C2 59110622 59110781
refGene_gn=BTLA;refGene_tx=XM_003991628.4;refGene_gn=BTLA;refGene_tx=XM_006936039.3;refGene_gn=BTLA;refGene_tx=XM_023260133.1;refGene_gn=BTLA;refGene_tx=XM_023260134.1;refGene_gn=BTLA;refGene_tx=XM_023260135.1
C2 59111938 59111974
refGene_gn=BTLA;refGene_tx=XM_003991628.4;refGene_gn=BTLA;refGene_tx=XM_006936039.3;refGene_gn=BTLA;refGene_tx=XM_023260133.1;refGene_gn=BTLA;refGene_tx=XM_023260134.1;refGene_gn=BTLA;refGene_tx=XM_023260135.1
C2 59118958 59119282
refGene_gn=BTLA;refGene_tx=XM_003991628.4;refGene_gn=BTLA;refGene_tx=XM_006936039.3;refGene_gn=BTLA;refGene_tx=XM_023260133.1;refGene_gn=BTLA;refGene_tx=XM_023260134.1;refGene_gn=BTLA;refGene_tx=XM_023260135.1
C2 59136200 59136288
refGene_gn=BTLA;refGene_tx=XM_003991628.4;refGene_gn=BTLA;refGene_tx=XM_006936039.3;refGene_gn=BTLA;refGene_tx=XM_023260133.1;refGene_gn=BTLA;refGene_tx=XM_023260134.1;refGene_gn=BTLA;refGene_tx=XM_023260135.1
C2 59168894 59168976 refGene_gn=ATG3;refGene_tx=XM_003991630.5
C2 59170549 59170618 refGene_gn=ATG3;refGene_tx=XM_003991630.5
C2 59172281 59172409 refGene_gn=ATG3;refGene_tx=XM_003991630.5
C2 59173395 59173551 refGene_gn=ATG3;refGene_tx=XM_003991630.5
C2 59173801 59173836 refGene_gn=ATG3;refGene_tx=XM_003991630.5
C2 59177410 59177492 refGene_gn=ATG3;refGene_tx=XM_003991630.5
C2 59179776 59179826 refGene_gn=ATG3;refGene_tx=XM_003991630.5
C2 59184524 59184632 refGene_gn=ATG3;refGene_tx=XM_003991630.5
C2 59186212 59186283 refGene_gn=ATG3;refGene_tx=XM_003991630.5
C2 59188968 59189018 refGene_gn=ATG3;refGene_tx=XM_003991630.5
C2 59192794 59192836 refGene_gn=ATG3;refGene_tx=XM_003991630.5
C2 59195551 59195623 refGene_gn=ATG3;refGene_tx=XM_003991630.5
C2 59196323 59196394
refGene_gn=SLC35A5;refGene_tx=XM_006936041.4;refGene_gn=SLC35A5;refGen

e_tx=XM_006936043.4;refGene_gn=SLC35A5;refGene_tx=XM_019839716.2;refGene_gn=SLC35A5;refGene_tx=XM_019839717.2;refGene_gn=SLC35A5;refGene_tx=XM_019839718.2
C2 59197166 59197343 refGene_gn=SLC35A5;refGene_tx=XM_006936040.4;refGene_gn=SLC35A5;refGene_tx=XM_006936041.4;refGene_gn=SLC35A5;refGene_tx=XM_006936042.4;refGene_gn=SLC35A5;refGene_tx=XM_006936043.4;refGene_gn=SLC35A5;refGene_tx=XM_019839716.2;refGene_gn=SLC35A5;refGene_tx=XM_019839717.2;refGene_gn=SLC35A5;refGene_tx=XM_019839718.2;refGene_gn=SLC35A5;refGene_tx=XM_019839719.2
C2 59201208 59201307 refGene_gn=SLC35A5;refGene_tx=XM_006936040.4;refGene_gn=SLC35A5;refGene_tx=XM_006936041.4;refGene_gn=SLC35A5;refGene_tx=XM_006936042.4;refGene_gn=SLC35A5;refGene_tx=XM_006936043.4;refGene_gn=SLC35A5;refGene_tx=XM_019839716.2;refGene_gn=SLC35A5;refGene_tx=XM_019839717.2;refGene_gn=SLC35A5;refGene_tx=XM_019839718.2;refGene_gn=SLC35A5;refGene_tx=XM_019839719.2
C2 59202846 59202980 refGene_gn=SLC35A5;refGene_tx=XM_006936040.4;refGene_gn=SLC35A5;refGene_tx=XM_006936041.4;refGene_gn=SLC35A5;refGene_tx=XM_006936042.4;refGene_gn=SLC35A5;refGene_tx=XM_006936043.4;refGene_gn=SLC35A5;refGene_tx=XM_019839716.2;refGene_gn=SLC35A5;refGene_tx=XM_019839717.2;refGene_gn=SLC35A5;refGene_tx=XM_019839718.2;refGene_gn=SLC35A5;refGene_tx=XM_019839719.2
C2 59205582 59205650 refGene_gn=SLC35A5;refGene_tx=XM_006936040.4;refGene_gn=SLC35A5;refGene_tx=XM_006936041.4;refGene_gn=SLC35A5;refGene_tx=XM_006936042.4;refGene_gn=SLC35A5;refGene_tx=XM_006936043.4;refGene_gn=SLC35A5;refGene_tx=XM_019839716.2;refGene_gn=SLC35A5;refGene_tx=XM_019839717.2;refGene_gn=SLC35A5;refGene_tx=XM_019839718.2;refGene_gn=SLC35A5;refGene_tx=XM_019839719.2
C2 59213438 59214219 refGene_gn=SLC35A5;refGene_tx=XM_006936040.4;refGene_gn=SLC35A5;refGene_tx=XM_006936041.4;refGene_gn=SLC35A5;refGene_tx=XM_006936042.4;refGene_gn=SLC35A5;refGene_tx=XM_006936043.4;refGene_gn=SLC35A5;refGene_tx=XM_019839716.2;refGene_gn=SLC35A5;refGene_tx=XM_019839717.2;refGene_gn=SLC35A5;refGene_tx=XM_019839718.2;refGene_gn=SLC35A5;refGene_tx=XM_019839719.2
C2 59216067 59216136 refGene_gn=SLC35A5;refGene_tx=XM_006936040.4;refGene_gn=SLC35A5;refGene_tx=XM_006936041.4;refGene_gn=SLC35A5;refGene_tx=XM_006936042.4;refGene_gn=SLC35A5;refGene_tx=XM_006936043.4;refGene_gn=SLC35A5;refGene_tx=XM_019839716.2;refGene_gn=SLC35A5;refGene_tx=XM_019839717.2;refGene_gn=SLC35A5;refGene_tx=XM_019839718.2;refGene_gn=SLC35A5;refGene_tx=XM_019839719.2
C2 59229993 59230340 refGene_gn=CCDC80;refGene_tx=XM_003991632.5
C2 59231891 59231972 refGene_gn=CCDC80;refGene_tx=XM_003991632.5
C2 59234658 59234762 refGene_gn=CCDC80;refGene_tx=XM_003991632.5
C2 59240824 59240973 refGene_gn=CCDC80;refGene_tx=XM_003991632.5
C2 59242652 59242789 refGene_gn=CCDC80;refGene_tx=XM_003991632.5

C2 59255013 59255170 refGene_gn=CCDC80;refGene_tx=XM_003991632.5
C2 59263855 59265751 refGene_gn=CCDC80;refGene_tx=XM_003991632.5
C2 59409326 59409333
refGene_gn=LOC101096593;refGene_tx=XM_019839721.1
C2 59409982 59410082
refGene_gn=LOC101096593;refGene_tx=XM_023260136.1;refGene_gn=LOC101096593;refGene_tx=XM_023260137.1
C2 59410207 59410331
refGene_gn=LOC101096593;refGene_tx=XM_019839721.1;refGene_gn=LOC101096593;refGene_tx=XM_023260136.1;refGene_gn=LOC101096593;refGene_tx=XM_023260137.1
C2 59414678 59414927
refGene_gn=LOC101096593;refGene_tx=XM_019839721.1;refGene_gn=LOC101096593;refGene_tx=XM_023260136.1;refGene_gn=LOC101096593;refGene_tx=XM_023260137.1
C2 59415045 59415366
refGene_gn=LOC101096593;refGene_tx=XM_019839721.1;refGene_gn=LOC101096593;refGene_tx=XM_023260136.1;refGene_gn=LOC101096593;refGene_tx=XM_023260137.1
C2 59418333 59418393
refGene_gn=LOC101096593;refGene_tx=XM_019839721.1;refGene_gn=LOC101096593;refGene_tx=XM_023260136.1;refGene_gn=LOC101096593;refGene_tx=XM_023260137.1
C2 59421846 59421930
refGene_gn=LOC101096593;refGene_tx=XM_019839721.1;refGene_gn=LOC101096593;refGene_tx=XM_023260136.1;refGene_gn=LOC101096593;refGene_tx=XM_023260137.1
C2 59434139 59434185
refGene_gn=LOC101096593;refGene_tx=XM_019839721.1;refGene_gn=LOC101096593;refGene_tx=XM_023260136.1
C2 59450087 59450431
refGene_gn=LOC111562227;refGene_tx=XM_023260165.1
C2 59450549 59450864
refGene_gn=LOC111562227;refGene_tx=XM_023260165.1
C2 59453505 59453644
refGene_gn=LOC111562227;refGene_tx=XM_023260165.1
C2 59490343 59490597
refGene_gn=LOC101095726;refGene_tx=XM_023260138.1
C2 59490715 59491036
refGene_gn=LOC101095726;refGene_tx=XM_023260138.1
C2 59494263 59494323
refGene_gn=LOC101095726;refGene_tx=XM_023260138.1
C2 59503812 59503876
refGene_gn=LOC101095726;refGene_tx=XM_023260138.1
C2 59529243 59529384
refGene_gn=LOC101097085;refGene_tx=XM_011285955.3
C2 59530049 59530095
refGene_gn=LOC101097085;refGene_tx=XM_011285955.3
C2 59530631 59530737
refGene_gn=LOC101097085;refGene_tx=XM_011285955.3

C2 59535943 59536192
refGene_gn=LOC101097085;refGene_tx=XM_011285955.3
C2 59536310 59536471
refGene_gn=LOC101097085;refGene_tx=XM_011285955.3
C2 59536472 59536629
refGene_gn=LOC101097085;refGene_tx=XM_011285955.3
C2 59539595 59539655
refGene_gn=LOC101097085;refGene_tx=XM_011285955.3
C2 59542879 59542948
refGene_gn=LOC101097085;refGene_tx=XM_011285955.3
C2 59558883 59558950
refGene_gn=LOC101097085;refGene_tx=XM_011285955.3
C2 59579357 59579693
refGene_gn=GTPBP8;refGene_tx=XM_003991656.5;refGene_gn=GTPBP8;refGene_
tx=XM_006936045.4
C2 59581478 59581577 refGene_gn=GTPBP8;refGene_tx=XM_003991656.5
C2 59584372 59584503
refGene_gn=GTPBP8;refGene_tx=XM_003991656.5;refGene_gn=GTPBP8;refGene_
tx=XM_006936045.4
C2 59585478 59585578
refGene_gn=GTPBP8;refGene_tx=XM_003991656.5;refGene_gn=GTPBP8;refGene_
tx=XM_006936045.4
C2 59587746 59587865
refGene_gn=GTPBP8;refGene_tx=XM_003991656.5;refGene_gn=GTPBP8;refGene_
tx=XM_006936045.4
C2 59592911 59592993
refGene_gn=GTPBP8;refGene_tx=XM_003991656.5;refGene_gn=GTPBP8;refGene_
tx=XM_006936045.4
C2 59598092 59598555 refGene_gn=NEPRO;refGene_tx=XM_003991633.3
C2 59601094 59601348 refGene_gn=NEPRO;refGene_tx=XM_003991633.3
C2 59603411 59603473 refGene_gn=NEPRO;refGene_tx=XM_003991633.3
C2 59603617 59603828 refGene_gn=NEPRO;refGene_tx=XM_003991633.3
C2 59603829 59603980 refGene_gn=NEPRO;refGene_tx=XM_003991633.3
C2 59604655 59604733 refGene_gn=NEPRO;refGene_tx=XM_003991633.3
C2 59605385 59605517 refGene_gn=NEPRO;refGene_tx=XM_003991633.3
C2 59605901 59605981 refGene_gn=NEPRO;refGene_tx=XM_003991633.3
C2 59609514 59609669 refGene_gn=NEPRO;refGene_tx=XM_003991633.3
C2 59611774 59611877 refGene_gn=NEPRO;refGene_tx=XM_003991633.3
C2 59772681 59772738
refGene_gn=LOC111562228;refGene_tx=XM_023260166.1
C2 59774356 59774424
refGene_gn=LOC111562228;refGene_tx=XM_023260166.1
C2 59779649 59779773
refGene_gn=LOC111562228;refGene_tx=XM_023260166.1
C2 59779954 59780693
refGene_gn=LOC111562228;refGene_tx=XM_023260166.1
C2 59783988 59784017
refGene_gn=LOC111562228;refGene_tx=XM_023260166.1
C2 59816577 59816722 refGene_gn=BOC;refGene_tx=XM_023260140.1
C2 59817401 59817680 refGene_gn=BOC;refGene_tx=XM_023260140.1

C2 59833520 59833667 refGene_gn=BOC;refGene_tx=XM_023260140.1
C2 59835474 59835618 refGene_gn=BOC;refGene_tx=XM_023260140.1
C2 59837517 59837811 refGene_gn=BOC;refGene_tx=XM_023260140.1
C2 59838194 59838467 refGene_gn=BOC;refGene_tx=XM_023260140.1
C2 59839581 59839889 refGene_gn=BOC;refGene_tx=XM_023260140.1
C2 59843201 59843361 refGene_gn=BOC;refGene_tx=XM_023260140.1
C2 59843796 59843907 refGene_gn=BOC;refGene_tx=XM_023260140.1
C2 59844421 59844628 refGene_gn=BOC;refGene_tx=XM_023260140.1
C2 59844964 59845146 refGene_gn=BOC;refGene_tx=XM_023260140.1
C2 59845628 59845734 refGene_gn=BOC;refGene_tx=XM_023260140.1
C2 59846060 59846183 refGene_gn=BOC;refGene_tx=XM_023260140.1
C2 59848481 59848703 refGene_gn=BOC;refGene_tx=XM_023260140.1
C2 59849392 59849628 refGene_gn=BOC;refGene_tx=XM_023260140.1
C2 59849839 59849916 refGene_gn=BOC;refGene_tx=XM_023260140.1
C2 59850377 59850571 refGene_gn=BOC;refGene_tx=XM_023260140.1
C2 59851582 59851770 refGene_gn=BOC;refGene_tx=XM_023260140.1
C2 59856116 59856308
refGene_gn=CFAP44;refGene_tx=XM_003991635.4;refGene_gn=CFAP44;refGene_tx=XM_019839725.2;refGene_gn=CFAP44;refGene_tx=XM_023260139.1
C2 59859470 59859605
refGene_gn=CFAP44;refGene_tx=XM_003991635.4;refGene_gn=CFAP44;refGene_tx=XM_019839725.2;refGene_gn=CFAP44;refGene_tx=XM_023260139.1
C2 59860810 59860971
refGene_gn=CFAP44;refGene_tx=XM_003991635.4;refGene_gn=CFAP44;refGene_tx=XM_019839725.2;refGene_gn=CFAP44;refGene_tx=XM_023260139.1
C2 59864979 59865181
refGene_gn=CFAP44;refGene_tx=XM_003991635.4;refGene_gn=CFAP44;refGene_tx=XM_019839725.2;refGene_gn=CFAP44;refGene_tx=XM_023260139.1
C2 59866190 59866307
refGene_gn=CFAP44;refGene_tx=XM_003991635.4;refGene_gn=CFAP44;refGene_tx=XM_019839725.2;refGene_gn=CFAP44;refGene_tx=XM_023260139.1
C2 59867587 59867718
refGene_gn=CFAP44;refGene_tx=XM_003991635.4;refGene_gn=CFAP44;refGene_tx=XM_019839725.2;refGene_gn=CFAP44;refGene_tx=XM_023260139.1
C2 59869320 59869431
refGene_gn=CFAP44;refGene_tx=XM_003991635.4;refGene_gn=CFAP44;refGene_tx=XM_019839725.2;refGene_gn=CFAP44;refGene_tx=XM_023260139.1
C2 59879155 59879351
refGene_gn=CFAP44;refGene_tx=XM_003991635.4;refGene_gn=CFAP44;refGene_tx=XM_019839725.2;refGene_gn=CFAP44;refGene_tx=XM_023260139.1
C2 59879589 59879793
refGene_gn=CFAP44;refGene_tx=XM_003991635.4;refGene_gn=CFAP44;refGene_tx=XM_019839725.2;refGene_gn=CFAP44;refGene_tx=XM_023260139.1
C2 59882325 59882826
refGene_gn=CFAP44;refGene_tx=XM_003991635.4;refGene_gn=CFAP44;refGene_tx=XM_019839725.2;refGene_gn=CFAP44;refGene_tx=XM_023260139.1
C2 59895194 59895372
refGene_gn=CFAP44;refGene_tx=XM_003991635.4;refGene_gn=CFAP44;refGene_tx=XM_019839725.2;refGene_gn=CFAP44;refGene_tx=XM_023260139.1
C2 59917822 59917997

refGene_gn=CFAP44;refGene_tx=XM_003991635.4;refGene_gn=CFAP44;refGene_tx=XM_019839725.2;refGene_gn=CFAP44;refGene_tx=XM_023260139.1
C2 59919617 59919814
refGene_gn=CFAP44;refGene_tx=XM_003991635.4;refGene_gn=CFAP44;refGene_tx=XM_019839725.2;refGene_gn=CFAP44;refGene_tx=XM_023260139.1
C2 59930747 59930878
refGene_gn=CFAP44;refGene_tx=XM_003991635.4;refGene_gn=CFAP44;refGene_tx=XM_019839725.2;refGene_gn=CFAP44;refGene_tx=XM_023260139.1
C2 59932808 59932971
refGene_gn=CFAP44;refGene_tx=XM_003991635.4;refGene_gn=CFAP44;refGene_tx=XM_019839725.2;refGene_gn=CFAP44;refGene_tx=XM_023260139.1
C2 59933124 59933180
refGene_gn=CFAP44;refGene_tx=XM_003991635.4;refGene_gn=CFAP44;refGene_tx=XM_019839725.2;refGene_gn=CFAP44;refGene_tx=XM_023260139.1
C2 59934572 59934843
refGene_gn=CFAP44;refGene_tx=XM_003991635.4;refGene_gn=CFAP44;refGene_tx=XM_019839725.2;refGene_gn=CFAP44;refGene_tx=XM_023260139.1
C2 59935851 59935997
refGene_gn=CFAP44;refGene_tx=XM_003991635.4;refGene_gn=CFAP44;refGene_tx=XM_019839725.2;refGene_gn=CFAP44;refGene_tx=XM_023260139.1
C2 59941763 59942003
refGene_gn=CFAP44;refGene_tx=XM_003991635.4;refGene_gn=CFAP44;refGene_tx=XM_019839725.2;refGene_gn=CFAP44;refGene_tx=XM_023260139.1
C2 59958671 59958830
refGene_gn=CFAP44;refGene_tx=XM_003991635.4;refGene_gn=CFAP44;refGene_tx=XM_019839725.2;refGene_gn=CFAP44;refGene_tx=XM_023260139.1
C2 59963343 59963454
refGene_gn=CFAP44;refGene_tx=XM_003991635.4;refGene_gn=CFAP44;refGene_tx=XM_019839725.2;refGene_gn=CFAP44;refGene_tx=XM_023260139.1
C2 59964103 59964313
refGene_gn=CFAP44;refGene_tx=XM_003991635.4;refGene_gn=CFAP44;refGene_tx=XM_019839725.2;refGene_gn=CFAP44;refGene_tx=XM_023260139.1
C2 59967375 59967470
refGene_gn=CFAP44;refGene_tx=XM_003991635.4;refGene_gn=CFAP44;refGene_tx=XM_019839725.2;refGene_gn=CFAP44;refGene_tx=XM_023260139.1
C2 59968062 59968162
refGene_gn=CFAP44;refGene_tx=XM_003991635.4;refGene_gn=CFAP44;refGene_tx=XM_019839725.2;refGene_gn=CFAP44;refGene_tx=XM_023260139.1
C2 59968939 59968987
refGene_gn=CFAP44;refGene_tx=XM_003991635.4;refGene_gn=CFAP44;refGene_tx=XM_019839725.2;refGene_gn=CFAP44;refGene_tx=XM_023260139.1
C2 59969898 59970054
refGene_gn=CFAP44;refGene_tx=XM_003991635.4;refGene_gn=CFAP44;refGene_tx=XM_019839725.2;refGene_gn=CFAP44;refGene_tx=XM_023260139.1
C2 59971993 59972158
refGene_gn=CFAP44;refGene_tx=XM_003991635.4;refGene_gn=CFAP44;refGene_tx=XM_019839725.2;refGene_gn=CFAP44;refGene_tx=XM_023260139.1
C2 59975858 59975973
refGene_gn=CFAP44;refGene_tx=XM_003991635.4;refGene_gn=CFAP44;refGene_tx=XM_019839725.2;refGene_gn=CFAP44;refGene_tx=XM_023260139.1

C2 59978174 59978391
refGene_gn=CFAP44;refGene_tx=XM_003991635.4;refGene_gn=CFAP44;refGene_tx=XM_019839725.2;refGene_gn=CFAP44;refGene_tx=XM_023260139.1
C2 59983127 59983230
refGene_gn=CFAP44;refGene_tx=XM_003991635.4;refGene_gn=CFAP44;refGene_tx=XM_019839725.2;refGene_gn=CFAP44;refGene_tx=XM_023260139.1
C2 59987518 59987681
refGene_gn=CFAP44;refGene_tx=XM_003991635.4;refGene_gn=CFAP44;refGene_tx=XM_019839725.2;refGene_gn=CFAP44;refGene_tx=XM_023260139.1
C2 59991554 59991714
refGene_gn=CFAP44;refGene_tx=XM_003991635.4;refGene_gn=CFAP44;refGene_tx=XM_019839725.2;refGene_gn=CFAP44;refGene_tx=XM_023260139.1
C2 59992247 59992403
refGene_gn=CFAP44;refGene_tx=XM_003991635.4;refGene_gn=CFAP44;refGene_tx=XM_019839725.2;refGene_gn=CFAP44;refGene_tx=XM_023260139.1
C2 59997030 59997130
refGene_gn=CFAP44;refGene_tx=XM_003991635.4;refGene_gn=CFAP44;refGene_tx=XM_019839725.2
C2 60010079 60010133
refGene_gn=SPICE1;refGene_tx=XM_006936046.3;refGene_gn=SPICE1;refGene_tx=XM_011285943.3;refGene_gn=SPICE1;refGene_tx=XM_011285944.3;refGene_gn=SPICE1;refGene_tx=XM_011285945.3;refGene_gn=SPICE1;refGene_tx=XM_011285946.3;refGene_gn=SPICE1;refGene_tx=XM_011285947.3;refGene_gn=SPICE1;refGene_tx=XM_019839731.2;refGene_gn=SPICE1;refGene_tx=XM_023260141.1
C2 60011097 60011185
refGene_gn=SPICE1;refGene_tx=XM_006936046.3;refGene_gn=SPICE1;refGene_tx=XM_011285943.3;refGene_gn=SPICE1;refGene_tx=XM_011285944.3;refGene_gn=SPICE1;refGene_tx=XM_011285945.3;refGene_gn=SPICE1;refGene_tx=XM_011285946.3;refGene_gn=SPICE1;refGene_tx=XM_011285947.3;refGene_gn=SPICE1;refGene_tx=XM_019839731.2;refGene_gn=SPICE1;refGene_tx=XM_023260141.1
C2 60012174 60012277
refGene_gn=SPICE1;refGene_tx=XM_006936046.3;refGene_gn=SPICE1;refGene_tx=XM_011285943.3;refGene_gn=SPICE1;refGene_tx=XM_011285944.3;refGene_gn=SPICE1;refGene_tx=XM_011285945.3;refGene_gn=SPICE1;refGene_tx=XM_011285946.3;refGene_gn=SPICE1;refGene_tx=XM_011285947.3;refGene_gn=SPICE1;refGene_tx=XM_019839731.2;refGene_gn=SPICE1;refGene_tx=XM_023260141.1
C2 60012530 60012541 refGene_gn=SPICE1;refGene_tx=XM_011285948.2
C2 60015058 60015242
refGene_gn=SPICE1;refGene_tx=XM_006936046.3;refGene_gn=SPICE1;refGene_tx=XM_011285943.3;refGene_gn=SPICE1;refGene_tx=XM_011285944.3;refGene_gn=SPICE1;refGene_tx=XM_011285945.3;refGene_gn=SPICE1;refGene_tx=XM_011285946.3;refGene_gn=SPICE1;refGene_tx=XM_011285947.3;refGene_gn=SPICE1;refGene_tx=XM_011285948.2;refGene_gn=SPICE1;refGene_tx=XM_019839731.2;refGene_gn=SPICE1;refGene_tx=XM_023260141.1
C2 60018073 60018564
refGene_gn=SPICE1;refGene_tx=XM_006936046.3;refGene_gn=SPICE1;refGene_tx=XM_011285943.3;refGene_gn=SPICE1;refGene_tx=XM_011285944.3;refGene_

gn=SPICE1;refGene_tx=XM_011285945.3;refGene_gn=SPICE1;refGene_tx=XM_011285946.3;refGene_gn=SPICE1;refGene_tx=XM_011285947.3;refGene_gn=SPICE1;refGene_tx=XM_011285948.2;refGene_gn=SPICE1;refGene_tx=XM_019839731.2;refGene_gn=SPICE1;refGene_tx=XM_023260141.1
C2 60025564 60025789
refGene_gn=SPICE1;refGene_tx=XM_006936046.3;refGene_gn=SPICE1;refGene_tx=XM_011285943.3;refGene_gn=SPICE1;refGene_tx=XM_011285944.3;refGene_gn=SPICE1;refGene_tx=XM_011285945.3;refGene_gn=SPICE1;refGene_tx=XM_011285946.3;refGene_gn=SPICE1;refGene_tx=XM_011285947.3;refGene_gn=SPICE1;refGene_tx=XM_011285948.2;refGene_gn=SPICE1;refGene_tx=XM_019839731.2;refGene_gn=SPICE1;refGene_tx=XM_023260141.1
C2 60026679 60026827
refGene_gn=SPICE1;refGene_tx=XM_006936046.3;refGene_gn=SPICE1;refGene_tx=XM_011285943.3;refGene_gn=SPICE1;refGene_tx=XM_011285944.3;refGene_gn=SPICE1;refGene_tx=XM_011285945.3;refGene_gn=SPICE1;refGene_tx=XM_011285946.3;refGene_gn=SPICE1;refGene_tx=XM_011285947.3;refGene_gn=SPICE1;refGene_tx=XM_011285948.2;refGene_gn=SPICE1;refGene_tx=XM_019839731.2;refGene_gn=SPICE1;refGene_tx=XM_023260141.1
C2 60030380 60030512
refGene_gn=SPICE1;refGene_tx=XM_006936046.3;refGene_gn=SPICE1;refGene_tx=XM_011285943.3;refGene_gn=SPICE1;refGene_tx=XM_011285944.3;refGene_gn=SPICE1;refGene_tx=XM_011285945.3;refGene_gn=SPICE1;refGene_tx=XM_011285946.3;refGene_gn=SPICE1;refGene_tx=XM_011285947.3;refGene_gn=SPICE1;refGene_tx=XM_011285948.2;refGene_gn=SPICE1;refGene_tx=XM_019839731.2;refGene_gn=SPICE1;refGene_tx=XM_023260141.1
C2 60033132 60033398
refGene_gn=SPICE1;refGene_tx=XM_006936046.3;refGene_gn=SPICE1;refGene_tx=XM_011285943.3;refGene_gn=SPICE1;refGene_tx=XM_011285944.3;refGene_gn=SPICE1;refGene_tx=XM_011285945.3;refGene_gn=SPICE1;refGene_tx=XM_011285946.3;refGene_gn=SPICE1;refGene_tx=XM_011285947.3;refGene_gn=SPICE1;refGene_tx=XM_011285948.2;refGene_gn=SPICE1;refGene_tx=XM_019839731.2;refGene_gn=SPICE1;refGene_tx=XM_023260141.1
C2 60033969 60034107
refGene_gn=SPICE1;refGene_tx=XM_006936046.3;refGene_gn=SPICE1;refGene_tx=XM_011285943.3;refGene_gn=SPICE1;refGene_tx=XM_011285944.3;refGene_gn=SPICE1;refGene_tx=XM_011285945.3;refGene_gn=SPICE1;refGene_tx=XM_011285946.3;refGene_gn=SPICE1;refGene_tx=XM_011285947.3;refGene_gn=SPICE1;refGene_tx=XM_011285948.2;refGene_gn=SPICE1;refGene_tx=XM_019839731.2;refGene_gn=SPICE1;refGene_tx=XM_023260141.1
C2 60034266 60034457
refGene_gn=SPICE1;refGene_tx=XM_006936046.3;refGene_gn=SPICE1;refGene_tx=XM_011285943.3;refGene_gn=SPICE1;refGene_tx=XM_011285944.3;refGene_gn=SPICE1;refGene_tx=XM_011285945.3;refGene_gn=SPICE1;refGene_tx=XM_011285946.3;refGene_gn=SPICE1;refGene_tx=XM_011285947.3;refGene_gn=SPICE1;refGene_tx=XM_011285948.2;refGene_gn=SPICE1;refGene_tx=XM_019839731.2;refGene_gn=SPICE1;refGene_tx=XM_023260141.1
C2 60044688 60044804
refGene_gn=SPICE1;refGene_tx=XM_006936046.3;refGene_gn=SPICE1;refGene_tx=XM_011285943.3;refGene_gn=SPICE1;refGene_tx=XM_011285944.3;refGene_gn=SPICE1;refGene_tx=XM_011285945.3;refGene_gn=SPICE1;refGene_tx=XM_01

1285946.3;refGene_gn=SPICE1;refGene_tx=XM_011285947.3;refGene_gn=SPICE1;refGene_tx=XM_011285948.2;refGene_gn=SPICE1;refGene_tx=XM_019839731.2;refGene_gn=SPICE1;refGene_tx=XM_023260141.1
C2 60047665 60047772
refGene_gn=SPICE1;refGene_tx=XM_006936046.3;refGene_gn=SPICE1;refGene_tx=XM_011285943.3;refGene_gn=SPICE1;refGene_tx=XM_011285944.3;refGene_gn=SPICE1;refGene_tx=XM_011285945.3;refGene_gn=SPICE1;refGene_tx=XM_011285946.3;refGene_gn=SPICE1;refGene_tx=XM_011285947.3;refGene_gn=SPICE1;refGene_tx=XM_011285948.2;refGene_gn=SPICE1;refGene_tx=XM_019839731.2;refGene_gn=SPICE1;refGene_tx=XM_023260141.1
C2 60048832 60048935
refGene_gn=SPICE1;refGene_tx=XM_006936046.3;refGene_gn=SPICE1;refGene_tx=XM_011285943.3;refGene_gn=SPICE1;refGene_tx=XM_011285944.3;refGene_gn=SPICE1;refGene_tx=XM_011285945.3;refGene_gn=SPICE1;refGene_tx=XM_011285946.3;refGene_gn=SPICE1;refGene_tx=XM_011285947.3;refGene_gn=SPICE1;refGene_tx=XM_011285948.2;refGene_gn=SPICE1;refGene_tx=XM_019839731.2;refGene_gn=SPICE1;refGene_tx=XM_023260141.1
C2 60053106 60053250
refGene_gn=SPICE1;refGene_tx=XM_006936046.3;refGene_gn=SPICE1;refGene_tx=XM_011285943.3;refGene_gn=SPICE1;refGene_tx=XM_011285944.3;refGene_gn=SPICE1;refGene_tx=XM_011285945.3;refGene_gn=SPICE1;refGene_tx=XM_011285946.3;refGene_gn=SPICE1;refGene_tx=XM_011285947.3;refGene_gn=SPICE1;refGene_tx=XM_011285948.2;refGene_gn=SPICE1;refGene_tx=XM_019839731.2;refGene_gn=SPICE1;refGene_tx=XM_023260141.1
C2 60057450 60057498
refGene_gn=SPICE1;refGene_tx=XM_006936046.3;refGene_gn=SPICE1;refGene_tx=XM_011285943.3;refGene_gn=SPICE1;refGene_tx=XM_011285944.3;refGene_gn=SPICE1;refGene_tx=XM_011285945.3;refGene_gn=SPICE1;refGene_tx=XM_011285946.3;refGene_gn=SPICE1;refGene_tx=XM_011285947.3;refGene_gn=SPICE1;refGene_tx=XM_011285948.2;refGene_gn=SPICE1;refGene_tx=XM_019839731.2
C2 60060765 60060864
refGene_gn=SPICE1;refGene_tx=XM_006936046.3;refGene_gn=SPICE1;refGene_tx=XM_011285943.3;refGene_gn=SPICE1;refGene_tx=XM_011285944.3;refGene_gn=SPICE1;refGene_tx=XM_011285945.3;refGene_gn=SPICE1;refGene_tx=XM_011285946.3;refGene_gn=SPICE1;refGene_tx=XM_011285947.3;refGene_gn=SPICE1;refGene_tx=XM_011285948.2;refGene_gn=SPICE1;refGene_tx=XM_019839731.2
C2 60087299 60087509
refGene_gn=SIDT1;refGene_tx=XM_011285942.2;refGene_gn=SIDT1;refGene_tx=XM_019839732.1;refGene_gn=SIDT1;refGene_tx=XM_019839733.1;refGene_gn=SIDT1;refGene_tx=XM_023260142.1
C2 60129802 60129924
refGene_gn=SIDT1;refGene_tx=XM_011285942.2;refGene_gn=SIDT1;refGene_tx=XM_019839732.1;refGene_gn=SIDT1;refGene_tx=XM_019839733.1;refGene_gn=SIDT1;refGene_tx=XM_023260142.1
C2 60130944 60131115
refGene_gn=SIDT1;refGene_tx=XM_011285942.2;refGene_gn=SIDT1;refGene_tx=XM_019839732.1;refGene_gn=SIDT1;refGene_tx=XM_019839733.1;refGene_gn=SIDT1;refGene_tx=XM_023260142.1

C2 60140034 60140080
refGene_gn=SIDT1;refGene_tx=XM_011285942.2;refGene_gn=SIDT1;refGene_tx=XM_019839732.1;refGene_gn=SIDT1;refGene_tx=XM_019839733.1;refGene_gn=SIDT1;refGene_tx=XM_023260142.1
C2 60143695 60143797
refGene_gn=SIDT1;refGene_tx=XM_011285942.2;refGene_gn=SIDT1;refGene_tx=XM_019839732.1;refGene_gn=SIDT1;refGene_tx=XM_023260142.1
C2 60144414 60144498
refGene_gn=SIDT1;refGene_tx=XM_011285942.2;refGene_gn=SIDT1;refGene_tx=XM_019839732.1;refGene_gn=SIDT1;refGene_tx=XM_023260142.1
C2 60145977 60146065
refGene_gn=SIDT1;refGene_tx=XM_011285942.2;refGene_gn=SIDT1;refGene_tx=XM_019839732.1;refGene_gn=SIDT1;refGene_tx=XM_019839733.1;refGene_gn=SIDT1;refGene_tx=XM_023260142.1
C2 60147265 60147337
refGene_gn=SIDT1;refGene_tx=XM_011285942.2;refGene_gn=SIDT1;refGene_tx=XM_019839732.1;refGene_gn=SIDT1;refGene_tx=XM_019839733.1;refGene_gn=SIDT1;refGene_tx=XM_023260142.1
C2 60147754 60147848
refGene_gn=SIDT1;refGene_tx=XM_011285942.2;refGene_gn=SIDT1;refGene_tx=XM_019839732.1;refGene_gn=SIDT1;refGene_tx=XM_019839733.1;refGene_gn=SIDT1;refGene_tx=XM_023260142.1
C2 60159280 60159324
refGene_gn=SIDT1;refGene_tx=XM_011285942.2;refGene_gn=SIDT1;refGene_tx=XM_019839732.1;refGene_gn=SIDT1;refGene_tx=XM_019839733.1;refGene_gn=SIDT1;refGene_tx=XM_023260142.1
C2 60173739 60173811
refGene_gn=SIDT1;refGene_tx=XM_011285942.2;refGene_gn=SIDT1;refGene_tx=XM_019839732.1;refGene_gn=SIDT1;refGene_tx=XM_019839733.1
C2 60175086 60175232
refGene_gn=SIDT1;refGene_tx=XM_011285942.2;refGene_gn=SIDT1;refGene_tx=XM_019839732.1;refGene_gn=SIDT1;refGene_tx=XM_019839733.1;refGene_gn=SIDT1;refGene_tx=XM_023260142.1;refGene_gn=SIDT1;refGene_tx=XM_023260143.1
C2 60176079 60176153
refGene_gn=SIDT1;refGene_tx=XM_011285942.2;refGene_gn=SIDT1;refGene_tx=XM_019839732.1;refGene_gn=SIDT1;refGene_tx=XM_019839733.1;refGene_gn=SIDT1;refGene_tx=XM_023260142.1;refGene_gn=SIDT1;refGene_tx=XM_023260143.1
C2 60177398 60177465
refGene_gn=SIDT1;refGene_tx=XM_011285942.2;refGene_gn=SIDT1;refGene_tx=XM_019839732.1;refGene_gn=SIDT1;refGene_tx=XM_019839733.1;refGene_gn=SIDT1;refGene_tx=XM_023260142.1;refGene_gn=SIDT1;refGene_tx=XM_023260143.1
C2 60179029 60179103
refGene_gn=SIDT1;refGene_tx=XM_011285942.2;refGene_gn=SIDT1;refGene_tx=XM_019839732.1;refGene_gn=SIDT1;refGene_tx=XM_019839733.1;refGene_gn=SIDT1;refGene_tx=XM_023260142.1;refGene_gn=SIDT1;refGene_tx=XM_023260143.1
C2 60180271 60180395

refGene_gn=SIDT1;refGene_tx=XM_011285942.2;refGene_gn=SIDT1;refGene_tx=XM_019839732.1;refGene_gn=SIDT1;refGene_tx=XM_019839733.1;refGene_gn=SIDT1;refGene_tx=XM_023260142.1;refGene_gn=SIDT1;refGene_tx=XM_023260143.1
C2 60180604 60180722
refGene_gn=SIDT1;refGene_tx=XM_011285942.2;refGene_gn=SIDT1;refGene_tx=XM_019839732.1;refGene_gn=SIDT1;refGene_tx=XM_019839733.1;refGene_gn=SIDT1;refGene_tx=XM_023260142.1;refGene_gn=SIDT1;refGene_tx=XM_023260143.1
C2 60183691 60183828
refGene_gn=SIDT1;refGene_tx=XM_011285942.2;refGene_gn=SIDT1;refGene_tx=XM_019839732.1;refGene_gn=SIDT1;refGene_tx=XM_019839733.1;refGene_gn=SIDT1;refGene_tx=XM_023260142.1;refGene_gn=SIDT1;refGene_tx=XM_023260143.1
C2 60184508 60184617
refGene_gn=SIDT1;refGene_tx=XM_011285942.2;refGene_gn=SIDT1;refGene_tx=XM_019839732.1;refGene_gn=SIDT1;refGene_tx=XM_019839733.1;refGene_gn=SIDT1;refGene_tx=XM_023260142.1;refGene_gn=SIDT1;refGene_tx=XM_023260143.1
C2 60187441 60187456
refGene_gn=SIDT1;refGene_tx=XM_019839732.1;refGene_gn=SIDT1;refGene_tx=XM_019839733.1;refGene_gn=SIDT1;refGene_tx=XM_023260142.1;refGene_gn=SIDT1;refGene_tx=XM_023260143.1
C2 60188742 60188819
refGene_gn=SIDT1;refGene_tx=XM_011285942.2;refGene_gn=SIDT1;refGene_tx=XM_019839732.1;refGene_gn=SIDT1;refGene_tx=XM_019839733.1;refGene_gn=SIDT1;refGene_tx=XM_023260142.1;refGene_gn=SIDT1;refGene_tx=XM_023260143.1
C2 60191553 60191600
refGene_gn=SIDT1;refGene_tx=XM_011285942.2;refGene_gn=SIDT1;refGene_tx=XM_019839732.1;refGene_gn=SIDT1;refGene_tx=XM_019839733.1;refGene_gn=SIDT1;refGene_tx=XM_023260142.1;refGene_gn=SIDT1;refGene_tx=XM_023260143.1
C2 60194156 60194262
refGene_gn=SIDT1;refGene_tx=XM_011285942.2;refGene_gn=SIDT1;refGene_tx=XM_019839732.1;refGene_gn=SIDT1;refGene_tx=XM_019839733.1;refGene_gn=SIDT1;refGene_tx=XM_023260142.1;refGene_gn=SIDT1;refGene_tx=XM_023260143.1
C2 60194336 60194447
refGene_gn=SIDT1;refGene_tx=XM_011285942.2;refGene_gn=SIDT1;refGene_tx=XM_019839732.1;refGene_gn=SIDT1;refGene_tx=XM_019839733.1;refGene_gn=SIDT1;refGene_tx=XM_023260142.1;refGene_gn=SIDT1;refGene_tx=XM_023260143.1
C2 60195375 60195489
refGene_gn=SIDT1;refGene_tx=XM_011285942.2;refGene_gn=SIDT1;refGene_tx=XM_019839732.1;refGene_gn=SIDT1;refGene_tx=XM_019839733.1;refGene_gn=SIDT1;refGene_tx=XM_023260142.1;refGene_gn=SIDT1;refGene_tx=XM_023260143.1
C2 60196890 60196953
refGene_gn=SIDT1;refGene_tx=XM_011285942.2;refGene_gn=SIDT1;refGene_tx

=XM_019839732.1;refGene_gn=SIDT1;refGene_tx=XM_019839733.1;refGene_gn=SIDT1;refGene_tx=XM_023260142.1;refGene_gn=SIDT1;refGene_tx=XM_023260143.1
C2 60218524 60224994
refGene_gn=USF3;refGene_tx=XM_023260144.1;refGene_gn=USF3;refGene_tx=XM_023260145.1
C2 60227660 60227757
refGene_gn=USF3;refGene_tx=XM_023260144.1;refGene_gn=USF3;refGene_tx=XM_023260145.1
C2 60232531 60232614 refGene_gn=USF3;refGene_tx=XM_023260144.1
C2 60234492 60234521 refGene_gn=USF3;refGene_tx=XM_023260144.1
C2 60235646 60235693 refGene_gn=USF3;refGene_tx=XM_023260144.1
C2 60282821 60282999
refGene_gn=NAA50;refGene_tx=XM_003991640.5;refGene_gn=NAA50;refGene_tx=XM_006936052.4
C2 60283958 60284025
refGene_gn=NAA50;refGene_tx=XM_003991640.5;refGene_gn=NAA50;refGene_tx=XM_006936052.4
C2 60284455 60284575
refGene_gn=NAA50;refGene_tx=XM_003991640.5;refGene_gn=NAA50;refGene_tx=XM_006936052.4
C2 60285006 60285143
refGene_gn=NAA50;refGene_tx=XM_003991640.5;refGene_gn=NAA50;refGene_tx=XM_006936052.4
C2 60306212 60306220
refGene_gn=NAA50;refGene_tx=XM_003991640.5;refGene_gn=NAA50;refGene_tx=XM_006936052.4
C2 60336072 60336157
refGene_gn=ATP6V1A;refGene_tx=XM_006936054.4;refGene_gn=ATP6V1A;refGene_tx=XM_006936055.4;refGene_gn=ATP6V1A;refGene_tx=XM_023260146.1
C2 60339599 60339728
refGene_gn=ATP6V1A;refGene_tx=XM_006936054.4;refGene_gn=ATP6V1A;refGene_tx=XM_006936055.4;refGene_gn=ATP6V1A;refGene_tx=XM_023260146.1
C2 60342786 60343001
refGene_gn=ATP6V1A;refGene_tx=XM_006936054.4;refGene_gn=ATP6V1A;refGene_tx=XM_006936055.4;refGene_gn=ATP6V1A;refGene_tx=XM_023260146.1
C2 60343282 60343420
refGene_gn=ATP6V1A;refGene_tx=XM_006936054.4;refGene_gn=ATP6V1A;refGene_tx=XM_006936055.4;refGene_gn=ATP6V1A;refGene_tx=XM_023260146.1
C2 60345847 60345999
refGene_gn=ATP6V1A;refGene_tx=XM_006936054.4;refGene_gn=ATP6V1A;refGene_tx=XM_006936055.4;refGene_gn=ATP6V1A;refGene_tx=XM_023260146.1
C2 60349930 60350093
refGene_gn=ATP6V1A;refGene_tx=XM_006936054.4;refGene_gn=ATP6V1A;refGene_tx=XM_006936055.4;refGene_gn=ATP6V1A;refGene_tx=XM_023260146.1
C2 60350629 60350738
refGene_gn=ATP6V1A;refGene_tx=XM_006936054.4;refGene_gn=ATP6V1A;refGene_tx=XM_006936055.4;refGene_gn=ATP6V1A;refGene_tx=XM_023260146.1
C2 60356550 60356673
refGene_gn=ATP6V1A;refGene_tx=XM_006936054.4;refGene_gn=ATP6V1A;refGen

e_tx=XM_006936055.4;refGene_gn=ATP6V1A;refGene_tx=XM_023260146.1
C2 60357028 60357143
refGene_gn=ATP6V1A;refGene_tx=XM_006936054.4;refGene_gn=ATP6V1A;refGene_tx=XM_006936055.4;refGene_gn=ATP6V1A;refGene_tx=XM_023260146.1
C2 60357699 60357763
refGene_gn=ATP6V1A;refGene_tx=XM_006936054.4;refGene_gn=ATP6V1A;refGene_tx=XM_006936055.4;refGene_gn=ATP6V1A;refGene_tx=XM_023260146.1
C2 60359598 60359802
refGene_gn=ATP6V1A;refGene_tx=XM_006936054.4;refGene_gn=ATP6V1A;refGene_tx=XM_006936055.4;refGene_gn=ATP6V1A;refGene_tx=XM_023260146.1
C2 60364526 60364621
refGene_gn=ATP6V1A;refGene_tx=XM_006936054.4;refGene_gn=ATP6V1A;refGene_tx=XM_006936055.4;refGene_gn=ATP6V1A;refGene_tx=XM_023260146.1
C2 60366683 60366855
refGene_gn=ATP6V1A;refGene_tx=XM_006936054.4;refGene_gn=ATP6V1A;refGene_tx=XM_006936055.4;refGene_gn=ATP6V1A;refGene_tx=XM_023260146.1
C2 60370227 60370320
refGene_gn=ATP6V1A;refGene_tx=XM_006936054.4;refGene_gn=ATP6V1A;refGene_tx=XM_006936055.4;refGene_gn=ATP6V1A;refGene_tx=XM_023260146.1
C2 60406906 60406933 refGene_gn=GRAMD1C;refGene_tx=XM_003991642.5
C2 60412654 60412804 refGene_gn=GRAMD1C;refGene_tx=XM_003991642.5
C2 60428384 60428469 refGene_gn=GRAMD1C;refGene_tx=XM_003991642.5
C2 60436769 60436873 refGene_gn=GRAMD1C;refGene_tx=XM_003991642.5
C2 60437463 60437559 refGene_gn=GRAMD1C;refGene_tx=XM_003991642.5
C2 60450436 60450517 refGene_gn=GRAMD1C;refGene_tx=XM_003991642.5
C2 60475536 60475655
refGene_gn=GRAMD1C;refGene_tx=XM_003991642.5;refGene_gn=GRAMD1C;refGene_tx=XM_019839738.2
C2 60479461 60479594
refGene_gn=GRAMD1C;refGene_tx=XM_003991642.5;refGene_gn=GRAMD1C;refGene_tx=XM_019839738.2
C2 60482626 60482789
refGene_gn=GRAMD1C;refGene_tx=XM_003991642.5;refGene_gn=GRAMD1C;refGene_tx=XM_019839738.2
C2 60489517 60489655
refGene_gn=GRAMD1C;refGene_tx=XM_003991642.5;refGene_gn=GRAMD1C;refGene_tx=XM_019839738.2
C2 60495404 60495523
refGene_gn=GRAMD1C;refGene_tx=XM_003991642.5;refGene_gn=GRAMD1C;refGene_tx=XM_019839738.2
C2 60497101 60497244
refGene_gn=GRAMD1C;refGene_tx=XM_003991642.5;refGene_gn=GRAMD1C;refGene_tx=XM_019839738.2
C2 60498036 60498140
refGene_gn=GRAMD1C;refGene_tx=XM_003991642.5;refGene_gn=GRAMD1C;refGene_tx=XM_019839738.2
C2 60499685 60499862
refGene_gn=GRAMD1C;refGene_tx=XM_003991642.5;refGene_gn=GRAMD1C;refGene_tx=XM_019839738.2
C2 60500952 60501010

refGene_gn=GRAMD1C;refGene_tx=XM_003991642.5;refGene_gn=GRAMD1C;refGene_tx=XM_019839738.2
C2 60502975 60503086
refGene_gn=GRAMD1C;refGene_tx=XM_003991642.5;refGene_gn=GRAMD1C;refGene_tx=XM_019839738.2
C2 60503312 60503418
refGene_gn=GRAMD1C;refGene_tx=XM_003991642.5;refGene_gn=GRAMD1C;refGene_tx=XM_019839738.2
C2 60509580 60509661
refGene_gn=GRAMD1C;refGene_tx=XM_003991642.5;refGene_gn=GRAMD1C;refGene_tx=XM_019839738.2
C2 60512913 60513065
refGene_gn=ZDHHC23;refGene_tx=XM_023260147.1;refGene_gn=ZDHHC23;refGene_tx=XM_023260148.1;refGene_gn=ZDHHC23;refGene_tx=XM_023260149.1
C2 60515055 60515766
refGene_gn=ZDHHC23;refGene_tx=XM_023260147.1;refGene_gn=ZDHHC23;refGene_tx=XM_023260148.1;refGene_gn=ZDHHC23;refGene_tx=XM_023260149.1
C2 60517506 60517674
refGene_gn=ZDHHC23;refGene_tx=XM_023260147.1;refGene_gn=ZDHHC23;refGene_tx=XM_023260148.1;refGene_gn=ZDHHC23;refGene_tx=XM_023260149.1
C2 60519774 60520042
refGene_gn=ZDHHC23;refGene_tx=XM_023260147.1;refGene_gn=ZDHHC23;refGene_tx=XM_023260148.1;refGene_gn=ZDHHC23;refGene_tx=XM_023260149.1
C2 60526173 60526318
refGene_gn=CCDC191;refGene_tx=XM_003991658.3;refGene_gn=CCDC191;refGene_tx=XM_019839735.1;refGene_gn=CCDC191;refGene_tx=XM_019839736.1;refGene_gn=CCDC191;refGene_tx=XM_019839737.1
C2 60540754 60540900
refGene_gn=CCDC191;refGene_tx=XM_003991658.3;refGene_gn=CCDC191;refGene_tx=XM_019839735.1;refGene_gn=CCDC191;refGene_tx=XM_019839736.1;refGene_gn=CCDC191;refGene_tx=XM_019839737.1
C2 60541414 60541567
refGene_gn=CCDC191;refGene_tx=XM_003991658.3;refGene_gn=CCDC191;refGene_tx=XM_019839735.1;refGene_gn=CCDC191;refGene_tx=XM_019839736.1;refGene_gn=CCDC191;refGene_tx=XM_019839737.1
C2 60546750 60546894
refGene_gn=CCDC191;refGene_tx=XM_003991658.3;refGene_gn=CCDC191;refGene_tx=XM_019839735.1;refGene_gn=CCDC191;refGene_tx=XM_019839736.1;refGene_gn=CCDC191;refGene_tx=XM_019839737.1
C2 60557420 60557522
refGene_gn=CCDC191;refGene_tx=XM_003991658.3;refGene_gn=CCDC191;refGene_tx=XM_019839735.1;refGene_gn=CCDC191;refGene_tx=XM_019839736.1;refGene_gn=CCDC191;refGene_tx=XM_019839737.1
C2 60558259 60558342
refGene_gn=CCDC191;refGene_tx=XM_003991658.3;refGene_gn=CCDC191;refGene_tx=XM_019839735.1;refGene_gn=CCDC191;refGene_tx=XM_019839736.1;refGene_gn=CCDC191;refGene_tx=XM_019839737.1
C2 60560454 60560567
refGene_gn=CCDC191;refGene_tx=XM_003991658.3;refGene_gn=CCDC191;refGene_tx=XM_019839735.1;refGene_gn=CCDC191;refGene_tx=XM_019839736.1;refGe

ne_gn=CCDC191;refGene_tx=XM_019839737.1
C2 60561336 60561794
refGene_gn=CCDC191;refGene_tx=XM_003991658.3;refGene_gn=CCDC191;refGene_tx=XM_019839735.1;refGene_gn=CCDC191;refGene_tx=XM_019839736.1;refGene_gn=CCDC191;refGene_tx=XM_019839737.1
C2 60568361 60568629
refGene_gn=CCDC191;refGene_tx=XM_003991658.3;refGene_gn=CCDC191;refGene_tx=XM_019839735.1;refGene_gn=CCDC191;refGene_tx=XM_019839736.1;refGene_gn=CCDC191;refGene_tx=XM_019839737.1
C2 60580290 60580481
refGene_gn=CCDC191;refGene_tx=XM_003991658.3;refGene_gn=CCDC191;refGene_tx=XM_019839735.1;refGene_gn=CCDC191;refGene_tx=XM_019839736.1;refGene_gn=CCDC191;refGene_tx=XM_019839737.1
C2 60592814 60592965
refGene_gn=CCDC191;refGene_tx=XM_003991658.3;refGene_gn=CCDC191;refGene_tx=XM_019839735.1;refGene_gn=CCDC191;refGene_tx=XM_019839736.1;refGene_gn=CCDC191;refGene_tx=XM_019839737.1
C2 60595963 60596187
refGene_gn=CCDC191;refGene_tx=XM_003991658.3;refGene_gn=CCDC191;refGene_tx=XM_019839735.1;refGene_gn=CCDC191;refGene_tx=XM_019839736.1;refGene_gn=CCDC191;refGene_tx=XM_019839737.1
C2 60597857 60598036
refGene_gn=CCDC191;refGene_tx=XM_003991658.3;refGene_gn=CCDC191;refGene_tx=XM_019839735.1;refGene_gn=CCDC191;refGene_tx=XM_019839736.1;refGene_gn=CCDC191;refGene_tx=XM_019839737.1
C2 60602009 60602150
refGene_gn=CCDC191;refGene_tx=XM_003991658.3;refGene_gn=CCDC191;refGene_tx=XM_019839735.1;refGene_gn=CCDC191;refGene_tx=XM_019839737.1
C2 60605705 60605847
refGene_gn=CCDC191;refGene_tx=XM_003991658.3;refGene_gn=CCDC191;refGene_tx=XM_019839735.1;refGene_gn=CCDC191;refGene_tx=XM_019839736.1;refGene_gn=CCDC191;refGene_tx=XM_019839737.1
C2 60612592 60612631
refGene_gn=CCDC191;refGene_tx=XM_003991658.3;refGene_gn=CCDC191;refGene_tx=XM_019839735.1;refGene_gn=CCDC191;refGene_tx=XM_019839736.1
C2 60614976 60615066
refGene_gn=CCDC191;refGene_tx=XM_003991658.3;refGene_gn=CCDC191;refGene_tx=XM_019839735.1;refGene_gn=CCDC191;refGene_tx=XM_019839736.1
C2 60620235 60620435
refGene_gn=QTRT2;refGene_tx=XM_006936056.4;refGene_gn=QTRT2;refGene_tx=XM_019839739.2;refGene_gn=QTRT2;refGene_tx=XM_023260150.1
C2 60622140 60622196
refGene_gn=QTRT2;refGene_tx=XM_006936056.4;refGene_gn=QTRT2;refGene_tx=XM_019839739.2;refGene_gn=QTRT2;refGene_tx=XM_023260150.1
C2 60624394 60624471
refGene_gn=QTRT2;refGene_tx=XM_006936056.4;refGene_gn=QTRT2;refGene_tx=XM_006936057.4;refGene_gn=QTRT2;refGene_tx=XM_019839739.2;refGene_gn=QTRT2;refGene_tx=XM_023260150.1
C2 60626329 60626542
refGene_gn=QTRT2;refGene_tx=XM_006936056.4;refGene_gn=QTRT2;refGene_tx

=XM_006936057.4;refGene_gn=QTRT2;refGene_tx=XM_019839739.2;refGene_gn=QTRT2;refGene_tx=XM_023260150.1
C2 60631968 60632168
refGene_gn=QTRT2;refGene_tx=XM_006936056.4;refGene_gn=QTRT2;refGene_tx=XM_006936057.4;refGene_gn=QTRT2;refGene_tx=XM_019839739.2;refGene_gn=QTRT2;refGene_tx=XM_023260150.1
C2 60634290 60634442
refGene_gn=QTRT2;refGene_tx=XM_006936056.4;refGene_gn=QTRT2;refGene_tx=XM_006936057.4;refGene_gn=QTRT2;refGene_tx=XM_019839739.2;refGene_gn=QTRT2;refGene_tx=XM_023260150.1
C2 60637016 60637134
refGene_gn=QTRT2;refGene_tx=XM_006936056.4;refGene_gn=QTRT2;refGene_tx=XM_006936057.4;refGene_gn=QTRT2;refGene_tx=XM_019839739.2;refGene_gn=QTRT2;refGene_tx=XM_023260150.1
C2 60638725 60638957
refGene_gn=QTRT2;refGene_tx=XM_006936056.4;refGene_gn=QTRT2;refGene_tx=XM_006936057.4;refGene_gn=QTRT2;refGene_tx=XM_019839739.2;refGene_gn=QTRT2;refGene_tx=XM_023260150.1
C2 60681679 60681876 refGene_gn=DRD3;refGene_tx=XM_019839742.2
C2 60683021 60683367 refGene_gn=DRD3;refGene_tx=XM_019839742.2
C2 60686681 60686759 refGene_gn=DRD3;refGene_tx=XM_019839742.2
C2 60687834 60688031 refGene_gn=DRD3;refGene_tx=XM_019839742.2
C2 60693598 60693741 refGene_gn=DRD3;refGene_tx=XM_019839742.2
C2 60702736 60702849 refGene_gn=DRD3;refGene_tx=XM_019839742.2
C2 60714521 60714791 refGene_gn=DRD3;refGene_tx=XM_019839742.2
C2 60797579 60797643
refGene_gn=TIGIT;refGene_tx=XM_019839745.1;refGene_gn=TIGIT;refGene_tx=XM_019839746.1;refGene_gn=TIGIT;refGene_tx=XM_019839748.1;refGene_gn=TIGIT;refGene_tx=XM_019839749.1
C2 60798645 60798975
refGene_gn=TIGIT;refGene_tx=XM_019839745.1;refGene_gn=TIGIT;refGene_tx=XM_019839746.1;refGene_gn=TIGIT;refGene_tx=XM_019839748.1;refGene_gn=TIGIT;refGene_tx=XM_019839749.1
C2 60801976 60802083
refGene_gn=TIGIT;refGene_tx=XM_019839745.1;refGene_gn=TIGIT;refGene_tx=XM_019839746.1;refGene_gn=TIGIT;refGene_tx=XM_019839748.1;refGene_gn=TIGIT;refGene_tx=XM_019839749.1
C2 60810741 60810993
refGene_gn=TIGIT;refGene_tx=XM_019839745.1;refGene_gn=TIGIT;refGene_tx=XM_019839746.1;refGene_gn=TIGIT;refGene_tx=XM_019839748.1;refGene_gn=TIGIT;refGene_tx=XM_019839749.1
C2 60841266 60841688
refGene_gn=ZBTB20;refGene_tx=XM_019839752.2;refGene_gn=ZBTB20;refGene_tx=XM_019839757.2;refGene_gn=ZBTB20;refGene_tx=XM_019839758.2;refGene_gn=ZBTB20;refGene_tx=XM_019839761.2;refGene_gn=ZBTB20;refGene_tx=XM_019839762.2;refGene_gn=ZBTB20;refGene_tx=XM_019839766.2;refGene_gn=ZBTB20;refGene_tx=XM_019839769.2;refGene_gn=ZBTB20;refGene_tx=XM_023260830.1;refGene_gn=ZBTB20;refGene_tx=XM_023260831.1;refGene_gn=ZBTB20;refGene_tx=XM_023260832.1;refGene_gn=ZBTB20;refGene_tx=XM_023260833.1;refGene_gn=ZBTB20;refGene_tx=XM_023260834.1;refGene_gn=ZBTB20;refGene_tx=XM_

023260835.1;refGene_gn=ZBTB20;refGene_tx=XM_023260836.1;refGene_gn=ZBTB20;refGene_tx=XM_023260837.1;refGene_gn=ZBTB20;refGene_tx=XM_023260838.1;refGene_gn=ZBTB20;refGene_tx=XM_023260840.1;refGene_gn=ZBTB20;refGene_tx=XM_023260841.1
C2 60852755 60854360
refGene_gn=ZBTB20;refGene_tx=XM_019839752.2;refGene_gn=ZBTB20;refGene_tx=XM_019839757.2;refGene_gn=ZBTB20;refGene_tx=XM_019839758.2;refGene_gn=ZBTB20;refGene_tx=XM_019839761.2;refGene_gn=ZBTB20;refGene_tx=XM_019839762.2;refGene_gn=ZBTB20;refGene_tx=XM_019839766.2;refGene_gn=ZBTB20;refGene_tx=XM_019839769.2;refGene_gn=ZBTB20;refGene_tx=XM_023260830.1;refGene_gn=ZBTB20;refGene_tx=XM_023260831.1;refGene_gn=ZBTB20;refGene_tx=XM_023260832.1;refGene_gn=ZBTB20;refGene_tx=XM_023260833.1;refGene_gn=ZBTB20;refGene_tx=XM_023260834.1;refGene_gn=ZBTB20;refGene_tx=XM_023260835.1;refGene_gn=ZBTB20;refGene_tx=XM_023260836.1;refGene_gn=ZBTB20;refGene_tx=XM_023260837.1;refGene_gn=ZBTB20;refGene_tx=XM_023260838.1;refGene_gn=ZBTB20;refGene_tx=XM_023260840.1;refGene_gn=ZBTB20;refGene_tx=XM_023260841.1
C2 60883916 60884104
refGene_gn=ZBTB20;refGene_tx=XM_019839752.2;refGene_gn=ZBTB20;refGene_tx=XM_019839757.2;refGene_gn=ZBTB20;refGene_tx=XM_019839758.2;refGene_gn=ZBTB20;refGene_tx=XM_019839761.2;refGene_gn=ZBTB20;refGene_tx=XM_019839762.2;refGene_gn=ZBTB20;refGene_tx=XM_019839766.2;refGene_gn=ZBTB20;refGene_tx=XM_019839769.2;refGene_gn=ZBTB20;refGene_tx=XM_023260830.1;refGene_gn=ZBTB20;refGene_tx=XM_023260831.1;refGene_gn=ZBTB20;refGene_tx=XM_023260832.1;refGene_gn=ZBTB20;refGene_tx=XM_023260833.1;refGene_gn=ZBTB20;refGene_tx=XM_023260834.1;refGene_gn=ZBTB20;refGene_tx=XM_023260835.1;refGene_gn=ZBTB20;refGene_tx=XM_023260836.1;refGene_gn=ZBTB20;refGene_tx=XM_023260837.1;refGene_gn=ZBTB20;refGene_tx=XM_023260838.1;refGene_gn=ZBTB20;refGene_tx=XM_023260840.1
C2 60884474 60884485
refGene_gn=ZBTB20;refGene_tx=XM_019839752.2;refGene_gn=ZBTB20;refGene_tx=XM_019839757.2;refGene_gn=ZBTB20;refGene_tx=XM_019839758.2;refGene_gn=ZBTB20;refGene_tx=XM_019839761.2;refGene_gn=ZBTB20;refGene_tx=XM_019839762.2;refGene_gn=ZBTB20;refGene_tx=XM_019839766.2;refGene_gn=ZBTB20;refGene_tx=XM_019839769.2;refGene_gn=ZBTB20;refGene_tx=XM_023260830.1;refGene_gn=ZBTB20;refGene_tx=XM_023260831.1;refGene_gn=ZBTB20;refGene_tx=XM_023260832.1;refGene_gn=ZBTB20;refGene_tx=XM_023260833.1;refGene_gn=ZBTB20;refGene_tx=XM_023260834.1;refGene_gn=ZBTB20;refGene_tx=XM_023260835.1;refGene_gn=ZBTB20;refGene_tx=XM_023260836.1;refGene_gn=ZBTB20;refGene_tx=XM_023260837.1;refGene_gn=ZBTB20;refGene_tx=XM_023260838.1;refGene_gn=ZBTB20;refGene_tx=XM_023260840.1
C2 62021938 62021959 refGene_gn=GAP43;refGene_tx=XM_019839229.1
C2 62047739 62047769 refGene_gn=GAP43;refGene_tx=NM_001009303.1
C2 62105263 62105873
refGene_gn=GAP43;refGene_tx=NM_001009303.1;refGene_gn=GAP43;refGene_tx=XM_019839229.1
C2 62149492 62149581
refGene_gn=GAP43;refGene_tx=NM_001009303.1;refGene_gn=GAP43;refGene_tx=XM_019839229.1
C2 62220816 62220914

refGene_gn=LSAMP;refGene_tx=XM_003991647.5;refGene_gn=LSAMP;refGene_tx=XM_006936060.4;refGene_gn=LSAMP;refGene_tx=XM_023260151.1
C2 62227033 62227066 refGene_gn=LSAMP;refGene_tx=XM_006936060.4
C2 62241868 62241948 refGene_gn=LSAMP;refGene_tx=XM_023260152.1
C2 62241975 62242016 refGene_gn=LSAMP;refGene_tx=XM_023260153.1
C2 62244920 62244956
refGene_gn=LSAMP;refGene_tx=XM_006936060.4;refGene_gn=LSAMP;refGene_tx=XM_023260151.1;refGene_gn=LSAMP;refGene_tx=XM_023260152.1;refGene_gn=LSAMP;refGene_tx=XM_023260153.1
C2 62251920 62252069
refGene_gn=LSAMP;refGene_tx=XM_003991647.5;refGene_gn=LSAMP;refGene_tx=XM_006936060.4;refGene_gn=LSAMP;refGene_tx=XM_023260151.1;refGene_gn=LSAMP;refGene_tx=XM_023260152.1;refGene_gn=LSAMP;refGene_tx=XM_023260153.1
C2 62252558 62252679
refGene_gn=LSAMP;refGene_tx=XM_003991647.5;refGene_gn=LSAMP;refGene_tx=XM_006936060.4;refGene_gn=LSAMP;refGene_tx=XM_023260151.1;refGene_gn=LSAMP;refGene_tx=XM_023260152.1;refGene_gn=LSAMP;refGene_tx=XM_023260153.1
C2 62263003 62263138
refGene_gn=LSAMP;refGene_tx=XM_003991647.5;refGene_gn=LSAMP;refGene_tx=XM_006936060.4;refGene_gn=LSAMP;refGene_tx=XM_023260151.1;refGene_gn=LSAMP;refGene_tx=XM_023260152.1;refGene_gn=LSAMP;refGene_tx=XM_023260153.1
C2 62449462 62449588
refGene_gn=LSAMP;refGene_tx=XM_003991647.5;refGene_gn=LSAMP;refGene_tx=XM_006936060.4;refGene_gn=LSAMP;refGene_tx=XM_023260151.1;refGene_gn=LSAMP;refGene_tx=XM_023260152.1;refGene_gn=LSAMP;refGene_tx=XM_023260153.1
C2 62524426 62524659
refGene_gn=LSAMP;refGene_tx=XM_003991647.5;refGene_gn=LSAMP;refGene_tx=XM_006936060.4;refGene_gn=LSAMP;refGene_tx=XM_023260151.1;refGene_gn=LSAMP;refGene_tx=XM_023260152.1;refGene_gn=LSAMP;refGene_tx=XM_023260153.1
C2 62870620 62870775
refGene_gn=LSAMP;refGene_tx=XM_003991647.5;refGene_gn=LSAMP;refGene_tx=XM_006936060.4;refGene_gn=LSAMP;refGene_tx=XM_023260151.1;refGene_gn=LSAMP;refGene_tx=XM_023260152.1;refGene_gn=LSAMP;refGene_tx=XM_023260153.1
C2 62904644 62904797
refGene_gn=LOC101098816;refGene_tx=XM_003991660.2
C2 62904881 62904992
refGene_gn=LOC101098816;refGene_tx=XM_003991660.2
C2 62970103 62970256
refGene_gn=LOC101098816;refGene_tx=XM_003991660.2
C2 65169322 65169752
refGene_gn=IGSF11;refGene_tx=XM_023260192.1;refGene_gn=IGSF11;refGene_tx=XM_023260193.1;refGene_gn=IGSF11;refGene_tx=XM_023260194.1
C2 65171586 65171737
refGene_gn=IGSF11;refGene_tx=XM_023260192.1;refGene_gn=IGSF11;refGene_

tx=XM_023260193.1;refGene_gn=IGSF11;refGene_tx=XM_023260194.1
C2	65172566 65172689
refGene_gn=IGSF11;refGene_tx=XM_023260192.1;refGene_gn=IGSF11;refGene_tx=XM_023260193.1;refGene_gn=IGSF11;refGene_tx=XM_023260194.1
C2	65187946 65188102
refGene_gn=IGSF11;refGene_tx=XM_023260192.1;refGene_gn=IGSF11;refGene_tx=XM_023260194.1
C2	65190795 65191003
refGene_gn=IGSF11;refGene_tx=XM_023260192.1;refGene_gn=IGSF11;refGene_tx=XM_023260193.1;refGene_gn=IGSF11;refGene_tx=XM_023260194.1
C2	65192412 65192451
refGene_gn=IGSF11;refGene_tx=XM_023260192.1;refGene_gn=IGSF11;refGene_tx=XM_023260193.1;refGene_gn=IGSF11;refGene_tx=XM_023260194.1
C2	65300191 65300273
refGene_gn=LOC111562229;refGene_tx=XM_023260167.1
C2	65311471 65311613
refGene_gn=LOC111562229;refGene_tx=XM_023260167.1
C2	65314996 65315055
refGene_gn=LOC111562229;refGene_tx=XM_023260167.1
C2	65315230 65315511
refGene_gn=LOC111562229;refGene_tx=XM_023260167.1
C2	65322912 65323029
refGene_gn=LOC111562229;refGene_tx=XM_023260167.1
C2	65386873 65388076
refGene_gn=CC2H3orf30;refGene_tx=XM_019839777.1;refGene_gn=CC2H3orf30;refGene_tx=XM_019839779.1;refGene_gn=CC2H3orf30;refGene_tx=XM_019839780.1;refGene_gn=CC2H3orf30;refGene_tx=XM_019839781.1;refGene_gn=CC2H3orf30;refGene_tx=XM_019839782.1;refGene_gn=CC2H3orf30;refGene_tx=XM_019839783.1;refGene_gn=CC2H3orf30;refGene_tx=XM_019839784.1;refGene_gn=CC2H3orf30;refGene_tx=XM_019839787.1
C2	65388638 65388803
refGene_gn=CC2H3orf30;refGene_tx=XM_019839777.1;refGene_gn=CC2H3orf30;refGene_tx=XM_019839779.1;refGene_gn=CC2H3orf30;refGene_tx=XM_019839780.1;refGene_gn=CC2H3orf30;refGene_tx=XM_019839781.1;refGene_gn=CC2H3orf30;refGene_tx=XM_019839782.1;refGene_gn=CC2H3orf30;refGene_tx=XM_019839783.1;refGene_gn=CC2H3orf30;refGene_tx=XM_019839784.1;refGene_gn=CC2H3orf30;refGene_tx=XM_019839787.1
C2	65390597 65390660
refGene_gn=CC2H3orf30;refGene_tx=XM_019839777.1;refGene_gn=CC2H3orf30;refGene_tx=XM_019839779.1;refGene_gn=CC2H3orf30;refGene_tx=XM_019839780.1;refGene_gn=CC2H3orf30;refGene_tx=XM_019839781.1;refGene_gn=CC2H3orf30;refGene_tx=XM_019839782.1;refGene_gn=CC2H3orf30;refGene_tx=XM_019839783.1;refGene_gn=CC2H3orf30;refGene_tx=XM_019839784.1
C2	65396099 65396198
refGene_gn=CC2H3orf30;refGene_tx=XM_019839777.1;refGene_gn=CC2H3orf30;refGene_tx=XM_019839779.1;refGene_gn=CC2H3orf30;refGene_tx=XM_019839780.1
C2	65403112 65403163
refGene_gn=CC2H3orf30;refGene_tx=XM_019839777.1;refGene_gn=CC2H3orf30;refGene_tx=XM_019839779.1;refGene_gn=CC2H3orf30;refGene_tx=XM_01983978

0.1;refGene_gn=CC2H3orf30;refGene_tx=XM_019839781.1;refGene_gn=CC2H3orf30;refGene_tx=XM_019839782.1;refGene_gn=CC2H3orf30;refGene_tx=XM_019839787.1
C2 65419300 65419348 refGene_gn=CC2H3orf30;refGene_tx=XM_019839783.1;refGene_gn=CC2H3orf30;refGene_tx=XM_019839784.1
C2 65430330 65430399 refGene_gn=UPK1B;refGene_tx=XM_003991661.4
C2 65431536 65431737 refGene_gn=UPK1B;refGene_tx=XM_003991661.4
C2 65434092 65434167 refGene_gn=UPK1B;refGene_tx=XM_003991661.4
C2 65435103 65435226 refGene_gn=UPK1B;refGene_tx=XM_003991661.4
C2 65437315 65437495 refGene_gn=UPK1B;refGene_tx=XM_003991661.4
C2 65444202 65444286 refGene_gn=UPK1B;refGene_tx=XM_003991661.4
C2 65447793 65447844 refGene_gn=UPK1B;refGene_tx=XM_003991661.4
C2 65455771 65455904 refGene_gn=B4GALT4;refGene_tx=XM_023260195.1;refGene_gn=B4GALT4;refGene_tx=XM_023260196.1
C2 65456202 65456301 refGene_gn=B4GALT4;refGene_tx=XM_023260195.1;refGene_gn=B4GALT4;refGene_tx=XM_023260198.1
C2 65456312 65456367 refGene_gn=B4GALT4;refGene_tx=XM_023260199.1
C2 65456626 65456741 refGene_gn=B4GALT4;refGene_tx=XM_023260197.1
C2 65459168 65459273 refGene_gn=B4GALT4;refGene_tx=XM_023260195.1;refGene_gn=B4GALT4;refGene_tx=XM_023260196.1;refGene_gn=B4GALT4;refGene_tx=XM_023260197.1;refGene_gn=B4GALT4;refGene_tx=XM_023260198.1;refGene_gn=B4GALT4;refGene_tx=XM_023260199.1
C2 65461338 65461461 refGene_gn=B4GALT4;refGene_tx=XM_023260195.1;refGene_gn=B4GALT4;refGene_tx=XM_023260196.1;refGene_gn=B4GALT4;refGene_tx=XM_023260197.1;refGene_gn=B4GALT4;refGene_tx=XM_023260198.1;refGene_gn=B4GALT4;refGene_tx=XM_023260199.1
C2 65466286 65466474 refGene_gn=B4GALT4;refGene_tx=XM_023260195.1;refGene_gn=B4GALT4;refGene_tx=XM_023260196.1;refGene_gn=B4GALT4;refGene_tx=XM_023260197.1;refGene_gn=B4GALT4;refGene_tx=XM_023260198.1;refGene_gn=B4GALT4;refGene_tx=XM_023260199.1
C2 65468748 65468981 refGene_gn=B4GALT4;refGene_tx=XM_023260195.1;refGene_gn=B4GALT4;refGene_tx=XM_023260196.1;refGene_gn=B4GALT4;refGene_tx=XM_023260197.1;refGene_gn=B4GALT4;refGene_tx=XM_023260198.1;refGene_gn=B4GALT4;refGene_tx=XM_023260199.1
C2 65472012 65472265 refGene_gn=B4GALT4;refGene_tx=XM_023260195.1;refGene_gn=B4GALT4;refGene_tx=XM_023260196.1;refGene_gn=B4GALT4;refGene_tx=XM_023260197.1;refGene_gn=B4GALT4;refGene_tx=XM_023260198.1;refGene_gn=B4GALT4;refGene_tx=XM_023260199.1
C2 65519921 65520021 refGene_gn=ARHGAP31;refGene_tx=XM_003991663.5
C2 65585131 65585234 refGene_gn=ARHGAP31;refGene_tx=XM_003991663.5
C2 65588152 65588297 refGene_gn=ARHGAP31;refGene_tx=XM_003991663.5
C2 65601136 65601219 refGene_gn=ARHGAP31;refGene_tx=XM_003991663.5

C2 65602167 65602275 refGene_gn=ARHGAP31;refGene_tx=XM_003991663.5
C2 65603383 65603526 refGene_gn=ARHGAP31;refGene_tx=XM_003991663.5
C2 65608013 65608215 refGene_gn=ARHGAP31;refGene_tx=XM_003991663.5
C2 65610261 65610386 refGene_gn=ARHGAP31;refGene_tx=XM_003991663.5
C2 65614701 65614764 refGene_gn=ARHGAP31;refGene_tx=XM_003991663.5
C2 65616822 65617398 refGene_gn=ARHGAP31;refGene_tx=XM_003991663.5
C2 65620422 65620703 refGene_gn=ARHGAP31;refGene_tx=XM_003991663.5
C2 65624834 65627258 refGene_gn=ARHGAP31;refGene_tx=XM_003991663.5
C2 65638129 65638363 refGene_gn=TMEM39A;refGene_tx=XM_003991664.4
C2 65641508 65641629 refGene_gn=TMEM39A;refGene_tx=XM_003991664.4
C2 65643557 65643745 refGene_gn=TMEM39A;refGene_tx=XM_003991664.4
C2 65644520 65644869 refGene_gn=TMEM39A;refGene_tx=XM_003991664.4
C2 65651900 65652055 refGene_gn=TMEM39A;refGene_tx=XM_003991664.4
C2 65655838 65655922 refGene_gn=TMEM39A;refGene_tx=XM_003991664.4
C2 65662119 65662342 refGene_gn=TMEM39A;refGene_tx=XM_003991664.4
C2 65667356 65667469 refGene_gn=TMEM39A;refGene_tx=XM_003991664.4
C2 65673073 65673158 refGene_gn=POGLUT1;refGene_tx=XM_011285936.3
C2 65673948 65674039 refGene_gn=POGLUT1;refGene_tx=XM_011285936.3
C2 65675140 65675284 refGene_gn=POGLUT1;refGene_tx=XM_011285936.3
C2 65684285 65684421 refGene_gn=POGLUT1;refGene_tx=XM_011285936.3
C2 65686227 65686349 refGene_gn=POGLUT1;refGene_tx=XM_011285936.3
C2 65691928 65691988 refGene_gn=POGLUT1;refGene_tx=XM_011285936.3
C2 65693420 65693520 refGene_gn=POGLUT1;refGene_tx=XM_011285936.3
C2 65695344 65695403 refGene_gn=POGLUT1;refGene_tx=XM_011285936.3
C2 65696435 65696603 refGene_gn=POGLUT1;refGene_tx=XM_011285936.3
C2 65697751 65697808 refGene_gn=POGLUT1;refGene_tx=XM_011285936.3
C2 65698500 65698657 refGene_gn=POGLUT1;refGene_tx=XM_011285936.3
C2 65704669 65704851 refGene_gn=TIMMDC1;refGene_tx=XM_003991665.4
C2 65707253 65707419 refGene_gn=TIMMDC1;refGene_tx=XM_003991665.4
C2 65711914 65712003 refGene_gn=TIMMDC1;refGene_tx=XM_003991665.4
C2 65712442 65712510 refGene_gn=TIMMDC1;refGene_tx=XM_003991665.4
C2 65723052 65723131 refGene_gn=TIMMDC1;refGene_tx=XM_003991665.4
C2 65727956 65728067 refGene_gn=TIMMDC1;refGene_tx=XM_003991665.4
C2 65730688 65730839 refGene_gn=TIMMDC1;refGene_tx=XM_003991665.4
C2 65736206 65736280 refGene_gn=CD80;refGene_tx=XM_019839227.2
C2 65736848 65737216
refGene_gn=CD80;refGene_tx=XM_019839223.2;refGene_gn=CD80;refGene_tx=XM_019839224.2;refGene_gn=CD80;refGene_tx=XM_019839227.2
C2 65741828 65742110
refGene_gn=CD80;refGene_tx=XM_019839223.2;refGene_gn=CD80;refGene_tx=XM_019839224.2;refGene_gn=CD80;refGene_tx=XM_019839227.2
C2 65746872 65747190
refGene_gn=CD80;refGene_tx=NM_001009253.1;refGene_gn=CD80;refGene_tx=XM_019839223.2;refGene_gn=CD80;refGene_tx=XM_019839224.2;refGene_gn=CD80;refGene_tx=XM_019839227.2
C2 65756526 65756626
refGene_gn=CD80;refGene_tx=NM_001009253.1;refGene_gn=CD80;refGene_tx=XM_019839223.2;refGene_gn=CD80;refGene_tx=XM_019839224.2;refGene_gn=CD80;refGene_tx=XM_019839227.2
C2 65777809 65778122

refGene_gn=ADPRH;refGene_tx=XM_019839795.2;refGene_gn=ADPRH;refGene_tx=XM_019839796.2;refGene_gn=ADPRH;refGene_tx=XM_019839797.2;refGene_gn=ADPRH;refGene_tx=XM_019839798.2;refGene_gn=ADPRH;refGene_tx=XM_019839801.2;refGene_gn=ADPRH;refGene_tx=XM_023260200.1
C2 65779203 65779564
refGene_gn=ADPRH;refGene_tx=XM_006936075.3;refGene_gn=ADPRH;refGene_tx=XM_019839795.2;refGene_gn=ADPRH;refGene_tx=XM_019839796.2;refGene_gn=ADPRH;refGene_tx=XM_019839797.2;refGene_gn=ADPRH;refGene_tx=XM_019839798.2;refGene_gn=ADPRH;refGene_tx=XM_019839801.2;refGene_gn=ADPRH;refGene_tx=XM_023260200.1
C2 65781002 65781417 refGene_gn=ADPRH;refGene_tx=XM_019839801.2
C2 65789470 65789583
refGene_gn=ADPRH;refGene_tx=XM_003991667.4;refGene_gn=ADPRH;refGene_tx=XM_006936075.3;refGene_gn=ADPRH;refGene_tx=XM_019839795.2;refGene_gn=ADPRH;refGene_tx=XM_019839796.2;refGene_gn=ADPRH;refGene_tx=XM_019839797.2;refGene_gn=ADPRH;refGene_tx=XM_019839798.2;refGene_gn=ADPRH;refGene_tx=XM_019839799.2;refGene_gn=ADPRH;refGene_tx=XM_019839800.1;refGene_gn=ADPRH;refGene_tx=XM_023260200.1
C2 65794130 65794332
refGene_gn=ADPRH;refGene_tx=XM_003991667.4;refGene_gn=ADPRH;refGene_tx=XM_006936075.3;refGene_gn=ADPRH;refGene_tx=XM_019839795.2;refGene_gn=ADPRH;refGene_tx=XM_019839796.2;refGene_gn=ADPRH;refGene_tx=XM_019839797.2;refGene_gn=ADPRH;refGene_tx=XM_019839798.2;refGene_gn=ADPRH;refGene_tx=XM_019839799.2;refGene_gn=ADPRH;refGene_tx=XM_019839800.1;refGene_gn=ADPRH;refGene_tx=XM_023260200.1
C2 65796351 65796529
refGene_gn=ADPRH;refGene_tx=XM_003991667.4;refGene_gn=ADPRH;refGene_tx=XM_006936075.3;refGene_gn=ADPRH;refGene_tx=XM_019839795.2;refGene_gn=ADPRH;refGene_tx=XM_019839796.2;refGene_gn=ADPRH;refGene_tx=XM_019839797.2;refGene_gn=ADPRH;refGene_tx=XM_019839798.2;refGene_gn=ADPRH;refGene_tx=XM_019839799.2;refGene_gn=ADPRH;refGene_tx=XM_019839800.1;refGene_gn=ADPRH;refGene_tx=XM_023260200.1
C2 65797028 65797137
refGene_gn=ADPRH;refGene_tx=XM_003991667.4;refGene_gn=ADPRH;refGene_tx=XM_006936075.3;refGene_gn=ADPRH;refGene_tx=XM_019839795.2;refGene_gn=ADPRH;refGene_tx=XM_019839796.2;refGene_gn=ADPRH;refGene_tx=XM_019839797.2;refGene_gn=ADPRH;refGene_tx=XM_019839798.2;refGene_gn=ADPRH;refGene_tx=XM_019839799.2;refGene_gn=ADPRH;refGene_tx=XM_019839800.1;refGene_gn=ADPRH;refGene_tx=XM_023260200.1
C2 65798140 65798242
refGene_gn=ADPRH;refGene_tx=XM_003991667.4;refGene_gn=ADPRH;refGene_tx=XM_006936075.3;refGene_gn=ADPRH;refGene_tx=XM_019839795.2;refGene_gn=ADPRH;refGene_tx=XM_019839796.2;refGene_gn=ADPRH;refGene_tx=XM_019839797.2;refGene_gn=ADPRH;refGene_tx=XM_019839798.2;refGene_gn=ADPRH;refGene_tx=XM_019839799.2;refGene_gn=ADPRH;refGene_tx=XM_019839800.1;refGene_gn=ADPRH;refGene_tx=XM_023260200.1
C2 65801073 65801163
refGene_gn=ADPRH;refGene_tx=XM_003991667.4;refGene_gn=ADPRH;refGene_tx=XM_006936075.3;refGene_gn=ADPRH;refGene_tx=XM_019839795.2;refGene_gn=ADPRH;refGene_tx=XM_019839796.2;refGene_gn=ADPRH;refGene_tx=XM_0198397

97.2;refGene_gn=ADPRH;refGene_tx=XM_019839798.2;refGene_gn=ADPRH;refGene_tx=XM_019839799.2;refGene_gn=ADPRH;refGene_tx=XM_019839800.1;refGene_gn=ADPRH;refGene_tx=XM_023260200.1
C2 65803755 65803923
refGene_gn=ADPRH;refGene_tx=XM_003991667.4;refGene_gn=ADPRH;refGene_tx=XM_006936075.3;refGene_gn=ADPRH;refGene_tx=XM_019839795.2;refGene_gn=ADPRH;refGene_tx=XM_019839796.2;refGene_gn=ADPRH;refGene_tx=XM_019839797.2;refGene_gn=ADPRH;refGene_tx=XM_019839798.2;refGene_gn=ADPRH;refGene_tx=XM_019839799.2;refGene_gn=ADPRH;refGene_tx=XM_019839800.1;refGene_gn=ADPRH;refGene_tx=XM_023260200.1
C2 65805262 65805352
refGene_gn=ADPRH;refGene_tx=XM_003991667.4;refGene_gn=ADPRH;refGene_tx=XM_006936075.3;refGene_gn=ADPRH;refGene_tx=XM_019839795.2;refGene_gn=ADPRH;refGene_tx=XM_019839796.2;refGene_gn=ADPRH;refGene_tx=XM_019839797.2;refGene_gn=ADPRH;refGene_tx=XM_019839798.2;refGene_gn=ADPRH;refGene_tx=XM_019839799.2;refGene_gn=ADPRH;refGene_tx=XM_019839800.1;refGene_gn=ADPRH;refGene_tx=XM_023260200.1
C2 65808918 65809031
refGene_gn=ADPRH;refGene_tx=XM_003991667.4;refGene_gn=ADPRH;refGene_tx=XM_006936075.3;refGene_gn=ADPRH;refGene_tx=XM_019839795.2;refGene_gn=ADPRH;refGene_tx=XM_019839796.2;refGene_gn=ADPRH;refGene_tx=XM_019839797.2;refGene_gn=ADPRH;refGene_tx=XM_019839798.2;refGene_gn=ADPRH;refGene_tx=XM_019839799.2;refGene_gn=ADPRH;refGene_tx=XM_019839800.1;refGene_gn=ADPRH;refGene_tx=XM_023260200.1
C2 65812494 65812659
refGene_gn=ADPRH;refGene_tx=XM_003991667.4;refGene_gn=ADPRH;refGene_tx=XM_006936075.3;refGene_gn=ADPRH;refGene_tx=XM_019839795.2;refGene_gn=ADPRH;refGene_tx=XM_019839796.2;refGene_gn=ADPRH;refGene_tx=XM_019839797.2;refGene_gn=ADPRH;refGene_tx=XM_019839798.2;refGene_gn=ADPRH;refGene_tx=XM_019839799.2;refGene_gn=ADPRH;refGene_tx=XM_019839800.1;refGene_gn=ADPRH;refGene_tx=XM_023260200.1
C2 65813258 65813343
refGene_gn=ADPRH;refGene_tx=XM_003991667.4;refGene_gn=ADPRH;refGene_tx=XM_006936075.3;refGene_gn=ADPRH;refGene_tx=XM_019839795.2;refGene_gn=ADPRH;refGene_tx=XM_019839797.2;refGene_gn=ADPRH;refGene_tx=XM_019839799.2;refGene_gn=ADPRH;refGene_tx=XM_019839800.1;refGene_gn=ADPRH;refGene_tx=XM_023260200.1
C2 65827559 65827645 refGene_gn=POPDC2;refGene_tx=XM_003991669.4
C2 65828477 65828551 refGene_gn=POPDC2;refGene_tx=XM_019839810.2
C2 65833820 65834309
refGene_gn=POPDC2;refGene_tx=XM_003991669.4;refGene_gn=POPDC2;refGene_tx=XM_006936077.3;refGene_gn=POPDC2;refGene_tx=XM_019839810.2
C2 65840578 65840687
refGene_gn=POPDC2;refGene_tx=XM_003991669.4;refGene_gn=POPDC2;refGene_tx=XM_006936077.3;refGene_gn=POPDC2;refGene_tx=XM_019839810.2
C2 65845209 65845700
refGene_gn=POPDC2;refGene_tx=XM_003991669.4;refGene_gn=POPDC2;refGene_tx=XM_006936077.3;refGene_gn=POPDC2;refGene_tx=XM_019839810.2
C2 65859006 65859091
refGene_gn=LOC101080819;refGene_tx=XM_006936078.3

C2 65861286 65861393
refGene_gn=LOC101080819;refGene_tx=XM_006936078.3
C2 65874212 65874385
refGene_gn=MAATS1;refGene_tx=XM_006936079.3;refGene_gn=MAATS1;refGene_tx=XM_019839802.2;refGene_gn=MAATS1;refGene_tx=XM_019839803.2;refGene_gn=MAATS1;refGene_tx=XM_019839804.2
C2 65879526 65879603
refGene_gn=MAATS1;refGene_tx=XM_006936079.3;refGene_gn=MAATS1;refGene_tx=XM_019839802.2
C2 65880113 65880271
refGene_gn=MAATS1;refGene_tx=XM_006936079.3;refGene_gn=MAATS1;refGene_tx=XM_019839802.2;refGene_gn=MAATS1;refGene_tx=XM_019839803.2;refGene_gn=MAATS1;refGene_tx=XM_019839805.2
C2 65885317 65885401
refGene_gn=MAATS1;refGene_tx=XM_006936079.3;refGene_gn=MAATS1;refGene_tx=XM_019839802.2;refGene_gn=MAATS1;refGene_tx=XM_019839803.2;refGene_gn=MAATS1;refGene_tx=XM_019839804.2;refGene_gn=MAATS1;refGene_tx=XM_019839805.2;refGene_gn=MAATS1;refGene_tx=XM_019839807.2
C2 65886947 65887004
refGene_gn=MAATS1;refGene_tx=XM_006936079.3;refGene_gn=MAATS1;refGene_tx=XM_019839802.2;refGene_gn=MAATS1;refGene_tx=XM_019839803.2;refGene_gn=MAATS1;refGene_tx=XM_019839804.2;refGene_gn=MAATS1;refGene_tx=XM_019839805.2;refGene_gn=MAATS1;refGene_tx=XM_019839807.2
C2 65900276 65900458
refGene_gn=MAATS1;refGene_tx=XM_006936079.3;refGene_gn=MAATS1;refGene_tx=XM_019839802.2;refGene_gn=MAATS1;refGene_tx=XM_019839803.2;refGene_gn=MAATS1;refGene_tx=XM_019839804.2;refGene_gn=MAATS1;refGene_tx=XM_019839805.2;refGene_gn=MAATS1;refGene_tx=XM_019839807.2
C2 65914279 65914457
refGene_gn=MAATS1;refGene_tx=XM_006936079.3;refGene_gn=MAATS1;refGene_tx=XM_019839802.2;refGene_gn=MAATS1;refGene_tx=XM_019839803.2;refGene_gn=MAATS1;refGene_tx=XM_019839804.2;refGene_gn=MAATS1;refGene_tx=XM_019839805.2;refGene_gn=MAATS1;refGene_tx=XM_019839807.2
C2 65918972 65919130
refGene_gn=MAATS1;refGene_tx=XM_006936079.3;refGene_gn=MAATS1;refGene_tx=XM_019839802.2;refGene_gn=MAATS1;refGene_tx=XM_019839803.2;refGene_gn=MAATS1;refGene_tx=XM_019839804.2;refGene_gn=MAATS1;refGene_tx=XM_019839805.2;refGene_gn=MAATS1;refGene_tx=XM_019839807.2
C2 65921395 65921578
refGene_gn=MAATS1;refGene_tx=XM_006936079.3;refGene_gn=MAATS1;refGene_tx=XM_019839802.2;refGene_gn=MAATS1;refGene_tx=XM_019839803.2;refGene_gn=MAATS1;refGene_tx=XM_019839804.2;refGene_gn=MAATS1;refGene_tx=XM_019839805.2;refGene_gn=MAATS1;refGene_tx=XM_019839807.2
C2 65922458 65922601
refGene_gn=MAATS1;refGene_tx=XM_006936079.3;refGene_gn=MAATS1;refGene_tx=XM_019839802.2;refGene_gn=MAATS1;refGene_tx=XM_019839803.2;refGene_gn=MAATS1;refGene_tx=XM_019839804.2;refGene_gn=MAATS1;refGene_tx=XM_019839805.2;refGene_gn=MAATS1;refGene_tx=XM_019839807.2
C2 65928440 65928557
refGene_gn=MAATS1;refGene_tx=XM_006936079.3;refGene_gn=MAATS1;refGene_

tx=XM_019839802.2;refGene_gn=MAATS1;refGene_tx=XM_019839803.2;refGene_gn=MAATS1;refGene_tx=XM_019839804.2;refGene_gn=MAATS1;refGene_tx=XM_019839805.2;refGene_gn=MAATS1;refGene_tx=XM_019839807.2
C2 65929904 65929976
refGene_gn=MAATS1;refGene_tx=XM_006936079.3;refGene_gn=MAATS1;refGene_tx=XM_019839802.2;refGene_gn=MAATS1;refGene_tx=XM_019839803.2;refGene_gn=MAATS1;refGene_tx=XM_019839804.2;refGene_gn=MAATS1;refGene_tx=XM_019839805.2;refGene_gn=MAATS1;refGene_tx=XM_019839807.2
C2 65931513 65931660
refGene_gn=MAATS1;refGene_tx=XM_006936079.3;refGene_gn=MAATS1;refGene_tx=XM_019839802.2;refGene_gn=MAATS1;refGene_tx=XM_019839803.2;refGene_gn=MAATS1;refGene_tx=XM_019839804.2;refGene_gn=MAATS1;refGene_tx=XM_019839805.2;refGene_gn=MAATS1;refGene_tx=XM_019839807.2
C2 65933135 65933357
refGene_gn=MAATS1;refGene_tx=XM_006936079.3;refGene_gn=MAATS1;refGene_tx=XM_019839802.2;refGene_gn=MAATS1;refGene_tx=XM_019839803.2;refGene_gn=MAATS1;refGene_tx=XM_019839804.2;refGene_gn=MAATS1;refGene_tx=XM_019839805.2;refGene_gn=MAATS1;refGene_tx=XM_019839807.2
C2 65937731 65937880
refGene_gn=MAATS1;refGene_tx=XM_006936079.3;refGene_gn=MAATS1;refGene_tx=XM_019839802.2;refGene_gn=MAATS1;refGene_tx=XM_019839803.2;refGene_gn=MAATS1;refGene_tx=XM_019839804.2;refGene_gn=MAATS1;refGene_tx=XM_019839805.2;refGene_gn=MAATS1;refGene_tx=XM_019839807.2
C2 65938465 65938557
refGene_gn=MAATS1;refGene_tx=XM_006936079.3;refGene_gn=MAATS1;refGene_tx=XM_019839802.2;refGene_gn=MAATS1;refGene_tx=XM_019839803.2;refGene_gn=MAATS1;refGene_tx=XM_019839804.2;refGene_gn=MAATS1;refGene_tx=XM_019839805.2;refGene_gn=MAATS1;refGene_tx=XM_019839807.2
C2 65951404 65951565
refGene_gn=MAATS1;refGene_tx=XM_006936079.3;refGene_gn=MAATS1;refGene_tx=XM_019839802.2;refGene_gn=MAATS1;refGene_tx=XM_019839803.2;refGene_gn=MAATS1;refGene_tx=XM_019839804.2;refGene_gn=MAATS1;refGene_tx=XM_019839805.2;refGene_gn=MAATS1;refGene_tx=XM_019839807.2
C2 65991933 65992538
refGene_gn=NR1I2;refGene_tx=XM_019839808.2;refGene_gn=NR1I2;refGene_tx=XM_023260201.1
C2 65993620 65993754
refGene_gn=NR1I2;refGene_tx=XM_019839808.2;refGene_gn=NR1I2;refGene_tx=XM_023260201.1
C2 65995073 65995261
refGene_gn=NR1I2;refGene_tx=XM_019839808.2;refGene_gn=NR1I2;refGene_tx=XM_023260201.1
C2 65996373 65996648
refGene_gn=NR1I2;refGene_tx=XM_019839808.2;refGene_gn=NR1I2;refGene_tx=XM_023260201.1
C2 65998370 65998513
refGene_gn=NR1I2;refGene_tx=XM_019839808.2;refGene_gn=NR1I2;refGene_tx=XM_023260201.1
C2 65998681 65998798
refGene_gn=NR1I2;refGene_tx=XM_019839808.2;refGene_gn=NR1I2;refGene_tx

=XM_023260201.1
C2 65999181 65999287
refGene_gn=NR1I2;refGene_tx=XM_019839808.2;refGene_gn=NR1I2;refGene_tx
=XM_023260201.1
C2 66000669 66000814
refGene_gn=NR1I2;refGene_tx=XM_019839808.2;refGene_gn=NR1I2;refGene_tx
=XM_023260201.1
C2 66009439 66009507
refGene_gn=GSK3B;refGene_tx=XM_003991674.4;refGene_gn=GSK3B;refGene_tx
=XM_003991675.4
C2 66018121 66018126 refGene_gn=GSK3B;refGene_tx=XM_019839811.2
C2 66023504 66023603
refGene_gn=GSK3B;refGene_tx=XM_003991674.4;refGene_gn=GSK3B;refGene_tx
=XM_003991675.4;refGene_gn=GSK3B;refGene_tx=XM_019839811.2
C2 66035217 66035404
refGene_gn=GSK3B;refGene_tx=XM_003991674.4;refGene_gn=GSK3B;refGene_tx
=XM_003991675.4;refGene_gn=GSK3B;refGene_tx=XM_019839811.2
C2 66038070 66038109
refGene_gn=GSK3B;refGene_tx=XM_003991675.4;refGene_gn=GSK3B;refGene_tx
=XM_019839811.2
C2 66046553 66046649
refGene_gn=GSK3B;refGene_tx=XM_003991674.4;refGene_gn=GSK3B;refGene_tx
=XM_003991675.4;refGene_gn=GSK3B;refGene_tx=XM_019839811.2
C2 66070230 66070328
refGene_gn=GSK3B;refGene_tx=XM_003991674.4;refGene_gn=GSK3B;refGene_tx
=XM_003991675.4;refGene_gn=GSK3B;refGene_tx=XM_019839811.2
C2 66078886 66078993
refGene_gn=GSK3B;refGene_tx=XM_003991674.4;refGene_gn=GSK3B;refGene_tx
=XM_003991675.4;refGene_gn=GSK3B;refGene_tx=XM_019839811.2
C2 66082191 66082322
refGene_gn=GSK3B;refGene_tx=XM_003991674.4;refGene_gn=GSK3B;refGene_tx
=XM_003991675.4;refGene_gn=GSK3B;refGene_tx=XM_019839811.2
C2 66094721 66094832
refGene_gn=GSK3B;refGene_tx=XM_003991674.4;refGene_gn=GSK3B;refGene_tx
=XM_003991675.4;refGene_gn=GSK3B;refGene_tx=XM_019839811.2
C2 66113119 66113203
refGene_gn=GSK3B;refGene_tx=XM_003991674.4;refGene_gn=GSK3B;refGene_tx
=XM_003991675.4;refGene_gn=GSK3B;refGene_tx=XM_019839811.2
C2 66137424 66137618
refGene_gn=GSK3B;refGene_tx=XM_003991674.4;refGene_gn=GSK3B;refGene_tx
=XM_003991675.4;refGene_gn=GSK3B;refGene_tx=XM_019839811.2
C2 66202678 66202766
refGene_gn=GSK3B;refGene_tx=XM_003991674.4;refGene_gn=GSK3B;refGene_tx
=XM_003991675.4;refGene_gn=GSK3B;refGene_tx=XM_019839811.2
C2 66202948 66202972
refGene_gn=LOC109503463;refGene_tx=XM_023260168.1
C2 66203232 66203904
refGene_gn=LOC109503463;refGene_tx=XM_023260168.1
C2 66267767 66269102 refGene_gn=GPR156;refGene_tx=XM_019839812.2
C2 66273476 66273635 refGene_gn=GPR156;refGene_tx=XM_019839812.2

C2 66279877 66280114 refGene_gn=GPR156;refGene_tx=XM_019839812.2
C2 66284262 66284358 refGene_gn=GPR156;refGene_tx=XM_019839812.2
C2 66285330 66285456 refGene_gn=GPR156;refGene_tx=XM_019839812.2
C2 66290517 66290628 refGene_gn=GPR156;refGene_tx=XM_019839812.2
C2 66290877 66291013 refGene_gn=GPR156;refGene_tx=XM_019839812.2
C2 66331226 66331363 refGene_gn=GPR156;refGene_tx=XM_019839812.2
C2 66331604 66331697 refGene_gn=GPR156;refGene_tx=XM_019839812.2
C2 66362735 66362748 refGene_gn=LRRC58;refGene_tx=XM_023260203.1
C2 66392012 66392067 refGene_gn=LRRC58;refGene_tx=XM_023260203.1
C2 66420550 66420759 refGene_gn=LRRC58;refGene_tx=XM_023260202.1
C2 66423989 66424267
refGene_gn=LRRC58;refGene_tx=XM_023260202.1;refGene_gn=LRRC58;refGene_tx=XM_023260203.1
C2 66424914 66425043
refGene_gn=LRRC58;refGene_tx=XM_023260202.1;refGene_gn=LRRC58;refGene_tx=XM_023260203.1
C2 66440518 66441018
refGene_gn=LRRC58;refGene_tx=XM_023260202.1;refGene_gn=LRRC58;refGene_tx=XM_023260203.1
C2 66481186 66481231 refGene_gn=FSTL1;refGene_tx=XM_003991677.3
C2 66484350 66484427 refGene_gn=FSTL1;refGene_tx=XM_003991677.3
C2 66486795 66486906 refGene_gn=FSTL1;refGene_tx=XM_003991677.3
C2 66487243 66487356 refGene_gn=FSTL1;refGene_tx=XM_003991677.3
C2 66488708 66488827 refGene_gn=FSTL1;refGene_tx=XM_003991677.3
C2 66493469 66493600 refGene_gn=FSTL1;refGene_tx=XM_003991677.3
C2 66494923 66494956 refGene_gn=FSTL1;refGene_tx=XM_003991677.3
C2 66495796 66495926 refGene_gn=FSTL1;refGene_tx=XM_003991677.3
C2 66498181 66498286 refGene_gn=FSTL1;refGene_tx=XM_003991677.3
C2 66532440 66532497 refGene_gn=FSTL1;refGene_tx=XM_003991677.3
C2 66532658 66532661 refGene_gn=FSTL1;refGene_tx=XM_003991677.3
C2 66639760 66639940
refGene_gn=NDUFB4;refGene_tx=XM_011285932.3;refGene_gn=NDUFB4;refGene_tx=XM_023260204.1
C2 66644589 66644736
refGene_gn=NDUFB4;refGene_tx=XM_011285932.3;refGene_gn=NDUFB4;refGene_tx=XM_023260204.1
C2 66647037 66647100 refGene_gn=NDUFB4;refGene_tx=XM_011285932.3
C2 66648380 66648464 refGene_gn=NDUFB4;refGene_tx=XM_023260204.1
C2 66663534 66663684 refGene_gn=HGD;refGene_tx=XM_006936082.3
C2 66666769 66666951 refGene_gn=HGD;refGene_tx=XM_006936082.3
C2 66672223 66672350 refGene_gn=HGD;refGene_tx=XM_006936082.3
C2 66674842 66674947 refGene_gn=HGD;refGene_tx=XM_006936082.3
C2 66677361 66677486 refGene_gn=HGD;refGene_tx=XM_006936082.3
C2 66679573 66679673 refGene_gn=HGD;refGene_tx=XM_006936082.3
C2 66680492 66680572 refGene_gn=HGD;refGene_tx=XM_006936082.3
C2 66681383 66681418 refGene_gn=HGD;refGene_tx=XM_006936082.3
C2 66682563 66682655 refGene_gn=HGD;refGene_tx=XM_006936082.3
C2 66684345 66684405 refGene_gn=HGD;refGene_tx=XM_006936082.3
C2 66701234 66701340 refGene_gn=HGD;refGene_tx=XM_006936082.3
C2 66705317 66705406 refGene_gn=HGD;refGene_tx=XM_006936082.3

C2 66706223 66706295 refGene_gn=HGD;refGene_tx=XM_006936082.3
C2 66711769 66711784 refGene_gn=HGD;refGene_tx=XM_006936082.3
C2 66719978 66720044 refGene_gn=RABL3;refGene_tx=XM_003991680.5
C2 66720606 66720645 refGene_gn=RABL3;refGene_tx=XM_003991680.5
C2 66723722 66723794 refGene_gn=RABL3;refGene_tx=XM_003991680.5
C2 66727952 66728103 refGene_gn=RABL3;refGene_tx=XM_003991680.5
C2 66731392 66731507 refGene_gn=RABL3;refGene_tx=XM_003991680.5
C2 66735390 66735520 refGene_gn=RABL3;refGene_tx=XM_003991680.5
C2 66748978 66749070 refGene_gn=RABL3;refGene_tx=XM_003991680.5
C2 66753652 66753698 refGene_gn=RABL3;refGene_tx=XM_003991680.5
C2 66758753 66759201
refGene_gn=GTF2E1;refGene_tx=XM_006936083.3;refGene_gn=GTF2E1;refGene_tx=XM_006936084.4;refGene_gn=GTF2E1;refGene_tx=XM_006936085.3;refGene_gn=GTF2E1;refGene_tx=XM_006936086.4;refGene_gn=GTF2E1;refGene_tx=XM_019839817.2;refGene_gn=GTF2E1;refGene_tx=XM_019839818.1
C2 66776825 66777027
refGene_gn=GTF2E1;refGene_tx=XM_006936083.3;refGene_gn=GTF2E1;refGene_tx=XM_006936084.4;refGene_gn=GTF2E1;refGene_tx=XM_006936085.3;refGene_gn=GTF2E1;refGene_tx=XM_006936086.4;refGene_gn=GTF2E1;refGene_tx=XM_019839817.2;refGene_gn=GTF2E1;refGene_tx=XM_019839818.1
C2 66782788 66783027
refGene_gn=GTF2E1;refGene_tx=XM_006936083.3;refGene_gn=GTF2E1;refGene_tx=XM_006936084.4;refGene_gn=GTF2E1;refGene_tx=XM_006936085.3;refGene_gn=GTF2E1;refGene_tx=XM_006936086.4;refGene_gn=GTF2E1;refGene_tx=XM_019839817.2;refGene_gn=GTF2E1;refGene_tx=XM_019839818.1
C2 66788050 66788478
refGene_gn=GTF2E1;refGene_tx=XM_006936083.3;refGene_gn=GTF2E1;refGene_tx=XM_006936084.4;refGene_gn=GTF2E1;refGene_tx=XM_006936085.3;refGene_gn=GTF2E1;refGene_tx=XM_006936086.4;refGene_gn=GTF2E1;refGene_tx=XM_019839817.2;refGene_gn=GTF2E1;refGene_tx=XM_019839818.1
C2 66924590 66924779 refGene_gn=STXBP5L;refGene_tx=XM_011285929.3
C2 66975484 66975582 refGene_gn=STXBP5L;refGene_tx=XM_011285929.3
C2 67008593 67008675 refGene_gn=STXBP5L;refGene_tx=XM_011285929.3
C2 67013022 67013123 refGene_gn=STXBP5L;refGene_tx=XM_011285929.3
C2 67055922 67056057 refGene_gn=STXBP5L;refGene_tx=XM_011285929.3
C2 67062687 67062751 refGene_gn=STXBP5L;refGene_tx=XM_011285929.3
C2 67102161 67102245 refGene_gn=STXBP5L;refGene_tx=XM_011285929.3
C2 67107140 67107264 refGene_gn=STXBP5L;refGene_tx=XM_011285929.3
C2 67122815 67122894 refGene_gn=STXBP5L;refGene_tx=XM_011285929.3
C2 67129496 67129651 refGene_gn=STXBP5L;refGene_tx=XM_011285929.3
C2 67134827 67134900 refGene_gn=STXBP5L;refGene_tx=XM_011285929.3
C2 67152316 67152464 refGene_gn=STXBP5L;refGene_tx=XM_011285929.3
C2 67153339 67153407 refGene_gn=STXBP5L;refGene_tx=XM_011285929.3
C2 67182518 67182559 refGene_gn=STXBP5L;refGene_tx=XM_011285929.3
C2 67185925 67186063 refGene_gn=STXBP5L;refGene_tx=XM_011285929.3
C2 67186065 67186142 refGene_gn=STXBP5L;refGene_tx=XM_011285929.3
C2 67187862 67188032 refGene_gn=STXBP5L;refGene_tx=XM_011285929.3
C2 67189925 67190051 refGene_gn=STXBP5L;refGene_tx=XM_011285929.3
C2 67207755 67207907 refGene_gn=STXBP5L;refGene_tx=XM_011285929.3
C2 67209877 67209949 refGene_gn=STXBP5L;refGene_tx=XM_011285929.3

| | | | |
|---|---|---|---|
| C2 | 67232570 | 67232636 | refGene_gn=STXBP5L;refGene_tx=XM_011285929.3 |
| C2 | 67245970 | 67246063 | refGene_gn=STXBP5L;refGene_tx=XM_011285929.3 |
| C2 | 67289333 | 67289504 | refGene_gn=STXBP5L;refGene_tx=XM_011285929.3 |
| C2 | 67291598 | 67291838 | refGene_gn=STXBP5L;refGene_tx=XM_011285929.3 |
| C2 | 67300067 | 67300428 | refGene_gn=STXBP5L;refGene_tx=XM_011285929.3 |
| C2 | 67305617 | 67305783 | refGene_gn=STXBP5L;refGene_tx=XM_011285929.3 |
| C2 | 67308215 | 67308308 | refGene_gn=STXBP5L;refGene_tx=XM_011285929.3 |
| C2 | 67308309 | 67308328 | refGene_gn=STXBP5L;refGene_tx=XM_011285929.3 |
| C2 | 67310002 | 67310223 | refGene_gn=STXBP5L;refGene_tx=XM_011285929.3 |
| C2 | 67310787 | 67310829 | refGene_gn=STXBP5L;refGene_tx=XM_011285929.3 |
| C2 | 67338405 | 67338519 | refGene_gn=POLQ;refGene_tx=XM_023260169.1 |
| C2 | 67339236 | 67339352 | refGene_gn=POLQ;refGene_tx=XM_023260169.1 |
| C2 | 67342364 | 67342518 | refGene_gn=POLQ;refGene_tx=XM_023260169.1 |
| C2 | 67351637 | 67351762 | refGene_gn=POLQ;refGene_tx=XM_023260169.1 |
| C2 | 67358262 | 67358374 | refGene_gn=POLQ;refGene_tx=XM_023260169.1 |
| C2 | 67361644 | 67361829 | refGene_gn=POLQ;refGene_tx=XM_023260169.1 |
| C2 | 67367230 | 67367352 | refGene_gn=POLQ;refGene_tx=XM_023260169.1 |
| C2 | 67368252 | 67368379 | refGene_gn=POLQ;refGene_tx=XM_023260169.1 |
| C2 | 67370804 | 67370979 | refGene_gn=POLQ;refGene_tx=XM_023260169.1 |
| C2 | 67372423 | 67372561 | refGene_gn=POLQ;refGene_tx=XM_023260169.1 |
| C2 | 67375659 | 67375853 | refGene_gn=POLQ;refGene_tx=XM_023260169.1 |
| C2 | 67377057 | 67377298 | refGene_gn=POLQ;refGene_tx=XM_023260169.1 |
| C2 | 67385115 | 67385312 | refGene_gn=POLQ;refGene_tx=XM_023260169.1 |
| C2 | 67386578 | 67386722 | refGene_gn=POLQ;refGene_tx=XM_023260169.1 |
| C2 | 67389442 | 67392552 | refGene_gn=POLQ;refGene_tx=XM_023260169.1 |
| C2 | 67406220 | 67406411 | refGene_gn=POLQ;refGene_tx=XM_023260169.1 |
| C2 | 67408469 | 67408588 | refGene_gn=POLQ;refGene_tx=XM_023260169.1 |
| C2 | 67409074 | 67409199 | refGene_gn=POLQ;refGene_tx=XM_023260169.1 |
| C2 | 67410103 | 67410297 | refGene_gn=POLQ;refGene_tx=XM_023260169.1 |
| C2 | 67425330 | 67425473 | refGene_gn=POLQ;refGene_tx=XM_023260169.1 |
| C2 | 67425797 | 67426002 | refGene_gn=POLQ;refGene_tx=XM_023260169.1 |
| C2 | 67427764 | 67427907 | refGene_gn=POLQ;refGene_tx=XM_023260169.1 |
| C2 | 67431518 | 67431731 | refGene_gn=POLQ;refGene_tx=XM_023260169.1 |
| C2 | 67439206 | 67439353 | refGene_gn=POLQ;refGene_tx=XM_023260169.1 |
| C2 | 67443342 | 67443490 | refGene_gn=POLQ;refGene_tx=XM_023260169.1 |
| C2 | 67446073 | 67446293 | refGene_gn=POLQ;refGene_tx=XM_023260169.1 |
| C2 | 67449985 | 67450094 | refGene_gn=POLQ;refGene_tx=XM_023260169.1 |
| C2 | 67451985 | 67452142 | refGene_gn=POLQ;refGene_tx=XM_023260169.1 |
| C2 | 67454024 | 67454155 | refGene_gn=POLQ;refGene_tx=XM_023260169.1 |
| C2 | 67455426 | 67455606 | refGene_gn=POLQ;refGene_tx=XM_023260169.1 |
| C2 | 67456069 | 67456250 | refGene_gn=POLQ;refGene_tx=XM_023260169.1 |
| C2 | 67507266 | 67507269 | refGene_gn=FBXO40;refGene_tx=XM_006936088.3 |
| C2 | 67508327 | 67510247 | refGene_gn=FBXO40;refGene_tx=XM_006936088.3 |
| C2 | 67512930 | 67513146 | refGene_gn=FBXO40;refGene_tx=XM_006936088.3 |
| C2 | 67517930 | 67518067 | refGene_gn=HCLS1;refGene_tx=XM_003991685.4 |
| C2 | 67518267 | 67518336 | refGene_gn=HCLS1;refGene_tx=XM_003991685.4 |
| C2 | 67518488 | 67518759 | refGene_gn=HCLS1;refGene_tx=XM_003991685.4 |
| C2 | 67519245 | 67519350 | refGene_gn=HCLS1;refGene_tx=XM_003991685.4 |
| C2 | 67519746 | 67519958 | refGene_gn=HCLS1;refGene_tx=XM_003991685.4 |
| C2 | 67520570 | 67520640 | refGene_gn=HCLS1;refGene_tx=XM_003991685.4 |

C2 67521217 67521273 refGene_gn=HCLS1;refGene_tx=XM_003991685.4
C2 67521926 67522037 refGene_gn=HCLS1;refGene_tx=XM_003991685.4
C2 67526381 67526436 refGene_gn=HCLS1;refGene_tx=XM_003991685.4
C2 67527904 67528015 refGene_gn=HCLS1;refGene_tx=XM_003991685.4
C2 67529893 67530023 refGene_gn=HCLS1;refGene_tx=XM_003991685.4
C2 67540104 67540178 refGene_gn=HCLS1;refGene_tx=XM_003991685.4
C2 67540763 67540847 refGene_gn=HCLS1;refGene_tx=XM_003991685.4
C2 67547194 67547329
refGene_gn=GOLGB1;refGene_tx=XM_019839828.2;refGene_gn=GOLGB1;refGene_tx=XM_023260205.1;refGene_gn=GOLGB1;refGene_tx=XM_023260206.1;refGene_gn=GOLGB1;refGene_tx=XM_023260207.1;refGene_gn=GOLGB1;refGene_tx=XM_023260208.1
C2 67547611 67547717
refGene_gn=GOLGB1;refGene_tx=XM_019839828.2;refGene_gn=GOLGB1;refGene_tx=XM_023260205.1;refGene_gn=GOLGB1;refGene_tx=XM_023260206.1;refGene_gn=GOLGB1;refGene_tx=XM_023260207.1;refGene_gn=GOLGB1;refGene_tx=XM_023260208.1
C2 67549316 67549451
refGene_gn=GOLGB1;refGene_tx=XM_019839828.2;refGene_gn=GOLGB1;refGene_tx=XM_023260205.1;refGene_gn=GOLGB1;refGene_tx=XM_023260206.1;refGene_gn=GOLGB1;refGene_tx=XM_023260207.1;refGene_gn=GOLGB1;refGene_tx=XM_023260208.1
C2 67549961 67550059
refGene_gn=GOLGB1;refGene_tx=XM_019839828.2;refGene_gn=GOLGB1;refGene_tx=XM_023260205.1;refGene_gn=GOLGB1;refGene_tx=XM_023260206.1;refGene_gn=GOLGB1;refGene_tx=XM_023260207.1;refGene_gn=GOLGB1;refGene_tx=XM_023260208.1
C2 67550835 67550979
refGene_gn=GOLGB1;refGene_tx=XM_019839828.2;refGene_gn=GOLGB1;refGene_tx=XM_023260205.1;refGene_gn=GOLGB1;refGene_tx=XM_023260206.1;refGene_gn=GOLGB1;refGene_tx=XM_023260207.1;refGene_gn=GOLGB1;refGene_tx=XM_023260208.1
C2 67556841 67556979
refGene_gn=GOLGB1;refGene_tx=XM_019839828.2;refGene_gn=GOLGB1;refGene_tx=XM_023260205.1;refGene_gn=GOLGB1;refGene_tx=XM_023260206.1;refGene_gn=GOLGB1;refGene_tx=XM_023260207.1
C2 67557180 67557346
refGene_gn=GOLGB1;refGene_tx=XM_019839828.2;refGene_gn=GOLGB1;refGene_tx=XM_023260205.1;refGene_gn=GOLGB1;refGene_tx=XM_023260206.1;refGene_gn=GOLGB1;refGene_tx=XM_023260207.1;refGene_gn=GOLGB1;refGene_tx=XM_023260208.1
C2 67563974 67564153
refGene_gn=GOLGB1;refGene_tx=XM_019839828.2;refGene_gn=GOLGB1;refGene_tx=XM_023260205.1;refGene_gn=GOLGB1;refGene_tx=XM_023260206.1;refGene_gn=GOLGB1;refGene_tx=XM_023260207.1;refGene_gn=GOLGB1;refGene_tx=XM_023260208.1;refGene_gn=GOLGB1;refGene_tx=XM_023260209.1
C2 67565244 67565291 refGene_gn=GOLGB1;refGene_tx=XM_023260209.1
C2 67578219 67580134
refGene_gn=GOLGB1;refGene_tx=XM_019839828.2;refGene_gn=GOLGB1;refGene_tx=XM_023260205.1;refGene_gn=GOLGB1;refGene_tx=XM_023260206.1;refGene_

gn=GOLGB1;refGene_tx=XM_023260207.1;refGene_gn=GOLGB1;refGene_tx=XM_023260208.1;refGene_gn=GOLGB1;refGene_tx=XM_023260209.1
C2 67580961 67586147
refGene_gn=GOLGB1;refGene_tx=XM_019839828.2;refGene_gn=GOLGB1;refGene_tx=XM_023260205.1;refGene_gn=GOLGB1;refGene_tx=XM_023260206.1;refGene_gn=GOLGB1;refGene_tx=XM_023260207.1;refGene_gn=GOLGB1;refGene_tx=XM_023260208.1;refGene_gn=GOLGB1;refGene_tx=XM_023260209.1
C2 67589232 67589306
refGene_gn=GOLGB1;refGene_tx=XM_019839828.2;refGene_gn=GOLGB1;refGene_tx=XM_023260205.1;refGene_gn=GOLGB1;refGene_tx=XM_023260206.1;refGene_gn=GOLGB1;refGene_tx=XM_023260207.1;refGene_gn=GOLGB1;refGene_tx=XM_023260208.1;refGene_gn=GOLGB1;refGene_tx=XM_023260209.1
C2 67591740 67591855
refGene_gn=GOLGB1;refGene_tx=XM_019839828.2;refGene_gn=GOLGB1;refGene_tx=XM_023260205.1;refGene_gn=GOLGB1;refGene_tx=XM_023260206.1;refGene_gn=GOLGB1;refGene_tx=XM_023260207.1;refGene_gn=GOLGB1;refGene_tx=XM_023260208.1;refGene_gn=GOLGB1;refGene_tx=XM_023260209.1
C2 67598550 67598642
refGene_gn=GOLGB1;refGene_tx=XM_019839828.2;refGene_gn=GOLGB1;refGene_tx=XM_023260205.1;refGene_gn=GOLGB1;refGene_tx=XM_023260206.1;refGene_gn=GOLGB1;refGene_tx=XM_023260207.1;refGene_gn=GOLGB1;refGene_tx=XM_023260208.1;refGene_gn=GOLGB1;refGene_tx=XM_023260209.1
C2 67600892 67601295
refGene_gn=GOLGB1;refGene_tx=XM_019839828.2;refGene_gn=GOLGB1;refGene_tx=XM_023260205.1;refGene_gn=GOLGB1;refGene_tx=XM_023260206.1;refGene_gn=GOLGB1;refGene_tx=XM_023260207.1;refGene_gn=GOLGB1;refGene_tx=XM_023260208.1;refGene_gn=GOLGB1;refGene_tx=XM_023260209.1
C2 67603289 67603403
refGene_gn=GOLGB1;refGene_tx=XM_019839828.2;refGene_gn=GOLGB1;refGene_tx=XM_023260205.1;refGene_gn=GOLGB1;refGene_tx=XM_023260206.1;refGene_gn=GOLGB1;refGene_tx=XM_023260207.1;refGene_gn=GOLGB1;refGene_tx=XM_023260208.1;refGene_gn=GOLGB1;refGene_tx=XM_023260209.1
C2 67604505 67604628
refGene_gn=GOLGB1;refGene_tx=XM_019839828.2;refGene_gn=GOLGB1;refGene_tx=XM_023260205.1;refGene_gn=GOLGB1;refGene_tx=XM_023260206.1;refGene_gn=GOLGB1;refGene_tx=XM_023260208.1;refGene_gn=GOLGB1;refGene_tx=XM_023260209.1
C2 67607232 67607349
refGene_gn=GOLGB1;refGene_tx=XM_019839828.2;refGene_gn=GOLGB1;refGene_tx=XM_023260205.1;refGene_gn=GOLGB1;refGene_tx=XM_023260206.1;refGene_gn=GOLGB1;refGene_tx=XM_023260207.1;refGene_gn=GOLGB1;refGene_tx=XM_023260208.1;refGene_gn=GOLGB1;refGene_tx=XM_023260209.1
C2 67609973 67610102
refGene_gn=GOLGB1;refGene_tx=XM_019839828.2;refGene_gn=GOLGB1;refGene_tx=XM_023260205.1;refGene_gn=GOLGB1;refGene_tx=XM_023260206.1;refGene_gn=GOLGB1;refGene_tx=XM_023260207.1;refGene_gn=GOLGB1;refGene_tx=XM_023260208.1;refGene_gn=GOLGB1;refGene_tx=XM_023260209.1
C2 67612594 67612747
refGene_gn=GOLGB1;refGene_tx=XM_019839828.2;refGene_gn=GOLGB1;refGene_tx=XM_023260205.1;refGene_gn=GOLGB1;refGene_tx=XM_023260206.1;refGene_

gn=GOLGB1;refGene_tx=XM_023260207.1;refGene_gn=GOLGB1;refGene_tx=XM_02
3260208.1;refGene_gn=GOLGB1;refGene_tx=XM_023260209.1
C2	67612957 67613110
refGene_gn=GOLGB1;refGene_tx=XM_019839828.2;refGene_gn=GOLGB1;refGene_
tx=XM_023260205.1;refGene_gn=GOLGB1;refGene_tx=XM_023260206.1;refGene_
gn=GOLGB1;refGene_tx=XM_023260207.1;refGene_gn=GOLGB1;refGene_tx=XM_02
3260208.1;refGene_gn=GOLGB1;refGene_tx=XM_023260209.1
C2	67613911 67614004
refGene_gn=GOLGB1;refGene_tx=XM_019839828.2;refGene_gn=GOLGB1;refGene_
tx=XM_023260205.1;refGene_gn=GOLGB1;refGene_tx=XM_023260206.1;refGene_
gn=GOLGB1;refGene_tx=XM_023260207.1;refGene_gn=GOLGB1;refGene_tx=XM_02
3260208.1;refGene_gn=GOLGB1;refGene_tx=XM_023260209.1
C2	67671682 67671912
refGene_gn=IQCB1;refGene_tx=XM_006936090.3;refGene_gn=IQCB1;refGene_tx
=XM_006936091.3;refGene_gn=IQCB1;refGene_tx=XM_006936092.4;refGene_gn=
IQCB1;refGene_tx=XM_006936093.3;refGene_gn=IQCB1;refGene_tx=XM_0112859
22.3;refGene_gn=IQCB1;refGene_tx=XM_011285923.3;refGene_gn=IQCB1;refGe
ne_tx=XM_011285924.3;refGene_gn=IQCB1;refGene_tx=XM_011285925.3;refGen
e_gn=IQCB1;refGene_tx=XM_011285926.3;refGene_gn=IQCB1;refGene_tx=XM_01
9839830.2;refGene_gn=IQCB1;refGene_tx=XM_019839831.2
C2	67672986 67673143
refGene_gn=IQCB1;refGene_tx=XM_006936090.3;refGene_gn=IQCB1;refGene_tx
=XM_006936091.3;refGene_gn=IQCB1;refGene_tx=XM_006936092.4;refGene_gn=
IQCB1;refGene_tx=XM_006936093.3;refGene_gn=IQCB1;refGene_tx=XM_0112859
22.3;refGene_gn=IQCB1;refGene_tx=XM_011285923.3;refGene_gn=IQCB1;refGe
ne_tx=XM_011285924.3;refGene_gn=IQCB1;refGene_tx=XM_011285925.3;refGen
e_gn=IQCB1;refGene_tx=XM_011285926.3;refGene_gn=IQCB1;refGene_tx=XM_01
9839830.2;refGene_gn=IQCB1;refGene_tx=XM_019839831.2
C2	67683062 67683194
refGene_gn=IQCB1;refGene_tx=XM_006936090.3;refGene_gn=IQCB1;refGene_tx
=XM_006936091.3;refGene_gn=IQCB1;refGene_tx=XM_006936092.4;refGene_gn=
IQCB1;refGene_tx=XM_006936093.3;refGene_gn=IQCB1;refGene_tx=XM_0112859
22.3;refGene_gn=IQCB1;refGene_tx=XM_011285923.3;refGene_gn=IQCB1;refGe
ne_tx=XM_011285924.3;refGene_gn=IQCB1;refGene_tx=XM_011285925.3;refGen
e_gn=IQCB1;refGene_tx=XM_011285926.3;refGene_gn=IQCB1;refGene_tx=XM_01
9839830.2;refGene_gn=IQCB1;refGene_tx=XM_019839831.2
C2	67688640 67688789
refGene_gn=IQCB1;refGene_tx=XM_006936090.3;refGene_gn=IQCB1;refGene_tx
=XM_006936091.3;refGene_gn=IQCB1;refGene_tx=XM_006936092.4;refGene_gn=
IQCB1;refGene_tx=XM_006936093.3;refGene_gn=IQCB1;refGene_tx=XM_0112859
22.3;refGene_gn=IQCB1;refGene_tx=XM_011285923.3;refGene_gn=IQCB1;refGe
ne_tx=XM_011285924.3;refGene_gn=IQCB1;refGene_tx=XM_011285925.3;refGen
e_gn=IQCB1;refGene_tx=XM_011285926.3;refGene_gn=IQCB1;refGene_tx=XM_01
9839830.2;refGene_gn=IQCB1;refGene_tx=XM_019839831.2
C2	67690297 67690440
refGene_gn=IQCB1;refGene_tx=XM_006936090.3;refGene_gn=IQCB1;refGene_tx
=XM_006936091.3;refGene_gn=IQCB1;refGene_tx=XM_006936092.4;refGene_gn=
IQCB1;refGene_tx=XM_006936093.3;refGene_gn=IQCB1;refGene_tx=XM_0112859
22.3;refGene_gn=IQCB1;refGene_tx=XM_011285923.3;refGene_gn=IQCB1;refGe
ne_tx=XM_011285924.3;refGene_gn=IQCB1;refGene_tx=XM_011285925.3;refGen

e_gn=IQCB1;refGene_tx=XM_011285926.3;refGene_gn=IQCB1;refGene_tx=XM_019839830.2;refGene_gn=IQCB1;refGene_tx=XM_019839831.2
C2 67702631 67702741
refGene_gn=IQCB1;refGene_tx=XM_006936090.3;refGene_gn=IQCB1;refGene_tx=XM_006936091.3;refGene_gn=IQCB1;refGene_tx=XM_006936092.4;refGene_gn=IQCB1;refGene_tx=XM_006936093.3;refGene_gn=IQCB1;refGene_tx=XM_011285922.3;refGene_gn=IQCB1;refGene_tx=XM_011285923.3;refGene_gn=IQCB1;refGene_tx=XM_011285924.3;refGene_gn=IQCB1;refGene_tx=XM_011285925.3;refGene_gn=IQCB1;refGene_tx=XM_011285926.3;refGene_gn=IQCB1;refGene_tx=XM_019839830.2;refGene_gn=IQCB1;refGene_tx=XM_019839831.2
C2 67703352 67703462
refGene_gn=IQCB1;refGene_tx=XM_006936090.3;refGene_gn=IQCB1;refGene_tx=XM_006936091.3;refGene_gn=IQCB1;refGene_tx=XM_006936092.4;refGene_gn=IQCB1;refGene_tx=XM_006936093.3;refGene_gn=IQCB1;refGene_tx=XM_011285922.3;refGene_gn=IQCB1;refGene_tx=XM_011285923.3;refGene_gn=IQCB1;refGene_tx=XM_011285924.3;refGene_gn=IQCB1;refGene_tx=XM_011285925.3;refGene_gn=IQCB1;refGene_tx=XM_011285926.3;refGene_gn=IQCB1;refGene_tx=XM_019839830.2;refGene_gn=IQCB1;refGene_tx=XM_019839831.2
C2 67705996 67706175
refGene_gn=IQCB1;refGene_tx=XM_006936090.3;refGene_gn=IQCB1;refGene_tx=XM_006936091.3;refGene_gn=IQCB1;refGene_tx=XM_006936092.4;refGene_gn=IQCB1;refGene_tx=XM_006936093.3;refGene_gn=IQCB1;refGene_tx=XM_011285922.3;refGene_gn=IQCB1;refGene_tx=XM_011285923.3;refGene_gn=IQCB1;refGene_tx=XM_011285924.3;refGene_gn=IQCB1;refGene_tx=XM_011285925.3;refGene_gn=IQCB1;refGene_tx=XM_011285926.3;refGene_gn=IQCB1;refGene_tx=XM_019839830.2;refGene_gn=IQCB1;refGene_tx=XM_019839831.2
C2 67716337 67716437
refGene_gn=IQCB1;refGene_tx=XM_006936090.3;refGene_gn=IQCB1;refGene_tx=XM_006936091.3;refGene_gn=IQCB1;refGene_tx=XM_006936092.4;refGene_gn=IQCB1;refGene_tx=XM_006936093.3;refGene_gn=IQCB1;refGene_tx=XM_011285922.3;refGene_gn=IQCB1;refGene_tx=XM_011285923.3;refGene_gn=IQCB1;refGene_tx=XM_011285924.3;refGene_gn=IQCB1;refGene_tx=XM_011285925.3;refGene_gn=IQCB1;refGene_tx=XM_011285926.3;refGene_gn=IQCB1;refGene_tx=XM_019839830.2;refGene_gn=IQCB1;refGene_tx=XM_019839831.2
C2 67721476 67721570
refGene_gn=IQCB1;refGene_tx=XM_006936090.3;refGene_gn=IQCB1;refGene_tx=XM_006936091.3;refGene_gn=IQCB1;refGene_tx=XM_006936092.4;refGene_gn=IQCB1;refGene_tx=XM_006936093.3;refGene_gn=IQCB1;refGene_tx=XM_011285922.3;refGene_gn=IQCB1;refGene_tx=XM_011285923.3;refGene_gn=IQCB1;refGene_tx=XM_011285924.3;refGene_gn=IQCB1;refGene_tx=XM_011285925.3;refGene_gn=IQCB1;refGene_tx=XM_011285926.3;refGene_gn=IQCB1;refGene_tx=XM_019839830.2;refGene_gn=IQCB1;refGene_tx=XM_019839831.2
C2 67726116 67726246
refGene_gn=IQCB1;refGene_tx=XM_006936090.3;refGene_gn=IQCB1;refGene_tx=XM_006936091.3;refGene_gn=IQCB1;refGene_tx=XM_006936092.4;refGene_gn=IQCB1;refGene_tx=XM_006936093.3;refGene_gn=IQCB1;refGene_tx=XM_011285922.3;refGene_gn=IQCB1;refGene_tx=XM_011285923.3;refGene_gn=IQCB1;refGene_tx=XM_011285924.3;refGene_gn=IQCB1;refGene_tx=XM_011285925.3;refGene_gn=IQCB1;refGene_tx=XM_011285926.3;refGene_gn=IQCB1;refGene_tx=XM_019839830.2;refGene_gn=IQCB1;refGene_tx=XM_019839831.2

C2 67728225 67728388
refGene_gn=IQCB1;refGene_tx=XM_006936090.3;refGene_gn=IQCB1;refGene_tx=XM_006936091.3;refGene_gn=IQCB1;refGene_tx=XM_006936092.4;refGene_gn=IQCB1;refGene_tx=XM_006936093.3;refGene_gn=IQCB1;refGene_tx=XM_011285922.3;refGene_gn=IQCB1;refGene_tx=XM_011285923.3;refGene_gn=IQCB1;refGene_tx=XM_011285924.3;refGene_gn=IQCB1;refGene_tx=XM_011285925.3;refGene_gn=IQCB1;refGene_tx=XM_011285926.3;refGene_gn=IQCB1;refGene_tx=XM_019839830.2;refGene_gn=IQCB1;refGene_tx=XM_019839831.2
C2 67728620 67728720
refGene_gn=IQCB1;refGene_tx=XM_006936090.3;refGene_gn=IQCB1;refGene_tx=XM_006936091.3;refGene_gn=IQCB1;refGene_tx=XM_006936092.4;refGene_gn=IQCB1;refGene_tx=XM_006936093.3;refGene_gn=IQCB1;refGene_tx=XM_011285922.3;refGene_gn=IQCB1;refGene_tx=XM_011285923.3;refGene_gn=IQCB1;refGene_tx=XM_011285924.3;refGene_gn=IQCB1;refGene_tx=XM_011285925.3;refGene_gn=IQCB1;refGene_tx=XM_011285926.3;refGene_gn=IQCB1;refGene_tx=XM_019839830.2;refGene_gn=IQCB1;refGene_tx=XM_019839831.2
C2 67734237 67734274 refGene_gn=EAF2;refGene_tx=XM_023260210.1
C2 67734296 67734405 refGene_gn=EAF2;refGene_tx=XM_003991690.5
C2 67742137 67742232
refGene_gn=EAF2;refGene_tx=XM_003991690.5;refGene_gn=EAF2;refGene_tx=XM_023260210.1
C2 67749769 67749906
refGene_gn=EAF2;refGene_tx=XM_003991690.5;refGene_gn=EAF2;refGene_tx=XM_023260210.1
C2 67752696 67752842
refGene_gn=EAF2;refGene_tx=XM_003991690.5;refGene_gn=EAF2;refGene_tx=XM_023260210.1;refGene_gn=EAF2;refGene_tx=XM_023260211.1
C2 67763391 67763652
refGene_gn=EAF2;refGene_tx=XM_003991690.5;refGene_gn=EAF2;refGene_tx=XM_023260210.1;refGene_gn=EAF2;refGene_tx=XM_023260211.1
C2 67768493 67768540
refGene_gn=EAF2;refGene_tx=XM_003991690.5;refGene_gn=EAF2;refGene_tx=XM_023260210.1;refGene_gn=EAF2;refGene_tx=XM_023260211.1
C2 67778747 67778885
refGene_gn=SLC15A2;refGene_tx=XM_003991691.5;refGene_gn=SLC15A2;refGene_tx=XM_019839833.2;refGene_gn=SLC15A2;refGene_tx=XM_019839834.2;refGene_gn=SLC15A2;refGene_tx=XM_019839835.2
C2 67780914 67781002
refGene_gn=SLC15A2;refGene_tx=XM_003991691.5;refGene_gn=SLC15A2;refGene_tx=XM_019839833.2;refGene_gn=SLC15A2;refGene_tx=XM_019839834.2;refGene_gn=SLC15A2;refGene_tx=XM_019839835.2
C2 67781597 67781739
refGene_gn=SLC15A2;refGene_tx=XM_003991691.5;refGene_gn=SLC15A2;refGene_tx=XM_019839833.2;refGene_gn=SLC15A2;refGene_tx=XM_019839834.2;refGene_gn=SLC15A2;refGene_tx=XM_019839835.2
C2 67785201 67785294
refGene_gn=SLC15A2;refGene_tx=XM_003991691.5;refGene_gn=SLC15A2;refGene_tx=XM_019839833.2;refGene_gn=SLC15A2;refGene_tx=XM_019839834.2;refGene_gn=SLC15A2;refGene_tx=XM_019839835.2
C2 67786619 67786719

refGene_gn=SLC15A2;refGene_tx=XM_003991691.5;refGene_gn=SLC15A2;refGene_tx=XM_019839833.2;refGene_gn=SLC15A2;refGene_tx=XM_019839834.2;refGene_gn=SLC15A2;refGene_tx=XM_019839835.2
C2 67788566 67788657
refGene_gn=SLC15A2;refGene_tx=XM_003991691.5;refGene_gn=SLC15A2;refGene_tx=XM_019839833.2;refGene_gn=SLC15A2;refGene_tx=XM_019839834.2;refGene_gn=SLC15A2;refGene_tx=XM_019839835.2
C2 67788925 67789003
refGene_gn=SLC15A2;refGene_tx=XM_003991691.5;refGene_gn=SLC15A2;refGene_tx=XM_019839833.2;refGene_gn=SLC15A2;refGene_tx=XM_019839834.2;refGene_gn=SLC15A2;refGene_tx=XM_019839835.2
C2 67794054 67794137
refGene_gn=SLC15A2;refGene_tx=XM_003991691.5;refGene_gn=SLC15A2;refGene_tx=XM_019839833.2;refGene_gn=SLC15A2;refGene_tx=XM_019839834.2;refGene_gn=SLC15A2;refGene_tx=XM_019839835.2
C2 67794593 67794680
refGene_gn=SLC15A2;refGene_tx=XM_003991691.5;refGene_gn=SLC15A2;refGene_tx=XM_019839833.2;refGene_gn=SLC15A2;refGene_tx=XM_019839834.2;refGene_gn=SLC15A2;refGene_tx=XM_019839835.2
C2 67794851 67794941
refGene_gn=SLC15A2;refGene_tx=XM_003991691.5;refGene_gn=SLC15A2;refGene_tx=XM_019839833.2;refGene_gn=SLC15A2;refGene_tx=XM_019839834.2;refGene_gn=SLC15A2;refGene_tx=XM_019839835.2
C2 67795033 67795078
refGene_gn=SLC15A2;refGene_tx=XM_003991691.5;refGene_gn=SLC15A2;refGene_tx=XM_019839833.2;refGene_gn=SLC15A2;refGene_tx=XM_019839834.2;refGene_gn=SLC15A2;refGene_tx=XM_019839835.2
C2 67796113 67796146
refGene_gn=SLC15A2;refGene_tx=XM_003991691.5;refGene_gn=SLC15A2;refGene_tx=XM_019839833.2;refGene_gn=SLC15A2;refGene_tx=XM_019839834.2;refGene_gn=SLC15A2;refGene_tx=XM_019839835.2
C2 67796826 67796915
refGene_gn=SLC15A2;refGene_tx=XM_003991691.5;refGene_gn=SLC15A2;refGene_tx=XM_019839833.2;refGene_gn=SLC15A2;refGene_tx=XM_019839834.2;refGene_gn=SLC15A2;refGene_tx=XM_019839835.2
C2 67798103 67798185
refGene_gn=SLC15A2;refGene_tx=XM_003991691.5;refGene_gn=SLC15A2;refGene_tx=XM_019839833.2;refGene_gn=SLC15A2;refGene_tx=XM_019839834.2;refGene_gn=SLC15A2;refGene_tx=XM_019839835.2
C2 67798770 67798905
refGene_gn=SLC15A2;refGene_tx=XM_003991691.5;refGene_gn=SLC15A2;refGene_tx=XM_019839833.2;refGene_gn=SLC15A2;refGene_tx=XM_019839834.2;refGene_gn=SLC15A2;refGene_tx=XM_019839835.2
C2 67800293 67800458
refGene_gn=SLC15A2;refGene_tx=XM_003991691.5;refGene_gn=SLC15A2;refGene_tx=XM_019839833.2;refGene_gn=SLC15A2;refGene_tx=XM_019839834.2;refGene_gn=SLC15A2;refGene_tx=XM_019839835.2
C2 67800607 67800654
refGene_gn=SLC15A2;refGene_tx=XM_003991691.5;refGene_gn=SLC15A2;refGene_tx=XM_019839833.2;refGene_gn=SLC15A2;refGene_tx=XM_019839834.2;refGe

ne_gn=SLC15A2;refGene_tx=XM_019839835.2
C2 67801369 67801480
refGene_gn=SLC15A2;refGene_tx=XM_003991691.5;refGene_gn=SLC15A2;refGene_tx=XM_019839833.2;refGene_gn=SLC15A2;refGene_tx=XM_019839834.2;refGene_gn=SLC15A2;refGene_tx=XM_019839835.2
C2 67802080 67802177
refGene_gn=SLC15A2;refGene_tx=XM_003991691.5;refGene_gn=SLC15A2;refGene_tx=XM_019839833.2;refGene_gn=SLC15A2;refGene_tx=XM_019839834.2;refGene_gn=SLC15A2;refGene_tx=XM_019839835.2
C2 67812205 67812352
refGene_gn=SLC15A2;refGene_tx=XM_003991691.5;refGene_gn=SLC15A2;refGene_tx=XM_019839833.2;refGene_gn=SLC15A2;refGene_tx=XM_019839834.2;refGene_gn=SLC15A2;refGene_tx=XM_019839835.2
C2 67813189 67813294
refGene_gn=SLC15A2;refGene_tx=XM_003991691.5;refGene_gn=SLC15A2;refGene_tx=XM_019839833.2;refGene_gn=SLC15A2;refGene_tx=XM_019839834.2;refGene_gn=SLC15A2;refGene_tx=XM_019839835.2
C2 67813629 67813806
refGene_gn=SLC15A2;refGene_tx=XM_003991691.5;refGene_gn=SLC15A2;refGene_tx=XM_019839833.2;refGene_gn=SLC15A2;refGene_tx=XM_019839834.2
C2 67833487 67833529
refGene_gn=ILDR1;refGene_tx=XM_019839837.2;refGene_gn=ILDR1;refGene_tx=XM_019839838.2;refGene_gn=ILDR1;refGene_tx=XM_023260214.1
C2 67835910 67835937 refGene_gn=ILDR1;refGene_tx=XM_023260212.1
C2 67837559 67837580 refGene_gn=ILDR1;refGene_tx=XM_023260213.1
C2 67837696 67838517
refGene_gn=ILDR1;refGene_tx=XM_019839837.2;refGene_gn=ILDR1;refGene_tx=XM_019839838.2;refGene_gn=ILDR1;refGene_tx=XM_023260212.1;refGene_gn=ILDR1;refGene_tx=XM_023260213.1;refGene_gn=ILDR1;refGene_tx=XM_023260214.1
C2 67838731 67838863
refGene_gn=ILDR1;refGene_tx=XM_019839837.2;refGene_gn=ILDR1;refGene_tx=XM_019839838.2;refGene_gn=ILDR1;refGene_tx=XM_023260212.1;refGene_gn=ILDR1;refGene_tx=XM_023260213.1;refGene_gn=ILDR1;refGene_tx=XM_023260214.1
C2 67845195 67845342
refGene_gn=ILDR1;refGene_tx=XM_019839837.2;refGene_gn=ILDR1;refGene_tx=XM_019839838.2;refGene_gn=ILDR1;refGene_tx=XM_023260212.1;refGene_gn=ILDR1;refGene_tx=XM_023260213.1;refGene_gn=ILDR1;refGene_tx=XM_023260214.1
C2 67845644 67845764
refGene_gn=ILDR1;refGene_tx=XM_019839837.2;refGene_gn=ILDR1;refGene_tx=XM_019839838.2;refGene_gn=ILDR1;refGene_tx=XM_023260212.1;refGene_gn=ILDR1;refGene_tx=XM_023260213.1;refGene_gn=ILDR1;refGene_tx=XM_023260214.1
C2 67846125 67846143 refGene_gn=SLC15A2;refGene_tx=XM_019839835.2
C2 67849392 67849542
refGene_gn=ILDR1;refGene_tx=XM_019839837.2;refGene_gn=ILDR1;refGene_tx=XM_019839838.2;refGene_gn=ILDR1;refGene_tx=XM_023260212.1;refGene_gn=ILDR1;refGene_tx=XM_023260213.1;refGene_gn=ILDR1;refGene_tx=XM_0232602

14.1
C2	67850887	67851058
refGene_gn=ILDR1;refGene_tx=XM_019839837.2;refGene_gn=ILDR1;refGene_tx=XM_019839838.2;refGene_gn=ILDR1;refGene_tx=XM_023260212.1;refGene_gn=ILDR1;refGene_tx=XM_023260213.1;refGene_gn=ILDR1;refGene_tx=XM_023260214.1
C2	67862560	67862615
refGene_gn=ILDR1;refGene_tx=XM_019839837.2;refGene_gn=ILDR1;refGene_tx=XM_023260212.1;refGene_gn=ILDR1;refGene_tx=XM_023260213.1;refGene_gn=ILDR1;refGene_tx=XM_023260214.1
C2	67874254	67874264	refGene_gn=ILDR1;refGene_tx=XM_019839838.2
C2	67904896	67904913	refGene_gn=CD86;refGene_tx=NM_001009229.1
C2	67917061	67917111	refGene_gn=CD86;refGene_tx=NM_001009229.1
C2	67929107	67929443	refGene_gn=CD86;refGene_tx=NM_001009229.1
C2	67931741	67932047	refGene_gn=CD86;refGene_tx=NM_001009229.1
C2	67934989	67935139	refGene_gn=CD86;refGene_tx=NM_001009229.1
C2	67944070	67944116	refGene_gn=CD86;refGene_tx=NM_001009229.1
C2	67946052	67946146	refGene_gn=CD86;refGene_tx=NM_001009229.1
C2	68042714	68042899
refGene_gn=CASR;refGene_tx=NM_001164654.1;refGene_gn=CASR;refGene_tx=XM_019839214.1;refGene_gn=CASR;refGene_tx=XM_019839215.2;refGene_gn=CASR;refGene_tx=XM_019839216.2;refGene_gn=CASR;refGene_tx=XM_019839219.1
C2	68045460	68045767
refGene_gn=CASR;refGene_tx=NM_001164654.1;refGene_gn=CASR;refGene_tx=XM_019839214.1;refGene_gn=CASR;refGene_tx=XM_019839215.2;refGene_gn=CASR;refGene_tx=XM_019839216.2
C2	68048690	68049575
refGene_gn=CASR;refGene_tx=NM_001164654.1;refGene_gn=CASR;refGene_tx=XM_019839214.1;refGene_gn=CASR;refGene_tx=XM_019839215.2;refGene_gn=CASR;refGene_tx=XM_019839216.2;refGene_gn=CASR;refGene_tx=XM_019839219.1
C2	68064560	68064791
refGene_gn=CASR;refGene_tx=NM_001164654.1;refGene_gn=CASR;refGene_tx=XM_019839214.1;refGene_gn=CASR;refGene_tx=XM_019839215.2;refGene_gn=CASR;refGene_tx=XM_019839216.2;refGene_gn=CASR;refGene_tx=XM_019839219.1
C2	68068020	68068144
refGene_gn=CASR;refGene_tx=NM_001164654.1;refGene_gn=CASR;refGene_tx=XM_019839214.1;refGene_gn=CASR;refGene_tx=XM_019839215.2;refGene_gn=CASR;refGene_tx=XM_019839216.2;refGene_gn=CASR;refGene_tx=XM_019839219.1
C2	68069429	68070943
refGene_gn=CASR;refGene_tx=NM_001164654.1;refGene_gn=CASR;refGene_tx=XM_019839214.1;refGene_gn=CASR;refGene_tx=XM_019839215.2;refGene_gn=CASR;refGene_tx=XM_019839216.2;refGene_gn=CASR;refGene_tx=XM_019839219.1
C2	68113731	68114226
refGene_gn=LOC101086085;refGene_tx=XM_011285920.3
C2	68118156	68118222	refGene_gn=CSTA;refGene_tx=NM_001009864.1
C2	68126077	68126179	refGene_gn=CSTA;refGene_tx=NM_001009864.1
C2	68129005	68129134	refGene_gn=CSTA;refGene_tx=NM_001009864.1
C2	68142873	68142924	refGene_gn=CCDC58;refGene_tx=XM_003991692.4
C2	68146040	68146100	refGene_gn=CCDC58;refGene_tx=XM_003991692.4
C2	68151638	68151785	refGene_gn=CCDC58;refGene_tx=XM_003991692.4

C2	68157040 68157166 refGene_gn=CCDC58;refGene_tx=XM_003991692.4
C2	68168679 68168730 refGene_gn=CCDC58;refGene_tx=XM_003991692.4
C2	68169957 68169991 refGene_gn=FAM162A;refGene_tx=XM_006936096.4
C2	68188430 68188553 refGene_gn=FAM162A;refGene_tx=XM_006936096.4
C2	68190915 68191024 refGene_gn=FAM162A;refGene_tx=XM_006936096.4
C2	68194226 68194335 refGene_gn=FAM162A;refGene_tx=XM_006936096.4
C2	68196748 68196841 refGene_gn=FAM162A;refGene_tx=XM_006936096.4
C2	68199598 68200588
refGene_gn=WDR5B;refGene_tx=XM_003991693.5;refGene_gn=WDR5B;refGene_tx=XM_006936097.3
C2	68210417 68210605
refGene_gn=KPNA1;refGene_tx=XM_003991694.5;refGene_gn=KPNA1;refGene_tx=XM_019839839.1;refGene_gn=KPNA1;refGene_tx=XM_019839840.2;refGene_gn=KPNA1;refGene_tx=XM_023260215.1
C2	68212108 68212287
refGene_gn=KPNA1;refGene_tx=XM_003991694.5;refGene_gn=KPNA1;refGene_tx=XM_019839839.1;refGene_gn=KPNA1;refGene_tx=XM_019839840.2;refGene_gn=KPNA1;refGene_tx=XM_023260215.1
C2	68216765 68216893
refGene_gn=KPNA1;refGene_tx=XM_003991694.5;refGene_gn=KPNA1;refGene_tx=XM_019839839.1;refGene_gn=KPNA1;refGene_tx=XM_019839840.2;refGene_gn=KPNA1;refGene_tx=XM_023260215.1
C2	68218523 68218649
refGene_gn=KPNA1;refGene_tx=XM_003991694.5;refGene_gn=KPNA1;refGene_tx=XM_019839839.1;refGene_gn=KPNA1;refGene_tx=XM_019839840.2;refGene_gn=KPNA1;refGene_tx=XM_023260215.1
C2	68223396 68223475
refGene_gn=KPNA1;refGene_tx=XM_003991694.5;refGene_gn=KPNA1;refGene_tx=XM_019839839.1;refGene_gn=KPNA1;refGene_tx=XM_019839840.2;refGene_gn=KPNA1;refGene_tx=XM_023260215.1
C2	68224824 68224988
refGene_gn=KPNA1;refGene_tx=XM_003991694.5;refGene_gn=KPNA1;refGene_tx=XM_019839839.1;refGene_gn=KPNA1;refGene_tx=XM_019839840.2;refGene_gn=KPNA1;refGene_tx=XM_023260215.1
C2	68225532 68225632
refGene_gn=KPNA1;refGene_tx=XM_003991694.5;refGene_gn=KPNA1;refGene_tx=XM_019839839.1;refGene_gn=KPNA1;refGene_tx=XM_019839840.2;refGene_gn=KPNA1;refGene_tx=XM_023260215.1
C2	68225971 68226060
refGene_gn=KPNA1;refGene_tx=XM_003991694.5;refGene_gn=KPNA1;refGene_tx=XM_019839839.1;refGene_gn=KPNA1;refGene_tx=XM_019839840.2;refGene_gn=KPNA1;refGene_tx=XM_023260215.1
C2	68227182 68227314
refGene_gn=KPNA1;refGene_tx=XM_003991694.5;refGene_gn=KPNA1;refGene_tx=XM_019839839.1;refGene_gn=KPNA1;refGene_tx=XM_019839840.2;refGene_gn=KPNA1;refGene_tx=XM_023260215.1
C2	68233824 68233919
refGene_gn=KPNA1;refGene_tx=XM_003991694.5;refGene_gn=KPNA1;refGene_tx=XM_019839839.1;refGene_gn=KPNA1;refGene_tx=XM_019839840.2;refGene_gn=KPNA1;refGene_tx=XM_023260215.1

C2 68236877 68236977
refGene_gn=KPNA1;refGene_tx=XM_003991694.5;refGene_gn=KPNA1;refGene_tx=XM_019839839.1;refGene_gn=KPNA1;refGene_tx=XM_019839840.2;refGene_gn=KPNA1;refGene_tx=XM_023260215.1
C2 68240107 68240215
refGene_gn=KPNA1;refGene_tx=XM_003991694.5;refGene_gn=KPNA1;refGene_tx=XM_019839839.1;refGene_gn=KPNA1;refGene_tx=XM_019839840.2;refGene_gn=KPNA1;refGene_tx=XM_023260215.1
C2 68256271 68256405
refGene_gn=KPNA1;refGene_tx=XM_003991694.5;refGene_gn=KPNA1;refGene_tx=XM_019839839.1;refGene_gn=KPNA1;refGene_tx=XM_019839840.2;refGene_gn=KPNA1;refGene_tx=XM_023260215.1
C2 68260718 68260737 refGene_gn=KPNA1;refGene_tx=XM_019839839.1
C2 68294959 68295339
refGene_gn=PARP9;refGene_tx=XM_006936099.4;refGene_gn=PARP9;refGene_tx=XM_006936100.4;refGene_gn=PARP9;refGene_tx=XM_019839841.2
C2 68301646 68301821
refGene_gn=PARP9;refGene_tx=XM_006936099.4;refGene_gn=PARP9;refGene_tx=XM_006936100.4;refGene_gn=PARP9;refGene_tx=XM_019839841.2
C2 68302437 68302574
refGene_gn=PARP9;refGene_tx=XM_006936099.4;refGene_gn=PARP9;refGene_tx=XM_006936100.4;refGene_gn=PARP9;refGene_tx=XM_019839841.2
C2 68306627 68307008
refGene_gn=PARP9;refGene_tx=XM_006936099.4;refGene_gn=PARP9;refGene_tx=XM_006936100.4;refGene_gn=PARP9;refGene_tx=XM_019839841.2
C2 68310945 68311006
refGene_gn=PARP9;refGene_tx=XM_006936099.4;refGene_gn=PARP9;refGene_tx=XM_006936100.4;refGene_gn=PARP9;refGene_tx=XM_019839841.2
C2 68315860 68316079
refGene_gn=PARP9;refGene_tx=XM_006936099.4;refGene_gn=PARP9;refGene_tx=XM_006936100.4;refGene_gn=PARP9;refGene_tx=XM_019839841.2
C2 68317974 68318202
refGene_gn=PARP9;refGene_tx=XM_006936099.4;refGene_gn=PARP9;refGene_tx=XM_006936100.4;refGene_gn=PARP9;refGene_tx=XM_019839841.2
C2 68321961 68322809
refGene_gn=PARP9;refGene_tx=XM_006936099.4;refGene_gn=PARP9;refGene_tx=XM_006936100.4;refGene_gn=PARP9;refGene_tx=XM_019839841.2
C2 68324879 68324913
refGene_gn=PARP9;refGene_tx=XM_006936099.4;refGene_gn=PARP9;refGene_tx=XM_006936100.4;refGene_gn=PARP9;refGene_tx=XM_019839841.2
C2 68325861 68325885
refGene_gn=PARP9;refGene_tx=XM_006936099.4;refGene_gn=PARP9;refGene_tx=XM_006936100.4;refGene_gn=PARP9;refGene_tx=XM_019839841.2
C2 68329891 68330075 refGene_gn=DTX3L;refGene_tx=XM_003991695.5
C2 68331619 68331825 refGene_gn=DTX3L;refGene_tx=XM_003991695.5
C2 68333616 68335155 refGene_gn=DTX3L;refGene_tx=XM_003991695.5
C2 68335579 68335797 refGene_gn=DTX3L;refGene_tx=XM_003991695.5
C2 68338134 68338204 refGene_gn=DTX3L;refGene_tx=XM_003991695.5
C2 68346880 68347093
refGene_gn=PARP15;refGene_tx=XM_023260216.1;refGene_gn=PARP15;refGene_

tx=XM_023260217.1
C2	68354812 68354818 refGene_gn=PARP15;refGene_tx=XM_023260218.1
C2	68365691 68365808
refGene_gn=PARP15;refGene_tx=XM_023260216.1;refGene_gn=PARP15;refGene_tx=XM_023260217.1;refGene_gn=PARP15;refGene_tx=XM_023260218.1;refGene_gn=PARP15;refGene_tx=XM_023260219.1;refGene_gn=PARP15;refGene_tx=XM_023260220.1;refGene_gn=PARP15;refGene_tx=XM_023260221.1;refGene_gn=PARP15;refGene_tx=XM_023260222.1
C2	68369209 68369446
refGene_gn=PARP15;refGene_tx=XM_023260216.1;refGene_gn=PARP15;refGene_tx=XM_023260217.1;refGene_gn=PARP15;refGene_tx=XM_023260218.1;refGene_gn=PARP15;refGene_tx=XM_023260219.1;refGene_gn=PARP15;refGene_tx=XM_023260220.1;refGene_gn=PARP15;refGene_tx=XM_023260221.1;refGene_gn=PARP15;refGene_tx=XM_023260222.1
C2	68370424 68370646
refGene_gn=PARP15;refGene_tx=XM_023260216.1;refGene_gn=PARP15;refGene_tx=XM_023260217.1;refGene_gn=PARP15;refGene_tx=XM_023260218.1;refGene_gn=PARP15;refGene_tx=XM_023260219.1;refGene_gn=PARP15;refGene_tx=XM_023260220.1;refGene_gn=PARP15;refGene_tx=XM_023260221.1;refGene_gn=PARP15;refGene_tx=XM_023260222.1
C2	68371843 68371922
refGene_gn=PARP15;refGene_tx=XM_023260216.1;refGene_gn=PARP15;refGene_tx=XM_023260217.1;refGene_gn=PARP15;refGene_tx=XM_023260218.1;refGene_gn=PARP15;refGene_tx=XM_023260219.1;refGene_gn=PARP15;refGene_tx=XM_023260220.1;refGene_gn=PARP15;refGene_tx=XM_023260221.1;refGene_gn=PARP15;refGene_tx=XM_023260222.1
C2	68374526 68374676
refGene_gn=PARP15;refGene_tx=XM_023260216.1;refGene_gn=PARP15;refGene_tx=XM_023260217.1;refGene_gn=PARP15;refGene_tx=XM_023260218.1;refGene_gn=PARP15;refGene_tx=XM_023260219.1;refGene_gn=PARP15;refGene_tx=XM_023260220.1;refGene_gn=PARP15;refGene_tx=XM_023260221.1;refGene_gn=PARP15;refGene_tx=XM_023260222.1
C2	68380105 68380168
refGene_gn=PARP15;refGene_tx=XM_023260216.1;refGene_gn=PARP15;refGene_tx=XM_023260217.1;refGene_gn=PARP15;refGene_tx=XM_023260218.1;refGene_gn=PARP15;refGene_tx=XM_023260219.1;refGene_gn=PARP15;refGene_tx=XM_023260220.1;refGene_gn=PARP15;refGene_tx=XM_023260221.1;refGene_gn=PARP15;refGene_tx=XM_023260222.1
C2	68383254 68383422
refGene_gn=PARP15;refGene_tx=XM_023260216.1;refGene_gn=PARP15;refGene_tx=XM_023260217.1;refGene_gn=PARP15;refGene_tx=XM_023260218.1;refGene_gn=PARP15;refGene_tx=XM_023260219.1;refGene_gn=PARP15;refGene_tx=XM_023260220.1;refGene_gn=PARP15;refGene_tx=XM_023260221.1;refGene_gn=PARP15;refGene_tx=XM_023260222.1
C2	68386454 68386661
refGene_gn=PARP15;refGene_tx=XM_023260216.1;refGene_gn=PARP15;refGene_tx=XM_023260217.1;refGene_gn=PARP15;refGene_tx=XM_023260218.1;refGene_gn=PARP15;refGene_tx=XM_023260219.1;refGene_gn=PARP15;refGene_tx=XM_023260220.1;refGene_gn=PARP15;refGene_tx=XM_023260221.1;refGene_gn=PARP15;refGene_tx=XM_023260222.1

C2 68390783 68390917
refGene_gn=PARP15;refGene_tx=XM_023260216.1;refGene_gn=PARP15;refGene_tx=XM_023260217.1;refGene_gn=PARP15;refGene_tx=XM_023260218.1;refGene_gn=PARP15;refGene_tx=XM_023260219.1;refGene_gn=PARP15;refGene_tx=XM_023260220.1;refGene_gn=PARP15;refGene_tx=XM_023260221.1;refGene_gn=PARP15;refGene_tx=XM_023260222.1
C2 68394097 68394280
refGene_gn=PARP15;refGene_tx=XM_023260216.1;refGene_gn=PARP15;refGene_tx=XM_023260217.1;refGene_gn=PARP15;refGene_tx=XM_023260218.1;refGene_gn=PARP15;refGene_tx=XM_023260219.1;refGene_gn=PARP15;refGene_tx=XM_023260220.1;refGene_gn=PARP15;refGene_tx=XM_023260221.1;refGene_gn=PARP15;refGene_tx=XM_023260222.1
C2 68394583 68394873
refGene_gn=PARP15;refGene_tx=XM_023260216.1;refGene_gn=PARP15;refGene_tx=XM_023260217.1;refGene_gn=PARP15;refGene_tx=XM_023260218.1;refGene_gn=PARP15;refGene_tx=XM_023260219.1;refGene_gn=PARP15;refGene_tx=XM_023260220.1;refGene_gn=PARP15;refGene_tx=XM_023260221.1;refGene_gn=PARP15;refGene_tx=XM_023260222.1
C2 68424002 68424189
refGene_gn=PARP14;refGene_tx=XM_003991707.5;refGene_gn=PARP14;refGene_tx=XM_011285918.3
C2 68429175 68429309
refGene_gn=PARP14;refGene_tx=XM_003991707.5;refGene_gn=PARP14;refGene_tx=XM_011285918.3
C2 68430070 68430104
refGene_gn=PARP14;refGene_tx=XM_003991707.5;refGene_gn=PARP14;refGene_tx=XM_011285918.3
C2 68434488 68434731
refGene_gn=PARP14;refGene_tx=XM_003991707.5;refGene_gn=PARP14;refGene_tx=XM_011285918.3;refGene_gn=PARP14;refGene_tx=XM_019839847.1
C2 68437412 68437649
refGene_gn=PARP14;refGene_tx=XM_003991707.5;refGene_gn=PARP14;refGene_tx=XM_011285918.3;refGene_gn=PARP14;refGene_tx=XM_019839847.1
C2 68442402 68444651
refGene_gn=PARP14;refGene_tx=XM_003991707.5;refGene_gn=PARP14;refGene_tx=XM_011285918.3;refGene_gn=PARP14;refGene_tx=XM_019839847.1
C2 68445558 68445795
refGene_gn=PARP14;refGene_tx=XM_003991707.5;refGene_gn=PARP14;refGene_tx=XM_011285918.3;refGene_gn=PARP14;refGene_tx=XM_019839847.1
C2 68446335 68446557
refGene_gn=PARP14;refGene_tx=XM_003991707.5;refGene_gn=PARP14;refGene_tx=XM_011285918.3;refGene_gn=PARP14;refGene_tx=XM_019839847.1
C2 68448823 68448902
refGene_gn=PARP14;refGene_tx=XM_003991707.5;refGene_gn=PARP14;refGene_tx=XM_011285918.3;refGene_gn=PARP14;refGene_tx=XM_019839847.1
C2 68451567 68451717
refGene_gn=PARP14;refGene_tx=XM_003991707.5;refGene_gn=PARP14;refGene_tx=XM_011285918.3;refGene_gn=PARP14;refGene_tx=XM_019839847.1
C2 68452248 68452311
refGene_gn=PARP14;refGene_tx=XM_003991707.5;refGene_gn=PARP14;refGene_

tx=XM_011285918.3;refGene_gn=PARP14;refGene_tx=XM_019839847.1
C2	68453077	68453245
refGene_gn=PARP14;refGene_tx=XM_003991707.5;refGene_gn=PARP14;refGene_
tx=XM_011285918.3;refGene_gn=PARP14;refGene_tx=XM_019839847.1
C2	68456695	68456902
refGene_gn=PARP14;refGene_tx=XM_003991707.5;refGene_gn=PARP14;refGene_
tx=XM_011285918.3;refGene_gn=PARP14;refGene_tx=XM_019839847.1
C2	68459542	68460142
refGene_gn=PARP14;refGene_tx=XM_003991707.5;refGene_gn=PARP14;refGene_
tx=XM_011285918.3;refGene_gn=PARP14;refGene_tx=XM_019839847.1
C2	68461646	68461780
refGene_gn=PARP14;refGene_tx=XM_003991707.5;refGene_gn=PARP14;refGene_
tx=XM_019839847.1
C2	68465562	68465737
refGene_gn=PARP14;refGene_tx=XM_003991707.5;refGene_gn=PARP14;refGene_
tx=XM_011285918.3;refGene_gn=PARP14;refGene_tx=XM_019839847.1
C2	68466291	68466581
refGene_gn=PARP14;refGene_tx=XM_003991707.5;refGene_gn=PARP14;refGene_
tx=XM_019839847.1
C2	68471954	68472494
refGene_gn=HSPBAP1;refGene_tx=XM_011285916.2;refGene_gn=HSPBAP1;refGen
e_tx=XM_011285917.3
C2	68472621	68472732	refGene_gn=HSPBAP1;refGene_tx=XM_011285916.2
C2	68483444	68483528
refGene_gn=HSPBAP1;refGene_tx=XM_011285916.2;refGene_gn=HSPBAP1;refGen
e_tx=XM_011285917.3
C2	68486567	68486739
refGene_gn=HSPBAP1;refGene_tx=XM_011285916.2;refGene_gn=HSPBAP1;refGen
e_tx=XM_011285917.3
C2	68490167	68490304
refGene_gn=HSPBAP1;refGene_tx=XM_011285916.2;refGene_gn=HSPBAP1;refGen
e_tx=XM_011285917.3
C2	68505524	68505706
refGene_gn=HSPBAP1;refGene_tx=XM_011285916.2;refGene_gn=HSPBAP1;refGen
e_tx=XM_011285917.3
C2	68513239	68513425
refGene_gn=HSPBAP1;refGene_tx=XM_011285916.2;refGene_gn=HSPBAP1;refGen
e_tx=XM_011285917.3
C2	68530947	68531011
refGene_gn=HSPBAP1;refGene_tx=XM_011285916.2;refGene_gn=HSPBAP1;refGen
e_tx=XM_011285917.3
C2	68532522	68532874	refGene_gn=DIRC2;refGene_tx=XM_003991709.5
C2	68541319	68541413	refGene_gn=DIRC2;refGene_tx=XM_003991709.5
C2	68560121	68560387	refGene_gn=DIRC2;refGene_tx=XM_003991709.5
C2	68565174	68565304	refGene_gn=DIRC2;refGene_tx=XM_003991709.5
C2	68573918	68574027	refGene_gn=DIRC2;refGene_tx=XM_003991709.5
C2	68584719	68584787	refGene_gn=DIRC2;refGene_tx=XM_003991709.5
C2	68587881	68588009	refGene_gn=DIRC2;refGene_tx=XM_003991709.5
C2	68602245	68602428	refGene_gn=DIRC2;refGene_tx=XM_003991709.5
C2	68610532	68610648	refGene_gn=DIRC2;refGene_tx=XM_003991709.5

C2 68643224 68643383
refGene_gn=SEMA5B;refGene_tx=XM_006936108.4;refGene_gn=SEMA5B;refGene_tx=XM_019839850.2;refGene_gn=SEMA5B;refGene_tx=XM_023260223.1;refGene_gn=SEMA5B;refGene_tx=XM_023260224.1;refGene_gn=SEMA5B;refGene_tx=XM_023260225.1
C2 68643611 68643817
refGene_gn=SEMA5B;refGene_tx=XM_006936108.4;refGene_gn=SEMA5B;refGene_tx=XM_019839850.2;refGene_gn=SEMA5B;refGene_tx=XM_023260223.1;refGene_gn=SEMA5B;refGene_tx=XM_023260224.1;refGene_gn=SEMA5B;refGene_tx=XM_023260225.1
C2 68644257 68644302
refGene_gn=SEMA5B;refGene_tx=XM_006936108.4;refGene_gn=SEMA5B;refGene_tx=XM_019839850.2;refGene_gn=SEMA5B;refGene_tx=XM_023260224.1;refGene_gn=SEMA5B;refGene_tx=XM_023260225.1
C2 68644622 68644772
refGene_gn=SEMA5B;refGene_tx=XM_006936108.4;refGene_gn=SEMA5B;refGene_tx=XM_019839850.2;refGene_gn=SEMA5B;refGene_tx=XM_023260223.1;refGene_gn=SEMA5B;refGene_tx=XM_023260224.1;refGene_gn=SEMA5B;refGene_tx=XM_023260225.1
C2 68644875 68645046
refGene_gn=SEMA5B;refGene_tx=XM_006936108.4;refGene_gn=SEMA5B;refGene_tx=XM_019839850.2;refGene_gn=SEMA5B;refGene_tx=XM_023260223.1;refGene_gn=SEMA5B;refGene_tx=XM_023260224.1;refGene_gn=SEMA5B;refGene_tx=XM_023260225.1
C2 68645534 68645753
refGene_gn=SEMA5B;refGene_tx=XM_006936108.4;refGene_gn=SEMA5B;refGene_tx=XM_019839850.2;refGene_gn=SEMA5B;refGene_tx=XM_023260223.1;refGene_gn=SEMA5B;refGene_tx=XM_023260224.1;refGene_gn=SEMA5B;refGene_tx=XM_023260225.1
C2 68645949 68646175
refGene_gn=SEMA5B;refGene_tx=XM_006936108.4;refGene_gn=SEMA5B;refGene_tx=XM_019839850.2;refGene_gn=SEMA5B;refGene_tx=XM_023260223.1;refGene_gn=SEMA5B;refGene_tx=XM_023260224.1;refGene_gn=SEMA5B;refGene_tx=XM_023260225.1
C2 68646293 68646441
refGene_gn=SEMA5B;refGene_tx=XM_006936108.4;refGene_gn=SEMA5B;refGene_tx=XM_019839850.2;refGene_gn=SEMA5B;refGene_tx=XM_023260223.1;refGene_gn=SEMA5B;refGene_tx=XM_023260224.1;refGene_gn=SEMA5B;refGene_tx=XM_023260225.1
C2 68646644 68646788
refGene_gn=SEMA5B;refGene_tx=XM_006936108.4;refGene_gn=SEMA5B;refGene_tx=XM_019839850.2;refGene_gn=SEMA5B;refGene_tx=XM_023260223.1;refGene_gn=SEMA5B;refGene_tx=XM_023260224.1;refGene_gn=SEMA5B;refGene_tx=XM_023260225.1
C2 68647796 68647978
refGene_gn=SEMA5B;refGene_tx=XM_006936108.4;refGene_gn=SEMA5B;refGene_tx=XM_019839850.2;refGene_gn=SEMA5B;refGene_tx=XM_023260223.1;refGene_gn=SEMA5B;refGene_tx=XM_023260224.1;refGene_gn=SEMA5B;refGene_tx=XM_023260225.1
C2 68648119 68648237

refGene_gn=SEMA5B;refGene_tx=XM_006936108.4;refGene_gn=SEMA5B;refGene_tx=XM_019839850.2;refGene_gn=SEMA5B;refGene_tx=XM_023260223.1;refGene_gn=SEMA5B;refGene_tx=XM_023260224.1;refGene_gn=SEMA5B;refGene_tx=XM_023260225.1
C2 68654827 68655035 refGene_gn=SEMA5B;refGene_tx=XM_006936108.4;refGene_gn=SEMA5B;refGene_tx=XM_019839850.2;refGene_gn=SEMA5B;refGene_tx=XM_023260223.1;refGene_gn=SEMA5B;refGene_tx=XM_023260224.1;refGene_gn=SEMA5B;refGene_tx=XM_023260225.1
C2 68655132 68655340 refGene_gn=SEMA5B;refGene_tx=XM_006936108.4;refGene_gn=SEMA5B;refGene_tx=XM_019839850.2;refGene_gn=SEMA5B;refGene_tx=XM_023260223.1;refGene_gn=SEMA5B;refGene_tx=XM_023260224.1;refGene_gn=SEMA5B;refGene_tx=XM_023260225.1
C2 68656552 68656688 refGene_gn=SEMA5B;refGene_tx=XM_006936108.4;refGene_gn=SEMA5B;refGene_tx=XM_019839850.2;refGene_gn=SEMA5B;refGene_tx=XM_023260223.1;refGene_gn=SEMA5B;refGene_tx=XM_023260224.1;refGene_gn=SEMA5B;refGene_tx=XM_023260225.1
C2 68658995 68659281 refGene_gn=SEMA5B;refGene_tx=XM_006936108.4;refGene_gn=SEMA5B;refGene_tx=XM_019839850.2;refGene_gn=SEMA5B;refGene_tx=XM_023260223.1;refGene_gn=SEMA5B;refGene_tx=XM_023260224.1;refGene_gn=SEMA5B;refGene_tx=XM_023260225.1
C2 68659967 68660181 refGene_gn=SEMA5B;refGene_tx=XM_006936108.4;refGene_gn=SEMA5B;refGene_tx=XM_019839850.2;refGene_gn=SEMA5B;refGene_tx=XM_023260223.1;refGene_gn=SEMA5B;refGene_tx=XM_023260224.1;refGene_gn=SEMA5B;refGene_tx=XM_023260225.1
C2 68660708 68660807 refGene_gn=SEMA5B;refGene_tx=XM_006936108.4;refGene_gn=SEMA5B;refGene_tx=XM_019839850.2;refGene_gn=SEMA5B;refGene_tx=XM_023260223.1;refGene_gn=SEMA5B;refGene_tx=XM_023260224.1;refGene_gn=SEMA5B;refGene_tx=XM_023260225.1
C2 68661130 68661193 refGene_gn=SEMA5B;refGene_tx=XM_006936108.4;refGene_gn=SEMA5B;refGene_tx=XM_019839850.2;refGene_gn=SEMA5B;refGene_tx=XM_023260223.1;refGene_gn=SEMA5B;refGene_tx=XM_023260224.1;refGene_gn=SEMA5B;refGene_tx=XM_023260225.1
C2 68662425 68662503 refGene_gn=SEMA5B;refGene_tx=XM_019839850.2
C2 68671478 68671524 refGene_gn=SEMA5B;refGene_tx=XM_006936108.4;refGene_gn=SEMA5B;refGene_tx=XM_023260223.1;refGene_gn=SEMA5B;refGene_tx=XM_023260224.1;refGene_gn=SEMA5B;refGene_tx=XM_023260225.1
C2 68676414 68676514 refGene_gn=SEMA5B;refGene_tx=XM_006936108.4;refGene_gn=SEMA5B;refGene_tx=XM_023260223.1;refGene_gn=SEMA5B;refGene_tx=XM_023260224.1;refGene_gn=SEMA5B;refGene_tx=XM_023260225.1
C2 68681691 68681845 refGene_gn=SEMA5B;refGene_tx=XM_006936108.4;refGene_gn=SEMA5B;refGene_

tx=XM_023260223.1;refGene_gn=SEMA5B;refGene_tx=XM_023260224.1;refGene_gn=SEMA5B;refGene_tx=XM_023260225.1
C2 68800511 68800559
refGene_gn=PDIA5;refGene_tx=XM_003991711.5;refGene_gn=PDIA5;refGene_tx=XM_019839851.2
C2 68822067 68822194
refGene_gn=PDIA5;refGene_tx=XM_003991711.5;refGene_gn=PDIA5;refGene_tx=XM_019839851.2
C2 68825085 68825173
refGene_gn=PDIA5;refGene_tx=XM_003991711.5;refGene_gn=PDIA5;refGene_tx=XM_019839851.2
C2 68833418 68833502
refGene_gn=PDIA5;refGene_tx=XM_003991711.5;refGene_gn=PDIA5;refGene_tx=XM_019839851.2
C2 68833751 68833797
refGene_gn=PDIA5;refGene_tx=XM_003991711.5;refGene_gn=PDIA5;refGene_tx=XM_019839851.2
C2 68837827 68837920
refGene_gn=PDIA5;refGene_tx=XM_003991711.5;refGene_gn=PDIA5;refGene_tx=XM_019839851.2
C2 68842326 68842387
refGene_gn=PDIA5;refGene_tx=XM_003991711.5;refGene_gn=PDIA5;refGene_tx=XM_019839851.2
C2 68848932 68849000
refGene_gn=PDIA5;refGene_tx=XM_003991711.5;refGene_gn=PDIA5;refGene_tx=XM_019839851.2
C2 68855890 68855982
refGene_gn=PDIA5;refGene_tx=XM_003991711.5;refGene_gn=PDIA5;refGene_tx=XM_019839851.2
C2 68856075 68856147
refGene_gn=PDIA5;refGene_tx=XM_003991711.5;refGene_gn=PDIA5;refGene_tx=XM_019839851.2
C2 68862545 68862682
refGene_gn=PDIA5;refGene_tx=XM_003991711.5;refGene_gn=PDIA5;refGene_tx=XM_019839851.2
C2 68875179 68875250
refGene_gn=PDIA5;refGene_tx=XM_003991711.5;refGene_gn=PDIA5;refGene_tx=XM_019839851.2
C2 68875796 68875957
refGene_gn=PDIA5;refGene_tx=XM_003991711.5;refGene_gn=PDIA5;refGene_tx=XM_019839851.2
C2 68880087 68880218
refGene_gn=PDIA5;refGene_tx=XM_003991711.5;refGene_gn=PDIA5;refGene_tx=XM_019839851.2
C2 68880591 68880593 refGene_gn=PDIA5;refGene_tx=XM_019839851.2
C2 68884399 68884470 refGene_gn=PDIA5;refGene_tx=XM_003991711.5
C2 68890763 68890898 refGene_gn=PDIA5;refGene_tx=XM_003991711.5
C2 68891270 68891351 refGene_gn=PDIA5;refGene_tx=XM_003991711.5
C2 68932183 68932230 refGene_gn=SEC22A;refGene_tx=XM_003991713.4
C2 68938161 68938362

refGene_gn=SEC22A;refGene_tx=XM_003991713.4;refGene_gn=SEC22A;refGene_tx=XM_003991714.4;refGene_gn=SEC22A;refGene_tx=XM_023260227.1
C2 68951709 68951873
refGene_gn=SEC22A;refGene_tx=XM_003991713.4;refGene_gn=SEC22A;refGene_tx=XM_003991714.4;refGene_gn=SEC22A;refGene_tx=XM_023260227.1
C2 68952997 68953192
refGene_gn=SEC22A;refGene_tx=XM_003991713.4;refGene_gn=SEC22A;refGene_tx=XM_003991714.4;refGene_gn=SEC22A;refGene_tx=XM_023260227.1
C2 68979380 68979496
refGene_gn=SEC22A;refGene_tx=XM_003991713.4;refGene_gn=SEC22A;refGene_tx=XM_003991714.4;refGene_gn=SEC22A;refGene_tx=XM_023260227.1
C2 68995706 68995772
refGene_gn=SEC22A;refGene_tx=XM_003991713.4;refGene_gn=SEC22A;refGene_tx=XM_003991714.4;refGene_gn=SEC22A;refGene_tx=XM_023260227.1
C2 69004261 69004462
refGene_gn=SEC22A;refGene_tx=XM_003991713.4;refGene_gn=SEC22A;refGene_tx=XM_003991714.4;refGene_gn=SEC22A;refGene_tx=XM_023260227.1
C2 69017623 69017752
refGene_gn=ADCY5;refGene_tx=XM_019839852.2;refGene_gn=ADCY5;refGene_tx=XM_023260228.1
C2 69019552 69019677
refGene_gn=ADCY5;refGene_tx=XM_019839852.2;refGene_gn=ADCY5;refGene_tx=XM_023260228.1
C2 69020620 69020825
refGene_gn=ADCY5;refGene_tx=XM_019839852.2;refGene_gn=ADCY5;refGene_tx=XM_023260228.1
C2 69021789 69022053
refGene_gn=ADCY5;refGene_tx=XM_019839852.2;refGene_gn=ADCY5;refGene_tx=XM_023260228.1
C2 69025687 69025820
refGene_gn=ADCY5;refGene_tx=XM_019839852.2;refGene_gn=ADCY5;refGene_tx=XM_023260228.1
C2 69027111 69027141
refGene_gn=ADCY5;refGene_tx=XM_019839852.2;refGene_gn=ADCY5;refGene_tx=XM_023260228.1
C2 69028794 69028970
refGene_gn=ADCY5;refGene_tx=XM_019839852.2;refGene_gn=ADCY5;refGene_tx=XM_023260228.1
C2 69031714 69031879
refGene_gn=ADCY5;refGene_tx=XM_019839852.2;refGene_gn=ADCY5;refGene_tx=XM_023260228.1
C2 69032246 69032363
refGene_gn=ADCY5;refGene_tx=XM_019839852.2;refGene_gn=ADCY5;refGene_tx=XM_023260228.1
C2 69041543 69041631
refGene_gn=ADCY5;refGene_tx=XM_019839852.2;refGene_gn=ADCY5;refGene_tx=XM_023260228.1
C2 69044884 69044982
refGene_gn=ADCY5;refGene_tx=XM_019839852.2;refGene_gn=ADCY5;refGene_tx=XM_023260228.1

C2 69046161 69046306
refGene_gn=ADCY5;refGene_tx=XM_019839852.2;refGene_gn=ADCY5;refGene_tx=XM_023260228.1
C2 69047229 69047252
refGene_gn=ADCY5;refGene_tx=XM_019839852.2;refGene_gn=ADCY5;refGene_tx=XM_023260228.1
C2 69051360 69051501
refGene_gn=ADCY5;refGene_tx=XM_019839852.2;refGene_gn=ADCY5;refGene_tx=XM_023260228.1
C2 69053731 69053873
refGene_gn=ADCY5;refGene_tx=XM_019839852.2;refGene_gn=ADCY5;refGene_tx=XM_023260228.1
C2 69054741 69054966
refGene_gn=ADCY5;refGene_tx=XM_019839852.2;refGene_gn=ADCY5;refGene_tx=XM_023260228.1
C2 69056892 69057020
refGene_gn=ADCY5;refGene_tx=XM_019839852.2;refGene_gn=ADCY5;refGene_tx=XM_023260228.1
C2 69058624 69058736
refGene_gn=ADCY5;refGene_tx=XM_019839852.2;refGene_gn=ADCY5;refGene_tx=XM_023260228.1
C2 69072060 69072182
refGene_gn=ADCY5;refGene_tx=XM_019839852.2;refGene_gn=ADCY5;refGene_tx=XM_023260228.1
C2 69075439 69075589
refGene_gn=ADCY5;refGene_tx=XM_019839852.2;refGene_gn=ADCY5;refGene_tx=XM_023260228.1
C2 69161648 69162785
refGene_gn=ADCY5;refGene_tx=XM_019839852.2;refGene_gn=ADCY5;refGene_tx=XM_023260228.1
C2 69206368 69206451
refGene_gn=HACD2;refGene_tx=XM_023260229.1;refGene_gn=HACD2;refGene_tx=XM_023260230.1
C2 69211924 69212103 refGene_gn=HACD2;refGene_tx=XM_023260229.1
C2 69213991 69214113
refGene_gn=HACD2;refGene_tx=XM_023260229.1;refGene_gn=HACD2;refGene_tx=XM_023260230.1
C2 69238401 69238490
refGene_gn=HACD2;refGene_tx=XM_023260229.1;refGene_gn=HACD2;refGene_tx=XM_023260230.1
C2 69282960 69282979
refGene_gn=HACD2;refGene_tx=XM_023260229.1;refGene_gn=HACD2;refGene_tx=XM_023260230.1
C2 69306032 69306150
refGene_gn=HACD2;refGene_tx=XM_023260229.1;refGene_gn=HACD2;refGene_tx=XM_023260230.1
C2 69308805 69309230
refGene_gn=HACD2;refGene_tx=XM_023260229.1;refGene_gn=HACD2;refGene_tx=XM_023260230.1
C2 69330030 69330281

refGene_gn=MYLK;refGene_tx=NM_001009384.1;refGene_gn=MYLK;refGene_tx=XM_006936114.3;refGene_gn=MYLK;refGene_tx=XM_006936115.3;refGene_gn=MYLK;refGene_tx=XM_006936116.4;refGene_gn=MYLK;refGene_tx=XM_019839240.2;refGene_gn=MYLK;refGene_tx=XM_019839241.2;refGene_gn=MYLK;refGene_tx=XM_019839242.2
C2 69333723 69333855
refGene_gn=MYLK;refGene_tx=NM_001009384.1;refGene_gn=MYLK;refGene_tx=XM_006936114.3;refGene_gn=MYLK;refGene_tx=XM_006936115.3;refGene_gn=MYLK;refGene_tx=XM_006936116.4;refGene_gn=MYLK;refGene_tx=XM_019839240.2;refGene_gn=MYLK;refGene_tx=XM_019839241.2;refGene_gn=MYLK;refGene_tx=XM_019839242.2
C2 69335287 69335417
refGene_gn=MYLK;refGene_tx=NM_001009384.1;refGene_gn=MYLK;refGene_tx=XM_006936114.3;refGene_gn=MYLK;refGene_tx=XM_006936115.3;refGene_gn=MYLK;refGene_tx=XM_006936116.4;refGene_gn=MYLK;refGene_tx=XM_019839240.2;refGene_gn=MYLK;refGene_tx=XM_019839241.2;refGene_gn=MYLK;refGene_tx=XM_019839242.2
C2 69341295 69341419
refGene_gn=MYLK;refGene_tx=XM_006936114.3;refGene_gn=MYLK;refGene_tx=XM_006936115.3;refGene_gn=MYLK;refGene_tx=XM_006936116.4;refGene_gn=MYLK;refGene_tx=XM_019839240.2;refGene_gn=MYLK;refGene_tx=XM_019839241.2;refGene_gn=MYLK;refGene_tx=XM_019839242.2
C2 69344322 69344475
refGene_gn=MYLK;refGene_tx=XM_006936114.3;refGene_gn=MYLK;refGene_tx=XM_006936115.3;refGene_gn=MYLK;refGene_tx=XM_006936116.4;refGene_gn=MYLK;refGene_tx=XM_019839240.2;refGene_gn=MYLK;refGene_tx=XM_019839241.2;refGene_gn=MYLK;refGene_tx=XM_019839242.2
C2 69354000 69354124
refGene_gn=MYLK;refGene_tx=XM_006936114.3;refGene_gn=MYLK;refGene_tx=XM_006936115.3;refGene_gn=MYLK;refGene_tx=XM_006936116.4;refGene_gn=MYLK;refGene_tx=XM_019839240.2;refGene_gn=MYLK;refGene_tx=XM_019839241.2;refGene_gn=MYLK;refGene_tx=XM_019839242.2
C2 69356097 69356315
refGene_gn=MYLK;refGene_tx=XM_006936114.3;refGene_gn=MYLK;refGene_tx=XM_006936115.3;refGene_gn=MYLK;refGene_tx=XM_006936116.4;refGene_gn=MYLK;refGene_tx=XM_019839240.2;refGene_gn=MYLK;refGene_tx=XM_019839241.2;refGene_gn=MYLK;refGene_tx=XM_019839242.2
C2 69362126 69362330
refGene_gn=MYLK;refGene_tx=XM_006936114.3;refGene_gn=MYLK;refGene_tx=XM_006936115.3;refGene_gn=MYLK;refGene_tx=XM_006936116.4;refGene_gn=MYLK;refGene_tx=XM_019839240.2;refGene_gn=MYLK;refGene_tx=XM_019839241.2;refGene_gn=MYLK;refGene_tx=XM_019839242.2
C2 69363639 69363733
refGene_gn=MYLK;refGene_tx=XM_006936114.3;refGene_gn=MYLK;refGene_tx=XM_006936115.3;refGene_gn=MYLK;refGene_tx=XM_006936116.4;refGene_gn=MYLK;refGene_tx=XM_019839240.2;refGene_gn=MYLK;refGene_tx=XM_019839241.2;refGene_gn=MYLK;refGene_tx=XM_019839242.2
C2 69363829 69363862
refGene_gn=MYLK;refGene_tx=XM_006936114.3;refGene_gn=MYLK;refGene_tx=XM_006936115.3;refGene_gn=MYLK;refGene_tx=XM_006936116.4;refGene_gn=MYL

K;refGene_tx=XM_019839240.2;refGene_gn=MYLK;refGene_tx=XM_019839241.2;
refGene_gn=MYLK;refGene_tx=XM_019839242.2
C2 69371654 69371957
refGene_gn=MYLK;refGene_tx=XM_006936114.3;refGene_gn=MYLK;refGene_tx=X
M_006936115.3;refGene_gn=MYLK;refGene_tx=XM_006936116.4;refGene_gn=MYL
K;refGene_tx=XM_019839240.2;refGene_gn=MYLK;refGene_tx=XM_019839241.2;
refGene_gn=MYLK;refGene_tx=XM_019839242.2
C2 69377858 69378012
refGene_gn=MYLK;refGene_tx=XM_006936114.3;refGene_gn=MYLK;refGene_tx=X
M_006936115.3;refGene_gn=MYLK;refGene_tx=XM_006936116.4;refGene_gn=MYL
K;refGene_tx=XM_019839240.2;refGene_gn=MYLK;refGene_tx=XM_019839241.2;
refGene_gn=MYLK;refGene_tx=XM_019839242.2
C2 69379938 69380066
refGene_gn=MYLK;refGene_tx=XM_006936114.3;refGene_gn=MYLK;refGene_tx=X
M_006936115.3;refGene_gn=MYLK;refGene_tx=XM_006936116.4;refGene_gn=MYL
K;refGene_tx=XM_019839240.2;refGene_gn=MYLK;refGene_tx=XM_019839241.2;
refGene_gn=MYLK;refGene_tx=XM_019839242.2
C2 69380846 69380897
refGene_gn=MYLK;refGene_tx=XM_006936114.3;refGene_gn=MYLK;refGene_tx=X
M_006936115.3;refGene_gn=MYLK;refGene_tx=XM_019839240.2;refGene_gn=MYL
K;refGene_tx=XM_019839241.2;refGene_gn=MYLK;refGene_tx=XM_019839242.2
C2 69396038 69396125
refGene_gn=MYLK;refGene_tx=XM_006936114.3;refGene_gn=MYLK;refGene_tx=X
M_006936115.3;refGene_gn=MYLK;refGene_tx=XM_019839240.2;refGene_gn=MYL
K;refGene_tx=XM_019839241.2;refGene_gn=MYLK;refGene_tx=XM_019839242.2
C2 69406168 69406285
refGene_gn=MYLK;refGene_tx=XM_006936114.3;refGene_gn=MYLK;refGene_tx=X
M_019839240.2;refGene_gn=MYLK;refGene_tx=XM_019839241.2;refGene_gn=MYL
K;refGene_tx=XM_019839242.2
C2 69413396 69414706
refGene_gn=MYLK;refGene_tx=XM_006936114.3;refGene_gn=MYLK;refGene_tx=X
M_019839240.2;refGene_gn=MYLK;refGene_tx=XM_019839241.2;refGene_gn=MYL
K;refGene_tx=XM_019839242.2
C2 69415145 69415217
refGene_gn=MYLK;refGene_tx=XM_006936114.3;refGene_gn=MYLK;refGene_tx=X
M_019839240.2;refGene_gn=MYLK;refGene_tx=XM_019839241.2;refGene_gn=MYL
K;refGene_tx=XM_019839242.2
C2 69421274 69421524
refGene_gn=MYLK;refGene_tx=XM_006936114.3;refGene_gn=MYLK;refGene_tx=X
M_019839240.2;refGene_gn=MYLK;refGene_tx=XM_019839241.2;refGene_gn=MYL
K;refGene_tx=XM_019839242.2
C2 69422312 69422510
refGene_gn=MYLK;refGene_tx=XM_006936114.3;refGene_gn=MYLK;refGene_tx=X
M_019839240.2;refGene_gn=MYLK;refGene_tx=XM_019839241.2;refGene_gn=MYL
K;refGene_tx=XM_019839242.2
C2 69424111 69424249
refGene_gn=MYLK;refGene_tx=XM_006936114.3;refGene_gn=MYLK;refGene_tx=X
M_019839240.2;refGene_gn=MYLK;refGene_tx=XM_019839241.2;refGene_gn=MYL
K;refGene_tx=XM_019839242.2
C2 69437486 69437639

refGene_gn=MYLK;refGene_tx=XM_006936114.3;refGene_gn=MYLK;refGene_tx=XM_019839240.2;refGene_gn=MYLK;refGene_tx=XM_019839241.2;refGene_gn=MYLK;refGene_tx=XM_019839242.2
C2 69440688 69440823
refGene_gn=MYLK;refGene_tx=XM_006936114.3;refGene_gn=MYLK;refGene_tx=XM_019839240.2;refGene_gn=MYLK;refGene_tx=XM_019839241.2;refGene_gn=MYLK;refGene_tx=XM_019839242.2
C2 69445838 69446045
refGene_gn=MYLK;refGene_tx=XM_006936114.3;refGene_gn=MYLK;refGene_tx=XM_019839240.2;refGene_gn=MYLK;refGene_tx=XM_019839241.2;refGene_gn=MYLK;refGene_tx=XM_019839242.2
C2 69446570 69447106
refGene_gn=MYLK;refGene_tx=XM_006936114.3;refGene_gn=MYLK;refGene_tx=XM_019839240.2;refGene_gn=MYLK;refGene_tx=XM_019839241.2;refGene_gn=MYLK;refGene_tx=XM_019839242.2
C2 69448216 69448235
refGene_gn=MYLK;refGene_tx=XM_006936114.3;refGene_gn=MYLK;refGene_tx=XM_019839240.2;refGene_gn=MYLK;refGene_tx=XM_019839241.2;refGene_gn=MYLK;refGene_tx=XM_019839242.2
C2 69450318 69450484
refGene_gn=MYLK;refGene_tx=XM_006936114.3;refGene_gn=MYLK;refGene_tx=XM_019839240.2;refGene_gn=MYLK;refGene_tx=XM_019839241.2;refGene_gn=MYLK;refGene_tx=XM_019839242.2
C2 69451418 69451584
refGene_gn=MYLK;refGene_tx=XM_006936114.3;refGene_gn=MYLK;refGene_tx=XM_019839240.2;refGene_gn=MYLK;refGene_tx=XM_019839241.2;refGene_gn=MYLK;refGene_tx=XM_019839242.2
C2 69452486 69452535
refGene_gn=MYLK;refGene_tx=XM_006936114.3;refGene_gn=MYLK;refGene_tx=XM_019839240.2;refGene_gn=MYLK;refGene_tx=XM_019839241.2;refGene_gn=MYLK;refGene_tx=XM_019839242.2
C2 69467187 69467395
refGene_gn=MYLK;refGene_tx=XM_006936114.3;refGene_gn=MYLK;refGene_tx=XM_019839240.2;refGene_gn=MYLK;refGene_tx=XM_019839241.2;refGene_gn=MYLK;refGene_tx=XM_019839242.2
C2 69508456 69508624
refGene_gn=MYLK;refGene_tx=XM_006936114.3;refGene_gn=MYLK;refGene_tx=XM_019839240.2;refGene_gn=MYLK;refGene_tx=XM_019839241.2;refGene_gn=MYLK;refGene_tx=XM_019839242.2
C2 69514330 69514360
refGene_gn=MYLK;refGene_tx=XM_006936114.3;refGene_gn=MYLK;refGene_tx=XM_019839240.2
C2 69524405 69524408 refGene_gn=MYLK;refGene_tx=XM_019839241.2
C2 69631217 69632151
refGene_gn=CCDC14;refGene_tx=XM_006936118.4;refGene_gn=CCDC14;refGene_tx=XM_011285914.3
C2 69635798 69635931
refGene_gn=CCDC14;refGene_tx=XM_006936118.4;refGene_gn=CCDC14;refGene_tx=XM_011285914.3
C2 69636006 69636225

refGene_gn=CCDC14;refGene_tx=XM_006936118.4;refGene_gn=CCDC14;refGene_tx=XM_011285914.3
C2 69645432 69645515
refGene_gn=CCDC14;refGene_tx=XM_006936118.4;refGene_gn=CCDC14;refGene_tx=XM_011285914.3
C2 69657475 69657617
refGene_gn=CCDC14;refGene_tx=XM_006936118.4;refGene_gn=CCDC14;refGene_tx=XM_011285914.3
C2 69660022 69660521
refGene_gn=CCDC14;refGene_tx=XM_006936118.4;refGene_gn=CCDC14;refGene_tx=XM_011285914.3
C2 69661424 69661519
refGene_gn=CCDC14;refGene_tx=XM_006936118.4;refGene_gn=CCDC14;refGene_tx=XM_011285914.3
C2 69661629 69661866
refGene_gn=CCDC14;refGene_tx=XM_006936118.4;refGene_gn=CCDC14;refGene_tx=XM_011285914.3
C2 69669164 69669284
refGene_gn=CCDC14;refGene_tx=XM_006936118.4;refGene_gn=CCDC14;refGene_tx=XM_011285914.3
C2 69669377 69669447
refGene_gn=CCDC14;refGene_tx=XM_006936118.4;refGene_gn=CCDC14;refGene_tx=XM_011285914.3
C2 69669920 69669993
refGene_gn=CCDC14;refGene_tx=XM_006936118.4;refGene_gn=CCDC14;refGene_tx=XM_011285914.3
C2 69670311 69670367
refGene_gn=CCDC14;refGene_tx=XM_006936118.4;refGene_gn=CCDC14;refGene_tx=XM_011285914.3
C2 69679669 69679843
refGene_gn=CCDC14;refGene_tx=XM_006936118.4;refGene_gn=CCDC14;refGene_tx=XM_011285914.3
C2 69689159 69689226
refGene_gn=ROPN1;refGene_tx=XM_003991716.5;refGene_gn=ROPN1;refGene_tx=XM_006936119.4;refGene_gn=ROPN1;refGene_tx=XM_006936120.4;refGene_gn=ROPN1;refGene_tx=XM_006936121.4;refGene_gn=ROPN1;refGene_tx=XM_006936122.4;refGene_gn=ROPN1;refGene_tx=XM_006936123.4;refGene_gn=ROPN1;refGene_tx=XM_006936124.4;refGene_gn=ROPN1;refGene_tx=XM_019839857.2;refGene_gn=ROPN1;refGene_tx=XM_019839858.2
C2 69689716 69689892
refGene_gn=ROPN1;refGene_tx=XM_003991716.5;refGene_gn=ROPN1;refGene_tx=XM_006936119.4;refGene_gn=ROPN1;refGene_tx=XM_006936120.4;refGene_gn=ROPN1;refGene_tx=XM_006936121.4;refGene_gn=ROPN1;refGene_tx=XM_006936122.4;refGene_gn=ROPN1;refGene_tx=XM_006936123.4;refGene_gn=ROPN1;refGene_tx=XM_006936124.4;refGene_gn=ROPN1;refGene_tx=XM_019839857.2;refGene_gn=ROPN1;refGene_tx=XM_019839858.2
C2 69695691 69695853
refGene_gn=ROPN1;refGene_tx=XM_003991716.5;refGene_gn=ROPN1;refGene_tx=XM_006936119.4;refGene_gn=ROPN1;refGene_tx=XM_006936120.4;refGene_gn=ROPN1;refGene_tx=XM_006936121.4;refGene_gn=ROPN1;refGene_tx=XM_0069361

22.4;refGene_gn=ROPN1;refGene_tx=XM_006936123.4;refGene_gn=ROPN1;refGene_tx=XM_006936124.4;refGene_gn=ROPN1;refGene_tx=XM_019839857.2;refGene_gn=ROPN1;refGene_tx=XM_019839858.2
C2 69697154 69697272
refGene_gn=ROPN1;refGene_tx=XM_003991716.5;refGene_gn=ROPN1;refGene_tx=XM_006936119.4;refGene_gn=ROPN1;refGene_tx=XM_006936120.4;refGene_gn=ROPN1;refGene_tx=XM_006936121.4;refGene_gn=ROPN1;refGene_tx=XM_006936122.4;refGene_gn=ROPN1;refGene_tx=XM_006936123.4;refGene_gn=ROPN1;refGene_tx=XM_006936124.4;refGene_gn=ROPN1;refGene_tx=XM_019839857.2;refGene_gn=ROPN1;refGene_tx=XM_019839858.2
C2 69700939 69701055
refGene_gn=ROPN1;refGene_tx=XM_003991716.5;refGene_gn=ROPN1;refGene_tx=XM_006936119.4;refGene_gn=ROPN1;refGene_tx=XM_006936120.4;refGene_gn=ROPN1;refGene_tx=XM_006936121.4;refGene_gn=ROPN1;refGene_tx=XM_006936122.4;refGene_gn=ROPN1;refGene_tx=XM_006936123.4;refGene_gn=ROPN1;refGene_tx=XM_006936124.4;refGene_gn=ROPN1;refGene_tx=XM_019839857.2
C2 69756636 69756709
refGene_gn=KALRN;refGene_tx=XM_023260231.1;refGene_gn=KALRN;refGene_tx=XM_023260232.1;refGene_gn=KALRN;refGene_tx=XM_023260233.1;refGene_gn=KALRN;refGene_tx=XM_023260234.1;refGene_gn=KALRN;refGene_tx=XM_023260235.1;refGene_gn=KALRN;refGene_tx=XM_023260236.1;refGene_gn=KALRN;refGene_tx=XM_023260237.1;refGene_gn=KALRN;refGene_tx=XM_023260238.1;refGene_gn=KALRN;refGene_tx=XM_023260239.1;refGene_gn=KALRN;refGene_tx=XM_023260240.1;refGene_gn=KALRN;refGene_tx=XM_023260241.1;refGene_gn=KALRN;refGene_tx=XM_023260243.1;refGene_gn=KALRN;refGene_tx=XM_023260245.1;refGene_gn=KALRN;refGene_tx=XM_023260246.1
C2 69821607 69821674
refGene_gn=KALRN;refGene_tx=XM_019839863.2;refGene_gn=KALRN;refGene_tx=XM_019839865.2;refGene_gn=KALRN;refGene_tx=XM_019839867.2;refGene_gn=KALRN;refGene_tx=XM_023260242.1;refGene_gn=KALRN;refGene_tx=XM_023260244.1
C2 69949458 69949533
refGene_gn=KALRN;refGene_tx=XM_019839863.2;refGene_gn=KALRN;refGene_tx=XM_019839865.2;refGene_gn=KALRN;refGene_tx=XM_019839867.2;refGene_gn=KALRN;refGene_tx=XM_023260231.1;refGene_gn=KALRN;refGene_tx=XM_023260232.1;refGene_gn=KALRN;refGene_tx=XM_023260233.1;refGene_gn=KALRN;refGene_tx=XM_023260234.1;refGene_gn=KALRN;refGene_tx=XM_023260235.1;refGene_gn=KALRN;refGene_tx=XM_023260236.1;refGene_gn=KALRN;refGene_tx=XM_023260237.1;refGene_gn=KALRN;refGene_tx=XM_023260238.1;refGene_gn=KALRN;refGene_tx=XM_023260239.1;refGene_gn=KALRN;refGene_tx=XM_023260240.1;refGene_gn=KALRN;refGene_tx=XM_023260241.1;refGene_gn=KALRN;refGene_tx=XM_023260242.1;refGene_gn=KALRN;refGene_tx=XM_023260243.1;refGene_gn=KALRN;refGene_tx=XM_023260244.1;refGene_gn=KALRN;refGene_tx=XM_023260245.1;refGene_gn=KALRN;refGene_tx=XM_023260246.1
C2 69956700 69956815
refGene_gn=KALRN;refGene_tx=XM_019839863.2;refGene_gn=KALRN;refGene_tx=XM_019839865.2;refGene_gn=KALRN;refGene_tx=XM_019839867.2;refGene_gn=KALRN;refGene_tx=XM_023260231.1;refGene_gn=KALRN;refGene_tx=XM_023260232.1;refGene_gn=KALRN;refGene_tx=XM_023260233.1;refGene_gn=KALRN;refGene_tx=XM_023260234.1;refGene_gn=KALRN;refGene_tx=XM_023260235.1;refGen

e_gn=KALRN;refGene_tx=XM_023260236.1;refGene_gn=KALRN;refGene_tx=XM_02
3260237.1;refGene_gn=KALRN;refGene_tx=XM_023260238.1;refGene_gn=KALRN;
refGene_tx=XM_023260239.1;refGene_gn=KALRN;refGene_tx=XM_023260240.1;r
efGene_gn=KALRN;refGene_tx=XM_023260241.1;refGene_gn=KALRN;refGene_tx=
XM_023260242.1;refGene_gn=KALRN;refGene_tx=XM_023260243.1;refGene_gn=K
ALRN;refGene_tx=XM_023260244.1;refGene_gn=KALRN;refGene_tx=XM_02326024
5.1;refGene_gn=KALRN;refGene_tx=XM_023260246.1
C2 69984002 69984195
refGene_gn=KALRN;refGene_tx=XM_019839863.2;refGene_gn=KALRN;refGene_tx
=XM_019839865.2;refGene_gn=KALRN;refGene_tx=XM_019839867.2;refGene_gn=
KALRN;refGene_tx=XM_023260231.1;refGene_gn=KALRN;refGene_tx=XM_0232602
32.1;refGene_gn=KALRN;refGene_tx=XM_023260233.1;refGene_gn=KALRN;refGe
ne_tx=XM_023260234.1;refGene_gn=KALRN;refGene_tx=XM_023260235.1;refGen
e_gn=KALRN;refGene_tx=XM_023260236.1;refGene_gn=KALRN;refGene_tx=XM_02
3260237.1;refGene_gn=KALRN;refGene_tx=XM_023260238.1;refGene_gn=KALRN;
refGene_tx=XM_023260239.1;refGene_gn=KALRN;refGene_tx=XM_023260240.1;r
efGene_gn=KALRN;refGene_tx=XM_023260241.1;refGene_gn=KALRN;refGene_tx=
XM_023260242.1;refGene_gn=KALRN;refGene_tx=XM_023260243.1;refGene_gn=K
ALRN;refGene_tx=XM_023260244.1;refGene_gn=KALRN;refGene_tx=XM_02326024
5.1;refGene_gn=KALRN;refGene_tx=XM_023260246.1
C2 69984613 69985126
refGene_gn=KALRN;refGene_tx=XM_019839863.2;refGene_gn=KALRN;refGene_tx
=XM_019839865.2;refGene_gn=KALRN;refGene_tx=XM_019839867.2;refGene_gn=
KALRN;refGene_tx=XM_023260231.1;refGene_gn=KALRN;refGene_tx=XM_0232602
32.1;refGene_gn=KALRN;refGene_tx=XM_023260233.1;refGene_gn=KALRN;refGe
ne_tx=XM_023260234.1;refGene_gn=KALRN;refGene_tx=XM_023260235.1;refGen
e_gn=KALRN;refGene_tx=XM_023260236.1;refGene_gn=KALRN;refGene_tx=XM_02
3260237.1;refGene_gn=KALRN;refGene_tx=XM_023260238.1;refGene_gn=KALRN;
refGene_tx=XM_023260239.1;refGene_gn=KALRN;refGene_tx=XM_023260240.1;r
efGene_gn=KALRN;refGene_tx=XM_023260241.1;refGene_gn=KALRN;refGene_tx=
XM_023260242.1;refGene_gn=KALRN;refGene_tx=XM_023260243.1;refGene_gn=K
ALRN;refGene_tx=XM_023260244.1;refGene_gn=KALRN;refGene_tx=XM_02326024
5.1;refGene_gn=KALRN;refGene_tx=XM_023260246.1
C2 70011485 70011608
refGene_gn=KALRN;refGene_tx=XM_019839863.2;refGene_gn=KALRN;refGene_tx
=XM_019839865.2;refGene_gn=KALRN;refGene_tx=XM_019839867.2;refGene_gn=
KALRN;refGene_tx=XM_023260231.1;refGene_gn=KALRN;refGene_tx=XM_0232602
32.1;refGene_gn=KALRN;refGene_tx=XM_023260233.1;refGene_gn=KALRN;refGe
ne_tx=XM_023260234.1;refGene_gn=KALRN;refGene_tx=XM_023260235.1;refGen
e_gn=KALRN;refGene_tx=XM_023260236.1;refGene_gn=KALRN;refGene_tx=XM_02
3260237.1;refGene_gn=KALRN;refGene_tx=XM_023260238.1;refGene_gn=KALRN;
refGene_tx=XM_023260239.1;refGene_gn=KALRN;refGene_tx=XM_023260240.1;r
efGene_gn=KALRN;refGene_tx=XM_023260241.1;refGene_gn=KALRN;refGene_tx=
XM_023260242.1;refGene_gn=KALRN;refGene_tx=XM_023260243.1;refGene_gn=K
ALRN;refGene_tx=XM_023260244.1;refGene_gn=KALRN;refGene_tx=XM_02326024
5.1;refGene_gn=KALRN;refGene_tx=XM_023260246.1
C2 70038537 70038729
refGene_gn=KALRN;refGene_tx=XM_019839863.2;refGene_gn=KALRN;refGene_tx
=XM_019839865.2;refGene_gn=KALRN;refGene_tx=XM_019839867.2;refGene_gn=
KALRN;refGene_tx=XM_023260231.1;refGene_gn=KALRN;refGene_tx=XM_0232602

32.1;refGene_gn=KALRN;refGene_tx=XM_023260233.1;refGene_gn=KALRN;refGene_tx=XM_023260234.1;refGene_gn=KALRN;refGene_tx=XM_023260235.1;refGene_gn=KALRN;refGene_tx=XM_023260236.1;refGene_gn=KALRN;refGene_tx=XM_023260237.1;refGene_gn=KALRN;refGene_tx=XM_023260238.1;refGene_gn=KALRN;refGene_tx=XM_023260239.1;refGene_gn=KALRN;refGene_tx=XM_023260240.1;refGene_gn=KALRN;refGene_tx=XM_023260241.1;refGene_gn=KALRN;refGene_tx=XM_023260242.1;refGene_gn=KALRN;refGene_tx=XM_023260243.1;refGene_gn=KALRN;refGene_tx=XM_023260244.1;refGene_gn=KALRN;refGene_tx=XM_023260245.1;refGene_gn=KALRN;refGene_tx=XM_023260246.1
C2 70042412 70042544
refGene_gn=KALRN;refGene_tx=XM_019839863.2;refGene_gn=KALRN;refGene_tx=XM_019839865.2;refGene_gn=KALRN;refGene_tx=XM_019839867.2;refGene_gn=KALRN;refGene_tx=XM_023260231.1;refGene_gn=KALRN;refGene_tx=XM_023260232.1;refGene_gn=KALRN;refGene_tx=XM_023260233.1;refGene_gn=KALRN;refGene_tx=XM_023260234.1;refGene_gn=KALRN;refGene_tx=XM_023260235.1;refGene_gn=KALRN;refGene_tx=XM_023260236.1;refGene_gn=KALRN;refGene_tx=XM_023260237.1;refGene_gn=KALRN;refGene_tx=XM_023260238.1;refGene_gn=KALRN;refGene_tx=XM_023260239.1;refGene_gn=KALRN;refGene_tx=XM_023260240.1;refGene_gn=KALRN;refGene_tx=XM_023260241.1;refGene_gn=KALRN;refGene_tx=XM_023260242.1;refGene_gn=KALRN;refGene_tx=XM_023260243.1;refGene_gn=KALRN;refGene_tx=XM_023260244.1;refGene_gn=KALRN;refGene_tx=XM_023260245.1;refGene_gn=KALRN;refGene_tx=XM_023260246.1
C2 70045992 70046223
refGene_gn=KALRN;refGene_tx=XM_019839863.2;refGene_gn=KALRN;refGene_tx=XM_019839865.2;refGene_gn=KALRN;refGene_tx=XM_019839867.2;refGene_gn=KALRN;refGene_tx=XM_023260231.1;refGene_gn=KALRN;refGene_tx=XM_023260232.1;refGene_gn=KALRN;refGene_tx=XM_023260233.1;refGene_gn=KALRN;refGene_tx=XM_023260234.1;refGene_gn=KALRN;refGene_tx=XM_023260235.1;refGene_gn=KALRN;refGene_tx=XM_023260236.1;refGene_gn=KALRN;refGene_tx=XM_023260237.1;refGene_gn=KALRN;refGene_tx=XM_023260238.1;refGene_gn=KALRN;refGene_tx=XM_023260239.1;refGene_gn=KALRN;refGene_tx=XM_023260240.1;refGene_gn=KALRN;refGene_tx=XM_023260241.1;refGene_gn=KALRN;refGene_tx=XM_023260242.1;refGene_gn=KALRN;refGene_tx=XM_023260243.1;refGene_gn=KALRN;refGene_tx=XM_023260244.1;refGene_gn=KALRN;refGene_tx=XM_023260245.1;refGene_gn=KALRN;refGene_tx=XM_023260246.1
C2 70064704 70064827
refGene_gn=KALRN;refGene_tx=XM_019839863.2;refGene_gn=KALRN;refGene_tx=XM_019839865.2;refGene_gn=KALRN;refGene_tx=XM_019839867.2;refGene_gn=KALRN;refGene_tx=XM_023260231.1;refGene_gn=KALRN;refGene_tx=XM_023260232.1;refGene_gn=KALRN;refGene_tx=XM_023260233.1;refGene_gn=KALRN;refGene_tx=XM_023260234.1;refGene_gn=KALRN;refGene_tx=XM_023260235.1;refGene_gn=KALRN;refGene_tx=XM_023260236.1;refGene_gn=KALRN;refGene_tx=XM_023260237.1;refGene_gn=KALRN;refGene_tx=XM_023260238.1;refGene_gn=KALRN;refGene_tx=XM_023260239.1;refGene_gn=KALRN;refGene_tx=XM_023260240.1;refGene_gn=KALRN;refGene_tx=XM_023260241.1;refGene_gn=KALRN;refGene_tx=XM_023260242.1;refGene_gn=KALRN;refGene_tx=XM_023260243.1;refGene_gn=KALRN;refGene_tx=XM_023260244.1;refGene_gn=KALRN;refGene_tx=XM_023260245.1;refGene_gn=KALRN;refGene_tx=XM_023260246.1
C2 70100586 70100778
refGene_gn=KALRN;refGene_tx=XM_019839863.2;refGene_gn=KALRN;refGene_tx

=XM_019839865.2;refGene_gn=KALRN;refGene_tx=XM_019839867.2;refGene_gn=KALRN;refGene_tx=XM_023260231.1;refGene_gn=KALRN;refGene_tx=XM_023260232.1;refGene_gn=KALRN;refGene_tx=XM_023260233.1;refGene_gn=KALRN;refGene_tx=XM_023260234.1;refGene_gn=KALRN;refGene_tx=XM_023260235.1;refGene_gn=KALRN;refGene_tx=XM_023260236.1;refGene_gn=KALRN;refGene_tx=XM_023260237.1;refGene_gn=KALRN;refGene_tx=XM_023260238.1;refGene_gn=KALRN;refGene_tx=XM_023260239.1;refGene_gn=KALRN;refGene_tx=XM_023260240.1;refGene_gn=KALRN;refGene_tx=XM_023260241.1;refGene_gn=KALRN;refGene_tx=XM_023260242.1;refGene_gn=KALRN;refGene_tx=XM_023260243.1;refGene_gn=KALRN;refGene_tx=XM_023260244.1;refGene_gn=KALRN;refGene_tx=XM_023260245.1;refGene_gn=KALRN;refGene_tx=XM_023260246.1
C2 70110270 70110479
refGene_gn=KALRN;refGene_tx=XM_019839863.2;refGene_gn=KALRN;refGene_tx=XM_019839865.2;refGene_gn=KALRN;refGene_tx=XM_019839867.2;refGene_gn=KALRN;refGene_tx=XM_023260231.1;refGene_gn=KALRN;refGene_tx=XM_023260232.1;refGene_gn=KALRN;refGene_tx=XM_023260233.1;refGene_gn=KALRN;refGene_tx=XM_023260234.1;refGene_gn=KALRN;refGene_tx=XM_023260235.1;refGene_gn=KALRN;refGene_tx=XM_023260236.1;refGene_gn=KALRN;refGene_tx=XM_023260237.1;refGene_gn=KALRN;refGene_tx=XM_023260238.1;refGene_gn=KALRN;refGene_tx=XM_023260239.1;refGene_gn=KALRN;refGene_tx=XM_023260240.1;refGene_gn=KALRN;refGene_tx=XM_023260241.1;refGene_gn=KALRN;refGene_tx=XM_023260242.1;refGene_gn=KALRN;refGene_tx=XM_023260243.1;refGene_gn=KALRN;refGene_tx=XM_023260244.1;refGene_gn=KALRN;refGene_tx=XM_023260245.1;refGene_gn=KALRN;refGene_tx=XM_023260246.1
C2 70114026 70114201
refGene_gn=KALRN;refGene_tx=XM_019839863.2;refGene_gn=KALRN;refGene_tx=XM_019839865.2;refGene_gn=KALRN;refGene_tx=XM_019839867.2;refGene_gn=KALRN;refGene_tx=XM_023260231.1;refGene_gn=KALRN;refGene_tx=XM_023260232.1;refGene_gn=KALRN;refGene_tx=XM_023260233.1;refGene_gn=KALRN;refGene_tx=XM_023260234.1;refGene_gn=KALRN;refGene_tx=XM_023260235.1;refGene_gn=KALRN;refGene_tx=XM_023260236.1;refGene_gn=KALRN;refGene_tx=XM_023260237.1;refGene_gn=KALRN;refGene_tx=XM_023260238.1;refGene_gn=KALRN;refGene_tx=XM_023260239.1;refGene_gn=KALRN;refGene_tx=XM_023260240.1;refGene_gn=KALRN;refGene_tx=XM_023260241.1;refGene_gn=KALRN;refGene_tx=XM_023260242.1;refGene_gn=KALRN;refGene_tx=XM_023260243.1;refGene_gn=KALRN;refGene_tx=XM_023260244.1;refGene_gn=KALRN;refGene_tx=XM_023260245.1;refGene_gn=KALRN;refGene_tx=XM_023260246.1
C2 70128546 70128742
refGene_gn=KALRN;refGene_tx=XM_019839863.2;refGene_gn=KALRN;refGene_tx=XM_019839865.2;refGene_gn=KALRN;refGene_tx=XM_019839867.2;refGene_gn=KALRN;refGene_tx=XM_023260231.1;refGene_gn=KALRN;refGene_tx=XM_023260232.1;refGene_gn=KALRN;refGene_tx=XM_023260233.1;refGene_gn=KALRN;refGene_tx=XM_023260234.1;refGene_gn=KALRN;refGene_tx=XM_023260235.1;refGene_gn=KALRN;refGene_tx=XM_023260236.1;refGene_gn=KALRN;refGene_tx=XM_023260237.1;refGene_gn=KALRN;refGene_tx=XM_023260238.1;refGene_gn=KALRN;refGene_tx=XM_023260239.1;refGene_gn=KALRN;refGene_tx=XM_023260240.1;refGene_gn=KALRN;refGene_tx=XM_023260241.1;refGene_gn=KALRN;refGene_tx=XM_023260242.1;refGene_gn=KALRN;refGene_tx=XM_023260243.1;refGene_gn=KALRN;refGene_tx=XM_023260244.1;refGene_gn=KALRN;refGene_tx=XM_023260245.1;refGene_gn=KALRN;refGene_tx=XM_023260246.1

C2	70138038	70138205
refGene_gn=KALRN;refGene_tx=XM_019839863.2;refGene_gn=KALRN;refGene_tx=XM_019839865.2;refGene_gn=KALRN;refGene_tx=XM_019839867.2;refGene_gn=KALRN;refGene_tx=XM_023260231.1;refGene_gn=KALRN;refGene_tx=XM_023260232.1;refGene_gn=KALRN;refGene_tx=XM_023260233.1;refGene_gn=KALRN;refGene_tx=XM_023260234.1;refGene_gn=KALRN;refGene_tx=XM_023260235.1;refGene_gn=KALRN;refGene_tx=XM_023260236.1;refGene_gn=KALRN;refGene_tx=XM_023260237.1;refGene_gn=KALRN;refGene_tx=XM_023260238.1;refGene_gn=KALRN;refGene_tx=XM_023260239.1;refGene_gn=KALRN;refGene_tx=XM_023260240.1;refGene_gn=KALRN;refGene_tx=XM_023260241.1;refGene_gn=KALRN;refGene_tx=XM_023260242.1;refGene_gn=KALRN;refGene_tx=XM_023260243.1;refGene_gn=KALRN;refGene_tx=XM_023260244.1;refGene_gn=KALRN;refGene_tx=XM_023260245.1;refGene_gn=KALRN;refGene_tx=XM_023260246.1
C2	70145824	70145944
refGene_gn=KALRN;refGene_tx=XM_019839863.2;refGene_gn=KALRN;refGene_tx=XM_019839865.2;refGene_gn=KALRN;refGene_tx=XM_019839867.2;refGene_gn=KALRN;refGene_tx=XM_023260231.1;refGene_gn=KALRN;refGene_tx=XM_023260232.1;refGene_gn=KALRN;refGene_tx=XM_023260233.1;refGene_gn=KALRN;refGene_tx=XM_023260234.1;refGene_gn=KALRN;refGene_tx=XM_023260235.1;refGene_gn=KALRN;refGene_tx=XM_023260236.1;refGene_gn=KALRN;refGene_tx=XM_023260237.1;refGene_gn=KALRN;refGene_tx=XM_023260238.1;refGene_gn=KALRN;refGene_tx=XM_023260239.1;refGene_gn=KALRN;refGene_tx=XM_023260240.1;refGene_gn=KALRN;refGene_tx=XM_023260241.1;refGene_gn=KALRN;refGene_tx=XM_023260242.1;refGene_gn=KALRN;refGene_tx=XM_023260243.1;refGene_gn=KALRN;refGene_tx=XM_023260244.1;refGene_gn=KALRN;refGene_tx=XM_023260245.1;refGene_gn=KALRN;refGene_tx=XM_023260246.1
C2	70152111	70152330
refGene_gn=KALRN;refGene_tx=XM_019839863.2;refGene_gn=KALRN;refGene_tx=XM_019839865.2;refGene_gn=KALRN;refGene_tx=XM_019839867.2;refGene_gn=KALRN;refGene_tx=XM_023260231.1;refGene_gn=KALRN;refGene_tx=XM_023260232.1;refGene_gn=KALRN;refGene_tx=XM_023260233.1;refGene_gn=KALRN;refGene_tx=XM_023260234.1;refGene_gn=KALRN;refGene_tx=XM_023260235.1;refGene_gn=KALRN;refGene_tx=XM_023260236.1;refGene_gn=KALRN;refGene_tx=XM_023260237.1;refGene_gn=KALRN;refGene_tx=XM_023260238.1;refGene_gn=KALRN;refGene_tx=XM_023260239.1;refGene_gn=KALRN;refGene_tx=XM_023260240.1;refGene_gn=KALRN;refGene_tx=XM_023260241.1;refGene_gn=KALRN;refGene_tx=XM_023260242.1;refGene_gn=KALRN;refGene_tx=XM_023260243.1;refGene_gn=KALRN;refGene_tx=XM_023260244.1;refGene_gn=KALRN;refGene_tx=XM_023260245.1;refGene_gn=KALRN;refGene_tx=XM_023260246.1
C2	70155977	70156127
refGene_gn=KALRN;refGene_tx=XM_019839863.2;refGene_gn=KALRN;refGene_tx=XM_019839865.2;refGene_gn=KALRN;refGene_tx=XM_019839867.2;refGene_gn=KALRN;refGene_tx=XM_023260231.1;refGene_gn=KALRN;refGene_tx=XM_023260232.1;refGene_gn=KALRN;refGene_tx=XM_023260233.1;refGene_gn=KALRN;refGene_tx=XM_023260234.1;refGene_gn=KALRN;refGene_tx=XM_023260235.1;refGene_gn=KALRN;refGene_tx=XM_023260236.1;refGene_gn=KALRN;refGene_tx=XM_023260237.1;refGene_gn=KALRN;refGene_tx=XM_023260238.1;refGene_gn=KALRN;refGene_tx=XM_023260239.1;refGene_gn=KALRN;refGene_tx=XM_023260240.1;refGene_gn=KALRN;refGene_tx=XM_023260241.1;refGene_gn=KALRN;refGene_tx=XM_023260242.1;refGene_gn=KALRN;refGene_tx=XM_023260243.1;refGene_gn=K

ALRN;refGene_tx=XM_023260244.1;refGene_gn=KALRN;refGene_tx=XM_02326024
5.1;refGene_gn=KALRN;refGene_tx=XM_023260246.1
C2 70158944 70159059
refGene_gn=KALRN;refGene_tx=XM_019839863.2;refGene_gn=KALRN;refGene_tx
=XM_019839865.2;refGene_gn=KALRN;refGene_tx=XM_019839867.2;refGene_gn=
KALRN;refGene_tx=XM_023260231.1;refGene_gn=KALRN;refGene_tx=XM_0232602
32.1;refGene_gn=KALRN;refGene_tx=XM_023260233.1;refGene_gn=KALRN;refGe
ne_tx=XM_023260234.1;refGene_gn=KALRN;refGene_tx=XM_023260235.1;refGen
e_gn=KALRN;refGene_tx=XM_023260236.1;refGene_gn=KALRN;refGene_tx=XM_02
3260237.1;refGene_gn=KALRN;refGene_tx=XM_023260238.1;refGene_gn=KALRN;
refGene_tx=XM_023260239.1;refGene_gn=KALRN;refGene_tx=XM_023260240.1;r
efGene_gn=KALRN;refGene_tx=XM_023260241.1;refGene_gn=KALRN;refGene_tx=
XM_023260242.1;refGene_gn=KALRN;refGene_tx=XM_023260243.1;refGene_gn=K
ALRN;refGene_tx=XM_023260244.1;refGene_gn=KALRN;refGene_tx=XM_02326024
5.1;refGene_gn=KALRN;refGene_tx=XM_023260246.1
C2 70162833 70162949
refGene_gn=KALRN;refGene_tx=XM_019839863.2;refGene_gn=KALRN;refGene_tx
=XM_019839865.2;refGene_gn=KALRN;refGene_tx=XM_019839867.2;refGene_gn=
KALRN;refGene_tx=XM_023260231.1;refGene_gn=KALRN;refGene_tx=XM_0232602
32.1;refGene_gn=KALRN;refGene_tx=XM_023260233.1;refGene_gn=KALRN;refGe
ne_tx=XM_023260234.1;refGene_gn=KALRN;refGene_tx=XM_023260235.1;refGen
e_gn=KALRN;refGene_tx=XM_023260236.1;refGene_gn=KALRN;refGene_tx=XM_02
3260237.1;refGene_gn=KALRN;refGene_tx=XM_023260238.1;refGene_gn=KALRN;
refGene_tx=XM_023260239.1;refGene_gn=KALRN;refGene_tx=XM_023260240.1;r
efGene_gn=KALRN;refGene_tx=XM_023260241.1;refGene_gn=KALRN;refGene_tx=
XM_023260242.1;refGene_gn=KALRN;refGene_tx=XM_023260243.1;refGene_gn=K
ALRN;refGene_tx=XM_023260244.1;refGene_gn=KALRN;refGene_tx=XM_02326024
5.1;refGene_gn=KALRN;refGene_tx=XM_023260246.1
C2 70163411 70163534
refGene_gn=KALRN;refGene_tx=XM_019839863.2;refGene_gn=KALRN;refGene_tx
=XM_019839865.2;refGene_gn=KALRN;refGene_tx=XM_019839867.2;refGene_gn=
KALRN;refGene_tx=XM_023260231.1;refGene_gn=KALRN;refGene_tx=XM_0232602
32.1;refGene_gn=KALRN;refGene_tx=XM_023260233.1;refGene_gn=KALRN;refGe
ne_tx=XM_023260234.1;refGene_gn=KALRN;refGene_tx=XM_023260235.1;refGen
e_gn=KALRN;refGene_tx=XM_023260236.1;refGene_gn=KALRN;refGene_tx=XM_02
3260237.1;refGene_gn=KALRN;refGene_tx=XM_023260238.1;refGene_gn=KALRN;
refGene_tx=XM_023260239.1;refGene_gn=KALRN;refGene_tx=XM_023260240.1;r
efGene_gn=KALRN;refGene_tx=XM_023260241.1;refGene_gn=KALRN;refGene_tx=
XM_023260242.1;refGene_gn=KALRN;refGene_tx=XM_023260243.1;refGene_gn=K
ALRN;refGene_tx=XM_023260244.1;refGene_gn=KALRN;refGene_tx=XM_02326024
5.1;refGene_gn=KALRN;refGene_tx=XM_023260246.1
C2 70171247 70171430
refGene_gn=KALRN;refGene_tx=XM_019839863.2;refGene_gn=KALRN;refGene_tx
=XM_019839865.2;refGene_gn=KALRN;refGene_tx=XM_019839867.2;refGene_gn=
KALRN;refGene_tx=XM_023260231.1;refGene_gn=KALRN;refGene_tx=XM_0232602
32.1;refGene_gn=KALRN;refGene_tx=XM_023260233.1;refGene_gn=KALRN;refGe
ne_tx=XM_023260234.1;refGene_gn=KALRN;refGene_tx=XM_023260235.1;refGen
e_gn=KALRN;refGene_tx=XM_023260236.1;refGene_gn=KALRN;refGene_tx=XM_02
3260237.1;refGene_gn=KALRN;refGene_tx=XM_023260238.1;refGene_gn=KALRN;
refGene_tx=XM_023260239.1;refGene_gn=KALRN;refGene_tx=XM_023260240.1;r

efGene_gn=KALRN;refGene_tx=XM_023260241.1;refGene_gn=KALRN;refGene_tx=XM_023260242.1;refGene_gn=KALRN;refGene_tx=XM_023260243.1;refGene_gn=KALRN;refGene_tx=XM_023260244.1;refGene_gn=KALRN;refGene_tx=XM_023260245.1;refGene_gn=KALRN;refGene_tx=XM_023260246.1
C2 70172690 70172809
refGene_gn=KALRN;refGene_tx=XM_019839863.2;refGene_gn=KALRN;refGene_tx=XM_019839865.2;refGene_gn=KALRN;refGene_tx=XM_019839867.2;refGene_gn=KALRN;refGene_tx=XM_023260231.1;refGene_gn=KALRN;refGene_tx=XM_023260232.1;refGene_gn=KALRN;refGene_tx=XM_023260233.1;refGene_gn=KALRN;refGene_tx=XM_023260234.1;refGene_gn=KALRN;refGene_tx=XM_023260235.1;refGene_gn=KALRN;refGene_tx=XM_023260236.1;refGene_gn=KALRN;refGene_tx=XM_023260237.1;refGene_gn=KALRN;refGene_tx=XM_023260238.1;refGene_gn=KALRN;refGene_tx=XM_023260239.1;refGene_gn=KALRN;refGene_tx=XM_023260240.1;refGene_gn=KALRN;refGene_tx=XM_023260241.1;refGene_gn=KALRN;refGene_tx=XM_023260242.1;refGene_gn=KALRN;refGene_tx=XM_023260243.1;refGene_gn=KALRN;refGene_tx=XM_023260244.1;refGene_gn=KALRN;refGene_tx=XM_023260245.1;refGene_gn=KALRN;refGene_tx=XM_023260246.1
C2 70176124 70176191
refGene_gn=KALRN;refGene_tx=XM_019839863.2;refGene_gn=KALRN;refGene_tx=XM_019839865.2;refGene_gn=KALRN;refGene_tx=XM_019839867.2;refGene_gn=KALRN;refGene_tx=XM_023260231.1;refGene_gn=KALRN;refGene_tx=XM_023260232.1;refGene_gn=KALRN;refGene_tx=XM_023260233.1;refGene_gn=KALRN;refGene_tx=XM_023260234.1;refGene_gn=KALRN;refGene_tx=XM_023260235.1;refGene_gn=KALRN;refGene_tx=XM_023260236.1;refGene_gn=KALRN;refGene_tx=XM_023260237.1;refGene_gn=KALRN;refGene_tx=XM_023260238.1;refGene_gn=KALRN;refGene_tx=XM_023260239.1;refGene_gn=KALRN;refGene_tx=XM_023260240.1;refGene_gn=KALRN;refGene_tx=XM_023260241.1;refGene_gn=KALRN;refGene_tx=XM_023260242.1;refGene_gn=KALRN;refGene_tx=XM_023260243.1;refGene_gn=KALRN;refGene_tx=XM_023260244.1;refGene_gn=KALRN;refGene_tx=XM_023260245.1;refGene_gn=KALRN;refGene_tx=XM_023260246.1
C2 70176758 70176868
refGene_gn=KALRN;refGene_tx=XM_019839863.2;refGene_gn=KALRN;refGene_tx=XM_019839865.2;refGene_gn=KALRN;refGene_tx=XM_019839867.2;refGene_gn=KALRN;refGene_tx=XM_023260231.1;refGene_gn=KALRN;refGene_tx=XM_023260232.1;refGene_gn=KALRN;refGene_tx=XM_023260233.1;refGene_gn=KALRN;refGene_tx=XM_023260234.1;refGene_gn=KALRN;refGene_tx=XM_023260235.1;refGene_gn=KALRN;refGene_tx=XM_023260236.1;refGene_gn=KALRN;refGene_tx=XM_023260237.1;refGene_gn=KALRN;refGene_tx=XM_023260238.1;refGene_gn=KALRN;refGene_tx=XM_023260239.1;refGene_gn=KALRN;refGene_tx=XM_023260240.1;refGene_gn=KALRN;refGene_tx=XM_023260241.1;refGene_gn=KALRN;refGene_tx=XM_023260242.1;refGene_gn=KALRN;refGene_tx=XM_023260243.1;refGene_gn=KALRN;refGene_tx=XM_023260244.1;refGene_gn=KALRN;refGene_tx=XM_023260245.1;refGene_gn=KALRN;refGene_tx=XM_023260246.1
C2 70192657 70192727
refGene_gn=KALRN;refGene_tx=XM_019839863.2;refGene_gn=KALRN;refGene_tx=XM_019839865.2;refGene_gn=KALRN;refGene_tx=XM_019839867.2;refGene_gn=KALRN;refGene_tx=XM_023260231.1;refGene_gn=KALRN;refGene_tx=XM_023260232.1;refGene_gn=KALRN;refGene_tx=XM_023260233.1;refGene_gn=KALRN;refGene_tx=XM_023260234.1;refGene_gn=KALRN;refGene_tx=XM_023260235.1;refGene_gn=KALRN;refGene_tx=XM_023260236.1;refGene_gn=KALRN;refGene_tx=XM_02

3260237.1;refGene_gn=KALRN;refGene_tx=XM_023260238.1;refGene_gn=KALRN;
refGene_tx=XM_023260239.1;refGene_gn=KALRN;refGene_tx=XM_023260240.1;r
efGene_gn=KALRN;refGene_tx=XM_023260241.1;refGene_gn=KALRN;refGene_tx=
XM_023260242.1;refGene_gn=KALRN;refGene_tx=XM_023260243.1;refGene_gn=K
ALRN;refGene_tx=XM_023260244.1;refGene_gn=KALRN;refGene_tx=XM_02326024
5.1;refGene_gn=KALRN;refGene_tx=XM_023260246.1
C2 70195147 70195237
refGene_gn=KALRN;refGene_tx=XM_019839863.2;refGene_gn=KALRN;refGene_tx
=XM_019839865.2;refGene_gn=KALRN;refGene_tx=XM_019839867.2;refGene_gn=
KALRN;refGene_tx=XM_023260231.1;refGene_gn=KALRN;refGene_tx=XM_0232602
32.1;refGene_gn=KALRN;refGene_tx=XM_023260233.1;refGene_gn=KALRN;refGe
ne_tx=XM_023260234.1;refGene_gn=KALRN;refGene_tx=XM_023260235.1;refGen
e_gn=KALRN;refGene_tx=XM_023260236.1;refGene_gn=KALRN;refGene_tx=XM_02
3260237.1;refGene_gn=KALRN;refGene_tx=XM_023260238.1;refGene_gn=KALRN;
refGene_tx=XM_023260239.1;refGene_gn=KALRN;refGene_tx=XM_023260240.1;r
efGene_gn=KALRN;refGene_tx=XM_023260241.1;refGene_gn=KALRN;refGene_tx=
XM_023260242.1;refGene_gn=KALRN;refGene_tx=XM_023260243.1;refGene_gn=K
ALRN;refGene_tx=XM_023260244.1;refGene_gn=KALRN;refGene_tx=XM_02326024
5.1;refGene_gn=KALRN;refGene_tx=XM_023260246.1
C2 70200321 70200414
refGene_gn=KALRN;refGene_tx=XM_019839863.2;refGene_gn=KALRN;refGene_tx
=XM_019839865.2;refGene_gn=KALRN;refGene_tx=XM_019839867.2;refGene_gn=
KALRN;refGene_tx=XM_023260231.1;refGene_gn=KALRN;refGene_tx=XM_0232602
32.1;refGene_gn=KALRN;refGene_tx=XM_023260233.1;refGene_gn=KALRN;refGe
ne_tx=XM_023260234.1;refGene_gn=KALRN;refGene_tx=XM_023260235.1;refGen
e_gn=KALRN;refGene_tx=XM_023260236.1;refGene_gn=KALRN;refGene_tx=XM_02
3260237.1;refGene_gn=KALRN;refGene_tx=XM_023260238.1;refGene_gn=KALRN;
refGene_tx=XM_023260239.1;refGene_gn=KALRN;refGene_tx=XM_023260240.1;r
efGene_gn=KALRN;refGene_tx=XM_023260241.1;refGene_gn=KALRN;refGene_tx=
XM_023260242.1;refGene_gn=KALRN;refGene_tx=XM_023260243.1;refGene_gn=K
ALRN;refGene_tx=XM_023260244.1;refGene_gn=KALRN;refGene_tx=XM_02326024
5.1;refGene_gn=KALRN;refGene_tx=XM_023260246.1
C2 70204652 70204764
refGene_gn=KALRN;refGene_tx=XM_019839863.2;refGene_gn=KALRN;refGene_tx
=XM_019839865.2;refGene_gn=KALRN;refGene_tx=XM_019839867.2;refGene_gn=
KALRN;refGene_tx=XM_023260231.1;refGene_gn=KALRN;refGene_tx=XM_0232602
32.1;refGene_gn=KALRN;refGene_tx=XM_023260233.1;refGene_gn=KALRN;refGe
ne_tx=XM_023260234.1;refGene_gn=KALRN;refGene_tx=XM_023260235.1;refGen
e_gn=KALRN;refGene_tx=XM_023260236.1;refGene_gn=KALRN;refGene_tx=XM_02
3260237.1;refGene_gn=KALRN;refGene_tx=XM_023260238.1;refGene_gn=KALRN;
refGene_tx=XM_023260239.1;refGene_gn=KALRN;refGene_tx=XM_023260240.1;r
efGene_gn=KALRN;refGene_tx=XM_023260241.1;refGene_gn=KALRN;refGene_tx=
XM_023260242.1;refGene_gn=KALRN;refGene_tx=XM_023260243.1;refGene_gn=K
ALRN;refGene_tx=XM_023260244.1;refGene_gn=KALRN;refGene_tx=XM_02326024
5.1;refGene_gn=KALRN;refGene_tx=XM_023260246.1
C2 70206798 70206989
refGene_gn=KALRN;refGene_tx=XM_019839863.2;refGene_gn=KALRN;refGene_tx
=XM_019839865.2;refGene_gn=KALRN;refGene_tx=XM_019839867.2;refGene_gn=
KALRN;refGene_tx=XM_023260231.1;refGene_gn=KALRN;refGene_tx=XM_0232602
32.1;refGene_gn=KALRN;refGene_tx=XM_023260233.1;refGene_gn=KALRN;refGe

ne_tx=XM_023260234.1;refGene_gn=KALRN;refGene_tx=XM_023260235.1;refGene_gn=KALRN;refGene_tx=XM_023260236.1;refGene_gn=KALRN;refGene_tx=XM_023260237.1;refGene_gn=KALRN;refGene_tx=XM_023260238.1;refGene_gn=KALRN;refGene_tx=XM_023260239.1;refGene_gn=KALRN;refGene_tx=XM_023260240.1;refGene_gn=KALRN;refGene_tx=XM_023260241.1;refGene_gn=KALRN;refGene_tx=XM_023260242.1;refGene_gn=KALRN;refGene_tx=XM_023260243.1;refGene_gn=KALRN;refGene_tx=XM_023260244.1;refGene_gn=KALRN;refGene_tx=XM_023260245.1;refGene_gn=KALRN;refGene_tx=XM_023260246.1
C2 70207419 70207521
refGene_gn=KALRN;refGene_tx=XM_019839863.2;refGene_gn=KALRN;refGene_tx=XM_019839865.2;refGene_gn=KALRN;refGene_tx=XM_019839867.2;refGene_gn=KALRN;refGene_tx=XM_023260231.1;refGene_gn=KALRN;refGene_tx=XM_023260232.1;refGene_gn=KALRN;refGene_tx=XM_023260233.1;refGene_gn=KALRN;refGene_tx=XM_023260234.1;refGene_gn=KALRN;refGene_tx=XM_023260235.1;refGene_gn=KALRN;refGene_tx=XM_023260236.1;refGene_gn=KALRN;refGene_tx=XM_023260237.1;refGene_gn=KALRN;refGene_tx=XM_023260238.1;refGene_gn=KALRN;refGene_tx=XM_023260239.1;refGene_gn=KALRN;refGene_tx=XM_023260240.1;refGene_gn=KALRN;refGene_tx=XM_023260241.1;refGene_gn=KALRN;refGene_tx=XM_023260242.1;refGene_gn=KALRN;refGene_tx=XM_023260243.1;refGene_gn=KALRN;refGene_tx=XM_023260244.1;refGene_gn=KALRN;refGene_tx=XM_023260245.1;refGene_gn=KALRN;refGene_tx=XM_023260246.1
C2 70208836 70208979
refGene_gn=KALRN;refGene_tx=XM_019839863.2;refGene_gn=KALRN;refGene_tx=XM_019839865.2;refGene_gn=KALRN;refGene_tx=XM_019839867.2;refGene_gn=KALRN;refGene_tx=XM_023260231.1;refGene_gn=KALRN;refGene_tx=XM_023260232.1;refGene_gn=KALRN;refGene_tx=XM_023260233.1;refGene_gn=KALRN;refGene_tx=XM_023260234.1;refGene_gn=KALRN;refGene_tx=XM_023260235.1;refGene_gn=KALRN;refGene_tx=XM_023260236.1;refGene_gn=KALRN;refGene_tx=XM_023260237.1;refGene_gn=KALRN;refGene_tx=XM_023260238.1;refGene_gn=KALRN;refGene_tx=XM_023260239.1;refGene_gn=KALRN;refGene_tx=XM_023260240.1;refGene_gn=KALRN;refGene_tx=XM_023260241.1;refGene_gn=KALRN;refGene_tx=XM_023260242.1;refGene_gn=KALRN;refGene_tx=XM_023260243.1;refGene_gn=KALRN;refGene_tx=XM_023260244.1;refGene_gn=KALRN;refGene_tx=XM_023260245.1;refGene_gn=KALRN;refGene_tx=XM_023260246.1
C2 70211966 70212069
refGene_gn=KALRN;refGene_tx=XM_019839863.2;refGene_gn=KALRN;refGene_tx=XM_019839865.2;refGene_gn=KALRN;refGene_tx=XM_019839867.2;refGene_gn=KALRN;refGene_tx=XM_023260231.1;refGene_gn=KALRN;refGene_tx=XM_023260232.1;refGene_gn=KALRN;refGene_tx=XM_023260233.1;refGene_gn=KALRN;refGene_tx=XM_023260234.1;refGene_gn=KALRN;refGene_tx=XM_023260235.1;refGene_gn=KALRN;refGene_tx=XM_023260236.1;refGene_gn=KALRN;refGene_tx=XM_023260237.1;refGene_gn=KALRN;refGene_tx=XM_023260238.1;refGene_gn=KALRN;refGene_tx=XM_023260239.1;refGene_gn=KALRN;refGene_tx=XM_023260240.1;refGene_gn=KALRN;refGene_tx=XM_023260241.1;refGene_gn=KALRN;refGene_tx=XM_023260242.1;refGene_gn=KALRN;refGene_tx=XM_023260243.1;refGene_gn=KALRN;refGene_tx=XM_023260244.1;refGene_gn=KALRN;refGene_tx=XM_023260245.1;refGene_gn=KALRN;refGene_tx=XM_023260246.1
C2 70235968 70236031
refGene_gn=KALRN;refGene_tx=XM_023260241.1;refGene_gn=KALRN;refGene_tx=XM_023260242.1;refGene_gn=KALRN;refGene_tx=XM_023260243.1;refGene_gn=

KALRN;refGene_tx=XM_023260244.1;refGene_gn=KALRN;refGene_tx=XM_0232602
45.1;refGene_gn=KALRN;refGene_tx=XM_023260246.1
C2 70285691 70285938
refGene_gn=KALRN;refGene_tx=XM_019839863.2;refGene_gn=KALRN;refGene_tx
=XM_019839865.2;refGene_gn=KALRN;refGene_tx=XM_019839867.2;refGene_gn=
KALRN;refGene_tx=XM_023260231.1;refGene_gn=KALRN;refGene_tx=XM_0232602
32.1;refGene_gn=KALRN;refGene_tx=XM_023260233.1;refGene_gn=KALRN;refGe
ne_tx=XM_023260234.1;refGene_gn=KALRN;refGene_tx=XM_023260235.1;refGen
e_gn=KALRN;refGene_tx=XM_023260236.1;refGene_gn=KALRN;refGene_tx=XM_02
3260237.1;refGene_gn=KALRN;refGene_tx=XM_023260238.1;refGene_gn=KALRN;
refGene_tx=XM_023260239.1;refGene_gn=KALRN;refGene_tx=XM_023260240.1
C2 70305596 70305684
refGene_gn=KALRN;refGene_tx=XM_006936125.4;refGene_gn=KALRN;refGene_tx
=XM_019839878.2;refGene_gn=KALRN;refGene_tx=XM_019839879.2;refGene_gn=
KALRN;refGene_tx=XM_019839880.2;refGene_gn=KALRN;refGene_tx=XM_0198398
81.2;refGene_gn=KALRN;refGene_tx=XM_019839882.2;refGene_gn=KALRN;refGe
ne_tx=XM_019839883.2
C2 70348854 70349138
refGene_gn=KALRN;refGene_tx=XM_006936125.4;refGene_gn=KALRN;refGene_tx
=XM_019839863.2;refGene_gn=KALRN;refGene_tx=XM_019839865.2;refGene_gn=
KALRN;refGene_tx=XM_019839867.2;refGene_gn=KALRN;refGene_tx=XM_0198398
78.2;refGene_gn=KALRN;refGene_tx=XM_019839879.2;refGene_gn=KALRN;refGe
ne_tx=XM_019839880.2;refGene_gn=KALRN;refGene_tx=XM_019839881.2;refGen
e_gn=KALRN;refGene_tx=XM_019839882.2;refGene_gn=KALRN;refGene_tx=XM_01
9839883.2;refGene_gn=KALRN;refGene_tx=XM_023260231.1;refGene_gn=KALRN;
refGene_tx=XM_023260232.1;refGene_gn=KALRN;refGene_tx=XM_023260233.1;r
efGene_gn=KALRN;refGene_tx=XM_023260234.1;refGene_gn=KALRN;refGene_tx=
XM_023260235.1;refGene_gn=KALRN;refGene_tx=XM_023260236.1;refGene_gn=K
ALRN;refGene_tx=XM_023260237.1;refGene_gn=KALRN;refGene_tx=XM_02326023
8.1;refGene_gn=KALRN;refGene_tx=XM_023260239.1;refGene_gn=KALRN;refGen
e_tx=XM_023260240.1
C2 70350312 70350417
refGene_gn=KALRN;refGene_tx=XM_006936125.4;refGene_gn=KALRN;refGene_tx
=XM_019839863.2;refGene_gn=KALRN;refGene_tx=XM_019839865.2;refGene_gn=
KALRN;refGene_tx=XM_019839867.2;refGene_gn=KALRN;refGene_tx=XM_0198398
78.2;refGene_gn=KALRN;refGene_tx=XM_019839879.2;refGene_gn=KALRN;refGe
ne_tx=XM_019839880.2;refGene_gn=KALRN;refGene_tx=XM_019839881.2;refGen
e_gn=KALRN;refGene_tx=XM_019839882.2;refGene_gn=KALRN;refGene_tx=XM_01
9839883.2;refGene_gn=KALRN;refGene_tx=XM_023260231.1;refGene_gn=KALRN;
refGene_tx=XM_023260232.1;refGene_gn=KALRN;refGene_tx=XM_023260233.1;r
efGene_gn=KALRN;refGene_tx=XM_023260234.1;refGene_gn=KALRN;refGene_tx=
XM_023260235.1;refGene_gn=KALRN;refGene_tx=XM_023260236.1;refGene_gn=K
ALRN;refGene_tx=XM_023260237.1;refGene_gn=KALRN;refGene_tx=XM_02326023
8.1;refGene_gn=KALRN;refGene_tx=XM_023260239.1;refGene_gn=KALRN;refGen
e_tx=XM_023260240.1
C2 70353834 70353930
refGene_gn=KALRN;refGene_tx=XM_006936125.4;refGene_gn=KALRN;refGene_tx
=XM_019839863.2;refGene_gn=KALRN;refGene_tx=XM_019839865.2;refGene_gn=
KALRN;refGene_tx=XM_019839867.2;refGene_gn=KALRN;refGene_tx=XM_0198398
78.2;refGene_gn=KALRN;refGene_tx=XM_019839879.2;refGene_gn=KALRN;refGe

ne_tx=XM_023260231.1;refGene_gn=KALRN;refGene_tx=XM_023260232.1;refGen
e_gn=KALRN;refGene_tx=XM_023260233.1;refGene_gn=KALRN;refGene_tx=XM_02
3260234.1;refGene_gn=KALRN;refGene_tx=XM_023260235.1;refGene_gn=KALRN;
refGene_tx=XM_023260237.1;refGene_gn=KALRN;refGene_tx=XM_023260238.1;r
efGene_gn=KALRN;refGene_tx=XM_023260239.1;refGene_gn=KALRN;refGene_tx=
XM_023260240.1
C2 70363454 70363585
refGene_gn=KALRN;refGene_tx=XM_006936125.4;refGene_gn=KALRN;refGene_tx
=XM_019839863.2;refGene_gn=KALRN;refGene_tx=XM_019839865.2;refGene_gn=
KALRN;refGene_tx=XM_019839867.2;refGene_gn=KALRN;refGene_tx=XM_0198398
78.2;refGene_gn=KALRN;refGene_tx=XM_019839879.2;refGene_gn=KALRN;refGe
ne_tx=XM_019839880.2;refGene_gn=KALRN;refGene_tx=XM_019839881.2;refGen
e_gn=KALRN;refGene_tx=XM_019839882.2;refGene_gn=KALRN;refGene_tx=XM_01
9839883.2;refGene_gn=KALRN;refGene_tx=XM_023260231.1;refGene_gn=KALRN;
refGene_tx=XM_023260232.1;refGene_gn=KALRN;refGene_tx=XM_023260233.1;r
efGene_gn=KALRN;refGene_tx=XM_023260234.1;refGene_gn=KALRN;refGene_tx=
XM_023260235.1;refGene_gn=KALRN;refGene_tx=XM_023260236.1;refGene_gn=K
ALRN;refGene_tx=XM_023260237.1;refGene_gn=KALRN;refGene_tx=XM_02326023
8.1;refGene_gn=KALRN;refGene_tx=XM_023260239.1;refGene_gn=KALRN;refGen
e_tx=XM_023260240.1
C2 70367964 70368031
refGene_gn=KALRN;refGene_tx=XM_006936125.4;refGene_gn=KALRN;refGene_tx
=XM_019839863.2;refGene_gn=KALRN;refGene_tx=XM_019839865.2;refGene_gn=
KALRN;refGene_tx=XM_019839867.2;refGene_gn=KALRN;refGene_tx=XM_0198398
78.2;refGene_gn=KALRN;refGene_tx=XM_019839879.2;refGene_gn=KALRN;refGe
ne_tx=XM_019839880.2;refGene_gn=KALRN;refGene_tx=XM_019839881.2;refGen
e_gn=KALRN;refGene_tx=XM_019839882.2;refGene_gn=KALRN;refGene_tx=XM_01
9839883.2;refGene_gn=KALRN;refGene_tx=XM_023260231.1;refGene_gn=KALRN;
refGene_tx=XM_023260232.1;refGene_gn=KALRN;refGene_tx=XM_023260233.1;r
efGene_gn=KALRN;refGene_tx=XM_023260234.1;refGene_gn=KALRN;refGene_tx=
XM_023260235.1;refGene_gn=KALRN;refGene_tx=XM_023260236.1;refGene_gn=K
ALRN;refGene_tx=XM_023260237.1;refGene_gn=KALRN;refGene_tx=XM_02326023
8.1;refGene_gn=KALRN;refGene_tx=XM_023260239.1;refGene_gn=KALRN;refGen
e_tx=XM_023260240.1
C2 70368827 70368931
refGene_gn=KALRN;refGene_tx=XM_006936125.4;refGene_gn=KALRN;refGene_tx
=XM_019839863.2;refGene_gn=KALRN;refGene_tx=XM_019839865.2;refGene_gn=
KALRN;refGene_tx=XM_019839867.2;refGene_gn=KALRN;refGene_tx=XM_0198398
78.2;refGene_gn=KALRN;refGene_tx=XM_019839879.2;refGene_gn=KALRN;refGe
ne_tx=XM_019839880.2;refGene_gn=KALRN;refGene_tx=XM_019839881.2;refGen
e_gn=KALRN;refGene_tx=XM_019839882.2;refGene_gn=KALRN;refGene_tx=XM_01
9839883.2;refGene_gn=KALRN;refGene_tx=XM_023260231.1;refGene_gn=KALRN;
refGene_tx=XM_023260232.1;refGene_gn=KALRN;refGene_tx=XM_023260233.1;r
efGene_gn=KALRN;refGene_tx=XM_023260234.1;refGene_gn=KALRN;refGene_tx=
XM_023260235.1;refGene_gn=KALRN;refGene_tx=XM_023260236.1;refGene_gn=K
ALRN;refGene_tx=XM_023260237.1;refGene_gn=KALRN;refGene_tx=XM_02326023
8.1;refGene_gn=KALRN;refGene_tx=XM_023260239.1;refGene_gn=KALRN;refGen
e_tx=XM_023260240.1
C2 70369118 70369188
refGene_gn=KALRN;refGene_tx=XM_006936125.4;refGene_gn=KALRN;refGene_tx

=XM_019839863.2;refGene_gn=KALRN;refGene_tx=XM_019839865.2;refGene_gn=KALRN;refGene_tx=XM_019839867.2;refGene_gn=KALRN;refGene_tx=XM_019839878.2;refGene_gn=KALRN;refGene_tx=XM_019839879.2;refGene_gn=KALRN;refGene_tx=XM_019839880.2;refGene_gn=KALRN;refGene_tx=XM_019839881.2;refGene_gn=KALRN;refGene_tx=XM_019839882.2;refGene_gn=KALRN;refGene_tx=XM_019839883.2;refGene_gn=KALRN;refGene_tx=XM_023260231.1;refGene_gn=KALRN;refGene_tx=XM_023260232.1;refGene_gn=KALRN;refGene_tx=XM_023260233.1;refGene_gn=KALRN;refGene_tx=XM_023260234.1;refGene_gn=KALRN;refGene_tx=XM_023260235.1;refGene_gn=KALRN;refGene_tx=XM_023260236.1;refGene_gn=KALRN;refGene_tx=XM_023260237.1;refGene_gn=KALRN;refGene_tx=XM_023260238.1;refGene_gn=KALRN;refGene_tx=XM_023260239.1;refGene_gn=KALRN;refGene_tx=XM_023260240.1

C2 70369537 70369624

refGene_gn=KALRN;refGene_tx=XM_006936125.4;refGene_gn=KALRN;refGene_tx=XM_019839863.2;refGene_gn=KALRN;refGene_tx=XM_019839865.2;refGene_gn=KALRN;refGene_tx=XM_019839867.2;refGene_gn=KALRN;refGene_tx=XM_019839878.2;refGene_gn=KALRN;refGene_tx=XM_019839879.2;refGene_gn=KALRN;refGene_tx=XM_019839880.2;refGene_gn=KALRN;refGene_tx=XM_019839881.2;refGene_gn=KALRN;refGene_tx=XM_019839882.2;refGene_gn=KALRN;refGene_tx=XM_019839883.2;refGene_gn=KALRN;refGene_tx=XM_023260231.1;refGene_gn=KALRN;refGene_tx=XM_023260232.1;refGene_gn=KALRN;refGene_tx=XM_023260233.1;refGene_gn=KALRN;refGene_tx=XM_023260234.1;refGene_gn=KALRN;refGene_tx=XM_023260235.1;refGene_gn=KALRN;refGene_tx=XM_023260236.1;refGene_gn=KALRN;refGene_tx=XM_023260237.1;refGene_gn=KALRN;refGene_tx=XM_023260238.1;refGene_gn=KALRN;refGene_tx=XM_023260239.1;refGene_gn=KALRN;refGene_tx=XM_023260240.1

C2 70370464 70370557

refGene_gn=KALRN;refGene_tx=XM_006936125.4;refGene_gn=KALRN;refGene_tx=XM_019839863.2;refGene_gn=KALRN;refGene_tx=XM_019839865.2;refGene_gn=KALRN;refGene_tx=XM_019839867.2;refGene_gn=KALRN;refGene_tx=XM_019839878.2;refGene_gn=KALRN;refGene_tx=XM_019839879.2;refGene_gn=KALRN;refGene_tx=XM_019839880.2;refGene_gn=KALRN;refGene_tx=XM_019839881.2;refGene_gn=KALRN;refGene_tx=XM_019839882.2;refGene_gn=KALRN;refGene_tx=XM_019839883.2;refGene_gn=KALRN;refGene_tx=XM_023260231.1;refGene_gn=KALRN;refGene_tx=XM_023260232.1;refGene_gn=KALRN;refGene_tx=XM_023260233.1;refGene_gn=KALRN;refGene_tx=XM_023260234.1;refGene_gn=KALRN;refGene_tx=XM_023260235.1;refGene_gn=KALRN;refGene_tx=XM_023260236.1;refGene_gn=KALRN;refGene_tx=XM_023260237.1;refGene_gn=KALRN;refGene_tx=XM_023260238.1;refGene_gn=KALRN;refGene_tx=XM_023260239.1;refGene_gn=KALRN;refGene_tx=XM_023260240.1

C2 70371341 70371392

refGene_gn=KALRN;refGene_tx=XM_006936125.4;refGene_gn=KALRN;refGene_tx=XM_019839863.2;refGene_gn=KALRN;refGene_tx=XM_019839865.2;refGene_gn=KALRN;refGene_tx=XM_019839867.2;refGene_gn=KALRN;refGene_tx=XM_019839878.2;refGene_gn=KALRN;refGene_tx=XM_019839879.2;refGene_gn=KALRN;refGene_tx=XM_019839880.2;refGene_gn=KALRN;refGene_tx=XM_019839881.2;refGene_gn=KALRN;refGene_tx=XM_019839882.2;refGene_gn=KALRN;refGene_tx=XM_019839883.2;refGene_gn=KALRN;refGene_tx=XM_023260231.1;refGene_gn=KALRN;refGene_tx=XM_023260232.1;refGene_gn=KALRN;refGene_tx=XM_023260233.1;refGene_gn=KALRN;refGene_tx=XM_023260234.1;refGene_gn=KALRN;refGene_tx=

XM_023260235.1;refGene_gn=KALRN;refGene_tx=XM_023260236.1;refGene_gn=KALRN;refGene_tx=XM_023260237.1;refGene_gn=KALRN;refGene_tx=XM_023260238.1;refGene_gn=KALRN;refGene_tx=XM_023260239.1;refGene_gn=KALRN;refGene_tx=XM_023260240.1

C2 70372482 70372560

refGene_gn=KALRN;refGene_tx=XM_006936125.4;refGene_gn=KALRN;refGene_tx=XM_019839863.2;refGene_gn=KALRN;refGene_tx=XM_019839865.2;refGene_gn=KALRN;refGene_tx=XM_019839867.2;refGene_gn=KALRN;refGene_tx=XM_019839878.2;refGene_gn=KALRN;refGene_tx=XM_019839879.2;refGene_gn=KALRN;refGene_tx=XM_019839880.2;refGene_gn=KALRN;refGene_tx=XM_019839881.2;refGene_gn=KALRN;refGene_tx=XM_019839882.2;refGene_gn=KALRN;refGene_tx=XM_019839883.2;refGene_gn=KALRN;refGene_tx=XM_023260231.1;refGene_gn=KALRN;refGene_tx=XM_023260232.1;refGene_gn=KALRN;refGene_tx=XM_023260233.1;refGene_gn=KALRN;refGene_tx=XM_023260234.1;refGene_gn=KALRN;refGene_tx=XM_023260235.1;refGene_gn=KALRN;refGene_tx=XM_023260236.1;refGene_gn=KALRN;refGene_tx=XM_023260237.1;refGene_gn=KALRN;refGene_tx=XM_023260238.1;refGene_gn=KALRN;refGene_tx=XM_023260239.1;refGene_gn=KALRN;refGene_tx=XM_023260240.1

C2 70376095 70376281

refGene_gn=KALRN;refGene_tx=XM_006936125.4;refGene_gn=KALRN;refGene_tx=XM_019839863.2;refGene_gn=KALRN;refGene_tx=XM_019839865.2;refGene_gn=KALRN;refGene_tx=XM_019839867.2;refGene_gn=KALRN;refGene_tx=XM_019839878.2;refGene_gn=KALRN;refGene_tx=XM_019839879.2;refGene_gn=KALRN;refGene_tx=XM_019839880.2;refGene_gn=KALRN;refGene_tx=XM_019839881.2;refGene_gn=KALRN;refGene_tx=XM_019839882.2;refGene_gn=KALRN;refGene_tx=XM_019839883.2;refGene_gn=KALRN;refGene_tx=XM_023260231.1;refGene_gn=KALRN;refGene_tx=XM_023260232.1;refGene_gn=KALRN;refGene_tx=XM_023260233.1;refGene_gn=KALRN;refGene_tx=XM_023260234.1;refGene_gn=KALRN;refGene_tx=XM_023260235.1;refGene_gn=KALRN;refGene_tx=XM_023260236.1;refGene_gn=KALRN;refGene_tx=XM_023260237.1;refGene_gn=KALRN;refGene_tx=XM_023260238.1;refGene_gn=KALRN;refGene_tx=XM_023260239.1;refGene_gn=KALRN;refGene_tx=XM_023260240.1

C2 70376706 70376878

refGene_gn=KALRN;refGene_tx=XM_006936125.4;refGene_gn=KALRN;refGene_tx=XM_019839863.2;refGene_gn=KALRN;refGene_tx=XM_019839865.2;refGene_gn=KALRN;refGene_tx=XM_019839867.2;refGene_gn=KALRN;refGene_tx=XM_019839878.2;refGene_gn=KALRN;refGene_tx=XM_019839879.2;refGene_gn=KALRN;refGene_tx=XM_019839880.2;refGene_gn=KALRN;refGene_tx=XM_019839881.2;refGene_gn=KALRN;refGene_tx=XM_019839882.2;refGene_gn=KALRN;refGene_tx=XM_019839883.2;refGene_gn=KALRN;refGene_tx=XM_023260231.1;refGene_gn=KALRN;refGene_tx=XM_023260232.1;refGene_gn=KALRN;refGene_tx=XM_023260233.1;refGene_gn=KALRN;refGene_tx=XM_023260234.1;refGene_gn=KALRN;refGene_tx=XM_023260235.1;refGene_gn=KALRN;refGene_tx=XM_023260236.1;refGene_gn=KALRN;refGene_tx=XM_023260237.1;refGene_gn=KALRN;refGene_tx=XM_023260238.1;refGene_gn=KALRN;refGene_tx=XM_023260239.1;refGene_gn=KALRN;refGene_tx=XM_023260240.1

C2 70380329 70380568

refGene_gn=KALRN;refGene_tx=XM_006936125.4;refGene_gn=KALRN;refGene_tx=XM_019839863.2;refGene_gn=KALRN;refGene_tx=XM_019839865.2;refGene_gn=KALRN;refGene_tx=XM_019839867.2;refGene_gn=KALRN;refGene_tx=XM_0198398

78.2;refGene_gn=KALRN;refGene_tx=XM_019839879.2;refGene_gn=KALRN;refGene_tx=XM_019839880.2;refGene_gn=KALRN;refGene_tx=XM_019839881.2;refGene_gn=KALRN;refGene_tx=XM_019839882.2;refGene_gn=KALRN;refGene_tx=XM_019839883.2;refGene_gn=KALRN;refGene_tx=XM_023260231.1;refGene_gn=KALRN;refGene_tx=XM_023260232.1;refGene_gn=KALRN;refGene_tx=XM_023260233.1;refGene_gn=KALRN;refGene_tx=XM_023260234.1;refGene_gn=KALRN;refGene_tx=XM_023260235.1;refGene_gn=KALRN;refGene_tx=XM_023260236.1;refGene_gn=KALRN;refGene_tx=XM_023260237.1;refGene_gn=KALRN;refGene_tx=XM_023260238.1;refGene_gn=KALRN;refGene_tx=XM_023260239.1;refGene_gn=KALRN;refGene_tx=XM_023260240.1
C2 70382274 70382535
refGene_gn=KALRN;refGene_tx=XM_006936125.4;refGene_gn=KALRN;refGene_tx=XM_019839863.2;refGene_gn=KALRN;refGene_tx=XM_019839865.2;refGene_gn=KALRN;refGene_tx=XM_019839867.2;refGene_gn=KALRN;refGene_tx=XM_019839878.2;refGene_gn=KALRN;refGene_tx=XM_019839879.2;refGene_gn=KALRN;refGene_tx=XM_019839880.2;refGene_gn=KALRN;refGene_tx=XM_019839881.2;refGene_gn=KALRN;refGene_tx=XM_019839882.2;refGene_gn=KALRN;refGene_tx=XM_019839883.2;refGene_gn=KALRN;refGene_tx=XM_023260231.1;refGene_gn=KALRN;refGene_tx=XM_023260232.1;refGene_gn=KALRN;refGene_tx=XM_023260233.1;refGene_gn=KALRN;refGene_tx=XM_023260234.1;refGene_gn=KALRN;refGene_tx=XM_023260235.1;refGene_gn=KALRN;refGene_tx=XM_023260236.1;refGene_gn=KALRN;refGene_tx=XM_023260237.1;refGene_gn=KALRN;refGene_tx=XM_023260238.1;refGene_gn=KALRN;refGene_tx=XM_023260239.1;refGene_gn=KALRN;refGene_tx=XM_023260240.1
C2 70382815 70382831 refGene_gn=KALRN;refGene_tx=XM_023260239.1
C2 70385058 70385113 refGene_gn=KALRN;refGene_tx=XM_023260238.1
C2 70386106 70386230
refGene_gn=KALRN;refGene_tx=XM_006936125.4;refGene_gn=KALRN;refGene_tx=XM_019839863.2;refGene_gn=KALRN;refGene_tx=XM_019839865.2;refGene_gn=KALRN;refGene_tx=XM_019839867.2;refGene_gn=KALRN;refGene_tx=XM_019839878.2;refGene_gn=KALRN;refGene_tx=XM_023260231.1;refGene_gn=KALRN;refGene_tx=XM_023260232.1;refGene_gn=KALRN;refGene_tx=XM_023260233.1;refGene_gn=KALRN;refGene_tx=XM_023260234.1;refGene_gn=KALRN;refGene_tx=XM_023260235.1;refGene_gn=KALRN;refGene_tx=XM_023260236.1;refGene_gn=KALRN;refGene_tx=XM_023260237.1
C2 70387367 70387427
refGene_gn=KALRN;refGene_tx=XM_006936125.4;refGene_gn=KALRN;refGene_tx=XM_019839863.2;refGene_gn=KALRN;refGene_tx=XM_019839865.2;refGene_gn=KALRN;refGene_tx=XM_019839867.2;refGene_gn=KALRN;refGene_tx=XM_019839878.2;refGene_gn=KALRN;refGene_tx=XM_023260231.1;refGene_gn=KALRN;refGene_tx=XM_023260232.1;refGene_gn=KALRN;refGene_tx=XM_023260233.1;refGene_gn=KALRN;refGene_tx=XM_023260234.1;refGene_gn=KALRN;refGene_tx=XM_023260235.1;refGene_gn=KALRN;refGene_tx=XM_023260236.1;refGene_gn=KALRN;refGene_tx=XM_023260237.1
C2 70401486 70401514
refGene_gn=KALRN;refGene_tx=XM_006936125.4;refGene_gn=KALRN;refGene_tx=XM_019839863.2;refGene_gn=KALRN;refGene_tx=XM_019839865.2;refGene_gn=KALRN;refGene_tx=XM_019839867.2;refGene_gn=KALRN;refGene_tx=XM_019839878.2;refGene_gn=KALRN;refGene_tx=XM_023260231.1;refGene_gn=KALRN;refGene_tx=XM_023260232.1;refGene_gn=KALRN;refGene_tx=XM_023260233.1;refGen

e_gn=KALRN;refGene_tx=XM_023260234.1;refGene_gn=KALRN;refGene_tx=XM_02
3260235.1;refGene_gn=KALRN;refGene_tx=XM_023260236.1;refGene_gn=KALRN;
refGene_tx=XM_023260237.1
C2 70401995 70402167
refGene_gn=KALRN;refGene_tx=XM_006936125.4;refGene_gn=KALRN;refGene_tx
=XM_019839863.2;refGene_gn=KALRN;refGene_tx=XM_019839865.2;refGene_gn=
KALRN;refGene_tx=XM_019839867.2;refGene_gn=KALRN;refGene_tx=XM_0198398
78.2;refGene_gn=KALRN;refGene_tx=XM_023260231.1;refGene_gn=KALRN;refGe
ne_tx=XM_023260232.1;refGene_gn=KALRN;refGene_tx=XM_023260233.1;refGen
e_gn=KALRN;refGene_tx=XM_023260234.1;refGene_gn=KALRN;refGene_tx=XM_02
3260235.1;refGene_gn=KALRN;refGene_tx=XM_023260236.1;refGene_gn=KALRN;
refGene_tx=XM_023260237.1
C2 70403752 70403874
refGene_gn=KALRN;refGene_tx=XM_006936125.4;refGene_gn=KALRN;refGene_tx
=XM_019839863.2;refGene_gn=KALRN;refGene_tx=XM_019839865.2;refGene_gn=
KALRN;refGene_tx=XM_019839867.2;refGene_gn=KALRN;refGene_tx=XM_0198398
78.2;refGene_gn=KALRN;refGene_tx=XM_023260231.1;refGene_gn=KALRN;refGe
ne_tx=XM_023260232.1;refGene_gn=KALRN;refGene_tx=XM_023260233.1;refGen
e_gn=KALRN;refGene_tx=XM_023260234.1;refGene_gn=KALRN;refGene_tx=XM_02
3260235.1;refGene_gn=KALRN;refGene_tx=XM_023260236.1;refGene_gn=KALRN;
refGene_tx=XM_023260237.1
C2 70404537 70404669
refGene_gn=KALRN;refGene_tx=XM_006936125.4;refGene_gn=KALRN;refGene_tx
=XM_019839863.2;refGene_gn=KALRN;refGene_tx=XM_019839865.2;refGene_gn=
KALRN;refGene_tx=XM_019839867.2;refGene_gn=KALRN;refGene_tx=XM_0198398
78.2;refGene_gn=KALRN;refGene_tx=XM_023260231.1;refGene_gn=KALRN;refGe
ne_tx=XM_023260232.1;refGene_gn=KALRN;refGene_tx=XM_023260233.1;refGen
e_gn=KALRN;refGene_tx=XM_023260234.1;refGene_gn=KALRN;refGene_tx=XM_02
3260235.1;refGene_gn=KALRN;refGene_tx=XM_023260236.1;refGene_gn=KALRN;
refGene_tx=XM_023260237.1
C2 70406298 70406463
refGene_gn=KALRN;refGene_tx=XM_006936125.4;refGene_gn=KALRN;refGene_tx
=XM_019839863.2;refGene_gn=KALRN;refGene_tx=XM_019839865.2;refGene_gn=
KALRN;refGene_tx=XM_019839867.2;refGene_gn=KALRN;refGene_tx=XM_0198398
78.2;refGene_gn=KALRN;refGene_tx=XM_023260231.1;refGene_gn=KALRN;refGe
ne_tx=XM_023260232.1;refGene_gn=KALRN;refGene_tx=XM_023260233.1;refGen
e_gn=KALRN;refGene_tx=XM_023260234.1;refGene_gn=KALRN;refGene_tx=XM_02
3260235.1;refGene_gn=KALRN;refGene_tx=XM_023260236.1;refGene_gn=KALRN;
refGene_tx=XM_023260237.1
C2 70408200 70408279
refGene_gn=KALRN;refGene_tx=XM_006936125.4;refGene_gn=KALRN;refGene_tx
=XM_019839863.2;refGene_gn=KALRN;refGene_tx=XM_019839865.2;refGene_gn=
KALRN;refGene_tx=XM_019839867.2;refGene_gn=KALRN;refGene_tx=XM_0198398
78.2;refGene_gn=KALRN;refGene_tx=XM_023260231.1;refGene_gn=KALRN;refGe
ne_tx=XM_023260232.1;refGene_gn=KALRN;refGene_tx=XM_023260233.1;refGen
e_gn=KALRN;refGene_tx=XM_023260234.1;refGene_gn=KALRN;refGene_tx=XM_02
3260235.1;refGene_gn=KALRN;refGene_tx=XM_023260236.1;refGene_gn=KALRN;
refGene_tx=XM_023260237.1
C2 70415685 70415886
refGene_gn=KALRN;refGene_tx=XM_006936125.4;refGene_gn=KALRN;refGene_tx

=XM_019839863.2;refGene_gn=KALRN;refGene_tx=XM_019839865.2;refGene_gn=KALRN;refGene_tx=XM_019839867.2;refGene_gn=KALRN;refGene_tx=XM_019839878.2;refGene_gn=KALRN;refGene_tx=XM_023260231.1;refGene_gn=KALRN;refGene_tx=XM_023260232.1;refGene_gn=KALRN;refGene_tx=XM_023260233.1;refGene_gn=KALRN;refGene_tx=XM_023260234.1;refGene_gn=KALRN;refGene_tx=XM_023260235.1;refGene_gn=KALRN;refGene_tx=XM_023260236.1;refGene_gn=KALRN;refGene_tx=XM_023260237.1
C2 70418916 70419055
refGene_gn=KALRN;refGene_tx=XM_006936125.4;refGene_gn=KALRN;refGene_tx=XM_019839863.2;refGene_gn=KALRN;refGene_tx=XM_019839865.2;refGene_gn=KALRN;refGene_tx=XM_019839867.2;refGene_gn=KALRN;refGene_tx=XM_019839878.2;refGene_gn=KALRN;refGene_tx=XM_023260231.1;refGene_gn=KALRN;refGene_tx=XM_023260232.1;refGene_gn=KALRN;refGene_tx=XM_023260233.1;refGene_gn=KALRN;refGene_tx=XM_023260234.1;refGene_gn=KALRN;refGene_tx=XM_023260235.1;refGene_gn=KALRN;refGene_tx=XM_023260236.1;refGene_gn=KALRN;refGene_tx=XM_023260237.1
C2 70419632 70420178
refGene_gn=KALRN;refGene_tx=XM_006936125.4;refGene_gn=KALRN;refGene_tx=XM_019839863.2;refGene_gn=KALRN;refGene_tx=XM_019839865.2;refGene_gn=KALRN;refGene_tx=XM_019839867.2;refGene_gn=KALRN;refGene_tx=XM_019839878.2;refGene_gn=KALRN;refGene_tx=XM_023260231.1;refGene_gn=KALRN;refGene_tx=XM_023260232.1;refGene_gn=KALRN;refGene_tx=XM_023260233.1;refGene_gn=KALRN;refGene_tx=XM_023260234.1;refGene_gn=KALRN;refGene_tx=XM_023260235.1;refGene_gn=KALRN;refGene_tx=XM_023260236.1;refGene_gn=KALRN;refGene_tx=XM_023260237.1
C2 70428066 70428222
refGene_gn=UMPS;refGene_tx=XM_003991720.5;refGene_gn=UMPS;refGene_tx=XM_023260247.1
C2 70438030 70438184
refGene_gn=UMPS;refGene_tx=XM_003991720.5;refGene_gn=UMPS;refGene_tx=XM_023260247.1
C2 70439957 70440635
refGene_gn=UMPS;refGene_tx=XM_003991720.5;refGene_gn=UMPS;refGene_tx=XM_023260247.1
C2 70442763 70442939
refGene_gn=UMPS;refGene_tx=XM_003991720.5;refGene_gn=UMPS;refGene_tx=XM_023260247.1
C2 70444574 70444689
refGene_gn=UMPS;refGene_tx=XM_003991720.5;refGene_gn=UMPS;refGene_tx=XM_023260247.1
C2 70462013 70462183 refGene_gn=UMPS;refGene_tx=XM_003991720.5
C2 70472858 70472920 refGene_gn=UMPS;refGene_tx=XM_023260247.1
C2 70477390 70477486
refGene_gn=ITGB5;refGene_tx=XM_023260248.1;refGene_gn=ITGB5;refGene_tx=XM_023260249.1
C2 70478130 70478297
refGene_gn=ITGB5;refGene_tx=XM_023260248.1;refGene_gn=ITGB5;refGene_tx=XM_023260249.1
C2 70479689 70479809
refGene_gn=ITGB5;refGene_tx=XM_023260248.1;refGene_gn=ITGB5;refGene_tx

=XM_023260249.1
C2	70483323	70483427
refGene_gn=ITGB5;refGene_tx=XM_023260248.1;refGene_gn=ITGB5;refGene_tx
=XM_023260249.1
C2	70487835	70488058
refGene_gn=ITGB5;refGene_tx=XM_023260248.1;refGene_gn=ITGB5;refGene_tx
=XM_023260249.1
C2	70509786	70510216
refGene_gn=ITGB5;refGene_tx=XM_023260248.1;refGene_gn=ITGB5;refGene_tx
=XM_023260249.1
C2	70519395	70519530
refGene_gn=ITGB5;refGene_tx=XM_023260248.1;refGene_gn=ITGB5;refGene_tx
=XM_023260249.1
C2	70528295	70528385
refGene_gn=ITGB5;refGene_tx=XM_023260248.1;refGene_gn=ITGB5;refGene_tx
=XM_023260249.1
C2	70530240	70530336
refGene_gn=ITGB5;refGene_tx=XM_023260248.1;refGene_gn=ITGB5;refGene_tx
=XM_023260249.1
C2	70531982	70532144	refGene_gn=ITGB5;refGene_tx=XM_023260248.1
C2	70554723	70554892
refGene_gn=ITGB5;refGene_tx=XM_023260248.1;refGene_gn=ITGB5;refGene_tx
=XM_023260249.1
C2	70564855	70565105
refGene_gn=ITGB5;refGene_tx=XM_023260248.1;refGene_gn=ITGB5;refGene_tx
=XM_023260249.1
C2	70577458	70577663
refGene_gn=ITGB5;refGene_tx=XM_023260248.1;refGene_gn=ITGB5;refGene_tx
=XM_023260249.1
C2	70587515	70587601
refGene_gn=ITGB5;refGene_tx=XM_023260248.1;refGene_gn=ITGB5;refGene_tx
=XM_023260249.1
C2	70598459	70599225
refGene_gn=ITGB5;refGene_tx=XM_023260248.1;refGene_gn=ITGB5;refGene_tx
=XM_023260249.1;refGene_gn=LOC102900615;refGene_tx=XM_023260170.1
C2	70600766	70600869
refGene_gn=LOC102900615;refGene_tx=XM_023260170.1
C2	70613083	70613282	refGene_gn=MUC13;refGene_tx=XM_019839884.2
C2	70615527	70615612	refGene_gn=MUC13;refGene_tx=XM_019839884.2
C2	70617422	70617460	refGene_gn=MUC13;refGene_tx=XM_019839884.2
C2	70618114	70618247	refGene_gn=MUC13;refGene_tx=XM_019839884.2
C2	70618574	70618700	refGene_gn=MUC13;refGene_tx=XM_019839884.2
C2	70621125	70621289	refGene_gn=MUC13;refGene_tx=XM_019839884.2
C2	70623018	70623074	refGene_gn=MUC13;refGene_tx=XM_019839884.2
C2	70626986	70627093	refGene_gn=MUC13;refGene_tx=XM_019839884.2
C2	70628478	70628601	refGene_gn=MUC13;refGene_tx=XM_019839884.2
C2	70632642	70633278	refGene_gn=MUC13;refGene_tx=XM_019839884.2
C2	70635181	70635326	refGene_gn=MUC13;refGene_tx=XM_019839884.2
C2	70673777	70673927	refGene_gn=HEG1;refGene_tx=XM_023260172.1
C2	70676707	70676882	refGene_gn=HEG1;refGene_tx=XM_023260172.1

C2 70680211 70680299 refGene_gn=HEG1;refGene_tx=XM_023260172.1
C2 70692992 70693030 refGene_gn=HEG1;refGene_tx=XM_023260172.1
C2 70693163 70693206 refGene_gn=HEG1;refGene_tx=XM_023260172.1
C2 70698458 70698593 refGene_gn=HEG1;refGene_tx=XM_023260172.1
C2 70702972 70703133 refGene_gn=HEG1;refGene_tx=XM_023260172.1
C2 70703381 70703440 refGene_gn=HEG1;refGene_tx=XM_023260172.1
C2 70706564 70706668 refGene_gn=HEG1;refGene_tx=XM_023260172.1
C2 70710544 70710664 refGene_gn=HEG1;refGene_tx=XM_023260172.1
C2 70711289 70711406 refGene_gn=HEG1;refGene_tx=XM_023260172.1
C2 70713691 70714620 refGene_gn=HEG1;refGene_tx=XM_023260172.1
C2 70714675 70715084 refGene_gn=HEG1;refGene_tx=XM_023260172.1
C2 70715716 70715983 refGene_gn=HEG1;refGene_tx=XM_023260172.1
C2 70719649 70719997 refGene_gn=HEG1;refGene_tx=XM_023260172.1
C2 70720924 70721278 refGene_gn=HEG1;refGene_tx=XM_023260172.1
C2 70728558 70728840 refGene_gn=HEG1;refGene_tx=XM_023260172.1
C2 70730504 70730792 refGene_gn=HEG1;refGene_tx=XM_023260172.1
C2 70758770 70759019 refGene_gn=HEG1;refGene_tx=XM_023260172.1
C2 70760488 70760549 refGene_gn=HEG1;refGene_tx=XM_023260172.1
C2 70781742 70781905 refGene_gn=SLC12A8;refGene_tx=XM_019839892.2
C2 70785491 70785552 refGene_gn=SLC12A8;refGene_tx=XM_019839892.2
C2 70794533 70794651 refGene_gn=SLC12A8;refGene_tx=XM_019839892.2
C2 70795200 70795298 refGene_gn=SLC12A8;refGene_tx=XM_019839892.2
C2 70807170 70807816 refGene_gn=SLC12A8;refGene_tx=XM_019839892.2
C2 70814106 70814253 refGene_gn=SLC12A8;refGene_tx=XM_019839892.2
C2 70821761 70821849 refGene_gn=SLC12A8;refGene_tx=XM_019839892.2
C2 70823376 70823464 refGene_gn=SLC12A8;refGene_tx=XM_019839892.2
C2 70839522 70839636 refGene_gn=SLC12A8;refGene_tx=XM_019839892.2
C2 70885853 70886085 refGene_gn=SLC12A8;refGene_tx=XM_019839892.2
C2 70894483 70894675 refGene_gn=SLC12A8;refGene_tx=XM_019839892.2
C2 70899242 70899389 refGene_gn=SLC12A8;refGene_tx=XM_019839892.2
C2 70922562 70922604 refGene_gn=SLC12A8;refGene_tx=XM_019839892.2
C2 70939856 70941455
refGene_gn=ZNF148;refGene_tx=XM_003991722.5;refGene_gn=ZNF148;refGene_tx=XM_003991724.5;refGene_gn=ZNF148;refGene_tx=XM_006936126.4;refGene_gn=ZNF148;refGene_tx=XM_019839899.2;refGene_gn=ZNF148;refGene_tx=XM_019839900.2;refGene_gn=ZNF148;refGene_tx=XM_019839901.2
C2 70941776 70941895
refGene_gn=ZNF148;refGene_tx=XM_003991722.5;refGene_gn=ZNF148;refGene_tx=XM_003991724.5;refGene_gn=ZNF148;refGene_tx=XM_006936126.4;refGene_gn=ZNF148;refGene_tx=XM_019839899.2;refGene_gn=ZNF148;refGene_tx=XM_019839900.2;refGene_gn=ZNF148;refGene_tx=XM_019839901.2
C2 70968935 70969019
refGene_gn=ZNF148;refGene_tx=XM_003991722.5;refGene_gn=ZNF148;refGene_tx=XM_003991724.5;refGene_gn=ZNF148;refGene_tx=XM_006936126.4;refGene_gn=ZNF148;refGene_tx=XM_019839899.2;refGene_gn=ZNF148;refGene_tx=XM_019839900.2;refGene_gn=ZNF148;refGene_tx=XM_019839901.2
C2 70970420 70970544
refGene_gn=ZNF148;refGene_tx=XM_003991722.5;refGene_gn=ZNF148;refGene_tx=XM_003991724.5;refGene_gn=ZNF148;refGene_tx=XM_006936126.4;refGene_gn=ZNF148;refGene_tx=XM_019839899.2;refGene_gn=ZNF148;refGene_tx=XM_01

9839900.2;refGene_gn=ZNF148;refGene_tx=XM_019839901.2
C2 70980169 70980295
refGene_gn=ZNF148;refGene_tx=XM_003991722.5;refGene_gn=ZNF148;refGene_tx=XM_003991724.5;refGene_gn=ZNF148;refGene_tx=XM_006936126.4;refGene_gn=ZNF148;refGene_tx=XM_019839899.2;refGene_gn=ZNF148;refGene_tx=XM_019839900.2;refGene_gn=ZNF148;refGene_tx=XM_019839901.2
C2 71007546 71007879
refGene_gn=ZNF148;refGene_tx=XM_003991722.5;refGene_gn=ZNF148;refGene_tx=XM_003991724.5;refGene_gn=ZNF148;refGene_tx=XM_006936126.4;refGene_gn=ZNF148;refGene_tx=XM_019839899.2;refGene_gn=ZNF148;refGene_tx=XM_019839900.2;refGene_gn=ZNF148;refGene_tx=XM_019839901.2
C2 71125753 71125801 refGene_gn=SNX4;refGene_tx=XM_003991725.3
C2 71130531 71130646 refGene_gn=SNX4;refGene_tx=XM_003991725.3
C2 71132824 71132970 refGene_gn=SNX4;refGene_tx=XM_003991725.3
C2 71135507 71135607 refGene_gn=SNX4;refGene_tx=XM_003991725.3
C2 71137895 71137985 refGene_gn=SNX4;refGene_tx=XM_003991725.3
C2 71142502 71142568 refGene_gn=SNX4;refGene_tx=XM_003991725.3
C2 71153460 71153522 refGene_gn=SNX4;refGene_tx=XM_003991725.3
C2 71163408 71163481 refGene_gn=SNX4;refGene_tx=XM_003991725.3
C2 71168111 71168167 refGene_gn=SNX4;refGene_tx=XM_003991725.3
C2 71173619 71173667 refGene_gn=SNX4;refGene_tx=XM_003991725.3
C2 71174145 71174295 refGene_gn=SNX4;refGene_tx=XM_003991725.3
C2 71174372 71174508 refGene_gn=SNX4;refGene_tx=XM_003991725.3
C2 71181006 71181128 refGene_gn=SNX4;refGene_tx=XM_003991725.3
C2 71195122 71195263 refGene_gn=SNX4;refGene_tx=XM_003991725.3
C2 71221402 71221468 refGene_gn=OSBPL11;refGene_tx=XM_003991726.5
C2 71225543 71225697 refGene_gn=OSBPL11;refGene_tx=XM_003991726.5
C2 71233400 71233583 refGene_gn=OSBPL11;refGene_tx=XM_003991726.5
C2 71247058 71247245 refGene_gn=OSBPL11;refGene_tx=XM_003991726.5
C2 71250528 71251027 refGene_gn=OSBPL11;refGene_tx=XM_003991726.5
C2 71259872 71260013 refGene_gn=OSBPL11;refGene_tx=XM_003991726.5
C2 71265608 71265754 refGene_gn=OSBPL11;refGene_tx=XM_003991726.5
C2 71270445 71270647 refGene_gn=OSBPL11;refGene_tx=XM_003991726.5
C2 71273974 71274151 refGene_gn=OSBPL11;refGene_tx=XM_003991726.5
C2 71276691 71276771 refGene_gn=OSBPL11;refGene_tx=XM_003991726.5
C2 71277674 71277850 refGene_gn=OSBPL11;refGene_tx=XM_003991726.5
C2 71287376 71287445 refGene_gn=OSBPL11;refGene_tx=XM_003991726.5
C2 71307666 71307983 refGene_gn=OSBPL11;refGene_tx=XM_003991726.5
C2 71324227 71324422
refGene_gn=LOC109503486;refGene_tx=XM_019839920.2
C2 71375505 71375673
refGene_gn=LMLN;refGene_tx=XM_019839909.2;refGene_gn=LMLN;refGene_tx=XM_019839911.2
C2 71385224 71385435
refGene_gn=LMLN;refGene_tx=XM_019839909.2;refGene_gn=LMLN;refGene_tx=XM_019839911.2;refGene_gn=LMLN;refGene_tx=XM_019839913.2
C2 71392402 71392533
refGene_gn=LMLN;refGene_tx=XM_019839909.2;refGene_gn=LMLN;refGene_tx=XM_019839911.2
C2 71395847 71396007

refGene_gn=LMLN;refGene_tx=XM_019839909.2;refGene_gn=LMLN;refGene_tx=XM_019839911.2;refGene_gn=LMLN;refGene_tx=XM_019839913.2
C2 71397176 71397309
refGene_gn=LMLN;refGene_tx=XM_019839909.2;refGene_gn=LMLN;refGene_tx=XM_019839911.2;refGene_gn=LMLN;refGene_tx=XM_019839913.2
C2 71402105 71402216
refGene_gn=LMLN;refGene_tx=XM_019839909.2;refGene_gn=LMLN;refGene_tx=XM_019839911.2;refGene_gn=LMLN;refGene_tx=XM_019839913.2
C2 71416468 71416545
refGene_gn=LMLN;refGene_tx=XM_019839909.2;refGene_gn=LMLN;refGene_tx=XM_019839911.2;refGene_gn=LMLN;refGene_tx=XM_019839913.2
C2 71420547 71420655
refGene_gn=LMLN;refGene_tx=XM_019839909.2;refGene_gn=LMLN;refGene_tx=XM_019839911.2;refGene_gn=LMLN;refGene_tx=XM_019839913.2
C2 71422259 71422377
refGene_gn=LMLN;refGene_tx=XM_019839909.2;refGene_gn=LMLN;refGene_tx=XM_019839911.2;refGene_gn=LMLN;refGene_tx=XM_019839913.2
C2 71426655 71426750
refGene_gn=LMLN;refGene_tx=XM_019839909.2;refGene_gn=LMLN;refGene_tx=XM_019839911.2;refGene_gn=LMLN;refGene_tx=XM_019839913.2
C2 71430657 71430763
refGene_gn=LMLN;refGene_tx=XM_019839909.2;refGene_gn=LMLN;refGene_tx=XM_019839911.2;refGene_gn=LMLN;refGene_tx=XM_019839913.2
C2 71434200 71434379
refGene_gn=LMLN;refGene_tx=XM_019839909.2;refGene_gn=LMLN;refGene_tx=XM_019839911.2;refGene_gn=LMLN;refGene_tx=XM_019839913.2
C2 71437811 71437926
refGene_gn=LMLN;refGene_tx=XM_019839909.2;refGene_gn=LMLN;refGene_tx=XM_019839911.2;refGene_gn=LMLN;refGene_tx=XM_019839913.2
C2 71438976 71439059
refGene_gn=LMLN;refGene_tx=XM_019839909.2;refGene_gn=LMLN;refGene_tx=XM_019839911.2;refGene_gn=LMLN;refGene_tx=XM_019839913.2
C2 71440166 71440197
refGene_gn=LMLN;refGene_tx=XM_019839909.2;refGene_gn=LMLN;refGene_tx=XM_019839911.2;refGene_gn=LMLN;refGene_tx=XM_019839913.2
C2 71441514 71441612
refGene_gn=LMLN;refGene_tx=XM_019839909.2;refGene_gn=LMLN;refGene_tx=XM_019839911.2;refGene_gn=LMLN;refGene_tx=XM_019839913.2
C2 71457186 71457405
refGene_gn=LMLN;refGene_tx=XM_019839909.2;refGene_gn=LMLN;refGene_tx=XM_019839911.2;refGene_gn=LMLN;refGene_tx=XM_019839913.2
C2 71462551 71462696
refGene_gn=RPL35A;refGene_tx=XM_003991729.4;refGene_gn=RPL35A;refGene_tx=XM_006936129.4;refGene_gn=RPL35A;refGene_tx=XM_019839919.2
C2 71463715 71463868
refGene_gn=RPL35A;refGene_tx=XM_003991729.4;refGene_gn=RPL35A;refGene_tx=XM_006936129.4;refGene_gn=RPL35A;refGene_tx=XM_019839919.2
C2 71464069 71464114
refGene_gn=RPL35A;refGene_tx=XM_003991729.4;refGene_gn=RPL35A;refGene_tx=XM_006936129.4;refGene_gn=RPL35A;refGene_tx=XM_019839919.2

C2	71464854	71464946	refGene_gn=RPL35A;refGene_tx=XM_003991729.4
C2	71470390	71470425	refGene_gn=IQCG;refGene_tx=XM_006936130.3;refGene_gn=IQCG;refGene_tx=XM_019839914.2;refGene_gn=IQCG;refGene_tx=XM_019839918.2
C2	71471501	71471769	refGene_gn=IQCG;refGene_tx=XM_006936130.3;refGene_gn=IQCG;refGene_tx=XM_019839914.2;refGene_gn=IQCG;refGene_tx=XM_019839918.2
C2	71476149	71476385	refGene_gn=IQCG;refGene_tx=XM_006936130.3;refGene_gn=IQCG;refGene_tx=XM_019839914.2;refGene_gn=IQCG;refGene_tx=XM_019839918.2
C2	71479810	71479939	refGene_gn=IQCG;refGene_tx=XM_006936130.3;refGene_gn=IQCG;refGene_tx=XM_019839914.2;refGene_gn=IQCG;refGene_tx=XM_019839918.2
C2	71497223	71497305	refGene_gn=IQCG;refGene_tx=XM_006936130.3;refGene_gn=IQCG;refGene_tx=XM_019839914.2;refGene_gn=IQCG;refGene_tx=XM_019839918.2
C2	71505509	71505701	refGene_gn=IQCG;refGene_tx=XM_006936130.3;refGene_gn=IQCG;refGene_tx=XM_019839914.2;refGene_gn=IQCG;refGene_tx=XM_019839918.2
C2	71505794	71505830	refGene_gn=IQCG;refGene_tx=XM_019839918.2
C2	71506918	71506993	refGene_gn=IQCG;refGene_tx=XM_006936130.3;refGene_gn=IQCG;refGene_tx=XM_019839914.2
C2	71518194	71518209	refGene_gn=LRCH3;refGene_tx=XM_023260252.1
C2	71519019	71519151	refGene_gn=IQCG;refGene_tx=XM_006936130.3;refGene_gn=IQCG;refGene_tx=XM_019839914.2
C2	71519882	71519984	refGene_gn=IQCG;refGene_tx=XM_006936130.3;refGene_gn=IQCG;refGene_tx=XM_019839914.2
C2	71521443	71521461	refGene_gn=LRCH3;refGene_tx=XM_023260251.1
C2	71521526	71521664	refGene_gn=IQCG;refGene_tx=XM_006936130.3;refGene_gn=IQCG;refGene_tx=XM_019839914.2
C2	71525610	71525733	refGene_gn=LRCH3;refGene_tx=XM_011285906.2
C2	71528370	71528571	refGene_gn=LRCH3;refGene_tx=XM_019839906.2
C2	71529386	71529716	refGene_gn=LRCH3;refGene_tx=XM_019839902.2;refGene_gn=LRCH3;refGene_tx=XM_019839903.2;refGene_gn=LRCH3;refGene_tx=XM_019839904.2;refGene_gn=LRCH3;refGene_tx=XM_019839905.2;refGene_gn=LRCH3;refGene_tx=XM_019839907.2
C2	71531912	71531990	refGene_gn=LRCH3;refGene_tx=XM_011285906.2;refGene_gn=LRCH3;refGene_tx=XM_019839902.2;refGene_gn=LRCH3;refGene_tx=XM_019839903.2;refGene_gn=LRCH3;refGene_tx=XM_019839904.2;refGene_gn=LRCH3;refGene_tx=XM_019839905.2;refGene_gn=LRCH3;refGene_tx=XM_019839906.2;refGene_gn=LRCH3;refGene_tx=XM_019839907.2;refGene_gn=LRCH3;refGene_tx=XM_023260251.1;refGene_gn=LRCH3;refGene_tx=XM_023260252.1
C2	71542720	71542858	refGene_gn=LRCH3;refGene_tx=XM_011285906.2;refGene_gn=LRCH3;refGene_tx

=XM_019839902.2;refGene_gn=LRCH3;refGene_tx=XM_019839903.2;refGene_gn=LRCH3;refGene_tx=XM_019839904.2;refGene_gn=LRCH3;refGene_tx=XM_019839905.2;refGene_gn=LRCH3;refGene_tx=XM_019839906.2;refGene_gn=LRCH3;refGene_tx=XM_019839907.2;refGene_gn=LRCH3;refGene_tx=XM_023260251.1;refGene_gn=LRCH3;refGene_tx=XM_023260252.1
C2 71543640 71543759
refGene_gn=LRCH3;refGene_tx=XM_011285906.2;refGene_gn=LRCH3;refGene_tx=XM_019839902.2;refGene_gn=LRCH3;refGene_tx=XM_019839903.2;refGene_gn=LRCH3;refGene_tx=XM_019839904.2;refGene_gn=LRCH3;refGene_tx=XM_019839905.2;refGene_gn=LRCH3;refGene_tx=XM_019839906.2;refGene_gn=LRCH3;refGene_tx=XM_019839907.2;refGene_gn=LRCH3;refGene_tx=XM_023260251.1;refGene_gn=LRCH3;refGene_tx=XM_023260252.1
C2 71545561 71545669
refGene_gn=LRCH3;refGene_tx=XM_011285906.2;refGene_gn=LRCH3;refGene_tx=XM_019839902.2;refGene_gn=LRCH3;refGene_tx=XM_019839903.2;refGene_gn=LRCH3;refGene_tx=XM_019839904.2;refGene_gn=LRCH3;refGene_tx=XM_019839905.2;refGene_gn=LRCH3;refGene_tx=XM_019839906.2;refGene_gn=LRCH3;refGene_tx=XM_019839907.2;refGene_gn=LRCH3;refGene_tx=XM_023260251.1;refGene_gn=LRCH3;refGene_tx=XM_023260252.1
C2 71546171 71546220
refGene_gn=LRCH3;refGene_tx=XM_011285906.2;refGene_gn=LRCH3;refGene_tx=XM_019839902.2;refGene_gn=LRCH3;refGene_tx=XM_019839903.2;refGene_gn=LRCH3;refGene_tx=XM_019839904.2;refGene_gn=LRCH3;refGene_tx=XM_019839905.2;refGene_gn=LRCH3;refGene_tx=XM_019839906.2;refGene_gn=LRCH3;refGene_tx=XM_019839907.2;refGene_gn=LRCH3;refGene_tx=XM_023260251.1;refGene_gn=LRCH3;refGene_tx=XM_023260252.1
C2 71548709 71548781
refGene_gn=LRCH3;refGene_tx=XM_011285906.2;refGene_gn=LRCH3;refGene_tx=XM_019839902.2;refGene_gn=LRCH3;refGene_tx=XM_019839904.2;refGene_gn=LRCH3;refGene_tx=XM_019839906.2;refGene_gn=LRCH3;refGene_tx=XM_023260251.1;refGene_gn=LRCH3;refGene_tx=XM_023260252.1
C2 71553032 71553086
refGene_gn=LRCH3;refGene_tx=XM_011285906.2;refGene_gn=LRCH3;refGene_tx=XM_019839902.2;refGene_gn=LRCH3;refGene_tx=XM_019839903.2;refGene_gn=LRCH3;refGene_tx=XM_019839906.2;refGene_gn=LRCH3;refGene_tx=XM_023260251.1;refGene_gn=LRCH3;refGene_tx=XM_023260252.1
C2 71554592 71554643
refGene_gn=LRCH3;refGene_tx=XM_011285906.2;refGene_gn=LRCH3;refGene_tx=XM_019839902.2;refGene_gn=LRCH3;refGene_tx=XM_019839903.2;refGene_gn=LRCH3;refGene_tx=XM_019839905.2;refGene_gn=LRCH3;refGene_tx=XM_019839906.2;refGene_gn=LRCH3;refGene_tx=XM_023260251.1;refGene_gn=LRCH3;refGene_tx=XM_023260252.1
C2 71557998 71558148
refGene_gn=LRCH3;refGene_tx=XM_011285906.2;refGene_gn=LRCH3;refGene_tx=XM_019839902.2;refGene_gn=LRCH3;refGene_tx=XM_019839903.2;refGene_gn=LRCH3;refGene_tx=XM_019839904.2;refGene_gn=LRCH3;refGene_tx=XM_019839905.2;refGene_gn=LRCH3;refGene_tx=XM_019839906.2;refGene_gn=LRCH3;refGene_tx=XM_019839907.2;refGene_gn=LRCH3;refGene_tx=XM_023260251.1;refGene_gn=LRCH3;refGene_tx=XM_023260252.1
C2 71558569 71558621

refGene_gn=LRCH3;refGene_tx=XM_011285906.2;refGene_gn=LRCH3;refGene_tx=XM_019839902.2;refGene_gn=LRCH3;refGene_tx=XM_019839903.2;refGene_gn=LRCH3;refGene_tx=XM_019839904.2;refGene_gn=LRCH3;refGene_tx=XM_019839905.2;refGene_gn=LRCH3;refGene_tx=XM_019839906.2;refGene_gn=LRCH3;refGene_tx=XM_019839907.2;refGene_gn=LRCH3;refGene_tx=XM_023260251.1;refGene_gn=LRCH3;refGene_tx=XM_023260252.1
C2 71564203 71564280
refGene_gn=LRCH3;refGene_tx=XM_011285906.2;refGene_gn=LRCH3;refGene_tx=XM_019839902.2;refGene_gn=LRCH3;refGene_tx=XM_019839903.2;refGene_gn=LRCH3;refGene_tx=XM_019839904.2;refGene_gn=LRCH3;refGene_tx=XM_019839905.2;refGene_gn=LRCH3;refGene_tx=XM_019839906.2;refGene_gn=LRCH3;refGene_tx=XM_019839907.2;refGene_gn=LRCH3;refGene_tx=XM_023260251.1;refGene_gn=LRCH3;refGene_tx=XM_023260252.1
C2 71567017 71567166
refGene_gn=LRCH3;refGene_tx=XM_011285906.2;refGene_gn=LRCH3;refGene_tx=XM_019839902.2;refGene_gn=LRCH3;refGene_tx=XM_019839903.2;refGene_gn=LRCH3;refGene_tx=XM_019839904.2;refGene_gn=LRCH3;refGene_tx=XM_019839905.2;refGene_gn=LRCH3;refGene_tx=XM_019839906.2;refGene_gn=LRCH3;refGene_tx=XM_019839907.2;refGene_gn=LRCH3;refGene_tx=XM_023260251.1;refGene_gn=LRCH3;refGene_tx=XM_023260252.1
C2 71570363 71570484
refGene_gn=LRCH3;refGene_tx=XM_011285906.2;refGene_gn=LRCH3;refGene_tx=XM_019839902.2;refGene_gn=LRCH3;refGene_tx=XM_019839903.2;refGene_gn=LRCH3;refGene_tx=XM_019839904.2;refGene_gn=LRCH3;refGene_tx=XM_019839905.2;refGene_gn=LRCH3;refGene_tx=XM_019839906.2;refGene_gn=LRCH3;refGene_tx=XM_019839907.2;refGene_gn=LRCH3;refGene_tx=XM_023260251.1;refGene_gn=LRCH3;refGene_tx=XM_023260252.1
C2 71571709 71571803
refGene_gn=LRCH3;refGene_tx=XM_011285906.2;refGene_gn=LRCH3;refGene_tx=XM_019839902.2;refGene_gn=LRCH3;refGene_tx=XM_019839903.2;refGene_gn=LRCH3;refGene_tx=XM_019839904.2;refGene_gn=LRCH3;refGene_tx=XM_019839905.2;refGene_gn=LRCH3;refGene_tx=XM_019839906.2;refGene_gn=LRCH3;refGene_tx=XM_019839907.2;refGene_gn=LRCH3;refGene_tx=XM_023260251.1;refGene_gn=LRCH3;refGene_tx=XM_023260252.1
C2 71579807 71579917
refGene_gn=LRCH3;refGene_tx=XM_011285906.2;refGene_gn=LRCH3;refGene_tx=XM_019839902.2;refGene_gn=LRCH3;refGene_tx=XM_019839903.2;refGene_gn=LRCH3;refGene_tx=XM_019839904.2;refGene_gn=LRCH3;refGene_tx=XM_019839905.2;refGene_gn=LRCH3;refGene_tx=XM_019839906.2;refGene_gn=LRCH3;refGene_tx=XM_019839907.2;refGene_gn=LRCH3;refGene_tx=XM_023260251.1;refGene_gn=LRCH3;refGene_tx=XM_023260252.1
C2 71581104 71581241
refGene_gn=LRCH3;refGene_tx=XM_011285906.2;refGene_gn=LRCH3;refGene_tx=XM_019839902.2;refGene_gn=LRCH3;refGene_tx=XM_019839903.2;refGene_gn=LRCH3;refGene_tx=XM_019839904.2;refGene_gn=LRCH3;refGene_tx=XM_019839905.2;refGene_gn=LRCH3;refGene_tx=XM_019839906.2;refGene_gn=LRCH3;refGene_tx=XM_019839907.2;refGene_gn=LRCH3;refGene_tx=XM_023260251.1;refGene_gn=LRCH3;refGene_tx=XM_023260252.1
C2 71586016 71586122
refGene_gn=LRCH3;refGene_tx=XM_011285906.2;refGene_gn=LRCH3;refGene_tx

=XM_019839902.2;refGene_gn=LRCH3;refGene_tx=XM_019839903.2;refGene_gn=
LRCH3;refGene_tx=XM_019839904.2;refGene_gn=LRCH3;refGene_tx=XM_0198399
05.2;refGene_gn=LRCH3;refGene_tx=XM_019839906.2;refGene_gn=LRCH3;refGe
ne_tx=XM_019839907.2;refGene_gn=LRCH3;refGene_tx=XM_023260251.1;refGen
e_gn=LRCH3;refGene_tx=XM_023260252.1
C2 71588972 71589099
refGene_gn=LRCH3;refGene_tx=XM_011285906.2;refGene_gn=LRCH3;refGene_tx
=XM_019839902.2;refGene_gn=LRCH3;refGene_tx=XM_019839903.2;refGene_gn=
LRCH3;refGene_tx=XM_019839904.2;refGene_gn=LRCH3;refGene_tx=XM_0198399
05.2;refGene_gn=LRCH3;refGene_tx=XM_019839906.2;refGene_gn=LRCH3;refGe
ne_tx=XM_019839907.2;refGene_gn=LRCH3;refGene_tx=XM_023260251.1;refGen
e_gn=LRCH3;refGene_tx=XM_023260252.1
C2 71591102 71591247
refGene_gn=LRCH3;refGene_tx=XM_011285906.2;refGene_gn=LRCH3;refGene_tx
=XM_019839902.2;refGene_gn=LRCH3;refGene_tx=XM_019839903.2;refGene_gn=
LRCH3;refGene_tx=XM_019839904.2;refGene_gn=LRCH3;refGene_tx=XM_0198399
05.2;refGene_gn=LRCH3;refGene_tx=XM_019839906.2;refGene_gn=LRCH3;refGe
ne_tx=XM_019839907.2;refGene_gn=LRCH3;refGene_tx=XM_023260251.1;refGen
e_gn=LRCH3;refGene_tx=XM_023260252.1
C2 71642817 71643079
refGene_gn=LRCH3;refGene_tx=XM_011285906.2;refGene_gn=LRCH3;refGene_tx
=XM_019839902.2;refGene_gn=LRCH3;refGene_tx=XM_019839903.2;refGene_gn=
LRCH3;refGene_tx=XM_019839904.2;refGene_gn=LRCH3;refGene_tx=XM_0198399
05.2;refGene_gn=LRCH3;refGene_tx=XM_019839906.2;refGene_gn=LRCH3;refGe
ne_tx=XM_019839907.2;refGene_gn=LRCH3;refGene_tx=XM_023260251.1;refGen
e_gn=LRCH3;refGene_tx=XM_023260252.1
C2 71649986 71650085
refGene_gn=FYTTD1;refGene_tx=XM_006936131.4;refGene_gn=FYTTD1;refGene_
tx=XM_011285903.3;refGene_gn=FYTTD1;refGene_tx=XM_011285904.3;refGene_
gn=FYTTD1;refGene_tx=XM_023260254.1
C2 71651778 71651905
refGene_gn=FYTTD1;refGene_tx=XM_006936131.4;refGene_gn=FYTTD1;refGene_
tx=XM_011285903.3;refGene_gn=FYTTD1;refGene_tx=XM_011285904.3;refGene_
gn=FYTTD1;refGene_tx=XM_023260254.1
C2 71653221 71653296
refGene_gn=FYTTD1;refGene_tx=XM_006936131.4;refGene_gn=FYTTD1;refGene_
tx=XM_011285903.3;refGene_gn=FYTTD1;refGene_tx=XM_011285904.3;refGene_
gn=FYTTD1;refGene_tx=XM_023260254.1
C2 71654801 71654863
refGene_gn=FYTTD1;refGene_tx=XM_006936131.4;refGene_gn=FYTTD1;refGene_
tx=XM_011285903.3;refGene_gn=FYTTD1;refGene_tx=XM_011285904.3;refGene_
gn=FYTTD1;refGene_tx=XM_023260254.1
C2 71655274 71655371
refGene_gn=FYTTD1;refGene_tx=XM_006936131.4;refGene_gn=FYTTD1;refGene_
tx=XM_011285903.3;refGene_gn=FYTTD1;refGene_tx=XM_011285904.3;refGene_
gn=FYTTD1;refGene_tx=XM_023260254.1
C2 71656968 71657081
refGene_gn=FYTTD1;refGene_tx=XM_006936131.4;refGene_gn=FYTTD1;refGene_
tx=XM_011285903.3;refGene_gn=FYTTD1;refGene_tx=XM_011285904.3;refGene_
gn=FYTTD1;refGene_tx=XM_023260254.1

C2 71662379 71662528
refGene_gn=FYTTD1;refGene_tx=XM_006936131.4;refGene_gn=FYTTD1;refGene_tx=XM_011285903.3;refGene_gn=FYTTD1;refGene_tx=XM_011285904.3;refGene_gn=FYTTD1;refGene_tx=XM_023260254.1
C2 71671654 71671783
refGene_gn=FYTTD1;refGene_tx=XM_006936131.4;refGene_gn=FYTTD1;refGene_tx=XM_011285903.3;refGene_gn=FYTTD1;refGene_tx=XM_011285904.3;refGene_gn=FYTTD1;refGene_tx=XM_023260254.1
C2 71673527 71673570 refGene_gn=FYTTD1;refGene_tx=XM_023260254.1
C2 71675787 71675815 refGene_gn=FYTTD1;refGene_tx=XM_006936131.4
C2 71676377 71676576 refGene_gn=FYTTD1;refGene_tx=XM_011285903.3
C2 71676814 71676920 refGene_gn=FYTTD1;refGene_tx=XM_011285904.3
C2 71688645 71688710
refGene_gn=RUBCN;refGene_tx=XM_006936133.3;refGene_gn=RUBCN;refGene_tx=XM_011285902.3;refGene_gn=RUBCN;refGene_tx=XM_019839921.2;refGene_gn=RUBCN;refGene_tx=XM_019839922.2;refGene_gn=RUBCN;refGene_tx=XM_019839925.2;refGene_gn=RUBCN;refGene_tx=XM_019839926.2;refGene_gn=RUBCN;refGene_tx=XM_019839927.2;refGene_gn=RUBCN;refGene_tx=XM_019839928.2
C2 71705946 71706100
refGene_gn=RUBCN;refGene_tx=XM_006936133.3;refGene_gn=RUBCN;refGene_tx=XM_011285902.3;refGene_gn=RUBCN;refGene_tx=XM_019839921.2;refGene_gn=RUBCN;refGene_tx=XM_019839922.2;refGene_gn=RUBCN;refGene_tx=XM_019839925.2;refGene_gn=RUBCN;refGene_tx=XM_019839926.2;refGene_gn=RUBCN;refGene_tx=XM_019839927.2;refGene_gn=RUBCN;refGene_tx=XM_019839928.2
C2 71715646 71715733
refGene_gn=RUBCN;refGene_tx=XM_006936133.3;refGene_gn=RUBCN;refGene_tx=XM_011285902.3;refGene_gn=RUBCN;refGene_tx=XM_019839921.2;refGene_gn=RUBCN;refGene_tx=XM_019839922.2;refGene_gn=RUBCN;refGene_tx=XM_019839925.2;refGene_gn=RUBCN;refGene_tx=XM_019839926.2;refGene_gn=RUBCN;refGene_tx=XM_019839927.2;refGene_gn=RUBCN;refGene_tx=XM_019839928.2
C2 71717541 71717701
refGene_gn=RUBCN;refGene_tx=XM_006936133.3;refGene_gn=RUBCN;refGene_tx=XM_011285902.3;refGene_gn=RUBCN;refGene_tx=XM_019839921.2;refGene_gn=RUBCN;refGene_tx=XM_019839922.2;refGene_gn=RUBCN;refGene_tx=XM_019839925.2;refGene_gn=RUBCN;refGene_tx=XM_019839926.2;refGene_gn=RUBCN;refGene_tx=XM_019839927.2;refGene_gn=RUBCN;refGene_tx=XM_019839928.2
C2 71718362 71718469
refGene_gn=RUBCN;refGene_tx=XM_006936133.3;refGene_gn=RUBCN;refGene_tx=XM_011285902.3;refGene_gn=RUBCN;refGene_tx=XM_019839921.2;refGene_gn=RUBCN;refGene_tx=XM_019839922.2;refGene_gn=RUBCN;refGene_tx=XM_019839925.2;refGene_gn=RUBCN;refGene_tx=XM_019839926.2;refGene_gn=RUBCN;refGene_tx=XM_019839927.2;refGene_gn=RUBCN;refGene_tx=XM_019839928.2
C2 71721805 71721962
refGene_gn=RUBCN;refGene_tx=XM_006936133.3;refGene_gn=RUBCN;refGene_tx=XM_011285902.3;refGene_gn=RUBCN;refGene_tx=XM_019839921.2;refGene_gn=RUBCN;refGene_tx=XM_019839922.2;refGene_gn=RUBCN;refGene_tx=XM_019839925.2;refGene_gn=RUBCN;refGene_tx=XM_019839926.2;refGene_gn=RUBCN;refGene_tx=XM_019839927.2;refGene_gn=RUBCN;refGene_tx=XM_019839928.2
C2 71722791 71723325
refGene_gn=RUBCN;refGene_tx=XM_006936133.3;refGene_gn=RUBCN;refGene_tx

=XM_011285902.3;refGene_gn=RUBCN;refGene_tx=XM_019839921.2;refGene_gn=RUBCN;refGene_tx=XM_019839922.2;refGene_gn=RUBCN;refGene_tx=XM_019839925.2;refGene_gn=RUBCN;refGene_tx=XM_019839926.2;refGene_gn=RUBCN;refGene_tx=XM_019839927.2;refGene_gn=RUBCN;refGene_tx=XM_019839928.2
C2 71724675 71724720
refGene_gn=RUBCN;refGene_tx=XM_011285902.3;refGene_gn=RUBCN;refGene_tx=XM_019839921.2;refGene_gn=RUBCN;refGene_tx=XM_019839922.2;refGene_gn=RUBCN;refGene_tx=XM_019839926.2
C2 71726834 71726930
refGene_gn=RUBCN;refGene_tx=XM_006936133.3;refGene_gn=RUBCN;refGene_tx=XM_011285902.3;refGene_gn=RUBCN;refGene_tx=XM_019839921.2;refGene_gn=RUBCN;refGene_tx=XM_019839922.2;refGene_gn=RUBCN;refGene_tx=XM_019839925.2;refGene_gn=RUBCN;refGene_tx=XM_019839926.2
C2 71727631 71727747
refGene_gn=RUBCN;refGene_tx=XM_006936133.3;refGene_gn=RUBCN;refGene_tx=XM_011285902.3;refGene_gn=RUBCN;refGene_tx=XM_019839921.2;refGene_gn=RUBCN;refGene_tx=XM_019839922.2;refGene_gn=RUBCN;refGene_tx=XM_019839925.2;refGene_gn=RUBCN;refGene_tx=XM_019839926.2;refGene_gn=RUBCN;refGene_tx=XM_019839927.2;refGene_gn=RUBCN;refGene_tx=XM_019839928.2
C2 71728917 71729128
refGene_gn=RUBCN;refGene_tx=XM_006936133.3;refGene_gn=RUBCN;refGene_tx=XM_011285902.3;refGene_gn=RUBCN;refGene_tx=XM_019839921.2;refGene_gn=RUBCN;refGene_tx=XM_019839922.2;refGene_gn=RUBCN;refGene_tx=XM_019839925.2;refGene_gn=RUBCN;refGene_tx=XM_019839926.2;refGene_gn=RUBCN;refGene_tx=XM_019839927.2;refGene_gn=RUBCN;refGene_tx=XM_019839928.2
C2 71729618 71729723
refGene_gn=RUBCN;refGene_tx=XM_006936133.3;refGene_gn=RUBCN;refGene_tx=XM_011285902.3;refGene_gn=RUBCN;refGene_tx=XM_019839921.2;refGene_gn=RUBCN;refGene_tx=XM_019839922.2;refGene_gn=RUBCN;refGene_tx=XM_019839925.2;refGene_gn=RUBCN;refGene_tx=XM_019839926.2;refGene_gn=RUBCN;refGene_tx=XM_019839927.2;refGene_gn=RUBCN;refGene_tx=XM_019839928.2
C2 71732306 71732381
refGene_gn=RUBCN;refGene_tx=XM_019839921.2;refGene_gn=RUBCN;refGene_tx=XM_019839922.2;refGene_gn=RUBCN;refGene_tx=XM_019839925.2;refGene_gn=RUBCN;refGene_tx=XM_019839927.2
C2 71738741 71738802
refGene_gn=RUBCN;refGene_tx=XM_006936133.3;refGene_gn=RUBCN;refGene_tx=XM_011285902.3;refGene_gn=RUBCN;refGene_tx=XM_019839921.2;refGene_gn=RUBCN;refGene_tx=XM_019839922.2;refGene_gn=RUBCN;refGene_tx=XM_019839925.2;refGene_gn=RUBCN;refGene_tx=XM_019839926.2;refGene_gn=RUBCN;refGene_tx=XM_019839927.2;refGene_gn=RUBCN;refGene_tx=XM_019839928.2
C2 71739840 71739973
refGene_gn=RUBCN;refGene_tx=XM_006936133.3;refGene_gn=RUBCN;refGene_tx=XM_011285902.3;refGene_gn=RUBCN;refGene_tx=XM_019839921.2;refGene_gn=RUBCN;refGene_tx=XM_019839922.2;refGene_gn=RUBCN;refGene_tx=XM_019839925.2;refGene_gn=RUBCN;refGene_tx=XM_019839926.2;refGene_gn=RUBCN;refGene_tx=XM_019839927.2;refGene_gn=RUBCN;refGene_tx=XM_019839928.2
C2 71740632 71740778
refGene_gn=RUBCN;refGene_tx=XM_006936133.3;refGene_gn=RUBCN;refGene_tx=XM_011285902.3;refGene_gn=RUBCN;refGene_tx=XM_019839921.2;refGene_gn=

RUBCN;refGene_tx=XM_019839922.2;refGene_gn=RUBCN;refGene_tx=XM_0198399
25.2;refGene_gn=RUBCN;refGene_tx=XM_019839926.2;refGene_gn=RUBCN;refGe
ne_tx=XM_019839927.2;refGene_gn=RUBCN;refGene_tx=XM_019839928.2
C2 71741623 71741688
refGene_gn=RUBCN;refGene_tx=XM_006936133.3;refGene_gn=RUBCN;refGene_tx
=XM_011285902.3;refGene_gn=RUBCN;refGene_tx=XM_019839921.2;refGene_gn=
RUBCN;refGene_tx=XM_019839922.2;refGene_gn=RUBCN;refGene_tx=XM_0198399
25.2;refGene_gn=RUBCN;refGene_tx=XM_019839926.2;refGene_gn=RUBCN;refGe
ne_tx=XM_019839927.2;refGene_gn=RUBCN;refGene_tx=XM_019839928.2
C2 71742151 71742390
refGene_gn=RUBCN;refGene_tx=XM_006936133.3;refGene_gn=RUBCN;refGene_tx
=XM_011285902.3;refGene_gn=RUBCN;refGene_tx=XM_019839921.2;refGene_gn=
RUBCN;refGene_tx=XM_019839922.2;refGene_gn=RUBCN;refGene_tx=XM_0198399
25.2;refGene_gn=RUBCN;refGene_tx=XM_019839926.2;refGene_gn=RUBCN;refGe
ne_tx=XM_019839927.2;refGene_gn=RUBCN;refGene_tx=XM_019839928.2
C2 71742953 71743015
refGene_gn=RUBCN;refGene_tx=XM_006936133.3;refGene_gn=RUBCN;refGene_tx
=XM_011285902.3;refGene_gn=RUBCN;refGene_tx=XM_019839921.2;refGene_gn=
RUBCN;refGene_tx=XM_019839922.2;refGene_gn=RUBCN;refGene_tx=XM_0198399
25.2;refGene_gn=RUBCN;refGene_tx=XM_019839926.2;refGene_gn=RUBCN;refGe
ne_tx=XM_019839927.2;refGene_gn=RUBCN;refGene_tx=XM_019839928.2
C2 71743715 71743869
refGene_gn=RUBCN;refGene_tx=XM_006936133.3;refGene_gn=RUBCN;refGene_tx
=XM_011285902.3;refGene_gn=RUBCN;refGene_tx=XM_019839921.2;refGene_gn=
RUBCN;refGene_tx=XM_019839922.2;refGene_gn=RUBCN;refGene_tx=XM_0198399
25.2;refGene_gn=RUBCN;refGene_tx=XM_019839926.2;refGene_gn=RUBCN;refGe
ne_tx=XM_019839927.2;refGene_gn=RUBCN;refGene_tx=XM_019839928.2
C2 71744984 71745078
refGene_gn=RUBCN;refGene_tx=XM_006936133.3;refGene_gn=RUBCN;refGene_tx
=XM_011285902.3;refGene_gn=RUBCN;refGene_tx=XM_019839921.2;refGene_gn=
RUBCN;refGene_tx=XM_019839922.2;refGene_gn=RUBCN;refGene_tx=XM_0198399
25.2;refGene_gn=RUBCN;refGene_tx=XM_019839926.2;refGene_gn=RUBCN;refGe
ne_tx=XM_019839927.2;refGene_gn=RUBCN;refGene_tx=XM_019839928.2
C2 71745306 71745485
refGene_gn=RUBCN;refGene_tx=XM_006936133.3;refGene_gn=RUBCN;refGene_tx
=XM_011285902.3;refGene_gn=RUBCN;refGene_tx=XM_019839921.2;refGene_gn=
RUBCN;refGene_tx=XM_019839922.2;refGene_gn=RUBCN;refGene_tx=XM_0198399
25.2;refGene_gn=RUBCN;refGene_tx=XM_019839926.2;refGene_gn=RUBCN;refGe
ne_tx=XM_019839927.2;refGene_gn=RUBCN;refGene_tx=XM_019839928.2
C2 71779927 71780019 refGene_gn=MUC20;refGene_tx=XM_023260255.1
C2 71783290 71783496 refGene_gn=MUC20;refGene_tx=XM_023260255.1
C2 71783498 71784212 refGene_gn=MUC20;refGene_tx=XM_023260255.1
C2 71784214 71784511 refGene_gn=MUC20;refGene_tx=XM_023260255.1
C2 71784513 71784600 refGene_gn=MUC20;refGene_tx=XM_023260255.1
C2 71786425 71786517 refGene_gn=MUC20;refGene_tx=XM_023260255.1
C2 71788003 71788072 refGene_gn=MUC20;refGene_tx=XM_023260255.1
C2 71804396 71804592
refGene_gn=MUC4;refGene_tx=XM_019839933.2;refGene_gn=MUC4;refGene_tx=X
M_023260256.1
C2 71805919 71806082

refGene_gn=MUC4;refGene_tx=XM_019839933.2;refGene_gn=MUC4;refGene_tx=XM_023260256.1
C2	71808004 71808243
refGene_gn=MUC4;refGene_tx=XM_019839933.2;refGene_gn=MUC4;refGene_tx=XM_023260256.1
C2	71808326 71808391
refGene_gn=MUC4;refGene_tx=XM_019839933.2;refGene_gn=MUC4;refGene_tx=XM_023260256.1
C2	71809740 71809814
refGene_gn=MUC4;refGene_tx=XM_019839933.2;refGene_gn=MUC4;refGene_tx=XM_023260256.1
C2	71810253 71810433
refGene_gn=MUC4;refGene_tx=XM_019839933.2;refGene_gn=MUC4;refGene_tx=XM_023260256.1
C2	71811206 71811366
refGene_gn=MUC4;refGene_tx=XM_019839933.2;refGene_gn=MUC4;refGene_tx=XM_023260256.1
C2	71811729 71811911
refGene_gn=MUC4;refGene_tx=XM_019839933.2;refGene_gn=MUC4;refGene_tx=XM_023260256.1
C2	71813318 71813456
refGene_gn=MUC4;refGene_tx=XM_019839933.2;refGene_gn=MUC4;refGene_tx=XM_023260256.1
C2	71814764 71814995
refGene_gn=MUC4;refGene_tx=XM_019839933.2;refGene_gn=MUC4;refGene_tx=XM_023260256.1
C2	71815393 71815495
refGene_gn=MUC4;refGene_tx=XM_019839933.2;refGene_gn=MUC4;refGene_tx=XM_023260256.1
C2	71816021 71816189
refGene_gn=MUC4;refGene_tx=XM_019839933.2;refGene_gn=MUC4;refGene_tx=XM_023260256.1
C2	71816616 71816707
refGene_gn=MUC4;refGene_tx=XM_019839933.2;refGene_gn=MUC4;refGene_tx=XM_023260256.1
C2	71817121 71817330
refGene_gn=MUC4;refGene_tx=XM_019839933.2;refGene_gn=MUC4;refGene_tx=XM_023260256.1
C2	71817710 71817830
refGene_gn=MUC4;refGene_tx=XM_019839933.2;refGene_gn=MUC4;refGene_tx=XM_023260256.1
C2	71818586 71818712
refGene_gn=MUC4;refGene_tx=XM_019839933.2;refGene_gn=MUC4;refGene_tx=XM_023260256.1
C2	71818884 71819064
refGene_gn=MUC4;refGene_tx=XM_019839933.2;refGene_gn=MUC4;refGene_tx=XM_023260256.1
C2	71821025 71821114
refGene_gn=MUC4;refGene_tx=XM_019839933.2;refGene_gn=MUC4;refGene_tx=XM_023260256.1

C2	71822708	71822839
refGene_gn=MUC4;refGene_tx=XM_019839933.2;refGene_gn=MUC4;refGene_tx=X
M_023260256.1
C2	71823801	71823957
refGene_gn=MUC4;refGene_tx=XM_019839933.2;refGene_gn=MUC4;refGene_tx=X
M_023260256.1
C2	71824211	71824376
refGene_gn=MUC4;refGene_tx=XM_019839933.2;refGene_gn=MUC4;refGene_tx=X
M_023260256.1
C2	71825269	71825403
refGene_gn=MUC4;refGene_tx=XM_019839933.2;refGene_gn=MUC4;refGene_tx=X
M_023260256.1
C2	71826657	71826810
refGene_gn=MUC4;refGene_tx=XM_019839933.2;refGene_gn=MUC4;refGene_tx=X
M_023260256.1
C2	71827151	71834010	refGene_gn=MUC4;refGene_tx=XM_019839933.2
C2	71853390	71853472	refGene_gn=MUC4;refGene_tx=XM_023260256.1
C2	71908284	71908291
refGene_gn=TNK2;refGene_tx=XM_023260257.1;refGene_gn=TNK2;refGene_tx=X
M_023260259.1;refGene_gn=TNK2;refGene_tx=XM_023260261.1;refGene_gn=TNK
2;refGene_tx=XM_023260262.1;refGene_gn=TNK2;refGene_tx=XM_023260265.1;
refGene_gn=TNK2;refGene_tx=XM_023260266.1;refGene_gn=TNK2;refGene_tx=X
M_023260267.1;refGene_gn=TNK2;refGene_tx=XM_023260269.1
C2	71909467	71909599
refGene_gn=TNK2;refGene_tx=XM_019839934.2;refGene_gn=TNK2;refGene_tx=X
M_023260257.1;refGene_gn=TNK2;refGene_tx=XM_023260258.1;refGene_gn=TNK
2;refGene_tx=XM_023260259.1;refGene_gn=TNK2;refGene_tx=XM_023260260.1;
refGene_gn=TNK2;refGene_tx=XM_023260261.1;refGene_gn=TNK2;refGene_tx=X
M_023260262.1;refGene_gn=TNK2;refGene_tx=XM_023260263.1;refGene_gn=TNK
2;refGene_tx=XM_023260265.1;refGene_gn=TNK2;refGene_tx=XM_023260266.1;
refGene_gn=TNK2;refGene_tx=XM_023260267.1;refGene_gn=TNK2;refGene_tx=X
M_023260268.1;refGene_gn=TNK2;refGene_tx=XM_023260269.1
C2	71909729	71909738	refGene_gn=TNK2;refGene_tx=XM_023260264.1
C2	71909771	71909867
refGene_gn=TNK2;refGene_tx=XM_023260257.1;refGene_gn=TNK2;refGene_tx=X
M_023260258.1;refGene_gn=TNK2;refGene_tx=XM_023260259.1;refGene_gn=TNK
2;refGene_tx=XM_023260260.1;refGene_gn=TNK2;refGene_tx=XM_023260261.1;
refGene_gn=TNK2;refGene_tx=XM_023260265.1;refGene_gn=TNK2;refGene_tx=X
M_023260266.1;refGene_gn=TNK2;refGene_tx=XM_023260267.1;refGene_gn=TNK
2;refGene_tx=XM_023260269.1
C2	71909969	71911324
refGene_gn=TNK2;refGene_tx=XM_019839934.2;refGene_gn=TNK2;refGene_tx=X
M_023260257.1;refGene_gn=TNK2;refGene_tx=XM_023260258.1;refGene_gn=TNK
2;refGene_tx=XM_023260259.1;refGene_gn=TNK2;refGene_tx=XM_023260260.1;
refGene_gn=TNK2;refGene_tx=XM_023260261.1;refGene_gn=TNK2;refGene_tx=X
M_023260262.1;refGene_gn=TNK2;refGene_tx=XM_023260263.1;refGene_gn=TNK
2;refGene_tx=XM_023260264.1;refGene_gn=TNK2;refGene_tx=XM_023260265.1;
refGene_gn=TNK2;refGene_tx=XM_023260266.1;refGene_gn=TNK2;refGene_tx=X
M_023260267.1;refGene_gn=TNK2;refGene_tx=XM_023260268.1;refGene_gn=TNK
2;refGene_tx=XM_023260269.1

C2 71912056 71912101
refGene_gn=TNK2;refGene_tx=XM_019839934.2;refGene_gn=TNK2;refGene_tx=XM_023260257.1;refGene_gn=TNK2;refGene_tx=XM_023260258.1;refGene_gn=TNK2;refGene_tx=XM_023260261.1;refGene_gn=TNK2;refGene_tx=XM_023260262.1;refGene_gn=TNK2;refGene_tx=XM_023260264.1;refGene_gn=TNK2;refGene_tx=XM_023260265.1;refGene_gn=TNK2;refGene_tx=XM_023260266.1;refGene_gn=TNK2;refGene_tx=XM_023260267.1;refGene_gn=TNK2;refGene_tx=XM_023260269.1
C2 71912676 71912768
refGene_gn=TNK2;refGene_tx=XM_019839934.2;refGene_gn=TNK2;refGene_tx=XM_023260257.1;refGene_gn=TNK2;refGene_tx=XM_023260258.1;refGene_gn=TNK2;refGene_tx=XM_023260259.1;refGene_gn=TNK2;refGene_tx=XM_023260260.1;refGene_gn=TNK2;refGene_tx=XM_023260261.1;refGene_gn=TNK2;refGene_tx=XM_023260262.1;refGene_gn=TNK2;refGene_tx=XM_023260263.1;refGene_gn=TNK2;refGene_tx=XM_023260264.1;refGene_gn=TNK2;refGene_tx=XM_023260265.1;refGene_gn=TNK2;refGene_tx=XM_023260266.1;refGene_gn=TNK2;refGene_tx=XM_023260267.1;refGene_gn=TNK2;refGene_tx=XM_023260268.1;refGene_gn=TNK2;refGene_tx=XM_023260269.1
C2 71914168 71914363
refGene_gn=TNK2;refGene_tx=XM_019839934.2;refGene_gn=TNK2;refGene_tx=XM_023260257.1;refGene_gn=TNK2;refGene_tx=XM_023260258.1;refGene_gn=TNK2;refGene_tx=XM_023260259.1;refGene_gn=TNK2;refGene_tx=XM_023260260.1;refGene_gn=TNK2;refGene_tx=XM_023260261.1;refGene_gn=TNK2;refGene_tx=XM_023260262.1;refGene_gn=TNK2;refGene_tx=XM_023260263.1;refGene_gn=TNK2;refGene_tx=XM_023260264.1;refGene_gn=TNK2;refGene_tx=XM_023260265.1;refGene_gn=TNK2;refGene_tx=XM_023260266.1;refGene_gn=TNK2;refGene_tx=XM_023260267.1;refGene_gn=TNK2;refGene_tx=XM_023260268.1;refGene_gn=TNK2;refGene_tx=XM_023260269.1
C2 71919298 71919393
refGene_gn=TNK2;refGene_tx=XM_019839934.2;refGene_gn=TNK2;refGene_tx=XM_023260257.1;refGene_gn=TNK2;refGene_tx=XM_023260258.1;refGene_gn=TNK2;refGene_tx=XM_023260259.1;refGene_gn=TNK2;refGene_tx=XM_023260260.1;refGene_gn=TNK2;refGene_tx=XM_023260261.1;refGene_gn=TNK2;refGene_tx=XM_023260262.1;refGene_gn=TNK2;refGene_tx=XM_023260263.1;refGene_gn=TNK2;refGene_tx=XM_023260264.1;refGene_gn=TNK2;refGene_tx=XM_023260265.1;refGene_gn=TNK2;refGene_tx=XM_023260266.1;refGene_gn=TNK2;refGene_tx=XM_023260267.1;refGene_gn=TNK2;refGene_tx=XM_023260268.1;refGene_gn=TNK2;refGene_tx=XM_023260269.1
C2 71919477 71919624
refGene_gn=TNK2;refGene_tx=XM_019839934.2;refGene_gn=TNK2;refGene_tx=XM_023260257.1;refGene_gn=TNK2;refGene_tx=XM_023260258.1;refGene_gn=TNK2;refGene_tx=XM_023260259.1;refGene_gn=TNK2;refGene_tx=XM_023260260.1;refGene_gn=TNK2;refGene_tx=XM_023260261.1;refGene_gn=TNK2;refGene_tx=XM_023260262.1;refGene_gn=TNK2;refGene_tx=XM_023260263.1;refGene_gn=TNK2;refGene_tx=XM_023260264.1;refGene_gn=TNK2;refGene_tx=XM_023260265.1;refGene_gn=TNK2;refGene_tx=XM_023260266.1;refGene_gn=TNK2;refGene_tx=XM_023260267.1;refGene_gn=TNK2;refGene_tx=XM_023260268.1;refGene_gn=TNK2;refGene_tx=XM_023260269.1
C2 71920036 71920163
refGene_gn=TNK2;refGene_tx=XM_019839934.2;refGene_gn=TNK2;refGene_tx=XM_023260257.1;refGene_gn=TNK2;refGene_tx=XM_023260258.1;refGene_gn=TNK

2;refGene_tx=XM_023260259.1;refGene_gn=TNK2;refGene_tx=XM_023260260.1;
refGene_gn=TNK2;refGene_tx=XM_023260261.1;refGene_gn=TNK2;refGene_tx=X
M_023260262.1;refGene_gn=TNK2;refGene_tx=XM_023260263.1;refGene_gn=TNK
2;refGene_tx=XM_023260264.1;refGene_gn=TNK2;refGene_tx=XM_023260265.1;
refGene_gn=TNK2;refGene_tx=XM_023260266.1;refGene_gn=TNK2;refGene_tx=X
M_023260267.1;refGene_gn=TNK2;refGene_tx=XM_023260268.1;refGene_gn=TNK
2;refGene_tx=XM_023260269.1
C2 71921604 71921882
refGene_gn=TNK2;refGene_tx=XM_019839934.2;refGene_gn=TNK2;refGene_tx=X
M_023260257.1;refGene_gn=TNK2;refGene_tx=XM_023260258.1;refGene_gn=TNK
2;refGene_tx=XM_023260259.1;refGene_gn=TNK2;refGene_tx=XM_023260260.1;
refGene_gn=TNK2;refGene_tx=XM_023260261.1;refGene_gn=TNK2;refGene_tx=X
M_023260262.1;refGene_gn=TNK2;refGene_tx=XM_023260263.1;refGene_gn=TNK
2;refGene_tx=XM_023260264.1;refGene_gn=TNK2;refGene_tx=XM_023260265.1;
refGene_gn=TNK2;refGene_tx=XM_023260266.1;refGene_gn=TNK2;refGene_tx=X
M_023260267.1;refGene_gn=TNK2;refGene_tx=XM_023260268.1;refGene_gn=TNK
2;refGene_tx=XM_023260269.1
C2 71922398 71922551
refGene_gn=TNK2;refGene_tx=XM_019839934.2;refGene_gn=TNK2;refGene_tx=X
M_023260257.1;refGene_gn=TNK2;refGene_tx=XM_023260258.1;refGene_gn=TNK
2;refGene_tx=XM_023260259.1;refGene_gn=TNK2;refGene_tx=XM_023260260.1;
refGene_gn=TNK2;refGene_tx=XM_023260261.1;refGene_gn=TNK2;refGene_tx=X
M_023260262.1;refGene_gn=TNK2;refGene_tx=XM_023260263.1;refGene_gn=TNK
2;refGene_tx=XM_023260264.1;refGene_gn=TNK2;refGene_tx=XM_023260265.1;
refGene_gn=TNK2;refGene_tx=XM_023260266.1;refGene_gn=TNK2;refGene_tx=X
M_023260267.1;refGene_gn=TNK2;refGene_tx=XM_023260268.1;refGene_gn=TNK
2;refGene_tx=XM_023260269.1
C2 71923536 71923758
refGene_gn=TNK2;refGene_tx=XM_019839934.2;refGene_gn=TNK2;refGene_tx=X
M_023260257.1;refGene_gn=TNK2;refGene_tx=XM_023260258.1;refGene_gn=TNK
2;refGene_tx=XM_023260259.1;refGene_gn=TNK2;refGene_tx=XM_023260260.1;
refGene_gn=TNK2;refGene_tx=XM_023260261.1;refGene_gn=TNK2;refGene_tx=X
M_023260262.1;refGene_gn=TNK2;refGene_tx=XM_023260263.1;refGene_gn=TNK
2;refGene_tx=XM_023260264.1;refGene_gn=TNK2;refGene_tx=XM_023260265.1;
refGene_gn=TNK2;refGene_tx=XM_023260266.1;refGene_gn=TNK2;refGene_tx=X
M_023260267.1;refGene_gn=TNK2;refGene_tx=XM_023260268.1;refGene_gn=TNK
2;refGene_tx=XM_023260269.1
C2 71925693 71925764
refGene_gn=TNK2;refGene_tx=XM_019839934.2;refGene_gn=TNK2;refGene_tx=X
M_023260257.1;refGene_gn=TNK2;refGene_tx=XM_023260258.1;refGene_gn=TNK
2;refGene_tx=XM_023260259.1;refGene_gn=TNK2;refGene_tx=XM_023260260.1;
refGene_gn=TNK2;refGene_tx=XM_023260261.1;refGene_gn=TNK2;refGene_tx=X
M_023260262.1;refGene_gn=TNK2;refGene_tx=XM_023260263.1;refGene_gn=TNK
2;refGene_tx=XM_023260264.1;refGene_gn=TNK2;refGene_tx=XM_023260265.1;
refGene_gn=TNK2;refGene_tx=XM_023260266.1;refGene_gn=TNK2;refGene_tx=X
M_023260267.1;refGene_gn=TNK2;refGene_tx=XM_023260268.1;refGene_gn=TNK
2;refGene_tx=XM_023260269.1
C2 71926792 71926973
refGene_gn=TNK2;refGene_tx=XM_019839934.2;refGene_gn=TNK2;refGene_tx=X
M_023260257.1;refGene_gn=TNK2;refGene_tx=XM_023260258.1;refGene_gn=TNK

2;refGene_tx=XM_023260259.1;refGene_gn=TNK2;refGene_tx=XM_023260260.1;refGene_gn=TNK2;refGene_tx=XM_023260261.1;refGene_gn=TNK2;refGene_tx=XM_023260262.1;refGene_gn=TNK2;refGene_tx=XM_023260263.1;refGene_gn=TNK2;refGene_tx=XM_023260264.1;refGene_gn=TNK2;refGene_tx=XM_023260265.1;refGene_gn=TNK2;refGene_tx=XM_023260266.1;refGene_gn=TNK2;refGene_tx=XM_023260267.1;refGene_gn=TNK2;refGene_tx=XM_023260268.1
C2 71930610 71930688
refGene_gn=TNK2;refGene_tx=XM_023260265.1;refGene_gn=TNK2;refGene_tx=XM_023260268.1
C2 71933296 71933479
refGene_gn=TNK2;refGene_tx=XM_019839934.2;refGene_gn=TNK2;refGene_tx=XM_023260257.1;refGene_gn=TNK2;refGene_tx=XM_023260258.1;refGene_gn=TNK2;refGene_tx=XM_023260259.1;refGene_gn=TNK2;refGene_tx=XM_023260260.1;refGene_gn=TNK2;refGene_tx=XM_023260261.1;refGene_gn=TNK2;refGene_tx=XM_023260262.1;refGene_gn=TNK2;refGene_tx=XM_023260263.1;refGene_gn=TNK2;refGene_tx=XM_023260264.1
C2 71953170 71954662
refGene_gn=LOC111562231;refGene_tx=XM_023260173.1
C2 71958324 71958386
refGene_gn=LOC111562231;refGene_tx=XM_023260173.1
C2 71971945 71972188
refGene_gn=TFRC;refGene_tx=NM_001009312.1;refGene_gn=TFRC;refGene_tx=XM_019839232.2;refGene_gn=TFRC;refGene_tx=XM_019839233.2;refGene_gn=TFRC;refGene_tx=XM_019839234.1;refGene_gn=TFRC;refGene_tx=XM_023260086.1;refGene_gn=TFRC;refGene_tx=XM_023260087.1
C2 71972802 71972943
refGene_gn=TFRC;refGene_tx=NM_001009312.1;refGene_gn=TFRC;refGene_tx=XM_019839232.2;refGene_gn=TFRC;refGene_tx=XM_019839233.2;refGene_gn=TFRC;refGene_tx=XM_019839234.1;refGene_gn=TFRC;refGene_tx=XM_023260086.1;refGene_gn=TFRC;refGene_tx=XM_023260087.1
C2 71974068 71974290
refGene_gn=TFRC;refGene_tx=NM_001009312.1;refGene_gn=TFRC;refGene_tx=XM_019839232.2;refGene_gn=TFRC;refGene_tx=XM_019839233.2;refGene_gn=TFRC;refGene_tx=XM_019839234.1;refGene_gn=TFRC;refGene_tx=XM_023260086.1;refGene_gn=TFRC;refGene_tx=XM_023260087.1
C2 71975790 71975872
refGene_gn=TFRC;refGene_tx=NM_001009312.1;refGene_gn=TFRC;refGene_tx=XM_019839232.2;refGene_gn=TFRC;refGene_tx=XM_019839233.2;refGene_gn=TFRC;refGene_tx=XM_019839234.1;refGene_gn=TFRC;refGene_tx=XM_023260086.1;refGene_gn=TFRC;refGene_tx=XM_023260087.1
C2 71976084 71976143
refGene_gn=TFRC;refGene_tx=NM_001009312.1;refGene_gn=TFRC;refGene_tx=XM_019839232.2;refGene_gn=TFRC;refGene_tx=XM_019839233.2;refGene_gn=TFRC;refGene_tx=XM_019839234.1;refGene_gn=TFRC;refGene_tx=XM_023260086.1;refGene_gn=TFRC;refGene_tx=XM_023260087.1
C2 71977288 71977356
refGene_gn=TFRC;refGene_tx=NM_001009312.1;refGene_gn=TFRC;refGene_tx=XM_019839232.2;refGene_gn=TFRC;refGene_tx=XM_019839233.2;refGene_gn=TFRC;refGene_tx=XM_019839234.1;refGene_gn=TFRC;refGene_tx=XM_023260086.1;refGene_gn=TFRC;refGene_tx=XM_023260087.1

C2	71978992	71979056
refGene_gn=TFRC;refGene_tx=NM_001009312.1;refGene_gn=TFRC;refGene_tx=XM_019839232.2;refGene_gn=TFRC;refGene_tx=XM_019839233.2;refGene_gn=TFRC;refGene_tx=XM_019839234.1;refGene_gn=TFRC;refGene_tx=XM_023260086.1;refGene_gn=TFRC;refGene_tx=XM_023260087.1
C2	71979625	71979711
refGene_gn=TFRC;refGene_tx=NM_001009312.1;refGene_gn=TFRC;refGene_tx=XM_019839232.2;refGene_gn=TFRC;refGene_tx=XM_019839233.2;refGene_gn=TFRC;refGene_tx=XM_019839234.1;refGene_gn=TFRC;refGene_tx=XM_023260086.1;refGene_gn=TFRC;refGene_tx=XM_023260087.1
C2	71980793	71980913
refGene_gn=TFRC;refGene_tx=NM_001009312.1;refGene_gn=TFRC;refGene_tx=XM_019839232.2;refGene_gn=TFRC;refGene_tx=XM_019839233.2;refGene_gn=TFRC;refGene_tx=XM_019839234.1;refGene_gn=TFRC;refGene_tx=XM_023260086.1;refGene_gn=TFRC;refGene_tx=XM_023260087.1
C2	71981832	71981990
refGene_gn=TFRC;refGene_tx=NM_001009312.1;refGene_gn=TFRC;refGene_tx=XM_019839232.2;refGene_gn=TFRC;refGene_tx=XM_019839233.2;refGene_gn=TFRC;refGene_tx=XM_019839234.1;refGene_gn=TFRC;refGene_tx=XM_023260086.1;refGene_gn=TFRC;refGene_tx=XM_023260087.1
C2	71984346	71984486
refGene_gn=TFRC;refGene_tx=NM_001009312.1;refGene_gn=TFRC;refGene_tx=XM_019839232.2;refGene_gn=TFRC;refGene_tx=XM_019839233.2;refGene_gn=TFRC;refGene_tx=XM_019839234.1;refGene_gn=TFRC;refGene_tx=XM_023260086.1;refGene_gn=TFRC;refGene_tx=XM_023260087.1
C2	71984886	71984985
refGene_gn=TFRC;refGene_tx=NM_001009312.1;refGene_gn=TFRC;refGene_tx=XM_019839232.2;refGene_gn=TFRC;refGene_tx=XM_019839233.2;refGene_gn=TFRC;refGene_tx=XM_019839234.1;refGene_gn=TFRC;refGene_tx=XM_023260086.1;refGene_gn=TFRC;refGene_tx=XM_023260087.1
C2	71985915	71986029
refGene_gn=TFRC;refGene_tx=NM_001009312.1;refGene_gn=TFRC;refGene_tx=XM_019839232.2;refGene_gn=TFRC;refGene_tx=XM_019839233.2;refGene_gn=TFRC;refGene_tx=XM_019839234.1;refGene_gn=TFRC;refGene_tx=XM_023260086.1;refGene_gn=TFRC;refGene_tx=XM_023260087.1
C2	71987836	71987939
refGene_gn=TFRC;refGene_tx=NM_001009312.1;refGene_gn=TFRC;refGene_tx=XM_019839232.2;refGene_gn=TFRC;refGene_tx=XM_019839233.2;refGene_gn=TFRC;refGene_tx=XM_019839234.1;refGene_gn=TFRC;refGene_tx=XM_023260086.1;refGene_gn=TFRC;refGene_tx=XM_023260087.1
C2	71988114	71988264
refGene_gn=TFRC;refGene_tx=NM_001009312.1;refGene_gn=TFRC;refGene_tx=XM_019839232.2;refGene_gn=TFRC;refGene_tx=XM_019839233.2;refGene_gn=TFRC;refGene_tx=XM_019839234.1;refGene_gn=TFRC;refGene_tx=XM_023260086.1;refGene_gn=TFRC;refGene_tx=XM_023260087.1
C2	71989429	71989643
refGene_gn=TFRC;refGene_tx=NM_001009312.1;refGene_gn=TFRC;refGene_tx=XM_019839232.2;refGene_gn=TFRC;refGene_tx=XM_019839233.2;refGene_gn=TFRC;refGene_tx=XM_019839234.1;refGene_gn=TFRC;refGene_tx=XM_023260086.1;refGene_gn=TFRC;refGene_tx=XM_023260087.1

C2	71990846	71991057
refGene_gn=TFRC;refGene_tx=NM_001009312.1;refGene_gn=TFRC;refGene_tx=XM_019839232.2;refGene_gn=TFRC;refGene_tx=XM_019839233.2;refGene_gn=TFRC;refGene_tx=XM_019839234.1;refGene_gn=TFRC;refGene_tx=XM_023260086.1;refGene_gn=TFRC;refGene_tx=XM_023260087.1
C2	71991628	71991664
refGene_gn=TFRC;refGene_tx=NM_001009312.1;refGene_gn=TFRC;refGene_tx=XM_019839232.2;refGene_gn=TFRC;refGene_tx=XM_019839233.2;refGene_gn=TFRC;refGene_tx=XM_019839234.1;refGene_gn=TFRC;refGene_tx=XM_023260086.1;refGene_gn=TFRC;refGene_tx=XM_023260087.1
C2	72078589	72078609	refGene_gn=ZDHHC19;refGene_tx=XM_019839235.2
C2	72079550	72079720
refGene_gn=ZDHHC19;refGene_tx=XM_023260270.1;refGene_gn=ZDHHC19;refGene_tx=XM_023260271.1
C2	72080353	72080553
refGene_gn=ZDHHC19;refGene_tx=XM_019839235.2;refGene_gn=ZDHHC19;refGene_tx=XM_023260270.1;refGene_gn=ZDHHC19;refGene_tx=XM_023260271.1
C2	72080901	72080987
refGene_gn=ZDHHC19;refGene_tx=XM_019839235.2;refGene_gn=ZDHHC19;refGene_tx=XM_023260270.1;refGene_gn=ZDHHC19;refGene_tx=XM_023260271.1
C2	72087439	72087545
refGene_gn=ZDHHC19;refGene_tx=XM_019839235.2;refGene_gn=ZDHHC19;refGene_tx=XM_023260270.1;refGene_gn=ZDHHC19;refGene_tx=XM_023260271.1
C2	72087945	72088118
refGene_gn=ZDHHC19;refGene_tx=XM_019839235.2;refGene_gn=ZDHHC19;refGene_tx=XM_023260270.1;refGene_gn=ZDHHC19;refGene_tx=XM_023260271.1
C2	72088920	72089060
refGene_gn=ZDHHC19;refGene_tx=XM_019839235.2;refGene_gn=ZDHHC19;refGene_tx=XM_023260270.1;refGene_gn=ZDHHC19;refGene_tx=XM_023260271.1
C2	72089566	72089688
refGene_gn=ZDHHC19;refGene_tx=XM_019839235.2;refGene_gn=ZDHHC19;refGene_tx=XM_023260270.1;refGene_gn=ZDHHC19;refGene_tx=XM_023260271.1
C2	72090055	72090201
refGene_gn=ZDHHC19;refGene_tx=XM_019839235.2;refGene_gn=ZDHHC19;refGene_tx=XM_023260270.1;refGene_gn=ZDHHC19;refGene_tx=XM_023260271.1
C2	72100521	72100559	refGene_gn=SLC51A;refGene_tx=XM_003991737.3
C2	72101151	72101246	refGene_gn=SLC51A;refGene_tx=XM_003991737.3
C2	72114309	72114464	refGene_gn=SLC51A;refGene_tx=XM_003991737.3
C2	72115395	72115469	refGene_gn=SLC51A;refGene_tx=XM_003991737.3
C2	72115894	72116053	refGene_gn=SLC51A;refGene_tx=XM_003991737.3
C2	72116608	72116720	refGene_gn=SLC51A;refGene_tx=XM_003991737.3
C2	72117014	72117161	refGene_gn=SLC51A;refGene_tx=XM_003991737.3
C2	72119448	72119554	refGene_gn=SLC51A;refGene_tx=XM_003991737.3
C2	72120090	72120299	refGene_gn=SLC51A;refGene_tx=XM_003991737.3
C2	72125446	72125653
refGene_gn=PCYT1A;refGene_tx=XM_003991738.5;refGene_gn=PCYT1A;refGene_tx=XM_019839936.2
C2	72126523	72126712
refGene_gn=PCYT1A;refGene_tx=XM_003991738.5;refGene_gn=PCYT1A;refGene_tx=XM_019839936.2

C2 72128378 72128521
refGene_gn=PCYT1A;refGene_tx=XM_003991738.5;refGene_gn=PCYT1A;refGene_tx=XM_019839936.2
C2 72129239 72129318
refGene_gn=PCYT1A;refGene_tx=XM_003991738.5;refGene_gn=PCYT1A;refGene_tx=XM_019839936.2
C2 72131371 72131523
refGene_gn=PCYT1A;refGene_tx=XM_003991738.5;refGene_gn=PCYT1A;refGene_tx=XM_019839936.2
C2 72132541 72132658
refGene_gn=PCYT1A;refGene_tx=XM_003991738.5;refGene_gn=PCYT1A;refGene_tx=XM_019839936.2
C2 72137972 72138072
refGene_gn=PCYT1A;refGene_tx=XM_003991738.5;refGene_gn=PCYT1A;refGene_tx=XM_019839936.2
C2 72151812 72151929
refGene_gn=PCYT1A;refGene_tx=XM_003991738.5;refGene_gn=PCYT1A;refGene_tx=XM_019839936.2
C2 72171287 72171335 refGene_gn=TCTEX1D2;refGene_tx=XM_003991739.5
C2 72173117 72173181 refGene_gn=TCTEX1D2;refGene_tx=XM_003991739.5
C2 72183309 72183379 refGene_gn=TCTEX1D2;refGene_tx=XM_003991739.5
C2 72188036 72188170 refGene_gn=TCTEX1D2;refGene_tx=XM_003991739.5
C2 72191028 72191147 refGene_gn=TCTEX1D2;refGene_tx=XM_003991739.5
C2 72195614 72195795 refGene_gn=TM4SF19;refGene_tx=XM_003991740.4
C2 72196050 72196220 refGene_gn=TM4SF19;refGene_tx=XM_003991740.4
C2 72198797 72198875 refGene_gn=TM4SF19;refGene_tx=XM_003991740.4
C2 72199412 72199613 refGene_gn=TM4SF19;refGene_tx=XM_003991740.4
C2 72222480 72222642 refGene_gn=UBXN7;refGene_tx=XM_003991742.5
C2 72223977 72224057 refGene_gn=UBXN7;refGene_tx=XM_003991742.5
C2 72224181 72224575 refGene_gn=UBXN7;refGene_tx=XM_003991742.5
C2 72230455 72230583 refGene_gn=UBXN7;refGene_tx=XM_003991742.5
C2 72231458 72231549 refGene_gn=UBXN7;refGene_tx=XM_003991742.5
C2 72232801 72232948 refGene_gn=UBXN7;refGene_tx=XM_003991742.5
C2 72238537 72238650 refGene_gn=UBXN7;refGene_tx=XM_003991742.5
C2 72241979 72242045 refGene_gn=UBXN7;refGene_tx=XM_003991742.5
C2 72250323 72250391 refGene_gn=UBXN7;refGene_tx=XM_003991742.5
C2 72254376 72254524 refGene_gn=UBXN7;refGene_tx=XM_003991742.5
C2 72273991 72274154 refGene_gn=UBXN7;refGene_tx=XM_003991742.5
C2 72306709 72307645
refGene_gn=RNF168;refGene_tx=XM_003991771.5;refGene_gn=RNF168;refGene_tx=XM_019839937.2
C2 72310912 72310994
refGene_gn=RNF168;refGene_tx=XM_003991771.5;refGene_gn=RNF168;refGene_tx=XM_019839937.2
C2 72320393 72320515
refGene_gn=RNF168;refGene_tx=XM_003991771.5;refGene_gn=RNF168;refGene_tx=XM_019839937.2
C2 72323299 72323479
refGene_gn=RNF168;refGene_tx=XM_003991771.5;refGene_gn=RNF168;refGene_tx=XM_019839937.2

C2 72325865 72325942
refGene_gn=RNF168;refGene_tx=XM_003991771.5;refGene_gn=RNF168;refGene_tx=XM_019839937.2
C2 72342284 72342585
refGene_gn=RNF168;refGene_tx=XM_003991771.5;refGene_gn=RNF168;refGene_tx=XM_019839937.2
C2 72347367 72347812 refGene_gn=SMCO1;refGene_tx=XM_003991752.5
C2 72348252 72348402 refGene_gn=SMCO1;refGene_tx=XM_003991752.5
C2 72351773 72351823 refGene_gn=SMCO1;refGene_tx=XM_003991752.5
C2 72376843 72377440 refGene_gn=WDR53;refGene_tx=XM_003991772.5
C2 72383221 72383701 refGene_gn=WDR53;refGene_tx=XM_003991772.5
C2 72390388 72390703 refGene_gn=FBXO45;refGene_tx=XM_003991753.5
C2 72398196 72398553 refGene_gn=FBXO45;refGene_tx=XM_003991753.5
C2 72402808 72402994 refGene_gn=FBXO45;refGene_tx=XM_003991753.5
C2 72448408 72448516 refGene_gn=NRROS;refGene_tx=XM_006936156.3
C2 72469734 72469842 refGene_gn=NRROS;refGene_tx=XM_003991754.3
C2 72474172 72476143
refGene_gn=NRROS;refGene_tx=XM_003991754.3;refGene_gn=NRROS;refGene_tx=XM_006936156.3
C2 72514809 72515171
refGene_gn=CEP19;refGene_tx=XM_003991755.4;refGene_gn=CEP19;refGene_tx=XM_006936157.3;refGene_gn=CEP19;refGene_tx=XM_006936158.3;refGene_gn=CEP19;refGene_tx=XM_006936159.3
C2 72515782 72515912
refGene_gn=CEP19;refGene_tx=XM_003991755.4;refGene_gn=CEP19;refGene_tx=XM_006936157.3;refGene_gn=CEP19;refGene_tx=XM_006936158.3;refGene_gn=CEP19;refGene_tx=XM_006936159.3
C2 72519775 72519791 refGene_gn=PIGX;refGene_tx=XM_006936179.3
C2 72521664 72522099 refGene_gn=PIGX;refGene_tx=XM_006936179.3
C2 72524738 72524802 refGene_gn=PIGX;refGene_tx=XM_006936179.3
C2 72537373 72537515 refGene_gn=PIGX;refGene_tx=XM_006936179.3
C2 72546409 72546623 refGene_gn=PIGX;refGene_tx=XM_006936179.3
C2 72549546 72549647 refGene_gn=PIGX;refGene_tx=XM_006936179.3
C2 72553102 72553246 refGene_gn=PIGX;refGene_tx=XM_006936179.3
C2 72599610 72599797 refGene_gn=PAK2;refGene_tx=XM_003991756.5
C2 72618769 72618870 refGene_gn=PAK2;refGene_tx=XM_003991756.5
C2 72619227 72619375 refGene_gn=PAK2;refGene_tx=XM_003991756.5
C2 72622186 72622218 refGene_gn=PAK2;refGene_tx=XM_003991756.5
C2 72623000 72623108 refGene_gn=PAK2;refGene_tx=XM_003991756.5
C2 72624150 72624283 refGene_gn=PAK2;refGene_tx=XM_003991756.5
C2 72625655 72625719 refGene_gn=PAK2;refGene_tx=XM_003991756.5
C2 72629118 72629167 refGene_gn=PAK2;refGene_tx=XM_003991756.5
C2 72629653 72629766 refGene_gn=PAK2;refGene_tx=XM_003991756.5
C2 72630852 72630970 refGene_gn=PAK2;refGene_tx=XM_003991756.5
C2 72632536 72632636 refGene_gn=PAK2;refGene_tx=XM_003991756.5
C2 72635882 72636079 refGene_gn=PAK2;refGene_tx=XM_003991756.5
C2 72647348 72647486 refGene_gn=PAK2;refGene_tx=XM_003991756.5
C2 72647943 72648030 refGene_gn=PAK2;refGene_tx=XM_003991756.5
C2 72704313 72705811
refGene_gn=SENP5;refGene_tx=XM_023260272.1;refGene_gn=SENP5;refGene_tx

=XM_023260273.1;refGene_gn=SENP5;refGene_tx=XM_023260274.1;refGene_gn=SENP5;refGene_tx=XM_023260275.1;refGene_gn=SENP5;refGene_tx=XM_023260276.1;refGene_gn=SENP5;refGene_tx=XM_023260277.1;refGene_gn=SENP5;refGene_tx=XM_023260278.1;refGene_gn=SENP5;refGene_tx=XM_023260279.1
C2 72714062 72714168
refGene_gn=SENP5;refGene_tx=XM_023260272.1;refGene_gn=SENP5;refGene_tx=XM_023260273.1;refGene_gn=SENP5;refGene_tx=XM_023260274.1;refGene_gn=SENP5;refGene_tx=XM_023260275.1;refGene_gn=SENP5;refGene_tx=XM_023260276.1;refGene_gn=SENP5;refGene_tx=XM_023260277.1;refGene_gn=SENP5;refGene_tx=XM_023260278.1;refGene_gn=SENP5;refGene_tx=XM_023260279.1
C2 72714289 72714428
refGene_gn=SENP5;refGene_tx=XM_023260272.1;refGene_gn=SENP5;refGene_tx=XM_023260273.1;refGene_gn=SENP5;refGene_tx=XM_023260274.1;refGene_gn=SENP5;refGene_tx=XM_023260275.1;refGene_gn=SENP5;refGene_tx=XM_023260276.1;refGene_gn=SENP5;refGene_tx=XM_023260277.1;refGene_gn=SENP5;refGene_tx=XM_023260278.1;refGene_gn=SENP5;refGene_tx=XM_023260279.1
C2 72714681 72714729
refGene_gn=SENP5;refGene_tx=XM_023260272.1;refGene_gn=SENP5;refGene_tx=XM_023260273.1;refGene_gn=SENP5;refGene_tx=XM_023260274.1;refGene_gn=SENP5;refGene_tx=XM_023260275.1;refGene_gn=SENP5;refGene_tx=XM_023260276.1;refGene_gn=SENP5;refGene_tx=XM_023260277.1;refGene_gn=SENP5;refGene_tx=XM_023260278.1;refGene_gn=SENP5;refGene_tx=XM_023260279.1
C2 72717449 72717527
refGene_gn=SENP5;refGene_tx=XM_023260272.1;refGene_gn=SENP5;refGene_tx=XM_023260273.1;refGene_gn=SENP5;refGene_tx=XM_023260274.1;refGene_gn=SENP5;refGene_tx=XM_023260275.1;refGene_gn=SENP5;refGene_tx=XM_023260276.1;refGene_gn=SENP5;refGene_tx=XM_023260277.1;refGene_gn=SENP5;refGene_tx=XM_023260278.1;refGene_gn=SENP5;refGene_tx=XM_023260279.1
C2 72726188 72726326
refGene_gn=SENP5;refGene_tx=XM_023260272.1;refGene_gn=SENP5;refGene_tx=XM_023260273.1;refGene_gn=SENP5;refGene_tx=XM_023260274.1;refGene_gn=SENP5;refGene_tx=XM_023260275.1;refGene_gn=SENP5;refGene_tx=XM_023260276.1;refGene_gn=SENP5;refGene_tx=XM_023260279.1
C2 72729079 72729163
refGene_gn=SENP5;refGene_tx=XM_023260272.1;refGene_gn=SENP5;refGene_tx=XM_023260273.1;refGene_gn=SENP5;refGene_tx=XM_023260274.1;refGene_gn=SENP5;refGene_tx=XM_023260275.1;refGene_gn=SENP5;refGene_tx=XM_023260279.1
C2 72730998 72731049
refGene_gn=SENP5;refGene_tx=XM_023260272.1;refGene_gn=SENP5;refGene_tx=XM_023260273.1;refGene_gn=SENP5;refGene_tx=XM_023260274.1;refGene_gn=SENP5;refGene_tx=XM_023260275.1;refGene_gn=SENP5;refGene_tx=XM_023260276.1;refGene_gn=SENP5;refGene_tx=XM_023260277.1;refGene_gn=SENP5;refGene_tx=XM_023260279.1
C2 72731822 72731933
refGene_gn=SENP5;refGene_tx=XM_023260272.1;refGene_gn=SENP5;refGene_tx=XM_023260273.1;refGene_gn=SENP5;refGene_tx=XM_023260274.1;refGene_gn=SENP5;refGene_tx=XM_023260275.1;refGene_gn=SENP5;refGene_tx=XM_023260276.1;refGene_gn=SENP5;refGene_tx=XM_023260277.1;refGene_gn=SENP5;refGene_tx=XM_023260279.1

C2 72736670 72736742 refGene_gn=NCBP2;refGene_tx=XM_003991757.5
C2 72737261 72737400 refGene_gn=NCBP2;refGene_tx=XM_003991757.5
C2 72740279 72740461 refGene_gn=NCBP2;refGene_tx=XM_003991757.5
C2 72746071 72746149 refGene_gn=NCBP2;refGene_tx=XM_003991757.5
C2 72746378 72746678
refGene_gn=LOC111562251;refGene_tx=XM_023260281.1
C2 72751015 72752583 refGene_gn=PIGZ;refGene_tx=XM_023260280.1
C2 72755690 72755901 refGene_gn=PIGZ;refGene_tx=XM_023260280.1
C2 72759468 72759542 refGene_gn=PIGZ;refGene_tx=XM_023260280.1
C2 72764536 72764641 refGene_gn=PIGZ;refGene_tx=XM_023260280.1
C2 72767612 72767637 refGene_gn=PIGZ;refGene_tx=XM_023260280.1
C2 72802048 72802131 refGene_gn=MELTF;refGene_tx=XM_006936165.4
C2 72802603 72802781 refGene_gn=MELTF;refGene_tx=XM_006936165.4
C2 72802782 72802803 refGene_gn=MELTF;refGene_tx=XM_006936165.4
C2 72804437 72804625 refGene_gn=MELTF;refGene_tx=XM_006936165.4
C2 72806703 72806771 refGene_gn=MELTF;refGene_tx=XM_006936165.4
C2 72806859 72807016 refGene_gn=MELTF;refGene_tx=XM_006936165.4
C2 72808220 72808415 refGene_gn=MELTF;refGene_tx=XM_006936165.4
C2 72809592 72809689 refGene_gn=MELTF;refGene_tx=XM_006936165.4
C2 72811936 72812088 refGene_gn=MELTF;refGene_tx=XM_006936165.4
C2 72812796 72812977 refGene_gn=MELTF;refGene_tx=XM_006936165.4
C2 72813717 72813905 refGene_gn=MELTF;refGene_tx=XM_006936165.4
C2 72816518 72816586 refGene_gn=MELTF;refGene_tx=XM_006936165.4
C2 72817807 72817964 refGene_gn=MELTF;refGene_tx=XM_006936165.4
C2 72820197 72820380 refGene_gn=MELTF;refGene_tx=XM_006936165.4
C2 72821199 72821299 refGene_gn=MELTF;refGene_tx=XM_006936165.4
C2 72822642 72822797 refGene_gn=MELTF;refGene_tx=XM_006936165.4
C2 72824630 72824679 refGene_gn=MELTF;refGene_tx=XM_006936165.4
C2 72841910 72842017
refGene_gn=DLG1;refGene_tx=XM_006936166.3;refGene_gn=DLG1;refGene_tx=XM_006936171.3;refGene_gn=DLG1;refGene_tx=XM_006936173.3;refGene_gn=DLG1;refGene_tx=XM_011285884.2;refGene_gn=DLG1;refGene_tx=XM_011285885.3;refGene_gn=DLG1;refGene_tx=XM_011285886.2;refGene_gn=DLG1;refGene_tx=XM_011285887.3;refGene_gn=DLG1;refGene_tx=XM_011285888.3;refGene_gn=DLG1;refGene_tx=XM_011285889.3;refGene_gn=DLG1;refGene_tx=XM_011285890.3;refGene_gn=DLG1;refGene_tx=XM_011285892.3;refGene_gn=DLG1;refGene_tx=XM_019839942.1;refGene_gn=DLG1;refGene_tx=XM_019839943.2;refGene_gn=DLG1;refGene_tx=XM_019839944.1;refGene_gn=DLG1;refGene_tx=XM_019839945.1;refGene_gn=DLG1;refGene_tx=XM_019839946.1;refGene_gn=DLG1;refGene_tx=XM_019839947.1;refGene_gn=DLG1;refGene_tx=XM_019839948.1;refGene_gn=DLG1;refGene_tx=XM_019839949.1;refGene_gn=DLG1;refGene_tx=XM_019839950.2
C2 72842901 72842993
refGene_gn=DLG1;refGene_tx=XM_006936166.3;refGene_gn=DLG1;refGene_tx=XM_006936171.3;refGene_gn=DLG1;refGene_tx=XM_006936173.3;refGene_gn=DLG1;refGene_tx=XM_011285884.2;refGene_gn=DLG1;refGene_tx=XM_011285885.3;refGene_gn=DLG1;refGene_tx=XM_011285886.2;refGene_gn=DLG1;refGene_tx=XM_011285887.3;refGene_gn=DLG1;refGene_tx=XM_011285888.3;refGene_gn=DLG1;refGene_tx=XM_011285889.3;refGene_gn=DLG1;refGene_tx=XM_011285890.3;refGene_gn=DLG1;refGene_tx=XM_011285892.3;refGene_gn=DLG1;refGene_tx=XM_019839942.1;refGene_gn=DLG1;refGene_tx=XM_019839943.2;refGene_gn=DLG

1;refGene_tx=XM_019839944.1;refGene_gn=DLG1;refGene_tx=XM_019839945.1;
refGene_gn=DLG1;refGene_tx=XM_019839946.1;refGene_gn=DLG1;refGene_tx=X
M_019839947.1;refGene_gn=DLG1;refGene_tx=XM_019839948.1;refGene_gn=DLG
1;refGene_tx=XM_019839949.1;refGene_gn=DLG1;refGene_tx=XM_019839950.2
C2 72855010 72855120
refGene_gn=DLG1;refGene_tx=XM_006936166.3;refGene_gn=DLG1;refGene_tx=X
M_006936171.3;refGene_gn=DLG1;refGene_tx=XM_006936173.3;refGene_gn=DLG
1;refGene_tx=XM_011285884.2;refGene_gn=DLG1;refGene_tx=XM_011285885.3;
refGene_gn=DLG1;refGene_tx=XM_011285886.2;refGene_gn=DLG1;refGene_tx=X
M_011285887.3;refGene_gn=DLG1;refGene_tx=XM_011285888.3;refGene_gn=DLG
1;refGene_tx=XM_011285889.3;refGene_gn=DLG1;refGene_tx=XM_011285890.3;
refGene_gn=DLG1;refGene_tx=XM_011285892.3;refGene_gn=DLG1;refGene_tx=X
M_019839942.1;refGene_gn=DLG1;refGene_tx=XM_019839943.2;refGene_gn=DLG
1;refGene_tx=XM_019839944.1;refGene_gn=DLG1;refGene_tx=XM_019839945.1;
refGene_gn=DLG1;refGene_tx=XM_019839946.1;refGene_gn=DLG1;refGene_tx=X
M_019839947.1;refGene_gn=DLG1;refGene_tx=XM_019839948.1;refGene_gn=DLG
1;refGene_tx=XM_019839949.1;refGene_gn=DLG1;refGene_tx=XM_019839950.2
C2 72867276 72867449
refGene_gn=DLG1;refGene_tx=XM_006936166.3;refGene_gn=DLG1;refGene_tx=X
M_006936171.3;refGene_gn=DLG1;refGene_tx=XM_006936173.3;refGene_gn=DLG
1;refGene_tx=XM_011285884.2;refGene_gn=DLG1;refGene_tx=XM_011285885.3;
refGene_gn=DLG1;refGene_tx=XM_011285886.2;refGene_gn=DLG1;refGene_tx=X
M_011285887.3;refGene_gn=DLG1;refGene_tx=XM_011285888.3;refGene_gn=DLG
1;refGene_tx=XM_011285889.3;refGene_gn=DLG1;refGene_tx=XM_011285890.3;
refGene_gn=DLG1;refGene_tx=XM_011285892.3;refGene_gn=DLG1;refGene_tx=X
M_019839942.1;refGene_gn=DLG1;refGene_tx=XM_019839943.2;refGene_gn=DLG
1;refGene_tx=XM_019839944.1;refGene_gn=DLG1;refGene_tx=XM_019839945.1;
refGene_gn=DLG1;refGene_tx=XM_019839946.1;refGene_gn=DLG1;refGene_tx=X
M_019839947.1;refGene_gn=DLG1;refGene_tx=XM_019839948.1;refGene_gn=DLG
1;refGene_tx=XM_019839949.1;refGene_gn=DLG1;refGene_tx=XM_019839950.2
C2 72867693 72867795
refGene_gn=DLG1;refGene_tx=XM_006936166.3;refGene_gn=DLG1;refGene_tx=X
M_006936171.3;refGene_gn=DLG1;refGene_tx=XM_006936173.3;refGene_gn=DLG
1;refGene_tx=XM_011285884.2;refGene_gn=DLG1;refGene_tx=XM_011285885.3;
refGene_gn=DLG1;refGene_tx=XM_011285886.2;refGene_gn=DLG1;refGene_tx=X
M_011285887.3;refGene_gn=DLG1;refGene_tx=XM_011285888.3;refGene_gn=DLG
1;refGene_tx=XM_011285889.3;refGene_gn=DLG1;refGene_tx=XM_011285890.3;
refGene_gn=DLG1;refGene_tx=XM_011285892.3;refGene_gn=DLG1;refGene_tx=X
M_019839942.1;refGene_gn=DLG1;refGene_tx=XM_019839943.2;refGene_gn=DLG
1;refGene_tx=XM_019839944.1;refGene_gn=DLG1;refGene_tx=XM_019839945.1;
refGene_gn=DLG1;refGene_tx=XM_019839946.1;refGene_gn=DLG1;refGene_tx=X
M_019839947.1;refGene_gn=DLG1;refGene_tx=XM_019839948.1;refGene_gn=DLG
1;refGene_tx=XM_019839949.1;refGene_gn=DLG1;refGene_tx=XM_019839950.2
C2 72868681 72868732
refGene_gn=DLG1;refGene_tx=XM_006936166.3;refGene_gn=DLG1;refGene_tx=X
M_006936171.3;refGene_gn=DLG1;refGene_tx=XM_006936173.3;refGene_gn=DLG
1;refGene_tx=XM_011285884.2;refGene_gn=DLG1;refGene_tx=XM_011285885.3;
refGene_gn=DLG1;refGene_tx=XM_011285886.2;refGene_gn=DLG1;refGene_tx=X
M_011285887.3;refGene_gn=DLG1;refGene_tx=XM_011285888.3;refGene_gn=DLG
1;refGene_tx=XM_011285889.3;refGene_gn=DLG1;refGene_tx=XM_011285890.3;

refGene_gn=DLG1;refGene_tx=XM_011285892.3;refGene_gn=DLG1;refGene_tx=XM_019839942.1;refGene_gn=DLG1;refGene_tx=XM_019839943.2;refGene_gn=DLG1;refGene_tx=XM_019839944.1;refGene_gn=DLG1;refGene_tx=XM_019839945.1;refGene_gn=DLG1;refGene_tx=XM_019839946.1;refGene_gn=DLG1;refGene_tx=XM_019839947.1;refGene_gn=DLG1;refGene_tx=XM_019839948.1;refGene_gn=DLG1;refGene_tx=XM_019839949.1;refGene_gn=DLG1;refGene_tx=XM_019839950.2
C2 72870558 72870594
refGene_gn=DLG1;refGene_tx=XM_011285884.2;refGene_gn=DLG1;refGene_tx=XM_011285885.3;refGene_gn=DLG1;refGene_tx=XM_011285886.2;refGene_gn=DLG1;refGene_tx=XM_011285887.3;refGene_gn=DLG1;refGene_tx=XM_011285888.3;refGene_gn=DLG1;refGene_tx=XM_011285889.3;refGene_gn=DLG1;refGene_tx=XM_011285892.3;refGene_gn=DLG1;refGene_tx=XM_019839942.1;refGene_gn=DLG1;refGene_tx=XM_019839943.2;refGene_gn=DLG1;refGene_tx=XM_019839944.1;refGene_gn=DLG1;refGene_tx=XM_019839945.1;refGene_gn=DLG1;refGene_tx=XM_019839946.1;refGene_gn=DLG1;refGene_tx=XM_019839947.1;refGene_gn=DLG1;refGene_tx=XM_019839948.1;refGene_gn=DLG1;refGene_tx=XM_019839949.1;refGene_gn=DLG1;refGene_tx=XM_019839950.2
C2 72871297 72871339
refGene_gn=DLG1;refGene_tx=XM_006936166.3;refGene_gn=DLG1;refGene_tx=XM_006936171.3;refGene_gn=DLG1;refGene_tx=XM_006936173.3;refGene_gn=DLG1;refGene_tx=XM_011285884.2;refGene_gn=DLG1;refGene_tx=XM_011285885.3;refGene_gn=DLG1;refGene_tx=XM_011285886.2;refGene_gn=DLG1;refGene_tx=XM_011285887.3;refGene_gn=DLG1;refGene_tx=XM_011285888.3;refGene_gn=DLG1;refGene_tx=XM_011285889.3;refGene_gn=DLG1;refGene_tx=XM_011285890.3;refGene_gn=DLG1;refGene_tx=XM_011285892.3;refGene_gn=DLG1;refGene_tx=XM_019839942.1;refGene_gn=DLG1;refGene_tx=XM_019839943.2;refGene_gn=DLG1;refGene_tx=XM_019839944.1;refGene_gn=DLG1;refGene_tx=XM_019839945.1;refGene_gn=DLG1;refGene_tx=XM_019839946.1;refGene_gn=DLG1;refGene_tx=XM_019839947.1;refGene_gn=DLG1;refGene_tx=XM_019839948.1;refGene_gn=DLG1;refGene_tx=XM_019839949.1;refGene_gn=DLG1;refGene_tx=XM_019839950.2
C2 72878641 72878675
refGene_gn=DLG1;refGene_tx=XM_006936171.3;refGene_gn=DLG1;refGene_tx=XM_011285889.3;refGene_gn=DLG1;refGene_tx=XM_011285890.3;refGene_gn=DLG1;refGene_tx=XM_011285892.3;refGene_gn=DLG1;refGene_tx=XM_019839946.1;refGene_gn=DLG1;refGene_tx=XM_019839947.1
C2 72879364 72879464
refGene_gn=DLG1;refGene_tx=XM_006936166.3;refGene_gn=DLG1;refGene_tx=XM_006936173.3;refGene_gn=DLG1;refGene_tx=XM_011285884.2;refGene_gn=DLG1;refGene_tx=XM_011285885.3;refGene_gn=DLG1;refGene_tx=XM_011285886.2;refGene_gn=DLG1;refGene_tx=XM_011285887.3;refGene_gn=DLG1;refGene_tx=XM_011285888.3;refGene_gn=DLG1;refGene_tx=XM_019839942.1;refGene_gn=DLG1;refGene_tx=XM_019839943.2;refGene_gn=DLG1;refGene_tx=XM_019839944.1;refGene_gn=DLG1;refGene_tx=XM_019839945.1;refGene_gn=DLG1;refGene_tx=XM_019839948.1;refGene_gn=DLG1;refGene_tx=XM_019839949.1;refGene_gn=DLG1;refGene_tx=XM_019839950.2
C2 72883914 72883984
refGene_gn=DLG1;refGene_tx=XM_006936166.3;refGene_gn=DLG1;refGene_tx=XM_006936171.3;refGene_gn=DLG1;refGene_tx=XM_006936173.3;refGene_gn=DLG1;refGene_tx=XM_011285884.2;refGene_gn=DLG1;refGene_tx=XM_011285885.3;refGene_gn=DLG1;refGene_tx=XM_011285886.2;refGene_gn=DLG1;refGene_tx=X

M_011285887.3;refGene_gn=DLG1;refGene_tx=XM_011285888.3;refGene_gn=DLG1;refGene_tx=XM_011285889.3;refGene_gn=DLG1;refGene_tx=XM_011285890.3;refGene_gn=DLG1;refGene_tx=XM_011285892.3;refGene_gn=DLG1;refGene_tx=XM_019839942.1;refGene_gn=DLG1;refGene_tx=XM_019839943.2;refGene_gn=DLG1;refGene_tx=XM_019839944.1;refGene_gn=DLG1;refGene_tx=XM_019839945.1;refGene_gn=DLG1;refGene_tx=XM_019839946.1;refGene_gn=DLG1;refGene_tx=XM_019839947.1;refGene_gn=DLG1;refGene_tx=XM_019839948.1;refGene_gn=DLG1;refGene_tx=XM_019839949.1;refGene_gn=DLG1;refGene_tx=XM_019839950.2
C2 72886723 72886900
refGene_gn=DLG1;refGene_tx=XM_006936166.3;refGene_gn=DLG1;refGene_tx=XM_006936171.3;refGene_gn=DLG1;refGene_tx=XM_006936173.3;refGene_gn=DLG1;refGene_tx=XM_011285884.2;refGene_gn=DLG1;refGene_tx=XM_011285885.3;refGene_gn=DLG1;refGene_tx=XM_011285886.2;refGene_gn=DLG1;refGene_tx=XM_011285887.3;refGene_gn=DLG1;refGene_tx=XM_011285888.3;refGene_gn=DLG1;refGene_tx=XM_011285889.3;refGene_gn=DLG1;refGene_tx=XM_011285890.3;refGene_gn=DLG1;refGene_tx=XM_011285892.3;refGene_gn=DLG1;refGene_tx=XM_019839942.1;refGene_gn=DLG1;refGene_tx=XM_019839943.2;refGene_gn=DLG1;refGene_tx=XM_019839944.1;refGene_gn=DLG1;refGene_tx=XM_019839945.1;refGene_gn=DLG1;refGene_tx=XM_019839946.1;refGene_gn=DLG1;refGene_tx=XM_019839947.1;refGene_gn=DLG1;refGene_tx=XM_019839948.1;refGene_gn=DLG1;refGene_tx=XM_019839949.1;refGene_gn=DLG1;refGene_tx=XM_019839950.2
C2 72891781 72891896
refGene_gn=DLG1;refGene_tx=XM_006936166.3;refGene_gn=DLG1;refGene_tx=XM_006936171.3;refGene_gn=DLG1;refGene_tx=XM_006936173.3;refGene_gn=DLG1;refGene_tx=XM_011285884.2;refGene_gn=DLG1;refGene_tx=XM_011285885.3;refGene_gn=DLG1;refGene_tx=XM_011285886.2;refGene_gn=DLG1;refGene_tx=XM_011285887.3;refGene_gn=DLG1;refGene_tx=XM_011285888.3;refGene_gn=DLG1;refGene_tx=XM_011285889.3;refGene_gn=DLG1;refGene_tx=XM_011285890.3;refGene_gn=DLG1;refGene_tx=XM_011285892.3;refGene_gn=DLG1;refGene_tx=XM_019839942.1;refGene_gn=DLG1;refGene_tx=XM_019839943.2;refGene_gn=DLG1;refGene_tx=XM_019839944.1;refGene_gn=DLG1;refGene_tx=XM_019839945.1;refGene_gn=DLG1;refGene_tx=XM_019839946.1;refGene_gn=DLG1;refGene_tx=XM_019839947.1;refGene_gn=DLG1;refGene_tx=XM_019839948.1;refGene_gn=DLG1;refGene_tx=XM_019839949.1;refGene_gn=DLG1;refGene_tx=XM_019839950.2
C2 72909241 72909344
refGene_gn=DLG1;refGene_tx=XM_006936166.3;refGene_gn=DLG1;refGene_tx=XM_006936171.3;refGene_gn=DLG1;refGene_tx=XM_006936173.3;refGene_gn=DLG1;refGene_tx=XM_011285884.2;refGene_gn=DLG1;refGene_tx=XM_011285885.3;refGene_gn=DLG1;refGene_tx=XM_011285886.2;refGene_gn=DLG1;refGene_tx=XM_011285887.3;refGene_gn=DLG1;refGene_tx=XM_011285888.3;refGene_gn=DLG1;refGene_tx=XM_011285889.3;refGene_gn=DLG1;refGene_tx=XM_011285890.3;refGene_gn=DLG1;refGene_tx=XM_011285892.3;refGene_gn=DLG1;refGene_tx=XM_019839942.1;refGene_gn=DLG1;refGene_tx=XM_019839943.2;refGene_gn=DLG1;refGene_tx=XM_019839944.1;refGene_gn=DLG1;refGene_tx=XM_019839945.1;refGene_gn=DLG1;refGene_tx=XM_019839946.1;refGene_gn=DLG1;refGene_tx=XM_019839947.1;refGene_gn=DLG1;refGene_tx=XM_019839948.1;refGene_gn=DLG1;refGene_tx=XM_019839949.1;refGene_gn=DLG1;refGene_tx=XM_019839950.2
C2 72922765 72922922
refGene_gn=DLG1;refGene_tx=XM_006936166.3;refGene_gn=DLG1;refGene_tx=XM_006936171.3;refGene_gn=DLG1;refGene_tx=XM_006936173.3;refGene_gn=DLG

1;refGene_tx=XM_011285884.2;refGene_gn=DLG1;refGene_tx=XM_011285885.3;
refGene_gn=DLG1;refGene_tx=XM_011285886.2;refGene_gn=DLG1;refGene_tx=X
M_011285887.3;refGene_gn=DLG1;refGene_tx=XM_011285888.3;refGene_gn=DLG
1;refGene_tx=XM_011285889.3;refGene_gn=DLG1;refGene_tx=XM_011285890.3;
refGene_gn=DLG1;refGene_tx=XM_011285892.3;refGene_gn=DLG1;refGene_tx=X
M_019839942.1;refGene_gn=DLG1;refGene_tx=XM_019839943.2;refGene_gn=DLG
1;refGene_tx=XM_019839944.1;refGene_gn=DLG1;refGene_tx=XM_019839945.1;
refGene_gn=DLG1;refGene_tx=XM_019839946.1;refGene_gn=DLG1;refGene_tx=X
M_019839947.1;refGene_gn=DLG1;refGene_tx=XM_019839948.1;refGene_gn=DLG
1;refGene_tx=XM_019839949.1;refGene_gn=DLG1;refGene_tx=XM_019839950.2
C2 72928661 72928782
refGene_gn=DLG1;refGene_tx=XM_006936166.3;refGene_gn=DLG1;refGene_tx=X
M_006936171.3;refGene_gn=DLG1;refGene_tx=XM_006936173.3;refGene_gn=DLG
1;refGene_tx=XM_011285884.2;refGene_gn=DLG1;refGene_tx=XM_011285885.3;
refGene_gn=DLG1;refGene_tx=XM_011285886.2;refGene_gn=DLG1;refGene_tx=X
M_011285887.3;refGene_gn=DLG1;refGene_tx=XM_011285888.3;refGene_gn=DLG
1;refGene_tx=XM_011285889.3;refGene_gn=DLG1;refGene_tx=XM_011285890.3;
refGene_gn=DLG1;refGene_tx=XM_011285892.3;refGene_gn=DLG1;refGene_tx=X
M_019839942.1;refGene_gn=DLG1;refGene_tx=XM_019839943.2;refGene_gn=DLG
1;refGene_tx=XM_019839944.1;refGene_gn=DLG1;refGene_tx=XM_019839945.1;
refGene_gn=DLG1;refGene_tx=XM_019839946.1;refGene_gn=DLG1;refGene_tx=X
M_019839947.1;refGene_gn=DLG1;refGene_tx=XM_019839948.1;refGene_gn=DLG
1;refGene_tx=XM_019839949.1;refGene_gn=DLG1;refGene_tx=XM_019839950.2
C2 72931407 72931552
refGene_gn=DLG1;refGene_tx=XM_006936166.3;refGene_gn=DLG1;refGene_tx=X
M_006936171.3;refGene_gn=DLG1;refGene_tx=XM_006936173.3;refGene_gn=DLG
1;refGene_tx=XM_011285884.2;refGene_gn=DLG1;refGene_tx=XM_011285885.3;
refGene_gn=DLG1;refGene_tx=XM_011285886.2;refGene_gn=DLG1;refGene_tx=X
M_011285887.3;refGene_gn=DLG1;refGene_tx=XM_011285888.3;refGene_gn=DLG
1;refGene_tx=XM_011285889.3;refGene_gn=DLG1;refGene_tx=XM_011285890.3;
refGene_gn=DLG1;refGene_tx=XM_011285892.3;refGene_gn=DLG1;refGene_tx=X
M_019839942.1;refGene_gn=DLG1;refGene_tx=XM_019839943.2;refGene_gn=DLG
1;refGene_tx=XM_019839944.1;refGene_gn=DLG1;refGene_tx=XM_019839945.1;
refGene_gn=DLG1;refGene_tx=XM_019839946.1;refGene_gn=DLG1;refGene_tx=X
M_019839947.1;refGene_gn=DLG1;refGene_tx=XM_019839948.1;refGene_gn=DLG
1;refGene_tx=XM_019839949.1;refGene_gn=DLG1;refGene_tx=XM_019839950.2
C2 72937298 72937435
refGene_gn=DLG1;refGene_tx=XM_006936166.3;refGene_gn=DLG1;refGene_tx=X
M_006936171.3;refGene_gn=DLG1;refGene_tx=XM_006936173.3;refGene_gn=DLG
1;refGene_tx=XM_011285884.2;refGene_gn=DLG1;refGene_tx=XM_011285885.3;
refGene_gn=DLG1;refGene_tx=XM_011285886.2;refGene_gn=DLG1;refGene_tx=X
M_011285887.3;refGene_gn=DLG1;refGene_tx=XM_011285888.3;refGene_gn=DLG
1;refGene_tx=XM_011285889.3;refGene_gn=DLG1;refGene_tx=XM_011285890.3;
refGene_gn=DLG1;refGene_tx=XM_011285892.3;refGene_gn=DLG1;refGene_tx=X
M_019839942.1;refGene_gn=DLG1;refGene_tx=XM_019839943.2;refGene_gn=DLG
1;refGene_tx=XM_019839944.1;refGene_gn=DLG1;refGene_tx=XM_019839945.1;
refGene_gn=DLG1;refGene_tx=XM_019839946.1;refGene_gn=DLG1;refGene_tx=X
M_019839947.1;refGene_gn=DLG1;refGene_tx=XM_019839948.1;refGene_gn=DLG
1;refGene_tx=XM_019839949.1;refGene_gn=DLG1;refGene_tx=XM_019839950.2
C2 72938645 72938815

refGene_gn=DLG1;refGene_tx=XM_006936166.3;refGene_gn=DLG1;refGene_tx=XM_006936171.3;refGene_gn=DLG1;refGene_tx=XM_006936173.3;refGene_gn=DLG1;refGene_tx=XM_011285884.2;refGene_gn=DLG1;refGene_tx=XM_011285885.3;refGene_gn=DLG1;refGene_tx=XM_011285886.2;refGene_gn=DLG1;refGene_tx=XM_011285887.3;refGene_gn=DLG1;refGene_tx=XM_011285888.3;refGene_gn=DLG1;refGene_tx=XM_011285889.3;refGene_gn=DLG1;refGene_tx=XM_011285890.3;refGene_gn=DLG1;refGene_tx=XM_011285892.3;refGene_gn=DLG1;refGene_tx=XM_019839942.1;refGene_gn=DLG1;refGene_tx=XM_019839943.2;refGene_gn=DLG1;refGene_tx=XM_019839944.1;refGene_gn=DLG1;refGene_tx=XM_019839945.1;refGene_gn=DLG1;refGene_tx=XM_019839946.1;refGene_gn=DLG1;refGene_tx=XM_019839947.1;refGene_gn=DLG1;refGene_tx=XM_019839948.1;refGene_gn=DLG1;refGene_tx=XM_019839949.1;refGene_gn=DLG1;refGene_tx=XM_019839950.2
C2 72941061 72941186
refGene_gn=DLG1;refGene_tx=XM_006936166.3;refGene_gn=DLG1;refGene_tx=XM_006936171.3;refGene_gn=DLG1;refGene_tx=XM_006936173.3;refGene_gn=DLG1;refGene_tx=XM_011285884.2;refGene_gn=DLG1;refGene_tx=XM_011285885.3;refGene_gn=DLG1;refGene_tx=XM_011285886.2;refGene_gn=DLG1;refGene_tx=XM_011285887.3;refGene_gn=DLG1;refGene_tx=XM_011285888.3;refGene_gn=DLG1;refGene_tx=XM_011285889.3;refGene_gn=DLG1;refGene_tx=XM_011285890.3;refGene_gn=DLG1;refGene_tx=XM_011285892.3;refGene_gn=DLG1;refGene_tx=XM_019839942.1;refGene_gn=DLG1;refGene_tx=XM_019839943.2;refGene_gn=DLG1;refGene_tx=XM_019839944.1;refGene_gn=DLG1;refGene_tx=XM_019839945.1;refGene_gn=DLG1;refGene_tx=XM_019839946.1;refGene_gn=DLG1;refGene_tx=XM_019839947.1;refGene_gn=DLG1;refGene_tx=XM_019839948.1;refGene_gn=DLG1;refGene_tx=XM_019839949.1;refGene_gn=DLG1;refGene_tx=XM_019839950.2
C2 72944219 72944270
refGene_gn=DLG1;refGene_tx=XM_006936166.3;refGene_gn=DLG1;refGene_tx=XM_006936171.3;refGene_gn=DLG1;refGene_tx=XM_006936173.3;refGene_gn=DLG1;refGene_tx=XM_011285884.2;refGene_gn=DLG1;refGene_tx=XM_011285885.3;refGene_gn=DLG1;refGene_tx=XM_011285886.2;refGene_gn=DLG1;refGene_tx=XM_011285887.3;refGene_gn=DLG1;refGene_tx=XM_011285888.3;refGene_gn=DLG1;refGene_tx=XM_011285889.3;refGene_gn=DLG1;refGene_tx=XM_011285890.3;refGene_gn=DLG1;refGene_tx=XM_011285892.3;refGene_gn=DLG1;refGene_tx=XM_019839942.1;refGene_gn=DLG1;refGene_tx=XM_019839943.2;refGene_gn=DLG1;refGene_tx=XM_019839944.1;refGene_gn=DLG1;refGene_tx=XM_019839945.1;refGene_gn=DLG1;refGene_tx=XM_019839946.1;refGene_gn=DLG1;refGene_tx=XM_019839947.1;refGene_gn=DLG1;refGene_tx=XM_019839948.1;refGene_gn=DLG1;refGene_tx=XM_019839949.1;refGene_gn=DLG1;refGene_tx=XM_019839950.2
C2 72952316 72952370
refGene_gn=DLG1;refGene_tx=XM_006936166.3;refGene_gn=DLG1;refGene_tx=XM_006936171.3;refGene_gn=DLG1;refGene_tx=XM_006936173.3;refGene_gn=DLG1;refGene_tx=XM_011285884.2;refGene_gn=DLG1;refGene_tx=XM_011285885.3;refGene_gn=DLG1;refGene_tx=XM_011285886.2;refGene_gn=DLG1;refGene_tx=XM_011285887.3;refGene_gn=DLG1;refGene_tx=XM_011285888.3;refGene_gn=DLG1;refGene_tx=XM_011285889.3;refGene_gn=DLG1;refGene_tx=XM_011285890.3;refGene_gn=DLG1;refGene_tx=XM_011285892.3;refGene_gn=DLG1;refGene_tx=XM_019839942.1;refGene_gn=DLG1;refGene_tx=XM_019839943.2;refGene_gn=DLG1;refGene_tx=XM_019839944.1;refGene_gn=DLG1;refGene_tx=XM_019839946.1;refGene_gn=DLG1;refGene_tx=XM_019839947.1;refGene_gn=DLG1;refGene_tx=XM_019839948.1;refGene_gn=DLG1;refGene_tx=XM_019839950.2

C2 72964553 72964652
refGene_gn=DLG1;refGene_tx=XM_006936166.3;refGene_gn=DLG1;refGene_tx=XM_006936171.3;refGene_gn=DLG1;refGene_tx=XM_011285884.2;refGene_gn=DLG1;refGene_tx=XM_011285885.3;refGene_gn=DLG1;refGene_tx=XM_011285886.2;refGene_gn=DLG1;refGene_tx=XM_011285887.3;refGene_gn=DLG1;refGene_tx=XM_011285892.3;refGene_gn=DLG1;refGene_tx=XM_019839942.1;refGene_gn=DLG1;refGene_tx=XM_019839943.2;refGene_gn=DLG1;refGene_tx=XM_019839944.1;refGene_gn=DLG1;refGene_tx=XM_019839945.1;refGene_gn=DLG1;refGene_tx=XM_019839946.1;refGene_gn=DLG1;refGene_tx=XM_019839947.1;refGene_gn=DLG1;refGene_tx=XM_019839950.2
C2 72988920 72989154
refGene_gn=DLG1;refGene_tx=XM_011285888.3;refGene_gn=DLG1;refGene_tx=XM_011285889.3;refGene_gn=DLG1;refGene_tx=XM_011285890.3;refGene_gn=DLG1;refGene_tx=XM_011285892.3;refGene_gn=DLG1;refGene_tx=XM_019839950.2
C2 72999992 73000157
refGene_gn=DLG1;refGene_tx=XM_006936166.3;refGene_gn=DLG1;refGene_tx=XM_006936171.3;refGene_gn=DLG1;refGene_tx=XM_006936173.3;refGene_gn=DLG1;refGene_tx=XM_011285884.2;refGene_gn=DLG1;refGene_tx=XM_011285885.3;refGene_gn=DLG1;refGene_tx=XM_011285886.2;refGene_gn=DLG1;refGene_tx=XM_011285887.3;refGene_gn=DLG1;refGene_tx=XM_019839942.1;refGene_gn=DLG1;refGene_tx=XM_019839943.2;refGene_gn=DLG1;refGene_tx=XM_019839944.1;refGene_gn=DLG1;refGene_tx=XM_019839945.1;refGene_gn=DLG1;refGene_tx=XM_019839946.1;refGene_gn=DLG1;refGene_tx=XM_019839947.1;refGene_gn=DLG1;refGene_tx=XM_019839948.1;refGene_gn=DLG1;refGene_tx=XM_019839949.1
C2 73076669 73076699 refGene_gn=DLG1;refGene_tx=XM_011285887.3
C2 73098468 73098635
refGene_gn=DLG1;refGene_tx=XM_006936166.3;refGene_gn=DLG1;refGene_tx=XM_006936171.3;refGene_gn=DLG1;refGene_tx=XM_006936173.3;refGene_gn=DLG1;refGene_tx=XM_011285884.2;refGene_gn=DLG1;refGene_tx=XM_011285885.3;refGene_gn=DLG1;refGene_tx=XM_011285886.2;refGene_gn=DLG1;refGene_tx=XM_019839942.1;refGene_gn=DLG1;refGene_tx=XM_019839943.2;refGene_gn=DLG1;refGene_tx=XM_019839944.1;refGene_gn=DLG1;refGene_tx=XM_019839945.1;refGene_gn=DLG1;refGene_tx=XM_019839946.1;refGene_gn=DLG1;refGene_tx=XM_019839947.1;refGene_gn=DLG1;refGene_tx=XM_019839948.1;refGene_gn=DLG1;refGene_tx=XM_019839949.1
C2 73116629 73116761
refGene_gn=DLG1;refGene_tx=XM_006936166.3;refGene_gn=DLG1;refGene_tx=XM_006936171.3;refGene_gn=DLG1;refGene_tx=XM_006936173.3;refGene_gn=DLG1;refGene_tx=XM_011285884.2;refGene_gn=DLG1;refGene_tx=XM_011285885.3;refGene_gn=DLG1;refGene_tx=XM_011285886.2;refGene_gn=DLG1;refGene_tx=XM_019839942.1;refGene_gn=DLG1;refGene_tx=XM_019839943.2;refGene_gn=DLG1;refGene_tx=XM_019839944.1;refGene_gn=DLG1;refGene_tx=XM_019839945.1;refGene_gn=DLG1;refGene_tx=XM_019839946.1;refGene_gn=DLG1;refGene_tx=XM_019839947.1;refGene_gn=DLG1;refGene_tx=XM_019839948.1;refGene_gn=DLG1;refGene_tx=XM_019839949.1
C2 73117468 73117487
refGene_gn=DLG1;refGene_tx=XM_006936166.3;refGene_gn=DLG1;refGene_tx=XM_006936171.3;refGene_gn=DLG1;refGene_tx=XM_006936173.3;refGene_gn=DLG1;refGene_tx=XM_011285884.2;refGene_gn=DLG1;refGene_tx=XM_011285885.3;refGene_gn=DLG1;refGene_tx=XM_011285886.2;refGene_gn=DLG1;refGene_tx=X

M_019839942.1;refGene_gn=DLG1;refGene_tx=XM_019839943.2;refGene_gn=DLG1;refGene_tx=XM_019839944.1;refGene_gn=DLG1;refGene_tx=XM_019839945.1;refGene_gn=DLG1;refGene_tx=XM_019839946.1;refGene_gn=DLG1;refGene_tx=XM_019839947.1;refGene_gn=DLG1;refGene_tx=XM_019839948.1;refGene_gn=DLG1;refGene_tx=XM_019839949.1
C2 73280655 73281125
refGene_gn=BDH1;refGene_tx=XM_003991764.5;refGene_gn=BDH1;refGene_tx=XM_019839951.2;refGene_gn=BDH1;refGene_tx=XM_019839952.1;refGene_gn=BDH1;refGene_tx=XM_019839953.1;refGene_gn=BDH1;refGene_tx=XM_019839954.2
C2 73282369 73282522
refGene_gn=BDH1;refGene_tx=XM_003991764.5;refGene_gn=BDH1;refGene_tx=XM_019839951.2;refGene_gn=BDH1;refGene_tx=XM_019839952.1;refGene_gn=BDH1;refGene_tx=XM_019839953.1;refGene_gn=BDH1;refGene_tx=XM_019839954.2
C2 73289567 73289709
refGene_gn=BDH1;refGene_tx=XM_003991764.5;refGene_gn=BDH1;refGene_tx=XM_019839951.2;refGene_gn=BDH1;refGene_tx=XM_019839952.1;refGene_gn=BDH1;refGene_tx=XM_019839953.1;refGene_gn=BDH1;refGene_tx=XM_019839954.2
C2 73303982 73304093
refGene_gn=BDH1;refGene_tx=XM_003991764.5;refGene_gn=BDH1;refGene_tx=XM_019839951.2;refGene_gn=BDH1;refGene_tx=XM_019839952.1;refGene_gn=BDH1;refGene_tx=XM_019839953.1;refGene_gn=BDH1;refGene_tx=XM_019839954.2
C2 73304714 73304787
refGene_gn=BDH1;refGene_tx=XM_003991764.5;refGene_gn=BDH1;refGene_tx=XM_019839951.2;refGene_gn=BDH1;refGene_tx=XM_019839952.1;refGene_gn=BDH1;refGene_tx=XM_019839953.1;refGene_gn=BDH1;refGene_tx=XM_019839954.2
C2 73311986 73312113
refGene_gn=BDH1;refGene_tx=XM_003991764.5;refGene_gn=BDH1;refGene_tx=XM_019839951.2;refGene_gn=BDH1;refGene_tx=XM_019839952.1;refGene_gn=BDH1;refGene_tx=XM_019839953.1;refGene_gn=BDH1;refGene_tx=XM_019839954.2
C2 73313731 73313804 refGene_gn=BDH1;refGene_tx=XM_003991764.5
C2 73345329 73345348 refGene_gn=BDH1;refGene_tx=XM_019839951.2
C2 73393082 73393205
refGene_gn=APOD;refGene_tx=XM_006936175.2;refGene_gn=APOD;refGene_tx=XM_019839955.1
C2 73398542 73398664
refGene_gn=APOD;refGene_tx=XM_006936175.2;refGene_gn=APOD;refGene_tx=XM_019839955.1
C2 73401952 73402041
refGene_gn=APOD;refGene_tx=XM_006936175.2;refGene_gn=APOD;refGene_tx=XM_019839955.1
C2 73404148 73404280 refGene_gn=APOD;refGene_tx=XM_019839955.1
C2 73405395 73405631
refGene_gn=APOD;refGene_tx=XM_006936175.2;refGene_gn=APOD;refGene_tx=XM_019839955.1
C2 73427394 73427516 refGene_gn=PPP1R2;refGene_tx=XM_003991767.5
C2 73440154 73440262 refGene_gn=PPP1R2;refGene_tx=XM_003991767.5
C2 73442418 73442496 refGene_gn=PPP1R2;refGene_tx=XM_003991767.5
C2 73443153 73443248 refGene_gn=PPP1R2;refGene_tx=XM_003991767.5
C2 73446560 73446728 refGene_gn=PPP1R2;refGene_tx=XM_003991767.5
C2 73448689 73448736 refGene_gn=PPP1R2;refGene_tx=XM_003991767.5

C2 73490534 73490587
refGene_gn=ACAP2;refGene_tx=XM_003991768.5;refGene_gn=ACAP2;refGene_tx=XM_019839956.2;refGene_gn=ACAP2;refGene_tx=XM_019839957.2;refGene_gn=ACAP2;refGene_tx=XM_023260282.1;refGene_gn=ACAP2;refGene_tx=XM_023260283.1;refGene_gn=ACAP2;refGene_tx=XM_023260284.1;refGene_gn=ACAP2;refGene_tx=XM_023260287.1
C2 73548669 73548727
refGene_gn=ACAP2;refGene_tx=XM_003991768.5;refGene_gn=ACAP2;refGene_tx=XM_019839956.2;refGene_gn=ACAP2;refGene_tx=XM_019839957.2;refGene_gn=ACAP2;refGene_tx=XM_023260282.1;refGene_gn=ACAP2;refGene_tx=XM_023260283.1;refGene_gn=ACAP2;refGene_tx=XM_023260284.1;refGene_gn=ACAP2;refGene_tx=XM_023260287.1
C2 73560875 73560995
refGene_gn=ACAP2;refGene_tx=XM_003991768.5;refGene_gn=ACAP2;refGene_tx=XM_019839956.2;refGene_gn=ACAP2;refGene_tx=XM_019839957.2;refGene_gn=ACAP2;refGene_tx=XM_023260282.1;refGene_gn=ACAP2;refGene_tx=XM_023260283.1;refGene_gn=ACAP2;refGene_tx=XM_023260284.1;refGene_gn=ACAP2;refGene_tx=XM_023260285.1;refGene_gn=ACAP2;refGene_tx=XM_023260286.1;refGene_gn=ACAP2;refGene_tx=XM_023260287.1
C2 73561792 73561846
refGene_gn=ACAP2;refGene_tx=XM_003991768.5;refGene_gn=ACAP2;refGene_tx=XM_019839956.2;refGene_gn=ACAP2;refGene_tx=XM_019839957.2;refGene_gn=ACAP2;refGene_tx=XM_023260282.1;refGene_gn=ACAP2;refGene_tx=XM_023260283.1;refGene_gn=ACAP2;refGene_tx=XM_023260284.1;refGene_gn=ACAP2;refGene_tx=XM_023260285.1;refGene_gn=ACAP2;refGene_tx=XM_023260286.1;refGene_gn=ACAP2;refGene_tx=XM_023260287.1
C2 73582978 73583037
refGene_gn=ACAP2;refGene_tx=XM_003991768.5;refGene_gn=ACAP2;refGene_tx=XM_019839956.2;refGene_gn=ACAP2;refGene_tx=XM_019839957.2;refGene_gn=ACAP2;refGene_tx=XM_023260282.1;refGene_gn=ACAP2;refGene_tx=XM_023260283.1;refGene_gn=ACAP2;refGene_tx=XM_023260284.1;refGene_gn=ACAP2;refGene_tx=XM_023260285.1;refGene_gn=ACAP2;refGene_tx=XM_023260286.1;refGene_gn=ACAP2;refGene_tx=XM_023260287.1
C2 73584188 73584372
refGene_gn=ACAP2;refGene_tx=XM_003991768.5;refGene_gn=ACAP2;refGene_tx=XM_019839956.2;refGene_gn=ACAP2;refGene_tx=XM_019839957.2;refGene_gn=ACAP2;refGene_tx=XM_023260282.1;refGene_gn=ACAP2;refGene_tx=XM_023260283.1;refGene_gn=ACAP2;refGene_tx=XM_023260284.1;refGene_gn=ACAP2;refGene_tx=XM_023260285.1;refGene_gn=ACAP2;refGene_tx=XM_023260286.1;refGene_gn=ACAP2;refGene_tx=XM_023260287.1
C2 73588420 73588465
refGene_gn=ACAP2;refGene_tx=XM_003991768.5;refGene_gn=ACAP2;refGene_tx=XM_019839956.2;refGene_gn=ACAP2;refGene_tx=XM_019839957.2;refGene_gn=ACAP2;refGene_tx=XM_023260282.1;refGene_gn=ACAP2;refGene_tx=XM_023260283.1;refGene_gn=ACAP2;refGene_tx=XM_023260284.1;refGene_gn=ACAP2;refGene_tx=XM_023260285.1;refGene_gn=ACAP2;refGene_tx=XM_023260286.1;refGene_gn=ACAP2;refGene_tx=XM_023260287.1
C2 73592431 73592527
refGene_gn=ACAP2;refGene_tx=XM_003991768.5;refGene_gn=ACAP2;refGene_tx=XM_019839956.2;refGene_gn=ACAP2;refGene_tx=XM_019839957.2;refGene_gn=

ACAP2;refGene_tx=XM_023260282.1;refGene_gn=ACAP2;refGene_tx=XM_0232602
83.1;refGene_gn=ACAP2;refGene_tx=XM_023260284.1;refGene_gn=ACAP2;refGe
ne_tx=XM_023260285.1;refGene_gn=ACAP2;refGene_tx=XM_023260286.1;refGen
e_gn=ACAP2;refGene_tx=XM_023260287.1
C2 73599386 73599461
refGene_gn=ACAP2;refGene_tx=XM_003991768.5;refGene_gn=ACAP2;refGene_tx
=XM_019839956.2;refGene_gn=ACAP2;refGene_tx=XM_019839957.2;refGene_gn=
ACAP2;refGene_tx=XM_023260282.1;refGene_gn=ACAP2;refGene_tx=XM_0232602
83.1;refGene_gn=ACAP2;refGene_tx=XM_023260284.1;refGene_gn=ACAP2;refGe
ne_tx=XM_023260285.1;refGene_gn=ACAP2;refGene_tx=XM_023260286.1;refGen
e_gn=ACAP2;refGene_tx=XM_023260287.1
C2 73600993 73601005
refGene_gn=ACAP2;refGene_tx=XM_019839956.2;refGene_gn=ACAP2;refGene_tx
=XM_023260282.1;refGene_gn=ACAP2;refGene_tx=XM_023260284.1;refGene_gn=
ACAP2;refGene_tx=XM_023260285.1;refGene_gn=ACAP2;refGene_tx=XM_0232602
86.1;refGene_gn=ACAP2;refGene_tx=XM_023260287.1
C2 73606519 73606632
refGene_gn=ACAP2;refGene_tx=XM_003991768.5;refGene_gn=ACAP2;refGene_tx
=XM_019839956.2;refGene_gn=ACAP2;refGene_tx=XM_019839957.2;refGene_gn=
ACAP2;refGene_tx=XM_023260282.1;refGene_gn=ACAP2;refGene_tx=XM_0232602
83.1;refGene_gn=ACAP2;refGene_tx=XM_023260284.1;refGene_gn=ACAP2;refGe
ne_tx=XM_023260285.1;refGene_gn=ACAP2;refGene_tx=XM_023260286.1;refGen
e_gn=ACAP2;refGene_tx=XM_023260287.1
C2 73611317 73611338
refGene_gn=ACAP2;refGene_tx=XM_023260282.1;refGene_gn=ACAP2;refGene_tx
=XM_023260283.1;refGene_gn=ACAP2;refGene_tx=XM_023260284.1;refGene_gn=
ACAP2;refGene_tx=XM_023260285.1;refGene_gn=ACAP2;refGene_tx=XM_0232602
86.1;refGene_gn=ACAP2;refGene_tx=XM_023260287.1
C2 73614788 73614840
refGene_gn=ACAP2;refGene_tx=XM_003991768.5;refGene_gn=ACAP2;refGene_tx
=XM_019839956.2;refGene_gn=ACAP2;refGene_tx=XM_019839957.2;refGene_gn=
ACAP2;refGene_tx=XM_023260282.1;refGene_gn=ACAP2;refGene_tx=XM_0232602
83.1;refGene_gn=ACAP2;refGene_tx=XM_023260284.1;refGene_gn=ACAP2;refGe
ne_tx=XM_023260285.1;refGene_gn=ACAP2;refGene_tx=XM_023260286.1;refGen
e_gn=ACAP2;refGene_tx=XM_023260287.1
C2 73616145 73616246
refGene_gn=ACAP2;refGene_tx=XM_003991768.5;refGene_gn=ACAP2;refGene_tx
=XM_019839956.2;refGene_gn=ACAP2;refGene_tx=XM_019839957.2;refGene_gn=
ACAP2;refGene_tx=XM_023260282.1;refGene_gn=ACAP2;refGene_tx=XM_0232602
83.1;refGene_gn=ACAP2;refGene_tx=XM_023260284.1;refGene_gn=ACAP2;refGe
ne_tx=XM_023260285.1;refGene_gn=ACAP2;refGene_tx=XM_023260286.1;refGen
e_gn=ACAP2;refGene_tx=XM_023260287.1
C2 73616826 73616932
refGene_gn=ACAP2;refGene_tx=XM_003991768.5;refGene_gn=ACAP2;refGene_tx
=XM_019839956.2;refGene_gn=ACAP2;refGene_tx=XM_019839957.2;refGene_gn=
ACAP2;refGene_tx=XM_023260282.1;refGene_gn=ACAP2;refGene_tx=XM_0232602
83.1;refGene_gn=ACAP2;refGene_tx=XM_023260284.1;refGene_gn=ACAP2;refGe
ne_tx=XM_023260285.1;refGene_gn=ACAP2;refGene_tx=XM_023260286.1;refGen
e_gn=ACAP2;refGene_tx=XM_023260287.1
C2 73625491 73625700

refGene_gn=ACAP2;refGene_tx=XM_003991768.5;refGene_gn=ACAP2;refGene_tx=XM_019839956.2;refGene_gn=ACAP2;refGene_tx=XM_019839957.2;refGene_gn=ACAP2;refGene_tx=XM_023260282.1;refGene_gn=ACAP2;refGene_tx=XM_023260283.1;refGene_gn=ACAP2;refGene_tx=XM_023260284.1;refGene_gn=ACAP2;refGene_tx=XM_023260285.1;refGene_gn=ACAP2;refGene_tx=XM_023260286.1;refGene_gn=ACAP2;refGene_tx=XM_023260287.1
C2 73626035 73626105
refGene_gn=ACAP2;refGene_tx=XM_003991768.5;refGene_gn=ACAP2;refGene_tx=XM_019839956.2;refGene_gn=ACAP2;refGene_tx=XM_019839957.2;refGene_gn=ACAP2;refGene_tx=XM_023260282.1;refGene_gn=ACAP2;refGene_tx=XM_023260283.1;refGene_gn=ACAP2;refGene_tx=XM_023260284.1;refGene_gn=ACAP2;refGene_tx=XM_023260285.1;refGene_gn=ACAP2;refGene_tx=XM_023260286.1;refGene_gn=ACAP2;refGene_tx=XM_023260287.1
C2 73628531 73628623
refGene_gn=ACAP2;refGene_tx=XM_003991768.5;refGene_gn=ACAP2;refGene_tx=XM_019839956.2;refGene_gn=ACAP2;refGene_tx=XM_019839957.2;refGene_gn=ACAP2;refGene_tx=XM_023260282.1;refGene_gn=ACAP2;refGene_tx=XM_023260283.1;refGene_gn=ACAP2;refGene_tx=XM_023260284.1;refGene_gn=ACAP2;refGene_tx=XM_023260285.1;refGene_gn=ACAP2;refGene_tx=XM_023260286.1;refGene_gn=ACAP2;refGene_tx=XM_023260287.1
C2 73630199 73630384
refGene_gn=ACAP2;refGene_tx=XM_003991768.5;refGene_gn=ACAP2;refGene_tx=XM_019839956.2;refGene_gn=ACAP2;refGene_tx=XM_019839957.2;refGene_gn=ACAP2;refGene_tx=XM_023260282.1;refGene_gn=ACAP2;refGene_tx=XM_023260283.1;refGene_gn=ACAP2;refGene_tx=XM_023260284.1;refGene_gn=ACAP2;refGene_tx=XM_023260285.1;refGene_gn=ACAP2;refGene_tx=XM_023260286.1;refGene_gn=ACAP2;refGene_tx=XM_023260287.1
C2 73630774 73630879
refGene_gn=ACAP2;refGene_tx=XM_019839956.2;refGene_gn=ACAP2;refGene_tx=XM_019839957.2;refGene_gn=ACAP2;refGene_tx=XM_023260282.1;refGene_gn=ACAP2;refGene_tx=XM_023260283.1;refGene_gn=ACAP2;refGene_tx=XM_023260285.1;refGene_gn=ACAP2;refGene_tx=XM_023260286.1
C2 73631261 73631354
refGene_gn=ACAP2;refGene_tx=XM_003991768.5;refGene_gn=ACAP2;refGene_tx=XM_019839956.2;refGene_gn=ACAP2;refGene_tx=XM_019839957.2;refGene_gn=ACAP2;refGene_tx=XM_023260282.1;refGene_gn=ACAP2;refGene_tx=XM_023260283.1;refGene_gn=ACAP2;refGene_tx=XM_023260284.1;refGene_gn=ACAP2;refGene_tx=XM_023260285.1;refGene_gn=ACAP2;refGene_tx=XM_023260286.1
C2 73632693 73632881
refGene_gn=ACAP2;refGene_tx=XM_003991768.5;refGene_gn=ACAP2;refGene_tx=XM_019839956.2;refGene_gn=ACAP2;refGene_tx=XM_019839957.2;refGene_gn=ACAP2;refGene_tx=XM_023260282.1;refGene_gn=ACAP2;refGene_tx=XM_023260283.1;refGene_gn=ACAP2;refGene_tx=XM_023260284.1;refGene_gn=ACAP2;refGene_tx=XM_023260285.1;refGene_gn=ACAP2;refGene_tx=XM_023260286.1;refGene_gn=ACAP2;refGene_tx=XM_023260287.1
C2 73633495 73633605
refGene_gn=ACAP2;refGene_tx=XM_003991768.5;refGene_gn=ACAP2;refGene_tx=XM_019839956.2;refGene_gn=ACAP2;refGene_tx=XM_019839957.2;refGene_gn=ACAP2;refGene_tx=XM_023260282.1;refGene_gn=ACAP2;refGene_tx=XM_023260283.1;refGene_gn=ACAP2;refGene_tx=XM_023260284.1;refGene_gn=ACAP2;refGe

ne_tx=XM_023260285.1;refGene_gn=ACAP2;refGene_tx=XM_023260286.1;refGene_gn=ACAP2;refGene_tx=XM_023260287.1
C2	73634945	73635056
refGene_gn=ACAP2;refGene_tx=XM_003991768.5;refGene_gn=ACAP2;refGene_tx=XM_019839956.2;refGene_gn=ACAP2;refGene_tx=XM_019839957.2;refGene_gn=ACAP2;refGene_tx=XM_023260282.1;refGene_gn=ACAP2;refGene_tx=XM_023260283.1;refGene_gn=ACAP2;refGene_tx=XM_023260284.1;refGene_gn=ACAP2;refGene_tx=XM_023260285.1;refGene_gn=ACAP2;refGene_tx=XM_023260286.1;refGene_gn=ACAP2;refGene_tx=XM_023260287.1
C2	73637344	73637406
refGene_gn=ACAP2;refGene_tx=XM_003991768.5;refGene_gn=ACAP2;refGene_tx=XM_019839956.2;refGene_gn=ACAP2;refGene_tx=XM_019839957.2;refGene_gn=ACAP2;refGene_tx=XM_023260282.1;refGene_gn=ACAP2;refGene_tx=XM_023260283.1;refGene_gn=ACAP2;refGene_tx=XM_023260284.1;refGene_gn=ACAP2;refGene_tx=XM_023260285.1;refGene_gn=ACAP2;refGene_tx=XM_023260286.1;refGene_gn=ACAP2;refGene_tx=XM_023260287.1
C2	73643266	73643367
refGene_gn=ACAP2;refGene_tx=XM_003991768.5;refGene_gn=ACAP2;refGene_tx=XM_019839956.2;refGene_gn=ACAP2;refGene_tx=XM_019839957.2;refGene_gn=ACAP2;refGene_tx=XM_023260282.1;refGene_gn=ACAP2;refGene_tx=XM_023260283.1;refGene_gn=ACAP2;refGene_tx=XM_023260284.1;refGene_gn=ACAP2;refGene_tx=XM_023260285.1;refGene_gn=ACAP2;refGene_tx=XM_023260286.1;refGene_gn=ACAP2;refGene_tx=XM_023260287.1
C2	73653293	73653794	refGene_gn=XXYLT1;refGene_tx=XM_023260288.1
C2	73684601	73684749	refGene_gn=XXYLT1;refGene_tx=XM_023260288.1
C2	73749241	73749374	refGene_gn=XXYLT1;refGene_tx=XM_023260288.1
C2	73821852	73822249	refGene_gn=XXYLT1;refGene_tx=XM_023260288.1
C2	74152431	74153739	refGene_gn=FAM43A;refGene_tx=XM_011285881.3
C2	74167933	74168032	refGene_gn=LSG1;refGene_tx=XM_019839962.2
C2	74169819	74169946	refGene_gn=LSG1;refGene_tx=XM_019839962.2
C2	74176548	74176669	refGene_gn=LSG1;refGene_tx=XM_019839962.2
C2	74176845	74176932	refGene_gn=LSG1;refGene_tx=XM_019839962.2
C2	74177478	74177565	refGene_gn=LSG1;refGene_tx=XM_019839962.2
C2	74185595	74185656	refGene_gn=LSG1;refGene_tx=XM_019839962.2
C2	74186708	74186885	refGene_gn=LSG1;refGene_tx=XM_019839962.2
C2	74191551	74191968	refGene_gn=LSG1;refGene_tx=XM_019839962.2
C2	74194680	74194782	refGene_gn=LSG1;refGene_tx=XM_019839962.2
C2	74194919	74195063	refGene_gn=LSG1;refGene_tx=XM_019839962.2
C2	74196801	74196925	refGene_gn=LSG1;refGene_tx=XM_019839962.2
C2	74199209	74199289	refGene_gn=LSG1;refGene_tx=XM_019839962.2
C2	74200839	74201013	refGene_gn=LSG1;refGene_tx=XM_019839962.2
C2	74204179	74204359	refGene_gn=LSG1;refGene_tx=XM_019839962.2
C2	74216422	74216559
refGene_gn=TMEM44;refGene_tx=XM_023260289.1;refGene_gn=TMEM44;refGene_tx=XM_023260290.1;refGene_gn=TMEM44;refGene_tx=XM_023260291.1;refGene_gn=TMEM44;refGene_tx=XM_023260292.1
C2	74219589	74219716
refGene_gn=TMEM44;refGene_tx=XM_023260289.1;refGene_gn=TMEM44;refGene_tx=XM_023260290.1;refGene_gn=TMEM44;refGene_tx=XM_023260291.1;refGene_gn=TMEM44;refGene_tx=XM_023260292.1

C2	74221697 74221791
refGene_gn=TMEM44;refGene_tx=XM_023260289.1;refGene_gn=TMEM44;refGene_tx=XM_023260290.1;refGene_gn=TMEM44;refGene_tx=XM_023260291.1;refGene_gn=TMEM44;refGene_tx=XM_023260292.1
C2	74222842 74223009
refGene_gn=TMEM44;refGene_tx=XM_023260289.1;refGene_gn=TMEM44;refGene_tx=XM_023260290.1;refGene_gn=TMEM44;refGene_tx=XM_023260291.1;refGene_gn=TMEM44;refGene_tx=XM_023260292.1
C2	74223203 74223290
refGene_gn=TMEM44;refGene_tx=XM_023260289.1;refGene_gn=TMEM44;refGene_tx=XM_023260290.1;refGene_gn=TMEM44;refGene_tx=XM_023260291.1;refGene_gn=TMEM44;refGene_tx=XM_023260292.1
C2	74228073 74228244
refGene_gn=TMEM44;refGene_tx=XM_023260289.1;refGene_gn=TMEM44;refGene_tx=XM_023260290.1;refGene_gn=TMEM44;refGene_tx=XM_023260291.1
C2	74230140 74230309
refGene_gn=TMEM44;refGene_tx=XM_023260289.1;refGene_gn=TMEM44;refGene_tx=XM_023260290.1;refGene_gn=TMEM44;refGene_tx=XM_023260291.1
C2	74232134 74232239
refGene_gn=TMEM44;refGene_tx=XM_023260289.1;refGene_gn=TMEM44;refGene_tx=XM_023260290.1;refGene_gn=TMEM44;refGene_tx=XM_023260291.1;refGene_gn=TMEM44;refGene_tx=XM_023260292.1
C2	74237534 74237693
refGene_gn=TMEM44;refGene_tx=XM_023260289.1;refGene_gn=TMEM44;refGene_tx=XM_023260290.1;refGene_gn=TMEM44;refGene_tx=XM_023260291.1;refGene_gn=TMEM44;refGene_tx=XM_023260292.1
C2	74248414 74248447
refGene_gn=TMEM44;refGene_tx=XM_023260289.1;refGene_gn=TMEM44;refGene_tx=XM_023260292.1
C2	74251196 74251307
refGene_gn=TMEM44;refGene_tx=XM_023260289.1;refGene_gn=TMEM44;refGene_tx=XM_023260290.1;refGene_gn=TMEM44;refGene_tx=XM_023260292.1
C2	74341887 74341938
refGene_gn=ATP13A3;refGene_tx=XM_011285878.3;refGene_gn=ATP13A3;refGene_tx=XM_019839963.2;refGene_gn=ATP13A3;refGene_tx=XM_019839964.2;refGene_gn=ATP13A3;refGene_tx=XM_019839965.2;refGene_gn=ATP13A3;refGene_tx=XM_019839966.2
C2	74343366 74343540
refGene_gn=ATP13A3;refGene_tx=XM_011285878.3;refGene_gn=ATP13A3;refGene_tx=XM_019839963.2;refGene_gn=ATP13A3;refGene_tx=XM_019839964.2;refGene_gn=ATP13A3;refGene_tx=XM_019839965.2;refGene_gn=ATP13A3;refGene_tx=XM_019839966.2
C2	74344499 74344682
refGene_gn=ATP13A3;refGene_tx=XM_011285878.3;refGene_gn=ATP13A3;refGene_tx=XM_019839963.2;refGene_gn=ATP13A3;refGene_tx=XM_019839964.2;refGene_gn=ATP13A3;refGene_tx=XM_019839965.2;refGene_gn=ATP13A3;refGene_tx=XM_019839966.2
C2	74344972 74345043
refGene_gn=ATP13A3;refGene_tx=XM_011285878.3;refGene_gn=ATP13A3;refGene_tx=XM_019839963.2;refGene_gn=ATP13A3;refGene_tx=XM_019839964.2;refGe

ne_gn=ATP13A3;refGene_tx=XM_019839965.2;refGene_gn=ATP13A3;refGene_tx=XM_019839966.2
C2 74346703 74346784
refGene_gn=ATP13A3;refGene_tx=XM_011285878.3;refGene_gn=ATP13A3;refGene_tx=XM_019839963.2;refGene_gn=ATP13A3;refGene_tx=XM_019839964.2;refGene_gn=ATP13A3;refGene_tx=XM_019839966.2
C2 74347959 74348029
refGene_gn=ATP13A3;refGene_tx=XM_011285878.3;refGene_gn=ATP13A3;refGene_tx=XM_019839963.2;refGene_gn=ATP13A3;refGene_tx=XM_019839964.2;refGene_gn=ATP13A3;refGene_tx=XM_019839965.2;refGene_gn=ATP13A3;refGene_tx=XM_019839966.2
C2 74350248 74350383
refGene_gn=ATP13A3;refGene_tx=XM_011285878.3;refGene_gn=ATP13A3;refGene_tx=XM_019839963.2;refGene_gn=ATP13A3;refGene_tx=XM_019839964.2;refGene_gn=ATP13A3;refGene_tx=XM_019839965.2;refGene_gn=ATP13A3;refGene_tx=XM_019839966.2
C2 74350828 74350901
refGene_gn=ATP13A3;refGene_tx=XM_011285878.3;refGene_gn=ATP13A3;refGene_tx=XM_019839963.2;refGene_gn=ATP13A3;refGene_tx=XM_019839964.2;refGene_gn=ATP13A3;refGene_tx=XM_019839965.2;refGene_gn=ATP13A3;refGene_tx=XM_019839966.2
C2 74353860 74353992
refGene_gn=ATP13A3;refGene_tx=XM_011285878.3;refGene_gn=ATP13A3;refGene_tx=XM_019839963.2;refGene_gn=ATP13A3;refGene_tx=XM_019839964.2;refGene_gn=ATP13A3;refGene_tx=XM_019839965.2;refGene_gn=ATP13A3;refGene_tx=XM_019839966.2
C2 74355246 74355426
refGene_gn=ATP13A3;refGene_tx=XM_011285878.3;refGene_gn=ATP13A3;refGene_tx=XM_019839963.2;refGene_gn=ATP13A3;refGene_tx=XM_019839964.2;refGene_gn=ATP13A3;refGene_tx=XM_019839965.2;refGene_gn=ATP13A3;refGene_tx=XM_019839966.2
C2 74355563 74355721
refGene_gn=ATP13A3;refGene_tx=XM_011285878.3;refGene_gn=ATP13A3;refGene_tx=XM_019839963.2;refGene_gn=ATP13A3;refGene_tx=XM_019839964.2;refGene_gn=ATP13A3;refGene_tx=XM_019839965.2;refGene_gn=ATP13A3;refGene_tx=XM_019839966.2
C2 74356661 74356850
refGene_gn=ATP13A3;refGene_tx=XM_011285878.3;refGene_gn=ATP13A3;refGene_tx=XM_019839963.2;refGene_gn=ATP13A3;refGene_tx=XM_019839964.2;refGene_gn=ATP13A3;refGene_tx=XM_019839965.2;refGene_gn=ATP13A3;refGene_tx=XM_019839966.2
C2 74358546 74358608
refGene_gn=ATP13A3;refGene_tx=XM_011285878.3;refGene_gn=ATP13A3;refGene_tx=XM_019839963.2;refGene_gn=ATP13A3;refGene_tx=XM_019839964.2;refGene_gn=ATP13A3;refGene_tx=XM_019839965.2;refGene_gn=ATP13A3;refGene_tx=XM_019839966.2
C2 74362352 74362503
refGene_gn=ATP13A3;refGene_tx=XM_011285878.3;refGene_gn=ATP13A3;refGene_tx=XM_019839963.2;refGene_gn=ATP13A3;refGene_tx=XM_019839964.2;refGene_gn=ATP13A3;refGene_tx=XM_019839965.2;refGene_gn=ATP13A3;refGene_tx=

XM_019839966.2
C2 74365213 74365330
refGene_gn=ATP13A3;refGene_tx=XM_011285878.3;refGene_gn=ATP13A3;refGene_tx=XM_019839963.2;refGene_gn=ATP13A3;refGene_tx=XM_019839964.2;refGene_gn=ATP13A3;refGene_tx=XM_019839965.2;refGene_gn=ATP13A3;refGene_tx=XM_019839966.2
C2 74366202 74366220
refGene_gn=ATP13A3;refGene_tx=XM_011285878.3;refGene_gn=ATP13A3;refGene_tx=XM_019839963.2;refGene_gn=ATP13A3;refGene_tx=XM_019839964.2;refGene_gn=ATP13A3;refGene_tx=XM_019839965.2;refGene_gn=ATP13A3;refGene_tx=XM_019839966.2
C2 74366312 74366466
refGene_gn=ATP13A3;refGene_tx=XM_011285878.3;refGene_gn=ATP13A3;refGene_tx=XM_019839963.2;refGene_gn=ATP13A3;refGene_tx=XM_019839964.2;refGene_gn=ATP13A3;refGene_tx=XM_019839965.2;refGene_gn=ATP13A3;refGene_tx=XM_019839966.2
C2 74366560 74366681
refGene_gn=ATP13A3;refGene_tx=XM_011285878.3;refGene_gn=ATP13A3;refGene_tx=XM_019839963.2;refGene_gn=ATP13A3;refGene_tx=XM_019839964.2;refGene_gn=ATP13A3;refGene_tx=XM_019839965.2;refGene_gn=ATP13A3;refGene_tx=XM_019839966.2
C2 74375527 74375652
refGene_gn=ATP13A3;refGene_tx=XM_011285878.3;refGene_gn=ATP13A3;refGene_tx=XM_019839963.2;refGene_gn=ATP13A3;refGene_tx=XM_019839964.2;refGene_gn=ATP13A3;refGene_tx=XM_019839965.2;refGene_gn=ATP13A3;refGene_tx=XM_019839966.2
C2 74377218 74377394
refGene_gn=ATP13A3;refGene_tx=XM_011285878.3;refGene_gn=ATP13A3;refGene_tx=XM_019839963.2;refGene_gn=ATP13A3;refGene_tx=XM_019839964.2;refGene_gn=ATP13A3;refGene_tx=XM_019839965.2;refGene_gn=ATP13A3;refGene_tx=XM_019839966.2
C2 74378395 74378518
refGene_gn=ATP13A3;refGene_tx=XM_011285878.3;refGene_gn=ATP13A3;refGene_tx=XM_019839963.2;refGene_gn=ATP13A3;refGene_tx=XM_019839964.2;refGene_gn=ATP13A3;refGene_tx=XM_019839965.2;refGene_gn=ATP13A3;refGene_tx=XM_019839966.2
C2 74378601 74378681
refGene_gn=ATP13A3;refGene_tx=XM_011285878.3;refGene_gn=ATP13A3;refGene_tx=XM_019839963.2;refGene_gn=ATP13A3;refGene_tx=XM_019839964.2;refGene_gn=ATP13A3;refGene_tx=XM_019839965.2;refGene_gn=ATP13A3;refGene_tx=XM_019839966.2
C2 74379520 74379563
refGene_gn=ATP13A3;refGene_tx=XM_011285878.3;refGene_gn=ATP13A3;refGene_tx=XM_019839963.2;refGene_gn=ATP13A3;refGene_tx=XM_019839964.2;refGene_gn=ATP13A3;refGene_tx=XM_019839965.2;refGene_gn=ATP13A3;refGene_tx=XM_019839966.2
C2 74379650 74379760
refGene_gn=ATP13A3;refGene_tx=XM_011285878.3;refGene_gn=ATP13A3;refGene_tx=XM_019839963.2;refGene_gn=ATP13A3;refGene_tx=XM_019839964.2;refGene_gn=ATP13A3;refGene_tx=XM_019839965.2;refGene_gn=ATP13A3;refGene_tx=

XM_019839966.2
C2	74380043 74380140
refGene_gn=ATP13A3;refGene_tx=XM_011285878.3;refGene_gn=ATP13A3;refGene_tx=XM_019839963.2;refGene_gn=ATP13A3;refGene_tx=XM_019839964.2;refGene_gn=ATP13A3;refGene_tx=XM_019839965.2;refGene_gn=ATP13A3;refGene_tx=XM_019839966.2
C2	74381055 74381128
refGene_gn=ATP13A3;refGene_tx=XM_011285878.3;refGene_gn=ATP13A3;refGene_tx=XM_019839963.2;refGene_gn=ATP13A3;refGene_tx=XM_019839964.2;refGene_gn=ATP13A3;refGene_tx=XM_019839965.2;refGene_gn=ATP13A3;refGene_tx=XM_019839966.2
C2	74382938 74383116
refGene_gn=ATP13A3;refGene_tx=XM_011285878.3;refGene_gn=ATP13A3;refGene_tx=XM_019839963.2;refGene_gn=ATP13A3;refGene_tx=XM_019839964.2;refGene_gn=ATP13A3;refGene_tx=XM_019839965.2;refGene_gn=ATP13A3;refGene_tx=XM_019839966.2
C2	74384721 74384909
refGene_gn=ATP13A3;refGene_tx=XM_011285878.3;refGene_gn=ATP13A3;refGene_tx=XM_019839963.2;refGene_gn=ATP13A3;refGene_tx=XM_019839964.2;refGene_gn=ATP13A3;refGene_tx=XM_019839965.2;refGene_gn=ATP13A3;refGene_tx=XM_019839966.2
C2	74389816 74389905
refGene_gn=ATP13A3;refGene_tx=XM_011285878.3;refGene_gn=ATP13A3;refGene_tx=XM_019839963.2;refGene_gn=ATP13A3;refGene_tx=XM_019839964.2;refGene_gn=ATP13A3;refGene_tx=XM_019839965.2;refGene_gn=ATP13A3;refGene_tx=XM_019839966.2
C2	74393309 74393390
refGene_gn=ATP13A3;refGene_tx=XM_011285878.3;refGene_gn=ATP13A3;refGene_tx=XM_019839963.2;refGene_gn=ATP13A3;refGene_tx=XM_019839964.2;refGene_gn=ATP13A3;refGene_tx=XM_019839965.2;refGene_gn=ATP13A3;refGene_tx=XM_019839966.2
C2	74394882 74394972
refGene_gn=ATP13A3;refGene_tx=XM_011285878.3;refGene_gn=ATP13A3;refGene_tx=XM_019839963.2;refGene_gn=ATP13A3;refGene_tx=XM_019839964.2;refGene_gn=ATP13A3;refGene_tx=XM_019839965.2
C2	74399134 74399154 refGene_gn=ATP13A3;refGene_tx=XM_019839964.2
C2	74401615 74401813
refGene_gn=ATP13A3;refGene_tx=XM_011285878.3;refGene_gn=ATP13A3;refGene_tx=XM_019839963.2;refGene_gn=ATP13A3;refGene_tx=XM_019839964.2;refGene_gn=ATP13A3;refGene_tx=XM_019839965.2;refGene_gn=ATP13A3;refGene_tx=XM_019839966.2
C2	74410020 74411709
refGene_gn=GP5;refGene_tx=XM_023260293.1;refGene_gn=GP5;refGene_tx=XM_023260294.1
C2	74450904 74452659 refGene_gn=LRRC15;refGene_tx=XM_006936178.2
C2	74465810 74467454
refGene_gn=CPN2;refGene_tx=XM_003991783.3;refGene_gn=CPN2;refGene_tx=XM_019839967.1
C2	74648090 74648641 refGene_gn=HES1;refGene_tx=XM_011285877.3
C2	74649307 74649395 refGene_gn=HES1;refGene_tx=XM_011285877.3

C2	74649657	74649753	refGene_gn=HES1;refGene_tx=XM_011285877.3
C2	74649885	74649993	refGene_gn=HES1;refGene_tx=XM_011285877.3
C2	75036626	75036691
refGene_gn=OPA1;refGene_tx=XM_003991770.5;refGene_gn=OPA1;refGene_tx=XM_011285874.2;refGene_gn=OPA1;refGene_tx=XM_011285875.2;refGene_gn=OPA1;refGene_tx=XM_011285876.2
C2	75040396	75040443	refGene_gn=OPA1;refGene_tx=XM_019839970.2
C2	75058395	75058506
refGene_gn=OPA1;refGene_tx=XM_003991770.5;refGene_gn=OPA1;refGene_tx=XM_011285874.2;refGene_gn=OPA1;refGene_tx=XM_011285875.2;refGene_gn=OPA1;refGene_tx=XM_011285876.2;refGene_gn=OPA1;refGene_tx=XM_019839970.2
C2	75059013	75059107
refGene_gn=OPA1;refGene_tx=XM_003991770.5;refGene_gn=OPA1;refGene_tx=XM_011285874.2;refGene_gn=OPA1;refGene_tx=XM_011285875.2;refGene_gn=OPA1;refGene_tx=XM_011285876.2;refGene_gn=OPA1;refGene_tx=XM_019839970.2
C2	75060393	75060510
refGene_gn=OPA1;refGene_tx=XM_003991770.5;refGene_gn=OPA1;refGene_tx=XM_011285874.2;refGene_gn=OPA1;refGene_tx=XM_011285875.2;refGene_gn=OPA1;refGene_tx=XM_011285876.2;refGene_gn=OPA1;refGene_tx=XM_019839970.2
C2	75062746	75062887
refGene_gn=OPA1;refGene_tx=XM_003991770.5;refGene_gn=OPA1;refGene_tx=XM_011285874.2;refGene_gn=OPA1;refGene_tx=XM_011285875.2;refGene_gn=OPA1;refGene_tx=XM_011285876.2;refGene_gn=OPA1;refGene_tx=XM_019839970.2
C2	75066865	75066945
refGene_gn=OPA1;refGene_tx=XM_003991770.5;refGene_gn=OPA1;refGene_tx=XM_011285874.2;refGene_gn=OPA1;refGene_tx=XM_011285875.2;refGene_gn=OPA1;refGene_tx=XM_011285876.2;refGene_gn=OPA1;refGene_tx=XM_019839970.2
C2	75067477	75067586
refGene_gn=OPA1;refGene_tx=XM_003991770.5;refGene_gn=OPA1;refGene_tx=XM_011285874.2;refGene_gn=OPA1;refGene_tx=XM_011285875.2;refGene_gn=OPA1;refGene_tx=XM_011285876.2;refGene_gn=OPA1;refGene_tx=XM_019839970.2
C2	75069404	75069557
refGene_gn=OPA1;refGene_tx=XM_003991770.5;refGene_gn=OPA1;refGene_tx=XM_011285874.2;refGene_gn=OPA1;refGene_tx=XM_011285875.2;refGene_gn=OPA1;refGene_tx=XM_011285876.2;refGene_gn=OPA1;refGene_tx=XM_019839970.2
C2	75071533	75071699
refGene_gn=OPA1;refGene_tx=XM_003991770.5;refGene_gn=OPA1;refGene_tx=XM_011285874.2;refGene_gn=OPA1;refGene_tx=XM_011285875.2;refGene_gn=OPA1;refGene_tx=XM_011285876.2;refGene_gn=OPA1;refGene_tx=XM_019839970.2
C2	75077759	75077836
refGene_gn=OPA1;refGene_tx=XM_003991770.5;refGene_gn=OPA1;refGene_tx=XM_011285874.2;refGene_gn=OPA1;refGene_tx=XM_011285875.2;refGene_gn=OPA1;refGene_tx=XM_011285876.2;refGene_gn=OPA1;refGene_tx=XM_019839970.2
C2	75078623	75078688
refGene_gn=OPA1;refGene_tx=XM_003991770.5;refGene_gn=OPA1;refGene_tx=XM_011285874.2;refGene_gn=OPA1;refGene_tx=XM_011285875.2;refGene_gn=OPA1;refGene_tx=XM_011285876.2;refGene_gn=OPA1;refGene_tx=XM_019839970.2
C2	75079559	75079675
refGene_gn=OPA1;refGene_tx=XM_003991770.5;refGene_gn=OPA1;refGene_tx=XM_011285874.2;refGene_gn=OPA1;refGene_tx=XM_011285875.2;refGene_gn=OPA

1;refGene_tx=XM_011285876.2;refGene_gn=OPA1;refGene_tx=XM_019839970.2
C2 75080998 75081071
refGene_gn=OPA1;refGene_tx=XM_003991770.5;refGene_gn=OPA1;refGene_tx=XM_011285874.2;refGene_gn=OPA1;refGene_tx=XM_011285875.2;refGene_gn=OPA1;refGene_tx=XM_011285876.2;refGene_gn=OPA1;refGene_tx=XM_019839970.2
C2 75081165 75081238
refGene_gn=OPA1;refGene_tx=XM_003991770.5;refGene_gn=OPA1;refGene_tx=XM_011285874.2;refGene_gn=OPA1;refGene_tx=XM_011285875.2;refGene_gn=OPA1;refGene_tx=XM_011285876.2;refGene_gn=OPA1;refGene_tx=XM_019839970.2
C2 75083782 75083913
refGene_gn=OPA1;refGene_tx=XM_003991770.5;refGene_gn=OPA1;refGene_tx=XM_011285874.2;refGene_gn=OPA1;refGene_tx=XM_011285875.2;refGene_gn=OPA1;refGene_tx=XM_011285876.2;refGene_gn=OPA1;refGene_tx=XM_019839970.2
C2 75084228 75084328
refGene_gn=OPA1;refGene_tx=XM_003991770.5;refGene_gn=OPA1;refGene_tx=XM_011285874.2;refGene_gn=OPA1;refGene_tx=XM_011285875.2;refGene_gn=OPA1;refGene_tx=XM_011285876.2;refGene_gn=OPA1;refGene_tx=XM_019839970.2
C2 75084411 75084483
refGene_gn=OPA1;refGene_tx=XM_003991770.5;refGene_gn=OPA1;refGene_tx=XM_011285874.2;refGene_gn=OPA1;refGene_tx=XM_011285875.2;refGene_gn=OPA1;refGene_tx=XM_011285876.2;refGene_gn=OPA1;refGene_tx=XM_019839970.2
C2 75084798 75084873
refGene_gn=OPA1;refGene_tx=XM_003991770.5;refGene_gn=OPA1;refGene_tx=XM_011285874.2;refGene_gn=OPA1;refGene_tx=XM_011285875.2;refGene_gn=OPA1;refGene_tx=XM_011285876.2;refGene_gn=OPA1;refGene_tx=XM_019839970.2
C2 75084999 75085080
refGene_gn=OPA1;refGene_tx=XM_003991770.5;refGene_gn=OPA1;refGene_tx=XM_011285874.2;refGene_gn=OPA1;refGene_tx=XM_011285875.2;refGene_gn=OPA1;refGene_tx=XM_011285876.2;refGene_gn=OPA1;refGene_tx=XM_019839970.2
C2 75089042 75089156
refGene_gn=OPA1;refGene_tx=XM_003991770.5;refGene_gn=OPA1;refGene_tx=XM_011285874.2;refGene_gn=OPA1;refGene_tx=XM_011285875.2;refGene_gn=OPA1;refGene_tx=XM_011285876.2;refGene_gn=OPA1;refGene_tx=XM_019839970.2
C2 75090887 75090974
refGene_gn=OPA1;refGene_tx=XM_003991770.5;refGene_gn=OPA1;refGene_tx=XM_011285874.2;refGene_gn=OPA1;refGene_tx=XM_011285875.2;refGene_gn=OPA1;refGene_tx=XM_011285876.2;refGene_gn=OPA1;refGene_tx=XM_019839970.2
C2 75092658 75092763
refGene_gn=OPA1;refGene_tx=XM_003991770.5;refGene_gn=OPA1;refGene_tx=XM_011285874.2;refGene_gn=OPA1;refGene_tx=XM_011285875.2;refGene_gn=OPA1;refGene_tx=XM_011285876.2;refGene_gn=OPA1;refGene_tx=XM_019839970.2
C2 75096165 75096219
refGene_gn=OPA1;refGene_tx=XM_003991770.5;refGene_gn=OPA1;refGene_tx=XM_011285874.2;refGene_gn=OPA1;refGene_tx=XM_011285875.2;refGene_gn=OPA1;refGene_tx=XM_011285876.2;refGene_gn=OPA1;refGene_tx=XM_019839970.2
C2 75100938 75101049
refGene_gn=OPA1;refGene_tx=XM_003991770.5;refGene_gn=OPA1;refGene_tx=XM_011285874.2;refGene_gn=OPA1;refGene_tx=XM_019839970.2
C2 75109482 75109550
refGene_gn=OPA1;refGene_tx=XM_003991770.5;refGene_gn=OPA1;refGene_tx=X

M_011285874.2;refGene_gn=OPA1;refGene_tx=XM_011285875.2;refGene_gn=OPA1;refGene_tx=XM_011285876.2;refGene_gn=OPA1;refGene_tx=XM_019839970.2
C2 75110220 75110274
refGene_gn=OPA1;refGene_tx=XM_003991770.5;refGene_gn=OPA1;refGene_tx=XM_011285875.2;refGene_gn=OPA1;refGene_tx=XM_019839970.2
C2 75110772 75110880
refGene_gn=OPA1;refGene_tx=XM_003991770.5;refGene_gn=OPA1;refGene_tx=XM_011285874.2;refGene_gn=OPA1;refGene_tx=XM_011285875.2;refGene_gn=OPA1;refGene_tx=XM_011285876.2;refGene_gn=OPA1;refGene_tx=XM_019839970.2
C2 75112584 75112681
refGene_gn=OPA1;refGene_tx=XM_003991770.5;refGene_gn=OPA1;refGene_tx=XM_011285874.2;refGene_gn=OPA1;refGene_tx=XM_011285875.2;refGene_gn=OPA1;refGene_tx=XM_011285876.2;refGene_gn=OPA1;refGene_tx=XM_019839970.2
C2 75113325 75113644
refGene_gn=OPA1;refGene_tx=XM_003991770.5;refGene_gn=OPA1;refGene_tx=XM_011285874.2;refGene_gn=OPA1;refGene_tx=XM_011285875.2;refGene_gn=OPA1;refGene_tx=XM_011285876.2;refGene_gn=OPA1;refGene_tx=XM_019839970.2
C2 75119590 75119622
refGene_gn=OPA1;refGene_tx=XM_003991770.5;refGene_gn=OPA1;refGene_tx=XM_011285874.2;refGene_gn=OPA1;refGene_tx=XM_011285875.2;refGene_gn=OPA1;refGene_tx=XM_011285876.2;refGene_gn=OPA1;refGene_tx=XM_019839970.2
C2 75149239 75149302
refGene_gn=LOC101091472;refGene_tx=XM_023260295.1
C2 75182076 75182250
refGene_gn=LOC101091472;refGene_tx=XM_023260295.1
C2 75194403 75194550
refGene_gn=LOC101091472;refGene_tx=XM_023260295.1;refGene_gn=LOC101091472;refGene_tx=XM_023260296.1
C2 75198875 75198946
refGene_gn=LOC101091472;refGene_tx=XM_023260295.1;refGene_gn=LOC101091472;refGene_tx=XM_023260296.1
C2 75199034 75199115
refGene_gn=LOC101091472;refGene_tx=XM_023260295.1;refGene_gn=LOC101091472;refGene_tx=XM_023260296.1
C2 75200600 75200670
refGene_gn=LOC101091472;refGene_tx=XM_023260295.1;refGene_gn=LOC101091472;refGene_tx=XM_023260296.1
C2 75202166 75202301
refGene_gn=LOC101091472;refGene_tx=XM_023260295.1;refGene_gn=LOC101091472;refGene_tx=XM_023260296.1
C2 75206697 75206767
refGene_gn=LOC101091472;refGene_tx=XM_023260295.1;refGene_gn=LOC101091472;refGene_tx=XM_023260296.1
C2 75217449 75217584
refGene_gn=LOC101091472;refGene_tx=XM_023260295.1;refGene_gn=LOC101091472;refGene_tx=XM_023260296.1
C2 75220099 75220270
refGene_gn=LOC101091472;refGene_tx=XM_023260295.1;refGene_gn=LOC101091472;refGene_tx=XM_023260296.1
C2 75221332 75221490

refGene_gn=LOC101091472;refGene_tx=XM_023260295.1;refGene_gn=LOC101091472;refGene_tx=XM_023260296.1
C2 75222098 75222287
refGene_gn=LOC101091472;refGene_tx=XM_023260295.1;refGene_gn=LOC101091472;refGene_tx=XM_023260296.1
C2 75224802 75224868
refGene_gn=LOC101091472;refGene_tx=XM_023260295.1;refGene_gn=LOC101091472;refGene_tx=XM_023260296.1
C2 77228572 77228659
refGene_gn=LOC109491645;refGene_tx=XM_023260297.1
C2 77230287 77230374
refGene_gn=LOC109491645;refGene_tx=XM_023260297.1
C2 77230593 77230747
refGene_gn=LOC109491645;refGene_tx=XM_023260297.1
C2 77231895 77232007
refGene_gn=LOC109491645;refGene_tx=XM_023260297.1
C2 77237416 77237541
refGene_gn=LOC109491645;refGene_tx=XM_023260297.1
C2 77242035 77242196
refGene_gn=LOC109491645;refGene_tx=XM_023260297.1
C2 77243030 77243153
refGene_gn=LOC109491645;refGene_tx=XM_023260297.1
C2 77244377 77244457
refGene_gn=LOC109491645;refGene_tx=XM_023260297.1
C2 77245922 77245965
refGene_gn=LOC109491645;refGene_tx=XM_023260297.1
C2 77246354 77246464
refGene_gn=LOC109491645;refGene_tx=XM_023260297.1
C2 77248547 77248644
refGene_gn=LOC109491645;refGene_tx=XM_023260297.1
C2 77250159 77250232
refGene_gn=LOC109491645;refGene_tx=XM_023260297.1
C2 77257387 77257559
refGene_gn=LOC109491645;refGene_tx=XM_023260297.1
C2 77258555 77258749
refGene_gn=LOC109491645;refGene_tx=XM_023260297.1
C2 77260332 77260421
refGene_gn=LOC109491645;refGene_tx=XM_023260297.1
C2 77263776 77263857
refGene_gn=LOC109491645;refGene_tx=XM_023260297.1
C2 77269272 77269485
refGene_gn=LOC109491645;refGene_tx=XM_023260297.1
C2 77285037 77285100
refGene_gn=LOC101091724;refGene_tx=XM_023260298.1;refGene_gn=LOC101091724;refGene_tx=XM_023260299.1
C2 77308539 77308713
refGene_gn=LOC101091724;refGene_tx=XM_023260298.1;refGene_gn=LOC101091724;refGene_tx=XM_023260299.1
C2 77309412 77309559
refGene_gn=LOC101091724;refGene_tx=XM_023260298.1;refGene_gn=LOC101091

724;refGene_tx=XM_023260299.1
C2 77309826 77309897
refGene_gn=LOC101091724;refGene_tx=XM_023260298.1;refGene_gn=LOC101091724;refGene_tx=XM_023260299.1
C2 77309996 77310077
refGene_gn=LOC101091724;refGene_tx=XM_023260298.1;refGene_gn=LOC101091724;refGene_tx=XM_023260299.1
C2 77313851 77313921
refGene_gn=LOC101091724;refGene_tx=XM_023260298.1;refGene_gn=LOC101091724;refGene_tx=XM_023260299.1
C2 77324723 77324858
refGene_gn=LOC101091724;refGene_tx=XM_023260298.1;refGene_gn=LOC101091724;refGene_tx=XM_023260299.1
C2 77334039 77334112
refGene_gn=LOC101091724;refGene_tx=XM_023260298.1;refGene_gn=LOC101091724;refGene_tx=XM_023260299.1
C2 77334935 77335064
refGene_gn=LOC101091724;refGene_tx=XM_023260298.1;refGene_gn=LOC101091724;refGene_tx=XM_023260299.1
C2 77342375 77342546
refGene_gn=LOC101091724;refGene_tx=XM_023260298.1;refGene_gn=LOC101091724;refGene_tx=XM_023260299.1
C2 77344719 77344877
refGene_gn=LOC101091724;refGene_tx=XM_023260298.1;refGene_gn=LOC101091724;refGene_tx=XM_023260299.1
C2 77347610 77347799
refGene_gn=LOC101091724;refGene_tx=XM_023260298.1;refGene_gn=LOC101091724;refGene_tx=XM_023260299.1
C2 77351763 77351825
refGene_gn=LOC101091724;refGene_tx=XM_023260298.1;refGene_gn=LOC101091724;refGene_tx=XM_023260299.1
C2 77353417 77353568
refGene_gn=LOC101091724;refGene_tx=XM_023260298.1;refGene_gn=LOC101091724;refGene_tx=XM_023260299.1
C2 77356101 77356188
refGene_gn=LOC101091724;refGene_tx=XM_023260298.1;refGene_gn=LOC101091724;refGene_tx=XM_023260299.1
C2 77356776 77356933
refGene_gn=LOC101091724;refGene_tx=XM_023260298.1;refGene_gn=LOC101091724;refGene_tx=XM_023260299.1
C2 77358893 77359011
refGene_gn=LOC101091724;refGene_tx=XM_023260298.1;refGene_gn=LOC101091724;refGene_tx=XM_023260299.1
C2 77362141 77362266
refGene_gn=LOC101091724;refGene_tx=XM_023260298.1;refGene_gn=LOC101091724;refGene_tx=XM_023260299.1
C2 77363070 77363231
refGene_gn=LOC101091724;refGene_tx=XM_023260298.1;refGene_gn=LOC101091724;refGene_tx=XM_023260299.1
C2 77365650 77365776

refGene_gn=LOC101091724;refGene_tx=XM_023260298.1;refGene_gn=LOC101091724;refGene_tx=XM_023260299.1
C2 77367116 77367196
refGene_gn=LOC101091724;refGene_tx=XM_023260298.1;refGene_gn=LOC101091724;refGene_tx=XM_023260299.1
C2 77369399 77369442
refGene_gn=LOC101091724;refGene_tx=XM_023260298.1;refGene_gn=LOC101091724;refGene_tx=XM_023260299.1
C2 77371132 77371242
refGene_gn=LOC101091724;refGene_tx=XM_023260298.1;refGene_gn=LOC101091724;refGene_tx=XM_023260299.1
C2 77374598 77374695
refGene_gn=LOC101091724;refGene_tx=XM_023260298.1;refGene_gn=LOC101091724;refGene_tx=XM_023260299.1
C2 77376694 77376767
refGene_gn=LOC101091724;refGene_tx=XM_023260298.1;refGene_gn=LOC101091724;refGene_tx=XM_023260299.1
C2 77384819 77384998
refGene_gn=LOC101091724;refGene_tx=XM_023260298.1;refGene_gn=LOC101091724;refGene_tx=XM_023260299.1
C2 77387430 77387633
refGene_gn=LOC101091724;refGene_tx=XM_023260299.1
C2 77393227 77393316
refGene_gn=LOC101091724;refGene_tx=XM_023260299.1
C2 77395622 77395703
refGene_gn=LOC101091724;refGene_tx=XM_023260299.1
C2 77397602 77397863
refGene_gn=LOC101091724;refGene_tx=XM_023260299.1
C2 77401585 77401687
refGene_gn=HRASLS;refGene_tx=XM_023260300.1;refGene_gn=HRASLS;refGene_tx=XM_023260301.1;refGene_gn=HRASLS;refGene_tx=XM_023260302.1;refGene_gn=HRASLS;refGene_tx=XM_023260303.1
C2 77410257 77410526
refGene_gn=HRASLS;refGene_tx=XM_023260300.1;refGene_gn=HRASLS;refGene_tx=XM_023260301.1;refGene_gn=HRASLS;refGene_tx=XM_023260302.1;refGene_gn=HRASLS;refGene_tx=XM_023260303.1
C2 77415962 77416101
refGene_gn=HRASLS;refGene_tx=XM_023260300.1;refGene_gn=HRASLS;refGene_tx=XM_023260301.1;refGene_gn=HRASLS;refGene_tx=XM_023260302.1;refGene_gn=HRASLS;refGene_tx=XM_023260303.1
C2 77423259 77423310 refGene_gn=HRASLS;refGene_tx=XM_023260303.1
C2 77423993 77424112
refGene_gn=HRASLS;refGene_tx=XM_023260300.1;refGene_gn=HRASLS;refGene_tx=XM_023260301.1
C2 77424274 77424374 refGene_gn=HRASLS;refGene_tx=XM_023260300.1
C2 77424516 77424576 refGene_gn=HRASLS;refGene_tx=XM_023260302.1
C2 77424914 77424981 refGene_gn=HRASLS;refGene_tx=XM_023260301.1
C2 77680401 77680612 refGene_gn=MB21D2;refGene_tx=XM_003991787.5
C2 77792712 77793977 refGene_gn=MB21D2;refGene_tx=XM_003991787.5
C2 77853424 77853437 refGene_gn=FGF12;refGene_tx=XM_011286026.3

C2 78040218 78040309 refGene_gn=FGF12;refGene_tx=XM_019839972.2
C2 78150062 78150261 refGene_gn=FGF12;refGene_tx=XM_003991788.5
C2 78196612 78196723
refGene_gn=FGF12;refGene_tx=XM_003991788.5;refGene_gn=FGF12;refGene_tx=XM_011286026.3;refGene_gn=FGF12;refGene_tx=XM_019839972.2
C2 78225896 78226000
refGene_gn=FGF12;refGene_tx=XM_003991788.5;refGene_gn=FGF12;refGene_tx=XM_011286026.3;refGene_gn=FGF12;refGene_tx=XM_019839972.2
C2 78393251 78393450
refGene_gn=FGF12;refGene_tx=XM_003991788.5;refGene_gn=FGF12;refGene_tx=XM_011286026.3;refGene_gn=FGF12;refGene_tx=XM_019839972.2
C2 78409808 78409927
refGene_gn=FGF12;refGene_tx=XM_003991788.5;refGene_gn=FGF12;refGene_tx=XM_011286026.3;refGene_gn=FGF12;refGene_tx=XM_019839972.2
C2 79042463 79042483
refGene_gn=CCDC50;refGene_tx=XM_023260304.1;refGene_gn=CCDC50;refGene_tx=XM_023260305.1
C2 79043849 79043956
refGene_gn=CCDC50;refGene_tx=XM_023260304.1;refGene_gn=CCDC50;refGene_tx=XM_023260305.1
C2 79048656 79048736
refGene_gn=CCDC50;refGene_tx=XM_023260304.1;refGene_gn=CCDC50;refGene_tx=XM_023260305.1
C2 79050557 79050662
refGene_gn=CCDC50;refGene_tx=XM_023260304.1;refGene_gn=CCDC50;refGene_tx=XM_023260305.1
C2 79050771 79050816
refGene_gn=CCDC50;refGene_tx=XM_023260304.1;refGene_gn=CCDC50;refGene_tx=XM_023260305.1
C2 79051197 79051313
refGene_gn=CCDC50;refGene_tx=XM_023260304.1;refGene_gn=CCDC50;refGene_tx=XM_023260305.1
C2 79058322 79058850 refGene_gn=CCDC50;refGene_tx=XM_023260304.1
C2 79063497 79063615
refGene_gn=CCDC50;refGene_tx=XM_023260304.1;refGene_gn=CCDC50;refGene_tx=XM_023260305.1
C2 79069225 79069316
refGene_gn=CCDC50;refGene_tx=XM_023260304.1;refGene_gn=CCDC50;refGene_tx=XM_023260305.1
C2 79073579 79073706
refGene_gn=CCDC50;refGene_tx=XM_023260304.1;refGene_gn=CCDC50;refGene_tx=XM_023260305.1
C2 79074517 79074580
refGene_gn=CCDC50;refGene_tx=XM_023260304.1;refGene_gn=CCDC50;refGene_tx=XM_023260305.1
C2 79101315 79101364
refGene_gn=CCDC50;refGene_tx=XM_023260304.1;refGene_gn=CCDC50;refGene_tx=XM_023260305.1
C2 79130696 79130799 refGene_gn=UTS2B;refGene_tx=XM_023260863.1
C2 79135289 79135388 refGene_gn=UTS2B;refGene_tx=XM_023260863.1

C2 79136538 79136576 refGene_gn=UTS2B;refGene_tx=XM_023260863.1
C2 79137846 79137940 refGene_gn=UTS2B;refGene_tx=XM_023260863.1
C2 79148523 79148549 refGene_gn=UTS2B;refGene_tx=XM_023260863.1
C2 79157446 79157531
refGene_gn=OSTN;refGene_tx=XM_023260306.1;refGene_gn=OSTN;refGene_tx=XM_023260307.1
C2 79183289 79183504
refGene_gn=OSTN;refGene_tx=XM_023260306.1;refGene_gn=OSTN;refGene_tx=XM_023260307.1
C2 79186877 79186976
refGene_gn=OSTN;refGene_tx=XM_023260306.1;refGene_gn=OSTN;refGene_tx=XM_023260307.1
C2 79472630 79472639 refGene_gn=GMNC;refGene_tx=XM_023260175.1
C2 79473159 79473334 refGene_gn=GMNC;refGene_tx=XM_023260175.1
C2 79475493 79475582 refGene_gn=GMNC;refGene_tx=XM_023260175.1
C2 79476962 79477079 refGene_gn=GMNC;refGene_tx=XM_023260175.1
C2 79478727 79479348 refGene_gn=GMNC;refGene_tx=XM_023260175.1
C2 79661501 79662220 refGene_gn=IL1RAP;refGene_tx=XM_023260308.1
C2 79671607 79671751 refGene_gn=IL1RAP;refGene_tx=XM_023260308.1
C2 79673320 79673464 refGene_gn=IL1RAP;refGene_tx=XM_023260308.1
C2 79682183 79682332 refGene_gn=IL1RAP;refGene_tx=XM_023260308.1
C2 79685053 79685180 refGene_gn=IL1RAP;refGene_tx=XM_023260308.1
C2 79687196 79687268 refGene_gn=IL1RAP;refGene_tx=XM_023260308.1
C2 79689741 79689907 refGene_gn=IL1RAP;refGene_tx=XM_023260308.1
C2 79703205 79703392 refGene_gn=IL1RAP;refGene_tx=XM_023260308.1
C2 79707894 79708102 refGene_gn=IL1RAP;refGene_tx=XM_023260308.1
C2 79708104 79708219 refGene_gn=IL1RAP;refGene_tx=XM_023260308.1
C2 79855857 79855932 refGene_gn=TMEM207;refGene_tx=XM_003991829.2
C2 79857826 79857864 refGene_gn=TMEM207;refGene_tx=XM_003991829.2
C2 79864648 79864693 refGene_gn=TMEM207;refGene_tx=XM_003991829.2
C2 79865500 79865643 refGene_gn=TMEM207;refGene_tx=XM_003991829.2
C2 79874746 79874883 refGene_gn=TMEM207;refGene_tx=XM_003991829.2
C2 79887866 79888000 refGene_gn=CLDN16;refGene_tx=XM_003991793.4
C2 79888738 79888930 refGene_gn=CLDN16;refGene_tx=XM_003991793.4
C2 79892342 79892507 refGene_gn=CLDN16;refGene_tx=XM_003991793.4
C2 79895777 79895880 refGene_gn=CLDN16;refGene_tx=XM_003991793.4
C2 79909671 79909806 refGene_gn=CLDN16;refGene_tx=XM_003991793.4
C2 79909807 79909855 refGene_gn=CLDN16;refGene_tx=XM_003991793.4
C2 79909857 79910001 refGene_gn=CLDN16;refGene_tx=XM_003991793.4
C2 79989709 79989932 refGene_gn=CLDN1;refGene_tx=XM_003991794.3
C2 79998745 79998910 refGene_gn=CLDN1;refGene_tx=XM_003991794.3
C2 80001400 80001485 refGene_gn=CLDN1;refGene_tx=XM_003991794.3
C2 80003185 80003348 refGene_gn=CLDN1;refGene_tx=XM_003991794.3
C2 80129642 80130119 refGene_gn=P3H2;refGene_tx=XM_003991795.3
C2 80243877 80244030
refGene_gn=P3H2;refGene_tx=XM_003991795.3;refGene_gn=P3H2;refGene_tx=XM_006936186.3
C2 80244962 80245152
refGene_gn=P3H2;refGene_tx=XM_003991795.3;refGene_gn=P3H2;refGene_tx=XM_006936186.3

C2 80250722 80250854
refGene_gn=P3H2;refGene_tx=XM_003991795.3;refGene_gn=P3H2;refGene_tx=XM_006936186.3
C2 80253183 80253326
refGene_gn=P3H2;refGene_tx=XM_003991795.3;refGene_gn=P3H2;refGene_tx=XM_006936186.3
C2 80253663 80253753
refGene_gn=P3H2;refGene_tx=XM_003991795.3;refGene_gn=P3H2;refGene_tx=XM_006936186.3
C2 80256115 80256156
refGene_gn=P3H2;refGene_tx=XM_003991795.3;refGene_gn=P3H2;refGene_tx=XM_006936186.3
C2 80257404 80257499
refGene_gn=P3H2;refGene_tx=XM_003991795.3;refGene_gn=P3H2;refGene_tx=XM_006936186.3
C2 80263882 80264010
refGene_gn=P3H2;refGene_tx=XM_003991795.3;refGene_gn=P3H2;refGene_tx=XM_006936186.3
C2 80264571 80264667
refGene_gn=P3H2;refGene_tx=XM_003991795.3;refGene_gn=P3H2;refGene_tx=XM_006936186.3
C2 80265132 80265283
refGene_gn=P3H2;refGene_tx=XM_003991795.3;refGene_gn=P3H2;refGene_tx=XM_006936186.3
C2 80266157 80266275
refGene_gn=P3H2;refGene_tx=XM_003991795.3;refGene_gn=P3H2;refGene_tx=XM_006936186.3
C2 80267255 80267331
refGene_gn=P3H2;refGene_tx=XM_003991795.3;refGene_gn=P3H2;refGene_tx=XM_006936186.3
C2 80272967 80273108
refGene_gn=P3H2;refGene_tx=XM_003991795.3;refGene_gn=P3H2;refGene_tx=XM_006936186.3
C2 80277615 80277708
refGene_gn=P3H2;refGene_tx=XM_003991795.3;refGene_gn=P3H2;refGene_tx=XM_006936186.3
C2 80333902 80334199
refGene_gn=TP63;refGene_tx=XM_003991796.5;refGene_gn=TP63;refGene_tx=XM_019839985.2;refGene_gn=TP63;refGene_tx=XM_019839986.1;refGene_gn=TP63;refGene_tx=XM_019839987.1;refGene_gn=TP63;refGene_tx=XM_019839988.2;refGene_gn=TP63;refGene_tx=XM_019839989.1;refGene_gn=TP63;refGene_tx=XM_019839990.1;refGene_gn=TP63;refGene_tx=XM_019839991.2;refGene_gn=TP63;refGene_tx=XM_019839992.2
C2 80337213 80337307
refGene_gn=TP63;refGene_tx=XM_003991796.5;refGene_gn=TP63;refGene_tx=XM_019839985.2;refGene_gn=TP63;refGene_tx=XM_019839986.1;refGene_gn=TP63;refGene_tx=XM_019839987.1;refGene_gn=TP63;refGene_tx=XM_019839989.1;refGene_gn=TP63;refGene_tx=XM_019839990.1
C2 80338628 80338773
refGene_gn=TP63;refGene_tx=XM_003991796.5;refGene_gn=TP63;refGene_tx=X

M_019839985.2;refGene_gn=TP63;refGene_tx=XM_019839986.1;refGene_gn=TP63;refGene_tx=XM_019839987.1;refGene_gn=TP63;refGene_tx=XM_019839988.2;refGene_gn=TP63;refGene_tx=XM_019839989.1;refGene_gn=TP63;refGene_tx=XM_019839990.1
C2 80340900 80341058
refGene_gn=TP63;refGene_tx=XM_003991796.5;refGene_gn=TP63;refGene_tx=XM_019839985.2;refGene_gn=TP63;refGene_tx=XM_019839986.1;refGene_gn=TP63;refGene_tx=XM_019839987.1;refGene_gn=TP63;refGene_tx=XM_019839988.2;refGene_gn=TP63;refGene_tx=XM_019839989.1;refGene_gn=TP63;refGene_tx=XM_019839990.1;refGene_gn=TP63;refGene_tx=XM_019839991.2;refGene_gn=TP63;refGene_tx=XM_019839992.2
C2 80348395 80348510 refGene_gn=TP63;refGene_tx=XM_003991798.4
C2 80354734 80354871
refGene_gn=TP63;refGene_tx=XM_003991796.5;refGene_gn=TP63;refGene_tx=XM_003991798.4;refGene_gn=TP63;refGene_tx=XM_019839985.2;refGene_gn=TP63;refGene_tx=XM_019839986.1;refGene_gn=TP63;refGene_tx=XM_019839987.1;refGene_gn=TP63;refGene_tx=XM_019839988.2;refGene_gn=TP63;refGene_tx=XM_019839989.1;refGene_gn=TP63;refGene_tx=XM_019839990.1;refGene_gn=TP63;refGene_tx=XM_019839991.2;refGene_gn=TP63;refGene_tx=XM_019839992.2
C2 80357163 80357246
refGene_gn=TP63;refGene_tx=XM_003991796.5;refGene_gn=TP63;refGene_tx=XM_003991798.4;refGene_gn=TP63;refGene_tx=XM_019839985.2;refGene_gn=TP63;refGene_tx=XM_019839986.1;refGene_gn=TP63;refGene_tx=XM_019839987.1;refGene_gn=TP63;refGene_tx=XM_019839988.2;refGene_gn=TP63;refGene_tx=XM_019839989.1;refGene_gn=TP63;refGene_tx=XM_019839990.1;refGene_gn=TP63;refGene_tx=XM_019839991.2;refGene_gn=TP63;refGene_tx=XM_019839992.2
C2 80359033 80359170
refGene_gn=TP63;refGene_tx=XM_003991796.5;refGene_gn=TP63;refGene_tx=XM_003991798.4;refGene_gn=TP63;refGene_tx=XM_019839985.2;refGene_gn=TP63;refGene_tx=XM_019839986.1;refGene_gn=TP63;refGene_tx=XM_019839987.1;refGene_gn=TP63;refGene_tx=XM_019839988.2;refGene_gn=TP63;refGene_tx=XM_019839989.1;refGene_gn=TP63;refGene_tx=XM_019839990.1;refGene_gn=TP63;refGene_tx=XM_019839991.2;refGene_gn=TP63;refGene_tx=XM_019839992.2
C2 80359837 80359947
refGene_gn=TP63;refGene_tx=XM_003991796.5;refGene_gn=TP63;refGene_tx=XM_003991798.4;refGene_gn=TP63;refGene_tx=XM_019839985.2;refGene_gn=TP63;refGene_tx=XM_019839986.1;refGene_gn=TP63;refGene_tx=XM_019839987.1;refGene_gn=TP63;refGene_tx=XM_019839988.2;refGene_gn=TP63;refGene_tx=XM_019839989.1;refGene_gn=TP63;refGene_tx=XM_019839990.1;refGene_gn=TP63;refGene_tx=XM_019839991.2;refGene_gn=TP63;refGene_tx=XM_019839992.2
C2 80360718 80360834
refGene_gn=TP63;refGene_tx=XM_003991796.5;refGene_gn=TP63;refGene_tx=XM_003991798.4;refGene_gn=TP63;refGene_tx=XM_019839985.2;refGene_gn=TP63;refGene_tx=XM_019839986.1;refGene_gn=TP63;refGene_tx=XM_019839987.1;refGene_gn=TP63;refGene_tx=XM_019839988.2;refGene_gn=TP63;refGene_tx=XM_019839989.1;refGene_gn=TP63;refGene_tx=XM_019839990.1;refGene_gn=TP63;refGene_tx=XM_019839991.2;refGene_gn=TP63;refGene_tx=XM_019839992.2
C2 80363381 80363568
refGene_gn=TP63;refGene_tx=XM_003991796.5;refGene_gn=TP63;refGene_tx=XM_003991798.4;refGene_gn=TP63;refGene_tx=XM_019839985.2;refGene_gn=TP6

3;refGene_tx=XM_019839986.1;refGene_gn=TP63;refGene_tx=XM_019839987.1;
refGene_gn=TP63;refGene_tx=XM_019839988.2;refGene_gn=TP63;refGene_tx=X
M_019839989.1;refGene_gn=TP63;refGene_tx=XM_019839990.1;refGene_gn=TP6
3;refGene_tx=XM_019839991.2;refGene_gn=TP63;refGene_tx=XM_019839992.2
C2 80414336 80414591
refGene_gn=TP63;refGene_tx=XM_003991796.5;refGene_gn=TP63;refGene_tx=X
M_003991798.4;refGene_gn=TP63;refGene_tx=XM_019839985.2;refGene_gn=TP6
3;refGene_tx=XM_019839986.1;refGene_gn=TP63;refGene_tx=XM_019839987.1;
refGene_gn=TP63;refGene_tx=XM_019839988.2;refGene_gn=TP63;refGene_tx=X
M_019839989.1;refGene_gn=TP63;refGene_tx=XM_019839990.1;refGene_gn=TP6
3;refGene_tx=XM_019839991.2;refGene_gn=TP63;refGene_tx=XM_019839992.2
C2 80433288 80433330 refGene_gn=TP63;refGene_tx=XM_019839987.1
C2 80475453 80475586
refGene_gn=TP63;refGene_tx=XM_003991796.5;refGene_gn=TP63;refGene_tx=X
M_003991798.4;refGene_gn=TP63;refGene_tx=XM_019839985.2;refGene_gn=TP6
3;refGene_tx=XM_019839986.1;refGene_gn=TP63;refGene_tx=XM_019839988.2;
refGene_gn=TP63;refGene_tx=XM_019839991.2;refGene_gn=TP63;refGene_tx=X
M_019839992.2
C2 80476358 80476487
refGene_gn=TP63;refGene_tx=XM_003991796.5;refGene_gn=TP63;refGene_tx=X
M_003991798.4;refGene_gn=TP63;refGene_tx=XM_019839985.2;refGene_gn=TP6
3;refGene_tx=XM_019839986.1;refGene_gn=TP63;refGene_tx=XM_019839988.2;
refGene_gn=TP63;refGene_tx=XM_019839991.2;refGene_gn=TP63;refGene_tx=X
M_019839992.2
C2 80561408 80561521
refGene_gn=TP63;refGene_tx=XM_003991796.5;refGene_gn=TP63;refGene_tx=X
M_003991798.4;refGene_gn=TP63;refGene_tx=XM_019839985.2;refGene_gn=TP6
3;refGene_tx=XM_019839988.2;refGene_gn=TP63;refGene_tx=XM_019839991.2;
refGene_gn=TP63;refGene_tx=XM_019839992.2
C2 80561711 80561722 refGene_gn=TP63;refGene_tx=XM_019839986.1
C2 80817284 80817479
refGene_gn=TPRG1;refGene_tx=XM_003991801.4;refGene_gn=TPRG1;refGene_tx
=XM_011286027.3;refGene_gn=TPRG1;refGene_tx=XM_019839995.2;refGene_gn=
TPRG1;refGene_tx=XM_019839996.1
C2 80831463 80831617
refGene_gn=TPRG1;refGene_tx=XM_003991801.4;refGene_gn=TPRG1;refGene_tx
=XM_011286027.3;refGene_gn=TPRG1;refGene_tx=XM_019839995.2;refGene_gn=
TPRG1;refGene_tx=XM_019839996.1
C2 80892871 80893048
refGene_gn=TPRG1;refGene_tx=XM_003991801.4;refGene_gn=TPRG1;refGene_tx
=XM_011286027.3;refGene_gn=TPRG1;refGene_tx=XM_019839995.2;refGene_gn=
TPRG1;refGene_tx=XM_019839996.1
C2 80919062 80919154
refGene_gn=TPRG1;refGene_tx=XM_003991801.4;refGene_gn=TPRG1;refGene_tx
=XM_011286027.3;refGene_gn=TPRG1;refGene_tx=XM_019839995.2;refGene_gn=
TPRG1;refGene_tx=XM_019839996.1
C2 80929140 80929371
refGene_gn=TPRG1;refGene_tx=XM_003991801.4;refGene_gn=TPRG1;refGene_tx
=XM_011286027.3;refGene_gn=TPRG1;refGene_tx=XM_019839995.2;refGene_gn=
TPRG1;refGene_tx=XM_019839996.1

C2 80940900 80940936 refGene_gn=TPRG1;refGene_tx=XM_019839995.2
C2 81034690 81034702 refGene_gn=TPRG1;refGene_tx=XM_003991801.4
C2 81221325 81221454
refGene_gn=LPP;refGene_tx=XM_019839993.2;refGene_gn=LPP;refGene_tx=XM_019839994.2;refGene_gn=LPP;refGene_tx=XM_023260309.1;refGene_gn=LPP;refGene_tx=XM_023260310.1;refGene_gn=LPP;refGene_tx=XM_023260311.1
C2 81224559 81224680
refGene_gn=LPP;refGene_tx=XM_019839993.2;refGene_gn=LPP;refGene_tx=XM_019839994.2;refGene_gn=LPP;refGene_tx=XM_023260309.1;refGene_gn=LPP;refGene_tx=XM_023260310.1;refGene_gn=LPP;refGene_tx=XM_023260311.1
C2 81230359 81230538
refGene_gn=LPP;refGene_tx=XM_019839993.2;refGene_gn=LPP;refGene_tx=XM_019839994.2;refGene_gn=LPP;refGene_tx=XM_023260309.1;refGene_gn=LPP;refGene_tx=XM_023260310.1;refGene_gn=LPP;refGene_tx=XM_023260311.1
C2 81330807 81330977
refGene_gn=LPP;refGene_tx=XM_019839993.2;refGene_gn=LPP;refGene_tx=XM_019839994.2;refGene_gn=LPP;refGene_tx=XM_023260309.1;refGene_gn=LPP;refGene_tx=XM_023260310.1;refGene_gn=LPP;refGene_tx=XM_023260311.1
C2 81381190 81381317
refGene_gn=LPP;refGene_tx=XM_019839993.2;refGene_gn=LPP;refGene_tx=XM_019839994.2;refGene_gn=LPP;refGene_tx=XM_023260309.1;refGene_gn=LPP;refGene_tx=XM_023260310.1;refGene_gn=LPP;refGene_tx=XM_023260311.1
C2 81474318 81475008
refGene_gn=LPP;refGene_tx=XM_019839993.2;refGene_gn=LPP;refGene_tx=XM_019839994.2;refGene_gn=LPP;refGene_tx=XM_023260309.1;refGene_gn=LPP;refGene_tx=XM_023260310.1;refGene_gn=LPP;refGene_tx=XM_023260311.1
C2 81564216 81564342
refGene_gn=LPP;refGene_tx=XM_019839993.2;refGene_gn=LPP;refGene_tx=XM_019839994.2;refGene_gn=LPP;refGene_tx=XM_023260309.1;refGene_gn=LPP;refGene_tx=XM_023260310.1;refGene_gn=LPP;refGene_tx=XM_023260311.1
C2 81600116 81600229
refGene_gn=LPP;refGene_tx=XM_019839993.2;refGene_gn=LPP;refGene_tx=XM_019839994.2;refGene_gn=LPP;refGene_tx=XM_023260309.1;refGene_gn=LPP;refGene_tx=XM_023260310.1;refGene_gn=LPP;refGene_tx=XM_023260311.1
C2 81673005 81673195
refGene_gn=LPP;refGene_tx=XM_019839993.2;refGene_gn=LPP;refGene_tx=XM_019839994.2;refGene_gn=LPP;refGene_tx=XM_023260309.1;refGene_gn=LPP;refGene_tx=XM_023260310.1;refGene_gn=LPP;refGene_tx=XM_023260311.1
C2 82274089 82274250
refGene_gn=BCL6;refGene_tx=XM_003991804.4;refGene_gn=BCL6;refGene_tx=XM_006936189.4;refGene_gn=BCL6;refGene_tx=XM_006936190.4
C2 82276462 82276684
refGene_gn=BCL6;refGene_tx=XM_003991804.4;refGene_gn=BCL6;refGene_tx=XM_006936189.4;refGene_gn=BCL6;refGene_tx=XM_006936190.4
C2 82278348 82279320
refGene_gn=BCL6;refGene_tx=XM_003991804.4;refGene_gn=BCL6;refGene_tx=XM_006936189.4;refGene_gn=BCL6;refGene_tx=XM_006936190.4
C2 82279853 82280038
refGene_gn=BCL6;refGene_tx=XM_003991804.4;refGene_gn=BCL6;refGene_tx=XM_006936189.4;refGene_gn=BCL6;refGene_tx=XM_006936190.4

C2 82281464 82281632
refGene_gn=BCL6;refGene_tx=XM_003991804.4;refGene_gn=BCL6;refGene_tx=XM_006936189.4;refGene_gn=BCL6;refGene_tx=XM_006936190.4
C2 82282530 82282661
refGene_gn=BCL6;refGene_tx=XM_003991804.4;refGene_gn=BCL6;refGene_tx=XM_006936189.4;refGene_gn=BCL6;refGene_tx=XM_006936190.4
C2 82282778 82282916
refGene_gn=BCL6;refGene_tx=XM_003991804.4;refGene_gn=BCL6;refGene_tx=XM_006936189.4;refGene_gn=BCL6;refGene_tx=XM_006936190.4
C2 82284881 82285025
refGene_gn=BCL6;refGene_tx=XM_003991804.4;refGene_gn=BCL6;refGene_tx=XM_006936189.4;refGene_gn=BCL6;refGene_tx=XM_006936190.4
C2 82310194 82310361 refGene_gn=RTP2;refGene_tx=XM_011286029.2
C2 82313010 82313527 refGene_gn=RTP2;refGene_tx=XM_011286029.2
C2 82334221 82334359 refGene_gn=SST;refGene_tx=XM_003991805.4
C2 82335255 82335468 refGene_gn=SST;refGene_tx=XM_003991805.4
C2 82564777 82565792 refGene_gn=RTP4;refGene_tx=XM_011286030.3
C2 82566728 82566973 refGene_gn=RTP4;refGene_tx=XM_011286030.3
C2 82639679 82639684
refGene_gn=MASP1;refGene_tx=XM_003991806.4;refGene_gn=MASP1;refGene_tx=XM_011286032.3;refGene_gn=MASP1;refGene_tx=XM_019839998.2;refGene_gn=MASP1;refGene_tx=XM_019840002.2
C2 82646313 82646545
refGene_gn=MASP1;refGene_tx=XM_003991806.4;refGene_gn=MASP1;refGene_tx=XM_011286032.3;refGene_gn=MASP1;refGene_tx=XM_019839998.2;refGene_gn=MASP1;refGene_tx=XM_019840002.2
C2 82650197 82650215 refGene_gn=MASP1;refGene_tx=XM_019839997.1
C2 82653176 82653266 refGene_gn=MASP1;refGene_tx=XM_023260312.1
C2 82666611 82666789
refGene_gn=MASP1;refGene_tx=XM_003991806.4;refGene_gn=MASP1;refGene_tx=XM_011286032.3;refGene_gn=MASP1;refGene_tx=XM_019839997.1;refGene_gn=MASP1;refGene_tx=XM_019839998.2;refGene_gn=MASP1;refGene_tx=XM_019839999.2;refGene_gn=MASP1;refGene_tx=XM_019840000.2;refGene_gn=MASP1;refGene_tx=XM_019840002.2;refGene_gn=MASP1;refGene_tx=XM_023260312.1
C2 82668859 82668991
refGene_gn=MASP1;refGene_tx=XM_003991806.4;refGene_gn=MASP1;refGene_tx=XM_011286032.3;refGene_gn=MASP1;refGene_tx=XM_019839997.1;refGene_gn=MASP1;refGene_tx=XM_019839998.2;refGene_gn=MASP1;refGene_tx=XM_019839999.2;refGene_gn=MASP1;refGene_tx=XM_019840000.2;refGene_gn=MASP1;refGene_tx=XM_019840002.2;refGene_gn=MASP1;refGene_tx=XM_023260312.1
C2 82669097 82669257 refGene_gn=MASP1;refGene_tx=XM_019840001.1
C2 82671149 82671346
refGene_gn=MASP1;refGene_tx=XM_003991806.4;refGene_gn=MASP1;refGene_tx=XM_011286032.3;refGene_gn=MASP1;refGene_tx=XM_019839997.1;refGene_gn=MASP1;refGene_tx=XM_019839998.2;refGene_gn=MASP1;refGene_tx=XM_019839999.2;refGene_gn=MASP1;refGene_tx=XM_019840000.2;refGene_gn=MASP1;refGene_tx=XM_019840001.1;refGene_gn=MASP1;refGene_tx=XM_019840002.2;refGene_gn=MASP1;refGene_tx=XM_023260312.1
C2 82672830 82672978
refGene_gn=MASP1;refGene_tx=XM_003991806.4;refGene_gn=MASP1;refGene_tx

=XM_011286032.3;refGene_gn=MASP1;refGene_tx=XM_019839997.1;refGene_gn=MASP1;refGene_tx=XM_019839998.2;refGene_gn=MASP1;refGene_tx=XM_019839999.2;refGene_gn=MASP1;refGene_tx=XM_019840000.2;refGene_gn=MASP1;refGene_tx=XM_019840001.1;refGene_gn=MASP1;refGene_tx=XM_019840002.2;refGene_gn=MASP1;refGene_tx=XM_023260312.1
C2 82674306 82674425
refGene_gn=MASP1;refGene_tx=XM_003991806.4;refGene_gn=MASP1;refGene_tx=XM_011286032.3;refGene_gn=MASP1;refGene_tx=XM_019839997.1;refGene_gn=MASP1;refGene_tx=XM_019839998.2;refGene_gn=MASP1;refGene_tx=XM_019839999.2;refGene_gn=MASP1;refGene_tx=XM_019840000.2;refGene_gn=MASP1;refGene_tx=XM_019840001.1;refGene_gn=MASP1;refGene_tx=XM_019840002.2;refGene_gn=MASP1;refGene_tx=XM_023260312.1
C2 82675406 82675485
refGene_gn=MASP1;refGene_tx=XM_003991806.4;refGene_gn=MASP1;refGene_tx=XM_011286032.3;refGene_gn=MASP1;refGene_tx=XM_019839997.1;refGene_gn=MASP1;refGene_tx=XM_019839998.2;refGene_gn=MASP1;refGene_tx=XM_019839999.2;refGene_gn=MASP1;refGene_tx=XM_019840000.2;refGene_gn=MASP1;refGene_tx=XM_019840001.1;refGene_gn=MASP1;refGene_tx=XM_019840002.2;refGene_gn=MASP1;refGene_tx=XM_023260312.1
C2 82677952 82678005 refGene_gn=MASP1;refGene_tx=XM_019840002.2
C2 82681702 82681840
refGene_gn=MASP1;refGene_tx=XM_003991806.4;refGene_gn=MASP1;refGene_tx=XM_011286032.3;refGene_gn=MASP1;refGene_tx=XM_019839997.1;refGene_gn=MASP1;refGene_tx=XM_019839998.2;refGene_gn=MASP1;refGene_tx=XM_019839999.2;refGene_gn=MASP1;refGene_tx=XM_019840000.2;refGene_gn=MASP1;refGene_tx=XM_019840001.1;refGene_gn=MASP1;refGene_tx=XM_023260312.1
C2 82682647 82682722
refGene_gn=MASP1;refGene_tx=XM_003991806.4;refGene_gn=MASP1;refGene_tx=XM_011286032.3;refGene_gn=MASP1;refGene_tx=XM_019839997.1;refGene_gn=MASP1;refGene_tx=XM_019839998.2;refGene_gn=MASP1;refGene_tx=XM_019839999.2;refGene_gn=MASP1;refGene_tx=XM_019840000.2;refGene_gn=MASP1;refGene_tx=XM_019840001.1;refGene_gn=MASP1;refGene_tx=XM_023260312.1
C2 82686672 82687556
refGene_gn=MASP1;refGene_tx=XM_003991806.4;refGene_gn=MASP1;refGene_tx=XM_019839997.1;refGene_gn=MASP1;refGene_tx=XM_019839999.2;refGene_gn=MASP1;refGene_tx=XM_019840000.2;refGene_gn=MASP1;refGene_tx=XM_019840001.1;refGene_gn=MASP1;refGene_tx=XM_023260312.1
C2 82693047 82693185
refGene_gn=MASP1;refGene_tx=XM_011286032.3;refGene_gn=MASP1;refGene_tx=XM_019839998.2
C2 82694593 82694707
refGene_gn=MASP1;refGene_tx=XM_011286032.3;refGene_gn=MASP1;refGene_tx=XM_019839998.2
C2 82695596 82695782 refGene_gn=MASP1;refGene_tx=XM_011286032.3
C2 82697553 82697621
refGene_gn=MASP1;refGene_tx=XM_011286032.3;refGene_gn=MASP1;refGene_tx=XM_019839998.2
C2 82699583 82699683
refGene_gn=MASP1;refGene_tx=XM_011286032.3;refGene_gn=MASP1;refGene_tx=XM_019839998.2

C2 82700112 82700303
refGene_gn=MASP1;refGene_tx=XM_011286032.3;refGene_gn=MASP1;refGene_tx
=XM_019839998.2
C2 82713621 82714141 refGene_gn=RTP1;refGene_tx=XM_003991808.3
C2 82715342 82715614 refGene_gn=RTP1;refGene_tx=XM_003991808.3
C2 82810282 82810524
refGene_gn=ST6GAL1;refGene_tx=XM_019840003.2;refGene_gn=ST6GAL1;refGen
e_tx=XM_023260313.1;refGene_gn=ST6GAL1;refGene_tx=XM_023260314.1;refGe
ne_gn=ST6GAL1;refGene_tx=XM_023260315.1
C2 82812311 82812334 refGene_gn=ST6GAL1;refGene_tx=XM_023260316.1
C2 82812815 82812990
refGene_gn=ST6GAL1;refGene_tx=XM_019840003.2;refGene_gn=ST6GAL1;refGen
e_tx=XM_023260313.1;refGene_gn=ST6GAL1;refGene_tx=XM_023260314.1;refGe
ne_gn=ST6GAL1;refGene_tx=XM_023260315.1;refGene_gn=ST6GAL1;refGene_tx=
XM_023260316.1
C2 82814340 82814439
refGene_gn=ST6GAL1;refGene_tx=XM_019840003.2;refGene_gn=ST6GAL1;refGen
e_tx=XM_023260313.1;refGene_gn=ST6GAL1;refGene_tx=XM_023260314.1;refGe
ne_gn=ST6GAL1;refGene_tx=XM_023260315.1;refGene_gn=ST6GAL1;refGene_tx=
XM_023260316.1
C2 82828424 82828522
refGene_gn=ST6GAL1;refGene_tx=XM_019840003.2;refGene_gn=ST6GAL1;refGen
e_tx=XM_023260313.1;refGene_gn=ST6GAL1;refGene_tx=XM_023260314.1;refGe
ne_gn=ST6GAL1;refGene_tx=XM_023260315.1;refGene_gn=ST6GAL1;refGene_tx=
XM_023260316.1
C2 82836197 82836783
refGene_gn=ST6GAL1;refGene_tx=XM_019840003.2;refGene_gn=ST6GAL1;refGen
e_tx=XM_023260313.1;refGene_gn=ST6GAL1;refGene_tx=XM_023260314.1;refGe
ne_gn=ST6GAL1;refGene_tx=XM_023260315.1;refGene_gn=ST6GAL1;refGene_tx=
XM_023260316.1
C2 82988920 82989441
refGene_gn=ADIPOQ;refGene_tx=NM_001085438.1;refGene_gn=ADIPOQ;refGene_
tx=XM_006936183.3;refGene_gn=ADIPOQ;refGene_tx=XM_006936184.2
C2 82990313 82990562
refGene_gn=ADIPOQ;refGene_tx=NM_001085438.1;refGene_gn=ADIPOQ;refGene_
tx=XM_006936183.3;refGene_gn=ADIPOQ;refGene_tx=XM_006936184.2
C2 82999264 82999346
refGene_gn=ADIPOQ;refGene_tx=XM_006936183.3;refGene_gn=ADIPOQ;refGene_
tx=XM_006936184.2
C2 83011234 83011419 refGene_gn=RFC4;refGene_tx=XM_006936198.4
C2 83012067 83012150 refGene_gn=RFC4;refGene_tx=XM_006936199.4
C2 83012300 83012445
refGene_gn=RFC4;refGene_tx=XM_006936198.4;refGene_gn=RFC4;refGene_tx=X
M_006936199.4
C2 83027512 83027527 refGene_gn=RFC4;refGene_tx=XM_006936200.4
C2 83028934 83029003 refGene_gn=RFC4;refGene_tx=XM_006936202.4
C2 83029111 83029248
refGene_gn=RFC4;refGene_tx=XM_003991813.5;refGene_gn=RFC4;refGene_tx=X
M_006936198.4;refGene_gn=RFC4;refGene_tx=XM_006936199.4;refGene_gn=RFC
4;refGene_tx=XM_006936200.4;refGene_gn=RFC4;refGene_tx=XM_006936202.4

C2 83031062 83031141
refGene_gn=RFC4;refGene_tx=XM_003991813.5;refGene_gn=RFC4;refGene_tx=XM_006936198.4;refGene_gn=RFC4;refGene_tx=XM_006936199.4;refGene_gn=RFC4;refGene_tx=XM_006936200.4;refGene_gn=RFC4;refGene_tx=XM_006936202.4
C2 83036182 83036262
refGene_gn=RFC4;refGene_tx=XM_003991813.5;refGene_gn=RFC4;refGene_tx=XM_006936198.4;refGene_gn=RFC4;refGene_tx=XM_006936199.4;refGene_gn=RFC4;refGene_tx=XM_006936200.4;refGene_gn=RFC4;refGene_tx=XM_006936202.4
C2 83038519 83038639
refGene_gn=RFC4;refGene_tx=XM_003991813.5;refGene_gn=RFC4;refGene_tx=XM_006936198.4;refGene_gn=RFC4;refGene_tx=XM_006936199.4;refGene_gn=RFC4;refGene_tx=XM_006936200.4;refGene_gn=RFC4;refGene_tx=XM_006936202.4
C2 83042905 83043049
refGene_gn=RFC4;refGene_tx=XM_003991813.5;refGene_gn=RFC4;refGene_tx=XM_006936198.4;refGene_gn=RFC4;refGene_tx=XM_006936199.4;refGene_gn=RFC4;refGene_tx=XM_006936200.4;refGene_gn=RFC4;refGene_tx=XM_006936202.4
C2 83043240 83043361
refGene_gn=RFC4;refGene_tx=XM_003991813.5;refGene_gn=RFC4;refGene_tx=XM_006936198.4;refGene_gn=RFC4;refGene_tx=XM_006936199.4;refGene_gn=RFC4;refGene_tx=XM_006936200.4;refGene_gn=RFC4;refGene_tx=XM_006936202.4
C2 83044029 83044155
refGene_gn=RFC4;refGene_tx=XM_003991813.5;refGene_gn=RFC4;refGene_tx=XM_006936198.4;refGene_gn=RFC4;refGene_tx=XM_006936199.4;refGene_gn=RFC4;refGene_tx=XM_006936200.4;refGene_gn=RFC4;refGene_tx=XM_006936202.4
C2 83045364 83045445
refGene_gn=RFC4;refGene_tx=XM_003991813.5;refGene_gn=RFC4;refGene_tx=XM_006936198.4;refGene_gn=RFC4;refGene_tx=XM_006936199.4;refGene_gn=RFC4;refGene_tx=XM_006936200.4;refGene_gn=RFC4;refGene_tx=XM_006936202.4
C2 83045526 83045643
refGene_gn=RFC4;refGene_tx=XM_003991813.5;refGene_gn=RFC4;refGene_tx=XM_006936198.4;refGene_gn=RFC4;refGene_tx=XM_006936199.4;refGene_gn=RFC4;refGene_tx=XM_006936200.4;refGene_gn=RFC4;refGene_tx=XM_006936202.4
C2 83045723 83045819
refGene_gn=RFC4;refGene_tx=XM_003991813.5;refGene_gn=RFC4;refGene_tx=XM_006936198.4;refGene_gn=RFC4;refGene_tx=XM_006936199.4;refGene_gn=RFC4;refGene_tx=XM_006936200.4;refGene_gn=RFC4;refGene_tx=XM_006936202.4
C2 83046519 83046664 refGene_gn=EIF4A2;refGene_tx=XM_003991810.5
C2 83047916 83047996 refGene_gn=EIF4A2;refGene_tx=XM_003991810.5
C2 83048217 83048307 refGene_gn=EIF4A2;refGene_tx=XM_003991810.5
C2 83048553 83048691 refGene_gn=EIF4A2;refGene_tx=XM_003991810.5
C2 83048987 83049131 refGene_gn=EIF4A2;refGene_tx=XM_003991810.5
C2 83049331 83049441 refGene_gn=EIF4A2;refGene_tx=XM_003991810.5
C2 83049551 83049720 refGene_gn=EIF4A2;refGene_tx=XM_003991810.5
C2 83050516 83050656 refGene_gn=EIF4A2;refGene_tx=XM_003991810.5
C2 83050936 83051069 refGene_gn=EIF4A2;refGene_tx=XM_003991810.5
C2 83051155 83051201 refGene_gn=EIF4A2;refGene_tx=XM_003991810.5
C2 83052005 83052034 refGene_gn=EIF4A2;refGene_tx=XM_003991810.5
C2 83052151 83052197
refGene_gn=LOC111562256;refGene_tx=XM_023260317.1
C2 83052361 83052645

refGene_gn=LOC111562256;refGene_tx=XM_023260317.1
C2 83090316 83090397
refGene_gn=LOC101099143;refGene_tx=XM_003991814.4;refGene_gn=LOC101099143;refGene_tx=XM_019840006.2
C2 83090609 83090630
refGene_gn=LOC101099143;refGene_tx=XM_023260318.1
C2 83091655 83092456
refGene_gn=LOC111562257;refGene_tx=XM_023260319.1
C2 83092477 83092555
refGene_gn=LOC101099143;refGene_tx=XM_003991814.4;refGene_gn=LOC101099143;refGene_tx=XM_023260318.1
C2 83093991 83094078
refGene_gn=LOC101099143;refGene_tx=XM_003991814.4;refGene_gn=LOC101099143;refGene_tx=XM_019840006.2;refGene_gn=LOC101099143;refGene_tx=XM_023260318.1
C2 83094195 83094303
refGene_gn=LOC101099143;refGene_tx=XM_003991814.4;refGene_gn=LOC101099143;refGene_tx=XM_019840006.2;refGene_gn=LOC101099143;refGene_tx=XM_023260318.1
C2 83099314 83099487
refGene_gn=LOC101099143;refGene_tx=XM_003991814.4;refGene_gn=LOC101099143;refGene_tx=XM_019840006.2;refGene_gn=LOC101099143;refGene_tx=XM_023260318.1
C2 83100358 83100443
refGene_gn=LOC101099143;refGene_tx=XM_003991814.4;refGene_gn=LOC101099143;refGene_tx=XM_019840006.2;refGene_gn=LOC101099143;refGene_tx=XM_023260318.1
C2 83107555 83107663
refGene_gn=LOC101099143;refGene_tx=XM_003991814.4;refGene_gn=LOC101099143;refGene_tx=XM_019840006.2;refGene_gn=LOC101099143;refGene_tx=XM_023260318.1
C2 83108546 83108719
refGene_gn=LOC101099143;refGene_tx=XM_003991814.4;refGene_gn=LOC101099143;refGene_tx=XM_019840006.2;refGene_gn=LOC101099143;refGene_tx=XM_023260318.1
C2 83112807 83112892
refGene_gn=LOC101099143;refGene_tx=XM_003991814.4;refGene_gn=LOC101099143;refGene_tx=XM_019840006.2;refGene_gn=LOC101099143;refGene_tx=XM_023260318.1
C2 83113924 83114035
refGene_gn=LOC101099143;refGene_tx=XM_003991814.4;refGene_gn=LOC101099143;refGene_tx=XM_019840006.2;refGene_gn=LOC101099143;refGene_tx=XM_023260318.1
C2 83119295 83119490
refGene_gn=LOC101099143;refGene_tx=XM_003991814.4;refGene_gn=LOC101099143;refGene_tx=XM_019840006.2;refGene_gn=LOC101099143;refGene_tx=XM_023260318.1
C2 83122425 83123349
refGene_gn=HRG;refGene_tx=XM_003991832.2;refGene_gn=HRG;refGene_tx=XM_019840005.1

C2 83124927 83125029
refGene_gn=HRG;refGene_tx=XM_003991832.2;refGene_gn=HRG;refGene_tx=XM_019840005.1
C2 83127267 83127351
refGene_gn=HRG;refGene_tx=XM_003991832.2;refGene_gn=HRG;refGene_tx=XM_019840005.1
C2 83128485 83128652
refGene_gn=HRG;refGene_tx=XM_003991832.2;refGene_gn=HRG;refGene_tx=XM_019840005.1
C2 83129293 83129381
refGene_gn=HRG;refGene_tx=XM_003991832.2;refGene_gn=HRG;refGene_tx=XM_019840005.1
C2 83130229 83130346
refGene_gn=HRG;refGene_tx=XM_003991832.2;refGene_gn=HRG;refGene_tx=XM_019840005.1
C2 83132472 83132664
refGene_gn=HRG;refGene_tx=XM_003991832.2;refGene_gn=HRG;refGene_tx=XM_019840005.1
C2 83136142 83136520
refGene_gn=FETUB;refGene_tx=XM_019840007.1;refGene_gn=FETUB;refGene_tx=XM_019840008.2
C2 83137644 83137728 refGene_gn=FETUB;refGene_tx=XM_019840008.2
C2 83142699 83142801
refGene_gn=FETUB;refGene_tx=XM_019840007.1;refGene_gn=FETUB;refGene_tx=XM_019840008.2
C2 83143523 83143687
refGene_gn=FETUB;refGene_tx=XM_019840007.1;refGene_gn=FETUB;refGene_tx=XM_019840008.2
C2 83145872 83145960
refGene_gn=FETUB;refGene_tx=XM_019840007.1;refGene_gn=FETUB;refGene_tx=XM_019840008.2
C2 83147754 83147868
refGene_gn=FETUB;refGene_tx=XM_019840007.1;refGene_gn=FETUB;refGene_tx=XM_019840008.2
C2 83148197 83148413
refGene_gn=FETUB;refGene_tx=XM_019840007.1;refGene_gn=FETUB;refGene_tx=XM_019840008.2
C2 83178531 83178858 refGene_gn=AHSG;refGene_tx=XM_006936204.2
C2 83179510 83179594 refGene_gn=AHSG;refGene_tx=XM_006936204.2
C2 83180967 83181069 refGene_gn=AHSG;refGene_tx=XM_006936204.2
C2 83181915 83182079 refGene_gn=AHSG;refGene_tx=XM_006936204.2
C2 83182417 83182502 refGene_gn=AHSG;refGene_tx=XM_006936204.2
C2 83182874 83182985 refGene_gn=AHSG;refGene_tx=XM_006936204.2
C2 83185396 83185609 refGene_gn=AHSG;refGene_tx=XM_006936204.2
C2 83201841 83201906 refGene_gn=DNAJB11;refGene_tx=XM_003991818.5
C2 83202622 83202782 refGene_gn=DNAJB11;refGene_tx=XM_003991818.5
C2 83203027 83203139 refGene_gn=DNAJB11;refGene_tx=XM_003991818.5
C2 83209170 83209228 refGene_gn=DNAJB11;refGene_tx=XM_003991818.5
C2 83210132 83210215 refGene_gn=DNAJB11;refGene_tx=XM_003991818.5
C2 83210998 83211141 refGene_gn=DNAJB11;refGene_tx=XM_003991818.5

C2	83214605	83214738	refGene_gn=DNAJB11;refGene_tx=XM_003991818.5
C2	83216436	83216534	refGene_gn=DNAJB11;refGene_tx=XM_003991818.5
C2	83220964	83221121	refGene_gn=DNAJB11;refGene_tx=XM_003991818.5
C2	83223842	83223910	refGene_gn=DNAJB11;refGene_tx=XM_003991818.5
C2	83228931	83228939
refGene_gn=TBCCD1;refGene_tx=XM_019840011.2;refGene_gn=TBCCD1;refGene_tx=XM_019840012.2;refGene_gn=TBCCD1;refGene_tx=XM_023260321.1;refGene_gn=TBCCD1;refGene_tx=XM_023260322.1
C2	83229442	83229821
refGene_gn=TBCCD1;refGene_tx=XM_003991819.5;refGene_gn=TBCCD1;refGene_tx=XM_019840009.2;refGene_gn=TBCCD1;refGene_tx=XM_019840011.2;refGene_gn=TBCCD1;refGene_tx=XM_019840012.2;refGene_gn=TBCCD1;refGene_tx=XM_019840013.2;refGene_gn=TBCCD1;refGene_tx=XM_023260320.1;refGene_gn=TBCCD1;refGene_tx=XM_023260321.1;refGene_gn=TBCCD1;refGene_tx=XM_023260322.1;refGene_gn=TBCCD1;refGene_tx=XM_023260323.1
C2	83233390	83233546
refGene_gn=TBCCD1;refGene_tx=XM_003991819.5;refGene_gn=TBCCD1;refGene_tx=XM_019840009.2;refGene_gn=TBCCD1;refGene_tx=XM_019840011.2;refGene_gn=TBCCD1;refGene_tx=XM_019840012.2;refGene_gn=TBCCD1;refGene_tx=XM_019840013.2;refGene_gn=TBCCD1;refGene_tx=XM_023260320.1;refGene_gn=TBCCD1;refGene_tx=XM_023260321.1;refGene_gn=TBCCD1;refGene_tx=XM_023260322.1;refGene_gn=TBCCD1;refGene_tx=XM_023260323.1
C2	83237510	83237877
refGene_gn=TBCCD1;refGene_tx=XM_003991819.5;refGene_gn=TBCCD1;refGene_tx=XM_019840009.2;refGene_gn=TBCCD1;refGene_tx=XM_019840011.2;refGene_gn=TBCCD1;refGene_tx=XM_019840012.2;refGene_gn=TBCCD1;refGene_tx=XM_019840013.2;refGene_gn=TBCCD1;refGene_tx=XM_023260320.1;refGene_gn=TBCCD1;refGene_tx=XM_023260321.1;refGene_gn=TBCCD1;refGene_tx=XM_023260322.1;refGene_gn=TBCCD1;refGene_tx=XM_023260323.1
C2	83239965	83240152
refGene_gn=TBCCD1;refGene_tx=XM_003991819.5;refGene_gn=TBCCD1;refGene_tx=XM_019840009.2;refGene_gn=TBCCD1;refGene_tx=XM_019840011.2;refGene_gn=TBCCD1;refGene_tx=XM_019840012.2;refGene_gn=TBCCD1;refGene_tx=XM_019840013.2;refGene_gn=TBCCD1;refGene_tx=XM_023260320.1;refGene_gn=TBCCD1;refGene_tx=XM_023260321.1;refGene_gn=TBCCD1;refGene_tx=XM_023260322.1;refGene_gn=TBCCD1;refGene_tx=XM_023260323.1
C2	83240309	83240807
refGene_gn=TBCCD1;refGene_tx=XM_003991819.5;refGene_gn=TBCCD1;refGene_tx=XM_019840009.2;refGene_gn=TBCCD1;refGene_tx=XM_019840011.2;refGene_gn=TBCCD1;refGene_tx=XM_019840012.2;refGene_gn=TBCCD1;refGene_tx=XM_019840013.2;refGene_gn=TBCCD1;refGene_tx=XM_023260320.1;refGene_gn=TBCCD1;refGene_tx=XM_023260321.1;refGene_gn=TBCCD1;refGene_tx=XM_023260322.1;refGene_gn=TBCCD1;refGene_tx=XM_023260323.1
C2	83241925	83242055
refGene_gn=TBCCD1;refGene_tx=XM_003991819.5;refGene_gn=TBCCD1;refGene_tx=XM_019840009.2;refGene_gn=TBCCD1;refGene_tx=XM_019840011.2;refGene_gn=TBCCD1;refGene_tx=XM_019840012.2;refGene_gn=TBCCD1;refGene_tx=XM_019840013.2;refGene_gn=TBCCD1;refGene_tx=XM_023260320.1;refGene_gn=TBCCD1;refGene_tx=XM_023260321.1;refGene_gn=TBCCD1;refGene_tx=XM_023260322.1;refGene_gn=TBCCD1;refGene_tx=XM_023260323.1

C2 83247399 83247519 refGene_gn=CRYGS;refGene_tx=XM_003991820.3
C2 83251969 83251972 refGene_gn=CRYGS;refGene_tx=XM_019840014.1
C2 83252285 83252528
refGene_gn=CRYGS;refGene_tx=XM_003991820.3;refGene_gn=CRYGS;refGene_tx
=XM_019840014.1
C2 83252952 83253225
refGene_gn=CRYGS;refGene_tx=XM_003991820.3;refGene_gn=CRYGS;refGene_tx
=XM_019840014.1
C2 83451083 83451150 refGene_gn=DGKG;refGene_tx=XM_023260324.1
C2 83463498 83463575 refGene_gn=DGKG;refGene_tx=XM_023260324.1
C2 83471688 83471851 refGene_gn=DGKG;refGene_tx=XM_023260324.1
C2 83472441 83472504 refGene_gn=DGKG;refGene_tx=XM_023260324.1
C2 83480677 83480848 refGene_gn=DGKG;refGene_tx=XM_023260324.1
C2 83485559 83485609 refGene_gn=DGKG;refGene_tx=XM_023260324.1
C2 83489497 83489572 refGene_gn=DGKG;refGene_tx=XM_023260324.1
C2 83490230 83490353 refGene_gn=DGKG;refGene_tx=XM_023260324.1
C2 83494997 83495115 refGene_gn=DGKG;refGene_tx=XM_023260324.1
C2 83496653 83496742 refGene_gn=DGKG;refGene_tx=XM_023260324.1
C2 83499777 83499894 refGene_gn=DGKG;refGene_tx=XM_023260324.1
C2 83500883 83500976 refGene_gn=DGKG;refGene_tx=XM_023260324.1
C2 83502705 83502765 refGene_gn=DGKG;refGene_tx=XM_023260324.1
C2 83505164 83505244 refGene_gn=DGKG;refGene_tx=XM_023260324.1
C2 83506392 83506467 refGene_gn=DGKG;refGene_tx=XM_023260324.1
C2 83509430 83509516 refGene_gn=DGKG;refGene_tx=XM_023260324.1
C2 83513899 83513989 refGene_gn=DGKG;refGene_tx=XM_023260324.1
C2 83516292 83516453 refGene_gn=DGKG;refGene_tx=XM_023260324.1
C2 83525681 83525746 refGene_gn=DGKG;refGene_tx=XM_023260324.1
C2 83554576 83554667 refGene_gn=DGKG;refGene_tx=XM_023260324.1
C2 83575533 83575711 refGene_gn=DGKG;refGene_tx=XM_023260324.1
C2 83597070 83597191 refGene_gn=DGKG;refGene_tx=XM_023260324.1
C2 83599591 83599652 refGene_gn=DGKG;refGene_tx=XM_023260324.1
C2 83609863 83609962 refGene_gn=DGKG;refGene_tx=XM_023260324.1
C2 83647456 83647501 refGene_gn=ETV5;refGene_tx=XM_003991824.4
C2 83647636 83647724 refGene_gn=ETV5;refGene_tx=XM_003991824.4
C2 83647887 83647935 refGene_gn=ETV5;refGene_tx=XM_003991824.4
C2 83648029 83648080 refGene_gn=ETV5;refGene_tx=XM_003991824.4
C2 83672388 83672518 refGene_gn=ETV5;refGene_tx=XM_003991824.4
C2 83673434 83673722 refGene_gn=ETV5;refGene_tx=XM_003991824.4
C2 83685327 83685587 refGene_gn=ETV5;refGene_tx=XM_003991824.4
C2 83687608 83687668 refGene_gn=ETV5;refGene_tx=XM_003991824.4
C2 83692209 83692278 refGene_gn=ETV5;refGene_tx=XM_003991824.4
C2 83692441 83692611 refGene_gn=ETV5;refGene_tx=XM_003991824.4
C2 83696973 83697075 refGene_gn=ETV5;refGene_tx=XM_003991824.4
C2 83702070 83702292 refGene_gn=ETV5;refGene_tx=XM_003991824.4
C2 83775328 83775364
refGene_gn=TRA2B;refGene_tx=XM_006936207.2;refGene_gn=TRA2B;refGene_tx
=XM_023260325.1
C2 83786806 83786940
refGene_gn=TRA2B;refGene_tx=XM_006936207.2;refGene_gn=TRA2B;refGene_tx
=XM_023260325.1

C2	83787990	83788153
refGene_gn=TRA2B;refGene_tx=XM_006936207.2;refGene_gn=TRA2B;refGene_tx=XM_019840015.1;refGene_gn=TRA2B;refGene_tx=XM_023260325.1
C2	83788780	83788969
refGene_gn=TRA2B;refGene_tx=XM_006936207.2;refGene_gn=TRA2B;refGene_tx=XM_019840015.1;refGene_gn=TRA2B;refGene_tx=XM_023260325.1
C2	83789197	83789313
refGene_gn=TRA2B;refGene_tx=XM_006936207.2;refGene_gn=TRA2B;refGene_tx=XM_019840015.1;refGene_gn=TRA2B;refGene_tx=XM_023260325.1
C2	83790450	83790534
refGene_gn=TRA2B;refGene_tx=XM_006936207.2;refGene_gn=TRA2B;refGene_tx=XM_019840015.1;refGene_gn=TRA2B;refGene_tx=XM_023260325.1
C2	83792497	83792557
refGene_gn=TRA2B;refGene_tx=XM_006936207.2;refGene_gn=TRA2B;refGene_tx=XM_019840015.1;refGene_gn=TRA2B;refGene_tx=XM_023260325.1
C2	83793128	83793202
refGene_gn=TRA2B;refGene_tx=XM_006936207.2;refGene_gn=TRA2B;refGene_tx=XM_019840015.1;refGene_gn=TRA2B;refGene_tx=XM_023260325.1
C2	83793679	83793690
refGene_gn=TRA2B;refGene_tx=XM_006936207.2;refGene_gn=TRA2B;refGene_tx=XM_019840015.1;refGene_gn=TRA2B;refGene_tx=XM_023260325.1
C2	83905502	83905680
refGene_gn=IGF2BP2;refGene_tx=XM_011286035.3;refGene_gn=IGF2BP2;refGene_tx=XM_019840016.2;refGene_gn=IGF2BP2;refGene_tx=XM_019840017.2;refGene_gn=IGF2BP2;refGene_tx=XM_019840018.2;refGene_gn=IGF2BP2;refGene_tx=XM_019840019.2
C2	83907263	83907324
refGene_gn=IGF2BP2;refGene_tx=XM_011286035.3;refGene_gn=IGF2BP2;refGene_tx=XM_019840016.2;refGene_gn=IGF2BP2;refGene_tx=XM_019840017.2;refGene_gn=IGF2BP2;refGene_tx=XM_019840018.2;refGene_gn=IGF2BP2;refGene_tx=XM_019840019.2
C2	84020239	84020288
refGene_gn=IGF2BP2;refGene_tx=XM_011286035.3;refGene_gn=IGF2BP2;refGene_tx=XM_019840016.2;refGene_gn=IGF2BP2;refGene_tx=XM_019840017.2;refGene_gn=IGF2BP2;refGene_tx=XM_019840018.2;refGene_gn=IGF2BP2;refGene_tx=XM_019840019.2
C2	84021393	84021445
refGene_gn=IGF2BP2;refGene_tx=XM_011286035.3;refGene_gn=IGF2BP2;refGene_tx=XM_019840016.2;refGene_gn=IGF2BP2;refGene_tx=XM_019840017.2;refGene_gn=IGF2BP2;refGene_tx=XM_019840018.2;refGene_gn=IGF2BP2;refGene_tx=XM_019840019.2
C2	84024984	84025048
refGene_gn=IGF2BP2;refGene_tx=XM_011286035.3;refGene_gn=IGF2BP2;refGene_tx=XM_019840016.2;refGene_gn=IGF2BP2;refGene_tx=XM_019840017.2;refGene_gn=IGF2BP2;refGene_tx=XM_019840018.2;refGene_gn=IGF2BP2;refGene_tx=XM_019840019.2
C2	84027049	84027322
refGene_gn=IGF2BP2;refGene_tx=XM_011286035.3;refGene_gn=IGF2BP2;refGene_tx=XM_019840016.2;refGene_gn=IGF2BP2;refGene_tx=XM_019840017.2;refGene_gn=IGF2BP2;refGene_tx=XM_019840018.2;refGene_gn=IGF2BP2;refGene_tx=

XM_019840019.2
C2	84029152 84029287
refGene_gn=IGF2BP2;refGene_tx=XM_011286035.3;refGene_gn=IGF2BP2;refGene_tx=XM_019840016.2;refGene_gn=IGF2BP2;refGene_tx=XM_019840017.2;refGene_gn=IGF2BP2;refGene_tx=XM_019840018.2;refGene_gn=IGF2BP2;refGene_tx=XM_019840019.2
C2	84037998 84038121
refGene_gn=IGF2BP2;refGene_tx=XM_011286035.3;refGene_gn=IGF2BP2;refGene_tx=XM_019840016.2;refGene_gn=IGF2BP2;refGene_tx=XM_019840017.2;refGene_gn=IGF2BP2;refGene_tx=XM_019840018.2;refGene_gn=IGF2BP2;refGene_tx=XM_019840019.2
C2	84038476 84038612
refGene_gn=IGF2BP2;refGene_tx=XM_011286035.3;refGene_gn=IGF2BP2;refGene_tx=XM_019840016.2;refGene_gn=IGF2BP2;refGene_tx=XM_019840017.2;refGene_gn=IGF2BP2;refGene_tx=XM_019840018.2;refGene_gn=IGF2BP2;refGene_tx=XM_019840019.2
C2	84040904 84041033
refGene_gn=IGF2BP2;refGene_tx=XM_011286035.3;refGene_gn=IGF2BP2;refGene_tx=XM_019840016.2;refGene_gn=IGF2BP2;refGene_tx=XM_019840019.2
C2	84051961 84052030
refGene_gn=IGF2BP2;refGene_tx=XM_011286035.3;refGene_gn=IGF2BP2;refGene_tx=XM_019840016.2;refGene_gn=IGF2BP2;refGene_tx=XM_019840017.2;refGene_gn=IGF2BP2;refGene_tx=XM_019840018.2;refGene_gn=IGF2BP2;refGene_tx=XM_019840019.2
C2	84052878 84052995
refGene_gn=IGF2BP2;refGene_tx=XM_011286035.3;refGene_gn=IGF2BP2;refGene_tx=XM_019840016.2;refGene_gn=IGF2BP2;refGene_tx=XM_019840017.2;refGene_gn=IGF2BP2;refGene_tx=XM_019840018.2;refGene_gn=IGF2BP2;refGene_tx=XM_019840019.2
C2	84056351 84056426
refGene_gn=IGF2BP2;refGene_tx=XM_011286035.3;refGene_gn=IGF2BP2;refGene_tx=XM_019840016.2;refGene_gn=IGF2BP2;refGene_tx=XM_019840017.2;refGene_gn=IGF2BP2;refGene_tx=XM_019840018.2;refGene_gn=IGF2BP2;refGene_tx=XM_019840019.2
C2	84057022 84057025 refGene_gn=IGF2BP2;refGene_tx=XM_019840019.2
C2	84057802 84057934
refGene_gn=IGF2BP2;refGene_tx=XM_011286035.3;refGene_gn=IGF2BP2;refGene_tx=XM_019840016.2;refGene_gn=IGF2BP2;refGene_tx=XM_019840017.2;refGene_gn=IGF2BP2;refGene_tx=XM_019840018.2
C2	84059568 84059682
refGene_gn=IGF2BP2;refGene_tx=XM_011286035.3;refGene_gn=IGF2BP2;refGene_tx=XM_019840016.2;refGene_gn=IGF2BP2;refGene_tx=XM_019840017.2;refGene_gn=IGF2BP2;refGene_tx=XM_019840018.2
C2	84061314 84061407
refGene_gn=IGF2BP2;refGene_tx=XM_011286035.3;refGene_gn=IGF2BP2;refGene_tx=XM_019840018.2
C2	84062325 84062460
refGene_gn=IGF2BP2;refGene_tx=XM_019840016.2;refGene_gn=IGF2BP2;refGene_tx=XM_019840017.2
C2	84076543 84076606

refGene_gn=SENP2;refGene_tx=XM_003991836.5;refGene_gn=SENP2;refGene_tx=XM_019840020.2;refGene_gn=SENP2;refGene_tx=XM_019840022.2;refGene_gn=SENP2;refGene_tx=XM_019840023.2;refGene_gn=SENP2;refGene_tx=XM_023260326.1
C2 84078059 84078155
refGene_gn=SENP2;refGene_tx=XM_003991836.5;refGene_gn=SENP2;refGene_tx=XM_019840020.2;refGene_gn=SENP2;refGene_tx=XM_019840022.2;refGene_gn=SENP2;refGene_tx=XM_019840023.2;refGene_gn=SENP2;refGene_tx=XM_023260326.1
C2 84079973 84080058
refGene_gn=SENP2;refGene_tx=XM_003991836.5;refGene_gn=SENP2;refGene_tx=XM_019840020.2;refGene_gn=SENP2;refGene_tx=XM_019840022.2;refGene_gn=SENP2;refGene_tx=XM_019840023.2;refGene_gn=SENP2;refGene_tx=XM_023260326.1
C2 84082933 84083013
refGene_gn=SENP2;refGene_tx=XM_003991836.5;refGene_gn=SENP2;refGene_tx=XM_019840020.2;refGene_gn=SENP2;refGene_tx=XM_019840022.2;refGene_gn=SENP2;refGene_tx=XM_019840023.2;refGene_gn=SENP2;refGene_tx=XM_023260326.1
C2 84083989 84084193
refGene_gn=SENP2;refGene_tx=XM_003991836.5;refGene_gn=SENP2;refGene_tx=XM_019840020.2;refGene_gn=SENP2;refGene_tx=XM_019840022.2;refGene_gn=SENP2;refGene_tx=XM_019840023.2;refGene_gn=SENP2;refGene_tx=XM_023260326.1
C2 84085827 84085959
refGene_gn=SENP2;refGene_tx=XM_003991836.5;refGene_gn=SENP2;refGene_tx=XM_019840020.2;refGene_gn=SENP2;refGene_tx=XM_019840022.2;refGene_gn=SENP2;refGene_tx=XM_019840023.2;refGene_gn=SENP2;refGene_tx=XM_023260326.1
C2 84087195 84087372
refGene_gn=SENP2;refGene_tx=XM_003991836.5;refGene_gn=SENP2;refGene_tx=XM_019840020.2;refGene_gn=SENP2;refGene_tx=XM_019840022.2;refGene_gn=SENP2;refGene_tx=XM_019840023.2;refGene_gn=SENP2;refGene_tx=XM_023260326.1
C2 84089269 84089336
refGene_gn=SENP2;refGene_tx=XM_003991836.5;refGene_gn=SENP2;refGene_tx=XM_019840020.2;refGene_gn=SENP2;refGene_tx=XM_019840022.2;refGene_gn=SENP2;refGene_tx=XM_019840023.2;refGene_gn=SENP2;refGene_tx=XM_023260326.1
C2 84090373 84090425
refGene_gn=SENP2;refGene_tx=XM_003991836.5;refGene_gn=SENP2;refGene_tx=XM_019840020.2;refGene_gn=SENP2;refGene_tx=XM_019840022.2;refGene_gn=SENP2;refGene_tx=XM_019840023.2;refGene_gn=SENP2;refGene_tx=XM_023260326.1
C2 84090850 84090945
refGene_gn=SENP2;refGene_tx=XM_003991836.5;refGene_gn=SENP2;refGene_tx=XM_019840020.2;refGene_gn=SENP2;refGene_tx=XM_019840022.2;refGene_gn=SENP2;refGene_tx=XM_019840023.2;refGene_gn=SENP2;refGene_tx=XM_023260326.1
C2 84092253 84092357

refGene_gn=SENP2;refGene_tx=XM_003991836.5;refGene_gn=SENP2;refGene_tx=XM_019840020.2;refGene_gn=SENP2;refGene_tx=XM_019840022.2;refGene_gn=SENP2;refGene_tx=XM_019840023.2;refGene_gn=SENP2;refGene_tx=XM_023260326.1
C2 84094093 84094262 refGene_gn=SENP2;refGene_tx=XM_003991836.5;refGene_gn=SENP2;refGene_tx=XM_019840020.2;refGene_gn=SENP2;refGene_tx=XM_019840022.2;refGene_gn=SENP2;refGene_tx=XM_019840023.2;refGene_gn=SENP2;refGene_tx=XM_023260326.1
C2 84096796 84096887 refGene_gn=SENP2;refGene_tx=XM_003991836.5;refGene_gn=SENP2;refGene_tx=XM_019840020.2;refGene_gn=SENP2;refGene_tx=XM_019840022.2;refGene_gn=SENP2;refGene_tx=XM_019840023.2;refGene_gn=SENP2;refGene_tx=XM_023260326.1
C2 84098912 84098979 refGene_gn=SENP2;refGene_tx=XM_003991836.5;refGene_gn=SENP2;refGene_tx=XM_019840020.2;refGene_gn=SENP2;refGene_tx=XM_019840022.2;refGene_gn=SENP2;refGene_tx=XM_019840023.2;refGene_gn=SENP2;refGene_tx=XM_023260326.1
C2 84099586 84099720 refGene_gn=SENP2;refGene_tx=XM_003991836.5;refGene_gn=SENP2;refGene_tx=XM_019840020.2;refGene_gn=SENP2;refGene_tx=XM_019840022.2;refGene_gn=SENP2;refGene_tx=XM_019840023.2
C2 84104499 84104555 refGene_gn=SENP2;refGene_tx=XM_003991836.5;refGene_gn=SENP2;refGene_tx=XM_019840020.2
C2 84105303 84105368 refGene_gn=SENP2;refGene_tx=XM_019840020.2
C2 84106583 84106684 refGene_gn=SENP2;refGene_tx=XM_003991836.5
C2 84116499 84116548 refGene_gn=LIPH;refGene_tx=XM_019840031.2;refGene_gn=LIPH;refGene_tx=XM_023260327.1
C2 84134684 84135052 refGene_gn=LIPH;refGene_tx=XM_019840031.2;refGene_gn=LIPH;refGene_tx=XM_023260327.1
C2 84136281 84136390 refGene_gn=LIPH;refGene_tx=XM_019840031.2;refGene_gn=LIPH;refGene_tx=XM_023260327.1
C2 84142401 84142503 refGene_gn=LIPH;refGene_tx=XM_019840031.2;refGene_gn=LIPH;refGene_tx=XM_023260327.1
C2 84144147 84144237 refGene_gn=LIPH;refGene_tx=XM_019840031.2;refGene_gn=LIPH;refGene_tx=XM_023260327.1
C2 84147339 84147507 refGene_gn=LIPH;refGene_tx=XM_019840031.2;refGene_gn=LIPH;refGene_tx=XM_023260327.1
C2 84148830 84148926 refGene_gn=LIPH;refGene_tx=XM_019840031.2;refGene_gn=LIPH;refGene_tx=XM_023260327.1
C2 84149779 84149892 refGene_gn=LIPH;refGene_tx=XM_023260327.1

C2	84150712	84150824	refGene_gn=LIPH;refGene_tx=XM_019840031.2
C2	84159599	84159773	refGene_gn=LIPH;refGene_tx=XM_019840031.2
C2	84161187	84161275	refGene_gn=LIPH;refGene_tx=XM_019840031.2
C2	84169759	84169878
refGene_gn=TMEM41A;refGene_tx=XM_003991837.5;refGene_gn=TMEM41A;refGen
e_tx=XM_011286038.3
C2	84171391	84171545
refGene_gn=TMEM41A;refGene_tx=XM_003991837.5;refGene_gn=TMEM41A;refGen
e_tx=XM_011286038.3
C2	84174584	84174746
refGene_gn=TMEM41A;refGene_tx=XM_003991837.5;refGene_gn=TMEM41A;refGen
e_tx=XM_011286038.3
C2	84175363	84175502
refGene_gn=TMEM41A;refGene_tx=XM_003991837.5;refGene_gn=TMEM41A;refGen
e_tx=XM_011286038.3
C2	84176894	84177115	refGene_gn=TMEM41A;refGene_tx=XM_003991837.5
C2	84182190	84182291	refGene_gn=TMEM41A;refGene_tx=XM_011286038.3
C2	84185098	84185200
refGene_gn=MAP3K13;refGene_tx=XM_011286037.2;refGene_gn=MAP3K13;refGen
e_tx=XM_019840024.1;refGene_gn=MAP3K13;refGene_tx=XM_019840025.2;refGe
ne_gn=MAP3K13;refGene_tx=XM_019840026.1;refGene_gn=MAP3K13;refGene_tx=
XM_019840028.1;refGene_gn=MAP3K13;refGene_tx=XM_019840029.2;refGene_gn
=MAP3K13;refGene_tx=XM_019840030.2
C2	84186352	84186650
refGene_gn=MAP3K13;refGene_tx=XM_011286037.2;refGene_gn=MAP3K13;refGen
e_tx=XM_019840024.1;refGene_gn=MAP3K13;refGene_tx=XM_019840025.2;refGe
ne_gn=MAP3K13;refGene_tx=XM_019840026.1;refGene_gn=MAP3K13;refGene_tx=
XM_019840028.1;refGene_gn=MAP3K13;refGene_tx=XM_019840029.2;refGene_gn
=MAP3K13;refGene_tx=XM_019840030.2
C2	84189093	84189164
refGene_gn=MAP3K13;refGene_tx=XM_011286037.2;refGene_gn=MAP3K13;refGen
e_tx=XM_019840024.1;refGene_gn=MAP3K13;refGene_tx=XM_019840025.2;refGe
ne_gn=MAP3K13;refGene_tx=XM_019840026.1;refGene_gn=MAP3K13;refGene_tx=
XM_019840028.1;refGene_gn=MAP3K13;refGene_tx=XM_019840029.2;refGene_gn
=MAP3K13;refGene_tx=XM_019840030.2
C2	84193457	84194244
refGene_gn=MAP3K13;refGene_tx=XM_011286037.2;refGene_gn=MAP3K13;refGen
e_tx=XM_019840024.1;refGene_gn=MAP3K13;refGene_tx=XM_019840025.2;refGe
ne_gn=MAP3K13;refGene_tx=XM_019840026.1;refGene_gn=MAP3K13;refGene_tx=
XM_019840028.1;refGene_gn=MAP3K13;refGene_tx=XM_019840029.2;refGene_gn
=MAP3K13;refGene_tx=XM_019840030.2
C2	84199713	84199851
refGene_gn=MAP3K13;refGene_tx=XM_011286037.2;refGene_gn=MAP3K13;refGen
e_tx=XM_019840024.1;refGene_gn=MAP3K13;refGene_tx=XM_019840025.2;refGe
ne_gn=MAP3K13;refGene_tx=XM_019840026.1;refGene_gn=MAP3K13;refGene_tx=
XM_019840028.1;refGene_gn=MAP3K13;refGene_tx=XM_019840029.2;refGene_gn
=MAP3K13;refGene_tx=XM_019840030.2
C2	84200546	84200663
refGene_gn=MAP3K13;refGene_tx=XM_011286037.2;refGene_gn=MAP3K13;refGen
e_tx=XM_019840024.1;refGene_gn=MAP3K13;refGene_tx=XM_019840025.2;refGe

ne_gn=MAP3K13;refGene_tx=XM_019840026.1;refGene_gn=MAP3K13;refGene_tx=XM_019840028.1;refGene_gn=MAP3K13;refGene_tx=XM_019840029.2;refGene_gn=MAP3K13;refGene_tx=XM_019840030.2
C2 84201633 84201743
refGene_gn=MAP3K13;refGene_tx=XM_011286037.2;refGene_gn=MAP3K13;refGene_tx=XM_019840024.1;refGene_gn=MAP3K13;refGene_tx=XM_019840025.2;refGene_gn=MAP3K13;refGene_tx=XM_019840026.1;refGene_gn=MAP3K13;refGene_tx=XM_019840028.1;refGene_gn=MAP3K13;refGene_tx=XM_019840029.2;refGene_gn=MAP3K13;refGene_tx=XM_019840030.2
C2 84209541 84209650
refGene_gn=MAP3K13;refGene_tx=XM_011286037.2;refGene_gn=MAP3K13;refGene_tx=XM_019840024.1;refGene_gn=MAP3K13;refGene_tx=XM_019840025.2;refGene_gn=MAP3K13;refGene_tx=XM_019840026.1;refGene_gn=MAP3K13;refGene_tx=XM_019840028.1;refGene_gn=MAP3K13;refGene_tx=XM_019840029.2;refGene_gn=MAP3K13;refGene_tx=XM_019840030.2
C2 84210098 84210257
refGene_gn=MAP3K13;refGene_tx=XM_011286037.2;refGene_gn=MAP3K13;refGene_tx=XM_019840024.1;refGene_gn=MAP3K13;refGene_tx=XM_019840025.2;refGene_gn=MAP3K13;refGene_tx=XM_019840026.1;refGene_gn=MAP3K13;refGene_tx=XM_019840028.1;refGene_gn=MAP3K13;refGene_tx=XM_019840029.2;refGene_gn=MAP3K13;refGene_tx=XM_019840030.2
C2 84211909 84212068
refGene_gn=MAP3K13;refGene_tx=XM_011286037.2;refGene_gn=MAP3K13;refGene_tx=XM_019840024.1;refGene_gn=MAP3K13;refGene_tx=XM_019840025.2;refGene_gn=MAP3K13;refGene_tx=XM_019840026.1;refGene_gn=MAP3K13;refGene_tx=XM_019840028.1;refGene_gn=MAP3K13;refGene_tx=XM_019840029.2;refGene_gn=MAP3K13;refGene_tx=XM_019840030.2
C2 84215455 84215647
refGene_gn=MAP3K13;refGene_tx=XM_011286037.2;refGene_gn=MAP3K13;refGene_tx=XM_019840024.1;refGene_gn=MAP3K13;refGene_tx=XM_019840025.2;refGene_gn=MAP3K13;refGene_tx=XM_019840026.1;refGene_gn=MAP3K13;refGene_tx=XM_019840028.1;refGene_gn=MAP3K13;refGene_tx=XM_019840029.2;refGene_gn=MAP3K13;refGene_tx=XM_019840030.2
C2 84220823 84221007
refGene_gn=MAP3K13;refGene_tx=XM_011286037.2;refGene_gn=MAP3K13;refGene_tx=XM_019840024.1;refGene_gn=MAP3K13;refGene_tx=XM_019840025.2;refGene_gn=MAP3K13;refGene_tx=XM_019840026.1;refGene_gn=MAP3K13;refGene_tx=XM_019840028.1;refGene_gn=MAP3K13;refGene_tx=XM_019840029.2;refGene_gn=MAP3K13;refGene_tx=XM_019840030.2
C2 84229067 84229542
refGene_gn=MAP3K13;refGene_tx=XM_011286037.2;refGene_gn=MAP3K13;refGene_tx=XM_019840024.1;refGene_gn=MAP3K13;refGene_tx=XM_019840025.2;refGene_gn=MAP3K13;refGene_tx=XM_019840026.1;refGene_gn=MAP3K13;refGene_tx=XM_019840028.1;refGene_gn=MAP3K13;refGene_tx=XM_019840029.2;refGene_gn=MAP3K13;refGene_tx=XM_019840030.2
C2 84367083 84367157 refGene_gn=EHHADH;refGene_tx=XM_003991838.3
C2 84369494 84369598 refGene_gn=EHHADH;refGene_tx=XM_003991838.3
C2 84374987 84375160 refGene_gn=EHHADH;refGene_tx=XM_003991838.3
C2 84380962 84381074 refGene_gn=EHHADH;refGene_tx=XM_003991838.3
C2 84389869 84389974 refGene_gn=EHHADH;refGene_tx=XM_003991838.3

C2	84401577	84401919	refGene_gn=EHHADH;refGene_tx=XM_003991838.3
C2	84409268	84410530	refGene_gn=EHHADH;refGene_tx=XM_003991838.3
C2	84448716	84448912
refGene_gn=CC2H3orf70;refGene_tx=XM_011286039.3
C2	84533950	84534507
refGene_gn=CC2H3orf70;refGene_tx=XM_011286039.3
C2	84555275	84555425
refGene_gn=VPS8;refGene_tx=XM_011286040.2;refGene_gn=VPS8;refGene_tx=X
M_011286041.2;refGene_gn=VPS8;refGene_tx=XM_019840035.1;refGene_gn=VPS
8;refGene_tx=XM_019840036.1;refGene_gn=VPS8;refGene_tx=XM_019840037.1
C2	84558278	84558359
refGene_gn=VPS8;refGene_tx=XM_011286040.2;refGene_gn=VPS8;refGene_tx=X
M_011286041.2;refGene_gn=VPS8;refGene_tx=XM_019840035.1;refGene_gn=VPS
8;refGene_tx=XM_019840036.1;refGene_gn=VPS8;refGene_tx=XM_019840037.1
C2	84569587	84569668
refGene_gn=VPS8;refGene_tx=XM_019840035.1;refGene_gn=VPS8;refGene_tx=X
M_019840036.1;refGene_gn=VPS8;refGene_tx=XM_019840037.1
C2	84572682	84572697	refGene_gn=VPS8;refGene_tx=XM_019840038.2
C2	84592409	84592463
refGene_gn=VPS8;refGene_tx=XM_011286040.2;refGene_gn=VPS8;refGene_tx=X
M_011286041.2;refGene_gn=VPS8;refGene_tx=XM_019840035.1;refGene_gn=VPS
8;refGene_tx=XM_019840036.1;refGene_gn=VPS8;refGene_tx=XM_019840037.1;
refGene_gn=VPS8;refGene_tx=XM_019840038.2
C2	84611834	84612000
refGene_gn=VPS8;refGene_tx=XM_011286040.2;refGene_gn=VPS8;refGene_tx=X
M_011286041.2;refGene_gn=VPS8;refGene_tx=XM_019840035.1;refGene_gn=VPS
8;refGene_tx=XM_019840036.1;refGene_gn=VPS8;refGene_tx=XM_019840037.1;
refGene_gn=VPS8;refGene_tx=XM_019840038.2
C2	84614050	84614220
refGene_gn=VPS8;refGene_tx=XM_011286040.2;refGene_gn=VPS8;refGene_tx=X
M_011286041.2;refGene_gn=VPS8;refGene_tx=XM_019840035.1;refGene_gn=VPS
8;refGene_tx=XM_019840036.1;refGene_gn=VPS8;refGene_tx=XM_019840037.1;
refGene_gn=VPS8;refGene_tx=XM_019840038.2
C2	84616045	84616126
refGene_gn=VPS8;refGene_tx=XM_011286040.2;refGene_gn=VPS8;refGene_tx=X
M_011286041.2;refGene_gn=VPS8;refGene_tx=XM_019840035.1;refGene_gn=VPS
8;refGene_tx=XM_019840036.1;refGene_gn=VPS8;refGene_tx=XM_019840037.1;
refGene_gn=VPS8;refGene_tx=XM_019840038.2;refGene_gn=VPS8;refGene_tx=X
M_019840040.1
C2	84625342	84625441	refGene_gn=VPS8;refGene_tx=XM_019840039.1
C2	84626114	84626197
refGene_gn=VPS8;refGene_tx=XM_011286040.2;refGene_gn=VPS8;refGene_tx=X
M_011286041.2;refGene_gn=VPS8;refGene_tx=XM_019840035.1;refGene_gn=VPS
8;refGene_tx=XM_019840036.1;refGene_gn=VPS8;refGene_tx=XM_019840037.1;
refGene_gn=VPS8;refGene_tx=XM_019840038.2;refGene_gn=VPS8;refGene_tx=X
M_019840039.1
C2	84626519	84626601
refGene_gn=VPS8;refGene_tx=XM_011286040.2;refGene_gn=VPS8;refGene_tx=X
M_011286041.2;refGene_gn=VPS8;refGene_tx=XM_019840035.1;refGene_gn=VPS
8;refGene_tx=XM_019840036.1;refGene_gn=VPS8;refGene_tx=XM_019840037.1;

refGene_gn=VPS8;refGene_tx=XM_019840038.2;refGene_gn=VPS8;refGene_tx=XM_019840039.1;refGene_gn=VPS8;refGene_tx=XM_019840040.1
C2 84631852 84631956
refGene_gn=VPS8;refGene_tx=XM_011286040.2;refGene_gn=VPS8;refGene_tx=XM_011286041.2;refGene_gn=VPS8;refGene_tx=XM_019840035.1;refGene_gn=VPS8;refGene_tx=XM_019840036.1;refGene_gn=VPS8;refGene_tx=XM_019840037.1;refGene_gn=VPS8;refGene_tx=XM_019840038.2;refGene_gn=VPS8;refGene_tx=XM_019840039.1;refGene_gn=VPS8;refGene_tx=XM_019840040.1
C2 84636636 84636679
refGene_gn=VPS8;refGene_tx=XM_011286040.2;refGene_gn=VPS8;refGene_tx=XM_011286041.2;refGene_gn=VPS8;refGene_tx=XM_019840035.1;refGene_gn=VPS8;refGene_tx=XM_019840036.1;refGene_gn=VPS8;refGene_tx=XM_019840037.1;refGene_gn=VPS8;refGene_tx=XM_019840038.2;refGene_gn=VPS8;refGene_tx=XM_019840039.1;refGene_gn=VPS8;refGene_tx=XM_019840040.1
C2 84640077 84640167
refGene_gn=VPS8;refGene_tx=XM_011286040.2;refGene_gn=VPS8;refGene_tx=XM_011286041.2;refGene_gn=VPS8;refGene_tx=XM_019840035.1;refGene_gn=VPS8;refGene_tx=XM_019840036.1;refGene_gn=VPS8;refGene_tx=XM_019840037.1;refGene_gn=VPS8;refGene_tx=XM_019840038.2;refGene_gn=VPS8;refGene_tx=XM_019840039.1;refGene_gn=VPS8;refGene_tx=XM_019840040.1
C2 84645055 84645203
refGene_gn=VPS8;refGene_tx=XM_011286040.2;refGene_gn=VPS8;refGene_tx=XM_011286041.2;refGene_gn=VPS8;refGene_tx=XM_019840035.1;refGene_gn=VPS8;refGene_tx=XM_019840036.1;refGene_gn=VPS8;refGene_tx=XM_019840037.1;refGene_gn=VPS8;refGene_tx=XM_019840038.2;refGene_gn=VPS8;refGene_tx=XM_019840039.1;refGene_gn=VPS8;refGene_tx=XM_019840040.1
C2 84660413 84660460
refGene_gn=VPS8;refGene_tx=XM_011286040.2;refGene_gn=VPS8;refGene_tx=XM_011286041.2;refGene_gn=VPS8;refGene_tx=XM_019840035.1;refGene_gn=VPS8;refGene_tx=XM_019840036.1;refGene_gn=VPS8;refGene_tx=XM_019840037.1;refGene_gn=VPS8;refGene_tx=XM_019840038.2;refGene_gn=VPS8;refGene_tx=XM_019840039.1;refGene_gn=VPS8;refGene_tx=XM_019840040.1
C2 84664159 84664249
refGene_gn=VPS8;refGene_tx=XM_011286040.2;refGene_gn=VPS8;refGene_tx=XM_011286041.2;refGene_gn=VPS8;refGene_tx=XM_019840035.1;refGene_gn=VPS8;refGene_tx=XM_019840036.1;refGene_gn=VPS8;refGene_tx=XM_019840037.1;refGene_gn=VPS8;refGene_tx=XM_019840038.2;refGene_gn=VPS8;refGene_tx=XM_019840039.1;refGene_gn=VPS8;refGene_tx=XM_019840040.1
C2 84675514 84675613
refGene_gn=VPS8;refGene_tx=XM_011286040.2;refGene_gn=VPS8;refGene_tx=XM_011286041.2;refGene_gn=VPS8;refGene_tx=XM_019840035.1;refGene_gn=VPS8;refGene_tx=XM_019840036.1;refGene_gn=VPS8;refGene_tx=XM_019840037.1;refGene_gn=VPS8;refGene_tx=XM_019840038.2;refGene_gn=VPS8;refGene_tx=XM_019840039.1;refGene_gn=VPS8;refGene_tx=XM_019840040.1
C2 84676398 84676483
refGene_gn=VPS8;refGene_tx=XM_011286040.2;refGene_gn=VPS8;refGene_tx=XM_011286041.2;refGene_gn=VPS8;refGene_tx=XM_019840035.1;refGene_gn=VPS8;refGene_tx=XM_019840036.1;refGene_gn=VPS8;refGene_tx=XM_019840037.1;refGene_gn=VPS8;refGene_tx=XM_019840038.2;refGene_gn=VPS8;refGene_tx=XM_019840039.1;refGene_gn=VPS8;refGene_tx=XM_019840040.1

C2 84677227 84677310
refGene_gn=VPS8;refGene_tx=XM_011286040.2;refGene_gn=VPS8;refGene_tx=XM_011286041.2;refGene_gn=VPS8;refGene_tx=XM_019840035.1;refGene_gn=VPS8;refGene_tx=XM_019840036.1;refGene_gn=VPS8;refGene_tx=XM_019840037.1;refGene_gn=VPS8;refGene_tx=XM_019840038.2;refGene_gn=VPS8;refGene_tx=XM_019840039.1;refGene_gn=VPS8;refGene_tx=XM_019840040.1
C2 84681131 84681188
refGene_gn=VPS8;refGene_tx=XM_011286040.2;refGene_gn=VPS8;refGene_tx=XM_011286041.2;refGene_gn=VPS8;refGene_tx=XM_019840035.1;refGene_gn=VPS8;refGene_tx=XM_019840036.1;refGene_gn=VPS8;refGene_tx=XM_019840037.1;refGene_gn=VPS8;refGene_tx=XM_019840038.2;refGene_gn=VPS8;refGene_tx=XM_019840039.1;refGene_gn=VPS8;refGene_tx=XM_019840040.1
C2 84682502 84682622
refGene_gn=VPS8;refGene_tx=XM_011286040.2;refGene_gn=VPS8;refGene_tx=XM_011286041.2;refGene_gn=VPS8;refGene_tx=XM_019840035.1;refGene_gn=VPS8;refGene_tx=XM_019840036.1;refGene_gn=VPS8;refGene_tx=XM_019840037.1;refGene_gn=VPS8;refGene_tx=XM_019840038.2;refGene_gn=VPS8;refGene_tx=XM_019840039.1;refGene_gn=VPS8;refGene_tx=XM_019840040.1
C2 84688042 84688114
refGene_gn=VPS8;refGene_tx=XM_011286040.2;refGene_gn=VPS8;refGene_tx=XM_011286041.2;refGene_gn=VPS8;refGene_tx=XM_019840035.1;refGene_gn=VPS8;refGene_tx=XM_019840036.1;refGene_gn=VPS8;refGene_tx=XM_019840037.1;refGene_gn=VPS8;refGene_tx=XM_019840038.2;refGene_gn=VPS8;refGene_tx=XM_019840039.1;refGene_gn=VPS8;refGene_tx=XM_019840040.1
C2 84701951 84702071
refGene_gn=VPS8;refGene_tx=XM_011286040.2;refGene_gn=VPS8;refGene_tx=XM_011286041.2;refGene_gn=VPS8;refGene_tx=XM_019840035.1;refGene_gn=VPS8;refGene_tx=XM_019840036.1;refGene_gn=VPS8;refGene_tx=XM_019840037.1;refGene_gn=VPS8;refGene_tx=XM_019840038.2;refGene_gn=VPS8;refGene_tx=XM_019840039.1;refGene_gn=VPS8;refGene_tx=XM_019840040.1
C2 84702404 84702477
refGene_gn=VPS8;refGene_tx=XM_011286040.2;refGene_gn=VPS8;refGene_tx=XM_011286041.2;refGene_gn=VPS8;refGene_tx=XM_019840035.1;refGene_gn=VPS8;refGene_tx=XM_019840036.1;refGene_gn=VPS8;refGene_tx=XM_019840037.1;refGene_gn=VPS8;refGene_tx=XM_019840038.2;refGene_gn=VPS8;refGene_tx=XM_019840039.1;refGene_gn=VPS8;refGene_tx=XM_019840040.1
C2 84703919 84703962
refGene_gn=VPS8;refGene_tx=XM_011286040.2;refGene_gn=VPS8;refGene_tx=XM_011286041.2;refGene_gn=VPS8;refGene_tx=XM_019840035.1;refGene_gn=VPS8;refGene_tx=XM_019840036.1;refGene_gn=VPS8;refGene_tx=XM_019840037.1;refGene_gn=VPS8;refGene_tx=XM_019840038.2;refGene_gn=VPS8;refGene_tx=XM_019840039.1;refGene_gn=VPS8;refGene_tx=XM_019840040.1
C2 84713419 84713471
refGene_gn=VPS8;refGene_tx=XM_011286040.2;refGene_gn=VPS8;refGene_tx=XM_011286041.2;refGene_gn=VPS8;refGene_tx=XM_019840035.1;refGene_gn=VPS8;refGene_tx=XM_019840036.1;refGene_gn=VPS8;refGene_tx=XM_019840037.1;refGene_gn=VPS8;refGene_tx=XM_019840038.2;refGene_gn=VPS8;refGene_tx=XM_019840039.1;refGene_gn=VPS8;refGene_tx=XM_019840040.1
C2 84716420 84716510
refGene_gn=VPS8;refGene_tx=XM_011286040.2;refGene_gn=VPS8;refGene_tx=X

M_011286041.2;refGene_gn=VPS8;refGene_tx=XM_019840035.1;refGene_gn=VPS8;refGene_tx=XM_019840036.1;refGene_gn=VPS8;refGene_tx=XM_019840037.1;refGene_gn=VPS8;refGene_tx=XM_019840038.2;refGene_gn=VPS8;refGene_tx=XM_019840039.1;refGene_gn=VPS8;refGene_tx=XM_019840040.1
C2 84720689 84720912
refGene_gn=VPS8;refGene_tx=XM_011286040.2;refGene_gn=VPS8;refGene_tx=XM_011286041.2;refGene_gn=VPS8;refGene_tx=XM_019840035.1;refGene_gn=VPS8;refGene_tx=XM_019840036.1;refGene_gn=VPS8;refGene_tx=XM_019840037.1;refGene_gn=VPS8;refGene_tx=XM_019840038.2;refGene_gn=VPS8;refGene_tx=XM_019840039.1;refGene_gn=VPS8;refGene_tx=XM_019840040.1
C2 84727992 84728039
refGene_gn=VPS8;refGene_tx=XM_011286040.2;refGene_gn=VPS8;refGene_tx=XM_011286041.2;refGene_gn=VPS8;refGene_tx=XM_019840035.1;refGene_gn=VPS8;refGene_tx=XM_019840036.1;refGene_gn=VPS8;refGene_tx=XM_019840037.1;refGene_gn=VPS8;refGene_tx=XM_019840038.2;refGene_gn=VPS8;refGene_tx=XM_019840039.1;refGene_gn=VPS8;refGene_tx=XM_019840040.1
C2 84743418 84743508
refGene_gn=VPS8;refGene_tx=XM_011286040.2;refGene_gn=VPS8;refGene_tx=XM_011286041.2;refGene_gn=VPS8;refGene_tx=XM_019840035.1;refGene_gn=VPS8;refGene_tx=XM_019840036.1;refGene_gn=VPS8;refGene_tx=XM_019840037.1;refGene_gn=VPS8;refGene_tx=XM_019840038.2;refGene_gn=VPS8;refGene_tx=XM_019840039.1;refGene_gn=VPS8;refGene_tx=XM_019840040.1
C2 84745374 84745421
refGene_gn=VPS8;refGene_tx=XM_011286040.2;refGene_gn=VPS8;refGene_tx=XM_011286041.2;refGene_gn=VPS8;refGene_tx=XM_019840035.1;refGene_gn=VPS8;refGene_tx=XM_019840036.1;refGene_gn=VPS8;refGene_tx=XM_019840037.1;refGene_gn=VPS8;refGene_tx=XM_019840038.2;refGene_gn=VPS8;refGene_tx=XM_019840039.1;refGene_gn=VPS8;refGene_tx=XM_019840040.1
C2 84745813 84745904
refGene_gn=VPS8;refGene_tx=XM_011286040.2;refGene_gn=VPS8;refGene_tx=XM_011286041.2;refGene_gn=VPS8;refGene_tx=XM_019840035.1;refGene_gn=VPS8;refGene_tx=XM_019840036.1;refGene_gn=VPS8;refGene_tx=XM_019840037.1;refGene_gn=VPS8;refGene_tx=XM_019840038.2;refGene_gn=VPS8;refGene_tx=XM_019840039.1;refGene_gn=VPS8;refGene_tx=XM_019840040.1
C2 84747040 84747076
refGene_gn=VPS8;refGene_tx=XM_011286040.2;refGene_gn=VPS8;refGene_tx=XM_011286041.2;refGene_gn=VPS8;refGene_tx=XM_019840035.1;refGene_gn=VPS8;refGene_tx=XM_019840036.1;refGene_gn=VPS8;refGene_tx=XM_019840037.1;refGene_gn=VPS8;refGene_tx=XM_019840038.2;refGene_gn=VPS8;refGene_tx=XM_019840039.1;refGene_gn=VPS8;refGene_tx=XM_019840040.1
C2 84747955 84748030
refGene_gn=VPS8;refGene_tx=XM_011286040.2;refGene_gn=VPS8;refGene_tx=XM_011286041.2;refGene_gn=VPS8;refGene_tx=XM_019840035.1;refGene_gn=VPS8;refGene_tx=XM_019840036.1;refGene_gn=VPS8;refGene_tx=XM_019840037.1;refGene_gn=VPS8;refGene_tx=XM_019840038.2;refGene_gn=VPS8;refGene_tx=XM_019840039.1;refGene_gn=VPS8;refGene_tx=XM_019840040.1
C2 84756799 84756970
refGene_gn=VPS8;refGene_tx=XM_011286040.2;refGene_gn=VPS8;refGene_tx=XM_011286041.2;refGene_gn=VPS8;refGene_tx=XM_019840035.1;refGene_gn=VPS8;refGene_tx=XM_019840036.1;refGene_gn=VPS8;refGene_tx=XM_019840037.1;

refGene_gn=VPS8;refGene_tx=XM_019840038.2;refGene_gn=VPS8;refGene_tx=XM_019840039.1;refGene_gn=VPS8;refGene_tx=XM_019840040.1
C2 84760397 84760478
refGene_gn=VPS8;refGene_tx=XM_011286040.2;refGene_gn=VPS8;refGene_tx=XM_011286041.2;refGene_gn=VPS8;refGene_tx=XM_019840035.1;refGene_gn=VPS8;refGene_tx=XM_019840036.1;refGene_gn=VPS8;refGene_tx=XM_019840037.1;refGene_gn=VPS8;refGene_tx=XM_019840038.2;refGene_gn=VPS8;refGene_tx=XM_019840039.1;refGene_gn=VPS8;refGene_tx=XM_019840040.1
C2 84764110 84764218
refGene_gn=VPS8;refGene_tx=XM_011286040.2;refGene_gn=VPS8;refGene_tx=XM_011286041.2;refGene_gn=VPS8;refGene_tx=XM_019840035.1;refGene_gn=VPS8;refGene_tx=XM_019840036.1;refGene_gn=VPS8;refGene_tx=XM_019840038.2;refGene_gn=VPS8;refGene_tx=XM_019840039.1;refGene_gn=VPS8;refGene_tx=XM_019840040.1
C2 84765732 84765792
refGene_gn=VPS8;refGene_tx=XM_011286040.2;refGene_gn=VPS8;refGene_tx=XM_011286041.2;refGene_gn=VPS8;refGene_tx=XM_019840035.1;refGene_gn=VPS8;refGene_tx=XM_019840036.1;refGene_gn=VPS8;refGene_tx=XM_019840037.1;refGene_gn=VPS8;refGene_tx=XM_019840038.2;refGene_gn=VPS8;refGene_tx=XM_019840039.1;refGene_gn=VPS8;refGene_tx=XM_019840040.1
C2 84765889 84766043
refGene_gn=VPS8;refGene_tx=XM_011286040.2;refGene_gn=VPS8;refGene_tx=XM_011286041.2;refGene_gn=VPS8;refGene_tx=XM_019840035.1;refGene_gn=VPS8;refGene_tx=XM_019840036.1;refGene_gn=VPS8;refGene_tx=XM_019840037.1;refGene_gn=VPS8;refGene_tx=XM_019840038.2;refGene_gn=VPS8;refGene_tx=XM_019840039.1;refGene_gn=VPS8;refGene_tx=XM_019840040.1
C2 84767285 84767353
refGene_gn=VPS8;refGene_tx=XM_011286040.2;refGene_gn=VPS8;refGene_tx=XM_011286041.2;refGene_gn=VPS8;refGene_tx=XM_019840035.1;refGene_gn=VPS8;refGene_tx=XM_019840036.1;refGene_gn=VPS8;refGene_tx=XM_019840037.1;refGene_gn=VPS8;refGene_tx=XM_019840038.2;refGene_gn=VPS8;refGene_tx=XM_019840039.1;refGene_gn=VPS8;refGene_tx=XM_019840040.1
C2 84770141 84770228
refGene_gn=VPS8;refGene_tx=XM_011286040.2;refGene_gn=VPS8;refGene_tx=XM_011286041.2;refGene_gn=VPS8;refGene_tx=XM_019840035.1;refGene_gn=VPS8;refGene_tx=XM_019840036.1;refGene_gn=VPS8;refGene_tx=XM_019840037.1;refGene_gn=VPS8;refGene_tx=XM_019840038.2;refGene_gn=VPS8;refGene_tx=XM_019840039.1;refGene_gn=VPS8;refGene_tx=XM_019840040.1
C2 84770799 84770924
refGene_gn=VPS8;refGene_tx=XM_011286040.2;refGene_gn=VPS8;refGene_tx=XM_011286041.2;refGene_gn=VPS8;refGene_tx=XM_019840035.1;refGene_gn=VPS8;refGene_tx=XM_019840036.1;refGene_gn=VPS8;refGene_tx=XM_019840037.1;refGene_gn=VPS8;refGene_tx=XM_019840038.2;refGene_gn=VPS8;refGene_tx=XM_019840039.1;refGene_gn=VPS8;refGene_tx=XM_019840040.1
C2 84775763 84775769
refGene_gn=VPS8;refGene_tx=XM_011286040.2;refGene_gn=VPS8;refGene_tx=XM_019840035.1;refGene_gn=VPS8;refGene_tx=XM_019840037.1;refGene_gn=VPS8;refGene_tx=XM_019840038.2;refGene_gn=VPS8;refGene_tx=XM_019840039.1;refGene_gn=VPS8;refGene_tx=XM_019840040.1
C2 84779055 84779110

refGene_gn=VPS8;refGene_tx=XM_011286040.2;refGene_gn=VPS8;refGene_tx=XM_011286041.2;refGene_gn=VPS8;refGene_tx=XM_019840035.1;refGene_gn=VPS8;refGene_tx=XM_019840036.1;refGene_gn=VPS8;refGene_tx=XM_019840037.1;refGene_gn=VPS8;refGene_tx=XM_019840038.2;refGene_gn=VPS8;refGene_tx=XM_019840039.1;refGene_gn=VPS8;refGene_tx=XM_019840040.1
C2 84780051 84780084
refGene_gn=VPS8;refGene_tx=XM_011286040.2;refGene_gn=VPS8;refGene_tx=XM_011286041.2;refGene_gn=VPS8;refGene_tx=XM_019840035.1;refGene_gn=VPS8;refGene_tx=XM_019840036.1;refGene_gn=VPS8;refGene_tx=XM_019840037.1;refGene_gn=VPS8;refGene_tx=XM_019840038.2;refGene_gn=VPS8;refGene_tx=XM_019840039.1;refGene_gn=VPS8;refGene_tx=XM_019840040.1
C2 84784124 84784218
refGene_gn=VPS8;refGene_tx=XM_011286040.2;refGene_gn=VPS8;refGene_tx=XM_011286041.2;refGene_gn=VPS8;refGene_tx=XM_019840035.1;refGene_gn=VPS8;refGene_tx=XM_019840036.1;refGene_gn=VPS8;refGene_tx=XM_019840037.1;refGene_gn=VPS8;refGene_tx=XM_019840038.2;refGene_gn=VPS8;refGene_tx=XM_019840039.1;refGene_gn=VPS8;refGene_tx=XM_019840040.1
C2 84789740 84789871
refGene_gn=VPS8;refGene_tx=XM_011286040.2;refGene_gn=VPS8;refGene_tx=XM_011286041.2;refGene_gn=VPS8;refGene_tx=XM_019840035.1;refGene_gn=VPS8;refGene_tx=XM_019840036.1;refGene_gn=VPS8;refGene_tx=XM_019840037.1;refGene_gn=VPS8;refGene_tx=XM_019840038.2;refGene_gn=VPS8;refGene_tx=XM_019840039.1;refGene_gn=VPS8;refGene_tx=XM_019840040.1
C2 84796886 84796955
refGene_gn=VPS8;refGene_tx=XM_011286040.2;refGene_gn=VPS8;refGene_tx=XM_011286041.2;refGene_gn=VPS8;refGene_tx=XM_019840035.1;refGene_gn=VPS8;refGene_tx=XM_019840036.1;refGene_gn=VPS8;refGene_tx=XM_019840037.1;refGene_gn=VPS8;refGene_tx=XM_019840038.2;refGene_gn=VPS8;refGene_tx=XM_019840039.1;refGene_gn=VPS8;refGene_tx=XM_019840040.1
C2 84797777 84797930
refGene_gn=VPS8;refGene_tx=XM_011286040.2;refGene_gn=VPS8;refGene_tx=XM_011286041.2;refGene_gn=VPS8;refGene_tx=XM_019840035.1;refGene_gn=VPS8;refGene_tx=XM_019840036.1;refGene_gn=VPS8;refGene_tx=XM_019840037.1;refGene_gn=VPS8;refGene_tx=XM_019840038.2;refGene_gn=VPS8;refGene_tx=XM_019840039.1;refGene_gn=VPS8;refGene_tx=XM_019840040.1
C2 84894892 84895843 refGene_gn=MAGEF1;refGene_tx=XM_011286042.2
C2 85005807 85005916
refGene_gn=EPHB3;refGene_tx=XM_023260329.1;refGene_gn=EPHB3;refGene_tx=XM_023260330.1
C2 85006021 85006177
refGene_gn=EPHB3;refGene_tx=XM_023260329.1;refGene_gn=EPHB3;refGene_tx=XM_023260330.1
C2 85006280 85006474
refGene_gn=EPHB3;refGene_tx=XM_023260329.1;refGene_gn=EPHB3;refGene_tx=XM_023260330.1
C2 85006565 85006736
refGene_gn=EPHB3;refGene_tx=XM_023260329.1;refGene_gn=EPHB3;refGene_tx=XM_023260330.1
C2 85006826 85007042
refGene_gn=EPHB3;refGene_tx=XM_023260329.1;refGene_gn=EPHB3;refGene_tx

=XM_023260330.1
C2	85007422	85007670
refGene_gn=EPHB3;refGene_tx=XM_023260329.1;refGene_gn=EPHB3;refGene_tx
=XM_023260330.1
C2	85007792	85007915
refGene_gn=EPHB3;refGene_tx=XM_023260329.1;refGene_gn=EPHB3;refGene_tx
=XM_023260330.1
C2	85008981	85009034
refGene_gn=EPHB3;refGene_tx=XM_023260329.1;refGene_gn=EPHB3;refGene_tx
=XM_023260330.1
C2	85009391	85009500
refGene_gn=EPHB3;refGene_tx=XM_023260329.1;refGene_gn=EPHB3;refGene_tx
=XM_023260330.1
C2	85009579	85009739
refGene_gn=EPHB3;refGene_tx=XM_023260329.1;refGene_gn=EPHB3;refGene_tx
=XM_023260330.1
C2	85009917	85010042
refGene_gn=EPHB3;refGene_tx=XM_023260329.1;refGene_gn=EPHB3;refGene_tx
=XM_023260330.1
C2	85010214	85010565
refGene_gn=EPHB3;refGene_tx=XM_023260329.1;refGene_gn=EPHB3;refGene_tx
=XM_023260330.1
C2	85011342	85011498
refGene_gn=EPHB3;refGene_tx=XM_023260329.1;refGene_gn=EPHB3;refGene_tx
=XM_023260330.1
C2	85013802	85014475
refGene_gn=EPHB3;refGene_tx=XM_023260329.1;refGene_gn=EPHB3;refGene_tx
=XM_023260330.1
C2	85015837	85015902
refGene_gn=EPHB3;refGene_tx=XM_023260329.1;refGene_gn=EPHB3;refGene_tx
=XM_023260330.1
C2	85024391	85024506
refGene_gn=EPHB3;refGene_tx=XM_023260329.1;refGene_gn=EPHB3;refGene_tx
=XM_023260330.1
C2	85125976	85126144
refGene_gn=LOC101084135;refGene_tx=XM_019811151.1
C2	85126146	85126446
refGene_gn=LOC101084135;refGene_tx=XM_019811151.1
C2	85126447	85126495
refGene_gn=LOC101084135;refGene_tx=XM_019811151.1
C2	85132097	85132255
refGene_gn=LINC02054;refGene_tx=XM_023260176.1
C2	85132381	85132452
refGene_gn=LINC02054;refGene_tx=XM_023260176.1
C2	85132453	85132537
refGene_gn=LINC02054;refGene_tx=XM_023260176.1
C2	85132538	85132747
refGene_gn=LINC02054;refGene_tx=XM_023260176.1
C2	85132748	85132798
refGene_gn=LINC02054;refGene_tx=XM_023260176.1

C2 85159124 85159152
refGene_gn=LINC02054;refGene_tx=XM_023260176.1
C2 85172544 85172600
refGene_gn=CHRD;refGene_tx=XM_023260331.1;refGene_gn=CHRD;refGene_tx=XM_023260332.1;refGene_gn=CHRD;refGene_tx=XM_023260333.1
C2 85172825 85172925
refGene_gn=CHRD;refGene_tx=XM_023260331.1;refGene_gn=CHRD;refGene_tx=XM_023260332.1;refGene_gn=CHRD;refGene_tx=XM_023260333.1
C2 85173095 85173250
refGene_gn=CHRD;refGene_tx=XM_023260331.1;refGene_gn=CHRD;refGene_tx=XM_023260332.1;refGene_gn=CHRD;refGene_tx=XM_023260333.1
C2 85173662 85173765
refGene_gn=CHRD;refGene_tx=XM_023260331.1;refGene_gn=CHRD;refGene_tx=XM_023260332.1;refGene_gn=CHRD;refGene_tx=XM_023260333.1
C2 85174216 85174320
refGene_gn=CHRD;refGene_tx=XM_023260331.1;refGene_gn=CHRD;refGene_tx=XM_023260332.1;refGene_gn=CHRD;refGene_tx=XM_023260333.1
C2 85174610 85174667
refGene_gn=CHRD;refGene_tx=XM_023260331.1;refGene_gn=CHRD;refGene_tx=XM_023260332.1;refGene_gn=CHRD;refGene_tx=XM_023260333.1
C2 85174784 85174878
refGene_gn=CHRD;refGene_tx=XM_023260331.1;refGene_gn=CHRD;refGene_tx=XM_023260332.1;refGene_gn=CHRD;refGene_tx=XM_023260333.1
C2 85174961 85175225
refGene_gn=CHRD;refGene_tx=XM_023260331.1;refGene_gn=CHRD;refGene_tx=XM_023260332.1;refGene_gn=CHRD;refGene_tx=XM_023260333.1
C2 85175567 85175681
refGene_gn=CHRD;refGene_tx=XM_023260331.1;refGene_gn=CHRD;refGene_tx=XM_023260332.1;refGene_gn=CHRD;refGene_tx=XM_023260333.1
C2 85176130 85176351
refGene_gn=CHRD;refGene_tx=XM_023260331.1;refGene_gn=CHRD;refGene_tx=XM_023260332.1;refGene_gn=CHRD;refGene_tx=XM_023260333.1
C2 85176695 85176852
refGene_gn=CHRD;refGene_tx=XM_023260331.1;refGene_gn=CHRD;refGene_tx=XM_023260332.1;refGene_gn=CHRD;refGene_tx=XM_023260333.1
C2 85178066 85178186
refGene_gn=CHRD;refGene_tx=XM_023260331.1;refGene_gn=CHRD;refGene_tx=XM_023260332.1;refGene_gn=CHRD;refGene_tx=XM_023260333.1
C2 85178309 85178416
refGene_gn=CHRD;refGene_tx=XM_023260331.1;refGene_gn=CHRD;refGene_tx=XM_023260332.1;refGene_gn=CHRD;refGene_tx=XM_023260333.1
C2 85178564 85178712
refGene_gn=CHRD;refGene_tx=XM_023260331.1;refGene_gn=CHRD;refGene_tx=XM_023260332.1;refGene_gn=CHRD;refGene_tx=XM_023260333.1
C2 85178788 85178871
refGene_gn=CHRD;refGene_tx=XM_023260331.1;refGene_gn=CHRD;refGene_tx=XM_023260332.1;refGene_gn=CHRD;refGene_tx=XM_023260333.1
C2 85178954 85179095
refGene_gn=CHRD;refGene_tx=XM_023260331.1;refGene_gn=CHRD;refGene_tx=XM_023260332.1;refGene_gn=CHRD;refGene_tx=XM_023260333.1

C2 85179210 85179352
refGene_gn=CHRD;refGene_tx=XM_023260331.1;refGene_gn=CHRD;refGene_tx=XM_023260332.1;refGene_gn=CHRD;refGene_tx=XM_023260333.1
C2 85179712 85179800
refGene_gn=CHRD;refGene_tx=XM_023260331.1;refGene_gn=CHRD;refGene_tx=XM_023260332.1;refGene_gn=CHRD;refGene_tx=XM_023260333.1
C2 85179923 85180023
refGene_gn=CHRD;refGene_tx=XM_023260331.1;refGene_gn=CHRD;refGene_tx=XM_023260332.1;refGene_gn=CHRD;refGene_tx=XM_023260333.1
C2 85180112 85180241
refGene_gn=CHRD;refGene_tx=XM_023260331.1;refGene_gn=CHRD;refGene_tx=XM_023260332.1;refGene_gn=CHRD;refGene_tx=XM_023260333.1
C2 85180391 85180524
refGene_gn=CHRD;refGene_tx=XM_023260331.1;refGene_gn=CHRD;refGene_tx=XM_023260332.1;refGene_gn=CHRD;refGene_tx=XM_023260333.1
C2 85181007 85181111
refGene_gn=CHRD;refGene_tx=XM_023260331.1;refGene_gn=CHRD;refGene_tx=XM_023260332.1;refGene_gn=CHRD;refGene_tx=XM_023260333.1
C2 85181341 85181489
refGene_gn=CHRD;refGene_tx=XM_023260331.1;refGene_gn=CHRD;refGene_tx=XM_023260332.1;refGene_gn=CHRD;refGene_tx=XM_023260333.1
C2 85185054 85185238
refGene_gn=THPO;refGene_tx=NM_001163656.1;refGene_gn=THPO;refGene_tx=XM_019839212.2;refGene_gn=THPO;refGene_tx=XM_019839213.2;refGene_gn=THPO;refGene_tx=XM_023260080.1
C2 85185459 85185587
refGene_gn=THPO;refGene_tx=NM_001163656.1;refGene_gn=THPO;refGene_tx=XM_019839212.2;refGene_gn=THPO;refGene_tx=XM_019839213.2;refGene_gn=THPO;refGene_tx=XM_023260080.1
C2 85185887 85185974
refGene_gn=THPO;refGene_tx=NM_001163656.1;refGene_gn=THPO;refGene_tx=XM_019839212.2;refGene_gn=THPO;refGene_tx=XM_019839213.2;refGene_gn=THPO;refGene_tx=XM_023260080.1
C2 85187450 85187618
refGene_gn=THPO;refGene_tx=NM_001163656.1;refGene_gn=THPO;refGene_tx=XM_019839212.2;refGene_gn=THPO;refGene_tx=XM_019839213.2;refGene_gn=THPO;refGene_tx=XM_023260080.1
C2 85187861 85188524
refGene_gn=THPO;refGene_tx=NM_001163656.1;refGene_gn=THPO;refGene_tx=XM_019839212.2;refGene_gn=THPO;refGene_tx=XM_019839213.2;refGene_gn=THPO;refGene_tx=XM_023260080.1
C2 85192179 85192297
refGene_gn=POLR2H;refGene_tx=XM_003991839.5;refGene_gn=POLR2H;refGene_tx=XM_006936214.4
C2 85193437 85193521
refGene_gn=POLR2H;refGene_tx=XM_003991839.5;refGene_gn=POLR2H;refGene_tx=XM_006936214.4
C2 85194069 85194163
refGene_gn=POLR2H;refGene_tx=XM_003991839.5;refGene_gn=POLR2H;refGene_tx=XM_006936214.4

C2 85194246 85194330
refGene_gn=POLR2H;refGene_tx=XM_003991839.5;refGene_gn=POLR2H;refGene_tx=XM_006936214.4
C2 85195293 85195366 refGene_gn=POLR2H;refGene_tx=XM_003991839.5
C2 85197462 85197534
refGene_gn=CLCN2;refGene_tx=XM_006936220.4;refGene_gn=CLCN2;refGene_tx=XM_006936221.4;refGene_gn=CLCN2;refGene_tx=XM_019840047.2;refGene_gn=CLCN2;refGene_tx=XM_019840048.2
C2 85199806 85199963
refGene_gn=CLCN2;refGene_tx=XM_006936220.4;refGene_gn=CLCN2;refGene_tx=XM_006936221.4;refGene_gn=CLCN2;refGene_tx=XM_019840047.2;refGene_gn=CLCN2;refGene_tx=XM_019840048.2
C2 85200137 85200269
refGene_gn=CLCN2;refGene_tx=XM_006936220.4;refGene_gn=CLCN2;refGene_tx=XM_006936221.4;refGene_gn=CLCN2;refGene_tx=XM_019840048.2
C2 85200645 85200774
refGene_gn=CLCN2;refGene_tx=XM_006936220.4;refGene_gn=CLCN2;refGene_tx=XM_006936221.4;refGene_gn=CLCN2;refGene_tx=XM_019840047.2;refGene_gn=CLCN2;refGene_tx=XM_019840048.2
C2 85200861 85200995
refGene_gn=CLCN2;refGene_tx=XM_006936220.4;refGene_gn=CLCN2;refGene_tx=XM_006936221.4;refGene_gn=CLCN2;refGene_tx=XM_019840047.2;refGene_gn=CLCN2;refGene_tx=XM_019840048.2
C2 85201102 85201180
refGene_gn=CLCN2;refGene_tx=XM_006936220.4;refGene_gn=CLCN2;refGene_tx=XM_006936221.4;refGene_gn=CLCN2;refGene_tx=XM_019840047.2;refGene_gn=CLCN2;refGene_tx=XM_019840048.2
C2 85201263 85201342
refGene_gn=CLCN2;refGene_tx=XM_006936220.4;refGene_gn=CLCN2;refGene_tx=XM_006936221.4;refGene_gn=CLCN2;refGene_tx=XM_019840047.2;refGene_gn=CLCN2;refGene_tx=XM_019840048.2
C2 85201466 85201592
refGene_gn=CLCN2;refGene_tx=XM_006936220.4;refGene_gn=CLCN2;refGene_tx=XM_006936221.4;refGene_gn=CLCN2;refGene_tx=XM_019840047.2;refGene_gn=CLCN2;refGene_tx=XM_019840048.2
C2 85201692 85201777
refGene_gn=CLCN2;refGene_tx=XM_006936220.4;refGene_gn=CLCN2;refGene_tx=XM_006936221.4;refGene_gn=CLCN2;refGene_tx=XM_019840047.2;refGene_gn=CLCN2;refGene_tx=XM_019840048.2
C2 85201869 85201971
refGene_gn=CLCN2;refGene_tx=XM_006936220.4;refGene_gn=CLCN2;refGene_tx=XM_006936221.4;refGene_gn=CLCN2;refGene_tx=XM_019840047.2;refGene_gn=CLCN2;refGene_tx=XM_019840048.2
C2 85203492 85203577
refGene_gn=CLCN2;refGene_tx=XM_006936220.4;refGene_gn=CLCN2;refGene_tx=XM_006936221.4;refGene_gn=CLCN2;refGene_tx=XM_019840047.2;refGene_gn=CLCN2;refGene_tx=XM_019840048.2
C2 85203714 85203870
refGene_gn=CLCN2;refGene_tx=XM_006936220.4;refGene_gn=CLCN2;refGene_tx=XM_006936221.4;refGene_gn=CLCN2;refGene_tx=XM_019840047.2;refGene_gn=

CLCN2;refGene_tx=XM_019840048.2
C2	85204247	85204317
refGene_gn=CLCN2;refGene_tx=XM_006936220.4;refGene_gn=CLCN2;refGene_tx=XM_006936221.4;refGene_gn=CLCN2;refGene_tx=XM_019840047.2;refGene_gn=CLCN2;refGene_tx=XM_019840048.2
C2	85204547	85204658
refGene_gn=CLCN2;refGene_tx=XM_006936220.4;refGene_gn=CLCN2;refGene_tx=XM_006936221.4;refGene_gn=CLCN2;refGene_tx=XM_019840047.2;refGene_gn=CLCN2;refGene_tx=XM_019840048.2
C2	85204909	85205123
refGene_gn=CLCN2;refGene_tx=XM_006936220.4;refGene_gn=CLCN2;refGene_tx=XM_006936221.4;refGene_gn=CLCN2;refGene_tx=XM_019840047.2;refGene_gn=CLCN2;refGene_tx=XM_019840048.2
C2	85205433	85205567
refGene_gn=CLCN2;refGene_tx=XM_006936220.4;refGene_gn=CLCN2;refGene_tx=XM_006936221.4;refGene_gn=CLCN2;refGene_tx=XM_019840047.2;refGene_gn=CLCN2;refGene_tx=XM_019840048.2
C2	85205803	85205976
refGene_gn=CLCN2;refGene_tx=XM_006936220.4;refGene_gn=CLCN2;refGene_tx=XM_006936221.4;refGene_gn=CLCN2;refGene_tx=XM_019840047.2;refGene_gn=CLCN2;refGene_tx=XM_019840048.2
C2	85206090	85206205
refGene_gn=CLCN2;refGene_tx=XM_006936220.4;refGene_gn=CLCN2;refGene_tx=XM_006936221.4;refGene_gn=CLCN2;refGene_tx=XM_019840047.2;refGene_gn=CLCN2;refGene_tx=XM_019840048.2
C2	85206420	85206494
refGene_gn=CLCN2;refGene_tx=XM_006936220.4;refGene_gn=CLCN2;refGene_tx=XM_006936221.4;refGene_gn=CLCN2;refGene_tx=XM_019840047.2;refGene_gn=CLCN2;refGene_tx=XM_019840048.2
C2	85206610	85206661	refGene_gn=CLCN2;refGene_tx=XM_019840048.2
C2	85206715	85206775
refGene_gn=CLCN2;refGene_tx=XM_006936220.4;refGene_gn=CLCN2;refGene_tx=XM_019840047.2
C2	85206915	85206954
refGene_gn=CLCN2;refGene_tx=XM_006936220.4;refGene_gn=CLCN2;refGene_tx=XM_006936221.4;refGene_gn=CLCN2;refGene_tx=XM_019840047.2
C2	85207126	85207231
refGene_gn=CLCN2;refGene_tx=XM_006936220.4;refGene_gn=CLCN2;refGene_tx=XM_006936221.4;refGene_gn=CLCN2;refGene_tx=XM_019840047.2
C2	85211278	85211365
refGene_gn=CLCN2;refGene_tx=XM_006936220.4;refGene_gn=CLCN2;refGene_tx=XM_006936221.4;refGene_gn=CLCN2;refGene_tx=XM_019840047.2
C2	85211510	85211705
refGene_gn=CLCN2;refGene_tx=XM_006936220.4;refGene_gn=CLCN2;refGene_tx=XM_006936221.4;refGene_gn=CLCN2;refGene_tx=XM_019840047.2
C2	85213315	85213791
refGene_gn=FAM131A;refGene_tx=XM_003991844.5;refGene_gn=FAM131A;refGene_tx=XM_006936215.4;refGene_gn=FAM131A;refGene_tx=XM_006936218.3;refGene_gn=FAM131A;refGene_tx=XM_011286045.3;refGene_gn=FAM131A;refGene_tx=XM_019840049.2;refGene_gn=FAM131A;refGene_tx=XM_019840050.2

C2 85215103 85215220
refGene_gn=FAM131A;refGene_tx=XM_003991844.5;refGene_gn=FAM131A;refGene_tx=XM_006936215.4;refGene_gn=FAM131A;refGene_tx=XM_006936218.3;refGene_gn=FAM131A;refGene_tx=XM_011286045.3;refGene_gn=FAM131A;refGene_tx=XM_019840049.2;refGene_gn=FAM131A;refGene_tx=XM_019840050.2
C2 85215421 85215604
refGene_gn=FAM131A;refGene_tx=XM_003991844.5;refGene_gn=FAM131A;refGene_tx=XM_006936215.4;refGene_gn=FAM131A;refGene_tx=XM_006936218.3;refGene_gn=FAM131A;refGene_tx=XM_011286045.3;refGene_gn=FAM131A;refGene_tx=XM_019840050.2
C2 85215852 85215946
refGene_gn=FAM131A;refGene_tx=XM_003991844.5;refGene_gn=FAM131A;refGene_tx=XM_006936215.4;refGene_gn=FAM131A;refGene_tx=XM_006936218.3;refGene_gn=FAM131A;refGene_tx=XM_011286045.3;refGene_gn=FAM131A;refGene_tx=XM_019840049.2;refGene_gn=FAM131A;refGene_tx=XM_019840050.2
C2 85218120 85218263
refGene_gn=FAM131A;refGene_tx=XM_003991844.5;refGene_gn=FAM131A;refGene_tx=XM_006936215.4;refGene_gn=FAM131A;refGene_tx=XM_019840049.2
C2 85218925 85219013
refGene_gn=FAM131A;refGene_tx=XM_003991844.5;refGene_gn=FAM131A;refGene_tx=XM_019840049.2
C2 85220536 85220573 refGene_gn=FAM131A;refGene_tx=XM_006936215.4
C2 85221774 85221959
refGene_gn=EIF4G1;refGene_tx=XM_006936222.3;refGene_gn=EIF4G1;refGene_tx=XM_006936224.4;refGene_gn=EIF4G1;refGene_tx=XM_011286046.2;refGene_gn=EIF4G1;refGene_tx=XM_019840042.1;refGene_gn=EIF4G1;refGene_tx=XM_019840043.2;refGene_gn=EIF4G1;refGene_tx=XM_019840044.1;refGene_gn=EIF4G1;refGene_tx=XM_019840045.1;refGene_gn=EIF4G1;refGene_tx=XM_019840046.2
C2 85224595 85224736
refGene_gn=EIF4G1;refGene_tx=XM_006936222.3;refGene_gn=EIF4G1;refGene_tx=XM_006936224.4;refGene_gn=EIF4G1;refGene_tx=XM_011286046.2;refGene_gn=EIF4G1;refGene_tx=XM_019840042.1;refGene_gn=EIF4G1;refGene_tx=XM_019840043.2;refGene_gn=EIF4G1;refGene_tx=XM_019840044.1;refGene_gn=EIF4G1;refGene_tx=XM_019840045.1;refGene_gn=EIF4G1;refGene_tx=XM_019840046.2
C2 85224877 85224959
refGene_gn=EIF4G1;refGene_tx=XM_006936222.3;refGene_gn=EIF4G1;refGene_tx=XM_006936224.4;refGene_gn=EIF4G1;refGene_tx=XM_011286046.2;refGene_gn=EIF4G1;refGene_tx=XM_019840042.1;refGene_gn=EIF4G1;refGene_tx=XM_019840043.2;refGene_gn=EIF4G1;refGene_tx=XM_019840044.1;refGene_gn=EIF4G1;refGene_tx=XM_019840045.1;refGene_gn=EIF4G1;refGene_tx=XM_019840046.2
C2 85225069 85225204
refGene_gn=EIF4G1;refGene_tx=XM_006936222.3;refGene_gn=EIF4G1;refGene_tx=XM_006936224.4;refGene_gn=EIF4G1;refGene_tx=XM_011286046.2;refGene_gn=EIF4G1;refGene_tx=XM_019840042.1;refGene_gn=EIF4G1;refGene_tx=XM_019840043.2;refGene_gn=EIF4G1;refGene_tx=XM_019840044.1;refGene_gn=EIF4G1;refGene_tx=XM_019840045.1;refGene_gn=EIF4G1;refGene_tx=XM_019840046.2

C2 85225318 85225417
refGene_gn=EIF4G1;refGene_tx=XM_006936222.3;refGene_gn=EIF4G1;refGene_tx=XM_006936224.4;refGene_gn=EIF4G1;refGene_tx=XM_011286046.2;refGene_gn=EIF4G1;refGene_tx=XM_019840042.1;refGene_gn=EIF4G1;refGene_tx=XM_019840043.2;refGene_gn=EIF4G1;refGene_tx=XM_019840044.1;refGene_gn=EIF4G1;refGene_tx=XM_019840045.1;refGene_gn=EIF4G1;refGene_tx=XM_019840046.2
C2 85227693 85227775
refGene_gn=EIF4G1;refGene_tx=XM_006936222.3;refGene_gn=EIF4G1;refGene_tx=XM_006936224.4;refGene_gn=EIF4G1;refGene_tx=XM_011286046.2;refGene_gn=EIF4G1;refGene_tx=XM_019840042.1;refGene_gn=EIF4G1;refGene_tx=XM_019840043.2;refGene_gn=EIF4G1;refGene_tx=XM_019840044.1;refGene_gn=EIF4G1;refGene_tx=XM_019840045.1;refGene_gn=EIF4G1;refGene_tx=XM_019840046.2
C2 85227933 85228059
refGene_gn=EIF4G1;refGene_tx=XM_006936222.3;refGene_gn=EIF4G1;refGene_tx=XM_006936224.4;refGene_gn=EIF4G1;refGene_tx=XM_011286046.2;refGene_gn=EIF4G1;refGene_tx=XM_019840042.1;refGene_gn=EIF4G1;refGene_tx=XM_019840043.2;refGene_gn=EIF4G1;refGene_tx=XM_019840044.1;refGene_gn=EIF4G1;refGene_tx=XM_019840045.1;refGene_gn=EIF4G1;refGene_tx=XM_019840046.2
C2 85228360 85228533
refGene_gn=EIF4G1;refGene_tx=XM_006936222.3;refGene_gn=EIF4G1;refGene_tx=XM_006936224.4;refGene_gn=EIF4G1;refGene_tx=XM_011286046.2;refGene_gn=EIF4G1;refGene_tx=XM_019840042.1;refGene_gn=EIF4G1;refGene_tx=XM_019840043.2;refGene_gn=EIF4G1;refGene_tx=XM_019840044.1;refGene_gn=EIF4G1;refGene_tx=XM_019840045.1;refGene_gn=EIF4G1;refGene_tx=XM_019840046.2
C2 85228659 85228778
refGene_gn=EIF4G1;refGene_tx=XM_006936222.3;refGene_gn=EIF4G1;refGene_tx=XM_006936224.4;refGene_gn=EIF4G1;refGene_tx=XM_011286046.2;refGene_gn=EIF4G1;refGene_tx=XM_019840042.1;refGene_gn=EIF4G1;refGene_tx=XM_019840043.2;refGene_gn=EIF4G1;refGene_tx=XM_019840044.1;refGene_gn=EIF4G1;refGene_tx=XM_019840045.1;refGene_gn=EIF4G1;refGene_tx=XM_019840046.2
C2 85228898 85229131
refGene_gn=EIF4G1;refGene_tx=XM_006936222.3;refGene_gn=EIF4G1;refGene_tx=XM_006936224.4;refGene_gn=EIF4G1;refGene_tx=XM_011286046.2;refGene_gn=EIF4G1;refGene_tx=XM_019840042.1;refGene_gn=EIF4G1;refGene_tx=XM_019840043.2;refGene_gn=EIF4G1;refGene_tx=XM_019840044.1;refGene_gn=EIF4G1;refGene_tx=XM_019840045.1;refGene_gn=EIF4G1;refGene_tx=XM_019840046.2
C2 85229361 85229464
refGene_gn=EIF4G1;refGene_tx=XM_006936222.3;refGene_gn=EIF4G1;refGene_tx=XM_006936224.4;refGene_gn=EIF4G1;refGene_tx=XM_011286046.2;refGene_gn=EIF4G1;refGene_tx=XM_019840042.1;refGene_gn=EIF4G1;refGene_tx=XM_019840043.2;refGene_gn=EIF4G1;refGene_tx=XM_019840044.1;refGene_gn=EIF4G1;refGene_tx=XM_019840045.1;refGene_gn=EIF4G1;refGene_tx=XM_019840046.2
C2 85229694 85229797

refGene_gn=EIF4G1;refGene_tx=XM_006936222.3;refGene_gn=EIF4G1;refGene_tx=XM_006936224.4;refGene_gn=EIF4G1;refGene_tx=XM_011286046.2;refGene_gn=EIF4G1;refGene_tx=XM_019840042.1;refGene_gn=EIF4G1;refGene_tx=XM_019840043.2;refGene_gn=EIF4G1;refGene_tx=XM_019840044.1;refGene_gn=EIF4G1;refGene_tx=XM_019840045.1;refGene_gn=EIF4G1;refGene_tx=XM_019840046.2
C2 85230398 85230499
refGene_gn=EIF4G1;refGene_tx=XM_006936222.3;refGene_gn=EIF4G1;refGene_tx=XM_006936224.4;refGene_gn=EIF4G1;refGene_tx=XM_011286046.2;refGene_gn=EIF4G1;refGene_tx=XM_019840042.1;refGene_gn=EIF4G1;refGene_tx=XM_019840043.2;refGene_gn=EIF4G1;refGene_tx=XM_019840044.1;refGene_gn=EIF4G1;refGene_tx=XM_019840045.1;refGene_gn=EIF4G1;refGene_tx=XM_019840046.2
C2 85230701 85230861
refGene_gn=EIF4G1;refGene_tx=XM_006936222.3;refGene_gn=EIF4G1;refGene_tx=XM_006936224.4;refGene_gn=EIF4G1;refGene_tx=XM_011286046.2;refGene_gn=EIF4G1;refGene_tx=XM_019840042.1;refGene_gn=EIF4G1;refGene_tx=XM_019840043.2;refGene_gn=EIF4G1;refGene_tx=XM_019840044.1;refGene_gn=EIF4G1;refGene_tx=XM_019840045.1;refGene_gn=EIF4G1;refGene_tx=XM_019840046.2
C2 85230982 85231087
refGene_gn=EIF4G1;refGene_tx=XM_006936222.3;refGene_gn=EIF4G1;refGene_tx=XM_006936224.4;refGene_gn=EIF4G1;refGene_tx=XM_011286046.2;refGene_gn=EIF4G1;refGene_tx=XM_019840042.1;refGene_gn=EIF4G1;refGene_tx=XM_019840043.2;refGene_gn=EIF4G1;refGene_tx=XM_019840044.1;refGene_gn=EIF4G1;refGene_tx=XM_019840045.1;refGene_gn=EIF4G1;refGene_tx=XM_019840046.2
C2 85231251 85231488
refGene_gn=EIF4G1;refGene_tx=XM_006936222.3;refGene_gn=EIF4G1;refGene_tx=XM_006936224.4;refGene_gn=EIF4G1;refGene_tx=XM_011286046.2;refGene_gn=EIF4G1;refGene_tx=XM_019840042.1;refGene_gn=EIF4G1;refGene_tx=XM_019840043.2;refGene_gn=EIF4G1;refGene_tx=XM_019840044.1;refGene_gn=EIF4G1;refGene_tx=XM_019840045.1;refGene_gn=EIF4G1;refGene_tx=XM_019840046.2
C2 85231713 85231860
refGene_gn=EIF4G1;refGene_tx=XM_006936222.3;refGene_gn=EIF4G1;refGene_tx=XM_006936224.4;refGene_gn=EIF4G1;refGene_tx=XM_011286046.2;refGene_gn=EIF4G1;refGene_tx=XM_019840042.1;refGene_gn=EIF4G1;refGene_tx=XM_019840043.2;refGene_gn=EIF4G1;refGene_tx=XM_019840044.1;refGene_gn=EIF4G1;refGene_tx=XM_019840045.1;refGene_gn=EIF4G1;refGene_tx=XM_019840046.2
C2 85232083 85232281
refGene_gn=EIF4G1;refGene_tx=XM_006936222.3;refGene_gn=EIF4G1;refGene_tx=XM_006936224.4;refGene_gn=EIF4G1;refGene_tx=XM_011286046.2;refGene_gn=EIF4G1;refGene_tx=XM_019840042.1;refGene_gn=EIF4G1;refGene_tx=XM_019840043.2;refGene_gn=EIF4G1;refGene_tx=XM_019840044.1;refGene_gn=EIF4G1;refGene_tx=XM_019840045.1;refGene_gn=EIF4G1;refGene_tx=XM_019840046.2
C2 85232722 85232911
refGene_gn=EIF4G1;refGene_tx=XM_006936222.3;refGene_gn=EIF4G1;refGene_

tx=XM_006936224.4;refGene_gn=EIF4G1;refGene_tx=XM_011286046.2;refGene_gn=EIF4G1;refGene_tx=XM_019840042.1;refGene_gn=EIF4G1;refGene_tx=XM_019840043.2;refGene_gn=EIF4G1;refGene_tx=XM_019840044.1;refGene_gn=EIF4G1;refGene_tx=XM_019840045.1;refGene_gn=EIF4G1;refGene_tx=XM_019840046.2
C2 85233076 85233235
refGene_gn=EIF4G1;refGene_tx=XM_006936222.3;refGene_gn=EIF4G1;refGene_tx=XM_006936224.4;refGene_gn=EIF4G1;refGene_tx=XM_011286046.2;refGene_gn=EIF4G1;refGene_tx=XM_019840042.1;refGene_gn=EIF4G1;refGene_tx=XM_019840043.2;refGene_gn=EIF4G1;refGene_tx=XM_019840044.1;refGene_gn=EIF4G1;refGene_tx=XM_019840045.1;refGene_gn=EIF4G1;refGene_tx=XM_019840046.2
C2 85233359 85233493
refGene_gn=EIF4G1;refGene_tx=XM_006936222.3;refGene_gn=EIF4G1;refGene_tx=XM_006936224.4;refGene_gn=EIF4G1;refGene_tx=XM_011286046.2;refGene_gn=EIF4G1;refGene_tx=XM_019840042.1;refGene_gn=EIF4G1;refGene_tx=XM_019840043.2;refGene_gn=EIF4G1;refGene_tx=XM_019840044.1;refGene_gn=EIF4G1;refGene_tx=XM_019840045.1;refGene_gn=EIF4G1;refGene_tx=XM_019840046.2
C2 85233587 85233774
refGene_gn=EIF4G1;refGene_tx=XM_006936222.3;refGene_gn=EIF4G1;refGene_tx=XM_006936224.4;refGene_gn=EIF4G1;refGene_tx=XM_011286046.2;refGene_gn=EIF4G1;refGene_tx=XM_019840042.1;refGene_gn=EIF4G1;refGene_tx=XM_019840043.2;refGene_gn=EIF4G1;refGene_tx=XM_019840044.1;refGene_gn=EIF4G1;refGene_tx=XM_019840045.1;refGene_gn=EIF4G1;refGene_tx=XM_019840046.2
C2 85233869 85233958
refGene_gn=EIF4G1;refGene_tx=XM_006936222.3;refGene_gn=EIF4G1;refGene_tx=XM_006936224.4;refGene_gn=EIF4G1;refGene_tx=XM_011286046.2;refGene_gn=EIF4G1;refGene_tx=XM_019840042.1;refGene_gn=EIF4G1;refGene_tx=XM_019840043.2;refGene_gn=EIF4G1;refGene_tx=XM_019840044.1;refGene_gn=EIF4G1;refGene_tx=XM_019840045.1;refGene_gn=EIF4G1;refGene_tx=XM_019840046.2
C2 85234242 85235073
refGene_gn=EIF4G1;refGene_tx=XM_006936222.3;refGene_gn=EIF4G1;refGene_tx=XM_006936224.4;refGene_gn=EIF4G1;refGene_tx=XM_011286046.2;refGene_gn=EIF4G1;refGene_tx=XM_019840042.1;refGene_gn=EIF4G1;refGene_tx=XM_019840043.2;refGene_gn=EIF4G1;refGene_tx=XM_019840044.1;refGene_gn=EIF4G1;refGene_tx=XM_019840045.1;refGene_gn=EIF4G1;refGene_tx=XM_019840046.2
C2 85235353 85235420
refGene_gn=EIF4G1;refGene_tx=XM_006936222.3;refGene_gn=EIF4G1;refGene_tx=XM_006936224.4;refGene_gn=EIF4G1;refGene_tx=XM_011286046.2;refGene_gn=EIF4G1;refGene_tx=XM_019840042.1;refGene_gn=EIF4G1;refGene_tx=XM_019840043.2;refGene_gn=EIF4G1;refGene_tx=XM_019840044.1;refGene_gn=EIF4G1;refGene_tx=XM_019840045.1;refGene_gn=EIF4G1;refGene_tx=XM_019840046.2
C2 85235624 85235717
refGene_gn=EIF4G1;refGene_tx=XM_006936222.3;refGene_gn=EIF4G1;refGene_tx=XM_006936224.4;refGene_gn=EIF4G1;refGene_tx=XM_011286046.2;refGene_

gn=EIF4G1;refGene_tx=XM_019840042.1;refGene_gn=EIF4G1;refGene_tx=XM_019840043.2;refGene_gn=EIF4G1;refGene_tx=XM_019840044.1;refGene_gn=EIF4G1;refGene_tx=XM_019840045.1;refGene_gn=EIF4G1;refGene_tx=XM_019840046.2
C2 85236537 85236650
refGene_gn=EIF4G1;refGene_tx=XM_006936222.3;refGene_gn=EIF4G1;refGene_tx=XM_011286046.2;refGene_gn=EIF4G1;refGene_tx=XM_019840042.1;refGene_gn=EIF4G1;refGene_tx=XM_019840043.2;refGene_gn=EIF4G1;refGene_tx=XM_019840044.1;refGene_gn=EIF4G1;refGene_tx=XM_019840045.1;refGene_gn=EIF4G1;refGene_tx=XM_019840046.2
C2 85237749 85237849
refGene_gn=EIF4G1;refGene_tx=XM_006936222.3;refGene_gn=EIF4G1;refGene_tx=XM_011286046.2;refGene_gn=EIF4G1;refGene_tx=XM_019840042.1;refGene_gn=EIF4G1;refGene_tx=XM_019840043.2;refGene_gn=EIF4G1;refGene_tx=XM_019840044.1;refGene_gn=EIF4G1;refGene_tx=XM_019840045.1;refGene_gn=EIF4G1;refGene_tx=XM_019840046.2
C2 85238072 85238249
refGene_gn=EIF4G1;refGene_tx=XM_006936222.3;refGene_gn=EIF4G1;refGene_tx=XM_011286046.2;refGene_gn=EIF4G1;refGene_tx=XM_019840042.1;refGene_gn=EIF4G1;refGene_tx=XM_019840043.2;refGene_gn=EIF4G1;refGene_tx=XM_019840044.1;refGene_gn=EIF4G1;refGene_tx=XM_019840045.1;refGene_gn=EIF4G1;refGene_tx=XM_019840046.2
C2 85238825 85238846
refGene_gn=EIF4G1;refGene_tx=XM_019840042.1;refGene_gn=EIF4G1;refGene_tx=XM_019840043.2;refGene_gn=EIF4G1;refGene_tx=XM_019840044.1
C2 85239350 85239437
refGene_gn=EIF4G1;refGene_tx=XM_006936222.3;refGene_gn=EIF4G1;refGene_tx=XM_019840042.1;refGene_gn=EIF4G1;refGene_tx=XM_019840043.2;refGene_gn=EIF4G1;refGene_tx=XM_019840044.1;refGene_gn=EIF4G1;refGene_tx=XM_019840045.1
C2 85239735 85239795
refGene_gn=EIF4G1;refGene_tx=XM_006936222.3;refGene_gn=EIF4G1;refGene_tx=XM_011286046.2;refGene_gn=EIF4G1;refGene_tx=XM_019840042.1;refGene_gn=EIF4G1;refGene_tx=XM_019840043.2;refGene_gn=EIF4G1;refGene_tx=XM_019840044.1;refGene_gn=EIF4G1;refGene_tx=XM_019840045.1;refGene_gn=EIF4G1;refGene_tx=XM_019840046.2
C2 85246064 85246247 refGene_gn=PSMD2;refGene_tx=XM_006936226.3
C2 85246399 85246518 refGene_gn=PSMD2;refGene_tx=XM_006936226.3
C2 85246957 85247084 refGene_gn=PSMD2;refGene_tx=XM_006936226.3
C2 85247296 85247391 refGene_gn=PSMD2;refGene_tx=XM_006936226.3
C2 85247480 85247649 refGene_gn=PSMD2;refGene_tx=XM_006936226.3
C2 85247878 85247962 refGene_gn=PSMD2;refGene_tx=XM_006936226.3
C2 85248216 85248362 refGene_gn=PSMD2;refGene_tx=XM_006936226.3
C2 85248773 85248875 refGene_gn=PSMD2;refGene_tx=XM_006936226.3
C2 85248994 85249157 refGene_gn=PSMD2;refGene_tx=XM_006936226.3
C2 85249940 85250028 refGene_gn=PSMD2;refGene_tx=XM_006936226.3
C2 85250252 85250380 refGene_gn=PSMD2;refGene_tx=XM_006936226.3
C2 85250574 85250681 refGene_gn=PSMD2;refGene_tx=XM_006936226.3
C2 85250851 85250998 refGene_gn=PSMD2;refGene_tx=XM_006936226.3
C2 85251500 85251561 refGene_gn=PSMD2;refGene_tx=XM_006936226.3

C2 85251747 85251892 refGene_gn=PSMD2;refGene_tx=XM_006936226.3
C2 85252048 85252207 refGene_gn=PSMD2;refGene_tx=XM_006936226.3
C2 85252495 85252720 refGene_gn=PSMD2;refGene_tx=XM_006936226.3
C2 85252883 85253005 refGene_gn=PSMD2;refGene_tx=XM_006936226.3
C2 85253901 85254066 refGene_gn=PSMD2;refGene_tx=XM_006936226.3
C2 85254419 85254476 refGene_gn=PSMD2;refGene_tx=XM_006936226.3
C2 85254935 85255070 refGene_gn=PSMD2;refGene_tx=XM_006936226.3
C2 85261471 85261648
refGene_gn=ECE2;refGene_tx=XM_003991853.4;refGene_gn=ECE2;refGene_tx=XM_023260334.1;refGene_gn=ECE2;refGene_tx=XM_023260335.1
C2 85262366 85262462
refGene_gn=ECE2;refGene_tx=XM_003991853.4;refGene_gn=ECE2;refGene_tx=XM_023260334.1;refGene_gn=ECE2;refGene_tx=XM_023260335.1
C2 85262578 85262769
refGene_gn=ECE2;refGene_tx=XM_003991853.4;refGene_gn=ECE2;refGene_tx=XM_023260334.1;refGene_gn=ECE2;refGene_tx=XM_023260335.1
C2 85262931 85262999
refGene_gn=ECE2;refGene_tx=XM_003991853.4;refGene_gn=ECE2;refGene_tx=XM_023260334.1;refGene_gn=ECE2;refGene_tx=XM_023260335.1
C2 85263124 85263235
refGene_gn=ECE2;refGene_tx=XM_003991853.4;refGene_gn=ECE2;refGene_tx=XM_023260334.1;refGene_gn=ECE2;refGene_tx=XM_023260335.1
C2 85263427 85263531
refGene_gn=ECE2;refGene_tx=XM_003991853.4;refGene_gn=ECE2;refGene_tx=XM_023260334.1;refGene_gn=ECE2;refGene_tx=XM_023260335.1
C2 85264041 85264119
refGene_gn=ECE2;refGene_tx=XM_003991853.4;refGene_gn=ECE2;refGene_tx=XM_023260334.1;refGene_gn=ECE2;refGene_tx=XM_023260335.1
C2 85264221 85264320
refGene_gn=ECE2;refGene_tx=XM_003991853.4;refGene_gn=ECE2;refGene_tx=XM_023260334.1;refGene_gn=ECE2;refGene_tx=XM_023260335.1
C2 85265439 85265550
refGene_gn=ECE2;refGene_tx=XM_003991853.4;refGene_gn=ECE2;refGene_tx=XM_023260334.1;refGene_gn=ECE2;refGene_tx=XM_023260335.1
C2 85266950 85267065
refGene_gn=ECE2;refGene_tx=XM_003991853.4;refGene_gn=ECE2;refGene_tx=XM_023260334.1;refGene_gn=ECE2;refGene_tx=XM_023260335.1
C2 85267446 85267589
refGene_gn=ECE2;refGene_tx=XM_003991853.4;refGene_gn=ECE2;refGene_tx=XM_023260334.1;refGene_gn=ECE2;refGene_tx=XM_023260335.1
C2 85268294 85268483
refGene_gn=ECE2;refGene_tx=XM_003991853.4;refGene_gn=ECE2;refGene_tx=XM_023260334.1;refGene_gn=ECE2;refGene_tx=XM_023260335.1
C2 85272326 85272392
refGene_gn=ECE2;refGene_tx=XM_003991853.4;refGene_gn=ECE2;refGene_tx=XM_023260334.1;refGene_gn=ECE2;refGene_tx=XM_023260335.1
C2 85272554 85272701
refGene_gn=ECE2;refGene_tx=XM_003991853.4;refGene_gn=ECE2;refGene_tx=XM_023260334.1;refGene_gn=ECE2;refGene_tx=XM_023260335.1
C2 85272793 85272918

refGene_gn=ECE2;refGene_tx=XM_003991853.4;refGene_gn=ECE2;refGene_tx=XM_023260334.1;refGene_gn=ECE2;refGene_tx=XM_023260335.1
C2 85273383 85273599
refGene_gn=ECE2;refGene_tx=XM_003991853.4;refGene_gn=ECE2;refGene_tx=XM_023260334.1;refGene_gn=ECE2;refGene_tx=XM_023260335.1
C2 85273803 85273939
refGene_gn=ECE2;refGene_tx=XM_003991853.4;refGene_gn=ECE2;refGene_tx=XM_023260334.1;refGene_gn=ECE2;refGene_tx=XM_023260335.1
C2 85274285 85274372
refGene_gn=ECE2;refGene_tx=XM_003991853.4;refGene_gn=ECE2;refGene_tx=XM_023260334.1
C2 85274641 85274680
refGene_gn=ECE2;refGene_tx=XM_023260334.1;refGene_gn=ECE2;refGene_tx=XM_023260335.1
C2 85283505 85283674 refGene_gn=CAMK2N2;refGene_tx=XM_023260336.1
C2 85284541 85284612 refGene_gn=CAMK2N2;refGene_tx=XM_023260336.1
C2 85286252 85286540 refGene_gn=ECE2;refGene_tx=XM_003991855.4
C2 85287052 85287336
refGene_gn=ECE2;refGene_tx=XM_003991853.4;refGene_gn=ECE2;refGene_tx=XM_003991855.4
C2 85292279 85292475
refGene_gn=ECE2;refGene_tx=XM_003991853.4;refGene_gn=ECE2;refGene_tx=XM_003991855.4
C2 85293237 85293433
refGene_gn=ALG3;refGene_tx=XM_003991856.3;refGene_gn=ALG3;refGene_tx=XM_003991857.3
C2 85295372 85295472
refGene_gn=ALG3;refGene_tx=XM_003991856.3;refGene_gn=ALG3;refGene_tx=XM_003991857.3
C2 85295557 85295705
refGene_gn=ALG3;refGene_tx=XM_003991856.3;refGene_gn=ALG3;refGene_tx=XM_003991857.3
C2 85295812 85295973
refGene_gn=ALG3;refGene_tx=XM_003991856.3;refGene_gn=ALG3;refGene_tx=XM_003991857.3
C2 85296444 85296565
refGene_gn=ALG3;refGene_tx=XM_003991856.3;refGene_gn=ALG3;refGene_tx=XM_003991857.3
C2 85296994 85297200
refGene_gn=ALG3;refGene_tx=XM_003991856.3;refGene_gn=ALG3;refGene_tx=XM_003991857.3
C2 85297344 85297421
refGene_gn=ALG3;refGene_tx=XM_003991856.3;refGene_gn=ALG3;refGene_tx=XM_003991857.3
C2 85298055 85298200
refGene_gn=ALG3;refGene_tx=XM_003991856.3;refGene_gn=ALG3;refGene_tx=XM_003991857.3
C2 85298340 85298503
refGene_gn=ALG3;refGene_tx=XM_003991856.3;refGene_gn=ALG3;refGene_tx=XM_003991857.3

C2 85298947 85299496
refGene_gn=VWA5B2;refGene_tx=XM_023260337.1;refGene_gn=VWA5B2;refGene_tx=XM_023260338.1;refGene_gn=VWA5B2;refGene_tx=XM_023260339.1;refGene_gn=VWA5B2;refGene_tx=XM_023260340.1;refGene_gn=VWA5B2;refGene_tx=XM_023260341.1;refGene_gn=VWA5B2;refGene_tx=XM_023260342.1;refGene_gn=VWA5B2;refGene_tx=XM_023260343.1;refGene_gn=VWA5B2;refGene_tx=XM_023260344.1;refGene_gn=VWA5B2;refGene_tx=XM_023260345.1;refGene_gn=VWA5B2;refGene_tx=XM_023260346.1
C2 85299582 85299806
refGene_gn=VWA5B2;refGene_tx=XM_023260337.1;refGene_gn=VWA5B2;refGene_tx=XM_023260338.1;refGene_gn=VWA5B2;refGene_tx=XM_023260339.1;refGene_gn=VWA5B2;refGene_tx=XM_023260340.1;refGene_gn=VWA5B2;refGene_tx=XM_023260341.1;refGene_gn=VWA5B2;refGene_tx=XM_023260342.1;refGene_gn=VWA5B2;refGene_tx=XM_023260343.1;refGene_gn=VWA5B2;refGene_tx=XM_023260344.1;refGene_gn=VWA5B2;refGene_tx=XM_023260345.1;refGene_gn=VWA5B2;refGene_tx=XM_023260346.1
C2 85299885 85299966
refGene_gn=VWA5B2;refGene_tx=XM_023260337.1;refGene_gn=VWA5B2;refGene_tx=XM_023260338.1;refGene_gn=VWA5B2;refGene_tx=XM_023260339.1;refGene_gn=VWA5B2;refGene_tx=XM_023260340.1;refGene_gn=VWA5B2;refGene_tx=XM_023260341.1;refGene_gn=VWA5B2;refGene_tx=XM_023260342.1;refGene_gn=VWA5B2;refGene_tx=XM_023260343.1;refGene_gn=VWA5B2;refGene_tx=XM_023260344.1;refGene_gn=VWA5B2;refGene_tx=XM_023260345.1;refGene_gn=VWA5B2;refGene_tx=XM_023260346.1
C2 85300056 85300194
refGene_gn=VWA5B2;refGene_tx=XM_023260337.1;refGene_gn=VWA5B2;refGene_tx=XM_023260338.1;refGene_gn=VWA5B2;refGene_tx=XM_023260339.1;refGene_gn=VWA5B2;refGene_tx=XM_023260340.1;refGene_gn=VWA5B2;refGene_tx=XM_023260341.1;refGene_gn=VWA5B2;refGene_tx=XM_023260342.1;refGene_gn=VWA5B2;refGene_tx=XM_023260343.1;refGene_gn=VWA5B2;refGene_tx=XM_023260344.1;refGene_gn=VWA5B2;refGene_tx=XM_023260345.1;refGene_gn=VWA5B2;refGene_tx=XM_023260346.1
C2 85300971 85301325
refGene_gn=VWA5B2;refGene_tx=XM_023260337.1;refGene_gn=VWA5B2;refGene_tx=XM_023260338.1;refGene_gn=VWA5B2;refGene_tx=XM_023260339.1;refGene_gn=VWA5B2;refGene_tx=XM_023260340.1;refGene_gn=VWA5B2;refGene_tx=XM_023260341.1;refGene_gn=VWA5B2;refGene_tx=XM_023260342.1;refGene_gn=VWA5B2;refGene_tx=XM_023260343.1;refGene_gn=VWA5B2;refGene_tx=XM_023260344.1;refGene_gn=VWA5B2;refGene_tx=XM_023260345.1;refGene_gn=VWA5B2;refGene_tx=XM_023260346.1
C2 85301427 85301617
refGene_gn=VWA5B2;refGene_tx=XM_023260337.1;refGene_gn=VWA5B2;refGene_tx=XM_023260338.1;refGene_gn=VWA5B2;refGene_tx=XM_023260339.1;refGene_gn=VWA5B2;refGene_tx=XM_023260340.1;refGene_gn=VWA5B2;refGene_tx=XM_023260341.1;refGene_gn=VWA5B2;refGene_tx=XM_023260342.1;refGene_gn=VWA5B2;refGene_tx=XM_023260343.1;refGene_gn=VWA5B2;refGene_tx=XM_023260344.1;refGene_gn=VWA5B2;refGene_tx=XM_023260345.1;refGene_gn=VWA5B2;refGene_tx=XM_023260346.1
C2 85302112 85302429
refGene_gn=VWA5B2;refGene_tx=XM_023260337.1;refGene_gn=VWA5B2;refGene_

tx=XM_023260338.1;refGene_gn=VWA5B2;refGene_tx=XM_023260339.1;refGene_gn=VWA5B2;refGene_tx=XM_023260340.1;refGene_gn=VWA5B2;refGene_tx=XM_023260341.1;refGene_gn=VWA5B2;refGene_tx=XM_023260342.1;refGene_gn=VWA5B2;refGene_tx=XM_023260343.1;refGene_gn=VWA5B2;refGene_tx=XM_023260344.1;refGene_gn=VWA5B2;refGene_tx=XM_023260345.1;refGene_gn=VWA5B2;refGene_tx=XM_023260346.1
C2 85302519 85302691
refGene_gn=VWA5B2;refGene_tx=XM_023260337.1;refGene_gn=VWA5B2;refGene_tx=XM_023260338.1;refGene_gn=VWA5B2;refGene_tx=XM_023260339.1;refGene_gn=VWA5B2;refGene_tx=XM_023260340.1;refGene_gn=VWA5B2;refGene_tx=XM_023260341.1;refGene_gn=VWA5B2;refGene_tx=XM_023260342.1;refGene_gn=VWA5B2;refGene_tx=XM_023260343.1;refGene_gn=VWA5B2;refGene_tx=XM_023260344.1;refGene_gn=VWA5B2;refGene_tx=XM_023260345.1;refGene_gn=VWA5B2;refGene_tx=XM_023260346.1
C2 85303656 85303842
refGene_gn=VWA5B2;refGene_tx=XM_023260337.1;refGene_gn=VWA5B2;refGene_tx=XM_023260338.1;refGene_gn=VWA5B2;refGene_tx=XM_023260339.1;refGene_gn=VWA5B2;refGene_tx=XM_023260340.1;refGene_gn=VWA5B2;refGene_tx=XM_023260341.1;refGene_gn=VWA5B2;refGene_tx=XM_023260342.1;refGene_gn=VWA5B2;refGene_tx=XM_023260343.1;refGene_gn=VWA5B2;refGene_tx=XM_023260344.1;refGene_gn=VWA5B2;refGene_tx=XM_023260345.1;refGene_gn=VWA5B2;refGene_tx=XM_023260346.1
C2 85304347 85304459
refGene_gn=VWA5B2;refGene_tx=XM_023260337.1;refGene_gn=VWA5B2;refGene_tx=XM_023260338.1;refGene_gn=VWA5B2;refGene_tx=XM_023260339.1;refGene_gn=VWA5B2;refGene_tx=XM_023260340.1;refGene_gn=VWA5B2;refGene_tx=XM_023260341.1;refGene_gn=VWA5B2;refGene_tx=XM_023260342.1;refGene_gn=VWA5B2;refGene_tx=XM_023260343.1;refGene_gn=VWA5B2;refGene_tx=XM_023260344.1;refGene_gn=VWA5B2;refGene_tx=XM_023260345.1;refGene_gn=VWA5B2;refGene_tx=XM_023260346.1
C2 85304548 85304757
refGene_gn=VWA5B2;refGene_tx=XM_023260337.1;refGene_gn=VWA5B2;refGene_tx=XM_023260338.1;refGene_gn=VWA5B2;refGene_tx=XM_023260339.1;refGene_gn=VWA5B2;refGene_tx=XM_023260340.1;refGene_gn=VWA5B2;refGene_tx=XM_023260341.1;refGene_gn=VWA5B2;refGene_tx=XM_023260342.1;refGene_gn=VWA5B2;refGene_tx=XM_023260343.1;refGene_gn=VWA5B2;refGene_tx=XM_023260344.1;refGene_gn=VWA5B2;refGene_tx=XM_023260345.1;refGene_gn=VWA5B2;refGene_tx=XM_023260346.1
C2 85304835 85304946
refGene_gn=VWA5B2;refGene_tx=XM_023260337.1;refGene_gn=VWA5B2;refGene_tx=XM_023260338.1;refGene_gn=VWA5B2;refGene_tx=XM_023260339.1;refGene_gn=VWA5B2;refGene_tx=XM_023260340.1;refGene_gn=VWA5B2;refGene_tx=XM_023260341.1;refGene_gn=VWA5B2;refGene_tx=XM_023260342.1;refGene_gn=VWA5B2;refGene_tx=XM_023260343.1;refGene_gn=VWA5B2;refGene_tx=XM_023260344.1;refGene_gn=VWA5B2;refGene_tx=XM_023260345.1;refGene_gn=VWA5B2;refGene_tx=XM_023260346.1
C2 85305375 85305531
refGene_gn=VWA5B2;refGene_tx=XM_023260337.1;refGene_gn=VWA5B2;refGene_tx=XM_023260338.1;refGene_gn=VWA5B2;refGene_tx=XM_023260339.1;refGene_gn=VWA5B2;refGene_tx=XM_023260340.1;refGene_gn=VWA5B2;refGene_tx=XM_02

3260341.1;refGene_gn=VWA5B2;refGene_tx=XM_023260342.1;refGene_gn=VWA5B2;refGene_tx=XM_023260343.1;refGene_gn=VWA5B2;refGene_tx=XM_023260344.1;refGene_gn=VWA5B2;refGene_tx=XM_023260345.1;refGene_gn=VWA5B2;refGene_tx=XM_023260346.1
C2 85305934 85306059
refGene_gn=VWA5B2;refGene_tx=XM_023260337.1;refGene_gn=VWA5B2;refGene_tx=XM_023260338.1;refGene_gn=VWA5B2;refGene_tx=XM_023260339.1;refGene_gn=VWA5B2;refGene_tx=XM_023260340.1;refGene_gn=VWA5B2;refGene_tx=XM_023260341.1;refGene_gn=VWA5B2;refGene_tx=XM_023260342.1;refGene_gn=VWA5B2;refGene_tx=XM_023260343.1;refGene_gn=VWA5B2;refGene_tx=XM_023260344.1;refGene_gn=VWA5B2;refGene_tx=XM_023260345.1;refGene_gn=VWA5B2;refGene_tx=XM_023260346.1
C2 85306275 85306407
refGene_gn=VWA5B2;refGene_tx=XM_023260337.1;refGene_gn=VWA5B2;refGene_tx=XM_023260338.1;refGene_gn=VWA5B2;refGene_tx=XM_023260339.1;refGene_gn=VWA5B2;refGene_tx=XM_023260340.1;refGene_gn=VWA5B2;refGene_tx=XM_023260341.1;refGene_gn=VWA5B2;refGene_tx=XM_023260342.1;refGene_gn=VWA5B2;refGene_tx=XM_023260343.1;refGene_gn=VWA5B2;refGene_tx=XM_023260344.1;refGene_gn=VWA5B2;refGene_tx=XM_023260345.1;refGene_gn=VWA5B2;refGene_tx=XM_023260346.1
C2 85306933 85307091
refGene_gn=VWA5B2;refGene_tx=XM_023260337.1;refGene_gn=VWA5B2;refGene_tx=XM_023260338.1;refGene_gn=VWA5B2;refGene_tx=XM_023260339.1;refGene_gn=VWA5B2;refGene_tx=XM_023260340.1;refGene_gn=VWA5B2;refGene_tx=XM_023260341.1;refGene_gn=VWA5B2;refGene_tx=XM_023260342.1;refGene_gn=VWA5B2;refGene_tx=XM_023260343.1;refGene_gn=VWA5B2;refGene_tx=XM_023260344.1;refGene_gn=VWA5B2;refGene_tx=XM_023260345.1;refGene_gn=VWA5B2;refGene_tx=XM_023260346.1
C2 85307258 85307490
refGene_gn=VWA5B2;refGene_tx=XM_023260337.1;refGene_gn=VWA5B2;refGene_tx=XM_023260338.1;refGene_gn=VWA5B2;refGene_tx=XM_023260339.1;refGene_gn=VWA5B2;refGene_tx=XM_023260340.1;refGene_gn=VWA5B2;refGene_tx=XM_023260341.1;refGene_gn=VWA5B2;refGene_tx=XM_023260342.1;refGene_gn=VWA5B2;refGene_tx=XM_023260343.1;refGene_gn=VWA5B2;refGene_tx=XM_023260344.1;refGene_gn=VWA5B2;refGene_tx=XM_023260345.1;refGene_gn=VWA5B2;refGene_tx=XM_023260346.1
C2 85309382 85309553
refGene_gn=VWA5B2;refGene_tx=XM_023260337.1;refGene_gn=VWA5B2;refGene_tx=XM_023260338.1;refGene_gn=VWA5B2;refGene_tx=XM_023260339.1;refGene_gn=VWA5B2;refGene_tx=XM_023260340.1;refGene_gn=VWA5B2;refGene_tx=XM_023260341.1;refGene_gn=VWA5B2;refGene_tx=XM_023260342.1;refGene_gn=VWA5B2;refGene_tx=XM_023260343.1;refGene_gn=VWA5B2;refGene_tx=XM_023260344.1;refGene_gn=VWA5B2;refGene_tx=XM_023260345.1;refGene_gn=VWA5B2;refGene_tx=XM_023260346.1
C2 85309638 85309777
refGene_gn=VWA5B2;refGene_tx=XM_023260337.1;refGene_gn=VWA5B2;refGene_tx=XM_023260338.1;refGene_gn=VWA5B2;refGene_tx=XM_023260339.1;refGene_gn=VWA5B2;refGene_tx=XM_023260340.1;refGene_gn=VWA5B2;refGene_tx=XM_023260341.1;refGene_gn=VWA5B2;refGene_tx=XM_023260342.1;refGene_gn=VWA5B2;refGene_tx=XM_023260343.1;refGene_gn=VWA5B2;refGene_tx=XM_023260344.

1;refGene_gn=VWA5B2;refGene_tx=XM_023260345.1;refGene_gn=VWA5B2;refGene_tx=XM_023260346.1
C2 85344099 85344258 refGene_gn=ABCF3;refGene_tx=XM_003991859.5
C2 85344347 85344435 refGene_gn=ABCF3;refGene_tx=XM_003991859.5
C2 85344584 85344717 refGene_gn=ABCF3;refGene_tx=XM_003991859.5
C2 85344904 85344996 refGene_gn=ABCF3;refGene_tx=XM_003991859.5
C2 85345094 85345183 refGene_gn=ABCF3;refGene_tx=XM_003991859.5
C2 85345718 85345851 refGene_gn=ABCF3;refGene_tx=XM_003991859.5
C2 85345929 85345974 refGene_gn=ABCF3;refGene_tx=XM_003991859.5
C2 85347050 85347127 refGene_gn=ABCF3;refGene_tx=XM_003991859.5
C2 85347218 85347419 refGene_gn=ABCF3;refGene_tx=XM_003991859.5
C2 85347545 85347601 refGene_gn=ABCF3;refGene_tx=XM_003991859.5
C2 85347703 85347726 refGene_gn=ABCF3;refGene_tx=XM_003991859.5
C2 85347839 85347896 refGene_gn=ABCF3;refGene_tx=XM_003991859.5
C2 85347994 85348054 refGene_gn=ABCF3;refGene_tx=XM_003991859.5
C2 85348143 85348224 refGene_gn=ABCF3;refGene_tx=XM_003991859.5
C2 85348541 85348808 refGene_gn=ABCF3;refGene_tx=XM_003991859.5
C2 85348972 85349095 refGene_gn=ABCF3;refGene_tx=XM_003991859.5
C2 85349196 85349294 refGene_gn=ABCF3;refGene_tx=XM_003991859.5
C2 85349476 85349523 refGene_gn=ABCF3;refGene_tx=XM_003991859.5
C2 85350033 85350113 refGene_gn=ABCF3;refGene_tx=XM_003991859.5
C2 85350271 85350419 refGene_gn=ABCF3;refGene_tx=XM_003991859.5
C2 85350649 85350722 refGene_gn=ABCF3;refGene_tx=XM_003991859.5
C2 85353110 85353245
refGene_gn=AP2M1;refGene_tx=XM_003991861.4;refGene_gn=AP2M1;refGene_tx=XM_019840056.1;refGene_gn=AP2M1;refGene_tx=XM_019840057.1;refGene_gn=AP2M1;refGene_tx=XM_019840058.1
C2 85353902 85354014
refGene_gn=AP2M1;refGene_tx=XM_003991861.4;refGene_gn=AP2M1;refGene_tx=XM_019840056.1;refGene_gn=AP2M1;refGene_tx=XM_019840057.1;refGene_gn=AP2M1;refGene_tx=XM_019840058.1
C2 85354536 85354634
refGene_gn=AP2M1;refGene_tx=XM_003991861.4;refGene_gn=AP2M1;refGene_tx=XM_019840056.1;refGene_gn=AP2M1;refGene_tx=XM_019840057.1;refGene_gn=AP2M1;refGene_tx=XM_019840058.1
C2 85354734 85354870
refGene_gn=AP2M1;refGene_tx=XM_003991861.4;refGene_gn=AP2M1;refGene_tx=XM_019840056.1;refGene_gn=AP2M1;refGene_tx=XM_019840057.1;refGene_gn=AP2M1;refGene_tx=XM_019840058.1
C2 85354972 85355092
refGene_gn=AP2M1;refGene_tx=XM_003991861.4;refGene_gn=AP2M1;refGene_tx=XM_019840056.1;refGene_gn=AP2M1;refGene_tx=XM_019840057.1;refGene_gn=AP2M1;refGene_tx=XM_019840058.1
C2 85355565 85355707
refGene_gn=AP2M1;refGene_tx=XM_003991861.4;refGene_gn=AP2M1;refGene_tx=XM_019840056.1;refGene_gn=AP2M1;refGene_tx=XM_019840057.1;refGene_gn=AP2M1;refGene_tx=XM_019840058.1
C2 85355796 85355932
refGene_gn=AP2M1;refGene_tx=XM_003991861.4;refGene_gn=AP2M1;refGene_tx=XM_019840056.1;refGene_gn=AP2M1;refGene_tx=XM_019840057.1;refGene_gn=

AP2M1;refGene_tx=XM_019840058.1
C2 85356131 85356137
refGene_gn=AP2M1;refGene_tx=XM_003991861.4;refGene_gn=AP2M1;refGene_tx
=XM_019840056.1
C2 85356538 85356621
refGene_gn=AP2M1;refGene_tx=XM_003991861.4;refGene_gn=AP2M1;refGene_tx
=XM_019840056.1;refGene_gn=AP2M1;refGene_tx=XM_019840057.1;refGene_gn=
AP2M1;refGene_tx=XM_019840058.1
C2 85357318 85357584
refGene_gn=AP2M1;refGene_tx=XM_003991861.4;refGene_gn=AP2M1;refGene_tx
=XM_019840056.1;refGene_gn=AP2M1;refGene_tx=XM_019840057.1;refGene_gn=
AP2M1;refGene_tx=XM_019840058.1
C2 85358224 85358230
refGene_gn=AP2M1;refGene_tx=XM_019840056.1;refGene_gn=AP2M1;refGene_tx
=XM_019840057.1
C2 85359455 85359529
refGene_gn=AP2M1;refGene_tx=XM_003991861.4;refGene_gn=AP2M1;refGene_tx
=XM_019840056.1;refGene_gn=AP2M1;refGene_tx=XM_019840057.1;refGene_gn=
AP2M1;refGene_tx=XM_019840058.1
C2 85364494 85364526
refGene_gn=DVL3;refGene_tx=XM_023260347.1;refGene_gn=DVL3;refGene_tx=X
M_023260348.1
C2 85365355 85365792
refGene_gn=DVL3;refGene_tx=XM_011286047.3;refGene_gn=DVL3;refGene_tx=X
M_023260347.1;refGene_gn=DVL3;refGene_tx=XM_023260348.1
C2 85365890 85366106
refGene_gn=DVL3;refGene_tx=XM_011286047.3;refGene_gn=DVL3;refGene_tx=X
M_023260347.1
C2 85369091 85369259
refGene_gn=DVL3;refGene_tx=XM_011286047.3;refGene_gn=DVL3;refGene_tx=X
M_023260347.1;refGene_gn=DVL3;refGene_tx=XM_023260348.1
C2 85369444 85369576
refGene_gn=DVL3;refGene_tx=XM_011286047.3;refGene_gn=DVL3;refGene_tx=X
M_023260347.1;refGene_gn=DVL3;refGene_tx=XM_023260348.1
C2 85370137 85370287
refGene_gn=DVL3;refGene_tx=XM_011286047.3;refGene_gn=DVL3;refGene_tx=X
M_023260347.1;refGene_gn=DVL3;refGene_tx=XM_023260348.1
C2 85370442 85370510
refGene_gn=DVL3;refGene_tx=XM_011286047.3;refGene_gn=DVL3;refGene_tx=X
M_023260347.1;refGene_gn=DVL3;refGene_tx=XM_023260348.1
C2 85370625 85370702
refGene_gn=DVL3;refGene_tx=XM_011286047.3;refGene_gn=DVL3;refGene_tx=X
M_023260347.1;refGene_gn=DVL3;refGene_tx=XM_023260348.1
C2 85370885 85371025
refGene_gn=DVL3;refGene_tx=XM_011286047.3;refGene_gn=DVL3;refGene_tx=X
M_023260347.1;refGene_gn=DVL3;refGene_tx=XM_023260348.1
C2 85371644 85371714
refGene_gn=DVL3;refGene_tx=XM_011286047.3;refGene_gn=DVL3;refGene_tx=X
M_023260347.1;refGene_gn=DVL3;refGene_tx=XM_023260348.1
C2 85371928 85372022

refGene_gn=DVL3;refGene_tx=XM_011286047.3;refGene_gn=DVL3;refGene_tx=XM_023260347.1;refGene_gn=DVL3;refGene_tx=XM_023260348.1
C2 85372203 85372339
refGene_gn=DVL3;refGene_tx=XM_011286047.3;refGene_gn=DVL3;refGene_tx=XM_023260347.1;refGene_gn=DVL3;refGene_tx=XM_023260348.1
C2 85372507 85372617
refGene_gn=DVL3;refGene_tx=XM_011286047.3;refGene_gn=DVL3;refGene_tx=XM_023260347.1;refGene_gn=DVL3;refGene_tx=XM_023260348.1
C2 85372738 85372860
refGene_gn=DVL3;refGene_tx=XM_011286047.3;refGene_gn=DVL3;refGene_tx=XM_023260347.1;refGene_gn=DVL3;refGene_tx=XM_023260348.1
C2 85373371 85373441
refGene_gn=DVL3;refGene_tx=XM_011286047.3;refGene_gn=DVL3;refGene_tx=XM_023260347.1;refGene_gn=DVL3;refGene_tx=XM_023260348.1
C2 85378736 85378897
refGene_gn=DVL3;refGene_tx=XM_011286047.3;refGene_gn=DVL3;refGene_tx=XM_023260347.1;refGene_gn=DVL3;refGene_tx=XM_023260348.1
C2 85384517 85384577 refGene_gn=EIF2B5;refGene_tx=XM_003991863.4
C2 85384753 85384864 refGene_gn=EIF2B5;refGene_tx=XM_003991863.4
C2 85385280 85385406 refGene_gn=EIF2B5;refGene_tx=XM_003991863.4
C2 85385912 85386036 refGene_gn=EIF2B5;refGene_tx=XM_003991863.4
C2 85386344 85386435 refGene_gn=EIF2B5;refGene_tx=XM_003991863.4
C2 85386864 85386972 refGene_gn=EIF2B5;refGene_tx=XM_003991863.4
C2 85387172 85387274 refGene_gn=EIF2B5;refGene_tx=XM_003991863.4
C2 85387395 85387537 refGene_gn=EIF2B5;refGene_tx=XM_003991863.4
C2 85388129 85388275 refGene_gn=EIF2B5;refGene_tx=XM_003991863.4
C2 85389502 85389815 refGene_gn=EIF2B5;refGene_tx=XM_003991863.4
C2 85390093 85390171 refGene_gn=EIF2B5;refGene_tx=XM_003991863.4
C2 85390447 85390528 refGene_gn=EIF2B5;refGene_tx=XM_003991863.4
C2 85390615 85390793 refGene_gn=EIF2B5;refGene_tx=XM_003991863.4
C2 85390886 85391072 refGene_gn=EIF2B5;refGene_tx=XM_003991863.4
C2 85391917 85392042 refGene_gn=EIF2B5;refGene_tx=XM_003991863.4
C2 85393044 85393239 refGene_gn=EIF2B5;refGene_tx=XM_003991863.4
C2 85413930 85414160
refGene_gn=LOC101089387;refGene_tx=XM_003991865.2
C2 85414288 85414504
refGene_gn=LOC101089387;refGene_tx=XM_003991865.2
C2 85414681 85414886
refGene_gn=LOC101089387;refGene_tx=XM_003991865.2
C2 85415231 85415392
refGene_gn=LOC101089387;refGene_tx=XM_003991865.2
C2 85415716 85415886
refGene_gn=LOC101089387;refGene_tx=XM_003991865.2
C2 85416055 85416165
refGene_gn=LOC101089387;refGene_tx=XM_003991865.2
C2 85418811 85418978
refGene_gn=LOC101089387;refGene_tx=XM_003991865.2
C2 85420936 85421003
refGene_gn=LOC101089387;refGene_tx=XM_003991865.2
C2 85443943 85444164

refGene_gn=LOC101089793;refGene_tx=XM_023260177.1
C2 85444253 85444540
refGene_gn=LOC101089793;refGene_tx=XM_023260177.1
C2 85444632 85445048
refGene_gn=LOC101089793;refGene_tx=XM_023260177.1
C2 85445209 85445307
refGene_gn=LOC101089793;refGene_tx=XM_023260177.1
C2 85445706 85445876
refGene_gn=LOC101089793;refGene_tx=XM_023260177.1
C2 85446300 85446410
refGene_gn=LOC101089793;refGene_tx=XM_023260177.1
C2 85447879 85447975
refGene_gn=LOC101089793;refGene_tx=XM_023260177.1
C2 85448808 85448880
refGene_gn=LOC101089793;refGene_tx=XM_023260177.1
C2 85456361 85456591
refGene_gn=LOC101090040;refGene_tx=XM_003991866.2
C2 85456722 85456938
refGene_gn=LOC101090040;refGene_tx=XM_003991866.2
C2 85457121 85457326
refGene_gn=LOC101090040;refGene_tx=XM_003991866.2
C2 85457854 85458015
refGene_gn=LOC101090040;refGene_tx=XM_003991866.2
C2 85459639 85459809
refGene_gn=LOC101090040;refGene_tx=XM_003991866.2
C2 85460078 85460188
refGene_gn=LOC101090040;refGene_tx=XM_003991866.2
C2 85460657 85460702
refGene_gn=LOC101090040;refGene_tx=XM_003991866.2
C2 85461112 85461279
refGene_gn=LOC101090040;refGene_tx=XM_003991866.2
C2 85462587 85462654
refGene_gn=LOC101090040;refGene_tx=XM_003991866.2
C2 85469367 85469597
refGene_gn=LOC101090286;refGene_tx=XM_006936227.1;refGene_gn=LOC101090
286;refGene_tx=XM_023260349.1
C2 85469726 85469942
refGene_gn=LOC101090286;refGene_tx=XM_006936227.1;refGene_gn=LOC101090
286;refGene_tx=XM_023260349.1
C2 85470068 85470321
refGene_gn=LOC101090286;refGene_tx=XM_006936227.1;refGene_gn=LOC101090
286;refGene_tx=XM_023260349.1
C2 85471129 85471290
refGene_gn=LOC101090286;refGene_tx=XM_006936227.1
C2 85471480 85471650
refGene_gn=LOC101090286;refGene_tx=XM_006936227.1;refGene_gn=LOC101090
286;refGene_tx=XM_023260349.1
C2 85471915 85472025
refGene_gn=LOC101090286;refGene_tx=XM_006936227.1
C2 85472467 85472512

refGene_gn=LOC101090286;refGene_tx=XM_006936227.1
C2 85472895 85473062
refGene_gn=LOC101090286;refGene_tx=XM_006936227.1
C2 85474466 85474533
refGene_gn=LOC101090286;refGene_tx=XM_006936227.1
C2 85491411 85491540
refGene_gn=ABCC5;refGene_tx=XM_003991868.5;refGene_gn=ABCC5;refGene_tx=XM_023260350.1;refGene_gn=ABCC5;refGene_tx=XM_023260353.1;refGene_gn=ABCC5;refGene_tx=XM_023260354.1;refGene_gn=ABCC5;refGene_tx=XM_023260355.1;refGene_gn=ABCC5;refGene_tx=XM_023260356.1
C2 85512120 85512278
refGene_gn=ABCC5;refGene_tx=XM_003991868.5;refGene_gn=ABCC5;refGene_tx=XM_023260350.1;refGene_gn=ABCC5;refGene_tx=XM_023260353.1;refGene_gn=ABCC5;refGene_tx=XM_023260354.1;refGene_gn=ABCC5;refGene_tx=XM_023260355.1;refGene_gn=ABCC5;refGene_tx=XM_023260356.1
C2 85512469 85512625
refGene_gn=ABCC5;refGene_tx=XM_003991868.5;refGene_gn=ABCC5;refGene_tx=XM_023260350.1;refGene_gn=ABCC5;refGene_tx=XM_023260353.1;refGene_gn=ABCC5;refGene_tx=XM_023260354.1;refGene_gn=ABCC5;refGene_tx=XM_023260355.1;refGene_gn=ABCC5;refGene_tx=XM_023260356.1
C2 85513299 85513447
refGene_gn=ABCC5;refGene_tx=XM_003991868.5;refGene_gn=ABCC5;refGene_tx=XM_023260350.1;refGene_gn=ABCC5;refGene_tx=XM_023260353.1;refGene_gn=ABCC5;refGene_tx=XM_023260354.1;refGene_gn=ABCC5;refGene_tx=XM_023260355.1;refGene_gn=ABCC5;refGene_tx=XM_023260356.1
C2 85515922 85515958
refGene_gn=ABCC5;refGene_tx=XM_023260353.1;refGene_gn=ABCC5;refGene_tx=XM_023260354.1;refGene_gn=ABCC5;refGene_tx=XM_023260355.1;refGene_gn=ABCC5;refGene_tx=XM_023260356.1
C2 85517865 85517922 refGene_gn=ABCC5;refGene_tx=XM_023260351.1
C2 85518367 85518601
refGene_gn=ABCC5;refGene_tx=XM_003991868.5;refGene_gn=ABCC5;refGene_tx=XM_023260350.1;refGene_gn=ABCC5;refGene_tx=XM_023260351.1;refGene_gn=ABCC5;refGene_tx=XM_023260352.1
C2 85518761 85518935
refGene_gn=ABCC5;refGene_tx=XM_003991868.5;refGene_gn=ABCC5;refGene_tx=XM_023260350.1;refGene_gn=ABCC5;refGene_tx=XM_023260351.1;refGene_gn=ABCC5;refGene_tx=XM_023260352.1
C2 85519711 85519859
refGene_gn=ABCC5;refGene_tx=XM_003991868.5;refGene_gn=ABCC5;refGene_tx=XM_023260350.1;refGene_gn=ABCC5;refGene_tx=XM_023260351.1;refGene_gn=ABCC5;refGene_tx=XM_023260352.1
C2 85522820 85522969
refGene_gn=ABCC5;refGene_tx=XM_003991868.5;refGene_gn=ABCC5;refGene_tx=XM_023260350.1;refGene_gn=ABCC5;refGene_tx=XM_023260351.1;refGene_gn=ABCC5;refGene_tx=XM_023260352.1
C2 85523361 85523469
refGene_gn=ABCC5;refGene_tx=XM_003991868.5;refGene_gn=ABCC5;refGene_tx=XM_023260350.1;refGene_gn=ABCC5;refGene_tx=XM_023260351.1;refGene_gn=ABCC5;refGene_tx=XM_023260352.1

C2 85527526 85527883
refGene_gn=ABCC5;refGene_tx=XM_003991868.5;refGene_gn=ABCC5;refGene_tx=XM_023260350.1;refGene_gn=ABCC5;refGene_tx=XM_023260351.1;refGene_gn=ABCC5;refGene_tx=XM_023260352.1
C2 85531115 85531187
refGene_gn=ABCC5;refGene_tx=XM_003991868.5;refGene_gn=ABCC5;refGene_tx=XM_023260350.1;refGene_gn=ABCC5;refGene_tx=XM_023260351.1;refGene_gn=ABCC5;refGene_tx=XM_023260352.1
C2 85533336 85533461
refGene_gn=ABCC5;refGene_tx=XM_003991868.5;refGene_gn=ABCC5;refGene_tx=XM_023260350.1;refGene_gn=ABCC5;refGene_tx=XM_023260351.1;refGene_gn=ABCC5;refGene_tx=XM_023260352.1
C2 85533583 85533656
refGene_gn=ABCC5;refGene_tx=XM_003991868.5;refGene_gn=ABCC5;refGene_tx=XM_023260350.1;refGene_gn=ABCC5;refGene_tx=XM_023260351.1;refGene_gn=ABCC5;refGene_tx=XM_023260352.1
C2 85536287 85536491
refGene_gn=ABCC5;refGene_tx=XM_003991868.5;refGene_gn=ABCC5;refGene_tx=XM_023260350.1;refGene_gn=ABCC5;refGene_tx=XM_023260351.1;refGene_gn=ABCC5;refGene_tx=XM_023260352.1
C2 85537340 85537484
refGene_gn=ABCC5;refGene_tx=XM_003991868.5;refGene_gn=ABCC5;refGene_tx=XM_023260350.1;refGene_gn=ABCC5;refGene_tx=XM_023260351.1;refGene_gn=ABCC5;refGene_tx=XM_023260352.1
C2 85539843 85539946
refGene_gn=ABCC5;refGene_tx=XM_003991868.5;refGene_gn=ABCC5;refGene_tx=XM_023260350.1;refGene_gn=ABCC5;refGene_tx=XM_023260351.1;refGene_gn=ABCC5;refGene_tx=XM_023260352.1
C2 85542828 85543013
refGene_gn=ABCC5;refGene_tx=XM_003991868.5;refGene_gn=ABCC5;refGene_tx=XM_023260350.1;refGene_gn=ABCC5;refGene_tx=XM_023260351.1;refGene_gn=ABCC5;refGene_tx=XM_023260352.1
C2 85543396 85543543
refGene_gn=ABCC5;refGene_tx=XM_003991868.5;refGene_gn=ABCC5;refGene_tx=XM_023260350.1;refGene_gn=ABCC5;refGene_tx=XM_023260351.1;refGene_gn=ABCC5;refGene_tx=XM_023260352.1
C2 85543849 85543979
refGene_gn=ABCC5;refGene_tx=XM_003991868.5;refGene_gn=ABCC5;refGene_tx=XM_023260350.1;refGene_gn=ABCC5;refGene_tx=XM_023260351.1;refGene_gn=ABCC5;refGene_tx=XM_023260352.1
C2 85545403 85545557
refGene_gn=ABCC5;refGene_tx=XM_003991868.5;refGene_gn=ABCC5;refGene_tx=XM_023260350.1;refGene_gn=ABCC5;refGene_tx=XM_023260351.1;refGene_gn=ABCC5;refGene_tx=XM_023260352.1
C2 85545641 85545770
refGene_gn=ABCC5;refGene_tx=XM_003991868.5;refGene_gn=ABCC5;refGene_tx=XM_023260350.1;refGene_gn=ABCC5;refGene_tx=XM_023260351.1;refGene_gn=ABCC5;refGene_tx=XM_023260352.1
C2 85548257 85548444
refGene_gn=ABCC5;refGene_tx=XM_003991868.5;refGene_gn=ABCC5;refGene_tx

=XM_023260350.1;refGene_gn=ABCC5;refGene_tx=XM_023260351.1;refGene_gn=ABCC5;refGene_tx=XM_023260352.1
C2 85549979 85550069
refGene_gn=ABCC5;refGene_tx=XM_003991868.5;refGene_gn=ABCC5;refGene_tx=XM_023260350.1;refGene_gn=ABCC5;refGene_tx=XM_023260351.1;refGene_gn=ABCC5;refGene_tx=XM_023260352.1
C2 85552060 85552250
refGene_gn=ABCC5;refGene_tx=XM_003991868.5;refGene_gn=ABCC5;refGene_tx=XM_023260350.1;refGene_gn=ABCC5;refGene_tx=XM_023260351.1;refGene_gn=ABCC5;refGene_tx=XM_023260352.1
C2 85555991 85556151
refGene_gn=ABCC5;refGene_tx=XM_003991868.5;refGene_gn=ABCC5;refGene_tx=XM_023260350.1;refGene_gn=ABCC5;refGene_tx=XM_023260351.1;refGene_gn=ABCC5;refGene_tx=XM_023260352.1
C2 85564234 85564313
refGene_gn=ABCC5;refGene_tx=XM_003991868.5;refGene_gn=ABCC5;refGene_tx=XM_023260350.1;refGene_gn=ABCC5;refGene_tx=XM_023260351.1;refGene_gn=ABCC5;refGene_tx=XM_023260352.1
C2 85565540 85565654
refGene_gn=ABCC5;refGene_tx=XM_003991868.5;refGene_gn=ABCC5;refGene_tx=XM_023260350.1;refGene_gn=ABCC5;refGene_tx=XM_023260351.1;refGene_gn=ABCC5;refGene_tx=XM_023260352.1
C2 85566244 85566409
refGene_gn=ABCC5;refGene_tx=XM_003991868.5;refGene_gn=ABCC5;refGene_tx=XM_023260350.1;refGene_gn=ABCC5;refGene_tx=XM_023260351.1;refGene_gn=ABCC5;refGene_tx=XM_023260352.1
C2 85569026 85569128
refGene_gn=ABCC5;refGene_tx=XM_003991868.5;refGene_gn=ABCC5;refGene_tx=XM_023260350.1;refGene_gn=ABCC5;refGene_tx=XM_023260351.1;refGene_gn=ABCC5;refGene_tx=XM_023260352.1
C2 85588792 85588917
refGene_gn=PARL;refGene_tx=XM_004001479.5;refGene_gn=PARL;refGene_tx=XM_011286065.3
C2 85602071 85602267
refGene_gn=PARL;refGene_tx=XM_004001479.5;refGene_gn=PARL;refGene_tx=XM_011286065.3
C2 85602756 85602897
refGene_gn=PARL;refGene_tx=XM_004001479.5;refGene_gn=PARL;refGene_tx=XM_011286065.3
C2 85605516 85605565
refGene_gn=PARL;refGene_tx=XM_004001479.5;refGene_gn=PARL;refGene_tx=XM_011286065.3
C2 85617769 85617865
refGene_gn=PARL;refGene_tx=XM_004001479.5;refGene_gn=PARL;refGene_tx=XM_011286065.3
C2 85619135 85619285
refGene_gn=PARL;refGene_tx=XM_004001479.5;refGene_gn=PARL;refGene_tx=XM_011286065.3
C2 85620963 85621034 refGene_gn=PARL;refGene_tx=XM_004001479.5
C2 85635127 85635229

refGene_gn=PARL;refGene_tx=XM_004001479.5;refGene_gn=PARL;refGene_tx=XM_011286065.3
C2 85635368 85635466
refGene_gn=PARL;refGene_tx=XM_004001479.5;refGene_gn=PARL;refGene_tx=XM_011286065.3
C2 85638749 85638861 refGene_gn=PARL;refGene_tx=XM_004001479.5
C2 85642525 85642902 refGene_gn=MAP6D1;refGene_tx=XM_023260360.1
C2 85647761 85647879 refGene_gn=MAP6D1;refGene_tx=XM_023260360.1
C2 85648383 85648464 refGene_gn=MAP6D1;refGene_tx=XM_023260360.1
C2 85653213 85653322
refGene_gn=YEATS2;refGene_tx=XM_003991869.4;refGene_gn=YEATS2;refGene_tx=XM_023260357.1;refGene_gn=YEATS2;refGene_tx=XM_023260358.1;refGene_gn=YEATS2;refGene_tx=XM_023260359.1
C2 85654744 85654818
refGene_gn=YEATS2;refGene_tx=XM_003991869.4;refGene_gn=YEATS2;refGene_tx=XM_023260357.1;refGene_gn=YEATS2;refGene_tx=XM_023260358.1;refGene_gn=YEATS2;refGene_tx=XM_023260359.1
C2 85655789 85655864
refGene_gn=YEATS2;refGene_tx=XM_003991869.4;refGene_gn=YEATS2;refGene_tx=XM_023260357.1;refGene_gn=YEATS2;refGene_tx=XM_023260358.1;refGene_gn=YEATS2;refGene_tx=XM_023260359.1
C2 85656772 85656999
refGene_gn=YEATS2;refGene_tx=XM_003991869.4;refGene_gn=YEATS2;refGene_tx=XM_023260357.1;refGene_gn=YEATS2;refGene_tx=XM_023260358.1;refGene_gn=YEATS2;refGene_tx=XM_023260359.1
C2 85658538 85658740
refGene_gn=YEATS2;refGene_tx=XM_003991869.4;refGene_gn=YEATS2;refGene_tx=XM_023260357.1;refGene_gn=YEATS2;refGene_tx=XM_023260358.1;refGene_gn=YEATS2;refGene_tx=XM_023260359.1
C2 85659841 85659921
refGene_gn=YEATS2;refGene_tx=XM_003991869.4;refGene_gn=YEATS2;refGene_tx=XM_023260357.1;refGene_gn=YEATS2;refGene_tx=XM_023260358.1;refGene_gn=YEATS2;refGene_tx=XM_023260359.1
C2 85661695 85661769
refGene_gn=YEATS2;refGene_tx=XM_003991869.4;refGene_gn=YEATS2;refGene_tx=XM_023260357.1;refGene_gn=YEATS2;refGene_tx=XM_023260358.1;refGene_gn=YEATS2;refGene_tx=XM_023260359.1
C2 85662675 85662778
refGene_gn=YEATS2;refGene_tx=XM_003991869.4;refGene_gn=YEATS2;refGene_tx=XM_023260357.1;refGene_gn=YEATS2;refGene_tx=XM_023260358.1;refGene_gn=YEATS2;refGene_tx=XM_023260359.1
C2 85668488 85668587
refGene_gn=YEATS2;refGene_tx=XM_003991869.4;refGene_gn=YEATS2;refGene_tx=XM_023260357.1;refGene_gn=YEATS2;refGene_tx=XM_023260358.1;refGene_gn=YEATS2;refGene_tx=XM_023260359.1
C2 85669077 85669206
refGene_gn=YEATS2;refGene_tx=XM_003991869.4;refGene_gn=YEATS2;refGene_tx=XM_023260357.1;refGene_gn=YEATS2;refGene_tx=XM_023260358.1;refGene_gn=YEATS2;refGene_tx=XM_023260359.1
C2 85676626 85676810

refGene_gn=YEATS2;refGene_tx=XM_003991869.4;refGene_gn=YEATS2;refGene_tx=XM_023260357.1;refGene_gn=YEATS2;refGene_tx=XM_023260358.1;refGene_gn=YEATS2;refGene_tx=XM_023260359.1
C2 85679751 85679928
refGene_gn=YEATS2;refGene_tx=XM_003991869.4;refGene_gn=YEATS2;refGene_tx=XM_023260357.1;refGene_gn=YEATS2;refGene_tx=XM_023260358.1
C2 85683273 85683432
refGene_gn=YEATS2;refGene_tx=XM_003991869.4;refGene_gn=YEATS2;refGene_tx=XM_023260357.1;refGene_gn=YEATS2;refGene_tx=XM_023260358.1;refGene_gn=YEATS2;refGene_tx=XM_023260359.1
C2 85684212 85684385
refGene_gn=YEATS2;refGene_tx=XM_003991869.4;refGene_gn=YEATS2;refGene_tx=XM_023260357.1;refGene_gn=YEATS2;refGene_tx=XM_023260358.1;refGene_gn=YEATS2;refGene_tx=XM_023260359.1
C2 85686557 85686719
refGene_gn=YEATS2;refGene_tx=XM_003991869.4;refGene_gn=YEATS2;refGene_tx=XM_023260357.1;refGene_gn=YEATS2;refGene_tx=XM_023260358.1;refGene_gn=YEATS2;refGene_tx=XM_023260359.1
C2 85689485 85689744
refGene_gn=YEATS2;refGene_tx=XM_003991869.4;refGene_gn=YEATS2;refGene_tx=XM_023260357.1;refGene_gn=YEATS2;refGene_tx=XM_023260358.1;refGene_gn=YEATS2;refGene_tx=XM_023260359.1
C2 85697579 85697762
refGene_gn=YEATS2;refGene_tx=XM_003991869.4;refGene_gn=YEATS2;refGene_tx=XM_023260357.1;refGene_gn=YEATS2;refGene_tx=XM_023260358.1;refGene_gn=YEATS2;refGene_tx=XM_023260359.1
C2 85698284 85698392
refGene_gn=YEATS2;refGene_tx=XM_003991869.4;refGene_gn=YEATS2;refGene_tx=XM_023260357.1;refGene_gn=YEATS2;refGene_tx=XM_023260358.1;refGene_gn=YEATS2;refGene_tx=XM_023260359.1
C2 85700020 85700124
refGene_gn=YEATS2;refGene_tx=XM_003991869.4;refGene_gn=YEATS2;refGene_tx=XM_023260357.1;refGene_gn=YEATS2;refGene_tx=XM_023260358.1;refGene_gn=YEATS2;refGene_tx=XM_023260359.1
C2 85701750 85701909
refGene_gn=YEATS2;refGene_tx=XM_003991869.4;refGene_gn=YEATS2;refGene_tx=XM_023260357.1;refGene_gn=YEATS2;refGene_tx=XM_023260358.1;refGene_gn=YEATS2;refGene_tx=XM_023260359.1
C2 85703367 85703607
refGene_gn=YEATS2;refGene_tx=XM_003991869.4;refGene_gn=YEATS2;refGene_tx=XM_023260357.1;refGene_gn=YEATS2;refGene_tx=XM_023260358.1;refGene_gn=YEATS2;refGene_tx=XM_023260359.1
C2 85708678 85708859
refGene_gn=YEATS2;refGene_tx=XM_003991869.4;refGene_gn=YEATS2;refGene_tx=XM_023260357.1;refGene_gn=YEATS2;refGene_tx=XM_023260358.1;refGene_gn=YEATS2;refGene_tx=XM_023260359.1
C2 85711578 85711623
refGene_gn=YEATS2;refGene_tx=XM_003991869.4;refGene_gn=YEATS2;refGene_tx=XM_023260357.1;refGene_gn=YEATS2;refGene_tx=XM_023260358.1;refGene_gn=YEATS2;refGene_tx=XM_023260359.1

C2 85718796 85718908
refGene_gn=YEATS2;refGene_tx=XM_003991869.4;refGene_gn=YEATS2;refGene_tx=XM_023260357.1;refGene_gn=YEATS2;refGene_tx=XM_023260358.1;refGene_gn=YEATS2;refGene_tx=XM_023260359.1
C2 85724017 85724179
refGene_gn=YEATS2;refGene_tx=XM_003991869.4;refGene_gn=YEATS2;refGene_tx=XM_023260357.1;refGene_gn=YEATS2;refGene_tx=XM_023260358.1;refGene_gn=YEATS2;refGene_tx=XM_023260359.1
C2 85734719 85734835
refGene_gn=YEATS2;refGene_tx=XM_003991869.4;refGene_gn=YEATS2;refGene_tx=XM_023260357.1;refGene_gn=YEATS2;refGene_tx=XM_023260358.1;refGene_gn=YEATS2;refGene_tx=XM_023260359.1
C2 85737230 85737476
refGene_gn=YEATS2;refGene_tx=XM_003991869.4;refGene_gn=YEATS2;refGene_tx=XM_023260357.1;refGene_gn=YEATS2;refGene_tx=XM_023260358.1;refGene_gn=YEATS2;refGene_tx=XM_023260359.1
C2 85740753 85740846
refGene_gn=YEATS2;refGene_tx=XM_003991869.4;refGene_gn=YEATS2;refGene_tx=XM_023260357.1;refGene_gn=YEATS2;refGene_tx=XM_023260358.1;refGene_gn=YEATS2;refGene_tx=XM_023260359.1
C2 85741395 85741493
refGene_gn=YEATS2;refGene_tx=XM_003991869.4;refGene_gn=YEATS2;refGene_tx=XM_023260357.1;refGene_gn=YEATS2;refGene_tx=XM_023260358.1;refGene_gn=YEATS2;refGene_tx=XM_023260359.1
C2 85745086 85745189
refGene_gn=YEATS2;refGene_tx=XM_003991869.4;refGene_gn=YEATS2;refGene_tx=XM_023260357.1;refGene_gn=YEATS2;refGene_tx=XM_023260358.1;refGene_gn=YEATS2;refGene_tx=XM_023260359.1
C2 85794767 85794968
refGene_gn=KLHL24;refGene_tx=XM_006936229.4;refGene_gn=KLHL24;refGene_tx=XM_006936230.4;refGene_gn=KLHL24;refGene_tx=XM_006936232.4;refGene_gn=KLHL24;refGene_tx=XM_011286064.3;refGene_gn=KLHL24;refGene_tx=XM_019840061.2;refGene_gn=KLHL24;refGene_tx=XM_019840062.2;refGene_gn=KLHL24;refGene_tx=XM_023260361.1
C2 85799676 85799865
refGene_gn=KLHL24;refGene_tx=XM_006936229.4;refGene_gn=KLHL24;refGene_tx=XM_006936230.4;refGene_gn=KLHL24;refGene_tx=XM_006936232.4;refGene_gn=KLHL24;refGene_tx=XM_011286064.3;refGene_gn=KLHL24;refGene_tx=XM_019840061.2;refGene_gn=KLHL24;refGene_tx=XM_019840062.2;refGene_gn=KLHL24;refGene_tx=XM_023260361.1
C2 85801265 85801454
refGene_gn=KLHL24;refGene_tx=XM_006936229.4;refGene_gn=KLHL24;refGene_tx=XM_006936230.4;refGene_gn=KLHL24;refGene_tx=XM_006936232.4;refGene_gn=KLHL24;refGene_tx=XM_011286064.3;refGene_gn=KLHL24;refGene_tx=XM_019840061.2;refGene_gn=KLHL24;refGene_tx=XM_019840062.2;refGene_gn=KLHL24;refGene_tx=XM_023260361.1
C2 85805620 85805739
refGene_gn=KLHL24;refGene_tx=XM_006936229.4;refGene_gn=KLHL24;refGene_tx=XM_006936230.4;refGene_gn=KLHL24;refGene_tx=XM_006936232.4;refGene_gn=KLHL24;refGene_tx=XM_011286064.3;refGene_gn=KLHL24;refGene_tx=XM_01

9840061.2;refGene_gn=KLHL24;refGene_tx=XM_019840062.2;refGene_gn=KLHL24;refGene_tx=XM_023260361.1
C2 85806943 85807128
refGene_gn=KLHL24;refGene_tx=XM_006936229.4;refGene_gn=KLHL24;refGene_tx=XM_006936230.4;refGene_gn=KLHL24;refGene_tx=XM_006936232.4;refGene_gn=KLHL24;refGene_tx=XM_011286064.3;refGene_gn=KLHL24;refGene_tx=XM_019840061.2;refGene_gn=KLHL24;refGene_tx=XM_019840062.2;refGene_gn=KLHL24;refGene_tx=XM_023260361.1
C2 85814073 85814147 refGene_gn=KLHL24;refGene_tx=XM_023260361.1
C2 85816910 85817830
refGene_gn=KLHL24;refGene_tx=XM_006936229.4;refGene_gn=KLHL24;refGene_tx=XM_006936230.4;refGene_gn=KLHL24;refGene_tx=XM_006936232.4;refGene_gn=KLHL24;refGene_tx=XM_011286064.3;refGene_gn=KLHL24;refGene_tx=XM_019840061.2;refGene_gn=KLHL24;refGene_tx=XM_019840062.2
C2 85913382 85913675 refGene_gn=KLHL6;refGene_tx=XM_003991872.3
C2 85936581 85936747 refGene_gn=KLHL6;refGene_tx=XM_003991872.3
C2 85952096 85952546 refGene_gn=KLHL6;refGene_tx=XM_003991872.3
C2 85960525 85960763 refGene_gn=KLHL6;refGene_tx=XM_003991872.3
C2 85965880 85966083 refGene_gn=KLHL6;refGene_tx=XM_003991872.3
C2 85967669 85967883 refGene_gn=KLHL6;refGene_tx=XM_003991872.3
C2 85969472 85969774 refGene_gn=KLHL6;refGene_tx=XM_003991872.3
C2 86039268 86039751
refGene_gn=LOC101092708;refGene_tx=XM_011286063.3
C2 86043845 86043921 refGene_gn=MCF2L2;refGene_tx=XM_023260362.1
C2 86084334 86084418 refGene_gn=MCF2L2;refGene_tx=XM_023260362.1
C2 86093692 86093807 refGene_gn=MCF2L2;refGene_tx=XM_023260362.1
C2 86133731 86133822 refGene_gn=MCF2L2;refGene_tx=XM_023260362.1
C2 86136370 86136490 refGene_gn=MCF2L2;refGene_tx=XM_023260362.1
C2 86156420 86156537 refGene_gn=MCF2L2;refGene_tx=XM_023260362.1
C2 86160736 86160886 refGene_gn=MCF2L2;refGene_tx=XM_023260362.1
C2 86171445 86171570 refGene_gn=MCF2L2;refGene_tx=XM_023260362.1
C2 86172172 86172287 refGene_gn=MCF2L2;refGene_tx=XM_023260362.1
C2 86173012 86173132 refGene_gn=MCF2L2;refGene_tx=XM_023260362.1
C2 86181550 86181742 refGene_gn=MCF2L2;refGene_tx=XM_023260362.1
C2 86184132 86184324 refGene_gn=MCF2L2;refGene_tx=XM_023260362.1
C2 86190414 86190592 refGene_gn=MCF2L2;refGene_tx=XM_023260362.1
C2 86197149 86197226 refGene_gn=MCF2L2;refGene_tx=XM_023260362.1
C2 86212204 86212290 refGene_gn=MCF2L2;refGene_tx=XM_023260362.1
C2 86217224 86218358
refGene_gn=B3GNT5;refGene_tx=XM_006936240.3;refGene_gn=B3GNT5;refGene_tx=XM_019840071.1;refGene_gn=B3GNT5;refGene_tx=XM_019840072.1
C2 86251035 86251102 refGene_gn=MCF2L2;refGene_tx=XM_023260362.1
C2 86259141 86259257 refGene_gn=MCF2L2;refGene_tx=XM_023260362.1
C2 86260411 86260481 refGene_gn=MCF2L2;refGene_tx=XM_023260362.1
C2 86266110 86266203 refGene_gn=MCF2L2;refGene_tx=XM_023260362.1
C2 86267193 86267286 refGene_gn=MCF2L2;refGene_tx=XM_023260362.1
C2 86270884 86270956 refGene_gn=MCF2L2;refGene_tx=XM_023260362.1
C2 86275450 86275576 refGene_gn=MCF2L2;refGene_tx=XM_023260362.1
C2 86283036 86283252 refGene_gn=MCF2L2;refGene_tx=XM_023260362.1
C2 86284148 86284241 refGene_gn=MCF2L2;refGene_tx=XM_023260362.1

C2 86284415 86284494 refGene_gn=MCF2L2;refGene_tx=XM_023260362.1
C2 86312604 86312693 refGene_gn=MCF2L2;refGene_tx=XM_023260362.1
C2 86313089 86313203 refGene_gn=MCF2L2;refGene_tx=XM_023260362.1
C2 86322203 86322234 refGene_gn=LAMP3;refGene_tx=XM_019840069.2
C2 86330972 86331021
refGene_gn=LAMP3;refGene_tx=XM_003991887.4;refGene_gn=LAMP3;refGene_tx=XM_019840068.2;refGene_gn=LAMP3;refGene_tx=XM_019840070.2
C2 86338733 86339464
refGene_gn=LAMP3;refGene_tx=XM_003991887.4;refGene_gn=LAMP3;refGene_tx=XM_019840068.2;refGene_gn=LAMP3;refGene_tx=XM_019840069.2;refGene_gn=LAMP3;refGene_tx=XM_019840070.2
C2 86340543 86340672
refGene_gn=LAMP3;refGene_tx=XM_003991887.4;refGene_gn=LAMP3;refGene_tx=XM_019840068.2;refGene_gn=LAMP3;refGene_tx=XM_019840069.2;refGene_gn=LAMP3;refGene_tx=XM_019840070.2
C2 86350488 86350546
refGene_gn=LAMP3;refGene_tx=XM_003991887.4;refGene_gn=LAMP3;refGene_tx=XM_019840068.2;refGene_gn=LAMP3;refGene_tx=XM_019840069.2;refGene_gn=LAMP3;refGene_tx=XM_019840070.2
C2 86353915 86354086
refGene_gn=LAMP3;refGene_tx=XM_003991887.4;refGene_gn=LAMP3;refGene_tx=XM_019840068.2;refGene_gn=LAMP3;refGene_tx=XM_019840069.2;refGene_gn=LAMP3;refGene_tx=XM_019840070.2
C2 86356018 86356158 refGene_gn=LAMP3;refGene_tx=XM_019840070.2
C2 86357954 86358130
refGene_gn=LAMP3;refGene_tx=XM_019840068.2;refGene_gn=LAMP3;refGene_tx=XM_019840069.2
C2 86367305 86367439 refGene_gn=LAMP3;refGene_tx=XM_003991887.4
C2 86374419 86374508
refGene_gn=MCCC1;refGene_tx=XM_003991888.5;refGene_gn=MCCC1;refGene_tx=XM_006936244.3
C2 86378186 86378233
refGene_gn=MCCC1;refGene_tx=XM_003991888.5;refGene_gn=MCCC1;refGene_tx=XM_006936244.3
C2 86379719 86379859
refGene_gn=MCCC1;refGene_tx=XM_003991888.5;refGene_gn=MCCC1;refGene_tx=XM_006936244.3
C2 86383247 86383343
refGene_gn=MCCC1;refGene_tx=XM_003991888.5;refGene_gn=MCCC1;refGene_tx=XM_006936244.3
C2 86387132 86387254
refGene_gn=MCCC1;refGene_tx=XM_003991888.5;refGene_gn=MCCC1;refGene_tx=XM_006936244.3
C2 86387795 86387943
refGene_gn=MCCC1;refGene_tx=XM_003991888.5;refGene_gn=MCCC1;refGene_tx=XM_006936244.3
C2 86388031 86388153
refGene_gn=MCCC1;refGene_tx=XM_003991888.5;refGene_gn=MCCC1;refGene_tx=XM_006936244.3
C2 86394643 86394755

refGene_gn=MCCC1;refGene_tx=XM_003991888.5;refGene_gn=MCCC1;refGene_tx=XM_006936244.3
C2 86396525 86396607
refGene_gn=MCCC1;refGene_tx=XM_003991888.5;refGene_gn=MCCC1;refGene_tx=XM_006936244.3
C2 86403469 86403597
refGene_gn=MCCC1;refGene_tx=XM_003991888.5;refGene_gn=MCCC1;refGene_tx=XM_006936244.3
C2 86407233 86407417
refGene_gn=MCCC1;refGene_tx=XM_003991888.5;refGene_gn=MCCC1;refGene_tx=XM_006936244.3
C2 86409999 86410109
refGene_gn=MCCC1;refGene_tx=XM_003991888.5;refGene_gn=MCCC1;refGene_tx=XM_006936244.3
C2 86413811 86414028
refGene_gn=MCCC1;refGene_tx=XM_003991888.5;refGene_gn=MCCC1;refGene_tx=XM_006936244.3
C2 86416017 86416104
refGene_gn=MCCC1;refGene_tx=XM_003991888.5;refGene_gn=MCCC1;refGene_tx=XM_006936244.3
C2 86422256 86422306
refGene_gn=MCCC1;refGene_tx=XM_003991888.5;refGene_gn=MCCC1;refGene_tx=XM_006936244.3
C2 86425464 86425602
refGene_gn=MCCC1;refGene_tx=XM_003991888.5;refGene_gn=MCCC1;refGene_tx=XM_006936244.3
C2 86426916 86427024
refGene_gn=MCCC1;refGene_tx=XM_003991888.5;refGene_gn=MCCC1;refGene_tx=XM_006936244.3
C2 86429347 86429419
refGene_gn=MCCC1;refGene_tx=XM_003991888.5;refGene_gn=MCCC1;refGene_tx=XM_006936244.3
C2 86430194 86430323
refGene_gn=MCCC1;refGene_tx=XM_003991888.5;refGene_gn=MCCC1;refGene_tx=XM_006936244.3
C2 86454303 86454315 refGene_gn=DCUN1D1;refGene_tx=XM_006936245.4
C2 86458860 86458863 refGene_gn=DCUN1D1;refGene_tx=XM_023260363.1
C2 86470299 86470516
refGene_gn=DCUN1D1;refGene_tx=XM_006936245.4;refGene_gn=DCUN1D1;refGene_tx=XM_006936246.4;refGene_gn=DCUN1D1;refGene_tx=XM_023260363.1
C2 86471966 86472135
refGene_gn=DCUN1D1;refGene_tx=XM_006936245.4;refGene_gn=DCUN1D1;refGene_tx=XM_006936246.4;refGene_gn=DCUN1D1;refGene_tx=XM_023260363.1
C2 86474368 86474499
refGene_gn=DCUN1D1;refGene_tx=XM_006936245.4;refGene_gn=DCUN1D1;refGene_tx=XM_006936246.4;refGene_gn=DCUN1D1;refGene_tx=XM_023260363.1
C2 86484060 86484143
refGene_gn=DCUN1D1;refGene_tx=XM_006936245.4;refGene_gn=DCUN1D1;refGene_tx=XM_006936246.4;refGene_gn=DCUN1D1;refGene_tx=XM_023260363.1
C2 86484602 86484699

refGene_gn=DCUN1D1;refGene_tx=XM_006936245.4;refGene_gn=DCUN1D1;refGene_tx=XM_006936246.4;refGene_gn=DCUN1D1;refGene_tx=XM_023260363.1
C2	86485997	86486077
refGene_gn=DCUN1D1;refGene_tx=XM_006936245.4;refGene_gn=DCUN1D1;refGene_tx=XM_006936246.4;refGene_gn=DCUN1D1;refGene_tx=XM_023260363.1
C2	86532645	86532727
refGene_gn=ATP11B;refGene_tx=XM_023260364.1;refGene_gn=ATP11B;refGene_tx=XM_023260368.1;refGene_gn=ATP11B;refGene_tx=XM_023260369.1
C2	86537542	86537676
refGene_gn=ATP11B;refGene_tx=XM_023260364.1;refGene_gn=ATP11B;refGene_tx=XM_023260365.1;refGene_gn=ATP11B;refGene_tx=XM_023260368.1;refGene_gn=ATP11B;refGene_tx=XM_023260369.1
C2	86545576	86545702
refGene_gn=ATP11B;refGene_tx=XM_023260366.1;refGene_gn=ATP11B;refGene_tx=XM_023260367.1
C2	86548657	86548743
refGene_gn=ATP11B;refGene_tx=XM_023260365.1;refGene_gn=ATP11B;refGene_tx=XM_023260366.1
C2	86551178	86551344
refGene_gn=ATP11B;refGene_tx=XM_023260364.1;refGene_gn=ATP11B;refGene_tx=XM_023260365.1;refGene_gn=ATP11B;refGene_tx=XM_023260366.1;refGene_gn=ATP11B;refGene_tx=XM_023260367.1;refGene_gn=ATP11B;refGene_tx=XM_023260368.1;refGene_gn=ATP11B;refGene_tx=XM_023260369.1
C2	86552877	86552981
refGene_gn=ATP11B;refGene_tx=XM_023260364.1;refGene_gn=ATP11B;refGene_tx=XM_023260365.1;refGene_gn=ATP11B;refGene_tx=XM_023260366.1;refGene_gn=ATP11B;refGene_tx=XM_023260367.1;refGene_gn=ATP11B;refGene_tx=XM_023260368.1;refGene_gn=ATP11B;refGene_tx=XM_023260369.1
C2	86553515	86553581
refGene_gn=ATP11B;refGene_tx=XM_023260364.1;refGene_gn=ATP11B;refGene_tx=XM_023260365.1;refGene_gn=ATP11B;refGene_tx=XM_023260366.1;refGene_gn=ATP11B;refGene_tx=XM_023260367.1;refGene_gn=ATP11B;refGene_tx=XM_023260368.1;refGene_gn=ATP11B;refGene_tx=XM_023260369.1
C2	86556114	86556138
refGene_gn=ATP11B;refGene_tx=XM_023260365.1;refGene_gn=ATP11B;refGene_tx=XM_023260366.1;refGene_gn=ATP11B;refGene_tx=XM_023260367.1;refGene_gn=ATP11B;refGene_tx=XM_023260368.1;refGene_gn=ATP11B;refGene_tx=XM_023260369.1
C2	86563479	86563618
refGene_gn=ATP11B;refGene_tx=XM_023260364.1;refGene_gn=ATP11B;refGene_tx=XM_023260365.1;refGene_gn=ATP11B;refGene_tx=XM_023260366.1;refGene_gn=ATP11B;refGene_tx=XM_023260367.1;refGene_gn=ATP11B;refGene_tx=XM_023260368.1;refGene_gn=ATP11B;refGene_tx=XM_023260369.1
C2	86564679	86564807
refGene_gn=ATP11B;refGene_tx=XM_023260364.1;refGene_gn=ATP11B;refGene_tx=XM_023260365.1;refGene_gn=ATP11B;refGene_tx=XM_023260366.1;refGene_gn=ATP11B;refGene_tx=XM_023260367.1;refGene_gn=ATP11B;refGene_tx=XM_023260368.1;refGene_gn=ATP11B;refGene_tx=XM_023260369.1
C2	86566374	86566434
refGene_gn=ATP11B;refGene_tx=XM_023260364.1;refGene_gn=ATP11B;refGene_

tx=XM_023260365.1;refGene_gn=ATP11B;refGene_tx=XM_023260366.1;refGene_gn=ATP11B;refGene_tx=XM_023260367.1;refGene_gn=ATP11B;refGene_tx=XM_023260368.1;refGene_gn=ATP11B;refGene_tx=XM_023260369.1
C2 86568339 86568485
refGene_gn=ATP11B;refGene_tx=XM_023260364.1;refGene_gn=ATP11B;refGene_tx=XM_023260365.1;refGene_gn=ATP11B;refGene_tx=XM_023260366.1;refGene_gn=ATP11B;refGene_tx=XM_023260367.1;refGene_gn=ATP11B;refGene_tx=XM_023260368.1;refGene_gn=ATP11B;refGene_tx=XM_023260369.1
C2 86572447 86572550
refGene_gn=ATP11B;refGene_tx=XM_023260364.1;refGene_gn=ATP11B;refGene_tx=XM_023260365.1;refGene_gn=ATP11B;refGene_tx=XM_023260366.1;refGene_gn=ATP11B;refGene_tx=XM_023260367.1;refGene_gn=ATP11B;refGene_tx=XM_023260368.1;refGene_gn=ATP11B;refGene_tx=XM_023260369.1
C2 86573751 86573905
refGene_gn=ATP11B;refGene_tx=XM_023260364.1;refGene_gn=ATP11B;refGene_tx=XM_023260365.1;refGene_gn=ATP11B;refGene_tx=XM_023260366.1;refGene_gn=ATP11B;refGene_tx=XM_023260367.1;refGene_gn=ATP11B;refGene_tx=XM_023260368.1;refGene_gn=ATP11B;refGene_tx=XM_023260369.1
C2 86578737 86578941
refGene_gn=ATP11B;refGene_tx=XM_023260364.1;refGene_gn=ATP11B;refGene_tx=XM_023260365.1;refGene_gn=ATP11B;refGene_tx=XM_023260366.1;refGene_gn=ATP11B;refGene_tx=XM_023260367.1;refGene_gn=ATP11B;refGene_tx=XM_023260368.1;refGene_gn=ATP11B;refGene_tx=XM_023260369.1
C2 86580090 86580272
refGene_gn=ATP11B;refGene_tx=XM_023260364.1;refGene_gn=ATP11B;refGene_tx=XM_023260365.1;refGene_gn=ATP11B;refGene_tx=XM_023260366.1;refGene_gn=ATP11B;refGene_tx=XM_023260367.1;refGene_gn=ATP11B;refGene_tx=XM_023260368.1;refGene_gn=ATP11B;refGene_tx=XM_023260369.1
C2 86582608 86582712
refGene_gn=ATP11B;refGene_tx=XM_023260364.1;refGene_gn=ATP11B;refGene_tx=XM_023260365.1;refGene_gn=ATP11B;refGene_tx=XM_023260366.1;refGene_gn=ATP11B;refGene_tx=XM_023260367.1;refGene_gn=ATP11B;refGene_tx=XM_023260368.1;refGene_gn=ATP11B;refGene_tx=XM_023260369.1
C2 86582785 86582859
refGene_gn=ATP11B;refGene_tx=XM_023260364.1;refGene_gn=ATP11B;refGene_tx=XM_023260365.1;refGene_gn=ATP11B;refGene_tx=XM_023260366.1;refGene_gn=ATP11B;refGene_tx=XM_023260367.1;refGene_gn=ATP11B;refGene_tx=XM_023260368.1;refGene_gn=ATP11B;refGene_tx=XM_023260369.1
C2 86584127 86584196
refGene_gn=ATP11B;refGene_tx=XM_023260364.1;refGene_gn=ATP11B;refGene_tx=XM_023260365.1;refGene_gn=ATP11B;refGene_tx=XM_023260366.1;refGene_gn=ATP11B;refGene_tx=XM_023260367.1;refGene_gn=ATP11B;refGene_tx=XM_023260368.1;refGene_gn=ATP11B;refGene_tx=XM_023260369.1
C2 86585070 86585246
refGene_gn=ATP11B;refGene_tx=XM_023260364.1;refGene_gn=ATP11B;refGene_tx=XM_023260365.1;refGene_gn=ATP11B;refGene_tx=XM_023260366.1;refGene_gn=ATP11B;refGene_tx=XM_023260367.1;refGene_gn=ATP11B;refGene_tx=XM_023260368.1;refGene_gn=ATP11B;refGene_tx=XM_023260369.1
C2 86585825 86586068
refGene_gn=ATP11B;refGene_tx=XM_023260364.1;refGene_gn=ATP11B;refGene_

tx=XM_023260365.1;refGene_gn=ATP11B;refGene_tx=XM_023260366.1;refGene_gn=ATP11B;refGene_tx=XM_023260367.1;refGene_gn=ATP11B;refGene_tx=XM_023260368.1;refGene_gn=ATP11B;refGene_tx=XM_023260369.1
C2 86590883 86591081
refGene_gn=ATP11B;refGene_tx=XM_023260364.1;refGene_gn=ATP11B;refGene_tx=XM_023260365.1;refGene_gn=ATP11B;refGene_tx=XM_023260366.1;refGene_gn=ATP11B;refGene_tx=XM_023260367.1;refGene_gn=ATP11B;refGene_tx=XM_023260368.1;refGene_gn=ATP11B;refGene_tx=XM_023260369.1
C2 86592198 86592349
refGene_gn=ATP11B;refGene_tx=XM_023260364.1;refGene_gn=ATP11B;refGene_tx=XM_023260365.1;refGene_gn=ATP11B;refGene_tx=XM_023260366.1;refGene_gn=ATP11B;refGene_tx=XM_023260367.1;refGene_gn=ATP11B;refGene_tx=XM_023260368.1;refGene_gn=ATP11B;refGene_tx=XM_023260369.1
C2 86605463 86605545
refGene_gn=ATP11B;refGene_tx=XM_023260364.1;refGene_gn=ATP11B;refGene_tx=XM_023260365.1;refGene_gn=ATP11B;refGene_tx=XM_023260366.1;refGene_gn=ATP11B;refGene_tx=XM_023260367.1;refGene_gn=ATP11B;refGene_tx=XM_023260368.1;refGene_gn=ATP11B;refGene_tx=XM_023260369.1
C2 86608587 86608652
refGene_gn=ATP11B;refGene_tx=XM_023260364.1;refGene_gn=ATP11B;refGene_tx=XM_023260365.1;refGene_gn=ATP11B;refGene_tx=XM_023260366.1;refGene_gn=ATP11B;refGene_tx=XM_023260367.1;refGene_gn=ATP11B;refGene_tx=XM_023260368.1;refGene_gn=ATP11B;refGene_tx=XM_023260369.1
C2 86609165 86609167 refGene_gn=ATP11B;refGene_tx=XM_023260369.1
C2 86610835 86610883
refGene_gn=ATP11B;refGene_tx=XM_023260364.1;refGene_gn=ATP11B;refGene_tx=XM_023260365.1;refGene_gn=ATP11B;refGene_tx=XM_023260366.1;refGene_gn=ATP11B;refGene_tx=XM_023260367.1;refGene_gn=ATP11B;refGene_tx=XM_023260368.1
C2 86613796 86613900
refGene_gn=ATP11B;refGene_tx=XM_023260364.1;refGene_gn=ATP11B;refGene_tx=XM_023260365.1;refGene_gn=ATP11B;refGene_tx=XM_023260366.1;refGene_gn=ATP11B;refGene_tx=XM_023260367.1;refGene_gn=ATP11B;refGene_tx=XM_023260368.1
C2 86614745 86614874
refGene_gn=ATP11B;refGene_tx=XM_023260364.1;refGene_gn=ATP11B;refGene_tx=XM_023260365.1;refGene_gn=ATP11B;refGene_tx=XM_023260366.1;refGene_gn=ATP11B;refGene_tx=XM_023260367.1;refGene_gn=ATP11B;refGene_tx=XM_023260368.1
C2 86615064 86615172
refGene_gn=ATP11B;refGene_tx=XM_023260364.1;refGene_gn=ATP11B;refGene_tx=XM_023260365.1;refGene_gn=ATP11B;refGene_tx=XM_023260366.1;refGene_gn=ATP11B;refGene_tx=XM_023260367.1;refGene_gn=ATP11B;refGene_tx=XM_023260368.1
C2 86621649 86621730
refGene_gn=ATP11B;refGene_tx=XM_023260364.1;refGene_gn=ATP11B;refGene_tx=XM_023260365.1;refGene_gn=ATP11B;refGene_tx=XM_023260366.1;refGene_gn=ATP11B;refGene_tx=XM_023260367.1;refGene_gn=ATP11B;refGene_tx=XM_023260368.1
C2 86623339 86623429

refGene_gn=ATP11B;refGene_tx=XM_023260364.1;refGene_gn=ATP11B;refGene_tx=XM_023260365.1;refGene_gn=ATP11B;refGene_tx=XM_023260366.1;refGene_gn=ATP11B;refGene_tx=XM_023260367.1;refGene_gn=ATP11B;refGene_tx=XM_023260368.1
C2 86628279 86628396
refGene_gn=ATP11B;refGene_tx=XM_023260364.1;refGene_gn=ATP11B;refGene_tx=XM_023260365.1;refGene_gn=ATP11B;refGene_tx=XM_023260366.1;refGene_gn=ATP11B;refGene_tx=XM_023260367.1;refGene_gn=ATP11B;refGene_tx=XM_023260368.1
C2 86656913 86656940
refGene_gn=ATP11B;refGene_tx=XM_023260364.1;refGene_gn=ATP11B;refGene_tx=XM_023260365.1;refGene_gn=ATP11B;refGene_tx=XM_023260366.1;refGene_gn=ATP11B;refGene_tx=XM_023260367.1;refGene_gn=ATP11B;refGene_tx=XM_023260368.1
C2 87648708 87649674 refGene_gn=SOX2;refGene_tx=NM_001173447.1
C2 88344853 88344856 refGene_gn=DNAJC19;refGene_tx=XM_004001179.3
C2 88345387 88345462
refGene_gn=DNAJC19;refGene_tx=XM_019840077.1;refGene_gn=DNAJC19;refGene_tx=XM_019840078.1
C2 88345692 88345744
refGene_gn=DNAJC19;refGene_tx=XM_004001179.3;refGene_gn=DNAJC19;refGene_tx=XM_019840077.1;refGene_gn=DNAJC19;refGene_tx=XM_019840078.1
C2 88345823 88345897
refGene_gn=DNAJC19;refGene_tx=XM_004001179.3;refGene_gn=DNAJC19;refGene_tx=XM_019840077.1;refGene_gn=DNAJC19;refGene_tx=XM_019840078.1
C2 88346968 88347048
refGene_gn=DNAJC19;refGene_tx=XM_004001179.3;refGene_gn=DNAJC19;refGene_tx=XM_019840077.1;refGene_gn=DNAJC19;refGene_tx=XM_019840078.1
C2 88347594 88347665
refGene_gn=DNAJC19;refGene_tx=XM_004001179.3;refGene_gn=DNAJC19;refGene_tx=XM_019840077.1;refGene_gn=DNAJC19;refGene_tx=XM_019840078.1
C2 88349158 88349229
refGene_gn=DNAJC19;refGene_tx=XM_004001179.3;refGene_gn=DNAJC19;refGene_tx=XM_019840077.1;refGene_gn=DNAJC19;refGene_tx=XM_019840078.1
C2 88357807 88357978 refGene_gn=FXR1;refGene_tx=XM_006944239.4
C2 88358743 88358835 refGene_gn=FXR1;refGene_tx=XM_006944239.4
C2 88362352 88362433 refGene_gn=FXR1;refGene_tx=XM_006944239.4
C2 88363159 88363360 refGene_gn=FXR1;refGene_tx=XM_006944239.4
C2 88365136 88365340 refGene_gn=FXR1;refGene_tx=XM_006944239.4
C2 88370288 88370351 refGene_gn=FXR1;refGene_tx=XM_006944239.4
C2 88370438 88370496 refGene_gn=FXR1;refGene_tx=XM_006944239.4
C2 88371910 88371997 refGene_gn=FXR1;refGene_tx=XM_006944239.4
C2 88376780 88376890 refGene_gn=FXR1;refGene_tx=XM_006944239.4
C2 88380522 88380601 refGene_gn=FXR1;refGene_tx=XM_006944239.4
C2 88381869 88382040 refGene_gn=FXR1;refGene_tx=XM_006944239.4
C2 88383730 88383847 refGene_gn=FXR1;refGene_tx=XM_006944239.4
C2 88384273 88384311 refGene_gn=FXR1;refGene_tx=XM_006944239.4
C2 88384313 88384366 refGene_gn=FXR1;refGene_tx=XM_006944239.4
C2 88384592 88384741 refGene_gn=FXR1;refGene_tx=XM_006944239.4
C2 88385161 88385233 refGene_gn=FXR1;refGene_tx=XM_006944239.4

C2 88395990 88396084 refGene_gn=FXR1;refGene_tx=XM_006944239.4
C2 88397672 88397725 refGene_gn=FXR1;refGene_tx=XM_006944239.4
C2 88417442 88417493 refGene_gn=FXR1;refGene_tx=XM_006944239.4
C2 88592821 88592911 refGene_gn=CCDC39;refGene_tx=XM_004001176.5
C2 88600959 88601079 refGene_gn=CCDC39;refGene_tx=XM_004001176.5
C2 88602926 88603073 refGene_gn=CCDC39;refGene_tx=XM_004001176.5
C2 88604269 88604428 refGene_gn=CCDC39;refGene_tx=XM_004001176.5
C2 88605183 88605276 refGene_gn=CCDC39;refGene_tx=XM_004001176.5
C2 88605372 88605501 refGene_gn=CCDC39;refGene_tx=XM_004001176.5
C2 88609123 88609315 refGene_gn=CCDC39;refGene_tx=XM_004001176.5
C2 88612052 88612156 refGene_gn=CCDC39;refGene_tx=XM_004001176.5
C2 88612488 88612621 refGene_gn=CCDC39;refGene_tx=XM_004001176.5
C2 88615940 88616135 refGene_gn=CCDC39;refGene_tx=XM_004001176.5
C2 88617073 88617238 refGene_gn=CCDC39;refGene_tx=XM_004001176.5
C2 88619935 88620073 refGene_gn=CCDC39;refGene_tx=XM_004001176.5
C2 88622321 88622530 refGene_gn=CCDC39;refGene_tx=XM_004001176.5
C2 88629967 88630091 refGene_gn=CCDC39;refGene_tx=XM_004001176.5
C2 88633917 88634077 refGene_gn=CCDC39;refGene_tx=XM_004001176.5
C2 88634495 88634602 refGene_gn=CCDC39;refGene_tx=XM_004001176.5
C2 88635116 88635257 refGene_gn=CCDC39;refGene_tx=XM_004001176.5
C2 88635407 88635587 refGene_gn=CCDC39;refGene_tx=XM_004001176.5
C2 88635727 88635813 refGene_gn=CCDC39;refGene_tx=XM_004001176.5
C2 88637068 88637225 refGene_gn=CCDC39;refGene_tx=XM_004001176.5
C2 88640586 88641499
refGene_gn=TTC14;refGene_tx=XM_004001175.5;refGene_gn=TTC14;refGene_tx
=XM_019840079.2
C2 88642331 88642441
refGene_gn=TTC14;refGene_tx=XM_004001175.5;refGene_gn=TTC14;refGene_tx
=XM_019840079.2
C2 88643364 88643482
refGene_gn=TTC14;refGene_tx=XM_004001175.5;refGene_gn=TTC14;refGene_tx
=XM_019840079.2
C2 88645693 88645816
refGene_gn=TTC14;refGene_tx=XM_004001175.5;refGene_gn=TTC14;refGene_tx
=XM_019840079.2
C2 88645919 88646039
refGene_gn=TTC14;refGene_tx=XM_004001175.5;refGene_gn=TTC14;refGene_tx
=XM_019840079.2
C2 88646682 88646754
refGene_gn=TTC14;refGene_tx=XM_004001175.5;refGene_gn=TTC14;refGene_tx
=XM_019840079.2
C2 88647152 88647308
refGene_gn=TTC14;refGene_tx=XM_004001175.5;refGene_gn=TTC14;refGene_tx
=XM_019840079.2
C2 88647588 88647718
refGene_gn=TTC14;refGene_tx=XM_004001175.5;refGene_gn=TTC14;refGene_tx
=XM_019840079.2
C2 88647888 88647973
refGene_gn=TTC14;refGene_tx=XM_004001175.5;refGene_gn=TTC14;refGene_tx
=XM_019840079.2

C2 88648909 88649109
refGene_gn=TTC14;refGene_tx=XM_004001175.5;refGene_gn=TTC14;refGene_tx=XM_019840079.2
C2 88649214 88649339 refGene_gn=TTC14;refGene_tx=XM_004001175.5
C2 88649841 88650002 refGene_gn=TTC14;refGene_tx=XM_004001175.5
C2 89159250 89159271
refGene_gn=PEX5L;refGene_tx=XM_011286056.3;refGene_gn=PEX5L;refGene_tx=XM_011286057.3;refGene_gn=PEX5L;refGene_tx=XM_011286058.3;refGene_gn=PEX5L;refGene_tx=XM_019840081.2
C2 89217529 89217622
refGene_gn=PEX5L;refGene_tx=XM_006944231.4;refGene_gn=PEX5L;refGene_tx=XM_019840080.2;refGene_gn=PEX5L;refGene_tx=XM_023260370.1;refGene_gn=PEX5L;refGene_tx=XM_023260371.1
C2 89219127 89219199
refGene_gn=PEX5L;refGene_tx=XM_006944231.4;refGene_gn=PEX5L;refGene_tx=XM_011286056.3;refGene_gn=PEX5L;refGene_tx=XM_019840080.2;refGene_gn=PEX5L;refGene_tx=XM_019840081.2;refGene_gn=PEX5L;refGene_tx=XM_023260370.1;refGene_gn=PEX5L;refGene_tx=XM_023260371.1
C2 89283830 89283896 refGene_gn=PEX5L;refGene_tx=XM_023260371.1
C2 89286206 89286311
refGene_gn=PEX5L;refGene_tx=XM_006944231.4;refGene_gn=PEX5L;refGene_tx=XM_011286056.3;refGene_gn=PEX5L;refGene_tx=XM_011286057.3;refGene_gn=PEX5L;refGene_tx=XM_019840082.2;refGene_gn=PEX5L;refGene_tx=XM_023260371.1
C2 89294861 89294973
refGene_gn=PEX5L;refGene_tx=XM_006944231.4;refGene_gn=PEX5L;refGene_tx=XM_011286056.3;refGene_gn=PEX5L;refGene_tx=XM_011286057.3;refGene_gn=PEX5L;refGene_tx=XM_011286058.3;refGene_gn=PEX5L;refGene_tx=XM_019840080.2;refGene_gn=PEX5L;refGene_tx=XM_019840081.2;refGene_gn=PEX5L;refGene_tx=XM_019840082.2;refGene_gn=PEX5L;refGene_tx=XM_023260370.1;refGene_gn=PEX5L;refGene_tx=XM_023260371.1
C2 89302885 89303080
refGene_gn=PEX5L;refGene_tx=XM_006944231.4;refGene_gn=PEX5L;refGene_tx=XM_011286056.3;refGene_gn=PEX5L;refGene_tx=XM_011286057.3;refGene_gn=PEX5L;refGene_tx=XM_011286058.3;refGene_gn=PEX5L;refGene_tx=XM_019840080.2;refGene_gn=PEX5L;refGene_tx=XM_019840081.2;refGene_gn=PEX5L;refGene_tx=XM_019840082.2;refGene_gn=PEX5L;refGene_tx=XM_023260370.1;refGene_gn=PEX5L;refGene_tx=XM_023260371.1
C2 89306599 89306723
refGene_gn=PEX5L;refGene_tx=XM_006944231.4;refGene_gn=PEX5L;refGene_tx=XM_011286056.3;refGene_gn=PEX5L;refGene_tx=XM_011286057.3;refGene_gn=PEX5L;refGene_tx=XM_011286058.3;refGene_gn=PEX5L;refGene_tx=XM_019840080.2;refGene_gn=PEX5L;refGene_tx=XM_019840081.2;refGene_gn=PEX5L;refGene_tx=XM_019840082.2;refGene_gn=PEX5L;refGene_tx=XM_023260370.1;refGene_gn=PEX5L;refGene_tx=XM_023260371.1
C2 89307629 89307726
refGene_gn=PEX5L;refGene_tx=XM_006944231.4;refGene_gn=PEX5L;refGene_tx=XM_011286056.3;refGene_gn=PEX5L;refGene_tx=XM_011286057.3;refGene_gn=PEX5L;refGene_tx=XM_011286058.3;refGene_gn=PEX5L;refGene_tx=XM_019840080.2;refGene_gn=PEX5L;refGene_tx=XM_019840081.2;refGene_gn=PEX5L;refGe

ne_tx=XM_019840082.2;refGene_gn=PEX5L;refGene_tx=XM_023260370.1;refGene_gn=PEX5L;refGene_tx=XM_023260371.1
C2 89318429 89318525
refGene_gn=PEX5L;refGene_tx=XM_006944231.4;refGene_gn=PEX5L;refGene_tx=XM_011286056.3;refGene_gn=PEX5L;refGene_tx=XM_011286057.3;refGene_gn=PEX5L;refGene_tx=XM_011286058.3;refGene_gn=PEX5L;refGene_tx=XM_019840080.2;refGene_gn=PEX5L;refGene_tx=XM_019840081.2;refGene_gn=PEX5L;refGene_tx=XM_019840082.2;refGene_gn=PEX5L;refGene_tx=XM_023260370.1;refGene_gn=PEX5L;refGene_tx=XM_023260371.1
C2 89365945 89366062
refGene_gn=PEX5L;refGene_tx=XM_006944231.4;refGene_gn=PEX5L;refGene_tx=XM_011286056.3;refGene_gn=PEX5L;refGene_tx=XM_011286057.3;refGene_gn=PEX5L;refGene_tx=XM_011286058.3;refGene_gn=PEX5L;refGene_tx=XM_019840080.2;refGene_gn=PEX5L;refGene_tx=XM_019840081.2;refGene_gn=PEX5L;refGene_tx=XM_019840082.2;refGene_gn=PEX5L;refGene_tx=XM_023260370.1;refGene_gn=PEX5L;refGene_tx=XM_023260371.1
C2 89368551 89368695
refGene_gn=PEX5L;refGene_tx=XM_006944231.4;refGene_gn=PEX5L;refGene_tx=XM_011286056.3;refGene_gn=PEX5L;refGene_tx=XM_011286057.3;refGene_gn=PEX5L;refGene_tx=XM_011286058.3;refGene_gn=PEX5L;refGene_tx=XM_019840080.2;refGene_gn=PEX5L;refGene_tx=XM_019840081.2;refGene_gn=PEX5L;refGene_tx=XM_019840082.2;refGene_gn=PEX5L;refGene_tx=XM_023260370.1;refGene_gn=PEX5L;refGene_tx=XM_023260371.1
C2 89371553 89371624
refGene_gn=PEX5L;refGene_tx=XM_006944231.4;refGene_gn=PEX5L;refGene_tx=XM_011286056.3;refGene_gn=PEX5L;refGene_tx=XM_011286057.3;refGene_gn=PEX5L;refGene_tx=XM_011286058.3;refGene_gn=PEX5L;refGene_tx=XM_019840080.2;refGene_gn=PEX5L;refGene_tx=XM_019840081.2;refGene_gn=PEX5L;refGene_tx=XM_019840082.2;refGene_gn=PEX5L;refGene_tx=XM_023260370.1;refGene_gn=PEX5L;refGene_tx=XM_023260371.1
C2 89373150 89373348
refGene_gn=PEX5L;refGene_tx=XM_006944231.4;refGene_gn=PEX5L;refGene_tx=XM_011286056.3;refGene_gn=PEX5L;refGene_tx=XM_011286057.3;refGene_gn=PEX5L;refGene_tx=XM_011286058.3;refGene_gn=PEX5L;refGene_tx=XM_019840080.2;refGene_gn=PEX5L;refGene_tx=XM_019840081.2;refGene_gn=PEX5L;refGene_tx=XM_019840082.2;refGene_gn=PEX5L;refGene_tx=XM_023260370.1;refGene_gn=PEX5L;refGene_tx=XM_023260371.1
C2 89374012 89374178
refGene_gn=PEX5L;refGene_tx=XM_006944231.4;refGene_gn=PEX5L;refGene_tx=XM_011286056.3;refGene_gn=PEX5L;refGene_tx=XM_011286057.3;refGene_gn=PEX5L;refGene_tx=XM_011286058.3;refGene_gn=PEX5L;refGene_tx=XM_019840080.2;refGene_gn=PEX5L;refGene_tx=XM_019840081.2;refGene_gn=PEX5L;refGene_tx=XM_019840082.2;refGene_gn=PEX5L;refGene_tx=XM_023260370.1;refGene_gn=PEX5L;refGene_tx=XM_023260371.1
C2 89374567 89374729
refGene_gn=PEX5L;refGene_tx=XM_006944231.4;refGene_gn=PEX5L;refGene_tx=XM_011286056.3;refGene_gn=PEX5L;refGene_tx=XM_011286057.3;refGene_gn=PEX5L;refGene_tx=XM_011286058.3;refGene_gn=PEX5L;refGene_tx=XM_019840080.2;refGene_gn=PEX5L;refGene_tx=XM_019840081.2;refGene_gn=PEX5L;refGene_tx=XM_019840082.2;refGene_gn=PEX5L;refGene_tx=XM_023260370.1;refGen

e_gn=PEX5L;refGene_tx=XM_023260371.1
C2 89380836 89381041
refGene_gn=PEX5L;refGene_tx=XM_006944231.4;refGene_gn=PEX5L;refGene_tx=XM_011286056.3;refGene_gn=PEX5L;refGene_tx=XM_011286057.3;refGene_gn=PEX5L;refGene_tx=XM_011286058.3;refGene_gn=PEX5L;refGene_tx=XM_019840080.2;refGene_gn=PEX5L;refGene_tx=XM_019840081.2;refGene_gn=PEX5L;refGene_tx=XM_019840082.2;refGene_gn=PEX5L;refGene_tx=XM_023260371.1
C2 89398174 89398268 refGene_gn=USP13;refGene_tx=XM_023260372.1
C2 89399914 89399999 refGene_gn=USP13;refGene_tx=XM_023260372.1
C2 89413326 89413480 refGene_gn=USP13;refGene_tx=XM_023260372.1
C2 89415037 89415204 refGene_gn=USP13;refGene_tx=XM_023260372.1
C2 89418054 89418198 refGene_gn=USP13;refGene_tx=XM_023260372.1
C2 89421286 89421313 refGene_gn=USP13;refGene_tx=XM_023260372.1
C2 89422926 89423049 refGene_gn=USP13;refGene_tx=XM_023260372.1
C2 89425387 89425476 refGene_gn=USP13;refGene_tx=XM_023260372.1
C2 89431983 89432158 refGene_gn=USP13;refGene_tx=XM_023260372.1
C2 89434541 89434695 refGene_gn=USP13;refGene_tx=XM_023260372.1
C2 89436419 89436545 refGene_gn=USP13;refGene_tx=XM_023260372.1
C2 89446321 89446396 refGene_gn=USP13;refGene_tx=XM_023260372.1
C2 89446397 89446416 refGene_gn=USP13;refGene_tx=XM_023260372.1
C2 89447031 89447103 refGene_gn=USP13;refGene_tx=XM_023260372.1
C2 89451995 89452183 refGene_gn=USP13;refGene_tx=XM_023260372.1
C2 89453507 89453602 refGene_gn=USP13;refGene_tx=XM_023260372.1
C2 89460522 89460707 refGene_gn=USP13;refGene_tx=XM_023260372.1
C2 89462335 89462478 refGene_gn=USP13;refGene_tx=XM_023260372.1
C2 89468124 89468246 refGene_gn=USP13;refGene_tx=XM_023260372.1
C2 89477541 89477602 refGene_gn=USP13;refGene_tx=XM_023260372.1
C2 89489239 89489365 refGene_gn=USP13;refGene_tx=XM_023260372.1
C2 89510960 89511128 refGene_gn=USP13;refGene_tx=XM_023260372.1
C2 89538778 89538899 refGene_gn=NDUFB5;refGene_tx=XM_004001169.4
C2 89542472 89542579 refGene_gn=NDUFB5;refGene_tx=XM_004001169.4
C2 89543793 89543855 refGene_gn=NDUFB5;refGene_tx=XM_004001169.4
C2 89545191 89545258 refGene_gn=NDUFB5;refGene_tx=XM_004001169.4
C2 89546312 89546401 refGene_gn=NDUFB5;refGene_tx=XM_004001169.4
C2 89552759 89552883 refGene_gn=NDUFB5;refGene_tx=XM_004001169.4
C2 89553080 89553178 refGene_gn=MRPL47;refGene_tx=XM_006944236.3
C2 89555753 89555883 refGene_gn=MRPL47;refGene_tx=XM_006944236.3
C2 89555884 89555906 refGene_gn=MRPL47;refGene_tx=XM_006944236.3
C2 89556475 89556536 refGene_gn=MRPL47;refGene_tx=XM_006944236.3
C2 89575135 89575232 refGene_gn=MRPL47;refGene_tx=XM_006944236.3
C2 89580662 89580793 refGene_gn=MRPL47;refGene_tx=XM_006944236.3
C2 89583625 89583721 refGene_gn=MRPL47;refGene_tx=XM_006944236.3
C2 89587923 89588047 refGene_gn=MRPL47;refGene_tx=XM_006944236.3
C2 89588855 89588936
refGene_gn=ACTL6A;refGene_tx=XM_004001172.4;refGene_gn=ACTL6A;refGene_tx=XM_019840084.2;refGene_gn=ACTL6A;refGene_tx=XM_023260373.1
C2 89591042 89591129
refGene_gn=ACTL6A;refGene_tx=XM_004001172.4;refGene_gn=ACTL6A;refGene_tx=XM_019840084.2;refGene_gn=ACTL6A;refGene_tx=XM_023260373.1
C2 89594545 89594641

refGene_gn=ACTL6A;refGene_tx=XM_004001172.4;refGene_gn=ACTL6A;refGene_tx=XM_019840084.2;refGene_gn=ACTL6A;refGene_tx=XM_023260373.1
C2 89596363 89596444
refGene_gn=ACTL6A;refGene_tx=XM_004001172.4;refGene_gn=ACTL6A;refGene_tx=XM_019840084.2;refGene_gn=ACTL6A;refGene_tx=XM_023260373.1
C2 89596599 89596714
refGene_gn=ACTL6A;refGene_tx=XM_004001172.4;refGene_gn=ACTL6A;refGene_tx=XM_019840084.2;refGene_gn=ACTL6A;refGene_tx=XM_023260373.1
C2 89596990 89597052
refGene_gn=ACTL6A;refGene_tx=XM_004001172.4;refGene_gn=ACTL6A;refGene_tx=XM_019840084.2;refGene_gn=ACTL6A;refGene_tx=XM_023260373.1
C2 89598647 89598737
refGene_gn=ACTL6A;refGene_tx=XM_004001172.4;refGene_gn=ACTL6A;refGene_tx=XM_019840084.2;refGene_gn=ACTL6A;refGene_tx=XM_023260373.1
C2 89598834 89598941
refGene_gn=ACTL6A;refGene_tx=XM_004001172.4;refGene_gn=ACTL6A;refGene_tx=XM_019840084.2;refGene_gn=ACTL6A;refGene_tx=XM_023260373.1
C2 89599261 89599356
refGene_gn=ACTL6A;refGene_tx=XM_004001172.4;refGene_gn=ACTL6A;refGene_tx=XM_019840084.2;refGene_gn=ACTL6A;refGene_tx=XM_023260373.1
C2 89602384 89602482
refGene_gn=ACTL6A;refGene_tx=XM_004001172.4;refGene_gn=ACTL6A;refGene_tx=XM_019840084.2;refGene_gn=ACTL6A;refGene_tx=XM_023260373.1
C2 89603587 89603688
refGene_gn=ACTL6A;refGene_tx=XM_004001172.4;refGene_gn=ACTL6A;refGene_tx=XM_019840084.2;refGene_gn=ACTL6A;refGene_tx=XM_023260373.1
C2 89606625 89606800
refGene_gn=ACTL6A;refGene_tx=XM_004001172.4;refGene_gn=ACTL6A;refGene_tx=XM_019840084.2;refGene_gn=ACTL6A;refGene_tx=XM_023260373.1
C2 89606963 89607040 refGene_gn=ACTL6A;refGene_tx=XM_004001172.4
C2 89618357 89618382 refGene_gn=ACTL6A;refGene_tx=XM_004001172.4
C2 89720557 89720614
refGene_gn=GNB4;refGene_tx=XM_023260374.1;refGene_gn=GNB4;refGene_tx=XM_023260375.1
C2 89730202 89730241
refGene_gn=GNB4;refGene_tx=XM_023260374.1;refGene_gn=GNB4;refGene_tx=XM_023260375.1
C2 89730948 89731055
refGene_gn=GNB4;refGene_tx=XM_023260374.1;refGene_gn=GNB4;refGene_tx=XM_023260375.1
C2 89734000 89734064
refGene_gn=GNB4;refGene_tx=XM_023260374.1;refGene_gn=GNB4;refGene_tx=XM_023260375.1
C2 89736317 89736480
refGene_gn=GNB4;refGene_tx=XM_023260374.1;refGene_gn=GNB4;refGene_tx=XM_023260375.1
C2 89737440 89737507
refGene_gn=GNB4;refGene_tx=XM_023260374.1;refGene_gn=GNB4;refGene_tx=XM_023260375.1
C2 89737614 89737816

refGene_gn=GNB4;refGene_tx=XM_023260374.1;refGene_gn=GNB4;refGene_tx=XM_023260375.1
C2 89747052 89747269
refGene_gn=GNB4;refGene_tx=XM_023260374.1;refGene_gn=GNB4;refGene_tx=XM_023260375.1
C2 89751001 89751108
refGene_gn=GNB4;refGene_tx=XM_023260374.1;refGene_gn=GNB4;refGene_tx=XM_023260375.1
C2 89757893 89757975
refGene_gn=MFN1;refGene_tx=XM_004001165.5;refGene_gn=MFN1;refGene_tx=XM_011286054.3
C2 89759764 89759899
refGene_gn=MFN1;refGene_tx=XM_004001165.5;refGene_gn=MFN1;refGene_tx=XM_011286054.3
C2 89762231 89762428
refGene_gn=MFN1;refGene_tx=XM_004001165.5;refGene_gn=MFN1;refGene_tx=XM_011286054.3
C2 89763221 89763374
refGene_gn=MFN1;refGene_tx=XM_004001165.5;refGene_gn=MFN1;refGene_tx=XM_011286054.3
C2 89765237 89765467
refGene_gn=MFN1;refGene_tx=XM_004001165.5;refGene_gn=MFN1;refGene_tx=XM_011286054.3
C2 89765600 89765703
refGene_gn=MFN1;refGene_tx=XM_004001165.5;refGene_gn=MFN1;refGene_tx=XM_011286054.3
C2 89765980 89766085
refGene_gn=MFN1;refGene_tx=XM_004001165.5;refGene_gn=MFN1;refGene_tx=XM_011286054.3
C2 89766216 89766343
refGene_gn=MFN1;refGene_tx=XM_004001165.5;refGene_gn=MFN1;refGene_tx=XM_011286054.3
C2 89768013 89768135
refGene_gn=MFN1;refGene_tx=XM_004001165.5;refGene_gn=MFN1;refGene_tx=XM_011286054.3
C2 89774708 89774776
refGene_gn=MFN1;refGene_tx=XM_004001165.5;refGene_gn=MFN1;refGene_tx=XM_011286054.3
C2 89774891 89775045
refGene_gn=MFN1;refGene_tx=XM_004001165.5;refGene_gn=MFN1;refGene_tx=XM_011286054.3
C2 89778708 89778816
refGene_gn=MFN1;refGene_tx=XM_004001165.5;refGene_gn=MFN1;refGene_tx=XM_011286054.3
C2 89780255 89780364
refGene_gn=MFN1;refGene_tx=XM_004001165.5;refGene_gn=MFN1;refGene_tx=XM_011286054.3
C2 89781643 89781768
refGene_gn=MFN1;refGene_tx=XM_004001165.5;refGene_gn=MFN1;refGene_tx=XM_011286054.3

C2 89787999 89788162
refGene_gn=MFN1;refGene_tx=XM_004001165.5;refGene_gn=MFN1;refGene_tx=XM_011286054.3
C2 89789003 89789139
refGene_gn=MFN1;refGene_tx=XM_004001165.5;refGene_gn=MFN1;refGene_tx=XM_011286054.3
C2 89791721 89791833
refGene_gn=MFN1;refGene_tx=XM_004001165.5;refGene_gn=MFN1;refGene_tx=XM_011286054.3
C2 89807529 89808683
refGene_gn=ZNF639;refGene_tx=XM_011286053.3;refGene_gn=ZNF639;refGene_tx=XM_019840085.2;refGene_gn=ZNF639;refGene_tx=XM_023260377.1
C2 89808815 89808950
refGene_gn=ZNF639;refGene_tx=XM_011286053.3;refGene_gn=ZNF639;refGene_tx=XM_019840085.2;refGene_gn=ZNF639;refGene_tx=XM_023260377.1
C2 89811235 89811346
refGene_gn=ZNF639;refGene_tx=XM_011286053.3;refGene_gn=ZNF639;refGene_tx=XM_019840085.2;refGene_gn=ZNF639;refGene_tx=XM_023260377.1
C2 89812344 89812402
refGene_gn=ZNF639;refGene_tx=XM_011286053.3;refGene_gn=ZNF639;refGene_tx=XM_019840085.2;refGene_gn=ZNF639;refGene_tx=XM_023260377.1
C2 89878244 89878351 refGene_gn=KCNMB3;refGene_tx=XM_011286052.3
C2 89900450 89900642 refGene_gn=KCNMB3;refGene_tx=XM_011286052.3
C2 89904495 89904694 refGene_gn=KCNMB3;refGene_tx=XM_011286052.3
C2 89905857 89906256 refGene_gn=KCNMB3;refGene_tx=XM_011286052.3
C2 89915366 89915637
refGene_gn=PIK3CA;refGene_tx=XM_019840086.2;refGene_gn=PIK3CA;refGene_tx=XM_023260376.1
C2 89916436 89916588
refGene_gn=PIK3CA;refGene_tx=XM_019840086.2;refGene_gn=PIK3CA;refGene_tx=XM_023260376.1
C2 89916695 89916813
refGene_gn=PIK3CA;refGene_tx=XM_019840086.2;refGene_gn=PIK3CA;refGene_tx=XM_023260376.1
C2 89917384 89917555
refGene_gn=PIK3CA;refGene_tx=XM_019840086.2;refGene_gn=PIK3CA;refGene_tx=XM_023260376.1
C2 89921715 89921794
refGene_gn=PIK3CA;refGene_tx=XM_019840086.2;refGene_gn=PIK3CA;refGene_tx=XM_023260376.1
C2 89922622 89922744
refGene_gn=PIK3CA;refGene_tx=XM_019840086.2;refGene_gn=PIK3CA;refGene_tx=XM_023260376.1
C2 89923468 89923575
refGene_gn=PIK3CA;refGene_tx=XM_019840086.2;refGene_gn=PIK3CA;refGene_tx=XM_023260376.1
C2 89926517 89926689
refGene_gn=PIK3CA;refGene_tx=XM_019840086.2;refGene_gn=PIK3CA;refGene_tx=XM_023260376.1
C2 89928522 89928626

refGene_gn=PIK3CA;refGene_tx=XM_019840086.2;refGene_gn=PIK3CA;refGene_tx=XM_023260376.1
C2 89928848 89929013
refGene_gn=PIK3CA;refGene_tx=XM_019840086.2;refGene_gn=PIK3CA;refGene_tx=XM_023260376.1
C2 89929341 89929423
refGene_gn=PIK3CA;refGene_tx=XM_019840086.2;refGene_gn=PIK3CA;refGene_tx=XM_023260376.1
C2 89930779 89930904
refGene_gn=PIK3CA;refGene_tx=XM_019840086.2;refGene_gn=PIK3CA;refGene_tx=XM_023260376.1
C2 89935595 89935730
refGene_gn=PIK3CA;refGene_tx=XM_019840086.2;refGene_gn=PIK3CA;refGene_tx=XM_023260376.1
C2 89935816 89935969
refGene_gn=PIK3CA;refGene_tx=XM_019840086.2;refGene_gn=PIK3CA;refGene_tx=XM_023260376.1
C2 89936479 89936585
refGene_gn=PIK3CA;refGene_tx=XM_019840086.2;refGene_gn=PIK3CA;refGene_tx=XM_023260376.1
C2 89941939 89942025
refGene_gn=PIK3CA;refGene_tx=XM_019840086.2;refGene_gn=PIK3CA;refGene_tx=XM_023260376.1
C2 89942658 89942904
refGene_gn=PIK3CA;refGene_tx=XM_019840086.2;refGene_gn=PIK3CA;refGene_tx=XM_023260376.1
C2 89943439 89943690
refGene_gn=PIK3CA;refGene_tx=XM_019840086.2;refGene_gn=PIK3CA;refGene_tx=XM_023260376.1
C2 89945309 89945519
refGene_gn=PIK3CA;refGene_tx=XM_019840086.2;refGene_gn=PIK3CA;refGene_tx=XM_023260376.1
C2 89946069 89946421
refGene_gn=PIK3CA;refGene_tx=XM_019840086.2;refGene_gn=PIK3CA;refGene_tx=XM_023260376.1
C2 90039714 90039984
refGene_gn=ZMAT3;refGene_tx=XM_011286051.3;refGene_gn=ZMAT3;refGene_tx=XM_023260379.1;refGene_gn=ZMAT3;refGene_tx=XM_023260380.1;refGene_gn=ZMAT3;refGene_tx=XM_023260381.1
C2 90054956 90055076
refGene_gn=ZMAT3;refGene_tx=XM_011286051.3;refGene_gn=ZMAT3;refGene_tx=XM_023260379.1;refGene_gn=ZMAT3;refGene_tx=XM_023260380.1;refGene_gn=ZMAT3;refGene_tx=XM_023260381.1
C2 90057399 90057566
refGene_gn=ZMAT3;refGene_tx=XM_011286051.3;refGene_gn=ZMAT3;refGene_tx=XM_023260379.1;refGene_gn=ZMAT3;refGene_tx=XM_023260380.1;refGene_gn=ZMAT3;refGene_tx=XM_023260381.1
C2 90057685 90057786
refGene_gn=ZMAT3;refGene_tx=XM_011286051.3;refGene_gn=ZMAT3;refGene_tx=XM_023260379.1;refGene_gn=ZMAT3;refGene_tx=XM_023260380.1;refGene_gn=

ZMAT3;refGene_tx=XM_023260381.1
C2 90061371 90061583
refGene_gn=ZMAT3;refGene_tx=XM_011286051.3;refGene_gn=ZMAT3;refGene_tx
=XM_023260379.1;refGene_gn=ZMAT3;refGene_tx=XM_023260380.1;refGene_gn=
ZMAT3;refGene_tx=XM_023260381.1
C2 90184096 90184381 refGene_gn=KCNMB2;refGene_tx=XM_003991892.4
C2 90198883 90199079 refGene_gn=KCNMB2;refGene_tx=XM_003991892.4
C2 90201469 90201640 refGene_gn=KCNMB2;refGene_tx=XM_003991892.4
C2 90217862 90217918 refGene_gn=KCNMB2;refGene_tx=XM_003991892.4
C2 91768414 91768472
refGene_gn=TBL1XR1;refGene_tx=XM_011286112.3;refGene_gn=TBL1XR1;refGen
e_tx=XM_011286113.3
C2 91779752 91779898
refGene_gn=TBL1XR1;refGene_tx=XM_011286112.3;refGene_gn=TBL1XR1;refGen
e_tx=XM_011286113.3
C2 91782115 91782338
refGene_gn=TBL1XR1;refGene_tx=XM_011286112.3;refGene_gn=TBL1XR1;refGen
e_tx=XM_011286113.3;refGene_gn=TBL1XR1;refGene_tx=XM_023260383.1;refGe
ne_gn=TBL1XR1;refGene_tx=XM_023260384.1
C2 91783050 91783183
refGene_gn=TBL1XR1;refGene_tx=XM_011286112.3;refGene_gn=TBL1XR1;refGen
e_tx=XM_011286113.3;refGene_gn=TBL1XR1;refGene_tx=XM_023260383.1;refGe
ne_gn=TBL1XR1;refGene_tx=XM_023260384.1
C2 91783565 91783707
refGene_gn=TBL1XR1;refGene_tx=XM_011286112.3;refGene_gn=TBL1XR1;refGen
e_tx=XM_011286113.3;refGene_gn=TBL1XR1;refGene_tx=XM_023260383.1;refGe
ne_gn=TBL1XR1;refGene_tx=XM_023260384.1
C2 91786300 91786364
refGene_gn=TBL1XR1;refGene_tx=XM_011286112.3;refGene_gn=TBL1XR1;refGen
e_tx=XM_011286113.3;refGene_gn=TBL1XR1;refGene_tx=XM_023260383.1;refGe
ne_gn=TBL1XR1;refGene_tx=XM_023260384.1
C2 91786466 91786564
refGene_gn=TBL1XR1;refGene_tx=XM_011286112.3;refGene_gn=TBL1XR1;refGen
e_tx=XM_011286113.3;refGene_gn=TBL1XR1;refGene_tx=XM_023260383.1;refGe
ne_gn=TBL1XR1;refGene_tx=XM_023260384.1
C2 91788047 91788108
refGene_gn=TBL1XR1;refGene_tx=XM_011286112.3;refGene_gn=TBL1XR1;refGen
e_tx=XM_011286113.3;refGene_gn=TBL1XR1;refGene_tx=XM_023260383.1;refGe
ne_gn=TBL1XR1;refGene_tx=XM_023260384.1
C2 91795732 91795854
refGene_gn=TBL1XR1;refGene_tx=XM_011286112.3;refGene_gn=TBL1XR1;refGen
e_tx=XM_011286113.3;refGene_gn=TBL1XR1;refGene_tx=XM_023260383.1;refGe
ne_gn=TBL1XR1;refGene_tx=XM_023260384.1
C2 91795992 91796067
refGene_gn=TBL1XR1;refGene_tx=XM_011286112.3;refGene_gn=TBL1XR1;refGen
e_tx=XM_011286113.3;refGene_gn=TBL1XR1;refGene_tx=XM_023260383.1;refGe
ne_gn=TBL1XR1;refGene_tx=XM_023260384.1
C2 91799178 91799306
refGene_gn=TBL1XR1;refGene_tx=XM_011286112.3;refGene_gn=TBL1XR1;refGen
e_tx=XM_011286113.3;refGene_gn=TBL1XR1;refGene_tx=XM_023260383.1;refGe

ne_gn=TBL1XR1;refGene_tx=XM_023260384.1
C2 91800081 91800247
refGene_gn=TBL1XR1;refGene_tx=XM_011286112.3;refGene_gn=TBL1XR1;refGene_tx=XM_011286113.3;refGene_gn=TBL1XR1;refGene_tx=XM_023260383.1;refGene_gn=TBL1XR1;refGene_tx=XM_023260384.1
C2 91803541 91803643
refGene_gn=TBL1XR1;refGene_tx=XM_011286112.3;refGene_gn=TBL1XR1;refGene_tx=XM_011286113.3;refGene_gn=TBL1XR1;refGene_tx=XM_023260383.1;refGene_gn=TBL1XR1;refGene_tx=XM_023260384.1
C2 91804509 91804536
refGene_gn=TBL1XR1;refGene_tx=XM_011286112.3;refGene_gn=TBL1XR1;refGene_tx=XM_011286113.3;refGene_gn=TBL1XR1;refGene_tx=XM_023260383.1;refGene_gn=TBL1XR1;refGene_tx=XM_023260384.1
C2 92880833 92881032
refGene_gn=NAALADL2;refGene_tx=XM_023260386.1;refGene_gn=NAALADL2;refGene_tx=XM_023260387.1;refGene_gn=NAALADL2;refGene_tx=XM_023260388.1;refGene_gn=NAALADL2;refGene_tx=XM_023260389.1;refGene_gn=NAALADL2;refGene_tx=XM_023260390.1;refGene_gn=NAALADL2;refGene_tx=XM_023260391.1
C2 92926721 92926920
refGene_gn=NAALADL2;refGene_tx=XM_023260386.1;refGene_gn=NAALADL2;refGene_tx=XM_023260387.1;refGene_gn=NAALADL2;refGene_tx=XM_023260388.1;refGene_gn=NAALADL2;refGene_tx=XM_023260389.1;refGene_gn=NAALADL2;refGene_tx=XM_023260390.1;refGene_gn=NAALADL2;refGene_tx=XM_023260391.1
C2 92939699 92939793
refGene_gn=NAALADL2;refGene_tx=XM_023260386.1;refGene_gn=NAALADL2;refGene_tx=XM_023260387.1;refGene_gn=NAALADL2;refGene_tx=XM_023260388.1;refGene_gn=NAALADL2;refGene_tx=XM_023260389.1;refGene_gn=NAALADL2;refGene_tx=XM_023260390.1;refGene_gn=NAALADL2;refGene_tx=XM_023260391.1
C2 93044223 93044319
refGene_gn=NAALADL2;refGene_tx=XM_023260386.1;refGene_gn=NAALADL2;refGene_tx=XM_023260387.1;refGene_gn=NAALADL2;refGene_tx=XM_023260388.1;refGene_gn=NAALADL2;refGene_tx=XM_023260389.1;refGene_gn=NAALADL2;refGene_tx=XM_023260390.1;refGene_gn=NAALADL2;refGene_tx=XM_023260391.1
C2 93092498 93092645
refGene_gn=NAALADL2;refGene_tx=XM_023260386.1;refGene_gn=NAALADL2;refGene_tx=XM_023260387.1;refGene_gn=NAALADL2;refGene_tx=XM_023260388.1;refGene_gn=NAALADL2;refGene_tx=XM_023260389.1;refGene_gn=NAALADL2;refGene_tx=XM_023260390.1;refGene_gn=NAALADL2;refGene_tx=XM_023260391.1
C2 93196818 93196938
refGene_gn=NAALADL2;refGene_tx=XM_023260386.1;refGene_gn=NAALADL2;refGene_tx=XM_023260387.1;refGene_gn=NAALADL2;refGene_tx=XM_023260388.1;refGene_gn=NAALADL2;refGene_tx=XM_023260389.1;refGene_gn=NAALADL2;refGene_tx=XM_023260390.1;refGene_gn=NAALADL2;refGene_tx=XM_023260391.1
C2 93201575 93201781
refGene_gn=NAALADL2;refGene_tx=XM_023260386.1;refGene_gn=NAALADL2;refGene_tx=XM_023260387.1;refGene_gn=NAALADL2;refGene_tx=XM_023260388.1;refGene_gn=NAALADL2;refGene_tx=XM_023260389.1;refGene_gn=NAALADL2;refGene_tx=XM_023260390.1;refGene_gn=NAALADL2;refGene_tx=XM_023260391.1
C2 93211999 93212092
refGene_gn=NAALADL2;refGene_tx=XM_023260386.1;refGene_gn=NAALADL2;refG

ene_tx=XM_023260387.1;refGene_gn=NAALADL2;refGene_tx=XM_023260388.1;refGene_gn=NAALADL2;refGene_tx=XM_023260389.1;refGene_gn=NAALADL2;refGene_tx=XM_023260390.1;refGene_gn=NAALADL2;refGene_tx=XM_023260391.1
C2 93235894 93236038
refGene_gn=NAALADL2;refGene_tx=XM_023260386.1;refGene_gn=NAALADL2;refGene_tx=XM_023260387.1;refGene_gn=NAALADL2;refGene_tx=XM_023260388.1;refGene_gn=NAALADL2;refGene_tx=XM_023260389.1;refGene_gn=NAALADL2;refGene_tx=XM_023260390.1;refGene_gn=NAALADL2;refGene_tx=XM_023260391.1
C2 93360657 93360808
refGene_gn=NAALADL2;refGene_tx=XM_023260386.1;refGene_gn=NAALADL2;refGene_tx=XM_023260387.1;refGene_gn=NAALADL2;refGene_tx=XM_023260388.1;refGene_gn=NAALADL2;refGene_tx=XM_023260389.1;refGene_gn=NAALADL2;refGene_tx=XM_023260390.1;refGene_gn=NAALADL2;refGene_tx=XM_023260391.1
C2 93426154 93426274
refGene_gn=NAALADL2;refGene_tx=XM_023260386.1;refGene_gn=NAALADL2;refGene_tx=XM_023260387.1;refGene_gn=NAALADL2;refGene_tx=XM_023260388.1;refGene_gn=NAALADL2;refGene_tx=XM_023260389.1;refGene_gn=NAALADL2;refGene_tx=XM_023260390.1;refGene_gn=NAALADL2;refGene_tx=XM_023260391.1
C2 93437288 93437562
refGene_gn=NAALADL2;refGene_tx=XM_023260386.1;refGene_gn=NAALADL2;refGene_tx=XM_023260387.1;refGene_gn=NAALADL2;refGene_tx=XM_023260388.1;refGene_gn=NAALADL2;refGene_tx=XM_023260389.1;refGene_gn=NAALADL2;refGene_tx=XM_023260390.1;refGene_gn=NAALADL2;refGene_tx=XM_023260391.1
C2 93575529 93576031
refGene_gn=NAALADL2;refGene_tx=XM_023260386.1;refGene_gn=NAALADL2;refGene_tx=XM_023260387.1;refGene_gn=NAALADL2;refGene_tx=XM_023260388.1;refGene_gn=NAALADL2;refGene_tx=XM_023260389.1;refGene_gn=NAALADL2;refGene_tx=XM_023260390.1;refGene_gn=NAALADL2;refGene_tx=XM_023260391.1
C2 93626857 93626873 refGene_gn=NAALADL2;refGene_tx=XM_023260389.1
C2 93762956 93762993 refGene_gn=NAALADL2;refGene_tx=XM_023260386.1
C2 93797696 93797718 refGene_gn=NAALADL2;refGene_tx=XM_023260388.1
C2 93802698 93802741
refGene_gn=NAALADL2;refGene_tx=XM_023260386.1;refGene_gn=NAALADL2;refGene_tx=XM_023260387.1
C2 94372046 94372869
refGene_gn=NLGN1;refGene_tx=XM_011286111.2;refGene_gn=NLGN1;refGene_tx=XM_019840091.1;refGene_gn=NLGN1;refGene_tx=XM_019840092.2;refGene_gn=NLGN1;refGene_tx=XM_019840093.2;refGene_gn=NLGN1;refGene_tx=XM_019840094.2;refGene_gn=NLGN1;refGene_tx=XM_019840095.1;refGene_gn=NLGN1;refGene_tx=XM_019840096.1;refGene_gn=NLGN1;refGene_tx=XM_019840097.1;refGene_gn=NLGN1;refGene_tx=XM_019840098.1;refGene_gn=NLGN1;refGene_tx=XM_019840099.1
C2 94373748 94374538
refGene_gn=NLGN1;refGene_tx=XM_011286111.2;refGene_gn=NLGN1;refGene_tx=XM_019840091.1;refGene_gn=NLGN1;refGene_tx=XM_019840092.2;refGene_gn=NLGN1;refGene_tx=XM_019840093.2;refGene_gn=NLGN1;refGene_tx=XM_019840094.2;refGene_gn=NLGN1;refGene_tx=XM_019840095.1;refGene_gn=NLGN1;refGene_tx=XM_019840096.1;refGene_gn=NLGN1;refGene_tx=XM_019840097.1;refGene_gn=NLGN1;refGene_tx=XM_019840098.1;refGene_gn=NLGN1;refGene_tx=XM_019840099.1

C2 94377999 94378212
refGene_gn=NLGN1;refGene_tx=XM_011286111.2;refGene_gn=NLGN1;refGene_tx=XM_019840091.1;refGene_gn=NLGN1;refGene_tx=XM_019840092.2;refGene_gn=NLGN1;refGene_tx=XM_019840093.2;refGene_gn=NLGN1;refGene_tx=XM_019840094.2;refGene_gn=NLGN1;refGene_tx=XM_019840095.1;refGene_gn=NLGN1;refGene_tx=XM_019840096.1;refGene_gn=NLGN1;refGene_tx=XM_019840097.1;refGene_gn=NLGN1;refGene_tx=XM_019840098.1;refGene_gn=NLGN1;refGene_tx=XM_019840099.1
C2 94835211 94835364
refGene_gn=NLGN1;refGene_tx=XM_011286111.2;refGene_gn=NLGN1;refGene_tx=XM_019840091.1;refGene_gn=NLGN1;refGene_tx=XM_019840092.2;refGene_gn=NLGN1;refGene_tx=XM_019840093.2;refGene_gn=NLGN1;refGene_tx=XM_019840094.2;refGene_gn=NLGN1;refGene_tx=XM_019840095.1;refGene_gn=NLGN1;refGene_tx=XM_019840096.1;refGene_gn=NLGN1;refGene_tx=XM_019840097.1;refGene_gn=NLGN1;refGene_tx=XM_019840098.1;refGene_gn=NLGN1;refGene_tx=XM_019840099.1
C2 94841720 94841780
refGene_gn=NLGN1;refGene_tx=XM_011286111.2;refGene_gn=NLGN1;refGene_tx=XM_019840091.1;refGene_gn=NLGN1;refGene_tx=XM_019840092.2;refGene_gn=NLGN1;refGene_tx=XM_019840093.2;refGene_gn=NLGN1;refGene_tx=XM_019840094.2;refGene_gn=NLGN1;refGene_tx=XM_019840095.1;refGene_gn=NLGN1;refGene_tx=XM_019840097.1
C2 95037161 95037221
refGene_gn=NLGN1;refGene_tx=XM_019840091.1;refGene_gn=NLGN1;refGene_tx=XM_019840092.2;refGene_gn=NLGN1;refGene_tx=XM_019840093.2;refGene_gn=NLGN1;refGene_tx=XM_019840094.2;refGene_gn=NLGN1;refGene_tx=XM_019840095.1;refGene_gn=NLGN1;refGene_tx=XM_019840096.1
C2 95037663 95038156
refGene_gn=NLGN1;refGene_tx=XM_011286111.2;refGene_gn=NLGN1;refGene_tx=XM_019840091.1;refGene_gn=NLGN1;refGene_tx=XM_019840092.2;refGene_gn=NLGN1;refGene_tx=XM_019840093.2;refGene_gn=NLGN1;refGene_tx=XM_019840094.2;refGene_gn=NLGN1;refGene_tx=XM_019840095.1;refGene_gn=NLGN1;refGene_tx=XM_019840096.1;refGene_gn=NLGN1;refGene_tx=XM_019840097.1;refGene_gn=NLGN1;refGene_tx=XM_019840098.1;refGene_gn=NLGN1;refGene_tx=XM_019840099.1
C2 95408396 95409008
refGene_gn=SPATA16;refGene_tx=XM_003991900.4;refGene_gn=SPATA16;refGene_tx=XM_019840100.2
C2 95464014 95464160
refGene_gn=SPATA16;refGene_tx=XM_003991900.4;refGene_gn=SPATA16;refGene_tx=XM_019840100.2
C2 95506383 95506473
refGene_gn=SPATA16;refGene_tx=XM_003991900.4;refGene_gn=SPATA16;refGene_tx=XM_019840100.2
C2 95546031 95546116
refGene_gn=SPATA16;refGene_tx=XM_003991900.4;refGene_gn=SPATA16;refGene_tx=XM_019840100.2
C2 95567959 95568107
refGene_gn=SPATA16;refGene_tx=XM_003991900.4;refGene_gn=SPATA16;refGene_tx=XM_019840100.2

C2	95585006 95585153
refGene_gn=SPATA16;refGene_tx=XM_003991900.4;refGene_gn=SPATA16;refGene_tx=XM_019840100.2
C2	95586212 95586325
refGene_gn=SPATA16;refGene_tx=XM_003991900.4;refGene_gn=SPATA16;refGene_tx=XM_019840100.2
C2	95593492 95593657
refGene_gn=SPATA16;refGene_tx=XM_003991900.4;refGene_gn=SPATA16;refGene_tx=XM_019840100.2
C2	95596239 95596323
refGene_gn=SPATA16;refGene_tx=XM_003991900.4;refGene_gn=SPATA16;refGene_tx=XM_019840100.2
C2	95629876 95629956 refGene_gn=SPATA16;refGene_tx=XM_019840100.2
C2	95630051 95630174
refGene_gn=SPATA16;refGene_tx=XM_003991900.4;refGene_gn=SPATA16;refGene_tx=XM_019840100.2
C2	95696689 95696779
refGene_gn=ECT2;refGene_tx=XM_011286110.3;refGene_gn=ECT2;refGene_tx=XM_023260397.1;refGene_gn=ECT2;refGene_tx=XM_023260398.1
C2	95698213 95698318
refGene_gn=ECT2;refGene_tx=XM_023260394.1;refGene_gn=ECT2;refGene_tx=XM_023260395.1;refGene_gn=ECT2;refGene_tx=XM_023260396.1
C2	95700048 95700195
refGene_gn=ECT2;refGene_tx=XM_011286110.3;refGene_gn=ECT2;refGene_tx=XM_023260394.1;refGene_gn=ECT2;refGene_tx=XM_023260395.1;refGene_gn=ECT2;refGene_tx=XM_023260396.1;refGene_gn=ECT2;refGene_tx=XM_023260397.1;refGene_gn=ECT2;refGene_tx=XM_023260398.1
C2	95701174 95701282
refGene_gn=ECT2;refGene_tx=XM_011286110.3;refGene_gn=ECT2;refGene_tx=XM_023260394.1;refGene_gn=ECT2;refGene_tx=XM_023260395.1;refGene_gn=ECT2;refGene_tx=XM_023260396.1;refGene_gn=ECT2;refGene_tx=XM_023260397.1;refGene_gn=ECT2;refGene_tx=XM_023260398.1
C2	95713236 95713391
refGene_gn=ECT2;refGene_tx=XM_011286110.3;refGene_gn=ECT2;refGene_tx=XM_023260394.1;refGene_gn=ECT2;refGene_tx=XM_023260395.1;refGene_gn=ECT2;refGene_tx=XM_023260396.1;refGene_gn=ECT2;refGene_tx=XM_023260397.1;refGene_gn=ECT2;refGene_tx=XM_023260398.1
C2	95714701 95714840
refGene_gn=ECT2;refGene_tx=XM_011286110.3;refGene_gn=ECT2;refGene_tx=XM_023260394.1;refGene_gn=ECT2;refGene_tx=XM_023260395.1;refGene_gn=ECT2;refGene_tx=XM_023260396.1;refGene_gn=ECT2;refGene_tx=XM_023260397.1;refGene_gn=ECT2;refGene_tx=XM_023260398.1
C2	95718228 95718348
refGene_gn=ECT2;refGene_tx=XM_011286110.3;refGene_gn=ECT2;refGene_tx=XM_023260394.1;refGene_gn=ECT2;refGene_tx=XM_023260395.1;refGene_gn=ECT2;refGene_tx=XM_023260396.1;refGene_gn=ECT2;refGene_tx=XM_023260397.1;refGene_gn=ECT2;refGene_tx=XM_023260398.1
C2	95718511 95718590
refGene_gn=ECT2;refGene_tx=XM_011286110.3;refGene_gn=ECT2;refGene_tx=XM_023260394.1;refGene_gn=ECT2;refGene_tx=XM_023260395.1;refGene_gn=ECT

2;refGene_tx=XM_023260396.1;refGene_gn=ECT2;refGene_tx=XM_023260397.1;
refGene_gn=ECT2;refGene_tx=XM_023260398.1
C2 95724108 95724190
refGene_gn=ECT2;refGene_tx=XM_011286110.3;refGene_gn=ECT2;refGene_tx=XM_023260394.1;refGene_gn=ECT2;refGene_tx=XM_023260395.1;refGene_gn=ECT2;refGene_tx=XM_023260396.1;refGene_gn=ECT2;refGene_tx=XM_023260397.1;
refGene_gn=ECT2;refGene_tx=XM_023260398.1
C2 95726025 95726122
refGene_gn=ECT2;refGene_tx=XM_011286110.3;refGene_gn=ECT2;refGene_tx=XM_023260394.1;refGene_gn=ECT2;refGene_tx=XM_023260395.1;refGene_gn=ECT2;refGene_tx=XM_023260396.1;refGene_gn=ECT2;refGene_tx=XM_023260397.1;
refGene_gn=ECT2;refGene_tx=XM_023260398.1
C2 95727050 95727161
refGene_gn=ECT2;refGene_tx=XM_011286110.3;refGene_gn=ECT2;refGene_tx=XM_023260394.1;refGene_gn=ECT2;refGene_tx=XM_023260395.1;refGene_gn=ECT2;refGene_tx=XM_023260396.1;refGene_gn=ECT2;refGene_tx=XM_023260397.1;
refGene_gn=ECT2;refGene_tx=XM_023260398.1
C2 95728981 95729050
refGene_gn=ECT2;refGene_tx=XM_011286110.3;refGene_gn=ECT2;refGene_tx=XM_023260394.1;refGene_gn=ECT2;refGene_tx=XM_023260395.1;refGene_gn=ECT2;refGene_tx=XM_023260396.1;refGene_gn=ECT2;refGene_tx=XM_023260397.1;
refGene_gn=ECT2;refGene_tx=XM_023260398.1
C2 95733718 95733838
refGene_gn=ECT2;refGene_tx=XM_011286110.3;refGene_gn=ECT2;refGene_tx=XM_023260394.1;refGene_gn=ECT2;refGene_tx=XM_023260395.1;refGene_gn=ECT2;refGene_tx=XM_023260396.1;refGene_gn=ECT2;refGene_tx=XM_023260397.1;
refGene_gn=ECT2;refGene_tx=XM_023260398.1
C2 95738395 95738532
refGene_gn=ECT2;refGene_tx=XM_011286110.3;refGene_gn=ECT2;refGene_tx=XM_023260394.1;refGene_gn=ECT2;refGene_tx=XM_023260395.1;refGene_gn=ECT2;refGene_tx=XM_023260396.1;refGene_gn=ECT2;refGene_tx=XM_023260397.1;
refGene_gn=ECT2;refGene_tx=XM_023260398.1
C2 95743170 95743393
refGene_gn=ECT2;refGene_tx=XM_011286110.3;refGene_gn=ECT2;refGene_tx=XM_023260394.1;refGene_gn=ECT2;refGene_tx=XM_023260395.1;refGene_gn=ECT2;refGene_tx=XM_023260396.1;refGene_gn=ECT2;refGene_tx=XM_023260397.1;
refGene_gn=ECT2;refGene_tx=XM_023260398.1
C2 95744751 95744814
refGene_gn=ECT2;refGene_tx=XM_011286110.3;refGene_gn=ECT2;refGene_tx=XM_023260394.1;refGene_gn=ECT2;refGene_tx=XM_023260395.1;refGene_gn=ECT2;refGene_tx=XM_023260396.1;refGene_gn=ECT2;refGene_tx=XM_023260397.1;
refGene_gn=ECT2;refGene_tx=XM_023260398.1
C2 95745980 95746096
refGene_gn=ECT2;refGene_tx=XM_011286110.3;refGene_gn=ECT2;refGene_tx=XM_023260394.1;refGene_gn=ECT2;refGene_tx=XM_023260395.1;refGene_gn=ECT2;refGene_tx=XM_023260396.1;refGene_gn=ECT2;refGene_tx=XM_023260397.1;
refGene_gn=ECT2;refGene_tx=XM_023260398.1
C2 95746244 95746375
refGene_gn=ECT2;refGene_tx=XM_011286110.3;refGene_gn=ECT2;refGene_tx=XM_023260394.1;refGene_gn=ECT2;refGene_tx=XM_023260395.1;refGene_gn=ECT

2;refGene_tx=XM_023260396.1;refGene_gn=ECT2;refGene_tx=XM_023260397.1;
refGene_gn=ECT2;refGene_tx=XM_023260398.1
C2 95748620 95748694
refGene_gn=ECT2;refGene_tx=XM_011286110.3;refGene_gn=ECT2;refGene_tx=X
M_023260394.1;refGene_gn=ECT2;refGene_tx=XM_023260395.1;refGene_gn=ECT
2;refGene_tx=XM_023260396.1;refGene_gn=ECT2;refGene_tx=XM_023260397.1;
refGene_gn=ECT2;refGene_tx=XM_023260398.1
C2 95749628 95749736
refGene_gn=ECT2;refGene_tx=XM_011286110.3;refGene_gn=ECT2;refGene_tx=X
M_023260394.1;refGene_gn=ECT2;refGene_tx=XM_023260395.1;refGene_gn=ECT
2;refGene_tx=XM_023260396.1;refGene_gn=ECT2;refGene_tx=XM_023260397.1;
refGene_gn=ECT2;refGene_tx=XM_023260398.1
C2 95750582 95750672
refGene_gn=ECT2;refGene_tx=XM_011286110.3;refGene_gn=ECT2;refGene_tx=X
M_023260394.1;refGene_gn=ECT2;refGene_tx=XM_023260395.1;refGene_gn=ECT
2;refGene_tx=XM_023260396.1;refGene_gn=ECT2;refGene_tx=XM_023260397.1;
refGene_gn=ECT2;refGene_tx=XM_023260398.1
C2 95752597 95752780
refGene_gn=ECT2;refGene_tx=XM_011286110.3;refGene_gn=ECT2;refGene_tx=X
M_023260394.1;refGene_gn=ECT2;refGene_tx=XM_023260395.1;refGene_gn=ECT
2;refGene_tx=XM_023260396.1;refGene_gn=ECT2;refGene_tx=XM_023260397.1;
refGene_gn=ECT2;refGene_tx=XM_023260398.1
C2 95754812 95754905
refGene_gn=ECT2;refGene_tx=XM_011286110.3;refGene_gn=ECT2;refGene_tx=X
M_023260394.1;refGene_gn=ECT2;refGene_tx=XM_023260395.1
C2 95755009 95755089
refGene_gn=ECT2;refGene_tx=XM_011286110.3;refGene_gn=ECT2;refGene_tx=X
M_023260394.1;refGene_gn=ECT2;refGene_tx=XM_023260395.1;refGene_gn=ECT
2;refGene_tx=XM_023260396.1;refGene_gn=ECT2;refGene_tx=XM_023260397.1;
refGene_gn=ECT2;refGene_tx=XM_023260398.1
C2 95756442 95756572
refGene_gn=ECT2;refGene_tx=XM_011286110.3;refGene_gn=ECT2;refGene_tx=X
M_023260394.1;refGene_gn=ECT2;refGene_tx=XM_023260395.1;refGene_gn=ECT
2;refGene_tx=XM_023260396.1;refGene_gn=ECT2;refGene_tx=XM_023260397.1;
refGene_gn=ECT2;refGene_tx=XM_023260398.1
C2 95784495 95784633 refGene_gn=NCEH1;refGene_tx=XM_003991902.5
C2 95824982 95825211 refGene_gn=NCEH1;refGene_tx=XM_003991902.5
C2 95827344 95827414 refGene_gn=NCEH1;refGene_tx=XM_003991902.5
C2 95838778 95838950 refGene_gn=NCEH1;refGene_tx=XM_003991902.5
C2 95840829 95841447 refGene_gn=NCEH1;refGene_tx=XM_003991902.5
C2 95939224 95939347 refGene_gn=TNFSF10;refGene_tx=NM_001130844.1
C2 95944946 95945084 refGene_gn=TNFSF10;refGene_tx=NM_001130844.1
C2 95947673 95947713 refGene_gn=TNFSF10;refGene_tx=NM_001130844.1
C2 95953528 95953633 refGene_gn=TNFSF10;refGene_tx=NM_001130844.1
C2 95955284 95955718 refGene_gn=TNFSF10;refGene_tx=NM_001130844.1
C2 95994223 95995019 refGene_gn=GHSR;refGene_tx=XM_023260870.1
C2 95997175 95997480 refGene_gn=GHSR;refGene_tx=XM_023260870.1
C2 96068564 96068876 refGene_gn=FNDC3B;refGene_tx=XM_023260400.1
C2 96080915 96081043 refGene_gn=FNDC3B;refGene_tx=XM_023260400.1
C2 96084487 96084654 refGene_gn=FNDC3B;refGene_tx=XM_023260400.1

C2 96098407 96098620 refGene_gn=FNDC3B;refGene_tx=XM_023260400.1
C2 96111890 96112171 refGene_gn=FNDC3B;refGene_tx=XM_023260400.1
C2 96116068 96116218 refGene_gn=FNDC3B;refGene_tx=XM_023260400.1
C2 96116554 96116668 refGene_gn=FNDC3B;refGene_tx=XM_023260400.1
C2 96118674 96118847 refGene_gn=FNDC3B;refGene_tx=XM_023260400.1
C2 96119751 96119857 refGene_gn=FNDC3B;refGene_tx=XM_023260400.1
C2 96121554 96121673 refGene_gn=FNDC3B;refGene_tx=XM_023260400.1
C2 96123799 96123871 refGene_gn=FNDC3B;refGene_tx=XM_023260400.1
C2 96126521 96126582 refGene_gn=FNDC3B;refGene_tx=XM_023260400.1
C2 96126583 96126661 refGene_gn=FNDC3B;refGene_tx=XM_023260400.1
C2 96128052 96128139 refGene_gn=FNDC3B;refGene_tx=XM_023260400.1
C2 96129972 96130147 refGene_gn=FNDC3B;refGene_tx=XM_023260400.1
C2 96131626 96131751 refGene_gn=FNDC3B;refGene_tx=XM_023260400.1
C2 96146291 96146345 refGene_gn=FNDC3B;refGene_tx=XM_023260400.1
C2 96150089 96150228 refGene_gn=FNDC3B;refGene_tx=XM_023260400.1
C2 96157966 96158026 refGene_gn=FNDC3B;refGene_tx=XM_023260400.1
C2 96161105 96161257 refGene_gn=FNDC3B;refGene_tx=XM_023260400.1
C2 96168902 96168961 refGene_gn=FNDC3B;refGene_tx=XM_023260400.1
C2 96214047 96214338 refGene_gn=FNDC3B;refGene_tx=XM_023260400.1
C2 96218253 96218497 refGene_gn=FNDC3B;refGene_tx=XM_023260400.1
C2 96235821 96235898 refGene_gn=FNDC3B;refGene_tx=XM_023260400.1
C2 96327110 96327186 refGene_gn=FNDC3B;refGene_tx=XM_023260400.1
C2 96349979 96350090 refGene_gn=FNDC3B;refGene_tx=XM_023260400.1
C2 96564293 96564308 refGene_gn=TMEM212;refGene_tx=XM_019840108.1
C2 96565407 96565428 refGene_gn=TMEM212;refGene_tx=XM_006936264.3
C2 96575949 96576273
refGene_gn=TMEM212;refGene_tx=XM_006936264.3;refGene_gn=TMEM212;refGene_tx=XM_019840108.1
C2 96577667 96577727
refGene_gn=TMEM212;refGene_tx=XM_006936264.3;refGene_gn=TMEM212;refGene_tx=XM_019840108.1
C2 96587667 96587826
refGene_gn=TMEM212;refGene_tx=XM_006936264.3;refGene_gn=TMEM212;refGene_tx=XM_019840108.1
C2 96652470 96652532 refGene_gn=PLD1;refGene_tx=XM_019840109.2
C2 96697169 96697360
refGene_gn=PLD1;refGene_tx=XM_003991907.5;refGene_gn=PLD1;refGene_tx=XM_019840109.2;refGene_gn=PLD1;refGene_tx=XM_019840110.2;refGene_gn=PLD1;refGene_tx=XM_019840111.2;refGene_gn=PLD1;refGene_tx=XM_019840112.2;refGene_gn=PLD1;refGene_tx=XM_023260401.1
C2 96697620 96697748
refGene_gn=PLD1;refGene_tx=XM_003991907.5;refGene_gn=PLD1;refGene_tx=XM_019840109.2;refGene_gn=PLD1;refGene_tx=XM_019840110.2;refGene_gn=PLD1;refGene_tx=XM_019840111.2;refGene_gn=PLD1;refGene_tx=XM_019840112.2;refGene_gn=PLD1;refGene_tx=XM_023260401.1
C2 96702402 96702548
refGene_gn=PLD1;refGene_tx=XM_003991907.5;refGene_gn=PLD1;refGene_tx=XM_019840109.2;refGene_gn=PLD1;refGene_tx=XM_019840110.2;refGene_gn=PLD1;refGene_tx=XM_019840111.2;refGene_gn=PLD1;refGene_tx=XM_019840112.2;refGene_gn=PLD1;refGene_tx=XM_023260401.1

C2 96702811 96702917
refGene_gn=PLD1;refGene_tx=XM_003991907.5;refGene_gn=PLD1;refGene_tx=XM_019840109.2;refGene_gn=PLD1;refGene_tx=XM_019840110.2;refGene_gn=PLD1;refGene_tx=XM_019840111.2;refGene_gn=PLD1;refGene_tx=XM_019840112.2;refGene_gn=PLD1;refGene_tx=XM_023260401.1
C2 96704434 96704500
refGene_gn=PLD1;refGene_tx=XM_003991907.5;refGene_gn=PLD1;refGene_tx=XM_019840109.2;refGene_gn=PLD1;refGene_tx=XM_019840110.2;refGene_gn=PLD1;refGene_tx=XM_019840111.2;refGene_gn=PLD1;refGene_tx=XM_019840112.2;refGene_gn=PLD1;refGene_tx=XM_023260401.1
C2 96712072 96712131
refGene_gn=PLD1;refGene_tx=XM_003991907.5;refGene_gn=PLD1;refGene_tx=XM_019840109.2;refGene_gn=PLD1;refGene_tx=XM_019840110.2;refGene_gn=PLD1;refGene_tx=XM_019840111.2;refGene_gn=PLD1;refGene_tx=XM_019840112.2;refGene_gn=PLD1;refGene_tx=XM_023260401.1
C2 96713599 96713692
refGene_gn=PLD1;refGene_tx=XM_003991907.5;refGene_gn=PLD1;refGene_tx=XM_019840109.2;refGene_gn=PLD1;refGene_tx=XM_019840110.2;refGene_gn=PLD1;refGene_tx=XM_019840111.2;refGene_gn=PLD1;refGene_tx=XM_019840112.2;refGene_gn=PLD1;refGene_tx=XM_023260401.1
C2 96722148 96722301
refGene_gn=PLD1;refGene_tx=XM_003991907.5;refGene_gn=PLD1;refGene_tx=XM_019840109.2;refGene_gn=PLD1;refGene_tx=XM_019840110.2;refGene_gn=PLD1;refGene_tx=XM_019840111.2;refGene_gn=PLD1;refGene_tx=XM_019840112.2;refGene_gn=PLD1;refGene_tx=XM_023260401.1
C2 96723337 96723487
refGene_gn=PLD1;refGene_tx=XM_003991907.5;refGene_gn=PLD1;refGene_tx=XM_019840109.2;refGene_gn=PLD1;refGene_tx=XM_019840110.2;refGene_gn=PLD1;refGene_tx=XM_019840111.2;refGene_gn=PLD1;refGene_tx=XM_019840112.2;refGene_gn=PLD1;refGene_tx=XM_023260401.1
C2 96724189 96724273
refGene_gn=PLD1;refGene_tx=XM_003991907.5;refGene_gn=PLD1;refGene_tx=XM_019840109.2;refGene_gn=PLD1;refGene_tx=XM_019840110.2;refGene_gn=PLD1;refGene_tx=XM_019840111.2;refGene_gn=PLD1;refGene_tx=XM_019840112.2;refGene_gn=PLD1;refGene_tx=XM_023260401.1
C2 96738203 96738285
refGene_gn=PLD1;refGene_tx=XM_003991907.5;refGene_gn=PLD1;refGene_tx=XM_019840109.2;refGene_gn=PLD1;refGene_tx=XM_019840110.2;refGene_gn=PLD1;refGene_tx=XM_019840111.2;refGene_gn=PLD1;refGene_tx=XM_019840112.2;refGene_gn=PLD1;refGene_tx=XM_023260401.1
C2 96748205 96748316
refGene_gn=PLD1;refGene_tx=XM_003991907.5;refGene_gn=PLD1;refGene_tx=XM_019840109.2;refGene_gn=PLD1;refGene_tx=XM_019840110.2;refGene_gn=PLD1;refGene_tx=XM_019840111.2;refGene_gn=PLD1;refGene_tx=XM_019840112.2;refGene_gn=PLD1;refGene_tx=XM_023260401.1
C2 96750435 96750636
refGene_gn=PLD1;refGene_tx=XM_003991907.5;refGene_gn=PLD1;refGene_tx=XM_019840109.2;refGene_gn=PLD1;refGene_tx=XM_019840110.2;refGene_gn=PLD1;refGene_tx=XM_019840111.2;refGene_gn=PLD1;refGene_tx=XM_019840112.2;refGene_gn=PLD1;refGene_tx=XM_023260401.1

C2 96751632 96751846
refGene_gn=PLD1;refGene_tx=XM_003991907.5;refGene_gn=PLD1;refGene_tx=XM_019840109.2;refGene_gn=PLD1;refGene_tx=XM_019840110.2;refGene_gn=PLD1;refGene_tx=XM_019840111.2;refGene_gn=PLD1;refGene_tx=XM_019840112.2;refGene_gn=PLD1;refGene_tx=XM_023260401.1
C2 96752430 96752544
refGene_gn=PLD1;refGene_tx=XM_003991907.5;refGene_gn=PLD1;refGene_tx=XM_019840109.2;refGene_gn=PLD1;refGene_tx=XM_019840111.2;refGene_gn=PLD1;refGene_tx=XM_019840112.2;refGene_gn=PLD1;refGene_tx=XM_023260401.1
C2 96757609 96757738
refGene_gn=PLD1;refGene_tx=XM_003991907.5;refGene_gn=PLD1;refGene_tx=XM_019840109.2;refGene_gn=PLD1;refGene_tx=XM_019840110.2;refGene_gn=PLD1;refGene_tx=XM_019840111.2;refGene_gn=PLD1;refGene_tx=XM_019840112.2;refGene_gn=PLD1;refGene_tx=XM_023260401.1
C2 96758482 96758601
refGene_gn=PLD1;refGene_tx=XM_003991907.5;refGene_gn=PLD1;refGene_tx=XM_019840109.2;refGene_gn=PLD1;refGene_tx=XM_019840110.2;refGene_gn=PLD1;refGene_tx=XM_019840111.2;refGene_gn=PLD1;refGene_tx=XM_019840112.2;refGene_gn=PLD1;refGene_tx=XM_023260401.1
C2 96760029 96760143
refGene_gn=PLD1;refGene_tx=XM_003991907.5;refGene_gn=PLD1;refGene_tx=XM_019840109.2;refGene_gn=PLD1;refGene_tx=XM_019840110.2;refGene_gn=PLD1;refGene_tx=XM_019840111.2;refGene_gn=PLD1;refGene_tx=XM_019840112.2;refGene_gn=PLD1;refGene_tx=XM_023260401.1
C2 96770885 96770996
refGene_gn=PLD1;refGene_tx=XM_003991907.5;refGene_gn=PLD1;refGene_tx=XM_019840109.2;refGene_gn=PLD1;refGene_tx=XM_019840110.2;refGene_gn=PLD1;refGene_tx=XM_019840111.2;refGene_gn=PLD1;refGene_tx=XM_019840112.2;refGene_gn=PLD1;refGene_tx=XM_023260401.1
C2 96773273 96773362
refGene_gn=PLD1;refGene_tx=XM_003991907.5;refGene_gn=PLD1;refGene_tx=XM_019840109.2;refGene_gn=PLD1;refGene_tx=XM_019840110.2;refGene_gn=PLD1;refGene_tx=XM_019840111.2;refGene_gn=PLD1;refGene_tx=XM_019840112.2;refGene_gn=PLD1;refGene_tx=XM_023260401.1
C2 96785233 96785347
refGene_gn=PLD1;refGene_tx=XM_003991907.5;refGene_gn=PLD1;refGene_tx=XM_019840109.2;refGene_gn=PLD1;refGene_tx=XM_019840110.2;refGene_gn=PLD1;refGene_tx=XM_019840111.2;refGene_gn=PLD1;refGene_tx=XM_019840112.2;refGene_gn=PLD1;refGene_tx=XM_023260401.1
C2 96787580 96787630
refGene_gn=PLD1;refGene_tx=XM_003991907.5;refGene_gn=PLD1;refGene_tx=XM_019840109.2;refGene_gn=PLD1;refGene_tx=XM_019840110.2;refGene_gn=PLD1;refGene_tx=XM_019840111.2;refGene_gn=PLD1;refGene_tx=XM_019840112.2;refGene_gn=PLD1;refGene_tx=XM_023260401.1
C2 96805713 96805848
refGene_gn=PLD1;refGene_tx=XM_003991907.5;refGene_gn=PLD1;refGene_tx=XM_019840109.2;refGene_gn=PLD1;refGene_tx=XM_019840110.2;refGene_gn=PLD1;refGene_tx=XM_019840111.2;refGene_gn=PLD1;refGene_tx=XM_019840112.2;refGene_gn=PLD1;refGene_tx=XM_023260401.1
C2 96813385 96813539

refGene_gn=PLD1;refGene_tx=XM_003991907.5;refGene_gn=PLD1;refGene_tx=XM_019840109.2;refGene_gn=PLD1;refGene_tx=XM_019840110.2;refGene_gn=PLD1;refGene_tx=XM_019840111.2;refGene_gn=PLD1;refGene_tx=XM_019840112.2;refGene_gn=PLD1;refGene_tx=XM_023260401.1
C2 96821432 96821550
refGene_gn=PLD1;refGene_tx=XM_003991907.5;refGene_gn=PLD1;refGene_tx=XM_019840109.2;refGene_gn=PLD1;refGene_tx=XM_019840110.2;refGene_gn=PLD1;refGene_tx=XM_019840111.2;refGene_gn=PLD1;refGene_tx=XM_023260401.1
C2 96823681 96823906
refGene_gn=PLD1;refGene_tx=XM_003991907.5;refGene_gn=PLD1;refGene_tx=XM_019840109.2;refGene_gn=PLD1;refGene_tx=XM_019840110.2;refGene_gn=PLD1;refGene_tx=XM_019840111.2;refGene_gn=PLD1;refGene_tx=XM_019840112.2;refGene_gn=PLD1;refGene_tx=XM_023260401.1
C2 96948479 96948536
refGene_gn=TNIK;refGene_tx=XM_006936265.4;refGene_gn=TNIK;refGene_tx=XM_019840114.2;refGene_gn=TNIK;refGene_tx=XM_019840115.2;refGene_gn=TNIK;refGene_tx=XM_019840116.2;refGene_gn=TNIK;refGene_tx=XM_019840117.2;refGene_gn=TNIK;refGene_tx=XM_019840118.2;refGene_gn=TNIK;refGene_tx=XM_019840119.2;refGene_gn=TNIK;refGene_tx=XM_019840120.2;refGene_gn=TNIK;refGene_tx=XM_023260402.1;refGene_gn=TNIK;refGene_tx=XM_023260403.1
C2 97035503 97035569
refGene_gn=TNIK;refGene_tx=XM_006936265.4;refGene_gn=TNIK;refGene_tx=XM_019840114.2;refGene_gn=TNIK;refGene_tx=XM_019840115.2;refGene_gn=TNIK;refGene_tx=XM_019840116.2;refGene_gn=TNIK;refGene_tx=XM_019840117.2;refGene_gn=TNIK;refGene_tx=XM_019840118.2;refGene_gn=TNIK;refGene_tx=XM_019840119.2;refGene_gn=TNIK;refGene_tx=XM_019840120.2;refGene_gn=TNIK;refGene_tx=XM_023260402.1;refGene_gn=TNIK;refGene_tx=XM_023260403.1
C2 97180064 97180121
refGene_gn=TNIK;refGene_tx=XM_006936265.4;refGene_gn=TNIK;refGene_tx=XM_019840114.2;refGene_gn=TNIK;refGene_tx=XM_019840115.2;refGene_gn=TNIK;refGene_tx=XM_019840116.2;refGene_gn=TNIK;refGene_tx=XM_019840117.2;refGene_gn=TNIK;refGene_tx=XM_019840118.2;refGene_gn=TNIK;refGene_tx=XM_019840119.2;refGene_gn=TNIK;refGene_tx=XM_019840120.2;refGene_gn=TNIK;refGene_tx=XM_023260402.1;refGene_gn=TNIK;refGene_tx=XM_023260403.1
C2 97194880 97195006
refGene_gn=TNIK;refGene_tx=XM_006936265.4;refGene_gn=TNIK;refGene_tx=XM_019840114.2;refGene_gn=TNIK;refGene_tx=XM_019840115.2;refGene_gn=TNIK;refGene_tx=XM_019840116.2;refGene_gn=TNIK;refGene_tx=XM_019840117.2;refGene_gn=TNIK;refGene_tx=XM_019840118.2;refGene_gn=TNIK;refGene_tx=XM_019840119.2;refGene_gn=TNIK;refGene_tx=XM_019840120.2;refGene_gn=TNIK;refGene_tx=XM_023260402.1;refGene_gn=TNIK;refGene_tx=XM_023260403.1
C2 97211565 97211676
refGene_gn=TNIK;refGene_tx=XM_006936265.4;refGene_gn=TNIK;refGene_tx=XM_019840114.2;refGene_gn=TNIK;refGene_tx=XM_019840115.2;refGene_gn=TNIK;refGene_tx=XM_019840116.2;refGene_gn=TNIK;refGene_tx=XM_019840117.2;refGene_gn=TNIK;refGene_tx=XM_019840118.2;refGene_gn=TNIK;refGene_tx=XM_019840119.2;refGene_gn=TNIK;refGene_tx=XM_019840120.2;refGene_gn=TNIK;refGene_tx=XM_023260402.1;refGene_gn=TNIK;refGene_tx=XM_023260403.1
C2 97215436 97215527
refGene_gn=TNIK;refGene_tx=XM_006936265.4;refGene_gn=TNIK;refGene_tx=X

M_019840114.2;refGene_gn=TNIK;refGene_tx=XM_019840115.2;refGene_gn=TNIK;refGene_tx=XM_019840116.2;refGene_gn=TNIK;refGene_tx=XM_019840117.2;refGene_gn=TNIK;refGene_tx=XM_019840118.2;refGene_gn=TNIK;refGene_tx=XM_019840119.2;refGene_gn=TNIK;refGene_tx=XM_019840120.2;refGene_gn=TNIK;refGene_tx=XM_023260402.1;refGene_gn=TNIK;refGene_tx=XM_023260403.1
C2 97217712 97217843
refGene_gn=TNIK;refGene_tx=XM_006936265.4;refGene_gn=TNIK;refGene_tx=XM_019840114.2;refGene_gn=TNIK;refGene_tx=XM_019840115.2;refGene_gn=TNIK;refGene_tx=XM_019840116.2;refGene_gn=TNIK;refGene_tx=XM_019840117.2;refGene_gn=TNIK;refGene_tx=XM_019840118.2;refGene_gn=TNIK;refGene_tx=XM_019840119.2;refGene_gn=TNIK;refGene_tx=XM_019840120.2;refGene_gn=TNIK;refGene_tx=XM_023260402.1;refGene_gn=TNIK;refGene_tx=XM_023260403.1
C2 97228876 97228931
refGene_gn=TNIK;refGene_tx=XM_006936265.4;refGene_gn=TNIK;refGene_tx=XM_019840114.2;refGene_gn=TNIK;refGene_tx=XM_019840115.2;refGene_gn=TNIK;refGene_tx=XM_019840116.2;refGene_gn=TNIK;refGene_tx=XM_019840117.2;refGene_gn=TNIK;refGene_tx=XM_019840118.2;refGene_gn=TNIK;refGene_tx=XM_019840119.2;refGene_gn=TNIK;refGene_tx=XM_019840120.2;refGene_gn=TNIK;refGene_tx=XM_023260402.1;refGene_gn=TNIK;refGene_tx=XM_023260403.1
C2 97230906 97230985
refGene_gn=TNIK;refGene_tx=XM_006936265.4;refGene_gn=TNIK;refGene_tx=XM_019840114.2;refGene_gn=TNIK;refGene_tx=XM_019840115.2;refGene_gn=TNIK;refGene_tx=XM_019840116.2;refGene_gn=TNIK;refGene_tx=XM_019840117.2;refGene_gn=TNIK;refGene_tx=XM_019840118.2;refGene_gn=TNIK;refGene_tx=XM_019840119.2;refGene_gn=TNIK;refGene_tx=XM_019840120.2;refGene_gn=TNIK;refGene_tx=XM_023260402.1;refGene_gn=TNIK;refGene_tx=XM_023260403.1
C2 97238717 97238893
refGene_gn=TNIK;refGene_tx=XM_006936265.4;refGene_gn=TNIK;refGene_tx=XM_019840114.2;refGene_gn=TNIK;refGene_tx=XM_019840115.2;refGene_gn=TNIK;refGene_tx=XM_019840116.2;refGene_gn=TNIK;refGene_tx=XM_019840117.2;refGene_gn=TNIK;refGene_tx=XM_019840118.2;refGene_gn=TNIK;refGene_tx=XM_019840119.2;refGene_gn=TNIK;refGene_tx=XM_019840120.2;refGene_gn=TNIK;refGene_tx=XM_023260402.1;refGene_gn=TNIK;refGene_tx=XM_023260403.1
C2 97243732 97243799
refGene_gn=TNIK;refGene_tx=XM_006936265.4;refGene_gn=TNIK;refGene_tx=XM_019840114.2;refGene_gn=TNIK;refGene_tx=XM_019840115.2;refGene_gn=TNIK;refGene_tx=XM_019840116.2;refGene_gn=TNIK;refGene_tx=XM_019840117.2;refGene_gn=TNIK;refGene_tx=XM_019840118.2;refGene_gn=TNIK;refGene_tx=XM_019840119.2;refGene_gn=TNIK;refGene_tx=XM_019840120.2;refGene_gn=TNIK;refGene_tx=XM_023260402.1;refGene_gn=TNIK;refGene_tx=XM_023260403.1
C2 97247487 97247692
refGene_gn=TNIK;refGene_tx=XM_006936265.4;refGene_gn=TNIK;refGene_tx=XM_019840114.2;refGene_gn=TNIK;refGene_tx=XM_019840115.2;refGene_gn=TNIK;refGene_tx=XM_019840116.2;refGene_gn=TNIK;refGene_tx=XM_019840117.2;refGene_gn=TNIK;refGene_tx=XM_019840118.2;refGene_gn=TNIK;refGene_tx=XM_019840119.2;refGene_gn=TNIK;refGene_tx=XM_019840120.2;refGene_gn=TNIK;refGene_tx=XM_023260402.1;refGene_gn=TNIK;refGene_tx=XM_023260403.1
C2 97267982 97268093
refGene_gn=TNIK;refGene_tx=XM_006936265.4;refGene_gn=TNIK;refGene_tx=XM_019840114.2;refGene_gn=TNIK;refGene_tx=XM_019840115.2;refGene_gn=TNI

K;refGene_tx=XM_019840116.2;refGene_gn=TNIK;refGene_tx=XM_019840117.2;
refGene_gn=TNIK;refGene_tx=XM_019840118.2;refGene_gn=TNIK;refGene_tx=X
M_019840119.2;refGene_gn=TNIK;refGene_tx=XM_019840120.2;refGene_gn=TNI
K;refGene_tx=XM_023260402.1;refGene_gn=TNIK;refGene_tx=XM_023260403.1
C2 97268921 97269008
refGene_gn=TNIK;refGene_tx=XM_006936265.4;refGene_gn=TNIK;refGene_tx=X
M_019840114.2;refGene_gn=TNIK;refGene_tx=XM_019840116.2;refGene_gn=TNI
K;refGene_tx=XM_019840117.2;refGene_gn=TNIK;refGene_tx=XM_019840120.2;
refGene_gn=TNIK;refGene_tx=XM_023260402.1;refGene_gn=TNIK;refGene_tx=X
M_023260403.1
C2 97270014 97270203
refGene_gn=TNIK;refGene_tx=XM_006936265.4;refGene_gn=TNIK;refGene_tx=X
M_019840114.2;refGene_gn=TNIK;refGene_tx=XM_019840115.2;refGene_gn=TNI
K;refGene_tx=XM_019840116.2;refGene_gn=TNIK;refGene_tx=XM_019840117.2;
refGene_gn=TNIK;refGene_tx=XM_019840118.2;refGene_gn=TNIK;refGene_tx=X
M_019840119.2;refGene_gn=TNIK;refGene_tx=XM_019840120.2;refGene_gn=TNI
K;refGene_tx=XM_023260402.1;refGene_gn=TNIK;refGene_tx=XM_023260403.1
C2 97280125 97280290
refGene_gn=TNIK;refGene_tx=XM_006936265.4;refGene_gn=TNIK;refGene_tx=X
M_019840114.2;refGene_gn=TNIK;refGene_tx=XM_019840115.2;refGene_gn=TNI
K;refGene_tx=XM_019840116.2;refGene_gn=TNIK;refGene_tx=XM_019840117.2;
refGene_gn=TNIK;refGene_tx=XM_019840118.2;refGene_gn=TNIK;refGene_tx=X
M_019840119.2;refGene_gn=TNIK;refGene_tx=XM_019840120.2;refGene_gn=TNI
K;refGene_tx=XM_023260402.1;refGene_gn=TNIK;refGene_tx=XM_023260403.1
C2 97282502 97282742
refGene_gn=TNIK;refGene_tx=XM_006936265.4;refGene_gn=TNIK;refGene_tx=X
M_019840114.2;refGene_gn=TNIK;refGene_tx=XM_019840115.2;refGene_gn=TNI
K;refGene_tx=XM_019840116.2;refGene_gn=TNIK;refGene_tx=XM_019840117.2;
refGene_gn=TNIK;refGene_tx=XM_019840118.2;refGene_gn=TNIK;refGene_tx=X
M_019840119.2;refGene_gn=TNIK;refGene_tx=XM_019840120.2;refGene_gn=TNI
K;refGene_tx=XM_023260402.1;refGene_gn=TNIK;refGene_tx=XM_023260403.1
C2 97285416 97285523
refGene_gn=TNIK;refGene_tx=XM_006936265.4;refGene_gn=TNIK;refGene_tx=X
M_019840114.2;refGene_gn=TNIK;refGene_tx=XM_019840115.2;refGene_gn=TNI
K;refGene_tx=XM_019840116.2;refGene_gn=TNIK;refGene_tx=XM_019840117.2;
refGene_gn=TNIK;refGene_tx=XM_019840118.2;refGene_gn=TNIK;refGene_tx=X
M_019840119.2;refGene_gn=TNIK;refGene_tx=XM_019840120.2;refGene_gn=TNI
K;refGene_tx=XM_023260402.1;refGene_gn=TNIK;refGene_tx=XM_023260403.1
C2 97294458 97294622
refGene_gn=TNIK;refGene_tx=XM_006936265.4;refGene_gn=TNIK;refGene_tx=X
M_019840114.2;refGene_gn=TNIK;refGene_tx=XM_019840115.2;refGene_gn=TNI
K;refGene_tx=XM_019840116.2;refGene_gn=TNIK;refGene_tx=XM_019840117.2;
refGene_gn=TNIK;refGene_tx=XM_019840118.2;refGene_gn=TNIK;refGene_tx=X
M_019840119.2;refGene_gn=TNIK;refGene_tx=XM_019840120.2;refGene_gn=TNI
K;refGene_tx=XM_023260402.1;refGene_gn=TNIK;refGene_tx=XM_023260403.1
C2 97296652 97296750
refGene_gn=TNIK;refGene_tx=XM_006936265.4;refGene_gn=TNIK;refGene_tx=X
M_019840114.2;refGene_gn=TNIK;refGene_tx=XM_019840115.2;refGene_gn=TNI
K;refGene_tx=XM_019840116.2;refGene_gn=TNIK;refGene_tx=XM_019840117.2;
refGene_gn=TNIK;refGene_tx=XM_019840118.2;refGene_gn=TNIK;refGene_tx=X

M_019840119.2;refGene_gn=TNIK;refGene_tx=XM_019840120.2;refGene_gn=TNIK;refGene_tx=XM_023260402.1;refGene_gn=TNIK;refGene_tx=XM_023260403.1
C2 97297609 97297633
refGene_gn=TNIK;refGene_tx=XM_006936265.4;refGene_gn=TNIK;refGene_tx=XM_019840114.2;refGene_gn=TNIK;refGene_tx=XM_019840115.2;refGene_gn=TNIK;refGene_tx=XM_019840116.2;refGene_gn=TNIK;refGene_tx=XM_019840118.2;refGene_gn=TNIK;refGene_tx=XM_019840120.2;refGene_gn=TNIK;refGene_tx=XM_023260403.1
C2 97302214 97302223
refGene_gn=TNIK;refGene_tx=XM_019840114.2;refGene_gn=TNIK;refGene_tx=XM_019840115.2;refGene_gn=TNIK;refGene_tx=XM_019840120.2;refGene_gn=TNIK;refGene_tx=XM_023260403.1
C2 97303275 97303460
refGene_gn=TNIK;refGene_tx=XM_006936265.4;refGene_gn=TNIK;refGene_tx=XM_019840114.2;refGene_gn=TNIK;refGene_tx=XM_019840115.2;refGene_gn=TNIK;refGene_tx=XM_019840116.2;refGene_gn=TNIK;refGene_tx=XM_019840117.2;refGene_gn=TNIK;refGene_tx=XM_019840118.2;refGene_gn=TNIK;refGene_tx=XM_019840119.2;refGene_gn=TNIK;refGene_tx=XM_019840120.2;refGene_gn=TNIK;refGene_tx=XM_023260402.1;refGene_gn=TNIK;refGene_tx=XM_023260403.1
C2 97315083 97315213
refGene_gn=TNIK;refGene_tx=XM_006936265.4;refGene_gn=TNIK;refGene_tx=XM_019840114.2;refGene_gn=TNIK;refGene_tx=XM_019840115.2;refGene_gn=TNIK;refGene_tx=XM_019840116.2;refGene_gn=TNIK;refGene_tx=XM_019840117.2;refGene_gn=TNIK;refGene_tx=XM_019840118.2;refGene_gn=TNIK;refGene_tx=XM_019840119.2;refGene_gn=TNIK;refGene_tx=XM_019840120.2;refGene_gn=TNIK;refGene_tx=XM_023260402.1;refGene_gn=TNIK;refGene_tx=XM_023260403.1
C2 97320312 97320477
refGene_gn=TNIK;refGene_tx=XM_006936265.4;refGene_gn=TNIK;refGene_tx=XM_019840114.2;refGene_gn=TNIK;refGene_tx=XM_019840115.2;refGene_gn=TNIK;refGene_tx=XM_019840116.2;refGene_gn=TNIK;refGene_tx=XM_019840117.2;refGene_gn=TNIK;refGene_tx=XM_019840118.2;refGene_gn=TNIK;refGene_tx=XM_019840119.2;refGene_gn=TNIK;refGene_tx=XM_019840120.2;refGene_gn=TNIK;refGene_tx=XM_023260402.1;refGene_gn=TNIK;refGene_tx=XM_023260403.1
C2 97322511 97322623
refGene_gn=TNIK;refGene_tx=XM_006936265.4;refGene_gn=TNIK;refGene_tx=XM_019840114.2;refGene_gn=TNIK;refGene_tx=XM_019840115.2;refGene_gn=TNIK;refGene_tx=XM_019840116.2;refGene_gn=TNIK;refGene_tx=XM_019840117.2;refGene_gn=TNIK;refGene_tx=XM_019840118.2;refGene_gn=TNIK;refGene_tx=XM_019840119.2;refGene_gn=TNIK;refGene_tx=XM_019840120.2;refGene_gn=TNIK;refGene_tx=XM_023260402.1;refGene_gn=TNIK;refGene_tx=XM_023260403.1
C2 97322874 97322961
refGene_gn=TNIK;refGene_tx=XM_006936265.4;refGene_gn=TNIK;refGene_tx=XM_019840114.2;refGene_gn=TNIK;refGene_tx=XM_019840115.2;refGene_gn=TNIK;refGene_tx=XM_019840120.2;refGene_gn=TNIK;refGene_tx=XM_023260402.1
C2 97323454 97323625
refGene_gn=TNIK;refGene_tx=XM_006936265.4;refGene_gn=TNIK;refGene_tx=XM_019840114.2;refGene_gn=TNIK;refGene_tx=XM_019840115.2;refGene_gn=TNIK;refGene_tx=XM_019840116.2;refGene_gn=TNIK;refGene_tx=XM_019840117.2;refGene_gn=TNIK;refGene_tx=XM_019840118.2;refGene_gn=TNIK;refGene_tx=XM_019840119.2;refGene_gn=TNIK;refGene_tx=XM_019840120.2;refGene_gn=TNI

K;refGene_tx=XM_023260402.1;refGene_gn=TNIK;refGene_tx=XM_023260403.1
C2 97325559 97325703
refGene_gn=TNIK;refGene_tx=XM_006936265.4;refGene_gn=TNIK;refGene_tx=XM_019840114.2;refGene_gn=TNIK;refGene_tx=XM_019840115.2;refGene_gn=TNIK;refGene_tx=XM_019840116.2;refGene_gn=TNIK;refGene_tx=XM_019840117.2;refGene_gn=TNIK;refGene_tx=XM_019840118.2;refGene_gn=TNIK;refGene_tx=XM_019840119.2;refGene_gn=TNIK;refGene_tx=XM_019840120.2;refGene_gn=TNIK;refGene_tx=XM_023260402.1;refGene_gn=TNIK;refGene_tx=XM_023260403.1
C2 97328673 97328808
refGene_gn=TNIK;refGene_tx=XM_006936265.4;refGene_gn=TNIK;refGene_tx=XM_019840114.2;refGene_gn=TNIK;refGene_tx=XM_019840115.2;refGene_gn=TNIK;refGene_tx=XM_019840116.2;refGene_gn=TNIK;refGene_tx=XM_019840117.2;refGene_gn=TNIK;refGene_tx=XM_019840118.2;refGene_gn=TNIK;refGene_tx=XM_019840119.2;refGene_gn=TNIK;refGene_tx=XM_019840120.2;refGene_gn=TNIK;refGene_tx=XM_023260402.1;refGene_gn=TNIK;refGene_tx=XM_023260403.1
C2 97332987 97333088
refGene_gn=TNIK;refGene_tx=XM_006936265.4;refGene_gn=TNIK;refGene_tx=XM_019840114.2;refGene_gn=TNIK;refGene_tx=XM_019840115.2;refGene_gn=TNIK;refGene_tx=XM_019840116.2;refGene_gn=TNIK;refGene_tx=XM_019840117.2;refGene_gn=TNIK;refGene_tx=XM_019840118.2;refGene_gn=TNIK;refGene_tx=XM_019840119.2;refGene_gn=TNIK;refGene_tx=XM_019840120.2;refGene_gn=TNIK;refGene_tx=XM_023260402.1;refGene_gn=TNIK;refGene_tx=XM_023260403.1
C2 97335335 97335485
refGene_gn=TNIK;refGene_tx=XM_006936265.4;refGene_gn=TNIK;refGene_tx=XM_019840114.2;refGene_gn=TNIK;refGene_tx=XM_019840115.2;refGene_gn=TNIK;refGene_tx=XM_019840116.2;refGene_gn=TNIK;refGene_tx=XM_019840117.2;refGene_gn=TNIK;refGene_tx=XM_019840118.2;refGene_gn=TNIK;refGene_tx=XM_019840119.2;refGene_gn=TNIK;refGene_tx=XM_019840120.2;refGene_gn=TNIK;refGene_tx=XM_023260402.1;refGene_gn=TNIK;refGene_tx=XM_023260403.1
C2 97337205 97337365
refGene_gn=TNIK;refGene_tx=XM_006936265.4;refGene_gn=TNIK;refGene_tx=XM_019840114.2;refGene_gn=TNIK;refGene_tx=XM_019840115.2;refGene_gn=TNIK;refGene_tx=XM_019840116.2;refGene_gn=TNIK;refGene_tx=XM_019840117.2;refGene_gn=TNIK;refGene_tx=XM_019840118.2;refGene_gn=TNIK;refGene_tx=XM_019840119.2;refGene_gn=TNIK;refGene_tx=XM_019840120.2;refGene_gn=TNIK;refGene_tx=XM_023260402.1;refGene_gn=TNIK;refGene_tx=XM_023260403.1
C2 97337609 97337749
refGene_gn=TNIK;refGene_tx=XM_006936265.4;refGene_gn=TNIK;refGene_tx=XM_019840114.2;refGene_gn=TNIK;refGene_tx=XM_019840115.2;refGene_gn=TNIK;refGene_tx=XM_019840116.2;refGene_gn=TNIK;refGene_tx=XM_019840117.2;refGene_gn=TNIK;refGene_tx=XM_019840118.2;refGene_gn=TNIK;refGene_tx=XM_019840119.2;refGene_gn=TNIK;refGene_tx=XM_019840120.2;refGene_gn=TNIK;refGene_tx=XM_023260402.1;refGene_gn=TNIK;refGene_tx=XM_023260403.1
C2 97339882 97339966
refGene_gn=TNIK;refGene_tx=XM_006936265.4;refGene_gn=TNIK;refGene_tx=XM_019840114.2;refGene_gn=TNIK;refGene_tx=XM_019840115.2;refGene_gn=TNIK;refGene_tx=XM_019840116.2;refGene_gn=TNIK;refGene_tx=XM_019840117.2;refGene_gn=TNIK;refGene_tx=XM_019840118.2;refGene_gn=TNIK;refGene_tx=XM_019840119.2;refGene_gn=TNIK;refGene_tx=XM_019840120.2;refGene_gn=TNIK;refGene_tx=XM_023260402.1;refGene_gn=TNIK;refGene_tx=XM_023260403.1

C2 97371591 97371606 refGene_gn=SLC2A2;refGene_tx=XM_003991916.4
C2 97379587 97379680 refGene_gn=SLC2A2;refGene_tx=XM_003991916.4
C2 97382908 97383171 refGene_gn=SLC2A2;refGene_tx=XM_003991916.4
C2 97390110 97390235 refGene_gn=SLC2A2;refGene_tx=XM_003991916.4
C2 97392863 97392979 refGene_gn=SLC2A2;refGene_tx=XM_003991916.4
C2 97394086 97394249 refGene_gn=SLC2A2;refGene_tx=XM_003991916.4
C2 97394748 97394936 refGene_gn=SLC2A2;refGene_tx=XM_003991916.4
C2 97397185 97397290 refGene_gn=SLC2A2;refGene_tx=XM_003991916.4
C2 97400638 97400740 refGene_gn=SLC2A2;refGene_tx=XM_003991916.4
C2 97401436 97401640 refGene_gn=SLC2A2;refGene_tx=XM_003991916.4
C2 97401723 97401924 refGene_gn=SLC2A2;refGene_tx=XM_003991916.4
C2 97452376 97452541
refGene_gn=EIF5A2;refGene_tx=XM_003991917.5;refGene_gn=EIF5A2;refGene_tx=XM_006936268.4;refGene_gn=EIF5A2;refGene_tx=XM_023260404.1;refGene_gn=EIF5A2;refGene_tx=XM_023260405.1
C2 97453076 97453181
refGene_gn=EIF5A2;refGene_tx=XM_003991917.5;refGene_gn=EIF5A2;refGene_tx=XM_006936268.4;refGene_gn=EIF5A2;refGene_tx=XM_023260404.1;refGene_gn=EIF5A2;refGene_tx=XM_023260405.1
C2 97455931 97456007 refGene_gn=EIF5A2;refGene_tx=XM_023260405.1
C2 97460011 97460143
refGene_gn=EIF5A2;refGene_tx=XM_003991917.5;refGene_gn=EIF5A2;refGene_tx=XM_006936268.4;refGene_gn=EIF5A2;refGene_tx=XM_023260404.1;refGene_gn=EIF5A2;refGene_tx=XM_023260405.1
C2 97460686 97460746
refGene_gn=EIF5A2;refGene_tx=XM_003991917.5;refGene_gn=EIF5A2;refGene_tx=XM_006936268.4;refGene_gn=EIF5A2;refGene_tx=XM_023260404.1
C2 97474586 97474595 refGene_gn=RPL22L1;refGene_tx=XM_003991918.4
C2 97476201 97476294 refGene_gn=RPL22L1;refGene_tx=XM_003991918.4
C2 97476459 97476578 refGene_gn=RPL22L1;refGene_tx=XM_003991918.4
C2 97478067 97478212 refGene_gn=RPL22L1;refGene_tx=XM_003991918.4
C2 97764051 97764355 refGene_gn=SLC7A14;refGene_tx=XM_006936269.4
C2 97785525 97785762 refGene_gn=SLC7A14;refGene_tx=XM_006936269.4
C2 97787417 97787635 refGene_gn=SLC7A14;refGene_tx=XM_006936269.4
C2 97796510 97796657 refGene_gn=SLC7A14;refGene_tx=XM_006936269.4
C2 97799133 97799342 refGene_gn=SLC7A14;refGene_tx=XM_006936269.4
C2 97801172 97802050 refGene_gn=SLC7A14;refGene_tx=XM_006936269.4
C2 97814407 97814730 refGene_gn=SLC7A14;refGene_tx=XM_006936269.4
C2 97846326 97846559 refGene_gn=CLDN11;refGene_tx=XM_003991920.5
C2 97856256 97856421 refGene_gn=CLDN11;refGene_tx=XM_003991920.5
C2 97860162 97860388 refGene_gn=CLDN11;refGene_tx=XM_003991920.5
C2 97878463 97878622
refGene_gn=SKIL;refGene_tx=XM_023260406.1;refGene_gn=SKIL;refGene_tx=XM_023260407.1
C2 97879276 97879501
refGene_gn=SKIL;refGene_tx=XM_023260406.1;refGene_gn=SKIL;refGene_tx=XM_023260407.1
C2 97879817 97880059
refGene_gn=SKIL;refGene_tx=XM_023260406.1;refGene_gn=SKIL;refGene_tx=XM_023260407.1

C2 97883303 97883536
refGene_gn=SKIL;refGene_tx=XM_023260406.1;refGene_gn=SKIL;refGene_tx=XM_023260407.1
C2 97885523 97885621
refGene_gn=SKIL;refGene_tx=XM_023260406.1;refGene_gn=SKIL;refGene_tx=XM_023260407.1
C2 97900688 97901786
refGene_gn=SKIL;refGene_tx=XM_023260406.1;refGene_gn=SKIL;refGene_tx=XM_023260407.1
C2 97939965 97940053 refGene_gn=PRKCI;refGene_tx=XM_003991921.4
C2 97944563 97944679 refGene_gn=PRKCI;refGene_tx=XM_003991921.4
C2 97945060 97945150 refGene_gn=PRKCI;refGene_tx=XM_003991921.4
C2 97947276 97947356 refGene_gn=PRKCI;refGene_tx=XM_003991921.4
C2 97948719 97948845 refGene_gn=PRKCI;refGene_tx=XM_003991921.4
C2 97949856 97949944 refGene_gn=PRKCI;refGene_tx=XM_003991921.4
C2 97954657 97954793 refGene_gn=PRKCI;refGene_tx=XM_003991921.4
C2 97957065 97957152 refGene_gn=PRKCI;refGene_tx=XM_003991921.4
C2 97958181 97958279 refGene_gn=PRKCI;refGene_tx=XM_003991921.4
C2 97958699 97958876 refGene_gn=PRKCI;refGene_tx=XM_003991921.4
C2 97961779 97961838 refGene_gn=PRKCI;refGene_tx=XM_003991921.4
C2 97968843 97968898 refGene_gn=PRKCI;refGene_tx=XM_003991921.4
C2 97972129 97972270 refGene_gn=PRKCI;refGene_tx=XM_003991921.4
C2 97973790 97973876 refGene_gn=PRKCI;refGene_tx=XM_003991921.4
C2 97976863 97976914 refGene_gn=PRKCI;refGene_tx=XM_003991921.4
C2 97978792 97978882 refGene_gn=PRKCI;refGene_tx=XM_003991921.4
C2 98001779 98001901 refGene_gn=PRKCI;refGene_tx=XM_003991921.4
C2 98018376 98018477 refGene_gn=PRKCI;refGene_tx=XM_003991921.4
C2 98041080 98041094
refGene_gn=PHC3;refGene_tx=XM_023260408.1;refGene_gn=PHC3;refGene_tx=XM_023260411.1;refGene_gn=PHC3;refGene_tx=XM_023260412.1;refGene_gn=PHC3;refGene_tx=XM_023260413.1;refGene_gn=PHC3;refGene_tx=XM_023260414.1
C2 98044095 98044261
refGene_gn=PHC3;refGene_tx=XM_023260408.1;refGene_gn=PHC3;refGene_tx=XM_023260409.1;refGene_gn=PHC3;refGene_tx=XM_023260410.1;refGene_gn=PHC3;refGene_tx=XM_023260411.1;refGene_gn=PHC3;refGene_tx=XM_023260412.1;refGene_gn=PHC3;refGene_tx=XM_023260413.1;refGene_gn=PHC3;refGene_tx=XM_023260414.1
C2 98048677 98048833
refGene_gn=PHC3;refGene_tx=XM_023260408.1;refGene_gn=PHC3;refGene_tx=XM_023260409.1;refGene_gn=PHC3;refGene_tx=XM_023260410.1;refGene_gn=PHC3;refGene_tx=XM_023260411.1;refGene_gn=PHC3;refGene_tx=XM_023260412.1;refGene_gn=PHC3;refGene_tx=XM_023260413.1;refGene_gn=PHC3;refGene_tx=XM_023260414.1
C2 98049946 98050024
refGene_gn=PHC3;refGene_tx=XM_023260408.1;refGene_gn=PHC3;refGene_tx=XM_023260409.1;refGene_gn=PHC3;refGene_tx=XM_023260410.1;refGene_gn=PHC3;refGene_tx=XM_023260411.1;refGene_gn=PHC3;refGene_tx=XM_023260412.1;refGene_gn=PHC3;refGene_tx=XM_023260413.1;refGene_gn=PHC3;refGene_tx=XM_023260414.1
C2 98063692 98063851

refGene_gn=PHC3;refGene_tx=XM_023260408.1;refGene_gn=PHC3;refGene_tx=XM_023260409.1;refGene_gn=PHC3;refGene_tx=XM_023260410.1;refGene_gn=PHC3;refGene_tx=XM_023260411.1;refGene_gn=PHC3;refGene_tx=XM_023260413.1;refGene_gn=PHC3;refGene_tx=XM_023260414.1
C2 98065872 98065971
refGene_gn=PHC3;refGene_tx=XM_023260408.1;refGene_gn=PHC3;refGene_tx=XM_023260409.1;refGene_gn=PHC3;refGene_tx=XM_023260410.1;refGene_gn=PHC3;refGene_tx=XM_023260411.1;refGene_gn=PHC3;refGene_tx=XM_023260412.1;refGene_gn=PHC3;refGene_tx=XM_023260413.1;refGene_gn=PHC3;refGene_tx=XM_023260414.1
C2 98074203 98074450
refGene_gn=PHC3;refGene_tx=XM_023260408.1;refGene_gn=PHC3;refGene_tx=XM_023260409.1;refGene_gn=PHC3;refGene_tx=XM_023260410.1;refGene_gn=PHC3;refGene_tx=XM_023260411.1;refGene_gn=PHC3;refGene_tx=XM_023260412.1;refGene_gn=PHC3;refGene_tx=XM_023260413.1;refGene_gn=PHC3;refGene_tx=XM_023260414.1
C2 98082283 98083155
refGene_gn=PHC3;refGene_tx=XM_023260408.1;refGene_gn=PHC3;refGene_tx=XM_023260409.1;refGene_gn=PHC3;refGene_tx=XM_023260410.1;refGene_gn=PHC3;refGene_tx=XM_023260411.1;refGene_gn=PHC3;refGene_tx=XM_023260412.1;refGene_gn=PHC3;refGene_tx=XM_023260413.1;refGene_gn=PHC3;refGene_tx=XM_023260414.1
C2 98083446 98083494
refGene_gn=PHC3;refGene_tx=XM_023260408.1;refGene_gn=PHC3;refGene_tx=XM_023260409.1;refGene_gn=PHC3;refGene_tx=XM_023260410.1;refGene_gn=PHC3;refGene_tx=XM_023260412.1;refGene_gn=PHC3;refGene_tx=XM_023260413.1;refGene_gn=PHC3;refGene_tx=XM_023260414.1
C2 98088851 98089002
refGene_gn=PHC3;refGene_tx=XM_023260408.1;refGene_gn=PHC3;refGene_tx=XM_023260409.1;refGene_gn=PHC3;refGene_tx=XM_023260410.1;refGene_gn=PHC3;refGene_tx=XM_023260411.1;refGene_gn=PHC3;refGene_tx=XM_023260412.1;refGene_gn=PHC3;refGene_tx=XM_023260413.1;refGene_gn=PHC3;refGene_tx=XM_023260414.1
C2 98093850 98094101
refGene_gn=PHC3;refGene_tx=XM_023260408.1;refGene_gn=PHC3;refGene_tx=XM_023260409.1;refGene_gn=PHC3;refGene_tx=XM_023260410.1;refGene_gn=PHC3;refGene_tx=XM_023260411.1;refGene_gn=PHC3;refGene_tx=XM_023260412.1;refGene_gn=PHC3;refGene_tx=XM_023260413.1;refGene_gn=PHC3;refGene_tx=XM_023260414.1
C2 98097944 98098104
refGene_gn=PHC3;refGene_tx=XM_023260408.1;refGene_gn=PHC3;refGene_tx=XM_023260409.1;refGene_gn=PHC3;refGene_tx=XM_023260410.1;refGene_gn=PHC3;refGene_tx=XM_023260411.1;refGene_gn=PHC3;refGene_tx=XM_023260412.1;refGene_gn=PHC3;refGene_tx=XM_023260413.1;refGene_gn=PHC3;refGene_tx=XM_023260414.1
C2 98105722 98105837
refGene_gn=PHC3;refGene_tx=XM_023260408.1;refGene_gn=PHC3;refGene_tx=XM_023260409.1;refGene_gn=PHC3;refGene_tx=XM_023260410.1;refGene_gn=PHC3;refGene_tx=XM_023260411.1;refGene_gn=PHC3;refGene_tx=XM_023260412.1;refGene_gn=PHC3;refGene_tx=XM_023260413.1;refGene_gn=PHC3;refGene_tx=X

M_023260414.1
C2	98109634	98109767
refGene_gn=PHC3;refGene_tx=XM_023260408.1;refGene_gn=PHC3;refGene_tx=XM_023260409.1;refGene_gn=PHC3;refGene_tx=XM_023260410.1;refGene_gn=PHC3;refGene_tx=XM_023260411.1;refGene_gn=PHC3;refGene_tx=XM_023260412.1;refGene_gn=PHC3;refGene_tx=XM_023260413.1
C2	98109858	98110090
refGene_gn=PHC3;refGene_tx=XM_023260408.1;refGene_gn=PHC3;refGene_tx=XM_023260409.1;refGene_gn=PHC3;refGene_tx=XM_023260410.1;refGene_gn=PHC3;refGene_tx=XM_023260411.1;refGene_gn=PHC3;refGene_tx=XM_023260412.1;refGene_gn=PHC3;refGene_tx=XM_023260413.1;refGene_gn=PHC3;refGene_tx=XM_023260414.1
C2	98115318	98115473
refGene_gn=PHC3;refGene_tx=XM_023260408.1;refGene_gn=PHC3;refGene_tx=XM_023260409.1;refGene_gn=PHC3;refGene_tx=XM_023260410.1;refGene_gn=PHC3;refGene_tx=XM_023260411.1;refGene_gn=PHC3;refGene_tx=XM_023260412.1
C2	98125500	98126517
refGene_gn=GPR160;refGene_tx=XM_011286107.3;refGene_gn=GPR160;refGene_tx=XM_019810448.2;refGene_gn=GPR160;refGene_tx=XM_019810450.1
C2	98203161	98203410
refGene_gn=LOC101087774;refGene_tx=XM_019811153.2
C2	98203621	98204170
refGene_gn=LOC101087774;refGene_tx=XM_019811153.2
C2	98219443	98219916	refGene_gn=SEC62;refGene_tx=XM_003991924.5
C2	98224022	98224142	refGene_gn=SEC62;refGene_tx=XM_003991924.5
C2	98225066	98225127	refGene_gn=SEC62;refGene_tx=XM_003991924.5
C2	98227510	98227603	refGene_gn=SEC62;refGene_tx=XM_003991924.5
C2	98227858	98228063	refGene_gn=SEC62;refGene_tx=XM_003991924.5
C2	98233401	98233507	refGene_gn=SEC62;refGene_tx=XM_003991924.5
C2	98234442	98234551	refGene_gn=SEC62;refGene_tx=XM_003991924.5
C2	98242754	98242790	refGene_gn=SEC62;refGene_tx=XM_003991924.5
C2	98270015	98270183
refGene_gn=SAMD7;refGene_tx=XM_003991925.3;refGene_gn=SAMD7;refGene_tx=XM_019810451.1
C2	98273303	98273414
refGene_gn=SAMD7;refGene_tx=XM_003991925.3;refGene_gn=SAMD7;refGene_tx=XM_019810451.1
C2	98280279	98280401
refGene_gn=SAMD7;refGene_tx=XM_003991925.3;refGene_gn=SAMD7;refGene_tx=XM_019810451.1
C2	98281634	98282257
refGene_gn=SAMD7;refGene_tx=XM_003991925.3;refGene_gn=SAMD7;refGene_tx=XM_019810451.1
C2	98283384	98283463
refGene_gn=SAMD7;refGene_tx=XM_003991925.3;refGene_gn=SAMD7;refGene_tx=XM_019810451.1
C2	98287059	98287184
refGene_gn=SAMD7;refGene_tx=XM_003991925.3;refGene_gn=SAMD7;refGene_tx=XM_019810451.1
C2	98288442	98288569

refGene_gn=SAMD7;refGene_tx=XM_003991925.3;refGene_gn=SAMD7;refGene_tx=XM_019810451.1
C2 98291038 98291141 refGene_gn=SAMD7;refGene_tx=XM_019810451.1
C2 98343586 98343779 refGene_gn=LRRC31;refGene_tx=XM_003991956.3
C2 98351844 98351988 refGene_gn=LRRC31;refGene_tx=XM_003991956.3
C2 98354115 98354283 refGene_gn=LRRC31;refGene_tx=XM_003991956.3
C2 98356931 98357099 refGene_gn=LRRC31;refGene_tx=XM_003991956.3
C2 98357319 98357487 refGene_gn=LRRC31;refGene_tx=XM_003991956.3
C2 98358459 98358627 refGene_gn=LRRC31;refGene_tx=XM_003991956.3
C2 98361672 98361840 refGene_gn=LRRC31;refGene_tx=XM_003991956.3
C2 98364541 98364709 refGene_gn=LRRC31;refGene_tx=XM_003991956.3
C2 98370926 98371282 refGene_gn=LRRC31;refGene_tx=XM_003991956.3
C2 98373508 98373658
refGene_gn=LRRIQ4;refGene_tx=XM_003991957.3;refGene_gn=LRRIQ4;refGene_tx=XM_019810453.2;refGene_gn=LRRIQ4;refGene_tx=XM_023260415.1
C2 98378730 98378927
refGene_gn=LRRIQ4;refGene_tx=XM_003991957.3;refGene_gn=LRRIQ4;refGene_tx=XM_019810453.2;refGene_gn=LRRIQ4;refGene_tx=XM_023260415.1
C2 98379938 98380077
refGene_gn=LRRIQ4;refGene_tx=XM_003991957.3;refGene_gn=LRRIQ4;refGene_tx=XM_019810453.2;refGene_gn=LRRIQ4;refGene_tx=XM_023260415.1
C2 98381559 98381733
refGene_gn=LRRIQ4;refGene_tx=XM_003991957.3;refGene_gn=LRRIQ4;refGene_tx=XM_019810453.2;refGene_gn=LRRIQ4;refGene_tx=XM_023260415.1
C2 98385663 98386711
refGene_gn=LRRIQ4;refGene_tx=XM_003991957.3;refGene_gn=LRRIQ4;refGene_tx=XM_019810453.2;refGene_gn=LRRIQ4;refGene_tx=XM_023260415.1
C2 98392523 98392540 refGene_gn=LRRIQ4;refGene_tx=XM_023260415.1
C2 98393594 98393697
refGene_gn=LRRC34;refGene_tx=XM_006936272.4;refGene_gn=LRRC34;refGene_tx=XM_023260416.1
C2 98398223 98398344
refGene_gn=LRRC34;refGene_tx=XM_006936272.4;refGene_gn=LRRC34;refGene_tx=XM_023260416.1
C2 98398853 98398972
refGene_gn=LRRC34;refGene_tx=XM_006936272.4;refGene_gn=LRRC34;refGene_tx=XM_023260416.1
C2 98399072 98399137
refGene_gn=LRRC34;refGene_tx=XM_006936272.4;refGene_gn=LRRC34;refGene_tx=XM_023260416.1
C2 98400163 98400247
refGene_gn=LRRC34;refGene_tx=XM_006936272.4;refGene_gn=LRRC34;refGene_tx=XM_023260416.1
C2 98403376 98403505
refGene_gn=LRRC34;refGene_tx=XM_006936272.4;refGene_gn=LRRC34;refGene_tx=XM_023260416.1
C2 98405235 98405331
refGene_gn=LRRC34;refGene_tx=XM_006936272.4;refGene_gn=LRRC34;refGene_tx=XM_023260416.1
C2 98408126 98408281

refGene_gn=LRRC34;refGene_tx=XM_006936272.4;refGene_gn=LRRC34;refGene_tx=XM_023260416.1
C2 98408667 98408823
refGene_gn=LRRC34;refGene_tx=XM_006936272.4;refGene_gn=LRRC34;refGene_tx=XM_023260416.1
C2 98409802 98409929 refGene_gn=LRRC34;refGene_tx=XM_006936272.4
C2 98411641 98411845
refGene_gn=LRRC34;refGene_tx=XM_006936272.4;refGene_gn=LRRC34;refGene_tx=XM_023260416.1
C2 98429639 98429902
refGene_gn=MYNN;refGene_tx=XM_003991926.5;refGene_gn=MYNN;refGene_tx=XM_006936273.4
C2 98431806 98431893
refGene_gn=MYNN;refGene_tx=XM_003991926.5;refGene_gn=MYNN;refGene_tx=XM_006936273.4
C2 98432905 98432989 refGene_gn=MYNN;refGene_tx=XM_003991926.5
C2 98433865 98434044
refGene_gn=MYNN;refGene_tx=XM_003991926.5;refGene_gn=MYNN;refGene_tx=XM_006936273.4
C2 98436038 98436198
refGene_gn=MYNN;refGene_tx=XM_003991926.5;refGene_gn=MYNN;refGene_tx=XM_006936273.4
C2 98436960 98437754
refGene_gn=MYNN;refGene_tx=XM_003991926.5;refGene_gn=MYNN;refGene_tx=XM_006936273.4
C2 98442402 98442668
refGene_gn=MYNN;refGene_tx=XM_003991926.5;refGene_gn=MYNN;refGene_tx=XM_006936273.4
C2 98446800 98447003 refGene_gn=ACTRT3;refGene_tx=XM_003991959.4
C2 98447799 98448718 refGene_gn=ACTRT3;refGene_tx=XM_003991959.4
C2 98524975 98525012
refGene_gn=MECOM;refGene_tx=XM_003991927.3;refGene_gn=MECOM;refGene_tx=XM_019810458.1;refGene_gn=MECOM;refGene_tx=XM_019810459.1;refGene_gn=MECOM;refGene_tx=XM_019810460.1;refGene_gn=MECOM;refGene_tx=XM_023260417.1
C2 98799414 98799752
refGene_gn=MECOM;refGene_tx=XM_003991927.3;refGene_gn=MECOM;refGene_tx=XM_019810458.1;refGene_gn=MECOM;refGene_tx=XM_019810459.1;refGene_gn=MECOM;refGene_tx=XM_019810460.1;refGene_gn=MECOM;refGene_tx=XM_023260417.1
C2 99027294 99027429
refGene_gn=MECOM;refGene_tx=XM_003991927.3;refGene_gn=MECOM;refGene_tx=XM_019810458.1;refGene_gn=MECOM;refGene_tx=XM_019810459.1;refGene_gn=MECOM;refGene_tx=XM_019810460.1;refGene_gn=MECOM;refGene_tx=XM_023260417.1
C2 99039404 99039507
refGene_gn=MECOM;refGene_tx=XM_003991927.3;refGene_gn=MECOM;refGene_tx=XM_019810458.1;refGene_gn=MECOM;refGene_tx=XM_019810459.1;refGene_gn=MECOM;refGene_tx=XM_019810460.1;refGene_gn=MECOM;refGene_tx=XM_019810461.2;refGene_gn=MECOM;refGene_tx=XM_023260417.1

C2 99042882 99043099
refGene_gn=MECOM;refGene_tx=XM_003991927.3;refGene_gn=MECOM;refGene_tx=XM_019810458.1;refGene_gn=MECOM;refGene_tx=XM_019810459.1;refGene_gn=MECOM;refGene_tx=XM_019810460.1;refGene_gn=MECOM;refGene_tx=XM_019810461.2;refGene_gn=MECOM;refGene_tx=XM_023260417.1
C2 99048719 99048867
refGene_gn=MECOM;refGene_tx=XM_003991927.3;refGene_gn=MECOM;refGene_tx=XM_019810458.1;refGene_gn=MECOM;refGene_tx=XM_019810459.1;refGene_gn=MECOM;refGene_tx=XM_019810460.1;refGene_gn=MECOM;refGene_tx=XM_019810461.2;refGene_gn=MECOM;refGene_tx=XM_023260417.1
C2 99050217 99050374
refGene_gn=MECOM;refGene_tx=XM_003991927.3;refGene_gn=MECOM;refGene_tx=XM_019810458.1;refGene_gn=MECOM;refGene_tx=XM_019810459.1;refGene_gn=MECOM;refGene_tx=XM_019810460.1;refGene_gn=MECOM;refGene_tx=XM_019810461.2;refGene_gn=MECOM;refGene_tx=XM_023260417.1
C2 99054767 99056124
refGene_gn=MECOM;refGene_tx=XM_003991927.3;refGene_gn=MECOM;refGene_tx=XM_019810458.1;refGene_gn=MECOM;refGene_tx=XM_019810459.1;refGene_gn=MECOM;refGene_tx=XM_019810460.1;refGene_gn=MECOM;refGene_tx=XM_019810461.2;refGene_gn=MECOM;refGene_tx=XM_023260417.1
C2 99058573 99058661
refGene_gn=MECOM;refGene_tx=XM_003991927.3;refGene_gn=MECOM;refGene_tx=XM_019810458.1;refGene_gn=MECOM;refGene_tx=XM_019810459.1;refGene_gn=MECOM;refGene_tx=XM_019810460.1;refGene_gn=MECOM;refGene_tx=XM_019810461.2;refGene_gn=MECOM;refGene_tx=XM_023260417.1
C2 99063835 99063862
refGene_gn=MECOM;refGene_tx=XM_003991927.3;refGene_gn=MECOM;refGene_tx=XM_019810458.1;refGene_gn=MECOM;refGene_tx=XM_019810461.2
C2 99069534 99069701
refGene_gn=MECOM;refGene_tx=XM_003991927.3;refGene_gn=MECOM;refGene_tx=XM_019810458.1;refGene_gn=MECOM;refGene_tx=XM_019810459.1;refGene_gn=MECOM;refGene_tx=XM_019810460.1;refGene_gn=MECOM;refGene_tx=XM_019810461.2;refGene_gn=MECOM;refGene_tx=XM_023260417.1
C2 99070757 99070835
refGene_gn=MECOM;refGene_tx=XM_003991927.3;refGene_gn=MECOM;refGene_tx=XM_019810458.1;refGene_gn=MECOM;refGene_tx=XM_019810459.1;refGene_gn=MECOM;refGene_tx=XM_019810460.1;refGene_gn=MECOM;refGene_tx=XM_019810461.2;refGene_gn=MECOM;refGene_tx=XM_023260417.1
C2 99074597 99074767
refGene_gn=MECOM;refGene_tx=XM_003991927.3;refGene_gn=MECOM;refGene_tx=XM_019810458.1;refGene_gn=MECOM;refGene_tx=XM_019810459.1;refGene_gn=MECOM;refGene_tx=XM_019810460.1;refGene_gn=MECOM;refGene_tx=XM_019810461.2;refGene_gn=MECOM;refGene_tx=XM_023260417.1
C2 99076958 99077103
refGene_gn=MECOM;refGene_tx=XM_003991927.3;refGene_gn=MECOM;refGene_tx=XM_019810458.1;refGene_gn=MECOM;refGene_tx=XM_019810459.1;refGene_gn=MECOM;refGene_tx=XM_019810460.1;refGene_gn=MECOM;refGene_tx=XM_019810461.2;refGene_gn=MECOM;refGene_tx=XM_023260417.1
C2 99080592 99080829
refGene_gn=MECOM;refGene_tx=XM_003991927.3;refGene_gn=MECOM;refGene_tx

=XM_019810458.1;refGene_gn=MECOM;refGene_tx=XM_019810459.1;refGene_gn=MECOM;refGene_tx=XM_019810460.1;refGene_gn=MECOM;refGene_tx=XM_019810461.2;refGene_gn=MECOM;refGene_tx=XM_023260417.1
C2 99081662 99081846
refGene_gn=MECOM;refGene_tx=XM_003991927.3;refGene_gn=MECOM;refGene_tx=XM_019810458.1;refGene_gn=MECOM;refGene_tx=XM_019810459.1;refGene_gn=MECOM;refGene_tx=XM_019810460.1;refGene_gn=MECOM;refGene_tx=XM_019810461.2;refGene_gn=MECOM;refGene_tx=XM_023260417.1
C2 99085465 99085600
refGene_gn=MECOM;refGene_tx=XM_003991927.3;refGene_gn=MECOM;refGene_tx=XM_019810458.1;refGene_gn=MECOM;refGene_tx=XM_019810459.1;refGene_gn=MECOM;refGene_tx=XM_019810460.1;refGene_gn=MECOM;refGene_tx=XM_019810461.2;refGene_gn=MECOM;refGene_tx=XM_023260417.1
C2 99275598 99275864
refGene_gn=LOC109499785;refGene_tx=XM_023260418.1
C2 99277970 99278099
refGene_gn=LOC109499785;refGene_tx=XM_023260418.1
C2 99284193 99284325
refGene_gn=LOC109499785;refGene_tx=XM_023260418.1
C2 99293842 99293894
refGene_gn=LOC109499785;refGene_tx=XM_023260418.1
C2 100003219 100003406
refGene_gn=GOLIM4;refGene_tx=XM_003991933.5;refGene_gn=GOLIM4;refGene_tx=XM_006936274.4;refGene_gn=GOLIM4;refGene_tx=XM_006936275.4;refGene_gn=GOLIM4;refGene_tx=XM_019810464.2
C2 100048874 100048949
refGene_gn=GOLIM4;refGene_tx=XM_003991933.5;refGene_gn=GOLIM4;refGene_tx=XM_006936274.4;refGene_gn=GOLIM4;refGene_tx=XM_006936275.4;refGene_gn=GOLIM4;refGene_tx=XM_019810464.2
C2 100050271 100050321
refGene_gn=GOLIM4;refGene_tx=XM_003991933.5;refGene_gn=GOLIM4;refGene_tx=XM_006936274.4;refGene_gn=GOLIM4;refGene_tx=XM_006936275.4;refGene_gn=GOLIM4;refGene_tx=XM_019810464.2
C2 100052116 100052170
refGene_gn=GOLIM4;refGene_tx=XM_003991933.5;refGene_gn=GOLIM4;refGene_tx=XM_006936274.4;refGene_gn=GOLIM4;refGene_tx=XM_006936275.4;refGene_gn=GOLIM4;refGene_tx=XM_019810464.2
C2 100053514 100053665
refGene_gn=GOLIM4;refGene_tx=XM_003991933.5;refGene_gn=GOLIM4;refGene_tx=XM_006936274.4;refGene_gn=GOLIM4;refGene_tx=XM_006936275.4;refGene_gn=GOLIM4;refGene_tx=XM_019810464.2
C2 100057588 100057671
refGene_gn=GOLIM4;refGene_tx=XM_003991933.5;refGene_gn=GOLIM4;refGene_tx=XM_006936274.4;refGene_gn=GOLIM4;refGene_tx=XM_006936275.4;refGene_gn=GOLIM4;refGene_tx=XM_019810464.2
C2 100058239 100058323
refGene_gn=GOLIM4;refGene_tx=XM_003991933.5;refGene_gn=GOLIM4;refGene_tx=XM_006936274.4
C2 100061601 100061760
refGene_gn=GOLIM4;refGene_tx=XM_003991933.5;refGene_gn=GOLIM4;refGene_

tx=XM_006936274.4;refGene_gn=GOLIM4;refGene_tx=XM_006936275.4;refGene_gn=GOLIM4;refGene_tx=XM_019810464.2
C2	100065575	100065908
refGene_gn=GOLIM4;refGene_tx=XM_003991933.5;refGene_gn=GOLIM4;refGene_tx=XM_006936274.4;refGene_gn=GOLIM4;refGene_tx=XM_006936275.4;refGene_gn=GOLIM4;refGene_tx=XM_019810464.2
C2	100067564	100067812
refGene_gn=GOLIM4;refGene_tx=XM_003991933.5;refGene_gn=GOLIM4;refGene_tx=XM_006936274.4;refGene_gn=GOLIM4;refGene_tx=XM_006936275.4;refGene_gn=GOLIM4;refGene_tx=XM_019810464.2
C2	100068547	100068630
refGene_gn=GOLIM4;refGene_tx=XM_003991933.5;refGene_gn=GOLIM4;refGene_tx=XM_006936274.4;refGene_gn=GOLIM4;refGene_tx=XM_006936275.4;refGene_gn=GOLIM4;refGene_tx=XM_019810464.2
C2	100069974	100070084
refGene_gn=GOLIM4;refGene_tx=XM_003991933.5;refGene_gn=GOLIM4;refGene_tx=XM_006936274.4;refGene_gn=GOLIM4;refGene_tx=XM_006936275.4;refGene_gn=GOLIM4;refGene_tx=XM_019810464.2
C2	100072976	100073144
refGene_gn=GOLIM4;refGene_tx=XM_003991933.5;refGene_gn=GOLIM4;refGene_tx=XM_006936274.4;refGene_gn=GOLIM4;refGene_tx=XM_006936275.4;refGene_gn=GOLIM4;refGene_tx=XM_019810464.2
C2	100073484	100073553
refGene_gn=GOLIM4;refGene_tx=XM_003991933.5;refGene_gn=GOLIM4;refGene_tx=XM_006936274.4;refGene_gn=GOLIM4;refGene_tx=XM_006936275.4;refGene_gn=GOLIM4;refGene_tx=XM_019810464.2
C2	100080565	100080646
refGene_gn=GOLIM4;refGene_tx=XM_003991933.5;refGene_gn=GOLIM4;refGene_tx=XM_006936274.4;refGene_gn=GOLIM4;refGene_tx=XM_006936275.4;refGene_gn=GOLIM4;refGene_tx=XM_019810464.2
C2	100080915	100081065
refGene_gn=GOLIM4;refGene_tx=XM_003991933.5;refGene_gn=GOLIM4;refGene_tx=XM_006936274.4;refGene_gn=GOLIM4;refGene_tx=XM_006936275.4;refGene_gn=GOLIM4;refGene_tx=XM_019810464.2
C2	100163047	100163159
refGene_gn=LOC101099321;refGene_tx=XM_019810466.2;refGene_gn=LOC101099321;refGene_tx=XM_019810467.2
C2	100164313	100164362
refGene_gn=LOC101099321;refGene_tx=XM_006936276.3
C2	100166172	100166370
refGene_gn=LOC101099321;refGene_tx=XM_006936276.3;refGene_gn=LOC101099321;refGene_tx=XM_019810466.2;refGene_gn=LOC101099321;refGene_tx=XM_019810467.2
C2	100169476	100169619
refGene_gn=LOC101099321;refGene_tx=XM_006936276.3;refGene_gn=LOC101099321;refGene_tx=XM_019810466.2;refGene_gn=LOC101099321;refGene_tx=XM_019810467.2
C2	100171860	100171979
refGene_gn=LOC101099321;refGene_tx=XM_006936276.3;refGene_gn=LOC101099321;refGene_tx=XM_019810466.2;refGene_gn=LOC101099321;refGene_tx=XM_01

9810467.2
C2	100177106	100177200
refGene_gn=LOC101099321;refGene_tx=XM_006936276.3;refGene_gn=LOC101099321;refGene_tx=XM_019810466.2;refGene_gn=LOC101099321;refGene_tx=XM_019810467.2
C2	100181437	100181579
refGene_gn=LOC101099321;refGene_tx=XM_006936276.3;refGene_gn=LOC101099321;refGene_tx=XM_019810466.2;refGene_gn=LOC101099321;refGene_tx=XM_019810467.2
C2	100189839	100189865
refGene_gn=LOC101099321;refGene_tx=XM_006936276.3;refGene_gn=LOC101099321;refGene_tx=XM_019810466.2;refGene_gn=LOC101099321;refGene_tx=XM_019810467.2
C2	100269652	100269729
refGene_gn=SERPINI1;refGene_tx=XM_003991934.5
C2	100269963	100270053
refGene_gn=SERPINI1;refGene_tx=XM_003991934.5
C2	100271446	100271533
refGene_gn=SERPINI1;refGene_tx=XM_003991934.5
C2	100284039	100284137
refGene_gn=SERPINI1;refGene_tx=XM_003991934.5
C2	100289646	100289851
refGene_gn=SERPINI1;refGene_tx=XM_003991934.5
C2	100291534	100291729
refGene_gn=SERPINI1;refGene_tx=XM_003991934.5
C2	100293692	100293923
refGene_gn=SERPINI1;refGene_tx=XM_003991934.5
C2	100294960	100295210
refGene_gn=SERPINI1;refGene_tx=XM_003991934.5
C2	100360868	100360964
refGene_gn=PDCD10;refGene_tx=XM_011286104.3;refGene_gn=PDCD10;refGene_tx=XM_019810468.2
C2	100370564	100370618
refGene_gn=PDCD10;refGene_tx=XM_011286104.3;refGene_gn=PDCD10;refGene_tx=XM_019810468.2
C2	100377486	100377604
refGene_gn=PDCD10;refGene_tx=XM_011286104.3;refGene_gn=PDCD10;refGene_tx=XM_019810468.2
C2	100378249	100378376
refGene_gn=PDCD10;refGene_tx=XM_011286104.3;refGene_gn=PDCD10;refGene_tx=XM_019810468.2
C2	100385968	100386047
refGene_gn=PDCD10;refGene_tx=XM_011286104.3;refGene_gn=PDCD10;refGene_tx=XM_019810468.2
C2	100386351	100386434
refGene_gn=PDCD10;refGene_tx=XM_011286104.3;refGene_gn=PDCD10;refGene_tx=XM_019810468.2
C2	100389288	100389370
refGene_gn=PDCD10;refGene_tx=XM_011286104.3;refGene_gn=PDCD10;refGene_tx=XM_019810468.2

C2	100412261	100412423
refGene_gn=WDR49;refGene_tx=XM_019810469.2
C2	100433437	100433878
refGene_gn=WDR49;refGene_tx=XM_019810469.2
C2	100439369	100439546
refGene_gn=WDR49;refGene_tx=XM_019810469.2
C2	100440369	100440544
refGene_gn=WDR49;refGene_tx=XM_019810469.2
C2	100451501	100451669
refGene_gn=WDR49;refGene_tx=XM_019810469.2
C2	100454465	100454614
refGene_gn=WDR49;refGene_tx=XM_019810469.2
C2	100471641	100471875
refGene_gn=WDR49;refGene_tx=XM_019810469.2
C2	100492218	100492383
refGene_gn=WDR49;refGene_tx=XM_019810469.2
C2	100496026	100496142
refGene_gn=WDR49;refGene_tx=XM_019810469.2
C2	100507371	100507502
refGene_gn=WDR49;refGene_tx=XM_019810469.2
C2	100509623	100509722
refGene_gn=WDR49;refGene_tx=XM_019810469.2
C2	100511581	100511746
refGene_gn=WDR49;refGene_tx=XM_019810469.2
C2	100513792	100513980
refGene_gn=WDR49;refGene_tx=XM_019810469.2
C2	100514784	100514982
refGene_gn=WDR49;refGene_tx=XM_019810469.2
C2	100519610	100519792
refGene_gn=WDR49;refGene_tx=XM_019810469.2
C2	100529032	100529142
refGene_gn=WDR49;refGene_tx=XM_019810469.2
C2	100531619	100531766
refGene_gn=WDR49;refGene_tx=XM_019810469.2
C2	100551653	100551784
refGene_gn=WDR49;refGene_tx=XM_019810469.2
C2	100560521	100560768
refGene_gn=SERPINI2;refGene_tx=XM_003991936.3
C2	100566019	100566250
refGene_gn=SERPINI2;refGene_tx=XM_003991936.3
C2	100567667	100567862
refGene_gn=SERPINI2;refGene_tx=XM_003991936.3
C2	100567940	100568133
refGene_gn=SERPINI2;refGene_tx=XM_003991936.3
C2	100580353	100580451
refGene_gn=SERPINI2;refGene_tx=XM_003991936.3
C2	100584312	100584399
refGene_gn=SERPINI2;refGene_tx=XM_003991936.3
C2	100587450	100587540
refGene_gn=SERPINI2;refGene_tx=XM_003991936.3

C2 100592675 100592752
refGene_gn=SERPINI2;refGene_tx=XM_003991936.3
C2 100650123 100650191
refGene_gn=ZBBX;refGene_tx=XM_019810470.2;refGene_gn=ZBBX;refGene_tx=XM_019810471.2
C2 100654934 100655048
refGene_gn=ZBBX;refGene_tx=XM_019810470.2;refGene_gn=ZBBX;refGene_tx=XM_019810471.2
C2 100657394 100657485
refGene_gn=ZBBX;refGene_tx=XM_019810470.2;refGene_gn=ZBBX;refGene_tx=XM_019810471.2
C2 100662016 100662019
refGene_gn=ZBBX;refGene_tx=XM_019810473.2
C2 100664243 100664292
refGene_gn=ZBBX;refGene_tx=XM_019810470.2;refGene_gn=ZBBX;refGene_tx=XM_019810471.2;refGene_gn=ZBBX;refGene_tx=XM_019810473.2
C2 100665435 100665545
refGene_gn=ZBBX;refGene_tx=XM_019810470.2;refGene_gn=ZBBX;refGene_tx=XM_019810471.2;refGene_gn=ZBBX;refGene_tx=XM_019810473.2
C2 100673420 100673516
refGene_gn=ZBBX;refGene_tx=XM_019810470.2;refGene_gn=ZBBX;refGene_tx=XM_019810471.2;refGene_gn=ZBBX;refGene_tx=XM_019810473.2
C2 100704703 100704862
refGene_gn=ZBBX;refGene_tx=XM_019810470.2;refGene_gn=ZBBX;refGene_tx=XM_019810471.2;refGene_gn=ZBBX;refGene_tx=XM_019810473.2
C2 100710476 100710651
refGene_gn=ZBBX;refGene_tx=XM_019810470.2;refGene_gn=ZBBX;refGene_tx=XM_019810471.2;refGene_gn=ZBBX;refGene_tx=XM_019810473.2
C2 100715087 100715208
refGene_gn=ZBBX;refGene_tx=XM_019810470.2;refGene_gn=ZBBX;refGene_tx=XM_019810471.2;refGene_gn=ZBBX;refGene_tx=XM_019810473.2
C2 100722684 100722794
refGene_gn=ZBBX;refGene_tx=XM_019810470.2;refGene_gn=ZBBX;refGene_tx=XM_019810471.2;refGene_gn=ZBBX;refGene_tx=XM_019810473.2
C2 100723417 100723515
refGene_gn=ZBBX;refGene_tx=XM_019810470.2;refGene_gn=ZBBX;refGene_tx=XM_019810471.2;refGene_gn=ZBBX;refGene_tx=XM_019810473.2
C2 100724694 100724774
refGene_gn=ZBBX;refGene_tx=XM_019810470.2;refGene_gn=ZBBX;refGene_tx=XM_019810471.2;refGene_gn=ZBBX;refGene_tx=XM_019810473.2
C2 100726664 100726804
refGene_gn=ZBBX;refGene_tx=XM_019810470.2;refGene_gn=ZBBX;refGene_tx=XM_019810471.2;refGene_gn=ZBBX;refGene_tx=XM_019810473.2
C2 100732923 100733231
refGene_gn=ZBBX;refGene_tx=XM_019810470.2;refGene_gn=ZBBX;refGene_tx=XM_019810471.2;refGene_gn=ZBBX;refGene_tx=XM_019810473.2
C2 100736559 100736713
refGene_gn=ZBBX;refGene_tx=XM_019810470.2;refGene_gn=ZBBX;refGene_tx=XM_019810471.2;refGene_gn=ZBBX;refGene_tx=XM_019810473.2
C2 100737495 100737612

refGene_gn=ZBBX;refGene_tx=XM_019810470.2;refGene_gn=ZBBX;refGene_tx=XM_019810471.2;refGene_gn=ZBBX;refGene_tx=XM_019810473.2
C2 100745105 100745360
refGene_gn=ZBBX;refGene_tx=XM_019810470.2;refGene_gn=ZBBX;refGene_tx=XM_019810471.2;refGene_gn=ZBBX;refGene_tx=XM_019810473.2
C2 100758934 100759073
refGene_gn=ZBBX;refGene_tx=XM_019810470.2;refGene_gn=ZBBX;refGene_tx=XM_019810471.2;refGene_gn=ZBBX;refGene_tx=XM_019810473.2
C2 100760849 100760967
refGene_gn=ZBBX;refGene_tx=XM_019810470.2;refGene_gn=ZBBX;refGene_tx=XM_019810471.2;refGene_gn=ZBBX;refGene_tx=XM_019810473.2
C2 102087815 102089332
refGene_gn=BCHE;refGene_tx=NM_001009364.1
C2 102140376 102140543
refGene_gn=BCHE;refGene_tx=NM_001009364.1
C2 102147032 102147157
refGene_gn=BCHE;refGene_tx=NM_001009364.1
C2 102645657 102648201
refGene_gn=SLITRK3;refGene_tx=XM_019810478.2
C2 102648202 102648598
refGene_gn=SLITRK3;refGene_tx=XM_019810478.2
C2 102748184 102748302
refGene_gn=SI;refGene_tx=NM_001123332.1
C2 102749571 102749708
refGene_gn=SI;refGene_tx=NM_001123332.1
C2 102755947 102756065
refGene_gn=SI;refGene_tx=NM_001123332.1
C2 102756365 102756475
refGene_gn=SI;refGene_tx=NM_001123332.1
C2 102757441 102757593
refGene_gn=SI;refGene_tx=NM_001123332.1
C2 102759214 102759386
refGene_gn=SI;refGene_tx=NM_001123332.1
C2 102761040 102761140
refGene_gn=SI;refGene_tx=NM_001123332.1
C2 102762095 102762208
refGene_gn=SI;refGene_tx=NM_001123332.1
C2 102764355 102764481
refGene_gn=SI;refGene_tx=NM_001123332.1
C2 102765054 102765186
refGene_gn=SI;refGene_tx=NM_001123332.1
C2 102765273 102765393
refGene_gn=SI;refGene_tx=NM_001123332.1
C2 102767134 102767248
refGene_gn=SI;refGene_tx=NM_001123332.1
C2 102772410 102772495
refGene_gn=SI;refGene_tx=NM_001123332.1
C2 102773225 102773343
refGene_gn=SI;refGene_tx=NM_001123332.1
C2 102774680 102774852

refGene_gn=SI;refGene_tx=NM_001123332.1
C2	102778733	102778850
refGene_gn=SI;refGene_tx=NM_001123332.1
C2	102780850	102781005
refGene_gn=SI;refGene_tx=NM_001123332.1
C2	102781890	102781975
refGene_gn=SI;refGene_tx=NM_001123332.1
C2	102783004	102783061
refGene_gn=SI;refGene_tx=NM_001123332.1
C2	102790104	102790229
refGene_gn=SI;refGene_tx=NM_001123332.1
C2	102791998	102792087
refGene_gn=SI;refGene_tx=NM_001123332.1
C2	102795007	102795057
refGene_gn=SI;refGene_tx=NM_001123332.1
C2	102795793	102795964
refGene_gn=SI;refGene_tx=NM_001123332.1
C2	102798439	102798595
refGene_gn=SI;refGene_tx=NM_001123332.1
C2	102800045	102800252
refGene_gn=SI;refGene_tx=NM_001123332.1
C2	102802164	102802319
refGene_gn=SI;refGene_tx=NM_001123332.1
C2	102803969	102804138
refGene_gn=SI;refGene_tx=NM_001123332.1
C2	102805629	102805726
refGene_gn=SI;refGene_tx=NM_001123332.1
C2	102805851	102805964
refGene_gn=SI;refGene_tx=NM_001123332.1
C2	102806282	102806408
refGene_gn=SI;refGene_tx=NM_001123332.1
C2	102807945	102808074
refGene_gn=SI;refGene_tx=NM_001123332.1
C2	102808807	102808918
refGene_gn=SI;refGene_tx=NM_001123332.1
C2	102810842	102810905
refGene_gn=SI;refGene_tx=NM_001123332.1
C2	102815854	102815971
refGene_gn=SI;refGene_tx=NM_001123332.1
C2	102821090	102821178
refGene_gn=SI;refGene_tx=NM_001123332.1
C2	102822121	102822260
refGene_gn=SI;refGene_tx=NM_001123332.1
C2	102830527	102830602
refGene_gn=SI;refGene_tx=NM_001123332.1
C2	102830603	102830662
refGene_gn=SI;refGene_tx=NM_001123332.1
C2	102832276	102832311
refGene_gn=SI;refGene_tx=NM_001123332.1
C2	102832395	102832512

refGene_gn=SI;refGene_tx=NM_001123332.1
C2 102835351 102835500
refGene_gn=SI;refGene_tx=NM_001123332.1
C2 102837689 102837774
refGene_gn=SI;refGene_tx=NM_001123332.1
C2 102837858 102837915
refGene_gn=SI;refGene_tx=NM_001123332.1
C2 102838509 102838634
refGene_gn=SI;refGene_tx=NM_001123332.1
C2 102840579 102840668
refGene_gn=SI;refGene_tx=NM_001123332.1
C2 102844602 102844652
refGene_gn=SI;refGene_tx=NM_001123332.1
C2 102844901 102845069
refGene_gn=SI;refGene_tx=NM_001123332.1
C2 102847882 102847951
refGene_gn=SI;refGene_tx=NM_001123332.1
C2 105805527 105806426
refGene_gn=OTOL1;refGene_tx=XM_023238566.1
C2 105807041 105807113
refGene_gn=OTOL1;refGene_tx=XM_023238566.1
C2 105811330 105811420
refGene_gn=OTOL1;refGene_tx=XM_023238566.1
C2 105814621 105814985
refGene_gn=OTOL1;refGene_tx=XM_023238566.1
C2 105946974 105947205
refGene_gn=SPTSSB;refGene_tx=XM_019810480.2
C2 106032889 106032924
refGene_gn=NMD3;refGene_tx=XM_023260419.1;refGene_gn=NMD3;refGene_tx=XM_023260420.1
C2 106034028 106034162
refGene_gn=NMD3;refGene_tx=XM_011286101.3;refGene_gn=NMD3;refGene_tx=XM_023260419.1;refGene_gn=NMD3;refGene_tx=XM_023260420.1
C2 106034600 106034671
refGene_gn=NMD3;refGene_tx=XM_011286101.3;refGene_gn=NMD3;refGene_tx=XM_023260419.1;refGene_gn=NMD3;refGene_tx=XM_023260420.1
C2 106036046 106036153
refGene_gn=NMD3;refGene_tx=XM_011286101.3;refGene_gn=NMD3;refGene_tx=XM_023260419.1;refGene_gn=NMD3;refGene_tx=XM_023260420.1
C2 106037367 106037440
refGene_gn=NMD3;refGene_tx=XM_011286101.3;refGene_gn=NMD3;refGene_tx=XM_023260419.1;refGene_gn=NMD3;refGene_tx=XM_023260420.1
C2 106038511 106038624
refGene_gn=NMD3;refGene_tx=XM_011286101.3;refGene_gn=NMD3;refGene_tx=XM_023260419.1;refGene_gn=NMD3;refGene_tx=XM_023260420.1
C2 106041110 106041256
refGene_gn=NMD3;refGene_tx=XM_011286101.3;refGene_gn=NMD3;refGene_tx=XM_023260419.1;refGene_gn=NMD3;refGene_tx=XM_023260420.1
C2 106042509 106042627
refGene_gn=NMD3;refGene_tx=XM_011286101.3;refGene_gn=NMD3;refGene_tx=X

M_023260419.1;refGene_gn=NMD3;refGene_tx=XM_023260420.1
C2 106044415 106044512
refGene_gn=NMD3;refGene_tx=XM_011286101.3;refGene_gn=NMD3;refGene_tx=XM_023260419.1;refGene_gn=NMD3;refGene_tx=XM_023260420.1
C2 106045601 106045680
refGene_gn=NMD3;refGene_tx=XM_011286101.3;refGene_gn=NMD3;refGene_tx=XM_023260419.1;refGene_gn=NMD3;refGene_tx=XM_023260420.1
C2 106052823 106052914
refGene_gn=NMD3;refGene_tx=XM_011286101.3;refGene_gn=NMD3;refGene_tx=XM_023260419.1;refGene_gn=NMD3;refGene_tx=XM_023260420.1
C2 106053187 106053316
refGene_gn=NMD3;refGene_tx=XM_011286101.3;refGene_gn=NMD3;refGene_tx=XM_023260419.1;refGene_gn=NMD3;refGene_tx=XM_023260420.1
C2 106054353 106054434
refGene_gn=NMD3;refGene_tx=XM_011286101.3;refGene_gn=NMD3;refGene_tx=XM_023260419.1;refGene_gn=NMD3;refGene_tx=XM_023260420.1
C2 106061315 106061412
refGene_gn=NMD3;refGene_tx=XM_011286101.3;refGene_gn=NMD3;refGene_tx=XM_023260419.1;refGene_gn=NMD3;refGene_tx=XM_023260420.1
C2 106063128 106063263
refGene_gn=NMD3;refGene_tx=XM_011286101.3;refGene_gn=NMD3;refGene_tx=XM_023260419.1;refGene_gn=NMD3;refGene_tx=XM_023260420.1
C2 106064436 106064480
refGene_gn=NMD3;refGene_tx=XM_011286101.3;refGene_gn=NMD3;refGene_tx=XM_023260419.1;refGene_gn=NMD3;refGene_tx=XM_023260420.1
C2 106186367 106187363
refGene_gn=B3GALNT1;refGene_tx=XM_003991941.5;refGene_gn=B3GALNT1;refGene_tx=XM_006936281.3;refGene_gn=B3GALNT1;refGene_tx=XM_006936282.3;refGene_gn=B3GALNT1;refGene_tx=XM_011286096.2;refGene_gn=B3GALNT1;refGene_tx=XM_011286097.3;refGene_gn=B3GALNT1;refGene_tx=XM_019810482.2;refGene_gn=B3GALNT1;refGene_tx=XM_019810483.2;refGene_gn=B3GALNT1;refGene_tx=XM_019810484.2;refGene_gn=B3GALNT1;refGene_tx=XM_019810485.2;refGene_gn=B3GALNT1;refGene_tx=XM_019810486.2;refGene_gn=B3GALNT1;refGene_tx=XM_019810487.2;refGene_gn=B3GALNT1;refGene_tx=XM_019810488.2;refGene_gn=B3GALNT1;refGene_tx=XM_019810489.2;refGene_gn=B3GALNT1;refGene_tx=XM_023260421.1;refGene_gn=B3GALNT1;refGene_tx=XM_023260422.1;refGene_gn=B3GALNT1;refGene_tx=XM_023260423.1;refGene_gn=B3GALNT1;refGene_tx=XM_023260424.1;refGene_gn=B3GALNT1;refGene_tx=XM_023260425.1;refGene_gn=B3GALNT1;refGene_tx=XM_023260426.1;refGene_gn=B3GALNT1;refGene_tx=XM_023260427.1
C2 106209027 106209374
refGene_gn=PPM1L;refGene_tx=XM_003991943.5;refGene_gn=PPM1L;refGene_tx=XM_019810492.2
C2 106212970 106213132
refGene_gn=PPM1L;refGene_tx=XM_003991943.5;refGene_gn=PPM1L;refGene_tx=XM_019810492.2
C2 106304242 106304417
refGene_gn=PPM1L;refGene_tx=XM_003991943.5;refGene_gn=PPM1L;refGene_tx=XM_019810492.2
C2 106498404 106498803

refGene_gn=PPM1L;refGene_tx=XM_003991943.5
C2	106575440	106576019
refGene_gn=ARL14;refGene_tx=XM_003991944.2
C2	106676391	106676460
refGene_gn=KPNA4;refGene_tx=XM_023260428.1
C2	106701911	106701956
refGene_gn=KPNA4;refGene_tx=XM_023260428.1
C2	106702832	106702922
refGene_gn=KPNA4;refGene_tx=XM_023260428.1
C2	106703032	106703062
refGene_gn=KPNA4;refGene_tx=XM_023260428.1
C2	106703145	106703198
refGene_gn=KPNA4;refGene_tx=XM_023260428.1
C2	106706292	106706338
refGene_gn=KPNA4;refGene_tx=XM_023260428.1
C2	106706339	106706389
refGene_gn=KPNA4;refGene_tx=XM_023260428.1
C2	106706914	106707000
refGene_gn=KPNA4;refGene_tx=XM_023260428.1
C2	106716603	106716690
refGene_gn=KPNA4;refGene_tx=XM_023260428.1
C2	106719083	106719253
refGene_gn=KPNA4;refGene_tx=XM_023260428.1
C2	106719342	106719387
refGene_gn=KPNA4;refGene_tx=XM_023260428.1
C2	106722343	106722475
refGene_gn=KPNA4;refGene_tx=XM_023260428.1
C2	106728824	106728953
refGene_gn=KPNA4;refGene_tx=XM_023260428.1
C2	106730365	106730470
refGene_gn=KPNA4;refGene_tx=XM_023260428.1
C2	106735063	106735135
refGene_gn=KPNA4;refGene_tx=XM_023260428.1
C2	106736354	106736517
refGene_gn=KPNA4;refGene_tx=XM_023260428.1
C2	106739043	106739138
refGene_gn=KPNA4;refGene_tx=XM_023260428.1
C2	106743458	106743557
refGene_gn=KPNA4;refGene_tx=XM_023260428.1
C2	106810135	106811347
refGene_gn=TRIM59;refGene_tx=XM_006936293.4;refGene_gn=TRIM59;refGene_tx=XM_019810496.2;refGene_gn=TRIM59;refGene_tx=XM_019810497.2;refGene_gn=TRIM59;refGene_tx=XM_023260429.1
C2	106814633	106814786
refGene_gn=SMC4;refGene_tx=XM_003991946.5;refGene_gn=SMC4;refGene_tx=XM_019810495.2
C2	106815259	106815443
refGene_gn=SMC4;refGene_tx=XM_003991946.5;refGene_gn=SMC4;refGene_tx=XM_019810495.2
C2	106815969	106816202

refGene_gn=SMC4;refGene_tx=XM_003991946.5;refGene_gn=SMC4;refGene_tx=XM_019810495.2
C2 106816346 106816529
refGene_gn=SMC4;refGene_tx=XM_003991946.5;refGene_gn=SMC4;refGene_tx=XM_019810495.2
C2 106816965 106817139
refGene_gn=SMC4;refGene_tx=XM_003991946.5;refGene_gn=SMC4;refGene_tx=XM_019810495.2
C2 106817413 106817558
refGene_gn=SMC4;refGene_tx=XM_003991946.5;refGene_gn=SMC4;refGene_tx=XM_019810495.2
C2 106818830 106819020
refGene_gn=SMC4;refGene_tx=XM_003991946.5;refGene_gn=SMC4;refGene_tx=XM_019810495.2
C2 106823527 106823654
refGene_gn=SMC4;refGene_tx=XM_003991946.5;refGene_gn=SMC4;refGene_tx=XM_019810495.2
C2 106824850 106825003
refGene_gn=SMC4;refGene_tx=XM_003991946.5;refGene_gn=SMC4;refGene_tx=XM_019810495.2
C2 106825804 106825884
refGene_gn=SMC4;refGene_tx=XM_003991946.5;refGene_gn=SMC4;refGene_tx=XM_019810495.2
C2 106826175 106826401
refGene_gn=SMC4;refGene_tx=XM_003991946.5;refGene_gn=SMC4;refGene_tx=XM_019810495.2
C2 106831653 106831815
refGene_gn=SMC4;refGene_tx=XM_003991946.5;refGene_gn=SMC4;refGene_tx=XM_019810495.2
C2 106833639 106833825
refGene_gn=SMC4;refGene_tx=XM_003991946.5;refGene_gn=SMC4;refGene_tx=XM_019810495.2
C2 106834785 106835019
refGene_gn=SMC4;refGene_tx=XM_003991946.5;refGene_gn=SMC4;refGene_tx=XM_019810495.2
C2 106837568 106837733
refGene_gn=SMC4;refGene_tx=XM_003991946.5;refGene_gn=SMC4;refGene_tx=XM_019810495.2
C2 106839093 106839244
refGene_gn=SMC4;refGene_tx=XM_003991946.5;refGene_gn=SMC4;refGene_tx=XM_019810495.2
C2 106840019 106840160
refGene_gn=SMC4;refGene_tx=XM_003991946.5;refGene_gn=SMC4;refGene_tx=XM_019810495.2
C2 106846990 106847118
refGene_gn=SMC4;refGene_tx=XM_003991946.5;refGene_gn=SMC4;refGene_tx=XM_019810495.2
C2 106847382 106847547
refGene_gn=SMC4;refGene_tx=XM_003991946.5;refGene_gn=SMC4;refGene_tx=XM_019810495.2

C2	106854823	106855000
refGene_gn=SMC4;refGene_tx=XM_003991946.5;refGene_gn=SMC4;refGene_tx=XM_019810495.2
C2	106856477	106856669
refGene_gn=SMC4;refGene_tx=XM_003991946.5;refGene_gn=SMC4;refGene_tx=XM_019810495.2
C2	106857112	106857291
refGene_gn=SMC4;refGene_tx=XM_003991946.5;refGene_gn=SMC4;refGene_tx=XM_019810495.2
C2	106858173	106858312
refGene_gn=SMC4;refGene_tx=XM_003991946.5;refGene_gn=SMC4;refGene_tx=XM_019810495.2
C2	106874982	106875019
refGene_gn=IFT80;refGene_tx=XM_003991947.5;refGene_gn=IFT80;refGene_tx=XM_019810498.2
C2	106877493	106877715
refGene_gn=IFT80;refGene_tx=XM_003991947.5;refGene_gn=IFT80;refGene_tx=XM_019810498.2
C2	106887150	106887261
refGene_gn=IFT80;refGene_tx=XM_003991947.5
C2	106888747	106888816
refGene_gn=IFT80;refGene_tx=XM_003991947.5
C2	106893181	106893233
refGene_gn=IFT80;refGene_tx=XM_019810499.2
C2	106902246	106902356
refGene_gn=IFT80;refGene_tx=XM_003991947.5;refGene_gn=IFT80;refGene_tx=XM_019810498.2;refGene_gn=IFT80;refGene_tx=XM_019810499.2
C2	106905901	106905991
refGene_gn=IFT80;refGene_tx=XM_003991947.5;refGene_gn=IFT80;refGene_tx=XM_019810498.2;refGene_gn=IFT80;refGene_tx=XM_019810499.2
C2	106906734	106906872
refGene_gn=IFT80;refGene_tx=XM_003991947.5;refGene_gn=IFT80;refGene_tx=XM_019810498.2;refGene_gn=IFT80;refGene_tx=XM_019810499.2
C2	106940839	106941019
refGene_gn=IFT80;refGene_tx=XM_003991947.5;refGene_gn=IFT80;refGene_tx=XM_019810498.2;refGene_gn=IFT80;refGene_tx=XM_019810499.2
C2	106951649	106951768
refGene_gn=IFT80;refGene_tx=XM_003991947.5;refGene_gn=IFT80;refGene_tx=XM_019810498.2;refGene_gn=IFT80;refGene_tx=XM_019810499.2
C2	106955779	106955854
refGene_gn=IFT80;refGene_tx=XM_003991947.5;refGene_gn=IFT80;refGene_tx=XM_019810498.2;refGene_gn=IFT80;refGene_tx=XM_019810499.2
C2	106959245	106959409
refGene_gn=IFT80;refGene_tx=XM_003991947.5;refGene_gn=IFT80;refGene_tx=XM_019810498.2;refGene_gn=IFT80;refGene_tx=XM_019810499.2
C2	106975262	106975327
refGene_gn=IFT80;refGene_tx=XM_003991947.5;refGene_gn=IFT80;refGene_tx=XM_019810498.2;refGene_gn=IFT80;refGene_tx=XM_019810499.2
C2	106977671	106977807
refGene_gn=IFT80;refGene_tx=XM_003991947.5;refGene_gn=IFT80;refGene_tx

=XM_019810498.2;refGene_gn=IFT80;refGene_tx=XM_019810499.2
C2 106978098 106978246
refGene_gn=IFT80;refGene_tx=XM_003991947.5;refGene_gn=IFT80;refGene_tx
=XM_019810498.2;refGene_gn=IFT80;refGene_tx=XM_019810499.2
C2 106979342 106979514
refGene_gn=IFT80;refGene_tx=XM_003991947.5;refGene_gn=IFT80;refGene_tx
=XM_019810498.2;refGene_gn=IFT80;refGene_tx=XM_019810499.2
C2 106981492 106981582
refGene_gn=IFT80;refGene_tx=XM_003991947.5;refGene_gn=IFT80;refGene_tx
=XM_019810498.2;refGene_gn=IFT80;refGene_tx=XM_019810499.2
C2 106981665 106981838
refGene_gn=IFT80;refGene_tx=XM_003991947.5;refGene_gn=IFT80;refGene_tx
=XM_019810498.2;refGene_gn=IFT80;refGene_tx=XM_019810499.2
C2 106986455 106986579
refGene_gn=IFT80;refGene_tx=XM_003991947.5;refGene_gn=IFT80;refGene_tx
=XM_019810498.2;refGene_gn=IFT80;refGene_tx=XM_019810499.2
C2 106995493 106995604
refGene_gn=IFT80;refGene_tx=XM_003991947.5;refGene_gn=IFT80;refGene_tx
=XM_019810498.2;refGene_gn=IFT80;refGene_tx=XM_019810499.2
C2 107033814 107034555
refGene_gn=CC2H3orf80;refGene_tx=XM_023260431.1
C2 107168245 107168727
refGene_gn=LOC102901165;refGene_tx=XM_023260179.1
C2 107178742 107178777
refGene_gn=LOC102901165;refGene_tx=XM_023260179.1
C2 107178892 107178979
refGene_gn=LOC102901165;refGene_tx=XM_023260179.1
C2 107181461 107181518
refGene_gn=LOC102901165;refGene_tx=XM_023260179.1
C2 107181842 107181872
refGene_gn=LOC102901165;refGene_tx=XM_023260179.1
C2 107185513 107185558
refGene_gn=LOC102901165;refGene_tx=XM_023260179.1
C2 107185915 107185963
refGene_gn=LOC102901165;refGene_tx=XM_023260179.1
C2 107186327 107186374
refGene_gn=LOC102901165;refGene_tx=XM_023260179.1
C2 107186483 107186533
refGene_gn=LOC102901165;refGene_tx=XM_023260179.1
C2 107186807 107187025
refGene_gn=LOC102901165;refGene_tx=XM_023260179.1
C2 107187136 107187387
refGene_gn=LOC102901165;refGene_tx=XM_023260179.1
C2 107190444 107190579
refGene_gn=LOC102901165;refGene_tx=XM_023260179.1
C2 107193526 107193665
refGene_gn=LOC102901165;refGene_tx=XM_023260179.1
C2 107207400 107207454
refGene_gn=LOC102901165;refGene_tx=XM_023260179.1
C2 107207931 107208012

refGene_gn=LOC102901165;refGene_tx=XM_023260179.1
C2 107210773 107211043
refGene_gn=LOC102901165;refGene_tx=XM_023260179.1
C2 107212544 107212564
refGene_gn=LOC102901165;refGene_tx=XM_023260179.1
C2 107221313 107221469
refGene_gn=IL12A;refGene_tx=NM_001009833.1
C2 107223215 107223359
refGene_gn=IL12A;refGene_tx=NM_001009833.1
C2 107223462 107223504
refGene_gn=IL12A;refGene_tx=NM_001009833.1
C2 107223579 107223621
refGene_gn=IL12A;refGene_tx=NM_001009833.1
C2 107223960 107224074
refGene_gn=IL12A;refGene_tx=NM_001009833.1
C2 107227411 107227566
refGene_gn=IL12A;refGene_tx=NM_001009833.1
C2 107228195 107228211
refGene_gn=IL12A;refGene_tx=NM_001009833.1
C2 107317734 107317786
refGene_gn=LOC101094201;refGene_tx=XM_011286086.3;refGene_gn=LOC101094201;refGene_tx=XM_011286087.2;refGene_gn=LOC101094201;refGene_tx=XM_019810501.2;refGene_gn=LOC101094201;refGene_tx=XM_023260432.1;refGene_gn=LOC101094201;refGene_tx=XM_023260433.1;refGene_gn=LOC101094201;refGene_tx=XM_023260434.1;refGene_gn=LOC101094201;refGene_tx=XM_023260435.1
C2 107321805 107321899
refGene_gn=LOC101094201;refGene_tx=XM_011286086.3;refGene_gn=LOC101094201;refGene_tx=XM_011286087.2;refGene_gn=LOC101094201;refGene_tx=XM_019810501.2;refGene_gn=LOC101094201;refGene_tx=XM_023260432.1;refGene_gn=LOC101094201;refGene_tx=XM_023260433.1;refGene_gn=LOC101094201;refGene_tx=XM_023260434.1;refGene_gn=LOC101094201;refGene_tx=XM_023260435.1
C2 107324334 107324458
refGene_gn=LOC101094201;refGene_tx=XM_011286086.3;refGene_gn=LOC101094201;refGene_tx=XM_011286087.2;refGene_gn=LOC101094201;refGene_tx=XM_019810501.2;refGene_gn=LOC101094201;refGene_tx=XM_023260432.1;refGene_gn=LOC101094201;refGene_tx=XM_023260433.1;refGene_gn=LOC101094201;refGene_tx=XM_023260434.1;refGene_gn=LOC101094201;refGene_tx=XM_023260435.1
C2 107325362 107325560
refGene_gn=LOC101094201;refGene_tx=XM_011286086.3;refGene_gn=LOC101094201;refGene_tx=XM_011286087.2;refGene_gn=LOC101094201;refGene_tx=XM_019810501.2;refGene_gn=LOC101094201;refGene_tx=XM_023260432.1;refGene_gn=LOC101094201;refGene_tx=XM_023260433.1;refGene_gn=LOC101094201;refGene_tx=XM_023260434.1;refGene_gn=LOC101094201;refGene_tx=XM_023260435.1
C2 107326765 107326883
refGene_gn=LOC101094201;refGene_tx=XM_011286086.3;refGene_gn=LOC101094201;refGene_tx=XM_011286087.2;refGene_gn=LOC101094201;refGene_tx=XM_019810501.2;refGene_gn=LOC101094201;refGene_tx=XM_023260432.1;refGene_gn=LOC101094201;refGene_tx=XM_023260433.1;refGene_gn=LOC101094201;refGene_tx=XM_023260434.1;refGene_gn=LOC101094201;refGene_tx=XM_023260435.1
C2 107345639 107345758

refGene_gn=LOC101094201;refGene_tx=XM_011286086.3;refGene_gn=LOC101094201;refGene_tx=XM_011286087.2;refGene_gn=LOC101094201;refGene_tx=XM_019810501.2;refGene_gn=LOC101094201;refGene_tx=XM_023260432.1;refGene_gn=LOC101094201;refGene_tx=XM_023260433.1;refGene_gn=LOC101094201;refGene_tx=XM_023260434.1;refGene_gn=LOC101094201;refGene_tx=XM_023260435.1
C2 107358356 107358434
refGene_gn=LOC101094201;refGene_tx=XM_011286086.3
C2 107370172 107370202
refGene_gn=LOC101094201;refGene_tx=XM_011286087.2
C2 107398603 107399303
refGene_gn=LOC109491414;refGene_tx=XM_023260436.1
C2 107399511 107399753
refGene_gn=LOC109491414;refGene_tx=XM_023260436.1
C2 107439136 107439838
refGene_gn=LOC101094201;refGene_tx=XM_019810501.2;refGene_gn=LOC101094201;refGene_tx=XM_023260432.1;refGene_gn=LOC101094201;refGene_tx=XM_023260433.1;refGene_gn=LOC101094201;refGene_tx=XM_023260434.1;refGene_gn=LOC101094201;refGene_tx=XM_023260435.1
C2 107440088 107440097
refGene_gn=LOC101094201;refGene_tx=XM_023260435.1
C2 107868496 107868559
refGene_gn=LOC101094201;refGene_tx=XM_023260432.1
C2 107879075 107879211
refGene_gn=LOC101094201;refGene_tx=XM_019810501.2
C2 107892062 107892143
refGene_gn=LOC101094201;refGene_tx=XM_019810501.2
C2 107897415 107897480
refGene_gn=LOC101094201;refGene_tx=XM_019810501.2
C2 108072658 108072667
refGene_gn=LOC101094201;refGene_tx=XM_019810501.2
C2 108340489 108340493
refGene_gn=MFSD1;refGene_tx=XM_003991949.5
C2 108341801 108341863
refGene_gn=MFSD1;refGene_tx=XM_003991949.5;refGene_gn=MFSD1;refGene_tx=XM_023260437.1
C2 108342999 108343047
refGene_gn=MFSD1;refGene_tx=XM_003991949.5;refGene_gn=MFSD1;refGene_tx=XM_023260437.1
C2 108345148 108345261
refGene_gn=MFSD1;refGene_tx=XM_003991949.5;refGene_gn=MFSD1;refGene_tx=XM_023260437.1
C2 108345856 108345954
refGene_gn=MFSD1;refGene_tx=XM_003991949.5;refGene_gn=MFSD1;refGene_tx=XM_023260437.1
C2 108347385 108347542
refGene_gn=MFSD1;refGene_tx=XM_003991949.5;refGene_gn=MFSD1;refGene_tx=XM_023260437.1
C2 108347874 108347931
refGene_gn=MFSD1;refGene_tx=XM_003991949.5;refGene_gn=MFSD1;refGene_tx=XM_023260437.1

C2	108349739	108349851
refGene_gn=MFSD1;refGene_tx=XM_003991949.5;refGene_gn=MFSD1;refGene_tx=XM_023260437.1
C2	108350303	108350402
refGene_gn=MFSD1;refGene_tx=XM_003991949.5;refGene_gn=MFSD1;refGene_tx=XM_023260437.1
C2	108352366	108352469
refGene_gn=MFSD1;refGene_tx=XM_003991949.5;refGene_gn=MFSD1;refGene_tx=XM_023260437.1
C2	108358548	108358657
refGene_gn=MFSD1;refGene_tx=XM_003991949.5;refGene_gn=MFSD1;refGene_tx=XM_023260437.1
C2	108360183	108360251
refGene_gn=MFSD1;refGene_tx=XM_003991949.5;refGene_gn=MFSD1;refGene_tx=XM_023260437.1
C2	108360558	108360601
refGene_gn=MFSD1;refGene_tx=XM_003991949.5;refGene_gn=MFSD1;refGene_tx=XM_023260437.1
C2	108362510	108362623
refGene_gn=MFSD1;refGene_tx=XM_003991949.5;refGene_gn=MFSD1;refGene_tx=XM_023260437.1
C2	108364211	108364264
refGene_gn=MFSD1;refGene_tx=XM_003991949.5;refGene_gn=MFSD1;refGene_tx=XM_023260437.1
C2	108366312	108366484
refGene_gn=MFSD1;refGene_tx=XM_003991949.5;refGene_gn=MFSD1;refGene_tx=XM_023260437.1
C2	108420757	108420944
refGene_gn=RARRES1;refGene_tx=XM_023260180.1
C2	108429367	108430272
refGene_gn=RARRES1;refGene_tx=XM_023260180.1
C2	108448187	108448250
refGene_gn=RARRES1;refGene_tx=XM_023260180.1
C2	108450963	108451147
refGene_gn=RARRES1;refGene_tx=XM_023260180.1
C2	108455628	108455765
refGene_gn=RARRES1;refGene_tx=XM_023260180.1
C2	108486206	108486269
refGene_gn=RARRES1;refGene_tx=XM_023260180.1
C2	108486829	108486976
refGene_gn=RARRES1;refGene_tx=XM_023260180.1
C2	108501547	108501679
refGene_gn=GFM1;refGene_tx=XM_003991950.4
C2	108501830	108501884
refGene_gn=GFM1;refGene_tx=XM_003991950.4
C2	108502863	108503024
refGene_gn=GFM1;refGene_tx=XM_003991950.4
C2	108508363	108508508
refGene_gn=GFM1;refGene_tx=XM_003991950.4
C2	108510398	108510561

refGene_gn=GFM1;refGene_tx=XM_003991950.4
C2 108515339 108515468
refGene_gn=LXN;refGene_tx=XM_003991951.3
C2 108516759 108516822
refGene_gn=LXN;refGene_tx=XM_003991951.3
C2 108517887 108518065
refGene_gn=LXN;refGene_tx=XM_003991951.3
C2 108518358 108518495
refGene_gn=LXN;refGene_tx=XM_003991951.3
C2 108520496 108520559
refGene_gn=LXN;refGene_tx=XM_003991951.3
C2 108520820 108520919
refGene_gn=LXN;refGene_tx=XM_003991951.3
C2 108521175 108521258
refGene_gn=GFM1;refGene_tx=XM_003991950.4
C2 108522139 108522277
refGene_gn=GFM1;refGene_tx=XM_003991950.4
C2 108525119 108525176
refGene_gn=GFM1;refGene_tx=XM_003991950.4
C2 108526796 108526898
refGene_gn=GFM1;refGene_tx=XM_003991950.4
C2 108528936 108529074
refGene_gn=GFM1;refGene_tx=XM_003991950.4
C2 108532192 108532277
refGene_gn=GFM1;refGene_tx=XM_003991950.4
C2 108532858 108533016
refGene_gn=GFM1;refGene_tx=XM_003991950.4
C2 108536770 108536921
refGene_gn=GFM1;refGene_tx=XM_003991950.4
C2 108540166 108540283
refGene_gn=GFM1;refGene_tx=XM_003991950.4
C2 108542693 108542898
refGene_gn=GFM1;refGene_tx=XM_003991950.4
C2 108543335 108543468
refGene_gn=GFM1;refGene_tx=XM_003991950.4
C2 108545014 108545167
refGene_gn=GFM1;refGene_tx=XM_003991950.4
C2 108546404 108546494
refGene_gn=GFM1;refGene_tx=XM_003991950.4
C2 108554066 108554172
refGene_gn=MLF1;refGene_tx=XM_003991952.5;refGene_gn=MLF1;refGene_tx=XM_006936297.4;refGene_gn=MLF1;refGene_tx=XM_006936298.4;refGene_gn=MLF1;refGene_tx=XM_006936299.4
C2 108556143 108556282
refGene_gn=MLF1;refGene_tx=XM_003991952.5;refGene_gn=MLF1;refGene_tx=XM_006936297.4;refGene_gn=MLF1;refGene_tx=XM_006936298.4;refGene_gn=MLF1;refGene_tx=XM_006936299.4
C2 108558081 108558241
refGene_gn=MLF1;refGene_tx=XM_003991952.5;refGene_gn=MLF1;refGene_tx=XM_006936297.4;refGene_gn=MLF1;refGene_tx=XM_006936298.4;refGene_gn=MLF

1;refGene_tx=XM_006936299.4
C2 108559944 108560073
refGene_gn=MLF1;refGene_tx=XM_003991952.5;refGene_gn=MLF1;refGene_tx=XM_006936297.4;refGene_gn=MLF1;refGene_tx=XM_006936298.4;refGene_gn=MLF1;refGene_tx=XM_006936299.4
C2 108561390 108561474
refGene_gn=MLF1;refGene_tx=XM_003991952.5;refGene_gn=MLF1;refGene_tx=XM_006936297.4;refGene_gn=MLF1;refGene_tx=XM_006936298.4
C2 108565032 108565074
refGene_gn=MLF1;refGene_tx=XM_006936297.4
C2 108569503 108569651
refGene_gn=MLF1;refGene_tx=XM_003991952.5;refGene_gn=MLF1;refGene_tx=XM_006936297.4;refGene_gn=MLF1;refGene_tx=XM_006936298.4;refGene_gn=MLF1;refGene_tx=XM_006936299.4
C2 108592070 108592117
refGene_gn=MLF1;refGene_tx=XM_003991952.5;refGene_gn=MLF1;refGene_tx=XM_006936297.4;refGene_gn=MLF1;refGene_tx=XM_006936298.4;refGene_gn=MLF1;refGene_tx=XM_006936299.4
C2 108616603 108616696
refGene_gn=LOC101084215;refGene_tx=XM_019810512.2
C2 108617374 108617527
refGene_gn=LOC101084215;refGene_tx=XM_019810512.2
C2 108625473 108625605
refGene_gn=LOC101084215;refGene_tx=XM_019810512.2
C2 108781451 108781507
refGene_gn=RSRC1;refGene_tx=XM_023260181.1
C2 108792496 108792548
refGene_gn=RSRC1;refGene_tx=XM_023260181.1
C2 108836392 108836429
refGene_gn=RSRC1;refGene_tx=XM_023260181.1
C2 108927797 108927971
refGene_gn=RSRC1;refGene_tx=XM_023260181.1
C2 109025356 109025482
refGene_gn=RSRC1;refGene_tx=XM_023260181.1
C2 109026255 109026449
refGene_gn=RSRC1;refGene_tx=XM_023260181.1
C2 109040480 109040832
refGene_gn=SHOX2;refGene_tx=XM_003991968.5;refGene_gn=SHOX2;refGene_tx=XM_023260439.1
C2 109043783 109043992
refGene_gn=SHOX2;refGene_tx=XM_003991968.5;refGene_gn=SHOX2;refGene_tx=XM_023260439.1
C2 109046429 109046487
refGene_gn=SHOX2;refGene_tx=XM_003991968.5;refGene_gn=SHOX2;refGene_tx=XM_023260439.1
C2 109046796 109046885
refGene_gn=SHOX2;refGene_tx=XM_003991968.5;refGene_gn=SHOX2;refGene_tx=XM_023260439.1
C2 109048444 109048738
refGene_gn=SHOX2;refGene_tx=XM_003991968.5;refGene_gn=SHOX2;refGene_tx

=XM_023260439.1
C2 109550961 109551107
refGene_gn=PQLC2L;refGene_tx=XM_019811112.2
C2 109551748 109551776
refGene_gn=PQLC2L;refGene_tx=XM_019811112.2
C2 109551777 109551880
refGene_gn=PQLC2L;refGene_tx=XM_019811112.2
C2 109564936 109565178
refGene_gn=PQLC2L;refGene_tx=XM_019811112.2
C2 109625036 109625174
refGene_gn=VEPH1;refGene_tx=XM_006936309.3;refGene_gn=VEPH1;refGene_tx=XM_019810513.2;refGene_gn=VEPH1;refGene_tx=XM_019810515.2;refGene_gn=VEPH1;refGene_tx=XM_019810516.2;refGene_gn=VEPH1;refGene_tx=XM_019810519.2;refGene_gn=VEPH1;refGene_tx=XM_019810520.2;refGene_gn=VEPH1;refGene_tx=XM_019810521.2
C2 109646889 109647105
refGene_gn=VEPH1;refGene_tx=XM_006936309.3;refGene_gn=VEPH1;refGene_tx=XM_019810513.2;refGene_gn=VEPH1;refGene_tx=XM_019810515.2;refGene_gn=VEPH1;refGene_tx=XM_019810516.2;refGene_gn=VEPH1;refGene_tx=XM_019810520.2;refGene_gn=VEPH1;refGene_tx=XM_019810521.2
C2 109656112 109656287
refGene_gn=VEPH1;refGene_tx=XM_006936309.3;refGene_gn=VEPH1;refGene_tx=XM_019810513.2;refGene_gn=VEPH1;refGene_tx=XM_019810515.2;refGene_gn=VEPH1;refGene_tx=XM_019810516.2;refGene_gn=VEPH1;refGene_tx=XM_019810519.2;refGene_gn=VEPH1;refGene_tx=XM_019810520.2;refGene_gn=VEPH1;refGene_tx=XM_019810521.2
C2 109666556 109667170
refGene_gn=PTX3;refGene_tx=XM_023260440.1
C2 109671153 109671555
refGene_gn=PTX3;refGene_tx=XM_023260440.1
C2 109672009 109672139
refGene_gn=PTX3;refGene_tx=XM_023260440.1
C2 109680008 109680175
refGene_gn=VEPH1;refGene_tx=XM_006936309.3;refGene_gn=VEPH1;refGene_tx=XM_019810513.2;refGene_gn=VEPH1;refGene_tx=XM_019810515.2;refGene_gn=VEPH1;refGene_tx=XM_019810516.2;refGene_gn=VEPH1;refGene_tx=XM_019810519.2;refGene_gn=VEPH1;refGene_tx=XM_019810520.2;refGene_gn=VEPH1;refGene_tx=XM_019810521.2
C2 109693650 109693860
refGene_gn=VEPH1;refGene_tx=XM_006936309.3;refGene_gn=VEPH1;refGene_tx=XM_019810513.2;refGene_gn=VEPH1;refGene_tx=XM_019810515.2;refGene_gn=VEPH1;refGene_tx=XM_019810516.2;refGene_gn=VEPH1;refGene_tx=XM_019810519.2;refGene_gn=VEPH1;refGene_tx=XM_019810520.2;refGene_gn=VEPH1;refGene_tx=XM_019810521.2
C2 109727995 109728216
refGene_gn=VEPH1;refGene_tx=XM_006936309.3;refGene_gn=VEPH1;refGene_tx=XM_019810513.2;refGene_gn=VEPH1;refGene_tx=XM_019810515.2;refGene_gn=VEPH1;refGene_tx=XM_019810516.2;refGene_gn=VEPH1;refGene_tx=XM_019810519.2;refGene_gn=VEPH1;refGene_tx=XM_019810520.2
C2 109729294 109729325

refGene_gn=VEPH1;refGene_tx=XM_019810520.2
C2 109751840 109752050
refGene_gn=VEPH1;refGene_tx=XM_006936309.3;refGene_gn=VEPH1;refGene_tx=XM_019810513.2;refGene_gn=VEPH1;refGene_tx=XM_019810515.2;refGene_gn=VEPH1;refGene_tx=XM_019810516.2;refGene_gn=VEPH1;refGene_tx=XM_019810519.2;refGene_gn=VEPH1;refGene_tx=XM_019810521.2
C2 109753082 109753480
refGene_gn=VEPH1;refGene_tx=XM_006936309.3;refGene_gn=VEPH1;refGene_tx=XM_019810513.2;refGene_gn=VEPH1;refGene_tx=XM_019810515.2;refGene_gn=VEPH1;refGene_tx=XM_019810516.2;refGene_gn=VEPH1;refGene_tx=XM_019810519.2
C2 109801866 109802006
refGene_gn=VEPH1;refGene_tx=XM_006936309.3;refGene_gn=VEPH1;refGene_tx=XM_019810513.2;refGene_gn=VEPH1;refGene_tx=XM_019810515.2;refGene_gn=VEPH1;refGene_tx=XM_019810516.2;refGene_gn=VEPH1;refGene_tx=XM_019810519.2
C2 109806512 109806647
refGene_gn=VEPH1;refGene_tx=XM_006936309.3;refGene_gn=VEPH1;refGene_tx=XM_019810513.2;refGene_gn=VEPH1;refGene_tx=XM_019810515.2;refGene_gn=VEPH1;refGene_tx=XM_019810516.2;refGene_gn=VEPH1;refGene_tx=XM_019810519.2
C2 109835379 109835497
refGene_gn=VEPH1;refGene_tx=XM_006936309.3;refGene_gn=VEPH1;refGene_tx=XM_019810513.2;refGene_gn=VEPH1;refGene_tx=XM_019810515.2;refGene_gn=VEPH1;refGene_tx=XM_019810516.2;refGene_gn=VEPH1;refGene_tx=XM_019810519.2
C2 109858789 109858926
refGene_gn=VEPH1;refGene_tx=XM_006936309.3;refGene_gn=VEPH1;refGene_tx=XM_019810513.2;refGene_gn=VEPH1;refGene_tx=XM_019810515.2;refGene_gn=VEPH1;refGene_tx=XM_019810516.2;refGene_gn=VEPH1;refGene_tx=XM_019810519.2
C2 109863384 109863621
refGene_gn=VEPH1;refGene_tx=XM_006936309.3;refGene_gn=VEPH1;refGene_tx=XM_019810513.2;refGene_gn=VEPH1;refGene_tx=XM_019810515.2;refGene_gn=VEPH1;refGene_tx=XM_019810519.2
C2 109868639 109868768
refGene_gn=VEPH1;refGene_tx=XM_019810516.2
C2 110020144 110020447
refGene_gn=CCNL1;refGene_tx=XM_011286074.3;refGene_gn=CCNL1;refGene_tx=XM_011286077.3
C2 110020745 110020820
refGene_gn=CCNL1;refGene_tx=XM_011286074.3;refGene_gn=CCNL1;refGene_tx=XM_011286077.3
C2 110021261 110021371
refGene_gn=CCNL1;refGene_tx=XM_011286074.3;refGene_gn=CCNL1;refGene_tx=XM_011286077.3
C2 110026965 110027086
refGene_gn=CCNL1;refGene_tx=XM_011286074.3;refGene_gn=CCNL1;refGene_tx=XM_011286077.3
C2 110028175 110028240

refGene_gn=CCNL1;refGene_tx=XM_011286074.3;refGene_gn=CCNL1;refGene_tx=XM_011286077.3
C2 110028479 110028493
refGene_gn=CCNL1;refGene_tx=XM_011286077.3
C2 110029985 110030085
refGene_gn=CCNL1;refGene_tx=XM_011286074.3;refGene_gn=CCNL1;refGene_tx=XM_011286077.3
C2 110030189 110030294
refGene_gn=CCNL1;refGene_tx=XM_011286074.3
C2 110030389 110030531
refGene_gn=CCNL1;refGene_tx=XM_011286074.3
C2 110030766 110030878
refGene_gn=CCNL1;refGene_tx=XM_011286074.3
C2 110030997 110031096
refGene_gn=CCNL1;refGene_tx=XM_011286074.3
C2 110031801 110032150
refGene_gn=CCNL1;refGene_tx=XM_011286074.3
C2 110110435 110110518
refGene_gn=LEKR1;refGene_tx=XM_023260441.1;refGene_gn=LEKR1;refGene_tx=XM_023260442.1
C2 110122645 110122744
refGene_gn=LEKR1;refGene_tx=XM_019810522.2;refGene_gn=LEKR1;refGene_tx=XM_023260441.1;refGene_gn=LEKR1;refGene_tx=XM_023260442.1
C2 110136393 110136807
refGene_gn=LEKR1;refGene_tx=XM_023260441.1;refGene_gn=LEKR1;refGene_tx=XM_023260442.1;refGene_gn=LEKR1;refGene_tx=XM_023260443.1
C2 110150379 110150679
refGene_gn=LEKR1;refGene_tx=XM_019810522.2;refGene_gn=LEKR1;refGene_tx=XM_023260441.1;refGene_gn=LEKR1;refGene_tx=XM_023260442.1;refGene_gn=LEKR1;refGene_tx=XM_023260443.1
C2 110153369 110153534
refGene_gn=LEKR1;refGene_tx=XM_019810522.2;refGene_gn=LEKR1;refGene_tx=XM_023260441.1;refGene_gn=LEKR1;refGene_tx=XM_023260442.1;refGene_gn=LEKR1;refGene_tx=XM_023260443.1
C2 110165946 110166040
refGene_gn=LEKR1;refGene_tx=XM_019810522.2;refGene_gn=LEKR1;refGene_tx=XM_023260441.1;refGene_gn=LEKR1;refGene_tx=XM_023260442.1;refGene_gn=LEKR1;refGene_tx=XM_023260443.1
C2 110186667 110186871
refGene_gn=LEKR1;refGene_tx=XM_019810522.2;refGene_gn=LEKR1;refGene_tx=XM_023260441.1;refGene_gn=LEKR1;refGene_tx=XM_023260442.1;refGene_gn=LEKR1;refGene_tx=XM_023260443.1
C2 110187207 110187285
refGene_gn=LEKR1;refGene_tx=XM_019810522.2;refGene_gn=LEKR1;refGene_tx=XM_023260441.1;refGene_gn=LEKR1;refGene_tx=XM_023260442.1;refGene_gn=LEKR1;refGene_tx=XM_023260443.1
C2 110196264 110196346
refGene_gn=LEKR1;refGene_tx=XM_019810522.2;refGene_gn=LEKR1;refGene_tx=XM_023260441.1;refGene_gn=LEKR1;refGene_tx=XM_023260442.1;refGene_gn=LEKR1;refGene_tx=XM_023260443.1

C2 110230240 110230426
refGene_gn=LEKR1;refGene_tx=XM_019810522.2;refGene_gn=LEKR1;refGene_tx=XM_023260441.1;refGene_gn=LEKR1;refGene_tx=XM_023260442.1;refGene_gn=LEKR1;refGene_tx=XM_023260443.1
C2 110239297 110239473
refGene_gn=LEKR1;refGene_tx=XM_019810522.2;refGene_gn=LEKR1;refGene_tx=XM_023260441.1;refGene_gn=LEKR1;refGene_tx=XM_023260442.1;refGene_gn=LEKR1;refGene_tx=XM_023260443.1
C2 110244415 110244535
refGene_gn=LEKR1;refGene_tx=XM_019810522.2;refGene_gn=LEKR1;refGene_tx=XM_023260441.1;refGene_gn=LEKR1;refGene_tx=XM_023260442.1;refGene_gn=LEKR1;refGene_tx=XM_023260443.1
C2 110300976 110301191
refGene_gn=LEKR1;refGene_tx=XM_019810522.2;refGene_gn=LEKR1;refGene_tx=XM_023260441.1;refGene_gn=LEKR1;refGene_tx=XM_023260442.1;refGene_gn=LEKR1;refGene_tx=XM_023260443.1
C2 110314564 110314612
refGene_gn=LEKR1;refGene_tx=XM_019810522.2;refGene_gn=LEKR1;refGene_tx=XM_023260441.1;refGene_gn=LEKR1;refGene_tx=XM_023260442.1;refGene_gn=LEKR1;refGene_tx=XM_023260443.1
C2 110445309 110445757
refGene_gn=TIPARP;refGene_tx=XM_003991970.5
C2 110446438 110446717
refGene_gn=TIPARP;refGene_tx=XM_003991970.5
C2 110451882 110452043
refGene_gn=TIPARP;refGene_tx=XM_003991970.5
C2 110453359 110453528
refGene_gn=TIPARP;refGene_tx=XM_003991970.5
C2 110469717 110470634
refGene_gn=TIPARP;refGene_tx=XM_003991970.5
C2 110583494 110583627
refGene_gn=SSR3;refGene_tx=XM_003991971.5
C2 110584802 110584929
refGene_gn=SSR3;refGene_tx=XM_003991971.5
C2 110589562 110589661
refGene_gn=SSR3;refGene_tx=XM_003991971.5
C2 110593982 110594114
refGene_gn=SSR3;refGene_tx=XM_003991971.5
C2 110594841 110594908
refGene_gn=SSR3;refGene_tx=XM_003991971.5
C2 110600961 110601051
refGene_gn=KCNAB1;refGene_tx=XM_011286073.3;refGene_gn=KCNAB1;refGene_tx=XM_019810528.2;refGene_gn=KCNAB1;refGene_tx=XM_019810529.2;refGene_gn=KCNAB1;refGene_tx=XM_019810530.2;refGene_gn=KCNAB1;refGene_tx=XM_019810531.2
C2 110606003 110606092
refGene_gn=KCNAB1;refGene_tx=XM_011286073.3;refGene_gn=KCNAB1;refGene_tx=XM_019810528.2;refGene_gn=KCNAB1;refGene_tx=XM_019810529.2;refGene_gn=KCNAB1;refGene_tx=XM_019810530.2;refGene_gn=KCNAB1;refGene_tx=XM_019810531.2

C2	110611049	110611170
refGene_gn=KCNAB1;refGene_tx=XM_011286073.3;refGene_gn=KCNAB1;refGene_tx=XM_019810528.2;refGene_gn=KCNAB1;refGene_tx=XM_019810529.2;refGene_gn=KCNAB1;refGene_tx=XM_019810530.2;refGene_gn=KCNAB1;refGene_tx=XM_019810531.2
C2	110618925	110619020
refGene_gn=KCNAB1;refGene_tx=XM_011286073.3;refGene_gn=KCNAB1;refGene_tx=XM_019810528.2;refGene_gn=KCNAB1;refGene_tx=XM_019810529.2;refGene_gn=KCNAB1;refGene_tx=XM_019810530.2;refGene_gn=KCNAB1;refGene_tx=XM_019810531.2
C2	110620362	110620483
refGene_gn=KCNAB1;refGene_tx=XM_011286073.3;refGene_gn=KCNAB1;refGene_tx=XM_019810528.2;refGene_gn=KCNAB1;refGene_tx=XM_019810529.2;refGene_gn=KCNAB1;refGene_tx=XM_019810530.2;refGene_gn=KCNAB1;refGene_tx=XM_019810531.2
C2	110621141	110621227
refGene_gn=KCNAB1;refGene_tx=XM_011286073.3;refGene_gn=KCNAB1;refGene_tx=XM_019810528.2;refGene_gn=KCNAB1;refGene_tx=XM_019810529.2;refGene_gn=KCNAB1;refGene_tx=XM_019810530.2;refGene_gn=KCNAB1;refGene_tx=XM_019810531.2
C2	110661289	110661376
refGene_gn=KCNAB1;refGene_tx=XM_011286073.3;refGene_gn=KCNAB1;refGene_tx=XM_019810528.2;refGene_gn=KCNAB1;refGene_tx=XM_019810529.2
C2	110672970	110673014
refGene_gn=KCNAB1;refGene_tx=XM_011286073.3;refGene_gn=KCNAB1;refGene_tx=XM_019810528.2;refGene_gn=KCNAB1;refGene_tx=XM_019810529.2;refGene_gn=KCNAB1;refGene_tx=XM_019810530.2;refGene_gn=KCNAB1;refGene_tx=XM_019810531.2
C2	110675031	110675076
refGene_gn=KCNAB1;refGene_tx=XM_011286073.3;refGene_gn=KCNAB1;refGene_tx=XM_019810528.2;refGene_gn=KCNAB1;refGene_tx=XM_019810529.2;refGene_gn=KCNAB1;refGene_tx=XM_019810530.2;refGene_gn=KCNAB1;refGene_tx=XM_019810531.2
C2	110678857	110678902
refGene_gn=KCNAB1;refGene_tx=XM_011286073.3;refGene_gn=KCNAB1;refGene_tx=XM_019810528.2;refGene_gn=KCNAB1;refGene_tx=XM_019810529.2;refGene_gn=KCNAB1;refGene_tx=XM_019810530.2;refGene_gn=KCNAB1;refGene_tx=XM_019810531.2
C2	110681459	110681539
refGene_gn=KCNAB1;refGene_tx=XM_011286073.3;refGene_gn=KCNAB1;refGene_tx=XM_019810528.2;refGene_gn=KCNAB1;refGene_tx=XM_019810529.2;refGene_gn=KCNAB1;refGene_tx=XM_019810530.2;refGene_gn=KCNAB1;refGene_tx=XM_019810531.2
C2	110685946	110685984
refGene_gn=KCNAB1;refGene_tx=XM_011286073.3;refGene_gn=KCNAB1;refGene_tx=XM_019810528.2;refGene_gn=KCNAB1;refGene_tx=XM_019810529.2;refGene_gn=KCNAB1;refGene_tx=XM_019810530.2;refGene_gn=KCNAB1;refGene_tx=XM_019810531.2
C2	110726816	110726860
refGene_gn=KCNAB1;refGene_tx=XM_011286073.3;refGene_gn=KCNAB1;refGene_

tx=XM_019810528.2;refGene_gn=KCNAB1;refGene_tx=XM_019810529.2;refGene_gn=KCNAB1;refGene_tx=XM_019810530.2;refGene_gn=KCNAB1;refGene_tx=XM_019810531.2
C2	110874868	110875089	refGene_gn=KCNAB1;refGene_tx=XM_019810529.2;refGene_gn=KCNAB1;refGene_tx=XM_019810531.2
C2	110997039	110997281	refGene_gn=KCNAB1;refGene_tx=XM_011286073.3
C2	111022660	111022935	refGene_gn=KCNAB1;refGene_tx=XM_019810528.2;refGene_gn=KCNAB1;refGene_tx=XM_019810530.2
C2	111155057	111155159	refGene_gn=GMPS;refGene_tx=XM_023260444.1;refGene_gn=GMPS;refGene_tx=XM_023260445.1
C2	111156211	111156384	refGene_gn=GMPS;refGene_tx=XM_023260444.1;refGene_gn=GMPS;refGene_tx=XM_023260445.1
C2	111157319	111157450	refGene_gn=GMPS;refGene_tx=XM_023260444.1;refGene_gn=GMPS;refGene_tx=XM_023260445.1
C2	111162157	111162273	refGene_gn=GMPS;refGene_tx=XM_023260444.1;refGene_gn=GMPS;refGene_tx=XM_023260445.1
C2	111173048	111173174	refGene_gn=GMPS;refGene_tx=XM_023260444.1;refGene_gn=GMPS;refGene_tx=XM_023260445.1
C2	111175314	111175430	refGene_gn=GMPS;refGene_tx=XM_023260444.1;refGene_gn=GMPS;refGene_tx=XM_023260445.1
C2	111177560	111177666	refGene_gn=GMPS;refGene_tx=XM_023260444.1;refGene_gn=GMPS;refGene_tx=XM_023260445.1
C2	111179944	111180118	refGene_gn=GMPS;refGene_tx=XM_023260444.1;refGene_gn=GMPS;refGene_tx=XM_023260445.1
C2	111180595	111180747	refGene_gn=GMPS;refGene_tx=XM_023260444.1;refGene_gn=GMPS;refGene_tx=XM_023260445.1
C2	111188268	111188434	refGene_gn=GMPS;refGene_tx=XM_023260444.1;refGene_gn=GMPS;refGene_tx=XM_023260445.1
C2	111188669	111188863	refGene_gn=GMPS;refGene_tx=XM_023260444.1;refGene_gn=GMPS;refGene_tx=XM_023260445.1
C2	111194270	111194374	refGene_gn=GMPS;refGene_tx=XM_023260444.1;refGene_gn=GMPS;refGene_tx=XM_023260445.1
C2	111198473	111198571	refGene_gn=GMPS;refGene_tx=XM_023260444.1;refGene_gn=GMPS;refGene_tx=XM_023260445.1

C2	111205573	111205688
refGene_gn=GMPS;refGene_tx=XM_023260444.1;refGene_gn=GMPS;refGene_tx=XM_023260445.1
C2	111209155	111209337
refGene_gn=GMPS;refGene_tx=XM_023260444.1;refGene_gn=GMPS;refGene_tx=XM_023260445.1
C2	111232906	111232930
refGene_gn=GMPS;refGene_tx=XM_023260445.1
C2	111234629	111234674
refGene_gn=GMPS;refGene_tx=XM_023260444.1
C2	111255361	111256136
refGene_gn=SLC33A1;refGene_tx=XM_011286071.3;refGene_gn=SLC33A1;refGene_tx=XM_019810533.2;refGene_gn=SLC33A1;refGene_tx=XM_023260446.1
C2	111268008	111268199
refGene_gn=SLC33A1;refGene_tx=XM_011286071.3;refGene_gn=SLC33A1;refGene_tx=XM_019810533.2;refGene_gn=SLC33A1;refGene_tx=XM_023260446.1
C2	111273200	111273385
refGene_gn=SLC33A1;refGene_tx=XM_011286071.3;refGene_gn=SLC33A1;refGene_tx=XM_019810533.2;refGene_gn=SLC33A1;refGene_tx=XM_023260446.1
C2	111273658	111273776
refGene_gn=SLC33A1;refGene_tx=XM_011286071.3;refGene_gn=SLC33A1;refGene_tx=XM_019810533.2;refGene_gn=SLC33A1;refGene_tx=XM_023260446.1
C2	111276284	111276500
refGene_gn=SLC33A1;refGene_tx=XM_011286071.3;refGene_gn=SLC33A1;refGene_tx=XM_023260446.1
C2	111277583	111277751
refGene_gn=SLC33A1;refGene_tx=XM_011286071.3;refGene_gn=SLC33A1;refGene_tx=XM_019810533.2;refGene_gn=SLC33A1;refGene_tx=XM_023260446.1
C2	111290093	111290207
refGene_gn=CC2H3orf33;refGene_tx=XM_023260447.1;refGene_gn=CC2H3orf33;refGene_tx=XM_023260448.1
C2	111291595	111291616
refGene_gn=CC2H3orf33;refGene_tx=XM_023260447.1
C2	111292563	111292623
refGene_gn=CC2H3orf33;refGene_tx=XM_023260447.1;refGene_gn=CC2H3orf33;refGene_tx=XM_023260448.1;refGene_gn=CC2H3orf33;refGene_tx=XM_023260449.1
C2	111296997	111297145
refGene_gn=CC2H3orf33;refGene_tx=XM_023260447.1;refGene_gn=CC2H3orf33;refGene_tx=XM_023260448.1;refGene_gn=CC2H3orf33;refGene_tx=XM_023260449.1
C2	111300450	111300611
refGene_gn=CC2H3orf33;refGene_tx=XM_023260447.1;refGene_gn=CC2H3orf33;refGene_tx=XM_023260448.1;refGene_gn=CC2H3orf33;refGene_tx=XM_023260449.1
C2	111304177	111304579
refGene_gn=CC2H3orf33;refGene_tx=XM_023260447.1;refGene_gn=CC2H3orf33;refGene_tx=XM_023260448.1;refGene_gn=CC2H3orf33;refGene_tx=XM_023260449.1
C2	111351632	111351711

refGene_gn=PLCH1;refGene_tx=XM_003991977.5;refGene_gn=PLCH1;refGene_tx=XM_006936314.4;refGene_gn=PLCH1;refGene_tx=XM_019810535.2;refGene_gn=PLCH1;refGene_tx=XM_019810536.2;refGene_gn=PLCH1;refGene_tx=XM_019810537.2;refGene_gn=PLCH1;refGene_tx=XM_019810538.2;refGene_gn=PLCH1;refGene_tx=XM_019810540.2;refGene_gn=PLCH1;refGene_tx=XM_023260450.1
C2 111382655 111382698
refGene_gn=PLCH1;refGene_tx=XM_019810539.2
C2 111450232 111450379
refGene_gn=PLCH1;refGene_tx=XM_003991977.5;refGene_gn=PLCH1;refGene_tx=XM_006936314.4;refGene_gn=PLCH1;refGene_tx=XM_019810535.2;refGene_gn=PLCH1;refGene_tx=XM_019810536.2;refGene_gn=PLCH1;refGene_tx=XM_019810537.2;refGene_gn=PLCH1;refGene_tx=XM_019810538.2;refGene_gn=PLCH1;refGene_tx=XM_019810539.2;refGene_gn=PLCH1;refGene_tx=XM_019810540.2;refGene_gn=PLCH1;refGene_tx=XM_023260450.1
C2 111452545 111452789
refGene_gn=PLCH1;refGene_tx=XM_003991977.5;refGene_gn=PLCH1;refGene_tx=XM_006936314.4;refGene_gn=PLCH1;refGene_tx=XM_019810535.2;refGene_gn=PLCH1;refGene_tx=XM_019810536.2;refGene_gn=PLCH1;refGene_tx=XM_019810537.2;refGene_gn=PLCH1;refGene_tx=XM_019810538.2;refGene_gn=PLCH1;refGene_tx=XM_019810539.2;refGene_gn=PLCH1;refGene_tx=XM_019810540.2;refGene_gn=PLCH1;refGene_tx=XM_023260450.1
C2 111463449 111463579
refGene_gn=PLCH1;refGene_tx=XM_003991977.5;refGene_gn=PLCH1;refGene_tx=XM_006936314.4;refGene_gn=PLCH1;refGene_tx=XM_019810535.2;refGene_gn=PLCH1;refGene_tx=XM_019810536.2;refGene_gn=PLCH1;refGene_tx=XM_019810537.2;refGene_gn=PLCH1;refGene_tx=XM_019810538.2;refGene_gn=PLCH1;refGene_tx=XM_019810539.2;refGene_gn=PLCH1;refGene_tx=XM_019810540.2;refGene_gn=PLCH1;refGene_tx=XM_023260450.1
C2 111467023 111467194
refGene_gn=PLCH1;refGene_tx=XM_003991977.5;refGene_gn=PLCH1;refGene_tx=XM_006936314.4;refGene_gn=PLCH1;refGene_tx=XM_019810535.2;refGene_gn=PLCH1;refGene_tx=XM_019810536.2;refGene_gn=PLCH1;refGene_tx=XM_019810537.2;refGene_gn=PLCH1;refGene_tx=XM_019810538.2;refGene_gn=PLCH1;refGene_tx=XM_019810539.2;refGene_gn=PLCH1;refGene_tx=XM_019810540.2;refGene_gn=PLCH1;refGene_tx=XM_023260450.1
C2 111479569 111479663
refGene_gn=PLCH1;refGene_tx=XM_003991977.5;refGene_gn=PLCH1;refGene_tx=XM_006936314.4;refGene_gn=PLCH1;refGene_tx=XM_019810535.2;refGene_gn=PLCH1;refGene_tx=XM_019810536.2;refGene_gn=PLCH1;refGene_tx=XM_019810537.2;refGene_gn=PLCH1;refGene_tx=XM_019810538.2;refGene_gn=PLCH1;refGene_tx=XM_019810539.2;refGene_gn=PLCH1;refGene_tx=XM_019810540.2;refGene_gn=PLCH1;refGene_tx=XM_023260450.1
C2 111482143 111482347
refGene_gn=PLCH1;refGene_tx=XM_003991977.5;refGene_gn=PLCH1;refGene_tx=XM_006936314.4;refGene_gn=PLCH1;refGene_tx=XM_019810535.2;refGene_gn=PLCH1;refGene_tx=XM_019810536.2;refGene_gn=PLCH1;refGene_tx=XM_019810537.2;refGene_gn=PLCH1;refGene_tx=XM_019810538.2;refGene_gn=PLCH1;refGene_tx=XM_019810539.2;refGene_gn=PLCH1;refGene_tx=XM_019810540.2;refGene_gn=PLCH1;refGene_tx=XM_023260450.1
C2 111491813 111491934

refGene_gn=PLCH1;refGene_tx=XM_003991977.5;refGene_gn=PLCH1;refGene_tx=XM_006936314.4;refGene_gn=PLCH1;refGene_tx=XM_019810535.2;refGene_gn=PLCH1;refGene_tx=XM_019810536.2;refGene_gn=PLCH1;refGene_tx=XM_019810537.2;refGene_gn=PLCH1;refGene_tx=XM_019810538.2;refGene_gn=PLCH1;refGene_tx=XM_019810539.2;refGene_gn=PLCH1;refGene_tx=XM_019810540.2;refGene_gn=PLCH1;refGene_tx=XM_023260450.1
C2 111497539 111497711
refGene_gn=PLCH1;refGene_tx=XM_003991977.5;refGene_gn=PLCH1;refGene_tx=XM_006936314.4;refGene_gn=PLCH1;refGene_tx=XM_019810535.2;refGene_gn=PLCH1;refGene_tx=XM_019810536.2;refGene_gn=PLCH1;refGene_tx=XM_019810537.2;refGene_gn=PLCH1;refGene_tx=XM_019810538.2;refGene_gn=PLCH1;refGene_tx=XM_019810539.2;refGene_gn=PLCH1;refGene_tx=XM_019810540.2;refGene_gn=PLCH1;refGene_tx=XM_023260450.1
C2 111501810 111501918
refGene_gn=PLCH1;refGene_tx=XM_003991977.5;refGene_gn=PLCH1;refGene_tx=XM_006936314.4;refGene_gn=PLCH1;refGene_tx=XM_019810535.2;refGene_gn=PLCH1;refGene_tx=XM_019810536.2;refGene_gn=PLCH1;refGene_tx=XM_019810537.2;refGene_gn=PLCH1;refGene_tx=XM_019810538.2;refGene_gn=PLCH1;refGene_tx=XM_019810539.2;refGene_gn=PLCH1;refGene_tx=XM_019810540.2;refGene_gn=PLCH1;refGene_tx=XM_023260450.1
C2 111511082 111511244
refGene_gn=PLCH1;refGene_tx=XM_003991977.5;refGene_gn=PLCH1;refGene_tx=XM_006936314.4;refGene_gn=PLCH1;refGene_tx=XM_019810535.2;refGene_gn=PLCH1;refGene_tx=XM_019810536.2;refGene_gn=PLCH1;refGene_tx=XM_019810537.2;refGene_gn=PLCH1;refGene_tx=XM_019810538.2;refGene_gn=PLCH1;refGene_tx=XM_019810539.2;refGene_gn=PLCH1;refGene_tx=XM_019810540.2;refGene_gn=PLCH1;refGene_tx=XM_023260450.1
C2 111516347 111516419
refGene_gn=PLCH1;refGene_tx=XM_003991977.5;refGene_gn=PLCH1;refGene_tx=XM_006936314.4;refGene_gn=PLCH1;refGene_tx=XM_019810535.2;refGene_gn=PLCH1;refGene_tx=XM_019810536.2;refGene_gn=PLCH1;refGene_tx=XM_019810537.2;refGene_gn=PLCH1;refGene_tx=XM_019810538.2;refGene_gn=PLCH1;refGene_tx=XM_019810539.2;refGene_gn=PLCH1;refGene_tx=XM_019810540.2;refGene_gn=PLCH1;refGene_tx=XM_023260450.1
C2 111519585 111519677
refGene_gn=PLCH1;refGene_tx=XM_003991977.5;refGene_gn=PLCH1;refGene_tx=XM_006936314.4;refGene_gn=PLCH1;refGene_tx=XM_019810535.2;refGene_gn=PLCH1;refGene_tx=XM_019810536.2;refGene_gn=PLCH1;refGene_tx=XM_019810537.2;refGene_gn=PLCH1;refGene_tx=XM_019810538.2;refGene_gn=PLCH1;refGene_tx=XM_019810539.2;refGene_gn=PLCH1;refGene_tx=XM_019810540.2;refGene_gn=PLCH1;refGene_tx=XM_023260450.1
C2 111520821 111520940
refGene_gn=PLCH1;refGene_tx=XM_003991977.5;refGene_gn=PLCH1;refGene_tx=XM_006936314.4;refGene_gn=PLCH1;refGene_tx=XM_019810535.2;refGene_gn=PLCH1;refGene_tx=XM_019810536.2;refGene_gn=PLCH1;refGene_tx=XM_019810537.2;refGene_gn=PLCH1;refGene_tx=XM_019810538.2;refGene_gn=PLCH1;refGene_tx=XM_019810539.2;refGene_gn=PLCH1;refGene_tx=XM_019810540.2;refGene_gn=PLCH1;refGene_tx=XM_023260450.1
C2 111524450 111524630
refGene_gn=PLCH1;refGene_tx=XM_003991977.5;refGene_gn=PLCH1;refGene_tx

=XM_006936314.4;refGene_gn=PLCH1;refGene_tx=XM_019810535.2;refGene_gn=PLCH1;refGene_tx=XM_019810536.2;refGene_gn=PLCH1;refGene_tx=XM_019810537.2;refGene_gn=PLCH1;refGene_tx=XM_019810538.2;refGene_gn=PLCH1;refGene_tx=XM_019810539.2;refGene_gn=PLCH1;refGene_tx=XM_019810540.2;refGene_gn=PLCH1;refGene_tx=XM_023260450.1
C2 111524722 111524830
refGene_gn=PLCH1;refGene_tx=XM_003991977.5;refGene_gn=PLCH1;refGene_tx=XM_006936314.4;refGene_gn=PLCH1;refGene_tx=XM_019810535.2;refGene_gn=PLCH1;refGene_tx=XM_019810536.2;refGene_gn=PLCH1;refGene_tx=XM_019810537.2;refGene_gn=PLCH1;refGene_tx=XM_019810538.2;refGene_gn=PLCH1;refGene_tx=XM_019810539.2;refGene_gn=PLCH1;refGene_tx=XM_019810540.2;refGene_gn=PLCH1;refGene_tx=XM_023260450.1
C2 111526314 111526439
refGene_gn=PLCH1;refGene_tx=XM_003991977.5;refGene_gn=PLCH1;refGene_tx=XM_006936314.4;refGene_gn=PLCH1;refGene_tx=XM_019810535.2;refGene_gn=PLCH1;refGene_tx=XM_019810536.2;refGene_gn=PLCH1;refGene_tx=XM_019810537.2;refGene_gn=PLCH1;refGene_tx=XM_019810538.2;refGene_gn=PLCH1;refGene_tx=XM_019810539.2;refGene_gn=PLCH1;refGene_tx=XM_019810540.2;refGene_gn=PLCH1;refGene_tx=XM_023260450.1
C2 111530674 111530759
refGene_gn=PLCH1;refGene_tx=XM_003991977.5;refGene_gn=PLCH1;refGene_tx=XM_006936314.4;refGene_gn=PLCH1;refGene_tx=XM_019810535.2;refGene_gn=PLCH1;refGene_tx=XM_019810536.2;refGene_gn=PLCH1;refGene_tx=XM_019810537.2;refGene_gn=PLCH1;refGene_tx=XM_019810538.2;refGene_gn=PLCH1;refGene_tx=XM_019810539.2;refGene_gn=PLCH1;refGene_tx=XM_019810540.2;refGene_gn=PLCH1;refGene_tx=XM_023260450.1
C2 111533091 111533238
refGene_gn=PLCH1;refGene_tx=XM_003991977.5;refGene_gn=PLCH1;refGene_tx=XM_006936314.4;refGene_gn=PLCH1;refGene_tx=XM_019810535.2;refGene_gn=PLCH1;refGene_tx=XM_019810536.2;refGene_gn=PLCH1;refGene_tx=XM_019810537.2;refGene_gn=PLCH1;refGene_tx=XM_019810538.2;refGene_gn=PLCH1;refGene_tx=XM_019810539.2;refGene_gn=PLCH1;refGene_tx=XM_019810540.2;refGene_gn=PLCH1;refGene_tx=XM_023260450.1
C2 111533485 111533565
refGene_gn=PLCH1;refGene_tx=XM_003991977.5;refGene_gn=PLCH1;refGene_tx=XM_006936314.4;refGene_gn=PLCH1;refGene_tx=XM_019810535.2;refGene_gn=PLCH1;refGene_tx=XM_019810536.2;refGene_gn=PLCH1;refGene_tx=XM_019810537.2;refGene_gn=PLCH1;refGene_tx=XM_019810538.2;refGene_gn=PLCH1;refGene_tx=XM_019810539.2;refGene_gn=PLCH1;refGene_tx=XM_019810540.2;refGene_gn=PLCH1;refGene_tx=XM_023260450.1
C2 111535101 111535161
refGene_gn=PLCH1;refGene_tx=XM_006936314.4;refGene_gn=PLCH1;refGene_tx=XM_019810535.2;refGene_gn=PLCH1;refGene_tx=XM_019810536.2;refGene_gn=PLCH1;refGene_tx=XM_019810537.2;refGene_gn=PLCH1;refGene_tx=XM_019810538.2;refGene_gn=PLCH1;refGene_tx=XM_019810539.2
C2 111535597 111535952
refGene_gn=PLCH1;refGene_tx=XM_003991977.5;refGene_gn=PLCH1;refGene_tx=XM_006936314.4;refGene_gn=PLCH1;refGene_tx=XM_019810535.2;refGene_gn=PLCH1;refGene_tx=XM_019810536.2;refGene_gn=PLCH1;refGene_tx=XM_019810537.2;refGene_gn=PLCH1;refGene_tx=XM_019810538.2;refGene_gn=PLCH1;refGe

ne_tx=XM_019810539.2;refGene_gn=PLCH1;refGene_tx=XM_019810540.2;refGen
e_gn=PLCH1;refGene_tx=XM_023260450.1
C2	111538281	111540362
refGene_gn=PLCH1;refGene_tx=XM_003991977.5;refGene_gn=PLCH1;refGene_tx
=XM_006936314.4;refGene_gn=PLCH1;refGene_tx=XM_019810535.2;refGene_gn=
PLCH1;refGene_tx=XM_019810536.2;refGene_gn=PLCH1;refGene_tx=XM_0198105
37.2;refGene_gn=PLCH1;refGene_tx=XM_019810538.2;refGene_gn=PLCH1;refGe
ne_tx=XM_019810539.2;refGene_gn=PLCH1;refGene_tx=XM_019810540.2;refGen
e_gn=PLCH1;refGene_tx=XM_023260450.1
C2	111720407	111720420
refGene_gn=LOC102902215;refGene_tx=XM_023260451.1
C2	111722917	111723009
refGene_gn=LOC102902215;refGene_tx=XM_023260451.1
C2	111806933	111807033
refGene_gn=MME;refGene_tx=XM_006936315.4;refGene_gn=MME;refGene_tx=XM_
019810542.2;refGene_gn=MME;refGene_tx=XM_023260452.1;refGene_gn=MME;re
fGene_tx=XM_023260453.1;refGene_gn=MME;refGene_tx=XM_023260454.1;refGe
ne_gn=MME;refGene_tx=XM_023260455.1
C2	111815851	111815928
refGene_gn=MME;refGene_tx=XM_006936315.4;refGene_gn=MME;refGene_tx=XM_
019810542.2;refGene_gn=MME;refGene_tx=XM_023260452.1;refGene_gn=MME;re
fGene_tx=XM_023260453.1;refGene_gn=MME;refGene_tx=XM_023260454.1;refGe
ne_gn=MME;refGene_tx=XM_023260455.1
C2	111816253	111816349
refGene_gn=MME;refGene_tx=XM_006936315.4;refGene_gn=MME;refGene_tx=XM_
019810542.2;refGene_gn=MME;refGene_tx=XM_023260452.1;refGene_gn=MME;re
fGene_tx=XM_023260453.1;refGene_gn=MME;refGene_tx=XM_023260454.1;refGe
ne_gn=MME;refGene_tx=XM_023260455.1
C2	111819256	111819322
refGene_gn=MME;refGene_tx=XM_006936315.4;refGene_gn=MME;refGene_tx=XM_
019810542.2;refGene_gn=MME;refGene_tx=XM_023260452.1;refGene_gn=MME;re
fGene_tx=XM_023260453.1;refGene_gn=MME;refGene_tx=XM_023260454.1;refGe
ne_gn=MME;refGene_tx=XM_023260455.1
C2	111820158	111820292
refGene_gn=MME;refGene_tx=XM_006936315.4;refGene_gn=MME;refGene_tx=XM_
019810542.2;refGene_gn=MME;refGene_tx=XM_023260452.1;refGene_gn=MME;re
fGene_tx=XM_023260453.1;refGene_gn=MME;refGene_tx=XM_023260454.1;refGe
ne_gn=MME;refGene_tx=XM_023260455.1
C2	111821322	111821442
refGene_gn=MME;refGene_tx=XM_006936315.4;refGene_gn=MME;refGene_tx=XM_
019810542.2;refGene_gn=MME;refGene_tx=XM_023260452.1;refGene_gn=MME;re
fGene_tx=XM_023260453.1;refGene_gn=MME;refGene_tx=XM_023260454.1;refGe
ne_gn=MME;refGene_tx=XM_023260455.1
C2	111827386	111827445
refGene_gn=MME;refGene_tx=XM_006936315.4;refGene_gn=MME;refGene_tx=XM_
019810542.2;refGene_gn=MME;refGene_tx=XM_023260452.1;refGene_gn=MME;re
fGene_tx=XM_023260453.1;refGene_gn=MME;refGene_tx=XM_023260454.1;refGe
ne_gn=MME;refGene_tx=XM_023260455.1
C2	111843986	111844090
refGene_gn=MME;refGene_tx=XM_006936315.4;refGene_gn=MME;refGene_tx=XM_

019810542.2;refGene_gn=MME;refGene_tx=XM_023260452.1;refGene_gn=MME;refGene_tx=XM_023260453.1;refGene_gn=MME;refGene_tx=XM_023260454.1;refGene_gn=MME;refGene_tx=XM_023260455.1
C2	111845371	111845452
refGene_gn=MME;refGene_tx=XM_006936315.4;refGene_gn=MME;refGene_tx=XM_019810542.2;refGene_gn=MME;refGene_tx=XM_023260452.1;refGene_gn=MME;refGene_tx=XM_023260453.1;refGene_gn=MME;refGene_tx=XM_023260454.1;refGene_gn=MME;refGene_tx=XM_023260455.1
C2	111848396	111848495
refGene_gn=MME;refGene_tx=XM_006936315.4;refGene_gn=MME;refGene_tx=XM_019810542.2;refGene_gn=MME;refGene_tx=XM_023260452.1;refGene_gn=MME;refGene_tx=XM_023260453.1;refGene_gn=MME;refGene_tx=XM_023260454.1;refGene_gn=MME;refGene_tx=XM_023260455.1
C2	111849466	111849595
refGene_gn=MME;refGene_tx=XM_006936315.4;refGene_gn=MME;refGene_tx=XM_019810542.2;refGene_gn=MME;refGene_tx=XM_023260452.1;refGene_gn=MME;refGene_tx=XM_023260453.1;refGene_gn=MME;refGene_tx=XM_023260454.1;refGene_gn=MME;refGene_tx=XM_023260455.1
C2	111850741	111850835
refGene_gn=MME;refGene_tx=XM_006936315.4;refGene_gn=MME;refGene_tx=XM_019810542.2;refGene_gn=MME;refGene_tx=XM_023260452.1;refGene_gn=MME;refGene_tx=XM_023260453.1;refGene_gn=MME;refGene_tx=XM_023260454.1;refGene_gn=MME;refGene_tx=XM_023260455.1
C2	111850944	111851081
refGene_gn=MME;refGene_tx=XM_006936315.4;refGene_gn=MME;refGene_tx=XM_019810542.2;refGene_gn=MME;refGene_tx=XM_023260452.1;refGene_gn=MME;refGene_tx=XM_023260453.1;refGene_gn=MME;refGene_tx=XM_023260454.1;refGene_gn=MME;refGene_tx=XM_023260455.1
C2	111852632	111852734
refGene_gn=MME;refGene_tx=XM_006936315.4;refGene_gn=MME;refGene_tx=XM_019810542.2;refGene_gn=MME;refGene_tx=XM_023260452.1;refGene_gn=MME;refGene_tx=XM_023260453.1;refGene_gn=MME;refGene_tx=XM_023260454.1;refGene_gn=MME;refGene_tx=XM_023260455.1
C2	111854925	111855060
refGene_gn=MME;refGene_tx=XM_006936315.4;refGene_gn=MME;refGene_tx=XM_019810542.2;refGene_gn=MME;refGene_tx=XM_023260452.1;refGene_gn=MME;refGene_tx=XM_023260453.1;refGene_gn=MME;refGene_tx=XM_023260454.1;refGene_gn=MME;refGene_tx=XM_023260455.1
C2	111866965	111867031
refGene_gn=MME;refGene_tx=XM_006936315.4;refGene_gn=MME;refGene_tx=XM_019810542.2;refGene_gn=MME;refGene_tx=XM_023260452.1;refGene_gn=MME;refGene_tx=XM_023260453.1;refGene_gn=MME;refGene_tx=XM_023260454.1;refGene_gn=MME;refGene_tx=XM_023260455.1
C2	111868813	111868932
refGene_gn=MME;refGene_tx=XM_006936315.4;refGene_gn=MME;refGene_tx=XM_019810542.2;refGene_gn=MME;refGene_tx=XM_023260452.1;refGene_gn=MME;refGene_tx=XM_023260453.1;refGene_gn=MME;refGene_tx=XM_023260454.1;refGene_gn=MME;refGene_tx=XM_023260455.1
C2	111869050	111869146
refGene_gn=MME;refGene_tx=XM_006936315.4;refGene_gn=MME;refGene_tx=XM_

019810542.2;refGene_gn=MME;refGene_tx=XM_023260452.1;refGene_gn=MME;refGene_tx=XM_023260453.1;refGene_gn=MME;refGene_tx=XM_023260454.1;refGene_gn=MME;refGene_tx=XM_023260455.1
C2 111869232 111869313
refGene_gn=MME;refGene_tx=XM_006936315.4;refGene_gn=MME;refGene_tx=XM_019810542.2;refGene_gn=MME;refGene_tx=XM_023260452.1;refGene_gn=MME;refGene_tx=XM_023260453.1;refGene_gn=MME;refGene_tx=XM_023260454.1;refGene_gn=MME;refGene_tx=XM_023260455.1
C2 111870481 111870643
refGene_gn=MME;refGene_tx=XM_006936315.4;refGene_gn=MME;refGene_tx=XM_019810542.2;refGene_gn=MME;refGene_tx=XM_023260452.1;refGene_gn=MME;refGene_tx=XM_023260453.1;refGene_gn=MME;refGene_tx=XM_023260454.1;refGene_gn=MME;refGene_tx=XM_023260455.1
C2 111900481 111900517
refGene_gn=MME;refGene_tx=XM_006936315.4;refGene_gn=MME;refGene_tx=XM_019810542.2;refGene_gn=MME;refGene_tx=XM_023260452.1;refGene_gn=MME;refGene_tx=XM_023260453.1;refGene_gn=MME;refGene_tx=XM_023260454.1;refGene_gn=MME;refGene_tx=XM_023260455.1
C2 111901247 111901404
refGene_gn=MME;refGene_tx=XM_006936315.4;refGene_gn=MME;refGene_tx=XM_019810542.2;refGene_gn=MME;refGene_tx=XM_023260452.1;refGene_gn=MME;refGene_tx=XM_023260453.1;refGene_gn=MME;refGene_tx=XM_023260454.1;refGene_gn=MME;refGene_tx=XM_023260455.1
C2 112385066 112386083
refGene_gn=GPR149;refGene_tx=XM_019810544.2
C2 112387204 112387397
refGene_gn=GPR149;refGene_tx=XM_019810544.2
C2 112392019 112392477
refGene_gn=GPR149;refGene_tx=XM_019810544.2
C2 112450152 112450728
refGene_gn=GPR149;refGene_tx=XM_019810544.2
C2 112461738 112461996
refGene_gn=DHX36;refGene_tx=XM_003991984.5
C2 112469463 112469588
refGene_gn=DHX36;refGene_tx=XM_003991984.5
C2 112470337 112470569
refGene_gn=DHX36;refGene_tx=XM_003991984.5
C2 112474379 112474418
refGene_gn=DHX36;refGene_tx=XM_003991984.5
C2 112475745 112475916
refGene_gn=DHX36;refGene_tx=XM_003991984.5
C2 112477576 112477656
refGene_gn=DHX36;refGene_tx=XM_003991984.5
C2 112478345 112478420
refGene_gn=DHX36;refGene_tx=XM_003991984.5
C2 112478570 112478737
refGene_gn=DHX36;refGene_tx=XM_003991984.5
C2 112480580 112480662
refGene_gn=DHX36;refGene_tx=XM_003991984.5
C2 112484458 112484599

refGene_gn=DHX36;refGene_tx=XM_003991984.5
C2 112484836 112484939
refGene_gn=DHX36;refGene_tx=XM_003991984.5
C2 112485543 112485631
refGene_gn=DHX36;refGene_tx=XM_003991984.5
C2 112488781 112488837
refGene_gn=DHX36;refGene_tx=XM_003991984.5
C2 112490527 112490592
refGene_gn=DHX36;refGene_tx=XM_003991984.5
C2 112491494 112491638
refGene_gn=DHX36;refGene_tx=XM_003991984.5
C2 112493527 112493645
refGene_gn=DHX36;refGene_tx=XM_003991984.5
C2 112494259 112494358
refGene_gn=DHX36;refGene_tx=XM_003991984.5
C2 112498509 112498683
refGene_gn=DHX36;refGene_tx=XM_003991984.5
C2 112498849 112498936
refGene_gn=DHX36;refGene_tx=XM_003991984.5
C2 112499982 112500066
refGene_gn=DHX36;refGene_tx=XM_003991984.5
C2 112502559 112502659
refGene_gn=DHX36;refGene_tx=XM_003991984.5
C2 112502751 112502842
refGene_gn=DHX36;refGene_tx=XM_003991984.5
C2 112507527 112507648
refGene_gn=DHX36;refGene_tx=XM_003991984.5
C2 112508528 112508681
refGene_gn=DHX36;refGene_tx=XM_003991984.5
C2 112509085 112509271
refGene_gn=DHX36;refGene_tx=XM_003991984.5
C2 112518764 112518907
refGene_gn=ARHGEF26;refGene_tx=XM_023260456.1;refGene_gn=ARHGEF26;refGene_tx=XM_023260457.1
C2 112519465 112519570
refGene_gn=ARHGEF26;refGene_tx=XM_023260456.1;refGene_gn=ARHGEF26;refGene_tx=XM_023260457.1
C2 112520873 112520941
refGene_gn=ARHGEF26;refGene_tx=XM_023260456.1;refGene_gn=ARHGEF26;refGene_tx=XM_023260457.1
C2 112531424 112531634
refGene_gn=ARHGEF26;refGene_tx=XM_023260456.1;refGene_gn=ARHGEF26;refGene_tx=XM_023260457.1
C2 112549541 112549696
refGene_gn=ARHGEF26;refGene_tx=XM_023260456.1;refGene_gn=ARHGEF26;refGene_tx=XM_023260457.1
C2 112558989 112559079
refGene_gn=ARHGEF26;refGene_tx=XM_023260456.1;refGene_gn=ARHGEF26;refGene_tx=XM_023260457.1
C2 112568464 112568539

refGene_gn=ARHGEF26;refGene_tx=XM_023260456.1;refGene_gn=ARHGEF26;refGene_tx=XM_023260457.1
C2	112572508	112572638
refGene_gn=ARHGEF26;refGene_tx=XM_023260456.1;refGene_gn=ARHGEF26;refGene_tx=XM_023260457.1
C2	112576218	112576371
refGene_gn=ARHGEF26;refGene_tx=XM_023260456.1
C2	112614539	112614700
refGene_gn=ARHGEF26;refGene_tx=XM_023260456.1;refGene_gn=ARHGEF26;refGene_tx=XM_023260457.1
C2	112618478	112618535
refGene_gn=ARHGEF26;refGene_tx=XM_023260456.1;refGene_gn=ARHGEF26;refGene_tx=XM_023260457.1
C2	112639856	112640002
refGene_gn=ARHGEF26;refGene_tx=XM_023260456.1;refGene_gn=ARHGEF26;refGene_tx=XM_023260457.1
C2	112643905	112643945
refGene_gn=ARHGEF26;refGene_tx=XM_023260456.1;refGene_gn=ARHGEF26;refGene_tx=XM_023260457.1
C2	112645307	112646387
refGene_gn=ARHGEF26;refGene_tx=XM_023260456.1;refGene_gn=ARHGEF26;refGene_tx=XM_023260457.1
C2	113247150	113247978
refGene_gn=LOC101095606;refGene_tx=XM_011286206.2
C2	113519049	113519601
refGene_gn=RAP2B;refGene_tx=XM_011286186.3
C2	113784318	113785443
refGene_gn=P2RY1;refGene_tx=XM_003991990.4;refGene_gn=P2RY1;refGene_tx=XM_023260458.1
C2	114119257	114119314
refGene_gn=MBNL1;refGene_tx=XM_006936319.4;refGene_gn=MBNL1;refGene_tx=XM_006936320.4;refGene_gn=MBNL1;refGene_tx=XM_006936322.4;refGene_gn=MBNL1;refGene_tx=XM_006936326.4;refGene_gn=MBNL1;refGene_tx=XM_019810547.2;refGene_gn=MBNL1;refGene_tx=XM_019810548.2;refGene_gn=MBNL1;refGene_tx=XM_019810549.2;refGene_gn=MBNL1;refGene_tx=XM_019810550.2;refGene_gn=MBNL1;refGene_tx=XM_019810551.2;refGene_gn=MBNL1;refGene_tx=XM_019810552.2;refGene_gn=MBNL1;refGene_tx=XM_019810554.2
C2	114122269	114122364
refGene_gn=MBNL1;refGene_tx=XM_006936319.4;refGene_gn=MBNL1;refGene_tx=XM_006936320.4;refGene_gn=MBNL1;refGene_tx=XM_006936322.4;refGene_gn=MBNL1;refGene_tx=XM_006936326.4;refGene_gn=MBNL1;refGene_tx=XM_019810547.2;refGene_gn=MBNL1;refGene_tx=XM_019810548.2;refGene_gn=MBNL1;refGene_tx=XM_019810549.2;refGene_gn=MBNL1;refGene_tx=XM_019810550.2;refGene_gn=MBNL1;refGene_tx=XM_019810552.2
C2	114123056	114123092
refGene_gn=MBNL1;refGene_tx=XM_006936319.4;refGene_gn=MBNL1;refGene_tx=XM_006936320.4;refGene_gn=MBNL1;refGene_tx=XM_006936322.4;refGene_gn=MBNL1;refGene_tx=XM_006936326.4;refGene_gn=MBNL1;refGene_tx=XM_019810547.2;refGene_gn=MBNL1;refGene_tx=XM_019810549.2;refGene_gn=MBNL1;refGene_tx=XM_019810551.2;refGene_gn=MBNL1;refGene_tx=XM_019810552.2

C2 114130696 114130850
refGene_gn=MBNL1;refGene_tx=XM_006936319.4;refGene_gn=MBNL1;refGene_tx=XM_006936320.4;refGene_gn=MBNL1;refGene_tx=XM_006936322.4;refGene_gn=MBNL1;refGene_tx=XM_006936326.4;refGene_gn=MBNL1;refGene_tx=XM_019810547.2;refGene_gn=MBNL1;refGene_tx=XM_019810548.2;refGene_gn=MBNL1;refGene_tx=XM_019810549.2;refGene_gn=MBNL1;refGene_tx=XM_019810550.2;refGene_gn=MBNL1;refGene_tx=XM_019810551.2;refGene_gn=MBNL1;refGene_tx=XM_019810552.2;refGene_gn=MBNL1;refGene_tx=XM_019810554.2
C2 114131762 114131816
refGene_gn=MBNL1;refGene_tx=XM_006936319.4;refGene_gn=MBNL1;refGene_tx=XM_006936320.4;refGene_gn=MBNL1;refGene_tx=XM_006936322.4;refGene_gn=MBNL1;refGene_tx=XM_006936326.4;refGene_gn=MBNL1;refGene_tx=XM_019810547.2;refGene_gn=MBNL1;refGene_tx=XM_019810548.2;refGene_gn=MBNL1;refGene_tx=XM_019810551.2;refGene_gn=MBNL1;refGene_tx=XM_019810552.2;refGene_gn=MBNL1;refGene_tx=XM_019810554.2
C2 114133006 114133264
refGene_gn=MBNL1;refGene_tx=XM_006936319.4;refGene_gn=MBNL1;refGene_tx=XM_006936320.4;refGene_gn=MBNL1;refGene_tx=XM_006936322.4;refGene_gn=MBNL1;refGene_tx=XM_006936326.4;refGene_gn=MBNL1;refGene_tx=XM_019810547.2;refGene_gn=MBNL1;refGene_tx=XM_019810548.2;refGene_gn=MBNL1;refGene_tx=XM_019810549.2;refGene_gn=MBNL1;refGene_tx=XM_019810550.2;refGene_gn=MBNL1;refGene_tx=XM_019810551.2;refGene_gn=MBNL1;refGene_tx=XM_019810552.2;refGene_gn=MBNL1;refGene_tx=XM_019810554.2
C2 114145564 114145765
refGene_gn=MBNL1;refGene_tx=XM_006936319.4;refGene_gn=MBNL1;refGene_tx=XM_006936320.4;refGene_gn=MBNL1;refGene_tx=XM_006936322.4;refGene_gn=MBNL1;refGene_tx=XM_006936326.4;refGene_gn=MBNL1;refGene_tx=XM_019810547.2;refGene_gn=MBNL1;refGene_tx=XM_019810548.2;refGene_gn=MBNL1;refGene_tx=XM_019810549.2;refGene_gn=MBNL1;refGene_tx=XM_019810550.2;refGene_gn=MBNL1;refGene_tx=XM_019810551.2;refGene_gn=MBNL1;refGene_tx=XM_019810552.2;refGene_gn=MBNL1;refGene_tx=XM_019810554.2
C2 114156025 114156196
refGene_gn=MBNL1;refGene_tx=XM_006936319.4;refGene_gn=MBNL1;refGene_tx=XM_006936320.4;refGene_gn=MBNL1;refGene_tx=XM_006936322.4;refGene_gn=MBNL1;refGene_tx=XM_006936326.4;refGene_gn=MBNL1;refGene_tx=XM_019810547.2;refGene_gn=MBNL1;refGene_tx=XM_019810548.2;refGene_gn=MBNL1;refGene_tx=XM_019810549.2;refGene_gn=MBNL1;refGene_tx=XM_019810550.2;refGene_gn=MBNL1;refGene_tx=XM_019810551.2;refGene_gn=MBNL1;refGene_tx=XM_019810552.2;refGene_gn=MBNL1;refGene_tx=XM_019810554.2
C2 114166135 114166231
refGene_gn=MBNL1;refGene_tx=XM_006936322.4
C2 114229691 114229697
refGene_gn=MBNL1;refGene_tx=XM_006936326.4
C2 114242451 114242481
refGene_gn=MBNL1;refGene_tx=XM_019810552.2
C2 114264229 114264403
refGene_gn=MBNL1;refGene_tx=XM_006936319.4;refGene_gn=MBNL1;refGene_tx=XM_006936320.4;refGene_gn=MBNL1;refGene_tx=XM_019810547.2;refGene_gn=MBNL1;refGene_tx=XM_019810548.2;refGene_gn=MBNL1;refGene_tx=XM_019810549.2;refGene_gn=MBNL1;refGene_tx=XM_019810550.2;refGene_gn=MBNL1;refGe

ne_tx=XM_019810551.2;refGene_gn=MBNL1;refGene_tx=XM_019810554.2
C2 114665948 114666902
refGene_gn=SUCNR1;refGene_tx=XM_019810555.2
C2 114672017 114672125
refGene_gn=SUCNR1;refGene_tx=XM_019810555.2
C2 114758265 114758427
refGene_gn=AADAC;refGene_tx=XM_003992010.3
C2 114758428 114758863
refGene_gn=AADAC;refGene_tx=XM_003992010.3
C2 114766581 114766721
refGene_gn=AADAC;refGene_tx=XM_003992010.3
C2 114766723 114766752
refGene_gn=AADAC;refGene_tx=XM_003992010.3
C2 114767847 114767917
refGene_gn=AADAC;refGene_tx=XM_003992010.3
C2 114770183 114770226
refGene_gn=AADAC;refGene_tx=XM_003992010.3
C2 114770227 114770407
refGene_gn=AADAC;refGene_tx=XM_003992010.3
C2 114776828 114776978
refGene_gn=AADAC;refGene_tx=XM_003992010.3
C2 114781706 114782119
refGene_gn=LOC101096595;refGene_tx=XM_023238569.1
C2 114782121 114782170
refGene_gn=LOC101096595;refGene_tx=XM_023238569.1
C2 114798518 114799121
refGene_gn=LOC101091891;refGene_tx=XM_023260460.1
C2 114812891 114812989
refGene_gn=LOC101091891;refGene_tx=XM_023260460.1
C2 114812991 114813062
refGene_gn=LOC101091891;refGene_tx=XM_023260460.1
C2 114815851 114815921
refGene_gn=LOC101091891;refGene_tx=XM_023260460.1
C2 114829935 114830158
refGene_gn=LOC101091891;refGene_tx=XM_023260460.1
C2 114834302 114834440
refGene_gn=LOC101091891;refGene_tx=XM_023260460.1
C2 114851619 114852222
refGene_gn=LOC101092136;refGene_tx=XM_003991994.3
C2 114854605 114854777
refGene_gn=LOC101092136;refGene_tx=XM_003991994.3
C2 114855698 114855768
refGene_gn=LOC101092136;refGene_tx=XM_003991994.3
C2 114858015 114858238
refGene_gn=LOC101092136;refGene_tx=XM_003991994.3
C2 114870843 114870981
refGene_gn=LOC101092136;refGene_tx=XM_003991994.3
C2 115140717 115140911
refGene_gn=IGSF10;refGene_tx=XM_003991995.5
C2 115150790 115150920

refGene_gn=IGSF10;refGene_tx=XM_003991995.5
C2	115196280	115196671
refGene_gn=IGSF10;refGene_tx=XM_003991995.5
C2	115207597	115211950
refGene_gn=IGSF10;refGene_tx=XM_003991995.5;refGene_gn=IGSF10;refGene_tx=XM_019810557.1
C2	115212449	115213350
refGene_gn=IGSF10;refGene_tx=XM_003991995.5;refGene_gn=IGSF10;refGene_tx=XM_019810557.1
C2	115216453	115218362
refGene_gn=IGSF10;refGene_tx=XM_003991995.5;refGene_gn=IGSF10;refGene_tx=XM_019810557.1
C2	115222183	115222236
refGene_gn=MED12L;refGene_tx=XM_019810565.1
C2	115224784	115224866
refGene_gn=MED12L;refGene_tx=XM_019810565.1
C2	115236270	115236357
refGene_gn=MED12L;refGene_tx=XM_019810565.1
C2	115239435	115239592
refGene_gn=MED12L;refGene_tx=XM_019810565.1
C2	115241780	115242010
refGene_gn=MED12L;refGene_tx=XM_019810565.1
C2	115243673	115243763
refGene_gn=MED12L;refGene_tx=XM_019810565.1
C2	115257293	115257505
refGene_gn=MED12L;refGene_tx=XM_019810565.1
C2	115261331	115261488
refGene_gn=MED12L;refGene_tx=XM_019810565.1
C2	115264356	115264719
refGene_gn=MED12L;refGene_tx=XM_019810565.1
C2	115267195	115267357
refGene_gn=MED12L;refGene_tx=XM_019810565.1
C2	115268163	115268299
refGene_gn=MED12L;refGene_tx=XM_019810565.1
C2	115268492	115268602
refGene_gn=MED12L;refGene_tx=XM_019810565.1
C2	115269652	115269742
refGene_gn=MED12L;refGene_tx=XM_019810565.1
C2	115271881	115271993
refGene_gn=MED12L;refGene_tx=XM_019810565.1
C2	115274304	115274466
refGene_gn=MED12L;refGene_tx=XM_019810565.1
C2	115275082	115275273
refGene_gn=MED12L;refGene_tx=XM_019810565.1
C2	115275386	115275461
refGene_gn=MED12L;refGene_tx=XM_019810565.1
C2	115276279	115276468
refGene_gn=MED12L;refGene_tx=XM_019810565.1
C2	115279733	115279933
refGene_gn=MED12L;refGene_tx=XM_019810565.1

C2	115283363	115283477
refGene_gn=MED12L;refGene_tx=XM_019810565.1
C2	115284176	115284278
refGene_gn=MED12L;refGene_tx=XM_019810565.1
C2	115284629	115284750
refGene_gn=MED12L;refGene_tx=XM_019810565.1
C2	115286372	115286514
refGene_gn=MED12L;refGene_tx=XM_019810565.1
C2	115287156	115287384
refGene_gn=MED12L;refGene_tx=XM_019810565.1
C2	115290625	115290757
refGene_gn=MED12L;refGene_tx=XM_019810565.1
C2	115293742	115293906
refGene_gn=MED12L;refGene_tx=XM_019810565.1
C2	115294976	115295120
refGene_gn=MED12L;refGene_tx=XM_019810565.1
C2	115296195	115296314
refGene_gn=MED12L;refGene_tx=XM_019810565.1
C2	115300541	115300689
refGene_gn=MED12L;refGene_tx=XM_019810565.1
C2	115312114	115313134
refGene_gn=P2RY12;refGene_tx=XM_003992001.5;refGene_gn=P2RY12;refGene_tx=XM_011286173.2;refGene_gn=P2RY12;refGene_tx=XM_011286175.2;refGene_gn=P2RY12;refGene_tx=XM_019810567.1;refGene_gn=P2RY12;refGene_tx=XM_019810568.1
C2	115320471	115320591
refGene_gn=P2RY13;refGene_tx=XM_006936330.2
C2	115321034	115322051
refGene_gn=P2RY13;refGene_tx=XM_006936330.2
C2	115350586	115350620
refGene_gn=GPR87;refGene_tx=XM_019810566.1
C2	115355089	115356132
refGene_gn=GPR87;refGene_tx=XM_019810566.1
C2	115426834	115427857
refGene_gn=P2RY14;refGene_tx=XM_006936329.4;refGene_gn=P2RY14;refGene_tx=XM_011286176.3;refGene_gn=P2RY14;refGene_tx=XM_011286178.3;refGene_gn=P2RY14;refGene_tx=XM_023260461.1
C2	115446040	115447000
refGene_gn=GPR171;refGene_tx=XM_019810569.2
C2	115451255	115451432
refGene_gn=MED12L;refGene_tx=XM_019810565.1
C2	115453972	115454187
refGene_gn=MED12L;refGene_tx=XM_019810565.1
C2	115456439	115456566
refGene_gn=MED12L;refGene_tx=XM_019810565.1
C2	115460228	115460360
refGene_gn=MED12L;refGene_tx=XM_019810565.1
C2	115477888	115478025
refGene_gn=MED12L;refGene_tx=XM_019810565.1
C2	115478342	115478442

refGene_gn=MED12L;refGene_tx=XM_019810565.1
C2 115479203 115479353
refGene_gn=MED12L;refGene_tx=XM_019810565.1
C2 115483254 115483524
refGene_gn=MED12L;refGene_tx=XM_019810565.1
C2 115484751 115484862
refGene_gn=MED12L;refGene_tx=XM_019810565.1
C2 115487246 115487312
refGene_gn=MED12L;refGene_tx=XM_019810565.1
C2 115487313 115487417
refGene_gn=MED12L;refGene_tx=XM_019810565.1
C2 115511361 115511521
refGene_gn=MED12L;refGene_tx=XM_019810565.1
C2 115516716 115516908
refGene_gn=MED12L;refGene_tx=XM_019810565.1
C2 115525761 115525866
refGene_gn=MED12L;refGene_tx=XM_019810565.1
C2 115556454 115556553
refGene_gn=MED12L;refGene_tx=XM_019810565.1
C2 115681476 115681729
refGene_gn=CLRN1;refGene_tx=XM_003992002.3
C2 115707386 115707566
refGene_gn=CLRN1;refGene_tx=XM_003992002.3
C2 115720604 115720870
refGene_gn=CLRN1;refGene_tx=XM_003992002.3
C2 115746768 115746875
refGene_gn=MINDY4B;refGene_tx=XM_023260182.1
C2 115747110 115747138
refGene_gn=MINDY4B;refGene_tx=XM_023260182.1
C2 115748793 115748961
refGene_gn=MINDY4B;refGene_tx=XM_023260182.1
C2 115749847 115749970
refGene_gn=MINDY4B;refGene_tx=XM_023260182.1
C2 115756874 115756994
refGene_gn=MINDY4B;refGene_tx=XM_023260182.1
C2 115757731 115757823
refGene_gn=MINDY4B;refGene_tx=XM_023260182.1
C2 115760621 115760787
refGene_gn=MINDY4B;refGene_tx=XM_023260182.1
C2 115761326 115761392
refGene_gn=MINDY4B;refGene_tx=XM_023260182.1
C2 115765743 115765814
refGene_gn=MINDY4B;refGene_tx=XM_023260182.1
C2 115768144 115768217
refGene_gn=MINDY4B;refGene_tx=XM_023260182.1
C2 115768812 115768974
refGene_gn=MINDY4B;refGene_tx=XM_023260182.1
C2 115780167 115780348
refGene_gn=MINDY4B;refGene_tx=XM_023260182.1
C2 115782973 115783116

refGene_gn=MINDY4B;refGene_tx=XM_023260182.1
C2 115859283 115859700
refGene_gn=SIAH2;refGene_tx=XM_019811114.2
C2 115877982 115878540
refGene_gn=SIAH2;refGene_tx=XM_019811114.2
C2 115979301 115979426
refGene_gn=SELENOT;refGene_tx=XM_003992003.5
C2 115981208 115981296
refGene_gn=SELENOT;refGene_tx=XM_003992003.5
C2 115983576 115983703
refGene_gn=SELENOT;refGene_tx=XM_003992003.5
C2 115984279 115984390
refGene_gn=SELENOT;refGene_tx=XM_003992003.5
C2 116001180 116001317
refGene_gn=SELENOT;refGene_tx=XM_003992003.5
C2 116015670 116015736
refGene_gn=EIF2A;refGene_tx=XM_003992004.5;refGene_gn=EIF2A;refGene_tx
=XM_019810570.2;refGene_gn=EIF2A;refGene_tx=XM_019810571.2
C2 116016337 116016403
refGene_gn=EIF2A;refGene_tx=XM_003992004.5;refGene_gn=EIF2A;refGene_tx
=XM_019810570.2;refGene_gn=EIF2A;refGene_tx=XM_019810571.2
C2 116017195 116017324
refGene_gn=EIF2A;refGene_tx=XM_003992004.5;refGene_gn=EIF2A;refGene_tx
=XM_019810570.2;refGene_gn=EIF2A;refGene_tx=XM_019810571.2
C2 116018587 116018701
refGene_gn=EIF2A;refGene_tx=XM_003992004.5;refGene_gn=EIF2A;refGene_tx
=XM_019810570.2;refGene_gn=EIF2A;refGene_tx=XM_019810571.2
C2 116022071 116022643
refGene_gn=EIF2A;refGene_tx=XM_003992004.5;refGene_gn=EIF2A;refGene_tx
=XM_019810570.2;refGene_gn=EIF2A;refGene_tx=XM_019810571.2
C2 116025008 116025125
refGene_gn=EIF2A;refGene_tx=XM_003992004.5;refGene_gn=EIF2A;refGene_tx
=XM_019810570.2;refGene_gn=EIF2A;refGene_tx=XM_019810571.2
C2 116025318 116025463
refGene_gn=EIF2A;refGene_tx=XM_003992004.5;refGene_gn=EIF2A;refGene_tx
=XM_019810570.2;refGene_gn=EIF2A;refGene_tx=XM_019810571.2
C2 116025599 116025673
refGene_gn=EIF2A;refGene_tx=XM_003992004.5;refGene_gn=EIF2A;refGene_tx
=XM_019810570.2;refGene_gn=EIF2A;refGene_tx=XM_019810571.2
C2 116027250 116027333
refGene_gn=EIF2A;refGene_tx=XM_003992004.5;refGene_gn=EIF2A;refGene_tx
=XM_019810570.2;refGene_gn=EIF2A;refGene_tx=XM_019810571.2
C2 116027691 116027791
refGene_gn=EIF2A;refGene_tx=XM_003992004.5;refGene_gn=EIF2A;refGene_tx
=XM_019810570.2;refGene_gn=EIF2A;refGene_tx=XM_019810571.2
C2 116028789 116028908
refGene_gn=EIF2A;refGene_tx=XM_003992004.5;refGene_gn=EIF2A;refGene_tx
=XM_019810570.2
C2 116032027 116032102
refGene_gn=EIF2A;refGene_tx=XM_003992004.5;refGene_gn=EIF2A;refGene_tx

=XM_019810570.2
C2	116034880	116034906
refGene_gn=EIF2A;refGene_tx=XM_019810570.2
C2	116035877	116035947
refGene_gn=EIF2A;refGene_tx=XM_003992004.5
C2	116045684	116045736
refGene_gn=EIF2A;refGene_tx=XM_003992004.5
C2	116046412	116046496
refGene_gn=SERP1;refGene_tx=XM_003992007.5
C2	116046778	116046854
refGene_gn=SERP1;refGene_tx=XM_003992007.5
C2	116048038	116048079
refGene_gn=SERP1;refGene_tx=XM_003992007.5
C2	116119488	116119749
refGene_gn=TSC22D2;refGene_tx=XM_019810572.2;refGene_gn=TSC22D2;refGen
e_tx=XM_019810573.2
C2	116120742	116120794
refGene_gn=TSC22D2;refGene_tx=XM_019810572.2;refGene_gn=TSC22D2;refGen
e_tx=XM_019810573.2
C2	116152076	116152148
refGene_gn=TSC22D2;refGene_tx=XM_019810572.2
C2	116162183	116162187
refGene_gn=TSC22D2;refGene_tx=XM_023260462.1
C2	116163490	116165469
refGene_gn=TSC22D2;refGene_tx=XM_019810572.2;refGene_gn=TSC22D2;refGen
e_tx=XM_019810573.2;refGene_gn=TSC22D2;refGene_tx=XM_023260462.1
C2	116582554	116582686
refGene_gn=PFN2;refGene_tx=XM_023260463.1;refGene_gn=PFN2;refGene_tx=X
M_023260464.1
C2	116584891	116585084
refGene_gn=PFN2;refGene_tx=XM_023260463.1;refGene_gn=PFN2;refGene_tx=X
M_023260464.1
C2	116586859	116586957
refGene_gn=PFN2;refGene_tx=XM_023260463.1
C2	116587182	116587280
refGene_gn=PFN2;refGene_tx=XM_023260464.1
C2	116592363	116592728
refGene_gn=RNF13;refGene_tx=XM_003992014.5;refGene_gn=RNF13;refGene_tx
=XM_006936334.4;refGene_gn=RNF13;refGene_tx=XM_019810574.1;refGene_gn=
RNF13;refGene_tx=XM_019810575.2
C2	116593017	116593098
refGene_gn=RNF13;refGene_tx=XM_003992014.5;refGene_gn=RNF13;refGene_tx
=XM_006936334.4;refGene_gn=RNF13;refGene_tx=XM_019810574.1;refGene_gn=
RNF13;refGene_tx=XM_019810575.2
C2	116629229	116629323
refGene_gn=RNF13;refGene_tx=XM_003992014.5;refGene_gn=RNF13;refGene_tx
=XM_006936334.4;refGene_gn=RNF13;refGene_tx=XM_019810574.1;refGene_gn=
RNF13;refGene_tx=XM_019810575.2
C2	116638209	116638315
refGene_gn=RNF13;refGene_tx=XM_003992014.5;refGene_gn=RNF13;refGene_tx

=XM_006936334.4;refGene_gn=RNF13;refGene_tx=XM_019810574.1;refGene_gn=RNF13;refGene_tx=XM_019810575.2
C2	116647994	116648085
refGene_gn=RNF13;refGene_tx=XM_003992014.5;refGene_gn=RNF13;refGene_tx=XM_006936334.4;refGene_gn=RNF13;refGene_tx=XM_019810574.1;refGene_gn=RNF13;refGene_tx=XM_019810575.2
C2	116655158	116655246
refGene_gn=RNF13;refGene_tx=XM_003992014.5;refGene_gn=RNF13;refGene_tx=XM_006936334.4;refGene_gn=RNF13;refGene_tx=XM_019810574.1;refGene_gn=RNF13;refGene_tx=XM_019810575.2
C2	116683605	116683731
refGene_gn=RNF13;refGene_tx=XM_003992014.5;refGene_gn=RNF13;refGene_tx=XM_006936334.4;refGene_gn=RNF13;refGene_tx=XM_019810574.1
C2	116704942	116705023
refGene_gn=RNF13;refGene_tx=XM_003992014.5;refGene_gn=RNF13;refGene_tx=XM_006936334.4;refGene_gn=RNF13;refGene_tx=XM_019810574.1;refGene_gn=RNF13;refGene_tx=XM_019810575.2
C2	116717402	116717516
refGene_gn=RNF13;refGene_tx=XM_003992014.5;refGene_gn=RNF13;refGene_tx=XM_006936334.4;refGene_gn=RNF13;refGene_tx=XM_019810574.1;refGene_gn=RNF13;refGene_tx=XM_019810575.2
C2	116765816	116765906
refGene_gn=ANKUB1;refGene_tx=XM_023260466.1
C2	116767744	116767888
refGene_gn=ANKUB1;refGene_tx=XM_023260466.1
C2	116777444	116777661
refGene_gn=ANKUB1;refGene_tx=XM_023260466.1
C2	116787080	116787195
refGene_gn=ANKUB1;refGene_tx=XM_023260466.1
C2	116791020	116791959
refGene_gn=ANKUB1;refGene_tx=XM_023260466.1
C2	116795778	116795908
refGene_gn=ANKUB1;refGene_tx=XM_023260466.1
C2	116803029	116803096
refGene_gn=COMMD2;refGene_tx=XM_003992016.5
C2	116803192	116803270
refGene_gn=COMMD2;refGene_tx=XM_003992016.5
C2	116803964	116804047
refGene_gn=COMMD2;refGene_tx=XM_003992016.5
C2	116804850	116805024
refGene_gn=COMMD2;refGene_tx=XM_003992016.5
C2	116809876	116810074
refGene_gn=COMMD2;refGene_tx=XM_003992016.5
C2	116876354	116876785
refGene_gn=WWTR1;refGene_tx=XM_011286170.3
C2	116956892	116957029
refGene_gn=WWTR1;refGene_tx=XM_011286170.3
C2	116986263	116986466
refGene_gn=WWTR1;refGene_tx=XM_011286170.3
C2	117000511	117000645

refGene_gn=WWTR1;refGene_tx=XM_011286170.3
C2 117001844 117001957
refGene_gn=WWTR1;refGene_tx=XM_011286170.3
C2 117008554 117008739
refGene_gn=WWTR1;refGene_tx=XM_011286170.3
C2 117031885 117031903
refGene_gn=TM4SF4;refGene_tx=XM_003992019.5
C2 117036164 117036354
refGene_gn=TM4SF4;refGene_tx=XM_003992019.5
C2 117047925 117048062
refGene_gn=TM4SF4;refGene_tx=XM_003992019.5
C2 117060441 117060531
refGene_gn=TM4SF4;refGene_tx=XM_003992019.5
C2 117061522 117061696
refGene_gn=TM4SF4;refGene_tx=XM_003992019.5
C2 117145023 117145200
refGene_gn=TM4SF1;refGene_tx=XM_003992020.4;refGene_gn=TM4SF1;refGene_tx=XM_003992021.4;refGene_gn=TM4SF1;refGene_tx=XM_011286168.3
C2 117146629 117146719
refGene_gn=TM4SF1;refGene_tx=XM_003992020.4;refGene_gn=TM4SF1;refGene_tx=XM_003992021.4;refGene_gn=TM4SF1;refGene_tx=XM_011286168.3
C2 117146807 117146953
refGene_gn=TM4SF1;refGene_tx=XM_003992020.4;refGene_gn=TM4SF1;refGene_tx=XM_003992021.4;refGene_gn=TM4SF1;refGene_tx=XM_011286168.3
C2 117149716 117149914
refGene_gn=TM4SF1;refGene_tx=XM_003992020.4;refGene_gn=TM4SF1;refGene_tx=XM_003992021.4;refGene_gn=TM4SF1;refGene_tx=XM_011286168.3
C2 117151796 117151902
refGene_gn=TM4SF1;refGene_tx=XM_003992020.4;refGene_gn=TM4SF1;refGene_tx=XM_003992021.4;refGene_gn=TM4SF1;refGene_tx=XM_011286168.3
C2 117207010 117207187
refGene_gn=TM4SF18;refGene_tx=XM_006936336.4;refGene_gn=TM4SF18;refGene_tx=XM_006936337.4;refGene_gn=TM4SF18;refGene_tx=XM_006936338.4;refGene_gn=TM4SF18;refGene_tx=XM_023260467.1
C2 117210221 117210311
refGene_gn=TM4SF18;refGene_tx=XM_006936336.4;refGene_gn=TM4SF18;refGene_tx=XM_006936337.4;refGene_gn=TM4SF18;refGene_tx=XM_023260467.1
C2 117225766 117225909
refGene_gn=TM4SF18;refGene_tx=XM_006936336.4;refGene_gn=TM4SF18;refGene_tx=XM_006936337.4;refGene_gn=TM4SF18;refGene_tx=XM_006936338.4;refGene_gn=TM4SF18;refGene_tx=XM_023260467.1
C2 117229945 117230141
refGene_gn=TM4SF18;refGene_tx=XM_006936336.4;refGene_gn=TM4SF18;refGene_tx=XM_006936337.4;refGene_gn=TM4SF18;refGene_tx=XM_006936338.4;refGene_gn=TM4SF18;refGene_tx=XM_023260467.1
C2 117230881 117230896
refGene_gn=TM4SF18;refGene_tx=XM_006936336.4;refGene_gn=TM4SF18;refGene_tx=XM_006936338.4
C2 117232653 117232665
refGene_gn=TM4SF18;refGene_tx=XM_006936337.4

C2 117251316 117251471
refGene_gn=LOC101080399;refGene_tx=XM_019810577.2
C2 117252588 117252839
refGene_gn=LOC101080399;refGene_tx=XM_019810577.2
C2 117254260 117254473
refGene_gn=LOC101080399;refGene_tx=XM_019810577.2
C2 117254976 117255147
refGene_gn=LOC101080399;refGene_tx=XM_019810577.2
C2 117255970 117256213
refGene_gn=LOC101080399;refGene_tx=XM_019810577.2
C2 117259381 117259547
refGene_gn=LOC101080399;refGene_tx=XM_019810577.2
C2 117261576 117261707
refGene_gn=LOC101080399;refGene_tx=XM_019810577.2
C2 117262060 117262201
refGene_gn=LOC101080399;refGene_tx=XM_019810577.2
C2 117262751 117262963
refGene_gn=LOC101080399;refGene_tx=XM_019810577.2
C2 117264647 117264798
refGene_gn=LOC101080399;refGene_tx=XM_019810577.2
C2 117265428 117265641
refGene_gn=LOC101080399;refGene_tx=XM_019810577.2
C2 117266765 117266973
refGene_gn=LOC101080399;refGene_tx=XM_019810577.2
C2 117267788 117267922
refGene_gn=LOC101080399;refGene_tx=XM_019810577.2
C2 117269535 117269664
refGene_gn=LOC101080399;refGene_tx=XM_019810577.2
C2 117271513 117271620
refGene_gn=LOC101080399;refGene_tx=XM_019810577.2
C2 117272570 117272772
refGene_gn=LOC101080399;refGene_tx=XM_019810577.2
C2 117275300 117275440
refGene_gn=LOC101080399;refGene_tx=XM_019810577.2
C2 117280412 117280571
refGene_gn=LOC101080399;refGene_tx=XM_019810577.2
C2 117295232 117295378
refGene_gn=LOC101081158;refGene_tx=XM_011286166.3;refGene_gn=LOC101081158;refGene_tx=XM_019810579.2;refGene_gn=LOC101081158;refGene_tx=XM_019810580.2;refGene_gn=LOC101081158;refGene_tx=XM_023260469.1
C2 117302948 117303196
refGene_gn=LOC101081158;refGene_tx=XM_011286166.3;refGene_gn=LOC101081158;refGene_tx=XM_019810579.2;refGene_gn=LOC101081158;refGene_tx=XM_019810580.2;refGene_gn=LOC101081158;refGene_tx=XM_023260469.1
C2 117304929 117305142
refGene_gn=LOC101081158;refGene_tx=XM_011286166.3;refGene_gn=LOC101081158;refGene_tx=XM_019810579.2;refGene_gn=LOC101081158;refGene_tx=XM_019810580.2;refGene_gn=LOC101081158;refGene_tx=XM_023260469.1
C2 117306022 117306196
refGene_gn=LOC101081158;refGene_tx=XM_011286166.3;refGene_gn=LOC101081

158;refGene_tx=XM_019810579.2;refGene_gn=LOC101081158;refGene_tx=XM_019810580.2;refGene_gn=LOC101081158;refGene_tx=XM_023260469.1
C2 117314544 117314799
refGene_gn=LOC101081158;refGene_tx=XM_011286166.3;refGene_gn=LOC101081158;refGene_tx=XM_019810579.2;refGene_gn=LOC101081158;refGene_tx=XM_019810580.2;refGene_gn=LOC101081158;refGene_tx=XM_023260469.1
C2 117315899 117316071
refGene_gn=LOC101081158;refGene_tx=XM_011286166.3;refGene_gn=LOC101081158;refGene_tx=XM_019810579.2;refGene_gn=LOC101081158;refGene_tx=XM_019810580.2;refGene_gn=LOC101081158;refGene_tx=XM_023260469.1
C2 117319494 117319634
refGene_gn=LOC101081158;refGene_tx=XM_011286166.3;refGene_gn=LOC101081158;refGene_tx=XM_019810579.2;refGene_gn=LOC101081158;refGene_tx=XM_019810580.2;refGene_gn=LOC101081158;refGene_tx=XM_023260469.1
C2 117320789 117320942
refGene_gn=LOC101081158;refGene_tx=XM_011286166.3;refGene_gn=LOC101081158;refGene_tx=XM_019810579.2;refGene_gn=LOC101081158;refGene_tx=XM_019810580.2;refGene_gn=LOC101081158;refGene_tx=XM_023260469.1
C2 117322437 117322649
refGene_gn=LOC101081158;refGene_tx=XM_011286166.3;refGene_gn=LOC101081158;refGene_tx=XM_019810579.2;refGene_gn=LOC101081158;refGene_tx=XM_019810580.2;refGene_gn=LOC101081158;refGene_tx=XM_023260469.1
C2 117335976 117336127
refGene_gn=LOC101081158;refGene_tx=XM_011286166.3;refGene_gn=LOC101081158;refGene_tx=XM_019810579.2;refGene_gn=LOC101081158;refGene_tx=XM_019810580.2;refGene_gn=LOC101081158;refGene_tx=XM_023260469.1
C2 117337426 117337639
refGene_gn=LOC101081158;refGene_tx=XM_011286166.3;refGene_gn=LOC101081158;refGene_tx=XM_019810579.2;refGene_gn=LOC101081158;refGene_tx=XM_019810580.2;refGene_gn=LOC101081158;refGene_tx=XM_023260469.1
C2 117338817 117339025
refGene_gn=LOC101081158;refGene_tx=XM_011286166.3;refGene_gn=LOC101081158;refGene_tx=XM_019810579.2;refGene_gn=LOC101081158;refGene_tx=XM_019810580.2;refGene_gn=LOC101081158;refGene_tx=XM_023260469.1
C2 117343085 117343225
refGene_gn=LOC101081158;refGene_tx=XM_011286166.3;refGene_gn=LOC101081158;refGene_tx=XM_019810579.2;refGene_gn=LOC101081158;refGene_tx=XM_019810580.2;refGene_gn=LOC101081158;refGene_tx=XM_023260469.1
C2 117344533 117344662
refGene_gn=LOC101081158;refGene_tx=XM_011286166.3;refGene_gn=LOC101081158;refGene_tx=XM_019810579.2;refGene_gn=LOC101081158;refGene_tx=XM_019810580.2;refGene_gn=LOC101081158;refGene_tx=XM_023260469.1
C2 117347307 117347414
refGene_gn=LOC101081158;refGene_tx=XM_011286166.3;refGene_gn=LOC101081158;refGene_tx=XM_019810579.2;refGene_gn=LOC101081158;refGene_tx=XM_019810580.2;refGene_gn=LOC101081158;refGene_tx=XM_023260469.1
C2 117348897 117349114
refGene_gn=LOC101081158;refGene_tx=XM_011286166.3;refGene_gn=LOC101081158;refGene_tx=XM_019810579.2;refGene_gn=LOC101081158;refGene_tx=XM_019810580.2;refGene_gn=LOC101081158;refGene_tx=XM_023260469.1

C2 117350004 117350144
refGene_gn=LOC101081158;refGene_tx=XM_011286166.3;refGene_gn=LOC101081
158;refGene_tx=XM_019810579.2;refGene_gn=LOC101081158;refGene_tx=XM_01
9810580.2;refGene_gn=LOC101081158;refGene_tx=XM_023260469.1
C2 117351286 117351449
refGene_gn=LOC101081158;refGene_tx=XM_011286166.3;refGene_gn=LOC101081
158;refGene_tx=XM_019810579.2;refGene_gn=LOC101081158;refGene_tx=XM_01
9810580.2
C2 117351848 117351916
refGene_gn=LOC101081158;refGene_tx=XM_019810579.2
C2 117355107 117355124
refGene_gn=LOC101081158;refGene_tx=XM_019810580.2
C2 117356852 117356980
refGene_gn=HPS3;refGene_tx=XM_023260470.1;refGene_gn=HPS3;refGene_tx=X
M_023260471.1;refGene_gn=HPS3;refGene_tx=XM_023260472.1;refGene_gn=HPS
3;refGene_tx=XM_023260473.1
C2 117360662 117360753
refGene_gn=HPS3;refGene_tx=XM_023260470.1;refGene_gn=HPS3;refGene_tx=X
M_023260471.1;refGene_gn=HPS3;refGene_tx=XM_023260472.1;refGene_gn=HPS
3;refGene_tx=XM_023260473.1
C2 117361631 117361838
refGene_gn=HPS3;refGene_tx=XM_023260470.1;refGene_gn=HPS3;refGene_tx=X
M_023260471.1;refGene_gn=HPS3;refGene_tx=XM_023260473.1
C2 117362839 117362943
refGene_gn=LOC101081158;refGene_tx=XM_011286166.3
C2 117365376 117365484
refGene_gn=HPS3;refGene_tx=XM_023260470.1;refGene_gn=HPS3;refGene_tx=X
M_023260472.1;refGene_gn=HPS3;refGene_tx=XM_023260473.1
C2 117366476 117366665
refGene_gn=HPS3;refGene_tx=XM_023260470.1;refGene_gn=HPS3;refGene_tx=X
M_023260471.1;refGene_gn=HPS3;refGene_tx=XM_023260472.1;refGene_gn=HPS
3;refGene_tx=XM_023260473.1
C2 117367012 117367198
refGene_gn=HPS3;refGene_tx=XM_023260470.1;refGene_gn=HPS3;refGene_tx=X
M_023260471.1;refGene_gn=HPS3;refGene_tx=XM_023260472.1;refGene_gn=HPS
3;refGene_tx=XM_023260473.1
C2 117367667 117367730
refGene_gn=LOC101081158;refGene_tx=XM_023260469.1
C2 117368246 117368480
refGene_gn=HPS3;refGene_tx=XM_023260470.1;refGene_gn=HPS3;refGene_tx=X
M_023260471.1;refGene_gn=HPS3;refGene_tx=XM_023260472.1;refGene_gn=HPS
3;refGene_tx=XM_023260473.1
C2 117369872 117370053
refGene_gn=HPS3;refGene_tx=XM_023260470.1;refGene_gn=HPS3;refGene_tx=X
M_023260471.1;refGene_gn=HPS3;refGene_tx=XM_023260472.1;refGene_gn=HPS
3;refGene_tx=XM_023260473.1
C2 117371153 117371335
refGene_gn=HPS3;refGene_tx=XM_023260470.1;refGene_gn=HPS3;refGene_tx=X
M_023260471.1;refGene_gn=HPS3;refGene_tx=XM_023260472.1;refGene_gn=HPS
3;refGene_tx=XM_023260473.1

C2 117373136 117373245
refGene_gn=HPS3;refGene_tx=XM_023260470.1;refGene_gn=HPS3;refGene_tx=XM_023260471.1;refGene_gn=HPS3;refGene_tx=XM_023260472.1;refGene_gn=HPS3;refGene_tx=XM_023260473.1
C2 117375339 117375494
refGene_gn=HPS3;refGene_tx=XM_023260470.1;refGene_gn=HPS3;refGene_tx=XM_023260471.1;refGene_gn=HPS3;refGene_tx=XM_023260472.1;refGene_gn=HPS3;refGene_tx=XM_023260473.1
C2 117378396 117378478
refGene_gn=HPS3;refGene_tx=XM_023260470.1;refGene_gn=HPS3;refGene_tx=XM_023260471.1;refGene_gn=HPS3;refGene_tx=XM_023260472.1;refGene_gn=HPS3;refGene_tx=XM_023260473.1
C2 117382889 117383082
refGene_gn=HPS3;refGene_tx=XM_023260470.1;refGene_gn=HPS3;refGene_tx=XM_023260471.1;refGene_gn=HPS3;refGene_tx=XM_023260472.1;refGene_gn=HPS3;refGene_tx=XM_023260473.1
C2 117385291 117385377
refGene_gn=HPS3;refGene_tx=XM_023260470.1;refGene_gn=HPS3;refGene_tx=XM_023260471.1;refGene_gn=HPS3;refGene_tx=XM_023260472.1;refGene_gn=HPS3;refGene_tx=XM_023260473.1
C2 117385482 117385654
refGene_gn=HPS3;refGene_tx=XM_023260470.1;refGene_gn=HPS3;refGene_tx=XM_023260471.1;refGene_gn=HPS3;refGene_tx=XM_023260472.1;refGene_gn=HPS3;refGene_tx=XM_023260473.1
C2 117386434 117386926
refGene_gn=HPS3;refGene_tx=XM_023260470.1;refGene_gn=HPS3;refGene_tx=XM_023260471.1;refGene_gn=HPS3;refGene_tx=XM_023260472.1;refGene_gn=HPS3;refGene_tx=XM_023260473.1
C2 117395309 117395526
refGene_gn=HPS3;refGene_tx=XM_023260470.1;refGene_gn=HPS3;refGene_tx=XM_023260471.1;refGene_gn=HPS3;refGene_tx=XM_023260472.1
C2 117402228 117403035
refGene_gn=LOC105259604;refGene_tx=XM_006936340.4
C2 117417951 117417971
refGene_gn=HLTF;refGene_tx=XM_019810583.2
C2 117419092 117419300
refGene_gn=HLTF;refGene_tx=XM_019810583.2
C2 117426720 117426887
refGene_gn=HLTF;refGene_tx=XM_019810583.2
C2 117428389 117428523
refGene_gn=HLTF;refGene_tx=XM_019810583.2
C2 117429295 117429393
refGene_gn=HLTF;refGene_tx=XM_019810583.2
C2 117431020 117431095
refGene_gn=HLTF;refGene_tx=XM_019810583.2
C2 117431254 117431303
refGene_gn=HLTF;refGene_tx=XM_019810583.2
C2 117431305 117431445
refGene_gn=HLTF;refGene_tx=XM_019810583.2
C2 117434074 117434170

refGene_gn=HLTF;refGene_tx=XM_019810583.2
C2	117437260	117437333
refGene_gn=HLTF;refGene_tx=XM_019810583.2
C2	117438662	117438753
refGene_gn=HLTF;refGene_tx=XM_019810583.2
C2	117440279	117440359
refGene_gn=HLTF;refGene_tx=XM_019810583.2
C2	117440444	117440486
refGene_gn=HLTF;refGene_tx=XM_019810583.2
C2	117441541	117441631
refGene_gn=HLTF;refGene_tx=XM_019810583.2
C2	117444488	117444586
refGene_gn=HLTF;refGene_tx=XM_019810583.2
C2	117448568	117448712
refGene_gn=HLTF;refGene_tx=XM_019810583.2
C2	117449830	117449969
refGene_gn=HLTF;refGene_tx=XM_019810583.2
C2	117450618	117450754
refGene_gn=HLTF;refGene_tx=XM_019810583.2
C2	117451880	117452060
refGene_gn=HLTF;refGene_tx=XM_019810583.2
C2	117457066	117457191
refGene_gn=HLTF;refGene_tx=XM_019810583.2
C2	117457657	117457836
refGene_gn=HLTF;refGene_tx=XM_019810583.2
C2	117459782	117459908
refGene_gn=HLTF;refGene_tx=XM_019810583.2
C2	117460229	117460342
refGene_gn=HLTF;refGene_tx=XM_019810583.2
C2	117460652	117460833
refGene_gn=HLTF;refGene_tx=XM_019810583.2
C2	117463716	117463797
refGene_gn=HLTF;refGene_tx=XM_019810583.2
C2	117466530	117466683
refGene_gn=HLTF;refGene_tx=XM_019810583.2
C2	117475869	117476043
refGene_gn=GYG1;refGene_tx=XM_019810587.2;refGene_gn=GYG1;refGene_tx=XM_023260474.1;refGene_gn=GYG1;refGene_tx=XM_023260475.1
C2	117476312	117476363
refGene_gn=GYG1;refGene_tx=XM_019810587.2;refGene_gn=GYG1;refGene_tx=XM_023260474.1
C2	117480064	117480284
refGene_gn=GYG1;refGene_tx=XM_019810587.2;refGene_gn=GYG1;refGene_tx=XM_023260474.1;refGene_gn=GYG1;refGene_tx=XM_023260475.1
C2	117494761	117494888
refGene_gn=GYG1;refGene_tx=XM_019810587.2;refGene_gn=GYG1;refGene_tx=XM_023260474.1;refGene_gn=GYG1;refGene_tx=XM_023260475.1
C2	117506366	117506529
refGene_gn=GYG1;refGene_tx=XM_019810587.2;refGene_gn=GYG1;refGene_tx=XM_023260474.1;refGene_gn=GYG1;refGene_tx=XM_023260475.1

C2 117506815 117506990
refGene_gn=GYG1;refGene_tx=XM_019810587.2;refGene_gn=GYG1;refGene_tx=XM_023260474.1;refGene_gn=GYG1;refGene_tx=XM_023260475.1
C2 117508836 117508972
refGene_gn=GYG1;refGene_tx=XM_019810587.2;refGene_gn=GYG1;refGene_tx=XM_023260474.1;refGene_gn=GYG1;refGene_tx=XM_023260475.1
C2 117511305 117511312
refGene_gn=GYG1;refGene_tx=XM_023260474.1
C2 117552054 117552106
refGene_gn=GYG1;refGene_tx=XM_019810587.2;refGene_gn=GYG1;refGene_tx=XM_023260475.1
C2 117610635 117610823
refGene_gn=CPA3;refGene_tx=XM_003992028.3
C2 117617694 117617779
refGene_gn=CPA3;refGene_tx=XM_003992028.3
C2 117619852 117620055
refGene_gn=CPA3;refGene_tx=XM_003992028.3
C2 117621068 117621159
refGene_gn=CPA3;refGene_tx=XM_003992028.3
C2 117622301 117622412
refGene_gn=CPA3;refGene_tx=XM_003992028.3
C2 117623707 117623809
refGene_gn=CPA3;refGene_tx=XM_003992028.3
C2 117625108 117625210
refGene_gn=CPA3;refGene_tx=XM_003992028.3
C2 117625314 117625417
refGene_gn=CPA3;refGene_tx=XM_003992028.3
C2 117634399 117634524
refGene_gn=CPA3;refGene_tx=XM_003992028.3
C2 117637527 117637603
refGene_gn=CPA3;refGene_tx=XM_003992028.3
C2 117637701 117637769
refGene_gn=CPA3;refGene_tx=XM_003992028.3
C2 117642373 117642561
refGene_gn=CPB1;refGene_tx=XM_019810588.1
C2 117644385 117644470
refGene_gn=CPB1;refGene_tx=XM_019810588.1
C2 117655564 117655767
refGene_gn=CPB1;refGene_tx=XM_019810588.1
C2 117656667 117656758
refGene_gn=CPB1;refGene_tx=XM_019810588.1
C2 117656847 117656958
refGene_gn=CPB1;refGene_tx=XM_019810588.1
C2 117659572 117659674
refGene_gn=CPB1;refGene_tx=XM_019810588.1
C2 117660521 117660623
refGene_gn=CPB1;refGene_tx=XM_019810588.1
C2 117660710 117660810
refGene_gn=CPB1;refGene_tx=XM_019810588.1
C2 117668726 117668851

refGene_gn=CPB1;refGene_tx=XM_019810588.1
C2 117675380 117675456
refGene_gn=CPB1;refGene_tx=XM_019810588.1
C2 117675568 117675678
refGene_gn=CPB1;refGene_tx=XM_019810588.1
C2 117739808 117740897
refGene_gn=AGTR1;refGene_tx=XM_003992030.5;refGene_gn=AGTR1;refGene_tx
=XM_019810589.2
C2 118895666 118895864
refGene_gn=ZIC1;refGene_tx=XM_003992031.4
C2 118896549 118896713
refGene_gn=ZIC1;refGene_tx=XM_003992031.4
C2 118898148 118899130
refGene_gn=ZIC1;refGene_tx=XM_003992031.4
C2 118903546 118903654
refGene_gn=ZIC4;refGene_tx=XM_006936348.3
C2 118905418 118905448
refGene_gn=ZIC4;refGene_tx=XM_019810591.2
C2 118906632 118906717
refGene_gn=ZIC4;refGene_tx=XM_006936348.3;refGene_gn=ZIC4;refGene_tx=X
M_019810591.2;refGene_gn=ZIC4;refGene_tx=XM_019810592.2;refGene_gn=ZIC
4;refGene_tx=XM_019810593.2;refGene_gn=ZIC4;refGene_tx=XM_019810594.2;
refGene_gn=ZIC4;refGene_tx=XM_019810595.2;refGene_gn=ZIC4;refGene_tx=X
M_019810597.2
C2 118912979 118913597
refGene_gn=ZIC4;refGene_tx=XM_006936348.3;refGene_gn=ZIC4;refGene_tx=X
M_019810591.2;refGene_gn=ZIC4;refGene_tx=XM_019810592.2;refGene_gn=ZIC
4;refGene_tx=XM_019810593.2;refGene_gn=ZIC4;refGene_tx=XM_019810594.2;
refGene_gn=ZIC4;refGene_tx=XM_019810595.2;refGene_gn=ZIC4;refGene_tx=X
M_019810597.2;refGene_gn=ZIC4;refGene_tx=XM_023260476.1
C2 118918360 118918679
refGene_gn=ZIC4;refGene_tx=XM_006936348.3;refGene_gn=ZIC4;refGene_tx=X
M_019810591.2;refGene_gn=ZIC4;refGene_tx=XM_019810592.2;refGene_gn=ZIC
4;refGene_tx=XM_019810593.2;refGene_gn=ZIC4;refGene_tx=XM_019810594.2;
refGene_gn=ZIC4;refGene_tx=XM_019810595.2;refGene_gn=ZIC4;refGene_tx=X
M_019810597.2;refGene_gn=ZIC4;refGene_tx=XM_023260476.1
C2 118920766 118920767
refGene_gn=ZIC4;refGene_tx=XM_006936348.3;refGene_gn=ZIC4;refGene_tx=X
M_019810591.2;refGene_gn=ZIC4;refGene_tx=XM_019810592.2;refGene_gn=ZIC
4;refGene_tx=XM_019810593.2;refGene_gn=ZIC4;refGene_tx=XM_019810594.2;
refGene_gn=ZIC4;refGene_tx=XM_019810595.2;refGene_gn=ZIC4;refGene_tx=X
M_019810597.2;refGene_gn=ZIC4;refGene_tx=XM_023260476.1
C2 119698362 119698375
refGene_gn=PLSCR5;refGene_tx=XM_019810598.2;refGene_gn=PLSCR5;refGene_
tx=XM_023260477.1
C2 119703145 119703309
refGene_gn=PLSCR5;refGene_tx=XM_019810598.2;refGene_gn=PLSCR5;refGene_
tx=XM_023260477.1
C2 119708305 119708348
refGene_gn=PLSCR5;refGene_tx=XM_019810598.2;refGene_gn=PLSCR5;refGene_

tx=XM_023260477.1
C2	119709584	119709805
refGene_gn=PLSCR5;refGene_tx=XM_019810598.2;refGene_gn=PLSCR5;refGene_tx=XM_023260477.1
C2	119712018	119712180
refGene_gn=PLSCR5;refGene_tx=XM_019810598.2;refGene_gn=PLSCR5;refGene_tx=XM_023260477.1
C2	119714288	119714450
refGene_gn=PLSCR5;refGene_tx=XM_019810598.2;refGene_gn=PLSCR5;refGene_tx=XM_023260477.1
C2	119717259	119717298
refGene_gn=PLSCR5;refGene_tx=XM_019810598.2;refGene_gn=PLSCR5;refGene_tx=XM_023260477.1
C2	119784977	119785037
refGene_gn=LOC111562293;refGene_tx=XM_023260478.1
C2	119786677	119786720
refGene_gn=LOC111562293;refGene_tx=XM_023260478.1
C2	119789835	119790056
refGene_gn=LOC111562293;refGene_tx=XM_023260478.1
C2	119791527	119791635
refGene_gn=LOC111562293;refGene_tx=XM_023260478.1
C2	119791636	119791690
refGene_gn=LOC111562293;refGene_tx=XM_023260478.1
C2	119798769	119798931
refGene_gn=LOC111562293;refGene_tx=XM_023260478.1
C2	119801789	119801828
refGene_gn=LOC111562293;refGene_tx=XM_023260478.1
C2	119826147	119826240
refGene_gn=LOC101086249;refGene_tx=XM_023260479.1
C2	119830423	119830466
refGene_gn=LOC101086249;refGene_tx=XM_023260479.1
C2	119833892	119834113
refGene_gn=LOC101086249;refGene_tx=XM_023260479.1
C2	119835990	119836152
refGene_gn=LOC101086249;refGene_tx=XM_023260479.1
C2	119840100	119840262
refGene_gn=LOC101086249;refGene_tx=XM_023260479.1
C2	119844666	119844696
refGene_gn=LOC101086249;refGene_tx=XM_023260479.1
C2	119995723	119995730
refGene_gn=PLSCR4;refGene_tx=XM_023260480.1;refGene_gn=PLSCR4;refGene_tx=XM_023260481.1;refGene_gn=PLSCR4;refGene_tx=XM_023260482.1;refGene_gn=PLSCR4;refGene_tx=XM_023260483.1
C2	119996258	119996265
refGene_gn=PLSCR4;refGene_tx=XM_019810602.2
C2	119996821	119996932
refGene_gn=PLSCR4;refGene_tx=XM_019810602.2;refGene_gn=PLSCR4;refGene_tx=XM_023260480.1;refGene_gn=PLSCR4;refGene_tx=XM_023260481.1;refGene_gn=PLSCR4;refGene_tx=XM_023260482.1;refGene_gn=PLSCR4;refGene_tx=XM_023260483.1

C2	120006018	120006254
refGene_gn=PLSCR4;refGene_tx=XM_019810602.2;refGene_gn=PLSCR4;refGene_tx=XM_023260480.1;refGene_gn=PLSCR4;refGene_tx=XM_023260481.1;refGene_gn=PLSCR4;refGene_tx=XM_023260482.1;refGene_gn=PLSCR4;refGene_tx=XM_023260483.1
C2	120010049	120010092
refGene_gn=PLSCR4;refGene_tx=XM_019810602.2;refGene_gn=PLSCR4;refGene_tx=XM_023260480.1;refGene_gn=PLSCR4;refGene_tx=XM_023260481.1;refGene_gn=PLSCR4;refGene_tx=XM_023260482.1;refGene_gn=PLSCR4;refGene_tx=XM_023260483.1
C2	120011001	120011228
refGene_gn=PLSCR4;refGene_tx=XM_019810602.2;refGene_gn=PLSCR4;refGene_tx=XM_023260480.1;refGene_gn=PLSCR4;refGene_tx=XM_023260481.1;refGene_gn=PLSCR4;refGene_tx=XM_023260482.1;refGene_gn=PLSCR4;refGene_tx=XM_023260483.1
C2	120013717	120013879
refGene_gn=PLSCR4;refGene_tx=XM_019810602.2;refGene_gn=PLSCR4;refGene_tx=XM_023260480.1;refGene_gn=PLSCR4;refGene_tx=XM_023260481.1;refGene_gn=PLSCR4;refGene_tx=XM_023260482.1;refGene_gn=PLSCR4;refGene_tx=XM_023260483.1
C2	120015714	120015873
refGene_gn=PLSCR4;refGene_tx=XM_019810602.2;refGene_gn=PLSCR4;refGene_tx=XM_023260480.1;refGene_gn=PLSCR4;refGene_tx=XM_023260481.1;refGene_gn=PLSCR4;refGene_tx=XM_023260482.1;refGene_gn=PLSCR4;refGene_tx=XM_023260483.1
C2	120016776	120016821
refGene_gn=PLSCR4;refGene_tx=XM_019810602.2;refGene_gn=PLSCR4;refGene_tx=XM_023260480.1;refGene_gn=PLSCR4;refGene_tx=XM_023260481.1;refGene_gn=PLSCR4;refGene_tx=XM_023260482.1;refGene_gn=PLSCR4;refGene_tx=XM_023260483.1
C2	120057102	120057217
refGene_gn=PLOD2;refGene_tx=XM_023260484.1
C2	120099309	120099401
refGene_gn=PLOD2;refGene_tx=XM_023260484.1
C2	120103176	120103313
refGene_gn=PLOD2;refGene_tx=XM_023260484.1
C2	120115897	120116061
refGene_gn=PLOD2;refGene_tx=XM_023260484.1
C2	120119213	120119326
refGene_gn=PLOD2;refGene_tx=XM_023260484.1
C2	120121630	120121694
refGene_gn=PLOD2;refGene_tx=XM_023260484.1
C2	120122929	120123027
refGene_gn=PLOD2;refGene_tx=XM_023260484.1
C2	120129800	120129902
refGene_gn=PLOD2;refGene_tx=XM_023260484.1
C2	120133222	120133348
refGene_gn=PLOD2;refGene_tx=XM_023260484.1
C2	120134997	120135119
refGene_gn=PLOD2;refGene_tx=XM_023260484.1

C2 120140537 120140569
refGene_gn=PLOD2;refGene_tx=XM_023260484.1
C2 120140571 120140641
refGene_gn=PLOD2;refGene_tx=XM_023260484.1
C2 120145803 120145929
refGene_gn=PLOD2;refGene_tx=XM_023260484.1
C2 120148224 120148366
refGene_gn=PLOD2;refGene_tx=XM_023260484.1
C2 120150584 120150698
refGene_gn=PLOD2;refGene_tx=XM_023260484.1
C2 120153480 120153546
refGene_gn=PLOD2;refGene_tx=XM_023260484.1
C2 120154610 120154715
refGene_gn=PLOD2;refGene_tx=XM_023260484.1
C2 120155851 120155998
refGene_gn=PLOD2;refGene_tx=XM_023260484.1
C2 120156109 120156235
refGene_gn=PLOD2;refGene_tx=XM_023260484.1
C2 120156402 120156558
refGene_gn=PLOD2;refGene_tx=XM_023260484.1
C2 122070687 122071019
refGene_gn=CC2H3orf58;refGene_tx=XM_003992038.5
C2 122076337 122076641
refGene_gn=CC2H3orf58;refGene_tx=XM_003992038.5
C2 122091774 122092431
refGene_gn=CC2H3orf58;refGene_tx=XM_003992038.5
C2 122115607 122115655
refGene_gn=SLC9A9;refGene_tx=XM_019810605.2;refGene_gn=SLC9A9;refGene_tx=XM_023260485.1;refGene_gn=SLC9A9;refGene_tx=XM_023260486.1
C2 122201293 122201357
refGene_gn=SLC9A9;refGene_tx=XM_019810605.2;refGene_gn=SLC9A9;refGene_tx=XM_023260485.1;refGene_gn=SLC9A9;refGene_tx=XM_023260486.1
C2 122216014 122216189
refGene_gn=SLC9A9;refGene_tx=XM_003992039.5
C2 122243333 122243536
refGene_gn=SLC9A9;refGene_tx=XM_003992039.5;refGene_gn=SLC9A9;refGene_tx=XM_019810605.2;refGene_gn=SLC9A9;refGene_tx=XM_019810608.2;refGene_gn=SLC9A9;refGene_tx=XM_019810610.2;refGene_gn=SLC9A9;refGene_tx=XM_023260485.1;refGene_gn=SLC9A9;refGene_tx=XM_023260486.1;refGene_gn=SLC9A9;refGene_tx=XM_023260487.1;refGene_gn=SLC9A9;refGene_tx=XM_023260489.1
C2 122285308 122285386
refGene_gn=SLC9A9;refGene_tx=XM_003992039.5;refGene_gn=SLC9A9;refGene_tx=XM_019810605.2;refGene_gn=SLC9A9;refGene_tx=XM_019810608.2;refGene_gn=SLC9A9;refGene_tx=XM_019810610.2;refGene_gn=SLC9A9;refGene_tx=XM_023260485.1;refGene_gn=SLC9A9;refGene_tx=XM_023260486.1;refGene_gn=SLC9A9;refGene_tx=XM_023260487.1;refGene_gn=SLC9A9;refGene_tx=XM_023260489.1
C2 122286935 122287012
refGene_gn=SLC9A9;refGene_tx=XM_003992039.5;refGene_gn=SLC9A9;refGene_

tx=XM_019810605.2;refGene_gn=SLC9A9;refGene_tx=XM_019810608.2;refGene_gn=SLC9A9;refGene_tx=XM_019810610.2;refGene_gn=SLC9A9;refGene_tx=XM_023260485.1;refGene_gn=SLC9A9;refGene_tx=XM_023260486.1;refGene_gn=SLC9A9;refGene_tx=XM_023260487.1;refGene_gn=SLC9A9;refGene_tx=XM_023260489.1
C2 122390876 122390992
refGene_gn=SLC9A9;refGene_tx=XM_003992039.5;refGene_gn=SLC9A9;refGene_tx=XM_019810605.2;refGene_gn=SLC9A9;refGene_tx=XM_019810608.2;refGene_gn=SLC9A9;refGene_tx=XM_019810610.2;refGene_gn=SLC9A9;refGene_tx=XM_023260485.1;refGene_gn=SLC9A9;refGene_tx=XM_023260486.1;refGene_gn=SLC9A9;refGene_tx=XM_023260487.1;refGene_gn=SLC9A9;refGene_tx=XM_023260489.1
C2 122423655 122423761
refGene_gn=SLC9A9;refGene_tx=XM_003992039.5;refGene_gn=SLC9A9;refGene_tx=XM_019810605.2;refGene_gn=SLC9A9;refGene_tx=XM_019810608.2;refGene_gn=SLC9A9;refGene_tx=XM_019810610.2;refGene_gn=SLC9A9;refGene_tx=XM_023260485.1;refGene_gn=SLC9A9;refGene_tx=XM_023260486.1;refGene_gn=SLC9A9;refGene_tx=XM_023260487.1;refGene_gn=SLC9A9;refGene_tx=XM_023260489.1
C2 122490917 122491056
refGene_gn=SLC9A9;refGene_tx=XM_003992039.5;refGene_gn=SLC9A9;refGene_tx=XM_019810605.2;refGene_gn=SLC9A9;refGene_tx=XM_019810608.2;refGene_gn=SLC9A9;refGene_tx=XM_019810610.2;refGene_gn=SLC9A9;refGene_tx=XM_023260485.1;refGene_gn=SLC9A9;refGene_tx=XM_023260486.1;refGene_gn=SLC9A9;refGene_tx=XM_023260487.1;refGene_gn=SLC9A9;refGene_tx=XM_023260489.1
C2 122495713 122495819
refGene_gn=SLC9A9;refGene_tx=XM_003992039.5;refGene_gn=SLC9A9;refGene_tx=XM_019810605.2;refGene_gn=SLC9A9;refGene_tx=XM_019810608.2;refGene_gn=SLC9A9;refGene_tx=XM_019810610.2;refGene_gn=SLC9A9;refGene_tx=XM_023260485.1;refGene_gn=SLC9A9;refGene_tx=XM_023260486.1;refGene_gn=SLC9A9;refGene_tx=XM_023260487.1;refGene_gn=SLC9A9;refGene_tx=XM_023260489.1
C2 122515979 122516068
refGene_gn=SLC9A9;refGene_tx=XM_003992039.5;refGene_gn=SLC9A9;refGene_tx=XM_019810605.2;refGene_gn=SLC9A9;refGene_tx=XM_019810608.2;refGene_gn=SLC9A9;refGene_tx=XM_019810610.2;refGene_gn=SLC9A9;refGene_tx=XM_023260485.1;refGene_gn=SLC9A9;refGene_tx=XM_023260486.1;refGene_gn=SLC9A9;refGene_tx=XM_023260487.1;refGene_gn=SLC9A9;refGene_tx=XM_023260489.1
C2 122559295 122559409
refGene_gn=SLC9A9;refGene_tx=XM_003992039.5;refGene_gn=SLC9A9;refGene_tx=XM_019810605.2;refGene_gn=SLC9A9;refGene_tx=XM_019810608.2;refGene_gn=SLC9A9;refGene_tx=XM_019810610.2;refGene_gn=SLC9A9;refGene_tx=XM_023260485.1;refGene_gn=SLC9A9;refGene_tx=XM_023260486.1;refGene_gn=SLC9A9;refGene_tx=XM_023260487.1;refGene_gn=SLC9A9;refGene_tx=XM_023260489.1
C2 122561026 122561138
refGene_gn=SLC9A9;refGene_tx=XM_003992039.5;refGene_gn=SLC9A9;refGene_tx=XM_019810605.2;refGene_gn=SLC9A9;refGene_tx=XM_019810608.2;refGene_

gn=SLC9A9;refGene_tx=XM_019810610.2;refGene_gn=SLC9A9;refGene_tx=XM_02
3260485.1;refGene_gn=SLC9A9;refGene_tx=XM_023260486.1;refGene_gn=SLC9A
9;refGene_tx=XM_023260487.1;refGene_gn=SLC9A9;refGene_tx=XM_023260489.
1
C2 122583369 122583523
refGene_gn=SLC9A9;refGene_tx=XM_003992039.5;refGene_gn=SLC9A9;refGene_
tx=XM_019810605.2;refGene_gn=SLC9A9;refGene_tx=XM_019810608.2;refGene_
gn=SLC9A9;refGene_tx=XM_019810610.2;refGene_gn=SLC9A9;refGene_tx=XM_02
3260485.1;refGene_gn=SLC9A9;refGene_tx=XM_023260486.1;refGene_gn=SLC9A
9;refGene_tx=XM_023260487.1;refGene_gn=SLC9A9;refGene_tx=XM_023260489.
1
C2 122672205 122672254
refGene_gn=SLC9A9;refGene_tx=XM_003992039.5;refGene_gn=SLC9A9;refGene_
tx=XM_019810605.2;refGene_gn=SLC9A9;refGene_tx=XM_019810608.2;refGene_
gn=SLC9A9;refGene_tx=XM_019810610.2;refGene_gn=SLC9A9;refGene_tx=XM_02
3260485.1;refGene_gn=SLC9A9;refGene_tx=XM_023260486.1;refGene_gn=SLC9A
9;refGene_tx=XM_023260487.1;refGene_gn=SLC9A9;refGene_tx=XM_023260489.
1
C2 122685754 122685834
refGene_gn=SLC9A9;refGene_tx=XM_003992039.5;refGene_gn=SLC9A9;refGene_
tx=XM_019810605.2;refGene_gn=SLC9A9;refGene_tx=XM_019810608.2;refGene_
gn=SLC9A9;refGene_tx=XM_019810610.2;refGene_gn=SLC9A9;refGene_tx=XM_02
3260485.1;refGene_gn=SLC9A9;refGene_tx=XM_023260486.1;refGene_gn=SLC9A
9;refGene_tx=XM_023260487.1;refGene_gn=SLC9A9;refGene_tx=XM_023260489.
1
C2 122785358 122785464
refGene_gn=SLC9A9;refGene_tx=XM_003992039.5;refGene_gn=SLC9A9;refGene_
tx=XM_019810605.2;refGene_gn=SLC9A9;refGene_tx=XM_019810608.2;refGene_
gn=SLC9A9;refGene_tx=XM_019810610.2;refGene_gn=SLC9A9;refGene_tx=XM_02
3260485.1;refGene_gn=SLC9A9;refGene_tx=XM_023260486.1;refGene_gn=SLC9A
9;refGene_tx=XM_023260487.1;refGene_gn=SLC9A9;refGene_tx=XM_023260489.
1
C2 122786810 122787038
refGene_gn=SLC9A9;refGene_tx=XM_003992039.5;refGene_gn=SLC9A9;refGene_
tx=XM_019810605.2;refGene_gn=SLC9A9;refGene_tx=XM_019810608.2;refGene_
gn=SLC9A9;refGene_tx=XM_019810610.2;refGene_gn=SLC9A9;refGene_tx=XM_02
3260485.1;refGene_gn=SLC9A9;refGene_tx=XM_023260486.1;refGene_gn=SLC9A
9;refGene_tx=XM_023260487.1;refGene_gn=SLC9A9;refGene_tx=XM_023260489.
1
C2 122917164 122918778
refGene_gn=CHST2;refGene_tx=XM_023260491.1
C2 122974715 122974854
refGene_gn=U2SURP;refGene_tx=XM_011286158.3;refGene_gn=U2SURP;refGene_
tx=XM_011286159.3;refGene_gn=U2SURP;refGene_tx=XM_019810612.2
C2 122976027 122976204
refGene_gn=U2SURP;refGene_tx=XM_011286158.3;refGene_gn=U2SURP;refGene_
tx=XM_011286159.3;refGene_gn=U2SURP;refGene_tx=XM_019810612.2
C2 122978178 122978297
refGene_gn=U2SURP;refGene_tx=XM_011286158.3;refGene_gn=U2SURP;refGene_
tx=XM_011286159.3;refGene_gn=U2SURP;refGene_tx=XM_019810612.2

C2	122980823	122980934
refGene_gn=U2SURP;refGene_tx=XM_011286158.3;refGene_gn=U2SURP;refGene_
tx=XM_011286159.3;refGene_gn=U2SURP;refGene_tx=XM_019810612.2
C2	122983615	122983775
refGene_gn=U2SURP;refGene_tx=XM_011286158.3;refGene_gn=U2SURP;refGene_
tx=XM_011286159.3;refGene_gn=U2SURP;refGene_tx=XM_019810612.2
C2	122986584	122986651
refGene_gn=U2SURP;refGene_tx=XM_011286158.3;refGene_gn=U2SURP;refGene_
tx=XM_011286159.3;refGene_gn=U2SURP;refGene_tx=XM_019810612.2
C2	122987336	122987432
refGene_gn=U2SURP;refGene_tx=XM_011286158.3;refGene_gn=U2SURP;refGene_
tx=XM_011286159.3;refGene_gn=U2SURP;refGene_tx=XM_019810612.2
C2	122988150	122988307
refGene_gn=U2SURP;refGene_tx=XM_011286158.3;refGene_gn=U2SURP;refGene_
tx=XM_011286159.3;refGene_gn=U2SURP;refGene_tx=XM_019810612.2
C2	122992773	122992896
refGene_gn=U2SURP;refGene_tx=XM_011286158.3;refGene_gn=U2SURP;refGene_
tx=XM_011286159.3;refGene_gn=U2SURP;refGene_tx=XM_019810612.2
C2	122994128	122994216
refGene_gn=U2SURP;refGene_tx=XM_011286158.3;refGene_gn=U2SURP;refGene_
tx=XM_011286159.3;refGene_gn=U2SURP;refGene_tx=XM_019810612.2
C2	122994889	122994969
refGene_gn=U2SURP;refGene_tx=XM_011286158.3;refGene_gn=U2SURP;refGene_
tx=XM_011286159.3;refGene_gn=U2SURP;refGene_tx=XM_019810612.2
C2	122995376	122995539
refGene_gn=U2SURP;refGene_tx=XM_011286158.3;refGene_gn=U2SURP;refGene_
tx=XM_011286159.3;refGene_gn=U2SURP;refGene_tx=XM_019810612.2
C2	122996875	122997039
refGene_gn=U2SURP;refGene_tx=XM_011286158.3;refGene_gn=U2SURP;refGene_
tx=XM_011286159.3;refGene_gn=U2SURP;refGene_tx=XM_019810612.2
C2	122997116	122997183
refGene_gn=U2SURP;refGene_tx=XM_011286158.3;refGene_gn=U2SURP;refGene_
tx=XM_011286159.3;refGene_gn=U2SURP;refGene_tx=XM_019810612.2
C2	122998367	122998472
refGene_gn=U2SURP;refGene_tx=XM_011286158.3;refGene_gn=U2SURP;refGene_
tx=XM_011286159.3;refGene_gn=U2SURP;refGene_tx=XM_019810612.2
C2	123001564	123001608
refGene_gn=U2SURP;refGene_tx=XM_011286158.3;refGene_gn=U2SURP;refGene_
tx=XM_011286159.3
C2	123002544	123002756
refGene_gn=U2SURP;refGene_tx=XM_011286158.3;refGene_gn=U2SURP;refGene_
tx=XM_011286159.3;refGene_gn=U2SURP;refGene_tx=XM_019810612.2
C2	123002938	123003104
refGene_gn=U2SURP;refGene_tx=XM_011286158.3;refGene_gn=U2SURP;refGene_
tx=XM_011286159.3
C2	123004612	123004695
refGene_gn=U2SURP;refGene_tx=XM_011286158.3;refGene_gn=U2SURP;refGene_
tx=XM_011286159.3
C2	123004786	123004822
refGene_gn=U2SURP;refGene_tx=XM_011286158.3;refGene_gn=U2SURP;refGene_

tx=XM_011286159.3
C2	123004904	123004999
refGene_gn=U2SURP;refGene_tx=XM_011286158.3;refGene_gn=U2SURP;refGene_
tx=XM_011286159.3
C2	123005576	123005644
refGene_gn=U2SURP;refGene_tx=XM_011286158.3;refGene_gn=U2SURP;refGene_
tx=XM_011286159.3
C2	123008279	123008413
refGene_gn=U2SURP;refGene_tx=XM_011286158.3;refGene_gn=U2SURP;refGene_
tx=XM_011286159.3
C2	123008882	123008997
refGene_gn=U2SURP;refGene_tx=XM_011286158.3;refGene_gn=U2SURP;refGene_
tx=XM_011286159.3
C2	123011504	123011603
refGene_gn=U2SURP;refGene_tx=XM_011286158.3;refGene_gn=U2SURP;refGene_
tx=XM_011286159.3
C2	123013922	123014054
refGene_gn=U2SURP;refGene_tx=XM_011286158.3;refGene_gn=U2SURP;refGene_
tx=XM_011286159.3
C2	123015502	123015547
refGene_gn=U2SURP;refGene_tx=XM_011286158.3;refGene_gn=U2SURP;refGene_
tx=XM_011286159.3
C2	123024119	123024164
refGene_gn=U2SURP;refGene_tx=XM_011286158.3;refGene_gn=U2SURP;refGene_
tx=XM_011286159.3
C2	123073627	123074761
refGene_gn=PAQR9;refGene_tx=XM_023260492.1;refGene_gn=PAQR9;refGene_tx
=XM_023260493.1;refGene_gn=PAQR9;refGene_tx=XM_023260495.1
C2	123132774	123132857
refGene_gn=PCOLCE2;refGene_tx=XM_023260496.1
C2	123133799	123133908
refGene_gn=PCOLCE2;refGene_tx=XM_023260496.1
C2	123171081	123171337
refGene_gn=PCOLCE2;refGene_tx=XM_023260496.1
C2	123175446	123175571
refGene_gn=PCOLCE2;refGene_tx=XM_023260496.1
C2	123178466	123178603
refGene_gn=PCOLCE2;refGene_tx=XM_023260496.1
C2	123185784	123185939
refGene_gn=PCOLCE2;refGene_tx=XM_023260496.1
C2	123189933	123190017
refGene_gn=PCOLCE2;refGene_tx=XM_023260496.1
C2	123192110	123192278
refGene_gn=PCOLCE2;refGene_tx=XM_023260496.1
C2	123194534	123194665
refGene_gn=PCOLCE2;refGene_tx=XM_023260496.1
C2	123207173	123207401
refGene_gn=TRPC1;refGene_tx=XM_006936359.4;refGene_gn=TRPC1;refGene_tx
=XM_023260497.1;refGene_gn=TRPC1;refGene_tx=XM_023260498.1;refGene_gn=
TRPC1;refGene_tx=XM_023260499.1;refGene_gn=TRPC1;refGene_tx=XM_0232605

00.1;refGene_gn=TRPC1;refGene_tx=XM_023260503.1
C2 123208388 123208593
refGene_gn=TRPC1;refGene_tx=XM_006936359.4;refGene_gn=TRPC1;refGene_tx=XM_023260497.1;refGene_gn=TRPC1;refGene_tx=XM_023260498.1;refGene_gn=TRPC1;refGene_tx=XM_023260499.1;refGene_gn=TRPC1;refGene_tx=XM_023260500.1;refGene_gn=TRPC1;refGene_tx=XM_023260502.1;refGene_gn=TRPC1;refGene_tx=XM_023260503.1
C2 123209183 123209385
refGene_gn=TRPC1;refGene_tx=XM_006936359.4;refGene_gn=TRPC1;refGene_tx=XM_023260497.1;refGene_gn=TRPC1;refGene_tx=XM_023260498.1;refGene_gn=TRPC1;refGene_tx=XM_023260499.1;refGene_gn=TRPC1;refGene_tx=XM_023260500.1;refGene_gn=TRPC1;refGene_tx=XM_023260502.1;refGene_gn=TRPC1;refGene_tx=XM_023260503.1
C2 123211044 123211220
refGene_gn=TRPC1;refGene_tx=XM_006936359.4;refGene_gn=TRPC1;refGene_tx=XM_023260497.1;refGene_gn=TRPC1;refGene_tx=XM_023260498.1;refGene_gn=TRPC1;refGene_tx=XM_023260499.1;refGene_gn=TRPC1;refGene_tx=XM_023260500.1;refGene_gn=TRPC1;refGene_tx=XM_023260502.1;refGene_gn=TRPC1;refGene_tx=XM_023260503.1
C2 123221089 123221233
refGene_gn=TRPC1;refGene_tx=XM_006936359.4;refGene_gn=TRPC1;refGene_tx=XM_023260497.1;refGene_gn=TRPC1;refGene_tx=XM_023260498.1;refGene_gn=TRPC1;refGene_tx=XM_023260499.1;refGene_gn=TRPC1;refGene_tx=XM_023260500.1;refGene_gn=TRPC1;refGene_tx=XM_023260502.1;refGene_gn=TRPC1;refGene_tx=XM_023260503.1
C2 123222902 123223042
refGene_gn=TRPC1;refGene_tx=XM_006936359.4;refGene_gn=TRPC1;refGene_tx=XM_023260497.1;refGene_gn=TRPC1;refGene_tx=XM_023260498.1;refGene_gn=TRPC1;refGene_tx=XM_023260499.1;refGene_gn=TRPC1;refGene_tx=XM_023260500.1;refGene_gn=TRPC1;refGene_tx=XM_023260502.1;refGene_gn=TRPC1;refGene_tx=XM_023260503.1
C2 123224843 123225180
refGene_gn=TRPC1;refGene_tx=XM_006936359.4;refGene_gn=TRPC1;refGene_tx=XM_023260497.1;refGene_gn=TRPC1;refGene_tx=XM_023260498.1;refGene_gn=TRPC1;refGene_tx=XM_023260499.1;refGene_gn=TRPC1;refGene_tx=XM_023260500.1;refGene_gn=TRPC1;refGene_tx=XM_023260502.1;refGene_gn=TRPC1;refGene_tx=XM_023260503.1
C2 123229064 123229260
refGene_gn=TRPC1;refGene_tx=XM_006936359.4;refGene_gn=TRPC1;refGene_tx=XM_023260497.1;refGene_gn=TRPC1;refGene_tx=XM_023260498.1;refGene_gn=TRPC1;refGene_tx=XM_023260499.1;refGene_gn=TRPC1;refGene_tx=XM_023260500.1;refGene_gn=TRPC1;refGene_tx=XM_023260502.1;refGene_gn=TRPC1;refGene_tx=XM_023260503.1
C2 123232033 123232165
refGene_gn=TRPC1;refGene_tx=XM_006936359.4;refGene_gn=TRPC1;refGene_tx=XM_023260498.1;refGene_gn=TRPC1;refGene_tx=XM_023260499.1;refGene_gn=TRPC1;refGene_tx=XM_023260500.1;refGene_gn=TRPC1;refGene_tx=XM_023260502.1;refGene_gn=TRPC1;refGene_tx=XM_023260503.1
C2 123246507 123246710
refGene_gn=TRPC1;refGene_tx=XM_006936359.4;refGene_gn=TRPC1;refGene_tx

=XM_023260497.1;refGene_gn=TRPC1;refGene_tx=XM_023260498.1;refGene_gn=TRPC1;refGene_tx=XM_023260499.1;refGene_gn=TRPC1;refGene_tx=XM_023260500.1;refGene_gn=TRPC1;refGene_tx=XM_023260502.1;refGene_gn=TRPC1;refGene_tx=XM_023260503.1
C2 123251096 123251198
refGene_gn=TRPC1;refGene_tx=XM_006936359.4;refGene_gn=TRPC1;refGene_tx=XM_023260497.1;refGene_gn=TRPC1;refGene_tx=XM_023260498.1;refGene_gn=TRPC1;refGene_tx=XM_023260502.1
C2 123257473 123257628
refGene_gn=TRPC1;refGene_tx=XM_006936359.4;refGene_gn=TRPC1;refGene_tx=XM_023260497.1;refGene_gn=TRPC1;refGene_tx=XM_023260499.1;refGene_gn=TRPC1;refGene_tx=XM_023260502.1
C2 123271465 123271640
refGene_gn=TRPC1;refGene_tx=XM_006936359.4;refGene_gn=TRPC1;refGene_tx=XM_023260497.1;refGene_gn=TRPC1;refGene_tx=XM_023260498.1;refGene_gn=TRPC1;refGene_tx=XM_023260499.1;refGene_gn=TRPC1;refGene_tx=XM_023260500.1;refGene_gn=TRPC1;refGene_tx=XM_023260502.1
C2 123276557 123276693
refGene_gn=PLS1;refGene_tx=XM_006936361.2;refGene_gn=PLS1;refGene_tx=XM_019810617.1
C2 123277162 123277287
refGene_gn=PLS1;refGene_tx=XM_006936361.2;refGene_gn=PLS1;refGene_tx=XM_019810617.1
C2 123283828 123283952
refGene_gn=PLS1;refGene_tx=XM_006936361.2;refGene_gn=PLS1;refGene_tx=XM_019810617.1
C2 123285354 123285488
refGene_gn=PLS1;refGene_tx=XM_006936361.2;refGene_gn=PLS1;refGene_tx=XM_019810617.1
C2 123291063 123291178
refGene_gn=PLS1;refGene_tx=XM_006936361.2;refGene_gn=PLS1;refGene_tx=XM_019810617.1
C2 123293768 123293847
refGene_gn=PLS1;refGene_tx=XM_006936361.2;refGene_gn=PLS1;refGene_tx=XM_019810617.1
C2 123298710 123298906
refGene_gn=PLS1;refGene_tx=XM_006936361.2;refGene_gn=PLS1;refGene_tx=XM_019810617.1
C2 123301488 123301584
refGene_gn=PLS1;refGene_tx=XM_006936361.2;refGene_gn=PLS1;refGene_tx=XM_019810617.1
C2 123304067 123304210
refGene_gn=PLS1;refGene_tx=XM_006936361.2;refGene_gn=PLS1;refGene_tx=XM_019810617.1
C2 123304291 123304457
refGene_gn=PLS1;refGene_tx=XM_006936361.2;refGene_gn=PLS1;refGene_tx=XM_019810617.1
C2 123307473 123307555
refGene_gn=PLS1;refGene_tx=XM_006936361.2;refGene_gn=PLS1;refGene_tx=XM_019810617.1

C2	123309347	123309480
refGene_gn=PLS1;refGene_tx=XM_006936361.2;refGene_gn=PLS1;refGene_tx=XM_019810617.1
C2	123313191	123313321
refGene_gn=PLS1;refGene_tx=XM_006936361.2;refGene_gn=PLS1;refGene_tx=XM_019810617.1
C2	123315389	123315553
refGene_gn=PLS1;refGene_tx=XM_006936361.2;refGene_gn=PLS1;refGene_tx=XM_019810617.1
C2	123319144	123319214
refGene_gn=PLS1;refGene_tx=XM_006936361.2;refGene_gn=PLS1;refGene_tx=XM_019810617.1
C2	123402289	123402348
refGene_gn=ATR;refGene_tx=XM_006936362.4
C2	123411858	123411950
refGene_gn=ATR;refGene_tx=XM_006936362.4
C2	123413316	123413463
refGene_gn=ATR;refGene_tx=XM_006936362.4
C2	123415204	123416082
refGene_gn=ATR;refGene_tx=XM_006936362.4
C2	123416879	123417058
refGene_gn=ATR;refGene_tx=XM_006936362.4
C2	123417060	123417074
refGene_gn=ATR;refGene_tx=XM_019810614.2
C2	123417535	123417727
refGene_gn=ATR;refGene_tx=XM_006936362.4;refGene_gn=ATR;refGene_tx=XM_019810614.2
C2	123418925	123419125
refGene_gn=ATR;refGene_tx=XM_006936362.4;refGene_gn=ATR;refGene_tx=XM_019810614.2
C2	123419433	123419586
refGene_gn=ATR;refGene_tx=XM_006936362.4;refGene_gn=ATR;refGene_tx=XM_019810614.2
C2	123420232	123420425
refGene_gn=ATR;refGene_tx=XM_006936362.4;refGene_gn=ATR;refGene_tx=XM_019810614.2
C2	123420658	123420921
refGene_gn=ATR;refGene_tx=XM_006936362.4;refGene_gn=ATR;refGene_tx=XM_019810614.2
C2	123422655	123422846
refGene_gn=ATR;refGene_tx=XM_006936362.4;refGene_gn=ATR;refGene_tx=XM_019810614.2
C2	123422929	123423030
refGene_gn=ATR;refGene_tx=XM_006936362.4;refGene_gn=ATR;refGene_tx=XM_019810614.2
C2	123423196	123423368
refGene_gn=ATR;refGene_tx=XM_006936362.4;refGene_gn=ATR;refGene_tx=XM_019810614.2
C2	123426626	123426797
refGene_gn=ATR;refGene_tx=XM_006936362.4;refGene_gn=ATR;refGene_tx=XM_

019810614.2
C2 123428388 123428583
refGene_gn=ATR;refGene_tx=XM_006936362.4;refGene_gn=ATR;refGene_tx=XM_
019810614.2
C2 123431063 123431249
refGene_gn=ATR;refGene_tx=XM_006936362.4;refGene_gn=ATR;refGene_tx=XM_
019810614.2
C2 123435916 123436009
refGene_gn=ATR;refGene_tx=XM_006936362.4;refGene_gn=ATR;refGene_tx=XM_
019810614.2
C2 123437863 123437994
refGene_gn=ATR;refGene_tx=XM_006936362.4;refGene_gn=ATR;refGene_tx=XM_
019810614.2
C2 123439341 123439485
refGene_gn=ATR;refGene_tx=XM_006936362.4;refGene_gn=ATR;refGene_tx=XM_
019810614.2
C2 123441610 123441704
refGene_gn=ATR;refGene_tx=XM_006936362.4;refGene_gn=ATR;refGene_tx=XM_
019810614.2
C2 123442155 123442281
refGene_gn=ATR;refGene_tx=XM_006936362.4;refGene_gn=ATR;refGene_tx=XM_
019810614.2
C2 123447877 123448084
refGene_gn=ATR;refGene_tx=XM_006936362.4;refGene_gn=ATR;refGene_tx=XM_
019810614.2
C2 123449443 123449557
refGene_gn=ATR;refGene_tx=XM_006936362.4;refGene_gn=ATR;refGene_tx=XM_
019810614.2
C2 123452731 123452847
refGene_gn=ATR;refGene_tx=XM_006936362.4;refGene_gn=ATR;refGene_tx=XM_
019810614.2
C2 123454498 123454619
refGene_gn=ATR;refGene_tx=XM_006936362.4;refGene_gn=ATR;refGene_tx=XM_
019810614.2
C2 123455926 123456064
refGene_gn=ATR;refGene_tx=XM_006936362.4;refGene_gn=ATR;refGene_tx=XM_
019810614.2
C2 123458624 123458835
refGene_gn=ATR;refGene_tx=XM_006936362.4;refGene_gn=ATR;refGene_tx=XM_
019810614.2
C2 123461958 123462137
refGene_gn=ATR;refGene_tx=XM_006936362.4;refGene_gn=ATR;refGene_tx=XM_
019810614.2
C2 123465796 123465961
refGene_gn=ATR;refGene_tx=XM_006936362.4;refGene_gn=ATR;refGene_tx=XM_
019810614.2
C2 123467002 123467094
refGene_gn=ATR;refGene_tx=XM_006936362.4;refGene_gn=ATR;refGene_tx=XM_
019810614.2
C2 123472515 123472607

refGene_gn=ATR;refGene_tx=XM_006936362.4;refGene_gn=ATR;refGene_tx=XM_019810614.2
C2 123472833 123473011
refGene_gn=ATR;refGene_tx=XM_006936362.4;refGene_gn=ATR;refGene_tx=XM_019810614.2
C2 123474283 123474463
refGene_gn=ATR;refGene_tx=XM_006936362.4;refGene_gn=ATR;refGene_tx=XM_019810614.2
C2 123474904 123475064
refGene_gn=ATR;refGene_tx=XM_006936362.4;refGene_gn=ATR;refGene_tx=XM_019810614.2
C2 123478782 123478962
refGene_gn=ATR;refGene_tx=XM_006936362.4;refGene_gn=ATR;refGene_tx=XM_019810614.2
C2 123480720 123480863
refGene_gn=ATR;refGene_tx=XM_006936362.4;refGene_gn=ATR;refGene_tx=XM_019810614.2
C2 123481664 123481762
refGene_gn=ATR;refGene_tx=XM_006936362.4;refGene_gn=ATR;refGene_tx=XM_019810614.2
C2 123482200 123482433
refGene_gn=ATR;refGene_tx=XM_006936362.4;refGene_gn=ATR;refGene_tx=XM_019810614.2
C2 123483370 123483505
refGene_gn=ATR;refGene_tx=XM_006936362.4;refGene_gn=ATR;refGene_tx=XM_019810614.2
C2 123485000 123485201
refGene_gn=ATR;refGene_tx=XM_006936362.4;refGene_gn=ATR;refGene_tx=XM_019810614.2
C2 123487749 123487893
refGene_gn=ATR;refGene_tx=XM_006936362.4;refGene_gn=ATR;refGene_tx=XM_019810614.2
C2 123490241 123490392
refGene_gn=ATR;refGene_tx=XM_006936362.4;refGene_gn=ATR;refGene_tx=XM_019810614.2
C2 123493386 123493543
refGene_gn=ATR;refGene_tx=XM_006936362.4;refGene_gn=ATR;refGene_tx=XM_019810614.2
C2 123493781 123493935
refGene_gn=ATR;refGene_tx=XM_006936362.4;refGene_gn=ATR;refGene_tx=XM_019810614.2
C2 123495109 123495261
refGene_gn=ATR;refGene_tx=XM_006936362.4;refGene_gn=ATR;refGene_tx=XM_019810614.2
C2 123502525 123502631
refGene_gn=ATR;refGene_tx=XM_006936362.4;refGene_gn=ATR;refGene_tx=XM_019810614.2
C2 123503519 123503693
refGene_gn=ATR;refGene_tx=XM_006936362.4;refGene_gn=ATR;refGene_tx=XM_019810614.2

C2 123505620 123505695
refGene_gn=XRN1;refGene_tx=XM_023260504.1
C2 123515680 123515913
refGene_gn=XRN1;refGene_tx=XM_023260504.1
C2 123520959 123521057
refGene_gn=XRN1;refGene_tx=XM_023260504.1
C2 123522238 123522348
refGene_gn=XRN1;refGene_tx=XM_023260504.1
C2 123522437 123522548
refGene_gn=XRN1;refGene_tx=XM_023260504.1
C2 123524239 123524322
refGene_gn=XRN1;refGene_tx=XM_023260504.1
C2 123526968 123527056
refGene_gn=XRN1;refGene_tx=XM_023260504.1
C2 123528237 123528406
refGene_gn=XRN1;refGene_tx=XM_023260504.1
C2 123529446 123529514
refGene_gn=XRN1;refGene_tx=XM_023260504.1
C2 123529826 123529964
refGene_gn=XRN1;refGene_tx=XM_023260504.1
C2 123534032 123534102
refGene_gn=XRN1;refGene_tx=XM_023260504.1
C2 123534313 123534372
refGene_gn=XRN1;refGene_tx=XM_023260504.1
C2 123534373 123534420
refGene_gn=XRN1;refGene_tx=XM_023260504.1
C2 123537187 123537277
refGene_gn=XRN1;refGene_tx=XM_023260504.1
C2 123542241 123542398
refGene_gn=XRN1;refGene_tx=XM_023260504.1
C2 123543980 123544100
refGene_gn=XRN1;refGene_tx=XM_023260504.1
C2 123548140 123548310
refGene_gn=XRN1;refGene_tx=XM_023260504.1
C2 123549975 123550096
refGene_gn=XRN1;refGene_tx=XM_023260504.1
C2 123550192 123550291
refGene_gn=XRN1;refGene_tx=XM_023260504.1
C2 123553949 123554053
refGene_gn=XRN1;refGene_tx=XM_023260504.1
C2 123557193 123557325
refGene_gn=XRN1;refGene_tx=XM_023260504.1
C2 123571317 123571480
refGene_gn=XRN1;refGene_tx=XM_023260504.1
C2 123574926 123575040
refGene_gn=XRN1;refGene_tx=XM_023260504.1
C2 123576733 123576832
refGene_gn=XRN1;refGene_tx=XM_023260504.1
C2 123579236 123579352
refGene_gn=XRN1;refGene_tx=XM_023260504.1

C2 123579876 123580023
refGene_gn=XRN1;refGene_tx=XM_023260504.1
C2 123582043 123582050
refGene_gn=XRN1;refGene_tx=XM_023260504.1
C2 123584826 123584909
refGene_gn=XRN1;refGene_tx=XM_023260504.1
C2 123585448 123585584
refGene_gn=XRN1;refGene_tx=XM_023260504.1
C2 123588929 123588986
refGene_gn=XRN1;refGene_tx=XM_023260504.1
C2 123589119 123589252
refGene_gn=XRN1;refGene_tx=XM_023260504.1
C2 123597200 123597270
refGene_gn=XRN1;refGene_tx=XM_023260504.1
C2 123599605 123599813
refGene_gn=XRN1;refGene_tx=XM_023260504.1
C2 123601109 123601205
refGene_gn=XRN1;refGene_tx=XM_023260504.1
C2 123606446 123606555
refGene_gn=XRN1;refGene_tx=XM_023260504.1
C2 123611435 123611497
refGene_gn=XRN1;refGene_tx=XM_023260504.1
C2 123612991 123613102
refGene_gn=XRN1;refGene_tx=XM_023260504.1
C2 123613257 123613380
refGene_gn=XRN1;refGene_tx=XM_023260504.1
C2 123613840 123614000
refGene_gn=XRN1;refGene_tx=XM_023260504.1
C2 123616146 123616328
refGene_gn=XRN1;refGene_tx=XM_023260504.1
C2 123617933 123618047
refGene_gn=XRN1;refGene_tx=XM_023260504.1
C2 123618141 123618244
refGene_gn=XRN1;refGene_tx=XM_023260504.1
C2 123621885 123621924
refGene_gn=XRN1;refGene_tx=XM_023260504.1
C2 123622273 123622434
refGene_gn=XRN1;refGene_tx=XM_023260504.1
C2 123623425 123623728
refGene_gn=XRN1;refGene_tx=XM_023260504.1
C2 123662705 123662867
refGene_gn=GK5;refGene_tx=XM_023260505.1
C2 123670196 123670290
refGene_gn=GK5;refGene_tx=XM_023260505.1
C2 123673115 123673191
refGene_gn=GK5;refGene_tx=XM_019810621.2;refGene_gn=GK5;refGene_tx=XM_023260505.1;refGene_gn=GK5;refGene_tx=XM_023260506.1;refGene_gn=GK5;refGene_tx=XM_023260507.1
C2 123686153 123686247
refGene_gn=GK5;refGene_tx=XM_019810621.2;refGene_gn=GK5;refGene_tx=XM_

023260505.1;refGene_gn=GK5;refGene_tx=XM_023260506.1;refGene_gn=GK5;refGene_tx=XM_023260507.1
C2 123693623 123693755
refGene_gn=GK5;refGene_tx=XM_019810621.2;refGene_gn=GK5;refGene_tx=XM_023260505.1;refGene_gn=GK5;refGene_tx=XM_023260506.1;refGene_gn=GK5;refGene_tx=XM_023260507.1
C2 123700305 123700381
refGene_gn=GK5;refGene_tx=XM_019810621.2;refGene_gn=GK5;refGene_tx=XM_023260505.1;refGene_gn=GK5;refGene_tx=XM_023260506.1;refGene_gn=GK5;refGene_tx=XM_023260507.1
C2 123701507 123701569
refGene_gn=GK5;refGene_tx=XM_019810621.2;refGene_gn=GK5;refGene_tx=XM_023260505.1;refGene_gn=GK5;refGene_tx=XM_023260506.1;refGene_gn=GK5;refGene_tx=XM_023260507.1
C2 123701651 123701725
refGene_gn=GK5;refGene_tx=XM_019810621.2;refGene_gn=GK5;refGene_tx=XM_023260505.1;refGene_gn=GK5;refGene_tx=XM_023260506.1;refGene_gn=GK5;refGene_tx=XM_023260507.1
C2 123701913 123701974
refGene_gn=GK5;refGene_tx=XM_019810621.2;refGene_gn=GK5;refGene_tx=XM_023260505.1;refGene_gn=GK5;refGene_tx=XM_023260506.1;refGene_gn=GK5;refGene_tx=XM_023260507.1
C2 123713613 123713740
refGene_gn=GK5;refGene_tx=XM_019810621.2;refGene_gn=GK5;refGene_tx=XM_023260505.1;refGene_gn=GK5;refGene_tx=XM_023260506.1;refGene_gn=GK5;refGene_tx=XM_023260507.1
C2 123715222 123715327
refGene_gn=GK5;refGene_tx=XM_019810621.2;refGene_gn=GK5;refGene_tx=XM_023260505.1;refGene_gn=GK5;refGene_tx=XM_023260506.1;refGene_gn=GK5;refGene_tx=XM_023260507.1
C2 123727615 123727710
refGene_gn=GK5;refGene_tx=XM_019810621.2;refGene_gn=GK5;refGene_tx=XM_023260505.1;refGene_gn=GK5;refGene_tx=XM_023260506.1;refGene_gn=GK5;refGene_tx=XM_023260507.1
C2 123732343 123732447
refGene_gn=GK5;refGene_tx=XM_019810621.2;refGene_gn=GK5;refGene_tx=XM_023260505.1;refGene_gn=GK5;refGene_tx=XM_023260506.1;refGene_gn=GK5;refGene_tx=XM_023260507.1
C2 123733799 123733859
refGene_gn=GK5;refGene_tx=XM_019810621.2;refGene_gn=GK5;refGene_tx=XM_023260505.1;refGene_gn=GK5;refGene_tx=XM_023260506.1;refGene_gn=GK5;refGene_tx=XM_023260507.1
C2 123734607 123734741
refGene_gn=GK5;refGene_tx=XM_019810621.2;refGene_gn=GK5;refGene_tx=XM_023260505.1;refGene_gn=GK5;refGene_tx=XM_023260506.1;refGene_gn=GK5;refGene_tx=XM_023260507.1
C2 123737472 123737627
refGene_gn=GK5;refGene_tx=XM_019810621.2;refGene_gn=GK5;refGene_tx=XM_023260505.1;refGene_gn=GK5;refGene_tx=XM_023260506.1;refGene_gn=GK5;refGene_tx=XM_023260507.1

C2	123801811	123801826
refGene_gn=TFDP2;refGene_tx=XM_003992044.4
C2	123809418	123809485
refGene_gn=TFDP2;refGene_tx=XM_003992044.4
C2	123873947	123874051
refGene_gn=TFDP2;refGene_tx=XM_003992044.4
C2	123881073	123881195
refGene_gn=TFDP2;refGene_tx=XM_003992044.4
C2	123882617	123882665
refGene_gn=TFDP2;refGene_tx=XM_003992044.4
C2	123900425	123900456
refGene_gn=TFDP2;refGene_tx=XM_003992044.4
C2	123900457	123900589
refGene_gn=TFDP2;refGene_tx=XM_003992044.4
C2	123903707	123903854
refGene_gn=TFDP2;refGene_tx=XM_003992044.4
C2	123907134	123907203
refGene_gn=TFDP2;refGene_tx=XM_003992044.4
C2	123917387	123918023
refGene_gn=ATP5S;refGene_tx=XM_023260508.1
C2	123919559	123919711
refGene_gn=TFDP2;refGene_tx=XM_003992044.4
C2	123923302	123923469
refGene_gn=TFDP2;refGene_tx=XM_003992044.4
C2	123927990	123928096
refGene_gn=TFDP2;refGene_tx=XM_003992044.4
C2	123928307	123928491
refGene_gn=TFDP2;refGene_tx=XM_003992044.4
C2	123976717	123976891
refGene_gn=ATP1B3;refGene_tx=XM_023260509.1
C2	123978453	123978540
refGene_gn=ATP1B3;refGene_tx=XM_023260509.1
C2	123982214	123982265
refGene_gn=ATP1B3;refGene_tx=XM_023260509.1
C2	123984356	123984541
refGene_gn=ATP1B3;refGene_tx=XM_023260509.1
C2	123990239	123990347
refGene_gn=ATP1B3;refGene_tx=XM_023260509.1
C2	123995377	123995506
refGene_gn=ATP1B3;refGene_tx=XM_023260509.1
C2	124022031	124022140
refGene_gn=ATP1B3;refGene_tx=XM_023260509.1
C2	124071275	124071304
refGene_gn=GRK7;refGene_tx=XM_023260510.1
C2	124071321	124071658
refGene_gn=GRK7;refGene_tx=XM_003992047.5;refGene_gn=GRK7;refGene_tx=XM_023260510.1
C2	124077484	124077759
refGene_gn=GRK7;refGene_tx=XM_003992047.5;refGene_gn=GRK7;refGene_tx=XM_023260510.1

C2 124095988 124096426
refGene_gn=GRK7;refGene_tx=XM_003992047.5;refGene_gn=GRK7;refGene_tx=XM_023260510.1
C2 124097758 124098370
refGene_gn=GRK7;refGene_tx=XM_003992047.5;refGene_gn=GRK7;refGene_tx=XM_023260510.1
C2 124123163 124123282
refGene_gn=RNF7;refGene_tx=XM_011286152.3;refGene_gn=RNF7;refGene_tx=XM_011286153.3;refGene_gn=RNF7;refGene_tx=XM_011286154.3
C2 124123458 124123582
refGene_gn=RNF7;refGene_tx=XM_011286152.3
C2 124124821 124124869
refGene_gn=RNF7;refGene_tx=XM_011286152.3;refGene_gn=RNF7;refGene_tx=XM_011286153.3;refGene_gn=RNF7;refGene_tx=XM_011286154.3
C2 124128454 124128629
refGene_gn=RNF7;refGene_tx=XM_011286152.3;refGene_gn=RNF7;refGene_tx=XM_011286153.3;refGene_gn=RNF7;refGene_tx=XM_011286154.3
C2 124133499 124133511
refGene_gn=LOC101098082;refGene_tx=XM_023260511.1
C2 124133941 124134054
refGene_gn=LOC101098082;refGene_tx=XM_023260511.1
C2 124145779 124145886
refGene_gn=LOC101098082;refGene_tx=XM_023260511.1
C2 124150263 124150390
refGene_gn=LOC101098082;refGene_tx=XM_023260511.1
C2 124159644 124159740
refGene_gn=LOC101098082;refGene_tx=XM_023260511.1
C2 124168294 124168425
refGene_gn=LOC101098082;refGene_tx=XM_023260511.1
C2 124168822 124168963
refGene_gn=LOC101098082;refGene_tx=XM_023260511.1
C2 124171680 124171829
refGene_gn=LOC101098082;refGene_tx=XM_023260511.1
C2 124178833 124178983
refGene_gn=LOC101098082;refGene_tx=XM_023260511.1
C2 124210860 124210891
refGene_gn=RASA2;refGene_tx=XM_023260512.1;refGene_gn=RASA2;refGene_tx=XM_023260513.1;refGene_gn=RASA2;refGene_tx=XM_023260514.1;refGene_gn=RASA2;refGene_tx=XM_023260515.1
C2 124213196 124213386
refGene_gn=RASA2;refGene_tx=XM_023260512.1;refGene_gn=RASA2;refGene_tx=XM_023260513.1;refGene_gn=RASA2;refGene_tx=XM_023260514.1;refGene_gn=RASA2;refGene_tx=XM_023260515.1
C2 124213738 124213842
refGene_gn=RASA2;refGene_tx=XM_023260512.1;refGene_gn=RASA2;refGene_tx=XM_023260513.1;refGene_gn=RASA2;refGene_tx=XM_023260514.1;refGene_gn=RASA2;refGene_tx=XM_023260515.1
C2 124214537 124214746
refGene_gn=RASA2;refGene_tx=XM_023260512.1;refGene_gn=RASA2;refGene_tx=XM_023260513.1;refGene_gn=RASA2;refGene_tx=XM_023260514.1;refGene_gn=

RASA2;refGene_tx=XM_023260515.1
C2	124215958	124216041
refGene_gn=RASA2;refGene_tx=XM_023260512.1;refGene_gn=RASA2;refGene_tx=XM_023260513.1;refGene_gn=RASA2;refGene_tx=XM_023260514.1;refGene_gn=RASA2;refGene_tx=XM_023260515.1
C2	124230273	124230285
refGene_gn=RASA2;refGene_tx=XM_023260512.1;refGene_gn=RASA2;refGene_tx=XM_023260513.1
C2	124232636	124232746
refGene_gn=RASA2;refGene_tx=XM_023260512.1;refGene_gn=RASA2;refGene_tx=XM_023260513.1;refGene_gn=RASA2;refGene_tx=XM_023260514.1;refGene_gn=RASA2;refGene_tx=XM_023260515.1
C2	124233284	124233358
refGene_gn=RASA2;refGene_tx=XM_023260512.1;refGene_gn=RASA2;refGene_tx=XM_023260513.1;refGene_gn=RASA2;refGene_tx=XM_023260514.1;refGene_gn=RASA2;refGene_tx=XM_023260515.1
C2	124236952	124237030
refGene_gn=RASA2;refGene_tx=XM_023260512.1;refGene_gn=RASA2;refGene_tx=XM_023260513.1;refGene_gn=RASA2;refGene_tx=XM_023260514.1;refGene_gn=RASA2;refGene_tx=XM_023260515.1
C2	124237675	124237759
refGene_gn=RASA2;refGene_tx=XM_023260512.1;refGene_gn=RASA2;refGene_tx=XM_023260513.1;refGene_gn=RASA2;refGene_tx=XM_023260514.1;refGene_gn=RASA2;refGene_tx=XM_023260515.1
C2	124241208	124241315
refGene_gn=RASA2;refGene_tx=XM_023260512.1;refGene_gn=RASA2;refGene_tx=XM_023260513.1;refGene_gn=RASA2;refGene_tx=XM_023260514.1;refGene_gn=RASA2;refGene_tx=XM_023260515.1
C2	124243675	124243799
refGene_gn=RASA2;refGene_tx=XM_023260512.1;refGene_gn=RASA2;refGene_tx=XM_023260513.1;refGene_gn=RASA2;refGene_tx=XM_023260514.1;refGene_gn=RASA2;refGene_tx=XM_023260515.1
C2	124244498	124244573
refGene_gn=RASA2;refGene_tx=XM_023260512.1;refGene_gn=RASA2;refGene_tx=XM_023260513.1;refGene_gn=RASA2;refGene_tx=XM_023260514.1;refGene_gn=RASA2;refGene_tx=XM_023260515.1
C2	124245004	124245119
refGene_gn=RASA2;refGene_tx=XM_023260512.1;refGene_gn=RASA2;refGene_tx=XM_023260513.1;refGene_gn=RASA2;refGene_tx=XM_023260514.1;refGene_gn=RASA2;refGene_tx=XM_023260515.1
C2	124245756	124245905
refGene_gn=RASA2;refGene_tx=XM_023260512.1;refGene_gn=RASA2;refGene_tx=XM_023260513.1;refGene_gn=RASA2;refGene_tx=XM_023260514.1;refGene_gn=RASA2;refGene_tx=XM_023260515.1
C2	124246213	124246370
refGene_gn=RASA2;refGene_tx=XM_023260512.1;refGene_gn=RASA2;refGene_tx=XM_023260513.1;refGene_gn=RASA2;refGene_tx=XM_023260514.1;refGene_gn=RASA2;refGene_tx=XM_023260515.1
C2	124252268	124252370
refGene_gn=RASA2;refGene_tx=XM_023260512.1;refGene_gn=RASA2;refGene_tx

=XM_023260513.1;refGene_gn=RASA2;refGene_tx=XM_023260514.1;refGene_gn=RASA2;refGene_tx=XM_023260515.1
C2	124253324	124253401
refGene_gn=RASA2;refGene_tx=XM_023260512.1;refGene_gn=RASA2;refGene_tx=XM_023260513.1;refGene_gn=RASA2;refGene_tx=XM_023260514.1;refGene_gn=RASA2;refGene_tx=XM_023260515.1
C2	124256884	124256957
refGene_gn=RASA2;refGene_tx=XM_023260512.1;refGene_gn=RASA2;refGene_tx=XM_023260513.1;refGene_gn=RASA2;refGene_tx=XM_023260514.1;refGene_gn=RASA2;refGene_tx=XM_023260515.1
C2	124259373	124259457
refGene_gn=RASA2;refGene_tx=XM_023260512.1;refGene_gn=RASA2;refGene_tx=XM_023260513.1;refGene_gn=RASA2;refGene_tx=XM_023260514.1;refGene_gn=RASA2;refGene_tx=XM_023260515.1
C2	124275944	124276021
refGene_gn=RASA2;refGene_tx=XM_023260512.1;refGene_gn=RASA2;refGene_tx=XM_023260513.1;refGene_gn=RASA2;refGene_tx=XM_023260514.1;refGene_gn=RASA2;refGene_tx=XM_023260515.1
C2	124287394	124287489
refGene_gn=RASA2;refGene_tx=XM_023260512.1;refGene_gn=RASA2;refGene_tx=XM_023260513.1;refGene_gn=RASA2;refGene_tx=XM_023260514.1;refGene_gn=RASA2;refGene_tx=XM_023260515.1
C2	124296619	124296723
refGene_gn=RASA2;refGene_tx=XM_023260512.1;refGene_gn=RASA2;refGene_tx=XM_023260513.1;refGene_gn=RASA2;refGene_tx=XM_023260514.1;refGene_gn=RASA2;refGene_tx=XM_023260515.1
C2	124305552	124305670
refGene_gn=RASA2;refGene_tx=XM_023260512.1;refGene_gn=RASA2;refGene_tx=XM_023260513.1;refGene_gn=RASA2;refGene_tx=XM_023260514.1;refGene_gn=RASA2;refGene_tx=XM_023260515.1
C2	124333602	124333735
refGene_gn=RASA2;refGene_tx=XM_023260512.1;refGene_gn=RASA2;refGene_tx=XM_023260513.1;refGene_gn=RASA2;refGene_tx=XM_023260514.1;refGene_gn=RASA2;refGene_tx=XM_023260515.1
C2	124380862	124384447
refGene_gn=ZBTB38;refGene_tx=XM_006936364.4;refGene_gn=ZBTB38;refGene_tx=XM_006936366.4;refGene_gn=ZBTB38;refGene_tx=XM_006936367.4;refGene_gn=ZBTB38;refGene_tx=XM_011286149.3;refGene_gn=ZBTB38;refGene_tx=XM_019810627.2;refGene_gn=ZBTB38;refGene_tx=XM_019810628.2;refGene_gn=ZBTB38;refGene_tx=XM_019810629.2;refGene_gn=ZBTB38;refGene_tx=XM_019810630.2;refGene_gn=ZBTB38;refGene_tx=XM_019810631.2
C2	124429275	124429320
refGene_gn=ZBTB38;refGene_tx=XM_019810627.2
C2	124530011	124530952
refGene_gn=PXYLP1;refGene_tx=XM_003992050.5;refGene_gn=PXYLP1;refGene_tx=XM_006936369.4;refGene_gn=PXYLP1;refGene_tx=XM_019810635.2;refGene_gn=PXYLP1;refGene_tx=XM_019810637.2;refGene_gn=PXYLP1;refGene_tx=XM_023260516.1;refGene_gn=PXYLP1;refGene_tx=XM_023260517.1;refGene_gn=PXYLP1;refGene_tx=XM_023260518.1
C2	124536395	124536535

refGene_gn=PXYLP1;refGene_tx=XM_003992050.5;refGene_gn=PXYLP1;refGene_tx=XM_006936369.4;refGene_gn=PXYLP1;refGene_tx=XM_019810635.2;refGene_gn=PXYLP1;refGene_tx=XM_019810637.2;refGene_gn=PXYLP1;refGene_tx=XM_023260516.1;refGene_gn=PXYLP1;refGene_tx=XM_023260517.1;refGene_gn=PXYLP1;refGene_tx=XM_023260518.1
C2 124546656 124546783
refGene_gn=PXYLP1;refGene_tx=XM_003992050.5;refGene_gn=PXYLP1;refGene_tx=XM_006936369.4;refGene_gn=PXYLP1;refGene_tx=XM_019810635.2;refGene_gn=PXYLP1;refGene_tx=XM_019810637.2;refGene_gn=PXYLP1;refGene_tx=XM_023260516.1;refGene_gn=PXYLP1;refGene_tx=XM_023260517.1;refGene_gn=PXYLP1;refGene_tx=XM_023260518.1
C2 124547634 124547793
refGene_gn=PXYLP1;refGene_tx=XM_003992050.5;refGene_gn=PXYLP1;refGene_tx=XM_006936369.4;refGene_gn=PXYLP1;refGene_tx=XM_019810635.2;refGene_gn=PXYLP1;refGene_tx=XM_019810637.2;refGene_gn=PXYLP1;refGene_tx=XM_023260516.1;refGene_gn=PXYLP1;refGene_tx=XM_023260517.1;refGene_gn=PXYLP1;refGene_tx=XM_023260518.1
C2 124562236 124562267
refGene_gn=PXYLP1;refGene_tx=XM_019810635.2
C2 124564622 124564701
refGene_gn=PXYLP1;refGene_tx=XM_003992050.5;refGene_gn=PXYLP1;refGene_tx=XM_006936369.4;refGene_gn=PXYLP1;refGene_tx=XM_023260516.1;refGene_gn=PXYLP1;refGene_tx=XM_023260517.1;refGene_gn=PXYLP1;refGene_tx=XM_023260518.1
C2 124660008 124660136
refGene_gn=SPSB4;refGene_tx=XM_006936390.4
C2 124723202 124723896
refGene_gn=SPSB4;refGene_tx=XM_006936390.4
C2 124797949 124798143
refGene_gn=SLC25A36;refGene_tx=XM_023260519.1
C2 124800686 124800976
refGene_gn=SLC25A36;refGene_tx=XM_023260519.1
C2 124803869 124803936
refGene_gn=SLC25A36;refGene_tx=XM_023260519.1
C2 124811679 124811780
refGene_gn=SLC25A36;refGene_tx=XM_023260519.1
C2 124815096 124815174
refGene_gn=SLC25A36;refGene_tx=XM_023260519.1
C2 124818629 124818794
refGene_gn=SLC25A36;refGene_tx=XM_023260519.1
C2 124833335 124833376
refGene_gn=SLC25A36;refGene_tx=XM_023260519.1
C2 125009750 125009837
refGene_gn=TRIM42;refGene_tx=XM_003992051.4
C2 125018680 125018905
refGene_gn=TRIM42;refGene_tx=XM_003992051.4
C2 125021875 125022699
refGene_gn=TRIM42;refGene_tx=XM_003992051.4
C2 125028570 125029268
refGene_gn=TRIM42;refGene_tx=XM_003992051.4

C2 125033081 125033422
refGene_gn=TRIM42;refGene_tx=XM_003992051.4
C2 125132480 125132684
refGene_gn=CLSTN2;refGene_tx=XM_003992052.5;refGene_gn=CLSTN2;refGene_tx=XM_019810639.2;refGene_gn=CLSTN2;refGene_tx=XM_019810642.2
C2 125134534 125134719
refGene_gn=CLSTN2;refGene_tx=XM_003992052.5;refGene_gn=CLSTN2;refGene_tx=XM_019810639.2;refGene_gn=CLSTN2;refGene_tx=XM_019810642.2
C2 125135394 125135518
refGene_gn=CLSTN2;refGene_tx=XM_003992052.5;refGene_gn=CLSTN2;refGene_tx=XM_019810639.2;refGene_gn=CLSTN2;refGene_tx=XM_019810642.2
C2 125135625 125135771
refGene_gn=CLSTN2;refGene_tx=XM_003992052.5;refGene_gn=CLSTN2;refGene_tx=XM_019810639.2;refGene_gn=CLSTN2;refGene_tx=XM_019810642.2
C2 125136244 125136415
refGene_gn=CLSTN2;refGene_tx=XM_003992052.5;refGene_gn=CLSTN2;refGene_tx=XM_019810639.2;refGene_gn=CLSTN2;refGene_tx=XM_019810642.2
C2 125146464 125146682
refGene_gn=CLSTN2;refGene_tx=XM_003992052.5;refGene_gn=CLSTN2;refGene_tx=XM_019810639.2;refGene_gn=CLSTN2;refGene_tx=XM_019810642.2
C2 125148593 125148742
refGene_gn=CLSTN2;refGene_tx=XM_003992052.5;refGene_gn=CLSTN2;refGene_tx=XM_019810639.2;refGene_gn=CLSTN2;refGene_tx=XM_019810642.2
C2 125157140 125157307
refGene_gn=CLSTN2;refGene_tx=XM_003992052.5;refGene_gn=CLSTN2;refGene_tx=XM_019810639.2;refGene_gn=CLSTN2;refGene_tx=XM_019810642.2
C2 125174236 125174399
refGene_gn=CLSTN2;refGene_tx=XM_003992052.5;refGene_gn=CLSTN2;refGene_tx=XM_019810639.2;refGene_gn=CLSTN2;refGene_tx=XM_019810642.2
C2 125223670 125223792
refGene_gn=CLSTN2;refGene_tx=XM_003992052.5;refGene_gn=CLSTN2;refGene_tx=XM_019810639.2;refGene_gn=CLSTN2;refGene_tx=XM_019810642.2
C2 125228951 125229200
refGene_gn=CLSTN2;refGene_tx=XM_003992052.5;refGene_gn=CLSTN2;refGene_tx=XM_019810639.2;refGene_gn=CLSTN2;refGene_tx=XM_019810642.2
C2 125239231 125239417
refGene_gn=CLSTN2;refGene_tx=XM_003992052.5;refGene_gn=CLSTN2;refGene_tx=XM_019810639.2;refGene_gn=CLSTN2;refGene_tx=XM_019810642.2
C2 125267429 125267579
refGene_gn=CLSTN2;refGene_tx=XM_003992052.5;refGene_gn=CLSTN2;refGene_tx=XM_019810639.2;refGene_gn=CLSTN2;refGene_tx=XM_019810642.2
C2 125274196 125274405
refGene_gn=CLSTN2;refGene_tx=XM_003992052.5;refGene_gn=CLSTN2;refGene_tx=XM_019810639.2;refGene_gn=CLSTN2;refGene_tx=XM_019810642.2
C2 125275146 125275342
refGene_gn=CLSTN2;refGene_tx=XM_003992052.5;refGene_gn=CLSTN2;refGene_tx=XM_019810639.2;refGene_gn=CLSTN2;refGene_tx=XM_019810642.2
C2 125294889 125294893
refGene_gn=CLSTN2;refGene_tx=XM_019810642.2
C2 125516987 125517110

refGene_gn=CLSTN2;refGene_tx=XM_003992052.5;refGene_gn=CLSTN2;refGene_tx=XM_019810639.2
C2 125536560 125536663
refGene_gn=CLSTN2;refGene_tx=XM_019810639.2
C2 125740655 125740764
refGene_gn=CLSTN2;refGene_tx=XM_003992052.5
C2 125999597 125999706
refGene_gn=NMNAT3;refGene_tx=XM_011286147.2;refGene_gn=NMNAT3;refGene_tx=XM_011286148.3
C2 126050893 126051077
refGene_gn=NMNAT3;refGene_tx=XM_011286147.2;refGene_gn=NMNAT3;refGene_tx=XM_011286148.3
C2 126055958 126056041
refGene_gn=NMNAT3;refGene_tx=XM_011286147.2;refGene_gn=NMNAT3;refGene_tx=XM_011286148.3
C2 126059569 126059835
refGene_gn=NMNAT3;refGene_tx=XM_011286148.3
C2 126062901 126063242
refGene_gn=NMNAT3;refGene_tx=XM_011286147.2
C2 126079649 126079665
refGene_gn=RBP1;refGene_tx=XM_023260183.1
C2 126081196 126081646
refGene_gn=RBP1;refGene_tx=XM_023260183.1
C2 126082099 126082278
refGene_gn=RBP1;refGene_tx=XM_023260183.1
C2 126103866 126103968
refGene_gn=RBP1;refGene_tx=XM_023260183.1
C2 126105813 126105867
refGene_gn=RBP1;refGene_tx=XM_023260183.1
C2 126151960 126152033
refGene_gn=RBP2;refGene_tx=XM_003992053.4;refGene_gn=RBP2;refGene_tx=XM_019810643.2;refGene_gn=RBP2;refGene_tx=XM_019810644.2;refGene_gn=RBP2;refGene_tx=XM_019810645.2
C2 126165045 126165224
refGene_gn=RBP2;refGene_tx=XM_003992053.4;refGene_gn=RBP2;refGene_tx=XM_019810643.2;refGene_gn=RBP2;refGene_tx=XM_019810644.2;refGene_gn=RBP2;refGene_tx=XM_019810645.2
C2 126172871 126172973
refGene_gn=RBP2;refGene_tx=XM_003992053.4;refGene_gn=RBP2;refGene_tx=XM_019810643.2;refGene_gn=RBP2;refGene_tx=XM_019810644.2;refGene_gn=RBP2;refGene_tx=XM_019810645.2
C2 126174546 126174597
refGene_gn=RBP2;refGene_tx=XM_003992053.4;refGene_gn=RBP2;refGene_tx=XM_019810643.2;refGene_gn=RBP2;refGene_tx=XM_019810644.2;refGene_gn=RBP2;refGene_tx=XM_019810645.2
C2 126244933 126244936
refGene_gn=COPB2;refGene_tx=XM_003992054.4
C2 126251477 126251615
refGene_gn=COPB2;refGene_tx=XM_003992054.4
C2 126255080 126255167

refGene_gn=COPB2;refGene_tx=XM_003992054.4
C2 126255602 126255729
refGene_gn=COPB2;refGene_tx=XM_003992054.4
C2 126256573 126256722
refGene_gn=COPB2;refGene_tx=XM_003992054.4
C2 126258814 126258961
refGene_gn=COPB2;refGene_tx=XM_003992054.4
C2 126260035 126260135
refGene_gn=COPB2;refGene_tx=XM_003992054.4
C2 126260756 126260899
refGene_gn=COPB2;refGene_tx=XM_003992054.4
C2 126261414 126261614
refGene_gn=COPB2;refGene_tx=XM_003992054.4
C2 126263256 126263367
refGene_gn=COPB2;refGene_tx=XM_003992054.4
C2 126265085 126265174
refGene_gn=COPB2;refGene_tx=XM_003992054.4
C2 126265258 126265365
refGene_gn=COPB2;refGene_tx=XM_003992054.4
C2 126266515 126266659
refGene_gn=COPB2;refGene_tx=XM_003992054.4
C2 126267299 126267430
refGene_gn=COPB2;refGene_tx=XM_003992054.4
C2 126267729 126267937
refGene_gn=COPB2;refGene_tx=XM_003992054.4
C2 126271749 126271860
refGene_gn=COPB2;refGene_tx=XM_003992054.4
C2 126273033 126273248
refGene_gn=COPB2;refGene_tx=XM_003992054.4
C2 126274654 126274747
refGene_gn=COPB2;refGene_tx=XM_003992054.4
C2 126274840 126275021
refGene_gn=COPB2;refGene_tx=XM_003992054.4
C2 126275168 126275237
refGene_gn=COPB2;refGene_tx=XM_003992054.4
C2 126275388 126275460
refGene_gn=COPB2;refGene_tx=XM_003992054.4
C2 126275687 126275780
refGene_gn=COPB2;refGene_tx=XM_003992054.4
C2 126276594 126276690
refGene_gn=MRPS22;refGene_tx=XM_003992056.5
C2 126277651 126277760
refGene_gn=MRPS22;refGene_tx=XM_003992056.5
C2 126281309 126281455
refGene_gn=MRPS22;refGene_tx=XM_003992056.5
C2 126282710 126282794
refGene_gn=MRPS22;refGene_tx=XM_003992056.5
C2 126283680 126283824
refGene_gn=MRPS22;refGene_tx=XM_003992056.5
C2 126285618 126285783

refGene_gn=MRPS22;refGene_tx=XM_003992056.5
C2	126286854	126287018
refGene_gn=MRPS22;refGene_tx=XM_003992056.5
C2	126297722	126298146
refGene_gn=MRPS22;refGene_tx=XM_003992056.5
C2	126599620	126600373
refGene_gn=LOC101100079;refGene_tx=XM_023238570.1
C2	126610753	126611506
refGene_gn=LOC101091647;refGene_tx=XM_003992057.3
C2	126656431	126657123
refGene_gn=FOXL2NB;refGene_tx=XM_006944357.3
C2	126659472	126659572
refGene_gn=FOXL2NB;refGene_tx=XM_006944357.3
C2	126660221	126660236
refGene_gn=FOXL2;refGene_tx=XM_023260523.1
C2	126660237	126661362
refGene_gn=FOXL2;refGene_tx=XM_023260523.1
C2	126706432	126706438
refGene_gn=LOC101100587;refGene_tx=XM_023260184.1
C2	126712598	126712725
refGene_gn=LOC101100587;refGene_tx=XM_023260184.1
C2	126712727	126712820
refGene_gn=LOC101100587;refGene_tx=XM_023260184.1
C2	126712822	126713229
refGene_gn=LOC101100587;refGene_tx=XM_023260184.1
C2	126840829	126841000
refGene_gn=PIK3CB;refGene_tx=XM_011286143.3
C2	126846345	126846571
refGene_gn=PIK3CB;refGene_tx=XM_011286143.3
C2	126852830	126853054
refGene_gn=PIK3CB;refGene_tx=XM_011286143.3
C2	126863183	126863363
refGene_gn=PIK3CB;refGene_tx=XM_011286143.3
C2	126866620	126866791
refGene_gn=PIK3CB;refGene_tx=XM_011286143.3
C2	126868220	126868298
refGene_gn=PIK3CB;refGene_tx=XM_011286143.3
C2	126879413	126879665
refGene_gn=PIK3CB;refGene_tx=XM_011286143.3
C2	126881461	126881558
refGene_gn=PIK3CB;refGene_tx=XM_011286143.3
C2	126884477	126884608
refGene_gn=PIK3CB;refGene_tx=XM_011286143.3
C2	126888578	126888629
refGene_gn=PIK3CB;refGene_tx=XM_011286143.3
C2	126895345	126895534
refGene_gn=PIK3CB;refGene_tx=XM_011286143.3
C2	126900964	126901086
refGene_gn=PIK3CB;refGene_tx=XM_011286143.3
C2	126904931	126905075

refGene_gn=PIK3CB;refGene_tx=XM_011286143.3
C2 126906280 126906296
refGene_gn=PIK3CB;refGene_tx=XM_011286143.3
C2 126906297 126906381
refGene_gn=PIK3CB;refGene_tx=XM_011286143.3
C2 126921786 126921965
refGene_gn=PIK3CB;refGene_tx=XM_011286143.3
C2 126922850 126922960
refGene_gn=PIK3CB;refGene_tx=XM_011286143.3
C2 126924430 126924509
refGene_gn=PIK3CB;refGene_tx=XM_011286143.3
C2 126940279 126940447
refGene_gn=PIK3CB;refGene_tx=XM_011286143.3
C2 126941478 126941602
refGene_gn=PIK3CB;refGene_tx=XM_011286143.3
C2 126942700 126942846
refGene_gn=PIK3CB;refGene_tx=XM_011286143.3
C2 126944098 126944231
refGene_gn=PIK3CB;refGene_tx=XM_011286143.3
C2 126944506 126944644
refGene_gn=PIK3CB;refGene_tx=XM_011286143.3
C2 126961960 126962110
refGene_gn=FAIM;refGene_tx=XM_006936373.4;refGene_gn=FAIM;refGene_tx=XM_006936374.4;refGene_gn=FAIM;refGene_tx=XM_019810648.2;refGene_gn=FAIM;refGene_tx=XM_019810649.2;refGene_gn=FAIM;refGene_tx=XM_019810651.2;refGene_gn=FAIM;refGene_tx=XM_023260524.1;refGene_gn=FAIM;refGene_tx=XM_023260525.1
C2 126965153 126965203
refGene_gn=FAIM;refGene_tx=XM_006936373.4;refGene_gn=FAIM;refGene_tx=XM_006936374.4;refGene_gn=FAIM;refGene_tx=XM_019810648.2;refGene_gn=FAIM;refGene_tx=XM_019810649.2;refGene_gn=FAIM;refGene_tx=XM_019810651.2;refGene_gn=FAIM;refGene_tx=XM_023260524.1;refGene_gn=FAIM;refGene_tx=XM_023260525.1
C2 126970242 126970471
refGene_gn=FAIM;refGene_tx=XM_006936373.4;refGene_gn=FAIM;refGene_tx=XM_006936374.4;refGene_gn=FAIM;refGene_tx=XM_019810648.2;refGene_gn=FAIM;refGene_tx=XM_019810649.2;refGene_gn=FAIM;refGene_tx=XM_019810651.2;refGene_gn=FAIM;refGene_tx=XM_023260524.1;refGene_gn=FAIM;refGene_tx=XM_023260525.1
C2 126971006 126971139
refGene_gn=FAIM;refGene_tx=XM_006936373.4;refGene_gn=FAIM;refGene_tx=XM_006936374.4;refGene_gn=FAIM;refGene_tx=XM_019810648.2;refGene_gn=FAIM;refGene_tx=XM_019810649.2;refGene_gn=FAIM;refGene_tx=XM_019810651.2;refGene_gn=FAIM;refGene_tx=XM_023260524.1;refGene_gn=FAIM;refGene_tx=XM_023260525.1
C2 126972224 126972283
refGene_gn=FAIM;refGene_tx=XM_019810648.2;refGene_gn=FAIM;refGene_tx=XM_019810649.2;refGene_gn=FAIM;refGene_tx=XM_023260524.1
C2 126975917 126975956
refGene_gn=FAIM;refGene_tx=XM_019810648.2

C2	127007013	127007028
refGene_gn=CEP70;refGene_tx=XM_023260526.1
C2	127009428	127009519
refGene_gn=CEP70;refGene_tx=XM_023260526.1
C2	127009642	127009764
refGene_gn=CEP70;refGene_tx=XM_023260526.1
C2	127010112	127010202
refGene_gn=CEP70;refGene_tx=XM_023260526.1
C2	127010203	127010296
refGene_gn=CEP70;refGene_tx=XM_023260526.1
C2	127019640	127019810
refGene_gn=CEP70;refGene_tx=XM_023260526.1
C2	127024168	127024225
refGene_gn=CEP70;refGene_tx=XM_023260526.1
C2	127034705	127034793
refGene_gn=CEP70;refGene_tx=XM_023260526.1
C2	127034880	127034969
refGene_gn=CEP70;refGene_tx=XM_023260526.1
C2	127036026	127036101
refGene_gn=CEP70;refGene_tx=XM_023260526.1
C2	127042059	127042165
refGene_gn=CEP70;refGene_tx=XM_023260526.1
C2	127049544	127049715
refGene_gn=CEP70;refGene_tx=XM_023260526.1
C2	127053243	127053390
refGene_gn=CEP70;refGene_tx=XM_023260526.1
C2	127053563	127053732
refGene_gn=CEP70;refGene_tx=XM_023260526.1
C2	127054012	127054127
refGene_gn=CEP70;refGene_tx=XM_023260526.1
C2	127056097	127056177
refGene_gn=CEP70;refGene_tx=XM_023260526.1
C2	127059498	127059560
refGene_gn=CEP70;refGene_tx=XM_023260526.1
C2	127079541	127079578
refGene_gn=ESYT3;refGene_tx=XM_003992061.5;refGene_gn=ESYT3;refGene_tx=XM_023260527.1;refGene_gn=ESYT3;refGene_tx=XM_023260528.1;refGene_gn=ESYT3;refGene_tx=XM_023260530.1
C2	127079924	127079974
refGene_gn=ESYT3;refGene_tx=XM_003992061.5;refGene_gn=ESYT3;refGene_tx=XM_023260527.1;refGene_gn=ESYT3;refGene_tx=XM_023260528.1;refGene_gn=ESYT3;refGene_tx=XM_023260530.1
C2	127081001	127081107
refGene_gn=ESYT3;refGene_tx=XM_003992061.5;refGene_gn=ESYT3;refGene_tx=XM_023260527.1;refGene_gn=ESYT3;refGene_tx=XM_023260528.1;refGene_gn=ESYT3;refGene_tx=XM_023260530.1
C2	127088310	127088442
refGene_gn=ESYT3;refGene_tx=XM_003992061.5;refGene_gn=ESYT3;refGene_tx=XM_023260528.1;refGene_gn=ESYT3;refGene_tx=XM_023260530.1
C2	127088980	127089083

refGene_gn=ESYT3;refGene_tx=XM_003992061.5;refGene_gn=ESYT3;refGene_tx=XM_023260527.1;refGene_gn=ESYT3;refGene_tx=XM_023260528.1;refGene_gn=ESYT3;refGene_tx=XM_023260529.1;refGene_gn=ESYT3;refGene_tx=XM_023260530.1
C2 127089721 127090218
refGene_gn=ESYT3;refGene_tx=XM_003992061.5;refGene_gn=ESYT3;refGene_tx=XM_023260527.1;refGene_gn=ESYT3;refGene_tx=XM_023260528.1;refGene_gn=ESYT3;refGene_tx=XM_023260529.1;refGene_gn=ESYT3;refGene_tx=XM_023260530.1
C2 127091808 127091958
refGene_gn=ESYT3;refGene_tx=XM_003992061.5;refGene_gn=ESYT3;refGene_tx=XM_023260527.1;refGene_gn=ESYT3;refGene_tx=XM_023260529.1;refGene_gn=ESYT3;refGene_tx=XM_023260530.1
C2 127092649 127092736
refGene_gn=ESYT3;refGene_tx=XM_003992061.5;refGene_gn=ESYT3;refGene_tx=XM_023260527.1;refGene_gn=ESYT3;refGene_tx=XM_023260528.1;refGene_gn=ESYT3;refGene_tx=XM_023260529.1;refGene_gn=ESYT3;refGene_tx=XM_023260530.1
C2 127093734 127093803
refGene_gn=ESYT3;refGene_tx=XM_003992061.5;refGene_gn=ESYT3;refGene_tx=XM_023260527.1;refGene_gn=ESYT3;refGene_tx=XM_023260528.1;refGene_gn=ESYT3;refGene_tx=XM_023260529.1;refGene_gn=ESYT3;refGene_tx=XM_023260530.1
C2 127094365 127094428
refGene_gn=ESYT3;refGene_tx=XM_003992061.5;refGene_gn=ESYT3;refGene_tx=XM_023260527.1;refGene_gn=ESYT3;refGene_tx=XM_023260528.1;refGene_gn=ESYT3;refGene_tx=XM_023260529.1;refGene_gn=ESYT3;refGene_tx=XM_023260530.1
C2 127094527 127094590
refGene_gn=ESYT3;refGene_tx=XM_003992061.5;refGene_gn=ESYT3;refGene_tx=XM_023260527.1;refGene_gn=ESYT3;refGene_tx=XM_023260528.1;refGene_gn=ESYT3;refGene_tx=XM_023260529.1;refGene_gn=ESYT3;refGene_tx=XM_023260530.1
C2 127095077 127095167
refGene_gn=ESYT3;refGene_tx=XM_003992061.5;refGene_gn=ESYT3;refGene_tx=XM_023260527.1;refGene_gn=ESYT3;refGene_tx=XM_023260528.1;refGene_gn=ESYT3;refGene_tx=XM_023260529.1;refGene_gn=ESYT3;refGene_tx=XM_023260530.1
C2 127095438 127095487
refGene_gn=ESYT3;refGene_tx=XM_003992061.5;refGene_gn=ESYT3;refGene_tx=XM_023260527.1;refGene_gn=ESYT3;refGene_tx=XM_023260528.1;refGene_gn=ESYT3;refGene_tx=XM_023260529.1;refGene_gn=ESYT3;refGene_tx=XM_023260530.1
C2 127097322 127097405
refGene_gn=ESYT3;refGene_tx=XM_003992061.5;refGene_gn=ESYT3;refGene_tx=XM_023260527.1;refGene_gn=ESYT3;refGene_tx=XM_023260528.1;refGene_gn=ESYT3;refGene_tx=XM_023260529.1;refGene_gn=ESYT3;refGene_tx=XM_023260530.1
C2 127098079 127098250
refGene_gn=ESYT3;refGene_tx=XM_003992061.5;refGene_gn=ESYT3;refGene_tx

=XM_023260527.1;refGene_gn=ESYT3;refGene_tx=XM_023260528.1;refGene_gn=ESYT3;refGene_tx=XM_023260529.1;refGene_gn=ESYT3;refGene_tx=XM_023260530.1
C2	127100353	127100474
refGene_gn=ESYT3;refGene_tx=XM_003992061.5;refGene_gn=ESYT3;refGene_tx=XM_023260527.1;refGene_gn=ESYT3;refGene_tx=XM_023260528.1;refGene_gn=ESYT3;refGene_tx=XM_023260529.1;refGene_gn=ESYT3;refGene_tx=XM_023260530.1
C2	127101760	127101816
refGene_gn=ESYT3;refGene_tx=XM_003992061.5;refGene_gn=ESYT3;refGene_tx=XM_023260527.1;refGene_gn=ESYT3;refGene_tx=XM_023260528.1;refGene_gn=ESYT3;refGene_tx=XM_023260529.1;refGene_gn=ESYT3;refGene_tx=XM_023260530.1
C2	127102339	127102429
refGene_gn=ESYT3;refGene_tx=XM_003992061.5;refGene_gn=ESYT3;refGene_tx=XM_023260527.1;refGene_gn=ESYT3;refGene_tx=XM_023260528.1;refGene_gn=ESYT3;refGene_tx=XM_023260529.1;refGene_gn=ESYT3;refGene_tx=XM_023260530.1
C2	127103063	127103130
refGene_gn=ESYT3;refGene_tx=XM_003992061.5;refGene_gn=ESYT3;refGene_tx=XM_023260527.1;refGene_gn=ESYT3;refGene_tx=XM_023260528.1;refGene_gn=ESYT3;refGene_tx=XM_023260529.1;refGene_gn=ESYT3;refGene_tx=XM_023260530.1
C2	127104624	127104701
refGene_gn=ESYT3;refGene_tx=XM_003992061.5;refGene_gn=ESYT3;refGene_tx=XM_023260527.1;refGene_gn=ESYT3;refGene_tx=XM_023260528.1;refGene_gn=ESYT3;refGene_tx=XM_023260529.1;refGene_gn=ESYT3;refGene_tx=XM_023260530.1
C2	127107891	127108026
refGene_gn=ESYT3;refGene_tx=XM_003992061.5;refGene_gn=ESYT3;refGene_tx=XM_023260527.1;refGene_gn=ESYT3;refGene_tx=XM_023260528.1;refGene_gn=ESYT3;refGene_tx=XM_023260529.1;refGene_gn=ESYT3;refGene_tx=XM_023260530.1
C2	127110216	127110258
refGene_gn=ESYT3;refGene_tx=XM_003992061.5;refGene_gn=ESYT3;refGene_tx=XM_023260527.1;refGene_gn=ESYT3;refGene_tx=XM_023260528.1;refGene_gn=ESYT3;refGene_tx=XM_023260529.1
C2	127127314	127127632
refGene_gn=ESYT3;refGene_tx=XM_003992061.5;refGene_gn=ESYT3;refGene_tx=XM_023260527.1;refGene_gn=ESYT3;refGene_tx=XM_023260528.1;refGene_gn=ESYT3;refGene_tx=XM_023260529.1
C2	127161318	127161418
refGene_gn=MRAS;refGene_tx=XM_006936375.2;refGene_gn=MRAS;refGene_tx=XM_006936376.3;refGene_gn=MRAS;refGene_tx=XM_019810658.2;refGene_gn=MRAS;refGene_tx=XM_019810659.2;refGene_gn=MRAS;refGene_tx=XM_023260531.1
C2	127162740	127162820
refGene_gn=MRAS;refGene_tx=XM_006936375.2;refGene_gn=MRAS;refGene_tx=XM_006936376.3;refGene_gn=MRAS;refGene_tx=XM_019810658.2;refGene_gn=MRAS;refGene_tx=XM_019810659.2;refGene_gn=MRAS;refGene_tx=XM_023260531.1
C2	127167169	127167269

refGene_gn=MRAS;refGene_tx=XM_006936375.2;refGene_gn=MRAS;refGene_tx=XM_006936376.3;refGene_gn=MRAS;refGene_tx=XM_019810658.2;refGene_gn=MRAS;refGene_tx=XM_019810659.2;refGene_gn=MRAS;refGene_tx=XM_023260531.1
C2 127168840 127168994
refGene_gn=MRAS;refGene_tx=XM_006936375.2;refGene_gn=MRAS;refGene_tx=XM_006936376.3;refGene_gn=MRAS;refGene_tx=XM_019810658.2;refGene_gn=MRAS;refGene_tx=XM_019810659.2;refGene_gn=MRAS;refGene_tx=XM_023260531.1
C2 127188754 127188947
refGene_gn=MRAS;refGene_tx=XM_019810658.2;refGene_gn=MRAS;refGene_tx=XM_019810659.2;refGene_gn=MRAS;refGene_tx=XM_023260531.1
C2 127230248 127230281
refGene_gn=NME9;refGene_tx=XM_006936377.3
C2 127236619 127236677
refGene_gn=NME9;refGene_tx=XM_006936377.3;refGene_gn=NME9;refGene_tx=XM_023260532.1
C2 127241125 127241229
refGene_gn=NME9;refGene_tx=XM_006936377.3;refGene_gn=NME9;refGene_tx=XM_023260532.1;refGene_gn=NME9;refGene_tx=XM_023260533.1
C2 127242275 127242347
refGene_gn=NME9;refGene_tx=XM_006936377.3;refGene_gn=NME9;refGene_tx=XM_023260532.1;refGene_gn=NME9;refGene_tx=XM_023260533.1
C2 127244710 127244827
refGene_gn=NME9;refGene_tx=XM_006936377.3;refGene_gn=NME9;refGene_tx=XM_023260532.1;refGene_gn=NME9;refGene_tx=XM_023260533.1
C2 127246085 127246161
refGene_gn=NME9;refGene_tx=XM_006936377.3;refGene_gn=NME9;refGene_tx=XM_023260532.1;refGene_gn=NME9;refGene_tx=XM_023260533.1
C2 127248762 127248845
refGene_gn=NME9;refGene_tx=XM_006936377.3;refGene_gn=NME9;refGene_tx=XM_023260532.1;refGene_gn=NME9;refGene_tx=XM_023260533.1
C2 127249136 127249229
refGene_gn=NME9;refGene_tx=XM_006936377.3;refGene_gn=NME9;refGene_tx=XM_023260532.1;refGene_gn=NME9;refGene_tx=XM_023260533.1
C2 127249950 127250105
refGene_gn=NME9;refGene_tx=XM_006936377.3;refGene_gn=NME9;refGene_tx=XM_023260532.1;refGene_gn=NME9;refGene_tx=XM_023260533.1
C2 127251420 127251557
refGene_gn=NME9;refGene_tx=XM_006936377.3;refGene_gn=NME9;refGene_tx=XM_023260532.1;refGene_gn=NME9;refGene_tx=XM_023260533.1
C2 127253555 127253620
refGene_gn=NME9;refGene_tx=XM_006936377.3;refGene_gn=NME9;refGene_tx=XM_023260532.1;refGene_gn=NME9;refGene_tx=XM_023260533.1
C2 127258652 127258686
refGene_gn=ARMC8;refGene_tx=XM_003992063.5;refGene_gn=ARMC8;refGene_tx=XM_019810660.2;refGene_gn=ARMC8;refGene_tx=XM_019810661.2
C2 127263034 127263128
refGene_gn=ARMC8;refGene_tx=XM_003992063.5;refGene_gn=ARMC8;refGene_tx=XM_019810660.2;refGene_gn=ARMC8;refGene_tx=XM_019810661.2
C2 127264491 127264564
refGene_gn=ARMC8;refGene_tx=XM_003992063.5;refGene_gn=ARMC8;refGene_tx

=XM_019810660.2;refGene_gn=ARMC8;refGene_tx=XM_019810661.2
C2	127269707	127269803
refGene_gn=ARMC8;refGene_tx=XM_003992063.5;refGene_gn=ARMC8;refGene_tx=XM_019810660.2;refGene_gn=ARMC8;refGene_tx=XM_019810661.2
C2	127269885	127269972
refGene_gn=ARMC8;refGene_tx=XM_019810660.2;refGene_gn=ARMC8;refGene_tx=XM_019810661.2
C2	127276042	127276138
refGene_gn=ARMC8;refGene_tx=XM_003992063.5;refGene_gn=ARMC8;refGene_tx=XM_019810660.2;refGene_gn=ARMC8;refGene_tx=XM_019810661.2
C2	127277196	127277346
refGene_gn=ARMC8;refGene_tx=XM_003992063.5;refGene_gn=ARMC8;refGene_tx=XM_019810660.2;refGene_gn=ARMC8;refGene_tx=XM_019810661.2
C2	127280166	127280259
refGene_gn=ARMC8;refGene_tx=XM_003992063.5;refGene_gn=ARMC8;refGene_tx=XM_019810660.2;refGene_gn=ARMC8;refGene_tx=XM_019810661.2
C2	127282924	127283011
refGene_gn=ARMC8;refGene_tx=XM_003992063.5;refGene_gn=ARMC8;refGene_tx=XM_019810660.2;refGene_gn=ARMC8;refGene_tx=XM_019810661.2
C2	127286730	127286812
refGene_gn=ARMC8;refGene_tx=XM_003992063.5;refGene_gn=ARMC8;refGene_tx=XM_019810660.2;refGene_gn=ARMC8;refGene_tx=XM_019810661.2
C2	127287124	127287207
refGene_gn=ARMC8;refGene_tx=XM_003992063.5;refGene_gn=ARMC8;refGene_tx=XM_019810660.2;refGene_gn=ARMC8;refGene_tx=XM_019810661.2
C2	127304965	127305061
refGene_gn=ARMC8;refGene_tx=XM_003992063.5;refGene_gn=ARMC8;refGene_tx=XM_019810660.2;refGene_gn=ARMC8;refGene_tx=XM_019810661.2
C2	127307692	127307893
refGene_gn=ARMC8;refGene_tx=XM_003992063.5;refGene_gn=ARMC8;refGene_tx=XM_019810660.2;refGene_gn=ARMC8;refGene_tx=XM_019810661.2
C2	127310353	127310414
refGene_gn=ARMC8;refGene_tx=XM_003992063.5;refGene_gn=ARMC8;refGene_tx=XM_019810660.2;refGene_gn=ARMC8;refGene_tx=XM_019810661.2
C2	127312048	127312139
refGene_gn=ARMC8;refGene_tx=XM_003992063.5;refGene_gn=ARMC8;refGene_tx=XM_019810660.2;refGene_gn=ARMC8;refGene_tx=XM_019810661.2
C2	127312235	127312311
refGene_gn=ARMC8;refGene_tx=XM_003992063.5;refGene_gn=ARMC8;refGene_tx=XM_019810660.2;refGene_gn=ARMC8;refGene_tx=XM_019810661.2
C2	127314196	127314277
refGene_gn=ARMC8;refGene_tx=XM_003992063.5;refGene_gn=ARMC8;refGene_tx=XM_019810660.2;refGene_gn=ARMC8;refGene_tx=XM_019810661.2
C2	127319911	127320004
refGene_gn=ARMC8;refGene_tx=XM_003992063.5;refGene_gn=ARMC8;refGene_tx=XM_019810660.2;refGene_gn=ARMC8;refGene_tx=XM_019810661.2
C2	127323745	127323843
refGene_gn=ARMC8;refGene_tx=XM_003992063.5;refGene_gn=ARMC8;refGene_tx=XM_019810660.2;refGene_gn=ARMC8;refGene_tx=XM_019810661.2
C2	127323947	127324090

refGene_gn=ARMC8;refGene_tx=XM_003992063.5;refGene_gn=ARMC8;refGene_tx
=XM_019810660.2;refGene_gn=ARMC8;refGene_tx=XM_019810661.2
C2	127325504	127325576
refGene_gn=ARMC8;refGene_tx=XM_003992063.5;refGene_gn=ARMC8;refGene_tx
=XM_019810660.2;refGene_gn=ARMC8;refGene_tx=XM_019810661.2
C2	127336852	127336929
refGene_gn=ARMC8;refGene_tx=XM_003992063.5;refGene_gn=ARMC8;refGene_tx
=XM_019810660.2
C2	127364948	127364993
refGene_gn=ARMC8;refGene_tx=XM_003992063.5;refGene_gn=ARMC8;refGene_tx
=XM_019810660.2
C2	127378632	127378829
refGene_gn=DBR1;refGene_tx=XM_011286137.3
C2	127379444	127379569
refGene_gn=DBR1;refGene_tx=XM_011286137.3
C2	127380916	127380997
refGene_gn=DBR1;refGene_tx=XM_011286137.3
C2	127383154	127383240
refGene_gn=DBR1;refGene_tx=XM_011286137.3
C2	127385028	127385253
refGene_gn=DBR1;refGene_tx=XM_011286137.3
C2	127388034	127388115
refGene_gn=DBR1;refGene_tx=XM_011286137.3
C2	127388363	127388509
refGene_gn=DBR1;refGene_tx=XM_011286137.3
C2	127389227	127389921
refGene_gn=DBR1;refGene_tx=XM_011286137.3
C2	127424777	127425188
refGene_gn=A4GNT;refGene_tx=XM_023260534.1
C2	127431749	127432364
refGene_gn=A4GNT;refGene_tx=XM_023260534.1
C2	127447703	127448204
refGene_gn=DZIP1L;refGene_tx=XM_006936380.3;refGene_gn=DZIP1L;refGene_
tx=XM_011286138.2;refGene_gn=DZIP1L;refGene_tx=XM_011286139.3;refGene_
gn=DZIP1L;refGene_tx=XM_011286140.2;refGene_gn=DZIP1L;refGene_tx=XM_01
9810663.2
C2	127450631	127450716
refGene_gn=DZIP1L;refGene_tx=XM_006936380.3;refGene_gn=DZIP1L;refGene_
tx=XM_011286138.2;refGene_gn=DZIP1L;refGene_tx=XM_011286139.3;refGene_
gn=DZIP1L;refGene_tx=XM_011286140.2;refGene_gn=DZIP1L;refGene_tx=XM_01
9810663.2
C2	127454486	127454608
refGene_gn=DZIP1L;refGene_tx=XM_006936380.3;refGene_gn=DZIP1L;refGene_
tx=XM_011286138.2;refGene_gn=DZIP1L;refGene_tx=XM_011286139.3;refGene_
gn=DZIP1L;refGene_tx=XM_011286140.2;refGene_gn=DZIP1L;refGene_tx=XM_01
9810663.2
C2	127455291	127455453
refGene_gn=DZIP1L;refGene_tx=XM_006936380.3;refGene_gn=DZIP1L;refGene_
tx=XM_011286138.2;refGene_gn=DZIP1L;refGene_tx=XM_011286139.3;refGene_
gn=DZIP1L;refGene_tx=XM_011286140.2;refGene_gn=DZIP1L;refGene_tx=XM_01

9810663.2
C2	127458760	127458889
refGene_gn=DZIP1L;refGene_tx=XM_006936380.3;refGene_gn=DZIP1L;refGene_tx=XM_011286138.2;refGene_gn=DZIP1L;refGene_tx=XM_011286139.3;refGene_gn=DZIP1L;refGene_tx=XM_011286140.2;refGene_gn=DZIP1L;refGene_tx=XM_019810663.2
C2	127463603	127463666
refGene_gn=DZIP1L;refGene_tx=XM_006936380.3;refGene_gn=DZIP1L;refGene_tx=XM_011286138.2;refGene_gn=DZIP1L;refGene_tx=XM_011286139.3;refGene_gn=DZIP1L;refGene_tx=XM_011286140.2;refGene_gn=DZIP1L;refGene_tx=XM_019810663.2
C2	127464888	127465029
refGene_gn=DZIP1L;refGene_tx=XM_006936380.3;refGene_gn=DZIP1L;refGene_tx=XM_011286138.2;refGene_gn=DZIP1L;refGene_tx=XM_011286139.3;refGene_gn=DZIP1L;refGene_tx=XM_011286140.2;refGene_gn=DZIP1L;refGene_tx=XM_019810663.2
C2	127467314	127467345
refGene_gn=DZIP1L;refGene_tx=XM_006936380.3;refGene_gn=DZIP1L;refGene_tx=XM_011286138.2;refGene_gn=DZIP1L;refGene_tx=XM_011286139.3;refGene_gn=DZIP1L;refGene_tx=XM_011286140.2;refGene_gn=DZIP1L;refGene_tx=XM_019810663.2
C2	127468382	127468436
refGene_gn=DZIP1L;refGene_tx=XM_006936380.3;refGene_gn=DZIP1L;refGene_tx=XM_011286138.2;refGene_gn=DZIP1L;refGene_tx=XM_011286139.3;refGene_gn=DZIP1L;refGene_tx=XM_011286140.2;refGene_gn=DZIP1L;refGene_tx=XM_019810663.2
C2	127468778	127468912
refGene_gn=DZIP1L;refGene_tx=XM_006936380.3;refGene_gn=DZIP1L;refGene_tx=XM_011286138.2;refGene_gn=DZIP1L;refGene_tx=XM_011286139.3;refGene_gn=DZIP1L;refGene_tx=XM_011286140.2;refGene_gn=DZIP1L;refGene_tx=XM_019810663.2
C2	127470658	127470851
refGene_gn=DZIP1L;refGene_tx=XM_006936380.3;refGene_gn=DZIP1L;refGene_tx=XM_011286138.2;refGene_gn=DZIP1L;refGene_tx=XM_011286139.3;refGene_gn=DZIP1L;refGene_tx=XM_011286140.2;refGene_gn=DZIP1L;refGene_tx=XM_019810663.2
C2	127474132	127474349
refGene_gn=DZIP1L;refGene_tx=XM_006936380.3;refGene_gn=DZIP1L;refGene_tx=XM_011286138.2;refGene_gn=DZIP1L;refGene_tx=XM_011286139.3;refGene_gn=DZIP1L;refGene_tx=XM_011286140.2;refGene_gn=DZIP1L;refGene_tx=XM_019810663.2
C2	127474791	127474979
refGene_gn=DZIP1L;refGene_tx=XM_006936380.3;refGene_gn=DZIP1L;refGene_tx=XM_011286138.2;refGene_gn=DZIP1L;refGene_tx=XM_011286139.3;refGene_gn=DZIP1L;refGene_tx=XM_011286140.2;refGene_gn=DZIP1L;refGene_tx=XM_019810663.2
C2	127478415	127478555
refGene_gn=DZIP1L;refGene_tx=XM_006936380.3;refGene_gn=DZIP1L;refGene_tx=XM_011286138.2;refGene_gn=DZIP1L;refGene_tx=XM_011286139.3;refGene_gn=DZIP1L;refGene_tx=XM_011286140.2;refGene_gn=DZIP1L;refGene_tx=XM_01

9810663.2
C2	127480128	127480290
refGene_gn=DZIP1L;refGene_tx=XM_006936380.3;refGene_gn=DZIP1L;refGene_tx=XM_011286138.2;refGene_gn=DZIP1L;refGene_tx=XM_011286139.3;refGene_gn=DZIP1L;refGene_tx=XM_011286140.2;refGene_gn=DZIP1L;refGene_tx=XM_019810663.2
C2	127520503	127520675
refGene_gn=CLDN18;refGene_tx=XM_003992066.3
C2	127521661	127521772
refGene_gn=CLDN18;refGene_tx=XM_003992066.3
C2	127525917	127526035
refGene_gn=CLDN18;refGene_tx=XM_003992066.3
C2	127526666	127526831
refGene_gn=CLDN18;refGene_tx=XM_003992066.3
C2	127537162	127537382
refGene_gn=CLDN18;refGene_tx=XM_003992066.3
C2	127724184	127724907
refGene_gn=SOX14;refGene_tx=XM_003992068.3
C2	128432694	128432802
refGene_gn=IL20RB;refGene_tx=XM_006936381.3;refGene_gn=IL20RB;refGene_tx=XM_019810665.1
C2	128439378	128439521
refGene_gn=IL20RB;refGene_tx=XM_006936381.3;refGene_gn=IL20RB;refGene_tx=XM_019810665.1
C2	128439987	128440142
refGene_gn=IL20RB;refGene_tx=XM_023260535.1
C2	128444006	128444157
refGene_gn=IL20RB;refGene_tx=XM_006936381.3;refGene_gn=IL20RB;refGene_tx=XM_019810665.1;refGene_gn=IL20RB;refGene_tx=XM_023260535.1
C2	128445922	128446047
refGene_gn=IL20RB;refGene_tx=XM_006936381.3;refGene_gn=IL20RB;refGene_tx=XM_019810665.1;refGene_gn=IL20RB;refGene_tx=XM_023260535.1
C2	128450110	128450301
refGene_gn=IL20RB;refGene_tx=XM_006936381.3;refGene_gn=IL20RB;refGene_tx=XM_019810665.1;refGene_gn=IL20RB;refGene_tx=XM_023260535.1
C2	128451778	128451905
refGene_gn=IL20RB;refGene_tx=XM_006936381.3;refGene_gn=IL20RB;refGene_tx=XM_019810665.1;refGene_gn=IL20RB;refGene_tx=XM_023260535.1
C2	128465090	128465130
refGene_gn=IL20RB;refGene_tx=XM_006936381.3;refGene_gn=IL20RB;refGene_tx=XM_023260535.1
C2	128476041	128476236
refGene_gn=NCK1;refGene_tx=XM_003992069.5;refGene_gn=NCK1;refGene_tx=XM_003992070.4;refGene_gn=NCK1;refGene_tx=XM_019810666.2;refGene_gn=NCK1;refGene_tx=XM_023260536.1;refGene_gn=NCK1;refGene_tx=XM_023260537.1;refGene_gn=NCK1;refGene_tx=XM_023260538.1
C2	128478215	128478928
refGene_gn=NCK1;refGene_tx=XM_003992069.5;refGene_gn=NCK1;refGene_tx=XM_003992070.4;refGene_gn=NCK1;refGene_tx=XM_019810666.2;refGene_gn=NCK1;refGene_tx=XM_023260536.1;refGene_gn=NCK1;refGene_tx=XM_023260537.1;

refGene_gn=NCK1;refGene_tx=XM_023260538.1
C2 128492312 128492346
refGene_gn=NCK1;refGene_tx=XM_003992070.4
C2 128495307 128495533
refGene_gn=NCK1;refGene_tx=XM_003992069.5;refGene_gn=NCK1;refGene_tx=XM_019810666.2;refGene_gn=NCK1;refGene_tx=XM_023260536.1;refGene_gn=NCK1;refGene_tx=XM_023260537.1;refGene_gn=NCK1;refGene_tx=XM_023260538.1
C2 128524855 128526211
refGene_gn=LOC101081405;refGene_tx=XM_011286193.3
C2 128569416 128570655
refGene_gn=SLC35G2;refGene_tx=XM_003992071.5;refGene_gn=SLC35G2;refGene_tx=XM_019810667.2
C2 128797831 128797860
refGene_gn=STAG1;refGene_tx=XM_023260539.1
C2 128808230 128808333
refGene_gn=STAG1;refGene_tx=XM_023260539.1
C2 128822688 128822853
refGene_gn=STAG1;refGene_tx=XM_019810669.1;refGene_gn=STAG1;refGene_tx=XM_023260539.1
C2 128860780 128860877
refGene_gn=STAG1;refGene_tx=XM_019810669.1;refGene_gn=STAG1;refGene_tx=XM_023260539.1
C2 128910959 128911036
refGene_gn=STAG1;refGene_tx=XM_019810669.1;refGene_gn=STAG1;refGene_tx=XM_023260539.1
C2 128926898 128927103
refGene_gn=STAG1;refGene_tx=XM_019810669.1;refGene_gn=STAG1;refGene_tx=XM_023260539.1
C2 128947039 128947191
refGene_gn=STAG1;refGene_tx=XM_019810669.1;refGene_gn=STAG1;refGene_tx=XM_019810670.2;refGene_gn=STAG1;refGene_tx=XM_023260539.1
C2 128949393 128949467
refGene_gn=STAG1;refGene_tx=XM_019810669.1;refGene_gn=STAG1;refGene_tx=XM_019810670.2;refGene_gn=STAG1;refGene_tx=XM_023260539.1
C2 128967664 128967788
refGene_gn=STAG1;refGene_tx=XM_019810669.1;refGene_gn=STAG1;refGene_tx=XM_019810670.2;refGene_gn=STAG1;refGene_tx=XM_023260539.1
C2 128983472 128983571
refGene_gn=STAG1;refGene_tx=XM_019810669.1;refGene_gn=STAG1;refGene_tx=XM_019810670.2;refGene_gn=STAG1;refGene_tx=XM_023260539.1
C2 128984081 128984161
refGene_gn=STAG1;refGene_tx=XM_019810669.1;refGene_gn=STAG1;refGene_tx=XM_019810670.2;refGene_gn=STAG1;refGene_tx=XM_023260539.1
C2 128990595 128990703
refGene_gn=STAG1;refGene_tx=XM_019810669.1;refGene_gn=STAG1;refGene_tx=XM_019810670.2;refGene_gn=STAG1;refGene_tx=XM_023260539.1
C2 128993686 128993801
refGene_gn=STAG1;refGene_tx=XM_019810669.1;refGene_gn=STAG1;refGene_tx=XM_019810670.2;refGene_gn=STAG1;refGene_tx=XM_023260539.1
C2 129001763 129001881

refGene_gn=STAG1;refGene_tx=XM_019810669.1;refGene_gn=STAG1;refGene_tx=XM_019810670.2;refGene_gn=STAG1;refGene_tx=XM_023260539.1
C2 129009699 129009803
refGene_gn=STAG1;refGene_tx=XM_019810669.1;refGene_gn=STAG1;refGene_tx=XM_019810670.2;refGene_gn=STAG1;refGene_tx=XM_023260539.1
C2 129017423 129017516
refGene_gn=STAG1;refGene_tx=XM_019810669.1;refGene_gn=STAG1;refGene_tx=XM_019810670.2;refGene_gn=STAG1;refGene_tx=XM_023260539.1
C2 129017621 129017711
refGene_gn=STAG1;refGene_tx=XM_019810669.1;refGene_gn=STAG1;refGene_tx=XM_019810670.2;refGene_gn=STAG1;refGene_tx=XM_023260539.1
C2 129017855 129018059
refGene_gn=STAG1;refGene_tx=XM_019810669.1;refGene_gn=STAG1;refGene_tx=XM_019810670.2;refGene_gn=STAG1;refGene_tx=XM_023260539.1
C2 129019242 129019313
refGene_gn=STAG1;refGene_tx=XM_019810669.1;refGene_gn=STAG1;refGene_tx=XM_019810670.2;refGene_gn=STAG1;refGene_tx=XM_023260539.1
C2 129021080 129021168
refGene_gn=STAG1;refGene_tx=XM_019810669.1;refGene_gn=STAG1;refGene_tx=XM_019810670.2;refGene_gn=STAG1;refGene_tx=XM_023260539.1
C2 129039200 129039281
refGene_gn=STAG1;refGene_tx=XM_019810669.1;refGene_gn=STAG1;refGene_tx=XM_019810670.2;refGene_gn=STAG1;refGene_tx=XM_023260539.1
C2 129053322 129053415
refGene_gn=STAG1;refGene_tx=XM_019810669.1;refGene_gn=STAG1;refGene_tx=XM_019810670.2;refGene_gn=STAG1;refGene_tx=XM_023260539.1
C2 129055707 129055882
refGene_gn=STAG1;refGene_tx=XM_019810669.1;refGene_gn=STAG1;refGene_tx=XM_019810670.2;refGene_gn=STAG1;refGene_tx=XM_023260539.1
C2 129057793 129057933
refGene_gn=STAG1;refGene_tx=XM_019810669.1;refGene_gn=STAG1;refGene_tx=XM_019810670.2;refGene_gn=STAG1;refGene_tx=XM_023260539.1
C2 129060601 129060703
refGene_gn=STAG1;refGene_tx=XM_019810669.1;refGene_gn=STAG1;refGene_tx=XM_019810670.2;refGene_gn=STAG1;refGene_tx=XM_023260539.1
C2 129064663 129064812
refGene_gn=STAG1;refGene_tx=XM_019810669.1;refGene_gn=STAG1;refGene_tx=XM_019810670.2;refGene_gn=STAG1;refGene_tx=XM_023260539.1
C2 129065529 129065658
refGene_gn=STAG1;refGene_tx=XM_019810669.1;refGene_gn=STAG1;refGene_tx=XM_019810670.2;refGene_gn=STAG1;refGene_tx=XM_023260539.1
C2 129081176 129081382
refGene_gn=STAG1;refGene_tx=XM_019810669.1;refGene_gn=STAG1;refGene_tx=XM_019810670.2;refGene_gn=STAG1;refGene_tx=XM_023260539.1
C2 129084764 129084939
refGene_gn=STAG1;refGene_tx=XM_019810669.1;refGene_gn=STAG1;refGene_tx=XM_019810670.2;refGene_gn=STAG1;refGene_tx=XM_023260539.1
C2 129086443 129086554
refGene_gn=STAG1;refGene_tx=XM_019810669.1;refGene_gn=STAG1;refGene_tx=XM_019810670.2;refGene_gn=STAG1;refGene_tx=XM_023260539.1

C2 129087112 129087227
refGene_gn=STAG1;refGene_tx=XM_019810669.1;refGene_gn=STAG1;refGene_tx=XM_019810670.2;refGene_gn=STAG1;refGene_tx=XM_023260539.1
C2 129089276 129089357
refGene_gn=STAG1;refGene_tx=XM_019810669.1;refGene_gn=STAG1;refGene_tx=XM_019810670.2;refGene_gn=STAG1;refGene_tx=XM_023260539.1
C2 129089439 129089463
refGene_gn=STAG1;refGene_tx=XM_019810669.1;refGene_gn=STAG1;refGene_tx=XM_019810670.2;refGene_gn=STAG1;refGene_tx=XM_023260539.1
C2 129096700 129096822
refGene_gn=PCCB;refGene_tx=XM_011286134.3;refGene_gn=PCCB;refGene_tx=XM_011286135.3
C2 129097912 129098012
refGene_gn=PCCB;refGene_tx=XM_011286134.3;refGene_gn=PCCB;refGene_tx=XM_011286135.3
C2 129098796 129098895
refGene_gn=PCCB;refGene_tx=XM_011286134.3;refGene_gn=PCCB;refGene_tx=XM_011286135.3
C2 129099325 129099426
refGene_gn=PCCB;refGene_tx=XM_011286134.3;refGene_gn=PCCB;refGene_tx=XM_011286135.3
C2 129099678 129099786
refGene_gn=PCCB;refGene_tx=XM_011286134.3;refGene_gn=PCCB;refGene_tx=XM_011286135.3
C2 129113386 129113510
refGene_gn=PCCB;refGene_tx=XM_011286134.3;refGene_gn=PCCB;refGene_tx=XM_011286135.3
C2 129132594 129132676
refGene_gn=PCCB;refGene_tx=XM_011286134.3;refGene_gn=PCCB;refGene_tx=XM_011286135.3
C2 129134703 129134824
refGene_gn=PCCB;refGene_tx=XM_011286134.3;refGene_gn=PCCB;refGene_tx=XM_011286135.3
C2 129137937 129138046
refGene_gn=PCCB;refGene_tx=XM_011286134.3;refGene_gn=PCCB;refGene_tx=XM_011286135.3
C2 129157809 129157920
refGene_gn=PCCB;refGene_tx=XM_011286134.3;refGene_gn=PCCB;refGene_tx=XM_011286135.3
C2 129185753 129185867
refGene_gn=PCCB;refGene_tx=XM_011286134.3
C2 129188380 129188437
refGene_gn=PCCB;refGene_tx=XM_011286134.3;refGene_gn=PCCB;refGene_tx=XM_011286135.3
C2 129191330 129191399
refGene_gn=PCCB;refGene_tx=XM_011286134.3;refGene_gn=PCCB;refGene_tx=XM_011286135.3
C2 129192399 129192519
refGene_gn=PCCB;refGene_tx=XM_011286134.3;refGene_gn=PCCB;refGene_tx=XM_011286135.3

C2 129194899 129195082
refGene_gn=PCCB;refGene_tx=XM_011286134.3;refGene_gn=PCCB;refGene_tx=XM_011286135.3
C2 129221822 129221964
refGene_gn=MSL2;refGene_tx=XM_003992096.5
C2 129252239 129252373
refGene_gn=MSL2;refGene_tx=XM_003992096.5
C2 129252374 129253832
refGene_gn=MSL2;refGene_tx=XM_003992096.5
C2 129259470 129259603
refGene_gn=PPP2R3A;refGene_tx=XM_003992098.5;refGene_gn=PPP2R3A;refGene_tx=XM_011286133.3;refGene_gn=PPP2R3A;refGene_tx=XM_023260540.1;refGene_gn=PPP2R3A;refGene_tx=XM_023260541.1
C2 129282882 129282989
refGene_gn=PPP2R3A;refGene_tx=XM_003992098.5;refGene_gn=PPP2R3A;refGene_tx=XM_011286133.3;refGene_gn=PPP2R3A;refGene_tx=XM_023260540.1;refGene_gn=PPP2R3A;refGene_tx=XM_023260541.1
C2 129285528 129285647
refGene_gn=PPP2R3A;refGene_tx=XM_003992098.5;refGene_gn=PPP2R3A;refGene_tx=XM_011286133.3;refGene_gn=PPP2R3A;refGene_tx=XM_023260540.1;refGene_gn=PPP2R3A;refGene_tx=XM_023260541.1
C2 129286162 129286338
refGene_gn=PPP2R3A;refGene_tx=XM_003992098.5;refGene_gn=PPP2R3A;refGene_tx=XM_011286133.3;refGene_gn=PPP2R3A;refGene_tx=XM_023260540.1;refGene_gn=PPP2R3A;refGene_tx=XM_023260541.1
C2 129322043 129322133
refGene_gn=PPP2R3A;refGene_tx=XM_003992098.5;refGene_gn=PPP2R3A;refGene_tx=XM_011286133.3;refGene_gn=PPP2R3A;refGene_tx=XM_023260540.1
C2 129322603 129322652
refGene_gn=PPP2R3A;refGene_tx=XM_003992098.5;refGene_gn=PPP2R3A;refGene_tx=XM_011286133.3;refGene_gn=PPP2R3A;refGene_tx=XM_023260540.1;refGene_gn=PPP2R3A;refGene_tx=XM_023260541.1
C2 129325334 129325491
refGene_gn=PPP2R3A;refGene_tx=XM_003992098.5;refGene_gn=PPP2R3A;refGene_tx=XM_011286133.3;refGene_gn=PPP2R3A;refGene_tx=XM_023260540.1;refGene_gn=PPP2R3A;refGene_tx=XM_023260541.1
C2 129329762 129329849
refGene_gn=PPP2R3A;refGene_tx=XM_003992098.5;refGene_gn=PPP2R3A;refGene_tx=XM_011286133.3;refGene_gn=PPP2R3A;refGene_tx=XM_023260540.1;refGene_gn=PPP2R3A;refGene_tx=XM_023260541.1
C2 129338248 129338323
refGene_gn=PPP2R3A;refGene_tx=XM_003992098.5;refGene_gn=PPP2R3A;refGene_tx=XM_011286133.3;refGene_gn=PPP2R3A;refGene_tx=XM_023260540.1;refGene_gn=PPP2R3A;refGene_tx=XM_023260541.1
C2 129378925 129379028
refGene_gn=PPP2R3A;refGene_tx=XM_003992098.5;refGene_gn=PPP2R3A;refGene_tx=XM_011286133.3;refGene_gn=PPP2R3A;refGene_tx=XM_023260540.1;refGene_gn=PPP2R3A;refGene_tx=XM_023260541.1
C2 129385848 129385952
refGene_gn=PPP2R3A;refGene_tx=XM_003992098.5;refGene_gn=PPP2R3A;refGen

e_tx=XM_011286133.3;refGene_gn=PPP2R3A;refGene_tx=XM_023260540.1;refGene_gn=PPP2R3A;refGene_tx=XM_023260541.1
C2 129392448 129392715
refGene_gn=PPP2R3A;refGene_tx=XM_003992098.5;refGene_gn=PPP2R3A;refGene_tx=XM_011286133.3;refGene_gn=PPP2R3A;refGene_tx=XM_023260540.1;refGene_gn=PPP2R3A;refGene_tx=XM_023260541.1
C2 129396305 129396437
refGene_gn=PPP2R3A;refGene_tx=XM_003992098.5
C2 129421696 129423682
refGene_gn=PPP2R3A;refGene_tx=XM_011286133.3;refGene_gn=PPP2R3A;refGene_tx=XM_023260540.1;refGene_gn=PPP2R3A;refGene_tx=XM_023260541.1
C2 130018031 130018140
refGene_gn=EPHB1;refGene_tx=XM_006936394.4;refGene_gn=EPHB1;refGene_tx=XM_023260542.1
C2 130021853 130022009
refGene_gn=EPHB1;refGene_tx=XM_006936394.4;refGene_gn=EPHB1;refGene_tx=XM_023260542.1
C2 130023871 130024065
refGene_gn=EPHB1;refGene_tx=XM_006936394.4;refGene_gn=EPHB1;refGene_tx=XM_023260542.1
C2 130031893 130032043
refGene_gn=EPHB1;refGene_tx=XM_006936394.4;refGene_gn=EPHB1;refGene_tx=XM_023260542.1
C2 130068969 130069185
refGene_gn=EPHB1;refGene_tx=XM_006936394.4;refGene_gn=EPHB1;refGene_tx=XM_023260542.1
C2 130077043 130077291
refGene_gn=EPHB1;refGene_tx=XM_006936394.4;refGene_gn=EPHB1;refGene_tx=XM_023260542.1
C2 130090416 130090539
refGene_gn=EPHB1;refGene_tx=XM_006936394.4;refGene_gn=EPHB1;refGene_tx=XM_023260542.1
C2 130102829 130102894
refGene_gn=EPHB1;refGene_tx=XM_006936394.4;refGene_gn=EPHB1;refGene_tx=XM_023260542.1
C2 130103700 130103809
refGene_gn=EPHB1;refGene_tx=XM_006936394.4;refGene_gn=EPHB1;refGene_tx=XM_023260542.1
C2 130107802 130107965
refGene_gn=EPHB1;refGene_tx=XM_006936394.4;refGene_gn=EPHB1;refGene_tx=XM_023260542.1
C2 130115895 130116020
refGene_gn=EPHB1;refGene_tx=XM_006936394.4;refGene_gn=EPHB1;refGene_tx=XM_023260542.1
C2 130137956 130138292
refGene_gn=EPHB1;refGene_tx=XM_006936394.4;refGene_gn=EPHB1;refGene_tx=XM_023260542.1
C2 130161169 130161325
refGene_gn=EPHB1;refGene_tx=XM_006936394.4;refGene_gn=EPHB1;refGene_tx=XM_023260542.1

C2	130286146	130286828
refGene_gn=EPHB1;refGene_tx=XM_006936394.4;refGene_gn=EPHB1;refGene_tx=XM_023260542.1
C2	130312200	130312265
refGene_gn=EPHB1;refGene_tx=XM_006936394.4;refGene_gn=EPHB1;refGene_tx=XM_023260542.1
C2	130447024	130447082
refGene_gn=EPHB1;refGene_tx=XM_023260542.1
C2	130562798	130562934
refGene_gn=KY;refGene_tx=XM_011286132.3
C2	130565982	130566045
refGene_gn=KY;refGene_tx=XM_011286132.3
C2	130569861	130569924
refGene_gn=KY;refGene_tx=XM_011286132.3
C2	130579893	130579968
refGene_gn=KY;refGene_tx=XM_011286132.3
C2	130581697	130581760
refGene_gn=KY;refGene_tx=XM_011286132.3
C2	130584707	130584790
refGene_gn=KY;refGene_tx=XM_011286132.3
C2	130589149	130589258
refGene_gn=KY;refGene_tx=XM_011286132.3
C2	130590758	130590876
refGene_gn=KY;refGene_tx=XM_011286132.3
C2	130599407	130599596
refGene_gn=KY;refGene_tx=XM_011286132.3
C2	130600862	130601053
refGene_gn=KY;refGene_tx=XM_011286132.3
C2	130604748	130605644
refGene_gn=KY;refGene_tx=XM_011286132.3
C2	130630906	130631065
refGene_gn=CEP63;refGene_tx=XM_023260543.1
C2	130633244	130633524
refGene_gn=CEP63;refGene_tx=XM_023260543.1
C2	130634294	130634500
refGene_gn=CEP63;refGene_tx=XM_023260543.1
C2	130638427	130638514
refGene_gn=CEP63;refGene_tx=XM_023260543.1
C2	130647374	130647572
refGene_gn=CEP63;refGene_tx=XM_023260543.1
C2	130648434	130648549
refGene_gn=CEP63;refGene_tx=XM_023260543.1
C2	130650303	130650441
refGene_gn=CEP63;refGene_tx=XM_023260543.1
C2	130651262	130651402
refGene_gn=CEP63;refGene_tx=XM_023260543.1
C2	130651840	130652074
refGene_gn=CEP63;refGene_tx=XM_023260543.1
C2	130666292	130666406
refGene_gn=CEP63;refGene_tx=XM_023260543.1

C2 130670830 130670953
refGene_gn=CEP63;refGene_tx=XM_023260543.1
C2 130671927 130672023
refGene_gn=CEP63;refGene_tx=XM_023260543.1
C2 130682888 130682948
refGene_gn=CEP63;refGene_tx=XM_023260543.1
C2 130682949 130683067
refGene_gn=CEP63;refGene_tx=XM_023260543.1
C2 130683996 130684065
refGene_gn=CEP63;refGene_tx=XM_023260543.1
C2 130693489 130693650
refGene_gn=CEP63;refGene_tx=XM_023260543.1
C2 130696502 130696601
refGene_gn=ANAPC13;refGene_tx=XM_003992102.5
C2 130701036 130701162
refGene_gn=ANAPC13;refGene_tx=XM_003992102.5
C2 130786507 130787241
refGene_gn=AMOTL2;refGene_tx=XM_006936400.4;refGene_gn=AMOTL2;refGene_tx=XM_006936401.4;refGene_gn=AMOTL2;refGene_tx=XM_006936402.4;refGene_gn=AMOTL2;refGene_tx=XM_019810678.1
C2 130789952 130790259
refGene_gn=AMOTL2;refGene_tx=XM_006936400.4;refGene_gn=AMOTL2;refGene_tx=XM_006936401.4;refGene_gn=AMOTL2;refGene_tx=XM_006936402.4;refGene_gn=AMOTL2;refGene_tx=XM_019810678.1
C2 130791267 130791412
refGene_gn=AMOTL2;refGene_tx=XM_006936400.4;refGene_gn=AMOTL2;refGene_tx=XM_006936401.4;refGene_gn=AMOTL2;refGene_tx=XM_006936402.4;refGene_gn=AMOTL2;refGene_tx=XM_019810678.1
C2 130791805 130791898
refGene_gn=AMOTL2;refGene_tx=XM_006936400.4;refGene_gn=AMOTL2;refGene_tx=XM_006936401.4;refGene_gn=AMOTL2;refGene_tx=XM_006936402.4;refGene_gn=AMOTL2;refGene_tx=XM_019810678.1
C2 130795765 130796061
refGene_gn=AMOTL2;refGene_tx=XM_006936400.4;refGene_gn=AMOTL2;refGene_tx=XM_006936401.4;refGene_gn=AMOTL2;refGene_tx=XM_006936402.4;refGene_gn=AMOTL2;refGene_tx=XM_019810678.1
C2 130796833 130797141
refGene_gn=AMOTL2;refGene_tx=XM_006936400.4;refGene_gn=AMOTL2;refGene_tx=XM_006936401.4;refGene_gn=AMOTL2;refGene_tx=XM_006936402.4;refGene_gn=AMOTL2;refGene_tx=XM_019810678.1
C2 130797918 130798136
refGene_gn=AMOTL2;refGene_tx=XM_006936400.4;refGene_gn=AMOTL2;refGene_tx=XM_006936401.4;refGene_gn=AMOTL2;refGene_tx=XM_006936402.4;refGene_gn=AMOTL2;refGene_tx=XM_019810678.1
C2 130798708 130798882
refGene_gn=AMOTL2;refGene_tx=XM_006936400.4;refGene_gn=AMOTL2;refGene_tx=XM_006936401.4;refGene_gn=AMOTL2;refGene_tx=XM_006936402.4;refGene_gn=AMOTL2;refGene_tx=XM_019810678.1
C2 130799894 130799953
refGene_gn=AMOTL2;refGene_tx=XM_006936400.4;refGene_gn=AMOTL2;refGene_

tx=XM_006936401.4;refGene_gn=AMOTL2;refGene_tx=XM_006936402.4;refGene_gn=AMOTL2;refGene_tx=XM_019810678.1
C2 130889701 130889930
refGene_gn=RYK;refGene_tx=XM_023260544.1;refGene_gn=RYK;refGene_tx=XM_023260545.1;refGene_gn=RYK;refGene_tx=XM_023260546.1
C2 130925725 130925847
refGene_gn=RYK;refGene_tx=XM_023260544.1;refGene_gn=RYK;refGene_tx=XM_023260545.1;refGene_gn=RYK;refGene_tx=XM_023260546.1
C2 130939417 130939517
refGene_gn=RYK;refGene_tx=XM_023260544.1;refGene_gn=RYK;refGene_tx=XM_023260545.1;refGene_gn=RYK;refGene_tx=XM_023260546.1
C2 130941678 130941813
refGene_gn=RYK;refGene_tx=XM_023260544.1;refGene_gn=RYK;refGene_tx=XM_023260545.1
C2 130943961 130944015
refGene_gn=RYK;refGene_tx=XM_023260544.1;refGene_gn=RYK;refGene_tx=XM_023260545.1
C2 130949309 130949454
refGene_gn=RYK;refGene_tx=XM_023260544.1;refGene_gn=RYK;refGene_tx=XM_023260545.1;refGene_gn=RYK;refGene_tx=XM_023260546.1
C2 130953410 130953511
refGene_gn=RYK;refGene_tx=XM_023260544.1;refGene_gn=RYK;refGene_tx=XM_023260545.1;refGene_gn=RYK;refGene_tx=XM_023260546.1
C2 130957737 130957872
refGene_gn=RYK;refGene_tx=XM_023260544.1;refGene_gn=RYK;refGene_tx=XM_023260545.1;refGene_gn=RYK;refGene_tx=XM_023260546.1
C2 130959484 130959571
refGene_gn=RYK;refGene_tx=XM_023260544.1;refGene_gn=RYK;refGene_tx=XM_023260545.1;refGene_gn=RYK;refGene_tx=XM_023260546.1
C2 130965517 130965587
refGene_gn=RYK;refGene_tx=XM_023260544.1;refGene_gn=RYK;refGene_tx=XM_023260545.1;refGene_gn=RYK;refGene_tx=XM_023260546.1
C2 130971718 130971851
refGene_gn=RYK;refGene_tx=XM_023260544.1;refGene_gn=RYK;refGene_tx=XM_023260545.1;refGene_gn=RYK;refGene_tx=XM_023260546.1
C2 130973103 130973213
refGene_gn=RYK;refGene_tx=XM_023260544.1;refGene_gn=RYK;refGene_tx=XM_023260545.1;refGene_gn=RYK;refGene_tx=XM_023260546.1
C2 130973367 130973527
refGene_gn=RYK;refGene_tx=XM_023260544.1;refGene_gn=RYK;refGene_tx=XM_023260545.1;refGene_gn=RYK;refGene_tx=XM_023260546.1
C2 130992493 130992630
refGene_gn=RYK;refGene_tx=XM_023260544.1;refGene_gn=RYK;refGene_tx=XM_023260545.1;refGene_gn=RYK;refGene_tx=XM_023260546.1
C2 130993544 130993656
refGene_gn=RYK;refGene_tx=XM_023260544.1;refGene_gn=RYK;refGene_tx=XM_023260545.1;refGene_gn=RYK;refGene_tx=XM_023260546.1
C2 131094138 131094234
refGene_gn=SLCO2A1;refGene_tx=XM_023260547.1
C2 131130276 131130414

refGene_gn=SLCO2A1;refGene_tx=XM_023260547.1
C2 131135815 131135978
refGene_gn=SLCO2A1;refGene_tx=XM_023260547.1
C2 131155218 131155446
refGene_gn=SLCO2A1;refGene_tx=XM_023260547.1
C2 131156307 131156406
refGene_gn=SLCO2A1;refGene_tx=XM_023260547.1
C2 131158946 131159083
refGene_gn=SLCO2A1;refGene_tx=XM_023260547.1
C2 131160872 131160951
refGene_gn=SLCO2A1;refGene_tx=XM_023260547.1
C2 131161506 131161671
refGene_gn=SLCO2A1;refGene_tx=XM_023260547.1
C2 131162717 131162907
refGene_gn=SLCO2A1;refGene_tx=XM_023260547.1
C2 131164587 131164753
refGene_gn=SLCO2A1;refGene_tx=XM_023260547.1
C2 131167149 131167313
refGene_gn=SLCO2A1;refGene_tx=XM_023260547.1
C2 131170951 131171016
refGene_gn=SLCO2A1;refGene_tx=XM_023260547.1
C2 131173450 131173574
refGene_gn=SLCO2A1;refGene_tx=XM_023260547.1
C2 131174525 131174646
refGene_gn=SLCO2A1;refGene_tx=XM_023260547.1
C2 131204133 131204203
refGene_gn=RAB6B;refGene_tx=XM_023260548.1
C2 131231961 131232020
refGene_gn=RAB6B;refGene_tx=XM_023260548.1
C2 131251976 131252030
refGene_gn=RAB6B;refGene_tx=XM_023260548.1
C2 131252225 131252331
refGene_gn=RAB6B;refGene_tx=XM_023260548.1
C2 131253902 131254014
refGene_gn=RAB6B;refGene_tx=XM_023260548.1
C2 131255078 131255172
refGene_gn=RAB6B;refGene_tx=XM_023260548.1
C2 131258018 131258085
refGene_gn=RAB6B;refGene_tx=XM_023260548.1
C2 131259654 131259719
refGene_gn=RAB6B;refGene_tx=XM_023260548.1
C2 131268558 131268772
refGene_gn=SRPRB;refGene_tx=XM_023260185.1
C2 131277957 131278012
refGene_gn=SRPRB;refGene_tx=XM_023260185.1
C2 131279152 131279289
refGene_gn=SRPRB;refGene_tx=XM_023260185.1
C2 131291331 131291414
refGene_gn=SRPRB;refGene_tx=XM_023260185.1
C2 131295972 131296067

refGene_gn=SRPRB;refGene_tx=XM_023260185.1
C2	131296713	131297038
refGene_gn=SRPRB;refGene_tx=XM_023260185.1
C2	131302224	131302259
refGene_gn=LOC101085148;refGene_tx=XM_019810683.1
C2	131303795	131303985
refGene_gn=LOC101085148;refGene_tx=XM_019810683.1
C2	131305179	131305364
refGene_gn=LOC101085148;refGene_tx=XM_019810683.1
C2	131308321	131308386
refGene_gn=LOC101085148;refGene_tx=XM_019810683.1
C2	131309543	131309688
refGene_gn=LOC101085148;refGene_tx=XM_019810683.1
C2	131312268	131312427
refGene_gn=LOC101085148;refGene_tx=XM_019810683.1
C2	131314114	131314153
refGene_gn=LOC101085148;refGene_tx=XM_019810683.1
C2	131314734	131314825
refGene_gn=LOC101085148;refGene_tx=XM_019810683.1
C2	131316184	131316339
refGene_gn=LOC101085148;refGene_tx=XM_019810683.1
C2	131317796	131317974
refGene_gn=LOC101085148;refGene_tx=XM_019810683.1
C2	131318690	131318869
refGene_gn=LOC101085148;refGene_tx=XM_019810683.1
C2	131319413	131319469
refGene_gn=LOC101085148;refGene_tx=XM_019810683.1
C2	131320325	131320473
refGene_gn=LOC101085148;refGene_tx=XM_019810683.1
C2	131321230	131321407
refGene_gn=LOC101085148;refGene_tx=XM_019810683.1
C2	131322207	131322316
refGene_gn=LOC101085148;refGene_tx=XM_019810683.1
C2	131324927	131325100
refGene_gn=LOC101085148;refGene_tx=XM_019810683.1
C2	131327120	131327163
refGene_gn=LOC101085148;refGene_tx=XM_019810683.1
C2	131350320	131350355
refGene_gn=LOC101087599;refGene_tx=XM_006936404.4
C2	131351187	131351374
refGene_gn=LOC101087599;refGene_tx=XM_006936404.4
C2	131352713	131352898
refGene_gn=LOC101087599;refGene_tx=XM_006936404.4
C2	131353405	131353470
refGene_gn=LOC101087599;refGene_tx=XM_006936404.4
C2	131359044	131359186
refGene_gn=LOC101087599;refGene_tx=XM_006936404.4
C2	131362080	131362236
refGene_gn=LOC101087599;refGene_tx=XM_006936404.4
C2	131365915	131365957

refGene_gn=LOC101087599;refGene_tx=XM_006936404.4
C2 131367132 131367223
refGene_gn=LOC101087599;refGene_tx=XM_006936404.4
C2 131369380 131369523
refGene_gn=LOC101087599;refGene_tx=XM_006936404.4
C2 131370426 131370598
refGene_gn=LOC101087599;refGene_tx=XM_006936404.4
C2 131371967 131372146
refGene_gn=LOC101087599;refGene_tx=XM_006936404.4
C2 131375440 131375496
refGene_gn=LOC101087599;refGene_tx=XM_006936404.4
C2 131376318 131376475
refGene_gn=LOC101087599;refGene_tx=XM_006936404.4
C2 131377680 131377842
refGene_gn=LOC101087599;refGene_tx=XM_006936404.4
C2 131378761 131378870
refGene_gn=LOC101087599;refGene_tx=XM_006936404.4
C2 131390920 131391093
refGene_gn=LOC101087599;refGene_tx=XM_006936404.4
C2 131408170 131408213
refGene_gn=LOC101087599;refGene_tx=XM_006936404.4
C2 131428679 131428763
refGene_gn=TOPBP1;refGene_tx=XM_006936405.4
C2 131430440 131430575
refGene_gn=TOPBP1;refGene_tx=XM_006936405.4;refGene_gn=TOPBP1;refGene_tx=XM_006936406.4
C2 131431191 131431218
refGene_gn=TOPBP1;refGene_tx=XM_006936407.2
C2 131432027 131432171
refGene_gn=TOPBP1;refGene_tx=XM_006936405.4;refGene_gn=TOPBP1;refGene_tx=XM_006936406.4;refGene_gn=TOPBP1;refGene_tx=XM_006936407.2;refGene_gn=TOPBP1;refGene_tx=XM_019810684.1
C2 131432818 131433000
refGene_gn=TOPBP1;refGene_tx=XM_006936405.4;refGene_gn=TOPBP1;refGene_tx=XM_006936406.4;refGene_gn=TOPBP1;refGene_tx=XM_006936407.2;refGene_gn=TOPBP1;refGene_tx=XM_019810684.1
C2 131434117 131434314
refGene_gn=TOPBP1;refGene_tx=XM_006936405.4;refGene_gn=TOPBP1;refGene_tx=XM_006936406.4;refGene_gn=TOPBP1;refGene_tx=XM_006936407.2;refGene_gn=TOPBP1;refGene_tx=XM_019810684.1
C2 131435433 131435613
refGene_gn=TOPBP1;refGene_tx=XM_006936405.4;refGene_gn=TOPBP1;refGene_tx=XM_006936406.4;refGene_gn=TOPBP1;refGene_tx=XM_006936407.2;refGene_gn=TOPBP1;refGene_tx=XM_019810684.1
C2 131436299 131436469
refGene_gn=TOPBP1;refGene_tx=XM_006936405.4;refGene_gn=TOPBP1;refGene_tx=XM_006936406.4;refGene_gn=TOPBP1;refGene_tx=XM_006936407.2;refGene_gn=TOPBP1;refGene_tx=XM_019810684.1
C2 131438794 131438958
refGene_gn=TOPBP1;refGene_tx=XM_006936405.4;refGene_gn=TOPBP1;refGene_

tx=XM_006936406.4;refGene_gn=TOPBP1;refGene_tx=XM_006936407.2;refGene_
gn=TOPBP1;refGene_tx=XM_019810684.1
C2	131439107	131439355
refGene_gn=TOPBP1;refGene_tx=XM_006936405.4;refGene_gn=TOPBP1;refGene_
tx=XM_006936406.4;refGene_gn=TOPBP1;refGene_tx=XM_006936407.2;refGene_
gn=TOPBP1;refGene_tx=XM_019810684.1
C2	131448675	131448989
refGene_gn=TOPBP1;refGene_tx=XM_006936405.4;refGene_gn=TOPBP1;refGene_
tx=XM_006936406.4;refGene_gn=TOPBP1;refGene_tx=XM_006936407.2;refGene_
gn=TOPBP1;refGene_tx=XM_019810684.1
C2	131449655	131449828
refGene_gn=TOPBP1;refGene_tx=XM_006936405.4;refGene_gn=TOPBP1;refGene_
tx=XM_006936406.4;refGene_gn=TOPBP1;refGene_tx=XM_006936407.2;refGene_
gn=TOPBP1;refGene_tx=XM_019810684.1
C2	131453732	131453944
refGene_gn=TOPBP1;refGene_tx=XM_006936405.4;refGene_gn=TOPBP1;refGene_
tx=XM_006936406.4;refGene_gn=TOPBP1;refGene_tx=XM_006936407.2;refGene_
gn=TOPBP1;refGene_tx=XM_019810684.1
C2	131455431	131455718
refGene_gn=TOPBP1;refGene_tx=XM_006936405.4;refGene_gn=TOPBP1;refGene_
tx=XM_006936406.4;refGene_gn=TOPBP1;refGene_tx=XM_006936407.2;refGene_
gn=TOPBP1;refGene_tx=XM_019810684.1
C2	131462202	131462376
refGene_gn=TOPBP1;refGene_tx=XM_006936405.4;refGene_gn=TOPBP1;refGene_
tx=XM_006936406.4;refGene_gn=TOPBP1;refGene_tx=XM_006936407.2;refGene_
gn=TOPBP1;refGene_tx=XM_019810684.1
C2	131462460	131462576
refGene_gn=TOPBP1;refGene_tx=XM_006936405.4;refGene_gn=TOPBP1;refGene_
tx=XM_006936406.4;refGene_gn=TOPBP1;refGene_tx=XM_006936407.2;refGene_
gn=TOPBP1;refGene_tx=XM_019810684.1
C2	131466654	131466778
refGene_gn=TOPBP1;refGene_tx=XM_006936405.4;refGene_gn=TOPBP1;refGene_
tx=XM_006936406.4;refGene_gn=TOPBP1;refGene_tx=XM_006936407.2;refGene_
gn=TOPBP1;refGene_tx=XM_019810684.1
C2	131467597	131467744
refGene_gn=TOPBP1;refGene_tx=XM_006936405.4;refGene_gn=TOPBP1;refGene_
tx=XM_006936406.4;refGene_gn=TOPBP1;refGene_tx=XM_006936407.2;refGene_
gn=TOPBP1;refGene_tx=XM_019810684.1
C2	131467862	131467965
refGene_gn=TOPBP1;refGene_tx=XM_006936405.4;refGene_gn=TOPBP1;refGene_
tx=XM_006936406.4;refGene_gn=TOPBP1;refGene_tx=XM_006936407.2;refGene_
gn=TOPBP1;refGene_tx=XM_019810684.1
C2	131469910	131470103
refGene_gn=TOPBP1;refGene_tx=XM_006936405.4;refGene_gn=TOPBP1;refGene_
tx=XM_006936406.4;refGene_gn=TOPBP1;refGene_tx=XM_006936407.2;refGene_
gn=TOPBP1;refGene_tx=XM_019810684.1
C2	131471880	131472101
refGene_gn=TOPBP1;refGene_tx=XM_006936405.4;refGene_gn=TOPBP1;refGene_
tx=XM_006936406.4;refGene_gn=TOPBP1;refGene_tx=XM_006936407.2;refGene_
gn=TOPBP1;refGene_tx=XM_019810684.1

C2 131472908 131473075
refGene_gn=TOPBP1;refGene_tx=XM_006936405.4;refGene_gn=TOPBP1;refGene_tx=XM_006936406.4;refGene_gn=TOPBP1;refGene_tx=XM_006936407.2;refGene_gn=TOPBP1;refGene_tx=XM_019810684.1
C2 131473316 131473428
refGene_gn=TOPBP1;refGene_tx=XM_006936405.4;refGene_gn=TOPBP1;refGene_tx=XM_006936406.4;refGene_gn=TOPBP1;refGene_tx=XM_006936407.2;refGene_gn=TOPBP1;refGene_tx=XM_019810684.1
C2 131476742 131476906
refGene_gn=TOPBP1;refGene_tx=XM_006936405.4;refGene_gn=TOPBP1;refGene_tx=XM_006936406.4;refGene_gn=TOPBP1;refGene_tx=XM_006936407.2;refGene_gn=TOPBP1;refGene_tx=XM_019810684.1
C2 131477912 131478050
refGene_gn=TOPBP1;refGene_tx=XM_006936405.4;refGene_gn=TOPBP1;refGene_tx=XM_006936406.4;refGene_gn=TOPBP1;refGene_tx=XM_006936407.2;refGene_gn=TOPBP1;refGene_tx=XM_019810684.1
C2 131479625 131479715
refGene_gn=TOPBP1;refGene_tx=XM_006936405.4;refGene_gn=TOPBP1;refGene_tx=XM_006936406.4;refGene_gn=TOPBP1;refGene_tx=XM_006936407.2;refGene_gn=TOPBP1;refGene_tx=XM_019810684.1
C2 131479884 131480046
refGene_gn=TOPBP1;refGene_tx=XM_006936405.4;refGene_gn=TOPBP1;refGene_tx=XM_006936406.4;refGene_gn=TOPBP1;refGene_tx=XM_006936407.2;refGene_gn=TOPBP1;refGene_tx=XM_019810684.1
C2 131488663 131488804
refGene_gn=TOPBP1;refGene_tx=XM_006936405.4;refGene_gn=TOPBP1;refGene_tx=XM_006936406.4;refGene_gn=TOPBP1;refGene_tx=XM_006936407.2;refGene_gn=TOPBP1;refGene_tx=XM_019810684.1
C2 131522686 131522703
refGene_gn=CDV3;refGene_tx=XM_023260551.1;refGene_gn=CDV3;refGene_tx=XM_023260554.1
C2 131522970 131522978
refGene_gn=CDV3;refGene_tx=XM_023260552.1;refGene_gn=CDV3;refGene_tx=XM_023260555.1
C2 131523005 131523156
refGene_gn=CDV3;refGene_tx=XM_006936415.4;refGene_gn=CDV3;refGene_tx=XM_019810687.2;refGene_gn=CDV3;refGene_tx=XM_023260549.1;refGene_gn=CDV3;refGene_tx=XM_023260550.1;refGene_gn=CDV3;refGene_tx=XM_023260551.1;refGene_gn=CDV3;refGene_tx=XM_023260552.1;refGene_gn=CDV3;refGene_tx=XM_023260556.1
C2 131524081 131524097
refGene_gn=CDV3;refGene_tx=XM_023260553.1
C2 131524515 131524675
refGene_gn=CDV3;refGene_tx=XM_006936415.4;refGene_gn=CDV3;refGene_tx=XM_019810687.2;refGene_gn=CDV3;refGene_tx=XM_023260549.1;refGene_gn=CDV3;refGene_tx=XM_023260550.1;refGene_gn=CDV3;refGene_tx=XM_023260551.1;refGene_gn=CDV3;refGene_tx=XM_023260552.1;refGene_gn=CDV3;refGene_tx=XM_023260553.1;refGene_gn=CDV3;refGene_tx=XM_023260554.1;refGene_gn=CDV3;refGene_tx=XM_023260555.1;refGene_gn=CDV3;refGene_tx=XM_023260556.1
C2 131527468 131527614

refGene_gn=CDV3;refGene_tx=XM_006936415.4;refGene_gn=CDV3;refGene_tx=XM_019810687.2;refGene_gn=CDV3;refGene_tx=XM_023260549.1;refGene_gn=CDV3;refGene_tx=XM_023260550.1;refGene_gn=CDV3;refGene_tx=XM_023260551.1;refGene_gn=CDV3;refGene_tx=XM_023260552.1;refGene_gn=CDV3;refGene_tx=XM_023260553.1;refGene_gn=CDV3;refGene_tx=XM_023260554.1;refGene_gn=CDV3;refGene_tx=XM_023260555.1;refGene_gn=CDV3;refGene_tx=XM_023260556.1
C2 131536726 131536803
refGene_gn=CDV3;refGene_tx=XM_006936415.4;refGene_gn=CDV3;refGene_tx=XM_019810687.2;refGene_gn=CDV3;refGene_tx=XM_023260549.1;refGene_gn=CDV3;refGene_tx=XM_023260550.1;refGene_gn=CDV3;refGene_tx=XM_023260551.1;refGene_gn=CDV3;refGene_tx=XM_023260552.1;refGene_gn=CDV3;refGene_tx=XM_023260553.1;refGene_gn=CDV3;refGene_tx=XM_023260554.1;refGene_gn=CDV3;refGene_tx=XM_023260555.1;refGene_gn=CDV3;refGene_tx=XM_023260556.1
C2 131537283 131537328
refGene_gn=CDV3;refGene_tx=XM_006936415.4
C2 131537582 131537822
refGene_gn=CDV3;refGene_tx=XM_023260549.1;refGene_gn=CDV3;refGene_tx=XM_023260550.1;refGene_gn=CDV3;refGene_tx=XM_023260551.1;refGene_gn=CDV3;refGene_tx=XM_023260552.1;refGene_gn=CDV3;refGene_tx=XM_023260553.1;refGene_gn=CDV3;refGene_tx=XM_023260554.1;refGene_gn=CDV3;refGene_tx=XM_023260555.1
C2 131605828 131605832
refGene_gn=BFSP2;refGene_tx=XM_003992114.3
C2 131608714 131608935
refGene_gn=BFSP2;refGene_tx=XM_003992114.3
C2 131615436 131615568
refGene_gn=BFSP2;refGene_tx=XM_003992114.3
C2 131631975 131632137
refGene_gn=BFSP2;refGene_tx=XM_003992114.3
C2 131633193 131633350
refGene_gn=BFSP2;refGene_tx=XM_003992114.3
C2 131639039 131639122
refGene_gn=BFSP2;refGene_tx=XM_003992114.3
C2 131673638 131674130
refGene_gn=BFSP2;refGene_tx=XM_003992114.3
C2 131678198 131678321
refGene_gn=TMEM108;refGene_tx=XM_003992115.3;refGene_gn=TMEM108;refGene_tx=XM_019810691.2;refGene_gn=TMEM108;refGene_tx=XM_019810692.1;refGene_gn=TMEM108;refGene_tx=XM_019810693.2;refGene_gn=TMEM108;refGene_tx=XM_019810694.2;refGene_gn=TMEM108;refGene_tx=XM_019810695.2
C2 131682811 131682966
refGene_gn=TMEM108;refGene_tx=XM_003992115.3;refGene_gn=TMEM108;refGene_tx=XM_019810691.2;refGene_gn=TMEM108;refGene_tx=XM_019810692.1;refGene_gn=TMEM108;refGene_tx=XM_019810693.2;refGene_gn=TMEM108;refGene_tx=XM_019810694.2;refGene_gn=TMEM108;refGene_tx=XM_019810695.2
C2 131691726 131693154
refGene_gn=TMEM108;refGene_tx=XM_003992115.3;refGene_gn=TMEM108;refGene_tx=XM_019810691.2;refGene_gn=TMEM108;refGene_tx=XM_019810692.1;refGene_gn=TMEM108;refGene_tx=XM_019810693.2;refGene_gn=TMEM108;refGene_tx=XM_019810694.2;refGene_gn=TMEM108;refGene_tx=XM_019810695.2

C2 131756553 131756602
refGene_gn=TMEM108;refGene_tx=XM_019810691.2
C2 131764370 131764392
refGene_gn=TMEM108;refGene_tx=XM_019810695.2
C2 131849992 131850032
refGene_gn=TMEM108;refGene_tx=XM_003992115.3;refGene_gn=TMEM108;refGen
e_tx=XM_019810692.1;refGene_gn=TMEM108;refGene_tx=XM_019810693.2;refGe
ne_gn=TMEM108;refGene_tx=XM_019810694.2
C2 132326374 132326770
refGene_gn=NPHP3;refGene_tx=XM_023260557.1;refGene_gn=NPHP3;refGene_tx
=XM_023260558.1;refGene_gn=NPHP3;refGene_tx=XM_023260559.1
C2 132329007 132329133
refGene_gn=NPHP3;refGene_tx=XM_023260557.1;refGene_gn=NPHP3;refGene_tx
=XM_023260558.1;refGene_gn=NPHP3;refGene_tx=XM_023260559.1
C2 132329666 132329817
refGene_gn=NPHP3;refGene_tx=XM_023260557.1;refGene_gn=NPHP3;refGene_tx
=XM_023260558.1;refGene_gn=NPHP3;refGene_tx=XM_023260559.1
C2 132331800 132331953
refGene_gn=NPHP3;refGene_tx=XM_023260557.1;refGene_gn=NPHP3;refGene_tx
=XM_023260558.1;refGene_gn=NPHP3;refGene_tx=XM_023260559.1
C2 132332984 132333118
refGene_gn=NPHP3;refGene_tx=XM_023260557.1;refGene_gn=NPHP3;refGene_tx
=XM_023260558.1;refGene_gn=NPHP3;refGene_tx=XM_023260559.1
C2 132334635 132334796
refGene_gn=NPHP3;refGene_tx=XM_023260557.1;refGene_gn=NPHP3;refGene_tx
=XM_023260558.1;refGene_gn=NPHP3;refGene_tx=XM_023260559.1
C2 132337978 132338135
refGene_gn=NPHP3;refGene_tx=XM_023260557.1;refGene_gn=NPHP3;refGene_tx
=XM_023260558.1;refGene_gn=NPHP3;refGene_tx=XM_023260559.1
C2 132341721 132341796
refGene_gn=NPHP3;refGene_tx=XM_023260557.1;refGene_gn=NPHP3;refGene_tx
=XM_023260558.1;refGene_gn=NPHP3;refGene_tx=XM_023260559.1
C2 132344134 132344308
refGene_gn=NPHP3;refGene_tx=XM_023260557.1;refGene_gn=NPHP3;refGene_tx
=XM_023260558.1;refGene_gn=NPHP3;refGene_tx=XM_023260559.1
C2 132346281 132346385
refGene_gn=NPHP3;refGene_tx=XM_023260557.1;refGene_gn=NPHP3;refGene_tx
=XM_023260558.1;refGene_gn=NPHP3;refGene_tx=XM_023260559.1
C2 132347700 132347815
refGene_gn=NPHP3;refGene_tx=XM_023260557.1;refGene_gn=NPHP3;refGene_tx
=XM_023260558.1;refGene_gn=NPHP3;refGene_tx=XM_023260559.1
C2 132348263 132348407
refGene_gn=NPHP3;refGene_tx=XM_023260557.1;refGene_gn=NPHP3;refGene_tx
=XM_023260558.1;refGene_gn=NPHP3;refGene_tx=XM_023260559.1
C2 132348880 132348978
refGene_gn=NPHP3;refGene_tx=XM_023260557.1;refGene_gn=NPHP3;refGene_tx
=XM_023260558.1;refGene_gn=NPHP3;refGene_tx=XM_023260559.1
C2 132350318 132350421
refGene_gn=NPHP3;refGene_tx=XM_023260557.1;refGene_gn=NPHP3;refGene_tx
=XM_023260558.1;refGene_gn=NPHP3;refGene_tx=XM_023260559.1

C2	132350829	132350912
refGene_gn=NPHP3;refGene_tx=XM_023260557.1;refGene_gn=NPHP3;refGene_tx=XM_023260558.1;refGene_gn=NPHP3;refGene_tx=XM_023260559.1
C2	132353417	132353556
refGene_gn=NPHP3;refGene_tx=XM_023260557.1;refGene_gn=NPHP3;refGene_tx=XM_023260558.1;refGene_gn=NPHP3;refGene_tx=XM_023260559.1
C2	132355204	132355369
refGene_gn=NPHP3;refGene_tx=XM_023260557.1;refGene_gn=NPHP3;refGene_tx=XM_023260558.1;refGene_gn=NPHP3;refGene_tx=XM_023260559.1
C2	132356750	132356845
refGene_gn=NPHP3;refGene_tx=XM_023260557.1;refGene_gn=NPHP3;refGene_tx=XM_023260558.1;refGene_gn=NPHP3;refGene_tx=XM_023260559.1
C2	132358066	132358189
refGene_gn=NPHP3;refGene_tx=XM_023260557.1;refGene_gn=NPHP3;refGene_tx=XM_023260558.1;refGene_gn=NPHP3;refGene_tx=XM_023260559.1
C2	132361498	132361688
refGene_gn=NPHP3;refGene_tx=XM_023260557.1;refGene_gn=NPHP3;refGene_tx=XM_023260558.1;refGene_gn=NPHP3;refGene_tx=XM_023260559.1
C2	132361848	132362090
refGene_gn=NPHP3;refGene_tx=XM_023260557.1;refGene_gn=NPHP3;refGene_tx=XM_023260558.1;refGene_gn=NPHP3;refGene_tx=XM_023260559.1
C2	132363140	132363216
refGene_gn=NPHP3;refGene_tx=XM_023260557.1;refGene_gn=NPHP3;refGene_tx=XM_023260558.1;refGene_gn=NPHP3;refGene_tx=XM_023260559.1
C2	132365271	132365399
refGene_gn=NPHP3;refGene_tx=XM_023260557.1;refGene_gn=NPHP3;refGene_tx=XM_023260558.1;refGene_gn=NPHP3;refGene_tx=XM_023260559.1
C2	132366313	132366554
refGene_gn=NPHP3;refGene_tx=XM_023260557.1;refGene_gn=NPHP3;refGene_tx=XM_023260558.1
C2	132367801	132367947
refGene_gn=NPHP3;refGene_tx=XM_023260557.1;refGene_gn=NPHP3;refGene_tx=XM_023260558.1;refGene_gn=NPHP3;refGene_tx=XM_023260559.1
C2	132368749	132368865
refGene_gn=NPHP3;refGene_tx=XM_023260557.1
C2	132369474	132369655
refGene_gn=NPHP3;refGene_tx=XM_023260557.1
C2	132374348	132374432
refGene_gn=UBA5;refGene_tx=XM_003992116.5;refGene_gn=UBA5;refGene_tx=XM_019810699.2;refGene_gn=UBA5;refGene_tx=XM_019810701.2;refGene_gn=UBA5;refGene_tx=XM_019810702.2;refGene_gn=UBA5;refGene_tx=XM_019810703.2
C2	132374964	132375071
refGene_gn=UBA5;refGene_tx=XM_003992116.5;refGene_gn=UBA5;refGene_tx=XM_019810699.2;refGene_gn=UBA5;refGene_tx=XM_019810701.2;refGene_gn=UBA5;refGene_tx=XM_019810702.2;refGene_gn=UBA5;refGene_tx=XM_019810703.2
C2	132375216	132375292
refGene_gn=UBA5;refGene_tx=XM_003992116.5;refGene_gn=UBA5;refGene_tx=XM_019810699.2;refGene_gn=UBA5;refGene_tx=XM_019810701.2;refGene_gn=UBA5;refGene_tx=XM_019810702.2;refGene_gn=UBA5;refGene_tx=XM_019810703.2
C2	132375582	132375718

refGene_gn=UBA5;refGene_tx=XM_003992116.5;refGene_gn=UBA5;refGene_tx=XM_019810699.2;refGene_gn=UBA5;refGene_tx=XM_019810701.2;refGene_gn=UBA5;refGene_tx=XM_019810702.2;refGene_gn=UBA5;refGene_tx=XM_019810703.2
C2 132377390 132377518
refGene_gn=UBA5;refGene_tx=XM_003992116.5;refGene_gn=UBA5;refGene_tx=XM_019810699.2;refGene_gn=UBA5;refGene_tx=XM_019810701.2;refGene_gn=UBA5;refGene_tx=XM_019810702.2;refGene_gn=UBA5;refGene_tx=XM_019810703.2
C2 132377687 132377792
refGene_gn=UBA5;refGene_tx=XM_003992116.5;refGene_gn=UBA5;refGene_tx=XM_019810699.2;refGene_gn=UBA5;refGene_tx=XM_019810701.2;refGene_gn=UBA5;refGene_tx=XM_019810702.2;refGene_gn=UBA5;refGene_tx=XM_019810703.2
C2 132378937 132379022
refGene_gn=UBA5;refGene_tx=XM_003992116.5;refGene_gn=UBA5;refGene_tx=XM_019810699.2;refGene_gn=UBA5;refGene_tx=XM_019810701.2;refGene_gn=UBA5;refGene_tx=XM_019810702.2;refGene_gn=UBA5;refGene_tx=XM_019810703.2
C2 132379882 132379969
refGene_gn=UBA5;refGene_tx=XM_003992116.5;refGene_gn=UBA5;refGene_tx=XM_019810699.2;refGene_gn=UBA5;refGene_tx=XM_019810701.2;refGene_gn=UBA5;refGene_tx=XM_019810702.2;refGene_gn=UBA5;refGene_tx=XM_019810703.2
C2 132380784 132380894
refGene_gn=UBA5;refGene_tx=XM_003992116.5;refGene_gn=UBA5;refGene_tx=XM_019810699.2;refGene_gn=UBA5;refGene_tx=XM_019810701.2;refGene_gn=UBA5;refGene_tx=XM_019810702.2;refGene_gn=UBA5;refGene_tx=XM_019810703.2
C2 132385422 132385569
refGene_gn=UBA5;refGene_tx=XM_003992116.5;refGene_gn=UBA5;refGene_tx=XM_019810699.2
C2 132385678 132385724
refGene_gn=UBA5;refGene_tx=XM_003992116.5;refGene_gn=UBA5;refGene_tx=XM_019810699.2
C2 132387048 132387209
refGene_gn=UBA5;refGene_tx=XM_003992116.5;refGene_gn=UBA5;refGene_tx=XM_019810699.2
C2 132388012 132388161
refGene_gn=ACAD11;refGene_tx=XM_003992118.5
C2 132395506 132395606
refGene_gn=ACAD11;refGene_tx=XM_003992118.5
C2 132398289 132398415
refGene_gn=ACAD11;refGene_tx=XM_003992118.5
C2 132399023 132399185
refGene_gn=ACAD11;refGene_tx=XM_003992118.5
C2 132400729 132400894
refGene_gn=ACAD11;refGene_tx=XM_003992118.5
C2 132415545 132415687
refGene_gn=ACAD11;refGene_tx=XM_003992118.5
C2 132416047 132416169
refGene_gn=ACAD11;refGene_tx=XM_003992118.5
C2 132418095 132418202
refGene_gn=ACAD11;refGene_tx=XM_003992118.5
C2 132419644 132419771
refGene_gn=ACAD11;refGene_tx=XM_003992118.5

C2	132426949	132427027
refGene_gn=ACAD11;refGene_tx=XM_003992118.5
C2	132427778	132427917
refGene_gn=ACAD11;refGene_tx=XM_003992118.5
C2	132432702	132432810
refGene_gn=ACAD11;refGene_tx=XM_003992118.5
C2	132434608	132434707
refGene_gn=ACAD11;refGene_tx=XM_003992118.5
C2	132435910	132436963
refGene_gn=ACKR4;refGene_tx=XM_003992117.5
C2	132461192	132461259
refGene_gn=ACAD11;refGene_tx=XM_003992118.5
C2	132461895	132461981
refGene_gn=ACAD11;refGene_tx=XM_003992118.5
C2	132464318	132464390
refGene_gn=ACAD11;refGene_tx=XM_003992118.5
C2	132465871	132466026
refGene_gn=ACAD11;refGene_tx=XM_003992118.5
C2	132476710	132476827
refGene_gn=ACAD11;refGene_tx=XM_003992118.5
C2	132477792	132477902
refGene_gn=ACAD11;refGene_tx=XM_003992118.5
C2	132478742	132478857
refGene_gn=ACAD11;refGene_tx=XM_003992118.5
C2	132501748	132501855
refGene_gn=DNAJC13;refGene_tx=XM_023260560.1;refGene_gn=DNAJC13;refGen
e_tx=XM_023260561.1
C2	132507845	132507945
refGene_gn=DNAJC13;refGene_tx=XM_023260560.1;refGene_gn=DNAJC13;refGen
e_tx=XM_023260561.1
C2	132510348	132510492
refGene_gn=DNAJC13;refGene_tx=XM_023260560.1;refGene_gn=DNAJC13;refGen
e_tx=XM_023260561.1
C2	132512305	132512446
refGene_gn=DNAJC13;refGene_tx=XM_023260560.1;refGene_gn=DNAJC13;refGen
e_tx=XM_023260561.1
C2	132512726	132512906
refGene_gn=DNAJC13;refGene_tx=XM_023260560.1;refGene_gn=DNAJC13;refGen
e_tx=XM_023260561.1
C2	132514682	132514856
refGene_gn=DNAJC13;refGene_tx=XM_023260560.1;refGene_gn=DNAJC13;refGen
e_tx=XM_023260561.1
C2	132515196	132515239
refGene_gn=DNAJC13;refGene_tx=XM_023260560.1;refGene_gn=DNAJC13;refGen
e_tx=XM_023260561.1
C2	132515387	132515557
refGene_gn=DNAJC13;refGene_tx=XM_023260560.1;refGene_gn=DNAJC13;refGen
e_tx=XM_023260561.1
C2	132522151	132522264
refGene_gn=DNAJC13;refGene_tx=XM_023260560.1;refGene_gn=DNAJC13;refGen

e_tx=XM_023260561.1
C2	132522489	132522564
refGene_gn=DNAJC13;refGene_tx=XM_023260560.1;refGene_gn=DNAJC13;refGene_tx=XM_023260561.1
C2	132524064	132524164
refGene_gn=DNAJC13;refGene_tx=XM_023260560.1;refGene_gn=DNAJC13;refGene_tx=XM_023260561.1
C2	132525856	132525948
refGene_gn=DNAJC13;refGene_tx=XM_023260560.1;refGene_gn=DNAJC13;refGene_tx=XM_023260561.1
C2	132527359	132527537
refGene_gn=DNAJC13;refGene_tx=XM_023260560.1;refGene_gn=DNAJC13;refGene_tx=XM_023260561.1
C2	132528827	132528944
refGene_gn=DNAJC13;refGene_tx=XM_023260560.1;refGene_gn=DNAJC13;refGene_tx=XM_023260561.1
C2	132530266	132530380
refGene_gn=DNAJC13;refGene_tx=XM_023260560.1;refGene_gn=DNAJC13;refGene_tx=XM_023260561.1
C2	132532443	132532611
refGene_gn=DNAJC13;refGene_tx=XM_023260560.1;refGene_gn=DNAJC13;refGene_tx=XM_023260561.1
C2	132533310	132533490
refGene_gn=DNAJC13;refGene_tx=XM_023260560.1;refGene_gn=DNAJC13;refGene_tx=XM_023260561.1
C2	132534451	132534571
refGene_gn=DNAJC13;refGene_tx=XM_023260560.1;refGene_gn=DNAJC13;refGene_tx=XM_023260561.1
C2	132536257	132536332
refGene_gn=DNAJC13;refGene_tx=XM_023260560.1;refGene_gn=DNAJC13;refGene_tx=XM_023260561.1
C2	132536773	132536958
refGene_gn=DNAJC13;refGene_tx=XM_023260560.1;refGene_gn=DNAJC13;refGene_tx=XM_023260561.1
C2	132538378	132538514
refGene_gn=DNAJC13;refGene_tx=XM_023260560.1;refGene_gn=DNAJC13;refGene_tx=XM_023260561.1
C2	132540140	132540219
refGene_gn=DNAJC13;refGene_tx=XM_023260560.1;refGene_gn=DNAJC13;refGene_tx=XM_023260561.1
C2	132541497	132541613
refGene_gn=DNAJC13;refGene_tx=XM_023260560.1;refGene_gn=DNAJC13;refGene_tx=XM_023260561.1
C2	132542914	132543116
refGene_gn=DNAJC13;refGene_tx=XM_023260560.1;refGene_gn=DNAJC13;refGene_tx=XM_023260561.1
C2	132544460	132544615
refGene_gn=DNAJC13;refGene_tx=XM_023260560.1;refGene_gn=DNAJC13;refGene_tx=XM_023260561.1
C2	132546675	132546721

refGene_gn=DNAJC13;refGene_tx=XM_023260560.1;refGene_gn=DNAJC13;refGene_tx=XM_023260561.1
C2	132547175	132547330
refGene_gn=DNAJC13;refGene_tx=XM_023260560.1;refGene_gn=DNAJC13;refGene_tx=XM_023260561.1
C2	132550020	132550105
refGene_gn=DNAJC13;refGene_tx=XM_023260560.1;refGene_gn=DNAJC13;refGene_tx=XM_023260561.1
C2	132551189	132551392
refGene_gn=DNAJC13;refGene_tx=XM_023260560.1;refGene_gn=DNAJC13;refGene_tx=XM_023260561.1
C2	132552211	132552316
refGene_gn=DNAJC13;refGene_tx=XM_023260560.1;refGene_gn=DNAJC13;refGene_tx=XM_023260561.1
C2	132554250	132554352
refGene_gn=DNAJC13;refGene_tx=XM_023260560.1;refGene_gn=DNAJC13;refGene_tx=XM_023260561.1
C2	132555405	132555468
refGene_gn=DNAJC13;refGene_tx=XM_023260560.1;refGene_gn=DNAJC13;refGene_tx=XM_023260561.1
C2	132557917	132558077
refGene_gn=DNAJC13;refGene_tx=XM_023260560.1;refGene_gn=DNAJC13;refGene_tx=XM_023260561.1
C2	132558164	132558268
refGene_gn=DNAJC13;refGene_tx=XM_023260560.1;refGene_gn=DNAJC13;refGene_tx=XM_023260561.1
C2	132561473	132561627
refGene_gn=DNAJC13;refGene_tx=XM_023260560.1;refGene_gn=DNAJC13;refGene_tx=XM_023260561.1
C2	132563006	132563089
refGene_gn=DNAJC13;refGene_tx=XM_023260560.1;refGene_gn=DNAJC13;refGene_tx=XM_023260561.1
C2	132563668	132563683
refGene_gn=DNAJC13;refGene_tx=XM_023260560.1
C2	132567281	132567425
refGene_gn=DNAJC13;refGene_tx=XM_023260560.1;refGene_gn=DNAJC13;refGene_tx=XM_023260561.1
C2	132568230	132568326
refGene_gn=DNAJC13;refGene_tx=XM_023260560.1;refGene_gn=DNAJC13;refGene_tx=XM_023260561.1
C2	132568582	132568658
refGene_gn=DNAJC13;refGene_tx=XM_023260560.1;refGene_gn=DNAJC13;refGene_tx=XM_023260561.1
C2	132571388	132571510
refGene_gn=DNAJC13;refGene_tx=XM_023260560.1;refGene_gn=DNAJC13;refGene_tx=XM_023260561.1
C2	132573533	132573590
refGene_gn=DNAJC13;refGene_tx=XM_023260560.1;refGene_gn=DNAJC13;refGene_tx=XM_023260561.1
C2	132574300	132574456

refGene_gn=DNAJC13;refGene_tx=XM_023260560.1;refGene_gn=DNAJC13;refGene_tx=XM_023260561.1
C2	132575318	132575426
refGene_gn=DNAJC13;refGene_tx=XM_023260560.1;refGene_gn=DNAJC13;refGene_tx=XM_023260561.1
C2	132577185	132577285
refGene_gn=DNAJC13;refGene_tx=XM_023260560.1;refGene_gn=DNAJC13;refGene_tx=XM_023260561.1
C2	132578156	132578326
refGene_gn=DNAJC13;refGene_tx=XM_023260560.1;refGene_gn=DNAJC13;refGene_tx=XM_023260561.1
C2	132578409	132578489
refGene_gn=DNAJC13;refGene_tx=XM_023260560.1;refGene_gn=DNAJC13;refGene_tx=XM_023260561.1
C2	132578593	132578760
refGene_gn=DNAJC13;refGene_tx=XM_023260560.1;refGene_gn=DNAJC13;refGene_tx=XM_023260561.1
C2	132583472	132583564
refGene_gn=DNAJC13;refGene_tx=XM_023260560.1;refGene_gn=DNAJC13;refGene_tx=XM_023260561.1
C2	132583934	132584030
refGene_gn=DNAJC13;refGene_tx=XM_023260560.1;refGene_gn=DNAJC13;refGene_tx=XM_023260561.1
C2	132584121	132584328
refGene_gn=DNAJC13;refGene_tx=XM_023260560.1;refGene_gn=DNAJC13;refGene_tx=XM_023260561.1
C2	132586850	132587051
refGene_gn=DNAJC13;refGene_tx=XM_023260560.1;refGene_gn=DNAJC13;refGene_tx=XM_023260561.1
C2	132590448	132590490
refGene_gn=DNAJC13;refGene_tx=XM_023260560.1;refGene_gn=DNAJC13;refGene_tx=XM_023260561.1
C2	132590920	132591070
refGene_gn=DNAJC13;refGene_tx=XM_023260560.1;refGene_gn=DNAJC13;refGene_tx=XM_023260561.1
C2	132591865	132591941
refGene_gn=DNAJC13;refGene_tx=XM_023260560.1;refGene_gn=DNAJC13;refGene_tx=XM_023260561.1
C2	132609170	132609238
refGene_gn=DNAJC13;refGene_tx=XM_023260560.1;refGene_gn=DNAJC13;refGene_tx=XM_023260561.1
C2	132682292	132682411
refGene_gn=ACPP;refGene_tx=XM_003992120.3
C2	132684925	132685026
refGene_gn=ACPP;refGene_tx=XM_019810704.1
C2	132691888	132691914
refGene_gn=ACPP;refGene_tx=XM_019810705.1
C2	132692583	132692753
refGene_gn=ACPP;refGene_tx=XM_003992120.3;refGene_gn=ACPP;refGene_tx=XM_019810704.1;refGene_gn=ACPP;refGene_tx=XM_019810705.1

C2	132697178	132697282
refGene_gn=ACPP;refGene_tx=XM_003992120.3;refGene_gn=ACPP;refGene_tx=XM_019810704.1;refGene_gn=ACPP;refGene_tx=XM_019810705.1
C2	132699123	132699206
refGene_gn=ACPP;refGene_tx=XM_003992120.3;refGene_gn=ACPP;refGene_tx=XM_019810704.1;refGene_gn=ACPP;refGene_tx=XM_019810705.1
C2	132703559	132703692
refGene_gn=ACPP;refGene_tx=XM_003992120.3;refGene_gn=ACPP;refGene_tx=XM_019810704.1;refGene_gn=ACPP;refGene_tx=XM_019810705.1
C2	132705301	132705394
refGene_gn=ACPP;refGene_tx=XM_003992120.3;refGene_gn=ACPP;refGene_tx=XM_019810704.1;refGene_gn=ACPP;refGene_tx=XM_019810705.1
C2	132709786	132709885
refGene_gn=ACPP;refGene_tx=XM_003992120.3;refGene_gn=ACPP;refGene_tx=XM_019810704.1;refGene_gn=ACPP;refGene_tx=XM_019810705.1
C2	132714567	132714720
refGene_gn=ACPP;refGene_tx=XM_003992120.3;refGene_gn=ACPP;refGene_tx=XM_019810704.1;refGene_gn=ACPP;refGene_tx=XM_019810705.1
C2	132715388	132715475
refGene_gn=ACPP;refGene_tx=XM_003992120.3;refGene_gn=ACPP;refGene_tx=XM_019810704.1;refGene_gn=ACPP;refGene_tx=XM_019810705.1
C2	132719533	132719629
refGene_gn=ACPP;refGene_tx=XM_003992120.3;refGene_gn=ACPP;refGene_tx=XM_019810704.1;refGene_gn=ACPP;refGene_tx=XM_019810705.1
C2	132727318	132727438
refGene_gn=ACPP;refGene_tx=XM_003992120.3;refGene_gn=ACPP;refGene_tx=XM_019810704.1;refGene_gn=ACPP;refGene_tx=XM_019810705.1
C2	132978299	132978352
refGene_gn=CPNE4;refGene_tx=XM_019810707.1
C2	133117954	133118135
refGene_gn=CPNE4;refGene_tx=XM_003992123.4;refGene_gn=CPNE4;refGene_tx=XM_019810707.1
C2	133309907	133310087
refGene_gn=CPNE4;refGene_tx=XM_003992123.4;refGene_gn=CPNE4;refGene_tx=XM_019810707.1
C2	133341853	133341925
refGene_gn=CPNE4;refGene_tx=XM_003992123.4;refGene_gn=CPNE4;refGene_tx=XM_019810707.1
C2	133346590	133346665
refGene_gn=CPNE4;refGene_tx=XM_003992123.4;refGene_gn=CPNE4;refGene_tx=XM_019810707.1
C2	133356398	133356482
refGene_gn=CPNE4;refGene_tx=XM_003992123.4;refGene_gn=CPNE4;refGene_tx=XM_019810707.1
C2	133359611	133359701
refGene_gn=CPNE4;refGene_tx=XM_003992123.4;refGene_gn=CPNE4;refGene_tx=XM_019810707.1
C2	133427680	133427779
refGene_gn=CPNE4;refGene_tx=XM_003992123.4;refGene_gn=CPNE4;refGene_tx=XM_019810707.1

C2 133433990 133434077
refGene_gn=CPNE4;refGene_tx=XM_003992123.4;refGene_gn=CPNE4;refGene_tx
=XM_019810707.1
C2 133437363 133437423
refGene_gn=CPNE4;refGene_tx=XM_003992123.4;refGene_gn=CPNE4;refGene_tx
=XM_019810707.1
C2 133449177 133449311
refGene_gn=CPNE4;refGene_tx=XM_003992123.4;refGene_gn=CPNE4;refGene_tx
=XM_019810707.1
C2 133457648 133457703
refGene_gn=CPNE4;refGene_tx=XM_003992123.4;refGene_gn=CPNE4;refGene_tx
=XM_019810707.1
C2 133460611 133460663
refGene_gn=CPNE4;refGene_tx=XM_003992123.4;refGene_gn=CPNE4;refGene_tx
=XM_019810707.1
C2 133462258 133462392
refGene_gn=CPNE4;refGene_tx=XM_003992123.4;refGene_gn=CPNE4;refGene_tx
=XM_019810707.1
C2 133468454 133468691
refGene_gn=CPNE4;refGene_tx=XM_003992123.4;refGene_gn=CPNE4;refGene_tx
=XM_019810707.1
C2 133474497 133474632
refGene_gn=CPNE4;refGene_tx=XM_003992123.4;refGene_gn=CPNE4;refGene_tx
=XM_019810707.1
C2 133501762 133501854
refGene_gn=MRPL3;refGene_tx=XM_003992124.5;refGene_gn=MRPL3;refGene_tx
=XM_019810708.1
C2 133503097 133503282
refGene_gn=MRPL3;refGene_tx=XM_003992124.5;refGene_gn=MRPL3;refGene_tx
=XM_019810708.1
C2 133504542 133504634
refGene_gn=MRPL3;refGene_tx=XM_003992124.5;refGene_gn=MRPL3;refGene_tx
=XM_019810708.1
C2 133513174 133513273
refGene_gn=MRPL3;refGene_tx=XM_003992124.5;refGene_gn=MRPL3;refGene_tx
=XM_019810708.1
C2 133520456 133520556
refGene_gn=MRPL3;refGene_tx=XM_003992124.5;refGene_gn=MRPL3;refGene_tx
=XM_019810708.1
C2 133522593 133522654
refGene_gn=MRPL3;refGene_tx=XM_003992124.5;refGene_gn=MRPL3;refGene_tx
=XM_019810708.1
C2 133536149 133536258
refGene_gn=MRPL3;refGene_tx=XM_003992124.5;refGene_gn=MRPL3;refGene_tx
=XM_019810708.1
C2 133539018 133539096
refGene_gn=MRPL3;refGene_tx=XM_003992124.5;refGene_gn=MRPL3;refGene_tx
=XM_019810708.1
C2 133544861 133544939
refGene_gn=MRPL3;refGene_tx=XM_003992124.5;refGene_gn=MRPL3;refGene_tx

=XM_019810708.1
C2 133555112 133555287
refGene_gn=MRPL3;refGene_tx=XM_003992124.5;refGene_gn=MRPL3;refGene_tx=XM_019810708.1
C2 133639573 133639753
refGene_gn=NUDT16;refGene_tx=XM_023260563.1
C2 133640502 133640772
refGene_gn=NUDT16;refGene_tx=XM_023260563.1
C2 133640870 133641008
refGene_gn=NUDT16;refGene_tx=XM_023260563.1
C2 133654976 133655247
refGene_gn=NEK11;refGene_tx=XM_023260564.1;refGene_gn=NEK11;refGene_tx=XM_023260565.1;refGene_gn=NEK11;refGene_tx=XM_023260566.1;refGene_gn=NEK11;refGene_tx=XM_023260567.1;refGene_gn=NEK11;refGene_tx=XM_023260568.1;refGene_gn=NEK11;refGene_tx=XM_023260569.1;refGene_gn=NEK11;refGene_tx=XM_023260570.1;refGene_gn=NEK11;refGene_tx=XM_023260571.1;refGene_gn=NEK11;refGene_tx=XM_023260572.1
C2 133710541 133710638
refGene_gn=NEK11;refGene_tx=XM_023260564.1;refGene_gn=NEK11;refGene_tx=XM_023260565.1;refGene_gn=NEK11;refGene_tx=XM_023260566.1;refGene_gn=NEK11;refGene_tx=XM_023260567.1;refGene_gn=NEK11;refGene_tx=XM_023260568.1;refGene_gn=NEK11;refGene_tx=XM_023260569.1;refGene_gn=NEK11;refGene_tx=XM_023260573.1
C2 133739175 133739236
refGene_gn=NEK11;refGene_tx=XM_023260564.1;refGene_gn=NEK11;refGene_tx=XM_023260565.1;refGene_gn=NEK11;refGene_tx=XM_023260566.1;refGene_gn=NEK11;refGene_tx=XM_023260567.1;refGene_gn=NEK11;refGene_tx=XM_023260568.1;refGene_gn=NEK11;refGene_tx=XM_023260569.1;refGene_gn=NEK11;refGene_tx=XM_023260570.1;refGene_gn=NEK11;refGene_tx=XM_023260571.1;refGene_gn=NEK11;refGene_tx=XM_023260572.1
C2 133744870 133744915
refGene_gn=NEK11;refGene_tx=XM_023260574.1
C2 133762715 133762876
refGene_gn=NEK11;refGene_tx=XM_023260564.1;refGene_gn=NEK11;refGene_tx=XM_023260565.1;refGene_gn=NEK11;refGene_tx=XM_023260566.1;refGene_gn=NEK11;refGene_tx=XM_023260567.1;refGene_gn=NEK11;refGene_tx=XM_023260568.1;refGene_gn=NEK11;refGene_tx=XM_023260569.1;refGene_gn=NEK11;refGene_tx=XM_023260570.1;refGene_gn=NEK11;refGene_tx=XM_023260571.1;refGene_gn=NEK11;refGene_tx=XM_023260572.1;refGene_gn=NEK11;refGene_tx=XM_023260573.1;refGene_gn=NEK11;refGene_tx=XM_023260574.1
C2 133805566 133805681
refGene_gn=NEK11;refGene_tx=XM_023260564.1;refGene_gn=NEK11;refGene_tx=XM_023260565.1;refGene_gn=NEK11;refGene_tx=XM_023260566.1;refGene_gn=NEK11;refGene_tx=XM_023260567.1;refGene_gn=NEK11;refGene_tx=XM_023260568.1;refGene_gn=NEK11;refGene_tx=XM_023260569.1;refGene_gn=NEK11;refGene_tx=XM_023260570.1;refGene_gn=NEK11;refGene_tx=XM_023260571.1;refGene_gn=NEK11;refGene_tx=XM_023260572.1;refGene_gn=NEK11;refGene_tx=XM_023260573.1;refGene_gn=NEK11;refGene_tx=XM_023260574.1
C2 133811485 133811579
refGene_gn=NEK11;refGene_tx=XM_023260564.1;refGene_gn=NEK11;refGene_tx

=XM_023260565.1;refGene_gn=NEK11;refGene_tx=XM_023260566.1;refGene_gn=NEK11;refGene_tx=XM_023260567.1;refGene_gn=NEK11;refGene_tx=XM_023260568.1;refGene_gn=NEK11;refGene_tx=XM_023260569.1;refGene_gn=NEK11;refGene_tx=XM_023260570.1;refGene_gn=NEK11;refGene_tx=XM_023260571.1;refGene_gn=NEK11;refGene_tx=XM_023260572.1;refGene_gn=NEK11;refGene_tx=XM_023260573.1;refGene_gn=NEK11;refGene_tx=XM_023260574.1
C2 133813963 133814083
refGene_gn=NEK11;refGene_tx=XM_023260564.1;refGene_gn=NEK11;refGene_tx=XM_023260565.1;refGene_gn=NEK11;refGene_tx=XM_023260566.1;refGene_gn=NEK11;refGene_tx=XM_023260567.1;refGene_gn=NEK11;refGene_tx=XM_023260568.1;refGene_gn=NEK11;refGene_tx=XM_023260569.1;refGene_gn=NEK11;refGene_tx=XM_023260570.1;refGene_gn=NEK11;refGene_tx=XM_023260571.1;refGene_gn=NEK11;refGene_tx=XM_023260572.1;refGene_gn=NEK11;refGene_tx=XM_023260573.1;refGene_gn=NEK11;refGene_tx=XM_023260574.1
C2 133817133 133817219
refGene_gn=NEK11;refGene_tx=XM_023260564.1;refGene_gn=NEK11;refGene_tx=XM_023260565.1;refGene_gn=NEK11;refGene_tx=XM_023260566.1;refGene_gn=NEK11;refGene_tx=XM_023260567.1;refGene_gn=NEK11;refGene_tx=XM_023260568.1;refGene_gn=NEK11;refGene_tx=XM_023260569.1;refGene_gn=NEK11;refGene_tx=XM_023260570.1;refGene_gn=NEK11;refGene_tx=XM_023260571.1;refGene_gn=NEK11;refGene_tx=XM_023260572.1;refGene_gn=NEK11;refGene_tx=XM_023260573.1;refGene_gn=NEK11;refGene_tx=XM_023260574.1
C2 133819548 133819584
refGene_gn=NEK11;refGene_tx=XM_023260564.1;refGene_gn=NEK11;refGene_tx=XM_023260565.1;refGene_gn=NEK11;refGene_tx=XM_023260566.1;refGene_gn=NEK11;refGene_tx=XM_023260569.1;refGene_gn=NEK11;refGene_tx=XM_023260570.1;refGene_gn=NEK11;refGene_tx=XM_023260571.1;refGene_gn=NEK11;refGene_tx=XM_023260573.1;refGene_gn=NEK11;refGene_tx=XM_023260574.1
C2 133820008 133820087
refGene_gn=NEK11;refGene_tx=XM_023260564.1;refGene_gn=NEK11;refGene_tx=XM_023260565.1;refGene_gn=NEK11;refGene_tx=XM_023260566.1;refGene_gn=NEK11;refGene_tx=XM_023260567.1;refGene_gn=NEK11;refGene_tx=XM_023260568.1;refGene_gn=NEK11;refGene_tx=XM_023260569.1;refGene_gn=NEK11;refGene_tx=XM_023260570.1;refGene_gn=NEK11;refGene_tx=XM_023260571.1;refGene_gn=NEK11;refGene_tx=XM_023260572.1;refGene_gn=NEK11;refGene_tx=XM_023260573.1;refGene_gn=NEK11;refGene_tx=XM_023260574.1
C2 133820223 133820373
refGene_gn=NEK11;refGene_tx=XM_023260564.1;refGene_gn=NEK11;refGene_tx=XM_023260565.1;refGene_gn=NEK11;refGene_tx=XM_023260566.1;refGene_gn=NEK11;refGene_tx=XM_023260567.1;refGene_gn=NEK11;refGene_tx=XM_023260568.1;refGene_gn=NEK11;refGene_tx=XM_023260569.1;refGene_gn=NEK11;refGene_tx=XM_023260570.1;refGene_gn=NEK11;refGene_tx=XM_023260571.1;refGene_gn=NEK11;refGene_tx=XM_023260572.1;refGene_gn=NEK11;refGene_tx=XM_023260573.1;refGene_gn=NEK11;refGene_tx=XM_023260574.1
C2 133839756 133839883
refGene_gn=NEK11;refGene_tx=XM_023260564.1;refGene_gn=NEK11;refGene_tx=XM_023260565.1;refGene_gn=NEK11;refGene_tx=XM_023260566.1;refGene_gn=NEK11;refGene_tx=XM_023260567.1;refGene_gn=NEK11;refGene_tx=XM_023260568.1;refGene_gn=NEK11;refGene_tx=XM_023260569.1;refGene_gn=NEK11;refGene_tx=XM_023260570.1;refGene_gn=NEK11;refGene_tx=XM_023260571.1;refGen

e_gn=NEK11;refGene_tx=XM_023260572.1;refGene_gn=NEK11;refGene_tx=XM_02
3260573.1;refGene_gn=NEK11;refGene_tx=XM_023260574.1
C2 133841086 133841151
refGene_gn=NEK11;refGene_tx=XM_023260564.1;refGene_gn=NEK11;refGene_tx
=XM_023260565.1;refGene_gn=NEK11;refGene_tx=XM_023260566.1;refGene_gn=
NEK11;refGene_tx=XM_023260567.1;refGene_gn=NEK11;refGene_tx=XM_0232605
68.1;refGene_gn=NEK11;refGene_tx=XM_023260569.1;refGene_gn=NEK11;refGe
ne_tx=XM_023260570.1;refGene_gn=NEK11;refGene_tx=XM_023260571.1;refGen
e_gn=NEK11;refGene_tx=XM_023260572.1;refGene_gn=NEK11;refGene_tx=XM_02
3260573.1;refGene_gn=NEK11;refGene_tx=XM_023260574.1
C2 133864448 133864567
refGene_gn=NEK11;refGene_tx=XM_023260564.1;refGene_gn=NEK11;refGene_tx
=XM_023260565.1;refGene_gn=NEK11;refGene_tx=XM_023260566.1;refGene_gn=
NEK11;refGene_tx=XM_023260567.1;refGene_gn=NEK11;refGene_tx=XM_0232605
68.1;refGene_gn=NEK11;refGene_tx=XM_023260569.1;refGene_gn=NEK11;refGe
ne_tx=XM_023260570.1;refGene_gn=NEK11;refGene_tx=XM_023260571.1;refGen
e_gn=NEK11;refGene_tx=XM_023260572.1;refGene_gn=NEK11;refGene_tx=XM_02
3260573.1;refGene_gn=NEK11;refGene_tx=XM_023260574.1
C2 133878795 133878961
refGene_gn=NEK11;refGene_tx=XM_023260564.1;refGene_gn=NEK11;refGene_tx
=XM_023260565.1;refGene_gn=NEK11;refGene_tx=XM_023260566.1;refGene_gn=
NEK11;refGene_tx=XM_023260567.1;refGene_gn=NEK11;refGene_tx=XM_0232605
68.1;refGene_gn=NEK11;refGene_tx=XM_023260569.1;refGene_gn=NEK11;refGe
ne_tx=XM_023260570.1;refGene_gn=NEK11;refGene_tx=XM_023260571.1;refGen
e_gn=NEK11;refGene_tx=XM_023260572.1;refGene_gn=NEK11;refGene_tx=XM_02
3260573.1;refGene_gn=NEK11;refGene_tx=XM_023260574.1
C2 133884916 133884984
refGene_gn=NEK11;refGene_tx=XM_023260569.1
C2 133921905 133922075
refGene_gn=NEK11;refGene_tx=XM_023260564.1;refGene_gn=NEK11;refGene_tx
=XM_023260566.1;refGene_gn=NEK11;refGene_tx=XM_023260567.1;refGene_gn=
NEK11;refGene_tx=XM_023260568.1;refGene_gn=NEK11;refGene_tx=XM_0232605
70.1;refGene_gn=NEK11;refGene_tx=XM_023260571.1;refGene_gn=NEK11;refGe
ne_tx=XM_023260572.1;refGene_gn=NEK11;refGene_tx=XM_023260573.1;refGen
e_gn=NEK11;refGene_tx=XM_023260574.1
C2 133925175 133925345
refGene_gn=NEK11;refGene_tx=XM_023260565.1
C2 133926925 133928230
refGene_gn=ASTE1;refGene_tx=XM_011286207.3;refGene_gn=ASTE1;refGene_tx
=XM_019810712.2;refGene_gn=ASTE1;refGene_tx=XM_019810713.2;refGene_gn=
ASTE1;refGene_tx=XM_019810714.2
C2 133934585 133934796
refGene_gn=ASTE1;refGene_tx=XM_011286207.3;refGene_gn=ASTE1;refGene_tx
=XM_019810712.2;refGene_gn=ASTE1;refGene_tx=XM_019810713.2;refGene_gn=
ASTE1;refGene_tx=XM_019810714.2
C2 133936632 133936858
refGene_gn=ASTE1;refGene_tx=XM_011286207.3;refGene_gn=ASTE1;refGene_tx
=XM_019810712.2;refGene_gn=ASTE1;refGene_tx=XM_019810713.2;refGene_gn=
ASTE1;refGene_tx=XM_019810714.2;refGene_gn=ATP2C1;refGene_tx=XM_011286
227.3;refGene_gn=ATP2C1;refGene_tx=XM_019810711.2

C2	133938594	133938925
refGene_gn=ASTE1;refGene_tx=XM_011286207.3;refGene_gn=ASTE1;refGene_tx=XM_019810712.2;refGene_gn=ASTE1;refGene_tx=XM_019810713.2;refGene_gn=ASTE1;refGene_tx=XM_019810714.2
C2	133952809	133952934
refGene_gn=ATP2C1;refGene_tx=XM_011286227.3;refGene_gn=ATP2C1;refGene_tx=XM_019810711.2
C2	133954174	133954316
refGene_gn=ATP2C1;refGene_tx=XM_011286227.3;refGene_gn=ATP2C1;refGene_tx=XM_019810711.2
C2	133957500	133957596
refGene_gn=ATP2C1;refGene_tx=XM_011286227.3;refGene_gn=ATP2C1;refGene_tx=XM_019810711.2
C2	133958134	133958282
refGene_gn=ATP2C1;refGene_tx=XM_011286227.3;refGene_gn=ATP2C1;refGene_tx=XM_019810711.2
C2	133959115	133959232
refGene_gn=ATP2C1;refGene_tx=XM_011286227.3;refGene_gn=ATP2C1;refGene_tx=XM_019810711.2
C2	133959863	133959932
refGene_gn=ATP2C1;refGene_tx=XM_011286227.3;refGene_gn=ATP2C1;refGene_tx=XM_019810711.2
C2	133961298	133961465
refGene_gn=ATP2C1;refGene_tx=XM_011286227.3;refGene_gn=ATP2C1;refGene_tx=XM_019810711.2
C2	133962387	133962438
refGene_gn=ATP2C1;refGene_tx=XM_011286227.3;refGene_gn=ATP2C1;refGene_tx=XM_019810711.2
C2	133971912	133972010
refGene_gn=ATP2C1;refGene_tx=XM_011286227.3;refGene_gn=ATP2C1;refGene_tx=XM_019810711.2
C2	133973597	133973768
refGene_gn=ATP2C1;refGene_tx=XM_011286227.3;refGene_gn=ATP2C1;refGene_tx=XM_019810711.2
C2	133976930	133977087
refGene_gn=ATP2C1;refGene_tx=XM_011286227.3;refGene_gn=ATP2C1;refGene_tx=XM_019810711.2
C2	133981309	133981414
refGene_gn=ATP2C1;refGene_tx=XM_011286227.3;refGene_gn=ATP2C1;refGene_tx=XM_019810711.2
C2	133983258	133983348
refGene_gn=ATP2C1;refGene_tx=XM_011286227.3;refGene_gn=ATP2C1;refGene_tx=XM_019810711.2
C2	133983435	133983531
refGene_gn=ATP2C1;refGene_tx=XM_011286227.3;refGene_gn=ATP2C1;refGene_tx=XM_019810711.2
C2	133985486	133985584
refGene_gn=ATP2C1;refGene_tx=XM_011286227.3;refGene_gn=ATP2C1;refGene_tx=XM_019810711.2
C2	133986466	133986591

refGene_gn=ATP2C1;refGene_tx=XM_011286227.3;refGene_gn=ATP2C1;refGene_tx=XM_019810711.2
C2 133991814 133991881
refGene_gn=ATP2C1;refGene_tx=XM_011286227.3;refGene_gn=ATP2C1;refGene_tx=XM_019810711.2
C2 133996417 133996493
refGene_gn=ATP2C1;refGene_tx=XM_011286227.3;refGene_gn=ATP2C1;refGene_tx=XM_019810711.2
C2 133997449 133997518
refGene_gn=ATP2C1;refGene_tx=XM_011286227.3;refGene_gn=ATP2C1;refGene_tx=XM_019810711.2
C2 133998923 133999079
refGene_gn=ATP2C1;refGene_tx=XM_011286227.3;refGene_gn=ATP2C1;refGene_tx=XM_019810711.2
C2 134016881 134016990
refGene_gn=ATP2C1;refGene_tx=XM_011286227.3;refGene_gn=ATP2C1;refGene_tx=XM_019810711.2
C2 134021785 134021847
refGene_gn=ATP2C1;refGene_tx=XM_011286227.3;refGene_gn=ATP2C1;refGene_tx=XM_019810711.2
C2 134024683 134024719
refGene_gn=ATP2C1;refGene_tx=XM_011286227.3;refGene_gn=ATP2C1;refGene_tx=XM_019810711.2
C2 134028832 134028922
refGene_gn=ATP2C1;refGene_tx=XM_011286227.3;refGene_gn=ATP2C1;refGene_tx=XM_019810711.2
C2 134030214 134030331
refGene_gn=ATP2C1;refGene_tx=XM_011286227.3;refGene_gn=ATP2C1;refGene_tx=XM_019810711.2
C2 134032708 134032819
refGene_gn=ATP2C1;refGene_tx=XM_011286227.3;refGene_gn=ATP2C1;refGene_tx=XM_019810711.2
C2 134110516 134110624
refGene_gn=ATP2C1;refGene_tx=XM_011286227.3;refGene_gn=ATP2C1;refGene_tx=XM_019810711.2
C2 134204849 134205582
refGene_gn=PIK3R4;refGene_tx=XM_003992127.4;refGene_gn=PIK3R4;refGene_tx=XM_019810710.2
C2 134214190 134214324
refGene_gn=PIK3R4;refGene_tx=XM_003992127.4;refGene_gn=PIK3R4;refGene_tx=XM_019810710.2
C2 134216582 134217165
refGene_gn=PIK3R4;refGene_tx=XM_003992127.4;refGene_gn=PIK3R4;refGene_tx=XM_019810710.2
C2 134220025 134220160
refGene_gn=PIK3R4;refGene_tx=XM_003992127.4;refGene_gn=PIK3R4;refGene_tx=XM_019810710.2
C2 134221748 134221970
refGene_gn=PIK3R4;refGene_tx=XM_003992127.4;refGene_gn=PIK3R4;refGene_tx=XM_019810710.2

C2	134228506	134228680
refGene_gn=PIK3R4;refGene_tx=XM_003992127.4;refGene_gn=PIK3R4;refGene_
tx=XM_019810710.2
C2	134233034	134233180
refGene_gn=PIK3R4;refGene_tx=XM_003992127.4;refGene_gn=PIK3R4;refGene_
tx=XM_019810710.2
C2	134235610	134235814
refGene_gn=PIK3R4;refGene_tx=XM_003992127.4;refGene_gn=PIK3R4;refGene_
tx=XM_019810710.2
C2	134245070	134245272
refGene_gn=PIK3R4;refGene_tx=XM_003992127.4;refGene_gn=PIK3R4;refGene_
tx=XM_019810710.2
C2	134247338	134247526
refGene_gn=PIK3R4;refGene_tx=XM_003992127.4;refGene_gn=PIK3R4;refGene_
tx=XM_019810710.2
C2	134248515	134248726
refGene_gn=PIK3R4;refGene_tx=XM_003992127.4;refGene_gn=PIK3R4;refGene_
tx=XM_019810710.2
C2	134251507	134251673
refGene_gn=PIK3R4;refGene_tx=XM_003992127.4;refGene_gn=PIK3R4;refGene_
tx=XM_019810710.2
C2	134267281	134267446
refGene_gn=PIK3R4;refGene_tx=XM_003992127.4;refGene_gn=PIK3R4;refGene_
tx=XM_019810710.2
C2	134271576	134271788
refGene_gn=PIK3R4;refGene_tx=XM_003992127.4;refGene_gn=PIK3R4;refGene_
tx=XM_019810710.2
C2	134273452	134273584
refGene_gn=PIK3R4;refGene_tx=XM_003992127.4;refGene_gn=PIK3R4;refGene_
tx=XM_019810710.2
C2	134276454	134276555
refGene_gn=PIK3R4;refGene_tx=XM_003992127.4;refGene_gn=PIK3R4;refGene_
tx=XM_019810710.2
C2	134277083	134277172
refGene_gn=PIK3R4;refGene_tx=XM_003992127.4;refGene_gn=PIK3R4;refGene_
tx=XM_019810710.2
C2	134277482	134277591
refGene_gn=PIK3R4;refGene_tx=XM_003992127.4;refGene_gn=PIK3R4;refGene_
tx=XM_019810710.2
C2	134278305	134278476
refGene_gn=PIK3R4;refGene_tx=XM_003992127.4;refGene_gn=PIK3R4;refGene_
tx=XM_019810710.2
C2	134282299	134282492
refGene_gn=COL6A6;refGene_tx=XM_011286221.3
C2	134299137	134299231
refGene_gn=COL6A6;refGene_tx=XM_011286221.3
C2	134302116	134302788
refGene_gn=COL6A6;refGene_tx=XM_011286221.3
C2	134306050	134306147
refGene_gn=COL6A6;refGene_tx=XM_011286221.3

C2	134313060	134313554
refGene_gn=COL6A6;refGene_tx=XM_011286221.3
C2	134316980	134316992
refGene_gn=COL6A6;refGene_tx=XM_011286221.3
C2	134318885	134318922
refGene_gn=COL6A6;refGene_tx=XM_011286221.3
C2	134319037	134319073
refGene_gn=COL6A6;refGene_tx=XM_011286221.3
C2	134320142	134320205
refGene_gn=COL6A6;refGene_tx=XM_011286221.3
C2	134331719	134331782
refGene_gn=COL6A6;refGene_tx=XM_011286221.3
C2	134332507	134332543
refGene_gn=COL6A6;refGene_tx=XM_011286221.3
C2	134334645	134334696
refGene_gn=COL6A6;refGene_tx=XM_011286221.3
C2	134335486	134335549
refGene_gn=COL6A6;refGene_tx=XM_011286221.3
C2	134339920	134339983
refGene_gn=COL6A6;refGene_tx=XM_011286221.3
C2	134347177	134347240
refGene_gn=COL6A6;refGene_tx=XM_011286221.3
C2	134348848	134348911
refGene_gn=COL6A6;refGene_tx=XM_011286221.3
C2	134350476	134350512
refGene_gn=COL6A6;refGene_tx=XM_011286221.3
C2	134360380	134360434
refGene_gn=COL6A6;refGene_tx=XM_011286221.3
C2	134361653	134361719
refGene_gn=COL6A6;refGene_tx=XM_011286221.3
C2	134367855	134367918
refGene_gn=COL6A6;refGene_tx=XM_011286221.3
C2	134368958	134369012
refGene_gn=COL6A6;refGene_tx=XM_011286221.3
C2	134369099	134369144
refGene_gn=COL6A6;refGene_tx=XM_011286221.3
C2	134369699	134369726
refGene_gn=COL6A6;refGene_tx=XM_011286221.3
C2	134369812	134369884
refGene_gn=COL6A6;refGene_tx=XM_011286221.3
C2	134371613	134371667
refGene_gn=COL6A6;refGene_tx=XM_011286221.3
C2	134373060	134373153
refGene_gn=COL6A6;refGene_tx=XM_011286221.3
C2	134375189	134375344
refGene_gn=COL6A6;refGene_tx=XM_011286221.3
C2	134379285	134379364
refGene_gn=COL6A6;refGene_tx=XM_011286221.3
C2	134379447	134379791
refGene_gn=COL6A6;refGene_tx=XM_011286221.3

C2 134387532 134388102
refGene_gn=COL6A6;refGene_tx=XM_011286221.3
C2 134389998 134390574
refGene_gn=COL6A6;refGene_tx=XM_011286221.3
C2 134392748 134393306
refGene_gn=COL6A6;refGene_tx=XM_011286221.3
C2 134394071 134394632
refGene_gn=COL6A6;refGene_tx=XM_011286221.3
C2 134395894 134396515
refGene_gn=COL6A6;refGene_tx=XM_011286221.3
C2 134397810 134398410
refGene_gn=COL6A6;refGene_tx=XM_011286221.3
C2 134400672 134400733
refGene_gn=COL6A6;refGene_tx=XM_011286221.3
C2 134476293 134476313
refGene_gn=COL6A5;refGene_tx=XM_023260575.1
C2 134485367 134485612
refGene_gn=COL6A5;refGene_tx=XM_023260575.1
C2 134486369 134486466
refGene_gn=COL6A5;refGene_tx=XM_023260575.1
C2 134487814 134488504
refGene_gn=COL6A5;refGene_tx=XM_023260575.1
C2 134501658 134501846
refGene_gn=COL6A5;refGene_tx=XM_023260575.1
C2 134504567 134504658
refGene_gn=COL6A5;refGene_tx=XM_023260575.1
C2 134507563 134508223
refGene_gn=COL6A5;refGene_tx=XM_023260575.1
C2 134508772 134508860
refGene_gn=COL6A5;refGene_tx=XM_023260575.1
C2 134511709 134512197
refGene_gn=COL6A5;refGene_tx=XM_023260575.1
C2 134514211 134514223
refGene_gn=COL6A5;refGene_tx=XM_023260575.1
C2 134517571 134517608
refGene_gn=COL6A5;refGene_tx=XM_023260575.1
C2 134517725 134517761
refGene_gn=COL6A5;refGene_tx=XM_023260575.1
C2 134520119 134520182
refGene_gn=COL6A5;refGene_tx=XM_023260575.1
C2 134521448 134521511
refGene_gn=COL6A5;refGene_tx=XM_023260575.1
C2 134522905 134522941
refGene_gn=COL6A5;refGene_tx=XM_023260575.1
C2 134523070 134523121
refGene_gn=COL6A5;refGene_tx=XM_023260575.1
C2 134524552 134524615
refGene_gn=COL6A5;refGene_tx=XM_023260575.1
C2 134526419 134526482
refGene_gn=COL6A5;refGene_tx=XM_023260575.1

C2 134527532 134527595
refGene_gn=COL6A5;refGene_tx=XM_023260575.1
C2 134529036 134529099
refGene_gn=COL6A5;refGene_tx=XM_023260575.1
C2 134529610 134529646
refGene_gn=COL6A5;refGene_tx=XM_023260575.1
C2 134533313 134533367
refGene_gn=COL6A5;refGene_tx=XM_023260575.1
C2 134533820 134533886
refGene_gn=COL6A5;refGene_tx=XM_023260575.1
C2 134535003 134535066
refGene_gn=COL6A5;refGene_tx=XM_023260575.1
C2 134536951 134537005
refGene_gn=COL6A5;refGene_tx=XM_023260575.1
C2 134537094 134537139
refGene_gn=COL6A5;refGene_tx=XM_023260575.1
C2 134537249 134537276
refGene_gn=COL6A5;refGene_tx=XM_023260575.1
C2 134537664 134537736
refGene_gn=COL6A5;refGene_tx=XM_023260575.1
C2 134539483 134539537
refGene_gn=COL6A5;refGene_tx=XM_023260575.1
C2 134540887 134540980
refGene_gn=COL6A5;refGene_tx=XM_023260575.1
C2 134541788 134541931
refGene_gn=COL6A5;refGene_tx=XM_023260575.1
C2 134544072 134544154
refGene_gn=COL6A5;refGene_tx=XM_023260575.1
C2 134544267 134544608
refGene_gn=COL6A5;refGene_tx=XM_023260575.1
C2 134546696 134547272
refGene_gn=COL6A5;refGene_tx=XM_023260575.1
C2 134549910 134550486
refGene_gn=COL6A5;refGene_tx=XM_023260575.1
C2 134552878 134553433
refGene_gn=COL6A5;refGene_tx=XM_023260575.1
C2 134556509 134557070
refGene_gn=COL6A5;refGene_tx=XM_023260575.1
C2 134562959 134563580
refGene_gn=COL6A5;refGene_tx=XM_023260575.1
C2 134567635 134568238
refGene_gn=COL6A5;refGene_tx=XM_023260575.1
C2 134569433 134569497
refGene_gn=COL6A5;refGene_tx=XM_023260575.1
C2 134677989 134678165
refGene_gn=LOC101096163;refGene_tx=XM_019810719.2
C2 134686546 134686758
refGene_gn=LOC101096163;refGene_tx=XM_019810719.2
C2 134687243 134687340
refGene_gn=LOC101096163;refGene_tx=XM_019810719.2

C2 134689272 134689956
refGene_gn=LOC101096163;refGene_tx=XM_019810719.2
C2 134691416 134691507
refGene_gn=LOC101096163;refGene_tx=XM_019810719.2
C2 134711789 134712283
refGene_gn=LOC101096163;refGene_tx=XM_019810719.2
C2 134712378 134712390
refGene_gn=LOC101096163;refGene_tx=XM_019810719.2
C2 134714708 134714748
refGene_gn=LOC101096163;refGene_tx=XM_019810719.2
C2 134715715 134715751
refGene_gn=LOC101096163;refGene_tx=XM_019810719.2
C2 134716923 134716986
refGene_gn=LOC101096163;refGene_tx=XM_019810719.2
C2 134718245 134718308
refGene_gn=LOC101096163;refGene_tx=XM_019810719.2
C2 134718412 134718448
refGene_gn=LOC101096163;refGene_tx=XM_019810719.2
C2 134723538 134723589
refGene_gn=LOC101096163;refGene_tx=XM_019810719.2
C2 134725551 134725614
refGene_gn=LOC101096163;refGene_tx=XM_019810719.2
C2 134726158 134726221
refGene_gn=LOC101096163;refGene_tx=XM_019810719.2
C2 134727285 134727348
refGene_gn=LOC101096163;refGene_tx=XM_019810719.2
C2 134729102 134729165
refGene_gn=LOC101096163;refGene_tx=XM_019810719.2
C2 134733800 134733836
refGene_gn=LOC101096163;refGene_tx=XM_019810719.2
C2 134733953 134734007
refGene_gn=LOC101096163;refGene_tx=XM_019810719.2
C2 134737781 134737847
refGene_gn=LOC101096163;refGene_tx=XM_019810719.2
C2 134738167 134738230
refGene_gn=LOC101096163;refGene_tx=XM_019810719.2
C2 134741409 134741463
refGene_gn=LOC101096163;refGene_tx=XM_019810719.2
C2 134743683 134743728
refGene_gn=LOC101096163;refGene_tx=XM_019810719.2
C2 134746432 134746459
refGene_gn=LOC101096163;refGene_tx=XM_019810719.2
C2 134747525 134747597
refGene_gn=LOC101096163;refGene_tx=XM_019810719.2
C2 134754685 134754739
refGene_gn=LOC101096163;refGene_tx=XM_019810719.2
C2 134756588 134756648
refGene_gn=LOC101096163;refGene_tx=XM_019810719.2
C2 134756649 134756682
refGene_gn=LOC101096163;refGene_tx=XM_019810719.2

C2 134757065 134757211
refGene_gn=LOC101096163;refGene_tx=XM_019810719.2
C2 134758974 134759053
refGene_gn=LOC101096163;refGene_tx=XM_019810719.2
C2 134760040 134760375
refGene_gn=LOC101096163;refGene_tx=XM_019810719.2
C2 134763720 134764207
refGene_gn=LOC101096163;refGene_tx=XM_019810719.2
C2 134764208 134764276
refGene_gn=LOC101096163;refGene_tx=XM_019810719.2
C2 134764622 134765168
refGene_gn=LOC101096163;refGene_tx=XM_019810719.2
C2 134769415 134770045
refGene_gn=LOC101096163;refGene_tx=XM_019810719.2
C2 134774229 134774835
refGene_gn=LOC101096163;refGene_tx=XM_019810719.2
C2 134778212 134778815
refGene_gn=LOC101096163;refGene_tx=XM_019810719.2
C2 134782223 134782299
refGene_gn=LOC101096163;refGene_tx=XM_019810719.2
C2 134854369 134854471
refGene_gn=CAPN7;refGene_tx=XM_006936421.4
C2 134857373 134857482
refGene_gn=CAPN7;refGene_tx=XM_006936421.4
C2 134862149 134862307
refGene_gn=CAPN7;refGene_tx=XM_006936421.4
C2 134863211 134863279
refGene_gn=CAPN7;refGene_tx=XM_006936421.4
C2 134863819 134863893
refGene_gn=CAPN7;refGene_tx=XM_023260576.1
C2 134865183 134865384
refGene_gn=CAPN7;refGene_tx=XM_006936421.4;refGene_gn=CAPN7;refGene_tx=XM_023260576.1
C2 134867105 134867192
refGene_gn=CAPN7;refGene_tx=XM_006936421.4;refGene_gn=CAPN7;refGene_tx=XM_023260576.1
C2 134870352 134870479
refGene_gn=CAPN7;refGene_tx=XM_006936421.4;refGene_gn=CAPN7;refGene_tx=XM_023260576.1
C2 134871265 134871351
refGene_gn=CAPN7;refGene_tx=XM_006936421.4;refGene_gn=CAPN7;refGene_tx=XM_023260576.1
C2 134872551 134872645
refGene_gn=CAPN7;refGene_tx=XM_006936421.4;refGene_gn=CAPN7;refGene_tx=XM_023260576.1
C2 134874073 134874220
refGene_gn=CAPN7;refGene_tx=XM_006936421.4;refGene_gn=CAPN7;refGene_tx=XM_023260576.1
C2 134875431 134875538
refGene_gn=CAPN7;refGene_tx=XM_006936421.4;refGene_gn=CAPN7;refGene_tx

=XM_023260576.1
C2	134876439	134876560
refGene_gn=CAPN7;refGene_tx=XM_006936421.4;refGene_gn=CAPN7;refGene_tx
=XM_023260576.1
C2	134880837	134880982
refGene_gn=CAPN7;refGene_tx=XM_006936421.4;refGene_gn=CAPN7;refGene_tx
=XM_023260576.1
C2	134881106	134881206
refGene_gn=CAPN7;refGene_tx=XM_006936421.4;refGene_gn=CAPN7;refGene_tx
=XM_023260576.1
C2	134881523	134881659
refGene_gn=CAPN7;refGene_tx=XM_006936421.4;refGene_gn=CAPN7;refGene_tx
=XM_023260576.1
C2	134882196	134882272
refGene_gn=CAPN7;refGene_tx=XM_006936421.4;refGene_gn=CAPN7;refGene_tx
=XM_023260576.1
C2	134885749	134885895
refGene_gn=CAPN7;refGene_tx=XM_006936421.4;refGene_gn=CAPN7;refGene_tx
=XM_023260576.1
C2	134886653	134886716
refGene_gn=CAPN7;refGene_tx=XM_006936421.4;refGene_gn=CAPN7;refGene_tx
=XM_023260576.1
C2	134892342	134892473
refGene_gn=CAPN7;refGene_tx=XM_006936421.4;refGene_gn=CAPN7;refGene_tx
=XM_023260576.1
C2	134894640	134894733
refGene_gn=CAPN7;refGene_tx=XM_006936421.4;refGene_gn=CAPN7;refGene_tx
=XM_023260576.1
C2	134894825	134894970
refGene_gn=CAPN7;refGene_tx=XM_006936421.4;refGene_gn=CAPN7;refGene_tx
=XM_023260576.1
C2	134899125	134899328
refGene_gn=SH3BP5;refGene_tx=XM_003992129.4
C2	134899819	134900080
refGene_gn=SH3BP5;refGene_tx=XM_003992129.4
C2	134902855	134903075
refGene_gn=SH3BP5;refGene_tx=XM_003992129.4
C2	134903781	134903824
refGene_gn=SH3BP5;refGene_tx=XM_003992129.4
C2	134906055	134906186
refGene_gn=SH3BP5;refGene_tx=XM_003992129.4
C2	134914363	134914528
refGene_gn=SH3BP5;refGene_tx=XM_003992129.4
C2	134947584	134947713
refGene_gn=SH3BP5;refGene_tx=XM_003992129.4
C2	134973018	134973081
refGene_gn=SH3BP5;refGene_tx=XM_003992129.4
C2	134974779	134974917
refGene_gn=SH3BP5;refGene_tx=XM_003992129.4
C2	135024913	135025095

refGene_gn=METTL6;refGene_tx=XM_023260582.1;refGene_gn=METTL6;refGene_tx=XM_023260583.1;refGene_gn=METTL6;refGene_tx=XM_023260584.1
C2 135026970 135027112
refGene_gn=METTL6;refGene_tx=XM_023260582.1;refGene_gn=METTL6;refGene_tx=XM_023260583.1;refGene_gn=METTL6;refGene_tx=XM_023260584.1
C2 135033677 135033848
refGene_gn=METTL6;refGene_tx=XM_023260582.1;refGene_gn=METTL6;refGene_tx=XM_023260583.1;refGene_gn=METTL6;refGene_tx=XM_023260584.1
C2 135039213 135039348
refGene_gn=METTL6;refGene_tx=XM_023260582.1;refGene_gn=METTL6;refGene_tx=XM_023260583.1
C2 135040788 135041013
refGene_gn=METTL6;refGene_tx=XM_023260582.1;refGene_gn=METTL6;refGene_tx=XM_023260583.1;refGene_gn=METTL6;refGene_tx=XM_023260584.1
C2 135041118 135041165
refGene_gn=EAF1;refGene_tx=XM_023260581.1
C2 135042124 135042630
refGene_gn=EAF1;refGene_tx=XM_023260581.1
C2 135044509 135044604
refGene_gn=EAF1;refGene_tx=XM_023260581.1
C2 135045926 135046062
refGene_gn=COLQ;refGene_tx=XM_006936431.4;refGene_gn=COLQ;refGene_tx=XM_006936432.4
C2 135046455 135046592
refGene_gn=EAF1;refGene_tx=XM_023260581.1
C2 135048761 135048952
refGene_gn=EAF1;refGene_tx=XM_023260581.1
C2 135050612 135050846
refGene_gn=EAF1;refGene_tx=XM_023260581.1
C2 135053424 135053471
refGene_gn=EAF1;refGene_tx=XM_023260581.1
C2 135064812 135064882
refGene_gn=COLQ;refGene_tx=XM_003992132.3;refGene_gn=COLQ;refGene_tx=XM_003992134.4;refGene_gn=COLQ;refGene_tx=XM_023260580.1
C2 135066813 135066916
refGene_gn=COLQ;refGene_tx=XM_003992132.3;refGene_gn=COLQ;refGene_tx=XM_003992134.4;refGene_gn=COLQ;refGene_tx=XM_006936431.4;refGene_gn=COLQ;refGene_tx=XM_006936432.4;refGene_gn=COLQ;refGene_tx=XM_023260580.1
C2 135068281 135068402
refGene_gn=COLQ;refGene_tx=XM_003992132.3;refGene_gn=COLQ;refGene_tx=XM_003992134.4;refGene_gn=COLQ;refGene_tx=XM_006936431.4;refGene_gn=COLQ;refGene_tx=XM_006936432.4;refGene_gn=COLQ;refGene_tx=XM_023260580.1
C2 135068784 135068904
refGene_gn=COLQ;refGene_tx=XM_003992132.3;refGene_gn=COLQ;refGene_tx=XM_003992134.4;refGene_gn=COLQ;refGene_tx=XM_006936431.4;refGene_gn=COLQ;refGene_tx=XM_006936432.4;refGene_gn=COLQ;refGene_tx=XM_023260580.1
C2 135069796 135069936
refGene_gn=COLQ;refGene_tx=XM_003992132.3;refGene_gn=COLQ;refGene_tx=XM_003992134.4;refGene_gn=COLQ;refGene_tx=XM_006936431.4;refGene_gn=COLQ;refGene_tx=XM_006936432.4;refGene_gn=COLQ;refGene_tx=XM_023260580.1

C2 135075726 135075832
refGene_gn=COLQ;refGene_tx=XM_003992132.3;refGene_gn=COLQ;refGene_tx=XM_003992134.4;refGene_gn=COLQ;refGene_tx=XM_006936431.4;refGene_gn=COLQ;refGene_tx=XM_006936432.4;refGene_gn=COLQ;refGene_tx=XM_023260580.1
C2 135080700 135080781
refGene_gn=COLQ;refGene_tx=XM_003992132.3;refGene_gn=COLQ;refGene_tx=XM_003992134.4;refGene_gn=COLQ;refGene_tx=XM_006936431.4;refGene_gn=COLQ;refGene_tx=XM_006936432.4;refGene_gn=COLQ;refGene_tx=XM_023260580.1
C2 135083985 135084021
refGene_gn=COLQ;refGene_tx=XM_003992132.3;refGene_gn=COLQ;refGene_tx=XM_003992134.4;refGene_gn=COLQ;refGene_tx=XM_006936431.4;refGene_gn=COLQ;refGene_tx=XM_006936432.4;refGene_gn=COLQ;refGene_tx=XM_023260580.1
C2 135084229 135084274
refGene_gn=COLQ;refGene_tx=XM_003992132.3;refGene_gn=COLQ;refGene_tx=XM_003992134.4;refGene_gn=COLQ;refGene_tx=XM_006936431.4;refGene_gn=COLQ;refGene_tx=XM_006936432.4;refGene_gn=COLQ;refGene_tx=XM_023260580.1
C2 135084924 135084951
refGene_gn=COLQ;refGene_tx=XM_003992132.3;refGene_gn=COLQ;refGene_tx=XM_003992134.4;refGene_gn=COLQ;refGene_tx=XM_006936431.4;refGene_gn=COLQ;refGene_tx=XM_006936432.4;refGene_gn=COLQ;refGene_tx=XM_023260580.1
C2 135085350 135085413
refGene_gn=COLQ;refGene_tx=XM_003992132.3;refGene_gn=COLQ;refGene_tx=XM_003992134.4;refGene_gn=COLQ;refGene_tx=XM_006936431.4;refGene_gn=COLQ;refGene_tx=XM_006936432.4;refGene_gn=COLQ;refGene_tx=XM_023260580.1
C2 135087129 135087192
refGene_gn=COLQ;refGene_tx=XM_003992132.3;refGene_gn=COLQ;refGene_tx=XM_003992134.4;refGene_gn=COLQ;refGene_tx=XM_006936431.4;refGene_gn=COLQ;refGene_tx=XM_006936432.4;refGene_gn=COLQ;refGene_tx=XM_023260580.1
C2 135089280 135089307
refGene_gn=COLQ;refGene_tx=XM_003992132.3;refGene_gn=COLQ;refGene_tx=XM_003992134.4;refGene_gn=COLQ;refGene_tx=XM_006936431.4;refGene_gn=COLQ;refGene_tx=XM_006936432.4;refGene_gn=COLQ;refGene_tx=XM_023260580.1
C2 135089640 135089685
refGene_gn=COLQ;refGene_tx=XM_003992132.3;refGene_gn=COLQ;refGene_tx=XM_003992134.4;refGene_gn=COLQ;refGene_tx=XM_006936431.4;refGene_gn=COLQ;refGene_tx=XM_006936432.4;refGene_gn=COLQ;refGene_tx=XM_023260580.1
C2 135093588 135093690
refGene_gn=COLQ;refGene_tx=XM_003992132.3;refGene_gn=COLQ;refGene_tx=XM_003992134.4;refGene_gn=COLQ;refGene_tx=XM_006936431.4;refGene_gn=COLQ;refGene_tx=XM_006936432.4;refGene_gn=COLQ;refGene_tx=XM_023260580.1
C2 135094798 135094911
refGene_gn=COLQ;refGene_tx=XM_003992132.3;refGene_gn=COLQ;refGene_tx=XM_003992134.4;refGene_gn=COLQ;refGene_tx=XM_006936431.4;refGene_gn=COLQ;refGene_tx=XM_006936432.4;refGene_gn=COLQ;refGene_tx=XM_023260580.1
C2 135100790 135100866
refGene_gn=COLQ;refGene_tx=XM_003992134.4;refGene_gn=COLQ;refGene_tx=XM_006936432.4
C2 135122311 135122417
refGene_gn=COLQ;refGene_tx=XM_003992132.3;refGene_gn=COLQ;refGene_tx=XM_006936431.4

C2 135141497 135141633
refGene_gn=COLQ;refGene_tx=XM_023260580.1
C2 135148306 135148339
refGene_gn=HACL1;refGene_tx=XM_003992135.5;refGene_gn=HACL1;refGene_tx=XM_023260577.1;refGene_gn=HACL1;refGene_tx=XM_023260578.1;refGene_gn=HACL1;refGene_tx=XM_023260579.1
C2 135151822 135152009
refGene_gn=HACL1;refGene_tx=XM_003992135.5;refGene_gn=HACL1;refGene_tx=XM_023260577.1;refGene_gn=HACL1;refGene_tx=XM_023260578.1;refGene_gn=HACL1;refGene_tx=XM_023260579.1
C2 135152937 135153045
refGene_gn=HACL1;refGene_tx=XM_003992135.5;refGene_gn=HACL1;refGene_tx=XM_023260577.1;refGene_gn=HACL1;refGene_tx=XM_023260578.1;refGene_gn=HACL1;refGene_tx=XM_023260579.1
C2 135154774 135154933
refGene_gn=HACL1;refGene_tx=XM_003992135.5;refGene_gn=HACL1;refGene_tx=XM_023260577.1;refGene_gn=HACL1;refGene_tx=XM_023260578.1;refGene_gn=HACL1;refGene_tx=XM_023260579.1
C2 135155350 135155505
refGene_gn=HACL1;refGene_tx=XM_003992135.5;refGene_gn=HACL1;refGene_tx=XM_023260577.1;refGene_gn=HACL1;refGene_tx=XM_023260578.1;refGene_gn=HACL1;refGene_tx=XM_023260579.1
C2 135157552 135157654
refGene_gn=HACL1;refGene_tx=XM_003992135.5;refGene_gn=HACL1;refGene_tx=XM_023260577.1;refGene_gn=HACL1;refGene_tx=XM_023260578.1;refGene_gn=HACL1;refGene_tx=XM_023260579.1
C2 135158591 135158675
refGene_gn=HACL1;refGene_tx=XM_003992135.5;refGene_gn=HACL1;refGene_tx=XM_023260577.1;refGene_gn=HACL1;refGene_tx=XM_023260578.1;refGene_gn=HACL1;refGene_tx=XM_023260579.1
C2 135160443 135160549
refGene_gn=HACL1;refGene_tx=XM_003992135.5;refGene_gn=HACL1;refGene_tx=XM_023260577.1;refGene_gn=HACL1;refGene_tx=XM_023260578.1;refGene_gn=HACL1;refGene_tx=XM_023260579.1
C2 135168524 135168660
refGene_gn=HACL1;refGene_tx=XM_003992135.5;refGene_gn=HACL1;refGene_tx=XM_023260577.1;refGene_gn=HACL1;refGene_tx=XM_023260578.1;refGene_gn=HACL1;refGene_tx=XM_023260579.1
C2 135169379 135169492
refGene_gn=HACL1;refGene_tx=XM_003992135.5;refGene_gn=HACL1;refGene_tx=XM_023260577.1;refGene_gn=HACL1;refGene_tx=XM_023260578.1;refGene_gn=HACL1;refGene_tx=XM_023260579.1
C2 135171272 135171367
refGene_gn=HACL1;refGene_tx=XM_003992135.5;refGene_gn=HACL1;refGene_tx=XM_023260577.1;refGene_gn=HACL1;refGene_tx=XM_023260578.1;refGene_gn=HACL1;refGene_tx=XM_023260579.1
C2 135172413 135172491
refGene_gn=HACL1;refGene_tx=XM_003992135.5;refGene_gn=HACL1;refGene_tx=XM_023260577.1;refGene_gn=HACL1;refGene_tx=XM_023260578.1;refGene_gn=HACL1;refGene_tx=XM_023260579.1

C2 135174662 135174735
refGene_gn=HACL1;refGene_tx=XM_003992135.5;refGene_gn=HACL1;refGene_tx=XM_023260577.1;refGene_gn=HACL1;refGene_tx=XM_023260578.1;refGene_gn=HACL1;refGene_tx=XM_023260579.1
C2 135176870 135176951
refGene_gn=HACL1;refGene_tx=XM_003992135.5;refGene_gn=HACL1;refGene_tx=XM_023260577.1;refGene_gn=HACL1;refGene_tx=XM_023260578.1
C2 135177575 135177616
refGene_gn=HACL1;refGene_tx=XM_003992135.5;refGene_gn=HACL1;refGene_tx=XM_023260577.1;refGene_gn=HACL1;refGene_tx=XM_023260578.1;refGene_gn=HACL1;refGene_tx=XM_023260579.1
C2 135183028 135183133
refGene_gn=HACL1;refGene_tx=XM_003992135.5;refGene_gn=HACL1;refGene_tx=XM_023260577.1;refGene_gn=HACL1;refGene_tx=XM_023260578.1;refGene_gn=HACL1;refGene_tx=XM_023260579.1
C2 135183310 135183400
refGene_gn=HACL1;refGene_tx=XM_003992135.5;refGene_gn=HACL1;refGene_tx=XM_023260577.1;refGene_gn=HACL1;refGene_tx=XM_023260578.1;refGene_gn=HACL1;refGene_tx=XM_023260579.1
C2 135183709 135183720
refGene_gn=BTD;refGene_tx=XM_003992138.5
C2 135185881 135185892
refGene_gn=BTD;refGene_tx=XM_006936433.4
C2 135205451 135205716
refGene_gn=BTD;refGene_tx=XM_003992138.5;refGene_gn=BTD;refGene_tx=XM_006936433.4
C2 135211015 135211165
refGene_gn=BTD;refGene_tx=XM_003992138.5;refGene_gn=BTD;refGene_tx=XM_006936433.4
C2 135212145 135213318
refGene_gn=BTD;refGene_tx=XM_003992138.5;refGene_gn=BTD;refGene_tx=XM_006936433.4
C2 135237036 135237323
refGene_gn=ANKRD28;refGene_tx=XM_006936434.4
C2 135244082 135244174
refGene_gn=ANKRD28;refGene_tx=XM_006936434.4
C2 135245240 135245323
refGene_gn=ANKRD28;refGene_tx=XM_006936434.4
C2 135245784 135245867
refGene_gn=ANKRD28;refGene_tx=XM_006936434.4
C2 135246470 135246616
refGene_gn=ANKRD28;refGene_tx=XM_006936434.4
C2 135247234 135247318
refGene_gn=ANKRD28;refGene_tx=XM_006936434.4
C2 135247406 135247494
refGene_gn=ANKRD28;refGene_tx=XM_006936434.4
C2 135253034 135253254
refGene_gn=ANKRD28;refGene_tx=XM_006936434.4
C2 135253821 135253939
refGene_gn=ANKRD28;refGene_tx=XM_006936434.4

C2 135254170 135254258
refGene_gn=ANKRD28;refGene_tx=XM_006936434.4
C2 135257811 135258013
refGene_gn=ANKRD28;refGene_tx=XM_006936434.4
C2 135261561 135261636
refGene_gn=ANKRD28;refGene_tx=XM_006936434.4
C2 135262057 135262084
refGene_gn=ANKRD28;refGene_tx=XM_006936434.4
C2 135263105 135263217
refGene_gn=ANKRD28;refGene_tx=XM_006936434.4
C2 135269791 135269932
refGene_gn=ANKRD28;refGene_tx=XM_006936434.4
C2 135272088 135272157
refGene_gn=ANKRD28;refGene_tx=XM_006936434.4
C2 135273595 135273659
refGene_gn=ANKRD28;refGene_tx=XM_006936434.4
C2 135275486 135275569
refGene_gn=ANKRD28;refGene_tx=XM_006936434.4
C2 135276827 135276942
refGene_gn=ANKRD28;refGene_tx=XM_006936434.4
C2 135277859 135277938
refGene_gn=ANKRD28;refGene_tx=XM_006936434.4
C2 135284736 135284949
refGene_gn=ANKRD28;refGene_tx=XM_006936434.4
C2 135288257 135288400
refGene_gn=ANKRD28;refGene_tx=XM_006936434.4
C2 135293866 135293954
refGene_gn=ANKRD28;refGene_tx=XM_006936434.4
C2 135295513 135295624
refGene_gn=ANKRD28;refGene_tx=XM_006936434.4
C2 135295625 135295715
refGene_gn=ANKRD28;refGene_tx=XM_006936434.4
C2 135302007 135302078
refGene_gn=ANKRD28;refGene_tx=XM_006936434.4
C2 135320232 135320311
refGene_gn=ANKRD28;refGene_tx=XM_006936434.4
C2 135355727 135355811
refGene_gn=ANKRD28;refGene_tx=XM_006936434.4
C2 135356936 135357056
refGene_gn=ANKRD28;refGene_tx=XM_006936434.4
C2 135680596 135681141
refGene_gn=GALNT15;refGene_tx=XM_003992147.4
C2 135699577 135699744
refGene_gn=GALNT15;refGene_tx=XM_003992147.4
C2 135702385 135702590
refGene_gn=GALNT15;refGene_tx=XM_003992147.4;refGene_gn=GALNT15;refGene_tx=XM_019810732.1
C2 135709872 135710040
refGene_gn=GALNT15;refGene_tx=XM_003992147.4;refGene_gn=GALNT15;refGene_tx=XM_019810732.1

C2	135712634	135712752
refGene_gn=GALNT15;refGene_tx=XM_003992147.4;refGene_gn=GALNT15;refGene_tx=XM_019810732.1
C2	135714502	135714697
refGene_gn=GALNT15;refGene_tx=XM_003992147.4;refGene_gn=GALNT15;refGene_tx=XM_019810732.1
C2	135717206	135717338
refGene_gn=GALNT15;refGene_tx=XM_003992147.4;refGene_gn=GALNT15;refGene_tx=XM_019810732.1
C2	135717722	135717827
refGene_gn=GALNT15;refGene_tx=XM_003992147.4;refGene_gn=GALNT15;refGene_tx=XM_019810732.1
C2	135722727	135722871
refGene_gn=GALNT15;refGene_tx=XM_003992147.4;refGene_gn=GALNT15;refGene_tx=XM_019810732.1
C2	135725266	135725410
refGene_gn=GALNT15;refGene_tx=XM_003992147.4;refGene_gn=GALNT15;refGene_tx=XM_019810732.1
C2	135751387	135753436
refGene_gn=LOC109491518;refGene_tx=XM_019810731.2
C2	135764815	135764881
refGene_gn=DPH3;refGene_tx=XM_003992148.4;refGene_gn=DPH3;refGene_tx=XM_019810736.2
C2	135769259	135769334
refGene_gn=DPH3;refGene_tx=XM_003992148.4
C2	135769867	135769975
refGene_gn=DPH3;refGene_tx=XM_003992148.4;refGene_gn=DPH3;refGene_tx=XM_019810736.2
C2	135777836	135777955
refGene_gn=OXNAD1;refGene_tx=XM_003992150.5;refGene_gn=OXNAD1;refGene_tx=XM_006936438.4;refGene_gn=OXNAD1;refGene_tx=XM_006936439.4;refGene_gn=OXNAD1;refGene_tx=XM_006936440.4;refGene_gn=OXNAD1;refGene_tx=XM_019810735.2
C2	135778231	135778295
refGene_gn=OXNAD1;refGene_tx=XM_003992150.5;refGene_gn=OXNAD1;refGene_tx=XM_006936438.4
C2	135779197	135779482
refGene_gn=OXNAD1;refGene_tx=XM_011286218.3;refGene_gn=OXNAD1;refGene_tx=XM_019810733.2;refGene_gn=OXNAD1;refGene_tx=XM_019810734.2
C2	135793605	135793712
refGene_gn=OXNAD1;refGene_tx=XM_003992150.5;refGene_gn=OXNAD1;refGene_tx=XM_006936438.4;refGene_gn=OXNAD1;refGene_tx=XM_006936439.4;refGene_gn=OXNAD1;refGene_tx=XM_006936440.4;refGene_gn=OXNAD1;refGene_tx=XM_011286218.3;refGene_gn=OXNAD1;refGene_tx=XM_019810733.2;refGene_gn=OXNAD1;refGene_tx=XM_019810734.2
C2	135802211	135802353
refGene_gn=OXNAD1;refGene_tx=XM_003992150.5;refGene_gn=OXNAD1;refGene_tx=XM_006936438.4;refGene_gn=OXNAD1;refGene_tx=XM_006936439.4;refGene_gn=OXNAD1;refGene_tx=XM_006936440.4;refGene_gn=OXNAD1;refGene_tx=XM_011286218.3;refGene_gn=OXNAD1;refGene_tx=XM_019810733.2;refGene_gn=OXNAD

1;refGene_tx=XM_019810734.2;refGene_gn=OXNAD1;refGene_tx=XM_019810735.
2
C2 135808711 135808954
refGene_gn=OXNAD1;refGene_tx=XM_003992150.5;refGene_gn=OXNAD1;refGene_
tx=XM_006936438.4;refGene_gn=OXNAD1;refGene_tx=XM_006936439.4;refGene_
gn=OXNAD1;refGene_tx=XM_006936440.4;refGene_gn=OXNAD1;refGene_tx=XM_01
1286218.3;refGene_gn=OXNAD1;refGene_tx=XM_019810733.2;refGene_gn=OXNAD
1;refGene_tx=XM_019810734.2;refGene_gn=OXNAD1;refGene_tx=XM_019810735.
2
C2 135809736 135809845
refGene_gn=OXNAD1;refGene_tx=XM_003992150.5;refGene_gn=OXNAD1;refGene_
tx=XM_006936438.4;refGene_gn=OXNAD1;refGene_tx=XM_006936439.4;refGene_
gn=OXNAD1;refGene_tx=XM_006936440.4;refGene_gn=OXNAD1;refGene_tx=XM_01
1286218.3;refGene_gn=OXNAD1;refGene_tx=XM_019810733.2;refGene_gn=OXNAD
1;refGene_tx=XM_019810734.2;refGene_gn=OXNAD1;refGene_tx=XM_019810735.
2
C2 135810541 135810696
refGene_gn=OXNAD1;refGene_tx=XM_003992150.5;refGene_gn=OXNAD1;refGene_
tx=XM_006936438.4;refGene_gn=OXNAD1;refGene_tx=XM_006936439.4;refGene_
gn=OXNAD1;refGene_tx=XM_006936440.4;refGene_gn=OXNAD1;refGene_tx=XM_01
1286218.3;refGene_gn=OXNAD1;refGene_tx=XM_019810733.2;refGene_gn=OXNAD
1;refGene_tx=XM_019810734.2;refGene_gn=OXNAD1;refGene_tx=XM_019810735.
2
C2 135825998 135826379
refGene_gn=RFTN1;refGene_tx=XM_006936442.3;refGene_gn=RFTN1;refGene_tx
=XM_006936444.4
C2 135836157 135836239
refGene_gn=RFTN1;refGene_tx=XM_006936442.3;refGene_gn=RFTN1;refGene_tx
=XM_006936444.4
C2 135838766 135838870
refGene_gn=RFTN1;refGene_tx=XM_006936442.3;refGene_gn=RFTN1;refGene_tx
=XM_006936444.4
C2 135870132 135870248
refGene_gn=RFTN1;refGene_tx=XM_006936442.3;refGene_gn=RFTN1;refGene_tx
=XM_006936444.4
C2 135881521 135881725
refGene_gn=RFTN1;refGene_tx=XM_006936442.3;refGene_gn=RFTN1;refGene_tx
=XM_006936444.4
C2 135889991 135890331
refGene_gn=RFTN1;refGene_tx=XM_006936442.3;refGene_gn=RFTN1;refGene_tx
=XM_006936444.4
C2 135923640 135923749
refGene_gn=RFTN1;refGene_tx=XM_006936442.3;refGene_gn=RFTN1;refGene_tx
=XM_006936444.4
C2 135951798 135951985
refGene_gn=RFTN1;refGene_tx=XM_006936442.3;refGene_gn=RFTN1;refGene_tx
=XM_006936444.4
C2 136013792 136013937
refGene_gn=RFTN1;refGene_tx=XM_006936442.3;refGene_gn=RFTN1;refGene_tx
=XM_006936444.4

C2 136085479 136085533
refGene_gn=DAZL;refGene_tx=XM_006936445.4
C2 136089516 136089615
refGene_gn=DAZL;refGene_tx=XM_006936445.4
C2 136090767 136090881
refGene_gn=DAZL;refGene_tx=XM_006936445.4
C2 136091924 136091975
refGene_gn=DAZL;refGene_tx=XM_006936445.4
C2 136092698 136092770
refGene_gn=DAZL;refGene_tx=XM_006936445.4
C2 136094144 136094284
refGene_gn=DAZL;refGene_tx=XM_006936445.4
C2 136094374 136094438
refGene_gn=DAZL;refGene_tx=XM_006936445.4
C2 136094849 136094901
refGene_gn=DAZL;refGene_tx=XM_006936445.4
C2 136095480 136095572
refGene_gn=DAZL;refGene_tx=XM_006936445.4
C2 136095846 136095993
refGene_gn=DAZL;refGene_tx=XM_006936445.4
C2 136102931 136102934
refGene_gn=DAZL;refGene_tx=XM_006936445.4
C2 136297092 136297099
refGene_gn=LOC111562241;refGene_tx=XM_023260186.1
C2 136301569 136301846
refGene_gn=LOC111562241;refGene_tx=XM_023260186.1
C2 136301884 136302295
refGene_gn=LOC111562241;refGene_tx=XM_023260186.1
C2 136302614 136302687
refGene_gn=LOC111562241;refGene_tx=XM_023260186.1
C2 136302853 136303204
refGene_gn=PLCL2;refGene_tx=XM_003992151.5;refGene_gn=PLCL2;refGene_tx=XM_023260585.1
C2 136349664 136349742
refGene_gn=PLCL2;refGene_tx=XM_019810738.2
C2 136418714 136421201
refGene_gn=PLCL2;refGene_tx=XM_003992151.5;refGene_gn=PLCL2;refGene_tx=XM_019810738.2;refGene_gn=PLCL2;refGene_tx=XM_023260585.1
C2 136424029 136424233
refGene_gn=PLCL2;refGene_tx=XM_003992151.5;refGene_gn=PLCL2;refGene_tx=XM_019810738.2;refGene_gn=PLCL2;refGene_tx=XM_023260585.1
C2 136450448 136450524
refGene_gn=PLCL2;refGene_tx=XM_003992151.5;refGene_gn=PLCL2;refGene_tx=XM_019810738.2;refGene_gn=PLCL2;refGene_tx=XM_023260585.1
C2 136457130 136457228
refGene_gn=PLCL2;refGene_tx=XM_023260585.1
C2 136474923 136475033
refGene_gn=PLCL2;refGene_tx=XM_003992151.5;refGene_gn=PLCL2;refGene_tx=XM_019810738.2
C2 136493237 136493417

refGene_gn=PLCL2;refGene_tx=XM_003992151.5;refGene_gn=PLCL2;refGene_tx=XM_019810738.2
C2 136571055 136571358
refGene_gn=TBC1D5;refGene_tx=XM_011286217.3;refGene_gn=TBC1D5;refGene_tx=XM_019810740.2;refGene_gn=TBC1D5;refGene_tx=XM_019810741.2;refGene_gn=TBC1D5;refGene_tx=XM_019810742.2;refGene_gn=TBC1D5;refGene_tx=XM_019810743.2;refGene_gn=TBC1D5;refGene_tx=XM_019810744.2;refGene_gn=TBC1D5;refGene_tx=XM_019810745.2;refGene_gn=TBC1D5;refGene_tx=XM_019810746.1;refGene_gn=TBC1D5;refGene_tx=XM_019810747.1;refGene_gn=TBC1D5;refGene_tx=XM_019810750.2;refGene_gn=TBC1D5;refGene_tx=XM_023260586.1
C2 136577055 136577211
refGene_gn=TBC1D5;refGene_tx=XM_011286217.3;refGene_gn=TBC1D5;refGene_tx=XM_019810740.2;refGene_gn=TBC1D5;refGene_tx=XM_019810741.2;refGene_gn=TBC1D5;refGene_tx=XM_019810742.2;refGene_gn=TBC1D5;refGene_tx=XM_019810743.2;refGene_gn=TBC1D5;refGene_tx=XM_019810744.2;refGene_gn=TBC1D5;refGene_tx=XM_019810745.2;refGene_gn=TBC1D5;refGene_tx=XM_019810746.1;refGene_gn=TBC1D5;refGene_tx=XM_019810747.1;refGene_gn=TBC1D5;refGene_tx=XM_019810750.2;refGene_gn=TBC1D5;refGene_tx=XM_023260586.1
C2 136578034 136578114
refGene_gn=TBC1D5;refGene_tx=XM_011286217.3;refGene_gn=TBC1D5;refGene_tx=XM_019810740.2;refGene_gn=TBC1D5;refGene_tx=XM_019810741.2;refGene_gn=TBC1D5;refGene_tx=XM_019810742.2;refGene_gn=TBC1D5;refGene_tx=XM_019810743.2;refGene_gn=TBC1D5;refGene_tx=XM_019810744.2;refGene_gn=TBC1D5;refGene_tx=XM_019810745.2;refGene_gn=TBC1D5;refGene_tx=XM_019810746.1;refGene_gn=TBC1D5;refGene_tx=XM_019810747.1;refGene_gn=TBC1D5;refGene_tx=XM_019810750.2;refGene_gn=TBC1D5;refGene_tx=XM_023260586.1
C2 136592102 136592202
refGene_gn=TBC1D5;refGene_tx=XM_011286217.3;refGene_gn=TBC1D5;refGene_tx=XM_019810740.2;refGene_gn=TBC1D5;refGene_tx=XM_019810741.2;refGene_gn=TBC1D5;refGene_tx=XM_019810742.2;refGene_gn=TBC1D5;refGene_tx=XM_019810743.2;refGene_gn=TBC1D5;refGene_tx=XM_019810744.2;refGene_gn=TBC1D5;refGene_tx=XM_019810745.2;refGene_gn=TBC1D5;refGene_tx=XM_019810746.1;refGene_gn=TBC1D5;refGene_tx=XM_019810747.1;refGene_gn=TBC1D5;refGene_tx=XM_019810750.2;refGene_gn=TBC1D5;refGene_tx=XM_023260586.1
C2 136612026 136612137
refGene_gn=TBC1D5;refGene_tx=XM_023260587.1
C2 136624629 136624793
refGene_gn=TBC1D5;refGene_tx=XM_011286217.3;refGene_gn=TBC1D5;refGene_tx=XM_019810740.2;refGene_gn=TBC1D5;refGene_tx=XM_019810741.2;refGene_gn=TBC1D5;refGene_tx=XM_019810742.2;refGene_gn=TBC1D5;refGene_tx=XM_019810743.2;refGene_gn=TBC1D5;refGene_tx=XM_019810744.2;refGene_gn=TBC1D5;refGene_tx=XM_019810745.2;refGene_gn=TBC1D5;refGene_tx=XM_019810746.1;refGene_gn=TBC1D5;refGene_tx=XM_019810747.1;refGene_gn=TBC1D5;refGene_tx=XM_019810750.2;refGene_gn=TBC1D5;refGene_tx=XM_023260586.1;refGene_gn=TBC1D5;refGene_tx=XM_023260587.1
C2 136633476 136633542
refGene_gn=TBC1D5;refGene_tx=XM_019810740.2;refGene_gn=TBC1D5;refGene_tx=XM_019810741.2;refGene_gn=TBC1D5;refGene_tx=XM_019810742.2;refGene_gn=TBC1D5;refGene_tx=XM_019810743.2;refGene_gn=TBC1D5;refGene_tx=XM_019810744.2;refGene_gn=TBC1D5;refGene_tx=XM_019810745.2;refGene_gn=TBC1D

5;refGene_tx=XM_019810746.1;refGene_gn=TBC1D5;refGene_tx=XM_019810747.1;refGene_gn=TBC1D5;refGene_tx=XM_019810750.2;refGene_gn=TBC1D5;refGene_tx=XM_023260587.1
C2 136637651 136637935
refGene_gn=TBC1D5;refGene_tx=XM_011286217.3;refGene_gn=TBC1D5;refGene_tx=XM_019810740.2;refGene_gn=TBC1D5;refGene_tx=XM_019810741.2;refGene_gn=TBC1D5;refGene_tx=XM_019810742.2;refGene_gn=TBC1D5;refGene_tx=XM_019810743.2;refGene_gn=TBC1D5;refGene_tx=XM_019810744.2;refGene_gn=TBC1D5;refGene_tx=XM_019810745.2;refGene_gn=TBC1D5;refGene_tx=XM_019810746.1;refGene_gn=TBC1D5;refGene_tx=XM_019810747.1;refGene_gn=TBC1D5;refGene_tx=XM_019810750.2;refGene_gn=TBC1D5;refGene_tx=XM_023260586.1;refGene_gn=TBC1D5;refGene_tx=XM_023260587.1
C2 136654770 136654856
refGene_gn=TBC1D5;refGene_tx=XM_011286217.3;refGene_gn=TBC1D5;refGene_tx=XM_019810740.2;refGene_gn=TBC1D5;refGene_tx=XM_019810741.2;refGene_gn=TBC1D5;refGene_tx=XM_019810742.2;refGene_gn=TBC1D5;refGene_tx=XM_019810743.2;refGene_gn=TBC1D5;refGene_tx=XM_019810744.2;refGene_gn=TBC1D5;refGene_tx=XM_019810745.2;refGene_gn=TBC1D5;refGene_tx=XM_019810746.1;refGene_gn=TBC1D5;refGene_tx=XM_019810747.1;refGene_gn=TBC1D5;refGene_tx=XM_019810750.2;refGene_gn=TBC1D5;refGene_tx=XM_023260586.1;refGene_gn=TBC1D5;refGene_tx=XM_023260587.1
C2 136686611 136686718
refGene_gn=TBC1D5;refGene_tx=XM_011286217.3;refGene_gn=TBC1D5;refGene_tx=XM_019810740.2;refGene_gn=TBC1D5;refGene_tx=XM_019810741.2;refGene_gn=TBC1D5;refGene_tx=XM_019810742.2;refGene_gn=TBC1D5;refGene_tx=XM_019810743.2;refGene_gn=TBC1D5;refGene_tx=XM_019810744.2;refGene_gn=TBC1D5;refGene_tx=XM_019810745.2;refGene_gn=TBC1D5;refGene_tx=XM_019810746.1;refGene_gn=TBC1D5;refGene_tx=XM_019810747.1;refGene_gn=TBC1D5;refGene_tx=XM_019810750.2;refGene_gn=TBC1D5;refGene_tx=XM_023260586.1;refGene_gn=TBC1D5;refGene_tx=XM_023260587.1
C2 136708259 136708402
refGene_gn=TBC1D5;refGene_tx=XM_011286217.3;refGene_gn=TBC1D5;refGene_tx=XM_019810740.2;refGene_gn=TBC1D5;refGene_tx=XM_019810741.2;refGene_gn=TBC1D5;refGene_tx=XM_019810742.2;refGene_gn=TBC1D5;refGene_tx=XM_019810743.2;refGene_gn=TBC1D5;refGene_tx=XM_019810744.2;refGene_gn=TBC1D5;refGene_tx=XM_019810745.2;refGene_gn=TBC1D5;refGene_tx=XM_019810746.1;refGene_gn=TBC1D5;refGene_tx=XM_019810747.1;refGene_gn=TBC1D5;refGene_tx=XM_019810750.2;refGene_gn=TBC1D5;refGene_tx=XM_023260586.1;refGene_gn=TBC1D5;refGene_tx=XM_023260587.1
C2 136773765 136773938
refGene_gn=TBC1D5;refGene_tx=XM_011286217.3;refGene_gn=TBC1D5;refGene_tx=XM_019810740.2;refGene_gn=TBC1D5;refGene_tx=XM_019810741.2;refGene_gn=TBC1D5;refGene_tx=XM_019810742.2;refGene_gn=TBC1D5;refGene_tx=XM_019810743.2;refGene_gn=TBC1D5;refGene_tx=XM_019810744.2;refGene_gn=TBC1D5;refGene_tx=XM_019810745.2;refGene_gn=TBC1D5;refGene_tx=XM_019810746.1;refGene_gn=TBC1D5;refGene_tx=XM_019810747.1;refGene_gn=TBC1D5;refGene_tx=XM_019810750.2;refGene_gn=TBC1D5;refGene_tx=XM_023260586.1;refGene_gn=TBC1D5;refGene_tx=XM_023260587.1
C2 136779710 136779780
refGene_gn=TBC1D5;refGene_tx=XM_011286217.3;refGene_gn=TBC1D5;refGene_

tx=XM_019810740.2;refGene_gn=TBC1D5;refGene_tx=XM_019810741.2;refGene_gn=TBC1D5;refGene_tx=XM_019810742.2;refGene_gn=TBC1D5;refGene_tx=XM_019810743.2;refGene_gn=TBC1D5;refGene_tx=XM_019810744.2;refGene_gn=TBC1D5;refGene_tx=XM_019810745.2;refGene_gn=TBC1D5;refGene_tx=XM_019810746.1;refGene_gn=TBC1D5;refGene_tx=XM_019810747.1;refGene_gn=TBC1D5;refGene_tx=XM_019810750.2;refGene_gn=TBC1D5;refGene_tx=XM_023260586.1;refGene_gn=TBC1D5;refGene_tx=XM_023260587.1
C2 136779876 136779927
refGene_gn=TBC1D5;refGene_tx=XM_011286217.3;refGene_gn=TBC1D5;refGene_tx=XM_019810740.2;refGene_gn=TBC1D5;refGene_tx=XM_019810741.2;refGene_gn=TBC1D5;refGene_tx=XM_019810742.2;refGene_gn=TBC1D5;refGene_tx=XM_019810743.2;refGene_gn=TBC1D5;refGene_tx=XM_019810744.2;refGene_gn=TBC1D5;refGene_tx=XM_019810745.2;refGene_gn=TBC1D5;refGene_tx=XM_019810746.1;refGene_gn=TBC1D5;refGene_tx=XM_019810747.1;refGene_gn=TBC1D5;refGene_tx=XM_019810750.2;refGene_gn=TBC1D5;refGene_tx=XM_023260586.1;refGene_gn=TBC1D5;refGene_tx=XM_023260587.1
C2 136781464 136781553
refGene_gn=TBC1D5;refGene_tx=XM_011286217.3;refGene_gn=TBC1D5;refGene_tx=XM_019810740.2;refGene_gn=TBC1D5;refGene_tx=XM_019810741.2;refGene_gn=TBC1D5;refGene_tx=XM_019810742.2;refGene_gn=TBC1D5;refGene_tx=XM_019810743.2;refGene_gn=TBC1D5;refGene_tx=XM_019810744.2;refGene_gn=TBC1D5;refGene_tx=XM_019810745.2;refGene_gn=TBC1D5;refGene_tx=XM_019810746.1;refGene_gn=TBC1D5;refGene_tx=XM_019810747.1;refGene_gn=TBC1D5;refGene_tx=XM_019810750.2;refGene_gn=TBC1D5;refGene_tx=XM_023260586.1;refGene_gn=TBC1D5;refGene_tx=XM_023260587.1
C2 136788622 136788725
refGene_gn=TBC1D5;refGene_tx=XM_011286217.3;refGene_gn=TBC1D5;refGene_tx=XM_019810740.2;refGene_gn=TBC1D5;refGene_tx=XM_019810741.2;refGene_gn=TBC1D5;refGene_tx=XM_019810742.2;refGene_gn=TBC1D5;refGene_tx=XM_019810743.2;refGene_gn=TBC1D5;refGene_tx=XM_019810744.2;refGene_gn=TBC1D5;refGene_tx=XM_019810745.2;refGene_gn=TBC1D5;refGene_tx=XM_019810746.1;refGene_gn=TBC1D5;refGene_tx=XM_019810747.1;refGene_gn=TBC1D5;refGene_tx=XM_019810750.2;refGene_gn=TBC1D5;refGene_tx=XM_023260586.1;refGene_gn=TBC1D5;refGene_tx=XM_023260587.1
C2 136798346 136798382
refGene_gn=TBC1D5;refGene_tx=XM_019810740.2;refGene_gn=TBC1D5;refGene_tx=XM_019810741.2;refGene_gn=TBC1D5;refGene_tx=XM_019810742.2;refGene_gn=TBC1D5;refGene_tx=XM_019810743.2;refGene_gn=TBC1D5;refGene_tx=XM_019810744.2;refGene_gn=TBC1D5;refGene_tx=XM_019810745.2;refGene_gn=TBC1D5;refGene_tx=XM_019810746.1;refGene_gn=TBC1D5;refGene_tx=XM_019810747.1;refGene_gn=TBC1D5;refGene_tx=XM_023260586.1;refGene_gn=TBC1D5;refGene_tx=XM_023260587.1
C2 136809652 136809720
refGene_gn=TBC1D5;refGene_tx=XM_011286217.3;refGene_gn=TBC1D5;refGene_tx=XM_019810740.2;refGene_gn=TBC1D5;refGene_tx=XM_019810741.2;refGene_gn=TBC1D5;refGene_tx=XM_019810742.2;refGene_gn=TBC1D5;refGene_tx=XM_019810743.2;refGene_gn=TBC1D5;refGene_tx=XM_019810744.2;refGene_gn=TBC1D5;refGene_tx=XM_019810745.2;refGene_gn=TBC1D5;refGene_tx=XM_019810746.1;refGene_gn=TBC1D5;refGene_tx=XM_019810747.1;refGene_gn=TBC1D5;refGene_tx=XM_019810750.2;refGene_gn=TBC1D5;refGene_tx=XM_023260586.1;refGen

e_gn=TBC1D5;refGene_tx=XM_023260587.1
C2	136811086	136811161
refGene_gn=TBC1D5;refGene_tx=XM_011286217.3;refGene_gn=TBC1D5;refGene_tx=XM_019810740.2;refGene_gn=TBC1D5;refGene_tx=XM_019810741.2;refGene_gn=TBC1D5;refGene_tx=XM_019810742.2;refGene_gn=TBC1D5;refGene_tx=XM_019810743.2;refGene_gn=TBC1D5;refGene_tx=XM_019810744.2;refGene_gn=TBC1D5;refGene_tx=XM_019810745.2;refGene_gn=TBC1D5;refGene_tx=XM_019810746.1;refGene_gn=TBC1D5;refGene_tx=XM_019810747.1;refGene_gn=TBC1D5;refGene_tx=XM_019810750.2;refGene_gn=TBC1D5;refGene_tx=XM_023260586.1;refGene_gn=TBC1D5;refGene_tx=XM_023260587.1
C2	136811261	136811351
refGene_gn=TBC1D5;refGene_tx=XM_011286217.3;refGene_gn=TBC1D5;refGene_tx=XM_019810740.2;refGene_gn=TBC1D5;refGene_tx=XM_019810741.2;refGene_gn=TBC1D5;refGene_tx=XM_019810742.2;refGene_gn=TBC1D5;refGene_tx=XM_019810743.2;refGene_gn=TBC1D5;refGene_tx=XM_019810744.2;refGene_gn=TBC1D5;refGene_tx=XM_019810745.2;refGene_gn=TBC1D5;refGene_tx=XM_019810746.1;refGene_gn=TBC1D5;refGene_tx=XM_019810747.1;refGene_gn=TBC1D5;refGene_tx=XM_019810750.2;refGene_gn=TBC1D5;refGene_tx=XM_023260586.1;refGene_gn=TBC1D5;refGene_tx=XM_023260587.1
C2	136813087	136813196
refGene_gn=TBC1D5;refGene_tx=XM_011286217.3;refGene_gn=TBC1D5;refGene_tx=XM_019810740.2;refGene_gn=TBC1D5;refGene_tx=XM_019810741.2;refGene_gn=TBC1D5;refGene_tx=XM_019810742.2;refGene_gn=TBC1D5;refGene_tx=XM_019810743.2;refGene_gn=TBC1D5;refGene_tx=XM_019810744.2;refGene_gn=TBC1D5;refGene_tx=XM_019810745.2;refGene_gn=TBC1D5;refGene_tx=XM_019810746.1;refGene_gn=TBC1D5;refGene_tx=XM_019810747.1;refGene_gn=TBC1D5;refGene_tx=XM_019810750.2;refGene_gn=TBC1D5;refGene_tx=XM_023260586.1;refGene_gn=TBC1D5;refGene_tx=XM_023260587.1
C2	136832930	136832997
refGene_gn=TBC1D5;refGene_tx=XM_011286217.3;refGene_gn=TBC1D5;refGene_tx=XM_019810740.2;refGene_gn=TBC1D5;refGene_tx=XM_019810741.2;refGene_gn=TBC1D5;refGene_tx=XM_019810742.2;refGene_gn=TBC1D5;refGene_tx=XM_019810743.2;refGene_gn=TBC1D5;refGene_tx=XM_019810744.2;refGene_gn=TBC1D5;refGene_tx=XM_019810745.2;refGene_gn=TBC1D5;refGene_tx=XM_019810746.1;refGene_gn=TBC1D5;refGene_tx=XM_019810747.1;refGene_gn=TBC1D5;refGene_tx=XM_019810750.2;refGene_gn=TBC1D5;refGene_tx=XM_023260586.1;refGene_gn=TBC1D5;refGene_tx=XM_023260587.1
C2	136917578	136917675
refGene_gn=TBC1D5;refGene_tx=XM_011286217.3;refGene_gn=TBC1D5;refGene_tx=XM_019810740.2;refGene_gn=TBC1D5;refGene_tx=XM_019810741.2;refGene_gn=TBC1D5;refGene_tx=XM_019810742.2;refGene_gn=TBC1D5;refGene_tx=XM_019810743.2;refGene_gn=TBC1D5;refGene_tx=XM_019810744.2;refGene_gn=TBC1D5;refGene_tx=XM_019810745.2;refGene_gn=TBC1D5;refGene_tx=XM_019810746.1;refGene_gn=TBC1D5;refGene_tx=XM_019810747.1;refGene_gn=TBC1D5;refGene_tx=XM_019810750.2;refGene_gn=TBC1D5;refGene_tx=XM_023260586.1;refGene_gn=TBC1D5;refGene_tx=XM_023260587.1
C2	137666804	137667308
refGene_gn=SATB1;refGene_tx=XM_011286212.3;refGene_gn=SATB1;refGene_tx=XM_011286213.3;refGene_gn=SATB1;refGene_tx=XM_011286214.3;refGene_gn=SATB1;refGene_tx=XM_019810753.2;refGene_gn=SATB1;refGene_tx=XM_0198107

54.2;refGene_gn=SATB1;refGene_tx=XM_019810755.2;refGene_gn=SATB1;refGene_tx=XM_019810756.2;refGene_gn=SATB1;refGene_tx=XM_023260588.1;refGene_gn=SATB1;refGene_tx=XM_023260589.1
C2 137668990 137669086
refGene_gn=SATB1;refGene_tx=XM_011286212.3;refGene_gn=SATB1;refGene_tx=XM_019810753.2;refGene_gn=SATB1;refGene_tx=XM_019810754.2;refGene_gn=SATB1;refGene_tx=XM_019810755.2;refGene_gn=SATB1;refGene_tx=XM_019810756.2;refGene_gn=SATB1;refGene_tx=XM_023260588.1;refGene_gn=SATB1;refGene_tx=XM_023260589.1
C2 137669679 137669883
refGene_gn=SATB1;refGene_tx=XM_011286212.3;refGene_gn=SATB1;refGene_tx=XM_011286213.3;refGene_gn=SATB1;refGene_tx=XM_011286214.3;refGene_gn=SATB1;refGene_tx=XM_019810753.2;refGene_gn=SATB1;refGene_tx=XM_019810754.2;refGene_gn=SATB1;refGene_tx=XM_019810755.2;refGene_gn=SATB1;refGene_tx=XM_019810756.2;refGene_gn=SATB1;refGene_tx=XM_023260588.1;refGene_gn=SATB1;refGene_tx=XM_023260589.1
C2 137697654 137697810
refGene_gn=SATB1;refGene_tx=XM_011286212.3;refGene_gn=SATB1;refGene_tx=XM_011286213.3;refGene_gn=SATB1;refGene_tx=XM_011286214.3;refGene_gn=SATB1;refGene_tx=XM_019810753.2;refGene_gn=SATB1;refGene_tx=XM_019810754.2;refGene_gn=SATB1;refGene_tx=XM_019810755.2;refGene_gn=SATB1;refGene_tx=XM_019810756.2;refGene_gn=SATB1;refGene_tx=XM_023260588.1;refGene_gn=SATB1;refGene_tx=XM_023260589.1
C2 137706377 137706590
refGene_gn=SATB1;refGene_tx=XM_011286212.3;refGene_gn=SATB1;refGene_tx=XM_011286213.3;refGene_gn=SATB1;refGene_tx=XM_011286214.3;refGene_gn=SATB1;refGene_tx=XM_019810753.2;refGene_gn=SATB1;refGene_tx=XM_019810754.2;refGene_gn=SATB1;refGene_tx=XM_019810755.2;refGene_gn=SATB1;refGene_tx=XM_019810756.2;refGene_gn=SATB1;refGene_tx=XM_023260588.1;refGene_gn=SATB1;refGene_tx=XM_023260589.1
C2 137714300 137714755
refGene_gn=SATB1;refGene_tx=XM_011286212.3;refGene_gn=SATB1;refGene_tx=XM_011286213.3;refGene_gn=SATB1;refGene_tx=XM_011286214.3;refGene_gn=SATB1;refGene_tx=XM_019810753.2;refGene_gn=SATB1;refGene_tx=XM_019810754.2;refGene_gn=SATB1;refGene_tx=XM_019810755.2;refGene_gn=SATB1;refGene_tx=XM_019810756.2;refGene_gn=SATB1;refGene_tx=XM_023260588.1;refGene_gn=SATB1;refGene_tx=XM_023260589.1
C2 137717057 137717169
refGene_gn=SATB1;refGene_tx=XM_011286212.3;refGene_gn=SATB1;refGene_tx=XM_011286213.3;refGene_gn=SATB1;refGene_tx=XM_011286214.3;refGene_gn=SATB1;refGene_tx=XM_019810753.2;refGene_gn=SATB1;refGene_tx=XM_019810754.2;refGene_gn=SATB1;refGene_tx=XM_019810755.2;refGene_gn=SATB1;refGene_tx=XM_019810756.2;refGene_gn=SATB1;refGene_tx=XM_023260588.1;refGene_gn=SATB1;refGene_tx=XM_023260589.1
C2 137736587 137736711
refGene_gn=SATB1;refGene_tx=XM_011286212.3;refGene_gn=SATB1;refGene_tx=XM_011286213.3;refGene_gn=SATB1;refGene_tx=XM_011286214.3;refGene_gn=SATB1;refGene_tx=XM_019810753.2;refGene_gn=SATB1;refGene_tx=XM_019810754.2;refGene_gn=SATB1;refGene_tx=XM_019810755.2;refGene_gn=SATB1;refGene_tx=XM_019810756.2;refGene_gn=SATB1;refGene_tx=XM_023260588.1;refGen

e_gn=SATB1;refGene_tx=XM_023260589.1
C2 137737476 137737603
refGene_gn=SATB1;refGene_tx=XM_011286212.3;refGene_gn=SATB1;refGene_tx=XM_011286213.3;refGene_gn=SATB1;refGene_tx=XM_011286214.3;refGene_gn=SATB1;refGene_tx=XM_019810753.2;refGene_gn=SATB1;refGene_tx=XM_019810754.2;refGene_gn=SATB1;refGene_tx=XM_019810755.2;refGene_gn=SATB1;refGene_tx=XM_019810756.2;refGene_gn=SATB1;refGene_tx=XM_023260588.1;refGene_gn=SATB1;refGene_tx=XM_023260589.1
C2 137738380 137738557
refGene_gn=SATB1;refGene_tx=XM_011286212.3;refGene_gn=SATB1;refGene_tx=XM_011286213.3;refGene_gn=SATB1;refGene_tx=XM_011286214.3;refGene_gn=SATB1;refGene_tx=XM_019810753.2;refGene_gn=SATB1;refGene_tx=XM_019810754.2;refGene_gn=SATB1;refGene_tx=XM_019810755.2;refGene_gn=SATB1;refGene_tx=XM_019810756.2;refGene_gn=SATB1;refGene_tx=XM_023260588.1;refGene_gn=SATB1;refGene_tx=XM_023260589.1
C2 137742257 137742468
refGene_gn=SATB1;refGene_tx=XM_011286212.3;refGene_gn=SATB1;refGene_tx=XM_011286213.3;refGene_gn=SATB1;refGene_tx=XM_011286214.3;refGene_gn=SATB1;refGene_tx=XM_019810753.2;refGene_gn=SATB1;refGene_tx=XM_019810754.2;refGene_gn=SATB1;refGene_tx=XM_019810755.2;refGene_gn=SATB1;refGene_tx=XM_019810756.2;refGene_gn=SATB1;refGene_tx=XM_023260588.1;refGene_gn=SATB1;refGene_tx=XM_023260589.1
C2 138467994 138468133
refGene_gn=KCNH8;refGene_tx=XM_023260187.1
C2 138478566 138478686
refGene_gn=KCNH8;refGene_tx=XM_023260187.1
C2 138522084 138522318
refGene_gn=KCNH8;refGene_tx=XM_023260187.1
C2 138557330 138557462
refGene_gn=KCNH8;refGene_tx=XM_023260187.1
C2 138610702 138610830
refGene_gn=KCNH8;refGene_tx=XM_023260187.1
C2 138616045 138616299
refGene_gn=KCNH8;refGene_tx=XM_023260187.1
C2 138616596 138616694
refGene_gn=KCNH8;refGene_tx=XM_023260187.1
C2 138617038 138617215
refGene_gn=KCNH8;refGene_tx=XM_023260187.1
C2 138618125 138618229
refGene_gn=KCNH8;refGene_tx=XM_023260187.1
C2 138629744 138629846
refGene_gn=KCNH8;refGene_tx=XM_023260187.1
C2 138630142 138630359
refGene_gn=KCNH8;refGene_tx=XM_023260187.1
C2 138645130 138645291
refGene_gn=KCNH8;refGene_tx=XM_023260187.1
C2 138649300 138649508
refGene_gn=KCNH8;refGene_tx=XM_023260187.1
C2 138687325 138687523
refGene_gn=KCNH8;refGene_tx=XM_023260187.1

C2 138698410 138698610
refGene_gn=KCNH8;refGene_tx=XM_023260187.1
C2 138699489 138699739
refGene_gn=KCNH8;refGene_tx=XM_023260187.1
C2 138705432 138705647
refGene_gn=KCNH8;refGene_tx=XM_023260187.1
C2 138728108 138728147
refGene_gn=KCNH8;refGene_tx=XM_023260187.1
C2 138732500 138732859
refGene_gn=KCNH8;refGene_tx=XM_023260187.1
C2 138734672 138734779
refGene_gn=KCNH8;refGene_tx=XM_023260187.1
C2 138737507 138737584
refGene_gn=KCNH8;refGene_tx=XM_023260187.1
C2 138759260 138759965
refGene_gn=KCNH8;refGene_tx=XM_023260187.1
C2 138989009 138989183
refGene_gn=EFHB;refGene_tx=XM_003992159.4;refGene_gn=EFHB;refGene_tx=XM_023260590.1
C2 138991681 138991863
refGene_gn=EFHB;refGene_tx=XM_003992159.4;refGene_gn=EFHB;refGene_tx=XM_023260590.1
C2 138993881 138994094
refGene_gn=EFHB;refGene_tx=XM_003992159.4;refGene_gn=EFHB;refGene_tx=XM_023260590.1
C2 138997077 138997285
refGene_gn=EFHB;refGene_tx=XM_003992159.4;refGene_gn=EFHB;refGene_tx=XM_023260590.1
C2 139002801 139002956
refGene_gn=EFHB;refGene_tx=XM_003992159.4;refGene_gn=EFHB;refGene_tx=XM_023260590.1
C2 139006646 139006714
refGene_gn=EFHB;refGene_tx=XM_003992159.4;refGene_gn=EFHB;refGene_tx=XM_023260590.1
C2 139007292 139007376
refGene_gn=EFHB;refGene_tx=XM_003992159.4;refGene_gn=EFHB;refGene_tx=XM_023260590.1
C2 139011807 139011937
refGene_gn=EFHB;refGene_tx=XM_003992159.4;refGene_gn=EFHB;refGene_tx=XM_023260590.1
C2 139015737 139015848
refGene_gn=EFHB;refGene_tx=XM_003992159.4
C2 139017517 139017698
refGene_gn=EFHB;refGene_tx=XM_003992159.4;refGene_gn=EFHB;refGene_tx=XM_023260590.1
C2 139018388 139018532
refGene_gn=EFHB;refGene_tx=XM_003992159.4;refGene_gn=EFHB;refGene_tx=XM_023260590.1
C2 139018983 139019046
refGene_gn=EFHB;refGene_tx=XM_003992159.4;refGene_gn=EFHB;refGene_tx=X

M_023260590.1
C2 139037170 139038067
refGene_gn=EFHB;refGene_tx=XM_003992159.4;refGene_gn=EFHB;refGene_tx=X
M_023260590.1
C2 139046719 139046882
refGene_gn=RAB5A;refGene_tx=XM_003992157.5;refGene_gn=RAB5A;refGene_tx
=XM_019810758.1
C2 139065795 139065947
refGene_gn=RAB5A;refGene_tx=XM_003992157.5;refGene_gn=RAB5A;refGene_tx
=XM_019810758.1
C2 139066259 139066382
refGene_gn=RAB5A;refGene_tx=XM_003992157.5;refGene_gn=RAB5A;refGene_tx
=XM_019810758.1
C2 139068093 139068187
refGene_gn=RAB5A;refGene_tx=XM_003992157.5;refGene_gn=RAB5A;refGene_tx
=XM_019810758.1
C2 139072532 139072648
refGene_gn=RAB5A;refGene_tx=XM_003992157.5;refGene_gn=RAB5A;refGene_tx
=XM_019810758.1
C2 139074173 139074976
refGene_gn=PP2D1;refGene_tx=XM_006936446.4;refGene_gn=PP2D1;refGene_tx
=XM_006936447.4;refGene_gn=PP2D1;refGene_tx=XM_006936448.4;refGene_gn=
PP2D1;refGene_tx=XM_019810757.2
C2 139085843 139086907
refGene_gn=PP2D1;refGene_tx=XM_006936446.4;refGene_gn=PP2D1;refGene_tx
=XM_006936447.4;refGene_gn=PP2D1;refGene_tx=XM_006936448.4;refGene_gn=
PP2D1;refGene_tx=XM_019810757.2
C2 139093266 139093289
refGene_gn=PP2D1;refGene_tx=XM_006936446.4;refGene_gn=PP2D1;refGene_tx
=XM_006936447.4;refGene_gn=PP2D1;refGene_tx=XM_019810757.2
C2 139116116 139116425
refGene_gn=KAT2B;refGene_tx=XM_023260591.1
C2 139117524 139117569
refGene_gn=KAT2B;refGene_tx=XM_006936449.4;refGene_gn=KAT2B;refGene_tx
=XM_023260592.1;refGene_gn=KAT2B;refGene_tx=XM_023260593.1
C2 139145123 139145378
refGene_gn=LOC109491531;refGene_tx=XM_023260594.1
C2 139148444 139148571
refGene_gn=KAT2B;refGene_tx=XM_006936449.4;refGene_gn=KAT2B;refGene_tx
=XM_023260591.1;refGene_gn=KAT2B;refGene_tx=XM_023260592.1;refGene_gn=
KAT2B;refGene_tx=XM_023260593.1
C2 139163057 139163203
refGene_gn=KAT2B;refGene_tx=XM_006936449.4;refGene_gn=KAT2B;refGene_tx
=XM_023260591.1;refGene_gn=KAT2B;refGene_tx=XM_023260592.1;refGene_gn=
KAT2B;refGene_tx=XM_023260593.1
C2 139167594 139167687
refGene_gn=KAT2B;refGene_tx=XM_006936449.4;refGene_gn=KAT2B;refGene_tx
=XM_023260591.1;refGene_gn=KAT2B;refGene_tx=XM_023260592.1;refGene_gn=
KAT2B;refGene_tx=XM_023260593.1
C2 139169020 139169202

refGene_gn=KAT2B;refGene_tx=XM_006936449.4;refGene_gn=KAT2B;refGene_tx=XM_023260591.1;refGene_gn=KAT2B;refGene_tx=XM_023260592.1;refGene_gn=KAT2B;refGene_tx=XM_023260593.1
C2 139176642 139176834
refGene_gn=KAT2B;refGene_tx=XM_006936449.4;refGene_gn=KAT2B;refGene_tx=XM_023260591.1;refGene_gn=KAT2B;refGene_tx=XM_023260592.1
C2 139182624 139182731
refGene_gn=KAT2B;refGene_tx=XM_006936449.4;refGene_gn=KAT2B;refGene_tx=XM_023260591.1;refGene_gn=KAT2B;refGene_tx=XM_023260592.1;refGene_gn=KAT2B;refGene_tx=XM_023260593.1
C2 139186645 139186771
refGene_gn=KAT2B;refGene_tx=XM_006936449.4;refGene_gn=KAT2B;refGene_tx=XM_023260591.1;refGene_gn=KAT2B;refGene_tx=XM_023260593.1
C2 139189533 139189670
refGene_gn=KAT2B;refGene_tx=XM_006936449.4;refGene_gn=KAT2B;refGene_tx=XM_023260591.1;refGene_gn=KAT2B;refGene_tx=XM_023260592.1;refGene_gn=KAT2B;refGene_tx=XM_023260593.1
C2 139192171 139192380
refGene_gn=KAT2B;refGene_tx=XM_006936449.4;refGene_gn=KAT2B;refGene_tx=XM_023260591.1;refGene_gn=KAT2B;refGene_tx=XM_023260592.1;refGene_gn=KAT2B;refGene_tx=XM_023260593.1
C2 139193543 139193670
refGene_gn=KAT2B;refGene_tx=XM_006936449.4;refGene_gn=KAT2B;refGene_tx=XM_023260591.1;refGene_gn=KAT2B;refGene_tx=XM_023260592.1;refGene_gn=KAT2B;refGene_tx=XM_023260593.1
C2 139202307 139202418
refGene_gn=KAT2B;refGene_tx=XM_006936449.4;refGene_gn=KAT2B;refGene_tx=XM_023260591.1;refGene_gn=KAT2B;refGene_tx=XM_023260592.1;refGene_gn=KAT2B;refGene_tx=XM_023260593.1
C2 139205613 139205757
refGene_gn=KAT2B;refGene_tx=XM_006936449.4;refGene_gn=KAT2B;refGene_tx=XM_023260591.1;refGene_gn=KAT2B;refGene_tx=XM_023260592.1;refGene_gn=KAT2B;refGene_tx=XM_023260593.1
C2 139210955 139211070
refGene_gn=KAT2B;refGene_tx=XM_006936449.4;refGene_gn=KAT2B;refGene_tx=XM_023260591.1;refGene_gn=KAT2B;refGene_tx=XM_023260592.1;refGene_gn=KAT2B;refGene_tx=XM_023260593.1
C2 139212580 139212617
refGene_gn=KAT2B;refGene_tx=XM_006936449.4;refGene_gn=KAT2B;refGene_tx=XM_023260591.1;refGene_gn=KAT2B;refGene_tx=XM_023260592.1;refGene_gn=KAT2B;refGene_tx=XM_023260593.1
C2 139212859 139212923
refGene_gn=KAT2B;refGene_tx=XM_006936449.4;refGene_gn=KAT2B;refGene_tx=XM_023260591.1;refGene_gn=KAT2B;refGene_tx=XM_023260592.1;refGene_gn=KAT2B;refGene_tx=XM_023260593.1
C2 139213021 139213106
refGene_gn=KAT2B;refGene_tx=XM_006936449.4;refGene_gn=KAT2B;refGene_tx=XM_023260591.1;refGene_gn=KAT2B;refGene_tx=XM_023260592.1;refGene_gn=KAT2B;refGene_tx=XM_023260593.1
C2 139217514 139217708

refGene_gn=KAT2B;refGene_tx=XM_006936449.4;refGene_gn=KAT2B;refGene_tx=XM_023260591.1;refGene_gn=KAT2B;refGene_tx=XM_023260592.1;refGene_gn=KAT2B;refGene_tx=XM_023260593.1
C2 139228409 139228512
refGene_gn=SGO1;refGene_tx=XM_023260596.1;refGene_gn=SGO1;refGene_tx=XM_023260597.1
C2 139228736 139228926
refGene_gn=SGO1;refGene_tx=XM_023260596.1;refGene_gn=SGO1;refGene_tx=XM_023260597.1
C2 139230942 139231755
refGene_gn=SGO1;refGene_tx=XM_023260596.1;refGene_gn=SGO1;refGene_tx=XM_023260597.1
C2 139232707 139232766
refGene_gn=SGO1;refGene_tx=XM_023260596.1;refGene_gn=SGO1;refGene_tx=XM_023260597.1
C2 139234394 139234468
refGene_gn=SGO1;refGene_tx=XM_023260596.1;refGene_gn=SGO1;refGene_tx=XM_023260597.1
C2 139238569 139238766
refGene_gn=SGO1;refGene_tx=XM_023260596.1;refGene_gn=SGO1;refGene_tx=XM_023260597.1
C2 139238850 139238992
refGene_gn=SGO1;refGene_tx=XM_023260596.1;refGene_gn=SGO1;refGene_tx=XM_023260597.1
C2 140352375 140352609
refGene_gn=ZNF385D;refGene_tx=XM_003992162.5;refGene_gn=ZNF385D;refGene_tx=XM_006936450.4;refGene_gn=ZNF385D;refGene_tx=XM_019810770.2;refGene_gn=ZNF385D;refGene_tx=XM_019810771.2;refGene_gn=ZNF385D;refGene_tx=XM_019810772.2
C2 140354657 140354759
refGene_gn=ZNF385D;refGene_tx=XM_003992162.5;refGene_gn=ZNF385D;refGene_tx=XM_006936450.4;refGene_gn=ZNF385D;refGene_tx=XM_019810770.2;refGene_gn=ZNF385D;refGene_tx=XM_019810771.2;refGene_gn=ZNF385D;refGene_tx=XM_019810772.2
C2 140355827 140356006
refGene_gn=ZNF385D;refGene_tx=XM_003992162.5;refGene_gn=ZNF385D;refGene_tx=XM_006936450.4;refGene_gn=ZNF385D;refGene_tx=XM_019810770.2;refGene_gn=ZNF385D;refGene_tx=XM_019810771.2;refGene_gn=ZNF385D;refGene_tx=XM_019810772.2
C2 140362981 140363212
refGene_gn=ZNF385D;refGene_tx=XM_003992162.5;refGene_gn=ZNF385D;refGene_tx=XM_006936450.4;refGene_gn=ZNF385D;refGene_tx=XM_019810770.2;refGene_gn=ZNF385D;refGene_tx=XM_019810771.2;refGene_gn=ZNF385D;refGene_tx=XM_019810772.2
C2 140432840 140433003
refGene_gn=ZNF385D;refGene_tx=XM_003992162.5;refGene_gn=ZNF385D;refGene_tx=XM_006936450.4;refGene_gn=ZNF385D;refGene_tx=XM_019810770.2;refGene_gn=ZNF385D;refGene_tx=XM_019810771.2;refGene_gn=ZNF385D;refGene_tx=XM_019810772.2
C2 140470344 140470455

refGene_gn=ZNF385D;refGene_tx=XM_003992162.5;refGene_gn=ZNF385D;refGene_tx=XM_006936450.4;refGene_gn=ZNF385D;refGene_tx=XM_019810770.2;refGene_gn=ZNF385D;refGene_tx=XM_019810771.2;refGene_gn=ZNF385D;refGene_tx=XM_019810772.2
C2 140572914 140573054
refGene_gn=ZNF385D;refGene_tx=XM_003992162.5;refGene_gn=ZNF385D;refGene_tx=XM_006936450.4;refGene_gn=ZNF385D;refGene_tx=XM_019810770.2;refGene_gn=ZNF385D;refGene_tx=XM_019810771.2;refGene_gn=ZNF385D;refGene_tx=XM_019810772.2
C2 140647866 140647882
refGene_gn=ZNF385D;refGene_tx=XM_019810770.2
C2 140648923 140648936
refGene_gn=ZNF385D;refGene_tx=XM_019810772.2
C2 140649760 140649782
refGene_gn=ZNF385D;refGene_tx=XM_003992162.5
C2 141018054 141018273
refGene_gn=ZNF385D;refGene_tx=XM_006936450.4;refGene_gn=ZNF385D;refGene_tx=XM_019810771.2
C2 141218018 141218049
refGene_gn=ZNF385D;refGene_tx=XM_006936450.4
C2 141240821 141240927
refGene_gn=ZNF385D;refGene_tx=XM_019810771.2
C2 141961684 141961860
refGene_gn=LOC101082513;refGene_tx=XM_023260599.1;refGene_gn=LOC101082513;refGene_tx=XM_023260600.1;refGene_gn=LOC101082513;refGene_tx=XM_023260601.1;refGene_gn=LOC101082513;refGene_tx=XM_023260602.1;refGene_gn=LOC101082513;refGene_tx=XM_023260603.1
C2 141970836 141970887
refGene_gn=LOC101082513;refGene_tx=XM_023260599.1;refGene_gn=LOC101082513;refGene_tx=XM_023260600.1;refGene_gn=LOC101082513;refGene_tx=XM_023260601.1;refGene_gn=LOC101082513;refGene_tx=XM_023260602.1;refGene_gn=LOC101082513;refGene_tx=XM_023260603.1
C2 142182657 142182739
refGene_gn=LOC101082513;refGene_tx=XM_023260599.1
C2 142183551 142183681
refGene_gn=LOC101082513;refGene_tx=XM_023260599.1;refGene_gn=LOC101082513;refGene_tx=XM_023260603.1
C2 142237677 142237810
refGene_gn=LOC101082513;refGene_tx=XM_023260600.1;refGene_gn=LOC101082513;refGene_tx=XM_023260601.1;refGene_gn=LOC101082513;refGene_tx=XM_023260602.1
C2 142279335 142279483
refGene_gn=LOC101082513;refGene_tx=XM_023260599.1;refGene_gn=LOC101082513;refGene_tx=XM_023260600.1;refGene_gn=LOC101082513;refGene_tx=XM_023260601.1;refGene_gn=LOC101082513;refGene_tx=XM_023260602.1;refGene_gn=LOC101082513;refGene_tx=XM_023260603.1
C2 142282368 142282465
refGene_gn=LOC101082513;refGene_tx=XM_023260599.1;refGene_gn=LOC101082513;refGene_tx=XM_023260600.1;refGene_gn=LOC101082513;refGene_tx=XM_023260601.1;refGene_gn=LOC101082513;refGene_tx=XM_023260603.1

C2	142326705	142326803
refGene_gn=LOC101082513;refGene_tx=XM_023260599.1;refGene_gn=LOC101082513;refGene_tx=XM_023260600.1;refGene_gn=LOC101082513;refGene_tx=XM_023260601.1;refGene_gn=LOC101082513;refGene_tx=XM_023260602.1;refGene_gn=LOC101082513;refGene_tx=XM_023260603.1
C2	142525530	142525682
refGene_gn=UBE2E1;refGene_tx=XM_023260604.1;refGene_gn=UBE2E1;refGene_tx=XM_023260605.1
C2	142529541	142529585
refGene_gn=UBE2E1;refGene_tx=XM_006936459.4
C2	142529780	142529831
refGene_gn=UBE2E1;refGene_tx=XM_006936459.4;refGene_gn=UBE2E1;refGene_tx=XM_023260604.1
C2	142602292	142602304
refGene_gn=NKIRAS1;refGene_tx=XM_023260611.1
C2	142603253	142603386
refGene_gn=UBE2E1;refGene_tx=XM_006936459.4;refGene_gn=UBE2E1;refGene_tx=XM_023260604.1;refGene_gn=UBE2E1;refGene_tx=XM_023260605.1
C2	142604375	142604523
refGene_gn=UBE2E1;refGene_tx=XM_006936459.4;refGene_gn=UBE2E1;refGene_tx=XM_023260604.1;refGene_gn=UBE2E1;refGene_tx=XM_023260605.1
C2	142605632	142605668
refGene_gn=NKIRAS1;refGene_tx=XM_023260610.1
C2	142605960	142606058
refGene_gn=UBE2E1;refGene_tx=XM_006936459.4;refGene_gn=UBE2E1;refGene_tx=XM_023260604.1;refGene_gn=UBE2E1;refGene_tx=XM_023260605.1
C2	142608813	142609056
refGene_gn=NKIRAS1;refGene_tx=XM_006936455.3;refGene_gn=NKIRAS1;refGene_tx=XM_023260606.1;refGene_gn=NKIRAS1;refGene_tx=XM_023260607.1;refGene_gn=NKIRAS1;refGene_tx=XM_023260608.1;refGene_gn=NKIRAS1;refGene_tx=XM_023260609.1;refGene_gn=NKIRAS1;refGene_tx=XM_023260610.1;refGene_gn=NKIRAS1;refGene_tx=XM_023260611.1
C2	142616290	142616532
refGene_gn=NKIRAS1;refGene_tx=XM_006936455.3;refGene_gn=NKIRAS1;refGene_tx=XM_023260606.1;refGene_gn=NKIRAS1;refGene_tx=XM_023260607.1;refGene_gn=NKIRAS1;refGene_tx=XM_023260608.1;refGene_gn=NKIRAS1;refGene_tx=XM_023260609.1;refGene_gn=NKIRAS1;refGene_tx=XM_023260610.1;refGene_gn=NKIRAS1;refGene_tx=XM_023260611.1
C2	142639508	142639602
refGene_gn=NKIRAS1;refGene_tx=XM_023260606.1;refGene_gn=NKIRAS1;refGene_tx=XM_023260607.1;refGene_gn=NKIRAS1;refGene_tx=XM_023260608.1;refGene_gn=NKIRAS1;refGene_tx=XM_023260610.1;refGene_gn=NKIRAS1;refGene_tx=XM_023260611.1
C2	142648806	142648978
refGene_gn=RPL15;refGene_tx=XM_006936460.3
C2	142649454	142649591
refGene_gn=RPL15;refGene_tx=XM_006936460.3
C2	142650400	142650706
refGene_gn=RPL15;refGene_tx=XM_006936460.3
C2	142679563	142679579

refGene_gn=NR1D2;refGene_tx=XM_023260612.1
C2	142687502	142687769
refGene_gn=NR1D2;refGene_tx=XM_023260612.1
C2	142688871	142688960
refGene_gn=NR1D2;refGene_tx=XM_023260612.1
C2	142690828	142690973
refGene_gn=NR1D2;refGene_tx=XM_023260612.1
C2	142692755	142693375
refGene_gn=NR1D2;refGene_tx=XM_023260612.1
C2	142696285	142696471
refGene_gn=NR1D2;refGene_tx=XM_023260612.1
C2	142698914	142699125
refGene_gn=NR1D2;refGene_tx=XM_023260612.1
C2	142705294	142705491
refGene_gn=NR1D2;refGene_tx=XM_023260612.1
C2	142828423	142828665
refGene_gn=THRB;refGene_tx=XM_011286224.3;refGene_gn=THRB;refGene_tx=XM_023260613.1;refGene_gn=THRB;refGene_tx=XM_023260614.1;refGene_gn=THRB;refGene_tx=XM_023260615.1;refGene_gn=THRB;refGene_tx=XM_023260616.1;refGene_gn=THRB;refGene_tx=XM_023260617.1;refGene_gn=THRB;refGene_tx=XM_023260618.1;refGene_gn=THRB;refGene_tx=XM_023260619.1
C2	142836084	142836343
refGene_gn=THRB;refGene_tx=XM_011286224.3;refGene_gn=THRB;refGene_tx=XM_023260613.1;refGene_gn=THRB;refGene_tx=XM_023260614.1;refGene_gn=THRB;refGene_tx=XM_023260615.1;refGene_gn=THRB;refGene_tx=XM_023260616.1;refGene_gn=THRB;refGene_tx=XM_023260617.1;refGene_gn=THRB;refGene_tx=XM_023260618.1;refGene_gn=THRB;refGene_tx=XM_023260619.1
C2	142843713	142843860
refGene_gn=THRB;refGene_tx=XM_011286224.3;refGene_gn=THRB;refGene_tx=XM_023260613.1;refGene_gn=THRB;refGene_tx=XM_023260614.1;refGene_gn=THRB;refGene_tx=XM_023260615.1;refGene_gn=THRB;refGene_tx=XM_023260616.1;refGene_gn=THRB;refGene_tx=XM_023260617.1;refGene_gn=THRB;refGene_tx=XM_023260618.1;refGene_gn=THRB;refGene_tx=XM_023260619.1
C2	142853901	142854107
refGene_gn=THRB;refGene_tx=XM_011286224.3;refGene_gn=THRB;refGene_tx=XM_023260613.1;refGene_gn=THRB;refGene_tx=XM_023260614.1;refGene_gn=THRB;refGene_tx=XM_023260615.1;refGene_gn=THRB;refGene_tx=XM_023260616.1;refGene_gn=THRB;refGene_tx=XM_023260617.1;refGene_gn=THRB;refGene_tx=XM_023260618.1;refGene_gn=THRB;refGene_tx=XM_023260619.1
C2	142857232	142857380
refGene_gn=THRB;refGene_tx=XM_011286224.3;refGene_gn=THRB;refGene_tx=XM_023260613.1;refGene_gn=THRB;refGene_tx=XM_023260614.1;refGene_gn=THRB;refGene_tx=XM_023260615.1;refGene_gn=THRB;refGene_tx=XM_023260616.1;refGene_gn=THRB;refGene_tx=XM_023260617.1;refGene_gn=THRB;refGene_tx=XM_023260618.1;refGene_gn=THRB;refGene_tx=XM_023260619.1
C2	142862866	142862967
refGene_gn=THRB;refGene_tx=XM_011286224.3;refGene_gn=THRB;refGene_tx=XM_023260613.1;refGene_gn=THRB;refGene_tx=XM_023260614.1;refGene_gn=THRB;refGene_tx=XM_023260615.1;refGene_gn=THRB;refGene_tx=XM_023260616.1;refGene_gn=THRB;refGene_tx=XM_023260617.1;refGene_gn=THRB;refGene_tx=X

M_023260618.1;refGene_gn=THRB;refGene_tx=XM_023260619.1
C2 142879633 142879961
refGene_gn=THRB;refGene_tx=XM_011286224.3
C2 142902135 142902396
refGene_gn=THRB;refGene_tx=XM_023260613.1;refGene_gn=THRB;refGene_tx=XM_023260614.1;refGene_gn=THRB;refGene_tx=XM_023260615.1;refGene_gn=THRB;refGene_tx=XM_023260616.1;refGene_gn=THRB;refGene_tx=XM_023260617.1;refGene_gn=THRB;refGene_tx=XM_023260618.1
C2 142934332 142934354
refGene_gn=THRB;refGene_tx=XM_023260613.1;refGene_gn=THRB;refGene_tx=XM_023260614.1;refGene_gn=THRB;refGene_tx=XM_023260615.1;refGene_gn=THRB;refGene_tx=XM_023260616.1;refGene_gn=THRB;refGene_tx=XM_023260617.1;refGene_gn=THRB;refGene_tx=XM_023260619.1
C2 143213065 143213397
refGene_gn=LOC111562308;refGene_tx=XM_023260620.1;refGene_gn=LOC111562308;refGene_tx=XM_023260623.1
C2 143214171 143214529
refGene_gn=LOC111562308;refGene_tx=XM_023260621.1;refGene_gn=LOC111562308;refGene_tx=XM_023260622.1
C2 143215358 143215788
refGene_gn=LOC111562308;refGene_tx=XM_023260620.1;refGene_gn=LOC111562308;refGene_tx=XM_023260621.1;refGene_gn=LOC111562308;refGene_tx=XM_023260622.1;refGene_gn=LOC111562308;refGene_tx=XM_023260623.1
C2 143956134 143956312
refGene_gn=RARB;refGene_tx=XM_011286272.3;refGene_gn=RARB;refGene_tx=XM_019810788.2;refGene_gn=RARB;refGene_tx=XM_019810789.2;refGene_gn=RARB;refGene_tx=XM_019810790.2;refGene_gn=RARB;refGene_tx=XM_023260626.1
C2 144189608 144189765
refGene_gn=RARB;refGene_tx=XM_003992173.4
C2 144199224 144199267
refGene_gn=RARB;refGene_tx=XM_019810791.2
C2 144218608 144218757
refGene_gn=RARB;refGene_tx=XM_003992173.4;refGene_gn=RARB;refGene_tx=XM_011286272.3;refGene_gn=RARB;refGene_tx=XM_019810788.2;refGene_gn=RARB;refGene_tx=XM_019810789.2;refGene_gn=RARB;refGene_tx=XM_019810790.2;refGene_gn=RARB;refGene_tx=XM_019810791.2;refGene_gn=RARB;refGene_tx=XM_023260626.1
C2 144259690 144259832
refGene_gn=RARB;refGene_tx=XM_003992173.4;refGene_gn=RARB;refGene_tx=XM_011286272.3;refGene_gn=RARB;refGene_tx=XM_019810788.2;refGene_gn=RARB;refGene_tx=XM_019810789.2;refGene_gn=RARB;refGene_tx=XM_019810790.2;refGene_gn=RARB;refGene_tx=XM_019810791.2;refGene_gn=RARB;refGene_tx=XM_019810792.2;refGene_gn=RARB;refGene_tx=XM_019810793.2;refGene_gn=RARB;refGene_tx=XM_023260626.1
C2 144335208 144335369
refGene_gn=RARB;refGene_tx=XM_003992173.4;refGene_gn=RARB;refGene_tx=XM_011286272.3;refGene_gn=RARB;refGene_tx=XM_019810788.2;refGene_gn=RARB;refGene_tx=XM_019810789.2;refGene_gn=RARB;refGene_tx=XM_019810790.2;refGene_gn=RARB;refGene_tx=XM_019810791.2;refGene_gn=RARB;refGene_tx=XM_019810792.2;refGene_gn=RARB;refGene_tx=XM_019810793.2;refGene_gn=RAR

B;refGene_tx=XM_023260626.1
C2	144347783	144347960
refGene_gn=RARB;refGene_tx=XM_003992173.4;refGene_gn=RARB;refGene_tx=XM_011286272.3;refGene_gn=RARB;refGene_tx=XM_019810788.2;refGene_gn=RARB;refGene_tx=XM_019810789.2;refGene_gn=RARB;refGene_tx=XM_019810790.2;refGene_gn=RARB;refGene_tx=XM_019810791.2;refGene_gn=RARB;refGene_tx=XM_019810792.2;refGene_gn=RARB;refGene_tx=XM_019810793.2;refGene_gn=RARB;refGene_tx=XM_023260626.1
C2	144359510	144359715
refGene_gn=RARB;refGene_tx=XM_003992173.4;refGene_gn=RARB;refGene_tx=XM_011286272.3;refGene_gn=RARB;refGene_tx=XM_019810788.2;refGene_gn=RARB;refGene_tx=XM_019810789.2;refGene_gn=RARB;refGene_tx=XM_019810790.2;refGene_gn=RARB;refGene_tx=XM_019810791.2;refGene_gn=RARB;refGene_tx=XM_019810792.2;refGene_gn=RARB;refGene_tx=XM_019810793.2;refGene_gn=RARB;refGene_tx=XM_023260626.1
C2	144360546	144360705
refGene_gn=RARB;refGene_tx=XM_003992173.4;refGene_gn=RARB;refGene_tx=XM_011286272.3;refGene_gn=RARB;refGene_tx=XM_019810788.2;refGene_gn=RARB;refGene_tx=XM_019810789.2;refGene_gn=RARB;refGene_tx=XM_019810790.2;refGene_gn=RARB;refGene_tx=XM_019810791.2;refGene_gn=RARB;refGene_tx=XM_019810792.2;refGene_gn=RARB;refGene_tx=XM_019810793.2;refGene_gn=RARB;refGene_tx=XM_023260626.1
C2	144362259	144362456
refGene_gn=RARB;refGene_tx=XM_003992173.4;refGene_gn=RARB;refGene_tx=XM_011286272.3;refGene_gn=RARB;refGene_tx=XM_019810788.2;refGene_gn=RARB;refGene_tx=XM_019810789.2;refGene_gn=RARB;refGene_tx=XM_019810790.2;refGene_gn=RARB;refGene_tx=XM_019810791.2;refGene_gn=RARB;refGene_tx=XM_019810792.2;refGene_gn=RARB;refGene_tx=XM_019810793.2;refGene_gn=RARB;refGene_tx=XM_023260626.1
C2	144364112	144364280
refGene_gn=TOP2B;refGene_tx=XM_023260624.1;refGene_gn=TOP2B;refGene_tx=XM_023260625.1
C2	144365157	144365252
refGene_gn=TOP2B;refGene_tx=XM_023260624.1;refGene_gn=TOP2B;refGene_tx=XM_023260625.1
C2	144366726	144366849
refGene_gn=TOP2B;refGene_tx=XM_023260624.1;refGene_gn=TOP2B;refGene_tx=XM_023260625.1
C2	144369420	144369531
refGene_gn=TOP2B;refGene_tx=XM_023260624.1;refGene_gn=TOP2B;refGene_tx=XM_023260625.1
C2	144370953	144371033
refGene_gn=TOP2B;refGene_tx=XM_023260624.1;refGene_gn=TOP2B;refGene_tx=XM_023260625.1
C2	144371979	144372181
refGene_gn=TOP2B;refGene_tx=XM_023260624.1;refGene_gn=TOP2B;refGene_tx=XM_023260625.1
C2	144373882	144374044
refGene_gn=TOP2B;refGene_tx=XM_023260624.1;refGene_gn=TOP2B;refGene_tx=XM_023260625.1

C2 144374511 144374656
refGene_gn=TOP2B;refGene_tx=XM_023260624.1;refGene_gn=TOP2B;refGene_tx=XM_023260625.1
C2 144376835 144377030
refGene_gn=TOP2B;refGene_tx=XM_023260624.1;refGene_gn=TOP2B;refGene_tx=XM_023260625.1
C2 144378860 144378944
refGene_gn=TOP2B;refGene_tx=XM_023260624.1;refGene_gn=TOP2B;refGene_tx=XM_023260625.1
C2 144379093 144379249
refGene_gn=TOP2B;refGene_tx=XM_023260624.1;refGene_gn=TOP2B;refGene_tx=XM_023260625.1
C2 144382048 144382140
refGene_gn=TOP2B;refGene_tx=XM_023260624.1;refGene_gn=TOP2B;refGene_tx=XM_023260625.1
C2 144382284 144382480
refGene_gn=TOP2B;refGene_tx=XM_023260624.1;refGene_gn=TOP2B;refGene_tx=XM_023260625.1
C2 144383478 144383679
refGene_gn=TOP2B;refGene_tx=XM_023260624.1;refGene_gn=TOP2B;refGene_tx=XM_023260625.1
C2 144384250 144384385
refGene_gn=TOP2B;refGene_tx=XM_023260624.1;refGene_gn=TOP2B;refGene_tx=XM_023260625.1
C2 144386258 144386490
refGene_gn=TOP2B;refGene_tx=XM_023260624.1;refGene_gn=TOP2B;refGene_tx=XM_023260625.1
C2 144387029 144387178
refGene_gn=TOP2B;refGene_tx=XM_023260624.1;refGene_gn=TOP2B;refGene_tx=XM_023260625.1
C2 144387377 144387499
refGene_gn=TOP2B;refGene_tx=XM_023260624.1;refGene_gn=TOP2B;refGene_tx=XM_023260625.1
C2 144389538 144389653
refGene_gn=TOP2B;refGene_tx=XM_023260624.1;refGene_gn=TOP2B;refGene_tx=XM_023260625.1
C2 144389745 144389838
refGene_gn=TOP2B;refGene_tx=XM_023260624.1;refGene_gn=TOP2B;refGene_tx=XM_023260625.1
C2 144390187 144390297
refGene_gn=TOP2B;refGene_tx=XM_023260624.1;refGene_gn=TOP2B;refGene_tx=XM_023260625.1
C2 144392061 144392167
refGene_gn=TOP2B;refGene_tx=XM_023260624.1;refGene_gn=TOP2B;refGene_tx=XM_023260625.1
C2 144392248 144392359
refGene_gn=TOP2B;refGene_tx=XM_023260624.1;refGene_gn=TOP2B;refGene_tx=XM_023260625.1
C2 144398063 144398189
refGene_gn=TOP2B;refGene_tx=XM_023260624.1;refGene_gn=TOP2B;refGene_tx

=XM_023260625.1
C2	144398438	144398596
refGene_gn=TOP2B;refGene_tx=XM_023260624.1;refGene_gn=TOP2B;refGene_tx
=XM_023260625.1
C2	144398944	144399083
refGene_gn=TOP2B;refGene_tx=XM_023260624.1;refGene_gn=TOP2B;refGene_tx
=XM_023260625.1
C2	144400689	144400827
refGene_gn=TOP2B;refGene_tx=XM_023260624.1;refGene_gn=TOP2B;refGene_tx
=XM_023260625.1
C2	144400910	144401012
refGene_gn=TOP2B;refGene_tx=XM_023260624.1;refGene_gn=TOP2B;refGene_tx
=XM_023260625.1
C2	144402731	144402905
refGene_gn=TOP2B;refGene_tx=XM_023260624.1;refGene_gn=TOP2B;refGene_tx
=XM_023260625.1
C2	144404886	144405099
refGene_gn=TOP2B;refGene_tx=XM_023260624.1;refGene_gn=TOP2B;refGene_tx
=XM_023260625.1
C2	144406233	144406331
refGene_gn=TOP2B;refGene_tx=XM_023260624.1;refGene_gn=TOP2B;refGene_tx
=XM_023260625.1
C2	144407198	144407344
refGene_gn=TOP2B;refGene_tx=XM_023260624.1;refGene_gn=TOP2B;refGene_tx
=XM_023260625.1
C2	144412241	144412305
refGene_gn=TOP2B;refGene_tx=XM_023260624.1;refGene_gn=TOP2B;refGene_tx
=XM_023260625.1
C2	144413653	144413744
refGene_gn=TOP2B;refGene_tx=XM_023260624.1;refGene_gn=TOP2B;refGene_tx
=XM_023260625.1
C2	144414617	144414788
refGene_gn=TOP2B;refGene_tx=XM_023260624.1;refGene_gn=TOP2B;refGene_tx
=XM_023260625.1
C2	144428570	144428633
refGene_gn=TOP2B;refGene_tx=XM_023260624.1;refGene_gn=TOP2B;refGene_tx
=XM_023260625.1
C2	144479133	144479309
refGene_gn=NGLY1;refGene_tx=XM_003992175.5;refGene_gn=NGLY1;refGene_tx
=XM_003992176.5;refGene_gn=NGLY1;refGene_tx=XM_006936470.4;refGene_gn=
NGLY1;refGene_tx=XM_006936471.4
C2	144479677	144479855
refGene_gn=NGLY1;refGene_tx=XM_003992175.5;refGene_gn=NGLY1;refGene_tx
=XM_006936470.4;refGene_gn=NGLY1;refGene_tx=XM_006936471.4
C2	144487679	144487865
refGene_gn=NGLY1;refGene_tx=XM_003992175.5;refGene_gn=NGLY1;refGene_tx
=XM_003992176.5;refGene_gn=NGLY1;refGene_tx=XM_006936471.4
C2	144491908	144492073
refGene_gn=NGLY1;refGene_tx=XM_003992175.5;refGene_gn=NGLY1;refGene_tx
=XM_003992176.5;refGene_gn=NGLY1;refGene_tx=XM_006936470.4;refGene_gn=

NGLY1;refGene_tx=XM_006936471.4
C2 144493350 144493461
refGene_gn=NGLY1;refGene_tx=XM_003992175.5;refGene_gn=NGLY1;refGene_tx=XM_003992176.5;refGene_gn=NGLY1;refGene_tx=XM_006936470.4;refGene_gn=NGLY1;refGene_tx=XM_006936471.4
C2 144495834 144495980
refGene_gn=NGLY1;refGene_tx=XM_003992175.5;refGene_gn=NGLY1;refGene_tx=XM_003992176.5;refGene_gn=NGLY1;refGene_tx=XM_006936470.4;refGene_gn=NGLY1;refGene_tx=XM_006936471.4
C2 144496957 144497079
refGene_gn=NGLY1;refGene_tx=XM_003992175.5;refGene_gn=NGLY1;refGene_tx=XM_003992176.5;refGene_gn=NGLY1;refGene_tx=XM_006936470.4;refGene_gn=NGLY1;refGene_tx=XM_006936471.4
C2 144500975 144501198
refGene_gn=NGLY1;refGene_tx=XM_003992175.5;refGene_gn=NGLY1;refGene_tx=XM_003992176.5;refGene_gn=NGLY1;refGene_tx=XM_006936470.4;refGene_gn=NGLY1;refGene_tx=XM_006936471.4
C2 144511231 144511397
refGene_gn=NGLY1;refGene_tx=XM_003992175.5;refGene_gn=NGLY1;refGene_tx=XM_003992176.5;refGene_gn=NGLY1;refGene_tx=XM_006936470.4;refGene_gn=NGLY1;refGene_tx=XM_006936471.4
C2 144525691 144525937
refGene_gn=NGLY1;refGene_tx=XM_003992175.5;refGene_gn=NGLY1;refGene_tx=XM_003992176.5;refGene_gn=NGLY1;refGene_tx=XM_006936470.4
C2 144530429 144530544
refGene_gn=NGLY1;refGene_tx=XM_003992175.5;refGene_gn=NGLY1;refGene_tx=XM_003992176.5;refGene_gn=NGLY1;refGene_tx=XM_006936470.4;refGene_gn=NGLY1;refGene_tx=XM_006936471.4
C2 144540752 144540883
refGene_gn=NGLY1;refGene_tx=XM_003992175.5;refGene_gn=NGLY1;refGene_tx=XM_003992176.5;refGene_gn=NGLY1;refGene_tx=XM_006936470.4;refGene_gn=NGLY1;refGene_tx=XM_006936471.4
C2 144548326 144549306
refGene_gn=OXSM;refGene_tx=XM_006936465.3;refGene_gn=OXSM;refGene_tx=XM_006936466.4;refGene_gn=OXSM;refGene_tx=XM_006936467.4;refGene_gn=OXSM;refGene_tx=XM_006936468.4;refGene_gn=OXSM;refGene_tx=XM_006936469.3
C2 144552275 144552678
refGene_gn=OXSM;refGene_tx=XM_006936465.3;refGene_gn=OXSM;refGene_tx=XM_006936466.4;refGene_gn=OXSM;refGene_tx=XM_006936467.4;refGene_gn=OXSM;refGene_tx=XM_006936468.4;refGene_gn=OXSM;refGene_tx=XM_006936469.3
C2 145289390 145290170
refGene_gn=LRRC3B;refGene_tx=XM_011286269.3;refGene_gn=LRRC3B;refGene_tx=XM_011286270.3;refGene_gn=LRRC3B;refGene_tx=XM_019810794.2;refGene_gn=LRRC3B;refGene_tx=XM_019810795.1
C2 145522066 145522562
refGene_gn=LOC101086839;refGene_tx=XM_023238572.1
C2 145522796 145522839
refGene_gn=LOC101086839;refGene_tx=XM_023238572.1
C2 145522840 145523177
refGene_gn=LOC101086839;refGene_tx=XM_023238572.1

C2	145600475	145600524
refGene_gn=NEK10;refGene_tx=XM_003992179.5;refGene_gn=NEK10;refGene_tx=XM_011286267.3;refGene_gn=NEK10;refGene_tx=XM_019810797.2;refGene_gn=NEK10;refGene_tx=XM_019810798.2;refGene_gn=NEK10;refGene_tx=XM_019810799.2;refGene_gn=NEK10;refGene_tx=XM_019810800.2;refGene_gn=NEK10;refGene_tx=XM_019810801.2;refGene_gn=NEK10;refGene_tx=XM_019810803.2;refGene_gn=NEK10;refGene_tx=XM_019810804.2;refGene_gn=NEK10;refGene_tx=XM_019810806.2;refGene_gn=NEK10;refGene_tx=XM_019810809.2;refGene_gn=NEK10;refGene_tx=XM_023260629.1
C2	145604618	145604674
refGene_gn=NEK10;refGene_tx=XM_003992179.5;refGene_gn=NEK10;refGene_tx=XM_011286267.3;refGene_gn=NEK10;refGene_tx=XM_019810797.2;refGene_gn=NEK10;refGene_tx=XM_019810798.2;refGene_gn=NEK10;refGene_tx=XM_019810799.2;refGene_gn=NEK10;refGene_tx=XM_019810800.2;refGene_gn=NEK10;refGene_tx=XM_019810801.2;refGene_gn=NEK10;refGene_tx=XM_019810803.2;refGene_gn=NEK10;refGene_tx=XM_019810804.2;refGene_gn=NEK10;refGene_tx=XM_019810806.2;refGene_gn=NEK10;refGene_tx=XM_019810809.2;refGene_gn=NEK10;refGene_tx=XM_023260629.1
C2	145604762	145604815
refGene_gn=NEK10;refGene_tx=XM_003992179.5;refGene_gn=NEK10;refGene_tx=XM_011286267.3;refGene_gn=NEK10;refGene_tx=XM_019810797.2;refGene_gn=NEK10;refGene_tx=XM_019810798.2;refGene_gn=NEK10;refGene_tx=XM_019810799.2;refGene_gn=NEK10;refGene_tx=XM_019810800.2;refGene_gn=NEK10;refGene_tx=XM_019810801.2;refGene_gn=NEK10;refGene_tx=XM_019810803.2;refGene_gn=NEK10;refGene_tx=XM_019810804.2;refGene_gn=NEK10;refGene_tx=XM_019810806.2;refGene_gn=NEK10;refGene_tx=XM_019810809.2;refGene_gn=NEK10;refGene_tx=XM_023260629.1
C2	145606746	145606855
refGene_gn=NEK10;refGene_tx=XM_003992179.5;refGene_gn=NEK10;refGene_tx=XM_011286267.3;refGene_gn=NEK10;refGene_tx=XM_019810797.2;refGene_gn=NEK10;refGene_tx=XM_019810798.2;refGene_gn=NEK10;refGene_tx=XM_019810799.2;refGene_gn=NEK10;refGene_tx=XM_019810800.2;refGene_gn=NEK10;refGene_tx=XM_019810801.2;refGene_gn=NEK10;refGene_tx=XM_019810803.2;refGene_gn=NEK10;refGene_tx=XM_019810804.2;refGene_gn=NEK10;refGene_tx=XM_019810806.2;refGene_gn=NEK10;refGene_tx=XM_019810809.2;refGene_gn=NEK10;refGene_tx=XM_023260629.1
C2	145612543	145612552
refGene_gn=NEK10;refGene_tx=XM_019810802.2
C2	145617148	145617259
refGene_gn=NEK10;refGene_tx=XM_003992179.5;refGene_gn=NEK10;refGene_tx=XM_011286267.3;refGene_gn=NEK10;refGene_tx=XM_019810797.2;refGene_gn=NEK10;refGene_tx=XM_019810798.2;refGene_gn=NEK10;refGene_tx=XM_019810799.2;refGene_gn=NEK10;refGene_tx=XM_019810800.2;refGene_gn=NEK10;refGene_tx=XM_019810801.2;refGene_gn=NEK10;refGene_tx=XM_019810802.2;refGene_gn=NEK10;refGene_tx=XM_019810803.2;refGene_gn=NEK10;refGene_tx=XM_019810804.2;refGene_gn=NEK10;refGene_tx=XM_019810806.2;refGene_gn=NEK10;refGene_tx=XM_019810809.2;refGene_gn=NEK10;refGene_tx=XM_023260629.1
C2	145627095	145627196
refGene_gn=NEK10;refGene_tx=XM_003992179.5;refGene_gn=NEK10;refGene_tx=XM_011286267.3;refGene_gn=NEK10;refGene_tx=XM_019810797.2;refGene_gn=

NEK10;refGene_tx=XM_019810798.2;refGene_gn=NEK10;refGene_tx=XM_0198107
99.2;refGene_gn=NEK10;refGene_tx=XM_019810800.2;refGene_gn=NEK10;refGe
ne_tx=XM_019810801.2;refGene_gn=NEK10;refGene_tx=XM_019810802.2;refGen
e_gn=NEK10;refGene_tx=XM_019810803.2;refGene_gn=NEK10;refGene_tx=XM_01
9810804.2;refGene_gn=NEK10;refGene_tx=XM_019810806.2;refGene_gn=NEK10;
refGene_tx=XM_019810809.2;refGene_gn=NEK10;refGene_tx=XM_023260629.1
C2 145629036 145629086
refGene_gn=NEK10;refGene_tx=XM_003992179.5;refGene_gn=NEK10;refGene_tx
=XM_019810797.2;refGene_gn=NEK10;refGene_tx=XM_019810798.2;refGene_gn=
NEK10;refGene_tx=XM_019810799.2;refGene_gn=NEK10;refGene_tx=XM_0198108
01.2;refGene_gn=NEK10;refGene_tx=XM_019810802.2;refGene_gn=NEK10;refGe
ne_tx=XM_019810803.2;refGene_gn=NEK10;refGene_tx=XM_019810804.2;refGen
e_gn=NEK10;refGene_tx=XM_019810806.2;refGene_gn=NEK10;refGene_tx=XM_01
9810807.2;refGene_gn=NEK10;refGene_tx=XM_019810809.2;refGene_gn=NEK10;
refGene_tx=XM_023260629.1
C2 145647752 145647893
refGene_gn=NEK10;refGene_tx=XM_003992179.5;refGene_gn=NEK10;refGene_tx
=XM_011286267.3;refGene_gn=NEK10;refGene_tx=XM_019810797.2;refGene_gn=
NEK10;refGene_tx=XM_019810798.2;refGene_gn=NEK10;refGene_tx=XM_0198108
01.2;refGene_gn=NEK10;refGene_tx=XM_019810802.2;refGene_gn=NEK10;refGe
ne_tx=XM_019810803.2;refGene_gn=NEK10;refGene_tx=XM_019810804.2;refGen
e_gn=NEK10;refGene_tx=XM_019810806.2;refGene_gn=NEK10;refGene_tx=XM_01
9810807.2;refGene_gn=NEK10;refGene_tx=XM_019810809.2;refGene_gn=NEK10;
refGene_tx=XM_023260629.1
C2 145647992 145648030
refGene_gn=NEK10;refGene_tx=XM_003992179.5;refGene_gn=NEK10;refGene_tx
=XM_011286267.3;refGene_gn=NEK10;refGene_tx=XM_019810797.2;refGene_gn=
NEK10;refGene_tx=XM_019810798.2;refGene_gn=NEK10;refGene_tx=XM_0198107
99.2;refGene_gn=NEK10;refGene_tx=XM_019810800.2;refGene_gn=NEK10;refGe
ne_tx=XM_019810801.2;refGene_gn=NEK10;refGene_tx=XM_019810802.2;refGen
e_gn=NEK10;refGene_tx=XM_019810803.2;refGene_gn=NEK10;refGene_tx=XM_01
9810804.2;refGene_gn=NEK10;refGene_tx=XM_019810806.2;refGene_gn=NEK10;
refGene_tx=XM_019810807.2;refGene_gn=NEK10;refGene_tx=XM_019810809.2;r
efGene_gn=NEK10;refGene_tx=XM_023260629.1
C2 145654953 145655002
refGene_gn=NEK10;refGene_tx=XM_011286268.3
C2 145658685 145658740
refGene_gn=NEK10;refGene_tx=XM_003992179.5;refGene_gn=NEK10;refGene_tx
=XM_011286267.3;refGene_gn=NEK10;refGene_tx=XM_011286268.3;refGene_gn=
NEK10;refGene_tx=XM_019810797.2;refGene_gn=NEK10;refGene_tx=XM_0198107
98.2;refGene_gn=NEK10;refGene_tx=XM_019810799.2;refGene_gn=NEK10;refGe
ne_tx=XM_019810800.2;refGene_gn=NEK10;refGene_tx=XM_019810801.2;refGen
e_gn=NEK10;refGene_tx=XM_019810802.2;refGene_gn=NEK10;refGene_tx=XM_01
9810803.2;refGene_gn=NEK10;refGene_tx=XM_019810804.2;refGene_gn=NEK10;
refGene_tx=XM_019810806.2;refGene_gn=NEK10;refGene_tx=XM_019810807.2;r
efGene_gn=NEK10;refGene_tx=XM_019810809.2;refGene_gn=NEK10;refGene_tx=
XM_023260629.1
C2 145660675 145660762
refGene_gn=NEK10;refGene_tx=XM_003992179.5;refGene_gn=NEK10;refGene_tx
=XM_011286267.3;refGene_gn=NEK10;refGene_tx=XM_011286268.3;refGene_gn=

NEK10;refGene_tx=XM_019810797.2;refGene_gn=NEK10;refGene_tx=XM_0198107
98.2;refGene_gn=NEK10;refGene_tx=XM_019810799.2;refGene_gn=NEK10;refGe
ne_tx=XM_019810800.2;refGene_gn=NEK10;refGene_tx=XM_019810801.2;refGen
e_gn=NEK10;refGene_tx=XM_019810802.2;refGene_gn=NEK10;refGene_tx=XM_01
9810803.2;refGene_gn=NEK10;refGene_tx=XM_019810804.2;refGene_gn=NEK10;
refGene_tx=XM_019810806.2;refGene_gn=NEK10;refGene_tx=XM_019810807.2;r
efGene_gn=NEK10;refGene_tx=XM_019810809.2;refGene_gn=NEK10;refGene_tx=
XM_023260629.1
C2 145660869 145661053
refGene_gn=NEK10;refGene_tx=XM_003992179.5;refGene_gn=NEK10;refGene_tx
=XM_011286267.3;refGene_gn=NEK10;refGene_tx=XM_011286268.3;refGene_gn=
NEK10;refGene_tx=XM_019810797.2;refGene_gn=NEK10;refGene_tx=XM_0198107
98.2;refGene_gn=NEK10;refGene_tx=XM_019810799.2;refGene_gn=NEK10;refGe
ne_tx=XM_019810800.2;refGene_gn=NEK10;refGene_tx=XM_019810801.2;refGen
e_gn=NEK10;refGene_tx=XM_019810802.2;refGene_gn=NEK10;refGene_tx=XM_01
9810803.2;refGene_gn=NEK10;refGene_tx=XM_019810804.2;refGene_gn=NEK10;
refGene_tx=XM_019810806.2;refGene_gn=NEK10;refGene_tx=XM_019810807.2;r
efGene_gn=NEK10;refGene_tx=XM_019810809.2;refGene_gn=NEK10;refGene_tx=
XM_023260629.1
C2 145677877 145678091
refGene_gn=NEK10;refGene_tx=XM_003992179.5;refGene_gn=NEK10;refGene_tx
=XM_011286267.3;refGene_gn=NEK10;refGene_tx=XM_011286268.3;refGene_gn=
NEK10;refGene_tx=XM_019810797.2;refGene_gn=NEK10;refGene_tx=XM_0198107
98.2;refGene_gn=NEK10;refGene_tx=XM_019810799.2;refGene_gn=NEK10;refGe
ne_tx=XM_019810800.2;refGene_gn=NEK10;refGene_tx=XM_019810801.2;refGen
e_gn=NEK10;refGene_tx=XM_019810802.2;refGene_gn=NEK10;refGene_tx=XM_01
9810803.2;refGene_gn=NEK10;refGene_tx=XM_019810804.2;refGene_gn=NEK10;
refGene_tx=XM_019810806.2;refGene_gn=NEK10;refGene_tx=XM_019810807.2;r
efGene_gn=NEK10;refGene_tx=XM_019810809.2;refGene_gn=NEK10;refGene_tx=
XM_023260629.1
C2 145689298 145689369
refGene_gn=NEK10;refGene_tx=XM_003992179.5;refGene_gn=NEK10;refGene_tx
=XM_011286267.3;refGene_gn=NEK10;refGene_tx=XM_011286268.3;refGene_gn=
NEK10;refGene_tx=XM_019810797.2;refGene_gn=NEK10;refGene_tx=XM_0198107
98.2;refGene_gn=NEK10;refGene_tx=XM_019810799.2;refGene_gn=NEK10;refGe
ne_tx=XM_019810800.2;refGene_gn=NEK10;refGene_tx=XM_019810801.2;refGen
e_gn=NEK10;refGene_tx=XM_019810802.2;refGene_gn=NEK10;refGene_tx=XM_01
9810803.2;refGene_gn=NEK10;refGene_tx=XM_019810804.2;refGene_gn=NEK10;
refGene_tx=XM_019810806.2;refGene_gn=NEK10;refGene_tx=XM_019810807.2;r
efGene_gn=NEK10;refGene_tx=XM_019810809.2;refGene_gn=NEK10;refGene_tx=
XM_023260629.1
C2 145689912 145690042
refGene_gn=NEK10;refGene_tx=XM_003992179.5;refGene_gn=NEK10;refGene_tx
=XM_011286267.3;refGene_gn=NEK10;refGene_tx=XM_011286268.3;refGene_gn=
NEK10;refGene_tx=XM_019810797.2;refGene_gn=NEK10;refGene_tx=XM_0198107
98.2;refGene_gn=NEK10;refGene_tx=XM_019810799.2;refGene_gn=NEK10;refGe
ne_tx=XM_019810800.2;refGene_gn=NEK10;refGene_tx=XM_019810801.2;refGen
e_gn=NEK10;refGene_tx=XM_019810802.2;refGene_gn=NEK10;refGene_tx=XM_01
9810803.2;refGene_gn=NEK10;refGene_tx=XM_019810804.2;refGene_gn=NEK10;
refGene_tx=XM_019810806.2;refGene_gn=NEK10;refGene_tx=XM_019810807.2;r

efGene_gn=NEK10;refGene_tx=XM_019810809.2;refGene_gn=NEK10;refGene_tx=XM_023260629.1
C2 145728316 145728320
refGene_gn=NEK10;refGene_tx=XM_006936474.3
C2 145730954 145731030
refGene_gn=NEK10;refGene_tx=XM_003992179.5;refGene_gn=NEK10;refGene_tx=XM_006936474.3;refGene_gn=NEK10;refGene_tx=XM_011286267.3;refGene_gn=NEK10;refGene_tx=XM_011286268.3;refGene_gn=NEK10;refGene_tx=XM_019810797.2;refGene_gn=NEK10;refGene_tx=XM_019810798.2;refGene_gn=NEK10;refGene_tx=XM_019810799.2;refGene_gn=NEK10;refGene_tx=XM_019810800.2;refGene_gn=NEK10;refGene_tx=XM_019810801.2;refGene_gn=NEK10;refGene_tx=XM_019810802.2;refGene_gn=NEK10;refGene_tx=XM_019810803.2;refGene_gn=NEK10;refGene_tx=XM_019810804.2;refGene_gn=NEK10;refGene_tx=XM_019810806.2;refGene_gn=NEK10;refGene_tx=XM_019810807.2;refGene_gn=NEK10;refGene_tx=XM_019810809.2;refGene_gn=NEK10;refGene_tx=XM_023260629.1
C2 145752737 145752840
refGene_gn=NEK10;refGene_tx=XM_003992179.5;refGene_gn=NEK10;refGene_tx=XM_006936474.3;refGene_gn=NEK10;refGene_tx=XM_011286267.3;refGene_gn=NEK10;refGene_tx=XM_011286268.3;refGene_gn=NEK10;refGene_tx=XM_019810797.2;refGene_gn=NEK10;refGene_tx=XM_019810798.2;refGene_gn=NEK10;refGene_tx=XM_019810799.2;refGene_gn=NEK10;refGene_tx=XM_019810800.2;refGene_gn=NEK10;refGene_tx=XM_019810801.2;refGene_gn=NEK10;refGene_tx=XM_019810802.2;refGene_gn=NEK10;refGene_tx=XM_019810803.2;refGene_gn=NEK10;refGene_tx=XM_019810804.2;refGene_gn=NEK10;refGene_tx=XM_019810806.2;refGene_gn=NEK10;refGene_tx=XM_019810807.2;refGene_gn=NEK10;refGene_tx=XM_019810809.2;refGene_gn=NEK10;refGene_tx=XM_023260629.1
C2 145752977 145753099
refGene_gn=NEK10;refGene_tx=XM_003992179.5;refGene_gn=NEK10;refGene_tx=XM_006936474.3;refGene_gn=NEK10;refGene_tx=XM_011286267.3;refGene_gn=NEK10;refGene_tx=XM_011286268.3;refGene_gn=NEK10;refGene_tx=XM_019810797.2;refGene_gn=NEK10;refGene_tx=XM_019810798.2;refGene_gn=NEK10;refGene_tx=XM_019810799.2;refGene_gn=NEK10;refGene_tx=XM_019810800.2;refGene_gn=NEK10;refGene_tx=XM_019810801.2;refGene_gn=NEK10;refGene_tx=XM_019810802.2;refGene_gn=NEK10;refGene_tx=XM_019810803.2;refGene_gn=NEK10;refGene_tx=XM_019810804.2;refGene_gn=NEK10;refGene_tx=XM_019810806.2;refGene_gn=NEK10;refGene_tx=XM_019810807.2;refGene_gn=NEK10;refGene_tx=XM_019810809.2;refGene_gn=NEK10;refGene_tx=XM_023260629.1
C2 145755821 145755867
refGene_gn=NEK10;refGene_tx=XM_003992179.5;refGene_gn=NEK10;refGene_tx=XM_006936474.3;refGene_gn=NEK10;refGene_tx=XM_011286267.3;refGene_gn=NEK10;refGene_tx=XM_011286268.3;refGene_gn=NEK10;refGene_tx=XM_019810797.2;refGene_gn=NEK10;refGene_tx=XM_019810798.2;refGene_gn=NEK10;refGene_tx=XM_019810799.2;refGene_gn=NEK10;refGene_tx=XM_019810800.2;refGene_gn=NEK10;refGene_tx=XM_019810801.2;refGene_gn=NEK10;refGene_tx=XM_019810802.2;refGene_gn=NEK10;refGene_tx=XM_019810803.2;refGene_gn=NEK10;refGene_tx=XM_019810804.2;refGene_gn=NEK10;refGene_tx=XM_019810806.2;refGene_gn=NEK10;refGene_tx=XM_019810807.2;refGene_gn=NEK10;refGene_tx=XM_019810809.2;refGene_gn=NEK10;refGene_tx=XM_023260629.1
C2 145759214 145759352
refGene_gn=NEK10;refGene_tx=XM_003992179.5;refGene_gn=NEK10;refGene_tx

=XM_006936474.3;refGene_gn=NEK10;refGene_tx=XM_011286267.3;refGene_gn=NEK10;refGene_tx=XM_011286268.3;refGene_gn=NEK10;refGene_tx=XM_019810797.2;refGene_gn=NEK10;refGene_tx=XM_019810798.2;refGene_gn=NEK10;refGene_tx=XM_019810799.2;refGene_gn=NEK10;refGene_tx=XM_019810800.2;refGene_gn=NEK10;refGene_tx=XM_019810801.2;refGene_gn=NEK10;refGene_tx=XM_019810802.2;refGene_gn=NEK10;refGene_tx=XM_019810803.2;refGene_gn=NEK10;refGene_tx=XM_019810804.2;refGene_gn=NEK10;refGene_tx=XM_019810806.2;refGene_gn=NEK10;refGene_tx=XM_019810807.2;refGene_gn=NEK10;refGene_tx=XM_019810809.2;refGene_gn=NEK10;refGene_tx=XM_023260629.1
C2 145759941 145760070
refGene_gn=NEK10;refGene_tx=XM_003992179.5;refGene_gn=NEK10;refGene_tx=XM_006936474.3;refGene_gn=NEK10;refGene_tx=XM_011286267.3;refGene_gn=NEK10;refGene_tx=XM_011286268.3;refGene_gn=NEK10;refGene_tx=XM_019810797.2;refGene_gn=NEK10;refGene_tx=XM_019810799.2;refGene_gn=NEK10;refGene_tx=XM_019810800.2;refGene_gn=NEK10;refGene_tx=XM_019810801.2;refGene_gn=NEK10;refGene_tx=XM_019810802.2;refGene_gn=NEK10;refGene_tx=XM_019810803.2;refGene_gn=NEK10;refGene_tx=XM_019810804.2;refGene_gn=NEK10;refGene_tx=XM_019810806.2;refGene_gn=NEK10;refGene_tx=XM_019810807.2;refGene_gn=NEK10;refGene_tx=XM_019810809.2;refGene_gn=NEK10;refGene_tx=XM_023260629.1
C2 145760180 145760283
refGene_gn=NEK10;refGene_tx=XM_003992179.5;refGene_gn=NEK10;refGene_tx=XM_006936474.3;refGene_gn=NEK10;refGene_tx=XM_011286267.3;refGene_gn=NEK10;refGene_tx=XM_011286268.3;refGene_gn=NEK10;refGene_tx=XM_019810797.2;refGene_gn=NEK10;refGene_tx=XM_019810798.2;refGene_gn=NEK10;refGene_tx=XM_019810799.2;refGene_gn=NEK10;refGene_tx=XM_019810800.2;refGene_gn=NEK10;refGene_tx=XM_019810801.2;refGene_gn=NEK10;refGene_tx=XM_019810802.2;refGene_gn=NEK10;refGene_tx=XM_019810803.2;refGene_gn=NEK10;refGene_tx=XM_019810804.2;refGene_gn=NEK10;refGene_tx=XM_019810806.2;refGene_gn=NEK10;refGene_tx=XM_019810807.2;refGene_gn=NEK10;refGene_tx=XM_019810809.2;refGene_gn=NEK10;refGene_tx=XM_023260629.1
C2 145764084 145764149
refGene_gn=NEK10;refGene_tx=XM_003992179.5;refGene_gn=NEK10;refGene_tx=XM_006936474.3;refGene_gn=NEK10;refGene_tx=XM_011286267.3;refGene_gn=NEK10;refGene_tx=XM_011286268.3;refGene_gn=NEK10;refGene_tx=XM_019810797.2;refGene_gn=NEK10;refGene_tx=XM_019810798.2;refGene_gn=NEK10;refGene_tx=XM_019810799.2;refGene_gn=NEK10;refGene_tx=XM_019810800.2;refGene_gn=NEK10;refGene_tx=XM_019810801.2;refGene_gn=NEK10;refGene_tx=XM_019810802.2;refGene_gn=NEK10;refGene_tx=XM_019810803.2;refGene_gn=NEK10;refGene_tx=XM_019810804.2;refGene_gn=NEK10;refGene_tx=XM_019810806.2;refGene_gn=NEK10;refGene_tx=XM_019810807.2;refGene_gn=NEK10;refGene_tx=XM_019810809.2;refGene_gn=NEK10;refGene_tx=XM_023260629.1
C2 145765967 145766045
refGene_gn=NEK10;refGene_tx=XM_003992179.5;refGene_gn=NEK10;refGene_tx=XM_006936474.3;refGene_gn=NEK10;refGene_tx=XM_011286267.3;refGene_gn=NEK10;refGene_tx=XM_011286268.3;refGene_gn=NEK10;refGene_tx=XM_019810797.2;refGene_gn=NEK10;refGene_tx=XM_019810798.2;refGene_gn=NEK10;refGene_tx=XM_019810799.2;refGene_gn=NEK10;refGene_tx=XM_019810800.2;refGene_gn=NEK10;refGene_tx=XM_019810801.2;refGene_gn=NEK10;refGene_tx=XM_019810802.2;refGene_gn=NEK10;refGene_tx=XM_019810803.2;refGene_gn=NEK10;

refGene_tx=XM_019810804.2;refGene_gn=NEK10;refGene_tx=XM_019810806.2;r
efGene_gn=NEK10;refGene_tx=XM_019810807.2;refGene_gn=NEK10;refGene_tx=
XM_023260629.1
C2	145768080	145768142
refGene_gn=NEK10;refGene_tx=XM_003992179.5;refGene_gn=NEK10;refGene_tx
=XM_006936474.3;refGene_gn=NEK10;refGene_tx=XM_011286267.3;refGene_gn=
NEK10;refGene_tx=XM_011286268.3;refGene_gn=NEK10;refGene_tx=XM_0198107
97.2;refGene_gn=NEK10;refGene_tx=XM_019810798.2;refGene_gn=NEK10;refGe
ne_tx=XM_019810799.2;refGene_gn=NEK10;refGene_tx=XM_019810800.2;refGen
e_gn=NEK10;refGene_tx=XM_019810801.2;refGene_gn=NEK10;refGene_tx=XM_01
9810802.2;refGene_gn=NEK10;refGene_tx=XM_019810803.2;refGene_gn=NEK10;
refGene_tx=XM_019810804.2;refGene_gn=NEK10;refGene_tx=XM_019810806.2;r
efGene_gn=NEK10;refGene_tx=XM_019810807.2;refGene_gn=NEK10;refGene_tx=
XM_023260629.1
C2	145770386	145770526
refGene_gn=NEK10;refGene_tx=XM_003992179.5;refGene_gn=NEK10;refGene_tx
=XM_006936474.3;refGene_gn=NEK10;refGene_tx=XM_011286267.3;refGene_gn=
NEK10;refGene_tx=XM_011286268.3;refGene_gn=NEK10;refGene_tx=XM_0198107
97.2;refGene_gn=NEK10;refGene_tx=XM_019810798.2;refGene_gn=NEK10;refGe
ne_tx=XM_019810799.2;refGene_gn=NEK10;refGene_tx=XM_019810800.2;refGen
e_gn=NEK10;refGene_tx=XM_019810801.2;refGene_gn=NEK10;refGene_tx=XM_01
9810802.2;refGene_gn=NEK10;refGene_tx=XM_019810803.2;refGene_gn=NEK10;
refGene_tx=XM_019810804.2;refGene_gn=NEK10;refGene_tx=XM_019810806.2;r
efGene_gn=NEK10;refGene_tx=XM_019810807.2;refGene_gn=NEK10;refGene_tx=
XM_023260629.1
C2	145773725	145773950
refGene_gn=NEK10;refGene_tx=XM_003992179.5;refGene_gn=NEK10;refGene_tx
=XM_006936474.3;refGene_gn=NEK10;refGene_tx=XM_011286267.3;refGene_gn=
NEK10;refGene_tx=XM_011286268.3;refGene_gn=NEK10;refGene_tx=XM_0198107
97.2;refGene_gn=NEK10;refGene_tx=XM_019810798.2;refGene_gn=NEK10;refGe
ne_tx=XM_019810799.2;refGene_gn=NEK10;refGene_tx=XM_019810800.2;refGen
e_gn=NEK10;refGene_tx=XM_019810801.2;refGene_gn=NEK10;refGene_tx=XM_01
9810802.2;refGene_gn=NEK10;refGene_tx=XM_019810803.2;refGene_gn=NEK10;
refGene_tx=XM_019810804.2;refGene_gn=NEK10;refGene_tx=XM_019810806.2;r
efGene_gn=NEK10;refGene_tx=XM_019810807.2;refGene_gn=NEK10;refGene_tx=
XM_023260629.1
C2	145776655	145776742
refGene_gn=NEK10;refGene_tx=XM_003992179.5;refGene_gn=NEK10;refGene_tx
=XM_006936474.3;refGene_gn=NEK10;refGene_tx=XM_011286267.3;refGene_gn=
NEK10;refGene_tx=XM_011286268.3;refGene_gn=NEK10;refGene_tx=XM_0198107
97.2;refGene_gn=NEK10;refGene_tx=XM_019810798.2;refGene_gn=NEK10;refGe
ne_tx=XM_019810799.2;refGene_gn=NEK10;refGene_tx=XM_019810800.2;refGen
e_gn=NEK10;refGene_tx=XM_019810801.2;refGene_gn=NEK10;refGene_tx=XM_01
9810802.2;refGene_gn=NEK10;refGene_tx=XM_019810803.2;refGene_gn=NEK10;
refGene_tx=XM_019810804.2;refGene_gn=NEK10;refGene_tx=XM_019810806.2;r
efGene_gn=NEK10;refGene_tx=XM_019810807.2;refGene_gn=NEK10;refGene_tx=
XM_023260629.1
C2	145778172	145778252
refGene_gn=NEK10;refGene_tx=XM_003992179.5;refGene_gn=NEK10;refGene_tx
=XM_006936474.3;refGene_gn=NEK10;refGene_tx=XM_011286267.3;refGene_gn=

NEK10;refGene_tx=XM_011286268.3;refGene_gn=NEK10;refGene_tx=XM_0198107
97.2;refGene_gn=NEK10;refGene_tx=XM_019810798.2;refGene_gn=NEK10;refGe
ne_tx=XM_019810799.2;refGene_gn=NEK10;refGene_tx=XM_019810800.2;refGen
e_gn=NEK10;refGene_tx=XM_019810801.2;refGene_gn=NEK10;refGene_tx=XM_01
9810802.2;refGene_gn=NEK10;refGene_tx=XM_019810803.2;refGene_gn=NEK10;
refGene_tx=XM_019810804.2;refGene_gn=NEK10;refGene_tx=XM_019810806.2;r
efGene_gn=NEK10;refGene_tx=XM_019810807.2;refGene_gn=NEK10;refGene_tx=
XM_023260629.1
C2 145779927 145779995
refGene_gn=NEK10;refGene_tx=XM_003992179.5;refGene_gn=NEK10;refGene_tx
=XM_006936474.3;refGene_gn=NEK10;refGene_tx=XM_011286267.3;refGene_gn=
NEK10;refGene_tx=XM_011286268.3;refGene_gn=NEK10;refGene_tx=XM_0198107
97.2;refGene_gn=NEK10;refGene_tx=XM_019810798.2;refGene_gn=NEK10;refGe
ne_tx=XM_019810799.2;refGene_gn=NEK10;refGene_tx=XM_019810800.2;refGen
e_gn=NEK10;refGene_tx=XM_019810801.2;refGene_gn=NEK10;refGene_tx=XM_01
9810802.2;refGene_gn=NEK10;refGene_tx=XM_019810803.2;refGene_gn=NEK10;
refGene_tx=XM_019810804.2;refGene_gn=NEK10;refGene_tx=XM_019810806.2;r
efGene_gn=NEK10;refGene_tx=XM_019810807.2;refGene_gn=NEK10;refGene_tx=
XM_023260629.1
C2 145781119 145781198
refGene_gn=NEK10;refGene_tx=XM_003992179.5;refGene_gn=NEK10;refGene_tx
=XM_006936474.3;refGene_gn=NEK10;refGene_tx=XM_011286267.3;refGene_gn=
NEK10;refGene_tx=XM_011286268.3;refGene_gn=NEK10;refGene_tx=XM_0198107
97.2;refGene_gn=NEK10;refGene_tx=XM_019810798.2;refGene_gn=NEK10;refGe
ne_tx=XM_019810799.2;refGene_gn=NEK10;refGene_tx=XM_019810800.2;refGen
e_gn=NEK10;refGene_tx=XM_019810801.2;refGene_gn=NEK10;refGene_tx=XM_01
9810802.2;refGene_gn=NEK10;refGene_tx=XM_019810803.2;refGene_gn=NEK10;
refGene_tx=XM_019810804.2;refGene_gn=NEK10;refGene_tx=XM_019810806.2;r
efGene_gn=NEK10;refGene_tx=XM_019810807.2;refGene_gn=NEK10;refGene_tx=
XM_023260629.1
C2 145782531 145782573
refGene_gn=NEK10;refGene_tx=XM_003992179.5;refGene_gn=NEK10;refGene_tx
=XM_006936474.3;refGene_gn=NEK10;refGene_tx=XM_011286267.3;refGene_gn=
NEK10;refGene_tx=XM_011286268.3;refGene_gn=NEK10;refGene_tx=XM_0198107
97.2;refGene_gn=NEK10;refGene_tx=XM_019810798.2;refGene_gn=NEK10;refGe
ne_tx=XM_019810799.2;refGene_gn=NEK10;refGene_tx=XM_019810800.2;refGen
e_gn=NEK10;refGene_tx=XM_019810801.2;refGene_gn=NEK10;refGene_tx=XM_01
9810802.2;refGene_gn=NEK10;refGene_tx=XM_019810803.2;refGene_gn=NEK10;
refGene_tx=XM_019810804.2;refGene_gn=NEK10;refGene_tx=XM_019810806.2;r
efGene_gn=NEK10;refGene_tx=XM_019810807.2;refGene_gn=NEK10;refGene_tx=
XM_023260629.1
C2 145790502 145790587
refGene_gn=NEK10;refGene_tx=XM_003992179.5;refGene_gn=NEK10;refGene_tx
=XM_006936474.3;refGene_gn=NEK10;refGene_tx=XM_011286267.3;refGene_gn=
NEK10;refGene_tx=XM_011286268.3;refGene_gn=NEK10;refGene_tx=XM_0198107
97.2;refGene_gn=NEK10;refGene_tx=XM_019810798.2;refGene_gn=NEK10;refGe
ne_tx=XM_019810799.2;refGene_gn=NEK10;refGene_tx=XM_019810800.2;refGen
e_gn=NEK10;refGene_tx=XM_019810801.2;refGene_gn=NEK10;refGene_tx=XM_01
9810802.2;refGene_gn=NEK10;refGene_tx=XM_019810804.2;refGene_gn=NEK10;
refGene_tx=XM_019810807.2;refGene_gn=NEK10;refGene_tx=XM_023260629.1

C2	145799765	145799864
refGene_gn=NEK10;refGene_tx=XM_003992179.5;refGene_gn=NEK10;refGene_tx=XM_006936474.3;refGene_gn=NEK10;refGene_tx=XM_011286267.3;refGene_gn=NEK10;refGene_tx=XM_011286268.3;refGene_gn=NEK10;refGene_tx=XM_019810797.2;refGene_gn=NEK10;refGene_tx=XM_019810798.2;refGene_gn=NEK10;refGene_tx=XM_019810799.2;refGene_gn=NEK10;refGene_tx=XM_019810800.2;refGene_gn=NEK10;refGene_tx=XM_019810801.2;refGene_gn=NEK10;refGene_tx=XM_019810802.2;refGene_gn=NEK10;refGene_tx=XM_019810806.2;refGene_gn=NEK10;refGene_tx=XM_019810807.2;refGene_gn=NEK10;refGene_tx=XM_023260629.1
C2	145801389	145801520
refGene_gn=NEK10;refGene_tx=XM_003992179.5;refGene_gn=NEK10;refGene_tx=XM_006936474.3;refGene_gn=NEK10;refGene_tx=XM_011286267.3;refGene_gn=NEK10;refGene_tx=XM_011286268.3;refGene_gn=NEK10;refGene_tx=XM_019810797.2;refGene_gn=NEK10;refGene_tx=XM_019810798.2;refGene_gn=NEK10;refGene_tx=XM_019810799.2;refGene_gn=NEK10;refGene_tx=XM_019810800.2;refGene_gn=NEK10;refGene_tx=XM_019810801.2;refGene_gn=NEK10;refGene_tx=XM_019810802.2;refGene_gn=NEK10;refGene_tx=XM_019810807.2;refGene_gn=NEK10;refGene_tx=XM_023260629.1
C2	145807126	145807187
refGene_gn=NEK10;refGene_tx=XM_003992179.5;refGene_gn=NEK10;refGene_tx=XM_006936474.3;refGene_gn=NEK10;refGene_tx=XM_011286267.3;refGene_gn=NEK10;refGene_tx=XM_011286268.3;refGene_gn=NEK10;refGene_tx=XM_019810797.2;refGene_gn=NEK10;refGene_tx=XM_019810798.2;refGene_gn=NEK10;refGene_tx=XM_019810799.2;refGene_gn=NEK10;refGene_tx=XM_019810800.2;refGene_gn=NEK10;refGene_tx=XM_019810802.2;refGene_gn=NEK10;refGene_tx=XM_019810803.2;refGene_gn=NEK10;refGene_tx=XM_019810807.2
C2	145807410	145807484
refGene_gn=NEK10;refGene_tx=XM_003992179.5;refGene_gn=NEK10;refGene_tx=XM_006936474.3;refGene_gn=NEK10;refGene_tx=XM_011286267.3;refGene_gn=NEK10;refGene_tx=XM_011286268.3;refGene_gn=NEK10;refGene_tx=XM_019810797.2;refGene_gn=NEK10;refGene_tx=XM_019810798.2;refGene_gn=NEK10;refGene_tx=XM_019810799.2;refGene_gn=NEK10;refGene_tx=XM_019810800.2;refGene_gn=NEK10;refGene_tx=XM_019810802.2;refGene_gn=NEK10;refGene_tx=XM_019810803.2;refGene_gn=NEK10;refGene_tx=XM_019810807.2
C2	145830834	145830916
refGene_gn=SLC4A7;refGene_tx=XM_019810815.2;refGene_gn=SLC4A7;refGene_tx=XM_019810819.2;refGene_gn=SLC4A7;refGene_tx=XM_019810820.2;refGene_gn=SLC4A7;refGene_tx=XM_019810821.2;refGene_gn=SLC4A7;refGene_tx=XM_023260631.1;refGene_gn=SLC4A7;refGene_tx=XM_023260632.1;refGene_gn=SLC4A7;refGene_tx=XM_023260633.1;refGene_gn=SLC4A7;refGene_tx=XM_023260634.1;refGene_gn=SLC4A7;refGene_tx=XM_023260635.1
C2	145832749	145832857
refGene_gn=SLC4A7;refGene_tx=XM_019810819.2;refGene_gn=SLC4A7;refGene_tx=XM_019810820.2;refGene_gn=SLC4A7;refGene_tx=XM_023260631.1;refGene_gn=SLC4A7;refGene_tx=XM_023260632.1;refGene_gn=SLC4A7;refGene_tx=XM_023260633.1;refGene_gn=SLC4A7;refGene_tx=XM_023260635.1
C2	145835762	145835860
refGene_gn=SLC4A7;refGene_tx=XM_019810815.2;refGene_gn=SLC4A7;refGene_tx=XM_019810819.2;refGene_gn=SLC4A7;refGene_tx=XM_019810820.2;refGene_gn=SLC4A7;refGene_tx=XM_019810821.2;refGene_gn=SLC4A7;refGene_tx=XM_02

3260631.1;refGene_gn=SLC4A7;refGene_tx=XM_023260632.1;refGene_gn=SLC4A
7;refGene_tx=XM_023260633.1;refGene_gn=SLC4A7;refGene_tx=XM_023260634.
1;refGene_gn=SLC4A7;refGene_tx=XM_023260635.1
C2 145838347 145838479
refGene_gn=SLC4A7;refGene_tx=XM_019810815.2;refGene_gn=SLC4A7;refGene_
tx=XM_019810819.2;refGene_gn=SLC4A7;refGene_tx=XM_019810820.2;refGene_
gn=SLC4A7;refGene_tx=XM_019810821.2;refGene_gn=SLC4A7;refGene_tx=XM_02
3260631.1;refGene_gn=SLC4A7;refGene_tx=XM_023260632.1;refGene_gn=SLC4A
7;refGene_tx=XM_023260633.1;refGene_gn=SLC4A7;refGene_tx=XM_023260634.
1;refGene_gn=SLC4A7;refGene_tx=XM_023260635.1
C2 145843491 145843665
refGene_gn=SLC4A7;refGene_tx=XM_019810815.2;refGene_gn=SLC4A7;refGene_
tx=XM_019810819.2;refGene_gn=SLC4A7;refGene_tx=XM_019810820.2;refGene_
gn=SLC4A7;refGene_tx=XM_019810821.2;refGene_gn=SLC4A7;refGene_tx=XM_02
3260631.1;refGene_gn=SLC4A7;refGene_tx=XM_023260632.1;refGene_gn=SLC4A
7;refGene_tx=XM_023260633.1;refGene_gn=SLC4A7;refGene_tx=XM_023260634.
1;refGene_gn=SLC4A7;refGene_tx=XM_023260635.1
C2 145844961 145845030
refGene_gn=SLC4A7;refGene_tx=XM_019810815.2;refGene_gn=SLC4A7;refGene_
tx=XM_019810819.2;refGene_gn=SLC4A7;refGene_tx=XM_019810820.2;refGene_
gn=SLC4A7;refGene_tx=XM_019810821.2;refGene_gn=SLC4A7;refGene_tx=XM_02
3260631.1;refGene_gn=SLC4A7;refGene_tx=XM_023260632.1;refGene_gn=SLC4A
7;refGene_tx=XM_023260633.1;refGene_gn=SLC4A7;refGene_tx=XM_023260634.
1;refGene_gn=SLC4A7;refGene_tx=XM_023260635.1
C2 145848372 145848624
refGene_gn=SLC4A7;refGene_tx=XM_019810815.2;refGene_gn=SLC4A7;refGene_
tx=XM_019810819.2;refGene_gn=SLC4A7;refGene_tx=XM_019810820.2;refGene_
gn=SLC4A7;refGene_tx=XM_019810821.2;refGene_gn=SLC4A7;refGene_tx=XM_02
3260631.1;refGene_gn=SLC4A7;refGene_tx=XM_023260632.1;refGene_gn=SLC4A
7;refGene_tx=XM_023260633.1;refGene_gn=SLC4A7;refGene_tx=XM_023260634.
1;refGene_gn=SLC4A7;refGene_tx=XM_023260635.1
C2 145848807 145848969
refGene_gn=SLC4A7;refGene_tx=XM_019810815.2;refGene_gn=SLC4A7;refGene_
tx=XM_019810819.2;refGene_gn=SLC4A7;refGene_tx=XM_019810820.2;refGene_
gn=SLC4A7;refGene_tx=XM_019810821.2;refGene_gn=SLC4A7;refGene_tx=XM_02
3260631.1;refGene_gn=SLC4A7;refGene_tx=XM_023260632.1;refGene_gn=SLC4A
7;refGene_tx=XM_023260633.1;refGene_gn=SLC4A7;refGene_tx=XM_023260634.
1;refGene_gn=SLC4A7;refGene_tx=XM_023260635.1
C2 145851847 145851961
refGene_gn=SLC4A7;refGene_tx=XM_019810815.2;refGene_gn=SLC4A7;refGene_
tx=XM_019810819.2;refGene_gn=SLC4A7;refGene_tx=XM_019810820.2;refGene_
gn=SLC4A7;refGene_tx=XM_019810821.2;refGene_gn=SLC4A7;refGene_tx=XM_02
3260631.1;refGene_gn=SLC4A7;refGene_tx=XM_023260632.1;refGene_gn=SLC4A
7;refGene_tx=XM_023260633.1;refGene_gn=SLC4A7;refGene_tx=XM_023260634.
1;refGene_gn=SLC4A7;refGene_tx=XM_023260635.1
C2 145852346 145852508
refGene_gn=SLC4A7;refGene_tx=XM_019810815.2;refGene_gn=SLC4A7;refGene_
tx=XM_019810819.2;refGene_gn=SLC4A7;refGene_tx=XM_019810820.2;refGene_
gn=SLC4A7;refGene_tx=XM_019810821.2;refGene_gn=SLC4A7;refGene_tx=XM_02
3260631.1;refGene_gn=SLC4A7;refGene_tx=XM_023260632.1;refGene_gn=SLC4A

7;refGene_tx=XM_023260633.1;refGene_gn=SLC4A7;refGene_tx=XM_023260634.
1;refGene_gn=SLC4A7;refGene_tx=XM_023260635.1
C2 145854623 145854729
refGene_gn=SLC4A7;refGene_tx=XM_019810815.2;refGene_gn=SLC4A7;refGene_
tx=XM_019810819.2;refGene_gn=SLC4A7;refGene_tx=XM_019810820.2;refGene_
gn=SLC4A7;refGene_tx=XM_019810821.2;refGene_gn=SLC4A7;refGene_tx=XM_02
3260631.1;refGene_gn=SLC4A7;refGene_tx=XM_023260632.1;refGene_gn=SLC4A
7;refGene_tx=XM_023260633.1;refGene_gn=SLC4A7;refGene_tx=XM_023260634.
1;refGene_gn=SLC4A7;refGene_tx=XM_023260635.1
C2 145856368 145856614
refGene_gn=SLC4A7;refGene_tx=XM_019810815.2;refGene_gn=SLC4A7;refGene_
tx=XM_019810819.2;refGene_gn=SLC4A7;refGene_tx=XM_019810820.2;refGene_
gn=SLC4A7;refGene_tx=XM_019810821.2;refGene_gn=SLC4A7;refGene_tx=XM_02
3260631.1;refGene_gn=SLC4A7;refGene_tx=XM_023260632.1;refGene_gn=SLC4A
7;refGene_tx=XM_023260633.1;refGene_gn=SLC4A7;refGene_tx=XM_023260634.
1;refGene_gn=SLC4A7;refGene_tx=XM_023260635.1
C2 145857776 145857910
refGene_gn=SLC4A7;refGene_tx=XM_019810815.2;refGene_gn=SLC4A7;refGene_
tx=XM_019810819.2;refGene_gn=SLC4A7;refGene_tx=XM_019810820.2;refGene_
gn=SLC4A7;refGene_tx=XM_019810821.2;refGene_gn=SLC4A7;refGene_tx=XM_02
3260631.1;refGene_gn=SLC4A7;refGene_tx=XM_023260632.1;refGene_gn=SLC4A
7;refGene_tx=XM_023260633.1;refGene_gn=SLC4A7;refGene_tx=XM_023260634.
1;refGene_gn=SLC4A7;refGene_tx=XM_023260635.1
C2 145861496 145861671
refGene_gn=SLC4A7;refGene_tx=XM_019810815.2;refGene_gn=SLC4A7;refGene_
tx=XM_019810819.2;refGene_gn=SLC4A7;refGene_tx=XM_019810820.2;refGene_
gn=SLC4A7;refGene_tx=XM_019810821.2;refGene_gn=SLC4A7;refGene_tx=XM_02
3260631.1;refGene_gn=SLC4A7;refGene_tx=XM_023260632.1;refGene_gn=SLC4A
7;refGene_tx=XM_023260633.1;refGene_gn=SLC4A7;refGene_tx=XM_023260634.
1;refGene_gn=SLC4A7;refGene_tx=XM_023260635.1
C2 145863856 145863963
refGene_gn=SLC4A7;refGene_tx=XM_019810815.2;refGene_gn=SLC4A7;refGene_
tx=XM_019810819.2;refGene_gn=SLC4A7;refGene_tx=XM_019810820.2;refGene_
gn=SLC4A7;refGene_tx=XM_019810821.2;refGene_gn=SLC4A7;refGene_tx=XM_02
3260631.1;refGene_gn=SLC4A7;refGene_tx=XM_023260632.1;refGene_gn=SLC4A
7;refGene_tx=XM_023260633.1;refGene_gn=SLC4A7;refGene_tx=XM_023260634.
1;refGene_gn=SLC4A7;refGene_tx=XM_023260635.1
C2 145867615 145867762
refGene_gn=SLC4A7;refGene_tx=XM_019810815.2;refGene_gn=SLC4A7;refGene_
tx=XM_019810819.2;refGene_gn=SLC4A7;refGene_tx=XM_019810820.2;refGene_
gn=SLC4A7;refGene_tx=XM_019810821.2;refGene_gn=SLC4A7;refGene_tx=XM_02
3260631.1;refGene_gn=SLC4A7;refGene_tx=XM_023260632.1;refGene_gn=SLC4A
7;refGene_tx=XM_023260633.1;refGene_gn=SLC4A7;refGene_tx=XM_023260634.
1;refGene_gn=SLC4A7;refGene_tx=XM_023260635.1
C2 145869497 145869585
refGene_gn=SLC4A7;refGene_tx=XM_019810815.2;refGene_gn=SLC4A7;refGene_
tx=XM_019810819.2;refGene_gn=SLC4A7;refGene_tx=XM_019810820.2;refGene_
gn=SLC4A7;refGene_tx=XM_019810821.2;refGene_gn=SLC4A7;refGene_tx=XM_02
3260631.1;refGene_gn=SLC4A7;refGene_tx=XM_023260632.1;refGene_gn=SLC4A
7;refGene_tx=XM_023260633.1;refGene_gn=SLC4A7;refGene_tx=XM_023260634.

1;refGene_gn=SLC4A7;refGene_tx=XM_023260635.1
C2 145871194 145871352
refGene_gn=SLC4A7;refGene_tx=XM_019810815.2;refGene_gn=SLC4A7;refGene_tx=XM_019810819.2;refGene_gn=SLC4A7;refGene_tx=XM_019810820.2;refGene_gn=SLC4A7;refGene_tx=XM_019810821.2;refGene_gn=SLC4A7;refGene_tx=XM_023260631.1;refGene_gn=SLC4A7;refGene_tx=XM_023260632.1;refGene_gn=SLC4A7;refGene_tx=XM_023260633.1;refGene_gn=SLC4A7;refGene_tx=XM_023260634.1;refGene_gn=SLC4A7;refGene_tx=XM_023260635.1
C2 145873873 145873989
refGene_gn=SLC4A7;refGene_tx=XM_019810815.2;refGene_gn=SLC4A7;refGene_tx=XM_019810819.2;refGene_gn=SLC4A7;refGene_tx=XM_019810820.2;refGene_gn=SLC4A7;refGene_tx=XM_019810821.2;refGene_gn=SLC4A7;refGene_tx=XM_023260631.1;refGene_gn=SLC4A7;refGene_tx=XM_023260632.1;refGene_gn=SLC4A7;refGene_tx=XM_023260633.1;refGene_gn=SLC4A7;refGene_tx=XM_023260634.1;refGene_gn=SLC4A7;refGene_tx=XM_023260635.1
C2 145879838 145880207
refGene_gn=SLC4A7;refGene_tx=XM_019810815.2;refGene_gn=SLC4A7;refGene_tx=XM_023260635.1
C2 145882775 145882964
refGene_gn=SLC4A7;refGene_tx=XM_019810815.2;refGene_gn=SLC4A7;refGene_tx=XM_019810819.2;refGene_gn=SLC4A7;refGene_tx=XM_019810820.2;refGene_gn=SLC4A7;refGene_tx=XM_019810821.2;refGene_gn=SLC4A7;refGene_tx=XM_023260631.1;refGene_gn=SLC4A7;refGene_tx=XM_023260632.1;refGene_gn=SLC4A7;refGene_tx=XM_023260633.1;refGene_gn=SLC4A7;refGene_tx=XM_023260634.1;refGene_gn=SLC4A7;refGene_tx=XM_023260635.1
C2 145884435 145884596
refGene_gn=SLC4A7;refGene_tx=XM_019810815.2;refGene_gn=SLC4A7;refGene_tx=XM_019810819.2;refGene_gn=SLC4A7;refGene_tx=XM_019810820.2;refGene_gn=SLC4A7;refGene_tx=XM_019810821.2;refGene_gn=SLC4A7;refGene_tx=XM_023260631.1;refGene_gn=SLC4A7;refGene_tx=XM_023260632.1;refGene_gn=SLC4A7;refGene_tx=XM_023260633.1;refGene_gn=SLC4A7;refGene_tx=XM_023260634.1;refGene_gn=SLC4A7;refGene_tx=XM_023260635.1
C2 145885168 145885307
refGene_gn=SLC4A7;refGene_tx=XM_019810815.2;refGene_gn=SLC4A7;refGene_tx=XM_019810819.2;refGene_gn=SLC4A7;refGene_tx=XM_019810820.2;refGene_gn=SLC4A7;refGene_tx=XM_019810821.2;refGene_gn=SLC4A7;refGene_tx=XM_023260631.1;refGene_gn=SLC4A7;refGene_tx=XM_023260632.1;refGene_gn=SLC4A7;refGene_tx=XM_023260633.1;refGene_gn=SLC4A7;refGene_tx=XM_023260634.1;refGene_gn=SLC4A7;refGene_tx=XM_023260635.1
C2 145892042 145892189
refGene_gn=SLC4A7;refGene_tx=XM_019810815.2;refGene_gn=SLC4A7;refGene_tx=XM_019810819.2;refGene_gn=SLC4A7;refGene_tx=XM_019810820.2;refGene_gn=SLC4A7;refGene_tx=XM_019810821.2;refGene_gn=SLC4A7;refGene_tx=XM_023260631.1;refGene_gn=SLC4A7;refGene_tx=XM_023260632.1;refGene_gn=SLC4A7;refGene_tx=XM_023260633.1;refGene_gn=SLC4A7;refGene_tx=XM_023260634.1;refGene_gn=SLC4A7;refGene_tx=XM_023260635.1
C2 145896419 145896501
refGene_gn=SLC4A7;refGene_tx=XM_019810815.2;refGene_gn=SLC4A7;refGene_tx=XM_019810819.2;refGene_gn=SLC4A7;refGene_tx=XM_019810820.2;refGene_gn=SLC4A7;refGene_tx=XM_019810821.2;refGene_gn=SLC4A7;refGene_tx=XM_02

3260631.1;refGene_gn=SLC4A7;refGene_tx=XM_023260632.1;refGene_gn=SLC4A7;refGene_tx=XM_023260633.1;refGene_gn=SLC4A7;refGene_tx=XM_023260634.1;refGene_gn=SLC4A7;refGene_tx=XM_023260635.1
C2 145905393 145905426
refGene_gn=SLC4A7;refGene_tx=XM_019810815.2;refGene_gn=SLC4A7;refGene_tx=XM_019810819.2;refGene_gn=SLC4A7;refGene_tx=XM_019810820.2;refGene_gn=SLC4A7;refGene_tx=XM_019810821.2
C2 145933716 145933749
refGene_gn=SLC4A7;refGene_tx=XM_023260632.1
C2 145938350 145938398
refGene_gn=SLC4A7;refGene_tx=XM_023260631.1;refGene_gn=SLC4A7;refGene_tx=XM_023260633.1;refGene_gn=SLC4A7;refGene_tx=XM_023260634.1;refGene_gn=SLC4A7;refGene_tx=XM_023260635.1
C2 146118872 146119611
refGene_gn=EOMES;refGene_tx=XM_019810822.2;refGene_gn=EOMES;refGene_tx=XM_019810823.2
C2 146120377 146120439
refGene_gn=EOMES;refGene_tx=XM_019810822.2;refGene_gn=EOMES;refGene_tx=XM_019810823.2
C2 146120525 146120684
refGene_gn=EOMES;refGene_tx=XM_019810822.2;refGene_gn=EOMES;refGene_tx=XM_019810823.2
C2 146121129 146121251
refGene_gn=EOMES;refGene_tx=XM_019810822.2;refGene_gn=EOMES;refGene_tx=XM_019810823.2
C2 146121926 146122081
refGene_gn=EOMES;refGene_tx=XM_019810822.2;refGene_gn=EOMES;refGene_tx=XM_019810823.2
C2 146123083 146123988
refGene_gn=EOMES;refGene_tx=XM_019810822.2;refGene_gn=EOMES;refGene_tx=XM_019810823.2
C2 146599787 146599806
refGene_gn=CMC1;refGene_tx=XM_023260636.1
C2 146626458 146626548
refGene_gn=CMC1;refGene_tx=XM_023260636.1
C2 146684867 146684958
refGene_gn=CMC1;refGene_tx=XM_023260636.1
C2 146688043 146688164
refGene_gn=CMC1;refGene_tx=XM_023260636.1
C2 146692096 146692509
refGene_gn=AZI2;refGene_tx=XM_003992184.5;refGene_gn=AZI2;refGene_tx=XM_019810825.2;refGene_gn=AZI2;refGene_tx=XM_019810826.2
C2 146694695 146694814
refGene_gn=AZI2;refGene_tx=XM_003992184.5;refGene_gn=AZI2;refGene_tx=XM_019810825.2;refGene_gn=AZI2;refGene_tx=XM_019810826.2
C2 146700035 146700094
refGene_gn=AZI2;refGene_tx=XM_003992184.5;refGene_gn=AZI2;refGene_tx=XM_019810825.2;refGene_gn=AZI2;refGene_tx=XM_019810826.2
C2 146709105 146709254
refGene_gn=AZI2;refGene_tx=XM_003992184.5;refGene_gn=AZI2;refGene_tx=X

M_019810825.2;refGene_gn=AZI2;refGene_tx=XM_019810826.2
C2 146710368 146710468
refGene_gn=AZI2;refGene_tx=XM_003992184.5;refGene_gn=AZI2;refGene_tx=XM_019810825.2;refGene_gn=AZI2;refGene_tx=XM_019810826.2
C2 146710938 146711064
refGene_gn=AZI2;refGene_tx=XM_003992184.5;refGene_gn=AZI2;refGene_tx=XM_019810825.2;refGene_gn=AZI2;refGene_tx=XM_019810826.2
C2 146712566 146712787
refGene_gn=AZI2;refGene_tx=XM_003992184.5;refGene_gn=AZI2;refGene_tx=XM_019810825.2;refGene_gn=AZI2;refGene_tx=XM_019810826.2
C2 146713859 146713872
refGene_gn=AZI2;refGene_tx=XM_019810825.2
C2 146783651 146783992
refGene_gn=ZCWPW2;refGene_tx=XM_006936475.3;refGene_gn=ZCWPW2;refGene_tx=XM_019810827.1;refGene_gn=ZCWPW2;refGene_tx=XM_019810828.2;refGene_gn=ZCWPW2;refGene_tx=XM_019810831.2
C2 146795185 146795345
refGene_gn=ZCWPW2;refGene_tx=XM_006936475.3;refGene_gn=ZCWPW2;refGene_tx=XM_019810827.1;refGene_gn=ZCWPW2;refGene_tx=XM_019810828.2;refGene_gn=ZCWPW2;refGene_tx=XM_019810831.2
C2 146820037 146820152
refGene_gn=ZCWPW2;refGene_tx=XM_006936475.3;refGene_gn=ZCWPW2;refGene_tx=XM_019810827.1;refGene_gn=ZCWPW2;refGene_tx=XM_019810828.2;refGene_gn=ZCWPW2;refGene_tx=XM_019810831.2
C2 146832732 146832779
refGene_gn=ZCWPW2;refGene_tx=XM_006936475.3;refGene_gn=ZCWPW2;refGene_tx=XM_019810827.1;refGene_gn=ZCWPW2;refGene_tx=XM_019810828.2;refGene_gn=ZCWPW2;refGene_tx=XM_019810831.2
C2 146856199 146856258
refGene_gn=ZCWPW2;refGene_tx=XM_006936475.3;refGene_gn=ZCWPW2;refGene_tx=XM_019810827.1;refGene_gn=ZCWPW2;refGene_tx=XM_019810828.2;refGene_gn=ZCWPW2;refGene_tx=XM_019810831.2
C2 146857345 146857413
refGene_gn=ZCWPW2;refGene_tx=XM_006936475.3;refGene_gn=ZCWPW2;refGene_tx=XM_019810827.1;refGene_gn=ZCWPW2;refGene_tx=XM_019810828.2;refGene_gn=ZCWPW2;refGene_tx=XM_019810831.2
C2 146863571 146863696
refGene_gn=ZCWPW2;refGene_tx=XM_006936475.3;refGene_gn=ZCWPW2;refGene_tx=XM_019810827.1;refGene_gn=ZCWPW2;refGene_tx=XM_019810828.2;refGene_gn=ZCWPW2;refGene_tx=XM_019810831.2
C2 146866999 146867161
refGene_gn=ZCWPW2;refGene_tx=XM_006936475.3;refGene_gn=ZCWPW2;refGene_tx=XM_019810827.1;refGene_gn=ZCWPW2;refGene_tx=XM_019810828.2;refGene_gn=ZCWPW2;refGene_tx=XM_019810831.2
C2 146910564 146910727
refGene_gn=RBMS3;refGene_tx=XM_023260637.1;refGene_gn=RBMS3;refGene_tx=XM_023260638.1;refGene_gn=RBMS3;refGene_tx=XM_023260639.1;refGene_gn=RBMS3;refGene_tx=XM_023260640.1;refGene_gn=RBMS3;refGene_tx=XM_023260641.1;refGene_gn=RBMS3;refGene_tx=XM_023260642.1;refGene_gn=RBMS3;refGene_tx=XM_023260643.1;refGene_gn=RBMS3;refGene_tx=XM_023260644.1

C2	146911640	146911921
refGene_gn=RBMS3;refGene_tx=XM_023260637.1;refGene_gn=RBMS3;refGene_tx=XM_023260638.1;refGene_gn=RBMS3;refGene_tx=XM_023260639.1;refGene_gn=RBMS3;refGene_tx=XM_023260640.1;refGene_gn=RBMS3;refGene_tx=XM_023260641.1;refGene_gn=RBMS3;refGene_tx=XM_023260642.1;refGene_gn=RBMS3;refGene_tx=XM_023260643.1;refGene_gn=RBMS3;refGene_tx=XM_023260644.1;refGene_gn=RBMS3;refGene_tx=XM_023260645.1
C2	147166355	147166424
refGene_gn=RBMS3;refGene_tx=XM_023260637.1;refGene_gn=RBMS3;refGene_tx=XM_023260638.1;refGene_gn=RBMS3;refGene_tx=XM_023260639.1;refGene_gn=RBMS3;refGene_tx=XM_023260640.1;refGene_gn=RBMS3;refGene_tx=XM_023260641.1;refGene_gn=RBMS3;refGene_tx=XM_023260642.1;refGene_gn=RBMS3;refGene_tx=XM_023260643.1;refGene_gn=RBMS3;refGene_tx=XM_023260644.1;refGene_gn=RBMS3;refGene_tx=XM_023260645.1
C2	147433205	147433379
refGene_gn=RBMS3;refGene_tx=XM_023260637.1;refGene_gn=RBMS3;refGene_tx=XM_023260638.1;refGene_gn=RBMS3;refGene_tx=XM_023260639.1;refGene_gn=RBMS3;refGene_tx=XM_023260640.1;refGene_gn=RBMS3;refGene_tx=XM_023260641.1;refGene_gn=RBMS3;refGene_tx=XM_023260642.1;refGene_gn=RBMS3;refGene_tx=XM_023260643.1;refGene_gn=RBMS3;refGene_tx=XM_023260644.1;refGene_gn=RBMS3;refGene_tx=XM_023260645.1
C2	147523824	147523899
refGene_gn=RBMS3;refGene_tx=XM_023260646.1
C2	147669868	147670041
refGene_gn=RBMS3;refGene_tx=XM_023260637.1;refGene_gn=RBMS3;refGene_tx=XM_023260638.1;refGene_gn=RBMS3;refGene_tx=XM_023260639.1;refGene_gn=RBMS3;refGene_tx=XM_023260640.1;refGene_gn=RBMS3;refGene_tx=XM_023260641.1;refGene_gn=RBMS3;refGene_tx=XM_023260642.1;refGene_gn=RBMS3;refGene_tx=XM_023260643.1;refGene_gn=RBMS3;refGene_tx=XM_023260644.1;refGene_gn=RBMS3;refGene_tx=XM_023260645.1;refGene_gn=RBMS3;refGene_tx=XM_023260646.1
C2	147728388	147728447
refGene_gn=RBMS3;refGene_tx=XM_023260637.1;refGene_gn=RBMS3;refGene_tx=XM_023260638.1;refGene_gn=RBMS3;refGene_tx=XM_023260639.1;refGene_gn=RBMS3;refGene_tx=XM_023260640.1;refGene_gn=RBMS3;refGene_tx=XM_023260641.1;refGene_gn=RBMS3;refGene_tx=XM_023260642.1;refGene_gn=RBMS3;refGene_tx=XM_023260643.1;refGene_gn=RBMS3;refGene_tx=XM_023260644.1;refGene_gn=RBMS3;refGene_tx=XM_023260645.1;refGene_gn=RBMS3;refGene_tx=XM_023260646.1
C2	147813584	147813676
refGene_gn=RBMS3;refGene_tx=XM_023260637.1;refGene_gn=RBMS3;refGene_tx=XM_023260638.1;refGene_gn=RBMS3;refGene_tx=XM_023260639.1;refGene_gn=RBMS3;refGene_tx=XM_023260640.1;refGene_gn=RBMS3;refGene_tx=XM_023260641.1;refGene_gn=RBMS3;refGene_tx=XM_023260642.1;refGene_gn=RBMS3;refGene_tx=XM_023260643.1;refGene_gn=RBMS3;refGene_tx=XM_023260644.1;refGene_gn=RBMS3;refGene_tx=XM_023260645.1;refGene_gn=RBMS3;refGene_tx=XM_023260646.1
C2	147971327	147971485
refGene_gn=RBMS3;refGene_tx=XM_023260637.1;refGene_gn=RBMS3;refGene_tx=XM_023260638.1;refGene_gn=RBMS3;refGene_tx=XM_023260639.1;refGene_gn=

RBMS3;refGene_tx=XM_023260640.1;refGene_gn=RBMS3;refGene_tx=XM_0232606
41.1;refGene_gn=RBMS3;refGene_tx=XM_023260642.1;refGene_gn=RBMS3;refGe
ne_tx=XM_023260643.1;refGene_gn=RBMS3;refGene_tx=XM_023260644.1;refGen
e_gn=RBMS3;refGene_tx=XM_023260645.1;refGene_gn=RBMS3;refGene_tx=XM_02
3260646.1
C2 147986470 147986550
refGene_gn=RBMS3;refGene_tx=XM_023260637.1;refGene_gn=RBMS3;refGene_tx
=XM_023260638.1;refGene_gn=RBMS3;refGene_tx=XM_023260639.1;refGene_gn=
RBMS3;refGene_tx=XM_023260640.1;refGene_gn=RBMS3;refGene_tx=XM_0232606
41.1;refGene_gn=RBMS3;refGene_tx=XM_023260642.1;refGene_gn=RBMS3;refGe
ne_tx=XM_023260643.1;refGene_gn=RBMS3;refGene_tx=XM_023260644.1;refGen
e_gn=RBMS3;refGene_tx=XM_023260645.1;refGene_gn=RBMS3;refGene_tx=XM_02
3260646.1
C2 148089077 148089184
refGene_gn=RBMS3;refGene_tx=XM_023260637.1;refGene_gn=RBMS3;refGene_tx
=XM_023260638.1;refGene_gn=RBMS3;refGene_tx=XM_023260639.1;refGene_gn=
RBMS3;refGene_tx=XM_023260640.1;refGene_gn=RBMS3;refGene_tx=XM_0232606
41.1;refGene_gn=RBMS3;refGene_tx=XM_023260642.1;refGene_gn=RBMS3;refGe
ne_tx=XM_023260643.1;refGene_gn=RBMS3;refGene_tx=XM_023260644.1;refGen
e_gn=RBMS3;refGene_tx=XM_023260645.1;refGene_gn=RBMS3;refGene_tx=XM_02
3260646.1
C2 148109070 148109117
refGene_gn=RBMS3;refGene_tx=XM_023260637.1;refGene_gn=RBMS3;refGene_tx
=XM_023260638.1;refGene_gn=RBMS3;refGene_tx=XM_023260639.1;refGene_gn=
RBMS3;refGene_tx=XM_023260640.1;refGene_gn=RBMS3;refGene_tx=XM_0232606
41.1;refGene_gn=RBMS3;refGene_tx=XM_023260642.1;refGene_gn=RBMS3;refGe
ne_tx=XM_023260643.1;refGene_gn=RBMS3;refGene_tx=XM_023260644.1;refGen
e_gn=RBMS3;refGene_tx=XM_023260645.1;refGene_gn=RBMS3;refGene_tx=XM_02
3260646.1
C2 148123045 148123142
refGene_gn=RBMS3;refGene_tx=XM_023260637.1;refGene_gn=RBMS3;refGene_tx
=XM_023260638.1;refGene_gn=RBMS3;refGene_tx=XM_023260639.1;refGene_gn=
RBMS3;refGene_tx=XM_023260640.1;refGene_gn=RBMS3;refGene_tx=XM_0232606
41.1;refGene_gn=RBMS3;refGene_tx=XM_023260642.1;refGene_gn=RBMS3;refGe
ne_tx=XM_023260643.1;refGene_gn=RBMS3;refGene_tx=XM_023260644.1;refGen
e_gn=RBMS3;refGene_tx=XM_023260645.1;refGene_gn=RBMS3;refGene_tx=XM_02
3260646.1
C2 148125380 148125431
refGene_gn=RBMS3;refGene_tx=XM_023260637.1;refGene_gn=RBMS3;refGene_tx
=XM_023260638.1;refGene_gn=RBMS3;refGene_tx=XM_023260639.1;refGene_gn=
RBMS3;refGene_tx=XM_023260640.1;refGene_gn=RBMS3;refGene_tx=XM_0232606
41.1;refGene_gn=RBMS3;refGene_tx=XM_023260645.1;refGene_gn=RBMS3;refGe
ne_tx=XM_023260646.1
C2 148169471 148169582
refGene_gn=RBMS3;refGene_tx=XM_023260637.1;refGene_gn=RBMS3;refGene_tx
=XM_023260638.1;refGene_gn=RBMS3;refGene_tx=XM_023260639.1;refGene_gn=
RBMS3;refGene_tx=XM_023260640.1;refGene_gn=RBMS3;refGene_tx=XM_0232606
41.1;refGene_gn=RBMS3;refGene_tx=XM_023260642.1;refGene_gn=RBMS3;refGe
ne_tx=XM_023260643.1;refGene_gn=RBMS3;refGene_tx=XM_023260644.1;refGen
e_gn=RBMS3;refGene_tx=XM_023260645.1;refGene_gn=RBMS3;refGene_tx=XM_02

3260646.1
C2 148177002 148177050
refGene_gn=RBMS3;refGene_tx=XM_023260637.1;refGene_gn=RBMS3;refGene_tx=XM_023260638.1;refGene_gn=RBMS3;refGene_tx=XM_023260639.1;refGene_gn=RBMS3;refGene_tx=XM_023260642.1;refGene_gn=RBMS3;refGene_tx=XM_023260643.1;refGene_gn=RBMS3;refGene_tx=XM_023260644.1;refGene_gn=RBMS3;refGene_tx=XM_023260645.1
C2 148217754 148217835
refGene_gn=RBMS3;refGene_tx=XM_023260637.1;refGene_gn=RBMS3;refGene_tx=XM_023260638.1;refGene_gn=RBMS3;refGene_tx=XM_023260639.1;refGene_gn=RBMS3;refGene_tx=XM_023260640.1;refGene_gn=RBMS3;refGene_tx=XM_023260641.1;refGene_gn=RBMS3;refGene_tx=XM_023260642.1;refGene_gn=RBMS3;refGene_tx=XM_023260643.1;refGene_gn=RBMS3;refGene_tx=XM_023260644.1;refGene_gn=RBMS3;refGene_tx=XM_023260645.1;refGene_gn=RBMS3;refGene_tx=XM_023260646.1
C2 148220959 148221091
refGene_gn=RBMS3;refGene_tx=XM_023260637.1;refGene_gn=RBMS3;refGene_tx=XM_023260638.1;refGene_gn=RBMS3;refGene_tx=XM_023260639.1;refGene_gn=RBMS3;refGene_tx=XM_023260640.1;refGene_gn=RBMS3;refGene_tx=XM_023260641.1;refGene_gn=RBMS3;refGene_tx=XM_023260642.1;refGene_gn=RBMS3;refGene_tx=XM_023260643.1;refGene_gn=RBMS3;refGene_tx=XM_023260644.1;refGene_gn=RBMS3;refGene_tx=XM_023260645.1;refGene_gn=RBMS3;refGene_tx=XM_023260646.1
C2 148236019 148236026
refGene_gn=RBMS3;refGene_tx=XM_023260637.1;refGene_gn=RBMS3;refGene_tx=XM_023260638.1;refGene_gn=RBMS3;refGene_tx=XM_023260640.1;refGene_gn=RBMS3;refGene_tx=XM_023260642.1;refGene_gn=RBMS3;refGene_tx=XM_023260645.1
C2 148749955 148750049
refGene_gn=TGFBR2;refGene_tx=XM_023260647.1
C2 148787319 148787488
refGene_gn=TGFBR2;refGene_tx=XM_023260647.1
C2 148793158 148793349
refGene_gn=TGFBR2;refGene_tx=XM_023260647.1
C2 148820345 148821145
refGene_gn=TGFBR2;refGene_tx=XM_023260647.1
C2 148822838 148822980
refGene_gn=TGFBR2;refGene_tx=XM_023260647.1
C2 148834656 148834784
refGene_gn=TGFBR2;refGene_tx=XM_023260647.1
C2 148837791 148837971
refGene_gn=TGFBR2;refGene_tx=XM_023260647.1
C2 148873415 148873589
refGene_gn=GADL1;refGene_tx=XM_003992189.4
C2 148928139 148928229
refGene_gn=GADL1;refGene_tx=XM_003992189.4
C2 148937611 148937663
refGene_gn=GADL1;refGene_tx=XM_003992189.4
C2 148947584 148947784
refGene_gn=GADL1;refGene_tx=XM_003992189.4

C2 148979144 148979226
refGene_gn=GADL1;refGene_tx=XM_003992189.4
C2 148979496 148979561
refGene_gn=GADL1;refGene_tx=XM_003992189.4
C2 148985954 148986071
refGene_gn=GADL1;refGene_tx=XM_003992189.4
C2 148990833 148990888
refGene_gn=GADL1;refGene_tx=XM_003992189.4
C2 148991010 148991090
refGene_gn=GADL1;refGene_tx=XM_003992189.4
C2 148996811 148996927
refGene_gn=GADL1;refGene_tx=XM_003992189.4
C2 148997631 148997738
refGene_gn=GADL1;refGene_tx=XM_003992189.4
C2 149001205 149001296
refGene_gn=GADL1;refGene_tx=XM_003992189.4
C2 149003767 149003894
refGene_gn=GADL1;refGene_tx=XM_003992189.4
C2 149011552 149011725
refGene_gn=GADL1;refGene_tx=XM_003992189.4
C2 149041551 149041588
refGene_gn=GADL1;refGene_tx=XM_003992189.4
C2 149626062 149626376
refGene_gn=STT3B;refGene_tx=XM_023260648.1
C2 149673314 149673423
refGene_gn=STT3B;refGene_tx=XM_023260648.1
C2 149677021 149677309
refGene_gn=STT3B;refGene_tx=XM_023260648.1
C2 149696163 149696229
refGene_gn=STT3B;refGene_tx=XM_023260648.1
C2 149697949 149698049
refGene_gn=STT3B;refGene_tx=XM_023260648.1
C2 149712755 149712854
refGene_gn=STT3B;refGene_tx=XM_023260648.1
C2 149714934 149715081
refGene_gn=STT3B;refGene_tx=XM_023260648.1
C2 149715911 149715960
refGene_gn=STT3B;refGene_tx=XM_023260648.1
C2 149717331 149717486
refGene_gn=STT3B;refGene_tx=XM_023260648.1
C2 149719841 149720053
refGene_gn=STT3B;refGene_tx=XM_023260648.1
C2 149721565 149721753
refGene_gn=STT3B;refGene_tx=XM_023260648.1
C2 149722162 149722334
refGene_gn=STT3B;refGene_tx=XM_023260648.1
C2 149723207 149723381
refGene_gn=STT3B;refGene_tx=XM_023260648.1
C2 149725259 149725373
refGene_gn=STT3B;refGene_tx=XM_023260648.1

C2	149727627	149727840
refGene_gn=STT3B;refGene_tx=XM_023260648.1
C2	149730157	149730238
refGene_gn=STT3B;refGene_tx=XM_023260648.1
C2	149752381	149752426
refGene_gn=OSBPL10;refGene_tx=XM_003992193.4;refGene_gn=OSBPL10;refGene_tx=XM_019810843.2
C2	149754762	149754916
refGene_gn=OSBPL10;refGene_tx=XM_003992193.4;refGene_gn=OSBPL10;refGene_tx=XM_019810843.2
C2	149757656	149757839
refGene_gn=OSBPL10;refGene_tx=XM_003992193.4;refGene_gn=OSBPL10;refGene_tx=XM_019810843.2
C2	149759581	149759768
refGene_gn=OSBPL10;refGene_tx=XM_003992193.4;refGene_gn=OSBPL10;refGene_tx=XM_019810843.2
C2	149772055	149772533
refGene_gn=OSBPL10;refGene_tx=XM_003992193.4;refGene_gn=OSBPL10;refGene_tx=XM_019810843.2
C2	149787262	149787412
refGene_gn=OSBPL10;refGene_tx=XM_003992193.4
C2	149818881	149819036
refGene_gn=OSBPL10;refGene_tx=XM_003992193.4;refGene_gn=OSBPL10;refGene_tx=XM_019810843.2
C2	149832590	149832801
refGene_gn=OSBPL10;refGene_tx=XM_003992193.4;refGene_gn=OSBPL10;refGene_tx=XM_019810843.2
C2	149907693	149907885
refGene_gn=OSBPL10;refGene_tx=XM_003992193.4;refGene_gn=OSBPL10;refGene_tx=XM_019810843.2
C2	149952872	149952952
refGene_gn=OSBPL10;refGene_tx=XM_003992193.4;refGene_gn=OSBPL10;refGene_tx=XM_019810843.2
C2	149955929	149956105
refGene_gn=OSBPL10;refGene_tx=XM_003992193.4;refGene_gn=OSBPL10;refGene_tx=XM_019810843.2
C2	150069688	150069969
refGene_gn=OSBPL10;refGene_tx=XM_003992193.4;refGene_gn=OSBPL10;refGene_tx=XM_019810843.2
C2	150166748	150166795
refGene_gn=GPD1L;refGene_tx=XM_023260649.1
C2	150179889	150180067
refGene_gn=GPD1L;refGene_tx=XM_023260649.1
C2	150188297	150188438
refGene_gn=GPD1L;refGene_tx=XM_023260649.1
C2	150189945	150190084
refGene_gn=GPD1L;refGene_tx=XM_023260649.1
C2	150203206	150203319
refGene_gn=GPD1L;refGene_tx=XM_023260649.1
C2	150213545	150213779

refGene_gn=GPD1L;refGene_tx=XM_023260649.1
C2 150214065 150214172
refGene_gn=GPD1L;refGene_tx=XM_023260649.1
C2 150221973 150222070
refGene_gn=GPD1L;refGene_tx=XM_023260649.1
C2 150293327 150293474
refGene_gn=CMTM8;refGene_tx=XM_023260650.1
C2 150392881 150393055
refGene_gn=CMTM8;refGene_tx=XM_023260650.1
C2 150403268 150403385
refGene_gn=CMTM8;refGene_tx=XM_023260650.1
C2 150404959 150405043
refGene_gn=CMTM8;refGene_tx=XM_023260650.1
C2 150421316 150421505
refGene_gn=CMTM7;refGene_tx=XM_023260651.1;refGene_gn=CMTM7;refGene_tx
=XM_023260652.1
C2 150458992 150459166
refGene_gn=CMTM7;refGene_tx=XM_023260651.1;refGene_gn=CMTM7;refGene_tx
=XM_023260652.1
C2 150465574 150465673
refGene_gn=CMTM7;refGene_tx=XM_023260651.1
C2 150468567 150468649
refGene_gn=CMTM7;refGene_tx=XM_023260651.1;refGene_gn=CMTM7;refGene_tx
=XM_023260652.1
C2 150471622 150471636
refGene_gn=CMTM7;refGene_tx=XM_023260651.1;refGene_gn=CMTM7;refGene_tx
=XM_023260652.1
C2 150484361 150484496
refGene_gn=CMTM6;refGene_tx=XM_023260653.1
C2 150487496 150487595
refGene_gn=CMTM6;refGene_tx=XM_023260653.1
C2 150492093 150492270
refGene_gn=CMTM6;refGene_tx=XM_023260653.1
C2 150504047 150504185
refGene_gn=CMTM6;refGene_tx=XM_023260653.1
C2 150522193 150522306
refGene_gn=DYNC1LI1;refGene_tx=XM_003992198.5
C2 150524879 150525035
refGene_gn=DYNC1LI1;refGene_tx=XM_003992198.5
C2 150525968 150526089
refGene_gn=DYNC1LI1;refGene_tx=XM_003992198.5
C2 150526761 150526806
refGene_gn=DYNC1LI1;refGene_tx=XM_003992198.5
C2 150527012 150527072
refGene_gn=DYNC1LI1;refGene_tx=XM_003992198.5
C2 150529035 150529147
refGene_gn=DYNC1LI1;refGene_tx=XM_003992198.5
C2 150529760 150529896
refGene_gn=DYNC1LI1;refGene_tx=XM_003992198.5
C2 150532547 150532641

refGene_gn=DYNC1LI1;refGene_tx=XM_003992198.5
C2	150533626	150533796
refGene_gn=DYNC1LI1;refGene_tx=XM_003992198.5
C2	150536152	150536183
refGene_gn=DYNC1LI1;refGene_tx=XM_003992198.5
C2	150536184	150536384
refGene_gn=DYNC1LI1;refGene_tx=XM_003992198.5
C2	150537110	150537227
refGene_gn=DYNC1LI1;refGene_tx=XM_003992198.5
C2	150557590	150557664
refGene_gn=DYNC1LI1;refGene_tx=XM_003992198.5
C2	150557868	150558014
refGene_gn=DYNC1LI1;refGene_tx=XM_003992198.5
C2	150704754	150704841
refGene_gn=POMGNT2;refGene_tx=XM_023260654.1
C2	150705478	150707326
refGene_gn=POMGNT2;refGene_tx=XM_003992200.4;refGene_gn=POMGNT2;refGene_tx=XM_023260654.1;refGene_gn=POMGNT2;refGene_tx=XM_023260655.1;refGene_gn=POMGNT2;refGene_tx=XM_023260656.1;refGene_gn=POMGNT2;refGene_tx=XM_023260657.1;refGene_gn=POMGNT2;refGene_tx=XM_023260658.1;refGene_gn=POMGNT2;refGene_tx=XM_023260659.1;refGene_gn=POMGNT2;refGene_tx=XM_023260660.1;refGene_gn=POMGNT2;refGene_tx=XM_023260661.1;refGene_gn=POMGNT2;refGene_tx=XM_023260662.1;refGene_gn=POMGNT2;refGene_tx=XM_023260663.1;refGene_gn=POMGNT2;refGene_tx=XM_023260664.1;refGene_gn=POMGNT2;refGene_tx=XM_023260665.1;refGene_gn=POMGNT2;refGene_tx=XM_023260667.1
C2	150725332	150725543
refGene_gn=FAM198A;refGene_tx=XM_006936487.3
C2	150726209	150726313
refGene_gn=FAM198A;refGene_tx=XM_006936487.3
C2	150728058	150728181
refGene_gn=FAM198A;refGene_tx=XM_006936487.3
C2	150744280	150745549
refGene_gn=FAM198A;refGene_tx=XM_006936487.3
C2	150781652	150781655
refGene_gn=FAM198A;refGene_tx=XM_006936487.3
C2	150823094	150824125
refGene_gn=ZNF662;refGene_tx=XM_006936491.4;refGene_gn=ZNF662;refGene_tx=XM_006936492.4;refGene_gn=ZNF662;refGene_tx=XM_019810851.2;refGene_gn=ZNF662;refGene_tx=XM_019810852.2
C2	150825359	150825461
refGene_gn=ZNF662;refGene_tx=XM_006936491.4;refGene_gn=ZNF662;refGene_tx=XM_006936492.4;refGene_gn=ZNF662;refGene_tx=XM_019810851.2;refGene_gn=ZNF662;refGene_tx=XM_019810852.2
C2	150827511	150827625
refGene_gn=ZNF662;refGene_tx=XM_006936491.4;refGene_gn=ZNF662;refGene_tx=XM_006936492.4;refGene_gn=ZNF662;refGene_tx=XM_019810851.2;refGene_gn=ZNF662;refGene_tx=XM_019810852.2
C2	150828275	150828402
refGene_gn=ZNF662;refGene_tx=XM_006936491.4;refGene_gn=ZNF662;refGene_tx=XM_006936492.4;refGene_gn=ZNF662;refGene_tx=XM_019810851.2;refGene_

gn=ZNF662;refGene_tx=XM_019810852.2
C2 150829694 150829859
refGene_gn=ZNF662;refGene_tx=XM_006936491.4
C2 150830004 150830094
refGene_gn=ZNF662;refGene_tx=XM_006936492.4
C2 150841276 150842782
refGene_gn=LOC101082004;refGene_tx=XM_003992237.5
C2 150847704 150848894
refGene_gn=ACKR2;refGene_tx=XM_006936494.4;refGene_gn=ACKR2;refGene_tx=XM_006936495.4;refGene_gn=ACKR2;refGene_tx=XM_023260668.1;refGene_gn=ACKR2;refGene_tx=XM_023260669.1
C2 150858994 150859148
refGene_gn=ACKR2;refGene_tx=XM_006936494.4
C2 150910151 150910248
refGene_gn=HIGD1A;refGene_tx=XM_006936497.3
C2 150910873 150911008
refGene_gn=HIGD1A;refGene_tx=XM_006936497.3
C2 150911682 150911732
refGene_gn=HIGD1A;refGene_tx=XM_006936497.3
C2 150924815 150924872
refGene_gn=CCDC13;refGene_tx=XM_019810857.2
C2 150934133 150934360
refGene_gn=CCDC13;refGene_tx=XM_019810857.2;refGene_gn=CCDC13;refGene_tx=XM_023260670.1;refGene_gn=CCDC13;refGene_tx=XM_023260671.1
C2 150936601 150936750
refGene_gn=CCDC13;refGene_tx=XM_019810857.2;refGene_gn=CCDC13;refGene_tx=XM_023260670.1;refGene_gn=CCDC13;refGene_tx=XM_023260671.1
C2 150940497 150940640
refGene_gn=CCDC13;refGene_tx=XM_019810857.2;refGene_gn=CCDC13;refGene_tx=XM_023260670.1;refGene_gn=CCDC13;refGene_tx=XM_023260671.1
C2 150942177 150942267
refGene_gn=CCDC13;refGene_tx=XM_019810857.2;refGene_gn=CCDC13;refGene_tx=XM_023260670.1;refGene_gn=CCDC13;refGene_tx=XM_023260671.1
C2 150946421 150946538
refGene_gn=CCDC13;refGene_tx=XM_019810857.2;refGene_gn=CCDC13;refGene_tx=XM_023260670.1;refGene_gn=CCDC13;refGene_tx=XM_023260671.1
C2 150947755 150947860
refGene_gn=CCDC13;refGene_tx=XM_019810857.2;refGene_gn=CCDC13;refGene_tx=XM_023260670.1;refGene_gn=CCDC13;refGene_tx=XM_023260671.1
C2 150949990 150950152
refGene_gn=CCDC13;refGene_tx=XM_019810857.2;refGene_gn=CCDC13;refGene_tx=XM_023260670.1;refGene_gn=CCDC13;refGene_tx=XM_023260671.1
C2 150952904 150953081
refGene_gn=CCDC13;refGene_tx=XM_019810857.2;refGene_gn=CCDC13;refGene_tx=XM_023260670.1;refGene_gn=CCDC13;refGene_tx=XM_023260671.1
C2 150954683 150954890
refGene_gn=CCDC13;refGene_tx=XM_019810857.2;refGene_gn=CCDC13;refGene_tx=XM_023260670.1;refGene_gn=CCDC13;refGene_tx=XM_023260671.1
C2 150957305 150957445
refGene_gn=CCDC13;refGene_tx=XM_019810857.2;refGene_gn=CCDC13;refGene_

tx=XM_023260670.1;refGene_gn=CCDC13;refGene_tx=XM_023260671.1
C2 150957960 150958044
refGene_gn=CCDC13;refGene_tx=XM_019810857.2;refGene_gn=CCDC13;refGene_tx=XM_023260670.1;refGene_gn=CCDC13;refGene_tx=XM_023260671.1
C2 150960304 150960427
refGene_gn=CCDC13;refGene_tx=XM_019810857.2;refGene_gn=CCDC13;refGene_tx=XM_023260670.1;refGene_gn=CCDC13;refGene_tx=XM_023260671.1
C2 150969877 150970032
refGene_gn=CCDC13;refGene_tx=XM_019810857.2;refGene_gn=CCDC13;refGene_tx=XM_023260670.1;refGene_gn=CCDC13;refGene_tx=XM_023260671.1
C2 150972667 150972782
refGene_gn=CCDC13;refGene_tx=XM_019810857.2;refGene_gn=CCDC13;refGene_tx=XM_023260670.1;refGene_gn=CCDC13;refGene_tx=XM_023260671.1
C2 150973343 150973503
refGene_gn=CCDC13;refGene_tx=XM_019810857.2;refGene_gn=CCDC13;refGene_tx=XM_023260670.1;refGene_gn=CCDC13;refGene_tx=XM_023260671.1
C2 150979528 150979851
refGene_gn=HHATL;refGene_tx=XM_003992205.5;refGene_gn=HHATL;refGene_tx=XM_006936500.4;refGene_gn=HHATL;refGene_tx=XM_019810861.2;refGene_gn=HHATL;refGene_tx=XM_023260674.1;refGene_gn=HHATL;refGene_tx=XM_023260675.1
C2 150981581 150981700
refGene_gn=HHATL;refGene_tx=XM_003992205.5;refGene_gn=HHATL;refGene_tx=XM_006936498.4;refGene_gn=HHATL;refGene_tx=XM_006936499.4;refGene_gn=HHATL;refGene_tx=XM_006936500.4;refGene_gn=HHATL;refGene_tx=XM_019810859.2;refGene_gn=HHATL;refGene_tx=XM_019810860.2;refGene_gn=HHATL;refGene_tx=XM_019810861.2;refGene_gn=HHATL;refGene_tx=XM_023260674.1;refGene_gn=HHATL;refGene_tx=XM_023260675.1
C2 150982108 150982176
refGene_gn=HHATL;refGene_tx=XM_003992205.5;refGene_gn=HHATL;refGene_tx=XM_006936498.4;refGene_gn=HHATL;refGene_tx=XM_006936499.4;refGene_gn=HHATL;refGene_tx=XM_006936500.4;refGene_gn=HHATL;refGene_tx=XM_019810859.2;refGene_gn=HHATL;refGene_tx=XM_019810860.2;refGene_gn=HHATL;refGene_tx=XM_019810861.2;refGene_gn=HHATL;refGene_tx=XM_023260674.1;refGene_gn=HHATL;refGene_tx=XM_023260675.1
C2 150982703 150982817
refGene_gn=HHATL;refGene_tx=XM_003992205.5;refGene_gn=HHATL;refGene_tx=XM_006936498.4;refGene_gn=HHATL;refGene_tx=XM_006936499.4;refGene_gn=HHATL;refGene_tx=XM_006936500.4;refGene_gn=HHATL;refGene_tx=XM_019810859.2;refGene_gn=HHATL;refGene_tx=XM_019810860.2;refGene_gn=HHATL;refGene_tx=XM_019810861.2;refGene_gn=HHATL;refGene_tx=XM_023260674.1;refGene_gn=HHATL;refGene_tx=XM_023260675.1
C2 150982931 150983126
refGene_gn=HHATL;refGene_tx=XM_003992205.5;refGene_gn=HHATL;refGene_tx=XM_006936498.4;refGene_gn=HHATL;refGene_tx=XM_006936499.4;refGene_gn=HHATL;refGene_tx=XM_006936500.4;refGene_gn=HHATL;refGene_tx=XM_019810859.2;refGene_gn=HHATL;refGene_tx=XM_019810860.2;refGene_gn=HHATL;refGene_tx=XM_023260674.1;refGene_gn=HHATL;refGene_tx=XM_023260675.1
C2 150983487 150983697
refGene_gn=HHATL;refGene_tx=XM_003992205.5;refGene_gn=HHATL;refGene_tx

=XM_006936498.4;refGene_gn=HHATL;refGene_tx=XM_006936499.4;refGene_gn=HHATL;refGene_tx=XM_006936500.4;refGene_gn=HHATL;refGene_tx=XM_019810859.2;refGene_gn=HHATL;refGene_tx=XM_019810860.2;refGene_gn=HHATL;refGene_tx=XM_019810861.2;refGene_gn=HHATL;refGene_tx=XM_023260674.1;refGene_gn=HHATL;refGene_tx=XM_023260675.1
C2 150984042 150984214
refGene_gn=HHATL;refGene_tx=XM_003992205.5;refGene_gn=HHATL;refGene_tx=XM_006936498.4;refGene_gn=HHATL;refGene_tx=XM_006936499.4;refGene_gn=HHATL;refGene_tx=XM_006936500.4;refGene_gn=HHATL;refGene_tx=XM_019810859.2;refGene_gn=HHATL;refGene_tx=XM_019810860.2;refGene_gn=HHATL;refGene_tx=XM_019810861.2;refGene_gn=HHATL;refGene_tx=XM_023260674.1;refGene_gn=HHATL;refGene_tx=XM_023260675.1
C2 150984674 150984819
refGene_gn=HHATL;refGene_tx=XM_003992205.5;refGene_gn=HHATL;refGene_tx=XM_006936498.4;refGene_gn=HHATL;refGene_tx=XM_006936499.4;refGene_gn=HHATL;refGene_tx=XM_006936500.4;refGene_gn=HHATL;refGene_tx=XM_019810859.2;refGene_gn=HHATL;refGene_tx=XM_019810860.2;refGene_gn=HHATL;refGene_tx=XM_019810861.2;refGene_gn=HHATL;refGene_tx=XM_023260674.1;refGene_gn=HHATL;refGene_tx=XM_023260675.1
C2 150984937 150984973
refGene_gn=HHATL;refGene_tx=XM_003992205.5;refGene_gn=HHATL;refGene_tx=XM_006936498.4;refGene_gn=HHATL;refGene_tx=XM_006936499.4;refGene_gn=HHATL;refGene_tx=XM_006936500.4;refGene_gn=HHATL;refGene_tx=XM_019810859.2;refGene_gn=HHATL;refGene_tx=XM_019810860.2;refGene_gn=HHATL;refGene_tx=XM_019810861.2;refGene_gn=HHATL;refGene_tx=XM_023260674.1;refGene_gn=HHATL;refGene_tx=XM_023260675.1
C2 150986617 150986690
refGene_gn=HHATL;refGene_tx=XM_023260674.1;refGene_gn=HHATL;refGene_tx=XM_023260675.1
C2 150987756 150987958
refGene_gn=HHATL;refGene_tx=XM_003992205.5;refGene_gn=HHATL;refGene_tx=XM_006936498.4;refGene_gn=HHATL;refGene_tx=XM_006936499.4;refGene_gn=HHATL;refGene_tx=XM_006936500.4;refGene_gn=HHATL;refGene_tx=XM_019810859.2;refGene_gn=HHATL;refGene_tx=XM_019810860.2;refGene_gn=HHATL;refGene_tx=XM_019810861.2
C2 150988573 150988715
refGene_gn=HHATL;refGene_tx=XM_003992205.5;refGene_gn=HHATL;refGene_tx=XM_006936498.4;refGene_gn=HHATL;refGene_tx=XM_006936499.4;refGene_gn=HHATL;refGene_tx=XM_019810859.2;refGene_gn=HHATL;refGene_tx=XM_019810860.2;refGene_gn=HHATL;refGene_tx=XM_019810861.2
C2 150988919 150989044
refGene_gn=HHATL;refGene_tx=XM_003992205.5;refGene_gn=HHATL;refGene_tx=XM_006936498.4;refGene_gn=HHATL;refGene_tx=XM_006936499.4;refGene_gn=HHATL;refGene_tx=XM_006936500.4;refGene_gn=HHATL;refGene_tx=XM_019810859.2;refGene_gn=HHATL;refGene_tx=XM_019810860.2;refGene_gn=HHATL;refGene_tx=XM_019810861.2
C2 150989834 150989946
refGene_gn=KLHL40;refGene_tx=XM_023260672.1;refGene_gn=KLHL40;refGene_tx=XM_023260673.1
C2 150990773 150990920

refGene_gn=KLHL40;refGene_tx=XM_023260672.1;refGene_gn=KLHL40;refGene_tx=XM_023260673.1
C2 150992688 150992874
refGene_gn=KLHL40;refGene_tx=XM_023260672.1;refGene_gn=KLHL40;refGene_tx=XM_023260673.1
C2 150993078 150993186
refGene_gn=KLHL40;refGene_tx=XM_023260672.1;refGene_gn=KLHL40;refGene_tx=XM_023260673.1
C2 150993447 150993608
refGene_gn=KLHL40;refGene_tx=XM_023260672.1;refGene_gn=KLHL40;refGene_tx=XM_023260673.1
C2 150994937 150996101
refGene_gn=KLHL40;refGene_tx=XM_023260672.1;refGene_gn=KLHL40;refGene_tx=XM_023260673.1
C2 151013729 151014091
refGene_gn=ZBTB47;refGene_tx=XM_011286258.3
C2 151014357 151014502
refGene_gn=ZBTB47;refGene_tx=XM_011286258.3
C2 151015138 151015254
refGene_gn=ZBTB47;refGene_tx=XM_011286258.3
C2 151016547 151016695
refGene_gn=ZBTB47;refGene_tx=XM_011286258.3
C2 151018240 151018669
refGene_gn=ZBTB47;refGene_tx=XM_011286258.3
C2 151018671 151019808
refGene_gn=ZBTB47;refGene_tx=XM_011286258.3
C2 151023437 151023440
refGene_gn=ZBTB47;refGene_tx=XM_011286258.3
C2 151031343 151031431
refGene_gn=NKTR;refGene_tx=XM_003992206.5;refGene_gn=NKTR;refGene_tx=XM_019810862.2;refGene_gn=NKTR;refGene_tx=XM_019810863.2;refGene_gn=NKTR;refGene_tx=XM_019810864.2
C2 151033150 151033255
refGene_gn=NKTR;refGene_tx=XM_003992206.5;refGene_gn=NKTR;refGene_tx=XM_019810862.2;refGene_gn=NKTR;refGene_tx=XM_019810864.2
C2 151033776 151033833
refGene_gn=NKTR;refGene_tx=XM_003992206.5;refGene_gn=NKTR;refGene_tx=XM_019810862.2;refGene_gn=NKTR;refGene_tx=XM_019810863.2;refGene_gn=NKTR;refGene_tx=XM_019810864.2
C2 151034580 151034676
refGene_gn=NKTR;refGene_tx=XM_003992206.5;refGene_gn=NKTR;refGene_tx=XM_019810862.2;refGene_gn=NKTR;refGene_tx=XM_019810863.2;refGene_gn=NKTR;refGene_tx=XM_019810864.2
C2 151037482 151040365
refGene_gn=NKTR;refGene_tx=XM_003992206.5;refGene_gn=NKTR;refGene_tx=XM_019810862.2;refGene_gn=NKTR;refGene_tx=XM_019810863.2;refGene_gn=NKTR;refGene_tx=XM_019810864.2
C2 151041660 151041806
refGene_gn=NKTR;refGene_tx=XM_003992206.5;refGene_gn=NKTR;refGene_tx=XM_019810862.2;refGene_gn=NKTR;refGene_tx=XM_019810863.2;refGene_gn=NKT

R;refGene_tx=XM_019810864.2
C2	151042265	151042353
refGene_gn=NKTR;refGene_tx=XM_003992206.5;refGene_gn=NKTR;refGene_tx=XM_019810862.2;refGene_gn=NKTR;refGene_tx=XM_019810863.2;refGene_gn=NKTR;refGene_tx=XM_019810864.2
C2	151048674	151048830
refGene_gn=NKTR;refGene_tx=XM_003992206.5;refGene_gn=NKTR;refGene_tx=XM_019810862.2;refGene_gn=NKTR;refGene_tx=XM_019810863.2;refGene_gn=NKTR;refGene_tx=XM_019810864.2
C2	151049523	151049746
refGene_gn=NKTR;refGene_tx=XM_003992206.5;refGene_gn=NKTR;refGene_tx=XM_019810862.2;refGene_gn=NKTR;refGene_tx=XM_019810863.2;refGene_gn=NKTR;refGene_tx=XM_019810864.2
C2	151050666	151050812
refGene_gn=NKTR;refGene_tx=XM_003992206.5;refGene_gn=NKTR;refGene_tx=XM_019810862.2;refGene_gn=NKTR;refGene_tx=XM_019810863.2;refGene_gn=NKTR;refGene_tx=XM_019810864.2;refGene_gn=NKTR;refGene_tx=XM_019810865.2
C2	151051333	151051363
refGene_gn=NKTR;refGene_tx=XM_003992206.5;refGene_gn=NKTR;refGene_tx=XM_019810862.2;refGene_gn=NKTR;refGene_tx=XM_019810863.2;refGene_gn=NKTR;refGene_tx=XM_019810864.2;refGene_gn=NKTR;refGene_tx=XM_019810865.2
C2	151054259	151054287
refGene_gn=NKTR;refGene_tx=XM_019810865.2
C2	151059905	151059993
refGene_gn=NKTR;refGene_tx=XM_003992206.5;refGene_gn=NKTR;refGene_tx=XM_011286257.3;refGene_gn=NKTR;refGene_tx=XM_019810862.2;refGene_gn=NKTR;refGene_tx=XM_019810863.2;refGene_gn=NKTR;refGene_tx=XM_019810864.2;refGene_gn=NKTR;refGene_tx=XM_019810865.2
C2	151061406	151061434
refGene_gn=NKTR;refGene_tx=XM_011286257.3
C2	151061741	151061786
refGene_gn=NKTR;refGene_tx=XM_003992206.5;refGene_gn=NKTR;refGene_tx=XM_011286257.3;refGene_gn=NKTR;refGene_tx=XM_019810862.2;refGene_gn=NKTR;refGene_tx=XM_019810863.2;refGene_gn=NKTR;refGene_tx=XM_019810865.2
C2	151062551	151062659
refGene_gn=NKTR;refGene_tx=XM_003992206.5;refGene_gn=NKTR;refGene_tx=XM_011286257.3;refGene_gn=NKTR;refGene_tx=XM_019810862.2;refGene_gn=NKTR;refGene_tx=XM_019810863.2;refGene_gn=NKTR;refGene_tx=XM_019810865.2
C2	151063637	151063712
refGene_gn=NKTR;refGene_tx=XM_003992206.5;refGene_gn=NKTR;refGene_tx=XM_011286257.3;refGene_gn=NKTR;refGene_tx=XM_019810862.2;refGene_gn=NKTR;refGene_tx=XM_019810863.2;refGene_gn=NKTR;refGene_tx=XM_019810865.2
C2	151087506	151087564
refGene_gn=NKTR;refGene_tx=XM_003992206.5;refGene_gn=NKTR;refGene_tx=XM_011286257.3;refGene_gn=NKTR;refGene_tx=XM_019810862.2;refGene_gn=NKTR;refGene_tx=XM_019810863.2;refGene_gn=NKTR;refGene_tx=XM_019810865.2
C2	151099632	151099720
refGene_gn=SS18L2;refGene_tx=XM_019810872.2
C2	151101556	151101633
refGene_gn=SS18L2;refGene_tx=XM_019810872.2

C2 151101899 151101968
refGene_gn=SS18L2;refGene_tx=XM_019810872.2
C2 151121687 151121869
refGene_gn=SEC22C;refGene_tx=XM_019810869.2;refGene_gn=SEC22C;refGene_tx=XM_023260676.1
C2 151125521 151125685
refGene_gn=SEC22C;refGene_tx=XM_019810869.2;refGene_gn=SEC22C;refGene_tx=XM_023260676.1
C2 151127602 151127782
refGene_gn=SEC22C;refGene_tx=XM_019810869.2;refGene_gn=SEC22C;refGene_tx=XM_023260676.1
C2 151130253 151130372
refGene_gn=SEC22C;refGene_tx=XM_019810869.2;refGene_gn=SEC22C;refGene_tx=XM_023260676.1
C2 151131570 151131636
refGene_gn=SEC22C;refGene_tx=XM_019810869.2;refGene_gn=SEC22C;refGene_tx=XM_023260676.1
C2 151133821 151134022
refGene_gn=SEC22C;refGene_tx=XM_019810869.2;refGene_gn=SEC22C;refGene_tx=XM_023260676.1
C2 151150525 151150717
refGene_gn=VIPR1;refGene_tx=XM_011286255.3;refGene_gn=VIPR1;refGene_tx=XM_019810877.2;refGene_gn=VIPR1;refGene_tx=XM_019810878.2;refGene_gn=VIPR1;refGene_tx=XM_023260677.1;refGene_gn=VIPR1;refGene_tx=XM_023260678.1;refGene_gn=VIPR1;refGene_tx=XM_023260679.1;refGene_gn=VIPR1;refGene_tx=XM_023260680.1
C2 151151439 151151481
refGene_gn=VIPR1;refGene_tx=XM_011286255.3;refGene_gn=VIPR1;refGene_tx=XM_019810877.2;refGene_gn=VIPR1;refGene_tx=XM_019810878.2;refGene_gn=VIPR1;refGene_tx=XM_023260677.1;refGene_gn=VIPR1;refGene_tx=XM_023260678.1;refGene_gn=VIPR1;refGene_tx=XM_023260679.1;refGene_gn=VIPR1;refGene_tx=XM_023260680.1
C2 151151726 151151856
refGene_gn=VIPR1;refGene_tx=XM_011286255.3;refGene_gn=VIPR1;refGene_tx=XM_019810877.2;refGene_gn=VIPR1;refGene_tx=XM_019810878.2;refGene_gn=VIPR1;refGene_tx=XM_023260677.1;refGene_gn=VIPR1;refGene_tx=XM_023260678.1;refGene_gn=VIPR1;refGene_tx=XM_023260679.1;refGene_gn=VIPR1;refGene_tx=XM_023260680.1
C2 151154290 151154382
refGene_gn=VIPR1;refGene_tx=XM_011286255.3;refGene_gn=VIPR1;refGene_tx=XM_019810877.2;refGene_gn=VIPR1;refGene_tx=XM_019810878.2;refGene_gn=VIPR1;refGene_tx=XM_023260677.1;refGene_gn=VIPR1;refGene_tx=XM_023260678.1;refGene_gn=VIPR1;refGene_tx=XM_023260679.1;refGene_gn=VIPR1;refGene_tx=XM_023260680.1
C2 151154755 151154822
refGene_gn=VIPR1;refGene_tx=XM_011286255.3;refGene_gn=VIPR1;refGene_tx=XM_019810877.2;refGene_gn=VIPR1;refGene_tx=XM_019810878.2;refGene_gn=VIPR1;refGene_tx=XM_023260677.1;refGene_gn=VIPR1;refGene_tx=XM_023260678.1;refGene_gn=VIPR1;refGene_tx=XM_023260679.1;refGene_gn=VIPR1;refGene_tx=XM_023260680.1

C2	151155023	151155084
refGene_gn=VIPR1;refGene_tx=XM_011286255.3;refGene_gn=VIPR1;refGene_tx=XM_019810877.2;refGene_gn=VIPR1;refGene_tx=XM_019810878.2;refGene_gn=VIPR1;refGene_tx=XM_023260677.1;refGene_gn=VIPR1;refGene_tx=XM_023260678.1;refGene_gn=VIPR1;refGene_tx=XM_023260679.1;refGene_gn=VIPR1;refGene_tx=XM_023260680.1
C2	151155606	151155760
refGene_gn=VIPR1;refGene_tx=XM_011286255.3;refGene_gn=VIPR1;refGene_tx=XM_019810877.2;refGene_gn=VIPR1;refGene_tx=XM_019810878.2;refGene_gn=VIPR1;refGene_tx=XM_023260677.1;refGene_gn=VIPR1;refGene_tx=XM_023260678.1;refGene_gn=VIPR1;refGene_tx=XM_023260679.1;refGene_gn=VIPR1;refGene_tx=XM_023260680.1
C2	151157859	151157992
refGene_gn=VIPR1;refGene_tx=XM_011286255.3;refGene_gn=VIPR1;refGene_tx=XM_019810877.2;refGene_gn=VIPR1;refGene_tx=XM_019810878.2;refGene_gn=VIPR1;refGene_tx=XM_023260677.1;refGene_gn=VIPR1;refGene_tx=XM_023260678.1;refGene_gn=VIPR1;refGene_tx=XM_023260679.1;refGene_gn=VIPR1;refGene_tx=XM_023260680.1
C2	151158424	151158531
refGene_gn=VIPR1;refGene_tx=XM_011286255.3;refGene_gn=VIPR1;refGene_tx=XM_019810877.2;refGene_gn=VIPR1;refGene_tx=XM_019810878.2;refGene_gn=VIPR1;refGene_tx=XM_023260677.1;refGene_gn=VIPR1;refGene_tx=XM_023260678.1;refGene_gn=VIPR1;refGene_tx=XM_023260679.1;refGene_gn=VIPR1;refGene_tx=XM_023260680.1
C2	151159953	151160060
refGene_gn=VIPR1;refGene_tx=XM_011286255.3;refGene_gn=VIPR1;refGene_tx=XM_019810877.2;refGene_gn=VIPR1;refGene_tx=XM_019810878.2;refGene_gn=VIPR1;refGene_tx=XM_023260677.1;refGene_gn=VIPR1;refGene_tx=XM_023260678.1;refGene_gn=VIPR1;refGene_tx=XM_023260679.1;refGene_gn=VIPR1;refGene_tx=XM_023260680.1
C2	151161372	151161559
refGene_gn=VIPR1;refGene_tx=XM_019810878.2
C2	151166685	151166793
refGene_gn=VIPR1;refGene_tx=XM_011286255.3;refGene_gn=VIPR1;refGene_tx=XM_019810877.2;refGene_gn=VIPR1;refGene_tx=XM_023260677.1;refGene_gn=VIPR1;refGene_tx=XM_023260678.1;refGene_gn=VIPR1;refGene_tx=XM_023260679.1;refGene_gn=VIPR1;refGene_tx=XM_023260680.1
C2	151171446	151171555
refGene_gn=VIPR1;refGene_tx=XM_019810877.2;refGene_gn=VIPR1;refGene_tx=XM_023260677.1;refGene_gn=VIPR1;refGene_tx=XM_023260678.1;refGene_gn=VIPR1;refGene_tx=XM_023260679.1
C2	151174553	151174556
refGene_gn=VIPR1;refGene_tx=XM_019810877.2
C2	151182625	151182703
refGene_gn=VIPR1;refGene_tx=XM_023260677.1;refGene_gn=VIPR1;refGene_tx=XM_023260678.1;refGene_gn=VIPR1;refGene_tx=XM_023260679.1;refGene_gn=VIPR1;refGene_tx=XM_023260680.1
C2	151284979	151285118
refGene_gn=LYZL4;refGene_tx=XM_003992209.5;refGene_gn=LYZL4;refGene_tx=XM_006936505.3;refGene_gn=LYZL4;refGene_tx=XM_023260681.1

C2	151285230	151285380
refGene_gn=LYZL4;refGene_tx=XM_003992209.5;refGene_gn=LYZL4;refGene_tx=XM_006936505.3;refGene_gn=LYZL4;refGene_tx=XM_023260681.1
C2	151288919	151288998
refGene_gn=LYZL4;refGene_tx=XM_003992209.5;refGene_gn=LYZL4;refGene_tx=XM_006936505.3;refGene_gn=LYZL4;refGene_tx=XM_023260681.1
C2	151295441	151295511
refGene_gn=LYZL4;refGene_tx=XM_003992209.5;refGene_gn=LYZL4;refGene_tx=XM_006936505.3;refGene_gn=LYZL4;refGene_tx=XM_023260681.1
C2	151425333	151425547
refGene_gn=CCK;refGene_tx=XM_003992210.5
C2	151429491	151429625
refGene_gn=CCK;refGene_tx=XM_003992210.5
C2	151471330	151472129
refGene_gn=TRAK1;refGene_tx=XM_011286249.3;refGene_gn=TRAK1;refGene_tx=XM_011286250.3;refGene_gn=TRAK1;refGene_tx=XM_011286252.3;refGene_gn=TRAK1;refGene_tx=XM_019810882.2;refGene_gn=TRAK1;refGene_tx=XM_019810884.2;refGene_gn=TRAK1;refGene_tx=XM_023260682.1;refGene_gn=TRAK1;refGene_tx=XM_023260683.1;refGene_gn=TRAK1;refGene_tx=XM_023260684.1;refGene_gn=TRAK1;refGene_tx=XM_023260685.1
C2	151474237	151474328
refGene_gn=TRAK1;refGene_tx=XM_023260685.1
C2	151475085	151475188
refGene_gn=TRAK1;refGene_tx=XM_011286249.3;refGene_gn=TRAK1;refGene_tx=XM_011286250.3;refGene_gn=TRAK1;refGene_tx=XM_011286252.3;refGene_gn=TRAK1;refGene_tx=XM_019810882.2;refGene_gn=TRAK1;refGene_tx=XM_019810884.2;refGene_gn=TRAK1;refGene_tx=XM_023260682.1;refGene_gn=TRAK1;refGene_tx=XM_023260683.1;refGene_gn=TRAK1;refGene_tx=XM_023260684.1;refGene_gn=TRAK1;refGene_tx=XM_023260685.1
C2	151484186	151484680
refGene_gn=TRAK1;refGene_tx=XM_011286249.3;refGene_gn=TRAK1;refGene_tx=XM_011286250.3;refGene_gn=TRAK1;refGene_tx=XM_011286252.3;refGene_gn=TRAK1;refGene_tx=XM_019810882.2;refGene_gn=TRAK1;refGene_tx=XM_019810884.2;refGene_gn=TRAK1;refGene_tx=XM_019810885.2;refGene_gn=TRAK1;refGene_tx=XM_023260682.1;refGene_gn=TRAK1;refGene_tx=XM_023260683.1;refGene_gn=TRAK1;refGene_tx=XM_023260684.1;refGene_gn=TRAK1;refGene_tx=XM_023260685.1
C2	151491190	151491656
refGene_gn=TRAK1;refGene_tx=XM_011286249.3;refGene_gn=TRAK1;refGene_tx=XM_011286250.3;refGene_gn=TRAK1;refGene_tx=XM_011286252.3;refGene_gn=TRAK1;refGene_tx=XM_019810882.2;refGene_gn=TRAK1;refGene_tx=XM_019810884.2;refGene_gn=TRAK1;refGene_tx=XM_019810885.2;refGene_gn=TRAK1;refGene_tx=XM_023260682.1;refGene_gn=TRAK1;refGene_tx=XM_023260683.1;refGene_gn=TRAK1;refGene_tx=XM_023260684.1;refGene_gn=TRAK1;refGene_tx=XM_023260685.1;refGene_gn=TRAK1;refGene_tx=XM_023260686.1
C2	151492670	151492907
refGene_gn=TRAK1;refGene_tx=XM_011286249.3;refGene_gn=TRAK1;refGene_tx=XM_011286250.3;refGene_gn=TRAK1;refGene_tx=XM_011286252.3;refGene_gn=TRAK1;refGene_tx=XM_019810882.2;refGene_gn=TRAK1;refGene_tx=XM_019810884.2;refGene_gn=TRAK1;refGene_tx=XM_019810885.2;refGene_gn=TRAK1;refGe

ne_tx=XM_023260682.1;refGene_gn=TRAK1;refGene_tx=XM_023260683.1;refGen
e_gn=TRAK1;refGene_tx=XM_023260684.1;refGene_gn=TRAK1;refGene_tx=XM_02
3260685.1;refGene_gn=TRAK1;refGene_tx=XM_023260686.1
C2 151494150 151494227
refGene_gn=TRAK1;refGene_tx=XM_011286249.3;refGene_gn=TRAK1;refGene_tx
=XM_011286250.3;refGene_gn=TRAK1;refGene_tx=XM_011286252.3;refGene_gn=
TRAK1;refGene_tx=XM_019810882.2;refGene_gn=TRAK1;refGene_tx=XM_0198108
84.2;refGene_gn=TRAK1;refGene_tx=XM_019810885.2;refGene_gn=TRAK1;refGe
ne_tx=XM_023260682.1;refGene_gn=TRAK1;refGene_tx=XM_023260683.1;refGen
e_gn=TRAK1;refGene_tx=XM_023260684.1;refGene_gn=TRAK1;refGene_tx=XM_02
3260685.1;refGene_gn=TRAK1;refGene_tx=XM_023260686.1
C2 151499772 151499910
refGene_gn=TRAK1;refGene_tx=XM_011286249.3;refGene_gn=TRAK1;refGene_tx
=XM_011286250.3;refGene_gn=TRAK1;refGene_tx=XM_011286252.3;refGene_gn=
TRAK1;refGene_tx=XM_019810882.2;refGene_gn=TRAK1;refGene_tx=XM_0198108
84.2;refGene_gn=TRAK1;refGene_tx=XM_019810885.2;refGene_gn=TRAK1;refGe
ne_tx=XM_023260682.1;refGene_gn=TRAK1;refGene_tx=XM_023260683.1;refGen
e_gn=TRAK1;refGene_tx=XM_023260684.1;refGene_gn=TRAK1;refGene_tx=XM_02
3260685.1;refGene_gn=TRAK1;refGene_tx=XM_023260686.1
C2 151500631 151500706
refGene_gn=TRAK1;refGene_tx=XM_011286249.3;refGene_gn=TRAK1;refGene_tx
=XM_011286250.3;refGene_gn=TRAK1;refGene_tx=XM_011286252.3;refGene_gn=
TRAK1;refGene_tx=XM_019810882.2;refGene_gn=TRAK1;refGene_tx=XM_0198108
84.2;refGene_gn=TRAK1;refGene_tx=XM_019810885.2;refGene_gn=TRAK1;refGe
ne_tx=XM_023260682.1;refGene_gn=TRAK1;refGene_tx=XM_023260683.1;refGen
e_gn=TRAK1;refGene_tx=XM_023260684.1;refGene_gn=TRAK1;refGene_tx=XM_02
3260685.1;refGene_gn=TRAK1;refGene_tx=XM_023260686.1
C2 151501580 151501711
refGene_gn=TRAK1;refGene_tx=XM_011286249.3;refGene_gn=TRAK1;refGene_tx
=XM_011286250.3;refGene_gn=TRAK1;refGene_tx=XM_011286252.3;refGene_gn=
TRAK1;refGene_tx=XM_019810882.2;refGene_gn=TRAK1;refGene_tx=XM_0198108
84.2;refGene_gn=TRAK1;refGene_tx=XM_019810885.2;refGene_gn=TRAK1;refGe
ne_tx=XM_023260682.1;refGene_gn=TRAK1;refGene_tx=XM_023260683.1;refGen
e_gn=TRAK1;refGene_tx=XM_023260684.1;refGene_gn=TRAK1;refGene_tx=XM_02
3260685.1;refGene_gn=TRAK1;refGene_tx=XM_023260686.1
C2 151502662 151502741
refGene_gn=TRAK1;refGene_tx=XM_011286249.3;refGene_gn=TRAK1;refGene_tx
=XM_011286250.3;refGene_gn=TRAK1;refGene_tx=XM_011286252.3;refGene_gn=
TRAK1;refGene_tx=XM_019810882.2;refGene_gn=TRAK1;refGene_tx=XM_0198108
84.2;refGene_gn=TRAK1;refGene_tx=XM_019810885.2;refGene_gn=TRAK1;refGe
ne_tx=XM_023260682.1;refGene_gn=TRAK1;refGene_tx=XM_023260683.1;refGen
e_gn=TRAK1;refGene_tx=XM_023260684.1;refGene_gn=TRAK1;refGene_tx=XM_02
3260685.1;refGene_gn=TRAK1;refGene_tx=XM_023260686.1
C2 151504568 151504677
refGene_gn=TRAK1;refGene_tx=XM_011286249.3;refGene_gn=TRAK1;refGene_tx
=XM_011286250.3;refGene_gn=TRAK1;refGene_tx=XM_011286252.3;refGene_gn=
TRAK1;refGene_tx=XM_019810882.2;refGene_gn=TRAK1;refGene_tx=XM_0198108
84.2;refGene_gn=TRAK1;refGene_tx=XM_019810885.2;refGene_gn=TRAK1;refGe
ne_tx=XM_023260682.1;refGene_gn=TRAK1;refGene_tx=XM_023260683.1;refGen
e_gn=TRAK1;refGene_tx=XM_023260684.1;refGene_gn=TRAK1;refGene_tx=XM_02

3260685.1;refGene_gn=TRAK1;refGene_tx=XM_023260686.1
C2 151505470 151505571
refGene_gn=TRAK1;refGene_tx=XM_011286249.3;refGene_gn=TRAK1;refGene_tx=XM_011286250.3;refGene_gn=TRAK1;refGene_tx=XM_011286252.3;refGene_gn=TRAK1;refGene_tx=XM_019810882.2;refGene_gn=TRAK1;refGene_tx=XM_019810884.2;refGene_gn=TRAK1;refGene_tx=XM_019810885.2;refGene_gn=TRAK1;refGene_tx=XM_023260682.1;refGene_gn=TRAK1;refGene_tx=XM_023260683.1;refGene_gn=TRAK1;refGene_tx=XM_023260684.1;refGene_gn=TRAK1;refGene_tx=XM_023260685.1;refGene_gn=TRAK1;refGene_tx=XM_023260686.1
C2 151508944 151509061
refGene_gn=TRAK1;refGene_tx=XM_011286249.3;refGene_gn=TRAK1;refGene_tx=XM_011286250.3;refGene_gn=TRAK1;refGene_tx=XM_011286252.3;refGene_gn=TRAK1;refGene_tx=XM_019810882.2;refGene_gn=TRAK1;refGene_tx=XM_019810884.2;refGene_gn=TRAK1;refGene_tx=XM_019810885.2;refGene_gn=TRAK1;refGene_tx=XM_023260682.1;refGene_gn=TRAK1;refGene_tx=XM_023260683.1;refGene_gn=TRAK1;refGene_tx=XM_023260684.1;refGene_gn=TRAK1;refGene_tx=XM_023260685.1;refGene_gn=TRAK1;refGene_tx=XM_023260686.1
C2 151515153 151515230
refGene_gn=TRAK1;refGene_tx=XM_011286249.3;refGene_gn=TRAK1;refGene_tx=XM_011286250.3;refGene_gn=TRAK1;refGene_tx=XM_011286252.3;refGene_gn=TRAK1;refGene_tx=XM_019810882.2;refGene_gn=TRAK1;refGene_tx=XM_019810884.2;refGene_gn=TRAK1;refGene_tx=XM_019810885.2;refGene_gn=TRAK1;refGene_tx=XM_023260682.1;refGene_gn=TRAK1;refGene_tx=XM_023260683.1;refGene_gn=TRAK1;refGene_tx=XM_023260684.1;refGene_gn=TRAK1;refGene_tx=XM_023260685.1;refGene_gn=TRAK1;refGene_tx=XM_023260686.1
C2 151530774 151530888
refGene_gn=TRAK1;refGene_tx=XM_019810882.2;refGene_gn=TRAK1;refGene_tx=XM_019810885.2
C2 151539578 151539642
refGene_gn=TRAK1;refGene_tx=XM_011286252.3;refGene_gn=TRAK1;refGene_tx=XM_019810882.2;refGene_gn=TRAK1;refGene_tx=XM_019810885.2;refGene_gn=TRAK1;refGene_tx=XM_023260682.1
C2 151561329 151561524
refGene_gn=TRAK1;refGene_tx=XM_011286249.3;refGene_gn=TRAK1;refGene_tx=XM_011286250.3;refGene_gn=TRAK1;refGene_tx=XM_023260685.1
C2 151591799 151591890
refGene_gn=TRAK1;refGene_tx=XM_011286249.3;refGene_gn=TRAK1;refGene_tx=XM_011286250.3;refGene_gn=TRAK1;refGene_tx=XM_023260685.1
C2 151667515 151668436
refGene_gn=LOC102902041;refGene_tx=XM_006936537.4
C2 151668525 151668606
refGene_gn=LOC102902041;refGene_tx=XM_006936537.4
C2 151668889 151668928
refGene_gn=LOC102902041;refGene_tx=XM_006936537.4
C2 151708256 151708394
refGene_gn=ULK4;refGene_tx=XM_023260687.1;refGene_gn=ULK4;refGene_tx=XM_023260688.1;refGene_gn=ULK4;refGene_tx=XM_023260689.1;refGene_gn=ULK4;refGene_tx=XM_023260690.1;refGene_gn=ULK4;refGene_tx=XM_023260691.1;refGene_gn=ULK4;refGene_tx=XM_023260692.1
C2 151714095 151714195

refGene_gn=ULK4;refGene_tx=XM_023260687.1;refGene_gn=ULK4;refGene_tx=XM_023260688.1;refGene_gn=ULK4;refGene_tx=XM_023260689.1;refGene_gn=ULK4;refGene_tx=XM_023260690.1;refGene_gn=ULK4;refGene_tx=XM_023260691.1;refGene_gn=ULK4;refGene_tx=XM_023260692.1
C2 151716212 151716352
refGene_gn=ULK4;refGene_tx=XM_023260687.1;refGene_gn=ULK4;refGene_tx=XM_023260688.1;refGene_gn=ULK4;refGene_tx=XM_023260689.1;refGene_gn=ULK4;refGene_tx=XM_023260690.1;refGene_gn=ULK4;refGene_tx=XM_023260691.1;refGene_gn=ULK4;refGene_tx=XM_023260692.1
C2 151718697 151718860
refGene_gn=ULK4;refGene_tx=XM_023260687.1;refGene_gn=ULK4;refGene_tx=XM_023260688.1;refGene_gn=ULK4;refGene_tx=XM_023260689.1;refGene_gn=ULK4;refGene_tx=XM_023260690.1;refGene_gn=ULK4;refGene_tx=XM_023260691.1;refGene_gn=ULK4;refGene_tx=XM_023260692.1
C2 151721480 151721582
refGene_gn=ULK4;refGene_tx=XM_023260687.1;refGene_gn=ULK4;refGene_tx=XM_023260688.1;refGene_gn=ULK4;refGene_tx=XM_023260689.1;refGene_gn=ULK4;refGene_tx=XM_023260690.1;refGene_gn=ULK4;refGene_tx=XM_023260692.1
C2 151722174 151722258
refGene_gn=ULK4;refGene_tx=XM_023260687.1;refGene_gn=ULK4;refGene_tx=XM_023260688.1;refGene_gn=ULK4;refGene_tx=XM_023260689.1;refGene_gn=ULK4;refGene_tx=XM_023260690.1;refGene_gn=ULK4;refGene_tx=XM_023260692.1
C2 151724028 151724104
refGene_gn=ULK4;refGene_tx=XM_023260687.1;refGene_gn=ULK4;refGene_tx=XM_023260688.1;refGene_gn=ULK4;refGene_tx=XM_023260689.1;refGene_gn=ULK4;refGene_tx=XM_023260690.1;refGene_gn=ULK4;refGene_tx=XM_023260691.1;refGene_gn=ULK4;refGene_tx=XM_023260692.1
C2 151727896 151727989
refGene_gn=ULK4;refGene_tx=XM_023260687.1;refGene_gn=ULK4;refGene_tx=XM_023260688.1;refGene_gn=ULK4;refGene_tx=XM_023260689.1;refGene_gn=ULK4;refGene_tx=XM_023260690.1;refGene_gn=ULK4;refGene_tx=XM_023260691.1;refGene_gn=ULK4;refGene_tx=XM_023260692.1
C2 151730229 151730342
refGene_gn=ULK4;refGene_tx=XM_023260687.1;refGene_gn=ULK4;refGene_tx=XM_023260688.1;refGene_gn=ULK4;refGene_tx=XM_023260689.1;refGene_gn=ULK4;refGene_tx=XM_023260690.1;refGene_gn=ULK4;refGene_tx=XM_023260691.1;refGene_gn=ULK4;refGene_tx=XM_023260692.1
C2 151730498 151730568
refGene_gn=ULK4;refGene_tx=XM_023260687.1;refGene_gn=ULK4;refGene_tx=XM_023260688.1;refGene_gn=ULK4;refGene_tx=XM_023260689.1;refGene_gn=ULK4;refGene_tx=XM_023260690.1;refGene_gn=ULK4;refGene_tx=XM_023260691.1;refGene_gn=ULK4;refGene_tx=XM_023260692.1
C2 151732729 151732826
refGene_gn=ULK4;refGene_tx=XM_023260687.1;refGene_gn=ULK4;refGene_tx=XM_023260688.1;refGene_gn=ULK4;refGene_tx=XM_023260689.1;refGene_gn=ULK4;refGene_tx=XM_023260690.1;refGene_gn=ULK4;refGene_tx=XM_023260691.1;refGene_gn=ULK4;refGene_tx=XM_023260692.1
C2 151738023 151738128
refGene_gn=ULK4;refGene_tx=XM_023260687.1;refGene_gn=ULK4;refGene_tx=XM_023260688.1;refGene_gn=ULK4;refGene_tx=XM_023260689.1;refGene_gn=ULK

4;refGene_tx=XM_023260690.1;refGene_gn=ULK4;refGene_tx=XM_023260691.1;
refGene_gn=ULK4;refGene_tx=XM_023260692.1
C2	151741176	151741237
refGene_gn=ULK4;refGene_tx=XM_023260687.1;refGene_gn=ULK4;refGene_tx=X
M_023260688.1;refGene_gn=ULK4;refGene_tx=XM_023260689.1;refGene_gn=ULK
4;refGene_tx=XM_023260690.1;refGene_gn=ULK4;refGene_tx=XM_023260691.1;
refGene_gn=ULK4;refGene_tx=XM_023260692.1
C2	151742797	151742979
refGene_gn=ULK4;refGene_tx=XM_023260687.1;refGene_gn=ULK4;refGene_tx=X
M_023260688.1;refGene_gn=ULK4;refGene_tx=XM_023260689.1;refGene_gn=ULK
4;refGene_tx=XM_023260690.1;refGene_gn=ULK4;refGene_tx=XM_023260691.1;
refGene_gn=ULK4;refGene_tx=XM_023260692.1
C2	151743880	151743927
refGene_gn=ULK4;refGene_tx=XM_023260687.1;refGene_gn=ULK4;refGene_tx=X
M_023260688.1;refGene_gn=ULK4;refGene_tx=XM_023260689.1;refGene_gn=ULK
4;refGene_tx=XM_023260690.1;refGene_gn=ULK4;refGene_tx=XM_023260691.1;
refGene_gn=ULK4;refGene_tx=XM_023260692.1
C2	151761856	151761935
refGene_gn=ULK4;refGene_tx=XM_023260687.1;refGene_gn=ULK4;refGene_tx=X
M_023260688.1;refGene_gn=ULK4;refGene_tx=XM_023260689.1;refGene_gn=ULK
4;refGene_tx=XM_023260690.1;refGene_gn=ULK4;refGene_tx=XM_023260691.1;
refGene_gn=ULK4;refGene_tx=XM_023260692.1
C2	151799182	151799290
refGene_gn=ULK4;refGene_tx=XM_023260687.1;refGene_gn=ULK4;refGene_tx=X
M_023260688.1;refGene_gn=ULK4;refGene_tx=XM_023260689.1;refGene_gn=ULK
4;refGene_tx=XM_023260690.1;refGene_gn=ULK4;refGene_tx=XM_023260691.1;
refGene_gn=ULK4;refGene_tx=XM_023260692.1
C2	151803545	151803629
refGene_gn=ULK4;refGene_tx=XM_023260687.1;refGene_gn=ULK4;refGene_tx=X
M_023260689.1;refGene_gn=ULK4;refGene_tx=XM_023260690.1;refGene_gn=ULK
4;refGene_tx=XM_023260691.1;refGene_gn=ULK4;refGene_tx=XM_023260692.1
C2	151821515	151821677
refGene_gn=ULK4;refGene_tx=XM_023260687.1;refGene_gn=ULK4;refGene_tx=X
M_023260688.1;refGene_gn=ULK4;refGene_tx=XM_023260689.1;refGene_gn=ULK
4;refGene_tx=XM_023260690.1;refGene_gn=ULK4;refGene_tx=XM_023260691.1;
refGene_gn=ULK4;refGene_tx=XM_023260692.1
C2	151831853	151832036
refGene_gn=ULK4;refGene_tx=XM_023260687.1;refGene_gn=ULK4;refGene_tx=X
M_023260688.1;refGene_gn=ULK4;refGene_tx=XM_023260689.1;refGene_gn=ULK
4;refGene_tx=XM_023260691.1;refGene_gn=ULK4;refGene_tx=XM_023260692.1
C2	151860203	151860331
refGene_gn=ULK4;refGene_tx=XM_023260687.1;refGene_gn=ULK4;refGene_tx=X
M_023260688.1;refGene_gn=ULK4;refGene_tx=XM_023260689.1;refGene_gn=ULK
4;refGene_tx=XM_023260690.1;refGene_gn=ULK4;refGene_tx=XM_023260691.1;
refGene_gn=ULK4;refGene_tx=XM_023260692.1
C2	151899620	151899754
refGene_gn=ULK4;refGene_tx=XM_023260687.1;refGene_gn=ULK4;refGene_tx=X
M_023260688.1;refGene_gn=ULK4;refGene_tx=XM_023260689.1;refGene_gn=ULK
4;refGene_tx=XM_023260690.1;refGene_gn=ULK4;refGene_tx=XM_023260691.1;
refGene_gn=ULK4;refGene_tx=XM_023260692.1

C2	151901661	151901783
refGene_gn=ULK4;refGene_tx=XM_023260687.1;refGene_gn=ULK4;refGene_tx=XM_023260688.1;refGene_gn=ULK4;refGene_tx=XM_023260689.1;refGene_gn=ULK4;refGene_tx=XM_023260690.1;refGene_gn=ULK4;refGene_tx=XM_023260691.1;refGene_gn=ULK4;refGene_tx=XM_023260692.1
C2	151901923	151901980
refGene_gn=ULK4;refGene_tx=XM_023260687.1;refGene_gn=ULK4;refGene_tx=XM_023260688.1;refGene_gn=ULK4;refGene_tx=XM_023260689.1;refGene_gn=ULK4;refGene_tx=XM_023260690.1;refGene_gn=ULK4;refGene_tx=XM_023260691.1;refGene_gn=ULK4;refGene_tx=XM_023260692.1
C2	151908907	151908959
refGene_gn=ULK4;refGene_tx=XM_023260687.1;refGene_gn=ULK4;refGene_tx=XM_023260688.1;refGene_gn=ULK4;refGene_tx=XM_023260690.1;refGene_gn=ULK4;refGene_tx=XM_023260691.1;refGene_gn=ULK4;refGene_tx=XM_023260692.1
C2	151909061	151909156
refGene_gn=ULK4;refGene_tx=XM_023260687.1;refGene_gn=ULK4;refGene_tx=XM_023260688.1;refGene_gn=ULK4;refGene_tx=XM_023260690.1;refGene_gn=ULK4;refGene_tx=XM_023260691.1;refGene_gn=ULK4;refGene_tx=XM_023260692.1
C2	151928172	151928224
refGene_gn=ULK4;refGene_tx=XM_023260687.1;refGene_gn=ULK4;refGene_tx=XM_023260688.1;refGene_gn=ULK4;refGene_tx=XM_023260689.1;refGene_gn=ULK4;refGene_tx=XM_023260690.1;refGene_gn=ULK4;refGene_tx=XM_023260691.1;refGene_gn=ULK4;refGene_tx=XM_023260692.1
C2	151928319	151928464
refGene_gn=ULK4;refGene_tx=XM_023260687.1;refGene_gn=ULK4;refGene_tx=XM_023260688.1;refGene_gn=ULK4;refGene_tx=XM_023260689.1;refGene_gn=ULK4;refGene_tx=XM_023260690.1;refGene_gn=ULK4;refGene_tx=XM_023260691.1;refGene_gn=ULK4;refGene_tx=XM_023260692.1
C2	151944384	151944477
refGene_gn=ULK4;refGene_tx=XM_023260687.1;refGene_gn=ULK4;refGene_tx=XM_023260688.1;refGene_gn=ULK4;refGene_tx=XM_023260689.1;refGene_gn=ULK4;refGene_tx=XM_023260690.1;refGene_gn=ULK4;refGene_tx=XM_023260691.1;refGene_gn=ULK4;refGene_tx=XM_023260692.1
C2	151977084	151977133
refGene_gn=ULK4;refGene_tx=XM_023260687.1;refGene_gn=ULK4;refGene_tx=XM_023260688.1;refGene_gn=ULK4;refGene_tx=XM_023260689.1;refGene_gn=ULK4;refGene_tx=XM_023260690.1;refGene_gn=ULK4;refGene_tx=XM_023260691.1;refGene_gn=ULK4;refGene_tx=XM_023260692.1
C2	151977550	151977580
refGene_gn=ULK4;refGene_tx=XM_023260692.1
C2	152037526	152037632
refGene_gn=ULK4;refGene_tx=XM_023260687.1;refGene_gn=ULK4;refGene_tx=XM_023260688.1;refGene_gn=ULK4;refGene_tx=XM_023260689.1;refGene_gn=ULK4;refGene_tx=XM_023260690.1;refGene_gn=ULK4;refGene_tx=XM_023260691.1
C2	152103562	152103729
refGene_gn=ULK4;refGene_tx=XM_023260687.1;refGene_gn=ULK4;refGene_tx=XM_023260688.1;refGene_gn=ULK4;refGene_tx=XM_023260689.1;refGene_gn=ULK4;refGene_tx=XM_023260690.1;refGene_gn=ULK4;refGene_tx=XM_023260691.1
C2	152114568	152114667
refGene_gn=ULK4;refGene_tx=XM_023260687.1;refGene_gn=ULK4;refGene_tx=X

M_023260688.1;refGene_gn=ULK4;refGene_tx=XM_023260689.1;refGene_gn=ULK4;refGene_tx=XM_023260690.1;refGene_gn=ULK4;refGene_tx=XM_023260691.1
C2 152165822 152166008
refGene_gn=ULK4;refGene_tx=XM_023260687.1;refGene_gn=ULK4;refGene_tx=XM_023260688.1;refGene_gn=ULK4;refGene_tx=XM_023260689.1;refGene_gn=ULK4;refGene_tx=XM_023260690.1;refGene_gn=ULK4;refGene_tx=XM_023260691.1
C2 152283514 152283600
refGene_gn=ULK4;refGene_tx=XM_023260687.1;refGene_gn=ULK4;refGene_tx=XM_023260688.1;refGene_gn=ULK4;refGene_tx=XM_023260689.1;refGene_gn=ULK4;refGene_tx=XM_023260690.1;refGene_gn=ULK4;refGene_tx=XM_023260691.1
C2 152285680 152285759
refGene_gn=ULK4;refGene_tx=XM_023260687.1;refGene_gn=ULK4;refGene_tx=XM_023260688.1;refGene_gn=ULK4;refGene_tx=XM_023260689.1;refGene_gn=ULK4;refGene_tx=XM_023260690.1;refGene_gn=ULK4;refGene_tx=XM_023260691.1
C2 152291588 152291797
refGene_gn=CTNNB1;refGene_tx=XM_003992212.4;refGene_gn=CTNNB1;refGene_tx=XM_011286247.2;refGene_gn=CTNNB1;refGene_tx=XM_011286248.2;refGene_gn=CTNNB1;refGene_tx=XM_019810887.2
C2 152292828 152292889
refGene_gn=CTNNB1;refGene_tx=XM_003992212.4;refGene_gn=CTNNB1;refGene_tx=XM_011286247.2;refGene_gn=CTNNB1;refGene_tx=XM_011286248.2;refGene_gn=CTNNB1;refGene_tx=XM_019810887.2
C2 152293833 152293955
refGene_gn=CTNNB1;refGene_tx=XM_003992212.4;refGene_gn=CTNNB1;refGene_tx=XM_011286247.2;refGene_gn=CTNNB1;refGene_tx=XM_011286248.2;refGene_gn=CTNNB1;refGene_tx=XM_019810887.2
C2 152294040 152294191
refGene_gn=CTNNB1;refGene_tx=XM_003992212.4;refGene_gn=CTNNB1;refGene_tx=XM_011286247.2;refGene_gn=CTNNB1;refGene_tx=XM_011286248.2;refGene_gn=CTNNB1;refGene_tx=XM_019810887.2
C2 152294791 152294911
refGene_gn=CTNNB1;refGene_tx=XM_003992212.4;refGene_gn=CTNNB1;refGene_tx=XM_011286247.2;refGene_gn=CTNNB1;refGene_tx=XM_011286248.2;refGene_gn=CTNNB1;refGene_tx=XM_019810887.2
C2 152296382 152296541
refGene_gn=CTNNB1;refGene_tx=XM_003992212.4;refGene_gn=CTNNB1;refGene_tx=XM_011286247.2;refGene_gn=CTNNB1;refGene_tx=XM_011286248.2;refGene_gn=CTNNB1;refGene_tx=XM_019810887.2
C2 152296806 152297145
refGene_gn=CTNNB1;refGene_tx=XM_003992212.4;refGene_gn=CTNNB1;refGene_tx=XM_011286247.2;refGene_gn=CTNNB1;refGene_tx=XM_011286248.2;refGene_gn=CTNNB1;refGene_tx=XM_019810887.2
C2 152297223 152297327
refGene_gn=CTNNB1;refGene_tx=XM_003992212.4;refGene_gn=CTNNB1;refGene_tx=XM_011286247.2;refGene_gn=CTNNB1;refGene_tx=XM_011286248.2;refGene_gn=CTNNB1;refGene_tx=XM_019810887.2
C2 152300383 152300528
refGene_gn=CTNNB1;refGene_tx=XM_003992212.4;refGene_gn=CTNNB1;refGene_tx=XM_011286247.2;refGene_gn=CTNNB1;refGene_tx=XM_011286248.2;refGene_gn=CTNNB1;refGene_tx=XM_019810887.2

C2 152301821 152302023
refGene_gn=CTNNB1;refGene_tx=XM_003992212.4;refGene_gn=CTNNB1;refGene_tx=XM_011286247.2;refGene_gn=CTNNB1;refGene_tx=XM_011286248.2;refGene_gn=CTNNB1;refGene_tx=XM_019810887.2
C2 152302111 152302350
refGene_gn=CTNNB1;refGene_tx=XM_003992212.4;refGene_gn=CTNNB1;refGene_tx=XM_011286247.2;refGene_gn=CTNNB1;refGene_tx=XM_011286248.2;refGene_gn=CTNNB1;refGene_tx=XM_019810887.2
C2 152302475 152302729
refGene_gn=CTNNB1;refGene_tx=XM_003992212.4;refGene_gn=CTNNB1;refGene_tx=XM_011286247.2;refGene_gn=CTNNB1;refGene_tx=XM_011286248.2;refGene_gn=CTNNB1;refGene_tx=XM_019810887.2
C2 152302953 152303181
refGene_gn=CTNNB1;refGene_tx=XM_003992212.4;refGene_gn=CTNNB1;refGene_tx=XM_011286247.2;refGene_gn=CTNNB1;refGene_tx=XM_011286248.2;refGene_gn=CTNNB1;refGene_tx=XM_019810887.2
C2 152303613 152303626
refGene_gn=CTNNB1;refGene_tx=XM_003992212.4;refGene_gn=CTNNB1;refGene_tx=XM_011286247.2;refGene_gn=CTNNB1;refGene_tx=XM_011286248.2
C2 152888808 152889869
refGene_gn=ZNF621;refGene_tx=XM_006936510.4;refGene_gn=ZNF621;refGene_tx=XM_006936511.3;refGene_gn=ZNF621;refGene_tx=XM_006936512.4;refGene_gn=ZNF621;refGene_tx=XM_006936513.3;refGene_gn=ZNF621;refGene_tx=XM_006936514.4;refGene_gn=ZNF621;refGene_tx=XM_006936515.4;refGene_gn=ZNF621;refGene_tx=XM_019810888.2;refGene_gn=ZNF621;refGene_tx=XM_019810889.2;refGene_gn=ZNF621;refGene_tx=XM_019810890.2;refGene_gn=ZNF621;refGene_tx=XM_019810891.2;refGene_gn=ZNF621;refGene_tx=XM_019810892.2;refGene_gn=ZNF621;refGene_tx=XM_019810893.2;refGene_gn=ZNF621;refGene_tx=XM_019810895.2
C2 152891348 152891453
refGene_gn=ZNF621;refGene_tx=XM_006936510.4;refGene_gn=ZNF621;refGene_tx=XM_006936511.3;refGene_gn=ZNF621;refGene_tx=XM_006936512.4;refGene_gn=ZNF621;refGene_tx=XM_006936513.3;refGene_gn=ZNF621;refGene_tx=XM_006936514.4;refGene_gn=ZNF621;refGene_tx=XM_006936515.4;refGene_gn=ZNF621;refGene_tx=XM_019810888.2;refGene_gn=ZNF621;refGene_tx=XM_019810889.2;refGene_gn=ZNF621;refGene_tx=XM_019810890.2;refGene_gn=ZNF621;refGene_tx=XM_019810891.2;refGene_gn=ZNF621;refGene_tx=XM_019810892.2;refGene_gn=ZNF621;refGene_tx=XM_019810893.2
C2 152892024 152892178
refGene_gn=ZNF621;refGene_tx=XM_006936510.4;refGene_gn=ZNF621;refGene_tx=XM_006936511.3;refGene_gn=ZNF621;refGene_tx=XM_006936512.4;refGene_gn=ZNF621;refGene_tx=XM_006936513.3;refGene_gn=ZNF621;refGene_tx=XM_006936514.4;refGene_gn=ZNF621;refGene_tx=XM_006936515.4;refGene_gn=ZNF621;refGene_tx=XM_019810888.2;refGene_gn=ZNF621;refGene_tx=XM_019810889.2;refGene_gn=ZNF621;refGene_tx=XM_019810890.2;refGene_gn=ZNF621;refGene_tx=XM_019810891.2;refGene_gn=ZNF621;refGene_tx=XM_019810892.2;refGene_gn=ZNF621;refGene_tx=XM_019810893.2;refGene_gn=ZNF621;refGene_tx=XM_019810895.2
C2 152901160 152901246
refGene_gn=ZNF621;refGene_tx=XM_006936510.4;refGene_gn=ZNF621;refGene_

tx=XM_006936511.3;refGene_gn=ZNF621;refGene_tx=XM_006936512.4;refGene_gn=ZNF621;refGene_tx=XM_006936513.3;refGene_gn=ZNF621;refGene_tx=XM_006936514.4;refGene_gn=ZNF621;refGene_tx=XM_006936515.4;refGene_gn=ZNF621;refGene_tx=XM_019810888.2;refGene_gn=ZNF621;refGene_tx=XM_019810889.2;refGene_gn=ZNF621;refGene_tx=XM_019810890.2;refGene_gn=ZNF621;refGene_tx=XM_019810891.2;refGene_gn=ZNF621;refGene_tx=XM_019810892.2;refGene_gn=ZNF621;refGene_tx=XM_019810893.2;refGene_gn=ZNF621;refGene_tx=XM_019810895.2
C2 152901518 152901543
refGene_gn=ZNF621;refGene_tx=XM_006936511.3;refGene_gn=ZNF621;refGene_tx=XM_019810889.2
C2 152902793 152902824
refGene_gn=ZNF621;refGene_tx=XM_006936510.4;refGene_gn=ZNF621;refGene_tx=XM_019810888.2;refGene_gn=ZNF621;refGene_tx=XM_019810895.2
C2 152909212 152910219
refGene_gn=ZNF620;refGene_tx=XM_006936516.4;refGene_gn=ZNF620;refGene_tx=XM_011286244.3;refGene_gn=ZNF620;refGene_tx=XM_011286245.3;refGene_gn=ZNF620;refGene_tx=XM_011286246.3;refGene_gn=ZNF620;refGene_tx=XM_019810897.2
C2 152913289 152913403
refGene_gn=ZNF620;refGene_tx=XM_006936516.4;refGene_gn=ZNF620;refGene_tx=XM_011286244.3;refGene_gn=ZNF620;refGene_tx=XM_011286245.3;refGene_gn=ZNF620;refGene_tx=XM_011286246.3;refGene_gn=ZNF620;refGene_tx=XM_019810897.2
C2 152913986 152914140
refGene_gn=ZNF620;refGene_tx=XM_006936516.4;refGene_gn=ZNF620;refGene_tx=XM_011286244.3;refGene_gn=ZNF620;refGene_tx=XM_011286245.3;refGene_gn=ZNF620;refGene_tx=XM_011286246.3;refGene_gn=ZNF620;refGene_tx=XM_019810897.2
C2 152920022 152920046
refGene_gn=ZNF620;refGene_tx=XM_006936516.4
C2 152938534 152939910
refGene_gn=ZNF619;refGene_tx=XM_023260188.1
C2 152943261 152943378
refGene_gn=ZNF619;refGene_tx=XM_023260188.1
C2 152945070 152945197
refGene_gn=ZNF619;refGene_tx=XM_023260188.1
C2 152949534 152949558
refGene_gn=ZNF619;refGene_tx=XM_023260188.1
C2 152954519 152954798
refGene_gn=RPL14;refGene_tx=XM_006936517.2
C2 152955227 152955281
refGene_gn=RPL14;refGene_tx=XM_006936517.2
C2 152955375 152955475
refGene_gn=RPL14;refGene_tx=XM_006936517.2
C2 152956709 152956804
refGene_gn=RPL14;refGene_tx=XM_006936517.2
C2 152957238 152957340
refGene_gn=RPL14;refGene_tx=XM_006936517.2
C2 152957855 152958109

refGene_gn=RPL14;refGene_tx=XM_006936517.2
C2 152959881 152959888
refGene_gn=RPL14;refGene_tx=XM_006936517.2
C2 152968178 152968184
refGene_gn=ENTPD3;refGene_tx=XM_023260698.1
C2 152975246 152975483
refGene_gn=ENTPD3;refGene_tx=XM_023260696.1;refGene_gn=ENTPD3;refGene_tx=XM_023260697.1
C2 152977939 152978077
refGene_gn=ENTPD3;refGene_tx=XM_023260696.1;refGene_gn=ENTPD3;refGene_tx=XM_023260697.1;refGene_gn=ENTPD3;refGene_tx=XM_023260698.1
C2 152978974 152979085
refGene_gn=ENTPD3;refGene_tx=XM_023260696.1;refGene_gn=ENTPD3;refGene_tx=XM_023260697.1;refGene_gn=ENTPD3;refGene_tx=XM_023260698.1
C2 152979259 152979532
refGene_gn=ENTPD3;refGene_tx=XM_023260696.1;refGene_gn=ENTPD3;refGene_tx=XM_023260697.1;refGene_gn=ENTPD3;refGene_tx=XM_023260698.1
C2 152986025 152986259
refGene_gn=ENTPD3;refGene_tx=XM_023260696.1;refGene_gn=ENTPD3;refGene_tx=XM_023260697.1;refGene_gn=ENTPD3;refGene_tx=XM_023260698.1
C2 152987271 152987431
refGene_gn=ENTPD3;refGene_tx=XM_023260696.1;refGene_gn=ENTPD3;refGene_tx=XM_023260697.1;refGene_gn=ENTPD3;refGene_tx=XM_023260698.1
C2 152990021 152990172
refGene_gn=ENTPD3;refGene_tx=XM_023260696.1;refGene_gn=ENTPD3;refGene_tx=XM_023260697.1;refGene_gn=ENTPD3;refGene_tx=XM_023260698.1
C2 152997789 152997907
refGene_gn=ENTPD3;refGene_tx=XM_023260696.1;refGene_gn=ENTPD3;refGene_tx=XM_023260697.1;refGene_gn=ENTPD3;refGene_tx=XM_023260698.1
C2 153005575 153005703
refGene_gn=ENTPD3;refGene_tx=XM_023260696.1;refGene_gn=ENTPD3;refGene_tx=XM_023260697.1;refGene_gn=ENTPD3;refGene_tx=XM_023260698.1
C2 153011355 153011395
refGene_gn=ENTPD3;refGene_tx=XM_023260696.1;refGene_gn=ENTPD3;refGene_tx=XM_023260697.1;refGene_gn=ENTPD3;refGene_tx=XM_023260698.1
C2 153073996 153074041
refGene_gn=EIF1B;refGene_tx=XM_023260700.1
C2 153074439 153074541
refGene_gn=EIF1B;refGene_tx=XM_023260700.1
C2 153074949 153075113
refGene_gn=EIF1B;refGene_tx=XM_023260700.1
C2 153076071 153076102
refGene_gn=EIF1B;refGene_tx=XM_023260700.1
C2 153107679 153107712
refGene_gn=MYRIP;refGene_tx=XM_011286243.3;refGene_gn=MYRIP;refGene_tx=XM_019810899.2
C2 153110252 153110258
refGene_gn=MYRIP;refGene_tx=XM_019810900.2
C2 153114802 153114808
refGene_gn=MYRIP;refGene_tx=XM_019810901.2

C2 153115173 153115292
refGene_gn=MYRIP;refGene_tx=XM_011286243.3;refGene_gn=MYRIP;refGene_tx=XM_019810899.2;refGene_gn=MYRIP;refGene_tx=XM_019810900.2;refGene_gn=MYRIP;refGene_tx=XM_019810901.2
C2 153116902 153116963
refGene_gn=MYRIP;refGene_tx=XM_011286243.3;refGene_gn=MYRIP;refGene_tx=XM_019810899.2;refGene_gn=MYRIP;refGene_tx=XM_019810900.2;refGene_gn=MYRIP;refGene_tx=XM_019810901.2
C2 153117075 153117180
refGene_gn=MYRIP;refGene_tx=XM_011286243.3;refGene_gn=MYRIP;refGene_tx=XM_019810899.2;refGene_gn=MYRIP;refGene_tx=XM_019810900.2;refGene_gn=MYRIP;refGene_tx=XM_019810901.2
C2 153121191 153121353
refGene_gn=MYRIP;refGene_tx=XM_011286243.3;refGene_gn=MYRIP;refGene_tx=XM_019810899.2;refGene_gn=MYRIP;refGene_tx=XM_019810900.2;refGene_gn=MYRIP;refGene_tx=XM_019810901.2
C2 153129051 153129246
refGene_gn=MYRIP;refGene_tx=XM_011286243.3;refGene_gn=MYRIP;refGene_tx=XM_019810899.2;refGene_gn=MYRIP;refGene_tx=XM_019810900.2;refGene_gn=MYRIP;refGene_tx=XM_019810901.2
C2 153156693 153156933
refGene_gn=MYRIP;refGene_tx=XM_011286243.3;refGene_gn=MYRIP;refGene_tx=XM_019810899.2;refGene_gn=MYRIP;refGene_tx=XM_019810900.2;refGene_gn=MYRIP;refGene_tx=XM_019810901.2
C2 153173870 153174508
refGene_gn=MYRIP;refGene_tx=XM_011286243.3;refGene_gn=MYRIP;refGene_tx=XM_019810899.2;refGene_gn=MYRIP;refGene_tx=XM_019810900.2;refGene_gn=MYRIP;refGene_tx=XM_019810901.2
C2 153182402 153182556
refGene_gn=MYRIP;refGene_tx=XM_011286243.3;refGene_gn=MYRIP;refGene_tx=XM_019810899.2;refGene_gn=MYRIP;refGene_tx=XM_019810900.2;refGene_gn=MYRIP;refGene_tx=XM_019810901.2
C2 153192727 153192871
refGene_gn=MYRIP;refGene_tx=XM_011286243.3;refGene_gn=MYRIP;refGene_tx=XM_019810899.2;refGene_gn=MYRIP;refGene_tx=XM_019810900.2;refGene_gn=MYRIP;refGene_tx=XM_019810901.2
C2 153195353 153195434
refGene_gn=MYRIP;refGene_tx=XM_011286243.3;refGene_gn=MYRIP;refGene_tx=XM_019810899.2;refGene_gn=MYRIP;refGene_tx=XM_019810900.2;refGene_gn=MYRIP;refGene_tx=XM_019810901.2
C2 153195652 153195750
refGene_gn=MYRIP;refGene_tx=XM_011286243.3;refGene_gn=MYRIP;refGene_tx=XM_019810899.2;refGene_gn=MYRIP;refGene_tx=XM_019810900.2;refGene_gn=MYRIP;refGene_tx=XM_019810901.2
C2 153199557 153199638
refGene_gn=MYRIP;refGene_tx=XM_011286243.3;refGene_gn=MYRIP;refGene_tx=XM_019810899.2;refGene_gn=MYRIP;refGene_tx=XM_019810900.2;refGene_gn=MYRIP;refGene_tx=XM_019810901.2
C2 153205549 153205686
refGene_gn=MYRIP;refGene_tx=XM_011286243.3;refGene_gn=MYRIP;refGene_tx

=XM_019810899.2;refGene_gn=MYRIP;refGene_tx=XM_019810900.2;refGene_gn=MYRIP;refGene_tx=XM_019810901.2
C2	153323521	153323743
refGene_gn=MYRIP;refGene_tx=XM_011286243.3;refGene_gn=MYRIP;refGene_tx=XM_019810899.2;refGene_gn=MYRIP;refGene_tx=XM_019810900.2;refGene_gn=MYRIP;refGene_tx=XM_019810901.2
C2	153423272	153423382
refGene_gn=MYRIP;refGene_tx=XM_011286243.3;refGene_gn=MYRIP;refGene_tx=XM_019810899.2;refGene_gn=MYRIP;refGene_tx=XM_019810900.2;refGene_gn=MYRIP;refGene_tx=XM_019810901.2
C2	153702509	153702549
refGene_gn=MOBP;refGene_tx=XM_006936519.4;refGene_gn=MOBP;refGene_tx=XM_019810906.2;refGene_gn=MOBP;refGene_tx=XM_023260701.1
C2	153714396	153714648
refGene_gn=MOBP;refGene_tx=XM_006936519.4;refGene_gn=MOBP;refGene_tx=XM_019810906.2;refGene_gn=MOBP;refGene_tx=XM_023260701.1
C2	153714909	153715115
refGene_gn=MOBP;refGene_tx=XM_006936519.4;refGene_gn=MOBP;refGene_tx=XM_019810906.2;refGene_gn=MOBP;refGene_tx=XM_023260701.1
C2	153776448	153776543
refGene_gn=RPSA;refGene_tx=XM_003992219.4;refGene_gn=RPSA;refGene_tx=XM_023260710.1
C2	153776769	153776935
refGene_gn=RPSA;refGene_tx=XM_003992219.4;refGene_gn=RPSA;refGene_tx=XM_023260710.1
C2	153777041	153777170
refGene_gn=RPSA;refGene_tx=XM_003992219.4;refGene_gn=RPSA;refGene_tx=XM_023260710.1
C2	153777897	153778143
refGene_gn=RPSA;refGene_tx=XM_003992219.4;refGene_gn=RPSA;refGene_tx=XM_023260710.1
C2	153779725	153779844
refGene_gn=RPSA;refGene_tx=XM_003992219.4;refGene_gn=RPSA;refGene_tx=XM_023260710.1
C2	153780884	153781017
refGene_gn=RPSA;refGene_tx=XM_003992219.4;refGene_gn=RPSA;refGene_tx=XM_023260710.1
C2	153786595	153786697
refGene_gn=LOC101084312;refGene_tx=XM_023260704.1
C2	153787344	153787647
refGene_gn=LOC101084312;refGene_tx=XM_023260702.1;refGene_gn=LOC101084312;refGene_tx=XM_023260703.1;refGene_gn=LOC101084312;refGene_tx=XM_023260706.1
C2	153787789	153787795
refGene_gn=LOC101084312;refGene_tx=XM_023260705.1
C2	153788274	153788384
refGene_gn=LOC101084312;refGene_tx=XM_023260702.1;refGene_gn=LOC101084312;refGene_tx=XM_023260703.1;refGene_gn=LOC101084312;refGene_tx=XM_023260704.1;refGene_gn=LOC101084312;refGene_tx=XM_023260705.1;refGene_gn=LOC101084312;refGene_tx=XM_023260706.1

C2 153792670 153792681
refGene_gn=LOC101084312;refGene_tx=XM_023260707.1;refGene_gn=LOC101084312;refGene_tx=XM_023260709.1
C2 153796764 153796881
refGene_gn=LOC101084312;refGene_tx=XM_023260702.1;refGene_gn=LOC101084312;refGene_tx=XM_023260704.1;refGene_gn=LOC101084312;refGene_tx=XM_023260705.1;refGene_gn=LOC101084312;refGene_tx=XM_023260706.1;refGene_gn=LOC101084312;refGene_tx=XM_023260707.1;refGene_gn=LOC101084312;refGene_tx=XM_023260709.1
C2 153797355 153797581
refGene_gn=LOC101084312;refGene_tx=XM_023260702.1;refGene_gn=LOC101084312;refGene_tx=XM_023260703.1;refGene_gn=LOC101084312;refGene_tx=XM_023260704.1;refGene_gn=LOC101084312;refGene_tx=XM_023260705.1;refGene_gn=LOC101084312;refGene_tx=XM_023260706.1;refGene_gn=LOC101084312;refGene_tx=XM_023260707.1;refGene_gn=LOC101084312;refGene_tx=XM_023260709.1
C2 153800599 153800749
refGene_gn=LOC101084312;refGene_tx=XM_023260702.1;refGene_gn=LOC101084312;refGene_tx=XM_023260703.1;refGene_gn=LOC101084312;refGene_tx=XM_023260704.1;refGene_gn=LOC101084312;refGene_tx=XM_023260705.1;refGene_gn=LOC101084312;refGene_tx=XM_023260706.1;refGene_gn=LOC101084312;refGene_tx=XM_023260707.1;refGene_gn=LOC101084312;refGene_tx=XM_023260709.1
C2 153801953 153802006
refGene_gn=LOC101084312;refGene_tx=XM_023260702.1;refGene_gn=LOC101084312;refGene_tx=XM_023260703.1;refGene_gn=LOC101084312;refGene_tx=XM_023260704.1;refGene_gn=LOC101084312;refGene_tx=XM_023260705.1;refGene_gn=LOC101084312;refGene_tx=XM_023260706.1;refGene_gn=LOC101084312;refGene_tx=XM_023260707.1;refGene_gn=LOC101084312;refGene_tx=XM_023260709.1
C2 153802768 153802802
refGene_gn=LOC101084312;refGene_tx=XM_023260706.1
C2 153803552 153803913
refGene_gn=LOC101084312;refGene_tx=XM_023260702.1;refGene_gn=LOC101084312;refGene_tx=XM_023260703.1;refGene_gn=LOC101084312;refGene_tx=XM_023260704.1;refGene_gn=LOC101084312;refGene_tx=XM_023260705.1;refGene_gn=LOC101084312;refGene_tx=XM_023260707.1;refGene_gn=LOC101084312;refGene_tx=XM_023260709.1
C2 153808603 153808878
refGene_gn=SLC25A38;refGene_tx=XM_003992221.5;refGene_gn=SLC25A38;refGene_tx=XM_023260711.1
C2 153810041 153810208
refGene_gn=SLC25A38;refGene_tx=XM_003992221.5
C2 153811862 153812031
refGene_gn=SLC25A38;refGene_tx=XM_003992221.5;refGene_gn=SLC25A38;refGene_tx=XM_023260711.1
C2 153812265 153812445
refGene_gn=SLC25A38;refGene_tx=XM_003992221.5;refGene_gn=SLC25A38;refGene_tx=XM_023260711.1
C2 153813152 153813237
refGene_gn=SLC25A38;refGene_tx=XM_003992221.5;refGene_gn=SLC25A38;refGene_tx=XM_023260711.1
C2 153813801 153813917

refGene_gn=SLC25A38;refGene_tx=XM_003992221.5;refGene_gn=SLC25A38;refGene_tx=XM_023260711.1
C2 153818941 153819010
refGene_gn=SLC25A38;refGene_tx=XM_003992221.5;refGene_gn=SLC25A38;refGene_tx=XM_023260711.1
C2 153837643 153838711
refGene_gn=CCR8;refGene_tx=XM_023238573.1
C2 153918539 153918659
refGene_gn=CX3CR1;refGene_tx=XM_019810907.2
C2 153930686 153931766
refGene_gn=CX3CR1;refGene_tx=XM_003992223.5;refGene_gn=CX3CR1;refGene_tx=XM_019810907.2;refGene_gn=CX3CR1;refGene_tx=XM_019810908.2
C2 154007890 154011214
refGene_gn=XIRP1;refGene_tx=XM_006936525.2
C2 154011629 154011638
refGene_gn=XIRP1;refGene_tx=XM_006936525.2
C2 154042108 154042319
refGene_gn=CSRNP1;refGene_tx=XM_023260716.1
C2 154044613 154044670
refGene_gn=CSRNP1;refGene_tx=XM_023260716.1
C2 154044672 154044872
refGene_gn=CSRNP1;refGene_tx=XM_023260716.1
C2 154045403 154045718
refGene_gn=CSRNP1;refGene_tx=XM_023260716.1
C2 154045801 154046782
refGene_gn=CSRNP1;refGene_tx=XM_023260716.1
C2 154051152 154051230
refGene_gn=TTC21A;refGene_tx=XM_023260713.1;refGene_gn=TTC21A;refGene_tx=XM_023260714.1;refGene_gn=TTC21A;refGene_tx=XM_023260715.1
C2 154051333 154051401
refGene_gn=TTC21A;refGene_tx=XM_023260713.1;refGene_gn=TTC21A;refGene_tx=XM_023260714.1;refGene_gn=TTC21A;refGene_tx=XM_023260715.1
C2 154051517 154051638
refGene_gn=TTC21A;refGene_tx=XM_023260713.1;refGene_gn=TTC21A;refGene_tx=XM_023260714.1;refGene_gn=TTC21A;refGene_tx=XM_023260715.1
C2 154052041 154052266
refGene_gn=TTC21A;refGene_tx=XM_023260713.1;refGene_gn=TTC21A;refGene_tx=XM_023260714.1;refGene_gn=TTC21A;refGene_tx=XM_023260715.1
C2 154052354 154052547
refGene_gn=TTC21A;refGene_tx=XM_023260713.1;refGene_gn=TTC21A;refGene_tx=XM_023260714.1;refGene_gn=TTC21A;refGene_tx=XM_023260715.1
C2 154052711 154052873
refGene_gn=TTC21A;refGene_tx=XM_023260713.1;refGene_gn=TTC21A;refGene_tx=XM_023260714.1;refGene_gn=TTC21A;refGene_tx=XM_023260715.1
C2 154053126 154053277
refGene_gn=TTC21A;refGene_tx=XM_023260713.1;refGene_gn=TTC21A;refGene_tx=XM_023260714.1;refGene_gn=TTC21A;refGene_tx=XM_023260715.1
C2 154054088 154054170
refGene_gn=TTC21A;refGene_tx=XM_023260713.1;refGene_gn=TTC21A;refGene_tx=XM_023260714.1;refGene_gn=TTC21A;refGene_tx=XM_023260715.1

C2 154054880 154054991
refGene_gn=TTC21A;refGene_tx=XM_023260713.1;refGene_gn=TTC21A;refGene_tx=XM_023260714.1;refGene_gn=TTC21A;refGene_tx=XM_023260715.1
C2 154055891 154056080
refGene_gn=TTC21A;refGene_tx=XM_023260713.1;refGene_gn=TTC21A;refGene_tx=XM_023260714.1;refGene_gn=TTC21A;refGene_tx=XM_023260715.1
C2 154057304 154057408
refGene_gn=TTC21A;refGene_tx=XM_023260713.1;refGene_gn=TTC21A;refGene_tx=XM_023260714.1;refGene_gn=TTC21A;refGene_tx=XM_023260715.1
C2 154057538 154057677
refGene_gn=TTC21A;refGene_tx=XM_023260713.1;refGene_gn=TTC21A;refGene_tx=XM_023260714.1;refGene_gn=TTC21A;refGene_tx=XM_023260715.1
C2 154058042 154058153
refGene_gn=TTC21A;refGene_tx=XM_023260713.1;refGene_gn=TTC21A;refGene_tx=XM_023260714.1;refGene_gn=TTC21A;refGene_tx=XM_023260715.1
C2 154058253 154058326
refGene_gn=TTC21A;refGene_tx=XM_023260713.1;refGene_gn=TTC21A;refGene_tx=XM_023260714.1;refGene_gn=TTC21A;refGene_tx=XM_023260715.1
C2 154059114 154059353
refGene_gn=TTC21A;refGene_tx=XM_023260713.1;refGene_gn=TTC21A;refGene_tx=XM_023260714.1;refGene_gn=TTC21A;refGene_tx=XM_023260715.1
C2 154063443 154063659
refGene_gn=TTC21A;refGene_tx=XM_023260713.1;refGene_gn=TTC21A;refGene_tx=XM_023260714.1;refGene_gn=TTC21A;refGene_tx=XM_023260715.1
C2 154063886 154064044
refGene_gn=TTC21A;refGene_tx=XM_023260713.1;refGene_gn=TTC21A;refGene_tx=XM_023260714.1;refGene_gn=TTC21A;refGene_tx=XM_023260715.1
C2 154065749 154065879
refGene_gn=TTC21A;refGene_tx=XM_023260713.1;refGene_gn=TTC21A;refGene_tx=XM_023260714.1;refGene_gn=TTC21A;refGene_tx=XM_023260715.1
C2 154066688 154066889
refGene_gn=TTC21A;refGene_tx=XM_023260713.1;refGene_gn=TTC21A;refGene_tx=XM_023260715.1
C2 154067056 154067154
refGene_gn=TTC21A;refGene_tx=XM_023260713.1;refGene_gn=TTC21A;refGene_tx=XM_023260714.1;refGene_gn=TTC21A;refGene_tx=XM_023260715.1
C2 154070823 154071016
refGene_gn=TTC21A;refGene_tx=XM_023260713.1;refGene_gn=TTC21A;refGene_tx=XM_023260714.1;refGene_gn=TTC21A;refGene_tx=XM_023260715.1
C2 154071995 154072094
refGene_gn=TTC21A;refGene_tx=XM_023260713.1;refGene_gn=TTC21A;refGene_tx=XM_023260714.1;refGene_gn=TTC21A;refGene_tx=XM_023260715.1
C2 154074409 154074494
refGene_gn=TTC21A;refGene_tx=XM_023260713.1;refGene_gn=TTC21A;refGene_tx=XM_023260714.1;refGene_gn=TTC21A;refGene_tx=XM_023260715.1
C2 154079252 154079410
refGene_gn=TTC21A;refGene_tx=XM_023260713.1;refGene_gn=TTC21A;refGene_tx=XM_023260714.1;refGene_gn=TTC21A;refGene_tx=XM_023260715.1
C2 154081757 154081880
refGene_gn=TTC21A;refGene_tx=XM_023260713.1;refGene_gn=TTC21A;refGene_

tx=XM_023260714.1;refGene_gn=TTC21A;refGene_tx=XM_023260715.1
C2 154083253 154083420
refGene_gn=TTC21A;refGene_tx=XM_023260713.1;refGene_gn=TTC21A;refGene_tx=XM_023260714.1;refGene_gn=TTC21A;refGene_tx=XM_023260715.1
C2 154084027 154084138
refGene_gn=TTC21A;refGene_tx=XM_023260713.1;refGene_gn=TTC21A;refGene_tx=XM_023260714.1;refGene_gn=TTC21A;refGene_tx=XM_023260715.1
C2 154084672 154084802
refGene_gn=TTC21A;refGene_tx=XM_023260713.1;refGene_gn=TTC21A;refGene_tx=XM_023260714.1
C2 154086098 154086125
refGene_gn=TTC21A;refGene_tx=XM_023260713.1;refGene_gn=TTC21A;refGene_tx=XM_023260714.1
C2 154086401 154086464
refGene_gn=GORASP1;refGene_tx=XM_023260717.1
C2 154087253 154087313
refGene_gn=GORASP1;refGene_tx=XM_006936527.4
C2 154090095 154090176
refGene_gn=GORASP1;refGene_tx=XM_006936527.4;refGene_gn=GORASP1;refGene_tx=XM_023260717.1
C2 154090711 154090915
refGene_gn=GORASP1;refGene_tx=XM_006936527.4;refGene_gn=GORASP1;refGene_tx=XM_023260717.1;refGene_gn=GORASP1;refGene_tx=XM_023260718.1
C2 154092159 154092246
refGene_gn=GORASP1;refGene_tx=XM_006936527.4;refGene_gn=GORASP1;refGene_tx=XM_023260717.1;refGene_gn=GORASP1;refGene_tx=XM_023260718.1
C2 154092395 154092526
refGene_gn=GORASP1;refGene_tx=XM_006936527.4;refGene_gn=GORASP1;refGene_tx=XM_023260717.1;refGene_gn=GORASP1;refGene_tx=XM_023260718.1
C2 154092755 154092963
refGene_gn=GORASP1;refGene_tx=XM_006936527.4;refGene_gn=GORASP1;refGene_tx=XM_023260717.1;refGene_gn=GORASP1;refGene_tx=XM_023260718.1
C2 154093720 154093871
refGene_gn=GORASP1;refGene_tx=XM_006936527.4;refGene_gn=GORASP1;refGene_tx=XM_023260717.1;refGene_gn=GORASP1;refGene_tx=XM_023260718.1
C2 154094342 154094495
refGene_gn=GORASP1;refGene_tx=XM_006936527.4;refGene_gn=GORASP1;refGene_tx=XM_023260717.1;refGene_gn=GORASP1;refGene_tx=XM_023260718.1
C2 154094729 154094992
refGene_gn=GORASP1;refGene_tx=XM_006936527.4;refGene_gn=GORASP1;refGene_tx=XM_023260717.1;refGene_gn=GORASP1;refGene_tx=XM_023260718.1
C2 154098773 154098869
refGene_gn=WDR48;refGene_tx=XM_003992225.5;refGene_gn=WDR48;refGene_tx=XM_019810913.2;refGene_gn=WDR48;refGene_tx=XM_019810914.2
C2 154099505 154099698
refGene_gn=WDR48;refGene_tx=XM_003992225.5;refGene_gn=WDR48;refGene_tx=XM_019810913.2;refGene_gn=WDR48;refGene_tx=XM_019810914.2
C2 154100847 154100924
refGene_gn=WDR48;refGene_tx=XM_003992225.5;refGene_gn=WDR48;refGene_tx=XM_019810913.2;refGene_gn=WDR48;refGene_tx=XM_019810914.2

C2	154103227	154103315
refGene_gn=WDR48;refGene_tx=XM_003992225.5;refGene_gn=WDR48;refGene_tx=XM_019810913.2;refGene_gn=WDR48;refGene_tx=XM_019810914.2
C2	154104441	154104547
refGene_gn=WDR48;refGene_tx=XM_003992225.5;refGene_gn=WDR48;refGene_tx=XM_019810913.2;refGene_gn=WDR48;refGene_tx=XM_019810914.2
C2	154106774	154106870
refGene_gn=WDR48;refGene_tx=XM_003992225.5;refGene_gn=WDR48;refGene_tx=XM_019810913.2;refGene_gn=WDR48;refGene_tx=XM_019810914.2
C2	154109905	154110059
refGene_gn=WDR48;refGene_tx=XM_003992225.5;refGene_gn=WDR48;refGene_tx=XM_019810913.2;refGene_gn=WDR48;refGene_tx=XM_019810914.2
C2	154110371	154110479
refGene_gn=WDR48;refGene_tx=XM_003992225.5;refGene_gn=WDR48;refGene_tx=XM_019810913.2;refGene_gn=WDR48;refGene_tx=XM_019810914.2
C2	154111618	154111716
refGene_gn=WDR48;refGene_tx=XM_003992225.5;refGene_gn=WDR48;refGene_tx=XM_019810913.2;refGene_gn=WDR48;refGene_tx=XM_019810914.2
C2	154113221	154113324
refGene_gn=WDR48;refGene_tx=XM_003992225.5;refGene_gn=WDR48;refGene_tx=XM_019810913.2;refGene_gn=WDR48;refGene_tx=XM_019810914.2
C2	154114419	154114494
refGene_gn=WDR48;refGene_tx=XM_003992225.5;refGene_gn=WDR48;refGene_tx=XM_019810913.2;refGene_gn=WDR48;refGene_tx=XM_019810914.2
C2	154116791	154117016
refGene_gn=WDR48;refGene_tx=XM_003992225.5;refGene_gn=WDR48;refGene_tx=XM_019810913.2;refGene_gn=WDR48;refGene_tx=XM_019810914.2
C2	154121384	154121486
refGene_gn=WDR48;refGene_tx=XM_003992225.5;refGene_gn=WDR48;refGene_tx=XM_019810913.2;refGene_gn=WDR48;refGene_tx=XM_019810914.2
C2	154121865	154121954
refGene_gn=WDR48;refGene_tx=XM_003992225.5;refGene_gn=WDR48;refGene_tx=XM_019810913.2;refGene_gn=WDR48;refGene_tx=XM_019810914.2
C2	154123711	154123825
refGene_gn=WDR48;refGene_tx=XM_019810914.2
C2	154123906	154124036
refGene_gn=WDR48;refGene_tx=XM_003992225.5;refGene_gn=WDR48;refGene_tx=XM_019810913.2;refGene_gn=WDR48;refGene_tx=XM_019810914.2
C2	154124149	154124232
refGene_gn=WDR48;refGene_tx=XM_003992225.5;refGene_gn=WDR48;refGene_tx=XM_019810913.2;refGene_gn=WDR48;refGene_tx=XM_019810914.2
C2	154124915	154124994
refGene_gn=WDR48;refGene_tx=XM_003992225.5;refGene_gn=WDR48;refGene_tx=XM_019810913.2;refGene_gn=WDR48;refGene_tx=XM_019810914.2
C2	154128612	154128753
refGene_gn=WDR48;refGene_tx=XM_003992225.5;refGene_gn=WDR48;refGene_tx=XM_019810913.2;refGene_gn=WDR48;refGene_tx=XM_019810914.2
C2	154140294	154140342
refGene_gn=WDR48;refGene_tx=XM_003992225.5;refGene_gn=WDR48;refGene_tx=XM_019810913.2;refGene_gn=WDR48;refGene_tx=XM_019810914.2

C2 154204411 154204684
refGene_gn=SCN11A;refGene_tx=XM_023260719.1
C2 154207771 154207890
refGene_gn=SCN11A;refGene_tx=XM_023260719.1
C2 154209677 154209779
refGene_gn=SCN11A;refGene_tx=XM_023260719.1
C2 154218395 154218524
refGene_gn=SCN11A;refGene_tx=XM_023260719.1
C2 154219823 154219927
refGene_gn=SCN11A;refGene_tx=XM_023260719.1
C2 154224012 154224186
refGene_gn=SCN11A;refGene_tx=XM_023260719.1
C2 154225147 154225211
refGene_gn=SCN11A;refGene_tx=XM_023260719.1
C2 154230624 154230766
refGene_gn=SCN11A;refGene_tx=XM_023260719.1
C2 154231332 154231530
refGene_gn=SCN11A;refGene_tx=XM_023260719.1
C2 154232365 154232539
refGene_gn=SCN11A;refGene_tx=XM_023260719.1
C2 154235537 154235667
refGene_gn=SCN11A;refGene_tx=XM_023260719.1
C2 154236750 154236989
refGene_gn=SCN11A;refGene_tx=XM_023260719.1
C2 154241177 154241360
refGene_gn=SCN11A;refGene_tx=XM_023260719.1
C2 154243791 154244160
refGene_gn=SCN11A;refGene_tx=XM_023260719.1
C2 154246299 154246731
refGene_gn=SCN11A;refGene_tx=XM_023260719.1
C2 154257273 154257387
refGene_gn=SCN11A;refGene_tx=XM_023260719.1
C2 154258121 154258236
refGene_gn=SCN11A;refGene_tx=XM_023260719.1
C2 154260226 154260381
refGene_gn=SCN11A;refGene_tx=XM_023260719.1
C2 154263321 154263495
refGene_gn=SCN11A;refGene_tx=XM_023260719.1
C2 154269195 154269297
refGene_gn=SCN11A;refGene_tx=XM_023260719.1
C2 154269750 154270020
refGene_gn=SCN11A;refGene_tx=XM_023260719.1
C2 154270702 154270756
refGene_gn=SCN11A;refGene_tx=XM_023260719.1
C2 154273397 154273535
refGene_gn=SCN11A;refGene_tx=XM_023260719.1
C2 154281496 154281601
refGene_gn=SCN11A;refGene_tx=XM_023260719.1
C2 154286847 154287118
refGene_gn=SCN11A;refGene_tx=XM_023260719.1

C2 154290047 154291078
refGene_gn=SCN11A;refGene_tx=XM_023260719.1
C2 154333811 154334081
refGene_gn=SCN10A;refGene_tx=XM_023260720.1
C2 154335325 154335453
refGene_gn=SCN10A;refGene_tx=XM_023260720.1
C2 154338487 154338568
refGene_gn=SCN10A;refGene_tx=XM_023260720.1
C2 154358976 154359105
refGene_gn=SCN10A;refGene_tx=XM_023260720.1
C2 154359685 154359777
refGene_gn=SCN10A;refGene_tx=XM_023260720.1
C2 154361785 154362007
refGene_gn=SCN10A;refGene_tx=XM_023260720.1
C2 154362404 154362468
refGene_gn=SCN10A;refGene_tx=XM_023260720.1
C2 154365895 154366037
refGene_gn=SCN10A;refGene_tx=XM_023260720.1
C2 154366181 154366379
refGene_gn=SCN10A;refGene_tx=XM_023260720.1
C2 154367087 154367258
refGene_gn=SCN10A;refGene_tx=XM_023260720.1
C2 154370519 154370813
refGene_gn=SCN10A;refGene_tx=XM_023260720.1
C2 154373226 154373338
refGene_gn=SCN10A;refGene_tx=XM_023260720.1
C2 154383328 154383567
refGene_gn=SCN10A;refGene_tx=XM_023260720.1
C2 154386062 154386236
refGene_gn=SCN10A;refGene_tx=XM_023260720.1
C2 154397637 154397997
refGene_gn=SCN10A;refGene_tx=XM_023260720.1
C2 154399507 154399981
refGene_gn=SCN10A;refGene_tx=XM_023260720.1
C2 154401362 154401497
refGene_gn=SCN10A;refGene_tx=XM_023260720.1
C2 154403133 154403257
refGene_gn=SCN10A;refGene_tx=XM_023260720.1
C2 154404297 154404452
refGene_gn=SCN10A;refGene_tx=XM_023260720.1
C2 154407527 154407701
refGene_gn=SCN10A;refGene_tx=XM_023260720.1
C2 154411655 154411778
refGene_gn=SCN10A;refGene_tx=XM_023260720.1
C2 154413312 154413597
refGene_gn=SCN10A;refGene_tx=XM_023260720.1
C2 154414339 154414393
refGene_gn=SCN10A;refGene_tx=XM_023260720.1
C2 154415258 154415396
refGene_gn=SCN10A;refGene_tx=XM_023260720.1

C2	154417504	154417609
refGene_gn=SCN10A;refGene_tx=XM_023260720.1
C2	154422890	154423161
refGene_gn=SCN10A;refGene_tx=XM_023260720.1
C2	154426916	154428133
refGene_gn=SCN10A;refGene_tx=XM_023260720.1
C2	154497727	154498000
refGene_gn=SCN5A;refGene_tx=XM_023260721.1;refGene_gn=SCN5A;refGene_tx=XM_023260722.1;refGene_gn=SCN5A;refGene_tx=XM_023260723.1;refGene_gn=SCN5A;refGene_tx=XM_023260724.1;refGene_gn=SCN5A;refGene_tx=XM_023260725.1;refGene_gn=SCN5A;refGene_tx=XM_023260726.1;refGene_gn=SCN5A;refGene_tx=XM_023260727.1;refGene_gn=SCN5A;refGene_tx=XM_023260728.1;refGene_gn=SCN5A;refGene_tx=XM_023260729.1;refGene_gn=SCN5A;refGene_tx=XM_023260730.1
C2	154500761	154500880
refGene_gn=SCN5A;refGene_tx=XM_023260721.1;refGene_gn=SCN5A;refGene_tx=XM_023260722.1;refGene_gn=SCN5A;refGene_tx=XM_023260723.1;refGene_gn=SCN5A;refGene_tx=XM_023260724.1;refGene_gn=SCN5A;refGene_tx=XM_023260725.1;refGene_gn=SCN5A;refGene_tx=XM_023260726.1;refGene_gn=SCN5A;refGene_tx=XM_023260727.1;refGene_gn=SCN5A;refGene_tx=XM_023260728.1;refGene_gn=SCN5A;refGene_tx=XM_023260729.1;refGene_gn=SCN5A;refGene_tx=XM_023260730.1
C2	154507957	154508047
refGene_gn=SCN5A;refGene_tx=XM_023260721.1;refGene_gn=SCN5A;refGene_tx=XM_023260722.1;refGene_gn=SCN5A;refGene_tx=XM_023260723.1;refGene_gn=SCN5A;refGene_tx=XM_023260724.1;refGene_gn=SCN5A;refGene_tx=XM_023260725.1;refGene_gn=SCN5A;refGene_tx=XM_023260726.1;refGene_gn=SCN5A;refGene_tx=XM_023260727.1;refGene_gn=SCN5A;refGene_tx=XM_023260728.1;refGene_gn=SCN5A;refGene_tx=XM_023260729.1;refGene_gn=SCN5A;refGene_tx=XM_023260730.1
C2	154509374	154509503
refGene_gn=SCN5A;refGene_tx=XM_023260721.1;refGene_gn=SCN5A;refGene_tx=XM_023260722.1;refGene_gn=SCN5A;refGene_tx=XM_023260723.1;refGene_gn=SCN5A;refGene_tx=XM_023260724.1;refGene_gn=SCN5A;refGene_tx=XM_023260725.1;refGene_gn=SCN5A;refGene_tx=XM_023260726.1;refGene_gn=SCN5A;refGene_tx=XM_023260727.1;refGene_gn=SCN5A;refGene_tx=XM_023260728.1;refGene_gn=SCN5A;refGene_tx=XM_023260729.1;refGene_gn=SCN5A;refGene_tx=XM_023260730.1
C2	154516092	154516184
refGene_gn=SCN5A;refGene_tx=XM_023260722.1;refGene_gn=SCN5A;refGene_tx=XM_023260724.1;refGene_gn=SCN5A;refGene_tx=XM_023260726.1;refGene_gn=SCN5A;refGene_tx=XM_023260728.1
C2	154516318	154516410
refGene_gn=SCN5A;refGene_tx=XM_023260721.1;refGene_gn=SCN5A;refGene_tx=XM_023260723.1;refGene_gn=SCN5A;refGene_tx=XM_023260725.1;refGene_gn=SCN5A;refGene_tx=XM_023260727.1;refGene_gn=SCN5A;refGene_tx=XM_023260729.1;refGene_gn=SCN5A;refGene_tx=XM_023260730.1
C2	154519539	154519770
refGene_gn=SCN5A;refGene_tx=XM_023260721.1;refGene_gn=SCN5A;refGene_tx=XM_023260722.1;refGene_gn=SCN5A;refGene_tx=XM_023260723.1;refGene_gn=

SCN5A;refGene_tx=XM_023260724.1;refGene_gn=SCN5A;refGene_tx=XM_0232607
25.1;refGene_gn=SCN5A;refGene_tx=XM_023260726.1;refGene_gn=SCN5A;refGe
ne_tx=XM_023260727.1;refGene_gn=SCN5A;refGene_tx=XM_023260728.1;refGen
e_gn=SCN5A;refGene_tx=XM_023260729.1;refGene_gn=SCN5A;refGene_tx=XM_02
3260730.1
C2 154521439 154521503
refGene_gn=SCN5A;refGene_tx=XM_023260721.1;refGene_gn=SCN5A;refGene_tx
=XM_023260722.1;refGene_gn=SCN5A;refGene_tx=XM_023260723.1;refGene_gn=
SCN5A;refGene_tx=XM_023260724.1;refGene_gn=SCN5A;refGene_tx=XM_0232607
25.1;refGene_gn=SCN5A;refGene_tx=XM_023260726.1;refGene_gn=SCN5A;refGe
ne_tx=XM_023260727.1;refGene_gn=SCN5A;refGene_tx=XM_023260728.1;refGen
e_gn=SCN5A;refGene_tx=XM_023260729.1;refGene_gn=SCN5A;refGene_tx=XM_02
3260730.1
C2 154522887 154523029
refGene_gn=SCN5A;refGene_tx=XM_023260721.1;refGene_gn=SCN5A;refGene_tx
=XM_023260722.1;refGene_gn=SCN5A;refGene_tx=XM_023260723.1;refGene_gn=
SCN5A;refGene_tx=XM_023260724.1;refGene_gn=SCN5A;refGene_tx=XM_0232607
25.1;refGene_gn=SCN5A;refGene_tx=XM_023260726.1;refGene_gn=SCN5A;refGe
ne_tx=XM_023260727.1;refGene_gn=SCN5A;refGene_tx=XM_023260728.1;refGen
e_gn=SCN5A;refGene_tx=XM_023260729.1;refGene_gn=SCN5A;refGene_tx=XM_02
3260730.1
C2 154523559 154523757
refGene_gn=SCN5A;refGene_tx=XM_023260721.1;refGene_gn=SCN5A;refGene_tx
=XM_023260722.1;refGene_gn=SCN5A;refGene_tx=XM_023260723.1;refGene_gn=
SCN5A;refGene_tx=XM_023260724.1;refGene_gn=SCN5A;refGene_tx=XM_0232607
25.1;refGene_gn=SCN5A;refGene_tx=XM_023260726.1;refGene_gn=SCN5A;refGe
ne_tx=XM_023260727.1;refGene_gn=SCN5A;refGene_tx=XM_023260728.1;refGen
e_gn=SCN5A;refGene_tx=XM_023260729.1;refGene_gn=SCN5A;refGene_tx=XM_02
3260730.1
C2 154525209 154525386
refGene_gn=SCN5A;refGene_tx=XM_023260721.1;refGene_gn=SCN5A;refGene_tx
=XM_023260722.1;refGene_gn=SCN5A;refGene_tx=XM_023260723.1;refGene_gn=
SCN5A;refGene_tx=XM_023260724.1;refGene_gn=SCN5A;refGene_tx=XM_0232607
25.1;refGene_gn=SCN5A;refGene_tx=XM_023260726.1;refGene_gn=SCN5A;refGe
ne_tx=XM_023260727.1;refGene_gn=SCN5A;refGene_tx=XM_023260728.1;refGen
e_gn=SCN5A;refGene_tx=XM_023260729.1;refGene_gn=SCN5A;refGene_tx=XM_02
3260730.1
C2 154526038 154526410
refGene_gn=SCN5A;refGene_tx=XM_023260721.1;refGene_gn=SCN5A;refGene_tx
=XM_023260722.1;refGene_gn=SCN5A;refGene_tx=XM_023260723.1;refGene_gn=
SCN5A;refGene_tx=XM_023260724.1;refGene_gn=SCN5A;refGene_tx=XM_0232607
25.1;refGene_gn=SCN5A;refGene_tx=XM_023260726.1;refGene_gn=SCN5A;refGe
ne_tx=XM_023260727.1;refGene_gn=SCN5A;refGene_tx=XM_023260728.1;refGen
e_gn=SCN5A;refGene_tx=XM_023260729.1;refGene_gn=SCN5A;refGene_tx=XM_02
3260730.1
C2 154531284 154531417
refGene_gn=SCN5A;refGene_tx=XM_023260721.1;refGene_gn=SCN5A;refGene_tx
=XM_023260722.1;refGene_gn=SCN5A;refGene_tx=XM_023260723.1;refGene_gn=
SCN5A;refGene_tx=XM_023260724.1;refGene_gn=SCN5A;refGene_tx=XM_0232607
25.1;refGene_gn=SCN5A;refGene_tx=XM_023260726.1;refGene_gn=SCN5A;refGe

ne_tx=XM_023260727.1;refGene_gn=SCN5A;refGene_tx=XM_023260728.1;refGene_gn=SCN5A;refGene_tx=XM_023260730.1
C2 154532287 154532526
refGene_gn=SCN5A;refGene_tx=XM_023260721.1;refGene_gn=SCN5A;refGene_tx=XM_023260722.1;refGene_gn=SCN5A;refGene_tx=XM_023260723.1;refGene_gn=SCN5A;refGene_tx=XM_023260724.1;refGene_gn=SCN5A;refGene_tx=XM_023260725.1;refGene_gn=SCN5A;refGene_tx=XM_023260726.1;refGene_gn=SCN5A;refGene_tx=XM_023260727.1;refGene_gn=SCN5A;refGene_tx=XM_023260728.1;refGene_gn=SCN5A;refGene_tx=XM_023260730.1
C2 154540517 154540691
refGene_gn=SCN5A;refGene_tx=XM_023260721.1;refGene_gn=SCN5A;refGene_tx=XM_023260722.1;refGene_gn=SCN5A;refGene_tx=XM_023260723.1;refGene_gn=SCN5A;refGene_tx=XM_023260724.1;refGene_gn=SCN5A;refGene_tx=XM_023260725.1;refGene_gn=SCN5A;refGene_tx=XM_023260726.1;refGene_gn=SCN5A;refGene_tx=XM_023260727.1;refGene_gn=SCN5A;refGene_tx=XM_023260728.1;refGene_gn=SCN5A;refGene_tx=XM_023260729.1;refGene_gn=SCN5A;refGene_tx=XM_023260730.1
C2 154541951 154542308
refGene_gn=SCN5A;refGene_tx=XM_023260721.1;refGene_gn=SCN5A;refGene_tx=XM_023260722.1;refGene_gn=SCN5A;refGene_tx=XM_023260723.1;refGene_gn=SCN5A;refGene_tx=XM_023260724.1;refGene_gn=SCN5A;refGene_tx=XM_023260725.1;refGene_gn=SCN5A;refGene_tx=XM_023260726.1;refGene_gn=SCN5A;refGene_tx=XM_023260727.1;refGene_gn=SCN5A;refGene_tx=XM_023260728.1;refGene_gn=SCN5A;refGene_tx=XM_023260729.1;refGene_gn=SCN5A;refGene_tx=XM_023260730.1
C2 154545857 154546304
refGene_gn=SCN5A;refGene_tx=XM_023260721.1;refGene_gn=SCN5A;refGene_tx=XM_023260722.1;refGene_gn=SCN5A;refGene_tx=XM_023260723.1;refGene_gn=SCN5A;refGene_tx=XM_023260724.1;refGene_gn=SCN5A;refGene_tx=XM_023260725.1;refGene_gn=SCN5A;refGene_tx=XM_023260726.1;refGene_gn=SCN5A;refGene_tx=XM_023260727.1;refGene_gn=SCN5A;refGene_tx=XM_023260728.1;refGene_gn=SCN5A;refGene_tx=XM_023260729.1;refGene_gn=SCN5A;refGene_tx=XM_023260730.1
C2 154547586 154547748
refGene_gn=SCN5A;refGene_tx=XM_023260721.1;refGene_gn=SCN5A;refGene_tx=XM_023260722.1;refGene_gn=SCN5A;refGene_tx=XM_023260723.1;refGene_gn=SCN5A;refGene_tx=XM_023260724.1;refGene_gn=SCN5A;refGene_tx=XM_023260725.1;refGene_gn=SCN5A;refGene_tx=XM_023260726.1;refGene_gn=SCN5A;refGene_tx=XM_023260729.1;refGene_gn=SCN5A;refGene_tx=XM_023260730.1
C2 154550265 154550386
refGene_gn=SCN5A;refGene_tx=XM_023260721.1;refGene_gn=SCN5A;refGene_tx=XM_023260722.1;refGene_gn=SCN5A;refGene_tx=XM_023260723.1;refGene_gn=SCN5A;refGene_tx=XM_023260724.1;refGene_gn=SCN5A;refGene_tx=XM_023260725.1;refGene_gn=SCN5A;refGene_tx=XM_023260726.1;refGene_gn=SCN5A;refGene_tx=XM_023260727.1;refGene_gn=SCN5A;refGene_tx=XM_023260728.1;refGene_gn=SCN5A;refGene_tx=XM_023260729.1;refGene_gn=SCN5A;refGene_tx=XM_023260730.1
C2 154551305 154551460
refGene_gn=SCN5A;refGene_tx=XM_023260721.1;refGene_gn=SCN5A;refGene_tx=XM_023260722.1;refGene_gn=SCN5A;refGene_tx=XM_023260723.1;refGene_gn=

SCN5A;refGene_tx=XM_023260724.1;refGene_gn=SCN5A;refGene_tx=XM_0232607
25.1;refGene_gn=SCN5A;refGene_tx=XM_023260726.1;refGene_gn=SCN5A;refGe
ne_tx=XM_023260727.1;refGene_gn=SCN5A;refGene_tx=XM_023260728.1;refGen
e_gn=SCN5A;refGene_tx=XM_023260729.1;refGene_gn=SCN5A;refGene_tx=XM_02
3260730.1
C2 154559798 154559972
refGene_gn=SCN5A;refGene_tx=XM_023260721.1;refGene_gn=SCN5A;refGene_tx
=XM_023260722.1;refGene_gn=SCN5A;refGene_tx=XM_023260723.1;refGene_gn=
SCN5A;refGene_tx=XM_023260724.1;refGene_gn=SCN5A;refGene_tx=XM_0232607
25.1;refGene_gn=SCN5A;refGene_tx=XM_023260726.1;refGene_gn=SCN5A;refGe
ne_tx=XM_023260727.1;refGene_gn=SCN5A;refGene_tx=XM_023260728.1;refGen
e_gn=SCN5A;refGene_tx=XM_023260729.1;refGene_gn=SCN5A;refGene_tx=XM_02
3260730.1
C2 154562923 154563046
refGene_gn=SCN5A;refGene_tx=XM_023260721.1;refGene_gn=SCN5A;refGene_tx
=XM_023260722.1;refGene_gn=SCN5A;refGene_tx=XM_023260723.1;refGene_gn=
SCN5A;refGene_tx=XM_023260724.1;refGene_gn=SCN5A;refGene_tx=XM_0232607
25.1;refGene_gn=SCN5A;refGene_tx=XM_023260726.1;refGene_gn=SCN5A;refGe
ne_tx=XM_023260727.1;refGene_gn=SCN5A;refGene_tx=XM_023260728.1;refGen
e_gn=SCN5A;refGene_tx=XM_023260729.1;refGene_gn=SCN5A;refGene_tx=XM_02
3260730.1
C2 154565540 154565822
refGene_gn=SCN5A;refGene_tx=XM_023260721.1;refGene_gn=SCN5A;refGene_tx
=XM_023260722.1;refGene_gn=SCN5A;refGene_tx=XM_023260723.1;refGene_gn=
SCN5A;refGene_tx=XM_023260724.1;refGene_gn=SCN5A;refGene_tx=XM_0232607
25.1;refGene_gn=SCN5A;refGene_tx=XM_023260726.1;refGene_gn=SCN5A;refGe
ne_tx=XM_023260727.1;refGene_gn=SCN5A;refGene_tx=XM_023260728.1;refGen
e_gn=SCN5A;refGene_tx=XM_023260729.1;refGene_gn=SCN5A;refGene_tx=XM_02
3260730.1
C2 154567789 154567843
refGene_gn=SCN5A;refGene_tx=XM_023260721.1;refGene_gn=SCN5A;refGene_tx
=XM_023260722.1;refGene_gn=SCN5A;refGene_tx=XM_023260723.1;refGene_gn=
SCN5A;refGene_tx=XM_023260724.1;refGene_gn=SCN5A;refGene_tx=XM_0232607
25.1;refGene_gn=SCN5A;refGene_tx=XM_023260726.1;refGene_gn=SCN5A;refGe
ne_tx=XM_023260727.1;refGene_gn=SCN5A;refGene_tx=XM_023260728.1;refGen
e_gn=SCN5A;refGene_tx=XM_023260729.1;refGene_gn=SCN5A;refGene_tx=XM_02
3260730.1
C2 154568452 154568590
refGene_gn=SCN5A;refGene_tx=XM_023260721.1;refGene_gn=SCN5A;refGene_tx
=XM_023260722.1;refGene_gn=SCN5A;refGene_tx=XM_023260723.1;refGene_gn=
SCN5A;refGene_tx=XM_023260724.1;refGene_gn=SCN5A;refGene_tx=XM_0232607
25.1;refGene_gn=SCN5A;refGene_tx=XM_023260726.1;refGene_gn=SCN5A;refGe
ne_tx=XM_023260727.1;refGene_gn=SCN5A;refGene_tx=XM_023260728.1;refGen
e_gn=SCN5A;refGene_tx=XM_023260729.1;refGene_gn=SCN5A;refGene_tx=XM_02
3260730.1
C2 154569225 154569330
refGene_gn=SCN5A;refGene_tx=XM_023260721.1;refGene_gn=SCN5A;refGene_tx
=XM_023260722.1;refGene_gn=SCN5A;refGene_tx=XM_023260723.1;refGene_gn=
SCN5A;refGene_tx=XM_023260724.1;refGene_gn=SCN5A;refGene_tx=XM_0232607
25.1;refGene_gn=SCN5A;refGene_tx=XM_023260726.1;refGene_gn=SCN5A;refGe

ne_tx=XM_023260727.1;refGene_gn=SCN5A;refGene_tx=XM_023260728.1;refGene_gn=SCN5A;refGene_tx=XM_023260729.1;refGene_gn=SCN5A;refGene_tx=XM_023260730.1
C2 154570861 154571132
refGene_gn=SCN5A;refGene_tx=XM_023260721.1;refGene_gn=SCN5A;refGene_tx=XM_023260722.1;refGene_gn=SCN5A;refGene_tx=XM_023260723.1;refGene_gn=SCN5A;refGene_tx=XM_023260724.1;refGene_gn=SCN5A;refGene_tx=XM_023260725.1;refGene_gn=SCN5A;refGene_tx=XM_023260726.1;refGene_gn=SCN5A;refGene_tx=XM_023260727.1;refGene_gn=SCN5A;refGene_tx=XM_023260728.1;refGene_gn=SCN5A;refGene_tx=XM_023260729.1;refGene_gn=SCN5A;refGene_tx=XM_023260730.1
C2 154573900 154575132
refGene_gn=SCN5A;refGene_tx=XM_023260721.1;refGene_gn=SCN5A;refGene_tx=XM_023260722.1;refGene_gn=SCN5A;refGene_tx=XM_023260723.1;refGene_gn=SCN5A;refGene_tx=XM_023260724.1;refGene_gn=SCN5A;refGene_tx=XM_023260725.1;refGene_gn=SCN5A;refGene_tx=XM_023260726.1;refGene_gn=SCN5A;refGene_tx=XM_023260727.1;refGene_gn=SCN5A;refGene_tx=XM_023260728.1;refGene_gn=SCN5A;refGene_tx=XM_023260729.1
C2 154577115 154577153
refGene_gn=SCN5A;refGene_tx=XM_023260730.1
C2 154598007 154598469
refGene_gn=EXOG;refGene_tx=XM_003992229.5
C2 154612975 154613090
refGene_gn=EXOG;refGene_tx=XM_003992229.5
C2 154615578 154615655
refGene_gn=EXOG;refGene_tx=XM_003992229.5
C2 154617392 154617532
refGene_gn=EXOG;refGene_tx=XM_003992229.5
C2 154620986 154621136
refGene_gn=EXOG;refGene_tx=XM_003992229.5
C2 154623586 154623749
refGene_gn=EXOG;refGene_tx=XM_003992229.5
C2 154637367 154637562
refGene_gn=ACVR2B;refGene_tx=XM_006936529.4;refGene_gn=ACVR2B;refGene_tx=XM_006936530.4;refGene_gn=ACVR2B;refGene_tx=XM_023260731.1
C2 154639165 154639296
refGene_gn=ACVR2B;refGene_tx=XM_006936529.4;refGene_gn=ACVR2B;refGene_tx=XM_006936530.4;refGene_gn=ACVR2B;refGene_tx=XM_023260731.1
C2 154639391 154639530
refGene_gn=ACVR2B;refGene_tx=XM_006936529.4;refGene_gn=ACVR2B;refGene_tx=XM_006936530.4;refGene_gn=ACVR2B;refGene_tx=XM_023260731.1
C2 154640243 154640358
refGene_gn=ACVR2B;refGene_tx=XM_006936529.4;refGene_gn=ACVR2B;refGene_tx=XM_006936530.4;refGene_gn=ACVR2B;refGene_tx=XM_023260731.1
C2 154642427 154642576
refGene_gn=ACVR2B;refGene_tx=XM_006936529.4;refGene_gn=ACVR2B;refGene_tx=XM_006936530.4;refGene_gn=ACVR2B;refGene_tx=XM_023260731.1
C2 154642859 154643003
refGene_gn=ACVR2B;refGene_tx=XM_006936529.4;refGene_gn=ACVR2B;refGene_tx=XM_006936530.4;refGene_gn=ACVR2B;refGene_tx=XM_023260731.1

C2 154643543 154643687
refGene_gn=ACVR2B;refGene_tx=XM_006936529.4;refGene_gn=ACVR2B;refGene_tx=XM_006936530.4;refGene_gn=ACVR2B;refGene_tx=XM_023260731.1
C2 154643775 154643927
refGene_gn=ACVR2B;refGene_tx=XM_006936529.4;refGene_gn=ACVR2B;refGene_tx=XM_006936530.4;refGene_gn=ACVR2B;refGene_tx=XM_023260731.1
C2 154644088 154644198
refGene_gn=ACVR2B;refGene_tx=XM_006936529.4;refGene_gn=ACVR2B;refGene_tx=XM_006936530.4;refGene_gn=ACVR2B;refGene_tx=XM_023260731.1
C2 154644598 154644806
refGene_gn=ACVR2B;refGene_tx=XM_006936529.4;refGene_gn=ACVR2B;refGene_tx=XM_006936530.4;refGene_gn=ACVR2B;refGene_tx=XM_023260731.1
C2 154661783 154661868
refGene_gn=ACVR2B;refGene_tx=XM_006936529.4;refGene_gn=ACVR2B;refGene_tx=XM_006936530.4
C2 154667115 154667167
refGene_gn=ACVR2B;refGene_tx=XM_023260731.1
C2 154683775 154683904
refGene_gn=XYLB;refGene_tx=XM_023260733.1
C2 154688198 154688293
refGene_gn=XYLB;refGene_tx=XM_023260733.1
C2 154691115 154691203
refGene_gn=XYLB;refGene_tx=XM_023260733.1
C2 154692687 154692746
refGene_gn=XYLB;refGene_tx=XM_023260733.1
C2 154707007 154707104
refGene_gn=XYLB;refGene_tx=XM_023260733.1
C2 154708604 154708678
refGene_gn=XYLB;refGene_tx=XM_023260733.1
C2 154709339 154709455
refGene_gn=XYLB;refGene_tx=XM_023260733.1
C2 154710189 154710305
refGene_gn=XYLB;refGene_tx=XM_023260733.1
C2 154710838 154710879
refGene_gn=XYLB;refGene_tx=XM_023260733.1
C2 154714353 154714435
refGene_gn=XYLB;refGene_tx=XM_023260733.1
C2 154718210 154718329
refGene_gn=XYLB;refGene_tx=XM_023260733.1
C2 154720130 154720203
refGene_gn=XYLB;refGene_tx=XM_023260733.1
C2 154721527 154721593
refGene_gn=XYLB;refGene_tx=XM_023260733.1
C2 154722368 154722497
refGene_gn=XYLB;refGene_tx=XM_023260733.1
C2 154723124 154723211
refGene_gn=XYLB;refGene_tx=XM_023260733.1
C2 154726301 154726382
refGene_gn=XYLB;refGene_tx=XM_023260733.1
C2 154728131 154728201

refGene_gn=XYLB;refGene_tx=XM_023260733.1
C2 154739854 154739937
refGene_gn=XYLB;refGene_tx=XM_023260733.1
C2 154740637 154740694
refGene_gn=XYLB;refGene_tx=XM_023260733.1
C2 154746355 154746384
refGene_gn=LOC111562244;refGene_tx=XM_023260189.1
C2 154754991 154755493
refGene_gn=LOC111562244;refGene_tx=XM_023260189.1
C2 154756262 154756424
refGene_gn=LOC111562244;refGene_tx=XM_023260189.1
C2 154760785 154760821
refGene_gn=LOC111562244;refGene_tx=XM_023260189.1
C2 154761253 154761422
refGene_gn=SLC22A14;refGene_tx=XM_019810930.2;refGene_gn=SLC22A14;refGene_tx=XM_019810931.2;refGene_gn=SLC22A14;refGene_tx=XM_023260734.1;refGene_gn=SLC22A14;refGene_tx=XM_023260735.1;refGene_gn=SLC22A14;refGene_tx=XM_023260736.1;refGene_gn=SLC22A14;refGene_tx=XM_023260737.1;refGene_gn=SLC22A14;refGene_tx=XM_023260738.1;refGene_gn=SLC22A14;refGene_tx=XM_023260739.1
C2 154762746 154762950
refGene_gn=SLC22A14;refGene_tx=XM_019810930.2;refGene_gn=SLC22A14;refGene_tx=XM_019810931.2;refGene_gn=SLC22A14;refGene_tx=XM_023260734.1;refGene_gn=SLC22A14;refGene_tx=XM_023260735.1;refGene_gn=SLC22A14;refGene_tx=XM_023260736.1;refGene_gn=SLC22A14;refGene_tx=XM_023260737.1;refGene_gn=SLC22A14;refGene_tx=XM_023260738.1;refGene_gn=SLC22A14;refGene_tx=XM_023260739.1
C2 154763381 154763406
refGene_gn=SLC22A14;refGene_tx=XM_019810933.2
C2 154763552 154763661
refGene_gn=SLC22A14;refGene_tx=XM_019810930.2;refGene_gn=SLC22A14;refGene_tx=XM_019810931.2;refGene_gn=SLC22A14;refGene_tx=XM_019810933.2;refGene_gn=SLC22A14;refGene_tx=XM_023260734.1;refGene_gn=SLC22A14;refGene_tx=XM_023260735.1;refGene_gn=SLC22A14;refGene_tx=XM_023260736.1;refGene_gn=SLC22A14;refGene_tx=XM_023260737.1;refGene_gn=SLC22A14;refGene_tx=XM_023260738.1;refGene_gn=SLC22A14;refGene_tx=XM_023260739.1
C2 154765832 154766047
refGene_gn=SLC22A14;refGene_tx=XM_019810930.2;refGene_gn=SLC22A14;refGene_tx=XM_019810931.2;refGene_gn=SLC22A14;refGene_tx=XM_019810933.2;refGene_gn=SLC22A14;refGene_tx=XM_023260734.1;refGene_gn=SLC22A14;refGene_tx=XM_023260735.1;refGene_gn=SLC22A14;refGene_tx=XM_023260736.1;refGene_gn=SLC22A14;refGene_tx=XM_023260737.1;refGene_gn=SLC22A14;refGene_tx=XM_023260738.1;refGene_gn=SLC22A14;refGene_tx=XM_023260739.1
C2 154766514 154766612
refGene_gn=SLC22A14;refGene_tx=XM_019810930.2;refGene_gn=SLC22A14;refGene_tx=XM_019810931.2;refGene_gn=SLC22A14;refGene_tx=XM_019810933.2;refGene_gn=SLC22A14;refGene_tx=XM_023260734.1;refGene_gn=SLC22A14;refGene_tx=XM_023260735.1;refGene_gn=SLC22A14;refGene_tx=XM_023260736.1;refGene_gn=SLC22A14;refGene_tx=XM_023260737.1;refGene_gn=SLC22A14;refGene_tx=XM_023260738.1;refGene_gn=SLC22A14;refGene_tx=XM_023260739.1

C2	154766893	154767014
refGene_gn=SLC22A14;refGene_tx=XM_019810930.2;refGene_gn=SLC22A14;refGene_tx=XM_019810931.2;refGene_gn=SLC22A14;refGene_tx=XM_019810933.2;refGene_gn=SLC22A14;refGene_tx=XM_023260734.1;refGene_gn=SLC22A14;refGene_tx=XM_023260735.1;refGene_gn=SLC22A14;refGene_tx=XM_023260736.1;refGene_gn=SLC22A14;refGene_tx=XM_023260737.1;refGene_gn=SLC22A14;refGene_tx=XM_023260738.1;refGene_gn=SLC22A14;refGene_tx=XM_023260739.1
C2	154770538	154770707
refGene_gn=SLC22A14;refGene_tx=XM_019810930.2;refGene_gn=SLC22A14;refGene_tx=XM_019810931.2;refGene_gn=SLC22A14;refGene_tx=XM_019810933.2;refGene_gn=SLC22A14;refGene_tx=XM_023260734.1;refGene_gn=SLC22A14;refGene_tx=XM_023260735.1;refGene_gn=SLC22A14;refGene_tx=XM_023260736.1;refGene_gn=SLC22A14;refGene_tx=XM_023260737.1;refGene_gn=SLC22A14;refGene_tx=XM_023260738.1;refGene_gn=SLC22A14;refGene_tx=XM_023260739.1
C2	154771733	154771888
refGene_gn=SLC22A14;refGene_tx=XM_019810930.2;refGene_gn=SLC22A14;refGene_tx=XM_019810931.2;refGene_gn=SLC22A14;refGene_tx=XM_019810933.2;refGene_gn=SLC22A14;refGene_tx=XM_023260734.1;refGene_gn=SLC22A14;refGene_tx=XM_023260735.1;refGene_gn=SLC22A14;refGene_tx=XM_023260736.1;refGene_gn=SLC22A14;refGene_tx=XM_023260737.1;refGene_gn=SLC22A14;refGene_tx=XM_023260738.1;refGene_gn=SLC22A14;refGene_tx=XM_023260739.1
C2	154772029	154772133
refGene_gn=SLC22A14;refGene_tx=XM_019810930.2;refGene_gn=SLC22A14;refGene_tx=XM_019810931.2;refGene_gn=SLC22A14;refGene_tx=XM_019810933.2;refGene_gn=SLC22A14;refGene_tx=XM_023260734.1;refGene_gn=SLC22A14;refGene_tx=XM_023260735.1;refGene_gn=SLC22A14;refGene_tx=XM_023260736.1;refGene_gn=SLC22A14;refGene_tx=XM_023260737.1;refGene_gn=SLC22A14;refGene_tx=XM_023260738.1;refGene_gn=SLC22A14;refGene_tx=XM_023260739.1
C2	154772873	154773389
refGene_gn=SLC22A14;refGene_tx=XM_019810930.2;refGene_gn=SLC22A14;refGene_tx=XM_019810931.2;refGene_gn=SLC22A14;refGene_tx=XM_019810933.2;refGene_gn=SLC22A14;refGene_tx=XM_023260734.1;refGene_gn=SLC22A14;refGene_tx=XM_023260735.1;refGene_gn=SLC22A14;refGene_tx=XM_023260736.1;refGene_gn=SLC22A14;refGene_tx=XM_023260737.1;refGene_gn=SLC22A14;refGene_tx=XM_023260738.1;refGene_gn=SLC22A14;refGene_tx=XM_023260739.1
C2	154799002	154799096
refGene_gn=SLC22A13;refGene_tx=XM_003992231.2
C2	154799299	154799515
refGene_gn=SLC22A13;refGene_tx=XM_003992231.2
C2	154799766	154799902
refGene_gn=SLC22A13;refGene_tx=XM_019810934.2
C2	154800056	154800165
refGene_gn=SLC22A13;refGene_tx=XM_003992231.2;refGene_gn=SLC22A13;refGene_tx=XM_019810934.2
C2	154800341	154800556
refGene_gn=SLC22A13;refGene_tx=XM_003992231.2;refGene_gn=SLC22A13;refGene_tx=XM_019810934.2
C2	154800792	154800887
refGene_gn=SLC22A13;refGene_tx=XM_003992231.2;refGene_gn=SLC22A13;refGene_tx=XM_019810934.2

C2 154800978 154801099
refGene_gn=SLC22A13;refGene_tx=XM_003992231.2;refGene_gn=SLC22A13;refGene_tx=XM_019810934.2
C2 154801323 154801492
refGene_gn=SLC22A13;refGene_tx=XM_003992231.2;refGene_gn=SLC22A13;refGene_tx=XM_019810934.2
C2 154801748 154801903
refGene_gn=SLC22A13;refGene_tx=XM_003992231.2;refGene_gn=SLC22A13;refGene_tx=XM_019810934.2
C2 154802126 154802230
refGene_gn=SLC22A13;refGene_tx=XM_003992231.2;refGene_gn=SLC22A13;refGene_tx=XM_019810934.2
C2 154807544 154807922
refGene_gn=SLC22A13;refGene_tx=XM_003992231.2;refGene_gn=SLC22A13;refGene_tx=XM_019810934.2
C2 154840973 154841048
refGene_gn=OXSR1;refGene_tx=XM_011286239.3;refGene_gn=OXSR1;refGene_tx=XM_019810939.2
C2 154841686 154841751
refGene_gn=OXSR1;refGene_tx=XM_011286239.3;refGene_gn=OXSR1;refGene_tx=XM_019810939.2
C2 154842620 154842689
refGene_gn=OXSR1;refGene_tx=XM_011286239.3;refGene_gn=OXSR1;refGene_tx=XM_019810939.2
C2 154844238 154844291
refGene_gn=OXSR1;refGene_tx=XM_011286239.3;refGene_gn=OXSR1;refGene_tx=XM_019810939.2
C2 154845431 154845496
refGene_gn=OXSR1;refGene_tx=XM_011286239.3;refGene_gn=OXSR1;refGene_tx=XM_019810939.2
C2 154847141 154847288
refGene_gn=OXSR1;refGene_tx=XM_011286239.3;refGene_gn=OXSR1;refGene_tx=XM_019810939.2
C2 154850848 154850884
refGene_gn=OXSR1;refGene_tx=XM_011286239.3;refGene_gn=OXSR1;refGene_tx=XM_019810939.2
C2 154855481 154855604
refGene_gn=OXSR1;refGene_tx=XM_011286239.3;refGene_gn=OXSR1;refGene_tx=XM_019810939.2
C2 154858431 154858497
refGene_gn=OXSR1;refGene_tx=XM_011286239.3;refGene_gn=OXSR1;refGene_tx=XM_019810939.2
C2 154859166 154859215
refGene_gn=OXSR1;refGene_tx=XM_011286239.3;refGene_gn=OXSR1;refGene_tx=XM_019810939.2
C2 154865838 154865972
refGene_gn=OXSR1;refGene_tx=XM_011286239.3;refGene_gn=OXSR1;refGene_tx=XM_019810939.2
C2 154868737 154868839
refGene_gn=OXSR1;refGene_tx=XM_011286239.3;refGene_gn=OXSR1;refGene_tx

=XM_019810939.2
C2	154872942	154873052
refGene_gn=OXSR1;refGene_tx=XM_011286239.3;refGene_gn=OXSR1;refGene_tx
=XM_019810939.2
C2	154887184	154887240
refGene_gn=OXSR1;refGene_tx=XM_011286239.3;refGene_gn=OXSR1;refGene_tx
=XM_019810939.2
C2	154900528	154900670
refGene_gn=OXSR1;refGene_tx=XM_011286239.3;refGene_gn=OXSR1;refGene_tx
=XM_019810939.2
C2	154907168	154907259
refGene_gn=OXSR1;refGene_tx=XM_019810939.2
C2	154909634	154909743
refGene_gn=OXSR1;refGene_tx=XM_011286239.3
C2	154920410	154920523
refGene_gn=OXSR1;refGene_tx=XM_011286239.3
C2	154944713	154944783
refGene_gn=OXSR1;refGene_tx=XM_011286239.3
C2	154972855	154973010
refGene_gn=MYD88;refGene_tx=XM_003992253.4
C2	154973286	154973378
refGene_gn=MYD88;refGene_tx=XM_003992253.4
C2	154973558	154973739
refGene_gn=MYD88;refGene_tx=XM_003992253.4
C2	154974082	154974217
refGene_gn=MYD88;refGene_tx=XM_003992253.4
C2	154975048	154975376
refGene_gn=MYD88;refGene_tx=XM_003992253.4
C2	154977080	154977257
refGene_gn=ACAA1;refGene_tx=XM_003992254.4
C2	154977441	154977535
refGene_gn=ACAA1;refGene_tx=XM_003992254.4
C2	154978510	154978568
refGene_gn=ACAA1;refGene_tx=XM_003992254.4
C2	154980620	154980700
refGene_gn=ACAA1;refGene_tx=XM_003992254.4
C2	154981002	154981045
refGene_gn=ACAA1;refGene_tx=XM_003992254.4
C2	154983105	154983204
refGene_gn=ACAA1;refGene_tx=XM_003992254.4
C2	154984048	154984129
refGene_gn=ACAA1;refGene_tx=XM_003992254.4
C2	154985276	154985467
refGene_gn=ACAA1;refGene_tx=XM_003992254.4
C2	154985611	154985791
refGene_gn=ACAA1;refGene_tx=XM_003992254.4
C2	154986077	154986133
refGene_gn=ACAA1;refGene_tx=XM_003992254.4
C2	154986245	154986391
refGene_gn=ACAA1;refGene_tx=XM_003992254.4

C2	154986660	154986736
refGene_gn=ACAA1;refGene_tx=XM_003992254.4
C2	154987193	154987317
refGene_gn=DLEC1;refGene_tx=XM_019810935.2;refGene_gn=DLEC1;refGene_tx=XM_019810936.2;refGene_gn=DLEC1;refGene_tx=XM_023260740.1;refGene_gn=DLEC1;refGene_tx=XM_023260741.1
C2	154987402	154987526
refGene_gn=DLEC1;refGene_tx=XM_019810935.2;refGene_gn=DLEC1;refGene_tx=XM_019810936.2;refGene_gn=DLEC1;refGene_tx=XM_023260740.1;refGene_gn=DLEC1;refGene_tx=XM_023260741.1
C2	154987702	154987856
refGene_gn=DLEC1;refGene_tx=XM_019810935.2;refGene_gn=DLEC1;refGene_tx=XM_019810936.2;refGene_gn=DLEC1;refGene_tx=XM_023260740.1;refGene_gn=DLEC1;refGene_tx=XM_023260741.1
C2	154988589	154988751
refGene_gn=DLEC1;refGene_tx=XM_019810935.2;refGene_gn=DLEC1;refGene_tx=XM_019810936.2;refGene_gn=DLEC1;refGene_tx=XM_023260740.1;refGene_gn=DLEC1;refGene_tx=XM_023260741.1
C2	154989917	154990130
refGene_gn=DLEC1;refGene_tx=XM_019810935.2;refGene_gn=DLEC1;refGene_tx=XM_019810936.2;refGene_gn=DLEC1;refGene_tx=XM_023260740.1;refGene_gn=DLEC1;refGene_tx=XM_023260741.1
C2	154990294	154990379
refGene_gn=DLEC1;refGene_tx=XM_019810935.2;refGene_gn=DLEC1;refGene_tx=XM_019810936.2;refGene_gn=DLEC1;refGene_tx=XM_023260740.1;refGene_gn=DLEC1;refGene_tx=XM_023260741.1
C2	154990529	154990624
refGene_gn=DLEC1;refGene_tx=XM_019810935.2;refGene_gn=DLEC1;refGene_tx=XM_019810936.2;refGene_gn=DLEC1;refGene_tx=XM_023260740.1;refGene_gn=DLEC1;refGene_tx=XM_023260741.1
C2	154990707	154990833
refGene_gn=DLEC1;refGene_tx=XM_019810935.2;refGene_gn=DLEC1;refGene_tx=XM_019810936.2;refGene_gn=DLEC1;refGene_tx=XM_023260740.1;refGene_gn=DLEC1;refGene_tx=XM_023260741.1
C2	154990922	154991039
refGene_gn=DLEC1;refGene_tx=XM_019810935.2;refGene_gn=DLEC1;refGene_tx=XM_019810936.2;refGene_gn=DLEC1;refGene_tx=XM_023260740.1;refGene_gn=DLEC1;refGene_tx=XM_023260741.1
C2	154991152	154991358
refGene_gn=DLEC1;refGene_tx=XM_019810935.2;refGene_gn=DLEC1;refGene_tx=XM_019810936.2;refGene_gn=DLEC1;refGene_tx=XM_023260740.1;refGene_gn=DLEC1;refGene_tx=XM_023260741.1
C2	154993672	154993743
refGene_gn=DLEC1;refGene_tx=XM_019810935.2;refGene_gn=DLEC1;refGene_tx=XM_019810936.2;refGene_gn=DLEC1;refGene_tx=XM_023260740.1;refGene_gn=DLEC1;refGene_tx=XM_023260741.1
C2	154994273	154994392
refGene_gn=DLEC1;refGene_tx=XM_019810935.2;refGene_gn=DLEC1;refGene_tx=XM_019810936.2;refGene_gn=DLEC1;refGene_tx=XM_023260740.1;refGene_gn=DLEC1;refGene_tx=XM_023260741.1

C2 154995010 154995162
refGene_gn=DLEC1;refGene_tx=XM_019810935.2;refGene_gn=DLEC1;refGene_tx=XM_019810936.2;refGene_gn=DLEC1;refGene_tx=XM_023260740.1;refGene_gn=DLEC1;refGene_tx=XM_023260741.1
C2 154995598 154995669
refGene_gn=DLEC1;refGene_tx=XM_019810935.2;refGene_gn=DLEC1;refGene_tx=XM_019810936.2;refGene_gn=DLEC1;refGene_tx=XM_023260740.1;refGene_gn=DLEC1;refGene_tx=XM_023260741.1
C2 154996853 154997045
refGene_gn=DLEC1;refGene_tx=XM_019810935.2;refGene_gn=DLEC1;refGene_tx=XM_019810936.2;refGene_gn=DLEC1;refGene_tx=XM_023260740.1;refGene_gn=DLEC1;refGene_tx=XM_023260741.1
C2 154997529 154997660
refGene_gn=DLEC1;refGene_tx=XM_019810935.2;refGene_gn=DLEC1;refGene_tx=XM_019810936.2;refGene_gn=DLEC1;refGene_tx=XM_023260740.1;refGene_gn=DLEC1;refGene_tx=XM_023260741.1
C2 154998519 154998630
refGene_gn=DLEC1;refGene_tx=XM_019810935.2;refGene_gn=DLEC1;refGene_tx=XM_019810936.2;refGene_gn=DLEC1;refGene_tx=XM_023260740.1;refGene_gn=DLEC1;refGene_tx=XM_023260741.1
C2 154999022 154999176
refGene_gn=DLEC1;refGene_tx=XM_019810935.2;refGene_gn=DLEC1;refGene_tx=XM_019810936.2;refGene_gn=DLEC1;refGene_tx=XM_023260740.1;refGene_gn=DLEC1;refGene_tx=XM_023260741.1
C2 155012509 155012649
refGene_gn=DLEC1;refGene_tx=XM_019810935.2;refGene_gn=DLEC1;refGene_tx=XM_019810936.2;refGene_gn=DLEC1;refGene_tx=XM_023260740.1;refGene_gn=DLEC1;refGene_tx=XM_023260741.1
C2 155015097 155015256
refGene_gn=DLEC1;refGene_tx=XM_019810935.2;refGene_gn=DLEC1;refGene_tx=XM_019810936.2;refGene_gn=DLEC1;refGene_tx=XM_023260740.1
C2 155015365 155015496
refGene_gn=DLEC1;refGene_tx=XM_019810935.2;refGene_gn=DLEC1;refGene_tx=XM_019810936.2;refGene_gn=DLEC1;refGene_tx=XM_023260740.1
C2 155015707 155015801
refGene_gn=DLEC1;refGene_tx=XM_019810935.2;refGene_gn=DLEC1;refGene_tx=XM_019810936.2;refGene_gn=DLEC1;refGene_tx=XM_023260740.1
C2 155016173 155016342
refGene_gn=DLEC1;refGene_tx=XM_019810935.2;refGene_gn=DLEC1;refGene_tx=XM_019810936.2;refGene_gn=DLEC1;refGene_tx=XM_023260740.1;refGene_gn=DLEC1;refGene_tx=XM_023260741.1
C2 155016956 155017015
refGene_gn=DLEC1;refGene_tx=XM_019810935.2;refGene_gn=DLEC1;refGene_tx=XM_019810936.2;refGene_gn=DLEC1;refGene_tx=XM_023260740.1;refGene_gn=DLEC1;refGene_tx=XM_023260741.1
C2 155017802 155017995
refGene_gn=DLEC1;refGene_tx=XM_019810935.2;refGene_gn=DLEC1;refGene_tx=XM_019810936.2;refGene_gn=DLEC1;refGene_tx=XM_023260740.1;refGene_gn=DLEC1;refGene_tx=XM_023260741.1
C2 155018905 155019068

refGene_gn=DLEC1;refGene_tx=XM_019810935.2;refGene_gn=DLEC1;refGene_tx
=XM_019810936.2;refGene_gn=DLEC1;refGene_tx=XM_023260740.1;refGene_gn=
DLEC1;refGene_tx=XM_023260741.1
C2 155019620 155019711
refGene_gn=DLEC1;refGene_tx=XM_019810935.2;refGene_gn=DLEC1;refGene_tx
=XM_019810936.2;refGene_gn=DLEC1;refGene_tx=XM_023260740.1;refGene_gn=
DLEC1;refGene_tx=XM_023260741.1
C2 155023119 155023212
refGene_gn=DLEC1;refGene_tx=XM_019810935.2;refGene_gn=DLEC1;refGene_tx
=XM_019810936.2;refGene_gn=DLEC1;refGene_tx=XM_023260740.1;refGene_gn=
DLEC1;refGene_tx=XM_023260741.1
C2 155024617 155024754
refGene_gn=DLEC1;refGene_tx=XM_019810935.2;refGene_gn=DLEC1;refGene_tx
=XM_019810936.2;refGene_gn=DLEC1;refGene_tx=XM_023260740.1;refGene_gn=
DLEC1;refGene_tx=XM_023260741.1
C2 155025394 155025568
refGene_gn=DLEC1;refGene_tx=XM_019810935.2;refGene_gn=DLEC1;refGene_tx
=XM_019810936.2;refGene_gn=DLEC1;refGene_tx=XM_023260740.1;refGene_gn=
DLEC1;refGene_tx=XM_023260741.1
C2 155026308 155026396
refGene_gn=DLEC1;refGene_tx=XM_019810935.2;refGene_gn=DLEC1;refGene_tx
=XM_019810936.2;refGene_gn=DLEC1;refGene_tx=XM_023260740.1;refGene_gn=
DLEC1;refGene_tx=XM_023260741.1
C2 155050719 155050798
refGene_gn=DLEC1;refGene_tx=XM_019810935.2;refGene_gn=DLEC1;refGene_tx
=XM_019810936.2;refGene_gn=DLEC1;refGene_tx=XM_023260740.1;refGene_gn=
DLEC1;refGene_tx=XM_023260741.1
C2 155051867 155052088
refGene_gn=DLEC1;refGene_tx=XM_019810935.2;refGene_gn=DLEC1;refGene_tx
=XM_019810936.2;refGene_gn=DLEC1;refGene_tx=XM_023260740.1;refGene_gn=
DLEC1;refGene_tx=XM_023260741.1
C2 155052315 155052506
refGene_gn=DLEC1;refGene_tx=XM_019810935.2;refGene_gn=DLEC1;refGene_tx
=XM_019810936.2;refGene_gn=DLEC1;refGene_tx=XM_023260740.1;refGene_gn=
DLEC1;refGene_tx=XM_023260741.1
C2 155054057 155054168
refGene_gn=DLEC1;refGene_tx=XM_019810935.2;refGene_gn=DLEC1;refGene_tx
=XM_019810936.2;refGene_gn=DLEC1;refGene_tx=XM_023260740.1;refGene_gn=
DLEC1;refGene_tx=XM_023260741.1
C2 155077203 155077354
refGene_gn=DLEC1;refGene_tx=XM_019810935.2;refGene_gn=DLEC1;refGene_tx
=XM_019810936.2;refGene_gn=DLEC1;refGene_tx=XM_023260740.1;refGene_gn=
DLEC1;refGene_tx=XM_023260741.1
C2 155078381 155078780
refGene_gn=DLEC1;refGene_tx=XM_019810935.2;refGene_gn=DLEC1;refGene_tx
=XM_023260740.1;refGene_gn=DLEC1;refGene_tx=XM_023260741.1
C2 155087409 155087443
refGene_gn=PLCD1;refGene_tx=XM_011286238.3;refGene_gn=PLCD1;refGene_tx
=XM_023260743.1
C2 155092046 155092143

refGene_gn=PLCD1;refGene_tx=XM_011286235.3
C2 155095564 155095729
refGene_gn=PLCD1;refGene_tx=XM_011286235.3;refGene_gn=PLCD1;refGene_tx=XM_011286238.3;refGene_gn=PLCD1;refGene_tx=XM_023260743.1
C2 155099136 155099365
refGene_gn=PLCD1;refGene_tx=XM_011286235.3;refGene_gn=PLCD1;refGene_tx=XM_011286238.3;refGene_gn=PLCD1;refGene_tx=XM_019810946.2;refGene_gn=PLCD1;refGene_tx=XM_023260743.1
C2 155102867 155102894
refGene_gn=PLCD1;refGene_tx=XM_011286237.3;refGene_gn=PLCD1;refGene_tx=XM_019810947.2
C2 155104791 155104934
refGene_gn=PLCD1;refGene_tx=XM_011286237.3;refGene_gn=PLCD1;refGene_tx=XM_019810947.2
C2 155105958 155106088
refGene_gn=PLCD1;refGene_tx=XM_011286235.3;refGene_gn=PLCD1;refGene_tx=XM_011286237.3;refGene_gn=PLCD1;refGene_tx=XM_011286238.3;refGene_gn=PLCD1;refGene_tx=XM_019810946.2;refGene_gn=PLCD1;refGene_tx=XM_019810947.2;refGene_gn=PLCD1;refGene_tx=XM_023260743.1
C2 155106181 155106413
refGene_gn=PLCD1;refGene_tx=XM_011286235.3;refGene_gn=PLCD1;refGene_tx=XM_011286237.3;refGene_gn=PLCD1;refGene_tx=XM_011286238.3;refGene_gn=PLCD1;refGene_tx=XM_019810946.2;refGene_gn=PLCD1;refGene_tx=XM_019810947.2;refGene_gn=PLCD1;refGene_tx=XM_023260743.1
C2 155106959 155107161
refGene_gn=PLCD1;refGene_tx=XM_011286235.3;refGene_gn=PLCD1;refGene_tx=XM_011286237.3;refGene_gn=PLCD1;refGene_tx=XM_011286238.3;refGene_gn=PLCD1;refGene_tx=XM_019810946.2;refGene_gn=PLCD1;refGene_tx=XM_019810947.2;refGene_gn=PLCD1;refGene_tx=XM_023260743.1
C2 155107270 155107415
refGene_gn=PLCD1;refGene_tx=XM_011286235.3;refGene_gn=PLCD1;refGene_tx=XM_011286237.3;refGene_gn=PLCD1;refGene_tx=XM_011286238.3;refGene_gn=PLCD1;refGene_tx=XM_019810946.2;refGene_gn=PLCD1;refGene_tx=XM_019810947.2;refGene_gn=PLCD1;refGene_tx=XM_023260743.1
C2 155107492 155107642
refGene_gn=PLCD1;refGene_tx=XM_011286235.3;refGene_gn=PLCD1;refGene_tx=XM_011286237.3;refGene_gn=PLCD1;refGene_tx=XM_011286238.3;refGene_gn=PLCD1;refGene_tx=XM_019810946.2;refGene_gn=PLCD1;refGene_tx=XM_019810947.2;refGene_gn=PLCD1;refGene_tx=XM_023260743.1
C2 155107740 155107899
refGene_gn=PLCD1;refGene_tx=XM_011286235.3;refGene_gn=PLCD1;refGene_tx=XM_011286237.3;refGene_gn=PLCD1;refGene_tx=XM_011286238.3;refGene_gn=PLCD1;refGene_tx=XM_019810946.2;refGene_gn=PLCD1;refGene_tx=XM_019810947.2;refGene_gn=PLCD1;refGene_tx=XM_023260743.1
C2 155108113 155108273
refGene_gn=PLCD1;refGene_tx=XM_011286235.3;refGene_gn=PLCD1;refGene_tx=XM_011286237.3;refGene_gn=PLCD1;refGene_tx=XM_011286238.3;refGene_gn=PLCD1;refGene_tx=XM_019810946.2;refGene_gn=PLCD1;refGene_tx=XM_019810947.2;refGene_gn=PLCD1;refGene_tx=XM_023260743.1
C2 155108464 155108581

refGene_gn=PLCD1;refGene_tx=XM_011286235.3;refGene_gn=PLCD1;refGene_tx=XM_011286237.3;refGene_gn=PLCD1;refGene_tx=XM_011286238.3;refGene_gn=PLCD1;refGene_tx=XM_019810946.2;refGene_gn=PLCD1;refGene_tx=XM_019810947.2;refGene_gn=PLCD1;refGene_tx=XM_023260743.1
C2	155108939	155109118
refGene_gn=PLCD1;refGene_tx=XM_011286235.3;refGene_gn=PLCD1;refGene_tx=XM_011286237.3;refGene_gn=PLCD1;refGene_tx=XM_011286238.3;refGene_gn=PLCD1;refGene_tx=XM_019810946.2;refGene_gn=PLCD1;refGene_tx=XM_019810947.2;refGene_gn=PLCD1;refGene_tx=XM_023260743.1
C2	155109212	155109570
refGene_gn=PLCD1;refGene_tx=XM_011286235.3;refGene_gn=PLCD1;refGene_tx=XM_011286237.3;refGene_gn=PLCD1;refGene_tx=XM_011286238.3;refGene_gn=PLCD1;refGene_tx=XM_019810946.2;refGene_gn=PLCD1;refGene_tx=XM_019810947.2;refGene_gn=PLCD1;refGene_tx=XM_023260743.1
C2	155109714	155109800
refGene_gn=PLCD1;refGene_tx=XM_011286235.3;refGene_gn=PLCD1;refGene_tx=XM_011286237.3;refGene_gn=PLCD1;refGene_tx=XM_011286238.3;refGene_gn=PLCD1;refGene_tx=XM_019810946.2;refGene_gn=PLCD1;refGene_tx=XM_019810947.2;refGene_gn=PLCD1;refGene_tx=XM_023260743.1
C2	155110531	155110645
refGene_gn=VILL;refGene_tx=XM_003992257.5;refGene_gn=VILL;refGene_tx=XM_019810941.2;refGene_gn=VILL;refGene_tx=XM_019810942.2;refGene_gn=VILL;refGene_tx=XM_023260742.1
C2	155110902	155111154
refGene_gn=VILL;refGene_tx=XM_003992257.5;refGene_gn=VILL;refGene_tx=XM_019810941.2;refGene_gn=VILL;refGene_tx=XM_019810942.2;refGene_gn=VILL;refGene_tx=XM_023260742.1
C2	155111369	155111441
refGene_gn=VILL;refGene_tx=XM_003992257.5;refGene_gn=VILL;refGene_tx=XM_019810941.2;refGene_gn=VILL;refGene_tx=XM_019810942.2;refGene_gn=VILL;refGene_tx=XM_023260742.1
C2	155111610	155111796
refGene_gn=VILL;refGene_tx=XM_003992257.5;refGene_gn=VILL;refGene_tx=XM_019810941.2;refGene_gn=VILL;refGene_tx=XM_019810942.2;refGene_gn=VILL;refGene_tx=XM_023260742.1
C2	155113427	155113572
refGene_gn=VILL;refGene_tx=XM_003992257.5;refGene_gn=VILL;refGene_tx=XM_019810941.2;refGene_gn=VILL;refGene_tx=XM_019810942.2;refGene_gn=VILL;refGene_tx=XM_023260742.1
C2	155114571	155114717
refGene_gn=VILL;refGene_tx=XM_003992257.5;refGene_gn=VILL;refGene_tx=XM_019810941.2;refGene_gn=VILL;refGene_tx=XM_019810942.2;refGene_gn=VILL;refGene_tx=XM_023260742.1
C2	155115259	155115439
refGene_gn=VILL;refGene_tx=XM_003992257.5;refGene_gn=VILL;refGene_tx=XM_019810941.2;refGene_gn=VILL;refGene_tx=XM_019810942.2;refGene_gn=VILL;refGene_tx=XM_023260742.1
C2	155115952	155116111
refGene_gn=VILL;refGene_tx=XM_003992257.5;refGene_gn=VILL;refGene_tx=XM_019810941.2;refGene_gn=VILL;refGene_tx=XM_019810942.2;refGene_gn=VIL

L;refGene_tx=XM_023260742.1
C2 155116216 155116354
refGene_gn=VILL;refGene_tx=XM_003992257.5;refGene_gn=VILL;refGene_tx=XM_019810941.2;refGene_gn=VILL;refGene_tx=XM_019810942.2
C2 155118219 155118320
refGene_gn=VILL;refGene_tx=XM_003992257.5;refGene_gn=VILL;refGene_tx=XM_019810941.2;refGene_gn=VILL;refGene_tx=XM_019810942.2;refGene_gn=VILL;refGene_tx=XM_023260742.1
C2 155118758 155118897
refGene_gn=VILL;refGene_tx=XM_003992257.5;refGene_gn=VILL;refGene_tx=XM_019810941.2;refGene_gn=VILL;refGene_tx=XM_019810942.2;refGene_gn=VILL;refGene_tx=XM_023260742.1
C2 155119260 155119359
refGene_gn=VILL;refGene_tx=XM_003992257.5;refGene_gn=VILL;refGene_tx=XM_019810941.2;refGene_gn=VILL;refGene_tx=XM_019810942.2;refGene_gn=VILL;refGene_tx=XM_023260742.1
C2 155119459 155119538
refGene_gn=VILL;refGene_tx=XM_003992257.5;refGene_gn=VILL;refGene_tx=XM_019810941.2;refGene_gn=VILL;refGene_tx=XM_019810942.2;refGene_gn=VILL;refGene_tx=XM_023260742.1
C2 155119912 155120112
refGene_gn=VILL;refGene_tx=XM_003992257.5;refGene_gn=VILL;refGene_tx=XM_019810941.2;refGene_gn=VILL;refGene_tx=XM_019810942.2;refGene_gn=VILL;refGene_tx=XM_023260742.1
C2 155120368 155120479
refGene_gn=VILL;refGene_tx=XM_003992257.5;refGene_gn=VILL;refGene_tx=XM_019810941.2;refGene_gn=VILL;refGene_tx=XM_019810942.2;refGene_gn=VILL;refGene_tx=XM_023260742.1
C2 155121491 155121600
refGene_gn=VILL;refGene_tx=XM_003992257.5;refGene_gn=VILL;refGene_tx=XM_019810941.2;refGene_gn=VILL;refGene_tx=XM_019810942.2;refGene_gn=VILL;refGene_tx=XM_023260742.1
C2 155123189 155123395
refGene_gn=VILL;refGene_tx=XM_003992257.5;refGene_gn=VILL;refGene_tx=XM_019810941.2;refGene_gn=VILL;refGene_tx=XM_019810942.2;refGene_gn=VILL;refGene_tx=XM_023260742.1
C2 155123675 155123750
refGene_gn=VILL;refGene_tx=XM_003992257.5;refGene_gn=VILL;refGene_tx=XM_019810941.2;refGene_gn=VILL;refGene_tx=XM_019810942.2;refGene_gn=VILL;refGene_tx=XM_023260742.1
C2 155123932 155123999
refGene_gn=VILL;refGene_tx=XM_003992257.5;refGene_gn=VILL;refGene_tx=XM_019810941.2;refGene_gn=VILL;refGene_tx=XM_019810942.2;refGene_gn=VILL;refGene_tx=XM_023260742.1
C2 155124091 155124279
refGene_gn=VILL;refGene_tx=XM_003992257.5;refGene_gn=VILL;refGene_tx=XM_023260742.1
C2 155126432 155126461
refGene_gn=VILL;refGene_tx=XM_019810942.2
C2 155128337 155128405

refGene_gn=VILL;refGene_tx=XM_019810941.2
C2 155135809 155135876
refGene_gn=CTDSPL;refGene_tx=XM_019810950.2;refGene_gn=CTDSPL;refGene_tx=XM_023260744.1;refGene_gn=CTDSPL;refGene_tx=XM_023260745.1
C2 155136838 155136964
refGene_gn=CTDSPL;refGene_tx=XM_019810950.2;refGene_gn=CTDSPL;refGene_tx=XM_023260744.1;refGene_gn=CTDSPL;refGene_tx=XM_023260745.1;refGene_gn=CTDSPL;refGene_tx=XM_023260746.1;refGene_gn=CTDSPL;refGene_tx=XM_023260747.1
C2 155137124 155137145
refGene_gn=CTDSPL;refGene_tx=XM_023260748.1
C2 155140794 155140980
refGene_gn=CTDSPL;refGene_tx=XM_019810950.2;refGene_gn=CTDSPL;refGene_tx=XM_023260744.1;refGene_gn=CTDSPL;refGene_tx=XM_023260745.1;refGene_gn=CTDSPL;refGene_tx=XM_023260746.1;refGene_gn=CTDSPL;refGene_tx=XM_023260747.1;refGene_gn=CTDSPL;refGene_tx=XM_023260748.1
C2 155144771 155144864
refGene_gn=CTDSPL;refGene_tx=XM_019810950.2;refGene_gn=CTDSPL;refGene_tx=XM_023260744.1;refGene_gn=CTDSPL;refGene_tx=XM_023260745.1;refGene_gn=CTDSPL;refGene_tx=XM_023260746.1;refGene_gn=CTDSPL;refGene_tx=XM_023260747.1;refGene_gn=CTDSPL;refGene_tx=XM_023260748.1
C2 155148299 155148356
refGene_gn=CTDSPL;refGene_tx=XM_019810950.2;refGene_gn=CTDSPL;refGene_tx=XM_023260744.1;refGene_gn=CTDSPL;refGene_tx=XM_023260745.1;refGene_gn=CTDSPL;refGene_tx=XM_023260746.1;refGene_gn=CTDSPL;refGene_tx=XM_023260747.1;refGene_gn=CTDSPL;refGene_tx=XM_023260748.1
C2 155154213 155154315
refGene_gn=CTDSPL;refGene_tx=XM_019810950.2;refGene_gn=CTDSPL;refGene_tx=XM_023260744.1;refGene_gn=CTDSPL;refGene_tx=XM_023260745.1;refGene_gn=CTDSPL;refGene_tx=XM_023260746.1;refGene_gn=CTDSPL;refGene_tx=XM_023260747.1;refGene_gn=CTDSPL;refGene_tx=XM_023260748.1
C2 155160811 155160844
refGene_gn=CTDSPL;refGene_tx=XM_019810950.2;refGene_gn=CTDSPL;refGene_tx=XM_023260744.1;refGene_gn=CTDSPL;refGene_tx=XM_023260746.1;refGene_gn=CTDSPL;refGene_tx=XM_023260748.1
C2 155169596 155169751
refGene_gn=CTDSPL;refGene_tx=XM_019810950.2;refGene_gn=CTDSPL;refGene_tx=XM_023260744.1;refGene_gn=CTDSPL;refGene_tx=XM_023260745.1;refGene_gn=CTDSPL;refGene_tx=XM_023260746.1;refGene_gn=CTDSPL;refGene_tx=XM_023260747.1;refGene_gn=CTDSPL;refGene_tx=XM_023260748.1
C2 155229519 155229526
refGene_gn=CTDSPL;refGene_tx=XM_019810950.2
C2 155254173 155254252
refGene_gn=CTDSPL;refGene_tx=XM_023260744.1;refGene_gn=CTDSPL;refGene_tx=XM_023260745.1;refGene_gn=CTDSPL;refGene_tx=XM_023260746.1;refGene_gn=CTDSPL;refGene_tx=XM_023260747.1;refGene_gn=CTDSPL;refGene_tx=XM_023260748.1
C2 155267509 155267647
refGene_gn=ITGA9;refGene_tx=XM_023260749.1;refGene_gn=ITGA9;refGene_tx=XM_023260752.1

C2 155289247 155289346
refGene_gn=ITGA9;refGene_tx=XM_023260750.1;refGene_gn=ITGA9;refGene_tx=XM_023260753.1
C2 155289626 155289713
refGene_gn=ITGA9;refGene_tx=XM_023260751.1
C2 155302412 155302532
refGene_gn=ITGA9;refGene_tx=XM_023260749.1;refGene_gn=ITGA9;refGene_tx=XM_023260750.1;refGene_gn=ITGA9;refGene_tx=XM_023260751.1;refGene_gn=ITGA9;refGene_tx=XM_023260752.1;refGene_gn=ITGA9;refGene_tx=XM_023260753.1
C2 155317517 155317619
refGene_gn=ITGA9;refGene_tx=XM_023260749.1;refGene_gn=ITGA9;refGene_tx=XM_023260750.1;refGene_gn=ITGA9;refGene_tx=XM_023260751.1;refGene_gn=ITGA9;refGene_tx=XM_023260752.1;refGene_gn=ITGA9;refGene_tx=XM_023260753.1
C2 155322377 155322497
refGene_gn=ITGA9;refGene_tx=XM_023260749.1;refGene_gn=ITGA9;refGene_tx=XM_023260750.1;refGene_gn=ITGA9;refGene_tx=XM_023260751.1;refGene_gn=ITGA9;refGene_tx=XM_023260752.1;refGene_gn=ITGA9;refGene_tx=XM_023260753.1
C2 155324198 155324324
refGene_gn=ITGA9;refGene_tx=XM_023260749.1;refGene_gn=ITGA9;refGene_tx=XM_023260750.1;refGene_gn=ITGA9;refGene_tx=XM_023260751.1;refGene_gn=ITGA9;refGene_tx=XM_023260752.1;refGene_gn=ITGA9;refGene_tx=XM_023260753.1
C2 155331052 155331157
refGene_gn=ITGA9;refGene_tx=XM_023260749.1;refGene_gn=ITGA9;refGene_tx=XM_023260750.1;refGene_gn=ITGA9;refGene_tx=XM_023260751.1
C2 155347899 155348007
refGene_gn=ITGA9;refGene_tx=XM_023260749.1;refGene_gn=ITGA9;refGene_tx=XM_023260750.1;refGene_gn=ITGA9;refGene_tx=XM_023260751.1;refGene_gn=ITGA9;refGene_tx=XM_023260752.1;refGene_gn=ITGA9;refGene_tx=XM_023260753.1
C2 155352916 155353025
refGene_gn=ITGA9;refGene_tx=XM_023260749.1;refGene_gn=ITGA9;refGene_tx=XM_023260750.1;refGene_gn=ITGA9;refGene_tx=XM_023260751.1;refGene_gn=ITGA9;refGene_tx=XM_023260752.1;refGene_gn=ITGA9;refGene_tx=XM_023260753.1
C2 155355188 155355278
refGene_gn=ITGA9;refGene_tx=XM_023260749.1;refGene_gn=ITGA9;refGene_tx=XM_023260750.1;refGene_gn=ITGA9;refGene_tx=XM_023260751.1;refGene_gn=ITGA9;refGene_tx=XM_023260752.1;refGene_gn=ITGA9;refGene_tx=XM_023260753.1
C2 155362614 155362694
refGene_gn=ITGA9;refGene_tx=XM_023260749.1;refGene_gn=ITGA9;refGene_tx=XM_023260750.1;refGene_gn=ITGA9;refGene_tx=XM_023260751.1;refGene_gn=ITGA9;refGene_tx=XM_023260752.1;refGene_gn=ITGA9;refGene_tx=XM_023260753.1
C2 155366408 155366495
refGene_gn=ITGA9;refGene_tx=XM_023260749.1;refGene_gn=ITGA9;refGene_tx

=XM_023260750.1;refGene_gn=ITGA9;refGene_tx=XM_023260751.1;refGene_gn=ITGA9;refGene_tx=XM_023260752.1;refGene_gn=ITGA9;refGene_tx=XM_023260753.1
C2	155414876	155415027
refGene_gn=ITGA9;refGene_tx=XM_023260749.1;refGene_gn=ITGA9;refGene_tx=XM_023260750.1;refGene_gn=ITGA9;refGene_tx=XM_023260751.1;refGene_gn=ITGA9;refGene_tx=XM_023260752.1;refGene_gn=ITGA9;refGene_tx=XM_023260753.1
C2	155438724	155438801
refGene_gn=ITGA9;refGene_tx=XM_023260749.1;refGene_gn=ITGA9;refGene_tx=XM_023260750.1;refGene_gn=ITGA9;refGene_tx=XM_023260751.1;refGene_gn=ITGA9;refGene_tx=XM_023260752.1;refGene_gn=ITGA9;refGene_tx=XM_023260753.1
C2	155454029	155454179
refGene_gn=ITGA9;refGene_tx=XM_023260749.1;refGene_gn=ITGA9;refGene_tx=XM_023260750.1;refGene_gn=ITGA9;refGene_tx=XM_023260751.1;refGene_gn=ITGA9;refGene_tx=XM_023260752.1;refGene_gn=ITGA9;refGene_tx=XM_023260753.1
C2	155539877	155540038
refGene_gn=ITGA9;refGene_tx=XM_023260749.1;refGene_gn=ITGA9;refGene_tx=XM_023260750.1;refGene_gn=ITGA9;refGene_tx=XM_023260751.1;refGene_gn=ITGA9;refGene_tx=XM_023260752.1;refGene_gn=ITGA9;refGene_tx=XM_023260753.1
C2	155551137	155551292
refGene_gn=ITGA9;refGene_tx=XM_023260749.1;refGene_gn=ITGA9;refGene_tx=XM_023260750.1;refGene_gn=ITGA9;refGene_tx=XM_023260751.1;refGene_gn=ITGA9;refGene_tx=XM_023260752.1;refGene_gn=ITGA9;refGene_tx=XM_023260753.1
C2	155558337	155558383
refGene_gn=ITGA9;refGene_tx=XM_023260749.1;refGene_gn=ITGA9;refGene_tx=XM_023260750.1;refGene_gn=ITGA9;refGene_tx=XM_023260751.1;refGene_gn=ITGA9;refGene_tx=XM_023260752.1;refGene_gn=ITGA9;refGene_tx=XM_023260753.1
C2	155561228	155561319
refGene_gn=ITGA9;refGene_tx=XM_023260749.1;refGene_gn=ITGA9;refGene_tx=XM_023260750.1;refGene_gn=ITGA9;refGene_tx=XM_023260751.1;refGene_gn=ITGA9;refGene_tx=XM_023260752.1;refGene_gn=ITGA9;refGene_tx=XM_023260753.1
C2	155566013	155566108
refGene_gn=ITGA9;refGene_tx=XM_023260749.1;refGene_gn=ITGA9;refGene_tx=XM_023260750.1;refGene_gn=ITGA9;refGene_tx=XM_023260751.1;refGene_gn=ITGA9;refGene_tx=XM_023260752.1;refGene_gn=ITGA9;refGene_tx=XM_023260753.1
C2	155567240	155567346
refGene_gn=ITGA9;refGene_tx=XM_023260749.1;refGene_gn=ITGA9;refGene_tx=XM_023260750.1;refGene_gn=ITGA9;refGene_tx=XM_023260751.1;refGene_gn=ITGA9;refGene_tx=XM_023260752.1;refGene_gn=ITGA9;refGene_tx=XM_023260753.1
C2	155570621	155570759
refGene_gn=ITGA9;refGene_tx=XM_023260749.1;refGene_gn=ITGA9;refGene_tx

=XM_023260750.1;refGene_gn=ITGA9;refGene_tx=XM_023260751.1;refGene_gn=ITGA9;refGene_tx=XM_023260752.1;refGene_gn=ITGA9;refGene_tx=XM_023260753.1
C2 155573685 155573754
refGene_gn=ITGA9;refGene_tx=XM_023260749.1;refGene_gn=ITGA9;refGene_tx=XM_023260750.1;refGene_gn=ITGA9;refGene_tx=XM_023260751.1;refGene_gn=ITGA9;refGene_tx=XM_023260752.1;refGene_gn=ITGA9;refGene_tx=XM_023260753.1
C2 155576466 155576552
refGene_gn=ITGA9;refGene_tx=XM_023260749.1;refGene_gn=ITGA9;refGene_tx=XM_023260750.1;refGene_gn=ITGA9;refGene_tx=XM_023260751.1;refGene_gn=ITGA9;refGene_tx=XM_023260752.1;refGene_gn=ITGA9;refGene_tx=XM_023260753.1
C2 155579445 155579575
refGene_gn=ITGA9;refGene_tx=XM_023260749.1;refGene_gn=ITGA9;refGene_tx=XM_023260750.1;refGene_gn=ITGA9;refGene_tx=XM_023260751.1;refGene_gn=ITGA9;refGene_tx=XM_023260752.1;refGene_gn=ITGA9;refGene_tx=XM_023260753.1
C2 155587520 155587588
refGene_gn=ITGA9;refGene_tx=XM_023260749.1;refGene_gn=ITGA9;refGene_tx=XM_023260750.1;refGene_gn=ITGA9;refGene_tx=XM_023260751.1;refGene_gn=ITGA9;refGene_tx=XM_023260752.1;refGene_gn=ITGA9;refGene_tx=XM_023260753.1
C2 155600931 155601055
refGene_gn=ITGA9;refGene_tx=XM_023260749.1;refGene_gn=ITGA9;refGene_tx=XM_023260750.1;refGene_gn=ITGA9;refGene_tx=XM_023260751.1;refGene_gn=ITGA9;refGene_tx=XM_023260752.1;refGene_gn=ITGA9;refGene_tx=XM_023260753.1
C2 155610394 155610501
refGene_gn=ITGA9;refGene_tx=XM_023260749.1;refGene_gn=ITGA9;refGene_tx=XM_023260750.1;refGene_gn=ITGA9;refGene_tx=XM_023260751.1;refGene_gn=ITGA9;refGene_tx=XM_023260752.1;refGene_gn=ITGA9;refGene_tx=XM_023260753.1
C2 155613095 155613223
refGene_gn=ITGA9;refGene_tx=XM_023260749.1;refGene_gn=ITGA9;refGene_tx=XM_023260750.1;refGene_gn=ITGA9;refGene_tx=XM_023260751.1;refGene_gn=ITGA9;refGene_tx=XM_023260752.1;refGene_gn=ITGA9;refGene_tx=XM_023260753.1
C2 155630681 155630866
refGene_gn=ITGA9;refGene_tx=XM_023260749.1;refGene_gn=ITGA9;refGene_tx=XM_023260750.1;refGene_gn=ITGA9;refGene_tx=XM_023260751.1;refGene_gn=ITGA9;refGene_tx=XM_023260752.1;refGene_gn=ITGA9;refGene_tx=XM_023260753.1
C2 155648911 155648986
refGene_gn=CC2H3orf35;refGene_tx=XM_006944352.3
C2 155648987 155649260
refGene_gn=CC2H3orf35;refGene_tx=XM_006944352.3
C2 155672615 155672639
refGene_gn=GOLGA4;refGene_tx=XM_023260755.1;refGene_gn=GOLGA4;refGene_tx=XM_023260760.1

C2	155675369	155675399
refGene_gn=GOLGA4;refGene_tx=XM_023260754.1;refGene_gn=GOLGA4;refGene_tx=XM_023260756.1;refGene_gn=GOLGA4;refGene_tx=XM_023260757.1;refGene_gn=GOLGA4;refGene_tx=XM_023260758.1;refGene_gn=GOLGA4;refGene_tx=XM_023260759.1
C2	155681815	155681902
refGene_gn=GOLGA4;refGene_tx=XM_023260754.1;refGene_gn=GOLGA4;refGene_tx=XM_023260755.1;refGene_gn=GOLGA4;refGene_tx=XM_023260756.1;refGene_gn=GOLGA4;refGene_tx=XM_023260757.1;refGene_gn=GOLGA4;refGene_tx=XM_023260758.1;refGene_gn=GOLGA4;refGene_tx=XM_023260759.1;refGene_gn=GOLGA4;refGene_tx=XM_023260760.1
C2	155686728	155686832
refGene_gn=GOLGA4;refGene_tx=XM_023260754.1;refGene_gn=GOLGA4;refGene_tx=XM_023260755.1;refGene_gn=GOLGA4;refGene_tx=XM_023260756.1;refGene_gn=GOLGA4;refGene_tx=XM_023260757.1;refGene_gn=GOLGA4;refGene_tx=XM_023260758.1;refGene_gn=GOLGA4;refGene_tx=XM_023260759.1;refGene_gn=GOLGA4;refGene_tx=XM_023260760.1
C2	155694372	155694448
refGene_gn=GOLGA4;refGene_tx=XM_023260754.1;refGene_gn=GOLGA4;refGene_tx=XM_023260755.1;refGene_gn=GOLGA4;refGene_tx=XM_023260756.1;refGene_gn=GOLGA4;refGene_tx=XM_023260757.1;refGene_gn=GOLGA4;refGene_tx=XM_023260758.1;refGene_gn=GOLGA4;refGene_tx=XM_023260759.1;refGene_gn=GOLGA4;refGene_tx=XM_023260760.1
C2	155697326	155697395
refGene_gn=GOLGA4;refGene_tx=XM_023260754.1;refGene_gn=GOLGA4;refGene_tx=XM_023260755.1;refGene_gn=GOLGA4;refGene_tx=XM_023260756.1;refGene_gn=GOLGA4;refGene_tx=XM_023260757.1;refGene_gn=GOLGA4;refGene_tx=XM_023260758.1;refGene_gn=GOLGA4;refGene_tx=XM_023260759.1;refGene_gn=GOLGA4;refGene_tx=XM_023260760.1
C2	155697603	155697624
refGene_gn=GOLGA4;refGene_tx=XM_023260754.1;refGene_gn=GOLGA4;refGene_tx=XM_023260755.1;refGene_gn=GOLGA4;refGene_tx=XM_023260756.1;refGene_gn=GOLGA4;refGene_tx=XM_023260757.1;refGene_gn=GOLGA4;refGene_tx=XM_023260758.1;refGene_gn=GOLGA4;refGene_tx=XM_023260759.1;refGene_gn=GOLGA4;refGene_tx=XM_023260760.1
C2	155700349	155700463
refGene_gn=GOLGA4;refGene_tx=XM_023260754.1;refGene_gn=GOLGA4;refGene_tx=XM_023260755.1;refGene_gn=GOLGA4;refGene_tx=XM_023260756.1;refGene_gn=GOLGA4;refGene_tx=XM_023260757.1;refGene_gn=GOLGA4;refGene_tx=XM_023260758.1;refGene_gn=GOLGA4;refGene_tx=XM_023260759.1;refGene_gn=GOLGA4;refGene_tx=XM_023260760.1
C2	155715545	155715676
refGene_gn=GOLGA4;refGene_tx=XM_023260754.1;refGene_gn=GOLGA4;refGene_tx=XM_023260755.1;refGene_gn=GOLGA4;refGene_tx=XM_023260756.1;refGene_gn=GOLGA4;refGene_tx=XM_023260757.1;refGene_gn=GOLGA4;refGene_tx=XM_023260758.1;refGene_gn=GOLGA4;refGene_tx=XM_023260759.1;refGene_gn=GOLGA4;refGene_tx=XM_023260760.1
C2	155716054	155716176
refGene_gn=GOLGA4;refGene_tx=XM_023260754.1;refGene_gn=GOLGA4;refGene_tx=XM_023260755.1;refGene_gn=GOLGA4;refGene_tx=XM_023260756.1;refGene_

gn=GOLGA4;refGene_tx=XM_023260757.1;refGene_gn=GOLGA4;refGene_tx=XM_023260758.1;refGene_gn=GOLGA4;refGene_tx=XM_023260759.1;refGene_gn=GOLGA4;refGene_tx=XM_023260760.1
C2	155716971	155721212
refGene_gn=GOLGA4;refGene_tx=XM_023260754.1;refGene_gn=GOLGA4;refGene_tx=XM_023260755.1;refGene_gn=GOLGA4;refGene_tx=XM_023260756.1;refGene_gn=GOLGA4;refGene_tx=XM_023260757.1;refGene_gn=GOLGA4;refGene_tx=XM_023260758.1;refGene_gn=GOLGA4;refGene_tx=XM_023260759.1;refGene_gn=GOLGA4;refGene_tx=XM_023260760.1
C2	155723106	155723262
refGene_gn=GOLGA4;refGene_tx=XM_023260754.1;refGene_gn=GOLGA4;refGene_tx=XM_023260755.1;refGene_gn=GOLGA4;refGene_tx=XM_023260756.1;refGene_gn=GOLGA4;refGene_tx=XM_023260757.1;refGene_gn=GOLGA4;refGene_tx=XM_023260758.1;refGene_gn=GOLGA4;refGene_tx=XM_023260759.1;refGene_gn=GOLGA4;refGene_tx=XM_023260760.1
C2	155726868	155726997
refGene_gn=GOLGA4;refGene_tx=XM_023260754.1;refGene_gn=GOLGA4;refGene_tx=XM_023260755.1;refGene_gn=GOLGA4;refGene_tx=XM_023260756.1;refGene_gn=GOLGA4;refGene_tx=XM_023260757.1;refGene_gn=GOLGA4;refGene_tx=XM_023260758.1;refGene_gn=GOLGA4;refGene_tx=XM_023260759.1;refGene_gn=GOLGA4;refGene_tx=XM_023260760.1
C2	155729294	155729473
refGene_gn=GOLGA4;refGene_tx=XM_023260754.1;refGene_gn=GOLGA4;refGene_tx=XM_023260755.1;refGene_gn=GOLGA4;refGene_tx=XM_023260756.1;refGene_gn=GOLGA4;refGene_tx=XM_023260757.1;refGene_gn=GOLGA4;refGene_tx=XM_023260758.1;refGene_gn=GOLGA4;refGene_tx=XM_023260759.1;refGene_gn=GOLGA4;refGene_tx=XM_023260760.1
C2	155741109	155741257
refGene_gn=GOLGA4;refGene_tx=XM_023260754.1;refGene_gn=GOLGA4;refGene_tx=XM_023260755.1;refGene_gn=GOLGA4;refGene_tx=XM_023260756.1;refGene_gn=GOLGA4;refGene_tx=XM_023260757.1;refGene_gn=GOLGA4;refGene_tx=XM_023260758.1;refGene_gn=GOLGA4;refGene_tx=XM_023260759.1;refGene_gn=GOLGA4;refGene_tx=XM_023260760.1
C2	155743373	155743457
refGene_gn=GOLGA4;refGene_tx=XM_023260754.1;refGene_gn=GOLGA4;refGene_tx=XM_023260755.1;refGene_gn=GOLGA4;refGene_tx=XM_023260756.1;refGene_gn=GOLGA4;refGene_tx=XM_023260758.1;refGene_gn=GOLGA4;refGene_tx=XM_023260759.1;refGene_gn=GOLGA4;refGene_tx=XM_023260760.1
C2	155743747	155743929
refGene_gn=GOLGA4;refGene_tx=XM_023260754.1;refGene_gn=GOLGA4;refGene_tx=XM_023260755.1;refGene_gn=GOLGA4;refGene_tx=XM_023260756.1;refGene_gn=GOLGA4;refGene_tx=XM_023260757.1;refGene_gn=GOLGA4;refGene_tx=XM_023260758.1;refGene_gn=GOLGA4;refGene_tx=XM_023260759.1;refGene_gn=GOLGA4;refGene_tx=XM_023260760.1
C2	155745281	155745411
refGene_gn=GOLGA4;refGene_tx=XM_023260754.1;refGene_gn=GOLGA4;refGene_tx=XM_023260755.1;refGene_gn=GOLGA4;refGene_tx=XM_023260756.1;refGene_gn=GOLGA4;refGene_tx=XM_023260757.1;refGene_gn=GOLGA4;refGene_tx=XM_023260758.1;refGene_gn=GOLGA4;refGene_tx=XM_023260759.1;refGene_gn=GOLGA4;refGene_tx=XM_023260760.1

C2 155746163 155746262
refGene_gn=GOLGA4;refGene_tx=XM_023260754.1;refGene_gn=GOLGA4;refGene_tx=XM_023260755.1;refGene_gn=GOLGA4;refGene_tx=XM_023260756.1;refGene_gn=GOLGA4;refGene_tx=XM_023260757.1;refGene_gn=GOLGA4;refGene_tx=XM_023260758.1;refGene_gn=GOLGA4;refGene_tx=XM_023260759.1;refGene_gn=GOLGA4;refGene_tx=XM_023260760.1
C2 155752984 155753041
refGene_gn=GOLGA4;refGene_tx=XM_023260754.1;refGene_gn=GOLGA4;refGene_tx=XM_023260755.1;refGene_gn=GOLGA4;refGene_tx=XM_023260756.1;refGene_gn=GOLGA4;refGene_tx=XM_023260757.1;refGene_gn=GOLGA4;refGene_tx=XM_023260758.1;refGene_gn=GOLGA4;refGene_tx=XM_023260759.1;refGene_gn=GOLGA4;refGene_tx=XM_023260760.1
C2 155754872 155754920
refGene_gn=GOLGA4;refGene_tx=XM_023260754.1;refGene_gn=GOLGA4;refGene_tx=XM_023260755.1;refGene_gn=GOLGA4;refGene_tx=XM_023260756.1;refGene_gn=GOLGA4;refGene_tx=XM_023260757.1;refGene_gn=GOLGA4;refGene_tx=XM_023260758.1;refGene_gn=GOLGA4;refGene_tx=XM_023260759.1;refGene_gn=GOLGA4;refGene_tx=XM_023260760.1
C2 155757541 155757856
refGene_gn=GOLGA4;refGene_tx=XM_023260754.1;refGene_gn=GOLGA4;refGene_tx=XM_023260755.1;refGene_gn=GOLGA4;refGene_tx=XM_023260756.1;refGene_gn=GOLGA4;refGene_tx=XM_023260757.1;refGene_gn=GOLGA4;refGene_tx=XM_023260758.1;refGene_gn=GOLGA4;refGene_tx=XM_023260759.1;refGene_gn=GOLGA4;refGene_tx=XM_023260760.1
C2 155767847 155767913
refGene_gn=GOLGA4;refGene_tx=XM_023260754.1;refGene_gn=GOLGA4;refGene_tx=XM_023260755.1;refGene_gn=GOLGA4;refGene_tx=XM_023260756.1;refGene_gn=GOLGA4;refGene_tx=XM_023260757.1;refGene_gn=GOLGA4;refGene_tx=XM_023260758.1;refGene_gn=GOLGA4;refGene_tx=XM_023260759.1;refGene_gn=GOLGA4;refGene_tx=XM_023260760.1
C2 155772292 155772391
refGene_gn=GOLGA4;refGene_tx=XM_023260754.1;refGene_gn=GOLGA4;refGene_tx=XM_023260755.1;refGene_gn=GOLGA4;refGene_tx=XM_023260756.1;refGene_gn=GOLGA4;refGene_tx=XM_023260757.1
C2 155785857 155785947
refGene_gn=GOLGA4;refGene_tx=XM_023260754.1;refGene_gn=GOLGA4;refGene_tx=XM_023260755.1;refGene_gn=GOLGA4;refGene_tx=XM_023260756.1;refGene_gn=GOLGA4;refGene_tx=XM_023260757.1;refGene_gn=GOLGA4;refGene_tx=XM_023260758.1;refGene_gn=GOLGA4;refGene_tx=XM_023260759.1;refGene_gn=GOLGA4;refGene_tx=XM_023260760.1
C2 155788021 155788081
refGene_gn=GOLGA4;refGene_tx=XM_023260754.1;refGene_gn=GOLGA4;refGene_tx=XM_023260755.1;refGene_gn=GOLGA4;refGene_tx=XM_023260757.1;refGene_gn=GOLGA4;refGene_tx=XM_023260758.1;refGene_gn=GOLGA4;refGene_tx=XM_023260760.1
C2 155798411 155798480
refGene_gn=GOLGA4;refGene_tx=XM_023260754.1;refGene_gn=GOLGA4;refGene_tx=XM_023260755.1;refGene_gn=GOLGA4;refGene_tx=XM_023260756.1;refGene_gn=GOLGA4;refGene_tx=XM_023260757.1;refGene_gn=GOLGA4;refGene_tx=XM_023260758.1;refGene_gn=GOLGA4;refGene_tx=XM_023260759.1;refGene_gn=GOLGA

4;refGene_tx=XM_023260760.1
C2 155872653 155872743
refGene_gn=LRRFIP2;refGene_tx=XM_003992258.5;refGene_gn=LRRFIP2;refGene_tx=XM_006936543.4;refGene_gn=LRRFIP2;refGene_tx=XM_019810956.1;refGene_gn=LRRFIP2;refGene_tx=XM_019810957.2;refGene_gn=LRRFIP2;refGene_tx=XM_023260762.1;refGene_gn=LRRFIP2;refGene_tx=XM_023260763.1;refGene_gn=LRRFIP2;refGene_tx=XM_023260764.1;refGene_gn=LRRFIP2;refGene_tx=XM_023260765.1;refGene_gn=LRRFIP2;refGene_tx=XM_023260766.1;refGene_gn=LRRFIP2;refGene_tx=XM_023260767.1;refGene_gn=LRRFIP2;refGene_tx=XM_023260768.1;refGene_gn=LRRFIP2;refGene_tx=XM_023260769.1;refGene_gn=LRRFIP2;refGene_tx=XM_023260770.1;refGene_gn=LRRFIP2;refGene_tx=XM_023260771.1;refGene_gn=LRRFIP2;refGene_tx=XM_023260772.1;refGene_gn=LRRFIP2;refGene_tx=XM_023260773.1;refGene_gn=LRRFIP2;refGene_tx=XM_023260774.1;refGene_gn=LRRFIP2;refGene_tx=XM_023260775.1;refGene_gn=LRRFIP2;refGene_tx=XM_023260776.1;refGene_gn=LRRFIP2;refGene_tx=XM_023260777.1;refGene_gn=LRRFIP2;refGene_tx=XM_023260778.1;refGene_gn=LRRFIP2;refGene_tx=XM_023260779.1
C2 155889700 155889787
refGene_gn=LRRFIP2;refGene_tx=XM_003992258.5;refGene_gn=LRRFIP2;refGene_tx=XM_006936543.4;refGene_gn=LRRFIP2;refGene_tx=XM_019810956.1;refGene_gn=LRRFIP2;refGene_tx=XM_019810957.2;refGene_gn=LRRFIP2;refGene_tx=XM_023260762.1;refGene_gn=LRRFIP2;refGene_tx=XM_023260763.1;refGene_gn=LRRFIP2;refGene_tx=XM_023260764.1;refGene_gn=LRRFIP2;refGene_tx=XM_023260765.1;refGene_gn=LRRFIP2;refGene_tx=XM_023260766.1;refGene_gn=LRRFIP2;refGene_tx=XM_023260767.1;refGene_gn=LRRFIP2;refGene_tx=XM_023260768.1;refGene_gn=LRRFIP2;refGene_tx=XM_023260769.1;refGene_gn=LRRFIP2;refGene_tx=XM_023260770.1;refGene_gn=LRRFIP2;refGene_tx=XM_023260771.1;refGene_gn=LRRFIP2;refGene_tx=XM_023260772.1;refGene_gn=LRRFIP2;refGene_tx=XM_023260773.1;refGene_gn=LRRFIP2;refGene_tx=XM_023260774.1;refGene_gn=LRRFIP2;refGene_tx=XM_023260775.1;refGene_gn=LRRFIP2;refGene_tx=XM_023260776.1;refGene_gn=LRRFIP2;refGene_tx=XM_023260777.1;refGene_gn=LRRFIP2;refGene_tx=XM_023260778.1;refGene_gn=LRRFIP2;refGene_tx=XM_023260779.1
C2 155891529 155891580
refGene_gn=LRRFIP2;refGene_tx=XM_003992258.5;refGene_gn=LRRFIP2;refGene_tx=XM_006936543.4;refGene_gn=LRRFIP2;refGene_tx=XM_019810956.1;refGene_gn=LRRFIP2;refGene_tx=XM_019810957.2;refGene_gn=LRRFIP2;refGene_tx=XM_023260762.1;refGene_gn=LRRFIP2;refGene_tx=XM_023260763.1;refGene_gn=LRRFIP2;refGene_tx=XM_023260764.1;refGene_gn=LRRFIP2;refGene_tx=XM_023260765.1;refGene_gn=LRRFIP2;refGene_tx=XM_023260766.1;refGene_gn=LRRFIP2;refGene_tx=XM_023260768.1;refGene_gn=LRRFIP2;refGene_tx=XM_023260769.1;refGene_gn=LRRFIP2;refGene_tx=XM_023260770.1;refGene_gn=LRRFIP2;refGene_tx=XM_023260771.1;refGene_gn=LRRFIP2;refGene_tx=XM_023260772.1;refGene_gn=LRRFIP2;refGene_tx=XM_023260774.1
C2 155896883 155896940
refGene_gn=LRRFIP2;refGene_tx=XM_003992258.5;refGene_gn=LRRFIP2;refGene_tx=XM_006936543.4;refGene_gn=LRRFIP2;refGene_tx=XM_019810956.1;refGene_gn=LRRFIP2;refGene_tx=XM_019810957.2;refGene_gn=LRRFIP2;refGene_tx=XM_023260762.1;refGene_gn=LRRFIP2;refGene_tx=XM_023260763.1;refGene_gn=LRRFIP2;refGene_tx=XM_023260764.1;refGene_gn=LRRFIP2;refGene_tx=XM_02

3260765.1;refGene_gn=LRRFIP2;refGene_tx=XM_023260766.1;refGene_gn=LRRF
IP2;refGene_tx=XM_023260768.1;refGene_gn=LRRFIP2;refGene_tx=XM_0232607
69.1;refGene_gn=LRRFIP2;refGene_tx=XM_023260770.1;refGene_gn=LRRFIP2;r
efGene_tx=XM_023260771.1;refGene_gn=LRRFIP2;refGene_tx=XM_023260772.1;
refGene_gn=LRRFIP2;refGene_tx=XM_023260774.1
C2 155897038 155897083
refGene_gn=LRRFIP2;refGene_tx=XM_003992258.5;refGene_gn=LRRFIP2;refGen
e_tx=XM_006936543.4;refGene_gn=LRRFIP2;refGene_tx=XM_019810956.1;refGe
ne_gn=LRRFIP2;refGene_tx=XM_019810957.2;refGene_gn=LRRFIP2;refGene_tx=
XM_023260762.1;refGene_gn=LRRFIP2;refGene_tx=XM_023260763.1;refGene_gn
=LRRFIP2;refGene_tx=XM_023260764.1;refGene_gn=LRRFIP2;refGene_tx=XM_02
3260765.1;refGene_gn=LRRFIP2;refGene_tx=XM_023260766.1;refGene_gn=LRRF
IP2;refGene_tx=XM_023260767.1;refGene_gn=LRRFIP2;refGene_tx=XM_0232607
68.1;refGene_gn=LRRFIP2;refGene_tx=XM_023260769.1;refGene_gn=LRRFIP2;r
efGene_tx=XM_023260770.1;refGene_gn=LRRFIP2;refGene_tx=XM_023260771.1;
refGene_gn=LRRFIP2;refGene_tx=XM_023260772.1;refGene_gn=LRRFIP2;refGen
e_tx=XM_023260774.1
C2 155901333 155901375
refGene_gn=LRRFIP2;refGene_tx=XM_003992258.5;refGene_gn=LRRFIP2;refGen
e_tx=XM_006936543.4;refGene_gn=LRRFIP2;refGene_tx=XM_019810956.1;refGe
ne_gn=LRRFIP2;refGene_tx=XM_019810957.2;refGene_gn=LRRFIP2;refGene_tx=
XM_023260762.1;refGene_gn=LRRFIP2;refGene_tx=XM_023260763.1;refGene_gn
=LRRFIP2;refGene_tx=XM_023260764.1;refGene_gn=LRRFIP2;refGene_tx=XM_02
3260765.1;refGene_gn=LRRFIP2;refGene_tx=XM_023260766.1;refGene_gn=LRRF
IP2;refGene_tx=XM_023260767.1;refGene_gn=LRRFIP2;refGene_tx=XM_0232607
68.1;refGene_gn=LRRFIP2;refGene_tx=XM_023260769.1;refGene_gn=LRRFIP2;r
efGene_tx=XM_023260770.1;refGene_gn=LRRFIP2;refGene_tx=XM_023260771.1
C2 155902818 155902884
refGene_gn=LRRFIP2;refGene_tx=XM_003992258.5;refGene_gn=LRRFIP2;refGen
e_tx=XM_006936543.4;refGene_gn=LRRFIP2;refGene_tx=XM_019810956.1;refGe
ne_gn=LRRFIP2;refGene_tx=XM_019810957.2;refGene_gn=LRRFIP2;refGene_tx=
XM_023260762.1;refGene_gn=LRRFIP2;refGene_tx=XM_023260763.1;refGene_gn
=LRRFIP2;refGene_tx=XM_023260764.1;refGene_gn=LRRFIP2;refGene_tx=XM_02
3260765.1;refGene_gn=LRRFIP2;refGene_tx=XM_023260766.1;refGene_gn=LRRF
IP2;refGene_tx=XM_023260767.1;refGene_gn=LRRFIP2;refGene_tx=XM_0232607
68.1;refGene_gn=LRRFIP2;refGene_tx=XM_023260769.1;refGene_gn=LRRFIP2;r
efGene_tx=XM_023260770.1;refGene_gn=LRRFIP2;refGene_tx=XM_023260771.1
C2 155904426 155904501
refGene_gn=LRRFIP2;refGene_tx=XM_003992258.5;refGene_gn=LRRFIP2;refGen
e_tx=XM_006936543.4;refGene_gn=LRRFIP2;refGene_tx=XM_019810956.1;refGe
ne_gn=LRRFIP2;refGene_tx=XM_019810957.2;refGene_gn=LRRFIP2;refGene_tx=
XM_023260762.1;refGene_gn=LRRFIP2;refGene_tx=XM_023260763.1;refGene_gn
=LRRFIP2;refGene_tx=XM_023260764.1;refGene_gn=LRRFIP2;refGene_tx=XM_02
3260765.1;refGene_gn=LRRFIP2;refGene_tx=XM_023260766.1;refGene_gn=LRRF
IP2;refGene_tx=XM_023260767.1;refGene_gn=LRRFIP2;refGene_tx=XM_0232607
68.1;refGene_gn=LRRFIP2;refGene_tx=XM_023260769.1;refGene_gn=LRRFIP2;r
efGene_tx=XM_023260770.1;refGene_gn=LRRFIP2;refGene_tx=XM_023260771.1
C2 155906281 155906329
refGene_gn=LRRFIP2;refGene_tx=XM_003992258.5;refGene_gn=LRRFIP2;refGen
e_tx=XM_006936543.4;refGene_gn=LRRFIP2;refGene_tx=XM_019810956.1;refGe

ne_gn=LRRFIP2;refGene_tx=XM_019810957.2;refGene_gn=LRRFIP2;refGene_tx=XM_023260762.1;refGene_gn=LRRFIP2;refGene_tx=XM_023260763.1;refGene_gn=LRRFIP2;refGene_tx=XM_023260764.1;refGene_gn=LRRFIP2;refGene_tx=XM_023260765.1;refGene_gn=LRRFIP2;refGene_tx=XM_023260766.1;refGene_gn=LRRFIP2;refGene_tx=XM_023260767.1;refGene_gn=LRRFIP2;refGene_tx=XM_023260768.1;refGene_gn=LRRFIP2;refGene_tx=XM_023260769.1;refGene_gn=LRRFIP2;refGene_tx=XM_023260770.1;refGene_gn=LRRFIP2;refGene_tx=XM_023260771.1
C2 155906410 155906455
refGene_gn=LRRFIP2;refGene_tx=XM_003992258.5;refGene_gn=LRRFIP2;refGene_tx=XM_006936543.4;refGene_gn=LRRFIP2;refGene_tx=XM_019810956.1;refGene_gn=LRRFIP2;refGene_tx=XM_019810957.2;refGene_gn=LRRFIP2;refGene_tx=XM_023260762.1;refGene_gn=LRRFIP2;refGene_tx=XM_023260764.1;refGene_gn=LRRFIP2;refGene_tx=XM_023260765.1;refGene_gn=LRRFIP2;refGene_tx=XM_023260766.1;refGene_gn=LRRFIP2;refGene_tx=XM_023260767.1;refGene_gn=LRRFIP2;refGene_tx=XM_023260768.1;refGene_gn=LRRFIP2;refGene_tx=XM_023260769.1;refGene_gn=LRRFIP2;refGene_tx=XM_023260771.1
C2 155907266 155907314
refGene_gn=LRRFIP2;refGene_tx=XM_003992258.5;refGene_gn=LRRFIP2;refGene_tx=XM_006936543.4;refGene_gn=LRRFIP2;refGene_tx=XM_019810956.1;refGene_gn=LRRFIP2;refGene_tx=XM_019810957.2;refGene_gn=LRRFIP2;refGene_tx=XM_023260762.1;refGene_gn=LRRFIP2;refGene_tx=XM_023260763.1;refGene_gn=LRRFIP2;refGene_tx=XM_023260764.1;refGene_gn=LRRFIP2;refGene_tx=XM_023260765.1;refGene_gn=LRRFIP2;refGene_tx=XM_023260766.1;refGene_gn=LRRFIP2;refGene_tx=XM_023260767.1;refGene_gn=LRRFIP2;refGene_tx=XM_023260768.1;refGene_gn=LRRFIP2;refGene_tx=XM_023260769.1;refGene_gn=LRRFIP2;refGene_tx=XM_023260770.1;refGene_gn=LRRFIP2;refGene_tx=XM_023260771.1
C2 155907780 155907837
refGene_gn=LRRFIP2;refGene_tx=XM_003992258.5;refGene_gn=LRRFIP2;refGene_tx=XM_006936543.4;refGene_gn=LRRFIP2;refGene_tx=XM_019810956.1;refGene_gn=LRRFIP2;refGene_tx=XM_019810957.2;refGene_gn=LRRFIP2;refGene_tx=XM_023260762.1;refGene_gn=LRRFIP2;refGene_tx=XM_023260763.1;refGene_gn=LRRFIP2;refGene_tx=XM_023260764.1;refGene_gn=LRRFIP2;refGene_tx=XM_023260765.1;refGene_gn=LRRFIP2;refGene_tx=XM_023260766.1;refGene_gn=LRRFIP2;refGene_tx=XM_023260767.1;refGene_gn=LRRFIP2;refGene_tx=XM_023260768.1;refGene_gn=LRRFIP2;refGene_tx=XM_023260769.1;refGene_gn=LRRFIP2;refGene_tx=XM_023260770.1;refGene_gn=LRRFIP2;refGene_tx=XM_023260771.1
C2 155911280 155911349
refGene_gn=LRRFIP2;refGene_tx=XM_003992258.5;refGene_gn=LRRFIP2;refGene_tx=XM_006936543.4;refGene_gn=LRRFIP2;refGene_tx=XM_019810956.1;refGene_gn=LRRFIP2;refGene_tx=XM_019810957.2;refGene_gn=LRRFIP2;refGene_tx=XM_023260762.1;refGene_gn=LRRFIP2;refGene_tx=XM_023260763.1;refGene_gn=LRRFIP2;refGene_tx=XM_023260764.1;refGene_gn=LRRFIP2;refGene_tx=XM_023260765.1;refGene_gn=LRRFIP2;refGene_tx=XM_023260766.1;refGene_gn=LRRFIP2;refGene_tx=XM_023260767.1;refGene_gn=LRRFIP2;refGene_tx=XM_023260768.1;refGene_gn=LRRFIP2;refGene_tx=XM_023260769.1;refGene_gn=LRRFIP2;refGene_tx=XM_023260770.1;refGene_gn=LRRFIP2;refGene_tx=XM_023260771.1
C2 155913565 155913655
refGene_gn=LRRFIP2;refGene_tx=XM_003992258.5;refGene_gn=LRRFIP2;refGene_tx=XM_006936543.4;refGene_gn=LRRFIP2;refGene_tx=XM_019810956.1;refGene_gn=LRRFIP2;refGene_tx=XM_019810957.2;refGene_gn=LRRFIP2;refGene_tx=

XM_023260762.1;refGene_gn=LRRFIP2;refGene_tx=XM_023260763.1;refGene_gn
=LRRFIP2;refGene_tx=XM_023260764.1;refGene_gn=LRRFIP2;refGene_tx=XM_02
3260765.1;refGene_gn=LRRFIP2;refGene_tx=XM_023260766.1;refGene_gn=LRRF
IP2;refGene_tx=XM_023260767.1;refGene_gn=LRRFIP2;refGene_tx=XM_0232607
68.1;refGene_gn=LRRFIP2;refGene_tx=XM_023260769.1;refGene_gn=LRRFIP2;r
efGene_tx=XM_023260770.1;refGene_gn=LRRFIP2;refGene_tx=XM_023260771.1
C2 155920433 155920478
refGene_gn=LRRFIP2;refGene_tx=XM_003992258.5;refGene_gn=LRRFIP2;refGen
e_tx=XM_006936543.4;refGene_gn=LRRFIP2;refGene_tx=XM_019810956.1;refGe
ne_gn=LRRFIP2;refGene_tx=XM_019810957.2;refGene_gn=LRRFIP2;refGene_tx=
XM_023260762.1;refGene_gn=LRRFIP2;refGene_tx=XM_023260763.1;refGene_gn
=LRRFIP2;refGene_tx=XM_023260764.1;refGene_gn=LRRFIP2;refGene_tx=XM_02
3260765.1;refGene_gn=LRRFIP2;refGene_tx=XM_023260766.1;refGene_gn=LRRF
IP2;refGene_tx=XM_023260767.1;refGene_gn=LRRFIP2;refGene_tx=XM_0232607
68.1;refGene_gn=LRRFIP2;refGene_tx=XM_023260769.1;refGene_gn=LRRFIP2;r
efGene_tx=XM_023260770.1;refGene_gn=LRRFIP2;refGene_tx=XM_023260771.1;
refGene_gn=LRRFIP2;refGene_tx=XM_023260772.1;refGene_gn=LRRFIP2;refGen
e_tx=XM_023260773.1;refGene_gn=LRRFIP2;refGene_tx=XM_023260774.1;refGe
ne_gn=LRRFIP2;refGene_tx=XM_023260775.1;refGene_gn=LRRFIP2;refGene_tx=
XM_023260776.1;refGene_gn=LRRFIP2;refGene_tx=XM_023260777.1;refGene_gn
=LRRFIP2;refGene_tx=XM_023260778.1;refGene_gn=LRRFIP2;refGene_tx=XM_02
3260779.1
C2 155922052 155922169
refGene_gn=LRRFIP2;refGene_tx=XM_003992258.5;refGene_gn=LRRFIP2;refGen
e_tx=XM_006936543.4;refGene_gn=LRRFIP2;refGene_tx=XM_019810956.1;refGe
ne_gn=LRRFIP2;refGene_tx=XM_019810957.2;refGene_gn=LRRFIP2;refGene_tx=
XM_023260762.1;refGene_gn=LRRFIP2;refGene_tx=XM_023260763.1;refGene_gn
=LRRFIP2;refGene_tx=XM_023260764.1;refGene_gn=LRRFIP2;refGene_tx=XM_02
3260765.1;refGene_gn=LRRFIP2;refGene_tx=XM_023260766.1;refGene_gn=LRRF
IP2;refGene_tx=XM_023260767.1;refGene_gn=LRRFIP2;refGene_tx=XM_0232607
68.1;refGene_gn=LRRFIP2;refGene_tx=XM_023260769.1;refGene_gn=LRRFIP2;r
efGene_tx=XM_023260770.1;refGene_gn=LRRFIP2;refGene_tx=XM_023260771.1;
refGene_gn=LRRFIP2;refGene_tx=XM_023260772.1;refGene_gn=LRRFIP2;refGen
e_tx=XM_023260773.1;refGene_gn=LRRFIP2;refGene_tx=XM_023260774.1;refGe
ne_gn=LRRFIP2;refGene_tx=XM_023260775.1;refGene_gn=LRRFIP2;refGene_tx=
XM_023260776.1;refGene_gn=LRRFIP2;refGene_tx=XM_023260777.1;refGene_gn
=LRRFIP2;refGene_tx=XM_023260778.1;refGene_gn=LRRFIP2;refGene_tx=XM_02
3260779.1
C2 155925010 155925082
refGene_gn=LRRFIP2;refGene_tx=XM_003992258.5;refGene_gn=LRRFIP2;refGen
e_tx=XM_006936543.4;refGene_gn=LRRFIP2;refGene_tx=XM_019810956.1;refGe
ne_gn=LRRFIP2;refGene_tx=XM_019810957.2;refGene_gn=LRRFIP2;refGene_tx=
XM_023260762.1;refGene_gn=LRRFIP2;refGene_tx=XM_023260763.1;refGene_gn
=LRRFIP2;refGene_tx=XM_023260765.1;refGene_gn=LRRFIP2;refGene_tx=XM_02
3260766.1;refGene_gn=LRRFIP2;refGene_tx=XM_023260767.1;refGene_gn=LRRF
IP2;refGene_tx=XM_023260769.1;refGene_gn=LRRFIP2;refGene_tx=XM_0232607
72.1;refGene_gn=LRRFIP2;refGene_tx=XM_023260773.1;refGene_gn=LRRFIP2;r
efGene_tx=XM_023260776.1;refGene_gn=LRRFIP2;refGene_tx=XM_023260778.1
C2 155930325 155930496
refGene_gn=LRRFIP2;refGene_tx=XM_003992258.5;refGene_gn=LRRFIP2;refGen

e_tx=XM_006936543.4;refGene_gn=LRRFIP2;refGene_tx=XM_019810956.1;refGene_gn=LRRFIP2;refGene_tx=XM_019810957.2;refGene_gn=LRRFIP2;refGene_tx=XM_023260762.1;refGene_gn=LRRFIP2;refGene_tx=XM_023260763.1;refGene_gn=LRRFIP2;refGene_tx=XM_023260764.1;refGene_gn=LRRFIP2;refGene_tx=XM_023260765.1;refGene_gn=LRRFIP2;refGene_tx=XM_023260766.1;refGene_gn=LRRFIP2;refGene_tx=XM_023260767.1;refGene_gn=LRRFIP2;refGene_tx=XM_023260768.1;refGene_gn=LRRFIP2;refGene_tx=XM_023260769.1;refGene_gn=LRRFIP2;refGene_tx=XM_023260770.1;refGene_gn=LRRFIP2;refGene_tx=XM_023260771.1;refGene_gn=LRRFIP2;refGene_tx=XM_023260772.1;refGene_gn=LRRFIP2;refGene_tx=XM_023260773.1;refGene_gn=LRRFIP2;refGene_tx=XM_023260774.1;refGene_gn=LRRFIP2;refGene_tx=XM_023260775.1;refGene_gn=LRRFIP2;refGene_tx=XM_023260776.1;refGene_gn=LRRFIP2;refGene_tx=XM_023260777.1;refGene_gn=LRRFIP2;refGene_tx=XM_023260778.1;refGene_gn=LRRFIP2;refGene_tx=XM_023260779.1

C2 155938738 155938831
refGene_gn=LRRFIP2;refGene_tx=XM_003992258.5;refGene_gn=LRRFIP2;refGene_tx=XM_006936543.4;refGene_gn=LRRFIP2;refGene_tx=XM_019810956.1;refGene_gn=LRRFIP2;refGene_tx=XM_019810957.2;refGene_gn=LRRFIP2;refGene_tx=XM_023260762.1;refGene_gn=LRRFIP2;refGene_tx=XM_023260763.1;refGene_gn=LRRFIP2;refGene_tx=XM_023260764.1;refGene_gn=LRRFIP2;refGene_tx=XM_023260765.1;refGene_gn=LRRFIP2;refGene_tx=XM_023260766.1;refGene_gn=LRRFIP2;refGene_tx=XM_023260767.1;refGene_gn=LRRFIP2;refGene_tx=XM_023260768.1;refGene_gn=LRRFIP2;refGene_tx=XM_023260769.1;refGene_gn=LRRFIP2;refGene_tx=XM_023260770.1;refGene_gn=LRRFIP2;refGene_tx=XM_023260771.1;refGene_gn=LRRFIP2;refGene_tx=XM_023260772.1;refGene_gn=LRRFIP2;refGene_tx=XM_023260773.1;refGene_gn=LRRFIP2;refGene_tx=XM_023260774.1;refGene_gn=LRRFIP2;refGene_tx=XM_023260775.1;refGene_gn=LRRFIP2;refGene_tx=XM_023260776.1;refGene_gn=LRRFIP2;refGene_tx=XM_023260777.1;refGene_gn=LRRFIP2;refGene_tx=XM_023260778.1;refGene_gn=LRRFIP2;refGene_tx=XM_023260779.1

C2 155941002 155941095
refGene_gn=LRRFIP2;refGene_tx=XM_003992258.5;refGene_gn=LRRFIP2;refGene_tx=XM_006936543.4;refGene_gn=LRRFIP2;refGene_tx=XM_019810956.1;refGene_gn=LRRFIP2;refGene_tx=XM_019810957.2;refGene_gn=LRRFIP2;refGene_tx=XM_023260762.1;refGene_gn=LRRFIP2;refGene_tx=XM_023260763.1;refGene_gn=LRRFIP2;refGene_tx=XM_023260764.1;refGene_gn=LRRFIP2;refGene_tx=XM_023260766.1;refGene_gn=LRRFIP2;refGene_tx=XM_023260767.1;refGene_gn=LRRFIP2;refGene_tx=XM_023260768.1;refGene_gn=LRRFIP2;refGene_tx=XM_023260773.1;refGene_gn=LRRFIP2;refGene_tx=XM_023260775.1

C2 155946646 155946748
refGene_gn=LRRFIP2;refGene_tx=XM_003992258.5;refGene_gn=LRRFIP2;refGene_tx=XM_006936543.4;refGene_gn=LRRFIP2;refGene_tx=XM_019810956.1;refGene_gn=LRRFIP2;refGene_tx=XM_019810957.2;refGene_gn=LRRFIP2;refGene_tx=XM_023260762.1;refGene_gn=LRRFIP2;refGene_tx=XM_023260763.1;refGene_gn=LRRFIP2;refGene_tx=XM_023260764.1;refGene_gn=LRRFIP2;refGene_tx=XM_023260765.1;refGene_gn=LRRFIP2;refGene_tx=XM_023260767.1;refGene_gn=LRRFIP2;refGene_tx=XM_023260770.1;refGene_gn=LRRFIP2;refGene_tx=XM_023260772.1;refGene_gn=LRRFIP2;refGene_tx=XM_023260773.1;refGene_gn=LRRFIP2;refGene_tx=XM_023260774.1;refGene_gn=LRRFIP2;refGene_tx=XM_023260775.1;refGene_gn=LRRFIP2;refGene_tx=XM_023260776.1;refGene_gn=LRRFIP2;refGen

e_tx=XM_023260777.1
C2	155947011	155947144
refGene_gn=LRRFIP2;refGene_tx=XM_003992258.5;refGene_gn=LRRFIP2;refGene_tx=XM_006936543.4;refGene_gn=LRRFIP2;refGene_tx=XM_019810956.1;refGene_gn=LRRFIP2;refGene_tx=XM_019810957.2;refGene_gn=LRRFIP2;refGene_tx=XM_023260762.1;refGene_gn=LRRFIP2;refGene_tx=XM_023260763.1;refGene_gn=LRRFIP2;refGene_tx=XM_023260764.1;refGene_gn=LRRFIP2;refGene_tx=XM_023260765.1;refGene_gn=LRRFIP2;refGene_tx=XM_023260766.1;refGene_gn=LRRFIP2;refGene_tx=XM_023260767.1;refGene_gn=LRRFIP2;refGene_tx=XM_023260768.1;refGene_gn=LRRFIP2;refGene_tx=XM_023260769.1;refGene_gn=LRRFIP2;refGene_tx=XM_023260770.1;refGene_gn=LRRFIP2;refGene_tx=XM_023260771.1;refGene_gn=LRRFIP2;refGene_tx=XM_023260772.1;refGene_gn=LRRFIP2;refGene_tx=XM_023260773.1;refGene_gn=LRRFIP2;refGene_tx=XM_023260774.1;refGene_gn=LRRFIP2;refGene_tx=XM_023260775.1;refGene_gn=LRRFIP2;refGene_tx=XM_023260776.1;refGene_gn=LRRFIP2;refGene_tx=XM_023260777.1;refGene_gn=LRRFIP2;refGene_tx=XM_023260778.1;refGene_gn=LRRFIP2;refGene_tx=XM_023260779.1
C2	155948901	155948951
refGene_gn=LRRFIP2;refGene_tx=XM_003992258.5;refGene_gn=LRRFIP2;refGene_tx=XM_006936543.4;refGene_gn=LRRFIP2;refGene_tx=XM_019810956.1;refGene_gn=LRRFIP2;refGene_tx=XM_019810957.2;refGene_gn=LRRFIP2;refGene_tx=XM_023260762.1;refGene_gn=LRRFIP2;refGene_tx=XM_023260763.1;refGene_gn=LRRFIP2;refGene_tx=XM_023260764.1;refGene_gn=LRRFIP2;refGene_tx=XM_023260765.1;refGene_gn=LRRFIP2;refGene_tx=XM_023260766.1;refGene_gn=LRRFIP2;refGene_tx=XM_023260767.1;refGene_gn=LRRFIP2;refGene_tx=XM_023260768.1;refGene_gn=LRRFIP2;refGene_tx=XM_023260769.1;refGene_gn=LRRFIP2;refGene_tx=XM_023260770.1;refGene_gn=LRRFIP2;refGene_tx=XM_023260771.1;refGene_gn=LRRFIP2;refGene_tx=XM_023260772.1;refGene_gn=LRRFIP2;refGene_tx=XM_023260773.1;refGene_gn=LRRFIP2;refGene_tx=XM_023260774.1;refGene_gn=LRRFIP2;refGene_tx=XM_023260775.1;refGene_gn=LRRFIP2;refGene_tx=XM_023260776.1;refGene_gn=LRRFIP2;refGene_tx=XM_023260777.1;refGene_gn=LRRFIP2;refGene_tx=XM_023260778.1;refGene_gn=LRRFIP2;refGene_tx=XM_023260779.1
C2	155954440	155954561
refGene_gn=LRRFIP2;refGene_tx=XM_003992258.5;refGene_gn=LRRFIP2;refGene_tx=XM_006936543.4;refGene_gn=LRRFIP2;refGene_tx=XM_019810956.1;refGene_gn=LRRFIP2;refGene_tx=XM_019810957.2;refGene_gn=LRRFIP2;refGene_tx=XM_023260762.1;refGene_gn=LRRFIP2;refGene_tx=XM_023260763.1;refGene_gn=LRRFIP2;refGene_tx=XM_023260764.1;refGene_gn=LRRFIP2;refGene_tx=XM_023260765.1;refGene_gn=LRRFIP2;refGene_tx=XM_023260766.1;refGene_gn=LRRFIP2;refGene_tx=XM_023260767.1;refGene_gn=LRRFIP2;refGene_tx=XM_023260768.1;refGene_gn=LRRFIP2;refGene_tx=XM_023260769.1;refGene_gn=LRRFIP2;refGene_tx=XM_023260770.1;refGene_gn=LRRFIP2;refGene_tx=XM_023260771.1;refGene_gn=LRRFIP2;refGene_tx=XM_023260772.1;refGene_gn=LRRFIP2;refGene_tx=XM_023260773.1;refGene_gn=LRRFIP2;refGene_tx=XM_023260774.1;refGene_gn=LRRFIP2;refGene_tx=XM_023260775.1;refGene_gn=LRRFIP2;refGene_tx=XM_023260776.1;refGene_gn=LRRFIP2;refGene_tx=XM_023260777.1;refGene_gn=LRRFIP2;refGene_tx=XM_023260778.1;refGene_gn=LRRFIP2;refGene_tx=XM_023260779.1
C2	155955436	155955516

refGene_gn=LRRFIP2;refGene_tx=XM_003992258.5;refGene_gn=LRRFIP2;refGene_tx=XM_006936543.4;refGene_gn=LRRFIP2;refGene_tx=XM_019810956.1;refGene_gn=LRRFIP2;refGene_tx=XM_019810957.2;refGene_gn=LRRFIP2;refGene_tx=XM_023260762.1;refGene_gn=LRRFIP2;refGene_tx=XM_023260763.1;refGene_gn=LRRFIP2;refGene_tx=XM_023260764.1;refGene_gn=LRRFIP2;refGene_tx=XM_023260765.1;refGene_gn=LRRFIP2;refGene_tx=XM_023260766.1;refGene_gn=LRRFIP2;refGene_tx=XM_023260767.1;refGene_gn=LRRFIP2;refGene_tx=XM_023260768.1;refGene_gn=LRRFIP2;refGene_tx=XM_023260769.1;refGene_gn=LRRFIP2;refGene_tx=XM_023260770.1;refGene_gn=LRRFIP2;refGene_tx=XM_023260771.1;refGene_gn=LRRFIP2;refGene_tx=XM_023260772.1;refGene_gn=LRRFIP2;refGene_tx=XM_023260773.1;refGene_gn=LRRFIP2;refGene_tx=XM_023260774.1;refGene_gn=LRRFIP2;refGene_tx=XM_023260775.1;refGene_gn=LRRFIP2;refGene_tx=XM_023260776.1;refGene_gn=LRRFIP2;refGene_tx=XM_023260777.1;refGene_gn=LRRFIP2;refGene_tx=XM_023260778.1;refGene_gn=LRRFIP2;refGene_tx=XM_023260779.1
C2 155956099 155956204
refGene_gn=LRRFIP2;refGene_tx=XM_003992258.5;refGene_gn=LRRFIP2;refGene_tx=XM_006936543.4;refGene_gn=LRRFIP2;refGene_tx=XM_019810956.1;refGene_gn=LRRFIP2;refGene_tx=XM_019810957.2;refGene_gn=LRRFIP2;refGene_tx=XM_023260762.1;refGene_gn=LRRFIP2;refGene_tx=XM_023260763.1;refGene_gn=LRRFIP2;refGene_tx=XM_023260764.1;refGene_gn=LRRFIP2;refGene_tx=XM_023260765.1;refGene_gn=LRRFIP2;refGene_tx=XM_023260766.1;refGene_gn=LRRFIP2;refGene_tx=XM_023260767.1;refGene_gn=LRRFIP2;refGene_tx=XM_023260768.1;refGene_gn=LRRFIP2;refGene_tx=XM_023260769.1;refGene_gn=LRRFIP2;refGene_tx=XM_023260770.1;refGene_gn=LRRFIP2;refGene_tx=XM_023260771.1;refGene_gn=LRRFIP2;refGene_tx=XM_023260772.1;refGene_gn=LRRFIP2;refGene_tx=XM_023260773.1;refGene_gn=LRRFIP2;refGene_tx=XM_023260774.1;refGene_gn=LRRFIP2;refGene_tx=XM_023260775.1;refGene_gn=LRRFIP2;refGene_tx=XM_023260776.1;refGene_gn=LRRFIP2;refGene_tx=XM_023260777.1;refGene_gn=LRRFIP2;refGene_tx=XM_023260778.1;refGene_gn=LRRFIP2;refGene_tx=XM_023260779.1
C2 155956576 155956687
refGene_gn=LRRFIP2;refGene_tx=XM_003992258.5;refGene_gn=LRRFIP2;refGene_tx=XM_006936543.4;refGene_gn=LRRFIP2;refGene_tx=XM_019810956.1;refGene_gn=LRRFIP2;refGene_tx=XM_019810957.2;refGene_gn=LRRFIP2;refGene_tx=XM_023260762.1;refGene_gn=LRRFIP2;refGene_tx=XM_023260763.1;refGene_gn=LRRFIP2;refGene_tx=XM_023260764.1;refGene_gn=LRRFIP2;refGene_tx=XM_023260765.1;refGene_gn=LRRFIP2;refGene_tx=XM_023260766.1;refGene_gn=LRRFIP2;refGene_tx=XM_023260767.1;refGene_gn=LRRFIP2;refGene_tx=XM_023260768.1;refGene_gn=LRRFIP2;refGene_tx=XM_023260769.1;refGene_gn=LRRFIP2;refGene_tx=XM_023260770.1;refGene_gn=LRRFIP2;refGene_tx=XM_023260771.1;refGene_gn=LRRFIP2;refGene_tx=XM_023260772.1;refGene_gn=LRRFIP2;refGene_tx=XM_023260773.1;refGene_gn=LRRFIP2;refGene_tx=XM_023260774.1;refGene_gn=LRRFIP2;refGene_tx=XM_023260775.1;refGene_gn=LRRFIP2;refGene_tx=XM_023260776.1;refGene_gn=LRRFIP2;refGene_tx=XM_023260777.1;refGene_gn=LRRFIP2;refGene_tx=XM_023260778.1;refGene_gn=LRRFIP2;refGene_tx=XM_023260779.1
C2 155959443 155959611
refGene_gn=MLH1;refGene_tx=XM_019810954.1;refGene_gn=MLH1;refGene_tx=XM_023260761.1

C2 155960345 155960459
refGene_gn=MLH1;refGene_tx=XM_019810954.1;refGene_gn=MLH1;refGene_tx=XM_023260761.1
C2 155960728 155960821
refGene_gn=MLH1;refGene_tx=XM_019810954.1;refGene_gn=MLH1;refGene_tx=XM_023260761.1
C2 155961359 155961524
refGene_gn=MLH1;refGene_tx=XM_019810954.1;refGene_gn=MLH1;refGene_tx=XM_023260761.1
C2 155965417 155965481
refGene_gn=MLH1;refGene_tx=XM_019810954.1;refGene_gn=MLH1;refGene_tx=XM_023260761.1
C2 155967381 155967490
refGene_gn=MLH1;refGene_tx=XM_019810954.1;refGene_gn=MLH1;refGene_tx=XM_023260761.1
C2 155974727 155974876
refGene_gn=MLH1;refGene_tx=XM_019810954.1;refGene_gn=MLH1;refGene_tx=XM_023260761.1
C2 155976462 155976839
refGene_gn=MLH1;refGene_tx=XM_019810954.1;refGene_gn=MLH1;refGene_tx=XM_023260761.1
C2 155982631 155982785
refGene_gn=MLH1;refGene_tx=XM_019810954.1;refGene_gn=MLH1;refGene_tx=XM_023260761.1
C2 155983608 155983702
refGene_gn=MLH1;refGene_tx=XM_019810954.1
C2 155986607 155986720
refGene_gn=MLH1;refGene_tx=XM_019810954.1
C2 155988618 155988707
refGene_gn=MLH1;refGene_tx=XM_019810954.1
C2 155988840 155988883
refGene_gn=MLH1;refGene_tx=XM_019810954.1
C2 155990330 155990422
refGene_gn=MLH1;refGene_tx=XM_019810954.1
C2 155991537 155991610
refGene_gn=MLH1;refGene_tx=XM_019810954.1
C2 155993679 155993753
refGene_gn=MLH1;refGene_tx=XM_019810954.1
C2 155997768 155997867
refGene_gn=MLH1;refGene_tx=XM_019810954.1
C2 156001105 156001196
refGene_gn=MLH1;refGene_tx=XM_019810954.1
C2 156003217 156003333
refGene_gn=MLH1;refGene_tx=XM_019810954.1
C2 156003814 156005641
refGene_gn=EPM2AIP1;refGene_tx=XM_003992266.5
C2 156026149 156026280
refGene_gn=TRANK1;refGene_tx=XM_003992272.5
C2 156029347 156029474
refGene_gn=TRANK1;refGene_tx=XM_003992272.5

C2 156032032 156032183
refGene_gn=TRANK1;refGene_tx=XM_003992272.5
C2 156034973 156035092
refGene_gn=TRANK1;refGene_tx=XM_003992272.5
C2 156037790 156037874
refGene_gn=TRANK1;refGene_tx=XM_003992272.5
C2 156038119 156038258
refGene_gn=TRANK1;refGene_tx=XM_003992272.5
C2 156040694 156040829
refGene_gn=TRANK1;refGene_tx=XM_003992272.5
C2 156052962 156053133
refGene_gn=TRANK1;refGene_tx=XM_003992272.5
C2 156062966 156063128
refGene_gn=TRANK1;refGene_tx=XM_003992272.5
C2 156065242 156065497
refGene_gn=TRANK1;refGene_tx=XM_003992272.5
C2 156067370 156067547
refGene_gn=TRANK1;refGene_tx=XM_003992272.5
C2 156068442 156071376
refGene_gn=TRANK1;refGene_tx=XM_003992272.5
C2 156076140 156076286
refGene_gn=TRANK1;refGene_tx=XM_003992272.5
C2 156076287 156076341
refGene_gn=TRANK1;refGene_tx=XM_003992272.5
C2 156076657 156076795
refGene_gn=TRANK1;refGene_tx=XM_003992272.5
C2 156081433 156081580
refGene_gn=TRANK1;refGene_tx=XM_003992272.5
C2 156083500 156083657
refGene_gn=TRANK1;refGene_tx=XM_003992272.5
C2 156087316 156087405
refGene_gn=TRANK1;refGene_tx=XM_003992272.5
C2 156092738 156092842
refGene_gn=TRANK1;refGene_tx=XM_003992272.5
C2 156092954 156093087
refGene_gn=TRANK1;refGene_tx=XM_003992272.5
C2 156097979 156098125
refGene_gn=TRANK1;refGene_tx=XM_003992272.5
C2 156101114 156104164
refGene_gn=TRANK1;refGene_tx=XM_003992272.5
C2 156105252 156105351
refGene_gn=TRANK1;refGene_tx=XM_003992272.5
C2 156106421 156106522
refGene_gn=TRANK1;refGene_tx=XM_003992272.5
C2 156181309 156182719
refGene_gn=DCLK3;refGene_tx=XM_023260780.1
C2 156200757 156200890
refGene_gn=DCLK3;refGene_tx=XM_023260780.1
C2 156205449 156205617
refGene_gn=DCLK3;refGene_tx=XM_023260780.1

C2 156207766 156207960
refGene_gn=DCLK3;refGene_tx=XM_023260780.1
C2 156311924 156312023
refGene_gn=STAC;refGene_tx=XM_011286318.2;refGene_gn=STAC;refGene_tx=XM_011286319.2
C2 156317366 156317378
refGene_gn=STAC;refGene_tx=XM_019810967.2
C2 156318957 156319095
refGene_gn=STAC;refGene_tx=XM_011286318.2;refGene_gn=STAC;refGene_tx=XM_011286319.2;refGene_gn=STAC;refGene_tx=XM_019810967.2
C2 156319211 156319263
refGene_gn=STAC;refGene_tx=XM_011286318.2;refGene_gn=STAC;refGene_tx=XM_011286319.2;refGene_gn=STAC;refGene_tx=XM_019810967.2
C2 156332284 156332489
refGene_gn=STAC;refGene_tx=XM_019810966.2
C2 156336419 156336508
refGene_gn=STAC;refGene_tx=XM_011286318.2;refGene_gn=STAC;refGene_tx=XM_011286319.2;refGene_gn=STAC;refGene_tx=XM_019810966.2;refGene_gn=STAC;refGene_tx=XM_019810967.2
C2 156338238 156338303
refGene_gn=STAC;refGene_tx=XM_011286318.2;refGene_gn=STAC;refGene_tx=XM_011286319.2;refGene_gn=STAC;refGene_tx=XM_019810966.2;refGene_gn=STAC;refGene_tx=XM_019810967.2
C2 156348533 156348612
refGene_gn=STAC;refGene_tx=XM_011286318.2;refGene_gn=STAC;refGene_tx=XM_011286319.2;refGene_gn=STAC;refGene_tx=XM_019810966.2;refGene_gn=STAC;refGene_tx=XM_019810967.2
C2 156355092 156355208
refGene_gn=STAC;refGene_tx=XM_011286318.2;refGene_gn=STAC;refGene_tx=XM_011286319.2;refGene_gn=STAC;refGene_tx=XM_019810966.2;refGene_gn=STAC;refGene_tx=XM_019810967.2
C2 156356363 156356445
refGene_gn=STAC;refGene_tx=XM_011286318.2;refGene_gn=STAC;refGene_tx=XM_011286319.2;refGene_gn=STAC;refGene_tx=XM_019810966.2;refGene_gn=STAC;refGene_tx=XM_019810967.2
C2 156358020 156358121
refGene_gn=STAC;refGene_tx=XM_011286318.2;refGene_gn=STAC;refGene_tx=XM_011286319.2;refGene_gn=STAC;refGene_tx=XM_019810966.2;refGene_gn=STAC;refGene_tx=XM_019810967.2
C2 156391587 156391867
refGene_gn=STAC;refGene_tx=XM_011286318.2;refGene_gn=STAC;refGene_tx=XM_011286319.2;refGene_gn=STAC;refGene_tx=XM_019810966.2;refGene_gn=STAC;refGene_tx=XM_019810967.2
C2 156449072 156449183
refGene_gn=STAC;refGene_tx=XM_011286318.2;refGene_gn=STAC;refGene_tx=XM_011286319.2;refGene_gn=STAC;refGene_tx=XM_019810966.2;refGene_gn=STAC;refGene_tx=XM_019810967.2
C2 156917467 156917725
refGene_gn=ARPP21;refGene_tx=XM_011286303.2;refGene_gn=ARPP21;refGene_tx=XM_011286304.3;refGene_gn=ARPP21;refGene_tx=XM_011286309.3;refGene_

gn=ARPP21;refGene_tx=XM_011286311.2;refGene_gn=ARPP21;refGene_tx=XM_01
1286312.2;refGene_gn=ARPP21;refGene_tx=XM_011286313.2;refGene_gn=ARPP2
1;refGene_tx=XM_011286314.2;refGene_gn=ARPP21;refGene_tx=XM_011286315.
2;refGene_gn=ARPP21;refGene_tx=XM_011286316.2;refGene_gn=ARPP21;refGen
e_tx=XM_011286317.2;refGene_gn=ARPP21;refGene_tx=XM_019810969.2;refGen
e_gn=ARPP21;refGene_tx=XM_019810970.2;refGene_gn=ARPP21;refGene_tx=XM_
019810971.2;refGene_gn=ARPP21;refGene_tx=XM_019810972.2;refGene_gn=ARP
P21;refGene_tx=XM_019810973.2;refGene_gn=ARPP21;refGene_tx=XM_01981097
4.2;refGene_gn=ARPP21;refGene_tx=XM_019810976.2;refGene_gn=ARPP21;refG
ene_tx=XM_019810977.2;refGene_gn=ARPP21;refGene_tx=XM_019810978.2;refG
ene_gn=ARPP21;refGene_tx=XM_019810979.2;refGene_gn=ARPP21;refGene_tx=X
M_019810980.2;refGene_gn=ARPP21;refGene_tx=XM_019810981.2;refGene_gn=A
RPP21;refGene_tx=XM_019810983.2;refGene_gn=ARPP21;refGene_tx=XM_019810
984.2;refGene_gn=ARPP21;refGene_tx=XM_019810985.2;refGene_gn=ARPP21;re
fGene_tx=XM_019810986.2;refGene_gn=ARPP21;refGene_tx=XM_019810987.2;re
fGene_gn=ARPP21;refGene_tx=XM_019810988.2;refGene_gn=ARPP21;refGene_tx
=XM_023260783.1;refGene_gn=ARPP21;refGene_tx=XM_023260786.1
C2 156918826 156918975
refGene_gn=ARPP21;refGene_tx=XM_011286303.2;refGene_gn=ARPP21;refGene_
tx=XM_011286304.3;refGene_gn=ARPP21;refGene_tx=XM_011286309.3;refGene_
gn=ARPP21;refGene_tx=XM_011286311.2;refGene_gn=ARPP21;refGene_tx=XM_01
1286312.2;refGene_gn=ARPP21;refGene_tx=XM_011286313.2;refGene_gn=ARPP2
1;refGene_tx=XM_011286314.2;refGene_gn=ARPP21;refGene_tx=XM_011286315.
2;refGene_gn=ARPP21;refGene_tx=XM_011286316.2;refGene_gn=ARPP21;refGen
e_tx=XM_011286317.2;refGene_gn=ARPP21;refGene_tx=XM_019810969.2;refGen
e_gn=ARPP21;refGene_tx=XM_019810970.2;refGene_gn=ARPP21;refGene_tx=XM_
019810971.2;refGene_gn=ARPP21;refGene_tx=XM_019810972.2;refGene_gn=ARP
P21;refGene_tx=XM_019810973.2;refGene_gn=ARPP21;refGene_tx=XM_01981097
4.2;refGene_gn=ARPP21;refGene_tx=XM_019810976.2;refGene_gn=ARPP21;refG
ene_tx=XM_019810977.2;refGene_gn=ARPP21;refGene_tx=XM_019810978.2;refG
ene_gn=ARPP21;refGene_tx=XM_019810979.2;refGene_gn=ARPP21;refGene_tx=X
M_019810980.2;refGene_gn=ARPP21;refGene_tx=XM_019810981.2;refGene_gn=A
RPP21;refGene_tx=XM_019810983.2;refGene_gn=ARPP21;refGene_tx=XM_019810
984.2;refGene_gn=ARPP21;refGene_tx=XM_019810985.2;refGene_gn=ARPP21;re
fGene_tx=XM_019810986.2;refGene_gn=ARPP21;refGene_tx=XM_019810987.2;re
fGene_gn=ARPP21;refGene_tx=XM_019810988.2;refGene_gn=ARPP21;refGene_tx
=XM_023260783.1;refGene_gn=ARPP21;refGene_tx=XM_023260786.1
C2 156969691 156969818
refGene_gn=ARPP21;refGene_tx=XM_011286303.2;refGene_gn=ARPP21;refGene_
tx=XM_011286304.3;refGene_gn=ARPP21;refGene_tx=XM_011286309.3;refGene_
gn=ARPP21;refGene_tx=XM_011286311.2;refGene_gn=ARPP21;refGene_tx=XM_01
1286312.2;refGene_gn=ARPP21;refGene_tx=XM_011286313.2;refGene_gn=ARPP2
1;refGene_tx=XM_011286314.2;refGene_gn=ARPP21;refGene_tx=XM_011286315.
2;refGene_gn=ARPP21;refGene_tx=XM_011286316.2;refGene_gn=ARPP21;refGen
e_tx=XM_011286317.2;refGene_gn=ARPP21;refGene_tx=XM_019810969.2;refGen
e_gn=ARPP21;refGene_tx=XM_019810970.2;refGene_gn=ARPP21;refGene_tx=XM_
019810971.2;refGene_gn=ARPP21;refGene_tx=XM_019810972.2;refGene_gn=ARP
P21;refGene_tx=XM_019810973.2;refGene_gn=ARPP21;refGene_tx=XM_01981097
4.2;refGene_gn=ARPP21;refGene_tx=XM_019810976.2;refGene_gn=ARPP21;refG
ene_tx=XM_019810977.2;refGene_gn=ARPP21;refGene_tx=XM_019810978.2;refG

ene_gn=ARPP21;refGene_tx=XM_019810979.2;refGene_gn=ARPP21;refGene_tx=XM_019810980.2;refGene_gn=ARPP21;refGene_tx=XM_019810981.2;refGene_gn=ARPP21;refGene_tx=XM_019810983.2;refGene_gn=ARPP21;refGene_tx=XM_019810984.2;refGene_gn=ARPP21;refGene_tx=XM_019810985.2;refGene_gn=ARPP21;refGene_tx=XM_019810986.2;refGene_gn=ARPP21;refGene_tx=XM_019810987.2;refGene_gn=ARPP21;refGene_tx=XM_019810988.2;refGene_gn=ARPP21;refGene_tx=XM_023260783.1;refGene_gn=ARPP21;refGene_tx=XM_023260786.1
C2 156974020 156974281
refGene_gn=ARPP21;refGene_tx=XM_011286303.2;refGene_gn=ARPP21;refGene_tx=XM_011286304.3;refGene_gn=ARPP21;refGene_tx=XM_011286309.3;refGene_gn=ARPP21;refGene_tx=XM_011286311.2;refGene_gn=ARPP21;refGene_tx=XM_011286312.2;refGene_gn=ARPP21;refGene_tx=XM_011286313.2;refGene_gn=ARPP21;refGene_tx=XM_011286314.2;refGene_gn=ARPP21;refGene_tx=XM_011286315.2;refGene_gn=ARPP21;refGene_tx=XM_011286316.2;refGene_gn=ARPP21;refGene_tx=XM_011286317.2;refGene_gn=ARPP21;refGene_tx=XM_019810969.2;refGene_gn=ARPP21;refGene_tx=XM_019810970.2;refGene_gn=ARPP21;refGene_tx=XM_019810971.2;refGene_gn=ARPP21;refGene_tx=XM_019810972.2;refGene_gn=ARPP21;refGene_tx=XM_019810973.2;refGene_gn=ARPP21;refGene_tx=XM_019810974.2;refGene_gn=ARPP21;refGene_tx=XM_019810976.2;refGene_gn=ARPP21;refGene_tx=XM_019810977.2;refGene_gn=ARPP21;refGene_tx=XM_019810978.2;refGene_gn=ARPP21;refGene_tx=XM_019810979.2;refGene_gn=ARPP21;refGene_tx=XM_019810980.2;refGene_gn=ARPP21;refGene_tx=XM_019810981.2;refGene_gn=ARPP21;refGene_tx=XM_019810983.2;refGene_gn=ARPP21;refGene_tx=XM_019810984.2;refGene_gn=ARPP21;refGene_tx=XM_019810985.2;refGene_gn=ARPP21;refGene_tx=XM_019810986.2;refGene_gn=ARPP21;refGene_tx=XM_019810987.2;refGene_gn=ARPP21;refGene_tx=XM_019810988.2;refGene_gn=ARPP21;refGene_tx=XM_023260783.1;refGene_gn=ARPP21;refGene_tx=XM_023260786.1
C2 156975235 156975340
refGene_gn=ARPP21;refGene_tx=XM_011286303.2;refGene_gn=ARPP21;refGene_tx=XM_011286304.3;refGene_gn=ARPP21;refGene_tx=XM_011286309.3;refGene_gn=ARPP21;refGene_tx=XM_011286311.2;refGene_gn=ARPP21;refGene_tx=XM_011286312.2;refGene_gn=ARPP21;refGene_tx=XM_011286313.2;refGene_gn=ARPP21;refGene_tx=XM_011286314.2;refGene_gn=ARPP21;refGene_tx=XM_011286315.2;refGene_gn=ARPP21;refGene_tx=XM_011286316.2;refGene_gn=ARPP21;refGene_tx=XM_011286317.2;refGene_gn=ARPP21;refGene_tx=XM_019810969.2;refGene_gn=ARPP21;refGene_tx=XM_019810970.2;refGene_gn=ARPP21;refGene_tx=XM_019810971.2;refGene_gn=ARPP21;refGene_tx=XM_019810972.2;refGene_gn=ARPP21;refGene_tx=XM_019810973.2;refGene_gn=ARPP21;refGene_tx=XM_019810974.2;refGene_gn=ARPP21;refGene_tx=XM_019810976.2;refGene_gn=ARPP21;refGene_tx=XM_019810979.2;refGene_gn=ARPP21;refGene_tx=XM_019810980.2;refGene_gn=ARPP21;refGene_tx=XM_019810981.2;refGene_gn=ARPP21;refGene_tx=XM_019810983.2;refGene_gn=ARPP21;refGene_tx=XM_019810984.2;refGene_gn=ARPP21;refGene_tx=XM_019810985.2;refGene_gn=ARPP21;refGene_tx=XM_019810986.2;refGene_gn=ARPP21;refGene_tx=XM_019810987.2;refGene_gn=ARPP21;refGene_tx=XM_019810988.2;refGene_gn=ARPP21;refGene_tx=XM_023260783.1;refGene_gn=ARPP21;refGene_tx=XM_023260786.1
C2 156976313 156976495
refGene_gn=ARPP21;refGene_tx=XM_011286303.2;refGene_gn=ARPP21;refGene_tx=XM_011286304.3;refGene_gn=ARPP21;refGene_tx=XM_011286309.3;refGene_gn=ARPP21;refGene_tx=XM_011286311.2;refGene_gn=ARPP21;refGene_tx=XM_01

1286312.2;refGene_gn=ARPP21;refGene_tx=XM_011286313.2;refGene_gn=ARPP2
1;refGene_tx=XM_011286314.2;refGene_gn=ARPP21;refGene_tx=XM_011286315.
2;refGene_gn=ARPP21;refGene_tx=XM_011286316.2;refGene_gn=ARPP21;refGen
e_tx=XM_011286317.2;refGene_gn=ARPP21;refGene_tx=XM_019810969.2;refGen
e_gn=ARPP21;refGene_tx=XM_019810970.2;refGene_gn=ARPP21;refGene_tx=XM_
019810971.2;refGene_gn=ARPP21;refGene_tx=XM_019810972.2;refGene_gn=ARP
P21;refGene_tx=XM_019810973.2;refGene_gn=ARPP21;refGene_tx=XM_01981097
4.2;refGene_gn=ARPP21;refGene_tx=XM_019810976.2;refGene_gn=ARPP21;refG
ene_tx=XM_019810977.2;refGene_gn=ARPP21;refGene_tx=XM_019810978.2;refG
ene_gn=ARPP21;refGene_tx=XM_019810979.2;refGene_gn=ARPP21;refGene_tx=X
M_019810980.2;refGene_gn=ARPP21;refGene_tx=XM_019810981.2;refGene_gn=A
RPP21;refGene_tx=XM_019810983.2;refGene_gn=ARPP21;refGene_tx=XM_019810
984.2;refGene_gn=ARPP21;refGene_tx=XM_019810985.2;refGene_gn=ARPP21;re
fGene_tx=XM_019810986.2;refGene_gn=ARPP21;refGene_tx=XM_019810987.2;re
fGene_gn=ARPP21;refGene_tx=XM_019810988.2;refGene_gn=ARPP21;refGene_tx
=XM_023260783.1;refGene_gn=ARPP21;refGene_tx=XM_023260786.1
C2 156984130 156984361
refGene_gn=ARPP21;refGene_tx=XM_011286303.2;refGene_gn=ARPP21;refGene_
tx=XM_011286304.3;refGene_gn=ARPP21;refGene_tx=XM_011286309.3;refGene_
gn=ARPP21;refGene_tx=XM_011286311.2;refGene_gn=ARPP21;refGene_tx=XM_01
1286312.2;refGene_gn=ARPP21;refGene_tx=XM_011286313.2;refGene_gn=ARPP2
1;refGene_tx=XM_011286314.2;refGene_gn=ARPP21;refGene_tx=XM_011286315.
2;refGene_gn=ARPP21;refGene_tx=XM_011286316.2;refGene_gn=ARPP21;refGen
e_tx=XM_011286317.2;refGene_gn=ARPP21;refGene_tx=XM_019810969.2;refGen
e_gn=ARPP21;refGene_tx=XM_019810970.2;refGene_gn=ARPP21;refGene_tx=XM_
019810971.2;refGene_gn=ARPP21;refGene_tx=XM_019810972.2;refGene_gn=ARP
P21;refGene_tx=XM_019810973.2;refGene_gn=ARPP21;refGene_tx=XM_01981097
4.2;refGene_gn=ARPP21;refGene_tx=XM_019810976.2;refGene_gn=ARPP21;refG
ene_tx=XM_019810977.2;refGene_gn=ARPP21;refGene_tx=XM_019810978.2;refG
ene_gn=ARPP21;refGene_tx=XM_019810979.2;refGene_gn=ARPP21;refGene_tx=X
M_019810980.2;refGene_gn=ARPP21;refGene_tx=XM_019810981.2;refGene_gn=A
RPP21;refGene_tx=XM_019810983.2;refGene_gn=ARPP21;refGene_tx=XM_019810
986.2;refGene_gn=ARPP21;refGene_tx=XM_019810987.2;refGene_gn=ARPP21;re
fGene_tx=XM_019810988.2;refGene_gn=ARPP21;refGene_tx=XM_023260783.1;re
fGene_gn=ARPP21;refGene_tx=XM_023260786.1
C2 156991002 156991235
refGene_gn=ARPP21;refGene_tx=XM_011286303.2;refGene_gn=ARPP21;refGene_
tx=XM_011286304.3;refGene_gn=ARPP21;refGene_tx=XM_011286309.3;refGene_
gn=ARPP21;refGene_tx=XM_011286311.2;refGene_gn=ARPP21;refGene_tx=XM_01
1286312.2;refGene_gn=ARPP21;refGene_tx=XM_011286313.2;refGene_gn=ARPP2
1;refGene_tx=XM_011286314.2;refGene_gn=ARPP21;refGene_tx=XM_011286315.
2;refGene_gn=ARPP21;refGene_tx=XM_011286316.2;refGene_gn=ARPP21;refGen
e_tx=XM_011286317.2;refGene_gn=ARPP21;refGene_tx=XM_019810969.2;refGen
e_gn=ARPP21;refGene_tx=XM_019810970.2;refGene_gn=ARPP21;refGene_tx=XM_
019810971.2;refGene_gn=ARPP21;refGene_tx=XM_019810972.2;refGene_gn=ARP
P21;refGene_tx=XM_019810973.2;refGene_gn=ARPP21;refGene_tx=XM_01981097
4.2;refGene_gn=ARPP21;refGene_tx=XM_019810976.2;refGene_gn=ARPP21;refG
ene_tx=XM_019810977.2;refGene_gn=ARPP21;refGene_tx=XM_019810978.2;refG
ene_gn=ARPP21;refGene_tx=XM_019810979.2;refGene_gn=ARPP21;refGene_tx=X
M_019810980.2;refGene_gn=ARPP21;refGene_tx=XM_019810981.2;refGene_gn=A

RPP21;refGene_tx=XM_019810983.2;refGene_gn=ARPP21;refGene_tx=XM_019810984.2;refGene_gn=ARPP21;refGene_tx=XM_019810985.2;refGene_gn=ARPP21;refGene_tx=XM_019810986.2;refGene_gn=ARPP21;refGene_tx=XM_019810987.2;refGene_gn=ARPP21;refGene_tx=XM_019810988.2;refGene_gn=ARPP21;refGene_tx=XM_023260783.1;refGene_gn=ARPP21;refGene_tx=XM_023260786.1
C2 156995292 156995356
refGene_gn=ARPP21;refGene_tx=XM_011286303.2;refGene_gn=ARPP21;refGene_tx=XM_011286304.3;refGene_gn=ARPP21;refGene_tx=XM_011286309.3;refGene_gn=ARPP21;refGene_tx=XM_011286311.2;refGene_gn=ARPP21;refGene_tx=XM_011286312.2;refGene_gn=ARPP21;refGene_tx=XM_011286313.2;refGene_gn=ARPP21;refGene_tx=XM_011286314.2;refGene_gn=ARPP21;refGene_tx=XM_011286315.2;refGene_gn=ARPP21;refGene_tx=XM_011286316.2;refGene_gn=ARPP21;refGene_tx=XM_011286317.2;refGene_gn=ARPP21;refGene_tx=XM_019810969.2;refGene_gn=ARPP21;refGene_tx=XM_019810970.2;refGene_gn=ARPP21;refGene_tx=XM_019810971.2;refGene_gn=ARPP21;refGene_tx=XM_019810972.2;refGene_gn=ARPP21;refGene_tx=XM_019810973.2;refGene_gn=ARPP21;refGene_tx=XM_019810974.2;refGene_gn=ARPP21;refGene_tx=XM_019810977.2;refGene_gn=ARPP21;refGene_tx=XM_019810978.2;refGene_gn=ARPP21;refGene_tx=XM_019810981.2;refGene_gn=ARPP21;refGene_tx=XM_019810983.2;refGene_gn=ARPP21;refGene_tx=XM_019810984.2;refGene_gn=ARPP21;refGene_tx=XM_019810985.2;refGene_gn=ARPP21;refGene_tx=XM_019810986.2;refGene_gn=ARPP21;refGene_tx=XM_019810988.2;refGene_gn=ARPP21;refGene_tx=XM_023260783.1;refGene_gn=ARPP21;refGene_tx=XM_023260786.1
C2 156997138 156997176
refGene_gn=ARPP21;refGene_tx=XM_011286303.2;refGene_gn=ARPP21;refGene_tx=XM_011286304.3;refGene_gn=ARPP21;refGene_tx=XM_011286309.3;refGene_gn=ARPP21;refGene_tx=XM_011286311.2;refGene_gn=ARPP21;refGene_tx=XM_011286312.2;refGene_gn=ARPP21;refGene_tx=XM_011286313.2;refGene_gn=ARPP21;refGene_tx=XM_011286314.2;refGene_gn=ARPP21;refGene_tx=XM_011286315.2;refGene_gn=ARPP21;refGene_tx=XM_011286316.2;refGene_gn=ARPP21;refGene_tx=XM_011286317.2;refGene_gn=ARPP21;refGene_tx=XM_019810969.2;refGene_gn=ARPP21;refGene_tx=XM_019810970.2;refGene_gn=ARPP21;refGene_tx=XM_019810971.2;refGene_gn=ARPP21;refGene_tx=XM_019810972.2;refGene_gn=ARPP21;refGene_tx=XM_019810973.2;refGene_gn=ARPP21;refGene_tx=XM_019810974.2;refGene_gn=ARPP21;refGene_tx=XM_019810976.2;refGene_gn=ARPP21;refGene_tx=XM_019810977.2;refGene_gn=ARPP21;refGene_tx=XM_019810978.2;refGene_gn=ARPP21;refGene_tx=XM_019810979.2;refGene_gn=ARPP21;refGene_tx=XM_019810980.2;refGene_gn=ARPP21;refGene_tx=XM_019810981.2;refGene_gn=ARPP21;refGene_tx=XM_019810983.2;refGene_gn=ARPP21;refGene_tx=XM_019810984.2;refGene_gn=ARPP21;refGene_tx=XM_019810985.2;refGene_gn=ARPP21;refGene_tx=XM_019810988.2;refGene_gn=ARPP21;refGene_tx=XM_023260783.1;refGene_gn=ARPP21;refGene_tx=XM_023260786.1
C2 157003440 157003542
refGene_gn=ARPP21;refGene_tx=XM_011286303.2;refGene_gn=ARPP21;refGene_tx=XM_011286304.3;refGene_gn=ARPP21;refGene_tx=XM_011286309.3;refGene_gn=ARPP21;refGene_tx=XM_011286311.2;refGene_gn=ARPP21;refGene_tx=XM_011286312.2;refGene_gn=ARPP21;refGene_tx=XM_011286313.2;refGene_gn=ARPP21;refGene_tx=XM_011286314.2;refGene_gn=ARPP21;refGene_tx=XM_019810969.2;refGene_gn=ARPP21;refGene_tx=XM_019810970.2;refGene_gn=ARPP21;refGene_tx=XM_019810971.2;refGene_gn=ARPP21;refGene_tx=XM_019810972.2;refGen

e_gn=ARPP21;refGene_tx=XM_019810976.2;refGene_gn=ARPP21;refGene_tx=XM_
019810977.2;refGene_gn=ARPP21;refGene_tx=XM_019810978.2;refGene_gn=ARP
P21;refGene_tx=XM_019810984.2;refGene_gn=ARPP21;refGene_tx=XM_01981098
5.2;refGene_gn=ARPP21;refGene_tx=XM_019810988.2;refGene_gn=ARPP21;refG
ene_tx=XM_023260783.1
C2 157005005 157005094
refGene_gn=ARPP21;refGene_tx=XM_019810973.2;refGene_gn=ARPP21;refGene_
tx=XM_019810974.2
C2 157005479 157005588
refGene_gn=ARPP21;refGene_tx=XM_011286303.2;refGene_gn=ARPP21;refGene_
tx=XM_011286304.3;refGene_gn=ARPP21;refGene_tx=XM_011286309.3;refGene_
gn=ARPP21;refGene_tx=XM_011286311.2;refGene_gn=ARPP21;refGene_tx=XM_01
1286312.2;refGene_gn=ARPP21;refGene_tx=XM_011286313.2;refGene_gn=ARPP2
1;refGene_tx=XM_011286314.2;refGene_gn=ARPP21;refGene_tx=XM_011286315.
2;refGene_gn=ARPP21;refGene_tx=XM_011286316.2;refGene_gn=ARPP21;refGen
e_tx=XM_011286317.2;refGene_gn=ARPP21;refGene_tx=XM_019810969.2;refGen
e_gn=ARPP21;refGene_tx=XM_019810970.2;refGene_gn=ARPP21;refGene_tx=XM_
019810971.2;refGene_gn=ARPP21;refGene_tx=XM_019810972.2;refGene_gn=ARP
P21;refGene_tx=XM_019810973.2;refGene_gn=ARPP21;refGene_tx=XM_01981097
4.2;refGene_gn=ARPP21;refGene_tx=XM_019810976.2;refGene_gn=ARPP21;refG
ene_tx=XM_019810977.2;refGene_gn=ARPP21;refGene_tx=XM_019810978.2;refG
ene_gn=ARPP21;refGene_tx=XM_019810979.2;refGene_gn=ARPP21;refGene_tx=X
M_019810980.2;refGene_gn=ARPP21;refGene_tx=XM_019810984.2;refGene_gn=A
RPP21;refGene_tx=XM_019810985.2;refGene_gn=ARPP21;refGene_tx=XM_019810
988.2;refGene_gn=ARPP21;refGene_tx=XM_023260783.1;refGene_gn=ARPP21;re
fGene_tx=XM_023260786.1
C2 157020101 157020242
refGene_gn=ARPP21;refGene_tx=XM_011286303.2;refGene_gn=ARPP21;refGene_
tx=XM_011286304.3;refGene_gn=ARPP21;refGene_tx=XM_011286309.3;refGene_
gn=ARPP21;refGene_tx=XM_011286311.2;refGene_gn=ARPP21;refGene_tx=XM_01
1286312.2;refGene_gn=ARPP21;refGene_tx=XM_011286313.2;refGene_gn=ARPP2
1;refGene_tx=XM_011286314.2;refGene_gn=ARPP21;refGene_tx=XM_011286315.
2;refGene_gn=ARPP21;refGene_tx=XM_011286316.2;refGene_gn=ARPP21;refGen
e_tx=XM_011286317.2;refGene_gn=ARPP21;refGene_tx=XM_019810969.2;refGen
e_gn=ARPP21;refGene_tx=XM_019810970.2;refGene_gn=ARPP21;refGene_tx=XM_
019810971.2;refGene_gn=ARPP21;refGene_tx=XM_019810972.2;refGene_gn=ARP
P21;refGene_tx=XM_019810973.2;refGene_gn=ARPP21;refGene_tx=XM_01981097
4.2;refGene_gn=ARPP21;refGene_tx=XM_019810976.2;refGene_gn=ARPP21;refG
ene_tx=XM_019810977.2;refGene_gn=ARPP21;refGene_tx=XM_019810978.2;refG
ene_gn=ARPP21;refGene_tx=XM_019810979.2;refGene_gn=ARPP21;refGene_tx=X
M_019810980.2;refGene_gn=ARPP21;refGene_tx=XM_019810981.2;refGene_gn=A
RPP21;refGene_tx=XM_019810983.2;refGene_gn=ARPP21;refGene_tx=XM_019810
984.2;refGene_gn=ARPP21;refGene_tx=XM_019810985.2;refGene_gn=ARPP21;re
fGene_tx=XM_019810986.2;refGene_gn=ARPP21;refGene_tx=XM_019810987.2;re
fGene_gn=ARPP21;refGene_tx=XM_019810988.2;refGene_gn=ARPP21;refGene_tx
=XM_023260783.1;refGene_gn=ARPP21;refGene_tx=XM_023260786.1
C2 157020974 157021034
refGene_gn=ARPP21;refGene_tx=XM_011286303.2;refGene_gn=ARPP21;refGene_
tx=XM_011286304.3;refGene_gn=ARPP21;refGene_tx=XM_011286309.3;refGene_
gn=ARPP21;refGene_tx=XM_011286311.2;refGene_gn=ARPP21;refGene_tx=XM_01

1286312.2;refGene_gn=ARPP21;refGene_tx=XM_011286313.2;refGene_gn=ARPP2
1;refGene_tx=XM_011286314.2;refGene_gn=ARPP21;refGene_tx=XM_011286315.
2;refGene_gn=ARPP21;refGene_tx=XM_011286316.2;refGene_gn=ARPP21;refGen
e_tx=XM_011286317.2;refGene_gn=ARPP21;refGene_tx=XM_019810969.2;refGen
e_gn=ARPP21;refGene_tx=XM_019810970.2;refGene_gn=ARPP21;refGene_tx=XM_
019810971.2;refGene_gn=ARPP21;refGene_tx=XM_019810972.2;refGene_gn=ARP
P21;refGene_tx=XM_019810973.2;refGene_gn=ARPP21;refGene_tx=XM_01981097
4.2;refGene_gn=ARPP21;refGene_tx=XM_019810976.2;refGene_gn=ARPP21;refG
ene_tx=XM_019810977.2;refGene_gn=ARPP21;refGene_tx=XM_019810978.2;refG
ene_gn=ARPP21;refGene_tx=XM_019810979.2;refGene_gn=ARPP21;refGene_tx=X
M_019810980.2;refGene_gn=ARPP21;refGene_tx=XM_019810981.2;refGene_gn=A
RPP21;refGene_tx=XM_019810983.2;refGene_gn=ARPP21;refGene_tx=XM_019810
984.2;refGene_gn=ARPP21;refGene_tx=XM_019810985.2;refGene_gn=ARPP21;re
fGene_tx=XM_019810986.2;refGene_gn=ARPP21;refGene_tx=XM_019810987.2;re
fGene_gn=ARPP21;refGene_tx=XM_019810988.2;refGene_gn=ARPP21;refGene_tx
=XM_023260783.1;refGene_gn=ARPP21;refGene_tx=XM_023260786.1
C2 157021734 157021813
refGene_gn=ARPP21;refGene_tx=XM_011286303.2;refGene_gn=ARPP21;refGene_
tx=XM_011286304.3;refGene_gn=ARPP21;refGene_tx=XM_011286309.3;refGene_
gn=ARPP21;refGene_tx=XM_011286311.2;refGene_gn=ARPP21;refGene_tx=XM_01
1286312.2;refGene_gn=ARPP21;refGene_tx=XM_011286313.2;refGene_gn=ARPP2
1;refGene_tx=XM_011286314.2;refGene_gn=ARPP21;refGene_tx=XM_011286315.
2;refGene_gn=ARPP21;refGene_tx=XM_011286316.2;refGene_gn=ARPP21;refGen
e_tx=XM_011286317.2;refGene_gn=ARPP21;refGene_tx=XM_019810969.2;refGen
e_gn=ARPP21;refGene_tx=XM_019810970.2;refGene_gn=ARPP21;refGene_tx=XM_
019810971.2;refGene_gn=ARPP21;refGene_tx=XM_019810972.2;refGene_gn=ARP
P21;refGene_tx=XM_019810973.2;refGene_gn=ARPP21;refGene_tx=XM_01981097
4.2;refGene_gn=ARPP21;refGene_tx=XM_019810976.2;refGene_gn=ARPP21;refG
ene_tx=XM_019810977.2;refGene_gn=ARPP21;refGene_tx=XM_019810978.2;refG
ene_gn=ARPP21;refGene_tx=XM_019810979.2;refGene_gn=ARPP21;refGene_tx=X
M_019810980.2;refGene_gn=ARPP21;refGene_tx=XM_019810981.2;refGene_gn=A
RPP21;refGene_tx=XM_019810983.2;refGene_gn=ARPP21;refGene_tx=XM_019810
984.2;refGene_gn=ARPP21;refGene_tx=XM_019810985.2;refGene_gn=ARPP21;re
fGene_tx=XM_019810986.2;refGene_gn=ARPP21;refGene_tx=XM_019810987.2;re
fGene_gn=ARPP21;refGene_tx=XM_023260783.1;refGene_gn=ARPP21;refGene_tx
=XM_023260786.1
C2 157023174 157023319
refGene_gn=ARPP21;refGene_tx=XM_011286303.2;refGene_gn=ARPP21;refGene_
tx=XM_011286304.3;refGene_gn=ARPP21;refGene_tx=XM_011286309.3;refGene_
gn=ARPP21;refGene_tx=XM_011286311.2;refGene_gn=ARPP21;refGene_tx=XM_01
1286312.2;refGene_gn=ARPP21;refGene_tx=XM_011286313.2;refGene_gn=ARPP2
1;refGene_tx=XM_011286314.2;refGene_gn=ARPP21;refGene_tx=XM_011286315.
2;refGene_gn=ARPP21;refGene_tx=XM_011286316.2;refGene_gn=ARPP21;refGen
e_tx=XM_011286317.2;refGene_gn=ARPP21;refGene_tx=XM_019810969.2;refGen
e_gn=ARPP21;refGene_tx=XM_019810970.2;refGene_gn=ARPP21;refGene_tx=XM_
019810971.2;refGene_gn=ARPP21;refGene_tx=XM_019810972.2;refGene_gn=ARP
P21;refGene_tx=XM_019810973.2;refGene_gn=ARPP21;refGene_tx=XM_01981097
4.2;refGene_gn=ARPP21;refGene_tx=XM_019810976.2;refGene_gn=ARPP21;refG
ene_tx=XM_019810977.2;refGene_gn=ARPP21;refGene_tx=XM_019810978.2;refG
ene_gn=ARPP21;refGene_tx=XM_019810979.2;refGene_gn=ARPP21;refGene_tx=X

M_019810980.2;refGene_gn=ARPP21;refGene_tx=XM_019810981.2;refGene_gn=ARPP21;refGene_tx=XM_019810983.2;refGene_gn=ARPP21;refGene_tx=XM_019810984.2;refGene_gn=ARPP21;refGene_tx=XM_019810985.2;refGene_gn=ARPP21;refGene_tx=XM_019810986.2;refGene_gn=ARPP21;refGene_tx=XM_019810987.2;refGene_gn=ARPP21;refGene_tx=XM_023260783.1;refGene_gn=ARPP21;refGene_tx=XM_023260786.1
C2 157027327 157027417
refGene_gn=ARPP21;refGene_tx=XM_011286303.2;refGene_gn=ARPP21;refGene_tx=XM_011286304.3;refGene_gn=ARPP21;refGene_tx=XM_011286309.3;refGene_gn=ARPP21;refGene_tx=XM_011286311.2;refGene_gn=ARPP21;refGene_tx=XM_011286312.2;refGene_gn=ARPP21;refGene_tx=XM_011286313.2;refGene_gn=ARPP21;refGene_tx=XM_011286314.2;refGene_gn=ARPP21;refGene_tx=XM_011286315.2;refGene_gn=ARPP21;refGene_tx=XM_011286316.2;refGene_gn=ARPP21;refGene_tx=XM_011286317.2;refGene_gn=ARPP21;refGene_tx=XM_019810969.2;refGene_gn=ARPP21;refGene_tx=XM_019810970.2;refGene_gn=ARPP21;refGene_tx=XM_019810971.2;refGene_gn=ARPP21;refGene_tx=XM_019810972.2;refGene_gn=ARPP21;refGene_tx=XM_019810973.2;refGene_gn=ARPP21;refGene_tx=XM_019810974.2;refGene_gn=ARPP21;refGene_tx=XM_019810976.2;refGene_gn=ARPP21;refGene_tx=XM_019810977.2;refGene_gn=ARPP21;refGene_tx=XM_019810978.2;refGene_gn=ARPP21;refGene_tx=XM_019810979.2;refGene_gn=ARPP21;refGene_tx=XM_019810980.2;refGene_gn=ARPP21;refGene_tx=XM_019810981.2;refGene_gn=ARPP21;refGene_tx=XM_019810983.2;refGene_gn=ARPP21;refGene_tx=XM_019810984.2;refGene_gn=ARPP21;refGene_tx=XM_019810985.2;refGene_gn=ARPP21;refGene_tx=XM_019810986.2;refGene_gn=ARPP21;refGene_tx=XM_019810987.2;refGene_gn=ARPP21;refGene_tx=XM_023260783.1;refGene_gn=ARPP21;refGene_tx=XM_023260786.1
C2 157028196 157028244
refGene_gn=ARPP21;refGene_tx=XM_011286303.2;refGene_gn=ARPP21;refGene_tx=XM_011286304.3;refGene_gn=ARPP21;refGene_tx=XM_011286309.3;refGene_gn=ARPP21;refGene_tx=XM_011286311.2;refGene_gn=ARPP21;refGene_tx=XM_011286312.2;refGene_gn=ARPP21;refGene_tx=XM_011286313.2;refGene_gn=ARPP21;refGene_tx=XM_011286314.2;refGene_gn=ARPP21;refGene_tx=XM_011286315.2;refGene_gn=ARPP21;refGene_tx=XM_011286316.2;refGene_gn=ARPP21;refGene_tx=XM_011286317.2;refGene_gn=ARPP21;refGene_tx=XM_019810969.2;refGene_gn=ARPP21;refGene_tx=XM_019810970.2;refGene_gn=ARPP21;refGene_tx=XM_019810971.2;refGene_gn=ARPP21;refGene_tx=XM_019810972.2;refGene_gn=ARPP21;refGene_tx=XM_019810973.2;refGene_gn=ARPP21;refGene_tx=XM_019810974.2;refGene_gn=ARPP21;refGene_tx=XM_019810976.2;refGene_gn=ARPP21;refGene_tx=XM_019810977.2;refGene_gn=ARPP21;refGene_tx=XM_019810978.2;refGene_gn=ARPP21;refGene_tx=XM_019810979.2;refGene_gn=ARPP21;refGene_tx=XM_019810980.2;refGene_gn=ARPP21;refGene_tx=XM_019810981.2;refGene_gn=ARPP21;refGene_tx=XM_019810983.2;refGene_gn=ARPP21;refGene_tx=XM_019810984.2;refGene_gn=ARPP21;refGene_tx=XM_019810985.2;refGene_gn=ARPP21;refGene_tx=XM_019810986.2;refGene_gn=ARPP21;refGene_tx=XM_019810987.2;refGene_gn=ARPP21;refGene_tx=XM_023260783.1;refGene_gn=ARPP21;refGene_tx=XM_023260786.1
C2 157029305 157029435
refGene_gn=ARPP21;refGene_tx=XM_011286303.2;refGene_gn=ARPP21;refGene_tx=XM_011286304.3;refGene_gn=ARPP21;refGene_tx=XM_011286309.3;refGene_gn=ARPP21;refGene_tx=XM_011286311.2;refGene_gn=ARPP21;refGene_tx=XM_01

1286312.2;refGene_gn=ARPP21;refGene_tx=XM_011286313.2;refGene_gn=ARPP2
1;refGene_tx=XM_011286314.2;refGene_gn=ARPP21;refGene_tx=XM_011286315.
2;refGene_gn=ARPP21;refGene_tx=XM_011286316.2;refGene_gn=ARPP21;refGen
e_tx=XM_011286317.2;refGene_gn=ARPP21;refGene_tx=XM_019810969.2;refGen
e_gn=ARPP21;refGene_tx=XM_019810970.2;refGene_gn=ARPP21;refGene_tx=XM_
019810971.2;refGene_gn=ARPP21;refGene_tx=XM_019810972.2;refGene_gn=ARP
P21;refGene_tx=XM_019810973.2;refGene_gn=ARPP21;refGene_tx=XM_01981097
4.2;refGene_gn=ARPP21;refGene_tx=XM_019810976.2;refGene_gn=ARPP21;refG
ene_tx=XM_019810977.2;refGene_gn=ARPP21;refGene_tx=XM_019810978.2;refG
ene_gn=ARPP21;refGene_tx=XM_019810979.2;refGene_gn=ARPP21;refGene_tx=X
M_019810980.2;refGene_gn=ARPP21;refGene_tx=XM_019810981.2;refGene_gn=A
RPP21;refGene_tx=XM_019810983.2;refGene_gn=ARPP21;refGene_tx=XM_019810
984.2;refGene_gn=ARPP21;refGene_tx=XM_019810985.2;refGene_gn=ARPP21;re
fGene_tx=XM_019810986.2;refGene_gn=ARPP21;refGene_tx=XM_019810987.2;re
fGene_gn=ARPP21;refGene_tx=XM_023260783.1;refGene_gn=ARPP21;refGene_tx
=XM_023260786.1
C2 158525589 158525921
refGene_gn=PDCD6IP;refGene_tx=XM_023260787.1;refGene_gn=PDCD6IP;refGen
e_tx=XM_023260788.1;refGene_gn=PDCD6IP;refGene_tx=XM_023260789.1;refGe
ne_gn=PDCD6IP;refGene_tx=XM_023260790.1
C2 158527004 158527219
refGene_gn=PDCD6IP;refGene_tx=XM_023260787.1;refGene_gn=PDCD6IP;refGen
e_tx=XM_023260788.1;refGene_gn=PDCD6IP;refGene_tx=XM_023260789.1;refGe
ne_gn=PDCD6IP;refGene_tx=XM_023260790.1
C2 158528353 158528477
refGene_gn=PDCD6IP;refGene_tx=XM_023260789.1;refGene_gn=PDCD6IP;refGen
e_tx=XM_023260790.1
C2 158538338 158538433
refGene_gn=PDCD6IP;refGene_tx=XM_023260787.1;refGene_gn=PDCD6IP;refGen
e_tx=XM_023260788.1;refGene_gn=PDCD6IP;refGene_tx=XM_023260789.1;refGe
ne_gn=PDCD6IP;refGene_tx=XM_023260790.1
C2 158539573 158539708
refGene_gn=PDCD6IP;refGene_tx=XM_023260787.1;refGene_gn=PDCD6IP;refGen
e_tx=XM_023260788.1;refGene_gn=PDCD6IP;refGene_tx=XM_023260789.1;refGe
ne_gn=PDCD6IP;refGene_tx=XM_023260790.1
C2 158540051 158540300
refGene_gn=PDCD6IP;refGene_tx=XM_023260787.1;refGene_gn=PDCD6IP;refGen
e_tx=XM_023260788.1;refGene_gn=PDCD6IP;refGene_tx=XM_023260789.1;refGe
ne_gn=PDCD6IP;refGene_tx=XM_023260790.1
C2 158549033 158549203
refGene_gn=PDCD6IP;refGene_tx=XM_023260787.1;refGene_gn=PDCD6IP;refGen
e_tx=XM_023260788.1;refGene_gn=PDCD6IP;refGene_tx=XM_023260789.1;refGe
ne_gn=PDCD6IP;refGene_tx=XM_023260790.1
C2 158550717 158550829
refGene_gn=PDCD6IP;refGene_tx=XM_023260787.1;refGene_gn=PDCD6IP;refGen
e_tx=XM_023260788.1;refGene_gn=PDCD6IP;refGene_tx=XM_023260789.1;refGe
ne_gn=PDCD6IP;refGene_tx=XM_023260790.1
C2 158555350 158555528
refGene_gn=PDCD6IP;refGene_tx=XM_023260787.1;refGene_gn=PDCD6IP;refGen
e_tx=XM_023260788.1;refGene_gn=PDCD6IP;refGene_tx=XM_023260789.1;refGe

ne_gn=PDCD6IP;refGene_tx=XM_023260790.1
C2	158557810	158557934
refGene_gn=PDCD6IP;refGene_tx=XM_023260787.1;refGene_gn=PDCD6IP;refGene_tx=XM_023260788.1;refGene_gn=PDCD6IP;refGene_tx=XM_023260789.1;refGene_gn=PDCD6IP;refGene_tx=XM_023260790.1
C2	158558948	158559171
refGene_gn=PDCD6IP;refGene_tx=XM_023260787.1;refGene_gn=PDCD6IP;refGene_tx=XM_023260788.1;refGene_gn=PDCD6IP;refGene_tx=XM_023260789.1;refGene_gn=PDCD6IP;refGene_tx=XM_023260790.1
C2	158570601	158570733
refGene_gn=PDCD6IP;refGene_tx=XM_023260787.1;refGene_gn=PDCD6IP;refGene_tx=XM_023260788.1;refGene_gn=PDCD6IP;refGene_tx=XM_023260789.1;refGene_gn=PDCD6IP;refGene_tx=XM_023260790.1
C2	158573134	158573235
refGene_gn=PDCD6IP;refGene_tx=XM_023260787.1;refGene_gn=PDCD6IP;refGene_tx=XM_023260788.1;refGene_gn=PDCD6IP;refGene_tx=XM_023260789.1;refGene_gn=PDCD6IP;refGene_tx=XM_023260790.1
C2	158573651	158573805
refGene_gn=PDCD6IP;refGene_tx=XM_023260787.1;refGene_gn=PDCD6IP;refGene_tx=XM_023260788.1;refGene_gn=PDCD6IP;refGene_tx=XM_023260789.1;refGene_gn=PDCD6IP;refGene_tx=XM_023260790.1
C2	158577068	158577196
refGene_gn=PDCD6IP;refGene_tx=XM_023260787.1;refGene_gn=PDCD6IP;refGene_tx=XM_023260788.1;refGene_gn=PDCD6IP;refGene_tx=XM_023260789.1;refGene_gn=PDCD6IP;refGene_tx=XM_023260790.1
C2	158581513	158581583
refGene_gn=PDCD6IP;refGene_tx=XM_023260787.1;refGene_gn=PDCD6IP;refGene_tx=XM_023260788.1;refGene_gn=PDCD6IP;refGene_tx=XM_023260789.1;refGene_gn=PDCD6IP;refGene_tx=XM_023260790.1
C2	158583271	158583326
refGene_gn=PDCD6IP;refGene_tx=XM_023260787.1;refGene_gn=PDCD6IP;refGene_tx=XM_023260788.1;refGene_gn=PDCD6IP;refGene_tx=XM_023260789.1;refGene_gn=PDCD6IP;refGene_tx=XM_023260790.1
C2	158599412	158599621
refGene_gn=PDCD6IP;refGene_tx=XM_023260787.1;refGene_gn=PDCD6IP;refGene_tx=XM_023260788.1;refGene_gn=PDCD6IP;refGene_tx=XM_023260789.1;refGene_gn=PDCD6IP;refGene_tx=XM_023260790.1
C2	158670700	158670722
refGene_gn=CNOT10;refGene_tx=XM_006936560.3
C2	158680981	158681183
refGene_gn=CNOT10;refGene_tx=XM_003992276.4;refGene_gn=CNOT10;refGene_tx=XM_006936558.3;refGene_gn=CNOT10;refGene_tx=XM_006936559.3;refGene_gn=CNOT10;refGene_tx=XM_019810995.2;refGene_gn=CNOT10;refGene_tx=XM_019810996.2;refGene_gn=CNOT10;refGene_tx=XM_019810997.2
C2	158688414	158688509
refGene_gn=CNOT10;refGene_tx=XM_003992276.4;refGene_gn=CNOT10;refGene_tx=XM_006936558.3;refGene_gn=CNOT10;refGene_tx=XM_006936559.3;refGene_gn=CNOT10;refGene_tx=XM_006936560.3;refGene_gn=CNOT10;refGene_tx=XM_019810995.2;refGene_gn=CNOT10;refGene_tx=XM_019810996.2;refGene_gn=CNOT10;refGene_tx=XM_019810997.2

C2 158689441 158689603
refGene_gn=CNOT10;refGene_tx=XM_003992276.4;refGene_gn=CNOT10;refGene_tx=XM_006936558.3;refGene_gn=CNOT10;refGene_tx=XM_006936559.3;refGene_gn=CNOT10;refGene_tx=XM_006936560.3;refGene_gn=CNOT10;refGene_tx=XM_019810995.2;refGene_gn=CNOT10;refGene_tx=XM_019810996.2;refGene_gn=CNOT10;refGene_tx=XM_019810997.2
C2 158694358 158694509
refGene_gn=CNOT10;refGene_tx=XM_003992276.4;refGene_gn=CNOT10;refGene_tx=XM_006936558.3;refGene_gn=CNOT10;refGene_tx=XM_006936559.3;refGene_gn=CNOT10;refGene_tx=XM_006936560.3;refGene_gn=CNOT10;refGene_tx=XM_019810995.2;refGene_gn=CNOT10;refGene_tx=XM_019810996.2;refGene_gn=CNOT10;refGene_tx=XM_019810997.2
C2 158698092 158698235
refGene_gn=CNOT10;refGene_tx=XM_003992276.4;refGene_gn=CNOT10;refGene_tx=XM_006936558.3;refGene_gn=CNOT10;refGene_tx=XM_006936559.3;refGene_gn=CNOT10;refGene_tx=XM_006936560.3;refGene_gn=CNOT10;refGene_tx=XM_019810995.2;refGene_gn=CNOT10;refGene_tx=XM_019810996.2;refGene_gn=CNOT10;refGene_tx=XM_019810997.2
C2 158701113 158701200
refGene_gn=CNOT10;refGene_tx=XM_003992276.4;refGene_gn=CNOT10;refGene_tx=XM_006936558.3;refGene_gn=CNOT10;refGene_tx=XM_006936559.3;refGene_gn=CNOT10;refGene_tx=XM_006936560.3;refGene_gn=CNOT10;refGene_tx=XM_019810995.2;refGene_gn=CNOT10;refGene_tx=XM_019810996.2;refGene_gn=CNOT10;refGene_tx=XM_019810997.2
C2 158702054 158702138
refGene_gn=CNOT10;refGene_tx=XM_003992276.4;refGene_gn=CNOT10;refGene_tx=XM_006936558.3;refGene_gn=CNOT10;refGene_tx=XM_006936559.3;refGene_gn=CNOT10;refGene_tx=XM_006936560.3;refGene_gn=CNOT10;refGene_tx=XM_019810995.2;refGene_gn=CNOT10;refGene_tx=XM_019810996.2;refGene_gn=CNOT10;refGene_tx=XM_019810997.2
C2 158703663 158703781
refGene_gn=CNOT10;refGene_tx=XM_003992276.4;refGene_gn=CNOT10;refGene_tx=XM_006936558.3;refGene_gn=CNOT10;refGene_tx=XM_006936559.3;refGene_gn=CNOT10;refGene_tx=XM_006936560.3;refGene_gn=CNOT10;refGene_tx=XM_019810995.2;refGene_gn=CNOT10;refGene_tx=XM_019810996.2;refGene_gn=CNOT10;refGene_tx=XM_019810997.2
C2 158708225 158708375
refGene_gn=CNOT10;refGene_tx=XM_003992276.4;refGene_gn=CNOT10;refGene_tx=XM_006936558.3;refGene_gn=CNOT10;refGene_tx=XM_006936559.3;refGene_gn=CNOT10;refGene_tx=XM_006936560.3;refGene_gn=CNOT10;refGene_tx=XM_019810995.2;refGene_gn=CNOT10;refGene_tx=XM_019810996.2;refGene_gn=CNOT10;refGene_tx=XM_019810997.2
C2 158710113 158710316
refGene_gn=CNOT10;refGene_tx=XM_003992276.4;refGene_gn=CNOT10;refGene_tx=XM_006936558.3;refGene_gn=CNOT10;refGene_tx=XM_006936559.3;refGene_gn=CNOT10;refGene_tx=XM_006936560.3;refGene_gn=CNOT10;refGene_tx=XM_019810995.2;refGene_gn=CNOT10;refGene_tx=XM_019810996.2;refGene_gn=CNOT10;refGene_tx=XM_019810997.2
C2 158714115 158714237
refGene_gn=CNOT10;refGene_tx=XM_003992276.4;refGene_gn=CNOT10;refGene_

tx=XM_006936558.3;refGene_gn=CNOT10;refGene_tx=XM_006936559.3;refGene_
gn=CNOT10;refGene_tx=XM_006936560.3;refGene_gn=CNOT10;refGene_tx=XM_01
9810995.2;refGene_gn=CNOT10;refGene_tx=XM_019810996.2;refGene_gn=CNOT1
0;refGene_tx=XM_019810997.2
C2 158716289 158716466
refGene_gn=CNOT10;refGene_tx=XM_003992276.4;refGene_gn=CNOT10;refGene_
tx=XM_006936558.3;refGene_gn=CNOT10;refGene_tx=XM_006936559.3;refGene_
gn=CNOT10;refGene_tx=XM_006936560.3;refGene_gn=CNOT10;refGene_tx=XM_01
9810995.2;refGene_gn=CNOT10;refGene_tx=XM_019810996.2;refGene_gn=CNOT1
0;refGene_tx=XM_019810997.2
C2 158717643 158717724
refGene_gn=CNOT10;refGene_tx=XM_003992276.4;refGene_gn=CNOT10;refGene_
tx=XM_006936558.3;refGene_gn=CNOT10;refGene_tx=XM_006936559.3;refGene_
gn=CNOT10;refGene_tx=XM_006936560.3;refGene_gn=CNOT10;refGene_tx=XM_01
9810995.2;refGene_gn=CNOT10;refGene_tx=XM_019810996.2;refGene_gn=CNOT1
0;refGene_tx=XM_019810997.2
C2 158732896 158733031
refGene_gn=CNOT10;refGene_tx=XM_003992276.4;refGene_gn=CNOT10;refGene_
tx=XM_006936558.3;refGene_gn=CNOT10;refGene_tx=XM_006936559.3;refGene_
gn=CNOT10;refGene_tx=XM_006936560.3;refGene_gn=CNOT10;refGene_tx=XM_01
9810995.2;refGene_gn=CNOT10;refGene_tx=XM_019810996.2;refGene_gn=CNOT1
0;refGene_tx=XM_019810997.2
C2 158736210 158736341
refGene_gn=CNOT10;refGene_tx=XM_003992276.4;refGene_gn=CNOT10;refGene_
tx=XM_006936558.3;refGene_gn=CNOT10;refGene_tx=XM_006936559.3;refGene_
gn=CNOT10;refGene_tx=XM_006936560.3;refGene_gn=CNOT10;refGene_tx=XM_01
9810995.2;refGene_gn=CNOT10;refGene_tx=XM_019810996.2;refGene_gn=CNOT1
0;refGene_tx=XM_019810997.2
C2 158737493 158737529
refGene_gn=CNOT10;refGene_tx=XM_003992276.4;refGene_gn=CNOT10;refGene_
tx=XM_006936558.3;refGene_gn=CNOT10;refGene_tx=XM_006936559.3;refGene_
gn=CNOT10;refGene_tx=XM_006936560.3;refGene_gn=CNOT10;refGene_tx=XM_01
9810995.2;refGene_gn=CNOT10;refGene_tx=XM_019810996.2;refGene_gn=CNOT1
0;refGene_tx=XM_019810997.2
C2 158737729 158737857
refGene_gn=CNOT10;refGene_tx=XM_003992276.4;refGene_gn=CNOT10;refGene_
tx=XM_006936558.3;refGene_gn=CNOT10;refGene_tx=XM_006936559.3;refGene_
gn=CNOT10;refGene_tx=XM_006936560.3;refGene_gn=CNOT10;refGene_tx=XM_01
9810995.2;refGene_gn=CNOT10;refGene_tx=XM_019810996.2;refGene_gn=CNOT1
0;refGene_tx=XM_019810997.2
C2 158738053 158738077
refGene_gn=CNOT10;refGene_tx=XM_019810997.2
C2 158741302 158741378
refGene_gn=CNOT10;refGene_tx=XM_003992276.4;refGene_gn=CNOT10;refGene_
tx=XM_006936558.3;refGene_gn=CNOT10;refGene_tx=XM_006936559.3;refGene_
gn=CNOT10;refGene_tx=XM_006936560.3;refGene_gn=CNOT10;refGene_tx=XM_01
9810995.2
C2 158742351 158742503
refGene_gn=CNOT10;refGene_tx=XM_003992276.4;refGene_gn=CNOT10;refGene_
tx=XM_006936558.3;refGene_gn=CNOT10;refGene_tx=XM_006936559.3;refGene_

gn=CNOT10;refGene_tx=XM_006936560.3;refGene_gn=CNOT10;refGene_tx=XM_01
9810995.2;refGene_gn=CNOT10;refGene_tx=XM_019810996.2
C2	158767327	158768221
refGene_gn=TRIM71;refGene_tx=XM_023260791.1
C2	158810245	158810413
refGene_gn=TRIM71;refGene_tx=XM_023260791.1
C2	158824678	158824813
refGene_gn=TRIM71;refGene_tx=XM_023260791.1
C2	158828028	158829480
refGene_gn=TRIM71;refGene_tx=XM_023260791.1
C2	158868254	158868307
refGene_gn=CCR4;refGene_tx=XM_019811003.2
C2	158879055	158880211
refGene_gn=CCR4;refGene_tx=XM_019811003.2;refGene_gn=CCR4;refGene_tx=X
M_019811004.2;refGene_gn=CCR4;refGene_tx=XM_019811008.2;refGene_gn=CCR
4;refGene_tx=XM_019811009.2;refGene_gn=CCR4;refGene_tx=XM_019811010.2;
refGene_gn=CCR4;refGene_tx=XM_019811012.2
C2	158910852	158910945
refGene_gn=GLB1;refGene_tx=XM_019839237.2
C2	158911053	158911068
refGene_gn=GLB1;refGene_tx=XM_019839238.2
C2	158912575	158912845
refGene_gn=GLB1;refGene_tx=NM_001009860.1
C2	158930212	158930470
refGene_gn=GLB1;refGene_tx=NM_001009860.1;refGene_gn=GLB1;refGene_tx=X
M_019839237.2;refGene_gn=GLB1;refGene_tx=XM_019839238.2
C2	158932132	158932264
refGene_gn=GLB1;refGene_tx=NM_001009860.1;refGene_gn=GLB1;refGene_tx=X
M_019839237.2;refGene_gn=GLB1;refGene_tx=XM_019839238.2
C2	158933729	158933843
refGene_gn=GLB1;refGene_tx=NM_001009860.1;refGene_gn=GLB1;refGene_tx=X
M_019839237.2;refGene_gn=GLB1;refGene_tx=XM_019839238.2
C2	158937501	158937591
refGene_gn=GLB1;refGene_tx=NM_001009860.1;refGene_gn=GLB1;refGene_tx=X
M_019839237.2;refGene_gn=GLB1;refGene_tx=XM_019839238.2
C2	158939763	158939838
refGene_gn=GLB1;refGene_tx=NM_001009860.1;refGene_gn=GLB1;refGene_tx=X
M_019839237.2;refGene_gn=GLB1;refGene_tx=XM_019839238.2
C2	158957078	158957191
refGene_gn=GLB1;refGene_tx=NM_001009860.1;refGene_gn=GLB1;refGene_tx=X
M_019839237.2;refGene_gn=GLB1;refGene_tx=XM_019839238.2
C2	158962270	158962311
refGene_gn=GLB1;refGene_tx=NM_001009860.1;refGene_gn=GLB1;refGene_tx=X
M_019839237.2;refGene_gn=GLB1;refGene_tx=XM_019839238.2
C2	158962396	158962518
refGene_gn=GLB1;refGene_tx=NM_001009860.1;refGene_gn=GLB1;refGene_tx=X
M_019839237.2;refGene_gn=GLB1;refGene_tx=XM_019839238.2
C2	158963776	158963835
refGene_gn=GLB1;refGene_tx=NM_001009860.1;refGene_gn=GLB1;refGene_tx=X
M_019839237.2;refGene_gn=GLB1;refGene_tx=XM_019839238.2

C2	158966518	158966699
refGene_gn=GLB1;refGene_tx=NM_001009860.1;refGene_gn=GLB1;refGene_tx=XM_019839237.2;refGene_gn=GLB1;refGene_tx=XM_019839238.2
C2	158970338	158970433
refGene_gn=GLB1;refGene_tx=NM_001009860.1;refGene_gn=GLB1;refGene_tx=XM_019839237.2;refGene_gn=GLB1;refGene_tx=XM_019839238.2
C2	158971492	158971553
refGene_gn=GLB1;refGene_tx=NM_001009860.1;refGene_gn=GLB1;refGene_tx=XM_019839237.2;refGene_gn=GLB1;refGene_tx=XM_019839238.2
C2	158972095	158972246
refGene_gn=GLB1;refGene_tx=NM_001009860.1;refGene_gn=GLB1;refGene_tx=XM_019839237.2;refGene_gn=GLB1;refGene_tx=XM_019839238.2
C2	158975358	158975528
refGene_gn=GLB1;refGene_tx=NM_001009860.1;refGene_gn=GLB1;refGene_tx=XM_019839237.2;refGene_gn=GLB1;refGene_tx=XM_019839238.2
C2	158995246	158996590
refGene_gn=TMPPE;refGene_tx=XM_023260792.1
C2	158998712	158998790
refGene_gn=GLB1;refGene_tx=NM_001009860.1;refGene_gn=GLB1;refGene_tx=XM_019839237.2;refGene_gn=GLB1;refGene_tx=XM_019839238.2
C2	159011923	159012394
refGene_gn=CRTAP;refGene_tx=XM_011286301.3;refGene_gn=CRTAP;refGene_tx=XM_019811013.2
C2	159016535	159016685
refGene_gn=CRTAP;refGene_tx=XM_011286301.3;refGene_gn=CRTAP;refGene_tx=XM_019811013.2
C2	159019137	159019309
refGene_gn=CRTAP;refGene_tx=XM_011286301.3;refGene_gn=CRTAP;refGene_tx=XM_019811013.2
C2	159022526	159022655
refGene_gn=CRTAP;refGene_tx=XM_011286301.3;refGene_gn=CRTAP;refGene_tx=XM_019811013.2
C2	159024330	159024528
refGene_gn=CRTAP;refGene_tx=XM_019811013.2
C2	159025517	159025663
refGene_gn=CRTAP;refGene_tx=XM_011286301.3;refGene_gn=CRTAP;refGene_tx=XM_019811013.2
C2	159027019	159027103
refGene_gn=CRTAP;refGene_tx=XM_011286301.3;refGene_gn=CRTAP;refGene_tx=XM_019811013.2
C2	159034738	159034792
refGene_gn=CRTAP;refGene_tx=XM_011286301.3;refGene_gn=CRTAP;refGene_tx=XM_019811013.2
C2	159042498	159043790
refGene_gn=SUSD5;refGene_tx=XM_019811014.2;refGene_gn=SUSD5;refGene_tx=XM_023260793.1
C2	159062971	159063160
refGene_gn=SUSD5;refGene_tx=XM_019811014.2;refGene_gn=SUSD5;refGene_tx=XM_023260793.1
C2	159076474	159076593

refGene_gn=SUSD5;refGene_tx=XM_019811014.2;refGene_gn=SUSD5;refGene_tx=XM_023260793.1
C2 159087074 159087252
refGene_gn=SUSD5;refGene_tx=XM_023260793.1
C2 159097336 159097439
refGene_gn=SUSD5;refGene_tx=XM_023260793.1
C2 159147097 159147169
refGene_gn=FBXL2;refGene_tx=XM_023260190.1
C2 159160614 159160676
refGene_gn=FBXL2;refGene_tx=XM_023260190.1
C2 159188777 159188832
refGene_gn=FBXL2;refGene_tx=XM_023260190.1
C2 159189196 159189271
refGene_gn=FBXL2;refGene_tx=XM_023260190.1
C2 159195294 159195389
refGene_gn=FBXL2;refGene_tx=XM_023260190.1
C2 159196112 159196182
refGene_gn=FBXL2;refGene_tx=XM_023260190.1
C2 159206025 159206274
refGene_gn=FBXL2;refGene_tx=XM_023260190.1
C2 159206396 159206471
refGene_gn=FBXL2;refGene_tx=XM_023260190.1
C2 159207968 159208099
refGene_gn=FBXL2;refGene_tx=XM_023260190.1
C2 159210109 159210170
refGene_gn=FBXL2;refGene_tx=XM_023260190.1
C2 159211016 159211061
refGene_gn=FBXL2;refGene_tx=XM_023260190.1
C2 159211516 159211573
refGene_gn=FBXL2;refGene_tx=XM_023260190.1
C2 159237627 159237840
refGene_gn=FBXL2;refGene_tx=XM_023260190.1
C2 159238501 159238609
refGene_gn=FBXL2;refGene_tx=XM_023260190.1
C2 159242459 159242497
refGene_gn=UBP1;refGene_tx=XM_023260794.1;refGene_gn=UBP1;refGene_tx=XM_023260795.1
C2 159244808 159244860
refGene_gn=UBP1;refGene_tx=XM_023260794.1;refGene_gn=UBP1;refGene_tx=XM_023260795.1
C2 159245582 159245725
refGene_gn=UBP1;refGene_tx=XM_023260794.1;refGene_gn=UBP1;refGene_tx=XM_023260795.1
C2 159248086 159248202
refGene_gn=UBP1;refGene_tx=XM_023260794.1;refGene_gn=UBP1;refGene_tx=XM_023260795.1
C2 159248959 159249050
refGene_gn=UBP1;refGene_tx=XM_023260794.1;refGene_gn=UBP1;refGene_tx=XM_023260795.1
C2 159252189 159252283

refGene_gn=UBP1;refGene_tx=XM_023260794.1;refGene_gn=UBP1;refGene_tx=XM_023260795.1
C2	159252535	159252590
refGene_gn=UBP1;refGene_tx=XM_023260794.1;refGene_gn=UBP1;refGene_tx=XM_023260795.1
C2	159254453	159254557
refGene_gn=UBP1;refGene_tx=XM_023260794.1;refGene_gn=UBP1;refGene_tx=XM_023260795.1
C2	159259732	159259840
refGene_gn=UBP1;refGene_tx=XM_023260794.1
C2	159260290	159260401
refGene_gn=UBP1;refGene_tx=XM_023260794.1;refGene_gn=UBP1;refGene_tx=XM_023260795.1
C2	159260505	159260658
refGene_gn=UBP1;refGene_tx=XM_023260794.1;refGene_gn=UBP1;refGene_tx=XM_023260795.1
C2	159262963	159263070
refGene_gn=UBP1;refGene_tx=XM_023260794.1;refGene_gn=UBP1;refGene_tx=XM_023260795.1
C2	159263522	159263628
refGene_gn=UBP1;refGene_tx=XM_023260794.1;refGene_gn=UBP1;refGene_tx=XM_023260795.1
C2	159268123	159268200
refGene_gn=UBP1;refGene_tx=XM_023260794.1;refGene_gn=UBP1;refGene_tx=XM_023260795.1
C2	159276920	159277072
refGene_gn=UBP1;refGene_tx=XM_023260794.1;refGene_gn=UBP1;refGene_tx=XM_023260795.1
C2	159291479	159291592
refGene_gn=UBP1;refGene_tx=XM_023260794.1;refGene_gn=UBP1;refGene_tx=XM_023260795.1
C2	159306532	159306619
refGene_gn=CLASP2;refGene_tx=XM_011286298.2;refGene_gn=CLASP2;refGene_tx=XM_011286299.3;refGene_gn=CLASP2;refGene_tx=XM_011286300.2;refGene_gn=CLASP2;refGene_tx=XM_019811031.1;refGene_gn=CLASP2;refGene_tx=XM_019811032.1;refGene_gn=CLASP2;refGene_tx=XM_019811033.1;refGene_gn=CLASP2;refGene_tx=XM_019811034.2;refGene_gn=CLASP2;refGene_tx=XM_023260796.1;refGene_gn=CLASP2;refGene_tx=XM_023260797.1;refGene_gn=CLASP2;refGene_tx=XM_023260798.1;refGene_gn=CLASP2;refGene_tx=XM_023260799.1;refGene_gn=CLASP2;refGene_tx=XM_023260800.1;refGene_gn=CLASP2;refGene_tx=XM_023260801.1;refGene_gn=CLASP2;refGene_tx=XM_023260802.1;refGene_gn=CLASP2;refGene_tx=XM_023260803.1;refGene_gn=CLASP2;refGene_tx=XM_023260804.1;refGene_gn=CLASP2;refGene_tx=XM_023260805.1;refGene_gn=CLASP2;refGene_tx=XM_023260806.1;refGene_gn=CLASP2;refGene_tx=XM_023260807.1;refGene_gn=CLASP2;refGene_tx=XM_023260808.1;refGene_gn=CLASP2;refGene_tx=XM_023260809.1
C2	159309230	159309347
refGene_gn=CLASP2;refGene_tx=XM_011286298.2;refGene_gn=CLASP2;refGene_tx=XM_011286299.3;refGene_gn=CLASP2;refGene_tx=XM_011286300.2;refGene_gn=CLASP2;refGene_tx=XM_019811031.1;refGene_gn=CLASP2;refGene_tx=XM_01

9811032.1;refGene_gn=CLASP2;refGene_tx=XM_019811033.1;refGene_gn=CLASP2;refGene_tx=XM_019811034.2;refGene_gn=CLASP2;refGene_tx=XM_023260796.1;refGene_gn=CLASP2;refGene_tx=XM_023260797.1;refGene_gn=CLASP2;refGene_tx=XM_023260798.1;refGene_gn=CLASP2;refGene_tx=XM_023260799.1;refGene_gn=CLASP2;refGene_tx=XM_023260800.1;refGene_gn=CLASP2;refGene_tx=XM_023260801.1;refGene_gn=CLASP2;refGene_tx=XM_023260802.1;refGene_gn=CLASP2;refGene_tx=XM_023260803.1;refGene_gn=CLASP2;refGene_tx=XM_023260804.1;refGene_gn=CLASP2;refGene_tx=XM_023260805.1;refGene_gn=CLASP2;refGene_tx=XM_023260806.1;refGene_gn=CLASP2;refGene_tx=XM_023260807.1;refGene_gn=CLASP2;refGene_tx=XM_023260808.1;refGene_gn=CLASP2;refGene_tx=XM_023260809.1
C2 159310876 159311083
refGene_gn=CLASP2;refGene_tx=XM_011286298.2;refGene_gn=CLASP2;refGene_tx=XM_011286299.3;refGene_gn=CLASP2;refGene_tx=XM_011286300.2;refGene_gn=CLASP2;refGene_tx=XM_019811031.1;refGene_gn=CLASP2;refGene_tx=XM_019811032.1;refGene_gn=CLASP2;refGene_tx=XM_019811033.1;refGene_gn=CLASP2;refGene_tx=XM_019811034.2;refGene_gn=CLASP2;refGene_tx=XM_023260796.1;refGene_gn=CLASP2;refGene_tx=XM_023260797.1;refGene_gn=CLASP2;refGene_tx=XM_023260798.1;refGene_gn=CLASP2;refGene_tx=XM_023260799.1;refGene_gn=CLASP2;refGene_tx=XM_023260800.1;refGene_gn=CLASP2;refGene_tx=XM_023260801.1;refGene_gn=CLASP2;refGene_tx=XM_023260802.1;refGene_gn=CLASP2;refGene_tx=XM_023260803.1;refGene_gn=CLASP2;refGene_tx=XM_023260804.1;refGene_gn=CLASP2;refGene_tx=XM_023260805.1;refGene_gn=CLASP2;refGene_tx=XM_023260806.1;refGene_gn=CLASP2;refGene_tx=XM_023260807.1;refGene_gn=CLASP2;refGene_tx=XM_023260808.1;refGene_gn=CLASP2;refGene_tx=XM_023260809.1
C2 159315942 159316071
refGene_gn=CLASP2;refGene_tx=XM_011286298.2;refGene_gn=CLASP2;refGene_tx=XM_011286299.3;refGene_gn=CLASP2;refGene_tx=XM_011286300.2;refGene_gn=CLASP2;refGene_tx=XM_019811031.1;refGene_gn=CLASP2;refGene_tx=XM_019811032.1;refGene_gn=CLASP2;refGene_tx=XM_019811033.1;refGene_gn=CLASP2;refGene_tx=XM_019811034.2;refGene_gn=CLASP2;refGene_tx=XM_023260796.1;refGene_gn=CLASP2;refGene_tx=XM_023260797.1;refGene_gn=CLASP2;refGene_tx=XM_023260798.1;refGene_gn=CLASP2;refGene_tx=XM_023260799.1;refGene_gn=CLASP2;refGene_tx=XM_023260800.1;refGene_gn=CLASP2;refGene_tx=XM_023260801.1;refGene_gn=CLASP2;refGene_tx=XM_023260802.1;refGene_gn=CLASP2;refGene_tx=XM_023260803.1;refGene_gn=CLASP2;refGene_tx=XM_023260804.1;refGene_gn=CLASP2;refGene_tx=XM_023260805.1;refGene_gn=CLASP2;refGene_tx=XM_023260806.1;refGene_gn=CLASP2;refGene_tx=XM_023260807.1;refGene_gn=CLASP2;refGene_tx=XM_023260808.1;refGene_gn=CLASP2;refGene_tx=XM_023260809.1
C2 159316578 159316772
refGene_gn=CLASP2;refGene_tx=XM_011286298.2;refGene_gn=CLASP2;refGene_tx=XM_011286299.3;refGene_gn=CLASP2;refGene_tx=XM_011286300.2;refGene_gn=CLASP2;refGene_tx=XM_019811031.1;refGene_gn=CLASP2;refGene_tx=XM_019811032.1;refGene_gn=CLASP2;refGene_tx=XM_019811033.1;refGene_gn=CLASP2;refGene_tx=XM_019811034.2;refGene_gn=CLASP2;refGene_tx=XM_023260796.1;refGene_gn=CLASP2;refGene_tx=XM_023260797.1;refGene_gn=CLASP2;refGene_tx=XM_023260798.1;refGene_gn=CLASP2;refGene_tx=XM_023260799.1;refGene_gn=CLASP2;refGene_tx=XM_023260800.1;refGene_gn=CLASP2;refGene_tx=XM_

023260801.1;refGene_gn=CLASP2;refGene_tx=XM_023260802.1;refGene_gn=CLASP2;refGene_tx=XM_023260803.1;refGene_gn=CLASP2;refGene_tx=XM_023260804.1;refGene_gn=CLASP2;refGene_tx=XM_023260805.1;refGene_gn=CLASP2;refGene_tx=XM_023260806.1;refGene_gn=CLASP2;refGene_tx=XM_023260807.1;refGene_gn=CLASP2;refGene_tx=XM_023260808.1;refGene_gn=CLASP2;refGene_tx=XM_023260809.1
C2 159323502 159323731
refGene_gn=CLASP2;refGene_tx=XM_011286298.2;refGene_gn=CLASP2;refGene_tx=XM_011286299.3;refGene_gn=CLASP2;refGene_tx=XM_011286300.2;refGene_gn=CLASP2;refGene_tx=XM_019811031.1;refGene_gn=CLASP2;refGene_tx=XM_019811032.1;refGene_gn=CLASP2;refGene_tx=XM_019811033.1;refGene_gn=CLASP2;refGene_tx=XM_019811034.2;refGene_gn=CLASP2;refGene_tx=XM_023260796.1;refGene_gn=CLASP2;refGene_tx=XM_023260797.1;refGene_gn=CLASP2;refGene_tx=XM_023260798.1;refGene_gn=CLASP2;refGene_tx=XM_023260799.1;refGene_gn=CLASP2;refGene_tx=XM_023260800.1;refGene_gn=CLASP2;refGene_tx=XM_023260801.1;refGene_gn=CLASP2;refGene_tx=XM_023260802.1;refGene_gn=CLASP2;refGene_tx=XM_023260803.1;refGene_gn=CLASP2;refGene_tx=XM_023260804.1;refGene_gn=CLASP2;refGene_tx=XM_023260805.1;refGene_gn=CLASP2;refGene_tx=XM_023260806.1;refGene_gn=CLASP2;refGene_tx=XM_023260807.1;refGene_gn=CLASP2;refGene_tx=XM_023260808.1;refGene_gn=CLASP2;refGene_tx=XM_023260809.1
C2 159327003 159327157
refGene_gn=CLASP2;refGene_tx=XM_011286298.2;refGene_gn=CLASP2;refGene_tx=XM_011286299.3;refGene_gn=CLASP2;refGene_tx=XM_011286300.2;refGene_gn=CLASP2;refGene_tx=XM_019811031.1;refGene_gn=CLASP2;refGene_tx=XM_019811032.1;refGene_gn=CLASP2;refGene_tx=XM_019811033.1;refGene_gn=CLASP2;refGene_tx=XM_019811034.2;refGene_gn=CLASP2;refGene_tx=XM_023260796.1;refGene_gn=CLASP2;refGene_tx=XM_023260797.1;refGene_gn=CLASP2;refGene_tx=XM_023260798.1;refGene_gn=CLASP2;refGene_tx=XM_023260799.1;refGene_gn=CLASP2;refGene_tx=XM_023260800.1;refGene_gn=CLASP2;refGene_tx=XM_023260801.1;refGene_gn=CLASP2;refGene_tx=XM_023260802.1;refGene_gn=CLASP2;refGene_tx=XM_023260803.1;refGene_gn=CLASP2;refGene_tx=XM_023260804.1;refGene_gn=CLASP2;refGene_tx=XM_023260805.1;refGene_gn=CLASP2;refGene_tx=XM_023260806.1;refGene_gn=CLASP2;refGene_tx=XM_023260807.1;refGene_gn=CLASP2;refGene_tx=XM_023260808.1;refGene_gn=CLASP2;refGene_tx=XM_023260809.1
C2 159328556 159328663
refGene_gn=CLASP2;refGene_tx=XM_011286298.2;refGene_gn=CLASP2;refGene_tx=XM_011286299.3;refGene_gn=CLASP2;refGene_tx=XM_011286300.2;refGene_gn=CLASP2;refGene_tx=XM_019811031.1;refGene_gn=CLASP2;refGene_tx=XM_019811032.1;refGene_gn=CLASP2;refGene_tx=XM_019811033.1;refGene_gn=CLASP2;refGene_tx=XM_019811034.2;refGene_gn=CLASP2;refGene_tx=XM_023260796.1;refGene_gn=CLASP2;refGene_tx=XM_023260797.1;refGene_gn=CLASP2;refGene_tx=XM_023260798.1;refGene_gn=CLASP2;refGene_tx=XM_023260799.1;refGene_gn=CLASP2;refGene_tx=XM_023260800.1;refGene_gn=CLASP2;refGene_tx=XM_023260801.1;refGene_gn=CLASP2;refGene_tx=XM_023260802.1;refGene_gn=CLASP2;refGene_tx=XM_023260803.1;refGene_gn=CLASP2;refGene_tx=XM_023260804.1;refGene_gn=CLASP2;refGene_tx=XM_023260805.1;refGene_gn=CLASP2;refGene_tx=XM_023260806.1;refGene_gn=CLASP2;refGene_tx=XM_023260807.1;refGene_gn=CLASP2;refGene_tx=XM_023260808.1;refGene_gn=CLASP2;refGene_tx=X

M_023260809.1
C2      159330376       159330520
refGene_gn=CLASP2;refGene_tx=XM_011286298.2;refGene_gn=CLASP2;refGene_tx=XM_011286299.3;refGene_gn=CLASP2;refGene_tx=XM_011286300.2;refGene_gn=CLASP2;refGene_tx=XM_019811031.1;refGene_gn=CLASP2;refGene_tx=XM_019811032.1;refGene_gn=CLASP2;refGene_tx=XM_019811033.1;refGene_gn=CLASP2;refGene_tx=XM_019811034.2;refGene_gn=CLASP2;refGene_tx=XM_023260796.1;refGene_gn=CLASP2;refGene_tx=XM_023260797.1;refGene_gn=CLASP2;refGene_tx=XM_023260798.1;refGene_gn=CLASP2;refGene_tx=XM_023260799.1;refGene_gn=CLASP2;refGene_tx=XM_023260800.1;refGene_gn=CLASP2;refGene_tx=XM_023260801.1;refGene_gn=CLASP2;refGene_tx=XM_023260802.1;refGene_gn=CLASP2;refGene_tx=XM_023260803.1;refGene_gn=CLASP2;refGene_tx=XM_023260804.1;refGene_gn=CLASP2;refGene_tx=XM_023260805.1;refGene_gn=CLASP2;refGene_tx=XM_023260806.1;refGene_gn=CLASP2;refGene_tx=XM_023260807.1;refGene_gn=CLASP2;refGene_tx=XM_023260808.1;refGene_gn=CLASP2;refGene_tx=XM_023260809.1
C2      159344432       159344576
refGene_gn=CLASP2;refGene_tx=XM_011286298.2;refGene_gn=CLASP2;refGene_tx=XM_011286299.3;refGene_gn=CLASP2;refGene_tx=XM_011286300.2;refGene_gn=CLASP2;refGene_tx=XM_019811031.1;refGene_gn=CLASP2;refGene_tx=XM_019811032.1;refGene_gn=CLASP2;refGene_tx=XM_019811033.1;refGene_gn=CLASP2;refGene_tx=XM_019811034.2;refGene_gn=CLASP2;refGene_tx=XM_023260796.1;refGene_gn=CLASP2;refGene_tx=XM_023260797.1;refGene_gn=CLASP2;refGene_tx=XM_023260798.1;refGene_gn=CLASP2;refGene_tx=XM_023260799.1;refGene_gn=CLASP2;refGene_tx=XM_023260800.1;refGene_gn=CLASP2;refGene_tx=XM_023260801.1;refGene_gn=CLASP2;refGene_tx=XM_023260802.1;refGene_gn=CLASP2;refGene_tx=XM_023260803.1;refGene_gn=CLASP2;refGene_tx=XM_023260804.1;refGene_gn=CLASP2;refGene_tx=XM_023260805.1;refGene_gn=CLASP2;refGene_tx=XM_023260806.1;refGene_gn=CLASP2;refGene_tx=XM_023260807.1;refGene_gn=CLASP2;refGene_tx=XM_023260808.1;refGene_gn=CLASP2;refGene_tx=XM_023260809.1
C2      159348293       159348344
refGene_gn=CLASP2;refGene_tx=XM_019811031.1;refGene_gn=CLASP2;refGene_tx=XM_019811032.1;refGene_gn=CLASP2;refGene_tx=XM_019811033.1;refGene_gn=CLASP2;refGene_tx=XM_023260796.1;refGene_gn=CLASP2;refGene_tx=XM_023260797.1;refGene_gn=CLASP2;refGene_tx=XM_023260798.1;refGene_gn=CLASP2;refGene_tx=XM_023260799.1;refGene_gn=CLASP2;refGene_tx=XM_023260800.1;refGene_gn=CLASP2;refGene_tx=XM_023260801.1;refGene_gn=CLASP2;refGene_tx=XM_023260802.1;refGene_gn=CLASP2;refGene_tx=XM_023260804.1;refGene_gn=CLASP2;refGene_tx=XM_023260805.1
C2      159348470       159348549
refGene_gn=CLASP2;refGene_tx=XM_011286298.2;refGene_gn=CLASP2;refGene_tx=XM_011286299.3;refGene_gn=CLASP2;refGene_tx=XM_011286300.2;refGene_gn=CLASP2;refGene_tx=XM_019811031.1;refGene_gn=CLASP2;refGene_tx=XM_019811032.1;refGene_gn=CLASP2;refGene_tx=XM_019811033.1;refGene_gn=CLASP2;refGene_tx=XM_019811034.2;refGene_gn=CLASP2;refGene_tx=XM_023260796.1;refGene_gn=CLASP2;refGene_tx=XM_023260797.1;refGene_gn=CLASP2;refGene_tx=XM_023260798.1;refGene_gn=CLASP2;refGene_tx=XM_023260799.1;refGene_gn=CLASP2;refGene_tx=XM_023260800.1;refGene_gn=CLASP2;refGene_tx=XM_023260801.1;refGene_gn=CLASP2;refGene_tx=XM_023260802.1;refGene_gn=CLA

SP2;refGene_tx=XM_023260803.1;refGene_gn=CLASP2;refGene_tx=XM_02326080
4.1;refGene_gn=CLASP2;refGene_tx=XM_023260805.1;refGene_gn=CLASP2;refG
ene_tx=XM_023260806.1;refGene_gn=CLASP2;refGene_tx=XM_023260807.1;refG
ene_gn=CLASP2;refGene_tx=XM_023260808.1;refGene_gn=CLASP2;refGene_tx=X
M_023260809.1
C2 159349300 159349464
refGene_gn=CLASP2;refGene_tx=XM_011286298.2;refGene_gn=CLASP2;refGene_
tx=XM_011286299.3;refGene_gn=CLASP2;refGene_tx=XM_011286300.2;refGene_
gn=CLASP2;refGene_tx=XM_019811031.1;refGene_gn=CLASP2;refGene_tx=XM_01
9811032.1;refGene_gn=CLASP2;refGene_tx=XM_019811033.1;refGene_gn=CLASP
2;refGene_tx=XM_019811034.2;refGene_gn=CLASP2;refGene_tx=XM_023260796.
1;refGene_gn=CLASP2;refGene_tx=XM_023260797.1;refGene_gn=CLASP2;refGen
e_tx=XM_023260798.1;refGene_gn=CLASP2;refGene_tx=XM_023260799.1;refGen
e_gn=CLASP2;refGene_tx=XM_023260800.1;refGene_gn=CLASP2;refGene_tx=XM_
023260801.1;refGene_gn=CLASP2;refGene_tx=XM_023260802.1;refGene_gn=CLA
SP2;refGene_tx=XM_023260803.1;refGene_gn=CLASP2;refGene_tx=XM_02326080
4.1;refGene_gn=CLASP2;refGene_tx=XM_023260805.1;refGene_gn=CLASP2;refG
ene_tx=XM_023260806.1;refGene_gn=CLASP2;refGene_tx=XM_023260807.1;refG
ene_gn=CLASP2;refGene_tx=XM_023260808.1;refGene_gn=CLASP2;refGene_tx=X
M_023260809.1
C2 159360078 159360142
refGene_gn=CLASP2;refGene_tx=XM_011286298.2;refGene_gn=CLASP2;refGene_
tx=XM_011286299.3;refGene_gn=CLASP2;refGene_tx=XM_011286300.2;refGene_
gn=CLASP2;refGene_tx=XM_019811031.1;refGene_gn=CLASP2;refGene_tx=XM_01
9811032.1;refGene_gn=CLASP2;refGene_tx=XM_019811033.1;refGene_gn=CLASP
2;refGene_tx=XM_019811034.2;refGene_gn=CLASP2;refGene_tx=XM_023260796.
1;refGene_gn=CLASP2;refGene_tx=XM_023260797.1;refGene_gn=CLASP2;refGen
e_tx=XM_023260798.1;refGene_gn=CLASP2;refGene_tx=XM_023260799.1;refGen
e_gn=CLASP2;refGene_tx=XM_023260800.1;refGene_gn=CLASP2;refGene_tx=XM_
023260801.1;refGene_gn=CLASP2;refGene_tx=XM_023260802.1;refGene_gn=CLA
SP2;refGene_tx=XM_023260803.1;refGene_gn=CLASP2;refGene_tx=XM_02326080
4.1;refGene_gn=CLASP2;refGene_tx=XM_023260805.1;refGene_gn=CLASP2;refG
ene_tx=XM_023260806.1;refGene_gn=CLASP2;refGene_tx=XM_023260807.1;refG
ene_gn=CLASP2;refGene_tx=XM_023260808.1;refGene_gn=CLASP2;refGene_tx=X
M_023260809.1
C2 159362502 159362747
refGene_gn=CLASP2;refGene_tx=XM_011286298.2;refGene_gn=CLASP2;refGene_
tx=XM_011286299.3;refGene_gn=CLASP2;refGene_tx=XM_011286300.2;refGene_
gn=CLASP2;refGene_tx=XM_019811031.1;refGene_gn=CLASP2;refGene_tx=XM_01
9811032.1;refGene_gn=CLASP2;refGene_tx=XM_019811033.1;refGene_gn=CLASP
2;refGene_tx=XM_019811034.2;refGene_gn=CLASP2;refGene_tx=XM_023260796.
1;refGene_gn=CLASP2;refGene_tx=XM_023260797.1;refGene_gn=CLASP2;refGen
e_tx=XM_023260798.1;refGene_gn=CLASP2;refGene_tx=XM_023260799.1;refGen
e_gn=CLASP2;refGene_tx=XM_023260800.1;refGene_gn=CLASP2;refGene_tx=XM_
023260801.1;refGene_gn=CLASP2;refGene_tx=XM_023260802.1;refGene_gn=CLA
SP2;refGene_tx=XM_023260803.1;refGene_gn=CLASP2;refGene_tx=XM_02326080
4.1;refGene_gn=CLASP2;refGene_tx=XM_023260805.1;refGene_gn=CLASP2;refG
ene_tx=XM_023260806.1;refGene_gn=CLASP2;refGene_tx=XM_023260807.1;refG
ene_gn=CLASP2;refGene_tx=XM_023260808.1;refGene_gn=CLASP2;refGene_tx=X
M_023260809.1

C2 159364142 159364166
refGene_gn=CLASP2;refGene_tx=XM_011286298.2;refGene_gn=CLASP2;refGene_tx=XM_011286299.3;refGene_gn=CLASP2;refGene_tx=XM_011286300.2;refGene_gn=CLASP2;refGene_tx=XM_019811031.1;refGene_gn=CLASP2;refGene_tx=XM_019811032.1;refGene_gn=CLASP2;refGene_tx=XM_019811033.1;refGene_gn=CLASP2;refGene_tx=XM_019811034.2;refGene_gn=CLASP2;refGene_tx=XM_023260796.1;refGene_gn=CLASP2;refGene_tx=XM_023260797.1;refGene_gn=CLASP2;refGene_tx=XM_023260798.1;refGene_gn=CLASP2;refGene_tx=XM_023260799.1;refGene_gn=CLASP2;refGene_tx=XM_023260801.1;refGene_gn=CLASP2;refGene_tx=XM_023260802.1;refGene_gn=CLASP2;refGene_tx=XM_023260803.1;refGene_gn=CLASP2;refGene_tx=XM_023260804.1;refGene_gn=CLASP2;refGene_tx=XM_023260805.1;refGene_gn=CLASP2;refGene_tx=XM_023260808.1;refGene_gn=CLASP2;refGene_tx=XM_023260809.1
C2 159365787 159365894
refGene_gn=CLASP2;refGene_tx=XM_011286298.2;refGene_gn=CLASP2;refGene_tx=XM_011286299.3;refGene_gn=CLASP2;refGene_tx=XM_011286300.2;refGene_gn=CLASP2;refGene_tx=XM_019811031.1;refGene_gn=CLASP2;refGene_tx=XM_019811032.1;refGene_gn=CLASP2;refGene_tx=XM_019811033.1;refGene_gn=CLASP2;refGene_tx=XM_019811034.2;refGene_gn=CLASP2;refGene_tx=XM_023260796.1;refGene_gn=CLASP2;refGene_tx=XM_023260797.1;refGene_gn=CLASP2;refGene_tx=XM_023260798.1;refGene_gn=CLASP2;refGene_tx=XM_023260799.1;refGene_gn=CLASP2;refGene_tx=XM_023260800.1;refGene_gn=CLASP2;refGene_tx=XM_023260801.1;refGene_gn=CLASP2;refGene_tx=XM_023260802.1;refGene_gn=CLASP2;refGene_tx=XM_023260803.1;refGene_gn=CLASP2;refGene_tx=XM_023260804.1;refGene_gn=CLASP2;refGene_tx=XM_023260805.1;refGene_gn=CLASP2;refGene_tx=XM_023260806.1;refGene_gn=CLASP2;refGene_tx=XM_023260807.1;refGene_gn=CLASP2;refGene_tx=XM_023260808.1;refGene_gn=CLASP2;refGene_tx=XM_023260809.1
C2 159366870 159366933
refGene_gn=CLASP2;refGene_tx=XM_011286298.2;refGene_gn=CLASP2;refGene_tx=XM_019811031.1;refGene_gn=CLASP2;refGene_tx=XM_019811032.1;refGene_gn=CLASP2;refGene_tx=XM_019811033.1;refGene_gn=CLASP2;refGene_tx=XM_023260796.1;refGene_gn=CLASP2;refGene_tx=XM_023260797.1;refGene_gn=CLASP2;refGene_tx=XM_023260798.1;refGene_gn=CLASP2;refGene_tx=XM_023260799.1;refGene_gn=CLASP2;refGene_tx=XM_023260800.1;refGene_gn=CLASP2;refGene_tx=XM_023260801.1;refGene_gn=CLASP2;refGene_tx=XM_023260802.1;refGene_gn=CLASP2;refGene_tx=XM_023260803.1;refGene_gn=CLASP2;refGene_tx=XM_023260804.1;refGene_gn=CLASP2;refGene_tx=XM_023260806.1;refGene_gn=CLASP2;refGene_tx=XM_023260807.1
C2 159370873 159370981
refGene_gn=CLASP2;refGene_tx=XM_011286298.2;refGene_gn=CLASP2;refGene_tx=XM_011286299.3;refGene_gn=CLASP2;refGene_tx=XM_011286300.2;refGene_gn=CLASP2;refGene_tx=XM_019811031.1;refGene_gn=CLASP2;refGene_tx=XM_019811032.1;refGene_gn=CLASP2;refGene_tx=XM_019811033.1;refGene_gn=CLASP2;refGene_tx=XM_019811034.2;refGene_gn=CLASP2;refGene_tx=XM_023260796.1;refGene_gn=CLASP2;refGene_tx=XM_023260797.1;refGene_gn=CLASP2;refGene_tx=XM_023260798.1;refGene_gn=CLASP2;refGene_tx=XM_023260799.1;refGene_gn=CLASP2;refGene_tx=XM_023260800.1;refGene_gn=CLASP2;refGene_tx=XM_023260801.1;refGene_gn=CLASP2;refGene_tx=XM_023260802.1;refGene_gn=CLASP2;refGene_tx=XM_023260803.1;refGene_gn=CLASP2;refGene_tx=XM_02326080

4.1;refGene_gn=CLASP2;refGene_tx=XM_023260805.1;refGene_gn=CLASP2;refGene_tx=XM_023260806.1;refGene_gn=CLASP2;refGene_tx=XM_023260807.1;refGene_gn=CLASP2;refGene_tx=XM_023260808.1;refGene_gn=CLASP2;refGene_tx=XM_023260809.1
C2 159371428 159371599
refGene_gn=CLASP2;refGene_tx=XM_011286298.2;refGene_gn=CLASP2;refGene_tx=XM_011286299.3;refGene_gn=CLASP2;refGene_tx=XM_011286300.2;refGene_gn=CLASP2;refGene_tx=XM_019811031.1;refGene_gn=CLASP2;refGene_tx=XM_019811032.1;refGene_gn=CLASP2;refGene_tx=XM_019811033.1;refGene_gn=CLASP2;refGene_tx=XM_019811034.2;refGene_gn=CLASP2;refGene_tx=XM_023260796.1;refGene_gn=CLASP2;refGene_tx=XM_023260797.1;refGene_gn=CLASP2;refGene_tx=XM_023260798.1;refGene_gn=CLASP2;refGene_tx=XM_023260799.1;refGene_gn=CLASP2;refGene_tx=XM_023260800.1;refGene_gn=CLASP2;refGene_tx=XM_023260801.1;refGene_gn=CLASP2;refGene_tx=XM_023260802.1;refGene_gn=CLASP2;refGene_tx=XM_023260803.1;refGene_gn=CLASP2;refGene_tx=XM_023260804.1;refGene_gn=CLASP2;refGene_tx=XM_023260805.1;refGene_gn=CLASP2;refGene_tx=XM_023260806.1;refGene_gn=CLASP2;refGene_tx=XM_023260807.1;refGene_gn=CLASP2;refGene_tx=XM_023260808.1;refGene_gn=CLASP2;refGene_tx=XM_023260809.1
C2 159374156 159374183
refGene_gn=CLASP2;refGene_tx=XM_019811031.1;refGene_gn=CLASP2;refGene_tx=XM_019811032.1;refGene_gn=CLASP2;refGene_tx=XM_019811033.1;refGene_gn=CLASP2;refGene_tx=XM_019811034.2;refGene_gn=CLASP2;refGene_tx=XM_023260796.1;refGene_gn=CLASP2;refGene_tx=XM_023260797.1;refGene_gn=CLASP2;refGene_tx=XM_023260798.1;refGene_gn=CLASP2;refGene_tx=XM_023260799.1;refGene_gn=CLASP2;refGene_tx=XM_023260800.1;refGene_gn=CLASP2;refGene_tx=XM_023260802.1;refGene_gn=CLASP2;refGene_tx=XM_023260803.1;refGene_gn=CLASP2;refGene_tx=XM_023260805.1;refGene_gn=CLASP2;refGene_tx=XM_023260806.1
C2 159377787 159377814
refGene_gn=CLASP2;refGene_tx=XM_019811031.1;refGene_gn=CLASP2;refGene_tx=XM_019811032.1;refGene_gn=CLASP2;refGene_tx=XM_019811033.1;refGene_gn=CLASP2;refGene_tx=XM_019811034.2;refGene_gn=CLASP2;refGene_tx=XM_023260796.1;refGene_gn=CLASP2;refGene_tx=XM_023260797.1;refGene_gn=CLASP2;refGene_tx=XM_023260798.1;refGene_gn=CLASP2;refGene_tx=XM_023260799.1;refGene_gn=CLASP2;refGene_tx=XM_023260800.1;refGene_gn=CLASP2;refGene_tx=XM_023260801.1;refGene_gn=CLASP2;refGene_tx=XM_023260802.1;refGene_gn=CLASP2;refGene_tx=XM_023260803.1;refGene_gn=CLASP2;refGene_tx=XM_023260805.1;refGene_gn=CLASP2;refGene_tx=XM_023260806.1
C2 159377970 159378075
refGene_gn=CLASP2;refGene_tx=XM_011286298.2;refGene_gn=CLASP2;refGene_tx=XM_011286299.3;refGene_gn=CLASP2;refGene_tx=XM_011286300.2;refGene_gn=CLASP2;refGene_tx=XM_019811031.1;refGene_gn=CLASP2;refGene_tx=XM_019811032.1;refGene_gn=CLASP2;refGene_tx=XM_019811033.1;refGene_gn=CLASP2;refGene_tx=XM_019811034.2;refGene_gn=CLASP2;refGene_tx=XM_023260796.1;refGene_gn=CLASP2;refGene_tx=XM_023260797.1;refGene_gn=CLASP2;refGene_tx=XM_023260798.1;refGene_gn=CLASP2;refGene_tx=XM_023260799.1;refGene_gn=CLASP2;refGene_tx=XM_023260800.1;refGene_gn=CLASP2;refGene_tx=XM_023260801.1;refGene_gn=CLASP2;refGene_tx=XM_023260802.1;refGene_gn=CLASP2;refGene_tx=XM_023260803.1;refGene_gn=CLASP2;refGene_tx=XM_02326080

4.1;refGene_gn=CLASP2;refGene_tx=XM_023260805.1;refGene_gn=CLASP2;refGene_tx=XM_023260806.1;refGene_gn=CLASP2;refGene_tx=XM_023260807.1;refGene_gn=CLASP2;refGene_tx=XM_023260808.1;refGene_gn=CLASP2;refGene_tx=XM_023260809.1
C2 159380099 159380117
refGene_gn=CLASP2;refGene_tx=XM_011286298.2;refGene_gn=CLASP2;refGene_tx=XM_011286299.3;refGene_gn=CLASP2;refGene_tx=XM_011286300.2;refGene_gn=CLASP2;refGene_tx=XM_019811031.1;refGene_gn=CLASP2;refGene_tx=XM_019811032.1;refGene_gn=CLASP2;refGene_tx=XM_019811033.1;refGene_gn=CLASP2;refGene_tx=XM_019811034.2;refGene_gn=CLASP2;refGene_tx=XM_023260796.1;refGene_gn=CLASP2;refGene_tx=XM_023260797.1;refGene_gn=CLASP2;refGene_tx=XM_023260798.1;refGene_gn=CLASP2;refGene_tx=XM_023260800.1;refGene_gn=CLASP2;refGene_tx=XM_023260801.1;refGene_gn=CLASP2;refGene_tx=XM_023260803.1;refGene_gn=CLASP2;refGene_tx=XM_023260804.1;refGene_gn=CLASP2;refGene_tx=XM_023260805.1;refGene_gn=CLASP2;refGene_tx=XM_023260806.1;refGene_gn=CLASP2;refGene_tx=XM_023260807.1;refGene_gn=CLASP2;refGene_tx=XM_023260808.1;refGene_gn=CLASP2;refGene_tx=XM_023260809.1
C2 159381823 159381847
refGene_gn=CLASP2;refGene_tx=XM_011286298.2;refGene_gn=CLASP2;refGene_tx=XM_011286299.3;refGene_gn=CLASP2;refGene_tx=XM_011286300.2;refGene_gn=CLASP2;refGene_tx=XM_019811031.1;refGene_gn=CLASP2;refGene_tx=XM_019811032.1;refGene_gn=CLASP2;refGene_tx=XM_019811033.1;refGene_gn=CLASP2;refGene_tx=XM_019811034.2;refGene_gn=CLASP2;refGene_tx=XM_023260796.1;refGene_gn=CLASP2;refGene_tx=XM_023260797.1;refGene_gn=CLASP2;refGene_tx=XM_023260798.1;refGene_gn=CLASP2;refGene_tx=XM_023260799.1;refGene_gn=CLASP2;refGene_tx=XM_023260800.1;refGene_gn=CLASP2;refGene_tx=XM_023260801.1;refGene_gn=CLASP2;refGene_tx=XM_023260803.1;refGene_gn=CLASP2;refGene_tx=XM_023260804.1;refGene_gn=CLASP2;refGene_tx=XM_023260805.1;refGene_gn=CLASP2;refGene_tx=XM_023260806.1;refGene_gn=CLASP2;refGene_tx=XM_023260807.1;refGene_gn=CLASP2;refGene_tx=XM_023260808.1;refGene_gn=CLASP2;refGene_tx=XM_023260809.1
C2 159388097 159388271
refGene_gn=CLASP2;refGene_tx=XM_011286298.2;refGene_gn=CLASP2;refGene_tx=XM_011286299.3;refGene_gn=CLASP2;refGene_tx=XM_011286300.2;refGene_gn=CLASP2;refGene_tx=XM_019811031.1;refGene_gn=CLASP2;refGene_tx=XM_019811032.1;refGene_gn=CLASP2;refGene_tx=XM_019811033.1;refGene_gn=CLASP2;refGene_tx=XM_019811034.2;refGene_gn=CLASP2;refGene_tx=XM_023260796.1;refGene_gn=CLASP2;refGene_tx=XM_023260797.1;refGene_gn=CLASP2;refGene_tx=XM_023260798.1;refGene_gn=CLASP2;refGene_tx=XM_023260799.1;refGene_gn=CLASP2;refGene_tx=XM_023260800.1;refGene_gn=CLASP2;refGene_tx=XM_023260801.1;refGene_gn=CLASP2;refGene_tx=XM_023260802.1;refGene_gn=CLASP2;refGene_tx=XM_023260803.1;refGene_gn=CLASP2;refGene_tx=XM_023260804.1;refGene_gn=CLASP2;refGene_tx=XM_023260805.1;refGene_gn=CLASP2;refGene_tx=XM_023260806.1;refGene_gn=CLASP2;refGene_tx=XM_023260807.1;refGene_gn=CLASP2;refGene_tx=XM_023260808.1;refGene_gn=CLASP2;refGene_tx=XM_023260809.1
C2 159389451 159389507
refGene_gn=CLASP2;refGene_tx=XM_011286298.2;refGene_gn=CLASP2;refGene_tx=XM_011286299.3;refGene_gn=CLASP2;refGene_tx=XM_011286300.2;refGene_gn=CLASP2;refGene_tx=XM_019811031.1;refGene_gn=CLASP2;refGene_tx=XM_01

9811032.1;refGene_gn=CLASP2;refGene_tx=XM_019811033.1;refGene_gn=CLASP
2;refGene_tx=XM_019811034.2;refGene_gn=CLASP2;refGene_tx=XM_023260796.
1;refGene_gn=CLASP2;refGene_tx=XM_023260797.1;refGene_gn=CLASP2;refGen
e_tx=XM_023260798.1;refGene_gn=CLASP2;refGene_tx=XM_023260799.1;refGen
e_gn=CLASP2;refGene_tx=XM_023260800.1;refGene_gn=CLASP2;refGene_tx=XM_
023260801.1;refGene_gn=CLASP2;refGene_tx=XM_023260802.1;refGene_gn=CLA
SP2;refGene_tx=XM_023260803.1;refGene_gn=CLASP2;refGene_tx=XM_02326080
4.1;refGene_gn=CLASP2;refGene_tx=XM_023260805.1;refGene_gn=CLASP2;refG
ene_tx=XM_023260806.1;refGene_gn=CLASP2;refGene_tx=XM_023260807.1;refG
ene_gn=CLASP2;refGene_tx=XM_023260808.1;refGene_gn=CLASP2;refGene_tx=X
M_023260809.1
C2 159391935 159392103
refGene_gn=CLASP2;refGene_tx=XM_011286298.2;refGene_gn=CLASP2;refGene_
tx=XM_011286299.3;refGene_gn=CLASP2;refGene_tx=XM_011286300.2;refGene_
gn=CLASP2;refGene_tx=XM_019811031.1;refGene_gn=CLASP2;refGene_tx=XM_01
9811032.1;refGene_gn=CLASP2;refGene_tx=XM_019811033.1;refGene_gn=CLASP
2;refGene_tx=XM_019811034.2;refGene_gn=CLASP2;refGene_tx=XM_023260796.
1;refGene_gn=CLASP2;refGene_tx=XM_023260797.1;refGene_gn=CLASP2;refGen
e_tx=XM_023260798.1;refGene_gn=CLASP2;refGene_tx=XM_023260799.1;refGen
e_gn=CLASP2;refGene_tx=XM_023260800.1;refGene_gn=CLASP2;refGene_tx=XM_
023260801.1;refGene_gn=CLASP2;refGene_tx=XM_023260802.1;refGene_gn=CLA
SP2;refGene_tx=XM_023260803.1;refGene_gn=CLASP2;refGene_tx=XM_02326080
4.1;refGene_gn=CLASP2;refGene_tx=XM_023260805.1;refGene_gn=CLASP2;refG
ene_tx=XM_023260806.1;refGene_gn=CLASP2;refGene_tx=XM_023260807.1;refG
ene_gn=CLASP2;refGene_tx=XM_023260808.1;refGene_gn=CLASP2;refGene_tx=X
M_023260809.1
C2 159392839 159392917
refGene_gn=CLASP2;refGene_tx=XM_011286298.2;refGene_gn=CLASP2;refGene_
tx=XM_011286299.3;refGene_gn=CLASP2;refGene_tx=XM_011286300.2;refGene_
gn=CLASP2;refGene_tx=XM_019811031.1;refGene_gn=CLASP2;refGene_tx=XM_01
9811032.1;refGene_gn=CLASP2;refGene_tx=XM_019811033.1;refGene_gn=CLASP
2;refGene_tx=XM_019811034.2;refGene_gn=CLASP2;refGene_tx=XM_023260796.
1;refGene_gn=CLASP2;refGene_tx=XM_023260797.1;refGene_gn=CLASP2;refGen
e_tx=XM_023260798.1;refGene_gn=CLASP2;refGene_tx=XM_023260799.1;refGen
e_gn=CLASP2;refGene_tx=XM_023260800.1;refGene_gn=CLASP2;refGene_tx=XM_
023260801.1;refGene_gn=CLASP2;refGene_tx=XM_023260802.1;refGene_gn=CLA
SP2;refGene_tx=XM_023260803.1;refGene_gn=CLASP2;refGene_tx=XM_02326080
4.1;refGene_gn=CLASP2;refGene_tx=XM_023260805.1;refGene_gn=CLASP2;refG
ene_tx=XM_023260806.1;refGene_gn=CLASP2;refGene_tx=XM_023260807.1;refG
ene_gn=CLASP2;refGene_tx=XM_023260808.1;refGene_gn=CLASP2;refGene_tx=X
M_023260809.1
C2 159394109 159394169
refGene_gn=CLASP2;refGene_tx=XM_011286298.2;refGene_gn=CLASP2;refGene_
tx=XM_011286299.3;refGene_gn=CLASP2;refGene_tx=XM_011286300.2;refGene_
gn=CLASP2;refGene_tx=XM_019811031.1;refGene_gn=CLASP2;refGene_tx=XM_01
9811032.1;refGene_gn=CLASP2;refGene_tx=XM_019811033.1;refGene_gn=CLASP
2;refGene_tx=XM_019811034.2;refGene_gn=CLASP2;refGene_tx=XM_023260796.
1;refGene_gn=CLASP2;refGene_tx=XM_023260797.1;refGene_gn=CLASP2;refGen
e_tx=XM_023260798.1;refGene_gn=CLASP2;refGene_tx=XM_023260799.1;refGen
e_gn=CLASP2;refGene_tx=XM_023260800.1;refGene_gn=CLASP2;refGene_tx=XM_

023260801.1;refGene_gn=CLASP2;refGene_tx=XM_023260802.1;refGene_gn=CLASP2;refGene_tx=XM_023260803.1;refGene_gn=CLASP2;refGene_tx=XM_023260804.1;refGene_gn=CLASP2;refGene_tx=XM_023260805.1;refGene_gn=CLASP2;refGene_tx=XM_023260806.1;refGene_gn=CLASP2;refGene_tx=XM_023260807.1;refGene_gn=CLASP2;refGene_tx=XM_023260808.1;refGene_gn=CLASP2;refGene_tx=XM_023260809.1
C2 159398352 159398423
refGene_gn=CLASP2;refGene_tx=XM_011286298.2;refGene_gn=CLASP2;refGene_tx=XM_011286299.3;refGene_gn=CLASP2;refGene_tx=XM_011286300.2;refGene_gn=CLASP2;refGene_tx=XM_019811031.1;refGene_gn=CLASP2;refGene_tx=XM_019811032.1;refGene_gn=CLASP2;refGene_tx=XM_019811033.1;refGene_gn=CLASP2;refGene_tx=XM_019811034.2;refGene_gn=CLASP2;refGene_tx=XM_023260796.1;refGene_gn=CLASP2;refGene_tx=XM_023260797.1;refGene_gn=CLASP2;refGene_tx=XM_023260798.1;refGene_gn=CLASP2;refGene_tx=XM_023260799.1;refGene_gn=CLASP2;refGene_tx=XM_023260800.1;refGene_gn=CLASP2;refGene_tx=XM_023260801.1;refGene_gn=CLASP2;refGene_tx=XM_023260802.1;refGene_gn=CLASP2;refGene_tx=XM_023260803.1;refGene_gn=CLASP2;refGene_tx=XM_023260804.1;refGene_gn=CLASP2;refGene_tx=XM_023260805.1;refGene_gn=CLASP2;refGene_tx=XM_023260806.1;refGene_gn=CLASP2;refGene_tx=XM_023260807.1;refGene_gn=CLASP2;refGene_tx=XM_023260808.1;refGene_gn=CLASP2;refGene_tx=XM_023260809.1
C2 159404087 159404223
refGene_gn=CLASP2;refGene_tx=XM_011286298.2;refGene_gn=CLASP2;refGene_tx=XM_011286299.3;refGene_gn=CLASP2;refGene_tx=XM_011286300.2;refGene_gn=CLASP2;refGene_tx=XM_019811031.1;refGene_gn=CLASP2;refGene_tx=XM_019811032.1;refGene_gn=CLASP2;refGene_tx=XM_019811033.1;refGene_gn=CLASP2;refGene_tx=XM_019811034.2;refGene_gn=CLASP2;refGene_tx=XM_023260796.1;refGene_gn=CLASP2;refGene_tx=XM_023260797.1;refGene_gn=CLASP2;refGene_tx=XM_023260798.1;refGene_gn=CLASP2;refGene_tx=XM_023260799.1;refGene_gn=CLASP2;refGene_tx=XM_023260800.1;refGene_gn=CLASP2;refGene_tx=XM_023260801.1;refGene_gn=CLASP2;refGene_tx=XM_023260802.1;refGene_gn=CLASP2;refGene_tx=XM_023260803.1;refGene_gn=CLASP2;refGene_tx=XM_023260804.1;refGene_gn=CLASP2;refGene_tx=XM_023260805.1;refGene_gn=CLASP2;refGene_tx=XM_023260806.1;refGene_gn=CLASP2;refGene_tx=XM_023260807.1;refGene_gn=CLASP2;refGene_tx=XM_023260808.1;refGene_gn=CLASP2;refGene_tx=XM_023260809.1
C2 159406838 159406984
refGene_gn=CLASP2;refGene_tx=XM_011286298.2;refGene_gn=CLASP2;refGene_tx=XM_011286299.3;refGene_gn=CLASP2;refGene_tx=XM_011286300.2;refGene_gn=CLASP2;refGene_tx=XM_019811031.1;refGene_gn=CLASP2;refGene_tx=XM_019811032.1;refGene_gn=CLASP2;refGene_tx=XM_019811033.1;refGene_gn=CLASP2;refGene_tx=XM_019811034.2;refGene_gn=CLASP2;refGene_tx=XM_023260796.1;refGene_gn=CLASP2;refGene_tx=XM_023260797.1;refGene_gn=CLASP2;refGene_tx=XM_023260798.1;refGene_gn=CLASP2;refGene_tx=XM_023260799.1;refGene_gn=CLASP2;refGene_tx=XM_023260800.1;refGene_gn=CLASP2;refGene_tx=XM_023260801.1;refGene_gn=CLASP2;refGene_tx=XM_023260802.1;refGene_gn=CLASP2;refGene_tx=XM_023260803.1;refGene_gn=CLASP2;refGene_tx=XM_023260804.1;refGene_gn=CLASP2;refGene_tx=XM_023260805.1;refGene_gn=CLASP2;refGene_tx=XM_023260806.1;refGene_gn=CLASP2;refGene_tx=XM_023260807.1;refGene_gn=CLASP2;refGene_tx=XM_023260808.1;refGene_gn=CLASP2;refGene_tx=X

M_023260809.1
C2	159410975	159411068
refGene_gn=CLASP2;refGene_tx=XM_011286298.2;refGene_gn=CLASP2;refGene_tx=XM_011286299.3;refGene_gn=CLASP2;refGene_tx=XM_011286300.2;refGene_gn=CLASP2;refGene_tx=XM_019811031.1;refGene_gn=CLASP2;refGene_tx=XM_019811032.1;refGene_gn=CLASP2;refGene_tx=XM_019811033.1;refGene_gn=CLASP2;refGene_tx=XM_019811034.2;refGene_gn=CLASP2;refGene_tx=XM_023260796.1;refGene_gn=CLASP2;refGene_tx=XM_023260797.1;refGene_gn=CLASP2;refGene_tx=XM_023260798.1;refGene_gn=CLASP2;refGene_tx=XM_023260799.1;refGene_gn=CLASP2;refGene_tx=XM_023260800.1;refGene_gn=CLASP2;refGene_tx=XM_023260801.1;refGene_gn=CLASP2;refGene_tx=XM_023260802.1;refGene_gn=CLASP2;refGene_tx=XM_023260803.1;refGene_gn=CLASP2;refGene_tx=XM_023260804.1;refGene_gn=CLASP2;refGene_tx=XM_023260805.1;refGene_gn=CLASP2;refGene_tx=XM_023260806.1;refGene_gn=CLASP2;refGene_tx=XM_023260807.1;refGene_gn=CLASP2;refGene_tx=XM_023260808.1;refGene_gn=CLASP2;refGene_tx=XM_023260809.1
C2	159414192	159414275
refGene_gn=CLASP2;refGene_tx=XM_011286298.2;refGene_gn=CLASP2;refGene_tx=XM_011286299.3;refGene_gn=CLASP2;refGene_tx=XM_011286300.2;refGene_gn=CLASP2;refGene_tx=XM_019811031.1;refGene_gn=CLASP2;refGene_tx=XM_019811032.1;refGene_gn=CLASP2;refGene_tx=XM_019811033.1;refGene_gn=CLASP2;refGene_tx=XM_019811034.2;refGene_gn=CLASP2;refGene_tx=XM_023260796.1;refGene_gn=CLASP2;refGene_tx=XM_023260797.1;refGene_gn=CLASP2;refGene_tx=XM_023260798.1;refGene_gn=CLASP2;refGene_tx=XM_023260799.1;refGene_gn=CLASP2;refGene_tx=XM_023260800.1;refGene_gn=CLASP2;refGene_tx=XM_023260801.1;refGene_gn=CLASP2;refGene_tx=XM_023260802.1;refGene_gn=CLASP2;refGene_tx=XM_023260803.1;refGene_gn=CLASP2;refGene_tx=XM_023260804.1;refGene_gn=CLASP2;refGene_tx=XM_023260805.1;refGene_gn=CLASP2;refGene_tx=XM_023260806.1;refGene_gn=CLASP2;refGene_tx=XM_023260807.1;refGene_gn=CLASP2;refGene_tx=XM_023260808.1;refGene_gn=CLASP2;refGene_tx=XM_023260809.1
C2	159423417	159423514
refGene_gn=CLASP2;refGene_tx=XM_019811033.1
C2	159429673	159429820
refGene_gn=CLASP2;refGene_tx=XM_011286298.2;refGene_gn=CLASP2;refGene_tx=XM_011286299.3;refGene_gn=CLASP2;refGene_tx=XM_011286300.2;refGene_gn=CLASP2;refGene_tx=XM_019811031.1;refGene_gn=CLASP2;refGene_tx=XM_019811032.1;refGene_gn=CLASP2;refGene_tx=XM_019811034.2;refGene_gn=CLASP2;refGene_tx=XM_023260796.1;refGene_gn=CLASP2;refGene_tx=XM_023260797.1;refGene_gn=CLASP2;refGene_tx=XM_023260798.1;refGene_gn=CLASP2;refGene_tx=XM_023260799.1;refGene_gn=CLASP2;refGene_tx=XM_023260800.1;refGene_gn=CLASP2;refGene_tx=XM_023260801.1;refGene_gn=CLASP2;refGene_tx=XM_023260802.1;refGene_gn=CLASP2;refGene_tx=XM_023260803.1;refGene_gn=CLASP2;refGene_tx=XM_023260804.1;refGene_gn=CLASP2;refGene_tx=XM_023260805.1;refGene_gn=CLASP2;refGene_tx=XM_023260806.1;refGene_gn=CLASP2;refGene_tx=XM_023260807.1;refGene_gn=CLASP2;refGene_tx=XM_023260808.1;refGene_gn=CLASP2;refGene_tx=XM_023260809.1
C2	159430202	159430236
refGene_gn=CLASP2;refGene_tx=XM_011286298.2;refGene_gn=CLASP2;refGene_tx=XM_011286299.3;refGene_gn=CLASP2;refGene_tx=XM_011286300.2;refGene_

gn=CLASP2;refGene_tx=XM_019811031.1;refGene_gn=CLASP2;refGene_tx=XM_019811034.2
C2 159446978 159446994
refGene_gn=CLASP2;refGene_tx=XM_019811032.1
C2 159452459 159452530
refGene_gn=CLASP2;refGene_tx=XM_023260796.1;refGene_gn=CLASP2;refGene_tx=XM_023260797.1;refGene_gn=CLASP2;refGene_tx=XM_023260798.1;refGene_gn=CLASP2;refGene_tx=XM_023260799.1;refGene_gn=CLASP2;refGene_tx=XM_023260800.1;refGene_gn=CLASP2;refGene_tx=XM_023260801.1;refGene_gn=CLASP2;refGene_tx=XM_023260802.1;refGene_gn=CLASP2;refGene_tx=XM_023260803.1;refGene_gn=CLASP2;refGene_tx=XM_023260804.1;refGene_gn=CLASP2;refGene_tx=XM_023260805.1;refGene_gn=CLASP2;refGene_tx=XM_023260806.1;refGene_gn=CLASP2;refGene_tx=XM_023260807.1;refGene_gn=CLASP2;refGene_tx=XM_023260808.1;refGene_gn=CLASP2;refGene_tx=XM_023260809.1
C2 159462290 159462388
refGene_gn=CLASP2;refGene_tx=XM_023260796.1;refGene_gn=CLASP2;refGene_tx=XM_023260797.1;refGene_gn=CLASP2;refGene_tx=XM_023260798.1;refGene_gn=CLASP2;refGene_tx=XM_023260799.1;refGene_gn=CLASP2;refGene_tx=XM_023260800.1;refGene_gn=CLASP2;refGene_tx=XM_023260801.1;refGene_gn=CLASP2;refGene_tx=XM_023260802.1;refGene_gn=CLASP2;refGene_tx=XM_023260803.1;refGene_gn=CLASP2;refGene_tx=XM_023260804.1;refGene_gn=CLASP2;refGene_tx=XM_023260805.1;refGene_gn=CLASP2;refGene_tx=XM_023260806.1;refGene_gn=CLASP2;refGene_tx=XM_023260807.1;refGene_gn=CLASP2;refGene_tx=XM_023260808.1;refGene_gn=CLASP2;refGene_tx=XM_023260809.1
C2 159464180 159464256
refGene_gn=CLASP2;refGene_tx=XM_023260796.1;refGene_gn=CLASP2;refGene_tx=XM_023260797.1;refGene_gn=CLASP2;refGene_tx=XM_023260798.1;refGene_gn=CLASP2;refGene_tx=XM_023260799.1;refGene_gn=CLASP2;refGene_tx=XM_023260800.1;refGene_gn=CLASP2;refGene_tx=XM_023260801.1;refGene_gn=CLASP2;refGene_tx=XM_023260802.1;refGene_gn=CLASP2;refGene_tx=XM_023260803.1;refGene_gn=CLASP2;refGene_tx=XM_023260804.1;refGene_gn=CLASP2;refGene_tx=XM_023260805.1;refGene_gn=CLASP2;refGene_tx=XM_023260806.1;refGene_gn=CLASP2;refGene_tx=XM_023260807.1;refGene_gn=CLASP2;refGene_tx=XM_023260808.1;refGene_gn=CLASP2;refGene_tx=XM_023260809.1
C2 159465421 159465513
refGene_gn=CLASP2;refGene_tx=XM_023260796.1;refGene_gn=CLASP2;refGene_tx=XM_023260797.1;refGene_gn=CLASP2;refGene_tx=XM_023260798.1;refGene_gn=CLASP2;refGene_tx=XM_023260799.1;refGene_gn=CLASP2;refGene_tx=XM_023260800.1;refGene_gn=CLASP2;refGene_tx=XM_023260801.1;refGene_gn=CLASP2;refGene_tx=XM_023260802.1;refGene_gn=CLASP2;refGene_tx=XM_023260803.1;refGene_gn=CLASP2;refGene_tx=XM_023260804.1;refGene_gn=CLASP2;refGene_tx=XM_023260805.1;refGene_gn=CLASP2;refGene_tx=XM_023260806.1;refGene_gn=CLASP2;refGene_tx=XM_023260807.1;refGene_gn=CLASP2;refGene_tx=XM_023260808.1;refGene_gn=CLASP2;refGene_tx=XM_023260809.1
C2 159467215 159467319
refGene_gn=CLASP2;refGene_tx=XM_023260796.1;refGene_gn=CLASP2;refGene_tx=XM_023260797.1;refGene_gn=CLASP2;refGene_tx=XM_023260798.1;refGene_gn=CLASP2;refGene_tx=XM_023260799.1;refGene_gn=CLASP2;refGene_tx=XM_023260800.1;refGene_gn=CLASP2;refGene_tx=XM_023260801.1;refGene_gn=CLASP2;refGene_tx=XM_023260802.1;refGene_gn=CLASP2;refGene_tx=XM_023260803.

1;refGene_gn=CLASP2;refGene_tx=XM_023260804.1;refGene_gn=CLASP2;refGene_tx=XM_023260805.1;refGene_gn=CLASP2;refGene_tx=XM_023260806.1;refGene_gn=CLASP2;refGene_tx=XM_023260807.1;refGene_gn=CLASP2;refGene_tx=XM_023260808.1;refGene_gn=CLASP2;refGene_tx=XM_023260809.1
C2 159473104 159473183
refGene_gn=CLASP2;refGene_tx=XM_023260796.1;refGene_gn=CLASP2;refGene_tx=XM_023260797.1;refGene_gn=CLASP2;refGene_tx=XM_023260798.1;refGene_gn=CLASP2;refGene_tx=XM_023260799.1;refGene_gn=CLASP2;refGene_tx=XM_023260800.1;refGene_gn=CLASP2;refGene_tx=XM_023260801.1;refGene_gn=CLASP2;refGene_tx=XM_023260802.1;refGene_gn=CLASP2;refGene_tx=XM_023260803.1;refGene_gn=CLASP2;refGene_tx=XM_023260804.1;refGene_gn=CLASP2;refGene_tx=XM_023260805.1;refGene_gn=CLASP2;refGene_tx=XM_023260806.1;refGene_gn=CLASP2;refGene_tx=XM_023260807.1;refGene_gn=CLASP2;refGene_tx=XM_023260808.1;refGene_gn=CLASP2;refGene_tx=XM_023260809.1
C2 159486927 159487122
refGene_gn=CLASP2;refGene_tx=XM_023260796.1;refGene_gn=CLASP2;refGene_tx=XM_023260797.1;refGene_gn=CLASP2;refGene_tx=XM_023260798.1;refGene_gn=CLASP2;refGene_tx=XM_023260799.1;refGene_gn=CLASP2;refGene_tx=XM_023260800.1;refGene_gn=CLASP2;refGene_tx=XM_023260801.1;refGene_gn=CLASP2;refGene_tx=XM_023260802.1;refGene_gn=CLASP2;refGene_tx=XM_023260803.1;refGene_gn=CLASP2;refGene_tx=XM_023260804.1;refGene_gn=CLASP2;refGene_tx=XM_023260805.1;refGene_gn=CLASP2;refGene_tx=XM_023260806.1;refGene_gn=CLASP2;refGene_tx=XM_023260807.1;refGene_gn=CLASP2;refGene_tx=XM_023260808.1;refGene_gn=CLASP2;refGene_tx=XM_023260809.1
C2 159780200 159780789
refGene_gn=SNRK;refGene_tx=XM_023260810.1
C2 159805989 159806131
refGene_gn=SNRK;refGene_tx=XM_023260810.1;refGene_gn=SNRK;refGene_tx=XM_023260811.1;refGene_gn=SNRK;refGene_tx=XM_023260812.1
C2 159811233 159811446
refGene_gn=SNRK;refGene_tx=XM_023260810.1;refGene_gn=SNRK;refGene_tx=XM_023260811.1;refGene_gn=SNRK;refGene_tx=XM_023260812.1
C2 159813496 159813592
refGene_gn=ANO10;refGene_tx=XM_023260817.1
C2 159813688 159813823
refGene_gn=SNRK;refGene_tx=XM_023260810.1;refGene_gn=SNRK;refGene_tx=XM_023260811.1;refGene_gn=SNRK;refGene_tx=XM_023260812.1
C2 159817728 159818920
refGene_gn=SNRK;refGene_tx=XM_023260810.1;refGene_gn=SNRK;refGene_tx=XM_023260811.1;refGene_gn=SNRK;refGene_tx=XM_023260812.1
C2 159833488 159833560
refGene_gn=ANO10;refGene_tx=XM_023260818.1
C2 159864827 159864998
refGene_gn=ANO10;refGene_tx=XM_023260815.1
C2 159866214 159866238
refGene_gn=ANO10;refGene_tx=XM_023260819.1
C2 159869551 159869674
refGene_gn=ANO10;refGene_tx=XM_023260816.1
C2 159879595 159879712
refGene_gn=ANO10;refGene_tx=XM_023260815.1;refGene_gn=ANO10;refGene_tx

=XM_023260816.1;refGene_gn=ANO10;refGene_tx=XM_023260817.1;refGene_gn=ANO10;refGene_tx=XM_023260818.1;refGene_gn=ANO10;refGene_tx=XM_023260819.1
C2 159920819 159921173
refGene_gn=ANO10;refGene_tx=XM_023260813.1;refGene_gn=ANO10;refGene_tx=XM_023260814.1
C2 159930511 159930622
refGene_gn=ANO10;refGene_tx=XM_023260820.1
C2 159984848 159984977
refGene_gn=ANO10;refGene_tx=XM_023260813.1;refGene_gn=ANO10;refGene_tx=XM_023260814.1;refGene_gn=ANO10;refGene_tx=XM_023260815.1;refGene_gn=ANO10;refGene_tx=XM_023260816.1;refGene_gn=ANO10;refGene_tx=XM_023260817.1;refGene_gn=ANO10;refGene_tx=XM_023260818.1;refGene_gn=ANO10;refGene_tx=XM_023260819.1;refGene_gn=ANO10;refGene_tx=XM_023260820.1
C2 159993322 159993514
refGene_gn=ANO10;refGene_tx=XM_023260813.1;refGene_gn=ANO10;refGene_tx=XM_023260814.1;refGene_gn=ANO10;refGene_tx=XM_023260815.1;refGene_gn=ANO10;refGene_tx=XM_023260816.1;refGene_gn=ANO10;refGene_tx=XM_023260817.1;refGene_gn=ANO10;refGene_tx=XM_023260818.1;refGene_gn=ANO10;refGene_tx=XM_023260819.1;refGene_gn=ANO10;refGene_tx=XM_023260820.1
C2 160005001 160005184
refGene_gn=ANO10;refGene_tx=XM_023260813.1;refGene_gn=ANO10;refGene_tx=XM_023260814.1;refGene_gn=ANO10;refGene_tx=XM_023260815.1;refGene_gn=ANO10;refGene_tx=XM_023260816.1;refGene_gn=ANO10;refGene_tx=XM_023260817.1;refGene_gn=ANO10;refGene_tx=XM_023260818.1;refGene_gn=ANO10;refGene_tx=XM_023260819.1;refGene_gn=ANO10;refGene_tx=XM_023260820.1
C2 160006610 160006685
refGene_gn=ANO10;refGene_tx=XM_023260813.1;refGene_gn=ANO10;refGene_tx=XM_023260814.1;refGene_gn=ANO10;refGene_tx=XM_023260815.1;refGene_gn=ANO10;refGene_tx=XM_023260816.1;refGene_gn=ANO10;refGene_tx=XM_023260817.1;refGene_gn=ANO10;refGene_tx=XM_023260818.1;refGene_gn=ANO10;refGene_tx=XM_023260819.1;refGene_gn=ANO10;refGene_tx=XM_023260820.1
C2 160008209 160008265
refGene_gn=ANO10;refGene_tx=XM_023260813.1;refGene_gn=ANO10;refGene_tx=XM_023260814.1;refGene_gn=ANO10;refGene_tx=XM_023260815.1;refGene_gn=ANO10;refGene_tx=XM_023260816.1;refGene_gn=ANO10;refGene_tx=XM_023260817.1;refGene_gn=ANO10;refGene_tx=XM_023260818.1;refGene_gn=ANO10;refGene_tx=XM_023260819.1;refGene_gn=ANO10;refGene_tx=XM_023260820.1
C2 160010078 160010648
refGene_gn=ANO10;refGene_tx=XM_023260813.1;refGene_gn=ANO10;refGene_tx=XM_023260814.1;refGene_gn=ANO10;refGene_tx=XM_023260815.1;refGene_gn=ANO10;refGene_tx=XM_023260816.1;refGene_gn=ANO10;refGene_tx=XM_023260817.1;refGene_gn=ANO10;refGene_tx=XM_023260818.1;refGene_gn=ANO10;refGene_tx=XM_023260819.1;refGene_gn=ANO10;refGene_tx=XM_023260820.1
C2 160011734 160011854
refGene_gn=ANO10;refGene_tx=XM_023260813.1;refGene_gn=ANO10;refGene_tx=XM_023260814.1;refGene_gn=ANO10;refGene_tx=XM_023260815.1;refGene_gn=ANO10;refGene_tx=XM_023260816.1;refGene_gn=ANO10;refGene_tx=XM_023260817.1;refGene_gn=ANO10;refGene_tx=XM_023260818.1;refGene_gn=ANO10;refGene_tx=XM_023260819.1;refGene_gn=ANO10;refGene_tx=XM_023260820.1

C2 160036543 160036678
refGene_gn=ANO10;refGene_tx=XM_023260813.1;refGene_gn=ANO10;refGene_tx
=XM_023260814.1;refGene_gn=ANO10;refGene_tx=XM_023260815.1;refGene_gn=
ANO10;refGene_tx=XM_023260816.1;refGene_gn=ANO10;refGene_tx=XM_0232608
17.1;refGene_gn=ANO10;refGene_tx=XM_023260818.1;refGene_gn=ANO10;refGe
ne_tx=XM_023260819.1;refGene_gn=ANO10;refGene_tx=XM_023260820.1
C2 160038281 160038479
refGene_gn=ANO10;refGene_tx=XM_023260813.1;refGene_gn=ANO10;refGene_tx
=XM_023260814.1;refGene_gn=ANO10;refGene_tx=XM_023260815.1;refGene_gn=
ANO10;refGene_tx=XM_023260816.1;refGene_gn=ANO10;refGene_tx=XM_0232608
17.1;refGene_gn=ANO10;refGene_tx=XM_023260818.1;refGene_gn=ANO10;refGe
ne_tx=XM_023260819.1;refGene_gn=ANO10;refGene_tx=XM_023260820.1
C2 160041621 160041760
refGene_gn=ANO10;refGene_tx=XM_023260813.1;refGene_gn=ANO10;refGene_tx
=XM_023260814.1;refGene_gn=ANO10;refGene_tx=XM_023260815.1;refGene_gn=
ANO10;refGene_tx=XM_023260816.1;refGene_gn=ANO10;refGene_tx=XM_0232608
17.1;refGene_gn=ANO10;refGene_tx=XM_023260818.1;refGene_gn=ANO10;refGe
ne_tx=XM_023260819.1;refGene_gn=ANO10;refGene_tx=XM_023260820.1
C2 160101219 160101266
refGene_gn=ABHD5;refGene_tx=XM_003992286.5
C2 160113238 160113324
refGene_gn=ABHD5;refGene_tx=XM_003992286.5
C2 160115719 160116092
refGene_gn=ABHD5;refGene_tx=XM_003992286.5
C2 160124466 160124621
refGene_gn=ABHD5;refGene_tx=XM_003992286.5
C2 160127007 160127119
refGene_gn=ABHD5;refGene_tx=XM_003992286.5
C2 160129567 160129754
refGene_gn=ABHD5;refGene_tx=XM_003992286.5
C2 160130287 160130377
refGene_gn=ABHD5;refGene_tx=XM_003992286.5
C2 160652056 160652261
refGene_gn=TOPAZ1;refGene_tx=XM_023260821.1
C2 160652832 160655296
refGene_gn=TOPAZ1;refGene_tx=XM_023260821.1
C2 160665520 160665582
refGene_gn=TOPAZ1;refGene_tx=XM_023260821.1
C2 160666276 160666385
refGene_gn=TOPAZ1;refGene_tx=XM_023260821.1
C2 160666487 160666506
refGene_gn=TOPAZ1;refGene_tx=XM_023260821.1
C2 160681578 160681643
refGene_gn=TOPAZ1;refGene_tx=XM_023260821.1
C2 160685585 160685722
refGene_gn=TOPAZ1;refGene_tx=XM_023260821.1
C2 160687118 160687204
refGene_gn=TOPAZ1;refGene_tx=XM_023260821.1
C2 160687929 160688055
refGene_gn=TOPAZ1;refGene_tx=XM_023260821.1

C2	160689575	160689639
refGene_gn=TOPAZ1;refGene_tx=XM_023260821.1
C2	160700327	160700479
refGene_gn=TOPAZ1;refGene_tx=XM_023260821.1
C2	160700686	160700779
refGene_gn=TOPAZ1;refGene_tx=XM_023260821.1
C2	160701823	160701939
refGene_gn=TOPAZ1;refGene_tx=XM_023260821.1
C2	160718892	160718959
refGene_gn=TOPAZ1;refGene_tx=XM_023260821.1
C2	160720261	160720436
refGene_gn=TOPAZ1;refGene_tx=XM_023260821.1
C2	160721755	160721856
refGene_gn=TOPAZ1;refGene_tx=XM_023260821.1
C2	160724318	160724484
refGene_gn=TOPAZ1;refGene_tx=XM_023260821.1
C2	160742215	160742380
refGene_gn=TOPAZ1;refGene_tx=XM_023260821.1
C2	160744083	160744287
refGene_gn=TOPAZ1;refGene_tx=XM_023260821.1
C2	160749902	160750086
refGene_gn=TOPAZ1;refGene_tx=XM_023260821.1
C2	160753689	160753909
refGene_gn=TOPAZ1;refGene_tx=XM_023260821.1
C2	160782999	160783028
refGene_gn=TCAIM;refGene_tx=XM_019811042.2;refGene_gn=TCAIM;refGene_tx
=XM_023260822.1;refGene_gn=TCAIM;refGene_tx=XM_023260823.1;refGene_gn=
TCAIM;refGene_tx=XM_023260824.1;refGene_gn=TCAIM;refGene_tx=XM_0232608
25.1
C2	160788672	160788808
refGene_gn=TCAIM;refGene_tx=XM_019811042.2;refGene_gn=TCAIM;refGene_tx
=XM_023260822.1;refGene_gn=TCAIM;refGene_tx=XM_023260823.1;refGene_gn=
TCAIM;refGene_tx=XM_023260824.1;refGene_gn=TCAIM;refGene_tx=XM_0232608
25.1
C2	160796817	160796971
refGene_gn=TCAIM;refGene_tx=XM_019811042.2;refGene_gn=TCAIM;refGene_tx
=XM_023260822.1;refGene_gn=TCAIM;refGene_tx=XM_023260823.1;refGene_gn=
TCAIM;refGene_tx=XM_023260824.1;refGene_gn=TCAIM;refGene_tx=XM_0232608
25.1
C2	160802929	160803182
refGene_gn=TCAIM;refGene_tx=XM_019811042.2;refGene_gn=TCAIM;refGene_tx
=XM_023260822.1;refGene_gn=TCAIM;refGene_tx=XM_023260823.1;refGene_gn=
TCAIM;refGene_tx=XM_023260824.1;refGene_gn=TCAIM;refGene_tx=XM_0232608
25.1
C2	160821074	160821197
refGene_gn=TCAIM;refGene_tx=XM_019811042.2;refGene_gn=TCAIM;refGene_tx
=XM_023260822.1;refGene_gn=TCAIM;refGene_tx=XM_023260823.1;refGene_gn=
TCAIM;refGene_tx=XM_023260824.1;refGene_gn=TCAIM;refGene_tx=XM_0232608
25.1
C2	160824495	160824593

refGene_gn=TCAIM;refGene_tx=XM_019811042.2;refGene_gn=TCAIM;refGene_tx=XM_023260822.1;refGene_gn=TCAIM;refGene_tx=XM_023260823.1;refGene_gn=TCAIM;refGene_tx=XM_023260824.1;refGene_gn=TCAIM;refGene_tx=XM_023260825.1
C2 160824872 160824964
refGene_gn=TCAIM;refGene_tx=XM_019811042.2;refGene_gn=TCAIM;refGene_tx=XM_023260822.1;refGene_gn=TCAIM;refGene_tx=XM_023260823.1;refGene_gn=TCAIM;refGene_tx=XM_023260824.1;refGene_gn=TCAIM;refGene_tx=XM_023260825.1
C2 160828204 160828437
refGene_gn=TCAIM;refGene_tx=XM_019811042.2;refGene_gn=TCAIM;refGene_tx=XM_023260822.1;refGene_gn=TCAIM;refGene_tx=XM_023260823.1;refGene_gn=TCAIM;refGene_tx=XM_023260824.1;refGene_gn=TCAIM;refGene_tx=XM_023260825.1
C2 160830415 160830547
refGene_gn=TCAIM;refGene_tx=XM_019811042.2;refGene_gn=TCAIM;refGene_tx=XM_023260822.1;refGene_gn=TCAIM;refGene_tx=XM_023260823.1;refGene_gn=TCAIM;refGene_tx=XM_023260824.1;refGene_gn=TCAIM;refGene_tx=XM_023260825.1
C2 160834391 160834632
refGene_gn=TCAIM;refGene_tx=XM_019811042.2;refGene_gn=TCAIM;refGene_tx=XM_023260822.1;refGene_gn=TCAIM;refGene_tx=XM_023260823.1;refGene_gn=TCAIM;refGene_tx=XM_023260824.1;refGene_gn=TCAIM;refGene_tx=XM_023260825.1
C2 160890370 160892529
refGene_gn=ZNF445;refGene_tx=XM_003992294.5;refGene_gn=ZNF445;refGene_tx=XM_006936568.4
C2 160893541 160893652
refGene_gn=ZNF445;refGene_tx=XM_003992294.5;refGene_gn=ZNF445;refGene_tx=XM_006936568.4
C2 160894380 160894507
refGene_gn=ZNF445;refGene_tx=XM_003992294.5;refGene_gn=ZNF445;refGene_tx=XM_006936568.4
C2 160894789 160894866
refGene_gn=ZNF445;refGene_tx=XM_003992294.5
C2 160895218 160895387
refGene_gn=ZNF445;refGene_tx=XM_003992294.5
C2 160897839 160898274
refGene_gn=ZNF445;refGene_tx=XM_003992294.5;refGene_gn=ZNF445;refGene_tx=XM_006936568.4
C2 160993934 160995376
refGene_gn=LOC105259958;refGene_tx=XM_023260826.1;refGene_gn=LOC105259958;refGene_tx=XM_023260827.1
C2 160997449 160997576
refGene_gn=LOC105259958;refGene_tx=XM_023260827.1
C2 160997865 160997880
refGene_gn=LOC105259958;refGene_tx=XM_023260827.1
C2 161024855 161025278
refGene_gn=LOC101099651;refGene_tx=XM_019811045.2
C2 161029388 161029557

refGene_gn=LOC101099651;refGene_tx=XM_019811045.2
C2 161031017 161031109
refGene_gn=LOC101099651;refGene_tx=XM_019811045.2
C2 161031398 161031525
refGene_gn=LOC101099651;refGene_tx=XM_019811045.2
C2 161033600 161035013
refGene_gn=LOC101099651;refGene_tx=XM_019811045.2
C2 161050145 161050252
refGene_gn=LOC101099651;refGene_tx=XM_019811045.2
C2 161050253 161050997
refGene_gn=LOC101099651;refGene_tx=XM_019811045.2
C2 161081752 161082151
refGene_gn=LOC101099396;refGene_tx=XM_003992295.5;refGene_gn=LOC101099396;refGene_tx=XM_019811046.2;refGene_gn=LOC101099396;refGene_tx=XM_019811047.2;refGene_gn=LOC101099396;refGene_tx=XM_019811048.2;refGene_gn=LOC101099396;refGene_tx=XM_019811049.2
C2 161083350 161083507
refGene_gn=LOC101099396;refGene_tx=XM_003992295.5;refGene_gn=LOC101099396;refGene_tx=XM_019811046.2;refGene_gn=LOC101099396;refGene_tx=XM_019811047.2;refGene_gn=LOC101099396;refGene_tx=XM_019811048.2;refGene_gn=LOC101099396;refGene_tx=XM_019811049.2
C2 161083902 161084012
refGene_gn=LOC101099396;refGene_tx=XM_003992295.5;refGene_gn=LOC101099396;refGene_tx=XM_019811046.2;refGene_gn=LOC101099396;refGene_tx=XM_019811047.2;refGene_gn=LOC101099396;refGene_tx=XM_019811048.2;refGene_gn=LOC101099396;refGene_tx=XM_019811049.2
C2 161084248 161084375
refGene_gn=LOC101099396;refGene_tx=XM_003992295.5;refGene_gn=LOC101099396;refGene_tx=XM_019811046.2;refGene_gn=LOC101099396;refGene_tx=XM_019811047.2;refGene_gn=LOC101099396;refGene_tx=XM_019811048.2;refGene_gn=LOC101099396;refGene_tx=XM_019811049.2
C2 161095456 161097774
refGene_gn=LOC101099396;refGene_tx=XM_003992295.5;refGene_gn=LOC101099396;refGene_tx=XM_019811046.2;refGene_gn=LOC101099396;refGene_tx=XM_019811047.2;refGene_gn=LOC101099396;refGene_tx=XM_019811048.2;refGene_gn=LOC101099396;refGene_tx=XM_019811049.2
C2 161104900 161105092
refGene_gn=ZNF35;refGene_tx=XM_011286320.3
C2 161106079 161106224
refGene_gn=ZNF35;refGene_tx=XM_011286320.3
C2 161111700 161111945
refGene_gn=ZNF35;refGene_tx=XM_011286320.3
C2 161111946 161112948
refGene_gn=ZNF35;refGene_tx=XM_011286320.3
D1 318321 318473
refGene_gn=TRPC6;refGene_tx=XM_019811185.2;refGene_gn=TRPC6;refGene_tx=XM_023239035.1;refGene_gn=TRPC6;refGene_tx=XM_023239036.1
D1 319233 319309
refGene_gn=TRPC6;refGene_tx=XM_019811185.2;refGene_gn=TRPC6;refGene_tx=XM_023239035.1;refGene_gn=TRPC6;refGene_tx=XM_023239036.1

D1	327452	327536
refGene_gn=TRPC6;refGene_tx=XM_019811185.2;refGene_gn=TRPC6;refGene_tx=XM_023239035.1;refGene_gn=TRPC6;refGene_tx=XM_023239036.1
D1	333467	333542
refGene_gn=TRPC6;refGene_tx=XM_019811185.2;refGene_gn=TRPC6;refGene_tx=XM_023239035.1;refGene_gn=TRPC6;refGene_tx=XM_023239036.1
D1	335579	335783
refGene_gn=TRPC6;refGene_tx=XM_019811185.2;refGene_gn=TRPC6;refGene_tx=XM_023239035.1;refGene_gn=TRPC6;refGene_tx=XM_023239036.1
D1	336535	336731
refGene_gn=TRPC6;refGene_tx=XM_019811185.2;refGene_gn=TRPC6;refGene_tx=XM_023239035.1;refGene_gn=TRPC6;refGene_tx=XM_023239036.1
D1	337962	338227
refGene_gn=TRPC6;refGene_tx=XM_019811185.2;refGene_gn=TRPC6;refGene_tx=XM_023239035.1;refGene_gn=TRPC6;refGene_tx=XM_023239036.1
D1	340107	340341
refGene_gn=TRPC6;refGene_tx=XM_019811185.2;refGene_gn=TRPC6;refGene_tx=XM_023239035.1;refGene_gn=TRPC6;refGene_tx=XM_023239036.1
D1	345485	345702
refGene_gn=TRPC6;refGene_tx=XM_019811185.2;refGene_gn=TRPC6;refGene_tx=XM_023239035.1;refGene_gn=TRPC6;refGene_tx=XM_023239036.1
D1	350929	351094
refGene_gn=TRPC6;refGene_tx=XM_019811185.2;refGene_gn=TRPC6;refGene_tx=XM_023239035.1
D1	353738	353924
refGene_gn=TRPC6;refGene_tx=XM_019811185.2;refGene_gn=TRPC6;refGene_tx=XM_023239035.1;refGene_gn=TRPC6;refGene_tx=XM_023239036.1
D1	358006	358781
refGene_gn=TRPC6;refGene_tx=XM_019811185.2;refGene_gn=TRPC6;refGene_tx=XM_023239035.1;refGene_gn=TRPC6;refGene_tx=XM_023239036.1
D1	380995	381087	refGene_gn=TRPC6;refGene_tx=XM_019811185.2
D1	451104	451274
refGene_gn=TRPC6;refGene_tx=XM_023239035.1;refGene_gn=TRPC6;refGene_tx=XM_023239036.1
D1	654804	655124	refGene_gn=ANGPTL5;refGene_tx=XM_003992300.4
D1	658190	658376	refGene_gn=ANGPTL5;refGene_tx=XM_003992300.4
D1	660553	660674	refGene_gn=ANGPTL5;refGene_tx=XM_003992300.4
D1	662902	663003	refGene_gn=ANGPTL5;refGene_tx=XM_003992300.4
D1	665107	665201	refGene_gn=ANGPTL5;refGene_tx=XM_003992300.4
D1	666351	666455	refGene_gn=ANGPTL5;refGene_tx=XM_003992300.4
D1	668522	668667	refGene_gn=ANGPTL5;refGene_tx=XM_003992300.4
D1	670256	670352	refGene_gn=ANGPTL5;refGene_tx=XM_003992300.4
D1	676114	676242
refGene_gn=CEP126;refGene_tx=XM_006936576.4;refGene_gn=CEP126;refGene_tx=XM_019811191.2;refGene_gn=CEP126;refGene_tx=XM_019811192.2;refGene_gn=CEP126;refGene_tx=XM_019811193.2;refGene_gn=CEP126;refGene_tx=XM_019811194.2;refGene_gn=CEP126;refGene_tx=XM_019811196.2
D1	679389	679509
refGene_gn=CEP126;refGene_tx=XM_006936576.4;refGene_gn=CEP126;refGene_tx=XM_019811191.2;refGene_gn=CEP126;refGene_tx=XM_019811192.2;refGene_

gn=CEP126;refGene_tx=XM_019811193.2;refGene_gn=CEP126;refGene_tx=XM_019811194.2;refGene_gn=CEP126;refGene_tx=XM_019811196.2
D1 713132 713278
refGene_gn=CEP126;refGene_tx=XM_006936576.4;refGene_gn=CEP126;refGene_tx=XM_019811191.2;refGene_gn=CEP126;refGene_tx=XM_019811192.2;refGene_gn=CEP126;refGene_tx=XM_019811193.2;refGene_gn=CEP126;refGene_tx=XM_019811194.2;refGene_gn=CEP126;refGene_tx=XM_019811196.2
D1 719575 719687
refGene_gn=CEP126;refGene_tx=XM_006936576.4;refGene_gn=CEP126;refGene_tx=XM_019811191.2;refGene_gn=CEP126;refGene_tx=XM_019811192.2;refGene_gn=CEP126;refGene_tx=XM_019811193.2;refGene_gn=CEP126;refGene_tx=XM_019811194.2;refGene_gn=CEP126;refGene_tx=XM_019811196.2
D1 730542 730741
refGene_gn=CEP126;refGene_tx=XM_006936576.4;refGene_gn=CEP126;refGene_tx=XM_011286327.2;refGene_gn=CEP126;refGene_tx=XM_019811191.2;refGene_gn=CEP126;refGene_tx=XM_019811192.2;refGene_gn=CEP126;refGene_tx=XM_019811193.2;refGene_gn=CEP126;refGene_tx=XM_019811194.2;refGene_gn=CEP126;refGene_tx=XM_019811196.2;refGene_gn=CEP126;refGene_tx=XM_019811197.1
D1 734455 736592
refGene_gn=CEP126;refGene_tx=XM_006936576.4;refGene_gn=CEP126;refGene_tx=XM_011286327.2;refGene_gn=CEP126;refGene_tx=XM_019811191.2;refGene_gn=CEP126;refGene_tx=XM_019811192.2;refGene_gn=CEP126;refGene_tx=XM_019811193.2;refGene_gn=CEP126;refGene_tx=XM_019811194.2;refGene_gn=CEP126;refGene_tx=XM_019811196.2;refGene_gn=CEP126;refGene_tx=XM_019811197.1
D1 747738 747791 refGene_gn=CEP126;refGene_tx=XM_019811196.2
D1 753044 753157
refGene_gn=CEP126;refGene_tx=XM_006936576.4;refGene_gn=CEP126;refGene_tx=XM_011286327.2;refGene_gn=CEP126;refGene_tx=XM_019811191.2;refGene_gn=CEP126;refGene_tx=XM_019811192.2;refGene_gn=CEP126;refGene_tx=XM_019811193.2;refGene_gn=CEP126;refGene_tx=XM_019811194.2;refGene_gn=CEP126;refGene_tx=XM_019811196.2;refGene_gn=CEP126;refGene_tx=XM_019811197.1
D1 755009 755162 refGene_gn=CEP126;refGene_tx=XM_019811194.2
D1 755837 755913
refGene_gn=CEP126;refGene_tx=XM_006936576.4;refGene_gn=CEP126;refGene_tx=XM_011286327.2;refGene_gn=CEP126;refGene_tx=XM_019811191.2;refGene_gn=CEP126;refGene_tx=XM_019811192.2;refGene_gn=CEP126;refGene_tx=XM_019811193.2;refGene_gn=CEP126;refGene_tx=XM_019811197.1
D1 758127 758337
refGene_gn=CEP126;refGene_tx=XM_006936576.4;refGene_gn=CEP126;refGene_tx=XM_011286327.2;refGene_gn=CEP126;refGene_tx=XM_019811191.2;refGene_gn=CEP126;refGene_tx=XM_019811192.2;refGene_gn=CEP126;refGene_tx=XM_019811197.1
D1 763520 763621 refGene_gn=CEP126;refGene_tx=XM_019811191.2
D1 769444 769509
refGene_gn=CEP126;refGene_tx=XM_006936576.4;refGene_gn=CEP126;refGene_tx=XM_011286327.2;refGene_gn=CEP126;refGene_tx=XM_019811193.2;refGene_gn=CEP126;refGene_tx=XM_019811197.1

D1 779456 779519
refGene_gn=CEP126;refGene_tx=XM_006936576.4;refGene_gn=CEP126;refGene_tx=XM_011286327.2;refGene_gn=CEP126;refGene_tx=XM_019811192.2;refGene_gn=CEP126;refGene_tx=XM_019811193.2;refGene_gn=CEP126;refGene_tx=XM_019811197.1
D1 812583 812693
refGene_gn=CD1H11orf70;refGene_tx=XM_023239037.1
D1 812778 812860
refGene_gn=CD1H11orf70;refGene_tx=XM_023239037.1
D1 816282 816358
refGene_gn=CD1H11orf70;refGene_tx=XM_023239037.1
D1 822440 822607
refGene_gn=CD1H11orf70;refGene_tx=XM_023239037.1
D1 827331 827504
refGene_gn=CD1H11orf70;refGene_tx=XM_023239037.1
D1 841307 841374
refGene_gn=CD1H11orf70;refGene_tx=XM_023239037.1
D1 842244 842373
refGene_gn=CD1H11orf70;refGene_tx=XM_023239037.1
D1 854216 854328
refGene_gn=LOC111556534;refGene_tx=XM_023239007.1
D1 855036 856257
refGene_gn=LOC111556534;refGene_tx=XM_023239007.1
D1 857303 857581
refGene_gn=LOC111556534;refGene_tx=XM_023239007.1
D1 857772 857973
refGene_gn=LOC111556534;refGene_tx=XM_023239007.1
D1 859659 859697 refGene_gn=YAP1;refGene_tx=XM_019811201.2
D1 912392 912508 refGene_gn=YAP1;refGene_tx=XM_019811201.2
D1 932337 932451 refGene_gn=YAP1;refGene_tx=XM_019811201.2
D1 946518 946694 refGene_gn=YAP1;refGene_tx=XM_019811201.2
D1 948866 948914 refGene_gn=YAP1;refGene_tx=XM_019811201.2
D1 958729 958860 refGene_gn=YAP1;refGene_tx=XM_019811201.2
D1 962560 962673 refGene_gn=YAP1;refGene_tx=XM_019811201.2
D1 964779 964839 refGene_gn=YAP1;refGene_tx=XM_019811201.2
D1 964841 965017 refGene_gn=YAP1;refGene_tx=XM_019811201.2
D1 1045105 1046018
refGene_gn=LOC101083444;refGene_tx=XM_019811205.2;refGene_gn=LOC101083444;refGene_tx=XM_023239038.1;refGene_gn=LOC101083444;refGene_tx=XM_023239039.1
D1 1046127 1046227
refGene_gn=LOC101083444;refGene_tx=XM_019811205.2;refGene_gn=LOC101083444;refGene_tx=XM_023239038.1;refGene_gn=LOC101083444;refGene_tx=XM_023239039.1
D1 1047602 1047681
refGene_gn=LOC101083444;refGene_tx=XM_019811205.2;refGene_gn=LOC101083444;refGene_tx=XM_023239038.1;refGene_gn=LOC101083444;refGene_tx=XM_023239039.1
D1 1050741 1050790
refGene_gn=LOC101083444;refGene_tx=XM_019811205.2;refGene_gn=LOC101083

444;refGene_tx=XM_023239038.1;refGene_gn=LOC101083444;refGene_tx=XM_023239039.1
D1 1052715 1052958 refGene_gn=LOC101083444;refGene_tx=XM_019811205.2;refGene_gn=LOC101083444;refGene_tx=XM_023239038.1;refGene_gn=LOC101083444;refGene_tx=XM_023239039.1
D1 1056460 1056715 refGene_gn=LOC101083444;refGene_tx=XM_019811205.2;refGene_gn=LOC101083444;refGene_tx=XM_023239038.1;refGene_gn=LOC101083444;refGene_tx=XM_023239039.1
D1 1057135 1057177 refGene_gn=LOC101083444;refGene_tx=XM_019811205.2;refGene_gn=LOC101083444;refGene_tx=XM_023239038.1;refGene_gn=LOC101083444;refGene_tx=XM_023239039.1
D1 1057521 1057715 refGene_gn=LOC101083444;refGene_tx=XM_019811205.2;refGene_gn=LOC101083444;refGene_tx=XM_023239038.1;refGene_gn=LOC101083444;refGene_tx=XM_023239039.1
D1 1070038 1070910 refGene_gn=LOC111556535;refGene_tx=XM_023239008.1
D1 1072810 1072820 refGene_gn=LOC111556535;refGene_tx=XM_023239008.1
D1 1073111 1074009 refGene_gn=BIRC2;refGene_tx=XM_023239040.1
D1 1074100 1074200 refGene_gn=BIRC2;refGene_tx=XM_023239040.1
D1 1077523 1077602 refGene_gn=BIRC2;refGene_tx=XM_023239040.1
D1 1078195 1078244 refGene_gn=BIRC2;refGene_tx=XM_023239040.1
D1 1093019 1093262 refGene_gn=BIRC2;refGene_tx=XM_023239040.1
D1 1095150 1095405 refGene_gn=BIRC2;refGene_tx=XM_023239040.1
D1 1095679 1095721 refGene_gn=BIRC2;refGene_tx=XM_023239040.1
D1 1095830 1096024 refGene_gn=BIRC2;refGene_tx=XM_023239040.1
D1 1103910 1103935 refGene_gn=TMEM123;refGene_tx=XM_019811208.1;refGene_gn=TMEM123;refGene_tx=XM_023239041.1
D1 1105461 1105615 refGene_gn=TMEM123;refGene_tx=XM_019811208.1;refGene_gn=TMEM123;refGene_tx=XM_023239041.1
D1 1106405 1106666 refGene_gn=TMEM123;refGene_tx=XM_019811208.1;refGene_gn=TMEM123;refGene_tx=XM_023239041.1
D1 1121897 1121997 refGene_gn=TMEM123;refGene_tx=XM_019811208.1
D1 1147972 1148029 refGene_gn=TMEM123;refGene_tx=XM_023239041.1
D1 1153064 1153164 refGene_gn=TMEM123;refGene_tx=XM_023239041.1
D1 1185587 1185616 refGene_gn=MMP7;refGene_tx=XM_003992303.2
D1 1187169 1187331 refGene_gn=MMP7;refGene_tx=XM_003992303.2
D1 1189123 1189252 refGene_gn=MMP7;refGene_tx=XM_003992303.2
D1 1190928 1191077 refGene_gn=MMP7;refGene_tx=XM_003992303.2
D1 1191163 1191390 refGene_gn=MMP7;refGene_tx=XM_003992303.2
D1 1193830 1193938 refGene_gn=MMP7;refGene_tx=XM_003992303.2
D1 1225191 1225292 refGene_gn=MMP20;refGene_tx=XM_003992304.2
D1 1226200 1226304 refGene_gn=MMP20;refGene_tx=XM_003992304.2

D1 1245747 1245904 refGene_gn=MMP20;refGene_tx=XM_003992304.2
D1 1247087 1247224 refGene_gn=MMP20;refGene_tx=XM_003992304.2
D1 1258488 1258630 refGene_gn=MMP20;refGene_tx=XM_003992304.2
D1 1261269 1261431 refGene_gn=MMP20;refGene_tx=XM_003992304.2
D1 1262198 1262324 refGene_gn=MMP20;refGene_tx=XM_003992304.2
D1 1263961 1264110 refGene_gn=MMP20;refGene_tx=XM_003992304.2
D1 1268060 1268308 refGene_gn=MMP20;refGene_tx=XM_003992304.2
D1 1276652 1276778 refGene_gn=MMP20;refGene_tx=XM_003992304.2
D1 1316072 1316308 refGene_gn=MMP27;refGene_tx=XM_006936578.3
D1 1317551 1317655 refGene_gn=MMP27;refGene_tx=XM_006936578.3
D1 1318679 1318839 refGene_gn=MMP27;refGene_tx=XM_006936578.3
D1 1319880 1320011 refGene_gn=MMP27;refGene_tx=XM_006936578.3
D1 1321282 1321403 refGene_gn=MMP27;refGene_tx=XM_006936578.3
D1 1321578 1321740 refGene_gn=MMP27;refGene_tx=XM_006936578.3
D1 1324696 1324825 refGene_gn=MMP27;refGene_tx=XM_006936578.3
D1 1324914 1325063 refGene_gn=MMP27;refGene_tx=XM_006936578.3
D1 1326142 1326381 refGene_gn=MMP27;refGene_tx=XM_006936578.3
D1 1327174 1327276 refGene_gn=MMP27;refGene_tx=XM_006936578.3
D1 1333003 1333107 refGene_gn=MMP8;refGene_tx=XM_003992305.4
D1 1333261 1333365 refGene_gn=MMP8;refGene_tx=XM_003992305.4
D1 1334271 1334425 refGene_gn=MMP8;refGene_tx=XM_003992305.4
D1 1334993 1335127 refGene_gn=MMP8;refGene_tx=XM_003992305.4
D1 1335869 1335987 refGene_gn=MMP8;refGene_tx=XM_003992305.4
D1 1337961 1338123 refGene_gn=MMP8;refGene_tx=XM_003992305.4
D1 1340661 1340787 refGene_gn=MMP8;refGene_tx=XM_003992305.4
D1 1340874 1341023 refGene_gn=MMP8;refGene_tx=XM_003992305.4
D1 1341579 1341824 refGene_gn=MMP8;refGene_tx=XM_003992305.4
D1 1343750 1343885 refGene_gn=MMP8;refGene_tx=XM_003992305.4
D1 1352104 1352376
refGene_gn=LOC101083957;refGene_tx=XM_023239009.1
D1 1361004 1361104
refGene_gn=LOC101083957;refGene_tx=XM_023239009.1
D1 1361260 1361372
refGene_gn=LOC101083957;refGene_tx=XM_023239009.1
D1 1361942 1362105
refGene_gn=LOC101083957;refGene_tx=XM_023239009.1
D1 1362610 1362732
refGene_gn=LOC101083957;refGene_tx=XM_023239009.1
D1 1364158 1364276
refGene_gn=LOC101083957;refGene_tx=XM_023239009.1
D1 1364659 1364815
refGene_gn=LOC101083957;refGene_tx=XM_023239009.1
D1 1365816 1365942
refGene_gn=LOC101083957;refGene_tx=XM_023239009.1
D1 1366392 1366541
refGene_gn=LOC101083957;refGene_tx=XM_023239009.1
D1 1366634 1366879
refGene_gn=LOC101083957;refGene_tx=XM_023239009.1
D1 1367304 1367403
refGene_gn=LOC101083957;refGene_tx=XM_023239009.1

D1 1381945 1382046 refGene_gn=MMP10;refGene_tx=XM_003992306.1
D1 1383066 1383170 refGene_gn=MMP10;refGene_tx=XM_003992306.1
D1 1383894 1384054 refGene_gn=MMP10;refGene_tx=XM_003992306.1
D1 1386846 1386980 refGene_gn=MMP10;refGene_tx=XM_003992306.1
D1 1388288 1388436 refGene_gn=MMP10;refGene_tx=XM_003992306.1
D1 1388621 1388786 refGene_gn=MMP10;refGene_tx=XM_003992306.1
D1 1390478 1390604 refGene_gn=MMP10;refGene_tx=XM_003992306.1
D1 1390711 1390860 refGene_gn=MMP10;refGene_tx=XM_003992306.1
D1 1390951 1391196 refGene_gn=MMP10;refGene_tx=XM_003992306.1
D1 1391823 1391928 refGene_gn=MMP10;refGene_tx=XM_003992306.1
D1 1402223 1402333 refGene_gn=MMP1;refGene_tx=XM_003992316.3
D1 1402513 1402617 refGene_gn=MMP1;refGene_tx=XM_003992316.3
D1 1403366 1403529 refGene_gn=MMP1;refGene_tx=XM_003992316.3
D1 1405027 1405161 refGene_gn=MMP1;refGene_tx=XM_003992316.3
D1 1407167 1407285 refGene_gn=MMP1;refGene_tx=XM_003992316.3
D1 1407415 1407571 refGene_gn=MMP1;refGene_tx=XM_003992316.3
D1 1407730 1407856 refGene_gn=MMP1;refGene_tx=XM_003992316.3
D1 1408084 1408233 refGene_gn=MMP1;refGene_tx=XM_003992316.3
D1 1408326 1408571 refGene_gn=MMP1;refGene_tx=XM_003992316.3
D1 1408991 1409102 refGene_gn=MMP1;refGene_tx=XM_003992316.3
D1 1427883 1427984
refGene_gn=LOC101081492;refGene_tx=XM_003992307.3
D1 1429253 1429357
refGene_gn=LOC101081492;refGene_tx=XM_003992307.3
D1 1430713 1430873
refGene_gn=LOC101081492;refGene_tx=XM_003992307.3
D1 1431296 1431430
refGene_gn=LOC101081492;refGene_tx=XM_003992307.3
D1 1432374 1432522
refGene_gn=LOC101081492;refGene_tx=XM_003992307.3
D1 1432707 1432872
refGene_gn=LOC101081492;refGene_tx=XM_003992307.3
D1 1434442 1434568
refGene_gn=LOC101081492;refGene_tx=XM_003992307.3
D1 1434692 1434841
refGene_gn=LOC101081492;refGene_tx=XM_003992307.3
D1 1434932 1435177
refGene_gn=LOC101081492;refGene_tx=XM_003992307.3
D1 1435939 1436044
refGene_gn=LOC101081492;refGene_tx=XM_003992307.3
D1 1451872 1451973
refGene_gn=MMP12;refGene_tx=XM_003992317.4;refGene_gn=MMP12;refGene_tx
=XM_019811212.2
D1 1452882 1452986
refGene_gn=MMP12;refGene_tx=XM_003992317.4;refGene_gn=MMP12;refGene_tx
=XM_019811212.2
D1 1453721 1453881
refGene_gn=MMP12;refGene_tx=XM_003992317.4;refGene_gn=MMP12;refGene_tx
=XM_019811212.2
D1 1454242 1454376

```
refGene_gn=MMP12;refGene_tx=XM_003992317.4;refGene_gn=MMP12;refGene_tx
=XM_019811212.2
D1	1458255	1458379
refGene_gn=MMP12;refGene_tx=XM_003992317.4;refGene_gn=MMP12;refGene_tx
=XM_019811212.2
D1	1458898	1459060
refGene_gn=MMP12;refGene_tx=XM_003992317.4;refGene_gn=MMP12;refGene_tx
=XM_019811212.2
D1	1461343	1461469
refGene_gn=MMP12;refGene_tx=XM_003992317.4;refGene_gn=MMP12;refGene_tx
=XM_019811212.2
D1	1461556	1461705
refGene_gn=MMP12;refGene_tx=XM_003992317.4;refGene_gn=MMP12;refGene_tx
=XM_019811212.2
D1	1462389	1462637
refGene_gn=MMP12;refGene_tx=XM_003992317.4;refGene_gn=MMP12;refGene_tx
=XM_019811212.2
D1	1464027	1464132	refGene_gn=MMP12;refGene_tx=XM_003992317.4
D1	1511408	1511509	refGene_gn=MMP13;refGene_tx=XM_023239043.1
D1	1512398	1512502	refGene_gn=MMP13;refGene_tx=XM_023239043.1
D1	1514319	1514479	refGene_gn=MMP13;refGene_tx=XM_023239043.1
D1	1515415	1515549	refGene_gn=MMP13;refGene_tx=XM_023239043.1
D1	1516558	1516676	refGene_gn=MMP13;refGene_tx=XM_023239043.1
D1	1518419	1518581	refGene_gn=MMP13;refGene_tx=XM_023239043.1
D1	1520506	1520632	refGene_gn=MMP13;refGene_tx=XM_023239043.1
D1	1520791	1520940	refGene_gn=MMP13;refGene_tx=XM_023239043.1
D1	1521868	1522110	refGene_gn=MMP13;refGene_tx=XM_023239043.1
D1	1522197	1522314	refGene_gn=MMP13;refGene_tx=XM_023239043.1
D1	1625269	1625325
refGene_gn=DCUN1D5;refGene_tx=XM_019811214.1;refGene_gn=DCUN1D5;refGen
e_tx=XM_023239045.1;refGene_gn=DCUN1D5;refGene_tx=XM_023239046.1;refGe
ne_gn=DCUN1D5;refGene_tx=XM_023239047.1
D1	1627212	1627315
refGene_gn=DCUN1D5;refGene_tx=XM_019811214.1;refGene_gn=DCUN1D5;refGen
e_tx=XM_023239045.1;refGene_gn=DCUN1D5;refGene_tx=XM_023239046.1;refGe
ne_gn=DCUN1D5;refGene_tx=XM_023239047.1
D1	1628898	1629003
refGene_gn=DCUN1D5;refGene_tx=XM_019811214.1;refGene_gn=DCUN1D5;refGen
e_tx=XM_023239045.1;refGene_gn=DCUN1D5;refGene_tx=XM_023239046.1;refGe
ne_gn=DCUN1D5;refGene_tx=XM_023239047.1
D1	1629088	1629197
refGene_gn=DCUN1D5;refGene_tx=XM_019811214.1;refGene_gn=DCUN1D5;refGen
e_tx=XM_023239045.1;refGene_gn=DCUN1D5;refGene_tx=XM_023239046.1;refGe
ne_gn=DCUN1D5;refGene_tx=XM_023239047.1
D1	1642939	1643031
refGene_gn=DCUN1D5;refGene_tx=XM_023239045.1;refGene_gn=DCUN1D5;refGen
e_tx=XM_023239046.1
D1	1643439	1643510
refGene_gn=DCUN1D5;refGene_tx=XM_019811214.1;refGene_gn=DCUN1D5;refGen
e_tx=XM_023239045.1;refGene_gn=DCUN1D5;refGene_tx=XM_023239046.1
```

D1 1648737 1648829 refGene_gn=DCUN1D5;refGene_tx=XM_023239045.1
D1 1650705 1650791
refGene_gn=DCUN1D5;refGene_tx=XM_023239045.1;refGene_gn=DCUN1D5;refGene_tx=XM_023239047.1
D1 1667113 1667308
refGene_gn=DYNC2H1;refGene_tx=XM_023239048.1;refGene_gn=DYNC2H1;refGene_tx=XM_023239049.1;refGene_gn=DYNC2H1;refGene_tx=XM_023239050.1;refGene_gn=DYNC2H1;refGene_tx=XM_023239051.1;refGene_gn=DYNC2H1;refGene_tx=XM_023239052.1;refGene_gn=DYNC2H1;refGene_tx=XM_023239053.1
D1 1673372 1673543
refGene_gn=DYNC2H1;refGene_tx=XM_023239048.1;refGene_gn=DYNC2H1;refGene_tx=XM_023239049.1;refGene_gn=DYNC2H1;refGene_tx=XM_023239050.1;refGene_gn=DYNC2H1;refGene_tx=XM_023239051.1;refGene_gn=DYNC2H1;refGene_tx=XM_023239052.1;refGene_gn=DYNC2H1;refGene_tx=XM_023239053.1
D1 1673968 1674104
refGene_gn=DYNC2H1;refGene_tx=XM_023239048.1;refGene_gn=DYNC2H1;refGene_tx=XM_023239049.1;refGene_gn=DYNC2H1;refGene_tx=XM_023239050.1;refGene_gn=DYNC2H1;refGene_tx=XM_023239051.1;refGene_gn=DYNC2H1;refGene_tx=XM_023239052.1;refGene_gn=DYNC2H1;refGene_tx=XM_023239053.1
D1 1675906 1676025
refGene_gn=DYNC2H1;refGene_tx=XM_023239048.1;refGene_gn=DYNC2H1;refGene_tx=XM_023239049.1;refGene_gn=DYNC2H1;refGene_tx=XM_023239050.1;refGene_gn=DYNC2H1;refGene_tx=XM_023239051.1;refGene_gn=DYNC2H1;refGene_tx=XM_023239052.1;refGene_gn=DYNC2H1;refGene_tx=XM_023239053.1
D1 1676989 1677134
refGene_gn=DYNC2H1;refGene_tx=XM_023239048.1;refGene_gn=DYNC2H1;refGene_tx=XM_023239049.1;refGene_gn=DYNC2H1;refGene_tx=XM_023239050.1;refGene_gn=DYNC2H1;refGene_tx=XM_023239051.1;refGene_gn=DYNC2H1;refGene_tx=XM_023239052.1;refGene_gn=DYNC2H1;refGene_tx=XM_023239053.1
D1 1678135 1678368
refGene_gn=DYNC2H1;refGene_tx=XM_023239048.1;refGene_gn=DYNC2H1;refGene_tx=XM_023239049.1;refGene_gn=DYNC2H1;refGene_tx=XM_023239050.1;refGene_gn=DYNC2H1;refGene_tx=XM_023239051.1;refGene_gn=DYNC2H1;refGene_tx=XM_023239052.1;refGene_gn=DYNC2H1;refGene_tx=XM_023239053.1
D1 1679906 1680041
refGene_gn=DYNC2H1;refGene_tx=XM_023239048.1;refGene_gn=DYNC2H1;refGene_tx=XM_023239049.1;refGene_gn=DYNC2H1;refGene_tx=XM_023239050.1;refGene_gn=DYNC2H1;refGene_tx=XM_023239051.1;refGene_gn=DYNC2H1;refGene_tx=XM_023239052.1;refGene_gn=DYNC2H1;refGene_tx=XM_023239053.1
D1 1680140 1680254
refGene_gn=DYNC2H1;refGene_tx=XM_023239048.1;refGene_gn=DYNC2H1;refGene_tx=XM_023239049.1;refGene_gn=DYNC2H1;refGene_tx=XM_023239050.1;refGene_gn=DYNC2H1;refGene_tx=XM_023239051.1;refGene_gn=DYNC2H1;refGene_tx=XM_023239052.1;refGene_gn=DYNC2H1;refGene_tx=XM_023239053.1
D1 1680358 1680470
refGene_gn=DYNC2H1;refGene_tx=XM_023239048.1;refGene_gn=DYNC2H1;refGene_tx=XM_023239049.1;refGene_gn=DYNC2H1;refGene_tx=XM_023239050.1;refGene_gn=DYNC2H1;refGene_tx=XM_023239051.1;refGene_gn=DYNC2H1;refGene_tx=XM_023239052.1;refGene_gn=DYNC2H1;refGene_tx=XM_023239053.1
D1 1680999 1681124

refGene_gn=DYNC2H1;refGene_tx=XM_023239048.1;refGene_gn=DYNC2H1;refGene_tx=XM_023239049.1;refGene_gn=DYNC2H1;refGene_tx=XM_023239050.1;refGene_gn=DYNC2H1;refGene_tx=XM_023239051.1;refGene_gn=DYNC2H1;refGene_tx=XM_023239052.1;refGene_gn=DYNC2H1;refGene_tx=XM_023239053.1
D1 1682254 1682430
refGene_gn=DYNC2H1;refGene_tx=XM_023239048.1;refGene_gn=DYNC2H1;refGene_tx=XM_023239049.1;refGene_gn=DYNC2H1;refGene_tx=XM_023239050.1;refGene_gn=DYNC2H1;refGene_tx=XM_023239051.1;refGene_gn=DYNC2H1;refGene_tx=XM_023239052.1;refGene_gn=DYNC2H1;refGene_tx=XM_023239053.1
D1 1685400 1685596
refGene_gn=DYNC2H1;refGene_tx=XM_023239048.1;refGene_gn=DYNC2H1;refGene_tx=XM_023239049.1;refGene_gn=DYNC2H1;refGene_tx=XM_023239050.1;refGene_gn=DYNC2H1;refGene_tx=XM_023239051.1;refGene_gn=DYNC2H1;refGene_tx=XM_023239052.1;refGene_gn=DYNC2H1;refGene_tx=XM_023239053.1
D1 1690173 1690269
refGene_gn=DYNC2H1;refGene_tx=XM_023239048.1;refGene_gn=DYNC2H1;refGene_tx=XM_023239049.1;refGene_gn=DYNC2H1;refGene_tx=XM_023239050.1;refGene_gn=DYNC2H1;refGene_tx=XM_023239051.1;refGene_gn=DYNC2H1;refGene_tx=XM_023239052.1;refGene_gn=DYNC2H1;refGene_tx=XM_023239053.1
D1 1693943 1694096
refGene_gn=DYNC2H1;refGene_tx=XM_023239048.1;refGene_gn=DYNC2H1;refGene_tx=XM_023239049.1;refGene_gn=DYNC2H1;refGene_tx=XM_023239050.1;refGene_gn=DYNC2H1;refGene_tx=XM_023239051.1;refGene_gn=DYNC2H1;refGene_tx=XM_023239052.1;refGene_gn=DYNC2H1;refGene_tx=XM_023239053.1
D1 1694945 1695044
refGene_gn=DYNC2H1;refGene_tx=XM_023239048.1;refGene_gn=DYNC2H1;refGene_tx=XM_023239049.1;refGene_gn=DYNC2H1;refGene_tx=XM_023239050.1;refGene_gn=DYNC2H1;refGene_tx=XM_023239051.1;refGene_gn=DYNC2H1;refGene_tx=XM_023239052.1;refGene_gn=DYNC2H1;refGene_tx=XM_023239053.1
D1 1697465 1697605
refGene_gn=DYNC2H1;refGene_tx=XM_023239048.1;refGene_gn=DYNC2H1;refGene_tx=XM_023239049.1;refGene_gn=DYNC2H1;refGene_tx=XM_023239050.1;refGene_gn=DYNC2H1;refGene_tx=XM_023239051.1;refGene_gn=DYNC2H1;refGene_tx=XM_023239052.1;refGene_gn=DYNC2H1;refGene_tx=XM_023239053.1
D1 1697687 1697916
refGene_gn=DYNC2H1;refGene_tx=XM_023239048.1;refGene_gn=DYNC2H1;refGene_tx=XM_023239049.1;refGene_gn=DYNC2H1;refGene_tx=XM_023239050.1;refGene_gn=DYNC2H1;refGene_tx=XM_023239051.1;refGene_gn=DYNC2H1;refGene_tx=XM_023239052.1;refGene_gn=DYNC2H1;refGene_tx=XM_023239053.1
D1 1698737 1698865
refGene_gn=DYNC2H1;refGene_tx=XM_023239048.1;refGene_gn=DYNC2H1;refGene_tx=XM_023239049.1;refGene_gn=DYNC2H1;refGene_tx=XM_023239050.1;refGene_gn=DYNC2H1;refGene_tx=XM_023239051.1;refGene_gn=DYNC2H1;refGene_tx=XM_023239052.1;refGene_gn=DYNC2H1;refGene_tx=XM_023239053.1
D1 1704168 1704284
refGene_gn=DYNC2H1;refGene_tx=XM_023239048.1;refGene_gn=DYNC2H1;refGene_tx=XM_023239049.1;refGene_gn=DYNC2H1;refGene_tx=XM_023239050.1;refGene_gn=DYNC2H1;refGene_tx=XM_023239051.1;refGene_gn=DYNC2H1;refGene_tx=XM_023239052.1;refGene_gn=DYNC2H1;refGene_tx=XM_023239053.1
D1 1704928 1705056

refGene_gn=DYNC2H1;refGene_tx=XM_023239048.1;refGene_gn=DYNC2H1;refGene_tx=XM_023239049.1;refGene_gn=DYNC2H1;refGene_tx=XM_023239050.1;refGene_gn=DYNC2H1;refGene_tx=XM_023239051.1;refGene_gn=DYNC2H1;refGene_tx=XM_023239052.1;refGene_gn=DYNC2H1;refGene_tx=XM_023239053.1
D1 1707423 1707573
refGene_gn=DYNC2H1;refGene_tx=XM_023239048.1;refGene_gn=DYNC2H1;refGene_tx=XM_023239049.1;refGene_gn=DYNC2H1;refGene_tx=XM_023239050.1;refGene_gn=DYNC2H1;refGene_tx=XM_023239051.1;refGene_gn=DYNC2H1;refGene_tx=XM_023239052.1;refGene_gn=DYNC2H1;refGene_tx=XM_023239053.1
D1 1708497 1708703
refGene_gn=DYNC2H1;refGene_tx=XM_023239048.1;refGene_gn=DYNC2H1;refGene_tx=XM_023239049.1;refGene_gn=DYNC2H1;refGene_tx=XM_023239050.1;refGene_gn=DYNC2H1;refGene_tx=XM_023239051.1;refGene_gn=DYNC2H1;refGene_tx=XM_023239052.1;refGene_gn=DYNC2H1;refGene_tx=XM_023239053.1
D1 1709593 1709749
refGene_gn=DYNC2H1;refGene_tx=XM_023239048.1;refGene_gn=DYNC2H1;refGene_tx=XM_023239049.1;refGene_gn=DYNC2H1;refGene_tx=XM_023239050.1;refGene_gn=DYNC2H1;refGene_tx=XM_023239051.1;refGene_gn=DYNC2H1;refGene_tx=XM_023239052.1;refGene_gn=DYNC2H1;refGene_tx=XM_023239053.1
D1 1709838 1709953
refGene_gn=DYNC2H1;refGene_tx=XM_023239048.1;refGene_gn=DYNC2H1;refGene_tx=XM_023239049.1;refGene_gn=DYNC2H1;refGene_tx=XM_023239050.1;refGene_gn=DYNC2H1;refGene_tx=XM_023239051.1;refGene_gn=DYNC2H1;refGene_tx=XM_023239052.1;refGene_gn=DYNC2H1;refGene_tx=XM_023239053.1
D1 1710489 1710660
refGene_gn=DYNC2H1;refGene_tx=XM_023239048.1;refGene_gn=DYNC2H1;refGene_tx=XM_023239049.1;refGene_gn=DYNC2H1;refGene_tx=XM_023239050.1;refGene_gn=DYNC2H1;refGene_tx=XM_023239051.1;refGene_gn=DYNC2H1;refGene_tx=XM_023239052.1;refGene_gn=DYNC2H1;refGene_tx=XM_023239053.1
D1 1711921 1712304
refGene_gn=DYNC2H1;refGene_tx=XM_023239048.1;refGene_gn=DYNC2H1;refGene_tx=XM_023239049.1;refGene_gn=DYNC2H1;refGene_tx=XM_023239050.1;refGene_gn=DYNC2H1;refGene_tx=XM_023239051.1;refGene_gn=DYNC2H1;refGene_tx=XM_023239052.1;refGene_gn=DYNC2H1;refGene_tx=XM_023239053.1
D1 1714005 1714138
refGene_gn=DYNC2H1;refGene_tx=XM_023239048.1;refGene_gn=DYNC2H1;refGene_tx=XM_023239049.1;refGene_gn=DYNC2H1;refGene_tx=XM_023239050.1;refGene_gn=DYNC2H1;refGene_tx=XM_023239051.1;refGene_gn=DYNC2H1;refGene_tx=XM_023239052.1;refGene_gn=DYNC2H1;refGene_tx=XM_023239053.1
D1 1714228 1714346
refGene_gn=DYNC2H1;refGene_tx=XM_023239048.1;refGene_gn=DYNC2H1;refGene_tx=XM_023239049.1;refGene_gn=DYNC2H1;refGene_tx=XM_023239050.1;refGene_gn=DYNC2H1;refGene_tx=XM_023239051.1;refGene_gn=DYNC2H1;refGene_tx=XM_023239052.1;refGene_gn=DYNC2H1;refGene_tx=XM_023239053.1
D1 1716535 1716648
refGene_gn=DYNC2H1;refGene_tx=XM_023239048.1;refGene_gn=DYNC2H1;refGene_tx=XM_023239049.1;refGene_gn=DYNC2H1;refGene_tx=XM_023239050.1;refGene_gn=DYNC2H1;refGene_tx=XM_023239051.1;refGene_gn=DYNC2H1;refGene_tx=XM_023239052.1;refGene_gn=DYNC2H1;refGene_tx=XM_023239053.1
D1 1718634 1718754

refGene_gn=DYNC2H1;refGene_tx=XM_023239048.1;refGene_gn=DYNC2H1;refGene_tx=XM_023239049.1;refGene_gn=DYNC2H1;refGene_tx=XM_023239050.1;refGene_gn=DYNC2H1;refGene_tx=XM_023239051.1;refGene_gn=DYNC2H1;refGene_tx=XM_023239052.1;refGene_gn=DYNC2H1;refGene_tx=XM_023239053.1
D1 1722811 1722962
refGene_gn=DYNC2H1;refGene_tx=XM_023239048.1;refGene_gn=DYNC2H1;refGene_tx=XM_023239049.1;refGene_gn=DYNC2H1;refGene_tx=XM_023239050.1;refGene_gn=DYNC2H1;refGene_tx=XM_023239051.1;refGene_gn=DYNC2H1;refGene_tx=XM_023239052.1;refGene_gn=DYNC2H1;refGene_tx=XM_023239053.1
D1 1724964 1725170
refGene_gn=DYNC2H1;refGene_tx=XM_023239048.1;refGene_gn=DYNC2H1;refGene_tx=XM_023239049.1;refGene_gn=DYNC2H1;refGene_tx=XM_023239050.1;refGene_gn=DYNC2H1;refGene_tx=XM_023239051.1;refGene_gn=DYNC2H1;refGene_tx=XM_023239052.1;refGene_gn=DYNC2H1;refGene_tx=XM_023239053.1
D1 1726230 1726413
refGene_gn=DYNC2H1;refGene_tx=XM_023239048.1;refGene_gn=DYNC2H1;refGene_tx=XM_023239049.1;refGene_gn=DYNC2H1;refGene_tx=XM_023239050.1;refGene_gn=DYNC2H1;refGene_tx=XM_023239051.1;refGene_gn=DYNC2H1;refGene_tx=XM_023239052.1;refGene_gn=DYNC2H1;refGene_tx=XM_023239053.1
D1 1727003 1727186
refGene_gn=DYNC2H1;refGene_tx=XM_023239048.1;refGene_gn=DYNC2H1;refGene_tx=XM_023239049.1;refGene_gn=DYNC2H1;refGene_tx=XM_023239050.1;refGene_gn=DYNC2H1;refGene_tx=XM_023239051.1;refGene_gn=DYNC2H1;refGene_tx=XM_023239052.1;refGene_gn=DYNC2H1;refGene_tx=XM_023239053.1
D1 1728994 1729218
refGene_gn=DYNC2H1;refGene_tx=XM_023239048.1;refGene_gn=DYNC2H1;refGene_tx=XM_023239049.1;refGene_gn=DYNC2H1;refGene_tx=XM_023239050.1;refGene_gn=DYNC2H1;refGene_tx=XM_023239051.1;refGene_gn=DYNC2H1;refGene_tx=XM_023239052.1;refGene_gn=DYNC2H1;refGene_tx=XM_023239053.1
D1 1729912 1730028
refGene_gn=DYNC2H1;refGene_tx=XM_023239048.1;refGene_gn=DYNC2H1;refGene_tx=XM_023239049.1;refGene_gn=DYNC2H1;refGene_tx=XM_023239050.1;refGene_gn=DYNC2H1;refGene_tx=XM_023239051.1;refGene_gn=DYNC2H1;refGene_tx=XM_023239052.1;refGene_gn=DYNC2H1;refGene_tx=XM_023239053.1
D1 1732713 1732913
refGene_gn=DYNC2H1;refGene_tx=XM_023239048.1;refGene_gn=DYNC2H1;refGene_tx=XM_023239049.1;refGene_gn=DYNC2H1;refGene_tx=XM_023239050.1;refGene_gn=DYNC2H1;refGene_tx=XM_023239051.1;refGene_gn=DYNC2H1;refGene_tx=XM_023239052.1;refGene_gn=DYNC2H1;refGene_tx=XM_023239053.1
D1 1734679 1734944
refGene_gn=DYNC2H1;refGene_tx=XM_023239048.1;refGene_gn=DYNC2H1;refGene_tx=XM_023239049.1;refGene_gn=DYNC2H1;refGene_tx=XM_023239050.1;refGene_gn=DYNC2H1;refGene_tx=XM_023239051.1;refGene_gn=DYNC2H1;refGene_tx=XM_023239052.1;refGene_gn=DYNC2H1;refGene_tx=XM_023239053.1
D1 1736267 1736475
refGene_gn=DYNC2H1;refGene_tx=XM_023239048.1;refGene_gn=DYNC2H1;refGene_tx=XM_023239049.1;refGene_gn=DYNC2H1;refGene_tx=XM_023239050.1;refGene_gn=DYNC2H1;refGene_tx=XM_023239051.1;refGene_gn=DYNC2H1;refGene_tx=XM_023239052.1;refGene_gn=DYNC2H1;refGene_tx=XM_023239053.1
D1 1738221 1738351

refGene_gn=DYNC2H1;refGene_tx=XM_023239048.1;refGene_gn=DYNC2H1;refGen
e_tx=XM_023239049.1;refGene_gn=DYNC2H1;refGene_tx=XM_023239050.1;refGe
ne_gn=DYNC2H1;refGene_tx=XM_023239051.1;refGene_gn=DYNC2H1;refGene_tx=
XM_023239052.1;refGene_gn=DYNC2H1;refGene_tx=XM_023239053.1
D1 1743892 1744048
refGene_gn=DYNC2H1;refGene_tx=XM_023239048.1;refGene_gn=DYNC2H1;refGen
e_tx=XM_023239049.1;refGene_gn=DYNC2H1;refGene_tx=XM_023239050.1;refGe
ne_gn=DYNC2H1;refGene_tx=XM_023239051.1;refGene_gn=DYNC2H1;refGene_tx=
XM_023239052.1;refGene_gn=DYNC2H1;refGene_tx=XM_023239053.1
D1 1744818 1745078
refGene_gn=DYNC2H1;refGene_tx=XM_023239048.1;refGene_gn=DYNC2H1;refGen
e_tx=XM_023239049.1;refGene_gn=DYNC2H1;refGene_tx=XM_023239050.1;refGe
ne_gn=DYNC2H1;refGene_tx=XM_023239051.1;refGene_gn=DYNC2H1;refGene_tx=
XM_023239052.1;refGene_gn=DYNC2H1;refGene_tx=XM_023239053.1
D1 1745903 1746150
refGene_gn=DYNC2H1;refGene_tx=XM_023239048.1;refGene_gn=DYNC2H1;refGen
e_tx=XM_023239049.1;refGene_gn=DYNC2H1;refGene_tx=XM_023239050.1;refGe
ne_gn=DYNC2H1;refGene_tx=XM_023239051.1;refGene_gn=DYNC2H1;refGene_tx=
XM_023239052.1;refGene_gn=DYNC2H1;refGene_tx=XM_023239053.1
D1 1747150 1747302
refGene_gn=DYNC2H1;refGene_tx=XM_023239048.1;refGene_gn=DYNC2H1;refGen
e_tx=XM_023239049.1;refGene_gn=DYNC2H1;refGene_tx=XM_023239050.1;refGe
ne_gn=DYNC2H1;refGene_tx=XM_023239051.1;refGene_gn=DYNC2H1;refGene_tx=
XM_023239052.1;refGene_gn=DYNC2H1;refGene_tx=XM_023239053.1
D1 1748282 1748427
refGene_gn=DYNC2H1;refGene_tx=XM_023239048.1;refGene_gn=DYNC2H1;refGen
e_tx=XM_023239049.1;refGene_gn=DYNC2H1;refGene_tx=XM_023239050.1;refGe
ne_gn=DYNC2H1;refGene_tx=XM_023239051.1;refGene_gn=DYNC2H1;refGene_tx=
XM_023239052.1;refGene_gn=DYNC2H1;refGene_tx=XM_023239053.1
D1 1749844 1749947
refGene_gn=DYNC2H1;refGene_tx=XM_023239048.1;refGene_gn=DYNC2H1;refGen
e_tx=XM_023239049.1;refGene_gn=DYNC2H1;refGene_tx=XM_023239050.1;refGe
ne_gn=DYNC2H1;refGene_tx=XM_023239051.1;refGene_gn=DYNC2H1;refGene_tx=
XM_023239052.1;refGene_gn=DYNC2H1;refGene_tx=XM_023239053.1
D1 1750507 1750675
refGene_gn=DYNC2H1;refGene_tx=XM_023239048.1;refGene_gn=DYNC2H1;refGen
e_tx=XM_023239049.1;refGene_gn=DYNC2H1;refGene_tx=XM_023239050.1;refGe
ne_gn=DYNC2H1;refGene_tx=XM_023239051.1;refGene_gn=DYNC2H1;refGene_tx=
XM_023239052.1;refGene_gn=DYNC2H1;refGene_tx=XM_023239053.1
D1 1756143 1756274
refGene_gn=DYNC2H1;refGene_tx=XM_023239048.1;refGene_gn=DYNC2H1;refGen
e_tx=XM_023239049.1;refGene_gn=DYNC2H1;refGene_tx=XM_023239050.1;refGe
ne_gn=DYNC2H1;refGene_tx=XM_023239051.1;refGene_gn=DYNC2H1;refGene_tx=
XM_023239052.1;refGene_gn=DYNC2H1;refGene_tx=XM_023239053.1
D1 1756678 1756927
refGene_gn=DYNC2H1;refGene_tx=XM_023239048.1;refGene_gn=DYNC2H1;refGen
e_tx=XM_023239049.1;refGene_gn=DYNC2H1;refGene_tx=XM_023239050.1;refGe
ne_gn=DYNC2H1;refGene_tx=XM_023239051.1;refGene_gn=DYNC2H1;refGene_tx=
XM_023239052.1;refGene_gn=DYNC2H1;refGene_tx=XM_023239053.1
D1 1757587 1757696

refGene_gn=DYNC2H1;refGene_tx=XM_023239048.1;refGene_gn=DYNC2H1;refGene_tx=XM_023239049.1;refGene_gn=DYNC2H1;refGene_tx=XM_023239050.1;refGene_gn=DYNC2H1;refGene_tx=XM_023239051.1;refGene_gn=DYNC2H1;refGene_tx=XM_023239052.1;refGene_gn=DYNC2H1;refGene_tx=XM_023239053.1
D1 1760487 1760601
refGene_gn=DYNC2H1;refGene_tx=XM_023239048.1;refGene_gn=DYNC2H1;refGene_tx=XM_023239049.1;refGene_gn=DYNC2H1;refGene_tx=XM_023239050.1;refGene_gn=DYNC2H1;refGene_tx=XM_023239051.1;refGene_gn=DYNC2H1;refGene_tx=XM_023239052.1;refGene_gn=DYNC2H1;refGene_tx=XM_023239053.1
D1 1761781 1761924
refGene_gn=DYNC2H1;refGene_tx=XM_023239048.1;refGene_gn=DYNC2H1;refGene_tx=XM_023239049.1;refGene_gn=DYNC2H1;refGene_tx=XM_023239050.1;refGene_gn=DYNC2H1;refGene_tx=XM_023239051.1;refGene_gn=DYNC2H1;refGene_tx=XM_023239052.1;refGene_gn=DYNC2H1;refGene_tx=XM_023239053.1
D1 1765118 1765203
refGene_gn=DYNC2H1;refGene_tx=XM_023239048.1;refGene_gn=DYNC2H1;refGene_tx=XM_023239049.1;refGene_gn=DYNC2H1;refGene_tx=XM_023239050.1;refGene_gn=DYNC2H1;refGene_tx=XM_023239051.1;refGene_gn=DYNC2H1;refGene_tx=XM_023239052.1;refGene_gn=DYNC2H1;refGene_tx=XM_023239053.1
D1 1766894 1767049
refGene_gn=DYNC2H1;refGene_tx=XM_023239048.1;refGene_gn=DYNC2H1;refGene_tx=XM_023239049.1;refGene_gn=DYNC2H1;refGene_tx=XM_023239050.1;refGene_gn=DYNC2H1;refGene_tx=XM_023239051.1;refGene_gn=DYNC2H1;refGene_tx=XM_023239052.1;refGene_gn=DYNC2H1;refGene_tx=XM_023239053.1
D1 1768816 1768954
refGene_gn=DYNC2H1;refGene_tx=XM_023239048.1;refGene_gn=DYNC2H1;refGene_tx=XM_023239049.1;refGene_gn=DYNC2H1;refGene_tx=XM_023239050.1;refGene_gn=DYNC2H1;refGene_tx=XM_023239051.1;refGene_gn=DYNC2H1;refGene_tx=XM_023239052.1;refGene_gn=DYNC2H1;refGene_tx=XM_023239053.1
D1 1778164 1778278
refGene_gn=DYNC2H1;refGene_tx=XM_023239048.1;refGene_gn=DYNC2H1;refGene_tx=XM_023239049.1;refGene_gn=DYNC2H1;refGene_tx=XM_023239050.1;refGene_gn=DYNC2H1;refGene_tx=XM_023239051.1;refGene_gn=DYNC2H1;refGene_tx=XM_023239052.1;refGene_gn=DYNC2H1;refGene_tx=XM_023239053.1
D1 1779996 1780157
refGene_gn=DYNC2H1;refGene_tx=XM_023239048.1;refGene_gn=DYNC2H1;refGene_tx=XM_023239049.1;refGene_gn=DYNC2H1;refGene_tx=XM_023239050.1;refGene_gn=DYNC2H1;refGene_tx=XM_023239051.1;refGene_gn=DYNC2H1;refGene_tx=XM_023239052.1;refGene_gn=DYNC2H1;refGene_tx=XM_023239053.1
D1 1781440 1781564
refGene_gn=DYNC2H1;refGene_tx=XM_023239048.1;refGene_gn=DYNC2H1;refGene_tx=XM_023239049.1;refGene_gn=DYNC2H1;refGene_tx=XM_023239050.1;refGene_gn=DYNC2H1;refGene_tx=XM_023239051.1;refGene_gn=DYNC2H1;refGene_tx=XM_023239052.1;refGene_gn=DYNC2H1;refGene_tx=XM_023239053.1
D1 1782383 1782505
refGene_gn=DYNC2H1;refGene_tx=XM_023239048.1;refGene_gn=DYNC2H1;refGene_tx=XM_023239049.1;refGene_gn=DYNC2H1;refGene_tx=XM_023239050.1;refGene_gn=DYNC2H1;refGene_tx=XM_023239051.1;refGene_gn=DYNC2H1;refGene_tx=XM_023239052.1;refGene_gn=DYNC2H1;refGene_tx=XM_023239053.1
D1 1785542 1785629

refGene_gn=DYNC2H1;refGene_tx=XM_023239048.1;refGene_gn=DYNC2H1;refGen
e_tx=XM_023239049.1;refGene_gn=DYNC2H1;refGene_tx=XM_023239050.1;refGe
ne_gn=DYNC2H1;refGene_tx=XM_023239051.1;refGene_gn=DYNC2H1;refGene_tx=
XM_023239052.1;refGene_gn=DYNC2H1;refGene_tx=XM_023239053.1
D1 1789300 1789427
refGene_gn=DYNC2H1;refGene_tx=XM_023239048.1;refGene_gn=DYNC2H1;refGen
e_tx=XM_023239049.1;refGene_gn=DYNC2H1;refGene_tx=XM_023239050.1;refGe
ne_gn=DYNC2H1;refGene_tx=XM_023239051.1;refGene_gn=DYNC2H1;refGene_tx=
XM_023239052.1;refGene_gn=DYNC2H1;refGene_tx=XM_023239053.1
D1 1797845 1797987
refGene_gn=DYNC2H1;refGene_tx=XM_023239048.1;refGene_gn=DYNC2H1;refGen
e_tx=XM_023239049.1;refGene_gn=DYNC2H1;refGene_tx=XM_023239050.1;refGe
ne_gn=DYNC2H1;refGene_tx=XM_023239051.1;refGene_gn=DYNC2H1;refGene_tx=
XM_023239052.1;refGene_gn=DYNC2H1;refGene_tx=XM_023239053.1
D1 1798524 1798634
refGene_gn=DYNC2H1;refGene_tx=XM_023239048.1;refGene_gn=DYNC2H1;refGen
e_tx=XM_023239049.1;refGene_gn=DYNC2H1;refGene_tx=XM_023239050.1;refGe
ne_gn=DYNC2H1;refGene_tx=XM_023239051.1;refGene_gn=DYNC2H1;refGene_tx=
XM_023239052.1;refGene_gn=DYNC2H1;refGene_tx=XM_023239053.1
D1 1808260 1808359
refGene_gn=DYNC2H1;refGene_tx=XM_023239048.1;refGene_gn=DYNC2H1;refGen
e_tx=XM_023239049.1;refGene_gn=DYNC2H1;refGene_tx=XM_023239050.1;refGe
ne_gn=DYNC2H1;refGene_tx=XM_023239051.1;refGene_gn=DYNC2H1;refGene_tx=
XM_023239052.1;refGene_gn=DYNC2H1;refGene_tx=XM_023239053.1
D1 1810235 1810359
refGene_gn=DYNC2H1;refGene_tx=XM_023239048.1;refGene_gn=DYNC2H1;refGen
e_tx=XM_023239049.1;refGene_gn=DYNC2H1;refGene_tx=XM_023239050.1;refGe
ne_gn=DYNC2H1;refGene_tx=XM_023239051.1;refGene_gn=DYNC2H1;refGene_tx=
XM_023239052.1;refGene_gn=DYNC2H1;refGene_tx=XM_023239053.1
D1 1818033 1818197
refGene_gn=DYNC2H1;refGene_tx=XM_023239048.1;refGene_gn=DYNC2H1;refGen
e_tx=XM_023239049.1;refGene_gn=DYNC2H1;refGene_tx=XM_023239050.1;refGe
ne_gn=DYNC2H1;refGene_tx=XM_023239051.1;refGene_gn=DYNC2H1;refGene_tx=
XM_023239052.1;refGene_gn=DYNC2H1;refGene_tx=XM_023239053.1
D1 1819204 1819282
refGene_gn=DYNC2H1;refGene_tx=XM_023239048.1;refGene_gn=DYNC2H1;refGen
e_tx=XM_023239049.1;refGene_gn=DYNC2H1;refGene_tx=XM_023239050.1
D1 1819691 1819811
refGene_gn=DYNC2H1;refGene_tx=XM_023239048.1;refGene_gn=DYNC2H1;refGen
e_tx=XM_023239049.1;refGene_gn=DYNC2H1;refGene_tx=XM_023239050.1;refGe
ne_gn=DYNC2H1;refGene_tx=XM_023239051.1;refGene_gn=DYNC2H1;refGene_tx=
XM_023239052.1;refGene_gn=DYNC2H1;refGene_tx=XM_023239053.1
D1 1820340 1820475
refGene_gn=DYNC2H1;refGene_tx=XM_023239048.1;refGene_gn=DYNC2H1;refGen
e_tx=XM_023239049.1;refGene_gn=DYNC2H1;refGene_tx=XM_023239050.1;refGe
ne_gn=DYNC2H1;refGene_tx=XM_023239051.1;refGene_gn=DYNC2H1;refGene_tx=
XM_023239052.1;refGene_gn=DYNC2H1;refGene_tx=XM_023239053.1
D1 1822062 1822206
refGene_gn=DYNC2H1;refGene_tx=XM_023239048.1;refGene_gn=DYNC2H1;refGen
e_tx=XM_023239049.1;refGene_gn=DYNC2H1;refGene_tx=XM_023239050.1;refGe

ne_gn=DYNC2H1;refGene_tx=XM_023239051.1;refGene_gn=DYNC2H1;refGene_tx=XM_023239052.1;refGene_gn=DYNC2H1;refGene_tx=XM_023239053.1
D1 1824789 1824900
refGene_gn=DYNC2H1;refGene_tx=XM_023239048.1;refGene_gn=DYNC2H1;refGene_tx=XM_023239049.1;refGene_gn=DYNC2H1;refGene_tx=XM_023239050.1;refGene_gn=DYNC2H1;refGene_tx=XM_023239051.1;refGene_gn=DYNC2H1;refGene_tx=XM_023239052.1;refGene_gn=DYNC2H1;refGene_tx=XM_023239053.1
D1 1832995 1833061
refGene_gn=DYNC2H1;refGene_tx=XM_023239048.1;refGene_gn=DYNC2H1;refGene_tx=XM_023239049.1;refGene_gn=DYNC2H1;refGene_tx=XM_023239050.1;refGene_gn=DYNC2H1;refGene_tx=XM_023239051.1;refGene_gn=DYNC2H1;refGene_tx=XM_023239052.1;refGene_gn=DYNC2H1;refGene_tx=XM_023239053.1
D1 1834757 1834808
refGene_gn=DYNC2H1;refGene_tx=XM_023239048.1;refGene_gn=DYNC2H1;refGene_tx=XM_023239049.1;refGene_gn=DYNC2H1;refGene_tx=XM_023239050.1;refGene_gn=DYNC2H1;refGene_tx=XM_023239051.1;refGene_gn=DYNC2H1;refGene_tx=XM_023239052.1;refGene_gn=DYNC2H1;refGene_tx=XM_023239053.1
D1 1835560 1835638
refGene_gn=DYNC2H1;refGene_tx=XM_023239048.1;refGene_gn=DYNC2H1;refGene_tx=XM_023239049.1;refGene_gn=DYNC2H1;refGene_tx=XM_023239050.1;refGene_gn=DYNC2H1;refGene_tx=XM_023239051.1;refGene_gn=DYNC2H1;refGene_tx=XM_023239052.1;refGene_gn=DYNC2H1;refGene_tx=XM_023239053.1
D1 1842005 1842137
refGene_gn=DYNC2H1;refGene_tx=XM_023239048.1;refGene_gn=DYNC2H1;refGene_tx=XM_023239049.1;refGene_gn=DYNC2H1;refGene_tx=XM_023239050.1;refGene_gn=DYNC2H1;refGene_tx=XM_023239051.1;refGene_gn=DYNC2H1;refGene_tx=XM_023239052.1;refGene_gn=DYNC2H1;refGene_tx=XM_023239053.1
D1 1843080 1843153
refGene_gn=DYNC2H1;refGene_tx=XM_023239048.1;refGene_gn=DYNC2H1;refGene_tx=XM_023239049.1;refGene_gn=DYNC2H1;refGene_tx=XM_023239050.1;refGene_gn=DYNC2H1;refGene_tx=XM_023239051.1;refGene_gn=DYNC2H1;refGene_tx=XM_023239052.1;refGene_gn=DYNC2H1;refGene_tx=XM_023239053.1
D1 1853193 1853354
refGene_gn=DYNC2H1;refGene_tx=XM_023239048.1;refGene_gn=DYNC2H1;refGene_tx=XM_023239049.1;refGene_gn=DYNC2H1;refGene_tx=XM_023239050.1;refGene_gn=DYNC2H1;refGene_tx=XM_023239051.1;refGene_gn=DYNC2H1;refGene_tx=XM_023239052.1;refGene_gn=DYNC2H1;refGene_tx=XM_023239053.1
D1 1855876 1856002
refGene_gn=DYNC2H1;refGene_tx=XM_023239048.1;refGene_gn=DYNC2H1;refGene_tx=XM_023239049.1;refGene_gn=DYNC2H1;refGene_tx=XM_023239050.1;refGene_gn=DYNC2H1;refGene_tx=XM_023239051.1;refGene_gn=DYNC2H1;refGene_tx=XM_023239052.1;refGene_gn=DYNC2H1;refGene_tx=XM_023239053.1
D1 1859721 1859832
refGene_gn=DYNC2H1;refGene_tx=XM_023239048.1;refGene_gn=DYNC2H1;refGene_tx=XM_023239049.1;refGene_gn=DYNC2H1;refGene_tx=XM_023239050.1;refGene_gn=DYNC2H1;refGene_tx=XM_023239051.1;refGene_gn=DYNC2H1;refGene_tx=XM_023239052.1;refGene_gn=DYNC2H1;refGene_tx=XM_023239053.1
D1 1864841 1864997
refGene_gn=DYNC2H1;refGene_tx=XM_023239048.1;refGene_gn=DYNC2H1;refGene_tx=XM_023239049.1;refGene_gn=DYNC2H1;refGene_tx=XM_023239050.1;refGe

ne_gn=DYNC2H1;refGene_tx=XM_023239051.1;refGene_gn=DYNC2H1;refGene_tx=XM_023239052.1;refGene_gn=DYNC2H1;refGene_tx=XM_023239053.1
D1 1875602 1875678
refGene_gn=DYNC2H1;refGene_tx=XM_023239048.1;refGene_gn=DYNC2H1;refGene_tx=XM_023239049.1;refGene_gn=DYNC2H1;refGene_tx=XM_023239050.1;refGene_gn=DYNC2H1;refGene_tx=XM_023239051.1;refGene_gn=DYNC2H1;refGene_tx=XM_023239052.1;refGene_gn=DYNC2H1;refGene_tx=XM_023239053.1
D1 1880867 1881076
refGene_gn=DYNC2H1;refGene_tx=XM_023239048.1;refGene_gn=DYNC2H1;refGene_tx=XM_023239049.1;refGene_gn=DYNC2H1;refGene_tx=XM_023239050.1;refGene_gn=DYNC2H1;refGene_tx=XM_023239051.1;refGene_gn=DYNC2H1;refGene_tx=XM_023239052.1;refGene_gn=DYNC2H1;refGene_tx=XM_023239053.1
D1 1883695 1883800
refGene_gn=DYNC2H1;refGene_tx=XM_023239048.1;refGene_gn=DYNC2H1;refGene_tx=XM_023239049.1;refGene_gn=DYNC2H1;refGene_tx=XM_023239050.1;refGene_gn=DYNC2H1;refGene_tx=XM_023239051.1;refGene_gn=DYNC2H1;refGene_tx=XM_023239052.1;refGene_gn=DYNC2H1;refGene_tx=XM_023239053.1
D1 1905918 1906035
refGene_gn=DYNC2H1;refGene_tx=XM_023239048.1;refGene_gn=DYNC2H1;refGene_tx=XM_023239049.1;refGene_gn=DYNC2H1;refGene_tx=XM_023239050.1;refGene_gn=DYNC2H1;refGene_tx=XM_023239051.1;refGene_gn=DYNC2H1;refGene_tx=XM_023239052.1;refGene_gn=DYNC2H1;refGene_tx=XM_023239053.1
D1 1926260 1926470
refGene_gn=DYNC2H1;refGene_tx=XM_023239048.1;refGene_gn=DYNC2H1;refGene_tx=XM_023239049.1;refGene_gn=DYNC2H1;refGene_tx=XM_023239050.1;refGene_gn=DYNC2H1;refGene_tx=XM_023239051.1;refGene_gn=DYNC2H1;refGene_tx=XM_023239052.1;refGene_gn=DYNC2H1;refGene_tx=XM_023239053.1
D1 1948627 1948717
refGene_gn=DYNC2H1;refGene_tx=XM_023239048.1;refGene_gn=DYNC2H1;refGene_tx=XM_023239049.1;refGene_gn=DYNC2H1;refGene_tx=XM_023239050.1;refGene_gn=DYNC2H1;refGene_tx=XM_023239051.1;refGene_gn=DYNC2H1;refGene_tx=XM_023239052.1;refGene_gn=DYNC2H1;refGene_tx=XM_023239053.1
D1 1964150 1964260
refGene_gn=DYNC2H1;refGene_tx=XM_023239048.1;refGene_gn=DYNC2H1;refGene_tx=XM_023239049.1;refGene_gn=DYNC2H1;refGene_tx=XM_023239050.1;refGene_gn=DYNC2H1;refGene_tx=XM_023239051.1;refGene_gn=DYNC2H1;refGene_tx=XM_023239052.1;refGene_gn=DYNC2H1;refGene_tx=XM_023239053.1
D1 1965236 1965318
refGene_gn=DYNC2H1;refGene_tx=XM_023239048.1;refGene_gn=DYNC2H1;refGene_tx=XM_023239049.1;refGene_gn=DYNC2H1;refGene_tx=XM_023239050.1;refGene_gn=DYNC2H1;refGene_tx=XM_023239051.1;refGene_gn=DYNC2H1;refGene_tx=XM_023239052.1;refGene_gn=DYNC2H1;refGene_tx=XM_023239053.1
D1 1983658 1983775
refGene_gn=DYNC2H1;refGene_tx=XM_023239048.1;refGene_gn=DYNC2H1;refGene_tx=XM_023239049.1;refGene_gn=DYNC2H1;refGene_tx=XM_023239050.1;refGene_gn=DYNC2H1;refGene_tx=XM_023239051.1;refGene_gn=DYNC2H1;refGene_tx=XM_023239052.1;refGene_gn=DYNC2H1;refGene_tx=XM_023239053.1
D1 2010396 2010555
refGene_gn=DYNC2H1;refGene_tx=XM_023239048.1;refGene_gn=DYNC2H1;refGene_tx=XM_023239049.1;refGene_gn=DYNC2H1;refGene_tx=XM_023239050.1;refGe

ne_gn=DYNC2H1;refGene_tx=XM_023239051.1;refGene_gn=DYNC2H1;refGene_tx=XM_023239052.1;refGene_gn=DYNC2H1;refGene_tx=XM_023239053.1
D1 2265310 2265436
refGene_gn=PDGFD;refGene_tx=XM_023239054.1;refGene_gn=PDGFD;refGene_tx=XM_023239055.1
D1 2275452 2275667
refGene_gn=PDGFD;refGene_tx=XM_023239054.1;refGene_gn=PDGFD;refGene_tx=XM_023239055.1
D1 2287436 2287635
refGene_gn=PDGFD;refGene_tx=XM_023239054.1;refGene_gn=PDGFD;refGene_tx=XM_023239055.1
D1 2291368 2291431
refGene_gn=PDGFD;refGene_tx=XM_023239054.1;refGene_gn=PDGFD;refGene_tx=XM_023239055.1
D1 2331914 2332095
refGene_gn=PDGFD;refGene_tx=XM_023239054.1;refGene_gn=PDGFD;refGene_tx=XM_023239055.1
D1 2334779 2334984
refGene_gn=PDGFD;refGene_tx=XM_023239054.1;refGene_gn=PDGFD;refGene_tx=XM_023239055.1
D1 2370312 2371512 refGene_gn=DDI1;refGene_tx=XM_019811218.2
D1 2483973 2484097
refGene_gn=PDGFD;refGene_tx=XM_023239054.1;refGene_gn=PDGFD;refGene_tx=XM_023239055.1
D1 3165342 3165441
refGene_gn=CASP12;refGene_tx=NM_001146147.1;refGene_gn=CASP12;refGene_tx=XM_006936581.4;refGene_gn=CASP12;refGene_tx=XM_006936583.3;refGene_gn=CASP12;refGene_tx=XM_019811158.2;refGene_gn=CASP12;refGene_tx=XM_019811159.1;refGene_gn=CASP12;refGene_tx=XM_023238906.1
D1 3165831 3165941
refGene_gn=CASP12;refGene_tx=NM_001146147.1;refGene_gn=CASP12;refGene_tx=XM_006936581.4;refGene_gn=CASP12;refGene_tx=XM_006936583.3;refGene_gn=CASP12;refGene_tx=XM_019811158.2;refGene_gn=CASP12;refGene_tx=XM_019811159.1;refGene_gn=CASP12;refGene_tx=XM_023238906.1
D1 3168743 3168890
refGene_gn=CASP12;refGene_tx=NM_001146147.1;refGene_gn=CASP12;refGene_tx=XM_006936581.4;refGene_gn=CASP12;refGene_tx=XM_006936583.3;refGene_gn=CASP12;refGene_tx=XM_019811158.2;refGene_gn=CASP12;refGene_tx=XM_019811159.1;refGene_gn=CASP12;refGene_tx=XM_023238906.1
D1 3169291 3169526
refGene_gn=CASP12;refGene_tx=NM_001146147.1;refGene_gn=CASP12;refGene_tx=XM_006936581.4;refGene_gn=CASP12;refGene_tx=XM_006936583.3;refGene_gn=CASP12;refGene_tx=XM_019811158.2;refGene_gn=CASP12;refGene_tx=XM_019811159.1;refGene_gn=CASP12;refGene_tx=XM_023238906.1
D1 3170401 3170575
refGene_gn=CASP12;refGene_tx=NM_001146147.1;refGene_gn=CASP12;refGene_tx=XM_006936581.4;refGene_gn=CASP12;refGene_tx=XM_006936583.3;refGene_gn=CASP12;refGene_tx=XM_019811158.2;refGene_gn=CASP12;refGene_tx=XM_019811159.1;refGene_gn=CASP12;refGene_tx=XM_023238906.1
D1 3170985 3171077

refGene_gn=CASP12;refGene_tx=NM_001146147.1;refGene_gn=CASP12;refGene_tx=XM_006936581.4;refGene_gn=CASP12;refGene_tx=XM_006936583.3;refGene_gn=CASP12;refGene_tx=XM_019811158.2;refGene_gn=CASP12;refGene_tx=XM_019811159.1;refGene_gn=CASP12;refGene_tx=XM_023238906.1
D1 3175101 3175164
refGene_gn=CASP12;refGene_tx=NM_001146147.1;refGene_gn=CASP12;refGene_tx=XM_006936581.4;refGene_gn=CASP12;refGene_tx=XM_006936583.3;refGene_gn=CASP12;refGene_tx=XM_019811158.2;refGene_gn=CASP12;refGene_tx=XM_023238906.1
D1 3176445 3176694
refGene_gn=CASP12;refGene_tx=NM_001146147.1;refGene_gn=CASP12;refGene_tx=XM_006936581.4;refGene_gn=CASP12;refGene_tx=XM_006936583.3;refGene_gn=CASP12;refGene_tx=XM_023238906.1
D1 3178254 3178398
refGene_gn=CASP12;refGene_tx=NM_001146147.1;refGene_gn=CASP12;refGene_tx=XM_006936581.4;refGene_gn=CASP12;refGene_tx=XM_006936583.3;refGene_gn=CASP12;refGene_tx=XM_019811158.2;refGene_gn=CASP12;refGene_tx=XM_023238906.1
D1 3178556 3178619
refGene_gn=CASP12;refGene_tx=NM_001146147.1;refGene_gn=CASP12;refGene_tx=XM_006936581.4;refGene_gn=CASP12;refGene_tx=XM_019811158.2;refGene_gn=CASP12;refGene_tx=XM_019811159.1;refGene_gn=CASP12;refGene_tx=XM_023238906.1
D1 3179898 3180106
refGene_gn=CASP12;refGene_tx=NM_001146147.1;refGene_gn=CASP12;refGene_tx=XM_006936581.4;refGene_gn=CASP12;refGene_tx=XM_019811158.2;refGene_gn=CASP12;refGene_tx=XM_019811159.1;refGene_gn=CASP12;refGene_tx=XM_023238906.1
D1 3190348 3190445 refGene_gn=CASP4;refGene_tx=NM_001009365.1
D1 3191925 3192035 refGene_gn=CASP4;refGene_tx=NM_001009365.1
D1 3193206 3193371 refGene_gn=CASP4;refGene_tx=NM_001009365.1
D1 3193905 3194143 refGene_gn=CASP4;refGene_tx=NM_001009365.1
D1 3195251 3195425 refGene_gn=CASP4;refGene_tx=NM_001009365.1
D1 3196260 3196370 refGene_gn=CASP4;refGene_tx=NM_001009365.1
D1 3201784 3201847 refGene_gn=CASP4;refGene_tx=NM_001009365.1
D1 3204159 3204426 refGene_gn=CASP4;refGene_tx=NM_001009365.1
D1 3727196 3727277
refGene_gn=LOC111556543;refGene_tx=XM_023239029.1
D1 3729200 3729545
refGene_gn=LOC111556543;refGene_tx=XM_023239029.1
D1 3729654 3729729
refGene_gn=LOC111556543;refGene_tx=XM_023239029.1
D1 3730062 3730150 refGene_gn=GRIA4;refGene_tx=XM_023239056.1
D1 3731339 3731498 refGene_gn=GRIA4;refGene_tx=XM_023239056.1
D1 3882985 3883225 refGene_gn=GRIA4;refGene_tx=XM_023239056.1
D1 4016034 4016219 refGene_gn=GRIA4;refGene_tx=XM_023239056.1
D1 4038615 4038669 refGene_gn=GRIA4;refGene_tx=XM_023239056.1
D1 4056034 4056193 refGene_gn=GRIA4;refGene_tx=XM_023239056.1
D1 4061840 4062008 refGene_gn=GRIA4;refGene_tx=XM_023239056.1
D1 4063190 4063295 refGene_gn=GRIA4;refGene_tx=XM_023239056.1

D1 4068440 4068551 refGene_gn=GRIA4;refGene_tx=XM_023239056.1
D1 4076497 4076704 refGene_gn=GRIA4;refGene_tx=XM_023239056.1
D1 4082185 4082556 refGene_gn=GRIA4;refGene_tx=XM_023239056.1
D1 4084623 4084645 refGene_gn=GRIA4;refGene_tx=XM_023239056.1
D1 4084646 4084823 refGene_gn=GRIA4;refGene_tx=XM_023239056.1
D1 4091320 4091568 refGene_gn=GRIA4;refGene_tx=XM_023239056.1
D1 4117531 4117646 refGene_gn=GRIA4;refGene_tx=XM_023239056.1
D1 4125957 4126092 refGene_gn=GRIA4;refGene_tx=XM_023239056.1
D1 4130426 4130591 refGene_gn=GRIA4;refGene_tx=XM_023239056.1
D1 4147638 4148166
refGene_gn=MSANTD4;refGene_tx=XM_019811230.2;refGene_gn=MSANTD4;refGene_tx=XM_023239057.1;refGene_gn=MSANTD4;refGene_tx=XM_023239058.1;refGene_gn=MSANTD4;refGene_tx=XM_023239059.1;refGene_gn=MSANTD4;refGene_tx=XM_023239060.1
D1 4148829 4149291
refGene_gn=MSANTD4;refGene_tx=XM_019811230.2;refGene_gn=MSANTD4;refGene_tx=XM_023239057.1;refGene_gn=MSANTD4;refGene_tx=XM_023239058.1;refGene_gn=MSANTD4;refGene_tx=XM_023239059.1;refGene_gn=MSANTD4;refGene_tx=XM_023239060.1
D1 4185062 4186665 refGene_gn=KBTBD3;refGene_tx=XM_011286329.3
D1 4191099 4191320 refGene_gn=KBTBD3;refGene_tx=XM_011286329.3
D1 4217940 4218123 refGene_gn=AASDHPPT;refGene_tx=XM_023239062.1
D1 4220504 4220730 refGene_gn=AASDHPPT;refGene_tx=XM_023239062.1
D1 4232921 4233043 refGene_gn=AASDHPPT;refGene_tx=XM_023239062.1
D1 4233671 4233833 refGene_gn=AASDHPPT;refGene_tx=XM_023239062.1
D1 4241359 4241431 refGene_gn=AASDHPPT;refGene_tx=XM_023239062.1
D1 4243082 4243247 refGene_gn=AASDHPPT;refGene_tx=XM_023239062.1
D1 5370812 5371020 refGene_gn=GUCY1A2;refGene_tx=XM_023239063.1
D1 5414855 5415010 refGene_gn=GUCY1A2;refGene_tx=XM_023239063.1
D1 5469443 5469587 refGene_gn=GUCY1A2;refGene_tx=XM_023239063.1
D1 5497936 5498422 refGene_gn=GUCY1A2;refGene_tx=XM_023239063.1
D1 5507823 5507862
refGene_gn=LOC101096214;refGene_tx=XM_019812911.1
D1 5508743 5508873
refGene_gn=LOC101096214;refGene_tx=XM_019812911.1
D1 5524496 5524597
refGene_gn=LOC101096214;refGene_tx=XM_019812911.1
D1 5524659 5524896
refGene_gn=LOC101096214;refGene_tx=XM_019812911.1
D1 5524897 5525032
refGene_gn=LOC101096214;refGene_tx=XM_019812911.1
D1 5614627 5615346 refGene_gn=GUCY1A2;refGene_tx=XM_023239063.1
D1 5652161 5652283 refGene_gn=GUCY1A2;refGene_tx=XM_023239063.1
D1 5661019 5661081 refGene_gn=GUCY1A2;refGene_tx=XM_023239063.1
D1 5692553 5692874 refGene_gn=GUCY1A2;refGene_tx=XM_023239063.1
D1 5757841 5758087
refGene_gn=LOC109492316;refGene_tx=XM_019813016.1;refGene_gn=LOC109492316;refGene_tx=XM_019813017.1
D1 5758297 5758441
refGene_gn=LOC109492316;refGene_tx=XM_019813016.1;refGene_gn=LOC109492

316;refGene_tx=XM_019813017.1
D1 5933854 5933998
refGene_gn=CWF19L2;refGene_tx=XM_003992324.4;refGene_gn=CWF19L2;refGene_tx=XM_019811233.2
D1 5937629 5937731
refGene_gn=CWF19L2;refGene_tx=XM_003992324.4;refGene_gn=CWF19L2;refGene_tx=XM_019811233.2
D1 5943234 5943315
refGene_gn=CWF19L2;refGene_tx=XM_003992324.4;refGene_gn=CWF19L2;refGene_tx=XM_019811233.2
D1 5945545 5945701
refGene_gn=CWF19L2;refGene_tx=XM_003992324.4;refGene_gn=CWF19L2;refGene_tx=XM_019811233.2
D1 5953054 5953171 refGene_gn=CWF19L2;refGene_tx=XM_003992324.4
D1 5956679 5956892 refGene_gn=CWF19L2;refGene_tx=XM_003992324.4
D1 5989240 5989378
refGene_gn=CWF19L2;refGene_tx=XM_003992324.4;refGene_gn=CWF19L2;refGene_tx=XM_019811233.2
D1 5990979 5991093
refGene_gn=CWF19L2;refGene_tx=XM_003992324.4;refGene_gn=CWF19L2;refGene_tx=XM_019811233.2
D1 6014741 6014834
refGene_gn=CWF19L2;refGene_tx=XM_003992324.4;refGene_gn=CWF19L2;refGene_tx=XM_019811233.2
D1 6016414 6016511
refGene_gn=CWF19L2;refGene_tx=XM_003992324.4;refGene_gn=CWF19L2;refGene_tx=XM_019811233.2
D1 6033481 6034143
refGene_gn=CWF19L2;refGene_tx=XM_003992324.4;refGene_gn=CWF19L2;refGene_tx=XM_019811233.2
D1 6050506 6050622
refGene_gn=CWF19L2;refGene_tx=XM_003992324.4;refGene_gn=CWF19L2;refGene_tx=XM_019811233.2
D1 6058985 6059079
refGene_gn=CWF19L2;refGene_tx=XM_003992324.4;refGene_gn=CWF19L2;refGene_tx=XM_019811233.2
D1 6065883 6066003
refGene_gn=CWF19L2;refGene_tx=XM_003992324.4;refGene_gn=CWF19L2;refGene_tx=XM_019811233.2
D1 6066725 6066836
refGene_gn=CWF19L2;refGene_tx=XM_003992324.4;refGene_gn=CWF19L2;refGene_tx=XM_019811233.2
D1 6078525 6078648
refGene_gn=CWF19L2;refGene_tx=XM_003992324.4;refGene_gn=CWF19L2;refGene_tx=XM_019811233.2
D1 6082708 6082819
refGene_gn=CWF19L2;refGene_tx=XM_003992324.4;refGene_gn=CWF19L2;refGene_tx=XM_019811233.2
D1 6084586 6084679
refGene_gn=CWF19L2;refGene_tx=XM_003992324.4;refGene_gn=CWF19L2;refGen

e_tx=XM_019811233.2
D1 6122065 6122623 refGene_gn=ALKBH8;refGene_tx=XM_023239031.1
D1 6123741 6123891 refGene_gn=ALKBH8;refGene_tx=XM_023239031.1
D1 6132943 6133200 refGene_gn=ALKBH8;refGene_tx=XM_023239031.1
D1 6135963 6136115 refGene_gn=ALKBH8;refGene_tx=XM_023239031.1
D1 6145552 6145659 refGene_gn=ALKBH8;refGene_tx=XM_023239031.1
D1 6167258 6167329 refGene_gn=ALKBH8;refGene_tx=XM_023239031.1
D1 6168436 6168541 refGene_gn=ALKBH8;refGene_tx=XM_023239031.1
D1 6169847 6169943 refGene_gn=ALKBH8;refGene_tx=XM_023239031.1
D1 6170601 6170733 refGene_gn=ALKBH8;refGene_tx=XM_023239031.1
D1 6173810 6174100 refGene_gn=ALKBH8;refGene_tx=XM_023239031.1
D1 6175384 6175464 refGene_gn=ALKBH8;refGene_tx=XM_023239031.1
D1 6235453 6235470 refGene_gn=ELMOD1;refGene_tx=XM_019811236.1
D1 6246085 6246231
refGene_gn=ELMOD1;refGene_tx=XM_019811234.2;refGene_gn=ELMOD1;refGene_tx=XM_019811235.2;refGene_gn=ELMOD1;refGene_tx=XM_019811236.1
D1 6246590 6246619
refGene_gn=ELMOD1;refGene_tx=XM_019811234.2;refGene_gn=ELMOD1;refGene_tx=XM_019811235.2;refGene_gn=ELMOD1;refGene_tx=XM_019811236.1
D1 6247579 6247677
refGene_gn=ELMOD1;refGene_tx=XM_019811234.2;refGene_gn=ELMOD1;refGene_tx=XM_019811235.2;refGene_gn=ELMOD1;refGene_tx=XM_019811236.1
D1 6251535 6251665
refGene_gn=ELMOD1;refGene_tx=XM_019811234.2;refGene_gn=ELMOD1;refGene_tx=XM_019811235.2;refGene_gn=ELMOD1;refGene_tx=XM_019811236.1
D1 6256767 6256901
refGene_gn=ELMOD1;refGene_tx=XM_019811234.2;refGene_gn=ELMOD1;refGene_tx=XM_019811235.2;refGene_gn=ELMOD1;refGene_tx=XM_019811236.1
D1 6259579 6259648
refGene_gn=ELMOD1;refGene_tx=XM_019811234.2;refGene_gn=ELMOD1;refGene_tx=XM_019811235.2;refGene_gn=ELMOD1;refGene_tx=XM_019811236.1
D1 6260156 6260180
refGene_gn=ELMOD1;refGene_tx=XM_019811234.2;refGene_gn=ELMOD1;refGene_tx=XM_019811236.1
D1 6263043 6263094
refGene_gn=ELMOD1;refGene_tx=XM_019811234.2;refGene_gn=ELMOD1;refGene_tx=XM_019811235.2;refGene_gn=ELMOD1;refGene_tx=XM_019811236.1
D1 6264212 6264346
refGene_gn=ELMOD1;refGene_tx=XM_019811234.2;refGene_gn=ELMOD1;refGene_tx=XM_019811235.2;refGene_gn=ELMOD1;refGene_tx=XM_019811236.1
D1 6271814 6271987
refGene_gn=ELMOD1;refGene_tx=XM_019811234.2;refGene_gn=ELMOD1;refGene_tx=XM_019811235.2;refGene_gn=ELMOD1;refGene_tx=XM_019811236.1
D1 6310497 6310593 refGene_gn=SLN;refGene_tx=XM_006936599.2
D1 6317369 6317375 refGene_gn=SLC35F2;refGene_tx=XM_023238905.1
D1 6358083 6358269
refGene_gn=SLC35F2;refGene_tx=NM_001101659.1;refGene_gn=SLC35F2;refGene_tx=XM_023238902.1;refGene_gn=SLC35F2;refGene_tx=XM_023238903.1;refGene_gn=SLC35F2;refGene_tx=XM_023238904.1
D1 6362498 6362653

refGene_gn=SLC35F2;refGene_tx=NM_001101659.1;refGene_gn=SLC35F2;refGene_tx=XM_023238902.1;refGene_gn=SLC35F2;refGene_tx=XM_023238903.1;refGene_gn=SLC35F2;refGene_tx=XM_023238904.1;refGene_gn=SLC35F2;refGene_tx=XM_023238905.1
D1 6363753 6363806
refGene_gn=SLC35F2;refGene_tx=NM_001101659.1;refGene_gn=SLC35F2;refGene_tx=XM_023238902.1;refGene_gn=SLC35F2;refGene_tx=XM_023238903.1;refGene_gn=SLC35F2;refGene_tx=XM_023238904.1;refGene_gn=SLC35F2;refGene_tx=XM_023238905.1
D1 6365069 6365226
refGene_gn=SLC35F2;refGene_tx=NM_001101659.1;refGene_gn=SLC35F2;refGene_tx=XM_023238902.1;refGene_gn=SLC35F2;refGene_tx=XM_023238903.1;refGene_gn=SLC35F2;refGene_tx=XM_023238904.1;refGene_gn=SLC35F2;refGene_tx=XM_023238905.1
D1 6366545 6366705
refGene_gn=SLC35F2;refGene_tx=NM_001101659.1;refGene_gn=SLC35F2;refGene_tx=XM_023238902.1;refGene_gn=SLC35F2;refGene_tx=XM_023238903.1;refGene_gn=SLC35F2;refGene_tx=XM_023238904.1;refGene_gn=SLC35F2;refGene_tx=XM_023238905.1
D1 6371000 6371128
refGene_gn=SLC35F2;refGene_tx=NM_001101659.1;refGene_gn=SLC35F2;refGene_tx=XM_023238902.1;refGene_gn=SLC35F2;refGene_tx=XM_023238903.1;refGene_gn=SLC35F2;refGene_tx=XM_023238904.1;refGene_gn=SLC35F2;refGene_tx=XM_023238905.1
D1 6374389 6374565
refGene_gn=SLC35F2;refGene_tx=NM_001101659.1;refGene_gn=SLC35F2;refGene_tx=XM_023238902.1;refGene_gn=SLC35F2;refGene_tx=XM_023238903.1;refGene_gn=SLC35F2;refGene_tx=XM_023238904.1;refGene_gn=SLC35F2;refGene_tx=XM_023238905.1
D1 6384575 6384652 refGene_gn=SLC35F2;refGene_tx=XM_023238902.1
D1 6414000 6414110
refGene_gn=SLC35F2;refGene_tx=NM_001101659.1;refGene_gn=SLC35F2;refGene_tx=XM_023238905.1
D1 6456858 6457085 refGene_gn=RAB39A;refGene_tx=XM_003992327.5
D1 6482995 6483422 refGene_gn=RAB39A;refGene_tx=XM_003992327.5
D1 6519819 6519843 refGene_gn=CUL5;refGene_tx=NM_001246321.1
D1 6540710 6540820 refGene_gn=CUL5;refGene_tx=NM_001246321.1
D1 6558521 6558621 refGene_gn=CUL5;refGene_tx=NM_001246321.1
D1 6567097 6567274 refGene_gn=CUL5;refGene_tx=NM_001246321.1
D1 6569090 6569232 refGene_gn=CUL5;refGene_tx=NM_001246321.1
D1 6571029 6571175 refGene_gn=CUL5;refGene_tx=NM_001246321.1
D1 6571258 6571339 refGene_gn=CUL5;refGene_tx=NM_001246321.1
D1 6590855 6590949 refGene_gn=CUL5;refGene_tx=NM_001246321.1
D1 6592287 6592418 refGene_gn=CUL5;refGene_tx=NM_001246321.1
D1 6592667 6592775 refGene_gn=CUL5;refGene_tx=NM_001246321.1
D1 6596200 6596265 refGene_gn=CUL5;refGene_tx=NM_001246321.1
D1 6602833 6602966 refGene_gn=CUL5;refGene_tx=NM_001246321.1
D1 6603845 6603977 refGene_gn=CUL5;refGene_tx=NM_001246321.1
D1 6606946 6607070 refGene_gn=CUL5;refGene_tx=NM_001246321.1
D1 6607626 6607802 refGene_gn=CUL5;refGene_tx=NM_001246321.1

D1	6608364	6608526	refGene_gn=CUL5;refGene_tx=NM_001246321.1
D1	6609468	6609587	refGene_gn=CUL5;refGene_tx=NM_001246321.1
D1	6610202	6610326	refGene_gn=CUL5;refGene_tx=NM_001246321.1
D1	6614952	6615147	refGene_gn=CUL5;refGene_tx=NM_001246321.1
D1	6630185	6630257	refGene_gn=ACAT1;refGene_tx=XM_003992328.4
D1	6630574	6630592	refGene_gn=ACAT1;refGene_tx=XM_023239065.1
D1	6637125	6637353	refGene_gn=ACAT1;refGene_tx=XM_019811238.1
D1	6637481	6637595	refGene_gn=ACAT1;refGene_tx=XM_019811238.1
D1	6638076	6638124
refGene_gn=ACAT1;refGene_tx=XM_003992328.4;refGene_gn=ACAT1;refGene_tx=XM_019811238.1;refGene_gn=ACAT1;refGene_tx=XM_023239065.1
D1	6639912	6640030
refGene_gn=ACAT1;refGene_tx=XM_003992328.4;refGene_gn=ACAT1;refGene_tx=XM_019811238.1;refGene_gn=ACAT1;refGene_tx=XM_023239065.1
D1	6641238	6641334
refGene_gn=ACAT1;refGene_tx=XM_003992328.4;refGene_gn=ACAT1;refGene_tx=XM_019811238.1;refGene_gn=ACAT1;refGene_tx=XM_023239065.1
D1	6641565	6641666
refGene_gn=ACAT1;refGene_tx=XM_003992328.4;refGene_gn=ACAT1;refGene_tx=XM_019811238.1;refGene_gn=ACAT1;refGene_tx=XM_023239065.1
D1	6642550	6642694
refGene_gn=ACAT1;refGene_tx=XM_003992328.4;refGene_gn=ACAT1;refGene_tx=XM_019811238.1;refGene_gn=ACAT1;refGene_tx=XM_023239065.1
D1	6643330	6643481
refGene_gn=ACAT1;refGene_tx=XM_003992328.4;refGene_gn=ACAT1;refGene_tx=XM_019811238.1;refGene_gn=ACAT1;refGene_tx=XM_023239065.1
D1	6644311	6644407
refGene_gn=ACAT1;refGene_tx=XM_003992328.4;refGene_gn=ACAT1;refGene_tx=XM_019811238.1;refGene_gn=ACAT1;refGene_tx=XM_023239065.1
D1	6645934	6646048
refGene_gn=ACAT1;refGene_tx=XM_003992328.4;refGene_gn=ACAT1;refGene_tx=XM_019811238.1;refGene_gn=ACAT1;refGene_tx=XM_023239065.1
D1	6646890	6646955
refGene_gn=ACAT1;refGene_tx=XM_003992328.4;refGene_gn=ACAT1;refGene_tx=XM_019811238.1;refGene_gn=ACAT1;refGene_tx=XM_023239065.1
D1	6648405	6648563
refGene_gn=ACAT1;refGene_tx=XM_003992328.4;refGene_gn=ACAT1;refGene_tx=XM_019811238.1;refGene_gn=ACAT1;refGene_tx=XM_023239065.1
D1	6649192	6649313
refGene_gn=ACAT1;refGene_tx=XM_003992328.4;refGene_gn=ACAT1;refGene_tx=XM_019811238.1;refGene_gn=ACAT1;refGene_tx=XM_023239065.1
D1	6665714	6665792
refGene_gn=LOC101089038;refGene_tx=XM_003992329.5;refGene_gn=LOC101089038;refGene_tx=XM_023239069.1
D1	6667026	6668185
refGene_gn=LOC101089038;refGene_tx=XM_003992329.5;refGene_gn=LOC101089038;refGene_tx=XM_023239069.1
D1	6668348	6668409
refGene_gn=LOC101089038;refGene_tx=XM_003992329.5;refGene_gn=LOC101089038;refGene_tx=XM_023239069.1

D1 6671085 6671194
refGene_gn=LOC101089038;refGene_tx=XM_003992329.5;refGene_gn=LOC101089038;refGene_tx=XM_023239069.1
D1 6671269 6671385
refGene_gn=LOC101089038;refGene_tx=XM_003992329.5;refGene_gn=LOC101089038;refGene_tx=XM_023239069.1
D1 6673530 6675201
refGene_gn=LOC101089038;refGene_tx=XM_003992329.5;refGene_gn=LOC101089038;refGene_tx=XM_023239069.1
D1 6676056 6676185
refGene_gn=LOC101089038;refGene_tx=XM_003992329.5;refGene_gn=LOC101089038;refGene_tx=XM_023239069.1
D1 6677025 6677122
refGene_gn=LOC101089038;refGene_tx=XM_003992329.5;refGene_gn=LOC101089038;refGene_tx=XM_023239069.1
D1 6683798 6683886
refGene_gn=LOC101089038;refGene_tx=XM_003992329.5;refGene_gn=LOC101089038;refGene_tx=XM_023239069.1
D1 6684201 6684293
refGene_gn=LOC101089038;refGene_tx=XM_003992329.5;refGene_gn=LOC101089038;refGene_tx=XM_023239069.1
D1 6684824 6684912
refGene_gn=LOC101089038;refGene_tx=XM_003992329.5;refGene_gn=LOC101089038;refGene_tx=XM_023239069.1
D1 6688037 6688119
refGene_gn=LOC101089038;refGene_tx=XM_003992329.5;refGene_gn=LOC101089038;refGene_tx=XM_023239069.1
D1 6689633 6689861
refGene_gn=LOC101089038;refGene_tx=XM_003992329.5;refGene_gn=LOC101089038;refGene_tx=XM_023239069.1
D1 6691134 6691175
refGene_gn=LOC101089038;refGene_tx=XM_003992329.5;refGene_gn=LOC101089038;refGene_tx=XM_023239069.1
D1 6692698 6692771
refGene_gn=LOC101089038;refGene_tx=XM_003992329.5;refGene_gn=LOC101089038;refGene_tx=XM_023239069.1
D1 6694624 6694685
refGene_gn=LOC101089038;refGene_tx=XM_003992329.5;refGene_gn=LOC101089038;refGene_tx=XM_023239069.1
D1 6698568 6698687
refGene_gn=LOC101089038;refGene_tx=XM_003992329.5
D1 6717262 6717299
refGene_gn=LOC101089038;refGene_tx=XM_003992329.5
D1 6723940 6724012
refGene_gn=ATM;refGene_tx=XM_019811239.2;refGene_gn=ATM;refGene_tx=XM_019811240.2;refGene_gn=ATM;refGene_tx=XM_019811241.2;refGene_gn=ATM;refGene_tx=XM_019811242.2;refGene_gn=ATM;refGene_tx=XM_019811243.2;refGene_gn=ATM;refGene_tx=XM_019811245.2;refGene_gn=ATM;refGene_tx=XM_019811246.2;refGene_gn=ATM;refGene_tx=XM_019811247.2;refGene_gn=ATM;refGene_tx=XM_023239067.1;refGene_gn=ATM;refGene_tx=XM_023239068.1

D1 6724094 6724207
refGene_gn=ATM;refGene_tx=XM_019811239.2;refGene_gn=ATM;refGene_tx=XM_019811240.2;refGene_gn=ATM;refGene_tx=XM_019811241.2;refGene_gn=ATM;refGene_tx=XM_019811242.2;refGene_gn=ATM;refGene_tx=XM_019811243.2;refGene_gn=ATM;refGene_tx=XM_019811245.2;refGene_gn=ATM;refGene_tx=XM_019811246.2;refGene_gn=ATM;refGene_tx=XM_019811247.2;refGene_gn=ATM;refGene_tx=XM_023239067.1;refGene_gn=ATM;refGene_tx=XM_023239068.1
D1 6726103 6726249
refGene_gn=ATM;refGene_tx=XM_019811239.2;refGene_gn=ATM;refGene_tx=XM_019811240.2;refGene_gn=ATM;refGene_tx=XM_019811241.2;refGene_gn=ATM;refGene_tx=XM_019811242.2;refGene_gn=ATM;refGene_tx=XM_019811243.2;refGene_gn=ATM;refGene_tx=XM_019811245.2;refGene_gn=ATM;refGene_tx=XM_019811246.2;refGene_gn=ATM;refGene_tx=XM_019811247.2;refGene_gn=ATM;refGene_tx=XM_023239067.1;refGene_gn=ATM;refGene_tx=XM_023239068.1
D1 6730586 6730751
refGene_gn=ATM;refGene_tx=XM_019811239.2;refGene_gn=ATM;refGene_tx=XM_019811240.2;refGene_gn=ATM;refGene_tx=XM_019811241.2;refGene_gn=ATM;refGene_tx=XM_019811242.2;refGene_gn=ATM;refGene_tx=XM_019811243.2;refGene_gn=ATM;refGene_tx=XM_019811245.2;refGene_gn=ATM;refGene_tx=XM_019811246.2;refGene_gn=ATM;refGene_tx=XM_019811247.2;refGene_gn=ATM;refGene_tx=XM_023239067.1;refGene_gn=ATM;refGene_tx=XM_023239068.1
D1 6731097 6731152 refGene_gn=ATM;refGene_tx=XM_023239066.1
D1 6739914 6740080
refGene_gn=ATM;refGene_tx=XM_019811239.2;refGene_gn=ATM;refGene_tx=XM_019811240.2;refGene_gn=ATM;refGene_tx=XM_019811241.2;refGene_gn=ATM;refGene_tx=XM_019811242.2;refGene_gn=ATM;refGene_tx=XM_019811243.2;refGene_gn=ATM;refGene_tx=XM_019811245.2;refGene_gn=ATM;refGene_tx=XM_019811246.2;refGene_gn=ATM;refGene_tx=XM_019811247.2;refGene_gn=ATM;refGene_tx=XM_023239066.1;refGene_gn=ATM;refGene_tx=XM_023239067.1;refGene_gn=ATM;refGene_tx=XM_023239068.1
D1 6740920 6741159
refGene_gn=ATM;refGene_tx=XM_019811239.2;refGene_gn=ATM;refGene_tx=XM_019811240.2;refGene_gn=ATM;refGene_tx=XM_019811241.2;refGene_gn=ATM;refGene_tx=XM_019811242.2;refGene_gn=ATM;refGene_tx=XM_019811243.2;refGene_gn=ATM;refGene_tx=XM_019811245.2;refGene_gn=ATM;refGene_tx=XM_019811246.2;refGene_gn=ATM;refGene_tx=XM_019811247.2;refGene_gn=ATM;refGene_tx=XM_023239066.1;refGene_gn=ATM;refGene_tx=XM_023239067.1;refGene_gn=ATM;refGene_tx=XM_023239068.1
D1 6742949 6743113
refGene_gn=ATM;refGene_tx=XM_019811239.2;refGene_gn=ATM;refGene_tx=XM_019811240.2;refGene_gn=ATM;refGene_tx=XM_019811241.2;refGene_gn=ATM;refGene_tx=XM_019811242.2;refGene_gn=ATM;refGene_tx=XM_019811243.2;refGene_gn=ATM;refGene_tx=XM_019811245.2;refGene_gn=ATM;refGene_tx=XM_019811246.2;refGene_gn=ATM;refGene_tx=XM_019811247.2;refGene_gn=ATM;refGene_tx=XM_023239066.1;refGene_gn=ATM;refGene_tx=XM_023239067.1;refGene_gn=ATM;refGene_tx=XM_023239068.1
D1 6744540 6744710
refGene_gn=ATM;refGene_tx=XM_019811239.2;refGene_gn=ATM;refGene_tx=XM_019811240.2;refGene_gn=ATM;refGene_tx=XM_019811241.2;refGene_gn=ATM;refGene_tx=XM_019811242.2;refGene_gn=ATM;refGene_tx=XM_019811243.2;refGe

ne_gn=ATM;refGene_tx=XM_019811245.2;refGene_gn=ATM;refGene_tx=XM_01981
1246.2;refGene_gn=ATM;refGene_tx=XM_019811247.2;refGene_gn=ATM;refGene
_tx=XM_023239066.1;refGene_gn=ATM;refGene_tx=XM_023239067.1;refGene_gn
=ATM;refGene_tx=XM_023239068.1
D1 6746495 6746867
refGene_gn=ATM;refGene_tx=XM_019811239.2;refGene_gn=ATM;refGene_tx=XM_
019811240.2;refGene_gn=ATM;refGene_tx=XM_019811241.2;refGene_gn=ATM;re
fGene_tx=XM_019811242.2;refGene_gn=ATM;refGene_tx=XM_019811243.2;refGe
ne_gn=ATM;refGene_tx=XM_019811245.2;refGene_gn=ATM;refGene_tx=XM_01981
1246.2;refGene_gn=ATM;refGene_tx=XM_019811247.2;refGene_gn=ATM;refGene
_tx=XM_023239066.1;refGene_gn=ATM;refGene_tx=XM_023239067.1;refGene_gn
=ATM;refGene_tx=XM_023239068.1
D1 6747522 6747717
refGene_gn=ATM;refGene_tx=XM_019811239.2;refGene_gn=ATM;refGene_tx=XM_
019811240.2;refGene_gn=ATM;refGene_tx=XM_019811241.2;refGene_gn=ATM;re
fGene_tx=XM_019811242.2;refGene_gn=ATM;refGene_tx=XM_019811243.2;refGe
ne_gn=ATM;refGene_tx=XM_019811245.2;refGene_gn=ATM;refGene_tx=XM_01981
1246.2;refGene_gn=ATM;refGene_tx=XM_019811247.2;refGene_gn=ATM;refGene
_tx=XM_023239066.1;refGene_gn=ATM;refGene_tx=XM_023239067.1;refGene_gn
=ATM;refGene_tx=XM_023239068.1
D1 6748496 6748592
refGene_gn=ATM;refGene_tx=XM_019811239.2;refGene_gn=ATM;refGene_tx=XM_
019811240.2;refGene_gn=ATM;refGene_tx=XM_019811241.2;refGene_gn=ATM;re
fGene_tx=XM_019811242.2;refGene_gn=ATM;refGene_tx=XM_019811243.2;refGe
ne_gn=ATM;refGene_tx=XM_019811245.2;refGene_gn=ATM;refGene_tx=XM_01981
1246.2;refGene_gn=ATM;refGene_tx=XM_019811247.2;refGene_gn=ATM;refGene
_tx=XM_023239066.1;refGene_gn=ATM;refGene_tx=XM_023239067.1;refGene_gn
=ATM;refGene_tx=XM_023239068.1
D1 6749444 6749687
refGene_gn=ATM;refGene_tx=XM_019811239.2;refGene_gn=ATM;refGene_tx=XM_
019811240.2;refGene_gn=ATM;refGene_tx=XM_019811241.2;refGene_gn=ATM;re
fGene_tx=XM_019811242.2;refGene_gn=ATM;refGene_tx=XM_019811243.2;refGe
ne_gn=ATM;refGene_tx=XM_019811245.2;refGene_gn=ATM;refGene_tx=XM_01981
1246.2;refGene_gn=ATM;refGene_tx=XM_019811247.2;refGene_gn=ATM;refGene
_tx=XM_023239066.1;refGene_gn=ATM;refGene_tx=XM_023239067.1;refGene_gn
=ATM;refGene_tx=XM_023239068.1
D1 6751733 6751859
refGene_gn=ATM;refGene_tx=XM_019811239.2;refGene_gn=ATM;refGene_tx=XM_
019811240.2;refGene_gn=ATM;refGene_tx=XM_019811241.2;refGene_gn=ATM;re
fGene_tx=XM_019811242.2;refGene_gn=ATM;refGene_tx=XM_019811243.2;refGe
ne_gn=ATM;refGene_tx=XM_019811245.2;refGene_gn=ATM;refGene_tx=XM_01981
1246.2;refGene_gn=ATM;refGene_tx=XM_019811247.2;refGene_gn=ATM;refGene
_tx=XM_023239066.1;refGene_gn=ATM;refGene_tx=XM_023239067.1
D1 6752426 6752552
refGene_gn=ATM;refGene_tx=XM_019811239.2;refGene_gn=ATM;refGene_tx=XM_
019811240.2;refGene_gn=ATM;refGene_tx=XM_019811241.2;refGene_gn=ATM;re
fGene_tx=XM_019811242.2;refGene_gn=ATM;refGene_tx=XM_019811243.2;refGe
ne_gn=ATM;refGene_tx=XM_019811245.2;refGene_gn=ATM;refGene_tx=XM_01981
1246.2;refGene_gn=ATM;refGene_tx=XM_019811247.2;refGene_gn=ATM;refGene
_tx=XM_023239066.1;refGene_gn=ATM;refGene_tx=XM_023239067.1

D1 6753497 6753587
refGene_gn=ATM;refGene_tx=XM_019811239.2;refGene_gn=ATM;refGene_tx=XM_019811240.2;refGene_gn=ATM;refGene_tx=XM_019811241.2;refGene_gn=ATM;refGene_tx=XM_019811242.2;refGene_gn=ATM;refGene_tx=XM_019811243.2;refGene_gn=ATM;refGene_tx=XM_019811245.2;refGene_gn=ATM;refGene_tx=XM_019811246.2;refGene_gn=ATM;refGene_tx=XM_019811247.2;refGene_gn=ATM;refGene_tx=XM_023239066.1;refGene_gn=ATM;refGene_tx=XM_023239067.1
D1 6754843 6755015
refGene_gn=ATM;refGene_tx=XM_019811239.2;refGene_gn=ATM;refGene_tx=XM_019811240.2;refGene_gn=ATM;refGene_tx=XM_019811241.2;refGene_gn=ATM;refGene_tx=XM_019811242.2;refGene_gn=ATM;refGene_tx=XM_019811243.2;refGene_gn=ATM;refGene_tx=XM_019811245.2;refGene_gn=ATM;refGene_tx=XM_019811246.2;refGene_gn=ATM;refGene_tx=XM_019811247.2;refGene_gn=ATM;refGene_tx=XM_023239066.1;refGene_gn=ATM;refGene_tx=XM_023239067.1
D1 6755969 6756169
refGene_gn=ATM;refGene_tx=XM_019811239.2;refGene_gn=ATM;refGene_tx=XM_019811240.2;refGene_gn=ATM;refGene_tx=XM_019811241.2;refGene_gn=ATM;refGene_tx=XM_019811242.2;refGene_gn=ATM;refGene_tx=XM_019811243.2;refGene_gn=ATM;refGene_tx=XM_019811245.2;refGene_gn=ATM;refGene_tx=XM_019811246.2;refGene_gn=ATM;refGene_tx=XM_019811247.2;refGene_gn=ATM;refGene_tx=XM_023239066.1;refGene_gn=ATM;refGene_tx=XM_023239067.1
D1 6761334 6761417
refGene_gn=ATM;refGene_tx=XM_019811239.2;refGene_gn=ATM;refGene_tx=XM_019811240.2;refGene_gn=ATM;refGene_tx=XM_019811241.2;refGene_gn=ATM;refGene_tx=XM_019811242.2;refGene_gn=ATM;refGene_tx=XM_019811243.2;refGene_gn=ATM;refGene_tx=XM_019811245.2;refGene_gn=ATM;refGene_tx=XM_019811246.2;refGene_gn=ATM;refGene_tx=XM_019811247.2;refGene_gn=ATM;refGene_tx=XM_023239066.1;refGene_gn=ATM;refGene_tx=XM_023239067.1
D1 6761513 6761669
refGene_gn=ATM;refGene_tx=XM_019811239.2;refGene_gn=ATM;refGene_tx=XM_019811240.2;refGene_gn=ATM;refGene_tx=XM_019811241.2;refGene_gn=ATM;refGene_tx=XM_019811242.2;refGene_gn=ATM;refGene_tx=XM_019811243.2;refGene_gn=ATM;refGene_tx=XM_019811245.2;refGene_gn=ATM;refGene_tx=XM_019811246.2;refGene_gn=ATM;refGene_tx=XM_019811247.2;refGene_gn=ATM;refGene_tx=XM_023239066.1;refGene_gn=ATM;refGene_tx=XM_023239067.1
D1 6762584 6762660
refGene_gn=ATM;refGene_tx=XM_019811239.2;refGene_gn=ATM;refGene_tx=XM_019811240.2;refGene_gn=ATM;refGene_tx=XM_019811241.2;refGene_gn=ATM;refGene_tx=XM_019811242.2;refGene_gn=ATM;refGene_tx=XM_019811243.2;refGene_gn=ATM;refGene_tx=XM_019811245.2;refGene_gn=ATM;refGene_tx=XM_019811246.2;refGene_gn=ATM;refGene_tx=XM_019811247.2;refGene_gn=ATM;refGene_tx=XM_023239066.1;refGene_gn=ATM;refGene_tx=XM_023239067.1
D1 6762788 6762919
refGene_gn=ATM;refGene_tx=XM_019811239.2;refGene_gn=ATM;refGene_tx=XM_019811240.2;refGene_gn=ATM;refGene_tx=XM_019811241.2;refGene_gn=ATM;refGene_tx=XM_019811242.2;refGene_gn=ATM;refGene_tx=XM_019811243.2;refGene_gn=ATM;refGene_tx=XM_019811245.2;refGene_gn=ATM;refGene_tx=XM_019811246.2;refGene_gn=ATM;refGene_tx=XM_019811247.2;refGene_gn=ATM;refGene_tx=XM_023239066.1;refGene_gn=ATM;refGene_tx=XM_023239067.1
D1 6763642 6763763

refGene_gn=ATM;refGene_tx=XM_019811239.2;refGene_gn=ATM;refGene_tx=XM_019811240.2;refGene_gn=ATM;refGene_tx=XM_019811241.2;refGene_gn=ATM;refGene_tx=XM_019811242.2;refGene_gn=ATM;refGene_tx=XM_019811243.2;refGene_gn=ATM;refGene_tx=XM_019811245.2;refGene_gn=ATM;refGene_tx=XM_019811246.2;refGene_gn=ATM;refGene_tx=XM_019811247.2;refGene_gn=ATM;refGene_tx=XM_023239066.1;refGene_gn=ATM;refGene_tx=XM_023239067.1
D1 6770429 6770603
refGene_gn=ATM;refGene_tx=XM_019811239.2;refGene_gn=ATM;refGene_tx=XM_019811240.2;refGene_gn=ATM;refGene_tx=XM_019811241.2;refGene_gn=ATM;refGene_tx=XM_019811242.2;refGene_gn=ATM;refGene_tx=XM_019811243.2;refGene_gn=ATM;refGene_tx=XM_019811245.2;refGene_gn=ATM;refGene_tx=XM_019811246.2;refGene_gn=ATM;refGene_tx=XM_019811247.2;refGene_gn=ATM;refGene_tx=XM_023239066.1;refGene_gn=ATM;refGene_tx=XM_023239067.1
D1 6771496 6771666
refGene_gn=ATM;refGene_tx=XM_019811239.2;refGene_gn=ATM;refGene_tx=XM_019811240.2;refGene_gn=ATM;refGene_tx=XM_019811241.2;refGene_gn=ATM;refGene_tx=XM_019811242.2;refGene_gn=ATM;refGene_tx=XM_019811243.2;refGene_gn=ATM;refGene_tx=XM_019811245.2;refGene_gn=ATM;refGene_tx=XM_019811246.2;refGene_gn=ATM;refGene_tx=XM_019811247.2;refGene_gn=ATM;refGene_tx=XM_023239066.1;refGene_gn=ATM;refGene_tx=XM_023239067.1
D1 6773141 6773388
refGene_gn=ATM;refGene_tx=XM_019811239.2;refGene_gn=ATM;refGene_tx=XM_019811240.2;refGene_gn=ATM;refGene_tx=XM_019811241.2;refGene_gn=ATM;refGene_tx=XM_019811242.2;refGene_gn=ATM;refGene_tx=XM_019811243.2;refGene_gn=ATM;refGene_tx=XM_019811245.2;refGene_gn=ATM;refGene_tx=XM_019811246.2;refGene_gn=ATM;refGene_tx=XM_019811247.2;refGene_gn=ATM;refGene_tx=XM_023239066.1;refGene_gn=ATM;refGene_tx=XM_023239067.1
D1 6777668 6777784
refGene_gn=ATM;refGene_tx=XM_019811239.2;refGene_gn=ATM;refGene_tx=XM_019811240.2;refGene_gn=ATM;refGene_tx=XM_019811241.2;refGene_gn=ATM;refGene_tx=XM_019811242.2;refGene_gn=ATM;refGene_tx=XM_019811243.2;refGene_gn=ATM;refGene_tx=XM_019811245.2;refGene_gn=ATM;refGene_tx=XM_019811246.2;refGene_gn=ATM;refGene_tx=XM_019811247.2;refGene_gn=ATM;refGene_tx=XM_023239066.1;refGene_gn=ATM;refGene_tx=XM_023239067.1
D1 6778664 6778791
refGene_gn=ATM;refGene_tx=XM_019811239.2;refGene_gn=ATM;refGene_tx=XM_019811240.2;refGene_gn=ATM;refGene_tx=XM_019811241.2;refGene_gn=ATM;refGene_tx=XM_019811242.2;refGene_gn=ATM;refGene_tx=XM_019811243.2;refGene_gn=ATM;refGene_tx=XM_019811245.2;refGene_gn=ATM;refGene_tx=XM_019811246.2;refGene_gn=ATM;refGene_tx=XM_019811247.2;refGene_gn=ATM;refGene_tx=XM_023239066.1;refGene_gn=ATM;refGene_tx=XM_023239067.1
D1 6779242 6779442
refGene_gn=ATM;refGene_tx=XM_019811239.2;refGene_gn=ATM;refGene_tx=XM_019811240.2;refGene_gn=ATM;refGene_tx=XM_019811241.2;refGene_gn=ATM;refGene_tx=XM_019811242.2;refGene_gn=ATM;refGene_tx=XM_019811243.2;refGene_gn=ATM;refGene_tx=XM_019811245.2;refGene_gn=ATM;refGene_tx=XM_019811246.2;refGene_gn=ATM;refGene_tx=XM_019811247.2;refGene_gn=ATM;refGene_tx=XM_023239066.1;refGene_gn=ATM;refGene_tx=XM_023239067.1
D1 6782552 6782727
refGene_gn=ATM;refGene_tx=XM_019811239.2;refGene_gn=ATM;refGene_tx=XM_

019811240.2;refGene_gn=ATM;refGene_tx=XM_019811241.2;refGene_gn=ATM;refGene_tx=XM_019811242.2;refGene_gn=ATM;refGene_tx=XM_019811243.2;refGene_gn=ATM;refGene_tx=XM_019811245.2;refGene_gn=ATM;refGene_tx=XM_019811246.2;refGene_gn=ATM;refGene_tx=XM_019811247.2;refGene_gn=ATM;refGene_tx=XM_023239066.1;refGene_gn=ATM;refGene_tx=XM_023239067.1
D1 6783503 6783668
refGene_gn=ATM;refGene_tx=XM_019811239.2;refGene_gn=ATM;refGene_tx=XM_019811240.2;refGene_gn=ATM;refGene_tx=XM_019811241.2;refGene_gn=ATM;refGene_tx=XM_019811242.2;refGene_gn=ATM;refGene_tx=XM_019811243.2;refGene_gn=ATM;refGene_tx=XM_019811245.2;refGene_gn=ATM;refGene_tx=XM_019811246.2;refGene_gn=ATM;refGene_tx=XM_019811247.2;refGene_gn=ATM;refGene_tx=XM_023239066.1;refGene_gn=ATM;refGene_tx=XM_023239067.1
D1 6784512 6784645
refGene_gn=ATM;refGene_tx=XM_019811239.2;refGene_gn=ATM;refGene_tx=XM_019811240.2;refGene_gn=ATM;refGene_tx=XM_019811241.2;refGene_gn=ATM;refGene_tx=XM_019811242.2;refGene_gn=ATM;refGene_tx=XM_019811243.2;refGene_gn=ATM;refGene_tx=XM_019811245.2;refGene_gn=ATM;refGene_tx=XM_019811246.2;refGene_gn=ATM;refGene_tx=XM_019811247.2;refGene_gn=ATM;refGene_tx=XM_023239066.1;refGene_gn=ATM;refGene_tx=XM_023239067.1
D1 6786267 6786374
refGene_gn=ATM;refGene_tx=XM_019811239.2;refGene_gn=ATM;refGene_tx=XM_019811240.2;refGene_gn=ATM;refGene_tx=XM_019811241.2;refGene_gn=ATM;refGene_tx=XM_019811242.2;refGene_gn=ATM;refGene_tx=XM_019811243.2;refGene_gn=ATM;refGene_tx=XM_019811245.2;refGene_gn=ATM;refGene_tx=XM_019811246.2;refGene_gn=ATM;refGene_tx=XM_019811247.2;refGene_gn=ATM;refGene_tx=XM_023239066.1;refGene_gn=ATM;refGene_tx=XM_023239067.1
D1 6788546 6788718
refGene_gn=ATM;refGene_tx=XM_019811239.2;refGene_gn=ATM;refGene_tx=XM_019811240.2;refGene_gn=ATM;refGene_tx=XM_019811241.2;refGene_gn=ATM;refGene_tx=XM_019811242.2;refGene_gn=ATM;refGene_tx=XM_019811243.2;refGene_gn=ATM;refGene_tx=XM_019811245.2;refGene_gn=ATM;refGene_tx=XM_019811246.2;refGene_gn=ATM;refGene_tx=XM_019811247.2;refGene_gn=ATM;refGene_tx=XM_023239066.1
D1 6791983 6792125
refGene_gn=ATM;refGene_tx=XM_019811239.2;refGene_gn=ATM;refGene_tx=XM_019811240.2;refGene_gn=ATM;refGene_tx=XM_019811241.2;refGene_gn=ATM;refGene_tx=XM_019811242.2;refGene_gn=ATM;refGene_tx=XM_019811243.2;refGene_gn=ATM;refGene_tx=XM_019811245.2;refGene_gn=ATM;refGene_tx=XM_019811246.2;refGene_gn=ATM;refGene_tx=XM_019811247.2;refGene_gn=ATM;refGene_tx=XM_023239066.1
D1 6795396 6795573
refGene_gn=ATM;refGene_tx=XM_019811239.2;refGene_gn=ATM;refGene_tx=XM_019811240.2;refGene_gn=ATM;refGene_tx=XM_019811241.2;refGene_gn=ATM;refGene_tx=XM_019811242.2;refGene_gn=ATM;refGene_tx=XM_019811243.2;refGene_gn=ATM;refGene_tx=XM_019811245.2;refGene_gn=ATM;refGene_tx=XM_019811246.2;refGene_gn=ATM;refGene_tx=XM_019811247.2;refGene_gn=ATM;refGene_tx=XM_023239066.1
D1 6797486 6797664
refGene_gn=ATM;refGene_tx=XM_019811239.2;refGene_gn=ATM;refGene_tx=XM_019811240.2;refGene_gn=ATM;refGene_tx=XM_019811241.2;refGene_gn=ATM;re

fGene_tx=XM_019811242.2;refGene_gn=ATM;refGene_tx=XM_019811243.2;refGene_gn=ATM;refGene_tx=XM_019811245.2;refGene_gn=ATM;refGene_tx=XM_019811246.2;refGene_gn=ATM;refGene_tx=XM_019811247.2;refGene_gn=ATM;refGene_tx=XM_023239066.1
D1 6800146 6800234
refGene_gn=ATM;refGene_tx=XM_019811239.2;refGene_gn=ATM;refGene_tx=XM_019811240.2;refGene_gn=ATM;refGene_tx=XM_019811241.2;refGene_gn=ATM;refGene_tx=XM_019811242.2;refGene_gn=ATM;refGene_tx=XM_019811243.2;refGene_gn=ATM;refGene_tx=XM_019811245.2;refGene_gn=ATM;refGene_tx=XM_019811246.2;refGene_gn=ATM;refGene_tx=XM_019811247.2;refGene_gn=ATM;refGene_tx=XM_023239066.1
D1 6802413 6802569
refGene_gn=ATM;refGene_tx=XM_019811239.2;refGene_gn=ATM;refGene_tx=XM_019811240.2;refGene_gn=ATM;refGene_tx=XM_019811241.2;refGene_gn=ATM;refGene_tx=XM_019811242.2;refGene_gn=ATM;refGene_tx=XM_019811243.2;refGene_gn=ATM;refGene_tx=XM_019811245.2;refGene_gn=ATM;refGene_tx=XM_019811246.2;refGene_gn=ATM;refGene_tx=XM_019811247.2;refGene_gn=ATM;refGene_tx=XM_023239066.1
D1 6805968 6806077
refGene_gn=ATM;refGene_tx=XM_019811239.2;refGene_gn=ATM;refGene_tx=XM_019811240.2;refGene_gn=ATM;refGene_tx=XM_019811241.2;refGene_gn=ATM;refGene_tx=XM_019811242.2;refGene_gn=ATM;refGene_tx=XM_019811243.2;refGene_gn=ATM;refGene_tx=XM_019811245.2;refGene_gn=ATM;refGene_tx=XM_019811246.2;refGene_gn=ATM;refGene_tx=XM_019811247.2;refGene_gn=ATM;refGene_tx=XM_023239066.1
D1 6808710 6808799
refGene_gn=ATM;refGene_tx=XM_019811239.2;refGene_gn=ATM;refGene_tx=XM_019811240.2;refGene_gn=ATM;refGene_tx=XM_019811241.2;refGene_gn=ATM;refGene_tx=XM_019811242.2;refGene_gn=ATM;refGene_tx=XM_019811243.2;refGene_gn=ATM;refGene_tx=XM_019811245.2;refGene_gn=ATM;refGene_tx=XM_019811246.2;refGene_gn=ATM;refGene_tx=XM_019811247.2;refGene_gn=ATM;refGene_tx=XM_023239066.1
D1 6808898 6809001
refGene_gn=ATM;refGene_tx=XM_019811239.2;refGene_gn=ATM;refGene_tx=XM_019811240.2;refGene_gn=ATM;refGene_tx=XM_019811241.2;refGene_gn=ATM;refGene_tx=XM_019811242.2;refGene_gn=ATM;refGene_tx=XM_019811243.2;refGene_gn=ATM;refGene_tx=XM_019811245.2;refGene_gn=ATM;refGene_tx=XM_019811246.2;refGene_gn=ATM;refGene_tx=XM_019811247.2;refGene_gn=ATM;refGene_tx=XM_023239066.1
D1 6809711 6809860
refGene_gn=ATM;refGene_tx=XM_019811239.2;refGene_gn=ATM;refGene_tx=XM_019811240.2;refGene_gn=ATM;refGene_tx=XM_019811241.2;refGene_gn=ATM;refGene_tx=XM_019811242.2;refGene_gn=ATM;refGene_tx=XM_019811243.2;refGene_gn=ATM;refGene_tx=XM_019811245.2;refGene_gn=ATM;refGene_tx=XM_019811246.2;refGene_gn=ATM;refGene_tx=XM_019811247.2;refGene_gn=ATM;refGene_tx=XM_023239066.1
D1 6811510 6811615
refGene_gn=ATM;refGene_tx=XM_019811239.2;refGene_gn=ATM;refGene_tx=XM_019811240.2;refGene_gn=ATM;refGene_tx=XM_019811241.2;refGene_gn=ATM;refGene_tx=XM_019811242.2;refGene_gn=ATM;refGene_tx=XM_019811243.2;refGe

ne_gn=ATM;refGene_tx=XM_019811245.2;refGene_gn=ATM;refGene_tx=XM_01981
1246.2;refGene_gn=ATM;refGene_tx=XM_019811247.2;refGene_gn=ATM;refGene
_tx=XM_023239066.1
D1 6812838 6812958
refGene_gn=ATM;refGene_tx=XM_019811239.2;refGene_gn=ATM;refGene_tx=XM_
019811240.2;refGene_gn=ATM;refGene_tx=XM_019811241.2;refGene_gn=ATM;re
fGene_tx=XM_019811242.2;refGene_gn=ATM;refGene_tx=XM_019811243.2;refGe
ne_gn=ATM;refGene_tx=XM_019811245.2;refGene_gn=ATM;refGene_tx=XM_01981
1246.2;refGene_gn=ATM;refGene_tx=XM_019811247.2;refGene_gn=ATM;refGene
_tx=XM_023239066.1
D1 6816651 6816886
refGene_gn=ATM;refGene_tx=XM_019811239.2;refGene_gn=ATM;refGene_tx=XM_
019811240.2;refGene_gn=ATM;refGene_tx=XM_019811241.2;refGene_gn=ATM;re
fGene_tx=XM_019811242.2;refGene_gn=ATM;refGene_tx=XM_019811243.2;refGe
ne_gn=ATM;refGene_tx=XM_019811245.2;refGene_gn=ATM;refGene_tx=XM_01981
1246.2;refGene_gn=ATM;refGene_tx=XM_019811247.2;refGene_gn=ATM;refGene
_tx=XM_023239066.1
D1 6817380 6817548
refGene_gn=ATM;refGene_tx=XM_019811239.2;refGene_gn=ATM;refGene_tx=XM_
019811240.2;refGene_gn=ATM;refGene_tx=XM_019811241.2;refGene_gn=ATM;re
fGene_tx=XM_019811242.2;refGene_gn=ATM;refGene_tx=XM_019811243.2;refGe
ne_gn=ATM;refGene_tx=XM_019811245.2;refGene_gn=ATM;refGene_tx=XM_01981
1246.2;refGene_gn=ATM;refGene_tx=XM_019811247.2;refGene_gn=ATM;refGene
_tx=XM_023239066.1
D1 6818903 6819017
refGene_gn=ATM;refGene_tx=XM_019811239.2;refGene_gn=ATM;refGene_tx=XM_
019811240.2;refGene_gn=ATM;refGene_tx=XM_019811241.2;refGene_gn=ATM;re
fGene_tx=XM_019811242.2;refGene_gn=ATM;refGene_tx=XM_019811243.2;refGe
ne_gn=ATM;refGene_tx=XM_019811245.2;refGene_gn=ATM;refGene_tx=XM_01981
1246.2;refGene_gn=ATM;refGene_tx=XM_019811247.2;refGene_gn=ATM;refGene
_tx=XM_023239066.1
D1 6819740 6819958
refGene_gn=ATM;refGene_tx=XM_019811239.2;refGene_gn=ATM;refGene_tx=XM_
019811240.2;refGene_gn=ATM;refGene_tx=XM_019811241.2;refGene_gn=ATM;re
fGene_tx=XM_019811242.2;refGene_gn=ATM;refGene_tx=XM_019811243.2;refGe
ne_gn=ATM;refGene_tx=XM_019811245.2;refGene_gn=ATM;refGene_tx=XM_01981
1246.2;refGene_gn=ATM;refGene_tx=XM_019811247.2;refGene_gn=ATM;refGene
_tx=XM_023239066.1
D1 6820698 6820906
refGene_gn=ATM;refGene_tx=XM_019811239.2;refGene_gn=ATM;refGene_tx=XM_
019811240.2;refGene_gn=ATM;refGene_tx=XM_019811241.2;refGene_gn=ATM;re
fGene_tx=XM_019811242.2;refGene_gn=ATM;refGene_tx=XM_019811243.2;refGe
ne_gn=ATM;refGene_tx=XM_019811245.2;refGene_gn=ATM;refGene_tx=XM_01981
1246.2;refGene_gn=ATM;refGene_tx=XM_019811247.2;refGene_gn=ATM;refGene
_tx=XM_023239066.1
D1 6821808 6821922
refGene_gn=ATM;refGene_tx=XM_019811239.2;refGene_gn=ATM;refGene_tx=XM_
019811240.2;refGene_gn=ATM;refGene_tx=XM_019811241.2;refGene_gn=ATM;re
fGene_tx=XM_019811242.2;refGene_gn=ATM;refGene_tx=XM_019811243.2;refGe
ne_gn=ATM;refGene_tx=XM_019811245.2;refGene_gn=ATM;refGene_tx=XM_01981

1246.2;refGene_gn=ATM;refGene_tx=XM_019811247.2;refGene_gn=ATM;refGene_tx=XM_023239066.1
D1 6822261 6822420
refGene_gn=ATM;refGene_tx=XM_019811239.2;refGene_gn=ATM;refGene_tx=XM_019811240.2;refGene_gn=ATM;refGene_tx=XM_019811241.2;refGene_gn=ATM;refGene_tx=XM_019811242.2;refGene_gn=ATM;refGene_tx=XM_019811243.2;refGene_gn=ATM;refGene_tx=XM_019811245.2;refGene_gn=ATM;refGene_tx=XM_019811246.2;refGene_gn=ATM;refGene_tx=XM_019811247.2;refGene_gn=ATM;refGene_tx=XM_023239066.1
D1 6822866 6823005
refGene_gn=ATM;refGene_tx=XM_019811239.2;refGene_gn=ATM;refGene_tx=XM_019811240.2;refGene_gn=ATM;refGene_tx=XM_019811241.2;refGene_gn=ATM;refGene_tx=XM_019811242.2;refGene_gn=ATM;refGene_tx=XM_019811243.2;refGene_gn=ATM;refGene_tx=XM_019811245.2;refGene_gn=ATM;refGene_tx=XM_019811246.2;refGene_gn=ATM;refGene_tx=XM_019811247.2;refGene_gn=ATM;refGene_tx=XM_023239066.1
D1 6823998 6824081
refGene_gn=ATM;refGene_tx=XM_019811239.2;refGene_gn=ATM;refGene_tx=XM_019811240.2;refGene_gn=ATM;refGene_tx=XM_019811241.2;refGene_gn=ATM;refGene_tx=XM_019811242.2;refGene_gn=ATM;refGene_tx=XM_019811243.2;refGene_gn=ATM;refGene_tx=XM_019811245.2;refGene_gn=ATM;refGene_tx=XM_019811246.2;refGene_gn=ATM;refGene_tx=XM_019811247.2;refGene_gn=ATM;refGene_tx=XM_023239066.1
D1 6825854 6825995
refGene_gn=ATM;refGene_tx=XM_019811239.2;refGene_gn=ATM;refGene_tx=XM_019811240.2;refGene_gn=ATM;refGene_tx=XM_019811241.2;refGene_gn=ATM;refGene_tx=XM_019811242.2;refGene_gn=ATM;refGene_tx=XM_019811243.2;refGene_gn=ATM;refGene_tx=XM_019811245.2;refGene_gn=ATM;refGene_tx=XM_019811246.2;refGene_gn=ATM;refGene_tx=XM_019811247.2;refGene_gn=ATM;refGene_tx=XM_023239066.1
D1 6826947 6827064
refGene_gn=ATM;refGene_tx=XM_019811239.2;refGene_gn=ATM;refGene_tx=XM_019811240.2;refGene_gn=ATM;refGene_tx=XM_019811241.2;refGene_gn=ATM;refGene_tx=XM_019811242.2;refGene_gn=ATM;refGene_tx=XM_019811243.2;refGene_gn=ATM;refGene_tx=XM_019811245.2;refGene_gn=ATM;refGene_tx=XM_019811246.2;refGene_gn=ATM;refGene_tx=XM_019811247.2;refGene_gn=ATM;refGene_tx=XM_023239066.1
D1 6830304 6830454
refGene_gn=ATM;refGene_tx=XM_019811239.2;refGene_gn=ATM;refGene_tx=XM_019811240.2;refGene_gn=ATM;refGene_tx=XM_019811241.2;refGene_gn=ATM;refGene_tx=XM_019811242.2;refGene_gn=ATM;refGene_tx=XM_019811243.2;refGene_gn=ATM;refGene_tx=XM_019811245.2;refGene_gn=ATM;refGene_tx=XM_019811246.2;refGene_gn=ATM;refGene_tx=XM_023239066.1
D1 6832231 6832255 refGene_gn=ATM;refGene_tx=XM_019811246.2
D1 6832312 6832478
refGene_gn=ATM;refGene_tx=XM_019811239.2;refGene_gn=ATM;refGene_tx=XM_019811240.2;refGene_gn=ATM;refGene_tx=XM_019811241.2;refGene_gn=ATM;refGene_tx=XM_019811242.2;refGene_gn=ATM;refGene_tx=XM_019811243.2;refGene_gn=ATM;refGene_tx=XM_019811245.2;refGene_gn=ATM;refGene_tx=XM_023239066.1

D1 6833834 6833921
refGene_gn=ATM;refGene_tx=XM_019811239.2;refGene_gn=ATM;refGene_tx=XM_
019811240.2;refGene_gn=ATM;refGene_tx=XM_019811241.2;refGene_gn=ATM;re
fGene_tx=XM_019811242.2;refGene_gn=ATM;refGene_tx=XM_019811243.2;refGe
ne_gn=ATM;refGene_tx=XM_019811245.2;refGene_gn=ATM;refGene_tx=XM_01981
1247.2;refGene_gn=ATM;refGene_tx=XM_023239066.1
D1 6835312 6835427
refGene_gn=ATM;refGene_tx=XM_019811239.2;refGene_gn=ATM;refGene_tx=XM_
019811240.2;refGene_gn=ATM;refGene_tx=XM_019811241.2;refGene_gn=ATM;re
fGene_tx=XM_019811242.2;refGene_gn=ATM;refGene_tx=XM_019811243.2;refGe
ne_gn=ATM;refGene_tx=XM_019811245.2;refGene_gn=ATM;refGene_tx=XM_02323
9066.1
D1 6836793 6836857
refGene_gn=ATM;refGene_tx=XM_019811239.2;refGene_gn=ATM;refGene_tx=XM_
019811240.2;refGene_gn=ATM;refGene_tx=XM_019811241.2;refGene_gn=ATM;re
fGene_tx=XM_019811242.2;refGene_gn=ATM;refGene_tx=XM_019811243.2;refGe
ne_gn=ATM;refGene_tx=XM_019811245.2;refGene_gn=ATM;refGene_tx=XM_02323
9066.1
D1 6837203 6837236 refGene_gn=ATM;refGene_tx=XM_019811245.2
D1 6843405 6843542
refGene_gn=ATM;refGene_tx=XM_019811239.2;refGene_gn=ATM;refGene_tx=XM_
019811240.2;refGene_gn=ATM;refGene_tx=XM_019811241.2;refGene_gn=ATM;re
fGene_tx=XM_019811242.2;refGene_gn=ATM;refGene_tx=XM_019811243.2;refGe
ne_gn=ATM;refGene_tx=XM_023239066.1
D1 6843631 6843815
refGene_gn=ATM;refGene_tx=XM_019811239.2;refGene_gn=ATM;refGene_tx=XM_
019811240.2;refGene_gn=ATM;refGene_tx=XM_019811241.2;refGene_gn=ATM;re
fGene_tx=XM_019811242.2;refGene_gn=ATM;refGene_tx=XM_019811243.2;refGe
ne_gn=ATM;refGene_tx=XM_023239066.1
D1 6859903 6860055
refGene_gn=CD1H11orf65;refGene_tx=XM_006936601.3;refGene_gn=CD1H11orf6
5;refGene_tx=XM_006936602.3;refGene_gn=CD1H11orf65;refGene_tx=XM_00693
6604.3;refGene_gn=CD1H11orf65;refGene_tx=XM_011286333.3;refGene_gn=CD1
H11orf65;refGene_tx=XM_019811249.2;refGene_gn=CD1H11orf65;refGene_tx=X
M_019811250.2
D1 6862390 6862446
refGene_gn=CD1H11orf65;refGene_tx=XM_006936601.3;refGene_gn=CD1H11orf6
5;refGene_tx=XM_006936602.3;refGene_gn=CD1H11orf65;refGene_tx=XM_00693
6604.3;refGene_gn=CD1H11orf65;refGene_tx=XM_011286333.3;refGene_gn=CD1
H11orf65;refGene_tx=XM_019811249.2;refGene_gn=CD1H11orf65;refGene_tx=X
M_019811250.2
D1 6870146 6870317
refGene_gn=CD1H11orf65;refGene_tx=XM_006936601.3;refGene_gn=CD1H11orf6
5;refGene_tx=XM_006936602.3;refGene_gn=CD1H11orf65;refGene_tx=XM_00693
6604.3;refGene_gn=CD1H11orf65;refGene_tx=XM_011286333.3;refGene_gn=CD1
H11orf65;refGene_tx=XM_019811249.2;refGene_gn=CD1H11orf65;refGene_tx=X
M_019811250.2
D1 6883093 6883224
refGene_gn=CD1H11orf65;refGene_tx=XM_006936601.3;refGene_gn=CD1H11orf6
5;refGene_tx=XM_006936602.3;refGene_gn=CD1H11orf65;refGene_tx=XM_00693

6604.3;refGene_gn=CD1H11orf65;refGene_tx=XM_011286333.3;refGene_gn=CD1H11orf65;refGene_tx=XM_019811249.2;refGene_gn=CD1H11orf65;refGene_tx=XM_019811250.2
D1 6884904 6885105
refGene_gn=CD1H11orf65;refGene_tx=XM_006936601.3;refGene_gn=CD1H11orf65;refGene_tx=XM_006936602.3;refGene_gn=CD1H11orf65;refGene_tx=XM_006936604.3;refGene_gn=CD1H11orf65;refGene_tx=XM_011286333.3;refGene_gn=CD1H11orf65;refGene_tx=XM_019811249.2;refGene_gn=CD1H11orf65;refGene_tx=XM_019811250.2
D1 6885233 6885287
refGene_gn=CD1H11orf65;refGene_tx=XM_006936601.3;refGene_gn=CD1H11orf65;refGene_tx=XM_006936602.3;refGene_gn=CD1H11orf65;refGene_tx=XM_011286333.3;refGene_gn=CD1H11orf65;refGene_tx=XM_019811249.2;refGene_gn=CD1H11orf65;refGene_tx=XM_019811250.2
D1 6891300 6891393
refGene_gn=CD1H11orf65;refGene_tx=XM_006936601.3;refGene_gn=CD1H11orf65;refGene_tx=XM_006936602.3;refGene_gn=CD1H11orf65;refGene_tx=XM_006936604.3;refGene_gn=CD1H11orf65;refGene_tx=XM_011286333.3;refGene_gn=CD1H11orf65;refGene_tx=XM_019811249.2
D1 6905940 6906021
refGene_gn=CD1H11orf65;refGene_tx=XM_006936601.3;refGene_gn=CD1H11orf65;refGene_tx=XM_006936602.3;refGene_gn=CD1H11orf65;refGene_tx=XM_006936604.3;refGene_gn=CD1H11orf65;refGene_tx=XM_011286333.3;refGene_gn=CD1H11orf65;refGene_tx=XM_019811249.2;refGene_gn=CD1H11orf65;refGene_tx=XM_019811250.2
D1 6924064 6924190 refGene_gn=KDELC2;refGene_tx=XM_023239070.1
D1 6926039 6926144 refGene_gn=KDELC2;refGene_tx=XM_023239070.1
D1 6927452 6927647 refGene_gn=KDELC2;refGene_tx=XM_023239070.1
D1 6929448 6929645 refGene_gn=KDELC2;refGene_tx=XM_023239070.1
D1 6931123 6931340 refGene_gn=KDELC2;refGene_tx=XM_023239070.1
D1 6934972 6935259 refGene_gn=KDELC2;refGene_tx=XM_023239070.1
D1 6938320 6938518 refGene_gn=KDELC2;refGene_tx=XM_023239070.1
D1 6950992 6951194 refGene_gn=KDELC2;refGene_tx=XM_023239070.1
D1 6960398 6965785
refGene_gn=EXPH5;refGene_tx=XM_019811252.2;refGene_gn=EXPH5;refGene_tx=XM_019811253.2;refGene_gn=EXPH5;refGene_tx=XM_019811254.1;refGene_gn=EXPH5;refGene_tx=XM_019811255.2;refGene_gn=EXPH5;refGene_tx=XM_019811256.2
D1 6970498 6970634
refGene_gn=EXPH5;refGene_tx=XM_019811252.2;refGene_gn=EXPH5;refGene_tx=XM_019811253.2;refGene_gn=EXPH5;refGene_tx=XM_019811254.1;refGene_gn=EXPH5;refGene_tx=XM_019811255.2;refGene_gn=EXPH5;refGene_tx=XM_019811256.2
D1 6983854 6983903
refGene_gn=EXPH5;refGene_tx=XM_019811252.2;refGene_gn=EXPH5;refGene_tx=XM_019811253.2;refGene_gn=EXPH5;refGene_tx=XM_019811254.1;refGene_gn=EXPH5;refGene_tx=XM_019811255.2
D1 6996727 6996890
refGene_gn=EXPH5;refGene_tx=XM_019811252.2;refGene_gn=EXPH5;refGene_tx=XM_019811253.2;refGene_gn=EXPH5;refGene_tx=XM_019811254.1;refGene_gn=

```
EXPH5;refGene_tx=XM_019811255.2
D1	6999010	6999171	
refGene_gn=EXPH5;refGene_tx=XM_019811252.2;refGene_gn=EXPH5;refGene_tx
=XM_019811253.2;refGene_gn=EXPH5;refGene_tx=XM_019811254.1;refGene_gn=
EXPH5;refGene_tx=XM_019811255.2
D1	7046358	7046477	
refGene_gn=EXPH5;refGene_tx=XM_019811252.2;refGene_gn=EXPH5;refGene_tx
=XM_019811253.2
D1	7115655	7115841	refGene_gn=DDX10;refGene_tx=XM_011286334.2
D1	7127231	7127292	refGene_gn=DDX10;refGene_tx=XM_011286334.2
D1	7128291	7128422	refGene_gn=DDX10;refGene_tx=XM_011286334.2
D1	7130154	7130313	refGene_gn=DDX10;refGene_tx=XM_011286334.2
D1	7137699	7137820	refGene_gn=DDX10;refGene_tx=XM_011286334.2
D1	7138446	7138636	refGene_gn=DDX10;refGene_tx=XM_011286334.2
D1	7142198	7142325	refGene_gn=DDX10;refGene_tx=XM_011286334.2
D1	7146579	7146742	refGene_gn=DDX10;refGene_tx=XM_011286334.2
D1	7148358	7148443	refGene_gn=DDX10;refGene_tx=XM_011286334.2
D1	7160114	7160216	refGene_gn=DDX10;refGene_tx=XM_011286334.2
D1	7161388	7161476	refGene_gn=DDX10;refGene_tx=XM_011286334.2
D1	7165146	7165235	refGene_gn=DDX10;refGene_tx=XM_011286334.2
D1	7168037	7168500	refGene_gn=DDX10;refGene_tx=XM_011286334.2
D1	7272992	7273112	refGene_gn=DDX10;refGene_tx=XM_011286334.2
D1	7276454	7276619	refGene_gn=DDX10;refGene_tx=XM_011286334.2
D1	7287547	7287604	refGene_gn=DDX10;refGene_tx=XM_011286334.2
D1	7358802	7358942	refGene_gn=DDX10;refGene_tx=XM_011286334.2
D1	7395302	7395480	refGene_gn=DDX10;refGene_tx=XM_011286334.2
D1	7804860	7805463	
refGene_gn=CD1H11orf87;refGene_tx=XM_011286437.3
D1	8367448	8367472	refGene_gn=ZC3H12C;refGene_tx=XM_023239072.1
D1	8367678	8367699	refGene_gn=ZC3H12C;refGene_tx=XM_023239073.1
D1	8400709	8401461	
refGene_gn=ZC3H12C;refGene_tx=XM_023239072.1;refGene_gn=ZC3H12C;refGen
e_tx=XM_023239073.1;refGene_gn=ZC3H12C;refGene_tx=XM_023239074.1;refGe
ne_gn=ZC3H12C;refGene_tx=XM_023239075.1
D1	8436299	8436439	
refGene_gn=ZC3H12C;refGene_tx=XM_023239072.1;refGene_gn=ZC3H12C;refGen
e_tx=XM_023239073.1;refGene_gn=ZC3H12C;refGene_tx=XM_023239074.1;refGe
ne_gn=ZC3H12C;refGene_tx=XM_023239075.1
D1	8441350	8441585	
refGene_gn=ZC3H12C;refGene_tx=XM_023239072.1;refGene_gn=ZC3H12C;refGen
e_tx=XM_023239073.1;refGene_gn=ZC3H12C;refGene_tx=XM_023239074.1;refGe
ne_gn=ZC3H12C;refGene_tx=XM_023239075.1
D1	8443426	8443533	
refGene_gn=ZC3H12C;refGene_tx=XM_023239072.1;refGene_gn=ZC3H12C;refGen
e_tx=XM_023239073.1;refGene_gn=ZC3H12C;refGene_tx=XM_023239074.1;refGe
ne_gn=ZC3H12C;refGene_tx=XM_023239075.1
D1	8444634	8446031	
refGene_gn=ZC3H12C;refGene_tx=XM_023239072.1;refGene_gn=ZC3H12C;refGen
e_tx=XM_023239073.1;refGene_gn=ZC3H12C;refGene_tx=XM_023239074.1;refGe
ne_gn=ZC3H12C;refGene_tx=XM_023239075.1
```

D1	8484293	8484360	refGene_gn=RDX;refGene_tx=XM_023239076.1
D1	8504061	8504071	refGene_gn=RDX;refGene_tx=XM_006936607.3
D1	8512165	8512326
refGene_gn=RDX;refGene_tx=XM_006936607.3;refGene_gn=RDX;refGene_tx=XM_023239076.1
D1	8513547	8513790
refGene_gn=RDX;refGene_tx=XM_006936607.3;refGene_gn=RDX;refGene_tx=XM_023239076.1
D1	8517955	8518048
refGene_gn=RDX;refGene_tx=XM_006936607.3;refGene_gn=RDX;refGene_tx=XM_023239076.1
D1	8518948	8519109
refGene_gn=RDX;refGene_tx=XM_006936607.3;refGene_gn=RDX;refGene_tx=XM_023239076.1
D1	8530353	8530484
refGene_gn=RDX;refGene_tx=XM_006936607.3;refGene_gn=RDX;refGene_tx=XM_023239076.1
D1	8535539	8535703
refGene_gn=RDX;refGene_tx=XM_006936607.3;refGene_gn=RDX;refGene_tx=XM_023239076.1
D1	8536172	8536269
refGene_gn=RDX;refGene_tx=XM_006936607.3;refGene_gn=RDX;refGene_tx=XM_023239076.1
D1	8539826	8539973
refGene_gn=RDX;refGene_tx=XM_006936607.3;refGene_gn=RDX;refGene_tx=XM_023239076.1
D1	8540166	8540250
refGene_gn=RDX;refGene_tx=XM_006936607.3;refGene_gn=RDX;refGene_tx=XM_023239076.1
D1	8545029	8545304
refGene_gn=RDX;refGene_tx=XM_006936607.3;refGene_gn=RDX;refGene_tx=XM_023239076.1
D1	8545726	8545822
refGene_gn=RDX;refGene_tx=XM_006936607.3;refGene_gn=RDX;refGene_tx=XM_023239076.1
D1	8551771	8551855
refGene_gn=RDX;refGene_tx=XM_006936607.3;refGene_gn=RDX;refGene_tx=XM_023239076.1
D1	8560509	8560521
refGene_gn=RDX;refGene_tx=XM_006936607.3;refGene_gn=RDX;refGene_tx=XM_023239076.1
D1	8647623	8647814	refGene_gn=FDX1;refGene_tx=XM_023239077.1
D1	8652027	8652152	refGene_gn=FDX1;refGene_tx=XM_023239077.1
D1	8677968	8678098	refGene_gn=FDX1;refGene_tx=XM_023239077.1
D1	8682935	8683050	refGene_gn=FDX1;refGene_tx=XM_023239077.1
D1	8772797	8774653
refGene_gn=ARHGAP20;refGene_tx=XM_023239078.1;refGene_gn=ARHGAP20;refGene_tx=XM_023239079.1;refGene_gn=ARHGAP20;refGene_tx=XM_023239080.1;refGene_gn=ARHGAP20;refGene_tx=XM_023239081.1;refGene_gn=ARHGAP20;refGene_tx=XM_023239082.1

D1	8775554	8775669
refGene_gn=ARHGAP20;refGene_tx=XM_023239078.1;refGene_gn=ARHGAP20;refGene_tx=XM_023239079.1;refGene_gn=ARHGAP20;refGene_tx=XM_023239080.1;refGene_gn=ARHGAP20;refGene_tx=XM_023239081.1;refGene_gn=ARHGAP20;refGene_tx=XM_023239082.1
D1	8776842	8777032
refGene_gn=ARHGAP20;refGene_tx=XM_023239078.1;refGene_gn=ARHGAP20;refGene_tx=XM_023239079.1;refGene_gn=ARHGAP20;refGene_tx=XM_023239080.1;refGene_gn=ARHGAP20;refGene_tx=XM_023239081.1;refGene_gn=ARHGAP20;refGene_tx=XM_023239082.1
D1	8779577	8779687
refGene_gn=ARHGAP20;refGene_tx=XM_023239078.1;refGene_gn=ARHGAP20;refGene_tx=XM_023239079.1;refGene_gn=ARHGAP20;refGene_tx=XM_023239080.1;refGene_gn=ARHGAP20;refGene_tx=XM_023239081.1;refGene_gn=ARHGAP20;refGene_tx=XM_023239082.1
D1	8783206	8783368
refGene_gn=ARHGAP20;refGene_tx=XM_023239078.1;refGene_gn=ARHGAP20;refGene_tx=XM_023239079.1;refGene_gn=ARHGAP20;refGene_tx=XM_023239080.1;refGene_gn=ARHGAP20;refGene_tx=XM_023239081.1;refGene_gn=ARHGAP20;refGene_tx=XM_023239082.1
D1	8784701	8784880
refGene_gn=ARHGAP20;refGene_tx=XM_023239078.1;refGene_gn=ARHGAP20;refGene_tx=XM_023239079.1;refGene_gn=ARHGAP20;refGene_tx=XM_023239080.1;refGene_gn=ARHGAP20;refGene_tx=XM_023239081.1;refGene_gn=ARHGAP20;refGene_tx=XM_023239082.1
D1	8796914	8797103
refGene_gn=ARHGAP20;refGene_tx=XM_023239078.1;refGene_gn=ARHGAP20;refGene_tx=XM_023239079.1;refGene_gn=ARHGAP20;refGene_tx=XM_023239080.1;refGene_gn=ARHGAP20;refGene_tx=XM_023239081.1;refGene_gn=ARHGAP20;refGene_tx=XM_023239082.1
D1	8799585	8799652
refGene_gn=ARHGAP20;refGene_tx=XM_023239078.1;refGene_gn=ARHGAP20;refGene_tx=XM_023239079.1;refGene_gn=ARHGAP20;refGene_tx=XM_023239080.1;refGene_gn=ARHGAP20;refGene_tx=XM_023239081.1;refGene_gn=ARHGAP20;refGene_tx=XM_023239082.1
D1	8802164	8802242
refGene_gn=ARHGAP20;refGene_tx=XM_023239078.1;refGene_gn=ARHGAP20;refGene_tx=XM_023239079.1;refGene_gn=ARHGAP20;refGene_tx=XM_023239080.1;refGene_gn=ARHGAP20;refGene_tx=XM_023239081.1;refGene_gn=ARHGAP20;refGene_tx=XM_023239082.1
D1	8805139	8805224
refGene_gn=ARHGAP20;refGene_tx=XM_023239078.1;refGene_gn=ARHGAP20;refGene_tx=XM_023239079.1;refGene_gn=ARHGAP20;refGene_tx=XM_023239080.1;refGene_gn=ARHGAP20;refGene_tx=XM_023239081.1;refGene_gn=ARHGAP20;refGene_tx=XM_023239082.1
D1	8806113	8806155
refGene_gn=ARHGAP20;refGene_tx=XM_023239078.1;refGene_gn=ARHGAP20;refGene_tx=XM_023239079.1;refGene_gn=ARHGAP20;refGene_tx=XM_023239080.1;refGene_gn=ARHGAP20;refGene_tx=XM_023239081.1;refGene_gn=ARHGAP20;refGene_tx=XM_023239082.1

D1 8808462 8808564
refGene_gn=ARHGAP20;refGene_tx=XM_023239078.1;refGene_gn=ARHGAP20;refGene_tx=XM_023239080.1;refGene_gn=ARHGAP20;refGene_tx=XM_023239081.1;refGene_gn=ARHGAP20;refGene_tx=XM_023239082.1
D1 8812217 8812367
refGene_gn=ARHGAP20;refGene_tx=XM_023239078.1;refGene_gn=ARHGAP20;refGene_tx=XM_023239079.1;refGene_gn=ARHGAP20;refGene_tx=XM_023239080.1;refGene_gn=ARHGAP20;refGene_tx=XM_023239081.1;refGene_gn=ARHGAP20;refGene_tx=XM_023239082.1
D1 8820675 8820840
refGene_gn=ARHGAP20;refGene_tx=XM_023239078.1;refGene_gn=ARHGAP20;refGene_tx=XM_023239079.1;refGene_gn=ARHGAP20;refGene_tx=XM_023239080.1;refGene_gn=ARHGAP20;refGene_tx=XM_023239081.1;refGene_gn=ARHGAP20;refGene_tx=XM_023239082.1
D1 8878041 8878124
refGene_gn=ARHGAP20;refGene_tx=XM_023239078.1;refGene_gn=ARHGAP20;refGene_tx=XM_023239079.1;refGene_gn=ARHGAP20;refGene_tx=XM_023239080.1;refGene_gn=ARHGAP20;refGene_tx=XM_023239081.1;refGene_gn=ARHGAP20;refGene_tx=XM_023239082.1
D1 8897237 8897342
refGene_gn=ARHGAP20;refGene_tx=XM_023239078.1;refGene_gn=ARHGAP20;refGene_tx=XM_023239079.1
D1 9397615 9397622 refGene_gn=COLCA2;refGene_tx=XM_019811258.2
D1 9399526 9399621 refGene_gn=COLCA2;refGene_tx=XM_019811258.2
D1 9402736 9402742
refGene_gn=COLCA2;refGene_tx=XM_019811259.2;refGene_gn=COLCA2;refGene_tx=XM_023239083.1
D1 9403733 9403788
refGene_gn=COLCA2;refGene_tx=XM_019811258.2;refGene_gn=COLCA2;refGene_tx=XM_019811259.2;refGene_gn=COLCA2;refGene_tx=XM_023239083.1
D1 9405059 9405289
refGene_gn=COLCA2;refGene_tx=XM_019811258.2;refGene_gn=COLCA2;refGene_tx=XM_019811259.2;refGene_gn=COLCA2;refGene_tx=XM_023239083.1
D1 9406678 9407050
refGene_gn=COLCA2;refGene_tx=XM_019811258.2;refGene_gn=COLCA2;refGene_tx=XM_019811259.2;refGene_gn=COLCA2;refGene_tx=XM_023239083.1
D1 9442957 9443272 refGene_gn=POU2AF1;refGene_tx=XM_003992337.4
D1 9445917 9446183 refGene_gn=POU2AF1;refGene_tx=XM_003992337.4
D1 9446267 9446310 refGene_gn=POU2AF1;refGene_tx=XM_003992337.4
D1 9446818 9446949 refGene_gn=POU2AF1;refGene_tx=XM_003992337.4
D1 9465450 9465466 refGene_gn=POU2AF1;refGene_tx=XM_003992337.4
D1 9554643 9554674 refGene_gn=BTG4;refGene_tx=XM_019811261.2
D1 9555098 9555250 refGene_gn=BTG4;refGene_tx=XM_019811261.2
D1 9557078 9557277 refGene_gn=BTG4;refGene_tx=XM_019811261.2
D1 9558096 9558234 refGene_gn=BTG4;refGene_tx=XM_019811261.2
D1 9558691 9558864 refGene_gn=BTG4;refGene_tx=XM_019811261.2
D1 9574305 9574534
refGene_gn=CD1H11orf88;refGene_tx=XM_006936608.3;refGene_gn=CD1H11orf88;refGene_tx=XM_019811262.2;refGene_gn=CD1H11orf88;refGene_tx=XM_019811264.2

D1 9575521 9575592
refGene_gn=CD1H11orf88;refGene_tx=XM_006936608.3;refGene_gn=CD1H11orf8
8;refGene_tx=XM_019811262.2;refGene_gn=CD1H11orf88;refGene_tx=XM_01981
1264.2
D1 9583593 9583692
refGene_gn=CD1H11orf88;refGene_tx=XM_019811262.2;refGene_gn=CD1H11orf8
8;refGene_tx=XM_019811264.2
D1 9594068 9594128
refGene_gn=CD1H11orf88;refGene_tx=XM_006936608.3;refGene_gn=CD1H11orf8
8;refGene_tx=XM_019811262.2;refGene_gn=CD1H11orf88;refGene_tx=XM_01981
1264.2
D1 9594713 9594766
refGene_gn=CD1H11orf88;refGene_tx=XM_006936608.3;refGene_gn=CD1H11orf8
8;refGene_tx=XM_019811262.2;refGene_gn=CD1H11orf88;refGene_tx=XM_01981
1264.2
D1 9597086 9597144
refGene_gn=CD1H11orf88;refGene_tx=XM_006936608.3;refGene_gn=CD1H11orf8
8;refGene_tx=XM_019811262.2;refGene_gn=CD1H11orf88;refGene_tx=XM_01981
1264.2
D1 9600884 9600969
refGene_gn=LAYN;refGene_tx=XM_023239084.1;refGene_gn=LAYN;refGene_tx=X
M_023239085.1
D1 9601617 9601641 refGene_gn=LAYN;refGene_tx=XM_023239084.1
D1 9603937 9604235
refGene_gn=LAYN;refGene_tx=XM_023239084.1;refGene_gn=LAYN;refGene_tx=X
M_023239085.1
D1 9610162 9610320
refGene_gn=LAYN;refGene_tx=XM_023239084.1;refGene_gn=LAYN;refGene_tx=X
M_023239085.1
D1 9614936 9614969
refGene_gn=LAYN;refGene_tx=XM_023239084.1;refGene_gn=LAYN;refGene_tx=X
M_023239085.1
D1 9615609 9615693
refGene_gn=LAYN;refGene_tx=XM_023239084.1;refGene_gn=LAYN;refGene_tx=X
M_023239085.1
D1 9617790 9617887
refGene_gn=LAYN;refGene_tx=XM_023239084.1;refGene_gn=LAYN;refGene_tx=X
M_023239085.1
D1 9620627 9620991
refGene_gn=LAYN;refGene_tx=XM_023239084.1;refGene_gn=LAYN;refGene_tx=X
M_023239085.1
D1 9657375 9657510 refGene_gn=SIK2;refGene_tx=XM_011286338.3
D1 9676189 9676306 refGene_gn=SIK2;refGene_tx=XM_011286338.3
D1 9678660 9678724 refGene_gn=SIK2;refGene_tx=XM_011286338.3
D1 9708411 9708466 refGene_gn=SIK2;refGene_tx=XM_023239086.1
D1 9739250 9739412
refGene_gn=SIK2;refGene_tx=XM_011286338.3;refGene_gn=SIK2;refGene_tx=X
M_023239086.1
D1 9752585 9752710
refGene_gn=SIK2;refGene_tx=XM_011286338.3;refGene_gn=SIK2;refGene_tx=X

M_023239086.1
D1 9753154 9753278
refGene_gn=SIK2;refGene_tx=XM_011286338.3;refGene_gn=SIK2;refGene_tx=X
M_023239086.1
D1 9754636 9754857
refGene_gn=SIK2;refGene_tx=XM_011286338.3;refGene_gn=SIK2;refGene_tx=X
M_023239086.1
D1 9756596 9756749
refGene_gn=SIK2;refGene_tx=XM_011286338.3;refGene_gn=SIK2;refGene_tx=X
M_023239086.1
D1 9763371 9763536
refGene_gn=SIK2;refGene_tx=XM_011286338.3;refGene_gn=SIK2;refGene_tx=X
M_023239086.1
D1 9770941 9771170
refGene_gn=SIK2;refGene_tx=XM_011286338.3;refGene_gn=SIK2;refGene_tx=X
M_023239086.1
D1 9771648 9771933
refGene_gn=SIK2;refGene_tx=XM_011286338.3;refGene_gn=SIK2;refGene_tx=X
M_023239086.1
D1 9772063 9772227
refGene_gn=SIK2;refGene_tx=XM_011286338.3;refGene_gn=SIK2;refGene_tx=X
M_023239086.1
D1 9773208 9773319
refGene_gn=SIK2;refGene_tx=XM_011286338.3;refGene_gn=SIK2;refGene_tx=X
M_023239086.1
D1 9774040 9774132
refGene_gn=SIK2;refGene_tx=XM_011286338.3;refGene_gn=SIK2;refGene_tx=X
M_023239086.1
D1 9774755 9775380
refGene_gn=SIK2;refGene_tx=XM_011286338.3;refGene_gn=SIK2;refGene_tx=X
M_023239086.1
D1 9777831 9777924 refGene_gn=PPP2R1B;refGene_tx=XM_011286439.3
D1 9778674 9778775
refGene_gn=PPP2R1B;refGene_tx=XM_011286439.3;refGene_gn=PPP2R1B;refGen
e_tx=XM_023239087.1;refGene_gn=PPP2R1B;refGene_tx=XM_023239088.1
D1 9785711 9785833
refGene_gn=PPP2R1B;refGene_tx=XM_011286439.3;refGene_gn=PPP2R1B;refGen
e_tx=XM_023239087.1;refGene_gn=PPP2R1B;refGene_tx=XM_023239088.1
D1 9789696 9789713 refGene_gn=PPP2R1B;refGene_tx=XM_023239089.1
D1 9790118 9790210
refGene_gn=PPP2R1B;refGene_tx=XM_011286439.3;refGene_gn=PPP2R1B;refGen
e_tx=XM_023239087.1;refGene_gn=PPP2R1B;refGene_tx=XM_023239088.1;refGe
ne_gn=PPP2R1B;refGene_tx=XM_023239089.1
D1 9790602 9790745
refGene_gn=PPP2R1B;refGene_tx=XM_011286439.3;refGene_gn=PPP2R1B;refGen
e_tx=XM_023239087.1;refGene_gn=PPP2R1B;refGene_tx=XM_023239088.1;refGe
ne_gn=PPP2R1B;refGene_tx=XM_023239089.1
D1 9791098 9791253
refGene_gn=PPP2R1B;refGene_tx=XM_011286439.3;refGene_gn=PPP2R1B;refGen
e_tx=XM_023239087.1;refGene_gn=PPP2R1B;refGene_tx=XM_023239088.1;refGe

ne_gn=PPP2R1B;refGene_tx=XM_023239089.1
D1 9795784 9795845
refGene_gn=PPP2R1B;refGene_tx=XM_011286439.3;refGene_gn=PPP2R1B;refGene_tx=XM_023239087.1;refGene_gn=PPP2R1B;refGene_tx=XM_023239088.1;refGene_gn=PPP2R1B;refGene_tx=XM_023239089.1
D1 9798455 9798629
refGene_gn=PPP2R1B;refGene_tx=XM_011286439.3;refGene_gn=PPP2R1B;refGene_tx=XM_023239087.1;refGene_gn=PPP2R1B;refGene_tx=XM_023239088.1;refGene_gn=PPP2R1B;refGene_tx=XM_023239089.1
D1 9799404 9799539
refGene_gn=PPP2R1B;refGene_tx=XM_011286439.3;refGene_gn=PPP2R1B;refGene_tx=XM_023239087.1;refGene_gn=PPP2R1B;refGene_tx=XM_023239088.1;refGene_gn=PPP2R1B;refGene_tx=XM_023239089.1
D1 9800136 9800207
refGene_gn=PPP2R1B;refGene_tx=XM_011286439.3;refGene_gn=PPP2R1B;refGene_tx=XM_023239087.1;refGene_gn=PPP2R1B;refGene_tx=XM_023239088.1;refGene_gn=PPP2R1B;refGene_tx=XM_023239089.1
D1 9800594 9800709
refGene_gn=PPP2R1B;refGene_tx=XM_011286439.3;refGene_gn=PPP2R1B;refGene_tx=XM_023239087.1;refGene_gn=PPP2R1B;refGene_tx=XM_023239088.1;refGene_gn=PPP2R1B;refGene_tx=XM_023239089.1
D1 9800912 9801068
refGene_gn=PPP2R1B;refGene_tx=XM_011286439.3;refGene_gn=PPP2R1B;refGene_tx=XM_023239087.1;refGene_gn=PPP2R1B;refGene_tx=XM_023239088.1;refGene_gn=PPP2R1B;refGene_tx=XM_023239089.1
D1 9805537 9805685
refGene_gn=PPP2R1B;refGene_tx=XM_011286439.3;refGene_gn=PPP2R1B;refGene_tx=XM_023239087.1;refGene_gn=PPP2R1B;refGene_tx=XM_023239088.1;refGene_gn=PPP2R1B;refGene_tx=XM_023239089.1
D1 9805998 9806231
refGene_gn=PPP2R1B;refGene_tx=XM_011286439.3;refGene_gn=PPP2R1B;refGene_tx=XM_023239087.1;refGene_gn=PPP2R1B;refGene_tx=XM_023239088.1;refGene_gn=PPP2R1B;refGene_tx=XM_023239089.1
D1 9809574 9809675
refGene_gn=PPP2R1B;refGene_tx=XM_011286439.3;refGene_gn=PPP2R1B;refGene_tx=XM_023239087.1;refGene_gn=PPP2R1B;refGene_tx=XM_023239088.1;refGene_gn=PPP2R1B;refGene_tx=XM_023239089.1
D1 9810066 9810157
refGene_gn=PPP2R1B;refGene_tx=XM_011286439.3;refGene_gn=PPP2R1B;refGene_tx=XM_023239087.1;refGene_gn=PPP2R1B;refGene_tx=XM_023239088.1;refGene_gn=PPP2R1B;refGene_tx=XM_023239089.1
D1 9810986 9811100
refGene_gn=PPP2R1B;refGene_tx=XM_011286439.3;refGene_gn=PPP2R1B;refGene_tx=XM_023239087.1;refGene_gn=PPP2R1B;refGene_tx=XM_023239088.1;refGene_gn=PPP2R1B;refGene_tx=XM_023239089.1
D1 9824975 9825096
refGene_gn=ALG9;refGene_tx=XM_019811268.2;refGene_gn=ALG9;refGene_tx=XM_023239092.1
D1 9826833 9827038
refGene_gn=ALG9;refGene_tx=XM_006936611.3;refGene_gn=ALG9;refGene_tx=X

M_023239090.1;refGene_gn=ALG9;refGene_tx=XM_023239091.1
D1 9827324 9827443
refGene_gn=ALG9;refGene_tx=XM_006936611.3;refGene_gn=ALG9;refGene_tx=XM_019811268.2;refGene_gn=ALG9;refGene_tx=XM_023239090.1;refGene_gn=ALG9;refGene_tx=XM_023239091.1;refGene_gn=ALG9;refGene_tx=XM_023239092.1
D1 9854290 9854421
refGene_gn=ALG9;refGene_tx=XM_006936611.3;refGene_gn=ALG9;refGene_tx=XM_019811268.2
D1 9881115 9881133 refGene_gn=ALG9;refGene_tx=XM_006936613.3
D1 9882563 9882656 refGene_gn=ALG9;refGene_tx=XM_006936612.4
D1 9884441 9884571
refGene_gn=ALG9;refGene_tx=XM_006936611.3;refGene_gn=ALG9;refGene_tx=XM_006936612.4;refGene_gn=ALG9;refGene_tx=XM_006936613.3;refGene_gn=ALG9;refGene_tx=XM_019811268.2;refGene_gn=ALG9;refGene_tx=XM_023239090.1;refGene_gn=ALG9;refGene_tx=XM_023239091.1;refGene_gn=ALG9;refGene_tx=XM_023239092.1
D1 9885698 9885846
refGene_gn=ALG9;refGene_tx=XM_006936611.3;refGene_gn=ALG9;refGene_tx=XM_006936612.4;refGene_gn=ALG9;refGene_tx=XM_006936613.3;refGene_gn=ALG9;refGene_tx=XM_019811268.2;refGene_gn=ALG9;refGene_tx=XM_023239090.1;refGene_gn=ALG9;refGene_tx=XM_023239091.1;refGene_gn=ALG9;refGene_tx=XM_023239092.1
D1 9886482 9886612
refGene_gn=ALG9;refGene_tx=XM_006936611.3;refGene_gn=ALG9;refGene_tx=XM_006936612.4;refGene_gn=ALG9;refGene_tx=XM_006936613.3;refGene_gn=ALG9;refGene_tx=XM_019811268.2;refGene_gn=ALG9;refGene_tx=XM_023239090.1;refGene_gn=ALG9;refGene_tx=XM_023239091.1;refGene_gn=ALG9;refGene_tx=XM_023239092.1
D1 9889741 9889896
refGene_gn=ALG9;refGene_tx=XM_006936611.3;refGene_gn=ALG9;refGene_tx=XM_006936612.4;refGene_gn=ALG9;refGene_tx=XM_006936613.3;refGene_gn=ALG9;refGene_tx=XM_019811268.2;refGene_gn=ALG9;refGene_tx=XM_023239090.1;refGene_gn=ALG9;refGene_tx=XM_023239091.1;refGene_gn=ALG9;refGene_tx=XM_023239092.1
D1 9892799 9892922
refGene_gn=ALG9;refGene_tx=XM_006936611.3;refGene_gn=ALG9;refGene_tx=XM_006936612.4;refGene_gn=ALG9;refGene_tx=XM_006936613.3;refGene_gn=ALG9;refGene_tx=XM_019811268.2;refGene_gn=ALG9;refGene_tx=XM_023239090.1;refGene_gn=ALG9;refGene_tx=XM_023239091.1;refGene_gn=ALG9;refGene_tx=XM_023239092.1
D1 9900067 9900173
refGene_gn=ALG9;refGene_tx=XM_006936611.3;refGene_gn=ALG9;refGene_tx=XM_006936612.4;refGene_gn=ALG9;refGene_tx=XM_006936613.3;refGene_gn=ALG9;refGene_tx=XM_019811268.2;refGene_gn=ALG9;refGene_tx=XM_023239090.1;refGene_gn=ALG9;refGene_tx=XM_023239091.1;refGene_gn=ALG9;refGene_tx=XM_023239092.1
D1 9900353 9900441
refGene_gn=ALG9;refGene_tx=XM_006936611.3;refGene_gn=ALG9;refGene_tx=XM_006936612.4;refGene_gn=ALG9;refGene_tx=XM_006936613.3;refGene_gn=ALG9;refGene_tx=XM_019811268.2;refGene_gn=ALG9;refGene_tx=XM_023239090.1;

refGene_gn=ALG9;refGene_tx=XM_023239091.1;refGene_gn=ALG9;refGene_tx=XM_023239092.1
D1 9904262 9904398
refGene_gn=ALG9;refGene_tx=XM_006936611.3;refGene_gn=ALG9;refGene_tx=XM_006936612.4;refGene_gn=ALG9;refGene_tx=XM_006936613.3;refGene_gn=ALG9;refGene_tx=XM_019811268.2;refGene_gn=ALG9;refGene_tx=XM_023239090.1;refGene_gn=ALG9;refGene_tx=XM_023239091.1;refGene_gn=ALG9;refGene_tx=XM_023239092.1
D1 9906412 9906501
refGene_gn=ALG9;refGene_tx=XM_006936611.3;refGene_gn=ALG9;refGene_tx=XM_006936612.4;refGene_gn=ALG9;refGene_tx=XM_006936613.3;refGene_gn=ALG9;refGene_tx=XM_019811268.2;refGene_gn=ALG9;refGene_tx=XM_023239090.1;refGene_gn=ALG9;refGene_tx=XM_023239091.1;refGene_gn=ALG9;refGene_tx=XM_023239092.1
D1 9911224 9911295
refGene_gn=ALG9;refGene_tx=XM_006936611.3;refGene_gn=ALG9;refGene_tx=XM_006936612.4;refGene_gn=ALG9;refGene_tx=XM_006936613.3;refGene_gn=ALG9;refGene_tx=XM_019811268.2;refGene_gn=ALG9;refGene_tx=XM_023239090.1;refGene_gn=ALG9;refGene_tx=XM_023239091.1;refGene_gn=ALG9;refGene_tx=XM_023239092.1
D1 9918304 9918439
refGene_gn=ALG9;refGene_tx=XM_006936611.3;refGene_gn=ALG9;refGene_tx=XM_006936612.4;refGene_gn=ALG9;refGene_tx=XM_006936613.3;refGene_gn=ALG9;refGene_tx=XM_019811268.2;refGene_gn=ALG9;refGene_tx=XM_023239090.1;refGene_gn=ALG9;refGene_tx=XM_023239091.1;refGene_gn=ALG9;refGene_tx=XM_023239092.1
D1 9919273 9919412
refGene_gn=ALG9;refGene_tx=XM_006936611.3;refGene_gn=ALG9;refGene_tx=XM_006936612.4;refGene_gn=ALG9;refGene_tx=XM_006936613.3;refGene_gn=ALG9;refGene_tx=XM_019811268.2;refGene_gn=ALG9;refGene_tx=XM_023239090.1;refGene_gn=ALG9;refGene_tx=XM_023239091.1;refGene_gn=ALG9;refGene_tx=XM_023239092.1
D1 9920466 9920597
refGene_gn=ALG9;refGene_tx=XM_006936611.3;refGene_gn=ALG9;refGene_tx=XM_006936612.4;refGene_gn=ALG9;refGene_tx=XM_006936613.3;refGene_gn=ALG9;refGene_tx=XM_019811268.2;refGene_gn=ALG9;refGene_tx=XM_023239090.1;refGene_gn=ALG9;refGene_tx=XM_023239091.1;refGene_gn=ALG9;refGene_tx=XM_023239092.1
D1 9924514 9925697 refGene_gn=FDXACB1;refGene_tx=XM_003992341.4
D1 9926075 9926234 refGene_gn=FDXACB1;refGene_tx=XM_003992341.4
D1 9926675 9926879 refGene_gn=FDXACB1;refGene_tx=XM_003992341.4
D1 9927807 9927964 refGene_gn=FDXACB1;refGene_tx=XM_003992341.4
D1 9928163 9928335 refGene_gn=FDXACB1;refGene_tx=XM_003992341.4
D1 9928787 9928829
refGene_gn=CD1H11orf1;refGene_tx=XM_011286343.3;refGene_gn=CD1H11orf1;refGene_tx=XM_019811271.2
D1 9929915 9930047
refGene_gn=CD1H11orf1;refGene_tx=XM_011286342.3;refGene_gn=CD1H11orf1;refGene_tx=XM_019811270.2
D1 9930825 9931056

refGene_gn=CD1H11orf1;refGene_tx=XM_011286342.3;refGene_gn=CD1H11orf1;
refGene_tx=XM_011286343.3;refGene_gn=CD1H11orf1;refGene_tx=XM_01981127
0.2;refGene_gn=CD1H11orf1;refGene_tx=XM_019811271.2
D1 9931550 9931617
refGene_gn=CD1H11orf1;refGene_tx=XM_011286342.3;refGene_gn=CD1H11orf1;
refGene_tx=XM_011286343.3
D1 9931871 9932047
refGene_gn=CD1H11orf1;refGene_tx=XM_011286342.3;refGene_gn=CD1H11orf1;
refGene_tx=XM_011286343.3;refGene_gn=CD1H11orf1;refGene_tx=XM_01981127
0.2;refGene_gn=CD1H11orf1;refGene_tx=XM_019811271.2
D1 9945526 9945730 refGene_gn=CRYAB;refGene_tx=XM_011286345.3
D1 9947007 9947130 refGene_gn=CRYAB;refGene_tx=XM_011286345.3
D1 9948231 9948432 refGene_gn=CRYAB;refGene_tx=XM_011286345.3
D1 9949495 9949589 refGene_gn=HSPB2;refGene_tx=XM_003992345.5
D1 9950101 9950556 refGene_gn=HSPB2;refGene_tx=XM_003992345.5
D1 9954682 9954773
refGene_gn=CD1H11orf52;refGene_tx=XM_003992346.5;refGene_gn=CD1H11orf5
2;refGene_tx=XM_019811272.2
D1 9959243 9959281
refGene_gn=CD1H11orf52;refGene_tx=XM_003992346.5
D1 9960264 9960320
refGene_gn=CD1H11orf52;refGene_tx=XM_003992346.5;refGene_gn=CD1H11orf5
2;refGene_tx=XM_019811272.2
D1 9960543 9960792
refGene_gn=CD1H11orf52;refGene_tx=XM_003992346.5;refGene_gn=CD1H11orf5
2;refGene_tx=XM_019811272.2
D1 9963126 9963183 refGene_gn=DIXDC1;refGene_tx=XM_019811274.2
D1 9970959 9971019
refGene_gn=DIXDC1;refGene_tx=XM_011286346.3;refGene_gn=DIXDC1;refGene_
tx=XM_011286347.3;refGene_gn=DIXDC1;refGene_tx=XM_019811273.2;refGene_
gn=DIXDC1;refGene_tx=XM_019811275.2;refGene_gn=DIXDC1;refGene_tx=XM_01
9811278.2;refGene_gn=DIXDC1;refGene_tx=XM_019811279.2;refGene_gn=DIXDC
1;refGene_tx=XM_019811280.2;refGene_gn=DIXDC1;refGene_tx=XM_019811281.
2
D1 9994428 9994558
refGene_gn=DIXDC1;refGene_tx=XM_011286346.3;refGene_gn=DIXDC1;refGene_
tx=XM_011286347.3;refGene_gn=DIXDC1;refGene_tx=XM_019811273.2;refGene_
gn=DIXDC1;refGene_tx=XM_019811274.2;refGene_gn=DIXDC1;refGene_tx=XM_01
9811275.2;refGene_gn=DIXDC1;refGene_tx=XM_019811278.2;refGene_gn=DIXDC
1;refGene_tx=XM_019811279.2;refGene_gn=DIXDC1;refGene_tx=XM_019811280.
2;refGene_gn=DIXDC1;refGene_tx=XM_019811281.2
D1 9997845 9997971
refGene_gn=DIXDC1;refGene_tx=XM_011286346.3;refGene_gn=DIXDC1;refGene_
tx=XM_011286347.3;refGene_gn=DIXDC1;refGene_tx=XM_019811273.2;refGene_
gn=DIXDC1;refGene_tx=XM_019811274.2;refGene_gn=DIXDC1;refGene_tx=XM_01
9811275.2;refGene_gn=DIXDC1;refGene_tx=XM_019811276.2;refGene_gn=DIXDC
1;refGene_tx=XM_019811277.2;refGene_gn=DIXDC1;refGene_tx=XM_019811278.
2;refGene_gn=DIXDC1;refGene_tx=XM_019811279.2;refGene_gn=DIXDC1;refGen
e_tx=XM_019811280.2;refGene_gn=DIXDC1;refGene_tx=XM_019811281.2
D1 10000752 10000984

refGene_gn=DIXDC1;refGene_tx=XM_011286346.3;refGene_gn=DIXDC1;refGene_tx=XM_011286347.3;refGene_gn=DIXDC1;refGene_tx=XM_019811273.2;refGene_gn=DIXDC1;refGene_tx=XM_019811274.2;refGene_gn=DIXDC1;refGene_tx=XM_019811275.2;refGene_gn=DIXDC1;refGene_tx=XM_019811276.2;refGene_gn=DIXDC1;refGene_tx=XM_019811277.2;refGene_gn=DIXDC1;refGene_tx=XM_019811278.2;refGene_gn=DIXDC1;refGene_tx=XM_019811279.2;refGene_gn=DIXDC1;refGene_tx=XM_019811280.2;refGene_gn=DIXDC1;refGene_tx=XM_019811281.2
D1 10002523 10002631
refGene_gn=DIXDC1;refGene_tx=XM_011286346.3;refGene_gn=DIXDC1;refGene_tx=XM_011286347.3;refGene_gn=DIXDC1;refGene_tx=XM_019811273.2;refGene_gn=DIXDC1;refGene_tx=XM_019811274.2;refGene_gn=DIXDC1;refGene_tx=XM_019811275.2;refGene_gn=DIXDC1;refGene_tx=XM_019811276.2;refGene_gn=DIXDC1;refGene_tx=XM_019811277.2;refGene_gn=DIXDC1;refGene_tx=XM_019811278.2;refGene_gn=DIXDC1;refGene_tx=XM_019811279.2;refGene_gn=DIXDC1;refGene_tx=XM_019811280.2;refGene_gn=DIXDC1;refGene_tx=XM_019811281.2
D1 10005327 10005350 refGene_gn=DIXDC1;refGene_tx=XM_003992348.5
D1 10008023 10008136
refGene_gn=DIXDC1;refGene_tx=XM_003992348.5;refGene_gn=DIXDC1;refGene_tx=XM_011286346.3;refGene_gn=DIXDC1;refGene_tx=XM_011286347.3;refGene_gn=DIXDC1;refGene_tx=XM_019811273.2;refGene_gn=DIXDC1;refGene_tx=XM_019811274.2;refGene_gn=DIXDC1;refGene_tx=XM_019811275.2;refGene_gn=DIXDC1;refGene_tx=XM_019811276.2;refGene_gn=DIXDC1;refGene_tx=XM_019811277.2;refGene_gn=DIXDC1;refGene_tx=XM_019811278.2;refGene_gn=DIXDC1;refGene_tx=XM_019811279.2;refGene_gn=DIXDC1;refGene_tx=XM_019811280.2;refGene_gn=DIXDC1;refGene_tx=XM_019811281.2
D1 10009624 10009773
refGene_gn=DIXDC1;refGene_tx=XM_003992348.5;refGene_gn=DIXDC1;refGene_tx=XM_011286346.3;refGene_gn=DIXDC1;refGene_tx=XM_011286347.3;refGene_gn=DIXDC1;refGene_tx=XM_019811273.2;refGene_gn=DIXDC1;refGene_tx=XM_019811274.2;refGene_gn=DIXDC1;refGene_tx=XM_019811275.2;refGene_gn=DIXDC1;refGene_tx=XM_019811276.2;refGene_gn=DIXDC1;refGene_tx=XM_019811277.2;refGene_gn=DIXDC1;refGene_tx=XM_019811278.2;refGene_gn=DIXDC1;refGene_tx=XM_019811279.2;refGene_gn=DIXDC1;refGene_tx=XM_019811280.2;refGene_gn=DIXDC1;refGene_tx=XM_019811281.2
D1 10015570 10015660
refGene_gn=DIXDC1;refGene_tx=XM_003992348.5;refGene_gn=DIXDC1;refGene_tx=XM_011286346.3;refGene_gn=DIXDC1;refGene_tx=XM_011286347.3;refGene_gn=DIXDC1;refGene_tx=XM_019811273.2;refGene_gn=DIXDC1;refGene_tx=XM_019811274.2;refGene_gn=DIXDC1;refGene_tx=XM_019811275.2;refGene_gn=DIXDC1;refGene_tx=XM_019811276.2;refGene_gn=DIXDC1;refGene_tx=XM_019811277.2;refGene_gn=DIXDC1;refGene_tx=XM_019811278.2;refGene_gn=DIXDC1;refGene_tx=XM_019811279.2;refGene_gn=DIXDC1;refGene_tx=XM_019811280.2;refGene_gn=DIXDC1;refGene_tx=XM_019811281.2
D1 10016818 10016872
refGene_gn=DIXDC1;refGene_tx=XM_003992348.5;refGene_gn=DIXDC1;refGene_tx=XM_011286346.3;refGene_gn=DIXDC1;refGene_tx=XM_011286347.3;refGene_gn=DIXDC1;refGene_tx=XM_019811273.2;refGene_gn=DIXDC1;refGene_tx=XM_019811274.2;refGene_gn=DIXDC1;refGene_tx=XM_019811275.2;refGene_gn=DIXDC1;refGene_tx=XM_019811276.2;refGene_gn=DIXDC1;refGene_tx=XM_019811277.2;refGene_gn=DIXDC1;refGene_tx=XM_019811278.2;refGene_gn=DIXDC1;refGen

e_tx=XM_019811279.2;refGene_gn=DIXDC1;refGene_tx=XM_019811280.2;refGene_gn=DIXDC1;refGene_tx=XM_019811281.2
D1 10018416 10018467
refGene_gn=DIXDC1;refGene_tx=XM_003992348.5;refGene_gn=DIXDC1;refGene_tx=XM_011286346.3;refGene_gn=DIXDC1;refGene_tx=XM_011286347.3;refGene_gn=DIXDC1;refGene_tx=XM_019811273.2;refGene_gn=DIXDC1;refGene_tx=XM_019811274.2;refGene_gn=DIXDC1;refGene_tx=XM_019811275.2;refGene_gn=DIXDC1;refGene_tx=XM_019811276.2;refGene_gn=DIXDC1;refGene_tx=XM_019811277.2;refGene_gn=DIXDC1;refGene_tx=XM_019811278.2;refGene_gn=DIXDC1;refGene_tx=XM_019811279.2;refGene_gn=DIXDC1;refGene_tx=XM_019811280.2;refGene_gn=DIXDC1;refGene_tx=XM_019811281.2
D1 10021293 10021398
refGene_gn=DIXDC1;refGene_tx=XM_003992348.5;refGene_gn=DIXDC1;refGene_tx=XM_011286346.3;refGene_gn=DIXDC1;refGene_tx=XM_011286347.3;refGene_gn=DIXDC1;refGene_tx=XM_019811273.2;refGene_gn=DIXDC1;refGene_tx=XM_019811274.2;refGene_gn=DIXDC1;refGene_tx=XM_019811275.2;refGene_gn=DIXDC1;refGene_tx=XM_019811276.2;refGene_gn=DIXDC1;refGene_tx=XM_019811277.2;refGene_gn=DIXDC1;refGene_tx=XM_019811278.2;refGene_gn=DIXDC1;refGene_tx=XM_019811279.2;refGene_gn=DIXDC1;refGene_tx=XM_019811280.2;refGene_gn=DIXDC1;refGene_tx=XM_019811281.2
D1 10021822 10021876
refGene_gn=DIXDC1;refGene_tx=XM_003992348.5;refGene_gn=DIXDC1;refGene_tx=XM_011286346.3;refGene_gn=DIXDC1;refGene_tx=XM_011286347.3;refGene_gn=DIXDC1;refGene_tx=XM_019811273.2;refGene_gn=DIXDC1;refGene_tx=XM_019811274.2;refGene_gn=DIXDC1;refGene_tx=XM_019811275.2;refGene_gn=DIXDC1;refGene_tx=XM_019811276.2;refGene_gn=DIXDC1;refGene_tx=XM_019811277.2;refGene_gn=DIXDC1;refGene_tx=XM_019811278.2;refGene_gn=DIXDC1;refGene_tx=XM_019811279.2;refGene_gn=DIXDC1;refGene_tx=XM_019811280.2;refGene_gn=DIXDC1;refGene_tx=XM_019811281.2
D1 10022396 10022489
refGene_gn=DIXDC1;refGene_tx=XM_003992348.5;refGene_gn=DIXDC1;refGene_tx=XM_011286346.3;refGene_gn=DIXDC1;refGene_tx=XM_011286347.3;refGene_gn=DIXDC1;refGene_tx=XM_019811273.2;refGene_gn=DIXDC1;refGene_tx=XM_019811274.2;refGene_gn=DIXDC1;refGene_tx=XM_019811275.2;refGene_gn=DIXDC1;refGene_tx=XM_019811276.2;refGene_gn=DIXDC1;refGene_tx=XM_019811277.2;refGene_gn=DIXDC1;refGene_tx=XM_019811278.2;refGene_gn=DIXDC1;refGene_tx=XM_019811279.2
D1 10022568 10022640
refGene_gn=DIXDC1;refGene_tx=XM_003992348.5;refGene_gn=DIXDC1;refGene_tx=XM_011286346.3;refGene_gn=DIXDC1;refGene_tx=XM_011286347.3;refGene_gn=DIXDC1;refGene_tx=XM_019811273.2;refGene_gn=DIXDC1;refGene_tx=XM_019811274.2;refGene_gn=DIXDC1;refGene_tx=XM_019811275.2;refGene_gn=DIXDC1;refGene_tx=XM_019811276.2;refGene_gn=DIXDC1;refGene_tx=XM_019811277.2;refGene_gn=DIXDC1;refGene_tx=XM_019811278.2;refGene_gn=DIXDC1;refGene_tx=XM_019811279.2
D1 10023433 10023523
refGene_gn=DIXDC1;refGene_tx=XM_003992348.5;refGene_gn=DIXDC1;refGene_tx=XM_011286346.3;refGene_gn=DIXDC1;refGene_tx=XM_011286347.3;refGene_gn=DIXDC1;refGene_tx=XM_019811273.2;refGene_gn=DIXDC1;refGene_tx=XM_019811274.2;refGene_gn=DIXDC1;refGene_tx=XM_019811275.2;refGene_gn=DIXDC

1;refGene_tx=XM_019811276.2;refGene_gn=DIXDC1;refGene_tx=XM_019811277.2;refGene_gn=DIXDC1;refGene_tx=XM_019811278.2;refGene_gn=DIXDC1;refGene_tx=XM_019811279.2
D1 10023846 10024008 refGene_gn=DIXDC1;refGene_tx=XM_003992348.5;refGene_gn=DIXDC1;refGene_tx=XM_011286346.3;refGene_gn=DIXDC1;refGene_tx=XM_011286347.3;refGene_gn=DIXDC1;refGene_tx=XM_019811273.2;refGene_gn=DIXDC1;refGene_tx=XM_019811274.2;refGene_gn=DIXDC1;refGene_tx=XM_019811275.2;refGene_gn=DIXDC1;refGene_tx=XM_019811276.2;refGene_gn=DIXDC1;refGene_tx=XM_019811277.2;refGene_gn=DIXDC1;refGene_tx=XM_019811278.2;refGene_gn=DIXDC1;refGene_tx=XM_019811279.2
D1 10024500 10024588 refGene_gn=DIXDC1;refGene_tx=XM_003992348.5;refGene_gn=DIXDC1;refGene_tx=XM_011286346.3;refGene_gn=DIXDC1;refGene_tx=XM_011286347.3;refGene_gn=DIXDC1;refGene_tx=XM_019811273.2;refGene_gn=DIXDC1;refGene_tx=XM_019811274.2;refGene_gn=DIXDC1;refGene_tx=XM_019811275.2;refGene_gn=DIXDC1;refGene_tx=XM_019811276.2;refGene_gn=DIXDC1;refGene_tx=XM_019811277.2;refGene_gn=DIXDC1;refGene_tx=XM_019811278.2;refGene_gn=DIXDC1;refGene_tx=XM_019811279.2;refGene_gn=DIXDC1;refGene_tx=XM_019811280.2;refGene_gn=DIXDC1;refGene_tx=XM_019811281.2
D1 10029579 10029685 refGene_gn=DIXDC1;refGene_tx=XM_003992348.5;refGene_gn=DIXDC1;refGene_tx=XM_011286346.3;refGene_gn=DIXDC1;refGene_tx=XM_011286347.3;refGene_gn=DIXDC1;refGene_tx=XM_019811273.2;refGene_gn=DIXDC1;refGene_tx=XM_019811274.2;refGene_gn=DIXDC1;refGene_tx=XM_019811275.2;refGene_gn=DIXDC1;refGene_tx=XM_019811276.2;refGene_gn=DIXDC1;refGene_tx=XM_019811277.2;refGene_gn=DIXDC1;refGene_tx=XM_019811280.2;refGene_gn=DIXDC1;refGene_tx=XM_019811281.2
D1 10030633 10030742 refGene_gn=DIXDC1;refGene_tx=XM_003992348.5;refGene_gn=DIXDC1;refGene_tx=XM_011286346.3;refGene_gn=DIXDC1;refGene_tx=XM_011286347.3;refGene_gn=DIXDC1;refGene_tx=XM_019811273.2;refGene_gn=DIXDC1;refGene_tx=XM_019811274.2;refGene_gn=DIXDC1;refGene_tx=XM_019811276.2;refGene_gn=DIXDC1;refGene_tx=XM_019811277.2;refGene_gn=DIXDC1;refGene_tx=XM_019811280.2;refGene_gn=DIXDC1;refGene_tx=XM_019811281.2
D1 10031837 10031918 refGene_gn=DIXDC1;refGene_tx=XM_003992348.5;refGene_gn=DIXDC1;refGene_tx=XM_011286346.3;refGene_gn=DIXDC1;refGene_tx=XM_011286347.3;refGene_gn=DIXDC1;refGene_tx=XM_019811273.2;refGene_gn=DIXDC1;refGene_tx=XM_019811274.2;refGene_gn=DIXDC1;refGene_tx=XM_019811275.2;refGene_gn=DIXDC1;refGene_tx=XM_019811276.2;refGene_gn=DIXDC1;refGene_tx=XM_019811277.2;refGene_gn=DIXDC1;refGene_tx=XM_019811278.2;refGene_gn=DIXDC1;refGene_tx=XM_019811279.2;refGene_gn=DIXDC1;refGene_tx=XM_019811280.2;refGene_gn=DIXDC1;refGene_tx=XM_019811281.2
D1 10037455 10037734 refGene_gn=DLAT;refGene_tx=XM_003992349.5
D1 10038209 10038311 refGene_gn=DLAT;refGene_tx=XM_003992349.5
D1 10038639 10038764 refGene_gn=DLAT;refGene_tx=XM_003992349.5
D1 10038922 10039076 refGene_gn=DLAT;refGene_tx=XM_003992349.5
D1 10041890 10042017 refGene_gn=DLAT;refGene_tx=XM_003992349.5
D1 10043765 10043953 refGene_gn=DLAT;refGene_tx=XM_003992349.5

D1 10045437 10045591 refGene_gn=DLAT;refGene_tx=XM_003992349.5
D1 10053369 10053437 refGene_gn=DLAT;refGene_tx=XM_003992349.5
D1 10054709 10054802 refGene_gn=DLAT;refGene_tx=XM_003992349.5
D1 10055156 10055264 refGene_gn=DLAT;refGene_tx=XM_003992349.5
D1 10059277 10059393 refGene_gn=DLAT;refGene_tx=XM_003992349.5
D1 10063513 10063676 refGene_gn=DLAT;refGene_tx=XM_003992349.5
D1 10064184 10064321 refGene_gn=DLAT;refGene_tx=XM_003992349.5
D1 10064888 10065018 refGene_gn=DLAT;refGene_tx=XM_003992349.5
D1 10068818 10068953
refGene_gn=PIH1D2;refGene_tx=XM_006936623.3;refGene_gn=PIH1D2;refGene_tx=XM_006936624.3;refGene_gn=PIH1D2;refGene_tx=XM_006936625.3
D1 10070182 10070442
refGene_gn=PIH1D2;refGene_tx=XM_006936623.3;refGene_gn=PIH1D2;refGene_tx=XM_006936624.3;refGene_gn=PIH1D2;refGene_tx=XM_006936625.3
D1 10070781 10071027
refGene_gn=PIH1D2;refGene_tx=XM_006936623.3;refGene_gn=PIH1D2;refGene_tx=XM_006936624.3;refGene_gn=PIH1D2;refGene_tx=XM_006936625.3
D1 10071390 10071514
refGene_gn=PIH1D2;refGene_tx=XM_006936623.3;refGene_gn=PIH1D2;refGene_tx=XM_006936624.3;refGene_gn=PIH1D2;refGene_tx=XM_006936625.3
D1 10072613 10072865
refGene_gn=PIH1D2;refGene_tx=XM_006936623.3;refGene_gn=PIH1D2;refGene_tx=XM_006936624.3;refGene_gn=PIH1D2;refGene_tx=XM_006936625.3
D1 10083170 10083338 refGene_gn=TIMM8B;refGene_tx=XM_003992353.3
D1 10084709 10084838 refGene_gn=TIMM8B;refGene_tx=XM_003992353.3
D1 10084979 10085031
refGene_gn=SDHD;refGene_tx=XM_003992354.4;refGene_gn=SDHD;refGene_tx=XM_023239096.1
D1 10088403 10088520
refGene_gn=SDHD;refGene_tx=XM_003992354.4;refGene_gn=SDHD;refGene_tx=XM_023239096.1
D1 10089948 10090093 refGene_gn=SDHD;refGene_tx=XM_003992354.4
D1 10095024 10095190
refGene_gn=SDHD;refGene_tx=XM_003992354.4;refGene_gn=SDHD;refGene_tx=XM_023239096.1
D1 10129151 10129373
refGene_gn=IL18;refGene_tx=NM_001009213.2;refGene_gn=IL18;refGene_tx=XM_006936584.3;refGene_gn=IL18;refGene_tx=XM_019811172.2;refGene_gn=IL18;refGene_tx=XM_019811173.1
D1 10132779 10132913
refGene_gn=IL18;refGene_tx=NM_001009213.2;refGene_gn=IL18;refGene_tx=XM_006936584.3;refGene_gn=IL18;refGene_tx=XM_019811172.2;refGene_gn=IL18;refGene_tx=XM_019811173.1
D1 10133723 10133858
refGene_gn=IL18;refGene_tx=NM_001009213.2;refGene_gn=IL18;refGene_tx=XM_006936584.3;refGene_gn=IL18;refGene_tx=XM_019811172.2;refGene_gn=IL18;refGene_tx=XM_019811173.1
D1 10136709 10136878 refGene_gn=IL18;refGene_tx=XM_019811172.2
D1 10137269 10137281
refGene_gn=IL18;refGene_tx=NM_001009213.2;refGene_gn=IL18;refGene_tx=X

M_006936584.3;refGene_gn=IL18;refGene_tx=XM_019811173.1
D1	10138103	10138179
refGene_gn=IL18;refGene_tx=NM_001009213.2;refGene_gn=IL18;refGene_tx=XM_006936584.3;refGene_gn=IL18;refGene_tx=XM_019811173.1
D1	10153013	10153076
refGene_gn=TEX12;refGene_tx=XM_003992355.4;refGene_gn=TEX12;refGene_tx=XM_006936629.4;refGene_gn=TEX12;refGene_tx=XM_019811286.2;refGene_gn=TEX12;refGene_tx=XM_019811287.2;refGene_gn=TEX12;refGene_tx=XM_019811288.2;refGene_gn=TEX12;refGene_tx=XM_023239097.1
D1	10153863	10153975
refGene_gn=TEX12;refGene_tx=XM_003992355.4;refGene_gn=TEX12;refGene_tx=XM_006936629.4;refGene_gn=TEX12;refGene_tx=XM_019811286.2;refGene_gn=TEX12;refGene_tx=XM_019811287.2;refGene_gn=TEX12;refGene_tx=XM_019811288.2;refGene_gn=TEX12;refGene_tx=XM_023239097.1
D1	10154066	10154118
refGene_gn=TEX12;refGene_tx=XM_003992355.4;refGene_gn=TEX12;refGene_tx=XM_006936629.4;refGene_gn=TEX12;refGene_tx=XM_019811286.2;refGene_gn=TEX12;refGene_tx=XM_019811287.2;refGene_gn=TEX12;refGene_tx=XM_019811288.2;refGene_gn=TEX12;refGene_tx=XM_023239097.1
D1	10154885	10155030
refGene_gn=TEX12;refGene_tx=XM_003992355.4;refGene_gn=TEX12;refGene_tx=XM_006936629.4;refGene_gn=TEX12;refGene_tx=XM_019811286.2;refGene_gn=TEX12;refGene_tx=XM_019811287.2;refGene_gn=TEX12;refGene_tx=XM_019811288.2;refGene_gn=TEX12;refGene_tx=XM_023239097.1
D1	10158350	10158438
refGene_gn=BCO2;refGene_tx=XM_003992357.5;refGene_gn=BCO2;refGene_tx=XM_023239094.1
D1	10163977	10164182
refGene_gn=BCO2;refGene_tx=XM_003992357.5;refGene_gn=BCO2;refGene_tx=XM_023239094.1
D1	10176138	10176350
refGene_gn=BCO2;refGene_tx=XM_003992357.5;refGene_gn=BCO2;refGene_tx=XM_023239094.1
D1	10176556	10176672
refGene_gn=BCO2;refGene_tx=XM_003992357.5;refGene_gn=BCO2;refGene_tx=XM_023239094.1
D1	10177575	10177678
refGene_gn=BCO2;refGene_tx=XM_003992357.5;refGene_gn=BCO2;refGene_tx=XM_023239094.1
D1	10183379	10183508
refGene_gn=BCO2;refGene_tx=XM_003992357.5;refGene_gn=BCO2;refGene_tx=XM_023239094.1
D1	10184483	10184644
refGene_gn=BCO2;refGene_tx=XM_003992357.5;refGene_gn=BCO2;refGene_tx=XM_023239094.1
D1	10185785	10185953
refGene_gn=BCO2;refGene_tx=XM_003992357.5;refGene_gn=BCO2;refGene_tx=XM_023239094.1
D1	10200630	10200768
refGene_gn=BCO2;refGene_tx=XM_003992357.5;refGene_gn=BCO2;refGene_tx=X

M_023239094.1
D1 10201336 10201519
refGene_gn=BCO2;refGene_tx=XM_003992357.5;refGene_gn=BCO2;refGene_tx=X
M_023239094.1
D1 10202323 10202434
refGene_gn=BCO2;refGene_tx=XM_003992357.5;refGene_gn=BCO2;refGene_tx=X
M_023239094.1
D1 10203967 10204081 refGene_gn=BCO2;refGene_tx=XM_003992357.5
D1 10207974 10208063 refGene_gn=PTS;refGene_tx=XM_003992359.5
D1 10210010 10210090 refGene_gn=PTS;refGene_tx=XM_003992359.5
D1 10211284 10211307 refGene_gn=PTS;refGene_tx=XM_003992359.5
D1 10211671 10211728 refGene_gn=PTS;refGene_tx=XM_003992359.5
D1 10213766 10213837 refGene_gn=PTS;refGene_tx=XM_003992359.5
D1 10214030 10214154 refGene_gn=PTS;refGene_tx=XM_003992359.5
D1 10227192 10227485 refGene_gn=PLET1;refGene_tx=XM_019811289.2
D1 10229425 10229487 refGene_gn=PLET1;refGene_tx=XM_019811289.2
D1 10232356 10232558 refGene_gn=PLET1;refGene_tx=XM_019811289.2
D1 10237827 10238107 refGene_gn=PLET1;refGene_tx=XM_019811289.2
D1 10908241 10908293
refGene_gn=NCAM1;refGene_tx=NM_001009869.1;refGene_gn=NCAM1;refGene_tx
=XM_006936585.4;refGene_gn=NCAM1;refGene_tx=XM_006936586.4;refGene_gn=
NCAM1;refGene_tx=XM_006936587.4;refGene_gn=NCAM1;refGene_tx=XM_0069365
88.4;refGene_gn=NCAM1;refGene_tx=XM_006936589.4;refGene_gn=NCAM1;refGe
ne_tx=XM_006936591.3;refGene_gn=NCAM1;refGene_tx=XM_006936592.3;refGen
e_gn=NCAM1;refGene_tx=XM_006936593.3;refGene_gn=NCAM1;refGene_tx=XM_00
6936594.4;refGene_gn=NCAM1;refGene_tx=XM_006936595.3;refGene_gn=NCAM1;
refGene_tx=XM_011286326.3;refGene_gn=NCAM1;refGene_tx=XM_023238914.1;r
efGene_gn=NCAM1;refGene_tx=XM_023238915.1;refGene_gn=NCAM1;refGene_tx=
XM_023238917.1;refGene_gn=NCAM1;refGene_tx=XM_023238918.1;refGene_gn=N
CAM1;refGene_tx=XM_023238919.1;refGene_gn=NCAM1;refGene_tx=XM_02323892
0.1;refGene_gn=NCAM1;refGene_tx=XM_023238921.1;refGene_gn=NCAM1;refGen
e_tx=XM_023238922.1;refGene_gn=NCAM1;refGene_tx=XM_023238923.1;refGene
_gn=NCAM1;refGene_tx=XM_023238924.1
D1 11155516 11155591
refGene_gn=NCAM1;refGene_tx=NM_001009869.1;refGene_gn=NCAM1;refGene_tx
=XM_006936585.4;refGene_gn=NCAM1;refGene_tx=XM_006936586.4;refGene_gn=
NCAM1;refGene_tx=XM_006936587.4;refGene_gn=NCAM1;refGene_tx=XM_0069365
88.4;refGene_gn=NCAM1;refGene_tx=XM_006936589.4;refGene_gn=NCAM1;refGe
ne_tx=XM_006936591.3;refGene_gn=NCAM1;refGene_tx=XM_006936592.3;refGen
e_gn=NCAM1;refGene_tx=XM_006936593.3;refGene_gn=NCAM1;refGene_tx=XM_00
6936594.4;refGene_gn=NCAM1;refGene_tx=XM_006936595.3;refGene_gn=NCAM1;
refGene_tx=XM_011286326.3;refGene_gn=NCAM1;refGene_tx=XM_023238914.1;r
efGene_gn=NCAM1;refGene_tx=XM_023238915.1;refGene_gn=NCAM1;refGene_tx=
XM_023238917.1;refGene_gn=NCAM1;refGene_tx=XM_023238918.1;refGene_gn=N
CAM1;refGene_tx=XM_023238919.1;refGene_gn=NCAM1;refGene_tx=XM_02323892
0.1;refGene_gn=NCAM1;refGene_tx=XM_023238921.1;refGene_gn=NCAM1;refGen
e_tx=XM_023238922.1;refGene_gn=NCAM1;refGene_tx=XM_023238923.1;refGene
_gn=NCAM1;refGene_tx=XM_023238924.1
D1 11157282 11157501
refGene_gn=NCAM1;refGene_tx=NM_001009869.1;refGene_gn=NCAM1;refGene_tx

=XM_006936585.4;refGene_gn=NCAM1;refGene_tx=XM_006936586.4;refGene_gn=NCAM1;refGene_tx=XM_006936587.4;refGene_gn=NCAM1;refGene_tx=XM_006936588.4;refGene_gn=NCAM1;refGene_tx=XM_006936589.4;refGene_gn=NCAM1;refGene_tx=XM_006936591.3;refGene_gn=NCAM1;refGene_tx=XM_006936592.3;refGene_gn=NCAM1;refGene_tx=XM_006936593.3;refGene_gn=NCAM1;refGene_tx=XM_006936594.4;refGene_gn=NCAM1;refGene_tx=XM_006936595.3;refGene_gn=NCAM1;refGene_tx=XM_011286326.3;refGene_gn=NCAM1;refGene_tx=XM_023238914.1;refGene_gn=NCAM1;refGene_tx=XM_023238915.1;refGene_gn=NCAM1;refGene_tx=XM_023238917.1;refGene_gn=NCAM1;refGene_tx=XM_023238918.1;refGene_gn=NCAM1;refGene_tx=XM_023238919.1;refGene_gn=NCAM1;refGene_tx=XM_023238920.1;refGene_gn=NCAM1;refGene_tx=XM_023238921.1;refGene_gn=NCAM1;refGene_tx=XM_023238922.1;refGene_gn=NCAM1;refGene_tx=XM_023238923.1;refGene_gn=NCAM1;refGene_tx=XM_023238924.1
D1 11158557 11158701
refGene_gn=NCAM1;refGene_tx=NM_001009869.1;refGene_gn=NCAM1;refGene_tx=XM_006936585.4;refGene_gn=NCAM1;refGene_tx=XM_006936586.4;refGene_gn=NCAM1;refGene_tx=XM_006936587.4;refGene_gn=NCAM1;refGene_tx=XM_006936588.4;refGene_gn=NCAM1;refGene_tx=XM_006936589.4;refGene_gn=NCAM1;refGene_tx=XM_006936591.3;refGene_gn=NCAM1;refGene_tx=XM_006936592.3;refGene_gn=NCAM1;refGene_tx=XM_006936593.3;refGene_gn=NCAM1;refGene_tx=XM_006936594.4;refGene_gn=NCAM1;refGene_tx=XM_006936595.3;refGene_gn=NCAM1;refGene_tx=XM_011286326.3;refGene_gn=NCAM1;refGene_tx=XM_023238914.1;refGene_gn=NCAM1;refGene_tx=XM_023238915.1;refGene_gn=NCAM1;refGene_tx=XM_023238917.1;refGene_gn=NCAM1;refGene_tx=XM_023238918.1;refGene_gn=NCAM1;refGene_tx=XM_023238919.1;refGene_gn=NCAM1;refGene_tx=XM_023238920.1;refGene_gn=NCAM1;refGene_tx=XM_023238921.1;refGene_gn=NCAM1;refGene_tx=XM_023238922.1;refGene_gn=NCAM1;refGene_tx=XM_023238923.1;refGene_gn=NCAM1;refGene_tx=XM_023238924.1
D1 11159088 11159226
refGene_gn=NCAM1;refGene_tx=NM_001009869.1;refGene_gn=NCAM1;refGene_tx=XM_006936585.4;refGene_gn=NCAM1;refGene_tx=XM_006936586.4;refGene_gn=NCAM1;refGene_tx=XM_006936587.4;refGene_gn=NCAM1;refGene_tx=XM_006936588.4;refGene_gn=NCAM1;refGene_tx=XM_006936589.4;refGene_gn=NCAM1;refGene_tx=XM_006936591.3;refGene_gn=NCAM1;refGene_tx=XM_006936592.3;refGene_gn=NCAM1;refGene_tx=XM_006936593.3;refGene_gn=NCAM1;refGene_tx=XM_006936594.4;refGene_gn=NCAM1;refGene_tx=XM_006936595.3;refGene_gn=NCAM1;refGene_tx=XM_011286326.3;refGene_gn=NCAM1;refGene_tx=XM_023238914.1;refGene_gn=NCAM1;refGene_tx=XM_023238915.1;refGene_gn=NCAM1;refGene_tx=XM_023238917.1;refGene_gn=NCAM1;refGene_tx=XM_023238918.1;refGene_gn=NCAM1;refGene_tx=XM_023238919.1;refGene_gn=NCAM1;refGene_tx=XM_023238920.1;refGene_gn=NCAM1;refGene_tx=XM_023238921.1;refGene_gn=NCAM1;refGene_tx=XM_023238922.1;refGene_gn=NCAM1;refGene_tx=XM_023238923.1;refGene_gn=NCAM1;refGene_tx=XM_023238924.1
D1 11160392 11160510
refGene_gn=NCAM1;refGene_tx=NM_001009869.1;refGene_gn=NCAM1;refGene_tx=XM_006936585.4;refGene_gn=NCAM1;refGene_tx=XM_006936586.4;refGene_gn=NCAM1;refGene_tx=XM_006936587.4;refGene_gn=NCAM1;refGene_tx=XM_006936588.4;refGene_gn=NCAM1;refGene_tx=XM_006936589.4;refGene_gn=NCAM1;refGene_tx=XM_006936591.3;refGene_gn=NCAM1;refGene_tx=XM_006936592.3;refGene_gn=NCAM1;refGene_tx=XM_006936593.3;refGene_gn=NCAM1;refGene_tx=XM_00

6936594.4;refGene_gn=NCAM1;refGene_tx=XM_006936595.3;refGene_gn=NCAM1;
refGene_tx=XM_011286326.3;refGene_gn=NCAM1;refGene_tx=XM_023238914.1;r
efGene_gn=NCAM1;refGene_tx=XM_023238915.1;refGene_gn=NCAM1;refGene_tx=
XM_023238917.1;refGene_gn=NCAM1;refGene_tx=XM_023238918.1;refGene_gn=N
CAM1;refGene_tx=XM_023238919.1;refGene_gn=NCAM1;refGene_tx=XM_02323892
0.1;refGene_gn=NCAM1;refGene_tx=XM_023238921.1;refGene_gn=NCAM1;refGen
e_tx=XM_023238922.1;refGene_gn=NCAM1;refGene_tx=XM_023238923.1;refGene
_gn=NCAM1;refGene_tx=XM_023238924.1
D1 11160955 11161122
refGene_gn=NCAM1;refGene_tx=NM_001009869.1;refGene_gn=NCAM1;refGene_tx
=XM_006936585.4;refGene_gn=NCAM1;refGene_tx=XM_006936586.4;refGene_gn=
NCAM1;refGene_tx=XM_006936587.4;refGene_gn=NCAM1;refGene_tx=XM_0069365
88.4;refGene_gn=NCAM1;refGene_tx=XM_006936589.4;refGene_gn=NCAM1;refGe
ne_tx=XM_006936591.3;refGene_gn=NCAM1;refGene_tx=XM_006936592.3;refGen
e_gn=NCAM1;refGene_tx=XM_006936593.3;refGene_gn=NCAM1;refGene_tx=XM_00
6936594.4;refGene_gn=NCAM1;refGene_tx=XM_006936595.3;refGene_gn=NCAM1;
refGene_tx=XM_011286326.3;refGene_gn=NCAM1;refGene_tx=XM_023238914.1;r
efGene_gn=NCAM1;refGene_tx=XM_023238915.1;refGene_gn=NCAM1;refGene_tx=
XM_023238917.1;refGene_gn=NCAM1;refGene_tx=XM_023238918.1;refGene_gn=N
CAM1;refGene_tx=XM_023238919.1;refGene_gn=NCAM1;refGene_tx=XM_02323892
0.1;refGene_gn=NCAM1;refGene_tx=XM_023238921.1;refGene_gn=NCAM1;refGen
e_tx=XM_023238922.1;refGene_gn=NCAM1;refGene_tx=XM_023238923.1;refGene
_gn=NCAM1;refGene_tx=XM_023238924.1
D1 11168562 11168705
refGene_gn=NCAM1;refGene_tx=NM_001009869.1;refGene_gn=NCAM1;refGene_tx
=XM_006936585.4;refGene_gn=NCAM1;refGene_tx=XM_006936586.4;refGene_gn=
NCAM1;refGene_tx=XM_006936587.4;refGene_gn=NCAM1;refGene_tx=XM_0069365
88.4;refGene_gn=NCAM1;refGene_tx=XM_006936589.4;refGene_gn=NCAM1;refGe
ne_tx=XM_006936591.3;refGene_gn=NCAM1;refGene_tx=XM_006936592.3;refGen
e_gn=NCAM1;refGene_tx=XM_006936593.3;refGene_gn=NCAM1;refGene_tx=XM_00
6936594.4;refGene_gn=NCAM1;refGene_tx=XM_006936595.3;refGene_gn=NCAM1;
refGene_tx=XM_011286326.3;refGene_gn=NCAM1;refGene_tx=XM_023238914.1;r
efGene_gn=NCAM1;refGene_tx=XM_023238915.1;refGene_gn=NCAM1;refGene_tx=
XM_023238917.1;refGene_gn=NCAM1;refGene_tx=XM_023238918.1;refGene_gn=N
CAM1;refGene_tx=XM_023238919.1;refGene_gn=NCAM1;refGene_tx=XM_02323892
0.1;refGene_gn=NCAM1;refGene_tx=XM_023238921.1;refGene_gn=NCAM1;refGen
e_tx=XM_023238922.1;refGene_gn=NCAM1;refGene_tx=XM_023238923.1;refGene
_gn=NCAM1;refGene_tx=XM_023238924.1
D1 11175155 11175185
refGene_gn=NCAM1;refGene_tx=XM_006936585.4;refGene_gn=NCAM1;refGene_tx
=XM_006936586.4;refGene_gn=NCAM1;refGene_tx=XM_006936587.4;refGene_gn=
NCAM1;refGene_tx=XM_006936588.4;refGene_gn=NCAM1;refGene_tx=XM_0069365
91.3;refGene_gn=NCAM1;refGene_tx=XM_006936592.3;refGene_gn=NCAM1;refGe
ne_tx=XM_023238914.1;refGene_gn=NCAM1;refGene_tx=XM_023238915.1;refGen
e_gn=NCAM1;refGene_tx=XM_023238917.1;refGene_gn=NCAM1;refGene_tx=XM_02
3238919.1;refGene_gn=NCAM1;refGene_tx=XM_023238920.1;refGene_gn=NCAM1;
refGene_tx=XM_023238921.1;refGene_gn=NCAM1;refGene_tx=XM_023238922.1;r
efGene_gn=NCAM1;refGene_tx=XM_023238923.1
D1 11179337 11179488
refGene_gn=NCAM1;refGene_tx=NM_001009869.1;refGene_gn=NCAM1;refGene_tx

=XM_006936585.4;refGene_gn=NCAM1;refGene_tx=XM_006936586.4;refGene_gn=NCAM1;refGene_tx=XM_006936587.4;refGene_gn=NCAM1;refGene_tx=XM_006936588.4;refGene_gn=NCAM1;refGene_tx=XM_006936589.4;refGene_gn=NCAM1;refGene_tx=XM_006936591.3;refGene_gn=NCAM1;refGene_tx=XM_006936592.3;refGene_gn=NCAM1;refGene_tx=XM_006936593.3;refGene_gn=NCAM1;refGene_tx=XM_006936594.4;refGene_gn=NCAM1;refGene_tx=XM_006936595.3;refGene_gn=NCAM1;refGene_tx=XM_011286326.3;refGene_gn=NCAM1;refGene_tx=XM_023238914.1;refGene_gn=NCAM1;refGene_tx=XM_023238915.1;refGene_gn=NCAM1;refGene_tx=XM_023238917.1;refGene_gn=NCAM1;refGene_tx=XM_023238918.1;refGene_gn=NCAM1;refGene_tx=XM_023238919.1;refGene_gn=NCAM1;refGene_tx=XM_023238920.1;refGene_gn=NCAM1;refGene_tx=XM_023238921.1;refGene_gn=NCAM1;refGene_tx=XM_023238922.1;refGene_gn=NCAM1;refGene_tx=XM_023238923.1;refGene_gn=NCAM1;refGene_tx=XM_023238924.1
D1 11179822 11180007
refGene_gn=NCAM1;refGene_tx=NM_001009869.1;refGene_gn=NCAM1;refGene_tx=XM_006936585.4;refGene_gn=NCAM1;refGene_tx=XM_006936586.4;refGene_gn=NCAM1;refGene_tx=XM_006936587.4;refGene_gn=NCAM1;refGene_tx=XM_006936588.4;refGene_gn=NCAM1;refGene_tx=XM_006936589.4;refGene_gn=NCAM1;refGene_tx=XM_006936591.3;refGene_gn=NCAM1;refGene_tx=XM_006936592.3;refGene_gn=NCAM1;refGene_tx=XM_006936593.3;refGene_gn=NCAM1;refGene_tx=XM_006936594.4;refGene_gn=NCAM1;refGene_tx=XM_006936595.3;refGene_gn=NCAM1;refGene_tx=XM_011286326.3;refGene_gn=NCAM1;refGene_tx=XM_023238914.1;refGene_gn=NCAM1;refGene_tx=XM_023238915.1;refGene_gn=NCAM1;refGene_tx=XM_023238917.1;refGene_gn=NCAM1;refGene_tx=XM_023238918.1;refGene_gn=NCAM1;refGene_tx=XM_023238919.1;refGene_gn=NCAM1;refGene_tx=XM_023238920.1;refGene_gn=NCAM1;refGene_tx=XM_023238921.1;refGene_gn=NCAM1;refGene_tx=XM_023238922.1;refGene_gn=NCAM1;refGene_tx=XM_023238923.1;refGene_gn=NCAM1;refGene_tx=XM_023238924.1
D1 11180376 11180473
refGene_gn=NCAM1;refGene_tx=NM_001009869.1;refGene_gn=NCAM1;refGene_tx=XM_006936585.4;refGene_gn=NCAM1;refGene_tx=XM_006936586.4;refGene_gn=NCAM1;refGene_tx=XM_006936587.4;refGene_gn=NCAM1;refGene_tx=XM_006936588.4;refGene_gn=NCAM1;refGene_tx=XM_006936589.4;refGene_gn=NCAM1;refGene_tx=XM_006936591.3;refGene_gn=NCAM1;refGene_tx=XM_006936592.3;refGene_gn=NCAM1;refGene_tx=XM_006936593.3;refGene_gn=NCAM1;refGene_tx=XM_006936594.4;refGene_gn=NCAM1;refGene_tx=XM_006936595.3;refGene_gn=NCAM1;refGene_tx=XM_011286326.3;refGene_gn=NCAM1;refGene_tx=XM_023238914.1;refGene_gn=NCAM1;refGene_tx=XM_023238915.1;refGene_gn=NCAM1;refGene_tx=XM_023238917.1;refGene_gn=NCAM1;refGene_tx=XM_023238918.1;refGene_gn=NCAM1;refGene_tx=XM_023238919.1;refGene_gn=NCAM1;refGene_tx=XM_023238920.1;refGene_gn=NCAM1;refGene_tx=XM_023238921.1;refGene_gn=NCAM1;refGene_tx=XM_023238922.1;refGene_gn=NCAM1;refGene_tx=XM_023238923.1;refGene_gn=NCAM1;refGene_tx=XM_023238924.1
D1 11180792 11180963
refGene_gn=NCAM1;refGene_tx=NM_001009869.1;refGene_gn=NCAM1;refGene_tx=XM_006936585.4;refGene_gn=NCAM1;refGene_tx=XM_006936586.4;refGene_gn=NCAM1;refGene_tx=XM_006936587.4;refGene_gn=NCAM1;refGene_tx=XM_006936588.4;refGene_gn=NCAM1;refGene_tx=XM_006936589.4;refGene_gn=NCAM1;refGene_tx=XM_006936591.3;refGene_gn=NCAM1;refGene_tx=XM_006936592.3;refGene_gn=NCAM1;refGene_tx=XM_006936593.3;refGene_gn=NCAM1;refGene_tx=XM_00

6936594.4;refGene_gn=NCAM1;refGene_tx=XM_006936595.3;refGene_gn=NCAM1;refGene_tx=XM_011286326.3;refGene_gn=NCAM1;refGene_tx=XM_023238914.1;refGene_gn=NCAM1;refGene_tx=XM_023238915.1;refGene_gn=NCAM1;refGene_tx=XM_023238917.1;refGene_gn=NCAM1;refGene_tx=XM_023238918.1;refGene_gn=NCAM1;refGene_tx=XM_023238919.1;refGene_gn=NCAM1;refGene_tx=XM_023238920.1;refGene_gn=NCAM1;refGene_tx=XM_023238921.1;refGene_gn=NCAM1;refGene_tx=XM_023238922.1;refGene_gn=NCAM1;refGene_tx=XM_023238923.1;refGene_gn=NCAM1;refGene_tx=XM_023238924.1

D1 11182895 11183027
refGene_gn=NCAM1;refGene_tx=NM_001009869.1;refGene_gn=NCAM1;refGene_tx=XM_006936585.4;refGene_gn=NCAM1;refGene_tx=XM_006936586.4;refGene_gn=NCAM1;refGene_tx=XM_006936587.4;refGene_gn=NCAM1;refGene_tx=XM_006936588.4;refGene_gn=NCAM1;refGene_tx=XM_006936589.4;refGene_gn=NCAM1;refGene_tx=XM_006936591.3;refGene_gn=NCAM1;refGene_tx=XM_006936592.3;refGene_gn=NCAM1;refGene_tx=XM_006936593.3;refGene_gn=NCAM1;refGene_tx=XM_006936594.4;refGene_gn=NCAM1;refGene_tx=XM_006936595.3;refGene_gn=NCAM1;refGene_tx=XM_011286326.3;refGene_gn=NCAM1;refGene_tx=XM_023238914.1;refGene_gn=NCAM1;refGene_tx=XM_023238915.1;refGene_gn=NCAM1;refGene_tx=XM_023238917.1;refGene_gn=NCAM1;refGene_tx=XM_023238918.1;refGene_gn=NCAM1;refGene_tx=XM_023238919.1;refGene_gn=NCAM1;refGene_tx=XM_023238920.1;refGene_gn=NCAM1;refGene_tx=XM_023238921.1;refGene_gn=NCAM1;refGene_tx=XM_023238922.1;refGene_gn=NCAM1;refGene_tx=XM_023238923.1;refGene_gn=NCAM1;refGene_tx=XM_023238924.1

D1 11184145 11184160
refGene_gn=NCAM1;refGene_tx=XM_006936585.4;refGene_gn=NCAM1;refGene_tx=XM_006936586.4;refGene_gn=NCAM1;refGene_tx=XM_006936591.3;refGene_gn=NCAM1;refGene_tx=XM_006936592.3;refGene_gn=NCAM1;refGene_tx=XM_006936593.3;refGene_gn=NCAM1;refGene_tx=XM_023238914.1;refGene_gn=NCAM1;refGene_tx=XM_023238915.1;refGene_gn=NCAM1;refGene_tx=XM_023238917.1;refGene_gn=NCAM1;refGene_tx=XM_023238918.1;refGene_gn=NCAM1;refGene_tx=XM_023238919.1;refGene_gn=NCAM1;refGene_tx=XM_023238920.1;refGene_gn=NCAM1;refGene_tx=XM_023238921.1

D1 11186465 11186468
refGene_gn=NCAM1;refGene_tx=XM_011286326.3;refGene_gn=NCAM1;refGene_tx=XM_023238920.1;refGene_gn=NCAM1;refGene_tx=XM_023238922.1

D1 11188377 11188425
refGene_gn=NCAM1;refGene_tx=XM_006936585.4;refGene_gn=NCAM1;refGene_tx=XM_006936586.4;refGene_gn=NCAM1;refGene_tx=XM_006936591.3;refGene_gn=NCAM1;refGene_tx=XM_006936592.3;refGene_gn=NCAM1;refGene_tx=XM_006936593.3;refGene_gn=NCAM1;refGene_tx=XM_023238914.1;refGene_gn=NCAM1;refGene_tx=XM_023238915.1;refGene_gn=NCAM1;refGene_tx=XM_023238917.1;refGene_gn=NCAM1;refGene_tx=XM_023238918.1;refGene_gn=NCAM1;refGene_tx=XM_023238919.1

D1 11190076 11190118
refGene_gn=NCAM1;refGene_tx=XM_006936585.4;refGene_gn=NCAM1;refGene_tx=XM_006936586.4;refGene_gn=NCAM1;refGene_tx=XM_006936591.3;refGene_gn=NCAM1;refGene_tx=XM_006936592.3;refGene_gn=NCAM1;refGene_tx=XM_006936593.3;refGene_gn=NCAM1;refGene_tx=XM_023238914.1;refGene_gn=NCAM1;refGene_tx=XM_023238915.1;refGene_gn=NCAM1;refGene_tx=XM_023238917.1;refGene_gn=NCAM1;refGene_tx=XM_023238918.1

D1 11193346 11193349
refGene_gn=NCAM1;refGene_tx=XM_006936585.4;refGene_gn=NCAM1;refGene_tx=XM_006936586.4;refGene_gn=NCAM1;refGene_tx=XM_006936591.3;refGene_gn=NCAM1;refGene_tx=XM_006936592.3;refGene_gn=NCAM1;refGene_tx=XM_006936593.3;refGene_gn=NCAM1;refGene_tx=XM_023238914.1;refGene_gn=NCAM1;refGene_tx=XM_023238915.1;refGene_gn=NCAM1;refGene_tx=XM_023238918.1;refGene_gn=NCAM1;refGene_tx=XM_023238919.1
D1 11201772 11201897
refGene_gn=NCAM1;refGene_tx=NM_001009869.1;refGene_gn=NCAM1;refGene_tx=XM_006936585.4;refGene_gn=NCAM1;refGene_tx=XM_006936586.4;refGene_gn=NCAM1;refGene_tx=XM_006936587.4;refGene_gn=NCAM1;refGene_tx=XM_006936588.4;refGene_gn=NCAM1;refGene_tx=XM_006936589.4;refGene_gn=NCAM1;refGene_tx=XM_006936591.3;refGene_gn=NCAM1;refGene_tx=XM_006936592.3;refGene_gn=NCAM1;refGene_tx=XM_006936593.3;refGene_gn=NCAM1;refGene_tx=XM_006936594.4;refGene_gn=NCAM1;refGene_tx=XM_006936595.3;refGene_gn=NCAM1;refGene_tx=XM_011286326.3;refGene_gn=NCAM1;refGene_tx=XM_023238914.1;refGene_gn=NCAM1;refGene_tx=XM_023238915.1;refGene_gn=NCAM1;refGene_tx=XM_023238917.1;refGene_gn=NCAM1;refGene_tx=XM_023238918.1;refGene_gn=NCAM1;refGene_tx=XM_023238919.1;refGene_gn=NCAM1;refGene_tx=XM_023238920.1;refGene_gn=NCAM1;refGene_tx=XM_023238921.1;refGene_gn=NCAM1;refGene_tx=XM_023238922.1;refGene_gn=NCAM1;refGene_tx=XM_023238923.1;refGene_gn=NCAM1;refGene_tx=XM_023238924.1
D1 11204714 11204895
refGene_gn=NCAM1;refGene_tx=NM_001009869.1;refGene_gn=NCAM1;refGene_tx=XM_006936585.4;refGene_gn=NCAM1;refGene_tx=XM_006936586.4;refGene_gn=NCAM1;refGene_tx=XM_006936587.4;refGene_gn=NCAM1;refGene_tx=XM_006936588.4;refGene_gn=NCAM1;refGene_tx=XM_006936589.4;refGene_gn=NCAM1;refGene_tx=XM_006936591.3;refGene_gn=NCAM1;refGene_tx=XM_006936592.3;refGene_gn=NCAM1;refGene_tx=XM_006936593.3;refGene_gn=NCAM1;refGene_tx=XM_006936594.4;refGene_gn=NCAM1;refGene_tx=XM_006936595.3;refGene_gn=NCAM1;refGene_tx=XM_011286326.3;refGene_gn=NCAM1;refGene_tx=XM_023238914.1;refGene_gn=NCAM1;refGene_tx=XM_023238915.1;refGene_gn=NCAM1;refGene_tx=XM_023238917.1;refGene_gn=NCAM1;refGene_tx=XM_023238918.1;refGene_gn=NCAM1;refGene_tx=XM_023238919.1;refGene_gn=NCAM1;refGene_tx=XM_023238920.1;refGene_gn=NCAM1;refGene_tx=XM_023238921.1;refGene_gn=NCAM1;refGene_tx=XM_023238922.1;refGene_gn=NCAM1;refGene_tx=XM_023238923.1;refGene_gn=NCAM1;refGene_tx=XM_023238924.1
D1 11207410 11207490
refGene_gn=NCAM1;refGene_tx=XM_006936591.3;refGene_gn=NCAM1;refGene_tx=XM_006936592.3;refGene_gn=NCAM1;refGene_tx=XM_006936593.3;refGene_gn=NCAM1;refGene_tx=XM_006936594.4;refGene_gn=NCAM1;refGene_tx=XM_006936595.3
D1 11214680 11214888
refGene_gn=NCAM1;refGene_tx=NM_001009869.1;refGene_gn=NCAM1;refGene_tx=XM_006936585.4;refGene_gn=NCAM1;refGene_tx=XM_006936586.4;refGene_gn=NCAM1;refGene_tx=XM_006936587.4;refGene_gn=NCAM1;refGene_tx=XM_006936588.4;refGene_gn=NCAM1;refGene_tx=XM_006936589.4;refGene_gn=NCAM1;refGene_tx=XM_011286326.3;refGene_gn=NCAM1;refGene_tx=XM_023238914.1;refGene_gn=NCAM1;refGene_tx=XM_023238915.1;refGene_gn=NCAM1;refGene_tx=XM_023238917.1;refGene_gn=NCAM1;refGene_tx=XM_023238918.1;refGene_gn=NCAM1;

refGene_tx=XM_023238919.1;refGene_gn=NCAM1;refGene_tx=XM_023238920.1;refGene_gn=NCAM1;refGene_tx=XM_023238921.1;refGene_gn=NCAM1;refGene_tx=XM_023238922.1;refGene_gn=NCAM1;refGene_tx=XM_023238923.1;refGene_gn=NCAM1;refGene_tx=XM_023238924.1
D1 11215858 11215975
refGene_gn=NCAM1;refGene_tx=NM_001009869.1;refGene_gn=NCAM1;refGene_tx=XM_006936585.4;refGene_gn=NCAM1;refGene_tx=XM_006936586.4;refGene_gn=NCAM1;refGene_tx=XM_006936587.4;refGene_gn=NCAM1;refGene_tx=XM_006936588.4;refGene_gn=NCAM1;refGene_tx=XM_006936589.4;refGene_gn=NCAM1;refGene_tx=XM_011286326.3;refGene_gn=NCAM1;refGene_tx=XM_023238914.1;refGene_gn=NCAM1;refGene_tx=XM_023238915.1;refGene_gn=NCAM1;refGene_tx=XM_023238917.1;refGene_gn=NCAM1;refGene_tx=XM_023238918.1;refGene_gn=NCAM1;refGene_tx=XM_023238919.1;refGene_gn=NCAM1;refGene_tx=XM_023238920.1;refGene_gn=NCAM1;refGene_tx=XM_023238921.1;refGene_gn=NCAM1;refGene_tx=XM_023238922.1;refGene_gn=NCAM1;refGene_tx=XM_023238923.1;refGene_gn=NCAM1;refGene_tx=XM_023238924.1
D1 11217004 11217811
refGene_gn=NCAM1;refGene_tx=XM_023238914.1;refGene_gn=NCAM1;refGene_tx=XM_023238915.1;refGene_gn=NCAM1;refGene_tx=XM_023238917.1;refGene_gn=NCAM1;refGene_tx=XM_023238918.1;refGene_gn=NCAM1;refGene_tx=XM_023238919.1;refGene_gn=NCAM1;refGene_tx=XM_023238920.1;refGene_gn=NCAM1;refGene_tx=XM_023238921.1;refGene_gn=NCAM1;refGene_tx=XM_023238922.1;refGene_gn=NCAM1;refGene_tx=XM_023238923.1;refGene_gn=NCAM1;refGene_tx=XM_023238924.1
D1 11219333 11219454
refGene_gn=NCAM1;refGene_tx=NM_001009869.1;refGene_gn=NCAM1;refGene_tx=XM_006936585.4;refGene_gn=NCAM1;refGene_tx=XM_006936586.4;refGene_gn=NCAM1;refGene_tx=XM_006936587.4;refGene_gn=NCAM1;refGene_tx=XM_006936588.4;refGene_gn=NCAM1;refGene_tx=XM_006936589.4;refGene_gn=NCAM1;refGene_tx=XM_011286326.3;refGene_gn=NCAM1;refGene_tx=XM_023238914.1;refGene_gn=NCAM1;refGene_tx=XM_023238915.1;refGene_gn=NCAM1;refGene_tx=XM_023238917.1;refGene_gn=NCAM1;refGene_tx=XM_023238918.1;refGene_gn=NCAM1;refGene_tx=XM_023238919.1;refGene_gn=NCAM1;refGene_tx=XM_023238920.1;refGene_gn=NCAM1;refGene_tx=XM_023238921.1;refGene_gn=NCAM1;refGene_tx=XM_023238922.1;refGene_gn=NCAM1;refGene_tx=XM_023238923.1;refGene_gn=NCAM1;refGene_tx=XM_023238924.1
D1 11272010 11272068
refGene_gn=TTC12;refGene_tx=XM_023239099.1;refGene_gn=TTC12;refGene_tx=XM_023239100.1;refGene_gn=TTC12;refGene_tx=XM_023239101.1;refGene_gn=TTC12;refGene_tx=XM_023239102.1;refGene_gn=TTC12;refGene_tx=XM_023239104.1;refGene_gn=TTC12;refGene_tx=XM_023239105.1
D1 11280489 11280653
refGene_gn=TTC12;refGene_tx=XM_023239099.1;refGene_gn=TTC12;refGene_tx=XM_023239100.1;refGene_gn=TTC12;refGene_tx=XM_023239101.1;refGene_gn=TTC12;refGene_tx=XM_023239102.1;refGene_gn=TTC12;refGene_tx=XM_023239103.1;refGene_gn=TTC12;refGene_tx=XM_023239104.1;refGene_gn=TTC12;refGene_tx=XM_023239105.1
D1 11281193 11281215
refGene_gn=TTC12;refGene_tx=XM_023239099.1;refGene_gn=TTC12;refGene_tx=XM_023239100.1;refGene_gn=TTC12;refGene_tx=XM_023239101.1;refGene_gn=

TTC12;refGene_tx=XM_023239102.1;refGene_gn=TTC12;refGene_tx=XM_0232391
03.1;refGene_gn=TTC12;refGene_tx=XM_023239104.1;refGene_gn=TTC12;refGe
ne_tx=XM_023239105.1
D1 11281858 11281936
refGene_gn=TTC12;refGene_tx=XM_023239099.1;refGene_gn=TTC12;refGene_tx
=XM_023239100.1;refGene_gn=TTC12;refGene_tx=XM_023239101.1;refGene_gn=
TTC12;refGene_tx=XM_023239102.1;refGene_gn=TTC12;refGene_tx=XM_0232391
03.1;refGene_gn=TTC12;refGene_tx=XM_023239104.1;refGene_gn=TTC12;refGe
ne_tx=XM_023239105.1
D1 11282965 11283087
refGene_gn=TTC12;refGene_tx=XM_023239099.1;refGene_gn=TTC12;refGene_tx
=XM_023239100.1;refGene_gn=TTC12;refGene_tx=XM_023239101.1;refGene_gn=
TTC12;refGene_tx=XM_023239102.1;refGene_gn=TTC12;refGene_tx=XM_0232391
03.1;refGene_gn=TTC12;refGene_tx=XM_023239104.1;refGene_gn=TTC12;refGe
ne_tx=XM_023239105.1
D1 11294615 11294675
refGene_gn=TTC12;refGene_tx=XM_023239099.1;refGene_gn=TTC12;refGene_tx
=XM_023239100.1;refGene_gn=TTC12;refGene_tx=XM_023239101.1;refGene_gn=
TTC12;refGene_tx=XM_023239102.1;refGene_gn=TTC12;refGene_tx=XM_0232391
03.1;refGene_gn=TTC12;refGene_tx=XM_023239104.1;refGene_gn=TTC12;refGe
ne_tx=XM_023239105.1
D1 11299681 11299753
refGene_gn=TTC12;refGene_tx=XM_023239099.1;refGene_gn=TTC12;refGene_tx
=XM_023239100.1;refGene_gn=TTC12;refGene_tx=XM_023239101.1;refGene_gn=
TTC12;refGene_tx=XM_023239102.1;refGene_gn=TTC12;refGene_tx=XM_0232391
03.1;refGene_gn=TTC12;refGene_tx=XM_023239104.1;refGene_gn=TTC12;refGe
ne_tx=XM_023239105.1
D1 11304035 11304096
refGene_gn=TTC12;refGene_tx=XM_023239099.1;refGene_gn=TTC12;refGene_tx
=XM_023239100.1;refGene_gn=TTC12;refGene_tx=XM_023239101.1;refGene_gn=
TTC12;refGene_tx=XM_023239102.1;refGene_gn=TTC12;refGene_tx=XM_0232391
03.1;refGene_gn=TTC12;refGene_tx=XM_023239104.1;refGene_gn=TTC12;refGe
ne_tx=XM_023239105.1
D1 11304533 11304722
refGene_gn=TTC12;refGene_tx=XM_023239099.1;refGene_gn=TTC12;refGene_tx
=XM_023239100.1;refGene_gn=TTC12;refGene_tx=XM_023239101.1;refGene_gn=
TTC12;refGene_tx=XM_023239102.1;refGene_gn=TTC12;refGene_tx=XM_0232391
03.1;refGene_gn=TTC12;refGene_tx=XM_023239104.1;refGene_gn=TTC12;refGe
ne_tx=XM_023239105.1
D1 11305374 11305444
refGene_gn=TTC12;refGene_tx=XM_023239099.1;refGene_gn=TTC12;refGene_tx
=XM_023239100.1;refGene_gn=TTC12;refGene_tx=XM_023239101.1;refGene_gn=
TTC12;refGene_tx=XM_023239102.1;refGene_gn=TTC12;refGene_tx=XM_0232391
03.1;refGene_gn=TTC12;refGene_tx=XM_023239104.1;refGene_gn=TTC12;refGe
ne_tx=XM_023239105.1
D1 11306287 11306376
refGene_gn=TTC12;refGene_tx=XM_023239099.1;refGene_gn=TTC12;refGene_tx
=XM_023239100.1;refGene_gn=TTC12;refGene_tx=XM_023239101.1;refGene_gn=
TTC12;refGene_tx=XM_023239102.1;refGene_gn=TTC12;refGene_tx=XM_0232391
03.1;refGene_gn=TTC12;refGene_tx=XM_023239104.1;refGene_gn=TTC12;refGe

ne_tx=XM_023239105.1
D1 11308431 11308600
refGene_gn=TTC12;refGene_tx=XM_023239099.1;refGene_gn=TTC12;refGene_tx=XM_023239100.1;refGene_gn=TTC12;refGene_tx=XM_023239101.1;refGene_gn=TTC12;refGene_tx=XM_023239102.1;refGene_gn=TTC12;refGene_tx=XM_023239103.1;refGene_gn=TTC12;refGene_tx=XM_023239104.1;refGene_gn=TTC12;refGene_tx=XM_023239105.1
D1 11314071 11314164
refGene_gn=TTC12;refGene_tx=XM_023239099.1;refGene_gn=TTC12;refGene_tx=XM_023239100.1;refGene_gn=TTC12;refGene_tx=XM_023239101.1;refGene_gn=TTC12;refGene_tx=XM_023239102.1;refGene_gn=TTC12;refGene_tx=XM_023239103.1;refGene_gn=TTC12;refGene_tx=XM_023239104.1;refGene_gn=TTC12;refGene_tx=XM_023239105.1
D1 11315143 11315204
refGene_gn=TTC12;refGene_tx=XM_023239099.1;refGene_gn=TTC12;refGene_tx=XM_023239100.1;refGene_gn=TTC12;refGene_tx=XM_023239101.1;refGene_gn=TTC12;refGene_tx=XM_023239102.1;refGene_gn=TTC12;refGene_tx=XM_023239103.1;refGene_gn=TTC12;refGene_tx=XM_023239104.1;refGene_gn=TTC12;refGene_tx=XM_023239105.1
D1 11315996 11316134
refGene_gn=TTC12;refGene_tx=XM_023239099.1;refGene_gn=TTC12;refGene_tx=XM_023239100.1;refGene_gn=TTC12;refGene_tx=XM_023239101.1;refGene_gn=TTC12;refGene_tx=XM_023239102.1;refGene_gn=TTC12;refGene_tx=XM_023239103.1;refGene_gn=TTC12;refGene_tx=XM_023239104.1
D1 11316603 11316702
refGene_gn=TTC12;refGene_tx=XM_023239099.1;refGene_gn=TTC12;refGene_tx=XM_023239100.1;refGene_gn=TTC12;refGene_tx=XM_023239101.1;refGene_gn=TTC12;refGene_tx=XM_023239102.1;refGene_gn=TTC12;refGene_tx=XM_023239103.1;refGene_gn=TTC12;refGene_tx=XM_023239104.1;refGene_gn=TTC12;refGene_tx=XM_023239105.1
D1 11317153 11317222
refGene_gn=TTC12;refGene_tx=XM_023239099.1;refGene_gn=TTC12;refGene_tx=XM_023239100.1;refGene_gn=TTC12;refGene_tx=XM_023239101.1;refGene_gn=TTC12;refGene_tx=XM_023239102.1;refGene_gn=TTC12;refGene_tx=XM_023239103.1;refGene_gn=TTC12;refGene_tx=XM_023239104.1;refGene_gn=TTC12;refGene_tx=XM_023239105.1
D1 11318628 11318712
refGene_gn=TTC12;refGene_tx=XM_023239099.1;refGene_gn=TTC12;refGene_tx=XM_023239100.1;refGene_gn=TTC12;refGene_tx=XM_023239101.1;refGene_gn=TTC12;refGene_tx=XM_023239102.1;refGene_gn=TTC12;refGene_tx=XM_023239103.1;refGene_gn=TTC12;refGene_tx=XM_023239105.1
D1 11319580 11319682
refGene_gn=TTC12;refGene_tx=XM_023239099.1;refGene_gn=TTC12;refGene_tx=XM_023239100.1;refGene_gn=TTC12;refGene_tx=XM_023239101.1;refGene_gn=TTC12;refGene_tx=XM_023239102.1;refGene_gn=TTC12;refGene_tx=XM_023239103.1;refGene_gn=TTC12;refGene_tx=XM_023239104.1;refGene_gn=TTC12;refGene_tx=XM_023239105.1
D1 11320930 11321030
refGene_gn=TTC12;refGene_tx=XM_023239099.1;refGene_gn=TTC12;refGene_tx=XM_023239100.1;refGene_gn=TTC12;refGene_tx=XM_023239101.1;refGene_gn=

TTC12;refGene_tx=XM_023239102.1;refGene_gn=TTC12;refGene_tx=XM_023239103.1;refGene_gn=TTC12;refGene_tx=XM_023239104.1;refGene_gn=TTC12;refGene_tx=XM_023239105.1
D1 11321966 11322192 refGene_gn=TTC12;refGene_tx=XM_023239099.1;refGene_gn=TTC12;refGene_tx=XM_023239100.1;refGene_gn=TTC12;refGene_tx=XM_023239101.1;refGene_gn=TTC12;refGene_tx=XM_023239102.1;refGene_gn=TTC12;refGene_tx=XM_023239103.1;refGene_gn=TTC12;refGene_tx=XM_023239104.1;refGene_gn=TTC12;refGene_tx=XM_023239105.1
D1 11323145 11323191 refGene_gn=TTC12;refGene_tx=XM_023239102.1
D1 11323222 11323298 refGene_gn=TTC12;refGene_tx=XM_023239099.1;refGene_gn=TTC12;refGene_tx=XM_023239100.1;refGene_gn=TTC12;refGene_tx=XM_023239101.1;refGene_gn=TTC12;refGene_tx=XM_023239103.1;refGene_gn=TTC12;refGene_tx=XM_023239104.1;refGene_gn=TTC12;refGene_tx=XM_023239105.1
D1 11347764 11347946 refGene_gn=ANKK1;refGene_tx=XM_006936637.3
D1 11353241 11353536 refGene_gn=ANKK1;refGene_tx=XM_006936637.3
D1 11354692 11354843 refGene_gn=ANKK1;refGene_tx=XM_006936637.3
D1 11355059 11355110 refGene_gn=ANKK1;refGene_tx=XM_006936637.3
D1 11355712 11355868 refGene_gn=ANKK1;refGene_tx=XM_006936637.3
D1 11356560 11356679 refGene_gn=ANKK1;refGene_tx=XM_006936637.3
D1 11357233 11357270 refGene_gn=ANKK1;refGene_tx=XM_006936637.3
D1 11358176 11359480 refGene_gn=ANKK1;refGene_tx=XM_006936637.3
D1 11367913 11368107 refGene_gn=DRD2;refGene_tx=XM_003992361.5;refGene_gn=DRD2;refGene_tx=XM_023239106.1
D1 11369898 11370226 refGene_gn=DRD2;refGene_tx=XM_003992361.5;refGene_gn=DRD2;refGene_tx=XM_023239106.1
D1 11371765 11371852 refGene_gn=DRD2;refGene_tx=XM_003992361.5
D1 11372765 11372956 refGene_gn=DRD2;refGene_tx=XM_003992361.5;refGene_gn=DRD2;refGene_tx=XM_023239106.1
D1 11374172 11374309 refGene_gn=DRD2;refGene_tx=XM_003992361.5;refGene_gn=DRD2;refGene_tx=XM_023239106.1
D1 11375299 11375409 refGene_gn=DRD2;refGene_tx=XM_003992361.5;refGene_gn=DRD2;refGene_tx=XM_023239106.1
D1 11379557 11379842 refGene_gn=DRD2;refGene_tx=XM_003992361.5;refGene_gn=DRD2;refGene_tx=XM_023239106.1
D1 11561790 11561827 refGene_gn=TMPRSS5;refGene_tx=XM_019811294.2;refGene_gn=TMPRSS5;refGene_tx=XM_023239110.1
D1 11614380 11614504 refGene_gn=TMPRSS5;refGene_tx=XM_003992380.5;refGene_gn=TMPRSS5;refGene_tx=XM_019811294.2
D1 11621628 11622075 refGene_gn=TMPRSS5;refGene_tx=XM_019811295.2;refGene_gn=TMPRSS5;refGen

e_tx=XM_019811297.2;refGene_gn=TMPRSS5;refGene_tx=XM_023239107.1;refGene_gn=TMPRSS5;refGene_tx=XM_023239108.1
D1 11623012 11623165 refGene_gn=TMPRSS5;refGene_tx=XM_003992380.5;refGene_gn=TMPRSS5;refGene_tx=XM_019811294.2;refGene_gn=TMPRSS5;refGene_tx=XM_019811295.2;refGene_gn=TMPRSS5;refGene_tx=XM_019811297.2;refGene_gn=TMPRSS5;refGene_tx=XM_023239107.1;refGene_gn=TMPRSS5;refGene_tx=XM_023239108.1;refGene_gn=TMPRSS5;refGene_tx=XM_023239110.1
D1 11623439 11623582 refGene_gn=TMPRSS5;refGene_tx=XM_003992380.5;refGene_gn=TMPRSS5;refGene_tx=XM_019811294.2;refGene_gn=TMPRSS5;refGene_tx=XM_019811295.2;refGene_gn=TMPRSS5;refGene_tx=XM_019811297.2;refGene_gn=TMPRSS5;refGene_tx=XM_023239107.1;refGene_gn=TMPRSS5;refGene_tx=XM_023239108.1;refGene_gn=TMPRSS5;refGene_tx=XM_023239110.1
D1 11624016 11624115 refGene_gn=TMPRSS5;refGene_tx=XM_003992380.5;refGene_gn=TMPRSS5;refGene_tx=XM_019811294.2;refGene_gn=TMPRSS5;refGene_tx=XM_019811295.2;refGene_gn=TMPRSS5;refGene_tx=XM_019811297.2;refGene_gn=TMPRSS5;refGene_tx=XM_023239107.1;refGene_gn=TMPRSS5;refGene_tx=XM_023239108.1;refGene_gn=TMPRSS5;refGene_tx=XM_023239110.1
D1 11626105 11626284 refGene_gn=TMPRSS5;refGene_tx=XM_003992380.5;refGene_gn=TMPRSS5;refGene_tx=XM_019811294.2;refGene_gn=TMPRSS5;refGene_tx=XM_019811295.2;refGene_gn=TMPRSS5;refGene_tx=XM_019811297.2;refGene_gn=TMPRSS5;refGene_tx=XM_023239107.1;refGene_gn=TMPRSS5;refGene_tx=XM_023239108.1;refGene_gn=TMPRSS5;refGene_tx=XM_023239110.1
D1 11627464 11627627 refGene_gn=TMPRSS5;refGene_tx=XM_003992380.5;refGene_gn=TMPRSS5;refGene_tx=XM_019811294.2;refGene_gn=TMPRSS5;refGene_tx=XM_019811295.2;refGene_gn=TMPRSS5;refGene_tx=XM_019811297.2;refGene_gn=TMPRSS5;refGene_tx=XM_023239107.1;refGene_gn=TMPRSS5;refGene_tx=XM_023239108.1;refGene_gn=TMPRSS5;refGene_tx=XM_023239110.1
D1 11628329 11628373 refGene_gn=TMPRSS5;refGene_tx=XM_003992380.5;refGene_gn=TMPRSS5;refGene_tx=XM_019811294.2;refGene_gn=TMPRSS5;refGene_tx=XM_019811295.2;refGene_gn=TMPRSS5;refGene_tx=XM_019811297.2;refGene_gn=TMPRSS5;refGene_tx=XM_023239107.1;refGene_gn=TMPRSS5;refGene_tx=XM_023239108.1;refGene_gn=TMPRSS5;refGene_tx=XM_023239110.1
D1 11629091 11629205 refGene_gn=TMPRSS5;refGene_tx=XM_003992380.5;refGene_gn=TMPRSS5;refGene_tx=XM_019811294.2;refGene_gn=TMPRSS5;refGene_tx=XM_019811295.2;refGene_gn=TMPRSS5;refGene_tx=XM_019811297.2;refGene_gn=TMPRSS5;refGene_tx=XM_023239107.1;refGene_gn=TMPRSS5;refGene_tx=XM_023239108.1;refGene_gn=TMPRSS5;refGene_tx=XM_023239110.1
D1 11629556 11629692 refGene_gn=TMPRSS5;refGene_tx=XM_003992380.5;refGene_gn=TMPRSS5;refGene_tx=XM_019811294.2;refGene_gn=TMPRSS5;refGene_tx=XM_019811295.2;refGene_gn=TMPRSS5;refGene_tx=XM_019811297.2;refGene_gn=TMPRSS5;refGene_tx=XM_023239107.1;refGene_gn=TMPRSS5;refGene_tx=XM_023239108.1;refGene_gn=TMPRSS5;refGene_tx=XM_023239110.1

D1 11630055 11630178
refGene_gn=TMPRSS5;refGene_tx=XM_003992380.5;refGene_gn=TMPRSS5;refGene_tx=XM_019811294.2;refGene_gn=TMPRSS5;refGene_tx=XM_019811295.2;refGene_gn=TMPRSS5;refGene_tx=XM_019811297.2;refGene_gn=TMPRSS5;refGene_tx=XM_023239107.1;refGene_gn=TMPRSS5;refGene_tx=XM_023239108.1;refGene_gn=TMPRSS5;refGene_tx=XM_023239110.1
D1 11630671 11630770
refGene_gn=TMPRSS5;refGene_tx=XM_003992380.5;refGene_gn=TMPRSS5;refGene_tx=XM_019811294.2;refGene_gn=TMPRSS5;refGene_tx=XM_023239107.1;refGene_gn=TMPRSS5;refGene_tx=XM_023239108.1;refGene_gn=TMPRSS5;refGene_tx=XM_023239110.1
D1 11631098 11631222
refGene_gn=TMPRSS5;refGene_tx=XM_003992380.5;refGene_gn=TMPRSS5;refGene_tx=XM_019811294.2;refGene_gn=TMPRSS5;refGene_tx=XM_023239107.1;refGene_gn=TMPRSS5;refGene_tx=XM_023239108.1;refGene_gn=TMPRSS5;refGene_tx=XM_023239110.1
D1 11632952 11632955 refGene_gn=TMPRSS5;refGene_tx=XM_023239108.1
D1 11653444 11653507
refGene_gn=TMPRSS5;refGene_tx=XM_003992380.5;refGene_gn=TMPRSS5;refGene_tx=XM_019811294.2;refGene_gn=TMPRSS5;refGene_tx=XM_023239107.1;refGene_gn=TMPRSS5;refGene_tx=XM_023239110.1
D1 11664037 11664074 refGene_gn=ZW10;refGene_tx=XM_019811299.2
D1 11664648 11664673 refGene_gn=ZW10;refGene_tx=XM_019811299.2
D1 11667892 11667978 refGene_gn=ZW10;refGene_tx=XM_019811298.2
D1 11674524 11674716
refGene_gn=ZW10;refGene_tx=XM_019811298.2;refGene_gn=ZW10;refGene_tx=XM_019811299.2
D1 11675189 11675342
refGene_gn=ZW10;refGene_tx=XM_019811298.2;refGene_gn=ZW10;refGene_tx=XM_019811299.2
D1 11676907 11677067
refGene_gn=ZW10;refGene_tx=XM_019811298.2;refGene_gn=ZW10;refGene_tx=XM_019811299.2
D1 11677220 11677298
refGene_gn=ZW10;refGene_tx=XM_019811298.2;refGene_gn=ZW10;refGene_tx=XM_019811299.2
D1 11677491 11677593
refGene_gn=ZW10;refGene_tx=XM_019811298.2;refGene_gn=ZW10;refGene_tx=XM_019811299.2
D1 11682314 11682449
refGene_gn=ZW10;refGene_tx=XM_019811298.2;refGene_gn=ZW10;refGene_tx=XM_019811299.2
D1 11685866 11685971
refGene_gn=ZW10;refGene_tx=XM_019811298.2;refGene_gn=ZW10;refGene_tx=XM_019811299.2
D1 11690944 11691631 refGene_gn=CLDN25;refGene_tx=XM_003992364.1
D1 11705922 11706098
refGene_gn=USP28;refGene_tx=XM_011286351.3;refGene_gn=USP28;refGene_tx=XM_019811300.2;refGene_gn=USP28;refGene_tx=XM_019811301.2;refGene_gn=USP28;refGene_tx=XM_023239111.1;refGene_gn=USP28;refGene_tx=XM_0232391

12.1;refGene_gn=USP28;refGene_tx=XM_023239113.1;refGene_gn=USP28;refGene_tx=XM_023239114.1;refGene_gn=USP28;refGene_tx=XM_023239115.1;refGene_gn=USP28;refGene_tx=XM_023239116.1
D1 11707723 11707919
refGene_gn=USP28;refGene_tx=XM_011286351.3;refGene_gn=USP28;refGene_tx=XM_019811300.2;refGene_gn=USP28;refGene_tx=XM_019811301.2;refGene_gn=USP28;refGene_tx=XM_023239111.1;refGene_gn=USP28;refGene_tx=XM_023239112.1;refGene_gn=USP28;refGene_tx=XM_023239113.1;refGene_gn=USP28;refGene_tx=XM_023239114.1;refGene_gn=USP28;refGene_tx=XM_023239115.1;refGene_gn=USP28;refGene_tx=XM_023239116.1
D1 11709847 11709971
refGene_gn=USP28;refGene_tx=XM_011286351.3;refGene_gn=USP28;refGene_tx=XM_019811300.2;refGene_gn=USP28;refGene_tx=XM_019811301.2;refGene_gn=USP28;refGene_tx=XM_023239111.1;refGene_gn=USP28;refGene_tx=XM_023239112.1;refGene_gn=USP28;refGene_tx=XM_023239113.1;refGene_gn=USP28;refGene_tx=XM_023239114.1;refGene_gn=USP28;refGene_tx=XM_023239115.1;refGene_gn=USP28;refGene_tx=XM_023239116.1
D1 11710515 11710595
refGene_gn=USP28;refGene_tx=XM_011286351.3;refGene_gn=USP28;refGene_tx=XM_019811300.2;refGene_gn=USP28;refGene_tx=XM_019811301.2;refGene_gn=USP28;refGene_tx=XM_023239111.1;refGene_gn=USP28;refGene_tx=XM_023239112.1;refGene_gn=USP28;refGene_tx=XM_023239113.1;refGene_gn=USP28;refGene_tx=XM_023239114.1;refGene_gn=USP28;refGene_tx=XM_023239115.1;refGene_gn=USP28;refGene_tx=XM_023239116.1
D1 11711419 11711498
refGene_gn=USP28;refGene_tx=XM_011286351.3;refGene_gn=USP28;refGene_tx=XM_019811300.2;refGene_gn=USP28;refGene_tx=XM_019811301.2;refGene_gn=USP28;refGene_tx=XM_023239111.1;refGene_gn=USP28;refGene_tx=XM_023239112.1;refGene_gn=USP28;refGene_tx=XM_023239113.1;refGene_gn=USP28;refGene_tx=XM_023239114.1;refGene_gn=USP28;refGene_tx=XM_023239115.1;refGene_gn=USP28;refGene_tx=XM_023239116.1
D1 11711608 11711787
refGene_gn=USP28;refGene_tx=XM_011286351.3;refGene_gn=USP28;refGene_tx=XM_019811300.2;refGene_gn=USP28;refGene_tx=XM_019811301.2;refGene_gn=USP28;refGene_tx=XM_023239111.1;refGene_gn=USP28;refGene_tx=XM_023239112.1;refGene_gn=USP28;refGene_tx=XM_023239113.1;refGene_gn=USP28;refGene_tx=XM_023239114.1;refGene_gn=USP28;refGene_tx=XM_023239115.1;refGene_gn=USP28;refGene_tx=XM_023239116.1
D1 11712475 11712571
refGene_gn=USP28;refGene_tx=XM_011286351.3;refGene_gn=USP28;refGene_tx=XM_019811300.2;refGene_gn=USP28;refGene_tx=XM_019811301.2;refGene_gn=USP28;refGene_tx=XM_023239111.1;refGene_gn=USP28;refGene_tx=XM_023239112.1;refGene_gn=USP28;refGene_tx=XM_023239113.1;refGene_gn=USP28;refGene_tx=XM_023239114.1
D1 11713596 11713782
refGene_gn=USP28;refGene_tx=XM_011286351.3;refGene_gn=USP28;refGene_tx=XM_019811300.2;refGene_gn=USP28;refGene_tx=XM_019811301.2;refGene_gn=USP28;refGene_tx=XM_023239111.1;refGene_gn=USP28;refGene_tx=XM_023239112.1
D1 11713927 11714067

refGene_gn=USP28;refGene_tx=XM_011286351.3;refGene_gn=USP28;refGene_tx=XM_019811300.2;refGene_gn=USP28;refGene_tx=XM_019811301.2;refGene_gn=USP28;refGene_tx=XM_023239111.1;refGene_gn=USP28;refGene_tx=XM_023239112.1;refGene_gn=USP28;refGene_tx=XM_023239113.1;refGene_gn=USP28;refGene_tx=XM_023239114.1;refGene_gn=USP28;refGene_tx=XM_023239115.1;refGene_gn=USP28;refGene_tx=XM_023239116.1
D1 11714595 11714790
refGene_gn=USP28;refGene_tx=XM_011286351.3;refGene_gn=USP28;refGene_tx=XM_019811300.2;refGene_gn=USP28;refGene_tx=XM_019811301.2;refGene_gn=USP28;refGene_tx=XM_023239111.1;refGene_gn=USP28;refGene_tx=XM_023239112.1;refGene_gn=USP28;refGene_tx=XM_023239113.1;refGene_gn=USP28;refGene_tx=XM_023239114.1;refGene_gn=USP28;refGene_tx=XM_023239115.1;refGene_gn=USP28;refGene_tx=XM_023239116.1
D1 11718091 11718320
refGene_gn=USP28;refGene_tx=XM_011286351.3;refGene_gn=USP28;refGene_tx=XM_019811300.2;refGene_gn=USP28;refGene_tx=XM_019811301.2;refGene_gn=USP28;refGene_tx=XM_023239111.1;refGene_gn=USP28;refGene_tx=XM_023239112.1;refGene_gn=USP28;refGene_tx=XM_023239113.1;refGene_gn=USP28;refGene_tx=XM_023239114.1;refGene_gn=USP28;refGene_tx=XM_023239115.1;refGene_gn=USP28;refGene_tx=XM_023239116.1
D1 11719665 11719745
refGene_gn=USP28;refGene_tx=XM_011286351.3;refGene_gn=USP28;refGene_tx=XM_019811300.2;refGene_gn=USP28;refGene_tx=XM_019811301.2;refGene_gn=USP28;refGene_tx=XM_023239111.1;refGene_gn=USP28;refGene_tx=XM_023239112.1;refGene_gn=USP28;refGene_tx=XM_023239113.1;refGene_gn=USP28;refGene_tx=XM_023239114.1;refGene_gn=USP28;refGene_tx=XM_023239115.1;refGene_gn=USP28;refGene_tx=XM_023239116.1;refGene_gn=USP28;refGene_tx=XM_023239117.1
D1 11721316 11721537
refGene_gn=USP28;refGene_tx=XM_011286351.3;refGene_gn=USP28;refGene_tx=XM_019811300.2;refGene_gn=USP28;refGene_tx=XM_019811301.2;refGene_gn=USP28;refGene_tx=XM_023239111.1;refGene_gn=USP28;refGene_tx=XM_023239112.1;refGene_gn=USP28;refGene_tx=XM_023239113.1;refGene_gn=USP28;refGene_tx=XM_023239114.1;refGene_gn=USP28;refGene_tx=XM_023239115.1;refGene_gn=USP28;refGene_tx=XM_023239116.1;refGene_gn=USP28;refGene_tx=XM_023239117.1
D1 11723381 11723558
refGene_gn=USP28;refGene_tx=XM_011286351.3;refGene_gn=USP28;refGene_tx=XM_019811300.2;refGene_gn=USP28;refGene_tx=XM_019811301.2;refGene_gn=USP28;refGene_tx=XM_023239111.1;refGene_gn=USP28;refGene_tx=XM_023239112.1;refGene_gn=USP28;refGene_tx=XM_023239113.1;refGene_gn=USP28;refGene_tx=XM_023239114.1;refGene_gn=USP28;refGene_tx=XM_023239115.1;refGene_gn=USP28;refGene_tx=XM_023239116.1;refGene_gn=USP28;refGene_tx=XM_023239117.1
D1 11727036 11727132
refGene_gn=USP28;refGene_tx=XM_011286351.3;refGene_gn=USP28;refGene_tx=XM_019811300.2;refGene_gn=USP28;refGene_tx=XM_019811301.2;refGene_gn=USP28;refGene_tx=XM_023239111.1;refGene_gn=USP28;refGene_tx=XM_023239112.1;refGene_gn=USP28;refGene_tx=XM_023239113.1;refGene_gn=USP28;refGene_tx=XM_023239114.1;refGene_gn=USP28;refGene_tx=XM_023239115.1;refGen

e_gn=USP28;refGene_tx=XM_023239116.1;refGene_gn=USP28;refGene_tx=XM_023239117.1
D1 11730230 11730358
refGene_gn=USP28;refGene_tx=XM_011286351.3;refGene_gn=USP28;refGene_tx=XM_019811300.2;refGene_gn=USP28;refGene_tx=XM_019811301.2;refGene_gn=USP28;refGene_tx=XM_023239111.1;refGene_gn=USP28;refGene_tx=XM_023239112.1;refGene_gn=USP28;refGene_tx=XM_023239113.1;refGene_gn=USP28;refGene_tx=XM_023239114.1;refGene_gn=USP28;refGene_tx=XM_023239115.1;refGene_gn=USP28;refGene_tx=XM_023239116.1;refGene_gn=USP28;refGene_tx=XM_023239117.1
D1 11731782 11731785 refGene_gn=USP28;refGene_tx=XM_019811301.2
D1 11732136 11732285
refGene_gn=USP28;refGene_tx=XM_011286351.3;refGene_gn=USP28;refGene_tx=XM_019811300.2;refGene_gn=USP28;refGene_tx=XM_023239111.1;refGene_gn=USP28;refGene_tx=XM_023239112.1;refGene_gn=USP28;refGene_tx=XM_023239113.1;refGene_gn=USP28;refGene_tx=XM_023239114.1;refGene_gn=USP28;refGene_tx=XM_023239115.1;refGene_gn=USP28;refGene_tx=XM_023239116.1;refGene_gn=USP28;refGene_tx=XM_023239117.1
D1 11733740 11733820
refGene_gn=USP28;refGene_tx=XM_011286351.3;refGene_gn=USP28;refGene_tx=XM_019811300.2;refGene_gn=USP28;refGene_tx=XM_023239111.1;refGene_gn=USP28;refGene_tx=XM_023239112.1;refGene_gn=USP28;refGene_tx=XM_023239113.1;refGene_gn=USP28;refGene_tx=XM_023239114.1;refGene_gn=USP28;refGene_tx=XM_023239115.1;refGene_gn=USP28;refGene_tx=XM_023239116.1;refGene_gn=USP28;refGene_tx=XM_023239117.1
D1 11734745 11734819
refGene_gn=USP28;refGene_tx=XM_011286351.3;refGene_gn=USP28;refGene_tx=XM_023239111.1;refGene_gn=USP28;refGene_tx=XM_023239112.1;refGene_gn=USP28;refGene_tx=XM_023239113.1;refGene_gn=USP28;refGene_tx=XM_023239114.1;refGene_gn=USP28;refGene_tx=XM_023239115.1;refGene_gn=USP28;refGene_tx=XM_023239116.1;refGene_gn=USP28;refGene_tx=XM_023239117.1
D1 11735676 11735814
refGene_gn=USP28;refGene_tx=XM_011286351.3;refGene_gn=USP28;refGene_tx=XM_023239111.1;refGene_gn=USP28;refGene_tx=XM_023239112.1;refGene_gn=USP28;refGene_tx=XM_023239113.1;refGene_gn=USP28;refGene_tx=XM_023239114.1;refGene_gn=USP28;refGene_tx=XM_023239115.1;refGene_gn=USP28;refGene_tx=XM_023239116.1;refGene_gn=USP28;refGene_tx=XM_023239117.1
D1 11736524 11736611
refGene_gn=USP28;refGene_tx=XM_011286351.3;refGene_gn=USP28;refGene_tx=XM_023239111.1;refGene_gn=USP28;refGene_tx=XM_023239112.1;refGene_gn=USP28;refGene_tx=XM_023239113.1;refGene_gn=USP28;refGene_tx=XM_023239114.1;refGene_gn=USP28;refGene_tx=XM_023239115.1;refGene_gn=USP28;refGene_tx=XM_023239116.1;refGene_gn=USP28;refGene_tx=XM_023239117.1
D1 11741399 11741559
refGene_gn=USP28;refGene_tx=XM_011286351.3;refGene_gn=USP28;refGene_tx=XM_023239111.1;refGene_gn=USP28;refGene_tx=XM_023239112.1;refGene_gn=USP28;refGene_tx=XM_023239113.1;refGene_gn=USP28;refGene_tx=XM_023239114.1;refGene_gn=USP28;refGene_tx=XM_023239115.1;refGene_gn=USP28;refGene_tx=XM_023239116.1;refGene_gn=USP28;refGene_tx=XM_023239117.1
D1 11742485 11742591

refGene_gn=USP28;refGene_tx=XM_011286351.3;refGene_gn=USP28;refGene_tx=XM_023239111.1;refGene_gn=USP28;refGene_tx=XM_023239112.1;refGene_gn=USP28;refGene_tx=XM_023239113.1;refGene_gn=USP28;refGene_tx=XM_023239114.1;refGene_gn=USP28;refGene_tx=XM_023239115.1;refGene_gn=USP28;refGene_tx=XM_023239116.1;refGene_gn=USP28;refGene_tx=XM_023239117.1
D1 11751904 11752037
refGene_gn=USP28;refGene_tx=XM_011286351.3;refGene_gn=USP28;refGene_tx=XM_023239111.1;refGene_gn=USP28;refGene_tx=XM_023239112.1;refGene_gn=USP28;refGene_tx=XM_023239113.1;refGene_gn=USP28;refGene_tx=XM_023239114.1;refGene_gn=USP28;refGene_tx=XM_023239115.1;refGene_gn=USP28;refGene_tx=XM_023239116.1;refGene_gn=USP28;refGene_tx=XM_023239117.1
D1 11753518 11753596
refGene_gn=USP28;refGene_tx=XM_011286351.3;refGene_gn=USP28;refGene_tx=XM_023239111.1;refGene_gn=USP28;refGene_tx=XM_023239112.1;refGene_gn=USP28;refGene_tx=XM_023239113.1;refGene_gn=USP28;refGene_tx=XM_023239114.1;refGene_gn=USP28;refGene_tx=XM_023239115.1;refGene_gn=USP28;refGene_tx=XM_023239116.1;refGene_gn=USP28;refGene_tx=XM_023239117.1
D1 11772724 11772781
refGene_gn=USP28;refGene_tx=XM_023239111.1;refGene_gn=USP28;refGene_tx=XM_023239112.1;refGene_gn=USP28;refGene_tx=XM_023239113.1;refGene_gn=USP28;refGene_tx=XM_023239114.1;refGene_gn=USP28;refGene_tx=XM_023239115.1;refGene_gn=USP28;refGene_tx=XM_023239116.1;refGene_gn=USP28;refGene_tx=XM_023239117.1
D1 11773659 11773662 refGene_gn=USP28;refGene_tx=XM_011286351.3
D1 11794357 11794424 refGene_gn=HTR3B;refGene_tx=XM_023239119.1
D1 11795664 11795778 refGene_gn=HTR3B;refGene_tx=XM_023239119.1
D1 11798754 11798915 refGene_gn=HTR3B;refGene_tx=XM_023239119.1
D1 11812481 11812526 refGene_gn=HTR3B;refGene_tx=XM_023239119.1
D1 11813076 11813186 refGene_gn=HTR3B;refGene_tx=XM_023239119.1
D1 11813613 11813722 refGene_gn=HTR3B;refGene_tx=XM_023239119.1
D1 11813723 11813784 refGene_gn=HTR3B;refGene_tx=XM_023239119.1
D1 11814318 11814476 refGene_gn=HTR3B;refGene_tx=XM_023239119.1
D1 11822845 11823056 refGene_gn=HTR3B;refGene_tx=XM_023239119.1
D1 11823199 11823382 refGene_gn=HTR3B;refGene_tx=XM_023239119.1
D1 11824617 11824853 refGene_gn=HTR3B;refGene_tx=XM_023239119.1
D1 11847970 11848085 refGene_gn=HTR3A;refGene_tx=XM_023239118.1
D1 11848182 11848287 refGene_gn=HTR3A;refGene_tx=XM_023239118.1
D1 11850189 11850444 refGene_gn=HTR3A;refGene_tx=XM_023239118.1
D1 11852671 11852838 refGene_gn=HTR3A;refGene_tx=XM_023239118.1
D1 11854404 11854449 refGene_gn=HTR3A;refGene_tx=XM_023239118.1
D1 11856842 11856952 refGene_gn=HTR3A;refGene_tx=XM_023239118.1
D1 11858228 11858398 refGene_gn=HTR3A;refGene_tx=XM_023239118.1
D1 11860365 11860526 refGene_gn=HTR3A;refGene_tx=XM_023239118.1
D1 11860890 11861101 refGene_gn=HTR3A;refGene_tx=XM_023239118.1
D1 11861217 11861439 refGene_gn=HTR3A;refGene_tx=XM_023239118.1
D1 11862915 11863214 refGene_gn=HTR3A;refGene_tx=XM_023239118.1
D1 11924059 11925327
refGene_gn=ZBTB16;refGene_tx=XM_011286356.2;refGene_gn=ZBTB16;refGene_tx=XM_011286357.3;refGene_gn=ZBTB16;refGene_tx=XM_011286359.3;refGene_gn=ZBTB16;refGene_tx=XM_011286360.3;refGene_gn=ZBTB16;refGene_tx=XM_02

3239120.1;refGene_gn=ZBTB16;refGene_tx=XM_023239121.1;refGene_gn=ZBTB1
6;refGene_tx=XM_023239122.1
D1 12018317 12018415
refGene_gn=ZBTB16;refGene_tx=XM_011286356.2;refGene_gn=ZBTB16;refGene_
tx=XM_011286357.3;refGene_gn=ZBTB16;refGene_tx=XM_011286359.3;refGene_
gn=ZBTB16;refGene_tx=XM_011286360.3;refGene_gn=ZBTB16;refGene_tx=XM_02
3239120.1;refGene_gn=ZBTB16;refGene_tx=XM_023239121.1;refGene_gn=ZBTB1
6;refGene_tx=XM_023239122.1
D1 12048927 12049014
refGene_gn=ZBTB16;refGene_tx=XM_011286356.2;refGene_gn=ZBTB16;refGene_
tx=XM_011286357.3;refGene_gn=ZBTB16;refGene_tx=XM_011286359.3;refGene_
gn=ZBTB16;refGene_tx=XM_011286360.3;refGene_gn=ZBTB16;refGene_tx=XM_02
3239120.1;refGene_gn=ZBTB16;refGene_tx=XM_023239121.1;refGene_gn=ZBTB1
6;refGene_tx=XM_023239122.1
D1 12099744 12099915
refGene_gn=ZBTB16;refGene_tx=XM_011286356.2;refGene_gn=ZBTB16;refGene_
tx=XM_011286357.3;refGene_gn=ZBTB16;refGene_tx=XM_011286359.3;refGene_
gn=ZBTB16;refGene_tx=XM_011286360.3;refGene_gn=ZBTB16;refGene_tx=XM_02
3239120.1;refGene_gn=ZBTB16;refGene_tx=XM_023239121.1;refGene_gn=ZBTB1
6;refGene_tx=XM_023239122.1
D1 12104652 12104820
refGene_gn=ZBTB16;refGene_tx=XM_011286356.2;refGene_gn=ZBTB16;refGene_
tx=XM_011286357.3;refGene_gn=ZBTB16;refGene_tx=XM_011286359.3;refGene_
gn=ZBTB16;refGene_tx=XM_011286360.3;refGene_gn=ZBTB16;refGene_tx=XM_02
3239120.1;refGene_gn=ZBTB16;refGene_tx=XM_023239121.1;refGene_gn=ZBTB1
6;refGene_tx=XM_023239122.1
D1 12106851 12107081
refGene_gn=ZBTB16;refGene_tx=XM_011286356.2;refGene_gn=ZBTB16;refGene_
tx=XM_011286357.3;refGene_gn=ZBTB16;refGene_tx=XM_011286359.3;refGene_
gn=ZBTB16;refGene_tx=XM_011286360.3;refGene_gn=ZBTB16;refGene_tx=XM_02
3239120.1;refGene_gn=ZBTB16;refGene_tx=XM_023239121.1;refGene_gn=ZBTB1
6;refGene_tx=XM_023239122.1
D1 12129174 12129325
refGene_gn=LOC101094108;refGene_tx=XM_003987756.4
D1 12130740 12130948
refGene_gn=LOC101094108;refGene_tx=XM_003987756.4
D1 12132114 12132547
refGene_gn=LOC101094108;refGene_tx=XM_003987756.4
D1 12149721 12149875
refGene_gn=LOC101083433;refGene_tx=XM_003987715.5
D1 12150703 12150911
refGene_gn=LOC101083433;refGene_tx=XM_003987715.5
D1 12166440 12166873
refGene_gn=LOC101083433;refGene_tx=XM_003987715.5
D1 12252821 12253052
refGene_gn=CD1H11orf71;refGene_tx=XM_003992451.5
D1 12253054 12253216
refGene_gn=CD1H11orf71;refGene_tx=XM_003992451.5
D1 12253551 12253647 refGene_gn=RBM7;refGene_tx=XM_023239123.1
D1 12254402 12254565 refGene_gn=RBM7;refGene_tx=XM_023239123.1

D1 12255431 12255519 refGene_gn=RBM7;refGene_tx=XM_023239123.1
D1 12258562 12258698 refGene_gn=RBM7;refGene_tx=XM_023239123.1
D1 12260297 12260660 refGene_gn=RBM7;refGene_tx=XM_023239123.1
D1 12284657 12284804 refGene_gn=REXO2;refGene_tx=XM_003992383.3
D1 12285839 12285923 refGene_gn=REXO2;refGene_tx=XM_003992383.3
D1 12288889 12288967 refGene_gn=REXO2;refGene_tx=XM_003992383.3
D1 12289578 12289690 refGene_gn=REXO2;refGene_tx=XM_003992383.3
D1 12291131 12291240 refGene_gn=REXO2;refGene_tx=XM_003992383.3
D1 12292612 12292666 refGene_gn=REXO2;refGene_tx=XM_003992383.3
D1 12294633 12294763 refGene_gn=REXO2;refGene_tx=XM_003992383.3
D1 12394037 12394174
refGene_gn=LOC101100996;refGene_tx=XM_023239033.1
D1 12394176 12394483
refGene_gn=LOC101100996;refGene_tx=XM_023239033.1
D1 12394484 12394562
refGene_gn=LOC101100996;refGene_tx=XM_023239033.1
D1 12394895 12395111
refGene_gn=LOC101100996;refGene_tx=XM_023239033.1
D1 12398884 12399098
refGene_gn=LOC101100996;refGene_tx=XM_023239033.1
D1 12400489 12400573
refGene_gn=LOC101100996;refGene_tx=XM_023239033.1
D1 12401521 12401753
refGene_gn=LOC101100996;refGene_tx=XM_023239033.1
D1 12401755 12401895
refGene_gn=LOC101100996;refGene_tx=XM_023239033.1
D1 12401896 12401964
refGene_gn=LOC101100996;refGene_tx=XM_023239033.1
D1 12481156 12481689 refGene_gn=NXPE4;refGene_tx=XM_019811309.2
D1 12484557 12484819 refGene_gn=NXPE4;refGene_tx=XM_019811309.2
D1 12566685 12566812 refGene_gn=NXPE2;refGene_tx=XM_023238980.1
D1 12613314 12613408 refGene_gn=NXPE2;refGene_tx=XM_023238980.1
D1 12613409 12613739 refGene_gn=NXPE2;refGene_tx=XM_023238980.1
D1 12613740 12613853 refGene_gn=NXPE2;refGene_tx=XM_023238980.1
D1 12613855 12613967 refGene_gn=NXPE2;refGene_tx=XM_023238980.1
D1 12613968 12614043 refGene_gn=NXPE2;refGene_tx=XM_023238980.1
D1 12620039 12620101 refGene_gn=NXPE2;refGene_tx=XM_023238980.1
D1 12635073 12635189 refGene_gn=NXPE2;refGene_tx=XM_023238980.1
D1 12635190 12635281 refGene_gn=NXPE2;refGene_tx=XM_023238980.1
D1 12638440 12638925 refGene_gn=NXPE2;refGene_tx=XM_023238980.1
D1 13067113 13067232
refGene_gn=CADM1;refGene_tx=XM_003992384.5;refGene_gn=CADM1;refGene_tx
=XM_003992385.5;refGene_gn=CADM1;refGene_tx=XM_003992386.5;refGene_gn=
CADM1;refGene_tx=XM_023239124.1;refGene_gn=CADM1;refGene_tx=XM_0232391
25.1;refGene_gn=CADM1;refGene_tx=XM_023239126.1
D1 13069298 13069430
refGene_gn=CADM1;refGene_tx=XM_003992384.5;refGene_gn=CADM1;refGene_tx
=XM_003992385.5;refGene_gn=CADM1;refGene_tx=XM_003992386.5;refGene_gn=
CADM1;refGene_tx=XM_023239124.1;refGene_gn=CADM1;refGene_tx=XM_0232391
25.1;refGene_gn=CADM1;refGene_tx=XM_023239126.1

D1 13080983 13081037
refGene_gn=CADM1;refGene_tx=XM_003992385.5;refGene_gn=CADM1;refGene_tx=XM_023239124.1;refGene_gn=CADM1;refGene_tx=XM_023239125.1
D1 13088294 13088327
refGene_gn=CADM1;refGene_tx=XM_003992385.5;refGene_gn=CADM1;refGene_tx=XM_003992386.5;refGene_gn=CADM1;refGene_tx=XM_023239124.1;refGene_gn=CADM1;refGene_tx=XM_023239125.1;refGene_gn=CADM1;refGene_tx=XM_023239126.1
D1 13099634 13099718
refGene_gn=CADM1;refGene_tx=XM_023239124.1;refGene_gn=CADM1;refGene_tx=XM_023239125.1;refGene_gn=CADM1;refGene_tx=XM_023239126.1
D1 13104324 13104497
refGene_gn=CADM1;refGene_tx=XM_003992384.5;refGene_gn=CADM1;refGene_tx=XM_003992385.5;refGene_gn=CADM1;refGene_tx=XM_003992386.5;refGene_gn=CADM1;refGene_tx=XM_023239124.1;refGene_gn=CADM1;refGene_tx=XM_023239125.1;refGene_gn=CADM1;refGene_tx=XM_023239126.1
D1 13108223 13108323
refGene_gn=CADM1;refGene_tx=XM_003992384.5;refGene_gn=CADM1;refGene_tx=XM_003992385.5;refGene_gn=CADM1;refGene_tx=XM_003992386.5;refGene_gn=CADM1;refGene_tx=XM_023239124.1;refGene_gn=CADM1;refGene_tx=XM_023239125.1;refGene_gn=CADM1;refGene_tx=XM_023239126.1
D1 13121565 13121724
refGene_gn=CADM1;refGene_tx=XM_003992384.5;refGene_gn=CADM1;refGene_tx=XM_003992385.5;refGene_gn=CADM1;refGene_tx=XM_003992386.5;refGene_gn=CADM1;refGene_tx=XM_023239124.1;refGene_gn=CADM1;refGene_tx=XM_023239125.1;refGene_gn=CADM1;refGene_tx=XM_023239126.1
D1 13123663 13123801
refGene_gn=CADM1;refGene_tx=XM_003992384.5;refGene_gn=CADM1;refGene_tx=XM_003992385.5;refGene_gn=CADM1;refGene_tx=XM_003992386.5;refGene_gn=CADM1;refGene_tx=XM_023239124.1;refGene_gn=CADM1;refGene_tx=XM_023239125.1;refGene_gn=CADM1;refGene_tx=XM_023239126.1
D1 13130225 13130378
refGene_gn=CADM1;refGene_tx=XM_003992384.5;refGene_gn=CADM1;refGene_tx=XM_003992385.5;refGene_gn=CADM1;refGene_tx=XM_003992386.5;refGene_gn=CADM1;refGene_tx=XM_023239124.1;refGene_gn=CADM1;refGene_tx=XM_023239125.1;refGene_gn=CADM1;refGene_tx=XM_023239126.1
D1 13131937 13132084
refGene_gn=CADM1;refGene_tx=XM_003992384.5;refGene_gn=CADM1;refGene_tx=XM_003992385.5;refGene_gn=CADM1;refGene_tx=XM_003992386.5;refGene_gn=CADM1;refGene_tx=XM_023239124.1;refGene_gn=CADM1;refGene_tx=XM_023239125.1;refGene_gn=CADM1;refGene_tx=XM_023239126.1
D1 13323549 13323634 refGene_gn=CADM1;refGene_tx=XM_023239125.1
D1 13394836 13394972
refGene_gn=CADM1;refGene_tx=XM_003992384.5;refGene_gn=CADM1;refGene_tx=XM_003992385.5;refGene_gn=CADM1;refGene_tx=XM_003992386.5;refGene_gn=CADM1;refGene_tx=XM_023239124.1;refGene_gn=CADM1;refGene_tx=XM_023239126.1
D1 14488295 14488498
refGene_gn=LOC102901651;refGene_tx=XM_006936657.3;refGene_gn=LOC102901651;refGene_tx=XM_019811317.2;refGene_gn=LOC102901651;refGene_tx=XM_01

9811318.2;refGene_gn=LOC102901651;refGene_tx=XM_023239128.1;refGene_gn=LOC102901651;refGene_tx=XM_023239129.1
D1 14489977 14490125
refGene_gn=LOC102901651;refGene_tx=XM_006936657.3;refGene_gn=LOC102901651;refGene_tx=XM_019811317.2;refGene_gn=LOC102901651;refGene_tx=XM_019811318.2;refGene_gn=LOC102901651;refGene_tx=XM_023239128.1;refGene_gn=LOC102901651;refGene_tx=XM_023239129.1
D1 14490455 14490644
refGene_gn=LOC102901651;refGene_tx=XM_006936657.3;refGene_gn=LOC102901651;refGene_tx=XM_019811317.2;refGene_gn=LOC102901651;refGene_tx=XM_019811318.2;refGene_gn=LOC102901651;refGene_tx=XM_023239128.1;refGene_gn=LOC102901651;refGene_tx=XM_023239129.1
D1 14490945 14491126
refGene_gn=LOC102901651;refGene_tx=XM_006936657.3;refGene_gn=LOC102901651;refGene_tx=XM_019811317.2;refGene_gn=LOC102901651;refGene_tx=XM_019811318.2;refGene_gn=LOC102901651;refGene_tx=XM_023239128.1;refGene_gn=LOC102901651;refGene_tx=XM_023239129.1
D1 14491267 14491371
refGene_gn=LOC102901651;refGene_tx=XM_006936657.3;refGene_gn=LOC102901651;refGene_tx=XM_019811317.2;refGene_gn=LOC102901651;refGene_tx=XM_019811318.2;refGene_gn=LOC102901651;refGene_tx=XM_023239128.1;refGene_gn=LOC102901651;refGene_tx=XM_023239129.1
D1 14497523 14497617
refGene_gn=BUD13;refGene_tx=XM_006936661.4;refGene_gn=BUD13;refGene_tx=XM_011286364.3
D1 14508392 14508474
refGene_gn=BUD13;refGene_tx=XM_006936661.4;refGene_gn=BUD13;refGene_tx=XM_011286364.3
D1 14508715 14508900
refGene_gn=BUD13;refGene_tx=XM_006936661.4;refGene_gn=BUD13;refGene_tx=XM_011286364.3
D1 14509213 14509352
refGene_gn=BUD13;refGene_tx=XM_006936661.4;refGene_gn=BUD13;refGene_tx=XM_011286364.3
D1 14509681 14509787
refGene_gn=BUD13;refGene_tx=XM_006936661.4;refGene_gn=BUD13;refGene_tx=XM_011286364.3
D1 14511330 14511635
refGene_gn=BUD13;refGene_tx=XM_006936661.4;refGene_gn=BUD13;refGene_tx=XM_011286364.3
D1 14513285 14514158
refGene_gn=BUD13;refGene_tx=XM_006936661.4;refGene_gn=BUD13;refGene_tx=XM_011286364.3
D1 14516857 14516942
refGene_gn=BUD13;refGene_tx=XM_006936661.4;refGene_gn=BUD13;refGene_tx=XM_011286364.3
D1 14520719 14520813
refGene_gn=BUD13;refGene_tx=XM_006936661.4;refGene_gn=BUD13;refGene_tx=XM_011286364.3
D1 14523406 14523549

refGene_gn=BUD13;refGene_tx=XM_006936661.4;refGene_gn=BUD13;refGene_tx=XM_011286364.3
D1 14525792 14525927 refGene_gn=ZPR1;refGene_tx=XM_003992390.4;refGene_gn=ZPR1;refGene_tx=XM_019811314.1
D1 14526658 14526724 refGene_gn=ZPR1;refGene_tx=XM_003992390.4;refGene_gn=ZPR1;refGene_tx=XM_019811314.1
D1 14527824 14527911 refGene_gn=ZPR1;refGene_tx=XM_003992390.4;refGene_gn=ZPR1;refGene_tx=XM_019811314.1
D1 14528552 14528663 refGene_gn=ZPR1;refGene_tx=XM_003992390.4
D1 14529145 14529235 refGene_gn=ZPR1;refGene_tx=XM_003992390.4;refGene_gn=ZPR1;refGene_tx=XM_019811314.1
D1 14529974 14530045 refGene_gn=ZPR1;refGene_tx=XM_003992390.4;refGene_gn=ZPR1;refGene_tx=XM_019811314.1
D1 14530460 14530527 refGene_gn=ZPR1;refGene_tx=XM_003992390.4;refGene_gn=ZPR1;refGene_tx=XM_019811314.1
D1 14530691 14530739 refGene_gn=ZPR1;refGene_tx=XM_003992390.4;refGene_gn=ZPR1;refGene_tx=XM_019811314.1
D1 14531347 14531470 refGene_gn=ZPR1;refGene_tx=XM_003992390.4;refGene_gn=ZPR1;refGene_tx=XM_019811314.1
D1 14531613 14531700 refGene_gn=ZPR1;refGene_tx=XM_003992390.4;refGene_gn=ZPR1;refGene_tx=XM_019811314.1
D1 14532440 14532511 refGene_gn=ZPR1;refGene_tx=XM_003992390.4;refGene_gn=ZPR1;refGene_tx=XM_019811314.1
D1 14532887 14532978 refGene_gn=ZPR1;refGene_tx=XM_003992390.4;refGene_gn=ZPR1;refGene_tx=XM_019811314.1
D1 14533367 14533529 refGene_gn=ZPR1;refGene_tx=XM_003992390.4;refGene_gn=ZPR1;refGene_tx=XM_019811314.1
D1 14533705 14533876 refGene_gn=ZPR1;refGene_tx=XM_003992390.4;refGene_gn=ZPR1;refGene_tx=XM_019811314.1
D1 14536191 14537143 refGene_gn=APOA5;refGene_tx=XM_003992391.4;refGene_gn=APOA5;refGene_tx=XM_019811319.2
D1 14537665 14537777 refGene_gn=APOA5;refGene_tx=XM_003992391.4;refGene_gn=APOA5;refGene_tx=XM_019811319.2
D1 14537890 14537939 refGene_gn=APOA5;refGene_tx=XM_003992391.4
D1 14542323 14542339 refGene_gn=APOA5;refGene_tx=XM_019811319.2

D1 14558162 14559135
refGene_gn=APOA4;refGene_tx=XM_011286365.3;refGene_gn=APOA4;refGene_tx=XM_019811320.2;refGene_gn=APOA4;refGene_tx=XM_019811321.2
D1 14559914 14560041
refGene_gn=APOA4;refGene_tx=XM_011286365.3;refGene_gn=APOA4;refGene_tx=XM_019811320.2;refGene_gn=APOA4;refGene_tx=XM_019811321.2
D1 14560406 14560455
refGene_gn=APOA4;refGene_tx=XM_011286365.3;refGene_gn=APOA4;refGene_tx=XM_019811320.2;refGene_gn=APOA4;refGene_tx=XM_019811321.2
D1 14566652 14566707
refGene_gn=APOC3;refGene_tx=XM_003992455.4;refGene_gn=APOC3;refGene_tx=XM_019811322.2;refGene_gn=APOC3;refGene_tx=XM_019811323.1
D1 14566873 14566997
refGene_gn=APOC3;refGene_tx=XM_003992455.4;refGene_gn=APOC3;refGene_tx=XM_019811322.2;refGene_gn=APOC3;refGene_tx=XM_019811323.1
D1 14568114 14568238
refGene_gn=APOC3;refGene_tx=XM_003992455.4;refGene_gn=APOC3;refGene_tx=XM_019811322.2;refGene_gn=APOC3;refGene_tx=XM_019811323.1
D1 14570874 14571478 refGene_gn=APOA1;refGene_tx=XM_011286366.3
D1 14572135 14572289 refGene_gn=APOA1;refGene_tx=XM_011286366.3
D1 14572484 14572527 refGene_gn=APOA1;refGene_tx=XM_011286366.3
D1 14580544 14580702
refGene_gn=SIK3;refGene_tx=XM_023239130.1;refGene_gn=SIK3;refGene_tx=XM_023239131.1;refGene_gn=SIK3;refGene_tx=XM_023239132.1;refGene_gn=SIK3;refGene_tx=XM_023239133.1;refGene_gn=SIK3;refGene_tx=XM_023239135.1;refGene_gn=SIK3;refGene_tx=XM_023239136.1;refGene_gn=SIK3;refGene_tx=XM_023239137.1
D1 14581586 14581719
refGene_gn=SIK3;refGene_tx=XM_023239130.1;refGene_gn=SIK3;refGene_tx=XM_023239131.1;refGene_gn=SIK3;refGene_tx=XM_023239132.1;refGene_gn=SIK3;refGene_tx=XM_023239133.1;refGene_gn=SIK3;refGene_tx=XM_023239135.1;refGene_gn=SIK3;refGene_tx=XM_023239136.1;refGene_gn=SIK3;refGene_tx=XM_023239137.1
D1 14582574 14582738
refGene_gn=SIK3;refGene_tx=XM_023239130.1;refGene_gn=SIK3;refGene_tx=XM_023239131.1;refGene_gn=SIK3;refGene_tx=XM_023239132.1;refGene_gn=SIK3;refGene_tx=XM_023239133.1;refGene_gn=SIK3;refGene_tx=XM_023239135.1;refGene_gn=SIK3;refGene_tx=XM_023239136.1;refGene_gn=SIK3;refGene_tx=XM_023239137.1
D1 14592160 14593050
refGene_gn=SIK3;refGene_tx=XM_023239130.1;refGene_gn=SIK3;refGene_tx=XM_023239131.1;refGene_gn=SIK3;refGene_tx=XM_023239132.1;refGene_gn=SIK3;refGene_tx=XM_023239133.1;refGene_gn=SIK3;refGene_tx=XM_023239135.1;refGene_gn=SIK3;refGene_tx=XM_023239136.1;refGene_gn=SIK3;refGene_tx=XM_023239137.1
D1 14593645 14593985
refGene_gn=SIK3;refGene_tx=XM_023239130.1;refGene_gn=SIK3;refGene_tx=XM_023239131.1;refGene_gn=SIK3;refGene_tx=XM_023239132.1;refGene_gn=SIK3;refGene_tx=XM_023239133.1;refGene_gn=SIK3;refGene_tx=XM_023239135.1;refGene_gn=SIK3;refGene_tx=XM_023239136.1;refGene_gn=SIK3;refGene_tx=X

M_023239137.1
D1 14594953 14595063
refGene_gn=SIK3;refGene_tx=XM_023239130.1;refGene_gn=SIK3;refGene_tx=XM_023239131.1;refGene_gn=SIK3;refGene_tx=XM_023239132.1;refGene_gn=SIK3;refGene_tx=XM_023239133.1;refGene_gn=SIK3;refGene_tx=XM_023239135.1;refGene_gn=SIK3;refGene_tx=XM_023239136.1;refGene_gn=SIK3;refGene_tx=XM_023239137.1
D1 14595578 14595664
refGene_gn=SIK3;refGene_tx=XM_023239130.1;refGene_gn=SIK3;refGene_tx=XM_023239131.1;refGene_gn=SIK3;refGene_tx=XM_023239132.1;refGene_gn=SIK3;refGene_tx=XM_023239133.1;refGene_gn=SIK3;refGene_tx=XM_023239135.1;refGene_gn=SIK3;refGene_tx=XM_023239136.1;refGene_gn=SIK3;refGene_tx=XM_023239137.1
D1 14595912 14596038
refGene_gn=SIK3;refGene_tx=XM_023239130.1;refGene_gn=SIK3;refGene_tx=XM_023239131.1;refGene_gn=SIK3;refGene_tx=XM_023239132.1;refGene_gn=SIK3;refGene_tx=XM_023239133.1;refGene_gn=SIK3;refGene_tx=XM_023239135.1;refGene_gn=SIK3;refGene_tx=XM_023239136.1;refGene_gn=SIK3;refGene_tx=XM_023239137.1
D1 14596780 14596931
refGene_gn=SIK3;refGene_tx=XM_023239130.1;refGene_gn=SIK3;refGene_tx=XM_023239131.1;refGene_gn=SIK3;refGene_tx=XM_023239132.1;refGene_gn=SIK3;refGene_tx=XM_023239133.1;refGene_gn=SIK3;refGene_tx=XM_023239135.1;refGene_gn=SIK3;refGene_tx=XM_023239136.1;refGene_gn=SIK3;refGene_tx=XM_023239137.1
D1 14600683 14600827
refGene_gn=SIK3;refGene_tx=XM_023239130.1;refGene_gn=SIK3;refGene_tx=XM_023239131.1;refGene_gn=SIK3;refGene_tx=XM_023239132.1;refGene_gn=SIK3;refGene_tx=XM_023239133.1;refGene_gn=SIK3;refGene_tx=XM_023239135.1;refGene_gn=SIK3;refGene_tx=XM_023239137.1
D1 14603097 14603168
refGene_gn=SIK3;refGene_tx=XM_023239130.1;refGene_gn=SIK3;refGene_tx=XM_023239131.1;refGene_gn=SIK3;refGene_tx=XM_023239132.1;refGene_gn=SIK3;refGene_tx=XM_023239133.1;refGene_gn=SIK3;refGene_tx=XM_023239135.1;refGene_gn=SIK3;refGene_tx=XM_023239136.1;refGene_gn=SIK3;refGene_tx=XM_023239137.1
D1 14606284 14606440
refGene_gn=SIK3;refGene_tx=XM_023239130.1;refGene_gn=SIK3;refGene_tx=XM_023239131.1;refGene_gn=SIK3;refGene_tx=XM_023239132.1;refGene_gn=SIK3;refGene_tx=XM_023239133.1;refGene_gn=SIK3;refGene_tx=XM_023239135.1;refGene_gn=SIK3;refGene_tx=XM_023239136.1;refGene_gn=SIK3;refGene_tx=XM_023239137.1
D1 14606701 14606855
refGene_gn=SIK3;refGene_tx=XM_023239130.1;refGene_gn=SIK3;refGene_tx=XM_023239131.1;refGene_gn=SIK3;refGene_tx=XM_023239132.1;refGene_gn=SIK3;refGene_tx=XM_023239133.1;refGene_gn=SIK3;refGene_tx=XM_023239135.1;refGene_gn=SIK3;refGene_tx=XM_023239136.1;refGene_gn=SIK3;refGene_tx=XM_023239137.1
D1 14608172 14608282
refGene_gn=SIK3;refGene_tx=XM_023239130.1;refGene_gn=SIK3;refGene_tx=X

M_023239131.1;refGene_gn=SIK3;refGene_tx=XM_023239132.1;refGene_gn=SIK3;refGene_tx=XM_023239133.1;refGene_gn=SIK3;refGene_tx=XM_023239135.1;refGene_gn=SIK3;refGene_tx=XM_023239136.1;refGene_gn=SIK3;refGene_tx=XM_023239137.1
D1 14608720 14608798
refGene_gn=SIK3;refGene_tx=XM_023239130.1;refGene_gn=SIK3;refGene_tx=XM_023239131.1;refGene_gn=SIK3;refGene_tx=XM_023239132.1;refGene_gn=SIK3;refGene_tx=XM_023239133.1;refGene_gn=SIK3;refGene_tx=XM_023239135.1;refGene_gn=SIK3;refGene_tx=XM_023239136.1
D1 14609226 14609370
refGene_gn=SIK3;refGene_tx=XM_023239130.1;refGene_gn=SIK3;refGene_tx=XM_023239131.1;refGene_gn=SIK3;refGene_tx=XM_023239132.1;refGene_gn=SIK3;refGene_tx=XM_023239133.1;refGene_gn=SIK3;refGene_tx=XM_023239136.1
D1 14609610 14609721
refGene_gn=SIK3;refGene_tx=XM_023239130.1;refGene_gn=SIK3;refGene_tx=XM_023239131.1;refGene_gn=SIK3;refGene_tx=XM_023239132.1;refGene_gn=SIK3;refGene_tx=XM_023239133.1;refGene_gn=SIK3;refGene_tx=XM_023239135.1;refGene_gn=SIK3;refGene_tx=XM_023239136.1;refGene_gn=SIK3;refGene_tx=XM_023239137.1
D1 14610269 14610388
refGene_gn=SIK3;refGene_tx=XM_023239130.1;refGene_gn=SIK3;refGene_tx=XM_023239131.1;refGene_gn=SIK3;refGene_tx=XM_023239132.1;refGene_gn=SIK3;refGene_tx=XM_023239133.1;refGene_gn=SIK3;refGene_tx=XM_023239135.1;refGene_gn=SIK3;refGene_tx=XM_023239136.1;refGene_gn=SIK3;refGene_tx=XM_023239137.1
D1 14628212 14628336
refGene_gn=SIK3;refGene_tx=XM_023239130.1;refGene_gn=SIK3;refGene_tx=XM_023239131.1;refGene_gn=SIK3;refGene_tx=XM_023239132.1;refGene_gn=SIK3;refGene_tx=XM_023239133.1;refGene_gn=SIK3;refGene_tx=XM_023239135.1;refGene_gn=SIK3;refGene_tx=XM_023239136.1;refGene_gn=SIK3;refGene_tx=XM_023239137.1
D1 14628895 14629020
refGene_gn=SIK3;refGene_tx=XM_023239130.1;refGene_gn=SIK3;refGene_tx=XM_023239131.1;refGene_gn=SIK3;refGene_tx=XM_023239132.1;refGene_gn=SIK3;refGene_tx=XM_023239133.1;refGene_gn=SIK3;refGene_tx=XM_023239135.1;refGene_gn=SIK3;refGene_tx=XM_023239136.1;refGene_gn=SIK3;refGene_tx=XM_023239137.1
D1 14656565 14656727
refGene_gn=SIK3;refGene_tx=XM_023239130.1;refGene_gn=SIK3;refGene_tx=XM_023239131.1;refGene_gn=SIK3;refGene_tx=XM_023239132.1;refGene_gn=SIK3;refGene_tx=XM_023239133.1;refGene_gn=SIK3;refGene_tx=XM_023239135.1;refGene_gn=SIK3;refGene_tx=XM_023239136.1;refGene_gn=SIK3;refGene_tx=XM_023239137.1
D1 14695903 14695967
refGene_gn=SIK3;refGene_tx=XM_023239130.1;refGene_gn=SIK3;refGene_tx=XM_023239131.1;refGene_gn=SIK3;refGene_tx=XM_023239132.1;refGene_gn=SIK3;refGene_tx=XM_023239133.1;refGene_gn=SIK3;refGene_tx=XM_023239135.1;refGene_gn=SIK3;refGene_tx=XM_023239136.1;refGene_gn=SIK3;refGene_tx=XM_023239137.1
D1 14699296 14699413

refGene_gn=SIK3;refGene_tx=XM_023239130.1;refGene_gn=SIK3;refGene_tx=XM_023239131.1;refGene_gn=SIK3;refGene_tx=XM_023239132.1;refGene_gn=SIK3;refGene_tx=XM_023239133.1;refGene_gn=SIK3;refGene_tx=XM_023239135.1;refGene_gn=SIK3;refGene_tx=XM_023239136.1;refGene_gn=SIK3;refGene_tx=XM_023239137.1
D1 14826590 14826863
refGene_gn=SIK3;refGene_tx=XM_023239130.1;refGene_gn=SIK3;refGene_tx=XM_023239131.1;refGene_gn=SIK3;refGene_tx=XM_023239132.1;refGene_gn=SIK3;refGene_tx=XM_023239133.1;refGene_gn=SIK3;refGene_tx=XM_023239135.1;refGene_gn=SIK3;refGene_tx=XM_023239136.1;refGene_gn=SIK3;refGene_tx=XM_023239137.1
D1 14878203 14878284
refGene_gn=PAFAH1B2;refGene_tx=XM_003992392.5;refGene_gn=PAFAH1B2;refGene_tx=XM_019811329.1
D1 14889650 14889740
refGene_gn=PAFAH1B2;refGene_tx=XM_003992392.5;refGene_gn=PAFAH1B2;refGene_tx=XM_019811328.2;refGene_gn=PAFAH1B2;refGene_tx=XM_019811329.1
D1 14890531 14890648
refGene_gn=PAFAH1B2;refGene_tx=XM_003992392.5;refGene_gn=PAFAH1B2;refGene_tx=XM_019811328.2;refGene_gn=PAFAH1B2;refGene_tx=XM_019811329.1
D1 14892135 14892258
refGene_gn=PAFAH1B2;refGene_tx=XM_003992392.5;refGene_gn=PAFAH1B2;refGene_tx=XM_019811328.2;refGene_gn=PAFAH1B2;refGene_tx=XM_019811329.1
D1 14892417 14892507 refGene_gn=PAFAH1B2;refGene_tx=XM_019811329.1
D1 14895921 14896200
refGene_gn=PAFAH1B2;refGene_tx=XM_003992392.5;refGene_gn=PAFAH1B2;refGene_tx=XM_019811328.2
D1 14907433 14907616
refGene_gn=SIDT2;refGene_tx=XM_011286371.3;refGene_gn=SIDT2;refGene_tx=XM_011286372.3;refGene_gn=SIDT2;refGene_tx=XM_019811330.2;refGene_gn=SIDT2;refGene_tx=XM_019811331.2
D1 14909301 14909423
refGene_gn=SIDT2;refGene_tx=XM_011286371.3;refGene_gn=SIDT2;refGene_tx=XM_011286372.3;refGene_gn=SIDT2;refGene_tx=XM_019811330.2;refGene_gn=SIDT2;refGene_tx=XM_019811331.2
D1 14909721 14909886
refGene_gn=SIDT2;refGene_tx=XM_011286371.3;refGene_gn=SIDT2;refGene_tx=XM_011286372.3;refGene_gn=SIDT2;refGene_tx=XM_019811330.2;refGene_gn=SIDT2;refGene_tx=XM_019811331.2;refGene_gn=SIDT2;refGene_tx=XM_019811332.1;refGene_gn=SIDT2;refGene_tx=XM_023239138.1
D1 14909985 14910031
refGene_gn=SIDT2;refGene_tx=XM_011286371.3;refGene_gn=SIDT2;refGene_tx=XM_011286372.3;refGene_gn=SIDT2;refGene_tx=XM_019811330.2;refGene_gn=SIDT2;refGene_tx=XM_019811331.2;refGene_gn=SIDT2;refGene_tx=XM_019811332.1;refGene_gn=SIDT2;refGene_tx=XM_023239138.1
D1 14910435 14910537
refGene_gn=SIDT2;refGene_tx=XM_011286371.3;refGene_gn=SIDT2;refGene_tx=XM_011286372.3;refGene_gn=SIDT2;refGene_tx=XM_019811330.2;refGene_gn=SIDT2;refGene_tx=XM_019811331.2;refGene_gn=SIDT2;refGene_tx=XM_019811332.1;refGene_gn=SIDT2;refGene_tx=XM_023239138.1

D1 14910645 14910729
refGene_gn=SIDT2;refGene_tx=XM_011286371.3;refGene_gn=SIDT2;refGene_tx=XM_011286372.3;refGene_gn=SIDT2;refGene_tx=XM_019811330.2;refGene_gn=SIDT2;refGene_tx=XM_019811331.2;refGene_gn=SIDT2;refGene_tx=XM_019811332.1;refGene_gn=SIDT2;refGene_tx=XM_023239138.1
D1 14911434 14911534
refGene_gn=SIDT2;refGene_tx=XM_011286371.3;refGene_gn=SIDT2;refGene_tx=XM_011286372.3;refGene_gn=SIDT2;refGene_tx=XM_019811330.2;refGene_gn=SIDT2;refGene_tx=XM_019811331.2;refGene_gn=SIDT2;refGene_tx=XM_019811332.1;refGene_gn=SIDT2;refGene_tx=XM_023239138.1
D1 14911701 14911767
refGene_gn=SIDT2;refGene_tx=XM_011286371.3;refGene_gn=SIDT2;refGene_tx=XM_011286372.3;refGene_gn=SIDT2;refGene_tx=XM_019811330.2;refGene_gn=SIDT2;refGene_tx=XM_019811331.2;refGene_gn=SIDT2;refGene_tx=XM_019811332.1;refGene_gn=SIDT2;refGene_tx=XM_023239138.1
D1 14912751 14912845
refGene_gn=SIDT2;refGene_tx=XM_011286371.3;refGene_gn=SIDT2;refGene_tx=XM_011286372.3;refGene_gn=SIDT2;refGene_tx=XM_019811330.2;refGene_gn=SIDT2;refGene_tx=XM_019811331.2;refGene_gn=SIDT2;refGene_tx=XM_019811332.1;refGene_gn=SIDT2;refGene_tx=XM_023239138.1
D1 14913180 14913233
refGene_gn=SIDT2;refGene_tx=XM_011286371.3;refGene_gn=SIDT2;refGene_tx=XM_011286372.3;refGene_gn=SIDT2;refGene_tx=XM_019811330.2;refGene_gn=SIDT2;refGene_tx=XM_019811331.2;refGene_gn=SIDT2;refGene_tx=XM_019811332.1;refGene_gn=SIDT2;refGene_tx=XM_023239138.1
D1 14913585 14913597
refGene_gn=SIDT2;refGene_tx=XM_011286371.3;refGene_gn=SIDT2;refGene_tx=XM_019811330.2;refGene_gn=SIDT2;refGene_tx=XM_019811332.1;refGene_gn=SIDT2;refGene_tx=XM_023239138.1
D1 14913971 14914043
refGene_gn=SIDT2;refGene_tx=XM_011286371.3;refGene_gn=SIDT2;refGene_tx=XM_011286372.3;refGene_gn=SIDT2;refGene_tx=XM_019811330.2;refGene_gn=SIDT2;refGene_tx=XM_019811331.2;refGene_gn=SIDT2;refGene_tx=XM_019811332.1;refGene_gn=SIDT2;refGene_tx=XM_023239138.1
D1 14914194 14914266
refGene_gn=SIDT2;refGene_tx=XM_011286371.3;refGene_gn=SIDT2;refGene_tx=XM_011286372.3;refGene_gn=SIDT2;refGene_tx=XM_019811330.2;refGene_gn=SIDT2;refGene_tx=XM_019811331.2;refGene_gn=SIDT2;refGene_tx=XM_019811332.1;refGene_gn=SIDT2;refGene_tx=XM_023239138.1
D1 14914456 14914519
refGene_gn=SIDT2;refGene_tx=XM_019811330.2;refGene_gn=SIDT2;refGene_tx=XM_019811331.2;refGene_gn=SIDT2;refGene_tx=XM_019811332.1;refGene_gn=SIDT2;refGene_tx=XM_023239138.1
D1 14915343 14915462
refGene_gn=SIDT2;refGene_tx=XM_011286371.3;refGene_gn=SIDT2;refGene_tx=XM_011286372.3;refGene_gn=SIDT2;refGene_tx=XM_019811330.2;refGene_gn=SIDT2;refGene_tx=XM_019811331.2;refGene_gn=SIDT2;refGene_tx=XM_019811332.1;refGene_gn=SIDT2;refGene_tx=XM_023239138.1
D1 14916005 14916079
refGene_gn=SIDT2;refGene_tx=XM_011286371.3;refGene_gn=SIDT2;refGene_tx

=XM_011286372.3;refGene_gn=SIDT2;refGene_tx=XM_019811330.2;refGene_gn=SIDT2;refGene_tx=XM_019811331.2;refGene_gn=SIDT2;refGene_tx=XM_019811332.1;refGene_gn=SIDT2;refGene_tx=XM_023239138.1
D1 14916185 14916252
refGene_gn=SIDT2;refGene_tx=XM_011286371.3;refGene_gn=SIDT2;refGene_tx=XM_011286372.3;refGene_gn=SIDT2;refGene_tx=XM_019811330.2;refGene_gn=SIDT2;refGene_tx=XM_019811331.2;refGene_gn=SIDT2;refGene_tx=XM_019811332.1;refGene_gn=SIDT2;refGene_tx=XM_023239138.1
D1 14916849 14916923
refGene_gn=SIDT2;refGene_tx=XM_011286371.3;refGene_gn=SIDT2;refGene_tx=XM_011286372.3;refGene_gn=SIDT2;refGene_tx=XM_019811330.2;refGene_gn=SIDT2;refGene_tx=XM_019811331.2;refGene_gn=SIDT2;refGene_tx=XM_019811332.1;refGene_gn=SIDT2;refGene_tx=XM_023239138.1
D1 14917041 14917165
refGene_gn=SIDT2;refGene_tx=XM_011286371.3;refGene_gn=SIDT2;refGene_tx=XM_011286372.3;refGene_gn=SIDT2;refGene_tx=XM_019811330.2;refGene_gn=SIDT2;refGene_tx=XM_019811331.2;refGene_gn=SIDT2;refGene_tx=XM_019811332.1;refGene_gn=SIDT2;refGene_tx=XM_023239138.1
D1 14917377 14917495
refGene_gn=SIDT2;refGene_tx=XM_011286371.3;refGene_gn=SIDT2;refGene_tx=XM_011286372.3;refGene_gn=SIDT2;refGene_tx=XM_019811330.2;refGene_gn=SIDT2;refGene_tx=XM_019811331.2;refGene_gn=SIDT2;refGene_tx=XM_019811332.1;refGene_gn=SIDT2;refGene_tx=XM_023239138.1
D1 14917966 14918103
refGene_gn=SIDT2;refGene_tx=XM_011286371.3;refGene_gn=SIDT2;refGene_tx=XM_011286372.3;refGene_gn=SIDT2;refGene_tx=XM_019811330.2;refGene_gn=SIDT2;refGene_tx=XM_019811331.2;refGene_gn=SIDT2;refGene_tx=XM_019811332.1;refGene_gn=SIDT2;refGene_tx=XM_023239138.1
D1 14918345 14918454
refGene_gn=SIDT2;refGene_tx=XM_011286371.3;refGene_gn=SIDT2;refGene_tx=XM_011286372.3;refGene_gn=SIDT2;refGene_tx=XM_019811330.2;refGene_gn=SIDT2;refGene_tx=XM_019811331.2;refGene_gn=SIDT2;refGene_tx=XM_019811332.1;refGene_gn=SIDT2;refGene_tx=XM_023239138.1
D1 14918722 14918799
refGene_gn=SIDT2;refGene_tx=XM_011286371.3;refGene_gn=SIDT2;refGene_tx=XM_011286372.3;refGene_gn=SIDT2;refGene_tx=XM_019811330.2;refGene_gn=SIDT2;refGene_tx=XM_019811331.2;refGene_gn=SIDT2;refGene_tx=XM_019811332.1;refGene_gn=SIDT2;refGene_tx=XM_023239138.1
D1 14918987 14919034
refGene_gn=SIDT2;refGene_tx=XM_011286371.3;refGene_gn=SIDT2;refGene_tx=XM_011286372.3;refGene_gn=SIDT2;refGene_tx=XM_019811330.2;refGene_gn=SIDT2;refGene_tx=XM_019811331.2;refGene_gn=SIDT2;refGene_tx=XM_019811332.1;refGene_gn=SIDT2;refGene_tx=XM_023239138.1
D1 14919307 14919413
refGene_gn=SIDT2;refGene_tx=XM_011286371.3;refGene_gn=SIDT2;refGene_tx=XM_011286372.3;refGene_gn=SIDT2;refGene_tx=XM_019811330.2;refGene_gn=SIDT2;refGene_tx=XM_019811331.2;refGene_gn=SIDT2;refGene_tx=XM_019811332.1;refGene_gn=SIDT2;refGene_tx=XM_023239138.1
D1 14920093 14920204
refGene_gn=SIDT2;refGene_tx=XM_011286371.3;refGene_gn=SIDT2;refGene_tx

=XM_011286372.3;refGene_gn=SIDT2;refGene_tx=XM_019811330.2;refGene_gn=SIDT2;refGene_tx=XM_019811331.2;refGene_gn=SIDT2;refGene_tx=XM_019811332.1;refGene_gn=SIDT2;refGene_tx=XM_023239138.1
D1 14921466 14921580 refGene_gn=SIDT2;refGene_tx=XM_011286371.3;refGene_gn=SIDT2;refGene_tx=XM_011286372.3;refGene_gn=SIDT2;refGene_tx=XM_019811330.2;refGene_gn=SIDT2;refGene_tx=XM_019811331.2;refGene_gn=SIDT2;refGene_tx=XM_019811332.1;refGene_gn=SIDT2;refGene_tx=XM_023239138.1
D1 14921663 14921726 refGene_gn=SIDT2;refGene_tx=XM_011286371.3;refGene_gn=SIDT2;refGene_tx=XM_011286372.3;refGene_gn=SIDT2;refGene_tx=XM_019811330.2;refGene_gn=SIDT2;refGene_tx=XM_019811331.2;refGene_gn=SIDT2;refGene_tx=XM_019811332.1;refGene_gn=SIDT2;refGene_tx=XM_023239138.1
D1 14928667 14928847 refGene_gn=TAGLN;refGene_tx=XM_003992393.3
D1 14928942 14929120 refGene_gn=TAGLN;refGene_tx=XM_003992393.3
D1 14929415 14929518 refGene_gn=TAGLN;refGene_tx=XM_003992393.3
D1 14929857 14930002 refGene_gn=TAGLN;refGene_tx=XM_003992393.3
D1 14931756 14932140 refGene_gn=PCSK7;refGene_tx=XM_023239142.1;refGene_gn=PCSK7;refGene_tx=XM_023239143.1;refGene_gn=PCSK7;refGene_tx=XM_023239144.1
D1 14932412 14932530 refGene_gn=PCSK7;refGene_tx=XM_023239142.1;refGene_gn=PCSK7;refGene_tx=XM_023239143.1;refGene_gn=PCSK7;refGene_tx=XM_023239144.1
D1 14932750 14932844 refGene_gn=PCSK7;refGene_tx=XM_023239142.1;refGene_gn=PCSK7;refGene_tx=XM_023239143.1;refGene_gn=PCSK7;refGene_tx=XM_023239144.1
D1 14933685 14933780 refGene_gn=PCSK7;refGene_tx=XM_023239142.1;refGene_gn=PCSK7;refGene_tx=XM_023239143.1;refGene_gn=PCSK7;refGene_tx=XM_023239144.1
D1 14934321 14934478 refGene_gn=PCSK7;refGene_tx=XM_023239142.1;refGene_gn=PCSK7;refGene_tx=XM_023239143.1;refGene_gn=PCSK7;refGene_tx=XM_023239144.1
D1 14943641 14943744 refGene_gn=PCSK7;refGene_tx=XM_023239142.1;refGene_gn=PCSK7;refGene_tx=XM_023239143.1;refGene_gn=PCSK7;refGene_tx=XM_023239144.1
D1 14944166 14944274 refGene_gn=PCSK7;refGene_tx=XM_023239142.1;refGene_gn=PCSK7;refGene_tx=XM_023239143.1;refGene_gn=PCSK7;refGene_tx=XM_023239144.1
D1 14945051 14945219 refGene_gn=PCSK7;refGene_tx=XM_023239142.1;refGene_gn=PCSK7;refGene_tx=XM_023239143.1;refGene_gn=PCSK7;refGene_tx=XM_023239144.1
D1 14947788 14947889 refGene_gn=PCSK7;refGene_tx=XM_023239142.1;refGene_gn=PCSK7;refGene_tx=XM_023239143.1;refGene_gn=PCSK7;refGene_tx=XM_023239144.1
D1 14948743 14948882 refGene_gn=PCSK7;refGene_tx=XM_023239142.1;refGene_gn=PCSK7;refGene_tx=XM_023239143.1;refGene_gn=PCSK7;refGene_tx=XM_023239144.1
D1 14949355 14949410 refGene_gn=PCSK7;refGene_tx=XM_023239142.1;refGene_gn=PCSK7;refGene_tx=XM_023239143.1;refGene_gn=PCSK7;refGene_tx=XM_023239144.1

D1 14950343 14950434
refGene_gn=PCSK7;refGene_tx=XM_023239142.1;refGene_gn=PCSK7;refGene_tx=XM_023239143.1;refGene_gn=PCSK7;refGene_tx=XM_023239144.1
D1 14951602 14951768
refGene_gn=PCSK7;refGene_tx=XM_023239142.1;refGene_gn=PCSK7;refGene_tx=XM_023239143.1;refGene_gn=PCSK7;refGene_tx=XM_023239144.1
D1 14952369 14952504
refGene_gn=PCSK7;refGene_tx=XM_023239142.1;refGene_gn=PCSK7;refGene_tx=XM_023239143.1;refGene_gn=PCSK7;refGene_tx=XM_023239144.1
D1 14953422 14953875
refGene_gn=PCSK7;refGene_tx=XM_023239142.1;refGene_gn=PCSK7;refGene_tx=XM_023239143.1;refGene_gn=PCSK7;refGene_tx=XM_023239144.1
D1 14956342 14956729
refGene_gn=RNF214;refGene_tx=XM_023239139.1;refGene_gn=RNF214;refGene_tx=XM_023239140.1;refGene_gn=RNF214;refGene_tx=XM_023239141.1
D1 14958509 14958634
refGene_gn=RNF214;refGene_tx=XM_023239139.1;refGene_gn=RNF214;refGene_tx=XM_023239140.1;refGene_gn=RNF214;refGene_tx=XM_023239141.1
D1 14960829 14961340
refGene_gn=RNF214;refGene_tx=XM_023239139.1;refGene_gn=RNF214;refGene_tx=XM_023239140.1;refGene_gn=RNF214;refGene_tx=XM_023239141.1
D1 14962033 14962093
refGene_gn=RNF214;refGene_tx=XM_023239139.1;refGene_gn=RNF214;refGene_tx=XM_023239140.1;refGene_gn=RNF214;refGene_tx=XM_023239141.1
D1 14967464 14967605
refGene_gn=RNF214;refGene_tx=XM_023239139.1;refGene_gn=RNF214;refGene_tx=XM_023239140.1;refGene_gn=RNF214;refGene_tx=XM_023239141.1
D1 14970559 14970699
refGene_gn=RNF214;refGene_tx=XM_023239139.1;refGene_gn=RNF214;refGene_tx=XM_023239140.1;refGene_gn=RNF214;refGene_tx=XM_023239141.1
D1 15001407 15001504
refGene_gn=RNF214;refGene_tx=XM_023239139.1;refGene_gn=RNF214;refGene_tx=XM_023239140.1;refGene_gn=RNF214;refGene_tx=XM_023239141.1
D1 15002043 15002132
refGene_gn=RNF214;refGene_tx=XM_023239139.1;refGene_gn=RNF214;refGene_tx=XM_023239140.1;refGene_gn=RNF214;refGene_tx=XM_023239141.1
D1 15002733 15002824
refGene_gn=RNF214;refGene_tx=XM_023239139.1;refGene_gn=RNF214;refGene_tx=XM_023239140.1;refGene_gn=RNF214;refGene_tx=XM_023239141.1
D1 15003024 15003123
refGene_gn=RNF214;refGene_tx=XM_023239139.1;refGene_gn=RNF214;refGene_tx=XM_023239140.1;refGene_gn=RNF214;refGene_tx=XM_023239141.1
D1 15003345 15003722
refGene_gn=RNF214;refGene_tx=XM_023239139.1;refGene_gn=RNF214;refGene_tx=XM_023239140.1;refGene_gn=RNF214;refGene_tx=XM_023239141.1
D1 15003851 15003984
refGene_gn=RNF214;refGene_tx=XM_023239139.1;refGene_gn=RNF214;refGene_tx=XM_023239140.1;refGene_gn=RNF214;refGene_tx=XM_023239141.1
D1 15004198 15004303
refGene_gn=RNF214;refGene_tx=XM_023239139.1;refGene_gn=RNF214;refGene_

tx=XM_023239140.1;refGene_gn=RNF214;refGene_tx=XM_023239141.1
D1 15004593 15004689
refGene_gn=RNF214;refGene_tx=XM_023239139.1;refGene_gn=RNF214;refGene_tx=XM_023239140.1;refGene_gn=RNF214;refGene_tx=XM_023239141.1
D1 15005574 15005640
refGene_gn=RNF214;refGene_tx=XM_023239139.1;refGene_gn=RNF214;refGene_tx=XM_023239140.1;refGene_gn=RNF214;refGene_tx=XM_023239141.1
D1 15010094 15010336 refGene_gn=BACE1;refGene_tx=XM_003992397.4
D1 15011005 15011177 refGene_gn=BACE1;refGene_tx=XM_003992397.4
D1 15011429 15011579 refGene_gn=BACE1;refGene_tx=XM_003992397.4
D1 15011940 15012042 refGene_gn=BACE1;refGene_tx=XM_003992397.4
D1 15013010 15013145 refGene_gn=BACE1;refGene_tx=XM_003992397.4
D1 15013875 15014013 refGene_gn=BACE1;refGene_tx=XM_003992397.4
D1 15014515 15014732 refGene_gn=BACE1;refGene_tx=XM_003992397.4
D1 15016227 15016316 refGene_gn=BACE1;refGene_tx=XM_003992397.4
D1 15029374 15029635 refGene_gn=BACE1;refGene_tx=XM_003992397.4
D1 15043725 15043807
refGene_gn=CEP164;refGene_tx=XM_011286373.3;refGene_gn=CEP164;refGene_tx=XM_011286374.3;refGene_gn=CEP164;refGene_tx=XM_011286375.3;refGene_gn=CEP164;refGene_tx=XM_019811336.2;refGene_gn=CEP164;refGene_tx=XM_019811337.2;refGene_gn=CEP164;refGene_tx=XM_019811338.2;refGene_gn=CEP164;refGene_tx=XM_023239145.1
D1 15048366 15048478
refGene_gn=CEP164;refGene_tx=XM_011286373.3;refGene_gn=CEP164;refGene_tx=XM_011286374.3;refGene_gn=CEP164;refGene_tx=XM_011286375.3;refGene_gn=CEP164;refGene_tx=XM_019811336.2;refGene_gn=CEP164;refGene_tx=XM_019811337.2;refGene_gn=CEP164;refGene_tx=XM_019811338.2;refGene_gn=CEP164;refGene_tx=XM_023239145.1
D1 15057971 15058176
refGene_gn=CEP164;refGene_tx=XM_011286373.3;refGene_gn=CEP164;refGene_tx=XM_011286374.3;refGene_gn=CEP164;refGene_tx=XM_011286375.3;refGene_gn=CEP164;refGene_tx=XM_019811336.2;refGene_gn=CEP164;refGene_tx=XM_019811337.2;refGene_gn=CEP164;refGene_tx=XM_019811338.2;refGene_gn=CEP164;refGene_tx=XM_023239145.1
D1 15060399 15062043
refGene_gn=CEP164;refGene_tx=XM_011286373.3;refGene_gn=CEP164;refGene_tx=XM_011286374.3;refGene_gn=CEP164;refGene_tx=XM_019811336.2;refGene_gn=CEP164;refGene_tx=XM_019811337.2;refGene_gn=CEP164;refGene_tx=XM_019811338.2;refGene_gn=CEP164;refGene_tx=XM_023239145.1
D1 15065145 15065301
refGene_gn=CEP164;refGene_tx=XM_011286373.3;refGene_gn=CEP164;refGene_tx=XM_011286374.3;refGene_gn=CEP164;refGene_tx=XM_011286375.3;refGene_gn=CEP164;refGene_tx=XM_019811336.2;refGene_gn=CEP164;refGene_tx=XM_019811337.2;refGene_gn=CEP164;refGene_tx=XM_019811338.2;refGene_gn=CEP164;refGene_tx=XM_023239145.1
D1 15065711 15065846
refGene_gn=CEP164;refGene_tx=XM_011286373.3;refGene_gn=CEP164;refGene_tx=XM_011286374.3;refGene_gn=CEP164;refGene_tx=XM_011286375.3;refGene_gn=CEP164;refGene_tx=XM_019811336.2;refGene_gn=CEP164;refGene_tx=XM_019811337.2;refGene_gn=CEP164;refGene_tx=XM_019811338.2;refGene_gn=CEP16

4;refGene_tx=XM_023239145.1
D1 15066468 15066546
refGene_gn=CEP164;refGene_tx=XM_011286373.3;refGene_gn=CEP164;refGene_tx=XM_011286374.3;refGene_gn=CEP164;refGene_tx=XM_011286375.3;refGene_gn=CEP164;refGene_tx=XM_019811336.2;refGene_gn=CEP164;refGene_tx=XM_019811337.2;refGene_gn=CEP164;refGene_tx=XM_019811338.2;refGene_gn=CEP164;refGene_tx=XM_023239145.1
D1 15075932 15076337
refGene_gn=CEP164;refGene_tx=XM_011286373.3;refGene_gn=CEP164;refGene_tx=XM_011286374.3;refGene_gn=CEP164;refGene_tx=XM_011286375.3;refGene_gn=CEP164;refGene_tx=XM_019811336.2;refGene_gn=CEP164;refGene_tx=XM_019811337.2;refGene_gn=CEP164;refGene_tx=XM_019811338.2;refGene_gn=CEP164;refGene_tx=XM_023239145.1
D1 15077303 15077384
refGene_gn=CEP164;refGene_tx=XM_011286373.3;refGene_gn=CEP164;refGene_tx=XM_011286374.3;refGene_gn=CEP164;refGene_tx=XM_011286375.3;refGene_gn=CEP164;refGene_tx=XM_019811336.2;refGene_gn=CEP164;refGene_tx=XM_019811337.2;refGene_gn=CEP164;refGene_tx=XM_019811338.2;refGene_gn=CEP164;refGene_tx=XM_023239145.1
D1 15079531 15079624
refGene_gn=CEP164;refGene_tx=XM_011286374.3;refGene_gn=CEP164;refGene_tx=XM_011286375.3;refGene_gn=CEP164;refGene_tx=XM_019811336.2;refGene_gn=CEP164;refGene_tx=XM_019811337.2;refGene_gn=CEP164;refGene_tx=XM_019811338.2;refGene_gn=CEP164;refGene_tx=XM_023239145.1
D1 15084345 15084431
refGene_gn=CEP164;refGene_tx=XM_011286373.3;refGene_gn=CEP164;refGene_tx=XM_011286374.3;refGene_gn=CEP164;refGene_tx=XM_011286375.3;refGene_gn=CEP164;refGene_tx=XM_019811336.2;refGene_gn=CEP164;refGene_tx=XM_019811337.2;refGene_gn=CEP164;refGene_tx=XM_019811338.2;refGene_gn=CEP164;refGene_tx=XM_023239145.1
D1 15085366 15085534
refGene_gn=CEP164;refGene_tx=XM_011286373.3;refGene_gn=CEP164;refGene_tx=XM_011286374.3;refGene_gn=CEP164;refGene_tx=XM_011286375.3;refGene_gn=CEP164;refGene_tx=XM_019811336.2;refGene_gn=CEP164;refGene_tx=XM_019811337.2;refGene_gn=CEP164;refGene_tx=XM_019811338.2;refGene_gn=CEP164;refGene_tx=XM_023239145.1
D1 15086399 15086513
refGene_gn=CEP164;refGene_tx=XM_011286373.3;refGene_gn=CEP164;refGene_tx=XM_011286375.3;refGene_gn=CEP164;refGene_tx=XM_019811336.2;refGene_gn=CEP164;refGene_tx=XM_019811337.2;refGene_gn=CEP164;refGene_tx=XM_019811338.2
D1 15088036 15088174
refGene_gn=CEP164;refGene_tx=XM_011286373.3;refGene_gn=CEP164;refGene_tx=XM_011286374.3;refGene_gn=CEP164;refGene_tx=XM_011286375.3;refGene_gn=CEP164;refGene_tx=XM_019811336.2;refGene_gn=CEP164;refGene_tx=XM_019811337.2;refGene_gn=CEP164;refGene_tx=XM_019811338.2
D1 15090800 15091001
refGene_gn=CEP164;refGene_tx=XM_011286373.3;refGene_gn=CEP164;refGene_tx=XM_011286374.3;refGene_gn=CEP164;refGene_tx=XM_011286375.3;refGene_gn=CEP164;refGene_tx=XM_019811336.2;refGene_gn=CEP164;refGene_tx=XM_01

9811337.2;refGene_gn=CEP164;refGene_tx=XM_019811338.2;refGene_gn=CEP164;refGene_tx=XM_023239145.1
D1 15094365 15094497
refGene_gn=CEP164;refGene_tx=XM_011286373.3;refGene_gn=CEP164;refGene_tx=XM_011286374.3;refGene_gn=CEP164;refGene_tx=XM_011286375.3;refGene_gn=CEP164;refGene_tx=XM_019811336.2;refGene_gn=CEP164;refGene_tx=XM_019811337.2;refGene_gn=CEP164;refGene_tx=XM_019811338.2;refGene_gn=CEP164;refGene_tx=XM_023239145.1
D1 15094578 15094795
refGene_gn=CEP164;refGene_tx=XM_011286373.3;refGene_gn=CEP164;refGene_tx=XM_011286374.3;refGene_gn=CEP164;refGene_tx=XM_011286375.3;refGene_gn=CEP164;refGene_tx=XM_019811336.2;refGene_gn=CEP164;refGene_tx=XM_019811337.2;refGene_gn=CEP164;refGene_tx=XM_019811338.2;refGene_gn=CEP164;refGene_tx=XM_023239145.1
D1 15095787 15095865
refGene_gn=CEP164;refGene_tx=XM_011286373.3;refGene_gn=CEP164;refGene_tx=XM_011286374.3;refGene_gn=CEP164;refGene_tx=XM_011286375.3;refGene_gn=CEP164;refGene_tx=XM_019811336.2;refGene_gn=CEP164;refGene_tx=XM_019811337.2;refGene_gn=CEP164;refGene_tx=XM_019811338.2;refGene_gn=CEP164;refGene_tx=XM_023239145.1
D1 15096098 15096230
refGene_gn=CEP164;refGene_tx=XM_011286373.3;refGene_gn=CEP164;refGene_tx=XM_011286374.3;refGene_gn=CEP164;refGene_tx=XM_011286375.3;refGene_gn=CEP164;refGene_tx=XM_019811336.2;refGene_gn=CEP164;refGene_tx=XM_019811337.2;refGene_gn=CEP164;refGene_tx=XM_019811338.2;refGene_gn=CEP164;refGene_tx=XM_023239145.1
D1 15096541 15096664
refGene_gn=CEP164;refGene_tx=XM_011286373.3;refGene_gn=CEP164;refGene_tx=XM_011286374.3;refGene_gn=CEP164;refGene_tx=XM_011286375.3;refGene_gn=CEP164;refGene_tx=XM_019811336.2;refGene_gn=CEP164;refGene_tx=XM_019811337.2;refGene_gn=CEP164;refGene_tx=XM_019811338.2;refGene_gn=CEP164;refGene_tx=XM_023239145.1
D1 15097628 15097772
refGene_gn=CEP164;refGene_tx=XM_011286373.3;refGene_gn=CEP164;refGene_tx=XM_011286374.3;refGene_gn=CEP164;refGene_tx=XM_011286375.3;refGene_gn=CEP164;refGene_tx=XM_019811336.2;refGene_gn=CEP164;refGene_tx=XM_019811337.2;refGene_gn=CEP164;refGene_tx=XM_019811338.2;refGene_gn=CEP164;refGene_tx=XM_023239145.1
D1 15098217 15098301
refGene_gn=CEP164;refGene_tx=XM_011286373.3;refGene_gn=CEP164;refGene_tx=XM_011286374.3;refGene_gn=CEP164;refGene_tx=XM_011286375.3;refGene_gn=CEP164;refGene_tx=XM_019811336.2;refGene_gn=CEP164;refGene_tx=XM_019811337.2;refGene_gn=CEP164;refGene_tx=XM_019811338.2;refGene_gn=CEP164;refGene_tx=XM_023239145.1
D1 15098446 15098515
refGene_gn=CEP164;refGene_tx=XM_011286373.3;refGene_gn=CEP164;refGene_tx=XM_011286374.3;refGene_gn=CEP164;refGene_tx=XM_011286375.3;refGene_gn=CEP164;refGene_tx=XM_019811336.2;refGene_gn=CEP164;refGene_tx=XM_019811337.2;refGene_gn=CEP164;refGene_tx=XM_019811338.2;refGene_gn=CEP164;refGene_tx=XM_023239145.1

D1 15098855 15099031
refGene_gn=CEP164;refGene_tx=XM_011286373.3;refGene_gn=CEP164;refGene_tx=XM_011286374.3;refGene_gn=CEP164;refGene_tx=XM_011286375.3;refGene_gn=CEP164;refGene_tx=XM_019811336.2;refGene_gn=CEP164;refGene_tx=XM_019811337.2;refGene_gn=CEP164;refGene_tx=XM_019811338.2;refGene_gn=CEP164;refGene_tx=XM_023239145.1
D1 15099408 15099535
refGene_gn=CEP164;refGene_tx=XM_011286373.3;refGene_gn=CEP164;refGene_tx=XM_011286374.3;refGene_gn=CEP164;refGene_tx=XM_011286375.3;refGene_gn=CEP164;refGene_tx=XM_019811336.2;refGene_gn=CEP164;refGene_tx=XM_019811337.2;refGene_gn=CEP164;refGene_tx=XM_019811338.2;refGene_gn=CEP164;refGene_tx=XM_023239145.1
D1 15099802 15099870
refGene_gn=CEP164;refGene_tx=XM_011286373.3;refGene_gn=CEP164;refGene_tx=XM_011286374.3;refGene_gn=CEP164;refGene_tx=XM_011286375.3;refGene_gn=CEP164;refGene_tx=XM_019811336.2;refGene_gn=CEP164;refGene_tx=XM_019811337.2;refGene_gn=CEP164;refGene_tx=XM_019811338.2;refGene_gn=CEP164;refGene_tx=XM_023239145.1
D1 15100315 15100523
refGene_gn=CEP164;refGene_tx=XM_011286373.3;refGene_gn=CEP164;refGene_tx=XM_011286374.3;refGene_gn=CEP164;refGene_tx=XM_011286375.3;refGene_gn=CEP164;refGene_tx=XM_019811336.2;refGene_gn=CEP164;refGene_tx=XM_019811337.2;refGene_gn=CEP164;refGene_tx=XM_019811338.2;refGene_gn=CEP164;refGene_tx=XM_023239145.1
D1 15101373 15101481
refGene_gn=CEP164;refGene_tx=XM_011286373.3;refGene_gn=CEP164;refGene_tx=XM_011286374.3;refGene_gn=CEP164;refGene_tx=XM_011286375.3;refGene_gn=CEP164;refGene_tx=XM_019811336.2;refGene_gn=CEP164;refGene_tx=XM_019811337.2;refGene_gn=CEP164;refGene_tx=XM_019811338.2;refGene_gn=CEP164;refGene_tx=XM_023239145.1
D1 15102060 15102175
refGene_gn=CEP164;refGene_tx=XM_011286373.3;refGene_gn=CEP164;refGene_tx=XM_011286374.3;refGene_gn=CEP164;refGene_tx=XM_011286375.3;refGene_gn=CEP164;refGene_tx=XM_019811336.2;refGene_gn=CEP164;refGene_tx=XM_019811337.2;refGene_gn=CEP164;refGene_tx=XM_019811338.2;refGene_gn=CEP164;refGene_tx=XM_023239145.1
D1 15102659 15103001
refGene_gn=CEP164;refGene_tx=XM_011286373.3;refGene_gn=CEP164;refGene_tx=XM_011286374.3;refGene_gn=CEP164;refGene_tx=XM_011286375.3;refGene_gn=CEP164;refGene_tx=XM_019811336.2;refGene_gn=CEP164;refGene_tx=XM_019811337.2;refGene_gn=CEP164;refGene_tx=XM_019811338.2;refGene_gn=CEP164;refGene_tx=XM_023239145.1
D1 15103874 15103941
refGene_gn=CEP164;refGene_tx=XM_011286373.3;refGene_gn=CEP164;refGene_tx=XM_011286374.3;refGene_gn=CEP164;refGene_tx=XM_011286375.3;refGene_gn=CEP164;refGene_tx=XM_019811336.2;refGene_gn=CEP164;refGene_tx=XM_019811337.2;refGene_gn=CEP164;refGene_tx=XM_019811338.2;refGene_gn=CEP164;refGene_tx=XM_023239145.1
D1 15104768 15104891
refGene_gn=CEP164;refGene_tx=XM_011286373.3;refGene_gn=CEP164;refGene_

tx=XM_011286374.3;refGene_gn=CEP164;refGene_tx=XM_011286375.3;refGene_gn=CEP164;refGene_tx=XM_019811336.2;refGene_gn=CEP164;refGene_tx=XM_019811337.2;refGene_gn=CEP164;refGene_tx=XM_019811338.2;refGene_gn=CEP164;refGene_tx=XM_023239145.1
D1 15105050 15105147 refGene_gn=CEP164;refGene_tx=XM_011286373.3;refGene_gn=CEP164;refGene_tx=XM_011286374.3;refGene_gn=CEP164;refGene_tx=XM_011286375.3;refGene_gn=CEP164;refGene_tx=XM_019811336.2;refGene_gn=CEP164;refGene_tx=XM_019811337.2;refGene_gn=CEP164;refGene_tx=XM_019811338.2;refGene_gn=CEP164;refGene_tx=XM_023239145.1
D1 15142425 15142901 refGene_gn=DSCAML1;refGene_tx=XM_023239146.1
D1 15144382 15144694 refGene_gn=DSCAML1;refGene_tx=XM_023239146.1
D1 15145314 15145509 refGene_gn=DSCAML1;refGene_tx=XM_023239146.1
D1 15146134 15146287 refGene_gn=DSCAML1;refGene_tx=XM_023239146.1
D1 15146569 15146688 refGene_gn=DSCAML1;refGene_tx=XM_023239146.1
D1 15146816 15146847 refGene_gn=DSCAML1;refGene_tx=XM_023239146.1
D1 15149089 15149245 refGene_gn=DSCAML1;refGene_tx=XM_023239146.1
D1 15150741 15151029 refGene_gn=DSCAML1;refGene_tx=XM_023239146.1
D1 15151477 15151666 refGene_gn=DSCAML1;refGene_tx=XM_023239146.1
D1 15152348 15152447 refGene_gn=DSCAML1;refGene_tx=XM_023239146.1
D1 15152695 15152859 refGene_gn=DSCAML1;refGene_tx=XM_023239146.1
D1 15153227 15153345 refGene_gn=DSCAML1;refGene_tx=XM_023239146.1
D1 15155693 15155847 refGene_gn=DSCAML1;refGene_tx=XM_023239146.1
D1 15161790 15161930 refGene_gn=DSCAML1;refGene_tx=XM_023239146.1
D1 15169144 15169300 refGene_gn=DSCAML1;refGene_tx=XM_023239146.1
D1 15171824 15171971 refGene_gn=DSCAML1;refGene_tx=XM_023239146.1
D1 15175159 15175400 refGene_gn=DSCAML1;refGene_tx=XM_023239146.1
D1 15180509 15180580 refGene_gn=DSCAML1;refGene_tx=XM_023239146.1
D1 15182318 15182486 refGene_gn=DSCAML1;refGene_tx=XM_023239146.1
D1 15198964 15199093 refGene_gn=DSCAML1;refGene_tx=XM_023239146.1
D1 15199732 15199829 refGene_gn=DSCAML1;refGene_tx=XM_023239146.1
D1 15200477 15200677 refGene_gn=DSCAML1;refGene_tx=XM_023239146.1
D1 15220544 15220721 refGene_gn=DSCAML1;refGene_tx=XM_023239146.1
D1 15221446 15221566 refGene_gn=DSCAML1;refGene_tx=XM_023239146.1
D1 15221969 15222248 refGene_gn=DSCAML1;refGene_tx=XM_023239146.1
D1 15231902 15232175 refGene_gn=DSCAML1;refGene_tx=XM_023239146.1
D1 15233833 15234130 refGene_gn=DSCAML1;refGene_tx=XM_023239146.1
D1 15236504 15236780 refGene_gn=DSCAML1;refGene_tx=XM_023239146.1
D1 15240380 15240659 refGene_gn=DSCAML1;refGene_tx=XM_023239146.1
D1 15246652 15246799 refGene_gn=DSCAML1;refGene_tx=XM_023239146.1
D1 15472678 15472825 refGene_gn=DSCAML1;refGene_tx=XM_023239146.1
D1 15477118 15477436 refGene_gn=DSCAML1;refGene_tx=XM_023239146.1
D1 15493386 15493609 refGene_gn=DSCAML1;refGene_tx=XM_023239146.1
D1 15512036 15512061 refGene_gn=FXYD2;refGene_tx=XM_003992398.2;refGene_gn=FXYD2;refGene_tx=XM_019811351.1
D1 15512249 15512286 refGene_gn=FXYD2;refGene_tx=XM_003992398.2;refGene_gn=FXYD2;refGene_tx=XM_019811351.1
D1 15513918 15513993

refGene_gn=FXYD2;refGene_tx=XM_003992398.2;refGene_gn=FXYD2;refGene_tx=XM_019811351.1
D1 15514176 15514215 refGene_gn=FXYD2;refGene_tx=XM_003992398.2;refGene_gn=FXYD2;refGene_tx=XM_019811351.1
D1 15516031 15516056 refGene_gn=FXYD2;refGene_tx=XM_019811351.1
D1 15518647 15518666 refGene_gn=FXYD2;refGene_tx=XM_003992398.2
D1 15529809 15529838 refGene_gn=LOC101087344;refGene_tx=XM_006936666.4;refGene_gn=LOC101087344;refGene_tx=XM_006936668.4;refGene_gn=LOC101087344;refGene_tx=XM_019811346.2;refGene_gn=LOC101087344;refGene_tx=XM_019811347.2;refGene_gn=LOC101087344;refGene_tx=XM_019811350.2;refGene_gn=LOC101087344;refGene_tx=XM_023239147.1;refGene_gn=LOC101087344;refGene_tx=XM_023239148.1;refGene_gn=LOC101087344;refGene_tx=XM_023239149.1
D1 15530355 15530405 refGene_gn=LOC101087344;refGene_tx=XM_006936666.4;refGene_gn=LOC101087344;refGene_tx=XM_006936668.4;refGene_gn=LOC101087344;refGene_tx=XM_019811346.2;refGene_gn=LOC101087344;refGene_tx=XM_019811347.2;refGene_gn=LOC101087344;refGene_tx=XM_019811350.2;refGene_gn=LOC101087344;refGene_tx=XM_023239147.1;refGene_gn=LOC101087344;refGene_tx=XM_023239148.1;refGene_gn=LOC101087344;refGene_tx=XM_023239149.1
D1 15531124 15531161 refGene_gn=LOC101087344;refGene_tx=XM_006936666.4;refGene_gn=LOC101087344;refGene_tx=XM_006936668.4;refGene_gn=LOC101087344;refGene_tx=XM_019811346.2;refGene_gn=LOC101087344;refGene_tx=XM_019811347.2;refGene_gn=LOC101087344;refGene_tx=XM_019811350.2;refGene_gn=LOC101087344;refGene_tx=XM_023239147.1;refGene_gn=LOC101087344;refGene_tx=XM_023239148.1;refGene_gn=LOC101087344;refGene_tx=XM_023239149.1
D1 15531978 15532053 refGene_gn=LOC101087344;refGene_tx=XM_006936666.4;refGene_gn=LOC101087344;refGene_tx=XM_006936668.4;refGene_gn=LOC101087344;refGene_tx=XM_019811346.2;refGene_gn=LOC101087344;refGene_tx=XM_019811347.2;refGene_gn=LOC101087344;refGene_tx=XM_019811350.2;refGene_gn=LOC101087344;refGene_tx=XM_023239147.1;refGene_gn=LOC101087344;refGene_tx=XM_023239148.1;refGene_gn=LOC101087344;refGene_tx=XM_023239149.1
D1 15532157 15532196 refGene_gn=LOC101087344;refGene_tx=XM_006936666.4;refGene_gn=LOC101087344;refGene_tx=XM_006936668.4;refGene_gn=LOC101087344;refGene_tx=XM_019811346.2;refGene_gn=LOC101087344;refGene_tx=XM_019811347.2;refGene_gn=LOC101087344;refGene_tx=XM_019811350.2;refGene_gn=LOC101087344;refGene_tx=XM_023239147.1;refGene_gn=LOC101087344;refGene_tx=XM_023239148.1;refGene_gn=LOC101087344;refGene_tx=XM_023239149.1
D1 15532803 15532866 refGene_gn=LOC101087344;refGene_tx=XM_006936666.4;refGene_gn=LOC101087344;refGene_tx=XM_006936668.4;refGene_gn=LOC101087344;refGene_tx=XM_019811346.2;refGene_gn=LOC101087344;refGene_tx=XM_019811347.2;refGene_gn=LOC101087344;refGene_tx=XM_019811350.2;refGene_gn=LOC101087344;refGene_tx=XM_023239147.1;refGene_gn=LOC101087344;refGene_tx=XM_023239148.1;refGene_gn=LOC101087344;refGene_tx=XM_023239149.1
D1 15546227 15546234

refGene_gn=LOC101087344;refGene_tx=XM_006936666.4;refGene_gn=LOC101087
344;refGene_tx=XM_006936668.4
D1 15552858 15552946
refGene_gn=LOC101087344;refGene_tx=XM_019811346.2;refGene_gn=LOC101087
344;refGene_tx=XM_019811347.2;refGene_gn=LOC101087344;refGene_tx=XM_02
3239147.1
D1 15579710 15579737
refGene_gn=TMPRSS13;refGene_tx=XM_006936671.3;refGene_gn=TMPRSS13;refG
ene_tx=XM_019811352.2
D1 15580431 15580584
refGene_gn=TMPRSS13;refGene_tx=XM_006936671.3;refGene_gn=TMPRSS13;refG
ene_tx=XM_019811352.2
D1 15580717 15580860
refGene_gn=TMPRSS13;refGene_tx=XM_006936671.3;refGene_gn=TMPRSS13;refG
ene_tx=XM_019811352.2
D1 15582705 15582804
refGene_gn=TMPRSS13;refGene_tx=XM_006936671.3;refGene_gn=TMPRSS13;refG
ene_tx=XM_019811352.2
D1 15585912 15586085
refGene_gn=TMPRSS13;refGene_tx=XM_006936671.3;refGene_gn=TMPRSS13;refG
ene_tx=XM_019811352.2
D1 15586914 15587077
refGene_gn=TMPRSS13;refGene_tx=XM_006936671.3;refGene_gn=TMPRSS13;refG
ene_tx=XM_019811352.2
D1 15588368 15588412
refGene_gn=TMPRSS13;refGene_tx=XM_006936671.3;refGene_gn=TMPRSS13;refG
ene_tx=XM_019811352.2
D1 15589451 15589544
refGene_gn=TMPRSS13;refGene_tx=XM_006936671.3;refGene_gn=TMPRSS13;refG
ene_tx=XM_019811352.2
D1 15591559 15591689
refGene_gn=TMPRSS13;refGene_tx=XM_006936671.3;refGene_gn=TMPRSS13;refG
ene_tx=XM_019811352.2
D1 15592605 15592728
refGene_gn=TMPRSS13;refGene_tx=XM_006936671.3;refGene_gn=TMPRSS13;refG
ene_tx=XM_019811352.2
D1 15592823 15592862 refGene_gn=TMPRSS13;refGene_tx=XM_019811352.2
D1 15595272 15595377
refGene_gn=TMPRSS13;refGene_tx=XM_006936671.3;refGene_gn=TMPRSS13;refG
ene_tx=XM_019811352.2
D1 15596383 15596594
refGene_gn=TMPRSS13;refGene_tx=XM_006936671.3;refGene_gn=TMPRSS13;refG
ene_tx=XM_019811352.2
D1 15606601 15606622
refGene_gn=TMPRSS13;refGene_tx=XM_006936671.3;refGene_gn=TMPRSS13;refG
ene_tx=XM_019811352.2
D1 15647731 15647798
refGene_gn=IL10RA;refGene_tx=XM_003992400.5;refGene_gn=IL10RA;refGene_
tx=XM_006936672.4
D1 15649819 15649940

refGene_gn=IL10RA;refGene_tx=XM_003992400.5;refGene_gn=IL10RA;refGene_tx=XM_006936672.4
D1 15650532 15650717
refGene_gn=IL10RA;refGene_tx=XM_003992400.5;refGene_gn=IL10RA;refGene_tx=XM_006936672.4
D1 15651609 15651776
refGene_gn=IL10RA;refGene_tx=XM_003992400.5;refGene_gn=IL10RA;refGene_tx=XM_006936672.4
D1 15652395 15652546
refGene_gn=IL10RA;refGene_tx=XM_003992400.5;refGene_gn=IL10RA;refGene_tx=XM_006936672.4
D1 15653391 15653513
refGene_gn=IL10RA;refGene_tx=XM_003992400.5;refGene_gn=IL10RA;refGene_tx=XM_006936672.4
D1 15656953 15657877
refGene_gn=IL10RA;refGene_tx=XM_003992400.5;refGene_gn=IL10RA;refGene_tx=XM_006936672.4
D1 15727375 15727399
refGene_gn=TMPRSS4;refGene_tx=XM_019811355.2;refGene_gn=TMPRSS4;refGene_tx=XM_019811356.2
D1 15742846 15742880
refGene_gn=TMPRSS4;refGene_tx=XM_019811355.2;refGene_gn=TMPRSS4;refGene_tx=XM_019811356.2
D1 15746429 15746543
refGene_gn=TMPRSS4;refGene_tx=XM_019811355.2;refGene_gn=TMPRSS4;refGene_tx=XM_019811356.2
D1 15749056 15749209
refGene_gn=TMPRSS4;refGene_tx=XM_019811355.2;refGene_gn=TMPRSS4;refGene_tx=XM_019811356.2
D1 15750309 15750439
refGene_gn=TMPRSS4;refGene_tx=XM_019811355.2;refGene_gn=TMPRSS4;refGene_tx=XM_019811356.2
D1 15752427 15752529
refGene_gn=TMPRSS4;refGene_tx=XM_019811355.2;refGene_gn=TMPRSS4;refGene_tx=XM_019811356.2
D1 15753444 15753485
refGene_gn=TMPRSS4;refGene_tx=XM_019811355.2;refGene_gn=TMPRSS4;refGene_tx=XM_019811356.2
D1 15755643 15755803
refGene_gn=TMPRSS4;refGene_tx=XM_019811355.2;refGene_gn=TMPRSS4;refGene_tx=XM_019811356.2
D1 15756554 15756721
refGene_gn=TMPRSS4;refGene_tx=XM_019811355.2;refGene_gn=TMPRSS4;refGene_tx=XM_019811356.2
D1 15757504 15757603
refGene_gn=TMPRSS4;refGene_tx=XM_019811355.2;refGene_gn=TMPRSS4;refGene_tx=XM_019811356.2
D1 15757789 15757932
refGene_gn=TMPRSS4;refGene_tx=XM_019811355.2;refGene_gn=TMPRSS4;refGene_tx=XM_019811356.2

D1 15758664 15758700 refGene_gn=TMPRSS4;refGene_tx=XM_019811356.2
D1 15759111 15759261 refGene_gn=TMPRSS4;refGene_tx=XM_019811355.2
D1 15759684 15759696 refGene_gn=TMPRSS4;refGene_tx=XM_019811355.2
D1 15775944 15776038
refGene_gn=SCN4B;refGene_tx=XM_003992401.3;refGene_gn=SCN4B;refGene_tx=XM_023239154.1
D1 15779564 15779694
refGene_gn=SCN4B;refGene_tx=XM_003992401.3;refGene_gn=SCN4B;refGene_tx=XM_023239154.1
D1 15782098 15782327
refGene_gn=SCN4B;refGene_tx=XM_003992401.3;refGene_gn=SCN4B;refGene_tx=XM_023239154.1
D1 15783312 15783485 refGene_gn=SCN4B;refGene_tx=XM_003992401.3
D1 15791995 15792056 refGene_gn=SCN4B;refGene_tx=XM_003992401.3
D1 15796691 15796696
refGene_gn=LOC102899230;refGene_tx=XM_006936673.3;refGene_gn=LOC102899230;refGene_tx=XM_006936674.3;refGene_gn=LOC102899230;refGene_tx=XM_011286376.3
D1 15805188 15805388
refGene_gn=SCN2B;refGene_tx=XM_003992403.5;refGene_gn=SCN2B;refGene_tx=XM_019811358.2
D1 15806044 15806255
refGene_gn=SCN2B;refGene_tx=XM_003992403.5;refGene_gn=SCN2B;refGene_tx=XM_019811358.2
D1 15806581 15806748
refGene_gn=SCN2B;refGene_tx=XM_003992403.5;refGene_gn=SCN2B;refGene_tx=XM_019811358.2
D1 15807008 15807244
refGene_gn=LOC102899230;refGene_tx=XM_006936673.3;refGene_gn=LOC102899230;refGene_tx=XM_006936674.3;refGene_gn=LOC102899230;refGene_tx=XM_011286376.3
D1 15808405 15808854
refGene_gn=LOC102899230;refGene_tx=XM_006936673.3;refGene_gn=LOC102899230;refGene_tx=XM_006936674.3
D1 15813542 15813683
refGene_gn=LOC102899230;refGene_tx=XM_011286376.3;refGene_gn=SCN2B;refGene_tx=XM_003992403.5
D1 15821709 15821802
refGene_gn=JAML;refGene_tx=XM_003992404.5;refGene_gn=JAML;refGene_tx=XM_006936675.4;refGene_gn=JAML;refGene_tx=XM_019811359.2;refGene_gn=JAML;refGene_tx=XM_019811360.2
D1 15823709 15823796
refGene_gn=JAML;refGene_tx=XM_003992404.5;refGene_gn=JAML;refGene_tx=XM_006936675.4;refGene_gn=JAML;refGene_tx=XM_019811359.2;refGene_gn=JAML;refGene_tx=XM_019811360.2
D1 15825327 15825388
refGene_gn=JAML;refGene_tx=XM_003992404.5;refGene_gn=JAML;refGene_tx=XM_006936675.4;refGene_gn=JAML;refGene_tx=XM_019811359.2;refGene_gn=JAML;refGene_tx=XM_019811360.2
D1 15827545 15827684

refGene_gn=JAML;refGene_tx=XM_003992404.5;refGene_gn=JAML;refGene_tx=XM_006936675.4;refGene_gn=JAML;refGene_tx=XM_019811359.2;refGene_gn=JAML;refGene_tx=XM_019811360.2
D1 15831151 15831389
refGene_gn=JAML;refGene_tx=XM_003992404.5;refGene_gn=JAML;refGene_tx=XM_006936675.4;refGene_gn=JAML;refGene_tx=XM_019811359.2;refGene_gn=JAML;refGene_tx=XM_019811360.2
D1 15833988 15834101
refGene_gn=JAML;refGene_tx=XM_003992404.5;refGene_gn=JAML;refGene_tx=XM_006936675.4;refGene_gn=JAML;refGene_tx=XM_019811359.2;refGene_gn=JAML;refGene_tx=XM_019811360.2
D1 15838171 15838397
refGene_gn=JAML;refGene_tx=XM_003992404.5;refGene_gn=JAML;refGene_tx=XM_006936675.4;refGene_gn=JAML;refGene_tx=XM_019811359.2;refGene_gn=JAML;refGene_tx=XM_019811360.2
D1 15839752 15839907
refGene_gn=JAML;refGene_tx=XM_003992404.5;refGene_gn=JAML;refGene_tx=XM_006936675.4;refGene_gn=JAML;refGene_tx=XM_019811359.2;refGene_gn=JAML;refGene_tx=XM_019811360.2
D1 15840573 15840673 refGene_gn=JAML;refGene_tx=XM_019811359.2
D1 15842750 15842793
refGene_gn=JAML;refGene_tx=XM_003992404.5;refGene_gn=JAML;refGene_tx=XM_006936675.4
D1 15852942 15852969 refGene_gn=MPZL3;refGene_tx=XM_003992405.5
D1 15856787 15856851 refGene_gn=MPZL3;refGene_tx=XM_003992405.5
D1 15858397 15858563 refGene_gn=MPZL3;refGene_tx=XM_003992405.5
D1 15860857 15861068 refGene_gn=MPZL3;refGene_tx=XM_003992405.5
D1 15864279 15864446 refGene_gn=MPZL3;refGene_tx=XM_003992405.5
D1 15873290 15873363 refGene_gn=MPZL3;refGene_tx=XM_003992405.5
D1 15879056 15879120 refGene_gn=MPZL2;refGene_tx=XM_003992406.5
D1 15882297 15882445 refGene_gn=MPZL2;refGene_tx=XM_003992406.5
D1 15884963 15885003 refGene_gn=MPZL2;refGene_tx=XM_003992406.5
D1 15885004 15885175 refGene_gn=MPZL2;refGene_tx=XM_003992406.5
D1 15885458 15885625 refGene_gn=MPZL2;refGene_tx=XM_003992406.5
D1 15886588 15886646 refGene_gn=MPZL2;refGene_tx=XM_003992406.5
D1 15915141 15915193
refGene_gn=CD3E;refGene_tx=NM_001009862.1;refGene_gn=CD3E;refGene_tx=XM_023238913.1
D1 15916862 15916950 refGene_gn=CD3E;refGene_tx=XM_023238912.1
D1 15917081 15917102
refGene_gn=CD3E;refGene_tx=NM_001009862.1;refGene_gn=CD3E;refGene_tx=XM_023238912.1;refGene_gn=CD3E;refGene_tx=XM_023238913.1
D1 15919750 15919768
refGene_gn=CD3E;refGene_tx=NM_001009862.1;refGene_gn=CD3E;refGene_tx=XM_023238912.1;refGene_gn=CD3E;refGene_tx=XM_023238913.1
D1 15923106 15923136
refGene_gn=CD3E;refGene_tx=NM_001009862.1;refGene_gn=CD3E;refGene_tx=XM_023238912.1;refGene_gn=CD3E;refGene_tx=XM_023238913.1
D1 15923591 15923810
refGene_gn=CD3E;refGene_tx=NM_001009862.1;refGene_gn=CD3E;refGene_tx=X

M_023238912.1;refGene_gn=CD3E;refGene_tx=XM_023238913.1
D1 15925090 15925258
refGene_gn=CD3E;refGene_tx=NM_001009862.1;refGene_gn=CD3E;refGene_tx=XM_023238912.1;refGene_gn=CD3E;refGene_tx=XM_023238913.1
D1 15926010 15926057
refGene_gn=CD3E;refGene_tx=NM_001009862.1;refGene_gn=CD3E;refGene_tx=XM_023238912.1;refGene_gn=CD3E;refGene_tx=XM_023238913.1
D1 15926939 15926996
refGene_gn=CD3E;refGene_tx=NM_001009862.1;refGene_gn=CD3E;refGene_tx=XM_023238912.1;refGene_gn=CD3E;refGene_tx=XM_023238913.1
D1 15933995 15934061
refGene_gn=CD3D;refGene_tx=XM_003992463.4;refGene_gn=CD3D;refGene_tx=XM_019811361.2
D1 15934281 15934325
refGene_gn=CD3D;refGene_tx=XM_003992463.4;refGene_gn=CD3D;refGene_tx=XM_019811361.2
D1 15934611 15934743 refGene_gn=CD3D;refGene_tx=XM_003992463.4
D1 15935210 15935426
refGene_gn=CD3D;refGene_tx=XM_003992463.4;refGene_gn=CD3D;refGene_tx=XM_019811361.2
D1 15936986 15937041
refGene_gn=CD3D;refGene_tx=XM_003992463.4;refGene_gn=CD3D;refGene_tx=XM_019811361.2
D1 15938692 15938747
refGene_gn=CD3G;refGene_tx=XM_003992407.4;refGene_gn=CD3G;refGene_tx=XM_006936676.3;refGene_gn=CD3G;refGene_tx=XM_006936677.3;refGene_gn=CD3G;refGene_tx=XM_006936678.3
D1 15943149 15943173
refGene_gn=CD3G;refGene_tx=XM_003992407.4;refGene_gn=CD3G;refGene_tx=XM_006936676.3;refGene_gn=CD3G;refGene_tx=XM_006936677.3;refGene_gn=CD3G;refGene_tx=XM_006936678.3
D1 15943635 15943863
refGene_gn=CD3G;refGene_tx=XM_003992407.4;refGene_gn=CD3G;refGene_tx=XM_006936676.3;refGene_gn=CD3G;refGene_tx=XM_006936677.3;refGene_gn=CD3G;refGene_tx=XM_006936678.3
D1 15944464 15944596
refGene_gn=CD3G;refGene_tx=XM_003992407.4;refGene_gn=CD3G;refGene_tx=XM_006936676.3;refGene_gn=CD3G;refGene_tx=XM_006936677.3;refGene_gn=CD3G;refGene_tx=XM_006936678.3
D1 15945226 15945270
refGene_gn=CD3G;refGene_tx=XM_003992407.4;refGene_gn=CD3G;refGene_tx=XM_006936676.3;refGene_gn=CD3G;refGene_tx=XM_006936677.3;refGene_gn=CD3G;refGene_tx=XM_006936678.3
D1 15945671 15945737
refGene_gn=CD3G;refGene_tx=XM_003992407.4;refGene_gn=CD3G;refGene_tx=XM_006936676.3;refGene_gn=CD3G;refGene_tx=XM_006936677.3;refGene_gn=CD3G;refGene_tx=XM_006936678.3
D1 15956525 15956646
refGene_gn=UBE4A;refGene_tx=XM_003992408.5;refGene_gn=UBE4A;refGene_tx=XM_019811362.2

D1 15961078 15961252
refGene_gn=UBE4A;refGene_tx=XM_003992408.5;refGene_gn=UBE4A;refGene_tx
=XM_019811362.2
D1 15962091 15962204
refGene_gn=UBE4A;refGene_tx=XM_003992408.5;refGene_gn=UBE4A;refGene_tx
=XM_019811362.2
D1 15963722 15963875
refGene_gn=UBE4A;refGene_tx=XM_003992408.5;refGene_gn=UBE4A;refGene_tx
=XM_019811362.2
D1 15965117 15965277
refGene_gn=UBE4A;refGene_tx=XM_003992408.5;refGene_gn=UBE4A;refGene_tx
=XM_019811362.2
D1 15965508 15965711
refGene_gn=UBE4A;refGene_tx=XM_003992408.5;refGene_gn=UBE4A;refGene_tx
=XM_019811362.2
D1 15965917 15966109
refGene_gn=UBE4A;refGene_tx=XM_003992408.5;refGene_gn=UBE4A;refGene_tx
=XM_019811362.2
D1 15967244 15967578
refGene_gn=UBE4A;refGene_tx=XM_003992408.5;refGene_gn=UBE4A;refGene_tx
=XM_019811362.2
D1 15968567 15968688
refGene_gn=UBE4A;refGene_tx=XM_003992408.5;refGene_gn=UBE4A;refGene_tx
=XM_019811362.2
D1 15970223 15970528
refGene_gn=UBE4A;refGene_tx=XM_003992408.5;refGene_gn=UBE4A;refGene_tx
=XM_019811362.2
D1 15971999 15972132
refGene_gn=UBE4A;refGene_tx=XM_003992408.5;refGene_gn=UBE4A;refGene_tx
=XM_019811362.2
D1 15973177 15973365
refGene_gn=UBE4A;refGene_tx=XM_003992408.5;refGene_gn=UBE4A;refGene_tx
=XM_019811362.2
D1 15974745 15974846
refGene_gn=UBE4A;refGene_tx=XM_003992408.5;refGene_gn=UBE4A;refGene_tx
=XM_019811362.2
D1 15974961 15975075
refGene_gn=UBE4A;refGene_tx=XM_003992408.5;refGene_gn=UBE4A;refGene_tx
=XM_019811362.2
D1 15976969 15977144
refGene_gn=UBE4A;refGene_tx=XM_003992408.5;refGene_gn=UBE4A;refGene_tx
=XM_019811362.2
D1 15980716 15980897
refGene_gn=UBE4A;refGene_tx=XM_003992408.5;refGene_gn=UBE4A;refGene_tx
=XM_019811362.2
D1 15981336 15981484
refGene_gn=UBE4A;refGene_tx=XM_003992408.5;refGene_gn=UBE4A;refGene_tx
=XM_019811362.2
D1 15981697 15981892 refGene_gn=UBE4A;refGene_tx=XM_019811362.2
D1 15982966 15983124 refGene_gn=UBE4A;refGene_tx=XM_003992408.5

D1 15984277 15984407 refGene_gn=UBE4A;refGene_tx=XM_003992408.5
D1 15989183 15989235 refGene_gn=ATP5MG;refGene_tx=XM_003992409.5
D1 15994005 15994166 refGene_gn=ATP5MG;refGene_tx=XM_003992409.5
D1 15995475 15995574 refGene_gn=ATP5MG;refGene_tx=XM_003992409.5
D1 16019992 16020424
refGene_gn=KMT2A;refGene_tx=XM_011286446.3;refGene_gn=KMT2A;refGene_tx=XM_023239150.1;refGene_gn=KMT2A;refGene_tx=XM_023239151.1;refGene_gn=KMT2A;refGene_tx=XM_023239152.1;refGene_gn=KMT2A;refGene_tx=XM_023239153.1
D1 16022510 16022609
refGene_gn=KMT2A;refGene_tx=XM_023239150.1;refGene_gn=KMT2A;refGene_tx=XM_023239151.1;refGene_gn=KMT2A;refGene_tx=XM_023239152.1
D1 16049388 16049458
refGene_gn=KMT2A;refGene_tx=XM_011286446.3;refGene_gn=KMT2A;refGene_tx=XM_023239150.1;refGene_gn=KMT2A;refGene_tx=XM_023239151.1;refGene_gn=KMT2A;refGene_tx=XM_023239152.1;refGene_gn=KMT2A;refGene_tx=XM_023239153.1
D1 16052129 16054783
refGene_gn=KMT2A;refGene_tx=XM_011286446.3;refGene_gn=KMT2A;refGene_tx=XM_023239150.1;refGene_gn=KMT2A;refGene_tx=XM_023239151.1;refGene_gn=KMT2A;refGene_tx=XM_023239152.1;refGene_gn=KMT2A;refGene_tx=XM_023239153.1
D1 16056327 16056505
refGene_gn=KMT2A;refGene_tx=XM_011286446.3;refGene_gn=KMT2A;refGene_tx=XM_023239150.1;refGene_gn=KMT2A;refGene_tx=XM_023239151.1;refGene_gn=KMT2A;refGene_tx=XM_023239152.1;refGene_gn=KMT2A;refGene_tx=XM_023239153.1
D1 16057378 16057613
refGene_gn=KMT2A;refGene_tx=XM_011286446.3;refGene_gn=KMT2A;refGene_tx=XM_023239150.1;refGene_gn=KMT2A;refGene_tx=XM_023239151.1;refGene_gn=KMT2A;refGene_tx=XM_023239152.1;refGene_gn=KMT2A;refGene_tx=XM_023239153.1
D1 16059290 16059355
refGene_gn=KMT2A;refGene_tx=XM_011286446.3;refGene_gn=KMT2A;refGene_tx=XM_023239150.1;refGene_gn=KMT2A;refGene_tx=XM_023239151.1;refGene_gn=KMT2A;refGene_tx=XM_023239152.1;refGene_gn=KMT2A;refGene_tx=XM_023239153.1
D1 16059767 16060145
refGene_gn=KMT2A;refGene_tx=XM_011286446.3;refGene_gn=KMT2A;refGene_tx=XM_023239150.1;refGene_gn=KMT2A;refGene_tx=XM_023239151.1;refGene_gn=KMT2A;refGene_tx=XM_023239152.1;refGene_gn=KMT2A;refGene_tx=XM_023239153.1
D1 16060467 16060541
refGene_gn=KMT2A;refGene_tx=XM_011286446.3;refGene_gn=KMT2A;refGene_tx=XM_023239150.1;refGene_gn=KMT2A;refGene_tx=XM_023239151.1;refGene_gn=KMT2A;refGene_tx=XM_023239152.1;refGene_gn=KMT2A;refGene_tx=XM_023239153.1
D1 16061525 16061657
refGene_gn=KMT2A;refGene_tx=XM_011286446.3;refGene_gn=KMT2A;refGene_tx=XM_023239150.1;refGene_gn=KMT2A;refGene_tx=XM_023239151.1;refGene_gn=

KMT2A;refGene_tx=XM_023239152.1;refGene_gn=KMT2A;refGene_tx=XM_0232391
53.1
D1 16062405 16062519
refGene_gn=KMT2A;refGene_tx=XM_011286446.3;refGene_gn=KMT2A;refGene_tx
=XM_023239150.1;refGene_gn=KMT2A;refGene_tx=XM_023239151.1;refGene_gn=
KMT2A;refGene_tx=XM_023239152.1;refGene_gn=KMT2A;refGene_tx=XM_0232391
53.1
D1 16066589 16066736
refGene_gn=KMT2A;refGene_tx=XM_011286446.3;refGene_gn=KMT2A;refGene_tx
=XM_023239150.1;refGene_gn=KMT2A;refGene_tx=XM_023239151.1;refGene_gn=
KMT2A;refGene_tx=XM_023239152.1;refGene_gn=KMT2A;refGene_tx=XM_0232391
53.1
D1 16067392 16067488
refGene_gn=KMT2A;refGene_tx=XM_011286446.3;refGene_gn=KMT2A;refGene_tx
=XM_023239150.1;refGene_gn=KMT2A;refGene_tx=XM_023239151.1;refGene_gn=
KMT2A;refGene_tx=XM_023239152.1;refGene_gn=KMT2A;refGene_tx=XM_0232391
53.1
D1 16067725 16067846
refGene_gn=KMT2A;refGene_tx=XM_011286446.3;refGene_gn=KMT2A;refGene_tx
=XM_023239150.1;refGene_gn=KMT2A;refGene_tx=XM_023239151.1;refGene_gn=
KMT2A;refGene_tx=XM_023239152.1;refGene_gn=KMT2A;refGene_tx=XM_0232391
53.1
D1 16068904 16069027
refGene_gn=KMT2A;refGene_tx=XM_011286446.3;refGene_gn=KMT2A;refGene_tx
=XM_023239150.1;refGene_gn=KMT2A;refGene_tx=XM_023239151.1;refGene_gn=
KMT2A;refGene_tx=XM_023239152.1;refGene_gn=KMT2A;refGene_tx=XM_0232391
53.1
D1 16069477 16069662
refGene_gn=KMT2A;refGene_tx=XM_011286446.3;refGene_gn=KMT2A;refGene_tx
=XM_023239150.1;refGene_gn=KMT2A;refGene_tx=XM_023239151.1;refGene_gn=
KMT2A;refGene_tx=XM_023239152.1;refGene_gn=KMT2A;refGene_tx=XM_0232391
53.1
D1 16070847 16071021
refGene_gn=KMT2A;refGene_tx=XM_011286446.3;refGene_gn=KMT2A;refGene_tx
=XM_023239150.1;refGene_gn=KMT2A;refGene_tx=XM_023239151.1;refGene_gn=
KMT2A;refGene_tx=XM_023239152.1;refGene_gn=KMT2A;refGene_tx=XM_0232391
53.1
D1 16071929 16072040
refGene_gn=KMT2A;refGene_tx=XM_011286446.3;refGene_gn=KMT2A;refGene_tx
=XM_023239150.1;refGene_gn=KMT2A;refGene_tx=XM_023239151.1;refGene_gn=
KMT2A;refGene_tx=XM_023239152.1;refGene_gn=KMT2A;refGene_tx=XM_0232391
53.1
D1 16072347 16072421
refGene_gn=KMT2A;refGene_tx=XM_011286446.3;refGene_gn=KMT2A;refGene_tx
=XM_023239150.1;refGene_gn=KMT2A;refGene_tx=XM_023239151.1;refGene_gn=
KMT2A;refGene_tx=XM_023239152.1;refGene_gn=KMT2A;refGene_tx=XM_0232391
53.1
D1 16073048 16073242
refGene_gn=KMT2A;refGene_tx=XM_011286446.3;refGene_gn=KMT2A;refGene_tx
=XM_023239150.1;refGene_gn=KMT2A;refGene_tx=XM_023239151.1;refGene_gn=

KMT2A;refGene_tx=XM_023239152.1;refGene_gn=KMT2A;refGene_tx=XM_0232391
53.1
D1 16073572 16073679
refGene_gn=KMT2A;refGene_tx=XM_011286446.3;refGene_gn=KMT2A;refGene_tx
=XM_023239150.1;refGene_gn=KMT2A;refGene_tx=XM_023239151.1;refGene_gn=
KMT2A;refGene_tx=XM_023239152.1;refGene_gn=KMT2A;refGene_tx=XM_0232391
53.1
D1 16074517 16074655
refGene_gn=KMT2A;refGene_tx=XM_011286446.3;refGene_gn=KMT2A;refGene_tx
=XM_023239150.1;refGene_gn=KMT2A;refGene_tx=XM_023239151.1;refGene_gn=
KMT2A;refGene_tx=XM_023239152.1;refGene_gn=KMT2A;refGene_tx=XM_0232391
53.1
D1 16075707 16075866
refGene_gn=KMT2A;refGene_tx=XM_011286446.3;refGene_gn=KMT2A;refGene_tx
=XM_023239150.1;refGene_gn=KMT2A;refGene_tx=XM_023239151.1;refGene_gn=
KMT2A;refGene_tx=XM_023239152.1;refGene_gn=KMT2A;refGene_tx=XM_0232391
53.1
D1 16076120 16076238
refGene_gn=KMT2A;refGene_tx=XM_011286446.3;refGene_gn=KMT2A;refGene_tx
=XM_023239150.1;refGene_gn=KMT2A;refGene_tx=XM_023239151.1;refGene_gn=
KMT2A;refGene_tx=XM_023239152.1;refGene_gn=KMT2A;refGene_tx=XM_0232391
53.1
D1 16076360 16076439
refGene_gn=KMT2A;refGene_tx=XM_011286446.3;refGene_gn=KMT2A;refGene_tx
=XM_023239150.1;refGene_gn=KMT2A;refGene_tx=XM_023239151.1;refGene_gn=
KMT2A;refGene_tx=XM_023239152.1;refGene_gn=KMT2A;refGene_tx=XM_0232391
53.1
D1 16077476 16077637
refGene_gn=KMT2A;refGene_tx=XM_011286446.3;refGene_gn=KMT2A;refGene_tx
=XM_023239150.1;refGene_gn=KMT2A;refGene_tx=XM_023239151.1;refGene_gn=
KMT2A;refGene_tx=XM_023239152.1;refGene_gn=KMT2A;refGene_tx=XM_0232391
53.1
D1 16078215 16078398
refGene_gn=KMT2A;refGene_tx=XM_011286446.3;refGene_gn=KMT2A;refGene_tx
=XM_023239150.1;refGene_gn=KMT2A;refGene_tx=XM_023239151.1;refGene_gn=
KMT2A;refGene_tx=XM_023239152.1;refGene_gn=KMT2A;refGene_tx=XM_0232391
53.1
D1 16078617 16082860
refGene_gn=KMT2A;refGene_tx=XM_011286446.3;refGene_gn=KMT2A;refGene_tx
=XM_023239150.1;refGene_gn=KMT2A;refGene_tx=XM_023239151.1;refGene_gn=
KMT2A;refGene_tx=XM_023239152.1;refGene_gn=KMT2A;refGene_tx=XM_0232391
53.1
D1 16083613 16083694
refGene_gn=KMT2A;refGene_tx=XM_011286446.3;refGene_gn=KMT2A;refGene_tx
=XM_023239150.1;refGene_gn=KMT2A;refGene_tx=XM_023239151.1;refGene_gn=
KMT2A;refGene_tx=XM_023239152.1;refGene_gn=KMT2A;refGene_tx=XM_0232391
53.1
D1 16084761 16084820
refGene_gn=KMT2A;refGene_tx=XM_011286446.3;refGene_gn=KMT2A;refGene_tx
=XM_023239150.1;refGene_gn=KMT2A;refGene_tx=XM_023239151.1;refGene_gn=

KMT2A;refGene_tx=XM_023239152.1;refGene_gn=KMT2A;refGene_tx=XM_0232391
53.1
D1 16085912 16086083
refGene_gn=KMT2A;refGene_tx=XM_011286446.3;refGene_gn=KMT2A;refGene_tx
=XM_023239150.1;refGene_gn=KMT2A;refGene_tx=XM_023239151.1;refGene_gn=
KMT2A;refGene_tx=XM_023239152.1;refGene_gn=KMT2A;refGene_tx=XM_0232391
53.1
D1 16087895 16087970
refGene_gn=KMT2A;refGene_tx=XM_011286446.3;refGene_gn=KMT2A;refGene_tx
=XM_023239150.1;refGene_gn=KMT2A;refGene_tx=XM_023239151.1;refGene_gn=
KMT2A;refGene_tx=XM_023239152.1;refGene_gn=KMT2A;refGene_tx=XM_0232391
53.1
D1 16098584 16098759
refGene_gn=KMT2A;refGene_tx=XM_011286446.3;refGene_gn=KMT2A;refGene_tx
=XM_023239150.1;refGene_gn=KMT2A;refGene_tx=XM_023239151.1;refGene_gn=
KMT2A;refGene_tx=XM_023239152.1;refGene_gn=KMT2A;refGene_tx=XM_0232391
53.1
D1 16098919 16099027
refGene_gn=KMT2A;refGene_tx=XM_011286446.3;refGene_gn=KMT2A;refGene_tx
=XM_023239150.1;refGene_gn=KMT2A;refGene_tx=XM_023239151.1;refGene_gn=
KMT2A;refGene_tx=XM_023239152.1;refGene_gn=KMT2A;refGene_tx=XM_0232391
53.1
D1 16099744 16099828
refGene_gn=KMT2A;refGene_tx=XM_011286446.3;refGene_gn=KMT2A;refGene_tx
=XM_023239150.1;refGene_gn=KMT2A;refGene_tx=XM_023239151.1;refGene_gn=
KMT2A;refGene_tx=XM_023239152.1;refGene_gn=KMT2A;refGene_tx=XM_0232391
53.1
D1 16100252 16100382
refGene_gn=KMT2A;refGene_tx=XM_011286446.3;refGene_gn=KMT2A;refGene_tx
=XM_023239150.1;refGene_gn=KMT2A;refGene_tx=XM_023239151.1;refGene_gn=
KMT2A;refGene_tx=XM_023239152.1;refGene_gn=KMT2A;refGene_tx=XM_0232391
53.1
D1 16100863 16101139
refGene_gn=KMT2A;refGene_tx=XM_011286446.3;refGene_gn=KMT2A;refGene_tx
=XM_023239150.1;refGene_gn=KMT2A;refGene_tx=XM_023239151.1;refGene_gn=
KMT2A;refGene_tx=XM_023239152.1;refGene_gn=KMT2A;refGene_tx=XM_0232391
53.1
D1 16106667 16106785 refGene_gn=TTC36;refGene_tx=XM_023239160.1
D1 16107329 16107518 refGene_gn=TTC36;refGene_tx=XM_023239160.1
D1 16108763 16108893 refGene_gn=TTC36;refGene_tx=XM_023239161.1
D1 16114034 16114297
refGene_gn=TTC36;refGene_tx=XM_023239160.1;refGene_gn=TTC36;refGene_tx
=XM_023239161.1
D1 16115093 16115163
refGene_gn=TMEM25;refGene_tx=XM_006936680.3;refGene_gn=TMEM25;refGene_
tx=XM_006936681.3;refGene_gn=TMEM25;refGene_tx=XM_006936682.3;refGene_
gn=TMEM25;refGene_tx=XM_006936683.3;refGene_gn=TMEM25;refGene_tx=XM_00
6936684.4;refGene_gn=TMEM25;refGene_tx=XM_006936685.4;refGene_gn=TMEM2
5;refGene_tx=XM_006936686.4;refGene_gn=TMEM25;refGene_tx=XM_023239157.
1;refGene_gn=TMEM25;refGene_tx=XM_023239158.1

D1 16115452 16115764
refGene_gn=TMEM25;refGene_tx=XM_006936680.3;refGene_gn=TMEM25;refGene_
tx=XM_006936681.3;refGene_gn=TMEM25;refGene_tx=XM_006936682.3;refGene_
gn=TMEM25;refGene_tx=XM_006936683.3;refGene_gn=TMEM25;refGene_tx=XM_00
6936684.4;refGene_gn=TMEM25;refGene_tx=XM_006936685.4;refGene_gn=TMEM2
5;refGene_tx=XM_006936686.4;refGene_gn=TMEM25;refGene_tx=XM_023239157.
1;refGene_gn=TMEM25;refGene_tx=XM_023239158.1
D1 16116195 16116523
refGene_gn=TMEM25;refGene_tx=XM_006936680.3;refGene_gn=TMEM25;refGene_
tx=XM_006936681.3;refGene_gn=TMEM25;refGene_tx=XM_006936682.3;refGene_
gn=TMEM25;refGene_tx=XM_006936683.3;refGene_gn=TMEM25;refGene_tx=XM_00
6936684.4;refGene_gn=TMEM25;refGene_tx=XM_006936685.4;refGene_gn=TMEM2
5;refGene_tx=XM_006936686.4;refGene_gn=TMEM25;refGene_tx=XM_023239157.
1;refGene_gn=TMEM25;refGene_tx=XM_023239158.1
D1 16116690 16116822
refGene_gn=TMEM25;refGene_tx=XM_006936680.3;refGene_gn=TMEM25;refGene_
tx=XM_006936681.3;refGene_gn=TMEM25;refGene_tx=XM_006936682.3;refGene_
gn=TMEM25;refGene_tx=XM_006936683.3;refGene_gn=TMEM25;refGene_tx=XM_00
6936684.4;refGene_gn=TMEM25;refGene_tx=XM_023239157.1
D1 16117150 16117181
refGene_gn=TMEM25;refGene_tx=XM_006936680.3;refGene_gn=TMEM25;refGene_
tx=XM_006936681.3;refGene_gn=TMEM25;refGene_tx=XM_006936682.3;refGene_
gn=TMEM25;refGene_tx=XM_006936683.3;refGene_gn=TMEM25;refGene_tx=XM_00
6936684.4;refGene_gn=TMEM25;refGene_tx=XM_006936685.4;refGene_gn=TMEM2
5;refGene_tx=XM_006936686.4;refGene_gn=TMEM25;refGene_tx=XM_023239157.
1
D1 16117336 16117437
refGene_gn=TMEM25;refGene_tx=XM_006936680.3;refGene_gn=TMEM25;refGene_
tx=XM_006936681.3;refGene_gn=TMEM25;refGene_tx=XM_006936682.3;refGene_
gn=TMEM25;refGene_tx=XM_006936683.3;refGene_gn=TMEM25;refGene_tx=XM_00
6936684.4;refGene_gn=TMEM25;refGene_tx=XM_006936685.4;refGene_gn=TMEM2
5;refGene_tx=XM_006936686.4;refGene_gn=TMEM25;refGene_tx=XM_023239157.
1;refGene_gn=TMEM25;refGene_tx=XM_023239158.1
D1 16117571 16117762
refGene_gn=TMEM25;refGene_tx=XM_006936680.3;refGene_gn=TMEM25;refGene_
tx=XM_006936681.3;refGene_gn=TMEM25;refGene_tx=XM_006936682.3;refGene_
gn=TMEM25;refGene_tx=XM_006936683.3;refGene_gn=TMEM25;refGene_tx=XM_00
6936684.4;refGene_gn=TMEM25;refGene_tx=XM_006936685.4;refGene_gn=TMEM2
5;refGene_tx=XM_006936686.4;refGene_gn=TMEM25;refGene_tx=XM_023239157.
1;refGene_gn=TMEM25;refGene_tx=XM_023239158.1
D1 16117788 16117862
refGene_gn=TMEM25;refGene_tx=XM_006936684.4;refGene_gn=TMEM25;refGene_
tx=XM_006936686.4
D1 16130791 16130887
refGene_gn=IFT46;refGene_tx=XM_003992416.5;refGene_gn=IFT46;refGene_tx
=XM_006936687.4
D1 16131376 16131462
refGene_gn=IFT46;refGene_tx=XM_003992416.5;refGene_gn=IFT46;refGene_tx
=XM_006936687.4
D1 16131748 16131809

refGene_gn=IFT46;refGene_tx=XM_003992416.5;refGene_gn=IFT46;refGene_tx=XM_006936687.4
D1	16134701	16134768	refGene_gn=IFT46;refGene_tx=XM_003992416.5;refGene_gn=IFT46;refGene_tx=XM_006936687.4
D1	16135024	16135146	refGene_gn=IFT46;refGene_tx=XM_003992416.5;refGene_gn=IFT46;refGene_tx=XM_006936687.4
D1	16136176	16136305	refGene_gn=IFT46;refGene_tx=XM_003992416.5;refGene_gn=IFT46;refGene_tx=XM_006936687.4
D1	16136738	16136832	refGene_gn=IFT46;refGene_tx=XM_003992416.5;refGene_gn=IFT46;refGene_tx=XM_006936687.4
D1	16136978	16137053	refGene_gn=IFT46;refGene_tx=XM_003992416.5;refGene_gn=IFT46;refGene_tx=XM_006936687.4
D1	16139678	16139812	refGene_gn=IFT46;refGene_tx=XM_003992416.5;refGene_gn=IFT46;refGene_tx=XM_006936687.4
D1	16144726	16144771	refGene_gn=IFT46;refGene_tx=XM_003992416.5;refGene_gn=IFT46;refGene_tx=XM_006936687.4
D1	16154793	16154796	refGene_gn=ARCN1;refGene_tx=XM_003992414.5
D1	16167049	16167313	refGene_gn=ARCN1;refGene_tx=XM_003992414.5
D1	16168117	16168297	refGene_gn=ARCN1;refGene_tx=XM_003992414.5
D1	16168407	16168613	refGene_gn=ARCN1;refGene_tx=XM_003992414.5
D1	16169275	16169440	refGene_gn=ARCN1;refGene_tx=XM_003992414.5
D1	16172145	16172311	refGene_gn=ARCN1;refGene_tx=XM_003992414.5
D1	16173708	16173856	refGene_gn=ARCN1;refGene_tx=XM_003992414.5
D1	16174176	16174285	refGene_gn=ARCN1;refGene_tx=XM_003992414.5
D1	16177470	16177675	refGene_gn=ARCN1;refGene_tx=XM_003992414.5
D1	16179365	16179455	refGene_gn=ARCN1;refGene_tx=XM_003992414.5
D1	16184077	16184095	refGene_gn=PHLDB1;refGene_tx=XM_019811367.2
D1	16184183	16184322	refGene_gn=PHLDB1;refGene_tx=XM_019811367.2
D1	16184516	16184578	refGene_gn=PHLDB1;refGene_tx=XM_019811367.2
D1	16187999	16188044	refGene_gn=PHLDB1;refGene_tx=XM_023239162.1;refGene_gn=PHLDB1;refGene_tx=XM_023239163.1;refGene_gn=PHLDB1;refGene_tx=XM_023239164.1;refGene_gn=PHLDB1;refGene_tx=XM_023239165.1;refGene_gn=PHLDB1;refGene_tx=XM_023239166.1;refGene_gn=PHLDB1;refGene_tx=XM_023239167.1;refGene_gn=PHLDB1;refGene_tx=XM_023239168.1;refGene_gn=PHLDB1;refGene_tx=XM_023239169.1;refGene_gn=PHLDB1;refGene_tx=XM_023239170.1;refGene_gn=PHLDB1;refGene_tx=XM_023239171.1;refGene_gn=PHLDB1;refGene_tx=XM_023239172.1
D1	16191177	16191258	refGene_gn=PHLDB1;refGene_tx=XM_019811364.2;refGene_gn=PHLDB1;refGene_tx=XM_019811367.2;refGene_gn=PHLDB1;refGene_tx=XM_023239162.1;refGene_gn=PHLDB1;refGene_tx=XM_023239163.1;refGene_gn=PHLDB1;refGene_tx=XM_023239164.1;refGene_gn=PHLDB1;refGene_tx=XM_023239165.1;refGene_gn=PHLDB1;refGene_tx=XM_023239166.1;refGene_gn=PHLDB1;refGene_tx=XM_023239167.

1;refGene_gn=PHLDB1;refGene_tx=XM_023239168.1;refGene_gn=PHLDB1;refGene_tx=XM_023239169.1;refGene_gn=PHLDB1;refGene_tx=XM_023239170.1;refGene_gn=PHLDB1;refGene_tx=XM_023239171.1;refGene_gn=PHLDB1;refGene_tx=XM_023239172.1
D1 16191907 16192031
refGene_gn=PHLDB1;refGene_tx=XM_019811364.2;refGene_gn=PHLDB1;refGene_tx=XM_019811367.2;refGene_gn=PHLDB1;refGene_tx=XM_023239162.1;refGene_gn=PHLDB1;refGene_tx=XM_023239163.1;refGene_gn=PHLDB1;refGene_tx=XM_023239164.1;refGene_gn=PHLDB1;refGene_tx=XM_023239165.1;refGene_gn=PHLDB1;refGene_tx=XM_023239166.1;refGene_gn=PHLDB1;refGene_tx=XM_023239167.1;refGene_gn=PHLDB1;refGene_tx=XM_023239168.1;refGene_gn=PHLDB1;refGene_tx=XM_023239169.1;refGene_gn=PHLDB1;refGene_tx=XM_023239170.1;refGene_gn=PHLDB1;refGene_tx=XM_023239171.1;refGene_gn=PHLDB1;refGene_tx=XM_023239172.1
D1 16192972 16193143
refGene_gn=PHLDB1;refGene_tx=XM_019811364.2;refGene_gn=PHLDB1;refGene_tx=XM_019811367.2;refGene_gn=PHLDB1;refGene_tx=XM_023239162.1;refGene_gn=PHLDB1;refGene_tx=XM_023239163.1;refGene_gn=PHLDB1;refGene_tx=XM_023239164.1;refGene_gn=PHLDB1;refGene_tx=XM_023239165.1;refGene_gn=PHLDB1;refGene_tx=XM_023239166.1;refGene_gn=PHLDB1;refGene_tx=XM_023239167.1;refGene_gn=PHLDB1;refGene_tx=XM_023239168.1;refGene_gn=PHLDB1;refGene_tx=XM_023239169.1;refGene_gn=PHLDB1;refGene_tx=XM_023239170.1;refGene_gn=PHLDB1;refGene_tx=XM_023239171.1;refGene_gn=PHLDB1;refGene_tx=XM_023239172.1
D1 16200405 16200531
refGene_gn=PHLDB1;refGene_tx=XM_019811364.2;refGene_gn=PHLDB1;refGene_tx=XM_019811367.2;refGene_gn=PHLDB1;refGene_tx=XM_023239162.1;refGene_gn=PHLDB1;refGene_tx=XM_023239163.1;refGene_gn=PHLDB1;refGene_tx=XM_023239164.1;refGene_gn=PHLDB1;refGene_tx=XM_023239165.1;refGene_gn=PHLDB1;refGene_tx=XM_023239166.1;refGene_gn=PHLDB1;refGene_tx=XM_023239167.1;refGene_gn=PHLDB1;refGene_tx=XM_023239168.1;refGene_gn=PHLDB1;refGene_tx=XM_023239169.1;refGene_gn=PHLDB1;refGene_tx=XM_023239170.1;refGene_gn=PHLDB1;refGene_tx=XM_023239171.1;refGene_gn=PHLDB1;refGene_tx=XM_023239172.1
D1 16201967 16203313
refGene_gn=PHLDB1;refGene_tx=XM_019811364.2;refGene_gn=PHLDB1;refGene_tx=XM_019811367.2;refGene_gn=PHLDB1;refGene_tx=XM_023239162.1;refGene_gn=PHLDB1;refGene_tx=XM_023239163.1;refGene_gn=PHLDB1;refGene_tx=XM_023239164.1;refGene_gn=PHLDB1;refGene_tx=XM_023239165.1;refGene_gn=PHLDB1;refGene_tx=XM_023239166.1;refGene_gn=PHLDB1;refGene_tx=XM_023239167.1;refGene_gn=PHLDB1;refGene_tx=XM_023239168.1;refGene_gn=PHLDB1;refGene_tx=XM_023239169.1;refGene_gn=PHLDB1;refGene_tx=XM_023239170.1;refGene_gn=PHLDB1;refGene_tx=XM_023239171.1;refGene_gn=PHLDB1;refGene_tx=XM_023239172.1
D1 16205931 16206213
refGene_gn=PHLDB1;refGene_tx=XM_019811364.2;refGene_gn=PHLDB1;refGene_tx=XM_019811367.2;refGene_gn=PHLDB1;refGene_tx=XM_023239162.1;refGene_gn=PHLDB1;refGene_tx=XM_023239163.1;refGene_gn=PHLDB1;refGene_tx=XM_023239164.1;refGene_gn=PHLDB1;refGene_tx=XM_023239165.1;refGene_gn=PHLDB1;refGene_tx=XM_023239166.1;refGene_gn=PHLDB1;refGene_tx=XM_023239167.

1;refGene_gn=PHLDB1;refGene_tx=XM_023239168.1;refGene_gn=PHLDB1;refGene_tx=XM_023239169.1;refGene_gn=PHLDB1;refGene_tx=XM_023239170.1;refGene_gn=PHLDB1;refGene_tx=XM_023239171.1;refGene_gn=PHLDB1;refGene_tx=XM_023239172.1
D1 16206634 16206775
refGene_gn=PHLDB1;refGene_tx=XM_019811364.2;refGene_gn=PHLDB1;refGene_tx=XM_019811367.2;refGene_gn=PHLDB1;refGene_tx=XM_023239162.1;refGene_gn=PHLDB1;refGene_tx=XM_023239163.1;refGene_gn=PHLDB1;refGene_tx=XM_023239164.1;refGene_gn=PHLDB1;refGene_tx=XM_023239165.1;refGene_gn=PHLDB1;refGene_tx=XM_023239166.1;refGene_gn=PHLDB1;refGene_tx=XM_023239167.1;refGene_gn=PHLDB1;refGene_tx=XM_023239168.1;refGene_gn=PHLDB1;refGene_tx=XM_023239169.1;refGene_gn=PHLDB1;refGene_tx=XM_023239170.1;refGene_gn=PHLDB1;refGene_tx=XM_023239171.1;refGene_gn=PHLDB1;refGene_tx=XM_023239172.1
D1 16206906 16207044
refGene_gn=PHLDB1;refGene_tx=XM_019811364.2;refGene_gn=PHLDB1;refGene_tx=XM_019811367.2;refGene_gn=PHLDB1;refGene_tx=XM_023239162.1;refGene_gn=PHLDB1;refGene_tx=XM_023239163.1;refGene_gn=PHLDB1;refGene_tx=XM_023239164.1;refGene_gn=PHLDB1;refGene_tx=XM_023239165.1;refGene_gn=PHLDB1;refGene_tx=XM_023239166.1;refGene_gn=PHLDB1;refGene_tx=XM_023239167.1;refGene_gn=PHLDB1;refGene_tx=XM_023239168.1;refGene_gn=PHLDB1;refGene_tx=XM_023239169.1;refGene_gn=PHLDB1;refGene_tx=XM_023239170.1;refGene_gn=PHLDB1;refGene_tx=XM_023239171.1;refGene_gn=PHLDB1;refGene_tx=XM_023239172.1
D1 16209628 16209757
refGene_gn=PHLDB1;refGene_tx=XM_019811364.2;refGene_gn=PHLDB1;refGene_tx=XM_019811367.2;refGene_gn=PHLDB1;refGene_tx=XM_023239162.1;refGene_gn=PHLDB1;refGene_tx=XM_023239163.1;refGene_gn=PHLDB1;refGene_tx=XM_023239165.1;refGene_gn=PHLDB1;refGene_tx=XM_023239166.1;refGene_gn=PHLDB1;refGene_tx=XM_023239167.1;refGene_gn=PHLDB1;refGene_tx=XM_023239168.1
D1 16210121 16210277
refGene_gn=PHLDB1;refGene_tx=XM_019811364.2;refGene_gn=PHLDB1;refGene_tx=XM_019811367.2;refGene_gn=PHLDB1;refGene_tx=XM_023239162.1;refGene_gn=PHLDB1;refGene_tx=XM_023239163.1;refGene_gn=PHLDB1;refGene_tx=XM_023239164.1;refGene_gn=PHLDB1;refGene_tx=XM_023239165.1;refGene_gn=PHLDB1;refGene_tx=XM_023239166.1;refGene_gn=PHLDB1;refGene_tx=XM_023239167.1;refGene_gn=PHLDB1;refGene_tx=XM_023239168.1;refGene_gn=PHLDB1;refGene_tx=XM_023239169.1;refGene_gn=PHLDB1;refGene_tx=XM_023239170.1;refGene_gn=PHLDB1;refGene_tx=XM_023239171.1;refGene_gn=PHLDB1;refGene_tx=XM_023239172.1
D1 16213350 16213461
refGene_gn=PHLDB1;refGene_tx=XM_019811364.2;refGene_gn=PHLDB1;refGene_tx=XM_019811367.2;refGene_gn=PHLDB1;refGene_tx=XM_023239162.1;refGene_gn=PHLDB1;refGene_tx=XM_023239163.1;refGene_gn=PHLDB1;refGene_tx=XM_023239164.1;refGene_gn=PHLDB1;refGene_tx=XM_023239165.1;refGene_gn=PHLDB1;refGene_tx=XM_023239166.1;refGene_gn=PHLDB1;refGene_tx=XM_023239167.1;refGene_gn=PHLDB1;refGene_tx=XM_023239168.1;refGene_gn=PHLDB1;refGene_tx=XM_023239169.1;refGene_gn=PHLDB1;refGene_tx=XM_023239170.1;refGene_gn=PHLDB1;refGene_tx=XM_023239171.1;refGene_gn=PHLDB1;refGene_tx=XM_

023239172.1
D1 16213636 16213726
refGene_gn=PHLDB1;refGene_tx=XM_019811364.2;refGene_gn=PHLDB1;refGene_tx=XM_019811367.2;refGene_gn=PHLDB1;refGene_tx=XM_023239162.1;refGene_gn=PHLDB1;refGene_tx=XM_023239163.1;refGene_gn=PHLDB1;refGene_tx=XM_023239164.1;refGene_gn=PHLDB1;refGene_tx=XM_023239165.1;refGene_gn=PHLDB1;refGene_tx=XM_023239166.1;refGene_gn=PHLDB1;refGene_tx=XM_023239167.1;refGene_gn=PHLDB1;refGene_tx=XM_023239168.1;refGene_gn=PHLDB1;refGene_tx=XM_023239169.1;refGene_gn=PHLDB1;refGene_tx=XM_023239170.1;refGene_gn=PHLDB1;refGene_tx=XM_023239171.1;refGene_gn=PHLDB1;refGene_tx=XM_023239172.1
D1 16214305 16214437
refGene_gn=PHLDB1;refGene_tx=XM_019811364.2;refGene_gn=PHLDB1;refGene_tx=XM_019811367.2;refGene_gn=PHLDB1;refGene_tx=XM_023239162.1;refGene_gn=PHLDB1;refGene_tx=XM_023239163.1;refGene_gn=PHLDB1;refGene_tx=XM_023239164.1;refGene_gn=PHLDB1;refGene_tx=XM_023239166.1
D1 16215548 16215653
refGene_gn=PHLDB1;refGene_tx=XM_019811364.2;refGene_gn=PHLDB1;refGene_tx=XM_019811367.2;refGene_gn=PHLDB1;refGene_tx=XM_023239162.1;refGene_gn=PHLDB1;refGene_tx=XM_023239164.1
D1 16216047 16216221
refGene_gn=PHLDB1;refGene_tx=XM_019811364.2;refGene_gn=PHLDB1;refGene_tx=XM_019811367.2;refGene_gn=PHLDB1;refGene_tx=XM_023239162.1;refGene_gn=PHLDB1;refGene_tx=XM_023239163.1;refGene_gn=PHLDB1;refGene_tx=XM_023239164.1;refGene_gn=PHLDB1;refGene_tx=XM_023239165.1
D1 16216715 16216856
refGene_gn=PHLDB1;refGene_tx=XM_019811364.2;refGene_gn=PHLDB1;refGene_tx=XM_019811367.2;refGene_gn=PHLDB1;refGene_tx=XM_023239162.1;refGene_gn=PHLDB1;refGene_tx=XM_023239163.1;refGene_gn=PHLDB1;refGene_tx=XM_023239164.1;refGene_gn=PHLDB1;refGene_tx=XM_023239165.1;refGene_gn=PHLDB1;refGene_tx=XM_023239166.1;refGene_gn=PHLDB1;refGene_tx=XM_023239167.1;refGene_gn=PHLDB1;refGene_tx=XM_023239169.1;refGene_gn=PHLDB1;refGene_tx=XM_023239170.1
D1 16218251 16218392
refGene_gn=PHLDB1;refGene_tx=XM_019811364.2;refGene_gn=PHLDB1;refGene_tx=XM_019811367.2;refGene_gn=PHLDB1;refGene_tx=XM_023239162.1;refGene_gn=PHLDB1;refGene_tx=XM_023239163.1;refGene_gn=PHLDB1;refGene_tx=XM_023239164.1;refGene_gn=PHLDB1;refGene_tx=XM_023239165.1;refGene_gn=PHLDB1;refGene_tx=XM_023239166.1;refGene_gn=PHLDB1;refGene_tx=XM_023239167.1;refGene_gn=PHLDB1;refGene_tx=XM_023239168.1;refGene_gn=PHLDB1;refGene_tx=XM_023239169.1;refGene_gn=PHLDB1;refGene_tx=XM_023239170.1;refGene_gn=PHLDB1;refGene_tx=XM_023239171.1;refGene_gn=PHLDB1;refGene_tx=XM_023239172.1
D1 16218561 16218664
refGene_gn=PHLDB1;refGene_tx=XM_019811364.2;refGene_gn=PHLDB1;refGene_tx=XM_019811367.2;refGene_gn=PHLDB1;refGene_tx=XM_023239162.1;refGene_gn=PHLDB1;refGene_tx=XM_023239163.1;refGene_gn=PHLDB1;refGene_tx=XM_023239164.1;refGene_gn=PHLDB1;refGene_tx=XM_023239165.1;refGene_gn=PHLDB1;refGene_tx=XM_023239166.1;refGene_gn=PHLDB1;refGene_tx=XM_023239167.1;refGene_gn=PHLDB1;refGene_tx=XM_023239168.1;refGene_gn=PHLDB1;refGen

e_tx=XM_023239169.1;refGene_gn=PHLDB1;refGene_tx=XM_023239170.1;refGene_gn=PHLDB1;refGene_tx=XM_023239171.1;refGene_gn=PHLDB1;refGene_tx=XM_023239172.1
D1 16219110 16219155
refGene_gn=PHLDB1;refGene_tx=XM_019811364.2;refGene_gn=PHLDB1;refGene_tx=XM_019811367.2;refGene_gn=PHLDB1;refGene_tx=XM_023239162.1;refGene_gn=PHLDB1;refGene_tx=XM_023239163.1;refGene_gn=PHLDB1;refGene_tx=XM_023239164.1;refGene_gn=PHLDB1;refGene_tx=XM_023239165.1;refGene_gn=PHLDB1;refGene_tx=XM_023239166.1;refGene_gn=PHLDB1;refGene_tx=XM_023239167.1;refGene_gn=PHLDB1;refGene_tx=XM_023239168.1;refGene_gn=PHLDB1;refGene_tx=XM_023239171.1
D1 16219711 16220006
refGene_gn=PHLDB1;refGene_tx=XM_019811364.2;refGene_gn=PHLDB1;refGene_tx=XM_019811367.2;refGene_gn=PHLDB1;refGene_tx=XM_023239162.1;refGene_gn=PHLDB1;refGene_tx=XM_023239163.1;refGene_gn=PHLDB1;refGene_tx=XM_023239164.1;refGene_gn=PHLDB1;refGene_tx=XM_023239165.1;refGene_gn=PHLDB1;refGene_tx=XM_023239166.1;refGene_gn=PHLDB1;refGene_tx=XM_023239167.1;refGene_gn=PHLDB1;refGene_tx=XM_023239168.1;refGene_gn=PHLDB1;refGene_tx=XM_023239169.1;refGene_gn=PHLDB1;refGene_tx=XM_023239170.1;refGene_gn=PHLDB1;refGene_tx=XM_023239171.1;refGene_gn=PHLDB1;refGene_tx=XM_023239172.1
D1 16220086 16220177
refGene_gn=PHLDB1;refGene_tx=XM_019811364.2;refGene_gn=PHLDB1;refGene_tx=XM_019811367.2;refGene_gn=PHLDB1;refGene_tx=XM_023239162.1;refGene_gn=PHLDB1;refGene_tx=XM_023239163.1;refGene_gn=PHLDB1;refGene_tx=XM_023239164.1;refGene_gn=PHLDB1;refGene_tx=XM_023239165.1;refGene_gn=PHLDB1;refGene_tx=XM_023239166.1;refGene_gn=PHLDB1;refGene_tx=XM_023239167.1;refGene_gn=PHLDB1;refGene_tx=XM_023239168.1;refGene_gn=PHLDB1;refGene_tx=XM_023239169.1;refGene_gn=PHLDB1;refGene_tx=XM_023239170.1;refGene_gn=PHLDB1;refGene_tx=XM_023239171.1;refGene_gn=PHLDB1;refGene_tx=XM_023239172.1
D1 16221766 16221913
refGene_gn=PHLDB1;refGene_tx=XM_019811364.2;refGene_gn=PHLDB1;refGene_tx=XM_019811367.2;refGene_gn=PHLDB1;refGene_tx=XM_023239162.1;refGene_gn=PHLDB1;refGene_tx=XM_023239163.1;refGene_gn=PHLDB1;refGene_tx=XM_023239164.1;refGene_gn=PHLDB1;refGene_tx=XM_023239165.1;refGene_gn=PHLDB1;refGene_tx=XM_023239166.1;refGene_gn=PHLDB1;refGene_tx=XM_023239167.1;refGene_gn=PHLDB1;refGene_tx=XM_023239168.1;refGene_gn=PHLDB1;refGene_tx=XM_023239169.1;refGene_gn=PHLDB1;refGene_tx=XM_023239170.1;refGene_gn=PHLDB1;refGene_tx=XM_023239171.1;refGene_gn=PHLDB1;refGene_tx=XM_023239172.1
D1 16223380 16223497
refGene_gn=PHLDB1;refGene_tx=XM_019811364.2;refGene_gn=PHLDB1;refGene_tx=XM_019811367.2;refGene_gn=PHLDB1;refGene_tx=XM_023239162.1;refGene_gn=PHLDB1;refGene_tx=XM_023239163.1;refGene_gn=PHLDB1;refGene_tx=XM_023239164.1;refGene_gn=PHLDB1;refGene_tx=XM_023239165.1;refGene_gn=PHLDB1;refGene_tx=XM_023239166.1;refGene_gn=PHLDB1;refGene_tx=XM_023239167.1;refGene_gn=PHLDB1;refGene_tx=XM_023239168.1;refGene_gn=PHLDB1;refGene_tx=XM_023239169.1;refGene_gn=PHLDB1;refGene_tx=XM_023239170.1;refGene_gn=PHLDB1;refGene_tx=XM_023239171.1;refGene_gn=PHLDB1;refGene_tx=XM_

023239172.1
D1 16223757 16223860
refGene_gn=PHLDB1;refGene_tx=XM_019811364.2;refGene_gn=PHLDB1;refGene_tx=XM_019811367.2;refGene_gn=PHLDB1;refGene_tx=XM_023239162.1;refGene_gn=PHLDB1;refGene_tx=XM_023239163.1;refGene_gn=PHLDB1;refGene_tx=XM_023239164.1;refGene_gn=PHLDB1;refGene_tx=XM_023239165.1;refGene_gn=PHLDB1;refGene_tx=XM_023239166.1;refGene_gn=PHLDB1;refGene_tx=XM_023239167.1;refGene_gn=PHLDB1;refGene_tx=XM_023239168.1;refGene_gn=PHLDB1;refGene_tx=XM_023239169.1;refGene_gn=PHLDB1;refGene_tx=XM_023239170.1;refGene_gn=PHLDB1;refGene_tx=XM_023239171.1;refGene_gn=PHLDB1;refGene_tx=XM_023239172.1
D1 16227541 16227627
refGene_gn=PHLDB1;refGene_tx=XM_019811364.2;refGene_gn=PHLDB1;refGene_tx=XM_019811367.2;refGene_gn=PHLDB1;refGene_tx=XM_023239162.1;refGene_gn=PHLDB1;refGene_tx=XM_023239163.1;refGene_gn=PHLDB1;refGene_tx=XM_023239164.1;refGene_gn=PHLDB1;refGene_tx=XM_023239165.1;refGene_gn=PHLDB1;refGene_tx=XM_023239166.1;refGene_gn=PHLDB1;refGene_tx=XM_023239167.1;refGene_gn=PHLDB1;refGene_tx=XM_023239168.1;refGene_gn=PHLDB1;refGene_tx=XM_023239169.1;refGene_gn=PHLDB1;refGene_tx=XM_023239170.1;refGene_gn=PHLDB1;refGene_tx=XM_023239171.1;refGene_gn=PHLDB1;refGene_tx=XM_023239172.1
D1 16227749 16227782
refGene_gn=PHLDB1;refGene_tx=XM_019811364.2;refGene_gn=PHLDB1;refGene_tx=XM_019811367.2;refGene_gn=PHLDB1;refGene_tx=XM_023239163.1;refGene_gn=PHLDB1;refGene_tx=XM_023239164.1;refGene_gn=PHLDB1;refGene_tx=XM_023239165.1;refGene_gn=PHLDB1;refGene_tx=XM_023239166.1;refGene_gn=PHLDB1;refGene_tx=XM_023239167.1;refGene_gn=PHLDB1;refGene_tx=XM_023239168.1;refGene_gn=PHLDB1;refGene_tx=XM_023239169.1;refGene_gn=PHLDB1;refGene_tx=XM_023239171.1
D1 16228503 16228644
refGene_gn=PHLDB1;refGene_tx=XM_019811364.2;refGene_gn=PHLDB1;refGene_tx=XM_019811367.2;refGene_gn=PHLDB1;refGene_tx=XM_023239162.1;refGene_gn=PHLDB1;refGene_tx=XM_023239163.1;refGene_gn=PHLDB1;refGene_tx=XM_023239164.1;refGene_gn=PHLDB1;refGene_tx=XM_023239165.1;refGene_gn=PHLDB1;refGene_tx=XM_023239166.1;refGene_gn=PHLDB1;refGene_tx=XM_023239167.1;refGene_gn=PHLDB1;refGene_tx=XM_023239168.1;refGene_gn=PHLDB1;refGene_tx=XM_023239169.1;refGene_gn=PHLDB1;refGene_tx=XM_023239170.1;refGene_gn=PHLDB1;refGene_tx=XM_023239171.1;refGene_gn=PHLDB1;refGene_tx=XM_023239172.1
D1 16230065 16230214 refGene_gn=TREH;refGene_tx=XM_023239034.1
D1 16230329 16230354 refGene_gn=TREH;refGene_tx=XM_023239034.1
D1 16230484 16230514 refGene_gn=TREH;refGene_tx=XM_023239034.1
D1 16230757 16230870 refGene_gn=TREH;refGene_tx=XM_023239034.1
D1 16231238 16231350 refGene_gn=TREH;refGene_tx=XM_023239034.1
D1 16231619 16231837 refGene_gn=TREH;refGene_tx=XM_023239034.1
D1 16232347 16232652 refGene_gn=TREH;refGene_tx=XM_023239034.1
D1 16232866 16232919 refGene_gn=TREH;refGene_tx=XM_023239034.1
D1 16232921 16233000 refGene_gn=TREH;refGene_tx=XM_023239034.1
D1 16233108 16233136 refGene_gn=TREH;refGene_tx=XM_023239034.1
D1 16233138 16233228 refGene_gn=TREH;refGene_tx=XM_023239034.1

D1 16233312 16233401 refGene_gn=TREH;refGene_tx=XM_023239034.1
D1 16233556 16233621 refGene_gn=TREH;refGene_tx=XM_023239034.1
D1 16233623 16233659 refGene_gn=TREH;refGene_tx=XM_023239034.1
D1 16235188 16235276 refGene_gn=TREH;refGene_tx=XM_023239034.1
D1 16235361 16235506 refGene_gn=TREH;refGene_tx=XM_023239034.1
D1 16235625 16235726 refGene_gn=TREH;refGene_tx=XM_023239034.1
D1 16242664 16242716 refGene_gn=TREH;refGene_tx=XM_023239034.1
D1 16242998 16243020 refGene_gn=TREH;refGene_tx=XM_023239034.1
D1 16285346 16285522
refGene_gn=DDX6;refGene_tx=XM_011286390.3;refGene_gn=DDX6;refGene_tx=XM_011286391.3;refGene_gn=DDX6;refGene_tx=XM_019811379.2;refGene_gn=DDX6;refGene_tx=XM_023239173.1
D1 16286278 16286380
refGene_gn=DDX6;refGene_tx=XM_011286390.3;refGene_gn=DDX6;refGene_tx=XM_011286391.3;refGene_gn=DDX6;refGene_tx=XM_019811379.2;refGene_gn=DDX6;refGene_tx=XM_023239173.1
D1 16289481 16289545
refGene_gn=DDX6;refGene_tx=XM_011286390.3;refGene_gn=DDX6;refGene_tx=XM_011286391.3;refGene_gn=DDX6;refGene_tx=XM_019811379.2;refGene_gn=DDX6;refGene_tx=XM_023239173.1
D1 16290927 16291044
refGene_gn=DDX6;refGene_tx=XM_011286390.3;refGene_gn=DDX6;refGene_tx=XM_011286391.3;refGene_gn=DDX6;refGene_tx=XM_019811379.2;refGene_gn=DDX6;refGene_tx=XM_023239173.1
D1 16292511 16292640
refGene_gn=DDX6;refGene_tx=XM_011286390.3;refGene_gn=DDX6;refGene_tx=XM_011286391.3;refGene_gn=DDX6;refGene_tx=XM_019811379.2;refGene_gn=DDX6;refGene_tx=XM_023239173.1
D1 16293125 16293248
refGene_gn=DDX6;refGene_tx=XM_011286390.3;refGene_gn=DDX6;refGene_tx=XM_011286391.3;refGene_gn=DDX6;refGene_tx=XM_019811379.2;refGene_gn=DDX6;refGene_tx=XM_023239173.1
D1 16295077 16295172
refGene_gn=DDX6;refGene_tx=XM_011286390.3;refGene_gn=DDX6;refGene_tx=XM_011286391.3;refGene_gn=DDX6;refGene_tx=XM_019811379.2;refGene_gn=DDX6;refGene_tx=XM_023239173.1
D1 16296281 16296428
refGene_gn=DDX6;refGene_tx=XM_011286390.3;refGene_gn=DDX6;refGene_tx=XM_011286391.3;refGene_gn=DDX6;refGene_tx=XM_019811379.2;refGene_gn=DDX6;refGene_tx=XM_023239173.1
D1 16298062 16298192
refGene_gn=DDX6;refGene_tx=XM_011286390.3;refGene_gn=DDX6;refGene_tx=XM_011286391.3;refGene_gn=DDX6;refGene_tx=XM_019811379.2;refGene_gn=DDX6;refGene_tx=XM_023239173.1
D1 16306399 16306504
refGene_gn=DDX6;refGene_tx=XM_011286390.3;refGene_gn=DDX6;refGene_tx=XM_011286391.3;refGene_gn=DDX6;refGene_tx=XM_019811379.2;refGene_gn=DDX6;refGene_tx=XM_023239173.1
D1 16307837 16307901
refGene_gn=DDX6;refGene_tx=XM_011286390.3;refGene_gn=DDX6;refGene_tx=X

M_011286391.3;refGene_gn=DDX6;refGene_tx=XM_019811379.2;refGene_gn=DDX6;refGene_tx=XM_023239173.1
D1 16312281 16312481 refGene_gn=DDX6;refGene_tx=XM_011286390.3;refGene_gn=DDX6;refGene_tx=XM_011286391.3;refGene_gn=DDX6;refGene_tx=XM_019811379.2;refGene_gn=DDX6;refGene_tx=XM_023239173.1
D1 16382410 16382467 refGene_gn=CXCR5;refGene_tx=XM_003992421.5
D1 16391221 16392289 refGene_gn=CXCR5;refGene_tx=XM_003992421.5
D1 16396140 16396616 refGene_gn=BCL9L;refGene_tx=XM_003992422.5
D1 16396618 16396916 refGene_gn=BCL9L;refGene_tx=XM_003992422.5
D1 16396917 16397231 refGene_gn=BCL9L;refGene_tx=XM_003992422.5
D1 16397666 16397826 refGene_gn=BCL9L;refGene_tx=XM_003992422.5
D1 16397827 16397949 refGene_gn=BCL9L;refGene_tx=XM_003992422.5
D1 16398407 16400694 refGene_gn=BCL9L;refGene_tx=XM_003992422.5
D1 16400777 16400862 refGene_gn=BCL9L;refGene_tx=XM_003992422.5
D1 16401024 16401235 refGene_gn=BCL9L;refGene_tx=XM_003992422.5
D1 16404073 16404193 refGene_gn=BCL9L;refGene_tx=XM_003992422.5
D1 16404902 16405288 refGene_gn=BCL9L;refGene_tx=XM_003992422.5
D1 16406552 16406578 refGene_gn=BCL9L;refGene_tx=XM_003992422.5
D1 16444956 16445032 refGene_gn=UPK2;refGene_tx=XM_019811382.2;refGene_gn=UPK2;refGene_tx=XM_019811384.2;refGene_gn=UPK2;refGene_tx=XM_023239174.1;refGene_gn=UPK2;refGene_tx=XM_023239175.1
D1 16445475 16445610 refGene_gn=UPK2;refGene_tx=XM_019811382.2;refGene_gn=UPK2;refGene_tx=XM_019811384.2;refGene_gn=UPK2;refGene_tx=XM_023239174.1;refGene_gn=UPK2;refGene_tx=XM_023239175.1
D1 16445798 16445937 refGene_gn=UPK2;refGene_tx=XM_019811382.2;refGene_gn=UPK2;refGene_tx=XM_019811384.2;refGene_gn=UPK2;refGene_tx=XM_023239174.1;refGene_gn=UPK2;refGene_tx=XM_023239175.1
D1 16446199 16446270 refGene_gn=UPK2;refGene_tx=XM_019811382.2;refGene_gn=UPK2;refGene_tx=XM_019811384.2;refGene_gn=UPK2;refGene_tx=XM_023239174.1;refGene_gn=UPK2;refGene_tx=XM_023239175.1
D1 16446639 16446776 refGene_gn=UPK2;refGene_tx=XM_019811382.2;refGene_gn=UPK2;refGene_tx=XM_019811384.2;refGene_gn=UPK2;refGene_tx=XM_023239174.1;refGene_gn=UPK2;refGene_tx=XM_023239175.1
D1 16454377 16454438 refGene_gn=FOXR1;refGene_tx=XM_023238981.1
D1 16463593 16463668 refGene_gn=FOXR1;refGene_tx=XM_023238981.1
D1 16463767 16464024 refGene_gn=FOXR1;refGene_tx=XM_023238981.1
D1 16464274 16464501 refGene_gn=FOXR1;refGene_tx=XM_023238981.1
D1 16465601 16465840 refGene_gn=FOXR1;refGene_tx=XM_023238981.1
D1 16466289 16466318 refGene_gn=FOXR1;refGene_tx=XM_023238981.1
D1 16471823 16471943 refGene_gn=CCDC84;refGene_tx=XM_003992426.4;refGene_gn=CCDC84;refGene_tx=XM_006936701.3;refGene_gn=CCDC84;refGene_tx=XM_006936702.3
D1 16472069 16472233 refGene_gn=CCDC84;refGene_tx=XM_003992426.4;refGene_gn=CCDC84;refGene_

tx=XM_006936701.3;refGene_gn=CCDC84;refGene_tx=XM_006936702.3
D1 16472642 16472741
refGene_gn=CCDC84;refGene_tx=XM_003992426.4;refGene_gn=CCDC84;refGene_tx=XM_006936701.3;refGene_gn=CCDC84;refGene_tx=XM_006936702.3
D1 16475079 16475211
refGene_gn=CCDC84;refGene_tx=XM_006936701.3;refGene_gn=CCDC84;refGene_tx=XM_006936702.3
D1 16475333 16475400
refGene_gn=CCDC84;refGene_tx=XM_003992426.4;refGene_gn=CCDC84;refGene_tx=XM_006936701.3;refGene_gn=CCDC84;refGene_tx=XM_006936702.3
D1 16475475 16475538
refGene_gn=CCDC84;refGene_tx=XM_003992426.4;refGene_gn=CCDC84;refGene_tx=XM_006936701.3;refGene_gn=CCDC84;refGene_tx=XM_006936702.3
D1 16475890 16475955
refGene_gn=CCDC84;refGene_tx=XM_003992426.4;refGene_gn=CCDC84;refGene_tx=XM_006936701.3;refGene_gn=CCDC84;refGene_tx=XM_006936702.3
D1 16476105 16476214
refGene_gn=CCDC84;refGene_tx=XM_003992426.4;refGene_gn=CCDC84;refGene_tx=XM_006936701.3;refGene_gn=CCDC84;refGene_tx=XM_006936702.3
D1 16477159 16477190
refGene_gn=CCDC84;refGene_tx=XM_003992426.4;refGene_gn=CCDC84;refGene_tx=XM_006936701.3
D1 16478186 16478276
refGene_gn=CCDC84;refGene_tx=XM_003992426.4;refGene_gn=CCDC84;refGene_tx=XM_006936701.3;refGene_gn=CCDC84;refGene_tx=XM_006936702.3
D1 16478350 16478483
refGene_gn=CCDC84;refGene_tx=XM_003992426.4;refGene_gn=CCDC84;refGene_tx=XM_006936701.3
D1 16478569 16478642
refGene_gn=CCDC84;refGene_tx=XM_003992426.4;refGene_gn=CCDC84;refGene_tx=XM_006936701.3
D1 16478898 16478993 refGene_gn=RPS25;refGene_tx=XM_003992425.5
D1 16479817 16480001 refGene_gn=RPS25;refGene_tx=XM_003992425.5
D1 16480545 16480641 refGene_gn=RPS25;refGene_tx=XM_003992425.5
D1 16480874 16480877 refGene_gn=RPS25;refGene_tx=XM_003992425.5
D1 16481362 16481537 refGene_gn=TRAPPC4;refGene_tx=XM_003992427.4
D1 16481700 16481875 refGene_gn=TRAPPC4;refGene_tx=XM_003992427.4
D1 16482875 16483035
refGene_gn=SLC37A4;refGene_tx=XM_006936706.3;refGene_gn=TRAPPC4;refGene_tx=XM_003992427.4
D1 16483594 16483721 refGene_gn=TRAPPC4;refGene_tx=XM_003992427.4
D1 16483999 16484078 refGene_gn=TRAPPC4;refGene_tx=XM_003992427.4
D1 16485771 16485938
refGene_gn=SLC37A4;refGene_tx=XM_003992428.4;refGene_gn=SLC37A4;refGene_tx=XM_006936703.3
D1 16486060 16486199
refGene_gn=SLC37A4;refGene_tx=XM_003992428.4;refGene_gn=SLC37A4;refGene_tx=XM_006936703.3;refGene_gn=SLC37A4;refGene_tx=XM_006936706.3
D1 16486536 16486602
refGene_gn=SLC37A4;refGene_tx=XM_006936703.3;refGene_gn=SLC37A4;refGen

e_tx=XM_006936706.3
D1 16486819 16486933
refGene_gn=SLC37A4;refGene_tx=XM_003992428.4;refGene_gn=SLC37A4;refGene_tx=XM_006936703.3;refGene_gn=SLC37A4;refGene_tx=XM_006936706.3
D1 16487457 16487543
refGene_gn=SLC37A4;refGene_tx=XM_003992428.4;refGene_gn=SLC37A4;refGene_tx=XM_006936703.3;refGene_gn=SLC37A4;refGene_tx=XM_006936706.3
D1 16487786 16487945
refGene_gn=SLC37A4;refGene_tx=XM_003992428.4;refGene_gn=SLC37A4;refGene_tx=XM_006936703.3;refGene_gn=SLC37A4;refGene_tx=XM_006936706.3
D1 16488458 16488702
refGene_gn=SLC37A4;refGene_tx=XM_003992428.4;refGene_gn=SLC37A4;refGene_tx=XM_006936703.3;refGene_gn=SLC37A4;refGene_tx=XM_006936706.3
D1 16489014 16489247
refGene_gn=SLC37A4;refGene_tx=XM_003992428.4;refGene_gn=SLC37A4;refGene_tx=XM_006936703.3;refGene_gn=SLC37A4;refGene_tx=XM_006936706.3
D1 16490115 16490263
refGene_gn=SLC37A4;refGene_tx=XM_003992428.4;refGene_gn=SLC37A4;refGene_tx=XM_006936703.3;refGene_gn=SLC37A4;refGene_tx=XM_006936706.3
D1 16506916 16506981
refGene_gn=HYOU1;refGene_tx=XM_006936709.4;refGene_gn=HYOU1;refGene_tx=XM_006936710.4;refGene_gn=HYOU1;refGene_tx=XM_006936711.4;refGene_gn=HYOU1;refGene_tx=XM_011286397.3;refGene_gn=HYOU1;refGene_tx=XM_011286398.3;refGene_gn=HYOU1;refGene_tx=XM_011286399.3;refGene_gn=HYOU1;refGene_tx=XM_019811387.2;refGene_gn=HYOU1;refGene_tx=XM_019811388.2
D1 16507104 16507182
refGene_gn=HYOU1;refGene_tx=XM_006936709.4;refGene_gn=HYOU1;refGene_tx=XM_006936710.4;refGene_gn=HYOU1;refGene_tx=XM_006936711.4;refGene_gn=HYOU1;refGene_tx=XM_011286397.3;refGene_gn=HYOU1;refGene_tx=XM_011286398.3;refGene_gn=HYOU1;refGene_tx=XM_011286399.3;refGene_gn=HYOU1;refGene_tx=XM_019811387.2;refGene_gn=HYOU1;refGene_tx=XM_019811388.2
D1 16507435 16507483
refGene_gn=HYOU1;refGene_tx=XM_006936709.4;refGene_gn=HYOU1;refGene_tx=XM_006936710.4;refGene_gn=HYOU1;refGene_tx=XM_006936711.4;refGene_gn=HYOU1;refGene_tx=XM_011286397.3;refGene_gn=HYOU1;refGene_tx=XM_011286398.3;refGene_gn=HYOU1;refGene_tx=XM_011286399.3;refGene_gn=HYOU1;refGene_tx=XM_019811387.2;refGene_gn=HYOU1;refGene_tx=XM_019811388.2
D1 16507577 16507818
refGene_gn=HYOU1;refGene_tx=XM_006936709.4;refGene_gn=HYOU1;refGene_tx=XM_006936710.4;refGene_gn=HYOU1;refGene_tx=XM_006936711.4;refGene_gn=HYOU1;refGene_tx=XM_011286397.3;refGene_gn=HYOU1;refGene_tx=XM_011286398.3;refGene_gn=HYOU1;refGene_tx=XM_011286399.3;refGene_gn=HYOU1;refGene_tx=XM_019811387.2;refGene_gn=HYOU1;refGene_tx=XM_019811388.2
D1 16508457 16508542
refGene_gn=HYOU1;refGene_tx=XM_006936709.4;refGene_gn=HYOU1;refGene_tx=XM_006936710.4;refGene_gn=HYOU1;refGene_tx=XM_006936711.4;refGene_gn=HYOU1;refGene_tx=XM_011286397.3;refGene_gn=HYOU1;refGene_tx=XM_011286398.3;refGene_gn=HYOU1;refGene_tx=XM_011286399.3;refGene_gn=HYOU1;refGene_tx=XM_019811387.2;refGene_gn=HYOU1;refGene_tx=XM_019811388.2
D1 16508674 16508808

refGene_gn=HYOU1;refGene_tx=XM_006936709.4;refGene_gn=HYOU1;refGene_tx=XM_006936710.4;refGene_gn=HYOU1;refGene_tx=XM_006936711.4;refGene_gn=HYOU1;refGene_tx=XM_011286397.3;refGene_gn=HYOU1;refGene_tx=XM_011286398.3;refGene_gn=HYOU1;refGene_tx=XM_011286399.3;refGene_gn=HYOU1;refGene_tx=XM_019811387.2;refGene_gn=HYOU1;refGene_tx=XM_019811388.2
D1 16508992 16509115
refGene_gn=HYOU1;refGene_tx=XM_006936709.4;refGene_gn=HYOU1;refGene_tx=XM_006936710.4;refGene_gn=HYOU1;refGene_tx=XM_006936711.4;refGene_gn=HYOU1;refGene_tx=XM_011286397.3;refGene_gn=HYOU1;refGene_tx=XM_011286398.3;refGene_gn=HYOU1;refGene_tx=XM_011286399.3;refGene_gn=HYOU1;refGene_tx=XM_019811387.2;refGene_gn=HYOU1;refGene_tx=XM_019811388.2
D1 16509229 16509317
refGene_gn=HYOU1;refGene_tx=XM_006936709.4;refGene_gn=HYOU1;refGene_tx=XM_006936710.4;refGene_gn=HYOU1;refGene_tx=XM_006936711.4;refGene_gn=HYOU1;refGene_tx=XM_011286397.3;refGene_gn=HYOU1;refGene_tx=XM_011286398.3;refGene_gn=HYOU1;refGene_tx=XM_011286399.3;refGene_gn=HYOU1;refGene_tx=XM_019811387.2;refGene_gn=HYOU1;refGene_tx=XM_019811388.2
D1 16509449 16509622
refGene_gn=HYOU1;refGene_tx=XM_006936709.4;refGene_gn=HYOU1;refGene_tx=XM_006936710.4;refGene_gn=HYOU1;refGene_tx=XM_006936711.4;refGene_gn=HYOU1;refGene_tx=XM_011286397.3;refGene_gn=HYOU1;refGene_tx=XM_011286398.3;refGene_gn=HYOU1;refGene_tx=XM_011286399.3;refGene_gn=HYOU1;refGene_tx=XM_019811387.2;refGene_gn=HYOU1;refGene_tx=XM_019811388.2
D1 16509713 16509899
refGene_gn=HYOU1;refGene_tx=XM_006936709.4;refGene_gn=HYOU1;refGene_tx=XM_006936710.4;refGene_gn=HYOU1;refGene_tx=XM_006936711.4;refGene_gn=HYOU1;refGene_tx=XM_011286397.3;refGene_gn=HYOU1;refGene_tx=XM_011286398.3;refGene_gn=HYOU1;refGene_tx=XM_011286399.3;refGene_gn=HYOU1;refGene_tx=XM_019811387.2;refGene_gn=HYOU1;refGene_tx=XM_019811388.2
D1 16510303 16510383
refGene_gn=HYOU1;refGene_tx=XM_006936709.4;refGene_gn=HYOU1;refGene_tx=XM_006936710.4;refGene_gn=HYOU1;refGene_tx=XM_006936711.4;refGene_gn=HYOU1;refGene_tx=XM_011286397.3;refGene_gn=HYOU1;refGene_tx=XM_011286398.3;refGene_gn=HYOU1;refGene_tx=XM_011286399.3;refGene_gn=HYOU1;refGene_tx=XM_019811387.2;refGene_gn=HYOU1;refGene_tx=XM_019811388.2
D1 16510557 16510618
refGene_gn=HYOU1;refGene_tx=XM_006936709.4;refGene_gn=HYOU1;refGene_tx=XM_006936710.4;refGene_gn=HYOU1;refGene_tx=XM_006936711.4;refGene_gn=HYOU1;refGene_tx=XM_011286397.3;refGene_gn=HYOU1;refGene_tx=XM_011286398.3;refGene_gn=HYOU1;refGene_tx=XM_011286399.3;refGene_gn=HYOU1;refGene_tx=XM_019811387.2;refGene_gn=HYOU1;refGene_tx=XM_019811388.2
D1 16511556 16511695
refGene_gn=HYOU1;refGene_tx=XM_006936709.4;refGene_gn=HYOU1;refGene_tx=XM_006936710.4;refGene_gn=HYOU1;refGene_tx=XM_006936711.4;refGene_gn=HYOU1;refGene_tx=XM_011286397.3;refGene_gn=HYOU1;refGene_tx=XM_011286398.3;refGene_gn=HYOU1;refGene_tx=XM_011286399.3;refGene_gn=HYOU1;refGene_tx=XM_019811387.2;refGene_gn=HYOU1;refGene_tx=XM_019811388.2
D1 16511897 16512085
refGene_gn=HYOU1;refGene_tx=XM_006936709.4;refGene_gn=HYOU1;refGene_tx=XM_006936710.4;refGene_gn=HYOU1;refGene_tx=XM_006936711.4;refGene_gn=

HYOU1;refGene_tx=XM_011286397.3;refGene_gn=HYOU1;refGene_tx=XM_0112863
98.3;refGene_gn=HYOU1;refGene_tx=XM_011286399.3;refGene_gn=HYOU1;refGe
ne_tx=XM_019811387.2;refGene_gn=HYOU1;refGene_tx=XM_019811388.2
D1 16512239 16512372
refGene_gn=HYOU1;refGene_tx=XM_006936709.4;refGene_gn=HYOU1;refGene_tx
=XM_006936710.4;refGene_gn=HYOU1;refGene_tx=XM_006936711.4;refGene_gn=
HYOU1;refGene_tx=XM_011286397.3;refGene_gn=HYOU1;refGene_tx=XM_0112863
98.3;refGene_gn=HYOU1;refGene_tx=XM_011286399.3;refGene_gn=HYOU1;refGe
ne_tx=XM_019811387.2;refGene_gn=HYOU1;refGene_tx=XM_019811388.2
D1 16512470 16512553
refGene_gn=HYOU1;refGene_tx=XM_006936709.4;refGene_gn=HYOU1;refGene_tx
=XM_006936710.4;refGene_gn=HYOU1;refGene_tx=XM_006936711.4;refGene_gn=
HYOU1;refGene_tx=XM_011286397.3;refGene_gn=HYOU1;refGene_tx=XM_0112863
98.3;refGene_gn=HYOU1;refGene_tx=XM_011286399.3;refGene_gn=HYOU1;refGe
ne_tx=XM_019811387.2;refGene_gn=HYOU1;refGene_tx=XM_019811388.2
D1 16512707 16512842
refGene_gn=HYOU1;refGene_tx=XM_006936709.4;refGene_gn=HYOU1;refGene_tx
=XM_006936710.4;refGene_gn=HYOU1;refGene_tx=XM_006936711.4;refGene_gn=
HYOU1;refGene_tx=XM_011286397.3;refGene_gn=HYOU1;refGene_tx=XM_0112863
98.3;refGene_gn=HYOU1;refGene_tx=XM_011286399.3;refGene_gn=HYOU1;refGe
ne_tx=XM_019811387.2;refGene_gn=HYOU1;refGene_tx=XM_019811388.2
D1 16513032 16513225
refGene_gn=HYOU1;refGene_tx=XM_006936709.4;refGene_gn=HYOU1;refGene_tx
=XM_006936710.4;refGene_gn=HYOU1;refGene_tx=XM_006936711.4;refGene_gn=
HYOU1;refGene_tx=XM_011286397.3;refGene_gn=HYOU1;refGene_tx=XM_0112863
98.3;refGene_gn=HYOU1;refGene_tx=XM_011286399.3;refGene_gn=HYOU1;refGe
ne_tx=XM_019811387.2;refGene_gn=HYOU1;refGene_tx=XM_019811388.2
D1 16514541 16514657
refGene_gn=HYOU1;refGene_tx=XM_006936709.4;refGene_gn=HYOU1;refGene_tx
=XM_006936710.4;refGene_gn=HYOU1;refGene_tx=XM_006936711.4;refGene_gn=
HYOU1;refGene_tx=XM_011286397.3;refGene_gn=HYOU1;refGene_tx=XM_0112863
98.3;refGene_gn=HYOU1;refGene_tx=XM_011286399.3;refGene_gn=HYOU1;refGe
ne_tx=XM_019811387.2;refGene_gn=HYOU1;refGene_tx=XM_019811388.2
D1 16514883 16515065
refGene_gn=HYOU1;refGene_tx=XM_006936709.4;refGene_gn=HYOU1;refGene_tx
=XM_006936710.4;refGene_gn=HYOU1;refGene_tx=XM_006936711.4;refGene_gn=
HYOU1;refGene_tx=XM_011286397.3;refGene_gn=HYOU1;refGene_tx=XM_0112863
98.3;refGene_gn=HYOU1;refGene_tx=XM_011286399.3;refGene_gn=HYOU1;refGe
ne_tx=XM_019811387.2;refGene_gn=HYOU1;refGene_tx=XM_019811388.2
D1 16515361 16515438
refGene_gn=HYOU1;refGene_tx=XM_006936709.4;refGene_gn=HYOU1;refGene_tx
=XM_006936710.4;refGene_gn=HYOU1;refGene_tx=XM_006936711.4;refGene_gn=
HYOU1;refGene_tx=XM_011286397.3;refGene_gn=HYOU1;refGene_tx=XM_0112863
98.3;refGene_gn=HYOU1;refGene_tx=XM_011286399.3;refGene_gn=HYOU1;refGe
ne_tx=XM_019811387.2;refGene_gn=HYOU1;refGene_tx=XM_019811388.2
D1 16515528 16515683
refGene_gn=HYOU1;refGene_tx=XM_006936709.4;refGene_gn=HYOU1;refGene_tx
=XM_006936710.4;refGene_gn=HYOU1;refGene_tx=XM_006936711.4;refGene_gn=
HYOU1;refGene_tx=XM_011286397.3;refGene_gn=HYOU1;refGene_tx=XM_0112863
98.3;refGene_gn=HYOU1;refGene_tx=XM_011286399.3;refGene_gn=HYOU1;refGe

ne_tx=XM_019811387.2;refGene_gn=HYOU1;refGene_tx=XM_019811388.2
D1 16515835 16515914
refGene_gn=HYOU1;refGene_tx=XM_006936709.4;refGene_gn=HYOU1;refGene_tx=XM_006936710.4;refGene_gn=HYOU1;refGene_tx=XM_006936711.4;refGene_gn=HYOU1;refGene_tx=XM_011286397.3;refGene_gn=HYOU1;refGene_tx=XM_011286398.3;refGene_gn=HYOU1;refGene_tx=XM_011286399.3;refGene_gn=HYOU1;refGene_tx=XM_019811387.2;refGene_gn=HYOU1;refGene_tx=XM_019811388.2
D1 16516092 16516186
refGene_gn=HYOU1;refGene_tx=XM_006936709.4;refGene_gn=HYOU1;refGene_tx=XM_006936710.4;refGene_gn=HYOU1;refGene_tx=XM_006936711.4;refGene_gn=HYOU1;refGene_tx=XM_011286397.3;refGene_gn=HYOU1;refGene_tx=XM_011286398.3;refGene_gn=HYOU1;refGene_tx=XM_011286399.3;refGene_gn=HYOU1;refGene_tx=XM_019811387.2;refGene_gn=HYOU1;refGene_tx=XM_019811388.2
D1 16516401 16516492
refGene_gn=HYOU1;refGene_tx=XM_006936709.4;refGene_gn=HYOU1;refGene_tx=XM_006936710.4;refGene_gn=HYOU1;refGene_tx=XM_006936711.4;refGene_gn=HYOU1;refGene_tx=XM_011286397.3;refGene_gn=HYOU1;refGene_tx=XM_011286398.3;refGene_gn=HYOU1;refGene_tx=XM_011286399.3;refGene_gn=HYOU1;refGene_tx=XM_019811387.2;refGene_gn=HYOU1;refGene_tx=XM_019811388.2
D1 16525448 16525716
refGene_gn=VPS11;refGene_tx=XM_003992430.3;refGene_gn=VPS11;refGene_tx=XM_019811389.2;refGene_gn=VPS11;refGene_tx=XM_019811390.2;refGene_gn=VPS11;refGene_tx=XM_019811391.2
D1 16526443 16526592
refGene_gn=VPS11;refGene_tx=XM_003992430.3;refGene_gn=VPS11;refGene_tx=XM_019811389.2;refGene_gn=VPS11;refGene_tx=XM_019811390.2;refGene_gn=VPS11;refGene_tx=XM_019811391.2
D1 16526684 16526820
refGene_gn=VPS11;refGene_tx=XM_003992430.3;refGene_gn=VPS11;refGene_tx=XM_019811389.2;refGene_gn=VPS11;refGene_tx=XM_019811390.2;refGene_gn=VPS11;refGene_tx=XM_019811391.2
D1 16527611 16527775
refGene_gn=VPS11;refGene_tx=XM_003992430.3;refGene_gn=VPS11;refGene_tx=XM_019811389.2;refGene_gn=VPS11;refGene_tx=XM_019811390.2;refGene_gn=VPS11;refGene_tx=XM_019811391.2
D1 16528286 16528534
refGene_gn=VPS11;refGene_tx=XM_003992430.3;refGene_gn=VPS11;refGene_tx=XM_019811389.2;refGene_gn=VPS11;refGene_tx=XM_019811390.2;refGene_gn=VPS11;refGene_tx=XM_019811391.2
D1 16529225 16529427
refGene_gn=VPS11;refGene_tx=XM_003992430.3;refGene_gn=VPS11;refGene_tx=XM_019811389.2;refGene_gn=VPS11;refGene_tx=XM_019811390.2;refGene_gn=VPS11;refGene_tx=XM_019811391.2
D1 16529836 16529988
refGene_gn=VPS11;refGene_tx=XM_003992430.3;refGene_gn=VPS11;refGene_tx=XM_019811389.2;refGene_gn=VPS11;refGene_tx=XM_019811390.2;refGene_gn=VPS11;refGene_tx=XM_019811391.2
D1 16531042 16531229
refGene_gn=VPS11;refGene_tx=XM_003992430.3;refGene_gn=VPS11;refGene_tx=XM_019811389.2;refGene_gn=VPS11;refGene_tx=XM_019811390.2;refGene_gn=

VPS11;refGene_tx=XM_019811391.2
D1 16531863 16532010
refGene_gn=VPS11;refGene_tx=XM_003992430.3;refGene_gn=VPS11;refGene_tx=XM_019811389.2;refGene_gn=VPS11;refGene_tx=XM_019811390.2;refGene_gn=VPS11;refGene_tx=XM_019811391.2
D1 16532133 16532322
refGene_gn=VPS11;refGene_tx=XM_003992430.3;refGene_gn=VPS11;refGene_tx=XM_019811389.2;refGene_gn=VPS11;refGene_tx=XM_019811390.2;refGene_gn=VPS11;refGene_tx=XM_019811391.2
D1 16532429 16532591
refGene_gn=VPS11;refGene_tx=XM_003992430.3;refGene_gn=VPS11;refGene_tx=XM_019811389.2;refGene_gn=VPS11;refGene_tx=XM_019811390.2;refGene_gn=VPS11;refGene_tx=XM_019811391.2
D1 16532817 16532957
refGene_gn=VPS11;refGene_tx=XM_003992430.3;refGene_gn=VPS11;refGene_tx=XM_019811389.2;refGene_gn=VPS11;refGene_tx=XM_019811390.2;refGene_gn=VPS11;refGene_tx=XM_019811391.2
D1 16533041 16533244
refGene_gn=VPS11;refGene_tx=XM_003992430.3;refGene_gn=VPS11;refGene_tx=XM_019811389.2;refGene_gn=VPS11;refGene_tx=XM_019811390.2;refGene_gn=VPS11;refGene_tx=XM_019811391.2
D1 16533367 16533539
refGene_gn=VPS11;refGene_tx=XM_003992430.3;refGene_gn=VPS11;refGene_tx=XM_019811389.2;refGene_gn=VPS11;refGene_tx=XM_019811390.2;refGene_gn=VPS11;refGene_tx=XM_019811391.2
D1 16534817 16535040
refGene_gn=VPS11;refGene_tx=XM_003992430.3;refGene_gn=VPS11;refGene_tx=XM_019811389.2;refGene_gn=VPS11;refGene_tx=XM_019811390.2;refGene_gn=VPS11;refGene_tx=XM_019811391.2
D1 16535166 16535331
refGene_gn=VPS11;refGene_tx=XM_003992430.3;refGene_gn=VPS11;refGene_tx=XM_019811389.2;refGene_gn=VPS11;refGene_tx=XM_019811390.2;refGene_gn=VPS11;refGene_tx=XM_019811391.2
D1 16537494 16537527 refGene_gn=HMBS;refGene_tx=NM_001177808.1
D1 16540526 16540580
refGene_gn=HMBS;refGene_tx=NM_001177808.1;refGene_gn=HMBS;refGene_tx=NM_001177809.1;refGene_gn=HMBS;refGene_tx=XM_006936655.3;refGene_gn=HMBS;refGene_tx=XM_006936656.1;refGene_gn=HMBS;refGene_tx=XM_019811157.1
D1 16540908 16540981
refGene_gn=HMBS;refGene_tx=NM_001177808.1;refGene_gn=HMBS;refGene_tx=NM_001177809.1;refGene_gn=HMBS;refGene_tx=XM_006936655.3;refGene_gn=HMBS;refGene_tx=XM_006936656.1;refGene_gn=HMBS;refGene_tx=XM_019811157.1
D1 16541439 16541489
refGene_gn=HMBS;refGene_tx=NM_001177808.1;refGene_gn=HMBS;refGene_tx=NM_001177809.1;refGene_gn=HMBS;refGene_tx=XM_006936655.3;refGene_gn=HMBS;refGene_tx=XM_006936656.1;refGene_gn=HMBS;refGene_tx=XM_019811157.1
D1 16541574 16541630
refGene_gn=HMBS;refGene_tx=NM_001177808.1;refGene_gn=HMBS;refGene_tx=NM_001177809.1;refGene_gn=HMBS;refGene_tx=XM_006936655.3;refGene_gn=HMBS;refGene_tx=XM_006936656.1;refGene_gn=HMBS;refGene_tx=XM_019811157.1

D1 16542011 16542089
refGene_gn=HMBS;refGene_tx=NM_001177808.1;refGene_gn=HMBS;refGene_tx=NM_001177809.1;refGene_gn=HMBS;refGene_tx=XM_006936655.3;refGene_gn=HMBS;refGene_tx=XM_006936656.1;refGene_gn=HMBS;refGene_tx=XM_019811157.1
D1 16542318 16542396
refGene_gn=HMBS;refGene_tx=NM_001177808.1;refGene_gn=HMBS;refGene_tx=NM_001177809.1;refGene_gn=HMBS;refGene_tx=XM_006936655.3;refGene_gn=HMBS;refGene_tx=XM_006936656.1;refGene_gn=HMBS;refGene_tx=XM_019811157.1
D1 16542518 16542594
refGene_gn=HMBS;refGene_tx=NM_001177808.1;refGene_gn=HMBS;refGene_tx=NM_001177809.1;refGene_gn=HMBS;refGene_tx=XM_006936655.3;refGene_gn=HMBS;refGene_tx=XM_006936656.1;refGene_gn=HMBS;refGene_tx=XM_019811157.1
D1 16543428 16543542
refGene_gn=HMBS;refGene_tx=NM_001177808.1;refGene_gn=HMBS;refGene_tx=NM_001177809.1;refGene_gn=HMBS;refGene_tx=XM_006936655.3;refGene_gn=HMBS;refGene_tx=XM_006936656.1;refGene_gn=HMBS;refGene_tx=XM_019811157.1
D1 16543793 16543832
refGene_gn=HMBS;refGene_tx=NM_001177808.1;refGene_gn=HMBS;refGene_tx=NM_001177809.1;refGene_gn=HMBS;refGene_tx=XM_006936655.3;refGene_gn=HMBS;refGene_tx=XM_006936656.1;refGene_gn=HMBS;refGene_tx=XM_019811157.1
D1 16544057 16544177
refGene_gn=HMBS;refGene_tx=NM_001177808.1;refGene_gn=HMBS;refGene_tx=NM_001177809.1;refGene_gn=HMBS;refGene_tx=XM_006936655.3;refGene_gn=HMBS;refGene_tx=XM_006936656.1;refGene_gn=HMBS;refGene_tx=XM_019811157.1
D1 16544415 16544469
refGene_gn=HMBS;refGene_tx=NM_001177808.1;refGene_gn=HMBS;refGene_tx=NM_001177809.1;refGene_gn=HMBS;refGene_tx=XM_006936655.3;refGene_gn=HMBS;refGene_tx=XM_006936656.1;refGene_gn=HMBS;refGene_tx=XM_019811157.1
D1 16544575 16544662
refGene_gn=HMBS;refGene_tx=NM_001177808.1;refGene_gn=HMBS;refGene_tx=NM_001177809.1;refGene_gn=HMBS;refGene_tx=XM_006936655.3;refGene_gn=HMBS;refGene_tx=XM_006936656.1;refGene_gn=HMBS;refGene_tx=XM_019811157.1
D1 16544759 16544933
refGene_gn=HMBS;refGene_tx=NM_001177808.1;refGene_gn=HMBS;refGene_tx=NM_001177809.1;refGene_gn=HMBS;refGene_tx=XM_006936655.3;refGene_gn=HMBS;refGene_tx=XM_006936656.1;refGene_gn=HMBS;refGene_tx=XM_019811157.1
D1 16546351 16546783
refGene_gn=LOC101095127;refGene_tx=XM_003992431.5
D1 16547988 16548100
refGene_gn=DPAGT1;refGene_tx=XM_003992432.4;refGene_gn=DPAGT1;refGene_tx=XM_019811392.2
D1 16548176 16548334
refGene_gn=DPAGT1;refGene_tx=XM_003992432.4;refGene_gn=DPAGT1;refGene_tx=XM_019811392.2
D1 16548493 16548581
refGene_gn=DPAGT1;refGene_tx=XM_003992432.4;refGene_gn=DPAGT1;refGene_tx=XM_019811392.2
D1 16548889 16549078
refGene_gn=DPAGT1;refGene_tx=XM_003992432.4;refGene_gn=DPAGT1;refGene_tx=XM_019811392.2

D1 16549441 16549526
refGene_gn=DPAGT1;refGene_tx=XM_003992432.4;refGene_gn=DPAGT1;refGene_tx=XM_019811392.2
D1 16551015 16551162
refGene_gn=DPAGT1;refGene_tx=XM_003992432.4;refGene_gn=DPAGT1;refGene_tx=XM_019811392.2
D1 16551374 16551588
refGene_gn=DPAGT1;refGene_tx=XM_003992432.4;refGene_gn=DPAGT1;refGene_tx=XM_019811392.2
D1 16551761 16551882
refGene_gn=DPAGT1;refGene_tx=XM_003992432.4;refGene_gn=DPAGT1;refGene_tx=XM_019811392.2
D1 16552213 16552374
refGene_gn=DPAGT1;refGene_tx=XM_003992432.4;refGene_gn=DPAGT1;refGene_tx=XM_019811392.2
D1 16557379 16557733
refGene_gn=C2CD2L;refGene_tx=XM_011286400.3;refGene_gn=C2CD2L;refGene_tx=XM_011286401.3
D1 16559772 16559868
refGene_gn=C2CD2L;refGene_tx=XM_011286400.3;refGene_gn=C2CD2L;refGene_tx=XM_011286401.3
D1 16560231 16560351
refGene_gn=C2CD2L;refGene_tx=XM_011286400.3;refGene_gn=C2CD2L;refGene_tx=XM_011286401.3
D1 16560517 16560628
refGene_gn=C2CD2L;refGene_tx=XM_011286400.3;refGene_gn=C2CD2L;refGene_tx=XM_011286401.3
D1 16560719 16560836
refGene_gn=C2CD2L;refGene_tx=XM_011286400.3;refGene_gn=C2CD2L;refGene_tx=XM_011286401.3
D1 16560930 16561042
refGene_gn=C2CD2L;refGene_tx=XM_011286400.3;refGene_gn=C2CD2L;refGene_tx=XM_011286401.3
D1 16561179 16561288
refGene_gn=C2CD2L;refGene_tx=XM_011286400.3;refGene_gn=C2CD2L;refGene_tx=XM_011286401.3
D1 16562182 16562247
refGene_gn=C2CD2L;refGene_tx=XM_011286400.3;refGene_gn=C2CD2L;refGene_tx=XM_011286401.3
D1 16562337 16562465
refGene_gn=C2CD2L;refGene_tx=XM_011286400.3;refGene_gn=C2CD2L;refGene_tx=XM_011286401.3
D1 16562556 16562731
refGene_gn=C2CD2L;refGene_tx=XM_011286400.3;refGene_gn=C2CD2L;refGene_tx=XM_011286401.3
D1 16563418 16563520
refGene_gn=C2CD2L;refGene_tx=XM_011286400.3;refGene_gn=C2CD2L;refGene_tx=XM_011286401.3
D1 16563659 16563796
refGene_gn=C2CD2L;refGene_tx=XM_011286400.3;refGene_gn=C2CD2L;refGene_

tx=XM_011286401.3
D1	16563883	16564169
refGene_gn=C2CD2L;refGene_tx=XM_011286400.3;refGene_gn=C2CD2L;refGene_tx=XM_011286401.3
D1	16565462	16565674
refGene_gn=C2CD2L;refGene_tx=XM_011286400.3;refGene_gn=C2CD2L;refGene_tx=XM_011286401.3
D1	16569908	16569955	refGene_gn=HINFP;refGene_tx=XM_003992433.4
D1	16575283	16575468
refGene_gn=HINFP;refGene_tx=XM_003992433.4;refGene_gn=HINFP;refGene_tx=XM_006936712.3;refGene_gn=HINFP;refGene_tx=XM_006936713.3;refGene_gn=HINFP;refGene_tx=XM_019811393.2;refGene_gn=HINFP;refGene_tx=XM_019811394.2;refGene_gn=HINFP;refGene_tx=XM_019811395.2;refGene_gn=HINFP;refGene_tx=XM_019811396.2
D1	16577588	16577818
refGene_gn=HINFP;refGene_tx=XM_003992433.4;refGene_gn=HINFP;refGene_tx=XM_006936712.3;refGene_gn=HINFP;refGene_tx=XM_006936713.3;refGene_gn=HINFP;refGene_tx=XM_019811393.2;refGene_gn=HINFP;refGene_tx=XM_019811394.2;refGene_gn=HINFP;refGene_tx=XM_019811395.2;refGene_gn=HINFP;refGene_tx=XM_019811396.2
D1	16578071	16578183
refGene_gn=HINFP;refGene_tx=XM_003992433.4;refGene_gn=HINFP;refGene_tx=XM_006936712.3;refGene_gn=HINFP;refGene_tx=XM_006936713.3;refGene_gn=HINFP;refGene_tx=XM_019811393.2;refGene_gn=HINFP;refGene_tx=XM_019811394.2;refGene_gn=HINFP;refGene_tx=XM_019811395.2;refGene_gn=HINFP;refGene_tx=XM_019811396.2
D1	16578351	16578504
refGene_gn=HINFP;refGene_tx=XM_003992433.4;refGene_gn=HINFP;refGene_tx=XM_006936712.3;refGene_gn=HINFP;refGene_tx=XM_006936713.3;refGene_gn=HINFP;refGene_tx=XM_019811393.2;refGene_gn=HINFP;refGene_tx=XM_019811394.2;refGene_gn=HINFP;refGene_tx=XM_019811395.2;refGene_gn=HINFP;refGene_tx=XM_019811396.2
D1	16579002	16579080
refGene_gn=HINFP;refGene_tx=XM_003992433.4;refGene_gn=HINFP;refGene_tx=XM_006936712.3;refGene_gn=HINFP;refGene_tx=XM_006936713.3;refGene_gn=HINFP;refGene_tx=XM_019811393.2;refGene_gn=HINFP;refGene_tx=XM_019811394.2;refGene_gn=HINFP;refGene_tx=XM_019811395.2;refGene_gn=HINFP;refGene_tx=XM_019811396.2
D1	16579167	16579288
refGene_gn=HINFP;refGene_tx=XM_003992433.4;refGene_gn=HINFP;refGene_tx=XM_006936712.3;refGene_gn=HINFP;refGene_tx=XM_006936713.3;refGene_gn=HINFP;refGene_tx=XM_019811393.2;refGene_gn=HINFP;refGene_tx=XM_019811394.2;refGene_gn=HINFP;refGene_tx=XM_019811395.2;refGene_gn=HINFP;refGene_tx=XM_019811396.2
D1	16579374	16579513
refGene_gn=HINFP;refGene_tx=XM_003992433.4;refGene_gn=HINFP;refGene_tx=XM_006936712.3;refGene_gn=HINFP;refGene_tx=XM_006936713.3;refGene_gn=HINFP;refGene_tx=XM_019811393.2;refGene_gn=HINFP;refGene_tx=XM_019811394.2;refGene_gn=HINFP;refGene_tx=XM_019811395.2;refGene_gn=HINFP;refGene_tx=XM_019811396.2

D1 16579605 16579730
refGene_gn=HINFP;refGene_tx=XM_003992433.4;refGene_gn=HINFP;refGene_tx=XM_006936712.3;refGene_gn=HINFP;refGene_tx=XM_006936713.3;refGene_gn=HINFP;refGene_tx=XM_019811393.2;refGene_gn=HINFP;refGene_tx=XM_019811394.2;refGene_gn=HINFP;refGene_tx=XM_019811395.2;refGene_gn=HINFP;refGene_tx=XM_019811396.2
D1 16580070 16580710
refGene_gn=HINFP;refGene_tx=XM_003992433.4;refGene_gn=HINFP;refGene_tx=XM_006936712.3;refGene_gn=HINFP;refGene_tx=XM_006936713.3;refGene_gn=HINFP;refGene_tx=XM_019811393.2;refGene_gn=HINFP;refGene_tx=XM_019811394.2;refGene_gn=HINFP;refGene_tx=XM_019811395.2;refGene_gn=HINFP;refGene_tx=XM_019811396.2
D1 16581056 16581068 refGene_gn=HINFP;refGene_tx=XM_019811394.2
D1 16581434 16581476 refGene_gn=HINFP;refGene_tx=XM_019811393.2
D1 16591293 16591531
refGene_gn=ABCG4;refGene_tx=XM_019811397.2;refGene_gn=ABCG4;refGene_tx=XM_019811399.2;refGene_gn=ABCG4;refGene_tx=XM_023239176.1;refGene_gn=ABCG4;refGene_tx=XM_023239177.1
D1 16595625 16595743
refGene_gn=ABCG4;refGene_tx=XM_019811397.2;refGene_gn=ABCG4;refGene_tx=XM_019811399.2;refGene_gn=ABCG4;refGene_tx=XM_019811401.2;refGene_gn=ABCG4;refGene_tx=XM_023239176.1;refGene_gn=ABCG4;refGene_tx=XM_023239177.1
D1 16595867 16596000
refGene_gn=ABCG4;refGene_tx=XM_019811397.2;refGene_gn=ABCG4;refGene_tx=XM_019811399.2;refGene_gn=ABCG4;refGene_tx=XM_019811401.2;refGene_gn=ABCG4;refGene_tx=XM_023239176.1;refGene_gn=ABCG4;refGene_tx=XM_023239177.1
D1 16596124 16596175
refGene_gn=ABCG4;refGene_tx=XM_019811397.2;refGene_gn=ABCG4;refGene_tx=XM_019811399.2;refGene_gn=ABCG4;refGene_tx=XM_019811401.2;refGene_gn=ABCG4;refGene_tx=XM_023239176.1;refGene_gn=ABCG4;refGene_tx=XM_023239177.1
D1 16596378 16596524
refGene_gn=ABCG4;refGene_tx=XM_019811397.2;refGene_gn=ABCG4;refGene_tx=XM_019811399.2;refGene_gn=ABCG4;refGene_tx=XM_019811401.2;refGene_gn=ABCG4;refGene_tx=XM_023239176.1;refGene_gn=ABCG4;refGene_tx=XM_023239177.1
D1 16597667 16597791
refGene_gn=ABCG4;refGene_tx=XM_019811397.2;refGene_gn=ABCG4;refGene_tx=XM_019811399.2;refGene_gn=ABCG4;refGene_tx=XM_019811401.2;refGene_gn=ABCG4;refGene_tx=XM_023239176.1;refGene_gn=ABCG4;refGene_tx=XM_023239177.1
D1 16597919 16598034
refGene_gn=ABCG4;refGene_tx=XM_019811397.2;refGene_gn=ABCG4;refGene_tx=XM_019811399.2;refGene_gn=ABCG4;refGene_tx=XM_019811401.2;refGene_gn=ABCG4;refGene_tx=XM_023239176.1
D1 16598204 16598347
refGene_gn=ABCG4;refGene_tx=XM_019811397.2;refGene_gn=ABCG4;refGene_tx=XM_019811399.2;refGene_gn=ABCG4;refGene_tx=XM_019811401.2;refGene_gn=

ABCG4;refGene_tx=XM_023239176.1
D1 16599600 16599699
refGene_gn=ABCG4;refGene_tx=XM_019811397.2;refGene_gn=ABCG4;refGene_tx=XM_019811399.2;refGene_gn=ABCG4;refGene_tx=XM_019811401.2;refGene_gn=ABCG4;refGene_tx=XM_023239176.1;refGene_gn=ABCG4;refGene_tx=XM_023239177.1
D1 16599925 16600094
refGene_gn=ABCG4;refGene_tx=XM_019811397.2;refGene_gn=ABCG4;refGene_tx=XM_019811399.2;refGene_gn=ABCG4;refGene_tx=XM_019811401.2;refGene_gn=ABCG4;refGene_tx=XM_023239176.1;refGene_gn=ABCG4;refGene_tx=XM_023239177.1
D1 16600201 16600302
refGene_gn=ABCG4;refGene_tx=XM_019811397.2;refGene_gn=ABCG4;refGene_tx=XM_019811399.2;refGene_gn=ABCG4;refGene_tx=XM_019811401.2;refGene_gn=ABCG4;refGene_tx=XM_023239176.1;refGene_gn=ABCG4;refGene_tx=XM_023239177.1
D1 16601644 16601803
refGene_gn=ABCG4;refGene_tx=XM_019811397.2;refGene_gn=ABCG4;refGene_tx=XM_019811399.2;refGene_gn=ABCG4;refGene_tx=XM_019811401.2;refGene_gn=ABCG4;refGene_tx=XM_023239176.1;refGene_gn=ABCG4;refGene_tx=XM_023239177.1
D1 16601970 16602089
refGene_gn=ABCG4;refGene_tx=XM_019811397.2;refGene_gn=ABCG4;refGene_tx=XM_019811399.2;refGene_gn=ABCG4;refGene_tx=XM_019811401.2;refGene_gn=ABCG4;refGene_tx=XM_023239176.1;refGene_gn=ABCG4;refGene_tx=XM_023239177.1
D1 16602252 16602559
refGene_gn=ABCG4;refGene_tx=XM_019811397.2;refGene_gn=ABCG4;refGene_tx=XM_019811399.2;refGene_gn=ABCG4;refGene_tx=XM_019811401.2;refGene_gn=ABCG4;refGene_tx=XM_023239176.1;refGene_gn=ABCG4;refGene_tx=XM_023239177.1
D1 16611260 16611330
refGene_gn=NLRX1;refGene_tx=XM_006936716.4;refGene_gn=NLRX1;refGene_tx=XM_006936717.4;refGene_gn=NLRX1;refGene_tx=XM_006936718.4;refGene_gn=NLRX1;refGene_tx=XM_019811402.2
D1 16611886 16611956
refGene_gn=NLRX1;refGene_tx=XM_006936716.4;refGene_gn=NLRX1;refGene_tx=XM_006936717.4;refGene_gn=NLRX1;refGene_tx=XM_006936718.4;refGene_gn=NLRX1;refGene_tx=XM_019811402.2
D1 16612439 16612528
refGene_gn=NLRX1;refGene_tx=XM_006936716.4;refGene_gn=NLRX1;refGene_tx=XM_006936717.4;refGene_gn=NLRX1;refGene_tx=XM_006936718.4;refGene_gn=NLRX1;refGene_tx=XM_019811402.2
D1 16613088 16613708
refGene_gn=NLRX1;refGene_tx=XM_006936716.4;refGene_gn=NLRX1;refGene_tx=XM_006936717.4;refGene_gn=NLRX1;refGene_tx=XM_006936718.4;refGene_gn=NLRX1;refGene_tx=XM_019811402.2
D1 16614290 16615112
refGene_gn=NLRX1;refGene_tx=XM_006936716.4;refGene_gn=NLRX1;refGene_tx=XM_006936717.4;refGene_gn=NLRX1;refGene_tx=XM_006936718.4;refGene_gn=

NLRX1;refGene_tx=XM_019811402.2
D1 16617949 16618545
refGene_gn=NLRX1;refGene_tx=XM_006936716.4;refGene_gn=NLRX1;refGene_tx=XM_006936717.4;refGene_gn=NLRX1;refGene_tx=XM_006936718.4;refGene_gn=NLRX1;refGene_tx=XM_019811402.2
D1 16618820 16618907
refGene_gn=NLRX1;refGene_tx=XM_006936716.4;refGene_gn=NLRX1;refGene_tx=XM_006936717.4;refGene_gn=NLRX1;refGene_tx=XM_006936718.4;refGene_gn=NLRX1;refGene_tx=XM_019811402.2
D1 16619207 16619235 refGene_gn=NLRX1;refGene_tx=XM_006936718.4
D1 16619642 16619894
refGene_gn=NLRX1;refGene_tx=XM_006936716.4;refGene_gn=NLRX1;refGene_tx=XM_006936717.4;refGene_gn=NLRX1;refGene_tx=XM_019811402.2
D1 16620302 16620624
refGene_gn=NLRX1;refGene_tx=XM_006936716.4;refGene_gn=NLRX1;refGene_tx=XM_006936717.4;refGene_gn=NLRX1;refGene_tx=XM_019811402.2
D1 16621653 16621669 refGene_gn=PDZD3;refGene_tx=XM_019811405.2
D1 16622055 16622083 refGene_gn=PDZD3;refGene_tx=XM_019811405.2
D1 16622226 16622300 refGene_gn=PDZD3;refGene_tx=XM_019811405.2
D1 16622636 16622848 refGene_gn=PDZD3;refGene_tx=XM_019811405.2
D1 16623123 16623313 refGene_gn=PDZD3;refGene_tx=XM_019811405.2
D1 16623428 16623565 refGene_gn=PDZD3;refGene_tx=XM_019811405.2
D1 16623817 16624022 refGene_gn=PDZD3;refGene_tx=XM_019811405.2
D1 16624244 16624327 refGene_gn=PDZD3;refGene_tx=XM_019811405.2
D1 16624328 16624442 refGene_gn=PDZD3;refGene_tx=XM_019811405.2
D1 16624547 16624775 refGene_gn=PDZD3;refGene_tx=XM_019811405.2
D1 16624915 16625117 refGene_gn=PDZD3;refGene_tx=XM_019811405.2
D1 16625349 16625438 refGene_gn=PDZD3;refGene_tx=XM_019811405.2
D1 16627241 16627370
refGene_gn=CCDC153;refGene_tx=XM_003992468.4;refGene_gn=CCDC153;refGene_tx=XM_023239179.1
D1 16627635 16627766
refGene_gn=CCDC153;refGene_tx=XM_003992468.4;refGene_gn=CCDC153;refGene_tx=XM_023239179.1
D1 16629389 16629482 refGene_gn=CCDC153;refGene_tx=XM_003992468.4
D1 16629992 16630112
refGene_gn=CCDC153;refGene_tx=XM_003992468.4;refGene_gn=CCDC153;refGene_tx=XM_023239179.1
D1 16630516 16630582
refGene_gn=CCDC153;refGene_tx=XM_003992468.4;refGene_gn=CCDC153;refGene_tx=XM_023239179.1
D1 16631040 16631110
refGene_gn=CCDC153;refGene_tx=XM_003992468.4;refGene_gn=CCDC153;refGene_tx=XM_023239179.1
D1 16636518 16636713
refGene_gn=LOC111556576;refGene_tx=XM_023239181.1
D1 16658054 16658306
refGene_gn=LOC111556576;refGene_tx=XM_023239181.1
D1 16659018 16659146
refGene_gn=LOC101096508;refGene_tx=XM_023239180.1

D1 16686845 16686992
refGene_gn=LOC101096508;refGene_tx=XM_023239180.1
D1 16690162 16690319
refGene_gn=LOC101096508;refGene_tx=XM_023239180.1
D1 16691103 16691225
refGene_gn=LOC101096508;refGene_tx=XM_023239180.1
D1 16691557 16691695
refGene_gn=LOC101096508;refGene_tx=XM_023239180.1
D1 16692523 16692611
refGene_gn=LOC101096508;refGene_tx=XM_023239180.1
D1 16693196 16693328
refGene_gn=LOC101096508;refGene_tx=XM_023239180.1
D1 16693541 16693745
refGene_gn=LOC101096508;refGene_tx=XM_023239180.1
D1 16701286 16701418
refGene_gn=LOC101096508;refGene_tx=XM_023239180.1
D1 16701564 16701942
refGene_gn=LOC101096508;refGene_tx=XM_023239180.1
D1 16703659 16703754
refGene_gn=LOC101096508;refGene_tx=XM_023239180.1
D1 16711533 16711674
refGene_gn=LOC101096508;refGene_tx=XM_023239180.1
D1 16712044 16712145
refGene_gn=LOC101096508;refGene_tx=XM_023239180.1
D1 16712854 16713037
refGene_gn=LOC101096508;refGene_tx=XM_023239180.1
D1 16714022 16714294
refGene_gn=LOC101096508;refGene_tx=XM_023239180.1
D1 16723587 16723617
refGene_gn=MCAM;refGene_tx=XM_006936722.4;refGene_gn=MCAM;refGene_tx=XM_006936723.4
D1 16724082 16724200 refGene_gn=MCAM;refGene_tx=XM_006936722.4
D1 16724532 16724680
refGene_gn=MCAM;refGene_tx=XM_006936722.4;refGene_gn=MCAM;refGene_tx=XM_006936723.4
D1 16724857 16724929
refGene_gn=MCAM;refGene_tx=XM_006936722.4;refGene_gn=MCAM;refGene_tx=XM_006936723.4
D1 16725021 16725163
refGene_gn=MCAM;refGene_tx=XM_006936722.4;refGene_gn=MCAM;refGene_tx=XM_006936723.4
D1 16725260 16725382
refGene_gn=MCAM;refGene_tx=XM_006936722.4;refGene_gn=MCAM;refGene_tx=XM_006936723.4
D1 16725553 16725695
refGene_gn=MCAM;refGene_tx=XM_006936722.4;refGene_gn=MCAM;refGene_tx=XM_006936723.4
D1 16725775 16725894
refGene_gn=MCAM;refGene_tx=XM_006936722.4;refGene_gn=MCAM;refGene_tx=XM_006936723.4

D1 16725982 16726145
refGene_gn=MCAM;refGene_tx=XM_006936722.4;refGene_gn=MCAM;refGene_tx=XM_006936723.4
D1 16726229 16726351
refGene_gn=MCAM;refGene_tx=XM_006936722.4;refGene_gn=MCAM;refGene_tx=XM_006936723.4
D1 16726468 16726648
refGene_gn=MCAM;refGene_tx=XM_006936722.4;refGene_gn=MCAM;refGene_tx=XM_006936723.4
D1 16727806 16727894
refGene_gn=MCAM;refGene_tx=XM_006936722.4;refGene_gn=MCAM;refGene_tx=XM_006936723.4
D1 16727971 16728042
refGene_gn=MCAM;refGene_tx=XM_006936722.4;refGene_gn=MCAM;refGene_tx=XM_006936723.4
D1 16728128 16728348
refGene_gn=MCAM;refGene_tx=XM_006936722.4;refGene_gn=MCAM;refGene_tx=XM_006936723.4
D1 16728440 16728565
refGene_gn=MCAM;refGene_tx=XM_006936722.4;refGene_gn=MCAM;refGene_tx=XM_006936723.4
D1 16730328 16730395
refGene_gn=MCAM;refGene_tx=XM_006936722.4;refGene_gn=MCAM;refGene_tx=XM_006936723.4
D1 16756936 16758469
refGene_gn=RNF26;refGene_tx=XM_019811422.2;refGene_gn=RNF26;refGene_tx=XM_019811423.2
D1 16761184 16761702 refGene_gn=C1QTNF5;refGene_tx=XM_023239188.1
D1 16762062 16762276 refGene_gn=C1QTNF5;refGene_tx=XM_023239188.1
D1 16763446 16763671
refGene_gn=MFRP;refGene_tx=XM_003992441.5;refGene_gn=MFRP;refGene_tx=XM_023239185.1;refGene_gn=MFRP;refGene_tx=XM_023239186.1;refGene_gn=MFRP;refGene_tx=XM_023239187.1
D1 16763781 16763909
refGene_gn=MFRP;refGene_tx=XM_003992441.5;refGene_gn=MFRP;refGene_tx=XM_023239185.1;refGene_gn=MFRP;refGene_tx=XM_023239186.1;refGene_gn=MFRP;refGene_tx=XM_023239187.1
D1 16765060 16765192
refGene_gn=MFRP;refGene_tx=XM_003992441.5;refGene_gn=MFRP;refGene_tx=XM_023239185.1;refGene_gn=MFRP;refGene_tx=XM_023239186.1;refGene_gn=MFRP;refGene_tx=XM_023239187.1
D1 16765339 16765470
refGene_gn=MFRP;refGene_tx=XM_003992441.5;refGene_gn=MFRP;refGene_tx=XM_023239185.1;refGene_gn=MFRP;refGene_tx=XM_023239186.1;refGene_gn=MFRP;refGene_tx=XM_023239187.1
D1 16765763 16765822 refGene_gn=MFRP;refGene_tx=XM_023239185.1
D1 16765902 16766051
refGene_gn=MFRP;refGene_tx=XM_003992441.5;refGene_gn=MFRP;refGene_tx=XM_023239186.1;refGene_gn=MFRP;refGene_tx=XM_023239187.1
D1 16766445 16766522

refGene_gn=MFRP;refGene_tx=XM_003992441.5;refGene_gn=MFRP;refGene_tx=XM_023239185.1;refGene_gn=MFRP;refGene_tx=XM_023239186.1;refGene_gn=MFRP;refGene_tx=XM_023239187.1
D1 16766755 16766881
refGene_gn=MFRP;refGene_tx=XM_003992441.5;refGene_gn=MFRP;refGene_tx=XM_023239185.1;refGene_gn=MFRP;refGene_tx=XM_023239187.1
D1 16767015 16767146
refGene_gn=MFRP;refGene_tx=XM_003992441.5;refGene_gn=MFRP;refGene_tx=XM_023239185.1;refGene_gn=MFRP;refGene_tx=XM_023239186.1;refGene_gn=MFRP;refGene_tx=XM_023239187.1
D1 16767545 16767759
refGene_gn=MFRP;refGene_tx=XM_003992441.5;refGene_gn=MFRP;refGene_tx=XM_023239185.1;refGene_gn=MFRP;refGene_tx=XM_023239186.1;refGene_gn=MFRP;refGene_tx=XM_023239187.1
D1 16767892 16768078
refGene_gn=MFRP;refGene_tx=XM_003992441.5;refGene_gn=MFRP;refGene_tx=XM_023239185.1;refGene_gn=MFRP;refGene_tx=XM_023239186.1
D1 16768200 16768314
refGene_gn=MFRP;refGene_tx=XM_003992441.5;refGene_gn=MFRP;refGene_tx=XM_023239185.1;refGene_gn=MFRP;refGene_tx=XM_023239186.1;refGene_gn=MFRP;refGene_tx=XM_023239187.1
D1 16768403 16768500
refGene_gn=MFRP;refGene_tx=XM_003992441.5;refGene_gn=MFRP;refGene_tx=XM_023239185.1;refGene_gn=MFRP;refGene_tx=XM_023239186.1;refGene_gn=MFRP;refGene_tx=XM_023239187.1
D1 16768625 16768679
refGene_gn=MFRP;refGene_tx=XM_003992441.5;refGene_gn=MFRP;refGene_tx=XM_023239185.1;refGene_gn=MFRP;refGene_tx=XM_023239186.1;refGene_gn=MFRP;refGene_tx=XM_023239187.1
D1 16775057 16775145
refGene_gn=USP2;refGene_tx=XM_006936724.4;refGene_gn=USP2;refGene_tx=XM_006936725.4;refGene_gn=USP2;refGene_tx=XM_006936727.4;refGene_gn=USP2;refGene_tx=XM_011286406.2;refGene_gn=USP2;refGene_tx=XM_019811412.2;refGene_gn=USP2;refGene_tx=XM_019811413.2;refGene_gn=USP2;refGene_tx=XM_019811414.2;refGene_gn=USP2;refGene_tx=XM_019811415.2;refGene_gn=USP2;refGene_tx=XM_023239183.1
D1 16775418 16775539
refGene_gn=USP2;refGene_tx=XM_006936724.4;refGene_gn=USP2;refGene_tx=XM_006936725.4;refGene_gn=USP2;refGene_tx=XM_006936727.4;refGene_gn=USP2;refGene_tx=XM_011286406.2;refGene_gn=USP2;refGene_tx=XM_019811412.2;refGene_gn=USP2;refGene_tx=XM_019811413.2;refGene_gn=USP2;refGene_tx=XM_019811414.2;refGene_gn=USP2;refGene_tx=XM_019811415.2;refGene_gn=USP2;refGene_tx=XM_023239183.1
D1 16775714 16775822
refGene_gn=USP2;refGene_tx=XM_006936724.4;refGene_gn=USP2;refGene_tx=XM_006936725.4;refGene_gn=USP2;refGene_tx=XM_006936727.4;refGene_gn=USP2;refGene_tx=XM_011286406.2;refGene_gn=USP2;refGene_tx=XM_019811412.2;refGene_gn=USP2;refGene_tx=XM_019811413.2;refGene_gn=USP2;refGene_tx=XM_019811414.2;refGene_gn=USP2;refGene_tx=XM_019811415.2;refGene_gn=USP2;refGene_tx=XM_023239183.1

D1 16776040 16776119
refGene_gn=USP2;refGene_tx=XM_006936724.4;refGene_gn=USP2;refGene_tx=XM_006936725.4;refGene_gn=USP2;refGene_tx=XM_006936727.4;refGene_gn=USP2;refGene_tx=XM_011286406.2;refGene_gn=USP2;refGene_tx=XM_019811412.2;refGene_gn=USP2;refGene_tx=XM_019811413.2;refGene_gn=USP2;refGene_tx=XM_019811414.2;refGene_gn=USP2;refGene_tx=XM_019811415.2;refGene_gn=USP2;refGene_tx=XM_023239183.1
D1 16776271 16776352
refGene_gn=USP2;refGene_tx=XM_006936724.4;refGene_gn=USP2;refGene_tx=XM_006936725.4;refGene_gn=USP2;refGene_tx=XM_006936727.4;refGene_gn=USP2;refGene_tx=XM_011286406.2;refGene_gn=USP2;refGene_tx=XM_019811412.2;refGene_gn=USP2;refGene_tx=XM_019811413.2;refGene_gn=USP2;refGene_tx=XM_019811414.2;refGene_gn=USP2;refGene_tx=XM_019811415.2;refGene_gn=USP2;refGene_tx=XM_023239183.1
D1 16776435 16776539
refGene_gn=USP2;refGene_tx=XM_006936724.4;refGene_gn=USP2;refGene_tx=XM_006936725.4;refGene_gn=USP2;refGene_tx=XM_006936727.4;refGene_gn=USP2;refGene_tx=XM_011286406.2;refGene_gn=USP2;refGene_tx=XM_019811412.2;refGene_gn=USP2;refGene_tx=XM_019811413.2;refGene_gn=USP2;refGene_tx=XM_019811414.2;refGene_gn=USP2;refGene_tx=XM_019811415.2;refGene_gn=USP2;refGene_tx=XM_023239183.1
D1 16776755 16776820
refGene_gn=USP2;refGene_tx=XM_006936724.4;refGene_gn=USP2;refGene_tx=XM_006936725.4;refGene_gn=USP2;refGene_tx=XM_006936727.4;refGene_gn=USP2;refGene_tx=XM_011286406.2;refGene_gn=USP2;refGene_tx=XM_019811412.2;refGene_gn=USP2;refGene_tx=XM_019811413.2;refGene_gn=USP2;refGene_tx=XM_019811414.2;refGene_gn=USP2;refGene_tx=XM_019811415.2;refGene_gn=USP2;refGene_tx=XM_023239183.1
D1 16776995 16777106
refGene_gn=USP2;refGene_tx=XM_006936724.4;refGene_gn=USP2;refGene_tx=XM_006936725.4;refGene_gn=USP2;refGene_tx=XM_006936727.4;refGene_gn=USP2;refGene_tx=XM_011286406.2;refGene_gn=USP2;refGene_tx=XM_019811412.2;refGene_gn=USP2;refGene_tx=XM_019811413.2;refGene_gn=USP2;refGene_tx=XM_019811414.2;refGene_gn=USP2;refGene_tx=XM_019811415.2;refGene_gn=USP2;refGene_tx=XM_023239183.1
D1 16777208 16777320
refGene_gn=USP2;refGene_tx=XM_006936724.4;refGene_gn=USP2;refGene_tx=XM_006936725.4;refGene_gn=USP2;refGene_tx=XM_006936727.4;refGene_gn=USP2;refGene_tx=XM_011286406.2;refGene_gn=USP2;refGene_tx=XM_019811412.2;refGene_gn=USP2;refGene_tx=XM_019811413.2;refGene_gn=USP2;refGene_tx=XM_019811414.2;refGene_gn=USP2;refGene_tx=XM_019811415.2;refGene_gn=USP2;refGene_tx=XM_023239183.1
D1 16777503 16777627
refGene_gn=USP2;refGene_tx=XM_006936724.4;refGene_gn=USP2;refGene_tx=XM_006936725.4;refGene_gn=USP2;refGene_tx=XM_006936727.4;refGene_gn=USP2;refGene_tx=XM_011286406.2;refGene_gn=USP2;refGene_tx=XM_019811412.2;refGene_gn=USP2;refGene_tx=XM_019811413.2;refGene_gn=USP2;refGene_tx=XM_019811414.2;refGene_gn=USP2;refGene_tx=XM_019811415.2;refGene_gn=USP2;refGene_tx=XM_023239183.1
D1 16778119 16778170

refGene_gn=USP2;refGene_tx=XM_006936724.4;refGene_gn=USP2;refGene_tx=XM_006936725.4;refGene_gn=USP2;refGene_tx=XM_006936727.4;refGene_gn=USP2;refGene_tx=XM_011286406.2;refGene_gn=USP2;refGene_tx=XM_019811412.2;refGene_gn=USP2;refGene_tx=XM_019811413.2;refGene_gn=USP2;refGene_tx=XM_019811414.2;refGene_gn=USP2;refGene_tx=XM_019811415.2;refGene_gn=USP2;refGene_tx=XM_023239183.1
D1 16781339 16781354 refGene_gn=USP2;refGene_tx=XM_011286406.2
D1 16781524 16781671 refGene_gn=USP2;refGene_tx=XM_023239183.1
D1 16789155 16789341
refGene_gn=USP2;refGene_tx=XM_006936724.4;refGene_gn=USP2;refGene_tx=XM_006936725.4;refGene_gn=USP2;refGene_tx=XM_006936727.4;refGene_gn=USP2;refGene_tx=XM_019811412.2;refGene_gn=USP2;refGene_tx=XM_019811413.2;refGene_gn=USP2;refGene_tx=XM_019811414.2
D1 16789569 16790346
refGene_gn=USP2;refGene_tx=XM_006936724.4;refGene_gn=USP2;refGene_tx=XM_006936725.4;refGene_gn=USP2;refGene_tx=XM_006936727.4;refGene_gn=USP2;refGene_tx=XM_019811412.2;refGene_gn=USP2;refGene_tx=XM_019811413.2;refGene_gn=USP2;refGene_tx=XM_019811414.2;refGene_gn=USP2;refGene_tx=XM_019811415.2
D1 16814773 16814886
refGene_gn=THY1;refGene_tx=XM_003992443.5;refGene_gn=THY1;refGene_tx=XM_006936729.4
D1 16815426 16815762
refGene_gn=THY1;refGene_tx=XM_003992443.5;refGene_gn=THY1;refGene_tx=XM_006936729.4
D1 16816254 16816291
refGene_gn=THY1;refGene_tx=XM_003992443.5;refGene_gn=THY1;refGene_tx=XM_006936729.4
D1 16996609 16996750 refGene_gn=NECTIN1;refGene_tx=XM_019811419.2
D1 17003998 17004325 refGene_gn=NECTIN1;refGene_tx=XM_019811418.2
D1 17004609 17004726
refGene_gn=NECTIN1;refGene_tx=XM_019811418.2;refGene_gn=NECTIN1;refGene_tx=XM_019811419.2
D1 17004879 17004978 refGene_gn=NECTIN1;refGene_tx=XM_019811420.2
D1 17005195 17005271
refGene_gn=NECTIN1;refGene_tx=XM_019811418.2;refGene_gn=NECTIN1;refGene_tx=XM_019811419.2;refGene_gn=NECTIN1;refGene_tx=XM_019811420.2
D1 17006488 17006636
refGene_gn=NECTIN1;refGene_tx=XM_019811418.2;refGene_gn=NECTIN1;refGene_tx=XM_019811419.2;refGene_gn=NECTIN1;refGene_tx=XM_019811420.2
D1 17030823 17031371 refGene_gn=NECTIN1;refGene_tx=XM_003992444.5
D1 17040019 17040171
refGene_gn=NECTIN1;refGene_tx=XM_003992444.5;refGene_gn=NECTIN1;refGene_tx=XM_019811418.2;refGene_gn=NECTIN1;refGene_tx=XM_019811419.2;refGene_gn=NECTIN1;refGene_tx=XM_019811420.2
D1 17041598 17041716
refGene_gn=NECTIN1;refGene_tx=XM_003992444.5;refGene_gn=NECTIN1;refGene_tx=XM_019811418.2;refGene_gn=NECTIN1;refGene_tx=XM_019811419.2;refGene_gn=NECTIN1;refGene_tx=XM_019811420.2
D1 17042129 17042432

refGene_gn=NECTIN1;refGene_tx=XM_003992444.5;refGene_gn=NECTIN1;refGene_tx=XM_019811418.2;refGene_gn=NECTIN1;refGene_tx=XM_019811419.2;refGene_gn=NECTIN1;refGene_tx=XM_019811420.2
D1 17042963 17043314
refGene_gn=NECTIN1;refGene_tx=XM_003992444.5;refGene_gn=NECTIN1;refGene_tx=XM_019811418.2;refGene_gn=NECTIN1;refGene_tx=XM_019811419.2;refGene_gn=NECTIN1;refGene_tx=XM_019811420.2
D1 17088818 17088897
refGene_gn=NECTIN1;refGene_tx=XM_003992444.5;refGene_gn=NECTIN1;refGene_tx=XM_019811418.2;refGene_gn=NECTIN1;refGene_tx=XM_019811419.2;refGene_gn=NECTIN1;refGene_tx=XM_019811420.2
D1 17444683 17444746 refGene_gn=TRIM29;refGene_tx=XM_003992445.5
D1 17447697 17447774 refGene_gn=TRIM29;refGene_tx=XM_003992445.5
D1 17450754 17450850 refGene_gn=TRIM29;refGene_tx=XM_003992445.5
D1 17452940 17453033 refGene_gn=TRIM29;refGene_tx=XM_003992445.5
D1 17455404 17455506 refGene_gn=TRIM29;refGene_tx=XM_003992445.5
D1 17458087 17458286 refGene_gn=TRIM29;refGene_tx=XM_003992445.5
D1 17459489 17459723 refGene_gn=TRIM29;refGene_tx=XM_003992445.5
D1 17460530 17460626 refGene_gn=TRIM29;refGene_tx=XM_003992445.5
D1 17468555 17469359 refGene_gn=TRIM29;refGene_tx=XM_003992445.5
D1 17530766 17531036 refGene_gn=OAF;refGene_tx=XM_019812919.2
D1 17544304 17544439 refGene_gn=OAF;refGene_tx=XM_019812919.2
D1 17545325 17545506 refGene_gn=OAF;refGene_tx=XM_019812919.2
D1 17547032 17547298 refGene_gn=OAF;refGene_tx=XM_019812919.2
D1 17558256 17558284 refGene_gn=POU2F3;refGene_tx=XM_019811427.2
D1 17565118 17565187 refGene_gn=POU2F3;refGene_tx=XM_019811427.2
D1 17589546 17589581 refGene_gn=POU2F3;refGene_tx=XM_019811427.2
D1 17617948 17618053 refGene_gn=POU2F3;refGene_tx=XM_019811427.2
D1 17619255 17619358 refGene_gn=POU2F3;refGene_tx=XM_019811427.2
D1 17622126 17622209 refGene_gn=POU2F3;refGene_tx=XM_019811427.2
D1 17624563 17624746 refGene_gn=POU2F3;refGene_tx=XM_019811427.2
D1 17625173 17625282 refGene_gn=POU2F3;refGene_tx=XM_019811427.2
D1 17625284 17625314 refGene_gn=POU2F3;refGene_tx=XM_019811427.2
D1 17627226 17627363 refGene_gn=POU2F3;refGene_tx=XM_019811427.2
D1 17629239 17629401 refGene_gn=POU2F3;refGene_tx=XM_019811427.2
D1 17634785 17634852 refGene_gn=POU2F3;refGene_tx=XM_019811427.2
D1 17636118 17636251 refGene_gn=POU2F3;refGene_tx=XM_019811427.2
D1 17637227 17637267 refGene_gn=POU2F3;refGene_tx=XM_019811427.2
D1 17645412 17645426 refGene_gn=TMEM136;refGene_tx=XM_011286408.3
D1 17647864 17648064
refGene_gn=TMEM136;refGene_tx=XM_011286408.3;refGene_gn=TMEM136;refGene_tx=XM_023239194.1
D1 17649712 17650251
refGene_gn=TMEM136;refGene_tx=XM_011286408.3;refGene_gn=TMEM136;refGene_tx=XM_019811428.2;refGene_gn=TMEM136;refGene_tx=XM_019811429.2;refGene_gn=TMEM136;refGene_tx=XM_023239194.1
D1 17656516 17656548
refGene_gn=ARHGEF12;refGene_tx=XM_006936730.4;refGene_gn=ARHGEF12;refGene_tx=XM_006936731.4;refGene_gn=ARHGEF12;refGene_tx=XM_019811430.2
D1 17728129 17728153

refGene_gn=ARHGEF12;refGene_tx=XM_006936730.4;refGene_gn=ARHGEF12;refGene_tx=XM_006936731.4;refGene_gn=ARHGEF12;refGene_tx=XM_019811430.2
D1 17731293 17731379
refGene_gn=ARHGEF12;refGene_tx=XM_006936730.4;refGene_gn=ARHGEF12;refGene_tx=XM_006936731.4;refGene_gn=ARHGEF12;refGene_tx=XM_019811430.2
D1 17732953 17733010
refGene_gn=ARHGEF12;refGene_tx=XM_006936730.4;refGene_gn=ARHGEF12;refGene_tx=XM_019811430.2
D1 17744947 17745046
refGene_gn=ARHGEF12;refGene_tx=XM_006936730.4;refGene_gn=ARHGEF12;refGene_tx=XM_006936731.4;refGene_gn=ARHGEF12;refGene_tx=XM_019811430.2
D1 17745673 17745723
refGene_gn=ARHGEF12;refGene_tx=XM_006936730.4;refGene_gn=ARHGEF12;refGene_tx=XM_006936731.4;refGene_gn=ARHGEF12;refGene_tx=XM_019811430.2;refGene_gn=ARHGEF12;refGene_tx=XM_023239192.1;refGene_gn=ARHGEF12;refGene_tx=XM_023239193.1
D1 17748136 17748194
refGene_gn=ARHGEF12;refGene_tx=XM_006936730.4;refGene_gn=ARHGEF12;refGene_tx=XM_006936731.4;refGene_gn=ARHGEF12;refGene_tx=XM_019811430.2;refGene_gn=ARHGEF12;refGene_tx=XM_023239192.1;refGene_gn=ARHGEF12;refGene_tx=XM_023239193.1
D1 17750726 17750905
refGene_gn=ARHGEF12;refGene_tx=XM_006936730.4;refGene_gn=ARHGEF12;refGene_tx=XM_006936731.4;refGene_gn=ARHGEF12;refGene_tx=XM_019811430.2;refGene_gn=ARHGEF12;refGene_tx=XM_023239192.1;refGene_gn=ARHGEF12;refGene_tx=XM_023239193.1
D1 17752097 17752175
refGene_gn=ARHGEF12;refGene_tx=XM_006936730.4;refGene_gn=ARHGEF12;refGene_tx=XM_006936731.4;refGene_gn=ARHGEF12;refGene_tx=XM_019811430.2;refGene_gn=ARHGEF12;refGene_tx=XM_023239192.1;refGene_gn=ARHGEF12;refGene_tx=XM_023239193.1
D1 17752367 17752487
refGene_gn=ARHGEF12;refGene_tx=XM_006936730.4;refGene_gn=ARHGEF12;refGene_tx=XM_006936731.4;refGene_gn=ARHGEF12;refGene_tx=XM_023239192.1;refGene_gn=ARHGEF12;refGene_tx=XM_023239193.1
D1 17754065 17754206
refGene_gn=ARHGEF12;refGene_tx=XM_006936730.4;refGene_gn=ARHGEF12;refGene_tx=XM_006936731.4;refGene_gn=ARHGEF12;refGene_tx=XM_019811430.2;refGene_gn=ARHGEF12;refGene_tx=XM_023239192.1;refGene_gn=ARHGEF12;refGene_tx=XM_023239193.1
D1 17759656 17759731
refGene_gn=ARHGEF12;refGene_tx=XM_006936730.4;refGene_gn=ARHGEF12;refGene_tx=XM_006936731.4;refGene_gn=ARHGEF12;refGene_tx=XM_019811430.2;refGene_gn=ARHGEF12;refGene_tx=XM_023239192.1;refGene_gn=ARHGEF12;refGene_tx=XM_023239193.1
D1 17762003 17762096
refGene_gn=ARHGEF12;refGene_tx=XM_006936730.4;refGene_gn=ARHGEF12;refGene_tx=XM_006936731.4;refGene_gn=ARHGEF12;refGene_tx=XM_019811430.2;refGene_gn=ARHGEF12;refGene_tx=XM_023239192.1;refGene_gn=ARHGEF12;refGene_tx=XM_023239193.1

D1 17764150 17764261
refGene_gn=ARHGEF12;refGene_tx=XM_006936730.4;refGene_gn=ARHGEF12;refGene_tx=XM_006936731.4;refGene_gn=ARHGEF12;refGene_tx=XM_019811430.2;refGene_gn=ARHGEF12;refGene_tx=XM_023239192.1;refGene_gn=ARHGEF12;refGene_tx=XM_023239193.1
D1 17764537 17764636
refGene_gn=ARHGEF12;refGene_tx=XM_006936730.4;refGene_gn=ARHGEF12;refGene_tx=XM_006936731.4;refGene_gn=ARHGEF12;refGene_tx=XM_019811430.2;refGene_gn=ARHGEF12;refGene_tx=XM_023239192.1;refGene_gn=ARHGEF12;refGene_tx=XM_023239193.1
D1 17770323 17770366
refGene_gn=ARHGEF12;refGene_tx=XM_006936730.4;refGene_gn=ARHGEF12;refGene_tx=XM_006936731.4;refGene_gn=ARHGEF12;refGene_tx=XM_019811430.2;refGene_gn=ARHGEF12;refGene_tx=XM_023239192.1;refGene_gn=ARHGEF12;refGene_tx=XM_023239193.1
D1 17770669 17770775
refGene_gn=ARHGEF12;refGene_tx=XM_006936730.4;refGene_gn=ARHGEF12;refGene_tx=XM_006936731.4;refGene_gn=ARHGEF12;refGene_tx=XM_019811430.2;refGene_gn=ARHGEF12;refGene_tx=XM_023239192.1;refGene_gn=ARHGEF12;refGene_tx=XM_023239193.1
D1 17771207 17771345
refGene_gn=ARHGEF12;refGene_tx=XM_006936730.4;refGene_gn=ARHGEF12;refGene_tx=XM_006936731.4;refGene_gn=ARHGEF12;refGene_tx=XM_019811430.2;refGene_gn=ARHGEF12;refGene_tx=XM_023239192.1;refGene_gn=ARHGEF12;refGene_tx=XM_023239193.1
D1 17772033 17772066
refGene_gn=ARHGEF12;refGene_tx=XM_006936730.4;refGene_gn=ARHGEF12;refGene_tx=XM_006936731.4;refGene_gn=ARHGEF12;refGene_tx=XM_019811430.2;refGene_gn=ARHGEF12;refGene_tx=XM_023239192.1;refGene_gn=ARHGEF12;refGene_tx=XM_023239193.1
D1 17772387 17772502
refGene_gn=ARHGEF12;refGene_tx=XM_006936730.4;refGene_gn=ARHGEF12;refGene_tx=XM_006936731.4;refGene_gn=ARHGEF12;refGene_tx=XM_019811430.2;refGene_gn=ARHGEF12;refGene_tx=XM_023239192.1;refGene_gn=ARHGEF12;refGene_tx=XM_023239193.1
D1 17773199 17773305
refGene_gn=ARHGEF12;refGene_tx=XM_006936730.4;refGene_gn=ARHGEF12;refGene_tx=XM_006936731.4;refGene_gn=ARHGEF12;refGene_tx=XM_019811430.2;refGene_gn=ARHGEF12;refGene_tx=XM_023239192.1;refGene_gn=ARHGEF12;refGene_tx=XM_023239193.1
D1 17775171 17775384
refGene_gn=ARHGEF12;refGene_tx=XM_006936730.4;refGene_gn=ARHGEF12;refGene_tx=XM_006936731.4;refGene_gn=ARHGEF12;refGene_tx=XM_019811430.2;refGene_gn=ARHGEF12;refGene_tx=XM_023239192.1;refGene_gn=ARHGEF12;refGene_tx=XM_023239193.1
D1 17780065 17780198
refGene_gn=ARHGEF12;refGene_tx=XM_006936730.4;refGene_gn=ARHGEF12;refGene_tx=XM_006936731.4;refGene_gn=ARHGEF12;refGene_tx=XM_019811430.2;refGene_gn=ARHGEF12;refGene_tx=XM_023239192.1;refGene_gn=ARHGEF12;refGene_tx=XM_023239193.1

D1 17781265 17781301
refGene_gn=ARHGEF12;refGene_tx=XM_006936730.4;refGene_gn=ARHGEF12;refGene_tx=XM_006936731.4;refGene_gn=ARHGEF12;refGene_tx=XM_019811430.2;refGene_gn=ARHGEF12;refGene_tx=XM_023239192.1;refGene_gn=ARHGEF12;refGene_tx=XM_023239193.1
D1 17781629 17781784
refGene_gn=ARHGEF12;refGene_tx=XM_006936730.4;refGene_gn=ARHGEF12;refGene_tx=XM_006936731.4;refGene_gn=ARHGEF12;refGene_tx=XM_019811430.2;refGene_gn=ARHGEF12;refGene_tx=XM_023239192.1;refGene_gn=ARHGEF12;refGene_tx=XM_023239193.1
D1 17782773 17782920
refGene_gn=ARHGEF12;refGene_tx=XM_006936730.4;refGene_gn=ARHGEF12;refGene_tx=XM_006936731.4;refGene_gn=ARHGEF12;refGene_tx=XM_019811430.2;refGene_gn=ARHGEF12;refGene_tx=XM_023239192.1;refGene_gn=ARHGEF12;refGene_tx=XM_023239193.1
D1 17783972 17784058
refGene_gn=ARHGEF12;refGene_tx=XM_006936730.4;refGene_gn=ARHGEF12;refGene_tx=XM_006936731.4;refGene_gn=ARHGEF12;refGene_tx=XM_019811430.2;refGene_gn=ARHGEF12;refGene_tx=XM_023239192.1;refGene_gn=ARHGEF12;refGene_tx=XM_023239193.1
D1 17787554 17787680
refGene_gn=ARHGEF12;refGene_tx=XM_006936730.4;refGene_gn=ARHGEF12;refGene_tx=XM_006936731.4;refGene_gn=ARHGEF12;refGene_tx=XM_019811430.2;refGene_gn=ARHGEF12;refGene_tx=XM_023239192.1;refGene_gn=ARHGEF12;refGene_tx=XM_023239193.1
D1 17789304 17789419
refGene_gn=ARHGEF12;refGene_tx=XM_006936730.4;refGene_gn=ARHGEF12;refGene_tx=XM_006936731.4;refGene_gn=ARHGEF12;refGene_tx=XM_019811430.2;refGene_gn=ARHGEF12;refGene_tx=XM_023239192.1;refGene_gn=ARHGEF12;refGene_tx=XM_023239193.1
D1 17790829 17790930
refGene_gn=ARHGEF12;refGene_tx=XM_006936730.4;refGene_gn=ARHGEF12;refGene_tx=XM_006936731.4;refGene_gn=ARHGEF12;refGene_tx=XM_019811430.2;refGene_gn=ARHGEF12;refGene_tx=XM_023239192.1;refGene_gn=ARHGEF12;refGene_tx=XM_023239193.1
D1 17794180 17794258
refGene_gn=ARHGEF12;refGene_tx=XM_006936730.4;refGene_gn=ARHGEF12;refGene_tx=XM_006936731.4;refGene_gn=ARHGEF12;refGene_tx=XM_019811430.2;refGene_gn=ARHGEF12;refGene_tx=XM_023239192.1;refGene_gn=ARHGEF12;refGene_tx=XM_023239193.1
D1 17795369 17795445
refGene_gn=ARHGEF12;refGene_tx=XM_006936730.4;refGene_gn=ARHGEF12;refGene_tx=XM_006936731.4;refGene_gn=ARHGEF12;refGene_tx=XM_019811430.2;refGene_gn=ARHGEF12;refGene_tx=XM_023239192.1;refGene_gn=ARHGEF12;refGene_tx=XM_023239193.1
D1 17796191 17796359
refGene_gn=ARHGEF12;refGene_tx=XM_006936730.4;refGene_gn=ARHGEF12;refGene_tx=XM_006936731.4;refGene_gn=ARHGEF12;refGene_tx=XM_019811430.2;refGene_gn=ARHGEF12;refGene_tx=XM_023239192.1;refGene_gn=ARHGEF12;refGene_tx=XM_023239193.1

D1 17797197 17797285
refGene_gn=ARHGEF12;refGene_tx=XM_006936730.4;refGene_gn=ARHGEF12;refGene_tx=XM_006936731.4;refGene_gn=ARHGEF12;refGene_tx=XM_019811430.2;refGene_gn=ARHGEF12;refGene_tx=XM_023239192.1;refGene_gn=ARHGEF12;refGene_tx=XM_023239193.1
D1 17797785 17797872
refGene_gn=ARHGEF12;refGene_tx=XM_006936730.4;refGene_gn=ARHGEF12;refGene_tx=XM_006936731.4;refGene_gn=ARHGEF12;refGene_tx=XM_019811430.2;refGene_gn=ARHGEF12;refGene_tx=XM_023239192.1;refGene_gn=ARHGEF12;refGene_tx=XM_023239193.1
D1 17798005 17798085
refGene_gn=ARHGEF12;refGene_tx=XM_006936730.4;refGene_gn=ARHGEF12;refGene_tx=XM_006936731.4;refGene_gn=ARHGEF12;refGene_tx=XM_019811430.2;refGene_gn=ARHGEF12;refGene_tx=XM_023239192.1;refGene_gn=ARHGEF12;refGene_tx=XM_023239193.1
D1 17798384 17798618
refGene_gn=ARHGEF12;refGene_tx=XM_006936730.4;refGene_gn=ARHGEF12;refGene_tx=XM_006936731.4;refGene_gn=ARHGEF12;refGene_tx=XM_019811430.2;refGene_gn=ARHGEF12;refGene_tx=XM_023239192.1;refGene_gn=ARHGEF12;refGene_tx=XM_023239193.1
D1 17800214 17800682
refGene_gn=ARHGEF12;refGene_tx=XM_006936730.4;refGene_gn=ARHGEF12;refGene_tx=XM_006936731.4;refGene_gn=ARHGEF12;refGene_tx=XM_019811430.2;refGene_gn=ARHGEF12;refGene_tx=XM_023239192.1;refGene_gn=ARHGEF12;refGene_tx=XM_023239193.1
D1 17801463 17801780
refGene_gn=ARHGEF12;refGene_tx=XM_006936730.4;refGene_gn=ARHGEF12;refGene_tx=XM_006936731.4;refGene_gn=ARHGEF12;refGene_tx=XM_019811430.2;refGene_gn=ARHGEF12;refGene_tx=XM_023239192.1;refGene_gn=ARHGEF12;refGene_tx=XM_023239193.1
D1 17803062 17803132
refGene_gn=ARHGEF12;refGene_tx=XM_006936730.4;refGene_gn=ARHGEF12;refGene_tx=XM_006936731.4;refGene_gn=ARHGEF12;refGene_tx=XM_019811430.2;refGene_gn=ARHGEF12;refGene_tx=XM_023239192.1;refGene_gn=ARHGEF12;refGene_tx=XM_023239193.1
D1 17803744 17803755
refGene_gn=ARHGEF12;refGene_tx=XM_006936730.4;refGene_gn=ARHGEF12;refGene_tx=XM_006936731.4;refGene_gn=ARHGEF12;refGene_tx=XM_019811430.2;refGene_gn=ARHGEF12;refGene_tx=XM_023239192.1;refGene_gn=ARHGEF12;refGene_tx=XM_023239193.1
D1 17958494 17958576
refGene_gn=GRIK4;refGene_tx=XM_023239195.1;refGene_gn=GRIK4;refGene_tx=XM_023239196.1;refGene_gn=GRIK4;refGene_tx=XM_023239197.1;refGene_gn=GRIK4;refGene_tx=XM_023239198.1
D1 18081154 18081319
refGene_gn=GRIK4;refGene_tx=XM_023239195.1;refGene_gn=GRIK4;refGene_tx=XM_023239196.1;refGene_gn=GRIK4;refGene_tx=XM_023239198.1
D1 18094256 18094354
refGene_gn=GRIK4;refGene_tx=XM_023239195.1;refGene_gn=GRIK4;refGene_tx=XM_023239196.1;refGene_gn=GRIK4;refGene_tx=XM_023239197.1;refGene_gn=

GRIK4;refGene_tx=XM_023239198.1
D1	18099086	18099252
refGene_gn=GRIK4;refGene_tx=XM_023239195.1;refGene_gn=GRIK4;refGene_tx=XM_023239196.1;refGene_gn=GRIK4;refGene_tx=XM_023239197.1;refGene_gn=GRIK4;refGene_tx=XM_023239198.1
D1	18109255	18109434
refGene_gn=GRIK4;refGene_tx=XM_023239195.1;refGene_gn=GRIK4;refGene_tx=XM_023239196.1;refGene_gn=GRIK4;refGene_tx=XM_023239197.1;refGene_gn=GRIK4;refGene_tx=XM_023239198.1
D1	18113749	18113803
refGene_gn=GRIK4;refGene_tx=XM_023239195.1;refGene_gn=GRIK4;refGene_tx=XM_023239196.1;refGene_gn=GRIK4;refGene_tx=XM_023239197.1;refGene_gn=GRIK4;refGene_tx=XM_023239198.1
D1	18141544	18141706
refGene_gn=GRIK4;refGene_tx=XM_023239195.1;refGene_gn=GRIK4;refGene_tx=XM_023239196.1;refGene_gn=GRIK4;refGene_tx=XM_023239197.1;refGene_gn=GRIK4;refGene_tx=XM_023239198.1
D1	18151751	18151904
refGene_gn=GRIK4;refGene_tx=XM_023239195.1;refGene_gn=GRIK4;refGene_tx=XM_023239196.1;refGene_gn=GRIK4;refGene_tx=XM_023239197.1;refGene_gn=GRIK4;refGene_tx=XM_023239198.1
D1	18152587	18152692
refGene_gn=GRIK4;refGene_tx=XM_023239195.1;refGene_gn=GRIK4;refGene_tx=XM_023239196.1;refGene_gn=GRIK4;refGene_tx=XM_023239197.1;refGene_gn=GRIK4;refGene_tx=XM_023239198.1
D1	18176033	18176141
refGene_gn=GRIK4;refGene_tx=XM_023239195.1;refGene_gn=GRIK4;refGene_tx=XM_023239196.1;refGene_gn=GRIK4;refGene_tx=XM_023239197.1;refGene_gn=GRIK4;refGene_tx=XM_023239198.1
D1	18182490	18182694
refGene_gn=GRIK4;refGene_tx=XM_023239195.1;refGene_gn=GRIK4;refGene_tx=XM_023239196.1;refGene_gn=GRIK4;refGene_tx=XM_023239197.1
D1	18214007	18214121
refGene_gn=GRIK4;refGene_tx=XM_023239195.1;refGene_gn=GRIK4;refGene_tx=XM_023239196.1;refGene_gn=GRIK4;refGene_tx=XM_023239197.1;refGene_gn=GRIK4;refGene_tx=XM_023239198.1
D1	18225589	18225699
refGene_gn=GRIK4;refGene_tx=XM_023239195.1;refGene_gn=GRIK4;refGene_tx=XM_023239196.1;refGene_gn=GRIK4;refGene_tx=XM_023239197.1;refGene_gn=GRIK4;refGene_tx=XM_023239198.1
D1	18229639	18229813
refGene_gn=GRIK4;refGene_tx=XM_023239195.1;refGene_gn=GRIK4;refGene_tx=XM_023239196.1;refGene_gn=GRIK4;refGene_tx=XM_023239197.1;refGene_gn=GRIK4;refGene_tx=XM_023239198.1
D1	18233359	18233525
refGene_gn=GRIK4;refGene_tx=XM_023239195.1;refGene_gn=GRIK4;refGene_tx=XM_023239196.1;refGene_gn=GRIK4;refGene_tx=XM_023239197.1;refGene_gn=GRIK4;refGene_tx=XM_023239198.1
D1	18235052	18235278
refGene_gn=GRIK4;refGene_tx=XM_023239195.1;refGene_gn=GRIK4;refGene_tx

=XM_023239196.1;refGene_gn=GRIK4;refGene_tx=XM_023239197.1;refGene_gn=GRIK4;refGene_tx=XM_023239198.1
D1 18239191 18239320 refGene_gn=GRIK4;refGene_tx=XM_023239195.1;refGene_gn=GRIK4;refGene_tx=XM_023239196.1;refGene_gn=GRIK4;refGene_tx=XM_023239197.1;refGene_gn=GRIK4;refGene_tx=XM_023239198.1
D1 18254061 18254180 refGene_gn=GRIK4;refGene_tx=XM_023239195.1;refGene_gn=GRIK4;refGene_tx=XM_023239196.1;refGene_gn=GRIK4;refGene_tx=XM_023239197.1;refGene_gn=GRIK4;refGene_tx=XM_023239198.1
D1 18258123 18258480 refGene_gn=GRIK4;refGene_tx=XM_023239195.1;refGene_gn=GRIK4;refGene_tx=XM_023239196.1;refGene_gn=GRIK4;refGene_tx=XM_023239197.1;refGene_gn=GRIK4;refGene_tx=XM_023239198.1
D1 18330666 18330799 refGene_gn=TBCEL;refGene_tx=XM_023239199.1;refGene_gn=TBCEL;refGene_tx=XM_023239200.1
D1 18332492 18332632 refGene_gn=TBCEL;refGene_tx=XM_023239199.1;refGene_gn=TBCEL;refGene_tx=XM_023239200.1
D1 18339226 18339408 refGene_gn=TBCEL;refGene_tx=XM_023239199.1;refGene_gn=TBCEL;refGene_tx=XM_023239200.1
D1 18340720 18340977 refGene_gn=TBCEL;refGene_tx=XM_023239199.1;refGene_gn=TBCEL;refGene_tx=XM_023239200.1
D1 18343545 18343672 refGene_gn=TBCEL;refGene_tx=XM_023239199.1;refGene_gn=TBCEL;refGene_tx=XM_023239200.1
D1 18345945 18346062 refGene_gn=TBCEL;refGene_tx=XM_023239199.1;refGene_gn=TBCEL;refGene_tx=XM_023239200.1
D1 18376145 18376464 refGene_gn=TBCEL;refGene_tx=XM_023239199.1;refGene_gn=TBCEL;refGene_tx=XM_023239200.1
D1 18397673 18397790 refGene_gn=TECTA;refGene_tx=XM_023239201.1
D1 18400618 18400906 refGene_gn=TECTA;refGene_tx=XM_023239201.1
D1 18404620 18404758 refGene_gn=TECTA;refGene_tx=XM_023239201.1
D1 18405102 18405268 refGene_gn=TECTA;refGene_tx=XM_023239201.1
D1 18409174 18409587 refGene_gn=TECTA;refGene_tx=XM_023239201.1
D1 18415266 18415837 refGene_gn=TECTA;refGene_tx=XM_023239201.1
D1 18417798 18418391 refGene_gn=TECTA;refGene_tx=XM_023239201.1
D1 18419708 18420282 refGene_gn=TECTA;refGene_tx=XM_023239201.1
D1 18425305 18425907 refGene_gn=TECTA;refGene_tx=XM_023239201.1
D1 18432980 18433542 refGene_gn=TECTA;refGene_tx=XM_023239201.1
D1 18442212 18442412 refGene_gn=TECTA;refGene_tx=XM_023239201.1
D1 18446501 18446885 refGene_gn=TECTA;refGene_tx=XM_023239201.1
D1 18448803 18449090 refGene_gn=TECTA;refGene_tx=XM_023239201.1
D1 18450841 18451122 refGene_gn=TECTA;refGene_tx=XM_023239201.1
D1 18454143 18454254 refGene_gn=TECTA;refGene_tx=XM_023239201.1

D1 18455486 18455689 refGene_gn=TECTA;refGene_tx=XM_023239201.1
D1 18457383 18457547 refGene_gn=TECTA;refGene_tx=XM_023239201.1
D1 18457981 18458230 refGene_gn=TECTA;refGene_tx=XM_023239201.1
D1 18461847 18462010 refGene_gn=TECTA;refGene_tx=XM_023239201.1
D1 18463154 18463242 refGene_gn=TECTA;refGene_tx=XM_023239201.1
D1 18463531 18463648 refGene_gn=TECTA;refGene_tx=XM_023239201.1
D1 18464246 18464347 refGene_gn=TECTA;refGene_tx=XM_023239201.1
D1 18584581 18584791 refGene_gn=SC5D;refGene_tx=XM_003992449.5
D1 18585574 18585707 refGene_gn=SC5D;refGene_tx=XM_003992449.5
D1 18587584 18587685 refGene_gn=SC5D;refGene_tx=XM_003992449.5
D1 18588034 18588490 refGene_gn=SC5D;refGene_tx=XM_003992449.5
D1 18719748 18720024
refGene_gn=SORL1;refGene_tx=XM_023239202.1;refGene_gn=SORL1;refGene_tx=XM_023239203.1;refGene_gn=SORL1;refGene_tx=XM_023239204.1
D1 18737586 18737703
refGene_gn=SORL1;refGene_tx=XM_023239202.1;refGene_gn=SORL1;refGene_tx=XM_023239203.1;refGene_gn=SORL1;refGene_tx=XM_023239204.1
D1 18744514 18744640
refGene_gn=SORL1;refGene_tx=XM_023239202.1;refGene_gn=SORL1;refGene_tx=XM_023239203.1;refGene_gn=SORL1;refGene_tx=XM_023239204.1
D1 18752510 18752672
refGene_gn=SORL1;refGene_tx=XM_023239202.1;refGene_gn=SORL1;refGene_tx=XM_023239203.1;refGene_gn=SORL1;refGene_tx=XM_023239204.1
D1 18754912 18754980
refGene_gn=SORL1;refGene_tx=XM_023239202.1;refGene_gn=SORL1;refGene_tx=XM_023239203.1;refGene_gn=SORL1;refGene_tx=XM_023239204.1
D1 18760541 18760722
refGene_gn=SORL1;refGene_tx=XM_023239202.1;refGene_gn=SORL1;refGene_tx=XM_023239203.1;refGene_gn=SORL1;refGene_tx=XM_023239204.1
D1 18773094 18773196
refGene_gn=SORL1;refGene_tx=XM_023239202.1;refGene_gn=SORL1;refGene_tx=XM_023239203.1;refGene_gn=SORL1;refGene_tx=XM_023239204.1
D1 18774728 18774898
refGene_gn=SORL1;refGene_tx=XM_023239202.1;refGene_gn=SORL1;refGene_tx=XM_023239203.1;refGene_gn=SORL1;refGene_tx=XM_023239204.1
D1 18780269 18780462
refGene_gn=SORL1;refGene_tx=XM_023239202.1;refGene_gn=SORL1;refGene_tx=XM_023239203.1;refGene_gn=SORL1;refGene_tx=XM_023239204.1;refGene_gn=SORL1;refGene_tx=XM_023239205.1
D1 18782852 18782970
refGene_gn=SORL1;refGene_tx=XM_023239202.1;refGene_gn=SORL1;refGene_tx=XM_023239203.1;refGene_gn=SORL1;refGene_tx=XM_023239204.1;refGene_gn=SORL1;refGene_tx=XM_023239205.1
D1 18783183 18783257
refGene_gn=SORL1;refGene_tx=XM_023239202.1;refGene_gn=SORL1;refGene_tx=XM_023239203.1;refGene_gn=SORL1;refGene_tx=XM_023239204.1;refGene_gn=SORL1;refGene_tx=XM_023239205.1
D1 18795886 18795975
refGene_gn=SORL1;refGene_tx=XM_023239202.1;refGene_gn=SORL1;refGene_tx=XM_023239203.1;refGene_gn=SORL1;refGene_tx=XM_023239204.1;refGene_gn=

SORL1;refGene_tx=XM_023239205.1
D1 18807674 18807853
refGene_gn=SORL1;refGene_tx=XM_023239202.1;refGene_gn=SORL1;refGene_tx=XM_023239203.1;refGene_gn=SORL1;refGene_tx=XM_023239204.1;refGene_gn=SORL1;refGene_tx=XM_023239205.1
D1 18809381 18809568
refGene_gn=SORL1;refGene_tx=XM_023239202.1;refGene_gn=SORL1;refGene_tx=XM_023239203.1;refGene_gn=SORL1;refGene_tx=XM_023239204.1;refGene_gn=SORL1;refGene_tx=XM_023239205.1
D1 18814836 18814965
refGene_gn=SORL1;refGene_tx=XM_023239202.1;refGene_gn=SORL1;refGene_tx=XM_023239203.1;refGene_gn=SORL1;refGene_tx=XM_023239204.1;refGene_gn=SORL1;refGene_tx=XM_023239205.1
D1 18815444 18815530
refGene_gn=SORL1;refGene_tx=XM_023239202.1;refGene_gn=SORL1;refGene_tx=XM_023239203.1;refGene_gn=SORL1;refGene_tx=XM_023239204.1;refGene_gn=SORL1;refGene_tx=XM_023239205.1
D1 18819308 18819481
refGene_gn=SORL1;refGene_tx=XM_023239202.1;refGene_gn=SORL1;refGene_tx=XM_023239203.1;refGene_gn=SORL1;refGene_tx=XM_023239204.1;refGene_gn=SORL1;refGene_tx=XM_023239205.1
D1 18820554 18820686
refGene_gn=SORL1;refGene_tx=XM_023239202.1;refGene_gn=SORL1;refGene_tx=XM_023239203.1;refGene_gn=SORL1;refGene_tx=XM_023239204.1;refGene_gn=SORL1;refGene_tx=XM_023239205.1
D1 18823225 18823317
refGene_gn=SORL1;refGene_tx=XM_023239202.1;refGene_gn=SORL1;refGene_tx=XM_023239203.1;refGene_gn=SORL1;refGene_tx=XM_023239204.1;refGene_gn=SORL1;refGene_tx=XM_023239205.1
D1 18824593 18824840
refGene_gn=SORL1;refGene_tx=XM_023239202.1;refGene_gn=SORL1;refGene_tx=XM_023239203.1;refGene_gn=SORL1;refGene_tx=XM_023239204.1;refGene_gn=SORL1;refGene_tx=XM_023239205.1
D1 18825524 18825663
refGene_gn=SORL1;refGene_tx=XM_023239202.1;refGene_gn=SORL1;refGene_tx=XM_023239203.1;refGene_gn=SORL1;refGene_tx=XM_023239204.1;refGene_gn=SORL1;refGene_tx=XM_023239205.1
D1 18831447 18831621
refGene_gn=SORL1;refGene_tx=XM_023239202.1;refGene_gn=SORL1;refGene_tx=XM_023239203.1;refGene_gn=SORL1;refGene_tx=XM_023239204.1;refGene_gn=SORL1;refGene_tx=XM_023239205.1
D1 18833619 18833733
refGene_gn=SORL1;refGene_tx=XM_023239202.1;refGene_gn=SORL1;refGene_tx=XM_023239203.1;refGene_gn=SORL1;refGene_tx=XM_023239204.1;refGene_gn=SORL1;refGene_tx=XM_023239205.1
D1 18837520 18837643
refGene_gn=SORL1;refGene_tx=XM_023239202.1;refGene_gn=SORL1;refGene_tx=XM_023239203.1;refGene_gn=SORL1;refGene_tx=XM_023239204.1;refGene_gn=SORL1;refGene_tx=XM_023239205.1
D1 18840787 18840907

refGene_gn=SORL1;refGene_tx=XM_023239202.1;refGene_gn=SORL1;refGene_tx=XM_023239203.1;refGene_gn=SORL1;refGene_tx=XM_023239204.1;refGene_gn=SORL1;refGene_tx=XM_023239205.1
D1 18846708 18846834
refGene_gn=SORL1;refGene_tx=XM_023239202.1;refGene_gn=SORL1;refGene_tx=XM_023239203.1;refGene_gn=SORL1;refGene_tx=XM_023239204.1;refGene_gn=SORL1;refGene_tx=XM_023239205.1
D1 18849829 18849937
refGene_gn=SORL1;refGene_tx=XM_023239202.1;refGene_gn=SORL1;refGene_tx=XM_023239203.1;refGene_gn=SORL1;refGene_tx=XM_023239204.1;refGene_gn=SORL1;refGene_tx=XM_023239205.1
D1 18852694 18852826
refGene_gn=SORL1;refGene_tx=XM_023239202.1;refGene_gn=SORL1;refGene_tx=XM_023239203.1;refGene_gn=SORL1;refGene_tx=XM_023239204.1;refGene_gn=SORL1;refGene_tx=XM_023239205.1
D1 18854382 18854514
refGene_gn=SORL1;refGene_tx=XM_023239202.1;refGene_gn=SORL1;refGene_tx=XM_023239203.1;refGene_gn=SORL1;refGene_tx=XM_023239204.1;refGene_gn=SORL1;refGene_tx=XM_023239205.1
D1 18855515 18855650
refGene_gn=SORL1;refGene_tx=XM_023239202.1;refGene_gn=SORL1;refGene_tx=XM_023239203.1;refGene_gn=SORL1;refGene_tx=XM_023239204.1;refGene_gn=SORL1;refGene_tx=XM_023239205.1
D1 18856422 18856578
refGene_gn=SORL1;refGene_tx=XM_023239202.1;refGene_gn=SORL1;refGene_tx=XM_023239203.1;refGene_gn=SORL1;refGene_tx=XM_023239204.1;refGene_gn=SORL1;refGene_tx=XM_023239205.1
D1 18861118 18861268
refGene_gn=SORL1;refGene_tx=XM_023239202.1;refGene_gn=SORL1;refGene_tx=XM_023239203.1;refGene_gn=SORL1;refGene_tx=XM_023239204.1;refGene_gn=SORL1;refGene_tx=XM_023239205.1
D1 18863790 18863922
refGene_gn=SORL1;refGene_tx=XM_023239202.1;refGene_gn=SORL1;refGene_tx=XM_023239203.1;refGene_gn=SORL1;refGene_tx=XM_023239204.1;refGene_gn=SORL1;refGene_tx=XM_023239205.1
D1 18864895 18865022
refGene_gn=SORL1;refGene_tx=XM_023239202.1;refGene_gn=SORL1;refGene_tx=XM_023239203.1;refGene_gn=SORL1;refGene_tx=XM_023239204.1;refGene_gn=SORL1;refGene_tx=XM_023239205.1
D1 18865187 18865357
refGene_gn=SORL1;refGene_tx=XM_023239202.1;refGene_gn=SORL1;refGene_tx=XM_023239203.1;refGene_gn=SORL1;refGene_tx=XM_023239204.1;refGene_gn=SORL1;refGene_tx=XM_023239205.1
D1 18867724 18867837
refGene_gn=SORL1;refGene_tx=XM_023239202.1;refGene_gn=SORL1;refGene_tx=XM_023239203.1;refGene_gn=SORL1;refGene_tx=XM_023239204.1;refGene_gn=SORL1;refGene_tx=XM_023239205.1
D1 18868063 18868168
refGene_gn=SORL1;refGene_tx=XM_023239202.1;refGene_gn=SORL1;refGene_tx=XM_023239203.1;refGene_gn=SORL1;refGene_tx=XM_023239204.1;refGene_gn=

SORL1;refGene_tx=XM_023239205.1
D1 18868951 18869024
refGene_gn=SORL1;refGene_tx=XM_023239202.1;refGene_gn=SORL1;refGene_tx
=XM_023239203.1;refGene_gn=SORL1;refGene_tx=XM_023239204.1;refGene_gn=
SORL1;refGene_tx=XM_023239205.1
D1 18871322 18871405
refGene_gn=SORL1;refGene_tx=XM_023239202.1;refGene_gn=SORL1;refGene_tx
=XM_023239203.1;refGene_gn=SORL1;refGene_tx=XM_023239204.1;refGene_gn=
SORL1;refGene_tx=XM_023239205.1
D1 18873353 18873450
refGene_gn=SORL1;refGene_tx=XM_023239202.1;refGene_gn=SORL1;refGene_tx
=XM_023239203.1;refGene_gn=SORL1;refGene_tx=XM_023239204.1;refGene_gn=
SORL1;refGene_tx=XM_023239205.1
D1 18876771 18876956
refGene_gn=SORL1;refGene_tx=XM_023239202.1;refGene_gn=SORL1;refGene_tx
=XM_023239203.1;refGene_gn=SORL1;refGene_tx=XM_023239204.1;refGene_gn=
SORL1;refGene_tx=XM_023239205.1
D1 18880082 18880202
refGene_gn=SORL1;refGene_tx=XM_023239202.1;refGene_gn=SORL1;refGene_tx
=XM_023239203.1;refGene_gn=SORL1;refGene_tx=XM_023239204.1;refGene_gn=
SORL1;refGene_tx=XM_023239205.1
D1 18881068 18881233
refGene_gn=SORL1;refGene_tx=XM_023239202.1;refGene_gn=SORL1;refGene_tx
=XM_023239203.1;refGene_gn=SORL1;refGene_tx=XM_023239204.1;refGene_gn=
SORL1;refGene_tx=XM_023239205.1
D1 18882798 18882973
refGene_gn=SORL1;refGene_tx=XM_023239202.1;refGene_gn=SORL1;refGene_tx
=XM_023239203.1;refGene_gn=SORL1;refGene_tx=XM_023239204.1;refGene_gn=
SORL1;refGene_tx=XM_023239205.1
D1 18883965 18884072
refGene_gn=SORL1;refGene_tx=XM_023239202.1;refGene_gn=SORL1;refGene_tx
=XM_023239203.1;refGene_gn=SORL1;refGene_tx=XM_023239204.1;refGene_gn=
SORL1;refGene_tx=XM_023239205.1
D1 18886342 18886535
refGene_gn=SORL1;refGene_tx=XM_023239202.1;refGene_gn=SORL1;refGene_tx
=XM_023239203.1;refGene_gn=SORL1;refGene_tx=XM_023239204.1;refGene_gn=
SORL1;refGene_tx=XM_023239205.1
D1 18889361 18889574
refGene_gn=SORL1;refGene_tx=XM_023239202.1;refGene_gn=SORL1;refGene_tx
=XM_023239203.1;refGene_gn=SORL1;refGene_tx=XM_023239204.1;refGene_gn=
SORL1;refGene_tx=XM_023239205.1
D1 18890788 18890856
refGene_gn=SORL1;refGene_tx=XM_023239202.1;refGene_gn=SORL1;refGene_tx
=XM_023239203.1;refGene_gn=SORL1;refGene_tx=XM_023239204.1;refGene_gn=
SORL1;refGene_tx=XM_023239205.1
D1 19786708 19786869 refGene_gn=UBASH3B;refGene_tx=XM_003992450.5
D1 19890556 19890610 refGene_gn=UBASH3B;refGene_tx=XM_003992450.5
D1 19891397 19891584 refGene_gn=UBASH3B;refGene_tx=XM_003992450.5
D1 19892890 19893089 refGene_gn=UBASH3B;refGene_tx=XM_003992450.5
D1 19896444 19896614 refGene_gn=UBASH3B;refGene_tx=XM_003992450.5

D1	19902181	19902390	refGene_gn=UBASH3B;refGene_tx=XM_003992450.5
D1	19906680	19906813	refGene_gn=UBASH3B;refGene_tx=XM_003992450.5
D1	19908125	19908246	refGene_gn=UBASH3B;refGene_tx=XM_003992450.5
D1	19908877	19909000	refGene_gn=UBASH3B;refGene_tx=XM_003992450.5
D1	19910248	19910341	refGene_gn=UBASH3B;refGene_tx=XM_003992450.5
D1	19912113	19912258	refGene_gn=UBASH3B;refGene_tx=XM_003992450.5
D1	19922112	19922219	refGene_gn=UBASH3B;refGene_tx=XM_003992450.5
D1	19924107	19924217	refGene_gn=UBASH3B;refGene_tx=XM_003992450.5
D1	19925915	19926053	refGene_gn=UBASH3B;refGene_tx=XM_003992450.5
D1	19941111	19941178	refGene_gn=CRTAM;refGene_tx=XM_003992475.2
D1	19949025	19949172	refGene_gn=CRTAM;refGene_tx=XM_003992475.2
D1	19952440	19952593	refGene_gn=CRTAM;refGene_tx=XM_003992475.2
D1	19954787	19954931	refGene_gn=CRTAM;refGene_tx=XM_003992475.2
D1	19955862	19956024	refGene_gn=CRTAM;refGene_tx=XM_003992475.2
D1	19960291	19960372	refGene_gn=CRTAM;refGene_tx=XM_003992475.2
D1	19962355	19962439	refGene_gn=CRTAM;refGene_tx=XM_003992475.2
D1	19963931	19964078	refGene_gn=CRTAM;refGene_tx=XM_003992475.2
D1	19966324	19966411	refGene_gn=CRTAM;refGene_tx=XM_003992475.2
D1	19968067	19968198	refGene_gn=CRTAM;refGene_tx=XM_003992475.2
D1	19992346	19992666
refGene_gn=JHY;refGene_tx=XM_003992476.4;refGene_gn=JHY;refGene_tx=XM_023239206.1;refGene_gn=JHY;refGene_tx=XM_023239207.1;refGene_gn=JHY;refGene_tx=XM_023239208.1;refGene_gn=JHY;refGene_tx=XM_023239210.1;refGene_gn=JHY;refGene_tx=XM_023239211.1
D1	20005912	20006429
refGene_gn=JHY;refGene_tx=XM_003992476.4;refGene_gn=JHY;refGene_tx=XM_023239206.1;refGene_gn=JHY;refGene_tx=XM_023239207.1;refGene_gn=JHY;refGene_tx=XM_023239208.1;refGene_gn=JHY;refGene_tx=XM_023239210.1;refGene_gn=JHY;refGene_tx=XM_023239211.1
D1	20026507	20026621
refGene_gn=JHY;refGene_tx=XM_003992476.4;refGene_gn=JHY;refGene_tx=XM_023239206.1;refGene_gn=JHY;refGene_tx=XM_023239207.1;refGene_gn=JHY;refGene_tx=XM_023239208.1;refGene_gn=JHY;refGene_tx=XM_023239211.1
D1	20033590	20034252
refGene_gn=JHY;refGene_tx=XM_003992476.4;refGene_gn=JHY;refGene_tx=XM_023239206.1;refGene_gn=JHY;refGene_tx=XM_023239207.1;refGene_gn=JHY;refGene_tx=XM_023239208.1;refGene_gn=JHY;refGene_tx=XM_023239210.1;refGene_gn=JHY;refGene_tx=XM_023239211.1
D1	20040135	20040427
refGene_gn=JHY;refGene_tx=XM_003992476.4;refGene_gn=JHY;refGene_tx=XM_023239206.1;refGene_gn=JHY;refGene_tx=XM_023239207.1;refGene_gn=JHY;refGene_tx=XM_023239208.1;refGene_gn=JHY;refGene_tx=XM_023239210.1;refGene_gn=JHY;refGene_tx=XM_023239211.1
D1	20048900	20048981
refGene_gn=JHY;refGene_tx=XM_003992476.4;refGene_gn=JHY;refGene_tx=XM_023239206.1;refGene_gn=JHY;refGene_tx=XM_023239207.1;refGene_gn=JHY;refGene_tx=XM_023239208.1;refGene_gn=JHY;refGene_tx=XM_023239210.1;refGene_gn=JHY;refGene_tx=XM_023239211.1
D1	20050268	20050397
refGene_gn=JHY;refGene_tx=XM_003992476.4;refGene_gn=JHY;refGene_tx=XM_

023239206.1;refGene_gn=JHY;refGene_tx=XM_023239207.1;refGene_gn=JHY;refGene_tx=XM_023239208.1;refGene_gn=JHY;refGene_tx=XM_023239210.1;refGene_gn=JHY;refGene_tx=XM_023239211.1
D1 20052938 20053136
refGene_gn=JHY;refGene_tx=XM_003992476.4;refGene_gn=JHY;refGene_tx=XM_023239206.1;refGene_gn=JHY;refGene_tx=XM_023239207.1;refGene_gn=JHY;refGene_tx=XM_023239208.1;refGene_gn=JHY;refGene_tx=XM_023239210.1;refGene_gn=JHY;refGene_tx=XM_023239211.1
D1 20071971 20072211 refGene_gn=BSX;refGene_tx=XM_011286409.2
D1 20073239 20073436 refGene_gn=BSX;refGene_tx=XM_011286409.2
D1 20075421 20075683 refGene_gn=BSX;refGene_tx=XM_011286409.2
D1 20145729 20145915 refGene_gn=HSPA8;refGene_tx=XM_003992478.3
D1 20146221 20146454 refGene_gn=HSPA8;refGene_tx=XM_003992478.3
D1 20146660 20146859 refGene_gn=HSPA8;refGene_tx=XM_003992478.3
D1 20147098 20147301 refGene_gn=HSPA8;refGene_tx=XM_003992478.3
D1 20147520 20148076 refGene_gn=HSPA8;refGene_tx=XM_003992478.3
D1 20148168 20148321 refGene_gn=HSPA8;refGene_tx=XM_003992478.3
D1 20148561 20148767 refGene_gn=HSPA8;refGene_tx=XM_003992478.3
D1 20149099 20149304 refGene_gn=HSPA8;refGene_tx=XM_003992478.3
D1 20156128 20156181 refGene_gn=CLMP;refGene_tx=XM_023239212.1
D1 20156183 20156428 refGene_gn=CLMP;refGene_tx=XM_023239212.1
D1 20157004 20157146 refGene_gn=CLMP;refGene_tx=XM_023239212.1
D1 20158155 20158278 refGene_gn=CLMP;refGene_tx=XM_023239212.1
D1 20158741 20158909 refGene_gn=CLMP;refGene_tx=XM_023239212.1
D1 20159900 20160102 refGene_gn=CLMP;refGene_tx=XM_023239212.1
D1 20177954 20178112 refGene_gn=CLMP;refGene_tx=XM_023239212.1
D1 20254548 20254579 refGene_gn=CLMP;refGene_tx=XM_023239212.1
D1 20404590 20404616 refGene_gn=GRAMD1B;refGene_tx=XM_023239225.1
D1 20479943 20480317
refGene_gn=GRAMD1B;refGene_tx=XM_023239213.1;refGene_gn=GRAMD1B;refGene_tx=XM_023239214.1;refGene_gn=GRAMD1B;refGene_tx=XM_023239215.1;refGene_gn=GRAMD1B;refGene_tx=XM_023239216.1;refGene_gn=GRAMD1B;refGene_tx=XM_023239217.1;refGene_gn=GRAMD1B;refGene_tx=XM_023239218.1;refGene_gn=GRAMD1B;refGene_tx=XM_023239219.1;refGene_gn=GRAMD1B;refGene_tx=XM_023239220.1;refGene_gn=GRAMD1B;refGene_tx=XM_023239222.1;refGene_gn=GRAMD1B;refGene_tx=XM_023239234.1;refGene_gn=GRAMD1B;refGene_tx=XM_023239235.1;refGene_gn=GRAMD1B;refGene_tx=XM_023239236.1
D1 20480600 20480694 refGene_gn=GRAMD1B;refGene_tx=XM_023239235.1
D1 20483226 20483251 refGene_gn=GRAMD1B;refGene_tx=XM_023239236.1
D1 20484088 20484203 refGene_gn=GRAMD1B;refGene_tx=XM_023239234.1
D1 20514983 20515061
refGene_gn=GRAMD1B;refGene_tx=XM_023239213.1;refGene_gn=GRAMD1B;refGene_tx=XM_023239214.1;refGene_gn=GRAMD1B;refGene_tx=XM_023239215.1;refGene_gn=GRAMD1B;refGene_tx=XM_023239216.1;refGene_gn=GRAMD1B;refGene_tx=XM_023239217.1;refGene_gn=GRAMD1B;refGene_tx=XM_023239218.1;refGene_gn=GRAMD1B;refGene_tx=XM_023239219.1;refGene_gn=GRAMD1B;refGene_tx=XM_023239220.1;refGene_gn=GRAMD1B;refGene_tx=XM_023239222.1;refGene_gn=GRAMD1B;refGene_tx=XM_023239225.1
D1 20525090 20525200 refGene_gn=GRAMD1B;refGene_tx=XM_023239224.1
D1 20555832 20555855 refGene_gn=GRAMD1B;refGene_tx=XM_023239227.1

D1 20574119 20574268 refGene_gn=GRAMD1B;refGene_tx=XM_023239223.1
D1 20588803 20588814 refGene_gn=GRAMD1B;refGene_tx=XM_023239228.1
D1 20605822 20606033
refGene_gn=GRAMD1B;refGene_tx=XM_023239213.1;refGene_gn=GRAMD1B;refGene_tx=XM_023239214.1;refGene_gn=GRAMD1B;refGene_tx=XM_023239215.1;refGene_gn=GRAMD1B;refGene_tx=XM_023239216.1;refGene_gn=GRAMD1B;refGene_tx=XM_023239217.1;refGene_gn=GRAMD1B;refGene_tx=XM_023239218.1;refGene_gn=GRAMD1B;refGene_tx=XM_023239219.1;refGene_gn=GRAMD1B;refGene_tx=XM_023239220.1;refGene_gn=GRAMD1B;refGene_tx=XM_023239222.1;refGene_gn=GRAMD1B;refGene_tx=XM_023239223.1;refGene_gn=GRAMD1B;refGene_tx=XM_023239224.1;refGene_gn=GRAMD1B;refGene_tx=XM_023239225.1;refGene_gn=GRAMD1B;refGene_tx=XM_023239226.1;refGene_gn=GRAMD1B;refGene_tx=XM_023239227.1;refGene_gn=GRAMD1B;refGene_tx=XM_023239228.1;refGene_gn=GRAMD1B;refGene_tx=XM_023239229.1;refGene_gn=GRAMD1B;refGene_tx=XM_023239230.1;refGene_gn=GRAMD1B;refGene_tx=XM_023239231.1;refGene_gn=GRAMD1B;refGene_tx=XM_023239232.1
D1 20606853 20606874
refGene_gn=GRAMD1B;refGene_tx=XM_023239213.1;refGene_gn=GRAMD1B;refGene_tx=XM_023239214.1;refGene_gn=GRAMD1B;refGene_tx=XM_023239215.1;refGene_gn=GRAMD1B;refGene_tx=XM_023239216.1;refGene_gn=GRAMD1B;refGene_tx=XM_023239218.1;refGene_gn=GRAMD1B;refGene_tx=XM_023239219.1;refGene_gn=GRAMD1B;refGene_tx=XM_023239223.1;refGene_gn=GRAMD1B;refGene_tx=XM_023239224.1;refGene_gn=GRAMD1B;refGene_tx=XM_023239225.1;refGene_gn=GRAMD1B;refGene_tx=XM_023239226.1;refGene_gn=GRAMD1B;refGene_tx=XM_023239227.1;refGene_gn=GRAMD1B;refGene_tx=XM_023239228.1;refGene_gn=GRAMD1B;refGene_tx=XM_023239229.1;refGene_gn=GRAMD1B;refGene_tx=XM_023239230.1;refGene_gn=GRAMD1B;refGene_tx=XM_023239231.1;refGene_gn=GRAMD1B;refGene_tx=XM_023239232.1
D1 20609593 20609617
refGene_gn=GRAMD1B;refGene_tx=XM_023239213.1;refGene_gn=GRAMD1B;refGene_tx=XM_023239214.1;refGene_gn=GRAMD1B;refGene_tx=XM_023239215.1;refGene_gn=GRAMD1B;refGene_tx=XM_023239218.1;refGene_gn=GRAMD1B;refGene_tx=XM_023239223.1;refGene_gn=GRAMD1B;refGene_tx=XM_023239224.1;refGene_gn=GRAMD1B;refGene_tx=XM_023239225.1;refGene_gn=GRAMD1B;refGene_tx=XM_023239226.1;refGene_gn=GRAMD1B;refGene_tx=XM_023239227.1;refGene_gn=GRAMD1B;refGene_tx=XM_023239228.1;refGene_gn=GRAMD1B;refGene_tx=XM_023239229.1;refGene_gn=GRAMD1B;refGene_tx=XM_023239230.1;refGene_gn=GRAMD1B;refGene_tx=XM_023239231.1;refGene_gn=GRAMD1B;refGene_tx=XM_023239232.1
D1 20612350 20612371
refGene_gn=GRAMD1B;refGene_tx=XM_023239213.1;refGene_gn=GRAMD1B;refGene_tx=XM_023239214.1;refGene_gn=GRAMD1B;refGene_tx=XM_023239215.1;refGene_gn=GRAMD1B;refGene_tx=XM_023239216.1;refGene_gn=GRAMD1B;refGene_tx=XM_023239217.1;refGene_gn=GRAMD1B;refGene_tx=XM_023239218.1;refGene_gn=GRAMD1B;refGene_tx=XM_023239219.1;refGene_gn=GRAMD1B;refGene_tx=XM_023239220.1;refGene_gn=GRAMD1B;refGene_tx=XM_023239222.1;refGene_gn=GRAMD1B;refGene_tx=XM_023239223.1;refGene_gn=GRAMD1B;refGene_tx=XM_023239224.1;refGene_gn=GRAMD1B;refGene_tx=XM_023239225.1;refGene_gn=GRAMD1B;refGene_tx=XM_023239226.1;refGene_gn=GRAMD1B;refGene_tx=XM_023239227.1;refGene_gn=GRAMD1B;refGene_tx=XM_023239228.1;refGene_gn=GRAMD1B;refGene_tx=XM_023239229.1;refGene_gn=GRAMD1B;refGene_tx=XM_023239230.1;refGe

ne_gn=GRAMD1B;refGene_tx=XM_023239231.1;refGene_gn=GRAMD1B;refGene_tx=XM_023239232.1
D1	20619430	20619577	refGene_gn=GRAMD1B;refGene_tx=XM_023239233.1
D1	20623112	20623197	refGene_gn=GRAMD1B;refGene_tx=XM_023239213.1;refGene_gn=GRAMD1B;refGene_tx=XM_023239214.1;refGene_gn=GRAMD1B;refGene_tx=XM_023239215.1;refGene_gn=GRAMD1B;refGene_tx=XM_023239216.1;refGene_gn=GRAMD1B;refGene_tx=XM_023239217.1;refGene_gn=GRAMD1B;refGene_tx=XM_023239218.1;refGene_gn=GRAMD1B;refGene_tx=XM_023239219.1;refGene_gn=GRAMD1B;refGene_tx=XM_023239220.1;refGene_gn=GRAMD1B;refGene_tx=XM_023239222.1;refGene_gn=GRAMD1B;refGene_tx=XM_023239223.1;refGene_gn=GRAMD1B;refGene_tx=XM_023239224.1;refGene_gn=GRAMD1B;refGene_tx=XM_023239225.1;refGene_gn=GRAMD1B;refGene_tx=XM_023239226.1;refGene_gn=GRAMD1B;refGene_tx=XM_023239227.1;refGene_gn=GRAMD1B;refGene_tx=XM_023239228.1;refGene_gn=GRAMD1B;refGene_tx=XM_023239229.1;refGene_gn=GRAMD1B;refGene_tx=XM_023239230.1;refGene_gn=GRAMD1B;refGene_tx=XM_023239231.1;refGene_gn=GRAMD1B;refGene_tx=XM_023239232.1;refGene_gn=GRAMD1B;refGene_tx=XM_023239233.1
D1	20623986	20624090	refGene_gn=GRAMD1B;refGene_tx=XM_023239213.1;refGene_gn=GRAMD1B;refGene_tx=XM_023239214.1;refGene_gn=GRAMD1B;refGene_tx=XM_023239215.1;refGene_gn=GRAMD1B;refGene_tx=XM_023239216.1;refGene_gn=GRAMD1B;refGene_tx=XM_023239217.1;refGene_gn=GRAMD1B;refGene_tx=XM_023239218.1;refGene_gn=GRAMD1B;refGene_tx=XM_023239219.1;refGene_gn=GRAMD1B;refGene_tx=XM_023239220.1;refGene_gn=GRAMD1B;refGene_tx=XM_023239222.1;refGene_gn=GRAMD1B;refGene_tx=XM_023239223.1;refGene_gn=GRAMD1B;refGene_tx=XM_023239224.1;refGene_gn=GRAMD1B;refGene_tx=XM_023239225.1;refGene_gn=GRAMD1B;refGene_tx=XM_023239226.1;refGene_gn=GRAMD1B;refGene_tx=XM_023239227.1;refGene_gn=GRAMD1B;refGene_tx=XM_023239228.1;refGene_gn=GRAMD1B;refGene_tx=XM_023239229.1;refGene_gn=GRAMD1B;refGene_tx=XM_023239230.1;refGene_gn=GRAMD1B;refGene_tx=XM_023239231.1;refGene_gn=GRAMD1B;refGene_tx=XM_023239232.1;refGene_gn=GRAMD1B;refGene_tx=XM_023239233.1
D1	20626502	20626598	refGene_gn=GRAMD1B;refGene_tx=XM_023239213.1;refGene_gn=GRAMD1B;refGene_tx=XM_023239214.1;refGene_gn=GRAMD1B;refGene_tx=XM_023239215.1;refGene_gn=GRAMD1B;refGene_tx=XM_023239216.1;refGene_gn=GRAMD1B;refGene_tx=XM_023239217.1;refGene_gn=GRAMD1B;refGene_tx=XM_023239218.1;refGene_gn=GRAMD1B;refGene_tx=XM_023239219.1;refGene_gn=GRAMD1B;refGene_tx=XM_023239220.1;refGene_gn=GRAMD1B;refGene_tx=XM_023239222.1;refGene_gn=GRAMD1B;refGene_tx=XM_023239223.1;refGene_gn=GRAMD1B;refGene_tx=XM_023239224.1;refGene_gn=GRAMD1B;refGene_tx=XM_023239225.1;refGene_gn=GRAMD1B;refGene_tx=XM_023239226.1;refGene_gn=GRAMD1B;refGene_tx=XM_023239227.1;refGene_gn=GRAMD1B;refGene_tx=XM_023239228.1;refGene_gn=GRAMD1B;refGene_tx=XM_023239229.1;refGene_gn=GRAMD1B;refGene_tx=XM_023239230.1;refGene_gn=GRAMD1B;refGene_tx=XM_023239231.1;refGene_gn=GRAMD1B;refGene_tx=XM_023239232.1;refGene_gn=GRAMD1B;refGene_tx=XM_023239233.1
D1	20628497	20628578	refGene_gn=GRAMD1B;refGene_tx=XM_023239213.1;refGene_gn=GRAMD1B;refGene_tx=XM_023239214.1;refGene_gn=GRAMD1B;refGene_tx=XM_023239215.1;refGene_gn=GRAMD1B;refGene_tx=XM_023239216.1;refGene_gn=GRAMD1B;refGene_tx=XM_023239217.1;refGene_gn=GRAMD1B;refGene_tx=XM_023239218.1;refGene_gn

=GRAMD1B;refGene_tx=XM_023239219.1;refGene_gn=GRAMD1B;refGene_tx=XM_023239220.1;refGene_gn=GRAMD1B;refGene_tx=XM_023239222.1;refGene_gn=GRAMD1B;refGene_tx=XM_023239223.1;refGene_gn=GRAMD1B;refGene_tx=XM_023239224.1;refGene_gn=GRAMD1B;refGene_tx=XM_023239225.1;refGene_gn=GRAMD1B;refGene_tx=XM_023239226.1;refGene_gn=GRAMD1B;refGene_tx=XM_023239227.1;refGene_gn=GRAMD1B;refGene_tx=XM_023239228.1;refGene_gn=GRAMD1B;refGene_tx=XM_023239229.1;refGene_gn=GRAMD1B;refGene_tx=XM_023239230.1;refGene_gn=GRAMD1B;refGene_tx=XM_023239231.1;refGene_gn=GRAMD1B;refGene_tx=XM_023239232.1;refGene_gn=GRAMD1B;refGene_tx=XM_023239233.1
D1 20632136 20632252
refGene_gn=GRAMD1B;refGene_tx=XM_023239213.1;refGene_gn=GRAMD1B;refGene_tx=XM_023239214.1;refGene_gn=GRAMD1B;refGene_tx=XM_023239215.1;refGene_gn=GRAMD1B;refGene_tx=XM_023239216.1;refGene_gn=GRAMD1B;refGene_tx=XM_023239217.1;refGene_gn=GRAMD1B;refGene_tx=XM_023239218.1;refGene_gn=GRAMD1B;refGene_tx=XM_023239219.1;refGene_gn=GRAMD1B;refGene_tx=XM_023239220.1;refGene_gn=GRAMD1B;refGene_tx=XM_023239222.1;refGene_gn=GRAMD1B;refGene_tx=XM_023239223.1;refGene_gn=GRAMD1B;refGene_tx=XM_023239224.1;refGene_gn=GRAMD1B;refGene_tx=XM_023239225.1;refGene_gn=GRAMD1B;refGene_tx=XM_023239226.1;refGene_gn=GRAMD1B;refGene_tx=XM_023239227.1;refGene_gn=GRAMD1B;refGene_tx=XM_023239228.1;refGene_gn=GRAMD1B;refGene_tx=XM_023239229.1;refGene_gn=GRAMD1B;refGene_tx=XM_023239230.1;refGene_gn=GRAMD1B;refGene_tx=XM_023239231.1;refGene_gn=GRAMD1B;refGene_tx=XM_023239232.1;refGene_gn=GRAMD1B;refGene_tx=XM_023239233.1
D1 20634053 20634210
refGene_gn=GRAMD1B;refGene_tx=XM_023239213.1;refGene_gn=GRAMD1B;refGene_tx=XM_023239214.1;refGene_gn=GRAMD1B;refGene_tx=XM_023239215.1;refGene_gn=GRAMD1B;refGene_tx=XM_023239216.1;refGene_gn=GRAMD1B;refGene_tx=XM_023239217.1;refGene_gn=GRAMD1B;refGene_tx=XM_023239218.1;refGene_gn=GRAMD1B;refGene_tx=XM_023239219.1;refGene_gn=GRAMD1B;refGene_tx=XM_023239220.1;refGene_gn=GRAMD1B;refGene_tx=XM_023239222.1;refGene_gn=GRAMD1B;refGene_tx=XM_023239223.1;refGene_gn=GRAMD1B;refGene_tx=XM_023239224.1;refGene_gn=GRAMD1B;refGene_tx=XM_023239225.1;refGene_gn=GRAMD1B;refGene_tx=XM_023239226.1;refGene_gn=GRAMD1B;refGene_tx=XM_023239227.1;refGene_gn=GRAMD1B;refGene_tx=XM_023239228.1;refGene_gn=GRAMD1B;refGene_tx=XM_023239229.1;refGene_gn=GRAMD1B;refGene_tx=XM_023239230.1;refGene_gn=GRAMD1B;refGene_tx=XM_023239231.1;refGene_gn=GRAMD1B;refGene_tx=XM_023239232.1;refGene_gn=GRAMD1B;refGene_tx=XM_023239233.1
D1 20635361 20635551
refGene_gn=GRAMD1B;refGene_tx=XM_023239213.1;refGene_gn=GRAMD1B;refGene_tx=XM_023239214.1;refGene_gn=GRAMD1B;refGene_tx=XM_023239215.1;refGene_gn=GRAMD1B;refGene_tx=XM_023239216.1;refGene_gn=GRAMD1B;refGene_tx=XM_023239217.1;refGene_gn=GRAMD1B;refGene_tx=XM_023239218.1;refGene_gn=GRAMD1B;refGene_tx=XM_023239219.1;refGene_gn=GRAMD1B;refGene_tx=XM_023239220.1;refGene_gn=GRAMD1B;refGene_tx=XM_023239222.1;refGene_gn=GRAMD1B;refGene_tx=XM_023239223.1;refGene_gn=GRAMD1B;refGene_tx=XM_023239224.1;refGene_gn=GRAMD1B;refGene_tx=XM_023239225.1;refGene_gn=GRAMD1B;refGene_tx=XM_023239226.1;refGene_gn=GRAMD1B;refGene_tx=XM_023239227.1;refGene_gn=GRAMD1B;refGene_tx=XM_023239228.1;refGene_gn=GRAMD1B;refGene_tx=XM_023239229.1;refGene_gn=GRAMD1B;refGene_tx=XM_023239230.1;refGene_gn=GRAMD1B;refGene_tx=XM_023239231.1;refGene_gn=GRAMD1B;refGene_tx=

XM_023239232.1;refGene_gn=GRAMD1B;refGene_tx=XM_023239233.1
D1 20637410 20637554
refGene_gn=GRAMD1B;refGene_tx=XM_023239213.1;refGene_gn=GRAMD1B;refGene_tx=XM_023239214.1;refGene_gn=GRAMD1B;refGene_tx=XM_023239215.1;refGene_gn=GRAMD1B;refGene_tx=XM_023239216.1;refGene_gn=GRAMD1B;refGene_tx=XM_023239217.1;refGene_gn=GRAMD1B;refGene_tx=XM_023239218.1;refGene_gn=GRAMD1B;refGene_tx=XM_023239219.1;refGene_gn=GRAMD1B;refGene_tx=XM_023239220.1;refGene_gn=GRAMD1B;refGene_tx=XM_023239222.1;refGene_gn=GRAMD1B;refGene_tx=XM_023239223.1;refGene_gn=GRAMD1B;refGene_tx=XM_023239224.1;refGene_gn=GRAMD1B;refGene_tx=XM_023239225.1;refGene_gn=GRAMD1B;refGene_tx=XM_023239226.1;refGene_gn=GRAMD1B;refGene_tx=XM_023239227.1;refGene_gn=GRAMD1B;refGene_tx=XM_023239228.1;refGene_gn=GRAMD1B;refGene_tx=XM_023239229.1;refGene_gn=GRAMD1B;refGene_tx=XM_023239230.1;refGene_gn=GRAMD1B;refGene_tx=XM_023239231.1;refGene_gn=GRAMD1B;refGene_tx=XM_023239232.1;refGene_gn=GRAMD1B;refGene_tx=XM_023239233.1
D1 20638505 20638624
refGene_gn=GRAMD1B;refGene_tx=XM_023239213.1;refGene_gn=GRAMD1B;refGene_tx=XM_023239214.1;refGene_gn=GRAMD1B;refGene_tx=XM_023239215.1;refGene_gn=GRAMD1B;refGene_tx=XM_023239216.1;refGene_gn=GRAMD1B;refGene_tx=XM_023239217.1;refGene_gn=GRAMD1B;refGene_tx=XM_023239218.1;refGene_gn=GRAMD1B;refGene_tx=XM_023239219.1;refGene_gn=GRAMD1B;refGene_tx=XM_023239220.1;refGene_gn=GRAMD1B;refGene_tx=XM_023239222.1;refGene_gn=GRAMD1B;refGene_tx=XM_023239223.1;refGene_gn=GRAMD1B;refGene_tx=XM_023239224.1;refGene_gn=GRAMD1B;refGene_tx=XM_023239225.1;refGene_gn=GRAMD1B;refGene_tx=XM_023239226.1;refGene_gn=GRAMD1B;refGene_tx=XM_023239227.1;refGene_gn=GRAMD1B;refGene_tx=XM_023239228.1;refGene_gn=GRAMD1B;refGene_tx=XM_023239229.1;refGene_gn=GRAMD1B;refGene_tx=XM_023239230.1;refGene_gn=GRAMD1B;refGene_tx=XM_023239231.1;refGene_gn=GRAMD1B;refGene_tx=XM_023239232.1;refGene_gn=GRAMD1B;refGene_tx=XM_023239233.1
D1 20638855 20638998
refGene_gn=GRAMD1B;refGene_tx=XM_023239213.1;refGene_gn=GRAMD1B;refGene_tx=XM_023239214.1;refGene_gn=GRAMD1B;refGene_tx=XM_023239215.1;refGene_gn=GRAMD1B;refGene_tx=XM_023239216.1;refGene_gn=GRAMD1B;refGene_tx=XM_023239217.1;refGene_gn=GRAMD1B;refGene_tx=XM_023239218.1;refGene_gn=GRAMD1B;refGene_tx=XM_023239219.1;refGene_gn=GRAMD1B;refGene_tx=XM_023239220.1;refGene_gn=GRAMD1B;refGene_tx=XM_023239222.1;refGene_gn=GRAMD1B;refGene_tx=XM_023239223.1;refGene_gn=GRAMD1B;refGene_tx=XM_023239224.1;refGene_gn=GRAMD1B;refGene_tx=XM_023239225.1;refGene_gn=GRAMD1B;refGene_tx=XM_023239226.1;refGene_gn=GRAMD1B;refGene_tx=XM_023239227.1;refGene_gn=GRAMD1B;refGene_tx=XM_023239228.1;refGene_gn=GRAMD1B;refGene_tx=XM_023239229.1;refGene_gn=GRAMD1B;refGene_tx=XM_023239230.1;refGene_gn=GRAMD1B;refGene_tx=XM_023239231.1;refGene_gn=GRAMD1B;refGene_tx=XM_023239232.1;refGene_gn=GRAMD1B;refGene_tx=XM_023239233.1
D1 20640633 20640737
refGene_gn=GRAMD1B;refGene_tx=XM_023239213.1;refGene_gn=GRAMD1B;refGene_tx=XM_023239214.1;refGene_gn=GRAMD1B;refGene_tx=XM_023239215.1;refGene_gn=GRAMD1B;refGene_tx=XM_023239216.1;refGene_gn=GRAMD1B;refGene_tx=XM_023239217.1;refGene_gn=GRAMD1B;refGene_tx=XM_023239218.1;refGene_gn=GRAMD1B;refGene_tx=XM_023239219.1;refGene_gn=GRAMD1B;refGene_tx=XM_023239220.1;refGene_gn=GRAMD1B;refGene_tx=XM_023239222.1;refGene_gn=GRAM

D1B;refGene_tx=XM_023239223.1;refGene_gn=GRAMD1B;refGene_tx=XM_0232392
24.1;refGene_gn=GRAMD1B;refGene_tx=XM_023239225.1;refGene_gn=GRAMD1B;r
efGene_tx=XM_023239226.1;refGene_gn=GRAMD1B;refGene_tx=XM_023239227.1;
refGene_gn=GRAMD1B;refGene_tx=XM_023239228.1;refGene_gn=GRAMD1B;refGen
e_tx=XM_023239229.1;refGene_gn=GRAMD1B;refGene_tx=XM_023239230.1;refGe
ne_gn=GRAMD1B;refGene_tx=XM_023239231.1;refGene_gn=GRAMD1B;refGene_tx=
XM_023239232.1;refGene_gn=GRAMD1B;refGene_tx=XM_023239233.1
D1 20641279 20641483
refGene_gn=GRAMD1B;refGene_tx=XM_023239213.1;refGene_gn=GRAMD1B;refGen
e_tx=XM_023239214.1;refGene_gn=GRAMD1B;refGene_tx=XM_023239215.1;refGe
ne_gn=GRAMD1B;refGene_tx=XM_023239216.1;refGene_gn=GRAMD1B;refGene_tx=
XM_023239217.1;refGene_gn=GRAMD1B;refGene_tx=XM_023239218.1;refGene_gn
=GRAMD1B;refGene_tx=XM_023239219.1;refGene_gn=GRAMD1B;refGene_tx=XM_02
3239220.1;refGene_gn=GRAMD1B;refGene_tx=XM_023239222.1;refGene_gn=GRAM
D1B;refGene_tx=XM_023239223.1;refGene_gn=GRAMD1B;refGene_tx=XM_0232392
24.1;refGene_gn=GRAMD1B;refGene_tx=XM_023239225.1;refGene_gn=GRAMD1B;r
efGene_tx=XM_023239226.1;refGene_gn=GRAMD1B;refGene_tx=XM_023239227.1;
refGene_gn=GRAMD1B;refGene_tx=XM_023239228.1;refGene_gn=GRAMD1B;refGen
e_tx=XM_023239229.1;refGene_gn=GRAMD1B;refGene_tx=XM_023239230.1;refGe
ne_gn=GRAMD1B;refGene_tx=XM_023239231.1;refGene_gn=GRAMD1B;refGene_tx=
XM_023239232.1;refGene_gn=GRAMD1B;refGene_tx=XM_023239233.1
D1 20642571 20642662
refGene_gn=GRAMD1B;refGene_tx=XM_023239213.1;refGene_gn=GRAMD1B;refGen
e_tx=XM_023239214.1;refGene_gn=GRAMD1B;refGene_tx=XM_023239215.1;refGe
ne_gn=GRAMD1B;refGene_tx=XM_023239216.1;refGene_gn=GRAMD1B;refGene_tx=
XM_023239217.1;refGene_gn=GRAMD1B;refGene_tx=XM_023239218.1;refGene_gn
=GRAMD1B;refGene_tx=XM_023239219.1;refGene_gn=GRAMD1B;refGene_tx=XM_02
3239220.1;refGene_gn=GRAMD1B;refGene_tx=XM_023239222.1;refGene_gn=GRAM
D1B;refGene_tx=XM_023239223.1;refGene_gn=GRAMD1B;refGene_tx=XM_0232392
24.1;refGene_gn=GRAMD1B;refGene_tx=XM_023239225.1;refGene_gn=GRAMD1B;r
efGene_tx=XM_023239226.1;refGene_gn=GRAMD1B;refGene_tx=XM_023239227.1;
refGene_gn=GRAMD1B;refGene_tx=XM_023239228.1;refGene_gn=GRAMD1B;refGen
e_tx=XM_023239229.1;refGene_gn=GRAMD1B;refGene_tx=XM_023239230.1;refGe
ne_gn=GRAMD1B;refGene_tx=XM_023239231.1;refGene_gn=GRAMD1B;refGene_tx=
XM_023239232.1;refGene_gn=GRAMD1B;refGene_tx=XM_023239233.1
D1 20646395 20646407
refGene_gn=GRAMD1B;refGene_tx=XM_023239213.1;refGene_gn=GRAMD1B;refGen
e_tx=XM_023239214.1;refGene_gn=GRAMD1B;refGene_tx=XM_023239216.1;refGe
ne_gn=GRAMD1B;refGene_tx=XM_023239217.1;refGene_gn=GRAMD1B;refGene_tx=
XM_023239218.1;refGene_gn=GRAMD1B;refGene_tx=XM_023239219.1;refGene_gn
=GRAMD1B;refGene_tx=XM_023239220.1;refGene_gn=GRAMD1B;refGene_tx=XM_02
3239223.1;refGene_gn=GRAMD1B;refGene_tx=XM_023239224.1;refGene_gn=GRAM
D1B;refGene_tx=XM_023239225.1;refGene_gn=GRAMD1B;refGene_tx=XM_0232392
26.1;refGene_gn=GRAMD1B;refGene_tx=XM_023239227.1;refGene_gn=GRAMD1B;r
efGene_tx=XM_023239228.1;refGene_gn=GRAMD1B;refGene_tx=XM_023239229.1;
refGene_gn=GRAMD1B;refGene_tx=XM_023239230.1;refGene_gn=GRAMD1B;refGen
e_tx=XM_023239231.1;refGene_gn=GRAMD1B;refGene_tx=XM_023239232.1;refGe
ne_gn=GRAMD1B;refGene_tx=XM_023239233.1
D1 20646864 20646972
refGene_gn=GRAMD1B;refGene_tx=XM_023239213.1;refGene_gn=GRAMD1B;refGen

e_tx=XM_023239214.1;refGene_gn=GRAMD1B;refGene_tx=XM_023239215.1;refGene_gn=GRAMD1B;refGene_tx=XM_023239216.1;refGene_gn=GRAMD1B;refGene_tx=XM_023239217.1;refGene_gn=GRAMD1B;refGene_tx=XM_023239218.1;refGene_gn=GRAMD1B;refGene_tx=XM_023239219.1;refGene_gn=GRAMD1B;refGene_tx=XM_023239220.1;refGene_gn=GRAMD1B;refGene_tx=XM_023239222.1;refGene_gn=GRAMD1B;refGene_tx=XM_023239223.1;refGene_gn=GRAMD1B;refGene_tx=XM_023239224.1;refGene_gn=GRAMD1B;refGene_tx=XM_023239225.1;refGene_gn=GRAMD1B;refGene_tx=XM_023239226.1;refGene_gn=GRAMD1B;refGene_tx=XM_023239227.1;refGene_gn=GRAMD1B;refGene_tx=XM_023239228.1;refGene_gn=GRAMD1B;refGene_tx=XM_023239229.1;refGene_gn=GRAMD1B;refGene_tx=XM_023239230.1;refGene_gn=GRAMD1B;refGene_tx=XM_023239231.1;refGene_gn=GRAMD1B;refGene_tx=XM_023239232.1;refGene_gn=GRAMD1B;refGene_tx=XM_023239233.1
D1 20647219 20647409
refGene_gn=GRAMD1B;refGene_tx=XM_023239213.1;refGene_gn=GRAMD1B;refGene_tx=XM_023239214.1;refGene_gn=GRAMD1B;refGene_tx=XM_023239215.1;refGene_gn=GRAMD1B;refGene_tx=XM_023239216.1;refGene_gn=GRAMD1B;refGene_tx=XM_023239217.1;refGene_gn=GRAMD1B;refGene_tx=XM_023239218.1;refGene_gn=GRAMD1B;refGene_tx=XM_023239219.1;refGene_gn=GRAMD1B;refGene_tx=XM_023239220.1;refGene_gn=GRAMD1B;refGene_tx=XM_023239222.1;refGene_gn=GRAMD1B;refGene_tx=XM_023239223.1;refGene_gn=GRAMD1B;refGene_tx=XM_023239224.1;refGene_gn=GRAMD1B;refGene_tx=XM_023239225.1;refGene_gn=GRAMD1B;refGene_tx=XM_023239226.1;refGene_gn=GRAMD1B;refGene_tx=XM_023239227.1;refGene_gn=GRAMD1B;refGene_tx=XM_023239228.1;refGene_gn=GRAMD1B;refGene_tx=XM_023239229.1;refGene_gn=GRAMD1B;refGene_tx=XM_023239230.1;refGene_gn=GRAMD1B;refGene_tx=XM_023239231.1;refGene_gn=GRAMD1B;refGene_tx=XM_023239232.1;refGene_gn=GRAMD1B;refGene_tx=XM_023239233.1
D1 20649959 20650049
refGene_gn=GRAMD1B;refGene_tx=XM_023239213.1;refGene_gn=GRAMD1B;refGene_tx=XM_023239214.1;refGene_gn=GRAMD1B;refGene_tx=XM_023239215.1;refGene_gn=GRAMD1B;refGene_tx=XM_023239216.1;refGene_gn=GRAMD1B;refGene_tx=XM_023239217.1;refGene_gn=GRAMD1B;refGene_tx=XM_023239218.1;refGene_gn=GRAMD1B;refGene_tx=XM_023239219.1;refGene_gn=GRAMD1B;refGene_tx=XM_023239220.1;refGene_gn=GRAMD1B;refGene_tx=XM_023239222.1;refGene_gn=GRAMD1B;refGene_tx=XM_023239223.1;refGene_gn=GRAMD1B;refGene_tx=XM_023239224.1;refGene_gn=GRAMD1B;refGene_tx=XM_023239225.1;refGene_gn=GRAMD1B;refGene_tx=XM_023239226.1;refGene_gn=GRAMD1B;refGene_tx=XM_023239227.1;refGene_gn=GRAMD1B;refGene_tx=XM_023239228.1;refGene_gn=GRAMD1B;refGene_tx=XM_023239229.1;refGene_gn=GRAMD1B;refGene_tx=XM_023239230.1;refGene_gn=GRAMD1B;refGene_tx=XM_023239231.1;refGene_gn=GRAMD1B;refGene_tx=XM_023239232.1;refGene_gn=GRAMD1B;refGene_tx=XM_023239233.1
D1 20660229 20660293
refGene_gn=SCN3B;refGene_tx=XM_006936745.3;refGene_gn=SCN3B;refGene_tx=XM_019811463.1
D1 20672292 20672435
refGene_gn=SCN3B;refGene_tx=XM_006936745.3;refGene_gn=SCN3B;refGene_tx=XM_019811463.1;refGene_gn=SCN3B;refGene_tx=XM_019811464.1
D1 20676497 20676723
refGene_gn=SCN3B;refGene_tx=XM_006936745.3;refGene_gn=SCN3B;refGene_tx=XM_019811463.1;refGene_gn=SCN3B;refGene_tx=XM_019811464.1
D1 20679306 20679470

refGene_gn=SCN3B;refGene_tx=XM_006936745.3;refGene_gn=SCN3B;refGene_tx=XM_019811463.1;refGene_gn=SCN3B;refGene_tx=XM_019811464.1
D1 20686350 20686405
refGene_gn=SCN3B;refGene_tx=XM_006936745.3;refGene_gn=SCN3B;refGene_tx=XM_019811463.1;refGene_gn=SCN3B;refGene_tx=XM_019811464.1
D1 20749847 20750842
refGene_gn=ZNF202;refGene_tx=XM_003992482.5;refGene_gn=ZNF202;refGene_tx=XM_006936746.4;refGene_gn=ZNF202;refGene_tx=XM_019811466.2;refGene_gn=ZNF202;refGene_tx=XM_019811467.1;refGene_gn=ZNF202;refGene_tx=XM_023239238.1
D1 20751371 20751491
refGene_gn=ZNF202;refGene_tx=XM_003992482.5;refGene_gn=ZNF202;refGene_tx=XM_006936746.4;refGene_gn=ZNF202;refGene_tx=XM_019811466.2;refGene_gn=ZNF202;refGene_tx=XM_019811467.1;refGene_gn=ZNF202;refGene_tx=XM_023239238.1
D1 20752042 20752172
refGene_gn=ZNF202;refGene_tx=XM_003992482.5;refGene_gn=ZNF202;refGene_tx=XM_006936746.4;refGene_gn=ZNF202;refGene_tx=XM_019811466.2;refGene_gn=ZNF202;refGene_tx=XM_019811467.1;refGene_gn=ZNF202;refGene_tx=XM_023239238.1
D1 20753094 20753183
refGene_gn=ZNF202;refGene_tx=XM_003992482.5;refGene_gn=ZNF202;refGene_tx=XM_006936746.4;refGene_gn=ZNF202;refGene_tx=XM_019811466.2;refGene_gn=ZNF202;refGene_tx=XM_019811467.1;refGene_gn=ZNF202;refGene_tx=XM_023239238.1
D1 20753531 20753742
refGene_gn=ZNF202;refGene_tx=XM_003992482.5;refGene_gn=ZNF202;refGene_tx=XM_006936746.4;refGene_gn=ZNF202;refGene_tx=XM_019811466.2;refGene_gn=ZNF202;refGene_tx=XM_019811467.1;refGene_gn=ZNF202;refGene_tx=XM_023239238.1
D1 20754453 20755101
refGene_gn=ZNF202;refGene_tx=XM_003992482.5;refGene_gn=ZNF202;refGene_tx=XM_006936746.4;refGene_gn=ZNF202;refGene_tx=XM_019811466.2;refGene_gn=ZNF202;refGene_tx=XM_019811467.1;refGene_gn=ZNF202;refGene_tx=XM_023239238.1
D1 20771837 20772501
refGene_gn=LOC101092538;refGene_tx=XM_019812921.1
D1 20772502 20772786
refGene_gn=LOC101092538;refGene_tx=XM_019812921.1
D1 20825074 20826022
refGene_gn=LOC101088780;refGene_tx=XM_003992483.2
D1 20846553 20847495
refGene_gn=LOC101089040;refGene_tx=XM_003992484.2
D1 20868292 20869240
refGene_gn=LOC105261380;refGene_tx=XM_019822585.2
D1 20879408 20880350
refGene_gn=LOC101093042;refGene_tx=XM_003992497.3
D1 20893326 20894262
refGene_gn=LOC101093294;refGene_tx=XM_023238983.1
D1 20913445 20913623

refGene_gn=TMEM225;refGene_tx=XM_003992499.4;refGene_gn=TMEM225;refGene_tx=XM_019811470.2
D1 20914237 20914384
refGene_gn=TMEM225;refGene_tx=XM_003992499.4;refGene_gn=TMEM225;refGene_tx=XM_019811470.2
D1 20914857 20914992 refGene_gn=TMEM225;refGene_tx=XM_003992499.4
D1 20915699 20915926
refGene_gn=TMEM225;refGene_tx=XM_003992499.4;refGene_gn=TMEM225;refGene_tx=XM_019811470.2
D1 20937240 20938185
refGene_gn=LOC101089293;refGene_tx=XM_003992485.1;refGene_gn=LOC101089293;refGene_tx=XM_019813022.1
D1 20941215 20942178
refGene_gn=LOC101093798;refGene_tx=XM_003992500.2
D1 20955142 20956075
refGene_gn=LOC101094045;refGene_tx=XM_006936748.3
D1 20964813 20965316
refGene_gn=LOC101094291;refGene_tx=XM_011286455.1
D1 20965318 20965541
refGene_gn=LOC101094291;refGene_tx=XM_011286455.1
D1 20965542 20966019
refGene_gn=LOC101094291;refGene_tx=XM_011286455.1
D1 20978225 20979161
refGene_gn=LOC101094536;refGene_tx=XM_011286415.3
D1 20994964 20995885
refGene_gn=LOC101094786;refGene_tx=XM_011286416.1
D1 21010036 21010966
refGene_gn=LOC101095042;refGene_tx=XM_003992504.1
D1 21023856 21024798
refGene_gn=LOC101095281;refGene_tx=XM_003992505.1
D1 21048113 21049049
refGene_gn=LOC101095517;refGene_tx=XM_003992506.4
D1 21060833 21060910
refGene_gn=LOC101095766;refGene_tx=XM_023239694.1
D1 21061154 21061619
refGene_gn=LOC101095766;refGene_tx=XM_023239694.1
D1 21061621 21062006
refGene_gn=LOC101095766;refGene_tx=XM_023239694.1
D1 21070047 21070090
refGene_gn=LOC101096008;refGene_tx=XM_023239240.1
D1 21070209 21070412
refGene_gn=LOC101096008;refGene_tx=XM_023239240.1
D1 21070715 21070938
refGene_gn=LOC101096008;refGene_tx=XM_023239240.1
D1 21071060 21071236
refGene_gn=LOC101096008;refGene_tx=XM_023239240.1
D1 21071516 21071588
refGene_gn=LOC101096008;refGene_tx=XM_023239240.1
D1 21071589 21071632
refGene_gn=LOC101096008;refGene_tx=XM_023239240.1

D1 21075227 21075262
refGene_gn=LOC101096008;refGene_tx=XM_023239240.1
D1 21075263 21075398
refGene_gn=LOC101096008;refGene_tx=XM_023239240.1
D1 21075650 21075739
refGene_gn=LOC101096008;refGene_tx=XM_023239240.1
D1 21075949 21076094
refGene_gn=LOC101096008;refGene_tx=XM_023239240.1
D1 21076556 21076640
refGene_gn=LOC101096008;refGene_tx=XM_023239240.1
D1 21085336 21085343
refGene_gn=LOC111556583;refGene_tx=XM_023239242.1
D1 21088959 21089197
refGene_gn=LOC111556583;refGene_tx=XM_023239241.1
D1 21096709 21096865
refGene_gn=LOC111556583;refGene_tx=XM_023239241.1;refGene_gn=LOC111556
583;refGene_tx=XM_023239242.1
D1 21098160 21098291
refGene_gn=LOC111556583;refGene_tx=XM_023239241.1;refGene_gn=LOC111556
583;refGene_tx=XM_023239242.1
D1 21101806 21101933
refGene_gn=LOC111556583;refGene_tx=XM_023239241.1;refGene_gn=LOC111556
583;refGene_tx=XM_023239242.1
D1 21102478 21102663
refGene_gn=LOC111556583;refGene_tx=XM_023239241.1;refGene_gn=LOC111556
583;refGene_tx=XM_023239242.1
D1 21114431 21115385
refGene_gn=LOC101096260;refGene_tx=XM_003992509.1
D1 21164854 21165814
refGene_gn=LOC111556641;refGene_tx=XM_023239695.1
D1 21171767 21172727
refGene_gn=LOC101096759;refGene_tx=XM_006936753.4
D1 21178678 21179614
refGene_gn=LOC101097015;refGene_tx=XM_006936754.4
D1 21196559 21197495
refGene_gn=LOC101097273;refGene_tx=XM_003992512.4
D1 21202379 21203312
refGene_gn=LOC101089797;refGene_tx=XM_003992486.1
D1 21219896 21220820
refGene_gn=LOC101097515;refGene_tx=XM_003992513.1
D1 21240542 21241469
refGene_gn=LOC101090043;refGene_tx=XM_011279812.2
D1 21256625 21257561
refGene_gn=LOC101090289;refGene_tx=XM_003992487.3
D1 21280427 21281363
refGene_gn=LOC101098006;refGene_tx=XM_003992514.3
D1 21288519 21289449
refGene_gn=LOC101098253;refGene_tx=XM_006936755.3
D1 21302849 21303776
refGene_gn=LOC105260951;refGene_tx=XM_019811480.2

D1 21324704 21325637
refGene_gn=LOC101098741;refGene_tx=XM_003992517.1
D1 21335438 21336371
refGene_gn=LOC101098984;refGene_tx=XM_006936756.1
D1 21353281 21354214
refGene_gn=LOC101090546;refGene_tx=XM_011286459.1
D1 21376944 21377066
refGene_gn=LOC101099242;refGene_tx=XM_011286460.3
D1 21377068 21377876
refGene_gn=LOC101099242;refGene_tx=XM_011286460.3
D1 21384476 21385437
refGene_gn=LOC101099496;refGene_tx=XM_023239696.1
D1 21387755 21387865
refGene_gn=LOC101099496;refGene_tx=XM_023239696.1
D1 21406330 21407263
refGene_gn=LOC105260952;refGene_tx=XM_011286461.2
D1 21419090 21420051
refGene_gn=LOC101099998;refGene_tx=XM_019812923.2
D1 21423331 21423474
refGene_gn=LOC101099998;refGene_tx=XM_019812923.2
D1 21438925 21438937
refGene_gn=LOC101100246;refGene_tx=XM_011286418.3
D1 21440992 21441928
refGene_gn=LOC101100246;refGene_tx=XM_011286418.3
D1 21466020 21466950
refGene_gn=LOC101090797;refGene_tx=XM_003992488.4
D1 21482422 21483367
refGene_gn=LOC101091045;refGene_tx=XM_019811473.2
D1 21517568 21518504
refGene_gn=LOC101091306;refGene_tx=XM_003992490.1
D1 21551253 21552189
refGene_gn=LOC101091557;refGene_tx=XM_003992491.1
D1 21569674 21570634
refGene_gn=LOC101100504;refGene_tx=XM_003992522.3
D1 21595941 21596874
refGene_gn=LOC101100752;refGene_tx=XM_006936758.4;refGene_gn=LOC101100752;refGene_tx=XM_023239243.1
D1 21611723 21612683
refGene_gn=LOC105260953;refGene_tx=XM_023239244.1
D1 21649299 21650253
refGene_gn=LOC101084218;refGene_tx=XM_006936772.4
D1 21686309 21686490 refGene_gn=PANX3;refGene_tx=XM_011286419.3
D1 21687713 21687856 refGene_gn=PANX3;refGene_tx=XM_011286419.3
D1 21690499 21690714 refGene_gn=PANX3;refGene_tx=XM_011286419.3
D1 21692201 21692841 refGene_gn=PANX3;refGene_tx=XM_011286419.3
D1 21697343 21697493
refGene_gn=TBRG1;refGene_tx=XM_003992525.4;refGene_gn=TBRG1;refGene_tx=XM_019811482.2
D1 21699185 21699256
refGene_gn=TBRG1;refGene_tx=XM_003992525.4;refGene_gn=TBRG1;refGene_tx

=XM_019811482.2
D1	21699903	21700136
refGene_gn=TBRG1;refGene_tx=XM_003992525.4;refGene_gn=TBRG1;refGene_tx=XM_019811482.2
D1	21700717	21700854
refGene_gn=TBRG1;refGene_tx=XM_003992525.4;refGene_gn=TBRG1;refGene_tx=XM_019811482.2
D1	21701148	21701295
refGene_gn=TBRG1;refGene_tx=XM_003992525.4;refGene_gn=TBRG1;refGene_tx=XM_019811482.2
D1	21706505	21706603
refGene_gn=TBRG1;refGene_tx=XM_003992525.4;refGene_gn=TBRG1;refGene_tx=XM_019811482.2
D1	21706863	21706974
refGene_gn=TBRG1;refGene_tx=XM_003992525.4;refGene_gn=TBRG1;refGene_tx=XM_019811482.2
D1	21707334	21707477
refGene_gn=TBRG1;refGene_tx=XM_003992525.4;refGene_gn=TBRG1;refGene_tx=XM_019811482.2
D1	21708161	21708307
refGene_gn=TBRG1;refGene_tx=XM_003992525.4;refGene_gn=TBRG1;refGene_tx=XM_019811482.2
D1	21712619	21712871
refGene_gn=SIAE;refGene_tx=XM_019811481.2;refGene_gn=SIAE;refGene_tx=XM_023239245.1;refGene_gn=SIAE;refGene_tx=XM_023239246.1;refGene_gn=SIAE;refGene_tx=XM_023239247.1;refGene_gn=SIAE;refGene_tx=XM_023239248.1
D1	21714223	21714413
refGene_gn=SIAE;refGene_tx=XM_019811481.2;refGene_gn=SIAE;refGene_tx=XM_023239245.1;refGene_gn=SIAE;refGene_tx=XM_023239246.1;refGene_gn=SIAE;refGene_tx=XM_023239247.1;refGene_gn=SIAE;refGene_tx=XM_023239248.1
D1	21715412	21715567
refGene_gn=SIAE;refGene_tx=XM_019811481.2;refGene_gn=SIAE;refGene_tx=XM_023239245.1;refGene_gn=SIAE;refGene_tx=XM_023239246.1;refGene_gn=SIAE;refGene_tx=XM_023239247.1;refGene_gn=SIAE;refGene_tx=XM_023239248.1
D1	21723622	21723756
refGene_gn=SIAE;refGene_tx=XM_019811481.2;refGene_gn=SIAE;refGene_tx=XM_023239245.1;refGene_gn=SIAE;refGene_tx=XM_023239246.1;refGene_gn=SIAE;refGene_tx=XM_023239247.1;refGene_gn=SIAE;refGene_tx=XM_023239248.1
D1	21724308	21724418
refGene_gn=SIAE;refGene_tx=XM_019811481.2;refGene_gn=SIAE;refGene_tx=XM_023239245.1;refGene_gn=SIAE;refGene_tx=XM_023239246.1;refGene_gn=SIAE;refGene_tx=XM_023239247.1;refGene_gn=SIAE;refGene_tx=XM_023239248.1
D1	21726051	21726229
refGene_gn=SIAE;refGene_tx=XM_019811481.2;refGene_gn=SIAE;refGene_tx=XM_023239245.1;refGene_gn=SIAE;refGene_tx=XM_023239246.1;refGene_gn=SIAE;refGene_tx=XM_023239247.1;refGene_gn=SIAE;refGene_tx=XM_023239248.1
D1	21728655	21728794
refGene_gn=SIAE;refGene_tx=XM_019811481.2;refGene_gn=SIAE;refGene_tx=XM_023239245.1;refGene_gn=SIAE;refGene_tx=XM_023239246.1;refGene_gn=SIAE;refGene_tx=XM_023239247.1;refGene_gn=SIAE;refGene_tx=XM_023239248.1

D1 21739053 21739229
refGene_gn=SIAE;refGene_tx=XM_019811481.2;refGene_gn=SIAE;refGene_tx=XM_023239245.1;refGene_gn=SIAE;refGene_tx=XM_023239246.1;refGene_gn=SIAE;refGene_tx=XM_023239247.1;refGene_gn=SIAE;refGene_tx=XM_023239248.1
D1 21746810 21746972
refGene_gn=SIAE;refGene_tx=XM_019811481.2;refGene_gn=SIAE;refGene_tx=XM_023239245.1;refGene_gn=SIAE;refGene_tx=XM_023239246.1;refGene_gn=SIAE;refGene_tx=XM_023239247.1
D1 21749889 21749949 refGene_gn=SIAE;refGene_tx=XM_019811481.2
D1 21750563 21750630
refGene_gn=SIAE;refGene_tx=XM_019811481.2;refGene_gn=SIAE;refGene_tx=XM_023239245.1;refGene_gn=SIAE;refGene_tx=XM_023239248.1
D1 21752238 21752392 refGene_gn=SPA17;refGene_tx=XM_003992527.5
D1 21756256 21756327 refGene_gn=SPA17;refGene_tx=XM_003992527.5
D1 21758872 21758959 refGene_gn=SPA17;refGene_tx=XM_003992527.5
D1 21762033 21762177 refGene_gn=SPA17;refGene_tx=XM_003992527.5
D1 21801572 21801587 refGene_gn=NRGN;refGene_tx=XM_003992528.4
D1 21806093 21806315 refGene_gn=NRGN;refGene_tx=XM_003992528.4
D1 21808136 21808269 refGene_gn=VSIG2;refGene_tx=XM_003992529.4
D1 21808963 21809108 refGene_gn=VSIG2;refGene_tx=XM_003992529.4
D1 21809224 21809344 refGene_gn=VSIG2;refGene_tx=XM_003992529.4
D1 21810235 21810394 refGene_gn=VSIG2;refGene_tx=XM_003992529.4
D1 21811243 21811451 refGene_gn=VSIG2;refGene_tx=XM_003992529.4
D1 21812261 21812425 refGene_gn=VSIG2;refGene_tx=XM_003992529.4
D1 21812984 21813042 refGene_gn=VSIG2;refGene_tx=XM_003992529.4
D1 21814592 21814688 refGene_gn=ESAM;refGene_tx=XM_003992530.5
D1 21814689 21814909 refGene_gn=ESAM;refGene_tx=XM_003992530.5
D1 21815144 21815271 refGene_gn=ESAM;refGene_tx=XM_003992530.5
D1 21815537 21815660 refGene_gn=ESAM;refGene_tx=XM_003992530.5
D1 21817723 21817879 refGene_gn=ESAM;refGene_tx=XM_003992530.5
D1 21818080 21818282 refGene_gn=ESAM;refGene_tx=XM_003992530.5
D1 21819348 21819536 refGene_gn=ESAM;refGene_tx=XM_003992530.5
D1 21822956 21823026 refGene_gn=ESAM;refGene_tx=XM_003992530.5
D1 21827881 21828734
refGene_gn=MSANTD2;refGene_tx=XM_023239249.1;refGene_gn=MSANTD2;refGene_tx=XM_023239250.1;refGene_gn=MSANTD2;refGene_tx=XM_023239251.1
D1 21833566 21833627
refGene_gn=MSANTD2;refGene_tx=XM_023239249.1;refGene_gn=MSANTD2;refGene_tx=XM_023239250.1;refGene_gn=MSANTD2;refGene_tx=XM_023239251.1
D1 21835263 21835519
refGene_gn=MSANTD2;refGene_tx=XM_023239249.1;refGene_gn=MSANTD2;refGene_tx=XM_023239250.1;refGene_gn=MSANTD2;refGene_tx=XM_023239251.1
D1 21839237 21839294
refGene_gn=MSANTD2;refGene_tx=XM_023239252.1;refGene_gn=MSANTD2;refGene_tx=XM_023239253.1
D1 21858206 21858716
refGene_gn=MSANTD2;refGene_tx=XM_023239249.1;refGene_gn=MSANTD2;refGene_tx=XM_023239250.1;refGene_gn=MSANTD2;refGene_tx=XM_023239252.1;refGene_gn=MSANTD2;refGene_tx=XM_023239253.1
D1 21905004 21905164

refGene_gn=ROBO3;refGene_tx=XM_011286422.3;refGene_gn=ROBO3;refGene_tx=XM_019811488.2;refGene_gn=ROBO3;refGene_tx=XM_019811489.2;refGene_gn=ROBO3;refGene_tx=XM_019811490.2;refGene_gn=ROBO3;refGene_tx=XM_019811491.2;refGene_gn=ROBO3;refGene_tx=XM_019811492.2
D1 21908345 21908672
refGene_gn=ROBO3;refGene_tx=XM_011286422.3;refGene_gn=ROBO3;refGene_tx=XM_019811488.2;refGene_gn=ROBO3;refGene_tx=XM_019811489.2;refGene_gn=ROBO3;refGene_tx=XM_019811490.2;refGene_gn=ROBO3;refGene_tx=XM_019811491.2;refGene_gn=ROBO3;refGene_tx=XM_019811492.2
D1 21909009 21909167
refGene_gn=ROBO3;refGene_tx=XM_011286422.3;refGene_gn=ROBO3;refGene_tx=XM_019811488.2;refGene_gn=ROBO3;refGene_tx=XM_019811489.2;refGene_gn=ROBO3;refGene_tx=XM_019811490.2;refGene_gn=ROBO3;refGene_tx=XM_019811491.2;refGene_gn=ROBO3;refGene_tx=XM_019811492.2
D1 21909645 21909766
refGene_gn=ROBO3;refGene_tx=XM_011286422.3;refGene_gn=ROBO3;refGene_tx=XM_019811488.2;refGene_gn=ROBO3;refGene_tx=XM_019811489.2;refGene_gn=ROBO3;refGene_tx=XM_019811490.2;refGene_gn=ROBO3;refGene_tx=XM_019811491.2;refGene_gn=ROBO3;refGene_tx=XM_019811492.2
D1 21909862 21910001
refGene_gn=ROBO3;refGene_tx=XM_011286422.3;refGene_gn=ROBO3;refGene_tx=XM_019811488.2;refGene_gn=ROBO3;refGene_tx=XM_019811489.2;refGene_gn=ROBO3;refGene_tx=XM_019811490.2;refGene_gn=ROBO3;refGene_tx=XM_019811491.2;refGene_gn=ROBO3;refGene_tx=XM_019811492.2
D1 21910301 21910429
refGene_gn=ROBO3;refGene_tx=XM_011286422.3;refGene_gn=ROBO3;refGene_tx=XM_019811488.2;refGene_gn=ROBO3;refGene_tx=XM_019811489.2;refGene_gn=ROBO3;refGene_tx=XM_019811490.2;refGene_gn=ROBO3;refGene_tx=XM_019811491.2;refGene_gn=ROBO3;refGene_tx=XM_019811492.2
D1 21910705 21910830
refGene_gn=ROBO3;refGene_tx=XM_011286422.3;refGene_gn=ROBO3;refGene_tx=XM_019811488.2;refGene_gn=ROBO3;refGene_tx=XM_019811489.2;refGene_gn=ROBO3;refGene_tx=XM_019811490.2;refGene_gn=ROBO3;refGene_tx=XM_019811491.2;refGene_gn=ROBO3;refGene_tx=XM_019811492.2
D1 21912038 21912210
refGene_gn=ROBO3;refGene_tx=XM_011286422.3;refGene_gn=ROBO3;refGene_tx=XM_019811488.2;refGene_gn=ROBO3;refGene_tx=XM_019811489.2;refGene_gn=ROBO3;refGene_tx=XM_019811490.2;refGene_gn=ROBO3;refGene_tx=XM_019811491.2;refGene_gn=ROBO3;refGene_tx=XM_019811492.2
D1 21912526 21912735
refGene_gn=ROBO3;refGene_tx=XM_011286422.3;refGene_gn=ROBO3;refGene_tx=XM_019811488.2;refGene_gn=ROBO3;refGene_tx=XM_019811489.2;refGene_gn=ROBO3;refGene_tx=XM_019811490.2;refGene_gn=ROBO3;refGene_tx=XM_019811491.2;refGene_gn=ROBO3;refGene_tx=XM_019811492.2
D1 21912950 21913032
refGene_gn=ROBO3;refGene_tx=XM_011286422.3;refGene_gn=ROBO3;refGene_tx=XM_019811488.2;refGene_gn=ROBO3;refGene_tx=XM_019811489.2;refGene_gn=ROBO3;refGene_tx=XM_019811490.2;refGene_gn=ROBO3;refGene_tx=XM_019811491.2;refGene_gn=ROBO3;refGene_tx=XM_019811492.2
D1 21913320 21913504

refGene_gn=ROBO3;refGene_tx=XM_011286422.3;refGene_gn=ROBO3;refGene_tx=XM_019811488.2;refGene_gn=ROBO3;refGene_tx=XM_019811489.2;refGene_gn=ROBO3;refGene_tx=XM_019811490.2;refGene_gn=ROBO3;refGene_tx=XM_019811491.2;refGene_gn=ROBO3;refGene_tx=XM_019811492.2
D1 21913694 21913861
refGene_gn=ROBO3;refGene_tx=XM_011286422.3;refGene_gn=ROBO3;refGene_tx=XM_019811488.2;refGene_gn=ROBO3;refGene_tx=XM_019811489.2;refGene_gn=ROBO3;refGene_tx=XM_019811490.2;refGene_gn=ROBO3;refGene_tx=XM_019811491.2;refGene_gn=ROBO3;refGene_tx=XM_019811492.2
D1 21914323 21914445
refGene_gn=ROBO3;refGene_tx=XM_011286422.3;refGene_gn=ROBO3;refGene_tx=XM_019811488.2;refGene_gn=ROBO3;refGene_tx=XM_019811489.2;refGene_gn=ROBO3;refGene_tx=XM_019811490.2;refGene_gn=ROBO3;refGene_tx=XM_019811491.2;refGene_gn=ROBO3;refGene_tx=XM_019811492.2
D1 21914635 21914861
refGene_gn=ROBO3;refGene_tx=XM_011286422.3;refGene_gn=ROBO3;refGene_tx=XM_019811488.2;refGene_gn=ROBO3;refGene_tx=XM_019811489.2;refGene_gn=ROBO3;refGene_tx=XM_019811490.2;refGene_gn=ROBO3;refGene_tx=XM_019811491.2;refGene_gn=ROBO3;refGene_tx=XM_019811492.2
D1 21915070 21915192
refGene_gn=ROBO3;refGene_tx=XM_011286422.3;refGene_gn=ROBO3;refGene_tx=XM_019811488.2;refGene_gn=ROBO3;refGene_tx=XM_019811489.2;refGene_gn=ROBO3;refGene_tx=XM_019811490.2;refGene_gn=ROBO3;refGene_tx=XM_019811491.2;refGene_gn=ROBO3;refGene_tx=XM_019811492.2
D1 21915443 21915615
refGene_gn=ROBO3;refGene_tx=XM_011286422.3;refGene_gn=ROBO3;refGene_tx=XM_019811488.2;refGene_gn=ROBO3;refGene_tx=XM_019811489.2;refGene_gn=ROBO3;refGene_tx=XM_019811490.2;refGene_gn=ROBO3;refGene_tx=XM_019811491.2;refGene_gn=ROBO3;refGene_tx=XM_019811492.2
D1 21915779 21915965
refGene_gn=ROBO3;refGene_tx=XM_011286422.3;refGene_gn=ROBO3;refGene_tx=XM_019811488.2;refGene_gn=ROBO3;refGene_tx=XM_019811489.2;refGene_gn=ROBO3;refGene_tx=XM_019811490.2;refGene_gn=ROBO3;refGene_tx=XM_019811491.2;refGene_gn=ROBO3;refGene_tx=XM_019811492.2
D1 21916727 21916875
refGene_gn=ROBO3;refGene_tx=XM_011286422.3;refGene_gn=ROBO3;refGene_tx=XM_019811488.2;refGene_gn=ROBO3;refGene_tx=XM_019811489.2;refGene_gn=ROBO3;refGene_tx=XM_019811490.2;refGene_gn=ROBO3;refGene_tx=XM_019811491.2;refGene_gn=ROBO3;refGene_tx=XM_019811492.2
D1 21917079 21917219
refGene_gn=ROBO3;refGene_tx=XM_011286422.3;refGene_gn=ROBO3;refGene_tx=XM_019811488.2;refGene_gn=ROBO3;refGene_tx=XM_019811489.2;refGene_gn=ROBO3;refGene_tx=XM_019811490.2;refGene_gn=ROBO3;refGene_tx=XM_019811491.2;refGene_gn=ROBO3;refGene_tx=XM_019811492.2
D1 21917536 21917731
refGene_gn=ROBO3;refGene_tx=XM_011286422.3;refGene_gn=ROBO3;refGene_tx=XM_019811488.2;refGene_gn=ROBO3;refGene_tx=XM_019811489.2;refGene_gn=ROBO3;refGene_tx=XM_019811490.2;refGene_gn=ROBO3;refGene_tx=XM_019811491.2;refGene_gn=ROBO3;refGene_tx=XM_019811492.2
D1 21917894 21918033

refGene_gn=ROBO3;refGene_tx=XM_011286422.3;refGene_gn=ROBO3;refGene_tx=XM_019811488.2;refGene_gn=ROBO3;refGene_tx=XM_019811489.2;refGene_gn=ROBO3;refGene_tx=XM_019811490.2;refGene_gn=ROBO3;refGene_tx=XM_019811491.2;refGene_gn=ROBO3;refGene_tx=XM_019811492.2
D1 21918189 21918402
refGene_gn=ROBO3;refGene_tx=XM_011286422.3;refGene_gn=ROBO3;refGene_tx=XM_019811488.2;refGene_gn=ROBO3;refGene_tx=XM_019811489.2;refGene_gn=ROBO3;refGene_tx=XM_019811490.2;refGene_gn=ROBO3;refGene_tx=XM_019811491.2
D1 21918792 21918944
refGene_gn=ROBO3;refGene_tx=XM_011286422.3;refGene_gn=ROBO3;refGene_tx=XM_019811488.2;refGene_gn=ROBO3;refGene_tx=XM_019811489.2;refGene_gn=ROBO3;refGene_tx=XM_019811490.2;refGene_gn=ROBO3;refGene_tx=XM_019811491.2;refGene_gn=ROBO3;refGene_tx=XM_019811492.2
D1 21919072 21919240
refGene_gn=ROBO3;refGene_tx=XM_011286422.3;refGene_gn=ROBO3;refGene_tx=XM_019811488.2;refGene_gn=ROBO3;refGene_tx=XM_019811489.2;refGene_gn=ROBO3;refGene_tx=XM_019811490.2;refGene_gn=ROBO3;refGene_tx=XM_019811491.2;refGene_gn=ROBO3;refGene_tx=XM_019811492.2
D1 21919400 21919568
refGene_gn=ROBO3;refGene_tx=XM_011286422.3;refGene_gn=ROBO3;refGene_tx=XM_019811488.2;refGene_gn=ROBO3;refGene_tx=XM_019811489.2;refGene_gn=ROBO3;refGene_tx=XM_019811490.2;refGene_gn=ROBO3;refGene_tx=XM_019811491.2;refGene_gn=ROBO3;refGene_tx=XM_019811492.2
D1 21920059 21920238
refGene_gn=ROBO3;refGene_tx=XM_011286422.3;refGene_gn=ROBO3;refGene_tx=XM_019811488.2;refGene_gn=ROBO3;refGene_tx=XM_019811489.2;refGene_gn=ROBO3;refGene_tx=XM_019811490.2;refGene_gn=ROBO3;refGene_tx=XM_019811491.2;refGene_gn=ROBO3;refGene_tx=XM_019811492.2
D1 21920556 21920568
refGene_gn=ROBO3;refGene_tx=XM_011286422.3;refGene_gn=ROBO3;refGene_tx=XM_019811488.2;refGene_gn=ROBO3;refGene_tx=XM_019811489.2;refGene_gn=ROBO3;refGene_tx=XM_019811490.2;refGene_gn=ROBO3;refGene_tx=XM_019811491.2;refGene_gn=ROBO3;refGene_tx=XM_019811492.2
D1 21923164 21923175
refGene_gn=ROBO4;refGene_tx=XM_006936762.4;refGene_gn=ROBO4;refGene_tx=XM_019811493.2;refGene_gn=ROBO4;refGene_tx=XM_019811494.2;refGene_gn=ROBO4;refGene_tx=XM_019811495.2;refGene_gn=ROBO4;refGene_tx=XM_019811496.2;refGene_gn=ROBO4;refGene_tx=XM_019811498.2
D1 21923297 21923504
refGene_gn=ROBO4;refGene_tx=XM_006936762.4;refGene_gn=ROBO4;refGene_tx=XM_019811493.2;refGene_gn=ROBO4;refGene_tx=XM_019811494.2;refGene_gn=ROBO4;refGene_tx=XM_019811495.2;refGene_gn=ROBO4;refGene_tx=XM_019811496.2;refGene_gn=ROBO4;refGene_tx=XM_019811498.2
D1 21924954 21925340
refGene_gn=ROBO4;refGene_tx=XM_006936762.4;refGene_gn=ROBO4;refGene_tx=XM_019811493.2;refGene_gn=ROBO4;refGene_tx=XM_019811494.2;refGene_gn=ROBO4;refGene_tx=XM_019811495.2;refGene_gn=ROBO4;refGene_tx=XM_019811496.2;refGene_gn=ROBO4;refGene_tx=XM_019811498.2
D1 21925449 21925695

refGene_gn=ROBO4;refGene_tx=XM_006936762.4;refGene_gn=ROBO4;refGene_tx=XM_019811493.2;refGene_gn=ROBO4;refGene_tx=XM_019811494.2;refGene_gn=ROBO4;refGene_tx=XM_019811495.2;refGene_gn=ROBO4;refGene_tx=XM_019811496.2;refGene_gn=ROBO4;refGene_tx=XM_019811498.2
D1 21925833 21925987
refGene_gn=ROBO4;refGene_tx=XM_006936762.4;refGene_gn=ROBO4;refGene_tx=XM_019811493.2;refGene_gn=ROBO4;refGene_tx=XM_019811494.2;refGene_gn=ROBO4;refGene_tx=XM_019811495.2;refGene_gn=ROBO4;refGene_tx=XM_019811496.2;refGene_gn=ROBO4;refGene_tx=XM_019811498.2
D1 21926113 21926221
refGene_gn=ROBO4;refGene_tx=XM_006936762.4;refGene_gn=ROBO4;refGene_tx=XM_019811493.2;refGene_gn=ROBO4;refGene_tx=XM_019811494.2;refGene_gn=ROBO4;refGene_tx=XM_019811495.2;refGene_gn=ROBO4;refGene_tx=XM_019811496.2;refGene_gn=ROBO4;refGene_tx=XM_019811498.2
D1 21929636 21929900
refGene_gn=ROBO4;refGene_tx=XM_006936762.4;refGene_gn=ROBO4;refGene_tx=XM_019811493.2;refGene_gn=ROBO4;refGene_tx=XM_019811494.2;refGene_gn=ROBO4;refGene_tx=XM_019811495.2;refGene_gn=ROBO4;refGene_tx=XM_019811496.2;refGene_gn=ROBO4;refGene_tx=XM_019811498.2
D1 21930017 21930154
refGene_gn=ROBO4;refGene_tx=XM_006936762.4;refGene_gn=ROBO4;refGene_tx=XM_019811493.2;refGene_gn=ROBO4;refGene_tx=XM_019811494.2;refGene_gn=ROBO4;refGene_tx=XM_019811495.2;refGene_gn=ROBO4;refGene_tx=XM_019811496.2;refGene_gn=ROBO4;refGene_tx=XM_019811498.2
D1 21932071 21932114
refGene_gn=ROBO4;refGene_tx=XM_006936762.4;refGene_gn=ROBO4;refGene_tx=XM_019811493.2;refGene_gn=ROBO4;refGene_tx=XM_019811494.2;refGene_gn=ROBO4;refGene_tx=XM_019811495.2;refGene_gn=ROBO4;refGene_tx=XM_019811496.2;refGene_gn=ROBO4;refGene_tx=XM_019811498.2
D1 21932224 21932407
refGene_gn=ROBO4;refGene_tx=XM_006936762.4;refGene_gn=ROBO4;refGene_tx=XM_019811493.2;refGene_gn=ROBO4;refGene_tx=XM_019811494.2;refGene_gn=ROBO4;refGene_tx=XM_019811495.2;refGene_gn=ROBO4;refGene_tx=XM_019811496.2;refGene_gn=ROBO4;refGene_tx=XM_019811498.2
D1 21932551 21932720
refGene_gn=ROBO4;refGene_tx=XM_006936762.4;refGene_gn=ROBO4;refGene_tx=XM_019811493.2;refGene_gn=ROBO4;refGene_tx=XM_019811494.2;refGene_gn=ROBO4;refGene_tx=XM_019811495.2;refGene_gn=ROBO4;refGene_tx=XM_019811496.2;refGene_gn=ROBO4;refGene_tx=XM_019811498.2
D1 21933446 21933559
refGene_gn=ROBO4;refGene_tx=XM_006936762.4;refGene_gn=ROBO4;refGene_tx=XM_019811493.2;refGene_gn=ROBO4;refGene_tx=XM_019811494.2;refGene_gn=ROBO4;refGene_tx=XM_019811495.2;refGene_gn=ROBO4;refGene_tx=XM_019811496.2;refGene_gn=ROBO4;refGene_tx=XM_019811498.2
D1 21933817 21934046
refGene_gn=ROBO4;refGene_tx=XM_006936762.4;refGene_gn=ROBO4;refGene_tx=XM_019811493.2;refGene_gn=ROBO4;refGene_tx=XM_019811494.2;refGene_gn=ROBO4;refGene_tx=XM_019811495.2;refGene_gn=ROBO4;refGene_tx=XM_019811496.2;refGene_gn=ROBO4;refGene_tx=XM_019811498.2
D1 21934145 21934273

refGene_gn=ROBO4;refGene_tx=XM_006936762.4;refGene_gn=ROBO4;refGene_tx=XM_019811493.2;refGene_gn=ROBO4;refGene_tx=XM_019811494.2;refGene_gn=ROBO4;refGene_tx=XM_019811495.2;refGene_gn=ROBO4;refGene_tx=XM_019811496.2;refGene_gn=ROBO4;refGene_tx=XM_019811498.2
D1 21934529 21934650
refGene_gn=ROBO4;refGene_tx=XM_006936762.4;refGene_gn=ROBO4;refGene_tx=XM_019811493.2;refGene_gn=ROBO4;refGene_tx=XM_019811494.2;refGene_gn=ROBO4;refGene_tx=XM_019811495.2;refGene_gn=ROBO4;refGene_tx=XM_019811496.2;refGene_gn=ROBO4;refGene_tx=XM_019811498.2
D1 21934849 21935007
refGene_gn=ROBO4;refGene_tx=XM_006936762.4;refGene_gn=ROBO4;refGene_tx=XM_019811493.2;refGene_gn=ROBO4;refGene_tx=XM_019811494.2;refGene_gn=ROBO4;refGene_tx=XM_019811495.2;refGene_gn=ROBO4;refGene_tx=XM_019811496.2;refGene_gn=ROBO4;refGene_tx=XM_019811498.2
D1 21935266 21935629
refGene_gn=ROBO4;refGene_tx=XM_006936762.4;refGene_gn=ROBO4;refGene_tx=XM_019811493.2;refGene_gn=ROBO4;refGene_tx=XM_019811494.2;refGene_gn=ROBO4;refGene_tx=XM_019811495.2;refGene_gn=ROBO4;refGene_tx=XM_019811496.2;refGene_gn=ROBO4;refGene_tx=XM_019811498.2
D1 21935925 21935995
refGene_gn=ROBO4;refGene_tx=XM_006936762.4;refGene_gn=ROBO4;refGene_tx=XM_019811493.2;refGene_gn=ROBO4;refGene_tx=XM_019811494.2;refGene_gn=ROBO4;refGene_tx=XM_019811495.2;refGene_gn=ROBO4;refGene_tx=XM_019811496.2;refGene_gn=ROBO4;refGene_tx=XM_019811498.2
D1 21948255 21948558
refGene_gn=HEPACAM;refGene_tx=XM_023239254.1;refGene_gn=HEPACAM;refGene_tx=XM_023239255.1;refGene_gn=HEPACAM;refGene_tx=XM_023239256.1;refGene_gn=HEPACAM;refGene_tx=XM_023239257.1;refGene_gn=HEPACAM;refGene_tx=XM_023239258.1;refGene_gn=HEPACAM;refGene_tx=XM_023239259.1
D1 21949086 21949157
refGene_gn=HEPACAM;refGene_tx=XM_023239254.1;refGene_gn=HEPACAM;refGene_tx=XM_023239255.1;refGene_gn=HEPACAM;refGene_tx=XM_023239256.1;refGene_gn=HEPACAM;refGene_tx=XM_023239257.1;refGene_gn=HEPACAM;refGene_tx=XM_023239258.1;refGene_gn=HEPACAM;refGene_tx=XM_023239259.1
D1 21949289 21949531
refGene_gn=HEPACAM;refGene_tx=XM_023239254.1;refGene_gn=HEPACAM;refGene_tx=XM_023239255.1;refGene_gn=HEPACAM;refGene_tx=XM_023239256.1;refGene_gn=HEPACAM;refGene_tx=XM_023239257.1;refGene_gn=HEPACAM;refGene_tx=XM_023239258.1;refGene_gn=HEPACAM;refGene_tx=XM_023239259.1
D1 21950074 21950174
refGene_gn=HEPACAM;refGene_tx=XM_023239254.1;refGene_gn=HEPACAM;refGene_tx=XM_023239255.1;refGene_gn=HEPACAM;refGene_tx=XM_023239256.1;refGene_gn=HEPACAM;refGene_tx=XM_023239257.1;refGene_gn=HEPACAM;refGene_tx=XM_023239258.1;refGene_gn=HEPACAM;refGene_tx=XM_023239259.1
D1 21950456 21950738
refGene_gn=HEPACAM;refGene_tx=XM_023239254.1;refGene_gn=HEPACAM;refGene_tx=XM_023239255.1;refGene_gn=HEPACAM;refGene_tx=XM_023239256.1;refGene_gn=HEPACAM;refGene_tx=XM_023239257.1;refGene_gn=HEPACAM;refGene_tx=XM_023239258.1;refGene_gn=HEPACAM;refGene_tx=XM_023239259.1
D1 21951453 21951795

refGene_gn=HEPACAM;refGene_tx=XM_023239254.1;refGene_gn=HEPACAM;refGene_tx=XM_023239255.1;refGene_gn=HEPACAM;refGene_tx=XM_023239256.1;refGene_gn=HEPACAM;refGene_tx=XM_023239257.1;refGene_gn=HEPACAM;refGene_tx=XM_023239258.1;refGene_gn=HEPACAM;refGene_tx=XM_023239259.1
D1 21961685 21962085
refGene_gn=HEPACAM;refGene_tx=XM_023239254.1;refGene_gn=HEPACAM;refGene_tx=XM_023239255.1;refGene_gn=HEPACAM;refGene_tx=XM_023239256.1;refGene_gn=HEPACAM;refGene_tx=XM_023239257.1;refGene_gn=HEPACAM;refGene_tx=XM_023239258.1;refGene_gn=HEPACAM;refGene_tx=XM_023239259.1
D1 21977479 21977659
refGene_gn=CCDC15;refGene_tx=XM_011286423.3;refGene_gn=CCDC15;refGene_tx=XM_023239260.1;refGene_gn=CCDC15;refGene_tx=XM_023239261.1;refGene_gn=CCDC15;refGene_tx=XM_023239262.1;refGene_gn=CCDC15;refGene_tx=XM_023239263.1;refGene_gn=CCDC15;refGene_tx=XM_023239264.1;refGene_gn=CCDC15;refGene_tx=XM_023239265.1
D1 21982741 21982891
refGene_gn=CCDC15;refGene_tx=XM_011286423.3;refGene_gn=CCDC15;refGene_tx=XM_023239260.1;refGene_gn=CCDC15;refGene_tx=XM_023239261.1;refGene_gn=CCDC15;refGene_tx=XM_023239262.1;refGene_gn=CCDC15;refGene_tx=XM_023239263.1;refGene_gn=CCDC15;refGene_tx=XM_023239264.1;refGene_gn=CCDC15;refGene_tx=XM_023239265.1
D1 21983305 21983497
refGene_gn=CCDC15;refGene_tx=XM_011286423.3;refGene_gn=CCDC15;refGene_tx=XM_023239260.1;refGene_gn=CCDC15;refGene_tx=XM_023239261.1;refGene_gn=CCDC15;refGene_tx=XM_023239262.1;refGene_gn=CCDC15;refGene_tx=XM_023239263.1;refGene_gn=CCDC15;refGene_tx=XM_023239264.1;refGene_gn=CCDC15;refGene_tx=XM_023239265.1
D1 21990371 21990485
refGene_gn=CCDC15;refGene_tx=XM_011286423.3;refGene_gn=CCDC15;refGene_tx=XM_023239260.1;refGene_gn=CCDC15;refGene_tx=XM_023239261.1;refGene_gn=CCDC15;refGene_tx=XM_023239262.1;refGene_gn=CCDC15;refGene_tx=XM_023239263.1;refGene_gn=CCDC15;refGene_tx=XM_023239265.1
D1 21995303 21995426
refGene_gn=CCDC15;refGene_tx=XM_011286423.3;refGene_gn=CCDC15;refGene_tx=XM_023239260.1;refGene_gn=CCDC15;refGene_tx=XM_023239261.1;refGene_gn=CCDC15;refGene_tx=XM_023239262.1;refGene_gn=CCDC15;refGene_tx=XM_023239263.1;refGene_gn=CCDC15;refGene_tx=XM_023239264.1;refGene_gn=CCDC15;refGene_tx=XM_023239265.1
D1 21995684 21996401
refGene_gn=CCDC15;refGene_tx=XM_011286423.3;refGene_gn=CCDC15;refGene_tx=XM_023239260.1;refGene_gn=CCDC15;refGene_tx=XM_023239261.1;refGene_gn=CCDC15;refGene_tx=XM_023239262.1;refGene_gn=CCDC15;refGene_tx=XM_023239263.1;refGene_gn=CCDC15;refGene_tx=XM_023239264.1;refGene_gn=CCDC15;refGene_tx=XM_023239265.1
D1 21996986 21997109
refGene_gn=CCDC15;refGene_tx=XM_011286423.3;refGene_gn=CCDC15;refGene_tx=XM_023239260.1;refGene_gn=CCDC15;refGene_tx=XM_023239261.1;refGene_gn=CCDC15;refGene_tx=XM_023239262.1;refGene_gn=CCDC15;refGene_tx=XM_023239263.1;refGene_gn=CCDC15;refGene_tx=XM_023239264.1;refGene_gn=CCDC15;refGene_tx=XM_023239265.1

D1 21997353 21997476
refGene_gn=CCDC15;refGene_tx=XM_011286423.3;refGene_gn=CCDC15;refGene_tx=XM_023239260.1;refGene_gn=CCDC15;refGene_tx=XM_023239261.1;refGene_gn=CCDC15;refGene_tx=XM_023239262.1;refGene_gn=CCDC15;refGene_tx=XM_023239263.1;refGene_gn=CCDC15;refGene_tx=XM_023239264.1;refGene_gn=CCDC15;refGene_tx=XM_023239265.1
D1 21999044 21999119
refGene_gn=CCDC15;refGene_tx=XM_011286423.3;refGene_gn=CCDC15;refGene_tx=XM_023239260.1;refGene_gn=CCDC15;refGene_tx=XM_023239261.1;refGene_gn=CCDC15;refGene_tx=XM_023239262.1;refGene_gn=CCDC15;refGene_tx=XM_023239263.1;refGene_gn=CCDC15;refGene_tx=XM_023239264.1;refGene_gn=CCDC15;refGene_tx=XM_023239265.1
D1 22000045 22000168
refGene_gn=CCDC15;refGene_tx=XM_011286423.3;refGene_gn=CCDC15;refGene_tx=XM_023239260.1;refGene_gn=CCDC15;refGene_tx=XM_023239261.1;refGene_gn=CCDC15;refGene_tx=XM_023239262.1;refGene_gn=CCDC15;refGene_tx=XM_023239263.1;refGene_gn=CCDC15;refGene_tx=XM_023239264.1;refGene_gn=CCDC15;refGene_tx=XM_023239265.1
D1 22005969 22006062
refGene_gn=CCDC15;refGene_tx=XM_011286423.3;refGene_gn=CCDC15;refGene_tx=XM_023239260.1;refGene_gn=CCDC15;refGene_tx=XM_023239261.1;refGene_gn=CCDC15;refGene_tx=XM_023239262.1;refGene_gn=CCDC15;refGene_tx=XM_023239263.1;refGene_gn=CCDC15;refGene_tx=XM_023239264.1;refGene_gn=CCDC15;refGene_tx=XM_023239265.1
D1 22007478 22007582
refGene_gn=CCDC15;refGene_tx=XM_011286423.3;refGene_gn=CCDC15;refGene_tx=XM_023239260.1;refGene_gn=CCDC15;refGene_tx=XM_023239261.1;refGene_gn=CCDC15;refGene_tx=XM_023239262.1;refGene_gn=CCDC15;refGene_tx=XM_023239263.1;refGene_gn=CCDC15;refGene_tx=XM_023239264.1;refGene_gn=CCDC15;refGene_tx=XM_023239265.1
D1 22042555 22042802 refGene_gn=CCDC15;refGene_tx=XM_023239265.1
D1 22060259 22060436
refGene_gn=CCDC15;refGene_tx=XM_011286423.3;refGene_gn=CCDC15;refGene_tx=XM_023239260.1;refGene_gn=CCDC15;refGene_tx=XM_023239261.1;refGene_gn=CCDC15;refGene_tx=XM_023239262.1;refGene_gn=CCDC15;refGene_tx=XM_023239263.1;refGene_gn=CCDC15;refGene_tx=XM_023239264.1
D1 22060640 22060789
refGene_gn=CCDC15;refGene_tx=XM_011286423.3;refGene_gn=CCDC15;refGene_tx=XM_023239260.1;refGene_gn=CCDC15;refGene_tx=XM_023239261.1;refGene_gn=CCDC15;refGene_tx=XM_023239262.1;refGene_gn=CCDC15;refGene_tx=XM_023239263.1;refGene_gn=CCDC15;refGene_tx=XM_023239264.1
D1 22062028 22062150
refGene_gn=CCDC15;refGene_tx=XM_011286423.3;refGene_gn=CCDC15;refGene_tx=XM_023239260.1;refGene_gn=CCDC15;refGene_tx=XM_023239261.1;refGene_gn=CCDC15;refGene_tx=XM_023239262.1;refGene_gn=CCDC15;refGene_tx=XM_023239263.1;refGene_gn=CCDC15;refGene_tx=XM_023239264.1
D1 22084291 22084350
refGene_gn=SLC37A2;refGene_tx=XM_023239266.1;refGene_gn=SLC37A2;refGene_tx=XM_023239267.1
D1 22097603 22097685

refGene_gn=SLC37A2;refGene_tx=XM_019811506.2;refGene_gn=SLC37A2;refGene_tx=XM_019811507.2;refGene_gn=SLC37A2;refGene_tx=XM_023239266.1;refGene_gn=SLC37A2;refGene_tx=XM_023239267.1
D1 22098139 22098233
refGene_gn=SLC37A2;refGene_tx=XM_019811506.2;refGene_gn=SLC37A2;refGene_tx=XM_019811507.2;refGene_gn=SLC37A2;refGene_tx=XM_023239266.1;refGene_gn=SLC37A2;refGene_tx=XM_023239267.1
D1 22098364 22098443
refGene_gn=SLC37A2;refGene_tx=XM_019811506.2;refGene_gn=SLC37A2;refGene_tx=XM_019811507.2;refGene_gn=SLC37A2;refGene_tx=XM_023239266.1;refGene_gn=SLC37A2;refGene_tx=XM_023239267.1
D1 22100144 22100280
refGene_gn=SLC37A2;refGene_tx=XM_019811506.2;refGene_gn=SLC37A2;refGene_tx=XM_019811507.2;refGene_gn=SLC37A2;refGene_tx=XM_023239266.1;refGene_gn=SLC37A2;refGene_tx=XM_023239267.1
D1 22100700 22100777
refGene_gn=SLC37A2;refGene_tx=XM_019811506.2;refGene_gn=SLC37A2;refGene_tx=XM_019811507.2;refGene_gn=SLC37A2;refGene_tx=XM_023239266.1;refGene_gn=SLC37A2;refGene_tx=XM_023239267.1
D1 22101670 22101837
refGene_gn=SLC37A2;refGene_tx=XM_019811506.2;refGene_gn=SLC37A2;refGene_tx=XM_019811507.2;refGene_gn=SLC37A2;refGene_tx=XM_023239266.1;refGene_gn=SLC37A2;refGene_tx=XM_023239267.1
D1 22102437 22102481
refGene_gn=SLC37A2;refGene_tx=XM_019811506.2;refGene_gn=SLC37A2;refGene_tx=XM_019811507.2;refGene_gn=SLC37A2;refGene_tx=XM_023239266.1;refGene_gn=SLC37A2;refGene_tx=XM_023239267.1
D1 22102790 22102943
refGene_gn=SLC37A2;refGene_tx=XM_019811506.2;refGene_gn=SLC37A2;refGene_tx=XM_019811507.2;refGene_gn=SLC37A2;refGene_tx=XM_023239266.1;refGene_gn=SLC37A2;refGene_tx=XM_023239267.1
D1 22103286 22103377
refGene_gn=SLC37A2;refGene_tx=XM_019811506.2;refGene_gn=SLC37A2;refGene_tx=XM_019811507.2;refGene_gn=SLC37A2;refGene_tx=XM_023239266.1;refGene_gn=SLC37A2;refGene_tx=XM_023239267.1
D1 22104712 22104775
refGene_gn=SLC37A2;refGene_tx=XM_019811506.2;refGene_gn=SLC37A2;refGene_tx=XM_019811507.2;refGene_gn=SLC37A2;refGene_tx=XM_023239266.1;refGene_gn=SLC37A2;refGene_tx=XM_023239267.1
D1 22105095 22105181
refGene_gn=SLC37A2;refGene_tx=XM_019811506.2;refGene_gn=SLC37A2;refGene_tx=XM_019811507.2;refGene_gn=SLC37A2;refGene_tx=XM_023239266.1;refGene_gn=SLC37A2;refGene_tx=XM_023239267.1
D1 22105693 22105742
refGene_gn=SLC37A2;refGene_tx=XM_019811506.2;refGene_gn=SLC37A2;refGene_tx=XM_019811507.2;refGene_gn=SLC37A2;refGene_tx=XM_023239266.1;refGene_gn=SLC37A2;refGene_tx=XM_023239267.1
D1 22105948 22106022
refGene_gn=SLC37A2;refGene_tx=XM_019811506.2;refGene_gn=SLC37A2;refGene_tx=XM_019811507.2;refGene_gn=SLC37A2;refGene_tx=XM_023239266.1;refGe

ne_gn=SLC37A2;refGene_tx=XM_023239267.1
D1 22106309 22106388
refGene_gn=SLC37A2;refGene_tx=XM_019811506.2;refGene_gn=SLC37A2;refGene_tx=XM_019811507.2;refGene_gn=SLC37A2;refGene_tx=XM_023239266.1;refGene_gn=SLC37A2;refGene_tx=XM_023239267.1
D1 22106531 22106629
refGene_gn=SLC37A2;refGene_tx=XM_019811506.2;refGene_gn=SLC37A2;refGene_tx=XM_019811507.2;refGene_gn=SLC37A2;refGene_tx=XM_023239266.1;refGene_gn=SLC37A2;refGene_tx=XM_023239267.1
D1 22106917 22106982
refGene_gn=SLC37A2;refGene_tx=XM_019811506.2;refGene_gn=SLC37A2;refGene_tx=XM_019811507.2;refGene_gn=SLC37A2;refGene_tx=XM_023239266.1;refGene_gn=SLC37A2;refGene_tx=XM_023239267.1
D1 22107183 22107214
refGene_gn=SLC37A2;refGene_tx=XM_019811506.2;refGene_gn=SLC37A2;refGene_tx=XM_019811507.2;refGene_gn=SLC37A2;refGene_tx=XM_023239266.1
D1 22108742 22108758 refGene_gn=SLC37A2;refGene_tx=XM_023239267.1
D1 22116648 22116783
refGene_gn=TMEM218;refGene_tx=XM_003992531.4;refGene_gn=TMEM218;refGene_tx=XM_003992532.3;refGene_gn=TMEM218;refGene_tx=XM_006936765.4;refGene_gn=TMEM218;refGene_tx=XM_006936766.4;refGene_gn=TMEM218;refGene_tx=XM_023239271.1;refGene_gn=TMEM218;refGene_tx=XM_023239272.1
D1 22120047 22120150
refGene_gn=TMEM218;refGene_tx=XM_003992531.4;refGene_gn=TMEM218;refGene_tx=XM_003992532.3;refGene_gn=TMEM218;refGene_tx=XM_006936765.4;refGene_gn=TMEM218;refGene_tx=XM_006936766.4;refGene_gn=TMEM218;refGene_tx=XM_023239271.1;refGene_gn=TMEM218;refGene_tx=XM_023239272.1
D1 22121016 22121126
refGene_gn=TMEM218;refGene_tx=XM_003992531.4;refGene_gn=TMEM218;refGene_tx=XM_003992532.3;refGene_gn=TMEM218;refGene_tx=XM_006936765.4;refGene_gn=TMEM218;refGene_tx=XM_006936766.4;refGene_gn=TMEM218;refGene_tx=XM_023239271.1;refGene_gn=TMEM218;refGene_tx=XM_023239272.1
D1 22370453 22370540
refGene_gn=PKNOX2;refGene_tx=XM_003992533.4;refGene_gn=PKNOX2;refGene_tx=XM_019811508.2;refGene_gn=PKNOX2;refGene_tx=XM_019811510.2;refGene_gn=PKNOX2;refGene_tx=XM_019811511.2;refGene_gn=PKNOX2;refGene_tx=XM_023239268.1;refGene_gn=PKNOX2;refGene_tx=XM_023239270.1
D1 22376116 22376119 refGene_gn=PKNOX2;refGene_tx=XM_006936768.3
D1 22386035 22386175
refGene_gn=PKNOX2;refGene_tx=XM_003992533.4;refGene_gn=PKNOX2;refGene_tx=XM_006936768.3;refGene_gn=PKNOX2;refGene_tx=XM_019811508.2;refGene_gn=PKNOX2;refGene_tx=XM_019811510.2;refGene_gn=PKNOX2;refGene_tx=XM_019811511.2;refGene_gn=PKNOX2;refGene_tx=XM_023239268.1;refGene_gn=PKNOX2;refGene_tx=XM_023239269.1;refGene_gn=PKNOX2;refGene_tx=XM_023239270.1
D1 22403383 22403555
refGene_gn=PKNOX2;refGene_tx=XM_003992533.4;refGene_gn=PKNOX2;refGene_tx=XM_006936768.3;refGene_gn=PKNOX2;refGene_tx=XM_019811508.2;refGene_gn=PKNOX2;refGene_tx=XM_019811510.2;refGene_gn=PKNOX2;refGene_tx=XM_019811511.2;refGene_gn=PKNOX2;refGene_tx=XM_023239268.1;refGene_gn=PKNOX

2;refGene_tx=XM_023239269.1;refGene_gn=PKNOX2;refGene_tx=XM_023239270.
1
D1 22415516 22415705
refGene_gn=PKNOX2;refGene_tx=XM_003992533.4;refGene_gn=PKNOX2;refGene_
tx=XM_006936768.3;refGene_gn=PKNOX2;refGene_tx=XM_019811508.2;refGene_
gn=PKNOX2;refGene_tx=XM_019811510.2;refGene_gn=PKNOX2;refGene_tx=XM_01
9811511.2;refGene_gn=PKNOX2;refGene_tx=XM_023239268.1;refGene_gn=PKNOX
2;refGene_tx=XM_023239269.1;refGene_gn=PKNOX2;refGene_tx=XM_023239270.
1
D1 22427126 22427256
refGene_gn=PKNOX2;refGene_tx=XM_003992533.4;refGene_gn=PKNOX2;refGene_
tx=XM_006936768.3;refGene_gn=PKNOX2;refGene_tx=XM_019811508.2;refGene_
gn=PKNOX2;refGene_tx=XM_019811510.2;refGene_gn=PKNOX2;refGene_tx=XM_01
9811511.2;refGene_gn=PKNOX2;refGene_tx=XM_023239268.1;refGene_gn=PKNOX
2;refGene_tx=XM_023239269.1
D1 22427709 22427807
refGene_gn=PKNOX2;refGene_tx=XM_003992533.4;refGene_gn=PKNOX2;refGene_
tx=XM_006936768.3;refGene_gn=PKNOX2;refGene_tx=XM_019811508.2;refGene_
gn=PKNOX2;refGene_tx=XM_019811510.2;refGene_gn=PKNOX2;refGene_tx=XM_01
9811511.2;refGene_gn=PKNOX2;refGene_tx=XM_023239268.1;refGene_gn=PKNOX
2;refGene_tx=XM_023239269.1
D1 22428617 22428737
refGene_gn=PKNOX2;refGene_tx=XM_003992533.4;refGene_gn=PKNOX2;refGene_
tx=XM_006936768.3;refGene_gn=PKNOX2;refGene_tx=XM_019811508.2;refGene_
gn=PKNOX2;refGene_tx=XM_019811510.2;refGene_gn=PKNOX2;refGene_tx=XM_01
9811511.2;refGene_gn=PKNOX2;refGene_tx=XM_023239268.1;refGene_gn=PKNOX
2;refGene_tx=XM_023239269.1;refGene_gn=PKNOX2;refGene_tx=XM_023239270.
1
D1 22445239 22445316
refGene_gn=PKNOX2;refGene_tx=XM_003992533.4;refGene_gn=PKNOX2;refGene_
tx=XM_006936768.3;refGene_gn=PKNOX2;refGene_tx=XM_019811508.2;refGene_
gn=PKNOX2;refGene_tx=XM_019811510.2;refGene_gn=PKNOX2;refGene_tx=XM_01
9811511.2;refGene_gn=PKNOX2;refGene_tx=XM_023239268.1;refGene_gn=PKNOX
2;refGene_tx=XM_023239269.1;refGene_gn=PKNOX2;refGene_tx=XM_023239270.
1
D1 22446221 22446400
refGene_gn=PKNOX2;refGene_tx=XM_003992533.4;refGene_gn=PKNOX2;refGene_
tx=XM_006936768.3;refGene_gn=PKNOX2;refGene_tx=XM_019811508.2;refGene_
gn=PKNOX2;refGene_tx=XM_019811510.2;refGene_gn=PKNOX2;refGene_tx=XM_01
9811511.2;refGene_gn=PKNOX2;refGene_tx=XM_023239268.1;refGene_gn=PKNOX
2;refGene_tx=XM_023239269.1;refGene_gn=PKNOX2;refGene_tx=XM_023239270.
1
D1 22447389 22447616
refGene_gn=PKNOX2;refGene_tx=XM_003992533.4;refGene_gn=PKNOX2;refGene_
tx=XM_006936768.3;refGene_gn=PKNOX2;refGene_tx=XM_019811508.2;refGene_
gn=PKNOX2;refGene_tx=XM_019811510.2;refGene_gn=PKNOX2;refGene_tx=XM_01
9811511.2;refGene_gn=PKNOX2;refGene_tx=XM_023239268.1;refGene_gn=PKNOX
2;refGene_tx=XM_023239269.1;refGene_gn=PKNOX2;refGene_tx=XM_023239270.
1
D1 22459563 22459580

refGene_gn=FEZ1;refGene_tx=XM_003992536.5;refGene_gn=FEZ1;refGene_tx=XM_023239273.1
D1 22462497 22462563
refGene_gn=FEZ1;refGene_tx=XM_003992536.5;refGene_gn=FEZ1;refGene_tx=XM_023239273.1
D1 22466372 22466448
refGene_gn=FEZ1;refGene_tx=XM_003992536.5;refGene_gn=FEZ1;refGene_tx=XM_023239273.1
D1 22467520 22467601
refGene_gn=FEZ1;refGene_tx=XM_003992536.5;refGene_gn=FEZ1;refGene_tx=XM_023239273.1
D1 22468532 22468804
refGene_gn=FEZ1;refGene_tx=XM_003992536.5;refGene_gn=FEZ1;refGene_tx=XM_023239273.1
D1 22471892 22472061
refGene_gn=FEZ1;refGene_tx=XM_003992536.5;refGene_gn=FEZ1;refGene_tx=XM_023239273.1
D1 22474369 22474456
refGene_gn=FEZ1;refGene_tx=XM_003992536.5;refGene_gn=FEZ1;refGene_tx=XM_023239273.1
D1 22490409 22490509
refGene_gn=FEZ1;refGene_tx=XM_003992536.5;refGene_gn=FEZ1;refGene_tx=XM_023239273.1
D1 22497434 22497745
refGene_gn=FEZ1;refGene_tx=XM_003992536.5;refGene_gn=FEZ1;refGene_tx=XM_023239273.1
D1 22539515 22539557 refGene_gn=EI24;refGene_tx=XM_023239274.1
D1 22542759 22542905 refGene_gn=EI24;refGene_tx=XM_023239274.1
D1 22543791 22543852 refGene_gn=EI24;refGene_tx=XM_023239274.1
D1 22544683 22544750 refGene_gn=EI24;refGene_tx=XM_023239274.1
D1 22545308 22545433 refGene_gn=EI24;refGene_tx=XM_023239274.1
D1 22545756 22545876 refGene_gn=EI24;refGene_tx=XM_023239274.1
D1 22546229 22546341 refGene_gn=EI24;refGene_tx=XM_023239274.1
D1 22547011 22547123 refGene_gn=EI24;refGene_tx=XM_023239274.1
D1 22547520 22547595 refGene_gn=EI24;refGene_tx=XM_023239274.1
D1 22548525 22548688 refGene_gn=EI24;refGene_tx=XM_023239274.1
D1 22557790 22557878 refGene_gn=STT3A;refGene_tx=XM_011286425.2
D1 22558683 22558744 refGene_gn=STT3A;refGene_tx=XM_011286425.2
D1 22561964 22562086 refGene_gn=STT3A;refGene_tx=XM_011286425.2
D1 22562488 22562634 refGene_gn=STT3A;refGene_tx=XM_011286425.2
D1 22564254 22564345 refGene_gn=STT3A;refGene_tx=XM_011286425.2
D1 22565715 22565822 refGene_gn=STT3A;refGene_tx=XM_011286425.2
D1 22566499 22566664 refGene_gn=STT3A;refGene_tx=XM_011286425.2
D1 22568203 22568384 refGene_gn=STT3A;refGene_tx=XM_011286425.2
D1 22569412 22569568 refGene_gn=STT3A;refGene_tx=XM_011286425.2
D1 22570949 22571041 refGene_gn=STT3A;refGene_tx=XM_011286425.2
D1 22571726 22571882 refGene_gn=STT3A;refGene_tx=XM_011286425.2
D1 22572111 22572300 refGene_gn=STT3A;refGene_tx=XM_011286425.2
D1 22573355 22573472 refGene_gn=STT3A;refGene_tx=XM_011286425.2
D1 22573591 22573694 refGene_gn=STT3A;refGene_tx=XM_011286425.2

D1 22576812 22577001 refGene_gn=STT3A;refGene_tx=XM_011286425.2
D1 22578299 22578415 refGene_gn=STT3A;refGene_tx=XM_011286425.2
D1 22579230 22579269 refGene_gn=STT3A;refGene_tx=XM_011286425.2
D1 22585669 22585734
refGene_gn=CHEK1;refGene_tx=XM_004001334.5;refGene_gn=CHEK1;refGene_tx=XM_011286426.3;refGene_gn=CHEK1;refGene_tx=XM_019811516.2
D1 22586812 22587036
refGene_gn=CHEK1;refGene_tx=XM_004001334.5;refGene_gn=CHEK1;refGene_tx=XM_011286426.3;refGene_gn=CHEK1;refGene_tx=XM_019811516.2
D1 22588237 22588302
refGene_gn=CHEK1;refGene_tx=XM_004001334.5;refGene_gn=CHEK1;refGene_tx=XM_011286426.3;refGene_gn=CHEK1;refGene_tx=XM_019811516.2
D1 22588391 22588461
refGene_gn=CHEK1;refGene_tx=XM_004001334.5;refGene_gn=CHEK1;refGene_tx=XM_011286426.3;refGene_gn=CHEK1;refGene_tx=XM_019811516.2
D1 22603560 22603749
refGene_gn=CHEK1;refGene_tx=XM_004001334.5;refGene_gn=CHEK1;refGene_tx=XM_011286426.3;refGene_gn=CHEK1;refGene_tx=XM_019811516.2
D1 22604189 22604294
refGene_gn=CHEK1;refGene_tx=XM_004001334.5;refGene_gn=CHEK1;refGene_tx=XM_011286426.3;refGene_gn=CHEK1;refGene_tx=XM_019811516.2
D1 22607302 22607398
refGene_gn=CHEK1;refGene_tx=XM_004001334.5;refGene_gn=CHEK1;refGene_tx=XM_011286426.3;refGene_gn=CHEK1;refGene_tx=XM_019811516.2
D1 22611364 22611473
refGene_gn=CHEK1;refGene_tx=XM_004001334.5;refGene_gn=CHEK1;refGene_tx=XM_011286426.3;refGene_gn=CHEK1;refGene_tx=XM_019811516.2
D1 22611663 22611841
refGene_gn=CHEK1;refGene_tx=XM_004001334.5;refGene_gn=CHEK1;refGene_tx=XM_011286426.3;refGene_gn=CHEK1;refGene_tx=XM_019811516.2
D1 22612110 22612242
refGene_gn=CHEK1;refGene_tx=XM_004001334.5;refGene_gn=CHEK1;refGene_tx=XM_011286426.3;refGene_gn=CHEK1;refGene_tx=XM_019811516.2
D1 22614957 22615059
refGene_gn=CHEK1;refGene_tx=XM_004001334.5;refGene_gn=CHEK1;refGene_tx=XM_011286426.3;refGene_gn=CHEK1;refGene_tx=XM_019811516.2
D1 22616814 22616910
refGene_gn=CHEK1;refGene_tx=XM_004001334.5;refGene_gn=CHEK1;refGene_tx=XM_011286426.3;refGene_gn=CHEK1;refGene_tx=XM_019811516.2
D1 22637256 22637381
refGene_gn=ACRV1;refGene_tx=XM_006936771.3;refGene_gn=ACRV1;refGene_tx=XM_023239275.1
D1 22638140 22638260
refGene_gn=ACRV1;refGene_tx=XM_006936771.3;refGene_gn=ACRV1;refGene_tx=XM_023239275.1
D1 22639629 22640220
refGene_gn=ACRV1;refGene_tx=XM_006936771.3;refGene_gn=ACRV1;refGene_tx=XM_023239275.1
D1 22642992 22643044 refGene_gn=ACRV1;refGene_tx=XM_006936771.3
D1 22645008 22645201

refGene_gn=LOC109491868;refGene_tx=XM_023239276.1;refGene_gn=LOC109491
868;refGene_tx=XM_023239277.1
D1 22647619 22647656
refGene_gn=LOC109491868;refGene_tx=XM_019811519.2
D1 22647854 22647909
refGene_gn=LOC109491868;refGene_tx=XM_023239278.1
D1 22648582 22648696
refGene_gn=LOC109491868;refGene_tx=XM_019811519.2;refGene_gn=LOC109491
868;refGene_tx=XM_023239276.1;refGene_gn=LOC109491868;refGene_tx=XM_02
3239277.1;refGene_gn=LOC109491868;refGene_tx=XM_023239278.1
D1 22649259 22649384
refGene_gn=LOC109491868;refGene_tx=XM_019811519.2;refGene_gn=LOC109491
868;refGene_tx=XM_023239276.1;refGene_gn=LOC109491868;refGene_tx=XM_02
3239278.1
D1 22651780 22651869
refGene_gn=LOC109491868;refGene_tx=XM_023239277.1
D1 22848530 22848667 refGene_gn=PATE2;refGene_tx=XM_006944244.4
D1 22848964 22849093 refGene_gn=PATE2;refGene_tx=XM_006944244.4
D1 22849714 22849766 refGene_gn=PATE2;refGene_tx=XM_006944244.4
D1 22864216 22864265 refGene_gn=PATE3;refGene_tx=XM_003992549.2
D1 22866095 22866218 refGene_gn=PATE3;refGene_tx=XM_003992549.2
D1 22866723 22866848 refGene_gn=PATE3;refGene_tx=XM_003992549.2
D1 22928545 22928856
refGene_gn=PUS3;refGene_tx=XM_006936780.4;refGene_gn=PUS3;refGene_tx=X
M_006936781.4;refGene_gn=PUS3;refGene_tx=XM_019811524.2
D1 22928997 22929188
refGene_gn=PUS3;refGene_tx=XM_006936780.4;refGene_gn=PUS3;refGene_tx=X
M_006936781.4;refGene_gn=PUS3;refGene_tx=XM_019811524.2
D1 22930090 22930656
refGene_gn=PUS3;refGene_tx=XM_006936780.4;refGene_gn=PUS3;refGene_tx=X
M_006936781.4;refGene_gn=PUS3;refGene_tx=XM_019811524.2
D1 22930768 22931143
refGene_gn=PUS3;refGene_tx=XM_006936780.4;refGene_gn=PUS3;refGene_tx=X
M_006936781.4;refGene_gn=PUS3;refGene_tx=XM_019811524.2
D1 22931820 22931834
refGene_gn=HYLS1;refGene_tx=XM_006936774.3;refGene_gn=HYLS1;refGene_tx
=XM_006936775.4;refGene_gn=HYLS1;refGene_tx=XM_006936777.4;refGene_gn=
HYLS1;refGene_tx=XM_006936778.4;refGene_gn=HYLS1;refGene_tx=XM_0069367
79.3;refGene_gn=HYLS1;refGene_tx=XM_023239280.1
D1 22934206 22935131
refGene_gn=HYLS1;refGene_tx=XM_006936774.3;refGene_gn=HYLS1;refGene_tx
=XM_006936775.4;refGene_gn=HYLS1;refGene_tx=XM_006936777.4;refGene_gn=
HYLS1;refGene_tx=XM_006936778.4;refGene_gn=HYLS1;refGene_tx=XM_0069367
79.3;refGene_gn=HYLS1;refGene_tx=XM_023239280.1
D1 22938966 22939029 refGene_gn=DDX25;refGene_tx=XM_023239279.1
D1 22939683 22939750 refGene_gn=DDX25;refGene_tx=XM_023239279.1
D1 22940005 22940047 refGene_gn=DDX25;refGene_tx=XM_023239279.1
D1 22940536 22940672 refGene_gn=DDX25;refGene_tx=XM_023239279.1
D1 22942300 22942393
refGene_gn=DDX25;refGene_tx=XM_003992552.5;refGene_gn=DDX25;refGene_tx

=XM_023239279.1
D1 22942481 22942584
refGene_gn=DDX25;refGene_tx=XM_003992552.5;refGene_gn=DDX25;refGene_tx=XM_023239279.1
D1 22944083 22944198
refGene_gn=DDX25;refGene_tx=XM_003992552.5;refGene_gn=DDX25;refGene_tx=XM_023239279.1
D1 22945056 22945234
refGene_gn=DDX25;refGene_tx=XM_003992552.5;refGene_gn=DDX25;refGene_tx=XM_023239279.1
D1 22952169 22952407
refGene_gn=DDX25;refGene_tx=XM_003992552.5;refGene_gn=DDX25;refGene_tx=XM_023239279.1
D1 22953500 22953663
refGene_gn=DDX25;refGene_tx=XM_003992552.5;refGene_gn=DDX25;refGene_tx=XM_023239279.1
D1 22955708 22955897
refGene_gn=DDX25;refGene_tx=XM_003992552.5;refGene_gn=DDX25;refGene_tx=XM_023239279.1
D1 22957362 22957424
refGene_gn=DDX25;refGene_tx=XM_003992552.5;refGene_gn=DDX25;refGene_tx=XM_023239279.1
D1 22981850 22981932 refGene_gn=VSIG10L2;refGene_tx=XM_023239697.1
D1 22983616 22983967 refGene_gn=VSIG10L2;refGene_tx=XM_023239697.1
D1 22984056 22984212 refGene_gn=VSIG10L2;refGene_tx=XM_023239697.1
D1 22984387 22984663 refGene_gn=VSIG10L2;refGene_tx=XM_023239697.1
D1 22985373 22985649 refGene_gn=VSIG10L2;refGene_tx=XM_023239697.1
D1 22986234 22986483 refGene_gn=VSIG10L2;refGene_tx=XM_023239697.1
D1 22987172 22987433 refGene_gn=VSIG10L2;refGene_tx=XM_023239697.1
D1 22988747 22989038 refGene_gn=VSIG10L2;refGene_tx=XM_023239697.1
D1 22989705 22990002 refGene_gn=VSIG10L2;refGene_tx=XM_023239697.1
D1 22990656 22990779 refGene_gn=VSIG10L2;refGene_tx=XM_023239697.1
D1 22991235 22991288 refGene_gn=VSIG10L2;refGene_tx=XM_023239697.1
D1 22991568 22991871 refGene_gn=VSIG10L2;refGene_tx=XM_023239697.1
D1 22996210 22996451
refGene_gn=CDON;refGene_tx=XM_003992553.4;refGene_gn=CDON;refGene_tx=XM_006936782.4;refGene_gn=CDON;refGene_tx=XM_006936783.3;refGene_gn=CDON;refGene_tx=XM_011286500.3;refGene_gn=CDON;refGene_tx=XM_019811527.1;refGene_gn=CDON;refGene_tx=XM_023239281.1
D1 22998201 22998476
refGene_gn=CDON;refGene_tx=XM_003992553.4;refGene_gn=CDON;refGene_tx=XM_006936782.4;refGene_gn=CDON;refGene_tx=XM_006936783.3;refGene_gn=CDON;refGene_tx=XM_011286500.3;refGene_gn=CDON;refGene_tx=XM_023239281.1
D1 23015177 23015257
refGene_gn=CDON;refGene_tx=XM_003992553.4;refGene_gn=CDON;refGene_tx=XM_006936782.4;refGene_gn=CDON;refGene_tx=XM_006936783.3;refGene_gn=CDON;refGene_tx=XM_011286500.3;refGene_gn=CDON;refGene_tx=XM_019811527.1;refGene_gn=CDON;refGene_tx=XM_023239281.1
D1 23018449 23018730
refGene_gn=CDON;refGene_tx=XM_003992553.4;refGene_gn=CDON;refGene_tx=X

M_006936782.4;refGene_gn=CDON;refGene_tx=XM_006936783.3;refGene_gn=CDON;refGene_tx=XM_011286500.3;refGene_gn=CDON;refGene_tx=XM_019811527.1;refGene_gn=CDON;refGene_tx=XM_023239281.1
D1 23021179 23021401
refGene_gn=CDON;refGene_tx=XM_003992553.4;refGene_gn=CDON;refGene_tx=XM_006936782.4;refGene_gn=CDON;refGene_tx=XM_006936783.3;refGene_gn=CDON;refGene_tx=XM_011286500.3;refGene_gn=CDON;refGene_tx=XM_019811527.1;refGene_gn=CDON;refGene_tx=XM_023239281.1
D1 23025887 23026010
refGene_gn=CDON;refGene_tx=XM_003992553.4;refGene_gn=CDON;refGene_tx=XM_006936782.4;refGene_gn=CDON;refGene_tx=XM_006936783.3;refGene_gn=CDON;refGene_tx=XM_011286500.3;refGene_gn=CDON;refGene_tx=XM_019811527.1;refGene_gn=CDON;refGene_tx=XM_023239281.1
D1 23030445 23030551
refGene_gn=CDON;refGene_tx=XM_003992553.4;refGene_gn=CDON;refGene_tx=XM_006936782.4;refGene_gn=CDON;refGene_tx=XM_006936783.3;refGene_gn=CDON;refGene_tx=XM_011286500.3;refGene_gn=CDON;refGene_tx=XM_019811527.1;refGene_gn=CDON;refGene_tx=XM_023239281.1
D1 23031217 23031399
refGene_gn=CDON;refGene_tx=XM_003992553.4;refGene_gn=CDON;refGene_tx=XM_006936782.4;refGene_gn=CDON;refGene_tx=XM_006936783.3;refGene_gn=CDON;refGene_tx=XM_011286500.3;refGene_gn=CDON;refGene_tx=XM_019811527.1;refGene_gn=CDON;refGene_tx=XM_023239281.1
D1 23032958 23033162
refGene_gn=CDON;refGene_tx=XM_003992553.4;refGene_gn=CDON;refGene_tx=XM_006936782.4;refGene_gn=CDON;refGene_tx=XM_006936783.3;refGene_gn=CDON;refGene_tx=XM_011286500.3;refGene_gn=CDON;refGene_tx=XM_019811527.1;refGene_gn=CDON;refGene_tx=XM_023239281.1
D1 23037168 23037297
refGene_gn=CDON;refGene_tx=XM_003992553.4;refGene_gn=CDON;refGene_tx=XM_006936782.4;refGene_gn=CDON;refGene_tx=XM_006936783.3;refGene_gn=CDON;refGene_tx=XM_011286500.3;refGene_gn=CDON;refGene_tx=XM_019811527.1;refGene_gn=CDON;refGene_tx=XM_023239281.1
D1 23039485 23039660
refGene_gn=CDON;refGene_tx=XM_003992553.4;refGene_gn=CDON;refGene_tx=XM_006936782.4;refGene_gn=CDON;refGene_tx=XM_006936783.3;refGene_gn=CDON;refGene_tx=XM_011286500.3;refGene_gn=CDON;refGene_tx=XM_019811527.1;refGene_gn=CDON;refGene_tx=XM_023239281.1
D1 23041230 23041520
refGene_gn=CDON;refGene_tx=XM_003992553.4;refGene_gn=CDON;refGene_tx=XM_006936782.4;refGene_gn=CDON;refGene_tx=XM_006936783.3;refGene_gn=CDON;refGene_tx=XM_011286500.3;refGene_gn=CDON;refGene_tx=XM_019811527.1;refGene_gn=CDON;refGene_tx=XM_023239281.1
D1 23048044 23048395
refGene_gn=CDON;refGene_tx=XM_003992553.4;refGene_gn=CDON;refGene_tx=XM_006936782.4;refGene_gn=CDON;refGene_tx=XM_006936783.3;refGene_gn=CDON;refGene_tx=XM_011286500.3;refGene_gn=CDON;refGene_tx=XM_019811527.1;refGene_gn=CDON;refGene_tx=XM_023239281.1
D1 23054045 23054315
refGene_gn=CDON;refGene_tx=XM_003992553.4;refGene_gn=CDON;refGene_tx=X

M_006936782.4;refGene_gn=CDON;refGene_tx=XM_006936783.3;refGene_gn=CDON;refGene_tx=XM_011286500.3;refGene_gn=CDON;refGene_tx=XM_019811527.1;refGene_gn=CDON;refGene_tx=XM_023239281.1
D1 23055580 23055868 refGene_gn=CDON;refGene_tx=XM_003992553.4;refGene_gn=CDON;refGene_tx=XM_006936782.4;refGene_gn=CDON;refGene_tx=XM_006936783.3;refGene_gn=CDON;refGene_tx=XM_011286500.3;refGene_gn=CDON;refGene_tx=XM_019811527.1;refGene_gn=CDON;refGene_tx=XM_023239281.1
D1 23056816 23056960 refGene_gn=CDON;refGene_tx=XM_003992553.4;refGene_gn=CDON;refGene_tx=XM_006936782.4;refGene_gn=CDON;refGene_tx=XM_006936783.3;refGene_gn=CDON;refGene_tx=XM_011286500.3;refGene_gn=CDON;refGene_tx=XM_019811527.1;refGene_gn=CDON;refGene_tx=XM_023239281.1
D1 23057796 23057943 refGene_gn=CDON;refGene_tx=XM_003992553.4;refGene_gn=CDON;refGene_tx=XM_006936782.4;refGene_gn=CDON;refGene_tx=XM_006936783.3;refGene_gn=CDON;refGene_tx=XM_011286500.3;refGene_gn=CDON;refGene_tx=XM_019811527.1;refGene_gn=CDON;refGene_tx=XM_023239281.1
D1 23059439 23059712 refGene_gn=CDON;refGene_tx=XM_003992553.4;refGene_gn=CDON;refGene_tx=XM_006936782.4;refGene_gn=CDON;refGene_tx=XM_006936783.3;refGene_gn=CDON;refGene_tx=XM_011286500.3;refGene_gn=CDON;refGene_tx=XM_019811527.1;refGene_gn=CDON;refGene_tx=XM_023239281.1
D1 23062119 23062198 refGene_gn=CDON;refGene_tx=XM_003992553.4;refGene_gn=CDON;refGene_tx=XM_006936782.4;refGene_gn=CDON;refGene_tx=XM_006936783.3;refGene_gn=CDON;refGene_tx=XM_011286500.3;refGene_gn=CDON;refGene_tx=XM_019811527.1;refGene_gn=CDON;refGene_tx=XM_023239281.1
D1 23229619 23229859 refGene_gn=RPUSD4;refGene_tx=XM_003992554.4
D1 23230377 23230475 refGene_gn=RPUSD4;refGene_tx=XM_003992554.4
D1 23231633 23231778 refGene_gn=RPUSD4;refGene_tx=XM_003992554.4
D1 23231853 23231947 refGene_gn=RPUSD4;refGene_tx=XM_003992554.4
D1 23235943 23236145 refGene_gn=RPUSD4;refGene_tx=XM_003992554.4
D1 23238163 23238329 refGene_gn=RPUSD4;refGene_tx=XM_003992554.4
D1 23238690 23238879 refGene_gn=RPUSD4;refGene_tx=XM_003992554.4
D1 23262996 23263082 refGene_gn=FAM118B;refGene_tx=XM_011286499.3
D1 23267397 23267600 refGene_gn=FAM118B;refGene_tx=XM_011286499.3
D1 23267602 23267649 refGene_gn=FAM118B;refGene_tx=XM_011286499.3
D1 23276690 23276918 refGene_gn=FAM118B;refGene_tx=XM_011286499.3
D1 23279034 23279163 refGene_gn=FAM118B;refGene_tx=XM_011286499.3
D1 23280480 23280766 refGene_gn=FAM118B;refGene_tx=XM_011286499.3
D1 23286893 23286950 refGene_gn=FAM118B;refGene_tx=XM_011286499.3
D1 23287213 23287227 refGene_gn=FAM118B;refGene_tx=XM_011286499.3
D1 23288976 23289105 refGene_gn=SRPRA;refGene_tx=XM_003992555.5
D1 23289271 23289370 refGene_gn=SRPRA;refGene_tx=XM_003992555.5
D1 23289457 23289621 refGene_gn=SRPRA;refGene_tx=XM_003992555.5
D1 23290038 23290252 refGene_gn=SRPRA;refGene_tx=XM_003992555.5
D1 23290338 23290511 refGene_gn=SRPRA;refGene_tx=XM_003992555.5
D1 23290794 23290881 refGene_gn=SRPRA;refGene_tx=XM_003992555.5
D1 23291014 23291133 refGene_gn=SRPRA;refGene_tx=XM_003992555.5

D1 23291238 23291330 refGene_gn=SRPRA;refGene_tx=XM_003992555.5
D1 23291554 23291708 refGene_gn=SRPRA;refGene_tx=XM_003992555.5
D1 23291842 23292002 refGene_gn=SRPRA;refGene_tx=XM_003992555.5
D1 23292244 23292408 refGene_gn=SRPRA;refGene_tx=XM_003992555.5
D1 23292588 23292752 refGene_gn=SRPRA;refGene_tx=XM_003992555.5
D1 23293022 23293106 refGene_gn=SRPRA;refGene_tx=XM_003992555.5
D1 23293737 23293854 refGene_gn=SRPRA;refGene_tx=XM_003992555.5
D1 23294257 23294342
refGene_gn=FOXRED1;refGene_tx=XM_003992558.4;refGene_gn=FOXRED1;refGene_tx=XM_019811532.1
D1 23296271 23296492
refGene_gn=FOXRED1;refGene_tx=XM_003992558.4;refGene_gn=FOXRED1;refGene_tx=XM_019811532.1
D1 23297191 23297302
refGene_gn=FOXRED1;refGene_tx=XM_003992558.4;refGene_gn=FOXRED1;refGene_tx=XM_019811532.1
D1 23297573 23297692
refGene_gn=FOXRED1;refGene_tx=XM_003992558.4;refGene_gn=FOXRED1;refGene_tx=XM_019811532.1
D1 23298838 23298933
refGene_gn=FOXRED1;refGene_tx=XM_003992558.4;refGene_gn=FOXRED1;refGene_tx=XM_019811532.1
D1 23299212 23299314
refGene_gn=FOXRED1;refGene_tx=XM_003992558.4;refGene_gn=FOXRED1;refGene_tx=XM_019811532.1
D1 23299668 23299745
refGene_gn=FOXRED1;refGene_tx=XM_003992558.4;refGene_gn=FOXRED1;refGene_tx=XM_019811532.1
D1 23299930 23300091
refGene_gn=FOXRED1;refGene_tx=XM_003992558.4;refGene_gn=FOXRED1;refGene_tx=XM_019811532.1
D1 23300201 23300331
refGene_gn=FOXRED1;refGene_tx=XM_003992558.4;refGene_gn=FOXRED1;refGene_tx=XM_019811532.1
D1 23300552 23300657
refGene_gn=FOXRED1;refGene_tx=XM_003992558.4;refGene_gn=FOXRED1;refGene_tx=XM_019811532.1
D1 23300880 23301135
refGene_gn=FOXRED1;refGene_tx=XM_003992558.4;refGene_gn=FOXRED1;refGene_tx=XM_019811532.1
D1 23317635 23317699
refGene_gn=TIRAP;refGene_tx=XM_011286494.3;refGene_gn=TIRAP;refGene_tx=XM_011286496.2;refGene_gn=TIRAP;refGene_tx=XM_019811537.1;refGene_gn=TIRAP;refGene_tx=XM_019811538.2;refGene_gn=TIRAP;refGene_tx=XM_023239283.1;refGene_gn=TIRAP;refGene_tx=XM_023239284.1;refGene_gn=TIRAP;refGene_tx=XM_023239285.1;refGene_gn=TIRAP;refGene_tx=XM_023239286.1;refGene_gn=TIRAP;refGene_tx=XM_023239287.1;refGene_gn=TIRAP;refGene_tx=XM_023239288.1
D1 23319365 23319967
refGene_gn=TIRAP;refGene_tx=XM_011286494.3;refGene_gn=TIRAP;refGene_tx

=XM_011286496.2;refGene_gn=TIRAP;refGene_tx=XM_019811537.1;refGene_gn=TIRAP;refGene_tx=XM_019811538.2;refGene_gn=TIRAP;refGene_tx=XM_023239283.1;refGene_gn=TIRAP;refGene_tx=XM_023239284.1;refGene_gn=TIRAP;refGene_tx=XM_023239285.1;refGene_gn=TIRAP;refGene_tx=XM_023239286.1;refGene_gn=TIRAP;refGene_tx=XM_023239287.1;refGene_gn=TIRAP;refGene_tx=XM_023239288.1
D1 23320528 23320603
refGene_gn=TIRAP;refGene_tx=XM_011286494.3;refGene_gn=TIRAP;refGene_tx=XM_011286496.2;refGene_gn=TIRAP;refGene_tx=XM_019811537.1;refGene_gn=TIRAP;refGene_tx=XM_019811538.2;refGene_gn=TIRAP;refGene_tx=XM_023239283.1;refGene_gn=TIRAP;refGene_tx=XM_023239284.1;refGene_gn=TIRAP;refGene_tx=XM_023239285.1;refGene_gn=TIRAP;refGene_tx=XM_023239286.1;refGene_gn=TIRAP;refGene_tx=XM_023239287.1;refGene_gn=TIRAP;refGene_tx=XM_023239288.1
D1 23328808 23329009
refGene_gn=DCPS;refGene_tx=XM_003992560.3;refGene_gn=DCPS;refGene_tx=XM_019811536.2;refGene_gn=DCPS;refGene_tx=XM_023239282.1
D1 23330378 23330553
refGene_gn=DCPS;refGene_tx=XM_003992560.3;refGene_gn=DCPS;refGene_tx=XM_019811536.2;refGene_gn=DCPS;refGene_tx=XM_023239282.1
D1 23362115 23362261
refGene_gn=DCPS;refGene_tx=XM_003992560.3;refGene_gn=DCPS;refGene_tx=XM_019811536.2;refGene_gn=DCPS;refGene_tx=XM_023239282.1
D1 23367438 23367474 refGene_gn=DCPS;refGene_tx=XM_023239282.1
D1 23368216 23368330
refGene_gn=DCPS;refGene_tx=XM_003992560.3;refGene_gn=DCPS;refGene_tx=XM_023239282.1
D1 23372781 23372892
refGene_gn=DCPS;refGene_tx=XM_003992560.3;refGene_gn=DCPS;refGene_tx=XM_019811536.2;refGene_gn=DCPS;refGene_tx=XM_023239282.1
D1 23374888 23375155
refGene_gn=DCPS;refGene_tx=XM_003992560.3;refGene_gn=DCPS;refGene_tx=XM_019811536.2;refGene_gn=DCPS;refGene_tx=XM_023239282.1
D1 23391758 23391803 refGene_gn=ST3GAL4;refGene_tx=XM_023239289.1
D1 23393033 23393084 refGene_gn=ST3GAL4;refGene_tx=XM_019811534.2
D1 23406949 23406958
refGene_gn=ST3GAL4;refGene_tx=XM_003992561.4;refGene_gn=ST3GAL4;refGene_tx=XM_019811533.2
D1 23407888 23407964
refGene_gn=ST3GAL4;refGene_tx=XM_003992561.4;refGene_gn=ST3GAL4;refGene_tx=XM_019811533.2;refGene_gn=ST3GAL4;refGene_tx=XM_019811534.2;refGene_gn=ST3GAL4;refGene_tx=XM_023239289.1;refGene_gn=ST3GAL4;refGene_tx=XM_023239290.1;refGene_gn=ST3GAL4;refGene_tx=XM_023239291.1
D1 23408269 23408354
refGene_gn=ST3GAL4;refGene_tx=XM_003992561.4;refGene_gn=ST3GAL4;refGene_tx=XM_019811533.2;refGene_gn=ST3GAL4;refGene_tx=XM_019811534.2;refGene_gn=ST3GAL4;refGene_tx=XM_023239289.1;refGene_gn=ST3GAL4;refGene_tx=XM_023239290.1;refGene_gn=ST3GAL4;refGene_tx=XM_023239291.1
D1 23408768 23408849
refGene_gn=ST3GAL4;refGene_tx=XM_003992561.4;refGene_gn=ST3GAL4;refGene_gen

e_tx=XM_019811533.2;refGene_gn=ST3GAL4;refGene_tx=XM_019811534.2;refGene_gn=ST3GAL4;refGene_tx=XM_023239289.1;refGene_gn=ST3GAL4;refGene_tx=XM_023239290.1;refGene_gn=ST3GAL4;refGene_tx=XM_023239291.1
D1 23409161 23409259
refGene_gn=ST3GAL4;refGene_tx=XM_003992561.4;refGene_gn=ST3GAL4;refGene_tx=XM_019811533.2;refGene_gn=ST3GAL4;refGene_tx=XM_019811534.2;refGene_gn=ST3GAL4;refGene_tx=XM_023239289.1;refGene_gn=ST3GAL4;refGene_tx=XM_023239290.1;refGene_gn=ST3GAL4;refGene_tx=XM_023239291.1
D1 23409482 23409543
refGene_gn=ST3GAL4;refGene_tx=XM_003992561.4;refGene_gn=ST3GAL4;refGene_tx=XM_019811533.2;refGene_gn=ST3GAL4;refGene_tx=XM_019811534.2;refGene_gn=ST3GAL4;refGene_tx=XM_023239289.1;refGene_gn=ST3GAL4;refGene_tx=XM_023239290.1;refGene_gn=ST3GAL4;refGene_tx=XM_023239291.1
D1 23410094 23410190
refGene_gn=ST3GAL4;refGene_tx=XM_003992561.4;refGene_gn=ST3GAL4;refGene_tx=XM_019811533.2;refGene_gn=ST3GAL4;refGene_tx=XM_019811534.2;refGene_gn=ST3GAL4;refGene_tx=XM_023239289.1;refGene_gn=ST3GAL4;refGene_tx=XM_023239290.1;refGene_gn=ST3GAL4;refGene_tx=XM_023239291.1
D1 23410286 23410476
refGene_gn=ST3GAL4;refGene_tx=XM_003992561.4;refGene_gn=ST3GAL4;refGene_tx=XM_019811533.2;refGene_gn=ST3GAL4;refGene_tx=XM_019811534.2;refGene_gn=ST3GAL4;refGene_tx=XM_023239289.1;refGene_gn=ST3GAL4;refGene_tx=XM_023239290.1;refGene_gn=ST3GAL4;refGene_tx=XM_023239291.1
D1 23411201 23411345
refGene_gn=ST3GAL4;refGene_tx=XM_003992561.4;refGene_gn=ST3GAL4;refGene_tx=XM_019811533.2;refGene_gn=ST3GAL4;refGene_tx=XM_019811534.2;refGene_gn=ST3GAL4;refGene_tx=XM_023239289.1;refGene_gn=ST3GAL4;refGene_tx=XM_023239290.1;refGene_gn=ST3GAL4;refGene_tx=XM_023239291.1
D1 23414665 23414809
refGene_gn=ST3GAL4;refGene_tx=XM_003992561.4;refGene_gn=ST3GAL4;refGene_tx=XM_019811533.2;refGene_gn=ST3GAL4;refGene_tx=XM_019811534.2;refGene_gn=ST3GAL4;refGene_tx=XM_023239289.1;refGene_gn=ST3GAL4;refGene_tx=XM_023239290.1;refGene_gn=ST3GAL4;refGene_tx=XM_023239291.1
D1 23415116 23415203
refGene_gn=ST3GAL4;refGene_tx=XM_003992561.4;refGene_gn=ST3GAL4;refGene_tx=XM_019811533.2;refGene_gn=ST3GAL4;refGene_tx=XM_019811534.2;refGene_gn=ST3GAL4;refGene_tx=XM_023239289.1;refGene_gn=ST3GAL4;refGene_tx=XM_023239290.1;refGene_gn=ST3GAL4;refGene_tx=XM_023239291.1
D1 23427603 23428047
refGene_gn=KIRREL3;refGene_tx=XM_023239292.1;refGene_gn=KIRREL3;refGene_tx=XM_023239293.1;refGene_gn=KIRREL3;refGene_tx=XM_023239294.1;refGene_gn=KIRREL3;refGene_tx=XM_023239295.1;refGene_gn=KIRREL3;refGene_tx=XM_023239296.1
D1 23428622 23428709
refGene_gn=KIRREL3;refGene_tx=XM_023239292.1;refGene_gn=KIRREL3;refGene_tx=XM_023239293.1;refGene_gn=KIRREL3;refGene_tx=XM_023239294.1;refGene_gn=KIRREL3;refGene_tx=XM_023239295.1;refGene_gn=KIRREL3;refGene_tx=XM_023239296.1
D1 23432089 23432199
refGene_gn=KIRREL3;refGene_tx=XM_023239292.1;refGene_gn=KIRREL3;refGen

e_tx=XM_023239293.1;refGene_gn=KIRREL3;refGene_tx=XM_023239294.1;refGene_gn=KIRREL3;refGene_tx=XM_023239295.1;refGene_gn=KIRREL3;refGene_tx=XM_023239296.1
D1 23433654 23433837
refGene_gn=KIRREL3;refGene_tx=XM_023239292.1;refGene_gn=KIRREL3;refGene_tx=XM_023239293.1;refGene_gn=KIRREL3;refGene_tx=XM_023239294.1;refGene_gn=KIRREL3;refGene_tx=XM_023239295.1;refGene_gn=KIRREL3;refGene_tx=XM_023239296.1
D1 23437325 23437361
refGene_gn=KIRREL3;refGene_tx=XM_023239292.1;refGene_gn=KIRREL3;refGene_tx=XM_023239293.1;refGene_gn=KIRREL3;refGene_tx=XM_023239295.1
D1 23438919 23439118
refGene_gn=KIRREL3;refGene_tx=XM_023239292.1;refGene_gn=KIRREL3;refGene_tx=XM_023239293.1;refGene_gn=KIRREL3;refGene_tx=XM_023239294.1;refGene_gn=KIRREL3;refGene_tx=XM_023239295.1;refGene_gn=KIRREL3;refGene_tx=XM_023239296.1
D1 23441217 23441318
refGene_gn=KIRREL3;refGene_tx=XM_023239292.1;refGene_gn=KIRREL3;refGene_tx=XM_023239293.1;refGene_gn=KIRREL3;refGene_tx=XM_023239294.1;refGene_gn=KIRREL3;refGene_tx=XM_023239295.1;refGene_gn=KIRREL3;refGene_tx=XM_023239296.1
D1 23445116 23445243
refGene_gn=KIRREL3;refGene_tx=XM_023239292.1;refGene_gn=KIRREL3;refGene_tx=XM_023239293.1;refGene_gn=KIRREL3;refGene_tx=XM_023239294.1;refGene_gn=KIRREL3;refGene_tx=XM_023239295.1;refGene_gn=KIRREL3;refGene_tx=XM_023239296.1
D1 23446730 23446858
refGene_gn=KIRREL3;refGene_tx=XM_023239292.1;refGene_gn=KIRREL3;refGene_tx=XM_023239293.1;refGene_gn=KIRREL3;refGene_tx=XM_023239294.1;refGene_gn=KIRREL3;refGene_tx=XM_023239295.1;refGene_gn=KIRREL3;refGene_tx=XM_023239296.1
D1 23448178 23448327
refGene_gn=KIRREL3;refGene_tx=XM_023239292.1;refGene_gn=KIRREL3;refGene_tx=XM_023239293.1;refGene_gn=KIRREL3;refGene_tx=XM_023239294.1;refGene_gn=KIRREL3;refGene_tx=XM_023239295.1;refGene_gn=KIRREL3;refGene_tx=XM_023239296.1
D1 23453949 23454055
refGene_gn=KIRREL3;refGene_tx=XM_023239292.1;refGene_gn=KIRREL3;refGene_tx=XM_023239293.1;refGene_gn=KIRREL3;refGene_tx=XM_023239294.1;refGene_gn=KIRREL3;refGene_tx=XM_023239295.1;refGene_gn=KIRREL3;refGene_tx=XM_023239296.1
D1 23460091 23460242
refGene_gn=KIRREL3;refGene_tx=XM_023239292.1;refGene_gn=KIRREL3;refGene_tx=XM_023239293.1;refGene_gn=KIRREL3;refGene_tx=XM_023239294.1;refGene_gn=KIRREL3;refGene_tx=XM_023239295.1;refGene_gn=KIRREL3;refGene_tx=XM_023239296.1
D1 23469847 23470005
refGene_gn=KIRREL3;refGene_tx=XM_023239292.1;refGene_gn=KIRREL3;refGene_tx=XM_023239293.1;refGene_gn=KIRREL3;refGene_tx=XM_023239294.1;refGene_gn=KIRREL3;refGene_tx=XM_023239295.1;refGene_gn=KIRREL3;refGene_tx=

XM_023239296.1
D1 23517079 23517229
refGene_gn=KIRREL3;refGene_tx=XM_023239292.1;refGene_gn=KIRREL3;refGene_tx=XM_023239293.1;refGene_gn=KIRREL3;refGene_tx=XM_023239294.1;refGene_gn=KIRREL3;refGene_tx=XM_023239295.1;refGene_gn=KIRREL3;refGene_tx=XM_023239296.1
D1 23521859 23522009
refGene_gn=KIRREL3;refGene_tx=XM_023239292.1;refGene_gn=KIRREL3;refGene_tx=XM_023239293.1;refGene_gn=KIRREL3;refGene_tx=XM_023239294.1;refGene_gn=KIRREL3;refGene_tx=XM_023239295.1;refGene_gn=KIRREL3;refGene_tx=XM_023239296.1
D1 23553450 23553528
refGene_gn=KIRREL3;refGene_tx=XM_023239292.1;refGene_gn=KIRREL3;refGene_tx=XM_023239293.1;refGene_gn=KIRREL3;refGene_tx=XM_023239294.1;refGene_gn=KIRREL3;refGene_tx=XM_023239295.1;refGene_gn=KIRREL3;refGene_tx=XM_023239296.1
D1 23977594 23977649
refGene_gn=KIRREL3;refGene_tx=XM_023239292.1;refGene_gn=KIRREL3;refGene_tx=XM_023239293.1;refGene_gn=KIRREL3;refGene_tx=XM_023239294.1;refGene_gn=KIRREL3;refGene_tx=XM_023239295.1;refGene_gn=KIRREL3;refGene_tx=XM_023239296.1
D1 25426482 25426698
refGene_gn=ETS1;refGene_tx=XM_011286487.3;refGene_gn=ETS1;refGene_tx=XM_011286488.3;refGene_gn=ETS1;refGene_tx=XM_019811545.2;refGene_gn=ETS1;refGene_tx=XM_023239297.1;refGene_gn=ETS1;refGene_tx=XM_023239298.1;refGene_gn=ETS1;refGene_tx=XM_023239299.1
D1 25427648 25427767
refGene_gn=ETS1;refGene_tx=XM_011286487.3;refGene_gn=ETS1;refGene_tx=XM_011286488.3;refGene_gn=ETS1;refGene_tx=XM_019811545.2;refGene_gn=ETS1;refGene_tx=XM_023239297.1;refGene_gn=ETS1;refGene_tx=XM_023239298.1;refGene_gn=ETS1;refGene_tx=XM_023239299.1
D1 25445595 25445856
refGene_gn=ETS1;refGene_tx=XM_011286487.3;refGene_gn=ETS1;refGene_tx=XM_011286488.3;refGene_gn=ETS1;refGene_tx=XM_023239297.1;refGene_gn=ETS1;refGene_tx=XM_023239298.1
D1 25450226 25450475
refGene_gn=ETS1;refGene_tx=XM_011286487.3;refGene_gn=ETS1;refGene_tx=XM_011286488.3;refGene_gn=ETS1;refGene_tx=XM_019811545.2;refGene_gn=ETS1;refGene_tx=XM_023239297.1;refGene_gn=ETS1;refGene_tx=XM_023239298.1;refGene_gn=ETS1;refGene_tx=XM_023239299.1
D1 25451351 25451429
refGene_gn=ETS1;refGene_tx=XM_011286487.3;refGene_gn=ETS1;refGene_tx=XM_011286488.3;refGene_gn=ETS1;refGene_tx=XM_019811545.2;refGene_gn=ETS1;refGene_tx=XM_023239297.1;refGene_gn=ETS1;refGene_tx=XM_023239298.1;refGene_gn=ETS1;refGene_tx=XM_023239299.1
D1 25454195 25454396
refGene_gn=ETS1;refGene_tx=XM_011286487.3;refGene_gn=ETS1;refGene_tx=XM_011286488.3;refGene_gn=ETS1;refGene_tx=XM_019811545.2;refGene_gn=ETS1;refGene_tx=XM_023239297.1;refGene_gn=ETS1;refGene_tx=XM_023239298.1;refGene_gn=ETS1;refGene_tx=XM_023239299.1

D1 25455919 25456039
refGene_gn=ETS1;refGene_tx=XM_011286487.3;refGene_gn=ETS1;refGene_tx=XM_011286488.3;refGene_gn=ETS1;refGene_tx=XM_019811545.2;refGene_gn=ETS1;refGene_tx=XM_023239297.1;refGene_gn=ETS1;refGene_tx=XM_023239298.1;refGene_gn=ETS1;refGene_tx=XM_023239299.1
D1 25464368 25464381 refGene_gn=ETS1;refGene_tx=XM_023239298.1
D1 25490194 25490276
refGene_gn=ETS1;refGene_tx=XM_011286488.3;refGene_gn=ETS1;refGene_tx=XM_019811545.2
D1 25524469 25524614
refGene_gn=ETS1;refGene_tx=XM_011286487.3;refGene_gn=ETS1;refGene_tx=XM_023239297.1;refGene_gn=ETS1;refGene_tx=XM_023239299.1
D1 25541192 25541261
refGene_gn=ETS1;refGene_tx=XM_011286487.3;refGene_gn=ETS1;refGene_tx=XM_023239297.1;refGene_gn=ETS1;refGene_tx=XM_023239299.1
D1 25648419 25648485
refGene_gn=FLI1;refGene_tx=XM_019811546.2;refGene_gn=FLI1;refGene_tx=XM_019811548.2
D1 25655931 25655990
refGene_gn=FLI1;refGene_tx=XM_011286485.3;refGene_gn=FLI1;refGene_tx=XM_011286486.3
D1 25720981 25721193
refGene_gn=FLI1;refGene_tx=XM_011286485.3;refGene_gn=FLI1;refGene_tx=XM_019811546.2;refGene_gn=FLI1;refGene_tx=XM_019811548.2;refGene_gn=FLI1;refGene_tx=XM_019811549.2
D1 25731811 25731966
refGene_gn=FLI1;refGene_tx=XM_011286485.3;refGene_gn=FLI1;refGene_tx=XM_011286486.3;refGene_gn=FLI1;refGene_tx=XM_019811546.2;refGene_gn=FLI1;refGene_tx=XM_019811548.2;refGene_gn=FLI1;refGene_tx=XM_019811549.2
D1 25736301 25736505
refGene_gn=FLI1;refGene_tx=XM_011286485.3;refGene_gn=FLI1;refGene_tx=XM_011286486.3;refGene_gn=FLI1;refGene_tx=XM_019811546.2;refGene_gn=FLI1;refGene_tx=XM_019811548.2;refGene_gn=FLI1;refGene_tx=XM_019811549.2
D1 25745576 25745642
refGene_gn=FLI1;refGene_tx=XM_011286485.3;refGene_gn=FLI1;refGene_tx=XM_011286486.3;refGene_gn=FLI1;refGene_tx=XM_019811546.2;refGene_gn=FLI1;refGene_tx=XM_019811548.2;refGene_gn=FLI1;refGene_tx=XM_019811549.2
D1 25769602 25769668
refGene_gn=FLI1;refGene_tx=XM_011286485.3;refGene_gn=FLI1;refGene_tx=XM_011286486.3;refGene_gn=FLI1;refGene_tx=XM_019811546.2;refGene_gn=FLI1;refGene_tx=XM_019811548.2;refGene_gn=FLI1;refGene_tx=XM_019811549.2
D1 25771348 25771408
refGene_gn=FLI1;refGene_tx=XM_011286485.3;refGene_gn=FLI1;refGene_tx=XM_011286486.3;refGene_gn=FLI1;refGene_tx=XM_019811546.2;refGene_gn=FLI1;refGene_tx=XM_019811548.2;refGene_gn=FLI1;refGene_tx=XM_019811549.2
D1 25773315 25773363
refGene_gn=FLI1;refGene_tx=XM_011286485.3;refGene_gn=FLI1;refGene_tx=XM_011286486.3;refGene_gn=FLI1;refGene_tx=XM_019811546.2;refGene_gn=FLI1;refGene_tx=XM_019811548.2;refGene_gn=FLI1;refGene_tx=XM_019811549.2
D1 25774623 25775153

refGene_gn=FLI1;refGene_tx=XM_011286485.3;refGene_gn=FLI1;refGene_tx=XM_011286486.3;refGene_gn=FLI1;refGene_tx=XM_019811546.2;refGene_gn=FLI1;refGene_tx=XM_019811548.2;refGene_gn=FLI1;refGene_tx=XM_019811549.2
D1 25802480 25803599
refGene_gn=KCNJ1;refGene_tx=XM_019811550.1;refGene_gn=KCNJ1;refGene_tx=XM_019811551.1;refGene_gn=KCNJ1;refGene_tx=XM_019811552.1
D1 25874379 25875316
refGene_gn=KCNJ5;refGene_tx=XM_006936792.3;refGene_gn=KCNJ5;refGene_tx=XM_019811553.2
D1 25878483 25878806
refGene_gn=KCNJ5;refGene_tx=XM_006936792.3;refGene_gn=KCNJ5;refGene_tx=XM_019811553.2
D1 25902450 25902585
refGene_gn=LOC109492330;refGene_tx=XM_023239699.1
D1 25903005 25903364
refGene_gn=LOC109492330;refGene_tx=XM_023239699.1
D1 25903903 25904058
refGene_gn=LOC109492330;refGene_tx=XM_023239699.1
D1 25904854 25905141
refGene_gn=LOC109492330;refGene_tx=XM_023239699.1
D1 25931074 25933327
refGene_gn=ARHGAP32;refGene_tx=XM_011286483.3;refGene_gn=ARHGAP32;refGene_tx=XM_019811554.2;refGene_gn=ARHGAP32;refGene_tx=XM_019811555.1;refGene_gn=ARHGAP32;refGene_tx=XM_019811556.2;refGene_gn=ARHGAP32;refGene_tx=XM_019811557.1;refGene_gn=ARHGAP32;refGene_tx=XM_023239300.1
D1 25934863 25935840
refGene_gn=ARHGAP32;refGene_tx=XM_011286483.3;refGene_gn=ARHGAP32;refGene_tx=XM_019811554.2;refGene_gn=ARHGAP32;refGene_tx=XM_019811555.1;refGene_gn=ARHGAP32;refGene_tx=XM_019811556.2;refGene_gn=ARHGAP32;refGene_tx=XM_019811557.1;refGene_gn=ARHGAP32;refGene_tx=XM_023239300.1
D1 25936564 25937443
refGene_gn=ARHGAP32;refGene_tx=XM_011286483.3;refGene_gn=ARHGAP32;refGene_tx=XM_019811554.2;refGene_gn=ARHGAP32;refGene_tx=XM_019811555.1;refGene_gn=ARHGAP32;refGene_tx=XM_019811556.2;refGene_gn=ARHGAP32;refGene_tx=XM_019811557.1;refGene_gn=ARHGAP32;refGene_tx=XM_023239300.1
D1 25939767 25939839
refGene_gn=ARHGAP32;refGene_tx=XM_011286483.3;refGene_gn=ARHGAP32;refGene_tx=XM_019811554.2;refGene_gn=ARHGAP32;refGene_tx=XM_019811555.1;refGene_gn=ARHGAP32;refGene_tx=XM_019811556.2;refGene_gn=ARHGAP32;refGene_tx=XM_019811557.1;refGene_gn=ARHGAP32;refGene_tx=XM_023239300.1
D1 25942498 25942644
refGene_gn=ARHGAP32;refGene_tx=XM_011286483.3;refGene_gn=ARHGAP32;refGene_tx=XM_019811554.2;refGene_gn=ARHGAP32;refGene_tx=XM_019811555.1;refGene_gn=ARHGAP32;refGene_tx=XM_019811556.2;refGene_gn=ARHGAP32;refGene_tx=XM_019811557.1;refGene_gn=ARHGAP32;refGene_tx=XM_023239300.1
D1 25948961 25949157
refGene_gn=ARHGAP32;refGene_tx=XM_011286483.3;refGene_gn=ARHGAP32;refGene_tx=XM_019811554.2;refGene_gn=ARHGAP32;refGene_tx=XM_019811555.1;refGene_gn=ARHGAP32;refGene_tx=XM_019811556.2;refGene_gn=ARHGAP32;refGene_tx=XM_019811557.1;refGene_gn=ARHGAP32;refGene_tx=XM_023239300.1

D1	25949767	25949913
refGene_gn=ARHGAP32;refGene_tx=XM_011286483.3;refGene_gn=ARHGAP32;refGene_tx=XM_019811554.2;refGene_gn=ARHGAP32;refGene_tx=XM_019811555.1;refGene_gn=ARHGAP32;refGene_tx=XM_019811556.2;refGene_gn=ARHGAP32;refGene_tx=XM_019811557.1;refGene_gn=ARHGAP32;refGene_tx=XM_023239300.1
D1	25950176	25950284
refGene_gn=ARHGAP32;refGene_tx=XM_011286483.3;refGene_gn=ARHGAP32;refGene_tx=XM_019811554.2;refGene_gn=ARHGAP32;refGene_tx=XM_019811555.1;refGene_gn=ARHGAP32;refGene_tx=XM_019811556.2;refGene_gn=ARHGAP32;refGene_tx=XM_019811557.1;refGene_gn=ARHGAP32;refGene_tx=XM_023239300.1
D1	25958379	25958462
refGene_gn=ARHGAP32;refGene_tx=XM_011286483.3;refGene_gn=ARHGAP32;refGene_tx=XM_019811554.2;refGene_gn=ARHGAP32;refGene_tx=XM_019811555.1;refGene_gn=ARHGAP32;refGene_tx=XM_019811556.2;refGene_gn=ARHGAP32;refGene_tx=XM_019811557.1;refGene_gn=ARHGAP32;refGene_tx=XM_023239300.1
D1	25958882	25959027
refGene_gn=ARHGAP32;refGene_tx=XM_011286483.3;refGene_gn=ARHGAP32;refGene_tx=XM_019811554.2;refGene_gn=ARHGAP32;refGene_tx=XM_019811555.1;refGene_gn=ARHGAP32;refGene_tx=XM_019811556.2;refGene_gn=ARHGAP32;refGene_tx=XM_019811557.1;refGene_gn=ARHGAP32;refGene_tx=XM_023239300.1
D1	25960837	25960940
refGene_gn=ARHGAP32;refGene_tx=XM_011286483.3;refGene_gn=ARHGAP32;refGene_tx=XM_019811554.2;refGene_gn=ARHGAP32;refGene_tx=XM_019811555.1;refGene_gn=ARHGAP32;refGene_tx=XM_019811556.2;refGene_gn=ARHGAP32;refGene_tx=XM_019811557.1;refGene_gn=ARHGAP32;refGene_tx=XM_023239300.1
D1	25970058	25970208
refGene_gn=ARHGAP32;refGene_tx=XM_011286483.3;refGene_gn=ARHGAP32;refGene_tx=XM_019811554.2;refGene_gn=ARHGAP32;refGene_tx=XM_019811555.1;refGene_gn=ARHGAP32;refGene_tx=XM_019811556.2;refGene_gn=ARHGAP32;refGene_tx=XM_019811557.1;refGene_gn=ARHGAP32;refGene_tx=XM_023239300.1
D1	26003074	26003156
refGene_gn=ARHGAP32;refGene_tx=XM_019811554.2;refGene_gn=ARHGAP32;refGene_tx=XM_019811555.1;refGene_gn=ARHGAP32;refGene_tx=XM_019811556.2;refGene_gn=ARHGAP32;refGene_tx=XM_019811557.1;refGene_gn=ARHGAP32;refGene_tx=XM_023239300.1
D1	26015591	26015669
refGene_gn=ARHGAP32;refGene_tx=XM_019811554.2;refGene_gn=ARHGAP32;refGene_tx=XM_019811555.1;refGene_gn=ARHGAP32;refGene_tx=XM_019811556.2;refGene_gn=ARHGAP32;refGene_tx=XM_019811557.1;refGene_gn=ARHGAP32;refGene_tx=XM_023239300.1
D1	26017270	26017393
refGene_gn=ARHGAP32;refGene_tx=XM_019811554.2;refGene_gn=ARHGAP32;refGene_tx=XM_019811555.1;refGene_gn=ARHGAP32;refGene_tx=XM_019811556.2;refGene_gn=ARHGAP32;refGene_tx=XM_019811557.1;refGene_gn=ARHGAP32;refGene_tx=XM_023239300.1
D1	26018733	26018826
refGene_gn=ARHGAP32;refGene_tx=XM_019811554.2;refGene_gn=ARHGAP32;refGene_tx=XM_019811555.1;refGene_gn=ARHGAP32;refGene_tx=XM_019811556.2;refGene_gn=ARHGAP32;refGene_tx=XM_019811557.1;refGene_gn=ARHGAP32;refGene_tx=XM_023239300.1

D1 26020975 26021113
refGene_gn=ARHGAP32;refGene_tx=XM_019811554.2;refGene_gn=ARHGAP32;refGene_tx=XM_019811555.1;refGene_gn=ARHGAP32;refGene_tx=XM_019811556.2;refGene_gn=ARHGAP32;refGene_tx=XM_019811557.1;refGene_gn=ARHGAP32;refGene_tx=XM_023239300.1
D1 26043396 26043483
refGene_gn=ARHGAP32;refGene_tx=XM_019811554.2;refGene_gn=ARHGAP32;refGene_tx=XM_019811555.1;refGene_gn=ARHGAP32;refGene_tx=XM_019811556.2;refGene_gn=ARHGAP32;refGene_tx=XM_019811557.1;refGene_gn=ARHGAP32;refGene_tx=XM_023239300.1
D1 26085584 26085669
refGene_gn=ARHGAP32;refGene_tx=XM_019811554.2;refGene_gn=ARHGAP32;refGene_tx=XM_019811555.1;refGene_gn=ARHGAP32;refGene_tx=XM_019811556.2;refGene_gn=ARHGAP32;refGene_tx=XM_019811557.1;refGene_gn=ARHGAP32;refGene_tx=XM_023239300.1
D1 26086045 26086087
refGene_gn=ARHGAP32;refGene_tx=XM_019811554.2;refGene_gn=ARHGAP32;refGene_tx=XM_019811556.2;refGene_gn=ARHGAP32;refGene_tx=XM_023239300.1
D1 26086713 26086805
refGene_gn=ARHGAP32;refGene_tx=XM_019811554.2;refGene_gn=ARHGAP32;refGene_tx=XM_019811555.1;refGene_gn=ARHGAP32;refGene_tx=XM_019811556.2;refGene_gn=ARHGAP32;refGene_tx=XM_019811557.1;refGene_gn=ARHGAP32;refGene_tx=XM_023239300.1
D1 26107028 26107137
refGene_gn=ARHGAP32;refGene_tx=XM_019811554.2;refGene_gn=ARHGAP32;refGene_tx=XM_019811555.1;refGene_gn=ARHGAP32;refGene_tx=XM_019811556.2;refGene_gn=ARHGAP32;refGene_tx=XM_023239300.1
D1 26124274 26124390
refGene_gn=ARHGAP32;refGene_tx=XM_019811554.2;refGene_gn=ARHGAP32;refGene_tx=XM_019811555.1
D1 26212283 26212481
refGene_gn=LOC111556643;refGene_tx=XM_023239700.1
D1 26226955 26227102
refGene_gn=LOC111556643;refGene_tx=XM_023239700.1
D1 26227445 26227682
refGene_gn=LOC111556643;refGene_tx=XM_023239700.1
D1 26315478 26315665 refGene_gn=BARX2;refGene_tx=XM_003992585.5
D1 26375937 26376238 refGene_gn=BARX2;refGene_tx=XM_003992585.5
D1 26380853 26380938 refGene_gn=BARX2;refGene_tx=XM_003992585.5
D1 26390570 26390837 refGene_gn=BARX2;refGene_tx=XM_003992585.5
D1 26682445 26682689
refGene_gn=TMEM45B;refGene_tx=XM_003992571.4;refGene_gn=TMEM45B;refGene_tx=XM_023239301.1
D1 26684642 26684849
refGene_gn=TMEM45B;refGene_tx=XM_003992571.4;refGene_gn=TMEM45B;refGene_tx=XM_023239301.1
D1 26685698 26685883
refGene_gn=TMEM45B;refGene_tx=XM_003992571.4;refGene_gn=TMEM45B;refGene_tx=XM_023239301.1
D1 26686443 26686589

refGene_gn=TMEM45B;refGene_tx=XM_003992571.4;refGene_gn=TMEM45B;refGene_tx=XM_023239301.1
D1 26687625 26687734
refGene_gn=TMEM45B;refGene_tx=XM_003992571.4;refGene_gn=TMEM45B;refGene_tx=XM_023239301.1
D1 26692198 26692324
refGene_gn=NFRKB;refGene_tx=XM_003992572.3;refGene_gn=NFRKB;refGene_tx=XM_006936796.2;refGene_gn=NFRKB;refGene_tx=XM_006936798.2
D1 26692439 26692575
refGene_gn=NFRKB;refGene_tx=XM_003992572.3;refGene_gn=NFRKB;refGene_tx=XM_006936796.2;refGene_gn=NFRKB;refGene_tx=XM_006936798.2
D1 26693388 26693495
refGene_gn=NFRKB;refGene_tx=XM_003992572.3;refGene_gn=NFRKB;refGene_tx=XM_006936796.2;refGene_gn=NFRKB;refGene_tx=XM_006936798.2
D1 26696303 26697071
refGene_gn=NFRKB;refGene_tx=XM_003992572.3;refGene_gn=NFRKB;refGene_tx=XM_006936796.2;refGene_gn=NFRKB;refGene_tx=XM_006936798.2
D1 26699657 26699867
refGene_gn=NFRKB;refGene_tx=XM_003992572.3;refGene_gn=NFRKB;refGene_tx=XM_006936796.2;refGene_gn=NFRKB;refGene_tx=XM_006936798.2
D1 26700486 26700757
refGene_gn=NFRKB;refGene_tx=XM_003992572.3;refGene_gn=NFRKB;refGene_tx=XM_006936796.2;refGene_gn=NFRKB;refGene_tx=XM_006936798.2
D1 26700867 26701088
refGene_gn=NFRKB;refGene_tx=XM_003992572.3;refGene_gn=NFRKB;refGene_tx=XM_006936796.2;refGene_gn=NFRKB;refGene_tx=XM_006936798.2
D1 26701223 26701303
refGene_gn=NFRKB;refGene_tx=XM_003992572.3;refGene_gn=NFRKB;refGene_tx=XM_006936796.2;refGene_gn=NFRKB;refGene_tx=XM_006936798.2
D1 26701515 26701639
refGene_gn=NFRKB;refGene_tx=XM_003992572.3;refGene_gn=NFRKB;refGene_tx=XM_006936796.2;refGene_gn=NFRKB;refGene_tx=XM_006936798.2
D1 26702008 26702115
refGene_gn=NFRKB;refGene_tx=XM_003992572.3;refGene_gn=NFRKB;refGene_tx=XM_006936796.2;refGene_gn=NFRKB;refGene_tx=XM_006936798.2
D1 26703882 26704057
refGene_gn=NFRKB;refGene_tx=XM_003992572.3;refGene_gn=NFRKB;refGene_tx=XM_006936796.2;refGene_gn=NFRKB;refGene_tx=XM_006936798.2
D1 26704475 26704536
refGene_gn=NFRKB;refGene_tx=XM_003992572.3;refGene_gn=NFRKB;refGene_tx=XM_006936796.2;refGene_gn=NFRKB;refGene_tx=XM_006936798.2
D1 26707093 26707140
refGene_gn=NFRKB;refGene_tx=XM_003992572.3;refGene_gn=NFRKB;refGene_tx=XM_006936796.2;refGene_gn=NFRKB;refGene_tx=XM_006936798.2
D1 26707466 26707546
refGene_gn=NFRKB;refGene_tx=XM_003992572.3;refGene_gn=NFRKB;refGene_tx=XM_006936796.2;refGene_gn=NFRKB;refGene_tx=XM_006936798.2
D1 26711924 26711990
refGene_gn=NFRKB;refGene_tx=XM_003992572.3;refGene_gn=NFRKB;refGene_tx=XM_006936796.2;refGene_gn=NFRKB;refGene_tx=XM_006936798.2

D1 26712239 26712366
refGene_gn=NFRKB;refGene_tx=XM_003992572.3;refGene_gn=NFRKB;refGene_tx=XM_006936796.2;refGene_gn=NFRKB;refGene_tx=XM_006936798.2
D1 26712585 26712694
refGene_gn=NFRKB;refGene_tx=XM_003992572.3;refGene_gn=NFRKB;refGene_tx=XM_006936796.2;refGene_gn=NFRKB;refGene_tx=XM_006936798.2
D1 26712994 26713175
refGene_gn=NFRKB;refGene_tx=XM_003992572.3;refGene_gn=NFRKB;refGene_tx=XM_006936796.2;refGene_gn=NFRKB;refGene_tx=XM_006936798.2
D1 26713578 26713663
refGene_gn=NFRKB;refGene_tx=XM_003992572.3;refGene_gn=NFRKB;refGene_tx=XM_006936796.2;refGene_gn=NFRKB;refGene_tx=XM_006936798.2
D1 26714442 26714516
refGene_gn=NFRKB;refGene_tx=XM_003992572.3;refGene_gn=NFRKB;refGene_tx=XM_006936796.2;refGene_gn=NFRKB;refGene_tx=XM_006936798.2
D1 26715145 26715283
refGene_gn=NFRKB;refGene_tx=XM_003992572.3;refGene_gn=NFRKB;refGene_tx=XM_006936796.2;refGene_gn=NFRKB;refGene_tx=XM_006936798.2
D1 26715832 26716007
refGene_gn=NFRKB;refGene_tx=XM_003992572.3;refGene_gn=NFRKB;refGene_tx=XM_006936796.2;refGene_gn=NFRKB;refGene_tx=XM_006936798.2
D1 26716722 26716850
refGene_gn=NFRKB;refGene_tx=XM_003992572.3;refGene_gn=NFRKB;refGene_tx=XM_006936796.2;refGene_gn=NFRKB;refGene_tx=XM_006936798.2
D1 26725960 26726162
refGene_gn=NFRKB;refGene_tx=XM_003992572.3;refGene_gn=NFRKB;refGene_tx=XM_006936796.2;refGene_gn=NFRKB;refGene_tx=XM_006936798.2
D1 26728688 26728823
refGene_gn=NFRKB;refGene_tx=XM_003992572.3;refGene_gn=NFRKB;refGene_tx=XM_006936796.2;refGene_gn=NFRKB;refGene_tx=XM_006936798.2
D1 26735064 26735265 refGene_gn=PRDM10;refGene_tx=XM_019811558.2
D1 26737897 26738001 refGene_gn=PRDM10;refGene_tx=XM_019811558.2
D1 26741897 26742069 refGene_gn=PRDM10;refGene_tx=XM_019811558.2
D1 26745025 26745166 refGene_gn=PRDM10;refGene_tx=XM_019811558.2
D1 26746876 26747047 refGene_gn=PRDM10;refGene_tx=XM_019811558.2
D1 26747435 26747579 refGene_gn=PRDM10;refGene_tx=XM_019811558.2
D1 26748258 26748459 refGene_gn=PRDM10;refGene_tx=XM_019811558.2
D1 26749415 26749526 refGene_gn=PRDM10;refGene_tx=XM_019811558.2
D1 26750450 26750630 refGene_gn=PRDM10;refGene_tx=XM_019811558.2
D1 26756570 26756726 refGene_gn=PRDM10;refGene_tx=XM_019811558.2
D1 26758182 26758530 refGene_gn=PRDM10;refGene_tx=XM_019811558.2
D1 26763031 26763043 refGene_gn=PRDM10;refGene_tx=XM_019811558.2
D1 26764650 26764893 refGene_gn=PRDM10;refGene_tx=XM_019811558.2
D1 26765850 26765980 refGene_gn=PRDM10;refGene_tx=XM_019811558.2
D1 26769308 26769426 refGene_gn=PRDM10;refGene_tx=XM_019811558.2
D1 26771511 26771584 refGene_gn=PRDM10;refGene_tx=XM_019811558.2
D1 26776987 26777191 refGene_gn=PRDM10;refGene_tx=XM_019811558.2
D1 26779454 26779696 refGene_gn=PRDM10;refGene_tx=XM_019811558.2
D1 26781522 26781748 refGene_gn=PRDM10;refGene_tx=XM_019811558.2
D1 26794159 26794219 refGene_gn=PRDM10;refGene_tx=XM_019811558.2

D1	26796227	26796392	refGene_gn=PRDM10;refGene_tx=XM_019811558.2
D1	26798656	26798671	refGene_gn=PRDM10;refGene_tx=XM_019811558.2
D1	26798673	26798730	refGene_gn=PRDM10;refGene_tx=XM_019811558.2
D1	26889151	26889256	
refGene_gn=APLP2;refGene_tx=XM_011286479.3;refGene_gn=APLP2;refGene_tx=XM_011286480.3;refGene_gn=APLP2;refGene_tx=XM_011286481.3;refGene_gn=APLP2;refGene_tx=XM_011286482.3
D1	26938480	26938654	
refGene_gn=APLP2;refGene_tx=XM_011286479.3;refGene_gn=APLP2;refGene_tx=XM_011286480.3;refGene_gn=APLP2;refGene_tx=XM_011286481.3;refGene_gn=APLP2;refGene_tx=XM_011286482.3
D1	26939747	26939871	
refGene_gn=APLP2;refGene_tx=XM_011286479.3;refGene_gn=APLP2;refGene_tx=XM_011286480.3;refGene_gn=APLP2;refGene_tx=XM_011286481.3;refGene_gn=APLP2;refGene_tx=XM_011286482.3
D1	26951742	26951855	
refGene_gn=APLP2;refGene_tx=XM_011286479.3;refGene_gn=APLP2;refGene_tx=XM_011286480.3;refGene_gn=APLP2;refGene_tx=XM_011286481.3;refGene_gn=APLP2;refGene_tx=XM_011286482.3
D1	26953283	26953480	
refGene_gn=APLP2;refGene_tx=XM_011286479.3;refGene_gn=APLP2;refGene_tx=XM_011286480.3;refGene_gn=APLP2;refGene_tx=XM_011286481.3;refGene_gn=APLP2;refGene_tx=XM_011286482.3
D1	26953877	26954086	
refGene_gn=APLP2;refGene_tx=XM_011286479.3;refGene_gn=APLP2;refGene_tx=XM_011286480.3;refGene_gn=APLP2;refGene_tx=XM_011286481.3;refGene_gn=APLP2;refGene_tx=XM_011286482.3
D1	26955482	26955650	
refGene_gn=APLP2;refGene_tx=XM_011286479.3;refGene_gn=APLP2;refGene_tx=XM_011286480.3
D1	26958653	26958784	
refGene_gn=APLP2;refGene_tx=XM_011286479.3;refGene_gn=APLP2;refGene_tx=XM_011286480.3;refGene_gn=APLP2;refGene_tx=XM_011286481.3;refGene_gn=APLP2;refGene_tx=XM_011286482.3
D1	26960148	26960223	
refGene_gn=APLP2;refGene_tx=XM_011286479.3;refGene_gn=APLP2;refGene_tx=XM_011286480.3;refGene_gn=APLP2;refGene_tx=XM_011286481.3;refGene_gn=APLP2;refGene_tx=XM_011286482.3
D1	26961440	26961599	
refGene_gn=APLP2;refGene_tx=XM_011286479.3;refGene_gn=APLP2;refGene_tx=XM_011286480.3;refGene_gn=APLP2;refGene_tx=XM_011286481.3;refGene_gn=APLP2;refGene_tx=XM_011286482.3
D1	26962669	26962798	
refGene_gn=APLP2;refGene_tx=XM_011286479.3;refGene_gn=APLP2;refGene_tx=XM_011286480.3;refGene_gn=APLP2;refGene_tx=XM_011286481.3;refGene_gn=APLP2;refGene_tx=XM_011286482.3
D1	26965411	26965511	
refGene_gn=APLP2;refGene_tx=XM_011286479.3;refGene_gn=APLP2;refGene_tx=XM_011286480.3;refGene_gn=APLP2;refGene_tx=XM_011286481.3;refGene_gn=APLP2;refGene_tx=XM_011286482.3

D1 26967044 26967197
refGene_gn=APLP2;refGene_tx=XM_011286479.3;refGene_gn=APLP2;refGene_tx=XM_011286480.3;refGene_gn=APLP2;refGene_tx=XM_011286481.3;refGene_gn=APLP2;refGene_tx=XM_011286482.3
D1 26968757 26968793
refGene_gn=APLP2;refGene_tx=XM_011286479.3;refGene_gn=APLP2;refGene_tx=XM_011286481.3
D1 26971861 26971947
refGene_gn=APLP2;refGene_tx=XM_011286479.3;refGene_gn=APLP2;refGene_tx=XM_011286480.3;refGene_gn=APLP2;refGene_tx=XM_011286481.3;refGene_gn=APLP2;refGene_tx=XM_011286482.3
D1 26973119 26973194
refGene_gn=APLP2;refGene_tx=XM_011286479.3;refGene_gn=APLP2;refGene_tx=XM_011286480.3;refGene_gn=APLP2;refGene_tx=XM_011286481.3;refGene_gn=APLP2;refGene_tx=XM_011286482.3
D1 26973569 26973725
refGene_gn=APLP2;refGene_tx=XM_011286479.3;refGene_gn=APLP2;refGene_tx=XM_011286480.3;refGene_gn=APLP2;refGene_tx=XM_011286481.3;refGene_gn=APLP2;refGene_tx=XM_011286482.3
D1 26975017 26975119
refGene_gn=APLP2;refGene_tx=XM_011286479.3;refGene_gn=APLP2;refGene_tx=XM_011286480.3;refGene_gn=APLP2;refGene_tx=XM_011286481.3;refGene_gn=APLP2;refGene_tx=XM_011286482.3
D1 26993180 26993261 refGene_gn=ST14;refGene_tx=XM_003992587.5
D1 27016582 27016742
refGene_gn=ST14;refGene_tx=XM_003992587.5;refGene_gn=ST14;refGene_tx=XM_019811561.2
D1 27017088 27017216
refGene_gn=ST14;refGene_tx=XM_003992587.5;refGene_gn=ST14;refGene_tx=XM_019811561.2
D1 27017422 27017493
refGene_gn=ST14;refGene_tx=XM_003992587.5;refGene_gn=ST14;refGene_tx=XM_019811561.2
D1 27018633 27018791
refGene_gn=ST14;refGene_tx=XM_003992587.5;refGene_gn=ST14;refGene_tx=XM_019811561.2
D1 27019008 27019044
refGene_gn=ST14;refGene_tx=XM_003992587.5;refGene_gn=ST14;refGene_tx=XM_019811561.2
D1 27019271 27019512
refGene_gn=ST14;refGene_tx=XM_003992587.5;refGene_gn=ST14;refGene_tx=XM_019811561.2
D1 27021593 27021733
refGene_gn=ST14;refGene_tx=XM_003992587.5;refGene_gn=ST14;refGene_tx=XM_019811561.2
D1 27022094 27022192
refGene_gn=ST14;refGene_tx=XM_003992587.5;refGene_gn=ST14;refGene_tx=XM_019811561.2
D1 27023187 27023297
refGene_gn=ST14;refGene_tx=XM_003992587.5;refGene_gn=ST14;refGene_tx=X

M_019811561.2
D1 27023387 27023518
refGene_gn=ST14;refGene_tx=XM_003992587.5;refGene_gn=ST14;refGene_tx=XM_019811561.2
D1 27024753 27024858
refGene_gn=ST14;refGene_tx=XM_003992587.5;refGene_gn=ST14;refGene_tx=XM_019811561.2
D1 27025183 27025294
refGene_gn=ST14;refGene_tx=XM_003992587.5;refGene_gn=ST14;refGene_tx=XM_019811561.2
D1 27025405 27025519
refGene_gn=ST14;refGene_tx=XM_003992587.5;refGene_gn=ST14;refGene_tx=XM_019811561.2
D1 27025837 27025960
refGene_gn=ST14;refGene_tx=XM_003992587.5;refGene_gn=ST14;refGene_tx=XM_019811561.2
D1 27026770 27026957
refGene_gn=ST14;refGene_tx=XM_003992587.5;refGene_gn=ST14;refGene_tx=XM_019811561.2
D1 27033473 27033748
refGene_gn=ST14;refGene_tx=XM_003992587.5;refGene_gn=ST14;refGene_tx=XM_019811561.2
D1 27034533 27034670
refGene_gn=ST14;refGene_tx=XM_003992587.5;refGene_gn=ST14;refGene_tx=XM_019811561.2
D1 27034773 27034935
refGene_gn=ST14;refGene_tx=XM_003992587.5;refGene_gn=ST14;refGene_tx=XM_019811561.2
D1 27059269 27059552
refGene_gn=ZBTB44;refGene_tx=XM_023239302.1;refGene_gn=ZBTB44;refGene_tx=XM_023239303.1
D1 27061635 27061936
refGene_gn=ZBTB44;refGene_tx=XM_023239302.1;refGene_gn=ZBTB44;refGene_tx=XM_023239303.1
D1 27063247 27063411
refGene_gn=ZBTB44;refGene_tx=XM_023239302.1;refGene_gn=ZBTB44;refGene_tx=XM_023239303.1
D1 27065302 27065387
refGene_gn=ZBTB44;refGene_tx=XM_023239302.1;refGene_gn=ZBTB44;refGene_tx=XM_023239303.1
D1 27082444 27083462
refGene_gn=ZBTB44;refGene_tx=XM_023239302.1;refGene_gn=ZBTB44;refGene_tx=XM_023239303.1
D1 27178225 27178796 refGene_gn=ADAMTS8;refGene_tx=XM_019812925.2
D1 27180930 27181106 refGene_gn=ADAMTS8;refGene_tx=XM_019812925.2
D1 27181227 27181400 refGene_gn=ADAMTS8;refGene_tx=XM_019812925.2
D1 27183808 27183992 refGene_gn=ADAMTS8;refGene_tx=XM_019812925.2
D1 27186306 27186608 refGene_gn=ADAMTS8;refGene_tx=XM_019812925.2
D1 27187584 27187752 refGene_gn=ADAMTS8;refGene_tx=XM_019812925.2
D1 27188420 27188556 refGene_gn=ADAMTS8;refGene_tx=XM_019812925.2

D1 27189361 27189601 refGene_gn=ADAMTS8;refGene_tx=XM_019812925.2
D1 27197575 27198766 refGene_gn=ADAMTS8;refGene_tx=XM_019812925.2
D1 27216796 27217753 refGene_gn=ADAMTS15;refGene_tx=XM_003992579.4
D1 27229645 27229778 refGene_gn=ADAMTS15;refGene_tx=XM_003992579.4
D1 27230291 27230459 refGene_gn=ADAMTS15;refGene_tx=XM_003992579.4
D1 27230692 27230976 refGene_gn=ADAMTS15;refGene_tx=XM_003992579.4
D1 27236527 27236705 refGene_gn=ADAMTS15;refGene_tx=XM_003992579.4
D1 27237746 27237928 refGene_gn=ADAMTS15;refGene_tx=XM_003992579.4
D1 27238033 27238209 refGene_gn=ADAMTS15;refGene_tx=XM_003992579.4
D1 27240202 27240977 refGene_gn=ADAMTS15;refGene_tx=XM_003992579.4
D1 27644372 27644505 refGene_gn=SNX19;refGene_tx=XM_003992580.5
D1 27645419 27645507 refGene_gn=SNX19;refGene_tx=XM_003992580.5
D1 27646609 27646794 refGene_gn=SNX19;refGene_tx=XM_003992580.5
D1 27669562 27669692 refGene_gn=SNX19;refGene_tx=XM_003992580.5
D1 27672205 27672386 refGene_gn=SNX19;refGene_tx=XM_003992580.5
D1 27673112 27673209 refGene_gn=SNX19;refGene_tx=XM_003992580.5
D1 27674454 27674585 refGene_gn=SNX19;refGene_tx=XM_003992580.5
D1 27676499 27676619 refGene_gn=SNX19;refGene_tx=XM_003992580.5
D1 27676735 27676836 refGene_gn=SNX19;refGene_tx=XM_003992580.5
D1 27678083 27678222 refGene_gn=SNX19;refGene_tx=XM_003992580.5
D1 27679961 27681281 refGene_gn=SNX19;refGene_tx=XM_003992580.5
D1 27681282 27681321 refGene_gn=SNX19;refGene_tx=XM_003992580.5
D1 27681323 27681662 refGene_gn=SNX19;refGene_tx=XM_003992580.5
D1 28108013 28108095 refGene_gn=NTM;refGene_tx=XM_006936805.4
D1 28386952 28387010 refGene_gn=NTM;refGene_tx=XM_019811567.1
D1 28649751 28649918
refGene_gn=NTM;refGene_tx=XM_003992582.5;refGene_gn=NTM;refGene_tx=XM_006936805.4;refGene_gn=NTM;refGene_tx=XM_006936806.4;refGene_gn=NTM;refGene_tx=XM_019811567.1;refGene_gn=NTM;refGene_tx=XM_019811568.2
D1 28878841 28879074
refGene_gn=NTM;refGene_tx=XM_003992582.5;refGene_gn=NTM;refGene_tx=XM_006936805.4;refGene_gn=NTM;refGene_tx=XM_006936806.4;refGene_gn=NTM;refGene_tx=XM_019811567.1;refGene_gn=NTM;refGene_tx=XM_019811568.2
D1 28940831 28940957
refGene_gn=NTM;refGene_tx=XM_003992582.5;refGene_gn=NTM;refGene_tx=XM_006936805.4;refGene_gn=NTM;refGene_tx=XM_006936806.4;refGene_gn=NTM;refGene_tx=XM_019811567.1;refGene_gn=NTM;refGene_tx=XM_019811568.2
D1 29026864 29026999
refGene_gn=NTM;refGene_tx=XM_003992582.5;refGene_gn=NTM;refGene_tx=XM_006936805.4;refGene_gn=NTM;refGene_tx=XM_006936806.4;refGene_gn=NTM;refGene_tx=XM_019811567.1;refGene_gn=NTM;refGene_tx=XM_019811568.2
D1 29028921 29029042
refGene_gn=NTM;refGene_tx=XM_003992582.5;refGene_gn=NTM;refGene_tx=XM_006936805.4;refGene_gn=NTM;refGene_tx=XM_006936806.4;refGene_gn=NTM;refGene_tx=XM_019811567.1;refGene_gn=NTM;refGene_tx=XM_019811568.2
D1 29033484 29033636
refGene_gn=NTM;refGene_tx=XM_003992582.5;refGene_gn=NTM;refGene_tx=XM_006936805.4;refGene_gn=NTM;refGene_tx=XM_006936806.4;refGene_gn=NTM;refGene_tx=XM_019811567.1;refGene_gn=NTM;refGene_tx=XM_019811568.2
D1 29036935 29036971

refGene_gn=NTM;refGene_tx=XM_006936805.4;refGene_gn=NTM;refGene_tx=XM_019811567.1;refGene_gn=NTM;refGene_tx=XM_019811568.2
D1 29042282 29042315
refGene_gn=NTM;refGene_tx=XM_003992582.5;refGene_gn=NTM;refGene_tx=XM_006936805.4;refGene_gn=NTM;refGene_tx=XM_019811567.1
D1 29047316 29047417
refGene_gn=NTM;refGene_tx=XM_003992582.5;refGene_gn=NTM;refGene_tx=XM_006936805.4;refGene_gn=NTM;refGene_tx=XM_006936806.4;refGene_gn=NTM;refGene_tx=XM_019811567.1;refGene_gn=NTM;refGene_tx=XM_019811568.2
D1 29121952 29122053
refGene_gn=OPCML;refGene_tx=XM_003992583.5;refGene_gn=OPCML;refGene_tx=XM_019811571.2;refGene_gn=OPCML;refGene_tx=XM_019811572.2;refGene_gn=OPCML;refGene_tx=XM_019811573.2;refGene_gn=OPCML;refGene_tx=XM_023239305.1;refGene_gn=OPCML;refGene_tx=XM_023239306.1
D1 29137524 29137551
refGene_gn=OPCML;refGene_tx=XM_019811571.2;refGene_gn=OPCML;refGene_tx=XM_019811572.2;refGene_gn=OPCML;refGene_tx=XM_023239305.1;refGene_gn=OPCML;refGene_tx=XM_023239306.1
D1 29138444 29138596
refGene_gn=OPCML;refGene_tx=XM_003992583.5;refGene_gn=OPCML;refGene_tx=XM_019811571.2;refGene_gn=OPCML;refGene_tx=XM_019811572.2;refGene_gn=OPCML;refGene_tx=XM_019811573.2;refGene_gn=OPCML;refGene_tx=XM_023239305.1;refGene_gn=OPCML;refGene_tx=XM_023239306.1
D1 29139077 29139198
refGene_gn=OPCML;refGene_tx=XM_003992583.5;refGene_gn=OPCML;refGene_tx=XM_019811571.2;refGene_gn=OPCML;refGene_tx=XM_019811572.2;refGene_gn=OPCML;refGene_tx=XM_019811573.2;refGene_gn=OPCML;refGene_tx=XM_023239305.1;refGene_gn=OPCML;refGene_tx=XM_023239306.1
D1 29139610 29139748
refGene_gn=OPCML;refGene_tx=XM_003992583.5;refGene_gn=OPCML;refGene_tx=XM_019811571.2;refGene_gn=OPCML;refGene_tx=XM_019811572.2;refGene_gn=OPCML;refGene_tx=XM_019811573.2;refGene_gn=OPCML;refGene_tx=XM_023239305.1;refGene_gn=OPCML;refGene_tx=XM_023239306.1
D1 29218577 29218703
refGene_gn=OPCML;refGene_tx=XM_003992583.5;refGene_gn=OPCML;refGene_tx=XM_019811571.2;refGene_gn=OPCML;refGene_tx=XM_019811572.2;refGene_gn=OPCML;refGene_tx=XM_019811573.2;refGene_gn=OPCML;refGene_tx=XM_023239305.1;refGene_gn=OPCML;refGene_tx=XM_023239306.1
D1 29333003 29333236
refGene_gn=OPCML;refGene_tx=XM_003992583.5;refGene_gn=OPCML;refGene_tx=XM_019811571.2;refGene_gn=OPCML;refGene_tx=XM_019811572.2;refGene_gn=OPCML;refGene_tx=XM_019811573.2;refGene_gn=OPCML;refGene_tx=XM_023239305.1;refGene_gn=OPCML;refGene_tx=XM_023239306.1
D1 29623749 29623916
refGene_gn=OPCML;refGene_tx=XM_003992583.5;refGene_gn=OPCML;refGene_tx=XM_019811571.2;refGene_gn=OPCML;refGene_tx=XM_019811572.2;refGene_gn=OPCML;refGene_tx=XM_019811573.2;refGene_gn=OPCML;refGene_tx=XM_023239305.1;refGene_gn=OPCML;refGene_tx=XM_023239306.1
D1 30190454 30190515 refGene_gn=OPCML;refGene_tx=XM_023239305.1
D1 30532601 30532698

refGene_gn=SPATA19;refGene_tx=XM_019813032.2;refGene_gn=SPATA19;refGene_tx=XM_023239307.1;refGene_gn=SPATA19;refGene_tx=XM_023239308.1
D1 30532970 30533048
refGene_gn=SPATA19;refGene_tx=XM_019813032.2;refGene_gn=SPATA19;refGene_tx=XM_023239307.1;refGene_gn=SPATA19;refGene_tx=XM_023239308.1
D1 30535489 30535581
refGene_gn=SPATA19;refGene_tx=XM_019813032.2;refGene_gn=SPATA19;refGene_tx=XM_023239307.1;refGene_gn=SPATA19;refGene_tx=XM_023239308.1
D1 30535757 30535850
refGene_gn=SPATA19;refGene_tx=XM_019813032.2;refGene_gn=SPATA19;refGene_tx=XM_023239307.1;refGene_gn=SPATA19;refGene_tx=XM_023239308.1
D1 30536312 30536369
refGene_gn=SPATA19;refGene_tx=XM_019813032.2;refGene_gn=SPATA19;refGene_tx=XM_023239307.1;refGene_gn=SPATA19;refGene_tx=XM_023239308.1
D1 30536540 30536618
refGene_gn=SPATA19;refGene_tx=XM_019813032.2;refGene_gn=SPATA19;refGene_tx=XM_023239307.1;refGene_gn=SPATA19;refGene_tx=XM_023239308.1
D1 30580719 30580928
refGene_gn=IGSF9B;refGene_tx=XM_011286471.3;refGene_gn=IGSF9B;refGene_tx=XM_019811574.2
D1 30583516 30583638
refGene_gn=IGSF9B;refGene_tx=XM_011286471.3;refGene_gn=IGSF9B;refGene_tx=XM_019811574.2
D1 30589752 30589798 refGene_gn=IGSF9B;refGene_tx=XM_019811575.2
D1 30591310 30592960
refGene_gn=IGSF9B;refGene_tx=XM_011286471.3;refGene_gn=IGSF9B;refGene_tx=XM_019811574.2;refGene_gn=IGSF9B;refGene_tx=XM_019811575.2
D1 30593532 30593578
refGene_gn=IGSF9B;refGene_tx=XM_011286471.3;refGene_gn=IGSF9B;refGene_tx=XM_019811574.2;refGene_gn=IGSF9B;refGene_tx=XM_019811575.2
D1 30593952 30594114
refGene_gn=IGSF9B;refGene_tx=XM_011286471.3;refGene_gn=IGSF9B;refGene_tx=XM_019811574.2;refGene_gn=IGSF9B;refGene_tx=XM_019811575.2
D1 30596304 30596389
refGene_gn=IGSF9B;refGene_tx=XM_011286471.3;refGene_gn=IGSF9B;refGene_tx=XM_019811574.2;refGene_gn=IGSF9B;refGene_tx=XM_019811575.2
D1 30597374 30597601
refGene_gn=IGSF9B;refGene_tx=XM_011286471.3;refGene_gn=IGSF9B;refGene_tx=XM_019811574.2;refGene_gn=IGSF9B;refGene_tx=XM_019811575.2
D1 30598922 30599098
refGene_gn=IGSF9B;refGene_tx=XM_011286471.3;refGene_gn=IGSF9B;refGene_tx=XM_019811574.2;refGene_gn=IGSF9B;refGene_tx=XM_019811575.2
D1 30601469 30601593
refGene_gn=IGSF9B;refGene_tx=XM_011286471.3;refGene_gn=IGSF9B;refGene_tx=XM_019811574.2;refGene_gn=IGSF9B;refGene_tx=XM_019811575.2
D1 30602680 30602831
refGene_gn=IGSF9B;refGene_tx=XM_011286471.3;refGene_gn=IGSF9B;refGene_tx=XM_019811574.2;refGene_gn=IGSF9B;refGene_tx=XM_019811575.2
D1 30603113 30603230
refGene_gn=IGSF9B;refGene_tx=XM_011286471.3;refGene_gn=IGSF9B;refGene_

tx=XM_019811574.2;refGene_gn=IGSF9B;refGene_tx=XM_019811575.2
D1	30603314	30603455
refGene_gn=IGSF9B;refGene_tx=XM_011286471.3;refGene_gn=IGSF9B;refGene_tx=XM_019811574.2;refGene_gn=IGSF9B;refGene_tx=XM_019811575.2
D1	30603770	30603913
refGene_gn=IGSF9B;refGene_tx=XM_011286471.3;refGene_gn=IGSF9B;refGene_tx=XM_019811574.2;refGene_gn=IGSF9B;refGene_tx=XM_019811575.2
D1	30606877	30607023
refGene_gn=IGSF9B;refGene_tx=XM_011286471.3;refGene_gn=IGSF9B;refGene_tx=XM_019811574.2;refGene_gn=IGSF9B;refGene_tx=XM_019811575.2
D1	30607328	30607470
refGene_gn=IGSF9B;refGene_tx=XM_011286471.3;refGene_gn=IGSF9B;refGene_tx=XM_019811574.2;refGene_gn=IGSF9B;refGene_tx=XM_019811575.2
D1	30608801	30608919
refGene_gn=IGSF9B;refGene_tx=XM_011286471.3;refGene_gn=IGSF9B;refGene_tx=XM_019811574.2;refGene_gn=IGSF9B;refGene_tx=XM_019811575.2
D1	30609189	30609341
refGene_gn=IGSF9B;refGene_tx=XM_011286471.3;refGene_gn=IGSF9B;refGene_tx=XM_019811574.2;refGene_gn=IGSF9B;refGene_tx=XM_019811575.2
D1	30614790	30614937
refGene_gn=IGSF9B;refGene_tx=XM_011286471.3;refGene_gn=IGSF9B;refGene_tx=XM_019811574.2;refGene_gn=IGSF9B;refGene_tx=XM_019811575.2
D1	30616691	30616889
refGene_gn=IGSF9B;refGene_tx=XM_011286471.3;refGene_gn=IGSF9B;refGene_tx=XM_019811574.2;refGene_gn=IGSF9B;refGene_tx=XM_019811575.2
D1	30627750	30627814
refGene_gn=IGSF9B;refGene_tx=XM_011286471.3;refGene_gn=IGSF9B;refGene_tx=XM_019811574.2;refGene_gn=IGSF9B;refGene_tx=XM_019811575.2
D1	30743241	30743278	refGene_gn=JAM3;refGene_tx=XM_019811576.2
D1	30819202	30819268	refGene_gn=JAM3;refGene_tx=XM_019811576.2
D1	30819782	30819896	refGene_gn=JAM3;refGene_tx=XM_019811576.2
D1	30823957	30824110	refGene_gn=JAM3;refGene_tx=XM_019811576.2
D1	30824577	30824780	refGene_gn=JAM3;refGene_tx=XM_019811576.2
D1	30825771	30825871	refGene_gn=JAM3;refGene_tx=XM_019811576.2
D1	30827621	30827751	refGene_gn=JAM3;refGene_tx=XM_019811576.2
D1	30827848	30827903	refGene_gn=JAM3;refGene_tx=XM_019811576.2
D1	30828191	30828227	refGene_gn=JAM3;refGene_tx=XM_019811576.2
D1	30832191	30832300
refGene_gn=NCAPD3;refGene_tx=XM_011286469.3;refGene_gn=NCAPD3;refGene_tx=XM_023239309.1;refGene_gn=NCAPD3;refGene_tx=XM_023239310.1
D1	30832390	30832451
refGene_gn=NCAPD3;refGene_tx=XM_011286469.3;refGene_gn=NCAPD3;refGene_tx=XM_023239309.1
D1	30832526	30832601
refGene_gn=NCAPD3;refGene_tx=XM_011286469.3;refGene_gn=NCAPD3;refGene_tx=XM_023239309.1;refGene_gn=NCAPD3;refGene_tx=XM_023239310.1
D1	30836681	30836759
refGene_gn=NCAPD3;refGene_tx=XM_011286469.3;refGene_gn=NCAPD3;refGene_tx=XM_023239309.1;refGene_gn=NCAPD3;refGene_tx=XM_023239310.1
D1	30838165	30838305

refGene_gn=NCAPD3;refGene_tx=XM_011286469.3;refGene_gn=NCAPD3;refGene_tx=XM_023239309.1;refGene_gn=NCAPD3;refGene_tx=XM_023239310.1
D1 30838523 30838690
refGene_gn=NCAPD3;refGene_tx=XM_011286469.3;refGene_gn=NCAPD3;refGene_tx=XM_023239309.1;refGene_gn=NCAPD3;refGene_tx=XM_023239310.1
D1 30839626 30839764
refGene_gn=NCAPD3;refGene_tx=XM_011286469.3;refGene_gn=NCAPD3;refGene_tx=XM_023239309.1;refGene_gn=NCAPD3;refGene_tx=XM_023239310.1
D1 30841646 30841757
refGene_gn=NCAPD3;refGene_tx=XM_011286469.3;refGene_gn=NCAPD3;refGene_tx=XM_023239309.1;refGene_gn=NCAPD3;refGene_tx=XM_023239310.1
D1 30843959 30844159
refGene_gn=NCAPD3;refGene_tx=XM_011286469.3;refGene_gn=NCAPD3;refGene_tx=XM_023239309.1;refGene_gn=NCAPD3;refGene_tx=XM_023239310.1
D1 30844408 30844542
refGene_gn=NCAPD3;refGene_tx=XM_011286469.3;refGene_gn=NCAPD3;refGene_tx=XM_023239309.1;refGene_gn=NCAPD3;refGene_tx=XM_023239310.1
D1 30844815 30844953
refGene_gn=NCAPD3;refGene_tx=XM_011286469.3;refGene_gn=NCAPD3;refGene_tx=XM_023239309.1;refGene_gn=NCAPD3;refGene_tx=XM_023239310.1
D1 30850750 30850830
refGene_gn=NCAPD3;refGene_tx=XM_011286469.3;refGene_gn=NCAPD3;refGene_tx=XM_023239309.1;refGene_gn=NCAPD3;refGene_tx=XM_023239310.1
D1 30851763 30852002
refGene_gn=NCAPD3;refGene_tx=XM_011286469.3;refGene_gn=NCAPD3;refGene_tx=XM_023239309.1;refGene_gn=NCAPD3;refGene_tx=XM_023239310.1
D1 30853121 30853229
refGene_gn=NCAPD3;refGene_tx=XM_011286469.3;refGene_gn=NCAPD3;refGene_tx=XM_023239309.1;refGene_gn=NCAPD3;refGene_tx=XM_023239310.1
D1 30853318 30853433
refGene_gn=NCAPD3;refGene_tx=XM_011286469.3;refGene_gn=NCAPD3;refGene_tx=XM_023239309.1;refGene_gn=NCAPD3;refGene_tx=XM_023239310.1
D1 30855214 30855322
refGene_gn=NCAPD3;refGene_tx=XM_011286469.3;refGene_gn=NCAPD3;refGene_tx=XM_023239309.1;refGene_gn=NCAPD3;refGene_tx=XM_023239310.1
D1 30859878 30859994
refGene_gn=NCAPD3;refGene_tx=XM_011286469.3;refGene_gn=NCAPD3;refGene_tx=XM_023239309.1;refGene_gn=NCAPD3;refGene_tx=XM_023239310.1
D1 30860088 30860186
refGene_gn=NCAPD3;refGene_tx=XM_011286469.3;refGene_gn=NCAPD3;refGene_tx=XM_023239309.1;refGene_gn=NCAPD3;refGene_tx=XM_023239310.1
D1 30860508 30860700
refGene_gn=NCAPD3;refGene_tx=XM_011286469.3;refGene_gn=NCAPD3;refGene_tx=XM_023239309.1;refGene_gn=NCAPD3;refGene_tx=XM_023239310.1
D1 30867792 30868013
refGene_gn=NCAPD3;refGene_tx=XM_011286469.3;refGene_gn=NCAPD3;refGene_tx=XM_023239309.1;refGene_gn=NCAPD3;refGene_tx=XM_023239310.1
D1 30869328 30869463
refGene_gn=NCAPD3;refGene_tx=XM_011286469.3;refGene_gn=NCAPD3;refGene_tx=XM_023239309.1;refGene_gn=NCAPD3;refGene_tx=XM_023239310.1

D1 30870253 30870327
refGene_gn=NCAPD3;refGene_tx=XM_011286469.3;refGene_gn=NCAPD3;refGene_tx=XM_023239309.1;refGene_gn=NCAPD3;refGene_tx=XM_023239310.1
D1 30875761 30875851
refGene_gn=NCAPD3;refGene_tx=XM_011286469.3;refGene_gn=NCAPD3;refGene_tx=XM_023239309.1;refGene_gn=NCAPD3;refGene_tx=XM_023239310.1
D1 30876107 30876164
refGene_gn=NCAPD3;refGene_tx=XM_011286469.3;refGene_gn=NCAPD3;refGene_tx=XM_023239309.1;refGene_gn=NCAPD3;refGene_tx=XM_023239310.1
D1 30877701 30877954
refGene_gn=NCAPD3;refGene_tx=XM_011286469.3;refGene_gn=NCAPD3;refGene_tx=XM_023239309.1;refGene_gn=NCAPD3;refGene_tx=XM_023239310.1
D1 30878086 30878212
refGene_gn=NCAPD3;refGene_tx=XM_011286469.3;refGene_gn=NCAPD3;refGene_tx=XM_023239309.1;refGene_gn=NCAPD3;refGene_tx=XM_023239310.1
D1 30879475 30879548
refGene_gn=NCAPD3;refGene_tx=XM_011286469.3;refGene_gn=NCAPD3;refGene_tx=XM_023239309.1;refGene_gn=NCAPD3;refGene_tx=XM_023239310.1
D1 30880951 30881085
refGene_gn=NCAPD3;refGene_tx=XM_011286469.3;refGene_gn=NCAPD3;refGene_tx=XM_023239309.1;refGene_gn=NCAPD3;refGene_tx=XM_023239310.1
D1 30882820 30882908
refGene_gn=NCAPD3;refGene_tx=XM_011286469.3;refGene_gn=NCAPD3;refGene_tx=XM_023239309.1;refGene_gn=NCAPD3;refGene_tx=XM_023239310.1
D1 30883103 30883165
refGene_gn=NCAPD3;refGene_tx=XM_011286469.3;refGene_gn=NCAPD3;refGene_tx=XM_023239309.1;refGene_gn=NCAPD3;refGene_tx=XM_023239310.1
D1 30883269 30883434
refGene_gn=NCAPD3;refGene_tx=XM_011286469.3;refGene_gn=NCAPD3;refGene_tx=XM_023239309.1;refGene_gn=NCAPD3;refGene_tx=XM_023239310.1
D1 30884079 30884264
refGene_gn=NCAPD3;refGene_tx=XM_011286469.3;refGene_gn=NCAPD3;refGene_tx=XM_023239309.1;refGene_gn=NCAPD3;refGene_tx=XM_023239310.1
D1 30892442 30892605
refGene_gn=NCAPD3;refGene_tx=XM_011286469.3;refGene_gn=NCAPD3;refGene_tx=XM_023239309.1;refGene_gn=NCAPD3;refGene_tx=XM_023239310.1
D1 30894092 30894247
refGene_gn=NCAPD3;refGene_tx=XM_011286469.3;refGene_gn=NCAPD3;refGene_tx=XM_023239309.1;refGene_gn=NCAPD3;refGene_tx=XM_023239310.1
D1 30896379 30896443
refGene_gn=NCAPD3;refGene_tx=XM_011286469.3;refGene_gn=NCAPD3;refGene_tx=XM_023239309.1;refGene_gn=NCAPD3;refGene_tx=XM_023239310.1
D1 30897605 30897828 refGene_gn=VPS26B;refGene_tx=XM_003992590.3
D1 30906833 30906990 refGene_gn=VPS26B;refGene_tx=XM_003992590.3
D1 30912271 30912436 refGene_gn=VPS26B;refGene_tx=XM_003992590.3
D1 30915257 30915433 refGene_gn=VPS26B;refGene_tx=XM_003992590.3
D1 30917109 30917252 refGene_gn=VPS26B;refGene_tx=XM_003992590.3
D1 30917670 30917817 refGene_gn=VPS26B;refGene_tx=XM_003992590.3
D1 30920628 30920675 refGene_gn=THYN1;refGene_tx=XM_006936810.3
D1 30920996 30921147 refGene_gn=THYN1;refGene_tx=XM_006936810.3

D1 30921400 30921496 refGene_gn=THYN1;refGene_tx=XM_006936810.3
D1 30922503 30922596 refGene_gn=THYN1;refGene_tx=XM_006936810.3
D1 30922903 30922972 refGene_gn=THYN1;refGene_tx=XM_006936810.3
D1 30923716 30923898 refGene_gn=THYN1;refGene_tx=XM_006936810.3
D1 30925634 30925686 refGene_gn=THYN1;refGene_tx=XM_006936810.3
D1 30926237 30926346 refGene_gn=ACAD8;refGene_tx=XM_003992595.4
D1 30928086 30928187 refGene_gn=ACAD8;refGene_tx=XM_003992595.4
D1 30928818 30928988 refGene_gn=ACAD8;refGene_tx=XM_003992595.4
D1 30929663 30929773 refGene_gn=ACAD8;refGene_tx=XM_003992595.4
D1 30930136 30930213 refGene_gn=ACAD8;refGene_tx=XM_003992595.4
D1 30930815 30930953 refGene_gn=ACAD8;refGene_tx=XM_003992595.4
D1 30935430 30935566 refGene_gn=ACAD8;refGene_tx=XM_003992595.4
D1 30935660 30935758 refGene_gn=ACAD8;refGene_tx=XM_003992595.4
D1 30936125 30936278 refGene_gn=ACAD8;refGene_tx=XM_003992595.4
D1 30936912 30937015 refGene_gn=ACAD8;refGene_tx=XM_003992595.4
D1 30942089 30942142 refGene_gn=ACAD8;refGene_tx=XM_003992595.4
D1 31020575 31020772
refGene_gn=LOC101098007;refGene_tx=XM_023239701.1
D1 31034405 31034474
refGene_gn=LOC101098007;refGene_tx=XM_023239701.1
D1 31037008 31037104
refGene_gn=LOC101098007;refGene_tx=XM_023239701.1
D1 31038272 31038381
refGene_gn=LOC101098007;refGene_tx=XM_023239701.1
D1 31041193 31041286
refGene_gn=LOC101098007;refGene_tx=XM_023239701.1
D1 31045161 31045243
refGene_gn=LOC101098007;refGene_tx=XM_023239701.1
D1 31046211 31046276
refGene_gn=LOC101098007;refGene_tx=XM_023239701.1
D1 31055086 31055171
refGene_gn=LOC101098007;refGene_tx=XM_023239701.1
D1 31055795 31055933
refGene_gn=LOC101098007;refGene_tx=XM_023239701.1
D1 31056994 31057075
refGene_gn=LOC101098007;refGene_tx=XM_023239701.1
D1 31057705 31057812
refGene_gn=LOC101098007;refGene_tx=XM_023239701.1
D1 31066302 31066449
refGene_gn=LOC101098007;refGene_tx=XM_023239701.1
D1 31068912 31068984
refGene_gn=LOC101098007;refGene_tx=XM_023239701.1
D1 31069496 31069584
refGene_gn=LOC101098007;refGene_tx=XM_023239701.1
D1 31070247 31070335
refGene_gn=LOC101098007;refGene_tx=XM_023239701.1
D1 31070696 31070808
refGene_gn=LOC101098007;refGene_tx=XM_023239701.1
D1 31071779 31071896
refGene_gn=LOC101098007;refGene_tx=XM_023239701.1

D1 31072004 31072068
refGene_gn=LOC101098007;refGene_tx=XM_023239701.1
D1 31073560 31074030
refGene_gn=LOC101098007;refGene_tx=XM_023239701.1
D1 31144877 31144945
refGene_gn=LOC101098254;refGene_tx=XM_019813033.2;refGene_gn=LOC101098254;refGene_tx=XM_023239311.1;refGene_gn=LOC101098254;refGene_tx=XM_023239312.1
D1 31145753 31145885
refGene_gn=LOC101098254;refGene_tx=XM_019813033.2;refGene_gn=LOC101098254;refGene_tx=XM_023239311.1;refGene_gn=LOC101098254;refGene_tx=XM_023239312.1
D1 31146031 31146247
refGene_gn=LOC101098254;refGene_tx=XM_019813033.2;refGene_gn=LOC101098254;refGene_tx=XM_023239311.1;refGene_gn=LOC101098254;refGene_tx=XM_023239312.1
D1 31151054 31151123
refGene_gn=LOC101098254;refGene_tx=XM_019813033.2;refGene_gn=LOC101098254;refGene_tx=XM_023239311.1;refGene_gn=LOC101098254;refGene_tx=XM_023239312.1
D1 31151683 31151779
refGene_gn=LOC101098254;refGene_tx=XM_019813033.2;refGene_gn=LOC101098254;refGene_tx=XM_023239311.1;refGene_gn=LOC101098254;refGene_tx=XM_023239312.1
D1 31152962 31153071
refGene_gn=LOC101098254;refGene_tx=XM_019813033.2;refGene_gn=LOC101098254;refGene_tx=XM_023239311.1;refGene_gn=LOC101098254;refGene_tx=XM_023239312.1
D1 31156301 31156394
refGene_gn=LOC101098254;refGene_tx=XM_019813033.2;refGene_gn=LOC101098254;refGene_tx=XM_023239311.1;refGene_gn=LOC101098254;refGene_tx=XM_023239312.1
D1 31157828 31157910
refGene_gn=LOC101098254;refGene_tx=XM_019813033.2;refGene_gn=LOC101098254;refGene_tx=XM_023239311.1;refGene_gn=LOC101098254;refGene_tx=XM_023239312.1
D1 31159302 31159367
refGene_gn=LOC101098254;refGene_tx=XM_019813033.2;refGene_gn=LOC101098254;refGene_tx=XM_023239311.1;refGene_gn=LOC101098254;refGene_tx=XM_023239312.1
D1 31187264 31187349
refGene_gn=LOC101098254;refGene_tx=XM_019813033.2;refGene_gn=LOC101098254;refGene_tx=XM_023239311.1;refGene_gn=LOC101098254;refGene_tx=XM_023239312.1
D1 31188902 31189040
refGene_gn=LOC101098254;refGene_tx=XM_019813033.2;refGene_gn=LOC101098254;refGene_tx=XM_023239311.1;refGene_gn=LOC101098254;refGene_tx=XM_023239312.1
D1 31189648 31189729
refGene_gn=LOC101098254;refGene_tx=XM_019813033.2;refGene_gn=LOC101098

254;refGene_tx=XM_023239311.1;refGene_gn=LOC101098254;refGene_tx=XM_023239312.1
D1 31190399 31190506
refGene_gn=LOC101098254;refGene_tx=XM_019813033.2;refGene_gn=LOC101098254;refGene_tx=XM_023239311.1;refGene_gn=LOC101098254;refGene_tx=XM_023239312.1
D1 31191358 31191499
refGene_gn=LOC101098254;refGene_tx=XM_019813033.2;refGene_gn=LOC101098254;refGene_tx=XM_023239311.1
D1 31191825 31191897
refGene_gn=LOC101098254;refGene_tx=XM_019813033.2;refGene_gn=LOC101098254;refGene_tx=XM_023239311.1;refGene_gn=LOC101098254;refGene_tx=XM_023239312.1
D1 31192526 31192605
refGene_gn=LOC101098254;refGene_tx=XM_019813033.2;refGene_gn=LOC101098254;refGene_tx=XM_023239311.1;refGene_gn=LOC101098254;refGene_tx=XM_023239312.1
D1 31193037 31193125
refGene_gn=LOC101098254;refGene_tx=XM_019813033.2;refGene_gn=LOC101098254;refGene_tx=XM_023239311.1;refGene_gn=LOC101098254;refGene_tx=XM_023239312.1
D1 31193412 31193524
refGene_gn=LOC101098254;refGene_tx=XM_019813033.2;refGene_gn=LOC101098254;refGene_tx=XM_023239311.1;refGene_gn=LOC101098254;refGene_tx=XM_023239312.1
D1 31211091 31211208
refGene_gn=LOC101098254;refGene_tx=XM_019813033.2;refGene_gn=LOC101098254;refGene_tx=XM_023239311.1;refGene_gn=LOC101098254;refGene_tx=XM_023239312.1
D1 31211348 31211414
refGene_gn=LOC101098254;refGene_tx=XM_019813033.2;refGene_gn=LOC101098254;refGene_tx=XM_023239311.1;refGene_gn=LOC101098254;refGene_tx=XM_023239312.1
D1 31231269 31231385
refGene_gn=LOC101098487;refGene_tx=XM_006936811.4;refGene_gn=LOC101098487;refGene_tx=XM_019811577.2
D1 31250665 31250713
refGene_gn=LOC101098487;refGene_tx=XM_019811577.2
D1 31251126 31251324
refGene_gn=LOC101098487;refGene_tx=XM_006936811.4;refGene_gn=LOC101098487;refGene_tx=XM_019811577.2;refGene_gn=LOC101098487;refGene_tx=XM_019811578.2
D1 31252981 31253050
refGene_gn=LOC101098487;refGene_tx=XM_006936811.4;refGene_gn=LOC101098487;refGene_tx=XM_019811577.2;refGene_gn=LOC101098487;refGene_tx=XM_019811578.2
D1 31253632 31253728
refGene_gn=LOC101098487;refGene_tx=XM_006936811.4;refGene_gn=LOC101098487;refGene_tx=XM_019811577.2;refGene_gn=LOC101098487;refGene_tx=XM_019811578.2

D1 31255508 31255617
refGene_gn=LOC101098487;refGene_tx=XM_006936811.4;refGene_gn=LOC101098
487;refGene_tx=XM_019811577.2;refGene_gn=LOC101098487;refGene_tx=XM_01
9811578.2
D1 31260270 31260363
refGene_gn=LOC101098487;refGene_tx=XM_006936811.4;refGene_gn=LOC101098
487;refGene_tx=XM_019811577.2;refGene_gn=LOC101098487;refGene_tx=XM_01
9811578.2
D1 31263193 31263275
refGene_gn=LOC101098487;refGene_tx=XM_006936811.4;refGene_gn=LOC101098
487;refGene_tx=XM_019811577.2;refGene_gn=LOC101098487;refGene_tx=XM_01
9811578.2
D1 31269654 31269725
refGene_gn=LOC101098487;refGene_tx=XM_006936811.4;refGene_gn=LOC101098
487;refGene_tx=XM_019811577.2;refGene_gn=LOC101098487;refGene_tx=XM_01
9811578.2
D1 31272811 31272896
refGene_gn=LOC101098487;refGene_tx=XM_006936811.4;refGene_gn=LOC101098
487;refGene_tx=XM_019811577.2;refGene_gn=LOC101098487;refGene_tx=XM_01
9811578.2
D1 31274722 31274860
refGene_gn=LOC101098487;refGene_tx=XM_006936811.4;refGene_gn=LOC101098
487;refGene_tx=XM_019811577.2;refGene_gn=LOC101098487;refGene_tx=XM_01
9811578.2
D1 31275421 31275502
refGene_gn=LOC101098487;refGene_tx=XM_006936811.4;refGene_gn=LOC101098
487;refGene_tx=XM_019811577.2;refGene_gn=LOC101098487;refGene_tx=XM_01
9811578.2
D1 31275902 31276009
refGene_gn=LOC101098487;refGene_tx=XM_006936811.4;refGene_gn=LOC101098
487;refGene_tx=XM_019811577.2;refGene_gn=LOC101098487;refGene_tx=XM_01
9811578.2
D1 31276661 31276802
refGene_gn=LOC101098487;refGene_tx=XM_006936811.4;refGene_gn=LOC101098
487;refGene_tx=XM_019811577.2;refGene_gn=LOC101098487;refGene_tx=XM_01
9811578.2
D1 31277097 31277169
refGene_gn=LOC101098487;refGene_tx=XM_006936811.4;refGene_gn=LOC101098
487;refGene_tx=XM_019811577.2;refGene_gn=LOC101098487;refGene_tx=XM_01
9811578.2
D1 31277433 31277512
refGene_gn=LOC101098487;refGene_tx=XM_006936811.4;refGene_gn=LOC101098
487;refGene_tx=XM_019811577.2;refGene_gn=LOC101098487;refGene_tx=XM_01
9811578.2
D1 31279680 31279768
refGene_gn=LOC101098487;refGene_tx=XM_006936811.4;refGene_gn=LOC101098
487;refGene_tx=XM_019811577.2;refGene_gn=LOC101098487;refGene_tx=XM_01
9811578.2
D1 31280048 31280160
refGene_gn=LOC101098487;refGene_tx=XM_006936811.4;refGene_gn=LOC101098

487;refGene_tx=XM_019811577.2;refGene_gn=LOC101098487;refGene_tx=XM_019811578.2
D1 31280598 31280715 refGene_gn=LOC101098487;refGene_tx=XM_006936811.4;refGene_gn=LOC101098487;refGene_tx=XM_019811577.2;refGene_gn=LOC101098487;refGene_tx=XM_019811578.2
D1 31280990 31281077 refGene_gn=LOC101098487;refGene_tx=XM_006936811.4;refGene_gn=LOC101098487;refGene_tx=XM_019811577.2;refGene_gn=LOC101098487;refGene_tx=XM_019811578.2
D1 31287450 31287537 refGene_gn=B3GAT1;refGene_tx=XM_011286467.2;refGene_gn=B3GAT1;refGene_tx=XM_011286468.3
D1 31288277 31288574 refGene_gn=B3GAT1;refGene_tx=XM_011286467.2;refGene_gn=B3GAT1;refGene_tx=XM_011286468.3
D1 31289215 31289724 refGene_gn=B3GAT1;refGene_tx=XM_011286467.2;refGene_gn=B3GAT1;refGene_tx=XM_011286468.3
D1 31293353 31293500 refGene_gn=B3GAT1;refGene_tx=XM_011286467.2;refGene_gn=B3GAT1;refGene_tx=XM_011286468.3
D1 31293665 31293669 refGene_gn=B3GAT1;refGene_tx=XM_011286467.2
D1 34185735 34185772 refGene_gn=LOC111556524;refGene_tx=XM_023238985.1
D1 34185976 34186031 refGene_gn=LOC111556524;refGene_tx=XM_023238984.1
D1 34186703 34186817 refGene_gn=LOC111556524;refGene_tx=XM_023238984.1;refGene_gn=LOC111556524;refGene_tx=XM_023238985.1
D1 34189917 34190006 refGene_gn=LOC111556524;refGene_tx=XM_023238984.1;refGene_gn=LOC111556524;refGene_tx=XM_023238985.1
D1 34217823 34217948 refGene_gn=LOC109492323;refGene_tx=XM_019813025.2
D1 34218706 34218995 refGene_gn=LOC109492323;refGene_tx=XM_019813025.2
D1 34541697 34542771 refGene_gn=LOC109491906;refGene_tx=XM_023239313.1
D1 34692124 34693758 refGene_gn=PGR;refGene_tx=XM_019811582.2
D1 34694995 34695147 refGene_gn=PGR;refGene_tx=XM_019811582.2
D1 34731487 34731604 refGene_gn=PGR;refGene_tx=XM_019811582.2
D1 34751965 34752052 refGene_gn=PGR;refGene_tx=XM_019811582.2
D1 34762362 34762574 refGene_gn=PGR;refGene_tx=XM_019811582.2
D1 34762575 34762669 refGene_gn=PGR;refGene_tx=XM_019811582.2
D1 34772978 34773123 refGene_gn=PGR;refGene_tx=XM_019811582.2
D1 34774538 34774669 refGene_gn=PGR;refGene_tx=XM_019811582.2
D1 34788922 34789080 refGene_gn=PGR;refGene_tx=XM_019811582.2
D1 34791549 34791705 refGene_gn=PGR;refGene_tx=XM_019811582.2
D1 34847451 34847540 refGene_gn=ARHGAP42;refGene_tx=XM_003992599.5

D1	34847969	34848049	refGene_gn=ARHGAP42;refGene_tx=XM_003992599.5
D1	34866000	34866063	refGene_gn=ARHGAP42;refGene_tx=XM_003992599.5
D1	34867990	34868147	refGene_gn=ARHGAP42;refGene_tx=XM_003992599.5
D1	34868514	34868895	refGene_gn=ARHGAP42;refGene_tx=XM_003992599.5
D1	34870128	34870273	refGene_gn=ARHGAP42;refGene_tx=XM_003992599.5
D1	34871453	34871613	refGene_gn=ARHGAP42;refGene_tx=XM_003992599.5
D1	34876801	34876907	refGene_gn=ARHGAP42;refGene_tx=XM_003992599.5
D1	34880380	34880439	refGene_gn=ARHGAP42;refGene_tx=XM_003992599.5
D1	34881052	34881137	refGene_gn=ARHGAP42;refGene_tx=XM_003992599.5
D1	34882080	34882155	refGene_gn=ARHGAP42;refGene_tx=XM_003992599.5
D1	34883098	34883161	refGene_gn=ARHGAP42;refGene_tx=XM_003992599.5
D1	34908292	34908332	refGene_gn=ARHGAP42;refGene_tx=XM_003992599.5
D1	34910168	34910201	refGene_gn=ARHGAP42;refGene_tx=XM_003992599.5
D1	34910202	34910248	refGene_gn=ARHGAP42;refGene_tx=XM_003992599.5
D1	34912856	34912966	refGene_gn=ARHGAP42;refGene_tx=XM_003992599.5
D1	34914889	34914990	refGene_gn=ARHGAP42;refGene_tx=XM_003992599.5
D1	34921815	34921945	refGene_gn=ARHGAP42;refGene_tx=XM_003992599.5
D1	34924465	34924570	refGene_gn=ARHGAP42;refGene_tx=XM_003992599.5
D1	34934426	34934537	refGene_gn=ARHGAP42;refGene_tx=XM_003992599.5
D1	34950876	34950978	refGene_gn=ARHGAP42;refGene_tx=XM_003992599.5
D1	35005698	35005770	refGene_gn=ARHGAP42;refGene_tx=XM_003992599.5
D1	35060171	35060233	refGene_gn=ARHGAP42;refGene_tx=XM_003992599.5
D1	35083458	35083554	refGene_gn=ARHGAP42;refGene_tx=XM_003992599.5
D1	35164528	35164682	refGene_gn=ARHGAP42;refGene_tx=XM_003992599.5
D1	35476076	35476180	refGene_gn=CNTN5;refGene_tx=XM_023239314.1
D1	35481056	35481225	refGene_gn=CNTN5;refGene_tx=XM_023239314.1
D1	35489143	35489256	refGene_gn=CNTN5;refGene_tx=XM_023239314.1
D1	35489697	35489884	refGene_gn=CNTN5;refGene_tx=XM_023239314.1
D1	35528134	35528244	refGene_gn=CNTN5;refGene_tx=XM_023239314.1
D1	35536827	35537062	refGene_gn=CNTN5;refGene_tx=XM_023239314.1
D1	35539362	35539433	refGene_gn=CNTN5;refGene_tx=XM_023239314.1
D1	35551788	35551938	refGene_gn=CNTN5;refGene_tx=XM_023239314.1
D1	35568369	35568528	refGene_gn=CNTN5;refGene_tx=XM_023239314.1
D1	35602539	35602660	refGene_gn=CNTN5;refGene_tx=XM_023239314.1
D1	35635580	35635756	refGene_gn=CNTN5;refGene_tx=XM_023239314.1
D1	35637966	35638094	refGene_gn=CNTN5;refGene_tx=XM_023239314.1
D1	35777849	35778000	refGene_gn=CNTN5;refGene_tx=XM_023239314.1
D1	35786474	35786604	refGene_gn=CNTN5;refGene_tx=XM_023239314.1
D1	35789091	35789228	refGene_gn=CNTN5;refGene_tx=XM_023239314.1
D1	35795871	35796053	refGene_gn=CNTN5;refGene_tx=XM_023239314.1
D1	35837296	35837399	refGene_gn=CNTN5;refGene_tx=XM_023239314.1
D1	35874501	35874705	refGene_gn=CNTN5;refGene_tx=XM_023239314.1
D1	35937536	35937632	refGene_gn=CNTN5;refGene_tx=XM_023239314.1
D1	36006473	36006649	refGene_gn=CNTN5;refGene_tx=XM_023239314.1
D1	36006776	36006900	refGene_gn=CNTN5;refGene_tx=XM_023239314.1
D1	36023578	36023800	refGene_gn=CNTN5;refGene_tx=XM_023239314.1
D1	36290821	36290876	refGene_gn=CNTN5;refGene_tx=XM_023239314.1
D1	38339767	38340183
refGene_gn=LOC101100411;refGene_tx=XM_023239317.1
D1	38340549	38342360

refGene_gn=LOC101100411;refGene_tx=XM_006936816.4;refGene_gn=LOC101100
411;refGene_tx=XM_006936818.2;refGene_gn=LOC101100411;refGene_tx=XM_02
3239315.1;refGene_gn=LOC101100411;refGene_tx=XM_023239316.1;refGene_gn
=LOC101100411;refGene_tx=XM_023239317.1
D1 39457874 39458579
refGene_gn=CCDC82;refGene_tx=XM_003992605.5;refGene_gn=CCDC82;refGene_
tx=XM_019811592.2;refGene_gn=CCDC82;refGene_tx=XM_019811593.2;refGene_
gn=CCDC82;refGene_tx=XM_019811594.2;refGene_gn=CCDC82;refGene_tx=XM_01
9811596.2;refGene_gn=CCDC82;refGene_tx=XM_023239318.1;refGene_gn=CCDC8
2;refGene_tx=XM_023239319.1
D1 39459057 39459262
refGene_gn=CCDC82;refGene_tx=XM_003992605.5;refGene_gn=CCDC82;refGene_
tx=XM_019811592.2;refGene_gn=CCDC82;refGene_tx=XM_019811593.2;refGene_
gn=CCDC82;refGene_tx=XM_019811594.2;refGene_gn=CCDC82;refGene_tx=XM_01
9811596.2;refGene_gn=CCDC82;refGene_tx=XM_023239318.1;refGene_gn=CCDC8
2;refGene_tx=XM_023239319.1
D1 39460633 39460722 refGene_gn=CCDC82;refGene_tx=XM_019811596.2
D1 39469441 39469534
refGene_gn=CCDC82;refGene_tx=XM_003992605.5;refGene_gn=CCDC82;refGene_
tx=XM_019811592.2;refGene_gn=CCDC82;refGene_tx=XM_019811593.2;refGene_
gn=CCDC82;refGene_tx=XM_019811594.2;refGene_gn=CCDC82;refGene_tx=XM_02
3239318.1;refGene_gn=CCDC82;refGene_tx=XM_023239319.1
D1 39471695 39471820
refGene_gn=CCDC82;refGene_tx=XM_003992605.5;refGene_gn=CCDC82;refGene_
tx=XM_019811592.2;refGene_gn=CCDC82;refGene_tx=XM_019811593.2;refGene_
gn=CCDC82;refGene_tx=XM_019811594.2;refGene_gn=CCDC82;refGene_tx=XM_02
3239318.1;refGene_gn=CCDC82;refGene_tx=XM_023239319.1
D1 39474883 39475054
refGene_gn=CCDC82;refGene_tx=XM_003992605.5;refGene_gn=CCDC82;refGene_
tx=XM_019811592.2;refGene_gn=CCDC82;refGene_tx=XM_019811593.2;refGene_
gn=CCDC82;refGene_tx=XM_019811594.2;refGene_gn=CCDC82;refGene_tx=XM_02
3239318.1;refGene_gn=CCDC82;refGene_tx=XM_023239319.1
D1 39480576 39480762
refGene_gn=CCDC82;refGene_tx=XM_003992605.5;refGene_gn=CCDC82;refGene_
tx=XM_019811592.2;refGene_gn=CCDC82;refGene_tx=XM_019811593.2;refGene_
gn=CCDC82;refGene_tx=XM_019811594.2;refGene_gn=CCDC82;refGene_tx=XM_02
3239318.1;refGene_gn=CCDC82;refGene_tx=XM_023239319.1
D1 39481138 39481186 refGene_gn=CCDC82;refGene_tx=XM_019811594.2
D1 39485009 39485078
refGene_gn=CCDC82;refGene_tx=XM_019811592.2;refGene_gn=CCDC82;refGene_
tx=XM_019811593.2;refGene_gn=CCDC82;refGene_tx=XM_023239318.1;refGene_
gn=CCDC82;refGene_tx=XM_023239319.1
D1 39486106 39486172 refGene_gn=CCDC82;refGene_tx=XM_003992605.5
D1 39491930 39492443
refGene_gn=MAML2;refGene_tx=XM_019811597.1;refGene_gn=MAML2;refGene_tx
=XM_019811598.1
D1 39725252 39726773
refGene_gn=MAML2;refGene_tx=XM_019811597.1;refGene_gn=MAML2;refGene_tx
=XM_019811598.1
D1 39824257 39824461 refGene_gn=MAML2;refGene_tx=XM_019811597.1

D1 39829581 39829693
refGene_gn=MAML2;refGene_tx=XM_019811597.1;refGene_gn=MAML2;refGene_tx
=XM_019811598.1
D1 39833306 39834322
refGene_gn=MAML2;refGene_tx=XM_019811597.1;refGene_gn=MAML2;refGene_tx
=XM_019811598.1
D1 39871310 39871390 refGene_gn=MTMR2;refGene_tx=XM_011286508.2
D1 39911308 39911414 refGene_gn=MTMR2;refGene_tx=XM_011286508.2
D1 39936538 39936614
refGene_gn=MTMR2;refGene_tx=XM_011286508.2;refGene_gn=MTMR2;refGene_tx
=XM_011286509.2;refGene_gn=MTMR2;refGene_tx=XM_019811599.2
D1 39941547 39941642
refGene_gn=MTMR2;refGene_tx=XM_011286508.2;refGene_gn=MTMR2;refGene_tx
=XM_011286509.2;refGene_gn=MTMR2;refGene_tx=XM_019811599.2
D1 39941793 39941904
refGene_gn=MTMR2;refGene_tx=XM_011286508.2;refGene_gn=MTMR2;refGene_tx
=XM_011286509.2;refGene_gn=MTMR2;refGene_tx=XM_019811599.2
D1 39945050 39945152
refGene_gn=MTMR2;refGene_tx=XM_011286508.2;refGene_gn=MTMR2;refGene_tx
=XM_011286509.2;refGene_gn=MTMR2;refGene_tx=XM_019811599.2
D1 39946498 39946582
refGene_gn=MTMR2;refGene_tx=XM_011286508.2;refGene_gn=MTMR2;refGene_tx
=XM_011286509.2;refGene_gn=MTMR2;refGene_tx=XM_019811599.2
D1 39958018 39958168
refGene_gn=MTMR2;refGene_tx=XM_011286508.2;refGene_gn=MTMR2;refGene_tx
=XM_011286509.2;refGene_gn=MTMR2;refGene_tx=XM_019811599.2
D1 39958918 39959107
refGene_gn=MTMR2;refGene_tx=XM_011286508.2;refGene_gn=MTMR2;refGene_tx
=XM_011286509.2;refGene_gn=MTMR2;refGene_tx=XM_019811599.2
D1 39974613 39974799
refGene_gn=MTMR2;refGene_tx=XM_011286508.2;refGene_gn=MTMR2;refGene_tx
=XM_011286509.2;refGene_gn=MTMR2;refGene_tx=XM_019811599.2
D1 39975952 39976159
refGene_gn=MTMR2;refGene_tx=XM_011286508.2;refGene_gn=MTMR2;refGene_tx
=XM_011286509.2;refGene_gn=MTMR2;refGene_tx=XM_019811599.2
D1 39982471 39982564
refGene_gn=MTMR2;refGene_tx=XM_011286508.2;refGene_gn=MTMR2;refGene_tx
=XM_011286509.2;refGene_gn=MTMR2;refGene_tx=XM_019811599.2
D1 39983759 39983873
refGene_gn=MTMR2;refGene_tx=XM_011286508.2;refGene_gn=MTMR2;refGene_tx
=XM_011286509.2;refGene_gn=MTMR2;refGene_tx=XM_019811599.2
D1 39985860 39986037
refGene_gn=MTMR2;refGene_tx=XM_011286508.2;refGene_gn=MTMR2;refGene_tx
=XM_011286509.2;refGene_gn=MTMR2;refGene_tx=XM_019811599.2
D1 39986749 39986911
refGene_gn=MTMR2;refGene_tx=XM_011286508.2;refGene_gn=MTMR2;refGene_tx
=XM_011286509.2;refGene_gn=MTMR2;refGene_tx=XM_019811599.2
D1 39998264 39998498
refGene_gn=CEP57;refGene_tx=XM_011286510.3;refGene_gn=CEP57;refGene_tx
=XM_023239321.1

D1 40000136 40000281
refGene_gn=CEP57;refGene_tx=XM_011286510.3;refGene_gn=CEP57;refGene_tx=XM_023239321.1
D1 40000862 40001104
refGene_gn=CEP57;refGene_tx=XM_011286510.3;refGene_gn=CEP57;refGene_tx=XM_023239321.1
D1 40007504 40007582
refGene_gn=CEP57;refGene_tx=XM_011286510.3;refGene_gn=CEP57;refGene_tx=XM_023239321.1
D1 40008227 40008335
refGene_gn=CEP57;refGene_tx=XM_011286510.3;refGene_gn=CEP57;refGene_tx=XM_023239321.1
D1 40011354 40011432
refGene_gn=CEP57;refGene_tx=XM_011286510.3;refGene_gn=CEP57;refGene_tx=XM_023239321.1
D1 40012018 40012135
refGene_gn=CEP57;refGene_tx=XM_011286510.3;refGene_gn=CEP57;refGene_tx=XM_023239321.1
D1 40013474 40013596
refGene_gn=CEP57;refGene_tx=XM_011286510.3;refGene_gn=CEP57;refGene_tx=XM_023239321.1
D1 40013976 40014156
refGene_gn=CEP57;refGene_tx=XM_011286510.3;refGene_gn=CEP57;refGene_tx=XM_023239321.1
D1 40023370 40023527
refGene_gn=CEP57;refGene_tx=XM_011286510.3;refGene_gn=CEP57;refGene_tx=XM_023239321.1
D1 40037049 40037094 refGene_gn=CEP57;refGene_tx=XM_023239321.1
D1 40037522 40037567 refGene_gn=CEP57;refGene_tx=XM_011286510.3
D1 40038796 40038883
refGene_gn=FAM76B;refGene_tx=XM_011286511.3;refGene_gn=FAM76B;refGene_tx=XM_011286513.3
D1 40039306 40039411
refGene_gn=FAM76B;refGene_tx=XM_019811602.2;refGene_gn=FAM76B;refGene_tx=XM_019811603.2
D1 40039712 40039777
refGene_gn=FAM76B;refGene_tx=XM_011286511.3;refGene_gn=FAM76B;refGene_tx=XM_011286513.3;refGene_gn=FAM76B;refGene_tx=XM_019811602.2;refGene_gn=FAM76B;refGene_tx=XM_019811603.2
D1 40040650 40040705
refGene_gn=FAM76B;refGene_tx=XM_011286511.3;refGene_gn=FAM76B;refGene_tx=XM_011286513.3;refGene_gn=FAM76B;refGene_tx=XM_019811602.2;refGene_gn=FAM76B;refGene_tx=XM_019811603.2
D1 40041736 40041892
refGene_gn=FAM76B;refGene_tx=XM_011286511.3;refGene_gn=FAM76B;refGene_tx=XM_011286513.3;refGene_gn=FAM76B;refGene_tx=XM_019811602.2;refGene_gn=FAM76B;refGene_tx=XM_019811603.2
D1 40043652 40043852
refGene_gn=FAM76B;refGene_tx=XM_011286511.3;refGene_gn=FAM76B;refGene_tx=XM_011286513.3;refGene_gn=FAM76B;refGene_tx=XM_019811602.2;refGene_

gn=FAM76B;refGene_tx=XM_019811603.2
D1 40047373 40047421
refGene_gn=FAM76B;refGene_tx=XM_011286511.3;refGene_gn=FAM76B;refGene_tx=XM_011286513.3;refGene_gn=FAM76B;refGene_tx=XM_019811602.2;refGene_gn=FAM76B;refGene_tx=XM_019811603.2
D1 40047611 40047692
refGene_gn=FAM76B;refGene_tx=XM_011286511.3;refGene_gn=FAM76B;refGene_tx=XM_011286513.3;refGene_gn=FAM76B;refGene_tx=XM_019811602.2;refGene_gn=FAM76B;refGene_tx=XM_019811603.2
D1 40048354 40048490
refGene_gn=FAM76B;refGene_tx=XM_011286511.3;refGene_gn=FAM76B;refGene_tx=XM_011286513.3;refGene_gn=FAM76B;refGene_tx=XM_019811602.2;refGene_gn=FAM76B;refGene_tx=XM_019811603.2
D1 40051182 40051284
refGene_gn=FAM76B;refGene_tx=XM_011286511.3;refGene_gn=FAM76B;refGene_tx=XM_011286513.3;refGene_gn=FAM76B;refGene_tx=XM_019811602.2;refGene_gn=FAM76B;refGene_tx=XM_019811603.2
D1 40055791 40055881
refGene_gn=FAM76B;refGene_tx=XM_011286511.3;refGene_gn=FAM76B;refGene_tx=XM_011286513.3;refGene_gn=FAM76B;refGene_tx=XM_019811602.2;refGene_gn=FAM76B;refGene_tx=XM_019811603.2
D1 40538895 40538973 refGene_gn=SESN3;refGene_tx=XM_023239322.1
D1 40583287 40583353 refGene_gn=SESN3;refGene_tx=XM_023239322.1
D1 40586186 40586384 refGene_gn=SESN3;refGene_tx=XM_023239322.1
D1 40587835 40588018 refGene_gn=SESN3;refGene_tx=XM_023239322.1
D1 40591894 40592131 refGene_gn=SESN3;refGene_tx=XM_023239322.1
D1 40592622 40592797 refGene_gn=SESN3;refGene_tx=XM_023239322.1
D1 40598575 40598694 refGene_gn=SESN3;refGene_tx=XM_023239322.1
D1 40599522 40599713 refGene_gn=SESN3;refGene_tx=XM_023239322.1
D1 40602487 40602632 refGene_gn=SESN3;refGene_tx=XM_023239322.1
D1 40604440 40604527 refGene_gn=SESN3;refGene_tx=XM_023239322.1
D1 40656873 40658076 refGene_gn=ENDOD1;refGene_tx=XM_023239323.1
D1 40680754 40681054 refGene_gn=ENDOD1;refGene_tx=XM_023239323.1
D1 40709019 40709685
refGene_gn=LOC111556644;refGene_tx=XM_023239702.1
D1 40717640 40719557
refGene_gn=LOC105259981;refGene_tx=XM_023239324.1
D1 40733729 40735307
refGene_gn=LOC101085402;refGene_tx=XM_023239325.1;refGene_gn=LOC101085402;refGene_tx=XM_023239326.1;refGene_gn=LOC101085402;refGene_tx=XM_023239327.1
D1 40769399 40769530
refGene_gn=CWC15;refGene_tx=XM_006936824.4;refGene_gn=CWC15;refGene_tx=XM_006936825.3;refGene_gn=CWC15;refGene_tx=XM_006936826.3;refGene_gn=CWC15;refGene_tx=XM_006936827.3;refGene_gn=CWC15;refGene_tx=XM_006936828.3
D1 40770011 40770124
refGene_gn=CWC15;refGene_tx=XM_006936824.4;refGene_gn=CWC15;refGene_tx=XM_006936825.3;refGene_gn=CWC15;refGene_tx=XM_006936826.3;refGene_gn=CWC15;refGene_tx=XM_006936827.3;refGene_gn=CWC15;refGene_tx=XM_0069368

28.3
D1	40770988	40771077
refGene_gn=CWC15;refGene_tx=XM_006936824.4;refGene_gn=CWC15;refGene_tx=XM_006936825.3;refGene_gn=CWC15;refGene_tx=XM_006936826.3;refGene_gn=CWC15;refGene_tx=XM_006936827.3;refGene_gn=CWC15;refGene_tx=XM_006936828.3
D1	40772445	40772559
refGene_gn=CWC15;refGene_tx=XM_006936824.4;refGene_gn=CWC15;refGene_tx=XM_006936825.3;refGene_gn=CWC15;refGene_tx=XM_006936826.3;refGene_gn=CWC15;refGene_tx=XM_006936827.3;refGene_gn=CWC15;refGene_tx=XM_006936828.3
D1	40776855	40776974
refGene_gn=CWC15;refGene_tx=XM_006936824.4;refGene_gn=CWC15;refGene_tx=XM_006936825.3;refGene_gn=CWC15;refGene_tx=XM_006936826.3;refGene_gn=CWC15;refGene_tx=XM_006936827.3;refGene_gn=CWC15;refGene_tx=XM_006936828.3
D1	40778185	40778315
refGene_gn=CWC15;refGene_tx=XM_006936824.4;refGene_gn=CWC15;refGene_tx=XM_006936825.3;refGene_gn=CWC15;refGene_tx=XM_006936826.3;refGene_gn=CWC15;refGene_tx=XM_006936827.3;refGene_gn=CWC15;refGene_tx=XM_006936828.3
D1	40853083	40853190
refGene_gn=AMOTL1;refGene_tx=XM_006936831.4;refGene_gn=AMOTL1;refGene_tx=XM_006936832.4;refGene_gn=AMOTL1;refGene_tx=XM_006936834.4;refGene_gn=AMOTL1;refGene_tx=XM_011286519.3;refGene_gn=AMOTL1;refGene_tx=XM_019811608.2;refGene_gn=AMOTL1;refGene_tx=XM_019811609.2;refGene_gn=AMOTL1;refGene_tx=XM_019811610.2;refGene_gn=AMOTL1;refGene_tx=XM_019811611.2;refGene_gn=AMOTL1;refGene_tx=XM_019811612.2;refGene_gn=AMOTL1;refGene_tx=XM_019811613.2;refGene_gn=AMOTL1;refGene_tx=XM_019811614.2;refGene_gn=AMOTL1;refGene_tx=XM_019811615.2;refGene_gn=AMOTL1;refGene_tx=XM_023239328.1;refGene_gn=AMOTL1;refGene_tx=XM_023239329.1
D1	40854244	40854505
refGene_gn=AMOTL1;refGene_tx=XM_006936831.4;refGene_gn=AMOTL1;refGene_tx=XM_006936832.4;refGene_gn=AMOTL1;refGene_tx=XM_006936834.4;refGene_gn=AMOTL1;refGene_tx=XM_011286519.3;refGene_gn=AMOTL1;refGene_tx=XM_019811608.2;refGene_gn=AMOTL1;refGene_tx=XM_019811609.2;refGene_gn=AMOTL1;refGene_tx=XM_019811610.2;refGene_gn=AMOTL1;refGene_tx=XM_019811611.2;refGene_gn=AMOTL1;refGene_tx=XM_019811612.2;refGene_gn=AMOTL1;refGene_tx=XM_019811613.2;refGene_gn=AMOTL1;refGene_tx=XM_019811614.2;refGene_gn=AMOTL1;refGene_tx=XM_019811615.2;refGene_gn=AMOTL1;refGene_tx=XM_023239328.1;refGene_gn=AMOTL1;refGene_tx=XM_023239329.1
D1	40857766	40857993
refGene_gn=AMOTL1;refGene_tx=XM_006936831.4;refGene_gn=AMOTL1;refGene_tx=XM_006936832.4;refGene_gn=AMOTL1;refGene_tx=XM_006936834.4;refGene_gn=AMOTL1;refGene_tx=XM_011286519.3;refGene_gn=AMOTL1;refGene_tx=XM_019811608.2;refGene_gn=AMOTL1;refGene_tx=XM_019811609.2;refGene_gn=AMOTL1;refGene_tx=XM_019811610.2;refGene_gn=AMOTL1;refGene_tx=XM_019811611.2;refGene_gn=AMOTL1;refGene_tx=XM_019811612.2;refGene_gn=AMOTL1;refGene_tx=XM_019811613.2;refGene_gn=AMOTL1;refGene_tx=XM_019811614.2;refGene_gn=AMOTL1;refGene_tx=XM_019811615.2;refGene_gn=AMOTL1;refGene_tx=XM_

023239328.1;refGene_gn=AMOTL1;refGene_tx=XM_023239329.1
D1 40858676 40858802
refGene_gn=AMOTL1;refGene_tx=XM_006936831.4;refGene_gn=AMOTL1;refGene_tx=XM_006936832.4;refGene_gn=AMOTL1;refGene_tx=XM_006936834.4;refGene_gn=AMOTL1;refGene_tx=XM_011286519.3;refGene_gn=AMOTL1;refGene_tx=XM_019811608.2;refGene_gn=AMOTL1;refGene_tx=XM_019811609.2;refGene_gn=AMOTL1;refGene_tx=XM_019811610.2;refGene_gn=AMOTL1;refGene_tx=XM_019811611.2;refGene_gn=AMOTL1;refGene_tx=XM_019811612.2;refGene_gn=AMOTL1;refGene_tx=XM_019811613.2;refGene_gn=AMOTL1;refGene_tx=XM_019811614.2;refGene_gn=AMOTL1;refGene_tx=XM_019811615.2;refGene_gn=AMOTL1;refGene_tx=XM_023239328.1;refGene_gn=AMOTL1;refGene_tx=XM_023239329.1
D1 40863528 40863719
refGene_gn=AMOTL1;refGene_tx=XM_006936831.4;refGene_gn=AMOTL1;refGene_tx=XM_006936832.4;refGene_gn=AMOTL1;refGene_tx=XM_006936834.4;refGene_gn=AMOTL1;refGene_tx=XM_011286519.3;refGene_gn=AMOTL1;refGene_tx=XM_019811608.2;refGene_gn=AMOTL1;refGene_tx=XM_019811609.2;refGene_gn=AMOTL1;refGene_tx=XM_019811610.2;refGene_gn=AMOTL1;refGene_tx=XM_019811611.2;refGene_gn=AMOTL1;refGene_tx=XM_019811612.2;refGene_gn=AMOTL1;refGene_tx=XM_019811613.2;refGene_gn=AMOTL1;refGene_tx=XM_019811614.2;refGene_gn=AMOTL1;refGene_tx=XM_019811615.2;refGene_gn=AMOTL1;refGene_tx=XM_023239328.1;refGene_gn=AMOTL1;refGene_tx=XM_023239329.1
D1 40867331 40867481
refGene_gn=AMOTL1;refGene_tx=XM_006936831.4;refGene_gn=AMOTL1;refGene_tx=XM_006936832.4;refGene_gn=AMOTL1;refGene_tx=XM_006936834.4;refGene_gn=AMOTL1;refGene_tx=XM_011286519.3;refGene_gn=AMOTL1;refGene_tx=XM_019811608.2;refGene_gn=AMOTL1;refGene_tx=XM_019811609.2;refGene_gn=AMOTL1;refGene_tx=XM_019811610.2;refGene_gn=AMOTL1;refGene_tx=XM_019811611.2;refGene_gn=AMOTL1;refGene_tx=XM_019811612.2;refGene_gn=AMOTL1;refGene_tx=XM_019811613.2;refGene_gn=AMOTL1;refGene_tx=XM_019811614.2;refGene_gn=AMOTL1;refGene_tx=XM_019811615.2;refGene_gn=AMOTL1;refGene_tx=XM_023239328.1;refGene_gn=AMOTL1;refGene_tx=XM_023239329.1
D1 40871853 40871999
refGene_gn=AMOTL1;refGene_tx=XM_006936831.4;refGene_gn=AMOTL1;refGene_tx=XM_006936832.4;refGene_gn=AMOTL1;refGene_tx=XM_006936834.4;refGene_gn=AMOTL1;refGene_tx=XM_011286519.3;refGene_gn=AMOTL1;refGene_tx=XM_019811608.2;refGene_gn=AMOTL1;refGene_tx=XM_019811609.2;refGene_gn=AMOTL1;refGene_tx=XM_019811610.2;refGene_gn=AMOTL1;refGene_tx=XM_019811611.2;refGene_gn=AMOTL1;refGene_tx=XM_019811612.2;refGene_gn=AMOTL1;refGene_tx=XM_019811613.2;refGene_gn=AMOTL1;refGene_tx=XM_019811614.2;refGene_gn=AMOTL1;refGene_tx=XM_019811615.2;refGene_gn=AMOTL1;refGene_tx=XM_023239328.1;refGene_gn=AMOTL1;refGene_tx=XM_023239329.1
D1 40891371 40891461
refGene_gn=AMOTL1;refGene_tx=XM_006936831.4;refGene_gn=AMOTL1;refGene_tx=XM_006936832.4;refGene_gn=AMOTL1;refGene_tx=XM_006936834.4;refGene_gn=AMOTL1;refGene_tx=XM_011286519.3;refGene_gn=AMOTL1;refGene_tx=XM_019811608.2;refGene_gn=AMOTL1;refGene_tx=XM_019811609.2;refGene_gn=AMOTL1;refGene_tx=XM_019811610.2;refGene_gn=AMOTL1;refGene_tx=XM_019811611.2;refGene_gn=AMOTL1;refGene_tx=XM_019811612.2;refGene_gn=AMOTL1;refGene_tx=XM_019811613.2;refGene_gn=AMOTL1;refGene_tx=XM_019811614.2;refGene_gn=AMOTL1;refGene_tx=XM_019811615.2;refGene_gn=AMOTL1;refGene_tx=XM_

023239328.1;refGene_gn=AMOTL1;refGene_tx=XM_023239329.1
D1 40892794 40892939
refGene_gn=AMOTL1;refGene_tx=XM_006936831.4;refGene_gn=AMOTL1;refGene_tx=XM_006936832.4;refGene_gn=AMOTL1;refGene_tx=XM_006936834.4;refGene_gn=AMOTL1;refGene_tx=XM_011286519.3;refGene_gn=AMOTL1;refGene_tx=XM_019811608.2;refGene_gn=AMOTL1;refGene_tx=XM_019811609.2;refGene_gn=AMOTL1;refGene_tx=XM_019811610.2;refGene_gn=AMOTL1;refGene_tx=XM_019811611.2;refGene_gn=AMOTL1;refGene_tx=XM_019811612.2;refGene_gn=AMOTL1;refGene_tx=XM_019811613.2;refGene_gn=AMOTL1;refGene_tx=XM_019811614.2;refGene_gn=AMOTL1;refGene_tx=XM_019811615.2;refGene_gn=AMOTL1;refGene_tx=XM_023239328.1;refGene_gn=AMOTL1;refGene_tx=XM_023239329.1
D1 40899507 40899799
refGene_gn=AMOTL1;refGene_tx=XM_006936831.4;refGene_gn=AMOTL1;refGene_tx=XM_006936832.4;refGene_gn=AMOTL1;refGene_tx=XM_006936834.4;refGene_gn=AMOTL1;refGene_tx=XM_011286519.3;refGene_gn=AMOTL1;refGene_tx=XM_019811608.2;refGene_gn=AMOTL1;refGene_tx=XM_019811609.2;refGene_gn=AMOTL1;refGene_tx=XM_019811610.2;refGene_gn=AMOTL1;refGene_tx=XM_019811611.2;refGene_gn=AMOTL1;refGene_tx=XM_019811612.2;refGene_gn=AMOTL1;refGene_tx=XM_019811613.2;refGene_gn=AMOTL1;refGene_tx=XM_019811614.2;refGene_gn=AMOTL1;refGene_tx=XM_019811615.2;refGene_gn=AMOTL1;refGene_tx=XM_023239328.1;refGene_gn=AMOTL1;refGene_tx=XM_023239329.1
D1 40914576 40915510
refGene_gn=AMOTL1;refGene_tx=XM_006936831.4;refGene_gn=AMOTL1;refGene_tx=XM_006936832.4;refGene_gn=AMOTL1;refGene_tx=XM_006936834.4;refGene_gn=AMOTL1;refGene_tx=XM_011286519.3;refGene_gn=AMOTL1;refGene_tx=XM_019811608.2;refGene_gn=AMOTL1;refGene_tx=XM_019811609.2;refGene_gn=AMOTL1;refGene_tx=XM_019811610.2;refGene_gn=AMOTL1;refGene_tx=XM_019811611.2;refGene_gn=AMOTL1;refGene_tx=XM_019811612.2;refGene_gn=AMOTL1;refGene_tx=XM_019811613.2;refGene_gn=AMOTL1;refGene_tx=XM_019811614.2;refGene_gn=AMOTL1;refGene_tx=XM_019811615.2;refGene_gn=AMOTL1;refGene_tx=XM_023239328.1;refGene_gn=AMOTL1;refGene_tx=XM_023239329.1
D1 40919750 40919900
refGene_gn=AMOTL1;refGene_tx=XM_006936831.4;refGene_gn=AMOTL1;refGene_tx=XM_006936832.4;refGene_gn=AMOTL1;refGene_tx=XM_006936834.4;refGene_gn=AMOTL1;refGene_tx=XM_011286519.3;refGene_gn=AMOTL1;refGene_tx=XM_019811608.2;refGene_gn=AMOTL1;refGene_tx=XM_019811609.2;refGene_gn=AMOTL1;refGene_tx=XM_019811610.2;refGene_gn=AMOTL1;refGene_tx=XM_019811611.2;refGene_gn=AMOTL1;refGene_tx=XM_023239328.1;refGene_gn=AMOTL1;refGene_tx=XM_023239329.1
D1 40936002 40936021 refGene_gn=AMOTL1;refGene_tx=XM_019811611.2
D1 40950993 40951042 refGene_gn=AMOTL1;refGene_tx=XM_011286519.3
D1 40972518 40972569
refGene_gn=AMOTL1;refGene_tx=XM_006936831.4;refGene_gn=AMOTL1;refGene_tx=XM_006936832.4;refGene_gn=AMOTL1;refGene_tx=XM_019811608.2;refGene_gn=AMOTL1;refGene_tx=XM_019811610.2;refGene_gn=AMOTL1;refGene_tx=XM_023239328.1;refGene_gn=AMOTL1;refGene_tx=XM_023239329.1
D1 40974085 40974161 refGene_gn=AMOTL1;refGene_tx=XM_019811608.2
D1 40983385 40983435
refGene_gn=AMOTL1;refGene_tx=XM_019811610.2;refGene_gn=AMOTL1;refGene_tx=XM_023239328.1

D1 40983761 40983966
refGene_gn=AMOTL1;refGene_tx=XM_006936831.4;refGene_gn=AMOTL1;refGene_tx=XM_006936832.4;refGene_gn=AMOTL1;refGene_tx=XM_006936834.4;refGene_gn=AMOTL1;refGene_tx=XM_019811609.2;refGene_gn=AMOTL1;refGene_tx=XM_019811610.2;refGene_gn=AMOTL1;refGene_tx=XM_023239328.1;refGene_gn=AMOTL1;refGene_tx=XM_023239329.1
D1 41003376 41003385
refGene_gn=AMOTL1;refGene_tx=XM_006936831.4;refGene_gn=AMOTL1;refGene_tx=XM_019811609.2
D1 41070261 41070378
refGene_gn=PIWIL4;refGene_tx=XM_019811619.1;refGene_gn=PIWIL4;refGene_tx=XM_019811620.2
D1 41071327 41071475
refGene_gn=PIWIL4;refGene_tx=XM_019811619.1;refGene_gn=PIWIL4;refGene_tx=XM_019811620.2
D1 41071586 41071712
refGene_gn=PIWIL4;refGene_tx=XM_019811619.1;refGene_gn=PIWIL4;refGene_tx=XM_019811620.2
D1 41073740 41073894
refGene_gn=PIWIL4;refGene_tx=XM_019811619.1;refGene_gn=PIWIL4;refGene_tx=XM_019811620.2
D1 41075269 41075340
refGene_gn=PIWIL4;refGene_tx=XM_019811619.1;refGene_gn=PIWIL4;refGene_tx=XM_019811620.2
D1 41078673 41078777
refGene_gn=PIWIL4;refGene_tx=XM_019811619.1;refGene_gn=PIWIL4;refGene_tx=XM_019811620.2
D1 41079959 41080160 refGene_gn=PIWIL4;refGene_tx=XM_019811619.1
D1 41084211 41084284
refGene_gn=PIWIL4;refGene_tx=XM_019811619.1;refGene_gn=PIWIL4;refGene_tx=XM_019811620.2
D1 41086456 41086641
refGene_gn=PIWIL4;refGene_tx=XM_019811619.1;refGene_gn=PIWIL4;refGene_tx=XM_019811620.2
D1 41089371 41089483
refGene_gn=PIWIL4;refGene_tx=XM_019811619.1;refGene_gn=PIWIL4;refGene_tx=XM_019811620.2
D1 41092022 41092140
refGene_gn=PIWIL4;refGene_tx=XM_019811619.1;refGene_gn=PIWIL4;refGene_tx=XM_019811620.2
D1 41094531 41094655
refGene_gn=PIWIL4;refGene_tx=XM_019811619.1;refGene_gn=PIWIL4;refGene_tx=XM_019811620.2
D1 41099027 41099139
refGene_gn=PIWIL4;refGene_tx=XM_019811619.1;refGene_gn=PIWIL4;refGene_tx=XM_019811620.2
D1 41100902 41101100
refGene_gn=PIWIL4;refGene_tx=XM_019811619.1;refGene_gn=PIWIL4;refGene_tx=XM_019811620.2
D1 41102601 41102682

refGene_gn=PIWIL4;refGene_tx=XM_019811619.1;refGene_gn=PIWIL4;refGene_tx=XM_019811620.2
D1 41104100 41104222
refGene_gn=PIWIL4;refGene_tx=XM_019811619.1;refGene_gn=PIWIL4;refGene_tx=XM_019811620.2
D1 41109036 41109251
refGene_gn=PIWIL4;refGene_tx=XM_019811619.1;refGene_gn=PIWIL4;refGene_tx=XM_019811620.2
D1 41111169 41111295
refGene_gn=PIWIL4;refGene_tx=XM_019811619.1;refGene_gn=PIWIL4;refGene_tx=XM_019811620.2
D1 41114021 41114100
refGene_gn=PIWIL4;refGene_tx=XM_019811619.1;refGene_gn=PIWIL4;refGene_tx=XM_019811620.2
D1 41115281 41115350
refGene_gn=PIWIL4;refGene_tx=XM_019811619.1;refGene_gn=PIWIL4;refGene_tx=XM_019811620.2
D1 41148921 41150495 refGene_gn=FUT4;refGene_tx=XM_011286559.3
D1 41150630 41150877 refGene_gn=FUT4;refGene_tx=XM_011286559.3
D1 41155013 41155088 refGene_gn=FUT4;refGene_tx=XM_011286559.3
D1 41155797 41155828 refGene_gn=FUT4;refGene_tx=XM_011286559.3
D1 41157665 41157719 refGene_gn=FUT4;refGene_tx=XM_011286559.3
D1 41158936 41158990 refGene_gn=FUT4;refGene_tx=XM_011286559.3
D1 41160655 41160672 refGene_gn=FUT4;refGene_tx=XM_011286559.3
D1 41167305 41167310
refGene_gn=CD1H11orf97;refGene_tx=XM_011286521.3
D1 41170576 41170755
refGene_gn=CD1H11orf97;refGene_tx=XM_011286520.3;refGene_gn=CD1H11orf97;refGene_tx=XM_011286521.3
D1 41182168 41182273
refGene_gn=CD1H11orf97;refGene_tx=XM_011286520.3;refGene_gn=CD1H11orf97;refGene_tx=XM_011286521.3
D1 41187219 41187367
refGene_gn=CD1H11orf97;refGene_tx=XM_011286520.3;refGene_gn=CD1H11orf97;refGene_tx=XM_011286521.3
D1 41207138 41207600
refGene_gn=ANKRD49;refGene_tx=XM_019811630.2;refGene_gn=ANKRD49;refGene_tx=XM_023239330.1
D1 41208781 41209039
refGene_gn=ANKRD49;refGene_tx=XM_019811630.2;refGene_gn=ANKRD49;refGene_tx=XM_023239330.1
D1 41213108 41213128
refGene_gn=MRE11;refGene_tx=XM_011286523.3;refGene_gn=MRE11;refGene_tx=XM_011286524.3;refGene_gn=MRE11;refGene_tx=XM_011286525.3;refGene_gn=MRE11;refGene_tx=XM_019811622.2;refGene_gn=MRE11;refGene_tx=XM_019811623.2;refGene_gn=MRE11;refGene_tx=XM_019811628.1
D1 41214213 41214221 refGene_gn=MRE11;refGene_tx=XM_019811624.2
D1 41214767 41214900
refGene_gn=MRE11;refGene_tx=XM_011286523.3;refGene_gn=MRE11;refGene_tx=XM_011286524.3;refGene_gn=MRE11;refGene_tx=XM_011286525.3;refGene_gn=

MRE11;refGene_tx=XM_019811622.2;refGene_gn=MRE11;refGene_tx=XM_0198116
23.2;refGene_gn=MRE11;refGene_tx=XM_019811624.2;refGene_gn=MRE11;refGe
ne_tx=XM_019811625.2;refGene_gn=MRE11;refGene_tx=XM_019811626.2;refGen
e_gn=MRE11;refGene_tx=XM_019811628.1
D1 41220625 41220786
refGene_gn=MRE11;refGene_tx=XM_011286523.3;refGene_gn=MRE11;refGene_tx
=XM_011286524.3;refGene_gn=MRE11;refGene_tx=XM_011286525.3;refGene_gn=
MRE11;refGene_tx=XM_019811622.2;refGene_gn=MRE11;refGene_tx=XM_0198116
23.2;refGene_gn=MRE11;refGene_tx=XM_019811624.2;refGene_gn=MRE11;refGe
ne_tx=XM_019811625.2;refGene_gn=MRE11;refGene_tx=XM_019811626.2;refGen
e_gn=MRE11;refGene_tx=XM_019811627.2;refGene_gn=MRE11;refGene_tx=XM_01
9811628.1
D1 41228493 41228581
refGene_gn=MRE11;refGene_tx=XM_011286523.3;refGene_gn=MRE11;refGene_tx
=XM_011286524.3;refGene_gn=MRE11;refGene_tx=XM_011286525.3;refGene_gn=
MRE11;refGene_tx=XM_019811622.2;refGene_gn=MRE11;refGene_tx=XM_0198116
23.2;refGene_gn=MRE11;refGene_tx=XM_019811624.2;refGene_gn=MRE11;refGe
ne_tx=XM_019811625.2;refGene_gn=MRE11;refGene_tx=XM_019811626.2;refGen
e_gn=MRE11;refGene_tx=XM_019811627.2;refGene_gn=MRE11;refGene_tx=XM_01
9811628.1
D1 41229363 41229505
refGene_gn=MRE11;refGene_tx=XM_011286523.3;refGene_gn=MRE11;refGene_tx
=XM_011286524.3;refGene_gn=MRE11;refGene_tx=XM_011286525.3;refGene_gn=
MRE11;refGene_tx=XM_019811622.2;refGene_gn=MRE11;refGene_tx=XM_0198116
23.2;refGene_gn=MRE11;refGene_tx=XM_019811624.2;refGene_gn=MRE11;refGe
ne_tx=XM_019811625.2;refGene_gn=MRE11;refGene_tx=XM_019811626.2;refGen
e_gn=MRE11;refGene_tx=XM_019811627.2;refGene_gn=MRE11;refGene_tx=XM_01
9811628.1
D1 41232751 41232866
refGene_gn=MRE11;refGene_tx=XM_011286523.3;refGene_gn=MRE11;refGene_tx
=XM_011286524.3;refGene_gn=MRE11;refGene_tx=XM_011286525.3;refGene_gn=
MRE11;refGene_tx=XM_019811622.2;refGene_gn=MRE11;refGene_tx=XM_0198116
23.2;refGene_gn=MRE11;refGene_tx=XM_019811624.2;refGene_gn=MRE11;refGe
ne_tx=XM_019811625.2;refGene_gn=MRE11;refGene_tx=XM_019811626.2;refGen
e_gn=MRE11;refGene_tx=XM_019811627.2;refGene_gn=MRE11;refGene_tx=XM_01
9811628.1
D1 41237481 41237667
refGene_gn=MRE11;refGene_tx=XM_011286523.3;refGene_gn=MRE11;refGene_tx
=XM_011286524.3;refGene_gn=MRE11;refGene_tx=XM_011286525.3;refGene_gn=
MRE11;refGene_tx=XM_019811622.2;refGene_gn=MRE11;refGene_tx=XM_0198116
23.2;refGene_gn=MRE11;refGene_tx=XM_019811624.2;refGene_gn=MRE11;refGe
ne_tx=XM_019811625.2;refGene_gn=MRE11;refGene_tx=XM_019811626.2;refGen
e_gn=MRE11;refGene_tx=XM_019811627.2;refGene_gn=MRE11;refGene_tx=XM_01
9811628.1
D1 41238566 41238738
refGene_gn=MRE11;refGene_tx=XM_011286523.3;refGene_gn=MRE11;refGene_tx
=XM_011286524.3;refGene_gn=MRE11;refGene_tx=XM_011286525.3;refGene_gn=
MRE11;refGene_tx=XM_019811622.2;refGene_gn=MRE11;refGene_tx=XM_0198116
23.2;refGene_gn=MRE11;refGene_tx=XM_019811624.2;refGene_gn=MRE11;refGe
ne_tx=XM_019811625.2;refGene_gn=MRE11;refGene_tx=XM_019811626.2;refGen

e_gn=MRE11;refGene_tx=XM_019811627.2;refGene_gn=MRE11;refGene_tx=XM_019811628.1
D1 41239977 41240058
refGene_gn=MRE11;refGene_tx=XM_011286523.3;refGene_gn=MRE11;refGene_tx=XM_011286524.3;refGene_gn=MRE11;refGene_tx=XM_011286525.3;refGene_gn=MRE11;refGene_tx=XM_019811622.2;refGene_gn=MRE11;refGene_tx=XM_019811623.2;refGene_gn=MRE11;refGene_tx=XM_019811624.2;refGene_gn=MRE11;refGene_tx=XM_019811625.2;refGene_gn=MRE11;refGene_tx=XM_019811626.2;refGene_gn=MRE11;refGene_tx=XM_019811627.2;refGene_gn=MRE11;refGene_tx=XM_019811628.1
D1 41243753 41243880
refGene_gn=MRE11;refGene_tx=XM_011286523.3;refGene_gn=MRE11;refGene_tx=XM_011286524.3;refGene_gn=MRE11;refGene_tx=XM_011286525.3;refGene_gn=MRE11;refGene_tx=XM_019811622.2;refGene_gn=MRE11;refGene_tx=XM_019811623.2;refGene_gn=MRE11;refGene_tx=XM_019811624.2;refGene_gn=MRE11;refGene_tx=XM_019811625.2;refGene_gn=MRE11;refGene_tx=XM_019811626.2;refGene_gn=MRE11;refGene_tx=XM_019811627.2;refGene_gn=MRE11;refGene_tx=XM_019811628.1
D1 41246801 41246902
refGene_gn=MRE11;refGene_tx=XM_011286523.3;refGene_gn=MRE11;refGene_tx=XM_011286524.3;refGene_gn=MRE11;refGene_tx=XM_011286525.3;refGene_gn=MRE11;refGene_tx=XM_019811622.2;refGene_gn=MRE11;refGene_tx=XM_019811623.2;refGene_gn=MRE11;refGene_tx=XM_019811624.2;refGene_gn=MRE11;refGene_tx=XM_019811625.2;refGene_gn=MRE11;refGene_tx=XM_019811626.2;refGene_gn=MRE11;refGene_tx=XM_019811627.2;refGene_gn=MRE11;refGene_tx=XM_019811628.1
D1 41248298 41248472
refGene_gn=MRE11;refGene_tx=XM_011286523.3;refGene_gn=MRE11;refGene_tx=XM_011286524.3;refGene_gn=MRE11;refGene_tx=XM_011286525.3;refGene_gn=MRE11;refGene_tx=XM_019811622.2;refGene_gn=MRE11;refGene_tx=XM_019811623.2;refGene_gn=MRE11;refGene_tx=XM_019811624.2;refGene_gn=MRE11;refGene_tx=XM_019811625.2;refGene_gn=MRE11;refGene_tx=XM_019811626.2;refGene_gn=MRE11;refGene_tx=XM_019811627.2;refGene_gn=MRE11;refGene_tx=XM_019811628.1
D1 41255439 41255502
refGene_gn=MRE11;refGene_tx=XM_011286523.3;refGene_gn=MRE11;refGene_tx=XM_011286524.3;refGene_gn=MRE11;refGene_tx=XM_011286525.3;refGene_gn=MRE11;refGene_tx=XM_019811622.2;refGene_gn=MRE11;refGene_tx=XM_019811623.2;refGene_gn=MRE11;refGene_tx=XM_019811624.2;refGene_gn=MRE11;refGene_tx=XM_019811625.2;refGene_gn=MRE11;refGene_tx=XM_019811626.2;refGene_gn=MRE11;refGene_tx=XM_019811627.2;refGene_gn=MRE11;refGene_tx=XM_019811628.1
D1 41263709 41263926
refGene_gn=MRE11;refGene_tx=XM_011286523.3;refGene_gn=MRE11;refGene_tx=XM_011286524.3;refGene_gn=MRE11;refGene_tx=XM_011286525.3;refGene_gn=MRE11;refGene_tx=XM_019811622.2;refGene_gn=MRE11;refGene_tx=XM_019811623.2;refGene_gn=MRE11;refGene_tx=XM_019811624.2;refGene_gn=MRE11;refGene_tx=XM_019811625.2;refGene_gn=MRE11;refGene_tx=XM_019811626.2;refGene_gn=MRE11;refGene_tx=XM_019811627.2;refGene_gn=MRE11;refGene_tx=XM_019811628.1

D1 41265597 41265681
refGene_gn=MRE11;refGene_tx=XM_011286523.3;refGene_gn=MRE11;refGene_tx
=XM_011286524.3;refGene_gn=MRE11;refGene_tx=XM_011286525.3;refGene_gn=
MRE11;refGene_tx=XM_019811622.2;refGene_gn=MRE11;refGene_tx=XM_0198116
23.2;refGene_gn=MRE11;refGene_tx=XM_019811624.2;refGene_gn=MRE11;refGe
ne_tx=XM_019811625.2;refGene_gn=MRE11;refGene_tx=XM_019811626.2;refGen
e_gn=MRE11;refGene_tx=XM_019811627.2;refGene_gn=MRE11;refGene_tx=XM_01
9811628.1
D1 41274496 41274555
refGene_gn=MRE11;refGene_tx=XM_011286523.3;refGene_gn=MRE11;refGene_tx
=XM_011286524.3;refGene_gn=MRE11;refGene_tx=XM_011286525.3;refGene_gn=
MRE11;refGene_tx=XM_019811622.2;refGene_gn=MRE11;refGene_tx=XM_0198116
23.2;refGene_gn=MRE11;refGene_tx=XM_019811624.2;refGene_gn=MRE11;refGe
ne_tx=XM_019811625.2;refGene_gn=MRE11;refGene_tx=XM_019811626.2;refGen
e_gn=MRE11;refGene_tx=XM_019811627.2
D1 41277628 41277696
refGene_gn=MRE11;refGene_tx=XM_011286523.3;refGene_gn=MRE11;refGene_tx
=XM_011286524.3;refGene_gn=MRE11;refGene_tx=XM_011286525.3;refGene_gn=
MRE11;refGene_tx=XM_019811622.2;refGene_gn=MRE11;refGene_tx=XM_0198116
23.2;refGene_gn=MRE11;refGene_tx=XM_019811624.2;refGene_gn=MRE11;refGe
ne_tx=XM_019811625.2;refGene_gn=MRE11;refGene_tx=XM_019811626.2;refGen
e_gn=MRE11;refGene_tx=XM_019811627.2
D1 41284333 41284415
refGene_gn=MRE11;refGene_tx=XM_011286523.3;refGene_gn=MRE11;refGene_tx
=XM_011286524.3;refGene_gn=MRE11;refGene_tx=XM_011286525.3;refGene_gn=
MRE11;refGene_tx=XM_019811622.2;refGene_gn=MRE11;refGene_tx=XM_0198116
23.2;refGene_gn=MRE11;refGene_tx=XM_019811624.2;refGene_gn=MRE11;refGe
ne_tx=XM_019811625.2;refGene_gn=MRE11;refGene_tx=XM_019811626.2;refGen
e_gn=MRE11;refGene_tx=XM_019811627.2
D1 41287018 41287075
refGene_gn=MRE11;refGene_tx=XM_011286523.3;refGene_gn=MRE11;refGene_tx
=XM_011286524.3;refGene_gn=MRE11;refGene_tx=XM_011286525.3;refGene_gn=
MRE11;refGene_tx=XM_019811622.2;refGene_gn=MRE11;refGene_tx=XM_0198116
23.2;refGene_gn=MRE11;refGene_tx=XM_019811624.2;refGene_gn=MRE11;refGe
ne_tx=XM_019811625.2;refGene_gn=MRE11;refGene_tx=XM_019811626.2;refGen
e_gn=MRE11;refGene_tx=XM_019811627.2
D1 41305857 41306241 refGene_gn=GPR83;refGene_tx=XM_023239331.1
D1 41310549 41310675 refGene_gn=GPR83;refGene_tx=XM_023239331.1
D1 41312934 41313068 refGene_gn=GPR83;refGene_tx=XM_023239331.1
D1 41316110 41316735 refGene_gn=GPR83;refGene_tx=XM_023239331.1
D1 41369477 41369746 refGene_gn=IZUMO1R;refGene_tx=XM_023238987.1
D1 41369887 41370023 refGene_gn=IZUMO1R;refGene_tx=XM_023238987.1
D1 41370907 41371096 refGene_gn=IZUMO1R;refGene_tx=XM_023238987.1
D1 41371857 41371995 refGene_gn=IZUMO1R;refGene_tx=XM_023238987.1
D1 41439899 41439979
refGene_gn=PANX1;refGene_tx=XM_003992612.4;refGene_gn=PANX1;refGene_tx
=XM_023239332.1
D1 41440528 41441187
refGene_gn=PANX1;refGene_tx=XM_003992612.4;refGene_gn=PANX1;refGene_tx
=XM_023239332.1

D1 41442280 41442504
refGene_gn=PANX1;refGene_tx=XM_003992612.4;refGene_gn=PANX1;refGene_tx
=XM_023239332.1
D1 41457460 41457553 refGene_gn=PANX1;refGene_tx=XM_023239332.1
D1 41458650 41458790
refGene_gn=PANX1;refGene_tx=XM_003992612.4;refGene_gn=PANX1;refGene_tx
=XM_023239332.1
D1 41487766 41487947
refGene_gn=PANX1;refGene_tx=XM_003992612.4;refGene_gn=PANX1;refGene_tx
=XM_023239332.1
D1 41494402 41494605 refGene_gn=HEPHL1;refGene_tx=XM_003992613.5
D1 41494701 41494764 refGene_gn=HEPHL1;refGene_tx=XM_003992613.5
D1 41495451 41495614 refGene_gn=HEPHL1;refGene_tx=XM_003992613.5
D1 41500371 41500511 refGene_gn=HEPHL1;refGene_tx=XM_003992613.5
D1 41502031 41502254 refGene_gn=HEPHL1;refGene_tx=XM_003992613.5
D1 41513604 41513711 refGene_gn=HEPHL1;refGene_tx=XM_003992613.5
D1 41517319 41517460 refGene_gn=HEPHL1;refGene_tx=XM_003992613.5
D1 41520875 41521015 refGene_gn=HEPHL1;refGene_tx=XM_003992613.5
D1 41526007 41526221 refGene_gn=HEPHL1;refGene_tx=XM_003992613.5
D1 41530884 41531097 refGene_gn=HEPHL1;refGene_tx=XM_003992613.5
D1 41534195 41534346 refGene_gn=HEPHL1;refGene_tx=XM_003992613.5
D1 41540507 41540719 refGene_gn=HEPHL1;refGene_tx=XM_003992613.5
D1 41542806 41542938 refGene_gn=HEPHL1;refGene_tx=XM_003992613.5
D1 41543091 41543231 refGene_gn=HEPHL1;refGene_tx=XM_003992613.5
D1 41545820 41545989 refGene_gn=HEPHL1;refGene_tx=XM_003992613.5
D1 41549520 41549775 refGene_gn=HEPHL1;refGene_tx=XM_003992613.5
D1 41552317 41552497 refGene_gn=HEPHL1;refGene_tx=XM_003992613.5
D1 41553176 41553389 refGene_gn=HEPHL1;refGene_tx=XM_003992613.5
D1 41560500 41560745 refGene_gn=HEPHL1;refGene_tx=XM_003992613.5
D1 41575336 41575506 refGene_gn=HEPHL1;refGene_tx=XM_003992613.5
D1 41720063 41720154
refGene_gn=VSTM5;refGene_tx=XM_023239333.1;refGene_gn=VSTM5;refGene_tx
=XM_023239334.1
D1 41746280 41746607
refGene_gn=VSTM5;refGene_tx=XM_023239333.1;refGene_gn=VSTM5;refGene_tx
=XM_023239334.1
D1 41746719 41746860
refGene_gn=VSTM5;refGene_tx=XM_023239333.1;refGene_gn=VSTM5;refGene_tx
=XM_023239334.1
D1 41747009 41747053
refGene_gn=VSTM5;refGene_tx=XM_023239333.1;refGene_gn=VSTM5;refGene_tx
=XM_023239334.1
D1 41757201 41757413
refGene_gn=MED17;refGene_tx=XM_003992615.4;refGene_gn=MED17;refGene_tx
=XM_019811638.2
D1 41758632 41758792
refGene_gn=MED17;refGene_tx=XM_003992615.4;refGene_gn=MED17;refGene_tx
=XM_019811638.2
D1 41760550 41760668
refGene_gn=MED17;refGene_tx=XM_003992615.4;refGene_gn=MED17;refGene_tx

=XM_019811638.2
D1 41765413 41765551
refGene_gn=MED17;refGene_tx=XM_003992615.4;refGene_gn=MED17;refGene_tx
=XM_019811638.2
D1 41766543 41766666 refGene_gn=MED17;refGene_tx=XM_019811638.2
D1 41769993 41770178
refGene_gn=MED17;refGene_tx=XM_003992615.4;refGene_gn=MED17;refGene_tx
=XM_019811638.2
D1 41770847 41770978
refGene_gn=MED17;refGene_tx=XM_003992615.4;refGene_gn=MED17;refGene_tx
=XM_019811638.2
D1 41772179 41772332
refGene_gn=MED17;refGene_tx=XM_003992615.4;refGene_gn=MED17;refGene_tx
=XM_019811638.2
D1 41772690 41772775
refGene_gn=MED17;refGene_tx=XM_003992615.4;refGene_gn=MED17;refGene_tx
=XM_019811638.2
D1 41773079 41773216
refGene_gn=MED17;refGene_tx=XM_003992615.4;refGene_gn=MED17;refGene_tx
=XM_019811638.2
D1 41775137 41775357
refGene_gn=MED17;refGene_tx=XM_003992615.4;refGene_gn=MED17;refGene_tx
=XM_019811638.2
D1 41777214 41777381
refGene_gn=MED17;refGene_tx=XM_003992615.4;refGene_gn=MED17;refGene_tx
=XM_019811638.2
D1 41780306 41780556
refGene_gn=MED17;refGene_tx=XM_003992615.4;refGene_gn=MED17;refGene_tx
=XM_019811638.2
D1 41786487 41786661
refGene_gn=CD1H11orf54;refGene_tx=XM_003992616.5;refGene_gn=CD1H11orf5
4;refGene_tx=XM_006936848.4;refGene_gn=CD1H11orf54;refGene_tx=XM_00693
6849.4;refGene_gn=CD1H11orf54;refGene_tx=XM_006936850.4;refGene_gn=CD1
H11orf54;refGene_tx=XM_019811639.2
D1 41787633 41787750
refGene_gn=CD1H11orf54;refGene_tx=XM_003992616.5;refGene_gn=CD1H11orf5
4;refGene_tx=XM_006936848.4;refGene_gn=CD1H11orf54;refGene_tx=XM_00693
6849.4;refGene_gn=CD1H11orf54;refGene_tx=XM_006936850.4;refGene_gn=CD1
H11orf54;refGene_tx=XM_019811639.2
D1 41793525 41793675
refGene_gn=CD1H11orf54;refGene_tx=XM_003992616.5;refGene_gn=CD1H11orf5
4;refGene_tx=XM_006936848.4;refGene_gn=CD1H11orf54;refGene_tx=XM_00693
6849.4;refGene_gn=CD1H11orf54;refGene_tx=XM_006936850.4;refGene_gn=CD1
H11orf54;refGene_tx=XM_019811639.2
D1 41794670 41794847
refGene_gn=CD1H11orf54;refGene_tx=XM_003992616.5;refGene_gn=CD1H11orf5
4;refGene_tx=XM_006936848.4;refGene_gn=CD1H11orf54;refGene_tx=XM_00693
6849.4;refGene_gn=CD1H11orf54;refGene_tx=XM_006936850.4;refGene_gn=CD1
H11orf54;refGene_tx=XM_019811639.2
D1 41795798 41795900

refGene_gn=CD1H11orf54;refGene_tx=XM_003992616.5;refGene_gn=CD1H11orf54;refGene_tx=XM_006936848.4;refGene_gn=CD1H11orf54;refGene_tx=XM_006936849.4;refGene_gn=CD1H11orf54;refGene_tx=XM_006936850.4;refGene_gn=CD1H11orf54;refGene_tx=XM_019811639.2
D1 41796079 41796153
refGene_gn=CD1H11orf54;refGene_tx=XM_003992616.5;refGene_gn=CD1H11orf54;refGene_tx=XM_006936848.4;refGene_gn=CD1H11orf54;refGene_tx=XM_006936849.4;refGene_gn=CD1H11orf54;refGene_tx=XM_006936850.4;refGene_gn=CD1H11orf54;refGene_tx=XM_019811639.2
D1 41799306 41799405
refGene_gn=CD1H11orf54;refGene_tx=XM_003992616.5;refGene_gn=CD1H11orf54;refGene_tx=XM_006936848.4;refGene_gn=CD1H11orf54;refGene_tx=XM_006936849.4;refGene_gn=CD1H11orf54;refGene_tx=XM_006936850.4;refGene_gn=CD1H11orf54;refGene_tx=XM_019811639.2
D1 41801807 41801862
refGene_gn=CD1H11orf54;refGene_tx=XM_003992616.5;refGene_gn=CD1H11orf54;refGene_tx=XM_006936848.4;refGene_gn=CD1H11orf54;refGene_tx=XM_006936849.4;refGene_gn=CD1H11orf54;refGene_tx=XM_006936850.4;refGene_gn=CD1H11orf54;refGene_tx=XM_019811639.2
D1 41808893 41808952
refGene_gn=TAF1D;refGene_tx=XM_019811640.2;refGene_gn=TAF1D;refGene_tx=XM_019811641.2
D1 41809657 41810036
refGene_gn=TAF1D;refGene_tx=XM_019811640.2;refGene_gn=TAF1D;refGene_tx=XM_019811641.2
D1 41811992 41812165
refGene_gn=TAF1D;refGene_tx=XM_019811640.2;refGene_gn=TAF1D;refGene_tx=XM_019811641.2
D1 41812460 41812521
refGene_gn=TAF1D;refGene_tx=XM_019811640.2;refGene_gn=TAF1D;refGene_tx=XM_019811641.2
D1 41812860 41813025
refGene_gn=TAF1D;refGene_tx=XM_019811640.2;refGene_gn=TAF1D;refGene_tx=XM_019811641.2
D1 41818342 41818381 refGene_gn=CEP295;refGene_tx=XM_006936838.4
D1 41818458 41818558 refGene_gn=CEP295;refGene_tx=XM_006936838.4
D1 41818648 41818748 refGene_gn=CEP295;refGene_tx=XM_006936838.4
D1 41818830 41818995 refGene_gn=CEP295;refGene_tx=XM_006936838.4
D1 41819124 41819221 refGene_gn=CEP295;refGene_tx=XM_006936838.4
D1 41820274 41820415 refGene_gn=CEP295;refGene_tx=XM_006936838.4
D1 41821280 41821921 refGene_gn=CEP295;refGene_tx=XM_006936838.4
D1 41822765 41822940 refGene_gn=CEP295;refGene_tx=XM_006936838.4
D1 41823843 41823924 refGene_gn=CEP295;refGene_tx=XM_006936838.4
D1 41823925 41823984 refGene_gn=CEP295;refGene_tx=XM_006936838.4
D1 41825102 41825354 refGene_gn=CEP295;refGene_tx=XM_006936838.4
D1 41825759 41825862 refGene_gn=CEP295;refGene_tx=XM_006936838.4
D1 41826408 41826506 refGene_gn=CEP295;refGene_tx=XM_006936838.4
D1 41830736 41830880 refGene_gn=CEP295;refGene_tx=XM_006936838.4
D1 41835171 41835315 refGene_gn=CEP295;refGene_tx=XM_006936838.4
D1 41835452 41835618 refGene_gn=CEP295;refGene_tx=XM_006936838.4

D1 41836632 41839981 refGene_gn=CEP295;refGene_tx=XM_006936838.4
D1 41840257 41840355 refGene_gn=CEP295;refGene_tx=XM_006936838.4
D1 41840785 41840917 refGene_gn=CEP295;refGene_tx=XM_006936838.4
D1 41842516 41842620 refGene_gn=CEP295;refGene_tx=XM_006936838.4
D1 41842765 41842858 refGene_gn=CEP295;refGene_tx=XM_006936838.4
D1 41847053 41847275 refGene_gn=CEP295;refGene_tx=XM_006936838.4
D1 41849861 41850029 refGene_gn=CEP295;refGene_tx=XM_006936838.4
D1 41850250 41850434 refGene_gn=CEP295;refGene_tx=XM_006936838.4
D1 41853468 41853609 refGene_gn=CEP295;refGene_tx=XM_006936838.4
D1 41859155 41859251 refGene_gn=CEP295;refGene_tx=XM_006936838.4
D1 41861374 41861468 refGene_gn=CEP295;refGene_tx=XM_006936838.4
D1 41862586 41862711 refGene_gn=CEP295;refGene_tx=XM_006936838.4
D1 41869277 41869478 refGene_gn=CEP295;refGene_tx=XM_006936838.4
D1 41870757 41870865 refGene_gn=CEP295;refGene_tx=XM_006936838.4
D1 41974283 41974507 refGene_gn=SMCO4;refGene_tx=XM_023239704.1
D1 41974600 41974786 refGene_gn=SMCO4;refGene_tx=XM_023239704.1
D1 41976533 41976600 refGene_gn=SMCO4;refGene_tx=XM_023239704.1
D1 41976729 41976940 refGene_gn=SMCO4;refGene_tx=XM_023239704.1
D1 42018995 42019072 refGene_gn=SMCO4;refGene_tx=XM_023239704.1
D1 42040015 42040429 refGene_gn=SMCO4;refGene_tx=XM_023239704.1
D1 42070824 42071001
refGene_gn=DEUP1;refGene_tx=XM_023239335.1;refGene_gn=DEUP1;refGene_tx=XM_023239336.1;refGene_gn=DEUP1;refGene_tx=XM_023239337.1;refGene_gn=DEUP1;refGene_tx=XM_023239338.1;refGene_gn=DEUP1;refGene_tx=XM_023239339.1;refGene_gn=DEUP1;refGene_tx=XM_023239340.1;refGene_gn=DEUP1;refGene_tx=XM_023239341.1;refGene_gn=DEUP1;refGene_tx=XM_023239342.1;refGene_gn=DEUP1;refGene_tx=XM_023239343.1
D1 42081173 42081288
refGene_gn=DEUP1;refGene_tx=XM_023239335.1;refGene_gn=DEUP1;refGene_tx=XM_023239336.1;refGene_gn=DEUP1;refGene_tx=XM_023239337.1;refGene_gn=DEUP1;refGene_tx=XM_023239338.1;refGene_gn=DEUP1;refGene_tx=XM_023239339.1;refGene_gn=DEUP1;refGene_tx=XM_023239340.1;refGene_gn=DEUP1;refGene_tx=XM_023239341.1;refGene_gn=DEUP1;refGene_tx=XM_023239342.1;refGene_gn=DEUP1;refGene_tx=XM_023239343.1
D1 42087646 42087843
refGene_gn=DEUP1;refGene_tx=XM_023239335.1;refGene_gn=DEUP1;refGene_tx=XM_023239336.1;refGene_gn=DEUP1;refGene_tx=XM_023239337.1;refGene_gn=DEUP1;refGene_tx=XM_023239338.1;refGene_gn=DEUP1;refGene_tx=XM_023239339.1;refGene_gn=DEUP1;refGene_tx=XM_023239340.1;refGene_gn=DEUP1;refGene_tx=XM_023239341.1;refGene_gn=DEUP1;refGene_tx=XM_023239342.1;refGene_gn=DEUP1;refGene_tx=XM_023239343.1
D1 42091562 42091586
refGene_gn=DEUP1;refGene_tx=XM_023239335.1;refGene_gn=DEUP1;refGene_tx=XM_023239336.1;refGene_gn=DEUP1;refGene_tx=XM_023239337.1;refGene_gn=DEUP1;refGene_tx=XM_023239338.1;refGene_gn=DEUP1;refGene_tx=XM_023239339.1;refGene_gn=DEUP1;refGene_tx=XM_023239340.1;refGene_gn=DEUP1;refGene_tx=XM_023239341.1;refGene_gn=DEUP1;refGene_tx=XM_023239342.1;refGene_gn=DEUP1;refGene_tx=XM_023239343.1
D1 42101938 42102025
refGene_gn=DEUP1;refGene_tx=XM_023239335.1;refGene_gn=DEUP1;refGene_tx

=XM_023239336.1;refGene_gn=DEUP1;refGene_tx=XM_023239337.1;refGene_gn=DEUP1;refGene_tx=XM_023239338.1;refGene_gn=DEUP1;refGene_tx=XM_023239339.1;refGene_gn=DEUP1;refGene_tx=XM_023239340.1;refGene_gn=DEUP1;refGene_tx=XM_023239341.1;refGene_gn=DEUP1;refGene_tx=XM_023239342.1;refGene_gn=DEUP1;refGene_tx=XM_023239343.1
D1 42105059 42105314
refGene_gn=DEUP1;refGene_tx=XM_023239335.1;refGene_gn=DEUP1;refGene_tx=XM_023239336.1;refGene_gn=DEUP1;refGene_tx=XM_023239337.1;refGene_gn=DEUP1;refGene_tx=XM_023239338.1;refGene_gn=DEUP1;refGene_tx=XM_023239339.1;refGene_gn=DEUP1;refGene_tx=XM_023239340.1;refGene_gn=DEUP1;refGene_tx=XM_023239341.1;refGene_gn=DEUP1;refGene_tx=XM_023239342.1
D1 42106814 42106865
refGene_gn=DEUP1;refGene_tx=XM_023239335.1;refGene_gn=DEUP1;refGene_tx=XM_023239336.1;refGene_gn=DEUP1;refGene_tx=XM_023239337.1;refGene_gn=DEUP1;refGene_tx=XM_023239338.1;refGene_gn=DEUP1;refGene_tx=XM_023239340.1;refGene_gn=DEUP1;refGene_tx=XM_023239342.1
D1 42109656 42109762
refGene_gn=DEUP1;refGene_tx=XM_023239335.1;refGene_gn=DEUP1;refGene_tx=XM_023239336.1;refGene_gn=DEUP1;refGene_tx=XM_023239337.1;refGene_gn=DEUP1;refGene_tx=XM_023239338.1;refGene_gn=DEUP1;refGene_tx=XM_023239339.1;refGene_gn=DEUP1;refGene_tx=XM_023239340.1;refGene_gn=DEUP1;refGene_tx=XM_023239341.1;refGene_gn=DEUP1;refGene_tx=XM_023239342.1;refGene_gn=DEUP1;refGene_tx=XM_023239343.1
D1 42113015 42113161
refGene_gn=DEUP1;refGene_tx=XM_023239335.1;refGene_gn=DEUP1;refGene_tx=XM_023239336.1;refGene_gn=DEUP1;refGene_tx=XM_023239337.1;refGene_gn=DEUP1;refGene_tx=XM_023239338.1;refGene_gn=DEUP1;refGene_tx=XM_023239339.1;refGene_gn=DEUP1;refGene_tx=XM_023239340.1;refGene_gn=DEUP1;refGene_tx=XM_023239341.1;refGene_gn=DEUP1;refGene_tx=XM_023239342.1;refGene_gn=DEUP1;refGene_tx=XM_023239343.1
D1 42119178 42119421
refGene_gn=DEUP1;refGene_tx=XM_023239335.1;refGene_gn=DEUP1;refGene_tx=XM_023239336.1;refGene_gn=DEUP1;refGene_tx=XM_023239337.1;refGene_gn=DEUP1;refGene_tx=XM_023239338.1;refGene_gn=DEUP1;refGene_tx=XM_023239339.1;refGene_gn=DEUP1;refGene_tx=XM_023239340.1;refGene_gn=DEUP1;refGene_tx=XM_023239341.1;refGene_gn=DEUP1;refGene_tx=XM_023239342.1;refGene_gn=DEUP1;refGene_tx=XM_023239343.1
D1 42121420 42121534
refGene_gn=DEUP1;refGene_tx=XM_023239335.1;refGene_gn=DEUP1;refGene_tx=XM_023239336.1;refGene_gn=DEUP1;refGene_tx=XM_023239337.1;refGene_gn=DEUP1;refGene_tx=XM_023239338.1;refGene_gn=DEUP1;refGene_tx=XM_023239339.1;refGene_gn=DEUP1;refGene_tx=XM_023239340.1;refGene_gn=DEUP1;refGene_tx=XM_023239341.1;refGene_gn=DEUP1;refGene_tx=XM_023239342.1;refGene_gn=DEUP1;refGene_tx=XM_023239343.1
D1 42124760 42124895
refGene_gn=DEUP1;refGene_tx=XM_023239335.1;refGene_gn=DEUP1;refGene_tx=XM_023239336.1;refGene_gn=DEUP1;refGene_tx=XM_023239337.1;refGene_gn=DEUP1;refGene_tx=XM_023239338.1;refGene_gn=DEUP1;refGene_tx=XM_023239339.1;refGene_gn=DEUP1;refGene_tx=XM_023239340.1;refGene_gn=DEUP1;refGene_tx=XM_023239341.1;refGene_gn=DEUP1;refGene_tx=XM_023239342.1;refGen

e_gn=DEUP1;refGene_tx=XM_023239343.1
D1 42132133 42132229
refGene_gn=DEUP1;refGene_tx=XM_023239335.1;refGene_gn=DEUP1;refGene_tx=XM_023239336.1;refGene_gn=DEUP1;refGene_tx=XM_023239337.1;refGene_gn=DEUP1;refGene_tx=XM_023239338.1;refGene_gn=DEUP1;refGene_tx=XM_023239339.1;refGene_gn=DEUP1;refGene_tx=XM_023239340.1;refGene_gn=DEUP1;refGene_tx=XM_023239341.1;refGene_gn=DEUP1;refGene_tx=XM_023239342.1;refGene_gn=DEUP1;refGene_tx=XM_023239343.1
D1 42133873 42134045
refGene_gn=DEUP1;refGene_tx=XM_023239335.1;refGene_gn=DEUP1;refGene_tx=XM_023239336.1;refGene_gn=DEUP1;refGene_tx=XM_023239337.1;refGene_gn=DEUP1;refGene_tx=XM_023239338.1;refGene_gn=DEUP1;refGene_tx=XM_023239339.1;refGene_gn=DEUP1;refGene_tx=XM_023239340.1;refGene_gn=DEUP1;refGene_tx=XM_023239341.1;refGene_gn=DEUP1;refGene_tx=XM_023239342.1;refGene_gn=DEUP1;refGene_tx=XM_023239343.1
D1 42140210 42140258
refGene_gn=DEUP1;refGene_tx=XM_023239335.1;refGene_gn=DEUP1;refGene_tx=XM_023239336.1;refGene_gn=DEUP1;refGene_tx=XM_023239339.1;refGene_gn=DEUP1;refGene_tx=XM_023239340.1;refGene_gn=DEUP1;refGene_tx=XM_023239341.1;refGene_gn=DEUP1;refGene_tx=XM_023239343.1
D1 42151127 42151156
refGene_gn=DEUP1;refGene_tx=XM_023239335.1;refGene_gn=DEUP1;refGene_tx=XM_023239336.1;refGene_gn=DEUP1;refGene_tx=XM_023239337.1;refGene_gn=DEUP1;refGene_tx=XM_023239338.1;refGene_gn=DEUP1;refGene_tx=XM_023239339.1;refGene_gn=DEUP1;refGene_tx=XM_023239340.1;refGene_gn=DEUP1;refGene_tx=XM_023239341.1;refGene_gn=DEUP1;refGene_tx=XM_023239343.1
D1 42281331 42281383 refGene_gn=SLC36A4;refGene_tx=XM_023239344.1
D1 42292915 42293039 refGene_gn=SLC36A4;refGene_tx=XM_023239344.1
D1 42294441 42294532 refGene_gn=SLC36A4;refGene_tx=XM_023239344.1
D1 42295734 42295823 refGene_gn=SLC36A4;refGene_tx=XM_023239344.1
D1 42296858 42296954
refGene_gn=SLC36A4;refGene_tx=XM_019811652.2;refGene_gn=SLC36A4;refGene_tx=XM_023239344.1
D1 42299284 42299369
refGene_gn=SLC36A4;refGene_tx=XM_019811652.2;refGene_gn=SLC36A4;refGene_tx=XM_023239344.1
D1 42304577 42304805
refGene_gn=SLC36A4;refGene_tx=XM_019811652.2;refGene_gn=SLC36A4;refGene_tx=XM_023239344.1
D1 42306724 42306823
refGene_gn=SLC36A4;refGene_tx=XM_019811652.2;refGene_gn=SLC36A4;refGene_tx=XM_023239344.1
D1 42313191 42313361
refGene_gn=SLC36A4;refGene_tx=XM_019811652.2;refGene_gn=SLC36A4;refGene_tx=XM_023239344.1
D1 42322809 42322979
refGene_gn=SLC36A4;refGene_tx=XM_019811652.2;refGene_gn=SLC36A4;refGene_tx=XM_023239344.1
D1 42328098 42328406
refGene_gn=SLC36A4;refGene_tx=XM_019811652.2;refGene_gn=SLC36A4;refGen

e_tx=XM_023239344.1
D1	42437567	42438439	refGene_gn=MTNR1B;refGene_tx=XM_003992620.4
D1	42449306	42449577	refGene_gn=MTNR1B;refGene_tx=XM_003992620.4
D1	42488195	42488339
refGene_gn=LOC101083959;refGene_tx=XM_003992621.3
D1	42488529	42488709
refGene_gn=LOC101083959;refGene_tx=XM_003992621.3
D1	42488971	42489268
refGene_gn=LOC101083959;refGene_tx=XM_003992621.3
D1	42489376	42489527
refGene_gn=LOC101083959;refGene_tx=XM_003992621.3
D1	42489845	42490102
refGene_gn=LOC101083959;refGene_tx=XM_003992621.3
D1	42490263	42490498
refGene_gn=LOC101083959;refGene_tx=XM_003992621.3
D1	42490623	42490742
refGene_gn=LOC101083959;refGene_tx=XM_003992621.3
D1	42512837	42513460
refGene_gn=FAT3;refGene_tx=XM_011286534.3;refGene_gn=FAT3;refGene_tx=XM_019811653.2;refGene_gn=FAT3;refGene_tx=XM_019811654.2;refGene_gn=FAT3;refGene_tx=XM_019811655.2
D1	42514074	42514110
refGene_gn=FAT3;refGene_tx=XM_019811653.2;refGene_gn=FAT3;refGene_tx=XM_019811654.2
D1	42514680	42514740
refGene_gn=FAT3;refGene_tx=XM_019811653.2;refGene_gn=FAT3;refGene_tx=XM_019811654.2
D1	42517035	42517149
refGene_gn=FAT3;refGene_tx=XM_011286534.3;refGene_gn=FAT3;refGene_tx=XM_019811653.2;refGene_gn=FAT3;refGene_tx=XM_019811654.2;refGene_gn=FAT3;refGene_tx=XM_019811655.2
D1	42520634	42521290
refGene_gn=FAT3;refGene_tx=XM_011286534.3;refGene_gn=FAT3;refGene_tx=XM_019811653.2;refGene_gn=FAT3;refGene_tx=XM_019811654.2;refGene_gn=FAT3;refGene_tx=XM_019811655.2
D1	42523228	42523382
refGene_gn=FAT3;refGene_tx=XM_011286534.3;refGene_gn=FAT3;refGene_tx=XM_019811653.2;refGene_gn=FAT3;refGene_tx=XM_019811654.2;refGene_gn=FAT3;refGene_tx=XM_019811655.2
D1	42537998	42538467
refGene_gn=FAT3;refGene_tx=XM_011286534.3;refGene_gn=FAT3;refGene_tx=XM_019811653.2;refGene_gn=FAT3;refGene_tx=XM_019811654.2;refGene_gn=FAT3;refGene_tx=XM_019811655.2
D1	42546113	42546271
refGene_gn=FAT3;refGene_tx=XM_011286534.3;refGene_gn=FAT3;refGene_tx=XM_019811653.2;refGene_gn=FAT3;refGene_tx=XM_019811654.2;refGene_gn=FAT3;refGene_tx=XM_019811655.2
D1	42548196	42548331
refGene_gn=FAT3;refGene_tx=XM_011286534.3;refGene_gn=FAT3;refGene_tx=XM_019811653.2;refGene_gn=FAT3;refGene_tx=XM_019811654.2;refGene_gn=FAT

3;refGene_tx=XM_019811655.2
D1	42560052	42560851
refGene_gn=FAT3;refGene_tx=XM_011286534.3;refGene_gn=FAT3;refGene_tx=XM_019811653.2;refGene_gn=FAT3;refGene_tx=XM_019811654.2;refGene_gn=FAT3;refGene_tx=XM_019811655.2
D1	42563933	42564131
refGene_gn=FAT3;refGene_tx=XM_011286534.3;refGene_gn=FAT3;refGene_tx=XM_019811653.2;refGene_gn=FAT3;refGene_tx=XM_019811654.2;refGene_gn=FAT3;refGene_tx=XM_019811655.2
D1	42567415	42567559
refGene_gn=FAT3;refGene_tx=XM_011286534.3;refGene_gn=FAT3;refGene_tx=XM_019811653.2;refGene_gn=FAT3;refGene_tx=XM_019811654.2;refGene_gn=FAT3;refGene_tx=XM_019811655.2
D1	42568764	42568902
refGene_gn=FAT3;refGene_tx=XM_011286534.3;refGene_gn=FAT3;refGene_tx=XM_019811653.2;refGene_gn=FAT3;refGene_tx=XM_019811654.2;refGene_gn=FAT3;refGene_tx=XM_019811655.2
D1	42570148	42570363
refGene_gn=FAT3;refGene_tx=XM_011286534.3;refGene_gn=FAT3;refGene_tx=XM_019811653.2;refGene_gn=FAT3;refGene_tx=XM_019811654.2;refGene_gn=FAT3;refGene_tx=XM_019811655.2
D1	42572891	42573281
refGene_gn=FAT3;refGene_tx=XM_011286534.3;refGene_gn=FAT3;refGene_tx=XM_019811653.2;refGene_gn=FAT3;refGene_tx=XM_019811654.2;refGene_gn=FAT3;refGene_tx=XM_019811655.2
D1	42593985	42594219
refGene_gn=FAT3;refGene_tx=XM_011286534.3;refGene_gn=FAT3;refGene_tx=XM_019811653.2;refGene_gn=FAT3;refGene_tx=XM_019811654.2;refGene_gn=FAT3;refGene_tx=XM_019811655.2
D1	42596465	42596619
refGene_gn=FAT3;refGene_tx=XM_011286534.3;refGene_gn=FAT3;refGene_tx=XM_019811653.2;refGene_gn=FAT3;refGene_tx=XM_019811654.2;refGene_gn=FAT3;refGene_tx=XM_019811655.2
D1	42597459	42597656
refGene_gn=FAT3;refGene_tx=XM_011286534.3;refGene_gn=FAT3;refGene_tx=XM_019811653.2;refGene_gn=FAT3;refGene_tx=XM_019811654.2;refGene_gn=FAT3;refGene_tx=XM_019811655.2
D1	42600939	42605013
refGene_gn=FAT3;refGene_tx=XM_011286534.3;refGene_gn=FAT3;refGene_tx=XM_019811653.2;refGene_gn=FAT3;refGene_tx=XM_019811654.2;refGene_gn=FAT3;refGene_tx=XM_019811655.2
D1	42609938	42610149
refGene_gn=FAT3;refGene_tx=XM_011286534.3;refGene_gn=FAT3;refGene_tx=XM_019811653.2;refGene_gn=FAT3;refGene_tx=XM_019811654.2;refGene_gn=FAT3;refGene_tx=XM_019811655.2
D1	42612619	42612895
refGene_gn=FAT3;refGene_tx=XM_011286534.3;refGene_gn=FAT3;refGene_tx=XM_019811653.2;refGene_gn=FAT3;refGene_tx=XM_019811654.2;refGene_gn=FAT3;refGene_tx=XM_019811655.2
D1	42626655	42626795

refGene_gn=FAT3;refGene_tx=XM_011286534.3;refGene_gn=FAT3;refGene_tx=XM_019811653.2;refGene_gn=FAT3;refGene_tx=XM_019811654.2;refGene_gn=FAT3;refGene_tx=XM_019811655.2
D1 42635213 42635282
refGene_gn=FAT3;refGene_tx=XM_019811653.2;refGene_gn=FAT3;refGene_tx=XM_019811655.2
D1 42635749 42635960
refGene_gn=FAT3;refGene_tx=XM_011286534.3;refGene_gn=FAT3;refGene_tx=XM_019811653.2;refGene_gn=FAT3;refGene_tx=XM_019811654.2;refGene_gn=FAT3;refGene_tx=XM_019811655.2
D1 42638286 42638601
refGene_gn=FAT3;refGene_tx=XM_011286534.3;refGene_gn=FAT3;refGene_tx=XM_019811653.2;refGene_gn=FAT3;refGene_tx=XM_019811654.2;refGene_gn=FAT3;refGene_tx=XM_019811655.2
D1 42698381 42698443
refGene_gn=FAT3;refGene_tx=XM_011286534.3;refGene_gn=FAT3;refGene_tx=XM_019811653.2;refGene_gn=FAT3;refGene_tx=XM_019811654.2;refGene_gn=FAT3;refGene_tx=XM_019811655.2
D1 42871477 42871792
refGene_gn=FAT3;refGene_tx=XM_011286534.3;refGene_gn=FAT3;refGene_tx=XM_019811653.2;refGene_gn=FAT3;refGene_tx=XM_019811654.2;refGene_gn=FAT3;refGene_tx=XM_019811655.2
D1 43029999 43033291
refGene_gn=FAT3;refGene_tx=XM_011286534.3;refGene_gn=FAT3;refGene_tx=XM_019811653.2;refGene_gn=FAT3;refGene_tx=XM_019811654.2;refGene_gn=FAT3;refGene_tx=XM_019811655.2
D1 45079588 45079652 refGene_gn=CHORDC1;refGene_tx=XM_003992634.5
D1 45083305 45083355 refGene_gn=CHORDC1;refGene_tx=XM_003992634.5
D1 45086399 45086456 refGene_gn=CHORDC1;refGene_tx=XM_003992634.5
D1 45088091 45088249 refGene_gn=CHORDC1;refGene_tx=XM_003992634.5
D1 45090775 45090879 refGene_gn=CHORDC1;refGene_tx=XM_003992634.5
D1 45091531 45091590 refGene_gn=CHORDC1;refGene_tx=XM_003992634.5
D1 45095365 45095436 refGene_gn=CHORDC1;refGene_tx=XM_003992634.5
D1 45096384 45096490 refGene_gn=CHORDC1;refGene_tx=XM_003992634.5
D1 45098410 45098530 refGene_gn=CHORDC1;refGene_tx=XM_003992634.5
D1 45099205 45099268 refGene_gn=CHORDC1;refGene_tx=XM_003992634.5
D1 45099947 45100094 refGene_gn=CHORDC1;refGene_tx=XM_003992634.5
D1 45114599 45114789
refGene_gn=NAALAD2;refGene_tx=XM_003992636.5;refGene_gn=NAALAD2;refGene_tx=XM_006936857.3
D1 45125479 45125572
refGene_gn=NAALAD2;refGene_tx=XM_003992636.5;refGene_gn=NAALAD2;refGene_tx=XM_006936857.3
D1 45126656 45126738
refGene_gn=NAALAD2;refGene_tx=XM_003992636.5;refGene_gn=NAALAD2;refGene_tx=XM_006936857.3
D1 45129491 45129756
refGene_gn=NAALAD2;refGene_tx=XM_003992636.5;refGene_gn=NAALAD2;refGene_tx=XM_006936857.3
D1 45131209 45131300

refGene_gn=NAALAD2;refGene_tx=XM_003992636.5;refGene_gn=NAALAD2;refGene_tx=XM_006936857.3
D1 45132495 45132587 refGene_gn=NAALAD2;refGene_tx=XM_003992636.5;refGene_gn=NAALAD2;refGene_tx=XM_006936857.3
D1 45135691 45135759 refGene_gn=NAALAD2;refGene_tx=XM_003992636.5;refGene_gn=NAALAD2;refGene_tx=XM_006936857.3
D1 45136453 45136517 refGene_gn=NAALAD2;refGene_tx=XM_003992636.5;refGene_gn=NAALAD2;refGene_tx=XM_006936857.3
D1 45146636 45146719 refGene_gn=NAALAD2;refGene_tx=XM_003992636.5;refGene_gn=NAALAD2;refGene_tx=XM_006936857.3
D1 45146824 45146944 refGene_gn=NAALAD2;refGene_tx=XM_003992636.5;refGene_gn=NAALAD2;refGene_tx=XM_006936857.3
D1 45147224 45147310 refGene_gn=NAALAD2;refGene_tx=XM_003992636.5;refGene_gn=NAALAD2;refGene_tx=XM_006936857.3
D1 45151812 45151911 refGene_gn=NAALAD2;refGene_tx=XM_003992636.5;refGene_gn=NAALAD2;refGene_tx=XM_006936857.3
D1 45153365 45153459 refGene_gn=NAALAD2;refGene_tx=XM_003992636.5;refGene_gn=NAALAD2;refGene_tx=XM_006936857.3
D1 45157303 45157490 refGene_gn=NAALAD2;refGene_tx=XM_003992636.5;refGene_gn=NAALAD2;refGene_tx=XM_006936857.3
D1 45159184 45159310 refGene_gn=NAALAD2;refGene_tx=XM_003992636.5;refGene_gn=NAALAD2;refGene_tx=XM_006936857.3
D1 45161875 45161977 refGene_gn=NAALAD2;refGene_tx=XM_003992636.5;refGene_gn=NAALAD2;refGene_tx=XM_006936857.3
D1 45163470 45163657 refGene_gn=NAALAD2;refGene_tx=XM_003992636.5;refGene_gn=NAALAD2;refGene_tx=XM_006936857.3
D1 45173596 45173705 refGene_gn=NAALAD2;refGene_tx=XM_003992636.5;refGene_gn=NAALAD2;refGene_tx=XM_006936857.3
D1 45174449 45174580 refGene_gn=NAALAD2;refGene_tx=XM_003992636.5;refGene_gn=NAALAD2;refGene_tx=XM_006936857.3
D1 45178838 45178974 refGene_gn=NAALAD2;refGene_tx=XM_006936857.3
D1 45181666 45181778 refGene_gn=NAALAD2;refGene_tx=XM_006936857.3
D1 45221934 45223170 refGene_gn=UBTFL1;refGene_tx=XM_006936859.3
D1 45244582 45244789 refGene_gn=LOC101090384;refGene_tx=XM_023239345.1
D1 45245251 45245274

refGene_gn=LOC101090384;refGene_tx=XM_023239345.1
D1 45246150 45246248
refGene_gn=LOC101090384;refGene_tx=XM_023239345.1
D1 45247260 45247754
refGene_gn=LOC101090384;refGene_tx=XM_023239345.1
D1 45256822 45257316
refGene_gn=LOC101090641;refGene_tx=XM_003992640.1
D1 45258799 45258897
refGene_gn=LOC101090641;refGene_tx=XM_003992640.1
D1 45259852 45259875
refGene_gn=LOC101090641;refGene_tx=XM_003992640.1
D1 45260568 45260799
refGene_gn=LOC101090641;refGene_tx=XM_003992640.1
D1 45263604 45263700
refGene_gn=LOC101090641;refGene_tx=XM_003992640.1
D1 45264358 45264922
refGene_gn=LOC101090641;refGene_tx=XM_003992640.1
D1 45319949 45320443
refGene_gn=LOC101091141;refGene_tx=XM_023239346.1
D1 45321455 45321553
refGene_gn=LOC101091141;refGene_tx=XM_023239346.1
D1 45322429 45322452
refGene_gn=LOC101091141;refGene_tx=XM_023239346.1
D1 45322918 45323125
refGene_gn=LOC101091141;refGene_tx=XM_023239346.1
D1 45358788 45359282
refGene_gn=LOC101091395;refGene_tx=XM_023239347.1
D1 45360409 45360507
refGene_gn=LOC101091395;refGene_tx=XM_023239347.1
D1 45361938 45361961
refGene_gn=LOC101091395;refGene_tx=XM_023239347.1
D1 45362737 45362968
refGene_gn=LOC101091395;refGene_tx=XM_023239347.1
D1 45364135 45364231
refGene_gn=LOC101091395;refGene_tx=XM_023239347.1
D1 45364793 45365216
refGene_gn=LOC101091395;refGene_tx=XM_023239347.1
D1 45382729 45382919
refGene_gn=FOLH1B;refGene_tx=XM_006936858.3;refGene_gn=FOLH1B;refGene_tx=XM_011286535.3;refGene_gn=FOLH1B;refGene_tx=XM_011286537.3;refGene_gn=FOLH1B;refGene_tx=XM_019811660.2;refGene_gn=FOLH1B;refGene_tx=XM_019811662.2
D1 45386629 45386722
refGene_gn=FOLH1B;refGene_tx=XM_006936858.3;refGene_gn=FOLH1B;refGene_tx=XM_011286535.3;refGene_gn=FOLH1B;refGene_tx=XM_011286537.3;refGene_gn=FOLH1B;refGene_tx=XM_019811660.2;refGene_gn=FOLH1B;refGene_tx=XM_019811662.2
D1 45391303 45391403
refGene_gn=FOLH1B;refGene_tx=XM_006936858.3;refGene_gn=FOLH1B;refGene_tx=XM_011286535.3;refGene_gn=FOLH1B;refGene_tx=XM_011286537.3;refGene_

gn=FOLH1B;refGene_tx=XM_019811660.2;refGene_gn=FOLH1B;refGene_tx=XM_019811662.2
D1 45391657 45391922
refGene_gn=FOLH1B;refGene_tx=XM_006936858.3;refGene_gn=FOLH1B;refGene_tx=XM_011286535.3;refGene_gn=FOLH1B;refGene_tx=XM_011286537.3;refGene_gn=FOLH1B;refGene_tx=XM_019811660.2;refGene_gn=FOLH1B;refGene_tx=XM_019811662.2
D1 45396895 45396986
refGene_gn=FOLH1B;refGene_tx=XM_006936858.3;refGene_gn=FOLH1B;refGene_tx=XM_011286535.3;refGene_gn=FOLH1B;refGene_tx=XM_011286537.3;refGene_gn=FOLH1B;refGene_tx=XM_019811660.2;refGene_gn=FOLH1B;refGene_tx=XM_019811662.2
D1 45398586 45398678
refGene_gn=FOLH1B;refGene_tx=XM_006936858.3;refGene_gn=FOLH1B;refGene_tx=XM_011286535.3;refGene_gn=FOLH1B;refGene_tx=XM_011286537.3;refGene_gn=FOLH1B;refGene_tx=XM_019811660.2;refGene_gn=FOLH1B;refGene_tx=XM_019811662.2
D1 45401292 45401360
refGene_gn=FOLH1B;refGene_tx=XM_006936858.3;refGene_gn=FOLH1B;refGene_tx=XM_011286535.3;refGene_gn=FOLH1B;refGene_tx=XM_011286537.3;refGene_gn=FOLH1B;refGene_tx=XM_019811660.2;refGene_gn=FOLH1B;refGene_tx=XM_019811662.2
D1 45405454 45405518
refGene_gn=FOLH1B;refGene_tx=XM_006936858.3;refGene_gn=FOLH1B;refGene_tx=XM_011286535.3;refGene_gn=FOLH1B;refGene_tx=XM_011286537.3;refGene_gn=FOLH1B;refGene_tx=XM_019811660.2;refGene_gn=FOLH1B;refGene_tx=XM_019811662.2
D1 45410608 45410691
refGene_gn=FOLH1B;refGene_tx=XM_006936858.3;refGene_gn=FOLH1B;refGene_tx=XM_011286535.3;refGene_gn=FOLH1B;refGene_tx=XM_011286537.3;refGene_gn=FOLH1B;refGene_tx=XM_019811660.2;refGene_gn=FOLH1B;refGene_tx=XM_019811662.2
D1 45413082 45413202
refGene_gn=FOLH1B;refGene_tx=XM_006936858.3;refGene_gn=FOLH1B;refGene_tx=XM_011286535.3;refGene_gn=FOLH1B;refGene_tx=XM_019811660.2;refGene_gn=FOLH1B;refGene_tx=XM_019811662.2
D1 45414719 45414805
refGene_gn=FOLH1B;refGene_tx=XM_006936858.3;refGene_gn=FOLH1B;refGene_tx=XM_011286535.3;refGene_gn=FOLH1B;refGene_tx=XM_011286537.3;refGene_gn=FOLH1B;refGene_tx=XM_019811660.2;refGene_gn=FOLH1B;refGene_tx=XM_019811662.2
D1 45415106 45415205
refGene_gn=FOLH1B;refGene_tx=XM_006936858.3;refGene_gn=FOLH1B;refGene_tx=XM_011286535.3;refGene_gn=FOLH1B;refGene_tx=XM_011286537.3;refGene_gn=FOLH1B;refGene_tx=XM_019811660.2;refGene_gn=FOLH1B;refGene_tx=XM_019811662.2
D1 45421781 45421875
refGene_gn=FOLH1B;refGene_tx=XM_006936858.3;refGene_gn=FOLH1B;refGene_tx=XM_011286535.3;refGene_gn=FOLH1B;refGene_tx=XM_011286537.3;refGene_gn=FOLH1B;refGene_tx=XM_019811660.2;refGene_gn=FOLH1B;refGene_tx=XM_01

9811662.2
D1 45423805 45423992
refGene_gn=FOLH1B;refGene_tx=XM_006936858.3;refGene_gn=FOLH1B;refGene_tx=XM_011286535.3;refGene_gn=FOLH1B;refGene_tx=XM_011286537.3;refGene_gn=FOLH1B;refGene_tx=XM_019811660.2;refGene_gn=FOLH1B;refGene_tx=XM_019811662.2
D1 45424775 45424901
refGene_gn=FOLH1B;refGene_tx=XM_006936858.3;refGene_gn=FOLH1B;refGene_tx=XM_011286535.3;refGene_gn=FOLH1B;refGene_tx=XM_011286537.3;refGene_gn=FOLH1B;refGene_tx=XM_019811660.2;refGene_gn=FOLH1B;refGene_tx=XM_019811662.2
D1 45435209 45435311
refGene_gn=FOLH1B;refGene_tx=XM_006936858.3;refGene_gn=FOLH1B;refGene_tx=XM_011286535.3;refGene_gn=FOLH1B;refGene_tx=XM_011286537.3;refGene_gn=FOLH1B;refGene_tx=XM_019811660.2;refGene_gn=FOLH1B;refGene_tx=XM_019811662.2
D1 45442219 45442406
refGene_gn=FOLH1B;refGene_tx=XM_006936858.3;refGene_gn=FOLH1B;refGene_tx=XM_011286535.3;refGene_gn=FOLH1B;refGene_tx=XM_011286537.3;refGene_gn=FOLH1B;refGene_tx=XM_019811660.2;refGene_gn=FOLH1B;refGene_tx=XM_019811662.2
D1 45446401 45446510
refGene_gn=FOLH1B;refGene_tx=XM_006936858.3;refGene_gn=FOLH1B;refGene_tx=XM_011286535.3;refGene_gn=FOLH1B;refGene_tx=XM_011286537.3;refGene_gn=FOLH1B;refGene_tx=XM_019811660.2
D1 45448947 45449065
refGene_gn=FOLH1B;refGene_tx=XM_006936858.3;refGene_gn=FOLH1B;refGene_tx=XM_011286535.3;refGene_gn=FOLH1B;refGene_tx=XM_011286537.3;refGene_gn=FOLH1B;refGene_tx=XM_019811660.2
D1 45601272 45601329 refGene_gn=NOX4;refGene_tx=XM_011286538.3
D1 45601982 45602078
refGene_gn=NOX4;refGene_tx=XM_011286538.3;refGene_gn=NOX4;refGene_tx=XM_019811663.2
D1 45623492 45623603
refGene_gn=NOX4;refGene_tx=XM_011286538.3;refGene_gn=NOX4;refGene_tx=XM_011286539.3;refGene_gn=NOX4;refGene_tx=XM_011286540.3;refGene_gn=NOX4;refGene_tx=XM_019811663.2;refGene_gn=NOX4;refGene_tx=XM_019811664.2
D1 45625431 45625516
refGene_gn=NOX4;refGene_tx=XM_011286538.3;refGene_gn=NOX4;refGene_tx=XM_011286539.3;refGene_gn=NOX4;refGene_tx=XM_011286540.3;refGene_gn=NOX4;refGene_tx=XM_019811663.2;refGene_gn=NOX4;refGene_tx=XM_019811664.2
D1 45628855 45628953
refGene_gn=NOX4;refGene_tx=XM_011286538.3;refGene_gn=NOX4;refGene_tx=XM_011286539.3;refGene_gn=NOX4;refGene_tx=XM_011286540.3;refGene_gn=NOX4;refGene_tx=XM_019811663.2;refGene_gn=NOX4;refGene_tx=XM_019811664.2
D1 45632808 45632836
refGene_gn=NOX4;refGene_tx=XM_011286538.3;refGene_gn=NOX4;refGene_tx=XM_011286539.3;refGene_gn=NOX4;refGene_tx=XM_011286540.3;refGene_gn=NOX4;refGene_tx=XM_019811663.2;refGene_gn=NOX4;refGene_tx=XM_019811664.2
D1 45639800 45639873

refGene_gn=NOX4;refGene_tx=XM_011286538.3;refGene_gn=NOX4;refGene_tx=XM_011286539.3;refGene_gn=NOX4;refGene_tx=XM_011286540.3;refGene_gn=NOX4;refGene_tx=XM_019811663.2;refGene_gn=NOX4;refGene_tx=XM_019811664.2
D1 45644169 45644250
refGene_gn=NOX4;refGene_tx=XM_011286538.3;refGene_gn=NOX4;refGene_tx=XM_011286539.3;refGene_gn=NOX4;refGene_tx=XM_011286540.3;refGene_gn=NOX4;refGene_tx=XM_019811663.2;refGene_gn=NOX4;refGene_tx=XM_019811664.2
D1 45663754 45663971
refGene_gn=NOX4;refGene_tx=XM_011286538.3;refGene_gn=NOX4;refGene_tx=XM_011286539.3;refGene_gn=NOX4;refGene_tx=XM_011286540.3;refGene_gn=NOX4;refGene_tx=XM_019811663.2;refGene_gn=NOX4;refGene_tx=XM_019811664.2
D1 45666302 45666467
refGene_gn=NOX4;refGene_tx=XM_011286538.3;refGene_gn=NOX4;refGene_tx=XM_011286539.3;refGene_gn=NOX4;refGene_tx=XM_011286540.3;refGene_gn=NOX4;refGene_tx=XM_019811663.2;refGene_gn=NOX4;refGene_tx=XM_019811664.2
D1 45666599 45666662
refGene_gn=NOX4;refGene_tx=XM_011286538.3;refGene_gn=NOX4;refGene_tx=XM_011286539.3;refGene_gn=NOX4;refGene_tx=XM_011286540.3;refGene_gn=NOX4;refGene_tx=XM_019811663.2;refGene_gn=NOX4;refGene_tx=XM_019811664.2
D1 45691485 45691546
refGene_gn=NOX4;refGene_tx=XM_011286538.3;refGene_gn=NOX4;refGene_tx=XM_011286539.3;refGene_gn=NOX4;refGene_tx=XM_011286540.3;refGene_gn=NOX4;refGene_tx=XM_019811663.2;refGene_gn=NOX4;refGene_tx=XM_019811664.2
D1 45717995 45718077
refGene_gn=NOX4;refGene_tx=XM_011286538.3;refGene_gn=NOX4;refGene_tx=XM_011286539.3;refGene_gn=NOX4;refGene_tx=XM_011286540.3;refGene_gn=NOX4;refGene_tx=XM_019811663.2;refGene_gn=NOX4;refGene_tx=XM_019811664.2
D1 45744710 45744830
refGene_gn=NOX4;refGene_tx=XM_011286538.3;refGene_gn=NOX4;refGene_tx=XM_011286539.3;refGene_gn=NOX4;refGene_tx=XM_011286540.3;refGene_gn=NOX4;refGene_tx=XM_019811663.2;refGene_gn=NOX4;refGene_tx=XM_019811664.2
D1 45747063 45747172
refGene_gn=NOX4;refGene_tx=XM_011286538.3;refGene_gn=NOX4;refGene_tx=XM_011286539.3;refGene_gn=NOX4;refGene_tx=XM_011286540.3;refGene_gn=NOX4;refGene_tx=XM_019811663.2;refGene_gn=NOX4;refGene_tx=XM_019811664.2
D1 45755108 45755177
refGene_gn=NOX4;refGene_tx=XM_011286538.3;refGene_gn=NOX4;refGene_tx=XM_011286539.3;refGene_gn=NOX4;refGene_tx=XM_011286540.3;refGene_gn=NOX4;refGene_tx=XM_019811663.2;refGene_gn=NOX4;refGene_tx=XM_019811664.2
D1 45756604 45756705
refGene_gn=NOX4;refGene_tx=XM_011286538.3;refGene_gn=NOX4;refGene_tx=XM_011286539.3;refGene_gn=NOX4;refGene_tx=XM_011286540.3;refGene_gn=NOX4;refGene_tx=XM_019811663.2;refGene_gn=NOX4;refGene_tx=XM_019811664.2
D1 45769114 45769235
refGene_gn=NOX4;refGene_tx=XM_011286538.3;refGene_gn=NOX4;refGene_tx=XM_011286539.3;refGene_gn=NOX4;refGene_tx=XM_011286540.3;refGene_gn=NOX4;refGene_tx=XM_019811663.2;refGene_gn=NOX4;refGene_tx=XM_019811664.2
D1 45803655 45803882 refGene_gn=TYR;refGene_tx=XM_003992642.4
D1 45812643 45812825 refGene_gn=TYR;refGene_tx=XM_003992642.4
D1 45862190 45862338 refGene_gn=TYR;refGene_tx=XM_003992642.4

D1 45898671 45898831 refGene_gn=TYR;refGene_tx=XM_003992642.4
D1 45898832 45898889 refGene_gn=TYR;refGene_tx=XM_003992642.4
D1 45907698 45908517 refGene_gn=TYR;refGene_tx=XM_003992642.4
D1 46006866 46007527
refGene_gn=GRM5;refGene_tx=XM_023239348.1;refGene_gn=GRM5;refGene_tx=XM_023239349.1;refGene_gn=GRM5;refGene_tx=XM_023239350.1;refGene_gn=GRM5;refGene_tx=XM_023239351.1
D1 46188268 46188518
refGene_gn=GRM5;refGene_tx=XM_023239348.1;refGene_gn=GRM5;refGene_tx=XM_023239349.1;refGene_gn=GRM5;refGene_tx=XM_023239350.1;refGene_gn=GRM5;refGene_tx=XM_023239351.1
D1 46370934 46371170
refGene_gn=GRM5;refGene_tx=XM_023239348.1;refGene_gn=GRM5;refGene_tx=XM_023239349.1;refGene_gn=GRM5;refGene_tx=XM_023239350.1;refGene_gn=GRM5;refGene_tx=XM_023239351.1
D1 46413841 46414088
refGene_gn=GRM5;refGene_tx=XM_023239348.1;refGene_gn=GRM5;refGene_tx=XM_023239349.1;refGene_gn=GRM5;refGene_tx=XM_023239350.1;refGene_gn=GRM5;refGene_tx=XM_023239351.1
D1 46427561 46427730
refGene_gn=GRM5;refGene_tx=XM_023239348.1;refGene_gn=GRM5;refGene_tx=XM_023239349.1;refGene_gn=GRM5;refGene_tx=XM_023239350.1;refGene_gn=GRM5;refGene_tx=XM_023239351.1
D1 46433333 46433460
refGene_gn=GRM5;refGene_tx=XM_023239348.1;refGene_gn=GRM5;refGene_tx=XM_023239349.1;refGene_gn=GRM5;refGene_tx=XM_023239350.1;refGene_gn=GRM5;refGene_tx=XM_023239351.1
D1 46458069 46459009
refGene_gn=GRM5;refGene_tx=XM_023239348.1;refGene_gn=GRM5;refGene_tx=XM_023239349.1;refGene_gn=GRM5;refGene_tx=XM_023239350.1;refGene_gn=GRM5;refGene_tx=XM_023239351.1
D1 46491697 46491793
refGene_gn=GRM5;refGene_tx=XM_023239348.1;refGene_gn=GRM5;refGene_tx=XM_023239349.1;refGene_gn=GRM5;refGene_tx=XM_023239350.1
D1 46510386 46511287
refGene_gn=GRM5;refGene_tx=XM_023239348.1;refGene_gn=GRM5;refGene_tx=XM_023239349.1;refGene_gn=GRM5;refGene_tx=XM_023239350.1;refGene_gn=GRM5;refGene_tx=XM_023239351.1
D1 46706547 46706719
refGene_gn=CTSC;refGene_tx=XM_003992644.5;refGene_gn=CTSC;refGene_tx=XM_003992645.5
D1 46708561 46708707
refGene_gn=CTSC;refGene_tx=XM_003992644.5;refGene_gn=CTSC;refGene_tx=XM_003992645.5
D1 46715587 46715672 refGene_gn=CTSC;refGene_tx=XM_003992645.5
D1 46717974 46717985 refGene_gn=CTSC;refGene_tx=XM_003992645.5
D1 46727535 46727702 refGene_gn=CTSC;refGene_tx=XM_003992644.5
D1 46732035 46732191 refGene_gn=CTSC;refGene_tx=XM_003992644.5
D1 46736850 46736966 refGene_gn=CTSC;refGene_tx=XM_003992644.5
D1 46742422 46742554 refGene_gn=CTSC;refGene_tx=XM_003992644.5

D1 46743848 46744351 refGene_gn=CTSC;refGene_tx=XM_003992644.5
D1 46823438 46823640 refGene_gn=RAB38;refGene_tx=XM_003992646.4
D1 46846016 46846297 refGene_gn=RAB38;refGene_tx=XM_003992646.4
D1 46879774 46879927 refGene_gn=RAB38;refGene_tx=XM_003992646.4
D1 47506036 47506043 refGene_gn=TMEM135;refGene_tx=XM_011286542.3
D1 47565595 47565728
refGene_gn=TMEM135;refGene_tx=XM_011286541.3;refGene_gn=TMEM135;refGen
e_tx=XM_019811666.2;refGene_gn=TMEM135;refGene_tx=XM_023239352.1
D1 47568167 47568235
refGene_gn=TMEM135;refGene_tx=XM_011286541.3;refGene_gn=TMEM135;refGen
e_tx=XM_011286542.3;refGene_gn=TMEM135;refGene_tx=XM_019811666.2;refGe
ne_gn=TMEM135;refGene_tx=XM_023239352.1
D1 47568992 47569091
refGene_gn=TMEM135;refGene_tx=XM_011286541.3;refGene_gn=TMEM135;refGen
e_tx=XM_011286542.3;refGene_gn=TMEM135;refGene_tx=XM_019811666.2;refGe
ne_gn=TMEM135;refGene_tx=XM_023239352.1
D1 47575236 47575313
refGene_gn=TMEM135;refGene_tx=XM_011286541.3;refGene_gn=TMEM135;refGen
e_tx=XM_011286542.3;refGene_gn=TMEM135;refGene_tx=XM_019811666.2;refGe
ne_gn=TMEM135;refGene_tx=XM_023239352.1
D1 47576297 47576361
refGene_gn=TMEM135;refGene_tx=XM_011286541.3;refGene_gn=TMEM135;refGen
e_tx=XM_011286542.3;refGene_gn=TMEM135;refGene_tx=XM_019811666.2;refGe
ne_gn=TMEM135;refGene_tx=XM_023239352.1
D1 47580148 47580316
refGene_gn=TMEM135;refGene_tx=XM_011286541.3;refGene_gn=TMEM135;refGen
e_tx=XM_011286542.3;refGene_gn=TMEM135;refGene_tx=XM_019811666.2;refGe
ne_gn=TMEM135;refGene_tx=XM_023239352.1
D1 47583452 47583522
refGene_gn=TMEM135;refGene_tx=XM_011286541.3;refGene_gn=TMEM135;refGen
e_tx=XM_011286542.3;refGene_gn=TMEM135;refGene_tx=XM_019811666.2;refGe
ne_gn=TMEM135;refGene_tx=XM_023239352.1
D1 47586696 47586843
refGene_gn=TMEM135;refGene_tx=XM_011286541.3;refGene_gn=TMEM135;refGen
e_tx=XM_011286542.3;refGene_gn=TMEM135;refGene_tx=XM_019811666.2;refGe
ne_gn=TMEM135;refGene_tx=XM_023239352.1
D1 47598145 47598187
refGene_gn=TMEM135;refGene_tx=XM_011286541.3;refGene_gn=TMEM135;refGen
e_tx=XM_011286542.3;refGene_gn=TMEM135;refGene_tx=XM_019811666.2;refGe
ne_gn=TMEM135;refGene_tx=XM_023239352.1
D1 47647764 47647811
refGene_gn=TMEM135;refGene_tx=XM_011286541.3;refGene_gn=TMEM135;refGen
e_tx=XM_011286542.3;refGene_gn=TMEM135;refGene_tx=XM_019811666.2;refGe
ne_gn=TMEM135;refGene_tx=XM_023239352.1
D1 47724523 47724589
refGene_gn=TMEM135;refGene_tx=XM_011286541.3;refGene_gn=TMEM135;refGen
e_tx=XM_011286542.3;refGene_gn=TMEM135;refGene_tx=XM_019811666.2;refGe
ne_gn=TMEM135;refGene_tx=XM_023239352.1
D1 47770872 47770906
refGene_gn=TMEM135;refGene_tx=XM_011286541.3;refGene_gn=TMEM135;refGen

e_tx=XM_011286542.3;refGene_gn=TMEM135;refGene_tx=XM_019811666.2;refGene_gn=TMEM135;refGene_tx=XM_023239352.1
D1 47791458 47791551
refGene_gn=TMEM135;refGene_tx=XM_011286541.3;refGene_gn=TMEM135;refGene_tx=XM_011286542.3;refGene_gn=TMEM135;refGene_tx=XM_019811666.2;refGene_gn=TMEM135;refGene_tx=XM_023239352.1
D1 47793880 47794008
refGene_gn=TMEM135;refGene_tx=XM_011286541.3;refGene_gn=TMEM135;refGene_tx=XM_011286542.3;refGene_gn=TMEM135;refGene_tx=XM_019811666.2;refGene_gn=TMEM135;refGene_tx=XM_023239352.1
D1 47820511 47820652
refGene_gn=TMEM135;refGene_tx=XM_011286541.3;refGene_gn=TMEM135;refGene_tx=XM_011286542.3
D1 47844634 47844691
refGene_gn=TMEM135;refGene_tx=XM_019811666.2;refGene_gn=TMEM135;refGene_tx=XM_023239352.1
D1 47844891 47844894
refGene_gn=TMEM135;refGene_tx=XM_019811666.2;refGene_gn=TMEM135;refGene_tx=XM_023239352.1
D1 47886528 47886822 refGene_gn=FZD4;refGene_tx=XM_003992649.4
D1 47888972 47890301
refGene_gn=FZD4;refGene_tx=XM_003992649.4;refGene_gn=FZD4;refGene_tx=XM_006936863.4
D1 47980440 47981571
refGene_gn=PRSS23;refGene_tx=XM_006936864.4;refGene_gn=PRSS23;refGene_tx=XM_023239353.1
D1 48114392 48114575
refGene_gn=ME3;refGene_tx=XM_011286564.2;refGene_gn=ME3;refGene_tx=XM_023239354.1
D1 48165841 48166081
refGene_gn=LOC101082433;refGene_tx=XM_023239006.1
D1 48166083 48166269
refGene_gn=LOC101082433;refGene_tx=XM_023239006.1
D1 48167317 48167431
refGene_gn=LOC101082433;refGene_tx=XM_023239006.1
D1 48172124 48172191
refGene_gn=LOC101082433;refGene_tx=XM_023239006.1
D1 48177486 48177681
refGene_gn=LOC101082433;refGene_tx=XM_023239006.1
D1 48177838 48177864
refGene_gn=LOC101082433;refGene_tx=XM_023239006.1
D1 48198510 48198644
refGene_gn=ME3;refGene_tx=XM_011286564.2;refGene_gn=ME3;refGene_tx=XM_023239354.1
D1 48201728 48201878
refGene_gn=ME3;refGene_tx=XM_011286564.2;refGene_gn=ME3;refGene_tx=XM_023239354.1
D1 48232539 48232615
refGene_gn=ME3;refGene_tx=XM_011286564.2;refGene_gn=ME3;refGene_tx=XM_023239354.1

D1 48243060 48243222 refGene_gn=ME3;refGene_tx=XM_011286564.2
D1 48253765 48253869
refGene_gn=ME3;refGene_tx=XM_011286564.2;refGene_gn=ME3;refGene_tx=XM_023239354.1
D1 48275427 48275537
refGene_gn=ME3;refGene_tx=XM_011286564.2;refGene_gn=ME3;refGene_tx=XM_023239354.1
D1 48289655 48289753
refGene_gn=ME3;refGene_tx=XM_011286564.2;refGene_gn=ME3;refGene_tx=XM_023239354.1
D1 48290023 48290137
refGene_gn=ME3;refGene_tx=XM_011286564.2;refGene_gn=ME3;refGene_tx=XM_023239354.1
D1 48292007 48292113
refGene_gn=ME3;refGene_tx=XM_011286564.2;refGene_gn=ME3;refGene_tx=XM_023239354.1
D1 48292718 48292861
refGene_gn=ME3;refGene_tx=XM_011286564.2;refGene_gn=ME3;refGene_tx=XM_023239354.1
D1 48293429 48293603
refGene_gn=ME3;refGene_tx=XM_011286564.2;refGene_gn=ME3;refGene_tx=XM_023239354.1
D1 48296797 48296896
refGene_gn=ME3;refGene_tx=XM_011286564.2;refGene_gn=ME3;refGene_tx=XM_023239354.1
D1 48298243 48298405
refGene_gn=ME3;refGene_tx=XM_011286564.2;refGene_gn=ME3;refGene_tx=XM_023239354.1
D1 48343267 48343409
refGene_gn=CCDC81;refGene_tx=XM_011286543.3;refGene_gn=CCDC81;refGene_tx=XM_019811670.2;refGene_gn=CCDC81;refGene_tx=XM_023239355.1;refGene_gn=CCDC81;refGene_tx=XM_023239356.1
D1 48345817 48345943
refGene_gn=CCDC81;refGene_tx=XM_011286543.3;refGene_gn=CCDC81;refGene_tx=XM_019811670.2;refGene_gn=CCDC81;refGene_tx=XM_023239355.1;refGene_gn=CCDC81;refGene_tx=XM_023239356.1
D1 48354106 48354327
refGene_gn=CCDC81;refGene_tx=XM_011286543.3;refGene_gn=CCDC81;refGene_tx=XM_019811670.2;refGene_gn=CCDC81;refGene_tx=XM_023239355.1;refGene_gn=CCDC81;refGene_tx=XM_023239356.1
D1 48354543 48354622
refGene_gn=CCDC81;refGene_tx=XM_011286543.3;refGene_gn=CCDC81;refGene_tx=XM_019811670.2;refGene_gn=CCDC81;refGene_tx=XM_023239355.1;refGene_gn=CCDC81;refGene_tx=XM_023239356.1
D1 48355857 48356030
refGene_gn=CCDC81;refGene_tx=XM_011286543.3;refGene_gn=CCDC81;refGene_tx=XM_019811670.2;refGene_gn=CCDC81;refGene_tx=XM_023239355.1;refGene_gn=CCDC81;refGene_tx=XM_023239356.1
D1 48356172 48356277
refGene_gn=CCDC81;refGene_tx=XM_011286543.3;refGene_gn=CCDC81;refGene_

tx=XM_019811670.2;refGene_gn=CCDC81;refGene_tx=XM_023239355.1;refGene_gn=CCDC81;refGene_tx=XM_023239356.1
D1 48357430 48357577
refGene_gn=CCDC81;refGene_tx=XM_011286543.3;refGene_gn=CCDC81;refGene_tx=XM_019811670.2;refGene_gn=CCDC81;refGene_tx=XM_023239355.1;refGene_gn=CCDC81;refGene_tx=XM_023239356.1
D1 48358181 48358275
refGene_gn=CCDC81;refGene_tx=XM_011286543.3;refGene_gn=CCDC81;refGene_tx=XM_019811670.2;refGene_gn=CCDC81;refGene_tx=XM_023239355.1;refGene_gn=CCDC81;refGene_tx=XM_023239356.1
D1 48361225 48361349
refGene_gn=CCDC81;refGene_tx=XM_011286543.3;refGene_gn=CCDC81;refGene_tx=XM_019811670.2;refGene_gn=CCDC81;refGene_tx=XM_023239355.1;refGene_gn=CCDC81;refGene_tx=XM_023239356.1
D1 48363248 48363373
refGene_gn=CCDC81;refGene_tx=XM_011286543.3;refGene_gn=CCDC81;refGene_tx=XM_019811670.2;refGene_gn=CCDC81;refGene_tx=XM_023239355.1;refGene_gn=CCDC81;refGene_tx=XM_023239356.1
D1 48365850 48365930
refGene_gn=CCDC81;refGene_tx=XM_011286543.3;refGene_gn=CCDC81;refGene_tx=XM_019811670.2;refGene_gn=CCDC81;refGene_tx=XM_023239355.1;refGene_gn=CCDC81;refGene_tx=XM_023239356.1
D1 48368784 48369041
refGene_gn=CCDC81;refGene_tx=XM_011286543.3;refGene_gn=CCDC81;refGene_tx=XM_019811670.2;refGene_gn=CCDC81;refGene_tx=XM_023239355.1;refGene_gn=CCDC81;refGene_tx=XM_023239356.1
D1 48372455 48372612
refGene_gn=CCDC81;refGene_tx=XM_011286543.3;refGene_gn=CCDC81;refGene_tx=XM_019811670.2;refGene_gn=CCDC81;refGene_tx=XM_023239355.1;refGene_gn=CCDC81;refGene_tx=XM_023239356.1
D1 48372973 48372976 refGene_gn=CCDC81;refGene_tx=XM_023239355.1
D1 48374009 48374071 refGene_gn=CCDC81;refGene_tx=XM_011286543.3
D1 48381003 48381148 refGene_gn=CCDC81;refGene_tx=XM_011286543.3
D1 48419009 48419064 refGene_gn=HIKESHI;refGene_tx=XM_003992652.4
D1 48419997 48420116 refGene_gn=HIKESHI;refGene_tx=XM_003992652.4
D1 48428372 48428524 refGene_gn=HIKESHI;refGene_tx=XM_003992652.4
D1 48457138 48457376 refGene_gn=HIKESHI;refGene_tx=XM_003992652.4
D1 48461894 48461924 refGene_gn=HIKESHI;refGene_tx=XM_003992652.4
D1 48480553 48480680
refGene_gn=EED;refGene_tx=XM_011286545.3;refGene_gn=EED;refGene_tx=XM_011286546.3;refGene_gn=EED;refGene_tx=XM_019811675.2;refGene_gn=EED;refGene_tx=XM_019811676.2;refGene_gn=EED;refGene_tx=XM_019811678.2;refGene_gn=EED;refGene_tx=XM_019811679.2;refGene_gn=EED;refGene_tx=XM_019811680.2;refGene_gn=EED;refGene_tx=XM_019811681.2;refGene_gn=EED;refGene_tx=XM_023239359.1
D1 48481046 48481120
refGene_gn=EED;refGene_tx=XM_011286545.3;refGene_gn=EED;refGene_tx=XM_011286546.3;refGene_gn=EED;refGene_tx=XM_019811675.2;refGene_gn=EED;refGene_tx=XM_019811676.2;refGene_gn=EED;refGene_tx=XM_019811678.2;refGene_gn=EED;refGene_tx=XM_019811679.2;refGene_gn=EED;refGene_tx=XM_01981

1680.2;refGene_gn=EED;refGene_tx=XM_019811681.2;refGene_gn=EED;refGene_tx=XM_023239359.1
D1 48482114 48482273
refGene_gn=EED;refGene_tx=XM_011286545.3;refGene_gn=EED;refGene_tx=XM_011286546.3;refGene_gn=EED;refGene_tx=XM_019811675.2;refGene_gn=EED;refGene_tx=XM_019811676.2;refGene_gn=EED;refGene_tx=XM_019811678.2;refGene_gn=EED;refGene_tx=XM_019811679.2;refGene_gn=EED;refGene_tx=XM_019811680.2;refGene_gn=EED;refGene_tx=XM_019811681.2;refGene_gn=EED;refGene_tx=XM_023239359.1
D1 48489385 48489460
refGene_gn=EED;refGene_tx=XM_011286545.3;refGene_gn=EED;refGene_tx=XM_019811675.2;refGene_gn=EED;refGene_tx=XM_019811676.2;refGene_gn=EED;refGene_tx=XM_019811678.2;refGene_gn=EED;refGene_tx=XM_019811679.2;refGene_gn=EED;refGene_tx=XM_019811680.2;refGene_gn=EED;refGene_tx=XM_019811681.2;refGene_gn=EED;refGene_tx=XM_023239359.1
D1 48490588 48490694
refGene_gn=EED;refGene_tx=XM_011286545.3;refGene_gn=EED;refGene_tx=XM_011286546.3;refGene_gn=EED;refGene_tx=XM_019811675.2;refGene_gn=EED;refGene_tx=XM_019811676.2;refGene_gn=EED;refGene_tx=XM_019811678.2;refGene_gn=EED;refGene_tx=XM_019811679.2;refGene_gn=EED;refGene_tx=XM_019811680.2;refGene_gn=EED;refGene_tx=XM_019811681.2;refGene_gn=EED;refGene_tx=XM_023239359.1
D1 48492661 48492795
refGene_gn=EED;refGene_tx=XM_011286545.3;refGene_gn=EED;refGene_tx=XM_011286546.3;refGene_gn=EED;refGene_tx=XM_019811675.2;refGene_gn=EED;refGene_tx=XM_019811676.2;refGene_gn=EED;refGene_tx=XM_019811678.2;refGene_gn=EED;refGene_tx=XM_019811679.2;refGene_gn=EED;refGene_tx=XM_019811680.2;refGene_gn=EED;refGene_tx=XM_019811681.2;refGene_gn=EED;refGene_tx=XM_023239359.1
D1 48495493 48495585
refGene_gn=EED;refGene_tx=XM_011286545.3;refGene_gn=EED;refGene_tx=XM_011286546.3;refGene_gn=EED;refGene_tx=XM_019811675.2;refGene_gn=EED;refGene_tx=XM_019811676.2;refGene_gn=EED;refGene_tx=XM_019811678.2;refGene_gn=EED;refGene_tx=XM_019811679.2;refGene_gn=EED;refGene_tx=XM_019811680.2;refGene_gn=EED;refGene_tx=XM_019811681.2;refGene_gn=EED;refGene_tx=XM_023239359.1
D1 48497776 48497858
refGene_gn=EED;refGene_tx=XM_011286545.3;refGene_gn=EED;refGene_tx=XM_011286546.3;refGene_gn=EED;refGene_tx=XM_019811675.2;refGene_gn=EED;refGene_tx=XM_019811676.2;refGene_gn=EED;refGene_tx=XM_019811678.2;refGene_gn=EED;refGene_tx=XM_019811679.2;refGene_gn=EED;refGene_tx=XM_019811680.2;refGene_gn=EED;refGene_tx=XM_019811681.2;refGene_gn=EED;refGene_tx=XM_023239359.1
D1 48498553 48498679
refGene_gn=EED;refGene_tx=XM_011286545.3;refGene_gn=EED;refGene_tx=XM_011286546.3;refGene_gn=EED;refGene_tx=XM_019811675.2;refGene_gn=EED;refGene_tx=XM_019811676.2;refGene_gn=EED;refGene_tx=XM_019811678.2;refGene_gn=EED;refGene_tx=XM_019811679.2;refGene_gn=EED;refGene_tx=XM_019811680.2;refGene_gn=EED;refGene_tx=XM_019811681.2;refGene_gn=EED;refGene_tx=XM_023239359.1

D1 48499642 48499708
refGene_gn=EED;refGene_tx=XM_011286545.3;refGene_gn=EED;refGene_tx=XM_011286546.3;refGene_gn=EED;refGene_tx=XM_019811675.2;refGene_gn=EED;refGene_tx=XM_019811676.2;refGene_gn=EED;refGene_tx=XM_019811678.2;refGene_gn=EED;refGene_tx=XM_019811679.2;refGene_gn=EED;refGene_tx=XM_019811680.2;refGene_gn=EED;refGene_tx=XM_019811681.2;refGene_gn=EED;refGene_tx=XM_023239359.1
D1 48502844 48502937
refGene_gn=EED;refGene_tx=XM_011286545.3;refGene_gn=EED;refGene_tx=XM_011286546.3;refGene_gn=EED;refGene_tx=XM_019811675.2;refGene_gn=EED;refGene_tx=XM_019811676.2;refGene_gn=EED;refGene_tx=XM_019811678.2;refGene_gn=EED;refGene_tx=XM_019811679.2;refGene_gn=EED;refGene_tx=XM_019811680.2;refGene_gn=EED;refGene_tx=XM_019811681.2;refGene_gn=EED;refGene_tx=XM_023239359.1
D1 48505226 48505379
refGene_gn=EED;refGene_tx=XM_011286545.3;refGene_gn=EED;refGene_tx=XM_011286546.3;refGene_gn=EED;refGene_tx=XM_019811675.2;refGene_gn=EED;refGene_tx=XM_019811676.2;refGene_gn=EED;refGene_tx=XM_019811678.2;refGene_gn=EED;refGene_tx=XM_019811679.2;refGene_gn=EED;refGene_tx=XM_019811680.2;refGene_gn=EED;refGene_tx=XM_023239359.1
D1 48510083 48510197
refGene_gn=EED;refGene_tx=XM_011286545.3;refGene_gn=EED;refGene_tx=XM_011286546.3;refGene_gn=EED;refGene_tx=XM_019811681.2
D1 48540094 48540097
refGene_gn=EED;refGene_tx=XM_019811675.2;refGene_gn=EED;refGene_tx=XM_019811676.2;refGene_gn=EED;refGene_tx=XM_019811678.2;refGene_gn=EED;refGene_tx=XM_019811679.2;refGene_gn=EED;refGene_tx=XM_019811680.2;refGene_gn=EED;refGene_tx=XM_023239359.1
D1 48672664 48672794
refGene_gn=PICALM;refGene_tx=XM_006936865.4;refGene_gn=PICALM;refGene_tx=XM_006936866.4;refGene_gn=PICALM;refGene_tx=XM_006936867.4;refGene_gn=PICALM;refGene_tx=XM_006936868.4;refGene_gn=PICALM;refGene_tx=XM_006936869.4;refGene_gn=PICALM;refGene_tx=XM_006936870.4;refGene_gn=PICALM;refGene_tx=XM_006936871.4;refGene_gn=PICALM;refGene_tx=XM_006936872.4;refGene_gn=PICALM;refGene_tx=XM_006936873.4;refGene_gn=PICALM;refGene_tx=XM_019811671.2;refGene_gn=PICALM;refGene_tx=XM_019811672.2;refGene_gn=PICALM;refGene_tx=XM_019811674.2
D1 48705878 48706021
refGene_gn=PICALM;refGene_tx=XM_006936865.4;refGene_gn=PICALM;refGene_tx=XM_006936866.4;refGene_gn=PICALM;refGene_tx=XM_006936867.4;refGene_gn=PICALM;refGene_tx=XM_006936868.4;refGene_gn=PICALM;refGene_tx=XM_006936869.4;refGene_gn=PICALM;refGene_tx=XM_006936870.4;refGene_gn=PICALM;refGene_tx=XM_006936871.4;refGene_gn=PICALM;refGene_tx=XM_006936872.4;refGene_gn=PICALM;refGene_tx=XM_006936873.4;refGene_gn=PICALM;refGene_tx=XM_019811671.2;refGene_gn=PICALM;refGene_tx=XM_019811672.2;refGene_gn=PICALM;refGene_tx=XM_019811674.2
D1 48709358 48709434
refGene_gn=PICALM;refGene_tx=XM_006936865.4;refGene_gn=PICALM;refGene_tx=XM_006936866.4;refGene_gn=PICALM;refGene_tx=XM_006936867.4;refGene_gn=PICALM;refGene_tx=XM_006936868.4;refGene_gn=PICALM;refGene_tx=XM_00

6936869.4;refGene_gn=PICALM;refGene_tx=XM_006936870.4;refGene_gn=PICALM;refGene_tx=XM_006936871.4;refGene_gn=PICALM;refGene_tx=XM_006936872.4;refGene_gn=PICALM;refGene_tx=XM_006936873.4;refGene_gn=PICALM;refGene_tx=XM_019811671.2;refGene_gn=PICALM;refGene_tx=XM_019811672.2;refGene_gn=PICALM;refGene_tx=XM_019811674.2

D1 48712499 48712602
refGene_gn=PICALM;refGene_tx=XM_006936865.4;refGene_gn=PICALM;refGene_tx=XM_006936866.4;refGene_gn=PICALM;refGene_tx=XM_006936867.4;refGene_gn=PICALM;refGene_tx=XM_006936868.4;refGene_gn=PICALM;refGene_tx=XM_006936869.4;refGene_gn=PICALM;refGene_tx=XM_006936870.4;refGene_gn=PICALM;refGene_tx=XM_006936871.4;refGene_gn=PICALM;refGene_tx=XM_006936872.4;refGene_gn=PICALM;refGene_tx=XM_006936873.4;refGene_gn=PICALM;refGene_tx=XM_019811671.2;refGene_gn=PICALM;refGene_tx=XM_019811672.2;refGene_gn=PICALM;refGene_tx=XM_019811674.2

D1 48719186 48719280
refGene_gn=PICALM;refGene_tx=XM_006936865.4;refGene_gn=PICALM;refGene_tx=XM_006936866.4;refGene_gn=PICALM;refGene_tx=XM_006936867.4;refGene_gn=PICALM;refGene_tx=XM_006936868.4;refGene_gn=PICALM;refGene_tx=XM_006936869.4;refGene_gn=PICALM;refGene_tx=XM_006936870.4;refGene_gn=PICALM;refGene_tx=XM_006936871.4;refGene_gn=PICALM;refGene_tx=XM_006936872.4;refGene_gn=PICALM;refGene_tx=XM_006936873.4;refGene_gn=PICALM;refGene_tx=XM_019811671.2;refGene_gn=PICALM;refGene_tx=XM_019811672.2;refGene_gn=PICALM;refGene_tx=XM_019811674.2

D1 48722442 48722554
refGene_gn=PICALM;refGene_tx=XM_006936865.4;refGene_gn=PICALM;refGene_tx=XM_006936866.4;refGene_gn=PICALM;refGene_tx=XM_006936867.4;refGene_gn=PICALM;refGene_tx=XM_006936868.4;refGene_gn=PICALM;refGene_tx=XM_006936869.4;refGene_gn=PICALM;refGene_tx=XM_006936870.4;refGene_gn=PICALM;refGene_tx=XM_006936871.4;refGene_gn=PICALM;refGene_tx=XM_006936872.4;refGene_gn=PICALM;refGene_tx=XM_006936873.4;refGene_gn=PICALM;refGene_tx=XM_019811671.2;refGene_gn=PICALM;refGene_tx=XM_019811672.2;refGene_gn=PICALM;refGene_tx=XM_019811674.2

D1 48723815 48723922
refGene_gn=PICALM;refGene_tx=XM_006936865.4;refGene_gn=PICALM;refGene_tx=XM_006936866.4;refGene_gn=PICALM;refGene_tx=XM_006936867.4;refGene_gn=PICALM;refGene_tx=XM_006936868.4;refGene_gn=PICALM;refGene_tx=XM_006936869.4;refGene_gn=PICALM;refGene_tx=XM_006936870.4;refGene_gn=PICALM;refGene_tx=XM_006936871.4;refGene_gn=PICALM;refGene_tx=XM_006936872.4;refGene_gn=PICALM;refGene_tx=XM_006936873.4;refGene_gn=PICALM;refGene_tx=XM_019811671.2;refGene_gn=PICALM;refGene_tx=XM_019811672.2;refGene_gn=PICALM;refGene_tx=XM_019811674.2

D1 48725906 48725948
refGene_gn=PICALM;refGene_tx=XM_006936865.4;refGene_gn=PICALM;refGene_tx=XM_006936866.4;refGene_gn=PICALM;refGene_tx=XM_006936867.4;refGene_gn=PICALM;refGene_tx=XM_006936868.4;refGene_gn=PICALM;refGene_tx=XM_006936869.4;refGene_gn=PICALM;refGene_tx=XM_006936870.4;refGene_gn=PICALM;refGene_tx=XM_006936871.4;refGene_gn=PICALM;refGene_tx=XM_006936872.4;refGene_gn=PICALM;refGene_tx=XM_006936873.4;refGene_gn=PICALM;refGene_tx=XM_019811671.2;refGene_gn=PICALM;refGene_tx=XM_019811672.2;refGene_gn=PICALM;refGene_tx=XM_019811674.2

D1 48728111 48728197
refGene_gn=PICALM;refGene_tx=XM_006936865.4;refGene_gn=PICALM;refGene_tx=XM_006936866.4;refGene_gn=PICALM;refGene_tx=XM_006936867.4;refGene_gn=PICALM;refGene_tx=XM_006936868.4;refGene_gn=PICALM;refGene_tx=XM_006936869.4;refGene_gn=PICALM;refGene_tx=XM_006936870.4;refGene_gn=PICALM;refGene_tx=XM_006936871.4;refGene_gn=PICALM;refGene_tx=XM_006936872.4;refGene_gn=PICALM;refGene_tx=XM_006936873.4;refGene_gn=PICALM;refGene_tx=XM_019811671.2;refGene_gn=PICALM;refGene_tx=XM_019811672.2;refGene_gn=PICALM;refGene_tx=XM_019811674.2
D1 48730298 48730422
refGene_gn=PICALM;refGene_tx=XM_006936865.4;refGene_gn=PICALM;refGene_tx=XM_006936866.4;refGene_gn=PICALM;refGene_tx=XM_006936867.4;refGene_gn=PICALM;refGene_tx=XM_006936868.4;refGene_gn=PICALM;refGene_tx=XM_006936869.4;refGene_gn=PICALM;refGene_tx=XM_006936870.4;refGene_gn=PICALM;refGene_tx=XM_006936871.4;refGene_gn=PICALM;refGene_tx=XM_006936872.4;refGene_gn=PICALM;refGene_tx=XM_006936873.4;refGene_gn=PICALM;refGene_tx=XM_019811671.2;refGene_gn=PICALM;refGene_tx=XM_019811672.2;refGene_gn=PICALM;refGene_tx=XM_019811674.2
D1 48730686 48730823
refGene_gn=PICALM;refGene_tx=XM_006936865.4;refGene_gn=PICALM;refGene_tx=XM_006936866.4;refGene_gn=PICALM;refGene_tx=XM_006936867.4;refGene_gn=PICALM;refGene_tx=XM_006936868.4;refGene_gn=PICALM;refGene_tx=XM_006936869.4;refGene_gn=PICALM;refGene_tx=XM_006936870.4;refGene_gn=PICALM;refGene_tx=XM_006936871.4;refGene_gn=PICALM;refGene_tx=XM_006936872.4;refGene_gn=PICALM;refGene_tx=XM_006936873.4;refGene_gn=PICALM;refGene_tx=XM_019811671.2;refGene_gn=PICALM;refGene_tx=XM_019811672.2;refGene_gn=PICALM;refGene_tx=XM_019811674.2
D1 48733423 48733527
refGene_gn=PICALM;refGene_tx=XM_006936865.4;refGene_gn=PICALM;refGene_tx=XM_006936866.4;refGene_gn=PICALM;refGene_tx=XM_006936867.4;refGene_gn=PICALM;refGene_tx=XM_006936868.4;refGene_gn=PICALM;refGene_tx=XM_006936869.4;refGene_gn=PICALM;refGene_tx=XM_006936870.4;refGene_gn=PICALM;refGene_tx=XM_006936871.4;refGene_gn=PICALM;refGene_tx=XM_006936872.4;refGene_gn=PICALM;refGene_tx=XM_006936873.4;refGene_gn=PICALM;refGene_tx=XM_019811671.2;refGene_gn=PICALM;refGene_tx=XM_019811672.2;refGene_gn=PICALM;refGene_tx=XM_019811674.2
D1 48739024 48739174
refGene_gn=PICALM;refGene_tx=XM_006936865.4;refGene_gn=PICALM;refGene_tx=XM_006936866.4;refGene_gn=PICALM;refGene_tx=XM_006936867.4;refGene_gn=PICALM;refGene_tx=XM_006936872.4;refGene_gn=PICALM;refGene_tx=XM_006936873.4;refGene_gn=PICALM;refGene_tx=XM_019811671.2;refGene_gn=PICALM;refGene_tx=XM_019811672.2;refGene_gn=PICALM;refGene_tx=XM_019811674.2
D1 48747107 48747215
refGene_gn=PICALM;refGene_tx=XM_006936865.4;refGene_gn=PICALM;refGene_tx=XM_006936866.4;refGene_gn=PICALM;refGene_tx=XM_006936867.4;refGene_gn=PICALM;refGene_tx=XM_006936868.4;refGene_gn=PICALM;refGene_tx=XM_006936869.4;refGene_gn=PICALM;refGene_tx=XM_006936870.4;refGene_gn=PICALM;refGene_tx=XM_006936871.4;refGene_gn=PICALM;refGene_tx=XM_006936872.4;refGene_gn=PICALM;refGene_tx=XM_006936873.4;refGene_gn=PICALM;refGen

e_tx=XM_019811671.2;refGene_gn=PICALM;refGene_tx=XM_019811672.2;refGene_gn=PICALM;refGene_tx=XM_019811674.2
D1 48749539 48749671
refGene_gn=PICALM;refGene_tx=XM_006936865.4;refGene_gn=PICALM;refGene_tx=XM_006936866.4;refGene_gn=PICALM;refGene_tx=XM_006936867.4;refGene_gn=PICALM;refGene_tx=XM_006936868.4;refGene_gn=PICALM;refGene_tx=XM_006936869.4;refGene_gn=PICALM;refGene_tx=XM_006936870.4;refGene_gn=PICALM;refGene_tx=XM_006936871.4;refGene_gn=PICALM;refGene_tx=XM_006936872.4;refGene_gn=PICALM;refGene_tx=XM_006936873.4;refGene_gn=PICALM;refGene_tx=XM_019811671.2;refGene_gn=PICALM;refGene_tx=XM_019811672.2;refGene_gn=PICALM;refGene_tx=XM_019811674.2
D1 48749765 48749796
refGene_gn=PICALM;refGene_tx=XM_006936865.4;refGene_gn=PICALM;refGene_tx=XM_006936866.4;refGene_gn=PICALM;refGene_tx=XM_006936867.4;refGene_gn=PICALM;refGene_tx=XM_006936868.4;refGene_gn=PICALM;refGene_tx=XM_006936869.4;refGene_gn=PICALM;refGene_tx=XM_006936870.4;refGene_gn=PICALM;refGene_tx=XM_006936871.4;refGene_gn=PICALM;refGene_tx=XM_006936872.4;refGene_gn=PICALM;refGene_tx=XM_006936873.4;refGene_gn=PICALM;refGene_tx=XM_019811671.2;refGene_gn=PICALM;refGene_tx=XM_019811672.2;refGene_gn=PICALM;refGene_tx=XM_019811674.2
D1 48749969 48750069
refGene_gn=PICALM;refGene_tx=XM_006936865.4;refGene_gn=PICALM;refGene_tx=XM_006936866.4;refGene_gn=PICALM;refGene_tx=XM_006936867.4;refGene_gn=PICALM;refGene_tx=XM_006936868.4;refGene_gn=PICALM;refGene_tx=XM_006936869.4;refGene_gn=PICALM;refGene_tx=XM_006936870.4;refGene_gn=PICALM;refGene_tx=XM_006936871.4;refGene_gn=PICALM;refGene_tx=XM_006936872.4;refGene_gn=PICALM;refGene_tx=XM_006936873.4;refGene_gn=PICALM;refGene_tx=XM_019811671.2;refGene_gn=PICALM;refGene_tx=XM_019811672.2;refGene_gn=PICALM;refGene_tx=XM_019811674.2
D1 48753622 48753646
refGene_gn=PICALM;refGene_tx=XM_006936865.4;refGene_gn=PICALM;refGene_tx=XM_006936867.4;refGene_gn=PICALM;refGene_tx=XM_006936868.4;refGene_gn=PICALM;refGene_tx=XM_006936869.4;refGene_gn=PICALM;refGene_tx=XM_006936872.4;refGene_gn=PICALM;refGene_tx=XM_019811671.2
D1 48755056 48755116
refGene_gn=PICALM;refGene_tx=XM_006936865.4;refGene_gn=PICALM;refGene_tx=XM_006936866.4;refGene_gn=PICALM;refGene_tx=XM_006936867.4;refGene_gn=PICALM;refGene_tx=XM_006936868.4;refGene_gn=PICALM;refGene_tx=XM_006936869.4;refGene_gn=PICALM;refGene_tx=XM_006936870.4;refGene_gn=PICALM;refGene_tx=XM_006936871.4;refGene_gn=PICALM;refGene_tx=XM_006936872.4;refGene_gn=PICALM;refGene_tx=XM_006936873.4;refGene_gn=PICALM;refGene_tx=XM_019811671.2;refGene_gn=PICALM;refGene_tx=XM_019811672.2;refGene_gn=PICALM;refGene_tx=XM_019811674.2
D1 48756632 48756737
refGene_gn=PICALM;refGene_tx=XM_006936865.4;refGene_gn=PICALM;refGene_tx=XM_006936866.4;refGene_gn=PICALM;refGene_tx=XM_006936867.4;refGene_gn=PICALM;refGene_tx=XM_006936868.4;refGene_gn=PICALM;refGene_tx=XM_006936869.4;refGene_gn=PICALM;refGene_tx=XM_006936870.4;refGene_gn=PICALM;refGene_tx=XM_006936871.4;refGene_gn=PICALM;refGene_tx=XM_006936872.4;refGene_gn=PICALM;refGene_tx=XM_006936873.4;refGene_gn=PICALM;refGen

e_tx=XM_019811671.2;refGene_gn=PICALM;refGene_tx=XM_019811672.2;refGene_gn=PICALM;refGene_tx=XM_019811674.2
D1 48769816 48769909
refGene_gn=PICALM;refGene_tx=XM_006936865.4;refGene_gn=PICALM;refGene_tx=XM_006936866.4;refGene_gn=PICALM;refGene_tx=XM_006936868.4;refGene_gn=PICALM;refGene_tx=XM_006936869.4;refGene_gn=PICALM;refGene_tx=XM_006936870.4;refGene_gn=PICALM;refGene_tx=XM_006936871.4;refGene_gn=PICALM;refGene_tx=XM_006936872.4;refGene_gn=PICALM;refGene_tx=XM_006936873.4;refGene_gn=PICALM;refGene_tx=XM_019811671.2;refGene_gn=PICALM;refGene_tx=XM_019811672.2;refGene_gn=PICALM;refGene_tx=XM_019811674.2
D1 48771188 48771203 refGene_gn=PICALM;refGene_tx=XM_006936867.4
D1 48788846 48789008
refGene_gn=CCDC83;refGene_tx=XM_019811699.2;refGene_gn=CCDC83;refGene_tx=XM_019811700.2;refGene_gn=CCDC83;refGene_tx=XM_019811702.2;refGene_gn=CCDC83;refGene_tx=XM_019811704.2;refGene_gn=CCDC83;refGene_tx=XM_019811705.2;refGene_gn=CCDC83;refGene_tx=XM_019811706.2;refGene_gn=CCDC83;refGene_tx=XM_019811707.1;refGene_gn=CCDC83;refGene_tx=XM_023239361.1;refGene_gn=CCDC83;refGene_tx=XM_023239362.1;refGene_gn=CCDC83;refGene_tx=XM_023239363.1
D1 48792065 48792268
refGene_gn=CCDC83;refGene_tx=XM_019811699.2;refGene_gn=CCDC83;refGene_tx=XM_019811700.2;refGene_gn=CCDC83;refGene_tx=XM_019811702.2;refGene_gn=CCDC83;refGene_tx=XM_019811704.2;refGene_gn=CCDC83;refGene_tx=XM_019811705.2;refGene_gn=CCDC83;refGene_tx=XM_019811706.2;refGene_gn=CCDC83;refGene_tx=XM_019811707.1;refGene_gn=CCDC83;refGene_tx=XM_023239361.1;refGene_gn=CCDC83;refGene_tx=XM_023239362.1;refGene_gn=CCDC83;refGene_tx=XM_023239363.1
D1 48792792 48792869
refGene_gn=CCDC83;refGene_tx=XM_019811699.2;refGene_gn=CCDC83;refGene_tx=XM_019811700.2;refGene_gn=CCDC83;refGene_tx=XM_019811702.2;refGene_gn=CCDC83;refGene_tx=XM_019811704.2;refGene_gn=CCDC83;refGene_tx=XM_019811705.2;refGene_gn=CCDC83;refGene_tx=XM_019811706.2;refGene_gn=CCDC83;refGene_tx=XM_019811707.1;refGene_gn=CCDC83;refGene_tx=XM_023239361.1;refGene_gn=CCDC83;refGene_tx=XM_023239362.1;refGene_gn=CCDC83;refGene_tx=XM_023239363.1
D1 48796900 48797022
refGene_gn=CCDC83;refGene_tx=XM_019811699.2;refGene_gn=CCDC83;refGene_tx=XM_019811700.2;refGene_gn=CCDC83;refGene_tx=XM_019811702.2;refGene_gn=CCDC83;refGene_tx=XM_019811704.2;refGene_gn=CCDC83;refGene_tx=XM_019811705.2;refGene_gn=CCDC83;refGene_tx=XM_019811706.2;refGene_gn=CCDC83;refGene_tx=XM_019811707.1;refGene_gn=CCDC83;refGene_tx=XM_023239361.1;refGene_gn=CCDC83;refGene_tx=XM_023239362.1;refGene_gn=CCDC83;refGene_tx=XM_023239363.1
D1 48802464 48802533
refGene_gn=CCDC83;refGene_tx=XM_019811699.2;refGene_gn=CCDC83;refGene_tx=XM_019811700.2;refGene_gn=CCDC83;refGene_tx=XM_019811702.2;refGene_gn=CCDC83;refGene_tx=XM_019811704.2;refGene_gn=CCDC83;refGene_tx=XM_019811705.2;refGene_gn=CCDC83;refGene_tx=XM_019811706.2;refGene_gn=CCDC83;refGene_tx=XM_019811707.1;refGene_gn=CCDC83;refGene_tx=XM_023239361.1;refGene_gn=CCDC83;refGene_tx=XM_023239362.1;refGene_gn=CCDC83;refGen

e_tx=XM_023239363.1
D1 48806295 48806387
refGene_gn=CCDC83;refGene_tx=XM_019811699.2;refGene_gn=CCDC83;refGene_tx=XM_019811700.2;refGene_gn=CCDC83;refGene_tx=XM_019811702.2;refGene_gn=CCDC83;refGene_tx=XM_019811704.2;refGene_gn=CCDC83;refGene_tx=XM_019811705.2;refGene_gn=CCDC83;refGene_tx=XM_019811706.2;refGene_gn=CCDC83;refGene_tx=XM_019811707.1;refGene_gn=CCDC83;refGene_tx=XM_023239361.1;refGene_gn=CCDC83;refGene_tx=XM_023239362.1;refGene_gn=CCDC83;refGene_tx=XM_023239363.1
D1 48814099 48814267
refGene_gn=CCDC83;refGene_tx=XM_019811699.2;refGene_gn=CCDC83;refGene_tx=XM_019811700.2;refGene_gn=CCDC83;refGene_tx=XM_019811702.2;refGene_gn=CCDC83;refGene_tx=XM_019811704.2;refGene_gn=CCDC83;refGene_tx=XM_019811705.2;refGene_gn=CCDC83;refGene_tx=XM_019811706.2;refGene_gn=CCDC83;refGene_tx=XM_019811707.1;refGene_gn=CCDC83;refGene_tx=XM_023239361.1;refGene_gn=CCDC83;refGene_tx=XM_023239362.1;refGene_gn=CCDC83;refGene_tx=XM_023239363.1
D1 48818171 48818334
refGene_gn=CCDC83;refGene_tx=XM_019811699.2;refGene_gn=CCDC83;refGene_tx=XM_019811700.2;refGene_gn=CCDC83;refGene_tx=XM_019811702.2;refGene_gn=CCDC83;refGene_tx=XM_019811704.2;refGene_gn=CCDC83;refGene_tx=XM_019811705.2;refGene_gn=CCDC83;refGene_tx=XM_019811706.2;refGene_gn=CCDC83;refGene_tx=XM_019811707.1;refGene_gn=CCDC83;refGene_tx=XM_023239361.1;refGene_gn=CCDC83;refGene_tx=XM_023239362.1;refGene_gn=CCDC83;refGene_tx=XM_023239363.1
D1 48823437 48823522
refGene_gn=CCDC83;refGene_tx=XM_019811699.2;refGene_gn=CCDC83;refGene_tx=XM_019811700.2;refGene_gn=CCDC83;refGene_tx=XM_019811702.2;refGene_gn=CCDC83;refGene_tx=XM_019811704.2;refGene_gn=CCDC83;refGene_tx=XM_019811705.2;refGene_gn=CCDC83;refGene_tx=XM_019811706.2;refGene_gn=CCDC83;refGene_tx=XM_019811707.1;refGene_gn=CCDC83;refGene_tx=XM_023239361.1;refGene_gn=CCDC83;refGene_tx=XM_023239362.1;refGene_gn=CCDC83;refGene_tx=XM_023239363.1
D1 48829636 48829731
refGene_gn=CCDC83;refGene_tx=XM_019811699.2;refGene_gn=CCDC83;refGene_tx=XM_019811700.2;refGene_gn=CCDC83;refGene_tx=XM_019811702.2;refGene_gn=CCDC83;refGene_tx=XM_019811704.2;refGene_gn=CCDC83;refGene_tx=XM_019811705.2;refGene_gn=CCDC83;refGene_tx=XM_019811706.2;refGene_gn=CCDC83;refGene_tx=XM_023239361.1;refGene_gn=CCDC83;refGene_tx=XM_023239362.1;refGene_gn=CCDC83;refGene_tx=XM_023239363.1
D1 48925434 48925547
refGene_gn=SYTL2;refGene_tx=XM_019811682.2;refGene_gn=SYTL2;refGene_tx=XM_019811683.2;refGene_gn=SYTL2;refGene_tx=XM_019811684.1;refGene_gn=SYTL2;refGene_tx=XM_019811685.2;refGene_gn=SYTL2;refGene_tx=XM_019811686.2;refGene_gn=SYTL2;refGene_tx=XM_019811687.2;refGene_gn=SYTL2;refGene_tx=XM_019811688.2;refGene_gn=SYTL2;refGene_tx=XM_019811689.2;refGene_gn=SYTL2;refGene_tx=XM_019811690.2;refGene_gn=SYTL2;refGene_tx=XM_019811691.2;refGene_gn=SYTL2;refGene_tx=XM_019811692.2;refGene_gn=SYTL2;refGene_tx=XM_019811693.2;refGene_gn=SYTL2;refGene_tx=XM_019811694.2;refGene_gn=SYTL2;refGene_tx=XM_019811695.2

D1 48938917 48939069
refGene_gn=SYTL2;refGene_tx=XM_019811682.2;refGene_gn=SYTL2;refGene_tx=XM_019811683.2;refGene_gn=SYTL2;refGene_tx=XM_019811684.1;refGene_gn=SYTL2;refGene_tx=XM_019811685.2;refGene_gn=SYTL2;refGene_tx=XM_019811686.2;refGene_gn=SYTL2;refGene_tx=XM_019811687.2;refGene_gn=SYTL2;refGene_tx=XM_019811688.2;refGene_gn=SYTL2;refGene_tx=XM_019811689.2;refGene_gn=SYTL2;refGene_tx=XM_019811690.2;refGene_gn=SYTL2;refGene_tx=XM_019811691.2;refGene_gn=SYTL2;refGene_tx=XM_019811692.2;refGene_gn=SYTL2;refGene_tx=XM_019811693.2;refGene_gn=SYTL2;refGene_tx=XM_019811694.2;refGene_gn=SYTL2;refGene_tx=XM_019811695.2
D1 48941361 48941494
refGene_gn=SYTL2;refGene_tx=XM_019811682.2;refGene_gn=SYTL2;refGene_tx=XM_019811683.2;refGene_gn=SYTL2;refGene_tx=XM_019811684.1;refGene_gn=SYTL2;refGene_tx=XM_019811685.2;refGene_gn=SYTL2;refGene_tx=XM_019811686.2;refGene_gn=SYTL2;refGene_tx=XM_019811687.2;refGene_gn=SYTL2;refGene_tx=XM_019811688.2;refGene_gn=SYTL2;refGene_tx=XM_019811689.2;refGene_gn=SYTL2;refGene_tx=XM_019811690.2;refGene_gn=SYTL2;refGene_tx=XM_019811691.2;refGene_gn=SYTL2;refGene_tx=XM_019811692.2;refGene_gn=SYTL2;refGene_tx=XM_019811693.2;refGene_gn=SYTL2;refGene_tx=XM_019811694.2;refGene_gn=SYTL2;refGene_tx=XM_019811695.2
D1 48952573 48952655
refGene_gn=SYTL2;refGene_tx=XM_019811682.2;refGene_gn=SYTL2;refGene_tx=XM_019811683.2;refGene_gn=SYTL2;refGene_tx=XM_019811684.1;refGene_gn=SYTL2;refGene_tx=XM_019811685.2;refGene_gn=SYTL2;refGene_tx=XM_019811686.2;refGene_gn=SYTL2;refGene_tx=XM_019811687.2;refGene_gn=SYTL2;refGene_tx=XM_019811688.2;refGene_gn=SYTL2;refGene_tx=XM_019811689.2;refGene_gn=SYTL2;refGene_tx=XM_019811690.2;refGene_gn=SYTL2;refGene_tx=XM_019811691.2;refGene_gn=SYTL2;refGene_tx=XM_019811692.2;refGene_gn=SYTL2;refGene_tx=XM_019811693.2;refGene_gn=SYTL2;refGene_tx=XM_019811694.2;refGene_gn=SYTL2;refGene_tx=XM_019811695.2
D1 48953576 48953691
refGene_gn=SYTL2;refGene_tx=XM_019811682.2;refGene_gn=SYTL2;refGene_tx=XM_019811683.2;refGene_gn=SYTL2;refGene_tx=XM_019811684.1;refGene_gn=SYTL2;refGene_tx=XM_019811685.2;refGene_gn=SYTL2;refGene_tx=XM_019811686.2;refGene_gn=SYTL2;refGene_tx=XM_019811687.2;refGene_gn=SYTL2;refGene_tx=XM_019811688.2;refGene_gn=SYTL2;refGene_tx=XM_019811689.2;refGene_gn=SYTL2;refGene_tx=XM_019811690.2;refGene_gn=SYTL2;refGene_tx=XM_019811691.2;refGene_gn=SYTL2;refGene_tx=XM_019811692.2;refGene_gn=SYTL2;refGene_tx=XM_019811693.2;refGene_gn=SYTL2;refGene_tx=XM_019811694.2;refGene_gn=SYTL2;refGene_tx=XM_019811695.2
D1 48955082 48955886
refGene_gn=SYTL2;refGene_tx=XM_019811682.2;refGene_gn=SYTL2;refGene_tx=XM_019811683.2;refGene_gn=SYTL2;refGene_tx=XM_019811684.1;refGene_gn=SYTL2;refGene_tx=XM_019811685.2;refGene_gn=SYTL2;refGene_tx=XM_019811686.2;refGene_gn=SYTL2;refGene_tx=XM_019811687.2;refGene_gn=SYTL2;refGene_tx=XM_019811688.2;refGene_gn=SYTL2;refGene_tx=XM_019811689.2;refGene_gn=SYTL2;refGene_tx=XM_019811690.2;refGene_gn=SYTL2;refGene_tx=XM_019811691.2;refGene_gn=SYTL2;refGene_tx=XM_019811692.2;refGene_gn=SYTL2;refGene_tx=XM_019811693.2;refGene_gn=SYTL2;refGene_tx=XM_019811694.2;refGene_gn=SYTL2;refGene_tx=XM_019811695.2

D1 48960226 48964120
refGene_gn=SYTL2;refGene_tx=XM_019811682.2;refGene_gn=SYTL2;refGene_tx=XM_019811683.2;refGene_gn=SYTL2;refGene_tx=XM_019811684.1;refGene_gn=SYTL2;refGene_tx=XM_019811685.2;refGene_gn=SYTL2;refGene_tx=XM_019811686.2;refGene_gn=SYTL2;refGene_tx=XM_019811687.2;refGene_gn=SYTL2;refGene_tx=XM_019811688.2;refGene_gn=SYTL2;refGene_tx=XM_019811689.2;refGene_gn=SYTL2;refGene_tx=XM_019811690.2;refGene_gn=SYTL2;refGene_tx=XM_019811691.2;refGene_gn=SYTL2;refGene_tx=XM_019811692.2;refGene_gn=SYTL2;refGene_tx=XM_019811693.2;refGene_gn=SYTL2;refGene_tx=XM_019811695.2
D1 48967218 48967320
refGene_gn=SYTL2;refGene_tx=XM_019811682.2;refGene_gn=SYTL2;refGene_tx=XM_019811683.2;refGene_gn=SYTL2;refGene_tx=XM_019811684.1;refGene_gn=SYTL2;refGene_tx=XM_019811685.2;refGene_gn=SYTL2;refGene_tx=XM_019811686.2;refGene_gn=SYTL2;refGene_tx=XM_019811687.2;refGene_gn=SYTL2;refGene_tx=XM_019811688.2;refGene_gn=SYTL2;refGene_tx=XM_019811689.2;refGene_gn=SYTL2;refGene_tx=XM_019811690.2;refGene_gn=SYTL2;refGene_tx=XM_019811691.2;refGene_gn=SYTL2;refGene_tx=XM_019811692.2;refGene_gn=SYTL2;refGene_tx=XM_019811693.2;refGene_gn=SYTL2;refGene_tx=XM_019811694.2;refGene_gn=SYTL2;refGene_tx=XM_019811695.2
D1 48969300 48969354
refGene_gn=SYTL2;refGene_tx=XM_019811682.2;refGene_gn=SYTL2;refGene_tx=XM_019811683.2;refGene_gn=SYTL2;refGene_tx=XM_019811684.1;refGene_gn=SYTL2;refGene_tx=XM_019811685.2;refGene_gn=SYTL2;refGene_tx=XM_019811686.2;refGene_gn=SYTL2;refGene_tx=XM_019811687.2;refGene_gn=SYTL2;refGene_tx=XM_019811688.2;refGene_gn=SYTL2;refGene_tx=XM_019811689.2;refGene_gn=SYTL2;refGene_tx=XM_019811690.2;refGene_gn=SYTL2;refGene_tx=XM_019811691.2;refGene_gn=SYTL2;refGene_tx=XM_019811692.2;refGene_gn=SYTL2;refGene_tx=XM_019811693.2;refGene_gn=SYTL2;refGene_tx=XM_019811694.2;refGene_gn=SYTL2;refGene_tx=XM_019811695.2
D1 48970597 48970648
refGene_gn=SYTL2;refGene_tx=XM_019811682.2;refGene_gn=SYTL2;refGene_tx=XM_019811683.2;refGene_gn=SYTL2;refGene_tx=XM_019811684.1;refGene_gn=SYTL2;refGene_tx=XM_019811685.2;refGene_gn=SYTL2;refGene_tx=XM_019811686.2;refGene_gn=SYTL2;refGene_tx=XM_019811687.2;refGene_gn=SYTL2;refGene_tx=XM_019811689.2;refGene_gn=SYTL2;refGene_tx=XM_019811690.2;refGene_gn=SYTL2;refGene_tx=XM_019811691.2;refGene_gn=SYTL2;refGene_tx=XM_019811692.2;refGene_gn=SYTL2;refGene_tx=XM_019811694.2;refGene_gn=SYTL2;refGene_tx=XM_019811695.2
D1 48973847 48973942
refGene_gn=SYTL2;refGene_tx=XM_019811682.2;refGene_gn=SYTL2;refGene_tx=XM_019811683.2;refGene_gn=SYTL2;refGene_tx=XM_019811684.1;refGene_gn=SYTL2;refGene_tx=XM_019811685.2;refGene_gn=SYTL2;refGene_tx=XM_019811686.2;refGene_gn=SYTL2;refGene_tx=XM_019811687.2;refGene_gn=SYTL2;refGene_tx=XM_019811688.2;refGene_gn=SYTL2;refGene_tx=XM_019811689.2;refGene_gn=SYTL2;refGene_tx=XM_019811690.2;refGene_gn=SYTL2;refGene_tx=XM_019811691.2;refGene_gn=SYTL2;refGene_tx=XM_019811692.2;refGene_gn=SYTL2;refGene_tx=XM_019811693.2;refGene_gn=SYTL2;refGene_tx=XM_019811694.2;refGene_gn=SYTL2;refGene_tx=XM_019811695.2;refGene_gn=SYTL2;refGene_tx=XM_019811696.1;refGene_gn=SYTL2;refGene_tx=XM_019811697.2;refGene_gn=SYTL2;refGene_tx=XM_019811698.2

D1	48977046 48977166
refGene_gn=SYTL2;refGene_tx=XM_019811682.2;refGene_gn=SYTL2;refGene_tx=XM_019811683.2;refGene_gn=SYTL2;refGene_tx=XM_019811684.1;refGene_gn=SYTL2;refGene_tx=XM_019811685.2;refGene_gn=SYTL2;refGene_tx=XM_019811686.2;refGene_gn=SYTL2;refGene_tx=XM_019811687.2;refGene_gn=SYTL2;refGene_tx=XM_019811688.2;refGene_gn=SYTL2;refGene_tx=XM_019811689.2;refGene_gn=SYTL2;refGene_tx=XM_019811691.2;refGene_gn=SYTL2;refGene_tx=XM_019811692.2;refGene_gn=SYTL2;refGene_tx=XM_019811693.2;refGene_gn=SYTL2;refGene_tx=XM_019811694.2;refGene_gn=SYTL2;refGene_tx=XM_019811696.1;refGene_gn=SYTL2;refGene_tx=XM_019811697.2;refGene_gn=SYTL2;refGene_tx=XM_019811698.2
D1	48978456 48978626
refGene_gn=SYTL2;refGene_tx=XM_019811682.2;refGene_gn=SYTL2;refGene_tx=XM_019811683.2;refGene_gn=SYTL2;refGene_tx=XM_019811684.1;refGene_gn=SYTL2;refGene_tx=XM_019811685.2;refGene_gn=SYTL2;refGene_tx=XM_019811686.2;refGene_gn=SYTL2;refGene_tx=XM_019811687.2;refGene_gn=SYTL2;refGene_tx=XM_019811688.2;refGene_gn=SYTL2;refGene_tx=XM_019811689.2;refGene_gn=SYTL2;refGene_tx=XM_019811690.2;refGene_gn=SYTL2;refGene_tx=XM_019811691.2;refGene_gn=SYTL2;refGene_tx=XM_019811692.2;refGene_gn=SYTL2;refGene_tx=XM_019811693.2;refGene_gn=SYTL2;refGene_tx=XM_019811694.2;refGene_gn=SYTL2;refGene_tx=XM_019811695.2;refGene_gn=SYTL2;refGene_tx=XM_019811696.1;refGene_gn=SYTL2;refGene_tx=XM_019811697.2;refGene_gn=SYTL2;refGene_tx=XM_019811698.2
D1	48979739 48979842
refGene_gn=SYTL2;refGene_tx=XM_019811682.2;refGene_gn=SYTL2;refGene_tx=XM_019811683.2;refGene_gn=SYTL2;refGene_tx=XM_019811684.1;refGene_gn=SYTL2;refGene_tx=XM_019811685.2;refGene_gn=SYTL2;refGene_tx=XM_019811686.2;refGene_gn=SYTL2;refGene_tx=XM_019811687.2;refGene_gn=SYTL2;refGene_tx=XM_019811688.2;refGene_gn=SYTL2;refGene_tx=XM_019811689.2;refGene_gn=SYTL2;refGene_tx=XM_019811690.2;refGene_gn=SYTL2;refGene_tx=XM_019811691.2;refGene_gn=SYTL2;refGene_tx=XM_019811692.2;refGene_gn=SYTL2;refGene_tx=XM_019811693.2;refGene_gn=SYTL2;refGene_tx=XM_019811694.2;refGene_gn=SYTL2;refGene_tx=XM_019811695.2;refGene_gn=SYTL2;refGene_tx=XM_019811696.1;refGene_gn=SYTL2;refGene_tx=XM_019811697.2;refGene_gn=SYTL2;refGene_tx=XM_019811698.2
D1	48982169 48982340
refGene_gn=SYTL2;refGene_tx=XM_019811682.2;refGene_gn=SYTL2;refGene_tx=XM_019811683.2;refGene_gn=SYTL2;refGene_tx=XM_019811684.1;refGene_gn=SYTL2;refGene_tx=XM_019811685.2;refGene_gn=SYTL2;refGene_tx=XM_019811686.2;refGene_gn=SYTL2;refGene_tx=XM_019811687.2;refGene_gn=SYTL2;refGene_tx=XM_019811688.2;refGene_gn=SYTL2;refGene_tx=XM_019811689.2;refGene_gn=SYTL2;refGene_tx=XM_019811690.2;refGene_gn=SYTL2;refGene_tx=XM_019811691.2;refGene_gn=SYTL2;refGene_tx=XM_019811692.2;refGene_gn=SYTL2;refGene_tx=XM_019811693.2;refGene_gn=SYTL2;refGene_tx=XM_019811694.2;refGene_gn=SYTL2;refGene_tx=XM_019811695.2;refGene_gn=SYTL2;refGene_tx=XM_019811696.1;refGene_gn=SYTL2;refGene_tx=XM_019811697.2;refGene_gn=SYTL2;refGene_tx=XM_019811698.2
D1	48984240 48984319
refGene_gn=SYTL2;refGene_tx=XM_019811682.2;refGene_gn=SYTL2;refGene_tx=XM_019811683.2;refGene_gn=SYTL2;refGene_tx=XM_019811684.1;refGene_gn=

SYTL2;refGene_tx=XM_019811685.2;refGene_gn=SYTL2;refGene_tx=XM_0198116
86.2;refGene_gn=SYTL2;refGene_tx=XM_019811687.2;refGene_gn=SYTL2;refGe
ne_tx=XM_019811688.2;refGene_gn=SYTL2;refGene_tx=XM_019811689.2;refGen
e_gn=SYTL2;refGene_tx=XM_019811690.2;refGene_gn=SYTL2;refGene_tx=XM_01
9811691.2;refGene_gn=SYTL2;refGene_tx=XM_019811692.2;refGene_gn=SYTL2;
refGene_tx=XM_019811693.2;refGene_gn=SYTL2;refGene_tx=XM_019811694.2;r
efGene_gn=SYTL2;refGene_tx=XM_019811695.2;refGene_gn=SYTL2;refGene_tx=
XM_019811696.1;refGene_gn=SYTL2;refGene_tx=XM_019811697.2;refGene_gn=S
YTL2;refGene_tx=XM_019811698.2
D1 48986567 48986667
refGene_gn=SYTL2;refGene_tx=XM_019811682.2;refGene_gn=SYTL2;refGene_tx
=XM_019811683.2;refGene_gn=SYTL2;refGene_tx=XM_019811684.1;refGene_gn=
SYTL2;refGene_tx=XM_019811685.2;refGene_gn=SYTL2;refGene_tx=XM_0198116
86.2;refGene_gn=SYTL2;refGene_tx=XM_019811687.2;refGene_gn=SYTL2;refGe
ne_tx=XM_019811688.2;refGene_gn=SYTL2;refGene_tx=XM_019811689.2;refGen
e_gn=SYTL2;refGene_tx=XM_019811690.2;refGene_gn=SYTL2;refGene_tx=XM_01
9811691.2;refGene_gn=SYTL2;refGene_tx=XM_019811692.2;refGene_gn=SYTL2;
refGene_tx=XM_019811693.2;refGene_gn=SYTL2;refGene_tx=XM_019811694.2;r
efGene_gn=SYTL2;refGene_tx=XM_019811695.2;refGene_gn=SYTL2;refGene_tx=
XM_019811696.1;refGene_gn=SYTL2;refGene_tx=XM_019811697.2;refGene_gn=S
YTL2;refGene_tx=XM_019811698.2
D1 48987781 48987987
refGene_gn=SYTL2;refGene_tx=XM_019811682.2;refGene_gn=SYTL2;refGene_tx
=XM_019811683.2;refGene_gn=SYTL2;refGene_tx=XM_019811684.1;refGene_gn=
SYTL2;refGene_tx=XM_019811685.2;refGene_gn=SYTL2;refGene_tx=XM_0198116
86.2;refGene_gn=SYTL2;refGene_tx=XM_019811687.2;refGene_gn=SYTL2;refGe
ne_tx=XM_019811688.2;refGene_gn=SYTL2;refGene_tx=XM_019811689.2;refGen
e_gn=SYTL2;refGene_tx=XM_019811690.2;refGene_gn=SYTL2;refGene_tx=XM_01
9811691.2;refGene_gn=SYTL2;refGene_tx=XM_019811692.2;refGene_gn=SYTL2;
refGene_tx=XM_019811693.2;refGene_gn=SYTL2;refGene_tx=XM_019811694.2;r
efGene_gn=SYTL2;refGene_tx=XM_019811695.2;refGene_gn=SYTL2;refGene_tx=
XM_019811696.1;refGene_gn=SYTL2;refGene_tx=XM_019811697.2;refGene_gn=S
YTL2;refGene_tx=XM_019811698.2
D1 48988789 48988935
refGene_gn=SYTL2;refGene_tx=XM_019811682.2;refGene_gn=SYTL2;refGene_tx
=XM_019811683.2;refGene_gn=SYTL2;refGene_tx=XM_019811684.1;refGene_gn=
SYTL2;refGene_tx=XM_019811685.2;refGene_gn=SYTL2;refGene_tx=XM_0198116
86.2;refGene_gn=SYTL2;refGene_tx=XM_019811687.2;refGene_gn=SYTL2;refGe
ne_tx=XM_019811688.2;refGene_gn=SYTL2;refGene_tx=XM_019811689.2;refGen
e_gn=SYTL2;refGene_tx=XM_019811690.2;refGene_gn=SYTL2;refGene_tx=XM_01
9811691.2;refGene_gn=SYTL2;refGene_tx=XM_019811692.2;refGene_gn=SYTL2;
refGene_tx=XM_019811693.2;refGene_gn=SYTL2;refGene_tx=XM_019811694.2;r
efGene_gn=SYTL2;refGene_tx=XM_019811695.2;refGene_gn=SYTL2;refGene_tx=
XM_019811696.1;refGene_gn=SYTL2;refGene_tx=XM_019811697.2;refGene_gn=S
YTL2;refGene_tx=XM_019811698.2
D1 48994274 48995387 refGene_gn=CCDC89;refGene_tx=XM_003992666.4
D1 49012408 49013473
refGene_gn=CREBZF;refGene_tx=XM_023239365.1;refGene_gn=CREBZF;refGene_
tx=XM_023239366.1;refGene_gn=CREBZF;refGene_tx=XM_023239367.1;refGene_
gn=CREBZF;refGene_tx=XM_023239368.1;refGene_gn=CREBZF;refGene_tx=XM_02

3239369.1;refGene_gn=CREBZF;refGene_tx=XM_023239370.1;refGene_gn=CREBZF;refGene_tx=XM_023239371.1;refGene_gn=CREBZF;refGene_tx=XM_023239372.1;refGene_gn=CREBZF;refGene_tx=XM_023239373.1
D1 49013678 49013694
refGene_gn=CREBZF;refGene_tx=XM_023239365.1;refGene_gn=CREBZF;refGene_tx=XM_023239366.1;refGene_gn=CREBZF;refGene_tx=XM_023239367.1;refGene_gn=CREBZF;refGene_tx=XM_023239368.1;refGene_gn=CREBZF;refGene_tx=XM_023239369.1;refGene_gn=CREBZF;refGene_tx=XM_023239370.1;refGene_gn=CREBZF;refGene_tx=XM_023239371.1;refGene_gn=CREBZF;refGene_tx=XM_023239372.1
D1 49021514 49021707 refGene_gn=TMEM126A;refGene_tx=XM_006936876.3
D1 49022195 49022310 refGene_gn=TMEM126A;refGene_tx=XM_006936876.3
D1 49024100 49024294 refGene_gn=TMEM126A;refGene_tx=XM_006936876.3
D1 49027358 49027450 refGene_gn=TMEM126A;refGene_tx=XM_006936876.3
D1 49048514 49048698
refGene_gn=TMEM126B;refGene_tx=XM_003992668.3;refGene_gn=TMEM126B;refGene_tx=XM_006936877.3;refGene_gn=TMEM126B;refGene_tx=XM_006936878.2
D1 49048984 49049096
refGene_gn=TMEM126B;refGene_tx=XM_003992668.3;refGene_gn=TMEM126B;refGene_tx=XM_006936877.3;refGene_gn=TMEM126B;refGene_tx=XM_006936878.2
D1 49050171 49050365
refGene_gn=TMEM126B;refGene_tx=XM_003992668.3;refGene_gn=TMEM126B;refGene_tx=XM_006936877.3;refGene_gn=TMEM126B;refGene_tx=XM_006936878.2
D1 49051969 49052091
refGene_gn=TMEM126B;refGene_tx=XM_003992668.3;refGene_gn=TMEM126B;refGene_tx=XM_006936877.3;refGene_gn=TMEM126B;refGene_tx=XM_006936878.2
D1 49055192 49055273
refGene_gn=TMEM126B;refGene_tx=XM_003992668.3;refGene_gn=TMEM126B;refGene_tx=XM_006936877.3
D1 49057260 49057279
refGene_gn=DLG2;refGene_tx=XM_023239374.1;refGene_gn=DLG2;refGene_tx=XM_023239375.1;refGene_gn=DLG2;refGene_tx=XM_023239376.1;refGene_gn=DLG2;refGene_tx=XM_023239377.1;refGene_gn=DLG2;refGene_tx=XM_023239378.1;refGene_gn=DLG2;refGene_tx=XM_023239379.1;refGene_gn=DLG2;refGene_tx=XM_023239380.1;refGene_gn=DLG2;refGene_tx=XM_023239381.1;refGene_gn=DLG2;refGene_tx=XM_023239382.1;refGene_gn=DLG2;refGene_tx=XM_023239388.1;refGene_gn=DLG2;refGene_tx=XM_023239393.1;refGene_gn=DLG2;refGene_tx=XM_023239394.1
D1 49072350 49072482
refGene_gn=DLG2;refGene_tx=XM_023239374.1;refGene_gn=DLG2;refGene_tx=XM_023239375.1;refGene_gn=DLG2;refGene_tx=XM_023239376.1;refGene_gn=DLG2;refGene_tx=XM_023239377.1;refGene_gn=DLG2;refGene_tx=XM_023239378.1;refGene_gn=DLG2;refGene_tx=XM_023239379.1;refGene_gn=DLG2;refGene_tx=XM_023239380.1;refGene_gn=DLG2;refGene_tx=XM_023239381.1;refGene_gn=DLG2;refGene_tx=XM_023239382.1;refGene_gn=DLG2;refGene_tx=XM_023239388.1;refGene_gn=DLG2;refGene_tx=XM_023239393.1;refGene_gn=DLG2;refGene_tx=XM_023239394.1
D1 49361364 49361510
refGene_gn=DLG2;refGene_tx=XM_023239374.1;refGene_gn=DLG2;refGene_tx=XM_023239375.1;refGene_gn=DLG2;refGene_tx=XM_023239376.1;refGene_gn=DLG

2;refGene_tx=XM_023239377.1;refGene_gn=DLG2;refGene_tx=XM_023239378.1;
refGene_gn=DLG2;refGene_tx=XM_023239379.1;refGene_gn=DLG2;refGene_tx=X
M_023239380.1;refGene_gn=DLG2;refGene_tx=XM_023239381.1;refGene_gn=DLG
2;refGene_tx=XM_023239382.1;refGene_gn=DLG2;refGene_tx=XM_023239388.1;
refGene_gn=DLG2;refGene_tx=XM_023239393.1;refGene_gn=DLG2;refGene_tx=X
M_023239394.1
D1 49519901 49519997
refGene_gn=DLG2;refGene_tx=XM_023239374.1;refGene_gn=DLG2;refGene_tx=X
M_023239375.1;refGene_gn=DLG2;refGene_tx=XM_023239376.1;refGene_gn=DLG
2;refGene_tx=XM_023239377.1;refGene_gn=DLG2;refGene_tx=XM_023239378.1;
refGene_gn=DLG2;refGene_tx=XM_023239379.1;refGene_gn=DLG2;refGene_tx=X
M_023239380.1;refGene_gn=DLG2;refGene_tx=XM_023239381.1;refGene_gn=DLG
2;refGene_tx=XM_023239382.1;refGene_gn=DLG2;refGene_tx=XM_023239388.1;
refGene_gn=DLG2;refGene_tx=XM_023239393.1;refGene_gn=DLG2;refGene_tx=X
M_023239394.1
D1 49532623 49532653
refGene_gn=DLG2;refGene_tx=XM_023239384.1;refGene_gn=DLG2;refGene_tx=X
M_023239386.1
D1 49556812 49556887
refGene_gn=DLG2;refGene_tx=XM_023239374.1;refGene_gn=DLG2;refGene_tx=X
M_023239375.1;refGene_gn=DLG2;refGene_tx=XM_023239376.1;refGene_gn=DLG
2;refGene_tx=XM_023239377.1;refGene_gn=DLG2;refGene_tx=XM_023239378.1;
refGene_gn=DLG2;refGene_tx=XM_023239379.1;refGene_gn=DLG2;refGene_tx=X
M_023239380.1;refGene_gn=DLG2;refGene_tx=XM_023239381.1;refGene_gn=DLG
2;refGene_tx=XM_023239382.1;refGene_gn=DLG2;refGene_tx=XM_023239384.1;
refGene_gn=DLG2;refGene_tx=XM_023239386.1;refGene_gn=DLG2;refGene_tx=X
M_023239388.1;refGene_gn=DLG2;refGene_tx=XM_023239393.1;refGene_gn=DLG
2;refGene_tx=XM_023239394.1
D1 49714163 49714205 refGene_gn=DLG2;refGene_tx=XM_023239389.1
D1 49902358 49902460
refGene_gn=DLG2;refGene_tx=XM_023239385.1;refGene_gn=DLG2;refGene_tx=X
M_023239387.1;refGene_gn=DLG2;refGene_tx=XM_023239392.1
D1 50102641 50102803
refGene_gn=DLG2;refGene_tx=XM_023239374.1;refGene_gn=DLG2;refGene_tx=X
M_023239375.1;refGene_gn=DLG2;refGene_tx=XM_023239376.1;refGene_gn=DLG
2;refGene_tx=XM_023239377.1;refGene_gn=DLG2;refGene_tx=XM_023239378.1;
refGene_gn=DLG2;refGene_tx=XM_023239379.1;refGene_gn=DLG2;refGene_tx=X
M_023239380.1;refGene_gn=DLG2;refGene_tx=XM_023239381.1;refGene_gn=DLG
2;refGene_tx=XM_023239382.1;refGene_gn=DLG2;refGene_tx=XM_023239384.1;
refGene_gn=DLG2;refGene_tx=XM_023239385.1;refGene_gn=DLG2;refGene_tx=X
M_023239386.1;refGene_gn=DLG2;refGene_tx=XM_023239387.1;refGene_gn=DLG
2;refGene_tx=XM_023239388.1;refGene_gn=DLG2;refGene_tx=XM_023239389.1;
refGene_gn=DLG2;refGene_tx=XM_023239392.1;refGene_gn=DLG2;refGene_tx=X
M_023239393.1;refGene_gn=DLG2;refGene_tx=XM_023239394.1
D1 50211845 50211923 refGene_gn=DLG2;refGene_tx=XM_023239391.1
D1 50331869 50332178
refGene_gn=DLG2;refGene_tx=XM_003992674.5;refGene_gn=DLG2;refGene_tx=X
M_023239383.1
D1 50368789 50368894
refGene_gn=DLG2;refGene_tx=XM_019811729.1;refGene_gn=DLG2;refGene_tx=X

M_023239390.1
D1 50389531 50389585
refGene_gn=DLG2;refGene_tx=XM_003992674.5;refGene_gn=DLG2;refGene_tx=XM_019811729.1;refGene_gn=DLG2;refGene_tx=XM_023239374.1;refGene_gn=DLG2;refGene_tx=XM_023239375.1;refGene_gn=DLG2;refGene_tx=XM_023239376.1;refGene_gn=DLG2;refGene_tx=XM_023239377.1;refGene_gn=DLG2;refGene_tx=XM_023239378.1;refGene_gn=DLG2;refGene_tx=XM_023239379.1;refGene_gn=DLG2;refGene_tx=XM_023239380.1;refGene_gn=DLG2;refGene_tx=XM_023239381.1;refGene_gn=DLG2;refGene_tx=XM_023239382.1;refGene_gn=DLG2;refGene_tx=XM_023239383.1;refGene_gn=DLG2;refGene_tx=XM_023239384.1;refGene_gn=DLG2;refGene_tx=XM_023239385.1;refGene_gn=DLG2;refGene_tx=XM_023239386.1;refGene_gn=DLG2;refGene_tx=XM_023239387.1;refGene_gn=DLG2;refGene_tx=XM_023239388.1;refGene_gn=DLG2;refGene_tx=XM_023239389.1;refGene_gn=DLG2;refGene_tx=XM_023239390.1;refGene_gn=DLG2;refGene_tx=XM_023239391.1;refGene_gn=DLG2;refGene_tx=XM_023239392.1;refGene_gn=DLG2;refGene_tx=XM_023239393.1;refGene_gn=DLG2;refGene_tx=XM_023239394.1
D1 50466144 50466195
refGene_gn=DLG2;refGene_tx=XM_003992674.5;refGene_gn=DLG2;refGene_tx=XM_019811729.1;refGene_gn=DLG2;refGene_tx=XM_023239374.1;refGene_gn=DLG2;refGene_tx=XM_023239375.1;refGene_gn=DLG2;refGene_tx=XM_023239376.1;refGene_gn=DLG2;refGene_tx=XM_023239377.1;refGene_gn=DLG2;refGene_tx=XM_023239378.1;refGene_gn=DLG2;refGene_tx=XM_023239379.1;refGene_gn=DLG2;refGene_tx=XM_023239380.1;refGene_gn=DLG2;refGene_tx=XM_023239381.1;refGene_gn=DLG2;refGene_tx=XM_023239382.1;refGene_gn=DLG2;refGene_tx=XM_023239383.1;refGene_gn=DLG2;refGene_tx=XM_023239384.1;refGene_gn=DLG2;refGene_tx=XM_023239385.1;refGene_gn=DLG2;refGene_tx=XM_023239386.1;refGene_gn=DLG2;refGene_tx=XM_023239387.1;refGene_gn=DLG2;refGene_tx=XM_023239388.1;refGene_gn=DLG2;refGene_tx=XM_023239389.1;refGene_gn=DLG2;refGene_tx=XM_023239390.1;refGene_gn=DLG2;refGene_tx=XM_023239391.1;refGene_gn=DLG2;refGene_tx=XM_023239392.1;refGene_gn=DLG2;refGene_tx=XM_023239393.1;refGene_gn=DLG2;refGene_tx=XM_023239394.1
D1 50523678 50523803
refGene_gn=DLG2;refGene_tx=XM_003992674.5;refGene_gn=DLG2;refGene_tx=XM_019811729.1;refGene_gn=DLG2;refGene_tx=XM_023239374.1;refGene_gn=DLG2;refGene_tx=XM_023239375.1;refGene_gn=DLG2;refGene_tx=XM_023239376.1;refGene_gn=DLG2;refGene_tx=XM_023239377.1;refGene_gn=DLG2;refGene_tx=XM_023239378.1;refGene_gn=DLG2;refGene_tx=XM_023239379.1;refGene_gn=DLG2;refGene_tx=XM_023239380.1;refGene_gn=DLG2;refGene_tx=XM_023239381.1;refGene_gn=DLG2;refGene_tx=XM_023239382.1;refGene_gn=DLG2;refGene_tx=XM_023239383.1;refGene_gn=DLG2;refGene_tx=XM_023239384.1;refGene_gn=DLG2;refGene_tx=XM_023239385.1;refGene_gn=DLG2;refGene_tx=XM_023239386.1;refGene_gn=DLG2;refGene_tx=XM_023239387.1;refGene_gn=DLG2;refGene_tx=XM_023239388.1;refGene_gn=DLG2;refGene_tx=XM_023239389.1;refGene_gn=DLG2;refGene_tx=XM_023239390.1;refGene_gn=DLG2;refGene_tx=XM_023239391.1;refGene_gn=DLG2;refGene_tx=XM_023239392.1;refGene_gn=DLG2;refGene_tx=XM_023239393.1;refGene_gn=DLG2;refGene_tx=XM_023239394.1
D1 50556964 50557134
refGene_gn=DLG2;refGene_tx=XM_003992674.5;refGene_gn=DLG2;refGene_tx=XM_019811729.1;refGene_gn=DLG2;refGene_tx=XM_023239374.1;refGene_gn=DLG2;refGene_tx=XM_023239375.1;refGene_gn=DLG2;refGene_tx=XM_023239376.1;

refGene_gn=DLG2;refGene_tx=XM_023239377.1;refGene_gn=DLG2;refGene_tx=XM_023239378.1;refGene_gn=DLG2;refGene_tx=XM_023239379.1;refGene_gn=DLG2;refGene_tx=XM_023239380.1;refGene_gn=DLG2;refGene_tx=XM_023239381.1;refGene_gn=DLG2;refGene_tx=XM_023239382.1;refGene_gn=DLG2;refGene_tx=XM_023239383.1;refGene_gn=DLG2;refGene_tx=XM_023239384.1;refGene_gn=DLG2;refGene_tx=XM_023239385.1;refGene_gn=DLG2;refGene_tx=XM_023239386.1;refGene_gn=DLG2;refGene_tx=XM_023239387.1;refGene_gn=DLG2;refGene_tx=XM_023239388.1;refGene_gn=DLG2;refGene_tx=XM_023239389.1;refGene_gn=DLG2;refGene_tx=XM_023239390.1;refGene_gn=DLG2;refGene_tx=XM_023239391.1;refGene_gn=DLG2;refGene_tx=XM_023239392.1;refGene_gn=DLG2;refGene_tx=XM_023239393.1;refGene_gn=DLG2;refGene_tx=XM_023239394.1

D1 50622877 50623014
refGene_gn=DLG2;refGene_tx=XM_003992674.5;refGene_gn=DLG2;refGene_tx=XM_019811729.1;refGene_gn=DLG2;refGene_tx=XM_023239374.1;refGene_gn=DLG2;refGene_tx=XM_023239375.1;refGene_gn=DLG2;refGene_tx=XM_023239376.1;refGene_gn=DLG2;refGene_tx=XM_023239377.1;refGene_gn=DLG2;refGene_tx=XM_023239378.1;refGene_gn=DLG2;refGene_tx=XM_023239379.1;refGene_gn=DLG2;refGene_tx=XM_023239380.1;refGene_gn=DLG2;refGene_tx=XM_023239381.1;refGene_gn=DLG2;refGene_tx=XM_023239382.1;refGene_gn=DLG2;refGene_tx=XM_023239383.1;refGene_gn=DLG2;refGene_tx=XM_023239384.1;refGene_gn=DLG2;refGene_tx=XM_023239385.1;refGene_gn=DLG2;refGene_tx=XM_023239386.1;refGene_gn=DLG2;refGene_tx=XM_023239387.1;refGene_gn=DLG2;refGene_tx=XM_023239388.1;refGene_gn=DLG2;refGene_tx=XM_023239389.1;refGene_gn=DLG2;refGene_tx=XM_023239390.1;refGene_gn=DLG2;refGene_tx=XM_023239391.1;refGene_gn=DLG2;refGene_tx=XM_023239392.1;refGene_gn=DLG2;refGene_tx=XM_023239393.1;refGene_gn=DLG2;refGene_tx=XM_023239394.1

D1 50635898 50636043
refGene_gn=DLG2;refGene_tx=XM_003992674.5;refGene_gn=DLG2;refGene_tx=XM_019811729.1;refGene_gn=DLG2;refGene_tx=XM_023239374.1;refGene_gn=DLG2;refGene_tx=XM_023239375.1;refGene_gn=DLG2;refGene_tx=XM_023239376.1;refGene_gn=DLG2;refGene_tx=XM_023239377.1;refGene_gn=DLG2;refGene_tx=XM_023239378.1;refGene_gn=DLG2;refGene_tx=XM_023239379.1;refGene_gn=DLG2;refGene_tx=XM_023239380.1;refGene_gn=DLG2;refGene_tx=XM_023239381.1;refGene_gn=DLG2;refGene_tx=XM_023239382.1;refGene_gn=DLG2;refGene_tx=XM_023239383.1;refGene_gn=DLG2;refGene_tx=XM_023239384.1;refGene_gn=DLG2;refGene_tx=XM_023239385.1;refGene_gn=DLG2;refGene_tx=XM_023239386.1;refGene_gn=DLG2;refGene_tx=XM_023239387.1;refGene_gn=DLG2;refGene_tx=XM_023239388.1;refGene_gn=DLG2;refGene_tx=XM_023239389.1;refGene_gn=DLG2;refGene_tx=XM_023239390.1;refGene_gn=DLG2;refGene_tx=XM_023239391.1;refGene_gn=DLG2;refGene_tx=XM_023239392.1;refGene_gn=DLG2;refGene_tx=XM_023239393.1;refGene_gn=DLG2;refGene_tx=XM_023239394.1

D1 50638296 50638435
refGene_gn=DLG2;refGene_tx=XM_003992674.5;refGene_gn=DLG2;refGene_tx=XM_019811729.1;refGene_gn=DLG2;refGene_tx=XM_023239374.1;refGene_gn=DLG2;refGene_tx=XM_023239375.1;refGene_gn=DLG2;refGene_tx=XM_023239376.1;refGene_gn=DLG2;refGene_tx=XM_023239377.1;refGene_gn=DLG2;refGene_tx=XM_023239378.1;refGene_gn=DLG2;refGene_tx=XM_023239379.1;refGene_gn=DLG2;refGene_tx=XM_023239380.1;refGene_gn=DLG2;refGene_tx=XM_023239381.1;refGene_gn=DLG2;refGene_tx=XM_023239382.1;refGene_gn=DLG2;refGene_tx=XM_023239383.1;refGene_gn=DLG2;refGene_tx=XM_023239384.1;refGene_gn=DLG

2;refGene_tx=XM_023239385.1;refGene_gn=DLG2;refGene_tx=XM_023239386.1;
refGene_gn=DLG2;refGene_tx=XM_023239387.1;refGene_gn=DLG2;refGene_tx=X
M_023239388.1;refGene_gn=DLG2;refGene_tx=XM_023239389.1;refGene_gn=DLG
2;refGene_tx=XM_023239390.1;refGene_gn=DLG2;refGene_tx=XM_023239391.1;
refGene_gn=DLG2;refGene_tx=XM_023239392.1;refGene_gn=DLG2;refGene_tx=X
M_023239393.1;refGene_gn=DLG2;refGene_tx=XM_023239394.1
D1 50672335 50672491
refGene_gn=DLG2;refGene_tx=XM_003992674.5;refGene_gn=DLG2;refGene_tx=X
M_023239374.1;refGene_gn=DLG2;refGene_tx=XM_023239375.1;refGene_gn=DLG
2;refGene_tx=XM_023239376.1;refGene_gn=DLG2;refGene_tx=XM_023239377.1;
refGene_gn=DLG2;refGene_tx=XM_023239378.1;refGene_gn=DLG2;refGene_tx=X
M_023239379.1;refGene_gn=DLG2;refGene_tx=XM_023239381.1;refGene_gn=DLG
2;refGene_tx=XM_023239382.1;refGene_gn=DLG2;refGene_tx=XM_023239383.1;
refGene_gn=DLG2;refGene_tx=XM_023239384.1;refGene_gn=DLG2;refGene_tx=X
M_023239385.1;refGene_gn=DLG2;refGene_tx=XM_023239386.1;refGene_gn=DLG
2;refGene_tx=XM_023239387.1;refGene_gn=DLG2;refGene_tx=XM_023239389.1;
refGene_gn=DLG2;refGene_tx=XM_023239390.1;refGene_gn=DLG2;refGene_tx=X
M_023239391.1;refGene_gn=DLG2;refGene_tx=XM_023239392.1;refGene_gn=DLG
2;refGene_tx=XM_023239393.1;refGene_gn=DLG2;refGene_tx=XM_023239394.1
D1 50727359 50727428
refGene_gn=DLG2;refGene_tx=XM_003992674.5;refGene_gn=DLG2;refGene_tx=X
M_019811729.1;refGene_gn=DLG2;refGene_tx=XM_023239374.1;refGene_gn=DLG
2;refGene_tx=XM_023239375.1;refGene_gn=DLG2;refGene_tx=XM_023239376.1;
refGene_gn=DLG2;refGene_tx=XM_023239377.1;refGene_gn=DLG2;refGene_tx=X
M_023239378.1;refGene_gn=DLG2;refGene_tx=XM_023239379.1;refGene_gn=DLG
2;refGene_tx=XM_023239380.1;refGene_gn=DLG2;refGene_tx=XM_023239381.1;
refGene_gn=DLG2;refGene_tx=XM_023239382.1;refGene_gn=DLG2;refGene_tx=X
M_023239383.1;refGene_gn=DLG2;refGene_tx=XM_023239384.1;refGene_gn=DLG
2;refGene_tx=XM_023239385.1;refGene_gn=DLG2;refGene_tx=XM_023239386.1;
refGene_gn=DLG2;refGene_tx=XM_023239387.1;refGene_gn=DLG2;refGene_tx=X
M_023239388.1;refGene_gn=DLG2;refGene_tx=XM_023239389.1;refGene_gn=DLG
2;refGene_tx=XM_023239390.1;refGene_gn=DLG2;refGene_tx=XM_023239391.1;
refGene_gn=DLG2;refGene_tx=XM_023239392.1;refGene_gn=DLG2;refGene_tx=X
M_023239393.1;refGene_gn=DLG2;refGene_tx=XM_023239394.1
D1 50767339 50767496
refGene_gn=DLG2;refGene_tx=XM_003992674.5;refGene_gn=DLG2;refGene_tx=X
M_019811729.1;refGene_gn=DLG2;refGene_tx=XM_023239374.1;refGene_gn=DLG
2;refGene_tx=XM_023239375.1;refGene_gn=DLG2;refGene_tx=XM_023239376.1;
refGene_gn=DLG2;refGene_tx=XM_023239377.1;refGene_gn=DLG2;refGene_tx=X
M_023239378.1;refGene_gn=DLG2;refGene_tx=XM_023239379.1;refGene_gn=DLG
2;refGene_tx=XM_023239380.1;refGene_gn=DLG2;refGene_tx=XM_023239381.1;
refGene_gn=DLG2;refGene_tx=XM_023239382.1;refGene_gn=DLG2;refGene_tx=X
M_023239383.1;refGene_gn=DLG2;refGene_tx=XM_023239384.1;refGene_gn=DLG
2;refGene_tx=XM_023239385.1;refGene_gn=DLG2;refGene_tx=XM_023239386.1;
refGene_gn=DLG2;refGene_tx=XM_023239387.1;refGene_gn=DLG2;refGene_tx=X
M_023239388.1;refGene_gn=DLG2;refGene_tx=XM_023239389.1;refGene_gn=DLG
2;refGene_tx=XM_023239390.1;refGene_gn=DLG2;refGene_tx=XM_023239391.1;
refGene_gn=DLG2;refGene_tx=XM_023239392.1;refGene_gn=DLG2;refGene_tx=X
M_023239393.1;refGene_gn=DLG2;refGene_tx=XM_023239394.1
D1 50813629 50813732

refGene_gn=DLG2;refGene_tx=XM_003992674.5;refGene_gn=DLG2;refGene_tx=XM_019811729.1;refGene_gn=DLG2;refGene_tx=XM_023239374.1;refGene_gn=DLG2;refGene_tx=XM_023239375.1;refGene_gn=DLG2;refGene_tx=XM_023239376.1;refGene_gn=DLG2;refGene_tx=XM_023239377.1;refGene_gn=DLG2;refGene_tx=XM_023239378.1;refGene_gn=DLG2;refGene_tx=XM_023239379.1;refGene_gn=DLG2;refGene_tx=XM_023239380.1;refGene_gn=DLG2;refGene_tx=XM_023239381.1;refGene_gn=DLG2;refGene_tx=XM_023239382.1;refGene_gn=DLG2;refGene_tx=XM_023239383.1;refGene_gn=DLG2;refGene_tx=XM_023239384.1;refGene_gn=DLG2;refGene_tx=XM_023239385.1;refGene_gn=DLG2;refGene_tx=XM_023239386.1;refGene_gn=DLG2;refGene_tx=XM_023239387.1;refGene_gn=DLG2;refGene_tx=XM_023239388.1;refGene_gn=DLG2;refGene_tx=XM_023239389.1;refGene_gn=DLG2;refGene_tx=XM_023239390.1;refGene_gn=DLG2;refGene_tx=XM_023239391.1;refGene_gn=DLG2;refGene_tx=XM_023239392.1;refGene_gn=DLG2;refGene_tx=XM_023239393.1;refGene_gn=DLG2;refGene_tx=XM_023239394.1
D1 50966476 50966591
refGene_gn=DLG2;refGene_tx=XM_003992674.5;refGene_gn=DLG2;refGene_tx=XM_019811729.1;refGene_gn=DLG2;refGene_tx=XM_023239374.1;refGene_gn=DLG2;refGene_tx=XM_023239375.1;refGene_gn=DLG2;refGene_tx=XM_023239376.1;refGene_gn=DLG2;refGene_tx=XM_023239377.1;refGene_gn=DLG2;refGene_tx=XM_023239378.1;refGene_gn=DLG2;refGene_tx=XM_023239379.1;refGene_gn=DLG2;refGene_tx=XM_023239380.1;refGene_gn=DLG2;refGene_tx=XM_023239381.1;refGene_gn=DLG2;refGene_tx=XM_023239382.1;refGene_gn=DLG2;refGene_tx=XM_023239383.1;refGene_gn=DLG2;refGene_tx=XM_023239384.1;refGene_gn=DLG2;refGene_tx=XM_023239385.1;refGene_gn=DLG2;refGene_tx=XM_023239386.1;refGene_gn=DLG2;refGene_tx=XM_023239387.1;refGene_gn=DLG2;refGene_tx=XM_023239388.1;refGene_gn=DLG2;refGene_tx=XM_023239389.1;refGene_gn=DLG2;refGene_tx=XM_023239390.1;refGene_gn=DLG2;refGene_tx=XM_023239391.1;refGene_gn=DLG2;refGene_tx=XM_023239392.1;refGene_gn=DLG2;refGene_tx=XM_023239393.1;refGene_gn=DLG2;refGene_tx=XM_023239394.1
D1 51047051 51047195
refGene_gn=DLG2;refGene_tx=XM_023239374.1;refGene_gn=DLG2;refGene_tx=XM_023239375.1;refGene_gn=DLG2;refGene_tx=XM_023239376.1;refGene_gn=DLG2;refGene_tx=XM_023239377.1;refGene_gn=DLG2;refGene_tx=XM_023239378.1;refGene_gn=DLG2;refGene_tx=XM_023239380.1;refGene_gn=DLG2;refGene_tx=XM_023239383.1;refGene_gn=DLG2;refGene_tx=XM_023239384.1;refGene_gn=DLG2;refGene_tx=XM_023239385.1;refGene_gn=DLG2;refGene_tx=XM_023239386.1;refGene_gn=DLG2;refGene_tx=XM_023239387.1;refGene_gn=DLG2;refGene_tx=XM_023239389.1;refGene_gn=DLG2;refGene_tx=XM_023239390.1;refGene_gn=DLG2;refGene_tx=XM_023239391.1;refGene_gn=DLG2;refGene_tx=XM_023239393.1;refGene_gn=DLG2;refGene_tx=XM_023239394.1
D1 51050142 51050319
refGene_gn=DLG2;refGene_tx=XM_003992674.5;refGene_gn=DLG2;refGene_tx=XM_019811729.1;refGene_gn=DLG2;refGene_tx=XM_023239374.1;refGene_gn=DLG2;refGene_tx=XM_023239375.1;refGene_gn=DLG2;refGene_tx=XM_023239376.1;refGene_gn=DLG2;refGene_tx=XM_023239377.1;refGene_gn=DLG2;refGene_tx=XM_023239378.1;refGene_gn=DLG2;refGene_tx=XM_023239379.1;refGene_gn=DLG2;refGene_tx=XM_023239380.1;refGene_gn=DLG2;refGene_tx=XM_023239381.1;refGene_gn=DLG2;refGene_tx=XM_023239382.1;refGene_gn=DLG2;refGene_tx=XM_023239383.1;refGene_gn=DLG2;refGene_tx=XM_023239384.1;refGene_gn=DLG2;refGene_tx=XM_023239385.1;refGene_gn=DLG2;refGene_tx=XM_023239386.1;

refGene_gn=DLG2;refGene_tx=XM_023239387.1;refGene_gn=DLG2;refGene_tx=XM_023239388.1;refGene_gn=DLG2;refGene_tx=XM_023239389.1;refGene_gn=DLG2;refGene_tx=XM_023239390.1;refGene_gn=DLG2;refGene_tx=XM_023239391.1;refGene_gn=DLG2;refGene_tx=XM_023239392.1;refGene_gn=DLG2;refGene_tx=XM_023239393.1;refGene_gn=DLG2;refGene_tx=XM_023239394.1
D1 51059055 51059131
refGene_gn=DLG2;refGene_tx=XM_003992674.5;refGene_gn=DLG2;refGene_tx=XM_019811729.1;refGene_gn=DLG2;refGene_tx=XM_023239374.1;refGene_gn=DLG2;refGene_tx=XM_023239375.1;refGene_gn=DLG2;refGene_tx=XM_023239376.1;refGene_gn=DLG2;refGene_tx=XM_023239377.1;refGene_gn=DLG2;refGene_tx=XM_023239378.1;refGene_gn=DLG2;refGene_tx=XM_023239379.1;refGene_gn=DLG2;refGene_tx=XM_023239380.1;refGene_gn=DLG2;refGene_tx=XM_023239381.1;refGene_gn=DLG2;refGene_tx=XM_023239382.1;refGene_gn=DLG2;refGene_tx=XM_023239383.1;refGene_gn=DLG2;refGene_tx=XM_023239384.1;refGene_gn=DLG2;refGene_tx=XM_023239385.1;refGene_gn=DLG2;refGene_tx=XM_023239386.1;refGene_gn=DLG2;refGene_tx=XM_023239387.1;refGene_gn=DLG2;refGene_tx=XM_023239388.1;refGene_gn=DLG2;refGene_tx=XM_023239389.1;refGene_gn=DLG2;refGene_tx=XM_023239390.1;refGene_gn=DLG2;refGene_tx=XM_023239391.1;refGene_gn=DLG2;refGene_tx=XM_023239392.1;refGene_gn=DLG2;refGene_tx=XM_023239393.1;refGene_gn=DLG2;refGene_tx=XM_023239394.1
D1 51084594 51084618 refGene_gn=DLG2;refGene_tx=XM_023239394.1
D1 51086519 51086619
refGene_gn=DLG2;refGene_tx=XM_003992674.5;refGene_gn=DLG2;refGene_tx=XM_023239374.1;refGene_gn=DLG2;refGene_tx=XM_023239375.1;refGene_gn=DLG2;refGene_tx=XM_023239376.1;refGene_gn=DLG2;refGene_tx=XM_023239379.1;refGene_gn=DLG2;refGene_tx=XM_023239380.1;refGene_gn=DLG2;refGene_tx=XM_023239381.1;refGene_gn=DLG2;refGene_tx=XM_023239383.1;refGene_gn=DLG2;refGene_tx=XM_023239384.1;refGene_gn=DLG2;refGene_tx=XM_023239385.1;refGene_gn=DLG2;refGene_tx=XM_023239386.1;refGene_gn=DLG2;refGene_tx=XM_023239387.1;refGene_gn=DLG2;refGene_tx=XM_023239389.1;refGene_gn=DLG2;refGene_tx=XM_023239390.1;refGene_gn=DLG2;refGene_tx=XM_023239391.1;refGene_gn=DLG2;refGene_tx=XM_023239393.1
D1 51087456 51087502
refGene_gn=DLG2;refGene_tx=XM_019811729.1;refGene_gn=DLG2;refGene_tx=XM_023239377.1;refGene_gn=DLG2;refGene_tx=XM_023239378.1;refGene_gn=DLG2;refGene_tx=XM_023239382.1;refGene_gn=DLG2;refGene_tx=XM_023239388.1;refGene_gn=DLG2;refGene_tx=XM_023239392.1
D1 51088089 51088115 refGene_gn=DLG2;refGene_tx=XM_023239393.1
D1 51090666 51090708
refGene_gn=DLG2;refGene_tx=XM_023239374.1;refGene_gn=DLG2;refGene_tx=XM_023239375.1;refGene_gn=DLG2;refGene_tx=XM_023239377.1;refGene_gn=DLG2;refGene_tx=XM_023239379.1;refGene_gn=DLG2;refGene_tx=XM_023239380.1;refGene_gn=DLG2;refGene_tx=XM_023239383.1;refGene_gn=DLG2;refGene_tx=XM_023239384.1;refGene_gn=DLG2;refGene_tx=XM_023239385.1;refGene_gn=DLG2;refGene_tx=XM_023239386.1;refGene_gn=DLG2;refGene_tx=XM_023239387.1;refGene_gn=DLG2;refGene_tx=XM_023239389.1;refGene_gn=DLG2;refGene_tx=XM_023239390.1;refGene_gn=DLG2;refGene_tx=XM_023239391.1
D1 51098051 51098102
refGene_gn=DLG2;refGene_tx=XM_003992674.5;refGene_gn=DLG2;refGene_tx=XM_019811729.1;refGene_gn=DLG2;refGene_tx=XM_023239374.1;refGene_gn=DLG

2;refGene_tx=XM_023239375.1;refGene_gn=DLG2;refGene_tx=XM_023239376.1;
refGene_gn=DLG2;refGene_tx=XM_023239377.1;refGene_gn=DLG2;refGene_tx=X
M_023239378.1;refGene_gn=DLG2;refGene_tx=XM_023239379.1;refGene_gn=DLG
2;refGene_tx=XM_023239380.1;refGene_gn=DLG2;refGene_tx=XM_023239381.1;
refGene_gn=DLG2;refGene_tx=XM_023239382.1;refGene_gn=DLG2;refGene_tx=X
M_023239383.1;refGene_gn=DLG2;refGene_tx=XM_023239384.1;refGene_gn=DLG
2;refGene_tx=XM_023239385.1;refGene_gn=DLG2;refGene_tx=XM_023239386.1;
refGene_gn=DLG2;refGene_tx=XM_023239387.1;refGene_gn=DLG2;refGene_tx=X
M_023239388.1;refGene_gn=DLG2;refGene_tx=XM_023239389.1;refGene_gn=DLG
2;refGene_tx=XM_023239390.1;refGene_gn=DLG2;refGene_tx=XM_023239391.1;
refGene_gn=DLG2;refGene_tx=XM_023239392.1
D1 51099329 51099431
refGene_gn=DLG2;refGene_tx=XM_003992674.5;refGene_gn=DLG2;refGene_tx=X
M_019811729.1;refGene_gn=DLG2;refGene_tx=XM_023239374.1;refGene_gn=DLG
2;refGene_tx=XM_023239375.1;refGene_gn=DLG2;refGene_tx=XM_023239376.1;
refGene_gn=DLG2;refGene_tx=XM_023239377.1;refGene_gn=DLG2;refGene_tx=X
M_023239378.1;refGene_gn=DLG2;refGene_tx=XM_023239379.1;refGene_gn=DLG
2;refGene_tx=XM_023239380.1;refGene_gn=DLG2;refGene_tx=XM_023239381.1;
refGene_gn=DLG2;refGene_tx=XM_023239382.1;refGene_gn=DLG2;refGene_tx=X
M_023239383.1;refGene_gn=DLG2;refGene_tx=XM_023239384.1;refGene_gn=DLG
2;refGene_tx=XM_023239385.1;refGene_gn=DLG2;refGene_tx=XM_023239386.1;
refGene_gn=DLG2;refGene_tx=XM_023239387.1;refGene_gn=DLG2;refGene_tx=X
M_023239388.1;refGene_gn=DLG2;refGene_tx=XM_023239389.1;refGene_gn=DLG
2;refGene_tx=XM_023239390.1;refGene_gn=DLG2;refGene_tx=XM_023239391.1;
refGene_gn=DLG2;refGene_tx=XM_023239392.1
D1 51101877 51102050
refGene_gn=DLG2;refGene_tx=XM_003992674.5;refGene_gn=DLG2;refGene_tx=X
M_019811729.1;refGene_gn=DLG2;refGene_tx=XM_023239374.1;refGene_gn=DLG
2;refGene_tx=XM_023239375.1;refGene_gn=DLG2;refGene_tx=XM_023239376.1;
refGene_gn=DLG2;refGene_tx=XM_023239377.1;refGene_gn=DLG2;refGene_tx=X
M_023239378.1;refGene_gn=DLG2;refGene_tx=XM_023239379.1;refGene_gn=DLG
2;refGene_tx=XM_023239380.1;refGene_gn=DLG2;refGene_tx=XM_023239381.1;
refGene_gn=DLG2;refGene_tx=XM_023239382.1;refGene_gn=DLG2;refGene_tx=X
M_023239383.1;refGene_gn=DLG2;refGene_tx=XM_023239384.1;refGene_gn=DLG
2;refGene_tx=XM_023239385.1;refGene_gn=DLG2;refGene_tx=XM_023239386.1;
refGene_gn=DLG2;refGene_tx=XM_023239387.1;refGene_gn=DLG2;refGene_tx=X
M_023239388.1;refGene_gn=DLG2;refGene_tx=XM_023239389.1;refGene_gn=DLG
2;refGene_tx=XM_023239390.1;refGene_gn=DLG2;refGene_tx=XM_023239391.1;
refGene_gn=DLG2;refGene_tx=XM_023239392.1
D1 51104420 51104530
refGene_gn=DLG2;refGene_tx=XM_003992674.5;refGene_gn=DLG2;refGene_tx=X
M_019811729.1;refGene_gn=DLG2;refGene_tx=XM_023239374.1;refGene_gn=DLG
2;refGene_tx=XM_023239375.1;refGene_gn=DLG2;refGene_tx=XM_023239376.1;
refGene_gn=DLG2;refGene_tx=XM_023239377.1;refGene_gn=DLG2;refGene_tx=X
M_023239378.1;refGene_gn=DLG2;refGene_tx=XM_023239379.1;refGene_gn=DLG
2;refGene_tx=XM_023239380.1;refGene_gn=DLG2;refGene_tx=XM_023239381.1;
refGene_gn=DLG2;refGene_tx=XM_023239382.1;refGene_gn=DLG2;refGene_tx=X
M_023239383.1;refGene_gn=DLG2;refGene_tx=XM_023239384.1;refGene_gn=DLG
2;refGene_tx=XM_023239385.1;refGene_gn=DLG2;refGene_tx=XM_023239386.1;
refGene_gn=DLG2;refGene_tx=XM_023239387.1;refGene_gn=DLG2;refGene_tx=X

M_023239388.1;refGene_gn=DLG2;refGene_tx=XM_023239389.1;refGene_gn=DLG2;refGene_tx=XM_023239390.1;refGene_gn=DLG2;refGene_tx=XM_023239391.1;refGene_gn=DLG2;refGene_tx=XM_023239392.1
D1 51109516 51109608
refGene_gn=DLG2;refGene_tx=XM_003992674.5;refGene_gn=DLG2;refGene_tx=XM_019811729.1;refGene_gn=DLG2;refGene_tx=XM_023239374.1;refGene_gn=DLG2;refGene_tx=XM_023239375.1;refGene_gn=DLG2;refGene_tx=XM_023239376.1;refGene_gn=DLG2;refGene_tx=XM_023239377.1;refGene_gn=DLG2;refGene_tx=XM_023239378.1;refGene_gn=DLG2;refGene_tx=XM_023239379.1;refGene_gn=DLG2;refGene_tx=XM_023239380.1;refGene_gn=DLG2;refGene_tx=XM_023239381.1;refGene_gn=DLG2;refGene_tx=XM_023239382.1;refGene_gn=DLG2;refGene_tx=XM_023239383.1;refGene_gn=DLG2;refGene_tx=XM_023239384.1;refGene_gn=DLG2;refGene_tx=XM_023239385.1;refGene_gn=DLG2;refGene_tx=XM_023239386.1;refGene_gn=DLG2;refGene_tx=XM_023239387.1;refGene_gn=DLG2;refGene_tx=XM_023239388.1;refGene_gn=DLG2;refGene_tx=XM_023239389.1;refGene_gn=DLG2;refGene_tx=XM_023239390.1;refGene_gn=DLG2;refGene_tx=XM_023239391.1;refGene_gn=DLG2;refGene_tx=XM_023239392.1
D1 51112394 51112501
refGene_gn=DLG2;refGene_tx=XM_003992674.5;refGene_gn=DLG2;refGene_tx=XM_019811729.1;refGene_gn=DLG2;refGene_tx=XM_023239374.1;refGene_gn=DLG2;refGene_tx=XM_023239375.1;refGene_gn=DLG2;refGene_tx=XM_023239376.1;refGene_gn=DLG2;refGene_tx=XM_023239377.1;refGene_gn=DLG2;refGene_tx=XM_023239378.1;refGene_gn=DLG2;refGene_tx=XM_023239379.1;refGene_gn=DLG2;refGene_tx=XM_023239380.1;refGene_gn=DLG2;refGene_tx=XM_023239381.1;refGene_gn=DLG2;refGene_tx=XM_023239382.1;refGene_gn=DLG2;refGene_tx=XM_023239383.1;refGene_gn=DLG2;refGene_tx=XM_023239384.1;refGene_gn=DLG2;refGene_tx=XM_023239385.1;refGene_gn=DLG2;refGene_tx=XM_023239386.1;refGene_gn=DLG2;refGene_tx=XM_023239387.1;refGene_gn=DLG2;refGene_tx=XM_023239388.1;refGene_gn=DLG2;refGene_tx=XM_023239389.1;refGene_gn=DLG2;refGene_tx=XM_023239390.1;refGene_gn=DLG2;refGene_tx=XM_023239391.1;refGene_gn=DLG2;refGene_tx=XM_023239392.1
D1 51214444 51214544
refGene_gn=CCDC90B;refGene_tx=XM_003992677.5;refGene_gn=CCDC90B;refGene_tx=XM_019811731.2;refGene_gn=CCDC90B;refGene_tx=XM_023239401.1;refGene_gn=CCDC90B;refGene_tx=XM_023239402.1
D1 51219718 51219838
refGene_gn=CCDC90B;refGene_tx=XM_003992677.5;refGene_gn=CCDC90B;refGene_tx=XM_019811731.2;refGene_gn=CCDC90B;refGene_tx=XM_023239401.1;refGene_gn=CCDC90B;refGene_tx=XM_023239402.1
D1 51221999 51222103
refGene_gn=CCDC90B;refGene_tx=XM_003992677.5;refGene_gn=CCDC90B;refGene_tx=XM_019811731.2;refGene_gn=CCDC90B;refGene_tx=XM_023239401.1;refGene_gn=CCDC90B;refGene_tx=XM_023239402.1
D1 51241632 51241734
refGene_gn=CCDC90B;refGene_tx=XM_003992677.5;refGene_gn=CCDC90B;refGene_tx=XM_019811731.2;refGene_gn=CCDC90B;refGene_tx=XM_023239401.1;refGene_gn=CCDC90B;refGene_tx=XM_023239402.1
D1 51242379 51242421
refGene_gn=CCDC90B;refGene_tx=XM_003992677.5;refGene_gn=CCDC90B;refGene_tx=XM_019811731.2;refGene_gn=CCDC90B;refGene_tx=XM_023239401.1;refGe

ne_gn=CCDC90B;refGene_tx=XM_023239402.1
D1 51242513 51242585
refGene_gn=CCDC90B;refGene_tx=XM_003992677.5;refGene_gn=CCDC90B;refGene_tx=XM_019811731.2;refGene_gn=CCDC90B;refGene_tx=XM_023239401.1;refGene_gn=CCDC90B;refGene_tx=XM_023239402.1
D1 51242666 51242720
refGene_gn=CCDC90B;refGene_tx=XM_003992677.5;refGene_gn=CCDC90B;refGene_tx=XM_019811731.2;refGene_gn=CCDC90B;refGene_tx=XM_023239401.1;refGene_gn=CCDC90B;refGene_tx=XM_023239402.1
D1 51244199 51244314
refGene_gn=CCDC90B;refGene_tx=XM_003992677.5;refGene_gn=CCDC90B;refGene_tx=XM_019811731.2;refGene_gn=CCDC90B;refGene_tx=XM_023239401.1;refGene_gn=CCDC90B;refGene_tx=XM_023239402.1
D1 51263379 51263435 refGene_gn=CCDC90B;refGene_tx=XM_003992677.5
D1 51268203 51268302
refGene_gn=ANKRD42;refGene_tx=XM_023239398.1;refGene_gn=ANKRD42;refGene_tx=XM_023239399.1
D1 51271859 51272001
refGene_gn=ANKRD42;refGene_tx=XM_023239398.1;refGene_gn=ANKRD42;refGene_tx=XM_023239399.1
D1 51275053 51275180
refGene_gn=ANKRD42;refGene_tx=XM_023239398.1;refGene_gn=ANKRD42;refGene_tx=XM_023239399.1
D1 51296705 51296881
refGene_gn=ANKRD42;refGene_tx=XM_023239398.1;refGene_gn=ANKRD42;refGene_tx=XM_023239399.1
D1 51300337 51300443
refGene_gn=ANKRD42;refGene_tx=XM_023239398.1;refGene_gn=ANKRD42;refGene_tx=XM_023239399.1
D1 51304740 51304866
refGene_gn=ANKRD42;refGene_tx=XM_023239398.1;refGene_gn=ANKRD42;refGene_tx=XM_023239399.1
D1 51308684 51308885
refGene_gn=ANKRD42;refGene_tx=XM_023239398.1;refGene_gn=ANKRD42;refGene_tx=XM_023239399.1
D1 51318217 51318353
refGene_gn=ANKRD42;refGene_tx=XM_023239398.1;refGene_gn=ANKRD42;refGene_tx=XM_023239399.1
D1 51319674 51319794
refGene_gn=ANKRD42;refGene_tx=XM_023239398.1;refGene_gn=ANKRD42;refGene_tx=XM_023239399.1
D1 51323329 51323389
refGene_gn=ANKRD42;refGene_tx=XM_023239398.1;refGene_gn=ANKRD42;refGene_tx=XM_023239399.1
D1 51348990 51349206
refGene_gn=PCF11;refGene_tx=XM_019811733.2;refGene_gn=PCF11;refGene_tx=XM_023239395.1;refGene_gn=PCF11;refGene_tx=XM_023239396.1;refGene_gn=PCF11;refGene_tx=XM_023239397.1
D1 51350325 51350361
refGene_gn=PCF11;refGene_tx=XM_019811733.2;refGene_gn=PCF11;refGene_tx

=XM_023239395.1;refGene_gn=PCF11;refGene_tx=XM_023239396.1;refGene_gn=PCF11;refGene_tx=XM_023239397.1
D1 51350873 51350966
refGene_gn=PCF11;refGene_tx=XM_019811733.2;refGene_gn=PCF11;refGene_tx=XM_023239395.1;refGene_gn=PCF11;refGene_tx=XM_023239396.1;refGene_gn=PCF11;refGene_tx=XM_023239397.1
D1 51351216 51351372
refGene_gn=PCF11;refGene_tx=XM_019811733.2;refGene_gn=PCF11;refGene_tx=XM_023239395.1;refGene_gn=PCF11;refGene_tx=XM_023239396.1;refGene_gn=PCF11;refGene_tx=XM_023239397.1
D1 51351883 51352067
refGene_gn=PCF11;refGene_tx=XM_019811733.2;refGene_gn=PCF11;refGene_tx=XM_023239395.1;refGene_gn=PCF11;refGene_tx=XM_023239396.1;refGene_gn=PCF11;refGene_tx=XM_023239397.1
D1 51353969 51354075
refGene_gn=PCF11;refGene_tx=XM_019811733.2;refGene_gn=PCF11;refGene_tx=XM_023239395.1;refGene_gn=PCF11;refGene_tx=XM_023239396.1;refGene_gn=PCF11;refGene_tx=XM_023239397.1
D1 51354581 51354701
refGene_gn=PCF11;refGene_tx=XM_019811733.2;refGene_gn=PCF11;refGene_tx=XM_023239395.1;refGene_gn=PCF11;refGene_tx=XM_023239396.1;refGene_gn=PCF11;refGene_tx=XM_023239397.1
D1 51357739 51357839
refGene_gn=PCF11;refGene_tx=XM_019811733.2;refGene_gn=PCF11;refGene_tx=XM_023239395.1;refGene_gn=PCF11;refGene_tx=XM_023239396.1;refGene_gn=PCF11;refGene_tx=XM_023239397.1
D1 51359420 51360985
refGene_gn=PCF11;refGene_tx=XM_019811733.2;refGene_gn=PCF11;refGene_tx=XM_023239395.1;refGene_gn=PCF11;refGene_tx=XM_023239396.1;refGene_gn=PCF11;refGene_tx=XM_023239397.1
D1 51361529 51362013
refGene_gn=PCF11;refGene_tx=XM_019811733.2;refGene_gn=PCF11;refGene_tx=XM_023239395.1;refGene_gn=PCF11;refGene_tx=XM_023239396.1;refGene_gn=PCF11;refGene_tx=XM_023239397.1
D1 51362119 51362300
refGene_gn=PCF11;refGene_tx=XM_019811733.2;refGene_gn=PCF11;refGene_tx=XM_023239395.1;refGene_gn=PCF11;refGene_tx=XM_023239396.1;refGene_gn=PCF11;refGene_tx=XM_023239397.1
D1 51362683 51363801
refGene_gn=PCF11;refGene_tx=XM_019811733.2;refGene_gn=PCF11;refGene_tx=XM_023239395.1;refGene_gn=PCF11;refGene_tx=XM_023239396.1;refGene_gn=PCF11;refGene_tx=XM_023239397.1
D1 51364746 51364941
refGene_gn=PCF11;refGene_tx=XM_019811733.2;refGene_gn=PCF11;refGene_tx=XM_023239395.1;refGene_gn=PCF11;refGene_tx=XM_023239396.1;refGene_gn=PCF11;refGene_tx=XM_023239397.1
D1 51365294 51365483
refGene_gn=PCF11;refGene_tx=XM_019811733.2;refGene_gn=PCF11;refGene_tx=XM_023239395.1;refGene_gn=PCF11;refGene_tx=XM_023239396.1;refGene_gn=PCF11;refGene_tx=XM_023239397.1

D1 51367387 51367513
refGene_gn=PCF11;refGene_tx=XM_019811733.2;refGene_gn=PCF11;refGene_tx=XM_023239395.1;refGene_gn=PCF11;refGene_tx=XM_023239396.1;refGene_gn=PCF11;refGene_tx=XM_023239397.1
D1 51371230 51371422
refGene_gn=PCF11;refGene_tx=XM_019811733.2;refGene_gn=PCF11;refGene_tx=XM_023239395.1;refGene_gn=PCF11;refGene_tx=XM_023239396.1;refGene_gn=PCF11;refGene_tx=XM_023239397.1
D1 51510421 51510514
refGene_gn=RAB30;refGene_tx=XM_003992679.5;refGene_gn=RAB30;refGene_tx=XM_006936880.4;refGene_gn=RAB30;refGene_tx=XM_019811734.2
D1 51513919 51514003
refGene_gn=RAB30;refGene_tx=XM_003992679.5;refGene_gn=RAB30;refGene_tx=XM_006936880.4;refGene_gn=RAB30;refGene_tx=XM_019811734.2
D1 51520878 51521062
refGene_gn=RAB30;refGene_tx=XM_003992679.5;refGene_gn=RAB30;refGene_tx=XM_006936880.4;refGene_gn=RAB30;refGene_tx=XM_019811734.2
D1 51525525 51525776
refGene_gn=RAB30;refGene_tx=XM_003992679.5;refGene_gn=RAB30;refGene_tx=XM_006936880.4;refGene_gn=RAB30;refGene_tx=XM_019811734.2
D1 51541570 51544147 refGene_gn=DDIAS;refGene_tx=XM_011286565.3
D1 51545476 51545594 refGene_gn=DDIAS;refGene_tx=XM_011286565.3
D1 51546492 51546654 refGene_gn=DDIAS;refGene_tx=XM_011286565.3
D1 51552997 51553110 refGene_gn=DDIAS;refGene_tx=XM_011286565.3
D1 51565188 51565207 refGene_gn=PRCP;refGene_tx=XM_003992681.5
D1 51565477 51565659 refGene_gn=PRCP;refGene_tx=XM_003992681.5
D1 51608442 51608583 refGene_gn=PRCP;refGene_tx=XM_003992681.5
D1 51611818 51611920 refGene_gn=PRCP;refGene_tx=XM_003992681.5
D1 51618418 51618600 refGene_gn=PRCP;refGene_tx=XM_003992681.5
D1 51619655 51619813 refGene_gn=PRCP;refGene_tx=XM_003992681.5
D1 51620384 51620557 refGene_gn=PRCP;refGene_tx=XM_003992681.5
D1 51631136 51631301 refGene_gn=PRCP;refGene_tx=XM_003992681.5
D1 51632018 51632206 refGene_gn=PRCP;refGene_tx=XM_003992681.5
D1 51640141 51640358 refGene_gn=PRCP;refGene_tx=XM_003992681.5
D1 55215656 55215666
refGene_gn=TENM4;refGene_tx=XM_023238927.1;refGene_gn=TENM4;refGene_tx=XM_023238928.1;refGene_gn=TENM4;refGene_tx=XM_023238929.1;refGene_gn=TENM4;refGene_tx=XM_023238930.1;refGene_gn=TENM4;refGene_tx=XM_023238931.1
D1 55342243 55342531
refGene_gn=TENM4;refGene_tx=XM_023238927.1;refGene_gn=TENM4;refGene_tx=XM_023238928.1;refGene_gn=TENM4;refGene_tx=XM_023238929.1;refGene_gn=TENM4;refGene_tx=XM_023238930.1;refGene_gn=TENM4;refGene_tx=XM_023238931.1
D1 55347118 55347388
refGene_gn=TENM4;refGene_tx=XM_023238927.1;refGene_gn=TENM4;refGene_tx=XM_023238928.1;refGene_gn=TENM4;refGene_tx=XM_023238929.1;refGene_gn=TENM4;refGene_tx=XM_023238930.1;refGene_gn=TENM4;refGene_tx=XM_023238931.1
D1 55440594 55440804

refGene_gn=TENM4;refGene_tx=XM_023238927.1;refGene_gn=TENM4;refGene_tx=XM_023238928.1;refGene_gn=TENM4;refGene_tx=XM_023238929.1;refGene_gn=TENM4;refGene_tx=XM_023238930.1
D1 55487388 55487523
refGene_gn=TENM4;refGene_tx=XM_023238927.1;refGene_gn=TENM4;refGene_tx=XM_023238928.1
D1 55496901 55497157
refGene_gn=TENM4;refGene_tx=XM_023238927.1;refGene_gn=TENM4;refGene_tx=XM_023238928.1;refGene_gn=TENM4;refGene_tx=XM_023238929.1;refGene_gn=TENM4;refGene_tx=XM_023238930.1;refGene_gn=TENM4;refGene_tx=XM_023238931.1
D1 55507355 55507454
refGene_gn=TENM4;refGene_tx=XM_023238927.1;refGene_gn=TENM4;refGene_tx=XM_023238928.1;refGene_gn=TENM4;refGene_tx=XM_023238929.1;refGene_gn=TENM4;refGene_tx=XM_023238931.1
D1 55508628 55508864
refGene_gn=TENM4;refGene_tx=XM_023238927.1;refGene_gn=TENM4;refGene_tx=XM_023238928.1;refGene_gn=TENM4;refGene_tx=XM_023238929.1;refGene_gn=TENM4;refGene_tx=XM_023238930.1;refGene_gn=TENM4;refGene_tx=XM_023238931.1
D1 55535280 55535451
refGene_gn=TENM4;refGene_tx=XM_023238927.1;refGene_gn=TENM4;refGene_tx=XM_023238928.1;refGene_gn=TENM4;refGene_tx=XM_023238929.1;refGene_gn=TENM4;refGene_tx=XM_023238930.1;refGene_gn=TENM4;refGene_tx=XM_023238931.1
D1 55543080 55543295
refGene_gn=TENM4;refGene_tx=XM_023238927.1;refGene_gn=TENM4;refGene_tx=XM_023238928.1;refGene_gn=TENM4;refGene_tx=XM_023238929.1;refGene_gn=TENM4;refGene_tx=XM_023238930.1;refGene_gn=TENM4;refGene_tx=XM_023238931.1
D1 55544929 55545140
refGene_gn=TENM4;refGene_tx=XM_023238927.1;refGene_gn=TENM4;refGene_tx=XM_023238928.1;refGene_gn=TENM4;refGene_tx=XM_023238929.1;refGene_gn=TENM4;refGene_tx=XM_023238930.1;refGene_gn=TENM4;refGene_tx=XM_023238931.1
D1 55580919 55581021
refGene_gn=TENM4;refGene_tx=XM_023238927.1;refGene_gn=TENM4;refGene_tx=XM_023238928.1;refGene_gn=TENM4;refGene_tx=XM_023238929.1;refGene_gn=TENM4;refGene_tx=XM_023238930.1;refGene_gn=TENM4;refGene_tx=XM_023238931.1
D1 55583036 55583231
refGene_gn=TENM4;refGene_tx=XM_023238927.1;refGene_gn=TENM4;refGene_tx=XM_023238928.1;refGene_gn=TENM4;refGene_tx=XM_023238929.1;refGene_gn=TENM4;refGene_tx=XM_023238930.1;refGene_gn=TENM4;refGene_tx=XM_023238931.1
D1 55589225 55589426
refGene_gn=TENM4;refGene_tx=XM_023238927.1;refGene_gn=TENM4;refGene_tx=XM_023238928.1;refGene_gn=TENM4;refGene_tx=XM_023238929.1;refGene_gn=TENM4;refGene_tx=XM_023238930.1;refGene_gn=TENM4;refGene_tx=XM_023238931.1

D1 55607381 55607567
refGene_gn=TENM4;refGene_tx=XM_023238927.1;refGene_gn=TENM4;refGene_tx=XM_023238928.1;refGene_gn=TENM4;refGene_tx=XM_023238929.1;refGene_gn=TENM4;refGene_tx=XM_023238930.1;refGene_gn=TENM4;refGene_tx=XM_023238931.1
D1 55615202 55615229
refGene_gn=TENM4;refGene_tx=XM_023238927.1;refGene_gn=TENM4;refGene_tx=XM_023238928.1;refGene_gn=TENM4;refGene_tx=XM_023238929.1;refGene_gn=TENM4;refGene_tx=XM_023238931.1
D1 55622675 55622822
refGene_gn=TENM4;refGene_tx=XM_023238927.1;refGene_gn=TENM4;refGene_tx=XM_023238928.1;refGene_gn=TENM4;refGene_tx=XM_023238929.1;refGene_gn=TENM4;refGene_tx=XM_023238930.1;refGene_gn=TENM4;refGene_tx=XM_023238931.1
D1 55635384 55635601
refGene_gn=TENM4;refGene_tx=XM_023238927.1;refGene_gn=TENM4;refGene_tx=XM_023238928.1;refGene_gn=TENM4;refGene_tx=XM_023238929.1;refGene_gn=TENM4;refGene_tx=XM_023238930.1;refGene_gn=TENM4;refGene_tx=XM_023238931.1
D1 55657770 55657890
refGene_gn=TENM4;refGene_tx=XM_023238927.1;refGene_gn=TENM4;refGene_tx=XM_023238928.1;refGene_gn=TENM4;refGene_tx=XM_023238929.1;refGene_gn=TENM4;refGene_tx=XM_023238930.1;refGene_gn=TENM4;refGene_tx=XM_023238931.1
D1 55662565 55662827
refGene_gn=TENM4;refGene_tx=XM_023238927.1;refGene_gn=TENM4;refGene_tx=XM_023238928.1;refGene_gn=TENM4;refGene_tx=XM_023238929.1;refGene_gn=TENM4;refGene_tx=XM_023238930.1;refGene_gn=TENM4;refGene_tx=XM_023238931.1
D1 55665462 55665730
refGene_gn=TENM4;refGene_tx=XM_023238927.1;refGene_gn=TENM4;refGene_tx=XM_023238928.1;refGene_gn=TENM4;refGene_tx=XM_023238929.1;refGene_gn=TENM4;refGene_tx=XM_023238930.1;refGene_gn=TENM4;refGene_tx=XM_023238931.1
D1 55668527 55668671
refGene_gn=TENM4;refGene_tx=XM_023238927.1;refGene_gn=TENM4;refGene_tx=XM_023238928.1;refGene_gn=TENM4;refGene_tx=XM_023238929.1;refGene_gn=TENM4;refGene_tx=XM_023238930.1;refGene_gn=TENM4;refGene_tx=XM_023238931.1
D1 55671680 55671930
refGene_gn=TENM4;refGene_tx=XM_023238927.1;refGene_gn=TENM4;refGene_tx=XM_023238928.1;refGene_gn=TENM4;refGene_tx=XM_023238929.1;refGene_gn=TENM4;refGene_tx=XM_023238930.1;refGene_gn=TENM4;refGene_tx=XM_023238931.1
D1 55674166 55674187
refGene_gn=TENM4;refGene_tx=XM_023238927.1;refGene_gn=TENM4;refGene_tx=XM_023238928.1;refGene_gn=TENM4;refGene_tx=XM_023238929.1;refGene_gn=TENM4;refGene_tx=XM_023238930.1;refGene_gn=TENM4;refGene_tx=XM_023238931.1
D1 55682141 55682374

refGene_gn=TENM4;refGene_tx=XM_023238927.1;refGene_gn=TENM4;refGene_tx=XM_023238928.1;refGene_gn=TENM4;refGene_tx=XM_023238929.1;refGene_gn=TENM4;refGene_tx=XM_023238930.1;refGene_gn=TENM4;refGene_tx=XM_023238931.1
D1 55686241 55686396
refGene_gn=TENM4;refGene_tx=XM_023238927.1;refGene_gn=TENM4;refGene_tx=XM_023238928.1;refGene_gn=TENM4;refGene_tx=XM_023238929.1;refGene_gn=TENM4;refGene_tx=XM_023238930.1;refGene_gn=TENM4;refGene_tx=XM_023238931.1
D1 55692396 55693274
refGene_gn=TENM4;refGene_tx=XM_023238927.1;refGene_gn=TENM4;refGene_tx=XM_023238928.1;refGene_gn=TENM4;refGene_tx=XM_023238929.1;refGene_gn=TENM4;refGene_tx=XM_023238930.1;refGene_gn=TENM4;refGene_tx=XM_023238931.1
D1 55706678 55706851
refGene_gn=TENM4;refGene_tx=XM_023238927.1;refGene_gn=TENM4;refGene_tx=XM_023238928.1;refGene_gn=TENM4;refGene_tx=XM_023238929.1;refGene_gn=TENM4;refGene_tx=XM_023238930.1;refGene_gn=TENM4;refGene_tx=XM_023238931.1
D1 55711694 55711930
refGene_gn=TENM4;refGene_tx=XM_023238927.1;refGene_gn=TENM4;refGene_tx=XM_023238928.1;refGene_gn=TENM4;refGene_tx=XM_023238929.1;refGene_gn=TENM4;refGene_tx=XM_023238930.1;refGene_gn=TENM4;refGene_tx=XM_023238931.1
D1 55715003 55715300
refGene_gn=TENM4;refGene_tx=XM_023238927.1;refGene_gn=TENM4;refGene_tx=XM_023238928.1;refGene_gn=TENM4;refGene_tx=XM_023238929.1;refGene_gn=TENM4;refGene_tx=XM_023238930.1;refGene_gn=TENM4;refGene_tx=XM_023238931.1
D1 55716750 55718365
refGene_gn=TENM4;refGene_tx=XM_023238927.1;refGene_gn=TENM4;refGene_tx=XM_023238928.1;refGene_gn=TENM4;refGene_tx=XM_023238929.1;refGene_gn=TENM4;refGene_tx=XM_023238930.1;refGene_gn=TENM4;refGene_tx=XM_023238931.1
D1 55723817 55723960
refGene_gn=TENM4;refGene_tx=XM_023238927.1;refGene_gn=TENM4;refGene_tx=XM_023238928.1;refGene_gn=TENM4;refGene_tx=XM_023238929.1;refGene_gn=TENM4;refGene_tx=XM_023238930.1;refGene_gn=TENM4;refGene_tx=XM_023238931.1
D1 55726563 55727322
refGene_gn=TENM4;refGene_tx=XM_023238927.1;refGene_gn=TENM4;refGene_tx=XM_023238928.1;refGene_gn=TENM4;refGene_tx=XM_023238929.1;refGene_gn=TENM4;refGene_tx=XM_023238930.1;refGene_gn=TENM4;refGene_tx=XM_023238931.1
D1 55764262 55764403
refGene_gn=NARS2;refGene_tx=XM_003992682.4;refGene_gn=NARS2;refGene_tx=XM_006936883.3;refGene_gn=NARS2;refGene_tx=XM_019811745.2;refGene_gn=NARS2;refGene_tx=XM_019811746.2
D1 55767672 55767782
refGene_gn=NARS2;refGene_tx=XM_003992682.4;refGene_gn=NARS2;refGene_tx

=XM_006936883.3;refGene_gn=NARS2;refGene_tx=XM_019811745.2;refGene_gn=NARS2;refGene_tx=XM_019811746.2
D1 55768530 55768651 refGene_gn=NARS2;refGene_tx=XM_003992682.4;refGene_gn=NARS2;refGene_tx=XM_006936883.3;refGene_gn=NARS2;refGene_tx=XM_019811745.2;refGene_gn=NARS2;refGene_tx=XM_019811746.2
D1 55770772 55770913 refGene_gn=NARS2;refGene_tx=XM_003992682.4;refGene_gn=NARS2;refGene_tx=XM_006936883.3;refGene_gn=NARS2;refGene_tx=XM_019811745.2
D1 55777123 55777204 refGene_gn=NARS2;refGene_tx=XM_003992682.4;refGene_gn=NARS2;refGene_tx=XM_006936883.3;refGene_gn=NARS2;refGene_tx=XM_019811745.2;refGene_gn=NARS2;refGene_tx=XM_019811746.2
D1 55822494 55822589 refGene_gn=NARS2;refGene_tx=XM_003992682.4;refGene_gn=NARS2;refGene_tx=XM_006936883.3;refGene_gn=NARS2;refGene_tx=XM_019811745.2;refGene_gn=NARS2;refGene_tx=XM_019811746.2
D1 55852157 55852290 refGene_gn=NARS2;refGene_tx=XM_003992682.4;refGene_gn=NARS2;refGene_tx=XM_006936883.3;refGene_gn=NARS2;refGene_tx=XM_019811745.2;refGene_gn=NARS2;refGene_tx=XM_019811746.2
D1 55856215 55856314 refGene_gn=NARS2;refGene_tx=XM_003992682.4;refGene_gn=NARS2;refGene_tx=XM_006936883.3;refGene_gn=NARS2;refGene_tx=XM_019811745.2;refGene_gn=NARS2;refGene_tx=XM_019811746.2
D1 55856422 55856460 refGene_gn=NARS2;refGene_tx=XM_003992682.4;refGene_gn=NARS2;refGene_tx=XM_006936883.3;refGene_gn=NARS2;refGene_tx=XM_019811745.2;refGene_gn=NARS2;refGene_tx=XM_019811746.2
D1 55865711 55865778 refGene_gn=NARS2;refGene_tx=XM_003992682.4;refGene_gn=NARS2;refGene_tx=XM_006936883.3;refGene_gn=NARS2;refGene_tx=XM_019811745.2;refGene_gn=NARS2;refGene_tx=XM_019811746.2
D1 55867095 55867233 refGene_gn=NARS2;refGene_tx=XM_003992682.4;refGene_gn=NARS2;refGene_tx=XM_006936883.3;refGene_gn=NARS2;refGene_tx=XM_019811745.2;refGene_gn=NARS2;refGene_tx=XM_019811746.2
D1 55874265 55874361 refGene_gn=NARS2;refGene_tx=XM_006936883.3
D1 55881006 55881104 refGene_gn=NARS2;refGene_tx=XM_003992682.4;refGene_gn=NARS2;refGene_tx=XM_019811745.2;refGene_gn=NARS2;refGene_tx=XM_019811746.2
D1 55883390 55883417 refGene_gn=NARS2;refGene_tx=XM_003992682.4;refGene_gn=NARS2;refGene_tx=XM_019811745.2;refGene_gn=NARS2;refGene_tx=XM_019811746.2
D1 55886687 55886832 refGene_gn=NARS2;refGene_tx=XM_003992682.4;refGene_gn=NARS2;refGene_tx=XM_019811745.2;refGene_gn=NARS2;refGene_tx=XM_019811746.2
D1 55903815 55903890 refGene_gn=GAB2;refGene_tx=XM_003992683.5;refGene_gn=GAB2;refGene_tx=XM_019811748.2

D1 56035116 56035416
refGene_gn=GAB2;refGene_tx=XM_003992683.5;refGene_gn=GAB2;refGene_tx=XM_019811748.2
D1 56064043 56064200 refGene_gn=GAB2;refGene_tx=XM_019811749.1
D1 56064339 56064583
refGene_gn=GAB2;refGene_tx=XM_003992683.5;refGene_gn=GAB2;refGene_tx=XM_019811748.2;refGene_gn=GAB2;refGene_tx=XM_019811749.1
D1 56082849 56082861
refGene_gn=GAB2;refGene_tx=XM_019811748.2;refGene_gn=GAB2;refGene_tx=XM_019811749.1
D1 56084588 56085181
refGene_gn=GAB2;refGene_tx=XM_003992683.5;refGene_gn=GAB2;refGene_tx=XM_019811748.2;refGene_gn=GAB2;refGene_tx=XM_019811749.1
D1 56086432 56086527
refGene_gn=GAB2;refGene_tx=XM_003992683.5;refGene_gn=GAB2;refGene_tx=XM_019811748.2;refGene_gn=GAB2;refGene_tx=XM_019811749.1
D1 56087507 56087772
refGene_gn=GAB2;refGene_tx=XM_003992683.5;refGene_gn=GAB2;refGene_tx=XM_019811748.2;refGene_gn=GAB2;refGene_tx=XM_019811749.1
D1 56088475 56088566
refGene_gn=GAB2;refGene_tx=XM_003992683.5;refGene_gn=GAB2;refGene_tx=XM_019811748.2;refGene_gn=GAB2;refGene_tx=XM_019811749.1
D1 56088836 56088939
refGene_gn=GAB2;refGene_tx=XM_003992683.5;refGene_gn=GAB2;refGene_tx=XM_019811748.2;refGene_gn=GAB2;refGene_tx=XM_019811749.1
D1 56090615 56090741
refGene_gn=GAB2;refGene_tx=XM_003992683.5;refGene_gn=GAB2;refGene_tx=XM_019811748.2;refGene_gn=GAB2;refGene_tx=XM_019811749.1
D1 56091422 56091566
refGene_gn=GAB2;refGene_tx=XM_003992683.5;refGene_gn=GAB2;refGene_tx=XM_019811748.2;refGene_gn=GAB2;refGene_tx=XM_019811749.1
D1 56096732 56096900 refGene_gn=USP35;refGene_tx=XM_006936884.4
D1 56098156 56099411 refGene_gn=USP35;refGene_tx=XM_006936884.4
D1 56099936 56100043 refGene_gn=USP35;refGene_tx=XM_006936884.4
D1 56100969 56101063 refGene_gn=USP35;refGene_tx=XM_006936884.4
D1 56136587 56136781 refGene_gn=USP35;refGene_tx=XM_006936884.4
D1 56141453 56141597 refGene_gn=USP35;refGene_tx=XM_006936884.4
D1 56141598 56141613 refGene_gn=USP35;refGene_tx=XM_006936884.4
D1 56143177 56143279 refGene_gn=USP35;refGene_tx=XM_006936884.4
D1 56143698 56143828 refGene_gn=USP35;refGene_tx=XM_006936884.4
D1 56145330 56145463 refGene_gn=USP35;refGene_tx=XM_006936884.4
D1 56146511 56147184 refGene_gn=USP35;refGene_tx=XM_006936884.4
D1 56166513 56167296
refGene_gn=KCTD21;refGene_tx=XM_003992684.4;refGene_gn=KCTD21;refGene_tx=XM_006936887.3;refGene_gn=KCTD21;refGene_tx=XM_019811750.2;refGene_gn=KCTD21;refGene_tx=XM_019811751.2;refGene_gn=KCTD21;refGene_tx=XM_019811752.2
D1 56213978 56214073
refGene_gn=ALG8;refGene_tx=XM_003992685.4;refGene_gn=ALG8;refGene_tx=XM_019811753.2

D1 56217737 56217816
refGene_gn=ALG8;refGene_tx=XM_003992685.4;refGene_gn=ALG8;refGene_tx=XM_019811753.2
D1 56222706 56222900
refGene_gn=ALG8;refGene_tx=XM_003992685.4;refGene_gn=ALG8;refGene_tx=XM_019811753.2;refGene_gn=ALG8;refGene_tx=XM_019811754.2
D1 56225839 56225949
refGene_gn=ALG8;refGene_tx=XM_003992685.4;refGene_gn=ALG8;refGene_tx=XM_019811753.2;refGene_gn=ALG8;refGene_tx=XM_019811754.2
D1 56227666 56227734
refGene_gn=ALG8;refGene_tx=XM_003992685.4;refGene_gn=ALG8;refGene_tx=XM_019811753.2;refGene_gn=ALG8;refGene_tx=XM_019811754.2
D1 56229993 56230120
refGene_gn=ALG8;refGene_tx=XM_003992685.4;refGene_gn=ALG8;refGene_tx=XM_019811753.2;refGene_gn=ALG8;refGene_tx=XM_019811754.2
D1 56230377 56230481
refGene_gn=ALG8;refGene_tx=XM_003992685.4;refGene_gn=ALG8;refGene_tx=XM_019811753.2;refGene_gn=ALG8;refGene_tx=XM_019811754.2
D1 56231178 56231299
refGene_gn=ALG8;refGene_tx=XM_003992685.4;refGene_gn=ALG8;refGene_tx=XM_019811753.2;refGene_gn=ALG8;refGene_tx=XM_019811754.2
D1 56234661 56234801
refGene_gn=ALG8;refGene_tx=XM_003992685.4;refGene_gn=ALG8;refGene_tx=XM_019811753.2;refGene_gn=ALG8;refGene_tx=XM_019811754.2
D1 56236362 56236502
refGene_gn=ALG8;refGene_tx=XM_003992685.4;refGene_gn=ALG8;refGene_tx=XM_019811753.2;refGene_gn=ALG8;refGene_tx=XM_019811754.2
D1 56237818 56237916
refGene_gn=ALG8;refGene_tx=XM_003992685.4;refGene_gn=ALG8;refGene_tx=XM_019811753.2;refGene_gn=ALG8;refGene_tx=XM_019811754.2
D1 56238231 56238304
refGene_gn=ALG8;refGene_tx=XM_003992685.4;refGene_gn=ALG8;refGene_tx=XM_019811753.2;refGene_gn=ALG8;refGene_tx=XM_019811754.2
D1 56240295 56240527
refGene_gn=ALG8;refGene_tx=XM_003992685.4;refGene_gn=ALG8;refGene_tx=XM_019811754.2
D1 56242907 56243049 refGene_gn=ALG8;refGene_tx=XM_019811753.2
D1 56256861 56257030
refGene_gn=LOC101080736;refGene_tx=XM_011286640.3
D1 56263296 56263440
refGene_gn=LOC101080736;refGene_tx=XM_011286640.3
D1 56265885 56265935
refGene_gn=LOC101080736;refGene_tx=XM_011286640.3
D1 56269749 56270220 refGene_gn=THRSP;refGene_tx=XM_003992699.5
D1 56299453 56299522 refGene_gn=KCTD14;refGene_tx=XM_019811756.1
D1 56302295 56302418 refGene_gn=KCTD14;refGene_tx=XM_019811755.1
D1 56304970 56305639
refGene_gn=KCTD14;refGene_tx=XM_019811755.1;refGene_gn=KCTD14;refGene_tx=XM_019811756.1
D1 56316426 56316480 refGene_gn=INTS4;refGene_tx=XM_006936891.4

D1 56321207 56321402 refGene_gn=INTS4;refGene_tx=XM_006936891.4
D1 56338417 56338535 refGene_gn=INTS4;refGene_tx=XM_006936891.4
D1 56344407 56344514 refGene_gn=INTS4;refGene_tx=XM_006936891.4
D1 56352468 56352480 refGene_gn=INTS4;refGene_tx=XM_019811758.1
D1 56366615 56366801
refGene_gn=INTS4;refGene_tx=XM_006936891.4;refGene_gn=INTS4;refGene_tx
=XM_019811758.1
D1 56367636 56367687
refGene_gn=INTS4;refGene_tx=XM_006936891.4;refGene_gn=INTS4;refGene_tx
=XM_019811758.1
D1 56371063 56371152
refGene_gn=INTS4;refGene_tx=XM_006936891.4;refGene_gn=INTS4;refGene_tx
=XM_019811758.1
D1 56372141 56372262
refGene_gn=INTS4;refGene_tx=XM_006936891.4;refGene_gn=INTS4;refGene_tx
=XM_019811758.1
D1 56378461 56378533
refGene_gn=INTS4;refGene_tx=XM_006936891.4;refGene_gn=INTS4;refGene_tx
=XM_019811758.1
D1 56380238 56380413
refGene_gn=INTS4;refGene_tx=XM_006936891.4;refGene_gn=INTS4;refGene_tx
=XM_019811758.1
D1 56389479 56389685
refGene_gn=INTS4;refGene_tx=XM_006936891.4;refGene_gn=INTS4;refGene_tx
=XM_019811758.1
D1 56396661 56396804
refGene_gn=INTS4;refGene_tx=XM_006936891.4;refGene_gn=INTS4;refGene_tx
=XM_019811758.1
D1 56399620 56399736
refGene_gn=INTS4;refGene_tx=XM_006936891.4;refGene_gn=INTS4;refGene_tx
=XM_019811758.1
D1 56402242 56402376
refGene_gn=INTS4;refGene_tx=XM_006936891.4;refGene_gn=INTS4;refGene_tx
=XM_019811758.1
D1 56403448 56403606
refGene_gn=INTS4;refGene_tx=XM_006936891.4;refGene_gn=INTS4;refGene_tx
=XM_019811758.1
D1 56413153 56413247
refGene_gn=INTS4;refGene_tx=XM_006936891.4;refGene_gn=INTS4;refGene_tx
=XM_019811758.1
D1 56415936 56416017
refGene_gn=INTS4;refGene_tx=XM_006936891.4;refGene_gn=INTS4;refGene_tx
=XM_019811758.1
D1 56418751 56418882
refGene_gn=INTS4;refGene_tx=XM_006936891.4;refGene_gn=INTS4;refGene_tx
=XM_019811758.1
D1 56424350 56424410
refGene_gn=INTS4;refGene_tx=XM_006936891.4;refGene_gn=INTS4;refGene_tx
=XM_019811758.1
D1 56426250 56426410

refGene_gn=INTS4;refGene_tx=XM_006936891.4;refGene_gn=INTS4;refGene_tx=XM_019811758.1
D1 56426615 56426759
refGene_gn=INTS4;refGene_tx=XM_006936891.4;refGene_gn=INTS4;refGene_tx=XM_019811758.1
D1 56430961 56431082
refGene_gn=INTS4;refGene_tx=XM_006936891.4;refGene_gn=INTS4;refGene_tx=XM_019811758.1
D1 56434752 56434931
refGene_gn=INTS4;refGene_tx=XM_006936891.4;refGene_gn=INTS4;refGene_tx=XM_019811758.1
D1 56441151 56441292 refGene_gn=AAMDC;refGene_tx=XM_003992689.4
D1 56443294 56443390 refGene_gn=AAMDC;refGene_tx=XM_003992689.4
D1 56463253 56463385 refGene_gn=AAMDC;refGene_tx=XM_003992689.4
D1 56471318 56471505
refGene_gn=RSF1;refGene_tx=XM_003992690.5;refGene_gn=RSF1;refGene_tx=XM_019811757.2
D1 56513750 56513842
refGene_gn=RSF1;refGene_tx=XM_003992690.5;refGene_gn=RSF1;refGene_tx=XM_019811757.2
D1 56558965 56559058
refGene_gn=RSF1;refGene_tx=XM_003992690.5;refGene_gn=RSF1;refGene_tx=XM_019811757.2
D1 56559499 56559705
refGene_gn=RSF1;refGene_tx=XM_003992690.5;refGene_gn=RSF1;refGene_tx=XM_019811757.2
D1 56571206 56571361
refGene_gn=RSF1;refGene_tx=XM_003992690.5;refGene_gn=RSF1;refGene_tx=XM_019811757.2
D1 56594315 56596081
refGene_gn=RSF1;refGene_tx=XM_003992690.5;refGene_gn=RSF1;refGene_tx=XM_019811757.2
D1 56597799 56598006
refGene_gn=RSF1;refGene_tx=XM_003992690.5;refGene_gn=RSF1;refGene_tx=XM_019811757.2
D1 56602979 56603084
refGene_gn=RSF1;refGene_tx=XM_003992690.5;refGene_gn=RSF1;refGene_tx=XM_019811757.2
D1 56604310 56604390
refGene_gn=RSF1;refGene_tx=XM_003992690.5;refGene_gn=RSF1;refGene_tx=XM_019811757.2
D1 56609370 56609425
refGene_gn=RSF1;refGene_tx=XM_003992690.5;refGene_gn=RSF1;refGene_tx=XM_019811757.2
D1 56610758 56610868
refGene_gn=RSF1;refGene_tx=XM_003992690.5;refGene_gn=RSF1;refGene_tx=XM_019811757.2
D1 56617211 56617279
refGene_gn=RSF1;refGene_tx=XM_003992690.5;refGene_gn=RSF1;refGene_tx=XM_019811757.2

D1 56618492 56618700
refGene_gn=RSF1;refGene_tx=XM_003992690.5;refGene_gn=RSF1;refGene_tx=X
M_019811757.2
D1 56620197 56620418
refGene_gn=RSF1;refGene_tx=XM_003992690.5;refGene_gn=RSF1;refGene_tx=X
M_019811757.2
D1 56623240 56623429
refGene_gn=RSF1;refGene_tx=XM_003992690.5;refGene_gn=RSF1;refGene_tx=X
M_019811757.2
D1 56626427 56626990
refGene_gn=RSF1;refGene_tx=XM_003992690.5;refGene_gn=RSF1;refGene_tx=X
M_019811757.2
D1 56644558 56644683 refGene_gn=CLNS1A;refGene_tx=XM_003992691.5
D1 56650306 56650443 refGene_gn=CLNS1A;refGene_tx=XM_003992691.5
D1 56653999 56654101 refGene_gn=CLNS1A;refGene_tx=XM_003992691.5
D1 56654749 56654857 refGene_gn=CLNS1A;refGene_tx=XM_003992691.5
D1 56656213 56656387 refGene_gn=CLNS1A;refGene_tx=XM_003992691.5
D1 56661298 56661366 refGene_gn=CLNS1A;refGene_tx=XM_003992691.5
D1 56676239 56676319 refGene_gn=AQP11;refGene_tx=XM_011286638.3
D1 56681769 56681886 refGene_gn=AQP11;refGene_tx=XM_011286638.3
D1 56689095 56689717 refGene_gn=AQP11;refGene_tx=XM_011286638.3
D1 56720974 56721378
refGene_gn=LOC102902403;refGene_tx=XM_023239403.1
D1 56726850 56726927
refGene_gn=LOC102902403;refGene_tx=XM_023239403.1
D1 56728028 56728126
refGene_gn=LOC102902403;refGene_tx=XM_023239403.1
D1 56859034 56859224
refGene_gn=PAK1;refGene_tx=XM_003992702.5;refGene_gn=PAK1;refGene_tx=X
M_006936899.4;refGene_gn=PAK1;refGene_tx=XM_023239404.1
D1 56879803 56879904
refGene_gn=PAK1;refGene_tx=XM_003992702.5;refGene_gn=PAK1;refGene_tx=X
M_006936899.4;refGene_gn=PAK1;refGene_tx=XM_023239404.1
D1 56880595 56880743
refGene_gn=PAK1;refGene_tx=XM_003992702.5;refGene_gn=PAK1;refGene_tx=X
M_006936899.4;refGene_gn=PAK1;refGene_tx=XM_023239404.1
D1 56884250 56884288
refGene_gn=PAK1;refGene_tx=XM_003992702.5;refGene_gn=PAK1;refGene_tx=X
M_006936899.4;refGene_gn=PAK1;refGene_tx=XM_023239404.1
D1 56897020 56897137
refGene_gn=PAK1;refGene_tx=XM_003992702.5;refGene_gn=PAK1;refGene_tx=X
M_006936899.4;refGene_gn=PAK1;refGene_tx=XM_023239404.1
D1 56898774 56898949
refGene_gn=PAK1;refGene_tx=XM_003992702.5;refGene_gn=PAK1;refGene_tx=X
M_006936899.4;refGene_gn=PAK1;refGene_tx=XM_023239404.1
D1 56901253 56901317
refGene_gn=PAK1;refGene_tx=XM_003992702.5;refGene_gn=PAK1;refGene_tx=X
M_006936899.4;refGene_gn=PAK1;refGene_tx=XM_023239404.1
D1 56905159 56905208
refGene_gn=PAK1;refGene_tx=XM_003992702.5;refGene_gn=PAK1;refGene_tx=X

M_006936899.4;refGene_gn=PAK1;refGene_tx=XM_023239404.1
D1	56910369	56910482
refGene_gn=PAK1;refGene_tx=XM_003992702.5;refGene_gn=PAK1;refGene_tx=XM_006936899.4;refGene_gn=PAK1;refGene_tx=XM_023239404.1
D1	56913784	56913902
refGene_gn=PAK1;refGene_tx=XM_003992702.5;refGene_gn=PAK1;refGene_tx=XM_006936899.4;refGene_gn=PAK1;refGene_tx=XM_023239404.1
D1	56920039	56920139
refGene_gn=PAK1;refGene_tx=XM_003992702.5;refGene_gn=PAK1;refGene_tx=XM_006936899.4;refGene_gn=PAK1;refGene_tx=XM_023239404.1
D1	56922198	56922395
refGene_gn=PAK1;refGene_tx=XM_003992702.5;refGene_gn=PAK1;refGene_tx=XM_006936899.4;refGene_gn=PAK1;refGene_tx=XM_023239404.1
D1	56925260	56925398
refGene_gn=PAK1;refGene_tx=XM_003992702.5;refGene_gn=PAK1;refGene_tx=XM_006936899.4;refGene_gn=PAK1;refGene_tx=XM_023239404.1
D1	56926900	56926987
refGene_gn=PAK1;refGene_tx=XM_003992702.5;refGene_gn=PAK1;refGene_tx=XM_006936899.4;refGene_gn=PAK1;refGene_tx=XM_023239404.1
D1	56960947	56961136
refGene_gn=GDPD4;refGene_tx=XM_019811760.1;refGene_gn=GDPD4;refGene_tx=XM_019811761.1;refGene_gn=GDPD4;refGene_tx=XM_019811762.2;refGene_gn=GDPD4;refGene_tx=XM_019811763.1
D1	56966072	56966176
refGene_gn=GDPD4;refGene_tx=XM_019811760.1;refGene_gn=GDPD4;refGene_tx=XM_019811761.1;refGene_gn=GDPD4;refGene_tx=XM_019811762.2;refGene_gn=GDPD4;refGene_tx=XM_019811763.1
D1	56989950	56990044
refGene_gn=GDPD4;refGene_tx=XM_019811760.1;refGene_gn=GDPD4;refGene_tx=XM_019811761.1;refGene_gn=GDPD4;refGene_tx=XM_019811762.2;refGene_gn=GDPD4;refGene_tx=XM_019811763.1
D1	56997825	56997885
refGene_gn=GDPD4;refGene_tx=XM_019811760.1;refGene_gn=GDPD4;refGene_tx=XM_019811761.1;refGene_gn=GDPD4;refGene_tx=XM_019811762.2;refGene_gn=GDPD4;refGene_tx=XM_019811763.1
D1	57004905	57005004
refGene_gn=GDPD4;refGene_tx=XM_019811760.1;refGene_gn=GDPD4;refGene_tx=XM_019811761.1;refGene_gn=GDPD4;refGene_tx=XM_019811762.2;refGene_gn=GDPD4;refGene_tx=XM_019811763.1
D1	57005075	57005169
refGene_gn=GDPD4;refGene_tx=XM_019811760.1;refGene_gn=GDPD4;refGene_tx=XM_019811761.1;refGene_gn=GDPD4;refGene_tx=XM_019811762.2;refGene_gn=GDPD4;refGene_tx=XM_019811763.1
D1	57006001	57006076
refGene_gn=GDPD4;refGene_tx=XM_019811760.1;refGene_gn=GDPD4;refGene_tx=XM_019811762.2;refGene_gn=GDPD4;refGene_tx=XM_019811763.1
D1	57006846	57006992
refGene_gn=GDPD4;refGene_tx=XM_019811760.1;refGene_gn=GDPD4;refGene_tx=XM_019811761.1;refGene_gn=GDPD4;refGene_tx=XM_019811762.2;refGene_gn=GDPD4;refGene_tx=XM_019811763.1

D1 57007364 57007447
refGene_gn=GDPD4;refGene_tx=XM_019811760.1;refGene_gn=GDPD4;refGene_tx=XM_019811761.1;refGene_gn=GDPD4;refGene_tx=XM_019811762.2;refGene_gn=GDPD4;refGene_tx=XM_019811763.1
D1 57011184 57011341
refGene_gn=GDPD4;refGene_tx=XM_019811760.1;refGene_gn=GDPD4;refGene_tx=XM_019811761.1;refGene_gn=GDPD4;refGene_tx=XM_019811762.2;refGene_gn=GDPD4;refGene_tx=XM_019811763.1
D1 57042559 57042781
refGene_gn=GDPD4;refGene_tx=XM_019811760.1;refGene_gn=GDPD4;refGene_tx=XM_019811761.1;refGene_gn=GDPD4;refGene_tx=XM_019811762.2
D1 57043699 57043854
refGene_gn=GDPD4;refGene_tx=XM_019811760.1;refGene_gn=GDPD4;refGene_tx=XM_019811761.1;refGene_gn=GDPD4;refGene_tx=XM_019811762.2;refGene_gn=GDPD4;refGene_tx=XM_019811763.1
D1 57065328 57065476
refGene_gn=GDPD4;refGene_tx=XM_019811760.1;refGene_gn=GDPD4;refGene_tx=XM_019811761.1;refGene_gn=GDPD4;refGene_tx=XM_019811762.2;refGene_gn=GDPD4;refGene_tx=XM_019811763.1
D1 57069455 57069538
refGene_gn=GDPD4;refGene_tx=XM_019811760.1;refGene_gn=GDPD4;refGene_tx=XM_019811761.1;refGene_gn=GDPD4;refGene_tx=XM_019811762.2;refGene_gn=GDPD4;refGene_tx=XM_019811763.1
D1 57070272 57070319
refGene_gn=GDPD4;refGene_tx=XM_019811760.1;refGene_gn=GDPD4;refGene_tx=XM_019811761.1;refGene_gn=GDPD4;refGene_tx=XM_019811762.2;refGene_gn=GDPD4;refGene_tx=XM_019811763.1
D1 57090909 57091403
refGene_gn=GDPD4;refGene_tx=XM_019811760.1;refGene_gn=GDPD4;refGene_tx=XM_019811761.1;refGene_gn=GDPD4;refGene_tx=XM_019811762.2;refGene_gn=GDPD4;refGene_tx=XM_019811763.1
D1 57119017 57119089 refGene_gn=GDPD4;refGene_tx=XM_019811762.2
D1 57130725 57130815
refGene_gn=MYO7A;refGene_tx=XM_019811764.2;refGene_gn=MYO7A;refGene_tx=XM_019811765.2;refGene_gn=MYO7A;refGene_tx=XM_019811766.1
D1 57131415 57131535
refGene_gn=MYO7A;refGene_tx=XM_019811764.2;refGene_gn=MYO7A;refGene_tx=XM_019811765.2;refGene_gn=MYO7A;refGene_tx=XM_019811766.1
D1 57132409 57132493
refGene_gn=MYO7A;refGene_tx=XM_019811764.2;refGene_gn=MYO7A;refGene_tx=XM_019811765.2;refGene_gn=MYO7A;refGene_tx=XM_019811766.1
D1 57133579 57133696
refGene_gn=MYO7A;refGene_tx=XM_019811764.2;refGene_gn=MYO7A;refGene_tx=XM_019811765.2;refGene_gn=MYO7A;refGene_tx=XM_019811766.1
D1 57134135 57134321
refGene_gn=MYO7A;refGene_tx=XM_019811764.2;refGene_gn=MYO7A;refGene_tx=XM_019811765.2;refGene_gn=MYO7A;refGene_tx=XM_019811766.1
D1 57136548 57136655
refGene_gn=MYO7A;refGene_tx=XM_019811764.2;refGene_gn=MYO7A;refGene_tx=XM_019811765.2;refGene_gn=MYO7A;refGene_tx=XM_019811766.1

D1 57136837 57136925
refGene_gn=MYO7A;refGene_tx=XM_019811764.2;refGene_gn=MYO7A;refGene_tx=XM_019811765.2;refGene_gn=MYO7A;refGene_tx=XM_019811766.1
D1 57137953 57138067
refGene_gn=MYO7A;refGene_tx=XM_019811764.2;refGene_gn=MYO7A;refGene_tx=XM_019811765.2;refGene_gn=MYO7A;refGene_tx=XM_019811766.1
D1 57139192 57139298
refGene_gn=MYO7A;refGene_tx=XM_019811764.2;refGene_gn=MYO7A;refGene_tx=XM_019811765.2;refGene_gn=MYO7A;refGene_tx=XM_019811766.1
D1 57139821 57139977
refGene_gn=MYO7A;refGene_tx=XM_019811764.2;refGene_gn=MYO7A;refGene_tx=XM_019811765.2;refGene_gn=MYO7A;refGene_tx=XM_019811766.1
D1 57141200 57141354
refGene_gn=MYO7A;refGene_tx=XM_019811764.2;refGene_gn=MYO7A;refGene_tx=XM_019811765.2;refGene_gn=MYO7A;refGene_tx=XM_019811766.1
D1 57142311 57142469
refGene_gn=MYO7A;refGene_tx=XM_019811764.2;refGene_gn=MYO7A;refGene_tx=XM_019811765.2;refGene_gn=MYO7A;refGene_tx=XM_019811766.1
D1 57143121 57143246
refGene_gn=MYO7A;refGene_tx=XM_019811764.2;refGene_gn=MYO7A;refGene_tx=XM_019811765.2;refGene_gn=MYO7A;refGene_tx=XM_019811766.1
D1 57143918 57144109
refGene_gn=MYO7A;refGene_tx=XM_019811764.2;refGene_gn=MYO7A;refGene_tx=XM_019811765.2;refGene_gn=MYO7A;refGene_tx=XM_019811766.1
D1 57145051 57145335
refGene_gn=MYO7A;refGene_tx=XM_019811764.2;refGene_gn=MYO7A;refGene_tx=XM_019811765.2;refGene_gn=MYO7A;refGene_tx=XM_019811766.1
D1 57146411 57146538
refGene_gn=MYO7A;refGene_tx=XM_019811764.2;refGene_gn=MYO7A;refGene_tx=XM_019811765.2;refGene_gn=MYO7A;refGene_tx=XM_019811766.1
D1 57147377 57147495
refGene_gn=MYO7A;refGene_tx=XM_019811764.2;refGene_gn=MYO7A;refGene_tx=XM_019811765.2;refGene_gn=MYO7A;refGene_tx=XM_019811766.1
D1 57149614 57149785
refGene_gn=MYO7A;refGene_tx=XM_019811764.2;refGene_gn=MYO7A;refGene_tx=XM_019811765.2;refGene_gn=MYO7A;refGene_tx=XM_019811766.1
D1 57151114 57151342
refGene_gn=MYO7A;refGene_tx=XM_019811764.2;refGene_gn=MYO7A;refGene_tx=XM_019811765.2;refGene_gn=MYO7A;refGene_tx=XM_019811766.1
D1 57152189 57152363
refGene_gn=MYO7A;refGene_tx=XM_019811764.2;refGene_gn=MYO7A;refGene_tx=XM_019811765.2;refGene_gn=MYO7A;refGene_tx=XM_019811766.1
D1 57153172 57153292
refGene_gn=MYO7A;refGene_tx=XM_019811764.2;refGene_gn=MYO7A;refGene_tx=XM_019811765.2;refGene_gn=MYO7A;refGene_tx=XM_019811766.1
D1 57154213 57154340
refGene_gn=MYO7A;refGene_tx=XM_019811764.2;refGene_gn=MYO7A;refGene_tx=XM_019811765.2;refGene_gn=MYO7A;refGene_tx=XM_019811766.1
D1 57156529 57156657
refGene_gn=MYO7A;refGene_tx=XM_019811764.2;refGene_gn=MYO7A;refGene_tx

=XM_019811765.2;refGene_gn=MYO7A;refGene_tx=XM_019811766.1
D1	57158019	57158109
refGene_gn=MYO7A;refGene_tx=XM_019811764.2;refGene_gn=MYO7A;refGene_tx
=XM_019811765.2;refGene_gn=MYO7A;refGene_tx=XM_019811766.1
D1	57158581	57158758
refGene_gn=MYO7A;refGene_tx=XM_019811764.2;refGene_gn=MYO7A;refGene_tx
=XM_019811765.2;refGene_gn=MYO7A;refGene_tx=XM_019811766.1
D1	57158948	57159152
refGene_gn=MYO7A;refGene_tx=XM_019811764.2;refGene_gn=MYO7A;refGene_tx
=XM_019811765.2;refGene_gn=MYO7A;refGene_tx=XM_019811766.1
D1	57159493	57159703
refGene_gn=MYO7A;refGene_tx=XM_019811764.2;refGene_gn=MYO7A;refGene_tx
=XM_019811765.2;refGene_gn=MYO7A;refGene_tx=XM_019811766.1
D1	57161249	57161357
refGene_gn=MYO7A;refGene_tx=XM_019811764.2;refGene_gn=MYO7A;refGene_tx
=XM_019811765.2;refGene_gn=MYO7A;refGene_tx=XM_019811766.1
D1	57161756	57161975
refGene_gn=MYO7A;refGene_tx=XM_019811764.2;refGene_gn=MYO7A;refGene_tx
=XM_019811765.2;refGene_gn=MYO7A;refGene_tx=XM_019811766.1
D1	57162572	57162657
refGene_gn=MYO7A;refGene_tx=XM_019811764.2;refGene_gn=MYO7A;refGene_tx
=XM_019811765.2;refGene_gn=MYO7A;refGene_tx=XM_019811766.1
D1	57163833	57163928
refGene_gn=MYO7A;refGene_tx=XM_019811764.2;refGene_gn=MYO7A;refGene_tx
=XM_019811765.2;refGene_gn=MYO7A;refGene_tx=XM_019811766.1
D1	57166127	57166220
refGene_gn=MYO7A;refGene_tx=XM_019811764.2;refGene_gn=MYO7A;refGene_tx
=XM_019811765.2;refGene_gn=MYO7A;refGene_tx=XM_019811766.1
D1	57166655	57166814
refGene_gn=MYO7A;refGene_tx=XM_019811764.2;refGene_gn=MYO7A;refGene_tx
=XM_019811765.2;refGene_gn=MYO7A;refGene_tx=XM_019811766.1
D1	57168549	57168687
refGene_gn=MYO7A;refGene_tx=XM_019811764.2;refGene_gn=MYO7A;refGene_tx
=XM_019811765.2;refGene_gn=MYO7A;refGene_tx=XM_019811766.1
D1	57174326	57174433
refGene_gn=MYO7A;refGene_tx=XM_019811764.2;refGene_gn=MYO7A;refGene_tx
=XM_019811765.2;refGene_gn=MYO7A;refGene_tx=XM_019811766.1
D1	57177699	57177835
refGene_gn=MYO7A;refGene_tx=XM_019811764.2;refGene_gn=MYO7A;refGene_tx
=XM_019811765.2;refGene_gn=MYO7A;refGene_tx=XM_019811766.1
D1	57178335	57178546
refGene_gn=MYO7A;refGene_tx=XM_019811764.2;refGene_gn=MYO7A;refGene_tx
=XM_019811765.2;refGene_gn=MYO7A;refGene_tx=XM_019811766.1
D1	57178945	57179088
refGene_gn=MYO7A;refGene_tx=XM_019811764.2;refGene_gn=MYO7A;refGene_tx
=XM_019811765.2;refGene_gn=MYO7A;refGene_tx=XM_019811766.1
D1	57179845	57179965
refGene_gn=MYO7A;refGene_tx=XM_019811764.2;refGene_gn=MYO7A;refGene_tx
=XM_019811765.2;refGene_gn=MYO7A;refGene_tx=XM_019811766.1
D1	57180564	57180641

refGene_gn=MYO7A;refGene_tx=XM_019811764.2;refGene_gn=MYO7A;refGene_tx
=XM_019811765.2;refGene_gn=MYO7A;refGene_tx=XM_019811766.1
D1 57181104 57181258
refGene_gn=MYO7A;refGene_tx=XM_019811764.2;refGene_gn=MYO7A;refGene_tx
=XM_019811765.2;refGene_gn=MYO7A;refGene_tx=XM_019811766.1
D1 57182139 57182253
refGene_gn=MYO7A;refGene_tx=XM_019811764.2;refGene_gn=MYO7A;refGene_tx
=XM_019811765.2;refGene_gn=MYO7A;refGene_tx=XM_019811766.1
D1 57182516 57182659
refGene_gn=MYO7A;refGene_tx=XM_019811764.2;refGene_gn=MYO7A;refGene_tx
=XM_019811765.2;refGene_gn=MYO7A;refGene_tx=XM_019811766.1
D1 57182739 57182861
refGene_gn=MYO7A;refGene_tx=XM_019811764.2;refGene_gn=MYO7A;refGene_tx
=XM_019811765.2;refGene_gn=MYO7A;refGene_tx=XM_019811766.1
D1 57183359 57183544
refGene_gn=MYO7A;refGene_tx=XM_019811764.2;refGene_gn=MYO7A;refGene_tx
=XM_019811765.2;refGene_gn=MYO7A;refGene_tx=XM_019811766.1
D1 57191151 57191304
refGene_gn=MYO7A;refGene_tx=XM_019811764.2;refGene_gn=MYO7A;refGene_tx
=XM_019811765.2;refGene_gn=MYO7A;refGene_tx=XM_019811766.1
D1 57198152 57198266
refGene_gn=MYO7A;refGene_tx=XM_019811764.2;refGene_gn=MYO7A;refGene_tx
=XM_019811765.2;refGene_gn=MYO7A;refGene_tx=XM_019811766.1
D1 57201863 57202013
refGene_gn=MYO7A;refGene_tx=XM_019811764.2;refGene_gn=MYO7A;refGene_tx
=XM_019811765.2
D1 57213486 57213504 refGene_gn=MYO7A;refGene_tx=XM_019811766.1
D1 57220388 57220571
refGene_gn=CAPN5;refGene_tx=XM_003992706.5;refGene_gn=CAPN5;refGene_tx
=XM_023239405.1;refGene_gn=CAPN5;refGene_tx=XM_023239406.1
D1 57221552 57221689
refGene_gn=CAPN5;refGene_tx=XM_003992706.5;refGene_gn=CAPN5;refGene_tx
=XM_023239405.1;refGene_gn=CAPN5;refGene_tx=XM_023239406.1
D1 57222260 57222376
refGene_gn=CAPN5;refGene_tx=XM_003992706.5;refGene_gn=CAPN5;refGene_tx
=XM_023239405.1;refGene_gn=CAPN5;refGene_tx=XM_023239406.1
D1 57223284 57223481
refGene_gn=CAPN5;refGene_tx=XM_003992706.5;refGene_gn=CAPN5;refGene_tx
=XM_023239405.1;refGene_gn=CAPN5;refGene_tx=XM_023239406.1
D1 57224594 57224717
refGene_gn=CAPN5;refGene_tx=XM_003992706.5;refGene_gn=CAPN5;refGene_tx
=XM_023239405.1;refGene_gn=CAPN5;refGene_tx=XM_023239406.1
D1 57225339 57225535
refGene_gn=CAPN5;refGene_tx=XM_003992706.5;refGene_gn=CAPN5;refGene_tx
=XM_023239405.1;refGene_gn=CAPN5;refGene_tx=XM_023239406.1
D1 57226363 57226441
refGene_gn=CAPN5;refGene_tx=XM_003992706.5;refGene_gn=CAPN5;refGene_tx
=XM_023239405.1;refGene_gn=CAPN5;refGene_tx=XM_023239406.1
D1 57227087 57227281
refGene_gn=CAPN5;refGene_tx=XM_003992706.5;refGene_gn=CAPN5;refGene_tx

=XM_023239405.1;refGene_gn=CAPN5;refGene_tx=XM_023239406.1
D1 57228022 57228215
refGene_gn=CAPN5;refGene_tx=XM_003992706.5;refGene_gn=CAPN5;refGene_tx=XM_023239405.1;refGene_gn=CAPN5;refGene_tx=XM_023239406.1
D1 57231447 57231656
refGene_gn=CAPN5;refGene_tx=XM_003992706.5;refGene_gn=CAPN5;refGene_tx=XM_023239405.1;refGene_gn=CAPN5;refGene_tx=XM_023239406.1
D1 57240800 57241292 refGene_gn=OMP;refGene_tx=XM_023239407.1
D1 57246922 57247054
refGene_gn=CAPN5;refGene_tx=XM_003992706.5;refGene_gn=CAPN5;refGene_tx=XM_023239405.1;refGene_gn=CAPN5;refGene_tx=XM_023239406.1
D1 57253687 57253852
refGene_gn=CAPN5;refGene_tx=XM_003992706.5;refGene_gn=CAPN5;refGene_tx=XM_023239405.1;refGene_gn=CAPN5;refGene_tx=XM_023239406.1
D1 57285563 57286739 refGene_gn=B3GNT6;refGene_tx=XM_023238988.1
D1 57305669 57305723
refGene_gn=ACER3;refGene_tx=XM_019811768.1;refGene_gn=ACER3;refGene_tx=XM_023239409.1;refGene_gn=ACER3;refGene_tx=XM_023239410.1
D1 57306244 57306290
refGene_gn=ACER3;refGene_tx=XM_019811768.1;refGene_gn=ACER3;refGene_tx=XM_023239409.1;refGene_gn=ACER3;refGene_tx=XM_023239410.1
D1 57315420 57315525
refGene_gn=ACER3;refGene_tx=XM_019811768.1;refGene_gn=ACER3;refGene_tx=XM_023239409.1;refGene_gn=ACER3;refGene_tx=XM_023239410.1
D1 57317892 57317994
refGene_gn=ACER3;refGene_tx=XM_019811768.1;refGene_gn=ACER3;refGene_tx=XM_023239409.1;refGene_gn=ACER3;refGene_tx=XM_023239410.1
D1 57334241 57334300
refGene_gn=ACER3;refGene_tx=XM_019811768.1;refGene_gn=ACER3;refGene_tx=XM_023239409.1;refGene_gn=ACER3;refGene_tx=XM_023239410.1
D1 57347099 57347135
refGene_gn=ACER3;refGene_tx=XM_019811768.1;refGene_gn=ACER3;refGene_tx=XM_023239409.1;refGene_gn=ACER3;refGene_tx=XM_023239410.1
D1 57351831 57351913
refGene_gn=ACER3;refGene_tx=XM_019811768.1;refGene_gn=ACER3;refGene_tx=XM_023239409.1;refGene_gn=ACER3;refGene_tx=XM_023239410.1
D1 57354107 57354160
refGene_gn=ACER3;refGene_tx=XM_019811768.1;refGene_gn=ACER3;refGene_tx=XM_023239409.1;refGene_gn=ACER3;refGene_tx=XM_023239410.1
D1 57371616 57371669
refGene_gn=ACER3;refGene_tx=XM_023239409.1;refGene_gn=ACER3;refGene_tx=XM_023239410.1
D1 57407676 57407787
refGene_gn=ACER3;refGene_tx=XM_023239409.1;refGene_gn=ACER3;refGene_tx=XM_023239410.1
D1 57470359 57470462 refGene_gn=ACER3;refGene_tx=XM_023239409.1
D1 57520354 57521436
refGene_gn=TSKU;refGene_tx=XM_006936903.4;refGene_gn=TSKU;refGene_tx=XM_019811769.2;refGene_gn=TSKU;refGene_tx=XM_019811770.2;refGene_gn=TSKU;refGene_tx=XM_023239411.1;refGene_gn=TSKU;refGene_tx=XM_023239412.1

D1 57521780 57521967 refGene_gn=TSKU;refGene_tx=XM_006936903.4
D1 57531749 57531822 refGene_gn=TSKU;refGene_tx=XM_019811769.2
D1 57599384 57600240
refGene_gn=LOC101099582;refGene_tx=XM_019812931.2
D1 57603017 57603313
refGene_gn=LOC101099582;refGene_tx=XM_019812931.2
D1 57605075 57605448
refGene_gn=LOC101099582;refGene_tx=XM_019812931.2
D1 57608997 57609082
refGene_gn=LOC101099582;refGene_tx=XM_019812931.2
D1 57609707 57609804
refGene_gn=LOC101099582;refGene_tx=XM_019812931.2
D1 57609887 57610016
refGene_gn=LOC101099582;refGene_tx=XM_019812931.2
D1 57611317 57611398
refGene_gn=LOC101099582;refGene_tx=XM_019812931.2
D1 57612705 57612882
refGene_gn=LOC101099582;refGene_tx=XM_019812931.2
D1 57613110 57613267
refGene_gn=LOC101099582;refGene_tx=XM_019812931.2
D1 57613560 57613710
refGene_gn=LOC101099582;refGene_tx=XM_019812931.2
D1 57617181 57617330
refGene_gn=LOC101099582;refGene_tx=XM_019812931.2
D1 57620342 57620506
refGene_gn=LOC101099582;refGene_tx=XM_019812931.2
D1 57620931 57621124
refGene_gn=LOC101099582;refGene_tx=XM_019812931.2
D1 57627360 57627535
refGene_gn=LOC101099582;refGene_tx=XM_019812931.2
D1 57628543 57628642
refGene_gn=LOC101099582;refGene_tx=XM_019812931.2
D1 57629478 57629573
refGene_gn=LOC101099582;refGene_tx=XM_019812931.2
D1 57629892 57629978
refGene_gn=LOC101099582;refGene_tx=XM_019812931.2
D1 57633806 57633915
refGene_gn=LOC101099582;refGene_tx=XM_019812931.2
D1 57650105 57650227 refGene_gn=LRRC32;refGene_tx=XM_003992711.4
D1 57654333 57654424
refGene_gn=LRRC32;refGene_tx=XM_003992711.4;refGene_gn=LRRC32;refGene_tx=XM_019811772.1;refGene_gn=LRRC32;refGene_tx=XM_023239415.1
D1 57659127 57661032
refGene_gn=LRRC32;refGene_tx=XM_003992711.4;refGene_gn=LRRC32;refGene_tx=XM_019811772.1;refGene_gn=LRRC32;refGene_tx=XM_023239415.1;refGene_gn=LRRC32;refGene_tx=XM_023239416.1
D1 57759104 57759299
refGene_gn=EMSY;refGene_tx=XM_003992713.5;refGene_gn=EMSY;refGene_tx=XM_006936908.4;refGene_gn=EMSY;refGene_tx=XM_006936909.4;refGene_gn=EMSY;refGene_tx=XM_011286634.3;refGene_gn=EMSY;refGene_tx=XM_019811773.2;

refGene_gn=EMSY;refGene_tx=XM_019811774.2;refGene_gn=EMSY;refGene_tx=X
M_019811775.2;refGene_gn=EMSY;refGene_tx=XM_019811777.2;refGene_gn=EMS
Y;refGene_tx=XM_019811778.2;refGene_gn=EMSY;refGene_tx=XM_019811779.2;
refGene_gn=EMSY;refGene_tx=XM_019811780.2;refGene_gn=EMSY;refGene_tx=X
M_019811781.2;refGene_gn=EMSY;refGene_tx=XM_019811782.2;refGene_gn=EMS
Y;refGene_tx=XM_019811783.2
D1 57763465 57763966
refGene_gn=EMSY;refGene_tx=XM_003992713.5;refGene_gn=EMSY;refGene_tx=X
M_006936908.4;refGene_gn=EMSY;refGene_tx=XM_006936909.4;refGene_gn=EMS
Y;refGene_tx=XM_011286634.3;refGene_gn=EMSY;refGene_tx=XM_019811773.2;
refGene_gn=EMSY;refGene_tx=XM_019811774.2;refGene_gn=EMSY;refGene_tx=X
M_019811775.2;refGene_gn=EMSY;refGene_tx=XM_019811777.2;refGene_gn=EMS
Y;refGene_tx=XM_019811778.2;refGene_gn=EMSY;refGene_tx=XM_019811779.2;
refGene_gn=EMSY;refGene_tx=XM_019811780.2;refGene_gn=EMSY;refGene_tx=X
M_019811781.2;refGene_gn=EMSY;refGene_tx=XM_019811782.2
D1 57764944 57765508
refGene_gn=EMSY;refGene_tx=XM_003992713.5;refGene_gn=EMSY;refGene_tx=X
M_006936908.4;refGene_gn=EMSY;refGene_tx=XM_006936909.4;refGene_gn=EMS
Y;refGene_tx=XM_011286634.3;refGene_gn=EMSY;refGene_tx=XM_019811773.2;
refGene_gn=EMSY;refGene_tx=XM_019811774.2;refGene_gn=EMSY;refGene_tx=X
M_019811775.2;refGene_gn=EMSY;refGene_tx=XM_019811777.2;refGene_gn=EMS
Y;refGene_tx=XM_019811778.2;refGene_gn=EMSY;refGene_tx=XM_019811779.2;
refGene_gn=EMSY;refGene_tx=XM_019811780.2;refGene_gn=EMSY;refGene_tx=X
M_019811781.2;refGene_gn=EMSY;refGene_tx=XM_019811782.2;refGene_gn=EMS
Y;refGene_tx=XM_019811783.2
D1 57767392 57767592
refGene_gn=EMSY;refGene_tx=XM_003992713.5;refGene_gn=EMSY;refGene_tx=X
M_006936908.4;refGene_gn=EMSY;refGene_tx=XM_006936909.4;refGene_gn=EMS
Y;refGene_tx=XM_011286634.3;refGene_gn=EMSY;refGene_tx=XM_019811773.2;
refGene_gn=EMSY;refGene_tx=XM_019811774.2;refGene_gn=EMSY;refGene_tx=X
M_019811775.2;refGene_gn=EMSY;refGene_tx=XM_019811777.2;refGene_gn=EMS
Y;refGene_tx=XM_019811778.2;refGene_gn=EMSY;refGene_tx=XM_019811779.2;
refGene_gn=EMSY;refGene_tx=XM_019811780.2;refGene_gn=EMSY;refGene_tx=X
M_019811781.2;refGene_gn=EMSY;refGene_tx=XM_019811782.2;refGene_gn=EMS
Y;refGene_tx=XM_019811783.2
D1 57769998 57770040
refGene_gn=EMSY;refGene_tx=XM_006936908.4;refGene_gn=EMSY;refGene_tx=X
M_011286634.3;refGene_gn=EMSY;refGene_tx=XM_019811773.2;refGene_gn=EMS
Y;refGene_tx=XM_019811774.2;refGene_gn=EMSY;refGene_tx=XM_019811775.2;
refGene_gn=EMSY;refGene_tx=XM_019811777.2;refGene_gn=EMSY;refGene_tx=X
M_019811778.2;refGene_gn=EMSY;refGene_tx=XM_019811779.2;refGene_gn=EMS
Y;refGene_tx=XM_019811780.2;refGene_gn=EMSY;refGene_tx=XM_019811781.2;
refGene_gn=EMSY;refGene_tx=XM_019811783.2
D1 57771085 57771241
refGene_gn=EMSY;refGene_tx=XM_003992713.5;refGene_gn=EMSY;refGene_tx=X
M_006936908.4;refGene_gn=EMSY;refGene_tx=XM_006936909.4;refGene_gn=EMS
Y;refGene_tx=XM_011286634.3;refGene_gn=EMSY;refGene_tx=XM_019811773.2;
refGene_gn=EMSY;refGene_tx=XM_019811774.2;refGene_gn=EMSY;refGene_tx=X
M_019811775.2;refGene_gn=EMSY;refGene_tx=XM_019811777.2;refGene_gn=EMS
Y;refGene_tx=XM_019811778.2;refGene_gn=EMSY;refGene_tx=XM_019811779.2;

refGene_gn=EMSY;refGene_tx=XM_019811780.2;refGene_gn=EMSY;refGene_tx=XM_019811781.2;refGene_gn=EMSY;refGene_tx=XM_019811783.2
D1 57772949 57773114
refGene_gn=EMSY;refGene_tx=XM_003992713.5;refGene_gn=EMSY;refGene_tx=XM_006936908.4;refGene_gn=EMSY;refGene_tx=XM_006936909.4;refGene_gn=EMSY;refGene_tx=XM_011286634.3;refGene_gn=EMSY;refGene_tx=XM_019811773.2;refGene_gn=EMSY;refGene_tx=XM_019811774.2;refGene_gn=EMSY;refGene_tx=XM_019811775.2;refGene_gn=EMSY;refGene_tx=XM_019811777.2;refGene_gn=EMSY;refGene_tx=XM_019811778.2;refGene_gn=EMSY;refGene_tx=XM_019811780.2;refGene_gn=EMSY;refGene_tx=XM_019811781.2;refGene_gn=EMSY;refGene_tx=XM_019811783.2
D1 57779719 57779918
refGene_gn=EMSY;refGene_tx=XM_003992713.5;refGene_gn=EMSY;refGene_tx=XM_006936908.4;refGene_gn=EMSY;refGene_tx=XM_006936909.4;refGene_gn=EMSY;refGene_tx=XM_011286634.3;refGene_gn=EMSY;refGene_tx=XM_019811773.2;refGene_gn=EMSY;refGene_tx=XM_019811774.2;refGene_gn=EMSY;refGene_tx=XM_019811775.2;refGene_gn=EMSY;refGene_tx=XM_019811777.2;refGene_gn=EMSY;refGene_tx=XM_019811778.2;refGene_gn=EMSY;refGene_tx=XM_019811779.2;refGene_gn=EMSY;refGene_tx=XM_019811780.2;refGene_gn=EMSY;refGene_tx=XM_019811781.2;refGene_gn=EMSY;refGene_tx=XM_019811782.2;refGene_gn=EMSY;refGene_tx=XM_019811783.2
D1 57781725 57781899
refGene_gn=EMSY;refGene_tx=XM_003992713.5;refGene_gn=EMSY;refGene_tx=XM_006936908.4;refGene_gn=EMSY;refGene_tx=XM_006936909.4;refGene_gn=EMSY;refGene_tx=XM_011286634.3;refGene_gn=EMSY;refGene_tx=XM_019811773.2;refGene_gn=EMSY;refGene_tx=XM_019811774.2;refGene_gn=EMSY;refGene_tx=XM_019811775.2;refGene_gn=EMSY;refGene_tx=XM_019811777.2;refGene_gn=EMSY;refGene_tx=XM_019811778.2;refGene_gn=EMSY;refGene_tx=XM_019811779.2;refGene_gn=EMSY;refGene_tx=XM_019811780.2;refGene_gn=EMSY;refGene_tx=XM_019811781.2;refGene_gn=EMSY;refGene_tx=XM_019811782.2;refGene_gn=EMSY;refGene_tx=XM_019811783.2
D1 57784294 57784431
refGene_gn=EMSY;refGene_tx=XM_003992713.5;refGene_gn=EMSY;refGene_tx=XM_006936908.4;refGene_gn=EMSY;refGene_tx=XM_006936909.4;refGene_gn=EMSY;refGene_tx=XM_011286634.3;refGene_gn=EMSY;refGene_tx=XM_019811773.2;refGene_gn=EMSY;refGene_tx=XM_019811774.2;refGene_gn=EMSY;refGene_tx=XM_019811775.2;refGene_gn=EMSY;refGene_tx=XM_019811777.2;refGene_gn=EMSY;refGene_tx=XM_019811778.2;refGene_gn=EMSY;refGene_tx=XM_019811779.2;refGene_gn=EMSY;refGene_tx=XM_019811780.2;refGene_gn=EMSY;refGene_tx=XM_019811781.2;refGene_gn=EMSY;refGene_tx=XM_019811782.2;refGene_gn=EMSY;refGene_tx=XM_019811783.2
D1 57790620 57790791
refGene_gn=EMSY;refGene_tx=XM_003992713.5;refGene_gn=EMSY;refGene_tx=XM_006936908.4;refGene_gn=EMSY;refGene_tx=XM_006936909.4;refGene_gn=EMSY;refGene_tx=XM_011286634.3;refGene_gn=EMSY;refGene_tx=XM_019811773.2;refGene_gn=EMSY;refGene_tx=XM_019811774.2;refGene_gn=EMSY;refGene_tx=XM_019811775.2;refGene_gn=EMSY;refGene_tx=XM_019811777.2;refGene_gn=EMSY;refGene_tx=XM_019811778.2;refGene_gn=EMSY;refGene_tx=XM_019811779.2;refGene_gn=EMSY;refGene_tx=XM_019811780.2;refGene_gn=EMSY;refGene_tx=XM_019811782.2;refGene_gn=EMSY;refGene_tx=XM_019811783.2

D1 57793359 57793509
refGene_gn=EMSY;refGene_tx=XM_003992713.5;refGene_gn=EMSY;refGene_tx=XM_006936908.4;refGene_gn=EMSY;refGene_tx=XM_006936909.4;refGene_gn=EMSY;refGene_tx=XM_011286634.3;refGene_gn=EMSY;refGene_tx=XM_019811773.2;refGene_gn=EMSY;refGene_tx=XM_019811774.2;refGene_gn=EMSY;refGene_tx=XM_019811775.2;refGene_gn=EMSY;refGene_tx=XM_019811777.2;refGene_gn=EMSY;refGene_tx=XM_019811778.2;refGene_gn=EMSY;refGene_tx=XM_019811779.2;refGene_gn=EMSY;refGene_tx=XM_019811780.2;refGene_gn=EMSY;refGene_tx=XM_019811782.2;refGene_gn=EMSY;refGene_tx=XM_019811783.2
D1 57804664 57804919
refGene_gn=EMSY;refGene_tx=XM_003992713.5;refGene_gn=EMSY;refGene_tx=XM_006936908.4;refGene_gn=EMSY;refGene_tx=XM_006936909.4;refGene_gn=EMSY;refGene_tx=XM_011286634.3;refGene_gn=EMSY;refGene_tx=XM_019811773.2;refGene_gn=EMSY;refGene_tx=XM_019811774.2;refGene_gn=EMSY;refGene_tx=XM_019811775.2;refGene_gn=EMSY;refGene_tx=XM_019811777.2;refGene_gn=EMSY;refGene_tx=XM_019811778.2;refGene_gn=EMSY;refGene_tx=XM_019811779.2;refGene_gn=EMSY;refGene_tx=XM_019811780.2;refGene_gn=EMSY;refGene_tx=XM_019811781.2;refGene_gn=EMSY;refGene_tx=XM_019811782.2;refGene_gn=EMSY;refGene_tx=XM_019811783.2
D1 57817102 57817379
refGene_gn=EMSY;refGene_tx=XM_003992713.5;refGene_gn=EMSY;refGene_tx=XM_006936908.4;refGene_gn=EMSY;refGene_tx=XM_006936909.4;refGene_gn=EMSY;refGene_tx=XM_011286634.3;refGene_gn=EMSY;refGene_tx=XM_019811773.2;refGene_gn=EMSY;refGene_tx=XM_019811774.2;refGene_gn=EMSY;refGene_tx=XM_019811775.2;refGene_gn=EMSY;refGene_tx=XM_019811777.2;refGene_gn=EMSY;refGene_tx=XM_019811778.2;refGene_gn=EMSY;refGene_tx=XM_019811779.2;refGene_gn=EMSY;refGene_tx=XM_019811780.2;refGene_gn=EMSY;refGene_tx=XM_019811781.2;refGene_gn=EMSY;refGene_tx=XM_019811782.2;refGene_gn=EMSY;refGene_tx=XM_019811783.2
D1 57826097 57826357
refGene_gn=EMSY;refGene_tx=XM_003992713.5;refGene_gn=EMSY;refGene_tx=XM_006936908.4;refGene_gn=EMSY;refGene_tx=XM_006936909.4;refGene_gn=EMSY;refGene_tx=XM_011286634.3;refGene_gn=EMSY;refGene_tx=XM_019811773.2;refGene_gn=EMSY;refGene_tx=XM_019811774.2;refGene_gn=EMSY;refGene_tx=XM_019811775.2;refGene_gn=EMSY;refGene_tx=XM_019811777.2;refGene_gn=EMSY;refGene_tx=XM_019811778.2;refGene_gn=EMSY;refGene_tx=XM_019811779.2;refGene_gn=EMSY;refGene_tx=XM_019811780.2;refGene_gn=EMSY;refGene_tx=XM_019811781.2;refGene_gn=EMSY;refGene_tx=XM_019811782.2;refGene_gn=EMSY;refGene_tx=XM_019811783.2
D1 57828999 57829152
refGene_gn=EMSY;refGene_tx=XM_006936908.4;refGene_gn=EMSY;refGene_tx=XM_006936909.4;refGene_gn=EMSY;refGene_tx=XM_011286634.3;refGene_gn=EMSY;refGene_tx=XM_019811773.2;refGene_gn=EMSY;refGene_tx=XM_019811774.2;refGene_gn=EMSY;refGene_tx=XM_019811775.2;refGene_gn=EMSY;refGene_tx=XM_019811777.2;refGene_gn=EMSY;refGene_tx=XM_019811779.2;refGene_gn=EMSY;refGene_tx=XM_019811780.2;refGene_gn=EMSY;refGene_tx=XM_019811781.2;refGene_gn=EMSY;refGene_tx=XM_019811782.2;refGene_gn=EMSY;refGene_tx=XM_019811783.2
D1 57830748 57830924
refGene_gn=EMSY;refGene_tx=XM_003992713.5;refGene_gn=EMSY;refGene_tx=X

M_006936908.4;refGene_gn=EMSY;refGene_tx=XM_006936909.4;refGene_gn=EMSY;refGene_tx=XM_011286634.3;refGene_gn=EMSY;refGene_tx=XM_019811773.2;refGene_gn=EMSY;refGene_tx=XM_019811774.2;refGene_gn=EMSY;refGene_tx=XM_019811775.2;refGene_gn=EMSY;refGene_tx=XM_019811777.2;refGene_gn=EMSY;refGene_tx=XM_019811778.2;refGene_gn=EMSY;refGene_tx=XM_019811779.2;refGene_gn=EMSY;refGene_tx=XM_019811780.2;refGene_gn=EMSY;refGene_tx=XM_019811781.2;refGene_gn=EMSY;refGene_tx=XM_019811782.2;refGene_gn=EMSY;refGene_tx=XM_019811783.2

D1 57834478 57834520 refGene_gn=EMSY;refGene_tx=XM_003992713.5;refGene_gn=EMSY;refGene_tx=XM_006936909.4;refGene_gn=EMSY;refGene_tx=XM_011286634.3;refGene_gn=EMSY;refGene_tx=XM_019811773.2;refGene_gn=EMSY;refGene_tx=XM_019811774.2;refGene_gn=EMSY;refGene_tx=XM_019811778.2;refGene_gn=EMSY;refGene_tx=XM_019811779.2;refGene_gn=EMSY;refGene_tx=XM_019811781.2;refGene_gn=EMSY;refGene_tx=XM_019811782.2;refGene_gn=EMSY;refGene_tx=XM_019811783.2

D1 57835881 57835956 refGene_gn=EMSY;refGene_tx=XM_003992713.5;refGene_gn=EMSY;refGene_tx=XM_006936908.4;refGene_gn=EMSY;refGene_tx=XM_006936909.4;refGene_gn=EMSY;refGene_tx=XM_011286634.3;refGene_gn=EMSY;refGene_tx=XM_019811773.2;refGene_gn=EMSY;refGene_tx=XM_019811774.2;refGene_gn=EMSY;refGene_tx=XM_019811775.2;refGene_gn=EMSY;refGene_tx=XM_019811777.2;refGene_gn=EMSY;refGene_tx=XM_019811778.2;refGene_gn=EMSY;refGene_tx=XM_019811779.2;refGene_gn=EMSY;refGene_tx=XM_019811781.2;refGene_gn=EMSY;refGene_tx=XM_019811782.2;refGene_gn=EMSY;refGene_tx=XM_019811783.2

D1 57837546 57837646 refGene_gn=EMSY;refGene_tx=XM_003992713.5;refGene_gn=EMSY;refGene_tx=XM_006936908.4;refGene_gn=EMSY;refGene_tx=XM_006936909.4;refGene_gn=EMSY;refGene_tx=XM_011286634.3;refGene_gn=EMSY;refGene_tx=XM_019811773.2;refGene_gn=EMSY;refGene_tx=XM_019811774.2;refGene_gn=EMSY;refGene_tx=XM_019811775.2;refGene_gn=EMSY;refGene_tx=XM_019811777.2;refGene_gn=EMSY;refGene_tx=XM_019811778.2;refGene_gn=EMSY;refGene_tx=XM_019811779.2;refGene_gn=EMSY;refGene_tx=XM_019811781.2;refGene_gn=EMSY;refGene_tx=XM_019811782.2;refGene_gn=EMSY;refGene_tx=XM_019811783.2

D1 57843360 57843430 refGene_gn=EMSY;refGene_tx=XM_003992713.5;refGene_gn=EMSY;refGene_tx=XM_006936908.4;refGene_gn=EMSY;refGene_tx=XM_006936909.4;refGene_gn=EMSY;refGene_tx=XM_011286634.3;refGene_gn=EMSY;refGene_tx=XM_019811773.2;refGene_gn=EMSY;refGene_tx=XM_019811774.2;refGene_gn=EMSY;refGene_tx=XM_019811775.2;refGene_gn=EMSY;refGene_tx=XM_019811777.2;refGene_gn=EMSY;refGene_tx=XM_019811778.2;refGene_gn=EMSY;refGene_tx=XM_019811779.2;refGene_gn=EMSY;refGene_tx=XM_019811780.2;refGene_gn=EMSY;refGene_tx=XM_019811781.2;refGene_gn=EMSY;refGene_tx=XM_019811782.2;refGene_gn=EMSY;refGene_tx=XM_019811783.2

D1 57902590 57903223 refGene_gn=GVQW3;refGene_tx=XM_019811785.2

D1 57904235 57904324 refGene_gn=THAP12;refGene_tx=XM_023239419.1;refGene_gn=THAP12;refGene_tx=XM_023239420.1;refGene_gn=THAP12;refGene_tx=XM_023239421.1

D1 57919747 57919868 refGene_gn=THAP12;refGene_tx=XM_023239419.1;refGene_gn=THAP12;refGene_tx=XM_023239420.1;refGene_gn=THAP12;refGene_tx=XM_023239421.1

D1 57923435 57923543
refGene_gn=THAP12;refGene_tx=XM_023239419.1;refGene_gn=THAP12;refGene_tx=XM_023239421.1
D1 57928067 57928104
refGene_gn=THAP12;refGene_tx=XM_023239419.1;refGene_gn=THAP12;refGene_tx=XM_023239420.1
D1 57929603 57931537
refGene_gn=THAP12;refGene_tx=XM_023239419.1;refGene_gn=THAP12;refGene_tx=XM_023239420.1;refGene_gn=THAP12;refGene_tx=XM_023239421.1
D1 58063049 58063132 refGene_gn=WNT11;refGene_tx=XM_023239422.1
D1 58070311 58070547 refGene_gn=WNT11;refGene_tx=XM_023239422.1
D1 58072167 58072445 refGene_gn=WNT11;refGene_tx=XM_023239422.1
D1 58075194 58075487 refGene_gn=WNT11;refGene_tx=XM_023239422.1
D1 58079465 58079640 refGene_gn=WNT11;refGene_tx=XM_023239422.1
D1 58126193 58126896 refGene_gn=UVRAG;refGene_tx=XM_023239423.1
D1 58151602 58151694 refGene_gn=UVRAG;refGene_tx=XM_023239423.1
D1 58205120 58205199 refGene_gn=UVRAG;refGene_tx=XM_023239423.1
D1 58246375 58246541 refGene_gn=UVRAG;refGene_tx=XM_023239423.1
D1 58254274 58254335 refGene_gn=UVRAG;refGene_tx=XM_023239423.1
D1 58256142 58256230 refGene_gn=UVRAG;refGene_tx=XM_023239423.1
D1 58258042 58258127 refGene_gn=UVRAG;refGene_tx=XM_023239423.1
D1 58275243 58275370 refGene_gn=UVRAG;refGene_tx=XM_023239423.1
D1 58289013 58289119 refGene_gn=UVRAG;refGene_tx=XM_023239423.1
D1 58339393 58339479 refGene_gn=UVRAG;refGene_tx=XM_023239423.1
D1 58354830 58354905 refGene_gn=UVRAG;refGene_tx=XM_023239423.1
D1 58363573 58363735 refGene_gn=UVRAG;refGene_tx=XM_023239423.1
D1 58381236 58381271 refGene_gn=UVRAG;refGene_tx=XM_023239423.1
D1 58388140 58388258 refGene_gn=UVRAG;refGene_tx=XM_023239423.1
D1 58419229 58419346 refGene_gn=UVRAG;refGene_tx=XM_023239423.1
D1 58436794 58436949 refGene_gn=DGAT2;refGene_tx=XM_011286632.2
D1 58438604 58438807 refGene_gn=DGAT2;refGene_tx=XM_011286632.2
D1 58439190 58439365 refGene_gn=DGAT2;refGene_tx=XM_011286632.2
D1 58439982 58440187 refGene_gn=DGAT2;refGene_tx=XM_011286632.2
D1 58446473 58446544 refGene_gn=DGAT2;refGene_tx=XM_011286632.2
D1 58446937 58447045 refGene_gn=DGAT2;refGene_tx=XM_011286632.2
D1 58452594 58452723 refGene_gn=DGAT2;refGene_tx=XM_011286632.2
D1 58467861 58467901 refGene_gn=DGAT2;refGene_tx=XM_011286632.2
D1 58501612 58501767 refGene_gn=MOGAT2;refGene_tx=XM_019811791.2
D1 58503972 58504172 refGene_gn=MOGAT2;refGene_tx=XM_019811791.2
D1 58504810 58504985 refGene_gn=MOGAT2;refGene_tx=XM_019811791.2
D1 58505591 58505796 refGene_gn=MOGAT2;refGene_tx=XM_019811791.2
D1 58514739 58514918 refGene_gn=MOGAT2;refGene_tx=XM_019811791.2
D1 58522030 58522121 refGene_gn=MOGAT2;refGene_tx=XM_019811791.2
D1 58555132 58556043 refGene_gn=MAP6;refGene_tx=XM_011286628.2
D1 58606777 58606991 refGene_gn=MAP6;refGene_tx=XM_011286628.2
D1 58608968 58609165 refGene_gn=MAP6;refGene_tx=XM_011286628.2
D1 58628325 58629316 refGene_gn=MAP6;refGene_tx=XM_011286628.2
D1 58641713 58642016
refGene_gn=SERPINH1;refGene_tx=XM_006936913.4;refGene_gn=SERPINH1;refGene_tx=XM_006936914.4

D1 58644803 58645036
refGene_gn=SERPINH1;refGene_tx=XM_006936913.4;refGene_gn=SERPINH1;refGene_tx=XM_006936914.4
D1 58645142 58645241
refGene_gn=SERPINH1;refGene_tx=XM_006936913.4;refGene_gn=SERPINH1;refGene_tx=XM_006936914.4
D1 58645657 58646279
refGene_gn=SERPINH1;refGene_tx=XM_006936913.4;refGene_gn=SERPINH1;refGene_tx=XM_006936914.4
D1 58727435 58727552
refGene_gn=GDPD5;refGene_tx=XM_003992719.5;refGene_gn=GDPD5;refGene_tx=XM_019811795.1;refGene_gn=GDPD5;refGene_tx=XM_019811796.1;refGene_gn=GDPD5;refGene_tx=XM_019811797.1;refGene_gn=GDPD5;refGene_tx=XM_019811798.1;refGene_gn=GDPD5;refGene_tx=XM_019811799.1;refGene_gn=GDPD5;refGene_tx=XM_019811800.1;refGene_gn=GDPD5;refGene_tx=XM_019811801.1
D1 58740366 58740369 refGene_gn=GDPD5;refGene_tx=XM_019811803.2
D1 58743131 58743235
refGene_gn=GDPD5;refGene_tx=XM_003992719.5;refGene_gn=GDPD5;refGene_tx=XM_019811795.1;refGene_gn=GDPD5;refGene_tx=XM_019811796.1;refGene_gn=GDPD5;refGene_tx=XM_019811797.1;refGene_gn=GDPD5;refGene_tx=XM_019811798.1;refGene_gn=GDPD5;refGene_tx=XM_019811799.1;refGene_gn=GDPD5;refGene_tx=XM_019811800.1;refGene_gn=GDPD5;refGene_tx=XM_019811801.1;refGene_gn=GDPD5;refGene_tx=XM_019811803.2
D1 58746327 58746421
refGene_gn=GDPD5;refGene_tx=XM_003992719.5;refGene_gn=GDPD5;refGene_tx=XM_019811795.1;refGene_gn=GDPD5;refGene_tx=XM_019811796.1;refGene_gn=GDPD5;refGene_tx=XM_019811797.1;refGene_gn=GDPD5;refGene_tx=XM_019811798.1;refGene_gn=GDPD5;refGene_tx=XM_019811799.1;refGene_gn=GDPD5;refGene_tx=XM_019811800.1;refGene_gn=GDPD5;refGene_tx=XM_019811801.1;refGene_gn=GDPD5;refGene_tx=XM_019811803.2;refGene_gn=GDPD5;refGene_tx=XM_019811804.2
D1 58747072 58747132
refGene_gn=GDPD5;refGene_tx=XM_003992719.5;refGene_gn=GDPD5;refGene_tx=XM_019811795.1;refGene_gn=GDPD5;refGene_tx=XM_019811796.1;refGene_gn=GDPD5;refGene_tx=XM_019811797.1;refGene_gn=GDPD5;refGene_tx=XM_019811798.1;refGene_gn=GDPD5;refGene_tx=XM_019811799.1;refGene_gn=GDPD5;refGene_tx=XM_019811800.1;refGene_gn=GDPD5;refGene_tx=XM_019811801.1;refGene_gn=GDPD5;refGene_tx=XM_019811803.2;refGene_gn=GDPD5;refGene_tx=XM_019811804.2
D1 58753945 58754044
refGene_gn=GDPD5;refGene_tx=XM_003992719.5;refGene_gn=GDPD5;refGene_tx=XM_019811795.1;refGene_gn=GDPD5;refGene_tx=XM_019811796.1;refGene_gn=GDPD5;refGene_tx=XM_019811797.1;refGene_gn=GDPD5;refGene_tx=XM_019811798.1;refGene_gn=GDPD5;refGene_tx=XM_019811799.1;refGene_gn=GDPD5;refGene_tx=XM_019811800.1;refGene_gn=GDPD5;refGene_tx=XM_019811801.1;refGene_gn=GDPD5;refGene_tx=XM_019811803.2;refGene_gn=GDPD5;refGene_tx=XM_019811804.2
D1 58754332 58754426
refGene_gn=GDPD5;refGene_tx=XM_003992719.5;refGene_gn=GDPD5;refGene_tx=XM_019811795.1;refGene_gn=GDPD5;refGene_tx=XM_019811796.1;refGene_gn=

GDPD5;refGene_tx=XM_019811797.1;refGene_gn=GDPD5;refGene_tx=XM_0198117
98.1;refGene_gn=GDPD5;refGene_tx=XM_019811799.1;refGene_gn=GDPD5;refGe
ne_tx=XM_019811800.1;refGene_gn=GDPD5;refGene_tx=XM_019811801.1;refGen
e_gn=GDPD5;refGene_tx=XM_019811803.2;refGene_gn=GDPD5;refGene_tx=XM_01
9811804.2
D1 58754770 58754916
refGene_gn=GDPD5;refGene_tx=XM_003992719.5;refGene_gn=GDPD5;refGene_tx
=XM_019811795.1;refGene_gn=GDPD5;refGene_tx=XM_019811796.1;refGene_gn=
GDPD5;refGene_tx=XM_019811797.1;refGene_gn=GDPD5;refGene_tx=XM_0198117
98.1;refGene_gn=GDPD5;refGene_tx=XM_019811799.1;refGene_gn=GDPD5;refGe
ne_tx=XM_019811800.1;refGene_gn=GDPD5;refGene_tx=XM_019811801.1;refGen
e_gn=GDPD5;refGene_tx=XM_019811803.2;refGene_gn=GDPD5;refGene_tx=XM_01
9811804.2
D1 58758746 58758829
refGene_gn=GDPD5;refGene_tx=XM_003992719.5;refGene_gn=GDPD5;refGene_tx
=XM_019811795.1;refGene_gn=GDPD5;refGene_tx=XM_019811796.1;refGene_gn=
GDPD5;refGene_tx=XM_019811797.1;refGene_gn=GDPD5;refGene_tx=XM_0198117
98.1;refGene_gn=GDPD5;refGene_tx=XM_019811799.1;refGene_gn=GDPD5;refGe
ne_tx=XM_019811800.1;refGene_gn=GDPD5;refGene_tx=XM_019811801.1;refGen
e_gn=GDPD5;refGene_tx=XM_019811803.2;refGene_gn=GDPD5;refGene_tx=XM_01
9811804.2
D1 58759993 58760144
refGene_gn=GDPD5;refGene_tx=XM_003992719.5;refGene_gn=GDPD5;refGene_tx
=XM_019811795.1;refGene_gn=GDPD5;refGene_tx=XM_019811796.1;refGene_gn=
GDPD5;refGene_tx=XM_019811797.1;refGene_gn=GDPD5;refGene_tx=XM_0198117
98.1;refGene_gn=GDPD5;refGene_tx=XM_019811799.1;refGene_gn=GDPD5;refGe
ne_tx=XM_019811800.1;refGene_gn=GDPD5;refGene_tx=XM_019811801.1;refGen
e_gn=GDPD5;refGene_tx=XM_019811803.2;refGene_gn=GDPD5;refGene_tx=XM_01
9811804.2
D1 58760702 58760921
refGene_gn=GDPD5;refGene_tx=XM_003992719.5;refGene_gn=GDPD5;refGene_tx
=XM_019811795.1;refGene_gn=GDPD5;refGene_tx=XM_019811796.1;refGene_gn=
GDPD5;refGene_tx=XM_019811797.1;refGene_gn=GDPD5;refGene_tx=XM_0198117
98.1;refGene_gn=GDPD5;refGene_tx=XM_019811799.1;refGene_gn=GDPD5;refGe
ne_tx=XM_019811800.1;refGene_gn=GDPD5;refGene_tx=XM_019811801.1;refGen
e_gn=GDPD5;refGene_tx=XM_019811803.2;refGene_gn=GDPD5;refGene_tx=XM_01
9811804.2
D1 58761491 58761649
refGene_gn=GDPD5;refGene_tx=XM_003992719.5;refGene_gn=GDPD5;refGene_tx
=XM_019811795.1;refGene_gn=GDPD5;refGene_tx=XM_019811796.1;refGene_gn=
GDPD5;refGene_tx=XM_019811797.1;refGene_gn=GDPD5;refGene_tx=XM_0198117
98.1;refGene_gn=GDPD5;refGene_tx=XM_019811799.1;refGene_gn=GDPD5;refGe
ne_tx=XM_019811800.1;refGene_gn=GDPD5;refGene_tx=XM_019811801.1;refGen
e_gn=GDPD5;refGene_tx=XM_019811803.2;refGene_gn=GDPD5;refGene_tx=XM_01
9811804.2
D1 58761899 58762047
refGene_gn=GDPD5;refGene_tx=XM_003992719.5;refGene_gn=GDPD5;refGene_tx
=XM_019811795.1;refGene_gn=GDPD5;refGene_tx=XM_019811796.1;refGene_gn=
GDPD5;refGene_tx=XM_019811797.1;refGene_gn=GDPD5;refGene_tx=XM_0198117
98.1;refGene_gn=GDPD5;refGene_tx=XM_019811799.1;refGene_gn=GDPD5;refGe

ne_tx=XM_019811800.1;refGene_gn=GDPD5;refGene_tx=XM_019811801.1;refGene_gn=GDPD5;refGene_tx=XM_019811803.2;refGene_gn=GDPD5;refGene_tx=XM_019811804.2
D1 58763511 58763594
refGene_gn=GDPD5;refGene_tx=XM_003992719.5;refGene_gn=GDPD5;refGene_tx=XM_019811795.1;refGene_gn=GDPD5;refGene_tx=XM_019811796.1;refGene_gn=GDPD5;refGene_tx=XM_019811797.1;refGene_gn=GDPD5;refGene_tx=XM_019811798.1;refGene_gn=GDPD5;refGene_tx=XM_019811799.1;refGene_gn=GDPD5;refGene_tx=XM_019811800.1;refGene_gn=GDPD5;refGene_tx=XM_019811801.1;refGene_gn=GDPD5;refGene_tx=XM_019811803.2;refGene_gn=GDPD5;refGene_tx=XM_019811804.2
D1 58764143 58764158
refGene_gn=GDPD5;refGene_tx=XM_019811795.1;refGene_gn=GDPD5;refGene_tx=XM_019811796.1;refGene_gn=GDPD5;refGene_tx=XM_019811797.1;refGene_gn=GDPD5;refGene_tx=XM_019811798.1;refGene_gn=GDPD5;refGene_tx=XM_019811799.1;refGene_gn=GDPD5;refGene_tx=XM_019811800.1;refGene_gn=GDPD5;refGene_tx=XM_019811803.2;refGene_gn=GDPD5;refGene_tx=XM_019811804.2
D1 58766379 58766492
refGene_gn=GDPD5;refGene_tx=XM_003992719.5;refGene_gn=GDPD5;refGene_tx=XM_019811795.1;refGene_gn=GDPD5;refGene_tx=XM_019811796.1;refGene_gn=GDPD5;refGene_tx=XM_019811797.1;refGene_gn=GDPD5;refGene_tx=XM_019811798.1;refGene_gn=GDPD5;refGene_tx=XM_019811799.1;refGene_gn=GDPD5;refGene_tx=XM_019811800.1;refGene_gn=GDPD5;refGene_tx=XM_019811801.1;refGene_gn=GDPD5;refGene_tx=XM_019811803.2;refGene_gn=GDPD5;refGene_tx=XM_019811804.2
D1 58769179 58769331
refGene_gn=GDPD5;refGene_tx=XM_003992719.5;refGene_gn=GDPD5;refGene_tx=XM_019811795.1;refGene_gn=GDPD5;refGene_tx=XM_019811796.1;refGene_gn=GDPD5;refGene_tx=XM_019811797.1;refGene_gn=GDPD5;refGene_tx=XM_019811798.1;refGene_gn=GDPD5;refGene_tx=XM_019811799.1;refGene_gn=GDPD5;refGene_tx=XM_019811800.1;refGene_gn=GDPD5;refGene_tx=XM_019811801.1;refGene_gn=GDPD5;refGene_tx=XM_019811803.2;refGene_gn=GDPD5;refGene_tx=XM_019811804.2
D1 58772794 58773159 refGene_gn=KLHL35;refGene_tx=XM_023239425.1
D1 58776044 58776926 refGene_gn=KLHL35;refGene_tx=XM_023239425.1
D1 58778108 58778293 refGene_gn=KLHL35;refGene_tx=XM_023239425.1
D1 58779953 58780072 refGene_gn=KLHL35;refGene_tx=XM_023239425.1
D1 58780724 58780913 refGene_gn=KLHL35;refGene_tx=XM_023239425.1
D1 58782335 58782524 refGene_gn=KLHL35;refGene_tx=XM_023239425.1
D1 58783394 58783583 refGene_gn=KLHL35;refGene_tx=XM_023239425.1
D1 58793191 58793385 refGene_gn=RPS3;refGene_tx=XM_003992721.4
D1 58793843 58794031 refGene_gn=RPS3;refGene_tx=XM_003992721.4
D1 58795452 58795547 refGene_gn=RPS3;refGene_tx=XM_003992721.4
D1 58796005 58796099 refGene_gn=RPS3;refGene_tx=XM_003992721.4
D1 58796369 58796500 refGene_gn=RPS3;refGene_tx=XM_003992721.4
D1 58797715 58797745 refGene_gn=RPS3;refGene_tx=XM_003992721.4
D1 58832529 58832549
refGene_gn=ARRB1;refGene_tx=XM_023239426.1;refGene_gn=ARRB1;refGene_tx=XM_023239427.1;refGene_gn=ARRB1;refGene_tx=XM_023239429.1
D1 58862243 58862296 refGene_gn=ARRB1;refGene_tx=XM_023239428.1

D1 58869594 58869599 refGene_gn=ARRB1;refGene_tx=XM_006936922.3
D1 58880319 58880375 refGene_gn=ARRB1;refGene_tx=XM_006936919.4
D1 58883328 58883359
refGene_gn=ARRB1;refGene_tx=XM_006936919.4;refGene_gn=ARRB1;refGene_tx=XM_006936922.3;refGene_gn=ARRB1;refGene_tx=XM_019811810.2;refGene_gn=ARRB1;refGene_tx=XM_023239426.1;refGene_gn=ARRB1;refGene_tx=XM_023239427.1;refGene_gn=ARRB1;refGene_tx=XM_023239428.1;refGene_gn=ARRB1;refGene_tx=XM_023239429.1;refGene_gn=ARRB1;refGene_tx=XM_023239430.1
D1 58884808 58884869
refGene_gn=ARRB1;refGene_tx=XM_006936919.4;refGene_gn=ARRB1;refGene_tx=XM_006936922.3;refGene_gn=ARRB1;refGene_tx=XM_019811810.2;refGene_gn=ARRB1;refGene_tx=XM_023239426.1;refGene_gn=ARRB1;refGene_tx=XM_023239427.1;refGene_gn=ARRB1;refGene_tx=XM_023239428.1;refGene_gn=ARRB1;refGene_tx=XM_023239429.1;refGene_gn=ARRB1;refGene_tx=XM_023239430.1
D1 58886348 58886393
refGene_gn=ARRB1;refGene_tx=XM_006936919.4;refGene_gn=ARRB1;refGene_tx=XM_006936922.3;refGene_gn=ARRB1;refGene_tx=XM_019811810.2;refGene_gn=ARRB1;refGene_tx=XM_023239426.1;refGene_gn=ARRB1;refGene_tx=XM_023239427.1;refGene_gn=ARRB1;refGene_tx=XM_023239428.1;refGene_gn=ARRB1;refGene_tx=XM_023239429.1;refGene_gn=ARRB1;refGene_tx=XM_023239430.1
D1 58887194 58887391
refGene_gn=ARRB1;refGene_tx=XM_006936919.4;refGene_gn=ARRB1;refGene_tx=XM_006936922.3;refGene_gn=ARRB1;refGene_tx=XM_023239426.1;refGene_gn=ARRB1;refGene_tx=XM_023239427.1;refGene_gn=ARRB1;refGene_tx=XM_023239428.1;refGene_gn=ARRB1;refGene_tx=XM_023239429.1
D1 58888669 58888729
refGene_gn=ARRB1;refGene_tx=XM_006936919.4;refGene_gn=ARRB1;refGene_tx=XM_006936922.3;refGene_gn=ARRB1;refGene_tx=XM_019811810.2;refGene_gn=ARRB1;refGene_tx=XM_023239426.1;refGene_gn=ARRB1;refGene_tx=XM_023239427.1;refGene_gn=ARRB1;refGene_tx=XM_023239428.1;refGene_gn=ARRB1;refGene_tx=XM_023239429.1;refGene_gn=ARRB1;refGene_tx=XM_023239430.1
D1 58889576 58889644
refGene_gn=ARRB1;refGene_tx=XM_006936919.4;refGene_gn=ARRB1;refGene_tx=XM_006936922.3;refGene_gn=ARRB1;refGene_tx=XM_019811810.2;refGene_gn=ARRB1;refGene_tx=XM_023239426.1;refGene_gn=ARRB1;refGene_tx=XM_023239427.1;refGene_gn=ARRB1;refGene_tx=XM_023239428.1;refGene_gn=ARRB1;refGene_tx=XM_023239429.1;refGene_gn=ARRB1;refGene_tx=XM_023239430.1
D1 58891897 58892033
refGene_gn=ARRB1;refGene_tx=XM_006936919.4;refGene_gn=ARRB1;refGene_tx=XM_006936922.3;refGene_gn=ARRB1;refGene_tx=XM_019811810.2;refGene_gn=ARRB1;refGene_tx=XM_023239426.1;refGene_gn=ARRB1;refGene_tx=XM_023239427.1;refGene_gn=ARRB1;refGene_tx=XM_023239428.1;refGene_gn=ARRB1;refGene_tx=XM_023239429.1;refGene_gn=ARRB1;refGene_tx=XM_023239430.1
D1 58892892 58892977
refGene_gn=ARRB1;refGene_tx=XM_006936919.4;refGene_gn=ARRB1;refGene_tx=XM_006936922.3;refGene_gn=ARRB1;refGene_tx=XM_019811810.2;refGene_gn=ARRB1;refGene_tx=XM_023239426.1;refGene_gn=ARRB1;refGene_tx=XM_023239427.1;refGene_gn=ARRB1;refGene_tx=XM_023239428.1;refGene_gn=ARRB1;refGene_tx=XM_023239429.1;refGene_gn=ARRB1;refGene_tx=XM_023239430.1
D1 58893416 58893489

refGene_gn=ARRB1;refGene_tx=XM_006936919.4;refGene_gn=ARRB1;refGene_tx=XM_006936922.3;refGene_gn=ARRB1;refGene_tx=XM_019811810.2;refGene_gn=ARRB1;refGene_tx=XM_023239426.1;refGene_gn=ARRB1;refGene_tx=XM_023239427.1;refGene_gn=ARRB1;refGene_tx=XM_023239428.1;refGene_gn=ARRB1;refGene_tx=XM_023239429.1;refGene_gn=ARRB1;refGene_tx=XM_023239430.1
D1 58894551 58894689
refGene_gn=ARRB1;refGene_tx=XM_006936919.4;refGene_gn=ARRB1;refGene_tx=XM_006936922.3;refGene_gn=ARRB1;refGene_tx=XM_019811810.2;refGene_gn=ARRB1;refGene_tx=XM_023239426.1;refGene_gn=ARRB1;refGene_tx=XM_023239427.1;refGene_gn=ARRB1;refGene_tx=XM_023239428.1;refGene_gn=ARRB1;refGene_tx=XM_023239429.1;refGene_gn=ARRB1;refGene_tx=XM_023239430.1
D1 58895871 58895955
refGene_gn=ARRB1;refGene_tx=XM_006936919.4;refGene_gn=ARRB1;refGene_tx=XM_006936922.3;refGene_gn=ARRB1;refGene_tx=XM_019811810.2;refGene_gn=ARRB1;refGene_tx=XM_023239426.1;refGene_gn=ARRB1;refGene_tx=XM_023239427.1;refGene_gn=ARRB1;refGene_tx=XM_023239428.1;refGene_gn=ARRB1;refGene_tx=XM_023239429.1;refGene_gn=ARRB1;refGene_tx=XM_023239430.1
D1 58897195 58897219
refGene_gn=ARRB1;refGene_tx=XM_006936919.4;refGene_gn=ARRB1;refGene_tx=XM_006936922.3;refGene_gn=ARRB1;refGene_tx=XM_019811810.2;refGene_gn=ARRB1;refGene_tx=XM_023239426.1;refGene_gn=ARRB1;refGene_tx=XM_023239428.1;refGene_gn=ARRB1;refGene_tx=XM_023239429.1;refGene_gn=ARRB1;refGene_tx=XM_023239430.1
D1 58899451 58899522
refGene_gn=ARRB1;refGene_tx=XM_006936919.4;refGene_gn=ARRB1;refGene_tx=XM_006936922.3;refGene_gn=ARRB1;refGene_tx=XM_019811810.2;refGene_gn=ARRB1;refGene_tx=XM_023239426.1;refGene_gn=ARRB1;refGene_tx=XM_023239427.1;refGene_gn=ARRB1;refGene_tx=XM_023239428.1;refGene_gn=ARRB1;refGene_tx=XM_023239429.1;refGene_gn=ARRB1;refGene_tx=XM_023239430.1
D1 58900273 58900325
refGene_gn=ARRB1;refGene_tx=XM_006936919.4;refGene_gn=ARRB1;refGene_tx=XM_006936922.3;refGene_gn=ARRB1;refGene_tx=XM_019811810.2;refGene_gn=ARRB1;refGene_tx=XM_023239426.1;refGene_gn=ARRB1;refGene_tx=XM_023239427.1;refGene_gn=ARRB1;refGene_tx=XM_023239428.1;refGene_gn=ARRB1;refGene_tx=XM_023239429.1;refGene_gn=ARRB1;refGene_tx=XM_023239430.1
D1 58900475 58900497 refGene_gn=ARRB1;refGene_tx=XM_023239429.1
D1 58901680 58901792
refGene_gn=ARRB1;refGene_tx=XM_006936919.4;refGene_gn=ARRB1;refGene_tx=XM_006936922.3;refGene_gn=ARRB1;refGene_tx=XM_019811810.2;refGene_gn=ARRB1;refGene_tx=XM_023239426.1;refGene_gn=ARRB1;refGene_tx=XM_023239427.1;refGene_gn=ARRB1;refGene_tx=XM_023239428.1;refGene_gn=ARRB1;refGene_tx=XM_023239430.1
D1 58921898 58923074 refGene_gn=TPBGL;refGene_tx=XM_023239432.1
D1 58955430 58955614
refGene_gn=SLCO2B1;refGene_tx=XM_006936923.4;refGene_gn=SLCO2B1;refGene_tx=XM_006936924.3;refGene_gn=SLCO2B1;refGene_tx=XM_019811812.2;refGene_gn=SLCO2B1;refGene_tx=XM_019811813.2;refGene_gn=SLCO2B1;refGene_tx=XM_023239431.1
D1 58956694 58956815
refGene_gn=SLCO2B1;refGene_tx=XM_006936923.4;refGene_gn=SLCO2B1;refGen

e_tx=XM_006936924.3;refGene_gn=SLCO2B1;refGene_tx=XM_019811812.2;refGene_gn=SLCO2B1;refGene_tx=XM_019811813.2;refGene_gn=SLCO2B1;refGene_tx=XM_023239431.1
D1 58957139 58957204
refGene_gn=SLCO2B1;refGene_tx=XM_006936923.4;refGene_gn=SLCO2B1;refGene_tx=XM_006936924.3;refGene_gn=SLCO2B1;refGene_tx=XM_019811812.2;refGene_gn=SLCO2B1;refGene_tx=XM_019811813.2;refGene_gn=SLCO2B1;refGene_tx=XM_023239431.1
D1 58958645 58958809
refGene_gn=SLCO2B1;refGene_tx=XM_006936923.4;refGene_gn=SLCO2B1;refGene_tx=XM_006936924.3;refGene_gn=SLCO2B1;refGene_tx=XM_019811812.2;refGene_gn=SLCO2B1;refGene_tx=XM_019811813.2;refGene_gn=SLCO2B1;refGene_tx=XM_023239431.1
D1 58963257 58963423
refGene_gn=SLCO2B1;refGene_tx=XM_006936923.4;refGene_gn=SLCO2B1;refGene_tx=XM_006936924.3;refGene_gn=SLCO2B1;refGene_tx=XM_019811812.2;refGene_gn=SLCO2B1;refGene_tx=XM_019811813.2;refGene_gn=SLCO2B1;refGene_tx=XM_023239431.1
D1 58966308 58966666
refGene_gn=SLCO2B1;refGene_tx=XM_006936923.4;refGene_gn=SLCO2B1;refGene_tx=XM_006936924.3;refGene_gn=SLCO2B1;refGene_tx=XM_019811812.2;refGene_gn=SLCO2B1;refGene_tx=XM_019811813.2;refGene_gn=SLCO2B1;refGene_tx=XM_023239431.1
D1 58970348 58970451
refGene_gn=SLCO2B1;refGene_tx=XM_006936923.4;refGene_gn=SLCO2B1;refGene_tx=XM_006936924.3;refGene_gn=SLCO2B1;refGene_tx=XM_019811812.2;refGene_gn=SLCO2B1;refGene_tx=XM_019811813.2;refGene_gn=SLCO2B1;refGene_tx=XM_023239431.1
D1 58979363 58979554
refGene_gn=SLCO2B1;refGene_tx=XM_006936923.4;refGene_gn=SLCO2B1;refGene_tx=XM_006936924.3;refGene_gn=SLCO2B1;refGene_tx=XM_019811812.2;refGene_gn=SLCO2B1;refGene_tx=XM_019811813.2;refGene_gn=SLCO2B1;refGene_tx=XM_023239431.1
D1 58981619 58981718
refGene_gn=SLCO2B1;refGene_tx=XM_006936923.4;refGene_gn=SLCO2B1;refGene_tx=XM_006936924.3;refGene_gn=SLCO2B1;refGene_tx=XM_019811812.2;refGene_gn=SLCO2B1;refGene_tx=XM_019811813.2;refGene_gn=SLCO2B1;refGene_tx=XM_023239431.1
D1 58981951 58982191
refGene_gn=SLCO2B1;refGene_tx=XM_006936923.4;refGene_gn=SLCO2B1;refGene_tx=XM_006936924.3;refGene_gn=SLCO2B1;refGene_tx=XM_019811812.2;refGene_gn=SLCO2B1;refGene_tx=XM_019811813.2;refGene_gn=SLCO2B1;refGene_tx=XM_023239431.1
D1 58984948 58985111
refGene_gn=SLCO2B1;refGene_tx=XM_006936923.4;refGene_gn=SLCO2B1;refGene_tx=XM_019811812.2;refGene_gn=SLCO2B1;refGene_tx=XM_019811813.2;refGene_gn=SLCO2B1;refGene_tx=XM_023239431.1
D1 58986575 58986713
refGene_gn=SLCO2B1;refGene_tx=XM_006936923.4;refGene_gn=SLCO2B1;refGene_tx=XM_019811812.2;refGene_gn=SLCO2B1;refGene_tx=XM_019811813.2;refGe

ne_gn=SLCO2B1;refGene_tx=XM_023239431.1
D1 58987811 58987942
refGene_gn=SLCO2B1;refGene_tx=XM_006936923.4;refGene_gn=SLCO2B1;refGene_tx=XM_019811812.2;refGene_gn=SLCO2B1;refGene_tx=XM_019811813.2;refGene_gn=SLCO2B1;refGene_tx=XM_023239431.1
D1 59000157 59000173 refGene_gn=SLCO2B1;refGene_tx=XM_006936923.4
D1 59036670 59037597
refGene_gn=LOC101092290;refGene_tx=XM_011286678.3
D1 59037599 59037635
refGene_gn=LOC101092290;refGene_tx=XM_011286678.3
D1 59068833 59069913
refGene_gn=NEU3;refGene_tx=XM_006936925.3;refGene_gn=NEU3;refGene_tx=XM_019811820.2;refGene_gn=NEU3;refGene_tx=XM_019811823.2;refGene_gn=NEU3;refGene_tx=XM_019811824.2;refGene_gn=NEU3;refGene_tx=XM_023239433.1;refGene_gn=NEU3;refGene_tx=XM_023239434.1;refGene_gn=NEU3;refGene_tx=XM_023239435.1
D1 59075245 59075457
refGene_gn=NEU3;refGene_tx=XM_006936925.3;refGene_gn=NEU3;refGene_tx=XM_019811820.2;refGene_gn=NEU3;refGene_tx=XM_023239433.1;refGene_gn=NEU3;refGene_tx=XM_023239434.1
D1 59079993 59080112
refGene_gn=NEU3;refGene_tx=XM_006936925.3;refGene_gn=NEU3;refGene_tx=XM_019811820.2;refGene_gn=NEU3;refGene_tx=XM_019811823.2;refGene_gn=NEU3;refGene_tx=XM_019811824.2;refGene_gn=NEU3;refGene_tx=XM_023239433.1
D1 59080479 59080546 refGene_gn=NEU3;refGene_tx=XM_019811824.2
D1 59088324 59088549
refGene_gn=NEU3;refGene_tx=XM_019811820.2;refGene_gn=NEU3;refGene_tx=XM_019811823.2;refGene_gn=NEU3;refGene_tx=XM_023239433.1;refGene_gn=NEU3;refGene_tx=XM_023239434.1;refGene_gn=NEU3;refGene_tx=XM_023239435.1
D1 59098112 59098299 refGene_gn=SPCS2;refGene_tx=XM_003992723.3
D1 59104677 59104812 refGene_gn=SPCS2;refGene_tx=XM_003992723.3
D1 59106174 59106335 refGene_gn=SPCS2;refGene_tx=XM_003992723.3
D1 59107164 59107248 refGene_gn=SPCS2;refGene_tx=XM_003992723.3
D1 59126198 59126312 refGene_gn=SPCS2;refGene_tx=XM_003992723.3
D1 59132531 59132625
refGene_gn=XRRA1;refGene_tx=XM_006936930.4;refGene_gn=XRRA1;refGene_tx=XM_006936933.4;refGene_gn=XRRA1;refGene_tx=XM_006936934.4;refGene_gn=XRRA1;refGene_tx=XM_006936935.4;refGene_gn=XRRA1;refGene_tx=XM_019811816.2;refGene_gn=XRRA1;refGene_tx=XM_023239436.1
D1 59137245 59137430
refGene_gn=XRRA1;refGene_tx=XM_006936930.4;refGene_gn=XRRA1;refGene_tx=XM_006936933.4;refGene_gn=XRRA1;refGene_tx=XM_006936934.4;refGene_gn=XRRA1;refGene_tx=XM_006936935.4;refGene_gn=XRRA1;refGene_tx=XM_019811816.2;refGene_gn=XRRA1;refGene_tx=XM_023239436.1
D1 59140701 59140773
refGene_gn=XRRA1;refGene_tx=XM_006936930.4;refGene_gn=XRRA1;refGene_tx=XM_006936933.4;refGene_gn=XRRA1;refGene_tx=XM_006936934.4;refGene_gn=XRRA1;refGene_tx=XM_006936935.4;refGene_gn=XRRA1;refGene_tx=XM_019811816.2;refGene_gn=XRRA1;refGene_tx=XM_023239436.1
D1 59142135 59142208

refGene_gn=XRRA1;refGene_tx=XM_006936933.4;refGene_gn=XRRA1;refGene_tx=XM_006936934.4;refGene_gn=XRRA1;refGene_tx=XM_006936935.4;refGene_gn=XRRA1;refGene_tx=XM_019811816.2;refGene_gn=XRRA1;refGene_tx=XM_023239436.1
D1 59144951 59145049
refGene_gn=XRRA1;refGene_tx=XM_006936933.4;refGene_gn=XRRA1;refGene_tx=XM_006936934.4;refGene_gn=XRRA1;refGene_tx=XM_006936935.4;refGene_gn=XRRA1;refGene_tx=XM_019811816.2;refGene_gn=XRRA1;refGene_tx=XM_023239436.1
D1 59150229 59150363
refGene_gn=XRRA1;refGene_tx=XM_006936930.4;refGene_gn=XRRA1;refGene_tx=XM_006936933.4;refGene_gn=XRRA1;refGene_tx=XM_006936934.4;refGene_gn=XRRA1;refGene_tx=XM_006936935.4;refGene_gn=XRRA1;refGene_tx=XM_019811816.2;refGene_gn=XRRA1;refGene_tx=XM_023239436.1
D1 59161362 59161491
refGene_gn=XRRA1;refGene_tx=XM_006936930.4;refGene_gn=XRRA1;refGene_tx=XM_006936933.4;refGene_gn=XRRA1;refGene_tx=XM_006936934.4;refGene_gn=XRRA1;refGene_tx=XM_006936935.4;refGene_gn=XRRA1;refGene_tx=XM_019811816.2;refGene_gn=XRRA1;refGene_tx=XM_023239436.1
D1 59162196 59162414
refGene_gn=XRRA1;refGene_tx=XM_006936930.4;refGene_gn=XRRA1;refGene_tx=XM_006936933.4;refGene_gn=XRRA1;refGene_tx=XM_006936934.4;refGene_gn=XRRA1;refGene_tx=XM_006936935.4;refGene_gn=XRRA1;refGene_tx=XM_019811816.2;refGene_gn=XRRA1;refGene_tx=XM_023239436.1
D1 59166086 59166127
refGene_gn=XRRA1;refGene_tx=XM_006936930.4;refGene_gn=XRRA1;refGene_tx=XM_006936933.4;refGene_gn=XRRA1;refGene_tx=XM_006936934.4;refGene_gn=XRRA1;refGene_tx=XM_006936935.4;refGene_gn=XRRA1;refGene_tx=XM_019811816.2;refGene_gn=XRRA1;refGene_tx=XM_023239436.1
D1 59168530 59168656
refGene_gn=XRRA1;refGene_tx=XM_006936930.4;refGene_gn=XRRA1;refGene_tx=XM_006936933.4;refGene_gn=XRRA1;refGene_tx=XM_006936934.4;refGene_gn=XRRA1;refGene_tx=XM_006936935.4;refGene_gn=XRRA1;refGene_tx=XM_019811816.2;refGene_gn=XRRA1;refGene_tx=XM_023239436.1
D1 59179959 59180053
refGene_gn=XRRA1;refGene_tx=XM_006936930.4;refGene_gn=XRRA1;refGene_tx=XM_006936933.4;refGene_gn=XRRA1;refGene_tx=XM_006936934.4;refGene_gn=XRRA1;refGene_tx=XM_006936935.4;refGene_gn=XRRA1;refGene_tx=XM_019811816.2;refGene_gn=XRRA1;refGene_tx=XM_023239436.1
D1 59180763 59180879
refGene_gn=XRRA1;refGene_tx=XM_006936930.4;refGene_gn=XRRA1;refGene_tx=XM_006936933.4;refGene_gn=XRRA1;refGene_tx=XM_006936934.4;refGene_gn=XRRA1;refGene_tx=XM_006936935.4;refGene_gn=XRRA1;refGene_tx=XM_019811816.2;refGene_gn=XRRA1;refGene_tx=XM_023239436.1
D1 59182538 59182886
refGene_gn=XRRA1;refGene_tx=XM_006936930.4;refGene_gn=XRRA1;refGene_tx=XM_006936933.4;refGene_gn=XRRA1;refGene_tx=XM_006936934.4;refGene_gn=XRRA1;refGene_tx=XM_019811816.2;refGene_gn=XRRA1;refGene_tx=XM_023239436.1
D1 59185833 59186032

refGene_gn=XRRA1;refGene_tx=XM_006936930.4;refGene_gn=XRRA1;refGene_tx=XM_006936933.4;refGene_gn=XRRA1;refGene_tx=XM_006936934.4;refGene_gn=XRRA1;refGene_tx=XM_006936935.4;refGene_gn=XRRA1;refGene_tx=XM_019811816.2;refGene_gn=XRRA1;refGene_tx=XM_023239436.1
D1 59186794 59186910
refGene_gn=XRRA1;refGene_tx=XM_006936930.4;refGene_gn=XRRA1;refGene_tx=XM_006936933.4;refGene_gn=XRRA1;refGene_tx=XM_006936934.4;refGene_gn=XRRA1;refGene_tx=XM_006936935.4;refGene_gn=XRRA1;refGene_tx=XM_019811816.2;refGene_gn=XRRA1;refGene_tx=XM_023239436.1
D1 59187123 59187229
refGene_gn=XRRA1;refGene_tx=XM_006936930.4;refGene_gn=XRRA1;refGene_tx=XM_006936934.4;refGene_gn=XRRA1;refGene_tx=XM_006936935.4;refGene_gn=XRRA1;refGene_tx=XM_019811816.2;refGene_gn=XRRA1;refGene_tx=XM_023239436.1
D1 59187621 59187875
refGene_gn=XRRA1;refGene_tx=XM_006936930.4;refGene_gn=XRRA1;refGene_tx=XM_006936933.4;refGene_gn=XRRA1;refGene_tx=XM_006936934.4;refGene_gn=XRRA1;refGene_tx=XM_006936935.4;refGene_gn=XRRA1;refGene_tx=XM_019811816.2;refGene_gn=XRRA1;refGene_tx=XM_023239436.1
D1 59196861 59198043 refGene_gn=RNF169;refGene_tx=XM_023239437.1
D1 59198614 59198714 refGene_gn=RNF169;refGene_tx=XM_023239437.1
D1 59225199 59225318 refGene_gn=RNF169;refGene_tx=XM_023239437.1
D1 59229577 59229724 refGene_gn=RNF169;refGene_tx=XM_023239437.1
D1 59265377 59265451 refGene_gn=RNF169;refGene_tx=XM_023239437.1
D1 59298900 59299402 refGene_gn=RNF169;refGene_tx=XM_023239437.1
D1 59314425 59314507
refGene_gn=CHRDL2;refGene_tx=XM_023239438.1;refGene_gn=CHRDL2;refGene_tx=XM_023239439.1;refGene_gn=CHRDL2;refGene_tx=XM_023239440.1;refGene_gn=CHRDL2;refGene_tx=XM_023239441.1
D1 59326074 59326187
refGene_gn=CHRDL2;refGene_tx=XM_023239438.1;refGene_gn=CHRDL2;refGene_tx=XM_023239439.1;refGene_gn=CHRDL2;refGene_tx=XM_023239440.1;refGene_gn=CHRDL2;refGene_tx=XM_023239441.1
D1 59329992 59330086
refGene_gn=CHRDL2;refGene_tx=XM_023239438.1;refGene_gn=CHRDL2;refGene_tx=XM_023239439.1;refGene_gn=CHRDL2;refGene_tx=XM_023239440.1;refGene_gn=CHRDL2;refGene_tx=XM_023239441.1
D1 59332107 59332250
refGene_gn=CHRDL2;refGene_tx=XM_023239438.1;refGene_gn=CHRDL2;refGene_tx=XM_023239439.1;refGene_gn=CHRDL2;refGene_tx=XM_023239440.1;refGene_gn=CHRDL2;refGene_tx=XM_023239441.1
D1 59334470 59334564
refGene_gn=CHRDL2;refGene_tx=XM_023239438.1;refGene_gn=CHRDL2;refGene_tx=XM_023239439.1;refGene_gn=CHRDL2;refGene_tx=XM_023239440.1;refGene_gn=CHRDL2;refGene_tx=XM_023239441.1
D1 59335768 59335824
refGene_gn=CHRDL2;refGene_tx=XM_023239438.1;refGene_gn=CHRDL2;refGene_tx=XM_023239439.1;refGene_gn=CHRDL2;refGene_tx=XM_023239440.1;refGene_gn=CHRDL2;refGene_tx=XM_023239441.1
D1 59337604 59337773

refGene_gn=CHRDL2;refGene_tx=XM_023239438.1;refGene_gn=CHRDL2;refGene_tx=XM_023239439.1;refGene_gn=CHRDL2;refGene_tx=XM_023239440.1;refGene_gn=CHRDL2;refGene_tx=XM_023239441.1
D1 59338707 59338902
refGene_gn=CHRDL2;refGene_tx=XM_023239438.1;refGene_gn=CHRDL2;refGene_tx=XM_023239439.1
D1 59341129 59341303
refGene_gn=CHRDL2;refGene_tx=XM_023239438.1;refGene_gn=CHRDL2;refGene_tx=XM_023239439.1;refGene_gn=CHRDL2;refGene_tx=XM_023239440.1;refGene_gn=CHRDL2;refGene_tx=XM_023239441.1
D1 59342170 59342225
refGene_gn=CHRDL2;refGene_tx=XM_023239439.1;refGene_gn=CHRDL2;refGene_tx=XM_023239441.1
D1 59345644 59345737
refGene_gn=CHRDL2;refGene_tx=XM_023239438.1;refGene_gn=CHRDL2;refGene_tx=XM_023239439.1;refGene_gn=CHRDL2;refGene_tx=XM_023239440.1;refGene_gn=CHRDL2;refGene_tx=XM_023239441.1
D1 59346463 59346543
refGene_gn=CHRDL2;refGene_tx=XM_023239438.1;refGene_gn=CHRDL2;refGene_tx=XM_023239440.1
D1 59398592 59398795
refGene_gn=POLD3;refGene_tx=XM_006936939.2;refGene_gn=POLD3;refGene_tx=XM_019811826.2
D1 59403323 59403399
refGene_gn=POLD3;refGene_tx=XM_006936939.2;refGene_gn=POLD3;refGene_tx=XM_019811826.2
D1 59404537 59404650
refGene_gn=POLD3;refGene_tx=XM_006936939.2;refGene_gn=POLD3;refGene_tx=XM_019811826.2
D1 59412086 59412193
refGene_gn=POLD3;refGene_tx=XM_006936939.2;refGene_gn=POLD3;refGene_tx=XM_019811826.2
D1 59414795 59414961
refGene_gn=POLD3;refGene_tx=XM_006936939.2;refGene_gn=POLD3;refGene_tx=XM_019811826.2
D1 59419633 59419706
refGene_gn=POLD3;refGene_tx=XM_006936939.2;refGene_gn=POLD3;refGene_tx=XM_019811826.2
D1 59421016 59421272
refGene_gn=POLD3;refGene_tx=XM_006936939.2;refGene_gn=POLD3;refGene_tx=XM_019811826.2
D1 59426985 59427118
refGene_gn=POLD3;refGene_tx=XM_006936939.2;refGene_gn=POLD3;refGene_tx=XM_019811826.2
D1 59428359 59428399
refGene_gn=POLD3;refGene_tx=XM_006936939.2;refGene_gn=POLD3;refGene_tx=XM_019811826.2
D1 59436547 59436650
refGene_gn=POLD3;refGene_tx=XM_006936939.2;refGene_gn=POLD3;refGene_tx=XM_019811826.2

D1 59447081 59447137 refGene_gn=POLD3;refGene_tx=XM_006936939.2
D1 59448435 59448495 refGene_gn=POLD3;refGene_tx=XM_006936939.2
D1 59508155 59508621 refGene_gn=LIPT2;refGene_tx=XM_003992728.4
D1 59509196 59509426 refGene_gn=LIPT2;refGene_tx=XM_003992728.4
D1 59535230 59535530
refGene_gn=KCNE3;refGene_tx=XM_003992729.5;refGene_gn=KCNE3;refGene_tx=XM_006936940.4;refGene_gn=KCNE3;refGene_tx=XM_006936941.4
D1 59582607 59582718 refGene_gn=PGM2L1;refGene_tx=XM_003992730.5
D1 59602546 59602714 refGene_gn=PGM2L1;refGene_tx=XM_003992730.5
D1 59605407 59605514 refGene_gn=PGM2L1;refGene_tx=XM_003992730.5
D1 59606231 59606316 refGene_gn=PGM2L1;refGene_tx=XM_003992730.5
D1 59606643 59606727 refGene_gn=PGM2L1;refGene_tx=XM_003992730.5
D1 59613597 59613791 refGene_gn=PGM2L1;refGene_tx=XM_003992730.5
D1 59614618 59614808 refGene_gn=PGM2L1;refGene_tx=XM_003992730.5
D1 59615307 59615405 refGene_gn=PGM2L1;refGene_tx=XM_003992730.5
D1 59616362 59616543 refGene_gn=PGM2L1;refGene_tx=XM_003992730.5
D1 59619045 59619139 refGene_gn=PGM2L1;refGene_tx=XM_003992730.5
D1 59619895 59620025 refGene_gn=PGM2L1;refGene_tx=XM_003992730.5
D1 59620259 59620449 refGene_gn=PGM2L1;refGene_tx=XM_003992730.5
D1 59623905 59624039 refGene_gn=PGM2L1;refGene_tx=XM_003992730.5
D1 59626184 59626287 refGene_gn=PGM2L1;refGene_tx=XM_003992730.5
D1 59654212 59654412
refGene_gn=P4HA3;refGene_tx=XM_006936977.4;refGene_gn=P4HA3;refGene_tx=XM_023239443.1;refGene_gn=P4HA3;refGene_tx=XM_023239444.1;refGene_gn=P4HA3;refGene_tx=XM_023239445.1
D1 59663441 59663584
refGene_gn=P4HA3;refGene_tx=XM_006936977.4;refGene_gn=P4HA3;refGene_tx=XM_023239443.1;refGene_gn=P4HA3;refGene_tx=XM_023239444.1;refGene_gn=P4HA3;refGene_tx=XM_023239445.1
D1 59665007 59665231
refGene_gn=P4HA3;refGene_tx=XM_006936977.4;refGene_gn=P4HA3;refGene_tx=XM_023239443.1;refGene_gn=P4HA3;refGene_tx=XM_023239444.1;refGene_gn=P4HA3;refGene_tx=XM_023239445.1
D1 59671242 59671392
refGene_gn=P4HA3;refGene_tx=XM_006936977.4;refGene_gn=P4HA3;refGene_tx=XM_023239443.1;refGene_gn=P4HA3;refGene_tx=XM_023239445.1
D1 59675903 59675955
refGene_gn=P4HA3;refGene_tx=XM_006936977.4;refGene_gn=P4HA3;refGene_tx=XM_023239443.1;refGene_gn=P4HA3;refGene_tx=XM_023239444.1;refGene_gn=P4HA3;refGene_tx=XM_023239445.1
D1 59678309 59678473
refGene_gn=P4HA3;refGene_tx=XM_006936977.4;refGene_gn=P4HA3;refGene_tx=XM_023239443.1;refGene_gn=P4HA3;refGene_tx=XM_023239444.1;refGene_gn=P4HA3;refGene_tx=XM_023239445.1
D1 59678714 59678891
refGene_gn=P4HA3;refGene_tx=XM_006936977.4;refGene_gn=P4HA3;refGene_tx=XM_023239443.1;refGene_gn=P4HA3;refGene_tx=XM_023239444.1;refGene_gn=P4HA3;refGene_tx=XM_023239445.1
D1 59684424 59684489
refGene_gn=P4HA3;refGene_tx=XM_006936977.4;refGene_gn=P4HA3;refGene_tx

=XM_023239443.1;refGene_gn=P4HA3;refGene_tx=XM_023239444.1;refGene_gn=P4HA3;refGene_tx=XM_023239445.1
D1 59685805 59685965
refGene_gn=P4HA3;refGene_tx=XM_006936977.4;refGene_gn=P4HA3;refGene_tx=XM_023239443.1;refGene_gn=P4HA3;refGene_tx=XM_023239444.1;refGene_gn=P4HA3;refGene_tx=XM_023239445.1
D1 59689970 59690033
refGene_gn=P4HA3;refGene_tx=XM_006936977.4;refGene_gn=P4HA3;refGene_tx=XM_023239443.1;refGene_gn=P4HA3;refGene_tx=XM_023239444.1;refGene_gn=P4HA3;refGene_tx=XM_023239445.1
D1 59692238 59692280 refGene_gn=P4HA3;refGene_tx=XM_023239445.1
D1 59693559 59693628
refGene_gn=P4HA3;refGene_tx=XM_006936977.4;refGene_gn=P4HA3;refGene_tx=XM_023239444.1
D1 59695021 59695118
refGene_gn=P4HA3;refGene_tx=XM_006936977.4;refGene_gn=P4HA3;refGene_tx=XM_023239443.1;refGene_gn=P4HA3;refGene_tx=XM_023239444.1
D1 59695945 59696016
refGene_gn=P4HA3;refGene_tx=XM_006936977.4;refGene_gn=P4HA3;refGene_tx=XM_023239443.1;refGene_gn=P4HA3;refGene_tx=XM_023239444.1
D1 59706118 59706137
refGene_gn=PPME1;refGene_tx=XM_003992731.5;refGene_gn=PPME1;refGene_tx=XM_006936942.3;refGene_gn=PPME1;refGene_tx=XM_019811830.1
D1 59707447 59707515
refGene_gn=PPME1;refGene_tx=XM_003992731.5;refGene_gn=PPME1;refGene_tx=XM_006936942.3;refGene_gn=PPME1;refGene_tx=XM_019811830.1
D1 59708124 59708189
refGene_gn=PPME1;refGene_tx=XM_003992731.5;refGene_gn=PPME1;refGene_tx=XM_006936942.3;refGene_gn=PPME1;refGene_tx=XM_019811830.1
D1 59712032 59712077
refGene_gn=PPME1;refGene_tx=XM_003992731.5;refGene_gn=PPME1;refGene_tx=XM_006936942.3;refGene_gn=PPME1;refGene_tx=XM_019811830.1
D1 59712966 59713096
refGene_gn=PPME1;refGene_tx=XM_003992731.5;refGene_gn=PPME1;refGene_tx=XM_006936942.3;refGene_gn=PPME1;refGene_tx=XM_019811830.1
D1 59723432 59723556
refGene_gn=PPME1;refGene_tx=XM_003992731.5;refGene_gn=PPME1;refGene_tx=XM_006936942.3;refGene_gn=PPME1;refGene_tx=XM_019811830.1
D1 59726731 59726797
refGene_gn=PPME1;refGene_tx=XM_003992731.5;refGene_gn=PPME1;refGene_tx=XM_006936942.3;refGene_gn=PPME1;refGene_tx=XM_019811830.1
D1 59731629 59731720
refGene_gn=PPME1;refGene_tx=XM_003992731.5;refGene_gn=PPME1;refGene_tx=XM_006936942.3;refGene_gn=PPME1;refGene_tx=XM_019811830.1
D1 59732595 59732750
refGene_gn=PPME1;refGene_tx=XM_003992731.5;refGene_gn=PPME1;refGene_tx=XM_006936942.3;refGene_gn=PPME1;refGene_tx=XM_019811830.1
D1 59738129 59738181
refGene_gn=PPME1;refGene_tx=XM_003992731.5;refGene_gn=PPME1;refGene_tx=XM_006936942.3;refGene_gn=PPME1;refGene_tx=XM_019811830.1

D1	59738645	59738703
refGene_gn=PPME1;refGene_tx=XM_003992731.5;refGene_gn=PPME1;refGene_tx=XM_006936942.3;refGene_gn=PPME1;refGene_tx=XM_019811830.1
D1	59756575	59756668
refGene_gn=PPME1;refGene_tx=XM_003992731.5;refGene_gn=PPME1;refGene_tx=XM_006936942.3;refGene_gn=PPME1;refGene_tx=XM_019811830.1
D1	59757213	59757307
refGene_gn=PPME1;refGene_tx=XM_003992731.5;refGene_gn=PPME1;refGene_tx=XM_006936942.3;refGene_gn=PPME1;refGene_tx=XM_019811830.1
D1	59761478	59761519	refGene_gn=PPME1;refGene_tx=XM_019811830.1
D1	59777311	59777316	refGene_gn=PPME1;refGene_tx=XM_006936942.3
D1	59792772	59792873	refGene_gn=PPME1;refGene_tx=XM_003992731.5
D1	59793494	59793549
refGene_gn=C2CD3;refGene_tx=XM_006936944.4;refGene_gn=C2CD3;refGene_tx=XM_019811829.2
D1	59797693	59797963
refGene_gn=C2CD3;refGene_tx=XM_006936944.4;refGene_gn=C2CD3;refGene_tx=XM_019811829.2
D1	59801947	59802105
refGene_gn=C2CD3;refGene_tx=XM_006936944.4;refGene_gn=C2CD3;refGene_tx=XM_019811829.2
D1	59814113	59814337
refGene_gn=C2CD3;refGene_tx=XM_006936944.4;refGene_gn=C2CD3;refGene_tx=XM_019811829.2
D1	59815129	59815377
refGene_gn=C2CD3;refGene_tx=XM_006936944.4;refGene_gn=C2CD3;refGene_tx=XM_019811829.2
D1	59821868	59822001
refGene_gn=C2CD3;refGene_tx=XM_006936944.4;refGene_gn=C2CD3;refGene_tx=XM_019811829.2
D1	59822420	59822597
refGene_gn=C2CD3;refGene_tx=XM_006936944.4;refGene_gn=C2CD3;refGene_tx=XM_019811829.2
D1	59829922	59830070
refGene_gn=C2CD3;refGene_tx=XM_006936944.4;refGene_gn=C2CD3;refGene_tx=XM_019811829.2
D1	59833989	59834138
refGene_gn=C2CD3;refGene_tx=XM_006936944.4;refGene_gn=C2CD3;refGene_tx=XM_019811829.2
D1	59835169	59835376
refGene_gn=C2CD3;refGene_tx=XM_006936944.4;refGene_gn=C2CD3;refGene_tx=XM_019811829.2
D1	59835999	59836112
refGene_gn=C2CD3;refGene_tx=XM_006936944.4;refGene_gn=C2CD3;refGene_tx=XM_019811829.2
D1	59842151	59842270
refGene_gn=C2CD3;refGene_tx=XM_006936944.4;refGene_gn=C2CD3;refGene_tx=XM_019811829.2
D1	59844458	59844581
refGene_gn=C2CD3;refGene_tx=XM_006936944.4;refGene_gn=C2CD3;refGene_tx

=XM_019811829.2
D1 59848721 59849216
refGene_gn=C2CD3;refGene_tx=XM_006936944.4;refGene_gn=C2CD3;refGene_tx
=XM_019811829.2
D1 59851901 59852053
refGene_gn=C2CD3;refGene_tx=XM_006936944.4;refGene_gn=C2CD3;refGene_tx
=XM_019811829.2
D1 59853844 59854091
refGene_gn=C2CD3;refGene_tx=XM_006936944.4;refGene_gn=C2CD3;refGene_tx
=XM_019811829.2
D1 59855208 59855389
refGene_gn=C2CD3;refGene_tx=XM_006936944.4;refGene_gn=C2CD3;refGene_tx
=XM_019811829.2
D1 59856789 59856973
refGene_gn=C2CD3;refGene_tx=XM_006936944.4;refGene_gn=C2CD3;refGene_tx
=XM_019811829.2
D1 59858610 59858783
refGene_gn=C2CD3;refGene_tx=XM_006936944.4;refGene_gn=C2CD3;refGene_tx
=XM_019811829.2
D1 59860440 59860564
refGene_gn=C2CD3;refGene_tx=XM_006936944.4;refGene_gn=C2CD3;refGene_tx
=XM_019811829.2
D1 59865501 59865770
refGene_gn=C2CD3;refGene_tx=XM_006936944.4;refGene_gn=C2CD3;refGene_tx
=XM_019811829.2
D1 59866147 59866237
refGene_gn=C2CD3;refGene_tx=XM_006936944.4;refGene_gn=C2CD3;refGene_tx
=XM_019811829.2
D1 59875382 59875985
refGene_gn=C2CD3;refGene_tx=XM_006936944.4;refGene_gn=C2CD3;refGene_tx
=XM_019811829.2
D1 59878425 59878764
refGene_gn=C2CD3;refGene_tx=XM_006936944.4;refGene_gn=C2CD3;refGene_tx
=XM_019811829.2
D1 59886947 59887086
refGene_gn=C2CD3;refGene_tx=XM_006936944.4;refGene_gn=C2CD3;refGene_tx
=XM_019811829.2
D1 59890068 59890133
refGene_gn=C2CD3;refGene_tx=XM_006936944.4;refGene_gn=C2CD3;refGene_tx
=XM_019811829.2
D1 59894132 59894338
refGene_gn=C2CD3;refGene_tx=XM_006936944.4;refGene_gn=C2CD3;refGene_tx
=XM_019811829.2
D1 59895667 59895801
refGene_gn=C2CD3;refGene_tx=XM_006936944.4;refGene_gn=C2CD3;refGene_tx
=XM_019811829.2
D1 59905993 59906158
refGene_gn=C2CD3;refGene_tx=XM_006936944.4;refGene_gn=C2CD3;refGene_tx
=XM_019811829.2
D1 59911190 59911408

refGene_gn=C2CD3;refGene_tx=XM_006936944.4;refGene_gn=C2CD3;refGene_tx=XM_019811829.2
D1 59914549 59915468
refGene_gn=C2CD3;refGene_tx=XM_006936944.4;refGene_gn=C2CD3;refGene_tx=XM_019811829.2
D1 59918465 59918577
refGene_gn=C2CD3;refGene_tx=XM_006936944.4;refGene_gn=C2CD3;refGene_tx=XM_019811829.2
D1 59934715 59934856
refGene_gn=C2CD3;refGene_tx=XM_006936944.4;refGene_gn=C2CD3;refGene_tx=XM_019811829.2
D1 59943685 59943811
refGene_gn=UCP3;refGene_tx=XM_003992734.5;refGene_gn=UCP3;refGene_tx=XM_006936948.4
D1 59944549 59944760
refGene_gn=UCP3;refGene_tx=XM_003992734.5;refGene_gn=UCP3;refGene_tx=XM_006936948.4
D1 59945043 59945244
refGene_gn=UCP3;refGene_tx=XM_003992734.5;refGene_gn=UCP3;refGene_tx=XM_006936948.4
D1 59946439 59946541
refGene_gn=UCP3;refGene_tx=XM_003992734.5;refGene_gn=UCP3;refGene_tx=XM_006936948.4
D1 59947094 59947275
refGene_gn=UCP3;refGene_tx=XM_003992734.5;refGene_gn=UCP3;refGene_tx=XM_006936948.4
D1 59949263 59949378
refGene_gn=UCP3;refGene_tx=XM_003992734.5;refGene_gn=UCP3;refGene_tx=XM_006936948.4
D1 59971588 59971714 refGene_gn=UCP2;refGene_tx=XM_003992735.4
D1 59971884 59972095 refGene_gn=UCP2;refGene_tx=XM_003992735.4
D1 59972917 59973112 refGene_gn=UCP2;refGene_tx=XM_003992735.4
D1 59973193 59973295 refGene_gn=UCP2;refGene_tx=XM_003992735.4
D1 59973638 59973819 refGene_gn=UCP2;refGene_tx=XM_003992735.4
D1 59974213 59974328 refGene_gn=UCP2;refGene_tx=XM_003992735.4
D1 59977930 59978084
refGene_gn=DNAJB13;refGene_tx=XM_023239446.1;refGene_gn=DNAJB13;refGene_tx=XM_023239447.1
D1 59978681 59978758
refGene_gn=DNAJB13;refGene_tx=XM_023239446.1;refGene_gn=DNAJB13;refGene_tx=XM_023239447.1
D1 59979197 59979287 refGene_gn=DNAJB13;refGene_tx=XM_019811835.2
D1 59979632 59979746
refGene_gn=DNAJB13;refGene_tx=XM_019811835.2;refGene_gn=DNAJB13;refGene_tx=XM_023239446.1;refGene_gn=DNAJB13;refGene_tx=XM_023239447.1
D1 59980859 59980973
refGene_gn=DNAJB13;refGene_tx=XM_019811835.2;refGene_gn=DNAJB13;refGene_tx=XM_023239446.1;refGene_gn=DNAJB13;refGene_tx=XM_023239447.1
D1 59981981 59982139
refGene_gn=DNAJB13;refGene_tx=XM_019811835.2;refGene_gn=DNAJB13;refGen

e_tx=XM_023239446.1;refGene_gn=DNAJB13;refGene_tx=XM_023239447.1
D1 59986119 59986281
refGene_gn=DNAJB13;refGene_tx=XM_019811835.2;refGene_gn=DNAJB13;refGen
e_tx=XM_023239446.1
D1 59987320 59987424
refGene_gn=DNAJB13;refGene_tx=XM_019811835.2;refGene_gn=DNAJB13;refGen
e_tx=XM_023239446.1;refGene_gn=DNAJB13;refGene_tx=XM_023239447.1
D1 59991779 59991847
refGene_gn=DNAJB13;refGene_tx=XM_019811835.2;refGene_gn=DNAJB13;refGen
e_tx=XM_023239446.1;refGene_gn=DNAJB13;refGene_tx=XM_023239447.1
D1 59996906 59996924 refGene_gn=PAAF1;refGene_tx=XM_019811833.2
D1 60000581 60000659
refGene_gn=PAAF1;refGene_tx=XM_003992737.4;refGene_gn=PAAF1;refGene_tx
=XM_006936950.3
D1 60005707 60005790
refGene_gn=PAAF1;refGene_tx=XM_003992737.4;refGene_gn=PAAF1;refGene_tx
=XM_006936950.3;refGene_gn=PAAF1;refGene_tx=XM_019811833.2
D1 60007749 60007832
refGene_gn=PAAF1;refGene_tx=XM_003992737.4;refGene_gn=PAAF1;refGene_tx
=XM_006936949.4;refGene_gn=PAAF1;refGene_tx=XM_006936950.3;refGene_gn=
PAAF1;refGene_tx=XM_019811833.2
D1 60009141 60009197 refGene_gn=PAAF1;refGene_tx=XM_006936949.4
D1 60009325 60009441
refGene_gn=PAAF1;refGene_tx=XM_003992737.4;refGene_gn=PAAF1;refGene_tx
=XM_006936949.4;refGene_gn=PAAF1;refGene_tx=XM_006936950.3;refGene_gn=
PAAF1;refGene_tx=XM_019811833.2
D1 60011257 60011349
refGene_gn=PAAF1;refGene_tx=XM_003992737.4;refGene_gn=PAAF1;refGene_tx
=XM_006936949.4;refGene_gn=PAAF1;refGene_tx=XM_006936950.3;refGene_gn=
PAAF1;refGene_tx=XM_019811833.2
D1 60013638 60013833
refGene_gn=PAAF1;refGene_tx=XM_003992737.4;refGene_gn=PAAF1;refGene_tx
=XM_006936949.4;refGene_gn=PAAF1;refGene_tx=XM_006936950.3;refGene_gn=
PAAF1;refGene_tx=XM_019811833.2
D1 60015317 60015468
refGene_gn=PAAF1;refGene_tx=XM_003992737.4;refGene_gn=PAAF1;refGene_tx
=XM_006936949.4;refGene_gn=PAAF1;refGene_tx=XM_006936950.3;refGene_gn=
PAAF1;refGene_tx=XM_019811833.2
D1 60016115 60016214
refGene_gn=PAAF1;refGene_tx=XM_003992737.4;refGene_gn=PAAF1;refGene_tx
=XM_006936949.4;refGene_gn=PAAF1;refGene_tx=XM_006936950.3;refGene_gn=
PAAF1;refGene_tx=XM_019811833.2
D1 60016750 60016783 refGene_gn=PAAF1;refGene_tx=XM_006936950.3
D1 60023218 60023308
refGene_gn=PAAF1;refGene_tx=XM_003992737.4;refGene_gn=PAAF1;refGene_tx
=XM_006936949.4;refGene_gn=PAAF1;refGene_tx=XM_019811833.2
D1 60027207 60027311
refGene_gn=PAAF1;refGene_tx=XM_003992737.4;refGene_gn=PAAF1;refGene_tx
=XM_006936949.4;refGene_gn=PAAF1;refGene_tx=XM_019811833.2
D1 60031280 60031321

refGene_gn=PAAF1;refGene_tx=XM_003992737.4;refGene_gn=PAAF1;refGene_tx=XM_006936949.4;refGene_gn=PAAF1;refGene_tx=XM_019811833.2
D1 60034829 60034876
refGene_gn=PAAF1;refGene_tx=XM_003992737.4;refGene_gn=PAAF1;refGene_tx=XM_006936949.4;refGene_gn=PAAF1;refGene_tx=XM_019811833.2
D1 60034983 60035142
refGene_gn=LOC101096761;refGene_tx=XM_003992739.4
D1 60035366 60035441
refGene_gn=LOC101096761;refGene_tx=XM_006936951.4
D1 60037015 60037282
refGene_gn=LOC101096761;refGene_tx=XM_003992739.4;refGene_gn=LOC101096761;refGene_tx=XM_006936951.4;refGene_gn=LOC101096761;refGene_tx=XM_006936952.2
D1 60045819 60045894
refGene_gn=MRPL48;refGene_tx=XM_006936953.3;refGene_gn=MRPL48;refGene_tx=XM_019811836.1;refGene_gn=MRPL48;refGene_tx=XM_019811837.2;refGene_gn=MRPL48;refGene_tx=XM_019811839.2;refGene_gn=MRPL48;refGene_tx=XM_019811840.2;refGene_gn=MRPL48;refGene_tx=XM_019811841.2;refGene_gn=MRPL48;refGene_tx=XM_023239449.1;refGene_gn=MRPL48;refGene_tx=XM_023239450.1
D1 60047061 60047151
refGene_gn=MRPL48;refGene_tx=XM_006936953.3;refGene_gn=MRPL48;refGene_tx=XM_019811836.1;refGene_gn=MRPL48;refGene_tx=XM_019811837.2;refGene_gn=MRPL48;refGene_tx=XM_019811839.2;refGene_gn=MRPL48;refGene_tx=XM_019811840.2;refGene_gn=MRPL48;refGene_tx=XM_019811841.2;refGene_gn=MRPL48;refGene_tx=XM_023239449.1;refGene_gn=MRPL48;refGene_tx=XM_023239450.1
D1 60049251 60049354
refGene_gn=MRPL48;refGene_tx=XM_006936953.3;refGene_gn=MRPL48;refGene_tx=XM_019811836.1;refGene_gn=MRPL48;refGene_tx=XM_019811837.2;refGene_gn=MRPL48;refGene_tx=XM_019811839.2;refGene_gn=MRPL48;refGene_tx=XM_019811840.2;refGene_gn=MRPL48;refGene_tx=XM_019811841.2;refGene_gn=MRPL48;refGene_tx=XM_023239449.1;refGene_gn=MRPL48;refGene_tx=XM_023239450.1
D1 60059301 60059471
refGene_gn=MRPL48;refGene_tx=XM_006936953.3;refGene_gn=MRPL48;refGene_tx=XM_019811836.1;refGene_gn=MRPL48;refGene_tx=XM_019811837.2;refGene_gn=MRPL48;refGene_tx=XM_019811839.2;refGene_gn=MRPL48;refGene_tx=XM_019811840.2;refGene_gn=MRPL48;refGene_tx=XM_019811841.2;refGene_gn=MRPL48;refGene_tx=XM_023239449.1;refGene_gn=MRPL48;refGene_tx=XM_023239450.1
D1 60081315 60081404
refGene_gn=MRPL48;refGene_tx=XM_006936953.3;refGene_gn=MRPL48;refGene_tx=XM_019811836.1;refGene_gn=MRPL48;refGene_tx=XM_019811837.2;refGene_gn=MRPL48;refGene_tx=XM_019811839.2;refGene_gn=MRPL48;refGene_tx=XM_019811840.2;refGene_gn=MRPL48;refGene_tx=XM_019811841.2;refGene_gn=MRPL48;refGene_tx=XM_023239449.1;refGene_gn=MRPL48;refGene_tx=XM_023239450.1
D1 60089817 60089871
refGene_gn=MRPL48;refGene_tx=XM_006936953.3;refGene_gn=MRPL48;refGene_

tx=XM_019811836.1;refGene_gn=MRPL48;refGene_tx=XM_019811839.2;refGene_gn=MRPL48;refGene_tx=XM_023239449.1
D1 60093776 60093814 refGene_gn=MRPL48;refGene_tx=XM_006936953.3;refGene_gn=MRPL48;refGene_tx=XM_019811836.1;refGene_gn=MRPL48;refGene_tx=XM_019811837.2;refGene_gn=MRPL48;refGene_tx=XM_019811839.2;refGene_gn=MRPL48;refGene_tx=XM_019811840.2;refGene_gn=MRPL48;refGene_tx=XM_019811841.2;refGene_gn=MRPL48;refGene_tx=XM_023239449.1;refGene_gn=MRPL48;refGene_tx=XM_023239450.1
D1 60096673 60096726 refGene_gn=MRPL48;refGene_tx=XM_006936953.3;refGene_gn=MRPL48;refGene_tx=XM_019811836.1;refGene_gn=MRPL48;refGene_tx=XM_019811837.2;refGene_gn=MRPL48;refGene_tx=XM_019811839.2;refGene_gn=MRPL48;refGene_tx=XM_019811840.2;refGene_gn=MRPL48;refGene_tx=XM_019811841.2;refGene_gn=MRPL48;refGene_tx=XM_023239449.1;refGene_gn=MRPL48;refGene_tx=XM_023239450.1
D1 60098989 60099082 refGene_gn=MRPL48;refGene_tx=XM_006936953.3;refGene_gn=MRPL48;refGene_tx=XM_019811837.2
D1 60099832 60099898 refGene_gn=MRPL48;refGene_tx=XM_019811836.1;refGene_gn=MRPL48;refGene_tx=XM_019811840.2
D1 60105617 60105629 refGene_gn=MRPL48;refGene_tx=XM_023239449.1;refGene_gn=MRPL48;refGene_tx=XM_023239450.1
D1 60106335 60106356 refGene_gn=MRPL48;refGene_tx=XM_019811839.2;refGene_gn=MRPL48;refGene_tx=XM_019811841.2
D1 60149499 60149569 refGene_gn=RAB6A;refGene_tx=XM_006936957.4;refGene_gn=RAB6A;refGene_tx=XM_019811842.2
D1 60194709 60194768 refGene_gn=RAB6A;refGene_tx=XM_006936957.4;refGene_gn=RAB6A;refGene_tx=XM_019811842.2
D1 60203906 60203960 refGene_gn=RAB6A;refGene_tx=XM_006936957.4;refGene_gn=RAB6A;refGene_tx=XM_019811842.2
D1 60205589 60205695 refGene_gn=RAB6A;refGene_tx=XM_019811842.2
D1 60205761 60205867 refGene_gn=RAB6A;refGene_tx=XM_006936957.4
D1 60207667 60207779 refGene_gn=RAB6A;refGene_tx=XM_006936957.4;refGene_gn=RAB6A;refGene_tx=XM_019811842.2
D1 60215776 60215870 refGene_gn=RAB6A;refGene_tx=XM_006936957.4;refGene_gn=RAB6A;refGene_tx=XM_019811842.2
D1 60232997 60233064 refGene_gn=RAB6A;refGene_tx=XM_006936957.4;refGene_gn=RAB6A;refGene_tx=XM_019811842.2
D1 60234523 60234588 refGene_gn=RAB6A;refGene_tx=XM_006936957.4;refGene_gn=RAB6A;refGene_tx

=XM_019811842.2
D1 60249283 60249420
refGene_gn=PLEKHB1;refGene_tx=XM_003992744.5;refGene_gn=PLEKHB1;refGene_tx=XM_019811845.2;refGene_gn=PLEKHB1;refGene_tx=XM_019811846.2;refGene_gn=PLEKHB1;refGene_tx=XM_019811847.2;refGene_gn=PLEKHB1;refGene_tx=XM_023239451.1
D1 60249991 60250091
refGene_gn=PLEKHB1;refGene_tx=XM_003992744.5;refGene_gn=PLEKHB1;refGene_tx=XM_019811845.2;refGene_gn=PLEKHB1;refGene_tx=XM_019811846.2;refGene_gn=PLEKHB1;refGene_tx=XM_019811847.2;refGene_gn=PLEKHB1;refGene_tx=XM_023239451.1
D1 60254595 60254700
refGene_gn=PLEKHB1;refGene_tx=XM_003992744.5;refGene_gn=PLEKHB1;refGene_tx=XM_019811845.2;refGene_gn=PLEKHB1;refGene_tx=XM_019811846.2
D1 60257305 60257345
refGene_gn=PLEKHB1;refGene_tx=XM_003992744.5;refGene_gn=PLEKHB1;refGene_tx=XM_019811845.2;refGene_gn=PLEKHB1;refGene_tx=XM_019811846.2;refGene_gn=PLEKHB1;refGene_tx=XM_019811847.2;refGene_gn=PLEKHB1;refGene_tx=XM_023239451.1
D1 60258441 60258544
refGene_gn=PLEKHB1;refGene_tx=XM_003992744.5;refGene_gn=PLEKHB1;refGene_tx=XM_019811845.2;refGene_gn=PLEKHB1;refGene_tx=XM_019811846.2;refGene_gn=PLEKHB1;refGene_tx=XM_019811847.2;refGene_gn=PLEKHB1;refGene_tx=XM_023239451.1
D1 60259124 60259277
refGene_gn=PLEKHB1;refGene_tx=XM_003992744.5;refGene_gn=PLEKHB1;refGene_tx=XM_019811845.2;refGene_gn=PLEKHB1;refGene_tx=XM_019811846.2;refGene_gn=PLEKHB1;refGene_tx=XM_019811847.2;refGene_gn=PLEKHB1;refGene_tx=XM_023239451.1
D1 60260418 60260494
refGene_gn=PLEKHB1;refGene_tx=XM_003992744.5;refGene_gn=PLEKHB1;refGene_tx=XM_019811845.2;refGene_gn=PLEKHB1;refGene_tx=XM_019811846.2;refGene_gn=PLEKHB1;refGene_tx=XM_019811847.2;refGene_gn=PLEKHB1;refGene_tx=XM_023239451.1
D1 60263209 60263227
refGene_gn=PLEKHB1;refGene_tx=XM_003992744.5;refGene_gn=PLEKHB1;refGene_tx=XM_019811847.2
D1 60428578 60428648
refGene_gn=FAM168A;refGene_tx=XM_019811850.2;refGene_gn=FAM168A;refGene_tx=XM_019811851.2;refGene_gn=FAM168A;refGene_tx=XM_023239452.1;refGene_gn=FAM168A;refGene_tx=XM_023239453.1;refGene_gn=FAM168A;refGene_tx=XM_023239454.1
D1 60468481 60468562
refGene_gn=FAM168A;refGene_tx=XM_019811850.2;refGene_gn=FAM168A;refGene_tx=XM_019811851.2;refGene_gn=FAM168A;refGene_tx=XM_023239452.1;refGene_gn=FAM168A;refGene_tx=XM_023239453.1;refGene_gn=FAM168A;refGene_tx=XM_023239454.1
D1 60473353 60473380
refGene_gn=FAM168A;refGene_tx=XM_019811850.2;refGene_gn=FAM168A;refGene_tx=XM_019811851.2;refGene_gn=FAM168A;refGene_tx=XM_023239452.1;refGe

ne_gn=FAM168A;refGene_tx=XM_023239454.1
D1 60477486 60477498
refGene_gn=FAM168A;refGene_tx=XM_019811850.2;refGene_gn=FAM168A;refGene_tx=XM_019811851.2
D1 60478355 60478481
refGene_gn=FAM168A;refGene_tx=XM_019811850.2;refGene_gn=FAM168A;refGene_tx=XM_019811851.2;refGene_gn=FAM168A;refGene_tx=XM_023239452.1;refGene_gn=FAM168A;refGene_tx=XM_023239453.1
D1 60485869 60486012
refGene_gn=FAM168A;refGene_tx=XM_019811850.2;refGene_gn=FAM168A;refGene_tx=XM_019811851.2;refGene_gn=FAM168A;refGene_tx=XM_023239452.1;refGene_gn=FAM168A;refGene_tx=XM_023239453.1;refGene_gn=FAM168A;refGene_tx=XM_023239454.1
D1 60487736 60487911
refGene_gn=FAM168A;refGene_tx=XM_019811850.2;refGene_gn=FAM168A;refGene_tx=XM_019811851.2;refGene_gn=FAM168A;refGene_tx=XM_023239452.1;refGene_gn=FAM168A;refGene_tx=XM_023239453.1;refGene_gn=FAM168A;refGene_tx=XM_023239454.1
D1 60489209 60489322
refGene_gn=FAM168A;refGene_tx=XM_019811850.2;refGene_gn=FAM168A;refGene_tx=XM_019811851.2;refGene_gn=FAM168A;refGene_tx=XM_023239452.1;refGene_gn=FAM168A;refGene_tx=XM_023239453.1;refGene_gn=FAM168A;refGene_tx=XM_023239454.1
D1 60501149 60501197
refGene_gn=RELT;refGene_tx=XM_003992748.5;refGene_gn=RELT;refGene_tx=XM_011286616.3;refGene_gn=RELT;refGene_tx=XM_019811843.2
D1 60501338 60501537
refGene_gn=RELT;refGene_tx=XM_003992748.5;refGene_gn=RELT;refGene_tx=XM_011286616.3;refGene_gn=RELT;refGene_tx=XM_019811843.2
D1 60501977 60502235
refGene_gn=RELT;refGene_tx=XM_003992748.5;refGene_gn=RELT;refGene_tx=XM_011286616.3;refGene_gn=RELT;refGene_tx=XM_019811843.2
D1 60502399 60502481
refGene_gn=RELT;refGene_tx=XM_003992748.5;refGene_gn=RELT;refGene_tx=XM_011286616.3;refGene_gn=RELT;refGene_tx=XM_019811843.2
D1 60502702 60502783
refGene_gn=RELT;refGene_tx=XM_003992748.5;refGene_gn=RELT;refGene_tx=XM_011286616.3;refGene_gn=RELT;refGene_tx=XM_019811843.2
D1 60504101 60504359
refGene_gn=RELT;refGene_tx=XM_003992748.5;refGene_gn=RELT;refGene_tx=XM_011286616.3
D1 60504998 60505078
refGene_gn=RELT;refGene_tx=XM_003992748.5;refGene_gn=RELT;refGene_tx=XM_011286616.3;refGene_gn=RELT;refGene_tx=XM_019811843.2
D1 60505278 60505445
refGene_gn=RELT;refGene_tx=XM_003992748.5;refGene_gn=RELT;refGene_tx=XM_011286616.3;refGene_gn=RELT;refGene_tx=XM_019811843.2
D1 60505539 60505614
refGene_gn=RELT;refGene_tx=XM_003992748.5;refGene_gn=RELT;refGene_tx=XM_011286616.3;refGene_gn=RELT;refGene_tx=XM_019811843.2

D1 60506646 60506715
refGene_gn=RELT;refGene_tx=XM_003992748.5;refGene_gn=RELT;refGene_tx=XM_011286616.3;refGene_gn=RELT;refGene_tx=XM_019811843.2
D1 60508677 60508779
refGene_gn=RELT;refGene_tx=XM_003992748.5;refGene_gn=RELT;refGene_tx=XM_019811843.2
D1 60527920 60528117
refGene_gn=ARHGEF17;refGene_tx=XM_003992762.5;refGene_gn=ARHGEF17;refGene_tx=XM_023239455.1
D1 60529020 60529290
refGene_gn=ARHGEF17;refGene_tx=XM_003992762.5;refGene_gn=ARHGEF17;refGene_tx=XM_023239455.1
D1 60529404 60529579
refGene_gn=ARHGEF17;refGene_tx=XM_003992762.5;refGene_gn=ARHGEF17;refGene_tx=XM_023239455.1
D1 60530024 60530173
refGene_gn=ARHGEF17;refGene_tx=XM_003992762.5;refGene_gn=ARHGEF17;refGene_tx=XM_023239455.1
D1 60530474 60530542
refGene_gn=ARHGEF17;refGene_tx=XM_003992762.5;refGene_gn=ARHGEF17;refGene_tx=XM_023239455.1
D1 60530858 60530945
refGene_gn=ARHGEF17;refGene_tx=XM_003992762.5;refGene_gn=ARHGEF17;refGene_tx=XM_023239455.1
D1 60531218 60531468
refGene_gn=ARHGEF17;refGene_tx=XM_003992762.5;refGene_gn=ARHGEF17;refGene_tx=XM_023239455.1
D1 60531994 60532293
refGene_gn=ARHGEF17;refGene_tx=XM_003992762.5;refGene_gn=ARHGEF17;refGene_tx=XM_023239455.1
D1 60532453 60532653
refGene_gn=ARHGEF17;refGene_tx=XM_003992762.5;refGene_gn=ARHGEF17;refGene_tx=XM_023239455.1
D1 60533238 60533312
refGene_gn=ARHGEF17;refGene_tx=XM_003992762.5;refGene_gn=ARHGEF17;refGene_tx=XM_023239455.1
D1 60533838 60534052
refGene_gn=ARHGEF17;refGene_tx=XM_003992762.5;refGene_gn=ARHGEF17;refGene_tx=XM_023239455.1
D1 60534432 60534551
refGene_gn=ARHGEF17;refGene_tx=XM_003992762.5;refGene_gn=ARHGEF17;refGene_tx=XM_023239455.1
D1 60536348 60536434
refGene_gn=ARHGEF17;refGene_tx=XM_003992762.5;refGene_gn=ARHGEF17;refGene_tx=XM_023239455.1
D1 60536539 60536649
refGene_gn=ARHGEF17;refGene_tx=XM_003992762.5;refGene_gn=ARHGEF17;refGene_tx=XM_023239455.1
D1 60536900 60536951
refGene_gn=ARHGEF17;refGene_tx=XM_003992762.5;refGene_gn=ARHGEF17;refG

ene_tx=XM_023239455.1
D1 60537281 60537458
refGene_gn=ARHGEF17;refGene_tx=XM_003992762.5;refGene_gn=ARHGEF17;refGene_tx=XM_023239455.1
D1 60537680 60537773
refGene_gn=ARHGEF17;refGene_tx=XM_003992762.5;refGene_gn=ARHGEF17;refGene_tx=XM_023239455.1
D1 60537999 60538116
refGene_gn=ARHGEF17;refGene_tx=XM_003992762.5;refGene_gn=ARHGEF17;refGene_tx=XM_023239455.1
D1 60540600 60540783
refGene_gn=ARHGEF17;refGene_tx=XM_003992762.5;refGene_gn=ARHGEF17;refGene_tx=XM_023239455.1
D1 60546454 60546532
refGene_gn=ARHGEF17;refGene_tx=XM_003992762.5;refGene_gn=ARHGEF17;refGene_tx=XM_023239455.1
D1 60581274 60584487
refGene_gn=ARHGEF17;refGene_tx=XM_003992762.5;refGene_gn=ARHGEF17;refGene_tx=XM_023239455.1
D1 60594156 60595177
refGene_gn=P2RY6;refGene_tx=XM_006936969.3;refGene_gn=P2RY6;refGene_tx=XM_006936970.3;refGene_gn=P2RY6;refGene_tx=XM_019811857.2;refGene_gn=P2RY6;refGene_tx=XM_019811858.1;refGene_gn=P2RY6;refGene_tx=XM_023239456.1
D1 60596010 60596150 refGene_gn=P2RY6;refGene_tx=XM_023239456.1
D1 60661398 60662532
refGene_gn=P2RY2;refGene_tx=XM_006936967.4;refGene_gn=P2RY2;refGene_tx=XM_019811864.2;refGene_gn=P2RY2;refGene_tx=XM_019811865.2;refGene_gn=P2RY2;refGene_tx=XM_019811866.2;refGene_gn=P2RY2;refGene_tx=XM_019811867.2
D1 60743551 60743566 refGene_gn=FCHSD2;refGene_tx=XM_019811869.1
D1 60754928 60754949
refGene_gn=FCHSD2;refGene_tx=XM_011286612.3;refGene_gn=FCHSD2;refGene_tx=XM_019811868.2
D1 60756671 60756769
refGene_gn=FCHSD2;refGene_tx=XM_011286612.3;refGene_gn=FCHSD2;refGene_tx=XM_019811868.2;refGene_gn=FCHSD2;refGene_tx=XM_019811869.1;refGene_gn=FCHSD2;refGene_tx=XM_019811870.2
D1 60801438 60801484
refGene_gn=FCHSD2;refGene_tx=XM_011286612.3;refGene_gn=FCHSD2;refGene_tx=XM_019811868.2;refGene_gn=FCHSD2;refGene_tx=XM_019811869.1;refGene_gn=FCHSD2;refGene_tx=XM_019811870.2
D1 60891694 60891771
refGene_gn=FCHSD2;refGene_tx=XM_011286612.3;refGene_gn=FCHSD2;refGene_tx=XM_019811868.2;refGene_gn=FCHSD2;refGene_tx=XM_019811869.1;refGene_gn=FCHSD2;refGene_tx=XM_019811870.2
D1 60901616 60901761
refGene_gn=FCHSD2;refGene_tx=XM_011286612.3;refGene_gn=FCHSD2;refGene_tx=XM_019811868.2;refGene_gn=FCHSD2;refGene_tx=XM_019811869.1;refGene_gn=FCHSD2;refGene_tx=XM_019811870.2;refGene_gn=FCHSD2;refGene_tx=XM_01

9811871.2
D1 60913472 60913606
refGene_gn=FCHSD2;refGene_tx=XM_011286612.3;refGene_gn=FCHSD2;refGene_tx=XM_019811868.2;refGene_gn=FCHSD2;refGene_tx=XM_019811869.1;refGene_gn=FCHSD2;refGene_tx=XM_019811870.2;refGene_gn=FCHSD2;refGene_tx=XM_019811871.2
D1 60918043 60918098
refGene_gn=FCHSD2;refGene_tx=XM_011286612.3;refGene_gn=FCHSD2;refGene_tx=XM_019811868.2;refGene_gn=FCHSD2;refGene_tx=XM_019811869.1;refGene_gn=FCHSD2;refGene_tx=XM_019811870.2;refGene_gn=FCHSD2;refGene_tx=XM_019811871.2
D1 60919086 60919215
refGene_gn=FCHSD2;refGene_tx=XM_011286612.3;refGene_gn=FCHSD2;refGene_tx=XM_019811868.2;refGene_gn=FCHSD2;refGene_tx=XM_019811869.1;refGene_gn=FCHSD2;refGene_tx=XM_019811870.2;refGene_gn=FCHSD2;refGene_tx=XM_019811871.2
D1 60962193 60962316
refGene_gn=FCHSD2;refGene_tx=XM_011286612.3;refGene_gn=FCHSD2;refGene_tx=XM_019811868.2;refGene_gn=FCHSD2;refGene_tx=XM_019811869.1;refGene_gn=FCHSD2;refGene_tx=XM_019811870.2;refGene_gn=FCHSD2;refGene_tx=XM_019811871.2
D1 60973609 60973705
refGene_gn=FCHSD2;refGene_tx=XM_011286612.3;refGene_gn=FCHSD2;refGene_tx=XM_019811868.2;refGene_gn=FCHSD2;refGene_tx=XM_019811869.1;refGene_gn=FCHSD2;refGene_tx=XM_019811870.2;refGene_gn=FCHSD2;refGene_tx=XM_019811871.2
D1 60974974 60975046
refGene_gn=FCHSD2;refGene_tx=XM_019811868.2;refGene_gn=FCHSD2;refGene_tx=XM_019811869.1;refGene_gn=FCHSD2;refGene_tx=XM_019811870.2;refGene_gn=FCHSD2;refGene_tx=XM_019811871.2
D1 60988021 60988138
refGene_gn=FCHSD2;refGene_tx=XM_011286612.3;refGene_gn=FCHSD2;refGene_tx=XM_019811868.2;refGene_gn=FCHSD2;refGene_tx=XM_019811869.1;refGene_gn=FCHSD2;refGene_tx=XM_019811870.2;refGene_gn=FCHSD2;refGene_tx=XM_019811871.2
D1 61005278 61005383
refGene_gn=FCHSD2;refGene_tx=XM_011286612.3;refGene_gn=FCHSD2;refGene_tx=XM_019811868.2;refGene_gn=FCHSD2;refGene_tx=XM_019811869.1;refGene_gn=FCHSD2;refGene_tx=XM_019811870.2;refGene_gn=FCHSD2;refGene_tx=XM_019811871.2
D1 61019774 61019936
refGene_gn=FCHSD2;refGene_tx=XM_011286612.3;refGene_gn=FCHSD2;refGene_tx=XM_019811868.2;refGene_gn=FCHSD2;refGene_tx=XM_019811869.1;refGene_gn=FCHSD2;refGene_tx=XM_019811870.2;refGene_gn=FCHSD2;refGene_tx=XM_019811871.2
D1 61032406 61032541
refGene_gn=FCHSD2;refGene_tx=XM_011286612.3;refGene_gn=FCHSD2;refGene_tx=XM_019811868.2;refGene_gn=FCHSD2;refGene_tx=XM_019811869.1;refGene_gn=FCHSD2;refGene_tx=XM_019811870.2;refGene_gn=FCHSD2;refGene_tx=XM_019811871.2

D1 61037830 61037914
refGene_gn=FCHSD2;refGene_tx=XM_011286612.3;refGene_gn=FCHSD2;refGene_tx=XM_019811868.2;refGene_gn=FCHSD2;refGene_tx=XM_019811869.1;refGene_gn=FCHSD2;refGene_tx=XM_019811870.2;refGene_gn=FCHSD2;refGene_tx=XM_019811871.2
D1 61038033 61038211
refGene_gn=FCHSD2;refGene_tx=XM_011286612.3;refGene_gn=FCHSD2;refGene_tx=XM_019811868.2;refGene_gn=FCHSD2;refGene_tx=XM_019811869.1;refGene_gn=FCHSD2;refGene_tx=XM_019811870.2;refGene_gn=FCHSD2;refGene_tx=XM_019811871.2
D1 61038510 61038731
refGene_gn=FCHSD2;refGene_tx=XM_011286612.3;refGene_gn=FCHSD2;refGene_tx=XM_019811868.2;refGene_gn=FCHSD2;refGene_tx=XM_019811869.1;refGene_gn=FCHSD2;refGene_tx=XM_019811870.2;refGene_gn=FCHSD2;refGene_tx=XM_019811871.2
D1 61039665 61039795
refGene_gn=FCHSD2;refGene_tx=XM_011286612.3;refGene_gn=FCHSD2;refGene_tx=XM_019811868.2;refGene_gn=FCHSD2;refGene_tx=XM_019811869.1;refGene_gn=FCHSD2;refGene_tx=XM_019811870.2;refGene_gn=FCHSD2;refGene_tx=XM_019811871.2
D1 61039937 61040020
refGene_gn=FCHSD2;refGene_tx=XM_011286612.3;refGene_gn=FCHSD2;refGene_tx=XM_019811868.2;refGene_gn=FCHSD2;refGene_tx=XM_019811869.1;refGene_gn=FCHSD2;refGene_tx=XM_019811870.2;refGene_gn=FCHSD2;refGene_tx=XM_019811871.2
D1 61042143 61042227
refGene_gn=FCHSD2;refGene_tx=XM_011286612.3;refGene_gn=FCHSD2;refGene_tx=XM_019811868.2;refGene_gn=FCHSD2;refGene_tx=XM_019811869.1;refGene_gn=FCHSD2;refGene_tx=XM_019811870.2;refGene_gn=FCHSD2;refGene_tx=XM_019811871.2
D1 61050301 61050389
refGene_gn=ATG16L2;refGene_tx=XM_003992764.4;refGene_gn=ATG16L2;refGene_tx=XM_006936987.3;refGene_gn=ATG16L2;refGene_tx=XM_011286611.2;refGene_gn=ATG16L2;refGene_tx=XM_019811872.2;refGene_gn=ATG16L2;refGene_tx=XM_019811873.1;refGene_gn=ATG16L2;refGene_tx=XM_019811874.1;refGene_gn=ATG16L2;refGene_tx=XM_019811875.1
D1 61050760 61050862
refGene_gn=ATG16L2;refGene_tx=XM_003992764.4;refGene_gn=ATG16L2;refGene_tx=XM_006936987.3;refGene_gn=ATG16L2;refGene_tx=XM_011286611.2;refGene_gn=ATG16L2;refGene_tx=XM_019811872.2;refGene_gn=ATG16L2;refGene_tx=XM_019811873.1;refGene_gn=ATG16L2;refGene_tx=XM_019811874.1;refGene_gn=ATG16L2;refGene_tx=XM_019811875.1
D1 61050977 61051025
refGene_gn=ATG16L2;refGene_tx=XM_003992764.4;refGene_gn=ATG16L2;refGene_tx=XM_006936987.3;refGene_gn=ATG16L2;refGene_tx=XM_011286611.2;refGene_gn=ATG16L2;refGene_tx=XM_019811872.2;refGene_gn=ATG16L2;refGene_tx=XM_019811873.1;refGene_gn=ATG16L2;refGene_tx=XM_019811874.1;refGene_gn=ATG16L2;refGene_tx=XM_019811875.1
D1 61051261 61051411
refGene_gn=ATG16L2;refGene_tx=XM_003992764.4;refGene_gn=ATG16L2;refGen

e_tx=XM_006936987.3;refGene_gn=ATG16L2;refGene_tx=XM_011286611.2;refGene_gn=ATG16L2;refGene_tx=XM_019811872.2;refGene_gn=ATG16L2;refGene_tx=XM_019811873.1;refGene_gn=ATG16L2;refGene_tx=XM_019811874.1;refGene_gn=ATG16L2;refGene_tx=XM_019811875.1
D1 61052069 61052175
refGene_gn=ATG16L2;refGene_tx=XM_003992764.4;refGene_gn=ATG16L2;refGene_tx=XM_006936987.3;refGene_gn=ATG16L2;refGene_tx=XM_011286611.2;refGene_gn=ATG16L2;refGene_tx=XM_019811872.2;refGene_gn=ATG16L2;refGene_tx=XM_019811873.1;refGene_gn=ATG16L2;refGene_tx=XM_019811874.1
D1 61052481 61052602
refGene_gn=ATG16L2;refGene_tx=XM_003992764.4;refGene_gn=ATG16L2;refGene_tx=XM_006936987.3;refGene_gn=ATG16L2;refGene_tx=XM_011286611.2;refGene_gn=ATG16L2;refGene_tx=XM_019811872.2;refGene_gn=ATG16L2;refGene_tx=XM_019811873.1;refGene_gn=ATG16L2;refGene_tx=XM_019811874.1
D1 61052716 61052788
refGene_gn=ATG16L2;refGene_tx=XM_003992764.4;refGene_gn=ATG16L2;refGene_tx=XM_006936987.3;refGene_gn=ATG16L2;refGene_tx=XM_011286611.2;refGene_gn=ATG16L2;refGene_tx=XM_019811872.2;refGene_gn=ATG16L2;refGene_tx=XM_019811873.1;refGene_gn=ATG16L2;refGene_tx=XM_019811874.1;refGene_gn=ATG16L2;refGene_tx=XM_019811875.1
D1 61053066 61053137
refGene_gn=ATG16L2;refGene_tx=XM_003992764.4;refGene_gn=ATG16L2;refGene_tx=XM_006936987.3;refGene_gn=ATG16L2;refGene_tx=XM_011286611.2;refGene_gn=ATG16L2;refGene_tx=XM_019811872.2;refGene_gn=ATG16L2;refGene_tx=XM_019811873.1;refGene_gn=ATG16L2;refGene_tx=XM_019811874.1;refGene_gn=ATG16L2;refGene_tx=XM_019811875.1
D1 61053791 61053897
refGene_gn=ATG16L2;refGene_tx=XM_003992764.4;refGene_gn=ATG16L2;refGene_tx=XM_006936987.3;refGene_gn=ATG16L2;refGene_tx=XM_011286611.2;refGene_gn=ATG16L2;refGene_tx=XM_019811872.2;refGene_gn=ATG16L2;refGene_tx=XM_019811873.1;refGene_gn=ATG16L2;refGene_tx=XM_019811874.1
D1 61054242 61054351
refGene_gn=ATG16L2;refGene_tx=XM_003992764.4;refGene_gn=ATG16L2;refGene_tx=XM_006936987.3;refGene_gn=ATG16L2;refGene_tx=XM_011286611.2;refGene_gn=ATG16L2;refGene_tx=XM_019811872.2;refGene_gn=ATG16L2;refGene_tx=XM_019811873.1;refGene_gn=ATG16L2;refGene_tx=XM_019811874.1;refGene_gn=ATG16L2;refGene_tx=XM_019811875.1
D1 61054931 61054994
refGene_gn=ATG16L2;refGene_tx=XM_003992764.4;refGene_gn=ATG16L2;refGene_tx=XM_006936987.3;refGene_gn=ATG16L2;refGene_tx=XM_011286611.2;refGene_gn=ATG16L2;refGene_tx=XM_019811872.2;refGene_gn=ATG16L2;refGene_tx=XM_019811874.1;refGene_gn=ATG16L2;refGene_tx=XM_019811875.1
D1 61056319 61056433
refGene_gn=ATG16L2;refGene_tx=XM_003992764.4;refGene_gn=ATG16L2;refGene_tx=XM_006936987.3;refGene_gn=ATG16L2;refGene_tx=XM_011286611.2;refGene_gn=ATG16L2;refGene_tx=XM_019811872.2;refGene_gn=ATG16L2;refGene_tx=XM_019811873.1;refGene_gn=ATG16L2;refGene_tx=XM_019811874.1;refGene_gn=ATG16L2;refGene_tx=XM_019811875.1
D1 61056760 61056826
refGene_gn=ATG16L2;refGene_tx=XM_003992764.4;refGene_gn=ATG16L2;refGen

e_tx=XM_006936987.3;refGene_gn=ATG16L2;refGene_tx=XM_011286611.2;refGene_gn=ATG16L2;refGene_tx=XM_019811872.2;refGene_gn=ATG16L2;refGene_tx=XM_019811873.1;refGene_gn=ATG16L2;refGene_tx=XM_019811874.1;refGene_gn=ATG16L2;refGene_tx=XM_019811875.1
D1 61056963 61057215
refGene_gn=ATG16L2;refGene_tx=XM_003992764.4;refGene_gn=ATG16L2;refGene_tx=XM_006936987.3;refGene_gn=ATG16L2;refGene_tx=XM_011286611.2;refGene_gn=ATG16L2;refGene_tx=XM_019811872.2;refGene_gn=ATG16L2;refGene_tx=XM_019811873.1;refGene_gn=ATG16L2;refGene_tx=XM_019811874.1;refGene_gn=ATG16L2;refGene_tx=XM_019811875.1
D1 61057495 61057569
refGene_gn=ATG16L2;refGene_tx=XM_003992764.4;refGene_gn=ATG16L2;refGene_tx=XM_006936987.3;refGene_gn=ATG16L2;refGene_tx=XM_011286611.2;refGene_gn=ATG16L2;refGene_tx=XM_019811872.2;refGene_gn=ATG16L2;refGene_tx=XM_019811873.1;refGene_gn=ATG16L2;refGene_tx=XM_019811874.1;refGene_gn=ATG16L2;refGene_tx=XM_019811875.1
D1 61061325 61061428
refGene_gn=ATG16L2;refGene_tx=XM_003992764.4;refGene_gn=ATG16L2;refGene_tx=XM_006936987.3;refGene_gn=ATG16L2;refGene_tx=XM_011286611.2;refGene_gn=ATG16L2;refGene_tx=XM_019811872.2;refGene_gn=ATG16L2;refGene_tx=XM_019811873.1;refGene_gn=ATG16L2;refGene_tx=XM_019811875.1
D1 61062287 61062387
refGene_gn=ATG16L2;refGene_tx=XM_003992764.4;refGene_gn=ATG16L2;refGene_tx=XM_006936987.3;refGene_gn=ATG16L2;refGene_tx=XM_011286611.2;refGene_gn=ATG16L2;refGene_tx=XM_019811872.2;refGene_gn=ATG16L2;refGene_tx=XM_019811873.1;refGene_gn=ATG16L2;refGene_tx=XM_019811875.1
D1 61064222 61064331
refGene_gn=ATG16L2;refGene_tx=XM_003992764.4;refGene_gn=ATG16L2;refGene_tx=XM_006936987.3;refGene_gn=ATG16L2;refGene_tx=XM_019811873.1;refGene_gn=ATG16L2;refGene_tx=XM_019811875.1
D1 61064547 61064653 refGene_gn=ATG16L2;refGene_tx=XM_019811872.2
D1 61087074 61087281
refGene_gn=STARD10;refGene_tx=XM_011286610.3;refGene_gn=STARD10;refGene_tx=XM_023239459.1
D1 61105237 61105385
refGene_gn=STARD10;refGene_tx=XM_011286610.3;refGene_gn=STARD10;refGene_tx=XM_023239459.1
D1 61105993 61106097
refGene_gn=STARD10;refGene_tx=XM_011286610.3;refGene_gn=STARD10;refGene_tx=XM_023239459.1
D1 61106816 61106934
refGene_gn=STARD10;refGene_tx=XM_011286610.3;refGene_gn=STARD10;refGene_tx=XM_023239459.1
D1 61108757 61108810
refGene_gn=STARD10;refGene_tx=XM_011286610.3;refGene_gn=STARD10;refGene_tx=XM_023239459.1
D1 61109060 61109306
refGene_gn=STARD10;refGene_tx=XM_011286610.3;refGene_gn=STARD10;refGene_tx=XM_023239459.1
D1 61136896 61137405

refGene_gn=ARAP1;refGene_tx=XM_011286606.3;refGene_gn=ARAP1;refGene_tx
=XM_011286608.3;refGene_gn=ARAP1;refGene_tx=XM_011286609.3;refGene_gn=
ARAP1;refGene_tx=XM_019811876.2;refGene_gn=ARAP1;refGene_tx=XM_0198118
78.2;refGene_gn=ARAP1;refGene_tx=XM_019811879.2
D1 61149428 61149598
refGene_gn=ARAP1;refGene_tx=XM_011286606.3;refGene_gn=ARAP1;refGene_tx
=XM_011286608.3;refGene_gn=ARAP1;refGene_tx=XM_011286609.3;refGene_gn=
ARAP1;refGene_tx=XM_019811876.2;refGene_gn=ARAP1;refGene_tx=XM_0198118
78.2;refGene_gn=ARAP1;refGene_tx=XM_019811879.2
D1 61150197 61150265
refGene_gn=ARAP1;refGene_tx=XM_011286606.3;refGene_gn=ARAP1;refGene_tx
=XM_011286608.3;refGene_gn=ARAP1;refGene_tx=XM_011286609.3;refGene_gn=
ARAP1;refGene_tx=XM_019811876.2;refGene_gn=ARAP1;refGene_tx=XM_0198118
78.2;refGene_gn=ARAP1;refGene_tx=XM_019811879.2;refGene_gn=ARAP1;refGe
ne_tx=XM_019811880.1;refGene_gn=ARAP1;refGene_tx=XM_019811881.1
D1 61150867 61151001
refGene_gn=ARAP1;refGene_tx=XM_011286606.3;refGene_gn=ARAP1;refGene_tx
=XM_011286608.3;refGene_gn=ARAP1;refGene_tx=XM_011286609.3;refGene_gn=
ARAP1;refGene_tx=XM_019811876.2;refGene_gn=ARAP1;refGene_tx=XM_0198118
78.2;refGene_gn=ARAP1;refGene_tx=XM_019811879.2;refGene_gn=ARAP1;refGe
ne_tx=XM_019811880.1;refGene_gn=ARAP1;refGene_tx=XM_019811881.1
D1 61151142 61151298
refGene_gn=ARAP1;refGene_tx=XM_011286606.3;refGene_gn=ARAP1;refGene_tx
=XM_011286608.3;refGene_gn=ARAP1;refGene_tx=XM_011286609.3;refGene_gn=
ARAP1;refGene_tx=XM_019811876.2;refGene_gn=ARAP1;refGene_tx=XM_0198118
78.2;refGene_gn=ARAP1;refGene_tx=XM_019811879.2;refGene_gn=ARAP1;refGe
ne_tx=XM_019811880.1;refGene_gn=ARAP1;refGene_tx=XM_019811881.1
D1 61152132 61152202
refGene_gn=ARAP1;refGene_tx=XM_011286606.3;refGene_gn=ARAP1;refGene_tx
=XM_011286608.3;refGene_gn=ARAP1;refGene_tx=XM_011286609.3;refGene_gn=
ARAP1;refGene_tx=XM_019811876.2;refGene_gn=ARAP1;refGene_tx=XM_0198118
78.2;refGene_gn=ARAP1;refGene_tx=XM_019811879.2;refGene_gn=ARAP1;refGe
ne_tx=XM_019811880.1;refGene_gn=ARAP1;refGene_tx=XM_019811881.1
D1 61152498 61152619
refGene_gn=ARAP1;refGene_tx=XM_011286606.3;refGene_gn=ARAP1;refGene_tx
=XM_011286608.3;refGene_gn=ARAP1;refGene_tx=XM_011286609.3;refGene_gn=
ARAP1;refGene_tx=XM_019811876.2;refGene_gn=ARAP1;refGene_tx=XM_0198118
78.2;refGene_gn=ARAP1;refGene_tx=XM_019811879.2;refGene_gn=ARAP1;refGe
ne_tx=XM_019811880.1;refGene_gn=ARAP1;refGene_tx=XM_019811881.1
D1 61153037 61153240
refGene_gn=ARAP1;refGene_tx=XM_011286606.3;refGene_gn=ARAP1;refGene_tx
=XM_011286608.3;refGene_gn=ARAP1;refGene_tx=XM_011286609.3;refGene_gn=
ARAP1;refGene_tx=XM_019811876.2;refGene_gn=ARAP1;refGene_tx=XM_0198118
78.2;refGene_gn=ARAP1;refGene_tx=XM_019811879.2;refGene_gn=ARAP1;refGe
ne_tx=XM_019811880.1;refGene_gn=ARAP1;refGene_tx=XM_019811881.1
D1 61154073 61154180
refGene_gn=ARAP1;refGene_tx=XM_011286606.3;refGene_gn=ARAP1;refGene_tx
=XM_011286608.3;refGene_gn=ARAP1;refGene_tx=XM_011286609.3;refGene_gn=
ARAP1;refGene_tx=XM_019811876.2;refGene_gn=ARAP1;refGene_tx=XM_0198118
78.2;refGene_gn=ARAP1;refGene_tx=XM_019811879.2;refGene_gn=ARAP1;refGe

ne_tx=XM_019811880.1;refGene_gn=ARAP1;refGene_tx=XM_019811881.1
D1 61156394 61156594
refGene_gn=ARAP1;refGene_tx=XM_011286606.3;refGene_gn=ARAP1;refGene_tx=XM_011286608.3;refGene_gn=ARAP1;refGene_tx=XM_011286609.3;refGene_gn=ARAP1;refGene_tx=XM_019811876.2;refGene_gn=ARAP1;refGene_tx=XM_019811878.2;refGene_gn=ARAP1;refGene_tx=XM_019811879.2;refGene_gn=ARAP1;refGene_tx=XM_019811880.1;refGene_gn=ARAP1;refGene_tx=XM_019811881.1
D1 61158658 61158744
refGene_gn=ARAP1;refGene_tx=XM_011286606.3;refGene_gn=ARAP1;refGene_tx=XM_011286608.3;refGene_gn=ARAP1;refGene_tx=XM_011286609.3;refGene_gn=ARAP1;refGene_tx=XM_019811876.2;refGene_gn=ARAP1;refGene_tx=XM_019811878.2;refGene_gn=ARAP1;refGene_tx=XM_019811879.2;refGene_gn=ARAP1;refGene_tx=XM_019811880.1;refGene_gn=ARAP1;refGene_tx=XM_019811881.1
D1 61159934 61160117
refGene_gn=ARAP1;refGene_tx=XM_011286606.3;refGene_gn=ARAP1;refGene_tx=XM_011286608.3;refGene_gn=ARAP1;refGene_tx=XM_011286609.3;refGene_gn=ARAP1;refGene_tx=XM_019811876.2;refGene_gn=ARAP1;refGene_tx=XM_019811878.2;refGene_gn=ARAP1;refGene_tx=XM_019811879.2;refGene_gn=ARAP1;refGene_tx=XM_019811880.1;refGene_gn=ARAP1;refGene_tx=XM_019811881.1
D1 61161212 61161387
refGene_gn=ARAP1;refGene_tx=XM_011286606.3;refGene_gn=ARAP1;refGene_tx=XM_011286608.3;refGene_gn=ARAP1;refGene_tx=XM_011286609.3;refGene_gn=ARAP1;refGene_tx=XM_019811876.2;refGene_gn=ARAP1;refGene_tx=XM_019811878.2;refGene_gn=ARAP1;refGene_tx=XM_019811879.2;refGene_gn=ARAP1;refGene_tx=XM_019811880.1;refGene_gn=ARAP1;refGene_tx=XM_019811881.1
D1 61162559 61162694
refGene_gn=ARAP1;refGene_tx=XM_011286606.3;refGene_gn=ARAP1;refGene_tx=XM_011286608.3;refGene_gn=ARAP1;refGene_tx=XM_011286609.3;refGene_gn=ARAP1;refGene_tx=XM_019811876.2;refGene_gn=ARAP1;refGene_tx=XM_019811878.2;refGene_gn=ARAP1;refGene_tx=XM_019811879.2;refGene_gn=ARAP1;refGene_tx=XM_019811880.1;refGene_gn=ARAP1;refGene_tx=XM_019811881.1
D1 61165005 61165141
refGene_gn=ARAP1;refGene_tx=XM_011286606.3;refGene_gn=ARAP1;refGene_tx=XM_011286608.3;refGene_gn=ARAP1;refGene_tx=XM_011286609.3;refGene_gn=ARAP1;refGene_tx=XM_019811876.2;refGene_gn=ARAP1;refGene_tx=XM_019811878.2;refGene_gn=ARAP1;refGene_tx=XM_019811879.2;refGene_gn=ARAP1;refGene_tx=XM_019811880.1;refGene_gn=ARAP1;refGene_tx=XM_019811881.1
D1 61165412 61165515
refGene_gn=ARAP1;refGene_tx=XM_011286606.3;refGene_gn=ARAP1;refGene_tx=XM_011286608.3;refGene_gn=ARAP1;refGene_tx=XM_011286609.3;refGene_gn=ARAP1;refGene_tx=XM_019811876.2;refGene_gn=ARAP1;refGene_tx=XM_019811878.2;refGene_gn=ARAP1;refGene_tx=XM_019811879.2;refGene_gn=ARAP1;refGene_tx=XM_019811880.1;refGene_gn=ARAP1;refGene_tx=XM_019811881.1
D1 61166058 61166254
refGene_gn=ARAP1;refGene_tx=XM_011286606.3;refGene_gn=ARAP1;refGene_tx=XM_011286608.3;refGene_gn=ARAP1;refGene_tx=XM_011286609.3;refGene_gn=ARAP1;refGene_tx=XM_019811876.2;refGene_gn=ARAP1;refGene_tx=XM_019811878.2;refGene_gn=ARAP1;refGene_tx=XM_019811879.2;refGene_gn=ARAP1;refGene_tx=XM_019811880.1;refGene_gn=ARAP1;refGene_tx=XM_019811881.1
D1 61166525 61166577

refGene_gn=ARAP1;refGene_tx=XM_011286606.3;refGene_gn=ARAP1;refGene_tx=XM_011286608.3;refGene_gn=ARAP1;refGene_tx=XM_011286609.3;refGene_gn=ARAP1;refGene_tx=XM_019811876.2;refGene_gn=ARAP1;refGene_tx=XM_019811878.2;refGene_gn=ARAP1;refGene_tx=XM_019811879.2;refGene_gn=ARAP1;refGene_tx=XM_019811880.1;refGene_gn=ARAP1;refGene_tx=XM_019811881.1
D1 61166689 61166853
refGene_gn=ARAP1;refGene_tx=XM_011286606.3;refGene_gn=ARAP1;refGene_tx=XM_011286608.3;refGene_gn=ARAP1;refGene_tx=XM_011286609.3;refGene_gn=ARAP1;refGene_tx=XM_019811876.2;refGene_gn=ARAP1;refGene_tx=XM_019811878.2;refGene_gn=ARAP1;refGene_tx=XM_019811879.2;refGene_gn=ARAP1;refGene_tx=XM_019811880.1;refGene_gn=ARAP1;refGene_tx=XM_019811881.1
D1 61167000 61167213
refGene_gn=ARAP1;refGene_tx=XM_011286606.3;refGene_gn=ARAP1;refGene_tx=XM_011286608.3;refGene_gn=ARAP1;refGene_tx=XM_011286609.3;refGene_gn=ARAP1;refGene_tx=XM_019811876.2;refGene_gn=ARAP1;refGene_tx=XM_019811878.2;refGene_gn=ARAP1;refGene_tx=XM_019811879.2;refGene_gn=ARAP1;refGene_tx=XM_019811880.1;refGene_gn=ARAP1;refGene_tx=XM_019811881.1
D1 61167493 61167599
refGene_gn=ARAP1;refGene_tx=XM_011286606.3;refGene_gn=ARAP1;refGene_tx=XM_011286608.3;refGene_gn=ARAP1;refGene_tx=XM_011286609.3;refGene_gn=ARAP1;refGene_tx=XM_019811876.2;refGene_gn=ARAP1;refGene_tx=XM_019811878.2;refGene_gn=ARAP1;refGene_tx=XM_019811879.2;refGene_gn=ARAP1;refGene_tx=XM_019811880.1;refGene_gn=ARAP1;refGene_tx=XM_019811881.1
D1 61168032 61168180
refGene_gn=ARAP1;refGene_tx=XM_011286606.3;refGene_gn=ARAP1;refGene_tx=XM_011286608.3;refGene_gn=ARAP1;refGene_tx=XM_011286609.3;refGene_gn=ARAP1;refGene_tx=XM_019811876.2;refGene_gn=ARAP1;refGene_tx=XM_019811878.2;refGene_gn=ARAP1;refGene_tx=XM_019811879.2;refGene_gn=ARAP1;refGene_tx=XM_019811880.1;refGene_gn=ARAP1;refGene_tx=XM_019811881.1
D1 61168268 61168355
refGene_gn=ARAP1;refGene_tx=XM_011286606.3;refGene_gn=ARAP1;refGene_tx=XM_011286608.3;refGene_gn=ARAP1;refGene_tx=XM_011286609.3;refGene_gn=ARAP1;refGene_tx=XM_019811876.2;refGene_gn=ARAP1;refGene_tx=XM_019811878.2;refGene_gn=ARAP1;refGene_tx=XM_019811879.2;refGene_gn=ARAP1;refGene_tx=XM_019811880.1;refGene_gn=ARAP1;refGene_tx=XM_019811881.1
D1 61168442 61168511
refGene_gn=ARAP1;refGene_tx=XM_011286606.3;refGene_gn=ARAP1;refGene_tx=XM_011286608.3;refGene_gn=ARAP1;refGene_tx=XM_011286609.3;refGene_gn=ARAP1;refGene_tx=XM_019811876.2;refGene_gn=ARAP1;refGene_tx=XM_019811878.2;refGene_gn=ARAP1;refGene_tx=XM_019811879.2;refGene_gn=ARAP1;refGene_tx=XM_019811880.1;refGene_gn=ARAP1;refGene_tx=XM_019811881.1
D1 61168816 61168934
refGene_gn=ARAP1;refGene_tx=XM_011286606.3;refGene_gn=ARAP1;refGene_tx=XM_011286608.3;refGene_gn=ARAP1;refGene_tx=XM_011286609.3;refGene_gn=ARAP1;refGene_tx=XM_019811876.2;refGene_gn=ARAP1;refGene_tx=XM_019811878.2;refGene_gn=ARAP1;refGene_tx=XM_019811879.2;refGene_gn=ARAP1;refGene_tx=XM_019811880.1;refGene_gn=ARAP1;refGene_tx=XM_019811881.1
D1 61170279 61170393
refGene_gn=ARAP1;refGene_tx=XM_011286606.3;refGene_gn=ARAP1;refGene_tx=XM_011286608.3;refGene_gn=ARAP1;refGene_tx=XM_011286609.3;refGene_gn=

ARAP1;refGene_tx=XM_019811876.2;refGene_gn=ARAP1;refGene_tx=XM_019811878.2;refGene_gn=ARAP1;refGene_tx=XM_019811879.2;refGene_gn=ARAP1;refGene_tx=XM_019811880.1;refGene_gn=ARAP1;refGene_tx=XM_019811881.1
D1 61170613 61170759
refGene_gn=ARAP1;refGene_tx=XM_011286606.3;refGene_gn=ARAP1;refGene_tx=XM_011286608.3;refGene_gn=ARAP1;refGene_tx=XM_011286609.3;refGene_gn=ARAP1;refGene_tx=XM_019811876.2;refGene_gn=ARAP1;refGene_tx=XM_019811878.2;refGene_gn=ARAP1;refGene_tx=XM_019811879.2;refGene_gn=ARAP1;refGene_tx=XM_019811880.1;refGene_gn=ARAP1;refGene_tx=XM_019811881.1
D1 61171285 61171318
refGene_gn=ARAP1;refGene_tx=XM_019811876.2;refGene_gn=ARAP1;refGene_tx=XM_019811878.2;refGene_gn=ARAP1;refGene_tx=XM_019811879.2;refGene_gn=ARAP1;refGene_tx=XM_019811880.1;refGene_gn=ARAP1;refGene_tx=XM_019811881.1
D1 61174861 61174944
refGene_gn=ARAP1;refGene_tx=XM_011286606.3;refGene_gn=ARAP1;refGene_tx=XM_011286608.3;refGene_gn=ARAP1;refGene_tx=XM_011286609.3;refGene_gn=ARAP1;refGene_tx=XM_019811876.2;refGene_gn=ARAP1;refGene_tx=XM_019811878.2;refGene_gn=ARAP1;refGene_tx=XM_019811879.2;refGene_gn=ARAP1;refGene_tx=XM_019811880.1;refGene_gn=ARAP1;refGene_tx=XM_019811881.1
D1 61175692 61175743
refGene_gn=ARAP1;refGene_tx=XM_011286606.3;refGene_gn=ARAP1;refGene_tx=XM_011286608.3;refGene_gn=ARAP1;refGene_tx=XM_011286609.3;refGene_gn=ARAP1;refGene_tx=XM_019811876.2;refGene_gn=ARAP1;refGene_tx=XM_019811878.2;refGene_gn=ARAP1;refGene_tx=XM_019811879.2;refGene_gn=ARAP1;refGene_tx=XM_019811880.1;refGene_gn=ARAP1;refGene_tx=XM_019811881.1
D1 61175904 61175968
refGene_gn=ARAP1;refGene_tx=XM_011286606.3;refGene_gn=ARAP1;refGene_tx=XM_011286608.3;refGene_gn=ARAP1;refGene_tx=XM_011286609.3;refGene_gn=ARAP1;refGene_tx=XM_019811876.2;refGene_gn=ARAP1;refGene_tx=XM_019811878.2;refGene_gn=ARAP1;refGene_tx=XM_019811879.2;refGene_gn=ARAP1;refGene_tx=XM_019811880.1;refGene_gn=ARAP1;refGene_tx=XM_019811881.1
D1 61177220 61177370
refGene_gn=ARAP1;refGene_tx=XM_011286606.3;refGene_gn=ARAP1;refGene_tx=XM_011286608.3;refGene_gn=ARAP1;refGene_tx=XM_011286609.3;refGene_gn=ARAP1;refGene_tx=XM_019811876.2;refGene_gn=ARAP1;refGene_tx=XM_019811878.2;refGene_gn=ARAP1;refGene_tx=XM_019811879.2;refGene_gn=ARAP1;refGene_tx=XM_019811880.1;refGene_gn=ARAP1;refGene_tx=XM_019811881.1
D1 61177508 61177553
refGene_gn=ARAP1;refGene_tx=XM_011286606.3;refGene_gn=ARAP1;refGene_tx=XM_019811876.2;refGene_gn=ARAP1;refGene_tx=XM_019811879.2;refGene_gn=ARAP1;refGene_tx=XM_019811880.1;refGene_gn=ARAP1;refGene_tx=XM_019811881.1
D1 61177670 61177688
refGene_gn=ARAP1;refGene_tx=XM_011286608.3;refGene_gn=ARAP1;refGene_tx=XM_011286609.3;refGene_gn=ARAP1;refGene_tx=XM_019811878.2
D1 61189852 61189923 refGene_gn=PDE2A;refGene_tx=XM_023239461.1
D1 61217728 61217801
refGene_gn=PDE2A;refGene_tx=XM_006936990.4;refGene_gn=PDE2A;refGene_tx=XM_023239461.1;refGene_gn=PDE2A;refGene_tx=XM_023239462.1

D1	61229519	61229581
refGene_gn=PDE2A;refGene_tx=XM_006936990.4;refGene_gn=PDE2A;refGene_tx=XM_023239462.1
D1	61251387	61251477
refGene_gn=PDE2A;refGene_tx=XM_006936990.4;refGene_gn=PDE2A;refGene_tx=XM_023239461.1;refGene_gn=PDE2A;refGene_tx=XM_023239462.1
D1	61255430	61255519
refGene_gn=PDE2A;refGene_tx=XM_006936990.4;refGene_gn=PDE2A;refGene_tx=XM_023239461.1;refGene_gn=PDE2A;refGene_tx=XM_023239462.1
D1	61265550	61265660
refGene_gn=PDE2A;refGene_tx=XM_006936990.4;refGene_gn=PDE2A;refGene_tx=XM_023239461.1;refGene_gn=PDE2A;refGene_tx=XM_023239462.1
D1	61266503	61266559
refGene_gn=PDE2A;refGene_tx=XM_006936990.4;refGene_gn=PDE2A;refGene_tx=XM_023239461.1;refGene_gn=PDE2A;refGene_tx=XM_023239462.1
D1	61272087	61272147
refGene_gn=PDE2A;refGene_tx=XM_006936990.4;refGene_gn=PDE2A;refGene_tx=XM_023239461.1;refGene_gn=PDE2A;refGene_tx=XM_023239462.1
D1	61272730	61272884
refGene_gn=PDE2A;refGene_tx=XM_006936990.4;refGene_gn=PDE2A;refGene_tx=XM_023239461.1;refGene_gn=PDE2A;refGene_tx=XM_023239462.1
D1	61273061	61273114
refGene_gn=PDE2A;refGene_tx=XM_006936990.4;refGene_gn=PDE2A;refGene_tx=XM_023239461.1;refGene_gn=PDE2A;refGene_tx=XM_023239462.1
D1	61273308	61273383
refGene_gn=PDE2A;refGene_tx=XM_006936990.4;refGene_gn=PDE2A;refGene_tx=XM_023239461.1;refGene_gn=PDE2A;refGene_tx=XM_023239462.1
D1	61273493	61273535
refGene_gn=PDE2A;refGene_tx=XM_006936990.4;refGene_gn=PDE2A;refGene_tx=XM_023239461.1;refGene_gn=PDE2A;refGene_tx=XM_023239462.1
D1	61274086	61274155
refGene_gn=PDE2A;refGene_tx=XM_006936990.4;refGene_gn=PDE2A;refGene_tx=XM_023239461.1;refGene_gn=PDE2A;refGene_tx=XM_023239462.1
D1	61274397	61274528
refGene_gn=PDE2A;refGene_tx=XM_006936990.4;refGene_gn=PDE2A;refGene_tx=XM_023239461.1;refGene_gn=PDE2A;refGene_tx=XM_023239462.1
D1	61277260	61277372
refGene_gn=PDE2A;refGene_tx=XM_006936990.4;refGene_gn=PDE2A;refGene_tx=XM_023239461.1;refGene_gn=PDE2A;refGene_tx=XM_023239462.1
D1	61277852	61277892
refGene_gn=PDE2A;refGene_tx=XM_006936990.4;refGene_gn=PDE2A;refGene_tx=XM_023239461.1;refGene_gn=PDE2A;refGene_tx=XM_023239462.1
D1	61278027	61278091
refGene_gn=PDE2A;refGene_tx=XM_006936990.4;refGene_gn=PDE2A;refGene_tx=XM_023239461.1;refGene_gn=PDE2A;refGene_tx=XM_023239462.1
D1	61278515	61278588
refGene_gn=PDE2A;refGene_tx=XM_006936990.4;refGene_gn=PDE2A;refGene_tx=XM_023239461.1;refGene_gn=PDE2A;refGene_tx=XM_023239462.1
D1	61278738	61278916
refGene_gn=PDE2A;refGene_tx=XM_006936990.4;refGene_gn=PDE2A;refGene_tx

=XM_023239461.1;refGene_gn=PDE2A;refGene_tx=XM_023239462.1
D1 61279152 61279265
refGene_gn=PDE2A;refGene_tx=XM_006936990.4;refGene_gn=PDE2A;refGene_tx=XM_023239461.1;refGene_gn=PDE2A;refGene_tx=XM_023239462.1
D1 61279936 61280014
refGene_gn=PDE2A;refGene_tx=XM_006936990.4;refGene_gn=PDE2A;refGene_tx=XM_023239461.1;refGene_gn=PDE2A;refGene_tx=XM_023239462.1
D1 61280884 61281007
refGene_gn=PDE2A;refGene_tx=XM_006936990.4;refGene_gn=PDE2A;refGene_tx=XM_023239461.1;refGene_gn=PDE2A;refGene_tx=XM_023239462.1
D1 61281497 61281568
refGene_gn=PDE2A;refGene_tx=XM_006936990.4;refGene_gn=PDE2A;refGene_tx=XM_023239461.1;refGene_gn=PDE2A;refGene_tx=XM_023239462.1
D1 61281848 61281971
refGene_gn=PDE2A;refGene_tx=XM_006936990.4;refGene_gn=PDE2A;refGene_tx=XM_023239461.1;refGene_gn=PDE2A;refGene_tx=XM_023239462.1
D1 61282325 61282413
refGene_gn=PDE2A;refGene_tx=XM_006936990.4;refGene_gn=PDE2A;refGene_tx=XM_023239461.1;refGene_gn=PDE2A;refGene_tx=XM_023239462.1
D1 61282677 61282725
refGene_gn=PDE2A;refGene_tx=XM_006936990.4;refGene_gn=PDE2A;refGene_tx=XM_023239461.1;refGene_gn=PDE2A;refGene_tx=XM_023239462.1
D1 61283767 61283842
refGene_gn=PDE2A;refGene_tx=XM_006936990.4;refGene_gn=PDE2A;refGene_tx=XM_023239461.1;refGene_gn=PDE2A;refGene_tx=XM_023239462.1
D1 61283980 61284080
refGene_gn=PDE2A;refGene_tx=XM_006936990.4;refGene_gn=PDE2A;refGene_tx=XM_023239461.1;refGene_gn=PDE2A;refGene_tx=XM_023239462.1
D1 61284319 61284432
refGene_gn=PDE2A;refGene_tx=XM_006936990.4;refGene_gn=PDE2A;refGene_tx=XM_023239461.1;refGene_gn=PDE2A;refGene_tx=XM_023239462.1
D1 61284830 61284869
refGene_gn=PDE2A;refGene_tx=XM_006936990.4;refGene_gn=PDE2A;refGene_tx=XM_023239461.1;refGene_gn=PDE2A;refGene_tx=XM_023239462.1
D1 61285006 61285113
refGene_gn=PDE2A;refGene_tx=XM_006936990.4;refGene_gn=PDE2A;refGene_tx=XM_023239461.1;refGene_gn=PDE2A;refGene_tx=XM_023239462.1
D1 61285353 61285564
refGene_gn=PDE2A;refGene_tx=XM_006936990.4;refGene_gn=PDE2A;refGene_tx=XM_023239461.1;refGene_gn=PDE2A;refGene_tx=XM_023239462.1
D1 61363869 61364269
refGene_gn=CLPB;refGene_tx=XM_011286604.3;refGene_gn=CLPB;refGene_tx=XM_011286605.3
D1 61367460 61367512
refGene_gn=CLPB;refGene_tx=XM_011286604.3;refGene_gn=CLPB;refGene_tx=XM_011286605.3
D1 61394029 61394116
refGene_gn=CLPB;refGene_tx=XM_011286604.3;refGene_gn=CLPB;refGene_tx=XM_011286605.3
D1 61416331 61416435

refGene_gn=CLPB;refGene_tx=XM_011286604.3;refGene_gn=CLPB;refGene_tx=XM_011286605.3
D1 61422714 61422804 refGene_gn=CLPB;refGene_tx=XM_011286604.3
D1 61437257 61437386
refGene_gn=CLPB;refGene_tx=XM_011286604.3;refGene_gn=CLPB;refGene_tx=XM_011286605.3
D1 61469864 61469962
refGene_gn=CLPB;refGene_tx=XM_011286604.3;refGene_gn=CLPB;refGene_tx=XM_011286605.3
D1 61482604 61482719
refGene_gn=CLPB;refGene_tx=XM_011286604.3;refGene_gn=CLPB;refGene_tx=XM_011286605.3
D1 61492716 61492794
refGene_gn=CLPB;refGene_tx=XM_011286604.3;refGene_gn=CLPB;refGene_tx=XM_011286605.3
D1 61494009 61494065
refGene_gn=CLPB;refGene_tx=XM_011286604.3;refGene_gn=CLPB;refGene_tx=XM_011286605.3
D1 61498873 61498918
refGene_gn=CLPB;refGene_tx=XM_011286604.3;refGene_gn=CLPB;refGene_tx=XM_011286605.3
D1 61499481 61499643
refGene_gn=CLPB;refGene_tx=XM_011286604.3;refGene_gn=CLPB;refGene_tx=XM_011286605.3
D1 61505536 61505693
refGene_gn=CLPB;refGene_tx=XM_011286604.3;refGene_gn=CLPB;refGene_tx=XM_011286605.3
D1 61506160 61506234
refGene_gn=CLPB;refGene_tx=XM_011286604.3;refGene_gn=CLPB;refGene_tx=XM_011286605.3
D1 61506379 61506499
refGene_gn=CLPB;refGene_tx=XM_011286604.3;refGene_gn=CLPB;refGene_tx=XM_011286605.3
D1 61506724 61506829
refGene_gn=CLPB;refGene_tx=XM_011286604.3;refGene_gn=CLPB;refGene_tx=XM_011286605.3
D1 61507237 61507486
refGene_gn=CLPB;refGene_tx=XM_011286604.3;refGene_gn=CLPB;refGene_tx=XM_011286605.3
D1 61542461 61542678 refGene_gn=PHOX2A;refGene_tx=XM_023239464.1
D1 61544849 61544866 refGene_gn=PHOX2A;refGene_tx=XM_023239464.1
D1 61544867 61545038 refGene_gn=PHOX2A;refGene_tx=XM_023239464.1
D1 61545951 61546401 refGene_gn=PHOX2A;refGene_tx=XM_023239464.1
D1 61547714 61547858
refGene_gn=INPPL1;refGene_tx=XM_003992812.5;refGene_gn=INPPL1;refGene_tx=XM_023239463.1
D1 61547962 61548096 refGene_gn=INPPL1;refGene_tx=XM_003992812.5
D1 61548339 61549012
refGene_gn=INPPL1;refGene_tx=XM_003992812.5;refGene_gn=INPPL1;refGene_tx=XM_023239463.1

D1 61550099 61550240
refGene_gn=INPPL1;refGene_tx=XM_003992812.5;refGene_gn=INPPL1;refGene_tx=XM_023239463.1
D1 61550332 61550411
refGene_gn=INPPL1;refGene_tx=XM_003992812.5;refGene_gn=INPPL1;refGene_tx=XM_023239463.1
D1 61550654 61550810
refGene_gn=INPPL1;refGene_tx=XM_003992812.5;refGene_gn=INPPL1;refGene_tx=XM_023239463.1
D1 61550916 61551004
refGene_gn=INPPL1;refGene_tx=XM_003992812.5;refGene_gn=INPPL1;refGene_tx=XM_023239463.1
D1 61551480 61551569
refGene_gn=INPPL1;refGene_tx=XM_003992812.5;refGene_gn=INPPL1;refGene_tx=XM_023239463.1
D1 61551707 61551821
refGene_gn=INPPL1;refGene_tx=XM_003992812.5;refGene_gn=INPPL1;refGene_tx=XM_023239463.1
D1 61552374 61552464
refGene_gn=INPPL1;refGene_tx=XM_003992812.5;refGene_gn=INPPL1;refGene_tx=XM_023239463.1
D1 61552588 61552670
refGene_gn=INPPL1;refGene_tx=XM_003992812.5;refGene_gn=INPPL1;refGene_tx=XM_023239463.1
D1 61552968 61553057
refGene_gn=INPPL1;refGene_tx=XM_003992812.5;refGene_gn=INPPL1;refGene_tx=XM_023239463.1
D1 61553157 61553257
refGene_gn=INPPL1;refGene_tx=XM_003992812.5;refGene_gn=INPPL1;refGene_tx=XM_023239463.1
D1 61553377 61553516
refGene_gn=INPPL1;refGene_tx=XM_003992812.5;refGene_gn=INPPL1;refGene_tx=XM_023239463.1
D1 61553804 61553901
refGene_gn=INPPL1;refGene_tx=XM_003992812.5;refGene_gn=INPPL1;refGene_tx=XM_023239463.1
D1 61554746 61554864
refGene_gn=INPPL1;refGene_tx=XM_003992812.5;refGene_gn=INPPL1;refGene_tx=XM_023239463.1
D1 61555185 61555382
refGene_gn=INPPL1;refGene_tx=XM_003992812.5;refGene_gn=INPPL1;refGene_tx=XM_023239463.1
D1 61555466 61555569
refGene_gn=INPPL1;refGene_tx=XM_003992812.5;refGene_gn=INPPL1;refGene_tx=XM_023239463.1
D1 61555866 61555973
refGene_gn=INPPL1;refGene_tx=XM_003992812.5;refGene_gn=INPPL1;refGene_tx=XM_023239463.1
D1 61556060 61556211
refGene_gn=INPPL1;refGene_tx=XM_003992812.5;refGene_gn=INPPL1;refGene_

tx=XM_023239463.1
D1 61556324 61556423
refGene_gn=INPPL1;refGene_tx=XM_003992812.5;refGene_gn=INPPL1;refGene_tx=XM_023239463.1
D1 61556605 61556695
refGene_gn=INPPL1;refGene_tx=XM_003992812.5;refGene_gn=INPPL1;refGene_tx=XM_023239463.1
D1 61556802 61556896
refGene_gn=INPPL1;refGene_tx=XM_003992812.5;refGene_gn=INPPL1;refGene_tx=XM_023239463.1
D1 61557138 61557279
refGene_gn=INPPL1;refGene_tx=XM_003992812.5;refGene_gn=INPPL1;refGene_tx=XM_023239463.1
D1 61557531 61557652
refGene_gn=INPPL1;refGene_tx=XM_003992812.5;refGene_gn=INPPL1;refGene_tx=XM_023239463.1
D1 61557885 61558036
refGene_gn=INPPL1;refGene_tx=XM_003992812.5;refGene_gn=INPPL1;refGene_tx=XM_023239463.1
D1 61558117 61558181
refGene_gn=INPPL1;refGene_tx=XM_003992812.5;refGene_gn=INPPL1;refGene_tx=XM_023239463.1
D1 61560983 61561180
refGene_gn=INPPL1;refGene_tx=XM_003992812.5;refGene_gn=INPPL1;refGene_tx=XM_023239463.1
D1 61564369 61564398
refGene_gn=LOC101095853;refGene_tx=XM_023239706.1
D1 61564516 61564626
refGene_gn=LOC101095853;refGene_tx=XM_023239706.1
D1 61564627 61564718
refGene_gn=LOC101095853;refGene_tx=XM_023239706.1
D1 61564850 61564967
refGene_gn=LOC101095853;refGene_tx=XM_023239706.1
D1 61564969 61564988
refGene_gn=LOC101095853;refGene_tx=XM_023239706.1
D1 61565113 61565302
refGene_gn=LOC101095853;refGene_tx=XM_023239706.1
D1 61567360 61567513
refGene_gn=LOC101095853;refGene_tx=XM_023239706.1
D1 61591970 61592257
refGene_gn=FOLR1;refGene_tx=XM_003992769.5;refGene_gn=FOLR1;refGene_tx=XM_006936993.4
D1 61592399 61592535
refGene_gn=FOLR1;refGene_tx=XM_003992769.5;refGene_gn=FOLR1;refGene_tx=XM_006936993.4
D1 61592683 61592872
refGene_gn=FOLR1;refGene_tx=XM_003992769.5;refGene_gn=FOLR1;refGene_tx=XM_006936993.4
D1 61594670 61594823
refGene_gn=FOLR1;refGene_tx=XM_003992769.5;refGene_gn=FOLR1;refGene_tx

=XM_006936993.4
D1	61602634	61602742
refGene_gn=LOC101084733;refGene_tx=XM_023239466.1;refGene_gn=LOC101084733;refGene_tx=XM_023239467.1
D1	61602789	61602968
refGene_gn=LOC101084733;refGene_tx=XM_023239466.1;refGene_gn=LOC101084733;refGene_tx=XM_023239467.1
D1	61603104	61603240
refGene_gn=LOC101084733;refGene_tx=XM_023239466.1;refGene_gn=LOC101084733;refGene_tx=XM_023239467.1
D1	61603361	61603550
refGene_gn=LOC101084733;refGene_tx=XM_023239466.1;refGene_gn=LOC101084733;refGene_tx=XM_023239467.1
D1	61606022	61606190
refGene_gn=LOC101084733;refGene_tx=XM_023239466.1;refGene_gn=LOC101084733;refGene_tx=XM_023239467.1
D1	61626164	61626256	refGene_gn=ANAPC15;refGene_tx=XM_006936996.3
D1	61627648	61627778
refGene_gn=ANAPC15;refGene_tx=XM_006936996.3;refGene_gn=ANAPC15;refGene_tx=XM_019811898.1
D1	61628265	61628325
refGene_gn=ANAPC15;refGene_tx=XM_006936996.3;refGene_gn=ANAPC15;refGene_tx=XM_019811898.1
D1	61628649	61628787
refGene_gn=ANAPC15;refGene_tx=XM_006936996.3;refGene_gn=ANAPC15;refGene_tx=XM_019811898.1
D1	61628938	61628986
refGene_gn=ANAPC15;refGene_tx=XM_006936996.3;refGene_gn=ANAPC15;refGene_tx=XM_019811898.1
D1	61629920	61630241
refGene_gn=TOMT;refGene_tx=NM_001282066.1;refGene_gn=TOMT;refGene_tx=XM_006936986.2;refGene_gn=TOMT;refGene_tx=XM_023238909.1
D1	61630695	61630892
refGene_gn=TOMT;refGene_tx=NM_001282066.1;refGene_gn=TOMT;refGene_tx=XM_006936986.2;refGene_gn=TOMT;refGene_tx=XM_023238909.1
D1	61631985	61632424
refGene_gn=TOMT;refGene_tx=NM_001282066.1;refGene_gn=TOMT;refGene_tx=XM_006936986.2;refGene_gn=TOMT;refGene_tx=XM_023238909.1
D1	61634656	61634698	refGene_gn=LAMTOR1;refGene_tx=XM_003992772.5
D1	61638455	61638601	refGene_gn=LAMTOR1;refGene_tx=XM_003992772.5
D1	61638851	61638929	refGene_gn=LAMTOR1;refGene_tx=XM_003992772.5
D1	61639286	61639413	refGene_gn=LAMTOR1;refGene_tx=XM_003992772.5
D1	61639762	61639855	refGene_gn=LAMTOR1;refGene_tx=XM_003992772.5
D1	61641204	61641346
refGene_gn=LRRC51;refGene_tx=NM_001282069.1;refGene_gn=LRRC51;refGene_tx=XM_006936980.4;refGene_gn=LRRC51;refGene_tx=XM_006936981.2;refGene_gn=LRRC51;refGene_tx=XM_006936983.4;refGene_gn=LRRC51;refGene_tx=XM_006936984.4;refGene_gn=LRRC51;refGene_tx=XM_011286568.3;refGene_gn=LRRC51;refGene_tx=XM_019811167.1
D1	61641613	61641762

refGene_gn=LRRC51;refGene_tx=NM_001282069.1;refGene_gn=LRRC51;refGene_tx=XM_006936980.4;refGene_gn=LRRC51;refGene_tx=XM_006936981.2;refGene_gn=LRRC51;refGene_tx=XM_006936983.4;refGene_gn=LRRC51;refGene_tx=XM_006936984.4;refGene_gn=LRRC51;refGene_tx=XM_011286568.3;refGene_gn=LRRC51;refGene_tx=XM_019811167.1
D1 61642738 61642944
refGene_gn=LRRC51;refGene_tx=NM_001282069.1;refGene_gn=LRRC51;refGene_tx=XM_006936980.4;refGene_gn=LRRC51;refGene_tx=XM_006936981.2;refGene_gn=LRRC51;refGene_tx=XM_006936983.4;refGene_gn=LRRC51;refGene_tx=XM_006936984.4;refGene_gn=LRRC51;refGene_tx=XM_011286568.3;refGene_gn=LRRC51;refGene_tx=XM_019811167.1
D1 61647533 61647615
refGene_gn=LRRC51;refGene_tx=NM_001282069.1;refGene_gn=LRRC51;refGene_tx=XM_006936980.4;refGene_gn=LRRC51;refGene_tx=XM_006936981.2;refGene_gn=LRRC51;refGene_tx=XM_006936983.4;refGene_gn=LRRC51;refGene_tx=XM_006936984.4;refGene_gn=LRRC51;refGene_tx=XM_011286568.3;refGene_gn=LRRC51;refGene_tx=XM_019811167.1
D1 61693373 61693415
refGene_gn=NUMA1;refGene_tx=XM_011286597.3;refGene_gn=NUMA1;refGene_tx=XM_011286598.3;refGene_gn=NUMA1;refGene_tx=XM_011286599.3;refGene_gn=NUMA1;refGene_tx=XM_011286600.3;refGene_gn=NUMA1;refGene_tx=XM_023239468.1;refGene_gn=NUMA1;refGene_tx=XM_023239469.1;refGene_gn=NUMA1;refGene_tx=XM_023239470.1
D1 61700481 61700567
refGene_gn=NUMA1;refGene_tx=XM_011286597.3;refGene_gn=NUMA1;refGene_tx=XM_011286598.3;refGene_gn=NUMA1;refGene_tx=XM_011286599.3;refGene_gn=NUMA1;refGene_tx=XM_011286600.3;refGene_gn=NUMA1;refGene_tx=XM_023239468.1;refGene_gn=NUMA1;refGene_tx=XM_023239469.1;refGene_gn=NUMA1;refGene_tx=XM_023239470.1
D1 61706299 61706379
refGene_gn=NUMA1;refGene_tx=XM_011286597.3;refGene_gn=NUMA1;refGene_tx=XM_011286598.3;refGene_gn=NUMA1;refGene_tx=XM_011286599.3;refGene_gn=NUMA1;refGene_tx=XM_011286600.3;refGene_gn=NUMA1;refGene_tx=XM_023239468.1;refGene_gn=NUMA1;refGene_tx=XM_023239469.1;refGene_gn=NUMA1;refGene_tx=XM_023239470.1
D1 61707479 61707562
refGene_gn=NUMA1;refGene_tx=XM_011286597.3;refGene_gn=NUMA1;refGene_tx=XM_011286598.3;refGene_gn=NUMA1;refGene_tx=XM_011286599.3;refGene_gn=NUMA1;refGene_tx=XM_011286600.3;refGene_gn=NUMA1;refGene_tx=XM_023239468.1;refGene_gn=NUMA1;refGene_tx=XM_023239469.1;refGene_gn=NUMA1;refGene_tx=XM_023239470.1
D1 61708128 61708209
refGene_gn=NUMA1;refGene_tx=XM_011286597.3;refGene_gn=NUMA1;refGene_tx=XM_011286598.3;refGene_gn=NUMA1;refGene_tx=XM_011286599.3;refGene_gn=NUMA1;refGene_tx=XM_011286600.3;refGene_gn=NUMA1;refGene_tx=XM_023239468.1;refGene_gn=NUMA1;refGene_tx=XM_023239469.1;refGene_gn=NUMA1;refGene_tx=XM_023239470.1
D1 61710239 61710327
refGene_gn=NUMA1;refGene_tx=XM_011286597.3;refGene_gn=NUMA1;refGene_tx=XM_011286598.3;refGene_gn=NUMA1;refGene_tx=XM_011286599.3;refGene_gn=

NUMA1;refGene_tx=XM_011286600.3;refGene_gn=NUMA1;refGene_tx=XM_0232394
68.1;refGene_gn=NUMA1;refGene_tx=XM_023239469.1;refGene_gn=NUMA1;refGe
ne_tx=XM_023239470.1
D1 61711868 61711989
refGene_gn=NUMA1;refGene_tx=XM_011286597.3;refGene_gn=NUMA1;refGene_tx
=XM_011286598.3;refGene_gn=NUMA1;refGene_tx=XM_011286599.3;refGene_gn=
NUMA1;refGene_tx=XM_011286600.3;refGene_gn=NUMA1;refGene_tx=XM_0232394
68.1;refGene_gn=NUMA1;refGene_tx=XM_023239469.1;refGene_gn=NUMA1;refGe
ne_tx=XM_023239470.1
D1 61712508 61712666
refGene_gn=NUMA1;refGene_tx=XM_011286597.3;refGene_gn=NUMA1;refGene_tx
=XM_011286598.3;refGene_gn=NUMA1;refGene_tx=XM_011286599.3;refGene_gn=
NUMA1;refGene_tx=XM_011286600.3;refGene_gn=NUMA1;refGene_tx=XM_0232394
68.1;refGene_gn=NUMA1;refGene_tx=XM_023239469.1;refGene_gn=NUMA1;refGe
ne_tx=XM_023239470.1
D1 61712953 61713071
refGene_gn=NUMA1;refGene_tx=XM_011286597.3;refGene_gn=NUMA1;refGene_tx
=XM_011286598.3;refGene_gn=NUMA1;refGene_tx=XM_011286599.3;refGene_gn=
NUMA1;refGene_tx=XM_011286600.3;refGene_gn=NUMA1;refGene_tx=XM_0232394
68.1;refGene_gn=NUMA1;refGene_tx=XM_023239469.1;refGene_gn=NUMA1;refGe
ne_tx=XM_023239470.1
D1 61713175 61713293
refGene_gn=NUMA1;refGene_tx=XM_011286597.3;refGene_gn=NUMA1;refGene_tx
=XM_011286598.3;refGene_gn=NUMA1;refGene_tx=XM_011286599.3;refGene_gn=
NUMA1;refGene_tx=XM_011286600.3;refGene_gn=NUMA1;refGene_tx=XM_0232394
68.1;refGene_gn=NUMA1;refGene_tx=XM_023239469.1;refGene_gn=NUMA1;refGe
ne_tx=XM_023239470.1
D1 61713642 61713783
refGene_gn=NUMA1;refGene_tx=XM_011286597.3;refGene_gn=NUMA1;refGene_tx
=XM_011286598.3;refGene_gn=NUMA1;refGene_tx=XM_011286599.3;refGene_gn=
NUMA1;refGene_tx=XM_011286600.3;refGene_gn=NUMA1;refGene_tx=XM_0232394
68.1;refGene_gn=NUMA1;refGene_tx=XM_023239469.1;refGene_gn=NUMA1;refGe
ne_tx=XM_023239470.1
D1 61715267 61715390
refGene_gn=NUMA1;refGene_tx=XM_011286597.3;refGene_gn=NUMA1;refGene_tx
=XM_011286598.3;refGene_gn=NUMA1;refGene_tx=XM_011286599.3;refGene_gn=
NUMA1;refGene_tx=XM_011286600.3;refGene_gn=NUMA1;refGene_tx=XM_0232394
68.1;refGene_gn=NUMA1;refGene_tx=XM_023239469.1;refGene_gn=NUMA1;refGe
ne_tx=XM_023239470.1
D1 61715541 61718940
refGene_gn=NUMA1;refGene_tx=XM_011286597.3;refGene_gn=NUMA1;refGene_tx
=XM_011286598.3;refGene_gn=NUMA1;refGene_tx=XM_011286599.3;refGene_gn=
NUMA1;refGene_tx=XM_011286600.3;refGene_gn=NUMA1;refGene_tx=XM_0232394
68.1;refGene_gn=NUMA1;refGene_tx=XM_023239469.1;refGene_gn=NUMA1;refGe
ne_tx=XM_023239470.1
D1 61719360 61719402
refGene_gn=NUMA1;refGene_tx=XM_011286597.3;refGene_gn=NUMA1;refGene_tx
=XM_011286598.3;refGene_gn=NUMA1;refGene_tx=XM_011286599.3;refGene_gn=
NUMA1;refGene_tx=XM_011286600.3;refGene_gn=NUMA1;refGene_tx=XM_0232394
68.1;refGene_gn=NUMA1;refGene_tx=XM_023239470.1

D1 61720877 61720946
refGene_gn=NUMA1;refGene_tx=XM_011286597.3;refGene_gn=NUMA1;refGene_tx=XM_011286598.3;refGene_gn=NUMA1;refGene_tx=XM_011286599.3;refGene_gn=NUMA1;refGene_tx=XM_011286600.3;refGene_gn=NUMA1;refGene_tx=XM_023239468.1;refGene_gn=NUMA1;refGene_tx=XM_023239469.1;refGene_gn=NUMA1;refGene_tx=XM_023239470.1
D1 61722319 61722430
refGene_gn=NUMA1;refGene_tx=XM_011286597.3;refGene_gn=NUMA1;refGene_tx=XM_011286598.3;refGene_gn=NUMA1;refGene_tx=XM_011286599.3;refGene_gn=NUMA1;refGene_tx=XM_011286600.3;refGene_gn=NUMA1;refGene_tx=XM_023239468.1;refGene_gn=NUMA1;refGene_tx=XM_023239469.1;refGene_gn=NUMA1;refGene_tx=XM_023239470.1
D1 61722536 61722755
refGene_gn=NUMA1;refGene_tx=XM_011286597.3;refGene_gn=NUMA1;refGene_tx=XM_011286598.3;refGene_gn=NUMA1;refGene_tx=XM_011286599.3;refGene_gn=NUMA1;refGene_tx=XM_011286600.3;refGene_gn=NUMA1;refGene_tx=XM_023239468.1;refGene_gn=NUMA1;refGene_tx=XM_023239469.1;refGene_gn=NUMA1;refGene_tx=XM_023239470.1
D1 61722870 61723028
refGene_gn=NUMA1;refGene_tx=XM_011286597.3;refGene_gn=NUMA1;refGene_tx=XM_011286598.3;refGene_gn=NUMA1;refGene_tx=XM_011286599.3;refGene_gn=NUMA1;refGene_tx=XM_011286600.3;refGene_gn=NUMA1;refGene_tx=XM_023239468.1;refGene_gn=NUMA1;refGene_tx=XM_023239469.1;refGene_gn=NUMA1;refGene_tx=XM_023239470.1
D1 61724380 61724627
refGene_gn=NUMA1;refGene_tx=XM_011286597.3;refGene_gn=NUMA1;refGene_tx=XM_011286598.3;refGene_gn=NUMA1;refGene_tx=XM_011286599.3;refGene_gn=NUMA1;refGene_tx=XM_011286600.3;refGene_gn=NUMA1;refGene_tx=XM_023239468.1;refGene_gn=NUMA1;refGene_tx=XM_023239469.1;refGene_gn=NUMA1;refGene_tx=XM_023239470.1
D1 61725482 61725711
refGene_gn=NUMA1;refGene_tx=XM_011286597.3;refGene_gn=NUMA1;refGene_tx=XM_011286598.3;refGene_gn=NUMA1;refGene_tx=XM_011286599.3;refGene_gn=NUMA1;refGene_tx=XM_011286600.3;refGene_gn=NUMA1;refGene_tx=XM_023239468.1;refGene_gn=NUMA1;refGene_tx=XM_023239469.1;refGene_gn=NUMA1;refGene_tx=XM_023239470.1
D1 61726246 61726383
refGene_gn=NUMA1;refGene_tx=XM_011286597.3;refGene_gn=NUMA1;refGene_tx=XM_011286598.3;refGene_gn=NUMA1;refGene_tx=XM_011286599.3;refGene_gn=NUMA1;refGene_tx=XM_011286600.3;refGene_gn=NUMA1;refGene_tx=XM_023239468.1;refGene_gn=NUMA1;refGene_tx=XM_023239469.1;refGene_gn=NUMA1;refGene_tx=XM_023239470.1
D1 61726748 61726925
refGene_gn=NUMA1;refGene_tx=XM_011286597.3;refGene_gn=NUMA1;refGene_tx=XM_011286598.3;refGene_gn=NUMA1;refGene_tx=XM_011286599.3;refGene_gn=NUMA1;refGene_tx=XM_011286600.3;refGene_gn=NUMA1;refGene_tx=XM_023239468.1;refGene_gn=NUMA1;refGene_tx=XM_023239469.1;refGene_gn=NUMA1;refGene_tx=XM_023239470.1
D1 61727101 61727215
refGene_gn=NUMA1;refGene_tx=XM_011286597.3;refGene_gn=NUMA1;refGene_tx

=XM_011286598.3;refGene_gn=NUMA1;refGene_tx=XM_011286599.3;refGene_gn=NUMA1;refGene_tx=XM_011286600.3;refGene_gn=NUMA1;refGene_tx=XM_023239468.1;refGene_gn=NUMA1;refGene_tx=XM_023239469.1;refGene_gn=NUMA1;refGene_tx=XM_023239470.1
D1 61727306 61727528
refGene_gn=NUMA1;refGene_tx=XM_011286597.3;refGene_gn=NUMA1;refGene_tx=XM_011286598.3;refGene_gn=NUMA1;refGene_tx=XM_011286599.3;refGene_gn=NUMA1;refGene_tx=XM_011286600.3;refGene_gn=NUMA1;refGene_tx=XM_023239468.1;refGene_gn=NUMA1;refGene_tx=XM_023239469.1;refGene_gn=NUMA1;refGene_tx=XM_023239470.1
D1 61727874 61727886
refGene_gn=NUMA1;refGene_tx=XM_011286597.3;refGene_gn=NUMA1;refGene_tx=XM_011286598.3;refGene_gn=NUMA1;refGene_tx=XM_011286599.3;refGene_gn=NUMA1;refGene_tx=XM_011286600.3;refGene_gn=NUMA1;refGene_tx=XM_023239468.1;refGene_gn=NUMA1;refGene_tx=XM_023239469.1;refGene_gn=NUMA1;refGene_tx=XM_023239470.1
D1 61729099 61729183
refGene_gn=IL18BP;refGene_tx=XM_003992816.4;refGene_gn=IL18BP;refGene_tx=XM_006937005.4;refGene_gn=IL18BP;refGene_tx=XM_019811897.2
D1 61729389 61729537
refGene_gn=IL18BP;refGene_tx=XM_003992816.4;refGene_gn=IL18BP;refGene_tx=XM_006937005.4;refGene_gn=IL18BP;refGene_tx=XM_019811897.2
D1 61729626 61729750
refGene_gn=IL18BP;refGene_tx=XM_003992816.4;refGene_gn=IL18BP;refGene_tx=XM_006937005.4;refGene_gn=IL18BP;refGene_tx=XM_019811897.2
D1 61730398 61730596
refGene_gn=IL18BP;refGene_tx=XM_003992816.4;refGene_gn=IL18BP;refGene_tx=XM_006937005.4;refGene_gn=IL18BP;refGene_tx=XM_019811897.2
D1 61730926 61730948
refGene_gn=IL18BP;refGene_tx=XM_003992816.4;refGene_gn=IL18BP;refGene_tx=XM_006937005.4;refGene_gn=IL18BP;refGene_tx=XM_019811897.2
D1 61734441 61734562
refGene_gn=RNF121;refGene_tx=XM_003992775.3;refGene_gn=RNF121;refGene_tx=XM_019811884.1;refGene_gn=RNF121;refGene_tx=XM_019811885.2
D1 61736403 61736505
refGene_gn=RNF121;refGene_tx=XM_003992775.3;refGene_gn=RNF121;refGene_tx=XM_019811884.1;refGene_gn=RNF121;refGene_tx=XM_019811885.2
D1 61737134 61737268
refGene_gn=RNF121;refGene_tx=XM_003992775.3;refGene_gn=RNF121;refGene_tx=XM_019811884.1;refGene_gn=RNF121;refGene_tx=XM_019811885.2
D1 61746894 61747015
refGene_gn=RNF121;refGene_tx=XM_003992775.3;refGene_gn=RNF121;refGene_tx=XM_019811884.1;refGene_gn=RNF121;refGene_tx=XM_019811885.2
D1 61750417 61750525
refGene_gn=RNF121;refGene_tx=XM_003992775.3;refGene_gn=RNF121;refGene_tx=XM_019811884.1;refGene_gn=RNF121;refGene_tx=XM_019811885.2
D1 61753246 61753401
refGene_gn=RNF121;refGene_tx=XM_003992775.3;refGene_gn=RNF121;refGene_tx=XM_019811884.1;refGene_gn=RNF121;refGene_tx=XM_019811885.2
D1 61782162 61782304

refGene_gn=RNF121;refGene_tx=XM_003992775.3;refGene_gn=RNF121;refGene_tx=XM_019811884.1;refGene_gn=RNF121;refGene_tx=XM_019811885.2
D1	61785420	61785458	refGene_gn=RNF121;refGene_tx=XM_003992775.3;refGene_gn=RNF121;refGene_tx=XM_019811884.1;refGene_gn=RNF121;refGene_tx=XM_019811885.2
D1	61814938	61815001	refGene_gn=RNF121;refGene_tx=XM_003992775.3
D1	61815479	61815500	refGene_gn=RNF121;refGene_tx=XM_019811884.1
D1	61816512	61816557	refGene_gn=LOC101086175;refGene_tx=XM_006937006.2
D1	61818568	61818710	refGene_gn=LOC101086175;refGene_tx=XM_006937006.2
D1	61819629	61819785	refGene_gn=LOC101086175;refGene_tx=XM_006937006.2
D1	61821849	61822028	refGene_gn=LOC101086175;refGene_tx=XM_006937006.2
D1	61826126	61826209	refGene_gn=LOC101086175;refGene_tx=XM_006937006.2
D1	61826453	61826626	refGene_gn=LOC101086175;refGene_tx=XM_006937006.2
D1	61829280	61829353	refGene_gn=LOC101086175;refGene_tx=XM_006937006.2
D1	61830935	61831063	refGene_gn=LOC101086175;refGene_tx=XM_006937006.2
D1	61831720	61831829	refGene_gn=LOC101086175;refGene_tx=XM_006937006.2
D1	61831906	61831911	refGene_gn=LOC101086175;refGene_tx=XM_006937006.2
D1	61832355	61832382	refGene_gn=LOC101086175;refGene_tx=XM_006937006.2
D1	61832717	61832734	refGene_gn=LOC101086175;refGene_tx=XM_006937006.2
D1	61834801	61835521	refGene_gn=LOC101086175;refGene_tx=XM_006937006.2
D1	61837377	61837590	refGene_gn=LOC101086175;refGene_tx=XM_006937006.2
D1	61838537	61838702	refGene_gn=LOC101086175;refGene_tx=XM_006937006.2
D1	61839721	61839896	refGene_gn=LOC101086175;refGene_tx=XM_006937006.2
D1	61842658	61842859	refGene_gn=LOC101086175;refGene_tx=XM_006937006.2
D1	61842943	61843106	refGene_gn=LOC101086175;refGene_tx=XM_006937006.2
D1	61844118	61844211	refGene_gn=LOC101086175;refGene_tx=XM_006937006.2
D1	61848387	61848580	refGene_gn=LOC101086175;refGene_tx=XM_006937006.2
D1	61849120	61849248	refGene_gn=LOC101086175;refGene_tx=XM_006937006.2
D1	61849370	61849449

refGene_gn=LOC101086175;refGene_tx=XM_006937006.2
D1 61851941 61852071
refGene_gn=LOC101086175;refGene_tx=XM_006937006.2
D1 61853198 61853578
refGene_gn=LOC101086175;refGene_tx=XM_006937006.2
D1 61854621 61854728
refGene_gn=ART5;refGene_tx=XM_006937007.4;refGene_gn=ART5;refGene_tx=XM_006937008.4;refGene_gn=ART5;refGene_tx=XM_019811889.2;refGene_gn=ART5;refGene_tx=XM_019811892.2;refGene_gn=ART5;refGene_tx=XM_019811894.2
D1 61854812 61854845
refGene_gn=ART5;refGene_tx=XM_006937007.4;refGene_gn=ART5;refGene_tx=XM_006937008.4;refGene_gn=ART5;refGene_tx=XM_019811889.2;refGene_gn=ART5;refGene_tx=XM_019811892.2;refGene_gn=ART5;refGene_tx=XM_019811894.2
D1 61855453 61856183
refGene_gn=ART5;refGene_tx=XM_006937007.4;refGene_gn=ART5;refGene_tx=XM_006937008.4;refGene_gn=ART5;refGene_tx=XM_019811889.2;refGene_gn=ART5;refGene_tx=XM_019811892.2;refGene_gn=ART5;refGene_tx=XM_019811894.2
D1 61857138 61857231
refGene_gn=ART5;refGene_tx=XM_006937007.4;refGene_gn=ART5;refGene_tx=XM_006937008.4;refGene_gn=ART5;refGene_tx=XM_019811889.2;refGene_gn=ART5;refGene_tx=XM_019811892.2;refGene_gn=ART5;refGene_tx=XM_019811894.2
D1 61865829 61865892
refGene_gn=ART1;refGene_tx=XM_003992777.3;refGene_gn=ART1;refGene_tx=XM_019811886.1;refGene_gn=ART1;refGene_tx=XM_019811887.1;refGene_gn=ART1;refGene_tx=XM_019811888.1
D1 61866181 61866962
refGene_gn=ART1;refGene_tx=XM_003992777.3;refGene_gn=ART1;refGene_tx=XM_019811886.1;refGene_gn=ART1;refGene_tx=XM_019811887.1;refGene_gn=ART1;refGene_tx=XM_019811888.1
D1 61868080 61868122
refGene_gn=ART1;refGene_tx=XM_003992777.3;refGene_gn=ART1;refGene_tx=XM_019811886.1;refGene_gn=ART1;refGene_tx=XM_019811887.1;refGene_gn=ART1;refGene_tx=XM_019811888.1
D1 61869687 61869767
refGene_gn=ART1;refGene_tx=XM_003992777.3;refGene_gn=ART1;refGene_tx=XM_019811886.1;refGene_gn=ART1;refGene_tx=XM_019811887.1;refGene_gn=ART1;refGene_tx=XM_019811888.1
D1 61871177 61871635 refGene_gn=CHRNA10;refGene_tx=XM_023239708.1
D1 61872294 61872827 refGene_gn=CHRNA10;refGene_tx=XM_023239708.1
D1 61873528 61873683 refGene_gn=CHRNA10;refGene_tx=XM_023239708.1
D1 61874116 61874664 refGene_gn=CHRNA10;refGene_tx=XM_023239708.1
D1 61875946 61876049 refGene_gn=CHRNA10;refGene_tx=XM_023239708.1
D1 61885903 61886121
refGene_gn=NUP98;refGene_tx=XM_011286589.3;refGene_gn=NUP98;refGene_tx=XM_011286590.3;refGene_gn=NUP98;refGene_tx=XM_011286591.3;refGene_gn=NUP98;refGene_tx=XM_011286592.3;refGene_gn=NUP98;refGene_tx=XM_011286593.3;refGene_gn=NUP98;refGene_tx=XM_019811901.2
D1 61886250 61886362
refGene_gn=NUP98;refGene_tx=XM_011286589.3;refGene_gn=NUP98;refGene_tx=XM_011286590.3;refGene_gn=NUP98;refGene_tx=XM_011286591.3;refGene_gn=

NUP98;refGene_tx=XM_011286592.3;refGene_gn=NUP98;refGene_tx=XM_0112865
93.3;refGene_gn=NUP98;refGene_tx=XM_019811901.2
D1 61892526 61892681
refGene_gn=NUP98;refGene_tx=XM_011286589.3;refGene_gn=NUP98;refGene_tx
=XM_011286590.3;refGene_gn=NUP98;refGene_tx=XM_011286591.3;refGene_gn=
NUP98;refGene_tx=XM_011286592.3;refGene_gn=NUP98;refGene_tx=XM_0112865
93.3;refGene_gn=NUP98;refGene_tx=XM_019811901.2
D1 61897804 61898046
refGene_gn=NUP98;refGene_tx=XM_011286589.3;refGene_gn=NUP98;refGene_tx
=XM_011286590.3;refGene_gn=NUP98;refGene_tx=XM_011286591.3;refGene_gn=
NUP98;refGene_tx=XM_011286592.3;refGene_gn=NUP98;refGene_tx=XM_0112865
93.3;refGene_gn=NUP98;refGene_tx=XM_019811901.2
D1 61900240 61900462
refGene_gn=NUP98;refGene_tx=XM_011286589.3;refGene_gn=NUP98;refGene_tx
=XM_011286590.3;refGene_gn=NUP98;refGene_tx=XM_011286591.3;refGene_gn=
NUP98;refGene_tx=XM_011286592.3;refGene_gn=NUP98;refGene_tx=XM_0112865
93.3;refGene_gn=NUP98;refGene_tx=XM_019811901.2
D1 61906004 61906147
refGene_gn=NUP98;refGene_tx=XM_011286589.3;refGene_gn=NUP98;refGene_tx
=XM_011286590.3;refGene_gn=NUP98;refGene_tx=XM_011286591.3;refGene_gn=
NUP98;refGene_tx=XM_011286592.3;refGene_gn=NUP98;refGene_tx=XM_0112865
93.3;refGene_gn=NUP98;refGene_tx=XM_019811901.2
D1 61908066 61908210
refGene_gn=NUP98;refGene_tx=XM_011286589.3;refGene_gn=NUP98;refGene_tx
=XM_011286590.3;refGene_gn=NUP98;refGene_tx=XM_011286591.3;refGene_gn=
NUP98;refGene_tx=XM_011286592.3;refGene_gn=NUP98;refGene_tx=XM_0112865
93.3;refGene_gn=NUP98;refGene_tx=XM_019811901.2
D1 61909134 61909292
refGene_gn=NUP98;refGene_tx=XM_011286589.3;refGene_gn=NUP98;refGene_tx
=XM_011286590.3;refGene_gn=NUP98;refGene_tx=XM_011286591.3;refGene_gn=
NUP98;refGene_tx=XM_011286592.3;refGene_gn=NUP98;refGene_tx=XM_0112865
93.3;refGene_gn=NUP98;refGene_tx=XM_019811901.2
D1 61913892 61914159
refGene_gn=NUP98;refGene_tx=XM_011286589.3;refGene_gn=NUP98;refGene_tx
=XM_011286590.3;refGene_gn=NUP98;refGene_tx=XM_011286591.3;refGene_gn=
NUP98;refGene_tx=XM_011286592.3;refGene_gn=NUP98;refGene_tx=XM_0112865
93.3;refGene_gn=NUP98;refGene_tx=XM_019811901.2
D1 61916390 61916617
refGene_gn=NUP98;refGene_tx=XM_011286589.3;refGene_gn=NUP98;refGene_tx
=XM_011286590.3;refGene_gn=NUP98;refGene_tx=XM_011286591.3;refGene_gn=
NUP98;refGene_tx=XM_011286592.3;refGene_gn=NUP98;refGene_tx=XM_0112865
93.3;refGene_gn=NUP98;refGene_tx=XM_019811901.2
D1 61918491 61918921
refGene_gn=NUP98;refGene_tx=XM_011286589.3;refGene_gn=NUP98;refGene_tx
=XM_011286590.3;refGene_gn=NUP98;refGene_tx=XM_011286591.3;refGene_gn=
NUP98;refGene_tx=XM_011286592.3;refGene_gn=NUP98;refGene_tx=XM_0112865
93.3;refGene_gn=NUP98;refGene_tx=XM_019811901.2
D1 61920292 61920449
refGene_gn=NUP98;refGene_tx=XM_011286589.3;refGene_gn=NUP98;refGene_tx
=XM_011286590.3;refGene_gn=NUP98;refGene_tx=XM_011286591.3;refGene_gn=

NUP98;refGene_tx=XM_011286592.3;refGene_gn=NUP98;refGene_tx=XM_0112865
93.3;refGene_gn=NUP98;refGene_tx=XM_019811901.2
D1 61921112 61921295
refGene_gn=NUP98;refGene_tx=XM_011286589.3;refGene_gn=NUP98;refGene_tx
=XM_011286590.3;refGene_gn=NUP98;refGene_tx=XM_011286591.3;refGene_gn=
NUP98;refGene_tx=XM_011286592.3;refGene_gn=NUP98;refGene_tx=XM_0112865
93.3;refGene_gn=NUP98;refGene_tx=XM_019811901.2
D1 61927818 61928001
refGene_gn=NUP98;refGene_tx=XM_011286589.3;refGene_gn=NUP98;refGene_tx
=XM_011286590.3;refGene_gn=NUP98;refGene_tx=XM_011286591.3;refGene_gn=
NUP98;refGene_tx=XM_011286592.3;refGene_gn=NUP98;refGene_tx=XM_0112865
93.3;refGene_gn=NUP98;refGene_tx=XM_019811901.2;refGene_gn=NUP98;refGe
ne_tx=XM_023239472.1
D1 61928880 61929058
refGene_gn=NUP98;refGene_tx=XM_011286589.3;refGene_gn=NUP98;refGene_tx
=XM_011286590.3;refGene_gn=NUP98;refGene_tx=XM_011286591.3;refGene_gn=
NUP98;refGene_tx=XM_011286592.3;refGene_gn=NUP98;refGene_tx=XM_0112865
93.3;refGene_gn=NUP98;refGene_tx=XM_019811901.2;refGene_gn=NUP98;refGe
ne_tx=XM_023239472.1
D1 61935347 61935486
refGene_gn=NUP98;refGene_tx=XM_011286589.3;refGene_gn=NUP98;refGene_tx
=XM_011286590.3;refGene_gn=NUP98;refGene_tx=XM_011286591.3;refGene_gn=
NUP98;refGene_tx=XM_011286592.3;refGene_gn=NUP98;refGene_tx=XM_0112865
93.3;refGene_gn=NUP98;refGene_tx=XM_019811901.2;refGene_gn=NUP98;refGe
ne_tx=XM_023239472.1
D1 61936304 61936418
refGene_gn=NUP98;refGene_tx=XM_011286589.3;refGene_gn=NUP98;refGene_tx
=XM_011286590.3;refGene_gn=NUP98;refGene_tx=XM_011286591.3;refGene_gn=
NUP98;refGene_tx=XM_011286592.3;refGene_gn=NUP98;refGene_tx=XM_0112865
93.3;refGene_gn=NUP98;refGene_tx=XM_019811901.2;refGene_gn=NUP98;refGe
ne_tx=XM_023239472.1
D1 61939837 61940136
refGene_gn=NUP98;refGene_tx=XM_011286589.3;refGene_gn=NUP98;refGene_tx
=XM_011286590.3;refGene_gn=NUP98;refGene_tx=XM_011286591.3;refGene_gn=
NUP98;refGene_tx=XM_011286592.3;refGene_gn=NUP98;refGene_tx=XM_0112865
93.3;refGene_gn=NUP98;refGene_tx=XM_019811901.2;refGene_gn=NUP98;refGe
ne_tx=XM_023239472.1
D1 61941371 61941488
refGene_gn=NUP98;refGene_tx=XM_011286589.3;refGene_gn=NUP98;refGene_tx
=XM_011286590.3;refGene_gn=NUP98;refGene_tx=XM_011286591.3;refGene_gn=
NUP98;refGene_tx=XM_011286592.3;refGene_gn=NUP98;refGene_tx=XM_0112865
93.3;refGene_gn=NUP98;refGene_tx=XM_019811901.2;refGene_gn=NUP98;refGe
ne_tx=XM_023239472.1
D1 61953047 61953235
refGene_gn=NUP98;refGene_tx=XM_011286589.3;refGene_gn=NUP98;refGene_tx
=XM_011286590.3;refGene_gn=NUP98;refGene_tx=XM_011286591.3;refGene_gn=
NUP98;refGene_tx=XM_011286592.3;refGene_gn=NUP98;refGene_tx=XM_0112865
93.3;refGene_gn=NUP98;refGene_tx=XM_019811901.2;refGene_gn=NUP98;refGe
ne_tx=XM_023239472.1
D1 61955721 61955855

refGene_gn=NUP98;refGene_tx=XM_011286589.3;refGene_gn=NUP98;refGene_tx=XM_011286590.3;refGene_gn=NUP98;refGene_tx=XM_011286591.3;refGene_gn=NUP98;refGene_tx=XM_011286592.3;refGene_gn=NUP98;refGene_tx=XM_011286593.3;refGene_gn=NUP98;refGene_tx=XM_019811901.2;refGene_gn=NUP98;refGene_tx=XM_023239472.1
D1 61962000 61962141
refGene_gn=NUP98;refGene_tx=XM_011286589.3;refGene_gn=NUP98;refGene_tx=XM_011286590.3;refGene_gn=NUP98;refGene_tx=XM_011286591.3;refGene_gn=NUP98;refGene_tx=XM_011286592.3;refGene_gn=NUP98;refGene_tx=XM_023239472.1
D1 61968021 61968114
refGene_gn=NUP98;refGene_tx=XM_011286589.3;refGene_gn=NUP98;refGene_tx=XM_011286590.3;refGene_gn=NUP98;refGene_tx=XM_011286591.3;refGene_gn=NUP98;refGene_tx=XM_011286592.3;refGene_gn=NUP98;refGene_tx=XM_011286593.3;refGene_gn=NUP98;refGene_tx=XM_019811901.2;refGene_gn=NUP98;refGene_tx=XM_023239472.1
D1 61974070 61974209
refGene_gn=NUP98;refGene_tx=XM_011286589.3;refGene_gn=NUP98;refGene_tx=XM_011286590.3;refGene_gn=NUP98;refGene_tx=XM_011286591.3;refGene_gn=NUP98;refGene_tx=XM_011286592.3;refGene_gn=NUP98;refGene_tx=XM_011286593.3;refGene_gn=NUP98;refGene_tx=XM_019811901.2;refGene_gn=NUP98;refGene_tx=XM_023239472.1
D1 61975624 61975762
refGene_gn=NUP98;refGene_tx=XM_011286589.3;refGene_gn=NUP98;refGene_tx=XM_011286590.3;refGene_gn=NUP98;refGene_tx=XM_011286591.3;refGene_gn=NUP98;refGene_tx=XM_011286592.3;refGene_gn=NUP98;refGene_tx=XM_011286593.3;refGene_gn=NUP98;refGene_tx=XM_019811901.2;refGene_gn=NUP98;refGene_tx=XM_023239472.1
D1 61979299 61979463
refGene_gn=NUP98;refGene_tx=XM_011286589.3;refGene_gn=NUP98;refGene_tx=XM_011286590.3;refGene_gn=NUP98;refGene_tx=XM_011286591.3;refGene_gn=NUP98;refGene_tx=XM_011286592.3;refGene_gn=NUP98;refGene_tx=XM_011286593.3;refGene_gn=NUP98;refGene_tx=XM_019811901.2;refGene_gn=NUP98;refGene_tx=XM_023239472.1
D1 61980946 61981127
refGene_gn=NUP98;refGene_tx=XM_011286589.3;refGene_gn=NUP98;refGene_tx=XM_011286590.3;refGene_gn=NUP98;refGene_tx=XM_011286591.3;refGene_gn=NUP98;refGene_tx=XM_011286592.3;refGene_gn=NUP98;refGene_tx=XM_011286593.3;refGene_gn=NUP98;refGene_tx=XM_019811901.2;refGene_gn=NUP98;refGene_tx=XM_023239472.1
D1 61982377 61982485
refGene_gn=NUP98;refGene_tx=XM_011286589.3;refGene_gn=NUP98;refGene_tx=XM_011286590.3;refGene_gn=NUP98;refGene_tx=XM_011286591.3;refGene_gn=NUP98;refGene_tx=XM_011286592.3;refGene_gn=NUP98;refGene_tx=XM_011286593.3;refGene_gn=NUP98;refGene_tx=XM_019811901.2;refGene_gn=NUP98;refGene_tx=XM_023239472.1
D1 61983780 61983920
refGene_gn=NUP98;refGene_tx=XM_011286589.3;refGene_gn=NUP98;refGene_tx=XM_011286590.3;refGene_gn=NUP98;refGene_tx=XM_011286591.3;refGene_gn=NUP98;refGene_tx=XM_011286592.3;refGene_gn=NUP98;refGene_tx=XM_0112865

93.3;refGene_gn=NUP98;refGene_tx=XM_019811901.2;refGene_gn=NUP98;refGene_tx=XM_023239472.1
D1 61985474 61985651
refGene_gn=NUP98;refGene_tx=XM_011286589.3;refGene_gn=NUP98;refGene_tx=XM_011286590.3;refGene_gn=NUP98;refGene_tx=XM_011286591.3;refGene_gn=NUP98;refGene_tx=XM_011286592.3;refGene_gn=NUP98;refGene_tx=XM_011286593.3;refGene_gn=NUP98;refGene_tx=XM_019811901.2;refGene_gn=NUP98;refGene_tx=XM_023239472.1
D1 61985761 61985863
refGene_gn=NUP98;refGene_tx=XM_011286589.3;refGene_gn=NUP98;refGene_tx=XM_011286590.3;refGene_gn=NUP98;refGene_tx=XM_011286591.3;refGene_gn=NUP98;refGene_tx=XM_011286592.3;refGene_gn=NUP98;refGene_tx=XM_011286593.3;refGene_gn=NUP98;refGene_tx=XM_019811901.2;refGene_gn=NUP98;refGene_tx=XM_023239472.1
D1 61990182 61990258
refGene_gn=NUP98;refGene_tx=XM_011286589.3;refGene_gn=NUP98;refGene_tx=XM_011286590.3;refGene_gn=NUP98;refGene_tx=XM_011286591.3;refGene_gn=NUP98;refGene_tx=XM_011286592.3;refGene_gn=NUP98;refGene_tx=XM_011286593.3;refGene_gn=NUP98;refGene_tx=XM_019811901.2;refGene_gn=NUP98;refGene_tx=XM_023239472.1
D1 62013877 62014113
refGene_gn=PGAP2;refGene_tx=XM_023239477.1;refGene_gn=PGAP2;refGene_tx=XM_023239480.1;refGene_gn=PGAP2;refGene_tx=XM_023239482.1
D1 62016668 62016844
refGene_gn=PGAP2;refGene_tx=XM_006937018.3;refGene_gn=PGAP2;refGene_tx=XM_019811905.2;refGene_gn=PGAP2;refGene_tx=XM_019811906.2;refGene_gn=PGAP2;refGene_tx=XM_019811907.2;refGene_gn=PGAP2;refGene_tx=XM_019811908.2;refGene_gn=PGAP2;refGene_tx=XM_023239473.1;refGene_gn=PGAP2;refGene_tx=XM_023239474.1;refGene_gn=PGAP2;refGene_tx=XM_023239475.1;refGene_gn=PGAP2;refGene_tx=XM_023239476.1;refGene_gn=PGAP2;refGene_tx=XM_023239478.1;refGene_gn=PGAP2;refGene_tx=XM_023239479.1;refGene_gn=PGAP2;refGene_tx=XM_023239480.1;refGene_gn=PGAP2;refGene_tx=XM_023239482.1;refGene_gn=PGAP2;refGene_tx=XM_023239483.1;refGene_gn=PGAP2;refGene_tx=XM_023239484.1;refGene_gn=PGAP2;refGene_tx=XM_023239485.1
D1 62020806 62020988
refGene_gn=PGAP2;refGene_tx=XM_006937018.3;refGene_gn=PGAP2;refGene_tx=XM_019811905.2;refGene_gn=PGAP2;refGene_tx=XM_019811906.2;refGene_gn=PGAP2;refGene_tx=XM_019811907.2;refGene_gn=PGAP2;refGene_tx=XM_019811908.2;refGene_gn=PGAP2;refGene_tx=XM_019811909.2;refGene_gn=PGAP2;refGene_tx=XM_023239473.1;refGene_gn=PGAP2;refGene_tx=XM_023239474.1;refGene_gn=PGAP2;refGene_tx=XM_023239475.1;refGene_gn=PGAP2;refGene_tx=XM_023239477.1;refGene_gn=PGAP2;refGene_tx=XM_023239480.1;refGene_gn=PGAP2;refGene_tx=XM_023239482.1
D1 62024627 62024880
refGene_gn=PGAP2;refGene_tx=XM_006937018.3;refGene_gn=PGAP2;refGene_tx=XM_019811905.2;refGene_gn=PGAP2;refGene_tx=XM_019811906.2;refGene_gn=PGAP2;refGene_tx=XM_019811907.2;refGene_gn=PGAP2;refGene_tx=XM_019811908.2;refGene_gn=PGAP2;refGene_tx=XM_019811909.2;refGene_gn=PGAP2;refGene_tx=XM_023239473.1;refGene_gn=PGAP2;refGene_tx=XM_023239474.1;refGene_gn=PGAP2;refGene_tx=XM_023239475.1;refGene_gn=PGAP2;refGene_tx=XM_02

3239476.1;refGene_gn=PGAP2;refGene_tx=XM_023239477.1;refGene_gn=PGAP2;
refGene_tx=XM_023239478.1;refGene_gn=PGAP2;refGene_tx=XM_023239479.1;r
efGene_gn=PGAP2;refGene_tx=XM_023239480.1;refGene_gn=PGAP2;refGene_tx=
XM_023239482.1;refGene_gn=PGAP2;refGene_tx=XM_023239483.1;refGene_gn=P
GAP2;refGene_tx=XM_023239484.1;refGene_gn=PGAP2;refGene_tx=XM_02323948
5.1
D1 62025022 62025129
refGene_gn=PGAP2;refGene_tx=XM_006937018.3;refGene_gn=PGAP2;refGene_tx
=XM_019811905.2;refGene_gn=PGAP2;refGene_tx=XM_019811906.2;refGene_gn=
PGAP2;refGene_tx=XM_019811907.2;refGene_gn=PGAP2;refGene_tx=XM_0198119
08.2;refGene_gn=PGAP2;refGene_tx=XM_019811909.2;refGene_gn=PGAP2;refGe
ne_tx=XM_023239473.1;refGene_gn=PGAP2;refGene_tx=XM_023239474.1;refGen
e_gn=PGAP2;refGene_tx=XM_023239475.1;refGene_gn=PGAP2;refGene_tx=XM_02
3239476.1;refGene_gn=PGAP2;refGene_tx=XM_023239477.1;refGene_gn=PGAP2;
refGene_tx=XM_023239478.1;refGene_gn=PGAP2;refGene_tx=XM_023239479.1;r
efGene_gn=PGAP2;refGene_tx=XM_023239480.1;refGene_gn=PGAP2;refGene_tx=
XM_023239482.1;refGene_gn=PGAP2;refGene_tx=XM_023239483.1;refGene_gn=P
GAP2;refGene_tx=XM_023239484.1;refGene_gn=PGAP2;refGene_tx=XM_02323948
5.1
D1 62025588 62025651
refGene_gn=PGAP2;refGene_tx=XM_019811905.2;refGene_gn=PGAP2;refGene_tx
=XM_019811906.2;refGene_gn=PGAP2;refGene_tx=XM_019811907.2;refGene_gn=
PGAP2;refGene_tx=XM_019811908.2;refGene_gn=PGAP2;refGene_tx=XM_0198119
09.2;refGene_gn=PGAP2;refGene_tx=XM_023239473.1;refGene_gn=PGAP2;refGe
ne_tx=XM_023239476.1;refGene_gn=PGAP2;refGene_tx=XM_023239477.1;refGen
e_gn=PGAP2;refGene_tx=XM_023239483.1
D1 62025712 62025724
refGene_gn=PGAP2;refGene_tx=XM_023239480.1;refGene_gn=PGAP2;refGene_tx
=XM_023239482.1
D1 62025744 62025853
refGene_gn=PGAP2;refGene_tx=XM_006937018.3;refGene_gn=PGAP2;refGene_tx
=XM_019811905.2;refGene_gn=PGAP2;refGene_tx=XM_019811906.2;refGene_gn=
PGAP2;refGene_tx=XM_019811907.2;refGene_gn=PGAP2;refGene_tx=XM_0198119
08.2;refGene_gn=PGAP2;refGene_tx=XM_019811909.2;refGene_gn=PGAP2;refGe
ne_tx=XM_023239473.1;refGene_gn=PGAP2;refGene_tx=XM_023239474.1;refGen
e_gn=PGAP2;refGene_tx=XM_023239475.1;refGene_gn=PGAP2;refGene_tx=XM_02
3239476.1;refGene_gn=PGAP2;refGene_tx=XM_023239477.1;refGene_gn=PGAP2;
refGene_tx=XM_023239478.1;refGene_gn=PGAP2;refGene_tx=XM_023239479.1;r
efGene_gn=PGAP2;refGene_tx=XM_023239483.1;refGene_gn=PGAP2;refGene_tx=
XM_023239484.1;refGene_gn=PGAP2;refGene_tx=XM_023239485.1
D1 62026059 62026190
refGene_gn=PGAP2;refGene_tx=XM_006937018.3;refGene_gn=PGAP2;refGene_tx
=XM_019811905.2;refGene_gn=PGAP2;refGene_tx=XM_019811906.2;refGene_gn=
PGAP2;refGene_tx=XM_019811907.2;refGene_gn=PGAP2;refGene_tx=XM_0198119
08.2;refGene_gn=PGAP2;refGene_tx=XM_019811909.2;refGene_gn=PGAP2;refGe
ne_tx=XM_023239473.1;refGene_gn=PGAP2;refGene_tx=XM_023239474.1;refGen
e_gn=PGAP2;refGene_tx=XM_023239475.1;refGene_gn=PGAP2;refGene_tx=XM_02
3239476.1;refGene_gn=PGAP2;refGene_tx=XM_023239477.1;refGene_gn=PGAP2;
refGene_tx=XM_023239478.1;refGene_gn=PGAP2;refGene_tx=XM_023239479.1;r
efGene_gn=PGAP2;refGene_tx=XM_023239483.1;refGene_gn=PGAP2;refGene_tx=

XM_023239484.1;refGene_gn=PGAP2;refGene_tx=XM_023239485.1
D1 62028377 62028953 refGene_gn=RHOG;refGene_tx=XM_023239487.1
D1 62050235 62050374
refGene_gn=STIM1;refGene_tx=XM_003992783.5;refGene_gn=STIM1;refGene_tx
=XM_006937019.4;refGene_gn=STIM1;refGene_tx=XM_006937020.4;refGene_gn=
STIM1;refGene_tx=XM_019811914.2
D1 62152045 62152176
refGene_gn=STIM1;refGene_tx=XM_003992783.5;refGene_gn=STIM1;refGene_tx
=XM_006937019.4;refGene_gn=STIM1;refGene_tx=XM_006937020.4;refGene_gn=
STIM1;refGene_tx=XM_019811914.2
D1 62163110 62163233 refGene_gn=STIM1;refGene_tx=XM_023239489.1
D1 62180445 62180560
refGene_gn=STIM1;refGene_tx=XM_003992783.5;refGene_gn=STIM1;refGene_tx
=XM_006937019.4;refGene_gn=STIM1;refGene_tx=XM_006937020.4;refGene_gn=
STIM1;refGene_tx=XM_019811914.2;refGene_gn=STIM1;refGene_tx=XM_0232394
89.1
D1 62200628 62200740
refGene_gn=STIM1;refGene_tx=XM_003992783.5;refGene_gn=STIM1;refGene_tx
=XM_006937019.4;refGene_gn=STIM1;refGene_tx=XM_006937020.4;refGene_gn=
STIM1;refGene_tx=XM_019811914.2;refGene_gn=STIM1;refGene_tx=XM_0232394
89.1
D1 62203858 62203974
refGene_gn=STIM1;refGene_tx=XM_003992783.5;refGene_gn=STIM1;refGene_tx
=XM_006937019.4;refGene_gn=STIM1;refGene_tx=XM_006937020.4;refGene_gn=
STIM1;refGene_tx=XM_019811914.2;refGene_gn=STIM1;refGene_tx=XM_0232394
89.1
D1 62212898 62213076
refGene_gn=STIM1;refGene_tx=XM_003992783.5;refGene_gn=STIM1;refGene_tx
=XM_006937019.4;refGene_gn=STIM1;refGene_tx=XM_006937020.4;refGene_gn=
STIM1;refGene_tx=XM_019811914.2;refGene_gn=STIM1;refGene_tx=XM_0232394
89.1
D1 62216881 62217059
refGene_gn=STIM1;refGene_tx=XM_003992783.5;refGene_gn=STIM1;refGene_tx
=XM_006937019.4;refGene_gn=STIM1;refGene_tx=XM_006937020.4;refGene_gn=
STIM1;refGene_tx=XM_019811914.2;refGene_gn=STIM1;refGene_tx=XM_0232394
89.1
D1 62224436 62224604
refGene_gn=STIM1;refGene_tx=XM_003992783.5;refGene_gn=STIM1;refGene_tx
=XM_006937019.4;refGene_gn=STIM1;refGene_tx=XM_006937020.4;refGene_gn=
STIM1;refGene_tx=XM_019811914.2;refGene_gn=STIM1;refGene_tx=XM_0232394
89.1
D1 62225122 62225223
refGene_gn=STIM1;refGene_tx=XM_003992783.5;refGene_gn=STIM1;refGene_tx
=XM_006937019.4;refGene_gn=STIM1;refGene_tx=XM_006937020.4;refGene_gn=
STIM1;refGene_tx=XM_019811914.2;refGene_gn=STIM1;refGene_tx=XM_0232394
89.1
D1 62225501 62225737
refGene_gn=STIM1;refGene_tx=XM_003992783.5;refGene_gn=STIM1;refGene_tx
=XM_006937019.4;refGene_gn=STIM1;refGene_tx=XM_006937020.4;refGene_gn=
STIM1;refGene_tx=XM_019811914.2;refGene_gn=STIM1;refGene_tx=XM_0232394

89.1
D1	62226868	62226961
refGene_gn=STIM1;refGene_tx=XM_006937020.4;refGene_gn=STIM1;refGene_tx=XM_023239489.1
D1	62228653	62229062
refGene_gn=STIM1;refGene_tx=XM_003992783.5;refGene_gn=STIM1;refGene_tx=XM_006937019.4;refGene_gn=STIM1;refGene_tx=XM_006937020.4;refGene_gn=STIM1;refGene_tx=XM_019811914.2;refGene_gn=STIM1;refGene_tx=XM_023239489.1
D1	62231074	62231111	refGene_gn=STIM1;refGene_tx=XM_019811914.2
D1	62233413	62233930
refGene_gn=STIM1;refGene_tx=XM_003992783.5;refGene_gn=STIM1;refGene_tx=XM_006937019.4;refGene_gn=STIM1;refGene_tx=XM_006937020.4;refGene_gn=STIM1;refGene_tx=XM_019811914.2;refGene_gn=STIM1;refGene_tx=XM_023239489.1
D1	62236694	62236713	refGene_gn=RRM1;refGene_tx=XM_011286586.3
D1	62244069	62244158	refGene_gn=RRM1;refGene_tx=XM_011286586.3
D1	62246921	62247099	refGene_gn=RRM1;refGene_tx=XM_011286586.3
D1	62248142	62248243	refGene_gn=RRM1;refGene_tx=XM_011286586.3
D1	62250192	62250252	refGene_gn=RRM1;refGene_tx=XM_011286586.3
D1	62251137	62251177	refGene_gn=RRM1;refGene_tx=XM_011286586.3
D1	62252513	62252676	refGene_gn=RRM1;refGene_tx=XM_011286586.3
D1	62256630	62256772	refGene_gn=RRM1;refGene_tx=XM_011286586.3
D1	62258102	62258186	refGene_gn=RRM1;refGene_tx=XM_011286586.3
D1	62259349	62259511	refGene_gn=RRM1;refGene_tx=XM_011286586.3
D1	62260120	62260200	refGene_gn=RRM1;refGene_tx=XM_011286586.3
D1	62260865	62261067	refGene_gn=RRM1;refGene_tx=XM_011286586.3
D1	62263266	62263416	refGene_gn=RRM1;refGene_tx=XM_011286586.3
D1	62263615	62263837	refGene_gn=RRM1;refGene_tx=XM_011286586.3
D1	62265935	62266012	refGene_gn=RRM1;refGene_tx=XM_011286586.3
D1	62269294	62269430	refGene_gn=RRM1;refGene_tx=XM_011286586.3
D1	62270222	62270318	refGene_gn=RRM1;refGene_tx=XM_011286586.3
D1	62271907	62272096	refGene_gn=RRM1;refGene_tx=XM_011286586.3
D1	62273105	62273294	refGene_gn=RRM1;refGene_tx=XM_011286586.3
D1	62278136	62279186
refGene_gn=LOC105260972;refGene_tx=XM_011286674.3
D1	62354138	62355083
refGene_gn=LOC101098084;refGene_tx=XM_003992821.1
D1	62388368	62389274
refGene_gn=LOC101088690;refGene_tx=XM_023238989.1
D1	62410153	62411092
refGene_gn=LOC101098332;refGene_tx=XM_019813037.1
D1	62422129	62422680
refGene_gn=TRIM21;refGene_tx=NM_001290243.1;refGene_gn=TRIM21;refGene_tx=XM_003992823.4
D1	62422962	62423063
refGene_gn=TRIM21;refGene_tx=NM_001290243.1;refGene_gn=TRIM21;refGene_tx=XM_003992823.4
D1	62423935	62423958
refGene_gn=TRIM21;refGene_tx=NM_001290243.1;refGene_gn=TRIM21;refGene_

tx=XM_003992823.4
D1	62425754	62425985
refGene_gn=TRIM21;refGene_tx=NM_001290243.1;refGene_gn=TRIM21;refGene_tx=XM_003992823.4
D1	62427079	62427175
refGene_gn=TRIM21;refGene_tx=NM_001290243.1;refGene_gn=TRIM21;refGene_tx=XM_003992823.4
D1	62427662	62428070
refGene_gn=TRIM21;refGene_tx=NM_001290243.1;refGene_gn=TRIM21;refGene_tx=XM_003992823.4
D1	62443133	62444087
refGene_gn=LOC101098821;refGene_tx=XM_003992824.1
D1	62458915	62459863
refGene_gn=LOC101099074;refGene_tx=XM_003992825.1
D1	62480191	62481142
refGene_gn=LOC101099324;refGene_tx=XM_003992826.2
D1	62490491	62491436
refGene_gn=LOC101088952;refGene_tx=XM_003992787.2
D1	62503740	62504694
refGene_gn=LOC101099583;refGene_tx=XM_011286673.2
D1	62544319	62545273
refGene_gn=LOC101089391;refGene_tx=XM_003992788.3
D1	62586864	62587839
refGene_gn=LOC101089643;refGene_tx=XM_006937021.2
D1	62596967	62597518
refGene_gn=TRIM68;refGene_tx=XM_003992790.5;refGene_gn=TRIM68;refGene_tx=XM_011286585.3;refGene_gn=TRIM68;refGene_tx=XM_019811917.2;refGene_gn=TRIM68;refGene_tx=XM_019811918.2;refGene_gn=TRIM68;refGene_tx=XM_019811919.1;refGene_gn=TRIM68;refGene_tx=XM_019811920.2;refGene_gn=TRIM68;refGene_tx=XM_019811921.1
D1	62597720	62597821
refGene_gn=TRIM68;refGene_tx=XM_003992790.5;refGene_gn=TRIM68;refGene_tx=XM_011286585.3;refGene_gn=TRIM68;refGene_tx=XM_019811917.2;refGene_gn=TRIM68;refGene_tx=XM_019811918.2;refGene_gn=TRIM68;refGene_tx=XM_019811919.1;refGene_gn=TRIM68;refGene_tx=XM_019811920.2;refGene_gn=TRIM68;refGene_tx=XM_019811921.1
D1	62598396	62598419
refGene_gn=TRIM68;refGene_tx=XM_003992790.5;refGene_gn=TRIM68;refGene_tx=XM_011286585.3;refGene_gn=TRIM68;refGene_tx=XM_019811917.2;refGene_gn=TRIM68;refGene_tx=XM_019811918.2;refGene_gn=TRIM68;refGene_tx=XM_019811919.1;refGene_gn=TRIM68;refGene_tx=XM_019811920.2;refGene_gn=TRIM68;refGene_tx=XM_019811921.1
D1	62599409	62599670
refGene_gn=TRIM68;refGene_tx=XM_003992790.5;refGene_gn=TRIM68;refGene_tx=XM_011286585.3;refGene_gn=TRIM68;refGene_tx=XM_019811917.2;refGene_gn=TRIM68;refGene_tx=XM_019811918.2;refGene_gn=TRIM68;refGene_tx=XM_019811919.1;refGene_gn=TRIM68;refGene_tx=XM_019811920.2;refGene_gn=TRIM68;refGene_tx=XM_019811921.1
D1	62600274	62600454	refGene_gn=TRIM68;refGene_tx=XM_019811917.2
D1	62600534	62600630

refGene_gn=TRIM68;refGene_tx=XM_003992790.5;refGene_gn=TRIM68;refGene_tx=XM_011286585.3;refGene_gn=TRIM68;refGene_tx=XM_019811918.2;refGene_gn=TRIM68;refGene_tx=XM_019811919.1;refGene_gn=TRIM68;refGene_tx=XM_019811920.2;refGene_gn=TRIM68;refGene_tx=XM_019811921.1
D1 62602282 62602402 refGene_gn=TRIM68;refGene_tx=XM_003992790.5
D1 62602436 62602739
refGene_gn=TRIM68;refGene_tx=XM_003992790.5;refGene_gn=TRIM68;refGene_tx=XM_011286585.3;refGene_gn=TRIM68;refGene_tx=XM_019811918.2
D1 62622654 62623725
refGene_gn=LOC101090137;refGene_tx=XM_003992791.4
D1 62639137 62640091
refGene_gn=LOC101090385;refGene_tx=XM_003992792.4
D1 62650389 62651352
refGene_gn=LOC101099832;refGene_tx=XM_003992828.2
D1 62659143 62660106
refGene_gn=LOC101090642;refGene_tx=XM_003992793.2
D1 62688220 62689183
refGene_gn=LOC101100330;refGene_tx=XM_023239490.1
D1 62697565 62698510
refGene_gn=LOC105260971;refGene_tx=XM_011286672.2
D1 62715270 62715769
refGene_gn=LOC101100589;refGene_tx=XM_011286671.3
D1 62715771 62716211
refGene_gn=LOC101100589;refGene_tx=XM_011286671.3
D1 62751960 62752905
refGene_gn=LOC101101079;refGene_tx=XM_019811924.2
D1 62764569 62765547
refGene_gn=LOC101101334;refGene_tx=XM_011286670.2
D1 62777407 62778352
refGene_gn=LOC101090892;refGene_tx=XM_003992794.3
D1 62801638 62802571
refGene_gn=LOC101080402;refGene_tx=XM_006937026.4
D1 62827410 62828358
refGene_gn=LOC101080657;refGene_tx=XM_011286584.3
D1 62837320 62838262
refGene_gn=LOC101091143;refGene_tx=XM_003992795.1
D1 62843048 62844017
refGene_gn=LOC101091397;refGene_tx=XM_003992796.2
D1 62868572 62869529
refGene_gn=LOC101091651;refGene_tx=XM_003992797.1
D1 62874944 62875898
refGene_gn=LOC101080913;refGene_tx=XM_003992833.2
D1 62901128 62902079
refGene_gn=LOC101081161;refGene_tx=XM_003992834.2
D1 62909628 62910612
refGene_gn=LOC101081408;refGene_tx=XM_023239491.1
D1 62916523 62917459
refGene_gn=LOC101091896;refGene_tx=XM_003992798.2
D1 62945643 62946585
refGene_gn=LOC101092141;refGene_tx=XM_003992799.2

D1 62954491 62955451
refGene_gn=LOC101081667;refGene_tx=XM_003992836.1
D1 62964622 62965567
refGene_gn=LOC101092395;refGene_tx=XM_003992800.4
D1 62973771 62973880
refGene_gn=LOC101081923;refGene_tx=XM_011286668.3
D1 62973881 62974723
refGene_gn=LOC101081923;refGene_tx=XM_011286668.3
D1 62980300 62981242
refGene_gn=LOC101082181;refGene_tx=XM_011286667.2
D1 62995750 62996698
refGene_gn=LOC101092641;refGene_tx=XM_003992801.1
D1 63009835 63010777
refGene_gn=LOC101092885;refGene_tx=XM_003992802.2
D1 63022907 63023846
refGene_gn=LOC101093137;refGene_tx=XM_003992803.2
D1 63033597 63034536
refGene_gn=LOC101093390;refGene_tx=XM_003992804.1
D1 63063478 63064429
refGene_gn=LOC101093639;refGene_tx=XM_011286582.2
D1 63078974 63079916
refGene_gn=LOC101082688;refGene_tx=XM_003992838.2
D1 63086196 63087147
refGene_gn=LOC101082941;refGene_tx=XM_006937028.3
D1 63096520 63097456
refGene_gn=LOC101083202;refGene_tx=XM_011286581.2
D1 63105716 63106652
refGene_gn=LOC101083448;refGene_tx=XM_003992840.2
D1 63131267 63132215
refGene_gn=LOC101083703;refGene_tx=XM_011286665.3
D1 63142059 63143004
refGene_gn=LOC101083961;refGene_tx=XM_003992841.1
D1 63149190 63150135
refGene_gn=LOC101084222;refGene_tx=XM_003992842.2
D1 63168590 63169541
refGene_gn=LOC101084734;refGene_tx=XM_023239709.1
D1 63180328 63181273
refGene_gn=LOC101084988;refGene_tx=XM_003992843.3
D1 63188540 63189485
refGene_gn=LOC101085238;refGene_tx=XM_003992844.3
D1 63203483 63204428
refGene_gn=LOC101085493;refGene_tx=XM_003992845.1
D1 63216898 63217849
refGene_gn=LOC101085745;refGene_tx=XM_003992846.1
D1 63231047 63231980
refGene_gn=LOC101086252;refGene_tx=XM_003992847.1
D1 63236155 63237115
refGene_gn=LOC101086501;refGene_tx=XM_023238990.1
D1 63245344 63245357
refGene_gn=LOC101086749;refGene_tx=XM_011286664.1

D1 63250845 63250977
refGene_gn=LOC101086749;refGene_tx=XM_011286664.1
D1 63251348 63251436
refGene_gn=LOC101086749;refGene_tx=XM_011286664.1
D1 63252525 63252823
refGene_gn=LOC101086749;refGene_tx=XM_011286664.1
D1 63252824 63253473
refGene_gn=LOC101086749;refGene_tx=XM_011286664.1
D1 63255326 63255412
refGene_gn=LOC101086749;refGene_tx=XM_011286664.1
D1 63261318 63262278
refGene_gn=LOC101087005;refGene_tx=XM_011286663.1
D1 63267208 63268228
refGene_gn=LOC101087266;refGene_tx=XM_023239710.1
D1 63305700 63306642
refGene_gn=LOC101088026;refGene_tx=XM_003992850.3
D1 63312602 63312697
refGene_gn=LOC101088271;refGene_tx=XM_023239711.1
D1 63312699 63312897
refGene_gn=LOC101088271;refGene_tx=XM_023239711.1
D1 63312899 63313527
refGene_gn=LOC101088271;refGene_tx=XM_023239711.1
D1 63321121 63321244
refGene_gn=LOC101088517;refGene_tx=XM_023239712.1
D1 63321246 63322012
refGene_gn=LOC101088517;refGene_tx=XM_023239712.1
D1 63322013 63322063
refGene_gn=LOC101088517;refGene_tx=XM_023239712.1
D1 63331416 63332411
refGene_gn=LOC101088781;refGene_tx=XM_011286579.3
D1 63335331 63335338
refGene_gn=LOC101088781;refGene_tx=XM_011286579.3
D1 63347237 63348194
refGene_gn=LOC101089041;refGene_tx=XM_019812937.2
D1 63368141 63369095
refGene_gn=LOC101094129;refGene_tx=XM_003992806.1
D1 63385844 63386801
refGene_gn=LOC101089545;refGene_tx=XM_011286662.2;refGene_gn=LOC101089545;refGene_tx=XM_019811928.1;refGene_gn=LOC101089545;refGene_tx=XM_019811929.2;refGene_gn=LOC101089545;refGene_tx=XM_019811930.2;refGene_gn=LOC101089545;refGene_tx=XM_023239492.1;refGene_gn=LOC101089545;refGene_tx=XM_023239493.1
D1 63397561 63398506
refGene_gn=LOC101094372;refGene_tx=XM_003992807.1
D1 63404605 63405547
refGene_gn=LOC105260970;refGene_tx=XM_023239713.1
D1 63415964 63416903
refGene_gn=LOC101089798;refGene_tx=XM_006937030.2
D1 63432735 63433686
refGene_gn=LOC105260969;refGene_tx=XM_011286661.2

D1 63440624 63441578
refGene_gn=LOC101090290;refGene_tx=XM_023238991.1
D1 63452823 63453777
refGene_gn=LOC111556281;refGene_tx=XM_023238992.1
D1 63473534 63473557
refGene_gn=LOC101097127;refGene_tx=XM_023239714.1
D1 63476346 63477292
refGene_gn=LOC101097127;refGene_tx=XM_023239714.1
D1 63485000 63485951
refGene_gn=LOC101090798;refGene_tx=XM_003992856.2
D1 63492755 63493712
refGene_gn=LOC105260968;refGene_tx=XM_023239715.1
D1 63504192 63505131
refGene_gn=LOC101090547;refGene_tx=XM_023238993.1
D1 63511426 63512374
refGene_gn=LOC105260967;refGene_tx=XM_011286657.1
D1 63529125 63529148
refGene_gn=LOC101086216;refGene_tx=XM_011286656.2
D1 63537175 63538121
refGene_gn=LOC101086216;refGene_tx=XM_011286656.2
D1 63545913 63546852
refGene_gn=LOC105260961;refGene_tx=XM_011286578.1
D1 63568560 63569499
refGene_gn=LOC101085192;refGene_tx=XM_023238994.1
D1 63598234 63599194
refGene_gn=LOC101084946;refGene_tx=XM_004001273.4;refGene_gn=LOC101084946;refGene_tx=XM_023238995.1
D1 63605997 63606960
refGene_gn=LOC101097913;refGene_tx=XM_023239716.1
D1 63608193 63608226
refGene_gn=LOC101097913;refGene_tx=XM_023239716.1
D1 63629886 63630015
refGene_gn=LOC101098159;refGene_tx=XM_003992882.4
D1 63630586 63630809
refGene_gn=LOC101098159;refGene_tx=XM_003992882.4
D1 63630940 63631032
refGene_gn=LOC101098159;refGene_tx=XM_003992882.4
D1 63633943 63634072
refGene_gn=LOC101091307;refGene_tx=XM_003992857.3
D1 63634831 63635054
refGene_gn=LOC101091307;refGene_tx=XM_003992857.3
D1 63635185 63635277
refGene_gn=LOC101091307;refGene_tx=XM_003992857.3
D1 63637584 63637713
refGene_gn=LOC101091558;refGene_tx=XM_003992858.2;refGene_gn=LOC101091558;refGene_tx=XM_019813043.1
D1 63638165 63638177
refGene_gn=LOC101091558;refGene_tx=XM_023239494.1
D1 63638581 63638804
refGene_gn=LOC101091558;refGene_tx=XM_003992858.2;refGene_gn=LOC101091

558;refGene_tx=XM_019813043.1;refGene_gn=LOC101091558;refGene_tx=XM_023239494.1
D1 63638916 63639008 refGene_gn=LOC101091558;refGene_tx=XM_003992858.2;refGene_gn=LOC101091558;refGene_tx=XM_019813043.1;refGene_gn=LOC101091558;refGene_tx=XM_023239494.1
D1 63649298 63649427 refGene_gn=HBE1;refGene_tx=XM_003992859.2
D1 63650228 63650451 refGene_gn=HBE1;refGene_tx=XM_003992859.2
D1 63650574 63650666 refGene_gn=HBE1;refGene_tx=XM_003992859.2
D1 63683036 63683722 refGene_gn=LOC101092048;refGene_tx=XM_011286652.1
D1 63683724 63683968 refGene_gn=LOC101092048;refGene_tx=XM_011286652.1
D1 63695756 63696692 refGene_gn=LOC101092291;refGene_tx=XM_003992860.1
D1 63713677 63714610 refGene_gn=LOC101098399;refGene_tx=XM_011286650.1
D1 63721280 63722249 refGene_gn=LOC111556648;refGene_tx=XM_023239717.1
D1 63739914 63740853 refGene_gn=LOC101092789;refGene_tx=XM_003992861.1
D1 63753844 63754792 refGene_gn=LOC101093045;refGene_tx=XM_003992862.1
D1 63776012 63776972 refGene_gn=LOC101099146;refGene_tx=XM_003992884.2
D1 63804980 63804998 refGene_gn=LOC101099654;refGene_tx=XM_006937035.4
D1 63807659 63808607 refGene_gn=LOC101099654;refGene_tx=XM_006937035.4
D1 63814718 63815705 refGene_gn=LOC101099902;refGene_tx=XM_003992887.4
D1 63820810 63821758 refGene_gn=LOC101100152;refGene_tx=XM_003992888.1
D1 63831527 63832472 refGene_gn=LOC101093295;refGene_tx=XM_003992863.2
D1 63852885 63853824 refGene_gn=LOC101100413;refGene_tx=XM_003992889.2
D1 63868259 63869198 refGene_gn=LOC101100666;refGene_tx=XM_011286649.1
D1 63878455 63879400 refGene_gn=LOC101100914;refGene_tx=XM_003992891.1
D1 63893070 63894015 refGene_gn=LOC101093544;refGene_tx=XM_003992864.1
D1 63906801 63907749 refGene_gn=LOC101101162;refGene_tx=XM_003992892.1
D1 63920008 63920965 refGene_gn=LOC101093799;refGene_tx=XM_003992865.5
D1 63932441 63933398 refGene_gn=LOC101101411;refGene_tx=XM_011286576.3
D1 63946749 63947706

refGene_gn=LOC101094046;refGene_tx=XM_003992866.4
D1 63957849 63958806
refGene_gn=LOC101101666;refGene_tx=XM_023239495.1
D1 63962644 63964264
refGene_gn=LOC102901394;refGene_tx=XM_011286574.3
D1 63975954 63977919
refGene_gn=LOC101094292;refGene_tx=XM_003992867.3
D1 63981966 63982110 refGene_gn=UBQLNL;refGene_tx=XM_003992894.4
D1 63982111 63983812 refGene_gn=UBQLNL;refGene_tx=XM_003992894.4
D1 64002770 64003815
refGene_gn=LOC111556649;refGene_tx=XM_023239719.1
D1 64004739 64004765
refGene_gn=LOC111556649;refGene_tx=XM_023239719.1
D1 64005992 64006115
refGene_gn=LOC111556649;refGene_tx=XM_023239719.1
D1 64013874 64014642
refGene_gn=LOC101094537;refGene_tx=XM_003992868.2
D1 64031673 64031924
refGene_gn=LOC101080998;refGene_tx=XM_019812939.1
D1 64031925 64032586
refGene_gn=LOC101080998;refGene_tx=XM_019812939.1
D1 64037410 64038358
refGene_gn=LOC101081249;refGene_tx=XM_003992895.4
D1 64049907 64050888
refGene_gn=LOC101081493;refGene_tx=XM_023238996.1
D1 64071310 64071733 refGene_gn=TRIM6;refGene_tx=XM_003992869.4
D1 64072211 64072307 refGene_gn=TRIM6;refGene_tx=XM_003992869.4
D1 64072988 64073219 refGene_gn=TRIM6;refGene_tx=XM_003992869.4
D1 64075955 64075978 refGene_gn=TRIM6;refGene_tx=XM_003992869.4
D1 64077303 64077404 refGene_gn=TRIM6;refGene_tx=XM_003992869.4
D1 64077690 64077717 refGene_gn=TRIM6;refGene_tx=XM_003992869.4
D1 64077939 64078502 refGene_gn=TRIM6;refGene_tx=XM_003992869.4
D1 64089155 64089222
refGene_gn=LOC101081753;refGene_tx=XM_019812940.1
D1 64091989 64092014
refGene_gn=LOC101081753;refGene_tx=XM_019812940.1
D1 64092942 64093906
refGene_gn=LOC101081753;refGene_tx=XM_019812940.1
D1 64115194 64115614
refGene_gn=TRIM5;refGene_tx=XM_006937032.3;refGene_gn=TRIM5;refGene_tx
=XM_023238908.1
D1 64116216 64116312
refGene_gn=TRIM5;refGene_tx=XM_006937032.3;refGene_gn=TRIM5;refGene_tx
=XM_023238908.1
D1 64117357 64117588
refGene_gn=TRIM5;refGene_tx=XM_006937032.3;refGene_gn=TRIM5;refGene_tx
=XM_023238908.1
D1 64123362 64123385
refGene_gn=TRIM5;refGene_tx=XM_006937032.3;refGene_gn=TRIM5;refGene_tx
=XM_023238908.1

D1 64124835 64124936
refGene_gn=TRIM5;refGene_tx=XM_006937032.3;refGene_gn=TRIM5;refGene_tx=XM_023238907.1;refGene_gn=TRIM5;refGene_tx=XM_023238908.1
D1 64125241 64125306 refGene_gn=TRIM5;refGene_tx=XM_023238908.1
D1 64137374 64137847
refGene_gn=TRIM5;refGene_tx=NM_001163659.1;refGene_gn=TRIM5;refGene_tx=XM_006937032.3;refGene_gn=TRIM5;refGene_tx=XM_019811165.2;refGene_gn=TRIM5;refGene_tx=XM_023238907.1
D1 64138493 64138589
refGene_gn=TRIM5;refGene_tx=NM_001163659.1;refGene_gn=TRIM5;refGene_tx=XM_006937032.3;refGene_gn=TRIM5;refGene_tx=XM_019811165.2;refGene_gn=TRIM5;refGene_tx=XM_023238907.1
D1 64139227 64139458
refGene_gn=TRIM5;refGene_tx=NM_001163659.1;refGene_gn=TRIM5;refGene_tx=XM_006937032.3;refGene_gn=TRIM5;refGene_tx=XM_019811165.2;refGene_gn=TRIM5;refGene_tx=XM_023238907.1
D1 64142305 64142328
refGene_gn=TRIM5;refGene_tx=NM_001163659.1;refGene_gn=TRIM5;refGene_tx=XM_006937032.3;refGene_gn=TRIM5;refGene_tx=XM_019811165.2;refGene_gn=TRIM5;refGene_tx=XM_023238907.1
D1 64143375 64143479
refGene_gn=TRIM5;refGene_tx=NM_001163659.1;refGene_gn=TRIM5;refGene_tx=XM_006937032.3;refGene_gn=TRIM5;refGene_tx=XM_019811165.2;refGene_gn=TRIM5;refGene_tx=XM_023238907.1
D1 64144086 64144106
refGene_gn=TRIM5;refGene_tx=NM_001163659.1;refGene_gn=TRIM5;refGene_tx=XM_006937032.3;refGene_gn=TRIM5;refGene_tx=XM_019811165.2;refGene_gn=TRIM5;refGene_tx=XM_023238907.1
D1 64158374 64158456 refGene_gn=TRIM22;refGene_tx=XM_019811931.1
D1 64158458 64158510 refGene_gn=TRIM22;refGene_tx=XM_019811931.1
D1 64158511 64158776 refGene_gn=TRIM22;refGene_tx=XM_019811931.1
D1 64159362 64159458 refGene_gn=TRIM22;refGene_tx=XM_019811931.1
D1 64160089 64160320 refGene_gn=TRIM22;refGene_tx=XM_019811931.1
D1 64163119 64163142 refGene_gn=TRIM22;refGene_tx=XM_019811931.1
D1 64164395 64164496 refGene_gn=TRIM22;refGene_tx=XM_019811931.1
D1 64165080 64165760 refGene_gn=TRIM22;refGene_tx=XM_019811931.1
D1 64183982 64184924
refGene_gn=LOC101082260;refGene_tx=XM_023238997.1
D1 64193951 64194137
refGene_gn=LOC101095434;refGene_tx=XM_011286572.2
D1 64194138 64194192
refGene_gn=LOC101095434;refGene_tx=XM_011286572.2
D1 64194193 64194901
refGene_gn=LOC101095434;refGene_tx=XM_011286572.2
D1 64208445 64209396
refGene_gn=LOC101095678;refGene_tx=XM_023238998.1
D1 64219646 64220624
refGene_gn=LOC105259998;refGene_tx=XM_011279821.2
D1 64239383 64240361
refGene_gn=LOC111556529;refGene_tx=XM_023238999.1

D1 64283781 64284771
refGene_gn=LOC101095915;refGene_tx=XM_003992874.3
D1 64291474 64291723
refGene_gn=LOC101096164;refGene_tx=XM_003992875.4
D1 64291724 64292441
refGene_gn=LOC101096164;refGene_tx=XM_003992875.4
D1 64303559 64304543
refGene_gn=LOC105259990;refGene_tx=XM_011279814.2
D1 64317705 64318689
refGene_gn=LOC101083291;refGene_tx=XM_003992902.3
D1 64328168 64328309
refGene_gn=LOC101096411;refGene_tx=XM_003992876.3
D1 64328311 64329136
refGene_gn=LOC101096411;refGene_tx=XM_003992876.3
D1 64340026 64340992
refGene_gn=LOC101083542;refGene_tx=XM_003992903.1
D1 64363305 64364214
refGene_gn=LOC101096667;refGene_tx=XM_011286571.1
D1 64377234 64377253
refGene_gn=LOC101083797;refGene_tx=XM_023239000.1
D1 64377430 64378387
refGene_gn=LOC101083797;refGene_tx=XM_003992904.1;refGene_gn=LOC101083
797;refGene_tx=XM_023239000.1
D1 64394798 64395758
refGene_gn=LOC111556652;refGene_tx=XM_023239721.1
D1 64403690 64404674
refGene_gn=LOC111556653;refGene_tx=XM_023239722.1
D1 64405858 64406842
refGene_gn=LOC101084313;refGene_tx=XM_003992906.2
D1 64414573 64415533
refGene_gn=LOC101084569;refGene_tx=XM_019812942.1
D1 64441614 64442577
refGene_gn=LOC101084823;refGene_tx=XM_011286644.1
D1 64458959 64459922
refGene_gn=LOC101096913;refGene_tx=XM_011286570.3
D1 64484013 64484258
refGene_gn=LOC101085068;refGene_tx=XM_011286643.1
D1 64484259 64484894
refGene_gn=LOC101085068;refGene_tx=XM_011286643.1
D1 64484896 64484973
refGene_gn=LOC101085068;refGene_tx=XM_011286643.1
D1 64491362 64492328
refGene_gn=LOC101085313;refGene_tx=XM_003992909.5
D1 64522984 64523923
refGene_gn=LOC101097169;refGene_tx=XM_003992878.1
D1 64534001 64534943
refGene_gn=LOC101097415;refGene_tx=XM_003992879.1
D1 64543647 64544553
refGene_gn=LOC101085828;refGene_tx=XM_003992910.5
D1 64544554 64544587

refGene_gn=LOC101085828;refGene_tx=XM_003992910.5
D1 64555884 64556439
refGene_gn=LOC101096456;refGene_tx=XM_004001181.3
D1 64556440 64556827
refGene_gn=LOC101096456;refGene_tx=XM_004001181.3
D1 64563175 64563541
refGene_gn=LOC101086088;refGene_tx=XM_003992911.5
D1 64563542 64563854
refGene_gn=LOC101086088;refGene_tx=XM_003992911.5
D1 64563855 64563939
refGene_gn=LOC101086088;refGene_tx=XM_003992911.5
D1 64563940 64564204
refGene_gn=LOC101086088;refGene_tx=XM_003992911.5
D1 64568509 64569073
refGene_gn=LOC101086342;refGene_tx=XM_019811942.2
D1 64579842 64580790
refGene_gn=LOC101086596;refGene_tx=XM_003992912.5
D1 64596460 64597402
refGene_gn=LOC101086841;refGene_tx=XM_003992913.1
D1 64622135 64623095
refGene_gn=LOC101097666;refGene_tx=XM_003992880.1
D1 64682374 64683322
refGene_gn=LOC101087345;refGene_tx=XM_011279808.2
D1 64696306 64697254
refGene_gn=LOC105260974;refGene_tx=XM_023239498.1
D1 64732475 64733435
refGene_gn=LOC101088105;refGene_tx=XM_003992915.4
D1 64744192 64745140
refGene_gn=LOC101096262;refGene_tx=XM_023239499.1
D1 64766824 64767766
refGene_gn=LOC101088358;refGene_tx=XM_003992916.1
D1 64773995 64774691
refGene_gn=LOC101096511;refGene_tx=XM_003992947.2
D1 64776577 64776826
refGene_gn=LOC101096511;refGene_tx=XM_003992947.2
D1 64799210 64800155
refGene_gn=LOC101096762;refGene_tx=XM_006937040.4
D1 64816539 64816961
refGene_gn=LOC101097018;refGene_tx=XM_011286695.3
D1 64816963 64816997
refGene_gn=LOC101097018;refGene_tx=XM_011286695.3
D1 64816998 64817219
refGene_gn=LOC101097018;refGene_tx=XM_011286695.3
D1 64817221 64817442
refGene_gn=LOC101097018;refGene_tx=XM_011286695.3
D1 64817443 64817481
refGene_gn=LOC101097018;refGene_tx=XM_011286695.3
D1 64853829 64854825
refGene_gn=LOC101088611;refGene_tx=XM_023239496.1;refGene_gn=LOC101088
611;refGene_tx=XM_023239497.1

D1 64871559 64872525
refGene_gn=LOC101097275;refGene_tx=XM_003992948.2
D1 64890202 64891198
refGene_gn=LOC101097517;refGene_tx=XM_003992949.3
D1 64898207 64899170
refGene_gn=LOC101097760;refGene_tx=XM_003992950.4
D1 64920636 64921354
refGene_gn=LOC101098008;refGene_tx=XM_011286696.1
D1 64921356 64921601
refGene_gn=LOC101098008;refGene_tx=XM_011286696.1
D1 64934547 64935513
refGene_gn=LOC101098255;refGene_tx=XM_003992951.3
D1 64969593 64970562
refGene_gn=LOC101098488;refGene_tx=XM_003992952.1
D1 64971431 64971643
refGene_gn=LOC111556613;refGene_tx=XM_023239500.1
D1 64971645 64972399
refGene_gn=LOC111556613;refGene_tx=XM_023239500.1
D1 65008790 65009753
refGene_gn=LOC102899190;refGene_tx=XM_019811938.2;refGene_gn=LOC102899190;refGene_tx=XM_019811941.2
D1 65014994 65015066
refGene_gn=CD1H11orf42;refGene_tx=XM_003992919.4;refGene_gn=CD1H11orf42;refGene_tx=XM_006937041.3
D1 65018582 65019381
refGene_gn=CD1H11orf42;refGene_tx=XM_003992919.4;refGene_gn=CD1H11orf42;refGene_tx=XM_006937041.3
D1 65019651 65019964
refGene_gn=CD1H11orf42;refGene_tx=XM_003992919.4;refGene_gn=CD1H11orf42;refGene_tx=XM_006937041.3
D1 65020255 65020617
refGene_gn=FAM160A2;refGene_tx=XM_006937042.3;refGene_gn=FAM160A2;refGene_tx=XM_006937043.3;refGene_gn=FAM160A2;refGene_tx=XM_006937044.3
D1 65023045 65023208
refGene_gn=FAM160A2;refGene_tx=XM_006937042.3;refGene_gn=FAM160A2;refGene_tx=XM_006937043.3;refGene_gn=FAM160A2;refGene_tx=XM_006937044.3;refGene_gn=FAM160A2;refGene_tx=XM_019811943.1
D1 65023342 65023544
refGene_gn=FAM160A2;refGene_tx=XM_006937042.3;refGene_gn=FAM160A2;refGene_tx=XM_006937043.3;refGene_gn=FAM160A2;refGene_tx=XM_006937044.3;refGene_gn=FAM160A2;refGene_tx=XM_019811943.1
D1 65025991 65026771
refGene_gn=FAM160A2;refGene_tx=XM_006937042.3;refGene_gn=FAM160A2;refGene_tx=XM_006937043.3;refGene_gn=FAM160A2;refGene_tx=XM_006937044.3;refGene_gn=FAM160A2;refGene_tx=XM_019811943.1
D1 65027155 65027361
refGene_gn=FAM160A2;refGene_tx=XM_006937042.3;refGene_gn=FAM160A2;refGene_tx=XM_006937043.3;refGene_gn=FAM160A2;refGene_tx=XM_006937044.3;refGene_gn=FAM160A2;refGene_tx=XM_019811943.1
D1 65027625 65027705

refGene_gn=FAM160A2;refGene_tx=XM_006937042.3;refGene_gn=FAM160A2;refGene_tx=XM_006937043.3;refGene_gn=FAM160A2;refGene_tx=XM_006937044.3;refGene_gn=FAM160A2;refGene_tx=XM_019811943.1
D1 65031252 65031420
refGene_gn=FAM160A2;refGene_tx=XM_006937042.3;refGene_gn=FAM160A2;refGene_tx=XM_006937043.3;refGene_gn=FAM160A2;refGene_tx=XM_006937044.3;refGene_gn=FAM160A2;refGene_tx=XM_019811943.1
D1 65031609 65031696
refGene_gn=FAM160A2;refGene_tx=XM_006937042.3;refGene_gn=FAM160A2;refGene_tx=XM_006937043.3;refGene_gn=FAM160A2;refGene_tx=XM_006937044.3;refGene_gn=FAM160A2;refGene_tx=XM_019811943.1
D1 65031885 65032044
refGene_gn=FAM160A2;refGene_tx=XM_006937042.3;refGene_gn=FAM160A2;refGene_tx=XM_006937043.3;refGene_gn=FAM160A2;refGene_tx=XM_006937044.3;refGene_gn=FAM160A2;refGene_tx=XM_019811943.1
D1 65032412 65033051
refGene_gn=FAM160A2;refGene_tx=XM_006937042.3;refGene_gn=FAM160A2;refGene_tx=XM_006937043.3;refGene_gn=FAM160A2;refGene_tx=XM_006937044.3;refGene_gn=FAM160A2;refGene_tx=XM_019811943.1
D1 65033178 65033316
refGene_gn=FAM160A2;refGene_tx=XM_006937042.3;refGene_gn=FAM160A2;refGene_tx=XM_006937043.3;refGene_gn=FAM160A2;refGene_tx=XM_006937044.3;refGene_gn=FAM160A2;refGene_tx=XM_019811943.1
D1 65048905 65048967 refGene_gn=CNGA4;refGene_tx=XM_003992922.3
D1 65049101 65049203
refGene_gn=CNGA4;refGene_tx=XM_003992922.3;refGene_gn=CNGA4;refGene_tx=XM_019811947.1
D1 65049391 65049498
refGene_gn=CNGA4;refGene_tx=XM_003992922.3;refGene_gn=CNGA4;refGene_tx=XM_019811947.1
D1 65049819 65050465
refGene_gn=CNGA4;refGene_tx=XM_003992922.3;refGene_gn=CNGA4;refGene_tx=XM_019811947.1
D1 65051236 65051586
refGene_gn=CNGA4;refGene_tx=XM_003992922.3;refGene_gn=CNGA4;refGene_tx=XM_019811947.1
D1 65053298 65053759
refGene_gn=CNGA4;refGene_tx=XM_003992922.3;refGene_gn=CNGA4;refGene_tx=XM_019811947.1
D1 65067951 65068102 refGene_gn=CCKBR;refGene_tx=XM_023239509.1
D1 65075831 65076083
refGene_gn=CCKBR;refGene_tx=XM_023239509.1;refGene_gn=CCKBR;refGene_tx=XM_023239510.1
D1 65076226 65076476
refGene_gn=CCKBR;refGene_tx=XM_023239509.1;refGene_gn=CCKBR;refGene_tx=XM_023239510.1
D1 65076770 65076943
refGene_gn=CCKBR;refGene_tx=XM_023239509.1;refGene_gn=CCKBR;refGene_tx=XM_023239510.1
D1 65077133 65077672

refGene_gn=CCKBR;refGene_tx=XM_023239509.1;refGene_gn=CCKBR;refGene_tx
=XM_023239510.1
D1 65133857 65134259 refGene_gn=CAVIN3;refGene_tx=XM_023239511.1
D1 65134803 65135187 refGene_gn=CAVIN3;refGene_tx=XM_023239511.1
D1 65188483 65188696 refGene_gn=SMPD1;refGene_tx=XM_006937046.3
D1 65188697 65188766 refGene_gn=SMPD1;refGene_tx=XM_006937046.3
D1 65189681 65190454 refGene_gn=SMPD1;refGene_tx=XM_006937046.3
D1 65190998 65191170 refGene_gn=SMPD1;refGene_tx=XM_006937046.3
D1 65191583 65191660 refGene_gn=SMPD1;refGene_tx=XM_006937046.3
D1 65191864 65192010 refGene_gn=SMPD1;refGene_tx=XM_006937046.3
D1 65192152 65192562 refGene_gn=SMPD1;refGene_tx=XM_006937046.3
D1 65193474 65193642
refGene_gn=APBB1;refGene_tx=XM_023239501.1;refGene_gn=APBB1;refGene_tx
=XM_023239502.1;refGene_gn=APBB1;refGene_tx=XM_023239503.1;refGene_gn=
APBB1;refGene_tx=XM_023239504.1;refGene_gn=APBB1;refGene_tx=XM_0232395
05.1;refGene_gn=APBB1;refGene_tx=XM_023239506.1;refGene_gn=APBB1;refGe
ne_tx=XM_023239508.1
D1 65193726 65193903
refGene_gn=APBB1;refGene_tx=XM_023239501.1;refGene_gn=APBB1;refGene_tx
=XM_023239502.1;refGene_gn=APBB1;refGene_tx=XM_023239503.1;refGene_gn=
APBB1;refGene_tx=XM_023239504.1;refGene_gn=APBB1;refGene_tx=XM_0232395
05.1;refGene_gn=APBB1;refGene_tx=XM_023239506.1;refGene_gn=APBB1;refGe
ne_tx=XM_023239508.1
D1 65194050 65194166
refGene_gn=APBB1;refGene_tx=XM_023239501.1;refGene_gn=APBB1;refGene_tx
=XM_023239502.1;refGene_gn=APBB1;refGene_tx=XM_023239503.1;refGene_gn=
APBB1;refGene_tx=XM_023239504.1;refGene_gn=APBB1;refGene_tx=XM_0232395
05.1;refGene_gn=APBB1;refGene_tx=XM_023239506.1;refGene_gn=APBB1;refGe
ne_tx=XM_023239508.1
D1 65195280 65195513 refGene_gn=APBB1;refGene_tx=XM_023239507.1
D1 65198907 65198991
refGene_gn=APBB1;refGene_tx=XM_023239501.1;refGene_gn=APBB1;refGene_tx
=XM_023239502.1;refGene_gn=APBB1;refGene_tx=XM_023239503.1;refGene_gn=
APBB1;refGene_tx=XM_023239504.1;refGene_gn=APBB1;refGene_tx=XM_0232395
05.1;refGene_gn=APBB1;refGene_tx=XM_023239506.1;refGene_gn=APBB1;refGe
ne_tx=XM_023239507.1;refGene_gn=APBB1;refGene_tx=XM_023239508.1
D1 65199271 65199356
refGene_gn=APBB1;refGene_tx=XM_023239501.1;refGene_gn=APBB1;refGene_tx
=XM_023239502.1;refGene_gn=APBB1;refGene_tx=XM_023239503.1;refGene_gn=
APBB1;refGene_tx=XM_023239504.1;refGene_gn=APBB1;refGene_tx=XM_0232395
05.1;refGene_gn=APBB1;refGene_tx=XM_023239506.1;refGene_gn=APBB1;refGe
ne_tx=XM_023239507.1;refGene_gn=APBB1;refGene_tx=XM_023239508.1
D1 65199505 65199620
refGene_gn=APBB1;refGene_tx=XM_023239501.1;refGene_gn=APBB1;refGene_tx
=XM_023239502.1;refGene_gn=APBB1;refGene_tx=XM_023239503.1;refGene_gn=
APBB1;refGene_tx=XM_023239504.1;refGene_gn=APBB1;refGene_tx=XM_0232395
05.1;refGene_gn=APBB1;refGene_tx=XM_023239506.1;refGene_gn=APBB1;refGe
ne_tx=XM_023239507.1;refGene_gn=APBB1;refGene_tx=XM_023239508.1
D1 65199915 65199921
refGene_gn=APBB1;refGene_tx=XM_023239501.1;refGene_gn=APBB1;refGene_tx

=XM_023239503.1;refGene_gn=APBB1;refGene_tx=XM_023239504.1;refGene_gn=APBB1;refGene_tx=XM_023239506.1;refGene_gn=APBB1;refGene_tx=XM_023239507.1;refGene_gn=APBB1;refGene_tx=XM_023239508.1
D1 65200027 65200155 refGene_gn=APBB1;refGene_tx=XM_023239501.1;refGene_gn=APBB1;refGene_tx=XM_023239502.1;refGene_gn=APBB1;refGene_tx=XM_023239503.1;refGene_gn=APBB1;refGene_tx=XM_023239504.1;refGene_gn=APBB1;refGene_tx=XM_023239505.1;refGene_gn=APBB1;refGene_tx=XM_023239506.1;refGene_gn=APBB1;refGene_tx=XM_023239507.1;refGene_gn=APBB1;refGene_tx=XM_023239508.1
D1 65200507 65200657 refGene_gn=APBB1;refGene_tx=XM_023239501.1;refGene_gn=APBB1;refGene_tx=XM_023239502.1;refGene_gn=APBB1;refGene_tx=XM_023239503.1;refGene_gn=APBB1;refGene_tx=XM_023239504.1;refGene_gn=APBB1;refGene_tx=XM_023239505.1;refGene_gn=APBB1;refGene_tx=XM_023239506.1;refGene_gn=APBB1;refGene_tx=XM_023239507.1;refGene_gn=APBB1;refGene_tx=XM_023239508.1
D1 65201104 65201168 refGene_gn=APBB1;refGene_tx=XM_023239501.1;refGene_gn=APBB1;refGene_tx=XM_023239502.1;refGene_gn=APBB1;refGene_tx=XM_023239503.1;refGene_gn=APBB1;refGene_tx=XM_023239504.1;refGene_gn=APBB1;refGene_tx=XM_023239505.1;refGene_gn=APBB1;refGene_tx=XM_023239506.1;refGene_gn=APBB1;refGene_tx=XM_023239507.1;refGene_gn=APBB1;refGene_tx=XM_023239508.1
D1 65201282 65201368 refGene_gn=APBB1;refGene_tx=XM_023239501.1;refGene_gn=APBB1;refGene_tx=XM_023239502.1;refGene_gn=APBB1;refGene_tx=XM_023239503.1;refGene_gn=APBB1;refGene_tx=XM_023239504.1;refGene_gn=APBB1;refGene_tx=XM_023239505.1;refGene_gn=APBB1;refGene_tx=XM_023239506.1;refGene_gn=APBB1;refGene_tx=XM_023239507.1;refGene_gn=APBB1;refGene_tx=XM_023239508.1
D1 65201447 65201504 refGene_gn=APBB1;refGene_tx=XM_023239501.1;refGene_gn=APBB1;refGene_tx=XM_023239502.1;refGene_gn=APBB1;refGene_tx=XM_023239503.1;refGene_gn=APBB1;refGene_tx=XM_023239504.1;refGene_gn=APBB1;refGene_tx=XM_023239505.1;refGene_gn=APBB1;refGene_tx=XM_023239506.1;refGene_gn=APBB1;refGene_tx=XM_023239507.1;refGene_gn=APBB1;refGene_tx=XM_023239508.1
D1 65201596 65201775 refGene_gn=APBB1;refGene_tx=XM_023239501.1;refGene_gn=APBB1;refGene_tx=XM_023239502.1;refGene_gn=APBB1;refGene_tx=XM_023239503.1;refGene_gn=APBB1;refGene_tx=XM_023239504.1;refGene_gn=APBB1;refGene_tx=XM_023239505.1;refGene_gn=APBB1;refGene_tx=XM_023239506.1;refGene_gn=APBB1;refGene_tx=XM_023239507.1;refGene_gn=APBB1;refGene_tx=XM_023239508.1
D1 65203301 65203415 refGene_gn=APBB1;refGene_tx=XM_023239501.1;refGene_gn=APBB1;refGene_tx=XM_023239502.1;refGene_gn=APBB1;refGene_tx=XM_023239506.1;refGene_gn=APBB1;refGene_tx=XM_023239507.1
D1 65207531 65208252 refGene_gn=APBB1;refGene_tx=XM_023239501.1;refGene_gn=APBB1;refGene_tx=XM_023239502.1;refGene_gn=APBB1;refGene_tx=XM_023239503.1;refGene_gn=APBB1;refGene_tx=XM_023239504.1;refGene_gn=APBB1;refGene_tx=XM_023239505.1;refGene_gn=APBB1;refGene_tx=XM_023239506.1;refGene_gn=APBB1;refGene_tx=XM_023239507.1
D1 65224318 65224578 refGene_gn=HPX;refGene_tx=XM_003992929.4

D1 65224716 65224879 refGene_gn=HPX;refGene_tx=XM_003992929.4
D1 65224959 65225090 refGene_gn=HPX;refGene_tx=XM_003992929.4
D1 65229307 65229439 refGene_gn=HPX;refGene_tx=XM_003992929.4
D1 65229696 65229909 refGene_gn=HPX;refGene_tx=XM_003992929.4
D1 65230586 65230740 refGene_gn=HPX;refGene_tx=XM_003992929.4
D1 65232384 65232506 refGene_gn=HPX;refGene_tx=XM_003992929.4
D1 65232673 65232745 refGene_gn=HPX;refGene_tx=XM_003992929.4
D1 65232842 65232907 refGene_gn=HPX;refGene_tx=XM_003992929.4
D1 65233054 65233137 refGene_gn=HPX;refGene_tx=XM_003992929.4
D1 65242585 65242738
refGene_gn=TRIM3;refGene_tx=XM_011286689.2;refGene_gn=TRIM3;refGene_tx
=XM_019811972.1;refGene_gn=TRIM3;refGene_tx=XM_019811973.1
D1 65242888 65243029
refGene_gn=TRIM3;refGene_tx=XM_011286689.2;refGene_gn=TRIM3;refGene_tx
=XM_019811972.1;refGene_gn=TRIM3;refGene_tx=XM_019811973.1
D1 65244086 65244157
refGene_gn=TRIM3;refGene_tx=XM_011286689.2;refGene_gn=TRIM3;refGene_tx
=XM_019811972.1;refGene_gn=TRIM3;refGene_tx=XM_019811973.1
D1 65244411 65244580
refGene_gn=TRIM3;refGene_tx=XM_011286689.2;refGene_gn=TRIM3;refGene_tx
=XM_019811972.1;refGene_gn=TRIM3;refGene_tx=XM_019811973.1
D1 65244761 65244929
refGene_gn=TRIM3;refGene_tx=XM_011286689.2;refGene_gn=TRIM3;refGene_tx
=XM_019811972.1;refGene_gn=TRIM3;refGene_tx=XM_019811973.1
D1 65250233 65250337
refGene_gn=TRIM3;refGene_tx=XM_011286689.2;refGene_gn=TRIM3;refGene_tx
=XM_019811972.1;refGene_gn=TRIM3;refGene_tx=XM_019811973.1
D1 65250459 65251192
refGene_gn=TRIM3;refGene_tx=XM_011286689.2;refGene_gn=TRIM3;refGene_tx
=XM_019811972.1;refGene_gn=TRIM3;refGene_tx=XM_019811973.1
D1 65251453 65251634
refGene_gn=TRIM3;refGene_tx=XM_011286689.2;refGene_gn=TRIM3;refGene_tx
=XM_019811972.1;refGene_gn=TRIM3;refGene_tx=XM_019811973.1
D1 65251866 65252018
refGene_gn=TRIM3;refGene_tx=XM_011286689.2;refGene_gn=TRIM3;refGene_tx
=XM_019811972.1;refGene_gn=TRIM3;refGene_tx=XM_019811973.1
D1 65252241 65252473
refGene_gn=TRIM3;refGene_tx=XM_011286689.2;refGene_gn=TRIM3;refGene_tx
=XM_019811972.1;refGene_gn=TRIM3;refGene_tx=XM_019811973.1
D1 65258062 65258193
refGene_gn=TRIM3;refGene_tx=XM_011286689.2;refGene_gn=TRIM3;refGene_tx
=XM_019811972.1;refGene_gn=TRIM3;refGene_tx=XM_019811973.1
D1 65269056 65269212
refGene_gn=ARFIP2;refGene_tx=XM_003992932.4;refGene_gn=ARFIP2;refGene_
tx=XM_011286690.1;refGene_gn=ARFIP2;refGene_tx=XM_019811974.1
D1 65269656 65269831
refGene_gn=ARFIP2;refGene_tx=XM_003992932.4;refGene_gn=ARFIP2;refGene_
tx=XM_011286690.1;refGene_gn=ARFIP2;refGene_tx=XM_019811974.1
D1 65269975 65270133
refGene_gn=ARFIP2;refGene_tx=XM_003992932.4;refGene_gn=ARFIP2;refGene_

tx=XM_011286690.1;refGene_gn=ARFIP2;refGene_tx=XM_019811974.1
D1 65270659 65270881
refGene_gn=ARFIP2;refGene_tx=XM_003992932.4;refGene_gn=ARFIP2;refGene_tx=XM_011286690.1;refGene_gn=ARFIP2;refGene_tx=XM_019811974.1
D1 65271068 65271187
refGene_gn=ARFIP2;refGene_tx=XM_003992932.4;refGene_gn=ARFIP2;refGene_tx=XM_019811974.1
D1 65271909 65272006
refGene_gn=ARFIP2;refGene_tx=XM_003992932.4;refGene_gn=ARFIP2;refGene_tx=XM_011286690.1;refGene_gn=ARFIP2;refGene_tx=XM_019811974.1
D1 65272259 65272358
refGene_gn=ARFIP2;refGene_tx=XM_003992932.4;refGene_gn=ARFIP2;refGene_tx=XM_019811974.1
D1 65273483 65273540 refGene_gn=TIMM10B;refGene_tx=XM_003992956.3
D1 65273716 65273812 refGene_gn=TIMM10B;refGene_tx=XM_003992956.3
D1 65273985 65274162 refGene_gn=TIMM10B;refGene_tx=XM_003992956.3
D1 65288420 65288484
refGene_gn=DNHD1;refGene_tx=XM_019811955.1;refGene_gn=DNHD1;refGene_tx=XM_019811967.1;refGene_gn=DNHD1;refGene_tx=XM_019811970.1;refGene_gn=DNHD1;refGene_tx=XM_019811971.1;refGene_gn=DNHD1;refGene_tx=XM_023239512.1;refGene_gn=DNHD1;refGene_tx=XM_023239513.1;refGene_gn=DNHD1;refGene_tx=XM_023239514.1;refGene_gn=DNHD1;refGene_tx=XM_023239516.1;refGene_gn=DNHD1;refGene_tx=XM_023239517.1
D1 65288754 65289589
refGene_gn=DNHD1;refGene_tx=XM_019811955.1;refGene_gn=DNHD1;refGene_tx=XM_019811967.1;refGene_gn=DNHD1;refGene_tx=XM_019811970.1;refGene_gn=DNHD1;refGene_tx=XM_019811971.1;refGene_gn=DNHD1;refGene_tx=XM_023239512.1;refGene_gn=DNHD1;refGene_tx=XM_023239513.1;refGene_gn=DNHD1;refGene_tx=XM_023239514.1;refGene_gn=DNHD1;refGene_tx=XM_023239516.1;refGene_gn=DNHD1;refGene_tx=XM_023239517.1
D1 65293699 65293873
refGene_gn=DNHD1;refGene_tx=XM_019811955.1;refGene_gn=DNHD1;refGene_tx=XM_019811967.1;refGene_gn=DNHD1;refGene_tx=XM_019811970.1;refGene_gn=DNHD1;refGene_tx=XM_019811971.1;refGene_gn=DNHD1;refGene_tx=XM_023239512.1;refGene_gn=DNHD1;refGene_tx=XM_023239513.1;refGene_gn=DNHD1;refGene_tx=XM_023239514.1;refGene_gn=DNHD1;refGene_tx=XM_023239516.1;refGene_gn=DNHD1;refGene_tx=XM_023239517.1
D1 65299160 65299364
refGene_gn=DNHD1;refGene_tx=XM_019811955.1;refGene_gn=DNHD1;refGene_tx=XM_019811967.1;refGene_gn=DNHD1;refGene_tx=XM_019811970.1;refGene_gn=DNHD1;refGene_tx=XM_019811971.1;refGene_gn=DNHD1;refGene_tx=XM_023239512.1;refGene_gn=DNHD1;refGene_tx=XM_023239513.1;refGene_gn=DNHD1;refGene_tx=XM_023239514.1;refGene_gn=DNHD1;refGene_tx=XM_023239516.1;refGene_gn=DNHD1;refGene_tx=XM_023239517.1
D1 65299443 65299554
refGene_gn=DNHD1;refGene_tx=XM_019811955.1;refGene_gn=DNHD1;refGene_tx=XM_019811967.1;refGene_gn=DNHD1;refGene_tx=XM_019811970.1;refGene_gn=DNHD1;refGene_tx=XM_019811971.1;refGene_gn=DNHD1;refGene_tx=XM_023239512.1;refGene_gn=DNHD1;refGene_tx=XM_023239513.1;refGene_gn=DNHD1;refGene_tx=XM_023239514.1;refGene_gn=DNHD1;refGene_tx=XM_023239516.1;refGen

e_gn=DNHD1;refGene_tx=XM_023239517.1
D1 65301108 65301265
refGene_gn=DNHD1;refGene_tx=XM_019811955.1;refGene_gn=DNHD1;refGene_tx=XM_019811967.1;refGene_gn=DNHD1;refGene_tx=XM_019811970.1;refGene_gn=DNHD1;refGene_tx=XM_019811971.1;refGene_gn=DNHD1;refGene_tx=XM_023239512.1;refGene_gn=DNHD1;refGene_tx=XM_023239513.1;refGene_gn=DNHD1;refGene_tx=XM_023239514.1;refGene_gn=DNHD1;refGene_tx=XM_023239516.1;refGene_gn=DNHD1;refGene_tx=XM_023239517.1
D1 65309989 65310244
refGene_gn=DNHD1;refGene_tx=XM_019811955.1;refGene_gn=DNHD1;refGene_tx=XM_019811967.1;refGene_gn=DNHD1;refGene_tx=XM_019811970.1;refGene_gn=DNHD1;refGene_tx=XM_019811971.1;refGene_gn=DNHD1;refGene_tx=XM_023239512.1;refGene_gn=DNHD1;refGene_tx=XM_023239513.1;refGene_gn=DNHD1;refGene_tx=XM_023239514.1;refGene_gn=DNHD1;refGene_tx=XM_023239516.1;refGene_gn=DNHD1;refGene_tx=XM_023239517.1
D1 65310353 65310491
refGene_gn=DNHD1;refGene_tx=XM_019811955.1;refGene_gn=DNHD1;refGene_tx=XM_019811967.1;refGene_gn=DNHD1;refGene_tx=XM_019811970.1;refGene_gn=DNHD1;refGene_tx=XM_019811971.1;refGene_gn=DNHD1;refGene_tx=XM_023239512.1;refGene_gn=DNHD1;refGene_tx=XM_023239513.1;refGene_gn=DNHD1;refGene_tx=XM_023239514.1;refGene_gn=DNHD1;refGene_tx=XM_023239516.1;refGene_gn=DNHD1;refGene_tx=XM_023239517.1
D1 65310629 65310681
refGene_gn=DNHD1;refGene_tx=XM_019811955.1;refGene_gn=DNHD1;refGene_tx=XM_019811967.1;refGene_gn=DNHD1;refGene_tx=XM_019811970.1;refGene_gn=DNHD1;refGene_tx=XM_019811971.1;refGene_gn=DNHD1;refGene_tx=XM_023239512.1;refGene_gn=DNHD1;refGene_tx=XM_023239513.1;refGene_gn=DNHD1;refGene_tx=XM_023239514.1;refGene_gn=DNHD1;refGene_tx=XM_023239516.1;refGene_gn=DNHD1;refGene_tx=XM_023239517.1
D1 65319919 65320185
refGene_gn=DNHD1;refGene_tx=XM_019811955.1;refGene_gn=DNHD1;refGene_tx=XM_019811967.1;refGene_gn=DNHD1;refGene_tx=XM_019811970.1;refGene_gn=DNHD1;refGene_tx=XM_019811971.1;refGene_gn=DNHD1;refGene_tx=XM_023239512.1;refGene_gn=DNHD1;refGene_tx=XM_023239513.1;refGene_gn=DNHD1;refGene_tx=XM_023239514.1;refGene_gn=DNHD1;refGene_tx=XM_023239516.1;refGene_gn=DNHD1;refGene_tx=XM_023239517.1
D1 65320275 65320519
refGene_gn=DNHD1;refGene_tx=XM_019811955.1;refGene_gn=DNHD1;refGene_tx=XM_019811967.1;refGene_gn=DNHD1;refGene_tx=XM_019811970.1;refGene_gn=DNHD1;refGene_tx=XM_019811971.1;refGene_gn=DNHD1;refGene_tx=XM_023239512.1;refGene_gn=DNHD1;refGene_tx=XM_023239513.1;refGene_gn=DNHD1;refGene_tx=XM_023239514.1;refGene_gn=DNHD1;refGene_tx=XM_023239516.1;refGene_gn=DNHD1;refGene_tx=XM_023239517.1
D1 65327723 65327881
refGene_gn=DNHD1;refGene_tx=XM_019811955.1;refGene_gn=DNHD1;refGene_tx=XM_019811967.1;refGene_gn=DNHD1;refGene_tx=XM_019811970.1;refGene_gn=DNHD1;refGene_tx=XM_019811971.1;refGene_gn=DNHD1;refGene_tx=XM_023239512.1;refGene_gn=DNHD1;refGene_tx=XM_023239513.1;refGene_gn=DNHD1;refGene_tx=XM_023239514.1;refGene_gn=DNHD1;refGene_tx=XM_023239516.1;refGene_gn=DNHD1;refGene_tx=XM_023239517.1

D1 65328331 65328834
refGene_gn=DNHD1;refGene_tx=XM_019811955.1;refGene_gn=DNHD1;refGene_tx=XM_019811967.1;refGene_gn=DNHD1;refGene_tx=XM_019811971.1;refGene_gn=DNHD1;refGene_tx=XM_023239512.1;refGene_gn=DNHD1;refGene_tx=XM_023239513.1;refGene_gn=DNHD1;refGene_tx=XM_023239514.1;refGene_gn=DNHD1;refGene_tx=XM_023239516.1;refGene_gn=DNHD1;refGene_tx=XM_023239518.1
D1 65334308 65334436
refGene_gn=DNHD1;refGene_tx=XM_019811955.1;refGene_gn=DNHD1;refGene_tx=XM_019811967.1;refGene_gn=DNHD1;refGene_tx=XM_019811970.1;refGene_gn=DNHD1;refGene_tx=XM_023239512.1;refGene_gn=DNHD1;refGene_tx=XM_023239513.1;refGene_gn=DNHD1;refGene_tx=XM_023239514.1;refGene_gn=DNHD1;refGene_tx=XM_023239516.1;refGene_gn=DNHD1;refGene_tx=XM_023239518.1
D1 65334536 65334735
refGene_gn=DNHD1;refGene_tx=XM_019811955.1;refGene_gn=DNHD1;refGene_tx=XM_019811967.1;refGene_gn=DNHD1;refGene_tx=XM_019811970.1;refGene_gn=DNHD1;refGene_tx=XM_023239512.1;refGene_gn=DNHD1;refGene_tx=XM_023239513.1;refGene_gn=DNHD1;refGene_tx=XM_023239514.1;refGene_gn=DNHD1;refGene_tx=XM_023239516.1;refGene_gn=DNHD1;refGene_tx=XM_023239517.1;refGene_gn=DNHD1;refGene_tx=XM_023239518.1
D1 65335137 65335232
refGene_gn=DNHD1;refGene_tx=XM_019811955.1;refGene_gn=DNHD1;refGene_tx=XM_019811967.1;refGene_gn=DNHD1;refGene_tx=XM_023239512.1;refGene_gn=DNHD1;refGene_tx=XM_023239513.1;refGene_gn=DNHD1;refGene_tx=XM_023239514.1;refGene_gn=DNHD1;refGene_tx=XM_023239516.1;refGene_gn=DNHD1;refGene_tx=XM_023239517.1;refGene_gn=DNHD1;refGene_tx=XM_023239518.1
D1 65335753 65335961
refGene_gn=DNHD1;refGene_tx=XM_019811955.1;refGene_gn=DNHD1;refGene_tx=XM_019811967.1;refGene_gn=DNHD1;refGene_tx=XM_023239512.1;refGene_gn=DNHD1;refGene_tx=XM_023239513.1;refGene_gn=DNHD1;refGene_tx=XM_023239514.1;refGene_gn=DNHD1;refGene_tx=XM_023239516.1;refGene_gn=DNHD1;refGene_tx=XM_023239517.1;refGene_gn=DNHD1;refGene_tx=XM_023239518.1
D1 65337494 65337620
refGene_gn=DNHD1;refGene_tx=XM_019811955.1;refGene_gn=DNHD1;refGene_tx=XM_019811967.1;refGene_gn=DNHD1;refGene_tx=XM_023239512.1;refGene_gn=DNHD1;refGene_tx=XM_023239513.1;refGene_gn=DNHD1;refGene_tx=XM_023239514.1;refGene_gn=DNHD1;refGene_tx=XM_023239516.1;refGene_gn=DNHD1;refGene_tx=XM_023239517.1;refGene_gn=DNHD1;refGene_tx=XM_023239518.1
D1 65337922 65338020
refGene_gn=DNHD1;refGene_tx=XM_019811955.1;refGene_gn=DNHD1;refGene_tx=XM_019811967.1;refGene_gn=DNHD1;refGene_tx=XM_023239512.1;refGene_gn=DNHD1;refGene_tx=XM_023239513.1;refGene_gn=DNHD1;refGene_tx=XM_023239514.1;refGene_gn=DNHD1;refGene_tx=XM_023239516.1;refGene_gn=DNHD1;refGene_tx=XM_023239517.1;refGene_gn=DNHD1;refGene_tx=XM_023239518.1
D1 65338121 65341005
refGene_gn=DNHD1;refGene_tx=XM_019811955.1;refGene_gn=DNHD1;refGene_tx=XM_019811967.1;refGene_gn=DNHD1;refGene_tx=XM_023239512.1;refGene_gn=DNHD1;refGene_tx=XM_023239513.1;refGene_gn=DNHD1;refGene_tx=XM_023239514.1;refGene_gn=DNHD1;refGene_tx=XM_023239516.1;refGene_gn=DNHD1;refGene_tx=XM_023239517.1;refGene_gn=DNHD1;refGene_tx=XM_023239518.1
D1 65341204 65341382

refGene_gn=DNHD1;refGene_tx=XM_019811955.1;refGene_gn=DNHD1;refGene_tx=XM_019811967.1;refGene_gn=DNHD1;refGene_tx=XM_023239512.1;refGene_gn=DNHD1;refGene_tx=XM_023239513.1;refGene_gn=DNHD1;refGene_tx=XM_023239514.1;refGene_gn=DNHD1;refGene_tx=XM_023239516.1;refGene_gn=DNHD1;refGene_tx=XM_023239517.1;refGene_gn=DNHD1;refGene_tx=XM_023239518.1
D1 65341553 65341746
refGene_gn=DNHD1;refGene_tx=XM_019811955.1;refGene_gn=DNHD1;refGene_tx=XM_019811967.1;refGene_gn=DNHD1;refGene_tx=XM_023239512.1;refGene_gn=DNHD1;refGene_tx=XM_023239513.1;refGene_gn=DNHD1;refGene_tx=XM_023239514.1;refGene_gn=DNHD1;refGene_tx=XM_023239516.1;refGene_gn=DNHD1;refGene_tx=XM_023239517.1;refGene_gn=DNHD1;refGene_tx=XM_023239518.1
D1 65341988 65342277
refGene_gn=DNHD1;refGene_tx=XM_019811955.1;refGene_gn=DNHD1;refGene_tx=XM_019811967.1;refGene_gn=DNHD1;refGene_tx=XM_023239512.1;refGene_gn=DNHD1;refGene_tx=XM_023239513.1;refGene_gn=DNHD1;refGene_tx=XM_023239514.1;refGene_gn=DNHD1;refGene_tx=XM_023239516.1;refGene_gn=DNHD1;refGene_tx=XM_023239517.1;refGene_gn=DNHD1;refGene_tx=XM_023239518.1
D1 65349538 65351147
refGene_gn=DNHD1;refGene_tx=XM_019811955.1;refGene_gn=DNHD1;refGene_tx=XM_019811967.1;refGene_gn=DNHD1;refGene_tx=XM_023239512.1;refGene_gn=DNHD1;refGene_tx=XM_023239513.1;refGene_gn=DNHD1;refGene_tx=XM_023239514.1;refGene_gn=DNHD1;refGene_tx=XM_023239516.1;refGene_gn=DNHD1;refGene_tx=XM_023239517.1;refGene_gn=DNHD1;refGene_tx=XM_023239518.1
D1 65351333 65351542
refGene_gn=DNHD1;refGene_tx=XM_019811955.1;refGene_gn=DNHD1;refGene_tx=XM_019811967.1;refGene_gn=DNHD1;refGene_tx=XM_023239512.1;refGene_gn=DNHD1;refGene_tx=XM_023239513.1;refGene_gn=DNHD1;refGene_tx=XM_023239514.1;refGene_gn=DNHD1;refGene_tx=XM_023239516.1;refGene_gn=DNHD1;refGene_tx=XM_023239517.1;refGene_gn=DNHD1;refGene_tx=XM_023239518.1
D1 65351742 65351947
refGene_gn=DNHD1;refGene_tx=XM_019811955.1;refGene_gn=DNHD1;refGene_tx=XM_019811967.1;refGene_gn=DNHD1;refGene_tx=XM_023239512.1;refGene_gn=DNHD1;refGene_tx=XM_023239513.1;refGene_gn=DNHD1;refGene_tx=XM_023239514.1;refGene_gn=DNHD1;refGene_tx=XM_023239516.1;refGene_gn=DNHD1;refGene_tx=XM_023239517.1;refGene_gn=DNHD1;refGene_tx=XM_023239518.1
D1 65352019 65352122
refGene_gn=DNHD1;refGene_tx=XM_019811955.1;refGene_gn=DNHD1;refGene_tx=XM_019811967.1;refGene_gn=DNHD1;refGene_tx=XM_023239512.1;refGene_gn=DNHD1;refGene_tx=XM_023239513.1;refGene_gn=DNHD1;refGene_tx=XM_023239514.1;refGene_gn=DNHD1;refGene_tx=XM_023239516.1;refGene_gn=DNHD1;refGene_tx=XM_023239517.1;refGene_gn=DNHD1;refGene_tx=XM_023239518.1
D1 65355993 65356143
refGene_gn=DNHD1;refGene_tx=XM_019811955.1;refGene_gn=DNHD1;refGene_tx=XM_019811967.1;refGene_gn=DNHD1;refGene_tx=XM_023239512.1;refGene_gn=DNHD1;refGene_tx=XM_023239513.1;refGene_gn=DNHD1;refGene_tx=XM_023239514.1;refGene_gn=DNHD1;refGene_tx=XM_023239516.1;refGene_gn=DNHD1;refGene_tx=XM_023239517.1;refGene_gn=DNHD1;refGene_tx=XM_023239518.1
D1 65356389 65356572
refGene_gn=DNHD1;refGene_tx=XM_019811955.1;refGene_gn=DNHD1;refGene_tx=XM_019811967.1;refGene_gn=DNHD1;refGene_tx=XM_023239512.1;refGene_gn=

DNHD1;refGene_tx=XM_023239513.1;refGene_gn=DNHD1;refGene_tx=XM_0232395
14.1;refGene_gn=DNHD1;refGene_tx=XM_023239516.1;refGene_gn=DNHD1;refGe
ne_tx=XM_023239517.1;refGene_gn=DNHD1;refGene_tx=XM_023239518.1
D1 65356701 65357133
refGene_gn=DNHD1;refGene_tx=XM_019811955.1;refGene_gn=DNHD1;refGene_tx
=XM_019811967.1;refGene_gn=DNHD1;refGene_tx=XM_023239512.1;refGene_gn=
DNHD1;refGene_tx=XM_023239513.1;refGene_gn=DNHD1;refGene_tx=XM_0232395
14.1;refGene_gn=DNHD1;refGene_tx=XM_023239516.1;refGene_gn=DNHD1;refGe
ne_tx=XM_023239517.1;refGene_gn=DNHD1;refGene_tx=XM_023239518.1
D1 65357352 65357812
refGene_gn=DNHD1;refGene_tx=XM_019811955.1;refGene_gn=DNHD1;refGene_tx
=XM_019811967.1;refGene_gn=DNHD1;refGene_tx=XM_023239512.1;refGene_gn=
DNHD1;refGene_tx=XM_023239513.1;refGene_gn=DNHD1;refGene_tx=XM_0232395
14.1;refGene_gn=DNHD1;refGene_tx=XM_023239516.1;refGene_gn=DNHD1;refGe
ne_tx=XM_023239517.1;refGene_gn=DNHD1;refGene_tx=XM_023239518.1
D1 65362098 65362431
refGene_gn=DNHD1;refGene_tx=XM_019811955.1;refGene_gn=DNHD1;refGene_tx
=XM_019811967.1;refGene_gn=DNHD1;refGene_tx=XM_023239512.1;refGene_gn=
DNHD1;refGene_tx=XM_023239513.1;refGene_gn=DNHD1;refGene_tx=XM_0232395
14.1;refGene_gn=DNHD1;refGene_tx=XM_023239516.1;refGene_gn=DNHD1;refGe
ne_tx=XM_023239517.1;refGene_gn=DNHD1;refGene_tx=XM_023239518.1
D1 65362658 65362811
refGene_gn=DNHD1;refGene_tx=XM_019811955.1;refGene_gn=DNHD1;refGene_tx
=XM_019811967.1;refGene_gn=DNHD1;refGene_tx=XM_023239512.1;refGene_gn=
DNHD1;refGene_tx=XM_023239513.1;refGene_gn=DNHD1;refGene_tx=XM_0232395
14.1;refGene_gn=DNHD1;refGene_tx=XM_023239516.1;refGene_gn=DNHD1;refGe
ne_tx=XM_023239517.1;refGene_gn=DNHD1;refGene_tx=XM_023239518.1
D1 65362988 65363167
refGene_gn=DNHD1;refGene_tx=XM_019811955.1;refGene_gn=DNHD1;refGene_tx
=XM_023239512.1;refGene_gn=DNHD1;refGene_tx=XM_023239513.1;refGene_gn=
DNHD1;refGene_tx=XM_023239514.1;refGene_gn=DNHD1;refGene_tx=XM_0232395
16.1;refGene_gn=DNHD1;refGene_tx=XM_023239517.1;refGene_gn=DNHD1;refGe
ne_tx=XM_023239518.1
D1 65363299 65364259
refGene_gn=DNHD1;refGene_tx=XM_019811955.1;refGene_gn=DNHD1;refGene_tx
=XM_019811967.1;refGene_gn=DNHD1;refGene_tx=XM_023239512.1;refGene_gn=
DNHD1;refGene_tx=XM_023239513.1;refGene_gn=DNHD1;refGene_tx=XM_0232395
14.1;refGene_gn=DNHD1;refGene_tx=XM_023239516.1;refGene_gn=DNHD1;refGe
ne_tx=XM_023239517.1;refGene_gn=DNHD1;refGene_tx=XM_023239518.1
D1 65364458 65364645
refGene_gn=DNHD1;refGene_tx=XM_019811955.1;refGene_gn=DNHD1;refGene_tx
=XM_023239512.1;refGene_gn=DNHD1;refGene_tx=XM_023239513.1;refGene_gn=
DNHD1;refGene_tx=XM_023239514.1;refGene_gn=DNHD1;refGene_tx=XM_0232395
16.1;refGene_gn=DNHD1;refGene_tx=XM_023239517.1;refGene_gn=DNHD1;refGe
ne_tx=XM_023239518.1
D1 65364849 65364972
refGene_gn=DNHD1;refGene_tx=XM_019811955.1;refGene_gn=DNHD1;refGene_tx
=XM_023239512.1;refGene_gn=DNHD1;refGene_tx=XM_023239513.1;refGene_gn=
DNHD1;refGene_tx=XM_023239514.1;refGene_gn=DNHD1;refGene_tx=XM_0232395
16.1;refGene_gn=DNHD1;refGene_tx=XM_023239517.1;refGene_gn=DNHD1;refGe

ne_tx=XM_023239518.1
D1	65365057	65365259
refGene_gn=DNHD1;refGene_tx=XM_019811955.1;refGene_gn=DNHD1;refGene_tx=XM_023239512.1;refGene_gn=DNHD1;refGene_tx=XM_023239513.1;refGene_gn=DNHD1;refGene_tx=XM_023239514.1;refGene_gn=DNHD1;refGene_tx=XM_023239516.1;refGene_gn=DNHD1;refGene_tx=XM_023239517.1;refGene_gn=DNHD1;refGene_tx=XM_023239518.1
D1	65366222	65366314
refGene_gn=DNHD1;refGene_tx=XM_019811955.1;refGene_gn=DNHD1;refGene_tx=XM_023239512.1;refGene_gn=DNHD1;refGene_tx=XM_023239513.1;refGene_gn=DNHD1;refGene_tx=XM_023239514.1;refGene_gn=DNHD1;refGene_tx=XM_023239516.1;refGene_gn=DNHD1;refGene_tx=XM_023239517.1;refGene_gn=DNHD1;refGene_tx=XM_023239518.1
D1	65366448	65366598
refGene_gn=DNHD1;refGene_tx=XM_019811955.1;refGene_gn=DNHD1;refGene_tx=XM_023239512.1;refGene_gn=DNHD1;refGene_tx=XM_023239513.1;refGene_gn=DNHD1;refGene_tx=XM_023239514.1;refGene_gn=DNHD1;refGene_tx=XM_023239516.1;refGene_gn=DNHD1;refGene_tx=XM_023239517.1;refGene_gn=DNHD1;refGene_tx=XM_023239518.1
D1	65366811	65367611
refGene_gn=DNHD1;refGene_tx=XM_019811955.1;refGene_gn=DNHD1;refGene_tx=XM_023239512.1;refGene_gn=DNHD1;refGene_tx=XM_023239513.1;refGene_gn=DNHD1;refGene_tx=XM_023239514.1;refGene_gn=DNHD1;refGene_tx=XM_023239516.1;refGene_gn=DNHD1;refGene_tx=XM_023239517.1;refGene_gn=DNHD1;refGene_tx=XM_023239518.1
D1	65367789	65368140
refGene_gn=DNHD1;refGene_tx=XM_019811955.1;refGene_gn=DNHD1;refGene_tx=XM_023239512.1;refGene_gn=DNHD1;refGene_tx=XM_023239513.1;refGene_gn=DNHD1;refGene_tx=XM_023239514.1;refGene_gn=DNHD1;refGene_tx=XM_023239516.1;refGene_gn=DNHD1;refGene_tx=XM_023239517.1;refGene_gn=DNHD1;refGene_tx=XM_023239518.1
D1	65392919	65393039	refGene_gn=RRP8;refGene_tx=XM_023239520.1
D1	65393264	65393361	refGene_gn=RRP8;refGene_tx=XM_023239520.1
D1	65393445	65393552	refGene_gn=RRP8;refGene_tx=XM_023239520.1
D1	65393729	65393859	refGene_gn=RRP8;refGene_tx=XM_023239520.1
D1	65393948	65394402	refGene_gn=RRP8;refGene_tx=XM_023239520.1
D1	65394628	65394995	refGene_gn=RRP8;refGene_tx=XM_023239520.1
D1	65396143	65396242	refGene_gn=RRP8;refGene_tx=XM_023239520.1
D1	65397021	65397110	refGene_gn=ILK;refGene_tx=XM_023239522.1
D1	65400520	65400686	refGene_gn=ILK;refGene_tx=XM_023239522.1
D1	65400829	65400925	refGene_gn=ILK;refGene_tx=XM_023239522.1
D1	65401126	65401223	refGene_gn=ILK;refGene_tx=XM_023239522.1
D1	65401323	65401407	refGene_gn=ILK;refGene_tx=XM_023239522.1
D1	65401513	65401599	refGene_gn=ILK;refGene_tx=XM_023239522.1
D1	65401740	65401860	refGene_gn=ILK;refGene_tx=XM_023239522.1
D1	65401958	65402085	refGene_gn=ILK;refGene_tx=XM_023239522.1
D1	65402166	65402288	refGene_gn=ILK;refGene_tx=XM_023239522.1
D1	65402386	65402486	refGene_gn=ILK;refGene_tx=XM_023239522.1
D1	65402593	65402724	refGene_gn=ILK;refGene_tx=XM_023239522.1
D1	65402908	65403058	refGene_gn=ILK;refGene_tx=XM_023239522.1

| | | | |
|---|---|---|---|
| D1 | 65403371 | 65403461 | refGene_gn=TAF10;refGene_tx=XM_023239521.1 |
| D1 | 65403643 | 65403758 | refGene_gn=TAF10;refGene_tx=XM_023239521.1 |
| D1 | 65403848 | 65403913 | refGene_gn=TAF10;refGene_tx=XM_023239521.1 |
| D1 | 65404148 | 65404303 | refGene_gn=TAF10;refGene_tx=XM_023239521.1 |
| D1 | 65404444 | 65404676 | refGene_gn=TAF10;refGene_tx=XM_023239521.1 |
| D1 | 65406999 | 65407140 | refGene_gn=TPP1;refGene_tx=XM_003992939.5 |
| D1 | 65407308 | 65407434 | refGene_gn=TPP1;refGene_tx=XM_003992939.5 |
| D1 | 65407606 | 65407765 | refGene_gn=TPP1;refGene_tx=XM_003992939.5 |
| D1 | 65407848 | 65407969 | refGene_gn=TPP1;refGene_tx=XM_003992939.5 |
| D1 | 65408409 | 65408479 | refGene_gn=TPP1;refGene_tx=XM_003992939.5 |
| D1 | 65408680 | 65408869 | refGene_gn=TPP1;refGene_tx=XM_003992939.5 |
| D1 | 65409015 | 65409214 | refGene_gn=TPP1;refGene_tx=XM_003992939.5 |
| D1 | 65409400 | 65409579 | refGene_gn=TPP1;refGene_tx=XM_003992939.5 |
| D1 | 65409729 | 65409857 | refGene_gn=TPP1;refGene_tx=XM_003992939.5 |
| D1 | 65410077 | 65410228 | refGene_gn=TPP1;refGene_tx=XM_003992939.5 |
| D1 | 65411256 | 65411337 | refGene_gn=TPP1;refGene_tx=XM_003992939.5 |
| D1 | 65411339 | 65411395 | refGene_gn=TPP1;refGene_tx=XM_003992939.5 |
| D1 | 65411657 | 65411729 | refGene_gn=TPP1;refGene_tx=XM_003992939.5 |
| D1 | 65411843 | 65411887 | refGene_gn=TPP1;refGene_tx=XM_003992939.5 |
| D1 | 65413107 | 65415278 | refGene_gn=DCHS1;refGene_tx=XM_023239523.1 |
| D1 | 65415280 | 65415355 | refGene_gn=DCHS1;refGene_tx=XM_023239523.1 |
| D1 | 65415357 | 65415519 | refGene_gn=DCHS1;refGene_tx=XM_023239523.1 |
| D1 | 65415521 | 65415552 | refGene_gn=DCHS1;refGene_tx=XM_023239523.1 |
| D1 | 65415553 | 65415720 | refGene_gn=DCHS1;refGene_tx=XM_023239523.1 |
| D1 | 65416052 | 65416104 | refGene_gn=DCHS1;refGene_tx=XM_023239523.1 |
| D1 | 65416105 | 65416192 | refGene_gn=DCHS1;refGene_tx=XM_023239523.1 |
| D1 | 65416554 | 65416838 | refGene_gn=DCHS1;refGene_tx=XM_023239523.1 |
| D1 | 65416953 | 65417084 | refGene_gn=DCHS1;refGene_tx=XM_023239523.1 |
| D1 | 65417281 | 65417436 | refGene_gn=DCHS1;refGene_tx=XM_023239523.1 |
| D1 | 65417524 | 65417736 | refGene_gn=DCHS1;refGene_tx=XM_023239523.1 |
| D1 | 65417902 | 65418016 | refGene_gn=DCHS1;refGene_tx=XM_023239523.1 |
| D1 | 65418142 | 65419021 | refGene_gn=DCHS1;refGene_tx=XM_023239523.1 |
| D1 | 65420275 | 65420485 | refGene_gn=DCHS1;refGene_tx=XM_023239523.1 |
| D1 | 65421113 | 65421239 | refGene_gn=DCHS1;refGene_tx=XM_023239523.1 |
| D1 | 65421331 | 65421571 | refGene_gn=DCHS1;refGene_tx=XM_023239523.1 |
| D1 | 65421663 | 65422528 | refGene_gn=DCHS1;refGene_tx=XM_023239523.1 |
| D1 | 65422727 | 65422849 | refGene_gn=DCHS1;refGene_tx=XM_023239523.1 |
| D1 | 65422851 | 65422884 | refGene_gn=DCHS1;refGene_tx=XM_023239523.1 |
| D1 | 65422985 | 65423082 | refGene_gn=DCHS1;refGene_tx=XM_023239523.1 |
| D1 | 65423349 | 65423543 | refGene_gn=DCHS1;refGene_tx=XM_023239523.1 |
| D1 | 65423751 | 65424705 | refGene_gn=DCHS1;refGene_tx=XM_023239523.1 |
| D1 | 65424707 | 65424776 | refGene_gn=DCHS1;refGene_tx=XM_023239523.1 |
| D1 | 65425051 | 65425288 | refGene_gn=DCHS1;refGene_tx=XM_023239523.1 |
| D1 | 65425427 | 65425659 | refGene_gn=DCHS1;refGene_tx=XM_023239523.1 |
| D1 | 65425755 | 65425944 | refGene_gn=DCHS1;refGene_tx=XM_023239523.1 |
| D1 | 65431664 | 65432113 | refGene_gn=DCHS1;refGene_tx=XM_023239523.1 |
| D1 | 65432115 | 65433457 | refGene_gn=DCHS1;refGene_tx=XM_023239523.1 |
| D1 | 65469479 | 65469755 | refGene_gn=MRPL17;refGene_tx=XM_003992941.5 |
| D1 | 65470106 | 65470175 | refGene_gn=MRPL17;refGene_tx=XM_003992941.5 |
| D1 | 65470474 | 65470735 | refGene_gn=MRPL17;refGene_tx=XM_003992941.5 |

D1 65515088 65522384
refGene_gn=LOC101094047;refGene_tx=XM_023239524.1
D1 65628516 65629440
refGene_gn=LOC101094538;refGene_tx=XM_003992942.3
D1 65639092 65640043
refGene_gn=LOC101097958;refGene_tx=XM_004001184.3
D1 65652776 65653709
refGene_gn=LOC101094789;refGene_tx=XM_011286698.2
D1 65660540 65661464
refGene_gn=LOC101095045;refGene_tx=XM_023239003.1
D1 65673868 65674792
refGene_gn=LOC101100247;refGene_tx=XM_003992958.3
D1 65687075 65688020
refGene_gn=LOC101100505;refGene_tx=XM_019811999.1
D1 65698787 65699744
refGene_gn=LOC101100753;refGene_tx=XM_003992960.1
D1 65723416 65724361
refGene_gn=LOC101100998;refGene_tx=XM_003992961.2
D1 65733547 65734531
refGene_gn=LOC101095283;refGene_tx=XM_011286699.3
D1 65780690 65781644
refGene_gn=LOC101095519;refGene_tx=XM_023239004.1
D1 65796883 65797837
refGene_gn=LOC101101252;refGene_tx=XM_006937059.3
D1 65809103 65810051
refGene_gn=LOC101095768;refGene_tx=XM_003992945.2
D1 65812393 65813341
refGene_gn=LOC101101503;refGene_tx=XM_003992963.1
D1 65849369 65850314
refGene_gn=LOC101080323;refGene_tx=XM_006937060.3
D1 65866139 65866539
refGene_gn=ZNF215;refGene_tx=XM_011286700.3;refGene_gn=ZNF215;refGene_tx=XM_011286701.3
D1 65874193 65874279
refGene_gn=ZNF215;refGene_tx=XM_011286700.3;refGene_gn=ZNF215;refGene_tx=XM_011286701.3
D1 65875669 65875802
refGene_gn=ZNF215;refGene_tx=XM_011286700.3;refGene_gn=ZNF215;refGene_tx=XM_011286701.3
D1 65876135 65876231 refGene_gn=ZNF215;refGene_tx=XM_011286701.3
D1 65884899 65885807
refGene_gn=ZNF215;refGene_tx=XM_011286700.3;refGene_gn=ZNF215;refGene_tx=XM_011286701.3
D1 65900529 65902226
refGene_gn=ZNF214;refGene_tx=XM_006937056.2;refGene_gn=ZNF214;refGene_tx=XM_006937057.2
D1 65903322 65903484
refGene_gn=ZNF214;refGene_tx=XM_006937056.2;refGene_gn=ZNF214;refGene_tx=XM_006937057.2
D1 65914813 65914902 refGene_gn=ZNF214;refGene_tx=XM_006937056.2

D1	65915492 65915509 refGene_gn=ZNF214;refGene_tx=XM_006937056.2
D1	65928161 65928453 refGene_gn=NLRP14;refGene_tx=XM_023239525.1
D1	65929223 65929295 refGene_gn=NLRP14;refGene_tx=XM_023239525.1
D1	65933207 65934801 refGene_gn=NLRP14;refGene_tx=XM_023239525.1
D1	65938403 65938574 refGene_gn=NLRP14;refGene_tx=XM_023239525.1
D1	65941002 65941170 refGene_gn=NLRP14;refGene_tx=XM_023239525.1
D1	65951361 65951532 refGene_gn=NLRP14;refGene_tx=XM_023239525.1
D1	65951919 65952090 refGene_gn=NLRP14;refGene_tx=XM_023239525.1
D1	65953879 65954050 refGene_gn=NLRP14;refGene_tx=XM_023239525.1
D1	65955934 65956105 refGene_gn=NLRP14;refGene_tx=XM_023239525.1
D1	65966127 65966298 refGene_gn=NLRP14;refGene_tx=XM_023239525.1
D1	65967699 65967835 refGene_gn=NLRP14;refGene_tx=XM_023239525.1
D1	65995461 65996646 refGene_gn=RBMXL2;refGene_tx=XM_011286755.3
D1	66154539 66154684
refGene_gn=SYT9;refGene_tx=XM_023239527.1;refGene_gn=SYT9;refGene_tx=XM_023239528.1;refGene_gn=SYT9;refGene_tx=XM_023239529.1
D1	66211967 66212319
refGene_gn=SYT9;refGene_tx=XM_023239527.1;refGene_gn=SYT9;refGene_tx=XM_023239528.1;refGene_gn=SYT9;refGene_tx=XM_023239529.1
D1	66222443 66222990
refGene_gn=SYT9;refGene_tx=XM_023239527.1;refGene_gn=SYT9;refGene_tx=XM_023239528.1;refGene_gn=SYT9;refGene_tx=XM_023239529.1
D1	66298740 66298861
refGene_gn=SYT9;refGene_tx=XM_023239527.1;refGene_gn=SYT9;refGene_tx=XM_023239528.1;refGene_gn=SYT9;refGene_tx=XM_023239529.1
D1	66300297 66300469
refGene_gn=SYT9;refGene_tx=XM_023239527.1;refGene_gn=SYT9;refGene_tx=XM_023239528.1;refGene_gn=SYT9;refGene_tx=XM_023239529.1
D1	66302248 66302348 refGene_gn=SYT9;refGene_tx=XM_023239528.1
D1	66302555 66302685
refGene_gn=SYT9;refGene_tx=XM_023239527.1;refGene_gn=SYT9;refGene_tx=XM_023239528.1;refGene_gn=SYT9;refGene_tx=XM_023239529.1
D1	66351418 66351427 refGene_gn=SYT9;refGene_tx=XM_023239527.1
D1	66370014 66370140 refGene_gn=OLFML1;refGene_tx=XM_003992967.4
D1	66372244 66372533 refGene_gn=OLFML1;refGene_tx=XM_003992967.4
D1	66394217 66395008 refGene_gn=OLFML1;refGene_tx=XM_003992967.4
D1	66430332 66430396
refGene_gn=PPFIBP2;refGene_tx=XM_006937066.4;refGene_gn=PPFIBP2;refGene_tx=XM_006937067.4;refGene_gn=PPFIBP2;refGene_tx=XM_019811984.1;refGene_gn=PPFIBP2;refGene_tx=XM_019811985.1
D1	66448665 66448868
refGene_gn=PPFIBP2;refGene_tx=XM_006937066.4;refGene_gn=PPFIBP2;refGene_tx=XM_006937067.4;refGene_gn=PPFIBP2;refGene_tx=XM_019811984.1;refGene_gn=PPFIBP2;refGene_tx=XM_019811985.1
D1	66462130 66462556
refGene_gn=PPFIBP2;refGene_tx=XM_019811979.2;refGene_gn=PPFIBP2;refGene_tx=XM_019811980.2;refGene_gn=PPFIBP2;refGene_tx=XM_019811982.2
D1	66462962 66463010 refGene_gn=PPFIBP2;refGene_tx=XM_019811987.2
D1	66478365 66478458
refGene_gn=PPFIBP2;refGene_tx=XM_006937066.4;refGene_gn=PPFIBP2;refGen

e_tx=XM_006937067.4;refGene_gn=PPFIBP2;refGene_tx=XM_019811979.2;refGene_gn=PPFIBP2;refGene_tx=XM_019811980.2;refGene_gn=PPFIBP2;refGene_tx=XM_019811982.2;refGene_gn=PPFIBP2;refGene_tx=XM_019811984.1;refGene_gn=PPFIBP2;refGene_tx=XM_019811985.1;refGene_gn=PPFIBP2;refGene_tx=XM_019811987.2
D1 66482903 66483017
refGene_gn=PPFIBP2;refGene_tx=XM_006937066.4;refGene_gn=PPFIBP2;refGene_tx=XM_006937067.4;refGene_gn=PPFIBP2;refGene_tx=XM_019811979.2;refGene_gn=PPFIBP2;refGene_tx=XM_019811980.2;refGene_gn=PPFIBP2;refGene_tx=XM_019811982.2;refGene_gn=PPFIBP2;refGene_tx=XM_019811984.1;refGene_gn=PPFIBP2;refGene_tx=XM_019811985.1;refGene_gn=PPFIBP2;refGene_tx=XM_019811987.2;refGene_gn=PPFIBP2;refGene_tx=XM_019811992.2;refGene_gn=PPFIBP2;refGene_tx=XM_023239526.1
D1 66488357 66488415
refGene_gn=PPFIBP2;refGene_tx=XM_019811992.2;refGene_gn=PPFIBP2;refGene_tx=XM_023239526.1
D1 66494479 66494611
refGene_gn=PPFIBP2;refGene_tx=XM_006937066.4;refGene_gn=PPFIBP2;refGene_tx=XM_006937067.4;refGene_gn=PPFIBP2;refGene_tx=XM_019811979.2;refGene_gn=PPFIBP2;refGene_tx=XM_019811980.2;refGene_gn=PPFIBP2;refGene_tx=XM_019811982.2;refGene_gn=PPFIBP2;refGene_tx=XM_019811984.1;refGene_gn=PPFIBP2;refGene_tx=XM_019811985.1;refGene_gn=PPFIBP2;refGene_tx=XM_019811987.2;refGene_gn=PPFIBP2;refGene_tx=XM_019811990.1;refGene_gn=PPFIBP2;refGene_tx=XM_019811991.2;refGene_gn=PPFIBP2;refGene_tx=XM_019811992.2;refGene_gn=PPFIBP2;refGene_tx=XM_023239526.1
D1 66508218 66508311
refGene_gn=PPFIBP2;refGene_tx=XM_006937066.4;refGene_gn=PPFIBP2;refGene_tx=XM_006937067.4;refGene_gn=PPFIBP2;refGene_tx=XM_019811979.2;refGene_gn=PPFIBP2;refGene_tx=XM_019811980.2;refGene_gn=PPFIBP2;refGene_tx=XM_019811982.2;refGene_gn=PPFIBP2;refGene_tx=XM_019811984.1;refGene_gn=PPFIBP2;refGene_tx=XM_019811985.1;refGene_gn=PPFIBP2;refGene_tx=XM_019811987.2;refGene_gn=PPFIBP2;refGene_tx=XM_019811990.1;refGene_gn=PPFIBP2;refGene_tx=XM_019811991.2;refGene_gn=PPFIBP2;refGene_tx=XM_019811992.2;refGene_gn=PPFIBP2;refGene_tx=XM_023239526.1
D1 66512880 66512992
refGene_gn=PPFIBP2;refGene_tx=XM_006937066.4;refGene_gn=PPFIBP2;refGene_tx=XM_006937067.4;refGene_gn=PPFIBP2;refGene_tx=XM_019811979.2;refGene_gn=PPFIBP2;refGene_tx=XM_019811980.2;refGene_gn=PPFIBP2;refGene_tx=XM_019811982.2;refGene_gn=PPFIBP2;refGene_tx=XM_019811984.1;refGene_gn=PPFIBP2;refGene_tx=XM_019811985.1;refGene_gn=PPFIBP2;refGene_tx=XM_019811987.2;refGene_gn=PPFIBP2;refGene_tx=XM_019811990.1;refGene_gn=PPFIBP2;refGene_tx=XM_019811991.2;refGene_gn=PPFIBP2;refGene_tx=XM_019811992.2;refGene_gn=PPFIBP2;refGene_tx=XM_023239526.1
D1 66515356 66515418
refGene_gn=PPFIBP2;refGene_tx=XM_006937066.4;refGene_gn=PPFIBP2;refGene_tx=XM_006937067.4;refGene_gn=PPFIBP2;refGene_tx=XM_019811979.2;refGene_gn=PPFIBP2;refGene_tx=XM_019811980.2;refGene_gn=PPFIBP2;refGene_tx=XM_019811982.2;refGene_gn=PPFIBP2;refGene_tx=XM_019811984.1;refGene_gn=PPFIBP2;refGene_tx=XM_019811985.1;refGene_gn=PPFIBP2;refGene_tx=XM_019811987.2;refGene_gn=PPFIBP2;refGene_tx=XM_019811990.1;refGene_gn=PPFI

BP2;refGene_tx=XM_019811991.2;refGene_gn=PPFIBP2;refGene_tx=XM_0198119
92.2;refGene_gn=PPFIBP2;refGene_tx=XM_023239526.1
D1 66516581 66516657
refGene_gn=PPFIBP2;refGene_tx=XM_006937066.4;refGene_gn=PPFIBP2;refGen
e_tx=XM_006937067.4;refGene_gn=PPFIBP2;refGene_tx=XM_019811979.2;refGe
ne_gn=PPFIBP2;refGene_tx=XM_019811980.2;refGene_gn=PPFIBP2;refGene_tx=
XM_019811982.2;refGene_gn=PPFIBP2;refGene_tx=XM_019811984.1;refGene_gn
=PPFIBP2;refGene_tx=XM_019811985.1;refGene_gn=PPFIBP2;refGene_tx=XM_01
9811987.2;refGene_gn=PPFIBP2;refGene_tx=XM_019811990.1;refGene_gn=PPFI
BP2;refGene_tx=XM_019811991.2;refGene_gn=PPFIBP2;refGene_tx=XM_0198119
92.2;refGene_gn=PPFIBP2;refGene_tx=XM_023239526.1
D1 66517692 66517796
refGene_gn=PPFIBP2;refGene_tx=XM_006937066.4;refGene_gn=PPFIBP2;refGen
e_tx=XM_006937067.4;refGene_gn=PPFIBP2;refGene_tx=XM_019811979.2;refGe
ne_gn=PPFIBP2;refGene_tx=XM_019811980.2;refGene_gn=PPFIBP2;refGene_tx=
XM_019811982.2;refGene_gn=PPFIBP2;refGene_tx=XM_019811984.1;refGene_gn
=PPFIBP2;refGene_tx=XM_019811985.1;refGene_gn=PPFIBP2;refGene_tx=XM_01
9811987.2;refGene_gn=PPFIBP2;refGene_tx=XM_019811990.1;refGene_gn=PPFI
BP2;refGene_tx=XM_019811991.2;refGene_gn=PPFIBP2;refGene_tx=XM_0198119
92.2;refGene_gn=PPFIBP2;refGene_tx=XM_023239526.1
D1 66519763 66519831
refGene_gn=PPFIBP2;refGene_tx=XM_006937066.4;refGene_gn=PPFIBP2;refGen
e_tx=XM_006937067.4;refGene_gn=PPFIBP2;refGene_tx=XM_019811979.2;refGe
ne_gn=PPFIBP2;refGene_tx=XM_019811980.2;refGene_gn=PPFIBP2;refGene_tx=
XM_019811982.2;refGene_gn=PPFIBP2;refGene_tx=XM_019811984.1;refGene_gn
=PPFIBP2;refGene_tx=XM_019811985.1;refGene_gn=PPFIBP2;refGene_tx=XM_01
9811987.2;refGene_gn=PPFIBP2;refGene_tx=XM_019811990.1;refGene_gn=PPFI
BP2;refGene_tx=XM_019811991.2;refGene_gn=PPFIBP2;refGene_tx=XM_0198119
92.2;refGene_gn=PPFIBP2;refGene_tx=XM_023239526.1
D1 66521423 66521481
refGene_gn=PPFIBP2;refGene_tx=XM_006937066.4;refGene_gn=PPFIBP2;refGen
e_tx=XM_006937067.4;refGene_gn=PPFIBP2;refGene_tx=XM_019811979.2;refGe
ne_gn=PPFIBP2;refGene_tx=XM_019811980.2;refGene_gn=PPFIBP2;refGene_tx=
XM_019811982.2;refGene_gn=PPFIBP2;refGene_tx=XM_019811984.1;refGene_gn
=PPFIBP2;refGene_tx=XM_019811985.1;refGene_gn=PPFIBP2;refGene_tx=XM_01
9811987.2;refGene_gn=PPFIBP2;refGene_tx=XM_019811990.1;refGene_gn=PPFI
BP2;refGene_tx=XM_019811991.2;refGene_gn=PPFIBP2;refGene_tx=XM_0198119
92.2;refGene_gn=PPFIBP2;refGene_tx=XM_023239526.1
D1 66522387 66522429
refGene_gn=PPFIBP2;refGene_tx=XM_006937066.4;refGene_gn=PPFIBP2;refGen
e_tx=XM_006937067.4;refGene_gn=PPFIBP2;refGene_tx=XM_019811979.2;refGe
ne_gn=PPFIBP2;refGene_tx=XM_019811980.2;refGene_gn=PPFIBP2;refGene_tx=
XM_019811982.2;refGene_gn=PPFIBP2;refGene_tx=XM_019811984.1;refGene_gn
=PPFIBP2;refGene_tx=XM_019811985.1;refGene_gn=PPFIBP2;refGene_tx=XM_01
9811987.2;refGene_gn=PPFIBP2;refGene_tx=XM_019811990.1;refGene_gn=PPFI
BP2;refGene_tx=XM_019811991.2;refGene_gn=PPFIBP2;refGene_tx=XM_0198119
92.2;refGene_gn=PPFIBP2;refGene_tx=XM_023239526.1
D1 66525842 66525981
refGene_gn=PPFIBP2;refGene_tx=XM_006937066.4;refGene_gn=PPFIBP2;refGen
e_tx=XM_006937067.4;refGene_gn=PPFIBP2;refGene_tx=XM_019811979.2;refGe

ne_gn=PPFIBP2;refGene_tx=XM_019811980.2;refGene_gn=PPFIBP2;refGene_tx=XM_019811982.2;refGene_gn=PPFIBP2;refGene_tx=XM_019811984.1;refGene_gn=PPFIBP2;refGene_tx=XM_019811985.1;refGene_gn=PPFIBP2;refGene_tx=XM_019811987.2;refGene_gn=PPFIBP2;refGene_tx=XM_019811990.1;refGene_gn=PPFIBP2;refGene_tx=XM_019811991.2;refGene_gn=PPFIBP2;refGene_tx=XM_019811992.2;refGene_gn=PPFIBP2;refGene_tx=XM_023239526.1
D1 66527180 66527213
refGene_gn=PPFIBP2;refGene_tx=XM_006937066.4;refGene_gn=PPFIBP2;refGene_tx=XM_019811979.2;refGene_gn=PPFIBP2;refGene_tx=XM_019811982.2;refGene_gn=PPFIBP2;refGene_tx=XM_019811984.1;refGene_gn=PPFIBP2;refGene_tx=XM_019811985.1;refGene_gn=PPFIBP2;refGene_tx=XM_019811987.2;refGene_gn=PPFIBP2;refGene_tx=XM_019811990.1;refGene_gn=PPFIBP2;refGene_tx=XM_019811991.2;refGene_gn=PPFIBP2;refGene_tx=XM_019811992.2;refGene_gn=PPFIBP2;refGene_tx=XM_023239526.1
D1 66527637 66527779
refGene_gn=PPFIBP2;refGene_tx=XM_006937066.4;refGene_gn=PPFIBP2;refGene_tx=XM_006937067.4;refGene_gn=PPFIBP2;refGene_tx=XM_019811979.2;refGene_gn=PPFIBP2;refGene_tx=XM_019811980.2;refGene_gn=PPFIBP2;refGene_tx=XM_019811982.2;refGene_gn=PPFIBP2;refGene_tx=XM_019811984.1;refGene_gn=PPFIBP2;refGene_tx=XM_019811985.1;refGene_gn=PPFIBP2;refGene_tx=XM_019811987.2;refGene_gn=PPFIBP2;refGene_tx=XM_019811990.1;refGene_gn=PPFIBP2;refGene_tx=XM_019811991.2;refGene_gn=PPFIBP2;refGene_tx=XM_019811992.2;refGene_gn=PPFIBP2;refGene_tx=XM_023239526.1
D1 66528424 66528553
refGene_gn=PPFIBP2;refGene_tx=XM_006937066.4;refGene_gn=PPFIBP2;refGene_tx=XM_006937067.4;refGene_gn=PPFIBP2;refGene_tx=XM_019811979.2;refGene_gn=PPFIBP2;refGene_tx=XM_019811980.2;refGene_gn=PPFIBP2;refGene_tx=XM_019811982.2;refGene_gn=PPFIBP2;refGene_tx=XM_019811984.1;refGene_gn=PPFIBP2;refGene_tx=XM_019811985.1;refGene_gn=PPFIBP2;refGene_tx=XM_019811987.2;refGene_gn=PPFIBP2;refGene_tx=XM_019811990.1;refGene_gn=PPFIBP2;refGene_tx=XM_019811991.2;refGene_gn=PPFIBP2;refGene_tx=XM_019811992.2;refGene_gn=PPFIBP2;refGene_tx=XM_023239526.1
D1 66534812 66534963
refGene_gn=PPFIBP2;refGene_tx=XM_006937066.4;refGene_gn=PPFIBP2;refGene_tx=XM_006937067.4;refGene_gn=PPFIBP2;refGene_tx=XM_019811979.2;refGene_gn=PPFIBP2;refGene_tx=XM_019811980.2;refGene_gn=PPFIBP2;refGene_tx=XM_019811982.2;refGene_gn=PPFIBP2;refGene_tx=XM_019811984.1;refGene_gn=PPFIBP2;refGene_tx=XM_019811985.1;refGene_gn=PPFIBP2;refGene_tx=XM_019811987.2;refGene_gn=PPFIBP2;refGene_tx=XM_019811990.1;refGene_gn=PPFIBP2;refGene_tx=XM_019811991.2;refGene_gn=PPFIBP2;refGene_tx=XM_019811992.2;refGene_gn=PPFIBP2;refGene_tx=XM_023239526.1
D1 66535248 66535360
refGene_gn=PPFIBP2;refGene_tx=XM_006937066.4;refGene_gn=PPFIBP2;refGene_tx=XM_006937067.4;refGene_gn=PPFIBP2;refGene_tx=XM_019811979.2;refGene_gn=PPFIBP2;refGene_tx=XM_019811980.2;refGene_gn=PPFIBP2;refGene_tx=XM_019811982.2;refGene_gn=PPFIBP2;refGene_tx=XM_019811984.1;refGene_gn=PPFIBP2;refGene_tx=XM_019811985.1;refGene_gn=PPFIBP2;refGene_tx=XM_019811987.2;refGene_gn=PPFIBP2;refGene_tx=XM_019811990.1;refGene_gn=PPFIBP2;refGene_tx=XM_019811991.2;refGene_gn=PPFIBP2;refGene_tx=XM_019811992.2;refGene_gn=PPFIBP2;refGene_tx=XM_023239526.1

D1 66535602 66535691
refGene_gn=PPFIBP2;refGene_tx=XM_006937066.4;refGene_gn=PPFIBP2;refGene_tx=XM_006937067.4;refGene_gn=PPFIBP2;refGene_tx=XM_019811979.2;refGene_gn=PPFIBP2;refGene_tx=XM_019811980.2;refGene_gn=PPFIBP2;refGene_tx=XM_019811982.2;refGene_gn=PPFIBP2;refGene_tx=XM_019811984.1;refGene_gn=PPFIBP2;refGene_tx=XM_019811985.1;refGene_gn=PPFIBP2;refGene_tx=XM_019811987.2;refGene_gn=PPFIBP2;refGene_tx=XM_019811990.1;refGene_gn=PPFIBP2;refGene_tx=XM_019811991.2;refGene_gn=PPFIBP2;refGene_tx=XM_019811992.2;refGene_gn=PPFIBP2;refGene_tx=XM_023239526.1
D1 66535998 66536121
refGene_gn=PPFIBP2;refGene_tx=XM_006937066.4;refGene_gn=PPFIBP2;refGene_tx=XM_006937067.4;refGene_gn=PPFIBP2;refGene_tx=XM_019811979.2;refGene_gn=PPFIBP2;refGene_tx=XM_019811980.2;refGene_gn=PPFIBP2;refGene_tx=XM_019811982.2;refGene_gn=PPFIBP2;refGene_tx=XM_019811984.1;refGene_gn=PPFIBP2;refGene_tx=XM_019811985.1;refGene_gn=PPFIBP2;refGene_tx=XM_019811987.2;refGene_gn=PPFIBP2;refGene_tx=XM_019811990.1;refGene_gn=PPFIBP2;refGene_tx=XM_019811991.2;refGene_gn=PPFIBP2;refGene_tx=XM_019811992.2;refGene_gn=PPFIBP2;refGene_tx=XM_023239526.1
D1 66537276 66537402
refGene_gn=PPFIBP2;refGene_tx=XM_006937066.4;refGene_gn=PPFIBP2;refGene_tx=XM_006937067.4;refGene_gn=PPFIBP2;refGene_tx=XM_019811979.2;refGene_gn=PPFIBP2;refGene_tx=XM_019811980.2;refGene_gn=PPFIBP2;refGene_tx=XM_019811982.2;refGene_gn=PPFIBP2;refGene_tx=XM_019811984.1;refGene_gn=PPFIBP2;refGene_tx=XM_019811985.1;refGene_gn=PPFIBP2;refGene_tx=XM_019811987.2;refGene_gn=PPFIBP2;refGene_tx=XM_019811990.1;refGene_gn=PPFIBP2;refGene_tx=XM_019811991.2;refGene_gn=PPFIBP2;refGene_tx=XM_019811992.2;refGene_gn=PPFIBP2;refGene_tx=XM_023239526.1
D1 66538063 66538252
refGene_gn=PPFIBP2;refGene_tx=XM_006937066.4;refGene_gn=PPFIBP2;refGene_tx=XM_006937067.4;refGene_gn=PPFIBP2;refGene_tx=XM_019811979.2;refGene_gn=PPFIBP2;refGene_tx=XM_019811980.2;refGene_gn=PPFIBP2;refGene_tx=XM_019811982.2;refGene_gn=PPFIBP2;refGene_tx=XM_019811984.1;refGene_gn=PPFIBP2;refGene_tx=XM_019811985.1;refGene_gn=PPFIBP2;refGene_tx=XM_019811987.2;refGene_gn=PPFIBP2;refGene_tx=XM_019811990.1;refGene_gn=PPFIBP2;refGene_tx=XM_019811991.2;refGene_gn=PPFIBP2;refGene_tx=XM_019811992.2;refGene_gn=PPFIBP2;refGene_tx=XM_023239526.1
D1 66539579 66539756
refGene_gn=PPFIBP2;refGene_tx=XM_006937066.4;refGene_gn=PPFIBP2;refGene_tx=XM_006937067.4;refGene_gn=PPFIBP2;refGene_tx=XM_019811979.2;refGene_gn=PPFIBP2;refGene_tx=XM_019811980.2;refGene_gn=PPFIBP2;refGene_tx=XM_019811982.2;refGene_gn=PPFIBP2;refGene_tx=XM_019811984.1;refGene_gn=PPFIBP2;refGene_tx=XM_019811985.1;refGene_gn=PPFIBP2;refGene_tx=XM_019811987.2;refGene_gn=PPFIBP2;refGene_tx=XM_019811990.1;refGene_gn=PPFIBP2;refGene_tx=XM_019811991.2;refGene_gn=PPFIBP2;refGene_tx=XM_019811992.2;refGene_gn=PPFIBP2;refGene_tx=XM_023239526.1
D1 66541968 66542034
refGene_gn=PPFIBP2;refGene_tx=XM_006937066.4;refGene_gn=PPFIBP2;refGene_tx=XM_006937067.4;refGene_gn=PPFIBP2;refGene_tx=XM_019811979.2;refGene_gn=PPFIBP2;refGene_tx=XM_019811980.2;refGene_gn=PPFIBP2;refGene_tx=XM_019811984.1;refGene_gn=PPFIBP2;refGene_tx=XM_019811985.1;refGene_gn

=PPFIBP2;refGene_tx=XM_019811987.2;refGene_gn=PPFIBP2;refGene_tx=XM_01
9811990.1;refGene_gn=PPFIBP2;refGene_tx=XM_019811991.2;refGene_gn=PPFI
BP2;refGene_tx=XM_019811992.2;refGene_gn=PPFIBP2;refGene_tx=XM_0232395
26.1
D1	66543299	66543371
refGene_gn=PPFIBP2;refGene_tx=XM_006937066.4;refGene_gn=PPFIBP2;refGen
e_tx=XM_006937067.4;refGene_gn=PPFIBP2;refGene_tx=XM_019811979.2;refGe
ne_gn=PPFIBP2;refGene_tx=XM_019811980.2;refGene_gn=PPFIBP2;refGene_tx=
XM_019811982.2;refGene_gn=PPFIBP2;refGene_tx=XM_019811984.1;refGene_gn
=PPFIBP2;refGene_tx=XM_019811985.1;refGene_gn=PPFIBP2;refGene_tx=XM_01
9811987.2;refGene_gn=PPFIBP2;refGene_tx=XM_019811990.1;refGene_gn=PPFI
BP2;refGene_tx=XM_019811991.2;refGene_gn=PPFIBP2;refGene_tx=XM_0198119
92.2;refGene_gn=PPFIBP2;refGene_tx=XM_023239526.1
D1	66546920	66547093
refGene_gn=LOC101082689;refGene_tx=XM_003992971.4;refGene_gn=LOC101082
689;refGene_tx=XM_023239530.1
D1	66547997	66548097
refGene_gn=LOC101082689;refGene_tx=XM_003992971.4;refGene_gn=LOC101082
689;refGene_tx=XM_023239530.1
D1	66549231	66549317
refGene_gn=LOC101082689;refGene_tx=XM_003992971.4;refGene_gn=LOC101082
689;refGene_tx=XM_023239530.1
D1	66550047	66550131
refGene_gn=LOC101082689;refGene_tx=XM_003992971.4;refGene_gn=LOC101082
689;refGene_tx=XM_023239530.1
D1	66550730	66550860
refGene_gn=LOC101082689;refGene_tx=XM_003992971.4;refGene_gn=LOC101082
689;refGene_tx=XM_023239530.1
D1	66551136	66551243
refGene_gn=LOC101082689;refGene_tx=XM_003992971.4;refGene_gn=LOC101082
689;refGene_tx=XM_023239530.1
D1	66553792	66553865
refGene_gn=LOC101082689;refGene_tx=XM_003992971.4;refGene_gn=LOC101082
689;refGene_tx=XM_023239530.1
D1	66554136	66554211
refGene_gn=LOC101082689;refGene_tx=XM_003992971.4;refGene_gn=LOC101082
689;refGene_tx=XM_023239530.1
D1	66569733	66569858	refGene_gn=OVCH2;refGene_tx=XM_019812007.2
D1	66571021	66571153	refGene_gn=OVCH2;refGene_tx=XM_019812007.2
D1	66571620	66571714	refGene_gn=OVCH2;refGene_tx=XM_019812007.2
D1	66574013	66574144	refGene_gn=OVCH2;refGene_tx=XM_019812007.2
D1	66574528	66574669	refGene_gn=OVCH2;refGene_tx=XM_019812007.2
D1	66575417	66575542	refGene_gn=OVCH2;refGene_tx=XM_019812007.2
D1	66575655	66575746	refGene_gn=OVCH2;refGene_tx=XM_019812007.2
D1	66577845	66577869	refGene_gn=OVCH2;refGene_tx=XM_019812007.2
D1	66578964	66579154	refGene_gn=OVCH2;refGene_tx=XM_019812007.2
D1	66579782	66579934	refGene_gn=OVCH2;refGene_tx=XM_019812007.2
D1	66580176	66580272	refGene_gn=OVCH2;refGene_tx=XM_019812007.2
D1	66580642	66580815	refGene_gn=OVCH2;refGene_tx=XM_019812007.2
D1	66582767	66582859	refGene_gn=OVCH2;refGene_tx=XM_019812007.2

D1 66584516 66584626 refGene_gn=OVCH2;refGene_tx=XM_019812007.2
D1 66586702 66586790 refGene_gn=OVCH2;refGene_tx=XM_019812007.2
D1 66587017 66587039
refGene_gn=LOC101087605;refGene_tx=XM_003993076.1
D1 66590541 66591248
refGene_gn=LOC101087605;refGene_tx=XM_003993076.1
D1 66602401 66603334
refGene_gn=LOC101087868;refGene_tx=XM_003993077.4;refGene_gn=LOC101087868;refGene_tx=XM_019812002.1;refGene_gn=LOC101087868;refGene_tx=XM_019812005.1
D1 66617041 66617104
refGene_gn=LOC101088107;refGene_tx=XM_003993078.2
D1 66620018 66620705
refGene_gn=LOC101088107;refGene_tx=XM_003993078.2
D1 66640961 66641954
refGene_gn=LOC101088359;refGene_tx=XM_003993079.3
D1 66651142 66651443
refGene_gn=LOC101088612;refGene_tx=XM_019812948.1
D1 66651444 66652018
refGene_gn=LOC101088612;refGene_tx=XM_019812948.1
D1 66652020 66652099
refGene_gn=LOC101088612;refGene_tx=XM_019812948.1
D1 66661852 66661867
refGene_gn=LOC101088612;refGene_tx=XM_019812948.1
D1 66671188 66672124
refGene_gn=LOC101088875;refGene_tx=XM_003993081.3
D1 66711034 66711979
refGene_gn=LOC101089140;refGene_tx=XM_011286757.1
D1 66736450 66737395
refGene_gn=LOC101089393;refGene_tx=XM_011286759.1
D1 66751506 66753143 refGene_gn=NLRP10;refGene_tx=XM_003993082.4
D1 66755003 66755286 refGene_gn=NLRP10;refGene_tx=XM_003993082.4
D1 66773132 66773532 refGene_gn=EIF3F;refGene_tx=XM_003992973.4
D1 66776784 66776855 refGene_gn=EIF3F;refGene_tx=XM_003992973.4
D1 66777095 66777175 refGene_gn=EIF3F;refGene_tx=XM_003992973.4
D1 66777579 66777717 refGene_gn=EIF3F;refGene_tx=XM_003992973.4
D1 66778884 66778976 refGene_gn=EIF3F;refGene_tx=XM_003992973.4
D1 66779392 66779529 refGene_gn=EIF3F;refGene_tx=XM_003992973.4
D1 66779664 66779778 refGene_gn=EIF3F;refGene_tx=XM_003992973.4
D1 66780353 66780431 refGene_gn=EIF3F;refGene_tx=XM_003992973.4
D1 66797201 66797257
refGene_gn=TUB;refGene_tx=XM_023239531.1;refGene_gn=TUB;refGene_tx=XM_023239532.1
D1 66860478 66860516 refGene_gn=TUB;refGene_tx=XM_011286702.3
D1 66870035 66870087
refGene_gn=TUB;refGene_tx=XM_011286702.3;refGene_gn=TUB;refGene_tx=XM_023239531.1;refGene_gn=TUB;refGene_tx=XM_023239532.1
D1 66870583 66870746
refGene_gn=TUB;refGene_tx=XM_011286702.3;refGene_gn=TUB;refGene_tx=XM_023239531.1;refGene_gn=TUB;refGene_tx=XM_023239532.1

D1 66872715 66872859
refGene_gn=TUB;refGene_tx=XM_011286702.3;refGene_gn=TUB;refGene_tx=XM_023239531.1;refGene_gn=TUB;refGene_tx=XM_023239532.1
D1 66874274 66874442
refGene_gn=TUB;refGene_tx=XM_011286702.3;refGene_gn=TUB;refGene_tx=XM_023239531.1
D1 66875462 66875584
refGene_gn=TUB;refGene_tx=XM_011286702.3;refGene_gn=TUB;refGene_tx=XM_023239531.1;refGene_gn=TUB;refGene_tx=XM_023239532.1
D1 66876047 66876245
refGene_gn=TUB;refGene_tx=XM_011286702.3;refGene_gn=TUB;refGene_tx=XM_023239531.1;refGene_gn=TUB;refGene_tx=XM_023239532.1
D1 66876529 66876642
refGene_gn=TUB;refGene_tx=XM_011286702.3;refGene_gn=TUB;refGene_tx=XM_023239531.1;refGene_gn=TUB;refGene_tx=XM_023239532.1
D1 66877516 66877634
refGene_gn=TUB;refGene_tx=XM_011286702.3;refGene_gn=TUB;refGene_tx=XM_023239531.1;refGene_gn=TUB;refGene_tx=XM_023239532.1
D1 66879164 66879263
refGene_gn=TUB;refGene_tx=XM_011286702.3;refGene_gn=TUB;refGene_tx=XM_023239531.1;refGene_gn=TUB;refGene_tx=XM_023239532.1
D1 66879495 66879667
refGene_gn=TUB;refGene_tx=XM_011286702.3;refGene_gn=TUB;refGene_tx=XM_023239531.1;refGene_gn=TUB;refGene_tx=XM_023239532.1
D1 66880103 66880237
refGene_gn=TUB;refGene_tx=XM_011286702.3;refGene_gn=TUB;refGene_tx=XM_023239531.1;refGene_gn=TUB;refGene_tx=XM_023239532.1
D1 66888217 66888690
refGene_gn=RIC3;refGene_tx=XM_003993084.4;refGene_gn=RIC3;refGene_tx=XM_006937071.4;refGene_gn=RIC3;refGene_tx=XM_006937072.4;refGene_gn=RIC3;refGene_tx=XM_019812011.2;refGene_gn=RIC3;refGene_tx=XM_019812014.2;refGene_gn=RIC3;refGene_tx=XM_019812015.2;refGene_gn=RIC3;refGene_tx=XM_019812017.2
D1 66888897 66888971
refGene_gn=RIC3;refGene_tx=XM_019812011.2;refGene_gn=RIC3;refGene_tx=XM_019812014.2
D1 66905518 66905606
refGene_gn=RIC3;refGene_tx=XM_019812015.2;refGene_gn=RIC3;refGene_tx=XM_019812017.2
D1 66906250 66906399
refGene_gn=RIC3;refGene_tx=XM_003993084.4;refGene_gn=RIC3;refGene_tx=XM_006937071.4;refGene_gn=RIC3;refGene_tx=XM_006937072.4;refGene_gn=RIC3;refGene_tx=XM_019812011.2;refGene_gn=RIC3;refGene_tx=XM_019812014.2;refGene_gn=RIC3;refGene_tx=XM_019812015.2;refGene_gn=RIC3;refGene_tx=XM_019812017.2
D1 66918948 66919042
refGene_gn=RIC3;refGene_tx=XM_003993084.4;refGene_gn=RIC3;refGene_tx=XM_006937071.4;refGene_gn=RIC3;refGene_tx=XM_006937072.4;refGene_gn=RIC3;refGene_tx=XM_019812011.2;refGene_gn=RIC3;refGene_tx=XM_019812014.2;refGene_gn=RIC3;refGene_tx=XM_019812015.2;refGene_gn=RIC3;refGene_tx=X

M_019812017.2
D1 66921051 66921127
refGene_gn=RIC3;refGene_tx=XM_003993084.4;refGene_gn=RIC3;refGene_tx=X
M_006937071.4;refGene_gn=RIC3;refGene_tx=XM_006937072.4;refGene_gn=RIC
3;refGene_tx=XM_019812011.2;refGene_gn=RIC3;refGene_tx=XM_019812014.2;
refGene_gn=RIC3;refGene_tx=XM_019812015.2;refGene_gn=RIC3;refGene_tx=X
M_019812017.2
D1 66922798 66923025
refGene_gn=RIC3;refGene_tx=XM_003993084.4;refGene_gn=RIC3;refGene_tx=X
M_006937071.4;refGene_gn=RIC3;refGene_tx=XM_006937072.4;refGene_gn=RIC
3;refGene_tx=XM_019812011.2;refGene_gn=RIC3;refGene_tx=XM_019812014.2;
refGene_gn=RIC3;refGene_tx=XM_019812015.2;refGene_gn=RIC3;refGene_tx=X
M_019812017.2
D1 66952944 66953068
refGene_gn=RIC3;refGene_tx=XM_003993084.4;refGene_gn=RIC3;refGene_tx=X
M_006937071.4;refGene_gn=RIC3;refGene_tx=XM_006937072.4;refGene_gn=RIC
3;refGene_tx=XM_019812011.2;refGene_gn=RIC3;refGene_tx=XM_019812014.2;
refGene_gn=RIC3;refGene_tx=XM_019812015.2;refGene_gn=RIC3;refGene_tx=X
M_019812017.2
D1 66996593 66996699
refGene_gn=LMO1;refGene_tx=XM_023239533.1;refGene_gn=LMO1;refGene_tx=X
M_023239534.1
D1 66998165 66998291
refGene_gn=LMO1;refGene_tx=XM_023239533.1;refGene_gn=LMO1;refGene_tx=X
M_023239534.1
D1 67001177 67001391
refGene_gn=LMO1;refGene_tx=XM_023239533.1;refGene_gn=LMO1;refGene_tx=X
M_023239534.1
D1 67032693 67032718 refGene_gn=LMO1;refGene_tx=XM_023239533.1
D1 67037478 67037500 refGene_gn=LMO1;refGene_tx=XM_023239534.1
D1 67145912 67146125
refGene_gn=STK33;refGene_tx=XM_023239535.1;refGene_gn=STK33;refGene_tx
=XM_023239536.1;refGene_gn=STK33;refGene_tx=XM_023239537.1;refGene_gn=
STK33;refGene_tx=XM_023239539.1
D1 67148586 67148607 refGene_gn=STK33;refGene_tx=XM_023239538.1
D1 67164470 67164677
refGene_gn=STK33;refGene_tx=XM_023239535.1;refGene_gn=STK33;refGene_tx
=XM_023239536.1;refGene_gn=STK33;refGene_tx=XM_023239537.1;refGene_gn=
STK33;refGene_tx=XM_023239538.1
D1 67182652 67182738
refGene_gn=STK33;refGene_tx=XM_023239535.1;refGene_gn=STK33;refGene_tx
=XM_023239536.1;refGene_gn=STK33;refGene_tx=XM_023239537.1;refGene_gn=
STK33;refGene_tx=XM_023239538.1;refGene_gn=STK33;refGene_tx=XM_0232395
39.1
D1 67183177 67183290
refGene_gn=STK33;refGene_tx=XM_023239535.1;refGene_gn=STK33;refGene_tx
=XM_023239536.1;refGene_gn=STK33;refGene_tx=XM_023239537.1;refGene_gn=
STK33;refGene_tx=XM_023239538.1;refGene_gn=STK33;refGene_tx=XM_0232395
39.1
D1 67187340 67187416

refGene_gn=STK33;refGene_tx=XM_023239535.1;refGene_gn=STK33;refGene_tx=XM_023239536.1;refGene_gn=STK33;refGene_tx=XM_023239537.1;refGene_gn=STK33;refGene_tx=XM_023239538.1;refGene_gn=STK33;refGene_tx=XM_023239539.1
D1 67194157 67194242
refGene_gn=STK33;refGene_tx=XM_023239535.1;refGene_gn=STK33;refGene_tx=XM_023239536.1;refGene_gn=STK33;refGene_tx=XM_023239537.1;refGene_gn=STK33;refGene_tx=XM_023239538.1;refGene_gn=STK33;refGene_tx=XM_023239539.1
D1 67196431 67196520
refGene_gn=STK33;refGene_tx=XM_023239535.1;refGene_gn=STK33;refGene_tx=XM_023239536.1;refGene_gn=STK33;refGene_tx=XM_023239537.1;refGene_gn=STK33;refGene_tx=XM_023239538.1;refGene_gn=STK33;refGene_tx=XM_023239539.1
D1 67198985 67199124
refGene_gn=STK33;refGene_tx=XM_023239535.1;refGene_gn=STK33;refGene_tx=XM_023239536.1;refGene_gn=STK33;refGene_tx=XM_023239537.1;refGene_gn=STK33;refGene_tx=XM_023239538.1;refGene_gn=STK33;refGene_tx=XM_023239539.1
D1 67204033 67204138
refGene_gn=STK33;refGene_tx=XM_023239535.1;refGene_gn=STK33;refGene_tx=XM_023239536.1;refGene_gn=STK33;refGene_tx=XM_023239537.1;refGene_gn=STK33;refGene_tx=XM_023239538.1;refGene_gn=STK33;refGene_tx=XM_023239539.1
D1 67206547 67206661
refGene_gn=STK33;refGene_tx=XM_023239535.1;refGene_gn=STK33;refGene_tx=XM_023239536.1;refGene_gn=STK33;refGene_tx=XM_023239537.1;refGene_gn=STK33;refGene_tx=XM_023239538.1;refGene_gn=STK33;refGene_tx=XM_023239539.1
D1 67211197 67211318
refGene_gn=STK33;refGene_tx=XM_023239535.1;refGene_gn=STK33;refGene_tx=XM_023239536.1;refGene_gn=STK33;refGene_tx=XM_023239537.1;refGene_gn=STK33;refGene_tx=XM_023239538.1;refGene_gn=STK33;refGene_tx=XM_023239539.1
D1 67212600 67212825
refGene_gn=STK33;refGene_tx=XM_023239535.1;refGene_gn=STK33;refGene_tx=XM_023239536.1;refGene_gn=STK33;refGene_tx=XM_023239537.1;refGene_gn=STK33;refGene_tx=XM_023239538.1;refGene_gn=STK33;refGene_tx=XM_023239539.1
D1 67331567 67331627
refGene_gn=TRIM66;refGene_tx=XM_006937075.3;refGene_gn=TRIM66;refGene_tx=XM_006937076.4;refGene_gn=TRIM66;refGene_tx=XM_019812024.2;refGene_gn=TRIM66;refGene_tx=XM_019812026.2;refGene_gn=TRIM66;refGene_tx=XM_019812027.2;refGene_gn=TRIM66;refGene_tx=XM_019812029.2;refGene_gn=TRIM66;refGene_tx=XM_023239540.1
D1 67332373 67332592
refGene_gn=TRIM66;refGene_tx=XM_006937075.3;refGene_gn=TRIM66;refGene_tx=XM_006937076.4;refGene_gn=TRIM66;refGene_tx=XM_019812024.2;refGene_gn=TRIM66;refGene_tx=XM_019812026.2;refGene_gn=TRIM66;refGene_tx=XM_019812027.2;refGene_gn=TRIM66;refGene_tx=XM_019812029.2;refGene_gn=TRIM6

6;refGene_tx=XM_023239540.1
D1 67333005 67333158
refGene_gn=TRIM66;refGene_tx=XM_006937075.3;refGene_gn=TRIM66;refGene_tx=XM_006937076.4;refGene_gn=TRIM66;refGene_tx=XM_019812024.2;refGene_gn=TRIM66;refGene_tx=XM_019812026.2;refGene_gn=TRIM66;refGene_tx=XM_019812027.2;refGene_gn=TRIM66;refGene_tx=XM_019812029.2;refGene_gn=TRIM66;refGene_tx=XM_023239540.1
D1 67333519 67333594
refGene_gn=TRIM66;refGene_tx=XM_006937075.3;refGene_gn=TRIM66;refGene_tx=XM_006937076.4;refGene_gn=TRIM66;refGene_tx=XM_019812024.2;refGene_gn=TRIM66;refGene_tx=XM_019812026.2;refGene_gn=TRIM66;refGene_tx=XM_019812027.2;refGene_gn=TRIM66;refGene_tx=XM_019812029.2;refGene_gn=TRIM66;refGene_tx=XM_023239540.1
D1 67333920 67334047
refGene_gn=TRIM66;refGene_tx=XM_006937075.3;refGene_gn=TRIM66;refGene_tx=XM_006937076.4;refGene_gn=TRIM66;refGene_tx=XM_019812024.2;refGene_gn=TRIM66;refGene_tx=XM_019812026.2;refGene_gn=TRIM66;refGene_tx=XM_019812027.2;refGene_gn=TRIM66;refGene_tx=XM_019812029.2;refGene_gn=TRIM66;refGene_tx=XM_023239540.1
D1 67334535 67334825
refGene_gn=TRIM66;refGene_tx=XM_006937075.3;refGene_gn=TRIM66;refGene_tx=XM_006937076.4;refGene_gn=TRIM66;refGene_tx=XM_019812024.2;refGene_gn=TRIM66;refGene_tx=XM_019812026.2;refGene_gn=TRIM66;refGene_tx=XM_019812027.2;refGene_gn=TRIM66;refGene_tx=XM_019812029.2;refGene_gn=TRIM66;refGene_tx=XM_023239540.1
D1 67335112 67335287
refGene_gn=TRIM66;refGene_tx=XM_006937075.3;refGene_gn=TRIM66;refGene_tx=XM_006937076.4;refGene_gn=TRIM66;refGene_tx=XM_019812024.2;refGene_gn=TRIM66;refGene_tx=XM_019812026.2;refGene_gn=TRIM66;refGene_tx=XM_019812027.2;refGene_gn=TRIM66;refGene_tx=XM_019812029.2;refGene_gn=TRIM66;refGene_tx=XM_023239540.1
D1 67335732 67335793
refGene_gn=TRIM66;refGene_tx=XM_006937075.3;refGene_gn=TRIM66;refGene_tx=XM_006937076.4;refGene_gn=TRIM66;refGene_tx=XM_019812024.2;refGene_gn=TRIM66;refGene_tx=XM_019812026.2;refGene_gn=TRIM66;refGene_tx=XM_019812027.2;refGene_gn=TRIM66;refGene_tx=XM_019812029.2;refGene_gn=TRIM66;refGene_tx=XM_023239540.1
D1 67337298 67337491
refGene_gn=TRIM66;refGene_tx=XM_006937075.3;refGene_gn=TRIM66;refGene_tx=XM_006937076.4;refGene_gn=TRIM66;refGene_tx=XM_019812024.2;refGene_gn=TRIM66;refGene_tx=XM_019812026.2;refGene_gn=TRIM66;refGene_tx=XM_019812027.2;refGene_gn=TRIM66;refGene_tx=XM_019812029.2;refGene_gn=TRIM66;refGene_tx=XM_023239540.1
D1 67337656 67338178
refGene_gn=TRIM66;refGene_tx=XM_006937075.3;refGene_gn=TRIM66;refGene_tx=XM_006937076.4;refGene_gn=TRIM66;refGene_tx=XM_019812024.2;refGene_gn=TRIM66;refGene_tx=XM_019812026.2;refGene_gn=TRIM66;refGene_tx=XM_019812027.2;refGene_gn=TRIM66;refGene_tx=XM_019812029.2;refGene_gn=TRIM66;refGene_tx=XM_023239540.1
D1 67349183 67349345

refGene_gn=TRIM66;refGene_tx=XM_006937075.3;refGene_gn=TRIM66;refGene_tx=XM_006937076.4;refGene_gn=TRIM66;refGene_tx=XM_019812024.2;refGene_gn=TRIM66;refGene_tx=XM_019812026.2;refGene_gn=TRIM66;refGene_tx=XM_019812027.2;refGene_gn=TRIM66;refGene_tx=XM_019812029.2;refGene_gn=TRIM66;refGene_tx=XM_023239540.1
D1 67350725 67351630
refGene_gn=TRIM66;refGene_tx=XM_006937075.3;refGene_gn=TRIM66;refGene_tx=XM_006937076.4;refGene_gn=TRIM66;refGene_tx=XM_019812024.2;refGene_gn=TRIM66;refGene_tx=XM_019812026.2;refGene_gn=TRIM66;refGene_tx=XM_019812027.2;refGene_gn=TRIM66;refGene_tx=XM_019812029.2;refGene_gn=TRIM66;refGene_tx=XM_023239540.1
D1 67353062 67353180
refGene_gn=TRIM66;refGene_tx=XM_006937075.3;refGene_gn=TRIM66;refGene_tx=XM_006937076.4;refGene_gn=TRIM66;refGene_tx=XM_019812024.2;refGene_gn=TRIM66;refGene_tx=XM_019812026.2;refGene_gn=TRIM66;refGene_tx=XM_019812027.2;refGene_gn=TRIM66;refGene_tx=XM_019812029.2;refGene_gn=TRIM66;refGene_tx=XM_023239540.1
D1 67355348 67355495
refGene_gn=TRIM66;refGene_tx=XM_006937075.3;refGene_gn=TRIM66;refGene_tx=XM_006937076.4;refGene_gn=TRIM66;refGene_tx=XM_019812024.2;refGene_gn=TRIM66;refGene_tx=XM_019812026.2;refGene_gn=TRIM66;refGene_tx=XM_019812027.2;refGene_gn=TRIM66;refGene_tx=XM_019812029.2;refGene_gn=TRIM66;refGene_tx=XM_023239540.1
D1 67356047 67356162
refGene_gn=TRIM66;refGene_tx=XM_006937075.3;refGene_gn=TRIM66;refGene_tx=XM_006937076.4;refGene_gn=TRIM66;refGene_tx=XM_019812024.2;refGene_gn=TRIM66;refGene_tx=XM_019812026.2;refGene_gn=TRIM66;refGene_tx=XM_019812027.2;refGene_gn=TRIM66;refGene_tx=XM_019812029.2;refGene_gn=TRIM66;refGene_tx=XM_023239540.1
D1 67357463 67357580
refGene_gn=TRIM66;refGene_tx=XM_006937075.3;refGene_gn=TRIM66;refGene_tx=XM_006937076.4;refGene_gn=TRIM66;refGene_tx=XM_019812024.2;refGene_gn=TRIM66;refGene_tx=XM_019812026.2;refGene_gn=TRIM66;refGene_tx=XM_019812027.2;refGene_gn=TRIM66;refGene_tx=XM_019812029.2;refGene_gn=TRIM66;refGene_tx=XM_023239540.1
D1 67357910 67358043
refGene_gn=TRIM66;refGene_tx=XM_006937075.3;refGene_gn=TRIM66;refGene_tx=XM_006937076.4;refGene_gn=TRIM66;refGene_tx=XM_019812024.2;refGene_gn=TRIM66;refGene_tx=XM_019812026.2;refGene_gn=TRIM66;refGene_tx=XM_019812027.2;refGene_gn=TRIM66;refGene_tx=XM_019812029.2;refGene_gn=TRIM66;refGene_tx=XM_023239540.1
D1 67358911 67359059
refGene_gn=TRIM66;refGene_tx=XM_006937075.3;refGene_gn=TRIM66;refGene_tx=XM_006937076.4;refGene_gn=TRIM66;refGene_tx=XM_019812024.2;refGene_gn=TRIM66;refGene_tx=XM_019812026.2;refGene_gn=TRIM66;refGene_tx=XM_019812027.2;refGene_gn=TRIM66;refGene_tx=XM_019812029.2;refGene_gn=TRIM66;refGene_tx=XM_023239540.1
D1 67360483 67360587
refGene_gn=TRIM66;refGene_tx=XM_006937075.3;refGene_gn=TRIM66;refGene_tx=XM_006937076.4;refGene_gn=TRIM66;refGene_tx=XM_019812024.2;refGene_

gn=TRIM66;refGene_tx=XM_019812026.2;refGene_gn=TRIM66;refGene_tx=XM_01
9812027.2;refGene_gn=TRIM66;refGene_tx=XM_019812029.2;refGene_gn=TRIM6
6;refGene_tx=XM_023239540.1
D1 67365750 67365772 refGene_gn=TRIM66;refGene_tx=XM_019812024.2
D1 67383791 67383843 refGene_gn=TRIM66;refGene_tx=XM_006937075.3
D1 67386551 67386554 refGene_gn=RPL27A;refGene_tx=XM_003992976.4
D1 67386945 67387009 refGene_gn=RPL27A;refGene_tx=XM_003992976.4
D1 67387451 67387527 refGene_gn=RPL27A;refGene_tx=XM_003992976.4
D1 67388204 67388379 refGene_gn=RPL27A;refGene_tx=XM_003992976.4
D1 67389559 67389688 refGene_gn=RPL27A;refGene_tx=XM_003992976.4
D1 67396932 67396967
refGene_gn=ST5;refGene_tx=XM_003992977.4;refGene_gn=ST5;refGene_tx=XM_
011286706.3;refGene_gn=ST5;refGene_tx=XM_011286708.3;refGene_gn=ST5;re
fGene_tx=XM_011286709.3;refGene_gn=ST5;refGene_tx=XM_019812031.2;refGe
ne_gn=ST5;refGene_tx=XM_019812032.2;refGene_gn=ST5;refGene_tx=XM_01981
2033.2;refGene_gn=ST5;refGene_tx=XM_019812034.2;refGene_gn=ST5;refGene
_tx=XM_023239541.1;refGene_gn=ST5;refGene_tx=XM_023239542.1
D1 67397591 67397678
refGene_gn=ST5;refGene_tx=XM_003992977.4;refGene_gn=ST5;refGene_tx=XM_
011286706.3;refGene_gn=ST5;refGene_tx=XM_011286708.3;refGene_gn=ST5;re
fGene_tx=XM_011286709.3;refGene_gn=ST5;refGene_tx=XM_019812031.2;refGe
ne_gn=ST5;refGene_tx=XM_019812032.2;refGene_gn=ST5;refGene_tx=XM_01981
2033.2;refGene_gn=ST5;refGene_tx=XM_019812034.2;refGene_gn=ST5;refGene
_tx=XM_023239541.1;refGene_gn=ST5;refGene_tx=XM_023239542.1
D1 67398557 67398797
refGene_gn=ST5;refGene_tx=XM_003992977.4;refGene_gn=ST5;refGene_tx=XM_
011286706.3;refGene_gn=ST5;refGene_tx=XM_011286708.3;refGene_gn=ST5;re
fGene_tx=XM_011286709.3;refGene_gn=ST5;refGene_tx=XM_019812031.2;refGe
ne_gn=ST5;refGene_tx=XM_019812032.2;refGene_gn=ST5;refGene_tx=XM_01981
2033.2;refGene_gn=ST5;refGene_tx=XM_019812034.2;refGene_gn=ST5;refGene
_tx=XM_023239541.1;refGene_gn=ST5;refGene_tx=XM_023239542.1
D1 67399537 67399649
refGene_gn=ST5;refGene_tx=XM_003992977.4;refGene_gn=ST5;refGene_tx=XM_
011286706.3;refGene_gn=ST5;refGene_tx=XM_011286708.3;refGene_gn=ST5;re
fGene_tx=XM_011286709.3;refGene_gn=ST5;refGene_tx=XM_019812031.2;refGe
ne_gn=ST5;refGene_tx=XM_019812032.2;refGene_gn=ST5;refGene_tx=XM_01981
2033.2;refGene_gn=ST5;refGene_tx=XM_019812034.2;refGene_gn=ST5;refGene
_tx=XM_023239541.1;refGene_gn=ST5;refGene_tx=XM_023239542.1
D1 67400533 67400575
refGene_gn=ST5;refGene_tx=XM_003992977.4;refGene_gn=ST5;refGene_tx=XM_
011286706.3;refGene_gn=ST5;refGene_tx=XM_011286708.3;refGene_gn=ST5;re
fGene_tx=XM_011286709.3;refGene_gn=ST5;refGene_tx=XM_019812031.2;refGe
ne_gn=ST5;refGene_tx=XM_019812032.2;refGene_gn=ST5;refGene_tx=XM_01981
2033.2;refGene_gn=ST5;refGene_tx=XM_019812034.2;refGene_gn=ST5;refGene
_tx=XM_023239541.1;refGene_gn=ST5;refGene_tx=XM_023239542.1
D1 67400875 67401053
refGene_gn=ST5;refGene_tx=XM_003992977.4;refGene_gn=ST5;refGene_tx=XM_
011286706.3;refGene_gn=ST5;refGene_tx=XM_011286708.3;refGene_gn=ST5;re
fGene_tx=XM_011286709.3;refGene_gn=ST5;refGene_tx=XM_019812031.2;refGe
ne_gn=ST5;refGene_tx=XM_019812032.2;refGene_gn=ST5;refGene_tx=XM_01981

2033.2;refGene_gn=ST5;refGene_tx=XM_019812034.2;refGene_gn=ST5;refGene_tx=XM_023239541.1;refGene_gn=ST5;refGene_tx=XM_023239542.1
D1 67404722 67404871
refGene_gn=ST5;refGene_tx=XM_003992977.4;refGene_gn=ST5;refGene_tx=XM_011286706.3;refGene_gn=ST5;refGene_tx=XM_011286708.3;refGene_gn=ST5;refGene_tx=XM_011286709.3;refGene_gn=ST5;refGene_tx=XM_019812031.2;refGene_gn=ST5;refGene_tx=XM_019812032.2;refGene_gn=ST5;refGene_tx=XM_019812033.2;refGene_gn=ST5;refGene_tx=XM_019812034.2;refGene_gn=ST5;refGene_tx=XM_023239541.1;refGene_gn=ST5;refGene_tx=XM_023239542.1
D1 67408544 67408685
refGene_gn=ST5;refGene_tx=XM_003992977.4;refGene_gn=ST5;refGene_tx=XM_011286706.3;refGene_gn=ST5;refGene_tx=XM_011286708.3;refGene_gn=ST5;refGene_tx=XM_011286709.3;refGene_gn=ST5;refGene_tx=XM_019812031.2;refGene_gn=ST5;refGene_tx=XM_019812032.2;refGene_gn=ST5;refGene_tx=XM_019812033.2;refGene_gn=ST5;refGene_tx=XM_019812034.2;refGene_gn=ST5;refGene_tx=XM_023239541.1;refGene_gn=ST5;refGene_tx=XM_023239542.1
D1 67409239 67409317
refGene_gn=ST5;refGene_tx=XM_003992977.4;refGene_gn=ST5;refGene_tx=XM_011286706.3;refGene_gn=ST5;refGene_tx=XM_011286708.3;refGene_gn=ST5;refGene_tx=XM_011286709.3;refGene_gn=ST5;refGene_tx=XM_019812031.2;refGene_gn=ST5;refGene_tx=XM_019812032.2;refGene_gn=ST5;refGene_tx=XM_019812033.2;refGene_gn=ST5;refGene_tx=XM_019812034.2;refGene_gn=ST5;refGene_tx=XM_023239541.1;refGene_gn=ST5;refGene_tx=XM_023239542.1
D1 67412070 67412140
refGene_gn=ST5;refGene_tx=XM_003992977.4;refGene_gn=ST5;refGene_tx=XM_011286706.3;refGene_gn=ST5;refGene_tx=XM_011286708.3;refGene_gn=ST5;refGene_tx=XM_011286709.3;refGene_gn=ST5;refGene_tx=XM_019812031.2;refGene_gn=ST5;refGene_tx=XM_019812032.2;refGene_gn=ST5;refGene_tx=XM_019812033.2;refGene_gn=ST5;refGene_tx=XM_019812034.2;refGene_gn=ST5;refGene_tx=XM_023239541.1;refGene_gn=ST5;refGene_tx=XM_023239542.1
D1 67412355 67412465
refGene_gn=ST5;refGene_tx=XM_003992977.4;refGene_gn=ST5;refGene_tx=XM_011286706.3;refGene_gn=ST5;refGene_tx=XM_011286708.3;refGene_gn=ST5;refGene_tx=XM_011286709.3;refGene_gn=ST5;refGene_tx=XM_019812031.2;refGene_gn=ST5;refGene_tx=XM_019812032.2;refGene_gn=ST5;refGene_tx=XM_019812033.2;refGene_gn=ST5;refGene_tx=XM_019812034.2;refGene_gn=ST5;refGene_tx=XM_023239541.1;refGene_gn=ST5;refGene_tx=XM_023239542.1
D1 67413536 67413721
refGene_gn=ST5;refGene_tx=XM_003992977.4;refGene_gn=ST5;refGene_tx=XM_011286706.3;refGene_gn=ST5;refGene_tx=XM_011286708.3;refGene_gn=ST5;refGene_tx=XM_011286709.3;refGene_gn=ST5;refGene_tx=XM_019812031.2;refGene_gn=ST5;refGene_tx=XM_019812032.2;refGene_gn=ST5;refGene_tx=XM_019812033.2;refGene_gn=ST5;refGene_tx=XM_019812034.2;refGene_gn=ST5;refGene_tx=XM_023239541.1;refGene_gn=ST5;refGene_tx=XM_023239542.1
D1 67414719 67414764
refGene_gn=ST5;refGene_tx=XM_003992977.4;refGene_gn=ST5;refGene_tx=XM_011286706.3;refGene_gn=ST5;refGene_tx=XM_011286708.3;refGene_gn=ST5;refGene_tx=XM_011286709.3;refGene_gn=ST5;refGene_tx=XM_019812031.2;refGene_gn=ST5;refGene_tx=XM_019812032.2;refGene_gn=ST5;refGene_tx=XM_019812034.2;refGene_gn=ST5;refGene_tx=XM_023239541.1;refGene_gn=ST5;refGene

_tx=XM_023239542.1
D1 67415360 67415457
refGene_gn=ST5;refGene_tx=XM_003992977.4;refGene_gn=ST5;refGene_tx=XM_011286706.3;refGene_gn=ST5;refGene_tx=XM_011286708.3;refGene_gn=ST5;refGene_tx=XM_011286709.3;refGene_gn=ST5;refGene_tx=XM_019812031.2;refGene_gn=ST5;refGene_tx=XM_019812032.2;refGene_gn=ST5;refGene_tx=XM_019812033.2;refGene_gn=ST5;refGene_tx=XM_019812034.2;refGene_gn=ST5;refGene_tx=XM_023239541.1;refGene_gn=ST5;refGene_tx=XM_023239542.1
D1 67416417 67416633
refGene_gn=ST5;refGene_tx=XM_003992977.4;refGene_gn=ST5;refGene_tx=XM_011286706.3;refGene_gn=ST5;refGene_tx=XM_011286708.3;refGene_gn=ST5;refGene_tx=XM_011286709.3;refGene_gn=ST5;refGene_tx=XM_019812031.2;refGene_gn=ST5;refGene_tx=XM_019812032.2;refGene_gn=ST5;refGene_tx=XM_019812033.2;refGene_gn=ST5;refGene_tx=XM_019812034.2;refGene_gn=ST5;refGene_tx=XM_023239541.1;refGene_gn=ST5;refGene_tx=XM_023239542.1
D1 67418145 67418297
refGene_gn=ST5;refGene_tx=XM_003992977.4;refGene_gn=ST5;refGene_tx=XM_011286706.3;refGene_gn=ST5;refGene_tx=XM_011286708.3;refGene_gn=ST5;refGene_tx=XM_011286709.3;refGene_gn=ST5;refGene_tx=XM_019812031.2;refGene_gn=ST5;refGene_tx=XM_019812032.2;refGene_gn=ST5;refGene_tx=XM_019812033.2;refGene_gn=ST5;refGene_tx=XM_019812034.2;refGene_gn=ST5;refGene_tx=XM_023239541.1;refGene_gn=ST5;refGene_tx=XM_023239542.1
D1 67425657 67425794
refGene_gn=ST5;refGene_tx=XM_003992977.4;refGene_gn=ST5;refGene_tx=XM_011286706.3;refGene_gn=ST5;refGene_tx=XM_011286708.3;refGene_gn=ST5;refGene_tx=XM_011286709.3;refGene_gn=ST5;refGene_tx=XM_019812031.2;refGene_gn=ST5;refGene_tx=XM_019812032.2;refGene_gn=ST5;refGene_tx=XM_019812033.2;refGene_gn=ST5;refGene_tx=XM_019812034.2;refGene_gn=ST5;refGene_tx=XM_023239541.1;refGene_gn=ST5;refGene_tx=XM_023239542.1
D1 67429244 67430489
refGene_gn=ST5;refGene_tx=XM_003992977.4;refGene_gn=ST5;refGene_tx=XM_011286706.3;refGene_gn=ST5;refGene_tx=XM_011286708.3;refGene_gn=ST5;refGene_tx=XM_011286709.3;refGene_gn=ST5;refGene_tx=XM_019812031.2;refGene_gn=ST5;refGene_tx=XM_019812032.2;refGene_gn=ST5;refGene_tx=XM_019812033.2;refGene_gn=ST5;refGene_tx=XM_023239541.1;refGene_gn=ST5;refGene_tx=XM_023239542.1
D1 67447176 67447256
refGene_gn=ST5;refGene_tx=XM_003992977.4;refGene_gn=ST5;refGene_tx=XM_011286706.3;refGene_gn=ST5;refGene_tx=XM_011286708.3;refGene_gn=ST5;refGene_tx=XM_011286709.3;refGene_gn=ST5;refGene_tx=XM_019812031.2;refGene_gn=ST5;refGene_tx=XM_019812032.2;refGene_gn=ST5;refGene_tx=XM_019812033.2;refGene_gn=ST5;refGene_tx=XM_019812034.2;refGene_gn=ST5;refGene_tx=XM_023239541.1;refGene_gn=ST5;refGene_tx=XM_023239542.1
D1 67576625 67576710
refGene_gn=AKIP1;refGene_tx=XM_006937080.4;refGene_gn=AKIP1;refGene_tx=XM_006937081.4;refGene_gn=AKIP1;refGene_tx=XM_006937082.4
D1 67576932 67577160
refGene_gn=AKIP1;refGene_tx=XM_006937080.4;refGene_gn=AKIP1;refGene_tx=XM_006937081.4;refGene_gn=AKIP1;refGene_tx=XM_006937082.4;refGene_gn=AKIP1;refGene_tx=XM_006937083.3

D1 67577624 67577705
refGene_gn=AKIP1;refGene_tx=XM_006937080.4;refGene_gn=AKIP1;refGene_tx=XM_006937081.4;refGene_gn=AKIP1;refGene_tx=XM_006937082.4;refGene_gn=AKIP1;refGene_tx=XM_006937083.3
D1 67579877 67579982
refGene_gn=AKIP1;refGene_tx=XM_006937080.4;refGene_gn=AKIP1;refGene_tx=XM_006937081.4;refGene_gn=AKIP1;refGene_tx=XM_006937082.4;refGene_gn=AKIP1;refGene_tx=XM_006937083.3
D1 67581561 67581636
refGene_gn=AKIP1;refGene_tx=XM_006937080.4;refGene_gn=AKIP1;refGene_tx=XM_006937081.4;refGene_gn=AKIP1;refGene_tx=XM_006937083.3
D1 67583825 67583966
refGene_gn=AKIP1;refGene_tx=XM_006937080.4;refGene_gn=AKIP1;refGene_tx=XM_006937081.4;refGene_gn=AKIP1;refGene_tx=XM_006937082.4;refGene_gn=AKIP1;refGene_tx=XM_006937083.3
D1 67597386 67597932 refGene_gn=ASCL3;refGene_tx=XM_003992979.3
D1 67606368 67606524
refGene_gn=TMEM9B;refGene_tx=XM_003992980.5;refGene_gn=TMEM9B;refGene_tx=XM_019812035.2
D1 67610831 67610966
refGene_gn=TMEM9B;refGene_tx=XM_003992980.5;refGene_gn=TMEM9B;refGene_tx=XM_019812035.2
D1 67612838 67612947
refGene_gn=TMEM9B;refGene_tx=XM_003992980.5;refGene_gn=TMEM9B;refGene_tx=XM_019812035.2
D1 67617061 67617153
refGene_gn=TMEM9B;refGene_tx=XM_003992980.5;refGene_gn=TMEM9B;refGene_tx=XM_019812035.2
D1 67619267 67619372
refGene_gn=TMEM9B;refGene_tx=XM_003992980.5;refGene_gn=TMEM9B;refGene_tx=XM_019812035.2
D1 67635046 67635062 refGene_gn=NRIP3;refGene_tx=XM_003992981.5
D1 67635345 67635440 refGene_gn=NRIP3;refGene_tx=XM_003992981.5
D1 67635566 67635619 refGene_gn=NRIP3;refGene_tx=XM_003992981.5
D1 67636796 67636936 refGene_gn=NRIP3;refGene_tx=XM_003992981.5
D1 67638848 67638931 refGene_gn=NRIP3;refGene_tx=XM_003992981.5
D1 67639433 67639595 refGene_gn=NRIP3;refGene_tx=XM_003992981.5
D1 67656447 67656621 refGene_gn=NRIP3;refGene_tx=XM_003992981.5
D1 67671245 67671398
refGene_gn=SCUBE2;refGene_tx=XM_006937084.4;refGene_gn=SCUBE2;refGene_tx=XM_023239543.1;refGene_gn=SCUBE2;refGene_tx=XM_023239544.1;refGene_gn=SCUBE2;refGene_tx=XM_023239545.1;refGene_gn=SCUBE2;refGene_tx=XM_023239546.1
D1 67672088 67672168
refGene_gn=SCUBE2;refGene_tx=XM_006937084.4;refGene_gn=SCUBE2;refGene_tx=XM_023239543.1;refGene_gn=SCUBE2;refGene_tx=XM_023239544.1;refGene_gn=SCUBE2;refGene_tx=XM_023239545.1;refGene_gn=SCUBE2;refGene_tx=XM_023239546.1
D1 67676136 67676289
refGene_gn=SCUBE2;refGene_tx=XM_006937084.4;refGene_gn=SCUBE2;refGene_

tx=XM_023239543.1;refGene_gn=SCUBE2;refGene_tx=XM_023239544.1;refGene_gn=SCUBE2;refGene_tx=XM_023239545.1;refGene_gn=SCUBE2;refGene_tx=XM_023239546.1
D1 67677484 67677682 refGene_gn=SCUBE2;refGene_tx=XM_006937084.4;refGene_gn=SCUBE2;refGene_tx=XM_023239543.1;refGene_gn=SCUBE2;refGene_tx=XM_023239544.1;refGene_gn=SCUBE2;refGene_tx=XM_023239545.1;refGene_gn=SCUBE2;refGene_tx=XM_023239546.1
D1 67679687 67679849 refGene_gn=SCUBE2;refGene_tx=XM_006937084.4;refGene_gn=SCUBE2;refGene_tx=XM_023239543.1;refGene_gn=SCUBE2;refGene_tx=XM_023239544.1;refGene_gn=SCUBE2;refGene_tx=XM_023239545.1;refGene_gn=SCUBE2;refGene_tx=XM_023239546.1
D1 67680556 67680724 refGene_gn=SCUBE2;refGene_tx=XM_006937084.4;refGene_gn=SCUBE2;refGene_tx=XM_023239543.1;refGene_gn=SCUBE2;refGene_tx=XM_023239544.1;refGene_gn=SCUBE2;refGene_tx=XM_023239545.1;refGene_gn=SCUBE2;refGene_tx=XM_023239546.1
D1 67682838 67683009 refGene_gn=SCUBE2;refGene_tx=XM_006937084.4;refGene_gn=SCUBE2;refGene_tx=XM_023239543.1;refGene_gn=SCUBE2;refGene_tx=XM_023239544.1;refGene_gn=SCUBE2;refGene_tx=XM_023239545.1;refGene_gn=SCUBE2;refGene_tx=XM_023239546.1
D1 67696395 67696602 refGene_gn=SCUBE2;refGene_tx=XM_006937084.4;refGene_gn=SCUBE2;refGene_tx=XM_023239543.1;refGene_gn=SCUBE2;refGene_tx=XM_023239544.1;refGene_gn=SCUBE2;refGene_tx=XM_023239545.1;refGene_gn=SCUBE2;refGene_tx=XM_023239546.1
D1 67697001 67697157 refGene_gn=SCUBE2;refGene_tx=XM_006937084.4;refGene_gn=SCUBE2;refGene_tx=XM_023239543.1;refGene_gn=SCUBE2;refGene_tx=XM_023239544.1
D1 67699767 67699872 refGene_gn=SCUBE2;refGene_tx=XM_006937084.4;refGene_gn=SCUBE2;refGene_tx=XM_023239543.1;refGene_gn=SCUBE2;refGene_tx=XM_023239544.1
D1 67701585 67701672 refGene_gn=SCUBE2;refGene_tx=XM_006937084.4;refGene_gn=SCUBE2;refGene_tx=XM_023239543.1;refGene_gn=SCUBE2;refGene_tx=XM_023239544.1
D1 67701932 67702049 refGene_gn=SCUBE2;refGene_tx=XM_006937084.4;refGene_gn=SCUBE2;refGene_tx=XM_023239543.1;refGene_gn=SCUBE2;refGene_tx=XM_023239544.1
D1 67702476 67702599 refGene_gn=SCUBE2;refGene_tx=XM_006937084.4;refGene_gn=SCUBE2;refGene_tx=XM_023239543.1;refGene_gn=SCUBE2;refGene_tx=XM_023239544.1;refGene_gn=SCUBE2;refGene_tx=XM_023239545.1;refGene_gn=SCUBE2;refGene_tx=XM_023239546.1
D1 67703876 67703993 refGene_gn=SCUBE2;refGene_tx=XM_006937084.4;refGene_gn=SCUBE2;refGene_tx=XM_023239543.1;refGene_gn=SCUBE2;refGene_tx=XM_023239544.1;refGene_gn=SCUBE2;refGene_tx=XM_023239545.1;refGene_gn=SCUBE2;refGene_tx=XM_023239546.1

D1 67706414 67706537
refGene_gn=SCUBE2;refGene_tx=XM_006937084.4;refGene_gn=SCUBE2;refGene_tx=XM_023239543.1;refGene_gn=SCUBE2;refGene_tx=XM_023239544.1;refGene_gn=SCUBE2;refGene_tx=XM_023239545.1;refGene_gn=SCUBE2;refGene_tx=XM_023239546.1
D1 67707490 67707607
refGene_gn=SCUBE2;refGene_tx=XM_006937084.4;refGene_gn=SCUBE2;refGene_tx=XM_023239543.1;refGene_gn=SCUBE2;refGene_tx=XM_023239544.1;refGene_gn=SCUBE2;refGene_tx=XM_023239545.1;refGene_gn=SCUBE2;refGene_tx=XM_023239546.1
D1 67712715 67712805
refGene_gn=SCUBE2;refGene_tx=XM_006937084.4;refGene_gn=SCUBE2;refGene_tx=XM_023239543.1;refGene_gn=SCUBE2;refGene_tx=XM_023239544.1;refGene_gn=SCUBE2;refGene_tx=XM_023239545.1;refGene_gn=SCUBE2;refGene_tx=XM_023239546.1
D1 67713549 67713666
refGene_gn=SCUBE2;refGene_tx=XM_006937084.4;refGene_gn=SCUBE2;refGene_tx=XM_023239543.1;refGene_gn=SCUBE2;refGene_tx=XM_023239544.1;refGene_gn=SCUBE2;refGene_tx=XM_023239545.1;refGene_gn=SCUBE2;refGene_tx=XM_023239546.1
D1 67716362 67716488
refGene_gn=SCUBE2;refGene_tx=XM_006937084.4;refGene_gn=SCUBE2;refGene_tx=XM_023239543.1;refGene_gn=SCUBE2;refGene_tx=XM_023239544.1;refGene_gn=SCUBE2;refGene_tx=XM_023239545.1;refGene_gn=SCUBE2;refGene_tx=XM_023239546.1
D1 67721691 67721826
refGene_gn=SCUBE2;refGene_tx=XM_006937084.4;refGene_gn=SCUBE2;refGene_tx=XM_023239543.1;refGene_gn=SCUBE2;refGene_tx=XM_023239544.1;refGene_gn=SCUBE2;refGene_tx=XM_023239545.1;refGene_gn=SCUBE2;refGene_tx=XM_023239546.1
D1 67726487 67726613
refGene_gn=SCUBE2;refGene_tx=XM_006937084.4;refGene_gn=SCUBE2;refGene_tx=XM_023239543.1;refGene_gn=SCUBE2;refGene_tx=XM_023239544.1;refGene_gn=SCUBE2;refGene_tx=XM_023239545.1;refGene_gn=SCUBE2;refGene_tx=XM_023239546.1
D1 67733535 67733658
refGene_gn=SCUBE2;refGene_tx=XM_006937084.4;refGene_gn=SCUBE2;refGene_tx=XM_023239543.1;refGene_gn=SCUBE2;refGene_tx=XM_023239544.1;refGene_gn=SCUBE2;refGene_tx=XM_023239545.1;refGene_gn=SCUBE2;refGene_tx=XM_023239546.1
D1 67733895 67733959 refGene_gn=SCUBE2;refGene_tx=XM_006937084.4
D1 67734504 67734775
refGene_gn=SCUBE2;refGene_tx=XM_023239543.1;refGene_gn=SCUBE2;refGene_tx=XM_023239545.1
D1 67735028 67735149
refGene_gn=SCUBE2;refGene_tx=XM_023239544.1;refGene_gn=SCUBE2;refGene_tx=XM_023239546.1
D1 67767526 67767710
refGene_gn=DENND5A;refGene_tx=XM_023239547.1;refGene_gn=DENND5A;refGene_tx=XM_023239548.1;refGene_gn=DENND5A;refGene_tx=XM_023239549.1;refGe

ne_gn=DENND5A;refGene_tx=XM_023239550.1;refGene_gn=DENND5A;refGene_tx=XM_023239551.1;refGene_gn=DENND5A;refGene_tx=XM_023239552.1
D1 67769213 67769382
refGene_gn=DENND5A;refGene_tx=XM_023239547.1;refGene_gn=DENND5A;refGene_tx=XM_023239548.1;refGene_gn=DENND5A;refGene_tx=XM_023239549.1;refGene_gn=DENND5A;refGene_tx=XM_023239550.1;refGene_gn=DENND5A;refGene_tx=XM_023239551.1;refGene_gn=DENND5A;refGene_tx=XM_023239552.1
D1 67769927 67770051
refGene_gn=DENND5A;refGene_tx=XM_023239547.1;refGene_gn=DENND5A;refGene_tx=XM_023239548.1;refGene_gn=DENND5A;refGene_tx=XM_023239549.1;refGene_gn=DENND5A;refGene_tx=XM_023239550.1;refGene_gn=DENND5A;refGene_tx=XM_023239551.1;refGene_gn=DENND5A;refGene_tx=XM_023239552.1
D1 67770544 67770627
refGene_gn=DENND5A;refGene_tx=XM_023239547.1;refGene_gn=DENND5A;refGene_tx=XM_023239548.1;refGene_gn=DENND5A;refGene_tx=XM_023239549.1;refGene_gn=DENND5A;refGene_tx=XM_023239550.1;refGene_gn=DENND5A;refGene_tx=XM_023239551.1;refGene_gn=DENND5A;refGene_tx=XM_023239552.1
D1 67771252 67771434
refGene_gn=DENND5A;refGene_tx=XM_023239547.1;refGene_gn=DENND5A;refGene_tx=XM_023239548.1;refGene_gn=DENND5A;refGene_tx=XM_023239549.1;refGene_gn=DENND5A;refGene_tx=XM_023239550.1;refGene_gn=DENND5A;refGene_tx=XM_023239551.1;refGene_gn=DENND5A;refGene_tx=XM_023239552.1
D1 67771844 67771963
refGene_gn=DENND5A;refGene_tx=XM_023239547.1;refGene_gn=DENND5A;refGene_tx=XM_023239548.1;refGene_gn=DENND5A;refGene_tx=XM_023239549.1;refGene_gn=DENND5A;refGene_tx=XM_023239550.1;refGene_gn=DENND5A;refGene_tx=XM_023239551.1;refGene_gn=DENND5A;refGene_tx=XM_023239552.1
D1 67772535 67772681
refGene_gn=DENND5A;refGene_tx=XM_023239547.1;refGene_gn=DENND5A;refGene_tx=XM_023239548.1;refGene_gn=DENND5A;refGene_tx=XM_023239549.1;refGene_gn=DENND5A;refGene_tx=XM_023239550.1;refGene_gn=DENND5A;refGene_tx=XM_023239551.1;refGene_gn=DENND5A;refGene_tx=XM_023239552.1
D1 67773892 67774014
refGene_gn=DENND5A;refGene_tx=XM_023239547.1;refGene_gn=DENND5A;refGene_tx=XM_023239548.1;refGene_gn=DENND5A;refGene_tx=XM_023239549.1;refGene_gn=DENND5A;refGene_tx=XM_023239550.1;refGene_gn=DENND5A;refGene_tx=XM_023239551.1;refGene_gn=DENND5A;refGene_tx=XM_023239552.1
D1 67778376 67778505
refGene_gn=DENND5A;refGene_tx=XM_023239547.1;refGene_gn=DENND5A;refGene_tx=XM_023239548.1;refGene_gn=DENND5A;refGene_tx=XM_023239549.1;refGene_gn=DENND5A;refGene_tx=XM_023239550.1;refGene_gn=DENND5A;refGene_tx=XM_023239551.1;refGene_gn=DENND5A;refGene_tx=XM_023239552.1
D1 67779186 67779271
refGene_gn=DENND5A;refGene_tx=XM_023239547.1;refGene_gn=DENND5A;refGene_tx=XM_023239548.1;refGene_gn=DENND5A;refGene_tx=XM_023239549.1;refGene_gn=DENND5A;refGene_tx=XM_023239550.1;refGene_gn=DENND5A;refGene_tx=XM_023239551.1;refGene_gn=DENND5A;refGene_tx=XM_023239552.1
D1 67782033 67782118
refGene_gn=DENND5A;refGene_tx=XM_023239547.1;refGene_gn=DENND5A;refGene_tx=XM_023239548.1;refGene_gn=DENND5A;refGene_tx=XM_023239549.1;refGe

ne_gn=DENND5A;refGene_tx=XM_023239550.1;refGene_gn=DENND5A;refGene_tx=XM_023239551.1;refGene_gn=DENND5A;refGene_tx=XM_023239552.1
D1 67787893 67788046
refGene_gn=DENND5A;refGene_tx=XM_023239547.1;refGene_gn=DENND5A;refGene_tx=XM_023239548.1;refGene_gn=DENND5A;refGene_tx=XM_023239549.1;refGene_gn=DENND5A;refGene_tx=XM_023239550.1;refGene_gn=DENND5A;refGene_tx=XM_023239551.1;refGene_gn=DENND5A;refGene_tx=XM_023239552.1
D1 67793374 67793506
refGene_gn=DENND5A;refGene_tx=XM_023239547.1;refGene_gn=DENND5A;refGene_tx=XM_023239548.1;refGene_gn=DENND5A;refGene_tx=XM_023239549.1;refGene_gn=DENND5A;refGene_tx=XM_023239550.1;refGene_gn=DENND5A;refGene_tx=XM_023239551.1;refGene_gn=DENND5A;refGene_tx=XM_023239552.1
D1 67796585 67796679
refGene_gn=DENND5A;refGene_tx=XM_023239547.1;refGene_gn=DENND5A;refGene_tx=XM_023239548.1;refGene_gn=DENND5A;refGene_tx=XM_023239549.1;refGene_gn=DENND5A;refGene_tx=XM_023239550.1;refGene_gn=DENND5A;refGene_tx=XM_023239551.1;refGene_gn=DENND5A;refGene_tx=XM_023239552.1
D1 67797351 67797502
refGene_gn=DENND5A;refGene_tx=XM_023239547.1;refGene_gn=DENND5A;refGene_tx=XM_023239548.1;refGene_gn=DENND5A;refGene_tx=XM_023239549.1;refGene_gn=DENND5A;refGene_tx=XM_023239550.1;refGene_gn=DENND5A;refGene_tx=XM_023239551.1;refGene_gn=DENND5A;refGene_tx=XM_023239552.1
D1 67811463 67811613
refGene_gn=DENND5A;refGene_tx=XM_023239547.1;refGene_gn=DENND5A;refGene_tx=XM_023239548.1;refGene_gn=DENND5A;refGene_tx=XM_023239551.1;refGene_gn=DENND5A;refGene_tx=XM_023239552.1
D1 67812026 67812261
refGene_gn=DENND5A;refGene_tx=XM_023239547.1;refGene_gn=DENND5A;refGene_tx=XM_023239548.1;refGene_gn=DENND5A;refGene_tx=XM_023239549.1;refGene_gn=DENND5A;refGene_tx=XM_023239550.1;refGene_gn=DENND5A;refGene_tx=XM_023239551.1;refGene_gn=DENND5A;refGene_tx=XM_023239552.1
D1 67812652 67812868
refGene_gn=DENND5A;refGene_tx=XM_023239547.1;refGene_gn=DENND5A;refGene_tx=XM_023239548.1;refGene_gn=DENND5A;refGene_tx=XM_023239549.1;refGene_gn=DENND5A;refGene_tx=XM_023239550.1;refGene_gn=DENND5A;refGene_tx=XM_023239551.1;refGene_gn=DENND5A;refGene_tx=XM_023239552.1
D1 67821315 67821633
refGene_gn=DENND5A;refGene_tx=XM_023239547.1;refGene_gn=DENND5A;refGene_tx=XM_023239548.1;refGene_gn=DENND5A;refGene_tx=XM_023239549.1;refGene_gn=DENND5A;refGene_tx=XM_023239550.1;refGene_gn=DENND5A;refGene_tx=XM_023239552.1
D1 67833222 67833410
refGene_gn=DENND5A;refGene_tx=XM_023239547.1;refGene_gn=DENND5A;refGene_tx=XM_023239548.1;refGene_gn=DENND5A;refGene_tx=XM_023239549.1;refGene_gn=DENND5A;refGene_tx=XM_023239550.1;refGene_gn=DENND5A;refGene_tx=XM_023239551.1;refGene_gn=DENND5A;refGene_tx=XM_023239552.1
D1 67842957 67843615
refGene_gn=DENND5A;refGene_tx=XM_023239547.1;refGene_gn=DENND5A;refGene_tx=XM_023239548.1;refGene_gn=DENND5A;refGene_tx=XM_023239549.1;refGene_gn=DENND5A;refGene_tx=XM_023239550.1;refGene_gn=DENND5A;refGene_tx=

XM_023239551.1;refGene_gn=DENND5A;refGene_tx=XM_023239552.1
D1 67845333 67845443
refGene_gn=DENND5A;refGene_tx=XM_023239547.1;refGene_gn=DENND5A;refGene_tx=XM_023239548.1;refGene_gn=DENND5A;refGene_tx=XM_023239549.1;refGene_gn=DENND5A;refGene_tx=XM_023239550.1;refGene_gn=DENND5A;refGene_tx=XM_023239551.1;refGene_gn=DENND5A;refGene_tx=XM_023239552.1
D1 67846210 67846282
refGene_gn=DENND5A;refGene_tx=XM_023239547.1;refGene_gn=DENND5A;refGene_tx=XM_023239549.1;refGene_gn=DENND5A;refGene_tx=XM_023239551.1;refGene_gn=DENND5A;refGene_tx=XM_023239552.1
D1 67884930 67885057
refGene_gn=DENND5A;refGene_tx=XM_023239547.1;refGene_gn=DENND5A;refGene_tx=XM_023239548.1;refGene_gn=DENND5A;refGene_tx=XM_023239549.1;refGene_gn=DENND5A;refGene_tx=XM_023239550.1;refGene_gn=DENND5A;refGene_tx=XM_023239551.1;refGene_gn=DENND5A;refGene_tx=XM_023239552.1
D1 67906575 67906745
refGene_gn=TMEM41B;refGene_tx=XM_006937086.3;refGene_gn=TMEM41B;refGene_tx=XM_023239553.1
D1 67908588 67908727
refGene_gn=TMEM41B;refGene_tx=XM_006937086.3;refGene_gn=TMEM41B;refGene_tx=XM_023239553.1
D1 67909838 67909943
refGene_gn=TMEM41B;refGene_tx=XM_006937086.3;refGene_gn=TMEM41B;refGene_tx=XM_023239553.1
D1 67910500 67910594
refGene_gn=TMEM41B;refGene_tx=XM_006937086.3;refGene_gn=TMEM41B;refGene_tx=XM_023239553.1
D1 67915561 67915690
refGene_gn=TMEM41B;refGene_tx=XM_006937086.3;refGene_gn=TMEM41B;refGene_tx=XM_023239553.1
D1 67917916 67918034
refGene_gn=TMEM41B;refGene_tx=XM_006937086.3;refGene_gn=TMEM41B;refGene_tx=XM_023239553.1
D1 67928902 67929023
refGene_gn=TMEM41B;refGene_tx=XM_006937086.3;refGene_gn=TMEM41B;refGene_tx=XM_023239553.1
D1 67992074 67992158 refGene_gn=IPO7;refGene_tx=XM_011286710.3
D1 68003252 68003334 refGene_gn=IPO7;refGene_tx=XM_011286710.3
D1 68006920 68007074 refGene_gn=IPO7;refGene_tx=XM_011286710.3
D1 68008538 68008697 refGene_gn=IPO7;refGene_tx=XM_011286710.3
D1 68012235 68012392 refGene_gn=IPO7;refGene_tx=XM_011286710.3
D1 68013899 68013989 refGene_gn=IPO7;refGene_tx=XM_011286710.3
D1 68016702 68016797 refGene_gn=IPO7;refGene_tx=XM_011286710.3
D1 68017175 68017260 refGene_gn=IPO7;refGene_tx=XM_011286710.3
D1 68019704 68019839 refGene_gn=IPO7;refGene_tx=XM_011286710.3
D1 68020505 68020605 refGene_gn=IPO7;refGene_tx=XM_011286710.3
D1 68022094 68022171 refGene_gn=IPO7;refGene_tx=XM_011286710.3
D1 68022325 68022442 refGene_gn=IPO7;refGene_tx=XM_011286710.3
D1 68024666 68024756 refGene_gn=IPO7;refGene_tx=XM_011286710.3
D1 68025167 68025333 refGene_gn=IPO7;refGene_tx=XM_011286710.3

D1	68025485	68025646	refGene_gn=IPO7;refGene_tx=XM_011286710.3
D1	68027240	68027369	refGene_gn=IPO7;refGene_tx=XM_011286710.3
D1	68029295	68029362	refGene_gn=IPO7;refGene_tx=XM_011286710.3
D1	68029441	68029567	refGene_gn=IPO7;refGene_tx=XM_011286710.3
D1	68029995	68030093	refGene_gn=IPO7;refGene_tx=XM_011286710.3
D1	68031827	68031923	refGene_gn=IPO7;refGene_tx=XM_011286710.3
D1	68032583	68032804	refGene_gn=IPO7;refGene_tx=XM_011286710.3
D1	68032893	68033099	refGene_gn=IPO7;refGene_tx=XM_011286710.3
D1	68034269	68034476	refGene_gn=IPO7;refGene_tx=XM_011286710.3
D1	68035783	68035900	refGene_gn=IPO7;refGene_tx=XM_011286710.3
D1	68036771	68036869	refGene_gn=IPO7;refGene_tx=XM_011286710.3
D1	68046859	68046901
refGene_gn=ZNF143;refGene_tx=XM_019812038.2;refGene_gn=ZNF143;refGene_tx=XM_019812039.2
D1	68064133	68064248
refGene_gn=ZNF143;refGene_tx=XM_003992984.3;refGene_gn=ZNF143;refGene_tx=XM_006937087.2;refGene_gn=ZNF143;refGene_tx=XM_006937088.3;refGene_gn=ZNF143;refGene_tx=XM_019812038.2;refGene_gn=ZNF143;refGene_tx=XM_019812039.2
D1	68064562	68064655
refGene_gn=ZNF143;refGene_tx=XM_003992984.3;refGene_gn=ZNF143;refGene_tx=XM_006937087.2;refGene_gn=ZNF143;refGene_tx=XM_006937088.3;refGene_gn=ZNF143;refGene_tx=XM_019812038.2;refGene_gn=ZNF143;refGene_tx=XM_019812039.2
D1	68065516	68065600
refGene_gn=ZNF143;refGene_tx=XM_003992984.3;refGene_gn=ZNF143;refGene_tx=XM_006937087.2;refGene_gn=ZNF143;refGene_tx=XM_006937088.3;refGene_gn=ZNF143;refGene_tx=XM_019812038.2;refGene_gn=ZNF143;refGene_tx=XM_019812039.2
D1	68066353	68066437
refGene_gn=ZNF143;refGene_tx=XM_003992984.3;refGene_gn=ZNF143;refGene_tx=XM_006937087.2;refGene_gn=ZNF143;refGene_tx=XM_006937088.3;refGene_gn=ZNF143;refGene_tx=XM_019812038.2;refGene_gn=ZNF143;refGene_tx=XM_019812039.2
D1	68067947	68068144
refGene_gn=ZNF143;refGene_tx=XM_003992984.3;refGene_gn=ZNF143;refGene_tx=XM_006937087.2;refGene_gn=ZNF143;refGene_tx=XM_006937088.3;refGene_gn=ZNF143;refGene_tx=XM_019812038.2;refGene_gn=ZNF143;refGene_tx=XM_019812039.2
D1	68068720	68068795
refGene_gn=ZNF143;refGene_tx=XM_003992984.3;refGene_gn=ZNF143;refGene_tx=XM_006937087.2;refGene_gn=ZNF143;refGene_tx=XM_006937088.3;refGene_gn=ZNF143;refGene_tx=XM_019812038.2;refGene_gn=ZNF143;refGene_tx=XM_019812039.2
D1	68080131	68080251
refGene_gn=ZNF143;refGene_tx=XM_003992984.3;refGene_gn=ZNF143;refGene_tx=XM_006937087.2;refGene_gn=ZNF143;refGene_tx=XM_006937088.3;refGene_gn=ZNF143;refGene_tx=XM_019812038.2;refGene_gn=ZNF143;refGene_tx=XM_019812039.2
D1	68082024	68082100

refGene_gn=ZNF143;refGene_tx=XM_003992984.3;refGene_gn=ZNF143;refGene_tx=XM_006937087.2;refGene_gn=ZNF143;refGene_tx=XM_006937088.3;refGene_gn=ZNF143;refGene_tx=XM_019812038.2;refGene_gn=ZNF143;refGene_tx=XM_019812039.2
D1 68082771 68082897
refGene_gn=ZNF143;refGene_tx=XM_003992984.3;refGene_gn=ZNF143;refGene_tx=XM_006937087.2;refGene_gn=ZNF143;refGene_tx=XM_006937088.3;refGene_gn=ZNF143;refGene_tx=XM_019812038.2;refGene_gn=ZNF143;refGene_tx=XM_019812039.2
D1 68085612 68085792
refGene_gn=ZNF143;refGene_tx=XM_003992984.3;refGene_gn=ZNF143;refGene_tx=XM_006937087.2;refGene_gn=ZNF143;refGene_tx=XM_006937088.3;refGene_gn=ZNF143;refGene_tx=XM_019812038.2;refGene_gn=ZNF143;refGene_tx=XM_019812039.2
D1 68088923 68089151
refGene_gn=ZNF143;refGene_tx=XM_003992984.3;refGene_gn=ZNF143;refGene_tx=XM_006937087.2;refGene_gn=ZNF143;refGene_tx=XM_006937088.3;refGene_gn=ZNF143;refGene_tx=XM_019812038.2;refGene_gn=ZNF143;refGene_tx=XM_019812039.2
D1 68092606 68092755
refGene_gn=ZNF143;refGene_tx=XM_003992984.3;refGene_gn=ZNF143;refGene_tx=XM_006937087.2;refGene_gn=ZNF143;refGene_tx=XM_006937088.3;refGene_gn=ZNF143;refGene_tx=XM_019812038.2;refGene_gn=ZNF143;refGene_tx=XM_019812039.2
D1 68095864 68096026
refGene_gn=ZNF143;refGene_tx=XM_003992984.3;refGene_gn=ZNF143;refGene_tx=XM_006937087.2;refGene_gn=ZNF143;refGene_tx=XM_006937088.3;refGene_gn=ZNF143;refGene_tx=XM_019812038.2;refGene_gn=ZNF143;refGene_tx=XM_019812039.2
D1 68101266 68101413
refGene_gn=ZNF143;refGene_tx=XM_003992984.3;refGene_gn=ZNF143;refGene_tx=XM_006937087.2;refGene_gn=ZNF143;refGene_tx=XM_006937088.3;refGene_gn=ZNF143;refGene_tx=XM_019812038.2;refGene_gn=ZNF143;refGene_tx=XM_019812039.2
D1 68102858 68102942
refGene_gn=ZNF143;refGene_tx=XM_003992984.3;refGene_gn=ZNF143;refGene_tx=XM_006937087.2;refGene_gn=ZNF143;refGene_tx=XM_006937088.3;refGene_gn=ZNF143;refGene_tx=XM_019812038.2;refGene_gn=ZNF143;refGene_tx=XM_019812039.2
D1 68127436 68128012 refGene_gn=WEE1;refGene_tx=XM_023239554.1
D1 68129479 68129685
refGene_gn=WEE1;refGene_tx=XM_023239554.1;refGene_gn=WEE1;refGene_tx=XM_023239555.1
D1 68130185 68130249
refGene_gn=WEE1;refGene_tx=XM_023239554.1;refGene_gn=WEE1;refGene_tx=XM_023239555.1
D1 68130437 68130610
refGene_gn=WEE1;refGene_tx=XM_023239554.1;refGene_gn=WEE1;refGene_tx=XM_023239555.1
D1 68131320 68131442

refGene_gn=WEE1;refGene_tx=XM_023239554.1;refGene_gn=WEE1;refGene_tx=XM_023239555.1
D1 68135105 68135252
refGene_gn=WEE1;refGene_tx=XM_023239554.1;refGene_gn=WEE1;refGene_tx=XM_023239555.1
D1 68138359 68138455
refGene_gn=WEE1;refGene_tx=XM_023239554.1;refGene_gn=WEE1;refGene_tx=XM_023239555.1
D1 68138542 68138628
refGene_gn=WEE1;refGene_tx=XM_023239554.1;refGene_gn=WEE1;refGene_tx=XM_023239555.1
D1 68139458 68139629
refGene_gn=WEE1;refGene_tx=XM_023239554.1;refGene_gn=WEE1;refGene_tx=XM_023239555.1
D1 68139728 68139874
refGene_gn=WEE1;refGene_tx=XM_023239554.1;refGene_gn=WEE1;refGene_tx=XM_023239555.1
D1 68140992 68141146
refGene_gn=WEE1;refGene_tx=XM_023239554.1;refGene_gn=WEE1;refGene_tx=XM_023239555.1
D1 68190685 68190784
refGene_gn=SWAP70;refGene_tx=XM_003992986.5;refGene_gn=SWAP70;refGene_tx=XM_023239556.1
D1 68216874 68217015
refGene_gn=SWAP70;refGene_tx=XM_003992986.5;refGene_gn=SWAP70;refGene_tx=XM_023239556.1
D1 68231590 68231764
refGene_gn=SWAP70;refGene_tx=XM_003992986.5;refGene_gn=SWAP70;refGene_tx=XM_023239556.1
D1 68246507 68246735
refGene_gn=SWAP70;refGene_tx=XM_003992986.5;refGene_gn=SWAP70;refGene_tx=XM_023239556.1
D1 68248894 68249041
refGene_gn=SWAP70;refGene_tx=XM_003992986.5;refGene_gn=SWAP70;refGene_tx=XM_023239556.1
D1 68250177 68250286
refGene_gn=SWAP70;refGene_tx=XM_003992986.5;refGene_gn=SWAP70;refGene_tx=XM_023239556.1
D1 68252444 68252626
refGene_gn=SWAP70;refGene_tx=XM_003992986.5;refGene_gn=SWAP70;refGene_tx=XM_023239556.1
D1 68258713 68258821 refGene_gn=SWAP70;refGene_tx=XM_003992986.5
D1 68260414 68260581
refGene_gn=SWAP70;refGene_tx=XM_003992986.5;refGene_gn=SWAP70;refGene_tx=XM_023239556.1
D1 68266409 68266608
refGene_gn=SWAP70;refGene_tx=XM_003992986.5;refGene_gn=SWAP70;refGene_tx=XM_023239556.1
D1 68267585 68267682
refGene_gn=SWAP70;refGene_tx=XM_003992986.5;refGene_gn=SWAP70;refGene_

tx=XM_023239556.1
D1 68268007 68268114
refGene_gn=SWAP70;refGene_tx=XM_003992986.5;refGene_gn=SWAP70;refGene_tx=XM_023239556.1
D1 68290504 68290603
refGene_gn=SBF2;refGene_tx=XM_023239557.1;refGene_gn=SBF2;refGene_tx=XM_023239558.1;refGene_gn=SBF2;refGene_tx=XM_023239559.1;refGene_gn=SBF2;refGene_tx=XM_023239560.1
D1 68291594 68291726
refGene_gn=SBF2;refGene_tx=XM_023239557.1;refGene_gn=SBF2;refGene_tx=XM_023239558.1;refGene_gn=SBF2;refGene_tx=XM_023239559.1;refGene_gn=SBF2;refGene_tx=XM_023239560.1
D1 68294391 68294479
refGene_gn=SBF2;refGene_tx=XM_023239557.1;refGene_gn=SBF2;refGene_tx=XM_023239558.1;refGene_gn=SBF2;refGene_tx=XM_023239559.1;refGene_gn=SBF2;refGene_tx=XM_023239560.1
D1 68295173 68295367
refGene_gn=SBF2;refGene_tx=XM_023239557.1;refGene_gn=SBF2;refGene_tx=XM_023239558.1;refGene_gn=SBF2;refGene_tx=XM_023239559.1;refGene_gn=SBF2;refGene_tx=XM_023239560.1
D1 68297507 68297612
refGene_gn=SBF2;refGene_tx=XM_023239557.1;refGene_gn=SBF2;refGene_tx=XM_023239558.1;refGene_gn=SBF2;refGene_tx=XM_023239559.1;refGene_gn=SBF2;refGene_tx=XM_023239560.1
D1 68298680 68298914
refGene_gn=SBF2;refGene_tx=XM_023239557.1;refGene_gn=SBF2;refGene_tx=XM_023239558.1;refGene_gn=SBF2;refGene_tx=XM_023239559.1;refGene_gn=SBF2;refGene_tx=XM_023239560.1
D1 68300154 68300282
refGene_gn=SBF2;refGene_tx=XM_023239557.1;refGene_gn=SBF2;refGene_tx=XM_023239558.1;refGene_gn=SBF2;refGene_tx=XM_023239559.1;refGene_gn=SBF2;refGene_tx=XM_023239560.1
D1 68303008 68303135
refGene_gn=SBF2;refGene_tx=XM_023239557.1;refGene_gn=SBF2;refGene_tx=XM_023239558.1;refGene_gn=SBF2;refGene_tx=XM_023239559.1;refGene_gn=SBF2;refGene_tx=XM_023239560.1
D1 68318945 68319131
refGene_gn=SBF2;refGene_tx=XM_023239557.1;refGene_gn=SBF2;refGene_tx=XM_023239558.1;refGene_gn=SBF2;refGene_tx=XM_023239559.1
D1 68319827 68319929
refGene_gn=SBF2;refGene_tx=XM_023239557.1;refGene_gn=SBF2;refGene_tx=XM_023239558.1;refGene_gn=SBF2;refGene_tx=XM_023239559.1;refGene_gn=SBF2;refGene_tx=XM_023239560.1
D1 68321955 68322132
refGene_gn=SBF2;refGene_tx=XM_023239557.1;refGene_gn=SBF2;refGene_tx=XM_023239558.1;refGene_gn=SBF2;refGene_tx=XM_023239559.1;refGene_gn=SBF2;refGene_tx=XM_023239560.1
D1 68325717 68325902
refGene_gn=SBF2;refGene_tx=XM_023239557.1;refGene_gn=SBF2;refGene_tx=XM_023239558.1;refGene_gn=SBF2;refGene_tx=XM_023239559.1;refGene_gn=SBF

2;refGene_tx=XM_023239560.1
D1 68327221 68327296
refGene_gn=SBF2;refGene_tx=XM_023239557.1;refGene_gn=SBF2;refGene_tx=XM_023239560.1
D1 68334192 68334288
refGene_gn=SBF2;refGene_tx=XM_023239557.1;refGene_gn=SBF2;refGene_tx=XM_023239558.1;refGene_gn=SBF2;refGene_tx=XM_023239560.1
D1 68335171 68335312
refGene_gn=SBF2;refGene_tx=XM_023239557.1;refGene_gn=SBF2;refGene_tx=XM_023239558.1;refGene_gn=SBF2;refGene_tx=XM_023239559.1;refGene_gn=SBF2;refGene_tx=XM_023239560.1
D1 68337653 68337850
refGene_gn=SBF2;refGene_tx=XM_023239557.1;refGene_gn=SBF2;refGene_tx=XM_023239558.1;refGene_gn=SBF2;refGene_tx=XM_023239559.1;refGene_gn=SBF2;refGene_tx=XM_023239560.1
D1 68345708 68345907
refGene_gn=SBF2;refGene_tx=XM_023239557.1;refGene_gn=SBF2;refGene_tx=XM_023239558.1;refGene_gn=SBF2;refGene_tx=XM_023239559.1;refGene_gn=SBF2;refGene_tx=XM_023239560.1
D1 68348433 68348579
refGene_gn=SBF2;refGene_tx=XM_023239557.1;refGene_gn=SBF2;refGene_tx=XM_023239558.1;refGene_gn=SBF2;refGene_tx=XM_023239559.1;refGene_gn=SBF2;refGene_tx=XM_023239560.1
D1 68350535 68350711
refGene_gn=SBF2;refGene_tx=XM_023239557.1;refGene_gn=SBF2;refGene_tx=XM_023239558.1;refGene_gn=SBF2;refGene_tx=XM_023239559.1;refGene_gn=SBF2;refGene_tx=XM_023239560.1
D1 68352389 68352517
refGene_gn=SBF2;refGene_tx=XM_023239557.1;refGene_gn=SBF2;refGene_tx=XM_023239558.1;refGene_gn=SBF2;refGene_tx=XM_023239559.1;refGene_gn=SBF2;refGene_tx=XM_023239560.1
D1 68362197 68362393
refGene_gn=SBF2;refGene_tx=XM_023239557.1;refGene_gn=SBF2;refGene_tx=XM_023239558.1;refGene_gn=SBF2;refGene_tx=XM_023239559.1;refGene_gn=SBF2;refGene_tx=XM_023239560.1
D1 68363849 68363923
refGene_gn=SBF2;refGene_tx=XM_023239557.1;refGene_gn=SBF2;refGene_tx=XM_023239558.1;refGene_gn=SBF2;refGene_tx=XM_023239559.1;refGene_gn=SBF2;refGene_tx=XM_023239560.1
D1 68364965 68365138
refGene_gn=SBF2;refGene_tx=XM_023239557.1;refGene_gn=SBF2;refGene_tx=XM_023239558.1;refGene_gn=SBF2;refGene_tx=XM_023239559.1;refGene_gn=SBF2;refGene_tx=XM_023239560.1
D1 68367534 68367797
refGene_gn=SBF2;refGene_tx=XM_023239557.1;refGene_gn=SBF2;refGene_tx=XM_023239558.1;refGene_gn=SBF2;refGene_tx=XM_023239559.1;refGene_gn=SBF2;refGene_tx=XM_023239560.1
D1 68375676 68375847
refGene_gn=SBF2;refGene_tx=XM_023239557.1;refGene_gn=SBF2;refGene_tx=XM_023239558.1;refGene_gn=SBF2;refGene_tx=XM_023239559.1;refGene_gn=SBF

2;refGene_tx=XM_023239560.1
D1 68419310 68419379
refGene_gn=SBF2;refGene_tx=XM_023239557.1;refGene_gn=SBF2;refGene_tx=XM_023239558.1;refGene_gn=SBF2;refGene_tx=XM_023239559.1;refGene_gn=SBF2;refGene_tx=XM_023239560.1
D1 68473188 68473338
refGene_gn=SBF2;refGene_tx=XM_023239557.1;refGene_gn=SBF2;refGene_tx=XM_023239558.1;refGene_gn=SBF2;refGene_tx=XM_023239559.1;refGene_gn=SBF2;refGene_tx=XM_023239560.1
D1 68475038 68475148
refGene_gn=SBF2;refGene_tx=XM_023239557.1;refGene_gn=SBF2;refGene_tx=XM_023239558.1;refGene_gn=SBF2;refGene_tx=XM_023239559.1;refGene_gn=SBF2;refGene_tx=XM_023239560.1
D1 68481838 68482043
refGene_gn=SBF2;refGene_tx=XM_023239557.1;refGene_gn=SBF2;refGene_tx=XM_023239558.1;refGene_gn=SBF2;refGene_tx=XM_023239559.1;refGene_gn=SBF2;refGene_tx=XM_023239560.1
D1 68493711 68493810
refGene_gn=SBF2;refGene_tx=XM_023239557.1;refGene_gn=SBF2;refGene_tx=XM_023239558.1;refGene_gn=SBF2;refGene_tx=XM_023239559.1;refGene_gn=SBF2;refGene_tx=XM_023239560.1
D1 68497165 68497294
refGene_gn=SBF2;refGene_tx=XM_023239557.1;refGene_gn=SBF2;refGene_tx=XM_023239558.1;refGene_gn=SBF2;refGene_tx=XM_023239559.1;refGene_gn=SBF2;refGene_tx=XM_023239560.1
D1 68497753 68497867
refGene_gn=SBF2;refGene_tx=XM_023239557.1;refGene_gn=SBF2;refGene_tx=XM_023239558.1;refGene_gn=SBF2;refGene_tx=XM_023239559.1;refGene_gn=SBF2;refGene_tx=XM_023239560.1
D1 68498982 68499060
refGene_gn=SBF2;refGene_tx=XM_023239557.1;refGene_gn=SBF2;refGene_tx=XM_023239558.1;refGene_gn=SBF2;refGene_tx=XM_023239559.1;refGene_gn=SBF2;refGene_tx=XM_023239560.1
D1 68501877 68501991
refGene_gn=SBF2;refGene_tx=XM_023239557.1;refGene_gn=SBF2;refGene_tx=XM_023239558.1;refGene_gn=SBF2;refGene_tx=XM_023239559.1;refGene_gn=SBF2;refGene_tx=XM_023239560.1
D1 68504124 68504233
refGene_gn=SBF2;refGene_tx=XM_023239557.1;refGene_gn=SBF2;refGene_tx=XM_023239558.1;refGene_gn=SBF2;refGene_tx=XM_023239559.1;refGene_gn=SBF2;refGene_tx=XM_023239560.1
D1 68505495 68505628
refGene_gn=SBF2;refGene_tx=XM_023239557.1;refGene_gn=SBF2;refGene_tx=XM_023239558.1;refGene_gn=SBF2;refGene_tx=XM_023239559.1;refGene_gn=SBF2;refGene_tx=XM_023239560.1
D1 68542647 68542753
refGene_gn=SBF2;refGene_tx=XM_023239557.1;refGene_gn=SBF2;refGene_tx=XM_023239558.1;refGene_gn=SBF2;refGene_tx=XM_023239559.1;refGene_gn=SBF2;refGene_tx=XM_023239560.1
D1 68545033 68545144

refGene_gn=SBF2;refGene_tx=XM_023239557.1;refGene_gn=SBF2;refGene_tx=XM_023239558.1;refGene_gn=SBF2;refGene_tx=XM_023239559.1;refGene_gn=SBF2;refGene_tx=XM_023239560.1
D1 68546892 68547015
refGene_gn=SBF2;refGene_tx=XM_023239557.1;refGene_gn=SBF2;refGene_tx=XM_023239558.1;refGene_gn=SBF2;refGene_tx=XM_023239559.1;refGene_gn=SBF2;refGene_tx=XM_023239560.1
D1 68563631 68563769
refGene_gn=SBF2;refGene_tx=XM_023239557.1;refGene_gn=SBF2;refGene_tx=XM_023239558.1;refGene_gn=SBF2;refGene_tx=XM_023239559.1;refGene_gn=SBF2;refGene_tx=XM_023239560.1
D1 68679652 68679738
refGene_gn=SBF2;refGene_tx=XM_023239557.1;refGene_gn=SBF2;refGene_tx=XM_023239558.1;refGene_gn=SBF2;refGene_tx=XM_023239559.1;refGene_gn=SBF2;refGene_tx=XM_023239560.1
D1 68736320 68736597
refGene_gn=LOC101086842;refGene_tx=XM_011286713.3
D1 68779332 68779387
refGene_gn=SBF2;refGene_tx=XM_023239557.1;refGene_gn=SBF2;refGene_tx=XM_023239558.1;refGene_gn=SBF2;refGene_tx=XM_023239559.1;refGene_gn=SBF2;refGene_tx=XM_023239560.1
D1 68789326 68789424
refGene_gn=ADM;refGene_tx=XM_006937091.3;refGene_gn=ADM;refGene_tx=XM_006937092.4;refGene_gn=ADM;refGene_tx=XM_019812049.1;refGene_gn=ADM;refGene_tx=XM_023239561.1
D1 68789575 68789725
refGene_gn=ADM;refGene_tx=XM_006937091.3;refGene_gn=ADM;refGene_tx=XM_006937092.4;refGene_gn=ADM;refGene_tx=XM_019812049.1;refGene_gn=ADM;refGene_tx=XM_023239561.1
D1 68789933 68790252
refGene_gn=ADM;refGene_tx=XM_006937091.3;refGene_gn=ADM;refGene_tx=XM_006937092.4;refGene_gn=ADM;refGene_tx=XM_019812049.1;refGene_gn=ADM;refGene_tx=XM_023239561.1
D1 68916772 68916881 refGene_gn=AMPD3;refGene_tx=XM_019812045.2
D1 68922605 68922630 refGene_gn=AMPD3;refGene_tx=XM_003992988.4
D1 68926526 68926752
refGene_gn=AMPD3;refGene_tx=XM_003992988.4;refGene_gn=AMPD3;refGene_tx=XM_006937093.3;refGene_gn=AMPD3;refGene_tx=XM_006937094.3;refGene_gn=AMPD3;refGene_tx=XM_006937095.3;refGene_gn=AMPD3;refGene_tx=XM_006937096.3;refGene_gn=AMPD3;refGene_tx=XM_019812045.2;refGene_gn=AMPD3;refGene_tx=XM_019812046.1;refGene_gn=AMPD3;refGene_tx=XM_019812047.2
D1 68937788 68937993
refGene_gn=AMPD3;refGene_tx=XM_003992988.4;refGene_gn=AMPD3;refGene_tx=XM_006937093.3;refGene_gn=AMPD3;refGene_tx=XM_006937094.3;refGene_gn=AMPD3;refGene_tx=XM_006937095.3;refGene_gn=AMPD3;refGene_tx=XM_006937096.3;refGene_gn=AMPD3;refGene_tx=XM_019812045.2;refGene_gn=AMPD3;refGene_tx=XM_019812046.1;refGene_gn=AMPD3;refGene_tx=XM_019812047.2
D1 68941610 68941773
refGene_gn=AMPD3;refGene_tx=XM_003992988.4;refGene_gn=AMPD3;refGene_tx=XM_006937093.3;refGene_gn=AMPD3;refGene_tx=XM_006937094.3;refGene_gn=

AMPD3;refGene_tx=XM_006937095.3;refGene_gn=AMPD3;refGene_tx=XM_0069370
96.3;refGene_gn=AMPD3;refGene_tx=XM_019812045.2;refGene_gn=AMPD3;refGe
ne_tx=XM_019812046.1;refGene_gn=AMPD3;refGene_tx=XM_019812047.2;refGen
e_gn=AMPD3;refGene_tx=XM_019812048.1
D1 68943892 68944112
refGene_gn=AMPD3;refGene_tx=XM_003992988.4;refGene_gn=AMPD3;refGene_tx
=XM_006937093.3;refGene_gn=AMPD3;refGene_tx=XM_006937094.3;refGene_gn=
AMPD3;refGene_tx=XM_006937095.3;refGene_gn=AMPD3;refGene_tx=XM_0069370
96.3;refGene_gn=AMPD3;refGene_tx=XM_019812045.2;refGene_gn=AMPD3;refGe
ne_tx=XM_019812046.1;refGene_gn=AMPD3;refGene_tx=XM_019812047.2;refGen
e_gn=AMPD3;refGene_tx=XM_019812048.1
D1 68946169 68946299
refGene_gn=AMPD3;refGene_tx=XM_003992988.4;refGene_gn=AMPD3;refGene_tx
=XM_006937093.3;refGene_gn=AMPD3;refGene_tx=XM_006937094.3;refGene_gn=
AMPD3;refGene_tx=XM_006937095.3;refGene_gn=AMPD3;refGene_tx=XM_0069370
96.3;refGene_gn=AMPD3;refGene_tx=XM_019812045.2;refGene_gn=AMPD3;refGe
ne_tx=XM_019812046.1;refGene_gn=AMPD3;refGene_tx=XM_019812047.2;refGen
e_gn=AMPD3;refGene_tx=XM_019812048.1
D1 68950555 68950750
refGene_gn=AMPD3;refGene_tx=XM_003992988.4;refGene_gn=AMPD3;refGene_tx
=XM_006937093.3;refGene_gn=AMPD3;refGene_tx=XM_006937094.3;refGene_gn=
AMPD3;refGene_tx=XM_006937095.3;refGene_gn=AMPD3;refGene_tx=XM_0069370
96.3;refGene_gn=AMPD3;refGene_tx=XM_019812045.2;refGene_gn=AMPD3;refGe
ne_tx=XM_019812046.1;refGene_gn=AMPD3;refGene_tx=XM_019812047.2;refGen
e_gn=AMPD3;refGene_tx=XM_019812048.1
D1 68952046 68952178
refGene_gn=AMPD3;refGene_tx=XM_003992988.4;refGene_gn=AMPD3;refGene_tx
=XM_006937093.3;refGene_gn=AMPD3;refGene_tx=XM_006937094.3;refGene_gn=
AMPD3;refGene_tx=XM_006937095.3;refGene_gn=AMPD3;refGene_tx=XM_0069370
96.3;refGene_gn=AMPD3;refGene_tx=XM_019812045.2;refGene_gn=AMPD3;refGe
ne_tx=XM_019812046.1;refGene_gn=AMPD3;refGene_tx=XM_019812047.2;refGen
e_gn=AMPD3;refGene_tx=XM_019812048.1
D1 68952710 68952874
refGene_gn=AMPD3;refGene_tx=XM_003992988.4;refGene_gn=AMPD3;refGene_tx
=XM_006937093.3;refGene_gn=AMPD3;refGene_tx=XM_006937094.3;refGene_gn=
AMPD3;refGene_tx=XM_006937095.3;refGene_gn=AMPD3;refGene_tx=XM_0069370
96.3;refGene_gn=AMPD3;refGene_tx=XM_019812045.2;refGene_gn=AMPD3;refGe
ne_tx=XM_019812046.1;refGene_gn=AMPD3;refGene_tx=XM_019812047.2;refGen
e_gn=AMPD3;refGene_tx=XM_019812048.1
D1 68953630 68953757
refGene_gn=AMPD3;refGene_tx=XM_003992988.4;refGene_gn=AMPD3;refGene_tx
=XM_006937093.3;refGene_gn=AMPD3;refGene_tx=XM_006937094.3;refGene_gn=
AMPD3;refGene_tx=XM_006937095.3;refGene_gn=AMPD3;refGene_tx=XM_0069370
96.3;refGene_gn=AMPD3;refGene_tx=XM_019812045.2;refGene_gn=AMPD3;refGe
ne_tx=XM_019812046.1;refGene_gn=AMPD3;refGene_tx=XM_019812047.2;refGen
e_gn=AMPD3;refGene_tx=XM_019812048.1
D1 68956460 68956624
refGene_gn=AMPD3;refGene_tx=XM_003992988.4;refGene_gn=AMPD3;refGene_tx
=XM_006937093.3;refGene_gn=AMPD3;refGene_tx=XM_006937094.3;refGene_gn=
AMPD3;refGene_tx=XM_006937095.3;refGene_gn=AMPD3;refGene_tx=XM_0069370

96.3;refGene_gn=AMPD3;refGene_tx=XM_019812045.2;refGene_gn=AMPD3;refGene_tx=XM_019812046.1;refGene_gn=AMPD3;refGene_tx=XM_019812047.2;refGene_gn=AMPD3;refGene_tx=XM_019812048.1
D1 68957756 68957877
refGene_gn=AMPD3;refGene_tx=XM_003992988.4;refGene_gn=AMPD3;refGene_tx=XM_006937093.3;refGene_gn=AMPD3;refGene_tx=XM_006937094.3;refGene_gn=AMPD3;refGene_tx=XM_006937095.3;refGene_gn=AMPD3;refGene_tx=XM_006937096.3;refGene_gn=AMPD3;refGene_tx=XM_019812045.2;refGene_gn=AMPD3;refGene_tx=XM_019812046.1;refGene_gn=AMPD3;refGene_tx=XM_019812047.2;refGene_gn=AMPD3;refGene_tx=XM_019812048.1
D1 68959066 68959240
refGene_gn=AMPD3;refGene_tx=XM_003992988.4;refGene_gn=AMPD3;refGene_tx=XM_006937093.3;refGene_gn=AMPD3;refGene_tx=XM_006937094.3;refGene_gn=AMPD3;refGene_tx=XM_006937095.3;refGene_gn=AMPD3;refGene_tx=XM_006937096.3;refGene_gn=AMPD3;refGene_tx=XM_019812045.2;refGene_gn=AMPD3;refGene_tx=XM_019812046.1;refGene_gn=AMPD3;refGene_tx=XM_019812047.2;refGene_gn=AMPD3;refGene_tx=XM_019812048.1
D1 68961093 68961204
refGene_gn=AMPD3;refGene_tx=XM_003992988.4;refGene_gn=AMPD3;refGene_tx=XM_006937093.3;refGene_gn=AMPD3;refGene_tx=XM_006937094.3;refGene_gn=AMPD3;refGene_tx=XM_006937095.3;refGene_gn=AMPD3;refGene_tx=XM_006937096.3;refGene_gn=AMPD3;refGene_tx=XM_019812045.2;refGene_gn=AMPD3;refGene_tx=XM_019812046.1;refGene_gn=AMPD3;refGene_tx=XM_019812047.2;refGene_gn=AMPD3;refGene_tx=XM_019812048.1
D1 68962239 68962416
refGene_gn=AMPD3;refGene_tx=XM_003992988.4;refGene_gn=AMPD3;refGene_tx=XM_006937093.3;refGene_gn=AMPD3;refGene_tx=XM_006937094.3;refGene_gn=AMPD3;refGene_tx=XM_006937095.3;refGene_gn=AMPD3;refGene_tx=XM_006937096.3;refGene_gn=AMPD3;refGene_tx=XM_019812045.2;refGene_gn=AMPD3;refGene_tx=XM_019812046.1;refGene_gn=AMPD3;refGene_tx=XM_019812047.2;refGene_gn=AMPD3;refGene_tx=XM_019812048.1
D1 68969211 68969362
refGene_gn=RNF141;refGene_tx=XM_003992989.3;refGene_gn=RNF141;refGene_tx=XM_006937097.2
D1 68974529 68974637
refGene_gn=RNF141;refGene_tx=XM_003992989.3;refGene_gn=RNF141;refGene_tx=XM_006937097.2
D1 68976305 68976487
refGene_gn=RNF141;refGene_tx=XM_003992989.3;refGene_gn=RNF141;refGene_tx=XM_006937097.2
D1 68978702 68978811
refGene_gn=RNF141;refGene_tx=XM_003992989.3;refGene_gn=RNF141;refGene_tx=XM_006937097.2
D1 68981742 68981885
refGene_gn=RNF141;refGene_tx=XM_003992989.3;refGene_gn=RNF141;refGene_tx=XM_006937097.2
D1 69014969 69015156 refGene_gn=LYVE1;refGene_tx=XM_003992990.4
D1 69015694 69015773 refGene_gn=LYVE1;refGene_tx=XM_003992990.4
D1 69016409 69016718 refGene_gn=LYVE1;refGene_tx=XM_003992990.4
D1 69023746 69023886 refGene_gn=LYVE1;refGene_tx=XM_003992990.4

D1 69023994 69024166 refGene_gn=LYVE1;refGene_tx=XM_003992990.4
D1 69027854 69027939 refGene_gn=LYVE1;refGene_tx=XM_003992990.4
D1 69040557 69040801
refGene_gn=MRVI1;refGene_tx=XM_003992992.3;refGene_gn=MRVI1;refGene_tx=XM_006937098.3;refGene_gn=MRVI1;refGene_tx=XM_006937099.3;refGene_gn=MRVI1;refGene_tx=XM_006937100.2;refGene_gn=MRVI1;refGene_tx=XM_006937101.2;refGene_gn=MRVI1;refGene_tx=XM_019812051.1;refGene_gn=MRVI1;refGene_tx=XM_019812052.1;refGene_gn=MRVI1;refGene_tx=XM_019812053.1;refGene_gn=MRVI1;refGene_tx=XM_019812054.1;refGene_gn=MRVI1;refGene_tx=XM_019812055.2;refGene_gn=MRVI1;refGene_tx=XM_019812056.2;refGene_gn=MRVI1;refGene_tx=XM_019812057.2
D1 69046011 69046146
refGene_gn=MRVI1;refGene_tx=XM_003992992.3;refGene_gn=MRVI1;refGene_tx=XM_006937098.3;refGene_gn=MRVI1;refGene_tx=XM_006937099.3;refGene_gn=MRVI1;refGene_tx=XM_006937100.2;refGene_gn=MRVI1;refGene_tx=XM_006937101.2;refGene_gn=MRVI1;refGene_tx=XM_019812051.1;refGene_gn=MRVI1;refGene_tx=XM_019812052.1;refGene_gn=MRVI1;refGene_tx=XM_019812053.1;refGene_gn=MRVI1;refGene_tx=XM_019812054.1;refGene_gn=MRVI1;refGene_tx=XM_019812055.2;refGene_gn=MRVI1;refGene_tx=XM_019812056.2;refGene_gn=MRVI1;refGene_tx=XM_019812057.2
D1 69047553 69047670
refGene_gn=MRVI1;refGene_tx=XM_003992992.3;refGene_gn=MRVI1;refGene_tx=XM_006937098.3;refGene_gn=MRVI1;refGene_tx=XM_006937099.3;refGene_gn=MRVI1;refGene_tx=XM_006937100.2;refGene_gn=MRVI1;refGene_tx=XM_006937101.2;refGene_gn=MRVI1;refGene_tx=XM_019812051.1;refGene_gn=MRVI1;refGene_tx=XM_019812052.1;refGene_gn=MRVI1;refGene_tx=XM_019812053.1;refGene_gn=MRVI1;refGene_tx=XM_019812054.1;refGene_gn=MRVI1;refGene_tx=XM_019812055.2;refGene_gn=MRVI1;refGene_tx=XM_019812056.2;refGene_gn=MRVI1;refGene_tx=XM_019812057.2
D1 69048335 69048413
refGene_gn=MRVI1;refGene_tx=XM_006937098.3;refGene_gn=MRVI1;refGene_tx=XM_006937099.3;refGene_gn=MRVI1;refGene_tx=XM_019812051.1;refGene_gn=MRVI1;refGene_tx=XM_019812052.1;refGene_gn=MRVI1;refGene_tx=XM_019812053.1;refGene_gn=MRVI1;refGene_tx=XM_019812054.1;refGene_gn=MRVI1;refGene_tx=XM_019812055.2;refGene_gn=MRVI1;refGene_tx=XM_019812056.2;refGene_gn=MRVI1;refGene_tx=XM_019812057.2
D1 69056422 69056487
refGene_gn=MRVI1;refGene_tx=XM_003992992.3;refGene_gn=MRVI1;refGene_tx=XM_006937098.3;refGene_gn=MRVI1;refGene_tx=XM_006937099.3;refGene_gn=MRVI1;refGene_tx=XM_006937100.2;refGene_gn=MRVI1;refGene_tx=XM_006937101.2;refGene_gn=MRVI1;refGene_tx=XM_019812051.1;refGene_gn=MRVI1;refGene_tx=XM_019812052.1;refGene_gn=MRVI1;refGene_tx=XM_019812053.1;refGene_gn=MRVI1;refGene_tx=XM_019812054.1;refGene_gn=MRVI1;refGene_tx=XM_019812055.2;refGene_gn=MRVI1;refGene_tx=XM_019812056.2;refGene_gn=MRVI1;refGene_tx=XM_019812057.2
D1 69058853 69058961
refGene_gn=MRVI1;refGene_tx=XM_003992992.3;refGene_gn=MRVI1;refGene_tx=XM_006937098.3;refGene_gn=MRVI1;refGene_tx=XM_006937099.3;refGene_gn=MRVI1;refGene_tx=XM_006937100.2;refGene_gn=MRVI1;refGene_tx=XM_006937101.2;refGene_gn=MRVI1;refGene_tx=XM_019812051.1;refGene_gn=MRVI1;refGe

ne_tx=XM_019812052.1;refGene_gn=MRVI1;refGene_tx=XM_019812053.1;refGene_gn=MRVI1;refGene_tx=XM_019812054.1;refGene_gn=MRVI1;refGene_tx=XM_019812055.2;refGene_gn=MRVI1;refGene_tx=XM_019812056.2;refGene_gn=MRVI1;refGene_tx=XM_019812057.2
D1 69059495 69059545
refGene_gn=MRVI1;refGene_tx=XM_003992992.3;refGene_gn=MRVI1;refGene_tx=XM_006937098.3;refGene_gn=MRVI1;refGene_tx=XM_006937099.3;refGene_gn=MRVI1;refGene_tx=XM_006937100.2;refGene_gn=MRVI1;refGene_tx=XM_006937101.2;refGene_gn=MRVI1;refGene_tx=XM_019812051.1;refGene_gn=MRVI1;refGene_tx=XM_019812052.1;refGene_gn=MRVI1;refGene_tx=XM_019812053.1;refGene_gn=MRVI1;refGene_tx=XM_019812054.1;refGene_gn=MRVI1;refGene_tx=XM_019812055.2;refGene_gn=MRVI1;refGene_tx=XM_019812056.2;refGene_gn=MRVI1;refGene_tx=XM_019812057.2
D1 69065314 69065456
refGene_gn=MRVI1;refGene_tx=XM_003992992.3;refGene_gn=MRVI1;refGene_tx=XM_006937098.3;refGene_gn=MRVI1;refGene_tx=XM_006937099.3;refGene_gn=MRVI1;refGene_tx=XM_006937100.2;refGene_gn=MRVI1;refGene_tx=XM_006937101.2;refGene_gn=MRVI1;refGene_tx=XM_019812051.1;refGene_gn=MRVI1;refGene_tx=XM_019812052.1;refGene_gn=MRVI1;refGene_tx=XM_019812053.1;refGene_gn=MRVI1;refGene_tx=XM_019812054.1;refGene_gn=MRVI1;refGene_tx=XM_019812055.2;refGene_gn=MRVI1;refGene_tx=XM_019812056.2;refGene_gn=MRVI1;refGene_tx=XM_019812057.2
D1 69067870 69068002
refGene_gn=MRVI1;refGene_tx=XM_003992992.3;refGene_gn=MRVI1;refGene_tx=XM_006937098.3;refGene_gn=MRVI1;refGene_tx=XM_006937099.3;refGene_gn=MRVI1;refGene_tx=XM_006937100.2;refGene_gn=MRVI1;refGene_tx=XM_006937101.2;refGene_gn=MRVI1;refGene_tx=XM_019812051.1;refGene_gn=MRVI1;refGene_tx=XM_019812052.1;refGene_gn=MRVI1;refGene_tx=XM_019812053.1;refGene_gn=MRVI1;refGene_tx=XM_019812054.1;refGene_gn=MRVI1;refGene_tx=XM_019812055.2;refGene_gn=MRVI1;refGene_tx=XM_019812056.2;refGene_gn=MRVI1;refGene_tx=XM_019812057.2
D1 69068490 69068631
refGene_gn=MRVI1;refGene_tx=XM_003992992.3;refGene_gn=MRVI1;refGene_tx=XM_006937098.3;refGene_gn=MRVI1;refGene_tx=XM_006937099.3;refGene_gn=MRVI1;refGene_tx=XM_006937100.2;refGene_gn=MRVI1;refGene_tx=XM_006937101.2;refGene_gn=MRVI1;refGene_tx=XM_019812051.1;refGene_gn=MRVI1;refGene_tx=XM_019812052.1;refGene_gn=MRVI1;refGene_tx=XM_019812053.1;refGene_gn=MRVI1;refGene_tx=XM_019812054.1;refGene_gn=MRVI1;refGene_tx=XM_019812055.2;refGene_gn=MRVI1;refGene_tx=XM_019812056.2;refGene_gn=MRVI1;refGene_tx=XM_019812057.2
D1 69070997 69071028
refGene_gn=MRVI1;refGene_tx=XM_003992992.3;refGene_gn=MRVI1;refGene_tx=XM_006937098.3;refGene_gn=MRVI1;refGene_tx=XM_006937099.3;refGene_gn=MRVI1;refGene_tx=XM_006937100.2;refGene_gn=MRVI1;refGene_tx=XM_006937101.2;refGene_gn=MRVI1;refGene_tx=XM_019812051.1;refGene_gn=MRVI1;refGene_tx=XM_019812052.1;refGene_gn=MRVI1;refGene_tx=XM_019812053.1;refGene_gn=MRVI1;refGene_tx=XM_019812054.1;refGene_gn=MRVI1;refGene_tx=XM_019812055.2;refGene_gn=MRVI1;refGene_tx=XM_019812056.2;refGene_gn=MRVI1;refGene_tx=XM_019812057.2
D1 69076660 69076784

refGene_gn=MRVI1;refGene_tx=XM_003992992.3;refGene_gn=MRVI1;refGene_tx=XM_006937098.3;refGene_gn=MRVI1;refGene_tx=XM_006937099.3;refGene_gn=MRVI1;refGene_tx=XM_006937100.2;refGene_gn=MRVI1;refGene_tx=XM_006937101.2;refGene_gn=MRVI1;refGene_tx=XM_019812051.1;refGene_gn=MRVI1;refGene_tx=XM_019812052.1;refGene_gn=MRVI1;refGene_tx=XM_019812053.1;refGene_gn=MRVI1;refGene_tx=XM_019812054.1;refGene_gn=MRVI1;refGene_tx=XM_019812055.2;refGene_gn=MRVI1;refGene_tx=XM_019812056.2;refGene_gn=MRVI1;refGene_tx=XM_019812057.2
D1 69087237 69087316
refGene_gn=MRVI1;refGene_tx=XM_003992992.3;refGene_gn=MRVI1;refGene_tx=XM_006937098.3;refGene_gn=MRVI1;refGene_tx=XM_006937099.3;refGene_gn=MRVI1;refGene_tx=XM_006937100.2;refGene_gn=MRVI1;refGene_tx=XM_006937101.2;refGene_gn=MRVI1;refGene_tx=XM_019812051.1;refGene_gn=MRVI1;refGene_tx=XM_019812052.1;refGene_gn=MRVI1;refGene_tx=XM_019812053.1;refGene_gn=MRVI1;refGene_tx=XM_019812054.1;refGene_gn=MRVI1;refGene_tx=XM_019812055.2;refGene_gn=MRVI1;refGene_tx=XM_019812056.2;refGene_gn=MRVI1;refGene_tx=XM_019812057.2
D1 69089154 69089775
refGene_gn=MRVI1;refGene_tx=XM_003992992.3;refGene_gn=MRVI1;refGene_tx=XM_006937098.3;refGene_gn=MRVI1;refGene_tx=XM_006937099.3;refGene_gn=MRVI1;refGene_tx=XM_006937100.2;refGene_gn=MRVI1;refGene_tx=XM_006937101.2;refGene_gn=MRVI1;refGene_tx=XM_019812051.1;refGene_gn=MRVI1;refGene_tx=XM_019812052.1;refGene_gn=MRVI1;refGene_tx=XM_019812053.1;refGene_gn=MRVI1;refGene_tx=XM_019812054.1;refGene_gn=MRVI1;refGene_tx=XM_019812055.2;refGene_gn=MRVI1;refGene_tx=XM_019812056.2;refGene_gn=MRVI1;refGene_tx=XM_019812057.2
D1 69090954 69091002
refGene_gn=MRVI1;refGene_tx=XM_003992992.3;refGene_gn=MRVI1;refGene_tx=XM_006937098.3;refGene_gn=MRVI1;refGene_tx=XM_006937099.3;refGene_gn=MRVI1;refGene_tx=XM_006937100.2;refGene_gn=MRVI1;refGene_tx=XM_006937101.2;refGene_gn=MRVI1;refGene_tx=XM_019812051.1;refGene_gn=MRVI1;refGene_tx=XM_019812052.1;refGene_gn=MRVI1;refGene_tx=XM_019812053.1;refGene_gn=MRVI1;refGene_tx=XM_019812054.1;refGene_gn=MRVI1;refGene_tx=XM_019812055.2;refGene_gn=MRVI1;refGene_tx=XM_019812056.2;refGene_gn=MRVI1;refGene_tx=XM_019812057.2
D1 69091205 69091258
refGene_gn=MRVI1;refGene_tx=XM_003992992.3;refGene_gn=MRVI1;refGene_tx=XM_006937098.3;refGene_gn=MRVI1;refGene_tx=XM_006937099.3;refGene_gn=MRVI1;refGene_tx=XM_006937100.2;refGene_gn=MRVI1;refGene_tx=XM_006937101.2;refGene_gn=MRVI1;refGene_tx=XM_019812051.1;refGene_gn=MRVI1;refGene_tx=XM_019812052.1;refGene_gn=MRVI1;refGene_tx=XM_019812053.1;refGene_gn=MRVI1;refGene_tx=XM_019812054.1;refGene_gn=MRVI1;refGene_tx=XM_019812055.2;refGene_gn=MRVI1;refGene_tx=XM_019812056.2;refGene_gn=MRVI1;refGene_tx=XM_019812057.2
D1 69092026 69092104
refGene_gn=MRVI1;refGene_tx=XM_003992992.3;refGene_gn=MRVI1;refGene_tx=XM_006937098.3;refGene_gn=MRVI1;refGene_tx=XM_006937099.3;refGene_gn=MRVI1;refGene_tx=XM_006937100.2;refGene_gn=MRVI1;refGene_tx=XM_006937101.2;refGene_gn=MRVI1;refGene_tx=XM_019812051.1;refGene_gn=MRVI1;refGene_tx=XM_019812052.1;refGene_gn=MRVI1;refGene_tx=XM_019812053.1;refGen

e_gn=MRVI1;refGene_tx=XM_019812054.1;refGene_gn=MRVI1;refGene_tx=XM_01
9812057.2
D1 69092668 69092842
refGene_gn=MRVI1;refGene_tx=XM_003992992.3;refGene_gn=MRVI1;refGene_tx
=XM_006937098.3;refGene_gn=MRVI1;refGene_tx=XM_006937099.3;refGene_gn=
MRVI1;refGene_tx=XM_006937100.2;refGene_gn=MRVI1;refGene_tx=XM_0069371
01.2;refGene_gn=MRVI1;refGene_tx=XM_019812051.1;refGene_gn=MRVI1;refGe
ne_tx=XM_019812052.1;refGene_gn=MRVI1;refGene_tx=XM_019812053.1;refGen
e_gn=MRVI1;refGene_tx=XM_019812054.1;refGene_gn=MRVI1;refGene_tx=XM_01
9812055.2;refGene_gn=MRVI1;refGene_tx=XM_019812056.2;refGene_gn=MRVI1;
refGene_tx=XM_019812057.2
D1 69095001 69095072
refGene_gn=MRVI1;refGene_tx=XM_003992992.3;refGene_gn=MRVI1;refGene_tx
=XM_006937098.3;refGene_gn=MRVI1;refGene_tx=XM_006937099.3;refGene_gn=
MRVI1;refGene_tx=XM_006937100.2;refGene_gn=MRVI1;refGene_tx=XM_0069371
01.2;refGene_gn=MRVI1;refGene_tx=XM_019812051.1;refGene_gn=MRVI1;refGe
ne_tx=XM_019812052.1;refGene_gn=MRVI1;refGene_tx=XM_019812053.1;refGen
e_gn=MRVI1;refGene_tx=XM_019812054.1;refGene_gn=MRVI1;refGene_tx=XM_01
9812055.2;refGene_gn=MRVI1;refGene_tx=XM_019812056.2;refGene_gn=MRVI1;
refGene_tx=XM_019812057.2
D1 69096163 69096264
refGene_gn=MRVI1;refGene_tx=XM_003992992.3;refGene_gn=MRVI1;refGene_tx
=XM_006937098.3;refGene_gn=MRVI1;refGene_tx=XM_006937099.3;refGene_gn=
MRVI1;refGene_tx=XM_006937100.2;refGene_gn=MRVI1;refGene_tx=XM_0069371
01.2;refGene_gn=MRVI1;refGene_tx=XM_019812051.1;refGene_gn=MRVI1;refGe
ne_tx=XM_019812052.1;refGene_gn=MRVI1;refGene_tx=XM_019812053.1;refGen
e_gn=MRVI1;refGene_tx=XM_019812054.1;refGene_gn=MRVI1;refGene_tx=XM_01
9812055.2;refGene_gn=MRVI1;refGene_tx=XM_019812056.2;refGene_gn=MRVI1;
refGene_tx=XM_019812057.2
D1 69113457 69113655
refGene_gn=MRVI1;refGene_tx=XM_003992992.3;refGene_gn=MRVI1;refGene_tx
=XM_006937098.3;refGene_gn=MRVI1;refGene_tx=XM_006937099.3;refGene_gn=
MRVI1;refGene_tx=XM_006937100.2;refGene_gn=MRVI1;refGene_tx=XM_0069371
01.2;refGene_gn=MRVI1;refGene_tx=XM_019812051.1;refGene_gn=MRVI1;refGe
ne_tx=XM_019812052.1;refGene_gn=MRVI1;refGene_tx=XM_019812053.1;refGen
e_gn=MRVI1;refGene_tx=XM_019812054.1;refGene_gn=MRVI1;refGene_tx=XM_01
9812055.2;refGene_gn=MRVI1;refGene_tx=XM_019812056.2;refGene_gn=MRVI1;
refGene_tx=XM_019812057.2
D1 69119382 69119401 refGene_gn=MRVI1;refGene_tx=XM_006937099.3
D1 69157505 69157572
refGene_gn=MRVI1;refGene_tx=XM_006937098.3;refGene_gn=MRVI1;refGene_tx
=XM_019812055.2;refGene_gn=MRVI1;refGene_tx=XM_019812056.2
D1 69202092 69202137 refGene_gn=CTR9;refGene_tx=XM_011286715.3
D1 69203365 69203464 refGene_gn=CTR9;refGene_tx=XM_011286715.3
D1 69205517 69205757 refGene_gn=CTR9;refGene_tx=XM_011286715.3
D1 69206274 69206392 refGene_gn=CTR9;refGene_tx=XM_011286715.3
D1 69207168 69207258 refGene_gn=CTR9;refGene_tx=XM_011286715.3
D1 69209568 69209717 refGene_gn=CTR9;refGene_tx=XM_011286715.3
D1 69211616 69211724 refGene_gn=CTR9;refGene_tx=XM_011286715.3
D1 69215156 69215264 refGene_gn=CTR9;refGene_tx=XM_011286715.3

D1	69215407	69215644	refGene_gn=CTR9;refGene_tx=XM_011286715.3
D1	69215877	69215967	refGene_gn=CTR9;refGene_tx=XM_011286715.3
D1	69216080	69216209	refGene_gn=CTR9;refGene_tx=XM_011286715.3
D1	69216317	69216501	refGene_gn=CTR9;refGene_tx=XM_011286715.3
D1	69217659	69217748	refGene_gn=CTR9;refGene_tx=XM_011286715.3
D1	69218241	69218427	refGene_gn=CTR9;refGene_tx=XM_011286715.3
D1	69218537	69218625	refGene_gn=CTR9;refGene_tx=XM_011286715.3
D1	69219253	69219402	refGene_gn=CTR9;refGene_tx=XM_011286715.3
D1	69221409	69221526	refGene_gn=CTR9;refGene_tx=XM_011286715.3
D1	69221683	69221829	refGene_gn=CTR9;refGene_tx=XM_011286715.3
D1	69222991	69223063	refGene_gn=CTR9;refGene_tx=XM_011286715.3
D1	69223527	69223663	refGene_gn=CTR9;refGene_tx=XM_011286715.3
D1	69223813	69223960	refGene_gn=CTR9;refGene_tx=XM_011286715.3
D1	69224340	69224498	refGene_gn=CTR9;refGene_tx=XM_011286715.3
D1	69225564	69225661	refGene_gn=CTR9;refGene_tx=XM_011286715.3
D1	69225892	69226005	refGene_gn=CTR9;refGene_tx=XM_011286715.3
D1	69228775	69229202	refGene_gn=CTR9;refGene_tx=XM_011286715.3
D1	69248944	69249010	refGene_gn=EIF4G2;refGene_tx=XM_011286761.1
D1	69250202	69250324	refGene_gn=EIF4G2;refGene_tx=XM_011286761.1
D1	69250418	69250630	refGene_gn=EIF4G2;refGene_tx=XM_011286761.1
D1	69250757	69250962	refGene_gn=EIF4G2;refGene_tx=XM_011286761.1
D1	69251275	69251534	refGene_gn=EIF4G2;refGene_tx=XM_011286761.1
D1	69251622	69251838	refGene_gn=EIF4G2;refGene_tx=XM_011286761.1
D1	69251926	69252031	refGene_gn=EIF4G2;refGene_tx=XM_011286761.1
D1	69252182	69252308	refGene_gn=EIF4G2;refGene_tx=XM_011286761.1
D1	69252867	69252981	refGene_gn=EIF4G2;refGene_tx=XM_011286761.1
D1	69253332	69253493	refGene_gn=EIF4G2;refGene_tx=XM_011286761.1
D1	69253595	69253737	refGene_gn=EIF4G2;refGene_tx=XM_011286761.1
D1	69253995	69254094	refGene_gn=EIF4G2;refGene_tx=XM_011286761.1
D1	69254184	69254268	refGene_gn=EIF4G2;refGene_tx=XM_011286761.1
D1	69254445	69254556	refGene_gn=EIF4G2;refGene_tx=XM_011286761.1
D1	69254811	69254963	refGene_gn=EIF4G2;refGene_tx=XM_011286761.1
D1	69255050	69255117	refGene_gn=EIF4G2;refGene_tx=XM_011286761.1
D1	69255205	69255337	refGene_gn=EIF4G2;refGene_tx=XM_011286761.1
D1	69256091	69256194	refGene_gn=EIF4G2;refGene_tx=XM_011286761.1
D1	69256730	69256871	refGene_gn=EIF4G2;refGene_tx=XM_011286761.1
D1	69257520	69257586	refGene_gn=EIF4G2;refGene_tx=XM_011286761.1
D1	69257913	69257978	refGene_gn=EIF4G2;refGene_tx=XM_011286761.1
D1	69259384	69259525	refGene_gn=EIF4G2;refGene_tx=XM_011286761.1
D1	69265106	69265202	refGene_gn=EIF4G2;refGene_tx=XM_011286761.1
D1	69290599	69292684
refGene_gn=ZBED5;refGene_tx=XM_011286716.3;refGene_gn=ZBED5;refGene_tx=XM_011286717.3;refGene_gn=ZBED5;refGene_tx=XM_011286718.3;refGene_gn=ZBED5;refGene_tx=XM_019812060.2;refGene_gn=ZBED5;refGene_tx=XM_019812061.1
D1	69643519	69643666	refGene_gn=GALNT18;refGene_tx=XM_003993000.4
D1	69659418	69659583	refGene_gn=GALNT18;refGene_tx=XM_003993000.4
D1	69686127	69686139	refGene_gn=GALNT18;refGene_tx=XM_023239562.1
D1	69690516	69690612
refGene_gn=GALNT18;refGene_tx=XM_003993000.4;refGene_gn=GALNT18;refGen

e_tx=XM_023239562.1
D1 69696218 69696356
refGene_gn=GALNT18;refGene_tx=XM_003993000.4;refGene_gn=GALNT18;refGene_tx=XM_023239562.1
D1 69704640 69704826
refGene_gn=GALNT18;refGene_tx=XM_003993000.4;refGene_gn=GALNT18;refGene_tx=XM_023239562.1
D1 69738039 69738154
refGene_gn=GALNT18;refGene_tx=XM_003993000.4;refGene_gn=GALNT18;refGene_tx=XM_023239562.1
D1 69743004 69743202
refGene_gn=GALNT18;refGene_tx=XM_003993000.4;refGene_gn=GALNT18;refGene_tx=XM_023239562.1
D1 69744716 69744900
refGene_gn=GALNT18;refGene_tx=XM_003993000.4;refGene_gn=GALNT18;refGene_tx=XM_023239562.1
D1 69802085 69802252
refGene_gn=GALNT18;refGene_tx=XM_003993000.4;refGene_gn=GALNT18;refGene_tx=XM_023239562.1
D1 69820308 69820501
refGene_gn=GALNT18;refGene_tx=XM_003993000.4;refGene_gn=GALNT18;refGene_tx=XM_023239562.1
D1 69996167 69996402
refGene_gn=GALNT18;refGene_tx=XM_003993000.4;refGene_gn=GALNT18;refGene_tx=XM_023239562.1
D1 70203717 70203756
refGene_gn=USP47;refGene_tx=XM_003993002.5;refGene_gn=USP47;refGene_tx=XM_006937103.4
D1 70237282 70237342 refGene_gn=USP47;refGene_tx=XM_006937103.4
D1 70243254 70243458
refGene_gn=USP47;refGene_tx=XM_003993002.5;refGene_gn=USP47;refGene_tx=XM_006937103.4
D1 70246756 70246870
refGene_gn=USP47;refGene_tx=XM_003993002.5;refGene_gn=USP47;refGene_tx=XM_006937103.4
D1 70248337 70248476
refGene_gn=USP47;refGene_tx=XM_003993002.5;refGene_gn=USP47;refGene_tx=XM_006937103.4
D1 70253507 70253604
refGene_gn=USP47;refGene_tx=XM_003993002.5;refGene_gn=USP47;refGene_tx=XM_006937103.4
D1 70255340 70255486
refGene_gn=USP47;refGene_tx=XM_003993002.5;refGene_gn=USP47;refGene_tx=XM_006937103.4
D1 70256035 70256115
refGene_gn=USP47;refGene_tx=XM_003993002.5;refGene_gn=USP47;refGene_tx=XM_006937103.4
D1 70258445 70258595
refGene_gn=USP47;refGene_tx=XM_003993002.5;refGene_gn=USP47;refGene_tx=XM_006937103.4

D1 70274997 70275093
refGene_gn=USP47;refGene_tx=XM_003993002.5;refGene_gn=USP47;refGene_tx
=XM_006937103.4
D1 70275168 70275321
refGene_gn=USP47;refGene_tx=XM_003993002.5;refGene_gn=USP47;refGene_tx
=XM_006937103.4
D1 70277976 70278144
refGene_gn=USP47;refGene_tx=XM_003993002.5;refGene_gn=USP47;refGene_tx
=XM_006937103.4
D1 70280181 70280313
refGene_gn=USP47;refGene_tx=XM_003993002.5;refGene_gn=USP47;refGene_tx
=XM_006937103.4
D1 70280983 70281060
refGene_gn=USP47;refGene_tx=XM_003993002.5;refGene_gn=USP47;refGene_tx
=XM_006937103.4
D1 70281610 70281666
refGene_gn=USP47;refGene_tx=XM_003993002.5;refGene_gn=USP47;refGene_tx
=XM_006937103.4
D1 70284719 70284832
refGene_gn=USP47;refGene_tx=XM_003993002.5;refGene_gn=USP47;refGene_tx
=XM_006937103.4
D1 70285550 70285655
refGene_gn=USP47;refGene_tx=XM_003993002.5;refGene_gn=USP47;refGene_tx
=XM_006937103.4
D1 70287697 70287905
refGene_gn=USP47;refGene_tx=XM_003993002.5;refGene_gn=USP47;refGene_tx
=XM_006937103.4
D1 70290607 70290723
refGene_gn=USP47;refGene_tx=XM_003993002.5;refGene_gn=USP47;refGene_tx
=XM_006937103.4
D1 70292711 70292831
refGene_gn=USP47;refGene_tx=XM_003993002.5;refGene_gn=USP47;refGene_tx
=XM_006937103.4
D1 70294492 70295273
refGene_gn=USP47;refGene_tx=XM_003993002.5;refGene_gn=USP47;refGene_tx
=XM_006937103.4
D1 70299960 70300136
refGene_gn=USP47;refGene_tx=XM_003993002.5;refGene_gn=USP47;refGene_tx
=XM_006937103.4
D1 70300486 70300567
refGene_gn=USP47;refGene_tx=XM_003993002.5;refGene_gn=USP47;refGene_tx
=XM_006937103.4
D1 70302600 70302716
refGene_gn=USP47;refGene_tx=XM_003993002.5;refGene_gn=USP47;refGene_tx
=XM_006937103.4
D1 70303302 70303421
refGene_gn=USP47;refGene_tx=XM_003993002.5;refGene_gn=USP47;refGene_tx
=XM_006937103.4
D1 70305491 70305622
refGene_gn=USP47;refGene_tx=XM_003993002.5;refGene_gn=USP47;refGene_tx

=XM_006937103.4
D1 70307614 70307662
refGene_gn=USP47;refGene_tx=XM_003993002.5;refGene_gn=USP47;refGene_tx
=XM_006937103.4
D1 70307753 70307884
refGene_gn=USP47;refGene_tx=XM_003993002.5;refGene_gn=USP47;refGene_tx
=XM_006937103.4
D1 70308712 70308887
refGene_gn=USP47;refGene_tx=XM_003993002.5;refGene_gn=USP47;refGene_tx
=XM_006937103.4
D1 70315311 70315531 refGene_gn=DKK3;refGene_tx=XM_019812064.2
D1 70316455 70316612 refGene_gn=DKK3;refGene_tx=XM_019812064.2
D1 70317553 70317698 refGene_gn=DKK3;refGene_tx=XM_019812064.2
D1 70318820 70318913 refGene_gn=DKK3;refGene_tx=XM_019812064.2
D1 70349116 70349200 refGene_gn=DKK3;refGene_tx=XM_019812064.2
D1 70352102 70352240 refGene_gn=DKK3;refGene_tx=XM_019812064.2
D1 70359934 70360147 refGene_gn=DKK3;refGene_tx=XM_019812064.2
D1 70485353 70485617
refGene_gn=LOC101091046;refGene_tx=XM_023239563.1;refGene_gn=LOC101091
046;refGene_tx=XM_023239564.1;refGene_gn=LOC101091046;refGene_tx=XM_02
3239565.1;refGene_gn=LOC101091046;refGene_tx=XM_023239566.1;refGene_gn
=LOC101091046;refGene_tx=XM_023239567.1;refGene_gn=LOC101091046;refGen
e_tx=XM_023239568.1;refGene_gn=LOC101091046;refGene_tx=XM_023239569.1;
refGene_gn=LOC101091046;refGene_tx=XM_023239570.1;refGene_gn=LOC101091
046;refGene_tx=XM_023239571.1
D1 70521492 70521700
refGene_gn=LOC101091046;refGene_tx=XM_023239563.1;refGene_gn=LOC101091
046;refGene_tx=XM_023239564.1;refGene_gn=LOC101091046;refGene_tx=XM_02
3239565.1;refGene_gn=LOC101091046;refGene_tx=XM_023239566.1;refGene_gn
=LOC101091046;refGene_tx=XM_023239567.1;refGene_gn=LOC101091046;refGen
e_tx=XM_023239568.1;refGene_gn=LOC101091046;refGene_tx=XM_023239569.1;
refGene_gn=LOC101091046;refGene_tx=XM_023239570.1;refGene_gn=LOC101091
046;refGene_tx=XM_023239571.1
D1 70525169 70525286
refGene_gn=LOC101091046;refGene_tx=XM_023239563.1;refGene_gn=LOC101091
046;refGene_tx=XM_023239564.1;refGene_gn=LOC101091046;refGene_tx=XM_02
3239565.1;refGene_gn=LOC101091046;refGene_tx=XM_023239566.1;refGene_gn
=LOC101091046;refGene_tx=XM_023239567.1;refGene_gn=LOC101091046;refGen
e_tx=XM_023239568.1;refGene_gn=LOC101091046;refGene_tx=XM_023239569.1;
refGene_gn=LOC101091046;refGene_tx=XM_023239570.1;refGene_gn=LOC101091
046;refGene_tx=XM_023239571.1
D1 70526596 70526698
refGene_gn=LOC101091046;refGene_tx=XM_023239563.1;refGene_gn=LOC101091
046;refGene_tx=XM_023239564.1;refGene_gn=LOC101091046;refGene_tx=XM_02
3239565.1;refGene_gn=LOC101091046;refGene_tx=XM_023239566.1;refGene_gn
=LOC101091046;refGene_tx=XM_023239567.1;refGene_gn=LOC101091046;refGen
e_tx=XM_023239568.1;refGene_gn=LOC101091046;refGene_tx=XM_023239569.1;
refGene_gn=LOC101091046;refGene_tx=XM_023239570.1;refGene_gn=LOC101091
046;refGene_tx=XM_023239571.1
D1 70529525 70529681

refGene_gn=LOC101091046;refGene_tx=XM_023239563.1;refGene_gn=LOC101091046;refGene_tx=XM_023239564.1;refGene_gn=LOC101091046;refGene_tx=XM_023239565.1;refGene_gn=LOC101091046;refGene_tx=XM_023239566.1;refGene_gn=LOC101091046;refGene_tx=XM_023239567.1;refGene_gn=LOC101091046;refGene_tx=XM_023239568.1;refGene_gn=LOC101091046;refGene_tx=XM_023239569.1;refGene_gn=LOC101091046;refGene_tx=XM_023239570.1;refGene_gn=LOC101091046;refGene_tx=XM_023239571.1
D1 70533473 70533574
refGene_gn=LOC101091046;refGene_tx=XM_023239563.1;refGene_gn=LOC101091046;refGene_tx=XM_023239564.1;refGene_gn=LOC101091046;refGene_tx=XM_023239565.1;refGene_gn=LOC101091046;refGene_tx=XM_023239566.1;refGene_gn=LOC101091046;refGene_tx=XM_023239567.1;refGene_gn=LOC101091046;refGene_tx=XM_023239568.1;refGene_gn=LOC101091046;refGene_tx=XM_023239569.1;refGene_gn=LOC101091046;refGene_tx=XM_023239570.1;refGene_gn=LOC101091046;refGene_tx=XM_023239571.1
D1 70537881 70538139
refGene_gn=LOC101091046;refGene_tx=XM_023239563.1;refGene_gn=LOC101091046;refGene_tx=XM_023239564.1;refGene_gn=LOC101091046;refGene_tx=XM_023239565.1;refGene_gn=LOC101091046;refGene_tx=XM_023239566.1;refGene_gn=LOC101091046;refGene_tx=XM_023239567.1;refGene_gn=LOC101091046;refGene_tx=XM_023239568.1;refGene_gn=LOC101091046;refGene_tx=XM_023239569.1;refGene_gn=LOC101091046;refGene_tx=XM_023239570.1;refGene_gn=LOC101091046;refGene_tx=XM_023239571.1
D1 70539478 70539594
refGene_gn=LOC101091046;refGene_tx=XM_023239563.1;refGene_gn=LOC101091046;refGene_tx=XM_023239564.1;refGene_gn=LOC101091046;refGene_tx=XM_023239565.1;refGene_gn=LOC101091046;refGene_tx=XM_023239566.1;refGene_gn=LOC101091046;refGene_tx=XM_023239567.1;refGene_gn=LOC101091046;refGene_tx=XM_023239568.1;refGene_gn=LOC101091046;refGene_tx=XM_023239569.1;refGene_gn=LOC101091046;refGene_tx=XM_023239570.1;refGene_gn=LOC101091046;refGene_tx=XM_023239571.1
D1 70540455 70540582
refGene_gn=LOC101091046;refGene_tx=XM_023239563.1;refGene_gn=LOC101091046;refGene_tx=XM_023239564.1;refGene_gn=LOC101091046;refGene_tx=XM_023239565.1;refGene_gn=LOC101091046;refGene_tx=XM_023239566.1;refGene_gn=LOC101091046;refGene_tx=XM_023239567.1;refGene_gn=LOC101091046;refGene_tx=XM_023239568.1;refGene_gn=LOC101091046;refGene_tx=XM_023239569.1;refGene_gn=LOC101091046;refGene_tx=XM_023239570.1;refGene_gn=LOC101091046;refGene_tx=XM_023239571.1
D1 70541383 70541474
refGene_gn=LOC101091046;refGene_tx=XM_023239563.1;refGene_gn=LOC101091046;refGene_tx=XM_023239564.1;refGene_gn=LOC101091046;refGene_tx=XM_023239565.1;refGene_gn=LOC101091046;refGene_tx=XM_023239566.1;refGene_gn=LOC101091046;refGene_tx=XM_023239567.1;refGene_gn=LOC101091046;refGene_tx=XM_023239568.1;refGene_gn=LOC101091046;refGene_tx=XM_023239569.1;refGene_gn=LOC101091046;refGene_tx=XM_023239570.1;refGene_gn=LOC101091046;refGene_tx=XM_023239571.1
D1 70542549 70542697
refGene_gn=LOC101091046;refGene_tx=XM_023239563.1;refGene_gn=LOC101091046;refGene_tx=XM_023239564.1;refGene_gn=LOC101091046;refGene_tx=XM_02

3239565.1;refGene_gn=LOC101091046;refGene_tx=XM_023239566.1;refGene_gn
=LOC101091046;refGene_tx=XM_023239567.1;refGene_gn=LOC101091046;refGen
e_tx=XM_023239568.1;refGene_gn=LOC101091046;refGene_tx=XM_023239569.1;
refGene_gn=LOC101091046;refGene_tx=XM_023239570.1;refGene_gn=LOC101091
046;refGene_tx=XM_023239571.1
D1 70543904 70544104
refGene_gn=LOC101091046;refGene_tx=XM_023239563.1;refGene_gn=LOC101091
046;refGene_tx=XM_023239564.1;refGene_gn=LOC101091046;refGene_tx=XM_02
3239565.1;refGene_gn=LOC101091046;refGene_tx=XM_023239566.1;refGene_gn
=LOC101091046;refGene_tx=XM_023239567.1;refGene_gn=LOC101091046;refGen
e_tx=XM_023239568.1;refGene_gn=LOC101091046;refGene_tx=XM_023239569.1;
refGene_gn=LOC101091046;refGene_tx=XM_023239570.1;refGene_gn=LOC101091
046;refGene_tx=XM_023239571.1
D1 70544503 70544610
refGene_gn=LOC101091046;refGene_tx=XM_023239563.1;refGene_gn=LOC101091
046;refGene_tx=XM_023239564.1;refGene_gn=LOC101091046;refGene_tx=XM_02
3239565.1;refGene_gn=LOC101091046;refGene_tx=XM_023239566.1;refGene_gn
=LOC101091046;refGene_tx=XM_023239567.1;refGene_gn=LOC101091046;refGen
e_tx=XM_023239568.1;refGene_gn=LOC101091046;refGene_tx=XM_023239569.1;
refGene_gn=LOC101091046;refGene_tx=XM_023239570.1;refGene_gn=LOC101091
046;refGene_tx=XM_023239571.1
D1 70552954 70553023
refGene_gn=LOC101091046;refGene_tx=XM_023239563.1;refGene_gn=LOC101091
046;refGene_tx=XM_023239564.1;refGene_gn=LOC101091046;refGene_tx=XM_02
3239565.1;refGene_gn=LOC101091046;refGene_tx=XM_023239566.1;refGene_gn
=LOC101091046;refGene_tx=XM_023239567.1;refGene_gn=LOC101091046;refGen
e_tx=XM_023239568.1;refGene_gn=LOC101091046;refGene_tx=XM_023239569.1;
refGene_gn=LOC101091046;refGene_tx=XM_023239570.1;refGene_gn=LOC101091
046;refGene_tx=XM_023239571.1
D1 70556435 70556585
refGene_gn=LOC101091046;refGene_tx=XM_023239563.1;refGene_gn=LOC101091
046;refGene_tx=XM_023239564.1;refGene_gn=LOC101091046;refGene_tx=XM_02
3239565.1;refGene_gn=LOC101091046;refGene_tx=XM_023239566.1;refGene_gn
=LOC101091046;refGene_tx=XM_023239567.1;refGene_gn=LOC101091046;refGen
e_tx=XM_023239568.1;refGene_gn=LOC101091046;refGene_tx=XM_023239569.1;
refGene_gn=LOC101091046;refGene_tx=XM_023239570.1;refGene_gn=LOC101091
046;refGene_tx=XM_023239571.1
D1 70557969 70558092
refGene_gn=LOC101091046;refGene_tx=XM_023239563.1;refGene_gn=LOC101091
046;refGene_tx=XM_023239565.1;refGene_gn=LOC101091046;refGene_tx=XM_02
3239568.1;refGene_gn=LOC101091046;refGene_tx=XM_023239569.1;refGene_gn
=LOC101091046;refGene_tx=XM_023239571.1
D1 70559347 70559566
refGene_gn=LOC101091046;refGene_tx=XM_023239563.1;refGene_gn=LOC101091
046;refGene_tx=XM_023239565.1;refGene_gn=LOC101091046;refGene_tx=XM_02
3239568.1;refGene_gn=LOC101091046;refGene_tx=XM_023239569.1;refGene_gn
=LOC101091046;refGene_tx=XM_023239571.1
D1 70559795 70559897
refGene_gn=LOC101091046;refGene_tx=XM_023239563.1;refGene_gn=LOC101091
046;refGene_tx=XM_023239565.1;refGene_gn=LOC101091046;refGene_tx=XM_02

3239568.1;refGene_gn=LOC101091046;refGene_tx=XM_023239569.1
D1	70561111	70561237
refGene_gn=LOC101091046;refGene_tx=XM_023239563.1;refGene_gn=LOC101091046;refGene_tx=XM_023239565.1;refGene_gn=LOC101091046;refGene_tx=XM_023239568.1;refGene_gn=LOC101091046;refGene_tx=XM_023239569.1
D1	70565727	70565790
refGene_gn=LOC101091046;refGene_tx=XM_023239563.1;refGene_gn=LOC101091046;refGene_tx=XM_023239564.1;refGene_gn=LOC101091046;refGene_tx=XM_023239565.1;refGene_gn=LOC101091046;refGene_tx=XM_023239566.1;refGene_gn=LOC101091046;refGene_tx=XM_023239568.1;refGene_gn=LOC101091046;refGene_tx=XM_023239569.1;refGene_gn=LOC101091046;refGene_tx=XM_023239570.1
D1	70570595	70570703
refGene_gn=LOC101091046;refGene_tx=XM_023239563.1;refGene_gn=LOC101091046;refGene_tx=XM_023239564.1;refGene_gn=LOC101091046;refGene_tx=XM_023239565.1;refGene_gn=LOC101091046;refGene_tx=XM_023239568.1;refGene_gn=LOC101091046;refGene_tx=XM_023239569.1;refGene_gn=LOC101091046;refGene_tx=XM_023239570.1
D1	70571753	70571940
refGene_gn=LOC101091046;refGene_tx=XM_023239563.1;refGene_gn=LOC101091046;refGene_tx=XM_023239564.1;refGene_gn=LOC101091046;refGene_tx=XM_023239565.1;refGene_gn=LOC101091046;refGene_tx=XM_023239566.1;refGene_gn=LOC101091046;refGene_tx=XM_023239567.1;refGene_gn=LOC101091046;refGene_tx=XM_023239568.1;refGene_gn=LOC101091046;refGene_tx=XM_023239569.1;refGene_gn=LOC101091046;refGene_tx=XM_023239570.1
D1	70573127	70573216
refGene_gn=LOC101091046;refGene_tx=XM_023239563.1;refGene_gn=LOC101091046;refGene_tx=XM_023239564.1;refGene_gn=LOC101091046;refGene_tx=XM_023239565.1;refGene_gn=LOC101091046;refGene_tx=XM_023239566.1;refGene_gn=LOC101091046;refGene_tx=XM_023239567.1;refGene_gn=LOC101091046;refGene_tx=XM_023239568.1;refGene_gn=LOC101091046;refGene_tx=XM_023239569.1;refGene_gn=LOC101091046;refGene_tx=XM_023239570.1
D1	70574950	70575053
refGene_gn=LOC101091046;refGene_tx=XM_023239563.1;refGene_gn=LOC101091046;refGene_tx=XM_023239564.1;refGene_gn=LOC101091046;refGene_tx=XM_023239565.1;refGene_gn=LOC101091046;refGene_tx=XM_023239566.1;refGene_gn=LOC101091046;refGene_tx=XM_023239567.1;refGene_gn=LOC101091046;refGene_tx=XM_023239568.1;refGene_gn=LOC101091046;refGene_tx=XM_023239569.1;refGene_gn=LOC101091046;refGene_tx=XM_023239570.1
D1	70577782	70577823
refGene_gn=LOC101091046;refGene_tx=XM_023239568.1;refGene_gn=LOC101091046;refGene_tx=XM_023239569.1;refGene_gn=LOC101091046;refGene_tx=XM_023239570.1
D1	70591309	70591493
refGene_gn=LOC101091046;refGene_tx=XM_023239563.1;refGene_gn=LOC101091046;refGene_tx=XM_023239564.1;refGene_gn=LOC101091046;refGene_tx=XM_023239565.1;refGene_gn=LOC101091046;refGene_tx=XM_023239566.1;refGene_gn=LOC101091046;refGene_tx=XM_023239567.1
D1	70595437	70595619
refGene_gn=LOC101091046;refGene_tx=XM_023239563.1;refGene_gn=LOC101091046;refGene_tx=XM_023239564.1;refGene_gn=LOC101091046;refGene_tx=XM_02

3239565.1;refGene_gn=LOC101091046;refGene_tx=XM_023239566.1;refGene_gn=LOC101091046;refGene_tx=XM_023239567.1
D1 70601425 70601500
refGene_gn=LOC101091046;refGene_tx=XM_023239563.1;refGene_gn=LOC101091046;refGene_tx=XM_023239564.1;refGene_gn=LOC101091046;refGene_tx=XM_023239565.1;refGene_gn=LOC101091046;refGene_tx=XM_023239566.1;refGene_gn=LOC101091046;refGene_tx=XM_023239567.1
D1 70605977 70606132
refGene_gn=LOC101091046;refGene_tx=XM_023239563.1;refGene_gn=LOC101091046;refGene_tx=XM_023239564.1;refGene_gn=LOC101091046;refGene_tx=XM_023239565.1;refGene_gn=LOC101091046;refGene_tx=XM_023239566.1;refGene_gn=LOC101091046;refGene_tx=XM_023239567.1
D1 70607378 70608681
refGene_gn=LOC101091046;refGene_tx=XM_023239563.1;refGene_gn=LOC101091046;refGene_tx=XM_023239564.1;refGene_gn=LOC101091046;refGene_tx=XM_023239565.1;refGene_gn=LOC101091046;refGene_tx=XM_023239566.1;refGene_gn=LOC101091046;refGene_tx=XM_023239567.1
D1 70614407 70614436
refGene_gn=LOC101091046;refGene_tx=XM_023239565.1
D1 70626020 70626136
refGene_gn=LOC101091046;refGene_tx=XM_023239563.1;refGene_gn=LOC101091046;refGene_tx=XM_023239564.1;refGene_gn=LOC101091046;refGene_tx=XM_023239566.1;refGene_gn=LOC101091046;refGene_tx=XM_023239567.1
D1 70631065 70631155
refGene_gn=LOC101091046;refGene_tx=XM_023239563.1;refGene_gn=LOC101091046;refGene_tx=XM_023239564.1;refGene_gn=LOC101091046;refGene_tx=XM_023239566.1;refGene_gn=LOC101091046;refGene_tx=XM_023239567.1
D1 70633897 70633991
refGene_gn=LOC101091046;refGene_tx=XM_023239563.1;refGene_gn=LOC101091046;refGene_tx=XM_023239564.1;refGene_gn=LOC101091046;refGene_tx=XM_023239566.1;refGene_gn=LOC101091046;refGene_tx=XM_023239567.1
D1 70651058 70651161
refGene_gn=LOC101091046;refGene_tx=XM_023239563.1;refGene_gn=LOC101091046;refGene_tx=XM_023239564.1;refGene_gn=LOC101091046;refGene_tx=XM_023239566.1;refGene_gn=LOC101091046;refGene_tx=XM_023239567.1
D1 70654474 70654548
refGene_gn=LOC101091046;refGene_tx=XM_023239563.1;refGene_gn=LOC101091046;refGene_tx=XM_023239564.1;refGene_gn=LOC101091046;refGene_tx=XM_023239566.1;refGene_gn=LOC101091046;refGene_tx=XM_023239567.1
D1 70658426 70658611
refGene_gn=LOC101091046;refGene_tx=XM_023239563.1;refGene_gn=LOC101091046;refGene_tx=XM_023239564.1;refGene_gn=LOC101091046;refGene_tx=XM_023239566.1;refGene_gn=LOC101091046;refGene_tx=XM_023239567.1
D1 70676832 70676968
refGene_gn=PARVA;refGene_tx=XM_006937152.2;refGene_gn=PARVA;refGene_tx=XM_023239572.1
D1 70779048 70779138
refGene_gn=PARVA;refGene_tx=XM_006937152.2;refGene_gn=PARVA;refGene_tx=XM_023239572.1
D1 70779217 70779288

refGene_gn=PARVA;refGene_tx=XM_006937152.2;refGene_gn=PARVA;refGene_tx=XM_023239572.1
D1 70781614 70781717
refGene_gn=PARVA;refGene_tx=XM_006937152.2;refGene_gn=PARVA;refGene_tx=XM_023239572.1
D1 70819914 70820055
refGene_gn=PARVA;refGene_tx=XM_006937152.2;refGene_gn=PARVA;refGene_tx=XM_023239572.1
D1 70826970 70827086
refGene_gn=PARVA;refGene_tx=XM_006937152.2;refGene_gn=PARVA;refGene_tx=XM_023239572.1
D1 70831288 70831347
refGene_gn=PARVA;refGene_tx=XM_006937152.2;refGene_gn=PARVA;refGene_tx=XM_023239572.1
D1 70833594 70833614
refGene_gn=PARVA;refGene_tx=XM_006937152.2;refGene_gn=PARVA;refGene_tx=XM_023239572.1
D1 70834969 70835031
refGene_gn=PARVA;refGene_tx=XM_006937152.2;refGene_gn=PARVA;refGene_tx=XM_023239572.1
D1 70835742 70835811
refGene_gn=PARVA;refGene_tx=XM_006937152.2;refGene_gn=PARVA;refGene_tx=XM_023239572.1
D1 70839400 70839502
refGene_gn=PARVA;refGene_tx=XM_006937152.2;refGene_gn=PARVA;refGene_tx=XM_023239572.1
D1 70840241 70840314
refGene_gn=PARVA;refGene_tx=XM_006937152.2;refGene_gn=PARVA;refGene_tx=XM_023239572.1
D1 70850009 70850086 refGene_gn=PARVA;refGene_tx=XM_006937152.2
D1 70867536 70867568 refGene_gn=PARVA;refGene_tx=XM_023239572.1
D1 71065678 71065835
refGene_gn=TEAD1;refGene_tx=XM_011286720.3;refGene_gn=TEAD1;refGene_tx=XM_011286722.3;refGene_gn=TEAD1;refGene_tx=XM_011286723.3;refGene_gn=TEAD1;refGene_tx=XM_011286724.3;refGene_gn=TEAD1;refGene_tx=XM_011286725.3;refGene_gn=TEAD1;refGene_tx=XM_011286726.3;refGene_gn=TEAD1;refGene_tx=XM_019812073.2;refGene_gn=TEAD1;refGene_tx=XM_019812074.2;refGene_gn=TEAD1;refGene_tx=XM_019812075.2;refGene_gn=TEAD1;refGene_tx=XM_023239574.1;refGene_gn=TEAD1;refGene_tx=XM_023239576.1
D1 71172299 71172364
refGene_gn=TEAD1;refGene_tx=XM_011286720.3;refGene_gn=TEAD1;refGene_tx=XM_011286722.3;refGene_gn=TEAD1;refGene_tx=XM_011286723.3;refGene_gn=TEAD1;refGene_tx=XM_011286724.3;refGene_gn=TEAD1;refGene_tx=XM_011286725.3;refGene_gn=TEAD1;refGene_tx=XM_011286726.3;refGene_gn=TEAD1;refGene_tx=XM_019812073.2;refGene_gn=TEAD1;refGene_tx=XM_019812074.2;refGene_gn=TEAD1;refGene_tx=XM_019812075.2;refGene_gn=TEAD1;refGene_tx=XM_023239574.1;refGene_gn=TEAD1;refGene_tx=XM_023239576.1
D1 71174750 71174813
refGene_gn=TEAD1;refGene_tx=XM_011286720.3;refGene_gn=TEAD1;refGene_tx=XM_011286722.3;refGene_gn=TEAD1;refGene_tx=XM_011286724.3;refGene_gn=

TEAD1;refGene_tx=XM_011286726.3;refGene_gn=TEAD1;refGene_tx=XM_0198120
73.2;refGene_gn=TEAD1;refGene_tx=XM_019812074.2;refGene_gn=TEAD1;refGe
ne_tx=XM_023239574.1
D1 71174902 71174965
refGene_gn=TEAD1;refGene_tx=XM_011286720.3;refGene_gn=TEAD1;refGene_tx
=XM_011286722.3;refGene_gn=TEAD1;refGene_tx=XM_011286723.3;refGene_gn=
TEAD1;refGene_tx=XM_011286725.3;refGene_gn=TEAD1;refGene_tx=XM_0112867
26.3;refGene_gn=TEAD1;refGene_tx=XM_019812073.2;refGene_gn=TEAD1;refGe
ne_tx=XM_023239574.1;refGene_gn=TEAD1;refGene_tx=XM_023239576.1
D1 71188423 71188435
refGene_gn=TEAD1;refGene_tx=XM_011286720.3;refGene_gn=TEAD1;refGene_tx
=XM_011286722.3;refGene_gn=TEAD1;refGene_tx=XM_011286723.3;refGene_gn=
TEAD1;refGene_tx=XM_011286724.3;refGene_gn=TEAD1;refGene_tx=XM_0112867
26.3;refGene_gn=TEAD1;refGene_tx=XM_019812073.2;refGene_gn=TEAD1;refGe
ne_tx=XM_023239576.1
D1 71189260 71189395
refGene_gn=TEAD1;refGene_tx=XM_011286720.3;refGene_gn=TEAD1;refGene_tx
=XM_011286722.3;refGene_gn=TEAD1;refGene_tx=XM_011286723.3;refGene_gn=
TEAD1;refGene_tx=XM_011286724.3;refGene_gn=TEAD1;refGene_tx=XM_0112867
25.3;refGene_gn=TEAD1;refGene_tx=XM_011286726.3;refGene_gn=TEAD1;refGe
ne_tx=XM_019812073.2;refGene_gn=TEAD1;refGene_tx=XM_019812074.2;refGen
e_gn=TEAD1;refGene_tx=XM_019812075.2;refGene_gn=TEAD1;refGene_tx=XM_02
3239574.1;refGene_gn=TEAD1;refGene_tx=XM_023239576.1
D1 71191033 71191080
refGene_gn=TEAD1;refGene_tx=XM_011286720.3;refGene_gn=TEAD1;refGene_tx
=XM_011286722.3;refGene_gn=TEAD1;refGene_tx=XM_011286723.3;refGene_gn=
TEAD1;refGene_tx=XM_011286724.3;refGene_gn=TEAD1;refGene_tx=XM_0112867
25.3;refGene_gn=TEAD1;refGene_tx=XM_011286726.3;refGene_gn=TEAD1;refGe
ne_tx=XM_019812073.2;refGene_gn=TEAD1;refGene_tx=XM_019812074.2;refGen
e_gn=TEAD1;refGene_tx=XM_019812075.2;refGene_gn=TEAD1;refGene_tx=XM_02
3239574.1;refGene_gn=TEAD1;refGene_tx=XM_023239576.1
D1 71191946 71192008
refGene_gn=TEAD1;refGene_tx=XM_011286720.3;refGene_gn=TEAD1;refGene_tx
=XM_011286722.3;refGene_gn=TEAD1;refGene_tx=XM_011286723.3;refGene_gn=
TEAD1;refGene_tx=XM_011286724.3;refGene_gn=TEAD1;refGene_tx=XM_0112867
25.3;refGene_gn=TEAD1;refGene_tx=XM_011286726.3;refGene_gn=TEAD1;refGe
ne_tx=XM_019812073.2;refGene_gn=TEAD1;refGene_tx=XM_019812074.2;refGen
e_gn=TEAD1;refGene_tx=XM_019812075.2;refGene_gn=TEAD1;refGene_tx=XM_02
3239574.1;refGene_gn=TEAD1;refGene_tx=XM_023239576.1
D1 71193641 71193766
refGene_gn=TEAD1;refGene_tx=XM_011286720.3;refGene_gn=TEAD1;refGene_tx
=XM_011286722.3;refGene_gn=TEAD1;refGene_tx=XM_011286723.3;refGene_gn=
TEAD1;refGene_tx=XM_011286724.3;refGene_gn=TEAD1;refGene_tx=XM_0112867
25.3;refGene_gn=TEAD1;refGene_tx=XM_011286726.3;refGene_gn=TEAD1;refGe
ne_tx=XM_019812073.2;refGene_gn=TEAD1;refGene_tx=XM_019812074.2;refGen
e_gn=TEAD1;refGene_tx=XM_019812075.2;refGene_gn=TEAD1;refGene_tx=XM_02
3239574.1;refGene_gn=TEAD1;refGene_tx=XM_023239576.1
D1 71209227 71209401
refGene_gn=TEAD1;refGene_tx=XM_011286720.3;refGene_gn=TEAD1;refGene_tx
=XM_011286722.3;refGene_gn=TEAD1;refGene_tx=XM_011286723.3;refGene_gn=

TEAD1;refGene_tx=XM_011286724.3;refGene_gn=TEAD1;refGene_tx=XM_011286725.3;refGene_gn=TEAD1;refGene_tx=XM_011286726.3;refGene_gn=TEAD1;refGene_tx=XM_019812073.2;refGene_gn=TEAD1;refGene_tx=XM_019812074.2;refGene_gn=TEAD1;refGene_tx=XM_019812075.2;refGene_gn=TEAD1;refGene_tx=XM_023239574.1

D1 71231900 71232041
refGene_gn=TEAD1;refGene_tx=XM_011286720.3;refGene_gn=TEAD1;refGene_tx=XM_011286722.3;refGene_gn=TEAD1;refGene_tx=XM_011286723.3;refGene_gn=TEAD1;refGene_tx=XM_011286724.3;refGene_gn=TEAD1;refGene_tx=XM_011286725.3;refGene_gn=TEAD1;refGene_tx=XM_011286726.3;refGene_gn=TEAD1;refGene_tx=XM_019812073.2;refGene_gn=TEAD1;refGene_tx=XM_019812074.2;refGene_gn=TEAD1;refGene_tx=XM_019812075.2;refGene_gn=TEAD1;refGene_tx=XM_023239574.1;refGene_gn=TEAD1;refGene_tx=XM_023239576.1

D1 71236399 71236552
refGene_gn=TEAD1;refGene_tx=XM_011286720.3;refGene_gn=TEAD1;refGene_tx=XM_011286722.3;refGene_gn=TEAD1;refGene_tx=XM_011286723.3;refGene_gn=TEAD1;refGene_tx=XM_011286724.3;refGene_gn=TEAD1;refGene_tx=XM_011286725.3;refGene_gn=TEAD1;refGene_tx=XM_011286726.3;refGene_gn=TEAD1;refGene_tx=XM_019812073.2;refGene_gn=TEAD1;refGene_tx=XM_019812074.2;refGene_gn=TEAD1;refGene_tx=XM_019812075.2;refGene_gn=TEAD1;refGene_tx=XM_023239574.1;refGene_gn=TEAD1;refGene_tx=XM_023239576.1

D1 71242481 71242595
refGene_gn=TEAD1;refGene_tx=XM_011286720.3;refGene_gn=TEAD1;refGene_tx=XM_011286722.3;refGene_gn=TEAD1;refGene_tx=XM_011286723.3;refGene_gn=TEAD1;refGene_tx=XM_011286724.3;refGene_gn=TEAD1;refGene_tx=XM_011286725.3;refGene_gn=TEAD1;refGene_tx=XM_011286726.3;refGene_gn=TEAD1;refGene_tx=XM_019812073.2;refGene_gn=TEAD1;refGene_tx=XM_019812074.2;refGene_gn=TEAD1;refGene_tx=XM_019812075.2;refGene_gn=TEAD1;refGene_tx=XM_023239574.1;refGene_gn=TEAD1;refGene_tx=XM_023239576.1

D1 71299331 71300846 refGene_gn=RASSF10;refGene_tx=XM_011286727.3

D1 71616714 71616854
refGene_gn=ARNTL;refGene_tx=XM_019812079.2;refGene_gn=ARNTL;refGene_tx=XM_019812080.2;refGene_gn=ARNTL;refGene_tx=XM_023239578.1;refGene_gn=ARNTL;refGene_tx=XM_023239579.1;refGene_gn=ARNTL;refGene_tx=XM_023239580.1;refGene_gn=ARNTL;refGene_tx=XM_023239581.1;refGene_gn=ARNTL;refGene_tx=XM_023239582.1;refGene_gn=ARNTL;refGene_tx=XM_023239583.1;refGene_gn=ARNTL;refGene_tx=XM_023239584.1;refGene_gn=ARNTL;refGene_tx=XM_023239586.1;refGene_gn=ARNTL;refGene_tx=XM_023239587.1;refGene_gn=ARNTL;refGene_tx=XM_023239588.1;refGene_gn=ARNTL;refGene_tx=XM_023239589.1

D1 71617412 71617433
refGene_gn=ARNTL;refGene_tx=XM_019812079.2;refGene_gn=ARNTL;refGene_tx=XM_019812080.2;refGene_gn=ARNTL;refGene_tx=XM_023239578.1;refGene_gn=ARNTL;refGene_tx=XM_023239579.1;refGene_gn=ARNTL;refGene_tx=XM_023239580.1;refGene_gn=ARNTL;refGene_tx=XM_023239581.1;refGene_gn=ARNTL;refGene_tx=XM_023239582.1;refGene_gn=ARNTL;refGene_tx=XM_023239583.1;refGene_gn=ARNTL;refGene_tx=XM_023239584.1

D1 71617632 71617721
refGene_gn=ARNTL;refGene_tx=XM_003993009.4;refGene_gn=ARNTL;refGene_tx=XM_006937112.3;refGene_gn=ARNTL;refGene_tx=XM_023239590.1;refGene_gn=ARNTL;refGene_tx=XM_023239591.1

D1 71619108 71619147
refGene_gn=ARNTL;refGene_tx=XM_003993009.4;refGene_gn=ARNTL;refGene_tx=XM_006937112.3;refGene_gn=ARNTL;refGene_tx=XM_019812079.2;refGene_gn=ARNTL;refGene_tx=XM_019812080.2;refGene_gn=ARNTL;refGene_tx=XM_019812084.1;refGene_gn=ARNTL;refGene_tx=XM_023239578.1;refGene_gn=ARNTL;refGene_tx=XM_023239579.1;refGene_gn=ARNTL;refGene_tx=XM_023239580.1;refGene_gn=ARNTL;refGene_tx=XM_023239581.1;refGene_gn=ARNTL;refGene_tx=XM_023239582.1;refGene_gn=ARNTL;refGene_tx=XM_023239583.1;refGene_gn=ARNTL;refGene_tx=XM_023239584.1;refGene_gn=ARNTL;refGene_tx=XM_023239586.1;refGene_gn=ARNTL;refGene_tx=XM_023239587.1;refGene_gn=ARNTL;refGene_tx=XM_023239588.1;refGene_gn=ARNTL;refGene_tx=XM_023239589.1;refGene_gn=ARNTL;refGene_tx=XM_023239590.1;refGene_gn=ARNTL;refGene_tx=XM_023239591.1
D1 71619478 71619520
refGene_gn=ARNTL;refGene_tx=XM_003993009.4;refGene_gn=ARNTL;refGene_tx=XM_006937112.3;refGene_gn=ARNTL;refGene_tx=XM_019812079.2;refGene_gn=ARNTL;refGene_tx=XM_019812080.2;refGene_gn=ARNTL;refGene_tx=XM_019812084.1;refGene_gn=ARNTL;refGene_tx=XM_023239578.1;refGene_gn=ARNTL;refGene_tx=XM_023239579.1;refGene_gn=ARNTL;refGene_tx=XM_023239580.1;refGene_gn=ARNTL;refGene_tx=XM_023239581.1;refGene_gn=ARNTL;refGene_tx=XM_023239582.1;refGene_gn=ARNTL;refGene_tx=XM_023239583.1;refGene_gn=ARNTL;refGene_tx=XM_023239584.1;refGene_gn=ARNTL;refGene_tx=XM_023239586.1;refGene_gn=ARNTL;refGene_tx=XM_023239587.1;refGene_gn=ARNTL;refGene_tx=XM_023239588.1;refGene_gn=ARNTL;refGene_tx=XM_023239589.1;refGene_gn=ARNTL;refGene_tx=XM_023239590.1;refGene_gn=ARNTL;refGene_tx=XM_023239591.1
D1 71621051 71621209
refGene_gn=ARNTL;refGene_tx=XM_003993009.4;refGene_gn=ARNTL;refGene_tx=XM_006937112.3;refGene_gn=ARNTL;refGene_tx=XM_019812079.2;refGene_gn=ARNTL;refGene_tx=XM_019812080.2;refGene_gn=ARNTL;refGene_tx=XM_019812084.1;refGene_gn=ARNTL;refGene_tx=XM_023239578.1;refGene_gn=ARNTL;refGene_tx=XM_023239579.1;refGene_gn=ARNTL;refGene_tx=XM_023239580.1;refGene_gn=ARNTL;refGene_tx=XM_023239581.1;refGene_gn=ARNTL;refGene_tx=XM_023239582.1;refGene_gn=ARNTL;refGene_tx=XM_023239583.1;refGene_gn=ARNTL;refGene_tx=XM_023239584.1;refGene_gn=ARNTL;refGene_tx=XM_023239586.1;refGene_gn=ARNTL;refGene_tx=XM_023239587.1;refGene_gn=ARNTL;refGene_tx=XM_023239588.1;refGene_gn=ARNTL;refGene_tx=XM_023239589.1;refGene_gn=ARNTL;refGene_tx=XM_023239590.1;refGene_gn=ARNTL;refGene_tx=XM_023239591.1
D1 71622826 71622906
refGene_gn=ARNTL;refGene_tx=XM_003993009.4;refGene_gn=ARNTL;refGene_tx=XM_006937112.3;refGene_gn=ARNTL;refGene_tx=XM_019812079.2;refGene_gn=ARNTL;refGene_tx=XM_019812080.2;refGene_gn=ARNTL;refGene_tx=XM_019812084.1;refGene_gn=ARNTL;refGene_tx=XM_023239578.1;refGene_gn=ARNTL;refGene_tx=XM_023239579.1;refGene_gn=ARNTL;refGene_tx=XM_023239580.1;refGene_gn=ARNTL;refGene_tx=XM_023239581.1;refGene_gn=ARNTL;refGene_tx=XM_023239582.1;refGene_gn=ARNTL;refGene_tx=XM_023239583.1;refGene_gn=ARNTL;refGene_tx=XM_023239584.1;refGene_gn=ARNTL;refGene_tx=XM_023239586.1;refGene_gn=ARNTL;refGene_tx=XM_023239587.1;refGene_gn=ARNTL;refGene_tx=XM_023239588.1;refGene_gn=ARNTL;refGene_tx=XM_023239589.1;refGene_gn=A

RNTL;refGene_tx=XM_023239590.1;refGene_gn=ARNTL;refGene_tx=XM_02323959
1.1
D1 71627267 71627360
refGene_gn=ARNTL;refGene_tx=XM_003993009.4;refGene_gn=ARNTL;refGene_tx
=XM_006937112.3;refGene_gn=ARNTL;refGene_tx=XM_019812079.2;refGene_gn=
ARNTL;refGene_tx=XM_019812080.2;refGene_gn=ARNTL;refGene_tx=XM_0198120
84.1;refGene_gn=ARNTL;refGene_tx=XM_023239578.1;refGene_gn=ARNTL;refGe
ne_tx=XM_023239579.1;refGene_gn=ARNTL;refGene_tx=XM_023239580.1;refGen
e_gn=ARNTL;refGene_tx=XM_023239581.1;refGene_gn=ARNTL;refGene_tx=XM_02
3239582.1;refGene_gn=ARNTL;refGene_tx=XM_023239583.1;refGene_gn=ARNTL;
refGene_tx=XM_023239584.1;refGene_gn=ARNTL;refGene_tx=XM_023239586.1;r
efGene_gn=ARNTL;refGene_tx=XM_023239587.1;refGene_gn=ARNTL;refGene_tx=
XM_023239588.1;refGene_gn=ARNTL;refGene_tx=XM_023239589.1;refGene_gn=A
RNTL;refGene_tx=XM_023239590.1;refGene_gn=ARNTL;refGene_tx=XM_02323959
1.1
D1 71628505 71628623
refGene_gn=ARNTL;refGene_tx=XM_003993009.4;refGene_gn=ARNTL;refGene_tx
=XM_006937112.3;refGene_gn=ARNTL;refGene_tx=XM_019812079.2;refGene_gn=
ARNTL;refGene_tx=XM_019812080.2;refGene_gn=ARNTL;refGene_tx=XM_0198120
84.1;refGene_gn=ARNTL;refGene_tx=XM_023239578.1;refGene_gn=ARNTL;refGe
ne_tx=XM_023239579.1;refGene_gn=ARNTL;refGene_tx=XM_023239580.1;refGen
e_gn=ARNTL;refGene_tx=XM_023239581.1;refGene_gn=ARNTL;refGene_tx=XM_02
3239582.1;refGene_gn=ARNTL;refGene_tx=XM_023239583.1;refGene_gn=ARNTL;
refGene_tx=XM_023239584.1;refGene_gn=ARNTL;refGene_tx=XM_023239586.1;r
efGene_gn=ARNTL;refGene_tx=XM_023239587.1;refGene_gn=ARNTL;refGene_tx=
XM_023239588.1;refGene_gn=ARNTL;refGene_tx=XM_023239589.1;refGene_gn=A
RNTL;refGene_tx=XM_023239590.1;refGene_gn=ARNTL;refGene_tx=XM_02323959
1.1
D1 71630852 71631002
refGene_gn=ARNTL;refGene_tx=XM_003993009.4;refGene_gn=ARNTL;refGene_tx
=XM_006937112.3;refGene_gn=ARNTL;refGene_tx=XM_019812079.2;refGene_gn=
ARNTL;refGene_tx=XM_019812080.2;refGene_gn=ARNTL;refGene_tx=XM_0198120
84.1;refGene_gn=ARNTL;refGene_tx=XM_023239578.1;refGene_gn=ARNTL;refGe
ne_tx=XM_023239579.1;refGene_gn=ARNTL;refGene_tx=XM_023239580.1;refGen
e_gn=ARNTL;refGene_tx=XM_023239581.1;refGene_gn=ARNTL;refGene_tx=XM_02
3239582.1;refGene_gn=ARNTL;refGene_tx=XM_023239583.1;refGene_gn=ARNTL;
refGene_tx=XM_023239584.1;refGene_gn=ARNTL;refGene_tx=XM_023239586.1;r
efGene_gn=ARNTL;refGene_tx=XM_023239587.1;refGene_gn=ARNTL;refGene_tx=
XM_023239588.1;refGene_gn=ARNTL;refGene_tx=XM_023239589.1;refGene_gn=A
RNTL;refGene_tx=XM_023239590.1;refGene_gn=ARNTL;refGene_tx=XM_02323959
1.1
D1 71633615 71633862
refGene_gn=ARNTL;refGene_tx=XM_003993009.4;refGene_gn=ARNTL;refGene_tx
=XM_006937112.3;refGene_gn=ARNTL;refGene_tx=XM_019812079.2;refGene_gn=
ARNTL;refGene_tx=XM_019812080.2;refGene_gn=ARNTL;refGene_tx=XM_0198120
84.1;refGene_gn=ARNTL;refGene_tx=XM_023239578.1;refGene_gn=ARNTL;refGe
ne_tx=XM_023239579.1;refGene_gn=ARNTL;refGene_tx=XM_023239580.1;refGen
e_gn=ARNTL;refGene_tx=XM_023239581.1;refGene_gn=ARNTL;refGene_tx=XM_02
3239582.1;refGene_gn=ARNTL;refGene_tx=XM_023239583.1;refGene_gn=ARNTL;
refGene_tx=XM_023239584.1;refGene_gn=ARNTL;refGene_tx=XM_023239586.1;r

efGene_gn=ARNTL;refGene_tx=XM_023239587.1;refGene_gn=ARNTL;refGene_tx=XM_023239588.1;refGene_gn=ARNTL;refGene_tx=XM_023239589.1;refGene_gn=ARNTL;refGene_tx=XM_023239590.1;refGene_gn=ARNTL;refGene_tx=XM_023239591.1
D1 71635383 71635490
refGene_gn=ARNTL;refGene_tx=XM_003993009.4;refGene_gn=ARNTL;refGene_tx=XM_006937112.3;refGene_gn=ARNTL;refGene_tx=XM_019812079.2;refGene_gn=ARNTL;refGene_tx=XM_019812080.2;refGene_gn=ARNTL;refGene_tx=XM_019812084.1;refGene_gn=ARNTL;refGene_tx=XM_023239578.1;refGene_gn=ARNTL;refGene_tx=XM_023239579.1;refGene_gn=ARNTL;refGene_tx=XM_023239580.1;refGene_gn=ARNTL;refGene_tx=XM_023239581.1;refGene_gn=ARNTL;refGene_tx=XM_023239582.1;refGene_gn=ARNTL;refGene_tx=XM_023239583.1;refGene_gn=ARNTL;refGene_tx=XM_023239584.1;refGene_gn=ARNTL;refGene_tx=XM_023239586.1;refGene_gn=ARNTL;refGene_tx=XM_023239587.1;refGene_gn=ARNTL;refGene_tx=XM_023239588.1;refGene_gn=ARNTL;refGene_tx=XM_023239589.1;refGene_gn=ARNTL;refGene_tx=XM_023239590.1;refGene_gn=ARNTL;refGene_tx=XM_023239591.1
D1 71636801 71636952
refGene_gn=ARNTL;refGene_tx=XM_003993009.4;refGene_gn=ARNTL;refGene_tx=XM_006937112.3;refGene_gn=ARNTL;refGene_tx=XM_019812079.2;refGene_gn=ARNTL;refGene_tx=XM_019812080.2;refGene_gn=ARNTL;refGene_tx=XM_019812084.1;refGene_gn=ARNTL;refGene_tx=XM_023239578.1;refGene_gn=ARNTL;refGene_tx=XM_023239579.1;refGene_gn=ARNTL;refGene_tx=XM_023239580.1;refGene_gn=ARNTL;refGene_tx=XM_023239581.1;refGene_gn=ARNTL;refGene_tx=XM_023239582.1;refGene_gn=ARNTL;refGene_tx=XM_023239583.1;refGene_gn=ARNTL;refGene_tx=XM_023239584.1;refGene_gn=ARNTL;refGene_tx=XM_023239586.1;refGene_gn=ARNTL;refGene_tx=XM_023239587.1;refGene_gn=ARNTL;refGene_tx=XM_023239588.1;refGene_gn=ARNTL;refGene_tx=XM_023239589.1;refGene_gn=ARNTL;refGene_tx=XM_023239590.1;refGene_gn=ARNTL;refGene_tx=XM_023239591.1
D1 71637801 71637890
refGene_gn=ARNTL;refGene_tx=XM_003993009.4;refGene_gn=ARNTL;refGene_tx=XM_006937112.3;refGene_gn=ARNTL;refGene_tx=XM_019812079.2;refGene_gn=ARNTL;refGene_tx=XM_019812080.2;refGene_gn=ARNTL;refGene_tx=XM_019812084.1;refGene_gn=ARNTL;refGene_tx=XM_023239578.1;refGene_gn=ARNTL;refGene_tx=XM_023239579.1;refGene_gn=ARNTL;refGene_tx=XM_023239580.1;refGene_gn=ARNTL;refGene_tx=XM_023239581.1;refGene_gn=ARNTL;refGene_tx=XM_023239582.1;refGene_gn=ARNTL;refGene_tx=XM_023239583.1;refGene_gn=ARNTL;refGene_tx=XM_023239584.1;refGene_gn=ARNTL;refGene_tx=XM_023239586.1;refGene_gn=ARNTL;refGene_tx=XM_023239587.1;refGene_gn=ARNTL;refGene_tx=XM_023239588.1;refGene_gn=ARNTL;refGene_tx=XM_023239589.1;refGene_gn=ARNTL;refGene_tx=XM_023239590.1;refGene_gn=ARNTL;refGene_tx=XM_023239591.1
D1 71639399 71639523
refGene_gn=ARNTL;refGene_tx=XM_003993009.4;refGene_gn=ARNTL;refGene_tx=XM_006937112.3;refGene_gn=ARNTL;refGene_tx=XM_019812079.2;refGene_gn=ARNTL;refGene_tx=XM_019812080.2;refGene_gn=ARNTL;refGene_tx=XM_019812084.1;refGene_gn=ARNTL;refGene_tx=XM_023239578.1;refGene_gn=ARNTL;refGene_tx=XM_023239579.1;refGene_gn=ARNTL;refGene_tx=XM_023239580.1;refGene_gn=ARNTL;refGene_tx=XM_023239581.1;refGene_gn=ARNTL;refGene_tx=XM_02

3239582.1;refGene_gn=ARNTL;refGene_tx=XM_023239583.1;refGene_gn=ARNTL;
refGene_tx=XM_023239584.1;refGene_gn=ARNTL;refGene_tx=XM_023239586.1;r
efGene_gn=ARNTL;refGene_tx=XM_023239587.1;refGene_gn=ARNTL;refGene_tx=
XM_023239588.1;refGene_gn=ARNTL;refGene_tx=XM_023239589.1;refGene_gn=A
RNTL;refGene_tx=XM_023239590.1;refGene_gn=ARNTL;refGene_tx=XM_02323959
1.1
D1 71642145 71642239
refGene_gn=ARNTL;refGene_tx=XM_003993009.4;refGene_gn=ARNTL;refGene_tx
=XM_006937112.3;refGene_gn=ARNTL;refGene_tx=XM_019812079.2;refGene_gn=
ARNTL;refGene_tx=XM_019812080.2;refGene_gn=ARNTL;refGene_tx=XM_0198120
84.1;refGene_gn=ARNTL;refGene_tx=XM_023239578.1;refGene_gn=ARNTL;refGe
ne_tx=XM_023239579.1;refGene_gn=ARNTL;refGene_tx=XM_023239580.1;refGen
e_gn=ARNTL;refGene_tx=XM_023239581.1;refGene_gn=ARNTL;refGene_tx=XM_02
3239582.1;refGene_gn=ARNTL;refGene_tx=XM_023239583.1;refGene_gn=ARNTL;
refGene_tx=XM_023239584.1;refGene_gn=ARNTL;refGene_tx=XM_023239586.1;r
efGene_gn=ARNTL;refGene_tx=XM_023239587.1;refGene_gn=ARNTL;refGene_tx=
XM_023239588.1;refGene_gn=ARNTL;refGene_tx=XM_023239589.1;refGene_gn=A
RNTL;refGene_tx=XM_023239590.1;refGene_gn=ARNTL;refGene_tx=XM_02323959
1.1
D1 71645842 71645945
refGene_gn=ARNTL;refGene_tx=XM_003993009.4;refGene_gn=ARNTL;refGene_tx
=XM_006937112.3;refGene_gn=ARNTL;refGene_tx=XM_019812079.2;refGene_gn=
ARNTL;refGene_tx=XM_019812080.2;refGene_gn=ARNTL;refGene_tx=XM_0198120
84.1;refGene_gn=ARNTL;refGene_tx=XM_023239578.1;refGene_gn=ARNTL;refGe
ne_tx=XM_023239579.1;refGene_gn=ARNTL;refGene_tx=XM_023239580.1;refGen
e_gn=ARNTL;refGene_tx=XM_023239581.1;refGene_gn=ARNTL;refGene_tx=XM_02
3239582.1;refGene_gn=ARNTL;refGene_tx=XM_023239583.1;refGene_gn=ARNTL;
refGene_tx=XM_023239584.1;refGene_gn=ARNTL;refGene_tx=XM_023239586.1;r
efGene_gn=ARNTL;refGene_tx=XM_023239587.1;refGene_gn=ARNTL;refGene_tx=
XM_023239588.1;refGene_gn=ARNTL;refGene_tx=XM_023239589.1;refGene_gn=A
RNTL;refGene_tx=XM_023239590.1;refGene_gn=ARNTL;refGene_tx=XM_02323959
1.1
D1 71646636 71646797
refGene_gn=ARNTL;refGene_tx=XM_003993009.4;refGene_gn=ARNTL;refGene_tx
=XM_006937112.3;refGene_gn=ARNTL;refGene_tx=XM_019812079.2;refGene_gn=
ARNTL;refGene_tx=XM_019812080.2;refGene_gn=ARNTL;refGene_tx=XM_0198120
84.1;refGene_gn=ARNTL;refGene_tx=XM_023239578.1;refGene_gn=ARNTL;refGe
ne_tx=XM_023239579.1;refGene_gn=ARNTL;refGene_tx=XM_023239580.1;refGen
e_gn=ARNTL;refGene_tx=XM_023239581.1;refGene_gn=ARNTL;refGene_tx=XM_02
3239582.1;refGene_gn=ARNTL;refGene_tx=XM_023239583.1;refGene_gn=ARNTL;
refGene_tx=XM_023239584.1;refGene_gn=ARNTL;refGene_tx=XM_023239586.1;r
efGene_gn=ARNTL;refGene_tx=XM_023239587.1;refGene_gn=ARNTL;refGene_tx=
XM_023239588.1;refGene_gn=ARNTL;refGene_tx=XM_023239589.1;refGene_gn=A
RNTL;refGene_tx=XM_023239590.1;refGene_gn=ARNTL;refGene_tx=XM_02323959
1.1
D1 71648791 71649102 refGene_gn=BTBD10;refGene_tx=XM_023239725.1
D1 71652078 71652189 refGene_gn=BTBD10;refGene_tx=XM_023239725.1
D1 71654617 71654815 refGene_gn=BTBD10;refGene_tx=XM_023239725.1
D1 71664407 71664528 refGene_gn=BTBD10;refGene_tx=XM_023239725.1
D1 71666315 71666418 refGene_gn=BTBD10;refGene_tx=XM_023239725.1

D1 71669974 71670300 refGene_gn=BTBD10;refGene_tx=XM_023239725.1
D1 71671234 71671248 refGene_gn=BTBD10;refGene_tx=XM_023239725.1
D1 71671533 71671619 refGene_gn=BTBD10;refGene_tx=XM_023239725.1
D1 71707012 71707113 refGene_gn=BTBD10;refGene_tx=XM_023239725.1
D1 71732693 71732965
refGene_gn=LOC111556624;refGene_tx=XM_023239594.1
D1 71744894 71745156
refGene_gn=PTH;refGene_tx=NM_001009210.1;refGene_gn=PTH;refGene_tx=XM_
019811171.2
D1 71745275 71745366
refGene_gn=PTH;refGene_tx=NM_001009210.1;refGene_gn=PTH;refGene_tx=XM_
019811171.2
D1 71747414 71747505 refGene_gn=PTH;refGene_tx=XM_019811171.2
D1 71753306 71753346
refGene_gn=LOC111556624;refGene_tx=XM_023239594.1
D1 71878880 71879534
refGene_gn=LOC111556623;refGene_tx=XM_023239593.1
D1 71921582 71921771
refGene_gn=FAR1;refGene_tx=XM_003993012.5;refGene_gn=FAR1;refGene_tx=X
M_006937119.4;refGene_gn=FAR1;refGene_tx=XM_011286730.3
D1 71925614 71925790
refGene_gn=FAR1;refGene_tx=XM_003993012.5;refGene_gn=FAR1;refGene_tx=X
M_006937119.4;refGene_gn=FAR1;refGene_tx=XM_011286730.3
D1 71932782 71932962
refGene_gn=FAR1;refGene_tx=XM_003993012.5;refGene_gn=FAR1;refGene_tx=X
M_006937119.4;refGene_gn=FAR1;refGene_tx=XM_011286730.3
D1 71935187 71935365
refGene_gn=FAR1;refGene_tx=XM_003993012.5;refGene_gn=FAR1;refGene_tx=X
M_006937119.4;refGene_gn=FAR1;refGene_tx=XM_011286730.3
D1 71936392 71936437
refGene_gn=FAR1;refGene_tx=XM_003993012.5;refGene_gn=FAR1;refGene_tx=X
M_006937119.4;refGene_gn=FAR1;refGene_tx=XM_011286730.3
D1 71936550 71936669
refGene_gn=FAR1;refGene_tx=XM_003993012.5;refGene_gn=FAR1;refGene_tx=X
M_006937119.4;refGene_gn=FAR1;refGene_tx=XM_011286730.3
D1 71937542 71937610
refGene_gn=FAR1;refGene_tx=XM_003993012.5;refGene_gn=FAR1;refGene_tx=X
M_006937119.4;refGene_gn=FAR1;refGene_tx=XM_011286730.3
D1 71939096 71939268
refGene_gn=FAR1;refGene_tx=XM_003993012.5;refGene_gn=FAR1;refGene_tx=X
M_011286730.3
D1 71939577 71939749 refGene_gn=FAR1;refGene_tx=XM_006937119.4
D1 71946954 71947084
refGene_gn=FAR1;refGene_tx=XM_003993012.5;refGene_gn=FAR1;refGene_tx=X
M_006937119.4;refGene_gn=FAR1;refGene_tx=XM_011286730.3
D1 71953154 71953282
refGene_gn=FAR1;refGene_tx=XM_003993012.5;refGene_gn=FAR1;refGene_tx=X
M_006937119.4;refGene_gn=FAR1;refGene_tx=XM_011286730.3
D1 71956032 71956195
refGene_gn=FAR1;refGene_tx=XM_003993012.5;refGene_gn=FAR1;refGene_tx=X

M_006937119.4;refGene_gn=FAR1;refGene_tx=XM_011286730.3
D1 72174089 72174327 refGene_gn=SPON1;refGene_tx=XM_023239596.1
D1 72194308 72194415 refGene_gn=SPON1;refGene_tx=XM_023239596.1
D1 72248109 72248243 refGene_gn=SPON1;refGene_tx=XM_023239596.1
D1 72282298 72282372 refGene_gn=SPON1;refGene_tx=XM_023239596.1
D1 72287605 72287728 refGene_gn=SPON1;refGene_tx=XM_023239596.1
D1 72321711 72321860 refGene_gn=SPON1;refGene_tx=XM_023239596.1
D1 72407086 72407151 refGene_gn=SPON1;refGene_tx=XM_023239596.1
D1 72420799 72421001 refGene_gn=SPON1;refGene_tx=XM_023239596.1
D1 72421963 72422104 refGene_gn=SPON1;refGene_tx=XM_023239596.1
D1 72422567 72422643 refGene_gn=SPON1;refGene_tx=XM_023239596.1
D1 72423626 72423809 refGene_gn=SPON1;refGene_tx=XM_023239596.1
D1 72428865 72429036 refGene_gn=SPON1;refGene_tx=XM_023239596.1
D1 72429128 72429296 refGene_gn=SPON1;refGene_tx=XM_023239596.1
D1 72430030 72430195 refGene_gn=SPON1;refGene_tx=XM_023239596.1
D1 72431924 72432188 refGene_gn=SPON1;refGene_tx=XM_023239596.1
D1 72433791 72433955 refGene_gn=SPON1;refGene_tx=XM_023239596.1
D1 72444653 72444741
refGene_gn=RRAS2;refGene_tx=XM_006937120.3;refGene_gn=RRAS2;refGene_tx
=XM_019812092.2;refGene_gn=RRAS2;refGene_tx=XM_023239597.1
D1 72446893 72447012
refGene_gn=RRAS2;refGene_tx=XM_006937120.3;refGene_gn=RRAS2;refGene_tx
=XM_019812092.2;refGene_gn=RRAS2;refGene_tx=XM_023239597.1
D1 72458114 72458223
refGene_gn=RRAS2;refGene_tx=XM_006937120.3;refGene_gn=RRAS2;refGene_tx
=XM_019812092.2;refGene_gn=RRAS2;refGene_tx=XM_023239597.1
D1 72458392 72458495
refGene_gn=RRAS2;refGene_tx=XM_006937120.3;refGene_gn=RRAS2;refGene_tx
=XM_019812092.2;refGene_gn=RRAS2;refGene_tx=XM_023239597.1
D1 72459445 72459533
refGene_gn=RRAS2;refGene_tx=XM_006937120.3;refGene_gn=RRAS2;refGene_tx
=XM_023239597.1
D1 72479581 72479617 refGene_gn=RRAS2;refGene_tx=XM_006937120.3
D1 72517644 72517752 refGene_gn=RRAS2;refGene_tx=XM_023239597.1
D1 72599816 72599876 refGene_gn=COPB1;refGene_tx=XM_003993013.4
D1 72600562 72600718 refGene_gn=COPB1;refGene_tx=XM_003993013.4
D1 72601869 72601959 refGene_gn=COPB1;refGene_tx=XM_003993013.4
D1 72602540 72602686 refGene_gn=COPB1;refGene_tx=XM_003993013.4
D1 72605363 72605483 refGene_gn=COPB1;refGene_tx=XM_003993013.4
D1 72606545 72606690 refGene_gn=COPB1;refGene_tx=XM_003993013.4
D1 72609882 72610062 refGene_gn=COPB1;refGene_tx=XM_003993013.4
D1 72610218 72610446 refGene_gn=COPB1;refGene_tx=XM_003993013.4
D1 72611696 72611817 refGene_gn=COPB1;refGene_tx=XM_003993013.4
D1 72612672 72612833 refGene_gn=COPB1;refGene_tx=XM_003993013.4
D1 72613913 72614010 refGene_gn=COPB1;refGene_tx=XM_003993013.4
D1 72615226 72615372 refGene_gn=COPB1;refGene_tx=XM_003993013.4
D1 72617017 72617164 refGene_gn=COPB1;refGene_tx=XM_003993013.4
D1 72617253 72617361 refGene_gn=COPB1;refGene_tx=XM_003993013.4
D1 72619186 72619306 refGene_gn=COPB1;refGene_tx=XM_003993013.4
D1 72621873 72622011 refGene_gn=COPB1;refGene_tx=XM_003993013.4

D1 72624867 72624960 refGene_gn=COPB1;refGene_tx=XM_003993013.4
D1 72626191 72626306 refGene_gn=COPB1;refGene_tx=XM_003993013.4
D1 72629101 72629271 refGene_gn=COPB1;refGene_tx=XM_003993013.4
D1 72630012 72630242 refGene_gn=COPB1;refGene_tx=XM_003993013.4
D1 72634892 72634983 refGene_gn=COPB1;refGene_tx=XM_003993013.4
D1 72640625 72640682
refGene_gn=PSMA1;refGene_tx=XM_004001597.4;refGene_gn=PSMA1;refGene_tx=XM_019812094.1
D1 72642694 72642805
refGene_gn=PSMA1;refGene_tx=XM_004001597.4;refGene_gn=PSMA1;refGene_tx=XM_019812094.1
D1 72646153 72646233
refGene_gn=PSMA1;refGene_tx=XM_004001597.4;refGene_gn=PSMA1;refGene_tx=XM_019812094.1
D1 72647148 72647278
refGene_gn=PSMA1;refGene_tx=XM_004001597.4;refGene_gn=PSMA1;refGene_tx=XM_019812094.1
D1 72647446 72647517
refGene_gn=PSMA1;refGene_tx=XM_004001597.4;refGene_gn=PSMA1;refGene_tx=XM_019812094.1
D1 72648380 72648469
refGene_gn=PSMA1;refGene_tx=XM_004001597.4;refGene_gn=PSMA1;refGene_tx=XM_019812094.1
D1 72650727 72650831
refGene_gn=PSMA1;refGene_tx=XM_004001597.4;refGene_gn=PSMA1;refGene_tx=XM_019812094.1
D1 72650951 72651053
refGene_gn=PSMA1;refGene_tx=XM_004001597.4;refGene_gn=PSMA1;refGene_tx=XM_019812094.1
D1 72651510 72651555
refGene_gn=PSMA1;refGene_tx=XM_004001597.4;refGene_gn=PSMA1;refGene_tx=XM_019812094.1
D1 72652781 72652784 refGene_gn=PSMA1;refGene_tx=XM_019812094.1
D1 72652839 72652842 refGene_gn=PSMA1;refGene_tx=XM_004001597.4
D1 72774343 72775321
refGene_gn=PDE3B;refGene_tx=XM_023239599.1;refGene_gn=PDE3B;refGene_tx=XM_023239600.1
D1 72881985 72882036
refGene_gn=PDE3B;refGene_tx=XM_023239599.1;refGene_gn=PDE3B;refGene_tx=XM_023239600.1
D1 72896113 72896362
refGene_gn=PDE3B;refGene_tx=XM_023239599.1;refGene_gn=PDE3B;refGene_tx=XM_023239600.1
D1 72899079 72899222
refGene_gn=PDE3B;refGene_tx=XM_023239599.1;refGene_gn=PDE3B;refGene_tx=XM_023239600.1
D1 72907150 72907257
refGene_gn=PDE3B;refGene_tx=XM_023239599.1;refGene_gn=PDE3B;refGene_tx=XM_023239600.1
D1 72918151 72918362

refGene_gn=PDE3B;refGene_tx=XM_023239599.1;refGene_gn=PDE3B;refGene_tx=XM_023239600.1
D1 72918922 72918999
refGene_gn=PDE3B;refGene_tx=XM_023239599.1;refGene_gn=PDE3B;refGene_tx=XM_023239600.1
D1 72925179 72925331
refGene_gn=PDE3B;refGene_tx=XM_023239599.1;refGene_gn=PDE3B;refGene_tx=XM_023239600.1
D1 72926971 72927109
refGene_gn=PDE3B;refGene_tx=XM_023239599.1;refGene_gn=PDE3B;refGene_tx=XM_023239600.1
D1 72928146 72928258
refGene_gn=PDE3B;refGene_tx=XM_023239599.1;refGene_gn=PDE3B;refGene_tx=XM_023239600.1
D1 72930928 72931042
refGene_gn=PDE3B;refGene_tx=XM_023239599.1;refGene_gn=PDE3B;refGene_tx=XM_023239600.1
D1 72938148 72938348
refGene_gn=PDE3B;refGene_tx=XM_023239599.1;refGene_gn=PDE3B;refGene_tx=XM_023239600.1
D1 72946254 72946458
refGene_gn=PDE3B;refGene_tx=XM_023239599.1;refGene_gn=PDE3B;refGene_tx=XM_023239600.1
D1 72948297 72948459
refGene_gn=PDE3B;refGene_tx=XM_023239599.1;refGene_gn=PDE3B;refGene_tx=XM_023239600.1
D1 72951293 72951549
refGene_gn=PDE3B;refGene_tx=XM_023239599.1;refGene_gn=PDE3B;refGene_tx=XM_023239600.1
D1 72952480 72952695
refGene_gn=PDE3B;refGene_tx=XM_023239599.1;refGene_gn=PDE3B;refGene_tx=XM_023239600.1
D1 72977210 72977386
refGene_gn=LOC101093889;refGene_tx=XM_003993015.5
D1 72978431 72978761
refGene_gn=LOC101093889;refGene_tx=XM_003993015.5
D1 72979388 72980021
refGene_gn=LOC101093889;refGene_tx=XM_003993015.5
D1 72983473 72983615
refGene_gn=LOC101093889;refGene_tx=XM_003993015.5
D1 72995663 72995888
refGene_gn=LOC101093889;refGene_tx=XM_003993015.5
D1 73057871 73057957
refGene_gn=LOC101095582;refGene_tx=XM_019812101.1;refGene_gn=LOC101095582;refGene_tx=XM_019812102.2;refGene_gn=LOC101095582;refGene_tx=XM_023239005.1
D1 73058273 73058411
refGene_gn=LOC101095582;refGene_tx=XM_019812101.1;refGene_gn=LOC101095582;refGene_tx=XM_019812102.2;refGene_gn=LOC101095582;refGene_tx=XM_023239005.1

D1 73059330 73059529
refGene_gn=LOC101095582;refGene_tx=XM_023239005.1
D1 73059631 73059728
refGene_gn=LOC101095582;refGene_tx=XM_019812102.2
D1 73060439 73060605
refGene_gn=LOC101095582;refGene_tx=XM_019812101.1
D1 73200087 73200307
refGene_gn=LOC111556655;refGene_tx=XM_023239726.1
D1 73200733 73200972
refGene_gn=LOC111556655;refGene_tx=XM_023239726.1
D1 73307306 73307312
refGene_gn=CALCB;refGene_tx=XM_019812103.2;refGene_gn=CALCB;refGene_tx=XM_019812104.2
D1 73307433 73307528
refGene_gn=CALCB;refGene_tx=XM_003993016.5;refGene_gn=CALCB;refGene_tx=XM_019812103.2;refGene_gn=CALCB;refGene_tx=XM_019812104.2
D1 73307788 73307935
refGene_gn=CALCB;refGene_tx=XM_003993016.5;refGene_gn=CALCB;refGene_tx=XM_019812103.2;refGene_gn=CALCB;refGene_tx=XM_019812104.2
D1 73308807 73308973
refGene_gn=CALCB;refGene_tx=XM_003993016.5;refGene_gn=CALCB;refGene_tx=XM_019812103.2
D1 73310055 73310215 refGene_gn=CALCB;refGene_tx=XM_019812104.2
D1 73379137 73379193 refGene_gn=INSC;refGene_tx=XM_011286732.3
D1 73403052 73403398 refGene_gn=INSC;refGene_tx=XM_011286732.3
D1 73404357 73404410 refGene_gn=INSC;refGene_tx=XM_011286732.3
D1 73406160 73406284 refGene_gn=INSC;refGene_tx=XM_011286732.3
D1 73421183 73421297 refGene_gn=INSC;refGene_tx=XM_011286732.3
D1 73426982 73427108 refGene_gn=INSC;refGene_tx=XM_011286732.3
D1 73445640 73445812 refGene_gn=INSC;refGene_tx=XM_011286732.3
D1 73449924 73450103 refGene_gn=INSC;refGene_tx=XM_011286732.3
D1 73458261 73458328 refGene_gn=INSC;refGene_tx=XM_011286732.3
D1 73462485 73462641 refGene_gn=INSC;refGene_tx=XM_011286732.3
D1 73464080 73464157 refGene_gn=INSC;refGene_tx=XM_011286732.3
D1 73470661 73470790 refGene_gn=INSC;refGene_tx=XM_011286732.3
D1 74174950 74175254
refGene_gn=SOX6;refGene_tx=XM_019812114.2;refGene_gn=SOX6;refGene_tx=XM_019812118.2;refGene_gn=SOX6;refGene_tx=XM_019812119.2;refGene_gn=SOX6;refGene_tx=XM_019812121.2;refGene_gn=SOX6;refGene_tx=XM_019812123.2;refGene_gn=SOX6;refGene_tx=XM_019812124.2;refGene_gn=SOX6;refGene_tx=XM_023239603.1;refGene_gn=SOX6;refGene_tx=XM_023239604.1;refGene_gn=SOX6;refGene_tx=XM_023239605.1;refGene_gn=SOX6;refGene_tx=XM_023239606.1;refGene_gn=SOX6;refGene_tx=XM_023239607.1;refGene_gn=SOX6;refGene_tx=XM_023239608.1;refGene_gn=SOX6;refGene_tx=XM_023239609.1;refGene_gn=SOX6;refGene_tx=XM_023239610.1;refGene_gn=SOX6;refGene_tx=XM_023239611.1;refGene_gn=SOX6;refGene_tx=XM_023239612.1;refGene_gn=SOX6;refGene_tx=XM_023239613.1
D1 74186543 74186760
refGene_gn=SOX6;refGene_tx=XM_019812114.2;refGene_gn=SOX6;refGene_tx=XM_019812118.2;refGene_gn=SOX6;refGene_tx=XM_019812119.2;refGene_gn=SOX

6;refGene_tx=XM_019812121.2;refGene_gn=SOX6;refGene_tx=XM_019812123.2;
refGene_gn=SOX6;refGene_tx=XM_019812124.2;refGene_gn=SOX6;refGene_tx=X
M_023239603.1;refGene_gn=SOX6;refGene_tx=XM_023239604.1;refGene_gn=SOX
6;refGene_tx=XM_023239605.1;refGene_gn=SOX6;refGene_tx=XM_023239606.1;
refGene_gn=SOX6;refGene_tx=XM_023239607.1;refGene_gn=SOX6;refGene_tx=X
M_023239608.1;refGene_gn=SOX6;refGene_tx=XM_023239609.1;refGene_gn=SOX
6;refGene_tx=XM_023239610.1;refGene_gn=SOX6;refGene_tx=XM_023239611.1;
refGene_gn=SOX6;refGene_tx=XM_023239612.1;refGene_gn=SOX6;refGene_tx=X
M_023239613.1
D1 74189472 74189706
refGene_gn=SOX6;refGene_tx=XM_019812114.2;refGene_gn=SOX6;refGene_tx=X
M_019812118.2;refGene_gn=SOX6;refGene_tx=XM_019812119.2;refGene_gn=SOX
6;refGene_tx=XM_019812121.2;refGene_gn=SOX6;refGene_tx=XM_019812123.2;
refGene_gn=SOX6;refGene_tx=XM_019812124.2;refGene_gn=SOX6;refGene_tx=X
M_023239603.1;refGene_gn=SOX6;refGene_tx=XM_023239604.1;refGene_gn=SOX
6;refGene_tx=XM_023239605.1;refGene_gn=SOX6;refGene_tx=XM_023239606.1;
refGene_gn=SOX6;refGene_tx=XM_023239607.1;refGene_gn=SOX6;refGene_tx=X
M_023239608.1;refGene_gn=SOX6;refGene_tx=XM_023239609.1;refGene_gn=SOX
6;refGene_tx=XM_023239610.1;refGene_gn=SOX6;refGene_tx=XM_023239611.1;
refGene_gn=SOX6;refGene_tx=XM_023239612.1;refGene_gn=SOX6;refGene_tx=X
M_023239613.1
D1 74214624 74214733
refGene_gn=SOX6;refGene_tx=XM_019812114.2;refGene_gn=SOX6;refGene_tx=X
M_019812118.2;refGene_gn=SOX6;refGene_tx=XM_019812119.2;refGene_gn=SOX
6;refGene_tx=XM_019812121.2;refGene_gn=SOX6;refGene_tx=XM_019812123.2;
refGene_gn=SOX6;refGene_tx=XM_019812124.2;refGene_gn=SOX6;refGene_tx=X
M_023239603.1;refGene_gn=SOX6;refGene_tx=XM_023239604.1;refGene_gn=SOX
6;refGene_tx=XM_023239605.1;refGene_gn=SOX6;refGene_tx=XM_023239606.1;
refGene_gn=SOX6;refGene_tx=XM_023239607.1;refGene_gn=SOX6;refGene_tx=X
M_023239608.1;refGene_gn=SOX6;refGene_tx=XM_023239609.1;refGene_gn=SOX
6;refGene_tx=XM_023239610.1;refGene_gn=SOX6;refGene_tx=XM_023239611.1;
refGene_gn=SOX6;refGene_tx=XM_023239612.1;refGene_gn=SOX6;refGene_tx=X
M_023239613.1
D1 74249264 74249452
refGene_gn=SOX6;refGene_tx=XM_019812114.2;refGene_gn=SOX6;refGene_tx=X
M_019812118.2;refGene_gn=SOX6;refGene_tx=XM_019812119.2;refGene_gn=SOX
6;refGene_tx=XM_019812121.2;refGene_gn=SOX6;refGene_tx=XM_019812123.2;
refGene_gn=SOX6;refGene_tx=XM_019812124.2;refGene_gn=SOX6;refGene_tx=X
M_023239603.1;refGene_gn=SOX6;refGene_tx=XM_023239604.1;refGene_gn=SOX
6;refGene_tx=XM_023239605.1;refGene_gn=SOX6;refGene_tx=XM_023239606.1;
refGene_gn=SOX6;refGene_tx=XM_023239607.1;refGene_gn=SOX6;refGene_tx=X
M_023239608.1;refGene_gn=SOX6;refGene_tx=XM_023239609.1;refGene_gn=SOX
6;refGene_tx=XM_023239610.1;refGene_gn=SOX6;refGene_tx=XM_023239611.1;
refGene_gn=SOX6;refGene_tx=XM_023239612.1;refGene_gn=SOX6;refGene_tx=X
M_023239613.1
D1 74252492 74252676
refGene_gn=SOX6;refGene_tx=XM_019812114.2;refGene_gn=SOX6;refGene_tx=X
M_019812118.2;refGene_gn=SOX6;refGene_tx=XM_019812119.2;refGene_gn=SOX
6;refGene_tx=XM_019812121.2;refGene_gn=SOX6;refGene_tx=XM_019812123.2;
refGene_gn=SOX6;refGene_tx=XM_019812124.2;refGene_gn=SOX6;refGene_tx=X

M_023239603.1;refGene_gn=SOX6;refGene_tx=XM_023239604.1;refGene_gn=SOX
6;refGene_tx=XM_023239605.1;refGene_gn=SOX6;refGene_tx=XM_023239606.1;
refGene_gn=SOX6;refGene_tx=XM_023239607.1;refGene_gn=SOX6;refGene_tx=X
M_023239608.1;refGene_gn=SOX6;refGene_tx=XM_023239609.1;refGene_gn=SOX
6;refGene_tx=XM_023239610.1;refGene_gn=SOX6;refGene_tx=XM_023239611.1;
refGene_gn=SOX6;refGene_tx=XM_023239612.1;refGene_gn=SOX6;refGene_tx=X
M_023239613.1
D1 74258317 74258467
refGene_gn=SOX6;refGene_tx=XM_019812114.2;refGene_gn=SOX6;refGene_tx=X
M_019812118.2;refGene_gn=SOX6;refGene_tx=XM_019812119.2;refGene_gn=SOX
6;refGene_tx=XM_019812121.2;refGene_gn=SOX6;refGene_tx=XM_019812123.2;
refGene_gn=SOX6;refGene_tx=XM_019812124.2;refGene_gn=SOX6;refGene_tx=X
M_023239603.1;refGene_gn=SOX6;refGene_tx=XM_023239604.1;refGene_gn=SOX
6;refGene_tx=XM_023239605.1;refGene_gn=SOX6;refGene_tx=XM_023239606.1;
refGene_gn=SOX6;refGene_tx=XM_023239607.1;refGene_gn=SOX6;refGene_tx=X
M_023239608.1;refGene_gn=SOX6;refGene_tx=XM_023239609.1;refGene_gn=SOX
6;refGene_tx=XM_023239610.1;refGene_gn=SOX6;refGene_tx=XM_023239611.1;
refGene_gn=SOX6;refGene_tx=XM_023239612.1;refGene_gn=SOX6;refGene_tx=X
M_023239613.1
D1 74286403 74286526
refGene_gn=SOX6;refGene_tx=XM_019812114.2;refGene_gn=SOX6;refGene_tx=X
M_019812118.2;refGene_gn=SOX6;refGene_tx=XM_019812119.2;refGene_gn=SOX
6;refGene_tx=XM_023239603.1;refGene_gn=SOX6;refGene_tx=XM_023239604.1;
refGene_gn=SOX6;refGene_tx=XM_023239605.1;refGene_gn=SOX6;refGene_tx=X
M_023239606.1;refGene_gn=SOX6;refGene_tx=XM_023239607.1;refGene_gn=SOX
6;refGene_tx=XM_023239608.1;refGene_gn=SOX6;refGene_tx=XM_023239609.1;
refGene_gn=SOX6;refGene_tx=XM_023239610.1;refGene_gn=SOX6;refGene_tx=X
M_023239611.1
D1 74288003 74288083
refGene_gn=SOX6;refGene_tx=XM_019812114.2;refGene_gn=SOX6;refGene_tx=X
M_019812118.2;refGene_gn=SOX6;refGene_tx=XM_019812119.2;refGene_gn=SOX
6;refGene_tx=XM_019812121.2;refGene_gn=SOX6;refGene_tx=XM_019812123.2;
refGene_gn=SOX6;refGene_tx=XM_019812124.2;refGene_gn=SOX6;refGene_tx=X
M_023239603.1;refGene_gn=SOX6;refGene_tx=XM_023239604.1;refGene_gn=SOX
6;refGene_tx=XM_023239605.1;refGene_gn=SOX6;refGene_tx=XM_023239606.1;
refGene_gn=SOX6;refGene_tx=XM_023239607.1;refGene_gn=SOX6;refGene_tx=X
M_023239608.1;refGene_gn=SOX6;refGene_tx=XM_023239609.1;refGene_gn=SOX
6;refGene_tx=XM_023239610.1;refGene_gn=SOX6;refGene_tx=XM_023239611.1;
refGene_gn=SOX6;refGene_tx=XM_023239612.1;refGene_gn=SOX6;refGene_tx=X
M_023239613.1
D1 74299135 74299259
refGene_gn=SOX6;refGene_tx=XM_019812114.2;refGene_gn=SOX6;refGene_tx=X
M_019812118.2;refGene_gn=SOX6;refGene_tx=XM_019812119.2;refGene_gn=SOX
6;refGene_tx=XM_019812121.2;refGene_gn=SOX6;refGene_tx=XM_019812123.2;
refGene_gn=SOX6;refGene_tx=XM_019812124.2;refGene_gn=SOX6;refGene_tx=X
M_023239603.1;refGene_gn=SOX6;refGene_tx=XM_023239604.1;refGene_gn=SOX
6;refGene_tx=XM_023239605.1;refGene_gn=SOX6;refGene_tx=XM_023239606.1;
refGene_gn=SOX6;refGene_tx=XM_023239607.1;refGene_gn=SOX6;refGene_tx=X
M_023239608.1;refGene_gn=SOX6;refGene_tx=XM_023239609.1;refGene_gn=SOX
6;refGene_tx=XM_023239610.1;refGene_gn=SOX6;refGene_tx=XM_023239611.1;

refGene_gn=SOX6;refGene_tx=XM_023239612.1;refGene_gn=SOX6;refGene_tx=XM_023239613.1
D1 74371067 74371136
refGene_gn=SOX6;refGene_tx=XM_019812114.2;refGene_gn=SOX6;refGene_tx=XM_019812118.2;refGene_gn=SOX6;refGene_tx=XM_019812119.2;refGene_gn=SOX6;refGene_tx=XM_019812121.2;refGene_gn=SOX6;refGene_tx=XM_019812123.2;refGene_gn=SOX6;refGene_tx=XM_019812124.2;refGene_gn=SOX6;refGene_tx=XM_023239603.1;refGene_gn=SOX6;refGene_tx=XM_023239604.1;refGene_gn=SOX6;refGene_tx=XM_023239605.1;refGene_gn=SOX6;refGene_tx=XM_023239606.1;refGene_gn=SOX6;refGene_tx=XM_023239607.1;refGene_gn=SOX6;refGene_tx=XM_023239608.1;refGene_gn=SOX6;refGene_tx=XM_023239609.1;refGene_gn=SOX6;refGene_tx=XM_023239610.1;refGene_gn=SOX6;refGene_tx=XM_023239611.1;refGene_gn=SOX6;refGene_tx=XM_023239612.1;refGene_gn=SOX6;refGene_tx=XM_023239613.1
D1 74373987 74374160
refGene_gn=SOX6;refGene_tx=XM_019812114.2;refGene_gn=SOX6;refGene_tx=XM_019812118.2;refGene_gn=SOX6;refGene_tx=XM_019812119.2;refGene_gn=SOX6;refGene_tx=XM_019812121.2;refGene_gn=SOX6;refGene_tx=XM_019812123.2;refGene_gn=SOX6;refGene_tx=XM_019812124.2;refGene_gn=SOX6;refGene_tx=XM_023239603.1;refGene_gn=SOX6;refGene_tx=XM_023239604.1;refGene_gn=SOX6;refGene_tx=XM_023239605.1;refGene_gn=SOX6;refGene_tx=XM_023239606.1;refGene_gn=SOX6;refGene_tx=XM_023239607.1;refGene_gn=SOX6;refGene_tx=XM_023239608.1;refGene_gn=SOX6;refGene_tx=XM_023239609.1;refGene_gn=SOX6;refGene_tx=XM_023239610.1;refGene_gn=SOX6;refGene_tx=XM_023239611.1;refGene_gn=SOX6;refGene_tx=XM_023239612.1;refGene_gn=SOX6;refGene_tx=XM_023239613.1
D1 74426911 74427001
refGene_gn=SOX6;refGene_tx=XM_019812114.2;refGene_gn=SOX6;refGene_tx=XM_019812118.2;refGene_gn=SOX6;refGene_tx=XM_019812119.2;refGene_gn=SOX6;refGene_tx=XM_019812121.2;refGene_gn=SOX6;refGene_tx=XM_019812123.2;refGene_gn=SOX6;refGene_tx=XM_019812124.2;refGene_gn=SOX6;refGene_tx=XM_023239603.1;refGene_gn=SOX6;refGene_tx=XM_023239604.1;refGene_gn=SOX6;refGene_tx=XM_023239605.1;refGene_gn=SOX6;refGene_tx=XM_023239606.1;refGene_gn=SOX6;refGene_tx=XM_023239607.1;refGene_gn=SOX6;refGene_tx=XM_023239608.1;refGene_gn=SOX6;refGene_tx=XM_023239609.1;refGene_gn=SOX6;refGene_tx=XM_023239610.1;refGene_gn=SOX6;refGene_tx=XM_023239611.1;refGene_gn=SOX6;refGene_tx=XM_023239612.1;refGene_gn=SOX6;refGene_tx=XM_023239613.1
D1 74507834 74508042
refGene_gn=SOX6;refGene_tx=XM_019812114.2;refGene_gn=SOX6;refGene_tx=XM_019812118.2;refGene_gn=SOX6;refGene_tx=XM_019812119.2;refGene_gn=SOX6;refGene_tx=XM_019812121.2;refGene_gn=SOX6;refGene_tx=XM_019812123.2;refGene_gn=SOX6;refGene_tx=XM_019812124.2;refGene_gn=SOX6;refGene_tx=XM_023239603.1;refGene_gn=SOX6;refGene_tx=XM_023239604.1;refGene_gn=SOX6;refGene_tx=XM_023239605.1;refGene_gn=SOX6;refGene_tx=XM_023239606.1;refGene_gn=SOX6;refGene_tx=XM_023239607.1;refGene_gn=SOX6;refGene_tx=XM_023239608.1;refGene_gn=SOX6;refGene_tx=XM_023239609.1;refGene_gn=SOX6;refGene_tx=XM_023239610.1;refGene_gn=SOX6;refGene_tx=XM_023239611.1;refGene_gn=SOX6;refGene_tx=XM_023239612.1;refGene_gn=SOX6;refGene_tx=XM_023239613.1

D1 74527659 74527900
refGene_gn=SOX6;refGene_tx=XM_019812114.2;refGene_gn=SOX6;refGene_tx=XM_019812118.2;refGene_gn=SOX6;refGene_tx=XM_019812119.2;refGene_gn=SOX6;refGene_tx=XM_019812121.2;refGene_gn=SOX6;refGene_tx=XM_019812123.2;refGene_gn=SOX6;refGene_tx=XM_019812124.2;refGene_gn=SOX6;refGene_tx=XM_023239603.1;refGene_gn=SOX6;refGene_tx=XM_023239604.1;refGene_gn=SOX6;refGene_tx=XM_023239605.1;refGene_gn=SOX6;refGene_tx=XM_023239606.1;refGene_gn=SOX6;refGene_tx=XM_023239607.1;refGene_gn=SOX6;refGene_tx=XM_023239608.1;refGene_gn=SOX6;refGene_tx=XM_023239609.1;refGene_gn=SOX6;refGene_tx=XM_023239610.1;refGene_gn=SOX6;refGene_tx=XM_023239611.1;refGene_gn=SOX6;refGene_tx=XM_023239612.1;refGene_gn=SOX6;refGene_tx=XM_023239613.1
D1 74543359 74543436
refGene_gn=SOX6;refGene_tx=XM_019812114.2;refGene_gn=SOX6;refGene_tx=XM_019812118.2;refGene_gn=SOX6;refGene_tx=XM_019812119.2;refGene_gn=SOX6;refGene_tx=XM_019812121.2;refGene_gn=SOX6;refGene_tx=XM_019812123.2;refGene_gn=SOX6;refGene_tx=XM_019812124.2;refGene_gn=SOX6;refGene_tx=XM_023239603.1;refGene_gn=SOX6;refGene_tx=XM_023239604.1;refGene_gn=SOX6;refGene_tx=XM_023239612.1;refGene_gn=SOX6;refGene_tx=XM_023239613.1
D1 74863648 74863711
refGene_gn=CD1H11orf58;refGene_tx=XM_003993022.5
D1 74868531 74868615
refGene_gn=CD1H11orf58;refGene_tx=XM_003993022.5
D1 74871778 74871839
refGene_gn=CD1H11orf58;refGene_tx=XM_003993022.5
D1 74873439 74873549
refGene_gn=CD1H11orf58;refGene_tx=XM_003993022.5
D1 74874961 74875192
refGene_gn=CD1H11orf58;refGene_tx=XM_003993022.5
D1 74896339 74896362
refGene_gn=PLEKHA7;refGene_tx=XM_023239616.1;refGene_gn=PLEKHA7;refGene_tx=XM_023239618.1;refGene_gn=PLEKHA7;refGene_tx=XM_023239619.1;refGene_gn=PLEKHA7;refGene_tx=XM_023239620.1;refGene_gn=PLEKHA7;refGene_tx=XM_023239621.1
D1 74898319 74898420
refGene_gn=PLEKHA7;refGene_tx=XM_019812131.2;refGene_gn=PLEKHA7;refGene_tx=XM_019812137.2;refGene_gn=PLEKHA7;refGene_tx=XM_023239614.1;refGene_gn=PLEKHA7;refGene_tx=XM_023239615.1;refGene_gn=PLEKHA7;refGene_tx=XM_023239617.1
D1 74899997 74900140
refGene_gn=PLEKHA7;refGene_tx=XM_019812131.2;refGene_gn=PLEKHA7;refGene_tx=XM_019812137.2;refGene_gn=PLEKHA7;refGene_tx=XM_023239614.1;refGene_gn=PLEKHA7;refGene_tx=XM_023239615.1;refGene_gn=PLEKHA7;refGene_tx=XM_023239616.1;refGene_gn=PLEKHA7;refGene_tx=XM_023239617.1;refGene_gn=PLEKHA7;refGene_tx=XM_023239618.1;refGene_gn=PLEKHA7;refGene_tx=XM_023239619.1
D1 74900921 74901055
refGene_gn=PLEKHA7;refGene_tx=XM_019812131.2;refGene_gn=PLEKHA7;refGene_tx=XM_019812137.2;refGene_gn=PLEKHA7;refGene_tx=XM_023239614.1;refGene_gn=PLEKHA7;refGene_tx=XM_023239615.1;refGene_gn=PLEKHA7;refGene_tx=

XM_023239616.1;refGene_gn=PLEKHA7;refGene_tx=XM_023239617.1;refGene_gn=PLEKHA7;refGene_tx=XM_023239618.1;refGene_gn=PLEKHA7;refGene_tx=XM_023239619.1
D1 74903282 74903441
refGene_gn=PLEKHA7;refGene_tx=XM_019812131.2;refGene_gn=PLEKHA7;refGene_tx=XM_019812137.2;refGene_gn=PLEKHA7;refGene_tx=XM_023239614.1;refGene_gn=PLEKHA7;refGene_tx=XM_023239615.1;refGene_gn=PLEKHA7;refGene_tx=XM_023239616.1;refGene_gn=PLEKHA7;refGene_tx=XM_023239617.1;refGene_gn=PLEKHA7;refGene_tx=XM_023239618.1;refGene_gn=PLEKHA7;refGene_tx=XM_023239619.1;refGene_gn=PLEKHA7;refGene_tx=XM_023239620.1;refGene_gn=PLEKHA7;refGene_tx=XM_023239621.1
D1 74906233 74906443
refGene_gn=PLEKHA7;refGene_tx=XM_019812131.2;refGene_gn=PLEKHA7;refGene_tx=XM_019812137.2;refGene_gn=PLEKHA7;refGene_tx=XM_023239614.1;refGene_gn=PLEKHA7;refGene_tx=XM_023239615.1;refGene_gn=PLEKHA7;refGene_tx=XM_023239616.1;refGene_gn=PLEKHA7;refGene_tx=XM_023239617.1;refGene_gn=PLEKHA7;refGene_tx=XM_023239618.1;refGene_gn=PLEKHA7;refGene_tx=XM_023239619.1;refGene_gn=PLEKHA7;refGene_tx=XM_023239620.1;refGene_gn=PLEKHA7;refGene_tx=XM_023239621.1;refGene_gn=PLEKHA7;refGene_tx=XM_023239622.1;refGene_gn=PLEKHA7;refGene_tx=XM_023239623.1
D1 74907051 74907155
refGene_gn=PLEKHA7;refGene_tx=XM_019812131.2;refGene_gn=PLEKHA7;refGene_tx=XM_019812137.2;refGene_gn=PLEKHA7;refGene_tx=XM_023239614.1;refGene_gn=PLEKHA7;refGene_tx=XM_023239615.1;refGene_gn=PLEKHA7;refGene_tx=XM_023239616.1;refGene_gn=PLEKHA7;refGene_tx=XM_023239617.1;refGene_gn=PLEKHA7;refGene_tx=XM_023239618.1;refGene_gn=PLEKHA7;refGene_tx=XM_023239619.1;refGene_gn=PLEKHA7;refGene_tx=XM_023239620.1;refGene_gn=PLEKHA7;refGene_tx=XM_023239621.1;refGene_gn=PLEKHA7;refGene_tx=XM_023239622.1;refGene_gn=PLEKHA7;refGene_tx=XM_023239623.1;refGene_gn=PLEKHA7;refGene_tx=XM_023239624.1
D1 74909147 74909265
refGene_gn=PLEKHA7;refGene_tx=XM_019812131.2;refGene_gn=PLEKHA7;refGene_tx=XM_019812137.2;refGene_gn=PLEKHA7;refGene_tx=XM_023239614.1;refGene_gn=PLEKHA7;refGene_tx=XM_023239615.1;refGene_gn=PLEKHA7;refGene_tx=XM_023239616.1;refGene_gn=PLEKHA7;refGene_tx=XM_023239617.1;refGene_gn=PLEKHA7;refGene_tx=XM_023239618.1;refGene_gn=PLEKHA7;refGene_tx=XM_023239619.1;refGene_gn=PLEKHA7;refGene_tx=XM_023239620.1;refGene_gn=PLEKHA7;refGene_tx=XM_023239621.1;refGene_gn=PLEKHA7;refGene_tx=XM_023239622.1
D1 74909364 74909553
refGene_gn=PLEKHA7;refGene_tx=XM_019812131.2;refGene_gn=PLEKHA7;refGene_tx=XM_019812137.2;refGene_gn=PLEKHA7;refGene_tx=XM_023239614.1;refGene_gn=PLEKHA7;refGene_tx=XM_023239615.1;refGene_gn=PLEKHA7;refGene_tx=XM_023239616.1;refGene_gn=PLEKHA7;refGene_tx=XM_023239617.1;refGene_gn=PLEKHA7;refGene_tx=XM_023239618.1;refGene_gn=PLEKHA7;refGene_tx=XM_023239619.1;refGene_gn=PLEKHA7;refGene_tx=XM_023239620.1;refGene_gn=PLEKHA7;refGene_tx=XM_023239621.1;refGene_gn=PLEKHA7;refGene_tx=XM_023239622.1;refGene_gn=PLEKHA7;refGene_tx=XM_023239623.1;refGene_gn=PLEKHA7;refGene_tx=XM_023239624.1
D1 74912535 74912762

refGene_gn=PLEKHA7;refGene_tx=XM_019812131.2;refGene_gn=PLEKHA7;refGene_tx=XM_019812137.2;refGene_gn=PLEKHA7;refGene_tx=XM_023239614.1;refGene_gn=PLEKHA7;refGene_tx=XM_023239615.1;refGene_gn=PLEKHA7;refGene_tx=XM_023239616.1;refGene_gn=PLEKHA7;refGene_tx=XM_023239617.1;refGene_gn=PLEKHA7;refGene_tx=XM_023239618.1;refGene_gn=PLEKHA7;refGene_tx=XM_023239619.1;refGene_gn=PLEKHA7;refGene_tx=XM_023239620.1;refGene_gn=PLEKHA7;refGene_tx=XM_023239621.1;refGene_gn=PLEKHA7;refGene_tx=XM_023239622.1;refGene_gn=PLEKHA7;refGene_tx=XM_023239623.1;refGene_gn=PLEKHA7;refGene_tx=XM_023239624.1
D1 74912927 74913036
refGene_gn=PLEKHA7;refGene_tx=XM_019812131.2;refGene_gn=PLEKHA7;refGene_tx=XM_019812137.2;refGene_gn=PLEKHA7;refGene_tx=XM_023239614.1;refGene_gn=PLEKHA7;refGene_tx=XM_023239615.1;refGene_gn=PLEKHA7;refGene_tx=XM_023239616.1;refGene_gn=PLEKHA7;refGene_tx=XM_023239617.1;refGene_gn=PLEKHA7;refGene_tx=XM_023239618.1;refGene_gn=PLEKHA7;refGene_tx=XM_023239619.1;refGene_gn=PLEKHA7;refGene_tx=XM_023239620.1;refGene_gn=PLEKHA7;refGene_tx=XM_023239621.1;refGene_gn=PLEKHA7;refGene_tx=XM_023239622.1;refGene_gn=PLEKHA7;refGene_tx=XM_023239623.1;refGene_gn=PLEKHA7;refGene_tx=XM_023239624.1
D1 74919812 74919914
refGene_gn=PLEKHA7;refGene_tx=XM_019812131.2;refGene_gn=PLEKHA7;refGene_tx=XM_019812137.2;refGene_gn=PLEKHA7;refGene_tx=XM_023239614.1;refGene_gn=PLEKHA7;refGene_tx=XM_023239615.1;refGene_gn=PLEKHA7;refGene_tx=XM_023239616.1;refGene_gn=PLEKHA7;refGene_tx=XM_023239617.1;refGene_gn=PLEKHA7;refGene_tx=XM_023239618.1;refGene_gn=PLEKHA7;refGene_tx=XM_023239619.1;refGene_gn=PLEKHA7;refGene_tx=XM_023239620.1;refGene_gn=PLEKHA7;refGene_tx=XM_023239621.1;refGene_gn=PLEKHA7;refGene_tx=XM_023239622.1;refGene_gn=PLEKHA7;refGene_tx=XM_023239623.1;refGene_gn=PLEKHA7;refGene_tx=XM_023239624.1
D1 74920453 74920603
refGene_gn=PLEKHA7;refGene_tx=XM_019812131.2;refGene_gn=PLEKHA7;refGene_tx=XM_019812137.2;refGene_gn=PLEKHA7;refGene_tx=XM_023239614.1;refGene_gn=PLEKHA7;refGene_tx=XM_023239615.1;refGene_gn=PLEKHA7;refGene_tx=XM_023239616.1;refGene_gn=PLEKHA7;refGene_tx=XM_023239617.1;refGene_gn=PLEKHA7;refGene_tx=XM_023239618.1;refGene_gn=PLEKHA7;refGene_tx=XM_023239619.1;refGene_gn=PLEKHA7;refGene_tx=XM_023239620.1;refGene_gn=PLEKHA7;refGene_tx=XM_023239621.1;refGene_gn=PLEKHA7;refGene_tx=XM_023239622.1;refGene_gn=PLEKHA7;refGene_tx=XM_023239623.1;refGene_gn=PLEKHA7;refGene_tx=XM_023239624.1
D1 74921492 74921573
refGene_gn=PLEKHA7;refGene_tx=XM_019812131.2;refGene_gn=PLEKHA7;refGene_tx=XM_019812137.2;refGene_gn=PLEKHA7;refGene_tx=XM_023239614.1;refGene_gn=PLEKHA7;refGene_tx=XM_023239615.1;refGene_gn=PLEKHA7;refGene_tx=XM_023239616.1;refGene_gn=PLEKHA7;refGene_tx=XM_023239617.1;refGene_gn=PLEKHA7;refGene_tx=XM_023239618.1;refGene_gn=PLEKHA7;refGene_tx=XM_023239619.1;refGene_gn=PLEKHA7;refGene_tx=XM_023239620.1;refGene_gn=PLEKHA7;refGene_tx=XM_023239621.1;refGene_gn=PLEKHA7;refGene_tx=XM_023239622.1;refGene_gn=PLEKHA7;refGene_tx=XM_023239623.1;refGene_gn=PLEKHA7;refGene_tx=XM_023239624.1
D1 74921753 74921822

refGene_gn=PLEKHA7;refGene_tx=XM_019812131.2;refGene_gn=PLEKHA7;refGen
e_tx=XM_019812137.2;refGene_gn=PLEKHA7;refGene_tx=XM_023239614.1;refGe
ne_gn=PLEKHA7;refGene_tx=XM_023239615.1;refGene_gn=PLEKHA7;refGene_tx=
XM_023239616.1;refGene_gn=PLEKHA7;refGene_tx=XM_023239617.1;refGene_gn
=PLEKHA7;refGene_tx=XM_023239618.1;refGene_gn=PLEKHA7;refGene_tx=XM_02
3239619.1;refGene_gn=PLEKHA7;refGene_tx=XM_023239620.1;refGene_gn=PLEK
HA7;refGene_tx=XM_023239621.1;refGene_gn=PLEKHA7;refGene_tx=XM_0232396
22.1;refGene_gn=PLEKHA7;refGene_tx=XM_023239623.1;refGene_gn=PLEKHA7;r
efGene_tx=XM_023239624.1
D1 74925692 74925773
refGene_gn=PLEKHA7;refGene_tx=XM_019812131.2;refGene_gn=PLEKHA7;refGen
e_tx=XM_019812137.2;refGene_gn=PLEKHA7;refGene_tx=XM_023239614.1;refGe
ne_gn=PLEKHA7;refGene_tx=XM_023239615.1;refGene_gn=PLEKHA7;refGene_tx=
XM_023239616.1;refGene_gn=PLEKHA7;refGene_tx=XM_023239618.1;refGene_gn
=PLEKHA7;refGene_tx=XM_023239620.1;refGene_gn=PLEKHA7;refGene_tx=XM_02
3239621.1;refGene_gn=PLEKHA7;refGene_tx=XM_023239622.1;refGene_gn=PLEK
HA7;refGene_tx=XM_023239623.1;refGene_gn=PLEKHA7;refGene_tx=XM_0232396
24.1
D1 74931434 74931488
refGene_gn=PLEKHA7;refGene_tx=XM_019812131.2;refGene_gn=PLEKHA7;refGen
e_tx=XM_019812137.2;refGene_gn=PLEKHA7;refGene_tx=XM_023239614.1;refGe
ne_gn=PLEKHA7;refGene_tx=XM_023239615.1;refGene_gn=PLEKHA7;refGene_tx=
XM_023239616.1;refGene_gn=PLEKHA7;refGene_tx=XM_023239617.1;refGene_gn
=PLEKHA7;refGene_tx=XM_023239618.1;refGene_gn=PLEKHA7;refGene_tx=XM_02
3239619.1;refGene_gn=PLEKHA7;refGene_tx=XM_023239620.1;refGene_gn=PLEK
HA7;refGene_tx=XM_023239621.1;refGene_gn=PLEKHA7;refGene_tx=XM_0232396
22.1;refGene_gn=PLEKHA7;refGene_tx=XM_023239623.1;refGene_gn=PLEKHA7;r
efGene_tx=XM_023239624.1
D1 74934400 74934487
refGene_gn=PLEKHA7;refGene_tx=XM_019812131.2;refGene_gn=PLEKHA7;refGen
e_tx=XM_019812137.2;refGene_gn=PLEKHA7;refGene_tx=XM_023239614.1;refGe
ne_gn=PLEKHA7;refGene_tx=XM_023239615.1;refGene_gn=PLEKHA7;refGene_tx=
XM_023239616.1;refGene_gn=PLEKHA7;refGene_tx=XM_023239617.1;refGene_gn
=PLEKHA7;refGene_tx=XM_023239618.1;refGene_gn=PLEKHA7;refGene_tx=XM_02
3239619.1;refGene_gn=PLEKHA7;refGene_tx=XM_023239620.1;refGene_gn=PLEK
HA7;refGene_tx=XM_023239621.1;refGene_gn=PLEKHA7;refGene_tx=XM_0232396
22.1;refGene_gn=PLEKHA7;refGene_tx=XM_023239623.1;refGene_gn=PLEKHA7;r
efGene_tx=XM_023239624.1
D1 74934983 74935506
refGene_gn=PLEKHA7;refGene_tx=XM_019812131.2;refGene_gn=PLEKHA7;refGen
e_tx=XM_019812137.2;refGene_gn=PLEKHA7;refGene_tx=XM_023239614.1;refGe
ne_gn=PLEKHA7;refGene_tx=XM_023239615.1;refGene_gn=PLEKHA7;refGene_tx=
XM_023239616.1;refGene_gn=PLEKHA7;refGene_tx=XM_023239617.1;refGene_gn
=PLEKHA7;refGene_tx=XM_023239618.1;refGene_gn=PLEKHA7;refGene_tx=XM_02
3239619.1;refGene_gn=PLEKHA7;refGene_tx=XM_023239620.1;refGene_gn=PLEK
HA7;refGene_tx=XM_023239621.1;refGene_gn=PLEKHA7;refGene_tx=XM_0232396
22.1;refGene_gn=PLEKHA7;refGene_tx=XM_023239623.1;refGene_gn=PLEKHA7;r
efGene_tx=XM_023239624.1
D1 74943541 74944012
refGene_gn=PLEKHA7;refGene_tx=XM_019812131.2;refGene_gn=PLEKHA7;refGen

e_tx=XM_019812137.2;refGene_gn=PLEKHA7;refGene_tx=XM_023239614.1;refGene_gn=PLEKHA7;refGene_tx=XM_023239615.1;refGene_gn=PLEKHA7;refGene_tx=XM_023239616.1;refGene_gn=PLEKHA7;refGene_tx=XM_023239617.1;refGene_gn=PLEKHA7;refGene_tx=XM_023239618.1;refGene_gn=PLEKHA7;refGene_tx=XM_023239619.1;refGene_gn=PLEKHA7;refGene_tx=XM_023239620.1;refGene_gn=PLEKHA7;refGene_tx=XM_023239621.1;refGene_gn=PLEKHA7;refGene_tx=XM_023239622.1;refGene_gn=PLEKHA7;refGene_tx=XM_023239623.1;refGene_gn=PLEKHA7;refGene_tx=XM_023239624.1
D1 74960430 74960606
refGene_gn=PLEKHA7;refGene_tx=XM_019812131.2;refGene_gn=PLEKHA7;refGene_tx=XM_019812137.2;refGene_gn=PLEKHA7;refGene_tx=XM_023239614.1;refGene_gn=PLEKHA7;refGene_tx=XM_023239615.1;refGene_gn=PLEKHA7;refGene_tx=XM_023239616.1;refGene_gn=PLEKHA7;refGene_tx=XM_023239617.1;refGene_gn=PLEKHA7;refGene_tx=XM_023239618.1;refGene_gn=PLEKHA7;refGene_tx=XM_023239619.1;refGene_gn=PLEKHA7;refGene_tx=XM_023239620.1;refGene_gn=PLEKHA7;refGene_tx=XM_023239621.1;refGene_gn=PLEKHA7;refGene_tx=XM_023239622.1;refGene_gn=PLEKHA7;refGene_tx=XM_023239623.1;refGene_gn=PLEKHA7;refGene_tx=XM_023239624.1
D1 74967822 74967923
refGene_gn=PLEKHA7;refGene_tx=XM_019812131.2;refGene_gn=PLEKHA7;refGene_tx=XM_019812137.2;refGene_gn=PLEKHA7;refGene_tx=XM_023239614.1;refGene_gn=PLEKHA7;refGene_tx=XM_023239615.1;refGene_gn=PLEKHA7;refGene_tx=XM_023239616.1;refGene_gn=PLEKHA7;refGene_tx=XM_023239617.1;refGene_gn=PLEKHA7;refGene_tx=XM_023239618.1;refGene_gn=PLEKHA7;refGene_tx=XM_023239619.1;refGene_gn=PLEKHA7;refGene_tx=XM_023239620.1;refGene_gn=PLEKHA7;refGene_tx=XM_023239621.1;refGene_gn=PLEKHA7;refGene_tx=XM_023239622.1;refGene_gn=PLEKHA7;refGene_tx=XM_023239623.1;refGene_gn=PLEKHA7;refGene_tx=XM_023239624.1
D1 74969334 74969407
refGene_gn=PLEKHA7;refGene_tx=XM_019812131.2;refGene_gn=PLEKHA7;refGene_tx=XM_019812137.2;refGene_gn=PLEKHA7;refGene_tx=XM_023239614.1;refGene_gn=PLEKHA7;refGene_tx=XM_023239615.1;refGene_gn=PLEKHA7;refGene_tx=XM_023239616.1;refGene_gn=PLEKHA7;refGene_tx=XM_023239617.1;refGene_gn=PLEKHA7;refGene_tx=XM_023239618.1;refGene_gn=PLEKHA7;refGene_tx=XM_023239619.1;refGene_gn=PLEKHA7;refGene_tx=XM_023239620.1;refGene_gn=PLEKHA7;refGene_tx=XM_023239621.1;refGene_gn=PLEKHA7;refGene_tx=XM_023239622.1;refGene_gn=PLEKHA7;refGene_tx=XM_023239623.1;refGene_gn=PLEKHA7;refGene_tx=XM_023239624.1
D1 74972024 74972129
refGene_gn=PLEKHA7;refGene_tx=XM_019812131.2;refGene_gn=PLEKHA7;refGene_tx=XM_019812137.2;refGene_gn=PLEKHA7;refGene_tx=XM_023239614.1;refGene_gn=PLEKHA7;refGene_tx=XM_023239615.1;refGene_gn=PLEKHA7;refGene_tx=XM_023239616.1;refGene_gn=PLEKHA7;refGene_tx=XM_023239617.1;refGene_gn=PLEKHA7;refGene_tx=XM_023239618.1;refGene_gn=PLEKHA7;refGene_tx=XM_023239619.1;refGene_gn=PLEKHA7;refGene_tx=XM_023239620.1;refGene_gn=PLEKHA7;refGene_tx=XM_023239621.1;refGene_gn=PLEKHA7;refGene_tx=XM_023239622.1;refGene_gn=PLEKHA7;refGene_tx=XM_023239623.1;refGene_gn=PLEKHA7;refGene_tx=XM_023239624.1
D1 74972868 74972980
refGene_gn=PLEKHA7;refGene_tx=XM_019812131.2;refGene_gn=PLEKHA7;refGen

e_tx=XM_019812137.2;refGene_gn=PLEKHA7;refGene_tx=XM_023239614.1;refGene_gn=PLEKHA7;refGene_tx=XM_023239615.1;refGene_gn=PLEKHA7;refGene_tx=XM_023239616.1;refGene_gn=PLEKHA7;refGene_tx=XM_023239617.1;refGene_gn=PLEKHA7;refGene_tx=XM_023239618.1;refGene_gn=PLEKHA7;refGene_tx=XM_023239619.1;refGene_gn=PLEKHA7;refGene_tx=XM_023239620.1;refGene_gn=PLEKHA7;refGene_tx=XM_023239621.1;refGene_gn=PLEKHA7;refGene_tx=XM_023239622.1;refGene_gn=PLEKHA7;refGene_tx=XM_023239623.1;refGene_gn=PLEKHA7;refGene_tx=XM_023239624.1
D1 74986645 74986729 refGene_gn=PLEKHA7;refGene_tx=XM_019812131.2;refGene_gn=PLEKHA7;refGene_tx=XM_023239614.1;refGene_gn=PLEKHA7;refGene_tx=XM_023239615.1;refGene_gn=PLEKHA7;refGene_tx=XM_023239616.1;refGene_gn=PLEKHA7;refGene_tx=XM_023239617.1;refGene_gn=PLEKHA7;refGene_tx=XM_023239618.1;refGene_gn=PLEKHA7;refGene_tx=XM_023239619.1;refGene_gn=PLEKHA7;refGene_tx=XM_023239620.1;refGene_gn=PLEKHA7;refGene_tx=XM_023239621.1;refGene_gn=PLEKHA7;refGene_tx=XM_023239622.1;refGene_gn=PLEKHA7;refGene_tx=XM_023239623.1;refGene_gn=PLEKHA7;refGene_tx=XM_023239624.1
D1 75037172 75037255 refGene_gn=PLEKHA7;refGene_tx=XM_019812131.2
D1 75117232 75117290 refGene_gn=PLEKHA7;refGene_tx=XM_023239614.1;refGene_gn=PLEKHA7;refGene_tx=XM_023239615.1;refGene_gn=PLEKHA7;refGene_tx=XM_023239616.1;refGene_gn=PLEKHA7;refGene_tx=XM_023239617.1;refGene_gn=PLEKHA7;refGene_tx=XM_023239618.1;refGene_gn=PLEKHA7;refGene_tx=XM_023239619.1;refGene_gn=PLEKHA7;refGene_tx=XM_023239620.1;refGene_gn=PLEKHA7;refGene_tx=XM_023239621.1;refGene_gn=PLEKHA7;refGene_tx=XM_023239622.1;refGene_gn=PLEKHA7;refGene_tx=XM_023239623.1;refGene_gn=PLEKHA7;refGene_tx=XM_023239624.1
D1 75117384 75117461 refGene_gn=PLEKHA7;refGene_tx=XM_023239614.1;refGene_gn=PLEKHA7;refGene_tx=XM_023239615.1;refGene_gn=PLEKHA7;refGene_tx=XM_023239616.1;refGene_gn=PLEKHA7;refGene_tx=XM_023239617.1;refGene_gn=PLEKHA7;refGene_tx=XM_023239618.1;refGene_gn=PLEKHA7;refGene_tx=XM_023239619.1;refGene_gn=PLEKHA7;refGene_tx=XM_023239620.1;refGene_gn=PLEKHA7;refGene_tx=XM_023239621.1;refGene_gn=PLEKHA7;refGene_tx=XM_023239622.1;refGene_gn=PLEKHA7;refGene_tx=XM_023239623.1;refGene_gn=PLEKHA7;refGene_tx=XM_023239624.1
D1 75117583 75117669 refGene_gn=PLEKHA7;refGene_tx=XM_023239614.1;refGene_gn=PLEKHA7;refGene_tx=XM_023239615.1;refGene_gn=PLEKHA7;refGene_tx=XM_023239616.1;refGene_gn=PLEKHA7;refGene_tx=XM_023239617.1;refGene_gn=PLEKHA7;refGene_tx=XM_023239618.1;refGene_gn=PLEKHA7;refGene_tx=XM_023239619.1;refGene_gn=PLEKHA7;refGene_tx=XM_023239620.1;refGene_gn=PLEKHA7;refGene_tx=XM_023239621.1;refGene_gn=PLEKHA7;refGene_tx=XM_023239622.1;refGene_gn=PLEKHA7;refGene_tx=XM_023239623.1;refGene_gn=PLEKHA7;refGene_tx=XM_023239624.1
D1 75148099 75148133 refGene_gn=RPS13;refGene_tx=XM_003993023.5
D1 75148760 75148861 refGene_gn=RPS13;refGene_tx=XM_003993023.5
D1 75149330 75149500 refGene_gn=RPS13;refGene_tx=XM_003993023.5
D1 75150315 75150394 refGene_gn=RPS13;refGene_tx=XM_003993023.5
D1 75150583 75150632 refGene_gn=RPS13;refGene_tx=XM_003993023.5

D1 75150791 75150814 refGene_gn=RPS13;refGene_tx=XM_003993023.5
D1 75157470 75157653 refGene_gn=PIK3C2A;refGene_tx=XM_006937126.4
D1 75159207 75159333 refGene_gn=PIK3C2A;refGene_tx=XM_006937126.4
D1 75159420 75159530 refGene_gn=PIK3C2A;refGene_tx=XM_006937126.4
D1 75159871 75159944 refGene_gn=PIK3C2A;refGene_tx=XM_006937126.4
D1 75160034 75160152 refGene_gn=PIK3C2A;refGene_tx=XM_006937126.4
D1 75161415 75161540 refGene_gn=PIK3C2A;refGene_tx=XM_006937126.4
D1 75162790 75162998 refGene_gn=PIK3C2A;refGene_tx=XM_006937126.4
D1 75165316 75165426 refGene_gn=PIK3C2A;refGene_tx=XM_006937126.4
D1 75166071 75166228 refGene_gn=PIK3C2A;refGene_tx=XM_006937126.4
D1 75167638 75167808 refGene_gn=PIK3C2A;refGene_tx=XM_006937126.4
D1 75170104 75170241 refGene_gn=PIK3C2A;refGene_tx=XM_006937126.4
D1 75175133 75175263 refGene_gn=PIK3C2A;refGene_tx=XM_006937126.4
D1 75176532 75176625 refGene_gn=PIK3C2A;refGene_tx=XM_006937126.4
D1 75178220 75178325 refGene_gn=PIK3C2A;refGene_tx=XM_006937126.4
D1 75182941 75183122 refGene_gn=PIK3C2A;refGene_tx=XM_006937126.4
D1 75183472 75183567 refGene_gn=PIK3C2A;refGene_tx=XM_006937126.4
D1 75184073 75184167 refGene_gn=PIK3C2A;refGene_tx=XM_006937126.4
D1 75185284 75185473 refGene_gn=PIK3C2A;refGene_tx=XM_006937126.4
D1 75187065 75187211 refGene_gn=PIK3C2A;refGene_tx=XM_006937126.4
D1 75187593 75187705 refGene_gn=PIK3C2A;refGene_tx=XM_006937126.4
D1 75193820 75193988 refGene_gn=PIK3C2A;refGene_tx=XM_006937126.4
D1 75195041 75195164 refGene_gn=PIK3C2A;refGene_tx=XM_006937126.4
D1 75197718 75197929 refGene_gn=PIK3C2A;refGene_tx=XM_006937126.4
D1 75198009 75198058 refGene_gn=PIK3C2A;refGene_tx=XM_006937126.4
D1 75199589 75199733 refGene_gn=PIK3C2A;refGene_tx=XM_006937126.4
D1 75206902 75206966 refGene_gn=PIK3C2A;refGene_tx=XM_006937126.4
D1 75207098 75207178 refGene_gn=PIK3C2A;refGene_tx=XM_006937126.4
D1 75208570 75208682 refGene_gn=PIK3C2A;refGene_tx=XM_006937126.4
D1 75209407 75209528 refGene_gn=PIK3C2A;refGene_tx=XM_006937126.4
D1 75211874 75212032 refGene_gn=PIK3C2A;refGene_tx=XM_006937126.4
D1 75221604 75221708 refGene_gn=PIK3C2A;refGene_tx=XM_006937126.4
D1 75249198 75250269 refGene_gn=PIK3C2A;refGene_tx=XM_006937126.4
D1 75275677 75275806
refGene_gn=LOC109492038;refGene_tx=XM_023239727.1
D1 75277011 75277424
refGene_gn=LOC109492038;refGene_tx=XM_023239727.1
D1 75278364 75278518
refGene_gn=LOC109492038;refGene_tx=XM_023239727.1
D1 75365520 75365664
refGene_gn=NUCB2;refGene_tx=XM_011286735.2;refGene_gn=NUCB2;refGene_tx
=XM_011286736.3
D1 75366445 75366553
refGene_gn=NUCB2;refGene_tx=XM_011286735.2;refGene_gn=NUCB2;refGene_tx
=XM_011286736.3
D1 75370450 75370577
refGene_gn=NUCB2;refGene_tx=XM_011286735.2;refGene_gn=NUCB2;refGene_tx
=XM_011286736.3
D1 75376914 75377018
refGene_gn=NUCB2;refGene_tx=XM_011286735.2;refGene_gn=NUCB2;refGene_tx

=XM_011286736.3
D1 75377824 75378010
refGene_gn=NUCB2;refGene_tx=XM_011286735.2;refGene_gn=NUCB2;refGene_tx
=XM_011286736.3
D1 75378199 75378290
refGene_gn=NUCB2;refGene_tx=XM_011286735.2;refGene_gn=NUCB2;refGene_tx
=XM_011286736.3
D1 75378881 75378940
refGene_gn=NUCB2;refGene_tx=XM_011286735.2;refGene_gn=NUCB2;refGene_tx
=XM_011286736.3
D1 75379036 75379129
refGene_gn=NUCB2;refGene_tx=XM_011286735.2;refGene_gn=NUCB2;refGene_tx
=XM_011286736.3
D1 75380158 75380248
refGene_gn=NUCB2;refGene_tx=XM_011286735.2;refGene_gn=NUCB2;refGene_tx
=XM_011286736.3
D1 75385584 75385755
refGene_gn=NUCB2;refGene_tx=XM_011286735.2;refGene_gn=NUCB2;refGene_tx
=XM_011286736.3
D1 75386695 75386770
refGene_gn=NUCB2;refGene_tx=XM_011286735.2;refGene_gn=NUCB2;refGene_tx
=XM_011286736.3
D1 75398375 75398487 refGene_gn=NCR3LG1;refGene_tx=XM_011286767.3
D1 75400721 75401069 refGene_gn=NCR3LG1;refGene_tx=XM_011286767.3
D1 75411506 75411821 refGene_gn=NCR3LG1;refGene_tx=XM_011286767.3
D1 75413421 75413519 refGene_gn=NCR3LG1;refGene_tx=XM_011286767.3
D1 75415804 75415897 refGene_gn=NCR3LG1;refGene_tx=XM_011286767.3
D1 75424734 75425907 refGene_gn=KCNJ11;refGene_tx=XM_003993026.4
D1 75430513 75430651 refGene_gn=ABCC8;refGene_tx=XM_003993103.5
D1 75431176 75431239 refGene_gn=ABCC8;refGene_tx=XM_003993103.5
D1 75431721 75431855 refGene_gn=ABCC8;refGene_tx=XM_003993103.5
D1 75432633 75432737 refGene_gn=ABCC8;refGene_tx=XM_003993103.5
D1 75433159 75433268 refGene_gn=ABCC8;refGene_tx=XM_003993103.5
D1 75433478 75433557 refGene_gn=ABCC8;refGene_tx=XM_003993103.5
D1 75436035 75436166 refGene_gn=ABCC8;refGene_tx=XM_003993103.5
D1 75436320 75436441 refGene_gn=ABCC8;refGene_tx=XM_003993103.5
D1 75436775 75436889 refGene_gn=ABCC8;refGene_tx=XM_003993103.5
D1 75437459 75437562 refGene_gn=ABCC8;refGene_tx=XM_003993103.5
D1 75441772 75441865 refGene_gn=ABCC8;refGene_tx=XM_003993103.5
D1 75443632 75443790 refGene_gn=ABCC8;refGene_tx=XM_003993103.5
D1 75444291 75444361 refGene_gn=ABCC8;refGene_tx=XM_003993103.5
D1 75445666 75445833 refGene_gn=ABCC8;refGene_tx=XM_003993103.5
D1 75445924 75446166 refGene_gn=ABCC8;refGene_tx=XM_003993103.5
D1 75446400 75446500 refGene_gn=ABCC8;refGene_tx=XM_003993103.5
D1 75447398 75447524 refGene_gn=ABCC8;refGene_tx=XM_003993103.5
D1 75448981 75449119 refGene_gn=ABCC8;refGene_tx=XM_003993103.5
D1 75450999 75451080 refGene_gn=ABCC8;refGene_tx=XM_003993103.5
D1 75451789 75451874 refGene_gn=ABCC8;refGene_tx=XM_003993103.5
D1 75452953 75453052 refGene_gn=ABCC8;refGene_tx=XM_003993103.5
D1 75453756 75453792 refGene_gn=ABCC8;refGene_tx=XM_003993103.5

D1 75455536 75455569 refGene_gn=ABCC8;refGene_tx=XM_003993103.5
D1 75463731 75463837 refGene_gn=ABCC8;refGene_tx=XM_003993103.5
D1 75464492 75464568 refGene_gn=ABCC8;refGene_tx=XM_003993103.5
D1 75464952 75465069 refGene_gn=ABCC8;refGene_tx=XM_003993103.5
D1 75465229 75465335 refGene_gn=ABCC8;refGene_tx=XM_003993103.5
D1 75467031 75467177 refGene_gn=ABCC8;refGene_tx=XM_003993103.5
D1 75468316 75468357 refGene_gn=ABCC8;refGene_tx=XM_003993103.5
D1 75478262 75478425 refGene_gn=ABCC8;refGene_tx=XM_003993103.5
D1 75478759 75478894 refGene_gn=ABCC8;refGene_tx=XM_003993103.5
D1 75483754 75483910 refGene_gn=ABCC8;refGene_tx=XM_003993103.5
D1 75486989 75487154 refGene_gn=ABCC8;refGene_tx=XM_003993103.5
D1 75494310 75494499 refGene_gn=ABCC8;refGene_tx=XM_003993103.5
D1 75495323 75495566 refGene_gn=ABCC8;refGene_tx=XM_003993103.5
D1 75497220 75497297 refGene_gn=ABCC8;refGene_tx=XM_003993103.5
D1 75497299 75497386 refGene_gn=ABCC8;refGene_tx=XM_003993103.5
D1 75503511 75503633 refGene_gn=ABCC8;refGene_tx=XM_003993103.5
D1 75508482 75508624 refGene_gn=ABCC8;refGene_tx=XM_003993103.5
D1 75509809 75509957 refGene_gn=ABCC8;refGene_tx=XM_003993103.5
D1 75529001 75529014
refGene_gn=USH1C;refGene_tx=XM_023239625.1;refGene_gn=USH1C;refGene_tx=XM_023239626.1;refGene_gn=USH1C;refGene_tx=XM_023239627.1;refGene_gn=USH1C;refGene_tx=XM_023239628.1
D1 75530147 75530271 refGene_gn=USH1C;refGene_tx=XM_023239629.1
D1 75531244 75531315
refGene_gn=USH1C;refGene_tx=XM_023239625.1;refGene_gn=USH1C;refGene_tx=XM_023239626.1;refGene_gn=USH1C;refGene_tx=XM_023239627.1;refGene_gn=USH1C;refGene_tx=XM_023239628.1;refGene_gn=USH1C;refGene_tx=XM_023239629.1
D1 75532580 75532687
refGene_gn=USH1C;refGene_tx=XM_023239625.1;refGene_gn=USH1C;refGene_tx=XM_023239626.1;refGene_gn=USH1C;refGene_tx=XM_023239627.1;refGene_gn=USH1C;refGene_tx=XM_023239628.1;refGene_gn=USH1C;refGene_tx=XM_023239629.1
D1 75537475 75537575
refGene_gn=USH1C;refGene_tx=XM_023239625.1;refGene_gn=USH1C;refGene_tx=XM_023239626.1;refGene_gn=USH1C;refGene_tx=XM_023239627.1;refGene_gn=USH1C;refGene_tx=XM_023239628.1;refGene_gn=USH1C;refGene_tx=XM_023239629.1
D1 75537902 75537956
refGene_gn=USH1C;refGene_tx=XM_023239625.1;refGene_gn=USH1C;refGene_tx=XM_023239626.1;refGene_gn=USH1C;refGene_tx=XM_023239627.1;refGene_gn=USH1C;refGene_tx=XM_023239628.1;refGene_gn=USH1C;refGene_tx=XM_023239629.1
D1 75538511 75538553
refGene_gn=USH1C;refGene_tx=XM_023239625.1;refGene_gn=USH1C;refGene_tx=XM_023239626.1;refGene_gn=USH1C;refGene_tx=XM_023239627.1;refGene_gn=USH1C;refGene_tx=XM_023239628.1;refGene_gn=USH1C;refGene_tx=XM_023239629.1
D1 75541263 75541314
refGene_gn=USH1C;refGene_tx=XM_023239625.1;refGene_gn=USH1C;refGene_tx

=XM_023239626.1;refGene_gn=USH1C;refGene_tx=XM_023239627.1;refGene_gn=USH1C;refGene_tx=XM_023239629.1
D1 75542396 75542516 refGene_gn=USH1C;refGene_tx=XM_023239625.1;refGene_gn=USH1C;refGene_tx=XM_023239626.1;refGene_gn=USH1C;refGene_tx=XM_023239627.1;refGene_gn=USH1C;refGene_tx=XM_023239629.1
D1 75546506 75546989 refGene_gn=USH1C;refGene_tx=XM_023239629.1
D1 75547515 75547632 refGene_gn=USH1C;refGene_tx=XM_023239629.1
D1 75548969 75549122 refGene_gn=USH1C;refGene_tx=XM_023239629.1
D1 75553026 75553076 refGene_gn=USH1C;refGene_tx=XM_023239629.1
D1 75554134 75554208 refGene_gn=USH1C;refGene_tx=XM_023239625.1;refGene_gn=USH1C;refGene_tx=XM_023239626.1;refGene_gn=USH1C;refGene_tx=XM_023239627.1;refGene_gn=USH1C;refGene_tx=XM_023239628.1
D1 75557276 75557401 refGene_gn=USH1C;refGene_tx=XM_023239625.1;refGene_gn=USH1C;refGene_tx=XM_023239626.1;refGene_gn=USH1C;refGene_tx=XM_023239627.1;refGene_gn=USH1C;refGene_tx=XM_023239628.1;refGene_gn=USH1C;refGene_tx=XM_023239629.1
D1 75557747 75557813 refGene_gn=USH1C;refGene_tx=XM_023239625.1;refGene_gn=USH1C;refGene_tx=XM_023239626.1;refGene_gn=USH1C;refGene_tx=XM_023239627.1;refGene_gn=USH1C;refGene_tx=XM_023239628.1;refGene_gn=USH1C;refGene_tx=XM_023239629.1
D1 75558855 75558998 refGene_gn=USH1C;refGene_tx=XM_023239625.1;refGene_gn=USH1C;refGene_tx=XM_023239626.1;refGene_gn=USH1C;refGene_tx=XM_023239627.1;refGene_gn=USH1C;refGene_tx=XM_023239628.1;refGene_gn=USH1C;refGene_tx=XM_023239629.1
D1 75559310 75559367 refGene_gn=USH1C;refGene_tx=XM_023239625.1;refGene_gn=USH1C;refGene_tx=XM_023239626.1;refGene_gn=USH1C;refGene_tx=XM_023239627.1;refGene_gn=USH1C;refGene_tx=XM_023239628.1;refGene_gn=USH1C;refGene_tx=XM_023239629.1
D1 75559535 75559595 refGene_gn=USH1C;refGene_tx=XM_023239625.1;refGene_gn=USH1C;refGene_tx=XM_023239626.1;refGene_gn=USH1C;refGene_tx=XM_023239627.1;refGene_gn=USH1C;refGene_tx=XM_023239628.1;refGene_gn=USH1C;refGene_tx=XM_023239629.1
D1 75560767 75560852 refGene_gn=USH1C;refGene_tx=XM_023239625.1;refGene_gn=USH1C;refGene_tx=XM_023239626.1;refGene_gn=USH1C;refGene_tx=XM_023239627.1;refGene_gn=USH1C;refGene_tx=XM_023239628.1;refGene_gn=USH1C;refGene_tx=XM_023239629.1
D1 75562442 75562537 refGene_gn=USH1C;refGene_tx=XM_023239625.1;refGene_gn=USH1C;refGene_tx=XM_023239626.1;refGene_gn=USH1C;refGene_tx=XM_023239627.1;refGene_gn=USH1C;refGene_tx=XM_023239628.1;refGene_gn=USH1C;refGene_tx=XM_023239629.1
D1 75562862 75562920

refGene_gn=USH1C;refGene_tx=XM_023239625.1;refGene_gn=USH1C;refGene_tx=XM_023239626.1;refGene_gn=USH1C;refGene_tx=XM_023239627.1;refGene_gn=USH1C;refGene_tx=XM_023239628.1;refGene_gn=USH1C;refGene_tx=XM_023239629.1
D1 75563125 75563150
refGene_gn=USH1C;refGene_tx=XM_023239625.1;refGene_gn=USH1C;refGene_tx=XM_023239626.1;refGene_gn=USH1C;refGene_tx=XM_023239627.1;refGene_gn=USH1C;refGene_tx=XM_023239628.1;refGene_gn=USH1C;refGene_tx=XM_023239629.1
D1 75563265 75563374
refGene_gn=USH1C;refGene_tx=XM_023239625.1;refGene_gn=USH1C;refGene_tx=XM_023239626.1;refGene_gn=USH1C;refGene_tx=XM_023239627.1;refGene_gn=USH1C;refGene_tx=XM_023239628.1;refGene_gn=USH1C;refGene_tx=XM_023239629.1
D1 75568026 75568165
refGene_gn=USH1C;refGene_tx=XM_023239625.1;refGene_gn=USH1C;refGene_tx=XM_023239626.1;refGene_gn=USH1C;refGene_tx=XM_023239627.1;refGene_gn=USH1C;refGene_tx=XM_023239628.1;refGene_gn=USH1C;refGene_tx=XM_023239629.1
D1 75568281 75568425
refGene_gn=USH1C;refGene_tx=XM_023239625.1;refGene_gn=USH1C;refGene_tx=XM_023239626.1;refGene_gn=USH1C;refGene_tx=XM_023239627.1;refGene_gn=USH1C;refGene_tx=XM_023239628.1;refGene_gn=USH1C;refGene_tx=XM_023239629.1
D1 75569834 75569902
refGene_gn=USH1C;refGene_tx=XM_023239625.1;refGene_gn=USH1C;refGene_tx=XM_023239626.1;refGene_gn=USH1C;refGene_tx=XM_023239627.1;refGene_gn=USH1C;refGene_tx=XM_023239628.1;refGene_gn=USH1C;refGene_tx=XM_023239629.1
D1 75577424 75577460
refGene_gn=USH1C;refGene_tx=XM_023239625.1;refGene_gn=USH1C;refGene_tx=XM_023239628.1;refGene_gn=USH1C;refGene_tx=XM_023239629.1
D1 75580308 75580420 refGene_gn=OTOG;refGene_tx=XM_023239728.1
D1 75580851 75580921 refGene_gn=OTOG;refGene_tx=XM_023239728.1
D1 75581089 75581150 refGene_gn=OTOG;refGene_tx=XM_023239728.1
D1 75584501 75584577 refGene_gn=OTOG;refGene_tx=XM_023239728.1
D1 75585508 75585601 refGene_gn=OTOG;refGene_tx=XM_023239728.1
D1 75585785 75585940 refGene_gn=OTOG;refGene_tx=XM_023239728.1
D1 75588277 75588396 refGene_gn=OTOG;refGene_tx=XM_023239728.1
D1 75589729 75589935 refGene_gn=OTOG;refGene_tx=XM_023239728.1
D1 75590768 75590899 refGene_gn=OTOG;refGene_tx=XM_023239728.1
D1 75591166 75591273 refGene_gn=OTOG;refGene_tx=XM_023239728.1
D1 75591699 75591809 refGene_gn=OTOG;refGene_tx=XM_023239728.1
D1 75592182 75592311 refGene_gn=OTOG;refGene_tx=XM_023239728.1
D1 75593207 75593316 refGene_gn=OTOG;refGene_tx=XM_023239728.1
D1 75593616 75593663 refGene_gn=OTOG;refGene_tx=XM_023239728.1
D1 75594150 75594296 refGene_gn=OTOG;refGene_tx=XM_023239728.1
D1 75601196 75601329 refGene_gn=OTOG;refGene_tx=XM_023239728.1
D1 75602194 75602372 refGene_gn=OTOG;refGene_tx=XM_023239728.1
D1 75604389 75604514 refGene_gn=OTOG;refGene_tx=XM_023239728.1

| | | | |
|---|---|---|---|
| D1 | 75605760 | 75605973 | refGene_gn=OTOG;refGene_tx=XM_023239728.1 |
| D1 | 75607355 | 75607548 | refGene_gn=OTOG;refGene_tx=XM_023239728.1 |
| D1 | 75609092 | 75609167 | refGene_gn=OTOG;refGene_tx=XM_023239728.1 |
| D1 | 75609418 | 75609462 | refGene_gn=OTOG;refGene_tx=XM_023239728.1 |
| D1 | 75610902 | 75611056 | refGene_gn=OTOG;refGene_tx=XM_023239728.1 |
| D1 | 75619411 | 75619519 | refGene_gn=OTOG;refGene_tx=XM_023239728.1 |
| D1 | 75619897 | 75620036 | refGene_gn=OTOG;refGene_tx=XM_023239728.1 |
| D1 | 75621654 | 75621789 | refGene_gn=OTOG;refGene_tx=XM_023239728.1 |
| D1 | 75622078 | 75622225 | refGene_gn=OTOG;refGene_tx=XM_023239728.1 |
| D1 | 75622530 | 75622650 | refGene_gn=OTOG;refGene_tx=XM_023239728.1 |
| D1 | 75624546 | 75624663 | refGene_gn=OTOG;refGene_tx=XM_023239728.1 |
| D1 | 75625714 | 75625871 | refGene_gn=OTOG;refGene_tx=XM_023239728.1 |
| D1 | 75628660 | 75628687 | refGene_gn=OTOG;refGene_tx=XM_023239728.1 |
| D1 | 75631164 | 75631332 | refGene_gn=OTOG;refGene_tx=XM_023239728.1 |
| D1 | 75635656 | 75635935 | refGene_gn=OTOG;refGene_tx=XM_023239728.1 |
| D1 | 75637867 | 75637985 | refGene_gn=OTOG;refGene_tx=XM_023239728.1 |
| D1 | 75638656 | 75638736 | refGene_gn=OTOG;refGene_tx=XM_023239728.1 |
| D1 | 75639301 | 75641097 | refGene_gn=OTOG;refGene_tx=XM_023239728.1 |
| D1 | 75642096 | 75642265 | refGene_gn=OTOG;refGene_tx=XM_023239728.1 |
| D1 | 75642575 | 75642721 | refGene_gn=OTOG;refGene_tx=XM_023239728.1 |
| D1 | 75643327 | 75643417 | refGene_gn=OTOG;refGene_tx=XM_023239728.1 |
| D1 | 75653111 | 75653295 | refGene_gn=OTOG;refGene_tx=XM_023239728.1 |
| D1 | 75655615 | 75655836 | refGene_gn=OTOG;refGene_tx=XM_023239728.1 |
| D1 | 75656046 | 75656185 | refGene_gn=OTOG;refGene_tx=XM_023239728.1 |
| D1 | 75657773 | 75657968 | refGene_gn=OTOG;refGene_tx=XM_023239728.1 |
| D1 | 75658180 | 75658393 | refGene_gn=OTOG;refGene_tx=XM_023239728.1 |
| D1 | 75658955 | 75659060 | refGene_gn=OTOG;refGene_tx=XM_023239728.1 |
| D1 | 75659227 | 75659335 | refGene_gn=OTOG;refGene_tx=XM_023239728.1 |
| D1 | 75659729 | 75659831 | refGene_gn=OTOG;refGene_tx=XM_023239728.1 |
| D1 | 75662668 | 75662767 | refGene_gn=OTOG;refGene_tx=XM_023239728.1 |
| D1 | 75663986 | 75664027 | refGene_gn=OTOG;refGene_tx=XM_023239728.1 |
| D1 | 75665006 | 75665082 | refGene_gn=OTOG;refGene_tx=XM_023239728.1 |
| D1 | 75665178 | 75665357 | refGene_gn=OTOG;refGene_tx=XM_023239728.1 |
| D1 | 75666115 | 75666220 | refGene_gn=OTOG;refGene_tx=XM_023239728.1 |
| D1 | 75666392 | 75666512 | refGene_gn=OTOG;refGene_tx=XM_023239728.1 |
| D1 | 75667653 | 75667699 | refGene_gn=OTOG;refGene_tx=XM_023239728.1 |
| D1 | 75670057 | 75670137 | refGene_gn=OTOG;refGene_tx=XM_023239728.1 |
| D1 | 75670219 | 75670420 | refGene_gn=OTOG;refGene_tx=XM_023239728.1 |
| D1 | 75741331 | 75741961 | refGene_gn=MYOD1;refGene_tx=XM_003993029.5 |
| D1 | 75742435 | 75742514 | refGene_gn=MYOD1;refGene_tx=XM_003993029.5 |
| D1 | 75742806 | 75743057 | refGene_gn=MYOD1;refGene_tx=XM_003993029.5 |
| D1 | 75757152 | 75757722 | refGene_gn=KCNC1;refGene_tx=XM_003993030.5 |
| D1 | 75789104 | 75790038 | refGene_gn=KCNC1;refGene_tx=XM_003993030.5 |
| D1 | 75798054 | 75798243 | refGene_gn=KCNC1;refGene_tx=XM_003993030.5 |
| D1 | 75800279 | 75800344 | refGene_gn=KCNC1;refGene_tx=XM_003993030.5 |
| D1 | 75806625 | 75806717 | refGene_gn=SERGEF;refGene_tx=XM_003993032.5 |
| D1 | 75806719 | 75806965 | refGene_gn=SERGEF;refGene_tx=XM_003993032.5 |
| D1 | 75897784 | 75897821 | refGene_gn=SERGEF;refGene_tx=XM_003993032.5 |
| D1 | 75993637 | 75993804 | refGene_gn=SERGEF;refGene_tx=XM_003993032.5 |
| D1 | 76020080 | 76020239 | refGene_gn=SERGEF;refGene_tx=XM_003993032.5 |

D1 76024860 76024923 refGene_gn=SERGEF;refGene_tx=XM_003993032.5
D1 76027686 76027800 refGene_gn=SERGEF;refGene_tx=XM_003993032.5
D1 76032321 76032382 refGene_gn=SERGEF;refGene_tx=XM_003993032.5
D1 76037008 76037103 refGene_gn=SERGEF;refGene_tx=XM_003993032.5
D1 76039056 76039212 refGene_gn=SERGEF;refGene_tx=XM_003993032.5
D1 76040400 76040536 refGene_gn=SERGEF;refGene_tx=XM_003993032.5
D1 76044032 76044092 refGene_gn=SERGEF;refGene_tx=XM_003993032.5
D1 76051762 76051937
refGene_gn=TPH1;refGene_tx=XM_003993033.4;refGene_gn=TPH1;refGene_tx=XM_023239630.1
D1 76053938 76054072
refGene_gn=TPH1;refGene_tx=XM_003993033.4;refGene_gn=TPH1;refGene_tx=XM_023239630.1
D1 76054873 76054969
refGene_gn=TPH1;refGene_tx=XM_003993033.4;refGene_gn=TPH1;refGene_tx=XM_023239630.1
D1 76059432 76059559
refGene_gn=TPH1;refGene_tx=XM_003993033.4;refGene_gn=TPH1;refGene_tx=XM_023239630.1
D1 76060257 76060393
refGene_gn=TPH1;refGene_tx=XM_003993033.4;refGene_gn=TPH1;refGene_tx=XM_023239630.1
D1 76063305 76063502
refGene_gn=TPH1;refGene_tx=XM_003993033.4;refGene_gn=TPH1;refGene_tx=XM_023239630.1
D1 76063628 76063696
refGene_gn=TPH1;refGene_tx=XM_003993033.4;refGene_gn=TPH1;refGene_tx=XM_023239630.1
D1 76067074 76067175
refGene_gn=TPH1;refGene_tx=XM_003993033.4;refGene_gn=TPH1;refGene_tx=XM_023239630.1
D1 76069363 76069547
refGene_gn=TPH1;refGene_tx=XM_003993033.4;refGene_gn=TPH1;refGene_tx=XM_023239630.1
D1 76076958 76077095
refGene_gn=TPH1;refGene_tx=XM_003993033.4;refGene_gn=TPH1;refGene_tx=XM_023239630.1
D1 76077218 76077324 refGene_gn=TPH1;refGene_tx=XM_023239630.1
D1 76110190 76110283
refGene_gn=SAAL1;refGene_tx=XM_003993034.4;refGene_gn=SAAL1;refGene_tx=XM_019812149.2
D1 76110972 76111065
refGene_gn=SAAL1;refGene_tx=XM_003993034.4;refGene_gn=SAAL1;refGene_tx=XM_019812149.2;refGene_gn=SAAL1;refGene_tx=XM_023239631.1
D1 76113196 76113390
refGene_gn=SAAL1;refGene_tx=XM_003993034.4;refGene_gn=SAAL1;refGene_tx=XM_019812149.2
D1 76116162 76116351
refGene_gn=SAAL1;refGene_tx=XM_003993034.4;refGene_gn=SAAL1;refGene_tx=XM_019812149.2;refGene_gn=SAAL1;refGene_tx=XM_023239631.1

D1 76116445 76116528
refGene_gn=SAAL1;refGene_tx=XM_003993034.4;refGene_gn=SAAL1;refGene_tx=XM_019812149.2;refGene_gn=SAAL1;refGene_tx=XM_023239631.1
D1 76119153 76119334
refGene_gn=SAAL1;refGene_tx=XM_003993034.4;refGene_gn=SAAL1;refGene_tx=XM_019812149.2;refGene_gn=SAAL1;refGene_tx=XM_023239631.1
D1 76120265 76120381
refGene_gn=SAAL1;refGene_tx=XM_003993034.4;refGene_gn=SAAL1;refGene_tx=XM_019812149.2;refGene_gn=SAAL1;refGene_tx=XM_023239631.1
D1 76120514 76120574
refGene_gn=SAAL1;refGene_tx=XM_003993034.4;refGene_gn=SAAL1;refGene_tx=XM_019812149.2;refGene_gn=SAAL1;refGene_tx=XM_023239631.1
D1 76122600 76122680
refGene_gn=SAAL1;refGene_tx=XM_003993034.4;refGene_gn=SAAL1;refGene_tx=XM_019812149.2;refGene_gn=SAAL1;refGene_tx=XM_023239631.1
D1 76126453 76126537
refGene_gn=SAAL1;refGene_tx=XM_003993034.4;refGene_gn=SAAL1;refGene_tx=XM_019812149.2;refGene_gn=SAAL1;refGene_tx=XM_023239631.1
D1 76134139 76134253
refGene_gn=SAAL1;refGene_tx=XM_003993034.4;refGene_gn=SAAL1;refGene_tx=XM_019812149.2;refGene_gn=SAAL1;refGene_tx=XM_023239631.1
D1 76136061 76136426
refGene_gn=SAAL1;refGene_tx=XM_003993034.4;refGene_gn=SAAL1;refGene_tx=XM_019812149.2;refGene_gn=SAAL1;refGene_tx=XM_023239631.1
D1 76140005 76140168
refGene_gn=LOC111556632;refGene_tx=XM_023239632.1
D1 76140572 76140618
refGene_gn=LOC111556632;refGene_tx=XM_023239632.1
D1 76140620 76140709
refGene_gn=LOC111556632;refGene_tx=XM_023239632.1
D1 76142657 76142752
refGene_gn=LOC111556632;refGene_tx=XM_023239632.1
D1 76145237 76145531
refGene_gn=LOC111556632;refGene_tx=XM_023239632.1
D1 76153544 76153707
refGene_gn=LOC101099498;refGene_tx=XM_023239633.1
D1 76154112 76154251
refGene_gn=LOC101099498;refGene_tx=XM_023239633.1
D1 76156033 76156121
refGene_gn=LOC101099498;refGene_tx=XM_023239633.1
D1 76171814 76171902
refGene_gn=LOC101099075;refGene_tx=NM_001309040.1;refGene_gn=LOC101099075;refGene_tx=XM_003993037.5;refGene_gn=LOC101099075;refGene_tx=XM_019811170.2
D1 76173684 76173823
refGene_gn=LOC101099075;refGene_tx=NM_001309040.1;refGene_gn=LOC101099075;refGene_tx=XM_003993037.5;refGene_gn=LOC101099075;refGene_tx=XM_019811170.2
D1 76174228 76174391
refGene_gn=LOC101099075;refGene_tx=NM_001309040.1;refGene_gn=LOC101099

075;refGene_tx=XM_003993037.5;refGene_gn=LOC101099075;refGene_tx=XM_019811170.2
D1 76185464 76185555
refGene_gn=LOC105260977;refGene_tx=XM_011286739.3
D1 76187467 76187606
refGene_gn=LOC105260977;refGene_tx=XM_011286739.3
D1 76188009 76188172
refGene_gn=LOC105260977;refGene_tx=XM_011286739.3
D1 76203721 76203782
refGene_gn=HPS5;refGene_tx=XM_003993038.4;refGene_gn=HPS5;refGene_tx=XM_006937131.3;refGene_gn=HPS5;refGene_tx=XM_019812150.2;refGene_gn=HPS5;refGene_tx=XM_019812151.2;refGene_gn=HPS5;refGene_tx=XM_023239634.1
D1 76204917 76205188
refGene_gn=HPS5;refGene_tx=XM_003993038.4;refGene_gn=HPS5;refGene_tx=XM_006937131.3;refGene_gn=HPS5;refGene_tx=XM_019812150.2;refGene_gn=HPS5;refGene_tx=XM_019812151.2;refGene_gn=HPS5;refGene_tx=XM_023239634.1
D1 76207775 76207882
refGene_gn=HPS5;refGene_tx=XM_003993038.4;refGene_gn=HPS5;refGene_tx=XM_006937131.3;refGene_gn=HPS5;refGene_tx=XM_019812150.2;refGene_gn=HPS5;refGene_tx=XM_019812151.2;refGene_gn=HPS5;refGene_tx=XM_023239634.1
D1 76209269 76209383
refGene_gn=HPS5;refGene_tx=XM_003993038.4;refGene_gn=HPS5;refGene_tx=XM_006937131.3;refGene_gn=HPS5;refGene_tx=XM_019812150.2;refGene_gn=HPS5;refGene_tx=XM_019812151.2;refGene_gn=HPS5;refGene_tx=XM_023239634.1
D1 76210223 76210343
refGene_gn=HPS5;refGene_tx=XM_003993038.4;refGene_gn=HPS5;refGene_tx=XM_006937131.3;refGene_gn=HPS5;refGene_tx=XM_019812150.2;refGene_gn=HPS5;refGene_tx=XM_019812151.2;refGene_gn=HPS5;refGene_tx=XM_023239634.1
D1 76211079 76211235
refGene_gn=HPS5;refGene_tx=XM_003993038.4;refGene_gn=HPS5;refGene_tx=XM_006937131.3;refGene_gn=HPS5;refGene_tx=XM_019812150.2;refGene_gn=HPS5;refGene_tx=XM_019812151.2;refGene_gn=HPS5;refGene_tx=XM_023239634.1
D1 76211659 76211780
refGene_gn=HPS5;refGene_tx=XM_003993038.4;refGene_gn=HPS5;refGene_tx=XM_006937131.3;refGene_gn=HPS5;refGene_tx=XM_019812150.2;refGene_gn=HPS5;refGene_tx=XM_019812151.2;refGene_gn=HPS5;refGene_tx=XM_023239634.1
D1 76212564 76213142
refGene_gn=HPS5;refGene_tx=XM_003993038.4;refGene_gn=HPS5;refGene_tx=XM_006937131.3;refGene_gn=HPS5;refGene_tx=XM_019812150.2;refGene_gn=HPS5;refGene_tx=XM_019812151.2;refGene_gn=HPS5;refGene_tx=XM_023239634.1
D1 76213674 76213752
refGene_gn=HPS5;refGene_tx=XM_003993038.4;refGene_gn=HPS5;refGene_tx=XM_006937131.3;refGene_gn=HPS5;refGene_tx=XM_019812150.2;refGene_gn=HPS5;refGene_tx=XM_019812151.2;refGene_gn=HPS5;refGene_tx=XM_023239634.1
D1 76215155 76215305
refGene_gn=HPS5;refGene_tx=XM_003993038.4;refGene_gn=HPS5;refGene_tx=XM_006937131.3;refGene_gn=HPS5;refGene_tx=XM_019812150.2;refGene_gn=HPS5;refGene_tx=XM_019812151.2;refGene_gn=HPS5;refGene_tx=XM_023239634.1
D1 76216256 76216380
refGene_gn=HPS5;refGene_tx=XM_003993038.4;refGene_gn=HPS5;refGene_tx=X

M_006937131.3;refGene_gn=HPS5;refGene_tx=XM_019812150.2;refGene_gn=HPS5;refGene_tx=XM_019812151.2;refGene_gn=HPS5;refGene_tx=XM_023239634.1
D1 76217053 76217249
refGene_gn=HPS5;refGene_tx=XM_003993038.4;refGene_gn=HPS5;refGene_tx=XM_006937131.3;refGene_gn=HPS5;refGene_tx=XM_019812150.2;refGene_gn=HPS5;refGene_tx=XM_019812151.2;refGene_gn=HPS5;refGene_tx=XM_023239634.1
D1 76217839 76217998
refGene_gn=HPS5;refGene_tx=XM_003993038.4;refGene_gn=HPS5;refGene_tx=XM_006937131.3;refGene_gn=HPS5;refGene_tx=XM_019812150.2;refGene_gn=HPS5;refGene_tx=XM_019812151.2;refGene_gn=HPS5;refGene_tx=XM_023239634.1
D1 76218439 76218618
refGene_gn=HPS5;refGene_tx=XM_003993038.4;refGene_gn=HPS5;refGene_tx=XM_006937131.3;refGene_gn=HPS5;refGene_tx=XM_019812150.2;refGene_gn=HPS5;refGene_tx=XM_019812151.2;refGene_gn=HPS5;refGene_tx=XM_023239634.1
D1 76220573 76220662
refGene_gn=HPS5;refGene_tx=XM_003993038.4;refGene_gn=HPS5;refGene_tx=XM_006937131.3;refGene_gn=HPS5;refGene_tx=XM_019812150.2;refGene_gn=HPS5;refGene_tx=XM_019812151.2;refGene_gn=HPS5;refGene_tx=XM_023239634.1
D1 76224113 76224185
refGene_gn=HPS5;refGene_tx=XM_003993038.4;refGene_gn=HPS5;refGene_tx=XM_006937131.3;refGene_gn=HPS5;refGene_tx=XM_019812150.2;refGene_gn=HPS5;refGene_tx=XM_019812151.2;refGene_gn=HPS5;refGene_tx=XM_023239634.1
D1 76224472 76224685
refGene_gn=HPS5;refGene_tx=XM_003993038.4;refGene_gn=HPS5;refGene_tx=XM_006937131.3;refGene_gn=HPS5;refGene_tx=XM_019812150.2;refGene_gn=HPS5;refGene_tx=XM_019812151.2;refGene_gn=HPS5;refGene_tx=XM_023239634.1
D1 76228130 76228264
refGene_gn=HPS5;refGene_tx=XM_003993038.4;refGene_gn=HPS5;refGene_tx=XM_006937131.3;refGene_gn=HPS5;refGene_tx=XM_019812150.2;refGene_gn=HPS5;refGene_tx=XM_019812151.2;refGene_gn=HPS5;refGene_tx=XM_023239634.1
D1 76229622 76229815
refGene_gn=HPS5;refGene_tx=XM_003993038.4;refGene_gn=HPS5;refGene_tx=XM_006937131.3;refGene_gn=HPS5;refGene_tx=XM_019812150.2;refGene_gn=HPS5;refGene_tx=XM_019812151.2;refGene_gn=HPS5;refGene_tx=XM_023239634.1
D1 76230273 76230338
refGene_gn=HPS5;refGene_tx=XM_003993038.4;refGene_gn=HPS5;refGene_tx=XM_006937131.3;refGene_gn=HPS5;refGene_tx=XM_019812150.2;refGene_gn=HPS5;refGene_tx=XM_019812151.2;refGene_gn=HPS5;refGene_tx=XM_023239634.1
D1 76230748 76230859
refGene_gn=HPS5;refGene_tx=XM_003993038.4;refGene_gn=HPS5;refGene_tx=XM_006937131.3;refGene_gn=HPS5;refGene_tx=XM_019812150.2;refGene_gn=HPS5;refGene_tx=XM_019812151.2;refGene_gn=HPS5;refGene_tx=XM_023239634.1
D1 76236208 76236316
refGene_gn=HPS5;refGene_tx=XM_003993038.4;refGene_gn=HPS5;refGene_tx=XM_006937131.3;refGene_gn=HPS5;refGene_tx=XM_019812150.2;refGene_gn=HPS5;refGene_tx=XM_019812151.2;refGene_gn=HPS5;refGene_tx=XM_023239634.1
D1 76251145 76251299 refGene_gn=GTF2H1;refGene_tx=XM_003993039.4
D1 76254016 76254209 refGene_gn=GTF2H1;refGene_tx=XM_003993039.4
D1 76254986 76255152 refGene_gn=GTF2H1;refGene_tx=XM_003993039.4
D1 76256815 76256909 refGene_gn=GTF2H1;refGene_tx=XM_003993039.4

D1 76258597 76258747 refGene_gn=GTF2H1;refGene_tx=XM_003993039.4
D1 76258880 76258960 refGene_gn=GTF2H1;refGene_tx=XM_003993039.4
D1 76263043 76263171 refGene_gn=GTF2H1;refGene_tx=XM_003993039.4
D1 76263280 76263368 refGene_gn=GTF2H1;refGene_tx=XM_003993039.4
D1 76267521 76267610 refGene_gn=GTF2H1;refGene_tx=XM_003993039.4
D1 76268161 76268279 refGene_gn=GTF2H1;refGene_tx=XM_003993039.4
D1 76271855 76271946 refGene_gn=GTF2H1;refGene_tx=XM_003993039.4
D1 76272415 76272531 refGene_gn=GTF2H1;refGene_tx=XM_003993039.4
D1 76273802 76273895 refGene_gn=GTF2H1;refGene_tx=XM_003993039.4
D1 76278720 76278807 refGene_gn=GTF2H1;refGene_tx=XM_003993039.4
D1 76301016 76301079 refGene_gn=LDHA;refGene_tx=XM_019812152.2
D1 76301289 76301439
refGene_gn=LDHA;refGene_tx=XM_006937133.4;refGene_gn=LDHA;refGene_tx=X
M_019812152.2
D1 76304265 76304383
refGene_gn=LDHA;refGene_tx=XM_006937133.4;refGene_gn=LDHA;refGene_tx=X
M_019812152.2
D1 76305312 76305486
refGene_gn=LDHA;refGene_tx=XM_006937133.4;refGene_gn=LDHA;refGene_tx=X
M_019812152.2
D1 76305930 76306104
refGene_gn=LDHA;refGene_tx=XM_006937133.4;refGene_gn=LDHA;refGene_tx=X
M_019812152.2
D1 76307412 76307530
refGene_gn=LDHA;refGene_tx=XM_006937133.4;refGene_gn=LDHA;refGene_tx=X
M_019812152.2
D1 76309063 76309187
refGene_gn=LDHA;refGene_tx=XM_006937133.4;refGene_gn=LDHA;refGene_tx=X
M_019812152.2
D1 76309737 76309902
refGene_gn=LDHA;refGene_tx=XM_006937133.4;refGene_gn=LDHA;refGene_tx=X
M_019812152.2
D1 76314977 76315103
refGene_gn=LDHC;refGene_tx=XM_011286740.3;refGene_gn=LDHC;refGene_tx=X
M_011286741.3
D1 76324055 76324173
refGene_gn=LDHC;refGene_tx=XM_011286740.3;refGene_gn=LDHC;refGene_tx=X
M_011286741.3
D1 76333142 76333316
refGene_gn=LDHC;refGene_tx=XM_011286740.3;refGene_gn=LDHC;refGene_tx=X
M_011286741.3
D1 76347241 76347415
refGene_gn=LDHC;refGene_tx=XM_011286740.3;refGene_gn=LDHC;refGene_tx=X
M_011286741.3
D1 76350105 76350223
refGene_gn=LDHC;refGene_tx=XM_011286740.3;refGene_gn=LDHC;refGene_tx=X
M_011286741.3
D1 76353702 76353826
refGene_gn=LDHC;refGene_tx=XM_011286740.3;refGene_gn=LDHC;refGene_tx=X
M_011286741.3

D1 76359883 76360048
refGene_gn=LDHC;refGene_tx=XM_011286740.3;refGene_gn=LDHC;refGene_tx=XM_011286741.3
D1 76363031 76363231 refGene_gn=LDHAL6A;refGene_tx=XM_023239729.1
D1 76363506 76363684 refGene_gn=LDHAL6A;refGene_tx=XM_023239729.1
D1 76368195 76368313 refGene_gn=LDHAL6A;refGene_tx=XM_023239729.1
D1 76369067 76369274 refGene_gn=LDHAL6A;refGene_tx=XM_023239729.1
D1 76375360 76375384 refGene_gn=LDHAL6A;refGene_tx=XM_023239729.1
D1 76375386 76375533 refGene_gn=LDHAL6A;refGene_tx=XM_023239729.1
D1 76376668 76376683 refGene_gn=LDHAL6A;refGene_tx=XM_023239729.1
D1 76376685 76376776 refGene_gn=LDHAL6A;refGene_tx=XM_023239729.1
D1 76377453 76377493 refGene_gn=LDHAL6A;refGene_tx=XM_023239729.1
D1 76377494 76377523 refGene_gn=LDHAL6A;refGene_tx=XM_023239729.1
D1 76377524 76377576 refGene_gn=LDHAL6A;refGene_tx=XM_023239729.1
D1 76378505 76378694 refGene_gn=LDHAL6A;refGene_tx=XM_023239729.1
D1 76380018 76380108
refGene_gn=TSG101;refGene_tx=XM_011286742.3;refGene_gn=TSG101;refGene_tx=XM_011286743.3
D1 76381356 76381596
refGene_gn=TSG101;refGene_tx=XM_011286742.3;refGene_gn=TSG101;refGene_tx=XM_011286743.3
D1 76382660 76382863
refGene_gn=TSG101;refGene_tx=XM_011286742.3;refGene_gn=TSG101;refGene_tx=XM_011286743.3
D1 76393392 76393484
refGene_gn=TSG101;refGene_tx=XM_011286742.3;refGene_gn=TSG101;refGene_tx=XM_011286743.3
D1 76397926 76397993
refGene_gn=TSG101;refGene_tx=XM_011286742.3;refGene_gn=TSG101;refGene_tx=XM_011286743.3
D1 76403213 76403240 refGene_gn=TSG101;refGene_tx=XM_011286742.3
D1 76403331 76403458
refGene_gn=TSG101;refGene_tx=XM_011286742.3;refGene_gn=TSG101;refGene_tx=XM_011286743.3
D1 76409981 76410145
refGene_gn=TSG101;refGene_tx=XM_011286742.3;refGene_gn=TSG101;refGene_tx=XM_011286743.3
D1 76412987 76413053
refGene_gn=TSG101;refGene_tx=XM_011286742.3;refGene_gn=TSG101;refGene_tx=XM_011286743.3
D1 76415293 76415378
refGene_gn=TSG101;refGene_tx=XM_011286742.3;refGene_gn=TSG101;refGene_tx=XM_011286743.3
D1 76419565 76419607
refGene_gn=TSG101;refGene_tx=XM_011286742.3;refGene_gn=TSG101;refGene_tx=XM_011286743.3
D1 76425008 76425119
refGene_gn=UEVLD;refGene_tx=XM_003993044.4;refGene_gn=UEVLD;refGene_tx=XM_019812156.2;refGene_gn=UEVLD;refGene_tx=XM_019812157.2;refGene_gn=UEVLD;refGene_tx=XM_019812159.2

D1 76427265 76427292 refGene_gn=UEVLD;refGene_tx=XM_019812158.2
D1 76428100 76428224
refGene_gn=UEVLD;refGene_tx=XM_003993044.4;refGene_gn=UEVLD;refGene_tx=XM_019812156.2;refGene_gn=UEVLD;refGene_tx=XM_019812157.2;refGene_gn=UEVLD;refGene_tx=XM_019812158.2;refGene_gn=UEVLD;refGene_tx=XM_019812159.2
D1 76430453 76430517
refGene_gn=UEVLD;refGene_tx=XM_003993044.4;refGene_gn=UEVLD;refGene_tx=XM_019812156.2;refGene_gn=UEVLD;refGene_tx=XM_019812157.2;refGene_gn=UEVLD;refGene_tx=XM_019812158.2;refGene_gn=UEVLD;refGene_tx=XM_019812159.2
D1 76436929 76437103
refGene_gn=UEVLD;refGene_tx=XM_003993044.4;refGene_gn=UEVLD;refGene_tx=XM_019812156.2;refGene_gn=UEVLD;refGene_tx=XM_019812157.2;refGene_gn=UEVLD;refGene_tx=XM_019812158.2
D1 76438569 76438740
refGene_gn=UEVLD;refGene_tx=XM_003993044.4;refGene_gn=UEVLD;refGene_tx=XM_019812156.2;refGene_gn=UEVLD;refGene_tx=XM_019812157.2;refGene_gn=UEVLD;refGene_tx=XM_019812158.2;refGene_gn=UEVLD;refGene_tx=XM_019812160.2
D1 76449321 76449424
refGene_gn=UEVLD;refGene_tx=XM_003993044.4;refGene_gn=UEVLD;refGene_tx=XM_019812156.2;refGene_gn=UEVLD;refGene_tx=XM_019812157.2;refGene_gn=UEVLD;refGene_tx=XM_019812158.2;refGene_gn=UEVLD;refGene_tx=XM_019812159.2;refGene_gn=UEVLD;refGene_tx=XM_019812160.2
D1 76452983 76453102
refGene_gn=UEVLD;refGene_tx=XM_003993044.4;refGene_gn=UEVLD;refGene_tx=XM_019812156.2;refGene_gn=UEVLD;refGene_tx=XM_019812157.2;refGene_gn=UEVLD;refGene_tx=XM_019812158.2;refGene_gn=UEVLD;refGene_tx=XM_019812159.2;refGene_gn=UEVLD;refGene_tx=XM_019812160.2
D1 76454272 76454408
refGene_gn=UEVLD;refGene_tx=XM_003993044.4;refGene_gn=UEVLD;refGene_tx=XM_019812156.2;refGene_gn=UEVLD;refGene_tx=XM_019812157.2;refGene_gn=UEVLD;refGene_tx=XM_019812158.2;refGene_gn=UEVLD;refGene_tx=XM_019812159.2;refGene_gn=UEVLD;refGene_tx=XM_019812160.2
D1 76457935 76458099
refGene_gn=UEVLD;refGene_tx=XM_003993044.4;refGene_gn=UEVLD;refGene_tx=XM_019812156.2;refGene_gn=UEVLD;refGene_tx=XM_019812157.2;refGene_gn=UEVLD;refGene_tx=XM_019812158.2;refGene_gn=UEVLD;refGene_tx=XM_019812159.2;refGene_gn=UEVLD;refGene_tx=XM_019812160.2
D1 76463876 76463942
refGene_gn=UEVLD;refGene_tx=XM_003993044.4;refGene_gn=UEVLD;refGene_tx=XM_019812156.2;refGene_gn=UEVLD;refGene_tx=XM_019812157.2;refGene_gn=UEVLD;refGene_tx=XM_019812158.2;refGene_gn=UEVLD;refGene_tx=XM_019812159.2;refGene_gn=UEVLD;refGene_tx=XM_019812160.2
D1 76464178 76464179 refGene_gn=UEVLD;refGene_tx=XM_019812157.2
D1 76468236 76468321
refGene_gn=UEVLD;refGene_tx=XM_003993044.4;refGene_gn=UEVLD;refGene_tx=XM_019812156.2;refGene_gn=UEVLD;refGene_tx=XM_019812158.2;refGene_gn=UEVLD;refGene_tx=XM_019812159.2;refGene_gn=UEVLD;refGene_tx=XM_0198121

60.2
D1	76476327	76476375
refGene_gn=UEVLD;refGene_tx=XM_003993044.4;refGene_gn=UEVLD;refGene_tx=XM_019812158.2;refGene_gn=UEVLD;refGene_tx=XM_019812159.2;refGene_gn=UEVLD;refGene_tx=XM_019812160.2
D1	76494252	76494346
refGene_gn=SPTY2D1;refGene_tx=XM_023239635.1;refGene_gn=SPTY2D1;refGene_tx=XM_023239636.1;refGene_gn=SPTY2D1;refGene_tx=XM_023239637.1;refGene_gn=SPTY2D1;refGene_tx=XM_023239638.1
D1	76496472	76496550
refGene_gn=SPTY2D1;refGene_tx=XM_023239635.1;refGene_gn=SPTY2D1;refGene_tx=XM_023239636.1;refGene_gn=SPTY2D1;refGene_tx=XM_023239637.1;refGene_gn=SPTY2D1;refGene_tx=XM_023239638.1
D1	76498112	76498284
refGene_gn=SPTY2D1;refGene_tx=XM_023239635.1;refGene_gn=SPTY2D1;refGene_tx=XM_023239636.1;refGene_gn=SPTY2D1;refGene_tx=XM_023239637.1;refGene_gn=SPTY2D1;refGene_tx=XM_023239638.1
D1	76499711	76501292
refGene_gn=SPTY2D1;refGene_tx=XM_023239635.1;refGene_gn=SPTY2D1;refGene_tx=XM_023239636.1;refGene_gn=SPTY2D1;refGene_tx=XM_023239637.1;refGene_gn=SPTY2D1;refGene_tx=XM_023239638.1
D1	76502054	76502169
refGene_gn=SPTY2D1;refGene_tx=XM_023239635.1;refGene_gn=SPTY2D1;refGene_tx=XM_023239636.1
D1	76513012	76513072
refGene_gn=SPTY2D1;refGene_tx=XM_023239635.1;refGene_gn=SPTY2D1;refGene_tx=XM_023239636.1
D1	76538425	76538728
refGene_gn=TMEM86A;refGene_tx=XM_019812161.2;refGene_gn=TMEM86A;refGene_tx=XM_023239639.1
D1	76538847	76538868	refGene_gn=TMEM86A;refGene_tx=XM_003993050.5
D1	76541003	76541268
refGene_gn=TMEM86A;refGene_tx=XM_003993050.5;refGene_gn=TMEM86A;refGene_tx=XM_019812161.2;refGene_gn=TMEM86A;refGene_tx=XM_023239639.1
D1	76541616	76542053
refGene_gn=TMEM86A;refGene_tx=XM_003993050.5;refGene_gn=TMEM86A;refGene_tx=XM_019812161.2
D1	76544509	76544580	refGene_gn=IGSF22;refGene_tx=XM_003993051.5
D1	76545839	76546169	refGene_gn=IGSF22;refGene_tx=XM_003993051.5
D1	76546364	76546522	refGene_gn=TMEM86A;refGene_tx=XM_023239639.1
D1	76546931	76547231	refGene_gn=IGSF22;refGene_tx=XM_003993051.5
D1	76547971	76548164	refGene_gn=IGSF22;refGene_tx=XM_003993051.5
D1	76548344	76548433	refGene_gn=IGSF22;refGene_tx=XM_003993051.5
D1	76548883	76549180	refGene_gn=IGSF22;refGene_tx=XM_003993051.5
D1	76550027	76550156	refGene_gn=IGSF22;refGene_tx=XM_003993051.5
D1	76550366	76550540	refGene_gn=IGSF22;refGene_tx=XM_003993051.5
D1	76551285	76551588	refGene_gn=IGSF22;refGene_tx=XM_003993051.5
D1	76552673	76552970	refGene_gn=IGSF22;refGene_tx=XM_003993051.5
D1	76553132	76553274	refGene_gn=IGSF22;refGene_tx=XM_003993051.5
D1	76553370	76553495	refGene_gn=IGSF22;refGene_tx=XM_003993051.5

D1 76554233 76554518 refGene_gn=IGSF22;refGene_tx=XM_003993051.5
D1 76555493 76555766 refGene_gn=IGSF22;refGene_tx=XM_003993051.5
D1 76556407 76556570 refGene_gn=IGSF22;refGene_tx=XM_003993051.5
D1 76557061 76557175 refGene_gn=IGSF22;refGene_tx=XM_003993051.5
D1 76557790 76557965 refGene_gn=IGSF22;refGene_tx=XM_003993051.5
D1 76558168 76558208 refGene_gn=IGSF22;refGene_tx=XM_003993051.5
D1 76558289 76558389 refGene_gn=IGSF22;refGene_tx=XM_003993051.5
D1 76559539 76559676 refGene_gn=IGSF22;refGene_tx=XM_003993051.5
D1 76559903 76560035 refGene_gn=IGSF22;refGene_tx=XM_003993051.5
D1 76562125 76562234 refGene_gn=IGSF22;refGene_tx=XM_003993051.5
D1 76567126 76567220
refGene_gn=PTPN5;refGene_tx=XM_019812154.2;refGene_gn=PTPN5;refGene_tx=XM_023239640.1;refGene_gn=PTPN5;refGene_tx=XM_023239641.1
D1 76567642 76567756
refGene_gn=PTPN5;refGene_tx=XM_019812154.2;refGene_gn=PTPN5;refGene_tx=XM_023239640.1;refGene_gn=PTPN5;refGene_tx=XM_023239641.1
D1 76567847 76568008
refGene_gn=PTPN5;refGene_tx=XM_019812154.2;refGene_gn=PTPN5;refGene_tx=XM_023239640.1;refGene_gn=PTPN5;refGene_tx=XM_023239641.1
D1 76569842 76569953
refGene_gn=PTPN5;refGene_tx=XM_019812154.2;refGene_gn=PTPN5;refGene_tx=XM_023239640.1;refGene_gn=PTPN5;refGene_tx=XM_023239641.1
D1 76570444 76570582
refGene_gn=PTPN5;refGene_tx=XM_019812154.2;refGene_gn=PTPN5;refGene_tx=XM_023239640.1;refGene_gn=PTPN5;refGene_tx=XM_023239641.1
D1 76570766 76570846
refGene_gn=PTPN5;refGene_tx=XM_019812154.2;refGene_gn=PTPN5;refGene_tx=XM_023239640.1;refGene_gn=PTPN5;refGene_tx=XM_023239641.1
D1 76574549 76574634
refGene_gn=PTPN5;refGene_tx=XM_019812154.2;refGene_gn=PTPN5;refGene_tx=XM_023239640.1;refGene_gn=PTPN5;refGene_tx=XM_023239641.1
D1 76576983 76577173
refGene_gn=PTPN5;refGene_tx=XM_019812154.2;refGene_gn=PTPN5;refGene_tx=XM_023239640.1;refGene_gn=PTPN5;refGene_tx=XM_023239641.1
D1 76578339 76578581
refGene_gn=PTPN5;refGene_tx=XM_019812154.2;refGene_gn=PTPN5;refGene_tx=XM_023239640.1;refGene_gn=PTPN5;refGene_tx=XM_023239641.1
D1 76579064 76579142
refGene_gn=PTPN5;refGene_tx=XM_023239640.1;refGene_gn=PTPN5;refGene_tx=XM_023239641.1
D1 76579389 76579497
refGene_gn=PTPN5;refGene_tx=XM_023239640.1;refGene_gn=PTPN5;refGene_tx=XM_023239641.1
D1 76580058 76580252
refGene_gn=PTPN5;refGene_tx=XM_019812154.2;refGene_gn=PTPN5;refGene_tx=XM_023239640.1;refGene_gn=PTPN5;refGene_tx=XM_023239641.1
D1 76599113 76599190
refGene_gn=PTPN5;refGene_tx=XM_019812154.2;refGene_gn=PTPN5;refGene_tx=XM_023239640.1;refGene_gn=PTPN5;refGene_tx=XM_023239641.1
D1 76605580 76605600

refGene_gn=PTPN5;refGene_tx=XM_019812154.2;refGene_gn=PTPN5;refGene_tx=XM_023239640.1
D1	76676190	76677208	refGene_gn=MRGPRX2;refGene_tx=XM_003993106.5
D1	76679038	76679049	refGene_gn=MRGPRX2;refGene_tx=XM_003993106.5
D1	76725333	76725360	refGene_gn=ZDHHC13;refGene_tx=XM_006937136.4;refGene_gn=ZDHHC13;refGene_tx=XM_019812164.2;refGene_gn=ZDHHC13;refGene_tx=XM_019812165.2;refGene_gn=ZDHHC13;refGene_tx=XM_023239642.1
D1	76739552	76739698	refGene_gn=ZDHHC13;refGene_tx=XM_006937136.4;refGene_gn=ZDHHC13;refGene_tx=XM_019812164.2;refGene_gn=ZDHHC13;refGene_tx=XM_019812165.2;refGene_gn=ZDHHC13;refGene_tx=XM_023239642.1
D1	76743208	76743331	refGene_gn=ZDHHC13;refGene_tx=XM_006937136.4;refGene_gn=ZDHHC13;refGene_tx=XM_019812164.2;refGene_gn=ZDHHC13;refGene_tx=XM_019812165.2;refGene_gn=ZDHHC13;refGene_tx=XM_023239642.1
D1	76744781	76744859	refGene_gn=ZDHHC13;refGene_tx=XM_006937136.4;refGene_gn=ZDHHC13;refGene_tx=XM_019812164.2;refGene_gn=ZDHHC13;refGene_tx=XM_019812165.2;refGene_gn=ZDHHC13;refGene_tx=XM_023239642.1
D1	76746752	76746897	refGene_gn=ZDHHC13;refGene_tx=XM_006937136.4;refGene_gn=ZDHHC13;refGene_tx=XM_019812164.2;refGene_gn=ZDHHC13;refGene_tx=XM_019812165.2;refGene_gn=ZDHHC13;refGene_tx=XM_019812167.2;refGene_gn=ZDHHC13;refGene_tx=XM_023239642.1
D1	76748320	76748385	refGene_gn=ZDHHC13;refGene_tx=XM_006937136.4;refGene_gn=ZDHHC13;refGene_tx=XM_019812164.2;refGene_gn=ZDHHC13;refGene_tx=XM_019812165.2;refGene_gn=ZDHHC13;refGene_tx=XM_019812167.2;refGene_gn=ZDHHC13;refGene_tx=XM_023239642.1
D1	76749788	76749951	refGene_gn=ZDHHC13;refGene_tx=XM_006937136.4;refGene_gn=ZDHHC13;refGene_tx=XM_019812164.2;refGene_gn=ZDHHC13;refGene_tx=XM_019812165.2;refGene_gn=ZDHHC13;refGene_tx=XM_019812167.2;refGene_gn=ZDHHC13;refGene_tx=XM_023239642.1
D1	76750205	76750331	refGene_gn=ZDHHC13;refGene_tx=XM_006937136.4;refGene_gn=ZDHHC13;refGene_tx=XM_019812164.2;refGene_gn=ZDHHC13;refGene_tx=XM_019812165.2;refGene_gn=ZDHHC13;refGene_tx=XM_019812167.2;refGene_gn=ZDHHC13;refGene_tx=XM_023239642.1
D1	76753882	76754016	refGene_gn=ZDHHC13;refGene_tx=XM_006937136.4;refGene_gn=ZDHHC13;refGene_tx=XM_019812164.2;refGene_gn=ZDHHC13;refGene_tx=XM_019812165.2;refGene_gn=ZDHHC13;refGene_tx=XM_019812167.2;refGene_gn=ZDHHC13;refGene_tx=XM_023239642.1
D1	76757422	76757523	refGene_gn=ZDHHC13;refGene_tx=XM_006937136.4;refGene_gn=ZDHHC13;refGene_tx=XM_019812164.2;refGene_gn=ZDHHC13;refGene_tx=XM_019812165.2;refGene_gn=ZDHHC13;refGene_tx=XM_019812167.2;refGene_gn=ZDHHC13;refGene_tx=XM_023239642.1

D1 76762515 76762640
refGene_gn=ZDHHC13;refGene_tx=XM_006937136.4;refGene_gn=ZDHHC13;refGene_tx=XM_019812164.2;refGene_gn=ZDHHC13;refGene_tx=XM_019812165.2;refGene_gn=ZDHHC13;refGene_tx=XM_019812167.2;refGene_gn=ZDHHC13;refGene_tx=XM_023239642.1
D1 76763483 76763546
refGene_gn=ZDHHC13;refGene_tx=XM_006937136.4;refGene_gn=ZDHHC13;refGene_tx=XM_019812164.2;refGene_gn=ZDHHC13;refGene_tx=XM_019812165.2;refGene_gn=ZDHHC13;refGene_tx=XM_019812167.2;refGene_gn=ZDHHC13;refGene_tx=XM_023239642.1
D1 76764243 76764337
refGene_gn=ZDHHC13;refGene_tx=XM_006937136.4;refGene_gn=ZDHHC13;refGene_tx=XM_019812164.2;refGene_gn=ZDHHC13;refGene_tx=XM_019812165.2;refGene_gn=ZDHHC13;refGene_tx=XM_019812167.2;refGene_gn=ZDHHC13;refGene_tx=XM_023239642.1
D1 76765903 76765987
refGene_gn=ZDHHC13;refGene_tx=XM_006937136.4;refGene_gn=ZDHHC13;refGene_tx=XM_019812164.2;refGene_gn=ZDHHC13;refGene_tx=XM_019812165.2;refGene_gn=ZDHHC13;refGene_tx=XM_019812167.2;refGene_gn=ZDHHC13;refGene_tx=XM_023239642.1
D1 76770018 76770176
refGene_gn=ZDHHC13;refGene_tx=XM_006937136.4;refGene_gn=ZDHHC13;refGene_tx=XM_019812164.2;refGene_gn=ZDHHC13;refGene_tx=XM_019812165.2;refGene_gn=ZDHHC13;refGene_tx=XM_019812167.2;refGene_gn=ZDHHC13;refGene_tx=XM_023239642.1
D1 76773448 76773546
refGene_gn=ZDHHC13;refGene_tx=XM_006937136.4;refGene_gn=ZDHHC13;refGene_tx=XM_019812164.2;refGene_gn=ZDHHC13;refGene_tx=XM_019812165.2;refGene_gn=ZDHHC13;refGene_tx=XM_019812167.2;refGene_gn=ZDHHC13;refGene_tx=XM_023239642.1
D1 76775972 76776111
refGene_gn=ZDHHC13;refGene_tx=XM_006937136.4;refGene_gn=ZDHHC13;refGene_tx=XM_019812167.2
D1 76777080 76777102 refGene_gn=ZDHHC13;refGene_tx=XM_023239642.1
D1 76782348 76782472
refGene_gn=ZDHHC13;refGene_tx=XM_019812164.2;refGene_gn=ZDHHC13;refGene_tx=XM_019812165.2
D1 76784714 76784715 refGene_gn=ZDHHC13;refGene_tx=XM_019812165.2
D1 76785202 76785279 refGene_gn=CSRP3;refGene_tx=XM_011286744.1
D1 76786862 76786956 refGene_gn=CSRP3;refGene_tx=XM_011286744.1
D1 76788118 76788251 refGene_gn=CSRP3;refGene_tx=XM_011286744.1
D1 76790188 76790357 refGene_gn=CSRP3;refGene_tx=XM_011286744.1
D1 76794953 76795065 refGene_gn=CSRP3;refGene_tx=XM_011286744.1
D1 76826118 76826301
refGene_gn=E2F8;refGene_tx=XM_006937137.3;refGene_gn=E2F8;refGene_tx=XM_019812163.2
D1 76826608 76826997
refGene_gn=E2F8;refGene_tx=XM_006937137.3;refGene_gn=E2F8;refGene_tx=XM_019812163.2
D1 76827111 76827244

refGene_gn=E2F8;refGene_tx=XM_006937137.3;refGene_gn=E2F8;refGene_tx=X
M_019812163.2
D1 76830616 76831148
refGene_gn=E2F8;refGene_tx=XM_006937137.3;refGene_gn=E2F8;refGene_tx=X
M_019812163.2
D1 76831398 76831486
refGene_gn=E2F8;refGene_tx=XM_006937137.3;refGene_gn=E2F8;refGene_tx=X
M_019812163.2
D1 76831798 76832002
refGene_gn=E2F8;refGene_tx=XM_006937137.3;refGene_gn=E2F8;refGene_tx=X
M_019812163.2
D1 76833745 76833883
refGene_gn=E2F8;refGene_tx=XM_006937137.3;refGene_gn=E2F8;refGene_tx=X
M_019812163.2
D1 76835470 76835632
refGene_gn=E2F8;refGene_tx=XM_006937137.3;refGene_gn=E2F8;refGene_tx=X
M_019812163.2
D1 76835857 76836172
refGene_gn=E2F8;refGene_tx=XM_006937137.3;refGene_gn=E2F8;refGene_tx=X
M_019812163.2
D1 76838562 76838719
refGene_gn=E2F8;refGene_tx=XM_006937137.3;refGene_gn=E2F8;refGene_tx=X
M_019812163.2
D1 76839117 76839396
refGene_gn=E2F8;refGene_tx=XM_006937137.3;refGene_gn=E2F8;refGene_tx=X
M_019812163.2
D1 76841157 76841172
refGene_gn=E2F8;refGene_tx=XM_006937137.3;refGene_gn=E2F8;refGene_tx=X
M_019812163.2
D1 76944531 76944606 refGene_gn=NAV2;refGene_tx=XM_023239646.1
D1 77284610 77284868
refGene_gn=NAV2;refGene_tx=XM_019812168.2;refGene_gn=NAV2;refGene_tx=X
M_019812169.2;refGene_gn=NAV2;refGene_tx=XM_019812170.2;refGene_gn=NAV
2;refGene_tx=XM_019812171.2;refGene_gn=NAV2;refGene_tx=XM_019812172.2;
refGene_gn=NAV2;refGene_tx=XM_019812173.2;refGene_gn=NAV2;refGene_tx=X
M_019812174.2;refGene_gn=NAV2;refGene_tx=XM_019812175.2;refGene_gn=NAV
2;refGene_tx=XM_019812176.2;refGene_gn=NAV2;refGene_tx=XM_019812177.2;
refGene_gn=NAV2;refGene_tx=XM_019812178.2;refGene_gn=NAV2;refGene_tx=X
M_019812183.2;refGene_gn=NAV2;refGene_tx=XM_023239644.1
D1 77352545 77352596 refGene_gn=NAV2;refGene_tx=XM_019812182.2
D1 77392753 77392825 refGene_gn=NAV2;refGene_tx=XM_019812181.2
D1 77411875 77411993
refGene_gn=NAV2;refGene_tx=XM_019812168.2;refGene_gn=NAV2;refGene_tx=X
M_019812169.2;refGene_gn=NAV2;refGene_tx=XM_019812170.2;refGene_gn=NAV
2;refGene_tx=XM_019812171.2;refGene_gn=NAV2;refGene_tx=XM_019812172.2;
refGene_gn=NAV2;refGene_tx=XM_019812173.2;refGene_gn=NAV2;refGene_tx=X
M_019812174.2;refGene_gn=NAV2;refGene_tx=XM_019812175.2;refGene_gn=NAV
2;refGene_tx=XM_019812176.2;refGene_gn=NAV2;refGene_tx=XM_019812177.2;
refGene_gn=NAV2;refGene_tx=XM_019812178.2;refGene_gn=NAV2;refGene_tx=X
M_019812181.2;refGene_gn=NAV2;refGene_tx=XM_019812182.2;refGene_gn=NAV

2;refGene_tx=XM_019812183.2;refGene_gn=NAV2;refGene_tx=XM_023239644.1;
refGene_gn=NAV2;refGene_tx=XM_023239646.1
D1 77422731 77422784
refGene_gn=NAV2;refGene_tx=XM_019812168.2;refGene_gn=NAV2;refGene_tx=X
M_019812169.2;refGene_gn=NAV2;refGene_tx=XM_019812170.2;refGene_gn=NAV
2;refGene_tx=XM_019812171.2;refGene_gn=NAV2;refGene_tx=XM_019812172.2;
refGene_gn=NAV2;refGene_tx=XM_019812173.2;refGene_gn=NAV2;refGene_tx=X
M_019812174.2;refGene_gn=NAV2;refGene_tx=XM_019812175.2;refGene_gn=NAV
2;refGene_tx=XM_019812176.2;refGene_gn=NAV2;refGene_tx=XM_019812177.2;
refGene_gn=NAV2;refGene_tx=XM_019812178.2;refGene_gn=NAV2;refGene_tx=X
M_019812181.2;refGene_gn=NAV2;refGene_tx=XM_019812182.2;refGene_gn=NAV
2;refGene_tx=XM_019812183.2;refGene_gn=NAV2;refGene_tx=XM_023239644.1;
refGene_gn=NAV2;refGene_tx=XM_023239646.1;refGene_gn=NAV2;refGene_tx=X
M_023239647.1
D1 77444747 77444820
refGene_gn=NAV2;refGene_tx=XM_019812168.2;refGene_gn=NAV2;refGene_tx=X
M_019812169.2;refGene_gn=NAV2;refGene_tx=XM_019812170.2;refGene_gn=NAV
2;refGene_tx=XM_019812171.2;refGene_gn=NAV2;refGene_tx=XM_019812172.2;
refGene_gn=NAV2;refGene_tx=XM_019812173.2;refGene_gn=NAV2;refGene_tx=X
M_019812174.2;refGene_gn=NAV2;refGene_tx=XM_019812175.2;refGene_gn=NAV
2;refGene_tx=XM_019812176.2;refGene_gn=NAV2;refGene_tx=XM_019812177.2;
refGene_gn=NAV2;refGene_tx=XM_019812178.2;refGene_gn=NAV2;refGene_tx=X
M_019812181.2;refGene_gn=NAV2;refGene_tx=XM_019812182.2;refGene_gn=NAV
2;refGene_tx=XM_019812183.2;refGene_gn=NAV2;refGene_tx=XM_023239644.1;
refGene_gn=NAV2;refGene_tx=XM_023239646.1;refGene_gn=NAV2;refGene_tx=X
M_023239647.1
D1 77454720 77454979
refGene_gn=NAV2;refGene_tx=XM_019812168.2;refGene_gn=NAV2;refGene_tx=X
M_019812169.2;refGene_gn=NAV2;refGene_tx=XM_019812170.2;refGene_gn=NAV
2;refGene_tx=XM_019812171.2;refGene_gn=NAV2;refGene_tx=XM_019812172.2;
refGene_gn=NAV2;refGene_tx=XM_019812173.2;refGene_gn=NAV2;refGene_tx=X
M_019812174.2;refGene_gn=NAV2;refGene_tx=XM_019812175.2;refGene_gn=NAV
2;refGene_tx=XM_019812176.2;refGene_gn=NAV2;refGene_tx=XM_019812177.2;
refGene_gn=NAV2;refGene_tx=XM_019812178.2;refGene_gn=NAV2;refGene_tx=X
M_019812181.2;refGene_gn=NAV2;refGene_tx=XM_019812182.2;refGene_gn=NAV
2;refGene_tx=XM_019812183.2;refGene_gn=NAV2;refGene_tx=XM_023239644.1;
refGene_gn=NAV2;refGene_tx=XM_023239646.1;refGene_gn=NAV2;refGene_tx=X
M_023239647.1
D1 77459671 77459740
refGene_gn=NAV2;refGene_tx=XM_019812168.2;refGene_gn=NAV2;refGene_tx=X
M_019812169.2;refGene_gn=NAV2;refGene_tx=XM_019812170.2;refGene_gn=NAV
2;refGene_tx=XM_019812171.2;refGene_gn=NAV2;refGene_tx=XM_019812173.2;
refGene_gn=NAV2;refGene_tx=XM_019812174.2;refGene_gn=NAV2;refGene_tx=X
M_019812175.2;refGene_gn=NAV2;refGene_tx=XM_019812177.2;refGene_gn=NAV
2;refGene_tx=XM_019812181.2;refGene_gn=NAV2;refGene_tx=XM_019812182.2;
refGene_gn=NAV2;refGene_tx=XM_019812183.2;refGene_gn=NAV2;refGene_tx=X
M_023239644.1;refGene_gn=NAV2;refGene_tx=XM_023239646.1;refGene_gn=NAV
2;refGene_tx=XM_023239647.1
D1 77468156 77468323
refGene_gn=NAV2;refGene_tx=XM_019812168.2;refGene_gn=NAV2;refGene_tx=X

M_019812169.2;refGene_gn=NAV2;refGene_tx=XM_019812170.2;refGene_gn=NAV2;refGene_tx=XM_019812171.2;refGene_gn=NAV2;refGene_tx=XM_019812172.2;refGene_gn=NAV2;refGene_tx=XM_019812173.2;refGene_gn=NAV2;refGene_tx=XM_019812174.2;refGene_gn=NAV2;refGene_tx=XM_019812175.2;refGene_gn=NAV2;refGene_tx=XM_019812176.2;refGene_gn=NAV2;refGene_tx=XM_019812177.2;refGene_gn=NAV2;refGene_tx=XM_019812178.2;refGene_gn=NAV2;refGene_tx=XM_019812181.2;refGene_gn=NAV2;refGene_tx=XM_019812182.2;refGene_gn=NAV2;refGene_tx=XM_019812183.2;refGene_gn=NAV2;refGene_tx=XM_023239644.1;refGene_gn=NAV2;refGene_tx=XM_023239646.1;refGene_gn=NAV2;refGene_tx=XM_023239647.1
D1 77513219 77514321
refGene_gn=NAV2;refGene_tx=XM_019812168.2;refGene_gn=NAV2;refGene_tx=XM_019812169.2;refGene_gn=NAV2;refGene_tx=XM_019812170.2;refGene_gn=NAV2;refGene_tx=XM_019812171.2;refGene_gn=NAV2;refGene_tx=XM_019812172.2;refGene_gn=NAV2;refGene_tx=XM_019812173.2;refGene_gn=NAV2;refGene_tx=XM_019812174.2;refGene_gn=NAV2;refGene_tx=XM_019812175.2;refGene_gn=NAV2;refGene_tx=XM_019812176.2;refGene_gn=NAV2;refGene_tx=XM_019812177.2;refGene_gn=NAV2;refGene_tx=XM_019812178.2;refGene_gn=NAV2;refGene_tx=XM_019812181.2;refGene_gn=NAV2;refGene_tx=XM_019812182.2;refGene_gn=NAV2;refGene_tx=XM_019812183.2;refGene_gn=NAV2;refGene_tx=XM_023239644.1;refGene_gn=NAV2;refGene_tx=XM_023239646.1;refGene_gn=NAV2;refGene_tx=XM_023239647.1
D1 77519345 77519458
refGene_gn=NAV2;refGene_tx=XM_019812168.2;refGene_gn=NAV2;refGene_tx=XM_019812169.2;refGene_gn=NAV2;refGene_tx=XM_019812170.2;refGene_gn=NAV2;refGene_tx=XM_019812171.2;refGene_gn=NAV2;refGene_tx=XM_019812172.2;refGene_gn=NAV2;refGene_tx=XM_019812173.2;refGene_gn=NAV2;refGene_tx=XM_019812174.2;refGene_gn=NAV2;refGene_tx=XM_019812175.2;refGene_gn=NAV2;refGene_tx=XM_019812176.2;refGene_gn=NAV2;refGene_tx=XM_019812177.2;refGene_gn=NAV2;refGene_tx=XM_019812178.2;refGene_gn=NAV2;refGene_tx=XM_019812181.2;refGene_gn=NAV2;refGene_tx=XM_019812182.2;refGene_gn=NAV2;refGene_tx=XM_019812183.2;refGene_gn=NAV2;refGene_tx=XM_023239644.1;refGene_gn=NAV2;refGene_tx=XM_023239646.1;refGene_gn=NAV2;refGene_tx=XM_023239647.1
D1 77524756 77524865
refGene_gn=NAV2;refGene_tx=XM_019812168.2;refGene_gn=NAV2;refGene_tx=XM_019812169.2;refGene_gn=NAV2;refGene_tx=XM_019812170.2;refGene_gn=NAV2;refGene_tx=XM_019812171.2;refGene_gn=NAV2;refGene_tx=XM_019812172.2;refGene_gn=NAV2;refGene_tx=XM_019812173.2;refGene_gn=NAV2;refGene_tx=XM_019812174.2;refGene_gn=NAV2;refGene_tx=XM_019812175.2;refGene_gn=NAV2;refGene_tx=XM_019812176.2;refGene_gn=NAV2;refGene_tx=XM_019812177.2;refGene_gn=NAV2;refGene_tx=XM_019812178.2;refGene_gn=NAV2;refGene_tx=XM_019812181.2;refGene_gn=NAV2;refGene_tx=XM_019812182.2;refGene_gn=NAV2;refGene_tx=XM_019812183.2;refGene_gn=NAV2;refGene_tx=XM_023239644.1;refGene_gn=NAV2;refGene_tx=XM_023239646.1;refGene_gn=NAV2;refGene_tx=XM_023239647.1
D1 77526773 77527163
refGene_gn=NAV2;refGene_tx=XM_019812168.2;refGene_gn=NAV2;refGene_tx=XM_019812169.2;refGene_gn=NAV2;refGene_tx=XM_019812170.2;refGene_gn=NAV2;refGene_tx=XM_019812171.2;refGene_gn=NAV2;refGene_tx=XM_019812172.2;

refGene_gn=NAV2;refGene_tx=XM_019812173.2;refGene_gn=NAV2;refGene_tx=XM_019812174.2;refGene_gn=NAV2;refGene_tx=XM_019812175.2;refGene_gn=NAV2;refGene_tx=XM_019812176.2;refGene_gn=NAV2;refGene_tx=XM_019812177.2;refGene_gn=NAV2;refGene_tx=XM_019812178.2;refGene_gn=NAV2;refGene_tx=XM_019812181.2;refGene_gn=NAV2;refGene_tx=XM_019812182.2;refGene_gn=NAV2;refGene_tx=XM_019812183.2;refGene_gn=NAV2;refGene_tx=XM_023239644.1;refGene_gn=NAV2;refGene_tx=XM_023239646.1;refGene_gn=NAV2;refGene_tx=XM_023239647.1
D1 77559402 77559525
refGene_gn=NAV2;refGene_tx=XM_019812168.2;refGene_gn=NAV2;refGene_tx=XM_019812169.2;refGene_gn=NAV2;refGene_tx=XM_019812170.2;refGene_gn=NAV2;refGene_tx=XM_019812171.2;refGene_gn=NAV2;refGene_tx=XM_019812172.2;refGene_gn=NAV2;refGene_tx=XM_019812173.2;refGene_gn=NAV2;refGene_tx=XM_019812174.2;refGene_gn=NAV2;refGene_tx=XM_019812175.2;refGene_gn=NAV2;refGene_tx=XM_019812176.2;refGene_gn=NAV2;refGene_tx=XM_019812177.2;refGene_gn=NAV2;refGene_tx=XM_019812178.2;refGene_gn=NAV2;refGene_tx=XM_019812181.2;refGene_gn=NAV2;refGene_tx=XM_019812182.2;refGene_gn=NAV2;refGene_tx=XM_019812183.2;refGene_gn=NAV2;refGene_tx=XM_023239644.1;refGene_gn=NAV2;refGene_tx=XM_023239646.1;refGene_gn=NAV2;refGene_tx=XM_023239647.1
D1 77608254 77608393
refGene_gn=NAV2;refGene_tx=XM_019812168.2;refGene_gn=NAV2;refGene_tx=XM_019812169.2;refGene_gn=NAV2;refGene_tx=XM_019812170.2;refGene_gn=NAV2;refGene_tx=XM_019812171.2;refGene_gn=NAV2;refGene_tx=XM_019812172.2;refGene_gn=NAV2;refGene_tx=XM_019812173.2;refGene_gn=NAV2;refGene_tx=XM_019812174.2;refGene_gn=NAV2;refGene_tx=XM_019812175.2;refGene_gn=NAV2;refGene_tx=XM_019812176.2;refGene_gn=NAV2;refGene_tx=XM_019812177.2;refGene_gn=NAV2;refGene_tx=XM_019812178.2;refGene_gn=NAV2;refGene_tx=XM_019812181.2;refGene_gn=NAV2;refGene_tx=XM_019812182.2;refGene_gn=NAV2;refGene_tx=XM_019812183.2;refGene_gn=NAV2;refGene_tx=XM_023239644.1;refGene_gn=NAV2;refGene_tx=XM_023239646.1;refGene_gn=NAV2;refGene_tx=XM_023239647.1
D1 77616860 77617152
refGene_gn=NAV2;refGene_tx=XM_019812168.2;refGene_gn=NAV2;refGene_tx=XM_019812169.2;refGene_gn=NAV2;refGene_tx=XM_019812170.2;refGene_gn=NAV2;refGene_tx=XM_019812171.2;refGene_gn=NAV2;refGene_tx=XM_019812172.2;refGene_gn=NAV2;refGene_tx=XM_019812173.2;refGene_gn=NAV2;refGene_tx=XM_019812174.2;refGene_gn=NAV2;refGene_tx=XM_019812175.2;refGene_gn=NAV2;refGene_tx=XM_019812176.2;refGene_gn=NAV2;refGene_tx=XM_019812177.2;refGene_gn=NAV2;refGene_tx=XM_019812178.2;refGene_gn=NAV2;refGene_tx=XM_019812181.2;refGene_gn=NAV2;refGene_tx=XM_019812182.2;refGene_gn=NAV2;refGene_tx=XM_019812183.2;refGene_gn=NAV2;refGene_tx=XM_023239644.1;refGene_gn=NAV2;refGene_tx=XM_023239646.1;refGene_gn=NAV2;refGene_tx=XM_023239647.1
D1 77617899 77618602
refGene_gn=NAV2;refGene_tx=XM_019812168.2;refGene_gn=NAV2;refGene_tx=XM_019812169.2;refGene_gn=NAV2;refGene_tx=XM_019812170.2;refGene_gn=NAV2;refGene_tx=XM_019812171.2;refGene_gn=NAV2;refGene_tx=XM_019812172.2;refGene_gn=NAV2;refGene_tx=XM_019812173.2;refGene_gn=NAV2;refGene_tx=XM_019812174.2;refGene_gn=NAV2;refGene_tx=XM_019812175.2;refGene_gn=NAV

2;refGene_tx=XM_019812176.2;refGene_gn=NAV2;refGene_tx=XM_019812177.2;
refGene_gn=NAV2;refGene_tx=XM_019812178.2;refGene_gn=NAV2;refGene_tx=X
M_019812181.2;refGene_gn=NAV2;refGene_tx=XM_019812182.2;refGene_gn=NAV
2;refGene_tx=XM_019812183.2;refGene_gn=NAV2;refGene_tx=XM_023239644.1;
refGene_gn=NAV2;refGene_tx=XM_023239646.1;refGene_gn=NAV2;refGene_tx=X
M_023239647.1
D1 77621259 77621727
refGene_gn=NAV2;refGene_tx=XM_019812168.2;refGene_gn=NAV2;refGene_tx=X
M_019812169.2;refGene_gn=NAV2;refGene_tx=XM_019812170.2;refGene_gn=NAV
2;refGene_tx=XM_019812171.2;refGene_gn=NAV2;refGene_tx=XM_019812172.2;
refGene_gn=NAV2;refGene_tx=XM_019812173.2;refGene_gn=NAV2;refGene_tx=X
M_019812174.2;refGene_gn=NAV2;refGene_tx=XM_019812175.2;refGene_gn=NAV
2;refGene_tx=XM_019812176.2;refGene_gn=NAV2;refGene_tx=XM_019812177.2;
refGene_gn=NAV2;refGene_tx=XM_019812178.2;refGene_gn=NAV2;refGene_tx=X
M_019812181.2;refGene_gn=NAV2;refGene_tx=XM_019812182.2;refGene_gn=NAV
2;refGene_tx=XM_019812183.2;refGene_gn=NAV2;refGene_tx=XM_023239644.1;
refGene_gn=NAV2;refGene_tx=XM_023239646.1;refGene_gn=NAV2;refGene_tx=X
M_023239647.1
D1 77622406 77622472
refGene_gn=NAV2;refGene_tx=XM_019812168.2;refGene_gn=NAV2;refGene_tx=X
M_019812169.2;refGene_gn=NAV2;refGene_tx=XM_019812170.2;refGene_gn=NAV
2;refGene_tx=XM_019812172.2;refGene_gn=NAV2;refGene_tx=XM_019812173.2;
refGene_gn=NAV2;refGene_tx=XM_019812175.2;refGene_gn=NAV2;refGene_tx=X
M_019812176.2;refGene_gn=NAV2;refGene_tx=XM_019812178.2;refGene_gn=NAV
2;refGene_tx=XM_019812181.2;refGene_gn=NAV2;refGene_tx=XM_019812182.2;
refGene_gn=NAV2;refGene_tx=XM_019812183.2;refGene_gn=NAV2;refGene_tx=X
M_023239644.1;refGene_gn=NAV2;refGene_tx=XM_023239646.1;refGene_gn=NAV
2;refGene_tx=XM_023239647.1
D1 77623852 77623897
refGene_gn=NAV2;refGene_tx=XM_019812168.2;refGene_gn=NAV2;refGene_tx=X
M_019812169.2;refGene_gn=NAV2;refGene_tx=XM_019812171.2;refGene_gn=NAV
2;refGene_tx=XM_019812172.2;refGene_gn=NAV2;refGene_tx=XM_019812173.2;
refGene_gn=NAV2;refGene_tx=XM_019812175.2;refGene_gn=NAV2;refGene_tx=X
M_019812176.2;refGene_gn=NAV2;refGene_tx=XM_019812177.2;refGene_gn=NAV
2;refGene_tx=XM_019812178.2;refGene_gn=NAV2;refGene_tx=XM_019812181.2;
refGene_gn=NAV2;refGene_tx=XM_019812182.2;refGene_gn=NAV2;refGene_tx=X
M_019812183.2;refGene_gn=NAV2;refGene_tx=XM_023239644.1;refGene_gn=NAV
2;refGene_tx=XM_023239646.1;refGene_gn=NAV2;refGene_tx=XM_023239647.1
D1 77626342 77626503
refGene_gn=NAV2;refGene_tx=XM_019812168.2;refGene_gn=NAV2;refGene_tx=X
M_019812169.2;refGene_gn=NAV2;refGene_tx=XM_019812170.2;refGene_gn=NAV
2;refGene_tx=XM_019812171.2;refGene_gn=NAV2;refGene_tx=XM_019812172.2;
refGene_gn=NAV2;refGene_tx=XM_019812173.2;refGene_gn=NAV2;refGene_tx=X
M_019812174.2;refGene_gn=NAV2;refGene_tx=XM_019812175.2;refGene_gn=NAV
2;refGene_tx=XM_019812176.2;refGene_gn=NAV2;refGene_tx=XM_019812177.2;
refGene_gn=NAV2;refGene_tx=XM_019812178.2;refGene_gn=NAV2;refGene_tx=X
M_019812181.2;refGene_gn=NAV2;refGene_tx=XM_019812182.2;refGene_gn=NAV
2;refGene_tx=XM_019812183.2;refGene_gn=NAV2;refGene_tx=XM_023239644.1;
refGene_gn=NAV2;refGene_tx=XM_023239646.1;refGene_gn=NAV2;refGene_tx=X
M_023239647.1

D1 77628304 77628493
refGene_gn=NAV2;refGene_tx=XM_019812168.2;refGene_gn=NAV2;refGene_tx=XM_019812169.2;refGene_gn=NAV2;refGene_tx=XM_019812170.2;refGene_gn=NAV2;refGene_tx=XM_019812171.2;refGene_gn=NAV2;refGene_tx=XM_019812172.2;refGene_gn=NAV2;refGene_tx=XM_019812173.2;refGene_gn=NAV2;refGene_tx=XM_019812174.2;refGene_gn=NAV2;refGene_tx=XM_019812175.2;refGene_gn=NAV2;refGene_tx=XM_019812176.2;refGene_gn=NAV2;refGene_tx=XM_019812177.2;refGene_gn=NAV2;refGene_tx=XM_019812178.2;refGene_gn=NAV2;refGene_tx=XM_019812181.2;refGene_gn=NAV2;refGene_tx=XM_019812182.2;refGene_gn=NAV2;refGene_tx=XM_019812183.2;refGene_gn=NAV2;refGene_tx=XM_023239644.1;refGene_gn=NAV2;refGene_tx=XM_023239646.1;refGene_gn=NAV2;refGene_tx=XM_023239647.1
D1 77634781 77634834
refGene_gn=NAV2;refGene_tx=XM_019812168.2;refGene_gn=NAV2;refGene_tx=XM_019812169.2;refGene_gn=NAV2;refGene_tx=XM_019812170.2;refGene_gn=NAV2;refGene_tx=XM_019812171.2;refGene_gn=NAV2;refGene_tx=XM_019812172.2;refGene_gn=NAV2;refGene_tx=XM_019812173.2;refGene_gn=NAV2;refGene_tx=XM_019812174.2;refGene_gn=NAV2;refGene_tx=XM_019812175.2;refGene_gn=NAV2;refGene_tx=XM_019812176.2;refGene_gn=NAV2;refGene_tx=XM_019812177.2;refGene_gn=NAV2;refGene_tx=XM_019812178.2;refGene_gn=NAV2;refGene_tx=XM_019812181.2;refGene_gn=NAV2;refGene_tx=XM_019812182.2;refGene_gn=NAV2;refGene_tx=XM_019812183.2;refGene_gn=NAV2;refGene_tx=XM_023239644.1;refGene_gn=NAV2;refGene_tx=XM_023239646.1;refGene_gn=NAV2;refGene_tx=XM_023239647.1
D1 77639767 77639791
refGene_gn=NAV2;refGene_tx=XM_019812168.2;refGene_gn=NAV2;refGene_tx=XM_019812169.2;refGene_gn=NAV2;refGene_tx=XM_019812170.2;refGene_gn=NAV2;refGene_tx=XM_019812171.2;refGene_gn=NAV2;refGene_tx=XM_019812172.2;refGene_gn=NAV2;refGene_tx=XM_019812173.2;refGene_gn=NAV2;refGene_tx=XM_019812174.2;refGene_gn=NAV2;refGene_tx=XM_019812175.2;refGene_gn=NAV2;refGene_tx=XM_019812176.2;refGene_gn=NAV2;refGene_tx=XM_019812177.2;refGene_gn=NAV2;refGene_tx=XM_019812178.2;refGene_gn=NAV2;refGene_tx=XM_019812181.2;refGene_gn=NAV2;refGene_tx=XM_019812182.2;refGene_gn=NAV2;refGene_tx=XM_019812183.2;refGene_gn=NAV2;refGene_tx=XM_023239644.1;refGene_gn=NAV2;refGene_tx=XM_023239646.1;refGene_gn=NAV2;refGene_tx=XM_023239647.1
D1 77639905 77639980
refGene_gn=NAV2;refGene_tx=XM_019812168.2;refGene_gn=NAV2;refGene_tx=XM_019812169.2;refGene_gn=NAV2;refGene_tx=XM_019812170.2;refGene_gn=NAV2;refGene_tx=XM_019812171.2;refGene_gn=NAV2;refGene_tx=XM_019812172.2;refGene_gn=NAV2;refGene_tx=XM_019812173.2;refGene_gn=NAV2;refGene_tx=XM_019812174.2;refGene_gn=NAV2;refGene_tx=XM_019812175.2;refGene_gn=NAV2;refGene_tx=XM_019812176.2;refGene_gn=NAV2;refGene_tx=XM_019812177.2;refGene_gn=NAV2;refGene_tx=XM_019812178.2;refGene_gn=NAV2;refGene_tx=XM_019812181.2;refGene_gn=NAV2;refGene_tx=XM_019812182.2;refGene_gn=NAV2;refGene_tx=XM_019812183.2;refGene_gn=NAV2;refGene_tx=XM_023239644.1;refGene_gn=NAV2;refGene_tx=XM_023239646.1;refGene_gn=NAV2;refGene_tx=XM_023239647.1
D1 77644521 77644605
refGene_gn=NAV2;refGene_tx=XM_019812168.2;refGene_gn=NAV2;refGene_tx=X

M_019812169.2;refGene_gn=NAV2;refGene_tx=XM_019812170.2;refGene_gn=NAV
2;refGene_tx=XM_019812171.2;refGene_gn=NAV2;refGene_tx=XM_019812172.2;
refGene_gn=NAV2;refGene_tx=XM_019812173.2;refGene_gn=NAV2;refGene_tx=X
M_019812174.2;refGene_gn=NAV2;refGene_tx=XM_019812175.2;refGene_gn=NAV
2;refGene_tx=XM_019812176.2;refGene_gn=NAV2;refGene_tx=XM_019812177.2;
refGene_gn=NAV2;refGene_tx=XM_019812178.2;refGene_gn=NAV2;refGene_tx=X
M_019812181.2;refGene_gn=NAV2;refGene_tx=XM_019812182.2;refGene_gn=NAV
2;refGene_tx=XM_019812183.2;refGene_gn=NAV2;refGene_tx=XM_023239644.1;
refGene_gn=NAV2;refGene_tx=XM_023239646.1;refGene_gn=NAV2;refGene_tx=X
M_023239647.1
D1 77644991 77645103
refGene_gn=NAV2;refGene_tx=XM_019812168.2;refGene_gn=NAV2;refGene_tx=X
M_019812169.2;refGene_gn=NAV2;refGene_tx=XM_019812170.2;refGene_gn=NAV
2;refGene_tx=XM_019812171.2;refGene_gn=NAV2;refGene_tx=XM_019812172.2;
refGene_gn=NAV2;refGene_tx=XM_019812173.2;refGene_gn=NAV2;refGene_tx=X
M_019812174.2;refGene_gn=NAV2;refGene_tx=XM_019812175.2;refGene_gn=NAV
2;refGene_tx=XM_019812176.2;refGene_gn=NAV2;refGene_tx=XM_019812177.2;
refGene_gn=NAV2;refGene_tx=XM_019812178.2;refGene_gn=NAV2;refGene_tx=X
M_019812181.2;refGene_gn=NAV2;refGene_tx=XM_019812182.2;refGene_gn=NAV
2;refGene_tx=XM_019812183.2;refGene_gn=NAV2;refGene_tx=XM_023239644.1;
refGene_gn=NAV2;refGene_tx=XM_023239646.1;refGene_gn=NAV2;refGene_tx=X
M_023239647.1
D1 77646353 77646574
refGene_gn=NAV2;refGene_tx=XM_019812168.2;refGene_gn=NAV2;refGene_tx=X
M_019812169.2;refGene_gn=NAV2;refGene_tx=XM_019812170.2;refGene_gn=NAV
2;refGene_tx=XM_019812171.2;refGene_gn=NAV2;refGene_tx=XM_019812172.2;
refGene_gn=NAV2;refGene_tx=XM_019812173.2;refGene_gn=NAV2;refGene_tx=X
M_019812174.2;refGene_gn=NAV2;refGene_tx=XM_019812175.2;refGene_gn=NAV
2;refGene_tx=XM_019812176.2;refGene_gn=NAV2;refGene_tx=XM_019812177.2;
refGene_gn=NAV2;refGene_tx=XM_019812178.2;refGene_gn=NAV2;refGene_tx=X
M_019812181.2;refGene_gn=NAV2;refGene_tx=XM_019812182.2;refGene_gn=NAV
2;refGene_tx=XM_019812183.2;refGene_gn=NAV2;refGene_tx=XM_023239644.1;
refGene_gn=NAV2;refGene_tx=XM_023239646.1;refGene_gn=NAV2;refGene_tx=X
M_023239647.1
D1 77649486 77649495
refGene_gn=NAV2;refGene_tx=XM_019812168.2;refGene_gn=NAV2;refGene_tx=X
M_019812170.2;refGene_gn=NAV2;refGene_tx=XM_019812171.2;refGene_gn=NAV
2;refGene_tx=XM_019812172.2;refGene_gn=NAV2;refGene_tx=XM_019812173.2;
refGene_gn=NAV2;refGene_tx=XM_019812174.2;refGene_gn=NAV2;refGene_tx=X
M_019812175.2;refGene_gn=NAV2;refGene_tx=XM_019812176.2;refGene_gn=NAV
2;refGene_tx=XM_019812181.2;refGene_gn=NAV2;refGene_tx=XM_019812182.2;
refGene_gn=NAV2;refGene_tx=XM_019812183.2;refGene_gn=NAV2;refGene_tx=X
M_023239646.1;refGene_gn=NAV2;refGene_tx=XM_023239647.1
D1 77649910 77650083
refGene_gn=NAV2;refGene_tx=XM_019812168.2;refGene_gn=NAV2;refGene_tx=X
M_019812169.2;refGene_gn=NAV2;refGene_tx=XM_019812170.2;refGene_gn=NAV
2;refGene_tx=XM_019812171.2;refGene_gn=NAV2;refGene_tx=XM_019812172.2;
refGene_gn=NAV2;refGene_tx=XM_019812173.2;refGene_gn=NAV2;refGene_tx=X
M_019812174.2;refGene_gn=NAV2;refGene_tx=XM_019812175.2;refGene_gn=NAV
2;refGene_tx=XM_019812176.2;refGene_gn=NAV2;refGene_tx=XM_019812177.2;

refGene_gn=NAV2;refGene_tx=XM_019812178.2;refGene_gn=NAV2;refGene_tx=XM_019812181.2;refGene_gn=NAV2;refGene_tx=XM_019812182.2;refGene_gn=NAV2;refGene_tx=XM_019812183.2;refGene_gn=NAV2;refGene_tx=XM_023239644.1;refGene_gn=NAV2;refGene_tx=XM_023239646.1;refGene_gn=NAV2;refGene_tx=XM_023239647.1
D1 77657923 77658077
refGene_gn=NAV2;refGene_tx=XM_019812168.2;refGene_gn=NAV2;refGene_tx=XM_019812169.2;refGene_gn=NAV2;refGene_tx=XM_019812170.2;refGene_gn=NAV2;refGene_tx=XM_019812171.2;refGene_gn=NAV2;refGene_tx=XM_019812172.2;refGene_gn=NAV2;refGene_tx=XM_019812173.2;refGene_gn=NAV2;refGene_tx=XM_019812174.2;refGene_gn=NAV2;refGene_tx=XM_019812175.2;refGene_gn=NAV2;refGene_tx=XM_019812176.2;refGene_gn=NAV2;refGene_tx=XM_019812177.2;refGene_gn=NAV2;refGene_tx=XM_019812178.2;refGene_gn=NAV2;refGene_tx=XM_019812181.2;refGene_gn=NAV2;refGene_tx=XM_019812182.2;refGene_gn=NAV2;refGene_tx=XM_019812183.2;refGene_gn=NAV2;refGene_tx=XM_023239644.1;refGene_gn=NAV2;refGene_tx=XM_023239646.1;refGene_gn=NAV2;refGene_tx=XM_023239647.1
D1 77659473 77659636
refGene_gn=NAV2;refGene_tx=XM_019812168.2;refGene_gn=NAV2;refGene_tx=XM_019812169.2;refGene_gn=NAV2;refGene_tx=XM_019812170.2;refGene_gn=NAV2;refGene_tx=XM_019812171.2;refGene_gn=NAV2;refGene_tx=XM_019812172.2;refGene_gn=NAV2;refGene_tx=XM_019812173.2;refGene_gn=NAV2;refGene_tx=XM_019812174.2;refGene_gn=NAV2;refGene_tx=XM_019812175.2;refGene_gn=NAV2;refGene_tx=XM_019812176.2;refGene_gn=NAV2;refGene_tx=XM_019812177.2;refGene_gn=NAV2;refGene_tx=XM_019812178.2;refGene_gn=NAV2;refGene_tx=XM_019812181.2;refGene_gn=NAV2;refGene_tx=XM_019812182.2;refGene_gn=NAV2;refGene_tx=XM_019812183.2;refGene_gn=NAV2;refGene_tx=XM_023239644.1;refGene_gn=NAV2;refGene_tx=XM_023239646.1;refGene_gn=NAV2;refGene_tx=XM_023239647.1
D1 77660359 77660460
refGene_gn=NAV2;refGene_tx=XM_019812168.2;refGene_gn=NAV2;refGene_tx=XM_019812169.2;refGene_gn=NAV2;refGene_tx=XM_019812170.2;refGene_gn=NAV2;refGene_tx=XM_019812171.2;refGene_gn=NAV2;refGene_tx=XM_019812172.2;refGene_gn=NAV2;refGene_tx=XM_019812173.2;refGene_gn=NAV2;refGene_tx=XM_019812174.2;refGene_gn=NAV2;refGene_tx=XM_019812175.2;refGene_gn=NAV2;refGene_tx=XM_019812176.2;refGene_gn=NAV2;refGene_tx=XM_019812177.2;refGene_gn=NAV2;refGene_tx=XM_019812178.2;refGene_gn=NAV2;refGene_tx=XM_019812181.2;refGene_gn=NAV2;refGene_tx=XM_019812182.2;refGene_gn=NAV2;refGene_tx=XM_023239644.1;refGene_gn=NAV2;refGene_tx=XM_023239646.1;refGene_gn=NAV2;refGene_tx=XM_023239647.1
D1 77662563 77662659
refGene_gn=NAV2;refGene_tx=XM_019812168.2;refGene_gn=NAV2;refGene_tx=XM_019812169.2;refGene_gn=NAV2;refGene_tx=XM_019812170.2;refGene_gn=NAV2;refGene_tx=XM_019812171.2;refGene_gn=NAV2;refGene_tx=XM_019812172.2;refGene_gn=NAV2;refGene_tx=XM_019812173.2;refGene_gn=NAV2;refGene_tx=XM_019812174.2;refGene_gn=NAV2;refGene_tx=XM_019812175.2;refGene_gn=NAV2;refGene_tx=XM_019812176.2;refGene_gn=NAV2;refGene_tx=XM_019812177.2;refGene_gn=NAV2;refGene_tx=XM_019812178.2;refGene_gn=NAV2;refGene_tx=XM_019812181.2;refGene_gn=NAV2;refGene_tx=XM_019812182.2;refGene_gn=NAV2;refGene_tx=XM_023239644.1;refGene_gn=NAV2;refGene_tx=XM_023239646.1;

refGene_gn=NAV2;refGene_tx=XM_023239647.1
D1 77664461 77664630
refGene_gn=NAV2;refGene_tx=XM_019812168.2;refGene_gn=NAV2;refGene_tx=XM_019812169.2;refGene_gn=NAV2;refGene_tx=XM_019812170.2;refGene_gn=NAV2;refGene_tx=XM_019812171.2;refGene_gn=NAV2;refGene_tx=XM_019812172.2;refGene_gn=NAV2;refGene_tx=XM_019812173.2;refGene_gn=NAV2;refGene_tx=XM_019812174.2;refGene_gn=NAV2;refGene_tx=XM_019812175.2;refGene_gn=NAV2;refGene_tx=XM_019812176.2;refGene_gn=NAV2;refGene_tx=XM_019812177.2;refGene_gn=NAV2;refGene_tx=XM_019812178.2;refGene_gn=NAV2;refGene_tx=XM_019812181.2;refGene_gn=NAV2;refGene_tx=XM_019812182.2;refGene_gn=NAV2;refGene_tx=XM_019812183.2;refGene_gn=NAV2;refGene_tx=XM_023239644.1;refGene_gn=NAV2;refGene_tx=XM_023239646.1;refGene_gn=NAV2;refGene_tx=XM_023239647.1
D1 77668190 77668426
refGene_gn=NAV2;refGene_tx=XM_019812168.2;refGene_gn=NAV2;refGene_tx=XM_019812169.2;refGene_gn=NAV2;refGene_tx=XM_019812170.2;refGene_gn=NAV2;refGene_tx=XM_019812171.2;refGene_gn=NAV2;refGene_tx=XM_019812172.2;refGene_gn=NAV2;refGene_tx=XM_019812173.2;refGene_gn=NAV2;refGene_tx=XM_019812174.2;refGene_gn=NAV2;refGene_tx=XM_019812175.2;refGene_gn=NAV2;refGene_tx=XM_019812176.2;refGene_gn=NAV2;refGene_tx=XM_019812177.2;refGene_gn=NAV2;refGene_tx=XM_019812178.2;refGene_gn=NAV2;refGene_tx=XM_019812181.2;refGene_gn=NAV2;refGene_tx=XM_019812182.2;refGene_gn=NAV2;refGene_tx=XM_023239644.1;refGene_gn=NAV2;refGene_tx=XM_023239646.1;refGene_gn=NAV2;refGene_tx=XM_023239647.1
D1 77670154 77670309
refGene_gn=NAV2;refGene_tx=XM_019812168.2;refGene_gn=NAV2;refGene_tx=XM_019812169.2;refGene_gn=NAV2;refGene_tx=XM_019812170.2;refGene_gn=NAV2;refGene_tx=XM_019812171.2;refGene_gn=NAV2;refGene_tx=XM_019812172.2;refGene_gn=NAV2;refGene_tx=XM_019812173.2;refGene_gn=NAV2;refGene_tx=XM_019812174.2;refGene_gn=NAV2;refGene_tx=XM_019812175.2;refGene_gn=NAV2;refGene_tx=XM_019812176.2;refGene_gn=NAV2;refGene_tx=XM_019812177.2;refGene_gn=NAV2;refGene_tx=XM_019812178.2;refGene_gn=NAV2;refGene_tx=XM_019812181.2;refGene_gn=NAV2;refGene_tx=XM_019812182.2;refGene_gn=NAV2;refGene_tx=XM_023239644.1;refGene_gn=NAV2;refGene_tx=XM_023239646.1;refGene_gn=NAV2;refGene_tx=XM_023239647.1
D1 77670525 77670597
refGene_gn=NAV2;refGene_tx=XM_019812168.2;refGene_gn=NAV2;refGene_tx=XM_019812169.2;refGene_gn=NAV2;refGene_tx=XM_019812170.2;refGene_gn=NAV2;refGene_tx=XM_019812171.2;refGene_gn=NAV2;refGene_tx=XM_019812172.2;refGene_gn=NAV2;refGene_tx=XM_019812173.2;refGene_gn=NAV2;refGene_tx=XM_019812174.2;refGene_gn=NAV2;refGene_tx=XM_019812175.2;refGene_gn=NAV2;refGene_tx=XM_019812176.2;refGene_gn=NAV2;refGene_tx=XM_019812177.2;refGene_gn=NAV2;refGene_tx=XM_019812178.2;refGene_gn=NAV2;refGene_tx=XM_019812181.2;refGene_gn=NAV2;refGene_tx=XM_019812182.2;refGene_gn=NAV2;refGene_tx=XM_023239644.1;refGene_gn=NAV2;refGene_tx=XM_023239646.1;refGene_gn=NAV2;refGene_tx=XM_023239647.1
D1 77672112 77672309
refGene_gn=NAV2;refGene_tx=XM_019812168.2;refGene_gn=NAV2;refGene_tx=XM_019812169.2;refGene_gn=NAV2;refGene_tx=XM_019812170.2;refGene_gn=NAV2;refGene_tx=XM_019812171.2;refGene_gn=NAV2;refGene_tx=XM_019812172.2;

refGene_gn=NAV2;refGene_tx=XM_019812173.2;refGene_gn=NAV2;refGene_tx=XM_019812174.2;refGene_gn=NAV2;refGene_tx=XM_019812175.2;refGene_gn=NAV2;refGene_tx=XM_019812176.2;refGene_gn=NAV2;refGene_tx=XM_019812177.2;refGene_gn=NAV2;refGene_tx=XM_019812178.2;refGene_gn=NAV2;refGene_tx=XM_019812181.2;refGene_gn=NAV2;refGene_tx=XM_019812182.2;refGene_gn=NAV2;refGene_tx=XM_023239644.1;refGene_gn=NAV2;refGene_tx=XM_023239646.1;refGene_gn=NAV2;refGene_tx=XM_023239647.1
D1 77673503 77673622
refGene_gn=NAV2;refGene_tx=XM_019812168.2;refGene_gn=NAV2;refGene_tx=XM_019812169.2;refGene_gn=NAV2;refGene_tx=XM_019812170.2;refGene_gn=NAV2;refGene_tx=XM_019812171.2;refGene_gn=NAV2;refGene_tx=XM_019812172.2;refGene_gn=NAV2;refGene_tx=XM_019812173.2;refGene_gn=NAV2;refGene_tx=XM_019812174.2;refGene_gn=NAV2;refGene_tx=XM_019812175.2;refGene_gn=NAV2;refGene_tx=XM_019812176.2;refGene_gn=NAV2;refGene_tx=XM_019812177.2;refGene_gn=NAV2;refGene_tx=XM_019812178.2;refGene_gn=NAV2;refGene_tx=XM_019812181.2;refGene_gn=NAV2;refGene_tx=XM_019812182.2;refGene_gn=NAV2;refGene_tx=XM_023239644.1;refGene_gn=NAV2;refGene_tx=XM_023239646.1;refGene_gn=NAV2;refGene_tx=XM_023239647.1
D1 77679748 77679952
refGene_gn=NAV2;refGene_tx=XM_019812168.2;refGene_gn=NAV2;refGene_tx=XM_019812169.2;refGene_gn=NAV2;refGene_tx=XM_019812170.2;refGene_gn=NAV2;refGene_tx=XM_019812171.2;refGene_gn=NAV2;refGene_tx=XM_019812172.2;refGene_gn=NAV2;refGene_tx=XM_019812173.2;refGene_gn=NAV2;refGene_tx=XM_019812174.2;refGene_gn=NAV2;refGene_tx=XM_019812175.2;refGene_gn=NAV2;refGene_tx=XM_019812176.2;refGene_gn=NAV2;refGene_tx=XM_019812177.2;refGene_gn=NAV2;refGene_tx=XM_019812178.2;refGene_gn=NAV2;refGene_tx=XM_019812181.2;refGene_gn=NAV2;refGene_tx=XM_019812182.2;refGene_gn=NAV2;refGene_tx=XM_023239644.1;refGene_gn=NAV2;refGene_tx=XM_023239646.1;refGene_gn=NAV2;refGene_tx=XM_023239647.1
D1 77682635 77682761
refGene_gn=NAV2;refGene_tx=XM_019812168.2;refGene_gn=NAV2;refGene_tx=XM_019812169.2;refGene_gn=NAV2;refGene_tx=XM_019812170.2;refGene_gn=NAV2;refGene_tx=XM_019812171.2;refGene_gn=NAV2;refGene_tx=XM_019812172.2;refGene_gn=NAV2;refGene_tx=XM_019812173.2;refGene_gn=NAV2;refGene_tx=XM_019812174.2;refGene_gn=NAV2;refGene_tx=XM_019812175.2;refGene_gn=NAV2;refGene_tx=XM_019812176.2;refGene_gn=NAV2;refGene_tx=XM_019812177.2;refGene_gn=NAV2;refGene_tx=XM_019812178.2;refGene_gn=NAV2;refGene_tx=XM_019812181.2;refGene_gn=NAV2;refGene_tx=XM_019812182.2;refGene_gn=NAV2;refGene_tx=XM_023239644.1;refGene_gn=NAV2;refGene_tx=XM_023239646.1;refGene_gn=NAV2;refGene_tx=XM_023239647.1
D1 77712823 77713183 refGene_gn=DBX1;refGene_tx=XM_003993062.2
D1 77713628 77713831 refGene_gn=DBX1;refGene_tx=XM_003993062.2
D1 77715781 77715883 refGene_gn=DBX1;refGene_tx=XM_003993062.2
D1 77716521 77716888 refGene_gn=DBX1;refGene_tx=XM_003993062.2
D1 77887761 77887956
refGene_gn=HTATIP2;refGene_tx=XM_006937140.4;refGene_gn=HTATIP2;refGene_tx=XM_023239648.1
D1 77890289 77890397
refGene_gn=HTATIP2;refGene_tx=XM_006937140.4;refGene_gn=HTATIP2;refGene_tx=XM_023239648.1

D1 77898089 77898227
refGene_gn=HTATIP2;refGene_tx=XM_006937140.4;refGene_gn=HTATIP2;refGene_tx=XM_023239648.1
D1 77900263 77900325
refGene_gn=HTATIP2;refGene_tx=XM_006937140.4;refGene_gn=HTATIP2;refGene_tx=XM_023239648.1
D1 77901043 77901269
refGene_gn=HTATIP2;refGene_tx=XM_006937140.4;refGene_gn=HTATIP2;refGene_tx=XM_023239648.1
D1 77904402 77904430 refGene_gn=PRMT3;refGene_tx=XM_003993064.4
D1 77904706 77904842 refGene_gn=PRMT3;refGene_tx=XM_003993064.4
D1 77906448 77906531 refGene_gn=PRMT3;refGene_tx=XM_003993064.4
D1 77908211 77908261 refGene_gn=PRMT3;refGene_tx=XM_003993064.4
D1 77908909 77909012 refGene_gn=PRMT3;refGene_tx=XM_003993064.4
D1 77916052 77916212 refGene_gn=PRMT3;refGene_tx=XM_003993064.4
D1 77917970 77918115 refGene_gn=PRMT3;refGene_tx=XM_003993064.4
D1 77921967 77922033 refGene_gn=PRMT3;refGene_tx=XM_003993064.4
D1 77933067 77933189 refGene_gn=PRMT3;refGene_tx=XM_003993064.4
D1 77953431 77953531 refGene_gn=PRMT3;refGene_tx=XM_003993064.4
D1 77984425 77984504 refGene_gn=PRMT3;refGene_tx=XM_003993064.4
D1 77998734 77998922 refGene_gn=PRMT3;refGene_tx=XM_003993064.4
D1 78000893 78000980 refGene_gn=PRMT3;refGene_tx=XM_003993064.4
D1 78030817 78030868 refGene_gn=PRMT3;refGene_tx=XM_003993064.4
D1 78031063 78031151 refGene_gn=PRMT3;refGene_tx=XM_003993064.4
D1 78044263 78044373 refGene_gn=PRMT3;refGene_tx=XM_003993064.4
D1 78107785 78107788 refGene_gn=SLC6A5;refGene_tx=XM_019812186.2
D1 78109202 78109754 refGene_gn=SLC6A5;refGene_tx=XM_019812186.2
D1 78112368 78112507 refGene_gn=SLC6A5;refGene_tx=XM_019812186.2
D1 78115205 78115337
refGene_gn=SLC6A5;refGene_tx=XM_019812186.2;refGene_gn=SLC6A5;refGene_tx=XM_019812187.2
D1 78115683 78115857
refGene_gn=SLC6A5;refGene_tx=XM_019812186.2;refGene_gn=SLC6A5;refGene_tx=XM_019812187.2
D1 78124256 78124398
refGene_gn=SLC6A5;refGene_tx=XM_019812186.2;refGene_gn=SLC6A5;refGene_tx=XM_019812187.2
D1 78127149 78127282
refGene_gn=SLC6A5;refGene_tx=XM_019812186.2;refGene_gn=SLC6A5;refGene_tx=XM_019812187.2
D1 78135288 78135423
refGene_gn=SLC6A5;refGene_tx=XM_019812186.2;refGene_gn=SLC6A5;refGene_tx=XM_019812187.2
D1 78137155 78137259
refGene_gn=SLC6A5;refGene_tx=XM_019812186.2;refGene_gn=SLC6A5;refGene_tx=XM_019812187.2
D1 78140184 78140309
refGene_gn=SLC6A5;refGene_tx=XM_019812186.2;refGene_gn=SLC6A5;refGene_tx=XM_019812187.2
D1 78145223 78145336

refGene_gn=SLC6A5;refGene_tx=XM_019812186.2;refGene_gn=SLC6A5;refGene_tx=XM_019812187.2
D1 78146084 78146216
refGene_gn=SLC6A5;refGene_tx=XM_019812186.2;refGene_gn=SLC6A5;refGene_tx=XM_019812187.2
D1 78147153 78147253
refGene_gn=SLC6A5;refGene_tx=XM_019812186.2;refGene_gn=SLC6A5;refGene_tx=XM_019812187.2
D1 78154673 78154774
refGene_gn=SLC6A5;refGene_tx=XM_019812186.2;refGene_gn=SLC6A5;refGene_tx=XM_019812187.2
D1 78159325 78159493
refGene_gn=SLC6A5;refGene_tx=XM_019812186.2;refGene_gn=SLC6A5;refGene_tx=XM_019812187.2
D1 78161683 78161839
refGene_gn=SLC6A5;refGene_tx=XM_019812186.2;refGene_gn=SLC6A5;refGene_tx=XM_019812187.2
D1 78178994 78179049
refGene_gn=NELL1;refGene_tx=XM_003993068.4;refGene_gn=NELL1;refGene_tx=XM_003993069.4
D1 78187288 78187417
refGene_gn=NELL1;refGene_tx=XM_003993068.4;refGene_gn=NELL1;refGene_tx=XM_003993069.4
D1 78262920 78263071
refGene_gn=NELL1;refGene_tx=XM_003993068.4;refGene_gn=NELL1;refGene_tx=XM_003993069.4
D1 78312742 78312913
refGene_gn=NELL1;refGene_tx=XM_003993068.4;refGene_gn=NELL1;refGene_tx=XM_003993069.4
D1 78352813 78352910
refGene_gn=NELL1;refGene_tx=XM_003993068.4;refGene_gn=NELL1;refGene_tx=XM_003993069.4
D1 78375190 78375263
refGene_gn=NELL1;refGene_tx=XM_003993068.4;refGene_gn=NELL1;refGene_tx=XM_003993069.4
D1 78375875 78375958
refGene_gn=NELL1;refGene_tx=XM_003993068.4;refGene_gn=NELL1;refGene_tx=XM_003993069.4
D1 78382620 78382755
refGene_gn=NELL1;refGene_tx=XM_003993068.4;refGene_gn=NELL1;refGene_tx=XM_003993069.4
D1 78383695 78383798
refGene_gn=NELL1;refGene_tx=XM_003993068.4;refGene_gn=NELL1;refGene_tx=XM_003993069.4
D1 78392982 78393056
refGene_gn=NELL1;refGene_tx=XM_003993068.4;refGene_gn=NELL1;refGene_tx=XM_003993069.4
D1 78399631 78399731
refGene_gn=NELL1;refGene_tx=XM_003993068.4;refGene_gn=NELL1;refGene_tx=XM_003993069.4

D1 78414571 78414700
refGene_gn=NELL1;refGene_tx=XM_003993068.4;refGene_gn=NELL1;refGene_tx
=XM_003993069.4
D1 78591040 78591166
refGene_gn=NELL1;refGene_tx=XM_003993068.4;refGene_gn=NELL1;refGene_tx
=XM_003993069.4
D1 78705009 78705132
refGene_gn=NELL1;refGene_tx=XM_003993068.4;refGene_gn=NELL1;refGene_tx
=XM_003993069.4
D1 78756924 78758091
refGene_gn=LOC101086177;refGene_tx=XM_011286745.3
D1 78829875 78829971
refGene_gn=NELL1;refGene_tx=XM_003993068.4;refGene_gn=NELL1;refGene_tx
=XM_003993069.4
D1 79020974 79021115 refGene_gn=NELL1;refGene_tx=XM_003993068.4
D1 79047731 79047925
refGene_gn=NELL1;refGene_tx=XM_003993068.4;refGene_gn=NELL1;refGene_tx
=XM_003993069.4
D1 79062563 79062740
refGene_gn=NELL1;refGene_tx=XM_003993068.4;refGene_gn=NELL1;refGene_tx
=XM_003993069.4
D1 79065116 79065341
refGene_gn=NELL1;refGene_tx=XM_003993068.4;refGene_gn=NELL1;refGene_tx
=XM_003993069.4
D1 79066793 79066844
refGene_gn=NELL1;refGene_tx=XM_003993068.4;refGene_gn=NELL1;refGene_tx
=XM_003993069.4
D1 79585499 79585539
refGene_gn=ANO5;refGene_tx=XM_023239650.1;refGene_gn=ANO5;refGene_tx=X
M_023239651.1
D1 79588299 79588387
refGene_gn=ANO5;refGene_tx=XM_006937143.4;refGene_gn=ANO5;refGene_tx=X
M_019812188.2;refGene_gn=ANO5;refGene_tx=XM_019812189.2;refGene_gn=ANO
5;refGene_tx=XM_019812190.2
D1 79598174 79598221
refGene_gn=ANO5;refGene_tx=XM_006937143.4;refGene_gn=ANO5;refGene_tx=X
M_019812188.2;refGene_gn=ANO5;refGene_tx=XM_019812189.2;refGene_gn=ANO
5;refGene_tx=XM_019812190.2;refGene_gn=ANO5;refGene_tx=XM_023239650.1;
refGene_gn=ANO5;refGene_tx=XM_023239651.1
D1 79601739 79601790
refGene_gn=ANO5;refGene_tx=XM_006937143.4;refGene_gn=ANO5;refGene_tx=X
M_019812188.2;refGene_gn=ANO5;refGene_tx=XM_019812189.2;refGene_gn=ANO
5;refGene_tx=XM_019812190.2;refGene_gn=ANO5;refGene_tx=XM_023239650.1;
refGene_gn=ANO5;refGene_tx=XM_023239651.1
D1 79609883 79609925
refGene_gn=ANO5;refGene_tx=XM_006937143.4;refGene_gn=ANO5;refGene_tx=X
M_019812188.2;refGene_gn=ANO5;refGene_tx=XM_023239650.1;refGene_gn=ANO
5;refGene_tx=XM_023239651.1
D1 79612866 79612980
refGene_gn=ANO5;refGene_tx=XM_006937143.4;refGene_gn=ANO5;refGene_tx=X

M_019812188.2;refGene_gn=ANO5;refGene_tx=XM_019812189.2;refGene_gn=ANO5;refGene_tx=XM_019812190.2;refGene_gn=ANO5;refGene_tx=XM_023239650.1;refGene_gn=ANO5;refGene_tx=XM_023239651.1
D1 79615573 79615642
refGene_gn=ANO5;refGene_tx=XM_006937143.4;refGene_gn=ANO5;refGene_tx=XM_019812188.2;refGene_gn=ANO5;refGene_tx=XM_019812189.2;refGene_gn=ANO5;refGene_tx=XM_019812190.2;refGene_gn=ANO5;refGene_tx=XM_023239650.1;refGene_gn=ANO5;refGene_tx=XM_023239651.1
D1 79616949 79617234
refGene_gn=ANO5;refGene_tx=XM_006937143.4;refGene_gn=ANO5;refGene_tx=XM_019812188.2;refGene_gn=ANO5;refGene_tx=XM_019812189.2;refGene_gn=ANO5;refGene_tx=XM_019812190.2;refGene_gn=ANO5;refGene_tx=XM_023239650.1;refGene_gn=ANO5;refGene_tx=XM_023239651.1
D1 79621863 79621977
refGene_gn=ANO5;refGene_tx=XM_006937143.4;refGene_gn=ANO5;refGene_tx=XM_019812188.2;refGene_gn=ANO5;refGene_tx=XM_019812189.2;refGene_gn=ANO5;refGene_tx=XM_019812190.2;refGene_gn=ANO5;refGene_tx=XM_023239650.1;refGene_gn=ANO5;refGene_tx=XM_023239651.1
D1 79625038 79625154
refGene_gn=ANO5;refGene_tx=XM_006937143.4;refGene_gn=ANO5;refGene_tx=XM_019812188.2;refGene_gn=ANO5;refGene_tx=XM_019812189.2;refGene_gn=ANO5;refGene_tx=XM_019812190.2;refGene_gn=ANO5;refGene_tx=XM_023239650.1;refGene_gn=ANO5;refGene_tx=XM_023239651.1
D1 79640304 79640439
refGene_gn=ANO5;refGene_tx=XM_006937143.4;refGene_gn=ANO5;refGene_tx=XM_019812188.2;refGene_gn=ANO5;refGene_tx=XM_019812189.2;refGene_gn=ANO5;refGene_tx=XM_019812190.2;refGene_gn=ANO5;refGene_tx=XM_023239650.1;refGene_gn=ANO5;refGene_tx=XM_023239651.1
D1 79640788 79640894
refGene_gn=ANO5;refGene_tx=XM_006937143.4;refGene_gn=ANO5;refGene_tx=XM_019812188.2;refGene_gn=ANO5;refGene_tx=XM_019812189.2;refGene_gn=ANO5;refGene_tx=XM_019812190.2;refGene_gn=ANO5;refGene_tx=XM_023239650.1;refGene_gn=ANO5;refGene_tx=XM_023239651.1
D1 79640991 79641052
refGene_gn=ANO5;refGene_tx=XM_006937143.4;refGene_gn=ANO5;refGene_tx=XM_019812188.2;refGene_gn=ANO5;refGene_tx=XM_019812189.2;refGene_gn=ANO5;refGene_tx=XM_019812190.2;refGene_gn=ANO5;refGene_tx=XM_023239650.1;refGene_gn=ANO5;refGene_tx=XM_023239651.1
D1 79654830 79654982
refGene_gn=ANO5;refGene_tx=XM_006937143.4;refGene_gn=ANO5;refGene_tx=XM_019812188.2;refGene_gn=ANO5;refGene_tx=XM_019812189.2;refGene_gn=ANO5;refGene_tx=XM_019812190.2;refGene_gn=ANO5;refGene_tx=XM_023239650.1;refGene_gn=ANO5;refGene_tx=XM_023239651.1
D1 79657195 79657270
refGene_gn=ANO5;refGene_tx=XM_006937143.4;refGene_gn=ANO5;refGene_tx=XM_019812188.2;refGene_gn=ANO5;refGene_tx=XM_019812189.2;refGene_gn=ANO5;refGene_tx=XM_019812190.2;refGene_gn=ANO5;refGene_tx=XM_023239650.1;refGene_gn=ANO5;refGene_tx=XM_023239651.1
D1 79659147 79659370
refGene_gn=ANO5;refGene_tx=XM_006937143.4;refGene_gn=ANO5;refGene_tx=X

M_019812188.2;refGene_gn=ANO5;refGene_tx=XM_019812189.2;refGene_gn=ANO5;refGene_tx=XM_019812190.2;refGene_gn=ANO5;refGene_tx=XM_023239650.1;refGene_gn=ANO5;refGene_tx=XM_023239651.1
D1 79661142 79661312 refGene_gn=ANO5;refGene_tx=XM_006937143.4;refGene_gn=ANO5;refGene_tx=XM_019812188.2;refGene_gn=ANO5;refGene_tx=XM_019812189.2;refGene_gn=ANO5;refGene_tx=XM_019812190.2;refGene_gn=ANO5;refGene_tx=XM_023239650.1;refGene_gn=ANO5;refGene_tx=XM_023239651.1
D1 79661984 79662082 refGene_gn=ANO5;refGene_tx=XM_006937143.4;refGene_gn=ANO5;refGene_tx=XM_019812188.2;refGene_gn=ANO5;refGene_tx=XM_019812189.2;refGene_gn=ANO5;refGene_tx=XM_019812190.2;refGene_gn=ANO5;refGene_tx=XM_023239650.1;refGene_gn=ANO5;refGene_tx=XM_023239651.1
D1 79665750 79665881 refGene_gn=ANO5;refGene_tx=XM_006937143.4;refGene_gn=ANO5;refGene_tx=XM_019812188.2;refGene_gn=ANO5;refGene_tx=XM_019812189.2;refGene_gn=ANO5;refGene_tx=XM_019812190.2;refGene_gn=ANO5;refGene_tx=XM_023239650.1;refGene_gn=ANO5;refGene_tx=XM_023239651.1
D1 79668115 79668321 refGene_gn=ANO5;refGene_tx=XM_006937143.4;refGene_gn=ANO5;refGene_tx=XM_019812188.2;refGene_gn=ANO5;refGene_tx=XM_019812189.2;refGene_gn=ANO5;refGene_tx=XM_019812190.2;refGene_gn=ANO5;refGene_tx=XM_023239650.1;refGene_gn=ANO5;refGene_tx=XM_023239651.1
D1 79670709 79670888 refGene_gn=ANO5;refGene_tx=XM_006937143.4;refGene_gn=ANO5;refGene_tx=XM_019812188.2;refGene_gn=ANO5;refGene_tx=XM_019812189.2;refGene_gn=ANO5;refGene_tx=XM_019812190.2;refGene_gn=ANO5;refGene_tx=XM_023239650.1;refGene_gn=ANO5;refGene_tx=XM_023239651.1
D1 79672429 79672535 refGene_gn=ANO5;refGene_tx=XM_006937143.4;refGene_gn=ANO5;refGene_tx=XM_019812188.2;refGene_gn=ANO5;refGene_tx=XM_019812189.2;refGene_gn=ANO5;refGene_tx=XM_019812190.2;refGene_gn=ANO5;refGene_tx=XM_023239650.1;refGene_gn=ANO5;refGene_tx=XM_023239651.1
D1 79675502 79675724 refGene_gn=ANO5;refGene_tx=XM_006937143.4;refGene_gn=ANO5;refGene_tx=XM_019812188.2;refGene_gn=ANO5;refGene_tx=XM_019812189.2;refGene_gn=ANO5;refGene_tx=XM_019812190.2;refGene_gn=ANO5;refGene_tx=XM_023239650.1;refGene_gn=ANO5;refGene_tx=XM_023239651.1
D1 79703027 79703113 refGene_gn=SLC17A6;refGene_tx=XM_011286746.3
D1 79705906 79706159 refGene_gn=SLC17A6;refGene_tx=XM_011286746.3
D1 79707695 79707814 refGene_gn=SLC17A6;refGene_tx=XM_011286746.3
D1 79724750 79724865 refGene_gn=SLC17A6;refGene_tx=XM_011286746.3
D1 79725844 79725932 refGene_gn=SLC17A6;refGene_tx=XM_011286746.3
D1 79726473 79726560 refGene_gn=SLC17A6;refGene_tx=XM_011286746.3
D1 79729288 79729431 refGene_gn=SLC17A6;refGene_tx=XM_011286746.3
D1 79733059 79733209 refGene_gn=SLC17A6;refGene_tx=XM_011286746.3
D1 79739549 79739682 refGene_gn=SLC17A6;refGene_tx=XM_011286746.3
D1 79740085 79740196 refGene_gn=SLC17A6;refGene_tx=XM_011286746.3
D1 79740712 79740840 refGene_gn=SLC17A6;refGene_tx=XM_011286746.3
D1 79741567 79741903 refGene_gn=SLC17A6;refGene_tx=XM_011286746.3

D1 79959404 79960484 refGene_gn=FANCF;refGene_tx=XM_004001521.5
D1 79993178 79993326
refGene_gn=GAS2;refGene_tx=XM_003993073.5;refGene_gn=GAS2;refGene_tx=XM_019812192.2;refGene_gn=GAS2;refGene_tx=XM_019812193.2;refGene_gn=GAS2;refGene_tx=XM_019812194.2;refGene_gn=GAS2;refGene_tx=XM_023239652.1
D1 80005858 80005980
refGene_gn=GAS2;refGene_tx=XM_003993073.5;refGene_gn=GAS2;refGene_tx=XM_019812192.2;refGene_gn=GAS2;refGene_tx=XM_019812193.2;refGene_gn=GAS2;refGene_tx=XM_019812194.2;refGene_gn=GAS2;refGene_tx=XM_023239652.1
D1 80037797 80037939
refGene_gn=GAS2;refGene_tx=XM_003993073.5;refGene_gn=GAS2;refGene_tx=XM_019812192.2;refGene_gn=GAS2;refGene_tx=XM_019812193.2;refGene_gn=GAS2;refGene_tx=XM_019812194.2;refGene_gn=GAS2;refGene_tx=XM_023239652.1
D1 80050504 80050568
refGene_gn=GAS2;refGene_tx=XM_003993073.5;refGene_gn=GAS2;refGene_tx=XM_019812192.2;refGene_gn=GAS2;refGene_tx=XM_019812193.2;refGene_gn=GAS2;refGene_tx=XM_019812194.2;refGene_gn=GAS2;refGene_tx=XM_023239652.1
D1 80060457 80060599
refGene_gn=GAS2;refGene_tx=XM_003993073.5;refGene_gn=GAS2;refGene_tx=XM_019812192.2;refGene_gn=GAS2;refGene_tx=XM_019812193.2;refGene_gn=GAS2;refGene_tx=XM_019812194.2;refGene_gn=GAS2;refGene_tx=XM_023239652.1
D1 80067644 80067752
refGene_gn=GAS2;refGene_tx=XM_003993073.5;refGene_gn=GAS2;refGene_tx=XM_019812192.2;refGene_gn=GAS2;refGene_tx=XM_019812193.2;refGene_gn=GAS2;refGene_tx=XM_019812194.2;refGene_gn=GAS2;refGene_tx=XM_023239652.1
D1 80116740 80116959
refGene_gn=GAS2;refGene_tx=XM_003993073.5;refGene_gn=GAS2;refGene_tx=XM_019812192.2;refGene_gn=GAS2;refGene_tx=XM_019812193.2;refGene_gn=GAS2;refGene_tx=XM_019812194.2;refGene_gn=GAS2;refGene_tx=XM_023239652.1
D1 80123694 80123709 refGene_gn=SVIP;refGene_tx=XM_011286748.3
D1 80128313 80128427 refGene_gn=SVIP;refGene_tx=XM_011286748.3
D1 80129140 80129191 refGene_gn=SVIP;refGene_tx=XM_011286748.3
D1 80131237 80131291 refGene_gn=SVIP;refGene_tx=XM_011286748.3
D1 80921557 80922441
refGene_gn=LOC101100590;refGene_tx=XM_011286771.3
D1 80922442 80922869
refGene_gn=LOC101100590;refGene_tx=XM_011286771.3
D1 81565748 81565810
refGene_gn=LUZP2;refGene_tx=XM_011286749.3;refGene_gn=LUZP2;refGene_tx=XM_023239654.1;refGene_gn=LUZP2;refGene_tx=XM_023239656.1;refGene_gn=LUZP2;refGene_tx=XM_023239657.1
D1 81800312 81800430
refGene_gn=LUZP2;refGene_tx=XM_011286749.3;refGene_gn=LUZP2;refGene_tx=XM_023239654.1;refGene_gn=LUZP2;refGene_tx=XM_023239656.1;refGene_gn=LUZP2;refGene_tx=XM_023239657.1
D1 81803946 81804017
refGene_gn=LUZP2;refGene_tx=XM_011286749.3;refGene_gn=LUZP2;refGene_tx=XM_023239654.1;refGene_gn=LUZP2;refGene_tx=XM_023239656.1;refGene_gn=LUZP2;refGene_tx=XM_023239657.1
D1 81808896 81808978

refGene_gn=LUZP2;refGene_tx=XM_011286749.3;refGene_gn=LUZP2;refGene_tx=XM_019812198.2;refGene_gn=LUZP2;refGene_tx=XM_023239654.1;refGene_gn=LUZP2;refGene_tx=XM_023239656.1;refGene_gn=LUZP2;refGene_tx=XM_023239657.1
D1 81814698 81814761
refGene_gn=LUZP2;refGene_tx=XM_011286749.3;refGene_gn=LUZP2;refGene_tx=XM_019812198.2;refGene_gn=LUZP2;refGene_tx=XM_023239654.1;refGene_gn=LUZP2;refGene_tx=XM_023239656.1;refGene_gn=LUZP2;refGene_tx=XM_023239657.1
D1 81924091 81924154
refGene_gn=LUZP2;refGene_tx=XM_011286749.3;refGene_gn=LUZP2;refGene_tx=XM_019812198.2;refGene_gn=LUZP2;refGene_tx=XM_023239654.1;refGene_gn=LUZP2;refGene_tx=XM_023239656.1;refGene_gn=LUZP2;refGene_tx=XM_023239657.1
D1 81929803 81929866
refGene_gn=LUZP2;refGene_tx=XM_011286749.3;refGene_gn=LUZP2;refGene_tx=XM_019812198.2;refGene_gn=LUZP2;refGene_tx=XM_023239654.1;refGene_gn=LUZP2;refGene_tx=XM_023239656.1;refGene_gn=LUZP2;refGene_tx=XM_023239657.1
D1 81977358 81977433
refGene_gn=LUZP2;refGene_tx=XM_011286749.3;refGene_gn=LUZP2;refGene_tx=XM_019812198.2;refGene_gn=LUZP2;refGene_tx=XM_023239654.1;refGene_gn=LUZP2;refGene_tx=XM_023239656.1;refGene_gn=LUZP2;refGene_tx=XM_023239657.1
D1 81986888 81987056
refGene_gn=LUZP2;refGene_tx=XM_011286749.3;refGene_gn=LUZP2;refGene_tx=XM_019812198.2;refGene_gn=LUZP2;refGene_tx=XM_023239654.1;refGene_gn=LUZP2;refGene_tx=XM_023239656.1;refGene_gn=LUZP2;refGene_tx=XM_023239657.1
D1 82033328 82033421
refGene_gn=LUZP2;refGene_tx=XM_011286749.3;refGene_gn=LUZP2;refGene_tx=XM_019812198.2;refGene_gn=LUZP2;refGene_tx=XM_023239654.1;refGene_gn=LUZP2;refGene_tx=XM_023239656.1
D1 82058554 82058641
refGene_gn=LUZP2;refGene_tx=XM_011286749.3;refGene_gn=LUZP2;refGene_tx=XM_019812198.2;refGene_gn=LUZP2;refGene_tx=XM_023239654.1;refGene_gn=LUZP2;refGene_tx=XM_023239656.1;refGene_gn=LUZP2;refGene_tx=XM_023239657.1
D1 82059400 82059505
refGene_gn=LUZP2;refGene_tx=XM_011286749.3;refGene_gn=LUZP2;refGene_tx=XM_019812198.2;refGene_gn=LUZP2;refGene_tx=XM_023239654.1;refGene_gn=LUZP2;refGene_tx=XM_023239656.1;refGene_gn=LUZP2;refGene_tx=XM_023239657.1
D1 82063559 82063725
refGene_gn=LUZP2;refGene_tx=XM_019812198.2;refGene_gn=LUZP2;refGene_tx=XM_023239654.1;refGene_gn=LUZP2;refGene_tx=XM_023239656.1;refGene_gn=LUZP2;refGene_tx=XM_023239657.1
D1 82064300 82064326 refGene_gn=LUZP2;refGene_tx=XM_023239656.1
D1 82982248 82982402 refGene_gn=ANO3;refGene_tx=XM_023239658.1
D1 83109342 83109417 refGene_gn=ANO3;refGene_tx=XM_023239658.1

D1 83131693 83131739 refGene_gn=ANO3;refGene_tx=XM_023239659.1
D1 83214584 83214779
refGene_gn=ANO3;refGene_tx=XM_023239658.1;refGene_gn=ANO3;refGene_tx=XM_023239659.1
D1 83216155 83216227
refGene_gn=ANO3;refGene_tx=XM_023239658.1;refGene_gn=ANO3;refGene_tx=XM_023239659.1
D1 83230995 83231114
refGene_gn=ANO3;refGene_tx=XM_023239658.1;refGene_gn=ANO3;refGene_tx=XM_023239659.1
D1 83275101 83275260
refGene_gn=ANO3;refGene_tx=XM_019812199.2;refGene_gn=ANO3;refGene_tx=XM_023239658.1;refGene_gn=ANO3;refGene_tx=XM_023239659.1
D1 83288076 83288177
refGene_gn=ANO3;refGene_tx=XM_019812199.2;refGene_gn=ANO3;refGene_tx=XM_023239658.1;refGene_gn=ANO3;refGene_tx=XM_023239659.1
D1 83296711 83296756
refGene_gn=ANO3;refGene_tx=XM_019812199.2;refGene_gn=ANO3;refGene_tx=XM_023239658.1;refGene_gn=ANO3;refGene_tx=XM_023239659.1
D1 83301767 83301899
refGene_gn=ANO3;refGene_tx=XM_019812199.2;refGene_gn=ANO3;refGene_tx=XM_023239658.1;refGene_gn=ANO3;refGene_tx=XM_023239659.1
D1 83304281 83304388
refGene_gn=ANO3;refGene_tx=XM_019812199.2;refGene_gn=ANO3;refGene_tx=XM_023239658.1;refGene_gn=ANO3;refGene_tx=XM_023239659.1
D1 83307366 83307422
refGene_gn=ANO3;refGene_tx=XM_019812199.2;refGene_gn=ANO3;refGene_tx=XM_023239658.1;refGene_gn=ANO3;refGene_tx=XM_023239659.1
D1 83311496 83311618
refGene_gn=ANO3;refGene_tx=XM_019812199.2;refGene_gn=ANO3;refGene_tx=XM_023239658.1;refGene_gn=ANO3;refGene_tx=XM_023239659.1
D1 83315317 83315452
refGene_gn=ANO3;refGene_tx=XM_019812199.2;refGene_gn=ANO3;refGene_tx=XM_023239658.1;refGene_gn=ANO3;refGene_tx=XM_023239659.1
D1 83322929 83323026
refGene_gn=ANO3;refGene_tx=XM_019812199.2;refGene_gn=ANO3;refGene_tx=XM_023239658.1;refGene_gn=ANO3;refGene_tx=XM_023239659.1
D1 83329785 83329846
refGene_gn=ANO3;refGene_tx=XM_019812199.2;refGene_gn=ANO3;refGene_tx=XM_023239658.1;refGene_gn=ANO3;refGene_tx=XM_023239659.1
D1 83331053 83331214
refGene_gn=MUC15;refGene_tx=XM_006937159.4;refGene_gn=MUC15;refGene_tx=XM_023239660.1
D1 83333293 83333443
refGene_gn=MUC15;refGene_tx=XM_006937159.4;refGene_gn=MUC15;refGene_tx=XM_023239660.1
D1 83335437 83336168
refGene_gn=MUC15;refGene_tx=XM_006937159.4;refGene_gn=MUC15;refGene_tx=XM_023239660.1
D1 83339722 83339742 refGene_gn=MUC15;refGene_tx=XM_023239660.1

D1 83360673 83360756
refGene_gn=ANO3;refGene_tx=XM_019812199.2;refGene_gn=ANO3;refGene_tx=XM_023239658.1;refGene_gn=ANO3;refGene_tx=XM_023239659.1
D1 83361179 83361320
refGene_gn=ANO3;refGene_tx=XM_019812199.2;refGene_gn=ANO3;refGene_tx=XM_023239658.1;refGene_gn=ANO3;refGene_tx=XM_023239659.1
D1 83361823 83361988
refGene_gn=ANO3;refGene_tx=XM_019812199.2;refGene_gn=ANO3;refGene_tx=XM_023239658.1;refGene_gn=ANO3;refGene_tx=XM_023239659.1
D1 83374346 83374383
refGene_gn=ANO3;refGene_tx=XM_019812199.2;refGene_gn=ANO3;refGene_tx=XM_023239658.1;refGene_gn=ANO3;refGene_tx=XM_023239659.1
D1 83377507 83377619
refGene_gn=ANO3;refGene_tx=XM_019812199.2;refGene_gn=ANO3;refGene_tx=XM_023239658.1;refGene_gn=ANO3;refGene_tx=XM_023239659.1
D1 83378376 83378434
refGene_gn=ANO3;refGene_tx=XM_019812199.2;refGene_gn=ANO3;refGene_tx=XM_023239658.1;refGene_gn=ANO3;refGene_tx=XM_023239659.1
D1 83383845 83383943
refGene_gn=ANO3;refGene_tx=XM_019812199.2;refGene_gn=ANO3;refGene_tx=XM_023239658.1;refGene_gn=ANO3;refGene_tx=XM_023239659.1
D1 83386508 83386642
refGene_gn=ANO3;refGene_tx=XM_019812199.2;refGene_gn=ANO3;refGene_tx=XM_023239658.1;refGene_gn=ANO3;refGene_tx=XM_023239659.1
D1 83387541 83387694
refGene_gn=ANO3;refGene_tx=XM_019812199.2;refGene_gn=ANO3;refGene_tx=XM_023239658.1;refGene_gn=ANO3;refGene_tx=XM_023239659.1
D1 83390919 83391067
refGene_gn=ANO3;refGene_tx=XM_019812199.2;refGene_gn=ANO3;refGene_tx=XM_023239658.1;refGene_gn=ANO3;refGene_tx=XM_023239659.1
D1 83395127 83395208
refGene_gn=ANO3;refGene_tx=XM_019812199.2;refGene_gn=ANO3;refGene_tx=XM_023239658.1;refGene_gn=ANO3;refGene_tx=XM_023239659.1
D1 83395384 83395490
refGene_gn=ANO3;refGene_tx=XM_019812199.2;refGene_gn=ANO3;refGene_tx=XM_023239658.1;refGene_gn=ANO3;refGene_tx=XM_023239659.1
D1 83400428 83400611
refGene_gn=ANO3;refGene_tx=XM_019812199.2;refGene_gn=ANO3;refGene_tx=XM_023239658.1;refGene_gn=ANO3;refGene_tx=XM_023239659.1
D1 83411968 83412121 refGene_gn=SLC5A12;refGene_tx=XM_003993114.3
D1 83414024 83414152 refGene_gn=SLC5A12;refGene_tx=XM_003993114.3
D1 83416732 83416836 refGene_gn=SLC5A12;refGene_tx=XM_003993114.3
D1 83418430 83418597 refGene_gn=SLC5A12;refGene_tx=XM_003993114.3
D1 83421294 83421381 refGene_gn=SLC5A12;refGene_tx=XM_003993114.3
D1 83426661 83426729 refGene_gn=SLC5A12;refGene_tx=XM_003993114.3
D1 83430606 83430719 refGene_gn=SLC5A12;refGene_tx=XM_003993114.3
D1 83435116 83435205 refGene_gn=SLC5A12;refGene_tx=XM_003993114.3
D1 83436418 83436548 refGene_gn=SLC5A12;refGene_tx=XM_003993114.3
D1 83442156 83442297 refGene_gn=SLC5A12;refGene_tx=XM_003993114.3
D1 83442415 83442570 refGene_gn=SLC5A12;refGene_tx=XM_003993114.3

D1	83447409	83447477	refGene_gn=SLC5A12;refGene_tx=XM_003993114.3
D1	83448322	83448374	refGene_gn=SLC5A12;refGene_tx=XM_003993114.3
D1	83449546	83449612	refGene_gn=SLC5A12;refGene_tx=XM_003993114.3
D1	83459175	83459514	refGene_gn=SLC5A12;refGene_tx=XM_003993114.3
D1	83677264	83677900	refGene_gn=FIBIN;refGene_tx=XM_006937160.4
D1	83723027	83723246
refGene_gn=BBOX1;refGene_tx=XM_003993116.5;refGene_gn=BBOX1;refGene_tx=XM_019812203.2;refGene_gn=BBOX1;refGene_tx=XM_019812204.2
D1	83724535	83724650
refGene_gn=BBOX1;refGene_tx=XM_003993116.5;refGene_gn=BBOX1;refGene_tx=XM_019812203.2;refGene_gn=BBOX1;refGene_tx=XM_019812204.2
D1	83742557	83742756
refGene_gn=BBOX1;refGene_tx=XM_003993116.5;refGene_gn=BBOX1;refGene_tx=XM_019812203.2;refGene_gn=BBOX1;refGene_tx=XM_019812204.2
D1	83758545	83758651
refGene_gn=BBOX1;refGene_tx=XM_003993116.5;refGene_gn=BBOX1;refGene_tx=XM_019812203.2;refGene_gn=BBOX1;refGene_tx=XM_019812204.2
D1	83764108	83764305
refGene_gn=BBOX1;refGene_tx=XM_003993116.5;refGene_gn=BBOX1;refGene_tx=XM_019812203.2;refGene_gn=BBOX1;refGene_tx=XM_019812204.2
D1	83772288	83772455
refGene_gn=BBOX1;refGene_tx=XM_003993116.5;refGene_gn=BBOX1;refGene_tx=XM_019812203.2;refGene_gn=BBOX1;refGene_tx=XM_019812204.2
D1	83775310	83775471
refGene_gn=BBOX1;refGene_tx=XM_003993116.5;refGene_gn=BBOX1;refGene_tx=XM_019812203.2;refGene_gn=BBOX1;refGene_tx=XM_019812204.2
D1	83973538	83973753
refGene_gn=CCDC34;refGene_tx=XM_011286751.3;refGene_gn=CCDC34;refGene_tx=XM_019812205.2;refGene_gn=CCDC34;refGene_tx=XM_019812206.2;refGene_gn=CCDC34;refGene_tx=XM_019812207.2;refGene_gn=CCDC34;refGene_tx=XM_019812208.2
D1	83975244	83975386
refGene_gn=CCDC34;refGene_tx=XM_011286751.3;refGene_gn=CCDC34;refGene_tx=XM_019812205.2;refGene_gn=CCDC34;refGene_tx=XM_019812206.2;refGene_gn=CCDC34;refGene_tx=XM_019812207.2;refGene_gn=CCDC34;refGene_tx=XM_019812208.2
D1	83976442	83976601	refGene_gn=CCDC34;refGene_tx=XM_011286751.3
D1	83984559	83984667
refGene_gn=CCDC34;refGene_tx=XM_011286751.3;refGene_gn=CCDC34;refGene_tx=XM_019812205.2;refGene_gn=CCDC34;refGene_tx=XM_019812206.2;refGene_gn=CCDC34;refGene_tx=XM_019812207.2;refGene_gn=CCDC34;refGene_tx=XM_019812208.2
D1	83991008	83991144
refGene_gn=CCDC34;refGene_tx=XM_011286751.3;refGene_gn=CCDC34;refGene_tx=XM_019812205.2;refGene_gn=CCDC34;refGene_tx=XM_019812206.2;refGene_gn=CCDC34;refGene_tx=XM_019812207.2;refGene_gn=CCDC34;refGene_tx=XM_019812208.2
D1	83996758	83997087
refGene_gn=CCDC34;refGene_tx=XM_011286751.3;refGene_gn=CCDC34;refGene_tx=XM_019812205.2;refGene_gn=CCDC34;refGene_tx=XM_019812206.2;refGene_

gn=CCDC34;refGene_tx=XM_019812207.2;refGene_gn=CCDC34;refGene_tx=XM_019812208.2
D1 84002141 84003418 refGene_gn=LGR4;refGene_tx=XM_023239663.1
D1 84006421 84006505 refGene_gn=LGR4;refGene_tx=XM_023239663.1
D1 84006760 84006876 refGene_gn=LGR4;refGene_tx=XM_023239663.1
D1 84007952 84008078 refGene_gn=LGR4;refGene_tx=XM_023239663.1
D1 84008373 84008445 refGene_gn=LGR4;refGene_tx=XM_023239663.1
D1 84010071 84010143 refGene_gn=LGR4;refGene_tx=XM_023239663.1
D1 84010861 84010927 refGene_gn=LGR4;refGene_tx=XM_023239663.1
D1 84012003 84012075 refGene_gn=LGR4;refGene_tx=XM_023239663.1
D1 84013594 84013663 refGene_gn=LGR4;refGene_tx=XM_023239663.1
D1 84014278 84014350 refGene_gn=LGR4;refGene_tx=XM_023239663.1
D1 84014536 84014608 refGene_gn=LGR4;refGene_tx=XM_023239663.1
D1 84015589 84015658 refGene_gn=LGR4;refGene_tx=XM_023239663.1
D1 84017388 84017460 refGene_gn=LGR4;refGene_tx=XM_023239663.1
D1 84018777 84018993 refGene_gn=LGR4;refGene_tx=XM_023239663.1
D1 84027424 84027496 refGene_gn=LGR4;refGene_tx=XM_023239663.1
D1 84028759 84028831 refGene_gn=LGR4;refGene_tx=XM_023239663.1
D1 84048701 84048773 refGene_gn=LGR4;refGene_tx=XM_023239663.1
D1 84103008 84103193 refGene_gn=LGR4;refGene_tx=XM_023239663.1
D1 84119611 84119767 refGene_gn=LIN7C;refGene_tx=XM_003993117.4
D1 84120353 84120563 refGene_gn=LIN7C;refGene_tx=XM_003993117.4
D1 84122813 84122885 refGene_gn=LIN7C;refGene_tx=XM_003993117.4
D1 84123912 84124031 refGene_gn=LIN7C;refGene_tx=XM_003993117.4
D1 84127170 84127207 refGene_gn=LIN7C;refGene_tx=XM_003993117.4
D1 84257818 84258562 refGene_gn=BDNF;refGene_tx=NM_001009828.1
D1 84580958 84581041 refGene_gn=KIF18A;refGene_tx=XM_011286752.3
D1 84583235 84583345 refGene_gn=KIF18A;refGene_tx=XM_011286752.3
D1 84589029 84589137 refGene_gn=KIF18A;refGene_tx=XM_011286752.3
D1 84589920 84590359 refGene_gn=KIF18A;refGene_tx=XM_011286752.3
D1 84617766 84618005 refGene_gn=KIF18A;refGene_tx=XM_011286752.3
D1 84624246 84624368 refGene_gn=KIF18A;refGene_tx=XM_011286752.3
D1 84628606 84628771 refGene_gn=KIF18A;refGene_tx=XM_011286752.3
D1 84633779 84633942 refGene_gn=KIF18A;refGene_tx=XM_011286752.3
D1 84637739 84637852 refGene_gn=KIF18A;refGene_tx=XM_011286752.3
D1 84638055 84638130 refGene_gn=KIF18A;refGene_tx=XM_011286752.3
D1 84639739 84639916 refGene_gn=KIF18A;refGene_tx=XM_011286752.3
D1 84642218 84642416 refGene_gn=KIF18A;refGene_tx=XM_011286752.3
D1 84644551 84644662 refGene_gn=KIF18A;refGene_tx=XM_011286752.3
D1 84645398 84645503 refGene_gn=KIF18A;refGene_tx=XM_011286752.3
D1 84648273 84648431 refGene_gn=KIF18A;refGene_tx=XM_011286752.3
D1 84650768 84651093 refGene_gn=KIF18A;refGene_tx=XM_011286752.3
D1 84666199 84666469 refGene_gn=METTL15;refGene_tx=XM_023239664.1;refGene_gn=METTL15;refGene_tx=XM_023239665.1;refGene_gn=METTL15;refGene_tx=XM_023239666.1;refGene_gn=METTL15;refGene_tx=XM_023239667.1;refGene_gn=METTL15;refGene_tx=XM_023239668.1
D1 84688084 84688161 refGene_gn=METTL15;refGene_tx=XM_023239664.1;refGene_gn=METTL15;refGene_tx=XM_023239665.1;refGene_gn=METTL15;refGene_tx=XM_023239666.1

D1 84689430 84689569
refGene_gn=METTL15;refGene_tx=XM_023239664.1;refGene_gn=METTL15;refGene_tx=XM_023239665.1;refGene_gn=METTL15;refGene_tx=XM_023239666.1;refGene_gn=METTL15;refGene_tx=XM_023239668.1
D1 84729893 84729929 refGene_gn=METTL15;refGene_tx=XM_023239669.1
D1 84749619 84749756
refGene_gn=METTL15;refGene_tx=XM_023239664.1;refGene_gn=METTL15;refGene_tx=XM_023239665.1;refGene_gn=METTL15;refGene_tx=XM_023239666.1;refGene_gn=METTL15;refGene_tx=XM_023239667.1;refGene_gn=METTL15;refGene_tx=XM_023239668.1;refGene_gn=METTL15;refGene_tx=XM_023239669.1
D1 84812256 84812448
refGene_gn=METTL15;refGene_tx=XM_023239664.1;refGene_gn=METTL15;refGene_tx=XM_023239665.1;refGene_gn=METTL15;refGene_tx=XM_023239666.1;refGene_gn=METTL15;refGene_tx=XM_023239667.1;refGene_gn=METTL15;refGene_tx=XM_023239668.1;refGene_gn=METTL15;refGene_tx=XM_023239669.1
D1 84816505 84816684
refGene_gn=METTL15;refGene_tx=XM_023239664.1;refGene_gn=METTL15;refGene_tx=XM_023239665.1;refGene_gn=METTL15;refGene_tx=XM_023239666.1;refGene_gn=METTL15;refGene_tx=XM_023239667.1;refGene_gn=METTL15;refGene_tx=XM_023239668.1;refGene_gn=METTL15;refGene_tx=XM_023239669.1
D1 84850990 84851436
refGene_gn=METTL15;refGene_tx=XM_023239664.1;refGene_gn=METTL15;refGene_tx=XM_023239665.1;refGene_gn=METTL15;refGene_tx=XM_023239666.1;refGene_gn=METTL15;refGene_tx=XM_023239667.1;refGene_gn=METTL15;refGene_tx=XM_023239668.1;refGene_gn=METTL15;refGene_tx=XM_023239669.1
D1 86358049 86360014 refGene_gn=KCNA4;refGene_tx=XM_006937163.4
D1 86364016 86364129
refGene_gn=LOC111556657;refGene_tx=XM_023239730.1
D1 86364323 86364752
refGene_gn=LOC111556657;refGene_tx=XM_023239730.1
D1 86366518 86366635
refGene_gn=LOC111556657;refGene_tx=XM_023239730.1
D1 86372901 86372932
refGene_gn=LOC111556657;refGene_tx=XM_023239730.1
D1 86563337 86563496 refGene_gn=FSHB;refGene_tx=XM_003993122.4
D1 86565415 86565646 refGene_gn=FSHB;refGene_tx=XM_003993122.4
D1 86651693 86652116 refGene_gn=ARL14EP;refGene_tx=XM_003993123.5
D1 86653983 86654111 refGene_gn=ARL14EP;refGene_tx=XM_003993123.5
D1 86655168 86655397 refGene_gn=ARL14EP;refGene_tx=XM_003993123.5
D1 86726631 86726750
refGene_gn=MPPED2;refGene_tx=XM_003993124.5;refGene_gn=MPPED2;refGene_tx=XM_019812217.2
D1 86729620 86729734
refGene_gn=MPPED2;refGene_tx=XM_003993124.5;refGene_gn=MPPED2;refGene_tx=XM_019812217.2
D1 86732903 86733019
refGene_gn=MPPED2;refGene_tx=XM_003993124.5;refGene_gn=MPPED2;refGene_tx=XM_019812217.2
D1 86813808 86814034
refGene_gn=MPPED2;refGene_tx=XM_003993124.5;refGene_gn=MPPED2;refGene_

tx=XM_019812217.2
D1 86852041 86852223
refGene_gn=MPPED2;refGene_tx=XM_003993124.5;refGene_gn=MPPED2;refGene_
tx=XM_019812217.2
D1 86894630 86894758
refGene_gn=MPPED2;refGene_tx=XM_003993124.5;refGene_gn=MPPED2;refGene_
tx=XM_019812217.2
D1 87138367 87138436
refGene_gn=LOC101086002;refGene_tx=XM_019812218.2
D1 87150069 87150179
refGene_gn=LOC101086002;refGene_tx=XM_019812218.2
D1 87152913 87153064
refGene_gn=LOC101086002;refGene_tx=XM_019812218.2
D1 87163748 87163928
refGene_gn=LOC101086002;refGene_tx=XM_019812218.2
D1 87165149 87165286
refGene_gn=LOC101086002;refGene_tx=XM_019812218.2
D1 87170002 87170104
refGene_gn=LOC101086002;refGene_tx=XM_019812218.2
D1 87170821 87170974
refGene_gn=LOC101086002;refGene_tx=XM_019812218.2
D1 87174163 87174365
refGene_gn=LOC101086002;refGene_tx=XM_019812218.2
D1 87175709 87175913
refGene_gn=LOC101086002;refGene_tx=XM_019812218.2
D1 87177772 87177958
refGene_gn=LOC101086002;refGene_tx=XM_019812218.2
D1 87184962 87185133
refGene_gn=LOC101086002;refGene_tx=XM_019812218.2
D1 87187508 87187602
refGene_gn=LOC101086002;refGene_tx=XM_019812218.2
D1 87192091 87192292
refGene_gn=LOC101086002;refGene_tx=XM_019812218.2
D1 87193413 87193572
refGene_gn=LOC101086002;refGene_tx=XM_019812218.2
D1 87203459 87203619
refGene_gn=LOC101086002;refGene_tx=XM_019812218.2
D1 87206471 87206607
refGene_gn=LOC101086002;refGene_tx=XM_019812218.2
D1 87208718 87208818
refGene_gn=LOC101086002;refGene_tx=XM_019812218.2
D1 87213131 87213230
refGene_gn=LOC101086002;refGene_tx=XM_019812218.2
D1 87217184 87217366
refGene_gn=LOC101086002;refGene_tx=XM_019812218.2
D1 87229998 87230122
refGene_gn=LOC101086002;refGene_tx=XM_019812218.2
D1 87326445 87326603
refGene_gn=LOC101086002;refGene_tx=XM_019812218.2
D1 87326992 87327121

refGene_gn=LOC101086002;refGene_tx=XM_019812218.2
D1 87336893 87336954
refGene_gn=LOC101086002;refGene_tx=XM_019812218.2
D1 87350265 87350384
refGene_gn=LOC101086002;refGene_tx=XM_019812218.2
D1 87354019 87354154
refGene_gn=LOC101086002;refGene_tx=XM_019812218.2
D1 87360302 87360408
refGene_gn=LOC101086002;refGene_tx=XM_019812218.2
D1 87363073 87363199
refGene_gn=LOC101086002;refGene_tx=XM_019812218.2
D1 87366249 87366413
refGene_gn=LOC101086002;refGene_tx=XM_019812218.2
D1 87369659 87369761
refGene_gn=LOC101086002;refGene_tx=XM_019812218.2
D1 87387694 87387865
refGene_gn=LOC101086002;refGene_tx=XM_019812218.2
D1 87395193 87395286
refGene_gn=LOC101086002;refGene_tx=XM_019812218.2
D1 87511394 87511561
refGene_gn=LOC101086002;refGene_tx=XM_019812218.2
D1 87529431 87529525
refGene_gn=LOC101086002;refGene_tx=XM_019812218.2
D1 87553554 87553760
refGene_gn=LOC101086002;refGene_tx=XM_019812218.2
D1 87573612 87573775
refGene_gn=LOC101086002;refGene_tx=XM_019812218.2
D1 87574265 87574422
refGene_gn=LOC101086002;refGene_tx=XM_019812218.2
D1 87575600 87575870
refGene_gn=LOC101086002;refGene_tx=XM_019812218.2
D1 87588414 87588584
refGene_gn=LOC101086002;refGene_tx=XM_019812218.2
D1 87609818 87609836
refGene_gn=LOC101086002;refGene_tx=XM_019812218.2
D1 87616281 87616392
refGene_gn=DNAJC24;refGene_tx=XM_003993126.4;refGene_gn=DNAJC24;refGene_tx=XM_006937171.3;refGene_gn=DNAJC24;refGene_tx=XM_006937172.3
D1 87655687 87655826
refGene_gn=DNAJC24;refGene_tx=XM_003993126.4;refGene_gn=DNAJC24;refGene_tx=XM_006937171.3;refGene_gn=DNAJC24;refGene_tx=XM_006937172.3
D1 87667512 87667581
refGene_gn=DNAJC24;refGene_tx=XM_003993126.4;refGene_gn=DNAJC24;refGene_tx=XM_006937171.3
D1 87671933 87672064
refGene_gn=DNAJC24;refGene_tx=XM_003993126.4;refGene_gn=DNAJC24;refGene_tx=XM_006937171.3;refGene_gn=DNAJC24;refGene_tx=XM_006937172.3
D1 87673492 87673561
refGene_gn=IMMP1L;refGene_tx=XM_006937165.3;refGene_gn=IMMP1L;refGene_tx=XM_006937166.3;refGene_gn=IMMP1L;refGene_tx=XM_006937168.4;refGene_

gn=IMMP1L;refGene_tx=XM_006937169.3;refGene_gn=IMMP1L;refGene_tx=XM_01
1286817.3;refGene_gn=IMMP1L;refGene_tx=XM_019812220.2;refGene_gn=IMMP1
L;refGene_tx=XM_019812221.2;refGene_gn=IMMP1L;refGene_tx=XM_023239670.
1;refGene_gn=IMMP1L;refGene_tx=XM_023239671.1
D1 87674411 87674522
refGene_gn=IMMP1L;refGene_tx=XM_006937165.3;refGene_gn=IMMP1L;refGene_
tx=XM_006937166.3;refGene_gn=IMMP1L;refGene_tx=XM_006937168.4;refGene_
gn=IMMP1L;refGene_tx=XM_006937169.3;refGene_gn=IMMP1L;refGene_tx=XM_01
1286817.3;refGene_gn=IMMP1L;refGene_tx=XM_019812220.2;refGene_gn=IMMP1
L;refGene_tx=XM_019812221.2;refGene_gn=IMMP1L;refGene_tx=XM_023239670.
1;refGene_gn=IMMP1L;refGene_tx=XM_023239671.1
D1 87693996 87694123
refGene_gn=IMMP1L;refGene_tx=XM_006937165.3;refGene_gn=IMMP1L;refGene_
tx=XM_006937166.3;refGene_gn=IMMP1L;refGene_tx=XM_006937168.4;refGene_
gn=IMMP1L;refGene_tx=XM_006937169.3;refGene_gn=IMMP1L;refGene_tx=XM_01
1286817.3;refGene_gn=IMMP1L;refGene_tx=XM_019812220.2;refGene_gn=IMMP1
L;refGene_tx=XM_019812221.2;refGene_gn=IMMP1L;refGene_tx=XM_023239670.
1;refGene_gn=IMMP1L;refGene_tx=XM_023239671.1
D1 87698099 87698188
refGene_gn=IMMP1L;refGene_tx=XM_006937165.3;refGene_gn=IMMP1L;refGene_
tx=XM_006937166.3;refGene_gn=IMMP1L;refGene_tx=XM_006937168.4;refGene_
gn=IMMP1L;refGene_tx=XM_006937169.3;refGene_gn=IMMP1L;refGene_tx=XM_01
1286817.3;refGene_gn=IMMP1L;refGene_tx=XM_019812220.2;refGene_gn=IMMP1
L;refGene_tx=XM_019812221.2;refGene_gn=IMMP1L;refGene_tx=XM_023239670.
1;refGene_gn=IMMP1L;refGene_tx=XM_023239671.1
D1 87699594 87699728
refGene_gn=IMMP1L;refGene_tx=XM_006937165.3;refGene_gn=IMMP1L;refGene_
tx=XM_006937166.3;refGene_gn=IMMP1L;refGene_tx=XM_006937168.4;refGene_
gn=IMMP1L;refGene_tx=XM_006937169.3;refGene_gn=IMMP1L;refGene_tx=XM_01
1286817.3;refGene_gn=IMMP1L;refGene_tx=XM_019812220.2;refGene_gn=IMMP1
L;refGene_tx=XM_019812221.2;refGene_gn=IMMP1L;refGene_tx=XM_023239670.
1;refGene_gn=IMMP1L;refGene_tx=XM_023239671.1
D1 87720862 87720884 refGene_gn=IMMP1L;refGene_tx=XM_023239671.1
D1 87755316 87755539
refGene_gn=ELP4;refGene_tx=XM_023239673.1;refGene_gn=ELP4;refGene_tx=X
M_023239674.1
D1 87763216 87763252
refGene_gn=ELP4;refGene_tx=XM_023239673.1;refGene_gn=ELP4;refGene_tx=X
M_023239674.1
D1 87786170 87786292
refGene_gn=ELP4;refGene_tx=XM_023239673.1;refGene_gn=ELP4;refGene_tx=X
M_023239674.1
D1 87813273 87813405
refGene_gn=ELP4;refGene_tx=XM_023239673.1;refGene_gn=ELP4;refGene_tx=X
M_023239674.1
D1 87820982 87821122
refGene_gn=ELP4;refGene_tx=XM_023239673.1;refGene_gn=ELP4;refGene_tx=X
M_023239674.1
D1 87843619 87843707
refGene_gn=ELP4;refGene_tx=XM_023239673.1;refGene_gn=ELP4;refGene_tx=X

M_023239674.1
D1 87849011 87849197
refGene_gn=ELP4;refGene_tx=XM_023239673.1;refGene_gn=ELP4;refGene_tx=XM_023239674.1
D1 87871114 87871223
refGene_gn=ELP4;refGene_tx=XM_023239673.1;refGene_gn=ELP4;refGene_tx=XM_023239674.1
D1 87874987 87875094
refGene_gn=ELP4;refGene_tx=XM_023239673.1;refGene_gn=ELP4;refGene_tx=XM_023239674.1
D1 88001513 88001645
refGene_gn=ELP4;refGene_tx=XM_023239673.1;refGene_gn=ELP4;refGene_tx=XM_023239674.1
D1 88008130 88008216 refGene_gn=PAX6;refGene_tx=XM_023239672.1
D1 88009173 88009324 refGene_gn=PAX6;refGene_tx=XM_023239672.1
D1 88011932 88012048 refGene_gn=PAX6;refGene_tx=XM_023239672.1
D1 88012142 88012293 refGene_gn=PAX6;refGene_tx=XM_023239672.1
D1 88012504 88012587 refGene_gn=PAX6;refGene_tx=XM_023239672.1
D1 88013115 88013274 refGene_gn=PAX6;refGene_tx=XM_023239672.1
D1 88019167 88019333 refGene_gn=PAX6;refGene_tx=XM_023239672.1
D1 88020155 88020371 refGene_gn=PAX6;refGene_tx=XM_023239672.1
D1 88020465 88020507 refGene_gn=PAX6;refGene_tx=XM_023239672.1
D1 88021310 88021346 refGene_gn=PAX6;refGene_tx=XM_023239672.1
D1 88021347 88021442 refGene_gn=PAX6;refGene_tx=XM_023239672.1
D1 88025200 88025210 refGene_gn=PAX6;refGene_tx=XM_023239672.1
D1 88029755 88029828
refGene_gn=LOC111556658;refGene_tx=XM_023239731.1
D1 88030992 88031641
refGene_gn=LOC111556658;refGene_tx=XM_023239731.1
D1 88031910 88032145
refGene_gn=LOC111556658;refGene_tx=XM_023239731.1
D1 88032416 88032435
refGene_gn=LOC111556659;refGene_tx=XM_023239732.1
D1 88032862 88033291
refGene_gn=LOC111556659;refGene_tx=XM_023239732.1
D1 88033406 88033756
refGene_gn=LOC111556659;refGene_tx=XM_023239732.1
D1 88296457 88296711 refGene_gn=RCN1;refGene_tx=XM_006937183.3
D1 88301130 88301324 refGene_gn=RCN1;refGene_tx=XM_006937183.3
D1 88302330 88302509 refGene_gn=RCN1;refGene_tx=XM_006937183.3
D1 88304229 88304290 refGene_gn=RCN1;refGene_tx=XM_006937183.3
D1 88307000 88307200 refGene_gn=RCN1;refGene_tx=XM_006937183.3
D1 88308307 88308415 refGene_gn=RCN1;refGene_tx=XM_006937183.3
D1 88545114 88545236
refGene_gn=WT1;refGene_tx=XM_019812232.2;refGene_gn=WT1;refGene_tx=XM_019812233.2;refGene_gn=WT1;refGene_tx=XM_019812234.2
D1 88547675 88547768
refGene_gn=WT1;refGene_tx=XM_019812232.2;refGene_gn=WT1;refGene_tx=XM_019812233.2;refGene_gn=WT1;refGene_tx=XM_019812234.2
D1 88548364 88548454

refGene_gn=WT1;refGene_tx=XM_019812232.2;refGene_gn=WT1;refGene_tx=XM_019812233.2;refGene_gn=WT1;refGene_tx=XM_019812234.2
D1 88551925 88552076
refGene_gn=WT1;refGene_tx=XM_019812232.2;refGene_gn=WT1;refGene_tx=XM_019812233.2;refGene_gn=WT1;refGene_tx=XM_019812234.2
D1 88555028 88555125
refGene_gn=WT1;refGene_tx=XM_019812232.2;refGene_gn=WT1;refGene_tx=XM_019812233.2;refGene_gn=WT1;refGene_tx=XM_019812234.2
D1 88574972 88575023
refGene_gn=WT1;refGene_tx=XM_019812232.2;refGene_gn=WT1;refGene_tx=XM_019812233.2
D1 88576103 88576181
refGene_gn=WT1;refGene_tx=XM_019812232.2;refGene_gn=WT1;refGene_tx=XM_019812233.2;refGene_gn=WT1;refGene_tx=XM_019812234.2
D1 88586586 88586689
refGene_gn=WT1;refGene_tx=XM_019812232.2;refGene_gn=WT1;refGene_tx=XM_019812233.2;refGene_gn=WT1;refGene_tx=XM_019812234.2
D1 88587163 88587286
refGene_gn=WT1;refGene_tx=XM_019812232.2;refGene_gn=WT1;refGene_tx=XM_019812233.2;refGene_gn=WT1;refGene_tx=XM_019812234.2
D1 88589058 88589068
refGene_gn=WT1;refGene_tx=XM_019812233.2;refGene_gn=WT1;refGene_tx=XM_019812234.2
D1 88592822 88593267 refGene_gn=WT1;refGene_tx=XM_019812232.2
D1 88727966 88728008 refGene_gn=EIF3M;refGene_tx=XM_003993139.5
D1 88730449 88730582 refGene_gn=EIF3M;refGene_tx=XM_003993139.5
D1 88731737 88731876 refGene_gn=EIF3M;refGene_tx=XM_003993139.5
D1 88732157 88732281 refGene_gn=EIF3M;refGene_tx=XM_003993139.5
D1 88732438 88732533 refGene_gn=EIF3M;refGene_tx=XM_003993139.5
D1 88743271 88743355 refGene_gn=EIF3M;refGene_tx=XM_003993139.5
D1 88744331 88744431 refGene_gn=EIF3M;refGene_tx=XM_003993139.5
D1 88745032 88745114 refGene_gn=EIF3M;refGene_tx=XM_003993139.5
D1 88749569 88749713 refGene_gn=EIF3M;refGene_tx=XM_003993139.5
D1 88750634 88750695 refGene_gn=EIF3M;refGene_tx=XM_003993139.5
D1 88751156 88751277 refGene_gn=EIF3M;refGene_tx=XM_003993139.5
D1 88751749 88751971 refGene_gn=CCDC73;refGene_tx=XM_023239733.1
D1 88755323 88755457 refGene_gn=CCDC73;refGene_tx=XM_023239733.1
D1 88757413 88758910 refGene_gn=CCDC73;refGene_tx=XM_023239733.1
D1 88760420 88760601 refGene_gn=CCDC73;refGene_tx=XM_023239733.1
D1 88770091 88770226 refGene_gn=CCDC73;refGene_tx=XM_023239733.1
D1 88774384 88774495 refGene_gn=CCDC73;refGene_tx=XM_023239733.1
D1 88783467 88783572 refGene_gn=CCDC73;refGene_tx=XM_023239733.1
D1 88784711 88784771 refGene_gn=CCDC73;refGene_tx=XM_023239733.1
D1 88785273 88785402 refGene_gn=CCDC73;refGene_tx=XM_023239733.1
D1 88809399 88809479 refGene_gn=CCDC73;refGene_tx=XM_023239733.1
D1 88809946 88810082 refGene_gn=CCDC73;refGene_tx=XM_023239733.1
D1 88820191 88820230 refGene_gn=CCDC73;refGene_tx=XM_023239733.1
D1 88835327 88835402 refGene_gn=CCDC73;refGene_tx=XM_023239733.1
D1 88836284 88836320 refGene_gn=CCDC73;refGene_tx=XM_023239733.1
D1 88838426 88838498 refGene_gn=CCDC73;refGene_tx=XM_023239733.1

D1 88849205 88849277 refGene_gn=CCDC73;refGene_tx=XM_023239733.1
D1 88878627 88878765 refGene_gn=CCDC73;refGene_tx=XM_023239733.1
D1 88949997 88950100
refGene_gn=PRRG4;refGene_tx=XM_003993140.3;refGene_gn=PRRG4;refGene_tx=XM_019812235.2
D1 88957437 88957601
refGene_gn=PRRG4;refGene_tx=XM_003993140.3;refGene_gn=PRRG4;refGene_tx=XM_019812235.2
D1 88959192 88959241
refGene_gn=PRRG4;refGene_tx=XM_003993140.3;refGene_gn=PRRG4;refGene_tx=XM_019812235.2
D1 88960425 88960558
refGene_gn=PRRG4;refGene_tx=XM_003993140.3;refGene_gn=PRRG4;refGene_tx=XM_019812235.2
D1 88965598 88965833
refGene_gn=PRRG4;refGene_tx=XM_003993140.3;refGene_gn=PRRG4;refGene_tx=XM_019812235.2
D1 88993949 88994140
refGene_gn=QSER1;refGene_tx=XM_023239677.1;refGene_gn=QSER1;refGene_tx=XM_023239678.1
D1 89031277 89031390
refGene_gn=QSER1;refGene_tx=XM_023239677.1;refGene_gn=QSER1;refGene_tx=XM_023239678.1
D1 89032480 89032642
refGene_gn=QSER1;refGene_tx=XM_006937176.4;refGene_gn=QSER1;refGene_tx=XM_023239677.1;refGene_gn=QSER1;refGene_tx=XM_023239678.1;refGene_gn=QSER1;refGene_tx=XM_023239679.1
D1 89035357 89039038
refGene_gn=QSER1;refGene_tx=XM_006937176.4;refGene_gn=QSER1;refGene_tx=XM_023239677.1;refGene_gn=QSER1;refGene_tx=XM_023239678.1;refGene_gn=QSER1;refGene_tx=XM_023239679.1
D1 89051297 89051620
refGene_gn=QSER1;refGene_tx=XM_006937176.4;refGene_gn=QSER1;refGene_tx=XM_023239677.1;refGene_gn=QSER1;refGene_tx=XM_023239678.1;refGene_gn=QSER1;refGene_tx=XM_023239679.1
D1 89052576 89052693
refGene_gn=QSER1;refGene_tx=XM_006937176.4;refGene_gn=QSER1;refGene_tx=XM_023239677.1;refGene_gn=QSER1;refGene_tx=XM_023239678.1;refGene_gn=QSER1;refGene_tx=XM_023239679.1
D1 89053789 89053923
refGene_gn=QSER1;refGene_tx=XM_006937176.4;refGene_gn=QSER1;refGene_tx=XM_023239677.1;refGene_gn=QSER1;refGene_tx=XM_023239678.1;refGene_gn=QSER1;refGene_tx=XM_023239679.1
D1 89055663 89055881
refGene_gn=QSER1;refGene_tx=XM_006937176.4;refGene_gn=QSER1;refGene_tx=XM_023239677.1;refGene_gn=QSER1;refGene_tx=XM_023239678.1;refGene_gn=QSER1;refGene_tx=XM_023239679.1
D1 89063882 89064020
refGene_gn=QSER1;refGene_tx=XM_006937176.4;refGene_gn=QSER1;refGene_tx=XM_023239677.1;refGene_gn=QSER1;refGene_tx=XM_023239678.1;refGene_gn=

QSER1;refGene_tx=XM_023239679.1
D1 89066402 89066500
refGene_gn=QSER1;refGene_tx=XM_006937176.4;refGene_gn=QSER1;refGene_tx=XM_023239677.1;refGene_gn=QSER1;refGene_tx=XM_023239678.1;refGene_gn=QSER1;refGene_tx=XM_023239679.1
D1 89071053 89071158 refGene_gn=QSER1;refGene_tx=XM_023239678.1
D1 89072320 89072473
refGene_gn=QSER1;refGene_tx=XM_006937176.4;refGene_gn=QSER1;refGene_tx=XM_023239677.1;refGene_gn=QSER1;refGene_tx=XM_023239679.1
D1 89073703 89073799
refGene_gn=QSER1;refGene_tx=XM_006937176.4;refGene_gn=QSER1;refGene_tx=XM_023239677.1;refGene_gn=QSER1;refGene_tx=XM_023239679.1
D1 89074776 89074917
refGene_gn=QSER1;refGene_tx=XM_006937176.4;refGene_gn=QSER1;refGene_tx=XM_023239677.1;refGene_gn=QSER1;refGene_tx=XM_023239679.1
D1 89100459 89100532 refGene_gn=DEPDC7;refGene_tx=XM_011286816.3
D1 89109218 89109609 refGene_gn=DEPDC7;refGene_tx=XM_011286816.3
D1 89111706 89111834 refGene_gn=DEPDC7;refGene_tx=XM_011286816.3
D1 89112662 89112852 refGene_gn=DEPDC7;refGene_tx=XM_011286816.3
D1 89114846 89115058 refGene_gn=DEPDC7;refGene_tx=XM_011286816.3
D1 89115823 89115966 refGene_gn=DEPDC7;refGene_tx=XM_011286816.3
D1 89116152 89116278 refGene_gn=DEPDC7;refGene_tx=XM_011286816.3
D1 89116394 89116473 refGene_gn=DEPDC7;refGene_tx=XM_011286816.3
D1 89117037 89117231 refGene_gn=DEPDC7;refGene_tx=XM_011286816.3
D1 89125252 89125415
refGene_gn=TCP11L1;refGene_tx=XM_003993142.5;refGene_gn=TCP11L1;refGene_tx=XM_006937179.4;refGene_gn=TCP11L1;refGene_tx=XM_006937180.4;refGene_gn=TCP11L1;refGene_tx=XM_019812236.2
D1 89136323 89136456
refGene_gn=TCP11L1;refGene_tx=XM_003993142.5;refGene_gn=TCP11L1;refGene_tx=XM_006937179.4;refGene_gn=TCP11L1;refGene_tx=XM_006937180.4;refGene_gn=TCP11L1;refGene_tx=XM_006937181.4;refGene_gn=TCP11L1;refGene_tx=XM_019812236.2
D1 89138050 89138171
refGene_gn=TCP11L1;refGene_tx=XM_003993142.5;refGene_gn=TCP11L1;refGene_tx=XM_006937179.4;refGene_gn=TCP11L1;refGene_tx=XM_006937180.4;refGene_gn=TCP11L1;refGene_tx=XM_006937181.4;refGene_gn=TCP11L1;refGene_tx=XM_019812236.2
D1 89139250 89139471
refGene_gn=TCP11L1;refGene_tx=XM_003993142.5;refGene_gn=TCP11L1;refGene_tx=XM_006937179.4;refGene_gn=TCP11L1;refGene_tx=XM_006937180.4;refGene_gn=TCP11L1;refGene_tx=XM_006937181.4;refGene_gn=TCP11L1;refGene_tx=XM_019812236.2
D1 89142210 89142347
refGene_gn=TCP11L1;refGene_tx=XM_003993142.5;refGene_gn=TCP11L1;refGene_tx=XM_006937179.4;refGene_gn=TCP11L1;refGene_tx=XM_006937180.4;refGene_gn=TCP11L1;refGene_tx=XM_006937181.4;refGene_gn=TCP11L1;refGene_tx=XM_019812236.2
D1 89144526 89144723
refGene_gn=TCP11L1;refGene_tx=XM_003993142.5;refGene_gn=TCP11L1;refGen

e_tx=XM_006937179.4;refGene_gn=TCP11L1;refGene_tx=XM_006937180.4;refGene_gn=TCP11L1;refGene_tx=XM_006937181.4;refGene_gn=TCP11L1;refGene_tx=XM_019812236.2
D1 89148173 89148355
refGene_gn=TCP11L1;refGene_tx=XM_003993142.5;refGene_gn=TCP11L1;refGene_tx=XM_006937179.4;refGene_gn=TCP11L1;refGene_tx=XM_006937180.4;refGene_gn=TCP11L1;refGene_tx=XM_006937181.4;refGene_gn=TCP11L1;refGene_tx=XM_019812236.2
D1 89150086 89150259
refGene_gn=TCP11L1;refGene_tx=XM_003993142.5;refGene_gn=TCP11L1;refGene_tx=XM_006937179.4;refGene_gn=TCP11L1;refGene_tx=XM_006937180.4;refGene_gn=TCP11L1;refGene_tx=XM_006937181.4;refGene_gn=TCP11L1;refGene_tx=XM_019812236.2
D1 89153366 89153569
refGene_gn=TCP11L1;refGene_tx=XM_003993142.5;refGene_gn=TCP11L1;refGene_tx=XM_006937179.4;refGene_gn=TCP11L1;refGene_tx=XM_006937180.4;refGene_gn=TCP11L1;refGene_tx=XM_006937181.4;refGene_gn=TCP11L1;refGene_tx=XM_019812236.2
D1 89161194 89161397 refGene_gn=CSTF3;refGene_tx=XM_003993143.5
D1 89161809 89161870 refGene_gn=CSTF3;refGene_tx=XM_003993143.5
D1 89162011 89162106 refGene_gn=CSTF3;refGene_tx=XM_003993143.5
D1 89162827 89162981 refGene_gn=CSTF3;refGene_tx=XM_003993143.5
D1 89165113 89165309 refGene_gn=CSTF3;refGene_tx=XM_003993143.5
D1 89166709 89166779 refGene_gn=CSTF3;refGene_tx=XM_003993143.5
D1 89171092 89171195 refGene_gn=CSTF3;refGene_tx=XM_003993143.5
D1 89171825 89171969 refGene_gn=CSTF3;refGene_tx=XM_003993143.5
D1 89173617 89173692 refGene_gn=CSTF3;refGene_tx=XM_003993143.5
D1 89173950 89174067 refGene_gn=CSTF3;refGene_tx=XM_003993143.5
D1 89174428 89174538 refGene_gn=CSTF3;refGene_tx=XM_003993143.5
D1 89176765 89176928 refGene_gn=CSTF3;refGene_tx=XM_003993143.5
D1 89178168 89178246 refGene_gn=CSTF3;refGene_tx=XM_003993143.5
D1 89180146 89180273 refGene_gn=CSTF3;refGene_tx=XM_003993143.5
D1 89180362 89180397 refGene_gn=CSTF3;refGene_tx=XM_003993143.5
D1 89180487 89180554 refGene_gn=CSTF3;refGene_tx=XM_003993143.5
D1 89181648 89181746 refGene_gn=CSTF3;refGene_tx=XM_003993143.5
D1 89182143 89182176 refGene_gn=CSTF3;refGene_tx=XM_003993143.5
D1 89217798 89217894 refGene_gn=CSTF3;refGene_tx=XM_003993143.5
D1 89218016 89218118 refGene_gn=CSTF3;refGene_tx=XM_003993143.5
D1 89232964 89232991 refGene_gn=CSTF3;refGene_tx=XM_003993143.5
D1 89351023 89352120
refGene_gn=HIPK3;refGene_tx=XM_019812241.2;refGene_gn=HIPK3;refGene_tx=XM_023239680.1;refGene_gn=HIPK3;refGene_tx=XM_023239681.1;refGene_gn=HIPK3;refGene_tx=XM_023239682.1;refGene_gn=HIPK3;refGene_tx=XM_023239683.1;refGene_gn=HIPK3;refGene_tx=XM_023239684.1;refGene_gn=HIPK3;refGene_tx=XM_023239685.1;refGene_gn=HIPK3;refGene_tx=XM_023239686.1
D1 89389552 89389676
refGene_gn=HIPK3;refGene_tx=XM_019812241.2;refGene_gn=HIPK3;refGene_tx=XM_023239680.1;refGene_gn=HIPK3;refGene_tx=XM_023239681.1;refGene_gn=HIPK3;refGene_tx=XM_023239682.1;refGene_gn=HIPK3;refGene_tx=XM_023239683.1;refGene_gn=HIPK3;refGene_tx=XM_023239684.1;refGene_gn=HIPK3;refGe

ne_tx=XM_023239685.1;refGene_gn=HIPK3;refGene_tx=XM_023239686.1
D1 89396843 89396963
refGene_gn=HIPK3;refGene_tx=XM_019812241.2;refGene_gn=HIPK3;refGene_tx=XM_023239680.1;refGene_gn=HIPK3;refGene_tx=XM_023239681.1;refGene_gn=HIPK3;refGene_tx=XM_023239682.1;refGene_gn=HIPK3;refGene_tx=XM_023239683.1;refGene_gn=HIPK3;refGene_tx=XM_023239684.1;refGene_gn=HIPK3;refGene_tx=XM_023239685.1;refGene_gn=HIPK3;refGene_tx=XM_023239686.1
D1 89397593 89397680
refGene_gn=HIPK3;refGene_tx=XM_019812241.2;refGene_gn=HIPK3;refGene_tx=XM_023239680.1;refGene_gn=HIPK3;refGene_tx=XM_023239681.1;refGene_gn=HIPK3;refGene_tx=XM_023239682.1;refGene_gn=HIPK3;refGene_tx=XM_023239683.1;refGene_gn=HIPK3;refGene_tx=XM_023239684.1;refGene_gn=HIPK3;refGene_tx=XM_023239685.1;refGene_gn=HIPK3;refGene_tx=XM_023239686.1
D1 89398187 89398372
refGene_gn=HIPK3;refGene_tx=XM_019812241.2;refGene_gn=HIPK3;refGene_tx=XM_023239680.1;refGene_gn=HIPK3;refGene_tx=XM_023239681.1;refGene_gn=HIPK3;refGene_tx=XM_023239682.1;refGene_gn=HIPK3;refGene_tx=XM_023239683.1;refGene_gn=HIPK3;refGene_tx=XM_023239684.1;refGene_gn=HIPK3;refGene_tx=XM_023239685.1;refGene_gn=HIPK3;refGene_tx=XM_023239686.1
D1 89400067 89400227
refGene_gn=HIPK3;refGene_tx=XM_019812241.2;refGene_gn=HIPK3;refGene_tx=XM_023239680.1;refGene_gn=HIPK3;refGene_tx=XM_023239681.1;refGene_gn=HIPK3;refGene_tx=XM_023239682.1;refGene_gn=HIPK3;refGene_tx=XM_023239683.1;refGene_gn=HIPK3;refGene_tx=XM_023239684.1;refGene_gn=HIPK3;refGene_tx=XM_023239685.1;refGene_gn=HIPK3;refGene_tx=XM_023239686.1
D1 89400616 89400740
refGene_gn=HIPK3;refGene_tx=XM_019812241.2;refGene_gn=HIPK3;refGene_tx=XM_023239680.1;refGene_gn=HIPK3;refGene_tx=XM_023239681.1;refGene_gn=HIPK3;refGene_tx=XM_023239682.1;refGene_gn=HIPK3;refGene_tx=XM_023239683.1;refGene_gn=HIPK3;refGene_tx=XM_023239684.1;refGene_gn=HIPK3;refGene_tx=XM_023239685.1;refGene_gn=HIPK3;refGene_tx=XM_023239686.1
D1 89404969 89405091
refGene_gn=HIPK3;refGene_tx=XM_019812241.2;refGene_gn=HIPK3;refGene_tx=XM_023239680.1;refGene_gn=HIPK3;refGene_tx=XM_023239681.1;refGene_gn=HIPK3;refGene_tx=XM_023239682.1;refGene_gn=HIPK3;refGene_tx=XM_023239683.1;refGene_gn=HIPK3;refGene_tx=XM_023239684.1;refGene_gn=HIPK3;refGene_tx=XM_023239685.1;refGene_gn=HIPK3;refGene_tx=XM_023239686.1
D1 89405344 89405472
refGene_gn=HIPK3;refGene_tx=XM_019812241.2;refGene_gn=HIPK3;refGene_tx=XM_023239680.1;refGene_gn=HIPK3;refGene_tx=XM_023239681.1;refGene_gn=HIPK3;refGene_tx=XM_023239682.1;refGene_gn=HIPK3;refGene_tx=XM_023239683.1;refGene_gn=HIPK3;refGene_tx=XM_023239684.1;refGene_gn=HIPK3;refGene_tx=XM_023239685.1;refGene_gn=HIPK3;refGene_tx=XM_023239686.1
D1 89405563 89405725
refGene_gn=HIPK3;refGene_tx=XM_019812241.2;refGene_gn=HIPK3;refGene_tx=XM_023239680.1;refGene_gn=HIPK3;refGene_tx=XM_023239681.1;refGene_gn=HIPK3;refGene_tx=XM_023239682.1;refGene_gn=HIPK3;refGene_tx=XM_023239683.1;refGene_gn=HIPK3;refGene_tx=XM_023239684.1;refGene_gn=HIPK3;refGene_tx=XM_023239685.1;refGene_gn=HIPK3;refGene_tx=XM_023239686.1
D1 89405881 89405944

refGene_gn=HIPK3;refGene_tx=XM_019812241.2;refGene_gn=HIPK3;refGene_tx=XM_023239680.1;refGene_gn=HIPK3;refGene_tx=XM_023239681.1;refGene_gn=HIPK3;refGene_tx=XM_023239682.1;refGene_gn=HIPK3;refGene_tx=XM_023239683.1
D1 89406229 89406526
refGene_gn=HIPK3;refGene_tx=XM_019812241.2;refGene_gn=HIPK3;refGene_tx=XM_023239680.1;refGene_gn=HIPK3;refGene_tx=XM_023239681.1;refGene_gn=HIPK3;refGene_tx=XM_023239682.1;refGene_gn=HIPK3;refGene_tx=XM_023239683.1;refGene_gn=HIPK3;refGene_tx=XM_023239684.1;refGene_gn=HIPK3;refGene_tx=XM_023239685.1;refGene_gn=HIPK3;refGene_tx=XM_023239686.1
D1 89406813 89406954
refGene_gn=HIPK3;refGene_tx=XM_019812241.2;refGene_gn=HIPK3;refGene_tx=XM_023239680.1;refGene_gn=HIPK3;refGene_tx=XM_023239681.1;refGene_gn=HIPK3;refGene_tx=XM_023239682.1;refGene_gn=HIPK3;refGene_tx=XM_023239683.1;refGene_gn=HIPK3;refGene_tx=XM_023239684.1;refGene_gn=HIPK3;refGene_tx=XM_023239685.1;refGene_gn=HIPK3;refGene_tx=XM_023239686.1
D1 89408893 89409141
refGene_gn=HIPK3;refGene_tx=XM_019812241.2;refGene_gn=HIPK3;refGene_tx=XM_023239680.1;refGene_gn=HIPK3;refGene_tx=XM_023239681.1;refGene_gn=HIPK3;refGene_tx=XM_023239682.1;refGene_gn=HIPK3;refGene_tx=XM_023239683.1;refGene_gn=HIPK3;refGene_tx=XM_023239684.1;refGene_gn=HIPK3;refGene_tx=XM_023239685.1;refGene_gn=HIPK3;refGene_tx=XM_023239686.1
D1 89409688 89409816
refGene_gn=HIPK3;refGene_tx=XM_019812241.2;refGene_gn=HIPK3;refGene_tx=XM_023239680.1;refGene_gn=HIPK3;refGene_tx=XM_023239681.1;refGene_gn=HIPK3;refGene_tx=XM_023239682.1;refGene_gn=HIPK3;refGene_tx=XM_023239683.1;refGene_gn=HIPK3;refGene_tx=XM_023239684.1;refGene_gn=HIPK3;refGene_tx=XM_023239685.1;refGene_gn=HIPK3;refGene_tx=XM_023239686.1
D1 89410665 89411142
refGene_gn=HIPK3;refGene_tx=XM_019812241.2;refGene_gn=HIPK3;refGene_tx=XM_023239680.1;refGene_gn=HIPK3;refGene_tx=XM_023239681.1;refGene_gn=HIPK3;refGene_tx=XM_023239682.1;refGene_gn=HIPK3;refGene_tx=XM_023239683.1;refGene_gn=HIPK3;refGene_tx=XM_023239684.1;refGene_gn=HIPK3;refGene_tx=XM_023239685.1;refGene_gn=HIPK3;refGene_tx=XM_023239686.1
D1 89582616 89582832
refGene_gn=KIAA1549L;refGene_tx=XM_019812248.2
D1 89582849 89585010
refGene_gn=KIAA1549L;refGene_tx=XM_019812245.2;refGene_gn=KIAA1549L;refGene_tx=XM_019812246.2;refGene_gn=KIAA1549L;refGene_tx=XM_019812247.2;refGene_gn=KIAA1549L;refGene_tx=XM_019812248.2;refGene_gn=KIAA1549L;refGene_tx=XM_023239688.1
D1 89585935 89586538
refGene_gn=KIAA1549L;refGene_tx=XM_019812245.2;refGene_gn=KIAA1549L;refGene_tx=XM_019812246.2;refGene_gn=KIAA1549L;refGene_tx=XM_019812247.2;refGene_gn=KIAA1549L;refGene_tx=XM_019812248.2;refGene_gn=KIAA1549L;refGene_tx=XM_023239688.1
D1 89588566 89588682
refGene_gn=KIAA1549L;refGene_tx=XM_019812245.2;refGene_gn=KIAA1549L;refGene_tx=XM_019812246.2;refGene_gn=KIAA1549L;refGene_tx=XM_019812247.2;refGene_gn=KIAA1549L;refGene_tx=XM_019812248.2;refGene_gn=KIAA1549L;r

efGene_tx=XM_023239688.1
D1 89591816 89592036
refGene_gn=KIAA1549L;refGene_tx=XM_019812245.2;refGene_gn=KIAA1549L;refGene_tx=XM_019812246.2;refGene_gn=KIAA1549L;refGene_tx=XM_019812247.2;refGene_gn=KIAA1549L;refGene_tx=XM_019812248.2;refGene_gn=KIAA1549L;refGene_tx=XM_023239688.1
D1 89592859 89592993
refGene_gn=KIAA1549L;refGene_tx=XM_019812245.2;refGene_gn=KIAA1549L;refGene_tx=XM_019812246.2;refGene_gn=KIAA1549L;refGene_tx=XM_019812247.2;refGene_gn=KIAA1549L;refGene_tx=XM_019812248.2;refGene_gn=KIAA1549L;refGene_tx=XM_023239688.1
D1 89600429 89600592
refGene_gn=KIAA1549L;refGene_tx=XM_019812245.2;refGene_gn=KIAA1549L;refGene_tx=XM_019812246.2;refGene_gn=KIAA1549L;refGene_tx=XM_019812247.2;refGene_gn=KIAA1549L;refGene_tx=XM_019812248.2;refGene_gn=KIAA1549L;refGene_tx=XM_023239688.1
D1 89602136 89602196
refGene_gn=KIAA1549L;refGene_tx=XM_019812245.2;refGene_gn=KIAA1549L;refGene_tx=XM_019812246.2;refGene_gn=KIAA1549L;refGene_tx=XM_019812247.2;refGene_gn=KIAA1549L;refGene_tx=XM_019812248.2;refGene_gn=KIAA1549L;refGene_tx=XM_023239688.1
D1 89608591 89608743
refGene_gn=KIAA1549L;refGene_tx=XM_019812245.2;refGene_gn=KIAA1549L;refGene_tx=XM_019812246.2;refGene_gn=KIAA1549L;refGene_tx=XM_019812247.2;refGene_gn=KIAA1549L;refGene_tx=XM_019812248.2;refGene_gn=KIAA1549L;refGene_tx=XM_023239688.1
D1 89614037 89614209
refGene_gn=KIAA1549L;refGene_tx=XM_019812245.2;refGene_gn=KIAA1549L;refGene_tx=XM_019812246.2;refGene_gn=KIAA1549L;refGene_tx=XM_019812247.2;refGene_gn=KIAA1549L;refGene_tx=XM_019812248.2;refGene_gn=KIAA1549L;refGene_tx=XM_023239688.1
D1 89624060 89624224
refGene_gn=KIAA1549L;refGene_tx=XM_019812245.2;refGene_gn=KIAA1549L;refGene_tx=XM_019812246.2;refGene_gn=KIAA1549L;refGene_tx=XM_019812247.2;refGene_gn=KIAA1549L;refGene_tx=XM_019812248.2;refGene_gn=KIAA1549L;refGene_tx=XM_023239688.1
D1 89631081 89631272
refGene_gn=KIAA1549L;refGene_tx=XM_019812245.2;refGene_gn=KIAA1549L;refGene_tx=XM_019812246.2;refGene_gn=KIAA1549L;refGene_tx=XM_019812247.2;refGene_gn=KIAA1549L;refGene_tx=XM_019812248.2;refGene_gn=KIAA1549L;refGene_tx=XM_023239688.1
D1 89638760 89638888
refGene_gn=KIAA1549L;refGene_tx=XM_019812245.2;refGene_gn=KIAA1549L;refGene_tx=XM_019812246.2;refGene_gn=KIAA1549L;refGene_tx=XM_019812247.2;refGene_gn=KIAA1549L;refGene_tx=XM_019812248.2;refGene_gn=KIAA1549L;refGene_tx=XM_023239688.1
D1 89644110 89644289
refGene_gn=KIAA1549L;refGene_tx=XM_019812245.2;refGene_gn=KIAA1549L;refGene_tx=XM_019812246.2;refGene_gn=KIAA1549L;refGene_tx=XM_019812247.2;refGene_gn=KIAA1549L;refGene_tx=XM_019812248.2;refGene_gn=KIAA1549L;r

efGene_tx=XM_023239688.1
D1 89650319 89650537
refGene_gn=KIAA1549L;refGene_tx=XM_019812245.2;refGene_gn=KIAA1549L;refGene_tx=XM_019812246.2;refGene_gn=KIAA1549L;refGene_tx=XM_019812247.2;refGene_gn=KIAA1549L;refGene_tx=XM_019812248.2;refGene_gn=KIAA1549L;refGene_tx=XM_023239688.1
D1 89657132 89657262
refGene_gn=KIAA1549L;refGene_tx=XM_019812245.2;refGene_gn=KIAA1549L;refGene_tx=XM_019812246.2;refGene_gn=KIAA1549L;refGene_tx=XM_019812247.2;refGene_gn=KIAA1549L;refGene_tx=XM_019812248.2;refGene_gn=KIAA1549L;refGene_tx=XM_023239688.1
D1 89677284 89677635
refGene_gn=KIAA1549L;refGene_tx=XM_019812245.2;refGene_gn=KIAA1549L;refGene_tx=XM_019812246.2;refGene_gn=KIAA1549L;refGene_tx=XM_019812247.2;refGene_gn=KIAA1549L;refGene_tx=XM_019812248.2;refGene_gn=KIAA1549L;refGene_tx=XM_023239688.1
D1 89688702 89688800
refGene_gn=KIAA1549L;refGene_tx=XM_019812245.2;refGene_gn=KIAA1549L;refGene_tx=XM_019812246.2;refGene_gn=KIAA1549L;refGene_tx=XM_019812248.2;refGene_gn=KIAA1549L;refGene_tx=XM_023239688.1
D1 89691224 89691373
refGene_gn=KIAA1549L;refGene_tx=XM_019812245.2;refGene_gn=KIAA1549L;refGene_tx=XM_019812246.2;refGene_gn=KIAA1549L;refGene_tx=XM_019812248.2;refGene_gn=KIAA1549L;refGene_tx=XM_023239688.1
D1 89692773 89692925
refGene_gn=KIAA1549L;refGene_tx=XM_019812245.2;refGene_gn=KIAA1549L;refGene_tx=XM_019812246.2;refGene_gn=KIAA1549L;refGene_tx=XM_019812248.2;refGene_gn=KIAA1549L;refGene_tx=XM_023239688.1
D1 89698299 89698581
refGene_gn=KIAA1549L;refGene_tx=XM_019812245.2;refGene_gn=KIAA1549L;refGene_tx=XM_019812246.2;refGene_gn=KIAA1549L;refGene_tx=XM_019812247.2;refGene_gn=KIAA1549L;refGene_tx=XM_019812248.2;refGene_gn=KIAA1549L;refGene_tx=XM_023239688.1
D1 89716447 89716533
refGene_gn=CD1H11orf91;refGene_tx=XM_003993149.4
D1 89717772 89718238
refGene_gn=CD1H11orf91;refGene_tx=XM_003993149.4
D1 89731224 89731427
refGene_gn=CD59;refGene_tx=XM_003993150.5;refGene_gn=CD59;refGene_tx=XM_006937186.2
D1 89738087 89738192
refGene_gn=CD59;refGene_tx=XM_003993150.5;refGene_gn=CD59;refGene_tx=XM_006937186.2
D1 89739960 89740033
refGene_gn=CD59;refGene_tx=XM_003993150.5;refGene_gn=CD59;refGene_tx=XM_006937186.2
D1 89755442 89755616 refGene_gn=FBXO3;refGene_tx=XM_003993151.4
D1 89760846 89761037 refGene_gn=FBXO3;refGene_tx=XM_003993151.4
D1 89761784 89761900 refGene_gn=FBXO3;refGene_tx=XM_003993151.4
D1 89763436 89763559 refGene_gn=FBXO3;refGene_tx=XM_003993151.4

| | | | |
|---|---|---|---|
| D1 | 89764398 | 89764483 | refGene_gn=FBXO3;refGene_tx=XM_003993151.4 |
| D1 | 89767295 | 89767341 | refGene_gn=FBXO3;refGene_tx=XM_003993151.4 |
| D1 | 89768455 | 89768660 | refGene_gn=FBXO3;refGene_tx=XM_003993151.4 |
| D1 | 89770410 | 89770525 | refGene_gn=FBXO3;refGene_tx=XM_003993151.4 |
| D1 | 89780203 | 89780367 | refGene_gn=FBXO3;refGene_tx=XM_003993151.4 |
| D1 | 89782231 | 89782321 | refGene_gn=FBXO3;refGene_tx=XM_003993151.4 |
| D1 | 89787587 | 89787691 | refGene_gn=FBXO3;refGene_tx=XM_003993151.4 |
| D1 | 89852741 | 89852961 | refGene_gn=LMO2;refGene_tx=XM_023239690.1 |
| D1 | 89857768 | 89857984 | refGene_gn=LMO2;refGene_tx=XM_023239690.1 |
| D1 | 89861673 | 89861714 | refGene_gn=LMO2;refGene_tx=XM_023239690.1 |
| D1 | 90006168 | 90006384 | refGene_gn=CAPRIN1;refGene_tx=XM_023239692.1 |
| D1 | 90020509 | 90020572 | refGene_gn=CAPRIN1;refGene_tx=XM_023239692.1 |
| D1 | 90020685 | 90020772 | refGene_gn=CAPRIN1;refGene_tx=XM_023239692.1 |
| D1 | 90024444 | 90024683 | refGene_gn=CAPRIN1;refGene_tx=XM_023239692.1 |
| D1 | 90024770 | 90024853 | refGene_gn=CAPRIN1;refGene_tx=XM_023239692.1 |
| D1 | 90028516 | 90028654 | refGene_gn=CAPRIN1;refGene_tx=XM_023239692.1 |
| D1 | 90029812 | 90029865 | refGene_gn=CAPRIN1;refGene_tx=XM_023239692.1 |
| D1 | 90029940 | 90030027 | refGene_gn=CAPRIN1;refGene_tx=XM_023239692.1 |
| D1 | 90032049 | 90032205 | refGene_gn=CAPRIN1;refGene_tx=XM_023239692.1 |
| D1 | 90032287 | 90032396 | refGene_gn=CAPRIN1;refGene_tx=XM_023239692.1 |
| D1 | 90033503 | 90033565 | refGene_gn=CAPRIN1;refGene_tx=XM_023239692.1 |
| D1 | 90034057 | 90034168 | refGene_gn=CAPRIN1;refGene_tx=XM_023239692.1 |
| D1 | 90034624 | 90034774 | refGene_gn=CAPRIN1;refGene_tx=XM_023239692.1 |
| D1 | 90035686 | 90035837 | refGene_gn=CAPRIN1;refGene_tx=XM_023239692.1 |
| D1 | 90038295 | 90038490 | refGene_gn=CAPRIN1;refGene_tx=XM_023239692.1 |
| D1 | 90038980 | 90039081 | refGene_gn=CAPRIN1;refGene_tx=XM_023239692.1 |
| D1 | 90040128 | 90040192 | refGene_gn=CAPRIN1;refGene_tx=XM_023239692.1 |
| D1 | 90041723 | 90041788 | refGene_gn=CAPRIN1;refGene_tx=XM_023239692.1 |
| D1 | 90049825 | 90049933 | refGene_gn=NAT10;refGene_tx=XM_006937187.4 |
| D1 | 90050245 | 90050337 | refGene_gn=NAT10;refGene_tx=XM_006937187.4 |
| D1 | 90051153 | 90051325 | refGene_gn=NAT10;refGene_tx=XM_006937187.4 |
| D1 | 90052809 | 90052932 | refGene_gn=NAT10;refGene_tx=XM_006937187.4 |
| D1 | 90054786 | 90054848 | refGene_gn=NAT10;refGene_tx=XM_006937187.4 |
| D1 | 90056254 | 90056369 | refGene_gn=NAT10;refGene_tx=XM_006937187.4 |
| D1 | 90056466 | 90056574 | refGene_gn=NAT10;refGene_tx=XM_006937187.4 |
| D1 | 90059679 | 90059813 | refGene_gn=NAT10;refGene_tx=XM_006937187.4 |
| D1 | 90061044 | 90061138 | refGene_gn=NAT10;refGene_tx=XM_006937187.4 |
| D1 | 90061280 | 90061379 | refGene_gn=NAT10;refGene_tx=XM_006937187.4 |
| D1 | 90064194 | 90064331 | refGene_gn=NAT10;refGene_tx=XM_006937187.4 |
| D1 | 90067078 | 90067203 | refGene_gn=NAT10;refGene_tx=XM_006937187.4 |
| D1 | 90067667 | 90067818 | refGene_gn=NAT10;refGene_tx=XM_006937187.4 |
| D1 | 90068092 | 90068189 | refGene_gn=NAT10;refGene_tx=XM_006937187.4 |
| D1 | 90069270 | 90069387 | refGene_gn=NAT10;refGene_tx=XM_006937187.4 |
| D1 | 90070233 | 90070335 | refGene_gn=NAT10;refGene_tx=XM_006937187.4 |
| D1 | 90070430 | 90070505 | refGene_gn=NAT10;refGene_tx=XM_006937187.4 |
| D1 | 90071313 | 90071430 | refGene_gn=NAT10;refGene_tx=XM_006937187.4 |
| D1 | 90072229 | 90072363 | refGene_gn=NAT10;refGene_tx=XM_006937187.4 |
| D1 | 90072575 | 90072624 | refGene_gn=NAT10;refGene_tx=XM_006937187.4 |
| D1 | 90075678 | 90075775 | refGene_gn=NAT10;refGene_tx=XM_006937187.4 |
| D1 | 90075853 | 90075964 | refGene_gn=NAT10;refGene_tx=XM_006937187.4 |

D1	90076856	90077029	refGene_gn=NAT10;refGene_tx=XM_006937187.4
D1	90077529	90077649	refGene_gn=NAT10;refGene_tx=XM_006937187.4
D1	90078013	90078112	refGene_gn=NAT10;refGene_tx=XM_006937187.4
D1	90078479	90078553	refGene_gn=NAT10;refGene_tx=XM_006937187.4
D1	90079722	90079806	refGene_gn=NAT10;refGene_tx=XM_006937187.4
D1	90082002	90082114	refGene_gn=NAT10;refGene_tx=XM_006937187.4
D1	90088316	90088514
refGene_gn=ABTB2;refGene_tx=XM_003993156.4;refGene_gn=ABTB2;refGene_tx=XM_019812251.2
D1	90089866	90089980
refGene_gn=ABTB2;refGene_tx=XM_003993156.4;refGene_gn=ABTB2;refGene_tx=XM_019812251.2
D1	90090613	90090682
refGene_gn=ABTB2;refGene_tx=XM_003993156.4;refGene_gn=ABTB2;refGene_tx=XM_019812251.2
D1	90094905	90094996
refGene_gn=ABTB2;refGene_tx=XM_003993156.4;refGene_gn=ABTB2;refGene_tx=XM_019812251.2
D1	90095487	90095590
refGene_gn=ABTB2;refGene_tx=XM_003993156.4;refGene_gn=ABTB2;refGene_tx=XM_019812251.2
D1	90095812	90095918
refGene_gn=ABTB2;refGene_tx=XM_003993156.4;refGene_gn=ABTB2;refGene_tx=XM_019812251.2
D1	90096297	90096476
refGene_gn=ABTB2;refGene_tx=XM_003993156.4;refGene_gn=ABTB2;refGene_tx=XM_019812251.2
D1	90097859	90098089
refGene_gn=ABTB2;refGene_tx=XM_003993156.4;refGene_gn=ABTB2;refGene_tx=XM_019812251.2
D1	90099808	90099944
refGene_gn=ABTB2;refGene_tx=XM_003993156.4;refGene_gn=ABTB2;refGene_tx=XM_019812251.2
D1	90100150	90100247
refGene_gn=ABTB2;refGene_tx=XM_003993156.4;refGene_gn=ABTB2;refGene_tx=XM_019812251.2
D1	90102619	90102721
refGene_gn=ABTB2;refGene_tx=XM_003993156.4;refGene_gn=ABTB2;refGene_tx=XM_019812251.2
D1	90103474	90103564
refGene_gn=ABTB2;refGene_tx=XM_003993156.4;refGene_gn=ABTB2;refGene_tx=XM_019812251.2
D1	90106393	90106559
refGene_gn=ABTB2;refGene_tx=XM_003993156.4;refGene_gn=ABTB2;refGene_tx=XM_019812251.2
D1	90107861	90108014
refGene_gn=ABTB2;refGene_tx=XM_003993156.4;refGene_gn=ABTB2;refGene_tx=XM_019812251.2
D1	90127214	90127428
refGene_gn=ABTB2;refGene_tx=XM_003993156.4;refGene_gn=ABTB2;refGene_tx

=XM_019812251.2
D1 90133586 90133733
refGene_gn=ABTB2;refGene_tx=XM_003993156.4;refGene_gn=ABTB2;refGene_tx
=XM_019812251.2
D1 90137475 90137494 refGene_gn=ABTB2;refGene_tx=XM_019812251.2
D1 90260329 90261212 refGene_gn=ABTB2;refGene_tx=XM_003993156.4
D1 90327005 90327071 refGene_gn=CAT;refGene_tx=XM_003993157.4
D1 90340695 90340867 refGene_gn=CAT;refGene_tx=XM_003993157.4
D1 90341936 90342047 refGene_gn=CAT;refGene_tx=XM_003993157.4
D1 90343243 90343374 refGene_gn=CAT;refGene_tx=XM_003993157.4
D1 90343870 90343975 refGene_gn=CAT;refGene_tx=XM_003993157.4
D1 90344571 90344697 refGene_gn=CAT;refGene_tx=XM_003993157.4
D1 90347696 90347888 refGene_gn=CAT;refGene_tx=XM_003993157.4
D1 90348319 90348472 refGene_gn=CAT;refGene_tx=XM_003993157.4
D1 90351729 90351868 refGene_gn=CAT;refGene_tx=XM_003993157.4
D1 90354718 90354849 refGene_gn=CAT;refGene_tx=XM_003993157.4
D1 90360574 90360682 refGene_gn=CAT;refGene_tx=XM_003993157.4
D1 90362668 90362752 refGene_gn=CAT;refGene_tx=XM_003993157.4
D1 90363068 90363134 refGene_gn=CAT;refGene_tx=XM_003993157.4
D1 90372188 90372285
refGene_gn=ELF5;refGene_tx=XM_006937190.3;refGene_gn=ELF5;refGene_tx=X
M_019812253.2;refGene_gn=ELF5;refGene_tx=XM_019812254.2
D1 90372740 90372936
refGene_gn=ELF5;refGene_tx=XM_006937190.3;refGene_gn=ELF5;refGene_tx=X
M_019812253.2;refGene_gn=ELF5;refGene_tx=XM_019812254.2
D1 90374130 90374199
refGene_gn=ELF5;refGene_tx=XM_006937190.3;refGene_gn=ELF5;refGene_tx=X
M_019812253.2;refGene_gn=ELF5;refGene_tx=XM_019812254.2
D1 90382032 90382083
refGene_gn=ELF5;refGene_tx=XM_006937190.3;refGene_gn=ELF5;refGene_tx=X
M_019812253.2;refGene_gn=ELF5;refGene_tx=XM_019812254.2
D1 90385286 90385520
refGene_gn=ELF5;refGene_tx=XM_006937190.3;refGene_gn=ELF5;refGene_tx=X
M_019812253.2;refGene_gn=ELF5;refGene_tx=XM_019812254.2
D1 90396170 90396295
refGene_gn=ELF5;refGene_tx=XM_006937190.3;refGene_gn=ELF5;refGene_tx=X
M_019812253.2;refGene_gn=ELF5;refGene_tx=XM_019812254.2
D1 90414765 90414821 refGene_gn=ELF5;refGene_tx=XM_019812253.2
D1 90536581 90536678
refGene_gn=EHF;refGene_tx=XM_019812258.2;refGene_gn=EHF;refGene_tx=XM_
023239693.1
D1 90540390 90540636
refGene_gn=EHF;refGene_tx=XM_019812258.2;refGene_gn=EHF;refGene_tx=XM_
023239693.1
D1 90542826 90542889
refGene_gn=EHF;refGene_tx=XM_019812258.2;refGene_gn=EHF;refGene_tx=XM_
023239693.1
D1 90545772 90545841
refGene_gn=EHF;refGene_tx=XM_019812258.2;refGene_gn=EHF;refGene_tx=XM_
023239693.1

D1 90545958 90546027
refGene_gn=EHF;refGene_tx=XM_019812258.2;refGene_gn=EHF;refGene_tx=XM_023239693.1
D1 90551089 90551152
refGene_gn=EHF;refGene_tx=XM_019812258.2;refGene_gn=EHF;refGene_tx=XM_023239693.1
D1 90552391 90552587
refGene_gn=EHF;refGene_tx=XM_019812258.2;refGene_gn=EHF;refGene_tx=XM_023239693.1
D1 90552676 90552776
refGene_gn=EHF;refGene_tx=XM_019812258.2;refGene_gn=EHF;refGene_tx=XM_023239693.1
D1 90708648 90708748 refGene_gn=APIP;refGene_tx=XM_003993163.5
D1 90709593 90709761 refGene_gn=APIP;refGene_tx=XM_003993163.5
D1 90713071 90713207 refGene_gn=APIP;refGene_tx=XM_003993163.5
D1 90714694 90714812 refGene_gn=APIP;refGene_tx=XM_003993163.5
D1 90715877 90715926 refGene_gn=APIP;refGene_tx=XM_003993163.5
D1 90721790 90721891 refGene_gn=APIP;refGene_tx=XM_003993163.5
D1 90732173 90732230 refGene_gn=APIP;refGene_tx=XM_003993163.5
D1 90732608 90732768
refGene_gn=PDHX;refGene_tx=XM_003993164.5;refGene_gn=PDHX;refGene_tx=XM_006937192.4
D1 90749149 90749230
refGene_gn=PDHX;refGene_tx=XM_003993164.5;refGene_gn=PDHX;refGene_tx=XM_006937192.4
D1 90761485 90761586
refGene_gn=PDHX;refGene_tx=XM_003993164.5;refGene_gn=PDHX;refGene_tx=XM_006937192.4
D1 90768548 90768748
refGene_gn=PDHX;refGene_tx=XM_003993164.5;refGene_gn=PDHX;refGene_tx=XM_006937192.4
D1 90772147 90772246
refGene_gn=PDHX;refGene_tx=XM_003993164.5;refGene_gn=PDHX;refGene_tx=XM_006937192.4
D1 90780407 90780582
refGene_gn=PDHX;refGene_tx=XM_003993164.5;refGene_gn=PDHX;refGene_tx=XM_006937192.4
D1 90782644 90782792
refGene_gn=PDHX;refGene_tx=XM_003993164.5;refGene_gn=PDHX;refGene_tx=XM_006937192.4
D1 90790375 90790434
refGene_gn=PDHX;refGene_tx=XM_003993164.5;refGene_gn=PDHX;refGene_tx=XM_006937192.4
D1 90793798 90793957
refGene_gn=PDHX;refGene_tx=XM_003993164.5;refGene_gn=PDHX;refGene_tx=XM_006937192.4
D1 90798302 90798367
refGene_gn=PDHX;refGene_tx=XM_003993164.5;refGene_gn=PDHX;refGene_tx=XM_006937192.4
D1 90801143 90801402

refGene_gn=PDHX;refGene_tx=XM_003993164.5;refGene_gn=PDHX;refGene_tx=XM_006937192.4
D1 90802565 90802710 refGene_gn=PDHX;refGene_tx=XM_006937192.4
D1 90920239 90920306
refGene_gn=CD44;refGene_tx=XM_011286808.3;refGene_gn=CD44;refGene_tx=XM_011286809.3;refGene_gn=CD44;refGene_tx=XM_011286810.3;refGene_gn=CD44;refGene_tx=XM_019812260.2;refGene_gn=CD44;refGene_tx=XM_019812261.2;refGene_gn=CD44;refGene_tx=XM_019812263.2;refGene_gn=CD44;refGene_tx=XM_019812264.2;refGene_gn=CD44;refGene_tx=XM_019812265.2;refGene_gn=CD44;refGene_tx=XM_019812266.2;refGene_gn=CD44;refGene_tx=XM_019812267.2;refGene_gn=CD44;refGene_tx=XM_019812268.2;refGene_gn=CD44;refGene_tx=XM_019812269.2;refGene_gn=CD44;refGene_tx=XM_019812270.2;refGene_gn=CD44;refGene_tx=XM_019812271.2;refGene_gn=CD44;refGene_tx=XM_019812273.2;refGene_gn=CD44;refGene_tx=XM_019812274.2
D1 90956358 90956524
refGene_gn=CD44;refGene_tx=XM_011286808.3;refGene_gn=CD44;refGene_tx=XM_011286809.3;refGene_gn=CD44;refGene_tx=XM_011286810.3;refGene_gn=CD44;refGene_tx=XM_019812260.2;refGene_gn=CD44;refGene_tx=XM_019812261.2;refGene_gn=CD44;refGene_tx=XM_019812263.2;refGene_gn=CD44;refGene_tx=XM_019812264.2;refGene_gn=CD44;refGene_tx=XM_019812265.2;refGene_gn=CD44;refGene_tx=XM_019812266.2;refGene_gn=CD44;refGene_tx=XM_019812267.2;refGene_gn=CD44;refGene_tx=XM_019812268.2;refGene_gn=CD44;refGene_tx=XM_019812269.2;refGene_gn=CD44;refGene_tx=XM_019812270.2;refGene_gn=CD44;refGene_tx=XM_019812271.2;refGene_gn=CD44;refGene_tx=XM_019812273.2;refGene_gn=CD44;refGene_tx=XM_019812274.2
D1 90960741 90960875
refGene_gn=CD44;refGene_tx=XM_011286808.3;refGene_gn=CD44;refGene_tx=XM_011286809.3;refGene_gn=CD44;refGene_tx=XM_011286810.3;refGene_gn=CD44;refGene_tx=XM_019812260.2;refGene_gn=CD44;refGene_tx=XM_019812261.2;refGene_gn=CD44;refGene_tx=XM_019812263.2;refGene_gn=CD44;refGene_tx=XM_019812264.2;refGene_gn=CD44;refGene_tx=XM_019812265.2;refGene_gn=CD44;refGene_tx=XM_019812266.2;refGene_gn=CD44;refGene_tx=XM_019812267.2;refGene_gn=CD44;refGene_tx=XM_019812268.2;refGene_gn=CD44;refGene_tx=XM_019812269.2;refGene_gn=CD44;refGene_tx=XM_019812270.2;refGene_gn=CD44;refGene_tx=XM_019812271.2;refGene_gn=CD44;refGene_tx=XM_019812273.2;refGene_gn=CD44;refGene_tx=XM_019812274.2
D1 90967893 90967962
refGene_gn=CD44;refGene_tx=XM_011286808.3;refGene_gn=CD44;refGene_tx=XM_011286809.3;refGene_gn=CD44;refGene_tx=XM_011286810.3;refGene_gn=CD44;refGene_tx=XM_019812260.2;refGene_gn=CD44;refGene_tx=XM_019812261.2;refGene_gn=CD44;refGene_tx=XM_019812263.2;refGene_gn=CD44;refGene_tx=XM_019812264.2;refGene_gn=CD44;refGene_tx=XM_019812265.2;refGene_gn=CD44;refGene_tx=XM_019812266.2;refGene_gn=CD44;refGene_tx=XM_019812267.2;refGene_gn=CD44;refGene_tx=XM_019812268.2;refGene_gn=CD44;refGene_tx=XM_019812269.2;refGene_gn=CD44;refGene_tx=XM_019812270.2;refGene_gn=CD44;refGene_tx=XM_019812271.2;refGene_gn=CD44;refGene_tx=XM_019812273.2;refGene_gn=CD44;refGene_tx=XM_019812274.2
D1 90970375 90970609
refGene_gn=CD44;refGene_tx=XM_011286808.3;refGene_gn=CD44;refGene_tx=XM_011286809.3;refGene_gn=CD44;refGene_tx=XM_011286810.3;refGene_gn=CD4

4;refGene_tx=XM_019812260.2;refGene_gn=CD44;refGene_tx=XM_019812261.2;
refGene_gn=CD44;refGene_tx=XM_019812263.2;refGene_gn=CD44;refGene_tx=X
M_019812264.2;refGene_gn=CD44;refGene_tx=XM_019812265.2;refGene_gn=CD4
4;refGene_tx=XM_019812266.2;refGene_gn=CD44;refGene_tx=XM_019812267.2;
refGene_gn=CD44;refGene_tx=XM_019812268.2;refGene_gn=CD44;refGene_tx=X
M_019812269.2;refGene_gn=CD44;refGene_tx=XM_019812270.2;refGene_gn=CD4
4;refGene_tx=XM_019812271.2;refGene_gn=CD44;refGene_tx=XM_019812273.2;
refGene_gn=CD44;refGene_tx=XM_019812274.2
D1 90975342 90975462
refGene_gn=CD44;refGene_tx=XM_019812260.2;refGene_gn=CD44;refGene_tx=X
M_019812261.2;refGene_gn=CD44;refGene_tx=XM_019812263.2;refGene_gn=CD4
4;refGene_tx=XM_019812264.2;refGene_gn=CD44;refGene_tx=XM_019812266.2;
refGene_gn=CD44;refGene_tx=XM_019812267.2;refGene_gn=CD44;refGene_tx=X
M_019812268.2;refGene_gn=CD44;refGene_tx=XM_019812269.2;refGene_gn=CD4
4;refGene_tx=XM_019812271.2;refGene_gn=CD44;refGene_tx=XM_019812273.2
D1 90977580 90977709
refGene_gn=CD44;refGene_tx=XM_019812260.2;refGene_gn=CD44;refGene_tx=X
M_019812261.2;refGene_gn=CD44;refGene_tx=XM_019812263.2;refGene_gn=CD4
4;refGene_tx=XM_019812264.2;refGene_gn=CD44;refGene_tx=XM_019812265.2;
refGene_gn=CD44;refGene_tx=XM_019812267.2;refGene_gn=CD44;refGene_tx=X
M_019812268.2;refGene_gn=CD44;refGene_tx=XM_019812269.2;refGene_gn=CD4
4;refGene_tx=XM_019812271.2
D1 90979022 90979148
refGene_gn=CD44;refGene_tx=XM_011286808.3;refGene_gn=CD44;refGene_tx=X
M_019812260.2;refGene_gn=CD44;refGene_tx=XM_019812261.2;refGene_gn=CD4
4;refGene_tx=XM_019812263.2;refGene_gn=CD44;refGene_tx=XM_019812264.2;
refGene_gn=CD44;refGene_tx=XM_019812265.2;refGene_gn=CD44;refGene_tx=X
M_019812266.2;refGene_gn=CD44;refGene_tx=XM_019812267.2;refGene_gn=CD4
4;refGene_tx=XM_019812268.2;refGene_gn=CD44;refGene_tx=XM_019812269.2;
refGene_gn=CD44;refGene_tx=XM_019812271.2
D1 90981508 90981622
refGene_gn=CD44;refGene_tx=XM_011286808.3;refGene_gn=CD44;refGene_tx=X
M_019812260.2;refGene_gn=CD44;refGene_tx=XM_019812261.2;refGene_gn=CD4
4;refGene_tx=XM_019812263.2;refGene_gn=CD44;refGene_tx=XM_019812264.2;
refGene_gn=CD44;refGene_tx=XM_019812265.2;refGene_gn=CD44;refGene_tx=X
M_019812266.2;refGene_gn=CD44;refGene_tx=XM_019812267.2;refGene_gn=CD4
4;refGene_tx=XM_019812269.2;refGene_gn=CD44;refGene_tx=XM_019812270.2;
refGene_gn=CD44;refGene_tx=XM_019812271.2
D1 90982080 90982197
refGene_gn=CD44;refGene_tx=XM_011286808.3;refGene_gn=CD44;refGene_tx=X
M_019812260.2;refGene_gn=CD44;refGene_tx=XM_019812261.2;refGene_gn=CD4
4;refGene_tx=XM_019812263.2;refGene_gn=CD44;refGene_tx=XM_019812265.2;
refGene_gn=CD44;refGene_tx=XM_019812266.2;refGene_gn=CD44;refGene_tx=X
M_019812267.2;refGene_gn=CD44;refGene_tx=XM_019812269.2;refGene_gn=CD4
4;refGene_tx=XM_019812270.2;refGene_gn=CD44;refGene_tx=XM_019812271.2
D1 90986558 90986687
refGene_gn=CD44;refGene_tx=XM_011286808.3;refGene_gn=CD44;refGene_tx=X
M_011286809.3;refGene_gn=CD44;refGene_tx=XM_019812260.2;refGene_gn=CD4
4;refGene_tx=XM_019812261.2;refGene_gn=CD44;refGene_tx=XM_019812263.2;
refGene_gn=CD44;refGene_tx=XM_019812264.2;refGene_gn=CD44;refGene_tx=X

M_019812265.2;refGene_gn=CD44;refGene_tx=XM_019812266.2;refGene_gn=CD4
4;refGene_tx=XM_019812267.2;refGene_gn=CD44;refGene_tx=XM_019812268.2;
refGene_gn=CD44;refGene_tx=XM_019812269.2;refGene_gn=CD44;refGene_tx=X
M_019812270.2;refGene_gn=CD44;refGene_tx=XM_019812271.2
D1 90989644 90989746
refGene_gn=CD44;refGene_tx=XM_011286808.3;refGene_gn=CD44;refGene_tx=X
M_011286809.3;refGene_gn=CD44;refGene_tx=XM_019812260.2;refGene_gn=CD4
4;refGene_tx=XM_019812261.2;refGene_gn=CD44;refGene_tx=XM_019812263.2;
refGene_gn=CD44;refGene_tx=XM_019812264.2;refGene_gn=CD44;refGene_tx=X
M_019812265.2;refGene_gn=CD44;refGene_tx=XM_019812266.2;refGene_gn=CD4
4;refGene_tx=XM_019812267.2;refGene_gn=CD44;refGene_tx=XM_019812268.2;
refGene_gn=CD44;refGene_tx=XM_019812269.2;refGene_gn=CD44;refGene_tx=X
M_019812270.2
D1 90991507 90991606
refGene_gn=CD44;refGene_tx=XM_011286808.3;refGene_gn=CD44;refGene_tx=X
M_011286809.3;refGene_gn=CD44;refGene_tx=XM_019812260.2;refGene_gn=CD4
4;refGene_tx=XM_019812261.2;refGene_gn=CD44;refGene_tx=XM_019812264.2;
refGene_gn=CD44;refGene_tx=XM_019812265.2;refGene_gn=CD44;refGene_tx=X
M_019812266.2;refGene_gn=CD44;refGene_tx=XM_019812267.2;refGene_gn=CD4
4;refGene_tx=XM_019812268.2;refGene_gn=CD44;refGene_tx=XM_019812270.2
D1 90992484 90992691
refGene_gn=CD44;refGene_tx=XM_011286808.3;refGene_gn=CD44;refGene_tx=X
M_011286809.3;refGene_gn=CD44;refGene_tx=XM_019812260.2;refGene_gn=CD4
4;refGene_tx=XM_019812261.2;refGene_gn=CD44;refGene_tx=XM_019812263.2;
refGene_gn=CD44;refGene_tx=XM_019812264.2;refGene_gn=CD44;refGene_tx=X
M_019812265.2;refGene_gn=CD44;refGene_tx=XM_019812266.2;refGene_gn=CD4
4;refGene_tx=XM_019812268.2;refGene_gn=CD44;refGene_tx=XM_019812270.2
D1 90995458 90995524
refGene_gn=CD44;refGene_tx=XM_011286808.3;refGene_gn=CD44;refGene_tx=X
M_011286809.3;refGene_gn=CD44;refGene_tx=XM_011286810.3;refGene_gn=CD4
4;refGene_tx=XM_019812260.2;refGene_gn=CD44;refGene_tx=XM_019812261.2;
refGene_gn=CD44;refGene_tx=XM_019812263.2;refGene_gn=CD44;refGene_tx=X
M_019812264.2;refGene_gn=CD44;refGene_tx=XM_019812265.2;refGene_gn=CD4
4;refGene_tx=XM_019812266.2;refGene_gn=CD44;refGene_tx=XM_019812267.2;
refGene_gn=CD44;refGene_tx=XM_019812268.2;refGene_gn=CD44;refGene_tx=X
M_019812269.2;refGene_gn=CD44;refGene_tx=XM_019812270.2;refGene_gn=CD4
4;refGene_tx=XM_019812271.2;refGene_gn=CD44;refGene_tx=XM_019812273.2
D1 90999467 90999539
refGene_gn=CD44;refGene_tx=XM_011286808.3;refGene_gn=CD44;refGene_tx=X
M_011286809.3;refGene_gn=CD44;refGene_tx=XM_011286810.3;refGene_gn=CD4
4;refGene_tx=XM_019812260.2;refGene_gn=CD44;refGene_tx=XM_019812261.2;
refGene_gn=CD44;refGene_tx=XM_019812263.2;refGene_gn=CD44;refGene_tx=X
M_019812264.2;refGene_gn=CD44;refGene_tx=XM_019812265.2;refGene_gn=CD4
4;refGene_tx=XM_019812266.2;refGene_gn=CD44;refGene_tx=XM_019812267.2;
refGene_gn=CD44;refGene_tx=XM_019812268.2;refGene_gn=CD44;refGene_tx=X
M_019812269.2;refGene_gn=CD44;refGene_tx=XM_019812270.2;refGene_gn=CD4
4;refGene_tx=XM_019812271.2;refGene_gn=CD44;refGene_tx=XM_019812273.2;
refGene_gn=CD44;refGene_tx=XM_019812274.2
D1 91001837 91001916
refGene_gn=CD44;refGene_tx=XM_011286808.3;refGene_gn=CD44;refGene_tx=X

M_011286809.3;refGene_gn=CD44;refGene_tx=XM_011286810.3;refGene_gn=CD4
4;refGene_tx=XM_019812260.2;refGene_gn=CD44;refGene_tx=XM_019812261.2;
refGene_gn=CD44;refGene_tx=XM_019812263.2;refGene_gn=CD44;refGene_tx=X
M_019812264.2;refGene_gn=CD44;refGene_tx=XM_019812265.2;refGene_gn=CD4
4;refGene_tx=XM_019812266.2;refGene_gn=CD44;refGene_tx=XM_019812267.2;
refGene_gn=CD44;refGene_tx=XM_019812268.2;refGene_gn=CD44;refGene_tx=X
M_019812269.2;refGene_gn=CD44;refGene_tx=XM_019812270.2;refGene_gn=CD4
4;refGene_tx=XM_019812271.2;refGene_gn=CD44;refGene_tx=XM_019812273.2;
refGene_gn=CD44;refGene_tx=XM_019812274.2
D1 91008326 91008531
refGene_gn=CD44;refGene_tx=XM_011286808.3;refGene_gn=CD44;refGene_tx=X
M_011286809.3;refGene_gn=CD44;refGene_tx=XM_011286810.3;refGene_gn=CD4
4;refGene_tx=XM_019812260.2;refGene_gn=CD44;refGene_tx=XM_019812261.2;
refGene_gn=CD44;refGene_tx=XM_019812263.2;refGene_gn=CD44;refGene_tx=X
M_019812264.2;refGene_gn=CD44;refGene_tx=XM_019812265.2;refGene_gn=CD4
4;refGene_tx=XM_019812266.2;refGene_gn=CD44;refGene_tx=XM_019812267.2;
refGene_gn=CD44;refGene_tx=XM_019812268.2;refGene_gn=CD44;refGene_tx=X
M_019812269.2;refGene_gn=CD44;refGene_tx=XM_019812270.2;refGene_gn=CD4
4;refGene_tx=XM_019812271.2;refGene_gn=CD44;refGene_tx=XM_019812273.2;
refGene_gn=CD44;refGene_tx=XM_019812274.2
D1 91039397 91039469 refGene_gn=SLC1A2;refGene_tx=NM_001009334.1
D1 91043425 91043657 refGene_gn=SLC1A2;refGene_tx=NM_001009334.1
D1 91056219 91056354 refGene_gn=SLC1A2;refGene_tx=NM_001009334.1
D1 91061146 91061341 refGene_gn=SLC1A2;refGene_tx=NM_001009334.1
D1 91066152 91066386 refGene_gn=SLC1A2;refGene_tx=NM_001009334.1
D1 91075052 91075179 refGene_gn=SLC1A2;refGene_tx=NM_001009334.1
D1 91079471 91079640 refGene_gn=SLC1A2;refGene_tx=NM_001009334.1
D1 91084761 91085012 refGene_gn=SLC1A2;refGene_tx=NM_001009334.1
D1 91087226 91087379 refGene_gn=SLC1A2;refGene_tx=NM_001009334.1
D1 91089286 91089426 refGene_gn=SLC1A2;refGene_tx=NM_001009334.1
D1 91193975 91194512 refGene_gn=PAMR1;refGene_tx=XM_006937193.4
D1 91195819 91196112 refGene_gn=PAMR1;refGene_tx=XM_006937193.4
D1 91196983 91197216 refGene_gn=PAMR1;refGene_tx=XM_006937193.4
D1 91200787 91200854 refGene_gn=PAMR1;refGene_tx=XM_006937193.4
D1 91202843 91203056 refGene_gn=PAMR1;refGene_tx=XM_006937193.4
D1 91220330 91220438 refGene_gn=PAMR1;refGene_tx=XM_006937193.4
D1 91222907 91223125 refGene_gn=PAMR1;refGene_tx=XM_006937193.4
D1 91227059 91227174 refGene_gn=PAMR1;refGene_tx=XM_006937193.4
D1 91243003 91243132 refGene_gn=PAMR1;refGene_tx=XM_006937193.4
D1 91248748 91248925 refGene_gn=PAMR1;refGene_tx=XM_006937193.4
D1 91268737 91268810 refGene_gn=PAMR1;refGene_tx=XM_006937193.4
D1 91351926 91353228 refGene_gn=FJX1;refGene_tx=XM_023239768.1
D1 91395858 91396527
refGene_gn=TRIM44;refGene_tx=XM_011286805.3;refGene_gn=TRIM44;refGene_
tx=XM_011286807.2
D1 91417058 91417136
refGene_gn=TRIM44;refGene_tx=XM_011286805.3;refGene_gn=TRIM44;refGene_
tx=XM_011286807.2
D1 91422074 91422188 refGene_gn=TRIM44;refGene_tx=XM_011286807.2
D1 91439910 91440150 refGene_gn=TRIM44;refGene_tx=XM_011286805.3

D1	91451653 91451673 refGene_gn=TRIM44;refGene_tx=XM_011286805.3
D1	91500191 91500219 refGene_gn=TRIM44;refGene_tx=XM_011286805.3
D1	91620036 91620082
refGene_gn=LDLRAD3;refGene_tx=XM_023239769.1;refGene_gn=LDLRAD3;refGen
e_tx=XM_023239770.1
D1	91691480 91691627
refGene_gn=LDLRAD3;refGene_tx=XM_011286804.3;refGene_gn=LDLRAD3;refGen
e_tx=XM_023239769.1
D1	91711919 91711965 refGene_gn=LDLRAD3;refGene_tx=XM_019812280.2
D1	91731029 91731189 refGene_gn=LDLRAD3;refGene_tx=XM_019812278.2
D1	91731375 91731501
refGene_gn=LDLRAD3;refGene_tx=XM_011286804.3;refGene_gn=LDLRAD3;refGen
e_tx=XM_019812278.2;refGene_gn=LDLRAD3;refGene_tx=XM_019812280.2;refGe
ne_gn=LDLRAD3;refGene_tx=XM_023239769.1;refGene_gn=LDLRAD3;refGene_tx=
XM_023239770.1
D1	91747284 91747419
refGene_gn=LDLRAD3;refGene_tx=XM_011286804.3;refGene_gn=LDLRAD3;refGen
e_tx=XM_019812278.2;refGene_gn=LDLRAD3;refGene_tx=XM_019812280.2;refGe
ne_gn=LDLRAD3;refGene_tx=XM_023239769.1;refGene_gn=LDLRAD3;refGene_tx=
XM_023239770.1
D1	91856837 91857183
refGene_gn=LDLRAD3;refGene_tx=XM_011286804.3;refGene_gn=LDLRAD3;refGen
e_tx=XM_019812278.2;refGene_gn=LDLRAD3;refGene_tx=XM_019812280.2;refGe
ne_gn=LDLRAD3;refGene_tx=XM_023239769.1;refGene_gn=LDLRAD3;refGene_tx=
XM_023239770.1
D1	91858857 91859095
refGene_gn=LDLRAD3;refGene_tx=XM_011286804.3;refGene_gn=LDLRAD3;refGen
e_tx=XM_019812278.2;refGene_gn=LDLRAD3;refGene_tx=XM_019812280.2;refGe
ne_gn=LDLRAD3;refGene_tx=XM_023239769.1;refGene_gn=LDLRAD3;refGene_tx=
XM_023239770.1
D1	91895605 91895746
refGene_gn=COMMD9;refGene_tx=XM_003993167.4;refGene_gn=COMMD9;refGene_
tx=XM_006937195.4
D1	91896815 91896919
refGene_gn=COMMD9;refGene_tx=XM_003993167.4;refGene_gn=COMMD9;refGene_
tx=XM_006937195.4
D1	91897806 91897841
refGene_gn=COMMD9;refGene_tx=XM_003993167.4;refGene_gn=COMMD9;refGene_
tx=XM_006937195.4
D1	91899210 91899350
refGene_gn=COMMD9;refGene_tx=XM_003993167.4;refGene_gn=COMMD9;refGene_
tx=XM_006937195.4
D1	91901074 91901200
refGene_gn=COMMD9;refGene_tx=XM_003993167.4;refGene_gn=COMMD9;refGene_
tx=XM_006937195.4
D1	91910251 91910368 refGene_gn=COMMD9;refGene_tx=XM_006937195.4
D1	91910538 91910589 refGene_gn=COMMD9;refGene_tx=XM_003993167.4
D1	92012231 92012395
refGene_gn=PRR5L;refGene_tx=XM_006937196.2;refGene_gn=PRR5L;refGene_tx
=XM_006937197.3;refGene_gn=PRR5L;refGene_tx=XM_011286799.2;refGene_gn=

PRR5L;refGene_tx=XM_011286800.2;refGene_gn=PRR5L;refGene_tx=XM_0112868
02.2;refGene_gn=PRR5L;refGene_tx=XM_019812283.2;refGene_gn=PRR5L;refGe
ne_tx=XM_019812284.1;refGene_gn=PRR5L;refGene_tx=XM_019812285.1;refGen
e_gn=PRR5L;refGene_tx=XM_023239771.1;refGene_gn=PRR5L;refGene_tx=XM_02
3239772.1
D1 92013727 92013808
refGene_gn=PRR5L;refGene_tx=XM_006937196.2;refGene_gn=PRR5L;refGene_tx
=XM_006937197.3;refGene_gn=PRR5L;refGene_tx=XM_011286799.2;refGene_gn=
PRR5L;refGene_tx=XM_011286800.2;refGene_gn=PRR5L;refGene_tx=XM_0112868
02.2;refGene_gn=PRR5L;refGene_tx=XM_019812283.2;refGene_gn=PRR5L;refGe
ne_tx=XM_019812284.1;refGene_gn=PRR5L;refGene_tx=XM_019812285.1;refGen
e_gn=PRR5L;refGene_tx=XM_023239771.1;refGene_gn=PRR5L;refGene_tx=XM_02
3239772.1
D1 92028858 92028907
refGene_gn=PRR5L;refGene_tx=XM_006937196.2;refGene_gn=PRR5L;refGene_tx
=XM_006937197.3;refGene_gn=PRR5L;refGene_tx=XM_011286799.2;refGene_gn=
PRR5L;refGene_tx=XM_011286800.2;refGene_gn=PRR5L;refGene_tx=XM_0198122
83.2;refGene_gn=PRR5L;refGene_tx=XM_019812284.1;refGene_gn=PRR5L;refGe
ne_tx=XM_019812285.1;refGene_gn=PRR5L;refGene_tx=XM_023239771.1;refGen
e_gn=PRR5L;refGene_tx=XM_023239772.1
D1 92041675 92041733
refGene_gn=PRR5L;refGene_tx=XM_006937196.2;refGene_gn=PRR5L;refGene_tx
=XM_006937197.3;refGene_gn=PRR5L;refGene_tx=XM_011286799.2;refGene_gn=
PRR5L;refGene_tx=XM_011286800.2;refGene_gn=PRR5L;refGene_tx=XM_0112868
02.2;refGene_gn=PRR5L;refGene_tx=XM_019812283.2;refGene_gn=PRR5L;refGe
ne_tx=XM_019812284.1;refGene_gn=PRR5L;refGene_tx=XM_019812285.1;refGen
e_gn=PRR5L;refGene_tx=XM_023239771.1;refGene_gn=PRR5L;refGene_tx=XM_02
3239772.1
D1 92046603 92046695
refGene_gn=PRR5L;refGene_tx=XM_006937196.2;refGene_gn=PRR5L;refGene_tx
=XM_006937197.3;refGene_gn=PRR5L;refGene_tx=XM_011286799.2;refGene_gn=
PRR5L;refGene_tx=XM_011286800.2;refGene_gn=PRR5L;refGene_tx=XM_0198122
83.2;refGene_gn=PRR5L;refGene_tx=XM_019812285.1;refGene_gn=PRR5L;refGe
ne_tx=XM_023239771.1;refGene_gn=PRR5L;refGene_tx=XM_023239772.1
D1 92049913 92050051
refGene_gn=PRR5L;refGene_tx=XM_006937196.2;refGene_gn=PRR5L;refGene_tx
=XM_006937197.3;refGene_gn=PRR5L;refGene_tx=XM_011286799.2;refGene_gn=
PRR5L;refGene_tx=XM_011286800.2;refGene_gn=PRR5L;refGene_tx=XM_0112868
02.2;refGene_gn=PRR5L;refGene_tx=XM_019812283.2;refGene_gn=PRR5L;refGe
ne_tx=XM_019812284.1;refGene_gn=PRR5L;refGene_tx=XM_019812285.1;refGen
e_gn=PRR5L;refGene_tx=XM_023239771.1;refGene_gn=PRR5L;refGene_tx=XM_02
3239772.1
D1 92054900 92055027
refGene_gn=PRR5L;refGene_tx=XM_006937196.2;refGene_gn=PRR5L;refGene_tx
=XM_006937197.3;refGene_gn=PRR5L;refGene_tx=XM_011286799.2;refGene_gn=
PRR5L;refGene_tx=XM_011286800.2;refGene_gn=PRR5L;refGene_tx=XM_0112868
02.2;refGene_gn=PRR5L;refGene_tx=XM_019812283.2;refGene_gn=PRR5L;refGe
ne_tx=XM_023239771.1;refGene_gn=PRR5L;refGene_tx=XM_023239772.1
D1 92057743 92057859 refGene_gn=PRR5L;refGene_tx=XM_023239772.1
D1 92063470 92063865

refGene_gn=PRR5L;refGene_tx=XM_006937196.2;refGene_gn=PRR5L;refGene_tx=XM_006937197.3;refGene_gn=PRR5L;refGene_tx=XM_011286799.2;refGene_gn=PRR5L;refGene_tx=XM_011286800.2;refGene_gn=PRR5L;refGene_tx=XM_011286802.2;refGene_gn=PRR5L;refGene_tx=XM_019812283.2;refGene_gn=PRR5L;refGene_tx=XM_019812284.1;refGene_gn=PRR5L;refGene_tx=XM_019812285.1;refGene_gn=PRR5L;refGene_tx=XM_023239771.1
D1 92086832 92087702 refGene_gn=TRAF6;refGene_tx=XM_011286798.2
D1 92089206 92089284 refGene_gn=TRAF6;refGene_tx=XM_011286798.2
D1 92091280 92091352 refGene_gn=TRAF6;refGene_tx=XM_011286798.2
D1 92092773 92092932 refGene_gn=TRAF6;refGene_tx=XM_011286798.2
D1 92096028 92096179 refGene_gn=TRAF6;refGene_tx=XM_011286798.2
D1 92100492 92100788 refGene_gn=TRAF6;refGene_tx=XM_011286798.2
D1 92143508 92146640 refGene_gn=RAG1;refGene_tx=XM_019812286.1
D1 92158367 92159951 refGene_gn=RAG2;refGene_tx=XM_004001424.5
D1 92177573 92177706
refGene_gn=CD1H11orf74;refGene_tx=XM_006937201.4;refGene_gn=CD1H11orf74;refGene_tx=XM_006937202.3;refGene_gn=CD1H11orf74;refGene_tx=XM_006937203.4;refGene_gn=CD1H11orf74;refGene_tx=XM_006937204.4;refGene_gn=CD1H11orf74;refGene_tx=XM_006937205.4;refGene_gn=CD1H11orf74;refGene_tx=XM_006937206.4;refGene_gn=CD1H11orf74;refGene_tx=XM_023239773.1
D1 92206145 92206219
refGene_gn=CD1H11orf74;refGene_tx=XM_006937201.4;refGene_gn=CD1H11orf74;refGene_tx=XM_006937202.3;refGene_gn=CD1H11orf74;refGene_tx=XM_006937203.4;refGene_gn=CD1H11orf74;refGene_tx=XM_006937204.4;refGene_gn=CD1H11orf74;refGene_tx=XM_006937206.4;refGene_gn=CD1H11orf74;refGene_tx=XM_023239773.1
D1 92207915 92207987
refGene_gn=CD1H11orf74;refGene_tx=XM_006937201.4;refGene_gn=CD1H11orf74;refGene_tx=XM_006937202.3;refGene_gn=CD1H11orf74;refGene_tx=XM_006937203.4;refGene_gn=CD1H11orf74;refGene_tx=XM_006937204.4;refGene_gn=CD1H11orf74;refGene_tx=XM_006937206.4;refGene_gn=CD1H11orf74;refGene_tx=XM_023239773.1
D1 92209370 92209437
refGene_gn=CD1H11orf74;refGene_tx=XM_006937201.4;refGene_gn=CD1H11orf74;refGene_tx=XM_006937202.3;refGene_gn=CD1H11orf74;refGene_tx=XM_006937203.4;refGene_gn=CD1H11orf74;refGene_tx=XM_006937204.4;refGene_gn=CD1H11orf74;refGene_tx=XM_006937206.4
D1 92220109 92220204
refGene_gn=CD1H11orf74;refGene_tx=XM_006937204.4
D1 92220526 92220666
refGene_gn=CD1H11orf74;refGene_tx=XM_006937201.4;refGene_gn=CD1H11orf74;refGene_tx=XM_006937202.3;refGene_gn=CD1H11orf74;refGene_tx=XM_006937203.4;refGene_gn=CD1H11orf74;refGene_tx=XM_006937204.4;refGene_gn=CD1H11orf74;refGene_tx=XM_006937205.4;refGene_gn=CD1H11orf74;refGene_tx=XM_023239773.1
D1 92229110 92229275
refGene_gn=CD1H11orf74;refGene_tx=XM_006937201.4;refGene_gn=CD1H11orf74;refGene_tx=XM_006937202.3;refGene_gn=CD1H11orf74;refGene_tx=XM_006937203.4;refGene_gn=CD1H11orf74;refGene_tx=XM_006937204.4;refGene_gn=CD1H11orf74;refGene_tx=XM_006937205.4;refGene_gn=CD1H11orf74;refGene_tx=X

M_006937206.4;refGene_gn=CD1H11orf74;refGene_tx=XM_023239773.1
D1 95071088 95073011
refGene_gn=LRRC4C;refGene_tx=XM_006937207.4;refGene_gn=LRRC4C;refGene_tx=XM_019812289.2;refGene_gn=LRRC4C;refGene_tx=XM_019812292.2;refGene_gn=LRRC4C;refGene_tx=XM_019812293.2;refGene_gn=LRRC4C;refGene_tx=XM_019812294.2;refGene_gn=LRRC4C;refGene_tx=XM_023239774.1;refGene_gn=LRRC4C;refGene_tx=XM_023239775.1;refGene_gn=LRRC4C;refGene_tx=XM_023239776.1
D1 97738322 97738391
refGene_gn=API5;refGene_tx=XM_003993181.4;refGene_gn=API5;refGene_tx=XM_006937211.3
D1 97745365 97745527
refGene_gn=API5;refGene_tx=XM_003993181.4;refGene_gn=API5;refGene_tx=XM_006937211.3
D1 97746664 97746758
refGene_gn=API5;refGene_tx=XM_003993181.4;refGene_gn=API5;refGene_tx=XM_006937211.3
D1 97747235 97747301
refGene_gn=API5;refGene_tx=XM_003993181.4;refGene_gn=API5;refGene_tx=XM_006937211.3
D1 97748473 97748625
refGene_gn=API5;refGene_tx=XM_003993181.4;refGene_gn=API5;refGene_tx=XM_006937211.3
D1 97750008 97750215
refGene_gn=API5;refGene_tx=XM_003993181.4;refGene_gn=API5;refGene_tx=XM_006937211.3
D1 97750913 97751018
refGene_gn=API5;refGene_tx=XM_003993181.4;refGene_gn=API5;refGene_tx=XM_006937211.3
D1 97751932 97752022
refGene_gn=API5;refGene_tx=XM_003993181.4;refGene_gn=API5;refGene_tx=XM_006937211.3
D1 97753332 97753514
refGene_gn=API5;refGene_tx=XM_003993181.4;refGene_gn=API5;refGene_tx=XM_006937211.3
D1 97754693 97754787
refGene_gn=API5;refGene_tx=XM_003993181.4;refGene_gn=API5;refGene_tx=XM_006937211.3
D1 97755742 97755799
refGene_gn=API5;refGene_tx=XM_003993181.4;refGene_gn=API5;refGene_tx=XM_006937211.3
D1 97757728 97757805
refGene_gn=API5;refGene_tx=XM_003993181.4;refGene_gn=API5;refGene_tx=XM_006937211.3
D1 97758311 97758448
refGene_gn=API5;refGene_tx=XM_003993181.4;refGene_gn=API5;refGene_tx=XM_006937211.3
D1 97762776 97762859
refGene_gn=API5;refGene_tx=XM_003993181.4;refGene_gn=API5;refGene_tx=XM_006937211.3

D1 97773702 97773861
refGene_gn=TTC17;refGene_tx=XM_003993184.5;refGene_gn=TTC17;refGene_tx=XM_006937212.4
D1 97797266 97797356
refGene_gn=TTC17;refGene_tx=XM_003993184.5;refGene_gn=TTC17;refGene_tx=XM_006937212.4
D1 97803108 97803278
refGene_gn=TTC17;refGene_tx=XM_003993184.5;refGene_gn=TTC17;refGene_tx=XM_006937212.4
D1 97803923 97804035
refGene_gn=TTC17;refGene_tx=XM_003993184.5;refGene_gn=TTC17;refGene_tx=XM_006937212.4
D1 97804527 97804659
refGene_gn=TTC17;refGene_tx=XM_003993184.5;refGene_gn=TTC17;refGene_tx=XM_006937212.4
D1 97806736 97806846
refGene_gn=TTC17;refGene_tx=XM_003993184.5;refGene_gn=TTC17;refGene_tx=XM_006937212.4
D1 97808100 97808245
refGene_gn=TTC17;refGene_tx=XM_003993184.5;refGene_gn=TTC17;refGene_tx=XM_006937212.4
D1 97808728 97808868
refGene_gn=TTC17;refGene_tx=XM_003993184.5;refGene_gn=TTC17;refGene_tx=XM_006937212.4
D1 97810274 97810435
refGene_gn=TTC17;refGene_tx=XM_003993184.5;refGene_gn=TTC17;refGene_tx=XM_006937212.4
D1 97812364 97812477
refGene_gn=TTC17;refGene_tx=XM_003993184.5;refGene_gn=TTC17;refGene_tx=XM_006937212.4
D1 97814274 97814421
refGene_gn=TTC17;refGene_tx=XM_003993184.5;refGene_gn=TTC17;refGene_tx=XM_006937212.4
D1 97816477 97816593
refGene_gn=TTC17;refGene_tx=XM_003993184.5;refGene_gn=TTC17;refGene_tx=XM_006937212.4
D1 97816749 97816915
refGene_gn=TTC17;refGene_tx=XM_003993184.5;refGene_gn=TTC17;refGene_tx=XM_006937212.4
D1 97818108 97818186
refGene_gn=TTC17;refGene_tx=XM_003993184.5;refGene_gn=TTC17;refGene_tx=XM_006937212.4
D1 97818321 97818546
refGene_gn=TTC17;refGene_tx=XM_003993184.5;refGene_gn=TTC17;refGene_tx=XM_006937212.4
D1 97828765 97828952
refGene_gn=TTC17;refGene_tx=XM_003993184.5;refGene_gn=TTC17;refGene_tx=XM_006937212.4
D1 97848545 97848716 refGene_gn=TTC17;refGene_tx=XM_006937212.4
D1 97859475 97859735

refGene_gn=TTC17;refGene_tx=XM_003993184.5;refGene_gn=TTC17;refGene_tx=XM_006937212.4
D1 97860179 97860333
refGene_gn=TTC17;refGene_tx=XM_003993184.5;refGene_gn=TTC17;refGene_tx=XM_006937212.4
D1 97867051 97867172
refGene_gn=TTC17;refGene_tx=XM_003993184.5;refGene_gn=TTC17;refGene_tx=XM_006937212.4
D1 97869337 97869497
refGene_gn=TTC17;refGene_tx=XM_003993184.5;refGene_gn=TTC17;refGene_tx=XM_006937212.4
D1 97870640 97870724
refGene_gn=TTC17;refGene_tx=XM_003993184.5;refGene_gn=TTC17;refGene_tx=XM_006937212.4
D1 97895041 97895161
refGene_gn=TTC17;refGene_tx=XM_003993184.5;refGene_gn=TTC17;refGene_tx=XM_006937212.4
D1 97896795 97896939
refGene_gn=TTC17;refGene_tx=XM_003993184.5;refGene_gn=TTC17;refGene_tx=XM_006937212.4
D1 97899052 97899184
refGene_gn=TTC17;refGene_tx=XM_003993184.5;refGene_gn=TTC17;refGene_tx=XM_006937212.4
D1 98061218 98061378 refGene_gn=HSD17B12;refGene_tx=XM_003993185.4
D1 98120768 98120815 refGene_gn=HSD17B12;refGene_tx=XM_003993185.4
D1 98124155 98124231 refGene_gn=HSD17B12;refGene_tx=XM_003993185.4
D1 98168651 98168759 refGene_gn=HSD17B12;refGene_tx=XM_003993185.4
D1 98184982 98185047 refGene_gn=HSD17B12;refGene_tx=XM_003993185.4
D1 98186375 98186420 refGene_gn=HSD17B12;refGene_tx=XM_003993185.4
D1 98203799 98203834 refGene_gn=HSD17B12;refGene_tx=XM_003993185.4
D1 98211481 98211563 refGene_gn=HSD17B12;refGene_tx=XM_003993185.4
D1 98213636 98213702 refGene_gn=HSD17B12;refGene_tx=XM_003993185.4
D1 98228081 98228231 refGene_gn=HSD17B12;refGene_tx=XM_003993185.4
D1 98228617 98228722 refGene_gn=HSD17B12;refGene_tx=XM_003993185.4
D1 98245573 98245652
refGene_gn=ALKBH3;refGene_tx=XM_006937213.3;refGene_gn=ALKBH3;refGene_tx=XM_019812296.2;refGene_gn=ALKBH3;refGene_tx=XM_019812297.2;refGene_gn=ALKBH3;refGene_tx=XM_019812298.2
D1 98245958 98246062
refGene_gn=ALKBH3;refGene_tx=XM_006937213.3;refGene_gn=ALKBH3;refGene_tx=XM_019812296.2;refGene_gn=ALKBH3;refGene_tx=XM_019812297.2;refGene_gn=ALKBH3;refGene_tx=XM_019812298.2
D1 98247679 98247714
refGene_gn=ALKBH3;refGene_tx=XM_006937213.3;refGene_gn=ALKBH3;refGene_tx=XM_019812296.2;refGene_gn=ALKBH3;refGene_tx=XM_019812297.2;refGene_gn=ALKBH3;refGene_tx=XM_019812298.2
D1 98250565 98250613
refGene_gn=ALKBH3;refGene_tx=XM_006937213.3;refGene_gn=ALKBH3;refGene_tx=XM_019812296.2;refGene_gn=ALKBH3;refGene_tx=XM_019812298.2
D1 98251987 98252091

refGene_gn=ALKBH3;refGene_tx=XM_006937213.3;refGene_gn=ALKBH3;refGene_tx=XM_019812296.2;refGene_gn=ALKBH3;refGene_tx=XM_019812297.2;refGene_gn=ALKBH3;refGene_tx=XM_019812298.2
D1 98253591 98253680 refGene_gn=ALKBH3;refGene_tx=XM_006937213.3;refGene_gn=ALKBH3;refGene_tx=XM_019812296.2;refGene_gn=ALKBH3;refGene_tx=XM_019812297.2;refGene_gn=ALKBH3;refGene_tx=XM_019812298.2
D1 98257405 98257462 refGene_gn=ALKBH3;refGene_tx=XM_019812296.2;refGene_gn=ALKBH3;refGene_tx=XM_019812297.2;refGene_gn=ALKBH3;refGene_tx=XM_019812298.2
D1 98258788 98258998 refGene_gn=ALKBH3;refGene_tx=XM_006937213.3;refGene_gn=ALKBH3;refGene_tx=XM_019812296.2;refGene_gn=ALKBH3;refGene_tx=XM_019812297.2;refGene_gn=ALKBH3;refGene_tx=XM_019812298.2
D1 98260242 98260350 refGene_gn=ALKBH3;refGene_tx=XM_019812298.2
D1 98276289 98276388 refGene_gn=ALKBH3;refGene_tx=XM_006937213.3;refGene_gn=ALKBH3;refGene_tx=XM_019812296.2;refGene_gn=ALKBH3;refGene_tx=XM_019812297.2
D1 98277121 98277214 refGene_gn=ALKBH3;refGene_tx=XM_006937213.3;refGene_gn=ALKBH3;refGene_tx=XM_019812296.2;refGene_gn=ALKBH3;refGene_tx=XM_019812297.2
D1 98298503 98298635 refGene_gn=CD1H11orf96;refGene_tx=XM_006937308.4
D1 98298637 98298877 refGene_gn=CD1H11orf96;refGene_tx=XM_006937308.4
D1 98404621 98405137 refGene_gn=ACCSL;refGene_tx=XM_023239778.1
D1 98406359 98406419 refGene_gn=ACCSL;refGene_tx=XM_023239778.1
D1 98406851 98406922 refGene_gn=ACCSL;refGene_tx=XM_023239778.1
D1 98407300 98407370 refGene_gn=ACCSL;refGene_tx=XM_023239778.1
D1 98407614 98407681 refGene_gn=ACCSL;refGene_tx=XM_023239778.1
D1 98408735 98408833 refGene_gn=ACCSL;refGene_tx=XM_023239778.1
D1 98409084 98409162 refGene_gn=ACCSL;refGene_tx=XM_023239778.1
D1 98409506 98409607 refGene_gn=ACCSL;refGene_tx=XM_023239778.1
D1 98410695 98410785 refGene_gn=ACCSL;refGene_tx=XM_023239778.1
D1 98412298 98412344 refGene_gn=ACCSL;refGene_tx=XM_023239778.1
D1 98412437 98412579 refGene_gn=ACCSL;refGene_tx=XM_023239778.1
D1 98413901 98414044 refGene_gn=ACCSL;refGene_tx=XM_023239778.1
D1 98414128 98414282 refGene_gn=ACCSL;refGene_tx=XM_023239778.1
D1 98415305 98415388 refGene_gn=ACCSL;refGene_tx=XM_023239778.1
D1 98421770 98422061 refGene_gn=ACCS;refGene_tx=XM_019812299.2;refGene_gn=ACCS;refGene_tx=XM_023239779.1;refGene_gn=ACCS;refGene_tx=XM_023239780.1
D1 98423160 98423220 refGene_gn=ACCS;refGene_tx=XM_019812299.2;refGene_gn=ACCS;refGene_tx=XM_023239779.1;refGene_gn=ACCS;refGene_tx=XM_023239780.1
D1 98425480 98425551 refGene_gn=ACCS;refGene_tx=XM_019812299.2;refGene_gn=ACCS;refGene_tx=XM_023239779.1;refGene_gn=ACCS;refGene_tx=XM_023239780.1
D1 98426712 98426782 refGene_gn=ACCS;refGene_tx=XM_019812299.2;refGene_gn=ACCS;refGene_tx=X

M_023239779.1;refGene_gn=ACCS;refGene_tx=XM_023239780.1
D1	98427278 98427345
refGene_gn=ACCS;refGene_tx=XM_019812299.2;refGene_gn=ACCS;refGene_tx=XM_023239779.1;refGene_gn=ACCS;refGene_tx=XM_023239780.1
D1	98429333 98429431
refGene_gn=ACCS;refGene_tx=XM_019812299.2;refGene_gn=ACCS;refGene_tx=XM_023239779.1;refGene_gn=ACCS;refGene_tx=XM_023239780.1
D1	98429962 98430040
refGene_gn=ACCS;refGene_tx=XM_019812299.2;refGene_gn=ACCS;refGene_tx=XM_023239779.1;refGene_gn=ACCS;refGene_tx=XM_023239780.1
D1	98430742 98430843
refGene_gn=ACCS;refGene_tx=XM_019812299.2;refGene_gn=ACCS;refGene_tx=XM_023239779.1;refGene_gn=ACCS;refGene_tx=XM_023239780.1
D1	98431464 98431554
refGene_gn=ACCS;refGene_tx=XM_019812299.2;refGene_gn=ACCS;refGene_tx=XM_023239779.1;refGene_gn=ACCS;refGene_tx=XM_023239780.1
D1	98433118 98433164
refGene_gn=ACCS;refGene_tx=XM_019812299.2;refGene_gn=ACCS;refGene_tx=XM_023239779.1;refGene_gn=ACCS;refGene_tx=XM_023239780.1
D1	98433273 98433415
refGene_gn=ACCS;refGene_tx=XM_019812299.2;refGene_gn=ACCS;refGene_tx=XM_023239779.1;refGene_gn=ACCS;refGene_tx=XM_023239780.1
D1	98434912 98435055
refGene_gn=ACCS;refGene_tx=XM_019812299.2;refGene_gn=ACCS;refGene_tx=XM_023239779.1;refGene_gn=ACCS;refGene_tx=XM_023239780.1
D1	98435179 98435333
refGene_gn=ACCS;refGene_tx=XM_019812299.2;refGene_gn=ACCS;refGene_tx=XM_023239779.1;refGene_gn=ACCS;refGene_tx=XM_023239780.1
D1	98435468 98435566
refGene_gn=ACCS;refGene_tx=XM_019812299.2;refGene_gn=ACCS;refGene_tx=XM_023239779.1;refGene_gn=ACCS;refGene_tx=XM_023239780.1
D1	98435717 98435748
refGene_gn=ACCS;refGene_tx=XM_019812299.2;refGene_gn=ACCS;refGene_tx=XM_023239779.1
D1	98453527 98453545
refGene_gn=EXT2;refGene_tx=XM_019812300.2;refGene_gn=EXT2;refGene_tx=XM_019812301.2
D1	98454775 98455341
refGene_gn=EXT2;refGene_tx=XM_019812300.2;refGene_gn=EXT2;refGene_tx=XM_019812301.2;refGene_gn=EXT2;refGene_tx=XM_023239782.1
D1	98456517 98456607
refGene_gn=EXT2;refGene_tx=XM_019812300.2;refGene_gn=EXT2;refGene_tx=XM_019812301.2;refGene_gn=EXT2;refGene_tx=XM_023239782.1
D1	98462675 98462792
refGene_gn=EXT2;refGene_tx=XM_019812300.2;refGene_gn=EXT2;refGene_tx=XM_019812301.2;refGene_gn=EXT2;refGene_tx=XM_023239782.1
D1	98471683 98471879
refGene_gn=EXT2;refGene_tx=XM_019812300.2;refGene_gn=EXT2;refGene_tx=XM_019812301.2;refGene_gn=EXT2;refGene_tx=XM_023239782.1
D1	98473712 98473852

refGene_gn=EXT2;refGene_tx=XM_019812300.2;refGene_gn=EXT2;refGene_tx=XM_019812301.2;refGene_gn=EXT2;refGene_tx=XM_023239782.1
D1 98476576 98476670
refGene_gn=EXT2;refGene_tx=XM_019812300.2;refGene_gn=EXT2;refGene_tx=XM_019812301.2;refGene_gn=EXT2;refGene_tx=XM_023239782.1
D1 98515095 98515227
refGene_gn=EXT2;refGene_tx=XM_019812300.2;refGene_gn=EXT2;refGene_tx=XM_019812301.2;refGene_gn=EXT2;refGene_tx=XM_023239782.1
D1 98542920 98543110
refGene_gn=EXT2;refGene_tx=XM_019812300.2;refGene_gn=EXT2;refGene_tx=XM_019812301.2;refGene_gn=EXT2;refGene_tx=XM_023239782.1
D1 98552319 98552486
refGene_gn=EXT2;refGene_tx=XM_019812300.2;refGene_gn=EXT2;refGene_tx=XM_019812301.2;refGene_gn=EXT2;refGene_tx=XM_023239782.1
D1 98577554 98577698
refGene_gn=EXT2;refGene_tx=XM_019812300.2;refGene_gn=EXT2;refGene_tx=XM_019812301.2;refGene_gn=EXT2;refGene_tx=XM_023239782.1
D1 98579799 98579928
refGene_gn=EXT2;refGene_tx=XM_019812300.2;refGene_gn=EXT2;refGene_tx=XM_019812301.2;refGene_gn=EXT2;refGene_tx=XM_023239782.1
D1 98581667 98581750
refGene_gn=EXT2;refGene_tx=XM_019812300.2;refGene_gn=EXT2;refGene_tx=XM_019812301.2;refGene_gn=EXT2;refGene_tx=XM_023239782.1
D1 98585424 98585512 refGene_gn=EXT2;refGene_tx=XM_019812301.2
D1 98588011 98588150
refGene_gn=EXT2;refGene_tx=XM_019812300.2;refGene_gn=EXT2;refGene_tx=XM_023239782.1
D1 98609820 98610150 refGene_gn=ALX4;refGene_tx=XM_003993190.4
D1 98612492 98612621 refGene_gn=ALX4;refGene_tx=XM_003993190.4
D1 98618386 98618697 refGene_gn=ALX4;refGene_tx=XM_003993190.4
D1 98651089 98651552 refGene_gn=ALX4;refGene_tx=XM_003993190.4
D1 98886617 98886680
refGene_gn=CD82;refGene_tx=XM_003993191.5;refGene_gn=CD82;refGene_tx=XM_006937217.4;refGene_gn=CD82;refGene_tx=XM_019812303.2;refGene_gn=CD82;refGene_tx=XM_019812304.2
D1 98891973 98892046
refGene_gn=CD82;refGene_tx=XM_003993191.5;refGene_gn=CD82;refGene_tx=XM_006937217.4;refGene_gn=CD82;refGene_tx=XM_019812303.2;refGene_gn=CD82;refGene_tx=XM_019812304.2
D1 98896774 98896899
refGene_gn=CD82;refGene_tx=XM_003993191.5;refGene_gn=CD82;refGene_tx=XM_006937217.4;refGene_gn=CD82;refGene_tx=XM_019812303.2;refGene_gn=CD82;refGene_tx=XM_019812304.2
D1 98896994 98897069
refGene_gn=CD82;refGene_tx=XM_003993191.5;refGene_gn=CD82;refGene_tx=XM_006937217.4;refGene_gn=CD82;refGene_tx=XM_019812303.2;refGene_gn=CD82;refGene_tx=XM_019812304.2
D1 98905462 98905564
refGene_gn=CD82;refGene_tx=XM_003993191.5;refGene_gn=CD82;refGene_tx=XM_006937217.4;refGene_gn=CD82;refGene_tx=XM_019812303.2;refGene_gn=CD8

2;refGene_tx=XM_019812304.2
D1 98907301 98907505
refGene_gn=CD82;refGene_tx=XM_003993191.5;refGene_gn=CD82;refGene_tx=XM_006937217.4;refGene_gn=CD82;refGene_tx=XM_019812303.2;refGene_gn=CD82;refGene_tx=XM_019812304.2
D1 98908176 98908260
refGene_gn=CD82;refGene_tx=XM_003993191.5;refGene_gn=CD82;refGene_tx=XM_006937217.4;refGene_gn=CD82;refGene_tx=XM_019812303.2;refGene_gn=CD82;refGene_tx=XM_019812304.2
D1 98908560 98908638
refGene_gn=CD82;refGene_tx=XM_003993191.5;refGene_gn=CD82;refGene_tx=XM_006937217.4;refGene_gn=CD82;refGene_tx=XM_019812303.2;refGene_gn=CD82;refGene_tx=XM_019812304.2
D1 99164551 99164614
refGene_gn=TSPAN18;refGene_tx=XM_003993193.5;refGene_gn=TSPAN18;refGene_tx=XM_019812305.2;refGene_gn=TSPAN18;refGene_tx=XM_019812306.2;refGene_gn=TSPAN18;refGene_tx=XM_019812307.2;refGene_gn=TSPAN18;refGene_tx=XM_019812308.2;refGene_gn=TSPAN18;refGene_tx=XM_019812309.2;refGene_gn=TSPAN18;refGene_tx=XM_023239784.1
D1 99166134 99166329
refGene_gn=TSPAN18;refGene_tx=XM_003993193.5;refGene_gn=TSPAN18;refGene_tx=XM_019812305.2;refGene_gn=TSPAN18;refGene_tx=XM_019812306.2;refGene_gn=TSPAN18;refGene_tx=XM_019812307.2;refGene_gn=TSPAN18;refGene_tx=XM_019812308.2;refGene_gn=TSPAN18;refGene_tx=XM_019812309.2;refGene_gn=TSPAN18;refGene_tx=XM_023239784.1
D1 99172700 99172775
refGene_gn=TSPAN18;refGene_tx=XM_003993193.5;refGene_gn=TSPAN18;refGene_tx=XM_019812305.2;refGene_gn=TSPAN18;refGene_tx=XM_019812306.2;refGene_gn=TSPAN18;refGene_tx=XM_019812307.2;refGene_gn=TSPAN18;refGene_tx=XM_019812308.2;refGene_gn=TSPAN18;refGene_tx=XM_019812309.2;refGene_gn=TSPAN18;refGene_tx=XM_023239784.1
D1 99173796 99173895
refGene_gn=TSPAN18;refGene_tx=XM_003993193.5;refGene_gn=TSPAN18;refGene_tx=XM_019812305.2;refGene_gn=TSPAN18;refGene_tx=XM_019812306.2;refGene_gn=TSPAN18;refGene_tx=XM_019812307.2;refGene_gn=TSPAN18;refGene_tx=XM_019812308.2;refGene_gn=TSPAN18;refGene_tx=XM_019812309.2;refGene_gn=TSPAN18;refGene_tx=XM_023239784.1
D1 99174432 99174615
refGene_gn=TSPAN18;refGene_tx=XM_003993193.5;refGene_gn=TSPAN18;refGene_tx=XM_019812305.2;refGene_gn=TSPAN18;refGene_tx=XM_019812306.2;refGene_gn=TSPAN18;refGene_tx=XM_019812307.2;refGene_gn=TSPAN18;refGene_tx=XM_019812308.2;refGene_gn=TSPAN18;refGene_tx=XM_019812309.2;refGene_gn=TSPAN18;refGene_tx=XM_023239784.1
D1 99177926 99178010
refGene_gn=TSPAN18;refGene_tx=XM_003993193.5;refGene_gn=TSPAN18;refGene_tx=XM_019812305.2;refGene_gn=TSPAN18;refGene_tx=XM_019812306.2;refGene_gn=TSPAN18;refGene_tx=XM_019812307.2;refGene_gn=TSPAN18;refGene_tx=XM_019812308.2;refGene_gn=TSPAN18;refGene_tx=XM_019812309.2;refGene_gn=TSPAN18;refGene_tx=XM_023239784.1
D1 99180118 99180166

refGene_gn=TSPAN18;refGene_tx=XM_003993193.5;refGene_gn=TSPAN18;refGene_tx=XM_019812305.2;refGene_gn=TSPAN18;refGene_tx=XM_019812306.2;refGene_gn=TSPAN18;refGene_tx=XM_019812307.2;refGene_gn=TSPAN18;refGene_tx=XM_019812308.2;refGene_gn=TSPAN18;refGene_tx=XM_019812309.2;refGene_gn=TSPAN18;refGene_tx=XM_023239784.1
D1 99185254 99185388
refGene_gn=TP53I11;refGene_tx=XM_003993194.5;refGene_gn=TP53I11;refGene_tx=XM_011286794.3;refGene_gn=TP53I11;refGene_tx=XM_011286795.3;refGene_gn=TP53I11;refGene_tx=XM_019812310.2;refGene_gn=TP53I11;refGene_tx=XM_019812311.2
D1 99186090 99186192
refGene_gn=TP53I11;refGene_tx=XM_003993194.5;refGene_gn=TP53I11;refGene_tx=XM_011286794.3;refGene_gn=TP53I11;refGene_tx=XM_011286795.3;refGene_gn=TP53I11;refGene_tx=XM_019812310.2;refGene_gn=TP53I11;refGene_tx=XM_019812311.2
D1 99187164 99187261
refGene_gn=TP53I11;refGene_tx=XM_003993194.5;refGene_gn=TP53I11;refGene_tx=XM_011286794.3;refGene_gn=TP53I11;refGene_tx=XM_011286795.3;refGene_gn=TP53I11;refGene_tx=XM_019812310.2;refGene_gn=TP53I11;refGene_tx=XM_019812311.2
D1 99187676 99187725
refGene_gn=TP53I11;refGene_tx=XM_003993194.5;refGene_gn=TP53I11;refGene_tx=XM_011286794.3;refGene_gn=TP53I11;refGene_tx=XM_011286795.3;refGene_gn=TP53I11;refGene_tx=XM_019812310.2;refGene_gn=TP53I11;refGene_tx=XM_019812311.2
D1 99187961 99188020
refGene_gn=TP53I11;refGene_tx=XM_003993194.5;refGene_gn=TP53I11;refGene_tx=XM_011286794.3;refGene_gn=TP53I11;refGene_tx=XM_011286795.3;refGene_gn=TP53I11;refGene_tx=XM_019812310.2;refGene_gn=TP53I11;refGene_tx=XM_019812311.2
D1 99188625 99188754
refGene_gn=TP53I11;refGene_tx=XM_003993194.5;refGene_gn=TP53I11;refGene_tx=XM_011286794.3;refGene_gn=TP53I11;refGene_tx=XM_011286795.3;refGene_gn=TP53I11;refGene_tx=XM_019812310.2;refGene_gn=TP53I11;refGene_tx=XM_019812311.2
D1 99398750 99398869
refGene_gn=PRDM11;refGene_tx=XM_003993196.4;refGene_gn=PRDM11;refGene_tx=XM_023239785.1;refGene_gn=PRDM11;refGene_tx=XM_023239786.1;refGene_gn=PRDM11;refGene_tx=XM_023239787.1;refGene_gn=PRDM11;refGene_tx=XM_023239788.1;refGene_gn=PRDM11;refGene_tx=XM_023239790.1
D1 99399240 99399344
refGene_gn=PRDM11;refGene_tx=XM_003993196.4;refGene_gn=PRDM11;refGene_tx=XM_023239785.1;refGene_gn=PRDM11;refGene_tx=XM_023239786.1;refGene_gn=PRDM11;refGene_tx=XM_023239787.1;refGene_gn=PRDM11;refGene_tx=XM_023239788.1;refGene_gn=PRDM11;refGene_tx=XM_023239790.1
D1 99399937 99400200
refGene_gn=PRDM11;refGene_tx=XM_003993196.4;refGene_gn=PRDM11;refGene_tx=XM_023239785.1;refGene_gn=PRDM11;refGene_tx=XM_023239786.1;refGene_gn=PRDM11;refGene_tx=XM_023239787.1;refGene_gn=PRDM11;refGene_tx=XM_023239788.1;refGene_gn=PRDM11;refGene_tx=XM_023239790.1

D1 99422604 99422672
refGene_gn=PRDM11;refGene_tx=XM_003993196.4;refGene_gn=PRDM11;refGene_tx=XM_019812317.2;refGene_gn=PRDM11;refGene_tx=XM_023239785.1;refGene_gn=PRDM11;refGene_tx=XM_023239786.1;refGene_gn=PRDM11;refGene_tx=XM_023239787.1;refGene_gn=PRDM11;refGene_tx=XM_023239788.1;refGene_gn=PRDM11;refGene_tx=XM_023239790.1
D1 99426576 99427291
refGene_gn=PRDM11;refGene_tx=XM_023239787.1;refGene_gn=PRDM11;refGene_tx=XM_023239790.1
D1 99429971 99430537 refGene_gn=PRDM11;refGene_tx=XM_023239787.1
D1 99437407 99437595
refGene_gn=PRDM11;refGene_tx=XM_003993196.4;refGene_gn=PRDM11;refGene_tx=XM_019812317.2;refGene_gn=PRDM11;refGene_tx=XM_023239785.1;refGene_gn=PRDM11;refGene_tx=XM_023239786.1;refGene_gn=PRDM11;refGene_tx=XM_023239788.1
D1 99441776 99442510
refGene_gn=PRDM11;refGene_tx=XM_003993196.4;refGene_gn=PRDM11;refGene_tx=XM_019812317.2;refGene_gn=PRDM11;refGene_tx=XM_023239785.1;refGene_gn=PRDM11;refGene_tx=XM_023239788.1
D1 99443571 99445736
refGene_gn=PRDM11;refGene_tx=XM_003993196.4;refGene_gn=PRDM11;refGene_tx=XM_019812317.2;refGene_gn=PRDM11;refGene_tx=XM_023239785.1;refGene_gn=PRDM11;refGene_tx=XM_023239786.1
D1 99461330 99461635 refGene_gn=SYT13;refGene_tx=XM_023239791.1
D1 99463289 99463419 refGene_gn=SYT13;refGene_tx=XM_023239791.1
D1 99468321 99468623 refGene_gn=SYT13;refGene_tx=XM_023239791.1
D1 99469795 99469930 refGene_gn=SYT13;refGene_tx=XM_023239791.1
D1 99472094 99472320 refGene_gn=SYT13;refGene_tx=XM_023239791.1
D1 99499383 99499563 refGene_gn=SYT13;refGene_tx=XM_023239791.1
D1 99829807 99831043
refGene_gn=CHST1;refGene_tx=XM_006937223.4;refGene_gn=CHST1;refGene_tx=XM_023239792.1;refGene_gn=CHST1;refGene_tx=XM_023239794.1;refGene_gn=CHST1;refGene_tx=XM_023239795.1;refGene_gn=CHST1;refGene_tx=XM_023239796.1
D1 99968741 99968773 refGene_gn=SLC35C1;refGene_tx=XM_003993198.5
D1 99970403 99970930
refGene_gn=SLC35C1;refGene_tx=XM_003993198.5;refGene_gn=SLC35C1;refGene_tx=XM_019812318.2
D1 99974212 99974772
refGene_gn=SLC35C1;refGene_tx=XM_003993198.5;refGene_gn=SLC35C1;refGene_tx=XM_019812318.2
D1 100006969 100007187
refGene_gn=CRY2;refGene_tx=XM_023239797.1
D1 100014790 100014899
refGene_gn=CRY2;refGene_tx=XM_023239797.1
D1 100017656 100017799
refGene_gn=CRY2;refGene_tx=XM_023239797.1
D1 100019659 100019844
refGene_gn=CRY2;refGene_tx=XM_023239797.1
D1 100020584 100020673

```
refGene_gn=CRY2;refGene_tx=XM_023239797.1
D1	100025473	100025614
refGene_gn=CRY2;refGene_tx=XM_023239797.1
D1	100025874	100026186
refGene_gn=CRY2;refGene_tx=XM_023239797.1
D1	100026430	100026582
refGene_gn=CRY2;refGene_tx=XM_023239797.1
D1	100026704	100026907
refGene_gn=CRY2;refGene_tx=XM_023239797.1
D1	100027241	100027334
refGene_gn=CRY2;refGene_tx=XM_023239797.1
D1	100028548	100028688
refGene_gn=CRY2;refGene_tx=XM_023239797.1
D1	100042924	100043025
refGene_gn=MAPK8IP1;refGene_tx=XM_023239798.1
D1	100052353	100052424
refGene_gn=MAPK8IP1;refGene_tx=XM_023239799.1
D1	100053502	100053608
refGene_gn=MAPK8IP1;refGene_tx=XM_023239798.1;refGene_gn=MAPK8IP1;refG
ene_tx=XM_023239799.1
D1	100055542	100055833
refGene_gn=MAPK8IP1;refGene_tx=XM_023239798.1;refGene_gn=MAPK8IP1;refG
ene_tx=XM_023239799.1
D1	100057418	100057500
refGene_gn=MAPK8IP1;refGene_tx=XM_023239798.1;refGene_gn=MAPK8IP1;refG
ene_tx=XM_023239799.1
D1	100057817	100058630
refGene_gn=MAPK8IP1;refGene_tx=XM_023239798.1;refGene_gn=MAPK8IP1;refG
ene_tx=XM_023239799.1
D1	100058808	100058884
refGene_gn=MAPK8IP1;refGene_tx=XM_023239798.1;refGene_gn=MAPK8IP1;refG
ene_tx=XM_023239799.1
D1	100059485	100059658
refGene_gn=MAPK8IP1;refGene_tx=XM_023239798.1;refGene_gn=MAPK8IP1;refG
ene_tx=XM_023239799.1
D1	100059861	100059971
refGene_gn=MAPK8IP1;refGene_tx=XM_023239798.1;refGene_gn=MAPK8IP1;refG
ene_tx=XM_023239799.1
D1	100060078	100060195
refGene_gn=MAPK8IP1;refGene_tx=XM_023239798.1;refGene_gn=MAPK8IP1;refG
ene_tx=XM_023239799.1
D1	100060288	100060359
refGene_gn=MAPK8IP1;refGene_tx=XM_023239798.1;refGene_gn=MAPK8IP1;refG
ene_tx=XM_023239799.1
D1	100060478	100060577
refGene_gn=MAPK8IP1;refGene_tx=XM_023239798.1;refGene_gn=MAPK8IP1;refG
ene_tx=XM_023239799.1
D1	100060903	100060976
refGene_gn=MAPK8IP1;refGene_tx=XM_023239798.1;refGene_gn=MAPK8IP1;refG
ene_tx=XM_023239799.1
```

D1 100061835 100061957
refGene_gn=CD1H11orf94;refGene_tx=XM_023239801.1
D1 100062140 100062239
refGene_gn=CD1H11orf94;refGene_tx=XM_023239801.1
D1 100062446 100062534
refGene_gn=CD1H11orf94;refGene_tx=XM_023239801.1
D1 100065878 100065937
refGene_gn=PEX16;refGene_tx=XM_023239802.1;refGene_gn=PEX16;refGene_tx=XM_023239803.1
D1 100066517 100066582
refGene_gn=PEX16;refGene_tx=XM_023239802.1;refGene_gn=PEX16;refGene_tx=XM_023239803.1
D1 100068456 100068576
refGene_gn=PEX16;refGene_tx=XM_023239802.1
D1 100068696 100068769
refGene_gn=PEX16;refGene_tx=XM_023239802.1;refGene_gn=PEX16;refGene_tx=XM_023239803.1
D1 100068870 100069023
refGene_gn=PEX16;refGene_tx=XM_023239802.1;refGene_gn=PEX16;refGene_tx=XM_023239803.1
D1 100069110 100069191
refGene_gn=PEX16;refGene_tx=XM_023239802.1;refGene_gn=PEX16;refGene_tx=XM_023239803.1
D1 100070203 100070304
refGene_gn=PEX16;refGene_tx=XM_023239802.1;refGene_gn=PEX16;refGene_tx=XM_023239803.1
D1 100070538 100070672
refGene_gn=PEX16;refGene_tx=XM_023239802.1;refGene_gn=PEX16;refGene_tx=XM_023239803.1
D1 100070969 100071046
refGene_gn=PEX16;refGene_tx=XM_023239802.1;refGene_gn=PEX16;refGene_tx=XM_023239803.1
D1 100071828 100071864
refGene_gn=PEX16;refGene_tx=XM_023239802.1;refGene_gn=PEX16;refGene_tx=XM_023239803.1
D1 100072068 100072180
refGene_gn=PEX16;refGene_tx=XM_023239802.1;refGene_gn=PEX16;refGene_tx=XM_023239803.1
D1 100076432 100076529
refGene_gn=LARGE2;refGene_tx=XM_023239804.1;refGene_gn=LARGE2;refGene_tx=XM_023239805.1;refGene_gn=LARGE2;refGene_tx=XM_023239806.1
D1 100076621 100076731
refGene_gn=LARGE2;refGene_tx=XM_023239804.1;refGene_gn=LARGE2;refGene_tx=XM_023239805.1;refGene_gn=LARGE2;refGene_tx=XM_023239806.1
D1 100077054 100077137
refGene_gn=LARGE2;refGene_tx=XM_023239804.1;refGene_gn=LARGE2;refGene_tx=XM_023239805.1;refGene_gn=LARGE2;refGene_tx=XM_023239806.1
D1 100077411 100077535
refGene_gn=LARGE2;refGene_tx=XM_023239804.1;refGene_gn=LARGE2;refGene_tx=XM_023239805.1;refGene_gn=LARGE2;refGene_tx=XM_023239806.1;refGene_

gn=LARGE2;refGene_tx=XM_023239807.1
D1 100077777 100077949
refGene_gn=LARGE2;refGene_tx=XM_023239804.1;refGene_gn=LARGE2;refGene_tx=XM_023239805.1;refGene_gn=LARGE2;refGene_tx=XM_023239806.1;refGene_gn=LARGE2;refGene_tx=XM_023239807.1
D1 100078076 100078181
refGene_gn=LARGE2;refGene_tx=XM_023239804.1;refGene_gn=LARGE2;refGene_tx=XM_023239805.1;refGene_gn=LARGE2;refGene_tx=XM_023239807.1
D1 100078660 100078773
refGene_gn=LARGE2;refGene_tx=XM_023239804.1;refGene_gn=LARGE2;refGene_tx=XM_023239805.1;refGene_gn=LARGE2;refGene_tx=XM_023239806.1;refGene_gn=LARGE2;refGene_tx=XM_023239807.1
D1 100078848 100078974
refGene_gn=LARGE2;refGene_tx=XM_023239804.1;refGene_gn=LARGE2;refGene_tx=XM_023239805.1;refGene_gn=LARGE2;refGene_tx=XM_023239806.1;refGene_gn=LARGE2;refGene_tx=XM_023239807.1
D1 100079051 100079207
refGene_gn=LARGE2;refGene_tx=XM_023239804.1;refGene_gn=LARGE2;refGene_tx=XM_023239805.1;refGene_gn=LARGE2;refGene_tx=XM_023239806.1;refGene_gn=LARGE2;refGene_tx=XM_023239807.1
D1 100079336 100079497
refGene_gn=LARGE2;refGene_tx=XM_023239804.1;refGene_gn=LARGE2;refGene_tx=XM_023239805.1;refGene_gn=LARGE2;refGene_tx=XM_023239806.1;refGene_gn=LARGE2;refGene_tx=XM_023239807.1
D1 100079923 100080202
refGene_gn=LARGE2;refGene_tx=XM_023239804.1;refGene_gn=LARGE2;refGene_tx=XM_023239805.1;refGene_gn=LARGE2;refGene_tx=XM_023239806.1;refGene_gn=LARGE2;refGene_tx=XM_023239807.1
D1 100080553 100080703
refGene_gn=LARGE2;refGene_tx=XM_023239804.1;refGene_gn=LARGE2;refGene_tx=XM_023239805.1;refGene_gn=LARGE2;refGene_tx=XM_023239806.1;refGene_gn=LARGE2;refGene_tx=XM_023239807.1
D1 100080825 100081021
refGene_gn=LARGE2;refGene_tx=XM_023239804.1;refGene_gn=LARGE2;refGene_tx=XM_023239805.1;refGene_gn=LARGE2;refGene_tx=XM_023239806.1;refGene_gn=LARGE2;refGene_tx=XM_023239807.1
D1 100081280 100081490
refGene_gn=LARGE2;refGene_tx=XM_023239804.1;refGene_gn=LARGE2;refGene_tx=XM_023239805.1;refGene_gn=LARGE2;refGene_tx=XM_023239806.1;refGene_gn=LARGE2;refGene_tx=XM_023239807.1
D1 100086547 100086805
refGene_gn=PHF21A;refGene_tx=XM_023239808.1
D1 100088192 100088251
refGene_gn=PHF21A;refGene_tx=XM_023239808.1
D1 100088253 100088295
refGene_gn=PHF21A;refGene_tx=XM_023239808.1
D1 100089150 100089226
refGene_gn=PHF21A;refGene_tx=XM_023239808.1
D1 100090924 100091080
refGene_gn=PHF21A;refGene_tx=XM_023239808.1

D1 100097407 100097571
refGene_gn=PHF21A;refGene_tx=XM_023239808.1
D1 100100163 100100224
refGene_gn=PHF21A;refGene_tx=XM_023239808.1
D1 100100688 100100768
refGene_gn=PHF21A;refGene_tx=XM_023239808.1
D1 100101551 100101603
refGene_gn=PHF21A;refGene_tx=XM_023239808.1
D1 100104134 100104233
refGene_gn=PHF21A;refGene_tx=XM_023239808.1
D1 100123272 100123566
refGene_gn=PHF21A;refGene_tx=XM_023239808.1
D1 100127582 100127672
refGene_gn=PHF21A;refGene_tx=XM_023239808.1
D1 100128901 100129153
refGene_gn=PHF21A;refGene_tx=XM_023239808.1
D1 100135790 100135994
refGene_gn=PHF21A;refGene_tx=XM_023239808.1
D1 100235357 100235423
refGene_gn=PHF21A;refGene_tx=XM_023239808.1
D1 100237743 100237776
refGene_gn=PHF21A;refGene_tx=XM_023239808.1
D1 100242903 100242957
refGene_gn=PHF21A;refGene_tx=XM_023239808.1
D1 100414244 100414346
refGene_gn=CREB3L1;refGene_tx=XM_003993204.4
D1 100432984 100433213
refGene_gn=CREB3L1;refGene_tx=XM_003993204.4
D1 100436469 100436654
refGene_gn=CREB3L1;refGene_tx=XM_003993204.4
D1 100439428 100439507
refGene_gn=CREB3L1;refGene_tx=XM_003993204.4
D1 100441205 100441363
refGene_gn=CREB3L1;refGene_tx=XM_003993204.4
D1 100442037 100442187
refGene_gn=CREB3L1;refGene_tx=XM_003993204.4
D1 100442339 100442398
refGene_gn=CREB3L1;refGene_tx=XM_003993204.4
D1 100442596 100442665
refGene_gn=CREB3L1;refGene_tx=XM_003993204.4
D1 100444468 100444568
refGene_gn=CREB3L1;refGene_tx=XM_003993204.4
D1 100445484 100445611
refGene_gn=CREB3L1;refGene_tx=XM_003993204.4
D1 100447908 100448176
refGene_gn=CREB3L1;refGene_tx=XM_003993204.4
D1 100448354 100448391
refGene_gn=CREB3L1;refGene_tx=XM_003993204.4
D1 100459837 100460049
refGene_gn=DGKZ;refGene_tx=XM_019812339.2;refGene_gn=DGKZ;refGene_tx=X

M_019812341.2
D1 100468990 100469166
refGene_gn=DGKZ;refGene_tx=XM_006937233.4;refGene_gn=DGKZ;refGene_tx=X
M_006937234.4
D1 100471284 100471445
refGene_gn=DGKZ;refGene_tx=XM_023239809.1;refGene_gn=DGKZ;refGene_tx=X
M_023239810.1;refGene_gn=DGKZ;refGene_tx=XM_023239811.1
D1 100486768 100486785
refGene_gn=DGKZ;refGene_tx=XM_019812342.2
D1 100489008 100489742
refGene_gn=DGKZ;refGene_tx=XM_003993206.5;refGene_gn=DGKZ;refGene_tx=X
M_006937232.4;refGene_gn=DGKZ;refGene_tx=XM_019812338.2
D1 100490085 100490194
refGene_gn=DGKZ;refGene_tx=XM_003993206.5;refGene_gn=DGKZ;refGene_tx=X
M_006937232.4;refGene_gn=DGKZ;refGene_tx=XM_006937233.4;refGene_gn=DGK
Z;refGene_tx=XM_006937234.4;refGene_gn=DGKZ;refGene_tx=XM_019812338.2;
refGene_gn=DGKZ;refGene_tx=XM_019812339.2;refGene_gn=DGKZ;refGene_tx=X
M_019812341.2;refGene_gn=DGKZ;refGene_tx=XM_019812342.2;refGene_gn=DGK
Z;refGene_tx=XM_023239809.1;refGene_gn=DGKZ;refGene_tx=XM_023239810.1;
refGene_gn=DGKZ;refGene_tx=XM_023239811.1
D1 100490650 100490746
refGene_gn=DGKZ;refGene_tx=XM_003993206.5;refGene_gn=DGKZ;refGene_tx=X
M_006937232.4;refGene_gn=DGKZ;refGene_tx=XM_006937233.4;refGene_gn=DGK
Z;refGene_tx=XM_006937234.4;refGene_gn=DGKZ;refGene_tx=XM_019812338.2;
refGene_gn=DGKZ;refGene_tx=XM_019812339.2;refGene_gn=DGKZ;refGene_tx=X
M_019812341.2;refGene_gn=DGKZ;refGene_tx=XM_019812342.2;refGene_gn=DGK
Z;refGene_tx=XM_023239809.1;refGene_gn=DGKZ;refGene_tx=XM_023239810.1;
refGene_gn=DGKZ;refGene_tx=XM_023239811.1
D1 100491035 100491113
refGene_gn=DGKZ;refGene_tx=XM_003993206.5;refGene_gn=DGKZ;refGene_tx=X
M_006937232.4;refGene_gn=DGKZ;refGene_tx=XM_006937233.4;refGene_gn=DGK
Z;refGene_tx=XM_006937234.4;refGene_gn=DGKZ;refGene_tx=XM_019812338.2;
refGene_gn=DGKZ;refGene_tx=XM_019812339.2;refGene_gn=DGKZ;refGene_tx=X
M_019812341.2;refGene_gn=DGKZ;refGene_tx=XM_019812342.2;refGene_gn=DGK
Z;refGene_tx=XM_023239809.1;refGene_gn=DGKZ;refGene_tx=XM_023239810.1;
refGene_gn=DGKZ;refGene_tx=XM_023239811.1
D1 100492178 100492235
refGene_gn=DGKZ;refGene_tx=XM_003993206.5;refGene_gn=DGKZ;refGene_tx=X
M_006937232.4;refGene_gn=DGKZ;refGene_tx=XM_006937233.4;refGene_gn=DGK
Z;refGene_tx=XM_006937234.4;refGene_gn=DGKZ;refGene_tx=XM_019812338.2;
refGene_gn=DGKZ;refGene_tx=XM_019812339.2;refGene_gn=DGKZ;refGene_tx=X
M_019812341.2;refGene_gn=DGKZ;refGene_tx=XM_019812342.2;refGene_gn=DGK
Z;refGene_tx=XM_023239809.1;refGene_gn=DGKZ;refGene_tx=XM_023239810.1;
refGene_gn=DGKZ;refGene_tx=XM_023239811.1
D1 100492601 100492670
refGene_gn=DGKZ;refGene_tx=XM_003993206.5;refGene_gn=DGKZ;refGene_tx=X
M_006937232.4;refGene_gn=DGKZ;refGene_tx=XM_006937233.4;refGene_gn=DGK
Z;refGene_tx=XM_006937234.4;refGene_gn=DGKZ;refGene_tx=XM_019812338.2;
refGene_gn=DGKZ;refGene_tx=XM_019812339.2;refGene_gn=DGKZ;refGene_tx=X
M_019812341.2;refGene_gn=DGKZ;refGene_tx=XM_019812342.2;refGene_gn=DGK

Z;refGene_tx=XM_023239809.1;refGene_gn=DGKZ;refGene_tx=XM_023239810.1;
refGene_gn=DGKZ;refGene_tx=XM_023239811.1
D1 100493892 100493964
refGene_gn=DGKZ;refGene_tx=XM_003993206.5;refGene_gn=DGKZ;refGene_tx=X
M_006937232.4;refGene_gn=DGKZ;refGene_tx=XM_006937233.4;refGene_gn=DGK
Z;refGene_tx=XM_006937234.4;refGene_gn=DGKZ;refGene_tx=XM_019812338.2;
refGene_gn=DGKZ;refGene_tx=XM_019812339.2;refGene_gn=DGKZ;refGene_tx=X
M_019812341.2;refGene_gn=DGKZ;refGene_tx=XM_019812342.2;refGene_gn=DGK
Z;refGene_tx=XM_023239809.1;refGene_gn=DGKZ;refGene_tx=XM_023239810.1;
refGene_gn=DGKZ;refGene_tx=XM_023239811.1
D1 100494069 100494186
refGene_gn=DGKZ;refGene_tx=XM_003993206.5;refGene_gn=DGKZ;refGene_tx=X
M_006937232.4;refGene_gn=DGKZ;refGene_tx=XM_006937233.4;refGene_gn=DGK
Z;refGene_tx=XM_006937234.4;refGene_gn=DGKZ;refGene_tx=XM_019812338.2;
refGene_gn=DGKZ;refGene_tx=XM_019812339.2;refGene_gn=DGKZ;refGene_tx=X
M_019812341.2;refGene_gn=DGKZ;refGene_tx=XM_019812342.2;refGene_gn=DGK
Z;refGene_tx=XM_023239809.1;refGene_gn=DGKZ;refGene_tx=XM_023239810.1;
refGene_gn=DGKZ;refGene_tx=XM_023239811.1
D1 100494286 100494358
refGene_gn=DGKZ;refGene_tx=XM_003993206.5;refGene_gn=DGKZ;refGene_tx=X
M_006937232.4;refGene_gn=DGKZ;refGene_tx=XM_006937233.4;refGene_gn=DGK
Z;refGene_tx=XM_006937234.4;refGene_gn=DGKZ;refGene_tx=XM_019812338.2;
refGene_gn=DGKZ;refGene_tx=XM_019812339.2;refGene_gn=DGKZ;refGene_tx=X
M_019812341.2;refGene_gn=DGKZ;refGene_tx=XM_019812342.2;refGene_gn=DGK
Z;refGene_tx=XM_023239809.1;refGene_gn=DGKZ;refGene_tx=XM_023239810.1;
refGene_gn=DGKZ;refGene_tx=XM_023239811.1
D1 100494627 100494723
refGene_gn=DGKZ;refGene_tx=XM_003993206.5;refGene_gn=DGKZ;refGene_tx=X
M_006937232.4;refGene_gn=DGKZ;refGene_tx=XM_006937233.4;refGene_gn=DGK
Z;refGene_tx=XM_006937234.4;refGene_gn=DGKZ;refGene_tx=XM_019812338.2;
refGene_gn=DGKZ;refGene_tx=XM_019812339.2;refGene_gn=DGKZ;refGene_tx=X
M_019812341.2;refGene_gn=DGKZ;refGene_tx=XM_019812342.2;refGene_gn=DGK
Z;refGene_tx=XM_023239809.1;refGene_gn=DGKZ;refGene_tx=XM_023239810.1;
refGene_gn=DGKZ;refGene_tx=XM_023239811.1
D1 100494985 100495068
refGene_gn=DGKZ;refGene_tx=XM_003993206.5;refGene_gn=DGKZ;refGene_tx=X
M_006937232.4;refGene_gn=DGKZ;refGene_tx=XM_006937233.4;refGene_gn=DGK
Z;refGene_tx=XM_006937234.4;refGene_gn=DGKZ;refGene_tx=XM_019812338.2;
refGene_gn=DGKZ;refGene_tx=XM_019812339.2;refGene_gn=DGKZ;refGene_tx=X
M_019812341.2;refGene_gn=DGKZ;refGene_tx=XM_019812342.2;refGene_gn=DGK
Z;refGene_tx=XM_023239809.1;refGene_gn=DGKZ;refGene_tx=XM_023239810.1;
refGene_gn=DGKZ;refGene_tx=XM_023239811.1
D1 100495169 100495230
refGene_gn=DGKZ;refGene_tx=XM_003993206.5;refGene_gn=DGKZ;refGene_tx=X
M_006937232.4;refGene_gn=DGKZ;refGene_tx=XM_006937233.4;refGene_gn=DGK
Z;refGene_tx=XM_006937234.4;refGene_gn=DGKZ;refGene_tx=XM_019812338.2;
refGene_gn=DGKZ;refGene_tx=XM_019812339.2;refGene_gn=DGKZ;refGene_tx=X
M_019812341.2;refGene_gn=DGKZ;refGene_tx=XM_019812342.2;refGene_gn=DGK
Z;refGene_tx=XM_023239809.1;refGene_gn=DGKZ;refGene_tx=XM_023239810.1;
refGene_gn=DGKZ;refGene_tx=XM_023239811.1

D1 100495332 100495446
refGene_gn=DGKZ;refGene_tx=XM_003993206.5;refGene_gn=DGKZ;refGene_tx=XM_006937232.4;refGene_gn=DGKZ;refGene_tx=XM_006937233.4;refGene_gn=DGKZ;refGene_tx=XM_006937234.4;refGene_gn=DGKZ;refGene_tx=XM_019812338.2;refGene_gn=DGKZ;refGene_tx=XM_019812339.2;refGene_gn=DGKZ;refGene_tx=XM_019812341.2;refGene_gn=DGKZ;refGene_tx=XM_019812342.2;refGene_gn=DGKZ;refGene_tx=XM_023239809.1;refGene_gn=DGKZ;refGene_tx=XM_023239810.1;refGene_gn=DGKZ;refGene_tx=XM_023239811.1
D1 100495547 100495688
refGene_gn=DGKZ;refGene_tx=XM_003993206.5;refGene_gn=DGKZ;refGene_tx=XM_006937232.4;refGene_gn=DGKZ;refGene_tx=XM_006937233.4;refGene_gn=DGKZ;refGene_tx=XM_006937234.4;refGene_gn=DGKZ;refGene_tx=XM_019812338.2;refGene_gn=DGKZ;refGene_tx=XM_019812339.2;refGene_gn=DGKZ;refGene_tx=XM_019812341.2;refGene_gn=DGKZ;refGene_tx=XM_019812342.2;refGene_gn=DGKZ;refGene_tx=XM_023239809.1;refGene_gn=DGKZ;refGene_tx=XM_023239810.1;refGene_gn=DGKZ;refGene_tx=XM_023239811.1
D1 100496281 100496360
refGene_gn=DGKZ;refGene_tx=XM_003993206.5;refGene_gn=DGKZ;refGene_tx=XM_006937232.4;refGene_gn=DGKZ;refGene_tx=XM_006937233.4;refGene_gn=DGKZ;refGene_tx=XM_006937234.4;refGene_gn=DGKZ;refGene_tx=XM_019812338.2;refGene_gn=DGKZ;refGene_tx=XM_019812339.2;refGene_gn=DGKZ;refGene_tx=XM_019812341.2;refGene_gn=DGKZ;refGene_tx=XM_019812342.2;refGene_gn=DGKZ;refGene_tx=XM_023239809.1;refGene_gn=DGKZ;refGene_tx=XM_023239810.1;refGene_gn=DGKZ;refGene_tx=XM_023239811.1
D1 100496555 100496611
refGene_gn=DGKZ;refGene_tx=XM_003993206.5;refGene_gn=DGKZ;refGene_tx=XM_006937232.4;refGene_gn=DGKZ;refGene_tx=XM_006937233.4;refGene_gn=DGKZ;refGene_tx=XM_006937234.4;refGene_gn=DGKZ;refGene_tx=XM_019812338.2;refGene_gn=DGKZ;refGene_tx=XM_019812339.2;refGene_gn=DGKZ;refGene_tx=XM_019812341.2;refGene_gn=DGKZ;refGene_tx=XM_019812342.2;refGene_gn=DGKZ;refGene_tx=XM_023239809.1;refGene_gn=DGKZ;refGene_tx=XM_023239810.1;refGene_gn=DGKZ;refGene_tx=XM_023239811.1
D1 100496769 100496832
refGene_gn=DGKZ;refGene_tx=XM_003993206.5;refGene_gn=DGKZ;refGene_tx=XM_006937232.4;refGene_gn=DGKZ;refGene_tx=XM_006937233.4;refGene_gn=DGKZ;refGene_tx=XM_006937234.4;refGene_gn=DGKZ;refGene_tx=XM_019812338.2;refGene_gn=DGKZ;refGene_tx=XM_019812339.2;refGene_gn=DGKZ;refGene_tx=XM_019812341.2;refGene_gn=DGKZ;refGene_tx=XM_019812342.2;refGene_gn=DGKZ;refGene_tx=XM_023239809.1;refGene_gn=DGKZ;refGene_tx=XM_023239810.1;refGene_gn=DGKZ;refGene_tx=XM_023239811.1
D1 100496935 100497009
refGene_gn=DGKZ;refGene_tx=XM_003993206.5;refGene_gn=DGKZ;refGene_tx=XM_006937232.4;refGene_gn=DGKZ;refGene_tx=XM_006937233.4;refGene_gn=DGKZ;refGene_tx=XM_006937234.4;refGene_gn=DGKZ;refGene_tx=XM_019812338.2;refGene_gn=DGKZ;refGene_tx=XM_019812339.2;refGene_gn=DGKZ;refGene_tx=XM_019812341.2;refGene_gn=DGKZ;refGene_tx=XM_019812342.2;refGene_gn=DGKZ;refGene_tx=XM_023239809.1;refGene_gn=DGKZ;refGene_tx=XM_023239810.1;refGene_gn=DGKZ;refGene_tx=XM_023239811.1
D1 100497097 100497209
refGene_gn=DGKZ;refGene_tx=XM_003993206.5;refGene_gn=DGKZ;refGene_tx=X

M_006937232.4;refGene_gn=DGKZ;refGene_tx=XM_006937233.4;refGene_gn=DGK
Z;refGene_tx=XM_006937234.4;refGene_gn=DGKZ;refGene_tx=XM_019812338.2;
refGene_gn=DGKZ;refGene_tx=XM_019812339.2;refGene_gn=DGKZ;refGene_tx=X
M_019812341.2;refGene_gn=DGKZ;refGene_tx=XM_019812342.2;refGene_gn=DGK
Z;refGene_tx=XM_023239809.1;refGene_gn=DGKZ;refGene_tx=XM_023239810.1;
refGene_gn=DGKZ;refGene_tx=XM_023239811.1
D1 100497577 100497777
refGene_gn=DGKZ;refGene_tx=XM_003993206.5;refGene_gn=DGKZ;refGene_tx=X
M_006937232.4;refGene_gn=DGKZ;refGene_tx=XM_006937233.4;refGene_gn=DGK
Z;refGene_tx=XM_006937234.4;refGene_gn=DGKZ;refGene_tx=XM_019812338.2;
refGene_gn=DGKZ;refGene_tx=XM_019812339.2;refGene_gn=DGKZ;refGene_tx=X
M_019812341.2;refGene_gn=DGKZ;refGene_tx=XM_019812342.2;refGene_gn=DGK
Z;refGene_tx=XM_023239809.1;refGene_gn=DGKZ;refGene_tx=XM_023239810.1;
refGene_gn=DGKZ;refGene_tx=XM_023239811.1
D1 100497980 100498081
refGene_gn=DGKZ;refGene_tx=XM_003993206.5;refGene_gn=DGKZ;refGene_tx=X
M_006937232.4;refGene_gn=DGKZ;refGene_tx=XM_006937233.4;refGene_gn=DGK
Z;refGene_tx=XM_006937234.4;refGene_gn=DGKZ;refGene_tx=XM_019812338.2;
refGene_gn=DGKZ;refGene_tx=XM_019812339.2;refGene_gn=DGKZ;refGene_tx=X
M_019812341.2;refGene_gn=DGKZ;refGene_tx=XM_019812342.2;refGene_gn=DGK
Z;refGene_tx=XM_023239809.1;refGene_gn=DGKZ;refGene_tx=XM_023239810.1;
refGene_gn=DGKZ;refGene_tx=XM_023239811.1
D1 100498182 100498262
refGene_gn=DGKZ;refGene_tx=XM_003993206.5;refGene_gn=DGKZ;refGene_tx=X
M_006937232.4;refGene_gn=DGKZ;refGene_tx=XM_006937233.4;refGene_gn=DGK
Z;refGene_tx=XM_006937234.4;refGene_gn=DGKZ;refGene_tx=XM_019812338.2;
refGene_gn=DGKZ;refGene_tx=XM_019812339.2;refGene_gn=DGKZ;refGene_tx=X
M_019812341.2;refGene_gn=DGKZ;refGene_tx=XM_019812342.2;refGene_gn=DGK
Z;refGene_tx=XM_023239809.1;refGene_gn=DGKZ;refGene_tx=XM_023239810.1;
refGene_gn=DGKZ;refGene_tx=XM_023239811.1
D1 100498351 100498424
refGene_gn=DGKZ;refGene_tx=XM_003993206.5;refGene_gn=DGKZ;refGene_tx=X
M_006937232.4;refGene_gn=DGKZ;refGene_tx=XM_006937233.4;refGene_gn=DGK
Z;refGene_tx=XM_006937234.4;refGene_gn=DGKZ;refGene_tx=XM_019812338.2;
refGene_gn=DGKZ;refGene_tx=XM_019812339.2;refGene_gn=DGKZ;refGene_tx=X
M_019812341.2;refGene_gn=DGKZ;refGene_tx=XM_019812342.2;refGene_gn=DGK
Z;refGene_tx=XM_023239809.1;refGene_gn=DGKZ;refGene_tx=XM_023239810.1;
refGene_gn=DGKZ;refGene_tx=XM_023239811.1
D1 100498567 100498608
refGene_gn=DGKZ;refGene_tx=XM_003993206.5;refGene_gn=DGKZ;refGene_tx=X
M_006937232.4;refGene_gn=DGKZ;refGene_tx=XM_006937233.4;refGene_gn=DGK
Z;refGene_tx=XM_006937234.4;refGene_gn=DGKZ;refGene_tx=XM_019812338.2;
refGene_gn=DGKZ;refGene_tx=XM_019812339.2;refGene_gn=DGKZ;refGene_tx=X
M_019812341.2;refGene_gn=DGKZ;refGene_tx=XM_019812342.2;refGene_gn=DGK
Z;refGene_tx=XM_023239809.1;refGene_gn=DGKZ;refGene_tx=XM_023239810.1;
refGene_gn=DGKZ;refGene_tx=XM_023239811.1
D1 100499115 100499255
refGene_gn=DGKZ;refGene_tx=XM_003993206.5;refGene_gn=DGKZ;refGene_tx=X
M_006937232.4;refGene_gn=DGKZ;refGene_tx=XM_006937233.4;refGene_gn=DGK
Z;refGene_tx=XM_006937234.4;refGene_gn=DGKZ;refGene_tx=XM_019812338.2;

refGene_gn=DGKZ;refGene_tx=XM_019812339.2;refGene_gn=DGKZ;refGene_tx=XM_019812341.2;refGene_gn=DGKZ;refGene_tx=XM_019812342.2;refGene_gn=DGKZ;refGene_tx=XM_023239809.1;refGene_gn=DGKZ;refGene_tx=XM_023239810.1;refGene_gn=DGKZ;refGene_tx=XM_023239811.1
D1 100500038 100500070
refGene_gn=DGKZ;refGene_tx=XM_003993206.5;refGene_gn=DGKZ;refGene_tx=XM_006937232.4;refGene_gn=DGKZ;refGene_tx=XM_006937233.4;refGene_gn=DGKZ;refGene_tx=XM_006937234.4;refGene_gn=DGKZ;refGene_tx=XM_019812338.2;refGene_gn=DGKZ;refGene_tx=XM_019812339.2;refGene_gn=DGKZ;refGene_tx=XM_019812341.2;refGene_gn=DGKZ;refGene_tx=XM_019812342.2;refGene_gn=DGKZ;refGene_tx=XM_023239809.1;refGene_gn=DGKZ;refGene_tx=XM_023239810.1;refGene_gn=DGKZ;refGene_tx=XM_023239811.1
D1 100500285 100500329
refGene_gn=DGKZ;refGene_tx=XM_003993206.5;refGene_gn=DGKZ;refGene_tx=XM_006937232.4;refGene_gn=DGKZ;refGene_tx=XM_006937233.4;refGene_gn=DGKZ;refGene_tx=XM_006937234.4;refGene_gn=DGKZ;refGene_tx=XM_019812338.2;refGene_gn=DGKZ;refGene_tx=XM_019812339.2;refGene_gn=DGKZ;refGene_tx=XM_019812341.2;refGene_gn=DGKZ;refGene_tx=XM_019812342.2;refGene_gn=DGKZ;refGene_tx=XM_023239809.1;refGene_gn=DGKZ;refGene_tx=XM_023239810.1;refGene_gn=DGKZ;refGene_tx=XM_023239811.1
D1 100500791 100500912
refGene_gn=DGKZ;refGene_tx=XM_003993206.5;refGene_gn=DGKZ;refGene_tx=XM_006937232.4;refGene_gn=DGKZ;refGene_tx=XM_006937233.4;refGene_gn=DGKZ;refGene_tx=XM_006937234.4;refGene_gn=DGKZ;refGene_tx=XM_019812338.2;refGene_gn=DGKZ;refGene_tx=XM_019812339.2;refGene_gn=DGKZ;refGene_tx=XM_019812341.2;refGene_gn=DGKZ;refGene_tx=XM_019812342.2;refGene_gn=DGKZ;refGene_tx=XM_023239809.1;refGene_gn=DGKZ;refGene_tx=XM_023239810.1;refGene_gn=DGKZ;refGene_tx=XM_023239811.1
D1 100501008 100501042
refGene_gn=DGKZ;refGene_tx=XM_003993206.5;refGene_gn=DGKZ;refGene_tx=XM_006937232.4;refGene_gn=DGKZ;refGene_tx=XM_006937233.4;refGene_gn=DGKZ;refGene_tx=XM_006937234.4;refGene_gn=DGKZ;refGene_tx=XM_019812338.2;refGene_gn=DGKZ;refGene_tx=XM_019812339.2;refGene_gn=DGKZ;refGene_tx=XM_019812341.2;refGene_gn=DGKZ;refGene_tx=XM_019812342.2;refGene_gn=DGKZ;refGene_tx=XM_023239809.1;refGene_gn=DGKZ;refGene_tx=XM_023239810.1;refGene_gn=DGKZ;refGene_tx=XM_023239811.1
D1 100501267 100501367
refGene_gn=DGKZ;refGene_tx=XM_003993206.5;refGene_gn=DGKZ;refGene_tx=XM_006937232.4;refGene_gn=DGKZ;refGene_tx=XM_006937233.4;refGene_gn=DGKZ;refGene_tx=XM_006937234.4;refGene_gn=DGKZ;refGene_tx=XM_019812338.2;refGene_gn=DGKZ;refGene_tx=XM_019812339.2;refGene_gn=DGKZ;refGene_tx=XM_019812341.2;refGene_gn=DGKZ;refGene_tx=XM_019812342.2;refGene_gn=DGKZ;refGene_tx=XM_023239809.1;refGene_gn=DGKZ;refGene_tx=XM_023239810.1;refGene_gn=DGKZ;refGene_tx=XM_023239811.1
D1 100501671 100501788
refGene_gn=DGKZ;refGene_tx=XM_003993206.5;refGene_gn=DGKZ;refGene_tx=XM_006937232.4;refGene_gn=DGKZ;refGene_tx=XM_006937233.4;refGene_gn=DGKZ;refGene_tx=XM_006937234.4;refGene_gn=DGKZ;refGene_tx=XM_019812338.2;refGene_gn=DGKZ;refGene_tx=XM_019812339.2;refGene_gn=DGKZ;refGene_tx=XM_019812341.2;refGene_gn=DGKZ;refGene_tx=XM_019812342.2;refGene_gn=DGK

Z;refGene_tx=XM_023239809.1;refGene_gn=DGKZ;refGene_tx=XM_023239810.1;
refGene_gn=DGKZ;refGene_tx=XM_023239811.1
D1 100504061 100504137
refGene_gn=MDK;refGene_tx=XM_023239812.1
D1 100504301 100504469
refGene_gn=MDK;refGene_tx=XM_023239812.1
D1 100504588 100504750
refGene_gn=MDK;refGene_tx=XM_023239812.1
D1 100505447 100505473
refGene_gn=MDK;refGene_tx=XM_023239812.1
D1 100506925 100508362
refGene_gn=CHRM4;refGene_tx=XM_003993210.4;refGene_gn=CHRM4;refGene_tx
=XM_019812344.2;refGene_gn=CHRM4;refGene_tx=XM_019812345.2
D1 100518003 100518506
refGene_gn=AMBRA1;refGene_tx=XM_011286774.3;refGene_gn=AMBRA1;refGene_
tx=XM_011286775.3;refGene_gn=AMBRA1;refGene_tx=XM_011286778.3;refGene_
gn=AMBRA1;refGene_tx=XM_011286779.3;refGene_gn=AMBRA1;refGene_tx=XM_01
1286780.3;refGene_gn=AMBRA1;refGene_tx=XM_011286781.3;refGene_gn=AMBRA
1;refGene_tx=XM_019812337.2
D1 100524348 100524542
refGene_gn=AMBRA1;refGene_tx=XM_011286774.3;refGene_gn=AMBRA1;refGene_
tx=XM_011286775.3;refGene_gn=AMBRA1;refGene_tx=XM_011286778.3;refGene_
gn=AMBRA1;refGene_tx=XM_011286779.3;refGene_gn=AMBRA1;refGene_tx=XM_01
1286780.3;refGene_gn=AMBRA1;refGene_tx=XM_011286781.3;refGene_gn=AMBRA
1;refGene_tx=XM_019812337.2
D1 100526087 100526180
refGene_gn=AMBRA1;refGene_tx=XM_011286774.3;refGene_gn=AMBRA1;refGene_
tx=XM_011286775.3;refGene_gn=AMBRA1;refGene_tx=XM_011286778.3;refGene_
gn=AMBRA1;refGene_tx=XM_011286779.3;refGene_gn=AMBRA1;refGene_tx=XM_01
1286780.3;refGene_gn=AMBRA1;refGene_tx=XM_011286781.3
D1 100533779 100533919
refGene_gn=AMBRA1;refGene_tx=XM_011286774.3;refGene_gn=AMBRA1;refGene_
tx=XM_011286775.3;refGene_gn=AMBRA1;refGene_tx=XM_011286778.3;refGene_
gn=AMBRA1;refGene_tx=XM_011286779.3;refGene_gn=AMBRA1;refGene_tx=XM_01
1286780.3;refGene_gn=AMBRA1;refGene_tx=XM_011286781.3;refGene_gn=AMBRA
1;refGene_tx=XM_019812337.2
D1 100547782 100547937
refGene_gn=AMBRA1;refGene_tx=XM_011286774.3;refGene_gn=AMBRA1;refGene_
tx=XM_011286775.3;refGene_gn=AMBRA1;refGene_tx=XM_011286778.3;refGene_
gn=AMBRA1;refGene_tx=XM_011286779.3;refGene_gn=AMBRA1;refGene_tx=XM_01
1286780.3;refGene_gn=AMBRA1;refGene_tx=XM_011286781.3;refGene_gn=AMBRA
1;refGene_tx=XM_019812337.2
D1 100548781 100548970
refGene_gn=AMBRA1;refGene_tx=XM_011286774.3;refGene_gn=AMBRA1;refGene_
tx=XM_011286775.3;refGene_gn=AMBRA1;refGene_tx=XM_011286778.3;refGene_
gn=AMBRA1;refGene_tx=XM_011286779.3;refGene_gn=AMBRA1;refGene_tx=XM_01
1286780.3;refGene_gn=AMBRA1;refGene_tx=XM_011286781.3;refGene_gn=AMBRA
1;refGene_tx=XM_019812337.2
D1 100558960 100559071
refGene_gn=AMBRA1;refGene_tx=XM_011286774.3;refGene_gn=AMBRA1;refGene_

tx=XM_011286775.3;refGene_gn=AMBRA1;refGene_tx=XM_011286778.3;refGene_gn=AMBRA1;refGene_tx=XM_011286779.3;refGene_gn=AMBRA1;refGene_tx=XM_011286780.3;refGene_gn=AMBRA1;refGene_tx=XM_011286781.3;refGene_gn=AMBRA1;refGene_tx=XM_019812337.2
D1 100610961 100611062
refGene_gn=AMBRA1;refGene_tx=XM_011286774.3;refGene_gn=AMBRA1;refGene_tx=XM_011286775.3;refGene_gn=AMBRA1;refGene_tx=XM_011286778.3;refGene_gn=AMBRA1;refGene_tx=XM_011286779.3;refGene_gn=AMBRA1;refGene_tx=XM_011286780.3;refGene_gn=AMBRA1;refGene_tx=XM_011286781.3;refGene_gn=AMBRA1;refGene_tx=XM_019812337.2
D1 100611487 100611568
refGene_gn=AMBRA1;refGene_tx=XM_011286774.3;refGene_gn=AMBRA1;refGene_tx=XM_011286775.3;refGene_gn=AMBRA1;refGene_tx=XM_011286778.3;refGene_gn=AMBRA1;refGene_tx=XM_011286779.3;refGene_gn=AMBRA1;refGene_tx=XM_011286780.3;refGene_gn=AMBRA1;refGene_tx=XM_011286781.3;refGene_gn=AMBRA1;refGene_tx=XM_019812337.2
D1 100623792 100623972
refGene_gn=AMBRA1;refGene_tx=XM_011286774.3;refGene_gn=AMBRA1;refGene_tx=XM_011286775.3;refGene_gn=AMBRA1;refGene_tx=XM_011286778.3;refGene_gn=AMBRA1;refGene_tx=XM_011286780.3;refGene_gn=AMBRA1;refGene_tx=XM_011286781.3;refGene_gn=AMBRA1;refGene_tx=XM_019812337.2
D1 100628184 100628271
refGene_gn=AMBRA1;refGene_tx=XM_011286774.3;refGene_gn=AMBRA1;refGene_tx=XM_011286775.3;refGene_gn=AMBRA1;refGene_tx=XM_011286780.3;refGene_gn=AMBRA1;refGene_tx=XM_019812337.2
D1 100659929 100661383
refGene_gn=AMBRA1;refGene_tx=XM_011286774.3;refGene_gn=AMBRA1;refGene_tx=XM_011286775.3;refGene_gn=AMBRA1;refGene_tx=XM_011286778.3;refGene_gn=AMBRA1;refGene_tx=XM_011286779.3;refGene_gn=AMBRA1;refGene_tx=XM_011286780.3;refGene_gn=AMBRA1;refGene_tx=XM_011286781.3;refGene_gn=AMBRA1;refGene_tx=XM_019812337.2
D1 100661937 100662004
refGene_gn=AMBRA1;refGene_tx=XM_011286774.3;refGene_gn=AMBRA1;refGene_tx=XM_011286775.3;refGene_gn=AMBRA1;refGene_tx=XM_011286778.3;refGene_gn=AMBRA1;refGene_tx=XM_011286779.3;refGene_gn=AMBRA1;refGene_tx=XM_011286780.3;refGene_gn=AMBRA1;refGene_tx=XM_011286781.3;refGene_gn=AMBRA1;refGene_tx=XM_019812337.2
D1 100662942 100663115
refGene_gn=AMBRA1;refGene_tx=XM_011286774.3;refGene_gn=AMBRA1;refGene_tx=XM_011286775.3;refGene_gn=AMBRA1;refGene_tx=XM_011286778.3;refGene_gn=AMBRA1;refGene_tx=XM_011286779.3;refGene_gn=AMBRA1;refGene_tx=XM_011286780.3;refGene_gn=AMBRA1;refGene_tx=XM_011286781.3;refGene_gn=AMBRA1;refGene_tx=XM_019812337.2
D1 100663748 100663932
refGene_gn=AMBRA1;refGene_tx=XM_011286774.3;refGene_gn=AMBRA1;refGene_tx=XM_011286775.3;refGene_gn=AMBRA1;refGene_tx=XM_011286778.3;refGene_gn=AMBRA1;refGene_tx=XM_011286779.3;refGene_gn=AMBRA1;refGene_tx=XM_011286780.3;refGene_gn=AMBRA1;refGene_tx=XM_011286781.3;refGene_gn=AMBRA1;refGene_tx=XM_019812337.2
D1 100664431 100664490

refGene_gn=AMBRA1;refGene_tx=XM_011286774.3;refGene_gn=AMBRA1;refGene_tx=XM_011286775.3;refGene_gn=AMBRA1;refGene_tx=XM_011286778.3;refGene_gn=AMBRA1;refGene_tx=XM_011286779.3;refGene_gn=AMBRA1;refGene_tx=XM_011286780.3;refGene_gn=AMBRA1;refGene_tx=XM_011286781.3;refGene_gn=AMBRA1;refGene_tx=XM_019812337.2
D1	100664894	100665029
refGene_gn=AMBRA1;refGene_tx=XM_011286774.3;refGene_gn=AMBRA1;refGene_tx=XM_011286775.3;refGene_gn=AMBRA1;refGene_tx=XM_011286778.3;refGene_gn=AMBRA1;refGene_tx=XM_011286779.3;refGene_gn=AMBRA1;refGene_tx=XM_011286780.3;refGene_gn=AMBRA1;refGene_tx=XM_011286781.3;refGene_gn=AMBRA1;refGene_tx=XM_019812337.2
D1	100705274	100705654
refGene_gn=HARBI1;refGene_tx=XM_019812353.2;refGene_gn=HARBI1;refGene_tx=XM_019812354.2
D1	100712322	100712992
refGene_gn=HARBI1;refGene_tx=XM_019812353.2;refGene_gn=HARBI1;refGene_tx=XM_019812354.2
D1	100742604	100742673
refGene_gn=ATG13;refGene_tx=XM_003993213.4;refGene_gn=ATG13;refGene_tx=XM_006937243.3;refGene_gn=ATG13;refGene_tx=XM_006937244.3;refGene_gn=ATG13;refGene_tx=XM_006937245.3;refGene_gn=ATG13;refGene_tx=XM_006937246.3;refGene_gn=ATG13;refGene_tx=XM_006937247.3;refGene_gn=ATG13;refGene_tx=XM_011286869.3;refGene_gn=ATG13;refGene_tx=XM_019812350.2;refGene_gn=ATG13;refGene_tx=XM_019812351.2
D1	100743637	100743718
refGene_gn=ATG13;refGene_tx=XM_003993213.4;refGene_gn=ATG13;refGene_tx=XM_006937243.3;refGene_gn=ATG13;refGene_tx=XM_006937244.3;refGene_gn=ATG13;refGene_tx=XM_006937245.3;refGene_gn=ATG13;refGene_tx=XM_006937246.3;refGene_gn=ATG13;refGene_tx=XM_006937247.3;refGene_gn=ATG13;refGene_tx=XM_011286869.3;refGene_gn=ATG13;refGene_tx=XM_019812350.2;refGene_gn=ATG13;refGene_tx=XM_019812351.2
D1	100744152	100744272
refGene_gn=ATG13;refGene_tx=XM_003993213.4;refGene_gn=ATG13;refGene_tx=XM_006937243.3;refGene_gn=ATG13;refGene_tx=XM_006937244.3;refGene_gn=ATG13;refGene_tx=XM_006937245.3;refGene_gn=ATG13;refGene_tx=XM_006937246.3;refGene_gn=ATG13;refGene_tx=XM_006937247.3;refGene_gn=ATG13;refGene_tx=XM_011286869.3;refGene_gn=ATG13;refGene_tx=XM_019812350.2;refGene_gn=ATG13;refGene_tx=XM_019812351.2
D1	100747352	100747399
refGene_gn=ATG13;refGene_tx=XM_003993213.4;refGene_gn=ATG13;refGene_tx=XM_006937243.3;refGene_gn=ATG13;refGene_tx=XM_006937244.3;refGene_gn=ATG13;refGene_tx=XM_006937245.3;refGene_gn=ATG13;refGene_tx=XM_006937246.3;refGene_gn=ATG13;refGene_tx=XM_006937247.3;refGene_gn=ATG13;refGene_tx=XM_011286869.3;refGene_gn=ATG13;refGene_tx=XM_019812350.2;refGene_gn=ATG13;refGene_tx=XM_019812351.2
D1	100748552	100748693
refGene_gn=ATG13;refGene_tx=XM_003993213.4;refGene_gn=ATG13;refGene_tx=XM_006937243.3;refGene_gn=ATG13;refGene_tx=XM_006937244.3;refGene_gn=ATG13;refGene_tx=XM_006937245.3;refGene_gn=ATG13;refGene_tx=XM_006937246.3;refGene_gn=ATG13;refGene_tx=XM_006937247.3;refGene_gn=ATG13;refGe

ne_tx=XM_011286869.3;refGene_gn=ATG13;refGene_tx=XM_019812350.2;refGen
e_gn=ATG13;refGene_tx=XM_019812351.2
D1 100753861 100753902
refGene_gn=ATG13;refGene_tx=XM_003993213.4;refGene_gn=ATG13;refGene_tx
=XM_006937243.3;refGene_gn=ATG13;refGene_tx=XM_006937244.3;refGene_gn=
ATG13;refGene_tx=XM_006937245.3;refGene_gn=ATG13;refGene_tx=XM_0069372
46.3;refGene_gn=ATG13;refGene_tx=XM_006937247.3;refGene_gn=ATG13;refGe
ne_tx=XM_011286869.3;refGene_gn=ATG13;refGene_tx=XM_019812350.2;refGen
e_gn=ATG13;refGene_tx=XM_019812351.2
D1 100754668 100754765
refGene_gn=ATG13;refGene_tx=XM_003993213.4;refGene_gn=ATG13;refGene_tx
=XM_006937243.3;refGene_gn=ATG13;refGene_tx=XM_006937244.3;refGene_gn=
ATG13;refGene_tx=XM_006937245.3;refGene_gn=ATG13;refGene_tx=XM_0069372
46.3;refGene_gn=ATG13;refGene_tx=XM_006937247.3;refGene_gn=ATG13;refGe
ne_tx=XM_011286869.3;refGene_gn=ATG13;refGene_tx=XM_019812350.2;refGen
e_gn=ATG13;refGene_tx=XM_019812351.2
D1 100755109 100755208
refGene_gn=ATG13;refGene_tx=XM_003993213.4;refGene_gn=ATG13;refGene_tx
=XM_006937243.3;refGene_gn=ATG13;refGene_tx=XM_006937244.3;refGene_gn=
ATG13;refGene_tx=XM_006937245.3;refGene_gn=ATG13;refGene_tx=XM_0069372
46.3;refGene_gn=ATG13;refGene_tx=XM_006937247.3;refGene_gn=ATG13;refGe
ne_tx=XM_011286869.3;refGene_gn=ATG13;refGene_tx=XM_019812350.2;refGen
e_gn=ATG13;refGene_tx=XM_019812351.2
D1 100755996 100756090
refGene_gn=ATG13;refGene_tx=XM_003993213.4;refGene_gn=ATG13;refGene_tx
=XM_006937243.3;refGene_gn=ATG13;refGene_tx=XM_006937244.3;refGene_gn=
ATG13;refGene_tx=XM_006937245.3;refGene_gn=ATG13;refGene_tx=XM_0069372
46.3;refGene_gn=ATG13;refGene_tx=XM_006937247.3;refGene_gn=ATG13;refGe
ne_tx=XM_011286869.3;refGene_gn=ATG13;refGene_tx=XM_019812350.2;refGen
e_gn=ATG13;refGene_tx=XM_019812351.2
D1 100759665 100759764
refGene_gn=ATG13;refGene_tx=XM_003993213.4;refGene_gn=ATG13;refGene_tx
=XM_006937243.3;refGene_gn=ATG13;refGene_tx=XM_006937244.3;refGene_gn=
ATG13;refGene_tx=XM_006937245.3;refGene_gn=ATG13;refGene_tx=XM_0069372
46.3
D1 100760736 100760847
refGene_gn=ATG13;refGene_tx=XM_003993213.4;refGene_gn=ATG13;refGene_tx
=XM_006937243.3;refGene_gn=ATG13;refGene_tx=XM_006937244.3;refGene_gn=
ATG13;refGene_tx=XM_006937245.3;refGene_gn=ATG13;refGene_tx=XM_0069372
46.3;refGene_gn=ATG13;refGene_tx=XM_011286869.3;refGene_gn=ATG13;refGe
ne_tx=XM_019812350.2
D1 100761259 100761396
refGene_gn=ATG13;refGene_tx=XM_003993213.4;refGene_gn=ATG13;refGene_tx
=XM_006937243.3;refGene_gn=ATG13;refGene_tx=XM_006937244.3;refGene_gn=
ATG13;refGene_tx=XM_006937245.3;refGene_gn=ATG13;refGene_tx=XM_0069372
46.3;refGene_gn=ATG13;refGene_tx=XM_006937247.3;refGene_gn=ATG13;refGe
ne_tx=XM_011286869.3;refGene_gn=ATG13;refGene_tx=XM_019812350.2;refGen
e_gn=ATG13;refGene_tx=XM_019812351.2
D1 100762685 100762800
refGene_gn=ATG13;refGene_tx=XM_003993213.4;refGene_gn=ATG13;refGene_tx

=XM_006937243.3;refGene_gn=ATG13;refGene_tx=XM_006937244.3;refGene_gn=ATG13;refGene_tx=XM_006937245.3;refGene_gn=ATG13;refGene_tx=XM_006937246.3;refGene_gn=ATG13;refGene_tx=XM_006937247.3;refGene_gn=ATG13;refGene_tx=XM_011286869.3;refGene_gn=ATG13;refGene_tx=XM_019812350.2;refGene_gn=ATG13;refGene_tx=XM_019812351.2
D1 100763357 100763435
refGene_gn=ATG13;refGene_tx=XM_003993213.4;refGene_gn=ATG13;refGene_tx=XM_006937243.3;refGene_gn=ATG13;refGene_tx=XM_006937244.3;refGene_gn=ATG13;refGene_tx=XM_006937245.3;refGene_gn=ATG13;refGene_tx=XM_006937246.3;refGene_gn=ATG13;refGene_tx=XM_006937247.3;refGene_gn=ATG13;refGene_tx=XM_011286869.3;refGene_gn=ATG13;refGene_tx=XM_019812350.2;refGene_gn=ATG13;refGene_tx=XM_019812351.2
D1 100763639 100763756
refGene_gn=ATG13;refGene_tx=XM_003993213.4;refGene_gn=ATG13;refGene_tx=XM_006937243.3;refGene_gn=ATG13;refGene_tx=XM_006937244.3;refGene_gn=ATG13;refGene_tx=XM_006937245.3;refGene_gn=ATG13;refGene_tx=XM_006937246.3;refGene_gn=ATG13;refGene_tx=XM_006937247.3;refGene_gn=ATG13;refGene_tx=XM_011286869.3;refGene_gn=ATG13;refGene_tx=XM_019812350.2;refGene_gn=ATG13;refGene_tx=XM_019812351.2
D1 100764310 100764439
refGene_gn=ATG13;refGene_tx=XM_003993213.4;refGene_gn=ATG13;refGene_tx=XM_006937243.3;refGene_gn=ATG13;refGene_tx=XM_006937244.3;refGene_gn=ATG13;refGene_tx=XM_006937245.3;refGene_gn=ATG13;refGene_tx=XM_006937246.3;refGene_gn=ATG13;refGene_tx=XM_006937247.3;refGene_gn=ATG13;refGene_tx=XM_011286869.3;refGene_gn=ATG13;refGene_tx=XM_019812350.2;refGene_gn=ATG13;refGene_tx=XM_019812351.2
D1 100766518 100766596
refGene_gn=ATG13;refGene_tx=XM_003993213.4;refGene_gn=ATG13;refGene_tx=XM_006937243.3;refGene_gn=ATG13;refGene_tx=XM_006937244.3;refGene_gn=ATG13;refGene_tx=XM_006937245.3;refGene_gn=ATG13;refGene_tx=XM_006937246.3;refGene_gn=ATG13;refGene_tx=XM_006937247.3;refGene_gn=ATG13;refGene_tx=XM_011286869.3;refGene_gn=ATG13;refGene_tx=XM_019812350.2;refGene_gn=ATG13;refGene_tx=XM_019812351.2
D1 100772889 100773078
refGene_gn=ARHGAP1;refGene_tx=XM_006937248.3;refGene_gn=ARHGAP1;refGene_tx=XM_019812352.2
D1 100774124 100774228
refGene_gn=ARHGAP1;refGene_tx=XM_006937248.3;refGene_gn=ARHGAP1;refGene_tx=XM_019812352.2
D1 100774408 100774537
refGene_gn=ARHGAP1;refGene_tx=XM_006937248.3;refGene_gn=ARHGAP1;refGene_tx=XM_019812352.2
D1 100775014 100775092
refGene_gn=ARHGAP1;refGene_tx=XM_006937248.3;refGene_gn=ARHGAP1;refGene_tx=XM_019812352.2
D1 100775304 100775381
refGene_gn=ARHGAP1;refGene_tx=XM_006937248.3;refGene_gn=ARHGAP1;refGene_tx=XM_019812352.2
D1 100775465 100775573
refGene_gn=ARHGAP1;refGene_tx=XM_006937248.3;refGene_gn=ARHGAP1;refGen

e_tx=XM_019812352.2
D1 100775939 100776038
refGene_gn=ARHGAP1;refGene_tx=XM_006937248.3;refGene_gn=ARHGAP1;refGen
e_tx=XM_019812352.2
D1 100776241 100776328
refGene_gn=ARHGAP1;refGene_tx=XM_006937248.3;refGene_gn=ARHGAP1;refGen
e_tx=XM_019812352.2
D1 100776938 100777070
refGene_gn=ARHGAP1;refGene_tx=XM_006937248.3;refGene_gn=ARHGAP1;refGen
e_tx=XM_019812352.2
D1 100781066 100781154
refGene_gn=ARHGAP1;refGene_tx=XM_006937248.3;refGene_gn=ARHGAP1;refGen
e_tx=XM_019812352.2
D1 100788540 100788636
refGene_gn=ARHGAP1;refGene_tx=XM_006937248.3;refGene_gn=ARHGAP1;refGen
e_tx=XM_019812352.2
D1 100788863 100788996
refGene_gn=ARHGAP1;refGene_tx=XM_006937248.3;refGene_gn=ARHGAP1;refGen
e_tx=XM_019812352.2
D1 100793092 100793141
refGene_gn=ZNF408;refGene_tx=XM_003993217.4
D1 100793441 100793719
refGene_gn=ZNF408;refGene_tx=XM_003993217.4
D1 100794363 100794425
refGene_gn=ZNF408;refGene_tx=XM_003993217.4
D1 100794665 100794925
refGene_gn=ZNF408;refGene_tx=XM_003993217.4
D1 100796005 100797534
refGene_gn=ZNF408;refGene_tx=XM_003993217.4
D1 100807922 100808001
refGene_gn=F2;refGene_tx=XM_003993218.4
D1 100808374 100808538
refGene_gn=F2;refGene_tx=XM_003993218.4
D1 100809245 100809270
refGene_gn=F2;refGene_tx=XM_003993218.4
D1 100809571 100809622
refGene_gn=F2;refGene_tx=XM_003993218.4
D1 100811060 100811166
refGene_gn=F2;refGene_tx=XM_003993218.4
D1 100811277 100811414
refGene_gn=F2;refGene_tx=XM_003993218.4
D1 100812781 100813096
refGene_gn=F2;refGene_tx=XM_003993218.4
D1 100813428 100813554
refGene_gn=F2;refGene_tx=XM_003993218.4
D1 100813634 100813761
refGene_gn=F2;refGene_tx=XM_003993218.4
D1 100814441 100814609
refGene_gn=F2;refGene_tx=XM_003993218.4
D1 100815138 100815312

refGene_gn=F2;refGene_tx=XM_003993218.4
D1	100815536	100815718
refGene_gn=F2;refGene_tx=XM_003993218.4
D1	100823981	100824052
refGene_gn=F2;refGene_tx=XM_003993218.4
D1	100824193	100824337
refGene_gn=F2;refGene_tx=XM_003993218.4
D1	100827597	100827843
refGene_gn=CKAP5;refGene_tx=XM_011286866.2;refGene_gn=CKAP5;refGene_tx=XM_019812357.1;refGene_gn=CKAP5;refGene_tx=XM_019812358.1;refGene_gn=CKAP5;refGene_tx=XM_019812359.2
D1	100827946	100828098
refGene_gn=CKAP5;refGene_tx=XM_011286866.2;refGene_gn=CKAP5;refGene_tx=XM_019812357.1;refGene_gn=CKAP5;refGene_tx=XM_019812358.1;refGene_gn=CKAP5;refGene_tx=XM_019812359.2
D1	100831165	100831325
refGene_gn=CKAP5;refGene_tx=XM_011286866.2;refGene_gn=CKAP5;refGene_tx=XM_019812357.1;refGene_gn=CKAP5;refGene_tx=XM_019812358.1;refGene_gn=CKAP5;refGene_tx=XM_019812359.2
D1	100831420	100831504
refGene_gn=CKAP5;refGene_tx=XM_011286866.2;refGene_gn=CKAP5;refGene_tx=XM_019812357.1;refGene_gn=CKAP5;refGene_tx=XM_019812358.1;refGene_gn=CKAP5;refGene_tx=XM_019812359.2
D1	100831999	100832137
refGene_gn=CKAP5;refGene_tx=XM_011286866.2;refGene_gn=CKAP5;refGene_tx=XM_019812357.1;refGene_gn=CKAP5;refGene_tx=XM_019812358.1;refGene_gn=CKAP5;refGene_tx=XM_019812359.2
D1	100832225	100832414
refGene_gn=CKAP5;refGene_tx=XM_011286866.2;refGene_gn=CKAP5;refGene_tx=XM_019812357.1;refGene_gn=CKAP5;refGene_tx=XM_019812358.1;refGene_gn=CKAP5;refGene_tx=XM_019812359.2
D1	100833122	100833198
refGene_gn=CKAP5;refGene_tx=XM_011286866.2;refGene_gn=CKAP5;refGene_tx=XM_019812357.1;refGene_gn=CKAP5;refGene_tx=XM_019812358.1;refGene_gn=CKAP5;refGene_tx=XM_019812359.2
D1	100833950	100834138
refGene_gn=CKAP5;refGene_tx=XM_011286866.2;refGene_gn=CKAP5;refGene_tx=XM_019812357.1;refGene_gn=CKAP5;refGene_tx=XM_019812358.1;refGene_gn=CKAP5;refGene_tx=XM_019812359.2
D1	100834618	100834798
refGene_gn=CKAP5;refGene_tx=XM_011286866.2;refGene_gn=CKAP5;refGene_tx=XM_019812357.1;refGene_gn=CKAP5;refGene_tx=XM_019812358.1;refGene_gn=CKAP5;refGene_tx=XM_019812359.2
D1	100847091	100847212
refGene_gn=CKAP5;refGene_tx=XM_011286866.2;refGene_gn=CKAP5;refGene_tx=XM_019812357.1;refGene_gn=CKAP5;refGene_tx=XM_019812358.1;refGene_gn=CKAP5;refGene_tx=XM_019812359.2
D1	100847434	100847608
refGene_gn=CKAP5;refGene_tx=XM_011286866.2;refGene_gn=CKAP5;refGene_tx=XM_019812357.1;refGene_gn=CKAP5;refGene_tx=XM_019812358.1;refGene_gn=

CKAP5;refGene_tx=XM_019812359.2
D1	100848543	100848567
refGene_gn=CKAP5;refGene_tx=XM_019812357.1;refGene_gn=CKAP5;refGene_tx=XM_019812358.1;refGene_gn=CKAP5;refGene_tx=XM_019812359.2
D1	100848803	100848976
refGene_gn=CKAP5;refGene_tx=XM_011286866.2;refGene_gn=CKAP5;refGene_tx=XM_019812357.1;refGene_gn=CKAP5;refGene_tx=XM_019812358.1;refGene_gn=CKAP5;refGene_tx=XM_019812359.2
D1	100850141	100850335
refGene_gn=CKAP5;refGene_tx=XM_011286866.2;refGene_gn=CKAP5;refGene_tx=XM_019812357.1;refGene_gn=CKAP5;refGene_tx=XM_019812358.1;refGene_gn=CKAP5;refGene_tx=XM_019812359.2
D1	100850777	100850913
refGene_gn=CKAP5;refGene_tx=XM_011286866.2;refGene_gn=CKAP5;refGene_tx=XM_019812357.1;refGene_gn=CKAP5;refGene_tx=XM_019812358.1;refGene_gn=CKAP5;refGene_tx=XM_019812359.2
D1	100851123	100851327
refGene_gn=CKAP5;refGene_tx=XM_011286866.2;refGene_gn=CKAP5;refGene_tx=XM_019812357.1;refGene_gn=CKAP5;refGene_tx=XM_019812358.1;refGene_gn=CKAP5;refGene_tx=XM_019812359.2
D1	100851605	100851755
refGene_gn=CKAP5;refGene_tx=XM_011286866.2;refGene_gn=CKAP5;refGene_tx=XM_019812357.1;refGene_gn=CKAP5;refGene_tx=XM_019812358.1;refGene_gn=CKAP5;refGene_tx=XM_019812359.2
D1	100853006	100853132
refGene_gn=CKAP5;refGene_tx=XM_011286866.2;refGene_gn=CKAP5;refGene_tx=XM_019812357.1;refGene_gn=CKAP5;refGene_tx=XM_019812358.1;refGene_gn=CKAP5;refGene_tx=XM_019812359.2
D1	100855602	100855691
refGene_gn=CKAP5;refGene_tx=XM_011286866.2;refGene_gn=CKAP5;refGene_tx=XM_019812357.1;refGene_gn=CKAP5;refGene_tx=XM_019812358.1;refGene_gn=CKAP5;refGene_tx=XM_019812359.2
D1	100857002	100857138
refGene_gn=CKAP5;refGene_tx=XM_011286866.2;refGene_gn=CKAP5;refGene_tx=XM_019812357.1;refGene_gn=CKAP5;refGene_tx=XM_019812358.1;refGene_gn=CKAP5;refGene_tx=XM_019812359.2
D1	100857511	100857706
refGene_gn=CKAP5;refGene_tx=XM_011286866.2;refGene_gn=CKAP5;refGene_tx=XM_019812357.1;refGene_gn=CKAP5;refGene_tx=XM_019812358.1;refGene_gn=CKAP5;refGene_tx=XM_019812359.2
D1	100862063	100862192
refGene_gn=CKAP5;refGene_tx=XM_011286866.2;refGene_gn=CKAP5;refGene_tx=XM_019812357.1;refGene_gn=CKAP5;refGene_tx=XM_019812358.1;refGene_gn=CKAP5;refGene_tx=XM_019812359.2
D1	100863231	100863345
refGene_gn=CKAP5;refGene_tx=XM_011286866.2;refGene_gn=CKAP5;refGene_tx=XM_019812357.1;refGene_gn=CKAP5;refGene_tx=XM_019812358.1;refGene_gn=CKAP5;refGene_tx=XM_019812359.2
D1	100864128	100864303
refGene_gn=CKAP5;refGene_tx=XM_011286866.2;refGene_gn=CKAP5;refGene_tx

=XM_019812357.1;refGene_gn=CKAP5;refGene_tx=XM_019812358.1;refGene_gn=CKAP5;refGene_tx=XM_019812359.2
D1	100864483	100864623
refGene_gn=CKAP5;refGene_tx=XM_011286866.2;refGene_gn=CKAP5;refGene_tx=XM_019812357.1;refGene_gn=CKAP5;refGene_tx=XM_019812358.1;refGene_gn=CKAP5;refGene_tx=XM_019812359.2
D1	100865240	100865366
refGene_gn=CKAP5;refGene_tx=XM_011286866.2;refGene_gn=CKAP5;refGene_tx=XM_019812357.1;refGene_gn=CKAP5;refGene_tx=XM_019812358.1;refGene_gn=CKAP5;refGene_tx=XM_019812359.2
D1	100865455	100865513
refGene_gn=CKAP5;refGene_tx=XM_011286866.2;refGene_gn=CKAP5;refGene_tx=XM_019812357.1;refGene_gn=CKAP5;refGene_tx=XM_019812358.1;refGene_gn=CKAP5;refGene_tx=XM_019812359.2
D1	100867981	100868076
refGene_gn=CKAP5;refGene_tx=XM_011286866.2;refGene_gn=CKAP5;refGene_tx=XM_019812357.1;refGene_gn=CKAP5;refGene_tx=XM_019812358.1;refGene_gn=CKAP5;refGene_tx=XM_019812359.2
D1	100869229	100869415
refGene_gn=CKAP5;refGene_tx=XM_011286866.2;refGene_gn=CKAP5;refGene_tx=XM_019812357.1;refGene_gn=CKAP5;refGene_tx=XM_019812358.1;refGene_gn=CKAP5;refGene_tx=XM_019812359.2
D1	100872592	100872685
refGene_gn=CKAP5;refGene_tx=XM_011286866.2;refGene_gn=CKAP5;refGene_tx=XM_019812357.1;refGene_gn=CKAP5;refGene_tx=XM_019812358.1;refGene_gn=CKAP5;refGene_tx=XM_019812359.2
D1	100873656	100873767
refGene_gn=CKAP5;refGene_tx=XM_011286866.2;refGene_gn=CKAP5;refGene_tx=XM_019812357.1;refGene_gn=CKAP5;refGene_tx=XM_019812358.1;refGene_gn=CKAP5;refGene_tx=XM_019812359.2
D1	100874057	100874171
refGene_gn=CKAP5;refGene_tx=XM_011286866.2;refGene_gn=CKAP5;refGene_tx=XM_019812357.1;refGene_gn=CKAP5;refGene_tx=XM_019812358.1;refGene_gn=CKAP5;refGene_tx=XM_019812359.2
D1	100876897	100877080
refGene_gn=CKAP5;refGene_tx=XM_011286866.2;refGene_gn=CKAP5;refGene_tx=XM_019812357.1;refGene_gn=CKAP5;refGene_tx=XM_019812358.1;refGene_gn=CKAP5;refGene_tx=XM_019812359.2
D1	100877634	100877763
refGene_gn=CKAP5;refGene_tx=XM_011286866.2;refGene_gn=CKAP5;refGene_tx=XM_019812357.1;refGene_gn=CKAP5;refGene_tx=XM_019812358.1;refGene_gn=CKAP5;refGene_tx=XM_019812359.2
D1	100878329	100878494
refGene_gn=CKAP5;refGene_tx=XM_011286866.2;refGene_gn=CKAP5;refGene_tx=XM_019812357.1;refGene_gn=CKAP5;refGene_tx=XM_019812358.1;refGene_gn=CKAP5;refGene_tx=XM_019812359.2
D1	100878606	100878696
refGene_gn=CKAP5;refGene_tx=XM_011286866.2;refGene_gn=CKAP5;refGene_tx=XM_019812357.1;refGene_gn=CKAP5;refGene_tx=XM_019812358.1;refGene_gn=CKAP5;refGene_tx=XM_019812359.2

D1 100881184 100881289
refGene_gn=CKAP5;refGene_tx=XM_011286866.2;refGene_gn=CKAP5;refGene_tx=XM_019812357.1;refGene_gn=CKAP5;refGene_tx=XM_019812358.1;refGene_gn=CKAP5;refGene_tx=XM_019812359.2
D1 100888323 100888437
refGene_gn=CKAP5;refGene_tx=XM_011286866.2;refGene_gn=CKAP5;refGene_tx=XM_019812357.1;refGene_gn=CKAP5;refGene_tx=XM_019812358.1;refGene_gn=CKAP5;refGene_tx=XM_019812359.2
D1 100890190 100890291
refGene_gn=CKAP5;refGene_tx=XM_011286866.2;refGene_gn=CKAP5;refGene_tx=XM_019812357.1;refGene_gn=CKAP5;refGene_tx=XM_019812358.1;refGene_gn=CKAP5;refGene_tx=XM_019812359.2
D1 100890523 100890656
refGene_gn=CKAP5;refGene_tx=XM_011286866.2;refGene_gn=CKAP5;refGene_tx=XM_019812357.1;refGene_gn=CKAP5;refGene_tx=XM_019812358.1;refGene_gn=CKAP5;refGene_tx=XM_019812359.2
D1 100891294 100891466
refGene_gn=CKAP5;refGene_tx=XM_011286866.2;refGene_gn=CKAP5;refGene_tx=XM_019812357.1;refGene_gn=CKAP5;refGene_tx=XM_019812358.1;refGene_gn=CKAP5;refGene_tx=XM_019812359.2
D1 100896861 100897068
refGene_gn=CKAP5;refGene_tx=XM_011286866.2;refGene_gn=CKAP5;refGene_tx=XM_019812357.1;refGene_gn=CKAP5;refGene_tx=XM_019812358.1;refGene_gn=CKAP5;refGene_tx=XM_019812359.2
D1 100898478 100898672
refGene_gn=CKAP5;refGene_tx=XM_011286866.2;refGene_gn=CKAP5;refGene_tx=XM_019812357.1;refGene_gn=CKAP5;refGene_tx=XM_019812358.1;refGene_gn=CKAP5;refGene_tx=XM_019812359.2
D1 100899168 100899225
refGene_gn=CKAP5;refGene_tx=XM_011286866.2;refGene_gn=CKAP5;refGene_tx=XM_019812357.1;refGene_gn=CKAP5;refGene_tx=XM_019812358.1;refGene_gn=CKAP5;refGene_tx=XM_019812359.2
D1 100949613 100949946
refGene_gn=LRP4;refGene_tx=XM_023239813.1
D1 100952382 100952524
refGene_gn=LRP4;refGene_tx=XM_023239813.1
D1 100954134 100954222
refGene_gn=LRP4;refGene_tx=XM_023239813.1
D1 100954788 100954856
refGene_gn=LRP4;refGene_tx=XM_023239813.1
D1 100958734 100958870
refGene_gn=LRP4;refGene_tx=XM_023239813.1
D1 100959329 100959443
refGene_gn=LRP4;refGene_tx=XM_023239813.1
D1 100959736 100959881
refGene_gn=LRP4;refGene_tx=XM_023239813.1
D1 100961745 100961854
refGene_gn=LRP4;refGene_tx=XM_023239813.1
D1 100962664 100962799
refGene_gn=LRP4;refGene_tx=XM_023239813.1

D1 100962967 100963186
refGene_gn=LRP4;refGene_tx=XM_023239813.1
D1 100964223 100964527
refGene_gn=LRP4;refGene_tx=XM_023239813.1
D1 100964876 100965102
refGene_gn=LRP4;refGene_tx=XM_023239813.1
D1 100965224 100965387
refGene_gn=LRP4;refGene_tx=XM_023239813.1
D1 100966188 100966360
refGene_gn=LRP4;refGene_tx=XM_023239813.1
D1 100966466 100966553
refGene_gn=LRP4;refGene_tx=XM_023239813.1
D1 100966897 100967038
refGene_gn=LRP4;refGene_tx=XM_023239813.1
D1 100968215 100968347
refGene_gn=LRP4;refGene_tx=XM_023239813.1
D1 100968435 100968625
refGene_gn=LRP4;refGene_tx=XM_023239813.1
D1 100970154 100970356
refGene_gn=LRP4;refGene_tx=XM_023239813.1
D1 100971703 100971809
refGene_gn=LRP4;refGene_tx=XM_023239813.1
D1 100972364 100972446
refGene_gn=LRP4;refGene_tx=XM_023239813.1
D1 100972638 100972847
refGene_gn=LRP4;refGene_tx=XM_023239813.1
D1 100974488 100974611
refGene_gn=LRP4;refGene_tx=XM_023239813.1
D1 100974990 100975167
refGene_gn=LRP4;refGene_tx=XM_023239813.1
D1 100975327 100975545
refGene_gn=LRP4;refGene_tx=XM_023239813.1
D1 100977749 100977906
refGene_gn=LRP4;refGene_tx=XM_023239813.1
D1 100978965 100979196
refGene_gn=LRP4;refGene_tx=XM_023239813.1
D1 100979531 100979657
refGene_gn=LRP4;refGene_tx=XM_023239813.1
D1 100980053 100980188
refGene_gn=LRP4;refGene_tx=XM_023239813.1
D1 100980379 100980505
refGene_gn=LRP4;refGene_tx=XM_023239813.1
D1 100981084 100981210
refGene_gn=LRP4;refGene_tx=XM_023239813.1
D1 100981617 100981737
refGene_gn=LRP4;refGene_tx=XM_023239813.1
D1 100981979 100982108
refGene_gn=LRP4;refGene_tx=XM_023239813.1
D1 100982424 100982541
refGene_gn=LRP4;refGene_tx=XM_023239813.1

D1 100982935 100983049
refGene_gn=LRP4;refGene_tx=XM_023239813.1
D1 100983325 100983442
refGene_gn=LRP4;refGene_tx=XM_023239813.1
D1 100984571 100984718
refGene_gn=LRP4;refGene_tx=XM_023239813.1
D1 100997306 100997358
refGene_gn=LRP4;refGene_tx=XM_023239813.1
D1 101007175 101007221
refGene_gn=CD1H11orf49;refGene_tx=XM_003993221.4;refGene_gn=CD1H11orf49;refGene_tx=XM_003993222.5;refGene_gn=CD1H11orf49;refGene_tx=XM_006937251.3;refGene_gn=CD1H11orf49;refGene_tx=XM_006937252.3
D1 101056380 101056483
refGene_gn=CD1H11orf49;refGene_tx=XM_003993221.4;refGene_gn=CD1H11orf49;refGene_tx=XM_003993222.5;refGene_gn=CD1H11orf49;refGene_tx=XM_006937251.3;refGene_gn=CD1H11orf49;refGene_tx=XM_006937252.3
D1 101092483 101092614
refGene_gn=CD1H11orf49;refGene_tx=XM_003993221.4;refGene_gn=CD1H11orf49;refGene_tx=XM_003993222.5;refGene_gn=CD1H11orf49;refGene_tx=XM_006937251.3;refGene_gn=CD1H11orf49;refGene_tx=XM_006937252.3
D1 101180442 101180527
refGene_gn=CD1H11orf49;refGene_tx=XM_003993221.4;refGene_gn=CD1H11orf49;refGene_tx=XM_003993222.5;refGene_gn=CD1H11orf49;refGene_tx=XM_006937251.3;refGene_gn=CD1H11orf49;refGene_tx=XM_006937252.3
D1 101199220 101199306
refGene_gn=CD1H11orf49;refGene_tx=XM_003993221.4;refGene_gn=CD1H11orf49;refGene_tx=XM_003993222.5;refGene_gn=CD1H11orf49;refGene_tx=XM_006937251.3;refGene_gn=CD1H11orf49;refGene_tx=XM_006937252.3
D1 101201320 101201516
refGene_gn=CD1H11orf49;refGene_tx=XM_003993221.4;refGene_gn=CD1H11orf49;refGene_tx=XM_003993222.5;refGene_gn=CD1H11orf49;refGene_tx=XM_006937251.3;refGene_gn=CD1H11orf49;refGene_tx=XM_006937252.3
D1 101202182 101202292
refGene_gn=CD1H11orf49;refGene_tx=XM_003993221.4;refGene_gn=CD1H11orf49;refGene_tx=XM_003993222.5;refGene_gn=CD1H11orf49;refGene_tx=XM_006937251.3;refGene_gn=CD1H11orf49;refGene_tx=XM_006937252.3
D1 101205361 101205429
refGene_gn=CD1H11orf49;refGene_tx=XM_003993221.4;refGene_gn=CD1H11orf49;refGene_tx=XM_003993222.5;refGene_gn=CD1H11orf49;refGene_tx=XM_006937251.3;refGene_gn=CD1H11orf49;refGene_tx=XM_006937252.3
D1 101205619 101205653
refGene_gn=CD1H11orf49;refGene_tx=XM_006937251.3
D1 101205697 101205872
refGene_gn=CD1H11orf49;refGene_tx=XM_003993222.5
D1 101207683 101207843
refGene_gn=CD1H11orf49;refGene_tx=XM_003993221.4
D1 101209091 101209112
refGene_gn=ARFGAP2;refGene_tx=XM_003993223.4;refGene_gn=ARFGAP2;refGene_tx=XM_006937249.3;refGene_gn=ARFGAP2;refGene_tx=XM_006937250.3
D1 101209562 101209681

refGene_gn=ARFGAP2;refGene_tx=XM_003993223.4;refGene_gn=ARFGAP2;refGene_tx=XM_006937249.3;refGene_gn=ARFGAP2;refGene_tx=XM_006937250.3;refGene_gn=ARFGAP2;refGene_tx=XM_019812385.1
D1 101209801 101209895
refGene_gn=ARFGAP2;refGene_tx=XM_003993223.4;refGene_gn=ARFGAP2;refGene_tx=XM_006937249.3;refGene_gn=ARFGAP2;refGene_tx=XM_006937250.3;refGene_gn=ARFGAP2;refGene_tx=XM_019812385.1
D1 101210055 101210182
refGene_gn=ARFGAP2;refGene_tx=XM_003993223.4;refGene_gn=ARFGAP2;refGene_tx=XM_006937249.3;refGene_gn=ARFGAP2;refGene_tx=XM_006937250.3
D1 101211158 101211293
refGene_gn=ARFGAP2;refGene_tx=XM_003993223.4;refGene_gn=ARFGAP2;refGene_tx=XM_006937249.3;refGene_gn=ARFGAP2;refGene_tx=XM_006937250.3;refGene_gn=ARFGAP2;refGene_tx=XM_019812385.1
D1 101211372 101211501
refGene_gn=ARFGAP2;refGene_tx=XM_003993223.4;refGene_gn=ARFGAP2;refGene_tx=XM_006937249.3;refGene_gn=ARFGAP2;refGene_tx=XM_006937250.3;refGene_gn=ARFGAP2;refGene_tx=XM_019812385.1
D1 101213433 101213565
refGene_gn=ARFGAP2;refGene_tx=XM_003993223.4;refGene_gn=ARFGAP2;refGene_tx=XM_006937249.3;refGene_gn=ARFGAP2;refGene_tx=XM_006937250.3;refGene_gn=ARFGAP2;refGene_tx=XM_019812385.1
D1 101213666 101213803
refGene_gn=ARFGAP2;refGene_tx=XM_003993223.4;refGene_gn=ARFGAP2;refGene_tx=XM_006937249.3;refGene_gn=ARFGAP2;refGene_tx=XM_006937250.3;refGene_gn=ARFGAP2;refGene_tx=XM_019812385.1
D1 101214236 101214289
refGene_gn=ARFGAP2;refGene_tx=XM_003993223.4;refGene_gn=ARFGAP2;refGene_tx=XM_006937249.3;refGene_gn=ARFGAP2;refGene_tx=XM_006937250.3;refGene_gn=ARFGAP2;refGene_tx=XM_019812385.1
D1 101214665 101214707
refGene_gn=ARFGAP2;refGene_tx=XM_006937249.3;refGene_gn=ARFGAP2;refGene_tx=XM_006937250.3;refGene_gn=ARFGAP2;refGene_tx=XM_019812385.1
D1 101215368 101215425
refGene_gn=ARFGAP2;refGene_tx=XM_003993223.4;refGene_gn=ARFGAP2;refGene_tx=XM_006937249.3;refGene_gn=ARFGAP2;refGene_tx=XM_006937250.3;refGene_gn=ARFGAP2;refGene_tx=XM_019812385.1
D1 101215735 101215817
refGene_gn=ARFGAP2;refGene_tx=XM_003993223.4;refGene_gn=ARFGAP2;refGene_tx=XM_006937249.3;refGene_gn=ARFGAP2;refGene_tx=XM_006937250.3;refGene_gn=ARFGAP2;refGene_tx=XM_019812385.1
D1 101216655 101216736
refGene_gn=ARFGAP2;refGene_tx=XM_003993223.4;refGene_gn=ARFGAP2;refGene_tx=XM_006937249.3;refGene_gn=ARFGAP2;refGene_tx=XM_006937250.3;refGene_gn=ARFGAP2;refGene_tx=XM_019812385.1
D1 101216823 101216955
refGene_gn=ARFGAP2;refGene_tx=XM_003993223.4;refGene_gn=ARFGAP2;refGene_tx=XM_006937249.3;refGene_gn=ARFGAP2;refGene_tx=XM_006937250.3;refGene_gn=ARFGAP2;refGene_tx=XM_019812385.1
D1 101217478 101217551

refGene_gn=ARFGAP2;refGene_tx=XM_003993223.4;refGene_gn=ARFGAP2;refGene_tx=XM_006937249.3;refGene_gn=ARFGAP2;refGene_tx=XM_006937250.3;refGene_gn=ARFGAP2;refGene_tx=XM_019812385.1
D1 101218115 101218234
refGene_gn=ARFGAP2;refGene_tx=XM_003993223.4;refGene_gn=ARFGAP2;refGene_tx=XM_006937249.3;refGene_gn=ARFGAP2;refGene_tx=XM_006937250.3;refGene_gn=ARFGAP2;refGene_tx=XM_019812385.1
D1 101218405 101218477
refGene_gn=ARFGAP2;refGene_tx=XM_003993223.4;refGene_gn=ARFGAP2;refGene_tx=XM_006937249.3;refGene_gn=ARFGAP2;refGene_tx=XM_006937250.3;refGene_gn=ARFGAP2;refGene_tx=XM_019812385.1
D1 101219620 101219736
refGene_gn=PACSIN3;refGene_tx=XM_006937253.4;refGene_gn=PACSIN3;refGene_tx=XM_011286864.3;refGene_gn=PACSIN3;refGene_tx=XM_019812392.1
D1 101220352 101220474
refGene_gn=PACSIN3;refGene_tx=XM_006937253.4;refGene_gn=PACSIN3;refGene_tx=XM_011286864.3;refGene_gn=PACSIN3;refGene_tx=XM_019812392.1
D1 101220892 101221029
refGene_gn=PACSIN3;refGene_tx=XM_006937253.4;refGene_gn=PACSIN3;refGene_tx=XM_011286864.3;refGene_gn=PACSIN3;refGene_tx=XM_019812392.1
D1 101221167 101221288
refGene_gn=PACSIN3;refGene_tx=XM_006937253.4;refGene_gn=PACSIN3;refGene_tx=XM_011286864.3;refGene_gn=PACSIN3;refGene_tx=XM_019812392.1
D1 101221431 101221607
refGene_gn=PACSIN3;refGene_tx=XM_006937253.4;refGene_gn=PACSIN3;refGene_tx=XM_011286864.3;refGene_gn=PACSIN3;refGene_tx=XM_019812392.1
D1 101222124 101222280
refGene_gn=PACSIN3;refGene_tx=XM_006937253.4;refGene_gn=PACSIN3;refGene_tx=XM_011286864.3;refGene_gn=PACSIN3;refGene_tx=XM_019812392.1
D1 101222378 101222614
refGene_gn=PACSIN3;refGene_tx=XM_006937253.4;refGene_gn=PACSIN3;refGene_tx=XM_011286864.3;refGene_gn=PACSIN3;refGene_tx=XM_019812392.1
D1 101223356 101223513
refGene_gn=PACSIN3;refGene_tx=XM_006937253.4;refGene_gn=PACSIN3;refGene_tx=XM_011286864.3;refGene_gn=PACSIN3;refGene_tx=XM_019812392.1
D1 101223653 101223707
refGene_gn=PACSIN3;refGene_tx=XM_006937253.4;refGene_gn=PACSIN3;refGene_tx=XM_011286864.3;refGene_gn=PACSIN3;refGene_tx=XM_019812392.1
D1 101227342 101227396
refGene_gn=DDB2;refGene_tx=XM_019812386.1;refGene_gn=DDB2;refGene_tx=XM_019812387.1;refGene_gn=DDB2;refGene_tx=XM_019812390.1
D1 101240553 101240710
refGene_gn=DDB2;refGene_tx=XM_019812386.1;refGene_gn=DDB2;refGene_tx=XM_019812387.1;refGene_gn=DDB2;refGene_tx=XM_019812388.2;refGene_gn=DDB2;refGene_tx=XM_019812389.2;refGene_gn=DDB2;refGene_tx=XM_019812390.1
D1 101241844 101241981
refGene_gn=DDB2;refGene_tx=XM_019812386.1;refGene_gn=DDB2;refGene_tx=XM_019812387.1;refGene_gn=DDB2;refGene_tx=XM_019812388.2;refGene_gn=DDB2;refGene_tx=XM_019812389.2;refGene_gn=DDB2;refGene_tx=XM_019812390.1
D1 101242368 101242560

refGene_gn=DDB2;refGene_tx=XM_019812386.1;refGene_gn=DDB2;refGene_tx=XM_019812387.1;refGene_gn=DDB2;refGene_tx=XM_019812388.2;refGene_gn=DDB2;refGene_tx=XM_019812389.2;refGene_gn=DDB2;refGene_tx=XM_019812390.1
D1 101256039 101256185
refGene_gn=DDB2;refGene_tx=XM_019812386.1;refGene_gn=DDB2;refGene_tx=XM_019812387.1;refGene_gn=DDB2;refGene_tx=XM_019812388.2;refGene_gn=DDB2;refGene_tx=XM_019812389.2;refGene_gn=DDB2;refGene_tx=XM_019812390.1
D1 101257496 101257596
refGene_gn=DDB2;refGene_tx=XM_019812386.1;refGene_gn=DDB2;refGene_tx=XM_019812387.1;refGene_gn=DDB2;refGene_tx=XM_019812388.2;refGene_gn=DDB2;refGene_tx=XM_019812389.2;refGene_gn=DDB2;refGene_tx=XM_019812390.1
D1 101257683 101257861
refGene_gn=DDB2;refGene_tx=XM_019812386.1;refGene_gn=DDB2;refGene_tx=XM_019812387.1;refGene_gn=DDB2;refGene_tx=XM_019812388.2;refGene_gn=DDB2;refGene_tx=XM_019812389.2;refGene_gn=DDB2;refGene_tx=XM_019812390.1
D1 101258186 101258329
refGene_gn=DDB2;refGene_tx=XM_019812386.1;refGene_gn=DDB2;refGene_tx=XM_019812387.1;refGene_gn=DDB2;refGene_tx=XM_019812388.2;refGene_gn=DDB2;refGene_tx=XM_019812389.2;refGene_gn=DDB2;refGene_tx=XM_019812390.1
D1 101258561 101258651
refGene_gn=DDB2;refGene_tx=XM_019812386.1;refGene_gn=DDB2;refGene_tx=XM_019812387.1;refGene_gn=DDB2;refGene_tx=XM_019812388.2;refGene_gn=DDB2;refGene_tx=XM_019812389.2
D1 101259742 101259907
refGene_gn=DDB2;refGene_tx=XM_019812386.1;refGene_gn=DDB2;refGene_tx=XM_019812387.1;refGene_gn=DDB2;refGene_tx=XM_019812388.2;refGene_gn=DDB2;refGene_tx=XM_019812389.2;refGene_gn=DDB2;refGene_tx=XM_019812390.1
D1 101260051 101260097
refGene_gn=DDB2;refGene_tx=XM_019812386.1;refGene_gn=DDB2;refGene_tx=XM_019812387.1;refGene_gn=DDB2;refGene_tx=XM_019812388.2;refGene_gn=DDB2;refGene_tx=XM_019812389.2;refGene_gn=DDB2;refGene_tx=XM_019812390.1
D1 101260406 101260453
refGene_gn=DDB2;refGene_tx=XM_019812386.1;refGene_gn=DDB2;refGene_tx=XM_019812387.1;refGene_gn=DDB2;refGene_tx=XM_019812388.2;refGene_gn=DDB2;refGene_tx=XM_019812389.2;refGene_gn=DDB2;refGene_tx=XM_019812390.1
D1 101261627 101261761
refGene_gn=ACP2;refGene_tx=XM_003993227.3;refGene_gn=ACP2;refGene_tx=XM_011286859.3;refGene_gn=ACP2;refGene_tx=XM_019812393.1;refGene_gn=ACP2;refGene_tx=XM_019812394.2;refGene_gn=ACP2;refGene_tx=XM_019812395.1
D1 101263721 101263897
refGene_gn=ACP2;refGene_tx=XM_003993227.3;refGene_gn=ACP2;refGene_tx=XM_011286859.3;refGene_gn=ACP2;refGene_tx=XM_019812393.1;refGene_gn=ACP2;refGene_tx=XM_019812394.2;refGene_gn=ACP2;refGene_tx=XM_019812395.1
D1 101264011 101264118
refGene_gn=ACP2;refGene_tx=XM_003993227.3;refGene_gn=ACP2;refGene_tx=XM_011286859.3;refGene_gn=ACP2;refGene_tx=XM_019812393.1;refGene_gn=ACP2;refGene_tx=XM_019812394.2;refGene_gn=ACP2;refGene_tx=XM_019812395.1
D1 101264249 101264332
refGene_gn=ACP2;refGene_tx=XM_003993227.3;refGene_gn=ACP2;refGene_tx=XM_011286859.3;refGene_gn=ACP2;refGene_tx=XM_019812393.1;refGene_gn=ACP

2;refGene_tx=XM_019812394.2;refGene_gn=ACP2;refGene_tx=XM_019812395.1
D1 101265118 101265251
refGene_gn=ACP2;refGene_tx=XM_003993227.3;refGene_gn=ACP2;refGene_tx=XM_011286859.3;refGene_gn=ACP2;refGene_tx=XM_019812393.1;refGene_gn=ACP2;refGene_tx=XM_019812394.2;refGene_gn=ACP2;refGene_tx=XM_019812395.1
D1 101265700 101265790
refGene_gn=ACP2;refGene_tx=XM_003993227.3;refGene_gn=ACP2;refGene_tx=XM_011286859.3;refGene_gn=ACP2;refGene_tx=XM_019812393.1;refGene_gn=ACP2;refGene_tx=XM_019812394.2;refGene_gn=ACP2;refGene_tx=XM_019812395.1
D1 101265868 101265967
refGene_gn=ACP2;refGene_tx=XM_003993227.3;refGene_gn=ACP2;refGene_tx=XM_011286859.3;refGene_gn=ACP2;refGene_tx=XM_019812393.1;refGene_gn=ACP2;refGene_tx=XM_019812394.2;refGene_gn=ACP2;refGene_tx=XM_019812395.1
D1 101266072 101266225
refGene_gn=ACP2;refGene_tx=XM_003993227.3;refGene_gn=ACP2;refGene_tx=XM_011286859.3;refGene_gn=ACP2;refGene_tx=XM_019812393.1;refGene_gn=ACP2;refGene_tx=XM_019812394.2;refGene_gn=ACP2;refGene_tx=XM_019812395.1
D1 101267300 101267387
refGene_gn=ACP2;refGene_tx=XM_003993227.3;refGene_gn=ACP2;refGene_tx=XM_011286859.3;refGene_gn=ACP2;refGene_tx=XM_019812393.1;refGene_gn=ACP2;refGene_tx=XM_019812394.2;refGene_gn=ACP2;refGene_tx=XM_019812395.1
D1 101267691 101267787
refGene_gn=ACP2;refGene_tx=XM_003993227.3;refGene_gn=ACP2;refGene_tx=XM_019812393.1;refGene_gn=ACP2;refGene_tx=XM_019812394.2
D1 101268178 101268208
refGene_gn=ACP2;refGene_tx=XM_019812394.2
D1 101268372 101268486
refGene_gn=ACP2;refGene_tx=XM_003993227.3;refGene_gn=ACP2;refGene_tx=XM_019812393.1
D1 101276255 101276298
refGene_gn=NR1H3;refGene_tx=XM_003993229.5;refGene_gn=NR1H3;refGene_tx=XM_006937254.4;refGene_gn=NR1H3;refGene_tx=XM_006937256.4;refGene_gn=NR1H3;refGene_tx=XM_006937257.3;refGene_gn=NR1H3;refGene_tx=XM_006937258.3;refGene_gn=NR1H3;refGene_tx=XM_006937260.4;refGene_gn=NR1H3;refGene_tx=XM_006937261.4;refGene_gn=NR1H3;refGene_tx=XM_011286860.3;refGene_gn=NR1H3;refGene_tx=XM_011286861.3;refGene_gn=NR1H3;refGene_tx=XM_011286862.3;refGene_gn=NR1H3;refGene_tx=XM_019812391.2
D1 101276809 101276998
refGene_gn=NR1H3;refGene_tx=XM_003993229.5;refGene_gn=NR1H3;refGene_tx=XM_006937254.4;refGene_gn=NR1H3;refGene_tx=XM_006937256.4;refGene_gn=NR1H3;refGene_tx=XM_006937257.3;refGene_gn=NR1H3;refGene_tx=XM_006937258.3;refGene_gn=NR1H3;refGene_tx=XM_006937260.4;refGene_gn=NR1H3;refGene_tx=XM_006937261.4;refGene_gn=NR1H3;refGene_tx=XM_011286860.3;refGene_gn=NR1H3;refGene_tx=XM_011286861.3;refGene_gn=NR1H3;refGene_tx=XM_011286862.3;refGene_gn=NR1H3;refGene_tx=XM_019812391.2
D1 101277536 101277803
refGene_gn=NR1H3;refGene_tx=XM_003993229.5;refGene_gn=NR1H3;refGene_tx=XM_006937254.4;refGene_gn=NR1H3;refGene_tx=XM_006937256.4;refGene_gn=NR1H3;refGene_tx=XM_006937257.3;refGene_gn=NR1H3;refGene_tx=XM_006937258.3;refGene_gn=NR1H3;refGene_tx=XM_006937260.4;refGene_gn=NR1H3;refGe

ne_tx=XM_006937261.4;refGene_gn=NR1H3;refGene_tx=XM_011286860.3;refGene_gn=NR1H3;refGene_tx=XM_011286861.3;refGene_gn=NR1H3;refGene_tx=XM_011286862.3
D1 101278046 101278255
refGene_gn=NR1H3;refGene_tx=XM_003993229.5;refGene_gn=NR1H3;refGene_tx=XM_006937254.4;refGene_gn=NR1H3;refGene_tx=XM_006937256.4;refGene_gn=NR1H3;refGene_tx=XM_006937257.3;refGene_gn=NR1H3;refGene_tx=XM_006937258.3;refGene_gn=NR1H3;refGene_tx=XM_006937260.4;refGene_gn=NR1H3;refGene_tx=XM_006937261.4;refGene_gn=NR1H3;refGene_tx=XM_011286860.3;refGene_gn=NR1H3;refGene_tx=XM_011286861.3;refGene_gn=NR1H3;refGene_tx=XM_011286862.3;refGene_gn=NR1H3;refGene_tx=XM_019812391.2
D1 101278360 101278540
refGene_gn=NR1H3;refGene_tx=XM_003993229.5;refGene_gn=NR1H3;refGene_tx=XM_006937254.4;refGene_gn=NR1H3;refGene_tx=XM_006937256.4;refGene_gn=NR1H3;refGene_tx=XM_006937257.3;refGene_gn=NR1H3;refGene_tx=XM_006937258.3;refGene_gn=NR1H3;refGene_tx=XM_006937260.4;refGene_gn=NR1H3;refGene_tx=XM_006937261.4;refGene_gn=NR1H3;refGene_tx=XM_011286860.3;refGene_gn=NR1H3;refGene_tx=XM_011286861.3;refGene_gn=NR1H3;refGene_tx=XM_011286862.3;refGene_gn=NR1H3;refGene_tx=XM_019812391.2
D1 101278687 101278787
refGene_gn=NR1H3;refGene_tx=XM_003993229.5;refGene_gn=NR1H3;refGene_tx=XM_006937254.4;refGene_gn=NR1H3;refGene_tx=XM_006937256.4;refGene_gn=NR1H3;refGene_tx=XM_006937257.3;refGene_gn=NR1H3;refGene_tx=XM_006937258.3;refGene_gn=NR1H3;refGene_tx=XM_006937260.4;refGene_gn=NR1H3;refGene_tx=XM_006937261.4;refGene_gn=NR1H3;refGene_tx=XM_011286860.3;refGene_gn=NR1H3;refGene_tx=XM_011286861.3;refGene_gn=NR1H3;refGene_tx=XM_011286862.3;refGene_gn=NR1H3;refGene_tx=XM_019812391.2
D1 101281422 101281536
refGene_gn=NR1H3;refGene_tx=XM_003993229.5;refGene_gn=NR1H3;refGene_tx=XM_006937254.4;refGene_gn=NR1H3;refGene_tx=XM_006937256.4;refGene_gn=NR1H3;refGene_tx=XM_006937257.3;refGene_gn=NR1H3;refGene_tx=XM_006937258.3;refGene_gn=NR1H3;refGene_tx=XM_006937260.4;refGene_gn=NR1H3;refGene_tx=XM_006937261.4;refGene_gn=NR1H3;refGene_tx=XM_011286860.3;refGene_gn=NR1H3;refGene_tx=XM_011286861.3;refGene_gn=NR1H3;refGene_tx=XM_011286862.3;refGene_gn=NR1H3;refGene_tx=XM_019812391.2
D1 101281750 101281845
refGene_gn=NR1H3;refGene_tx=XM_003993229.5;refGene_gn=NR1H3;refGene_tx=XM_006937254.4;refGene_gn=NR1H3;refGene_tx=XM_006937256.4;refGene_gn=NR1H3;refGene_tx=XM_006937257.3;refGene_gn=NR1H3;refGene_tx=XM_006937258.3;refGene_gn=NR1H3;refGene_tx=XM_006937261.4;refGene_gn=NR1H3;refGene_tx=XM_011286860.3;refGene_gn=NR1H3;refGene_tx=XM_011286861.3;refGene_gn=NR1H3;refGene_tx=XM_011286862.3;refGene_gn=NR1H3;refGene_tx=XM_019812391.2
D1 101282035 101282182
refGene_gn=NR1H3;refGene_tx=XM_003993229.5;refGene_gn=NR1H3;refGene_tx=XM_006937254.4;refGene_gn=NR1H3;refGene_tx=XM_006937256.4;refGene_gn=NR1H3;refGene_tx=XM_006937257.3;refGene_gn=NR1H3;refGene_tx=XM_006937258.3;refGene_gn=NR1H3;refGene_tx=XM_006937260.4;refGene_gn=NR1H3;refGene_tx=XM_006937261.4;refGene_gn=NR1H3;refGene_tx=XM_011286860.3;refGene_gn=NR1H3;refGene_tx=XM_011286861.3;refGene_gn=NR1H3;refGene_tx=XM_01

1286862.3;refGene_gn=NR1H3;refGene_tx=XM_019812391.2
D1 101287258 101287320
refGene_gn=MADD;refGene_tx=XM_003993232.5;refGene_gn=MADD;refGene_tx=XM_003993233.5;refGene_gn=MADD;refGene_tx=XM_003993234.5;refGene_gn=MADD;refGene_tx=XM_006937269.4;refGene_gn=MADD;refGene_tx=XM_006937270.4;refGene_gn=MADD;refGene_tx=XM_006937271.4;refGene_gn=MADD;refGene_tx=XM_006937278.4;refGene_gn=MADD;refGene_tx=XM_006937279.4;refGene_gn=MADD;refGene_tx=XM_006937280.4;refGene_gn=MADD;refGene_tx=XM_011286854.3;refGene_gn=MADD;refGene_tx=XM_011286856.3;refGene_gn=MADD;refGene_tx=XM_011286857.3;refGene_gn=MADD;refGene_tx=XM_011286858.3;refGene_gn=MADD;refGene_tx=XM_019812363.2;refGene_gn=MADD;refGene_tx=XM_019812364.2;refGene_gn=MADD;refGene_tx=XM_019812365.2;refGene_gn=MADD;refGene_tx=XM_019812366.1;refGene_gn=MADD;refGene_tx=XM_019812367.2;refGene_gn=MADD;refGene_tx=XM_019812368.2;refGene_gn=MADD;refGene_tx=XM_019812369.2;refGene_gn=MADD;refGene_tx=XM_019812370.2;refGene_gn=MADD;refGene_tx=XM_019812371.2;refGene_gn=MADD;refGene_tx=XM_019812372.2;refGene_gn=MADD;refGene_tx=XM_019812373.2;refGene_gn=MADD;refGene_tx=XM_019812374.2;refGene_gn=MADD;refGene_tx=XM_019812375.2;refGene_gn=MADD;refGene_tx=XM_019812376.2;refGene_gn=MADD;refGene_tx=XM_019812378.2;refGene_gn=MADD;refGene_tx=XM_019812379.2;refGene_gn=MADD;refGene_tx=XM_019812380.2;refGene_gn=MADD;refGene_tx=XM_019812381.2;refGene_gn=MADD;refGene_tx=XM_019812382.2;refGene_gn=MADD;refGene_tx=XM_019812383.2;refGene_gn=MADD;refGene_tx=XM_019812384.2
D1 101287963 101288560
refGene_gn=MADD;refGene_tx=XM_003993232.5;refGene_gn=MADD;refGene_tx=XM_003993233.5;refGene_gn=MADD;refGene_tx=XM_003993234.5;refGene_gn=MADD;refGene_tx=XM_006937269.4;refGene_gn=MADD;refGene_tx=XM_006937270.4;refGene_gn=MADD;refGene_tx=XM_006937271.4;refGene_gn=MADD;refGene_tx=XM_006937278.4;refGene_gn=MADD;refGene_tx=XM_006937279.4;refGene_gn=MADD;refGene_tx=XM_006937280.4;refGene_gn=MADD;refGene_tx=XM_011286854.3;refGene_gn=MADD;refGene_tx=XM_011286856.3;refGene_gn=MADD;refGene_tx=XM_011286857.3;refGene_gn=MADD;refGene_tx=XM_011286858.3;refGene_gn=MADD;refGene_tx=XM_019812363.2;refGene_gn=MADD;refGene_tx=XM_019812364.2;refGene_gn=MADD;refGene_tx=XM_019812365.2;refGene_gn=MADD;refGene_tx=XM_019812366.1;refGene_gn=MADD;refGene_tx=XM_019812367.2;refGene_gn=MADD;refGene_tx=XM_019812368.2;refGene_gn=MADD;refGene_tx=XM_019812369.2;refGene_gn=MADD;refGene_tx=XM_019812370.2;refGene_gn=MADD;refGene_tx=XM_019812371.2;refGene_gn=MADD;refGene_tx=XM_019812372.2;refGene_gn=MADD;refGene_tx=XM_019812373.2;refGene_gn=MADD;refGene_tx=XM_019812374.2;refGene_gn=MADD;refGene_tx=XM_019812375.2;refGene_gn=MADD;refGene_tx=XM_019812376.2;refGene_gn=MADD;refGene_tx=XM_019812378.2;refGene_gn=MADD;refGene_tx=XM_019812379.2;refGene_gn=MADD;refGene_tx=XM_019812380.2;refGene_gn=MADD;refGene_tx=XM_019812381.2;refGene_gn=MADD;refGene_tx=XM_019812382.2;refGene_gn=MADD;refGene_tx=XM_019812383.2;refGene_gn=MADD;refGene_tx=XM_019812384.2
D1 101289049 101289353
refGene_gn=MADD;refGene_tx=XM_003993232.5;refGene_gn=MADD;refGene_tx=XM_003993233.5;refGene_gn=MADD;refGene_tx=XM_003993234.5;refGene_gn=MADD;refGene_tx=XM_006937269.4;refGene_gn=MADD;refGene_tx=XM_006937270.4;refGene_gn=MADD;refGene_tx=XM_006937271.4;refGene_gn=MADD;refGene_tx=X

M_006937278.4;refGene_gn=MADD;refGene_tx=XM_006937279.4;refGene_gn=MAD
D;refGene_tx=XM_006937280.4;refGene_gn=MADD;refGene_tx=XM_011286854.3;
refGene_gn=MADD;refGene_tx=XM_011286856.3;refGene_gn=MADD;refGene_tx=X
M_011286857.3;refGene_gn=MADD;refGene_tx=XM_011286858.3;refGene_gn=MAD
D;refGene_tx=XM_019812363.2;refGene_gn=MADD;refGene_tx=XM_019812364.2;
refGene_gn=MADD;refGene_tx=XM_019812365.2;refGene_gn=MADD;refGene_tx=X
M_019812366.1;refGene_gn=MADD;refGene_tx=XM_019812367.2;refGene_gn=MAD
D;refGene_tx=XM_019812368.2;refGene_gn=MADD;refGene_tx=XM_019812369.2;
refGene_gn=MADD;refGene_tx=XM_019812370.2;refGene_gn=MADD;refGene_tx=X
M_019812371.2;refGene_gn=MADD;refGene_tx=XM_019812372.2;refGene_gn=MAD
D;refGene_tx=XM_019812373.2;refGene_gn=MADD;refGene_tx=XM_019812374.2;
refGene_gn=MADD;refGene_tx=XM_019812375.2;refGene_gn=MADD;refGene_tx=X
M_019812376.2;refGene_gn=MADD;refGene_tx=XM_019812378.2;refGene_gn=MAD
D;refGene_tx=XM_019812379.2;refGene_gn=MADD;refGene_tx=XM_019812380.2;
refGene_gn=MADD;refGene_tx=XM_019812381.2;refGene_gn=MADD;refGene_tx=X
M_019812382.2;refGene_gn=MADD;refGene_tx=XM_019812383.2;refGene_gn=MAD
D;refGene_tx=XM_019812384.2
D1 101289751 101289883
refGene_gn=MADD;refGene_tx=XM_003993232.5;refGene_gn=MADD;refGene_tx=X
M_003993233.5;refGene_gn=MADD;refGene_tx=XM_003993234.5;refGene_gn=MAD
D;refGene_tx=XM_006937269.4;refGene_gn=MADD;refGene_tx=XM_006937270.4;
refGene_gn=MADD;refGene_tx=XM_006937271.4;refGene_gn=MADD;refGene_tx=X
M_006937278.4;refGene_gn=MADD;refGene_tx=XM_006937279.4;refGene_gn=MAD
D;refGene_tx=XM_006937280.4;refGene_gn=MADD;refGene_tx=XM_011286854.3;
refGene_gn=MADD;refGene_tx=XM_011286856.3;refGene_gn=MADD;refGene_tx=X
M_011286857.3;refGene_gn=MADD;refGene_tx=XM_011286858.3;refGene_gn=MAD
D;refGene_tx=XM_019812363.2;refGene_gn=MADD;refGene_tx=XM_019812364.2;
refGene_gn=MADD;refGene_tx=XM_019812365.2;refGene_gn=MADD;refGene_tx=X
M_019812366.1;refGene_gn=MADD;refGene_tx=XM_019812367.2;refGene_gn=MAD
D;refGene_tx=XM_019812368.2;refGene_gn=MADD;refGene_tx=XM_019812369.2;
refGene_gn=MADD;refGene_tx=XM_019812370.2;refGene_gn=MADD;refGene_tx=X
M_019812371.2;refGene_gn=MADD;refGene_tx=XM_019812372.2;refGene_gn=MAD
D;refGene_tx=XM_019812373.2;refGene_gn=MADD;refGene_tx=XM_019812374.2;
refGene_gn=MADD;refGene_tx=XM_019812375.2;refGene_gn=MADD;refGene_tx=X
M_019812376.2;refGene_gn=MADD;refGene_tx=XM_019812378.2;refGene_gn=MAD
D;refGene_tx=XM_019812379.2;refGene_gn=MADD;refGene_tx=XM_019812380.2;
refGene_gn=MADD;refGene_tx=XM_019812381.2;refGene_gn=MADD;refGene_tx=X
M_019812382.2;refGene_gn=MADD;refGene_tx=XM_019812383.2;refGene_gn=MAD
D;refGene_tx=XM_019812384.2
D1 101290326 101290440
refGene_gn=MADD;refGene_tx=XM_003993232.5;refGene_gn=MADD;refGene_tx=X
M_003993233.5;refGene_gn=MADD;refGene_tx=XM_003993234.5;refGene_gn=MAD
D;refGene_tx=XM_006937269.4;refGene_gn=MADD;refGene_tx=XM_006937270.4;
refGene_gn=MADD;refGene_tx=XM_006937271.4;refGene_gn=MADD;refGene_tx=X
M_006937278.4;refGene_gn=MADD;refGene_tx=XM_006937279.4;refGene_gn=MAD
D;refGene_tx=XM_006937280.4;refGene_gn=MADD;refGene_tx=XM_011286854.3;
refGene_gn=MADD;refGene_tx=XM_011286856.3;refGene_gn=MADD;refGene_tx=X
M_011286857.3;refGene_gn=MADD;refGene_tx=XM_011286858.3;refGene_gn=MAD
D;refGene_tx=XM_019812363.2;refGene_gn=MADD;refGene_tx=XM_019812364.2;
refGene_gn=MADD;refGene_tx=XM_019812365.2;refGene_gn=MADD;refGene_tx=X

M_019812366.1;refGene_gn=MADD;refGene_tx=XM_019812367.2;refGene_gn=MAD
D;refGene_tx=XM_019812368.2;refGene_gn=MADD;refGene_tx=XM_019812369.2;
refGene_gn=MADD;refGene_tx=XM_019812370.2;refGene_gn=MADD;refGene_tx=X
M_019812371.2;refGene_gn=MADD;refGene_tx=XM_019812372.2;refGene_gn=MAD
D;refGene_tx=XM_019812373.2;refGene_gn=MADD;refGene_tx=XM_019812374.2;
refGene_gn=MADD;refGene_tx=XM_019812375.2;refGene_gn=MADD;refGene_tx=X
M_019812376.2;refGene_gn=MADD;refGene_tx=XM_019812378.2;refGene_gn=MAD
D;refGene_tx=XM_019812379.2;refGene_gn=MADD;refGene_tx=XM_019812380.2;
refGene_gn=MADD;refGene_tx=XM_019812381.2;refGene_gn=MADD;refGene_tx=X
M_019812382.2;refGene_gn=MADD;refGene_tx=XM_019812383.2;refGene_gn=MAD
D;refGene_tx=XM_019812384.2
D1 101291390 101291471
refGene_gn=MADD;refGene_tx=XM_003993232.5;refGene_gn=MADD;refGene_tx=X
M_003993233.5;refGene_gn=MADD;refGene_tx=XM_003993234.5;refGene_gn=MAD
D;refGene_tx=XM_006937269.4;refGene_gn=MADD;refGene_tx=XM_006937270.4;
refGene_gn=MADD;refGene_tx=XM_006937271.4;refGene_gn=MADD;refGene_tx=X
M_006937278.4;refGene_gn=MADD;refGene_tx=XM_006937279.4;refGene_gn=MAD
D;refGene_tx=XM_006937280.4;refGene_gn=MADD;refGene_tx=XM_011286854.3;
refGene_gn=MADD;refGene_tx=XM_011286856.3;refGene_gn=MADD;refGene_tx=X
M_011286857.3;refGene_gn=MADD;refGene_tx=XM_011286858.3;refGene_gn=MAD
D;refGene_tx=XM_019812363.2;refGene_gn=MADD;refGene_tx=XM_019812364.2;
refGene_gn=MADD;refGene_tx=XM_019812365.2;refGene_gn=MADD;refGene_tx=X
M_019812366.1;refGene_gn=MADD;refGene_tx=XM_019812367.2;refGene_gn=MAD
D;refGene_tx=XM_019812368.2;refGene_gn=MADD;refGene_tx=XM_019812369.2;
refGene_gn=MADD;refGene_tx=XM_019812370.2;refGene_gn=MADD;refGene_tx=X
M_019812371.2;refGene_gn=MADD;refGene_tx=XM_019812372.2;refGene_gn=MAD
D;refGene_tx=XM_019812373.2;refGene_gn=MADD;refGene_tx=XM_019812374.2;
refGene_gn=MADD;refGene_tx=XM_019812375.2;refGene_gn=MADD;refGene_tx=X
M_019812376.2;refGene_gn=MADD;refGene_tx=XM_019812378.2;refGene_gn=MAD
D;refGene_tx=XM_019812379.2;refGene_gn=MADD;refGene_tx=XM_019812380.2;
refGene_gn=MADD;refGene_tx=XM_019812381.2;refGene_gn=MADD;refGene_tx=X
M_019812382.2;refGene_gn=MADD;refGene_tx=XM_019812383.2;refGene_gn=MAD
D;refGene_tx=XM_019812384.2
D1 101291960 101291969
refGene_gn=MADD;refGene_tx=XM_019812363.2;refGene_gn=MADD;refGene_tx=X
M_019812364.2;refGene_gn=MADD;refGene_tx=XM_019812365.2;refGene_gn=MAD
D;refGene_tx=XM_019812366.1;refGene_gn=MADD;refGene_tx=XM_019812367.2;
refGene_gn=MADD;refGene_tx=XM_019812369.2;refGene_gn=MADD;refGene_tx=X
M_019812370.2;refGene_gn=MADD;refGene_tx=XM_019812371.2;refGene_gn=MAD
D;refGene_tx=XM_019812372.2;refGene_gn=MADD;refGene_tx=XM_019812373.2;
refGene_gn=MADD;refGene_tx=XM_019812374.2;refGene_gn=MADD;refGene_tx=X
M_019812376.2;refGene_gn=MADD;refGene_tx=XM_019812378.2;refGene_gn=MAD
D;refGene_tx=XM_019812379.2;refGene_gn=MADD;refGene_tx=XM_019812380.2;
refGene_gn=MADD;refGene_tx=XM_019812381.2;refGene_gn=MADD;refGene_tx=X
M_019812382.2;refGene_gn=MADD;refGene_tx=XM_019812383.2;refGene_gn=MAD
D;refGene_tx=XM_019812384.2
D1 101293004 101293183
refGene_gn=MADD;refGene_tx=XM_003993232.5;refGene_gn=MADD;refGene_tx=X
M_003993233.5;refGene_gn=MADD;refGene_tx=XM_003993234.5;refGene_gn=MAD
D;refGene_tx=XM_006937269.4;refGene_gn=MADD;refGene_tx=XM_006937270.4;

refGene_gn=MADD;refGene_tx=XM_006937271.4;refGene_gn=MADD;refGene_tx=X
M_006937278.4;refGene_gn=MADD;refGene_tx=XM_006937279.4;refGene_gn=MAD
D;refGene_tx=XM_006937280.4;refGene_gn=MADD;refGene_tx=XM_011286854.3;
refGene_gn=MADD;refGene_tx=XM_011286856.3;refGene_gn=MADD;refGene_tx=X
M_011286857.3;refGene_gn=MADD;refGene_tx=XM_011286858.3;refGene_gn=MAD
D;refGene_tx=XM_019812363.2;refGene_gn=MADD;refGene_tx=XM_019812364.2;
refGene_gn=MADD;refGene_tx=XM_019812365.2;refGene_gn=MADD;refGene_tx=X
M_019812366.1;refGene_gn=MADD;refGene_tx=XM_019812367.2;refGene_gn=MAD
D;refGene_tx=XM_019812368.2;refGene_gn=MADD;refGene_tx=XM_019812369.2;
refGene_gn=MADD;refGene_tx=XM_019812370.2;refGene_gn=MADD;refGene_tx=X
M_019812371.2;refGene_gn=MADD;refGene_tx=XM_019812372.2;refGene_gn=MAD
D;refGene_tx=XM_019812373.2;refGene_gn=MADD;refGene_tx=XM_019812374.2;
refGene_gn=MADD;refGene_tx=XM_019812375.2;refGene_gn=MADD;refGene_tx=X
M_019812376.2;refGene_gn=MADD;refGene_tx=XM_019812378.2;refGene_gn=MAD
D;refGene_tx=XM_019812379.2;refGene_gn=MADD;refGene_tx=XM_019812380.2;
refGene_gn=MADD;refGene_tx=XM_019812381.2;refGene_gn=MADD;refGene_tx=X
M_019812382.2;refGene_gn=MADD;refGene_tx=XM_019812383.2;refGene_gn=MAD
D;refGene_tx=XM_019812384.2
D1 101293474 101293710
refGene_gn=MADD;refGene_tx=XM_003993232.5;refGene_gn=MADD;refGene_tx=X
M_003993233.5;refGene_gn=MADD;refGene_tx=XM_003993234.5;refGene_gn=MAD
D;refGene_tx=XM_006937269.4;refGene_gn=MADD;refGene_tx=XM_006937270.4;
refGene_gn=MADD;refGene_tx=XM_006937271.4;refGene_gn=MADD;refGene_tx=X
M_006937278.4;refGene_gn=MADD;refGene_tx=XM_006937279.4;refGene_gn=MAD
D;refGene_tx=XM_006937280.4;refGene_gn=MADD;refGene_tx=XM_011286854.3;
refGene_gn=MADD;refGene_tx=XM_011286856.3;refGene_gn=MADD;refGene_tx=X
M_011286857.3;refGene_gn=MADD;refGene_tx=XM_011286858.3;refGene_gn=MAD
D;refGene_tx=XM_019812363.2;refGene_gn=MADD;refGene_tx=XM_019812364.2;
refGene_gn=MADD;refGene_tx=XM_019812365.2;refGene_gn=MADD;refGene_tx=X
M_019812366.1;refGene_gn=MADD;refGene_tx=XM_019812367.2;refGene_gn=MAD
D;refGene_tx=XM_019812368.2;refGene_gn=MADD;refGene_tx=XM_019812369.2;
refGene_gn=MADD;refGene_tx=XM_019812370.2;refGene_gn=MADD;refGene_tx=X
M_019812371.2;refGene_gn=MADD;refGene_tx=XM_019812372.2;refGene_gn=MAD
D;refGene_tx=XM_019812373.2;refGene_gn=MADD;refGene_tx=XM_019812374.2;
refGene_gn=MADD;refGene_tx=XM_019812375.2;refGene_gn=MADD;refGene_tx=X
M_019812376.2;refGene_gn=MADD;refGene_tx=XM_019812378.2;refGene_gn=MAD
D;refGene_tx=XM_019812379.2;refGene_gn=MADD;refGene_tx=XM_019812380.2;
refGene_gn=MADD;refGene_tx=XM_019812381.2;refGene_gn=MADD;refGene_tx=X
M_019812382.2;refGene_gn=MADD;refGene_tx=XM_019812383.2;refGene_gn=MAD
D;refGene_tx=XM_019812384.2
D1 101293898 101294055
refGene_gn=MADD;refGene_tx=XM_003993232.5;refGene_gn=MADD;refGene_tx=X
M_003993233.5;refGene_gn=MADD;refGene_tx=XM_003993234.5;refGene_gn=MAD
D;refGene_tx=XM_006937269.4;refGene_gn=MADD;refGene_tx=XM_006937270.4;
refGene_gn=MADD;refGene_tx=XM_006937271.4;refGene_gn=MADD;refGene_tx=X
M_006937278.4;refGene_gn=MADD;refGene_tx=XM_006937279.4;refGene_gn=MAD
D;refGene_tx=XM_006937280.4;refGene_gn=MADD;refGene_tx=XM_011286854.3;
refGene_gn=MADD;refGene_tx=XM_011286856.3;refGene_gn=MADD;refGene_tx=X
M_011286857.3;refGene_gn=MADD;refGene_tx=XM_011286858.3;refGene_gn=MAD
D;refGene_tx=XM_019812363.2;refGene_gn=MADD;refGene_tx=XM_019812364.2;

refGene_gn=MADD;refGene_tx=XM_019812365.2;refGene_gn=MADD;refGene_tx=X
M_019812366.1;refGene_gn=MADD;refGene_tx=XM_019812367.2;refGene_gn=MAD
D;refGene_tx=XM_019812368.2;refGene_gn=MADD;refGene_tx=XM_019812369.2;
refGene_gn=MADD;refGene_tx=XM_019812370.2;refGene_gn=MADD;refGene_tx=X
M_019812371.2;refGene_gn=MADD;refGene_tx=XM_019812372.2;refGene_gn=MAD
D;refGene_tx=XM_019812373.2;refGene_gn=MADD;refGene_tx=XM_019812374.2;
refGene_gn=MADD;refGene_tx=XM_019812375.2;refGene_gn=MADD;refGene_tx=X
M_019812376.2;refGene_gn=MADD;refGene_tx=XM_019812378.2;refGene_gn=MAD
D;refGene_tx=XM_019812379.2;refGene_gn=MADD;refGene_tx=XM_019812380.2;
refGene_gn=MADD;refGene_tx=XM_019812381.2;refGene_gn=MADD;refGene_tx=X
M_019812382.2;refGene_gn=MADD;refGene_tx=XM_019812383.2;refGene_gn=MAD
D;refGene_tx=XM_019812384.2
D1 101294764 101294868
refGene_gn=MADD;refGene_tx=XM_003993232.5;refGene_gn=MADD;refGene_tx=X
M_003993233.5;refGene_gn=MADD;refGene_tx=XM_003993234.5;refGene_gn=MAD
D;refGene_tx=XM_006937269.4;refGene_gn=MADD;refGene_tx=XM_006937270.4;
refGene_gn=MADD;refGene_tx=XM_006937271.4;refGene_gn=MADD;refGene_tx=X
M_006937278.4;refGene_gn=MADD;refGene_tx=XM_006937279.4;refGene_gn=MAD
D;refGene_tx=XM_006937280.4;refGene_gn=MADD;refGene_tx=XM_011286854.3;
refGene_gn=MADD;refGene_tx=XM_011286856.3;refGene_gn=MADD;refGene_tx=X
M_011286857.3;refGene_gn=MADD;refGene_tx=XM_011286858.3;refGene_gn=MAD
D;refGene_tx=XM_019812363.2;refGene_gn=MADD;refGene_tx=XM_019812364.2;
refGene_gn=MADD;refGene_tx=XM_019812365.2;refGene_gn=MADD;refGene_tx=X
M_019812366.1;refGene_gn=MADD;refGene_tx=XM_019812367.2;refGene_gn=MAD
D;refGene_tx=XM_019812368.2;refGene_gn=MADD;refGene_tx=XM_019812369.2;
refGene_gn=MADD;refGene_tx=XM_019812370.2;refGene_gn=MADD;refGene_tx=X
M_019812371.2;refGene_gn=MADD;refGene_tx=XM_019812372.2;refGene_gn=MAD
D;refGene_tx=XM_019812373.2;refGene_gn=MADD;refGene_tx=XM_019812374.2;
refGene_gn=MADD;refGene_tx=XM_019812375.2;refGene_gn=MADD;refGene_tx=X
M_019812376.2;refGene_gn=MADD;refGene_tx=XM_019812378.2;refGene_gn=MAD
D;refGene_tx=XM_019812379.2;refGene_gn=MADD;refGene_tx=XM_019812380.2;
refGene_gn=MADD;refGene_tx=XM_019812381.2;refGene_gn=MADD;refGene_tx=X
M_019812382.2;refGene_gn=MADD;refGene_tx=XM_019812383.2;refGene_gn=MAD
D;refGene_tx=XM_019812384.2
D1 101294958 101295149
refGene_gn=MADD;refGene_tx=XM_003993232.5;refGene_gn=MADD;refGene_tx=X
M_003993233.5;refGene_gn=MADD;refGene_tx=XM_003993234.5;refGene_gn=MAD
D;refGene_tx=XM_006937269.4;refGene_gn=MADD;refGene_tx=XM_006937270.4;
refGene_gn=MADD;refGene_tx=XM_006937271.4;refGene_gn=MADD;refGene_tx=X
M_006937278.4;refGene_gn=MADD;refGene_tx=XM_006937279.4;refGene_gn=MAD
D;refGene_tx=XM_006937280.4;refGene_gn=MADD;refGene_tx=XM_011286854.3;
refGene_gn=MADD;refGene_tx=XM_011286856.3;refGene_gn=MADD;refGene_tx=X
M_011286857.3;refGene_gn=MADD;refGene_tx=XM_011286858.3;refGene_gn=MAD
D;refGene_tx=XM_019812363.2;refGene_gn=MADD;refGene_tx=XM_019812364.2;
refGene_gn=MADD;refGene_tx=XM_019812365.2;refGene_gn=MADD;refGene_tx=X
M_019812366.1;refGene_gn=MADD;refGene_tx=XM_019812367.2;refGene_gn=MAD
D;refGene_tx=XM_019812368.2;refGene_gn=MADD;refGene_tx=XM_019812369.2;
refGene_gn=MADD;refGene_tx=XM_019812370.2;refGene_gn=MADD;refGene_tx=X
M_019812371.2;refGene_gn=MADD;refGene_tx=XM_019812372.2;refGene_gn=MAD
D;refGene_tx=XM_019812373.2;refGene_gn=MADD;refGene_tx=XM_019812374.2;

refGene_gn=MADD;refGene_tx=XM_019812375.2;refGene_gn=MADD;refGene_tx=XM_019812376.2;refGene_gn=MADD;refGene_tx=XM_019812378.2;refGene_gn=MADD;refGene_tx=XM_019812379.2;refGene_gn=MADD;refGene_tx=XM_019812380.2;refGene_gn=MADD;refGene_tx=XM_019812381.2;refGene_gn=MADD;refGene_tx=XM_019812382.2;refGene_gn=MADD;refGene_tx=XM_019812383.2;refGene_gn=MADD;refGene_tx=XM_019812384.2
D1 101295532 101295786
refGene_gn=MADD;refGene_tx=XM_003993232.5;refGene_gn=MADD;refGene_tx=XM_003993233.5;refGene_gn=MADD;refGene_tx=XM_003993234.5;refGene_gn=MADD;refGene_tx=XM_006937269.4;refGene_gn=MADD;refGene_tx=XM_006937270.4;refGene_gn=MADD;refGene_tx=XM_006937271.4;refGene_gn=MADD;refGene_tx=XM_006937278.4;refGene_gn=MADD;refGene_tx=XM_006937279.4;refGene_gn=MADD;refGene_tx=XM_006937280.4;refGene_gn=MADD;refGene_tx=XM_011286854.3;refGene_gn=MADD;refGene_tx=XM_011286856.3;refGene_gn=MADD;refGene_tx=XM_011286857.3;refGene_gn=MADD;refGene_tx=XM_011286858.3;refGene_gn=MADD;refGene_tx=XM_019812363.2;refGene_gn=MADD;refGene_tx=XM_019812364.2;refGene_gn=MADD;refGene_tx=XM_019812365.2;refGene_gn=MADD;refGene_tx=XM_019812366.1;refGene_gn=MADD;refGene_tx=XM_019812367.2;refGene_gn=MADD;refGene_tx=XM_019812368.2;refGene_gn=MADD;refGene_tx=XM_019812369.2;refGene_gn=MADD;refGene_tx=XM_019812370.2;refGene_gn=MADD;refGene_tx=XM_019812371.2;refGene_gn=MADD;refGene_tx=XM_019812372.2;refGene_gn=MADD;refGene_tx=XM_019812373.2;refGene_gn=MADD;refGene_tx=XM_019812374.2;refGene_gn=MADD;refGene_tx=XM_019812375.2;refGene_gn=MADD;refGene_tx=XM_019812376.2;refGene_gn=MADD;refGene_tx=XM_019812378.2;refGene_gn=MADD;refGene_tx=XM_019812379.2;refGene_gn=MADD;refGene_tx=XM_019812380.2;refGene_gn=MADD;refGene_tx=XM_019812381.2;refGene_gn=MADD;refGene_tx=XM_019812382.2;refGene_gn=MADD;refGene_tx=XM_019812383.2;refGene_gn=MADD;refGene_tx=XM_019812384.2
D1 101296030 101296170
refGene_gn=MADD;refGene_tx=XM_003993232.5;refGene_gn=MADD;refGene_tx=XM_003993233.5;refGene_gn=MADD;refGene_tx=XM_003993234.5;refGene_gn=MADD;refGene_tx=XM_006937269.4;refGene_gn=MADD;refGene_tx=XM_006937270.4;refGene_gn=MADD;refGene_tx=XM_006937271.4;refGene_gn=MADD;refGene_tx=XM_006937278.4;refGene_gn=MADD;refGene_tx=XM_006937279.4;refGene_gn=MADD;refGene_tx=XM_006937280.4;refGene_gn=MADD;refGene_tx=XM_011286854.3;refGene_gn=MADD;refGene_tx=XM_011286856.3;refGene_gn=MADD;refGene_tx=XM_011286857.3;refGene_gn=MADD;refGene_tx=XM_011286858.3;refGene_gn=MADD;refGene_tx=XM_019812363.2;refGene_gn=MADD;refGene_tx=XM_019812364.2;refGene_gn=MADD;refGene_tx=XM_019812365.2;refGene_gn=MADD;refGene_tx=XM_019812366.1;refGene_gn=MADD;refGene_tx=XM_019812367.2;refGene_gn=MADD;refGene_tx=XM_019812368.2;refGene_gn=MADD;refGene_tx=XM_019812369.2;refGene_gn=MADD;refGene_tx=XM_019812370.2;refGene_gn=MADD;refGene_tx=XM_019812371.2;refGene_gn=MADD;refGene_tx=XM_019812372.2;refGene_gn=MADD;refGene_tx=XM_019812373.2;refGene_gn=MADD;refGene_tx=XM_019812374.2;refGene_gn=MADD;refGene_tx=XM_019812375.2;refGene_gn=MADD;refGene_tx=XM_019812376.2;refGene_gn=MADD;refGene_tx=XM_019812378.2;refGene_gn=MADD;refGene_tx=XM_019812379.2;refGene_gn=MADD;refGene_tx=XM_019812380.2;refGene_gn=MADD;refGene_tx=XM_019812381.2;refGene_gn=MADD;refGene_tx=XM_019812382.2;refGene_gn=MADD;refGene_tx=XM_019812383.2;refGene_gn=MADD;refGene_tx=XM_019812384.2

D1	101296973	101297075
refGene_gn=MADD;refGene_tx=XM_003993232.5;refGene_gn=MADD;refGene_tx=XM_003993233.5;refGene_gn=MADD;refGene_tx=XM_003993234.5;refGene_gn=MADD;refGene_tx=XM_006937269.4;refGene_gn=MADD;refGene_tx=XM_006937270.4;refGene_gn=MADD;refGene_tx=XM_006937271.4;refGene_gn=MADD;refGene_tx=XM_006937278.4;refGene_gn=MADD;refGene_tx=XM_006937279.4;refGene_gn=MADD;refGene_tx=XM_006937280.4;refGene_gn=MADD;refGene_tx=XM_011286854.3;refGene_gn=MADD;refGene_tx=XM_011286856.3;refGene_gn=MADD;refGene_tx=XM_011286857.3;refGene_gn=MADD;refGene_tx=XM_011286858.3;refGene_gn=MADD;refGene_tx=XM_019812363.2;refGene_gn=MADD;refGene_tx=XM_019812364.2;refGene_gn=MADD;refGene_tx=XM_019812365.2;refGene_gn=MADD;refGene_tx=XM_019812366.1;refGene_gn=MADD;refGene_tx=XM_019812367.2;refGene_gn=MADD;refGene_tx=XM_019812368.2;refGene_gn=MADD;refGene_tx=XM_019812369.2;refGene_gn=MADD;refGene_tx=XM_019812370.2;refGene_gn=MADD;refGene_tx=XM_019812371.2;refGene_gn=MADD;refGene_tx=XM_019812372.2;refGene_gn=MADD;refGene_tx=XM_019812373.2;refGene_gn=MADD;refGene_tx=XM_019812374.2;refGene_gn=MADD;refGene_tx=XM_019812375.2;refGene_gn=MADD;refGene_tx=XM_019812376.2;refGene_gn=MADD;refGene_tx=XM_019812378.2;refGene_gn=MADD;refGene_tx=XM_019812379.2;refGene_gn=MADD;refGene_tx=XM_019812380.2;refGene_gn=MADD;refGene_tx=XM_019812381.2;refGene_gn=MADD;refGene_tx=XM_019812382.2;refGene_gn=MADD;refGene_tx=XM_019812383.2;refGene_gn=MADD;refGene_tx=XM_019812384.2
D1	101298383	101298443
refGene_gn=MADD;refGene_tx=XM_003993234.5;refGene_gn=MADD;refGene_tx=XM_006937269.4;refGene_gn=MADD;refGene_tx=XM_011286854.3;refGene_gn=MADD;refGene_tx=XM_011286856.3;refGene_gn=MADD;refGene_tx=XM_011286857.3;refGene_gn=MADD;refGene_tx=XM_019812363.2;refGene_gn=MADD;refGene_tx=XM_019812364.2;refGene_gn=MADD;refGene_tx=XM_019812365.2;refGene_gn=MADD;refGene_tx=XM_019812366.1;refGene_gn=MADD;refGene_tx=XM_019812367.2;refGene_gn=MADD;refGene_tx=XM_019812368.2;refGene_gn=MADD;refGene_tx=XM_019812369.2;refGene_gn=MADD;refGene_tx=XM_019812370.2;refGene_gn=MADD;refGene_tx=XM_019812371.2;refGene_gn=MADD;refGene_tx=XM_019812372.2;refGene_gn=MADD;refGene_tx=XM_019812374.2;refGene_gn=MADD;refGene_tx=XM_019812376.2;refGene_gn=MADD;refGene_tx=XM_019812378.2;refGene_gn=MADD;refGene_tx=XM_019812379.2;refGene_gn=MADD;refGene_tx=XM_019812380.2;refGene_gn=MADD;refGene_tx=XM_019812381.2;refGene_gn=MADD;refGene_tx=XM_019812382.2;refGene_gn=MADD;refGene_tx=XM_019812383.2;refGene_gn=MADD;refGene_tx=XM_019812384.2
D1	101298802	101298905
refGene_gn=MADD;refGene_tx=XM_003993232.5;refGene_gn=MADD;refGene_tx=XM_003993233.5;refGene_gn=MADD;refGene_tx=XM_003993234.5;refGene_gn=MADD;refGene_tx=XM_006937269.4;refGene_gn=MADD;refGene_tx=XM_006937270.4;refGene_gn=MADD;refGene_tx=XM_006937271.4;refGene_gn=MADD;refGene_tx=XM_006937278.4;refGene_gn=MADD;refGene_tx=XM_006937279.4;refGene_gn=MADD;refGene_tx=XM_006937280.4;refGene_gn=MADD;refGene_tx=XM_011286854.3;refGene_gn=MADD;refGene_tx=XM_011286856.3;refGene_gn=MADD;refGene_tx=XM_011286857.3;refGene_gn=MADD;refGene_tx=XM_011286858.3;refGene_gn=MADD;refGene_tx=XM_019812363.2;refGene_gn=MADD;refGene_tx=XM_019812364.2;refGene_gn=MADD;refGene_tx=XM_019812365.2;refGene_gn=MADD;refGene_tx=XM_019812366.1;refGene_gn=MADD;refGene_tx=XM_019812367.2;refGene_gn=MAD

D;refGene_tx=XM_019812368.2;refGene_gn=MADD;refGene_tx=XM_019812369.2;
refGene_gn=MADD;refGene_tx=XM_019812370.2;refGene_gn=MADD;refGene_tx=X
M_019812371.2;refGene_gn=MADD;refGene_tx=XM_019812372.2;refGene_gn=MAD
D;refGene_tx=XM_019812373.2;refGene_gn=MADD;refGene_tx=XM_019812374.2;
refGene_gn=MADD;refGene_tx=XM_019812375.2;refGene_gn=MADD;refGene_tx=X
M_019812376.2;refGene_gn=MADD;refGene_tx=XM_019812378.2;refGene_gn=MAD
D;refGene_tx=XM_019812379.2;refGene_gn=MADD;refGene_tx=XM_019812380.2;
refGene_gn=MADD;refGene_tx=XM_019812381.2;refGene_gn=MADD;refGene_tx=X
M_019812382.2;refGene_gn=MADD;refGene_tx=XM_019812383.2;refGene_gn=MAD
D;refGene_tx=XM_019812384.2
D1 101299266 101299453
refGene_gn=MADD;refGene_tx=XM_003993232.5;refGene_gn=MADD;refGene_tx=X
M_003993233.5;refGene_gn=MADD;refGene_tx=XM_003993234.5;refGene_gn=MAD
D;refGene_tx=XM_006937269.4;refGene_gn=MADD;refGene_tx=XM_006937270.4;
refGene_gn=MADD;refGene_tx=XM_006937271.4;refGene_gn=MADD;refGene_tx=X
M_006937278.4;refGene_gn=MADD;refGene_tx=XM_006937279.4;refGene_gn=MAD
D;refGene_tx=XM_006937280.4;refGene_gn=MADD;refGene_tx=XM_011286854.3;
refGene_gn=MADD;refGene_tx=XM_011286856.3;refGene_gn=MADD;refGene_tx=X
M_011286857.3;refGene_gn=MADD;refGene_tx=XM_011286858.3;refGene_gn=MAD
D;refGene_tx=XM_019812363.2;refGene_gn=MADD;refGene_tx=XM_019812364.2;
refGene_gn=MADD;refGene_tx=XM_019812365.2;refGene_gn=MADD;refGene_tx=X
M_019812366.1;refGene_gn=MADD;refGene_tx=XM_019812367.2;refGene_gn=MAD
D;refGene_tx=XM_019812368.2;refGene_gn=MADD;refGene_tx=XM_019812369.2;
refGene_gn=MADD;refGene_tx=XM_019812370.2;refGene_gn=MADD;refGene_tx=X
M_019812371.2;refGene_gn=MADD;refGene_tx=XM_019812372.2;refGene_gn=MAD
D;refGene_tx=XM_019812373.2;refGene_gn=MADD;refGene_tx=XM_019812374.2;
refGene_gn=MADD;refGene_tx=XM_019812375.2;refGene_gn=MADD;refGene_tx=X
M_019812376.2;refGene_gn=MADD;refGene_tx=XM_019812378.2;refGene_gn=MAD
D;refGene_tx=XM_019812379.2;refGene_gn=MADD;refGene_tx=XM_019812380.2;
refGene_gn=MADD;refGene_tx=XM_019812381.2;refGene_gn=MADD;refGene_tx=X
M_019812382.2;refGene_gn=MADD;refGene_tx=XM_019812383.2;refGene_gn=MAD
D;refGene_tx=XM_019812384.2
D1 101299549 101299700
refGene_gn=MADD;refGene_tx=XM_003993232.5;refGene_gn=MADD;refGene_tx=X
M_003993233.5;refGene_gn=MADD;refGene_tx=XM_003993234.5;refGene_gn=MAD
D;refGene_tx=XM_006937269.4;refGene_gn=MADD;refGene_tx=XM_006937270.4;
refGene_gn=MADD;refGene_tx=XM_006937271.4;refGene_gn=MADD;refGene_tx=X
M_006937278.4;refGene_gn=MADD;refGene_tx=XM_006937279.4;refGene_gn=MAD
D;refGene_tx=XM_006937280.4;refGene_gn=MADD;refGene_tx=XM_011286854.3;
refGene_gn=MADD;refGene_tx=XM_011286856.3;refGene_gn=MADD;refGene_tx=X
M_011286857.3;refGene_gn=MADD;refGene_tx=XM_011286858.3;refGene_gn=MAD
D;refGene_tx=XM_019812363.2;refGene_gn=MADD;refGene_tx=XM_019812364.2;
refGene_gn=MADD;refGene_tx=XM_019812365.2;refGene_gn=MADD;refGene_tx=X
M_019812366.1;refGene_gn=MADD;refGene_tx=XM_019812367.2;refGene_gn=MAD
D;refGene_tx=XM_019812368.2;refGene_gn=MADD;refGene_tx=XM_019812369.2;
refGene_gn=MADD;refGene_tx=XM_019812370.2;refGene_gn=MADD;refGene_tx=X
M_019812371.2;refGene_gn=MADD;refGene_tx=XM_019812372.2;refGene_gn=MAD
D;refGene_tx=XM_019812373.2;refGene_gn=MADD;refGene_tx=XM_019812374.2;
refGene_gn=MADD;refGene_tx=XM_019812375.2;refGene_gn=MADD;refGene_tx=X
M_019812376.2;refGene_gn=MADD;refGene_tx=XM_019812378.2;refGene_gn=MAD

D;refGene_tx=XM_019812379.2;refGene_gn=MADD;refGene_tx=XM_019812380.2;
refGene_gn=MADD;refGene_tx=XM_019812381.2;refGene_gn=MADD;refGene_tx=X
M_019812382.2;refGene_gn=MADD;refGene_tx=XM_019812383.2;refGene_gn=MAD
D;refGene_tx=XM_019812384.2
D1 101300001 101300208
refGene_gn=MADD;refGene_tx=XM_003993232.5;refGene_gn=MADD;refGene_tx=X
M_003993233.5;refGene_gn=MADD;refGene_tx=XM_003993234.5;refGene_gn=MAD
D;refGene_tx=XM_006937269.4;refGene_gn=MADD;refGene_tx=XM_006937270.4;
refGene_gn=MADD;refGene_tx=XM_006937271.4;refGene_gn=MADD;refGene_tx=X
M_006937278.4;refGene_gn=MADD;refGene_tx=XM_006937279.4;refGene_gn=MAD
D;refGene_tx=XM_006937280.4;refGene_gn=MADD;refGene_tx=XM_011286854.3;
refGene_gn=MADD;refGene_tx=XM_011286856.3;refGene_gn=MADD;refGene_tx=X
M_011286857.3;refGene_gn=MADD;refGene_tx=XM_011286858.3;refGene_gn=MAD
D;refGene_tx=XM_019812363.2;refGene_gn=MADD;refGene_tx=XM_019812364.2;
refGene_gn=MADD;refGene_tx=XM_019812365.2;refGene_gn=MADD;refGene_tx=X
M_019812366.1;refGene_gn=MADD;refGene_tx=XM_019812367.2;refGene_gn=MAD
D;refGene_tx=XM_019812368.2;refGene_gn=MADD;refGene_tx=XM_019812369.2;
refGene_gn=MADD;refGene_tx=XM_019812370.2;refGene_gn=MADD;refGene_tx=X
M_019812371.2;refGene_gn=MADD;refGene_tx=XM_019812372.2;refGene_gn=MAD
D;refGene_tx=XM_019812373.2;refGene_gn=MADD;refGene_tx=XM_019812374.2;
refGene_gn=MADD;refGene_tx=XM_019812375.2;refGene_gn=MADD;refGene_tx=X
M_019812376.2;refGene_gn=MADD;refGene_tx=XM_019812378.2;refGene_gn=MAD
D;refGene_tx=XM_019812379.2;refGene_gn=MADD;refGene_tx=XM_019812380.2;
refGene_gn=MADD;refGene_tx=XM_019812381.2;refGene_gn=MADD;refGene_tx=X
M_019812382.2;refGene_gn=MADD;refGene_tx=XM_019812383.2;refGene_gn=MAD
D;refGene_tx=XM_019812384.2
D1 101301918 101301972
refGene_gn=MADD;refGene_tx=XM_011286854.3;refGene_gn=MADD;refGene_tx=X
M_011286856.3;refGene_gn=MADD;refGene_tx=XM_011286857.3;refGene_gn=MAD
D;refGene_tx=XM_011286858.3;refGene_gn=MADD;refGene_tx=XM_019812363.2;
refGene_gn=MADD;refGene_tx=XM_019812364.2;refGene_gn=MADD;refGene_tx=X
M_019812365.2;refGene_gn=MADD;refGene_tx=XM_019812366.1;refGene_gn=MAD
D;refGene_tx=XM_019812367.2;refGene_gn=MADD;refGene_tx=XM_019812368.2;
refGene_gn=MADD;refGene_tx=XM_019812369.2;refGene_gn=MADD;refGene_tx=X
M_019812370.2;refGene_gn=MADD;refGene_tx=XM_019812371.2;refGene_gn=MAD
D;refGene_tx=XM_019812373.2;refGene_gn=MADD;refGene_tx=XM_019812374.2;
refGene_gn=MADD;refGene_tx=XM_019812375.2;refGene_gn=MADD;refGene_tx=X
M_019812376.2;refGene_gn=MADD;refGene_tx=XM_019812378.2;refGene_gn=MAD
D;refGene_tx=XM_019812379.2;refGene_gn=MADD;refGene_tx=XM_019812380.2;
refGene_gn=MADD;refGene_tx=XM_019812381.2;refGene_gn=MADD;refGene_tx=X
M_019812382.2;refGene_gn=MADD;refGene_tx=XM_019812383.2;refGene_gn=MAD
D;refGene_tx=XM_019812384.2
D1 101302955 101303056
refGene_gn=MADD;refGene_tx=XM_003993232.5;refGene_gn=MADD;refGene_tx=X
M_003993233.5;refGene_gn=MADD;refGene_tx=XM_003993234.5;refGene_gn=MAD
D;refGene_tx=XM_006937269.4;refGene_gn=MADD;refGene_tx=XM_006937270.4;
refGene_gn=MADD;refGene_tx=XM_006937271.4;refGene_gn=MADD;refGene_tx=X
M_006937278.4;refGene_gn=MADD;refGene_tx=XM_006937279.4;refGene_gn=MAD
D;refGene_tx=XM_006937280.4;refGene_gn=MADD;refGene_tx=XM_011286854.3;
refGene_gn=MADD;refGene_tx=XM_011286856.3;refGene_gn=MADD;refGene_tx=X

M_011286857.3;refGene_gn=MADD;refGene_tx=XM_011286858.3;refGene_gn=MAD
D;refGene_tx=XM_019812363.2;refGene_gn=MADD;refGene_tx=XM_019812364.2;
refGene_gn=MADD;refGene_tx=XM_019812365.2;refGene_gn=MADD;refGene_tx=X
M_019812366.1;refGene_gn=MADD;refGene_tx=XM_019812367.2;refGene_gn=MAD
D;refGene_tx=XM_019812368.2;refGene_gn=MADD;refGene_tx=XM_019812369.2;
refGene_gn=MADD;refGene_tx=XM_019812370.2;refGene_gn=MADD;refGene_tx=X
M_019812371.2;refGene_gn=MADD;refGene_tx=XM_019812372.2;refGene_gn=MAD
D;refGene_tx=XM_019812373.2;refGene_gn=MADD;refGene_tx=XM_019812374.2;
refGene_gn=MADD;refGene_tx=XM_019812375.2;refGene_gn=MADD;refGene_tx=X
M_019812376.2;refGene_gn=MADD;refGene_tx=XM_019812378.2;refGene_gn=MAD
D;refGene_tx=XM_019812379.2;refGene_gn=MADD;refGene_tx=XM_019812380.2;
refGene_gn=MADD;refGene_tx=XM_019812381.2;refGene_gn=MADD;refGene_tx=X
M_019812382.2;refGene_gn=MADD;refGene_tx=XM_019812383.2;refGene_gn=MAD
D;refGene_tx=XM_019812384.2
D1 101303642 101303726
refGene_gn=MADD;refGene_tx=XM_003993232.5;refGene_gn=MADD;refGene_tx=X
M_003993233.5;refGene_gn=MADD;refGene_tx=XM_003993234.5;refGene_gn=MAD
D;refGene_tx=XM_006937269.4;refGene_gn=MADD;refGene_tx=XM_006937270.4;
refGene_gn=MADD;refGene_tx=XM_006937271.4;refGene_gn=MADD;refGene_tx=X
M_006937278.4;refGene_gn=MADD;refGene_tx=XM_006937279.4;refGene_gn=MAD
D;refGene_tx=XM_006937280.4;refGene_gn=MADD;refGene_tx=XM_011286854.3;
refGene_gn=MADD;refGene_tx=XM_011286856.3;refGene_gn=MADD;refGene_tx=X
M_011286857.3;refGene_gn=MADD;refGene_tx=XM_011286858.3;refGene_gn=MAD
D;refGene_tx=XM_019812363.2;refGene_gn=MADD;refGene_tx=XM_019812364.2;
refGene_gn=MADD;refGene_tx=XM_019812365.2;refGene_gn=MADD;refGene_tx=X
M_019812366.1;refGene_gn=MADD;refGene_tx=XM_019812367.2;refGene_gn=MAD
D;refGene_tx=XM_019812368.2;refGene_gn=MADD;refGene_tx=XM_019812369.2;
refGene_gn=MADD;refGene_tx=XM_019812370.2;refGene_gn=MADD;refGene_tx=X
M_019812371.2;refGene_gn=MADD;refGene_tx=XM_019812372.2;refGene_gn=MAD
D;refGene_tx=XM_019812373.2;refGene_gn=MADD;refGene_tx=XM_019812374.2;
refGene_gn=MADD;refGene_tx=XM_019812375.2;refGene_gn=MADD;refGene_tx=X
M_019812376.2;refGene_gn=MADD;refGene_tx=XM_019812378.2;refGene_gn=MAD
D;refGene_tx=XM_019812379.2;refGene_gn=MADD;refGene_tx=XM_019812380.2;
refGene_gn=MADD;refGene_tx=XM_019812381.2;refGene_gn=MADD;refGene_tx=X
M_019812382.2;refGene_gn=MADD;refGene_tx=XM_019812383.2;refGene_gn=MAD
D;refGene_tx=XM_019812384.2
D1 101304057 101304216
refGene_gn=MADD;refGene_tx=XM_003993232.5;refGene_gn=MADD;refGene_tx=X
M_003993233.5;refGene_gn=MADD;refGene_tx=XM_003993234.5;refGene_gn=MAD
D;refGene_tx=XM_006937269.4;refGene_gn=MADD;refGene_tx=XM_006937270.4;
refGene_gn=MADD;refGene_tx=XM_006937271.4;refGene_gn=MADD;refGene_tx=X
M_006937278.4;refGene_gn=MADD;refGene_tx=XM_006937279.4;refGene_gn=MAD
D;refGene_tx=XM_006937280.4;refGene_gn=MADD;refGene_tx=XM_011286854.3;
refGene_gn=MADD;refGene_tx=XM_011286856.3;refGene_gn=MADD;refGene_tx=X
M_011286857.3;refGene_gn=MADD;refGene_tx=XM_011286858.3;refGene_gn=MAD
D;refGene_tx=XM_019812363.2;refGene_gn=MADD;refGene_tx=XM_019812364.2;
refGene_gn=MADD;refGene_tx=XM_019812365.2;refGene_gn=MADD;refGene_tx=X
M_019812366.1;refGene_gn=MADD;refGene_tx=XM_019812367.2;refGene_gn=MAD
D;refGene_tx=XM_019812368.2;refGene_gn=MADD;refGene_tx=XM_019812369.2;
refGene_gn=MADD;refGene_tx=XM_019812370.2;refGene_gn=MADD;refGene_tx=X

M_019812371.2;refGene_gn=MADD;refGene_tx=XM_019812372.2;refGene_gn=MAD
D;refGene_tx=XM_019812373.2;refGene_gn=MADD;refGene_tx=XM_019812374.2;
refGene_gn=MADD;refGene_tx=XM_019812375.2;refGene_gn=MADD;refGene_tx=X
M_019812376.2;refGene_gn=MADD;refGene_tx=XM_019812378.2;refGene_gn=MAD
D;refGene_tx=XM_019812379.2;refGene_gn=MADD;refGene_tx=XM_019812380.2;
refGene_gn=MADD;refGene_tx=XM_019812381.2;refGene_gn=MADD;refGene_tx=X
M_019812382.2;refGene_gn=MADD;refGene_tx=XM_019812383.2;refGene_gn=MAD
D;refGene_tx=XM_019812384.2
D1 101306536 101306645
refGene_gn=MADD;refGene_tx=XM_003993232.5;refGene_gn=MADD;refGene_tx=X
M_003993233.5;refGene_gn=MADD;refGene_tx=XM_003993234.5;refGene_gn=MAD
D;refGene_tx=XM_006937269.4;refGene_gn=MADD;refGene_tx=XM_006937270.4;
refGene_gn=MADD;refGene_tx=XM_006937271.4;refGene_gn=MADD;refGene_tx=X
M_006937278.4;refGene_gn=MADD;refGene_tx=XM_006937279.4;refGene_gn=MAD
D;refGene_tx=XM_006937280.4;refGene_gn=MADD;refGene_tx=XM_011286854.3;
refGene_gn=MADD;refGene_tx=XM_011286856.3;refGene_gn=MADD;refGene_tx=X
M_011286857.3;refGene_gn=MADD;refGene_tx=XM_011286858.3;refGene_gn=MAD
D;refGene_tx=XM_019812363.2;refGene_gn=MADD;refGene_tx=XM_019812364.2;
refGene_gn=MADD;refGene_tx=XM_019812365.2;refGene_gn=MADD;refGene_tx=X
M_019812366.1;refGene_gn=MADD;refGene_tx=XM_019812367.2;refGene_gn=MAD
D;refGene_tx=XM_019812368.2;refGene_gn=MADD;refGene_tx=XM_019812369.2;
refGene_gn=MADD;refGene_tx=XM_019812370.2;refGene_gn=MADD;refGene_tx=X
M_019812371.2;refGene_gn=MADD;refGene_tx=XM_019812372.2;refGene_gn=MAD
D;refGene_tx=XM_019812373.2;refGene_gn=MADD;refGene_tx=XM_019812374.2;
refGene_gn=MADD;refGene_tx=XM_019812375.2;refGene_gn=MADD;refGene_tx=X
M_019812376.2;refGene_gn=MADD;refGene_tx=XM_019812378.2;refGene_gn=MAD
D;refGene_tx=XM_019812379.2;refGene_gn=MADD;refGene_tx=XM_019812380.2;
refGene_gn=MADD;refGene_tx=XM_019812381.2;refGene_gn=MADD;refGene_tx=X
M_019812382.2;refGene_gn=MADD;refGene_tx=XM_019812383.2;refGene_gn=MAD
D;refGene_tx=XM_019812384.2
D1 101306927 101306990
refGene_gn=MADD;refGene_tx=XM_006937270.4;refGene_gn=MADD;refGene_tx=X
M_011286854.3;refGene_gn=MADD;refGene_tx=XM_011286856.3;refGene_gn=MAD
D;refGene_tx=XM_011286857.3;refGene_gn=MADD;refGene_tx=XM_011286858.3;
refGene_gn=MADD;refGene_tx=XM_019812363.2;refGene_gn=MADD;refGene_tx=X
M_019812364.2;refGene_gn=MADD;refGene_tx=XM_019812365.2;refGene_gn=MAD
D;refGene_tx=XM_019812366.1;refGene_gn=MADD;refGene_tx=XM_019812367.2;
refGene_gn=MADD;refGene_tx=XM_019812368.2;refGene_gn=MADD;refGene_tx=X
M_019812369.2;refGene_gn=MADD;refGene_tx=XM_019812370.2;refGene_gn=MAD
D;refGene_tx=XM_019812371.2;refGene_gn=MADD;refGene_tx=XM_019812372.2;
refGene_gn=MADD;refGene_tx=XM_019812373.2;refGene_gn=MADD;refGene_tx=X
M_019812375.2;refGene_gn=MADD;refGene_tx=XM_019812376.2;refGene_gn=MAD
D;refGene_tx=XM_019812378.2;refGene_gn=MADD;refGene_tx=XM_019812379.2;
refGene_gn=MADD;refGene_tx=XM_019812380.2;refGene_gn=MADD;refGene_tx=X
M_019812381.2;refGene_gn=MADD;refGene_tx=XM_019812382.2;refGene_gn=MAD
D;refGene_tx=XM_019812383.2;refGene_gn=MADD;refGene_tx=XM_019812384.2
D1 101307220 101307341
refGene_gn=MADD;refGene_tx=XM_003993232.5;refGene_gn=MADD;refGene_tx=X
M_003993233.5;refGene_gn=MADD;refGene_tx=XM_003993234.5;refGene_gn=MAD
D;refGene_tx=XM_006937269.4;refGene_gn=MADD;refGene_tx=XM_006937270.4;

refGene_gn=MADD;refGene_tx=XM_006937271.4;refGene_gn=MADD;refGene_tx=XM_006937278.4;refGene_gn=MADD;refGene_tx=XM_006937279.4;refGene_gn=MADD;refGene_tx=XM_006937280.4;refGene_gn=MADD;refGene_tx=XM_011286854.3;refGene_gn=MADD;refGene_tx=XM_011286856.3;refGene_gn=MADD;refGene_tx=XM_011286857.3;refGene_gn=MADD;refGene_tx=XM_011286858.3;refGene_gn=MADD;refGene_tx=XM_019812363.2;refGene_gn=MADD;refGene_tx=XM_019812364.2;refGene_gn=MADD;refGene_tx=XM_019812365.2;refGene_gn=MADD;refGene_tx=XM_019812366.1;refGene_gn=MADD;refGene_tx=XM_019812367.2;refGene_gn=MADD;refGene_tx=XM_019812368.2;refGene_gn=MADD;refGene_tx=XM_019812369.2;refGene_gn=MADD;refGene_tx=XM_019812370.2;refGene_gn=MADD;refGene_tx=XM_019812371.2;refGene_gn=MADD;refGene_tx=XM_019812372.2;refGene_gn=MADD;refGene_tx=XM_019812373.2;refGene_gn=MADD;refGene_tx=XM_019812374.2;refGene_gn=MADD;refGene_tx=XM_019812375.2;refGene_gn=MADD;refGene_tx=XM_019812376.2;refGene_gn=MADD;refGene_tx=XM_019812378.2;refGene_gn=MADD;refGene_tx=XM_019812379.2;refGene_gn=MADD;refGene_tx=XM_019812380.2;refGene_gn=MADD;refGene_tx=XM_019812381.2;refGene_gn=MADD;refGene_tx=XM_019812382.2;refGene_gn=MADD;refGene_tx=XM_019812383.2;refGene_gn=MADD;refGene_tx=XM_019812384.2
D1 101307440 101307546
refGene_gn=MADD;refGene_tx=XM_003993232.5;refGene_gn=MADD;refGene_tx=XM_003993233.5;refGene_gn=MADD;refGene_tx=XM_003993234.5;refGene_gn=MADD;refGene_tx=XM_006937269.4;refGene_gn=MADD;refGene_tx=XM_006937270.4;refGene_gn=MADD;refGene_tx=XM_006937271.4;refGene_gn=MADD;refGene_tx=XM_006937278.4;refGene_gn=MADD;refGene_tx=XM_006937279.4;refGene_gn=MADD;refGene_tx=XM_006937280.4;refGene_gn=MADD;refGene_tx=XM_011286854.3;refGene_gn=MADD;refGene_tx=XM_011286856.3;refGene_gn=MADD;refGene_tx=XM_011286857.3;refGene_gn=MADD;refGene_tx=XM_011286858.3;refGene_gn=MADD;refGene_tx=XM_019812363.2;refGene_gn=MADD;refGene_tx=XM_019812364.2;refGene_gn=MADD;refGene_tx=XM_019812365.2;refGene_gn=MADD;refGene_tx=XM_019812366.1;refGene_gn=MADD;refGene_tx=XM_019812367.2;refGene_gn=MADD;refGene_tx=XM_019812368.2;refGene_gn=MADD;refGene_tx=XM_019812369.2;refGene_gn=MADD;refGene_tx=XM_019812370.2;refGene_gn=MADD;refGene_tx=XM_019812371.2;refGene_gn=MADD;refGene_tx=XM_019812372.2;refGene_gn=MADD;refGene_tx=XM_019812373.2;refGene_gn=MADD;refGene_tx=XM_019812374.2;refGene_gn=MADD;refGene_tx=XM_019812375.2;refGene_gn=MADD;refGene_tx=XM_019812376.2;refGene_gn=MADD;refGene_tx=XM_019812378.2;refGene_gn=MADD;refGene_tx=XM_019812379.2;refGene_gn=MADD;refGene_tx=XM_019812380.2;refGene_gn=MADD;refGene_tx=XM_019812381.2;refGene_gn=MADD;refGene_tx=XM_019812382.2;refGene_gn=MADD;refGene_tx=XM_019812383.2;refGene_gn=MADD;refGene_tx=XM_019812384.2
D1 101308704 101308815
refGene_gn=MADD;refGene_tx=XM_003993232.5;refGene_gn=MADD;refGene_tx=XM_003993233.5;refGene_gn=MADD;refGene_tx=XM_003993234.5;refGene_gn=MADD;refGene_tx=XM_006937269.4;refGene_gn=MADD;refGene_tx=XM_006937270.4;refGene_gn=MADD;refGene_tx=XM_006937271.4;refGene_gn=MADD;refGene_tx=XM_006937278.4;refGene_gn=MADD;refGene_tx=XM_006937279.4;refGene_gn=MADD;refGene_tx=XM_006937280.4;refGene_gn=MADD;refGene_tx=XM_011286854.3;refGene_gn=MADD;refGene_tx=XM_011286856.3;refGene_gn=MADD;refGene_tx=XM_011286857.3;refGene_gn=MADD;refGene_tx=XM_011286858.3;refGene_gn=MADD;refGene_tx=XM_019812363.2;refGene_gn=MADD;refGene_tx=XM_019812364.2;

refGene_gn=MADD;refGene_tx=XM_019812365.2;refGene_gn=MADD;refGene_tx=XM_019812366.1;refGene_gn=MADD;refGene_tx=XM_019812367.2;refGene_gn=MADD;refGene_tx=XM_019812368.2;refGene_gn=MADD;refGene_tx=XM_019812369.2;refGene_gn=MADD;refGene_tx=XM_019812370.2;refGene_gn=MADD;refGene_tx=XM_019812371.2;refGene_gn=MADD;refGene_tx=XM_019812372.2;refGene_gn=MADD;refGene_tx=XM_019812373.2;refGene_gn=MADD;refGene_tx=XM_019812374.2;refGene_gn=MADD;refGene_tx=XM_019812375.2;refGene_gn=MADD;refGene_tx=XM_019812376.2;refGene_gn=MADD;refGene_tx=XM_019812378.2;refGene_gn=MADD;refGene_tx=XM_019812379.2;refGene_gn=MADD;refGene_tx=XM_019812380.2;refGene_gn=MADD;refGene_tx=XM_019812381.2;refGene_gn=MADD;refGene_tx=XM_019812382.2;refGene_gn=MADD;refGene_tx=XM_019812383.2;refGene_gn=MADD;refGene_tx=XM_019812384.2
D1 101311385 101311493
refGene_gn=MADD;refGene_tx=XM_003993232.5;refGene_gn=MADD;refGene_tx=XM_003993233.5;refGene_gn=MADD;refGene_tx=XM_003993234.5;refGene_gn=MADD;refGene_tx=XM_006937269.4;refGene_gn=MADD;refGene_tx=XM_006937270.4;refGene_gn=MADD;refGene_tx=XM_006937271.4;refGene_gn=MADD;refGene_tx=XM_006937278.4;refGene_gn=MADD;refGene_tx=XM_006937279.4;refGene_gn=MADD;refGene_tx=XM_006937280.4;refGene_gn=MADD;refGene_tx=XM_011286854.3;refGene_gn=MADD;refGene_tx=XM_011286856.3;refGene_gn=MADD;refGene_tx=XM_011286857.3;refGene_gn=MADD;refGene_tx=XM_011286858.3;refGene_gn=MADD;refGene_tx=XM_019812363.2;refGene_gn=MADD;refGene_tx=XM_019812364.2;refGene_gn=MADD;refGene_tx=XM_019812365.2;refGene_gn=MADD;refGene_tx=XM_019812366.1;refGene_gn=MADD;refGene_tx=XM_019812367.2;refGene_gn=MADD;refGene_tx=XM_019812368.2;refGene_gn=MADD;refGene_tx=XM_019812369.2;refGene_gn=MADD;refGene_tx=XM_019812370.2;refGene_gn=MADD;refGene_tx=XM_019812371.2;refGene_gn=MADD;refGene_tx=XM_019812372.2;refGene_gn=MADD;refGene_tx=XM_019812373.2;refGene_gn=MADD;refGene_tx=XM_019812374.2;refGene_gn=MADD;refGene_tx=XM_019812375.2;refGene_gn=MADD;refGene_tx=XM_019812376.2;refGene_gn=MADD;refGene_tx=XM_019812378.2;refGene_gn=MADD;refGene_tx=XM_019812379.2;refGene_gn=MADD;refGene_tx=XM_019812380.2;refGene_gn=MADD;refGene_tx=XM_019812381.2;refGene_gn=MADD;refGene_tx=XM_019812382.2;refGene_gn=MADD;refGene_tx=XM_019812383.2;refGene_gn=MADD;refGene_tx=XM_019812384.2
D1 101316934 101317099
refGene_gn=MADD;refGene_tx=XM_003993232.5;refGene_gn=MADD;refGene_tx=XM_003993233.5;refGene_gn=MADD;refGene_tx=XM_003993234.5;refGene_gn=MADD;refGene_tx=XM_006937269.4;refGene_gn=MADD;refGene_tx=XM_006937270.4;refGene_gn=MADD;refGene_tx=XM_006937271.4;refGene_gn=MADD;refGene_tx=XM_006937278.4;refGene_gn=MADD;refGene_tx=XM_006937279.4;refGene_gn=MADD;refGene_tx=XM_006937280.4;refGene_gn=MADD;refGene_tx=XM_011286854.3;refGene_gn=MADD;refGene_tx=XM_011286856.3;refGene_gn=MADD;refGene_tx=XM_011286857.3;refGene_gn=MADD;refGene_tx=XM_011286858.3;refGene_gn=MADD;refGene_tx=XM_019812363.2;refGene_gn=MADD;refGene_tx=XM_019812364.2;refGene_gn=MADD;refGene_tx=XM_019812365.2;refGene_gn=MADD;refGene_tx=XM_019812366.1;refGene_gn=MADD;refGene_tx=XM_019812367.2;refGene_gn=MADD;refGene_tx=XM_019812368.2;refGene_gn=MADD;refGene_tx=XM_019812369.2;refGene_gn=MADD;refGene_tx=XM_019812370.2;refGene_gn=MADD;refGene_tx=XM_019812371.2;refGene_gn=MADD;refGene_tx=XM_019812372.2;refGene_gn=MADD;refGene_tx=XM_019812373.2;refGene_gn=MADD;refGene_tx=XM_019812374.2;

refGene_gn=MADD;refGene_tx=XM_019812375.2;refGene_gn=MADD;refGene_tx=XM_019812376.2;refGene_gn=MADD;refGene_tx=XM_019812378.2;refGene_gn=MADD;refGene_tx=XM_019812379.2;refGene_gn=MADD;refGene_tx=XM_019812380.2;refGene_gn=MADD;refGene_tx=XM_019812381.2;refGene_gn=MADD;refGene_tx=XM_019812382.2;refGene_gn=MADD;refGene_tx=XM_019812383.2;refGene_gn=MADD;refGene_tx=XM_019812384.2
D1 101317347 101317420
refGene_gn=MADD;refGene_tx=XM_003993232.5;refGene_gn=MADD;refGene_tx=XM_003993233.5;refGene_gn=MADD;refGene_tx=XM_003993234.5;refGene_gn=MADD;refGene_tx=XM_006937269.4;refGene_gn=MADD;refGene_tx=XM_006937270.4;refGene_gn=MADD;refGene_tx=XM_006937271.4;refGene_gn=MADD;refGene_tx=XM_006937278.4;refGene_gn=MADD;refGene_tx=XM_006937279.4;refGene_gn=MADD;refGene_tx=XM_006937280.4;refGene_gn=MADD;refGene_tx=XM_011286854.3;refGene_gn=MADD;refGene_tx=XM_011286856.3;refGene_gn=MADD;refGene_tx=XM_011286857.3;refGene_gn=MADD;refGene_tx=XM_011286858.3;refGene_gn=MADD;refGene_tx=XM_019812363.2;refGene_gn=MADD;refGene_tx=XM_019812364.2;refGene_gn=MADD;refGene_tx=XM_019812365.2;refGene_gn=MADD;refGene_tx=XM_019812366.1;refGene_gn=MADD;refGene_tx=XM_019812367.2;refGene_gn=MADD;refGene_tx=XM_019812368.2;refGene_gn=MADD;refGene_tx=XM_019812369.2;refGene_gn=MADD;refGene_tx=XM_019812370.2;refGene_gn=MADD;refGene_tx=XM_019812371.2;refGene_gn=MADD;refGene_tx=XM_019812372.2;refGene_gn=MADD;refGene_tx=XM_019812373.2;refGene_gn=MADD;refGene_tx=XM_019812374.2;refGene_gn=MADD;refGene_tx=XM_019812375.2;refGene_gn=MADD;refGene_tx=XM_019812376.2;refGene_gn=MADD;refGene_tx=XM_019812378.2;refGene_gn=MADD;refGene_tx=XM_019812379.2;refGene_gn=MADD;refGene_tx=XM_019812380.2;refGene_gn=MADD;refGene_tx=XM_019812381.2;refGene_gn=MADD;refGene_tx=XM_019812382.2;refGene_gn=MADD;refGene_tx=XM_019812383.2;refGene_gn=MADD;refGene_tx=XM_019812384.2
D1 101317560 101317667
refGene_gn=MADD;refGene_tx=XM_003993232.5;refGene_gn=MADD;refGene_tx=XM_003993233.5;refGene_gn=MADD;refGene_tx=XM_003993234.5;refGene_gn=MADD;refGene_tx=XM_006937269.4;refGene_gn=MADD;refGene_tx=XM_006937270.4;refGene_gn=MADD;refGene_tx=XM_006937271.4;refGene_gn=MADD;refGene_tx=XM_006937278.4;refGene_gn=MADD;refGene_tx=XM_006937279.4;refGene_gn=MADD;refGene_tx=XM_006937280.4;refGene_gn=MADD;refGene_tx=XM_011286854.3;refGene_gn=MADD;refGene_tx=XM_011286856.3;refGene_gn=MADD;refGene_tx=XM_011286857.3;refGene_gn=MADD;refGene_tx=XM_011286858.3;refGene_gn=MADD;refGene_tx=XM_019812363.2;refGene_gn=MADD;refGene_tx=XM_019812364.2;refGene_gn=MADD;refGene_tx=XM_019812365.2;refGene_gn=MADD;refGene_tx=XM_019812366.1;refGene_gn=MADD;refGene_tx=XM_019812367.2;refGene_gn=MADD;refGene_tx=XM_019812368.2;refGene_gn=MADD;refGene_tx=XM_019812369.2;refGene_gn=MADD;refGene_tx=XM_019812370.2;refGene_gn=MADD;refGene_tx=XM_019812371.2;refGene_gn=MADD;refGene_tx=XM_019812372.2;refGene_gn=MADD;refGene_tx=XM_019812373.2;refGene_gn=MADD;refGene_tx=XM_019812374.2;refGene_gn=MADD;refGene_tx=XM_019812375.2;refGene_gn=MADD;refGene_tx=XM_019812376.2;refGene_gn=MADD;refGene_tx=XM_019812378.2;refGene_gn=MADD;refGene_tx=XM_019812379.2;refGene_gn=MADD;refGene_tx=XM_019812380.2;refGene_gn=MADD;refGene_tx=XM_019812381.2;refGene_gn=MADD;refGene_tx=XM_019812382.2;refGene_gn=MADD;refGene_tx=XM_019812383.2;refGene_gn=MADD;refGene_tx=XM_019812384.2

D1	101318345	101318348
refGene_gn=MADD;refGene_tx=XM_019812384.2
D1	101319640	101319649
refGene_gn=MADD;refGene_tx=XM_019812363.2;refGene_gn=MADD;refGene_tx=XM_019812364.2;refGene_gn=MADD;refGene_tx=XM_019812365.2;refGene_gn=MADD;refGene_tx=XM_019812366.1;refGene_gn=MADD;refGene_tx=XM_019812367.2;refGene_gn=MADD;refGene_tx=XM_019812368.2;refGene_gn=MADD;refGene_tx=XM_019812370.2;refGene_gn=MADD;refGene_tx=XM_019812371.2;refGene_gn=MADD;refGene_tx=XM_019812372.2;refGene_gn=MADD;refGene_tx=XM_019812373.2;refGene_gn=MADD;refGene_tx=XM_019812374.2;refGene_gn=MADD;refGene_tx=XM_019812375.2;refGene_gn=MADD;refGene_tx=XM_019812376.2;refGene_gn=MADD;refGene_tx=XM_019812378.2;refGene_gn=MADD;refGene_tx=XM_019812379.2;refGene_gn=MADD;refGene_tx=XM_019812380.2
D1	101319834	101319904
refGene_gn=MADD;refGene_tx=XM_003993232.5;refGene_gn=MADD;refGene_tx=XM_003993233.5;refGene_gn=MADD;refGene_tx=XM_003993234.5;refGene_gn=MADD;refGene_tx=XM_006937269.4;refGene_gn=MADD;refGene_tx=XM_006937270.4;refGene_gn=MADD;refGene_tx=XM_006937271.4;refGene_gn=MADD;refGene_tx=XM_006937278.4;refGene_gn=MADD;refGene_tx=XM_006937279.4;refGene_gn=MADD;refGene_tx=XM_011286854.3;refGene_gn=MADD;refGene_tx=XM_019812363.2;refGene_gn=MADD;refGene_tx=XM_019812364.2;refGene_gn=MADD;refGene_tx=XM_019812365.2;refGene_gn=MADD;refGene_tx=XM_019812366.1;refGene_gn=MADD;refGene_tx=XM_019812367.2;refGene_gn=MADD;refGene_tx=XM_019812368.2;refGene_gn=MADD;refGene_tx=XM_019812369.2;refGene_gn=MADD;refGene_tx=XM_019812370.2;refGene_gn=MADD;refGene_tx=XM_019812371.2;refGene_gn=MADD;refGene_tx=XM_019812372.2;refGene_gn=MADD;refGene_tx=XM_019812373.2;refGene_gn=MADD;refGene_tx=XM_019812374.2;refGene_gn=MADD;refGene_tx=XM_019812375.2;refGene_gn=MADD;refGene_tx=XM_019812376.2;refGene_gn=MADD;refGene_tx=XM_019812378.2
D1	101319928	101319982
refGene_gn=MADD;refGene_tx=XM_019812379.2;refGene_gn=MADD;refGene_tx=XM_019812380.2;refGene_gn=MADD;refGene_tx=XM_019812381.2;refGene_gn=MADD;refGene_tx=XM_019812382.2
D1	101321803	101321850
refGene_gn=MADD;refGene_tx=XM_003993232.5;refGene_gn=MADD;refGene_tx=XM_003993233.5;refGene_gn=MADD;refGene_tx=XM_003993234.5;refGene_gn=MADD;refGene_tx=XM_006937269.4;refGene_gn=MADD;refGene_tx=XM_006937270.4;refGene_gn=MADD;refGene_tx=XM_006937271.4;refGene_gn=MADD;refGene_tx=XM_006937278.4;refGene_gn=MADD;refGene_tx=XM_006937279.4;refGene_gn=MADD;refGene_tx=XM_006937280.4;refGene_gn=MADD;refGene_tx=XM_011286854.3;refGene_gn=MADD;refGene_tx=XM_011286856.3;refGene_gn=MADD;refGene_tx=XM_011286857.3;refGene_gn=MADD;refGene_tx=XM_011286858.3;refGene_gn=MADD;refGene_tx=XM_019812363.2;refGene_gn=MADD;refGene_tx=XM_019812364.2;refGene_gn=MADD;refGene_tx=XM_019812365.2;refGene_gn=MADD;refGene_tx=XM_019812366.1;refGene_gn=MADD;refGene_tx=XM_019812367.2;refGene_gn=MADD;refGene_tx=XM_019812368.2;refGene_gn=MADD;refGene_tx=XM_019812369.2;refGene_gn=MADD;refGene_tx=XM_019812370.2;refGene_gn=MADD;refGene_tx=XM_019812371.2;refGene_gn=MADD;refGene_tx=XM_019812372.2;refGene_gn=MADD;refGene_tx=XM_019812373.2;refGene_gn=MADD;refGene_tx=XM_019812374.2;refGene_gn=MADD;refGene_tx=XM_019812375.2;refGene_gn=MADD;refGene_tx=X

M_019812376.2;refGene_gn=MADD;refGene_tx=XM_019812378.2;refGene_gn=MAD
D;refGene_tx=XM_019812383.2
D1 101322183 101322288
refGene_gn=MADD;refGene_tx=XM_003993232.5;refGene_gn=MADD;refGene_tx=X
M_003993233.5;refGene_gn=MADD;refGene_tx=XM_003993234.5;refGene_gn=MAD
D;refGene_tx=XM_006937269.4;refGene_gn=MADD;refGene_tx=XM_006937270.4;
refGene_gn=MADD;refGene_tx=XM_006937271.4;refGene_gn=MADD;refGene_tx=X
M_006937278.4;refGene_gn=MADD;refGene_tx=XM_006937279.4;refGene_gn=MAD
D;refGene_tx=XM_011286854.3;refGene_gn=MADD;refGene_tx=XM_019812363.2;
refGene_gn=MADD;refGene_tx=XM_019812364.2;refGene_gn=MADD;refGene_tx=X
M_019812365.2;refGene_gn=MADD;refGene_tx=XM_019812366.1;refGene_gn=MAD
D;refGene_tx=XM_019812367.2;refGene_gn=MADD;refGene_tx=XM_019812368.2;
refGene_gn=MADD;refGene_tx=XM_019812369.2;refGene_gn=MADD;refGene_tx=X
M_019812370.2;refGene_gn=MADD;refGene_tx=XM_019812371.2;refGene_gn=MAD
D;refGene_tx=XM_019812372.2;refGene_gn=MADD;refGene_tx=XM_019812373.2;
refGene_gn=MADD;refGene_tx=XM_019812374.2;refGene_gn=MADD;refGene_tx=X
M_019812375.2;refGene_gn=MADD;refGene_tx=XM_019812376.2;refGene_gn=MAD
D;refGene_tx=XM_019812378.2
D1 101324988 101324999
refGene_gn=MYBPC3;refGene_tx=XM_019812396.1;refGene_gn=MYBPC3;refGene_
tx=XM_019812397.1
D1 101325187 101325374
refGene_gn=MYBPC3;refGene_tx=XM_019812396.1;refGene_gn=MYBPC3;refGene_
tx=XM_019812397.1
D1 101325648 101325785
refGene_gn=MYBPC3;refGene_tx=XM_019812396.1;refGene_gn=MYBPC3;refGene_
tx=XM_019812397.1
D1 101325898 101326058
refGene_gn=MYBPC3;refGene_tx=XM_019812396.1;refGene_gn=MYBPC3;refGene_
tx=XM_019812397.1
D1 101326283 101326423
refGene_gn=MYBPC3;refGene_tx=XM_019812396.1;refGene_gn=MYBPC3;refGene_
tx=XM_019812397.1
D1 101326581 101326777
refGene_gn=MYBPC3;refGene_tx=XM_019812396.1;refGene_gn=MYBPC3;refGene_
tx=XM_019812397.1
D1 101326911 101327000
refGene_gn=MYBPC3;refGene_tx=XM_019812396.1;refGene_gn=MYBPC3;refGene_
tx=XM_019812397.1
D1 101327693 101327861
refGene_gn=MYBPC3;refGene_tx=XM_019812396.1;refGene_gn=MYBPC3;refGene_
tx=XM_019812397.1
D1 101328043 101328178
refGene_gn=MYBPC3;refGene_tx=XM_019812396.1;refGene_gn=MYBPC3;refGene_
tx=XM_019812397.1
D1 101329501 101329690
refGene_gn=MYBPC3;refGene_tx=XM_019812396.1;refGene_gn=MYBPC3;refGene_
tx=XM_019812397.1
D1 101329779 101329884
refGene_gn=MYBPC3;refGene_tx=XM_019812396.1;refGene_gn=MYBPC3;refGene_

tx=XM_019812397.1
D1 101330153 101330313
refGene_gn=MYBPC3;refGene_tx=XM_019812396.1;refGene_gn=MYBPC3;refGene_tx=XM_019812397.1
D1 101331011 101331095
refGene_gn=MYBPC3;refGene_tx=XM_019812396.1;refGene_gn=MYBPC3;refGene_tx=XM_019812397.1
D1 101331369 101331509
refGene_gn=MYBPC3;refGene_tx=XM_019812396.1;refGene_gn=MYBPC3;refGene_tx=XM_019812397.1
D1 101333052 101333082
refGene_gn=MYBPC3;refGene_tx=XM_019812396.1;refGene_gn=MYBPC3;refGene_tx=XM_019812397.1
D1 101333171 101333278
refGene_gn=MYBPC3;refGene_tx=XM_019812396.1;refGene_gn=MYBPC3;refGene_tx=XM_019812397.1
D1 101333982 101334148
refGene_gn=MYBPC3;refGene_tx=XM_019812396.1;refGene_gn=MYBPC3;refGene_tx=XM_019812397.1
D1 101334578 101334745
refGene_gn=MYBPC3;refGene_tx=XM_019812396.1;refGene_gn=MYBPC3;refGene_tx=XM_019812397.1
D1 101334840 101334946
refGene_gn=MYBPC3;refGene_tx=XM_019812396.1;refGene_gn=MYBPC3;refGene_tx=XM_019812397.1
D1 101335034 101335159
refGene_gn=MYBPC3;refGene_tx=XM_019812396.1;refGene_gn=MYBPC3;refGene_tx=XM_019812397.1
D1 101335297 101335300
refGene_gn=MYBPC3;refGene_tx=XM_019812396.1;refGene_gn=MYBPC3;refGene_tx=XM_019812397.1
D1 101335575 101335708
refGene_gn=MYBPC3;refGene_tx=XM_019812396.1;refGene_gn=MYBPC3;refGene_tx=XM_019812397.1
D1 101336215 101336379
refGene_gn=MYBPC3;refGene_tx=XM_019812396.1;refGene_gn=MYBPC3;refGene_tx=XM_019812397.1
D1 101336602 101336620
refGene_gn=MYBPC3;refGene_tx=XM_019812396.1;refGene_gn=MYBPC3;refGene_tx=XM_019812397.1
D1 101336951 101336954
refGene_gn=MYBPC3;refGene_tx=XM_019812396.1;refGene_gn=MYBPC3;refGene_tx=XM_019812397.1
D1 101337567 101337621
refGene_gn=MYBPC3;refGene_tx=XM_019812396.1;refGene_gn=MYBPC3;refGene_tx=XM_019812397.1
D1 101337778 101337808
refGene_gn=MYBPC3;refGene_tx=XM_019812396.1;refGene_gn=MYBPC3;refGene_tx=XM_019812397.1
D1 101337986 101338035

refGene_gn=MYBPC3;refGene_tx=XM_019812396.1;refGene_gn=MYBPC3;refGene_tx=XM_019812397.1
D1 101338544 101338662
refGene_gn=MYBPC3;refGene_tx=XM_019812396.1;refGene_gn=MYBPC3;refGene_tx=XM_019812397.1
D1 101339492 101339641
refGene_gn=MYBPC3;refGene_tx=XM_019812396.1;refGene_gn=MYBPC3;refGene_tx=XM_019812397.1
D1 101339731 101339824
refGene_gn=MYBPC3;refGene_tx=XM_019812396.1;refGene_gn=MYBPC3;refGene_tx=XM_019812397.1
D1 101339913 101340027
refGene_gn=MYBPC3;refGene_tx=XM_019812396.1;refGene_gn=MYBPC3;refGene_tx=XM_019812397.1
D1 101340570 101340837
refGene_gn=MYBPC3;refGene_tx=XM_019812396.1;refGene_gn=MYBPC3;refGene_tx=XM_019812397.1
D1 101341928 101341953
refGene_gn=MYBPC3;refGene_tx=XM_019812396.1;refGene_gn=MYBPC3;refGene_tx=XM_019812397.1
D1 101344580 101344900
refGene_gn=SPI1;refGene_tx=XM_003993240.4;refGene_gn=SPI1;refGene_tx=XM_006937283.2;refGene_gn=SPI1;refGene_tx=XM_019812398.1;refGene_gn=SPI1;refGene_tx=XM_019812399.1
D1 101345325 101345488
refGene_gn=SPI1;refGene_tx=XM_003993240.4;refGene_gn=SPI1;refGene_tx=XM_006937283.2;refGene_gn=SPI1;refGene_tx=XM_019812398.1;refGene_gn=SPI1;refGene_tx=XM_019812399.1
D1 101346335 101346523
refGene_gn=SPI1;refGene_tx=XM_003993240.4;refGene_gn=SPI1;refGene_tx=XM_006937283.2;refGene_gn=SPI1;refGene_tx=XM_019812398.1;refGene_gn=SPI1;refGene_tx=XM_019812399.1
D1 101357139 101357239
refGene_gn=SPI1;refGene_tx=XM_003993240.4;refGene_gn=SPI1;refGene_tx=XM_006937283.2;refGene_gn=SPI1;refGene_tx=XM_019812398.1;refGene_gn=SPI1;refGene_tx=XM_019812399.1
D1 101360100 101360235
refGene_gn=SPI1;refGene_tx=XM_003993240.4;refGene_gn=SPI1;refGene_tx=XM_006937283.2;refGene_gn=SPI1;refGene_tx=XM_019812398.1;refGene_gn=SPI1;refGene_tx=XM_019812399.1
D1 101383482 101383526
refGene_gn=SLC39A13;refGene_tx=XM_023239820.1;refGene_gn=SLC39A13;refGene_tx=XM_023239823.1
D1 101385004 101385313
refGene_gn=SLC39A13;refGene_tx=XM_023239820.1;refGene_gn=SLC39A13;refGene_tx=XM_023239821.1;refGene_gn=SLC39A13;refGene_tx=XM_023239822.1;refGene_gn=SLC39A13;refGene_tx=XM_023239823.1;refGene_gn=SLC39A13;refGene_tx=XM_023239825.1
D1 101387213 101387347
refGene_gn=SLC39A13;refGene_tx=XM_023239820.1;refGene_gn=SLC39A13;refG

ene_tx=XM_023239821.1;refGene_gn=SLC39A13;refGene_tx=XM_023239822.1;refGene_gn=SLC39A13;refGene_tx=XM_023239823.1;refGene_gn=SLC39A13;refGene_tx=XM_023239825.1
D1 101387630 101387770
refGene_gn=SLC39A13;refGene_tx=XM_023239820.1;refGene_gn=SLC39A13;refGene_tx=XM_023239821.1;refGene_gn=SLC39A13;refGene_tx=XM_023239822.1;refGene_gn=SLC39A13;refGene_tx=XM_023239823.1;refGene_gn=SLC39A13;refGene_tx=XM_023239825.1
D1 101388168 101388273
refGene_gn=SLC39A13;refGene_tx=XM_023239820.1;refGene_gn=SLC39A13;refGene_tx=XM_023239821.1;refGene_gn=SLC39A13;refGene_tx=XM_023239822.1;refGene_gn=SLC39A13;refGene_tx=XM_023239825.1
D1 101388386 101388476
refGene_gn=SLC39A13;refGene_tx=XM_023239820.1;refGene_gn=SLC39A13;refGene_tx=XM_023239821.1;refGene_gn=SLC39A13;refGene_tx=XM_023239822.1;refGene_gn=SLC39A13;refGene_tx=XM_023239823.1;refGene_gn=SLC39A13;refGene_tx=XM_023239825.1
D1 101389221 101389272
refGene_gn=SLC39A13;refGene_tx=XM_023239820.1;refGene_gn=SLC39A13;refGene_tx=XM_023239821.1;refGene_gn=SLC39A13;refGene_tx=XM_023239822.1;refGene_gn=SLC39A13;refGene_tx=XM_023239823.1;refGene_gn=SLC39A13;refGene_tx=XM_023239825.1
D1 101389570 101389703
refGene_gn=SLC39A13;refGene_tx=XM_023239820.1;refGene_gn=SLC39A13;refGene_tx=XM_023239821.1;refGene_gn=SLC39A13;refGene_tx=XM_023239822.1;refGene_gn=SLC39A13;refGene_tx=XM_023239823.1;refGene_gn=SLC39A13;refGene_tx=XM_023239825.1
D1 101389881 101389981
refGene_gn=SLC39A13;refGene_tx=XM_023239820.1;refGene_gn=SLC39A13;refGene_tx=XM_023239821.1;refGene_gn=SLC39A13;refGene_tx=XM_023239822.1;refGene_gn=SLC39A13;refGene_tx=XM_023239823.1;refGene_gn=SLC39A13;refGene_tx=XM_023239825.1
D1 101390106 101390182
refGene_gn=SLC39A13;refGene_tx=XM_023239820.1;refGene_gn=SLC39A13;refGene_tx=XM_023239821.1;refGene_gn=SLC39A13;refGene_tx=XM_023239822.1;refGene_gn=SLC39A13;refGene_tx=XM_023239823.1
D1 101393620 101393731
refGene_gn=PSMC3;refGene_tx=XM_003993242.5
D1 101393910 101393992
refGene_gn=PSMC3;refGene_tx=XM_003993242.5
D1 101394855 101395001
refGene_gn=PSMC3;refGene_tx=XM_003993242.5
D1 101395219 101395316
refGene_gn=PSMC3;refGene_tx=XM_003993242.5
D1 101396132 101396281
refGene_gn=PSMC3;refGene_tx=XM_003993242.5
D1 101396402 101396546
refGene_gn=PSMC3;refGene_tx=XM_003993242.5
D1 101397111 101397249
refGene_gn=PSMC3;refGene_tx=XM_003993242.5

D1 101397513 101397576
refGene_gn=PSMC3;refGene_tx=XM_003993242.5
D1 101397689 101397794
refGene_gn=PSMC3;refGene_tx=XM_003993242.5
D1 101398325 101398451
refGene_gn=PSMC3;refGene_tx=XM_003993242.5
D1 101399027 101399111
refGene_gn=PSMC3;refGene_tx=XM_003993242.5
D1 101399367 101399442
refGene_gn=PSMC3;refGene_tx=XM_003993242.5
D1 101401028 101401101
refGene_gn=RAPSN;refGene_tx=XM_003993243.5;refGene_gn=RAPSN;refGene_tx=XM_006937289.3;refGene_gn=RAPSN;refGene_tx=XM_006937290.3
D1 101401558 101401758
refGene_gn=RAPSN;refGene_tx=XM_003993243.5;refGene_gn=RAPSN;refGene_tx=XM_006937289.3;refGene_gn=RAPSN;refGene_tx=XM_006937290.3
D1 101402979 101403033
refGene_gn=RAPSN;refGene_tx=XM_003993243.5;refGene_gn=RAPSN;refGene_tx=XM_006937289.3
D1 101403538 101403661
refGene_gn=RAPSN;refGene_tx=XM_003993243.5;refGene_gn=RAPSN;refGene_tx=XM_006937289.3;refGene_gn=RAPSN;refGene_tx=XM_006937290.3
D1 101403737 101403836
refGene_gn=RAPSN;refGene_tx=XM_003993243.5;refGene_gn=RAPSN;refGene_tx=XM_006937289.3;refGene_gn=RAPSN;refGene_tx=XM_006937290.3
D1 101404349 101404508
refGene_gn=RAPSN;refGene_tx=XM_003993243.5;refGene_gn=RAPSN;refGene_tx=XM_006937289.3;refGene_gn=RAPSN;refGene_tx=XM_006937290.3
D1 101410128 101410467
refGene_gn=RAPSN;refGene_tx=XM_003993243.5;refGene_gn=RAPSN;refGene_tx=XM_006937289.3;refGene_gn=RAPSN;refGene_tx=XM_006937290.3
D1 101410994 101411186
refGene_gn=RAPSN;refGene_tx=XM_003993243.5;refGene_gn=RAPSN;refGene_tx=XM_006937289.3;refGene_gn=RAPSN;refGene_tx=XM_006937290.3
D1 101430180 101430308
refGene_gn=CELF1;refGene_tx=XM_019812409.2;refGene_gn=CELF1;refGene_tx=XM_019812410.2;refGene_gn=CELF1;refGene_tx=XM_019812411.2;refGene_gn=CELF1;refGene_tx=XM_019812412.2;refGene_gn=CELF1;refGene_tx=XM_019812413.2;refGene_gn=CELF1;refGene_tx=XM_023239826.1;refGene_gn=CELF1;refGene_tx=XM_023239827.1;refGene_gn=CELF1;refGene_tx=XM_023239828.1;refGene_gn=CELF1;refGene_tx=XM_023239830.1;refGene_gn=CELF1;refGene_tx=XM_023239831.1
D1 101430699 101430843
refGene_gn=CELF1;refGene_tx=XM_019812409.2;refGene_gn=CELF1;refGene_tx=XM_019812410.2;refGene_gn=CELF1;refGene_tx=XM_019812411.2;refGene_gn=CELF1;refGene_tx=XM_019812412.2;refGene_gn=CELF1;refGene_tx=XM_019812413.2;refGene_gn=CELF1;refGene_tx=XM_023239826.1;refGene_gn=CELF1;refGene_tx=XM_023239827.1;refGene_gn=CELF1;refGene_tx=XM_023239828.1;refGene_gn=CELF1;refGene_tx=XM_023239830.1;refGene_gn=CELF1;refGene_tx=XM_023239831.1

D1 101433284 101433470
refGene_gn=CELF1;refGene_tx=XM_019812409.2;refGene_gn=CELF1;refGene_tx=XM_019812410.2;refGene_gn=CELF1;refGene_tx=XM_019812411.2;refGene_gn=CELF1;refGene_tx=XM_019812412.2;refGene_gn=CELF1;refGene_tx=XM_019812413.2;refGene_gn=CELF1;refGene_tx=XM_023239826.1;refGene_gn=CELF1;refGene_tx=XM_023239827.1;refGene_gn=CELF1;refGene_tx=XM_023239828.1;refGene_gn=CELF1;refGene_tx=XM_023239830.1;refGene_gn=CELF1;refGene_tx=XM_023239831.1
D1 101434137 101434248
refGene_gn=CELF1;refGene_tx=XM_019812409.2;refGene_gn=CELF1;refGene_tx=XM_019812410.2;refGene_gn=CELF1;refGene_tx=XM_019812411.2;refGene_gn=CELF1;refGene_tx=XM_019812412.2;refGene_gn=CELF1;refGene_tx=XM_019812413.2;refGene_gn=CELF1;refGene_tx=XM_023239826.1;refGene_gn=CELF1;refGene_tx=XM_023239827.1;refGene_gn=CELF1;refGene_tx=XM_023239828.1;refGene_gn=CELF1;refGene_tx=XM_023239830.1;refGene_gn=CELF1;refGene_tx=XM_023239831.1
D1 101434588 101434717
refGene_gn=CELF1;refGene_tx=XM_019812409.2;refGene_gn=CELF1;refGene_tx=XM_019812410.2;refGene_gn=CELF1;refGene_tx=XM_019812411.2;refGene_gn=CELF1;refGene_tx=XM_019812412.2;refGene_gn=CELF1;refGene_tx=XM_019812413.2;refGene_gn=CELF1;refGene_tx=XM_023239826.1;refGene_gn=CELF1;refGene_tx=XM_023239827.1;refGene_gn=CELF1;refGene_tx=XM_023239828.1;refGene_gn=CELF1;refGene_tx=XM_023239830.1;refGene_gn=CELF1;refGene_tx=XM_023239831.1
D1 101436223 101436299
refGene_gn=CELF1;refGene_tx=XM_019812409.2;refGene_gn=CELF1;refGene_tx=XM_019812410.2;refGene_gn=CELF1;refGene_tx=XM_019812411.2;refGene_gn=CELF1;refGene_tx=XM_019812412.2;refGene_gn=CELF1;refGene_tx=XM_019812413.2;refGene_gn=CELF1;refGene_tx=XM_023239826.1;refGene_gn=CELF1;refGene_tx=XM_023239827.1;refGene_gn=CELF1;refGene_tx=XM_023239828.1;refGene_gn=CELF1;refGene_tx=XM_023239830.1;refGene_gn=CELF1;refGene_tx=XM_023239831.1
D1 101438798 101438960
refGene_gn=CELF1;refGene_tx=XM_019812409.2;refGene_gn=CELF1;refGene_tx=XM_019812410.2;refGene_gn=CELF1;refGene_tx=XM_019812411.2;refGene_gn=CELF1;refGene_tx=XM_019812412.2;refGene_gn=CELF1;refGene_tx=XM_019812413.2;refGene_gn=CELF1;refGene_tx=XM_023239826.1;refGene_gn=CELF1;refGene_tx=XM_023239827.1;refGene_gn=CELF1;refGene_tx=XM_023239828.1;refGene_gn=CELF1;refGene_tx=XM_023239830.1;refGene_gn=CELF1;refGene_tx=XM_023239831.1
D1 101439282 101439362
refGene_gn=CELF1;refGene_tx=XM_019812409.2;refGene_gn=CELF1;refGene_tx=XM_019812410.2;refGene_gn=CELF1;refGene_tx=XM_019812411.2;refGene_gn=CELF1;refGene_tx=XM_019812412.2;refGene_gn=CELF1;refGene_tx=XM_019812413.2;refGene_gn=CELF1;refGene_tx=XM_023239826.1;refGene_gn=CELF1;refGene_tx=XM_023239827.1;refGene_gn=CELF1;refGene_tx=XM_023239828.1;refGene_gn=CELF1;refGene_tx=XM_023239830.1;refGene_gn=CELF1;refGene_tx=XM_023239831.1
D1 101439898 101440033
refGene_gn=CELF1;refGene_tx=XM_019812409.2;refGene_gn=CELF1;refGene_tx

=XM_019812410.2;refGene_gn=CELF1;refGene_tx=XM_019812411.2;refGene_gn=CELF1;refGene_tx=XM_019812412.2;refGene_gn=CELF1;refGene_tx=XM_019812413.2;refGene_gn=CELF1;refGene_tx=XM_023239826.1;refGene_gn=CELF1;refGene_tx=XM_023239827.1;refGene_gn=CELF1;refGene_tx=XM_023239828.1;refGene_gn=CELF1;refGene_tx=XM_023239830.1;refGene_gn=CELF1;refGene_tx=XM_023239831.1
D1 101443756 101443805
refGene_gn=CELF1;refGene_tx=XM_019812409.2;refGene_gn=CELF1;refGene_tx=XM_019812410.2;refGene_gn=CELF1;refGene_tx=XM_019812411.2;refGene_gn=CELF1;refGene_tx=XM_019812412.2;refGene_gn=CELF1;refGene_tx=XM_019812413.2;refGene_gn=CELF1;refGene_tx=XM_023239826.1;refGene_gn=CELF1;refGene_tx=XM_023239827.1;refGene_gn=CELF1;refGene_tx=XM_023239828.1;refGene_gn=CELF1;refGene_tx=XM_023239830.1;refGene_gn=CELF1;refGene_tx=XM_023239831.1
D1 101444181 101444264
refGene_gn=CELF1;refGene_tx=XM_019812409.2;refGene_gn=CELF1;refGene_tx=XM_019812410.2;refGene_gn=CELF1;refGene_tx=XM_019812411.2;refGene_gn=CELF1;refGene_tx=XM_019812412.2;refGene_gn=CELF1;refGene_tx=XM_019812413.2;refGene_gn=CELF1;refGene_tx=XM_023239826.1;refGene_gn=CELF1;refGene_tx=XM_023239827.1;refGene_gn=CELF1;refGene_tx=XM_023239828.1;refGene_gn=CELF1;refGene_tx=XM_023239830.1;refGene_gn=CELF1;refGene_tx=XM_023239831.1
D1 101446214 101446402
refGene_gn=CELF1;refGene_tx=XM_019812409.2;refGene_gn=CELF1;refGene_tx=XM_019812410.2;refGene_gn=CELF1;refGene_tx=XM_019812411.2;refGene_gn=CELF1;refGene_tx=XM_019812412.2;refGene_gn=CELF1;refGene_tx=XM_019812413.2;refGene_gn=CELF1;refGene_tx=XM_023239826.1;refGene_gn=CELF1;refGene_tx=XM_023239827.1;refGene_gn=CELF1;refGene_tx=XM_023239828.1;refGene_gn=CELF1;refGene_tx=XM_023239830.1;refGene_gn=CELF1;refGene_tx=XM_023239831.1
D1 101455628 101455699
refGene_gn=CELF1;refGene_tx=XM_019812409.2;refGene_gn=CELF1;refGene_tx=XM_023239826.1;refGene_gn=CELF1;refGene_tx=XM_023239827.1;refGene_gn=CELF1;refGene_tx=XM_023239828.1;refGene_gn=CELF1;refGene_tx=XM_023239830.1;refGene_gn=CELF1;refGene_tx=XM_023239831.1
D1 101517383 101517779
refGene_gn=PTPMT1;refGene_tx=XM_023239835.1
D1 101517879 101517960
refGene_gn=PTPMT1;refGene_tx=XM_023239835.1
D1 101525093 101525285
refGene_gn=PTPMT1;refGene_tx=XM_023239835.1
D1 101526605 101526764
refGene_gn=PTPMT1;refGene_tx=XM_023239835.1
D1 101528031 101528892
refGene_gn=KBTBD4;refGene_tx=XM_003993248.5;refGene_gn=KBTBD4;refGene_tx=XM_023239832.1
D1 101529339 101529446
refGene_gn=KBTBD4;refGene_tx=XM_003993248.5;refGene_gn=KBTBD4;refGene_tx=XM_023239832.1
D1 101531133 101531751

refGene_gn=KBTBD4;refGene_tx=XM_003993248.5;refGene_gn=KBTBD4;refGene_tx=XM_023239832.1
D1	101532668	101532687
refGene_gn=KBTBD4;refGene_tx=XM_003993248.5
D1	101532828	101532898
refGene_gn=NDUFS3;refGene_tx=XM_019812414.2
D1	101532997	101533063
refGene_gn=NDUFS3;refGene_tx=XM_019812414.2
D1	101534179	101534277
refGene_gn=NDUFS3;refGene_tx=XM_019812414.2
D1	101534486	101534636
refGene_gn=NDUFS3;refGene_tx=XM_019812414.2
D1	101535480	101535606
refGene_gn=NDUFS3;refGene_tx=XM_019812414.2
D1	101535738	101535858
refGene_gn=NDUFS3;refGene_tx=XM_019812414.2
D1	101537676	101537844
refGene_gn=NDUFS3;refGene_tx=XM_019812414.2
D1	101539145	101539218
refGene_gn=FAM180B;refGene_tx=XM_003993250.4
D1	101540050	101540121
refGene_gn=FAM180B;refGene_tx=XM_003993250.4
D1	101540336	101540732
refGene_gn=FAM180B;refGene_tx=XM_003993250.4
D1	101542025	101542991
refGene_gn=C1QTNF4;refGene_tx=XM_023239834.1
D1	101564902	101564989
refGene_gn=MTCH2;refGene_tx=XM_003993251.4;refGene_gn=MTCH2;refGene_tx=XM_006937299.3
D1	101566812	101566888
refGene_gn=MTCH2;refGene_tx=XM_003993251.4;refGene_gn=MTCH2;refGene_tx=XM_006937299.3
D1	101568177	101568245
refGene_gn=MTCH2;refGene_tx=XM_003993251.4;refGene_gn=MTCH2;refGene_tx=XM_006937299.3
D1	101568906	101568954
refGene_gn=MTCH2;refGene_tx=XM_003993251.4;refGene_gn=MTCH2;refGene_tx=XM_006937299.3
D1	101570004	101570098
refGene_gn=MTCH2;refGene_tx=XM_003993251.4;refGene_gn=MTCH2;refGene_tx=XM_006937299.3
D1	101572471	101572531
refGene_gn=MTCH2;refGene_tx=XM_003993251.4;refGene_gn=MTCH2;refGene_tx=XM_006937299.3
D1	101573073	101573125
refGene_gn=MTCH2;refGene_tx=XM_003993251.4;refGene_gn=MTCH2;refGene_tx=XM_006937299.3
D1	101573398	101573456
refGene_gn=MTCH2;refGene_tx=XM_003993251.4;refGene_gn=MTCH2;refGene_tx=XM_006937299.3

D1 101575547 101575610
refGene_gn=MTCH2;refGene_tx=XM_003993251.4;refGene_gn=MTCH2;refGene_tx
=XM_006937299.3
D1 101576013 101576040
refGene_gn=MTCH2;refGene_tx=XM_003993251.4
D1 101581491 101581598
refGene_gn=MTCH2;refGene_tx=XM_003993251.4;refGene_gn=MTCH2;refGene_tx
=XM_006937299.3
D1 101581756 101581841
refGene_gn=MTCH2;refGene_tx=XM_003993251.4;refGene_gn=MTCH2;refGene_tx
=XM_006937299.3
D1 101584175 101584262
refGene_gn=MTCH2;refGene_tx=XM_003993251.4;refGene_gn=MTCH2;refGene_tx
=XM_006937299.3
D1 101605018 101605192
refGene_gn=AGBL2;refGene_tx=XM_006937300.3;refGene_gn=AGBL2;refGene_tx
=XM_006937301.3;refGene_gn=AGBL2;refGene_tx=XM_006937302.3;refGene_gn=
AGBL2;refGene_tx=XM_019812416.1;refGene_gn=AGBL2;refGene_tx=XM_0198124
17.1
D1 101605526 101605532
refGene_gn=AGBL2;refGene_tx=XM_006937303.4;refGene_gn=AGBL2;refGene_tx
=XM_019812418.2
D1 101606734 101606821
refGene_gn=AGBL2;refGene_tx=XM_006937300.3;refGene_gn=AGBL2;refGene_tx
=XM_006937301.3;refGene_gn=AGBL2;refGene_tx=XM_006937302.3;refGene_gn=
AGBL2;refGene_tx=XM_006937303.4;refGene_gn=AGBL2;refGene_tx=XM_0198124
16.1;refGene_gn=AGBL2;refGene_tx=XM_019812417.1;refGene_gn=AGBL2;refGe
ne_tx=XM_019812418.2
D1 101609951 101610071
refGene_gn=AGBL2;refGene_tx=XM_006937300.3;refGene_gn=AGBL2;refGene_tx
=XM_006937301.3;refGene_gn=AGBL2;refGene_tx=XM_006937302.3;refGene_gn=
AGBL2;refGene_tx=XM_006937303.4;refGene_gn=AGBL2;refGene_tx=XM_0069373
04.3
D1 101610551 101610677
refGene_gn=AGBL2;refGene_tx=XM_006937300.3;refGene_gn=AGBL2;refGene_tx
=XM_006937301.3;refGene_gn=AGBL2;refGene_tx=XM_006937302.3;refGene_gn=
AGBL2;refGene_tx=XM_006937303.4;refGene_gn=AGBL2;refGene_tx=XM_0069373
04.3;refGene_gn=AGBL2;refGene_tx=XM_019812416.1;refGene_gn=AGBL2;refGe
ne_tx=XM_019812417.1;refGene_gn=AGBL2;refGene_tx=XM_019812418.2
D1 101613148 101613215
refGene_gn=AGBL2;refGene_tx=XM_006937300.3;refGene_gn=AGBL2;refGene_tx
=XM_006937301.3;refGene_gn=AGBL2;refGene_tx=XM_006937302.3;refGene_gn=
AGBL2;refGene_tx=XM_006937303.4;refGene_gn=AGBL2;refGene_tx=XM_0069373
04.3;refGene_gn=AGBL2;refGene_tx=XM_019812416.1;refGene_gn=AGBL2;refGe
ne_tx=XM_019812417.1;refGene_gn=AGBL2;refGene_tx=XM_019812418.2
D1 101616716 101616847
refGene_gn=AGBL2;refGene_tx=XM_006937300.3;refGene_gn=AGBL2;refGene_tx
=XM_006937301.3;refGene_gn=AGBL2;refGene_tx=XM_006937302.3;refGene_gn=
AGBL2;refGene_tx=XM_006937303.4;refGene_gn=AGBL2;refGene_tx=XM_0069373
04.3;refGene_gn=AGBL2;refGene_tx=XM_019812416.1;refGene_gn=AGBL2;refGe

ne_tx=XM_019812417.1;refGene_gn=AGBL2;refGene_tx=XM_019812418.2
D1	101618095	101618196
refGene_gn=AGBL2;refGene_tx=XM_006937300.3;refGene_gn=AGBL2;refGene_tx=XM_006937301.3;refGene_gn=AGBL2;refGene_tx=XM_006937302.3;refGene_gn=AGBL2;refGene_tx=XM_006937303.4;refGene_gn=AGBL2;refGene_tx=XM_006937304.3;refGene_gn=AGBL2;refGene_tx=XM_019812416.1;refGene_gn=AGBL2;refGene_tx=XM_019812417.1;refGene_gn=AGBL2;refGene_tx=XM_019812418.2
D1	101618792	101618919
refGene_gn=AGBL2;refGene_tx=XM_006937300.3;refGene_gn=AGBL2;refGene_tx=XM_006937301.3;refGene_gn=AGBL2;refGene_tx=XM_006937302.3;refGene_gn=AGBL2;refGene_tx=XM_006937303.4;refGene_gn=AGBL2;refGene_tx=XM_006937304.3;refGene_gn=AGBL2;refGene_tx=XM_019812416.1;refGene_gn=AGBL2;refGene_tx=XM_019812417.1;refGene_gn=AGBL2;refGene_tx=XM_019812418.2
D1	101623065	101623222
refGene_gn=AGBL2;refGene_tx=XM_006937300.3;refGene_gn=AGBL2;refGene_tx=XM_006937301.3;refGene_gn=AGBL2;refGene_tx=XM_006937302.3;refGene_gn=AGBL2;refGene_tx=XM_006937303.4;refGene_gn=AGBL2;refGene_tx=XM_006937304.3;refGene_gn=AGBL2;refGene_tx=XM_019812416.1;refGene_gn=AGBL2;refGene_tx=XM_019812417.1;refGene_gn=AGBL2;refGene_tx=XM_019812418.2
D1	101624761	101625544
refGene_gn=AGBL2;refGene_tx=XM_006937300.3;refGene_gn=AGBL2;refGene_tx=XM_006937301.3;refGene_gn=AGBL2;refGene_tx=XM_006937302.3;refGene_gn=AGBL2;refGene_tx=XM_006937303.4;refGene_gn=AGBL2;refGene_tx=XM_006937304.3;refGene_gn=AGBL2;refGene_tx=XM_019812416.1;refGene_gn=AGBL2;refGene_tx=XM_019812417.1;refGene_gn=AGBL2;refGene_tx=XM_019812418.2
D1	101628360	101628514
refGene_gn=AGBL2;refGene_tx=XM_006937300.3;refGene_gn=AGBL2;refGene_tx=XM_006937301.3;refGene_gn=AGBL2;refGene_tx=XM_006937302.3;refGene_gn=AGBL2;refGene_tx=XM_006937303.4;refGene_gn=AGBL2;refGene_tx=XM_006937304.3;refGene_gn=AGBL2;refGene_tx=XM_019812416.1;refGene_gn=AGBL2;refGene_tx=XM_019812417.1;refGene_gn=AGBL2;refGene_tx=XM_019812418.2
D1	101636157	101636265
refGene_gn=AGBL2;refGene_tx=XM_006937300.3;refGene_gn=AGBL2;refGene_tx=XM_006937301.3;refGene_gn=AGBL2;refGene_tx=XM_006937302.3;refGene_gn=AGBL2;refGene_tx=XM_006937303.4;refGene_gn=AGBL2;refGene_tx=XM_006937304.3;refGene_gn=AGBL2;refGene_tx=XM_019812416.1;refGene_gn=AGBL2;refGene_tx=XM_019812417.1;refGene_gn=AGBL2;refGene_tx=XM_019812418.2
D1	101644080	101644266
refGene_gn=AGBL2;refGene_tx=XM_006937300.3;refGene_gn=AGBL2;refGene_tx=XM_006937301.3;refGene_gn=AGBL2;refGene_tx=XM_006937302.3;refGene_gn=AGBL2;refGene_tx=XM_006937303.4;refGene_gn=AGBL2;refGene_tx=XM_006937304.3;refGene_gn=AGBL2;refGene_tx=XM_019812416.1;refGene_gn=AGBL2;refGene_tx=XM_019812417.1;refGene_gn=AGBL2;refGene_tx=XM_019812418.2
D1	101645003	101645057
refGene_gn=AGBL2;refGene_tx=XM_006937300.3;refGene_gn=AGBL2;refGene_tx=XM_006937301.3;refGene_gn=AGBL2;refGene_tx=XM_006937303.4;refGene_gn=AGBL2;refGene_tx=XM_006937304.3;refGene_gn=AGBL2;refGene_tx=XM_019812416.1;refGene_gn=AGBL2;refGene_tx=XM_019812418.2
D1	101649219	101649354
refGene_gn=AGBL2;refGene_tx=XM_006937300.3;refGene_gn=AGBL2;refGene_tx

=XM_006937301.3;refGene_gn=AGBL2;refGene_tx=XM_006937302.3;refGene_gn=AGBL2;refGene_tx=XM_006937303.4;refGene_gn=AGBL2;refGene_tx=XM_006937304.3;refGene_gn=AGBL2;refGene_tx=XM_019812416.1;refGene_gn=AGBL2;refGene_tx=XM_019812417.1;refGene_gn=AGBL2;refGene_tx=XM_019812418.2
D1 101650539 101650603
refGene_gn=AGBL2;refGene_tx=XM_006937300.3;refGene_gn=AGBL2;refGene_tx=XM_006937301.3;refGene_gn=AGBL2;refGene_tx=XM_006937302.3;refGene_gn=AGBL2;refGene_tx=XM_006937303.4;refGene_gn=AGBL2;refGene_tx=XM_006937304.3;refGene_gn=AGBL2;refGene_tx=XM_019812416.1;refGene_gn=AGBL2;refGene_tx=XM_019812417.1;refGene_gn=AGBL2;refGene_tx=XM_019812418.2
D1 101650864 101650900
refGene_gn=AGBL2;refGene_tx=XM_006937300.3;refGene_gn=AGBL2;refGene_tx=XM_006937301.3;refGene_gn=AGBL2;refGene_tx=XM_006937302.3;refGene_gn=AGBL2;refGene_tx=XM_006937303.4;refGene_gn=AGBL2;refGene_tx=XM_006937304.3;refGene_gn=AGBL2;refGene_tx=XM_019812416.1;refGene_gn=AGBL2;refGene_tx=XM_019812417.1;refGene_gn=AGBL2;refGene_tx=XM_019812418.2
D1 101653700 101653791
refGene_gn=FNBP4;refGene_tx=XM_011286845.2;refGene_gn=FNBP4;refGene_tx=XM_011286846.3
D1 101657029 101657187
refGene_gn=FNBP4;refGene_tx=XM_011286845.2;refGene_gn=FNBP4;refGene_tx=XM_011286846.3
D1 101665594 101665935
refGene_gn=FNBP4;refGene_tx=XM_011286845.2;refGene_gn=FNBP4;refGene_tx=XM_011286846.3
D1 101666461 101666606
refGene_gn=FNBP4;refGene_tx=XM_011286845.2;refGene_gn=FNBP4;refGene_tx=XM_011286846.3
D1 101666734 101667045
refGene_gn=FNBP4;refGene_tx=XM_011286845.2;refGene_gn=FNBP4;refGene_tx=XM_011286846.3
D1 101672476 101672664
refGene_gn=FNBP4;refGene_tx=XM_011286845.2;refGene_gn=FNBP4;refGene_tx=XM_011286846.3
D1 101673649 101673783
refGene_gn=FNBP4;refGene_tx=XM_011286845.2;refGene_gn=FNBP4;refGene_tx=XM_011286846.3
D1 101675107 101675212
refGene_gn=FNBP4;refGene_tx=XM_011286845.2;refGene_gn=FNBP4;refGene_tx=XM_011286846.3
D1 101677025 101677147
refGene_gn=FNBP4;refGene_tx=XM_011286845.2;refGene_gn=FNBP4;refGene_tx=XM_011286846.3
D1 101680447 101680658
refGene_gn=FNBP4;refGene_tx=XM_011286845.2;refGene_gn=FNBP4;refGene_tx=XM_011286846.3
D1 101681812 101682154
refGene_gn=FNBP4;refGene_tx=XM_011286845.2;refGene_gn=FNBP4;refGene_tx=XM_011286846.3
D1 101685893 101686014

refGene_gn=FNBP4;refGene_tx=XM_011286845.2;refGene_gn=FNBP4;refGene_tx=XM_011286846.3
D1 101686139 101686287
refGene_gn=FNBP4;refGene_tx=XM_011286845.2;refGene_gn=FNBP4;refGene_tx=XM_011286846.3
D1 101688540 101688727
refGene_gn=FNBP4;refGene_tx=XM_011286845.2;refGene_gn=FNBP4;refGene_tx=XM_011286846.3
D1 101689072 101689209
refGene_gn=FNBP4;refGene_tx=XM_011286845.2;refGene_gn=FNBP4;refGene_tx=XM_011286846.3
D1 101695147 101695240
refGene_gn=FNBP4;refGene_tx=XM_011286845.2;refGene_gn=FNBP4;refGene_tx=XM_011286846.3
D1 101696327 101696562
refGene_gn=FNBP4;refGene_tx=XM_011286845.2;refGene_gn=FNBP4;refGene_tx=XM_011286846.3
D1 101707849 101707939
refGene_gn=NUP160;refGene_tx=XM_011286844.2
D1 101708406 101708511
refGene_gn=NUP160;refGene_tx=XM_011286844.2
D1 101710277 101710403
refGene_gn=NUP160;refGene_tx=XM_011286844.2
D1 101711586 101711728
refGene_gn=NUP160;refGene_tx=XM_011286844.2
D1 101712350 101712452
refGene_gn=NUP160;refGene_tx=XM_011286844.2
D1 101713360 101713484
refGene_gn=NUP160;refGene_tx=XM_011286844.2
D1 101713674 101713785
refGene_gn=NUP160;refGene_tx=XM_011286844.2
D1 101714747 101714808
refGene_gn=NUP160;refGene_tx=XM_011286844.2
D1 101717404 101717565
refGene_gn=NUP160;refGene_tx=XM_011286844.2
D1 101722053 101722159
refGene_gn=NUP160;refGene_tx=XM_011286844.2
D1 101722246 101722343
refGene_gn=NUP160;refGene_tx=XM_011286844.2
D1 101722438 101722533
refGene_gn=NUP160;refGene_tx=XM_011286844.2
D1 101722631 101722727
refGene_gn=NUP160;refGene_tx=XM_011286844.2
D1 101727201 101727321
refGene_gn=NUP160;refGene_tx=XM_011286844.2
D1 101728243 101728342
refGene_gn=NUP160;refGene_tx=XM_011286844.2
D1 101729803 101729873
refGene_gn=NUP160;refGene_tx=XM_011286844.2
D1 101730474 101730634

refGene_gn=NUP160;refGene_tx=XM_011286844.2
D1 101731474 101731545
refGene_gn=NUP160;refGene_tx=XM_011286844.2
D1 101732242 101732376
refGene_gn=NUP160;refGene_tx=XM_011286844.2
D1 101734951 101735112
refGene_gn=NUP160;refGene_tx=XM_011286844.2
D1 101735201 101735329
refGene_gn=NUP160;refGene_tx=XM_011286844.2
D1 101735786 101735952
refGene_gn=NUP160;refGene_tx=XM_011286844.2
D1 101736223 101736323
refGene_gn=NUP160;refGene_tx=XM_011286844.2
D1 101737079 101737250
refGene_gn=NUP160;refGene_tx=XM_011286844.2
D1 101737606 101737690
refGene_gn=NUP160;refGene_tx=XM_011286844.2
D1 101740954 101741023
refGene_gn=NUP160;refGene_tx=XM_011286844.2
D1 101742593 101742678
refGene_gn=NUP160;refGene_tx=XM_011286844.2
D1 101744512 101744610
refGene_gn=NUP160;refGene_tx=XM_011286844.2
D1 101744866 101744944
refGene_gn=NUP160;refGene_tx=XM_011286844.2
D1 101750858 101751017
refGene_gn=NUP160;refGene_tx=XM_011286844.2
D1 101751507 101751622
refGene_gn=NUP160;refGene_tx=XM_011286844.2
D1 101752466 101752545
refGene_gn=NUP160;refGene_tx=XM_011286844.2
D1 101753851 101753894
refGene_gn=NUP160;refGene_tx=XM_011286844.2
D1 101753895 101754075
refGene_gn=NUP160;refGene_tx=XM_011286844.2
D1 101754386 101754597
refGene_gn=NUP160;refGene_tx=XM_011286844.2
D1 101759329 101759441
refGene_gn=NUP160;refGene_tx=XM_011286844.2
D1 101759709 101759926
refGene_gn=NUP160;refGene_tx=XM_011286844.2
D1 101983479 101983498
refGene_gn=PTPRJ;refGene_tx=XM_023239838.1
D1 101985586 101985790
refGene_gn=PTPRJ;refGene_tx=XM_023239838.1
D1 101987539 101987791
refGene_gn=PTPRJ;refGene_tx=XM_023239838.1
D1 101988904 101989129
refGene_gn=PTPRJ;refGene_tx=XM_023239838.1
D1 101992411 101992678

refGene_gn=PTPRJ;refGene_tx=XM_023239838.1
D1	101995637	101995873
refGene_gn=PTPRJ;refGene_tx=XM_023239838.1
D1	101995874	101995899
refGene_gn=PTPRJ;refGene_tx=XM_023239838.1
D1	102001020	102001278
refGene_gn=PTPRJ;refGene_tx=XM_023239838.1
D1	102002320	102002599
refGene_gn=PTPRJ;refGene_tx=XM_023239838.1
D1	102004611	102004902
refGene_gn=PTPRJ;refGene_tx=XM_023239838.1
D1	102008836	102008968
refGene_gn=PTPRJ;refGene_tx=XM_023239838.1
D1	102010411	102010622
refGene_gn=PTPRJ;refGene_tx=XM_023239838.1
D1	102010735	102010860
refGene_gn=PTPRJ;refGene_tx=XM_023239838.1
D1	102013090	102013178
refGene_gn=PTPRJ;refGene_tx=XM_023239838.1
D1	102015887	102015929
refGene_gn=PTPRJ;refGene_tx=XM_023239838.1
D1	102016913	102016922
refGene_gn=PTPRJ;refGene_tx=XM_023239838.1
D1	102017027	102017115
refGene_gn=PTPRJ;refGene_tx=XM_023239838.1
D1	102019215	102019306
refGene_gn=PTPRJ;refGene_tx=XM_023239838.1
D1	102021461	102021535
refGene_gn=PTPRJ;refGene_tx=XM_023239838.1
D1	102021638	102021773
refGene_gn=PTPRJ;refGene_tx=XM_023239838.1
D1	102024804	102024924
refGene_gn=PTPRJ;refGene_tx=XM_023239838.1
D1	102028791	102028952
refGene_gn=PTPRJ;refGene_tx=XM_023239838.1
D1	102030657	102030793
refGene_gn=PTPRJ;refGene_tx=XM_023239838.1
D1	102033380	102033539
refGene_gn=PTPRJ;refGene_tx=XM_023239838.1
D1	102049983	102051000
refGene_gn=LOC101098160;refGene_tx=XM_011286891.3
D1	102069752	102070688
refGene_gn=LOC101098400;refGene_tx=XM_003993255.2
D1	102090125	102091055
refGene_gn=LOC101098652;refGene_tx=XM_003993256.3
D1	102100205	102101135
refGene_gn=LOC101098890;refGene_tx=XM_023239010.1
D1	102110365	102111295
refGene_gn=LOC101086844;refGene_tx=XM_003993289.1
D1	102137544	102138474

refGene_gn=LOC101099147;refGene_tx=XM_003993258.2
D1 102154808 102155738
refGene_gn=LOC101087098;refGene_tx=XM_003993290.2
D1 102172999 102173908
refGene_gn=LOC101099399;refGene_tx=XM_003993259.1
D1 102389760 102389911
refGene_gn=LOC111556539;refGene_tx=XM_023239011.1
D1 102389913 102390677
refGene_gn=LOC111556539;refGene_tx=XM_023239011.1
D1 102412426 102413344
refGene_gn=LOC105260998;refGene_tx=XM_011286886.2
D1 102430654 102431584
refGene_gn=LOC101100414;refGene_tx=XM_003993263.2
D1 102442655 102443597
refGene_gn=LOC101089645;refGene_tx=XM_006937314.3
D1 102461245 102462199
refGene_gn=LOC101089895;refGene_tx=XM_003993298.4
D1 102463326 102463394
refGene_gn=LOC101090140;refGene_tx=XM_011286841.2
D1 102472760 102473700
refGene_gn=LOC101090140;refGene_tx=XM_011286841.2
D1 102479744 102480692
refGene_gn=LOC101090388;refGene_tx=XM_019812963.2
D1 102493223 102494231
refGene_gn=LOC101100668;refGene_tx=XM_023239013.1
D1 102504753 102505728
refGene_gn=LOC109492366;refGene_tx=XM_019813046.2
D1 102512853 102513768
refGene_gn=LOC109492367;refGene_tx=XM_019813047.1
D1 102526627 102527525
refGene_gn=LOC101091400;refGene_tx=XM_006937316.3
D1 102528107 102528292
refGene_gn=LOC101091400;refGene_tx=XM_006937316.3
D1 102541836 102542745
refGene_gn=LOC101091653;refGene_tx=XM_006937306.2
D1 102550082 102551024
refGene_gn=LOC101091898;refGene_tx=XM_003993304.3
D1 102578108 102579017
refGene_gn=LOC101092143;refGene_tx=XM_003993305.1
D1 102601990 102602923
refGene_gn=LOC101100916;refGene_tx=XM_003993265.1
D1 102618158 102619085
refGene_gn=LOC101092397;refGene_tx=XM_023239014.1
D1 102624756 102625692
refGene_gn=LOC101101164;refGene_tx=XM_003993266.2
D1 102638149 102639082
refGene_gn=LOC101092643;refGene_tx=XM_011286840.1
D1 102652941 102653871
refGene_gn=LOC101099903;refGene_tx=XM_003993261.2
D1 102668444 102669374

refGene_gn=LOC101092890;refGene_tx=XM_011286839.1
D1	102679074	102680031
refGene_gn=LOC105260996;refGene_tx=XM_011286884.1
D1	102688366	102689302
refGene_gn=LOC101093141;refGene_tx=XM_003993308.1
D1	102697430	102698360
refGene_gn=LOC101101667;refGene_tx=XM_003993268.1
D1	102709531	102710461
refGene_gn=LOC101093394;refGene_tx=XM_003993309.2
D1	102724075	102725011
refGene_gn=LOC101093643;refGene_tx=XM_023239015.1
D1	102744072	102745002
refGene_gn=LOC101094132;refGene_tx=XM_003993311.1
D1	102756133	102757069
refGene_gn=LOC101094374;refGene_tx=XM_003993312.1
D1	102768926	102769856
refGene_gn=LOC101094624;refGene_tx=XM_003993313.1
D1	102780480	102781419
refGene_gn=LOC101094878;refGene_tx=XM_003993314.2
D1	102786098	102787037
refGene_gn=LOC101095131;refGene_tx=XM_003993315.1
D1	102812912	102812929
refGene_gn=LOC101095613;refGene_tx=XM_023239736.1
D1	102816708	102817663
refGene_gn=LOC101095613;refGene_tx=XM_023239736.1
D1	102821710	102821837
refGene_gn=LOC101095855;refGene_tx=XM_003993317.1
D1	102829158	102830162
refGene_gn=LOC101095855;refGene_tx=XM_003993317.1
D1	102838104	102839055
refGene_gn=LOC101096103;refGene_tx=XM_003993318.1
D1	102851465	102851937
refGene_gn=LOC101096351;refGene_tx=XM_011286882.1
D1	102851939	102852406
refGene_gn=LOC101096351;refGene_tx=XM_011286882.1
D1	102859445	102860417
refGene_gn=LOC101096601;refGene_tx=XM_003993319.1
D1	102869497	102870433
refGene_gn=LOC101080481;refGene_tx=XM_003993269.1
D1	102882698	102883634
refGene_gn=LOC101080738;refGene_tx=XM_003993270.1
D1	102892892	102893843
refGene_gn=LOC101096841;refGene_tx=XM_011286881.3
D1	102897932	102898874
refGene_gn=LOC101097093;refGene_tx=XM_006937317.2
D1	102904121	102905072
refGene_gn=LOC101097345;refGene_tx=XM_019812967.1
D1	102917079	102918012
refGene_gn=LOC101097591;refGene_tx=XM_023239017.1
D1	102923271	102924255

refGene_gn=LOC101097839;refGene_tx=XM_003993322.4
D1	102947013	102947949
refGene_gn=LOC101098086;refGene_tx=XM_011286836.1
D1	102983530	102984463
refGene_gn=LOC101098824;refGene_tx=XM_011286835.2
D1	102990906	102991851
refGene_gn=LOC101081251;refGene_tx=XM_003993271.1
D1	103018271	103019222
refGene_gn=LOC101099077;refGene_tx=XM_003993324.5
D1	103027909	103028851
refGene_gn=LOC101099325;refGene_tx=XM_023239019.1
D1	103041937	103042888
refGene_gn=LOC101099585;refGene_tx=XM_003993326.4
D1	103057015	103057966
refGene_gn=LOC101099834;refGene_tx=XM_003993327.4
D1	103066509	103067250
refGene_gn=LOC101100083;refGene_tx=XM_003993328.5
D1	103067251	103067404
refGene_gn=LOC101100083;refGene_tx=XM_003993328.5
D1	103082788	103083781
refGene_gn=LOC101100332;refGene_tx=XM_003993329.2
D1	103091681	103092590
refGene_gn=LOC101100591;refGene_tx=XM_003993330.5
D1	103104503	103105436
refGene_gn=LOC101100835;refGene_tx=XM_023239737.1
D1	103112855	103112869
refGene_gn=LOC101100835;refGene_tx=XM_023239737.1
D1	103114181	103114770
refGene_gn=LOC101100835;refGene_tx=XM_023239737.1
D1	103132913	103133852
refGene_gn=LOC101101336;refGene_tx=XM_003993333.1
D1	103143331	103144270
refGene_gn=LOC101081495;refGene_tx=XM_003993272.1
D1	103158221	103158283
refGene_gn=LOC101101588;refGene_tx=XM_003993334.2
D1	103158284	103159227
refGene_gn=LOC101101588;refGene_tx=XM_003993334.2
D1	103166175	103167114
refGene_gn=LOC101080404;refGene_tx=XM_003993335.1
D1	103180662	103181595
refGene_gn=LOC101081755;refGene_tx=XM_003993273.2
D1	103190971	103191919
refGene_gn=LOC101080916;refGene_tx=XM_003993336.1
D1	103201099	103202032
refGene_gn=LOC101081163;refGene_tx=XM_011286833.3
D1	103212092	103213085
refGene_gn=LOC105260995;refGene_tx=XM_011286879.2
D1	103219230	103220250
refGene_gn=LOC101081670;refGene_tx=XM_011286832.2
D1	103238389	103239310

refGene_gn=LOC101082261;refGene_tx=XM_006937307.1
D1 103246103 103247051
refGene_gn=LOC101081925;refGene_tx=XM_003993339.2;refGene_gn=LOC101081
925;refGene_tx=XM_023239020.1
D1 103250212 103250223
refGene_gn=LOC101081925;refGene_tx=XM_003993339.2
D1 103256203 103257145
refGene_gn=LOC101082183;refGene_tx=XM_003993340.1
D1 103264046 103265036
refGene_gn=LOC101082436;refGene_tx=XM_003993341.1
D1 103282359 103283286
refGene_gn=LOC101082692;refGene_tx=XM_011286831.1
D1 103291009 103291951
refGene_gn=LOC101082945;refGene_tx=XM_003993343.1
D1 103307151 103308108
refGene_gn=LOC101082516;refGene_tx=XM_023239738.1
D1 103318641 103319580
refGene_gn=LOC101083205;refGene_tx=XM_003993344.1
D1 103334131 103335073
refGene_gn=LOC111556222;refGene_tx=XM_023239841.1
D1 103339311 103340271
refGene_gn=LOC101083451;refGene_tx=XM_023239839.1;refGene_gn=LOC101083
451;refGene_tx=XM_023239840.1
D1 103355880 103356828
refGene_gn=LOC101083031;refGene_tx=XM_003993276.1
D1 103367191 103368151
refGene_gn=LOC101083292;refGene_tx=XM_003993277.1
D1 103382205 103383147
refGene_gn=LOC111556662;refGene_tx=XM_023239739.1
D1 103385519 103386479
refGene_gn=LOC101083543;refGene_tx=XM_003993278.1
D1 103405251 103406208
refGene_gn=LOC101084226;refGene_tx=XM_003993346.1
D1 103413224 103414160
refGene_gn=LOC101084480;refGene_tx=XM_003993347.2
D1 103421129 103421729
refGene_gn=LOC101084480;refGene_tx=XM_003993347.2
D1 103437892 103438828
refGene_gn=LOC101083798;refGene_tx=XM_003993279.2
D1 103451023 103451980
refGene_gn=LOC101084990;refGene_tx=XM_003993349.2
D1 103465229 103466177
refGene_gn=LOC101097711;refGene_tx=XM_023239843.1
D1 103550437 103550962
refGene_gn=LOC101097711;refGene_tx=XM_023239843.1
D1 103550963 103551029
refGene_gn=LOC101097711;refGene_tx=XM_023239843.1
D1 103563873 103564812
refGene_gn=LOC105260993;refGene_tx=XM_023239845.1
D1 103574390 103575329

refGene_gn=LOC101084348;refGene_tx=XM_011279811.3
D1 103597966 103598914
refGene_gn=LOC101086382;refGene_tx=XM_004001251.4
D1 103611000 103611948
refGene_gn=LOC105260988;refGene_tx=XM_011286829.2
D1 103624126 103625071
refGene_gn=LOC111556682;refGene_tx=XM_023239844.1
D1 103641154 103642123
refGene_gn=LOC101085748;refGene_tx=XM_003993352.1
D1 103658165 103658591
refGene_gn=LOC101085240;refGene_tx=XM_003993350.3
D1 103658592 103659120
refGene_gn=LOC101085240;refGene_tx=XM_003993350.3
D1 103672170 103673103
refGene_gn=LOC101085496;refGene_tx=XM_003993351.2
D1 103677979 103678945
refGene_gn=LOC101086255;refGene_tx=XM_003993353.1
D1 103702279 103703197
refGene_gn=LOC101086504;refGene_tx=XM_023239021.1
D1 103715597 103716533
refGene_gn=LOC101086751;refGene_tx=XM_003993355.1
D1 103722714 103723653
refGene_gn=LOC101087007;refGene_tx=XM_011286874.3
D1 103729399 103730335
refGene_gn=LOC102899058;refGene_tx=XM_006937320.1
D1 103756862 103757804
refGene_gn=LOC101087267;refGene_tx=XM_003993357.1
D1 103771211 103772156
refGene_gn=LOC101087522;refGene_tx=XM_003993358.1
D1 103821432 103821514
refGene_gn=LOC109492144;refGene_tx=XM_023239740.1
D1 103822942 103823068
refGene_gn=LOC109492144;refGene_tx=XM_023239740.1
D1 103824393 103824473
refGene_gn=LOC109492144;refGene_tx=XM_023239740.1
D1 103826751 103826848
refGene_gn=LOC109492144;refGene_tx=XM_023239740.1
D1 103827826 103827924
refGene_gn=LOC109492144;refGene_tx=XM_023239740.1
D1 103828118 103828253
refGene_gn=LOC109492144;refGene_tx=XM_023239740.1
D1 103834349 103834496
refGene_gn=LOC109492144;refGene_tx=XM_023239740.1
D1 103838994 103839096
refGene_gn=LOC109492144;refGene_tx=XM_023239740.1
D1 103840598 103840796
refGene_gn=LOC109492144;refGene_tx=XM_023239740.1
D1 103842394 103842505
refGene_gn=LOC109492144;refGene_tx=XM_023239740.1
D1 103851086 103851413

refGene_gn=LOC109492144;refGene_tx=XM_023239740.1
D1 103888772 103889723
refGene_gn=LOC101085660;refGene_tx=XM_011286872.2
D1 103904383 103905334
refGene_gn=LOC111556663;refGene_tx=XM_023239741.1
D1 103938592 103939537
refGene_gn=LOC105259999;refGene_tx=XM_011286870.3
D1 103956585 103957530
refGene_gn=LOC101085153;refGene_tx=XM_019824703.2
D1 104142752 104143079
refGene_gn=LOC101085921;refGene_tx=XM_003993440.3
D1 104151799 104151910
refGene_gn=LOC101085921;refGene_tx=XM_003993440.3
D1 104154844 104155042
refGene_gn=LOC101085921;refGene_tx=XM_003993440.3
D1 104156522 104156624
refGene_gn=LOC101085921;refGene_tx=XM_003993440.3
D1 104161352 104161478
refGene_gn=LOC101085921;refGene_tx=XM_003993440.3
D1 104164081 104164142
refGene_gn=LOC101085921;refGene_tx=XM_003993440.3
D1 104168969 104169046
refGene_gn=LOC101085921;refGene_tx=XM_003993440.3
D1 104169241 104169339
refGene_gn=LOC101085921;refGene_tx=XM_003993440.3
D1 104170331 104170428
refGene_gn=LOC101085921;refGene_tx=XM_003993440.3
D1 104176958 104177038
refGene_gn=LOC101085921;refGene_tx=XM_003993440.3
D1 104178222 104178348
refGene_gn=LOC101085921;refGene_tx=XM_003993440.3
D1 104180009 104180061
refGene_gn=LOC101085921;refGene_tx=XM_003993440.3
D1 104204841 104205771
refGene_gn=LOC101086178;refGene_tx=XM_011286901.3
D1 104218422 104219364
refGene_gn=LOC101086420;refGene_tx=XM_003993441.3
D1 104316146 104316867
refGene_gn=LRRC55;refGene_tx=XM_019812422.2
D1 104320795 104321031
refGene_gn=LRRC55;refGene_tx=XM_019812422.2
D1 104372267 104373410
refGene_gn=APLNR;refGene_tx=XM_006937323.3
D1 104418925 104418986
refGene_gn=TNKS1BP1;refGene_tx=XM_006937325.3;refGene_gn=TNKS1BP1;refGene_tx=XM_011286903.2;refGene_gn=TNKS1BP1;refGene_tx=XM_011286904.1
D1 104419230 104419388
refGene_gn=TNKS1BP1;refGene_tx=XM_006937325.3;refGene_gn=TNKS1BP1;refGene_tx=XM_011286903.2;refGene_gn=TNKS1BP1;refGene_tx=XM_011286904.1
D1 104420136 104420273

refGene_gn=TNKS1BP1;refGene_tx=XM_006937325.3;refGene_gn=TNKS1BP1;refGene_tx=XM_011286903.2;refGene_gn=TNKS1BP1;refGene_tx=XM_011286904.1
D1 104420384 104420535
refGene_gn=TNKS1BP1;refGene_tx=XM_006937325.3;refGene_gn=TNKS1BP1;refGene_tx=XM_011286903.2;refGene_gn=TNKS1BP1;refGene_tx=XM_011286904.1
D1 104420793 104421160
refGene_gn=TNKS1BP1;refGene_tx=XM_006937325.3;refGene_gn=TNKS1BP1;refGene_tx=XM_011286903.2;refGene_gn=TNKS1BP1;refGene_tx=XM_011286904.1
D1 104426677 104428761
refGene_gn=TNKS1BP1;refGene_tx=XM_006937325.3;refGene_gn=TNKS1BP1;refGene_tx=XM_011286903.2;refGene_gn=TNKS1BP1;refGene_tx=XM_011286904.1
D1 104430728 104432132
refGene_gn=TNKS1BP1;refGene_tx=XM_006937325.3;refGene_gn=TNKS1BP1;refGene_tx=XM_011286903.2;refGene_gn=TNKS1BP1;refGene_tx=XM_011286904.1
D1 104436065 104436135
refGene_gn=TNKS1BP1;refGene_tx=XM_006937325.3;refGene_gn=TNKS1BP1;refGene_tx=XM_011286903.2;refGene_gn=TNKS1BP1;refGene_tx=XM_011286904.1
D1 104437547 104438181
refGene_gn=TNKS1BP1;refGene_tx=XM_006937325.3;refGene_gn=TNKS1BP1;refGene_tx=XM_011286903.2;refGene_gn=TNKS1BP1;refGene_tx=XM_011286904.1
D1 104439233 104439327
refGene_gn=TNKS1BP1;refGene_tx=XM_006937325.3;refGene_gn=TNKS1BP1;refGene_tx=XM_011286903.2;refGene_gn=TNKS1BP1;refGene_tx=XM_011286904.1
D1 104443926 104443998
refGene_gn=SSRP1;refGene_tx=XM_003993361.3;refGene_gn=SSRP1;refGene_tx=XM_006937326.2;refGene_gn=SSRP1;refGene_tx=XM_006937327.4
D1 104444229 104444416
refGene_gn=SSRP1;refGene_tx=XM_003993361.3;refGene_gn=SSRP1;refGene_tx=XM_006937326.2;refGene_gn=SSRP1;refGene_tx=XM_006937327.4
D1 104444947 104445036
refGene_gn=SSRP1;refGene_tx=XM_003993361.3;refGene_gn=SSRP1;refGene_tx=XM_006937326.2;refGene_gn=SSRP1;refGene_tx=XM_006937327.4
D1 104445192 104445363
refGene_gn=SSRP1;refGene_tx=XM_003993361.3;refGene_gn=SSRP1;refGene_tx=XM_006937326.2;refGene_gn=SSRP1;refGene_tx=XM_006937327.4
D1 104445789 104445919
refGene_gn=SSRP1;refGene_tx=XM_003993361.3;refGene_gn=SSRP1;refGene_tx=XM_006937326.2;refGene_gn=SSRP1;refGene_tx=XM_006937327.4
D1 104447350 104447396
refGene_gn=SSRP1;refGene_tx=XM_003993361.3;refGene_gn=SSRP1;refGene_tx=XM_006937326.2;refGene_gn=SSRP1;refGene_tx=XM_006937327.4
D1 104447525 104447664
refGene_gn=SSRP1;refGene_tx=XM_003993361.3;refGene_gn=SSRP1;refGene_tx=XM_006937326.2;refGene_gn=SSRP1;refGene_tx=XM_006937327.4
D1 104447957 104447966
refGene_gn=SSRP1;refGene_tx=XM_006937326.2;refGene_gn=SSRP1;refGene_tx=XM_006937327.4
D1 104448072 104448145
refGene_gn=SSRP1;refGene_tx=XM_003993361.3;refGene_gn=SSRP1;refGene_tx=XM_006937326.2;refGene_gn=SSRP1;refGene_tx=XM_006937327.4

D1 104448844 104449066
refGene_gn=SSRP1;refGene_tx=XM_003993361.3;refGene_gn=SSRP1;refGene_tx=XM_006937326.2;refGene_gn=SSRP1;refGene_tx=XM_006937327.4
D1 104449309 104449438
refGene_gn=SSRP1;refGene_tx=XM_003993361.3;refGene_gn=SSRP1;refGene_tx=XM_006937326.2;refGene_gn=SSRP1;refGene_tx=XM_006937327.4
D1 104449540 104449644
refGene_gn=SSRP1;refGene_tx=XM_003993361.3;refGene_gn=SSRP1;refGene_tx=XM_006937326.2;refGene_gn=SSRP1;refGene_tx=XM_006937327.4
D1 104449771 104450002
refGene_gn=SSRP1;refGene_tx=XM_003993361.3;refGene_gn=SSRP1;refGene_tx=XM_006937326.2;refGene_gn=SSRP1;refGene_tx=XM_006937327.4
D1 104450106 104450297
refGene_gn=SSRP1;refGene_tx=XM_003993361.3;refGene_gn=SSRP1;refGene_tx=XM_006937326.2;refGene_gn=SSRP1;refGene_tx=XM_006937327.4
D1 104450826 104450932
refGene_gn=SSRP1;refGene_tx=XM_003993361.3;refGene_gn=SSRP1;refGene_tx=XM_006937326.2;refGene_gn=SSRP1;refGene_tx=XM_006937327.4
D1 104451463 104451649
refGene_gn=SSRP1;refGene_tx=XM_003993361.3;refGene_gn=SSRP1;refGene_tx=XM_006937326.2;refGene_gn=SSRP1;refGene_tx=XM_006937327.4
D1 104452573 104452627
refGene_gn=SSRP1;refGene_tx=XM_003993361.3;refGene_gn=SSRP1;refGene_tx=XM_006937326.2;refGene_gn=SSRP1;refGene_tx=XM_006937327.4
D1 104455233 104455352
refGene_gn=P2RX3;refGene_tx=XM_003993362.3;refGene_gn=P2RX3;refGene_tx=XM_019812424.1
D1 104462374 104462510
refGene_gn=P2RX3;refGene_tx=XM_003993362.3;refGene_gn=P2RX3;refGene_tx=XM_019812424.1
D1 104462948 104463020
refGene_gn=P2RX3;refGene_tx=XM_003993362.3;refGene_gn=P2RX3;refGene_tx=XM_019812424.1
D1 104463256 104463320
refGene_gn=P2RX3;refGene_tx=XM_003993362.3;refGene_gn=P2RX3;refGene_tx=XM_019812424.1
D1 104464008 104464102
refGene_gn=P2RX3;refGene_tx=XM_003993362.3;refGene_gn=P2RX3;refGene_tx=XM_019812424.1
D1 104464406 104464484
refGene_gn=P2RX3;refGene_tx=XM_003993362.3;refGene_gn=P2RX3;refGene_tx=XM_019812424.1
D1 104465224 104465366
refGene_gn=P2RX3;refGene_tx=XM_003993362.3;refGene_gn=P2RX3;refGene_tx=XM_019812424.1
D1 104465832 104465969
refGene_gn=P2RX3;refGene_tx=XM_003993362.3;refGene_gn=P2RX3;refGene_tx=XM_019812424.1
D1 104466309 104466493
refGene_gn=P2RX3;refGene_tx=XM_019812424.1

D1	104479379	104479473
refGene_gn=P2RX3;refGene_tx=XM_003993362.3
D1	104479786	104479852
refGene_gn=P2RX3;refGene_tx=XM_003993362.3
D1	104480464	104480542
refGene_gn=P2RX3;refGene_tx=XM_003993362.3
D1	104480975	104481089
refGene_gn=P2RX3;refGene_tx=XM_003993362.3
D1	104486370	104486429
refGene_gn=LOC101086924;refGene_tx=XM_006937398.2
D1	104487229	104487341
refGene_gn=LOC101086924;refGene_tx=XM_006937398.2
D1	104488153	104488285
refGene_gn=LOC101086924;refGene_tx=XM_006937398.2
D1	104488967	104489281
refGene_gn=LOC101086924;refGene_tx=XM_006937398.2
D1	104489726	104489787
refGene_gn=LOC101086924;refGene_tx=XM_006937398.2
D1	104510933	104511035
refGene_gn=SLC43A3;refGene_tx=XM_023239849.1
D1	104512305	104512429
refGene_gn=SLC43A3;refGene_tx=XM_023239849.1
D1	104513052	104513239
refGene_gn=SLC43A3;refGene_tx=XM_023239849.1
D1	104516978	104517098
refGene_gn=SLC43A3;refGene_tx=XM_023239849.1
D1	104517347	104517539
refGene_gn=SLC43A3;refGene_tx=XM_023239849.1
D1	104518849	104518947
refGene_gn=SLC43A3;refGene_tx=XM_023239849.1
D1	104520235	104520375
refGene_gn=SLC43A3;refGene_tx=XM_023239849.1
D1	104522759	104522852
refGene_gn=SLC43A3;refGene_tx=XM_023239849.1
D1	104523073	104523150
refGene_gn=SLC43A3;refGene_tx=XM_023239849.1
D1	104524485	104524532
refGene_gn=SLC43A3;refGene_tx=XM_023239849.1
D1	104531077	104531154
refGene_gn=SLC43A3;refGene_tx=XM_023239849.1
D1	104556603	104556634
refGene_gn=RTN4RL2;refGene_tx=XM_023239850.1
D1	104562217	104562699
refGene_gn=RTN4RL2;refGene_tx=XM_023239850.1
D1	104569383	104570133
refGene_gn=RTN4RL2;refGene_tx=XM_023239850.1
D1	104579175	104579322
refGene_gn=SLC43A1;refGene_tx=XM_003993445.5;refGene_gn=SLC43A1;refGen
e_tx=XM_006937328.4;refGene_gn=SLC43A1;refGene_tx=XM_006937329.3;refGe
ne_gn=SLC43A1;refGene_tx=XM_006937330.3;refGene_gn=SLC43A1;refGene_tx=

XM_006937332.3;refGene_gn=SLC43A1;refGene_tx=XM_006937334.3;refGene_gn=SLC43A1;refGene_tx=XM_019812427.1;refGene_gn=SLC43A1;refGene_tx=XM_019812428.1;refGene_gn=SLC43A1;refGene_tx=XM_019812429.1;refGene_gn=SLC43A1;refGene_tx=XM_019812430.2;refGene_gn=SLC43A1;refGene_tx=XM_023239851.1;refGene_gn=SLC43A1;refGene_tx=XM_023239852.1;refGene_gn=SLC43A1;refGene_tx=XM_023239854.1
D1 104580396 104580520
refGene_gn=SLC43A1;refGene_tx=XM_003993445.5;refGene_gn=SLC43A1;refGene_tx=XM_006937328.4;refGene_gn=SLC43A1;refGene_tx=XM_006937329.3;refGene_gn=SLC43A1;refGene_tx=XM_006937330.3;refGene_gn=SLC43A1;refGene_tx=XM_006937332.3;refGene_gn=SLC43A1;refGene_tx=XM_006937334.3;refGene_gn=SLC43A1;refGene_tx=XM_019812427.1;refGene_gn=SLC43A1;refGene_tx=XM_019812428.1;refGene_gn=SLC43A1;refGene_tx=XM_019812429.1;refGene_gn=SLC43A1;refGene_tx=XM_019812430.2;refGene_gn=SLC43A1;refGene_tx=XM_023239851.1;refGene_gn=SLC43A1;refGene_tx=XM_023239852.1;refGene_gn=SLC43A1;refGene_tx=XM_023239854.1
D1 104582907 104582981
refGene_gn=SLC43A1;refGene_tx=XM_003993445.5;refGene_gn=SLC43A1;refGene_tx=XM_006937328.4;refGene_gn=SLC43A1;refGene_tx=XM_006937329.3;refGene_gn=SLC43A1;refGene_tx=XM_006937330.3;refGene_gn=SLC43A1;refGene_tx=XM_006937332.3;refGene_gn=SLC43A1;refGene_tx=XM_006937334.3;refGene_gn=SLC43A1;refGene_tx=XM_019812427.1;refGene_gn=SLC43A1;refGene_tx=XM_019812428.1;refGene_gn=SLC43A1;refGene_tx=XM_019812429.1;refGene_gn=SLC43A1;refGene_tx=XM_019812430.2;refGene_gn=SLC43A1;refGene_tx=XM_023239851.1;refGene_gn=SLC43A1;refGene_tx=XM_023239852.1;refGene_gn=SLC43A1;refGene_tx=XM_023239854.1
D1 104583248 104583390
refGene_gn=SLC43A1;refGene_tx=XM_003993445.5;refGene_gn=SLC43A1;refGene_tx=XM_006937328.4;refGene_gn=SLC43A1;refGene_tx=XM_006937329.3;refGene_gn=SLC43A1;refGene_tx=XM_006937330.3;refGene_gn=SLC43A1;refGene_tx=XM_006937332.3;refGene_gn=SLC43A1;refGene_tx=XM_006937334.3;refGene_gn=SLC43A1;refGene_tx=XM_019812427.1;refGene_gn=SLC43A1;refGene_tx=XM_019812428.1;refGene_gn=SLC43A1;refGene_tx=XM_019812429.1;refGene_gn=SLC43A1;refGene_tx=XM_019812430.2;refGene_gn=SLC43A1;refGene_tx=XM_023239851.1;refGene_gn=SLC43A1;refGene_tx=XM_023239852.1;refGene_gn=SLC43A1;refGene_tx=XM_023239854.1
D1 104585309 104585448
refGene_gn=SLC43A1;refGene_tx=XM_003993445.5;refGene_gn=SLC43A1;refGene_tx=XM_006937328.4;refGene_gn=SLC43A1;refGene_tx=XM_006937329.3;refGene_gn=SLC43A1;refGene_tx=XM_006937330.3;refGene_gn=SLC43A1;refGene_tx=XM_006937332.3;refGene_gn=SLC43A1;refGene_tx=XM_006937334.3;refGene_gn=SLC43A1;refGene_tx=XM_019812427.1;refGene_gn=SLC43A1;refGene_tx=XM_019812428.1;refGene_gn=SLC43A1;refGene_tx=XM_019812429.1;refGene_gn=SLC43A1;refGene_tx=XM_019812430.2;refGene_gn=SLC43A1;refGene_tx=XM_023239851.1;refGene_gn=SLC43A1;refGene_tx=XM_023239852.1;refGene_gn=SLC43A1;refGene_tx=XM_023239854.1
D1 104585685 104585721
refGene_gn=SLC43A1;refGene_tx=XM_003993445.5;refGene_gn=SLC43A1;refGene_tx=XM_006937328.4;refGene_gn=SLC43A1;refGene_tx=XM_006937329.3;refGene_gn=SLC43A1;refGene_tx=XM_006937330.3;refGene_gn=SLC43A1;refGene_tx=

XM_006937332.3;refGene_gn=SLC43A1;refGene_tx=XM_006937334.3;refGene_gn=SLC43A1;refGene_tx=XM_019812427.1;refGene_gn=SLC43A1;refGene_tx=XM_019812428.1;refGene_gn=SLC43A1;refGene_tx=XM_019812429.1;refGene_gn=SLC43A1;refGene_tx=XM_019812430.2;refGene_gn=SLC43A1;refGene_tx=XM_023239851.1;refGene_gn=SLC43A1;refGene_tx=XM_023239852.1;refGene_gn=SLC43A1;refGene_tx=XM_023239854.1
D1 104585816 104585963
refGene_gn=SLC43A1;refGene_tx=XM_003993445.5;refGene_gn=SLC43A1;refGene_tx=XM_006937328.4;refGene_gn=SLC43A1;refGene_tx=XM_006937329.3;refGene_gn=SLC43A1;refGene_tx=XM_006937330.3;refGene_gn=SLC43A1;refGene_tx=XM_006937332.3;refGene_gn=SLC43A1;refGene_tx=XM_006937334.3;refGene_gn=SLC43A1;refGene_tx=XM_019812427.1;refGene_gn=SLC43A1;refGene_tx=XM_019812428.1;refGene_gn=SLC43A1;refGene_tx=XM_019812429.1;refGene_gn=SLC43A1;refGene_tx=XM_019812430.2;refGene_gn=SLC43A1;refGene_tx=XM_023239851.1;refGene_gn=SLC43A1;refGene_tx=XM_023239852.1;refGene_gn=SLC43A1;refGene_tx=XM_023239854.1
D1 104587064 104587243
refGene_gn=SLC43A1;refGene_tx=XM_003993445.5;refGene_gn=SLC43A1;refGene_tx=XM_006937328.4;refGene_gn=SLC43A1;refGene_tx=XM_006937329.3;refGene_gn=SLC43A1;refGene_tx=XM_006937330.3;refGene_gn=SLC43A1;refGene_tx=XM_006937332.3;refGene_gn=SLC43A1;refGene_tx=XM_006937334.3;refGene_gn=SLC43A1;refGene_tx=XM_019812427.1;refGene_gn=SLC43A1;refGene_tx=XM_019812428.1;refGene_gn=SLC43A1;refGene_tx=XM_019812429.1;refGene_gn=SLC43A1;refGene_tx=XM_019812430.2;refGene_gn=SLC43A1;refGene_tx=XM_023239851.1;refGene_gn=SLC43A1;refGene_tx=XM_023239852.1;refGene_gn=SLC43A1;refGene_tx=XM_023239854.1
D1 104588818 104588952
refGene_gn=SLC43A1;refGene_tx=XM_003993445.5;refGene_gn=SLC43A1;refGene_tx=XM_006937328.4;refGene_gn=SLC43A1;refGene_tx=XM_006937329.3;refGene_gn=SLC43A1;refGene_tx=XM_006937330.3;refGene_gn=SLC43A1;refGene_tx=XM_006937332.3;refGene_gn=SLC43A1;refGene_tx=XM_006937334.3;refGene_gn=SLC43A1;refGene_tx=XM_019812427.1;refGene_gn=SLC43A1;refGene_tx=XM_019812428.1;refGene_gn=SLC43A1;refGene_tx=XM_019812429.1;refGene_gn=SLC43A1;refGene_tx=XM_019812430.2;refGene_gn=SLC43A1;refGene_tx=XM_023239851.1;refGene_gn=SLC43A1;refGene_tx=XM_023239852.1;refGene_gn=SLC43A1;refGene_tx=XM_023239854.1
D1 104590586 104590679
refGene_gn=SLC43A1;refGene_tx=XM_003993445.5;refGene_gn=SLC43A1;refGene_tx=XM_006937328.4;refGene_gn=SLC43A1;refGene_tx=XM_006937329.3;refGene_gn=SLC43A1;refGene_tx=XM_006937330.3;refGene_gn=SLC43A1;refGene_tx=XM_006937332.3;refGene_gn=SLC43A1;refGene_tx=XM_006937334.3;refGene_gn=SLC43A1;refGene_tx=XM_019812427.1;refGene_gn=SLC43A1;refGene_tx=XM_019812428.1;refGene_gn=SLC43A1;refGene_tx=XM_019812429.1;refGene_gn=SLC43A1;refGene_tx=XM_019812430.2;refGene_gn=SLC43A1;refGene_tx=XM_023239851.1;refGene_gn=SLC43A1;refGene_tx=XM_023239852.1;refGene_gn=SLC43A1;refGene_tx=XM_023239854.1
D1 104591651 104591915
refGene_gn=SLC43A1;refGene_tx=XM_023239852.1
D1 104592965 104593042
refGene_gn=SLC43A1;refGene_tx=XM_003993445.5;refGene_gn=SLC43A1;refGen

e_tx=XM_006937328.4;refGene_gn=SLC43A1;refGene_tx=XM_006937329.3;refGene_gn=SLC43A1;refGene_tx=XM_006937330.3;refGene_gn=SLC43A1;refGene_tx=XM_006937332.3;refGene_gn=SLC43A1;refGene_tx=XM_006937334.3;refGene_gn=SLC43A1;refGene_tx=XM_019812427.1;refGene_gn=SLC43A1;refGene_tx=XM_019812428.1;refGene_gn=SLC43A1;refGene_tx=XM_019812429.1;refGene_gn=SLC43A1;refGene_tx=XM_019812430.2;refGene_gn=SLC43A1;refGene_tx=XM_023239851.1;refGene_gn=SLC43A1;refGene_tx=XM_023239854.1
D1 104593177 104593233
refGene_gn=SLC43A1;refGene_tx=XM_003993445.5;refGene_gn=SLC43A1;refGene_tx=XM_006937328.4;refGene_gn=SLC43A1;refGene_tx=XM_006937329.3;refGene_gn=SLC43A1;refGene_tx=XM_006937330.3;refGene_gn=SLC43A1;refGene_tx=XM_006937332.3;refGene_gn=SLC43A1;refGene_tx=XM_006937334.3;refGene_gn=SLC43A1;refGene_tx=XM_019812427.1;refGene_gn=SLC43A1;refGene_tx=XM_019812428.1;refGene_gn=SLC43A1;refGene_tx=XM_019812429.1;refGene_gn=SLC43A1;refGene_tx=XM_019812430.2;refGene_gn=SLC43A1;refGene_tx=XM_023239851.1;refGene_gn=SLC43A1;refGene_tx=XM_023239854.1
D1 104593331 104593509
refGene_gn=SLC43A1;refGene_tx=XM_003993445.5;refGene_gn=SLC43A1;refGene_tx=XM_006937328.4;refGene_gn=SLC43A1;refGene_tx=XM_006937329.3;refGene_gn=SLC43A1;refGene_tx=XM_006937330.3;refGene_gn=SLC43A1;refGene_tx=XM_006937332.3;refGene_gn=SLC43A1;refGene_tx=XM_006937334.3;refGene_gn=SLC43A1;refGene_tx=XM_019812427.1;refGene_gn=SLC43A1;refGene_tx=XM_019812428.1;refGene_gn=SLC43A1;refGene_tx=XM_019812429.1;refGene_gn=SLC43A1;refGene_tx=XM_019812430.2;refGene_gn=SLC43A1;refGene_tx=XM_023239851.1;refGene_gn=SLC43A1;refGene_tx=XM_023239854.1
D1 104609900 104610054
refGene_gn=SLC43A1;refGene_tx=XM_003993445.5;refGene_gn=SLC43A1;refGene_tx=XM_006937328.4;refGene_gn=SLC43A1;refGene_tx=XM_023239851.1
D1 104621237 104621439
refGene_gn=TIMM10;refGene_tx=XM_003993364.4;refGene_gn=TIMM10;refGene_tx=XM_006937336.3
D1 104622659 104622730
refGene_gn=TIMM10;refGene_tx=XM_003993364.4;refGene_gn=TIMM10;refGene_tx=XM_006937336.3
D1 104629222 104629244
refGene_gn=SMTNL1;refGene_tx=XM_006937337.3
D1 104631362 104632042
refGene_gn=SMTNL1;refGene_tx=XM_006937337.3;refGene_gn=SMTNL1;refGene_tx=XM_006937338.2;refGene_gn=SMTNL1;refGene_tx=XM_006937339.3
D1 104632290 104632423
refGene_gn=SMTNL1;refGene_tx=XM_006937337.3;refGene_gn=SMTNL1;refGene_tx=XM_006937338.2;refGene_gn=SMTNL1;refGene_tx=XM_006937339.3
D1 104632542 104632597
refGene_gn=SMTNL1;refGene_tx=XM_006937337.3;refGene_gn=SMTNL1;refGene_tx=XM_006937338.2;refGene_gn=SMTNL1;refGene_tx=XM_006937339.3
D1 104633989 104634148
refGene_gn=SMTNL1;refGene_tx=XM_006937337.3;refGene_gn=SMTNL1;refGene_tx=XM_006937338.2;refGene_gn=SMTNL1;refGene_tx=XM_006937339.3
D1 104634311 104634426
refGene_gn=SMTNL1;refGene_tx=XM_006937337.3;refGene_gn=SMTNL1;refGene_

tx=XM_006937338.2;refGene_gn=SMTNL1;refGene_tx=XM_006937339.3
D1	104634552	104634704
refGene_gn=SMTNL1;refGene_tx=XM_006937337.3;refGene_gn=SMTNL1;refGene_tx=XM_006937338.2;refGene_gn=SMTNL1;refGene_tx=XM_006937339.3
D1	104639038	104639183
refGene_gn=SMTNL1;refGene_tx=XM_006937337.3;refGene_gn=SMTNL1;refGene_tx=XM_006937338.2;refGene_gn=SMTNL1;refGene_tx=XM_006937339.3
D1	104641531	104641683
refGene_gn=UBE2L6;refGene_tx=XM_003993366.5;refGene_gn=UBE2L6;refGene_tx=XM_006937340.4
D1	104643477	104643664
refGene_gn=UBE2L6;refGene_tx=XM_003993366.5;refGene_gn=UBE2L6;refGene_tx=XM_006937340.4
D1	104650637	104650733
refGene_gn=UBE2L6;refGene_tx=XM_003993366.5
D1	104654676	104654703
refGene_gn=UBE2L6;refGene_tx=XM_003993366.5;refGene_gn=UBE2L6;refGene_tx=XM_006937340.4
D1	104686136	104686187
refGene_gn=SERPING1;refGene_tx=XM_006937341.2
D1	104687762	104688294
refGene_gn=SERPING1;refGene_tx=XM_006937341.2
D1	104689822	104689957
refGene_gn=SERPING1;refGene_tx=XM_006937341.2
D1	104692035	104692239
refGene_gn=SERPING1;refGene_tx=XM_006937341.2
D1	104692432	104692572
refGene_gn=SERPING1;refGene_tx=XM_006937341.2
D1	104695973	104696193
refGene_gn=SERPING1;refGene_tx=XM_006937341.2
D1	104697447	104697701
refGene_gn=SERPING1;refGene_tx=XM_006937341.2
D1	104728177	104728267
refGene_gn=YPEL4;refGene_tx=XM_003993367.5;refGene_gn=YPEL4;refGene_tx=XM_023239856.1
D1	104728481	104728590
refGene_gn=YPEL4;refGene_tx=XM_003993367.5;refGene_gn=YPEL4;refGene_tx=XM_023239856.1
D1	104728928	104728972
refGene_gn=YPEL4;refGene_tx=XM_003993367.5;refGene_gn=YPEL4;refGene_tx=XM_023239856.1
D1	104729148	104729289
refGene_gn=YPEL4;refGene_tx=XM_003993367.5;refGene_gn=YPEL4;refGene_tx=XM_023239856.1
D1	104736853	104737459
refGene_gn=CLP1;refGene_tx=XM_003993368.4;refGene_gn=CLP1;refGene_tx=XM_019812440.1;refGene_gn=CLP1;refGene_tx=XM_019812441.1;refGene_gn=CLP1;refGene_tx=XM_023239858.1
D1	104737861	104738533
refGene_gn=CLP1;refGene_tx=XM_003993368.4;refGene_gn=CLP1;refGene_tx=X

M_019812440.1;refGene_gn=CLP1;refGene_tx=XM_019812441.1;refGene_gn=CLP1;refGene_tx=XM_023239858.1
D1 104748663 104748767
refGene_gn=ZDHHC5;refGene_tx=XM_023239860.1
D1 104756754 104756876
refGene_gn=ZDHHC5;refGene_tx=XM_023239860.1
D1 104760261 104760419
refGene_gn=ZDHHC5;refGene_tx=XM_023239860.1
D1 104761434 104761607
refGene_gn=ZDHHC5;refGene_tx=XM_023239860.1
D1 104761762 104761865
refGene_gn=ZDHHC5;refGene_tx=XM_023239860.1
D1 104762463 104762555
refGene_gn=ZDHHC5;refGene_tx=XM_023239860.1
D1 104762999 104763132
refGene_gn=ZDHHC5;refGene_tx=XM_023239860.1
D1 104763741 104763865
refGene_gn=ZDHHC5;refGene_tx=XM_023239860.1
D1 104764316 104764429
refGene_gn=ZDHHC5;refGene_tx=XM_023239860.1
D1 104765122 104765982
refGene_gn=ZDHHC5;refGene_tx=XM_023239860.1
D1 104766817 104766983
refGene_gn=ZDHHC5;refGene_tx=XM_023239860.1
D1 104769722 104769791
refGene_gn=MED19;refGene_tx=XM_003993371.5
D1 104769979 104770074
refGene_gn=MED19;refGene_tx=XM_003993371.5
D1 104770392 104770489
refGene_gn=MED19;refGene_tx=XM_003993371.5
D1 104770655 104770912
refGene_gn=MED19;refGene_tx=XM_003993371.5
D1 104776601 104776869
refGene_gn=MED19;refGene_tx=XM_003993371.5
D1 104777263 104777452
refGene_gn=TMX2;refGene_tx=XM_003993394.4;refGene_gn=TMX2;refGene_tx=XM_023239862.1
D1 104784442 104784503
refGene_gn=TMX2;refGene_tx=XM_003993394.4;refGene_gn=TMX2;refGene_tx=XM_023239862.1
D1 104784746 104784860
refGene_gn=TMX2;refGene_tx=XM_003993394.4;refGene_gn=TMX2;refGene_tx=XM_023239862.1
D1 104785123 104785200
refGene_gn=TMX2;refGene_tx=XM_003993394.4;refGene_gn=TMX2;refGene_tx=XM_023239862.1
D1 104785420 104785527
refGene_gn=TMX2;refGene_tx=XM_003993394.4;refGene_gn=TMX2;refGene_tx=XM_023239862.1
D1 104785949 104786015

refGene_gn=TMX2;refGene_tx=XM_003993394.4;refGene_gn=TMX2;refGene_tx=XM_023239862.1
D1 104786107 104786237
refGene_gn=TMX2;refGene_tx=XM_003993394.4;refGene_gn=TMX2;refGene_tx=XM_023239862.1
D1 104786737 104786884
refGene_gn=TMX2;refGene_tx=XM_003993394.4
D1 104788330 104788452
refGene_gn=SELENOH;refGene_tx=XM_003993396.5
D1 104788533 104788679
refGene_gn=SELENOH;refGene_tx=XM_003993396.5
D1 104788800 104788901
refGene_gn=SELENOH;refGene_tx=XM_003993396.5
D1 104790397 104792463
refGene_gn=BTBD18;refGene_tx=XM_023239859.1
D1 104795500 104795624
refGene_gn=BTBD18;refGene_tx=XM_023239859.1
D1 104827922 104828117
refGene_gn=CTNND1;refGene_tx=XM_019812432.2
D1 104830316 104830388
refGene_gn=CTNND1;refGene_tx=XM_019812432.2
D1 104831387 104831540
refGene_gn=CTNND1;refGene_tx=XM_019812432.2
D1 104832204 104832740
refGene_gn=CTNND1;refGene_tx=XM_019812432.2
D1 104836082 104836546
refGene_gn=CTNND1;refGene_tx=XM_019812432.2
D1 104837652 104837687
refGene_gn=CTNND1;refGene_tx=XM_019812432.2
D1 104837688 104837837
refGene_gn=CTNND1;refGene_tx=XM_019812432.2
D1 104839027 104839145
refGene_gn=CTNND1;refGene_tx=XM_019812432.2
D1 104839607 104839761
refGene_gn=CTNND1;refGene_tx=XM_019812432.2
D1 104840157 104840175
refGene_gn=CTNND1;refGene_tx=XM_019812432.2
D1 104840617 104840686
refGene_gn=CTNND1;refGene_tx=XM_019812432.2
D1 104841510 104841638
refGene_gn=CTNND1;refGene_tx=XM_019812432.2
D1 104841740 104841891
refGene_gn=CTNND1;refGene_tx=XM_019812432.2
D1 104842612 104842805
refGene_gn=CTNND1;refGene_tx=XM_019812432.2
D1 104843145 104843260
refGene_gn=CTNND1;refGene_tx=XM_019812432.2
D1 104843807 104843895
refGene_gn=CTNND1;refGene_tx=XM_019812432.2
D1 104846950 104847013

refGene_gn=CTNND1;refGene_tx=XM_019812432.2
D1 104847816 104847920
refGene_gn=CTNND1;refGene_tx=XM_019812432.2
D1 104848333 104848405
refGene_gn=CTNND1;refGene_tx=XM_019812432.2
D1 104848691 104848703
refGene_gn=CTNND1;refGene_tx=XM_019812432.2
D1 104894177 104895164
refGene_gn=LOC101088187;refGene_tx=XM_023239022.1
D1 104947242 104947448
refGene_gn=LOC101088432;refGene_tx=XM_023239742.1
D1 104947450 104947851
refGene_gn=LOC101088432;refGene_tx=XM_023239742.1
D1 104947852 104948187
refGene_gn=LOC101088432;refGene_tx=XM_023239742.1
D1 104995071 104996037
refGene_gn=LOC101096352;refGene_tx=XM_003993397.4
D1 105038531 105039662
refGene_gn=LOC101088954;refGene_tx=XM_003993449.4
D1 105064679 105065621
refGene_gn=LOC101089211;refGene_tx=XM_003993450.1
D1 105115911 105116082
refGene_gn=LOC101089463;refGene_tx=XM_003993451.1
D1 105116083 105116848
refGene_gn=LOC101089463;refGene_tx=XM_003993451.1
D1 105127637 105128582
refGene_gn=LOC101089717;refGene_tx=XM_003993452.2
D1 105137065 105138034
refGene_gn=LOC101089962;refGene_tx=XM_003993453.4
D1 105150141 105151101
refGene_gn=LOC101090211;refGene_tx=XM_023239743.1
D1 105167239 105168184
refGene_gn=LOC101096602;refGene_tx=XM_003993398.2
D1 105179967 105180927
refGene_gn=LOC101090464;refGene_tx=XM_003993455.1
D1 105193358 105194303
refGene_gn=LOC101090718;refGene_tx=XM_003993456.1
D1 105218528 105219479
refGene_gn=LOC101091225;refGene_tx=XM_003993457.4
D1 105232139 105233078
refGene_gn=LOC101091477;refGene_tx=XM_003993458.2
D1 105288301 105289252
refGene_gn=LOC101092212;refGene_tx=XM_011286947.1
D1 105303215 105304154
refGene_gn=LOC101092710;refGene_tx=XM_003993460.2
D1 105316956 105317901
refGene_gn=LOC101096842;refGene_tx=XM_003993399.4
D1 105338555 105339494
refGene_gn=LOC101093208;refGene_tx=XM_003993461.2
D1 105365519 105366464

refGene_gn=LOC101097094;refGene_tx=XM_003993400.2
D1 105393077 105394028
refGene_gn=LOC101093712;refGene_tx=XM_023239023.1
D1 105414410 105414680
refGene_gn=LPXN;refGene_tx=XM_003993401.4;refGene_gn=LPXN;refGene_tx=XM_023239864.1;refGene_gn=LPXN;refGene_tx=XM_023239865.1
D1 105414992 105415141
refGene_gn=LPXN;refGene_tx=XM_003993401.4;refGene_gn=LPXN;refGene_tx=XM_023239864.1;refGene_gn=LPXN;refGene_tx=XM_023239865.1
D1 105428829 105428911
refGene_gn=LPXN;refGene_tx=XM_003993401.4;refGene_gn=LPXN;refGene_tx=XM_023239864.1;refGene_gn=LPXN;refGene_tx=XM_023239865.1
D1 105428993 105429167
refGene_gn=LPXN;refGene_tx=XM_003993401.4;refGene_gn=LPXN;refGene_tx=XM_023239864.1;refGene_gn=LPXN;refGene_tx=XM_023239865.1
D1 105430087 105430255
refGene_gn=LPXN;refGene_tx=XM_003993401.4;refGene_gn=LPXN;refGene_tx=XM_023239864.1;refGene_gn=LPXN;refGene_tx=XM_023239865.1
D1 105433698 105433798
refGene_gn=LPXN;refGene_tx=XM_003993401.4;refGene_gn=LPXN;refGene_tx=XM_023239864.1;refGene_gn=LPXN;refGene_tx=XM_023239865.1
D1 105440663 105440710
refGene_gn=LPXN;refGene_tx=XM_003993401.4;refGene_gn=LPXN;refGene_tx=XM_023239864.1
D1 105446332 105446490
refGene_gn=LPXN;refGene_tx=XM_003993401.4;refGene_gn=LPXN;refGene_tx=XM_023239864.1;refGene_gn=LPXN;refGene_tx=XM_023239865.1
D1 105452099 105452112
refGene_gn=LPXN;refGene_tx=XM_003993401.4
D1 105454773 105454789
refGene_gn=LPXN;refGene_tx=XM_023239864.1
D1 105455768 105456118
refGene_gn=ZFP91;refGene_tx=XM_011286949.3;refGene_gn=ZFP91;refGene_tx=XM_023239863.1
D1 105461778 105461807
refGene_gn=ZFP91;refGene_tx=XM_011286949.3;refGene_gn=ZFP91;refGene_tx=XM_023239863.1
D1 105490105 105490318
refGene_gn=ZFP91;refGene_tx=XM_011286949.3;refGene_gn=ZFP91;refGene_tx=XM_023239863.1
D1 105490547 105490584
refGene_gn=ZFP91;refGene_tx=XM_011286949.3;refGene_gn=ZFP91;refGene_tx=XM_023239863.1
D1 105491205 105491310
refGene_gn=ZFP91;refGene_tx=XM_011286949.3;refGene_gn=ZFP91;refGene_tx=XM_023239863.1
D1 105491838 105491973
refGene_gn=ZFP91;refGene_tx=XM_011286949.3;refGene_gn=ZFP91;refGene_tx=XM_023239863.1
D1 105492741 105492792

refGene_gn=ZFP91;refGene_tx=XM_011286949.3;refGene_gn=ZFP91;refGene_tx=XM_023239863.1
D1 105493215 105493294
refGene_gn=ZFP91;refGene_tx=XM_011286949.3;refGene_gn=ZFP91;refGene_tx=XM_023239863.1
D1 105494645 105494760
refGene_gn=ZFP91;refGene_tx=XM_011286949.3;refGene_gn=ZFP91;refGene_tx=XM_023239863.1
D1 105498012 105498112
refGene_gn=ZFP91;refGene_tx=XM_011286949.3;refGene_gn=ZFP91;refGene_tx=XM_023239863.1
D1 105498461 105498972
refGene_gn=ZFP91;refGene_tx=XM_011286949.3;refGene_gn=ZFP91;refGene_tx=XM_023239863.1
D1 105504071 105504185
refGene_gn=CNTF;refGene_tx=XM_003993402.5
D1 105505400 105505889
refGene_gn=CNTF;refGene_tx=XM_003993402.5
D1 105555896 105556299
refGene_gn=GLYAT;refGene_tx=XM_011286912.3;refGene_gn=GLYAT;refGene_tx=XM_019812442.1
D1 105556712 105556888
refGene_gn=GLYAT;refGene_tx=XM_011286912.3;refGene_gn=GLYAT;refGene_tx=XM_019812442.1;refGene_gn=GLYAT;refGene_tx=XM_019812443.2
D1 105558248 105558375
refGene_gn=GLYAT;refGene_tx=XM_011286912.3;refGene_gn=GLYAT;refGene_tx=XM_019812442.1;refGene_gn=GLYAT;refGene_tx=XM_019812443.2
D1 105564416 105564524
refGene_gn=GLYAT;refGene_tx=XM_011286912.3;refGene_gn=GLYAT;refGene_tx=XM_019812442.1;refGene_gn=GLYAT;refGene_tx=XM_019812443.2
D1 105567868 105567964
refGene_gn=GLYAT;refGene_tx=XM_011286912.3;refGene_gn=GLYAT;refGene_tx=XM_019812442.1;refGene_gn=GLYAT;refGene_tx=XM_019812443.2
D1 105576019 105576199
refGene_gn=GLYAT;refGene_tx=XM_011286912.3;refGene_gn=GLYAT;refGene_tx=XM_019812443.2
D1 105595688 105596091
refGene_gn=LOC101098257;refGene_tx=XM_003993405.4
D1 105597689 105597861
refGene_gn=LOC101098257;refGene_tx=XM_003993405.4
D1 105598723 105598850
refGene_gn=LOC101098257;refGene_tx=XM_003993405.4
D1 105601441 105601549
refGene_gn=LOC101098257;refGene_tx=XM_003993405.4
D1 105604318 105604396
refGene_gn=LOC101098257;refGene_tx=XM_003993405.4
D1 105623089 105623501
refGene_gn=GLYATL2;refGene_tx=XM_023239024.1
D1 105626535 105626707
refGene_gn=GLYATL2;refGene_tx=XM_023239024.1

D1	105626800	105626927
refGene_gn=GLYATL2;refGene_tx=XM_023239024.1
D1	105628935	105629043
refGene_gn=GLYATL2;refGene_tx=XM_023239024.1
D1	105630170	105630248
refGene_gn=GLYATL2;refGene_tx=XM_023239024.1
D1	105702407	105702488
refGene_gn=FAM111A;refGene_tx=XM_003993406.4;refGene_gn=FAM111A;refGene_tx=XM_011286913.3;refGene_gn=FAM111A;refGene_tx=XM_011286914.3;refGene_gn=FAM111A;refGene_tx=XM_019812444.2;refGene_gn=FAM111A;refGene_tx=XM_019812445.2;refGene_gn=FAM111A;refGene_tx=XM_019812446.2;refGene_gn=FAM111A;refGene_tx=XM_019812447.2
D1	105705079	105706810
refGene_gn=FAM111A;refGene_tx=XM_003993406.4;refGene_gn=FAM111A;refGene_tx=XM_011286913.3;refGene_gn=FAM111A;refGene_tx=XM_011286914.3;refGene_gn=FAM111A;refGene_tx=XM_019812444.2;refGene_gn=FAM111A;refGene_tx=XM_019812445.2;refGene_gn=FAM111A;refGene_tx=XM_019812446.2;refGene_gn=FAM111A;refGene_tx=XM_019812447.2
D1	105720204	105720415
refGene_gn=DTX4;refGene_tx=XM_011286915.2
D1	105728486	105729213
refGene_gn=DTX4;refGene_tx=XM_011286915.2
D1	105735858	105735920
refGene_gn=DTX4;refGene_tx=XM_011286915.2
D1	105736241	105736403
refGene_gn=DTX4;refGene_tx=XM_011286915.2
D1	105738182	105738244
refGene_gn=DTX4;refGene_tx=XM_011286915.2
D1	105739705	105739858
refGene_gn=DTX4;refGene_tx=XM_011286915.2
D1	105742883	105743045
refGene_gn=DTX4;refGene_tx=XM_011286915.2
D1	105748331	105748421
refGene_gn=DTX4;refGene_tx=XM_011286915.2
D1	105752560	105752794
refGene_gn=DTX4;refGene_tx=XM_011286915.2
D1	105759074	105761228
refGene_gn=MPEG1;refGene_tx=XM_003993407.4
D1	105778490	105779453
refGene_gn=LOC101095681;refGene_tx=XM_003993466.4
D1	105804140	105805088
refGene_gn=LOC101095916;refGene_tx=XM_003993467.2
D1	105832504	105833449
refGene_gn=LOC101096165;refGene_tx=XM_003993468.3
D1	105862398	105863337
refGene_gn=LOC111556541;refGene_tx=XM_023239025.1
D1	105898114	105899053
refGene_gn=LOC101096915;refGene_tx=XM_003993470.1
D1	105926762	105927701
refGene_gn=LOC101097171;refGene_tx=XM_003993471.1

D1 105946547 105947513
refGene_gn=LOC101097669;refGene_tx=XM_011286950.1
D1 105964677 105965625
refGene_gn=LOC101098988;refGene_tx=XM_003993408.2
D1 105988790 105989756
refGene_gn=LOC101098161;refGene_tx=XM_003993472.1
D1 106003248 106004184
refGene_gn=LOC101098401;refGene_tx=XM_003993473.1
D1 106016956 106017892
refGene_gn=LOC101099246;refGene_tx=XM_003993409.2
D1 106026152 106026448
refGene_gn=LOC101099500;refGene_tx=XM_003993410.2
D1 106026449 106027086
refGene_gn=LOC101099500;refGene_tx=XM_003993410.2
D1 106036678 106037614
refGene_gn=LOC101099751;refGene_tx=XM_003993411.1
D1 106050707 106051643
refGene_gn=LOC101100002;refGene_tx=XM_003993412.1
D1 106077735 106077878
refGene_gn=OSBP;refGene_tx=XM_003993413.4
D1 106077976 106078197
refGene_gn=OSBP;refGene_tx=XM_003993413.4
D1 106079330 106079512
refGene_gn=OSBP;refGene_tx=XM_003993413.4
D1 106080272 106080368
refGene_gn=OSBP;refGene_tx=XM_003993413.4
D1 106081607 106081711
refGene_gn=OSBP;refGene_tx=XM_003993413.4
D1 106090049 106090170
refGene_gn=OSBP;refGene_tx=XM_003993413.4
D1 106090448 106090694
refGene_gn=OSBP;refGene_tx=XM_003993413.4
D1 106096217 106096349
refGene_gn=OSBP;refGene_tx=XM_003993413.4
D1 106096527 106096582
refGene_gn=OSBP;refGene_tx=XM_003993413.4
D1 106096904 106097007
refGene_gn=OSBP;refGene_tx=XM_003993413.4
D1 106097262 106097461
refGene_gn=OSBP;refGene_tx=XM_003993413.4
D1 106102470 106102721
refGene_gn=OSBP;refGene_tx=XM_003993413.4
D1 106105417 106105626
refGene_gn=OSBP;refGene_tx=XM_003993413.4
D1 106110197 106110562
refGene_gn=OSBP;refGene_tx=XM_003993413.4
D1 106134816 106134838
refGene_gn=PATL1;refGene_tx=XM_003993414.5
D1 106135148 106135298
refGene_gn=PATL1;refGene_tx=XM_003993414.5

D1 106135397 106135489
refGene_gn=PATL1;refGene_tx=XM_003993414.5
D1 106137421 106137577
refGene_gn=PATL1;refGene_tx=XM_003993414.5
D1 106141904 106142064
refGene_gn=PATL1;refGene_tx=XM_003993414.5
D1 106143340 106143489
refGene_gn=PATL1;refGene_tx=XM_003993414.5
D1 106144732 106144792
refGene_gn=PATL1;refGene_tx=XM_003993414.5
D1 106145574 106145649
refGene_gn=PATL1;refGene_tx=XM_003993414.5
D1 106145650 106145673
refGene_gn=PATL1;refGene_tx=XM_003993414.5
D1 106146133 106146257
refGene_gn=PATL1;refGene_tx=XM_003993414.5
D1 106147314 106147495
refGene_gn=PATL1;refGene_tx=XM_003993414.5
D1 106149868 106149958
refGene_gn=PATL1;refGene_tx=XM_003993414.5
D1 106151729 106151947
refGene_gn=PATL1;refGene_tx=XM_003993414.5
D1 106152161 106152251
refGene_gn=PATL1;refGene_tx=XM_003993414.5
D1 106153402 106153504
refGene_gn=PATL1;refGene_tx=XM_003993414.5
D1 106154440 106154635
refGene_gn=PATL1;refGene_tx=XM_003993414.5
D1 106155270 106155351
refGene_gn=PATL1;refGene_tx=XM_003993414.5
D1 106155643 106155861
refGene_gn=PATL1;refGene_tx=XM_003993414.5
D1 106163017 106163129
refGene_gn=PATL1;refGene_tx=XM_003993414.5
D1 106165036 106165051
refGene_gn=PATL1;refGene_tx=XM_003993414.5
D1 106205732 106206662
refGene_gn=LOC101100755;refGene_tx=XM_019812449.2
D1 106228833 106229856
refGene_gn=LOC111556270;refGene_tx=XM_023239026.1
D1 106241214 106242153
refGene_gn=LOC101101000;refGene_tx=XM_023239866.1
D1 106252234 106252264
refGene_gn=STX3;refGene_tx=XM_023239867.1;refGene_gn=STX3;refGene_tx=XM_023239868.1;refGene_gn=STX3;refGene_tx=XM_023239869.1
D1 106268034 106268118
refGene_gn=STX3;refGene_tx=XM_023239867.1;refGene_gn=STX3;refGene_tx=XM_023239868.1;refGene_gn=STX3;refGene_tx=XM_023239869.1
D1 106280125 106280225
refGene_gn=STX3;refGene_tx=XM_023239867.1;refGene_gn=STX3;refGene_tx=X

M_023239868.1;refGene_gn=STX3;refGene_tx=XM_023239869.1
D1 106281965 106282040
refGene_gn=STX3;refGene_tx=XM_023239867.1;refGene_gn=STX3;refGene_tx=XM_023239868.1;refGene_gn=STX3;refGene_tx=XM_023239869.1
D1 106283053 106283121
refGene_gn=STX3;refGene_tx=XM_023239867.1;refGene_gn=STX3;refGene_tx=XM_023239868.1;refGene_gn=STX3;refGene_tx=XM_023239869.1
D1 106285089 106285198
refGene_gn=STX3;refGene_tx=XM_023239867.1;refGene_gn=STX3;refGene_tx=XM_023239868.1;refGene_gn=STX3;refGene_tx=XM_023239869.1
D1 106286256 106286330
refGene_gn=STX3;refGene_tx=XM_023239867.1;refGene_gn=STX3;refGene_tx=XM_023239868.1;refGene_gn=STX3;refGene_tx=XM_023239869.1
D1 106286483 106286618
refGene_gn=STX3;refGene_tx=XM_023239867.1;refGene_gn=STX3;refGene_tx=XM_023239868.1;refGene_gn=STX3;refGene_tx=XM_023239869.1
D1 106288302 106288413
refGene_gn=STX3;refGene_tx=XM_023239867.1;refGene_gn=STX3;refGene_tx=XM_023239868.1;refGene_gn=STX3;refGene_tx=XM_023239869.1
D1 106289890 106289974
refGene_gn=STX3;refGene_tx=XM_023239868.1;refGene_gn=STX3;refGene_tx=XM_023239869.1
D1 106292315 106292438
refGene_gn=STX3;refGene_tx=XM_023239867.1
D1 106297761 106298247
refGene_gn=MRPL16;refGene_tx=XM_023239870.1
D1 106299281 106299430
refGene_gn=MRPL16;refGene_tx=XM_023239870.1
D1 106301831 106301897
refGene_gn=MRPL16;refGene_tx=XM_023239870.1
D1 106302250 106302305
refGene_gn=MRPL16;refGene_tx=XM_023239870.1
D1 106317419 106317481
refGene_gn=GIF;refGene_tx=XM_003993417.3;refGene_gn=GIF;refGene_tx=XM_019812451.2;refGene_gn=GIF;refGene_tx=XM_019812453.2
D1 106318592 106318711
refGene_gn=GIF;refGene_tx=XM_003993417.3;refGene_gn=GIF;refGene_tx=XM_019812451.2;refGene_gn=GIF;refGene_tx=XM_019812453.2
D1 106321348 106321394
refGene_gn=GIF;refGene_tx=XM_019812452.1
D1 106321502 106321704
refGene_gn=GIF;refGene_tx=XM_003993417.3;refGene_gn=GIF;refGene_tx=XM_019812451.2;refGene_gn=GIF;refGene_tx=XM_019812452.1;refGene_gn=GIF;refGene_tx=XM_019812453.2
D1 106323064 106323242
refGene_gn=GIF;refGene_tx=XM_003993417.3;refGene_gn=GIF;refGene_tx=XM_019812451.2;refGene_gn=GIF;refGene_tx=XM_019812452.1;refGene_gn=GIF;refGene_tx=XM_019812453.2
D1 106326982 106327164
refGene_gn=GIF;refGene_tx=XM_003993417.3;refGene_gn=GIF;refGene_tx=XM_

019812451.2;refGene_gn=GIF;refGene_tx=XM_019812452.1;refGene_gn=GIF;refGene_tx=XM_019812453.2
D1 106327529 106328047
refGene_gn=GIF;refGene_tx=XM_019812454.1
D1 106328976 106329117
refGene_gn=GIF;refGene_tx=XM_003993417.3;refGene_gn=GIF;refGene_tx=XM_019812451.2;refGene_gn=GIF;refGene_tx=XM_019812452.1;refGene_gn=GIF;refGene_tx=XM_019812453.2;refGene_gn=GIF;refGene_tx=XM_019812454.1
D1 106329549 106329663
refGene_gn=GIF;refGene_tx=XM_003993417.3;refGene_gn=GIF;refGene_tx=XM_019812451.2;refGene_gn=GIF;refGene_tx=XM_019812452.1;refGene_gn=GIF;refGene_tx=XM_019812453.2;refGene_gn=GIF;refGene_tx=XM_019812454.1
D1 106330718 106330895
refGene_gn=GIF;refGene_tx=XM_003993417.3;refGene_gn=GIF;refGene_tx=XM_019812451.2;refGene_gn=GIF;refGene_tx=XM_019812452.1;refGene_gn=GIF;refGene_tx=XM_019812454.1
D1 106331806 106331885
refGene_gn=GIF;refGene_tx=XM_003993417.3;refGene_gn=GIF;refGene_tx=XM_019812452.1;refGene_gn=GIF;refGene_tx=XM_019812453.2;refGene_gn=GIF;refGene_tx=XM_019812454.1
D1 106337793 106337855
refGene_gn=TCN1;refGene_tx=XM_003993476.2
D1 106338014 106338133
refGene_gn=TCN1;refGene_tx=XM_003993476.2
D1 106339632 106339819
refGene_gn=TCN1;refGene_tx=XM_003993476.2
D1 106340830 106341020
refGene_gn=TCN1;refGene_tx=XM_003993476.2
D1 106343346 106343534
refGene_gn=TCN1;refGene_tx=XM_003993476.2
D1 106344796 106344952
refGene_gn=TCN1;refGene_tx=XM_003993476.2
D1 106345843 106345984
refGene_gn=TCN1;refGene_tx=XM_003993476.2
D1 106346707 106346887
refGene_gn=TCN1;refGene_tx=XM_003993476.2
D1 106348855 106348934
refGene_gn=TCN1;refGene_tx=XM_003993476.2
D1 106421595 106421659
refGene_gn=OOSP2;refGene_tx=XM_003993477.3
D1 106425898 106426077
refGene_gn=OOSP2;refGene_tx=XM_003993477.3
D1 106426993 106427097
refGene_gn=OOSP2;refGene_tx=XM_003993477.3
D1 106428934 106429133
refGene_gn=OOSP2;refGene_tx=XM_003993477.3
D1 106450434 106450590
refGene_gn=MS4A3;refGene_tx=XM_011286920.3;refGene_gn=MS4A3;refGene_tx=XM_019812455.2;refGene_gn=MS4A3;refGene_tx=XM_019812456.2
D1 106451238 106451373

refGene_gn=MS4A3;refGene_tx=XM_011286920.3;refGene_gn=MS4A3;refGene_tx=XM_019812455.2
D1 106453829 106453886
refGene_gn=MS4A3;refGene_tx=XM_011286920.3;refGene_gn=MS4A3;refGene_tx=XM_019812455.2;refGene_gn=MS4A3;refGene_tx=XM_019812456.2
D1 106455801 106455960
refGene_gn=MS4A3;refGene_tx=XM_011286920.3;refGene_gn=MS4A3;refGene_tx=XM_019812455.2;refGene_gn=MS4A3;refGene_tx=XM_019812456.2
D1 106458184 106458286
refGene_gn=MS4A3;refGene_tx=XM_011286920.3;refGene_gn=MS4A3;refGene_tx=XM_019812455.2;refGene_gn=MS4A3;refGene_tx=XM_019812456.2
D1 106458905 106458935
refGene_gn=MS4A3;refGene_tx=XM_011286920.3;refGene_gn=MS4A3;refGene_tx=XM_019812455.2;refGene_gn=MS4A3;refGene_tx=XM_019812456.2
D1 106469255 106469317
refGene_gn=LOC101101505;refGene_tx=XM_023239872.1
D1 106470375 106470499
refGene_gn=LOC101101505;refGene_tx=XM_023239872.1
D1 106471034 106471169
refGene_gn=LOC101101505;refGene_tx=XM_023239872.1
D1 106473229 106473286
refGene_gn=LOC101101505;refGene_tx=XM_023239872.1
D1 106474065 106474221
refGene_gn=LOC101101505;refGene_tx=XM_023239872.1
D1 106474566 106474665
refGene_gn=LOC101101505;refGene_tx=XM_023239872.1
D1 106475527 106475626
refGene_gn=LOC101101505;refGene_tx=XM_023239872.1
D1 106488897 106488968
refGene_gn=LOC102901994;refGene_tx=XM_006937350.3
D1 106491922 106492061
refGene_gn=LOC102901994;refGene_tx=XM_006937350.3
D1 106492663 106492798
refGene_gn=LOC102901994;refGene_tx=XM_006937350.3
D1 106493007 106493064
refGene_gn=LOC102901994;refGene_tx=XM_006937350.3
D1 106493449 106493566
refGene_gn=LOC102901994;refGene_tx=XM_006937350.3
D1 106494344 106494446
refGene_gn=LOC102901994;refGene_tx=XM_006937350.3
D1 106494831 106494918
refGene_gn=LOC102901994;refGene_tx=XM_006937350.3
D1 106499465 106499606
refGene_gn=LOC102902042;refGene_tx=XM_019812458.1
D1 106503937 106504039
refGene_gn=LOC102902042;refGene_tx=XM_019812458.1
D1 106505405 106505549
refGene_gn=LOC102902042;refGene_tx=XM_019812458.1
D1 106506179 106506236
refGene_gn=LOC102902042;refGene_tx=XM_019812458.1

D1 106507249 106507381
refGene_gn=LOC102902042;refGene_tx=XM_019812458.1
D1 106507885 106508041
refGene_gn=LOC102902042;refGene_tx=XM_019812458.1
D1 106513728 106513818
refGene_gn=LOC102902084;refGene_tx=XM_006937354.4;refGene_gn=LOC102902
084;refGene_tx=XM_023239874.1;refGene_gn=LOC102902084;refGene_tx=XM_02
3239875.1
D1 106515060 106515162
refGene_gn=LOC102902084;refGene_tx=XM_006937354.4;refGene_gn=LOC102902
084;refGene_tx=XM_023239874.1;refGene_gn=LOC102902084;refGene_tx=XM_02
3239875.1
D1 106516434 106516566
refGene_gn=LOC102902084;refGene_tx=XM_006937354.4;refGene_gn=LOC102902
084;refGene_tx=XM_023239874.1;refGene_gn=LOC102902084;refGene_tx=XM_02
3239875.1
D1 106517154 106517211
refGene_gn=LOC102902084;refGene_tx=XM_006937354.4;refGene_gn=LOC102902
084;refGene_tx=XM_023239874.1;refGene_gn=LOC102902084;refGene_tx=XM_02
3239875.1
D1 106517391 106517526
refGene_gn=LOC102902084;refGene_tx=XM_006937354.4;refGene_gn=LOC102902
084;refGene_tx=XM_023239874.1;refGene_gn=LOC102902084;refGene_tx=XM_02
3239875.1
D1 106517866 106517986
refGene_gn=LOC102902084;refGene_tx=XM_006937354.4;refGene_gn=LOC102902
084;refGene_tx=XM_023239874.1;refGene_gn=LOC102902084;refGene_tx=XM_02
3239875.1
D1 106530557 106530677
refGene_gn=LOC102902189;refGene_tx=XM_023239876.1
D1 106533602 106533704
refGene_gn=LOC102902189;refGene_tx=XM_023239876.1
D1 106537271 106537415
refGene_gn=LOC102902189;refGene_tx=XM_023239876.1
D1 106538027 106538084
refGene_gn=LOC102902189;refGene_tx=XM_023239876.1
D1 106538862 106538994
refGene_gn=LOC102902189;refGene_tx=XM_023239876.1
D1 106539386 106539542
refGene_gn=LOC102902189;refGene_tx=XM_023239876.1
D1 106570881 106571031
refGene_gn=LOC102902224;refGene_tx=XM_023239027.1;refGene_gn=LOC102902
224;refGene_tx=XM_023239028.1
D1 106571225 106571282
refGene_gn=LOC102902224;refGene_tx=XM_023239027.1;refGene_gn=LOC102902
224;refGene_tx=XM_023239028.1
D1 106571682 106571814
refGene_gn=LOC102902224;refGene_tx=XM_023239027.1;refGene_gn=LOC102902
224;refGene_tx=XM_023239028.1
D1 106572574 106572673

refGene_gn=LOC102902224;refGene_tx=XM_023239027.1;refGene_gn=LOC102902
224;refGene_tx=XM_023239028.1
D1 106573107 106573182
refGene_gn=LOC102902224;refGene_tx=XM_023239027.1;refGene_gn=LOC102902
224;refGene_tx=XM_023239028.1
D1 106585462 106585588
refGene_gn=LOC102902259;refGene_tx=XM_011286924.2
D1 106586837 106586939
refGene_gn=LOC102902259;refGene_tx=XM_011286924.2
D1 106588141 106588285
refGene_gn=LOC102902259;refGene_tx=XM_011286924.2
D1 106589202 106589259
refGene_gn=LOC102902259;refGene_tx=XM_011286924.2
D1 106590138 106590270
refGene_gn=LOC102902259;refGene_tx=XM_011286924.2
D1 106590705 106590861
refGene_gn=LOC102902259;refGene_tx=XM_011286924.2
D1 106599510 106599606
refGene_gn=LOC102902259;refGene_tx=XM_023239877.1
D1 106600399 106600501
refGene_gn=LOC102902259;refGene_tx=XM_023239877.1
D1 106602390 106602600
refGene_gn=LOC102902259;refGene_tx=XM_023239877.1
D1 106605223 106605280
refGene_gn=LOC102902259;refGene_tx=XM_023239877.1
D1 106605604 106605739
refGene_gn=LOC102902259;refGene_tx=XM_023239877.1
D1 106607288 106607435
refGene_gn=LOC102902259;refGene_tx=XM_023239877.1
D1 106624168 106624240
refGene_gn=LOC101100669;refGene_tx=XM_006937359.4;refGene_gn=LOC101100
669;refGene_tx=XM_019812466.2;refGene_gn=LOC101100669;refGene_tx=XM_01
9812467.2;refGene_gn=LOC101100669;refGene_tx=XM_019812470.1
D1 106627427 106627529
refGene_gn=LOC101100669;refGene_tx=XM_006937359.4;refGene_gn=LOC101100
669;refGene_tx=XM_019812466.2;refGene_gn=LOC101100669;refGene_tx=XM_01
9812467.2;refGene_gn=LOC101100669;refGene_tx=XM_019812470.1
D1 106631880 106632039
refGene_gn=LOC101100669;refGene_tx=XM_006937359.4;refGene_gn=LOC101100
669;refGene_tx=XM_019812466.2;refGene_gn=LOC101100669;refGene_tx=XM_01
9812467.2;refGene_gn=LOC101100669;refGene_tx=XM_019812470.1
D1 106633241 106633298
refGene_gn=LOC101100669;refGene_tx=XM_006937359.4;refGene_gn=LOC101100
669;refGene_tx=XM_019812466.2;refGene_gn=LOC101100669;refGene_tx=XM_01
9812467.2;refGene_gn=LOC101100669;refGene_tx=XM_019812470.1
D1 106635233 106635362
refGene_gn=LOC101100669;refGene_tx=XM_006937359.4;refGene_gn=LOC101100
669;refGene_tx=XM_019812466.2;refGene_gn=LOC101100669;refGene_tx=XM_01
9812467.2;refGene_gn=LOC101100669;refGene_tx=XM_019812470.1
D1 106636457 106636617

refGene_gn=LOC101100669;refGene_tx=XM_006937359.4;refGene_gn=LOC101100669;refGene_tx=XM_019812466.2;refGene_gn=LOC101100669;refGene_tx=XM_019812467.2;refGene_gn=LOC101100669;refGene_tx=XM_019812470.1
D1 106677877 106677885
refGene_gn=MS4A7;refGene_tx=XM_003993481.4
D1 106687079 106687239
refGene_gn=MS4A7;refGene_tx=XM_003993481.4;refGene_gn=MS4A7;refGene_tx=XM_019812465.2
D1 106691294 106691429
refGene_gn=MS4A7;refGene_tx=XM_003993481.4;refGene_gn=MS4A7;refGene_tx=XM_019812465.2
D1 106691707 106691809
refGene_gn=LOC101100669;refGene_tx=XM_006937359.4;refGene_gn=LOC101100669;refGene_tx=XM_019812466.2;refGene_gn=LOC101100669;refGene_tx=XM_019812467.2
D1 106693139 106693196
refGene_gn=MS4A7;refGene_tx=XM_003993481.4;refGene_gn=MS4A7;refGene_tx=XM_019812465.2
D1 106694920 106695127
refGene_gn=MS4A7;refGene_tx=XM_003993481.4;refGene_gn=MS4A7;refGene_tx=XM_019812465.2
D1 106698610 106698712
refGene_gn=MS4A7;refGene_tx=XM_003993481.4;refGene_gn=MS4A7;refGene_tx=XM_019812465.2
D1 106699808 106699982
refGene_gn=MS4A7;refGene_tx=XM_003993481.4;refGene_gn=MS4A7;refGene_tx=XM_019812465.2
D1 106707678 106707813
refGene_gn=LOC101101165;refGene_tx=XM_006937358.3
D1 106709120 106709249
refGene_gn=LOC101101165;refGene_tx=XM_006937358.3
D1 106715405 106715456
refGene_gn=LOC101101165;refGene_tx=XM_006937358.3
D1 106717285 106717435
refGene_gn=LOC101101165;refGene_tx=XM_006937358.3
D1 106720060 106720159
refGene_gn=LOC101101165;refGene_tx=XM_006937358.3
D1 106723929 106723949
refGene_gn=LOC101100669;refGene_tx=XM_006937359.4;refGene_gn=LOC101100669;refGene_tx=XM_019812466.2;refGene_gn=LOC101100669;refGene_tx=XM_019812467.2
D1 106727440 106729660
refGene_gn=LOC101101165;refGene_tx=XM_006937358.3
D1 106748119 106748272
refGene_gn=MS4A5;refGene_tx=XM_003993483.4;refGene_gn=MS4A5;refGene_tx=XM_006937360.3
D1 106752183 106752312
refGene_gn=MS4A5;refGene_tx=XM_003993483.4;refGene_gn=MS4A5;refGene_tx=XM_006937360.3
D1 106754759 106754816

refGene_gn=MS4A5;refGene_tx=XM_003993483.4
D1	106756008	106756161
refGene_gn=MS4A5;refGene_tx=XM_003993483.4;refGene_gn=MS4A5;refGene_tx=XM_006937360.3
D1	106765357	106765459
refGene_gn=MS4A5;refGene_tx=XM_003993483.4;refGene_gn=MS4A5;refGene_tx=XM_006937360.3
D1	106776780	106776939
refGene_gn=MS4A1;refGene_tx=NM_001009388.2
D1	106777358	106777478
refGene_gn=MS4A1;refGene_tx=NM_001009388.2
D1	106778654	106778711
refGene_gn=MS4A1;refGene_tx=NM_001009388.2
D1	106780222	106780462
refGene_gn=MS4A1;refGene_tx=NM_001009388.2
D1	106781235	106781337
refGene_gn=MS4A1;refGene_tx=NM_001009388.2
D1	106782207	106782426
refGene_gn=MS4A1;refGene_tx=NM_001009388.2
D1	106809946	106810210
refGene_gn=MS4A12;refGene_tx=XM_023239745.1
D1	106814095	106814233
refGene_gn=MS4A12;refGene_tx=XM_023239745.1
D1	106816616	106816659
refGene_gn=MS4A12;refGene_tx=XM_023239745.1
D1	106816661	106816732
refGene_gn=MS4A12;refGene_tx=XM_023239745.1
D1	106820646	106820757
refGene_gn=MS4A12;refGene_tx=XM_023239745.1
D1	106820938	106821040
refGene_gn=MS4A12;refGene_tx=XM_023239745.1
D1	106833684	106833738
refGene_gn=MS4A13;refGene_tx=XM_019812471.2;refGene_gn=MS4A13;refGene_tx=XM_023239881.1
D1	106836282	106836423
refGene_gn=MS4A13;refGene_tx=XM_019812471.2;refGene_gn=MS4A13;refGene_tx=XM_023239881.1
D1	106839667	106839724
refGene_gn=MS4A13;refGene_tx=XM_019812471.2;refGene_gn=MS4A13;refGene_tx=XM_023239881.1
D1	106840662	106840782
refGene_gn=MS4A13;refGene_tx=XM_019812471.2;refGene_gn=MS4A13;refGene_tx=XM_023239881.1
D1	106844951	106845047
refGene_gn=MS4A13;refGene_tx=XM_019812471.2;refGene_gn=MS4A13;refGene_tx=XM_023239881.1
D1	106851664	106851724
refGene_gn=MS4A13;refGene_tx=XM_023239881.1
D1	106864172	106864220
refGene_gn=LOC102899106;refGene_tx=XM_006937367.4;refGene_gn=LOC102899

106;refGene_tx=XM_011286925.3;refGene_gn=LOC102899106;refGene_tx=XM_01
9812473.2;refGene_gn=LOC102899106;refGene_tx=XM_019812474.2;refGene_gn
=LOC102899106;refGene_tx=XM_019812475.2;refGene_gn=LOC102899106;refGen
e_tx=XM_019812476.2;refGene_gn=LOC102899106;refGene_tx=XM_019812477.2;
refGene_gn=LOC102899106;refGene_tx=XM_019812478.2;refGene_gn=LOC102899
106;refGene_tx=XM_019812479.2;refGene_gn=LOC102899106;refGene_tx=XM_02
3239882.1
D1 106869762 106869906
refGene_gn=LOC102899106;refGene_tx=XM_006937367.4;refGene_gn=LOC102899
106;refGene_tx=XM_019812473.2;refGene_gn=LOC102899106;refGene_tx=XM_01
9812474.2;refGene_gn=LOC102899106;refGene_tx=XM_019812475.2;refGene_gn
=LOC102899106;refGene_tx=XM_019812476.2;refGene_gn=LOC102899106;refGen
e_tx=XM_019812478.2;refGene_gn=LOC102899106;refGene_tx=XM_023239882.1
D1 106874845 106874902
refGene_gn=LOC102899106;refGene_tx=XM_006937367.4;refGene_gn=LOC102899
106;refGene_tx=XM_011286925.3;refGene_gn=LOC102899106;refGene_tx=XM_01
9812473.2;refGene_gn=LOC102899106;refGene_tx=XM_019812474.2;refGene_gn
=LOC102899106;refGene_tx=XM_019812475.2;refGene_gn=LOC102899106;refGen
e_tx=XM_019812476.2;refGene_gn=LOC102899106;refGene_tx=XM_019812477.2;
refGene_gn=LOC102899106;refGene_tx=XM_019812478.2;refGene_gn=LOC102899
106;refGene_tx=XM_019812479.2;refGene_gn=LOC102899106;refGene_tx=XM_02
3239882.1
D1 106885078 106885198
refGene_gn=LOC102899106;refGene_tx=XM_006937367.4;refGene_gn=LOC102899
106;refGene_tx=XM_011286925.3;refGene_gn=LOC102899106;refGene_tx=XM_01
9812473.2;refGene_gn=LOC102899106;refGene_tx=XM_019812474.2;refGene_gn
=LOC102899106;refGene_tx=XM_019812475.2;refGene_gn=LOC102899106;refGen
e_tx=XM_019812476.2;refGene_gn=LOC102899106;refGene_tx=XM_019812477.2;
refGene_gn=LOC102899106;refGene_tx=XM_019812478.2;refGene_gn=LOC102899
106;refGene_tx=XM_019812479.2;refGene_gn=LOC102899106;refGene_tx=XM_02
3239882.1
D1 106887840 106887897
refGene_gn=LOC102899106;refGene_tx=XM_006937367.4;refGene_gn=LOC102899
106;refGene_tx=XM_011286925.3;refGene_gn=LOC102899106;refGene_tx=XM_01
9812473.2;refGene_gn=LOC102899106;refGene_tx=XM_019812474.2;refGene_gn
=LOC102899106;refGene_tx=XM_019812475.2;refGene_gn=LOC102899106;refGen
e_tx=XM_019812476.2;refGene_gn=LOC102899106;refGene_tx=XM_019812477.2;
refGene_gn=LOC102899106;refGene_tx=XM_019812478.2;refGene_gn=LOC102899
106;refGene_tx=XM_019812479.2;refGene_gn=LOC102899106;refGene_tx=XM_02
3239882.1
D1 106889130 106889163
refGene_gn=LOC102899106;refGene_tx=XM_019812475.2;refGene_gn=LOC102899
106;refGene_tx=XM_019812478.2
D1 106890393 106890477
refGene_gn=LOC102899106;refGene_tx=XM_006937367.4;refGene_gn=LOC102899
106;refGene_tx=XM_011286925.3;refGene_gn=LOC102899106;refGene_tx=XM_01
9812473.2;refGene_gn=LOC102899106;refGene_tx=XM_019812474.2;refGene_gn
=LOC102899106;refGene_tx=XM_019812476.2;refGene_gn=LOC102899106;refGen
e_tx=XM_019812477.2;refGene_gn=LOC102899106;refGene_tx=XM_019812479.2;
refGene_gn=LOC102899106;refGene_tx=XM_023239882.1

D1 106898283 106898328
refGene_gn=LOC102899200;refGene_tx=XM_006937368.4
D1 106900736 106900880
refGene_gn=LOC102899200;refGene_tx=XM_006937368.4
D1 106904429 106904486
refGene_gn=LOC102899200;refGene_tx=XM_006937368.4
D1 106915941 106916061
refGene_gn=LOC102899200;refGene_tx=XM_006937368.4
D1 106919602 106919723
refGene_gn=LOC102899200;refGene_tx=XM_006937368.4
D1 106929729 106929764
refGene_gn=LOC102899200;refGene_tx=XM_006937368.4
D1 106947952 106948162
refGene_gn=LOC101081000;refGene_tx=XM_003993486.4
D1 106950078 106950201
refGene_gn=LOC101081000;refGene_tx=XM_003993486.4
D1 106953912 106953972
refGene_gn=LOC101081000;refGene_tx=XM_003993486.4
D1 106955177 106955315
refGene_gn=LOC101081000;refGene_tx=XM_003993486.4
D1 106957049 106957172
refGene_gn=LOC101081000;refGene_tx=XM_003993486.4
D1 106957343 106957448
refGene_gn=LOC101081000;refGene_tx=XM_003993486.4
D1 106982386 106982617
refGene_gn=MS4A15;refGene_tx=XM_003993487.4
D1 106985130 106985253
refGene_gn=MS4A15;refGene_tx=XM_003993487.4
D1 106986254 106986311
refGene_gn=MS4A15;refGene_tx=XM_003993487.4
D1 106988221 106988314
refGene_gn=MS4A15;refGene_tx=XM_003993487.4
D1 106988689 106988803
refGene_gn=MS4A15;refGene_tx=XM_003993487.4
D1 106989705 106989822
refGene_gn=MS4A15;refGene_tx=XM_003993487.4
D1 106994898 106994934
refGene_gn=MS4A10;refGene_tx=XM_023239746.1
D1 106998338 106998557
refGene_gn=MS4A10;refGene_tx=XM_023239746.1
D1 106998899 106999127
refGene_gn=MS4A10;refGene_tx=XM_023239746.1
D1 106999635 106999692
refGene_gn=MS4A10;refGene_tx=XM_023239746.1
D1 107001318 107001450
refGene_gn=MS4A10;refGene_tx=XM_023239746.1
D1 107002623 107002734
refGene_gn=MS4A10;refGene_tx=XM_023239746.1
D1 107004565 107004676
refGene_gn=MS4A10;refGene_tx=XM_023239746.1

D1 107037646 107038407
refGene_gn=CCDC86;refGene_tx=XM_003993419.4
D1 107041159 107041289
refGene_gn=CCDC86;refGene_tx=XM_003993419.4
D1 107042956 107043031
refGene_gn=CCDC86;refGene_tx=XM_003993419.4
D1 107043225 107043345
refGene_gn=CCDC86;refGene_tx=XM_003993419.4
D1 107045330 107046541
refGene_gn=PTGDR2;refGene_tx=XM_023239883.1;refGene_gn=PTGDR2;refGene_tx=XM_023239884.1;refGene_gn=PTGDR2;refGene_tx=XM_023239885.1
D1 107048611 107048774
refGene_gn=PTGDR2;refGene_tx=XM_023239883.1
D1 107055794 107056014
refGene_gn=ZP1;refGene_tx=NM_001284435.1
D1 107057303 107057425
refGene_gn=ZP1;refGene_tx=NM_001284435.1
D1 107057647 107057960
refGene_gn=ZP1;refGene_tx=NM_001284435.1
D1 107058125 107058269
refGene_gn=ZP1;refGene_tx=NM_001284435.1
D1 107059104 107059292
refGene_gn=ZP1;refGene_tx=NM_001284435.1
D1 107059371 107059469
refGene_gn=ZP1;refGene_tx=NM_001284435.1
D1 107060732 107060860
refGene_gn=ZP1;refGene_tx=NM_001284435.1
D1 107060942 107061132
refGene_gn=ZP1;refGene_tx=NM_001284435.1
D1 107061225 107061370
refGene_gn=ZP1;refGene_tx=NM_001284435.1
D1 107062456 107062538
refGene_gn=ZP1;refGene_tx=NM_001284435.1
D1 107062646 107062766
refGene_gn=ZP1;refGene_tx=NM_001284435.1
D1 107062973 107063080
refGene_gn=ZP1;refGene_tx=NM_001284435.1
D1 107063082 107063181
refGene_gn=ZP1;refGene_tx=NM_001284435.1
D1 107073425 107073523
refGene_gn=PRPF19;refGene_tx=XM_011286926.3
D1 107075455 107075561
refGene_gn=PRPF19;refGene_tx=XM_011286926.3
D1 107075704 107075875
refGene_gn=PRPF19;refGene_tx=XM_011286926.3
D1 107076223 107076309
refGene_gn=PRPF19;refGene_tx=XM_011286926.3
D1 107076539 107076609
refGene_gn=PRPF19;refGene_tx=XM_011286926.3
D1 107076733 107076889

refGene_gn=PRPF19;refGene_tx=XM_011286926.3
D1 107078207 107078317
refGene_gn=PRPF19;refGene_tx=XM_011286926.3
D1 107078478 107078554
refGene_gn=PRPF19;refGene_tx=XM_011286926.3
D1 107078882 107078957
refGene_gn=PRPF19;refGene_tx=XM_011286926.3
D1 107079081 107079123
refGene_gn=PRPF19;refGene_tx=XM_011286926.3
D1 107079987 107080050
refGene_gn=PRPF19;refGene_tx=XM_011286926.3
D1 107080169 107080243
refGene_gn=PRPF19;refGene_tx=XM_011286926.3
D1 107080326 107080468
refGene_gn=PRPF19;refGene_tx=XM_011286926.3
D1 107081199 107081276
refGene_gn=PRPF19;refGene_tx=XM_011286926.3
D1 107081464 107081614
refGene_gn=PRPF19;refGene_tx=XM_011286926.3
D1 107083304 107083323
refGene_gn=PRPF19;refGene_tx=XM_011286926.3
D1 107092926 107093151
refGene_gn=TMEM109;refGene_tx=XM_003993420.5;refGene_gn=TMEM109;refGen
e_tx=XM_019812481.2
D1 107094056 107094159
refGene_gn=TMEM109;refGene_tx=XM_003993420.5;refGene_gn=TMEM109;refGen
e_tx=XM_019812481.2
D1 107094611 107095003
refGene_gn=TMEM109;refGene_tx=XM_003993420.5;refGene_gn=TMEM109;refGen
e_tx=XM_019812481.2
D1 107097538 107097638
refGene_gn=TMEM132A;refGene_tx=XM_023239887.1;refGene_gn=TMEM132A;refG
ene_tx=XM_023239888.1
D1 107100028 107100243
refGene_gn=TMEM132A;refGene_tx=XM_023239887.1;refGene_gn=TMEM132A;refG
ene_tx=XM_023239888.1
D1 107100466 107100685
refGene_gn=TMEM132A;refGene_tx=XM_023239887.1;refGene_gn=TMEM132A;refG
ene_tx=XM_023239888.1
D1 107101452 107101784
refGene_gn=TMEM132A;refGene_tx=XM_023239887.1;refGene_gn=TMEM132A;refG
ene_tx=XM_023239888.1
D1 107103092 107103242
refGene_gn=TMEM132A;refGene_tx=XM_023239887.1;refGene_gn=TMEM132A;refG
ene_tx=XM_023239888.1
D1 107104538 107104737
refGene_gn=TMEM132A;refGene_tx=XM_023239887.1;refGene_gn=TMEM132A;refG
ene_tx=XM_023239888.1
D1 107104822 107104966
refGene_gn=TMEM132A;refGene_tx=XM_023239887.1;refGene_gn=TMEM132A;refG

ene_tx=XM_023239888.1
D1 107106060 107106263
refGene_gn=TMEM132A;refGene_tx=XM_023239887.1;refGene_gn=TMEM132A;refGene_tx=XM_023239888.1
D1 107106991 107107271
refGene_gn=TMEM132A;refGene_tx=XM_023239887.1;refGene_gn=TMEM132A;refGene_tx=XM_023239888.1
D1 107107630 107107822
refGene_gn=TMEM132A;refGene_tx=XM_023239887.1;refGene_gn=TMEM132A;refGene_tx=XM_023239888.1
D1 107108233 107109259
refGene_gn=TMEM132A;refGene_tx=XM_023239887.1;refGene_gn=TMEM132A;refGene_tx=XM_023239888.1
D1 107109576 107109731
refGene_gn=SLC15A3;refGene_tx=XM_019812480.2;refGene_gn=SLC15A3;refGene_tx=XM_023239889.1
D1 107110186 107110342
refGene_gn=SLC15A3;refGene_tx=XM_019812480.2;refGene_gn=SLC15A3;refGene_tx=XM_023239889.1
D1 107111578 107111737
refGene_gn=SLC15A3;refGene_tx=XM_019812480.2;refGene_gn=SLC15A3;refGene_tx=XM_023239889.1
D1 107114079 107114248
refGene_gn=SLC15A3;refGene_tx=XM_019812480.2;refGene_gn=SLC15A3;refGene_tx=XM_023239889.1
D1 107114948 107115059
refGene_gn=SLC15A3;refGene_tx=XM_019812480.2;refGene_gn=SLC15A3;refGene_tx=XM_023239889.1
D1 107116708 107116856
refGene_gn=SLC15A3;refGene_tx=XM_019812480.2;refGene_gn=SLC15A3;refGene_tx=XM_023239889.1
D1 107118718 107119011
refGene_gn=SLC15A3;refGene_tx=XM_019812480.2;refGene_gn=SLC15A3;refGene_tx=XM_023239889.1
D1 107122171 107122711
refGene_gn=SLC15A3;refGene_tx=XM_019812480.2;refGene_gn=SLC15A3;refGene_tx=XM_023239889.1
D1 107142335 107142396
refGene_gn=CD6;refGene_tx=XM_003993494.5;refGene_gn=CD6;refGene_tx=XM_006937371.4;refGene_gn=CD6;refGene_tx=XM_019812482.2
D1 107169789 107169858
refGene_gn=CD6;refGene_tx=XM_003993494.5;refGene_gn=CD6;refGene_tx=XM_006937371.4;refGene_gn=CD6;refGene_tx=XM_019812482.2
D1 107170945 107171296
refGene_gn=CD6;refGene_tx=XM_003993494.5;refGene_gn=CD6;refGene_tx=XM_006937371.4;refGene_gn=CD6;refGene_tx=XM_019812482.2
D1 107172084 107172396
refGene_gn=CD6;refGene_tx=XM_003993494.5;refGene_gn=CD6;refGene_tx=XM_006937371.4;refGene_gn=CD6;refGene_tx=XM_019812482.2
D1 107173121 107173424

refGene_gn=CD6;refGene_tx=XM_003993494.5;refGene_gn=CD6;refGene_tx=XM_006937371.4
D1 107174610 107174676
refGene_gn=CD6;refGene_tx=XM_003993494.5;refGene_gn=CD6;refGene_tx=XM_006937371.4;refGene_gn=CD6;refGene_tx=XM_019812482.2
D1 107176791 107176932
refGene_gn=CD6;refGene_tx=XM_003993494.5;refGene_gn=CD6;refGene_tx=XM_006937371.4;refGene_gn=CD6;refGene_tx=XM_019812482.2
D1 107177270 107177369
refGene_gn=CD6;refGene_tx=XM_003993494.5;refGene_gn=CD6;refGene_tx=XM_006937371.4;refGene_gn=CD6;refGene_tx=XM_019812482.2
D1 107178426 107178549
refGene_gn=CD6;refGene_tx=XM_003993494.5;refGene_gn=CD6;refGene_tx=XM_006937371.4;refGene_gn=CD6;refGene_tx=XM_019812482.2
D1 107180233 107180305
refGene_gn=CD6;refGene_tx=XM_003993494.5;refGene_gn=CD6;refGene_tx=XM_006937371.4;refGene_gn=CD6;refGene_tx=XM_019812482.2
D1 107180402 107180657
refGene_gn=CD6;refGene_tx=XM_003993494.5;refGene_gn=CD6;refGene_tx=XM_006937371.4;refGene_gn=CD6;refGene_tx=XM_019812482.2
D1 107180960 107181080
refGene_gn=CD6;refGene_tx=XM_003993494.5;refGene_gn=CD6;refGene_tx=XM_006937371.4;refGene_gn=CD6;refGene_tx=XM_019812482.2
D1 107181591 107181656
refGene_gn=CD6;refGene_tx=XM_003993494.5;refGene_gn=CD6;refGene_tx=XM_006937371.4;refGene_gn=CD6;refGene_tx=XM_019812482.2
D1 107249703 107249755
refGene_gn=CD5;refGene_tx=XM_006937372.3
D1 107259319 107259358
refGene_gn=CD5;refGene_tx=XM_006937372.3
D1 107261819 107262128
refGene_gn=CD5;refGene_tx=XM_006937372.3
D1 107262636 107262699
refGene_gn=CD5;refGene_tx=XM_006937372.3
D1 107262960 107263293
refGene_gn=CD5;refGene_tx=XM_006937372.3
D1 107265327 107265618
refGene_gn=CD5;refGene_tx=XM_006937372.3
D1 107266281 107266407
refGene_gn=CD5;refGene_tx=XM_006937372.3
D1 107267129 107267183
refGene_gn=CD5;refGene_tx=XM_006937372.3
D1 107268092 107268212
refGene_gn=CD5;refGene_tx=XM_006937372.3
D1 107268796 107268885
refGene_gn=CD5;refGene_tx=XM_006937372.3
D1 107275642 107276365
refGene_gn=VPS37C;refGene_tx=XM_023239891.1;refGene_gn=VPS37C;refGene_tx=XM_023239892.1;refGene_gn=VPS37C;refGene_tx=XM_023239893.1;refGene_gn=VPS37C;refGene_tx=XM_023239894.1;refGene_gn=VPS37C;refGene_tx=XM_02

3239895.1
D1 107277121 107277204
refGene_gn=VPS37C;refGene_tx=XM_023239891.1;refGene_gn=VPS37C;refGene_tx=XM_023239892.1;refGene_gn=VPS37C;refGene_tx=XM_023239893.1;refGene_gn=VPS37C;refGene_tx=XM_023239894.1;refGene_gn=VPS37C;refGene_tx=XM_023239895.1
D1 107277894 107278066
refGene_gn=VPS37C;refGene_tx=XM_023239891.1;refGene_gn=VPS37C;refGene_tx=XM_023239892.1;refGene_gn=VPS37C;refGene_tx=XM_023239893.1;refGene_gn=VPS37C;refGene_tx=XM_023239894.1;refGene_gn=VPS37C;refGene_tx=XM_023239895.1
D1 107281488 107281586
refGene_gn=VPS37C;refGene_tx=XM_023239891.1;refGene_gn=VPS37C;refGene_tx=XM_023239892.1;refGene_gn=VPS37C;refGene_tx=XM_023239893.1
D1 107294398 107294405
refGene_gn=VPS37C;refGene_tx=XM_023239891.1
D1 107322840 107322893
refGene_gn=LOC493954;refGene_tx=NM_001009868.1
D1 107323971 107324134
refGene_gn=LOC493954;refGene_tx=NM_001009868.1;refGene_gn=LOC493954;refGene_tx=XM_019811181.2
D1 107325004 107325122
refGene_gn=LOC493954;refGene_tx=NM_001009868.1;refGene_gn=LOC493954;refGene_tx=XM_019811181.2
D1 107325208 107325327
refGene_gn=LOC493954;refGene_tx=NM_001009868.1;refGene_gn=LOC493954;refGene_tx=XM_019811181.2
D1 107326112 107326312
refGene_gn=LOC493954;refGene_tx=NM_001009868.1;refGene_gn=LOC493954;refGene_tx=XM_019811181.2
D1 107329026 107329143
refGene_gn=LOC493954;refGene_tx=NM_001009868.1;refGene_gn=LOC493954;refGene_tx=XM_019811181.2
D1 107330043 107330188
refGene_gn=LOC493954;refGene_tx=NM_001009868.1;refGene_gn=LOC493954;refGene_tx=XM_019811181.2
D1 107330279 107330378
refGene_gn=LOC493954;refGene_tx=NM_001009868.1;refGene_gn=LOC493954;refGene_tx=XM_019811181.2
D1 107330734 107330887
refGene_gn=LOC493954;refGene_tx=NM_001009868.1;refGene_gn=LOC493954;refGene_tx=XM_019811181.2
D1 107349086 107349142
refGene_gn=LOC101080826;refGene_tx=XM_003993421.4
D1 107349566 107349726
refGene_gn=LOC101080826;refGene_tx=XM_003993421.4
D1 107351824 107351942
refGene_gn=LOC101080826;refGene_tx=XM_003993421.4
D1 107352482 107352601
refGene_gn=LOC101080826;refGene_tx=XM_003993421.4

D1 107353376 107353576
refGene_gn=LOC101080826;refGene_tx=XM_003993421.4
D1 107355012 107355129
refGene_gn=LOC101080826;refGene_tx=XM_003993421.4
D1 107356169 107356314
refGene_gn=LOC101080826;refGene_tx=XM_003993421.4
D1 107356412 107356511
refGene_gn=LOC101080826;refGene_tx=XM_003993421.4
D1 107357608 107357758
refGene_gn=LOC101080826;refGene_tx=XM_003993421.4
D1 107364462 107365058
refGene_gn=VWCE;refGene_tx=XM_023239896.1;refGene_gn=VWCE;refGene_tx=XM_023239897.1
D1 107367780 107367871
refGene_gn=VWCE;refGene_tx=XM_023239896.1;refGene_gn=VWCE;refGene_tx=XM_023239897.1
D1 107368172 107368255
refGene_gn=VWCE;refGene_tx=XM_023239896.1;refGene_gn=VWCE;refGene_tx=XM_023239897.1
D1 107368329 107368420
refGene_gn=VWCE;refGene_tx=XM_023239896.1;refGene_gn=VWCE;refGene_tx=XM_023239897.1
D1 107369529 107369612
refGene_gn=VWCE;refGene_tx=XM_023239896.1;refGene_gn=VWCE;refGene_tx=XM_023239897.1
D1 107370992 107371089
refGene_gn=VWCE;refGene_tx=XM_023239896.1;refGene_gn=VWCE;refGene_tx=XM_023239897.1
D1 107372442 107372528
refGene_gn=VWCE;refGene_tx=XM_023239896.1;refGene_gn=VWCE;refGene_tx=XM_023239897.1
D1 107373018 107373136
refGene_gn=VWCE;refGene_tx=XM_023239896.1;refGene_gn=VWCE;refGene_tx=XM_023239897.1
D1 107374365 107374451
refGene_gn=VWCE;refGene_tx=XM_023239896.1;refGene_gn=VWCE;refGene_tx=XM_023239897.1
D1 107376180 107376268
refGene_gn=VWCE;refGene_tx=XM_023239896.1;refGene_gn=VWCE;refGene_tx=XM_023239897.1
D1 107377969 107378052
refGene_gn=VWCE;refGene_tx=XM_023239896.1;refGene_gn=VWCE;refGene_tx=XM_023239897.1
D1 107379290 107379384
refGene_gn=VWCE;refGene_tx=XM_023239896.1;refGene_gn=VWCE;refGene_tx=XM_023239897.1
D1 107379466 107379879
refGene_gn=VWCE;refGene_tx=XM_023239896.1;refGene_gn=VWCE;refGene_tx=XM_023239897.1
D1 107380865 107380994

refGene_gn=VWCE;refGene_tx=XM_023239896.1;refGene_gn=VWCE;refGene_tx=XM_023239897.1
D1 107381890 107382007
refGene_gn=VWCE;refGene_tx=XM_023239896.1;refGene_gn=VWCE;refGene_tx=XM_023239897.1
D1 107384486 107384603
refGene_gn=VWCE;refGene_tx=XM_023239896.1;refGene_gn=VWCE;refGene_tx=XM_023239897.1
D1 107387080 107387209
refGene_gn=VWCE;refGene_tx=XM_023239896.1;refGene_gn=VWCE;refGene_tx=XM_023239897.1
D1 107387511 107387601
refGene_gn=VWCE;refGene_tx=XM_023239896.1;refGene_gn=VWCE;refGene_tx=XM_023239897.1
D1 107387724 107387819
refGene_gn=VWCE;refGene_tx=XM_023239896.1;refGene_gn=VWCE;refGene_tx=XM_023239897.1
D1 107390296 107390406
refGene_gn=VWCE;refGene_tx=XM_023239896.1;refGene_gn=VWCE;refGene_tx=XM_023239897.1
D1 107394970 107395054
refGene_gn=DDB1;refGene_tx=XM_003993422.5
D1 107395644 107395768
refGene_gn=DDB1;refGene_tx=XM_003993422.5
D1 107397404 107397507
refGene_gn=DDB1;refGene_tx=XM_003993422.5
D1 107397706 107397876
refGene_gn=DDB1;refGene_tx=XM_003993422.5
D1 107398152 107398262
refGene_gn=DDB1;refGene_tx=XM_003993422.5
D1 107398618 107398789
refGene_gn=DDB1;refGene_tx=XM_003993422.5
D1 107403369 107403464
refGene_gn=DDB1;refGene_tx=XM_003993422.5
D1 107404186 107404202
refGene_gn=DDB1;refGene_tx=XM_003993422.5
D1 107404203 107404352
refGene_gn=DDB1;refGene_tx=XM_003993422.5
D1 107404707 107404831
refGene_gn=DDB1;refGene_tx=XM_003993422.5
D1 107405783 107405895
refGene_gn=DDB1;refGene_tx=XM_003993422.5
D1 107405996 107406092
refGene_gn=DDB1;refGene_tx=XM_003993422.5
D1 107407047 107407255
refGene_gn=DDB1;refGene_tx=XM_003993422.5
D1 107407406 107407514
refGene_gn=DDB1;refGene_tx=XM_003993422.5
D1 107407597 107407761
refGene_gn=DDB1;refGene_tx=XM_003993422.5

D1	107407858	107408037
refGene_gn=DDB1;refGene_tx=XM_003993422.5
D1	107409974	107410083
refGene_gn=DDB1;refGene_tx=XM_003993422.5
D1	107410179	107410255
refGene_gn=DDB1;refGene_tx=XM_003993422.5
D1	107413780	107413883
refGene_gn=DDB1;refGene_tx=XM_003993422.5
D1	107414466	107414583
refGene_gn=DDB1;refGene_tx=XM_003993422.5
D1	107415166	107415250
refGene_gn=DDB1;refGene_tx=XM_003993422.5
D1	107415835	107415994
refGene_gn=DDB1;refGene_tx=XM_003993422.5
D1	107416795	107416893
refGene_gn=DDB1;refGene_tx=XM_003993422.5
D1	107417823	107417938
refGene_gn=DDB1;refGene_tx=XM_003993422.5
D1	107419968	107420190
refGene_gn=DDB1;refGene_tx=XM_003993422.5
D1	107420754	107420871
refGene_gn=DDB1;refGene_tx=XM_003993422.5
D1	107422191	107422340
refGene_gn=DDB1;refGene_tx=XM_003993422.5
D1	107423569	107423630
refGene_gn=DDB1;refGene_tx=XM_003993422.5
D1	107425438	107425441
refGene_gn=TKFC;refGene_tx=XM_003993423.3;refGene_gn=TKFC;refGene_tx=XM_006937375.3;refGene_gn=TKFC;refGene_tx=XM_006937376.2;refGene_gn=TKFC;refGene_tx=XM_006937377.4
D1	107427992	107428182
refGene_gn=TKFC;refGene_tx=XM_003993423.3;refGene_gn=TKFC;refGene_tx=XM_006937375.3;refGene_gn=TKFC;refGene_tx=XM_006937376.2;refGene_gn=TKFC;refGene_tx=XM_006937377.4
D1	107429290	107429401
refGene_gn=TKFC;refGene_tx=XM_003993423.3;refGene_gn=TKFC;refGene_tx=XM_006937375.3;refGene_gn=TKFC;refGene_tx=XM_006937376.2;refGene_gn=TKFC;refGene_tx=XM_006937377.4
D1	107429477	107429659
refGene_gn=TKFC;refGene_tx=XM_003993423.3;refGene_gn=TKFC;refGene_tx=XM_006937375.3;refGene_gn=TKFC;refGene_tx=XM_006937376.2;refGene_gn=TKFC;refGene_tx=XM_006937377.4
D1	107430413	107430492
refGene_gn=TKFC;refGene_tx=XM_003993423.3;refGene_gn=TKFC;refGene_tx=XM_006937375.3;refGene_gn=TKFC;refGene_tx=XM_006937376.2;refGene_gn=TKFC;refGene_tx=XM_006937377.4
D1	107430822	107430912
refGene_gn=TKFC;refGene_tx=XM_003993423.3;refGene_gn=TKFC;refGene_tx=XM_006937375.3;refGene_gn=TKFC;refGene_tx=XM_006937376.2;refGene_gn=TKFC;refGene_tx=XM_006937377.4

D1 107431662 107431697
refGene_gn=TKFC;refGene_tx=XM_003993423.3;refGene_gn=TKFC;refGene_tx=XM_006937375.3;refGene_gn=TKFC;refGene_tx=XM_006937376.2;refGene_gn=TKFC;refGene_tx=XM_006937377.4
D1 107431775 107431860
refGene_gn=TKFC;refGene_tx=XM_003993423.3;refGene_gn=TKFC;refGene_tx=XM_006937375.3;refGene_gn=TKFC;refGene_tx=XM_006937376.2;refGene_gn=TKFC;refGene_tx=XM_006937377.4
D1 107431950 107432040
refGene_gn=TKFC;refGene_tx=XM_003993423.3;refGene_gn=TKFC;refGene_tx=XM_006937375.3;refGene_gn=TKFC;refGene_tx=XM_006937376.2;refGene_gn=TKFC;refGene_tx=XM_006937377.4
D1 107432571 107432688
refGene_gn=TKFC;refGene_tx=XM_003993423.3;refGene_gn=TKFC;refGene_tx=XM_006937375.3;refGene_gn=TKFC;refGene_tx=XM_006937376.2;refGene_gn=TKFC;refGene_tx=XM_006937377.4
D1 107433004 107433124
refGene_gn=TKFC;refGene_tx=XM_003993423.3;refGene_gn=TKFC;refGene_tx=XM_006937375.3;refGene_gn=TKFC;refGene_tx=XM_006937376.2;refGene_gn=TKFC;refGene_tx=XM_006937377.4
D1 107433264 107433402
refGene_gn=TKFC;refGene_tx=XM_003993423.3;refGene_gn=TKFC;refGene_tx=XM_006937375.3;refGene_gn=TKFC;refGene_tx=XM_006937376.2;refGene_gn=TKFC;refGene_tx=XM_006937377.4
D1 107435271 107435378
refGene_gn=TKFC;refGene_tx=XM_003993423.3;refGene_gn=TKFC;refGene_tx=XM_006937375.3;refGene_gn=TKFC;refGene_tx=XM_006937376.2;refGene_gn=TKFC;refGene_tx=XM_006937377.4
D1 107435482 107435586
refGene_gn=TKFC;refGene_tx=XM_003993423.3;refGene_gn=TKFC;refGene_tx=XM_006937375.3;refGene_gn=TKFC;refGene_tx=XM_006937376.2;refGene_gn=TKFC;refGene_tx=XM_006937377.4
D1 107435723 107435757
refGene_gn=TKFC;refGene_tx=XM_003993423.3;refGene_gn=TKFC;refGene_tx=XM_006937375.3;refGene_gn=TKFC;refGene_tx=XM_006937376.2;refGene_gn=TKFC;refGene_tx=XM_006937377.4
D1 107435861 107435951
refGene_gn=TKFC;refGene_tx=XM_003993423.3;refGene_gn=TKFC;refGene_tx=XM_006937375.3;refGene_gn=TKFC;refGene_tx=XM_006937376.2;refGene_gn=TKFC;refGene_tx=XM_006937377.4
D1 107436321 107436483
refGene_gn=TKFC;refGene_tx=XM_003993423.3;refGene_gn=TKFC;refGene_tx=XM_006937375.3;refGene_gn=TKFC;refGene_tx=XM_006937376.2;refGene_gn=TKFC;refGene_tx=XM_006937377.4
D1 107439809 107439833
refGene_gn=LOC101084057;refGene_tx=XM_023239747.1
D1 107440476 107440633
refGene_gn=LOC101084057;refGene_tx=XM_023239747.1
D1 107441606 107441761
refGene_gn=LOC101084057;refGene_tx=XM_023239747.1

D1 107442359 107442625
refGene_gn=LOC101084057;refGene_tx=XM_023239747.1
D1 107443179 107443289
refGene_gn=LOC101084057;refGene_tx=XM_023239747.1
D1 107443723 107443914
refGene_gn=LOC101084057;refGene_tx=XM_023239747.1
D1 107451016 107451144
refGene_gn=TMEM138;refGene_tx=XM_003993424.5;refGene_gn=TMEM138;refGene_tx=XM_023239898.1
D1 107452250 107452422
refGene_gn=TMEM138;refGene_tx=XM_003993424.5;refGene_gn=TMEM138;refGene_tx=XM_023239898.1
D1 107453582 107453658
refGene_gn=TMEM138;refGene_tx=XM_003993424.5;refGene_gn=TMEM138;refGene_tx=XM_023239898.1
D1 107453930 107454043
refGene_gn=TMEM138;refGene_tx=XM_003993424.5;refGene_gn=TMEM138;refGene_tx=XM_023239898.1
D1 107470781 107470794
refGene_gn=TMEM216;refGene_tx=XM_006937379.3
D1 107471384 107471486
refGene_gn=TMEM216;refGene_tx=XM_006937379.3
D1 107472044 107472137
refGene_gn=TMEM216;refGene_tx=XM_003993425.4;refGene_gn=TMEM216;refGene_tx=XM_006937379.3;refGene_gn=TMEM216;refGene_tx=XM_006937380.4;refGene_gn=TMEM216;refGene_tx=XM_019812501.1
D1 107473772 107473974
refGene_gn=TMEM216;refGene_tx=XM_003993425.4;refGene_gn=TMEM216;refGene_tx=XM_006937379.3;refGene_gn=TMEM216;refGene_tx=XM_006937380.4;refGene_gn=TMEM216;refGene_tx=XM_019812501.1
D1 107474444 107474460
refGene_gn=TMEM216;refGene_tx=XM_003993425.4;refGene_gn=TMEM216;refGene_tx=XM_006937379.3;refGene_gn=TMEM216;refGene_tx=XM_006937380.4;refGene_gn=TMEM216;refGene_tx=XM_019812501.1
D1 107484761 107484924
refGene_gn=CPSF7;refGene_tx=XM_006937383.3;refGene_gn=CPSF7;refGene_tx=XM_006937384.4;refGene_gn=CPSF7;refGene_tx=XM_006937385.3;refGene_gn=CPSF7;refGene_tx=XM_019812497.2
D1 107485575 107485744
refGene_gn=CPSF7;refGene_tx=XM_006937383.3;refGene_gn=CPSF7;refGene_tx=XM_006937384.4;refGene_gn=CPSF7;refGene_tx=XM_006937385.3;refGene_gn=CPSF7;refGene_tx=XM_019812497.2
D1 107488072 107488191
refGene_gn=CPSF7;refGene_tx=XM_006937383.3;refGene_gn=CPSF7;refGene_tx=XM_006937384.4;refGene_gn=CPSF7;refGene_tx=XM_006937385.3;refGene_gn=CPSF7;refGene_tx=XM_019812497.2
D1 107488696 107489111
refGene_gn=CPSF7;refGene_tx=XM_006937383.3;refGene_gn=CPSF7;refGene_tx=XM_006937384.4;refGene_gn=CPSF7;refGene_tx=XM_006937385.3;refGene_gn=CPSF7;refGene_tx=XM_019812497.2

D1 107491931 107492104
refGene_gn=CPSF7;refGene_tx=XM_006937383.3;refGene_gn=CPSF7;refGene_tx
=XM_006937384.4;refGene_gn=CPSF7;refGene_tx=XM_006937385.3;refGene_gn=
CPSF7;refGene_tx=XM_019812497.2
D1 107492489 107492593
refGene_gn=CPSF7;refGene_tx=XM_006937383.3;refGene_gn=CPSF7;refGene_tx
=XM_006937384.4;refGene_gn=CPSF7;refGene_tx=XM_006937385.3;refGene_gn=
CPSF7;refGene_tx=XM_019812497.2
D1 107493417 107493636
refGene_gn=CPSF7;refGene_tx=XM_006937383.3;refGene_gn=CPSF7;refGene_tx
=XM_006937384.4;refGene_gn=CPSF7;refGene_tx=XM_006937385.3;refGene_gn=
CPSF7;refGene_tx=XM_019812497.2
D1 107500920 107501014
refGene_gn=CPSF7;refGene_tx=XM_006937383.3;refGene_gn=CPSF7;refGene_tx
=XM_006937384.4;refGene_gn=CPSF7;refGene_tx=XM_006937385.3;refGene_gn=
CPSF7;refGene_tx=XM_019812497.2;refGene_gn=SDHAF2;refGene_tx=XM_023239
900.1
D1 107501871 107501907
refGene_gn=SDHAF2;refGene_tx=XM_003993429.5;refGene_gn=SDHAF2;refGene_
tx=XM_019812498.2
D1 107505135 107505213
refGene_gn=SDHAF2;refGene_tx=XM_019812498.2
D1 107505549 107505773
refGene_gn=SDHAF2;refGene_tx=XM_003993429.5;refGene_gn=SDHAF2;refGene_
tx=XM_006937382.4;refGene_gn=SDHAF2;refGene_tx=XM_019812498.2;refGene_
gn=SDHAF2;refGene_tx=XM_019812499.2;refGene_gn=SDHAF2;refGene_tx=XM_02
3239900.1
D1 107505857 107505967
refGene_gn=SDHAF2;refGene_tx=XM_003993429.5;refGene_gn=SDHAF2;refGene_
tx=XM_006937382.4;refGene_gn=SDHAF2;refGene_tx=XM_019812498.2;refGene_
gn=SDHAF2;refGene_tx=XM_019812499.2;refGene_gn=SDHAF2;refGene_tx=XM_02
3239900.1
D1 107515271 107515402
refGene_gn=SDHAF2;refGene_tx=XM_003993429.5;refGene_gn=SDHAF2;refGene_
tx=XM_006937382.4;refGene_gn=SDHAF2;refGene_tx=XM_019812498.2;refGene_
gn=SDHAF2;refGene_tx=XM_019812499.2;refGene_gn=SDHAF2;refGene_tx=XM_02
3239900.1
D1 107538984 107539093
refGene_gn=PPP1R32;refGene_tx=XM_011286937.3;refGene_gn=PPP1R32;refGen
e_tx=XM_011286938.3;refGene_gn=PPP1R32;refGene_tx=XM_011286939.3;refGe
ne_gn=PPP1R32;refGene_tx=XM_011286940.3;refGene_gn=PPP1R32;refGene_tx=
XM_011286941.3;refGene_gn=PPP1R32;refGene_tx=XM_019812502.2
D1 107539599 107539798
refGene_gn=PPP1R32;refGene_tx=XM_011286937.3;refGene_gn=PPP1R32;refGen
e_tx=XM_011286938.3;refGene_gn=PPP1R32;refGene_tx=XM_011286939.3;refGe
ne_gn=PPP1R32;refGene_tx=XM_011286940.3;refGene_gn=PPP1R32;refGene_tx=
XM_011286941.3;refGene_gn=PPP1R32;refGene_tx=XM_019812502.2
D1 107540003 107540157
refGene_gn=PPP1R32;refGene_tx=XM_011286937.3;refGene_gn=PPP1R32;refGen
e_tx=XM_011286938.3;refGene_gn=PPP1R32;refGene_tx=XM_011286939.3;refGe

ne_gn=PPP1R32;refGene_tx=XM_011286940.3;refGene_gn=PPP1R32;refGene_tx=
XM_011286941.3;refGene_gn=PPP1R32;refGene_tx=XM_019812502.2
D1 107540884 107541013
refGene_gn=PPP1R32;refGene_tx=XM_011286937.3;refGene_gn=PPP1R32;refGen
e_tx=XM_011286938.3;refGene_gn=PPP1R32;refGene_tx=XM_011286939.3;refGe
ne_gn=PPP1R32;refGene_tx=XM_011286940.3;refGene_gn=PPP1R32;refGene_tx=
XM_019812502.2
D1 107541652 107541716
refGene_gn=PPP1R32;refGene_tx=XM_011286937.3;refGene_gn=PPP1R32;refGen
e_tx=XM_011286938.3;refGene_gn=PPP1R32;refGene_tx=XM_011286939.3;refGe
ne_gn=PPP1R32;refGene_tx=XM_011286940.3;refGene_gn=PPP1R32;refGene_tx=
XM_011286941.3;refGene_gn=PPP1R32;refGene_tx=XM_019812502.2
D1 107542168 107542281
refGene_gn=PPP1R32;refGene_tx=XM_011286937.3;refGene_gn=PPP1R32;refGen
e_tx=XM_011286938.3;refGene_gn=PPP1R32;refGene_tx=XM_011286939.3;refGe
ne_gn=PPP1R32;refGene_tx=XM_011286940.3;refGene_gn=PPP1R32;refGene_tx=
XM_011286941.3;refGene_gn=PPP1R32;refGene_tx=XM_019812502.2
D1 107542650 107542710
refGene_gn=PPP1R32;refGene_tx=XM_011286937.3;refGene_gn=PPP1R32;refGen
e_tx=XM_011286938.3;refGene_gn=PPP1R32;refGene_tx=XM_011286939.3;refGe
ne_gn=PPP1R32;refGene_tx=XM_011286940.3;refGene_gn=PPP1R32;refGene_tx=
XM_011286941.3;refGene_gn=PPP1R32;refGene_tx=XM_019812502.2
D1 107542846 107542933
refGene_gn=PPP1R32;refGene_tx=XM_011286937.3;refGene_gn=PPP1R32;refGen
e_tx=XM_011286938.3;refGene_gn=PPP1R32;refGene_tx=XM_011286940.3;refGe
ne_gn=PPP1R32;refGene_tx=XM_011286941.3;refGene_gn=PPP1R32;refGene_tx=
XM_019812502.2
D1 107543485 107543599
refGene_gn=PPP1R32;refGene_tx=XM_011286937.3;refGene_gn=PPP1R32;refGen
e_tx=XM_011286938.3;refGene_gn=PPP1R32;refGene_tx=XM_011286939.3;refGe
ne_gn=PPP1R32;refGene_tx=XM_011286941.3;refGene_gn=PPP1R32;refGene_tx=
XM_019812502.2
D1 107543680 107543775
refGene_gn=PPP1R32;refGene_tx=XM_011286937.3;refGene_gn=PPP1R32;refGen
e_tx=XM_011286938.3;refGene_gn=PPP1R32;refGene_tx=XM_011286939.3;refGe
ne_gn=PPP1R32;refGene_tx=XM_011286940.3;refGene_gn=PPP1R32;refGene_tx=
XM_011286941.3;refGene_gn=PPP1R32;refGene_tx=XM_019812502.2
D1 107545372 107545537
refGene_gn=PPP1R32;refGene_tx=XM_011286937.3;refGene_gn=PPP1R32;refGen
e_tx=XM_011286938.3;refGene_gn=PPP1R32;refGene_tx=XM_011286939.3;refGe
ne_gn=PPP1R32;refGene_tx=XM_011286940.3;refGene_gn=PPP1R32;refGene_tx=
XM_011286941.3;refGene_gn=PPP1R32;refGene_tx=XM_019812502.2
D1 107546004 107546074
refGene_gn=PPP1R32;refGene_tx=XM_011286937.3;refGene_gn=PPP1R32;refGen
e_tx=XM_011286938.3;refGene_gn=PPP1R32;refGene_tx=XM_011286939.3;refGe
ne_gn=PPP1R32;refGene_tx=XM_011286940.3;refGene_gn=PPP1R32;refGene_tx=
XM_011286941.3;refGene_gn=PPP1R32;refGene_tx=XM_019812502.2
D1 107560250 107561129
refGene_gn=LRRC10B;refGene_tx=XM_011286942.3;refGene_gn=LRRC10B;refGen
e_tx=XM_023239902.1

D1	107568458	107568563
refGene_gn=SYT7;refGene_tx=XM_023239904.1;refGene_gn=SYT7;refGene_tx=XM_023239905.1;refGene_gn=SYT7;refGene_tx=XM_023239906.1;refGene_gn=SYT7;refGene_tx=XM_023239907.1;refGene_gn=SYT7;refGene_tx=XM_023239908.1;refGene_gn=SYT7;refGene_tx=XM_023239909.1;refGene_gn=SYT7;refGene_tx=XM_023239910.1
D1	107570978	107571178
refGene_gn=SYT7;refGene_tx=XM_023239904.1;refGene_gn=SYT7;refGene_tx=XM_023239905.1;refGene_gn=SYT7;refGene_tx=XM_023239906.1;refGene_gn=SYT7;refGene_tx=XM_023239907.1;refGene_gn=SYT7;refGene_tx=XM_023239908.1;refGene_gn=SYT7;refGene_tx=XM_023239909.1;refGene_gn=SYT7;refGene_tx=XM_023239910.1
D1	107571710	107571825
refGene_gn=SYT7;refGene_tx=XM_023239904.1;refGene_gn=SYT7;refGene_tx=XM_023239905.1;refGene_gn=SYT7;refGene_tx=XM_023239906.1;refGene_gn=SYT7;refGene_tx=XM_023239907.1;refGene_gn=SYT7;refGene_tx=XM_023239908.1;refGene_gn=SYT7;refGene_tx=XM_023239909.1;refGene_gn=SYT7;refGene_tx=XM_023239910.1
D1	107572261	107572431
refGene_gn=SYT7;refGene_tx=XM_023239904.1;refGene_gn=SYT7;refGene_tx=XM_023239905.1;refGene_gn=SYT7;refGene_tx=XM_023239906.1;refGene_gn=SYT7;refGene_tx=XM_023239907.1;refGene_gn=SYT7;refGene_tx=XM_023239908.1;refGene_gn=SYT7;refGene_tx=XM_023239909.1;refGene_gn=SYT7;refGene_tx=XM_023239910.1
D1	107575778	107576049
refGene_gn=SYT7;refGene_tx=XM_023239904.1;refGene_gn=SYT7;refGene_tx=XM_023239905.1;refGene_gn=SYT7;refGene_tx=XM_023239906.1;refGene_gn=SYT7;refGene_tx=XM_023239907.1;refGene_gn=SYT7;refGene_tx=XM_023239908.1;refGene_gn=SYT7;refGene_tx=XM_023239909.1;refGene_gn=SYT7;refGene_tx=XM_023239910.1
D1	107580134	107580270
refGene_gn=SYT7;refGene_tx=XM_023239904.1;refGene_gn=SYT7;refGene_tx=XM_023239905.1;refGene_gn=SYT7;refGene_tx=XM_023239906.1;refGene_gn=SYT7;refGene_tx=XM_023239907.1;refGene_gn=SYT7;refGene_tx=XM_023239908.1;refGene_gn=SYT7;refGene_tx=XM_023239909.1;refGene_gn=SYT7;refGene_tx=XM_023239910.1
D1	107585097	107585220
refGene_gn=SYT7;refGene_tx=XM_023239904.1;refGene_gn=SYT7;refGene_tx=XM_023239905.1;refGene_gn=SYT7;refGene_tx=XM_023239906.1
D1	107589185	107589557
refGene_gn=SYT7;refGene_tx=XM_023239904.1;refGene_gn=SYT7;refGene_tx=XM_023239905.1;refGene_gn=SYT7;refGene_tx=XM_023239906.1
D1	107592946	107593171
refGene_gn=SYT7;refGene_tx=XM_023239904.1;refGene_gn=SYT7;refGene_tx=XM_023239905.1;refGene_gn=SYT7;refGene_tx=XM_023239907.1;refGene_gn=SYT7;refGene_tx=XM_023239908.1
D1	107594073	107594205
refGene_gn=SYT7;refGene_tx=XM_023239904.1;refGene_gn=SYT7;refGene_tx=XM_023239906.1;refGene_gn=SYT7;refGene_tx=XM_023239907.1;refGene_gn=SYT7;refGene_tx=XM_023239909.1

D1	107598193	107598273
refGene_gn=SYT7;refGene_tx=XM_023239904.1;refGene_gn=SYT7;refGene_tx=XM_023239905.1;refGene_gn=SYT7;refGene_tx=XM_023239906.1;refGene_gn=SYT7;refGene_tx=XM_023239907.1;refGene_gn=SYT7;refGene_tx=XM_023239908.1;refGene_gn=SYT7;refGene_tx=XM_023239909.1;refGene_gn=SYT7;refGene_tx=XM_023239910.1
D1	107602623	107602727
refGene_gn=SYT7;refGene_tx=XM_023239904.1;refGene_gn=SYT7;refGene_tx=XM_023239905.1;refGene_gn=SYT7;refGene_tx=XM_023239906.1;refGene_gn=SYT7;refGene_tx=XM_023239907.1;refGene_gn=SYT7;refGene_tx=XM_023239908.1;refGene_gn=SYT7;refGene_tx=XM_023239909.1;refGene_gn=SYT7;refGene_tx=XM_023239910.1
D1	107627644	107627675
refGene_gn=SYT7;refGene_tx=XM_023239904.1;refGene_gn=SYT7;refGene_tx=XM_023239905.1;refGene_gn=SYT7;refGene_tx=XM_023239906.1;refGene_gn=SYT7;refGene_tx=XM_023239907.1;refGene_gn=SYT7;refGene_tx=XM_023239908.1;refGene_gn=SYT7;refGene_tx=XM_023239909.1;refGene_gn=SYT7;refGene_tx=XM_023239910.1
D1	107736166	107736261
refGene_gn=DAGLA;refGene_tx=XM_019812513.2;refGene_gn=DAGLA;refGene_tx=XM_023239911.1;refGene_gn=DAGLA;refGene_tx=XM_023239913.1
D1	107736656	107736868
refGene_gn=DAGLA;refGene_tx=XM_019812513.2;refGene_gn=DAGLA;refGene_tx=XM_023239911.1;refGene_gn=DAGLA;refGene_tx=XM_023239913.1
D1	107737860	107737888
refGene_gn=DAGLA;refGene_tx=XM_023239914.1
D1	107738219	107738321
refGene_gn=DAGLA;refGene_tx=XM_019812513.2;refGene_gn=DAGLA;refGene_tx=XM_023239911.1;refGene_gn=DAGLA;refGene_tx=XM_023239913.1;refGene_gn=DAGLA;refGene_tx=XM_023239914.1
D1	107738753	107738892
refGene_gn=DAGLA;refGene_tx=XM_019812513.2;refGene_gn=DAGLA;refGene_tx=XM_023239911.1;refGene_gn=DAGLA;refGene_tx=XM_023239913.1;refGene_gn=DAGLA;refGene_tx=XM_023239914.1
D1	107742137	107742225
refGene_gn=DAGLA;refGene_tx=XM_019812513.2;refGene_gn=DAGLA;refGene_tx=XM_023239911.1;refGene_gn=DAGLA;refGene_tx=XM_023239913.1;refGene_gn=DAGLA;refGene_tx=XM_023239914.1
D1	107744083	107744218
refGene_gn=DAGLA;refGene_tx=XM_019812513.2;refGene_gn=DAGLA;refGene_tx=XM_023239911.1;refGene_gn=DAGLA;refGene_tx=XM_023239913.1;refGene_gn=DAGLA;refGene_tx=XM_023239914.1
D1	107744441	107744519
refGene_gn=DAGLA;refGene_tx=XM_019812513.2;refGene_gn=DAGLA;refGene_tx=XM_023239911.1;refGene_gn=DAGLA;refGene_tx=XM_023239913.1;refGene_gn=DAGLA;refGene_tx=XM_023239914.1
D1	107746425	107746550
refGene_gn=DAGLA;refGene_tx=XM_019812513.2;refGene_gn=DAGLA;refGene_tx=XM_023239911.1;refGene_gn=DAGLA;refGene_tx=XM_023239913.1;refGene_gn=DAGLA;refGene_tx=XM_023239914.1

D1 107749306 107749460
refGene_gn=DAGLA;refGene_tx=XM_019812513.2;refGene_gn=DAGLA;refGene_tx=XM_023239911.1;refGene_gn=DAGLA;refGene_tx=XM_023239913.1;refGene_gn=DAGLA;refGene_tx=XM_023239914.1
D1 107749819 107749903
refGene_gn=DAGLA;refGene_tx=XM_019812513.2;refGene_gn=DAGLA;refGene_tx=XM_023239911.1;refGene_gn=DAGLA;refGene_tx=XM_023239913.1;refGene_gn=DAGLA;refGene_tx=XM_023239914.1
D1 107750016 107750094
refGene_gn=DAGLA;refGene_tx=XM_019812513.2;refGene_gn=DAGLA;refGene_tx=XM_023239911.1;refGene_gn=DAGLA;refGene_tx=XM_023239913.1;refGene_gn=DAGLA;refGene_tx=XM_023239914.1
D1 107750653 107750734
refGene_gn=DAGLA;refGene_tx=XM_019812513.2;refGene_gn=DAGLA;refGene_tx=XM_023239911.1;refGene_gn=DAGLA;refGene_tx=XM_023239913.1;refGene_gn=DAGLA;refGene_tx=XM_023239914.1
D1 107751574 107751717
refGene_gn=DAGLA;refGene_tx=XM_019812513.2;refGene_gn=DAGLA;refGene_tx=XM_023239911.1;refGene_gn=DAGLA;refGene_tx=XM_023239913.1;refGene_gn=DAGLA;refGene_tx=XM_023239914.1
D1 107752044 107752113
refGene_gn=DAGLA;refGene_tx=XM_019812513.2;refGene_gn=DAGLA;refGene_tx=XM_023239911.1;refGene_gn=DAGLA;refGene_tx=XM_023239913.1;refGene_gn=DAGLA;refGene_tx=XM_023239914.1
D1 107752452 107752525
refGene_gn=DAGLA;refGene_tx=XM_019812513.2;refGene_gn=DAGLA;refGene_tx=XM_023239911.1;refGene_gn=DAGLA;refGene_tx=XM_023239913.1;refGene_gn=DAGLA;refGene_tx=XM_023239914.1
D1 107753806 107754003
refGene_gn=DAGLA;refGene_tx=XM_019812513.2;refGene_gn=DAGLA;refGene_tx=XM_023239911.1;refGene_gn=DAGLA;refGene_tx=XM_023239913.1;refGene_gn=DAGLA;refGene_tx=XM_023239914.1
D1 107754709 107754839
refGene_gn=DAGLA;refGene_tx=XM_019812513.2;refGene_gn=DAGLA;refGene_tx=XM_023239911.1;refGene_gn=DAGLA;refGene_tx=XM_023239913.1;refGene_gn=DAGLA;refGene_tx=XM_023239914.1
D1 107755361 107755549
refGene_gn=DAGLA;refGene_tx=XM_019812513.2;refGene_gn=DAGLA;refGene_tx=XM_023239911.1;refGene_gn=DAGLA;refGene_tx=XM_023239913.1;refGene_gn=DAGLA;refGene_tx=XM_023239914.1
D1 107757496 107758460
refGene_gn=DAGLA;refGene_tx=XM_019812513.2;refGene_gn=DAGLA;refGene_tx=XM_023239911.1;refGene_gn=DAGLA;refGene_tx=XM_023239913.1;refGene_gn=DAGLA;refGene_tx=XM_023239914.1
D1 107765905 107765951
refGene_gn=MYRF;refGene_tx=XM_023239915.1;refGene_gn=MYRF;refGene_tx=XM_023239916.1;refGene_gn=MYRF;refGene_tx=XM_023239917.1;refGene_gn=MYRF;refGene_tx=XM_023239918.1;refGene_gn=MYRF;refGene_tx=XM_023239919.1;refGene_gn=MYRF;refGene_tx=XM_023239921.1;refGene_gn=MYRF;refGene_tx=XM_023239922.1;refGene_gn=MYRF;refGene_tx=XM_023239923.1;refGene_gn=MYR

F;refGene_tx=XM_023239924.1
D1 107768774 107768814
refGene_gn=MYRF;refGene_tx=XM_019812514.1
D1 107778231 107778319
refGene_gn=MYRF;refGene_tx=XM_019812514.1;refGene_gn=MYRF;refGene_tx=XM_023239915.1;refGene_gn=MYRF;refGene_tx=XM_023239916.1;refGene_gn=MYRF;refGene_tx=XM_023239917.1;refGene_gn=MYRF;refGene_tx=XM_023239918.1;refGene_gn=MYRF;refGene_tx=XM_023239919.1;refGene_gn=MYRF;refGene_tx=XM_023239921.1;refGene_gn=MYRF;refGene_tx=XM_023239922.1;refGene_gn=MYRF;refGene_tx=XM_023239923.1;refGene_gn=MYRF;refGene_tx=XM_023239924.1
D1 107778504 107778768
refGene_gn=MYRF;refGene_tx=XM_019812514.1;refGene_gn=MYRF;refGene_tx=XM_023239915.1;refGene_gn=MYRF;refGene_tx=XM_023239916.1;refGene_gn=MYRF;refGene_tx=XM_023239917.1;refGene_gn=MYRF;refGene_tx=XM_023239918.1;refGene_gn=MYRF;refGene_tx=XM_023239919.1;refGene_gn=MYRF;refGene_tx=XM_023239921.1;refGene_gn=MYRF;refGene_tx=XM_023239922.1;refGene_gn=MYRF;refGene_tx=XM_023239923.1;refGene_gn=MYRF;refGene_tx=XM_023239924.1
D1 107781135 107781197
refGene_gn=MYRF;refGene_tx=XM_019812514.1;refGene_gn=MYRF;refGene_tx=XM_023239915.1;refGene_gn=MYRF;refGene_tx=XM_023239916.1;refGene_gn=MYRF;refGene_tx=XM_023239917.1;refGene_gn=MYRF;refGene_tx=XM_023239918.1;refGene_gn=MYRF;refGene_tx=XM_023239919.1;refGene_gn=MYRF;refGene_tx=XM_023239921.1;refGene_gn=MYRF;refGene_tx=XM_023239922.1;refGene_gn=MYRF;refGene_tx=XM_023239923.1;refGene_gn=MYRF;refGene_tx=XM_023239924.1
D1 107781441 107781460
refGene_gn=MYRF;refGene_tx=XM_019812522.1
D1 107782136 107782416
refGene_gn=MYRF;refGene_tx=XM_019812514.1;refGene_gn=MYRF;refGene_tx=XM_019812522.1;refGene_gn=MYRF;refGene_tx=XM_019812523.2;refGene_gn=MYRF;refGene_tx=XM_023239915.1;refGene_gn=MYRF;refGene_tx=XM_023239916.1;refGene_gn=MYRF;refGene_tx=XM_023239917.1;refGene_gn=MYRF;refGene_tx=XM_023239918.1;refGene_gn=MYRF;refGene_tx=XM_023239919.1;refGene_gn=MYRF;refGene_tx=XM_023239921.1;refGene_gn=MYRF;refGene_tx=XM_023239922.1;refGene_gn=MYRF;refGene_tx=XM_023239923.1;refGene_gn=MYRF;refGene_tx=XM_023239924.1
D1 107783390 107783641
refGene_gn=MYRF;refGene_tx=XM_019812514.1;refGene_gn=MYRF;refGene_tx=XM_019812522.1;refGene_gn=MYRF;refGene_tx=XM_019812523.2;refGene_gn=MYRF;refGene_tx=XM_023239915.1;refGene_gn=MYRF;refGene_tx=XM_023239916.1;refGene_gn=MYRF;refGene_tx=XM_023239917.1;refGene_gn=MYRF;refGene_tx=XM_023239918.1;refGene_gn=MYRF;refGene_tx=XM_023239919.1;refGene_gn=MYRF;refGene_tx=XM_023239921.1;refGene_gn=MYRF;refGene_tx=XM_023239922.1;refGene_gn=MYRF;refGene_tx=XM_023239923.1;refGene_gn=MYRF;refGene_tx=XM_023239924.1
D1 107783733 107783857
refGene_gn=MYRF;refGene_tx=XM_019812514.1;refGene_gn=MYRF;refGene_tx=XM_019812522.1;refGene_gn=MYRF;refGene_tx=XM_019812523.2;refGene_gn=MYRF;refGene_tx=XM_023239915.1;refGene_gn=MYRF;refGene_tx=XM_023239916.1;refGene_gn=MYRF;refGene_tx=XM_023239917.1;refGene_gn=MYRF;refGene_tx=XM_023239918.1;refGene_gn=MYRF;refGene_tx=XM_023239919.1;refGene_gn=MYR

F;refGene_tx=XM_023239921.1;refGene_gn=MYRF;refGene_tx=XM_023239922.1;
refGene_gn=MYRF;refGene_tx=XM_023239923.1;refGene_gn=MYRF;refGene_tx=X
M_023239924.1
D1 107785583 107785779
refGene_gn=MYRF;refGene_tx=XM_019812514.1;refGene_gn=MYRF;refGene_tx=X
M_019812522.1;refGene_gn=MYRF;refGene_tx=XM_019812523.2;refGene_gn=MYR
F;refGene_tx=XM_023239915.1;refGene_gn=MYRF;refGene_tx=XM_023239916.1;
refGene_gn=MYRF;refGene_tx=XM_023239917.1;refGene_gn=MYRF;refGene_tx=X
M_023239918.1;refGene_gn=MYRF;refGene_tx=XM_023239919.1;refGene_gn=MYR
F;refGene_tx=XM_023239921.1;refGene_gn=MYRF;refGene_tx=XM_023239922.1;
refGene_gn=MYRF;refGene_tx=XM_023239923.1;refGene_gn=MYRF;refGene_tx=X
M_023239924.1
D1 107786959 107787036
refGene_gn=MYRF;refGene_tx=XM_019812514.1;refGene_gn=MYRF;refGene_tx=X
M_019812522.1;refGene_gn=MYRF;refGene_tx=XM_019812523.2;refGene_gn=MYR
F;refGene_tx=XM_023239915.1;refGene_gn=MYRF;refGene_tx=XM_023239916.1;
refGene_gn=MYRF;refGene_tx=XM_023239917.1;refGene_gn=MYRF;refGene_tx=X
M_023239918.1;refGene_gn=MYRF;refGene_tx=XM_023239919.1;refGene_gn=MYR
F;refGene_tx=XM_023239921.1;refGene_gn=MYRF;refGene_tx=XM_023239922.1;
refGene_gn=MYRF;refGene_tx=XM_023239923.1;refGene_gn=MYRF;refGene_tx=X
M_023239924.1
D1 107787204 107787315
refGene_gn=MYRF;refGene_tx=XM_019812514.1;refGene_gn=MYRF;refGene_tx=X
M_019812522.1;refGene_gn=MYRF;refGene_tx=XM_019812523.2;refGene_gn=MYR
F;refGene_tx=XM_023239915.1;refGene_gn=MYRF;refGene_tx=XM_023239916.1;
refGene_gn=MYRF;refGene_tx=XM_023239917.1;refGene_gn=MYRF;refGene_tx=X
M_023239918.1;refGene_gn=MYRF;refGene_tx=XM_023239919.1;refGene_gn=MYR
F;refGene_tx=XM_023239921.1;refGene_gn=MYRF;refGene_tx=XM_023239922.1;
refGene_gn=MYRF;refGene_tx=XM_023239923.1;refGene_gn=MYRF;refGene_tx=X
M_023239924.1
D1 107787722 107787813
refGene_gn=MYRF;refGene_tx=XM_019812514.1;refGene_gn=MYRF;refGene_tx=X
M_019812522.1;refGene_gn=MYRF;refGene_tx=XM_019812523.2;refGene_gn=MYR
F;refGene_tx=XM_023239915.1;refGene_gn=MYRF;refGene_tx=XM_023239916.1;
refGene_gn=MYRF;refGene_tx=XM_023239917.1;refGene_gn=MYRF;refGene_tx=X
M_023239918.1;refGene_gn=MYRF;refGene_tx=XM_023239919.1;refGene_gn=MYR
F;refGene_tx=XM_023239921.1;refGene_gn=MYRF;refGene_tx=XM_023239922.1;
refGene_gn=MYRF;refGene_tx=XM_023239923.1;refGene_gn=MYRF;refGene_tx=X
M_023239924.1
D1 107788196 107788397
refGene_gn=MYRF;refGene_tx=XM_019812514.1;refGene_gn=MYRF;refGene_tx=X
M_019812522.1;refGene_gn=MYRF;refGene_tx=XM_019812523.2;refGene_gn=MYR
F;refGene_tx=XM_023239915.1;refGene_gn=MYRF;refGene_tx=XM_023239916.1;
refGene_gn=MYRF;refGene_tx=XM_023239917.1;refGene_gn=MYRF;refGene_tx=X
M_023239918.1;refGene_gn=MYRF;refGene_tx=XM_023239919.1;refGene_gn=MYR
F;refGene_tx=XM_023239921.1;refGene_gn=MYRF;refGene_tx=XM_023239922.1;
refGene_gn=MYRF;refGene_tx=XM_023239923.1;refGene_gn=MYRF;refGene_tx=X
M_023239924.1
D1 107788722 107788834
refGene_gn=MYRF;refGene_tx=XM_019812514.1;refGene_gn=MYRF;refGene_tx=X

M_019812522.1;refGene_gn=MYRF;refGene_tx=XM_019812523.2;refGene_gn=MYRF;refGene_tx=XM_023239915.1;refGene_gn=MYRF;refGene_tx=XM_023239916.1;refGene_gn=MYRF;refGene_tx=XM_023239917.1;refGene_gn=MYRF;refGene_tx=XM_023239918.1;refGene_gn=MYRF;refGene_tx=XM_023239919.1;refGene_gn=MYRF;refGene_tx=XM_023239921.1;refGene_gn=MYRF;refGene_tx=XM_023239922.1;refGene_gn=MYRF;refGene_tx=XM_023239923.1;refGene_gn=MYRF;refGene_tx=XM_023239924.1
D1 107789355 107789465 refGene_gn=MYRF;refGene_tx=XM_019812514.1;refGene_gn=MYRF;refGene_tx=XM_019812522.1;refGene_gn=MYRF;refGene_tx=XM_019812523.2;refGene_gn=MYRF;refGene_tx=XM_023239915.1;refGene_gn=MYRF;refGene_tx=XM_023239916.1;refGene_gn=MYRF;refGene_tx=XM_023239917.1;refGene_gn=MYRF;refGene_tx=XM_023239918.1;refGene_gn=MYRF;refGene_tx=XM_023239919.1;refGene_gn=MYRF;refGene_tx=XM_023239921.1;refGene_gn=MYRF;refGene_tx=XM_023239922.1;refGene_gn=MYRF;refGene_tx=XM_023239923.1;refGene_gn=MYRF;refGene_tx=XM_023239924.1
D1 107790167 107790328 refGene_gn=MYRF;refGene_tx=XM_019812514.1;refGene_gn=MYRF;refGene_tx=XM_019812522.1;refGene_gn=MYRF;refGene_tx=XM_019812523.2;refGene_gn=MYRF;refGene_tx=XM_023239915.1;refGene_gn=MYRF;refGene_tx=XM_023239916.1;refGene_gn=MYRF;refGene_tx=XM_023239917.1;refGene_gn=MYRF;refGene_tx=XM_023239918.1;refGene_gn=MYRF;refGene_tx=XM_023239919.1;refGene_gn=MYRF;refGene_tx=XM_023239921.1;refGene_gn=MYRF;refGene_tx=XM_023239922.1;refGene_gn=MYRF;refGene_tx=XM_023239923.1;refGene_gn=MYRF;refGene_tx=XM_023239924.1
D1 107790400 107790473 refGene_gn=MYRF;refGene_tx=XM_019812514.1;refGene_gn=MYRF;refGene_tx=XM_019812522.1;refGene_gn=MYRF;refGene_tx=XM_019812523.2;refGene_gn=MYRF;refGene_tx=XM_023239915.1;refGene_gn=MYRF;refGene_tx=XM_023239916.1;refGene_gn=MYRF;refGene_tx=XM_023239917.1;refGene_gn=MYRF;refGene_tx=XM_023239918.1;refGene_gn=MYRF;refGene_tx=XM_023239919.1;refGene_gn=MYRF;refGene_tx=XM_023239921.1;refGene_gn=MYRF;refGene_tx=XM_023239922.1;refGene_gn=MYRF;refGene_tx=XM_023239923.1;refGene_gn=MYRF;refGene_tx=XM_023239924.1
D1 107790739 107790828 refGene_gn=MYRF;refGene_tx=XM_019812514.1;refGene_gn=MYRF;refGene_tx=XM_019812522.1;refGene_gn=MYRF;refGene_tx=XM_019812523.2;refGene_gn=MYRF;refGene_tx=XM_023239915.1;refGene_gn=MYRF;refGene_tx=XM_023239916.1;refGene_gn=MYRF;refGene_tx=XM_023239917.1;refGene_gn=MYRF;refGene_tx=XM_023239918.1;refGene_gn=MYRF;refGene_tx=XM_023239919.1;refGene_gn=MYRF;refGene_tx=XM_023239921.1;refGene_gn=MYRF;refGene_tx=XM_023239922.1;refGene_gn=MYRF;refGene_tx=XM_023239923.1;refGene_gn=MYRF;refGene_tx=XM_023239924.1
D1 107791153 107791222 refGene_gn=MYRF;refGene_tx=XM_019812514.1;refGene_gn=MYRF;refGene_tx=XM_019812522.1;refGene_gn=MYRF;refGene_tx=XM_019812523.2;refGene_gn=MYRF;refGene_tx=XM_023239915.1;refGene_gn=MYRF;refGene_tx=XM_023239916.1;refGene_gn=MYRF;refGene_tx=XM_023239917.1;refGene_gn=MYRF;refGene_tx=XM_023239918.1;refGene_gn=MYRF;refGene_tx=XM_023239919.1;refGene_gn=MYRF;refGene_tx=XM_023239921.1;refGene_gn=MYRF;refGene_tx=XM_023239922.1;

refGene_gn=MYRF;refGene_tx=XM_023239923.1;refGene_gn=MYRF;refGene_tx=XM_023239924.1
D1	107791669	107791750
refGene_gn=MYRF;refGene_tx=XM_019812514.1;refGene_gn=MYRF;refGene_tx=XM_019812522.1;refGene_gn=MYRF;refGene_tx=XM_019812523.2;refGene_gn=MYRF;refGene_tx=XM_023239915.1;refGene_gn=MYRF;refGene_tx=XM_023239916.1;refGene_gn=MYRF;refGene_tx=XM_023239917.1;refGene_gn=MYRF;refGene_tx=XM_023239918.1;refGene_gn=MYRF;refGene_tx=XM_023239923.1;refGene_gn=MYRF;refGene_tx=XM_023239924.1
D1	107791918	107792007
refGene_gn=MYRF;refGene_tx=XM_019812514.1;refGene_gn=MYRF;refGene_tx=XM_019812522.1;refGene_gn=MYRF;refGene_tx=XM_019812523.2;refGene_gn=MYRF;refGene_tx=XM_023239915.1;refGene_gn=MYRF;refGene_tx=XM_023239916.1;refGene_gn=MYRF;refGene_tx=XM_023239917.1;refGene_gn=MYRF;refGene_tx=XM_023239918.1;refGene_gn=MYRF;refGene_tx=XM_023239919.1;refGene_gn=MYRF;refGene_tx=XM_023239921.1;refGene_gn=MYRF;refGene_tx=XM_023239922.1;refGene_gn=MYRF;refGene_tx=XM_023239923.1;refGene_gn=MYRF;refGene_tx=XM_023239924.1
D1	107792127	107792319
refGene_gn=MYRF;refGene_tx=XM_019812514.1;refGene_gn=MYRF;refGene_tx=XM_019812522.1;refGene_gn=MYRF;refGene_tx=XM_019812523.2;refGene_gn=MYRF;refGene_tx=XM_023239915.1;refGene_gn=MYRF;refGene_tx=XM_023239916.1;refGene_gn=MYRF;refGene_tx=XM_023239917.1;refGene_gn=MYRF;refGene_tx=XM_023239918.1;refGene_gn=MYRF;refGene_tx=XM_023239919.1;refGene_gn=MYRF;refGene_tx=XM_023239921.1;refGene_gn=MYRF;refGene_tx=XM_023239922.1;refGene_gn=MYRF;refGene_tx=XM_023239923.1;refGene_gn=MYRF;refGene_tx=XM_023239924.1
D1	107792571	107792823
refGene_gn=MYRF;refGene_tx=XM_019812514.1;refGene_gn=MYRF;refGene_tx=XM_019812522.1;refGene_gn=MYRF;refGene_tx=XM_019812523.2;refGene_gn=MYRF;refGene_tx=XM_023239915.1;refGene_gn=MYRF;refGene_tx=XM_023239916.1;refGene_gn=MYRF;refGene_tx=XM_023239917.1;refGene_gn=MYRF;refGene_tx=XM_023239918.1;refGene_gn=MYRF;refGene_tx=XM_023239919.1;refGene_gn=MYRF;refGene_tx=XM_023239921.1;refGene_gn=MYRF;refGene_tx=XM_023239922.1;refGene_gn=MYRF;refGene_tx=XM_023239924.1
D1	107794353	107794456
refGene_gn=MYRF;refGene_tx=XM_019812514.1;refGene_gn=MYRF;refGene_tx=XM_019812522.1;refGene_gn=MYRF;refGene_tx=XM_019812523.2;refGene_gn=MYRF;refGene_tx=XM_023239915.1;refGene_gn=MYRF;refGene_tx=XM_023239916.1;refGene_gn=MYRF;refGene_tx=XM_023239917.1;refGene_gn=MYRF;refGene_tx=XM_023239918.1;refGene_gn=MYRF;refGene_tx=XM_023239919.1;refGene_gn=MYRF;refGene_tx=XM_023239921.1;refGene_gn=MYRF;refGene_tx=XM_023239922.1;refGene_gn=MYRF;refGene_tx=XM_023239923.1;refGene_gn=MYRF;refGene_tx=XM_023239924.1
D1	107794684	107794759
refGene_gn=MYRF;refGene_tx=XM_019812514.1;refGene_gn=MYRF;refGene_tx=XM_019812522.1;refGene_gn=MYRF;refGene_tx=XM_019812523.2;refGene_gn=MYRF;refGene_tx=XM_023239915.1;refGene_gn=MYRF;refGene_tx=XM_023239916.1;refGene_gn=MYRF;refGene_tx=XM_023239917.1;refGene_gn=MYRF;refGene_tx=XM_023239918.1;refGene_gn=MYRF;refGene_tx=XM_023239919.1;refGene_gn=MYR

F;refGene_tx=XM_023239921.1;refGene_gn=MYRF;refGene_tx=XM_023239922.1;
refGene_gn=MYRF;refGene_tx=XM_023239923.1;refGene_gn=MYRF;refGene_tx=X
M_023239924.1
D1 107795108 107795214
refGene_gn=MYRF;refGene_tx=XM_019812514.1;refGene_gn=MYRF;refGene_tx=X
M_019812522.1;refGene_gn=MYRF;refGene_tx=XM_019812523.2;refGene_gn=MYR
F;refGene_tx=XM_023239915.1;refGene_gn=MYRF;refGene_tx=XM_023239916.1;
refGene_gn=MYRF;refGene_tx=XM_023239917.1;refGene_gn=MYRF;refGene_tx=X
M_023239918.1;refGene_gn=MYRF;refGene_tx=XM_023239919.1;refGene_gn=MYR
F;refGene_tx=XM_023239921.1;refGene_gn=MYRF;refGene_tx=XM_023239922.1;
refGene_gn=MYRF;refGene_tx=XM_023239923.1;refGene_gn=MYRF;refGene_tx=X
M_023239924.1
D1 107796720 107796795
refGene_gn=MYRF;refGene_tx=XM_019812514.1;refGene_gn=MYRF;refGene_tx=X
M_019812522.1;refGene_gn=MYRF;refGene_tx=XM_019812523.2;refGene_gn=MYR
F;refGene_tx=XM_023239915.1;refGene_gn=MYRF;refGene_tx=XM_023239916.1;
refGene_gn=MYRF;refGene_tx=XM_023239917.1;refGene_gn=MYRF;refGene_tx=X
M_023239918.1;refGene_gn=MYRF;refGene_tx=XM_023239919.1;refGene_gn=MYR
F;refGene_tx=XM_023239921.1;refGene_gn=MYRF;refGene_tx=XM_023239922.1;
refGene_gn=MYRF;refGene_tx=XM_023239923.1;refGene_gn=MYRF;refGene_tx=X
M_023239924.1
D1 107796996 107797077
refGene_gn=MYRF;refGene_tx=XM_019812514.1;refGene_gn=MYRF;refGene_tx=X
M_019812522.1;refGene_gn=MYRF;refGene_tx=XM_019812523.2;refGene_gn=MYR
F;refGene_tx=XM_023239915.1;refGene_gn=MYRF;refGene_tx=XM_023239916.1;
refGene_gn=MYRF;refGene_tx=XM_023239917.1;refGene_gn=MYRF;refGene_tx=X
M_023239918.1;refGene_gn=MYRF;refGene_tx=XM_023239919.1;refGene_gn=MYR
F;refGene_tx=XM_023239921.1;refGene_gn=MYRF;refGene_tx=XM_023239922.1;
refGene_gn=MYRF;refGene_tx=XM_023239923.1;refGene_gn=MYRF;refGene_tx=X
M_023239924.1
D1 107800892 107801019
refGene_gn=TMEM258;refGene_tx=XM_006937393.3
D1 107801604 107801714
refGene_gn=TMEM258;refGene_tx=XM_006937393.3
D1 107803236 107803677
refGene_gn=TMEM258;refGene_tx=XM_006937393.3
D1 107804237 107804291
refGene_gn=TMEM258;refGene_tx=XM_006937393.3
D1 107806729 107807872
refGene_gn=FEN1;refGene_tx=XM_003993433.5;refGene_gn=FEN1;refGene_tx=X
M_019812525.2
D1 107813441 107813493
refGene_gn=FADS1;refGene_tx=XM_003993434.5;refGene_gn=FADS1;refGene_tx
=XM_006937394.4
D1 107813849 107813975
refGene_gn=FADS1;refGene_tx=XM_003993434.5;refGene_gn=FADS1;refGene_tx
=XM_006937394.4
D1 107814085 107814165
refGene_gn=FADS1;refGene_tx=XM_003993434.5;refGene_gn=FADS1;refGene_tx
=XM_006937394.4

D1 107814445 107814542
refGene_gn=FADS1;refGene_tx=XM_003993434.5;refGene_gn=FADS1;refGene_tx=XM_006937394.4
D1 107814749 107814847
refGene_gn=FADS1;refGene_tx=XM_003993434.5;refGene_gn=FADS1;refGene_tx=XM_006937394.4
D1 107815765 107815842
refGene_gn=FADS1;refGene_tx=XM_003993434.5;refGene_gn=FADS1;refGene_tx=XM_006937394.4
D1 107817262 107817323
refGene_gn=FADS1;refGene_tx=XM_003993434.5;refGene_gn=FADS1;refGene_tx=XM_006937394.4
D1 107821444 107821573
refGene_gn=FADS1;refGene_tx=XM_003993434.5;refGene_gn=FADS1;refGene_tx=XM_006937394.4
D1 107821680 107821782
refGene_gn=FADS1;refGene_tx=XM_003993434.5;refGene_gn=FADS1;refGene_tx=XM_006937394.4
D1 107822493 107822691
refGene_gn=FADS1;refGene_tx=XM_003993434.5;refGene_gn=FADS1;refGene_tx=XM_006937394.4
D1 107823280 107823391
refGene_gn=FADS1;refGene_tx=XM_003993434.5;refGene_gn=FADS1;refGene_tx=XM_006937394.4
D1 107825396 107825588
refGene_gn=FADS1;refGene_tx=XM_003993434.5
D1 107894965 107895172
refGene_gn=FADS2;refGene_tx=XM_023239925.1;refGene_gn=FADS2;refGene_tx=XM_023239927.1
D1 107903926 107904037
refGene_gn=FADS2;refGene_tx=XM_023239925.1;refGene_gn=FADS2;refGene_tx=XM_023239927.1
D1 107905742 107905940
refGene_gn=FADS2;refGene_tx=XM_023239925.1;refGene_gn=FADS2;refGene_tx=XM_023239927.1
D1 107906065 107906167
refGene_gn=FADS2;refGene_tx=XM_023239925.1;refGene_gn=FADS2;refGene_tx=XM_023239927.1
D1 107911320 107911446
refGene_gn=FADS2;refGene_tx=XM_023239925.1;refGene_gn=FADS2;refGene_tx=XM_023239927.1
D1 107919418 107919479
refGene_gn=FADS2;refGene_tx=XM_023239925.1
D1 107919858 107919935
refGene_gn=FADS2;refGene_tx=XM_023239925.1
D1 107925441 107925539
refGene_gn=FADS2;refGene_tx=XM_023239925.1;refGene_gn=FADS2;refGene_tx=XM_023239927.1
D1 107925791 107925888
refGene_gn=FADS2;refGene_tx=XM_023239925.1;refGene_gn=FADS2;refGene_tx

=XM_023239927.1
D1	107926138	107926218
refGene_gn=FADS2;refGene_tx=XM_023239925.1;refGene_gn=FADS2;refGene_tx=XM_023239927.1
D1	107926603	107926729
refGene_gn=FADS2;refGene_tx=XM_023239925.1;refGene_gn=FADS2;refGene_tx=XM_023239927.1
D1	107926982	107927034
refGene_gn=FADS2;refGene_tx=XM_023239925.1;refGene_gn=FADS2;refGene_tx=XM_023239927.1
D1	107933686	107933738
refGene_gn=FADS3;refGene_tx=XM_003993435.5
D1	107934908	107935034
refGene_gn=FADS3;refGene_tx=XM_003993435.5
D1	107935252	107935332
refGene_gn=FADS3;refGene_tx=XM_003993435.5
D1	107935427	107935524
refGene_gn=FADS3;refGene_tx=XM_003993435.5
D1	107935924	107936022
refGene_gn=FADS3;refGene_tx=XM_003993435.5
D1	107936469	107936546
refGene_gn=FADS3;refGene_tx=XM_003993435.5
D1	107937090	107937151
refGene_gn=FADS3;refGene_tx=XM_003993435.5
D1	107937447	107937570
refGene_gn=FADS3;refGene_tx=XM_003993435.5
D1	107937688	107937790
refGene_gn=FADS3;refGene_tx=XM_003993435.5
D1	107938155	107938353
refGene_gn=FADS3;refGene_tx=XM_003993435.5
D1	107938918	107939029
refGene_gn=FADS3;refGene_tx=XM_003993435.5
D1	107946551	107946758
refGene_gn=FADS3;refGene_tx=XM_003993435.5
D1	107953269	107953352
refGene_gn=RAB3IL1;refGene_tx=XM_006937395.3;refGene_gn=RAB3IL1;refGene_tx=XM_006937396.4;refGene_gn=RAB3IL1;refGene_tx=XM_019812535.2;refGene_gn=RAB3IL1;refGene_tx=XM_023239929.1;refGene_gn=RAB3IL1;refGene_tx=XM_023239930.1;refGene_gn=RAB3IL1;refGene_tx=XM_023239931.1
D1	107954177	107954244
refGene_gn=RAB3IL1;refGene_tx=XM_006937395.3;refGene_gn=RAB3IL1;refGene_tx=XM_006937396.4;refGene_gn=RAB3IL1;refGene_tx=XM_019812535.2;refGene_gn=RAB3IL1;refGene_tx=XM_023239929.1;refGene_gn=RAB3IL1;refGene_tx=XM_023239930.1;refGene_gn=RAB3IL1;refGene_tx=XM_023239931.1
D1	107956813	107956913
refGene_gn=RAB3IL1;refGene_tx=XM_006937395.3;refGene_gn=RAB3IL1;refGene_tx=XM_006937396.4;refGene_gn=RAB3IL1;refGene_tx=XM_019812535.2;refGene_gn=RAB3IL1;refGene_tx=XM_023239929.1;refGene_gn=RAB3IL1;refGene_tx=XM_023239930.1;refGene_gn=RAB3IL1;refGene_tx=XM_023239931.1
D1	107958420	107958533

refGene_gn=RAB3IL1;refGene_tx=XM_006937395.3;refGene_gn=RAB3IL1;refGene_tx=XM_006937396.4;refGene_gn=RAB3IL1;refGene_tx=XM_019812535.2;refGene_gn=RAB3IL1;refGene_tx=XM_023239929.1;refGene_gn=RAB3IL1;refGene_tx=XM_023239930.1;refGene_gn=RAB3IL1;refGene_tx=XM_023239931.1
D1 107958627 107958756
refGene_gn=RAB3IL1;refGene_tx=XM_006937395.3;refGene_gn=RAB3IL1;refGene_tx=XM_006937396.4;refGene_gn=RAB3IL1;refGene_tx=XM_019812535.2;refGene_gn=RAB3IL1;refGene_tx=XM_023239929.1;refGene_gn=RAB3IL1;refGene_tx=XM_023239930.1;refGene_gn=RAB3IL1;refGene_tx=XM_023239931.1
D1 107959271 107959424
refGene_gn=RAB3IL1;refGene_tx=XM_006937395.3;refGene_gn=RAB3IL1;refGene_tx=XM_006937396.4;refGene_gn=RAB3IL1;refGene_tx=XM_019812535.2;refGene_gn=RAB3IL1;refGene_tx=XM_023239929.1
D1 107960356 107960575
refGene_gn=RAB3IL1;refGene_tx=XM_006937395.3;refGene_gn=RAB3IL1;refGene_tx=XM_006937396.4;refGene_gn=RAB3IL1;refGene_tx=XM_019812535.2;refGene_gn=RAB3IL1;refGene_tx=XM_023239929.1;refGene_gn=RAB3IL1;refGene_tx=XM_023239930.1;refGene_gn=RAB3IL1;refGene_tx=XM_023239931.1
D1 107961260 107961338
refGene_gn=RAB3IL1;refGene_tx=XM_006937395.3;refGene_gn=RAB3IL1;refGene_tx=XM_006937396.4;refGene_gn=RAB3IL1;refGene_tx=XM_019812535.2;refGene_gn=RAB3IL1;refGene_tx=XM_023239929.1;refGene_gn=RAB3IL1;refGene_tx=XM_023239930.1;refGene_gn=RAB3IL1;refGene_tx=XM_023239931.1
D1 107961447 107961543
refGene_gn=RAB3IL1;refGene_tx=XM_006937395.3;refGene_gn=RAB3IL1;refGene_tx=XM_006937396.4;refGene_gn=RAB3IL1;refGene_tx=XM_019812535.2;refGene_gn=RAB3IL1;refGene_tx=XM_023239929.1;refGene_gn=RAB3IL1;refGene_tx=XM_023239930.1;refGene_gn=RAB3IL1;refGene_tx=XM_023239931.1
D1 107961998 107962254
refGene_gn=RAB3IL1;refGene_tx=XM_006937395.3;refGene_gn=RAB3IL1;refGene_tx=XM_006937396.4;refGene_gn=RAB3IL1;refGene_tx=XM_019812535.2;refGene_gn=RAB3IL1;refGene_tx=XM_023239929.1;refGene_gn=RAB3IL1;refGene_tx=XM_023239930.1;refGene_gn=RAB3IL1;refGene_tx=XM_023239931.1
D1 107970065 107970076
refGene_gn=RAB3IL1;refGene_tx=XM_023239929.1;refGene_gn=RAB3IL1;refGene_tx=XM_023239931.1
D1 107972733 107972885
refGene_gn=RAB3IL1;refGene_tx=XM_006937395.3;refGene_gn=RAB3IL1;refGene_tx=XM_023239930.1
D1 107996496 107996618
refGene_gn=RAB3IL1;refGene_tx=XM_006937396.4
D1 108001386 108001538
refGene_gn=BEST1;refGene_tx=XM_003993438.5;refGene_gn=BEST1;refGene_tx=XM_019812527.2;refGene_gn=BEST1;refGene_tx=XM_019812528.2;refGene_gn=BEST1;refGene_tx=XM_019812529.2
D1 108002936 108003031
refGene_gn=BEST1;refGene_tx=XM_003993438.5;refGene_gn=BEST1;refGene_tx=XM_019812527.2;refGene_gn=BEST1;refGene_tx=XM_019812528.2;refGene_gn=BEST1;refGene_tx=XM_019812529.2
D1 108003517 108003751

refGene_gn=BEST1;refGene_tx=XM_003993438.5;refGene_gn=BEST1;refGene_tx=XM_019812527.2;refGene_gn=BEST1;refGene_tx=XM_019812528.2;refGene_gn=BEST1;refGene_tx=XM_019812529.2
D1 108004304 108004459
refGene_gn=BEST1;refGene_tx=XM_003993438.5;refGene_gn=BEST1;refGene_tx=XM_019812527.2;refGene_gn=BEST1;refGene_tx=XM_019812528.2;refGene_gn=BEST1;refGene_tx=XM_019812529.2
D1 108004817 108004895
refGene_gn=BEST1;refGene_tx=XM_003993438.5;refGene_gn=BEST1;refGene_tx=XM_019812527.2;refGene_gn=BEST1;refGene_tx=XM_019812528.2;refGene_gn=BEST1;refGene_tx=XM_019812529.2
D1 108005445 108005598
refGene_gn=BEST1;refGene_tx=XM_003993438.5;refGene_gn=BEST1;refGene_tx=XM_019812527.2;refGene_gn=BEST1;refGene_tx=XM_019812528.2;refGene_gn=BEST1;refGene_tx=XM_019812529.2
D1 108007183 108007264
refGene_gn=BEST1;refGene_tx=XM_003993438.5;refGene_gn=BEST1;refGene_tx=XM_019812527.2;refGene_gn=BEST1;refGene_tx=XM_019812528.2
D1 108007561 108007713
refGene_gn=BEST1;refGene_tx=XM_003993438.5;refGene_gn=BEST1;refGene_tx=XM_019812527.2;refGene_gn=BEST1;refGene_tx=XM_019812528.2;refGene_gn=BEST1;refGene_tx=XM_019812529.2
D1 108009945 108010590
refGene_gn=BEST1;refGene_tx=XM_003993438.5;refGene_gn=BEST1;refGene_tx=XM_019812527.2;refGene_gn=BEST1;refGene_tx=XM_019812528.2;refGene_gn=BEST1;refGene_tx=XM_019812529.2
D1 108011578 108011597
refGene_gn=BEST1;refGene_tx=XM_003993438.5;refGene_gn=BEST1;refGene_tx=XM_019812527.2;refGene_gn=BEST1;refGene_tx=XM_019812528.2;refGene_gn=BEST1;refGene_tx=XM_019812529.2
D1 108012218 108012383
refGene_gn=FTH1;refGene_tx=NM_001048151.1
D1 108012477 108012603
refGene_gn=FTH1;refGene_tx=NM_001048151.1
D1 108012907 108013054
refGene_gn=FTH1;refGene_tx=NM_001048151.1
D1 108014537 108014651
refGene_gn=FTH1;refGene_tx=NM_001048151.1
D1 108088118 108088258
refGene_gn=INCENP;refGene_tx=XM_023239932.1;refGene_gn=INCENP;refGene_tx=XM_023239934.1;refGene_gn=INCENP;refGene_tx=XM_023239935.1;refGene_gn=INCENP;refGene_tx=XM_023239936.1;refGene_gn=INCENP;refGene_tx=XM_023239937.1;refGene_gn=INCENP;refGene_tx=XM_023239938.1;refGene_gn=INCENP;refGene_tx=XM_023239939.1;refGene_gn=INCENP;refGene_tx=XM_023239940.1;refGene_gn=INCENP;refGene_tx=XM_023239941.1
D1 108088652 108088766
refGene_gn=INCENP;refGene_tx=XM_023239932.1;refGene_gn=INCENP;refGene_tx=XM_023239934.1;refGene_gn=INCENP;refGene_tx=XM_023239935.1;refGene_gn=INCENP;refGene_tx=XM_023239936.1;refGene_gn=INCENP;refGene_tx=XM_023239937.1;refGene_gn=INCENP;refGene_tx=XM_023239938.1;refGene_gn=INCEN

P;refGene_tx=XM_023239939.1;refGene_gn=INCENP;refGene_tx=XM_023239940.
1;refGene_gn=INCENP;refGene_tx=XM_023239941.1
D1	108089624	108090454
refGene_gn=INCENP;refGene_tx=XM_023239932.1;refGene_gn=INCENP;refGene_
tx=XM_023239934.1;refGene_gn=INCENP;refGene_tx=XM_023239935.1;refGene_
gn=INCENP;refGene_tx=XM_023239936.1;refGene_gn=INCENP;refGene_tx=XM_02
3239937.1;refGene_gn=INCENP;refGene_tx=XM_023239938.1;refGene_gn=INCEN
P;refGene_tx=XM_023239939.1;refGene_gn=INCENP;refGene_tx=XM_023239940.
1;refGene_gn=INCENP;refGene_tx=XM_023239941.1
D1	108096643	108096695
refGene_gn=INCENP;refGene_tx=XM_023239932.1;refGene_gn=INCENP;refGene_
tx=XM_023239934.1;refGene_gn=INCENP;refGene_tx=XM_023239935.1;refGene_
gn=INCENP;refGene_tx=XM_023239936.1;refGene_gn=INCENP;refGene_tx=XM_02
3239937.1;refGene_gn=INCENP;refGene_tx=XM_023239938.1;refGene_gn=INCEN
P;refGene_tx=XM_023239939.1;refGene_gn=INCENP;refGene_tx=XM_023239940.
1;refGene_gn=INCENP;refGene_tx=XM_023239941.1
D1	108097506	108097564
refGene_gn=INCENP;refGene_tx=XM_023239932.1;refGene_gn=INCENP;refGene_
tx=XM_023239934.1;refGene_gn=INCENP;refGene_tx=XM_023239935.1;refGene_
gn=INCENP;refGene_tx=XM_023239936.1;refGene_gn=INCENP;refGene_tx=XM_02
3239937.1;refGene_gn=INCENP;refGene_tx=XM_023239938.1;refGene_gn=INCEN
P;refGene_tx=XM_023239939.1;refGene_gn=INCENP;refGene_tx=XM_023239940.
1;refGene_gn=INCENP;refGene_tx=XM_023239941.1
D1	108097667	108097785
refGene_gn=INCENP;refGene_tx=XM_023239932.1;refGene_gn=INCENP;refGene_
tx=XM_023239934.1;refGene_gn=INCENP;refGene_tx=XM_023239935.1;refGene_
gn=INCENP;refGene_tx=XM_023239936.1;refGene_gn=INCENP;refGene_tx=XM_02
3239937.1;refGene_gn=INCENP;refGene_tx=XM_023239938.1;refGene_gn=INCEN
P;refGene_tx=XM_023239939.1;refGene_gn=INCENP;refGene_tx=XM_023239940.
1;refGene_gn=INCENP;refGene_tx=XM_023239941.1
D1	108098915	108098967
refGene_gn=INCENP;refGene_tx=XM_023239932.1;refGene_gn=INCENP;refGene_
tx=XM_023239934.1;refGene_gn=INCENP;refGene_tx=XM_023239935.1;refGene_
gn=INCENP;refGene_tx=XM_023239936.1;refGene_gn=INCENP;refGene_tx=XM_02
3239937.1;refGene_gn=INCENP;refGene_tx=XM_023239938.1;refGene_gn=INCEN
P;refGene_tx=XM_023239939.1;refGene_gn=INCENP;refGene_tx=XM_023239940.
1;refGene_gn=INCENP;refGene_tx=XM_023239941.1
D1	108099357	108099475
refGene_gn=INCENP;refGene_tx=XM_023239932.1;refGene_gn=INCENP;refGene_
tx=XM_023239934.1;refGene_gn=INCENP;refGene_tx=XM_023239935.1;refGene_
gn=INCENP;refGene_tx=XM_023239936.1;refGene_gn=INCENP;refGene_tx=XM_02
3239937.1;refGene_gn=INCENP;refGene_tx=XM_023239938.1;refGene_gn=INCEN
P;refGene_tx=XM_023239939.1;refGene_gn=INCENP;refGene_tx=XM_023239940.
1;refGene_gn=INCENP;refGene_tx=XM_023239941.1
D1	108099562	108099697
refGene_gn=INCENP;refGene_tx=XM_023239932.1;refGene_gn=INCENP;refGene_
tx=XM_023239934.1;refGene_gn=INCENP;refGene_tx=XM_023239935.1;refGene_
gn=INCENP;refGene_tx=XM_023239936.1;refGene_gn=INCENP;refGene_tx=XM_02
3239937.1;refGene_gn=INCENP;refGene_tx=XM_023239938.1;refGene_gn=INCEN
P;refGene_tx=XM_023239939.1;refGene_gn=INCENP;refGene_tx=XM_023239940.

1;refGene_gn=INCENP;refGene_tx=XM_023239941.1
D1	108100246	108100258
refGene_gn=INCENP;refGene_tx=XM_023239932.1;refGene_gn=INCENP;refGene_tx=XM_023239934.1;refGene_gn=INCENP;refGene_tx=XM_023239935.1;refGene_gn=INCENP;refGene_tx=XM_023239937.1;refGene_gn=INCENP;refGene_tx=XM_023239938.1;refGene_gn=INCENP;refGene_tx=XM_023239940.1
D1	108102314	108102452
refGene_gn=INCENP;refGene_tx=XM_023239932.1;refGene_gn=INCENP;refGene_tx=XM_023239934.1;refGene_gn=INCENP;refGene_tx=XM_023239935.1;refGene_gn=INCENP;refGene_tx=XM_023239936.1;refGene_gn=INCENP;refGene_tx=XM_023239937.1
D1	108104614	108104724
refGene_gn=INCENP;refGene_tx=XM_023239932.1;refGene_gn=INCENP;refGene_tx=XM_023239934.1;refGene_gn=INCENP;refGene_tx=XM_023239935.1;refGene_gn=INCENP;refGene_tx=XM_023239936.1;refGene_gn=INCENP;refGene_tx=XM_023239937.1;refGene_gn=INCENP;refGene_tx=XM_023239938.1;refGene_gn=INCENP;refGene_tx=XM_023239939.1;refGene_gn=INCENP;refGene_tx=XM_023239940.1;refGene_gn=INCENP;refGene_tx=XM_023239941.1
D1	108104804	108104925
refGene_gn=INCENP;refGene_tx=XM_023239932.1;refGene_gn=INCENP;refGene_tx=XM_023239934.1;refGene_gn=INCENP;refGene_tx=XM_023239935.1;refGene_gn=INCENP;refGene_tx=XM_023239936.1;refGene_gn=INCENP;refGene_tx=XM_023239937.1;refGene_gn=INCENP;refGene_tx=XM_023239938.1;refGene_gn=INCENP;refGene_tx=XM_023239939.1;refGene_gn=INCENP;refGene_tx=XM_023239940.1;refGene_gn=INCENP;refGene_tx=XM_023239941.1
D1	108105173	108105296
refGene_gn=INCENP;refGene_tx=XM_023239932.1;refGene_gn=INCENP;refGene_tx=XM_023239934.1;refGene_gn=INCENP;refGene_tx=XM_023239935.1;refGene_gn=INCENP;refGene_tx=XM_023239936.1;refGene_gn=INCENP;refGene_tx=XM_023239937.1;refGene_gn=INCENP;refGene_tx=XM_023239938.1;refGene_gn=INCENP;refGene_tx=XM_023239939.1;refGene_gn=INCENP;refGene_tx=XM_023239940.1;refGene_gn=INCENP;refGene_tx=XM_023239941.1
D1	108106377	108106556
refGene_gn=INCENP;refGene_tx=XM_023239932.1;refGene_gn=INCENP;refGene_tx=XM_023239934.1;refGene_gn=INCENP;refGene_tx=XM_023239935.1;refGene_gn=INCENP;refGene_tx=XM_023239936.1;refGene_gn=INCENP;refGene_tx=XM_023239937.1;refGene_gn=INCENP;refGene_tx=XM_023239938.1;refGene_gn=INCENP;refGene_tx=XM_023239939.1;refGene_gn=INCENP;refGene_tx=XM_023239940.1;refGene_gn=INCENP;refGene_tx=XM_023239941.1
D1	108106766	108106830
refGene_gn=INCENP;refGene_tx=XM_023239932.1;refGene_gn=INCENP;refGene_tx=XM_023239934.1;refGene_gn=INCENP;refGene_tx=XM_023239935.1;refGene_gn=INCENP;refGene_tx=XM_023239936.1;refGene_gn=INCENP;refGene_tx=XM_023239937.1;refGene_gn=INCENP;refGene_tx=XM_023239938.1;refGene_gn=INCENP;refGene_tx=XM_023239939.1;refGene_gn=INCENP;refGene_tx=XM_023239940.1;refGene_gn=INCENP;refGene_tx=XM_023239941.1
D1	108107013	108107121
refGene_gn=INCENP;refGene_tx=XM_023239932.1;refGene_gn=INCENP;refGene_tx=XM_023239934.1;refGene_gn=INCENP;refGene_tx=XM_023239935.1;refGene_gn=INCENP;refGene_tx=XM_023239936.1;refGene_gn=INCENP;refGene_tx=XM_02

3239937.1;refGene_gn=INCENP;refGene_tx=XM_023239938.1;refGene_gn=INCEN
P;refGene_tx=XM_023239939.1;refGene_gn=INCENP;refGene_tx=XM_023239940.
1;refGene_gn=INCENP;refGene_tx=XM_023239941.1
D1	108108568	108108719
refGene_gn=INCENP;refGene_tx=XM_023239932.1;refGene_gn=INCENP;refGene_
tx=XM_023239934.1;refGene_gn=INCENP;refGene_tx=XM_023239935.1;refGene_
gn=INCENP;refGene_tx=XM_023239936.1;refGene_gn=INCENP;refGene_tx=XM_02
3239937.1;refGene_gn=INCENP;refGene_tx=XM_023239938.1;refGene_gn=INCEN
P;refGene_tx=XM_023239939.1;refGene_gn=INCENP;refGene_tx=XM_023239940.
1;refGene_gn=INCENP;refGene_tx=XM_023239941.1
D1	108110432	108110647
refGene_gn=INCENP;refGene_tx=XM_023239932.1;refGene_gn=INCENP;refGene_
tx=XM_023239934.1;refGene_gn=INCENP;refGene_tx=XM_023239935.1;refGene_
gn=INCENP;refGene_tx=XM_023239936.1;refGene_gn=INCENP;refGene_tx=XM_02
3239937.1;refGene_gn=INCENP;refGene_tx=XM_023239938.1;refGene_gn=INCEN
P;refGene_tx=XM_023239939.1;refGene_gn=INCENP;refGene_tx=XM_023239940.
1;refGene_gn=INCENP;refGene_tx=XM_023239941.1
D1	108144179	108144234
refGene_gn=LOC101085573;refGene_tx=XM_003993504.3
D1	108145771	108145959
refGene_gn=LOC101085573;refGene_tx=XM_003993504.3
D1	108147136	108147166
refGene_gn=LOC101085573;refGene_tx=XM_003993504.3
D1	108161946	108162001
refGene_gn=LOC101085830;refGene_tx=XM_003993505.4
D1	108162385	108162645
refGene_gn=LOC101085830;refGene_tx=XM_003993505.4
D1	108173590	108173780
refGene_gn=ASRGL1;refGene_tx=XM_011286962.3
D1	108183303	108183446
refGene_gn=ASRGL1;refGene_tx=XM_011286962.3
D1	108184282	108184440
refGene_gn=ASRGL1;refGene_tx=XM_011286962.3
D1	108192091	108192210
refGene_gn=ASRGL1;refGene_tx=XM_011286962.3
D1	108192840	108192951
refGene_gn=ASRGL1;refGene_tx=XM_011286962.3
D1	108193595	108193801
refGene_gn=ASRGL1;refGene_tx=XM_011286962.3
D1	108226171	108226226
refGene_gn=SCGB1A1;refGene_tx=XM_011286963.2
D1	108228164	108228352
refGene_gn=SCGB1A1;refGene_tx=XM_011286963.2
D1	108229063	108229096
refGene_gn=SCGB1A1;refGene_tx=XM_011286963.2
D1	108306981	108315888
refGene_gn=AHNAK;refGene_tx=XM_023239943.1
D1	108316656	108320225
refGene_gn=AHNAK;refGene_tx=XM_023239943.1
D1	108320306	108321093

refGene_gn=AHNAK;refGene_tx=XM_023239943.1
D1	108321476	108326366
refGene_gn=AHNAK;refGene_tx=XM_023239943.1
D1	108327293	108327481
refGene_gn=AHNAK;refGene_tx=XM_023239943.1
D1	108327858	108328015
refGene_gn=AHNAK;refGene_tx=XM_023239943.1
D1	108349961	108350120
refGene_gn=EEF1G;refGene_tx=XM_003993509.4
D1	108350358	108350483
refGene_gn=EEF1G;refGene_tx=XM_003993509.4
D1	108350572	108350745
refGene_gn=EEF1G;refGene_tx=XM_003993509.4
D1	108354506	108354711
refGene_gn=EEF1G;refGene_tx=XM_003993509.4
D1	108355101	108355231
refGene_gn=EEF1G;refGene_tx=XM_003993509.4
D1	108357332	108357476
refGene_gn=EEF1G;refGene_tx=XM_003993509.4
D1	108358033	108358176
refGene_gn=EEF1G;refGene_tx=XM_003993509.4
D1	108358330	108358394
refGene_gn=EEF1G;refGene_tx=XM_003993509.4
D1	108359349	108359508
refGene_gn=EEF1G;refGene_tx=XM_003993509.4
D1	108360587	108360599
refGene_gn=EEF1G;refGene_tx=XM_003993509.4
D1	108361777	108362925
refGene_gn=TUT1;refGene_tx=XM_003993510.5
D1	108363342	108363435
refGene_gn=TUT1;refGene_tx=XM_003993510.5
D1	108363597	108363708
refGene_gn=TUT1;refGene_tx=XM_003993510.5
D1	108363857	108363967
refGene_gn=TUT1;refGene_tx=XM_003993510.5
D1	108364564	108365034
refGene_gn=TUT1;refGene_tx=XM_003993510.5
D1	108366437	108366538
refGene_gn=TUT1;refGene_tx=XM_003993510.5
D1	108366715	108367031
refGene_gn=TUT1;refGene_tx=XM_003993510.5
D1	108370849	108371040
refGene_gn=TUT1;refGene_tx=XM_003993510.5
D1	108372336	108372418
refGene_gn=TUT1;refGene_tx=XM_003993510.5
D1	108374794	108374960
refGene_gn=MTA2;refGene_tx=XM_023239949.1;refGene_gn=MTA2;refGene_tx=X
M_023239950.1
D1	108375179	108375328
refGene_gn=MTA2;refGene_tx=XM_023239949.1;refGene_gn=MTA2;refGene_tx=X

M_023239950.1
D1 108375437 108375556
refGene_gn=MTA2;refGene_tx=XM_023239949.1;refGene_gn=MTA2;refGene_tx=X
M_023239950.1
D1 108375845 108375935
refGene_gn=MTA2;refGene_tx=XM_023239949.1;refGene_gn=MTA2;refGene_tx=X
M_023239950.1
D1 108376138 108376367
refGene_gn=MTA2;refGene_tx=XM_023239949.1;refGene_gn=MTA2;refGene_tx=X
M_023239950.1
D1 108376626 108376766
refGene_gn=MTA2;refGene_tx=XM_023239949.1;refGene_gn=MTA2;refGene_tx=X
M_023239950.1
D1 108376885 108376983
refGene_gn=MTA2;refGene_tx=XM_023239949.1;refGene_gn=MTA2;refGene_tx=X
M_023239950.1
D1 108377209 108377268
refGene_gn=MTA2;refGene_tx=XM_023239949.1;refGene_gn=MTA2;refGene_tx=X
M_023239950.1
D1 108377393 108377468
refGene_gn=MTA2;refGene_tx=XM_023239949.1;refGene_gn=MTA2;refGene_tx=X
M_023239950.1
D1 108377571 108377760
refGene_gn=MTA2;refGene_tx=XM_023239949.1;refGene_gn=MTA2;refGene_tx=X
M_023239950.1
D1 108377898 108377998
refGene_gn=MTA2;refGene_tx=XM_023239949.1;refGene_gn=MTA2;refGene_tx=X
M_023239950.1
D1 108378175 108378278
refGene_gn=MTA2;refGene_tx=XM_023239949.1;refGene_gn=MTA2;refGene_tx=X
M_023239950.1
D1 108378576 108378694
refGene_gn=MTA2;refGene_tx=XM_023239949.1
D1 108378851 108378915
refGene_gn=MTA2;refGene_tx=XM_023239949.1
D1 108379052 108379170
refGene_gn=MTA2;refGene_tx=XM_023239949.1
D1 108380779 108380873
refGene_gn=MTA2;refGene_tx=XM_023239949.1
D1 108381569 108381637
refGene_gn=MTA2;refGene_tx=XM_023239949.1
D1 108382392 108382420
refGene_gn=MTA2;refGene_tx=XM_023239949.1
D1 108383280 108383633
refGene_gn=EML3;refGene_tx=XM_023239944.1;refGene_gn=EML3;refGene_tx=X
M_023239945.1;refGene_gn=EML3;refGene_tx=XM_023239946.1;refGene_gn=EML
3;refGene_tx=XM_023239948.1
D1 108383724 108383855
refGene_gn=EML3;refGene_tx=XM_023239944.1;refGene_gn=EML3;refGene_tx=X
M_023239945.1;refGene_gn=EML3;refGene_tx=XM_023239946.1;refGene_gn=EML

3;refGene_tx=XM_023239948.1
D1	108384148	108384247
refGene_gn=EML3;refGene_tx=XM_023239944.1;refGene_gn=EML3;refGene_tx=XM_023239945.1;refGene_gn=EML3;refGene_tx=XM_023239946.1;refGene_gn=EML3;refGene_tx=XM_023239948.1
D1	108384772	108384860
refGene_gn=EML3;refGene_tx=XM_023239944.1;refGene_gn=EML3;refGene_tx=XM_023239945.1;refGene_gn=EML3;refGene_tx=XM_023239946.1;refGene_gn=EML3;refGene_tx=XM_023239948.1
D1	108384989	108385087
refGene_gn=EML3;refGene_tx=XM_023239944.1;refGene_gn=EML3;refGene_tx=XM_023239945.1;refGene_gn=EML3;refGene_tx=XM_023239946.1;refGene_gn=EML3;refGene_tx=XM_023239948.1
D1	108385158	108385247
refGene_gn=EML3;refGene_tx=XM_023239944.1;refGene_gn=EML3;refGene_tx=XM_023239945.1;refGene_gn=EML3;refGene_tx=XM_023239946.1;refGene_gn=EML3;refGene_tx=XM_023239948.1
D1	108385737	108385805
refGene_gn=EML3;refGene_tx=XM_023239944.1;refGene_gn=EML3;refGene_tx=XM_023239945.1;refGene_gn=EML3;refGene_tx=XM_023239946.1;refGene_gn=EML3;refGene_tx=XM_023239948.1
D1	108386273	108386405
refGene_gn=EML3;refGene_tx=XM_023239944.1;refGene_gn=EML3;refGene_tx=XM_023239945.1;refGene_gn=EML3;refGene_tx=XM_023239946.1;refGene_gn=EML3;refGene_tx=XM_023239948.1
D1	108386488	108386614
refGene_gn=EML3;refGene_tx=XM_023239944.1;refGene_gn=EML3;refGene_tx=XM_023239945.1;refGene_gn=EML3;refGene_tx=XM_023239946.1;refGene_gn=EML3;refGene_tx=XM_023239948.1
D1	108386696	108386848
refGene_gn=EML3;refGene_tx=XM_023239944.1;refGene_gn=EML3;refGene_tx=XM_023239945.1;refGene_gn=EML3;refGene_tx=XM_023239946.1;refGene_gn=EML3;refGene_tx=XM_023239948.1
D1	108387271	108387413
refGene_gn=EML3;refGene_tx=XM_023239944.1;refGene_gn=EML3;refGene_tx=XM_023239945.1;refGene_gn=EML3;refGene_tx=XM_023239946.1;refGene_gn=EML3;refGene_tx=XM_023239948.1
D1	108387567	108387723
refGene_gn=EML3;refGene_tx=XM_023239944.1;refGene_gn=EML3;refGene_tx=XM_023239945.1;refGene_gn=EML3;refGene_tx=XM_023239946.1;refGene_gn=EML3;refGene_tx=XM_023239948.1
D1	108388106	108388202
refGene_gn=EML3;refGene_tx=XM_023239944.1;refGene_gn=EML3;refGene_tx=XM_023239945.1;refGene_gn=EML3;refGene_tx=XM_023239946.1;refGene_gn=EML3;refGene_tx=XM_023239948.1
D1	108388447	108388558
refGene_gn=EML3;refGene_tx=XM_023239944.1;refGene_gn=EML3;refGene_tx=XM_023239945.1;refGene_gn=EML3;refGene_tx=XM_023239946.1;refGene_gn=EML3;refGene_tx=XM_023239948.1
D1	108388645	108388715

refGene_gn=EML3;refGene_tx=XM_023239944.1;refGene_gn=EML3;refGene_tx=XM_023239945.1;refGene_gn=EML3;refGene_tx=XM_023239946.1;refGene_gn=EML3;refGene_tx=XM_023239948.1
D1 108388912 108389083
refGene_gn=EML3;refGene_tx=XM_023239944.1;refGene_gn=EML3;refGene_tx=XM_023239945.1;refGene_gn=EML3;refGene_tx=XM_023239946.1;refGene_gn=EML3;refGene_tx=XM_023239948.1
D1 108389303 108389427
refGene_gn=EML3;refGene_tx=XM_023239944.1;refGene_gn=EML3;refGene_tx=XM_023239945.1;refGene_gn=EML3;refGene_tx=XM_023239946.1;refGene_gn=EML3;refGene_tx=XM_023239948.1
D1 108389592 108389660
refGene_gn=EML3;refGene_tx=XM_023239944.1;refGene_gn=EML3;refGene_tx=XM_023239945.1;refGene_gn=EML3;refGene_tx=XM_023239946.1;refGene_gn=EML3;refGene_tx=XM_023239948.1
D1 108390703 108390817
refGene_gn=EML3;refGene_tx=XM_023239944.1;refGene_gn=EML3;refGene_tx=XM_023239945.1;refGene_gn=EML3;refGene_tx=XM_023239946.1;refGene_gn=EML3;refGene_tx=XM_023239948.1
D1 108390914 108391169
refGene_gn=EML3;refGene_tx=XM_023239944.1;refGene_gn=EML3;refGene_tx=XM_023239945.1;refGene_gn=EML3;refGene_tx=XM_023239946.1;refGene_gn=EML3;refGene_tx=XM_023239948.1
D1 108391240 108391412
refGene_gn=EML3;refGene_tx=XM_023239944.1;refGene_gn=EML3;refGene_tx=XM_023239945.1;refGene_gn=EML3;refGene_tx=XM_023239946.1;refGene_gn=EML3;refGene_tx=XM_023239948.1
D1 108392237 108392259
refGene_gn=EML3;refGene_tx=XM_023239944.1;refGene_gn=EML3;refGene_tx=XM_023239945.1;refGene_gn=EML3;refGene_tx=XM_023239946.1;refGene_gn=EML3;refGene_tx=XM_023239948.1
D1 108393077 108393667
refGene_gn=ROM1;refGene_tx=XM_003993512.5
D1 108394060 108394307
refGene_gn=ROM1;refGene_tx=XM_003993512.5
D1 108394422 108394641
refGene_gn=ROM1;refGene_tx=XM_003993512.5
D1 108395482 108395581
refGene_gn=B3GAT3;refGene_tx=XM_003993513.5
D1 108395924 108396215
refGene_gn=B3GAT3;refGene_tx=XM_003993513.5
D1 108396433 108396794
refGene_gn=B3GAT3;refGene_tx=XM_003993513.5
D1 108398253 108398428
refGene_gn=B3GAT3;refGene_tx=XM_003993513.5
D1 108399555 108399637
refGene_gn=B3GAT3;refGene_tx=XM_003993513.5
D1 108402362 108402472
refGene_gn=GANAB;refGene_tx=XM_003993515.4
D1 108402610 108402711

refGene_gn=GANAB;refGene_tx=XM_003993515.4
D1 108402874 108402987
refGene_gn=GANAB;refGene_tx=XM_003993515.4
D1 108403108 108403222
refGene_gn=GANAB;refGene_tx=XM_003993515.4
D1 108403414 108403489
refGene_gn=GANAB;refGene_tx=XM_003993515.4
D1 108403601 108403643
refGene_gn=GANAB;refGene_tx=XM_003993515.4
D1 108403644 108403679
refGene_gn=GANAB;refGene_tx=XM_003993515.4
D1 108403874 108403939
refGene_gn=GANAB;refGene_tx=XM_003993515.4
D1 108404769 108405013
refGene_gn=GANAB;refGene_tx=XM_003993515.4
D1 108405190 108405292
refGene_gn=GANAB;refGene_tx=XM_003993515.4
D1 108405620 108405717
refGene_gn=GANAB;refGene_tx=XM_003993515.4
D1 108405818 108405962
refGene_gn=GANAB;refGene_tx=XM_003993515.4
D1 108406192 108406272
refGene_gn=GANAB;refGene_tx=XM_003993515.4
D1 108406377 108406504
refGene_gn=GANAB;refGene_tx=XM_003993515.4
D1 108406586 108406822
refGene_gn=GANAB;refGene_tx=XM_003993515.4
D1 108407037 108407191
refGene_gn=GANAB;refGene_tx=XM_003993515.4
D1 108407788 108407969
refGene_gn=GANAB;refGene_tx=XM_003993515.4
D1 108408221 108408318
refGene_gn=GANAB;refGene_tx=XM_003993515.4
D1 108408405 108408493
refGene_gn=GANAB;refGene_tx=XM_003993515.4
D1 108408661 108408731
refGene_gn=GANAB;refGene_tx=XM_003993515.4
D1 108409535 108409601
refGene_gn=GANAB;refGene_tx=XM_003993515.4
D1 108410094 108410274
refGene_gn=GANAB;refGene_tx=XM_003993515.4
D1 108413765 108413893
refGene_gn=GANAB;refGene_tx=XM_003993515.4
D1 108414103 108414212
refGene_gn=GANAB;refGene_tx=XM_003993515.4
D1 108414356 108414461
refGene_gn=GANAB;refGene_tx=XM_003993515.4
D1 108417123 108417161
refGene_gn=GANAB;refGene_tx=XM_003993515.4
D1 108417598 108420578

refGene_gn=INTS5;refGene_tx=XM_003993519.5
D1 108421868 108421948
refGene_gn=INTS5;refGene_tx=XM_003993519.5
D1 108427764 108427795
refGene_gn=LBHD1;refGene_tx=XM_003993521.4;refGene_gn=LBHD1;refGene_tx=XM_019812559.2
D1 108427879 108427977
refGene_gn=LBHD1;refGene_tx=XM_003993521.4;refGene_gn=LBHD1;refGene_tx=XM_019812559.2
D1 108429285 108429410
refGene_gn=LBHD1;refGene_tx=XM_003993521.4;refGene_gn=LBHD1;refGene_tx=XM_019812559.2
D1 108430171 108430381
refGene_gn=CSKMT;refGene_tx=XM_006937414.3;refGene_gn=CSKMT;refGene_tx=XM_006937415.3;refGene_gn=CSKMT;refGene_tx=XM_006937416.4;refGene_gn=CSKMT;refGene_tx=XM_006937417.4;refGene_gn=CSKMT;refGene_tx=XM_006937418.3;refGene_gn=CSKMT;refGene_tx=XM_006937419.4;refGene_gn=CSKMT;refGene_tx=XM_019812557.2;refGene_gn=CSKMT;refGene_tx=XM_019812558.2
D1 108430506 108431217
refGene_gn=CSKMT;refGene_tx=XM_006937414.3;refGene_gn=CSKMT;refGene_tx=XM_006937415.3;refGene_gn=CSKMT;refGene_tx=XM_006937416.4;refGene_gn=CSKMT;refGene_tx=XM_006937417.4;refGene_gn=CSKMT;refGene_tx=XM_006937418.3;refGene_gn=CSKMT;refGene_tx=XM_006937419.4;refGene_gn=CSKMT;refGene_tx=XM_019812557.2;refGene_gn=CSKMT;refGene_tx=XM_019812558.2
D1 108431726 108431954
refGene_gn=LBHD1;refGene_tx=XM_003993521.4;refGene_gn=LBHD1;refGene_tx=XM_019812559.2
D1 108432864 108433024
refGene_gn=LBHD1;refGene_tx=XM_003993521.4;refGene_gn=LBHD1;refGene_tx=XM_019812559.2
D1 108433103 108433253
refGene_gn=LBHD1;refGene_tx=XM_003993521.4;refGene_gn=LBHD1;refGene_tx=XM_019812559.2
D1 108434478 108434598
refGene_gn=LOC101087173;refGene_tx=XM_003993600.4
D1 108434689 108434839
refGene_gn=LOC101087173;refGene_tx=XM_003993600.4
D1 108437362 108437412
refGene_gn=UBXN1;refGene_tx=XM_006937420.3
D1 108437578 108437771
refGene_gn=UBXN1;refGene_tx=XM_006937420.3
D1 108438275 108438389
refGene_gn=UBXN1;refGene_tx=XM_006937420.3
D1 108438530 108438582
refGene_gn=UBXN1;refGene_tx=XM_006937420.3
D1 108438676 108438868
refGene_gn=UBXN1;refGene_tx=XM_006937420.3
D1 108439110 108439180
refGene_gn=UBXN1;refGene_tx=XM_006937420.3
D1 108439263 108439370

refGene_gn=UBXN1;refGene_tx=XM_006937420.3
D1 108439458 108439512
refGene_gn=UBXN1;refGene_tx=XM_006937420.3
D1 108439651 108439710
refGene_gn=UBXN1;refGene_tx=XM_006937420.3
D1 108442684 108443407
refGene_gn=LRRN4CL;refGene_tx=XM_003993601.5
D1 108444937 108445104
refGene_gn=BSCL2;refGene_tx=XM_011286972.2;refGene_gn=BSCL2;refGene_tx=XM_019812554.2;refGene_gn=BSCL2;refGene_tx=XM_023239954.1;refGene_gn=BSCL2;refGene_tx=XM_023239955.1;refGene_gn=BSCL2;refGene_tx=XM_023239956.1;refGene_gn=BSCL2;refGene_tx=XM_023239957.1;refGene_gn=BSCL2;refGene_tx=XM_023239958.1;refGene_gn=BSCL2;refGene_tx=XM_023239959.1
D1 108445241 108445322
refGene_gn=BSCL2;refGene_tx=XM_011286972.2;refGene_gn=BSCL2;refGene_tx=XM_019812554.2;refGene_gn=BSCL2;refGene_tx=XM_023239954.1;refGene_gn=BSCL2;refGene_tx=XM_023239955.1;refGene_gn=BSCL2;refGene_tx=XM_023239956.1;refGene_gn=BSCL2;refGene_tx=XM_023239957.1;refGene_gn=BSCL2;refGene_tx=XM_023239958.1;refGene_gn=BSCL2;refGene_tx=XM_023239959.1
D1 108445422 108445515
refGene_gn=BSCL2;refGene_tx=XM_011286972.2;refGene_gn=BSCL2;refGene_tx=XM_019812554.2;refGene_gn=BSCL2;refGene_tx=XM_023239954.1;refGene_gn=BSCL2;refGene_tx=XM_023239955.1;refGene_gn=BSCL2;refGene_tx=XM_023239956.1;refGene_gn=BSCL2;refGene_tx=XM_023239957.1;refGene_gn=BSCL2;refGene_tx=XM_023239958.1;refGene_gn=BSCL2;refGene_tx=XM_023239959.1
D1 108445676 108445743
refGene_gn=BSCL2;refGene_tx=XM_011286972.2;refGene_gn=BSCL2;refGene_tx=XM_019812554.2;refGene_gn=BSCL2;refGene_tx=XM_023239954.1;refGene_gn=BSCL2;refGene_tx=XM_023239955.1;refGene_gn=BSCL2;refGene_tx=XM_023239956.1;refGene_gn=BSCL2;refGene_tx=XM_023239957.1;refGene_gn=BSCL2;refGene_tx=XM_023239958.1;refGene_gn=BSCL2;refGene_tx=XM_023239959.1
D1 108445899 108446041
refGene_gn=BSCL2;refGene_tx=XM_011286972.2;refGene_gn=BSCL2;refGene_tx=XM_019812554.2;refGene_gn=BSCL2;refGene_tx=XM_023239954.1;refGene_gn=BSCL2;refGene_tx=XM_023239955.1;refGene_gn=BSCL2;refGene_tx=XM_023239956.1;refGene_gn=BSCL2;refGene_tx=XM_023239957.1;refGene_gn=BSCL2;refGene_tx=XM_023239958.1;refGene_gn=BSCL2;refGene_tx=XM_023239959.1
D1 108446184 108446282
refGene_gn=BSCL2;refGene_tx=XM_011286972.2;refGene_gn=BSCL2;refGene_tx=XM_019812554.2;refGene_gn=BSCL2;refGene_tx=XM_023239954.1;refGene_gn=BSCL2;refGene_tx=XM_023239955.1;refGene_gn=BSCL2;refGene_tx=XM_023239956.1;refGene_gn=BSCL2;refGene_tx=XM_023239957.1;refGene_gn=BSCL2;refGene_tx=XM_023239958.1;refGene_gn=BSCL2;refGene_tx=XM_023239959.1
D1 108446463 108446598
refGene_gn=BSCL2;refGene_tx=XM_011286972.2;refGene_gn=BSCL2;refGene_tx=XM_019812554.2;refGene_gn=BSCL2;refGene_tx=XM_023239954.1;refGene_gn=BSCL2;refGene_tx=XM_023239955.1;refGene_gn=BSCL2;refGene_tx=XM_023239956.1;refGene_gn=BSCL2;refGene_tx=XM_023239957.1;refGene_gn=BSCL2;refGene_tx=XM_023239958.1;refGene_gn=BSCL2;refGene_tx=XM_023239959.1
D1 108447636 108447780

refGene_gn=BSCL2;refGene_tx=XM_011286972.2;refGene_gn=BSCL2;refGene_tx=XM_019812554.2;refGene_gn=BSCL2;refGene_tx=XM_023239954.1;refGene_gn=BSCL2;refGene_tx=XM_023239955.1;refGene_gn=BSCL2;refGene_tx=XM_023239956.1;refGene_gn=BSCL2;refGene_tx=XM_023239957.1;refGene_gn=BSCL2;refGene_tx=XM_023239958.1;refGene_gn=BSCL2;refGene_tx=XM_023239959.1
D1 108451119 108451201
refGene_gn=BSCL2;refGene_tx=XM_011286972.2;refGene_gn=BSCL2;refGene_tx=XM_019812554.2;refGene_gn=BSCL2;refGene_tx=XM_023239954.1;refGene_gn=BSCL2;refGene_tx=XM_023239955.1;refGene_gn=BSCL2;refGene_tx=XM_023239956.1;refGene_gn=BSCL2;refGene_tx=XM_023239957.1;refGene_gn=BSCL2;refGene_tx=XM_023239958.1;refGene_gn=BSCL2;refGene_tx=XM_023239959.1
D1 108452641 108452958
refGene_gn=BSCL2;refGene_tx=XM_011286972.2;refGene_gn=BSCL2;refGene_tx=XM_019812554.2;refGene_gn=BSCL2;refGene_tx=XM_023239954.1;refGene_gn=BSCL2;refGene_tx=XM_023239955.1;refGene_gn=BSCL2;refGene_tx=XM_023239956.1;refGene_gn=BSCL2;refGene_tx=XM_023239957.1;refGene_gn=BSCL2;refGene_tx=XM_023239958.1;refGene_gn=BSCL2;refGene_tx=XM_023239959.1
D1 108454378 108454486
refGene_gn=BSCL2;refGene_tx=XM_011286972.2;refGene_gn=BSCL2;refGene_tx=XM_019812554.2;refGene_gn=BSCL2;refGene_tx=XM_023239957.1;refGene_gn=BSCL2;refGene_tx=XM_023239958.1
D1 108455620 108455719
refGene_gn=GNG3;refGene_tx=XM_003993525.5;refGene_gn=GNG3;refGene_tx=XM_019812560.1
D1 108455994 108456123
refGene_gn=GNG3;refGene_tx=XM_003993525.5;refGene_gn=GNG3;refGene_tx=XM_019812560.1
D1 108461972 108462050
refGene_gn=HNRNPUL2;refGene_tx=XM_023239952.1
D1 108462165 108462273
refGene_gn=HNRNPUL2;refGene_tx=XM_023239952.1;refGene_gn=HNRNPUL2;refGene_tx=XM_023239953.1
D1 108462528 108462602
refGene_gn=HNRNPUL2;refGene_tx=XM_023239952.1
D1 108463705 108463906
refGene_gn=HNRNPUL2;refGene_tx=XM_023239952.1;refGene_gn=HNRNPUL2;refGene_tx=XM_023239953.1
D1 108465597 108465766
refGene_gn=HNRNPUL2;refGene_tx=XM_023239952.1;refGene_gn=HNRNPUL2;refGene_tx=XM_023239953.1
D1 108466306 108466435
refGene_gn=HNRNPUL2;refGene_tx=XM_023239952.1;refGene_gn=HNRNPUL2;refGene_tx=XM_023239953.1
D1 108466813 108466936
refGene_gn=HNRNPUL2;refGene_tx=XM_023239952.1;refGene_gn=HNRNPUL2;refGene_tx=XM_023239953.1
D1 108467098 108467362
refGene_gn=HNRNPUL2;refGene_tx=XM_023239952.1;refGene_gn=HNRNPUL2;refGene_tx=XM_023239953.1
D1 108467820 108467933

refGene_gn=HNRNPUL2;refGene_tx=XM_023239952.1;refGene_gn=HNRNPUL2;refGene_tx=XM_023239953.1
D1 108468030 108468121
refGene_gn=HNRNPUL2;refGene_tx=XM_023239952.1;refGene_gn=HNRNPUL2;refGene_tx=XM_023239953.1
D1 108468424 108468564
refGene_gn=HNRNPUL2;refGene_tx=XM_023239952.1;refGene_gn=HNRNPUL2;refGene_tx=XM_023239953.1
D1 108468763 108468840
refGene_gn=HNRNPUL2;refGene_tx=XM_023239952.1;refGene_gn=HNRNPUL2;refGene_tx=XM_023239953.1
D1 108469131 108469267
refGene_gn=HNRNPUL2;refGene_tx=XM_023239952.1;refGene_gn=HNRNPUL2;refGene_tx=XM_023239953.1
D1 108471313 108471854
refGene_gn=HNRNPUL2;refGene_tx=XM_023239952.1;refGene_gn=HNRNPUL2;refGene_tx=XM_023239953.1
D1 108473359 108473416
refGene_gn=TTC9C;refGene_tx=XM_006937427.3;refGene_gn=TTC9C;refGene_tx=XM_019812561.2
D1 108473550 108473845
refGene_gn=TTC9C;refGene_tx=XM_003993526.5;refGene_gn=TTC9C;refGene_tx=XM_006937427.3;refGene_gn=TTC9C;refGene_tx=XM_006937428.4;refGene_gn=TTC9C;refGene_tx=XM_006937429.3;refGene_gn=TTC9C;refGene_tx=XM_006937430.4;refGene_gn=TTC9C;refGene_tx=XM_019812561.2
D1 108476787 108476970
refGene_gn=TTC9C;refGene_tx=XM_003993526.5;refGene_gn=TTC9C;refGene_tx=XM_006937427.3;refGene_gn=TTC9C;refGene_tx=XM_006937428.4;refGene_gn=TTC9C;refGene_tx=XM_006937429.3;refGene_gn=TTC9C;refGene_tx=XM_006937430.4
D1 108479113 108479208
refGene_gn=TTC9C;refGene_tx=XM_003993526.5;refGene_gn=TTC9C;refGene_tx=XM_006937427.3;refGene_gn=TTC9C;refGene_tx=XM_006937428.4;refGene_gn=TTC9C;refGene_tx=XM_006937429.3;refGene_gn=TTC9C;refGene_tx=XM_006937430.4;refGene_gn=TTC9C;refGene_tx=XM_019812561.2
D1 108486476 108488024
refGene_gn=ZBTB3;refGene_tx=XM_006937432.4;refGene_gn=ZBTB3;refGene_tx=XM_006937433.4
D1 108498074 108498086
refGene_gn=POLR2G;refGene_tx=XM_003993527.3
D1 108498219 108498329
refGene_gn=POLR2G;refGene_tx=XM_003993527.3
D1 108499212 108499372
refGene_gn=POLR2G;refGene_tx=XM_003993527.3
D1 108500543 108500594
refGene_gn=POLR2G;refGene_tx=XM_003993527.3
D1 108500699 108500765
refGene_gn=POLR2G;refGene_tx=XM_003993527.3
D1 108500861 108500933
refGene_gn=POLR2G;refGene_tx=XM_003993527.3

D1 108501125 108501159
refGene_gn=POLR2G;refGene_tx=XM_003993527.3
D1 108501382 108501396
refGene_gn=POLR2G;refGene_tx=XM_003993527.3
D1 108506031 108506079
refGene_gn=TAF6L;refGene_tx=XM_023239960.1
D1 108506603 108506663
refGene_gn=TAF6L;refGene_tx=XM_023239960.1;refGene_gn=TAF6L;refGene_tx=XM_023239962.1;refGene_gn=TAF6L;refGene_tx=XM_023239963.1
D1 108508473 108508791
refGene_gn=TAF6L;refGene_tx=XM_023239960.1;refGene_gn=TAF6L;refGene_tx=XM_023239961.1;refGene_gn=TAF6L;refGene_tx=XM_023239962.1;refGene_gn=TAF6L;refGene_tx=XM_023239963.1;refGene_gn=TAF6L;refGene_tx=XM_023239964.1
D1 108509238 108509325
refGene_gn=TAF6L;refGene_tx=XM_023239960.1;refGene_gn=TAF6L;refGene_tx=XM_023239961.1;refGene_gn=TAF6L;refGene_tx=XM_023239962.1;refGene_gn=TAF6L;refGene_tx=XM_023239963.1;refGene_gn=TAF6L;refGene_tx=XM_023239964.1
D1 108510683 108510834
refGene_gn=TAF6L;refGene_tx=XM_023239960.1;refGene_gn=TAF6L;refGene_tx=XM_023239961.1;refGene_gn=TAF6L;refGene_tx=XM_023239962.1;refGene_gn=TAF6L;refGene_tx=XM_023239963.1;refGene_gn=TAF6L;refGene_tx=XM_023239964.1
D1 108510973 108511024
refGene_gn=TAF6L;refGene_tx=XM_023239960.1;refGene_gn=TAF6L;refGene_tx=XM_023239961.1;refGene_gn=TAF6L;refGene_tx=XM_023239962.1;refGene_gn=TAF6L;refGene_tx=XM_023239963.1;refGene_gn=TAF6L;refGene_tx=XM_023239964.1
D1 108511779 108511874
refGene_gn=TAF6L;refGene_tx=XM_023239960.1;refGene_gn=TAF6L;refGene_tx=XM_023239961.1;refGene_gn=TAF6L;refGene_tx=XM_023239962.1;refGene_gn=TAF6L;refGene_tx=XM_023239963.1;refGene_gn=TAF6L;refGene_tx=XM_023239964.1
D1 108513953 108514079
refGene_gn=TAF6L;refGene_tx=XM_023239960.1;refGene_gn=TAF6L;refGene_tx=XM_023239961.1;refGene_gn=TAF6L;refGene_tx=XM_023239962.1;refGene_gn=TAF6L;refGene_tx=XM_023239963.1;refGene_gn=TAF6L;refGene_tx=XM_023239964.1
D1 108514223 108514444
refGene_gn=TAF6L;refGene_tx=XM_023239960.1;refGene_gn=TAF6L;refGene_tx=XM_023239961.1;refGene_gn=TAF6L;refGene_tx=XM_023239962.1;refGene_gn=TAF6L;refGene_tx=XM_023239963.1;refGene_gn=TAF6L;refGene_tx=XM_023239964.1
D1 108514774 108514907
refGene_gn=TAF6L;refGene_tx=XM_023239960.1;refGene_gn=TAF6L;refGene_tx=XM_023239961.1;refGene_gn=TAF6L;refGene_tx=XM_023239962.1;refGene_gn=TAF6L;refGene_tx=XM_023239963.1;refGene_gn=TAF6L;refGene_tx=XM_023239964.1
D1 108516483 108516612

refGene_gn=TAF6L;refGene_tx=XM_023239960.1;refGene_gn=TAF6L;refGene_tx=XM_023239961.1;refGene_gn=TAF6L;refGene_tx=XM_023239962.1;refGene_gn=TAF6L;refGene_tx=XM_023239963.1;refGene_gn=TAF6L;refGene_tx=XM_023239964.1
D1 108516743 108517514
refGene_gn=TAF6L;refGene_tx=XM_023239960.1;refGene_gn=TAF6L;refGene_tx=XM_023239961.1;refGene_gn=TAF6L;refGene_tx=XM_023239962.1;refGene_gn=TAF6L;refGene_tx=XM_023239963.1;refGene_gn=TAF6L;refGene_tx=XM_023239964.1
D1 108517666 108517762
refGene_gn=TMEM179B;refGene_tx=XM_023239965.1
D1 108518663 108518851
refGene_gn=TMEM179B;refGene_tx=XM_023239965.1
D1 108518933 108519068
refGene_gn=TMEM179B;refGene_tx=XM_023239965.1
D1 108519211 108519290
refGene_gn=TMEM179B;refGene_tx=XM_023239965.1
D1 108519497 108519659
refGene_gn=TMEM179B;refGene_tx=XM_023239965.1
D1 108520046 108520339
refGene_gn=TMEM223;refGene_tx=XM_023239966.1
D1 108520740 108521056
refGene_gn=TMEM223;refGene_tx=XM_023239966.1
D1 108521577 108521616
refGene_gn=NXF1;refGene_tx=XM_003993528.5
D1 108521735 108521796
refGene_gn=NXF1;refGene_tx=XM_003993528.5
D1 108522519 108522702
refGene_gn=NXF1;refGene_tx=XM_003993528.5
D1 108523232 108523305
refGene_gn=NXF1;refGene_tx=XM_003993528.5
D1 108523892 108523935
refGene_gn=NXF1;refGene_tx=XM_003993528.5
D1 108524059 108524175
refGene_gn=NXF1;refGene_tx=XM_003993528.5
D1 108524279 108524338
refGene_gn=NXF1;refGene_tx=XM_003993528.5
D1 108524444 108524552
refGene_gn=NXF1;refGene_tx=XM_003993528.5
D1 108524687 108524743
refGene_gn=NXF1;refGene_tx=XM_003993528.5
D1 108524822 108524891
refGene_gn=NXF1;refGene_tx=XM_003993528.5
D1 108525200 108525237
refGene_gn=NXF1;refGene_tx=XM_003993528.5
D1 108527262 108527372
refGene_gn=NXF1;refGene_tx=XM_003993528.5
D1 108527503 108527611
refGene_gn=NXF1;refGene_tx=XM_003993528.5
D1 108527739 108527828

refGene_gn=NXF1;refGene_tx=XM_003993528.5
D1 108527975 108528045
refGene_gn=NXF1;refGene_tx=XM_003993528.5
D1 108528163 108528244
refGene_gn=NXF1;refGene_tx=XM_003993528.5
D1 108528340 108528445
refGene_gn=NXF1;refGene_tx=XM_003993528.5
D1 108528553 108528637
refGene_gn=NXF1;refGene_tx=XM_003993528.5
D1 108529219 108529373
refGene_gn=NXF1;refGene_tx=XM_003993528.5
D1 108529593 108529777
refGene_gn=NXF1;refGene_tx=XM_003993528.5
D1 108531593 108531621
refGene_gn=NXF1;refGene_tx=XM_003993528.5
D1 108533788 108533948
refGene_gn=STX5;refGene_tx=XM_003993529.5;refGene_gn=STX5;refGene_tx=XM_006937436.3
D1 108544572 108544694
refGene_gn=STX5;refGene_tx=XM_003993529.5;refGene_gn=STX5;refGene_tx=XM_006937436.3
D1 108544865 108544972
refGene_gn=STX5;refGene_tx=XM_003993529.5;refGene_gn=STX5;refGene_tx=XM_006937436.3
D1 108545682 108545764
refGene_gn=STX5;refGene_tx=XM_003993529.5;refGene_gn=STX5;refGene_tx=XM_006937436.3
D1 108545938 108545995
refGene_gn=STX5;refGene_tx=XM_003993529.5;refGene_gn=STX5;refGene_tx=XM_006937436.3
D1 108546081 108546198
refGene_gn=STX5;refGene_tx=XM_003993529.5;refGene_gn=STX5;refGene_tx=XM_006937436.3
D1 108550913 108550984
refGene_gn=STX5;refGene_tx=XM_003993529.5;refGene_gn=STX5;refGene_tx=XM_006937436.3
D1 108551098 108551154
refGene_gn=STX5;refGene_tx=XM_003993529.5;refGene_gn=STX5;refGene_tx=XM_006937436.3
D1 108551309 108551380
refGene_gn=STX5;refGene_tx=XM_003993529.5;refGene_gn=STX5;refGene_tx=XM_006937436.3
D1 108552364 108552589
refGene_gn=STX5;refGene_tx=XM_003993529.5;refGene_gn=STX5;refGene_tx=XM_006937436.3
D1 108554338 108554518
refGene_gn=WDR74;refGene_tx=XM_003993532.3
D1 108554819 108554876
refGene_gn=WDR74;refGene_tx=XM_003993532.3
D1 108554960 108555106

refGene_gn=WDR74;refGene_tx=XM_003993532.3
D1	108555436	108555492
refGene_gn=WDR74;refGene_tx=XM_003993532.3
D1	108555579	108555680
refGene_gn=WDR74;refGene_tx=XM_003993532.3
D1	108556887	108556989
refGene_gn=WDR74;refGene_tx=XM_003993532.3
D1	108557129	108557276
refGene_gn=WDR74;refGene_tx=XM_003993532.3
D1	108558062	108558138
refGene_gn=WDR74;refGene_tx=XM_003993532.3
D1	108559539	108559661
refGene_gn=WDR74;refGene_tx=XM_003993532.3
D1	108559753	108559860
refGene_gn=WDR74;refGene_tx=XM_003993532.3
D1	108559944	108560008
refGene_gn=WDR74;refGene_tx=XM_003993532.3
D1	108591664	108592097
refGene_gn=SLC3A2;refGene_tx=XM_003993534.5
D1	108592794	108592968
refGene_gn=SLC3A2;refGene_tx=XM_003993534.5
D1	108593570	108593665
refGene_gn=SLC3A2;refGene_tx=XM_003993534.5
D1	108595475	108595544
refGene_gn=SLC3A2;refGene_tx=XM_003993534.5
D1	108595648	108595707
refGene_gn=SLC3A2;refGene_tx=XM_003993534.5
D1	108595868	108596049
refGene_gn=SLC3A2;refGene_tx=XM_003993534.5
D1	108596162	108596306
refGene_gn=SLC3A2;refGene_tx=XM_003993534.5
D1	108597497	108597581
refGene_gn=SLC3A2;refGene_tx=XM_003993534.5
D1	108597693	108598041
refGene_gn=SLC3A2;refGene_tx=XM_003993534.5
D1	108615878	108617261
refGene_gn=CHRM1;refGene_tx=XM_006937437.4;refGene_gn=CHRM1;refGene_tx=XM_019812563.2
D1	108672641	108672729
refGene_gn=LOC101094048;refGene_tx=XM_019812564.2;refGene_gn=LOC101094048;refGene_tx=XM_019812566.1
D1	108672919	108673123
refGene_gn=LOC101094048;refGene_tx=XM_019812564.2
D1	108673998	108674122
refGene_gn=LOC101094048;refGene_tx=XM_019812565.2
D1	108674632	108674741
refGene_gn=LOC101094048;refGene_tx=XM_019812564.2;refGene_gn=LOC101094048;refGene_tx=XM_019812565.2;refGene_gn=LOC101094048;refGene_tx=XM_019812566.1
D1	108674841	108675056

refGene_gn=LOC101094048;refGene_tx=XM_019812564.2;refGene_gn=LOC101094048;refGene_tx=XM_019812565.2;refGene_gn=LOC101094048;refGene_tx=XM_019812566.1
D1 108676129 108676245
refGene_gn=LOC101094048;refGene_tx=XM_019812564.2;refGene_gn=LOC101094048;refGene_tx=XM_019812565.2;refGene_gn=LOC101094048;refGene_tx=XM_019812566.1
D1 108676402 108676526
refGene_gn=LOC101094048;refGene_tx=XM_019812564.2;refGene_gn=LOC101094048;refGene_tx=XM_019812565.2;refGene_gn=LOC101094048;refGene_tx=XM_019812566.1
D1 108676976 108677145
refGene_gn=LOC101094048;refGene_tx=XM_019812564.2;refGene_gn=LOC101094048;refGene_tx=XM_019812565.2;refGene_gn=LOC101094048;refGene_tx=XM_019812566.1
D1 108678394 108678549
refGene_gn=LOC101094048;refGene_tx=XM_019812564.2;refGene_gn=LOC101094048;refGene_tx=XM_019812565.2;refGene_gn=LOC101094048;refGene_tx=XM_019812566.1
D1 108678786 108678890
refGene_gn=LOC101094048;refGene_tx=XM_019812564.2;refGene_gn=LOC101094048;refGene_tx=XM_019812565.2;refGene_gn=LOC101094048;refGene_tx=XM_019812566.1
D1 108679267 108679636
refGene_gn=LOC101094048;refGene_tx=XM_019812564.2;refGene_gn=LOC101094048;refGene_tx=XM_019812565.2;refGene_gn=LOC101094048;refGene_tx=XM_019812566.1
D1 108687452 108687552
refGene_gn=LOC101088955;refGene_tx=XM_023239748.1
D1 108687640 108687844
refGene_gn=LOC101088955;refGene_tx=XM_023239748.1
D1 108687970 108688038
refGene_gn=LOC101088955;refGene_tx=XM_023239748.1
D1 108688039 108688080
refGene_gn=LOC101088955;refGene_tx=XM_023239748.1
D1 108688784 108688999
refGene_gn=LOC101088955;refGene_tx=XM_023239748.1
D1 108689935 108690051
refGene_gn=LOC101088955;refGene_tx=XM_023239748.1
D1 108690243 108690367
refGene_gn=LOC101088955;refGene_tx=XM_023239748.1
D1 108693028 108693197
refGene_gn=LOC101088955;refGene_tx=XM_023239748.1
D1 108694345 108694378
refGene_gn=LOC101088955;refGene_tx=XM_023239748.1
D1 108694380 108694505
refGene_gn=LOC101088955;refGene_tx=XM_023239748.1
D1 108695006 108695215
refGene_gn=LOC101088955;refGene_tx=XM_023239748.1
D1 108705370 108705471

refGene_gn=LOC101088955;refGene_tx=XM_023239748.1
D1 108705579 108705700
refGene_gn=LOC101088955;refGene_tx=XM_023239748.1
D1 108710966 108711073
refGene_gn=LOC101089212;refGene_tx=XM_004001518.2
D1 108769208 108769278
refGene_gn=LOC101089212;refGene_tx=XM_004001518.2
D1 108770787 108771189
refGene_gn=LOC101089212;refGene_tx=XM_004001518.2
D1 108790519 108790887
refGene_gn=LOC101089464;refGene_tx=XM_011286974.1
D1 108790888 108790913
refGene_gn=LOC101089464;refGene_tx=XM_011286974.1
D1 108791870 108791978
refGene_gn=LOC101089464;refGene_tx=XM_011286974.1
D1 108827274 108827676
refGene_gn=LOC101094449;refGene_tx=XM_003993538.3;refGene_gn=LOC101094449;refGene_tx=XM_019812568.2
D1 108828590 108828694
refGene_gn=LOC101094449;refGene_tx=XM_003993538.3;refGene_gn=LOC101094449;refGene_tx=XM_019812568.2
D1 108842585 108842740
refGene_gn=LOC101094449;refGene_tx=XM_003993538.3;refGene_gn=LOC101094449;refGene_tx=XM_019812568.2
D1 108842849 108843018
refGene_gn=LOC101094449;refGene_tx=XM_003993538.3;refGene_gn=LOC101094449;refGene_tx=XM_019812568.2
D1 108843770 108843894
refGene_gn=LOC101094449;refGene_tx=XM_003993538.3;refGene_gn=LOC101094449;refGene_tx=XM_019812568.2
D1 108856087 108856203
refGene_gn=LOC101094449;refGene_tx=XM_003993538.3;refGene_gn=LOC101094449;refGene_tx=XM_019812568.2
D1 108857254 108857469
refGene_gn=LOC101094449;refGene_tx=XM_003993538.3;refGene_gn=LOC101094449;refGene_tx=XM_019812568.2
D1 108859313 108859422
refGene_gn=LOC101094449;refGene_tx=XM_003993538.3;refGene_gn=LOC101094449;refGene_tx=XM_019812568.2
D1 108860348 108860552
refGene_gn=LOC101094449;refGene_tx=XM_003993538.3;refGene_gn=LOC101094449;refGene_tx=XM_019812568.2
D1 108861818 108861879
refGene_gn=LOC101094449;refGene_tx=XM_003993538.3;refGene_gn=LOC101094449;refGene_tx=XM_019812568.2
D1 108895297 108895390
refGene_gn=HRASLS5;refGene_tx=XM_006937438.4
D1 108897219 108897482
refGene_gn=HRASLS5;refGene_tx=XM_006937438.4
D1 108899513 108899622

refGene_gn=HRASLS5;refGene_tx=XM_006937438.4
D1	108910308	108910417
refGene_gn=HRASLS5;refGene_tx=XM_006937438.4
D1	108912296	108912381
refGene_gn=HRASLS5;refGene_tx=XM_006937438.4
D1	108913023	108913114
refGene_gn=HRASLS5;refGene_tx=XM_006937438.4
D1	108922231	108922300
refGene_gn=LGALS12;refGene_tx=XM_023239971.1
D1	108923516	108923667
refGene_gn=LGALS12;refGene_tx=XM_023239968.1;refGene_gn=LGALS12;refGene_tx=XM_023239969.1;refGene_gn=LGALS12;refGene_tx=XM_023239970.1
D1	108924289	108924378
refGene_gn=LGALS12;refGene_tx=XM_023239968.1;refGene_gn=LGALS12;refGene_tx=XM_023239969.1;refGene_gn=LGALS12;refGene_tx=XM_023239970.1;refGene_gn=LGALS12;refGene_tx=XM_023239971.1;refGene_gn=LGALS12;refGene_tx=XM_023239972.1;refGene_gn=LGALS12;refGene_tx=XM_023239973.1;refGene_gn=LGALS12;refGene_tx=XM_023239974.1;refGene_gn=LGALS12;refGene_tx=XM_023239975.1
D1	108924461	108924675
refGene_gn=LGALS12;refGene_tx=XM_023239968.1;refGene_gn=LGALS12;refGene_tx=XM_023239969.1;refGene_gn=LGALS12;refGene_tx=XM_023239970.1;refGene_gn=LGALS12;refGene_tx=XM_023239971.1;refGene_gn=LGALS12;refGene_tx=XM_023239972.1;refGene_gn=LGALS12;refGene_tx=XM_023239973.1;refGene_gn=LGALS12;refGene_tx=XM_023239974.1;refGene_gn=LGALS12;refGene_tx=XM_023239975.1
D1	108925570	108925690
refGene_gn=LGALS12;refGene_tx=XM_023239968.1;refGene_gn=LGALS12;refGene_tx=XM_023239969.1;refGene_gn=LGALS12;refGene_tx=XM_023239970.1;refGene_gn=LGALS12;refGene_tx=XM_023239971.1;refGene_gn=LGALS12;refGene_tx=XM_023239972.1;refGene_gn=LGALS12;refGene_tx=XM_023239973.1;refGene_gn=LGALS12;refGene_tx=XM_023239974.1;refGene_gn=LGALS12;refGene_tx=XM_023239975.1
D1	108926222	108926261
refGene_gn=LGALS12;refGene_tx=XM_023239968.1;refGene_gn=LGALS12;refGene_tx=XM_023239969.1;refGene_gn=LGALS12;refGene_tx=XM_023239970.1;refGene_gn=LGALS12;refGene_tx=XM_023239971.1;refGene_gn=LGALS12;refGene_tx=XM_023239972.1;refGene_gn=LGALS12;refGene_tx=XM_023239973.1;refGene_gn=LGALS12;refGene_tx=XM_023239974.1;refGene_gn=LGALS12;refGene_tx=XM_023239975.1
D1	108926848	108926875
refGene_gn=LGALS12;refGene_tx=XM_023239968.1;refGene_gn=LGALS12;refGene_tx=XM_023239969.1;refGene_gn=LGALS12;refGene_tx=XM_023239970.1;refGene_gn=LGALS12;refGene_tx=XM_023239971.1;refGene_gn=LGALS12;refGene_tx=XM_023239972.1;refGene_gn=LGALS12;refGene_tx=XM_023239973.1;refGene_gn=LGALS12;refGene_tx=XM_023239974.1;refGene_gn=LGALS12;refGene_tx=XM_023239975.1
D1	108927491	108927580
refGene_gn=LGALS12;refGene_tx=XM_023239968.1;refGene_gn=LGALS12;refGene_tx=XM_023239969.1;refGene_gn=LGALS12;refGene_tx=XM_023239970.1;refGe

ne_gn=LGALS12;refGene_tx=XM_023239971.1;refGene_gn=LGALS12;refGene_tx=
XM_023239972.1;refGene_gn=LGALS12;refGene_tx=XM_023239973.1;refGene_gn
=LGALS12;refGene_tx=XM_023239974.1;refGene_gn=LGALS12;refGene_tx=XM_02
3239975.1
D1 108932588 108932739
refGene_gn=LGALS12;refGene_tx=XM_023239968.1;refGene_gn=LGALS12;refGen
e_tx=XM_023239969.1;refGene_gn=LGALS12;refGene_tx=XM_023239970.1;refGe
ne_gn=LGALS12;refGene_tx=XM_023239971.1;refGene_gn=LGALS12;refGene_tx=
XM_023239972.1;refGene_gn=LGALS12;refGene_tx=XM_023239973.1;refGene_gn
=LGALS12;refGene_tx=XM_023239974.1;refGene_gn=LGALS12;refGene_tx=XM_02
3239975.1
D1 108933591 108933738
refGene_gn=LGALS12;refGene_tx=XM_023239968.1;refGene_gn=LGALS12;refGen
e_tx=XM_023239969.1;refGene_gn=LGALS12;refGene_tx=XM_023239970.1;refGe
ne_gn=LGALS12;refGene_tx=XM_023239971.1;refGene_gn=LGALS12;refGene_tx=
XM_023239972.1;refGene_gn=LGALS12;refGene_tx=XM_023239973.1;refGene_gn
=LGALS12;refGene_tx=XM_023239974.1;refGene_gn=LGALS12;refGene_tx=XM_02
3239975.1
D1 108956616 108956700
refGene_gn=LOC111556693;refGene_tx=XM_023239976.1
D1 108957745 108957781
refGene_gn=LOC111556693;refGene_tx=XM_023239977.1
D1 108957929 108958295
refGene_gn=LOC111556693;refGene_tx=XM_023239976.1;refGene_gn=LOC111556
693;refGene_tx=XM_023239977.1
D1 108961107 108961473
refGene_gn=LOC102899075;refGene_tx=XM_006937445.2;refGene_gn=LOC102899
075;refGene_tx=XM_019812576.1
D1 108961621 108961657
refGene_gn=LOC102899075;refGene_tx=XM_019812576.1
D1 108962702 108962786
refGene_gn=LOC102899075;refGene_tx=XM_006937445.2
D1 108986628 108986730
refGene_gn=LOC101090719;refGene_tx=XM_011286975.3;refGene_gn=LOC101090
719;refGene_tx=XM_011286976.2;refGene_gn=LOC101090719;refGene_tx=XM_01
1286977.3;refGene_gn=LOC101090719;refGene_tx=XM_019812577.1;refGene_gn
=LOC101090719;refGene_tx=XM_019812578.1
D1 108994951 108995220
refGene_gn=LOC101090719;refGene_tx=XM_011286975.3;refGene_gn=LOC101090
719;refGene_tx=XM_011286976.2;refGene_gn=LOC101090719;refGene_tx=XM_01
1286977.3;refGene_gn=LOC101090719;refGene_tx=XM_019812577.1;refGene_gn
=LOC101090719;refGene_tx=XM_019812578.1
D1 109002800 109002903
refGene_gn=LOC101090719;refGene_tx=XM_011286975.3;refGene_gn=LOC101090
719;refGene_tx=XM_011286976.2;refGene_gn=LOC101090719;refGene_tx=XM_01
1286977.3;refGene_gn=LOC101090719;refGene_tx=XM_019812577.1;refGene_gn
=LOC101090719;refGene_tx=XM_019812578.1
D1 109020676 109020745
refGene_gn=LOC101090719;refGene_tx=XM_019812577.1
D1 109024477 109024492

refGene_gn=LOC101090719;refGene_tx=XM_011286975.3;refGene_gn=LOC101090719;refGene_tx=XM_011286976.2;refGene_gn=LOC101090719;refGene_tx=XM_011286977.3;refGene_gn=LOC101090719;refGene_tx=XM_019812578.1
D1 109033356 109033443
refGene_gn=ATL3;refGene_tx=XM_003993539.4;refGene_gn=ATL3;refGene_tx=XM_023239979.1
D1 109034220 109034652
refGene_gn=ATL3;refGene_tx=XM_003993539.4;refGene_gn=ATL3;refGene_tx=XM_023239979.1
D1 109035040 109035112
refGene_gn=ATL3;refGene_tx=XM_003993539.4;refGene_gn=ATL3;refGene_tx=XM_023239979.1
D1 109036313 109036370
refGene_gn=ATL3;refGene_tx=XM_003993539.4;refGene_gn=ATL3;refGene_tx=XM_023239979.1
D1 109036998 109037126
refGene_gn=ATL3;refGene_tx=XM_003993539.4;refGene_gn=ATL3;refGene_tx=XM_023239979.1
D1 109045650 109045789
refGene_gn=ATL3;refGene_tx=XM_003993539.4;refGene_gn=ATL3;refGene_tx=XM_023239979.1
D1 109046490 109046583
refGene_gn=ATL3;refGene_tx=XM_003993539.4;refGene_gn=ATL3;refGene_tx=XM_023239979.1
D1 109048057 109048114
refGene_gn=ATL3;refGene_tx=XM_003993539.4;refGene_gn=ATL3;refGene_tx=XM_023239979.1
D1 109056800 109056851
refGene_gn=ATL3;refGene_tx=XM_003993539.4;refGene_gn=ATL3;refGene_tx=XM_023239979.1
D1 109057347 109057452
refGene_gn=ATL3;refGene_tx=XM_003993539.4;refGene_gn=ATL3;refGene_tx=XM_023239979.1
D1 109066015 109066159
refGene_gn=ATL3;refGene_tx=XM_003993539.4;refGene_gn=ATL3;refGene_tx=XM_023239979.1
D1 109066299 109066514
refGene_gn=ATL3;refGene_tx=XM_003993539.4;refGene_gn=ATL3;refGene_tx=XM_023239979.1
D1 109075383 109075429
refGene_gn=ATL3;refGene_tx=XM_023239979.1
D1 109082518 109082657
refGene_gn=RTN3;refGene_tx=XM_019812579.2;refGene_gn=RTN3;refGene_tx=XM_019812580.2;refGene_gn=RTN3;refGene_tx=XM_019812581.2;refGene_gn=RTN3;refGene_tx=XM_019812582.2
D1 109094721 109094778
refGene_gn=RTN3;refGene_tx=XM_019812579.2;refGene_gn=RTN3;refGene_tx=XM_019812581.2
D1 109100862 109100866
refGene_gn=RTN3;refGene_tx=XM_011287051.2

D1	109104851	109107209
refGene_gn=RTN3;refGene_tx=XM_011287051.2;refGene_gn=RTN3;refGene_tx=XM_019812579.2;refGene_gn=RTN3;refGene_tx=XM_019812580.2
D1	109134526	109134734
refGene_gn=RTN3;refGene_tx=XM_011287051.2;refGene_gn=RTN3;refGene_tx=XM_019812579.2;refGene_gn=RTN3;refGene_tx=XM_019812580.2;refGene_gn=RTN3;refGene_tx=XM_019812581.2;refGene_gn=RTN3;refGene_tx=XM_019812582.2
D1	109136611	109136750
refGene_gn=RTN3;refGene_tx=XM_011287051.2;refGene_gn=RTN3;refGene_tx=XM_019812579.2;refGene_gn=RTN3;refGene_tx=XM_019812580.2;refGene_gn=RTN3;refGene_tx=XM_019812581.2;refGene_gn=RTN3;refGene_tx=XM_019812582.2
D1	109137186	109137256
refGene_gn=RTN3;refGene_tx=XM_011287051.2;refGene_gn=RTN3;refGene_tx=XM_019812579.2;refGene_gn=RTN3;refGene_tx=XM_019812580.2;refGene_gn=RTN3;refGene_tx=XM_019812581.2;refGene_gn=RTN3;refGene_tx=XM_019812582.2
D1	109138479	109138526
refGene_gn=RTN3;refGene_tx=XM_011287051.2;refGene_gn=RTN3;refGene_tx=XM_019812579.2;refGene_gn=RTN3;refGene_tx=XM_019812580.2;refGene_gn=RTN3;refGene_tx=XM_019812581.2;refGene_gn=RTN3;refGene_tx=XM_019812582.2
D1	109139368	109139427
refGene_gn=RTN3;refGene_tx=XM_011287051.2;refGene_gn=RTN3;refGene_tx=XM_019812579.2;refGene_gn=RTN3;refGene_tx=XM_019812580.2;refGene_gn=RTN3;refGene_tx=XM_019812581.2;refGene_gn=RTN3;refGene_tx=XM_019812582.2
D1	109140874	109140920
refGene_gn=RTN3;refGene_tx=XM_011287051.2;refGene_gn=RTN3;refGene_tx=XM_019812579.2;refGene_gn=RTN3;refGene_tx=XM_019812580.2;refGene_gn=RTN3;refGene_tx=XM_019812581.2;refGene_gn=RTN3;refGene_tx=XM_019812582.2
D1	109146516	109146980
refGene_gn=CD1H11orf95;refGene_tx=XM_023239978.1
D1	109147218	109147785
refGene_gn=CD1H11orf95;refGene_tx=XM_023239978.1
D1	109148035	109148333
refGene_gn=CD1H11orf95;refGene_tx=XM_023239978.1
D1	109148334	109148423
refGene_gn=CD1H11orf95;refGene_tx=XM_023239978.1
D1	109148982	109149480
refGene_gn=CD1H11orf95;refGene_tx=XM_023239978.1
D1	109151685	109151824
refGene_gn=CD1H11orf95;refGene_tx=XM_023239978.1
D1	109209321	109209448
refGene_gn=CD1H11orf84;refGene_tx=XM_023239981.1;refGene_gn=CD1H11orf84;refGene_tx=XM_023239982.1;refGene_gn=CD1H11orf84;refGene_tx=XM_023239983.1;refGene_gn=CD1H11orf84;refGene_tx=XM_023239984.1
D1	109212906	109213236
refGene_gn=CD1H11orf84;refGene_tx=XM_023239981.1;refGene_gn=CD1H11orf84;refGene_tx=XM_023239982.1;refGene_gn=CD1H11orf84;refGene_tx=XM_023239983.1;refGene_gn=CD1H11orf84;refGene_tx=XM_023239984.1
D1	109213328	109213478
refGene_gn=CD1H11orf84;refGene_tx=XM_023239981.1;refGene_gn=CD1H11orf84;refGene_tx=XM_023239982.1;refGene_gn=CD1H11orf84;refGene_tx=XM_02323

9983.1;refGene_gn=CD1H11orf84;refGene_tx=XM_023239984.1
D1 109213739 109213948
refGene_gn=CD1H11orf84;refGene_tx=XM_023239981.1;refGene_gn=CD1H11orf8
4;refGene_tx=XM_023239982.1;refGene_gn=CD1H11orf84;refGene_tx=XM_02323
9983.1;refGene_gn=CD1H11orf84;refGene_tx=XM_023239984.1
D1 109214041 109214242
refGene_gn=CD1H11orf84;refGene_tx=XM_023239981.1;refGene_gn=CD1H11orf8
4;refGene_tx=XM_023239982.1;refGene_gn=CD1H11orf84;refGene_tx=XM_02323
9983.1;refGene_gn=CD1H11orf84;refGene_tx=XM_023239984.1
D1 109215804 109215815
refGene_gn=CD1H11orf84;refGene_tx=XM_023239983.1;refGene_gn=CD1H11orf8
4;refGene_tx=XM_023239984.1
D1 109220389 109220893
refGene_gn=CD1H11orf84;refGene_tx=XM_023239981.1;refGene_gn=CD1H11orf8
4;refGene_tx=XM_023239982.1
D1 109232918 109232972
refGene_gn=MARK2;refGene_tx=XM_006937449.4
D1 109274279 109274459
refGene_gn=MARK2;refGene_tx=XM_006937449.4;refGene_gn=MARK2;refGene_tx
=XM_019812587.1;refGene_gn=MARK2;refGene_tx=XM_019812590.2;refGene_gn=
MARK2;refGene_tx=XM_019812591.1;refGene_gn=MARK2;refGene_tx=XM_0198125
92.1
D1 109274685 109274739
refGene_gn=MARK2;refGene_tx=XM_006937449.4;refGene_gn=MARK2;refGene_tx
=XM_019812587.1;refGene_gn=MARK2;refGene_tx=XM_019812590.2;refGene_gn=
MARK2;refGene_tx=XM_019812591.1;refGene_gn=MARK2;refGene_tx=XM_0198125
92.1
D1 109277552 109277601
refGene_gn=MARK2;refGene_tx=XM_006937449.4;refGene_gn=MARK2;refGene_tx
=XM_019812587.1;refGene_gn=MARK2;refGene_tx=XM_019812590.2;refGene_gn=
MARK2;refGene_tx=XM_019812591.1;refGene_gn=MARK2;refGene_tx=XM_0198125
92.1
D1 109277923 109277989
refGene_gn=MARK2;refGene_tx=XM_006937449.4;refGene_gn=MARK2;refGene_tx
=XM_019812587.1;refGene_gn=MARK2;refGene_tx=XM_019812590.2;refGene_gn=
MARK2;refGene_tx=XM_019812591.1;refGene_gn=MARK2;refGene_tx=XM_0198125
92.1
D1 109278077 109278148
refGene_gn=MARK2;refGene_tx=XM_006937449.4;refGene_gn=MARK2;refGene_tx
=XM_019812587.1;refGene_gn=MARK2;refGene_tx=XM_019812590.2;refGene_gn=
MARK2;refGene_tx=XM_019812591.1;refGene_gn=MARK2;refGene_tx=XM_0198125
92.1
D1 109278358 109278415
refGene_gn=MARK2;refGene_tx=XM_006937449.4;refGene_gn=MARK2;refGene_tx
=XM_019812587.1;refGene_gn=MARK2;refGene_tx=XM_019812590.2;refGene_gn=
MARK2;refGene_tx=XM_019812591.1;refGene_gn=MARK2;refGene_tx=XM_0198125
92.1
D1 109279142 109279379
refGene_gn=MARK2;refGene_tx=XM_006937449.4;refGene_gn=MARK2;refGene_tx
=XM_019812587.1;refGene_gn=MARK2;refGene_tx=XM_019812590.2;refGene_gn=

MARK2;refGene_tx=XM_019812591.1;refGene_gn=MARK2;refGene_tx=XM_0198125
92.1
D1 109280381 109280501
refGene_gn=MARK2;refGene_tx=XM_006937449.4;refGene_gn=MARK2;refGene_tx
=XM_019812587.1;refGene_gn=MARK2;refGene_tx=XM_019812590.2;refGene_gn=
MARK2;refGene_tx=XM_019812591.1;refGene_gn=MARK2;refGene_tx=XM_0198125
92.1
D1 109280608 109280708
refGene_gn=MARK2;refGene_tx=XM_006937449.4;refGene_gn=MARK2;refGene_tx
=XM_019812587.1;refGene_gn=MARK2;refGene_tx=XM_019812590.2;refGene_gn=
MARK2;refGene_tx=XM_019812591.1;refGene_gn=MARK2;refGene_tx=XM_0198125
92.1
D1 109280785 109280898
refGene_gn=MARK2;refGene_tx=XM_006937449.4;refGene_gn=MARK2;refGene_tx
=XM_019812587.1;refGene_gn=MARK2;refGene_tx=XM_019812590.2;refGene_gn=
MARK2;refGene_tx=XM_019812591.1;refGene_gn=MARK2;refGene_tx=XM_0198125
92.1
D1 109281883 109282016
refGene_gn=MARK2;refGene_tx=XM_006937449.4;refGene_gn=MARK2;refGene_tx
=XM_019812587.1;refGene_gn=MARK2;refGene_tx=XM_019812590.2;refGene_gn=
MARK2;refGene_tx=XM_019812591.1;refGene_gn=MARK2;refGene_tx=XM_0198125
92.1
D1 109282317 109282496
refGene_gn=MARK2;refGene_tx=XM_006937449.4;refGene_gn=MARK2;refGene_tx
=XM_019812587.1;refGene_gn=MARK2;refGene_tx=XM_019812590.2;refGene_gn=
MARK2;refGene_tx=XM_019812591.1;refGene_gn=MARK2;refGene_tx=XM_0198125
92.1
D1 109282705 109282803
refGene_gn=MARK2;refGene_tx=XM_006937449.4;refGene_gn=MARK2;refGene_tx
=XM_019812587.1;refGene_gn=MARK2;refGene_tx=XM_019812590.2;refGene_gn=
MARK2;refGene_tx=XM_019812591.1;refGene_gn=MARK2;refGene_tx=XM_0198125
92.1
D1 109283816 109283978
refGene_gn=MARK2;refGene_tx=XM_006937449.4;refGene_gn=MARK2;refGene_tx
=XM_019812587.1;refGene_gn=MARK2;refGene_tx=XM_019812590.2
D1 109284632 109284890
refGene_gn=MARK2;refGene_tx=XM_006937449.4;refGene_gn=MARK2;refGene_tx
=XM_019812587.1;refGene_gn=MARK2;refGene_tx=XM_019812590.2;refGene_gn=
MARK2;refGene_tx=XM_019812591.1;refGene_gn=MARK2;refGene_tx=XM_0198125
92.1
D1 109288014 109288059
refGene_gn=MARK2;refGene_tx=XM_006937449.4;refGene_gn=MARK2;refGene_tx
=XM_019812587.1;refGene_gn=MARK2;refGene_tx=XM_019812590.2;refGene_gn=
MARK2;refGene_tx=XM_019812591.1;refGene_gn=MARK2;refGene_tx=XM_0198125
92.1
D1 109288711 109289072
refGene_gn=MARK2;refGene_tx=XM_006937449.4;refGene_gn=MARK2;refGene_tx
=XM_019812587.1;refGene_gn=MARK2;refGene_tx=XM_019812590.2;refGene_gn=
MARK2;refGene_tx=XM_019812591.1;refGene_gn=MARK2;refGene_tx=XM_0198125
92.1

D1 109291813 109292128
refGene_gn=RCOR2;refGene_tx=XM_023239986.1
D1 109292238 109292468
refGene_gn=RCOR2;refGene_tx=XM_023239986.1
D1 109292619 109292677
refGene_gn=RCOR2;refGene_tx=XM_023239986.1
D1 109292830 109292908
refGene_gn=RCOR2;refGene_tx=XM_023239986.1
D1 109293265 109293481
refGene_gn=RCOR2;refGene_tx=XM_023239986.1
D1 109293582 109293652
refGene_gn=RCOR2;refGene_tx=XM_023239986.1
D1 109293743 109293868
refGene_gn=RCOR2;refGene_tx=XM_023239986.1
D1 109293977 109294139
refGene_gn=RCOR2;refGene_tx=XM_023239986.1
D1 109294228 109294281
refGene_gn=RCOR2;refGene_tx=XM_023239986.1
D1 109294503 109294584
refGene_gn=RCOR2;refGene_tx=XM_023239986.1
D1 109294882 109294939
refGene_gn=RCOR2;refGene_tx=XM_023239986.1
D1 109295682 109295809
refGene_gn=RCOR2;refGene_tx=XM_023239986.1
D1 109316241 109316247
refGene_gn=NAA40;refGene_tx=XM_023239987.1
D1 109323363 109323459
refGene_gn=NAA40;refGene_tx=XM_023239987.1
D1 109324286 109324450
refGene_gn=NAA40;refGene_tx=XM_019812594.2;refGene_gn=NAA40;refGene_tx=XM_019812595.1;refGene_gn=NAA40;refGene_tx=XM_023239987.1
D1 109327634 109327730
refGene_gn=NAA40;refGene_tx=XM_019812594.2;refGene_gn=NAA40;refGene_tx=XM_019812595.1;refGene_gn=NAA40;refGene_tx=XM_023239987.1
D1 109327803 109327962
refGene_gn=NAA40;refGene_tx=XM_019812594.2;refGene_gn=NAA40;refGene_tx=XM_019812595.1;refGene_gn=NAA40;refGene_tx=XM_023239987.1
D1 109328134 109328218
refGene_gn=NAA40;refGene_tx=XM_019812594.2;refGene_gn=NAA40;refGene_tx=XM_019812595.1;refGene_gn=NAA40;refGene_tx=XM_023239987.1
D1 109328684 109328762
refGene_gn=NAA40;refGene_tx=XM_019812594.2;refGene_gn=NAA40;refGene_tx=XM_019812595.1;refGene_gn=NAA40;refGene_tx=XM_023239987.1
D1 109329042 109329184
refGene_gn=NAA40;refGene_tx=XM_019812594.2;refGene_gn=NAA40;refGene_tx=XM_019812595.1;refGene_gn=NAA40;refGene_tx=XM_023239987.1
D1 109340539 109340653
refGene_gn=LOC101096603;refGene_tx=XM_003993548.4
D1 109341974 109342070
refGene_gn=LOC101096603;refGene_tx=XM_003993548.4

D1 109347895 109347953
refGene_gn=OTUB1;refGene_tx=XM_011286981.3
D1 109349647 109349709
refGene_gn=OTUB1;refGene_tx=XM_011286981.3
D1 109349984 109350083
refGene_gn=OTUB1;refGene_tx=XM_011286981.3
D1 109353966 109354085
refGene_gn=OTUB1;refGene_tx=XM_011286981.3
D1 109354325 109354410
refGene_gn=OTUB1;refGene_tx=XM_011286981.3
D1 109354529 109354724
refGene_gn=OTUB1;refGene_tx=XM_011286981.3
D1 109354840 109355038
refGene_gn=OTUB1;refGene_tx=XM_011286981.3
D1 109356193 109356231
refGene_gn=MACROD1;refGene_tx=XM_023239988.1
D1 109356519 109356601
refGene_gn=MACROD1;refGene_tx=XM_023239988.1
D1 109356921 109356995
refGene_gn=MACROD1;refGene_tx=XM_023239988.1
D1 109357216 109357247
refGene_gn=MACROD1;refGene_tx=XM_023239988.1
D1 109357380 109357502
refGene_gn=MACROD1;refGene_tx=XM_023239988.1
D1 109357939 109358056
refGene_gn=MACROD1;refGene_tx=XM_023239988.1
D1 109370612 109370642
refGene_gn=MACROD1;refGene_tx=XM_023239988.1
D1 109456000 109458025
refGene_gn=FLRT1;refGene_tx=XM_003993550.5;refGene_gn=FLRT1;refGene_tx
=XM_019812597.2;refGene_gn=FLRT1;refGene_tx=XM_019812600.2
D1 109487219 109487336
refGene_gn=MACROD1;refGene_tx=XM_023239988.1
D1 109487986 109488088
refGene_gn=MACROD1;refGene_tx=XM_023239988.1
D1 109499421 109499509
refGene_gn=MACROD1;refGene_tx=XM_023239988.1
D1 109499511 109499595
refGene_gn=MACROD1;refGene_tx=XM_023239988.1
D1 109499596 109499725
refGene_gn=MACROD1;refGene_tx=XM_023239988.1
D1 109515653 109515662
refGene_gn=STIP1;refGene_tx=XM_003993553.3
D1 109518864 109519074
refGene_gn=STIP1;refGene_tx=XM_003993553.3
D1 109519563 109519705
refGene_gn=STIP1;refGene_tx=XM_003993553.3
D1 109520472 109520614
refGene_gn=STIP1;refGene_tx=XM_003993553.3
D1 109521571 109521740

refGene_gn=STIP1;refGene_tx=XM_003993553.3
D1 109522910 109523037
refGene_gn=STIP1;refGene_tx=XM_003993553.3
D1 109523129 109523232
refGene_gn=STIP1;refGene_tx=XM_003993553.3
D1 109523492 109523613
refGene_gn=STIP1;refGene_tx=XM_003993553.3
D1 109524283 109524380
refGene_gn=STIP1;refGene_tx=XM_003993553.3
D1 109524731 109524856
refGene_gn=STIP1;refGene_tx=XM_003993553.3
D1 109527052 109527089
refGene_gn=STIP1;refGene_tx=XM_003993553.3
D1 109527294 109527398
refGene_gn=STIP1;refGene_tx=XM_003993553.3
D1 109527607 109527780
refGene_gn=STIP1;refGene_tx=XM_003993553.3
D1 109528158 109528231
refGene_gn=STIP1;refGene_tx=XM_003993553.3
D1 109531688 109531848
refGene_gn=FERMT3;refGene_tx=XM_003993551.5
D1 109534677 109534911
refGene_gn=FERMT3;refGene_tx=XM_003993551.5
D1 109535140 109535260
refGene_gn=FERMT3;refGene_tx=XM_003993551.5
D1 109535347 109535516
refGene_gn=FERMT3;refGene_tx=XM_003993551.5
D1 109535739 109535842
refGene_gn=FERMT3;refGene_tx=XM_003993551.5
D1 109546170 109546278
refGene_gn=FERMT3;refGene_tx=XM_003993551.5
D1 109546439 109546574
refGene_gn=FERMT3;refGene_tx=XM_003993551.5
D1 109546661 109546711
refGene_gn=FERMT3;refGene_tx=XM_003993551.5
D1 109546803 109546928
refGene_gn=FERMT3;refGene_tx=XM_003993551.5
D1 109547128 109547235
refGene_gn=FERMT3;refGene_tx=XM_003993551.5
D1 109547361 109547595
refGene_gn=FERMT3;refGene_tx=XM_003993551.5
D1 109547918 109548043
refGene_gn=FERMT3;refGene_tx=XM_003993551.5
D1 109548575 109548717
refGene_gn=FERMT3;refGene_tx=XM_003993551.5
D1 109548875 109549055
refGene_gn=FERMT3;refGene_tx=XM_003993551.5
D1 109549419 109549511
refGene_gn=TRPT1;refGene_tx=XM_003993614.5;refGene_gn=TRPT1;refGene_tx
=XM_011286991.3;refGene_gn=TRPT1;refGene_tx=XM_019812617.2

D1	109549625	109549736
refGene_gn=TRPT1;refGene_tx=XM_003993614.5;refGene_gn=TRPT1;refGene_tx
=XM_011286991.3;refGene_gn=TRPT1;refGene_tx=XM_019812617.2
D1	109549811	109549868
refGene_gn=TRPT1;refGene_tx=XM_003993614.5;refGene_gn=TRPT1;refGene_tx
=XM_011286991.3;refGene_gn=TRPT1;refGene_tx=XM_019812617.2
D1	109550096	109550271
refGene_gn=TRPT1;refGene_tx=XM_003993614.5;refGene_gn=TRPT1;refGene_tx
=XM_011286991.3;refGene_gn=TRPT1;refGene_tx=XM_019812617.2
D1	109550357	109550527
refGene_gn=TRPT1;refGene_tx=XM_003993614.5;refGene_gn=TRPT1;refGene_tx
=XM_011286991.3;refGene_gn=TRPT1;refGene_tx=XM_019812617.2
D1	109551373	109551455
refGene_gn=TRPT1;refGene_tx=XM_003993614.5;refGene_gn=TRPT1;refGene_tx
=XM_011286991.3;refGene_gn=TRPT1;refGene_tx=XM_019812617.2
D1	109551708	109551783
refGene_gn=TRPT1;refGene_tx=XM_003993614.5;refGene_gn=TRPT1;refGene_tx
=XM_011286991.3
D1	109552642	109553122
refGene_gn=NUDT22;refGene_tx=XM_006937455.2;refGene_gn=NUDT22;refGene_
tx=XM_006937456.3;refGene_gn=NUDT22;refGene_tx=XM_006937458.2;refGene_
gn=NUDT22;refGene_tx=XM_006937459.1;refGene_gn=NUDT22;refGene_tx=XM_01
1279829.2;refGene_gn=NUDT22;refGene_tx=XM_019824699.2;refGene_gn=NUDT2
2;refGene_tx=XM_019824700.1;refGene_gn=NUDT22;refGene_tx=XM_023239989.
1
D1	109553480	109553579
refGene_gn=NUDT22;refGene_tx=XM_006937455.2;refGene_gn=NUDT22;refGene_
tx=XM_006937456.3;refGene_gn=NUDT22;refGene_tx=XM_006937458.2;refGene_
gn=NUDT22;refGene_tx=XM_006937459.1;refGene_gn=NUDT22;refGene_tx=XM_01
1279829.2;refGene_gn=NUDT22;refGene_tx=XM_019824699.2;refGene_gn=NUDT2
2;refGene_tx=XM_019824700.1;refGene_gn=NUDT22;refGene_tx=XM_023239989.
1
D1	109554076	109554174
refGene_gn=NUDT22;refGene_tx=XM_006937455.2;refGene_gn=NUDT22;refGene_
tx=XM_006937456.3;refGene_gn=NUDT22;refGene_tx=XM_006937458.2;refGene_
gn=NUDT22;refGene_tx=XM_006937459.1;refGene_gn=NUDT22;refGene_tx=XM_01
1279829.2;refGene_gn=NUDT22;refGene_tx=XM_019824699.2;refGene_gn=NUDT2
2;refGene_tx=XM_019824700.1;refGene_gn=NUDT22;refGene_tx=XM_023239989.
1
D1	109554312	109554406
refGene_gn=NUDT22;refGene_tx=XM_006937455.2;refGene_gn=NUDT22;refGene_
tx=XM_006937456.3;refGene_gn=NUDT22;refGene_tx=XM_006937458.2;refGene_
gn=NUDT22;refGene_tx=XM_006937459.1;refGene_gn=NUDT22;refGene_tx=XM_01
1279829.2;refGene_gn=NUDT22;refGene_tx=XM_019824699.2;refGene_gn=NUDT2
2;refGene_tx=XM_019824700.1;refGene_gn=NUDT22;refGene_tx=XM_023239989.
1
D1	109554721	109554832
refGene_gn=NUDT22;refGene_tx=XM_006937455.2;refGene_gn=NUDT22;refGene_
tx=XM_006937456.3;refGene_gn=NUDT22;refGene_tx=XM_006937458.2;refGene_
gn=NUDT22;refGene_tx=XM_006937459.1;refGene_gn=NUDT22;refGene_tx=XM_01

1279829.2;refGene_gn=NUDT22;refGene_tx=XM_019824699.2;refGene_gn=NUDT2
2;refGene_tx=XM_019824700.1;refGene_gn=NUDT22;refGene_tx=XM_023239989.
1
D1 109555689 109555814
refGene_gn=DNAJC4;refGene_tx=XM_011286993.2;refGene_gn=DNAJC4;refGene_
tx=XM_019824701.2
D1 109556706 109556800
refGene_gn=DNAJC4;refGene_tx=XM_011286993.2;refGene_gn=DNAJC4;refGene_
tx=XM_019824701.2
D1 109557294 109557476
refGene_gn=DNAJC4;refGene_tx=XM_011286993.2;refGene_gn=DNAJC4;refGene_
tx=XM_019824701.2
D1 109557577 109557739
refGene_gn=DNAJC4;refGene_tx=XM_011286993.2;refGene_gn=DNAJC4;refGene_
tx=XM_019824701.2
D1 109558468 109558555
refGene_gn=DNAJC4;refGene_tx=XM_011286993.2;refGene_gn=DNAJC4;refGene_
tx=XM_019824701.2
D1 109558648 109558778
refGene_gn=DNAJC4;refGene_tx=XM_011286993.2;refGene_gn=DNAJC4;refGene_
tx=XM_019824701.2
D1 109559446 109559506
refGene_gn=VEGFB;refGene_tx=XM_023239990.1;refGene_gn=VEGFB;refGene_tx
=XM_023239991.1;refGene_gn=VEGFB;refGene_tx=XM_023239992.1;refGene_gn=
VEGFB;refGene_tx=XM_023239993.1
D1 109559832 109559922
refGene_gn=VEGFB;refGene_tx=XM_023239990.1;refGene_gn=VEGFB;refGene_tx
=XM_023239991.1
D1 109560077 109560120
refGene_gn=VEGFB;refGene_tx=XM_023239990.1;refGene_gn=VEGFB;refGene_tx
=XM_023239991.1;refGene_gn=VEGFB;refGene_tx=XM_023239992.1;refGene_gn=
VEGFB;refGene_tx=XM_023239993.1
D1 109560428 109560625
refGene_gn=VEGFB;refGene_tx=XM_023239990.1;refGene_gn=VEGFB;refGene_tx
=XM_023239991.1;refGene_gn=VEGFB;refGene_tx=XM_023239992.1;refGene_gn=
VEGFB;refGene_tx=XM_023239993.1
D1 109560804 109560878
refGene_gn=VEGFB;refGene_tx=XM_023239990.1;refGene_gn=VEGFB;refGene_tx
=XM_023239991.1;refGene_gn=VEGFB;refGene_tx=XM_023239992.1;refGene_gn=
VEGFB;refGene_tx=XM_023239993.1
D1 109561073 109561109
refGene_gn=VEGFB;refGene_tx=XM_023239990.1;refGene_gn=VEGFB;refGene_tx
=XM_023239991.1;refGene_gn=VEGFB;refGene_tx=XM_023239992.1;refGene_gn=
VEGFB;refGene_tx=XM_023239993.1
D1 109561355 109561591
refGene_gn=VEGFB;refGene_tx=XM_023239990.1;refGene_gn=VEGFB;refGene_tx
=XM_023239991.1;refGene_gn=VEGFB;refGene_tx=XM_023239992.1;refGene_gn=
VEGFB;refGene_tx=XM_023239993.1
D1 109562155 109562177
refGene_gn=VEGFB;refGene_tx=XM_023239991.1;refGene_gn=VEGFB;refGene_tx

=XM_023239993.1
D1	109564490	109564552
refGene_gn=FKBP2;refGene_tx=XM_003993555.5
D1	109565334	109565462
refGene_gn=FKBP2;refGene_tx=XM_006937462.4
D1	109565799	109565968
refGene_gn=FKBP2;refGene_tx=XM_003993555.5;refGene_gn=FKBP2;refGene_tx
=XM_003993556.5;refGene_gn=FKBP2;refGene_tx=XM_006937461.4;refGene_gn=
FKBP2;refGene_tx=XM_006937462.4;refGene_gn=FKBP2;refGene_tx=XM_0069374
63.4
D1	109566297	109566410
refGene_gn=FKBP2;refGene_tx=XM_003993555.5;refGene_gn=FKBP2;refGene_tx
=XM_003993556.5;refGene_gn=FKBP2;refGene_tx=XM_006937461.4;refGene_gn=
FKBP2;refGene_tx=XM_006937462.4;refGene_gn=FKBP2;refGene_tx=XM_0069374
63.4
D1	109566531	109566578
refGene_gn=FKBP2;refGene_tx=XM_003993555.5;refGene_gn=FKBP2;refGene_tx
=XM_003993556.5;refGene_gn=FKBP2;refGene_tx=XM_006937461.4;refGene_gn=
FKBP2;refGene_tx=XM_006937462.4;refGene_gn=FKBP2;refGene_tx=XM_0069374
63.4
D1	109566921	109566957
refGene_gn=FKBP2;refGene_tx=XM_003993555.5;refGene_gn=FKBP2;refGene_tx
=XM_003993556.5;refGene_gn=FKBP2;refGene_tx=XM_006937461.4;refGene_gn=
FKBP2;refGene_tx=XM_006937462.4;refGene_gn=FKBP2;refGene_tx=XM_0069374
63.4
D1	109567046	109567108
refGene_gn=FKBP2;refGene_tx=XM_003993555.5;refGene_gn=FKBP2;refGene_tx
=XM_003993556.5;refGene_gn=FKBP2;refGene_tx=XM_006937461.4;refGene_gn=
FKBP2;refGene_tx=XM_006937462.4;refGene_gn=FKBP2;refGene_tx=XM_0069374
63.4
D1	109567865	109567934
refGene_gn=PPP1R14B;refGene_tx=XM_011287053.2
D1	109568029	109568062
refGene_gn=PPP1R14B;refGene_tx=XM_011287053.2
D1	109568382	109568397
refGene_gn=PPP1R14B;refGene_tx=XM_011287053.2
D1	109568398	109568467
refGene_gn=PPP1R14B;refGene_tx=XM_011287053.2
D1	109569573	109569855
refGene_gn=PPP1R14B;refGene_tx=XM_011287053.2
D1	109569856	109569960
refGene_gn=PPP1R14B;refGene_tx=XM_011287053.2
D1	109569962	109569999
refGene_gn=PPP1R14B;refGene_tx=XM_011287053.2
D1	109575328	109575427
refGene_gn=PLCB3;refGene_tx=XM_023239994.1;refGene_gn=PLCB3;refGene_tx
=XM_023239995.1
D1	109577918	109577996
refGene_gn=PLCB3;refGene_tx=XM_023239994.1;refGene_gn=PLCB3;refGene_tx
=XM_023239995.1

D1	109578265	109578334
refGene_gn=PLCB3;refGene_tx=XM_023239994.1;refGene_gn=PLCB3;refGene_tx=XM_023239995.1
D1	109578415	109578556
refGene_gn=PLCB3;refGene_tx=XM_023239994.1;refGene_gn=PLCB3;refGene_tx=XM_023239995.1
D1	109578791	109578871
refGene_gn=PLCB3;refGene_tx=XM_023239994.1;refGene_gn=PLCB3;refGene_tx=XM_023239995.1
D1	109578952	109579006
refGene_gn=PLCB3;refGene_tx=XM_023239994.1;refGene_gn=PLCB3;refGene_tx=XM_023239995.1
D1	109579095	109579171
refGene_gn=PLCB3;refGene_tx=XM_023239994.1;refGene_gn=PLCB3;refGene_tx=XM_023239995.1
D1	109579275	109579376
refGene_gn=PLCB3;refGene_tx=XM_023239994.1;refGene_gn=PLCB3;refGene_tx=XM_023239995.1
D1	109579893	109580060
refGene_gn=PLCB3;refGene_tx=XM_023239994.1;refGene_gn=PLCB3;refGene_tx=XM_023239995.1
D1	109580145	109580292
refGene_gn=PLCB3;refGene_tx=XM_023239994.1;refGene_gn=PLCB3;refGene_tx=XM_023239995.1
D1	109581020	109581261
refGene_gn=PLCB3;refGene_tx=XM_023239994.1;refGene_gn=PLCB3;refGene_tx=XM_023239995.1
D1	109581438	109581523
refGene_gn=PLCB3;refGene_tx=XM_023239994.1;refGene_gn=PLCB3;refGene_tx=XM_023239995.1
D1	109581601	109581788
refGene_gn=PLCB3;refGene_tx=XM_023239994.1;refGene_gn=PLCB3;refGene_tx=XM_023239995.1
D1	109582590	109582796
refGene_gn=PLCB3;refGene_tx=XM_023239994.1;refGene_gn=PLCB3;refGene_tx=XM_023239995.1
D1	109583669	109583766
refGene_gn=PLCB3;refGene_tx=XM_023239994.1;refGene_gn=PLCB3;refGene_tx=XM_023239995.1
D1	109583858	109583943
refGene_gn=PLCB3;refGene_tx=XM_023239994.1;refGene_gn=PLCB3;refGene_tx=XM_023239995.1
D1	109584215	109584340
refGene_gn=PLCB3;refGene_tx=XM_023239994.1;refGene_gn=PLCB3;refGene_tx=XM_023239995.1
D1	109584624	109584779
refGene_gn=PLCB3;refGene_tx=XM_023239994.1;refGene_gn=PLCB3;refGene_tx=XM_023239995.1
D1	109584859	109585021
refGene_gn=PLCB3;refGene_tx=XM_023239994.1;refGene_gn=PLCB3;refGene_tx

=XM_023239995.1
D1 109585661 109585761
refGene_gn=PLCB3;refGene_tx=XM_023239994.1;refGene_gn=PLCB3;refGene_tx=XM_023239995.1
D1 109585849 109585954
refGene_gn=PLCB3;refGene_tx=XM_023239994.1;refGene_gn=PLCB3;refGene_tx=XM_023239995.1
D1 109586175 109586267
refGene_gn=PLCB3;refGene_tx=XM_023239994.1;refGene_gn=PLCB3;refGene_tx=XM_023239995.1
D1 109587499 109587644
refGene_gn=PLCB3;refGene_tx=XM_023239994.1;refGene_gn=PLCB3;refGene_tx=XM_023239995.1
D1 109587734 109587770
refGene_gn=PLCB3;refGene_tx=XM_023239994.1;refGene_gn=PLCB3;refGene_tx=XM_023239995.1
D1 109587866 109588059
refGene_gn=PLCB3;refGene_tx=XM_023239994.1;refGene_gn=PLCB3;refGene_tx=XM_023239995.1
D1 109588357 109588565
refGene_gn=PLCB3;refGene_tx=XM_023239994.1;refGene_gn=PLCB3;refGene_tx=XM_023239995.1
D1 109588661 109588738
refGene_gn=PLCB3;refGene_tx=XM_023239994.1;refGene_gn=PLCB3;refGene_tx=XM_023239995.1
D1 109588856 109588946
refGene_gn=PLCB3;refGene_tx=XM_023239994.1;refGene_gn=PLCB3;refGene_tx=XM_023239995.1
D1 109589029 109589087
refGene_gn=PLCB3;refGene_tx=XM_023239994.1;refGene_gn=PLCB3;refGene_tx=XM_023239995.1
D1 109589459 109589546
refGene_gn=PLCB3;refGene_tx=XM_023239994.1;refGene_gn=PLCB3;refGene_tx=XM_023239995.1
D1 109589634 109589838
refGene_gn=PLCB3;refGene_tx=XM_023239994.1;refGene_gn=PLCB3;refGene_tx=XM_023239995.1
D1 109592779 109592905
refGene_gn=BAD;refGene_tx=XM_003993558.3;refGene_gn=BAD;refGene_tx=XM_006937465.3
D1 109593450 109593644
refGene_gn=BAD;refGene_tx=XM_003993558.3;refGene_gn=BAD;refGene_tx=XM_006937465.3
D1 109601891 109602078
refGene_gn=BAD;refGene_tx=XM_003993558.3;refGene_gn=BAD;refGene_tx=XM_006937465.3
D1 109604339 109604696
refGene_gn=GPR137;refGene_tx=XM_006937466.4;refGene_gn=GPR137;refGene_tx=XM_023239996.1
D1 109604786 109604836

refGene_gn=GPR137;refGene_tx=XM_006937466.4;refGene_gn=GPR137;refGene_tx=XM_023239996.1
D1 109605563 109605789
refGene_gn=GPR137;refGene_tx=XM_006937466.4;refGene_gn=GPR137;refGene_tx=XM_023239996.1
D1 109605908 109606058
refGene_gn=GPR137;refGene_tx=XM_006937466.4;refGene_gn=GPR137;refGene_tx=XM_023239996.1
D1 109606183 109606312
refGene_gn=GPR137;refGene_tx=XM_006937466.4;refGene_gn=GPR137;refGene_tx=XM_023239996.1
D1 109606418 109606537
refGene_gn=GPR137;refGene_tx=XM_006937466.4;refGene_gn=GPR137;refGene_tx=XM_023239996.1
D1 109606859 109607019
refGene_gn=GPR137;refGene_tx=XM_006937466.4;refGene_gn=GPR137;refGene_tx=XM_023239996.1
D1 109609187 109609188
refGene_gn=KCNK4;refGene_tx=XM_023239998.1
D1 109610346 109610727
refGene_gn=KCNK4;refGene_tx=XM_003993561.5;refGene_gn=KCNK4;refGene_tx=XM_019812622.2;refGene_gn=KCNK4;refGene_tx=XM_023239998.1
D1 109613626 109613750
refGene_gn=KCNK4;refGene_tx=XM_003993561.5;refGene_gn=KCNK4;refGene_tx=XM_019812622.2;refGene_gn=KCNK4;refGene_tx=XM_023239998.1
D1 109613898 109614059
refGene_gn=KCNK4;refGene_tx=XM_003993561.5;refGene_gn=KCNK4;refGene_tx=XM_019812622.2;refGene_gn=KCNK4;refGene_tx=XM_023239998.1
D1 109614257 109614444
refGene_gn=KCNK4;refGene_tx=XM_003993561.5;refGene_gn=KCNK4;refGene_tx=XM_019812622.2;refGene_gn=KCNK4;refGene_tx=XM_023239998.1
D1 109614970 109615110
refGene_gn=KCNK4;refGene_tx=XM_003993561.5;refGene_gn=KCNK4;refGene_tx=XM_019812622.2;refGene_gn=KCNK4;refGene_tx=XM_023239998.1
D1 109615814 109616219
refGene_gn=KCNK4;refGene_tx=XM_003993561.5;refGene_gn=KCNK4;refGene_tx=XM_019812622.2;refGene_gn=KCNK4;refGene_tx=XM_023239998.1
D1 109616977 109616998
refGene_gn=CATSPERZ;refGene_tx=XM_003993620.4
D1 109617244 109617587
refGene_gn=CATSPERZ;refGene_tx=XM_003993620.4
D1 109618319 109618399
refGene_gn=CATSPERZ;refGene_tx=XM_003993620.4
D1 109618615 109618682
refGene_gn=CATSPERZ;refGene_tx=XM_003993620.4
D1 109619435 109619545
refGene_gn=CATSPERZ;refGene_tx=XM_003993620.4
D1 109621540 109621600
refGene_gn=ESRRA;refGene_tx=XM_019812624.2
D1 109622091 109622428

refGene_gn=ESRRA;refGene_tx=XM_011286995.3;refGene_gn=ESRRA;refGene_tx=XM_019812624.2;refGene_gn=ESRRA;refGene_tx=XM_019812625.2;refGene_gn=ESRRA;refGene_tx=XM_023239997.1
D1 109625412 109625529
refGene_gn=ESRRA;refGene_tx=XM_011286995.3;refGene_gn=ESRRA;refGene_tx=XM_019812624.2;refGene_gn=ESRRA;refGene_tx=XM_019812625.2;refGene_gn=ESRRA;refGene_tx=XM_023239997.1
D1 109625680 109625809
refGene_gn=ESRRA;refGene_tx=XM_011286995.3;refGene_gn=ESRRA;refGene_tx=XM_019812624.2;refGene_gn=ESRRA;refGene_tx=XM_019812625.2;refGene_gn=ESRRA;refGene_tx=XM_023239997.1
D1 109626224 109626392
refGene_gn=ESRRA;refGene_tx=XM_011286995.3;refGene_gn=ESRRA;refGene_tx=XM_019812624.2;refGene_gn=ESRRA;refGene_tx=XM_019812625.2;refGene_gn=ESRRA;refGene_tx=XM_023239997.1
D1 109626491 109626761
refGene_gn=ESRRA;refGene_tx=XM_011286995.3;refGene_gn=ESRRA;refGene_tx=XM_019812624.2;refGene_gn=ESRRA;refGene_tx=XM_019812625.2;refGene_gn=ESRRA;refGene_tx=XM_023239997.1
D1 109627204 109627464
refGene_gn=ESRRA;refGene_tx=XM_011286995.3;refGene_gn=ESRRA;refGene_tx=XM_019812624.2;refGene_gn=ESRRA;refGene_tx=XM_019812625.2;refGene_gn=ESRRA;refGene_tx=XM_023239997.1
D1 109628355 109628463
refGene_gn=TRMT112;refGene_tx=XM_003993564.5
D1 109628544 109628634
refGene_gn=TRMT112;refGene_tx=XM_003993564.5
D1 109628760 109628862
refGene_gn=TRMT112;refGene_tx=XM_003993564.5
D1 109628945 109629023
refGene_gn=TRMT112;refGene_tx=XM_003993564.5
D1 109629731 109629923
refGene_gn=PRDX5;refGene_tx=XM_006937467.4
D1 109631396 109631531
refGene_gn=PRDX5;refGene_tx=XM_006937467.4
D1 109632274 109632406
refGene_gn=PRDX5;refGene_tx=XM_006937467.4
D1 109632488 109632527
refGene_gn=PRDX5;refGene_tx=XM_006937467.4
D1 109632698 109632760
refGene_gn=PRDX5;refGene_tx=XM_006937467.4
D1 109632869 109632975
refGene_gn=PRDX5;refGene_tx=XM_006937467.4
D1 109644324 109644384
refGene_gn=CCDC88B;refGene_tx=XM_019812630.2
D1 109644694 109644837
refGene_gn=CCDC88B;refGene_tx=XM_019812630.2
D1 109644983 109645095
refGene_gn=CCDC88B;refGene_tx=XM_019812630.2
D1 109645175 109645245

refGene_gn=CCDC88B;refGene_tx=XM_019812630.2
D1 109645321 109645380
refGene_gn=CCDC88B;refGene_tx=XM_019812630.2
D1 109645476 109645560
refGene_gn=CCDC88B;refGene_tx=XM_019812630.2
D1 109645658 109645802
refGene_gn=CCDC88B;refGene_tx=XM_019812630.2
D1 109646010 109646156
refGene_gn=CCDC88B;refGene_tx=XM_019812630.2
D1 109646286 109646368
refGene_gn=CCDC88B;refGene_tx=XM_019812630.2
D1 109646520 109646679
refGene_gn=CCDC88B;refGene_tx=XM_019812630.2
D1 109647142 109647289
refGene_gn=CCDC88B;refGene_tx=XM_019812630.2
D1 109647473 109647582
refGene_gn=CCDC88B;refGene_tx=XM_019812630.2
D1 109647782 109647919
refGene_gn=CCDC88B;refGene_tx=XM_019812630.2
D1 109648001 109649132
refGene_gn=CCDC88B;refGene_tx=XM_019812630.2
D1 109652500 109652628
refGene_gn=CCDC88B;refGene_tx=XM_019812630.2
D1 109652706 109652824
refGene_gn=CCDC88B;refGene_tx=XM_019812630.2
D1 109653526 109653622
refGene_gn=CCDC88B;refGene_tx=XM_019812630.2
D1 109653711 109653852
refGene_gn=CCDC88B;refGene_tx=XM_019812630.2
D1 109654457 109654714
refGene_gn=CCDC88B;refGene_tx=XM_019812630.2
D1 109655057 109655201
refGene_gn=CCDC88B;refGene_tx=XM_019812630.2
D1 109655394 109655581
refGene_gn=CCDC88B;refGene_tx=XM_019812630.2
D1 109655725 109655871
refGene_gn=CCDC88B;refGene_tx=XM_019812630.2
D1 109656139 109656238
refGene_gn=CCDC88B;refGene_tx=XM_019812630.2
D1 109656391 109656558
refGene_gn=CCDC88B;refGene_tx=XM_019812630.2
D1 109657116 109657169
refGene_gn=CCDC88B;refGene_tx=XM_019812630.2
D1 109657170 109657330
refGene_gn=CCDC88B;refGene_tx=XM_019812630.2
D1 109657474 109657543
refGene_gn=CCDC88B;refGene_tx=XM_019812630.2
D1 109658926 109658982
refGene_gn=CCDC88B;refGene_tx=XM_019812630.2
D1 109661151 109661206

refGene_gn=RPS6KA4;refGene_tx=XM_023239999.1
D1 109661324 109661396
refGene_gn=RPS6KA4;refGene_tx=XM_023239999.1
D1 109662111 109662330
refGene_gn=RPS6KA4;refGene_tx=XM_023239999.1
D1 109662435 109662551
refGene_gn=RPS6KA4;refGene_tx=XM_023239999.1
D1 109663094 109663202
refGene_gn=RPS6KA4;refGene_tx=XM_023239999.1
D1 109663429 109663510
refGene_gn=RPS6KA4;refGene_tx=XM_023239999.1
D1 109663589 109663693
refGene_gn=RPS6KA4;refGene_tx=XM_023239999.1
D1 109663785 109663936
refGene_gn=RPS6KA4;refGene_tx=XM_023239999.1
D1 109666632 109666797
refGene_gn=RPS6KA4;refGene_tx=XM_023239999.1
D1 109668488 109668617
refGene_gn=RPS6KA4;refGene_tx=XM_023239999.1
D1 109668788 109668922
refGene_gn=RPS6KA4;refGene_tx=XM_023239999.1
D1 109669019 109669113
refGene_gn=RPS6KA4;refGene_tx=XM_023239999.1
D1 109669583 109669757
refGene_gn=RPS6KA4;refGene_tx=XM_023239999.1
D1 109669838 109669967
refGene_gn=RPS6KA4;refGene_tx=XM_023239999.1
D1 109669968 109670034
refGene_gn=RPS6KA4;refGene_tx=XM_023239999.1
D1 109670389 109670549
refGene_gn=RPS6KA4;refGene_tx=XM_023239999.1
D1 109670774 109670938
refGene_gn=RPS6KA4;refGene_tx=XM_023239999.1
D1 109671192 109671387
refGene_gn=RPS6KA4;refGene_tx=XM_023239999.1
D1 109799071 109799473
refGene_gn=SLC22A11;refGene_tx=XM_003993623.3;refGene_gn=SLC22A11;refGene_tx=XM_019812632.1;refGene_gn=SLC22A11;refGene_tx=XM_019812633.1;refGene_gn=SLC22A11;refGene_tx=XM_019812634.1;refGene_gn=SLC22A11;refGene_tx=XM_019812635.1;refGene_gn=SLC22A11;refGene_tx=XM_019812636.1;refGene_gn=SLC22A11;refGene_tx=XM_019812637.1;refGene_gn=SLC22A11;refGene_tx=XM_019812638.2;refGene_gn=SLC22A11;refGene_tx=XM_019812639.1;refGene_gn=SLC22A11;refGene_tx=XM_019812640.1;refGene_gn=SLC22A11;refGene_tx=XM_019812641.1;refGene_gn=SLC22A11;refGene_tx=XM_019812642.1
D1 109800743 109800847
refGene_gn=SLC22A11;refGene_tx=XM_003993623.3;refGene_gn=SLC22A11;refGene_tx=XM_019812632.1;refGene_gn=SLC22A11;refGene_tx=XM_019812633.1;refGene_gn=SLC22A11;refGene_tx=XM_019812634.1;refGene_gn=SLC22A11;refGene_tx=XM_019812635.1;refGene_gn=SLC22A11;refGene_tx=XM_019812636.1;refGene_gn=SLC22A11;refGene_tx=XM_019812637.1;refGene_gn=SLC22A11;refGene_

tx=XM_019812638.2;refGene_gn=SLC22A11;refGene_tx=XM_019812639.1;refGene_gn=SLC22A11;refGene_tx=XM_019812640.1;refGene_gn=SLC22A11;refGene_tx=XM_019812641.1;refGene_gn=SLC22A11;refGene_tx=XM_019812642.1;refGene_gn=SLC22A11;refGene_tx=XM_019812643.1
D1 109805277 109805432
refGene_gn=SLC22A11;refGene_tx=XM_003993623.3;refGene_gn=SLC22A11;refGene_tx=XM_019812632.1;refGene_gn=SLC22A11;refGene_tx=XM_019812633.1;refGene_gn=SLC22A11;refGene_tx=XM_019812634.1;refGene_gn=SLC22A11;refGene_tx=XM_019812635.1;refGene_gn=SLC22A11;refGene_tx=XM_019812636.1;refGene_gn=SLC22A11;refGene_tx=XM_019812637.1;refGene_gn=SLC22A11;refGene_tx=XM_019812638.2;refGene_gn=SLC22A11;refGene_tx=XM_019812639.1;refGene_gn=SLC22A11;refGene_tx=XM_019812640.1;refGene_gn=SLC22A11;refGene_tx=XM_019812641.1;refGene_gn=SLC22A11;refGene_tx=XM_019812642.1;refGene_gn=SLC22A11;refGene_tx=XM_019812643.1
D1 109805532 109805701
refGene_gn=SLC22A11;refGene_tx=XM_003993623.3;refGene_gn=SLC22A11;refGene_tx=XM_019812632.1;refGene_gn=SLC22A11;refGene_tx=XM_019812633.1;refGene_gn=SLC22A11;refGene_tx=XM_019812634.1;refGene_gn=SLC22A11;refGene_tx=XM_019812635.1;refGene_gn=SLC22A11;refGene_tx=XM_019812636.1;refGene_gn=SLC22A11;refGene_tx=XM_019812637.1;refGene_gn=SLC22A11;refGene_tx=XM_019812638.2;refGene_gn=SLC22A11;refGene_tx=XM_019812639.1;refGene_gn=SLC22A11;refGene_tx=XM_019812640.1;refGene_gn=SLC22A11;refGene_tx=XM_019812641.1;refGene_gn=SLC22A11;refGene_tx=XM_019812642.1;refGene_gn=SLC22A11;refGene_tx=XM_019812643.1
D1 109806678 109806802
refGene_gn=SLC22A11;refGene_tx=XM_003993623.3;refGene_gn=SLC22A11;refGene_tx=XM_019812632.1;refGene_gn=SLC22A11;refGene_tx=XM_019812633.1;refGene_gn=SLC22A11;refGene_tx=XM_019812634.1;refGene_gn=SLC22A11;refGene_tx=XM_019812635.1;refGene_gn=SLC22A11;refGene_tx=XM_019812636.1;refGene_gn=SLC22A11;refGene_tx=XM_019812637.1;refGene_gn=SLC22A11;refGene_tx=XM_019812638.2;refGene_gn=SLC22A11;refGene_tx=XM_019812639.1;refGene_gn=SLC22A11;refGene_tx=XM_019812640.1;refGene_gn=SLC22A11;refGene_tx=XM_019812641.1;refGene_gn=SLC22A11;refGene_tx=XM_019812642.1;refGene_gn=SLC22A11;refGene_tx=XM_019812643.1
D1 109807745 109807861
refGene_gn=SLC22A11;refGene_tx=XM_003993623.3;refGene_gn=SLC22A11;refGene_tx=XM_019812632.1;refGene_gn=SLC22A11;refGene_tx=XM_019812633.1;refGene_gn=SLC22A11;refGene_tx=XM_019812634.1;refGene_gn=SLC22A11;refGene_tx=XM_019812635.1;refGene_gn=SLC22A11;refGene_tx=XM_019812636.1;refGene_gn=SLC22A11;refGene_tx=XM_019812637.1;refGene_gn=SLC22A11;refGene_tx=XM_019812638.2;refGene_gn=SLC22A11;refGene_tx=XM_019812639.1;refGene_gn=SLC22A11;refGene_tx=XM_019812640.1;refGene_gn=SLC22A11;refGene_tx=XM_019812641.1;refGene_gn=SLC22A11;refGene_tx=XM_019812642.1;refGene_gn=SLC22A11;refGene_tx=XM_019812643.1
D1 109811004 109811219
refGene_gn=SLC22A11;refGene_tx=XM_003993623.3;refGene_gn=SLC22A11;refGene_tx=XM_019812632.1;refGene_gn=SLC22A11;refGene_tx=XM_019812633.1;refGene_gn=SLC22A11;refGene_tx=XM_019812634.1;refGene_gn=SLC22A11;refGene_tx=XM_019812635.1;refGene_gn=SLC22A11;refGene_tx=XM_019812636.1;refGene_gn=SLC22A11;refGene_tx=XM_019812637.1;refGene_gn=SLC22A11;refGene_

tx=XM_019812638.2;refGene_gn=SLC22A11;refGene_tx=XM_019812639.1;refGen
e_gn=SLC22A11;refGene_tx=XM_019812640.1;refGene_gn=SLC22A11;refGene_tx
=XM_019812641.1;refGene_gn=SLC22A11;refGene_tx=XM_019812642.1;refGene_
gn=SLC22A11;refGene_tx=XM_019812643.1
D1 109812148 109812312
refGene_gn=SLC22A11;refGene_tx=XM_003993623.3;refGene_gn=SLC22A11;refG
ene_tx=XM_019812632.1;refGene_gn=SLC22A11;refGene_tx=XM_019812633.1;re
fGene_gn=SLC22A11;refGene_tx=XM_019812634.1;refGene_gn=SLC22A11;refGen
e_tx=XM_019812635.1;refGene_gn=SLC22A11;refGene_tx=XM_019812636.1;refG
ene_gn=SLC22A11;refGene_tx=XM_019812637.1;refGene_gn=SLC22A11;refGene_
tx=XM_019812638.2;refGene_gn=SLC22A11;refGene_tx=XM_019812639.1;refGen
e_gn=SLC22A11;refGene_tx=XM_019812640.1;refGene_gn=SLC22A11;refGene_tx
=XM_019812641.1;refGene_gn=SLC22A11;refGene_tx=XM_019812642.1;refGene_
gn=SLC22A11;refGene_tx=XM_019812643.1
D1 109812489 109812777
refGene_gn=SLC22A11;refGene_tx=XM_019812637.1;refGene_gn=SLC22A11;refG
ene_tx=XM_019812638.2;refGene_gn=SLC22A11;refGene_tx=XM_019812639.1;re
fGene_gn=SLC22A11;refGene_tx=XM_019812640.1;refGene_gn=SLC22A11;refGen
e_tx=XM_019812641.1
D1 109813309 109813516
refGene_gn=SLC22A11;refGene_tx=XM_003993623.3;refGene_gn=SLC22A11;refG
ene_tx=XM_019812632.1;refGene_gn=SLC22A11;refGene_tx=XM_019812633.1;re
fGene_gn=SLC22A11;refGene_tx=XM_019812634.1;refGene_gn=SLC22A11;refGen
e_tx=XM_019812635.1;refGene_gn=SLC22A11;refGene_tx=XM_019812636.1;refG
ene_gn=SLC22A11;refGene_tx=XM_019812642.1;refGene_gn=SLC22A11;refGene_
tx=XM_019812643.1
D1 109814056 109814219
refGene_gn=SLC22A11;refGene_tx=XM_019812632.1;refGene_gn=SLC22A11;refG
ene_tx=XM_019812633.1;refGene_gn=SLC22A11;refGene_tx=XM_019812634.1;re
fGene_gn=SLC22A11;refGene_tx=XM_019812635.1;refGene_gn=SLC22A11;refGen
e_tx=XM_019812643.1
D1 109814797 109814861
refGene_gn=SLC22A11;refGene_tx=XM_003993623.3;refGene_gn=SLC22A11;refG
ene_tx=XM_019812636.1
D1 109830321 109830723
refGene_gn=SLC22A12;refGene_tx=XM_003993566.4;refGene_gn=SLC22A12;refG
ene_tx=XM_019812644.2
D1 109831375 109831479
refGene_gn=SLC22A12;refGene_tx=XM_003993566.4;refGene_gn=SLC22A12;refG
ene_tx=XM_019812644.2;refGene_gn=SLC22A12;refGene_tx=XM_019812645.1
D1 109831882 109832037
refGene_gn=SLC22A12;refGene_tx=XM_003993566.4;refGene_gn=SLC22A12;refG
ene_tx=XM_019812644.2;refGene_gn=SLC22A12;refGene_tx=XM_019812645.1
D1 109832112 109832281
refGene_gn=SLC22A12;refGene_tx=XM_003993566.4;refGene_gn=SLC22A12;refG
ene_tx=XM_019812644.2;refGene_gn=SLC22A12;refGene_tx=XM_019812645.1
D1 109835186 109835310
refGene_gn=SLC22A12;refGene_tx=XM_003993566.4;refGene_gn=SLC22A12;refG
ene_tx=XM_019812644.2;refGene_gn=SLC22A12;refGene_tx=XM_019812645.1
D1 109835483 109835599

refGene_gn=SLC22A12;refGene_tx=XM_003993566.4;refGene_gn=SLC22A12;refGene_tx=XM_019812644.2;refGene_gn=SLC22A12;refGene_tx=XM_019812645.1
D1 109836291 109836506
refGene_gn=SLC22A12;refGene_tx=XM_003993566.4;refGene_gn=SLC22A12;refGene_tx=XM_019812644.2;refGene_gn=SLC22A12;refGene_tx=XM_019812645.1
D1 109836947 109837056
refGene_gn=SLC22A12;refGene_tx=XM_003993566.4;refGene_gn=SLC22A12;refGene_tx=XM_019812644.2;refGene_gn=SLC22A12;refGene_tx=XM_019812645.1
D1 109837301 109837505
refGene_gn=SLC22A12;refGene_tx=XM_003993566.4;refGene_gn=SLC22A12;refGene_tx=XM_019812644.2;refGene_gn=SLC22A12;refGene_tx=XM_019812645.1
D1 109838036 109838100
refGene_gn=SLC22A12;refGene_tx=XM_003993566.4;refGene_gn=SLC22A12;refGene_tx=XM_019812644.2;refGene_gn=SLC22A12;refGene_tx=XM_019812645.1
D1 109843338 109844225
refGene_gn=NRXN2;refGene_tx=XM_023240000.1;refGene_gn=NRXN2;refGene_tx=XM_023240001.1;refGene_gn=NRXN2;refGene_tx=XM_023240002.1;refGene_gn=NRXN2;refGene_tx=XM_023240003.1;refGene_gn=NRXN2;refGene_tx=XM_023240004.1;refGene_gn=NRXN2;refGene_tx=XM_023240005.1;refGene_gn=NRXN2;refGene_tx=XM_023240006.1;refGene_gn=NRXN2;refGene_tx=XM_023240007.1
D1 109855376 109855455
refGene_gn=NRXN2;refGene_tx=XM_023240000.1;refGene_gn=NRXN2;refGene_tx=XM_023240001.1;refGene_gn=NRXN2;refGene_tx=XM_023240002.1;refGene_gn=NRXN2;refGene_tx=XM_023240003.1;refGene_gn=NRXN2;refGene_tx=XM_023240004.1;refGene_gn=NRXN2;refGene_tx=XM_023240005.1;refGene_gn=NRXN2;refGene_tx=XM_023240006.1;refGene_gn=NRXN2;refGene_tx=XM_023240007.1
D1 109857791 109858117
refGene_gn=NRXN2;refGene_tx=XM_023240000.1;refGene_gn=NRXN2;refGene_tx=XM_023240001.1;refGene_gn=NRXN2;refGene_tx=XM_023240002.1;refGene_gn=NRXN2;refGene_tx=XM_023240003.1;refGene_gn=NRXN2;refGene_tx=XM_023240004.1;refGene_gn=NRXN2;refGene_tx=XM_023240005.1;refGene_gn=NRXN2;refGene_tx=XM_023240006.1;refGene_gn=NRXN2;refGene_tx=XM_023240007.1
D1 109861500 109861590
refGene_gn=NRXN2;refGene_tx=XM_023240000.1;refGene_gn=NRXN2;refGene_tx=XM_023240001.1;refGene_gn=NRXN2;refGene_tx=XM_023240002.1;refGene_gn=NRXN2;refGene_tx=XM_023240004.1;refGene_gn=NRXN2;refGene_tx=XM_023240006.1
D1 109865896 109866068
refGene_gn=NRXN2;refGene_tx=XM_023240000.1;refGene_gn=NRXN2;refGene_tx=XM_023240001.1;refGene_gn=NRXN2;refGene_tx=XM_023240002.1;refGene_gn=NRXN2;refGene_tx=XM_023240003.1;refGene_gn=NRXN2;refGene_tx=XM_023240004.1;refGene_gn=NRXN2;refGene_tx=XM_023240005.1;refGene_gn=NRXN2;refGene_tx=XM_023240006.1;refGene_gn=NRXN2;refGene_tx=XM_023240007.1
D1 109870802 109870984
refGene_gn=NRXN2;refGene_tx=XM_023240000.1;refGene_gn=NRXN2;refGene_tx=XM_023240001.1;refGene_gn=NRXN2;refGene_tx=XM_023240002.1;refGene_gn=NRXN2;refGene_tx=XM_023240003.1;refGene_gn=NRXN2;refGene_tx=XM_023240004.1;refGene_gn=NRXN2;refGene_tx=XM_023240005.1;refGene_gn=NRXN2;refGene_tx=XM_023240006.1;refGene_gn=NRXN2;refGene_tx=XM_023240007.1
D1 109878313 109878560

refGene_gn=NRXN2;refGene_tx=XM_023240004.1;refGene_gn=NRXN2;refGene_tx=XM_023240005.1;refGene_gn=NRXN2;refGene_tx=XM_023240006.1;refGene_gn=NRXN2;refGene_tx=XM_023240007.1
D1 109884262 109884382
refGene_gn=NRXN2;refGene_tx=XM_023240000.1;refGene_gn=NRXN2;refGene_tx=XM_023240001.1;refGene_gn=NRXN2;refGene_tx=XM_023240002.1;refGene_gn=NRXN2;refGene_tx=XM_023240003.1
D1 109884774 109884948
refGene_gn=NRXN2;refGene_tx=XM_023240000.1;refGene_gn=NRXN2;refGene_tx=XM_023240001.1;refGene_gn=NRXN2;refGene_tx=XM_023240002.1;refGene_gn=NRXN2;refGene_tx=XM_023240003.1
D1 109886585 109886776
refGene_gn=NRXN2;refGene_tx=XM_023240000.1;refGene_gn=NRXN2;refGene_tx=XM_023240001.1;refGene_gn=NRXN2;refGene_tx=XM_023240002.1;refGene_gn=NRXN2;refGene_tx=XM_023240003.1
D1 109887382 109887764
refGene_gn=NRXN2;refGene_tx=XM_023240000.1;refGene_gn=NRXN2;refGene_tx=XM_023240001.1;refGene_gn=NRXN2;refGene_tx=XM_023240002.1;refGene_gn=NRXN2;refGene_tx=XM_023240003.1
D1 109888131 109888260
refGene_gn=NRXN2;refGene_tx=XM_023240000.1;refGene_gn=NRXN2;refGene_tx=XM_023240001.1;refGene_gn=NRXN2;refGene_tx=XM_023240002.1;refGene_gn=NRXN2;refGene_tx=XM_023240003.1
D1 109889807 109889834
refGene_gn=NRXN2;refGene_tx=XM_023240000.1;refGene_gn=NRXN2;refGene_tx=XM_023240001.1;refGene_gn=NRXN2;refGene_tx=XM_023240002.1
D1 109895801 109896005
refGene_gn=NRXN2;refGene_tx=XM_023240000.1;refGene_gn=NRXN2;refGene_tx=XM_023240001.1;refGene_gn=NRXN2;refGene_tx=XM_023240002.1;refGene_gn=NRXN2;refGene_tx=XM_023240003.1
D1 109896183 109896570
refGene_gn=NRXN2;refGene_tx=XM_023240000.1;refGene_gn=NRXN2;refGene_tx=XM_023240001.1;refGene_gn=NRXN2;refGene_tx=XM_023240002.1;refGene_gn=NRXN2;refGene_tx=XM_023240003.1
D1 109899906 109900345
refGene_gn=NRXN2;refGene_tx=XM_023240000.1;refGene_gn=NRXN2;refGene_tx=XM_023240001.1;refGene_gn=NRXN2;refGene_tx=XM_023240002.1;refGene_gn=NRXN2;refGene_tx=XM_023240003.1
D1 109901033 109901195
refGene_gn=NRXN2;refGene_tx=XM_023240000.1;refGene_gn=NRXN2;refGene_tx=XM_023240001.1;refGene_gn=NRXN2;refGene_tx=XM_023240002.1;refGene_gn=NRXN2;refGene_tx=XM_023240003.1
D1 109909052 109909097
refGene_gn=NRXN2;refGene_tx=XM_023240000.1;refGene_gn=NRXN2;refGene_tx=XM_023240001.1;refGene_gn=NRXN2;refGene_tx=XM_023240002.1;refGene_gn=NRXN2;refGene_tx=XM_023240003.1
D1 109915186 109915488
refGene_gn=NRXN2;refGene_tx=XM_023240000.1;refGene_gn=NRXN2;refGene_tx=XM_023240001.1;refGene_gn=NRXN2;refGene_tx=XM_023240002.1;refGene_gn=NRXN2;refGene_tx=XM_023240003.1

D1 109919825 109919897
refGene_gn=NRXN2;refGene_tx=XM_023240000.1;refGene_gn=NRXN2;refGene_tx=XM_023240001.1;refGene_gn=NRXN2;refGene_tx=XM_023240002.1
D1 109922233 109922263
refGene_gn=NRXN2;refGene_tx=XM_023240000.1;refGene_gn=NRXN2;refGene_tx=XM_023240001.1;refGene_gn=NRXN2;refGene_tx=XM_023240002.1;refGene_gn=NRXN2;refGene_tx=XM_023240003.1
D1 109932002 109932020
refGene_gn=NRXN2;refGene_tx=XM_023240000.1;refGene_gn=NRXN2;refGene_tx=XM_023240001.1;refGene_gn=NRXN2;refGene_tx=XM_023240002.1;refGene_gn=NRXN2;refGene_tx=XM_023240003.1
D1 109945190 109945926
refGene_gn=NRXN2;refGene_tx=XM_023240000.1;refGene_gn=NRXN2;refGene_tx=XM_023240001.1;refGene_gn=NRXN2;refGene_tx=XM_023240002.1;refGene_gn=NRXN2;refGene_tx=XM_023240003.1
D1 109959136 109959195
refGene_gn=RASGRP2;refGene_tx=XM_023240008.1;refGene_gn=RASGRP2;refGene_tx=XM_023240009.1;refGene_gn=RASGRP2;refGene_tx=XM_023240010.1
D1 109960496 109960673
refGene_gn=RASGRP2;refGene_tx=XM_023240008.1;refGene_gn=RASGRP2;refGene_tx=XM_023240009.1;refGene_gn=RASGRP2;refGene_tx=XM_023240010.1
D1 109961036 109961073
refGene_gn=RASGRP2;refGene_tx=XM_023240008.1;refGene_gn=RASGRP2;refGene_tx=XM_023240009.1;refGene_gn=RASGRP2;refGene_tx=XM_023240010.1
D1 109961341 109961483
refGene_gn=RASGRP2;refGene_tx=XM_023240008.1;refGene_gn=RASGRP2;refGene_tx=XM_023240009.1;refGene_gn=RASGRP2;refGene_tx=XM_023240010.1
D1 109964689 109964805
refGene_gn=RASGRP2;refGene_tx=XM_023240008.1;refGene_gn=RASGRP2;refGene_tx=XM_023240009.1;refGene_gn=RASGRP2;refGene_tx=XM_023240010.1
D1 109965136 109965259
refGene_gn=RASGRP2;refGene_tx=XM_023240008.1;refGene_gn=RASGRP2;refGene_tx=XM_023240009.1;refGene_gn=RASGRP2;refGene_tx=XM_023240010.1
D1 109965456 109965534
refGene_gn=RASGRP2;refGene_tx=XM_023240008.1;refGene_gn=RASGRP2;refGene_tx=XM_023240009.1;refGene_gn=RASGRP2;refGene_tx=XM_023240010.1
D1 109966890 109967172
refGene_gn=RASGRP2;refGene_tx=XM_023240008.1;refGene_gn=RASGRP2;refGene_tx=XM_023240009.1;refGene_gn=RASGRP2;refGene_tx=XM_023240010.1
D1 109968501 109968618
refGene_gn=RASGRP2;refGene_tx=XM_023240008.1;refGene_gn=RASGRP2;refGene_tx=XM_023240009.1;refGene_gn=RASGRP2;refGene_tx=XM_023240010.1
D1 109968790 109968964
refGene_gn=RASGRP2;refGene_tx=XM_023240008.1;refGene_gn=RASGRP2;refGene_tx=XM_023240009.1;refGene_gn=RASGRP2;refGene_tx=XM_023240010.1
D1 109969150 109969301
refGene_gn=RASGRP2;refGene_tx=XM_023240008.1;refGene_gn=RASGRP2;refGene_tx=XM_023240009.1;refGene_gn=RASGRP2;refGene_tx=XM_023240010.1
D1 109970076 109970208
refGene_gn=RASGRP2;refGene_tx=XM_023240008.1;refGene_gn=RASGRP2;refGen

e_tx=XM_023240009.1;refGene_gn=RASGRP2;refGene_tx=XM_023240010.1
D1	109970553	109970616
refGene_gn=RASGRP2;refGene_tx=XM_023240008.1;refGene_gn=RASGRP2;refGen
e_tx=XM_023240009.1;refGene_gn=RASGRP2;refGene_tx=XM_023240010.1
D1	109971123	109971226
refGene_gn=RASGRP2;refGene_tx=XM_023240008.1;refGene_gn=RASGRP2;refGen
e_tx=XM_023240009.1;refGene_gn=RASGRP2;refGene_tx=XM_023240010.1
D1	109971550	109971644
refGene_gn=RASGRP2;refGene_tx=XM_023240009.1
D1	109971690	109971706
refGene_gn=RASGRP2;refGene_tx=XM_023240008.1
D1	109971885	109971958
refGene_gn=RASGRP2;refGene_tx=XM_023240010.1
D1	109975865	109976015
refGene_gn=PYGM;refGene_tx=XM_003993567.5
D1	109976128	109976195
refGene_gn=PYGM;refGene_tx=XM_003993567.5
D1	109976409	109976544
refGene_gn=PYGM;refGene_tx=XM_003993567.5
D1	109979047	109979255
refGene_gn=PYGM;refGene_tx=XM_003993567.5
D1	109979710	109979852
refGene_gn=PYGM;refGene_tx=XM_003993567.5
D1	109979979	109980038
refGene_gn=PYGM;refGene_tx=XM_003993567.5
D1	109980331	109980479
refGene_gn=PYGM;refGene_tx=XM_003993567.5
D1	109980826	109980928
refGene_gn=PYGM;refGene_tx=XM_003993567.5
D1	109981463	109981578
refGene_gn=PYGM;refGene_tx=XM_003993567.5
D1	109981853	109982017
refGene_gn=PYGM;refGene_tx=XM_003993567.5
D1	109982140	109982287
refGene_gn=PYGM;refGene_tx=XM_003993567.5
D1	109982534	109982627
refGene_gn=PYGM;refGene_tx=XM_003993567.5
D1	109982953	109983097
refGene_gn=PYGM;refGene_tx=XM_003993567.5
D1	109983683	109983766
refGene_gn=PYGM;refGene_tx=XM_003993567.5
D1	109983852	109983964
refGene_gn=PYGM;refGene_tx=XM_003993567.5
D1	109985223	109985355
refGene_gn=PYGM;refGene_tx=XM_003993567.5
D1	109985737	109985841
refGene_gn=PYGM;refGene_tx=XM_003993567.5
D1	109985928	109986007
refGene_gn=PYGM;refGene_tx=XM_003993567.5
D1	109986107	109986209

refGene_gn=PYGM;refGene_tx=XM_003993567.5
D1	109987251	109987494
refGene_gn=PYGM;refGene_tx=XM_003993567.5
D1	109992943	109993073
refGene_gn=SF1;refGene_tx=XM_023240011.1;refGene_gn=SF1;refGene_tx=XM_023240017.1;refGene_gn=SF1;refGene_tx=XM_023240018.1;refGene_gn=SF1;refGene_tx=XM_023240019.1;refGene_gn=SF1;refGene_tx=XM_023240020.1
D1	109993375	109993713
refGene_gn=SF1;refGene_tx=XM_023240012.1;refGene_gn=SF1;refGene_tx=XM_023240013.1;refGene_gn=SF1;refGene_tx=XM_023240014.1;refGene_gn=SF1;refGene_tx=XM_023240015.1;refGene_gn=SF1;refGene_tx=XM_023240016.1;refGene_gn=SF1;refGene_tx=XM_023240020.1
D1	109994476	109994656
refGene_gn=SF1;refGene_tx=XM_023240011.1;refGene_gn=SF1;refGene_tx=XM_023240012.1;refGene_gn=SF1;refGene_tx=XM_023240013.1;refGene_gn=SF1;refGene_tx=XM_023240014.1;refGene_gn=SF1;refGene_tx=XM_023240015.1;refGene_gn=SF1;refGene_tx=XM_023240016.1;refGene_gn=SF1;refGene_tx=XM_023240017.1;refGene_gn=SF1;refGene_tx=XM_023240018.1;refGene_gn=SF1;refGene_tx=XM_023240019.1;refGene_gn=SF1;refGene_tx=XM_023240020.1
D1	109994762	109994822
refGene_gn=SF1;refGene_tx=XM_023240011.1;refGene_gn=SF1;refGene_tx=XM_023240012.1;refGene_gn=SF1;refGene_tx=XM_023240013.1;refGene_gn=SF1;refGene_tx=XM_023240014.1;refGene_gn=SF1;refGene_tx=XM_023240015.1;refGene_gn=SF1;refGene_tx=XM_023240016.1;refGene_gn=SF1;refGene_tx=XM_023240017.1;refGene_gn=SF1;refGene_tx=XM_023240018.1;refGene_gn=SF1;refGene_tx=XM_023240019.1;refGene_gn=SF1;refGene_tx=XM_023240020.1
D1	109995153	109995431
refGene_gn=SF1;refGene_tx=XM_023240011.1;refGene_gn=SF1;refGene_tx=XM_023240012.1;refGene_gn=SF1;refGene_tx=XM_023240013.1;refGene_gn=SF1;refGene_tx=XM_023240014.1;refGene_gn=SF1;refGene_tx=XM_023240015.1;refGene_gn=SF1;refGene_tx=XM_023240016.1;refGene_gn=SF1;refGene_tx=XM_023240017.1;refGene_gn=SF1;refGene_tx=XM_023240018.1;refGene_gn=SF1;refGene_tx=XM_023240019.1;refGene_gn=SF1;refGene_tx=XM_023240020.1
D1	109995693	109995874
refGene_gn=SF1;refGene_tx=XM_023240011.1;refGene_gn=SF1;refGene_tx=XM_023240012.1;refGene_gn=SF1;refGene_tx=XM_023240013.1;refGene_gn=SF1;refGene_tx=XM_023240014.1;refGene_gn=SF1;refGene_tx=XM_023240015.1;refGene_gn=SF1;refGene_tx=XM_023240016.1;refGene_gn=SF1;refGene_tx=XM_023240017.1;refGene_gn=SF1;refGene_tx=XM_023240018.1;refGene_gn=SF1;refGene_tx=XM_023240019.1;refGene_gn=SF1;refGene_tx=XM_023240020.1
D1	109996560	109996668
refGene_gn=SF1;refGene_tx=XM_023240011.1;refGene_gn=SF1;refGene_tx=XM_023240012.1;refGene_gn=SF1;refGene_tx=XM_023240013.1;refGene_gn=SF1;refGene_tx=XM_023240014.1;refGene_gn=SF1;refGene_tx=XM_023240015.1;refGene_gn=SF1;refGene_tx=XM_023240016.1;refGene_gn=SF1;refGene_tx=XM_023240017.1;refGene_gn=SF1;refGene_tx=XM_023240018.1;refGene_gn=SF1;refGene_tx=XM_023240019.1;refGene_gn=SF1;refGene_tx=XM_023240020.1
D1	109996798	109996914
refGene_gn=SF1;refGene_tx=XM_023240011.1;refGene_gn=SF1;refGene_tx=XM_023240012.1;refGene_gn=SF1;refGene_tx=XM_023240013.1;refGene_gn=SF1;re

fGene_tx=XM_023240014.1;refGene_gn=SF1;refGene_tx=XM_023240015.1;refGene_gn=SF1;refGene_tx=XM_023240016.1;refGene_gn=SF1;refGene_tx=XM_023240017.1;refGene_gn=SF1;refGene_tx=XM_023240018.1;refGene_gn=SF1;refGene_tx=XM_023240019.1;refGene_gn=SF1;refGene_tx=XM_023240020.1
D1 109996999 109997183
refGene_gn=SF1;refGene_tx=XM_023240011.1;refGene_gn=SF1;refGene_tx=XM_023240012.1;refGene_gn=SF1;refGene_tx=XM_023240013.1;refGene_gn=SF1;refGene_tx=XM_023240014.1;refGene_gn=SF1;refGene_tx=XM_023240015.1;refGene_gn=SF1;refGene_tx=XM_023240016.1;refGene_gn=SF1;refGene_tx=XM_023240017.1;refGene_gn=SF1;refGene_tx=XM_023240018.1;refGene_gn=SF1;refGene_tx=XM_023240019.1;refGene_gn=SF1;refGene_tx=XM_023240020.1
D1 109997504 109997594
refGene_gn=SF1;refGene_tx=XM_023240011.1;refGene_gn=SF1;refGene_tx=XM_023240012.1;refGene_gn=SF1;refGene_tx=XM_023240013.1;refGene_gn=SF1;refGene_tx=XM_023240014.1;refGene_gn=SF1;refGene_tx=XM_023240015.1;refGene_gn=SF1;refGene_tx=XM_023240016.1;refGene_gn=SF1;refGene_tx=XM_023240017.1;refGene_gn=SF1;refGene_tx=XM_023240018.1;refGene_gn=SF1;refGene_tx=XM_023240019.1;refGene_gn=SF1;refGene_tx=XM_023240020.1
D1 109997808 109997961
refGene_gn=SF1;refGene_tx=XM_023240011.1;refGene_gn=SF1;refGene_tx=XM_023240012.1;refGene_gn=SF1;refGene_tx=XM_023240013.1;refGene_gn=SF1;refGene_tx=XM_023240014.1;refGene_gn=SF1;refGene_tx=XM_023240015.1;refGene_gn=SF1;refGene_tx=XM_023240016.1;refGene_gn=SF1;refGene_tx=XM_023240017.1;refGene_gn=SF1;refGene_tx=XM_023240018.1;refGene_gn=SF1;refGene_tx=XM_023240019.1;refGene_gn=SF1;refGene_tx=XM_023240020.1
D1 110001016 110001092
refGene_gn=SF1;refGene_tx=XM_023240011.1;refGene_gn=SF1;refGene_tx=XM_023240012.1;refGene_gn=SF1;refGene_tx=XM_023240013.1;refGene_gn=SF1;refGene_tx=XM_023240014.1;refGene_gn=SF1;refGene_tx=XM_023240017.1;refGene_gn=SF1;refGene_tx=XM_023240018.1;refGene_gn=SF1;refGene_tx=XM_023240019.1;refGene_gn=SF1;refGene_tx=XM_023240020.1
D1 110003814 110003943
refGene_gn=SF1;refGene_tx=XM_023240011.1;refGene_gn=SF1;refGene_tx=XM_023240012.1;refGene_gn=SF1;refGene_tx=XM_023240013.1;refGene_gn=SF1;refGene_tx=XM_023240014.1;refGene_gn=SF1;refGene_tx=XM_023240015.1;refGene_gn=SF1;refGene_tx=XM_023240016.1;refGene_gn=SF1;refGene_tx=XM_023240017.1;refGene_gn=SF1;refGene_tx=XM_023240018.1;refGene_gn=SF1;refGene_tx=XM_023240019.1;refGene_gn=SF1;refGene_tx=XM_023240020.1
D1 110005247 110005644
refGene_gn=SF1;refGene_tx=XM_023240011.1;refGene_gn=SF1;refGene_tx=XM_023240012.1;refGene_gn=SF1;refGene_tx=XM_023240013.1;refGene_gn=SF1;refGene_tx=XM_023240014.1;refGene_gn=SF1;refGene_tx=XM_023240017.1;refGene_gn=SF1;refGene_tx=XM_023240018.1;refGene_gn=SF1;refGene_tx=XM_023240019.1;refGene_gn=SF1;refGene_tx=XM_023240020.1
D1 110016228 110016316
refGene_gn=MAP4K2;refGene_tx=XM_011287057.3
D1 110016420 110016476
refGene_gn=MAP4K2;refGene_tx=XM_011287057.3
D1 110016575 110016646
refGene_gn=MAP4K2;refGene_tx=XM_011287057.3

D1 110016884 110016971
refGene_gn=MAP4K2;refGene_tx=XM_011287057.3
D1 110017107 110017176
refGene_gn=MAP4K2;refGene_tx=XM_011287057.3
D1 110017790 110017968
refGene_gn=MAP4K2;refGene_tx=XM_011287057.3
D1 110018069 110018173
refGene_gn=MAP4K2;refGene_tx=XM_011287057.3
D1 110018253 110018312
refGene_gn=MAP4K2;refGene_tx=XM_011287057.3
D1 110022189 110022307
refGene_gn=MAP4K2;refGene_tx=XM_011287057.3
D1 110022392 110022453
refGene_gn=MAP4K2;refGene_tx=XM_011287057.3
D1 110022558 110022638
refGene_gn=MAP4K2;refGene_tx=XM_011287057.3
D1 110022731 110022816
refGene_gn=MAP4K2;refGene_tx=XM_011287057.3
D1 110022913 110022955
refGene_gn=MAP4K2;refGene_tx=XM_011287057.3
D1 110023036 110023125
refGene_gn=MAP4K2;refGene_tx=XM_011287057.3
D1 110023205 110023311
refGene_gn=MAP4K2;refGene_tx=XM_011287057.3
D1 110023408 110023442
refGene_gn=MAP4K2;refGene_tx=XM_011287057.3
D1 110023538 110023577
refGene_gn=MAP4K2;refGene_tx=XM_011287057.3
D1 110023661 110023705
refGene_gn=MAP4K2;refGene_tx=XM_011287057.3
D1 110024207 110024266
refGene_gn=MAP4K2;refGene_tx=XM_011287057.3
D1 110024393 110024472
refGene_gn=MAP4K2;refGene_tx=XM_011287057.3
D1 110024875 110024983
refGene_gn=MAP4K2;refGene_tx=XM_011287057.3
D1 110025073 110025155
refGene_gn=MAP4K2;refGene_tx=XM_011287057.3
D1 110025512 110025575
refGene_gn=MAP4K2;refGene_tx=XM_011287057.3
D1 110025655 110025787
refGene_gn=MAP4K2;refGene_tx=XM_011287057.3
D1 110025883 110025956
refGene_gn=MAP4K2;refGene_tx=XM_011287057.3
D1 110026355 110026398
refGene_gn=MAP4K2;refGene_tx=XM_011287057.3
D1 110026484 110026532
refGene_gn=MAP4K2;refGene_tx=XM_011287057.3
D1 110026875 110026931
refGene_gn=MAP4K2;refGene_tx=XM_011287057.3

D1 110027237 110027302
refGene_gn=MAP4K2;refGene_tx=XM_011287057.3
D1 110027388 110027479
refGene_gn=MAP4K2;refGene_tx=XM_011287057.3
D1 110027683 110027741
refGene_gn=MAP4K2;refGene_tx=XM_011287057.3
D1 110027865 110027961
refGene_gn=MAP4K2;refGene_tx=XM_011287057.3
D1 110029242 110029725
refGene_gn=MEN1;refGene_tx=XM_006937482.4;refGene_gn=MEN1;refGene_tx=XM_006937483.3;refGene_gn=MEN1;refGene_tx=XM_019812647.2
D1 110029930 110030095
refGene_gn=MEN1;refGene_tx=XM_006937482.4;refGene_gn=MEN1;refGene_tx=XM_006937483.3;refGene_gn=MEN1;refGene_tx=XM_019812647.2
D1 110030570 110030706
refGene_gn=MEN1;refGene_tx=XM_006937482.4;refGene_gn=MEN1;refGene_tx=XM_006937483.3;refGene_gn=MEN1;refGene_tx=XM_019812647.2
D1 110031095 110031232
refGene_gn=MEN1;refGene_tx=XM_006937482.4;refGene_gn=MEN1;refGene_tx=XM_006937483.3;refGene_gn=MEN1;refGene_tx=XM_019812647.2
D1 110031936 110032024
refGene_gn=MEN1;refGene_tx=XM_006937482.4;refGene_gn=MEN1;refGene_tx=XM_006937483.3;refGene_gn=MEN1;refGene_tx=XM_019812647.2
D1 110032097 110032138
refGene_gn=MEN1;refGene_tx=XM_006937482.4;refGene_gn=MEN1;refGene_tx=XM_006937483.3;refGene_gn=MEN1;refGene_tx=XM_019812647.2
D1 110032411 110032540
refGene_gn=MEN1;refGene_tx=XM_006937482.4;refGene_gn=MEN1;refGene_tx=XM_006937483.3;refGene_gn=MEN1;refGene_tx=XM_019812647.2
D1 110032737 110032946
refGene_gn=MEN1;refGene_tx=XM_006937482.4;refGene_gn=MEN1;refGene_tx=XM_006937483.3;refGene_gn=MEN1;refGene_tx=XM_019812647.2
D1 110033915 110034376
refGene_gn=MEN1;refGene_tx=XM_006937482.4;refGene_gn=MEN1;refGene_tx=XM_006937483.3;refGene_gn=MEN1;refGene_tx=XM_019812647.2
D1 110034765 110034767
refGene_gn=MEN1;refGene_tx=XM_006937482.4
D1 110037483 110037489
refGene_gn=CDC42BPG;refGene_tx=XM_023240026.1
D1 110042588 110042645
refGene_gn=CDC42BPG;refGene_tx=XM_019812985.2;refGene_gn=CDC42BPG;refGene_tx=XM_023240022.1;refGene_gn=CDC42BPG;refGene_tx=XM_023240023.1;refGene_gn=CDC42BPG;refGene_tx=XM_023240024.1;refGene_gn=CDC42BPG;refGene_tx=XM_023240025.1;refGene_gn=CDC42BPG;refGene_tx=XM_023240027.1;refGene_gn=CDC42BPG;refGene_tx=XM_023240028.1;refGene_gn=CDC42BPG;refGene_tx=XM_023240029.1;refGene_gn=CDC42BPG;refGene_tx=XM_023240030.1
D1 110043660 110043746
refGene_gn=CDC42BPG;refGene_tx=XM_019812985.2;refGene_gn=CDC42BPG;refGene_tx=XM_023240022.1;refGene_gn=CDC42BPG;refGene_tx=XM_023240023.1;refGene_gn=CDC42BPG;refGene_tx=XM_023240024.1;refGene_gn=CDC42BPG;refGen

e_tx=XM_023240025.1;refGene_gn=CDC42BPG;refGene_tx=XM_023240026.1;refGene_gn=CDC42BPG;refGene_tx=XM_023240027.1;refGene_gn=CDC42BPG;refGene_tx=XM_023240028.1;refGene_gn=CDC42BPG;refGene_tx=XM_023240029.1;refGene_gn=CDC42BPG;refGene_tx=XM_023240030.1
D1 110043873 110044000 refGene_gn=CDC42BPG;refGene_tx=XM_019812985.2;refGene_gn=CDC42BPG;refGene_tx=XM_023240022.1;refGene_gn=CDC42BPG;refGene_tx=XM_023240023.1;refGene_gn=CDC42BPG;refGene_tx=XM_023240024.1;refGene_gn=CDC42BPG;refGene_tx=XM_023240025.1;refGene_gn=CDC42BPG;refGene_tx=XM_023240026.1;refGene_gn=CDC42BPG;refGene_tx=XM_023240027.1;refGene_gn=CDC42BPG;refGene_tx=XM_023240028.1;refGene_gn=CDC42BPG;refGene_tx=XM_023240029.1;refGene_gn=CDC42BPG;refGene_tx=XM_023240030.1
D1 110044170 110044288 refGene_gn=CDC42BPG;refGene_tx=XM_019812985.2;refGene_gn=CDC42BPG;refGene_tx=XM_023240022.1;refGene_gn=CDC42BPG;refGene_tx=XM_023240023.1;refGene_gn=CDC42BPG;refGene_tx=XM_023240024.1;refGene_gn=CDC42BPG;refGene_tx=XM_023240025.1;refGene_gn=CDC42BPG;refGene_tx=XM_023240026.1;refGene_gn=CDC42BPG;refGene_tx=XM_023240027.1;refGene_gn=CDC42BPG;refGene_tx=XM_023240028.1;refGene_gn=CDC42BPG;refGene_tx=XM_023240029.1;refGene_gn=CDC42BPG;refGene_tx=XM_023240030.1
D1 110044425 110044546 refGene_gn=CDC42BPG;refGene_tx=XM_019812985.2;refGene_gn=CDC42BPG;refGene_tx=XM_023240022.1;refGene_gn=CDC42BPG;refGene_tx=XM_023240023.1;refGene_gn=CDC42BPG;refGene_tx=XM_023240024.1;refGene_gn=CDC42BPG;refGene_tx=XM_023240025.1;refGene_gn=CDC42BPG;refGene_tx=XM_023240026.1;refGene_gn=CDC42BPG;refGene_tx=XM_023240027.1;refGene_gn=CDC42BPG;refGene_tx=XM_023240028.1;refGene_gn=CDC42BPG;refGene_tx=XM_023240029.1;refGene_gn=CDC42BPG;refGene_tx=XM_023240030.1
D1 110044675 110044760 refGene_gn=CDC42BPG;refGene_tx=XM_019812985.2;refGene_gn=CDC42BPG;refGene_tx=XM_023240022.1;refGene_gn=CDC42BPG;refGene_tx=XM_023240023.1;refGene_gn=CDC42BPG;refGene_tx=XM_023240024.1;refGene_gn=CDC42BPG;refGene_tx=XM_023240025.1;refGene_gn=CDC42BPG;refGene_tx=XM_023240026.1;refGene_gn=CDC42BPG;refGene_tx=XM_023240027.1;refGene_gn=CDC42BPG;refGene_tx=XM_023240028.1;refGene_gn=CDC42BPG;refGene_tx=XM_023240029.1;refGene_gn=CDC42BPG;refGene_tx=XM_023240030.1
D1 110044854 110045039 refGene_gn=CDC42BPG;refGene_tx=XM_019812985.2;refGene_gn=CDC42BPG;refGene_tx=XM_023240022.1;refGene_gn=CDC42BPG;refGene_tx=XM_023240023.1;refGene_gn=CDC42BPG;refGene_tx=XM_023240024.1;refGene_gn=CDC42BPG;refGene_tx=XM_023240025.1;refGene_gn=CDC42BPG;refGene_tx=XM_023240026.1;refGene_gn=CDC42BPG;refGene_tx=XM_023240027.1;refGene_gn=CDC42BPG;refGene_tx=XM_023240028.1;refGene_gn=CDC42BPG;refGene_tx=XM_023240029.1;refGene_gn=CDC42BPG;refGene_tx=XM_023240030.1
D1 110046048 110046648 refGene_gn=CDC42BPG;refGene_tx=XM_019812985.2;refGene_gn=CDC42BPG;refGene_tx=XM_023240022.1;refGene_gn=CDC42BPG;refGene_tx=XM_023240023.1;refGene_gn=CDC42BPG;refGene_tx=XM_023240024.1;refGene_gn=CDC42BPG;refGene_tx=XM_023240025.1;refGene_gn=CDC42BPG;refGene_tx=XM_023240026.1;refGene_gn=CDC42BPG;refGene_tx=XM_023240027.1;refGene_gn=CDC42BPG;refGene_

tx=XM_023240028.1;refGene_gn=CDC42BPG;refGene_tx=XM_023240029.1;refGene_gn=CDC42BPG;refGene_tx=XM_023240030.1
D1	110046760	110046823
refGene_gn=CDC42BPG;refGene_tx=XM_019812985.2;refGene_gn=CDC42BPG;refGene_tx=XM_023240022.1;refGene_gn=CDC42BPG;refGene_tx=XM_023240023.1;refGene_gn=CDC42BPG;refGene_tx=XM_023240024.1;refGene_gn=CDC42BPG;refGene_tx=XM_023240025.1;refGene_gn=CDC42BPG;refGene_tx=XM_023240026.1;refGene_gn=CDC42BPG;refGene_tx=XM_023240027.1;refGene_gn=CDC42BPG;refGene_tx=XM_023240028.1;refGene_gn=CDC42BPG;refGene_tx=XM_023240029.1;refGene_gn=CDC42BPG;refGene_tx=XM_023240030.1
D1	110047642	110047859
refGene_gn=CDC42BPG;refGene_tx=XM_019812985.2;refGene_gn=CDC42BPG;refGene_tx=XM_023240022.1;refGene_gn=CDC42BPG;refGene_tx=XM_023240023.1;refGene_gn=CDC42BPG;refGene_tx=XM_023240024.1;refGene_gn=CDC42BPG;refGene_tx=XM_023240025.1;refGene_gn=CDC42BPG;refGene_tx=XM_023240026.1;refGene_gn=CDC42BPG;refGene_tx=XM_023240027.1;refGene_gn=CDC42BPG;refGene_tx=XM_023240028.1;refGene_gn=CDC42BPG;refGene_tx=XM_023240029.1;refGene_gn=CDC42BPG;refGene_tx=XM_023240030.1
D1	110048515	110048597
refGene_gn=CDC42BPG;refGene_tx=XM_019812985.2;refGene_gn=CDC42BPG;refGene_tx=XM_023240022.1;refGene_gn=CDC42BPG;refGene_tx=XM_023240023.1;refGene_gn=CDC42BPG;refGene_tx=XM_023240024.1;refGene_gn=CDC42BPG;refGene_tx=XM_023240025.1;refGene_gn=CDC42BPG;refGene_tx=XM_023240026.1;refGene_gn=CDC42BPG;refGene_tx=XM_023240027.1;refGene_gn=CDC42BPG;refGene_tx=XM_023240028.1;refGene_gn=CDC42BPG;refGene_tx=XM_023240029.1;refGene_gn=CDC42BPG;refGene_tx=XM_023240030.1
D1	110048696	110048836
refGene_gn=CDC42BPG;refGene_tx=XM_019812985.2;refGene_gn=CDC42BPG;refGene_tx=XM_023240022.1;refGene_gn=CDC42BPG;refGene_tx=XM_023240023.1;refGene_gn=CDC42BPG;refGene_tx=XM_023240024.1;refGene_gn=CDC42BPG;refGene_tx=XM_023240025.1;refGene_gn=CDC42BPG;refGene_tx=XM_023240026.1;refGene_gn=CDC42BPG;refGene_tx=XM_023240027.1;refGene_gn=CDC42BPG;refGene_tx=XM_023240028.1;refGene_gn=CDC42BPG;refGene_tx=XM_023240029.1;refGene_gn=CDC42BPG;refGene_tx=XM_023240030.1
D1	110048919	110049053
refGene_gn=CDC42BPG;refGene_tx=XM_019812985.2;refGene_gn=CDC42BPG;refGene_tx=XM_023240022.1;refGene_gn=CDC42BPG;refGene_tx=XM_023240023.1;refGene_gn=CDC42BPG;refGene_tx=XM_023240024.1;refGene_gn=CDC42BPG;refGene_tx=XM_023240025.1;refGene_gn=CDC42BPG;refGene_tx=XM_023240026.1;refGene_gn=CDC42BPG;refGene_tx=XM_023240027.1;refGene_gn=CDC42BPG;refGene_tx=XM_023240028.1;refGene_gn=CDC42BPG;refGene_tx=XM_023240029.1;refGene_gn=CDC42BPG;refGene_tx=XM_023240030.1
D1	110049340	110049446
refGene_gn=CDC42BPG;refGene_tx=XM_019812985.2;refGene_gn=CDC42BPG;refGene_tx=XM_023240022.1;refGene_gn=CDC42BPG;refGene_tx=XM_023240023.1;refGene_gn=CDC42BPG;refGene_tx=XM_023240024.1;refGene_gn=CDC42BPG;refGene_tx=XM_023240025.1;refGene_gn=CDC42BPG;refGene_tx=XM_023240026.1;refGene_gn=CDC42BPG;refGene_tx=XM_023240027.1;refGene_gn=CDC42BPG;refGene_tx=XM_023240028.1;refGene_gn=CDC42BPG;refGene_tx=XM_023240029.1;refGene_gn=CDC42BPG;refGene_tx=XM_023240030.1

D1 110049740 110049800
refGene_gn=CDC42BPG;refGene_tx=XM_019812985.2;refGene_gn=CDC42BPG;refGene_tx=XM_023240022.1;refGene_gn=CDC42BPG;refGene_tx=XM_023240023.1;refGene_gn=CDC42BPG;refGene_tx=XM_023240024.1;refGene_gn=CDC42BPG;refGene_tx=XM_023240025.1;refGene_gn=CDC42BPG;refGene_tx=XM_023240026.1;refGene_gn=CDC42BPG;refGene_tx=XM_023240029.1;refGene_gn=CDC42BPG;refGene_tx=XM_023240030.1
D1 110049895 110049994
refGene_gn=CDC42BPG;refGene_tx=XM_019812985.2;refGene_gn=CDC42BPG;refGene_tx=XM_023240022.1;refGene_gn=CDC42BPG;refGene_tx=XM_023240023.1;refGene_gn=CDC42BPG;refGene_tx=XM_023240024.1;refGene_gn=CDC42BPG;refGene_tx=XM_023240025.1;refGene_gn=CDC42BPG;refGene_tx=XM_023240026.1;refGene_gn=CDC42BPG;refGene_tx=XM_023240027.1;refGene_gn=CDC42BPG;refGene_tx=XM_023240028.1;refGene_gn=CDC42BPG;refGene_tx=XM_023240029.1;refGene_gn=CDC42BPG;refGene_tx=XM_023240030.1
D1 110050079 110050132
refGene_gn=CDC42BPG;refGene_tx=XM_019812985.2;refGene_gn=CDC42BPG;refGene_tx=XM_023240022.1;refGene_gn=CDC42BPG;refGene_tx=XM_023240023.1;refGene_gn=CDC42BPG;refGene_tx=XM_023240024.1;refGene_gn=CDC42BPG;refGene_tx=XM_023240025.1;refGene_gn=CDC42BPG;refGene_tx=XM_023240026.1;refGene_gn=CDC42BPG;refGene_tx=XM_023240027.1;refGene_gn=CDC42BPG;refGene_tx=XM_023240028.1;refGene_gn=CDC42BPG;refGene_tx=XM_023240029.1;refGene_gn=CDC42BPG;refGene_tx=XM_023240030.1
D1 110050443 110050532
refGene_gn=CDC42BPG;refGene_tx=XM_019812985.2;refGene_gn=CDC42BPG;refGene_tx=XM_023240022.1;refGene_gn=CDC42BPG;refGene_tx=XM_023240023.1;refGene_gn=CDC42BPG;refGene_tx=XM_023240024.1;refGene_gn=CDC42BPG;refGene_tx=XM_023240025.1;refGene_gn=CDC42BPG;refGene_tx=XM_023240026.1;refGene_gn=CDC42BPG;refGene_tx=XM_023240027.1;refGene_gn=CDC42BPG;refGene_tx=XM_023240028.1;refGene_gn=CDC42BPG;refGene_tx=XM_023240029.1;refGene_gn=CDC42BPG;refGene_tx=XM_023240030.1
D1 110050624 110050773
refGene_gn=CDC42BPG;refGene_tx=XM_019812985.2;refGene_gn=CDC42BPG;refGene_tx=XM_023240022.1;refGene_gn=CDC42BPG;refGene_tx=XM_023240023.1;refGene_gn=CDC42BPG;refGene_tx=XM_023240024.1;refGene_gn=CDC42BPG;refGene_tx=XM_023240025.1;refGene_gn=CDC42BPG;refGene_tx=XM_023240026.1;refGene_gn=CDC42BPG;refGene_tx=XM_023240027.1;refGene_gn=CDC42BPG;refGene_tx=XM_023240028.1;refGene_gn=CDC42BPG;refGene_tx=XM_023240029.1;refGene_gn=CDC42BPG;refGene_tx=XM_023240030.1
D1 110051011 110051126
refGene_gn=CDC42BPG;refGene_tx=XM_019812985.2;refGene_gn=CDC42BPG;refGene_tx=XM_023240022.1;refGene_gn=CDC42BPG;refGene_tx=XM_023240023.1;refGene_gn=CDC42BPG;refGene_tx=XM_023240024.1;refGene_gn=CDC42BPG;refGene_tx=XM_023240025.1;refGene_gn=CDC42BPG;refGene_tx=XM_023240026.1;refGene_gn=CDC42BPG;refGene_tx=XM_023240027.1;refGene_gn=CDC42BPG;refGene_tx=XM_023240028.1;refGene_gn=CDC42BPG;refGene_tx=XM_023240029.1;refGene_gn=CDC42BPG;refGene_tx=XM_023240030.1
D1 110051213 110051320
refGene_gn=CDC42BPG;refGene_tx=XM_019812985.2;refGene_gn=CDC42BPG;refGene_tx=XM_023240022.1;refGene_gn=CDC42BPG;refGene_tx=XM_023240023.1;re

fGene_gn=CDC42BPG;refGene_tx=XM_023240024.1;refGene_gn=CDC42BPG;refGene_tx=XM_023240025.1;refGene_gn=CDC42BPG;refGene_tx=XM_023240026.1;refGene_gn=CDC42BPG;refGene_tx=XM_023240027.1;refGene_gn=CDC42BPG;refGene_tx=XM_023240028.1;refGene_gn=CDC42BPG;refGene_tx=XM_023240029.1;refGene_gn=CDC42BPG;refGene_tx=XM_023240030.1
D1	110051548	110051621	refGene_gn=CDC42BPG;refGene_tx=XM_019812985.2;refGene_gn=CDC42BPG;refGene_tx=XM_023240022.1;refGene_gn=CDC42BPG;refGene_tx=XM_023240023.1;refGene_gn=CDC42BPG;refGene_tx=XM_023240024.1;refGene_gn=CDC42BPG;refGene_tx=XM_023240025.1;refGene_gn=CDC42BPG;refGene_tx=XM_023240026.1;refGene_gn=CDC42BPG;refGene_tx=XM_023240027.1;refGene_gn=CDC42BPG;refGene_tx=XM_023240028.1;refGene_gn=CDC42BPG;refGene_tx=XM_023240029.1;refGene_gn=CDC42BPG;refGene_tx=XM_023240030.1
D1	110051697	110051819	refGene_gn=CDC42BPG;refGene_tx=XM_019812985.2;refGene_gn=CDC42BPG;refGene_tx=XM_023240022.1;refGene_gn=CDC42BPG;refGene_tx=XM_023240023.1;refGene_gn=CDC42BPG;refGene_tx=XM_023240024.1;refGene_gn=CDC42BPG;refGene_tx=XM_023240025.1;refGene_gn=CDC42BPG;refGene_tx=XM_023240026.1;refGene_gn=CDC42BPG;refGene_tx=XM_023240027.1;refGene_gn=CDC42BPG;refGene_tx=XM_023240028.1;refGene_gn=CDC42BPG;refGene_tx=XM_023240029.1;refGene_gn=CDC42BPG;refGene_tx=XM_023240030.1
D1	110051962	110052055	refGene_gn=CDC42BPG;refGene_tx=XM_019812985.2;refGene_gn=CDC42BPG;refGene_tx=XM_023240022.1;refGene_gn=CDC42BPG;refGene_tx=XM_023240023.1;refGene_gn=CDC42BPG;refGene_tx=XM_023240024.1;refGene_gn=CDC42BPG;refGene_tx=XM_023240025.1;refGene_gn=CDC42BPG;refGene_tx=XM_023240026.1;refGene_gn=CDC42BPG;refGene_tx=XM_023240027.1;refGene_gn=CDC42BPG;refGene_tx=XM_023240028.1;refGene_gn=CDC42BPG;refGene_tx=XM_023240029.1
D1	110052349	110052529	refGene_gn=CDC42BPG;refGene_tx=XM_019812985.2;refGene_gn=CDC42BPG;refGene_tx=XM_023240022.1;refGene_gn=CDC42BPG;refGene_tx=XM_023240023.1;refGene_gn=CDC42BPG;refGene_tx=XM_023240024.1;refGene_gn=CDC42BPG;refGene_tx=XM_023240025.1;refGene_gn=CDC42BPG;refGene_tx=XM_023240026.1;refGene_gn=CDC42BPG;refGene_tx=XM_023240027.1;refGene_gn=CDC42BPG;refGene_tx=XM_023240028.1;refGene_gn=CDC42BPG;refGene_tx=XM_023240029.1;refGene_gn=CDC42BPG;refGene_tx=XM_023240030.1
D1	110052620	110052718	refGene_gn=CDC42BPG;refGene_tx=XM_019812985.2;refGene_gn=CDC42BPG;refGene_tx=XM_023240022.1;refGene_gn=CDC42BPG;refGene_tx=XM_023240023.1;refGene_gn=CDC42BPG;refGene_tx=XM_023240024.1;refGene_gn=CDC42BPG;refGene_tx=XM_023240025.1;refGene_gn=CDC42BPG;refGene_tx=XM_023240026.1;refGene_gn=CDC42BPG;refGene_tx=XM_023240027.1;refGene_gn=CDC42BPG;refGene_tx=XM_023240028.1;refGene_gn=CDC42BPG;refGene_tx=XM_023240029.1;refGene_gn=CDC42BPG;refGene_tx=XM_023240030.1
D1	110053010	110053091	refGene_gn=CDC42BPG;refGene_tx=XM_019812985.2;refGene_gn=CDC42BPG;refGene_tx=XM_023240022.1;refGene_gn=CDC42BPG;refGene_tx=XM_023240023.1;refGene_gn=CDC42BPG;refGene_tx=XM_023240024.1;refGene_gn=CDC42BPG;refGene_tx=XM_023240025.1;refGene_gn=CDC42BPG;refGene_tx=XM_023240026.1;refGene_gn=CDC42BPG;refGene_tx=XM_023240027.1;refGene_gn=CDC42BPG;refGene_

tx=XM_023240028.1;refGene_gn=CDC42BPG;refGene_tx=XM_023240029.1;refGene_gn=CDC42BPG;refGene_tx=XM_023240030.1
D1 110053179 110053277
refGene_gn=CDC42BPG;refGene_tx=XM_019812985.2;refGene_gn=CDC42BPG;refGene_tx=XM_023240022.1;refGene_gn=CDC42BPG;refGene_tx=XM_023240023.1;refGene_gn=CDC42BPG;refGene_tx=XM_023240024.1;refGene_gn=CDC42BPG;refGene_tx=XM_023240025.1;refGene_gn=CDC42BPG;refGene_tx=XM_023240026.1;refGene_gn=CDC42BPG;refGene_tx=XM_023240027.1;refGene_gn=CDC42BPG;refGene_tx=XM_023240028.1;refGene_gn=CDC42BPG;refGene_tx=XM_023240029.1;refGene_gn=CDC42BPG;refGene_tx=XM_023240030.1
D1 110054096 110054176
refGene_gn=CDC42BPG;refGene_tx=XM_019812985.2;refGene_gn=CDC42BPG;refGene_tx=XM_023240022.1;refGene_gn=CDC42BPG;refGene_tx=XM_023240023.1;refGene_gn=CDC42BPG;refGene_tx=XM_023240024.1;refGene_gn=CDC42BPG;refGene_tx=XM_023240025.1;refGene_gn=CDC42BPG;refGene_tx=XM_023240026.1;refGene_gn=CDC42BPG;refGene_tx=XM_023240027.1;refGene_gn=CDC42BPG;refGene_tx=XM_023240028.1;refGene_gn=CDC42BPG;refGene_tx=XM_023240029.1;refGene_gn=CDC42BPG;refGene_tx=XM_023240030.1
D1 110054610 110054859
refGene_gn=CDC42BPG;refGene_tx=XM_019812985.2;refGene_gn=CDC42BPG;refGene_tx=XM_023240022.1;refGene_gn=CDC42BPG;refGene_tx=XM_023240023.1;refGene_gn=CDC42BPG;refGene_tx=XM_023240024.1;refGene_gn=CDC42BPG;refGene_tx=XM_023240025.1;refGene_gn=CDC42BPG;refGene_tx=XM_023240026.1;refGene_gn=CDC42BPG;refGene_tx=XM_023240027.1;refGene_gn=CDC42BPG;refGene_tx=XM_023240028.1;refGene_gn=CDC42BPG;refGene_tx=XM_023240029.1;refGene_gn=CDC42BPG;refGene_tx=XM_023240030.1
D1 110054996 110055197
refGene_gn=CDC42BPG;refGene_tx=XM_019812985.2;refGene_gn=CDC42BPG;refGene_tx=XM_023240022.1;refGene_gn=CDC42BPG;refGene_tx=XM_023240023.1;refGene_gn=CDC42BPG;refGene_tx=XM_023240024.1;refGene_gn=CDC42BPG;refGene_tx=XM_023240025.1;refGene_gn=CDC42BPG;refGene_tx=XM_023240026.1;refGene_gn=CDC42BPG;refGene_tx=XM_023240027.1;refGene_gn=CDC42BPG;refGene_tx=XM_023240028.1;refGene_gn=CDC42BPG;refGene_tx=XM_023240029.1;refGene_gn=CDC42BPG;refGene_tx=XM_023240030.1
D1 110055433 110055527
refGene_gn=CDC42BPG;refGene_tx=XM_019812985.2;refGene_gn=CDC42BPG;refGene_tx=XM_023240022.1;refGene_gn=CDC42BPG;refGene_tx=XM_023240023.1;refGene_gn=CDC42BPG;refGene_tx=XM_023240024.1;refGene_gn=CDC42BPG;refGene_tx=XM_023240025.1;refGene_gn=CDC42BPG;refGene_tx=XM_023240026.1;refGene_gn=CDC42BPG;refGene_tx=XM_023240027.1;refGene_gn=CDC42BPG;refGene_tx=XM_023240028.1;refGene_gn=CDC42BPG;refGene_tx=XM_023240029.1;refGene_gn=CDC42BPG;refGene_tx=XM_023240030.1
D1 110056084 110056233
refGene_gn=CDC42BPG;refGene_tx=XM_019812985.2;refGene_gn=CDC42BPG;refGene_tx=XM_023240022.1;refGene_gn=CDC42BPG;refGene_tx=XM_023240023.1;refGene_gn=CDC42BPG;refGene_tx=XM_023240024.1;refGene_gn=CDC42BPG;refGene_tx=XM_023240025.1;refGene_gn=CDC42BPG;refGene_tx=XM_023240026.1;refGene_gn=CDC42BPG;refGene_tx=XM_023240027.1;refGene_gn=CDC42BPG;refGene_tx=XM_023240028.1;refGene_gn=CDC42BPG;refGene_tx=XM_023240029.1;refGene_gn=CDC42BPG;refGene_tx=XM_023240030.1

D1 110056570 110056666
refGene_gn=CDC42BPG;refGene_tx=XM_019812985.2;refGene_gn=CDC42BPG;refGene_tx=XM_023240022.1;refGene_gn=CDC42BPG;refGene_tx=XM_023240023.1;refGene_gn=CDC42BPG;refGene_tx=XM_023240024.1;refGene_gn=CDC42BPG;refGene_tx=XM_023240025.1;refGene_gn=CDC42BPG;refGene_tx=XM_023240026.1;refGene_gn=CDC42BPG;refGene_tx=XM_023240027.1;refGene_gn=CDC42BPG;refGene_tx=XM_023240028.1;refGene_gn=CDC42BPG;refGene_tx=XM_023240029.1;refGene_gn=CDC42BPG;refGene_tx=XM_023240030.1
D1 110057035 110057119
refGene_gn=CDC42BPG;refGene_tx=XM_019812985.2;refGene_gn=CDC42BPG;refGene_tx=XM_023240022.1;refGene_gn=CDC42BPG;refGene_tx=XM_023240023.1;refGene_gn=CDC42BPG;refGene_tx=XM_023240024.1;refGene_gn=CDC42BPG;refGene_tx=XM_023240025.1;refGene_gn=CDC42BPG;refGene_tx=XM_023240026.1;refGene_gn=CDC42BPG;refGene_tx=XM_023240027.1;refGene_gn=CDC42BPG;refGene_tx=XM_023240028.1;refGene_gn=CDC42BPG;refGene_tx=XM_023240029.1;refGene_gn=CDC42BPG;refGene_tx=XM_023240030.1
D1 110057200 110057292
refGene_gn=CDC42BPG;refGene_tx=XM_019812985.2;refGene_gn=CDC42BPG;refGene_tx=XM_023240022.1;refGene_gn=CDC42BPG;refGene_tx=XM_023240023.1;refGene_gn=CDC42BPG;refGene_tx=XM_023240024.1;refGene_gn=CDC42BPG;refGene_tx=XM_023240025.1;refGene_gn=CDC42BPG;refGene_tx=XM_023240026.1;refGene_gn=CDC42BPG;refGene_tx=XM_023240027.1;refGene_gn=CDC42BPG;refGene_tx=XM_023240028.1;refGene_gn=CDC42BPG;refGene_tx=XM_023240029.1;refGene_gn=CDC42BPG;refGene_tx=XM_023240030.1
D1 110059930 110060090
refGene_gn=CDC42BPG;refGene_tx=XM_019812985.2;refGene_gn=CDC42BPG;refGene_tx=XM_023240022.1;refGene_gn=CDC42BPG;refGene_tx=XM_023240023.1;refGene_gn=CDC42BPG;refGene_tx=XM_023240024.1;refGene_gn=CDC42BPG;refGene_tx=XM_023240025.1;refGene_gn=CDC42BPG;refGene_tx=XM_023240026.1;refGene_gn=CDC42BPG;refGene_tx=XM_023240027.1;refGene_gn=CDC42BPG;refGene_tx=XM_023240028.1;refGene_gn=CDC42BPG;refGene_tx=XM_023240029.1;refGene_gn=CDC42BPG;refGene_tx=XM_023240030.1
D1 110068016 110068541
refGene_gn=EHD1;refGene_tx=XM_003993572.5
D1 110069017 110069182
refGene_gn=EHD1;refGene_tx=XM_003993572.5
D1 110071482 110071895
refGene_gn=EHD1;refGene_tx=XM_003993572.5
D1 110084481 110084579
refGene_gn=EHD1;refGene_tx=XM_003993572.5
D1 110088301 110088728
refGene_gn=EHD1;refGene_tx=XM_003993572.5
D1 110089918 110089937
refGene_gn=EHD1;refGene_tx=XM_003993572.5
D1 110103057 110103294
refGene_gn=ATG2A;refGene_tx=XM_023240031.1
D1 110103374 110103527
refGene_gn=ATG2A;refGene_tx=XM_023240031.1
D1 110104571 110104726
refGene_gn=ATG2A;refGene_tx=XM_023240031.1

D1 110104873 110104995
refGene_gn=ATG2A;refGene_tx=XM_023240031.1
D1 110105253 110105336
refGene_gn=ATG2A;refGene_tx=XM_023240031.1
D1 110105529 110105602
refGene_gn=ATG2A;refGene_tx=XM_023240031.1
D1 110105688 110105821
refGene_gn=ATG2A;refGene_tx=XM_023240031.1
D1 110105910 110105995
refGene_gn=ATG2A;refGene_tx=XM_023240031.1
D1 110106079 110106136
refGene_gn=ATG2A;refGene_tx=XM_023240031.1
D1 110106137 110106182
refGene_gn=ATG2A;refGene_tx=XM_023240031.1
D1 110106377 110106584
refGene_gn=ATG2A;refGene_tx=XM_023240031.1
D1 110107819 110107955
refGene_gn=ATG2A;refGene_tx=XM_023240031.1
D1 110108212 110108439
refGene_gn=ATG2A;refGene_tx=XM_023240031.1
D1 110108976 110109191
refGene_gn=ATG2A;refGene_tx=XM_023240031.1
D1 110109279 110109406
refGene_gn=ATG2A;refGene_tx=XM_023240031.1
D1 110109562 110109727
refGene_gn=ATG2A;refGene_tx=XM_023240031.1
D1 110110244 110110321
refGene_gn=ATG2A;refGene_tx=XM_023240031.1
D1 110110537 110110608
refGene_gn=ATG2A;refGene_tx=XM_023240031.1
D1 110111892 110111985
refGene_gn=ATG2A;refGene_tx=XM_023240031.1
D1 110112074 110112181
refGene_gn=ATG2A;refGene_tx=XM_023240031.1
D1 110112415 110112496
refGene_gn=ATG2A;refGene_tx=XM_023240031.1
D1 110112599 110112799
refGene_gn=ATG2A;refGene_tx=XM_023240031.1
D1 110112892 110113043
refGene_gn=ATG2A;refGene_tx=XM_023240031.1
D1 110113475 110113636
refGene_gn=ATG2A;refGene_tx=XM_023240031.1
D1 110113741 110113881
refGene_gn=ATG2A;refGene_tx=XM_023240031.1
D1 110113965 110114108
refGene_gn=ATG2A;refGene_tx=XM_023240031.1
D1 110114767 110114927
refGene_gn=ATG2A;refGene_tx=XM_023240031.1
D1 110115056 110115153
refGene_gn=ATG2A;refGene_tx=XM_023240031.1

D1 110115481 110115725
refGene_gn=ATG2A;refGene_tx=XM_023240031.1
D1 110115826 110115985
refGene_gn=ATG2A;refGene_tx=XM_023240031.1
D1 110116441 110116534
refGene_gn=ATG2A;refGene_tx=XM_023240031.1
D1 110116630 110116775
refGene_gn=ATG2A;refGene_tx=XM_023240031.1
D1 110116858 110117096
refGene_gn=ATG2A;refGene_tx=XM_023240031.1
D1 110117641 110117782
refGene_gn=ATG2A;refGene_tx=XM_023240031.1
D1 110117890 110118055
refGene_gn=ATG2A;refGene_tx=XM_023240031.1
D1 110118143 110118240
refGene_gn=ATG2A;refGene_tx=XM_023240031.1
D1 110118529 110118628
refGene_gn=ATG2A;refGene_tx=XM_023240031.1
D1 110118777 110118913
refGene_gn=ATG2A;refGene_tx=XM_023240031.1
D1 110119330 110119433
refGene_gn=ATG2A;refGene_tx=XM_023240031.1
D1 110119591 110119744
refGene_gn=ATG2A;refGene_tx=XM_023240031.1
D1 110119838 110120001
refGene_gn=ATG2A;refGene_tx=XM_023240031.1
D1 110122135 110122306
refGene_gn=ATG2A;refGene_tx=XM_023240031.1
D1 110128512 110128711
refGene_gn=PPP2R5B;refGene_tx=XM_003993575.5;refGene_gn=PPP2R5B;refGene_tx=XM_019812650.2
D1 110129467 110129664
refGene_gn=PPP2R5B;refGene_tx=XM_003993575.5;refGene_gn=PPP2R5B;refGene_tx=XM_019812650.2
D1 110130279 110130381
refGene_gn=PPP2R5B;refGene_tx=XM_003993575.5;refGene_gn=PPP2R5B;refGene_tx=XM_019812650.2
D1 110130514 110130607
refGene_gn=PPP2R5B;refGene_tx=XM_003993575.5;refGene_gn=PPP2R5B;refGene_tx=XM_019812650.2
D1 110130733 110130864
refGene_gn=PPP2R5B;refGene_tx=XM_003993575.5;refGene_gn=PPP2R5B;refGene_tx=XM_019812650.2
D1 110131959 110132019
refGene_gn=PPP2R5B;refGene_tx=XM_003993575.5;refGene_gn=PPP2R5B;refGene_tx=XM_019812650.2
D1 110132119 110132228
refGene_gn=PPP2R5B;refGene_tx=XM_003993575.5;refGene_gn=PPP2R5B;refGene_tx=XM_019812650.2
D1 110132778 110132832

refGene_gn=PPP2R5B;refGene_tx=XM_003993575.5;refGene_gn=PPP2R5B;refGene_tx=XM_019812650.2
D1 110132905 110132956
refGene_gn=PPP2R5B;refGene_tx=XM_003993575.5;refGene_gn=PPP2R5B;refGene_tx=XM_019812650.2
D1 110133085 110133205
refGene_gn=PPP2R5B;refGene_tx=XM_003993575.5;refGene_gn=PPP2R5B;refGene_tx=XM_019812650.2
D1 110134076 110134204
refGene_gn=PPP2R5B;refGene_tx=XM_003993575.5;refGene_gn=PPP2R5B;refGene_tx=XM_019812650.2
D1 110134438 110134540
refGene_gn=PPP2R5B;refGene_tx=XM_003993575.5;refGene_gn=PPP2R5B;refGene_tx=XM_019812650.2
D1 110134648 110134811
refGene_gn=PPP2R5B;refGene_tx=XM_019812650.2
D1 110134914 110135062
refGene_gn=PPP2R5B;refGene_tx=XM_003993575.5
D1 110135940 110136042
refGene_gn=GPHA2;refGene_tx=XM_011287010.2
D1 110136156 110136341
refGene_gn=GPHA2;refGene_tx=XM_011287010.2
D1 110136502 110136605
refGene_gn=GPHA2;refGene_tx=XM_011287010.2
D1 110136957 110137095
refGene_gn=GPHA2;refGene_tx=XM_011287010.2
D1 110137419 110137426
refGene_gn=MAJIN;refGene_tx=XM_023240034.1
D1 110139215 110139261
refGene_gn=MAJIN;refGene_tx=XM_019812651.2;refGene_gn=MAJIN;refGene_tx=XM_019812655.2;refGene_gn=MAJIN;refGene_tx=XM_019812656.2;refGene_gn=MAJIN;refGene_tx=XM_019812657.2;refGene_gn=MAJIN;refGene_tx=XM_019812659.2;refGene_gn=MAJIN;refGene_tx=XM_019812660.2;refGene_gn=MAJIN;refGene_tx=XM_019812665.2;refGene_gn=MAJIN;refGene_tx=XM_023240032.1;refGene_gn=MAJIN;refGene_tx=XM_023240033.1;refGene_gn=MAJIN;refGene_tx=XM_023240035.1
D1 110139868 110139986
refGene_gn=MAJIN;refGene_tx=XM_019812651.2;refGene_gn=MAJIN;refGene_tx=XM_019812655.2;refGene_gn=MAJIN;refGene_tx=XM_019812656.2;refGene_gn=MAJIN;refGene_tx=XM_019812657.2;refGene_gn=MAJIN;refGene_tx=XM_019812659.2;refGene_gn=MAJIN;refGene_tx=XM_019812660.2;refGene_gn=MAJIN;refGene_tx=XM_019812665.2;refGene_gn=MAJIN;refGene_tx=XM_023240032.1;refGene_gn=MAJIN;refGene_tx=XM_023240033.1;refGene_gn=MAJIN;refGene_tx=XM_023240034.1;refGene_gn=MAJIN;refGene_tx=XM_023240035.1
D1 110140751 110140824
refGene_gn=MAJIN;refGene_tx=XM_019812651.2;refGene_gn=MAJIN;refGene_tx=XM_019812655.2;refGene_gn=MAJIN;refGene_tx=XM_019812656.2;refGene_gn=MAJIN;refGene_tx=XM_019812657.2;refGene_gn=MAJIN;refGene_tx=XM_019812659.2;refGene_gn=MAJIN;refGene_tx=XM_019812660.2;refGene_gn=MAJIN;refGene_tx=XM_019812665.2;refGene_gn=MAJIN;refGene_tx=XM_023240032.1;refGen

e_gn=MAJIN;refGene_tx=XM_023240033.1;refGene_gn=MAJIN;refGene_tx=XM_023240034.1;refGene_gn=MAJIN;refGene_tx=XM_023240035.1
D1 110144660 110144724
refGene_gn=MAJIN;refGene_tx=XM_019812651.2;refGene_gn=MAJIN;refGene_tx=XM_019812655.2;refGene_gn=MAJIN;refGene_tx=XM_019812656.2;refGene_gn=MAJIN;refGene_tx=XM_019812657.2;refGene_gn=MAJIN;refGene_tx=XM_019812659.2;refGene_gn=MAJIN;refGene_tx=XM_019812660.2;refGene_gn=MAJIN;refGene_tx=XM_019812665.2;refGene_gn=MAJIN;refGene_tx=XM_023240032.1;refGene_gn=MAJIN;refGene_tx=XM_023240033.1;refGene_gn=MAJIN;refGene_tx=XM_023240034.1;refGene_gn=MAJIN;refGene_tx=XM_023240035.1
D1 110145560 110145732
refGene_gn=MAJIN;refGene_tx=XM_023240037.1;refGene_gn=MAJIN;refGene_tx=XM_023240038.1
D1 110145935 110146048
refGene_gn=MAJIN;refGene_tx=XM_019812651.2;refGene_gn=MAJIN;refGene_tx=XM_019812655.2;refGene_gn=MAJIN;refGene_tx=XM_019812656.2;refGene_gn=MAJIN;refGene_tx=XM_019812657.2;refGene_gn=MAJIN;refGene_tx=XM_019812659.2;refGene_gn=MAJIN;refGene_tx=XM_019812660.2;refGene_gn=MAJIN;refGene_tx=XM_019812665.2;refGene_gn=MAJIN;refGene_tx=XM_023240032.1;refGene_gn=MAJIN;refGene_tx=XM_023240033.1;refGene_gn=MAJIN;refGene_tx=XM_023240034.1;refGene_gn=MAJIN;refGene_tx=XM_023240035.1;refGene_gn=MAJIN;refGene_tx=XM_023240037.1;refGene_gn=MAJIN;refGene_tx=XM_023240038.1
D1 110146348 110146380
refGene_gn=MAJIN;refGene_tx=XM_019812651.2;refGene_gn=MAJIN;refGene_tx=XM_019812655.2;refGene_gn=MAJIN;refGene_tx=XM_019812656.2;refGene_gn=MAJIN;refGene_tx=XM_019812657.2;refGene_gn=MAJIN;refGene_tx=XM_019812659.2;refGene_gn=MAJIN;refGene_tx=XM_019812660.2;refGene_gn=MAJIN;refGene_tx=XM_019812665.2;refGene_gn=MAJIN;refGene_tx=XM_023240032.1;refGene_gn=MAJIN;refGene_tx=XM_023240033.1;refGene_gn=MAJIN;refGene_tx=XM_023240034.1;refGene_gn=MAJIN;refGene_tx=XM_023240035.1;refGene_gn=MAJIN;refGene_tx=XM_023240037.1;refGene_gn=MAJIN;refGene_tx=XM_023240038.1
D1 110147479 110147605
refGene_gn=MAJIN;refGene_tx=XM_019812651.2;refGene_gn=MAJIN;refGene_tx=XM_019812655.2;refGene_gn=MAJIN;refGene_tx=XM_019812656.2;refGene_gn=MAJIN;refGene_tx=XM_019812657.2;refGene_gn=MAJIN;refGene_tx=XM_019812659.2;refGene_gn=MAJIN;refGene_tx=XM_019812660.2;refGene_gn=MAJIN;refGene_tx=XM_019812665.2;refGene_gn=MAJIN;refGene_tx=XM_023240032.1;refGene_gn=MAJIN;refGene_tx=XM_023240033.1;refGene_gn=MAJIN;refGene_tx=XM_023240034.1;refGene_gn=MAJIN;refGene_tx=XM_023240035.1;refGene_gn=MAJIN;refGene_tx=XM_023240037.1;refGene_gn=MAJIN;refGene_tx=XM_023240038.1
D1 110147767 110147843
refGene_gn=MAJIN;refGene_tx=XM_019812651.2;refGene_gn=MAJIN;refGene_tx=XM_019812655.2;refGene_gn=MAJIN;refGene_tx=XM_019812656.2;refGene_gn=MAJIN;refGene_tx=XM_019812657.2;refGene_gn=MAJIN;refGene_tx=XM_019812659.2;refGene_gn=MAJIN;refGene_tx=XM_019812660.2;refGene_gn=MAJIN;refGene_tx=XM_023240032.1;refGene_gn=MAJIN;refGene_tx=XM_023240033.1;refGene_gn=MAJIN;refGene_tx=XM_023240034.1;refGene_gn=MAJIN;refGene_tx=XM_023240035.1;refGene_gn=MAJIN;refGene_tx=XM_023240037.1;refGene_gn=MAJIN;refGene_tx=XM_023240038.1
D1 110149504 110149550

refGene_gn=MAJIN;refGene_tx=XM_019812651.2;refGene_gn=MAJIN;refGene_tx=XM_019812655.2;refGene_gn=MAJIN;refGene_tx=XM_019812656.2;refGene_gn=MAJIN;refGene_tx=XM_019812657.2;refGene_gn=MAJIN;refGene_tx=XM_019812659.2;refGene_gn=MAJIN;refGene_tx=XM_019812660.2;refGene_gn=MAJIN;refGene_tx=XM_023240032.1;refGene_gn=MAJIN;refGene_tx=XM_023240033.1;refGene_gn=MAJIN;refGene_tx=XM_023240034.1;refGene_gn=MAJIN;refGene_tx=XM_023240035.1;refGene_gn=MAJIN;refGene_tx=XM_023240037.1;refGene_gn=MAJIN;refGene_tx=XM_023240038.1
D1 110156819 110156937
refGene_gn=MAJIN;refGene_tx=XM_019812651.2;refGene_gn=MAJIN;refGene_tx=XM_019812655.2;refGene_gn=MAJIN;refGene_tx=XM_019812656.2;refGene_gn=MAJIN;refGene_tx=XM_019812657.2;refGene_gn=MAJIN;refGene_tx=XM_019812659.2;refGene_gn=MAJIN;refGene_tx=XM_019812660.2;refGene_gn=MAJIN;refGene_tx=XM_023240032.1;refGene_gn=MAJIN;refGene_tx=XM_023240033.1;refGene_gn=MAJIN;refGene_tx=XM_023240034.1;refGene_gn=MAJIN;refGene_tx=XM_023240035.1;refGene_gn=MAJIN;refGene_tx=XM_023240037.1;refGene_gn=MAJIN;refGene_tx=XM_023240038.1
D1 110162301 110162338
refGene_gn=MAJIN;refGene_tx=XM_019812655.2;refGene_gn=MAJIN;refGene_tx=XM_023240033.1;refGene_gn=MAJIN;refGene_tx=XM_023240034.1;refGene_gn=MAJIN;refGene_tx=XM_023240037.1;refGene_gn=MAJIN;refGene_tx=XM_023240038.1
D1 110162433 110162455
refGene_gn=MAJIN;refGene_tx=XM_019812656.2
D1 110181903 110182590
refGene_gn=BATF2;refGene_tx=XM_023240039.1;refGene_gn=BATF2;refGene_tx=XM_023240040.1
D1 110185866 110185968
refGene_gn=BATF2;refGene_tx=XM_023240039.1;refGene_gn=BATF2;refGene_tx=XM_023240040.1
D1 110187292 110187331
refGene_gn=BATF2;refGene_tx=XM_023240039.1;refGene_gn=BATF2;refGene_tx=XM_023240040.1
D1 110202420 110202485
refGene_gn=ARL2;refGene_tx=XM_003993577.4
D1 110205587 110205698
refGene_gn=ARL2;refGene_tx=XM_003993577.4
D1 110205793 110205956
refGene_gn=ARL2;refGene_tx=XM_003993577.4
D1 110207448 110207529
refGene_gn=ARL2;refGene_tx=XM_003993577.4
D1 110208405 110208540
refGene_gn=ARL2;refGene_tx=XM_003993577.4
D1 110212262 110212361
refGene_gn=SNX15;refGene_tx=XM_003993579.3
D1 110217533 110217569
refGene_gn=SNX15;refGene_tx=XM_003993579.3
D1 110217777 110217898
refGene_gn=SNX15;refGene_tx=XM_003993579.3
D1 110219410 110219526

refGene_gn=SNX15;refGene_tx=XM_003993579.3
D1 110219626 110219774
refGene_gn=SNX15;refGene_tx=XM_003993579.3
D1 110220341 110220485
refGene_gn=SNX15;refGene_tx=XM_003993579.3
D1 110221465 110221723
refGene_gn=SNX15;refGene_tx=XM_003993579.3
D1 110222046 110222153
refGene_gn=SNX15;refGene_tx=XM_003993579.3
D1 110223024 110223128
refGene_gn=SAC3D1;refGene_tx=XM_003993581.4
D1 110223305 110223886
refGene_gn=SAC3D1;refGene_tx=XM_003993581.4
D1 110224996 110225499
refGene_gn=SAC3D1;refGene_tx=XM_003993581.4
D1 110226292 110226479
refGene_gn=NAALADL1;refGene_tx=XM_023240042.1
D1 110226824 110226917
refGene_gn=NAALADL1;refGene_tx=XM_023240042.1
D1 110227024 110227112
refGene_gn=NAALADL1;refGene_tx=XM_023240042.1
D1 110227187 110227361
refGene_gn=NAALADL1;refGene_tx=XM_023240042.1
D1 110227444 110227526
refGene_gn=NAALADL1;refGene_tx=XM_023240042.1
D1 110228817 110228908
refGene_gn=NAALADL1;refGene_tx=XM_023240042.1
D1 110228989 110229081
refGene_gn=NAALADL1;refGene_tx=XM_023240042.1
D1 110229307 110229358
refGene_gn=NAALADL1;refGene_tx=XM_023240042.1
D1 110229359 110229376
refGene_gn=NAALADL1;refGene_tx=XM_023240042.1
D1 110229460 110229524
refGene_gn=NAALADL1;refGene_tx=XM_023240042.1
D1 110229608 110229694
refGene_gn=NAALADL1;refGene_tx=XM_023240042.1
D1 110232337 110232457
refGene_gn=NAALADL1;refGene_tx=XM_023240042.1
D1 110232564 110232650
refGene_gn=NAALADL1;refGene_tx=XM_023240042.1
D1 110233357 110233462
refGene_gn=NAALADL1;refGene_tx=XM_023240042.1
D1 110233549 110233812
refGene_gn=NAALADL1;refGene_tx=XM_023240042.1
D1 110233813 110233834
refGene_gn=NAALADL1;refGene_tx=XM_023240042.1
D1 110235696 110235819
refGene_gn=NAALADL1;refGene_tx=XM_023240042.1
D1 110236179 110236301

refGene_gn=NAALADL1;refGene_tx=XM_023240042.1
D1 110236385 110236556
refGene_gn=NAALADL1;refGene_tx=XM_023240042.1
D1 110236664 110236866
refGene_gn=NAALADL1;refGene_tx=XM_023240042.1
D1 110243599 110243782
refGene_gn=CDCA5;refGene_tx=XM_023240043.1;refGene_gn=CDCA5;refGene_tx=XM_023240044.1
D1 110243931 110243998
refGene_gn=CDCA5;refGene_tx=XM_023240043.1;refGene_gn=CDCA5;refGene_tx=XM_023240044.1
D1 110244965 110245064
refGene_gn=CDCA5;refGene_tx=XM_023240043.1;refGene_gn=CDCA5;refGene_tx=XM_023240044.1
D1 110245247 110245354
refGene_gn=CDCA5;refGene_tx=XM_023240043.1;refGene_gn=CDCA5;refGene_tx=XM_023240044.1
D1 110245563 110245677
refGene_gn=CDCA5;refGene_tx=XM_023240043.1;refGene_gn=CDCA5;refGene_tx=XM_023240044.1
D1 110250683 110250764
refGene_gn=CDCA5;refGene_tx=XM_003993583.4
D1 110250949 110251384
refGene_gn=CDCA5;refGene_tx=XM_003993583.4;refGene_gn=CDCA5;refGene_tx=XM_023240043.1
D1 110254779 110254815
refGene_gn=CDCA5;refGene_tx=XM_003993583.4;refGene_gn=CDCA5;refGene_tx=XM_023240043.1;refGene_gn=CDCA5;refGene_tx=XM_023240044.1
D1 110254894 110254960
refGene_gn=CDCA5;refGene_tx=XM_003993583.4;refGene_gn=CDCA5;refGene_tx=XM_023240043.1;refGene_gn=CDCA5;refGene_tx=XM_023240044.1
D1 110255038 110255133
refGene_gn=CDCA5;refGene_tx=XM_003993583.4;refGene_gn=CDCA5;refGene_tx=XM_023240043.1;refGene_gn=CDCA5;refGene_tx=XM_023240044.1
D1 110255343 110255389
refGene_gn=CDCA5;refGene_tx=XM_003993583.4;refGene_gn=CDCA5;refGene_tx=XM_023240043.1;refGene_gn=CDCA5;refGene_tx=XM_023240044.1
D1 110256099 110256201
refGene_gn=ZFPL1;refGene_tx=XM_006937495.4
D1 110256493 110256605
refGene_gn=ZFPL1;refGene_tx=XM_006937495.4
D1 110258000 110258194
refGene_gn=ZFPL1;refGene_tx=XM_006937495.4
D1 110258302 110258375
refGene_gn=ZFPL1;refGene_tx=XM_006937495.4
D1 110258448 110258595
refGene_gn=ZFPL1;refGene_tx=XM_006937495.4
D1 110258844 110258968
refGene_gn=ZFPL1;refGene_tx=XM_006937495.4
D1 110259475 110259662

refGene_gn=ZFPL1;refGene_tx=XM_006937495.4
D1 110259847 110260052
refGene_gn=TMEM262;refGene_tx=XM_006937492.4;refGene_gn=TMEM262;refGene_tx=XM_006937493.4;refGene_gn=TMEM262;refGene_tx=XM_006937494.4
D1 110260158 110260258
refGene_gn=TMEM262;refGene_tx=XM_006937492.4;refGene_gn=TMEM262;refGene_tx=XM_006937493.4;refGene_gn=TMEM262;refGene_tx=XM_006937494.4
D1 110260365 110260539
refGene_gn=TMEM262;refGene_tx=XM_006937492.4;refGene_gn=TMEM262;refGene_tx=XM_006937493.4;refGene_gn=TMEM262;refGene_tx=XM_006937494.4
D1 110267075 110267300
refGene_gn=VPS51;refGene_tx=XM_003993629.4
D1 110267768 110267898
refGene_gn=VPS51;refGene_tx=XM_003993629.4
D1 110282744 110282891
refGene_gn=VPS51;refGene_tx=XM_003993629.4
D1 110282971 110283191
refGene_gn=VPS51;refGene_tx=XM_003993629.4
D1 110283324 110284042
refGene_gn=VPS51;refGene_tx=XM_003993629.4
D1 110284453 110284669
refGene_gn=VPS51;refGene_tx=XM_003993629.4
D1 110284858 110285077
refGene_gn=VPS51;refGene_tx=XM_003993629.4
D1 110285740 110285862
refGene_gn=VPS51;refGene_tx=XM_003993629.4
D1 110285951 110286039
refGene_gn=VPS51;refGene_tx=XM_003993629.4
D1 110286862 110287123
refGene_gn=VPS51;refGene_tx=XM_003993629.4
D1 110287567 110287619
refGene_gn=TM7SF2;refGene_tx=XM_003993585.4
D1 110288015 110288212
refGene_gn=TM7SF2;refGene_tx=XM_003993585.4
D1 110288295 110288350
refGene_gn=TM7SF2;refGene_tx=XM_003993585.4
D1 110288724 110288919
refGene_gn=TM7SF2;refGene_tx=XM_003993585.4
D1 110289002 110289106
refGene_gn=TM7SF2;refGene_tx=XM_003993585.4
D1 110290481 110290601
refGene_gn=TM7SF2;refGene_tx=XM_003993585.4
D1 110290688 110290857
refGene_gn=TM7SF2;refGene_tx=XM_003993585.4
D1 110291059 110291140
refGene_gn=TM7SF2;refGene_tx=XM_003993585.4
D1 110291234 110291357
refGene_gn=TM7SF2;refGene_tx=XM_003993585.4
D1 110291676 110291837
refGene_gn=TM7SF2;refGene_tx=XM_003993585.4

D1 110292267 110293464
refGene_gn=ZNHIT2;refGene_tx=XM_023240045.1
D1 110295911 110296037
refGene_gn=FAU;refGene_tx=XM_003993586.5
D1 110296213 110296269
refGene_gn=FAU;refGene_tx=XM_003993586.5
D1 110296534 110296679
refGene_gn=FAU;refGene_tx=XM_003993586.5
D1 110296789 110296864
refGene_gn=FAU;refGene_tx=XM_003993586.5
D1 110297400 110297499
refGene_gn=MRPL49;refGene_tx=XM_003993587.5;refGene_gn=MRPL49;refGene_tx=XM_023240046.1
D1 110298417 110298568
refGene_gn=MRPL49;refGene_tx=XM_003993587.5;refGene_gn=MRPL49;refGene_tx=XM_023240046.1
D1 110299358 110299483
refGene_gn=MRPL49;refGene_tx=XM_003993587.5;refGene_gn=MRPL49;refGene_tx=XM_023240046.1
D1 110299595 110299742
refGene_gn=MRPL49;refGene_tx=XM_003993587.5;refGene_gn=MRPL49;refGene_tx=XM_023240046.1
D1 110301855 110301962
refGene_gn=SYVN1;refGene_tx=XM_003993588.4;refGene_gn=SYVN1;refGene_tx=XM_006937497.4;refGene_gn=SYVN1;refGene_tx=XM_019812667.2;refGene_gn=SYVN1;refGene_tx=XM_019812668.2
D1 110302041 110302190
refGene_gn=SYVN1;refGene_tx=XM_003993588.4;refGene_gn=SYVN1;refGene_tx=XM_006937497.4;refGene_gn=SYVN1;refGene_tx=XM_019812667.2;refGene_gn=SYVN1;refGene_tx=XM_019812668.2
D1 110303182 110303369
refGene_gn=SYVN1;refGene_tx=XM_003993588.4;refGene_gn=SYVN1;refGene_tx=XM_006937497.4;refGene_gn=SYVN1;refGene_tx=XM_019812667.2;refGene_gn=SYVN1;refGene_tx=XM_019812668.2
D1 110303457 110303616
refGene_gn=SYVN1;refGene_tx=XM_003993588.4;refGene_gn=SYVN1;refGene_tx=XM_006937497.4;refGene_gn=SYVN1;refGene_tx=XM_019812667.2;refGene_gn=SYVN1;refGene_tx=XM_019812668.2
D1 110303711 110303840
refGene_gn=SYVN1;refGene_tx=XM_003993588.4;refGene_gn=SYVN1;refGene_tx=XM_006937497.4;refGene_gn=SYVN1;refGene_tx=XM_019812667.2
D1 110304123 110304287
refGene_gn=SYVN1;refGene_tx=XM_003993588.4;refGene_gn=SYVN1;refGene_tx=XM_006937497.4;refGene_gn=SYVN1;refGene_tx=XM_019812667.2;refGene_gn=SYVN1;refGene_tx=XM_019812668.2
D1 110304381 110304498
refGene_gn=SYVN1;refGene_tx=XM_003993588.4;refGene_gn=SYVN1;refGene_tx=XM_006937497.4;refGene_gn=SYVN1;refGene_tx=XM_019812667.2;refGene_gn=SYVN1;refGene_tx=XM_019812668.2
D1 110304593 110304659

refGene_gn=SYVN1;refGene_tx=XM_003993588.4;refGene_gn=SYVN1;refGene_tx=XM_006937497.4;refGene_gn=SYVN1;refGene_tx=XM_019812667.2;refGene_gn=SYVN1;refGene_tx=XM_019812668.2
D1 110304753 110305080
refGene_gn=SYVN1;refGene_tx=XM_003993588.4;refGene_gn=SYVN1;refGene_tx=XM_006937497.4;refGene_gn=SYVN1;refGene_tx=XM_019812667.2;refGene_gn=SYVN1;refGene_tx=XM_019812668.2
D1 110305714 110305818
refGene_gn=SYVN1;refGene_tx=XM_003993588.4;refGene_gn=SYVN1;refGene_tx=XM_006937497.4;refGene_gn=SYVN1;refGene_tx=XM_019812667.2;refGene_gn=SYVN1;refGene_tx=XM_019812668.2
D1 110306229 110306278
refGene_gn=SYVN1;refGene_tx=XM_003993588.4;refGene_gn=SYVN1;refGene_tx=XM_006937497.4;refGene_gn=SYVN1;refGene_tx=XM_019812667.2;refGene_gn=SYVN1;refGene_tx=XM_019812668.2
D1 110306418 110306571
refGene_gn=SYVN1;refGene_tx=XM_003993588.4;refGene_gn=SYVN1;refGene_tx=XM_006937497.4;refGene_gn=SYVN1;refGene_tx=XM_019812667.2;refGene_gn=SYVN1;refGene_tx=XM_019812668.2
D1 110306662 110306755
refGene_gn=SYVN1;refGene_tx=XM_003993588.4;refGene_gn=SYVN1;refGene_tx=XM_006937497.4;refGene_gn=SYVN1;refGene_tx=XM_019812667.2;refGene_gn=SYVN1;refGene_tx=XM_019812668.2
D1 110306982 110307114
refGene_gn=SYVN1;refGene_tx=XM_003993588.4;refGene_gn=SYVN1;refGene_tx=XM_006937497.4;refGene_gn=SYVN1;refGene_tx=XM_019812667.2;refGene_gn=SYVN1;refGene_tx=XM_019812668.2
D1 110345476 110345630
refGene_gn=SPDYC;refGene_tx=XM_023239750.1
D1 110346474 110346660
refGene_gn=SPDYC;refGene_tx=XM_023239750.1
D1 110346927 110346986
refGene_gn=SPDYC;refGene_tx=XM_023239750.1
D1 110347455 110347541
refGene_gn=SPDYC;refGene_tx=XM_023239750.1
D1 110347642 110347814
refGene_gn=SPDYC;refGene_tx=XM_023239750.1
D1 110347896 110348239
refGene_gn=SPDYC;refGene_tx=XM_023239750.1
D1 110352988 110353029
refGene_gn=SPDYC;refGene_tx=XM_023239750.1
D1 110357494 110357761
refGene_gn=CAPN1;refGene_tx=XM_023240047.1;refGene_gn=CAPN1;refGene_tx=XM_023240048.1;refGene_gn=CAPN1;refGene_tx=XM_023240049.1;refGene_gn=CAPN1;refGene_tx=XM_023240050.1
D1 110357922 110357992
refGene_gn=CAPN1;refGene_tx=XM_023240047.1;refGene_gn=CAPN1;refGene_tx=XM_023240048.1;refGene_gn=CAPN1;refGene_tx=XM_023240049.1;refGene_gn=CAPN1;refGene_tx=XM_023240050.1
D1 110358262 110358381

refGene_gn=CAPN1;refGene_tx=XM_023240047.1;refGene_gn=CAPN1;refGene_tx=XM_023240048.1;refGene_gn=CAPN1;refGene_tx=XM_023240049.1;refGene_gn=CAPN1;refGene_tx=XM_023240050.1
D1	110360121	110360255
refGene_gn=CAPN1;refGene_tx=XM_023240047.1;refGene_gn=CAPN1;refGene_tx=XM_023240048.1;refGene_gn=CAPN1;refGene_tx=XM_023240049.1;refGene_gn=CAPN1;refGene_tx=XM_023240050.1
D1	110360344	110360513
refGene_gn=CAPN1;refGene_tx=XM_023240047.1;refGene_gn=CAPN1;refGene_tx=XM_023240048.1;refGene_gn=CAPN1;refGene_tx=XM_023240049.1;refGene_gn=CAPN1;refGene_tx=XM_023240050.1
D1	110361284	110361368
refGene_gn=CAPN1;refGene_tx=XM_023240047.1;refGene_gn=CAPN1;refGene_tx=XM_023240048.1;refGene_gn=CAPN1;refGene_tx=XM_023240049.1;refGene_gn=CAPN1;refGene_tx=XM_023240050.1
D1	110361721	110361807
refGene_gn=CAPN1;refGene_tx=XM_023240047.1;refGene_gn=CAPN1;refGene_tx=XM_023240048.1;refGene_gn=CAPN1;refGene_tx=XM_023240049.1;refGene_gn=CAPN1;refGene_tx=XM_023240050.1
D1	110362158	110362233
refGene_gn=CAPN1;refGene_tx=XM_023240047.1;refGene_gn=CAPN1;refGene_tx=XM_023240048.1;refGene_gn=CAPN1;refGene_tx=XM_023240049.1;refGene_gn=CAPN1;refGene_tx=XM_023240050.1
D1	110362371	110362532
refGene_gn=CAPN1;refGene_tx=XM_023240047.1;refGene_gn=CAPN1;refGene_tx=XM_023240048.1;refGene_gn=CAPN1;refGene_tx=XM_023240049.1;refGene_gn=CAPN1;refGene_tx=XM_023240050.1
D1	110377838	110378020
refGene_gn=CAPN1;refGene_tx=XM_023240047.1;refGene_gn=CAPN1;refGene_tx=XM_023240048.1;refGene_gn=CAPN1;refGene_tx=XM_023240049.1;refGene_gn=CAPN1;refGene_tx=XM_023240050.1
D1	110378747	110378759
refGene_gn=CAPN1;refGene_tx=XM_023240047.1;refGene_gn=CAPN1;refGene_tx=XM_023240048.1;refGene_gn=CAPN1;refGene_tx=XM_023240049.1;refGene_gn=CAPN1;refGene_tx=XM_023240050.1
D1	110379554	110379766
refGene_gn=CAPN1;refGene_tx=XM_023240047.1;refGene_gn=CAPN1;refGene_tx=XM_023240048.1;refGene_gn=CAPN1;refGene_tx=XM_023240049.1;refGene_gn=CAPN1;refGene_tx=XM_023240050.1
D1	110379863	110379903
refGene_gn=CAPN1;refGene_tx=XM_023240047.1;refGene_gn=CAPN1;refGene_tx=XM_023240048.1;refGene_gn=CAPN1;refGene_tx=XM_023240049.1;refGene_gn=CAPN1;refGene_tx=XM_023240050.1
D1	110380477	110380543
refGene_gn=CAPN1;refGene_tx=XM_023240047.1;refGene_gn=CAPN1;refGene_tx=XM_023240048.1;refGene_gn=CAPN1;refGene_tx=XM_023240049.1;refGene_gn=CAPN1;refGene_tx=XM_023240050.1
D1	110380634	110380692
refGene_gn=CAPN1;refGene_tx=XM_023240047.1;refGene_gn=CAPN1;refGene_tx=XM_023240048.1;refGene_gn=CAPN1;refGene_tx=XM_023240049.1;refGene_gn=

CAPN1;refGene_tx=XM_023240050.1
D1 110381440 110381505
refGene_gn=CAPN1;refGene_tx=XM_023240047.1;refGene_gn=CAPN1;refGene_tx=XM_023240048.1;refGene_gn=CAPN1;refGene_tx=XM_023240049.1;refGene_gn=CAPN1;refGene_tx=XM_023240050.1
D1 110381910 110381979
refGene_gn=CAPN1;refGene_tx=XM_023240047.1;refGene_gn=CAPN1;refGene_tx=XM_023240048.1;refGene_gn=CAPN1;refGene_tx=XM_023240049.1;refGene_gn=CAPN1;refGene_tx=XM_023240050.1
D1 110382213 110382292
refGene_gn=CAPN1;refGene_tx=XM_023240047.1;refGene_gn=CAPN1;refGene_tx=XM_023240048.1;refGene_gn=CAPN1;refGene_tx=XM_023240049.1;refGene_gn=CAPN1;refGene_tx=XM_023240050.1
D1 110382880 110382997
refGene_gn=CAPN1;refGene_tx=XM_023240047.1;refGene_gn=CAPN1;refGene_tx=XM_023240048.1;refGene_gn=CAPN1;refGene_tx=XM_023240049.1;refGene_gn=CAPN1;refGene_tx=XM_023240050.1
D1 110383316 110383375
refGene_gn=CAPN1;refGene_tx=XM_023240047.1;refGene_gn=CAPN1;refGene_tx=XM_023240048.1;refGene_gn=CAPN1;refGene_tx=XM_023240049.1;refGene_gn=CAPN1;refGene_tx=XM_023240050.1
D1 110383711 110383738
refGene_gn=CAPN1;refGene_tx=XM_023240047.1;refGene_gn=CAPN1;refGene_tx=XM_023240048.1;refGene_gn=CAPN1;refGene_tx=XM_023240049.1;refGene_gn=CAPN1;refGene_tx=XM_023240050.1
D1 110386212 110386587
refGene_gn=SLC22A20P;refGene_tx=XM_003993589.3;refGene_gn=SLC22A20P;refGene_tx=XM_019812670.1;refGene_gn=SLC22A20P;refGene_tx=XM_019812671.1
D1 110386662 110386766
refGene_gn=SLC22A20P;refGene_tx=XM_003993589.3;refGene_gn=SLC22A20P;refGene_tx=XM_019812670.1;refGene_gn=SLC22A20P;refGene_tx=XM_019812671.1
D1 110388209 110388364
refGene_gn=SLC22A20P;refGene_tx=XM_003993589.3;refGene_gn=SLC22A20P;refGene_tx=XM_019812670.1;refGene_gn=SLC22A20P;refGene_tx=XM_019812671.1
D1 110390727 110390896
refGene_gn=SLC22A20P;refGene_tx=XM_003993589.3;refGene_gn=SLC22A20P;refGene_tx=XM_019812670.1;refGene_gn=SLC22A20P;refGene_tx=XM_019812671.1
D1 110391404 110391528
refGene_gn=SLC22A20P;refGene_tx=XM_003993589.3;refGene_gn=SLC22A20P;refGene_tx=XM_019812670.1;refGene_gn=SLC22A20P;refGene_tx=XM_019812671.1
D1 110392476 110392592
refGene_gn=SLC22A20P;refGene_tx=XM_003993589.3;refGene_gn=SLC22A20P;refGene_tx=XM_019812670.1;refGene_gn=SLC22A20P;refGene_tx=XM_019812671.1
D1 110398953 110399168
refGene_gn=SLC22A20P;refGene_tx=XM_003993589.3;refGene_gn=SLC22A20P;refGene_tx=XM_019812670.1;refGene_gn=SLC22A20P;refGene_tx=XM_019812671.1
D1 110399354 110399463
refGene_gn=SLC22A20P;refGene_tx=XM_003993589.3;refGene_gn=SLC22A20P;refGene_tx=XM_019812670.1;refGene_gn=SLC22A20P;refGene_tx=XM_019812671.1
D1 110399657 110399861

refGene_gn=SLC22A20P;refGene_tx=XM_003993589.3;refGene_gn=SLC22A20P;refGene_tx=XM_019812670.1;refGene_gn=SLC22A20P;refGene_tx=XM_019812671.1
D1 110406574 110406671
refGene_gn=SLC22A20P;refGene_tx=XM_003993589.3;refGene_gn=SLC22A20P;refGene_tx=XM_019812670.1
D1 110449830 110449909
refGene_gn=POLA2;refGene_tx=XM_003993590.5
D1 110453124 110453249
refGene_gn=POLA2;refGene_tx=XM_003993590.5
D1 110453737 110453829
refGene_gn=POLA2;refGene_tx=XM_003993590.5
D1 110454418 110454476
refGene_gn=POLA2;refGene_tx=XM_003993590.5
D1 110459017 110459124
refGene_gn=POLA2;refGene_tx=XM_003993590.5
D1 110461123 110461317
refGene_gn=POLA2;refGene_tx=XM_003993590.5
D1 110462195 110462284
refGene_gn=POLA2;refGene_tx=XM_003993590.5
D1 110463527 110463683
refGene_gn=POLA2;refGene_tx=XM_003993590.5
D1 110464550 110464613
refGene_gn=POLA2;refGene_tx=XM_003993590.5
D1 110465383 110465426
refGene_gn=POLA2;refGene_tx=XM_003993590.5
D1 110468763 110468888
refGene_gn=POLA2;refGene_tx=XM_003993590.5
D1 110469190 110469229
refGene_gn=POLA2;refGene_tx=XM_003993590.5
D1 110469959 110470033
refGene_gn=POLA2;refGene_tx=XM_003993590.5
D1 110473046 110473155
refGene_gn=POLA2;refGene_tx=XM_003993590.5
D1 110473652 110473759
refGene_gn=POLA2;refGene_tx=XM_003993590.5
D1 110474570 110474630
refGene_gn=POLA2;refGene_tx=XM_003993590.5
D1 110474885 110475012
refGene_gn=POLA2;refGene_tx=XM_003993590.5
D1 110475728 110475881
refGene_gn=POLA2;refGene_tx=XM_003993590.5
D1 110493390 110494035
refGene_gn=CDC42EP2;refGene_tx=XM_003993591.4
D1 110504650 110504682
refGene_gn=DPF2;refGene_tx=XM_003993592.2;refGene_gn=DPF2;refGene_tx=XM_006937501.2
D1 110509081 110509242
refGene_gn=DPF2;refGene_tx=XM_003993592.2;refGene_gn=DPF2;refGene_tx=XM_006937501.2;refGene_gn=DPF2;refGene_tx=XM_006937503.3
D1 110509671 110509779

refGene_gn=DPF2;refGene_tx=XM_003993592.2;refGene_gn=DPF2;refGene_tx=XM_006937501.2;refGene_gn=DPF2;refGene_tx=XM_006937503.3
D1 110510101 110510265
refGene_gn=DPF2;refGene_tx=XM_003993592.2;refGene_gn=DPF2;refGene_tx=XM_006937501.2;refGene_gn=DPF2;refGene_tx=XM_006937503.3
D1 110511304 110511397
refGene_gn=DPF2;refGene_tx=XM_003993592.2;refGene_gn=DPF2;refGene_tx=XM_006937501.2;refGene_gn=DPF2;refGene_tx=XM_006937503.3
D1 110511561 110511640
refGene_gn=DPF2;refGene_tx=XM_003993592.2;refGene_gn=DPF2;refGene_tx=XM_006937501.2;refGene_gn=DPF2;refGene_tx=XM_006937503.3
D1 110512129 110512171
refGene_gn=DPF2;refGene_tx=XM_006937501.2;refGene_gn=DPF2;refGene_tx=XM_006937503.3
D1 110513200 110513338
refGene_gn=DPF2;refGene_tx=XM_003993592.2;refGene_gn=DPF2;refGene_tx=XM_006937501.2;refGene_gn=DPF2;refGene_tx=XM_006937503.3
D1 110513454 110513583
refGene_gn=DPF2;refGene_tx=XM_003993592.2;refGene_gn=DPF2;refGene_tx=XM_006937501.2;refGene_gn=DPF2;refGene_tx=XM_006937503.3
D1 110513699 110513812
refGene_gn=DPF2;refGene_tx=XM_003993592.2;refGene_gn=DPF2;refGene_tx=XM_006937501.2;refGene_gn=DPF2;refGene_tx=XM_006937503.3
D1 110514931 110515013
refGene_gn=DPF2;refGene_tx=XM_003993592.2;refGene_gn=DPF2;refGene_tx=XM_006937501.2;refGene_gn=DPF2;refGene_tx=XM_006937503.3
D1 110517018 110517095
refGene_gn=DPF2;refGene_tx=XM_003993592.2;refGene_gn=DPF2;refGene_tx=XM_006937501.2;refGene_gn=DPF2;refGene_tx=XM_006937503.3
D1 110521555 110522971
refGene_gn=TIGD3;refGene_tx=XM_023240051.1
D1 110532809 110533078
refGene_gn=SLC25A45;refGene_tx=XM_011287017.3;refGene_gn=SLC25A45;refGene_tx=XM_023240053.1
D1 110533233 110533495
refGene_gn=SLC25A45;refGene_tx=XM_011287017.3;refGene_gn=SLC25A45;refGene_tx=XM_023240053.1
D1 110536225 110536411
refGene_gn=SLC25A45;refGene_tx=XM_011287017.3;refGene_gn=SLC25A45;refGene_tx=XM_023240053.1
D1 110536708 110536780
refGene_gn=SLC25A45;refGene_tx=XM_011287017.3;refGene_gn=SLC25A45;refGene_tx=XM_023240053.1
D1 110536966 110537010
refGene_gn=SLC25A45;refGene_tx=XM_011287017.3;refGene_gn=SLC25A45;refGene_tx=XM_023240053.1
D1 110537118 110537209
refGene_gn=SLC25A45;refGene_tx=XM_023240053.1
D1 110538040 110538077
refGene_gn=SLC25A45;refGene_tx=XM_011287017.3

D1 110543583 110543668
refGene_gn=FRMD8;refGene_tx=XM_003993593.4;refGene_gn=FRMD8;refGene_tx=XM_006937504.3;refGene_gn=FRMD8;refGene_tx=XM_019812680.2;refGene_gn=FRMD8;refGene_tx=XM_019812681.2
D1 110545095 110545263
refGene_gn=FRMD8;refGene_tx=XM_003993593.4;refGene_gn=FRMD8;refGene_tx=XM_019812680.2;refGene_gn=FRMD8;refGene_tx=XM_019812681.2
D1 110549191 110549293
refGene_gn=FRMD8;refGene_tx=XM_003993593.4;refGene_gn=FRMD8;refGene_tx=XM_006937504.3;refGene_gn=FRMD8;refGene_tx=XM_019812680.2;refGene_gn=FRMD8;refGene_tx=XM_019812681.2
D1 110549645 110549704
refGene_gn=FRMD8;refGene_tx=XM_003993593.4;refGene_gn=FRMD8;refGene_tx=XM_006937504.3;refGene_gn=FRMD8;refGene_tx=XM_019812680.2;refGene_gn=FRMD8;refGene_tx=XM_019812681.2
D1 110549834 110550001
refGene_gn=FRMD8;refGene_tx=XM_003993593.4;refGene_gn=FRMD8;refGene_tx=XM_006937504.3;refGene_gn=FRMD8;refGene_tx=XM_019812680.2;refGene_gn=FRMD8;refGene_tx=XM_019812681.2
D1 110552002 110552227
refGene_gn=FRMD8;refGene_tx=XM_003993593.4;refGene_gn=FRMD8;refGene_tx=XM_006937504.3;refGene_gn=FRMD8;refGene_tx=XM_019812680.2;refGene_gn=FRMD8;refGene_tx=XM_019812681.2
D1 110554390 110554514
refGene_gn=FRMD8;refGene_tx=XM_003993593.4;refGene_gn=FRMD8;refGene_tx=XM_006937504.3;refGene_gn=FRMD8;refGene_tx=XM_019812680.2;refGene_gn=FRMD8;refGene_tx=XM_019812681.2
D1 110555447 110555591
refGene_gn=FRMD8;refGene_tx=XM_003993593.4;refGene_gn=FRMD8;refGene_tx=XM_006937504.3;refGene_gn=FRMD8;refGene_tx=XM_019812680.2;refGene_gn=FRMD8;refGene_tx=XM_019812681.2
D1 110559420 110559696
refGene_gn=FRMD8;refGene_tx=XM_003993593.4;refGene_gn=FRMD8;refGene_tx=XM_006937504.3;refGene_gn=FRMD8;refGene_tx=XM_019812680.2;refGene_gn=FRMD8;refGene_tx=XM_019812681.2
D1 110562947 110563066
refGene_gn=FRMD8;refGene_tx=XM_003993593.4;refGene_gn=FRMD8;refGene_tx=XM_006937504.3
D1 110574792 110574866
refGene_gn=FRMD8;refGene_tx=XM_019812680.2
D1 110602659 110602851
refGene_gn=LOC102899282;refGene_tx=XM_006937505.4;refGene_gn=LOC102899282;refGene_tx=XM_019812684.1
D1 110602956 110603101
refGene_gn=LOC102899282;refGene_tx=XM_006937505.4;refGene_gn=LOC102899282;refGene_tx=XM_019812684.1
D1 110603505 110605307
refGene_gn=LOC102899282;refGene_tx=XM_006937505.4;refGene_gn=LOC102899282;refGene_tx=XM_019812684.1
D1 110662156 110662267

refGene_gn=SCYL1;refGene_tx=XM_006937506.3;refGene_gn=SCYL1;refGene_tx=XM_006937507.3
D1 110662616 110662757
refGene_gn=SCYL1;refGene_tx=XM_006937506.3;refGene_gn=SCYL1;refGene_tx=XM_006937507.3
D1 110662948 110663071
refGene_gn=SCYL1;refGene_tx=XM_006937506.3;refGene_gn=SCYL1;refGene_tx=XM_006937507.3
D1 110663150 110663377
refGene_gn=SCYL1;refGene_tx=XM_006937506.3;refGene_gn=SCYL1;refGene_tx=XM_006937507.3
D1 110663718 110663809
refGene_gn=SCYL1;refGene_tx=XM_006937506.3;refGene_gn=SCYL1;refGene_tx=XM_006937507.3
D1 110663885 110664041
refGene_gn=SCYL1;refGene_tx=XM_006937506.3;refGene_gn=SCYL1;refGene_tx=XM_006937507.3
D1 110666391 110666550
refGene_gn=SCYL1;refGene_tx=XM_006937506.3;refGene_gn=SCYL1;refGene_tx=XM_006937507.3
D1 110667414 110667522
refGene_gn=SCYL1;refGene_tx=XM_006937506.3;refGene_gn=SCYL1;refGene_tx=XM_006937507.3
D1 110668294 110668408
refGene_gn=SCYL1;refGene_tx=XM_006937506.3;refGene_gn=SCYL1;refGene_tx=XM_006937507.3
D1 110669895 110670051
refGene_gn=SCYL1;refGene_tx=XM_006937506.3;refGene_gn=SCYL1;refGene_tx=XM_006937507.3
D1 110670617 110670806
refGene_gn=SCYL1;refGene_tx=XM_006937506.3;refGene_gn=SCYL1;refGene_tx=XM_006937507.3
D1 110670928 110671004
refGene_gn=SCYL1;refGene_tx=XM_006937506.3;refGene_gn=SCYL1;refGene_tx=XM_006937507.3
D1 110671283 110671448
refGene_gn=SCYL1;refGene_tx=XM_006937506.3;refGene_gn=SCYL1;refGene_tx=XM_006937507.3
D1 110671687 110671830
refGene_gn=SCYL1;refGene_tx=XM_006937506.3;refGene_gn=SCYL1;refGene_tx=XM_006937507.3
D1 110672471 110672543
refGene_gn=SCYL1;refGene_tx=XM_006937506.3;refGene_gn=SCYL1;refGene_tx=XM_006937507.3
D1 110672639 110672849
refGene_gn=SCYL1;refGene_tx=XM_006937506.3;refGene_gn=SCYL1;refGene_tx=XM_006937507.3
D1 110672937 110672992
refGene_gn=SCYL1;refGene_tx=XM_006937506.3;refGene_gn=SCYL1;refGene_tx=XM_006937507.3

D1	110673123	110673248
refGene_gn=SCYL1;refGene_tx=XM_006937506.3;refGene_gn=SCYL1;refGene_tx=XM_006937507.3
D1	110673755	110673907
refGene_gn=LTBP3;refGene_tx=XM_023240055.1;refGene_gn=LTBP3;refGene_tx=XM_023240056.1;refGene_gn=LTBP3;refGene_tx=XM_023240057.1;refGene_gn=LTBP3;refGene_tx=XM_023240058.1
D1	110674073	110674205
refGene_gn=LTBP3;refGene_tx=XM_023240055.1;refGene_gn=LTBP3;refGene_tx=XM_023240056.1;refGene_gn=LTBP3;refGene_tx=XM_023240057.1;refGene_gn=LTBP3;refGene_tx=XM_023240058.1
D1	110674309	110674390
refGene_gn=LTBP3;refGene_tx=XM_023240055.1;refGene_gn=LTBP3;refGene_tx=XM_023240056.1;refGene_gn=LTBP3;refGene_tx=XM_023240057.1;refGene_gn=LTBP3;refGene_tx=XM_023240058.1
D1	110674477	110674639
refGene_gn=LTBP3;refGene_tx=XM_023240055.1;refGene_gn=LTBP3;refGene_tx=XM_023240056.1;refGene_gn=LTBP3;refGene_tx=XM_023240057.1;refGene_gn=LTBP3;refGene_tx=XM_023240058.1
D1	110674788	110674929
refGene_gn=LTBP3;refGene_tx=XM_023240055.1;refGene_gn=LTBP3;refGene_tx=XM_023240056.1;refGene_gn=LTBP3;refGene_tx=XM_023240057.1
D1	110675026	110675164
refGene_gn=LTBP3;refGene_tx=XM_023240055.1;refGene_gn=LTBP3;refGene_tx=XM_023240056.1;refGene_gn=LTBP3;refGene_tx=XM_023240057.1;refGene_gn=LTBP3;refGene_tx=XM_023240058.1
D1	110675268	110675397
refGene_gn=LTBP3;refGene_tx=XM_023240055.1;refGene_gn=LTBP3;refGene_tx=XM_023240056.1;refGene_gn=LTBP3;refGene_tx=XM_023240057.1;refGene_gn=LTBP3;refGene_tx=XM_023240058.1
D1	110675655	110675739
refGene_gn=LTBP3;refGene_tx=XM_023240055.1;refGene_gn=LTBP3;refGene_tx=XM_023240056.1;refGene_gn=LTBP3;refGene_tx=XM_023240057.1;refGene_gn=LTBP3;refGene_tx=XM_023240058.1
D1	110675916	110676084
refGene_gn=LTBP3;refGene_tx=XM_023240055.1;refGene_gn=LTBP3;refGene_tx=XM_023240056.1;refGene_gn=LTBP3;refGene_tx=XM_023240057.1;refGene_gn=LTBP3;refGene_tx=XM_023240058.1
D1	110676389	110676518
refGene_gn=LTBP3;refGene_tx=XM_023240055.1;refGene_gn=LTBP3;refGene_tx=XM_023240056.1;refGene_gn=LTBP3;refGene_tx=XM_023240057.1;refGene_gn=LTBP3;refGene_tx=XM_023240058.1
D1	110677409	110677529
refGene_gn=LTBP3;refGene_tx=XM_023240055.1;refGene_gn=LTBP3;refGene_tx=XM_023240056.1;refGene_gn=LTBP3;refGene_tx=XM_023240057.1;refGene_gn=LTBP3;refGene_tx=XM_023240058.1
D1	110677775	110677898
refGene_gn=LTBP3;refGene_tx=XM_023240055.1;refGene_gn=LTBP3;refGene_tx=XM_023240056.1;refGene_gn=LTBP3;refGene_tx=XM_023240057.1;refGene_gn=LTBP3;refGene_tx=XM_023240058.1

D1 110680551 110680674
refGene_gn=LTBP3;refGene_tx=XM_023240055.1;refGene_gn=LTBP3;refGene_tx=XM_023240056.1;refGene_gn=LTBP3;refGene_tx=XM_023240057.1;refGene_gn=LTBP3;refGene_tx=XM_023240058.1
D1 110680912 110681035
refGene_gn=LTBP3;refGene_tx=XM_023240055.1;refGene_gn=LTBP3;refGene_tx=XM_023240056.1;refGene_gn=LTBP3;refGene_tx=XM_023240057.1;refGene_gn=LTBP3;refGene_tx=XM_023240058.1
D1 110681273 110681402
refGene_gn=LTBP3;refGene_tx=XM_023240055.1;refGene_gn=LTBP3;refGene_tx=XM_023240056.1;refGene_gn=LTBP3;refGene_tx=XM_023240057.1;refGene_gn=LTBP3;refGene_tx=XM_023240058.1
D1 110681524 110681656
refGene_gn=LTBP3;refGene_tx=XM_023240055.1;refGene_gn=LTBP3;refGene_tx=XM_023240056.1;refGene_gn=LTBP3;refGene_tx=XM_023240057.1;refGene_gn=LTBP3;refGene_tx=XM_023240058.1
D1 110681746 110681872
refGene_gn=LTBP3;refGene_tx=XM_023240055.1;refGene_gn=LTBP3;refGene_tx=XM_023240056.1;refGene_gn=LTBP3;refGene_tx=XM_023240057.1;refGene_gn=LTBP3;refGene_tx=XM_023240058.1
D1 110683752 110683851
refGene_gn=LTBP3;refGene_tx=XM_023240055.1;refGene_gn=LTBP3;refGene_tx=XM_023240056.1;refGene_gn=LTBP3;refGene_tx=XM_023240057.1;refGene_gn=LTBP3;refGene_tx=XM_023240058.1
D1 110684028 110684102
refGene_gn=LTBP3;refGene_tx=XM_023240055.1;refGene_gn=LTBP3;refGene_tx=XM_023240056.1;refGene_gn=LTBP3;refGene_tx=XM_023240057.1;refGene_gn=LTBP3;refGene_tx=XM_023240058.1
D1 110684175 110684192
refGene_gn=LTBP3;refGene_tx=XM_023240055.1;refGene_gn=LTBP3;refGene_tx=XM_023240056.1;refGene_gn=LTBP3;refGene_tx=XM_023240058.1
D1 110684768 110684955
refGene_gn=LTBP3;refGene_tx=XM_023240055.1;refGene_gn=LTBP3;refGene_tx=XM_023240056.1;refGene_gn=LTBP3;refGene_tx=XM_023240057.1;refGene_gn=LTBP3;refGene_tx=XM_023240058.1
D1 110685041 110685200
refGene_gn=LTBP3;refGene_tx=XM_023240055.1;refGene_gn=LTBP3;refGene_tx=XM_023240056.1;refGene_gn=LTBP3;refGene_tx=XM_023240057.1;refGene_gn=LTBP3;refGene_tx=XM_023240058.1
D1 110685693 110685816
refGene_gn=LTBP3;refGene_tx=XM_023240055.1;refGene_gn=LTBP3;refGene_tx=XM_023240056.1;refGene_gn=LTBP3;refGene_tx=XM_023240057.1;refGene_gn=LTBP3;refGene_tx=XM_023240058.1
D1 110685991 110686084
refGene_gn=LTBP3;refGene_tx=XM_023240055.1;refGene_gn=LTBP3;refGene_tx=XM_023240056.1;refGene_gn=LTBP3;refGene_tx=XM_023240057.1;refGene_gn=LTBP3;refGene_tx=XM_023240058.1
D1 110686304 110686410
refGene_gn=LTBP3;refGene_tx=XM_023240055.1;refGene_gn=LTBP3;refGene_tx=XM_023240056.1;refGene_gn=LTBP3;refGene_tx=XM_023240057.1;refGene_gn=

LTBP3;refGene_tx=XM_023240058.1
D1 110686588 110686791
refGene_gn=LTBP3;refGene_tx=XM_023240055.1;refGene_gn=LTBP3;refGene_tx=XM_023240056.1;refGene_gn=LTBP3;refGene_tx=XM_023240057.1;refGene_gn=LTBP3;refGene_tx=XM_023240058.1
D1 110686924 110687254
refGene_gn=LTBP3;refGene_tx=XM_023240055.1;refGene_gn=LTBP3;refGene_tx=XM_023240056.1;refGene_gn=LTBP3;refGene_tx=XM_023240057.1;refGene_gn=LTBP3;refGene_tx=XM_023240058.1
D1 110690257 110690585
refGene_gn=LTBP3;refGene_tx=XM_023240055.1;refGene_gn=LTBP3;refGene_tx=XM_023240056.1;refGene_gn=LTBP3;refGene_tx=XM_023240057.1;refGene_gn=LTBP3;refGene_tx=XM_023240058.1
D1 110713179 110713198
refGene_gn=SSSCA1;refGene_tx=XM_003993597.3
D1 110713294 110713446
refGene_gn=SSSCA1;refGene_tx=XM_003993597.3
D1 110713575 110713660
refGene_gn=SSSCA1;refGene_tx=XM_003993597.3
D1 110714068 110714412
refGene_gn=SSSCA1;refGene_tx=XM_003993597.3
D1 110715317 110715617
refGene_gn=FAM89B;refGene_tx=XM_023240059.1
D1 110716032 110716311
refGene_gn=FAM89B;refGene_tx=XM_023240059.1
D1 110719323 110719427
refGene_gn=EHBP1L1;refGene_tx=XM_023240060.1;refGene_gn=EHBP1L1;refGene_tx=XM_023240061.1;refGene_gn=EHBP1L1;refGene_tx=XM_023240062.1;refGene_gn=EHBP1L1;refGene_tx=XM_023240063.1;refGene_gn=EHBP1L1;refGene_tx=XM_023240064.1;refGene_gn=EHBP1L1;refGene_tx=XM_023240065.1;refGene_gn=EHBP1L1;refGene_tx=XM_023240066.1
D1 110722151 110722209
refGene_gn=EHBP1L1;refGene_tx=XM_023240060.1;refGene_gn=EHBP1L1;refGene_tx=XM_023240061.1;refGene_gn=EHBP1L1;refGene_tx=XM_023240062.1;refGene_gn=EHBP1L1;refGene_tx=XM_023240063.1;refGene_gn=EHBP1L1;refGene_tx=XM_023240064.1;refGene_gn=EHBP1L1;refGene_tx=XM_023240065.1;refGene_gn=EHBP1L1;refGene_tx=XM_023240066.1
D1 110722854 110722950
refGene_gn=EHBP1L1;refGene_tx=XM_023240060.1;refGene_gn=EHBP1L1;refGene_tx=XM_023240061.1;refGene_gn=EHBP1L1;refGene_tx=XM_023240062.1;refGene_gn=EHBP1L1;refGene_tx=XM_023240063.1;refGene_gn=EHBP1L1;refGene_tx=XM_023240064.1;refGene_gn=EHBP1L1;refGene_tx=XM_023240065.1;refGene_gn=EHBP1L1;refGene_tx=XM_023240066.1
D1 110723243 110723297
refGene_gn=EHBP1L1;refGene_tx=XM_023240060.1;refGene_gn=EHBP1L1;refGene_tx=XM_023240061.1;refGene_gn=EHBP1L1;refGene_tx=XM_023240062.1;refGene_gn=EHBP1L1;refGene_tx=XM_023240063.1;refGene_gn=EHBP1L1;refGene_tx=XM_023240064.1;refGene_gn=EHBP1L1;refGene_tx=XM_023240065.1;refGene_gn=EHBP1L1;refGene_tx=XM_023240066.1
D1 110723388 110723567

refGene_gn=EHBP1L1;refGene_tx=XM_023240060.1;refGene_gn=EHBP1L1;refGen
e_tx=XM_023240061.1;refGene_gn=EHBP1L1;refGene_tx=XM_023240062.1;refGe
ne_gn=EHBP1L1;refGene_tx=XM_023240063.1;refGene_gn=EHBP1L1;refGene_tx=
XM_023240064.1;refGene_gn=EHBP1L1;refGene_tx=XM_023240065.1;refGene_gn
=EHBP1L1;refGene_tx=XM_023240066.1
D1	110723657	110723800
refGene_gn=EHBP1L1;refGene_tx=XM_023240060.1;refGene_gn=EHBP1L1;refGen
e_tx=XM_023240061.1;refGene_gn=EHBP1L1;refGene_tx=XM_023240062.1;refGe
ne_gn=EHBP1L1;refGene_tx=XM_023240063.1;refGene_gn=EHBP1L1;refGene_tx=
XM_023240064.1;refGene_gn=EHBP1L1;refGene_tx=XM_023240065.1;refGene_gn
=EHBP1L1;refGene_tx=XM_023240066.1
D1	110724259	110724295
refGene_gn=EHBP1L1;refGene_tx=XM_023240060.1;refGene_gn=EHBP1L1;refGen
e_tx=XM_023240062.1;refGene_gn=EHBP1L1;refGene_tx=XM_023240063.1;refGe
ne_gn=EHBP1L1;refGene_tx=XM_023240064.1;refGene_gn=EHBP1L1;refGene_tx=
XM_023240065.1
D1	110724384	110724453
refGene_gn=EHBP1L1;refGene_tx=XM_023240060.1;refGene_gn=EHBP1L1;refGen
e_tx=XM_023240061.1;refGene_gn=EHBP1L1;refGene_tx=XM_023240062.1;refGe
ne_gn=EHBP1L1;refGene_tx=XM_023240063.1;refGene_gn=EHBP1L1;refGene_tx=
XM_023240064.1;refGene_gn=EHBP1L1;refGene_tx=XM_023240065.1;refGene_gn
=EHBP1L1;refGene_tx=XM_023240066.1
D1	110724545	110724708
refGene_gn=EHBP1L1;refGene_tx=XM_023240060.1;refGene_gn=EHBP1L1;refGen
e_tx=XM_023240061.1;refGene_gn=EHBP1L1;refGene_tx=XM_023240062.1;refGe
ne_gn=EHBP1L1;refGene_tx=XM_023240063.1;refGene_gn=EHBP1L1;refGene_tx=
XM_023240064.1;refGene_gn=EHBP1L1;refGene_tx=XM_023240065.1;refGene_gn
=EHBP1L1;refGene_tx=XM_023240066.1
D1	110724841	110726843
refGene_gn=EHBP1L1;refGene_tx=XM_023240060.1;refGene_gn=EHBP1L1;refGen
e_tx=XM_023240061.1;refGene_gn=EHBP1L1;refGene_tx=XM_023240062.1;refGe
ne_gn=EHBP1L1;refGene_tx=XM_023240063.1;refGene_gn=EHBP1L1;refGene_tx=
XM_023240064.1
D1	110727303	110727461
refGene_gn=EHBP1L1;refGene_tx=XM_023240060.1;refGene_gn=EHBP1L1;refGen
e_tx=XM_023240061.1;refGene_gn=EHBP1L1;refGene_tx=XM_023240062.1;refGe
ne_gn=EHBP1L1;refGene_tx=XM_023240063.1;refGene_gn=EHBP1L1;refGene_tx=
XM_023240064.1;refGene_gn=EHBP1L1;refGene_tx=XM_023240065.1;refGene_gn
=EHBP1L1;refGene_tx=XM_023240066.1
D1	110727549	110727598
refGene_gn=EHBP1L1;refGene_tx=XM_023240060.1;refGene_gn=EHBP1L1;refGen
e_tx=XM_023240061.1;refGene_gn=EHBP1L1;refGene_tx=XM_023240062.1;refGe
ne_gn=EHBP1L1;refGene_tx=XM_023240063.1;refGene_gn=EHBP1L1;refGene_tx=
XM_023240064.1;refGene_gn=EHBP1L1;refGene_tx=XM_023240065.1;refGene_gn
=EHBP1L1;refGene_tx=XM_023240066.1
D1	110728029	110728692
refGene_gn=EHBP1L1;refGene_tx=XM_023240060.1;refGene_gn=EHBP1L1;refGen
e_tx=XM_023240061.1;refGene_gn=EHBP1L1;refGene_tx=XM_023240062.1;refGe
ne_gn=EHBP1L1;refGene_tx=XM_023240063.1;refGene_gn=EHBP1L1;refGene_tx=
XM_023240064.1;refGene_gn=EHBP1L1;refGene_tx=XM_023240065.1;refGene_gn

=EHBP1L1;refGene_tx=XM_023240066.1
D1 110732442 110732512
refGene_gn=EHBP1L1;refGene_tx=XM_023240060.1;refGene_gn=EHBP1L1;refGene_tx=XM_023240061.1;refGene_gn=EHBP1L1;refGene_tx=XM_023240062.1;refGene_gn=EHBP1L1;refGene_tx=XM_023240063.1;refGene_gn=EHBP1L1;refGene_tx=XM_023240064.1;refGene_gn=EHBP1L1;refGene_tx=XM_023240065.1;refGene_gn=EHBP1L1;refGene_tx=XM_023240066.1
D1 110732632 110732688
refGene_gn=EHBP1L1;refGene_tx=XM_023240060.1;refGene_gn=EHBP1L1;refGene_tx=XM_023240061.1;refGene_gn=EHBP1L1;refGene_tx=XM_023240062.1;refGene_gn=EHBP1L1;refGene_tx=XM_023240063.1;refGene_gn=EHBP1L1;refGene_tx=XM_023240064.1;refGene_gn=EHBP1L1;refGene_tx=XM_023240065.1;refGene_gn=EHBP1L1;refGene_tx=XM_023240066.1
D1 110732778 110732902
refGene_gn=EHBP1L1;refGene_tx=XM_023240060.1;refGene_gn=EHBP1L1;refGene_tx=XM_023240061.1;refGene_gn=EHBP1L1;refGene_tx=XM_023240062.1;refGene_gn=EHBP1L1;refGene_tx=XM_023240063.1;refGene_gn=EHBP1L1;refGene_tx=XM_023240064.1;refGene_gn=EHBP1L1;refGene_tx=XM_023240065.1;refGene_gn=EHBP1L1;refGene_tx=XM_023240066.1
D1 110733148 110733248
refGene_gn=EHBP1L1;refGene_tx=XM_023240060.1;refGene_gn=EHBP1L1;refGene_tx=XM_023240061.1;refGene_gn=EHBP1L1;refGene_tx=XM_023240062.1;refGene_gn=EHBP1L1;refGene_tx=XM_023240063.1;refGene_gn=EHBP1L1;refGene_tx=XM_023240064.1;refGene_gn=EHBP1L1;refGene_tx=XM_023240065.1;refGene_gn=EHBP1L1;refGene_tx=XM_023240066.1
D1 110734219 110734293
refGene_gn=EHBP1L1;refGene_tx=XM_023240060.1;refGene_gn=EHBP1L1;refGene_tx=XM_023240061.1;refGene_gn=EHBP1L1;refGene_tx=XM_023240062.1;refGene_gn=EHBP1L1;refGene_tx=XM_023240063.1;refGene_gn=EHBP1L1;refGene_tx=XM_023240064.1;refGene_gn=EHBP1L1;refGene_tx=XM_023240065.1;refGene_gn=EHBP1L1;refGene_tx=XM_023240066.1
D1 110734377 110734492
refGene_gn=EHBP1L1;refGene_tx=XM_023240060.1;refGene_gn=EHBP1L1;refGene_tx=XM_023240061.1;refGene_gn=EHBP1L1;refGene_tx=XM_023240062.1;refGene_gn=EHBP1L1;refGene_tx=XM_023240063.1;refGene_gn=EHBP1L1;refGene_tx=XM_023240064.1;refGene_gn=EHBP1L1;refGene_tx=XM_023240065.1;refGene_gn=EHBP1L1;refGene_tx=XM_023240066.1
D1 110734626 110734726
refGene_gn=EHBP1L1;refGene_tx=XM_023240060.1;refGene_gn=EHBP1L1;refGene_tx=XM_023240061.1;refGene_gn=EHBP1L1;refGene_tx=XM_023240062.1;refGene_gn=EHBP1L1;refGene_tx=XM_023240063.1;refGene_gn=EHBP1L1;refGene_tx=XM_023240064.1;refGene_gn=EHBP1L1;refGene_tx=XM_023240065.1;refGene_gn=EHBP1L1;refGene_tx=XM_023240066.1
D1 110735746 110735952
refGene_gn=KCNK7;refGene_tx=XM_003993719.3
D1 110736204 110736603
refGene_gn=KCNK7;refGene_tx=XM_003993719.3
D1 110737966 110738483
refGene_gn=KCNK7;refGene_tx=XM_003993719.3
D1 110740291 110740629

refGene_gn=MAP3K11;refGene_tx=XM_003993637.5;refGene_gn=MAP3K11;refGene_tx=XM_019812686.2;refGene_gn=MAP3K11;refGene_tx=XM_019812687.2;refGene_gn=MAP3K11;refGene_tx=XM_023240067.1
D1 110741277 110741703
refGene_gn=MAP3K11;refGene_tx=XM_003993637.5;refGene_gn=MAP3K11;refGene_tx=XM_019812686.2;refGene_gn=MAP3K11;refGene_tx=XM_019812687.2;refGene_gn=MAP3K11;refGene_tx=XM_023240067.1
D1 110747014 110747106
refGene_gn=MAP3K11;refGene_tx=XM_003993637.5;refGene_gn=MAP3K11;refGene_tx=XM_019812686.2;refGene_gn=MAP3K11;refGene_tx=XM_019812687.2;refGene_gn=MAP3K11;refGene_tx=XM_023240067.1
D1 110747194 110747330
refGene_gn=MAP3K11;refGene_tx=XM_003993637.5;refGene_gn=MAP3K11;refGene_tx=XM_019812686.2;refGene_gn=MAP3K11;refGene_tx=XM_019812687.2;refGene_gn=MAP3K11;refGene_tx=XM_023240067.1
D1 110747861 110747975
refGene_gn=MAP3K11;refGene_tx=XM_003993637.5;refGene_gn=MAP3K11;refGene_tx=XM_019812686.2;refGene_gn=MAP3K11;refGene_tx=XM_019812687.2;refGene_gn=MAP3K11;refGene_tx=XM_023240067.1
D1 110748456 110748700
refGene_gn=MAP3K11;refGene_tx=XM_003993637.5;refGene_gn=MAP3K11;refGene_tx=XM_019812686.2;refGene_gn=MAP3K11;refGene_tx=XM_019812687.2;refGene_gn=MAP3K11;refGene_tx=XM_023240067.1
D1 110748836 110749012
refGene_gn=MAP3K11;refGene_tx=XM_003993637.5;refGene_gn=MAP3K11;refGene_tx=XM_019812686.2;refGene_gn=MAP3K11;refGene_tx=XM_019812687.2;refGene_gn=MAP3K11;refGene_tx=XM_023240067.1
D1 110749116 110749265
refGene_gn=MAP3K11;refGene_tx=XM_003993637.5;refGene_gn=MAP3K11;refGene_tx=XM_019812686.2;refGene_gn=MAP3K11;refGene_tx=XM_019812687.2;refGene_gn=MAP3K11;refGene_tx=XM_023240067.1
D1 110749461 110749642
refGene_gn=MAP3K11;refGene_tx=XM_003993637.5;refGene_gn=MAP3K11;refGene_tx=XM_019812686.2;refGene_gn=MAP3K11;refGene_tx=XM_019812687.2;refGene_gn=MAP3K11;refGene_tx=XM_023240067.1
D1 110753878 110754620
refGene_gn=MAP3K11;refGene_tx=XM_003993637.5;refGene_gn=MAP3K11;refGene_tx=XM_019812686.2;refGene_gn=MAP3K11;refGene_tx=XM_019812687.2;refGene_gn=MAP3K11;refGene_tx=XM_023240067.1
D1 110756797 110756950
refGene_gn=PCNX3;refGene_tx=XM_003993638.3;refGene_gn=PCNX3;refGene_tx=XM_006937525.2;refGene_gn=PCNX3;refGene_tx=XM_006937526.2;refGene_gn=PCNX3;refGene_tx=XM_006937527.2
D1 110757326 110757514
refGene_gn=PCNX3;refGene_tx=XM_003993638.3;refGene_gn=PCNX3;refGene_tx=XM_006937525.2;refGene_gn=PCNX3;refGene_tx=XM_006937526.2;refGene_gn=PCNX3;refGene_tx=XM_006937527.2
D1 110757741 110757841
refGene_gn=PCNX3;refGene_tx=XM_003993638.3;refGene_gn=PCNX3;refGene_tx=XM_006937525.2;refGene_gn=PCNX3;refGene_tx=XM_006937526.2;refGene_gn=

PCNX3;refGene_tx=XM_006937527.2
D1 110757963 110758003
refGene_gn=PCNX3;refGene_tx=XM_003993638.3;refGene_gn=PCNX3;refGene_tx=XM_006937525.2;refGene_gn=PCNX3;refGene_tx=XM_006937526.2;refGene_gn=PCNX3;refGene_tx=XM_006937527.2
D1 110758102 110758198
refGene_gn=PCNX3;refGene_tx=XM_003993638.3;refGene_gn=PCNX3;refGene_tx=XM_006937525.2;refGene_gn=PCNX3;refGene_tx=XM_006937526.2;refGene_gn=PCNX3;refGene_tx=XM_006937527.2
D1 110758433 110759561
refGene_gn=PCNX3;refGene_tx=XM_003993638.3;refGene_gn=PCNX3;refGene_tx=XM_006937525.2;refGene_gn=PCNX3;refGene_tx=XM_006937526.2;refGene_gn=PCNX3;refGene_tx=XM_006937527.2
D1 110760066 110760190
refGene_gn=PCNX3;refGene_tx=XM_003993638.3;refGene_gn=PCNX3;refGene_tx=XM_006937525.2;refGene_gn=PCNX3;refGene_tx=XM_006937526.2;refGene_gn=PCNX3;refGene_tx=XM_006937527.2
D1 110760275 110760454
refGene_gn=PCNX3;refGene_tx=XM_003993638.3;refGene_gn=PCNX3;refGene_tx=XM_006937525.2;refGene_gn=PCNX3;refGene_tx=XM_006937526.2;refGene_gn=PCNX3;refGene_tx=XM_006937527.2
D1 110760918 110761009
refGene_gn=PCNX3;refGene_tx=XM_003993638.3;refGene_gn=PCNX3;refGene_tx=XM_006937525.2;refGene_gn=PCNX3;refGene_tx=XM_006937526.2;refGene_gn=PCNX3;refGene_tx=XM_006937527.2
D1 110761364 110761500
refGene_gn=PCNX3;refGene_tx=XM_003993638.3;refGene_gn=PCNX3;refGene_tx=XM_006937525.2;refGene_gn=PCNX3;refGene_tx=XM_006937526.2;refGene_gn=PCNX3;refGene_tx=XM_006937527.2
D1 110762750 110762911
refGene_gn=PCNX3;refGene_tx=XM_003993638.3;refGene_gn=PCNX3;refGene_tx=XM_006937525.2;refGene_gn=PCNX3;refGene_tx=XM_006937526.2;refGene_gn=PCNX3;refGene_tx=XM_006937527.2
D1 110763714 110763868
refGene_gn=PCNX3;refGene_tx=XM_003993638.3;refGene_gn=PCNX3;refGene_tx=XM_006937525.2;refGene_gn=PCNX3;refGene_tx=XM_006937526.2;refGene_gn=PCNX3;refGene_tx=XM_006937527.2
D1 110764153 110764186
refGene_gn=PCNX3;refGene_tx=XM_003993638.3;refGene_gn=PCNX3;refGene_tx=XM_006937525.2;refGene_gn=PCNX3;refGene_tx=XM_006937526.2;refGene_gn=PCNX3;refGene_tx=XM_006937527.2
D1 110764448 110764620
refGene_gn=PCNX3;refGene_tx=XM_003993638.3;refGene_gn=PCNX3;refGene_tx=XM_006937525.2;refGene_gn=PCNX3;refGene_tx=XM_006937526.2;refGene_gn=PCNX3;refGene_tx=XM_006937527.2
D1 110764728 110764839
refGene_gn=PCNX3;refGene_tx=XM_003993638.3;refGene_gn=PCNX3;refGene_tx=XM_006937525.2;refGene_gn=PCNX3;refGene_tx=XM_006937526.2;refGene_gn=PCNX3;refGene_tx=XM_006937527.2
D1 110765202 110765294

refGene_gn=PCNX3;refGene_tx=XM_003993638.3;refGene_gn=PCNX3;refGene_tx=XM_006937525.2;refGene_gn=PCNX3;refGene_tx=XM_006937526.2;refGene_gn=PCNX3;refGene_tx=XM_006937527.2
D1 110765468 110765578
refGene_gn=PCNX3;refGene_tx=XM_003993638.3;refGene_gn=PCNX3;refGene_tx=XM_006937525.2;refGene_gn=PCNX3;refGene_tx=XM_006937526.2;refGene_gn=PCNX3;refGene_tx=XM_006937527.2
D1 110765759 110765865
refGene_gn=PCNX3;refGene_tx=XM_003993638.3;refGene_gn=PCNX3;refGene_tx=XM_006937525.2;refGene_gn=PCNX3;refGene_tx=XM_006937526.2;refGene_gn=PCNX3;refGene_tx=XM_006937527.2
D1 110765971 110766064
refGene_gn=PCNX3;refGene_tx=XM_003993638.3;refGene_gn=PCNX3;refGene_tx=XM_006937525.2;refGene_gn=PCNX3;refGene_tx=XM_006937526.2;refGene_gn=PCNX3;refGene_tx=XM_006937527.2
D1 110766242 110766393
refGene_gn=PCNX3;refGene_tx=XM_003993638.3;refGene_gn=PCNX3;refGene_tx=XM_006937525.2;refGene_gn=PCNX3;refGene_tx=XM_006937526.2;refGene_gn=PCNX3;refGene_tx=XM_006937527.2
D1 110767186 110767331
refGene_gn=PCNX3;refGene_tx=XM_003993638.3;refGene_gn=PCNX3;refGene_tx=XM_006937525.2;refGene_gn=PCNX3;refGene_tx=XM_006937526.2;refGene_gn=PCNX3;refGene_tx=XM_006937527.2
D1 110767681 110767859
refGene_gn=PCNX3;refGene_tx=XM_003993638.3;refGene_gn=PCNX3;refGene_tx=XM_006937525.2;refGene_gn=PCNX3;refGene_tx=XM_006937526.2;refGene_gn=PCNX3;refGene_tx=XM_006937527.2
D1 110768890 110768999
refGene_gn=PCNX3;refGene_tx=XM_003993638.3;refGene_gn=PCNX3;refGene_tx=XM_006937525.2;refGene_gn=PCNX3;refGene_tx=XM_006937526.2;refGene_gn=PCNX3;refGene_tx=XM_006937527.2
D1 110769165 110769295
refGene_gn=PCNX3;refGene_tx=XM_003993638.3;refGene_gn=PCNX3;refGene_tx=XM_006937525.2;refGene_gn=PCNX3;refGene_tx=XM_006937526.2;refGene_gn=PCNX3;refGene_tx=XM_006937527.2
D1 110769686 110769745
refGene_gn=PCNX3;refGene_tx=XM_003993638.3;refGene_gn=PCNX3;refGene_tx=XM_006937525.2;refGene_gn=PCNX3;refGene_tx=XM_006937526.2;refGene_gn=PCNX3;refGene_tx=XM_006937527.2
D1 110769860 110770076
refGene_gn=PCNX3;refGene_tx=XM_003993638.3;refGene_gn=PCNX3;refGene_tx=XM_006937525.2;refGene_gn=PCNX3;refGene_tx=XM_006937526.2;refGene_gn=PCNX3;refGene_tx=XM_006937527.2
D1 110770725 110770979
refGene_gn=PCNX3;refGene_tx=XM_003993638.3;refGene_gn=PCNX3;refGene_tx=XM_006937525.2;refGene_gn=PCNX3;refGene_tx=XM_006937526.2;refGene_gn=PCNX3;refGene_tx=XM_006937527.2
D1 110773361 110773592
refGene_gn=PCNX3;refGene_tx=XM_003993638.3;refGene_gn=PCNX3;refGene_tx=XM_006937525.2;refGene_gn=PCNX3;refGene_tx=XM_006937526.2;refGene_gn=

PCNX3;refGene_tx=XM_006937527.2
D1 110773762 110773866
refGene_gn=PCNX3;refGene_tx=XM_003993638.3;refGene_gn=PCNX3;refGene_tx=XM_006937525.2;refGene_gn=PCNX3;refGene_tx=XM_006937526.2;refGene_gn=PCNX3;refGene_tx=XM_006937527.2
D1 110774172 110774320
refGene_gn=PCNX3;refGene_tx=XM_003993638.3;refGene_gn=PCNX3;refGene_tx=XM_006937525.2;refGene_gn=PCNX3;refGene_tx=XM_006937526.2;refGene_gn=PCNX3;refGene_tx=XM_006937527.2
D1 110774412 110774643
refGene_gn=PCNX3;refGene_tx=XM_003993638.3;refGene_gn=PCNX3;refGene_tx=XM_006937525.2;refGene_gn=PCNX3;refGene_tx=XM_006937526.2;refGene_gn=PCNX3;refGene_tx=XM_006937527.2
D1 110774727 110775002
refGene_gn=PCNX3;refGene_tx=XM_003993638.3;refGene_gn=PCNX3;refGene_tx=XM_006937525.2;refGene_gn=PCNX3;refGene_tx=XM_006937526.2;refGene_gn=PCNX3;refGene_tx=XM_006937527.2
D1 110775384 110775518
refGene_gn=PCNX3;refGene_tx=XM_003993638.3;refGene_gn=PCNX3;refGene_tx=XM_006937525.2;refGene_gn=PCNX3;refGene_tx=XM_006937526.2;refGene_gn=PCNX3;refGene_tx=XM_006937527.2
D1 110775601 110775900
refGene_gn=PCNX3;refGene_tx=XM_003993638.3;refGene_gn=PCNX3;refGene_tx=XM_006937525.2;refGene_gn=PCNX3;refGene_tx=XM_006937526.2;refGene_gn=PCNX3;refGene_tx=XM_006937527.2
D1 110775976 110776174
refGene_gn=PCNX3;refGene_tx=XM_003993638.3;refGene_gn=PCNX3;refGene_tx=XM_006937525.2;refGene_gn=PCNX3;refGene_tx=XM_006937526.2;refGene_gn=PCNX3;refGene_tx=XM_006937527.2
D1 110779843 110780516
refGene_gn=SIPA1;refGene_tx=XM_003993639.3;refGene_gn=SIPA1;refGene_tx=XM_006937528.2;refGene_gn=SIPA1;refGene_tx=XM_006937529.2;refGene_gn=SIPA1;refGene_tx=XM_006937530.3
D1 110781214 110781342
refGene_gn=SIPA1;refGene_tx=XM_003993639.3;refGene_gn=SIPA1;refGene_tx=XM_006937528.2;refGene_gn=SIPA1;refGene_tx=XM_006937529.2;refGene_gn=SIPA1;refGene_tx=XM_006937530.3
D1 110781427 110781604
refGene_gn=SIPA1;refGene_tx=XM_003993639.3;refGene_gn=SIPA1;refGene_tx=XM_006937528.2;refGene_gn=SIPA1;refGene_tx=XM_006937529.2;refGene_gn=SIPA1;refGene_tx=XM_006937530.3
D1 110783654 110783829
refGene_gn=SIPA1;refGene_tx=XM_003993639.3;refGene_gn=SIPA1;refGene_tx=XM_006937528.2;refGene_gn=SIPA1;refGene_tx=XM_006937529.2;refGene_gn=SIPA1;refGene_tx=XM_006937530.3
D1 110784536 110784640
refGene_gn=SIPA1;refGene_tx=XM_003993639.3;refGene_gn=SIPA1;refGene_tx=XM_006937528.2;refGene_gn=SIPA1;refGene_tx=XM_006937529.2;refGene_gn=SIPA1;refGene_tx=XM_006937530.3
D1 110784897 110785055

refGene_gn=SIPA1;refGene_tx=XM_003993639.3;refGene_gn=SIPA1;refGene_tx=XM_006937528.2;refGene_gn=SIPA1;refGene_tx=XM_006937529.2;refGene_gn=SIPA1;refGene_tx=XM_006937530.3
D1 110785135 110785745
refGene_gn=SIPA1;refGene_tx=XM_003993639.3;refGene_gn=SIPA1;refGene_tx=XM_006937528.2;refGene_gn=SIPA1;refGene_tx=XM_006937529.2;refGene_gn=SIPA1;refGene_tx=XM_006937530.3
D1 110786070 110786345
refGene_gn=SIPA1;refGene_tx=XM_003993639.3;refGene_gn=SIPA1;refGene_tx=XM_006937528.2;refGene_gn=SIPA1;refGene_tx=XM_006937529.2;refGene_gn=SIPA1;refGene_tx=XM_006937530.3
D1 110787251 110787473
refGene_gn=SIPA1;refGene_tx=XM_003993639.3;refGene_gn=SIPA1;refGene_tx=XM_006937528.2;refGene_gn=SIPA1;refGene_tx=XM_006937529.2;refGene_gn=SIPA1;refGene_tx=XM_006937530.3
D1 110787590 110787901
refGene_gn=SIPA1;refGene_tx=XM_003993639.3;refGene_gn=SIPA1;refGene_tx=XM_006937528.2;refGene_gn=SIPA1;refGene_tx=XM_006937529.2;refGene_gn=SIPA1;refGene_tx=XM_006937530.3
D1 110787999 110788101
refGene_gn=SIPA1;refGene_tx=XM_003993639.3;refGene_gn=SIPA1;refGene_tx=XM_006937528.2;refGene_gn=SIPA1;refGene_tx=XM_006937529.2;refGene_gn=SIPA1;refGene_tx=XM_006937530.3
D1 110788175 110788251
refGene_gn=SIPA1;refGene_tx=XM_003993639.3;refGene_gn=SIPA1;refGene_tx=XM_006937528.2;refGene_gn=SIPA1;refGene_tx=XM_006937529.2;refGene_gn=SIPA1;refGene_tx=XM_006937530.3
D1 110788473 110788552
refGene_gn=SIPA1;refGene_tx=XM_003993639.3;refGene_gn=SIPA1;refGene_tx=XM_006937528.2;refGene_gn=SIPA1;refGene_tx=XM_006937529.2;refGene_gn=SIPA1;refGene_tx=XM_006937530.3
D1 110788646 110788793
refGene_gn=SIPA1;refGene_tx=XM_003993639.3;refGene_gn=SIPA1;refGene_tx=XM_006937528.2;refGene_gn=SIPA1;refGene_tx=XM_006937529.2;refGene_gn=SIPA1;refGene_tx=XM_006937530.3
D1 110791656 110792291
refGene_gn=RELA;refGene_tx=XM_023240068.1
D1 110793041 110793116
refGene_gn=RELA;refGene_tx=XM_023240068.1
D1 110793204 110793285
refGene_gn=RELA;refGene_tx=XM_023240068.1
D1 110795430 110795643
refGene_gn=RELA;refGene_tx=XM_023240068.1
D1 110795864 110795969
refGene_gn=RELA;refGene_tx=XM_023240068.1
D1 110796805 110796937
refGene_gn=RELA;refGene_tx=XM_023240068.1
D1 110797300 110797392
refGene_gn=RELA;refGene_tx=XM_023240068.1
D1 110798045 110798194

refGene_gn=RELA;refGene_tx=XM_023240068.1
D1 110798343 110798495
refGene_gn=RELA;refGene_tx=XM_023240068.1
D1 110798597 110798624
refGene_gn=RELA;refGene_tx=XM_023240068.1
D1 110799254 110799261
refGene_gn=RELA;refGene_tx=XM_023240068.1
D1 110827629 110827807
refGene_gn=KAT5;refGene_tx=XM_003993641.5;refGene_gn=KAT5;refGene_tx=XM_003993642.4;refGene_gn=KAT5;refGene_tx=XM_006937531.4;refGene_gn=KAT5;refGene_tx=XM_006937532.4;refGene_gn=KAT5;refGene_tx=XM_019812689.2;refGene_gn=KAT5;refGene_tx=XM_019812690.2
D1 110828127 110828196
refGene_gn=KAT5;refGene_tx=XM_003993641.5;refGene_gn=KAT5;refGene_tx=XM_003993642.4;refGene_gn=KAT5;refGene_tx=XM_006937531.4;refGene_gn=KAT5;refGene_tx=XM_006937532.4;refGene_gn=KAT5;refGene_tx=XM_019812689.2;refGene_gn=KAT5;refGene_tx=XM_019812690.2
D1 110828287 110828424
refGene_gn=KAT5;refGene_tx=XM_003993641.5;refGene_gn=KAT5;refGene_tx=XM_003993642.4;refGene_gn=KAT5;refGene_tx=XM_006937531.4;refGene_gn=KAT5;refGene_tx=XM_006937532.4;refGene_gn=KAT5;refGene_tx=XM_019812689.2;refGene_gn=KAT5;refGene_tx=XM_019812690.2
D1 110828687 110828870
refGene_gn=KAT5;refGene_tx=XM_003993641.5;refGene_gn=KAT5;refGene_tx=XM_006937531.4;refGene_gn=KAT5;refGene_tx=XM_019812689.2;refGene_gn=KAT5;refGene_tx=XM_019812690.2
D1 110828959 110829034
refGene_gn=KAT5;refGene_tx=XM_003993641.5;refGene_gn=KAT5;refGene_tx=XM_003993642.4;refGene_gn=KAT5;refGene_tx=XM_006937531.4;refGene_gn=KAT5;refGene_tx=XM_006937532.4;refGene_gn=KAT5;refGene_tx=XM_019812689.2;refGene_gn=KAT5;refGene_tx=XM_019812690.2
D1 110829139 110829214
refGene_gn=KAT5;refGene_tx=XM_003993641.5;refGene_gn=KAT5;refGene_tx=XM_003993642.4;refGene_gn=KAT5;refGene_tx=XM_006937531.4;refGene_gn=KAT5;refGene_tx=XM_006937532.4;refGene_gn=KAT5;refGene_tx=XM_019812689.2;refGene_gn=KAT5;refGene_tx=XM_019812690.2
D1 110829782 110830031
refGene_gn=KAT5;refGene_tx=XM_003993641.5;refGene_gn=KAT5;refGene_tx=XM_003993642.4;refGene_gn=KAT5;refGene_tx=XM_006937531.4;refGene_gn=KAT5;refGene_tx=XM_006937532.4;refGene_gn=KAT5;refGene_tx=XM_019812689.2;refGene_gn=KAT5;refGene_tx=XM_019812690.2
D1 110830108 110830198
refGene_gn=KAT5;refGene_tx=XM_003993641.5;refGene_gn=KAT5;refGene_tx=XM_003993642.4;refGene_gn=KAT5;refGene_tx=XM_006937531.4;refGene_gn=KAT5;refGene_tx=XM_006937532.4;refGene_gn=KAT5;refGene_tx=XM_019812689.2;refGene_gn=KAT5;refGene_tx=XM_019812690.2
D1 110831554 110831695
refGene_gn=KAT5;refGene_tx=XM_003993641.5;refGene_gn=KAT5;refGene_tx=XM_003993642.4;refGene_gn=KAT5;refGene_tx=XM_006937531.4;refGene_gn=KAT5;refGene_tx=XM_006937532.4;refGene_gn=KAT5;refGene_tx=XM_019812689.2;

refGene_gn=KAT5;refGene_tx=XM_019812690.2
D1 110831774 110831868
refGene_gn=KAT5;refGene_tx=XM_003993641.5;refGene_gn=KAT5;refGene_tx=XM_003993642.4;refGene_gn=KAT5;refGene_tx=XM_006937531.4;refGene_gn=KAT5;refGene_tx=XM_006937532.4;refGene_gn=KAT5;refGene_tx=XM_019812689.2;refGene_gn=KAT5;refGene_tx=XM_019812690.2
D1 110832760 110832939
refGene_gn=KAT5;refGene_tx=XM_019812690.2
D1 110834137 110834297
refGene_gn=KAT5;refGene_tx=XM_003993641.5;refGene_gn=KAT5;refGene_tx=XM_003993642.4;refGene_gn=KAT5;refGene_tx=XM_006937531.4;refGene_gn=KAT5;refGene_tx=XM_006937532.4;refGene_gn=KAT5;refGene_tx=XM_019812689.2
D1 110834419 110834501
refGene_gn=KAT5;refGene_tx=XM_003993641.5;refGene_gn=KAT5;refGene_tx=XM_003993642.4;refGene_gn=KAT5;refGene_tx=XM_006937531.4;refGene_gn=KAT5;refGene_tx=XM_006937532.4;refGene_gn=KAT5;refGene_tx=XM_019812689.2
D1 110834590 110834725
refGene_gn=KAT5;refGene_tx=XM_003993641.5;refGene_gn=KAT5;refGene_tx=XM_003993642.4;refGene_gn=KAT5;refGene_tx=XM_006937531.4;refGene_gn=KAT5;refGene_tx=XM_006937532.4;refGene_gn=KAT5;refGene_tx=XM_019812689.2
D1 110835324 110835351
refGene_gn=RNASEH2C;refGene_tx=XM_003993643.4
D1 110835678 110835798
refGene_gn=RNASEH2C;refGene_tx=XM_003993643.4
D1 110835897 110836091
refGene_gn=RNASEH2C;refGene_tx=XM_003993643.4
D1 110836279 110836451
refGene_gn=RNASEH2C;refGene_tx=XM_003993643.4
D1 110856516 110859003
refGene_gn=AP5B1;refGene_tx=XM_023240069.1
D1 110859106 110859256
refGene_gn=AP5B1;refGene_tx=XM_023240069.1
D1 110868025 110868125
refGene_gn=OVOL1;refGene_tx=XM_023240070.1
D1 110869536 110869720
refGene_gn=OVOL1;refGene_tx=XM_006937533.4
D1 110875260 110875478
refGene_gn=OVOL1;refGene_tx=XM_006937533.4;refGene_gn=OVOL1;refGene_tx=XM_019812692.2;refGene_gn=OVOL1;refGene_tx=XM_019812693.2;refGene_gn=OVOL1;refGene_tx=XM_023240070.1;refGene_gn=OVOL1;refGene_tx=XM_023240071.1
D1 110877244 110877434
refGene_gn=OVOL1;refGene_tx=XM_006937533.4;refGene_gn=OVOL1;refGene_tx=XM_019812692.2;refGene_gn=OVOL1;refGene_tx=XM_019812693.2;refGene_gn=OVOL1;refGene_tx=XM_023240070.1;refGene_gn=OVOL1;refGene_tx=XM_023240071.1
D1 110877737 110878033
refGene_gn=OVOL1;refGene_tx=XM_006937533.4;refGene_gn=OVOL1;refGene_tx=XM_019812692.2;refGene_gn=OVOL1;refGene_tx=XM_019812693.2;refGene_gn=OVOL1;refGene_tx=XM_023240070.1;refGene_gn=OVOL1;refGene_tx=XM_0232400

71.1
D1 110910889 110911939
refGene_gn=LOC102901440;refGene_tx=XM_023240072.1;refGene_gn=LOC102901440;refGene_tx=XM_023240073.1;refGene_gn=LOC102901440;refGene_tx=XM_023240074.1;refGene_gn=LOC102901440;refGene_tx=XM_023240075.1;refGene_gn=LOC102901440;refGene_tx=XM_023240076.1
D1 110912571 110912607
refGene_gn=SNX32;refGene_tx=XM_003993723.4;refGene_gn=SNX32;refGene_tx=XM_019812694.2
D1 110921223 110921328
refGene_gn=SNX32;refGene_tx=XM_003993723.4
D1 110921728 110921839
refGene_gn=SNX32;refGene_tx=XM_003993723.4;refGene_gn=SNX32;refGene_tx=XM_019812694.2
D1 110921982 110922104
refGene_gn=SNX32;refGene_tx=XM_003993723.4;refGene_gn=SNX32;refGene_tx=XM_019812694.2
D1 110922281 110922405
refGene_gn=SNX32;refGene_tx=XM_003993723.4;refGene_gn=SNX32;refGene_tx=XM_019812694.2
D1 110922564 110922669
refGene_gn=SNX32;refGene_tx=XM_003993723.4;refGene_gn=SNX32;refGene_tx=XM_019812694.2
D1 110922863 110922969
refGene_gn=SNX32;refGene_tx=XM_003993723.4;refGene_gn=SNX32;refGene_tx=XM_019812694.2
D1 110923143 110923219
refGene_gn=SNX32;refGene_tx=XM_003993723.4;refGene_gn=SNX32;refGene_tx=XM_019812694.2
D1 110923481 110923521
refGene_gn=SNX32;refGene_tx=XM_003993723.4;refGene_gn=SNX32;refGene_tx=XM_019812694.2
D1 110924265 110924352
refGene_gn=SNX32;refGene_tx=XM_003993723.4;refGene_gn=SNX32;refGene_tx=XM_019812694.2
D1 110924465 110924625
refGene_gn=SNX32;refGene_tx=XM_003993723.4;refGene_gn=SNX32;refGene_tx=XM_019812694.2
D1 110924706 110924792
refGene_gn=SNX32;refGene_tx=XM_003993723.4;refGene_gn=SNX32;refGene_tx=XM_019812694.2
D1 110925125 110925179
refGene_gn=SNX32;refGene_tx=XM_003993723.4;refGene_gn=SNX32;refGene_tx=XM_019812694.2
D1 110927046 110927159
refGene_gn=CFL1;refGene_tx=XM_003993644.4;refGene_gn=CFL1;refGene_tx=XM_023240077.1
D1 110927375 110927452
refGene_gn=CFL1;refGene_tx=XM_003993644.4;refGene_gn=CFL1;refGene_tx=XM_023240077.1

D1 110927729 110928037
refGene_gn=CFL1;refGene_tx=XM_003993644.4;refGene_gn=CFL1;refGene_tx=X
M_023240077.1
D1 110929459 110929594
refGene_gn=CFL1;refGene_tx=XM_023240077.1
D1 110929958 110929961
refGene_gn=CFL1;refGene_tx=XM_003993644.4
D1 110932572 110932707
refGene_gn=MUS81;refGene_tx=XM_003993646.5;refGene_gn=MUS81;refGene_tx
=XM_006937537.4
D1 110932795 110932925
refGene_gn=MUS81;refGene_tx=XM_003993646.5;refGene_gn=MUS81;refGene_tx
=XM_006937537.4
D1 110933165 110933251
refGene_gn=MUS81;refGene_tx=XM_003993646.5;refGene_gn=MUS81;refGene_tx
=XM_006937537.4
D1 110933803 110933902
refGene_gn=MUS81;refGene_tx=XM_003993646.5;refGene_gn=MUS81;refGene_tx
=XM_006937537.4
D1 110934024 110934093
refGene_gn=MUS81;refGene_tx=XM_003993646.5;refGene_gn=MUS81;refGene_tx
=XM_006937537.4
D1 110934256 110934342
refGene_gn=MUS81;refGene_tx=XM_003993646.5;refGene_gn=MUS81;refGene_tx
=XM_006937537.4
D1 110934852 110934993
refGene_gn=MUS81;refGene_tx=XM_003993646.5;refGene_gn=MUS81;refGene_tx
=XM_006937537.4
D1 110935171 110935267
refGene_gn=MUS81;refGene_tx=XM_003993646.5;refGene_gn=MUS81;refGene_tx
=XM_006937537.4
D1 110935356 110935478
refGene_gn=MUS81;refGene_tx=XM_003993646.5;refGene_gn=MUS81;refGene_tx
=XM_006937537.4
D1 110935574 110935672
refGene_gn=MUS81;refGene_tx=XM_003993646.5;refGene_gn=MUS81;refGene_tx
=XM_006937537.4
D1 110936234 110936351
refGene_gn=MUS81;refGene_tx=XM_003993646.5;refGene_gn=MUS81;refGene_tx
=XM_006937537.4
D1 110936451 110936547
refGene_gn=MUS81;refGene_tx=XM_003993646.5;refGene_gn=MUS81;refGene_tx
=XM_006937537.4
D1 110936698 110936827
refGene_gn=MUS81;refGene_tx=XM_003993646.5;refGene_gn=MUS81;refGene_tx
=XM_006937537.4
D1 110936902 110937006
refGene_gn=MUS81;refGene_tx=XM_003993646.5;refGene_gn=MUS81;refGene_tx
=XM_006937537.4
D1 110937572 110937656

refGene_gn=MUS81;refGene_tx=XM_003993646.5
D1	110937748	110937815
refGene_gn=MUS81;refGene_tx=XM_003993646.5;refGene_gn=MUS81;refGene_tx=XM_006937537.4
D1	110938737	110938899
refGene_gn=EFEMP2;refGene_tx=XM_023240078.1
D1	110940013	110940209
refGene_gn=EFEMP2;refGene_tx=XM_023240078.1
D1	110940669	110940796
refGene_gn=EFEMP2;refGene_tx=XM_023240078.1
D1	110940883	110941003
refGene_gn=EFEMP2;refGene_tx=XM_023240078.1
D1	110942418	110942538
refGene_gn=EFEMP2;refGene_tx=XM_023240078.1
D1	110942677	110942794
refGene_gn=EFEMP2;refGene_tx=XM_023240078.1
D1	110943274	110943397
refGene_gn=EFEMP2;refGene_tx=XM_023240078.1
D1	110943986	110944193
refGene_gn=EFEMP2;refGene_tx=XM_023240078.1
D1	110944909	110944958
refGene_gn=EFEMP2;refGene_tx=XM_023240078.1
D1	110945188	110945299
refGene_gn=EFEMP2;refGene_tx=XM_023240078.1
D1	110949436	110949523
refGene_gn=CTSW;refGene_tx=NM_001122871.1;refGene_gn=CTSW;refGene_tx=XM_006937522.3;refGene_gn=CTSW;refGene_tx=XM_006937524.3;refGene_gn=CTSW;refGene_tx=XM_019811160.2;refGene_gn=CTSW;refGene_tx=XM_019811162.2
D1	110949790	110949875
refGene_gn=CTSW;refGene_tx=NM_001122871.1;refGene_gn=CTSW;refGene_tx=XM_006937522.3;refGene_gn=CTSW;refGene_tx=XM_006937524.3;refGene_gn=CTSW;refGene_tx=XM_019811160.2;refGene_gn=CTSW;refGene_tx=XM_019811162.2
D1	110950610	110950797
refGene_gn=CTSW;refGene_tx=NM_001122871.1;refGene_gn=CTSW;refGene_tx=XM_006937522.3;refGene_gn=CTSW;refGene_tx=XM_006937524.3;refGene_gn=CTSW;refGene_tx=XM_019811160.2;refGene_gn=CTSW;refGene_tx=XM_019811161.1;refGene_gn=CTSW;refGene_tx=XM_019811162.2
D1	110951182	110951527
refGene_gn=CTSW;refGene_tx=NM_001122871.1;refGene_gn=CTSW;refGene_tx=XM_006937522.3;refGene_gn=CTSW;refGene_tx=XM_006937524.3;refGene_gn=CTSW;refGene_tx=XM_019811160.2;refGene_gn=CTSW;refGene_tx=XM_019811161.1;refGene_gn=CTSW;refGene_tx=XM_019811162.2
D1	110951618	110951699
refGene_gn=CTSW;refGene_tx=NM_001122871.1;refGene_gn=CTSW;refGene_tx=XM_006937522.3;refGene_gn=CTSW;refGene_tx=XM_006937524.3;refGene_gn=CTSW;refGene_tx=XM_019811161.1
D1	110951805	110951931
refGene_gn=CTSW;refGene_tx=NM_001122871.1;refGene_gn=CTSW;refGene_tx=XM_006937522.3;refGene_gn=CTSW;refGene_tx=XM_006937524.3;refGene_gn=CTSW;refGene_tx=XM_019811160.2;refGene_gn=CTSW;refGene_tx=XM_019811161.1;

refGene_gn=CTSW;refGene_tx=XM_019811162.2
D1 110952116 110952181
refGene_gn=CTSW;refGene_tx=NM_001122871.1;refGene_gn=CTSW;refGene_tx=XM_006937522.3;refGene_gn=CTSW;refGene_tx=XM_006937524.3;refGene_gn=CTSW;refGene_tx=XM_019811160.2;refGene_gn=CTSW;refGene_tx=XM_019811161.1;refGene_gn=CTSW;refGene_tx=XM_019811162.2
D1 110952266 110952470
refGene_gn=CTSW;refGene_tx=NM_001122871.1;refGene_gn=CTSW;refGene_tx=XM_006937522.3;refGene_gn=CTSW;refGene_tx=XM_006937524.3;refGene_gn=CTSW;refGene_tx=XM_019811160.2;refGene_gn=CTSW;refGene_tx=XM_019811161.1;refGene_gn=CTSW;refGene_tx=XM_019811162.2
D1 110952552 110952663
refGene_gn=CTSW;refGene_tx=NM_001122871.1;refGene_gn=CTSW;refGene_tx=XM_006937522.3;refGene_gn=CTSW;refGene_tx=XM_006937524.3;refGene_gn=CTSW;refGene_tx=XM_019811160.2;refGene_gn=CTSW;refGene_tx=XM_019811161.1;refGene_gn=CTSW;refGene_tx=XM_019811162.2
D1 110953015 110953085
refGene_gn=FIBP;refGene_tx=XM_003993647.5
D1 110953426 110953524
refGene_gn=FIBP;refGene_tx=XM_003993647.5
D1 110953612 110953699
refGene_gn=FIBP;refGene_tx=XM_003993647.5
D1 110953976 110954040
refGene_gn=FIBP;refGene_tx=XM_003993647.5
D1 110954116 110954225
refGene_gn=FIBP;refGene_tx=XM_003993647.5
D1 110954559 110954693
refGene_gn=FIBP;refGene_tx=XM_003993647.5
D1 110955181 110955282
refGene_gn=FIBP;refGene_tx=XM_003993647.5
D1 110956143 110956270
refGene_gn=FIBP;refGene_tx=XM_003993647.5
D1 110956426 110956625
refGene_gn=FIBP;refGene_tx=XM_003993647.5
D1 110956821 110956906
refGene_gn=FIBP;refGene_tx=XM_003993647.5
D1 110959248 110959857
refGene_gn=CCDC85B;refGene_tx=XM_023240079.1
D1 110961358 110961781
refGene_gn=FOSL1;refGene_tx=XM_003993649.5;refGene_gn=FOSL1;refGene_tx=XM_006937538.3;refGene_gn=FOSL1;refGene_tx=XM_011287025.3
D1 110962202 110962310
refGene_gn=FOSL1;refGene_tx=XM_003993649.5;refGene_gn=FOSL1;refGene_tx=XM_006937538.3;refGene_gn=FOSL1;refGene_tx=XM_011287025.3
D1 110964261 110964663
refGene_gn=FOSL1;refGene_tx=XM_003993649.5;refGene_gn=FOSL1;refGene_tx=XM_006937538.3;refGene_gn=FOSL1;refGene_tx=XM_011287025.3
D1 110966970 110967060
refGene_gn=FOSL1;refGene_tx=XM_003993649.5;refGene_gn=FOSL1;refGene_tx=XM_011287025.3

D1 110981250 110982010
refGene_gn=CD1H11orf68;refGene_tx=XM_003993725.5;refGene_gn=CD1H11orf68;refGene_tx=XM_006937539.4;refGene_gn=CD1H11orf68;refGene_tx=XM_019812695.2
D1 110982701 110982829
refGene_gn=CD1H11orf68;refGene_tx=XM_003993725.5;refGene_gn=CD1H11orf68;refGene_tx=XM_006937539.4
D1 110983273 110983315
refGene_gn=DRAP1;refGene_tx=XM_006937540.4;refGene_gn=DRAP1;refGene_tx=XM_011287026.3
D1 110983560 110983633
refGene_gn=DRAP1;refGene_tx=XM_006937540.4;refGene_gn=DRAP1;refGene_tx=XM_011287026.3
D1 110983743 110983837
refGene_gn=DRAP1;refGene_tx=XM_006937540.4;refGene_gn=DRAP1;refGene_tx=XM_011287026.3
D1 110984119 110984239
refGene_gn=DRAP1;refGene_tx=XM_006937540.4;refGene_gn=DRAP1;refGene_tx=XM_011287026.3
D1 110984338 110984453
refGene_gn=DRAP1;refGene_tx=XM_006937540.4;refGene_gn=DRAP1;refGene_tx=XM_011287026.3
D1 110984669 110984758
refGene_gn=DRAP1;refGene_tx=XM_006937540.4;refGene_gn=DRAP1;refGene_tx=XM_011287026.3
D1 110985131 110985237
refGene_gn=DRAP1;refGene_tx=XM_006937540.4;refGene_gn=DRAP1;refGene_tx=XM_011287026.3
D1 111009780 111010011
refGene_gn=TSGA10IP;refGene_tx=XM_023239751.1
D1 111014418 111014738
refGene_gn=TSGA10IP;refGene_tx=XM_023239751.1
D1 111017296 111017396
refGene_gn=TSGA10IP;refGene_tx=XM_023239751.1
D1 111017597 111017722
refGene_gn=TSGA10IP;refGene_tx=XM_023239751.1
D1 111018434 111018543
refGene_gn=TSGA10IP;refGene_tx=XM_023239751.1
D1 111020041 111020354
refGene_gn=SART1;refGene_tx=XM_003993652.5;refGene_gn=SART1;refGene_tx=XM_006937541.3
D1 111021105 111021163
refGene_gn=SART1;refGene_tx=XM_003993652.5;refGene_gn=SART1;refGene_tx=XM_006937541.3
D1 111021509 111021565
refGene_gn=SART1;refGene_tx=XM_003993652.5;refGene_gn=SART1;refGene_tx=XM_006937541.3
D1 111022148 111022275
refGene_gn=SART1;refGene_tx=XM_003993652.5;refGene_gn=SART1;refGene_tx=XM_006937541.3

D1	111022397	111022503
refGene_gn=SART1;refGene_tx=XM_003993652.5;refGene_gn=SART1;refGene_tx=XM_006937541.3
D1	111022752	111022830
refGene_gn=SART1;refGene_tx=XM_003993652.5;refGene_gn=SART1;refGene_tx=XM_006937541.3
D1	111022953	111023053
refGene_gn=SART1;refGene_tx=XM_003993652.5;refGene_gn=SART1;refGene_tx=XM_006937541.3
D1	111023144	111023287
refGene_gn=SART1;refGene_tx=XM_003993652.5;refGene_gn=SART1;refGene_tx=XM_006937541.3
D1	111023376	111023583
refGene_gn=SART1;refGene_tx=XM_003993652.5;refGene_gn=SART1;refGene_tx=XM_006937541.3
D1	111024287	111024412
refGene_gn=SART1;refGene_tx=XM_003993652.5;refGene_gn=SART1;refGene_tx=XM_006937541.3
D1	111024493	111024627
refGene_gn=SART1;refGene_tx=XM_003993652.5;refGene_gn=SART1;refGene_tx=XM_006937541.3
D1	111024749	111024883
refGene_gn=SART1;refGene_tx=XM_003993652.5;refGene_gn=SART1;refGene_tx=XM_006937541.3
D1	111033331	111033505
refGene_gn=SART1;refGene_tx=XM_003993652.5;refGene_gn=SART1;refGene_tx=XM_006937541.3
D1	111033677	111033788
refGene_gn=SART1;refGene_tx=XM_003993652.5;refGene_gn=SART1;refGene_tx=XM_006937541.3
D1	111034084	111034172
refGene_gn=SART1;refGene_tx=XM_003993652.5;refGene_gn=SART1;refGene_tx=XM_006937541.3
D1	111034474	111034565
refGene_gn=SART1;refGene_tx=XM_003993652.5;refGene_gn=SART1;refGene_tx=XM_006937541.3
D1	111034678	111034814
refGene_gn=SART1;refGene_tx=XM_003993652.5;refGene_gn=SART1;refGene_tx=XM_006937541.3
D1	111035349	111035439
refGene_gn=SART1;refGene_tx=XM_003993652.5;refGene_gn=SART1;refGene_tx=XM_006937541.3
D1	111035547	111035669
refGene_gn=SART1;refGene_tx=XM_003993652.5;refGene_gn=SART1;refGene_tx=XM_006937541.3
D1	111035771	111035790
refGene_gn=SART1;refGene_tx=XM_003993652.5;refGene_gn=SART1;refGene_tx=XM_006937541.3
D1	111057127	111057272
refGene_gn=EIF1AD;refGene_tx=XM_003993653.5

D1 111057771 111057822
refGene_gn=EIF1AD;refGene_tx=XM_003993653.5
D1 111057990 111058099
refGene_gn=EIF1AD;refGene_tx=XM_003993653.5
D1 111058200 111058309
refGene_gn=EIF1AD;refGene_tx=XM_003993653.5
D1 111058455 111058542
refGene_gn=EIF1AD;refGene_tx=XM_003993653.5
D1 111061409 111061532
refGene_gn=BANF1;refGene_tx=XM_003993654.5
D1 111061765 111061912
refGene_gn=BANF1;refGene_tx=XM_003993654.5
D1 111068191 111068431
refGene_gn=CST6;refGene_tx=XM_003993727.4
D1 111068912 111069038
refGene_gn=CST6;refGene_tx=XM_003993727.4
D1 111069378 111069465
refGene_gn=CST6;refGene_tx=XM_003993727.4
D1 111071997 111072024
refGene_gn=CATSPER1;refGene_tx=XM_006937542.3
D1 111072184 111072299
refGene_gn=CATSPER1;refGene_tx=XM_006937542.3
D1 111074363 111074439
refGene_gn=CATSPER1;refGene_tx=XM_006937542.3
D1 111076110 111076171
refGene_gn=CATSPER1;refGene_tx=XM_006937542.3
D1 111076308 111076381
refGene_gn=CATSPER1;refGene_tx=XM_006937542.3
D1 111076576 111076640
refGene_gn=CATSPER1;refGene_tx=XM_006937542.3
D1 111076801 111076945
refGene_gn=CATSPER1;refGene_tx=XM_006937542.3
D1 111077071 111077163
refGene_gn=CATSPER1;refGene_tx=XM_006937542.3
D1 111077473 111077624
refGene_gn=CATSPER1;refGene_tx=XM_006937542.3
D1 111077809 111077923
refGene_gn=CATSPER1;refGene_tx=XM_006937542.3
D1 111079228 111079441
refGene_gn=CATSPER1;refGene_tx=XM_006937542.3
D1 111080424 111081781
refGene_gn=CATSPER1;refGene_tx=XM_006937542.3
D1 111102776 111103947
refGene_gn=GAL3ST3;refGene_tx=XM_006937543.4
D1 111106162 111106287
refGene_gn=GAL3ST3;refGene_tx=XM_006937543.4
D1 111113003 111113136
refGene_gn=SF3B2;refGene_tx=XM_003993658.5;refGene_gn=SF3B2;refGene_tx
=XM_019812699.2;refGene_gn=SF3B2;refGene_tx=XM_019812700.2
D1 111113312 111113362

refGene_gn=SF3B2;refGene_tx=XM_003993658.5;refGene_gn=SF3B2;refGene_tx=XM_019812699.2;refGene_gn=SF3B2;refGene_tx=XM_019812700.2
D1 111113900 111113978
refGene_gn=SF3B2;refGene_tx=XM_003993658.5;refGene_gn=SF3B2;refGene_tx=XM_019812699.2;refGene_gn=SF3B2;refGene_tx=XM_019812700.2
D1 111114739 111114988
refGene_gn=SF3B2;refGene_tx=XM_003993658.5;refGene_gn=SF3B2;refGene_tx=XM_019812699.2;refGene_gn=SF3B2;refGene_tx=XM_019812700.2
D1 111115205 111115256
refGene_gn=SF3B2;refGene_tx=XM_003993658.5;refGene_gn=SF3B2;refGene_tx=XM_019812699.2;refGene_gn=SF3B2;refGene_tx=XM_019812700.2
D1 111117414 111117532
refGene_gn=SF3B2;refGene_tx=XM_003993658.5;refGene_gn=SF3B2;refGene_tx=XM_019812699.2;refGene_gn=SF3B2;refGene_tx=XM_019812700.2
D1 111118039 111118140
refGene_gn=SF3B2;refGene_tx=XM_003993658.5;refGene_gn=SF3B2;refGene_tx=XM_019812699.2;refGene_gn=SF3B2;refGene_tx=XM_019812700.2
D1 111118447 111118544
refGene_gn=SF3B2;refGene_tx=XM_003993658.5;refGene_gn=SF3B2;refGene_tx=XM_019812699.2;refGene_gn=SF3B2;refGene_tx=XM_019812700.2
D1 111118725 111118817
refGene_gn=SF3B2;refGene_tx=XM_003993658.5;refGene_gn=SF3B2;refGene_tx=XM_019812699.2;refGene_gn=SF3B2;refGene_tx=XM_019812700.2
D1 111119204 111119420
refGene_gn=SF3B2;refGene_tx=XM_003993658.5;refGene_gn=SF3B2;refGene_tx=XM_019812699.2;refGene_gn=SF3B2;refGene_tx=XM_019812700.2
D1 111119569 111119707
refGene_gn=SF3B2;refGene_tx=XM_003993658.5;refGene_gn=SF3B2;refGene_tx=XM_019812699.2;refGene_gn=SF3B2;refGene_tx=XM_019812700.2
D1 111119872 111119953
refGene_gn=SF3B2;refGene_tx=XM_003993658.5;refGene_gn=SF3B2;refGene_tx=XM_019812699.2;refGene_gn=SF3B2;refGene_tx=XM_019812700.2
D1 111120170 111120398
refGene_gn=SF3B2;refGene_tx=XM_003993658.5;refGene_gn=SF3B2;refGene_tx=XM_019812699.2;refGene_gn=SF3B2;refGene_tx=XM_019812700.2
D1 111121178 111121328
refGene_gn=SF3B2;refGene_tx=XM_003993658.5;refGene_gn=SF3B2;refGene_tx=XM_019812699.2;refGene_gn=SF3B2;refGene_tx=XM_019812700.2
D1 111122306 111122396
refGene_gn=SF3B2;refGene_tx=XM_003993658.5;refGene_gn=SF3B2;refGene_tx=XM_019812699.2;refGene_gn=SF3B2;refGene_tx=XM_019812700.2
D1 111122508 111122616
refGene_gn=SF3B2;refGene_tx=XM_003993658.5;refGene_gn=SF3B2;refGene_tx=XM_019812699.2;refGene_gn=SF3B2;refGene_tx=XM_019812700.2
D1 111123865 111123973
refGene_gn=SF3B2;refGene_tx=XM_003993658.5;refGene_gn=SF3B2;refGene_tx=XM_019812699.2;refGene_gn=SF3B2;refGene_tx=XM_019812700.2
D1 111124248 111124391
refGene_gn=SF3B2;refGene_tx=XM_003993658.5;refGene_gn=SF3B2;refGene_tx=XM_019812699.2;refGene_gn=SF3B2;refGene_tx=XM_019812700.2

D1	111124474	111124576
refGene_gn=SF3B2;refGene_tx=XM_003993658.5;refGene_gn=SF3B2;refGene_tx
=XM_019812699.2;refGene_gn=SF3B2;refGene_tx=XM_019812700.2
D1	111128754	111128854
refGene_gn=SF3B2;refGene_tx=XM_003993658.5;refGene_gn=SF3B2;refGene_tx
=XM_019812699.2;refGene_gn=SF3B2;refGene_tx=XM_019812700.2
D1	111128943	111129129
refGene_gn=SF3B2;refGene_tx=XM_003993658.5;refGene_gn=SF3B2;refGene_tx
=XM_019812699.2;refGene_gn=SF3B2;refGene_tx=XM_019812700.2
D1	111129449	111129521
refGene_gn=SF3B2;refGene_tx=XM_003993658.5;refGene_gn=SF3B2;refGene_tx
=XM_019812699.2;refGene_gn=SF3B2;refGene_tx=XM_019812700.2
D1	111131561	111131911
refGene_gn=PACS1;refGene_tx=XM_023240080.1
D1	111245316	111245404
refGene_gn=PACS1;refGene_tx=XM_023240080.1
D1	111256163	111256253
refGene_gn=PACS1;refGene_tx=XM_023240080.1
D1	111256945	111257071
refGene_gn=PACS1;refGene_tx=XM_023240080.1
D1	111262198	111262343
refGene_gn=PACS1;refGene_tx=XM_023240080.1
D1	111262621	111262713
refGene_gn=PACS1;refGene_tx=XM_023240080.1
D1	111262809	111262890
refGene_gn=PACS1;refGene_tx=XM_023240080.1
D1	111266053	111266113
refGene_gn=PACS1;refGene_tx=XM_023240080.1
D1	111267098	111267259
refGene_gn=PACS1;refGene_tx=XM_023240080.1
D1	111267630	111267724
refGene_gn=PACS1;refGene_tx=XM_023240080.1
D1	111272425	111272506
refGene_gn=PACS1;refGene_tx=XM_023240080.1
D1	111273916	111274032
refGene_gn=PACS1;refGene_tx=XM_023240080.1
D1	111274209	111274345
refGene_gn=PACS1;refGene_tx=XM_023240080.1
D1	111275596	111275701
refGene_gn=PACS1;refGene_tx=XM_023240080.1
D1	111276432	111276539
refGene_gn=PACS1;refGene_tx=XM_023240080.1
D1	111277243	111277398
refGene_gn=PACS1;refGene_tx=XM_023240080.1
D1	111277536	111277647
refGene_gn=PACS1;refGene_tx=XM_023240080.1
D1	111278421	111278524
refGene_gn=PACS1;refGene_tx=XM_023240080.1
D1	111279246	111279289
refGene_gn=PACS1;refGene_tx=XM_023240080.1

D1 111281459 111281502
refGene_gn=PACS1;refGene_tx=XM_023240080.1
D1 111281760 111281896
refGene_gn=PACS1;refGene_tx=XM_023240080.1
D1 111284362 111284589
refGene_gn=PACS1;refGene_tx=XM_023240080.1
D1 111285802 111285922
refGene_gn=PACS1;refGene_tx=XM_023240080.1
D1 111286075 111286191
refGene_gn=PACS1;refGene_tx=XM_023240080.1
D1 111298653 111298881
refGene_gn=KLC2;refGene_tx=XM_003993660.5;refGene_gn=KLC2;refGene_tx=XM_003993661.5
D1 111301586 111301817
refGene_gn=KLC2;refGene_tx=XM_003993660.5;refGene_gn=KLC2;refGene_tx=XM_003993661.5
D1 111301949 111302019
refGene_gn=KLC2;refGene_tx=XM_003993660.5;refGene_gn=KLC2;refGene_tx=XM_003993661.5
D1 111302796 111303019
refGene_gn=KLC2;refGene_tx=XM_003993660.5;refGene_gn=KLC2;refGene_tx=XM_003993661.5
D1 111303726 111303814
refGene_gn=KLC2;refGene_tx=XM_003993660.5;refGene_gn=KLC2;refGene_tx=XM_003993661.5
D1 111303898 111304000
refGene_gn=KLC2;refGene_tx=XM_003993660.5;refGene_gn=KLC2;refGene_tx=XM_003993661.5
D1 111304092 111304266
refGene_gn=KLC2;refGene_tx=XM_003993660.5;refGene_gn=KLC2;refGene_tx=XM_003993661.5
D1 111304354 111304454
refGene_gn=KLC2;refGene_tx=XM_003993660.5;refGene_gn=KLC2;refGene_tx=XM_003993661.5
D1 111305165 111305215
refGene_gn=KLC2;refGene_tx=XM_003993660.5;refGene_gn=KLC2;refGene_tx=XM_003993661.5
D1 111305322 111305390
refGene_gn=KLC2;refGene_tx=XM_003993660.5;refGene_gn=KLC2;refGene_tx=XM_003993661.5
D1 111305823 111305932
refGene_gn=KLC2;refGene_tx=XM_003993660.5;refGene_gn=KLC2;refGene_tx=XM_003993661.5
D1 111306034 111306193
refGene_gn=KLC2;refGene_tx=XM_003993660.5;refGene_gn=KLC2;refGene_tx=XM_003993661.5
D1 111306267 111306392
refGene_gn=KLC2;refGene_tx=XM_003993660.5;refGene_gn=KLC2;refGene_tx=XM_003993661.5
D1 111306659 111306717

refGene_gn=KLC2;refGene_tx=XM_003993660.5;refGene_gn=KLC2;refGene_tx=XM_003993661.5
D1 111307089 111307173
refGene_gn=KLC2;refGene_tx=XM_003993660.5;refGene_gn=KLC2;refGene_tx=XM_003993661.5
D1 111308904 111308918
refGene_gn=RAB1B;refGene_tx=XM_003993664.5
D1 111310573 111310646
refGene_gn=RAB1B;refGene_tx=XM_003993664.5
D1 111310901 111310997
refGene_gn=RAB1B;refGene_tx=XM_003993664.5
D1 111311109 111311205
refGene_gn=RAB1B;refGene_tx=XM_003993664.5
D1 111314425 111314557
refGene_gn=RAB1B;refGene_tx=XM_003993664.5
D1 111314652 111314847
refGene_gn=RAB1B;refGene_tx=XM_003993664.5
D1 111317090 111317171
refGene_gn=CNIH2;refGene_tx=XM_023240082.1
D1 111320966 111321035
refGene_gn=CNIH2;refGene_tx=XM_023240082.1
D1 111321493 111321541
refGene_gn=CNIH2;refGene_tx=XM_023240082.1
D1 111321799 111321912
refGene_gn=CNIH2;refGene_tx=XM_023240082.1
D1 111322051 111322195
refGene_gn=CNIH2;refGene_tx=XM_023240082.1
D1 111322392 111322420
refGene_gn=CNIH2;refGene_tx=XM_023240082.1
D1 111323490 111323637
refGene_gn=YIF1A;refGene_tx=XM_003993666.3
D1 111323736 111323830
refGene_gn=YIF1A;refGene_tx=XM_003993666.3
D1 111324172 111324328
refGene_gn=YIF1A;refGene_tx=XM_003993666.3
D1 111324503 111324561
refGene_gn=YIF1A;refGene_tx=XM_003993666.3
D1 111326064 111326143
refGene_gn=YIF1A;refGene_tx=XM_003993666.3
D1 111326280 111326385
refGene_gn=YIF1A;refGene_tx=XM_003993666.3
D1 111326549 111326761
refGene_gn=YIF1A;refGene_tx=XM_003993666.3
D1 111327455 111327486
refGene_gn=YIF1A;refGene_tx=XM_003993666.3
D1 111330147 111330222
refGene_gn=TMEM151A;refGene_tx=XM_023240081.1
D1 111332490 111333816
refGene_gn=TMEM151A;refGene_tx=XM_023240081.1
D1 111348210 111350535

```
refGene_gn=CD248;refGene_tx=XM_003993667.4
D1	111360802	111361297	refGene_gn=RIN1;refGene_tx=XM_011287028.3
D1	111362277	111362429	refGene_gn=RIN1;refGene_tx=XM_011287028.3
D1	111362568	111362700	refGene_gn=RIN1;refGene_tx=XM_011287028.3
D1	111363011	111363317	refGene_gn=RIN1;refGene_tx=XM_011287028.3
D1	111363540	111364281	refGene_gn=RIN1;refGene_tx=XM_011287028.3
D1	111364438	111364530	refGene_gn=RIN1;refGene_tx=XM_011287028.3
D1	111364674	111364747	refGene_gn=RIN1;refGene_tx=XM_011287028.3
D1	111364846	111364961	refGene_gn=RIN1;refGene_tx=XM_011287028.3
D1	111365059	111365240	refGene_gn=RIN1;refGene_tx=XM_011287028.3
D1	111365373	111365459	refGene_gn=RIN1;refGene_tx=XM_011287028.3
D1	111367038	111367046	refGene_gn=BRMS1;refGene_tx=XM_006937545.3
D1	111367454	111367494	refGene_gn=BRMS1;refGene_tx=XM_006937545.3
D1	111367950	111368015	refGene_gn=BRMS1;refGene_tx=XM_006937545.3
D1	111368527	111368673	refGene_gn=BRMS1;refGene_tx=XM_019812706.1;refGene_gn=BRMS1;refGene_tx=XM_019812707.1;refGene_gn=BRMS1;refGene_tx=XM_019812708.1
D1	111369185	111369278	refGene_gn=BRMS1;refGene_tx=XM_006937545.3;refGene_gn=BRMS1;refGene_tx=XM_019812706.1;refGene_gn=BRMS1;refGene_tx=XM_019812707.1;refGene_gn=BRMS1;refGene_tx=XM_019812708.1
D1	111369860	111369957	refGene_gn=BRMS1;refGene_tx=XM_006937545.3;refGene_gn=BRMS1;refGene_tx=XM_019812706.1;refGene_gn=BRMS1;refGene_tx=XM_019812707.1;refGene_gn=BRMS1;refGene_tx=XM_019812708.1
D1	111370038	111370118	refGene_gn=BRMS1;refGene_tx=XM_006937545.3;refGene_gn=BRMS1;refGene_tx=XM_019812706.1;refGene_gn=BRMS1;refGene_tx=XM_019812707.1;refGene_gn=BRMS1;refGene_tx=XM_019812708.1
D1	111370283	111370411	refGene_gn=BRMS1;refGene_tx=XM_006937545.3;refGene_gn=BRMS1;refGene_tx=XM_019812706.1;refGene_gn=BRMS1;refGene_tx=XM_019812707.1;refGene_gn=BRMS1;refGene_tx=XM_019812708.1
D1	111370576	111370667	refGene_gn=BRMS1;refGene_tx=XM_006937545.3;refGene_gn=BRMS1;refGene_tx=XM_019812706.1;refGene_gn=BRMS1;refGene_tx=XM_019812707.1;refGene_gn=BRMS1;refGene_tx=XM_019812708.1
```

D1 111370741 111370880
refGene_gn=BRMS1;refGene_tx=XM_006937545.3;refGene_gn=BRMS1;refGene_tx=XM_019812706.1;refGene_gn=BRMS1;refGene_tx=XM_019812707.1;refGene_gn=BRMS1;refGene_tx=XM_019812708.1
D1 111374716 111374908
refGene_gn=B4GAT1;refGene_tx=XM_003993670.4
D1 111375153 111376209
refGene_gn=B4GAT1;refGene_tx=XM_003993670.4
D1 111385305 111385417
refGene_gn=SLC29A2;refGene_tx=XM_011287066.2
D1 111386068 111386268
refGene_gn=SLC29A2;refGene_tx=XM_011287066.2
D1 111387042 111387128
refGene_gn=SLC29A2;refGene_tx=XM_011287066.2
D1 111387242 111387348
refGene_gn=SLC29A2;refGene_tx=XM_011287066.2
D1 111387605 111387742
refGene_gn=SLC29A2;refGene_tx=XM_011287066.2
D1 111388293 111388378
refGene_gn=SLC29A2;refGene_tx=XM_011287066.2
D1 111388588 111388686
refGene_gn=SLC29A2;refGene_tx=XM_011287066.2
D1 111389333 111389468
refGene_gn=SLC29A2;refGene_tx=XM_011287066.2
D1 111389882 111390022
refGene_gn=SLC29A2;refGene_tx=XM_011287066.2
D1 111390244 111390405
refGene_gn=SLC29A2;refGene_tx=XM_011287066.2
D1 111391941 111392023
refGene_gn=SLC29A2;refGene_tx=XM_011287066.2
D1 111392282 111392311
refGene_gn=SLC29A2;refGene_tx=XM_011287066.2
D1 111428884 111429059
refGene_gn=NPAS4;refGene_tx=XM_003993672.5
D1 111429778 111429930
refGene_gn=NPAS4;refGene_tx=XM_003993672.5
D1 111430115 111430218
refGene_gn=NPAS4;refGene_tx=XM_003993672.5
D1 111430337 111430599
refGene_gn=NPAS4;refGene_tx=XM_003993672.5;refGene_gn=NPAS4;refGene_tx=XM_023240083.1
D1 111430780 111430890
refGene_gn=NPAS4;refGene_tx=XM_003993672.5
D1 111431238 111431374
refGene_gn=NPAS4;refGene_tx=XM_003993672.5;refGene_gn=NPAS4;refGene_tx=XM_023240083.1
D1 111431494 111432930
refGene_gn=NPAS4;refGene_tx=XM_003993672.5;refGene_gn=NPAS4;refGene_tx=XM_023240083.1
D1 111433652 111433681

refGene_gn=NPAS4;refGene_tx=XM_003993672.5;refGene_gn=NPAS4;refGene_tx=XM_023240083.1
D1 111443133 111443330
refGene_gn=MRPL11;refGene_tx=XM_003993673.5;refGene_gn=MRPL11;refGene_tx=XM_011287030.3;refGene_gn=MRPL11;refGene_tx=XM_019812711.2
D1 111444478 111444655
refGene_gn=MRPL11;refGene_tx=XM_003993673.5;refGene_gn=MRPL11;refGene_tx=XM_011287030.3;refGene_gn=MRPL11;refGene_tx=XM_019812711.2
D1 111444741 111444835
refGene_gn=MRPL11;refGene_tx=XM_003993673.5;refGene_gn=MRPL11;refGene_tx=XM_011287030.3;refGene_gn=MRPL11;refGene_tx=XM_019812711.2
D1 111445650 111445746
refGene_gn=MRPL11;refGene_tx=XM_003993673.5;refGene_gn=MRPL11;refGene_tx=XM_011287030.3
D1 111446119 111446242
refGene_gn=MRPL11;refGene_tx=XM_003993673.5;refGene_gn=MRPL11;refGene_tx=XM_011287030.3;refGene_gn=MRPL11;refGene_tx=XM_019812711.2
D1 111457163 111457315
refGene_gn=PELI3;refGene_tx=XM_023240084.1;refGene_gn=PELI3;refGene_tx=XM_023240085.1
D1 111457846 111457918
refGene_gn=PELI3;refGene_tx=XM_023240084.1
D1 111460328 111460458
refGene_gn=PELI3;refGene_tx=XM_023240084.1;refGene_gn=PELI3;refGene_tx=XM_023240085.1
D1 111461093 111461195
refGene_gn=PELI3;refGene_tx=XM_023240084.1;refGene_gn=PELI3;refGene_tx=XM_023240085.1
D1 111461999 111462194
refGene_gn=PELI3;refGene_tx=XM_023240084.1;refGene_gn=PELI3;refGene_tx=XM_023240085.1
D1 111462513 111462702
refGene_gn=PELI3;refGene_tx=XM_023240084.1;refGene_gn=PELI3;refGene_tx=XM_023240085.1
D1 111463984 111464554
refGene_gn=PELI3;refGene_tx=XM_023240084.1;refGene_gn=PELI3;refGene_tx=XM_023240085.1
D1 111468450 111468502
refGene_gn=DPP3;refGene_tx=XM_011287031.3;refGene_gn=DPP3;refGene_tx=XM_011287033.3
D1 111469602 111469880
refGene_gn=DPP3;refGene_tx=XM_011287031.3;refGene_gn=DPP3;refGene_tx=XM_011287032.3;refGene_gn=DPP3;refGene_tx=XM_011287033.3;refGene_gn=DPP3;refGene_tx=XM_011287034.2;refGene_gn=DPP3;refGene_tx=XM_011287035.2;refGene_gn=DPP3;refGene_tx=XM_011287036.3
D1 111479031 111479121
refGene_gn=DPP3;refGene_tx=XM_011287031.3;refGene_gn=DPP3;refGene_tx=XM_011287032.3;refGene_gn=DPP3;refGene_tx=XM_011287033.3;refGene_gn=DPP3;refGene_tx=XM_011287034.2;refGene_gn=DPP3;refGene_tx=XM_011287035.2;refGene_gn=DPP3;refGene_tx=XM_011287036.3

D1 111479726 111479864
refGene_gn=DPP3;refGene_tx=XM_011287031.3;refGene_gn=DPP3;refGene_tx=XM_011287032.3;refGene_gn=DPP3;refGene_tx=XM_011287033.3;refGene_gn=DPP3;refGene_tx=XM_011287034.2;refGene_gn=DPP3;refGene_tx=XM_011287035.2;refGene_gn=DPP3;refGene_tx=XM_011287036.3
D1 111480694 111480769
refGene_gn=DPP3;refGene_tx=XM_011287031.3;refGene_gn=DPP3;refGene_tx=XM_011287032.3;refGene_gn=DPP3;refGene_tx=XM_011287033.3;refGene_gn=DPP3;refGene_tx=XM_011287034.2;refGene_gn=DPP3;refGene_tx=XM_011287035.2;refGene_gn=DPP3;refGene_tx=XM_011287036.3
D1 111481396 111481490
refGene_gn=DPP3;refGene_tx=XM_011287031.3;refGene_gn=DPP3;refGene_tx=XM_011287032.3;refGene_gn=DPP3;refGene_tx=XM_011287033.3;refGene_gn=DPP3;refGene_tx=XM_011287034.2;refGene_gn=DPP3;refGene_tx=XM_011287035.2;refGene_gn=DPP3;refGene_tx=XM_011287036.3
D1 111485087 111485218
refGene_gn=DPP3;refGene_tx=XM_011287031.3;refGene_gn=DPP3;refGene_tx=XM_011287032.3;refGene_gn=DPP3;refGene_tx=XM_011287033.3;refGene_gn=DPP3;refGene_tx=XM_011287034.2;refGene_gn=DPP3;refGene_tx=XM_011287035.2;refGene_gn=DPP3;refGene_tx=XM_011287036.3
D1 111485328 111485459
refGene_gn=DPP3;refGene_tx=XM_011287031.3;refGene_gn=DPP3;refGene_tx=XM_011287032.3;refGene_gn=DPP3;refGene_tx=XM_011287033.3;refGene_gn=DPP3;refGene_tx=XM_011287034.2;refGene_gn=DPP3;refGene_tx=XM_011287035.2;refGene_gn=DPP3;refGene_tx=XM_011287036.3
D1 111485533 111485592
refGene_gn=DPP3;refGene_tx=XM_011287031.3;refGene_gn=DPP3;refGene_tx=XM_011287032.3;refGene_gn=DPP3;refGene_tx=XM_011287033.3;refGene_gn=DPP3;refGene_tx=XM_011287034.2;refGene_gn=DPP3;refGene_tx=XM_011287035.2;refGene_gn=DPP3;refGene_tx=XM_011287036.3
D1 111488560 111488755
refGene_gn=DPP3;refGene_tx=XM_011287031.3;refGene_gn=DPP3;refGene_tx=XM_011287032.3;refGene_gn=DPP3;refGene_tx=XM_011287033.3;refGene_gn=DPP3;refGene_tx=XM_011287034.2;refGene_gn=DPP3;refGene_tx=XM_011287035.2;refGene_gn=DPP3;refGene_tx=XM_011287036.3
D1 111488956 111489069
refGene_gn=DPP3;refGene_tx=XM_011287031.3;refGene_gn=DPP3;refGene_tx=XM_011287032.3;refGene_gn=DPP3;refGene_tx=XM_011287033.3;refGene_gn=DPP3;refGene_tx=XM_011287034.2;refGene_gn=DPP3;refGene_tx=XM_011287035.2;refGene_gn=DPP3;refGene_tx=XM_011287036.3
D1 111489402 111489495
refGene_gn=DPP3;refGene_tx=XM_011287031.3;refGene_gn=DPP3;refGene_tx=XM_011287032.3;refGene_gn=DPP3;refGene_tx=XM_011287033.3;refGene_gn=DPP3;refGene_tx=XM_011287034.2;refGene_gn=DPP3;refGene_tx=XM_011287035.2;refGene_gn=DPP3;refGene_tx=XM_011287036.3
D1 111490864 111490927
refGene_gn=DPP3;refGene_tx=XM_011287031.3;refGene_gn=DPP3;refGene_tx=XM_011287032.3;refGene_gn=DPP3;refGene_tx=XM_011287033.3;refGene_gn=DPP3;refGene_tx=XM_011287034.2;refGene_gn=DPP3;refGene_tx=XM_011287035.2;refGene_gn=DPP3;refGene_tx=XM_011287036.3

D1 111491022 111491147
refGene_gn=DPP3;refGene_tx=XM_011287031.3;refGene_gn=DPP3;refGene_tx=XM_011287032.3;refGene_gn=DPP3;refGene_tx=XM_011287033.3;refGene_gn=DPP3;refGene_tx=XM_011287034.2;refGene_gn=DPP3;refGene_tx=XM_011287035.2;refGene_gn=DPP3;refGene_tx=XM_011287036.3
D1 111491268 111491389
refGene_gn=DPP3;refGene_tx=XM_011287031.3;refGene_gn=DPP3;refGene_tx=XM_011287032.3;refGene_gn=DPP3;refGene_tx=XM_011287033.3;refGene_gn=DPP3;refGene_tx=XM_011287034.2;refGene_gn=DPP3;refGene_tx=XM_011287035.2;refGene_gn=DPP3;refGene_tx=XM_011287036.3
D1 111492464 111492644
refGene_gn=DPP3;refGene_tx=XM_011287031.3;refGene_gn=DPP3;refGene_tx=XM_011287032.3;refGene_gn=DPP3;refGene_tx=XM_011287033.3;refGene_gn=DPP3;refGene_tx=XM_011287034.2;refGene_gn=DPP3;refGene_tx=XM_011287035.2;refGene_gn=DPP3;refGene_tx=XM_011287036.3
D1 111500074 111500237
refGene_gn=DPP3;refGene_tx=XM_011287031.3;refGene_gn=DPP3;refGene_tx=XM_011287032.3;refGene_gn=DPP3;refGene_tx=XM_011287033.3;refGene_gn=DPP3;refGene_tx=XM_011287034.2;refGene_gn=DPP3;refGene_tx=XM_011287035.2;refGene_gn=DPP3;refGene_tx=XM_011287036.3
D1 111503216 111503389
refGene_gn=DPP3;refGene_tx=XM_011287031.3;refGene_gn=DPP3;refGene_tx=XM_011287032.3;refGene_gn=DPP3;refGene_tx=XM_011287033.3;refGene_gn=DPP3;refGene_tx=XM_011287034.2;refGene_gn=DPP3;refGene_tx=XM_011287035.2;refGene_gn=DPP3;refGene_tx=XM_011287036.3
D1 111504681 111504728
refGene_gn=BBS1;refGene_tx=XM_003993678.4;refGene_gn=BBS1;refGene_tx=XM_019812712.2
D1 111505034 111505138
refGene_gn=BBS1;refGene_tx=XM_003993678.4;refGene_gn=BBS1;refGene_tx=XM_019812712.2
D1 111505272 111505307
refGene_gn=BBS1;refGene_tx=XM_003993678.4;refGene_gn=BBS1;refGene_tx=XM_019812712.2
D1 111508648 111508921
refGene_gn=BBS1;refGene_tx=XM_003993678.4;refGene_gn=BBS1;refGene_tx=XM_019812712.2
D1 111509036 111509083
refGene_gn=BBS1;refGene_tx=XM_003993678.4;refGene_gn=BBS1;refGene_tx=XM_019812712.2
D1 111509225 111509264
refGene_gn=BBS1;refGene_tx=XM_003993678.4;refGene_gn=BBS1;refGene_tx=XM_019812712.2
D1 111509383 111509456
refGene_gn=BBS1;refGene_tx=XM_003993678.4;refGene_gn=BBS1;refGene_tx=XM_019812712.2
D1 111512088 111512220
refGene_gn=BBS1;refGene_tx=XM_003993678.4;refGene_gn=BBS1;refGene_tx=XM_019812712.2
D1 111512956 111513063

refGene_gn=BBS1;refGene_tx=XM_003993678.4;refGene_gn=BBS1;refGene_tx=XM_019812712.2
D1 111514416 111514537
refGene_gn=BBS1;refGene_tx=XM_003993678.4;refGene_gn=BBS1;refGene_tx=XM_019812712.2
D1 111514703 111514862
refGene_gn=BBS1;refGene_tx=XM_003993678.4;refGene_gn=BBS1;refGene_tx=XM_019812712.2
D1 111516270 111516340
refGene_gn=BBS1;refGene_tx=XM_003993678.4;refGene_gn=BBS1;refGene_tx=XM_019812712.2
D1 111516825 111516984
refGene_gn=BBS1;refGene_tx=XM_003993678.4;refGene_gn=BBS1;refGene_tx=XM_019812712.2
D1 111519607 111519741
refGene_gn=BBS1;refGene_tx=XM_003993678.4;refGene_gn=BBS1;refGene_tx=XM_019812712.2
D1 111520744 111520879
refGene_gn=BBS1;refGene_tx=XM_003993678.4;refGene_gn=BBS1;refGene_tx=XM_019812712.2
D1 111521496 111521583
refGene_gn=BBS1;refGene_tx=XM_003993678.4;refGene_gn=BBS1;refGene_tx=XM_019812712.2
D1 111521810 111521897
refGene_gn=BBS1;refGene_tx=XM_003993678.4;refGene_gn=BBS1;refGene_tx=XM_019812712.2
D1 111530251 111530547
refGene_gn=ZDHHC24;refGene_tx=XM_003993680.4;refGene_gn=ZDHHC24;refGene_tx=XM_019812713.2
D1 111533896 111534174
refGene_gn=ZDHHC24;refGene_tx=XM_003993680.4;refGene_gn=ZDHHC24;refGene_tx=XM_019812713.2
D1 111535057 111535188
refGene_gn=ZDHHC24;refGene_tx=XM_019812713.2
D1 111535266 111535547
refGene_gn=ZDHHC24;refGene_tx=XM_003993680.4
D1 111536464 111536611
refGene_gn=ACTN3;refGene_tx=XM_003993681.4
D1 111540259 111540374
refGene_gn=ACTN3;refGene_tx=XM_003993681.4
D1 111540549 111540669
refGene_gn=ACTN3;refGene_tx=XM_003993681.4
D1 111540984 111541071
refGene_gn=ACTN3;refGene_tx=XM_003993681.4
D1 111541153 111541241
refGene_gn=ACTN3;refGene_tx=XM_003993681.4
D1 111541697 111541776
refGene_gn=ACTN3;refGene_tx=XM_003993681.4
D1 111541859 111541941
refGene_gn=ACTN3;refGene_tx=XM_003993681.4

D1 111542707 111542793
refGene_gn=ACTN3;refGene_tx=XM_003993681.4
D1 111543083 111543176
refGene_gn=ACTN3;refGene_tx=XM_003993681.4
D1 111543614 111543845
refGene_gn=ACTN3;refGene_tx=XM_003993681.4
D1 111543969 111544117
refGene_gn=ACTN3;refGene_tx=XM_003993681.4
D1 111545268 111545419
refGene_gn=ACTN3;refGene_tx=XM_003993681.4
D1 111546019 111546128
refGene_gn=ACTN3;refGene_tx=XM_003993681.4
D1 111546223 111546364
refGene_gn=ACTN3;refGene_tx=XM_003993681.4
D1 111546627 111546810
refGene_gn=ACTN3;refGene_tx=XM_003993681.4
D1 111547248 111547383
refGene_gn=ACTN3;refGene_tx=XM_003993681.4
D1 111547471 111547651
refGene_gn=ACTN3;refGene_tx=XM_003993681.4
D1 111548075 111548222
refGene_gn=ACTN3;refGene_tx=XM_003993681.4
D1 111548314 111548380
refGene_gn=ACTN3;refGene_tx=XM_003993681.4
D1 111549082 111549241
refGene_gn=ACTN3;refGene_tx=XM_003993681.4
D1 111549324 111549483
refGene_gn=ACTN3;refGene_tx=XM_003993681.4
D1 111550256 111550331
refGene_gn=CTSF;refGene_tx=XM_023240086.1
D1 111550413 111550472
refGene_gn=CTSF;refGene_tx=XM_023240086.1
D1 111550858 111550949
refGene_gn=CTSF;refGene_tx=XM_023240086.1
D1 111551053 111551118
refGene_gn=CTSF;refGene_tx=XM_023240086.1
D1 111551197 111551317
refGene_gn=CTSF;refGene_tx=XM_023240086.1
D1 111552110 111552191
refGene_gn=CTSF;refGene_tx=XM_023240086.1
D1 111552270 111552367
refGene_gn=CTSF;refGene_tx=XM_023240086.1
D1 111552455 111552601
refGene_gn=CTSF;refGene_tx=XM_023240086.1
D1 111552720 111552834
refGene_gn=CTSF;refGene_tx=XM_023240086.1
D1 111553434 111553510
refGene_gn=CTSF;refGene_tx=XM_023240086.1
D1 111553643 111553705
refGene_gn=CTSF;refGene_tx=XM_023240086.1

D1	111553797	111553882
refGene_gn=CTSF;refGene_tx=XM_023240086.1
D1	111554176	111554275
refGene_gn=CTSF;refGene_tx=XM_023240086.1
D1	111554492	111554702
refGene_gn=CTSF;refGene_tx=XM_023240086.1
D1	111578756	111578780
refGene_gn=CCDC87;refGene_tx=XM_004001447.4
D1	111578781	111581310
refGene_gn=CCDC87;refGene_tx=XM_004001447.4
D1	111581587	111581626
refGene_gn=CCS;refGene_tx=XM_003993684.3
D1	111581958	111582031
refGene_gn=CCS;refGene_tx=XM_003993684.3
D1	111587765	111587903
refGene_gn=CCS;refGene_tx=XM_003993684.3
D1	111588110	111588288
refGene_gn=CCS;refGene_tx=XM_003993684.3
D1	111589053	111589114
refGene_gn=CCS;refGene_tx=XM_003993684.3
D1	111593163	111593241
refGene_gn=CCS;refGene_tx=XM_003993684.3
D1	111593311	111593415
refGene_gn=CCS;refGene_tx=XM_003993684.3
D1	111593529	111593683
refGene_gn=CCS;refGene_tx=XM_003993684.3
D1	111611644	111611981
refGene_gn=RBM14;refGene_tx=XM_011287038.3;refGene_gn=RBM14;refGene_tx
=XM_011287039.3;refGene_gn=RBM14;refGene_tx=XM_011287041.3
D1	111617517	111618982
refGene_gn=RBM14;refGene_tx=XM_011287038.3;refGene_gn=RBM14;refGene_tx
=XM_011287039.3;refGene_gn=RBM14;refGene_tx=XM_019812715.2
D1	111619758	111619966
refGene_gn=RBM14;refGene_tx=XM_011287038.3;refGene_gn=RBM14;refGene_tx
=XM_011287039.3;refGene_gn=RBM14;refGene_tx=XM_019812715.2
D1	111627557	111627577
refGene_gn=RBM14;refGene_tx=XM_011287041.3
D1	111628732	111629144
refGene_gn=LOC101088520;refGene_tx=XM_003993687.4;refGene_gn=LOC101088
520;refGene_tx=XM_006937552.3
D1	111632344	111633030
refGene_gn=LOC101088520;refGene_tx=XM_003993687.4
D1	111634894	111634914
refGene_gn=LOC101088520;refGene_tx=XM_006937552.3
D1	111657250	111657918
refGene_gn=RBM4B;refGene_tx=XM_003993688.5;refGene_gn=RBM4B;refGene_tx
=XM_006937553.4;refGene_gn=RBM4B;refGene_tx=XM_019812736.2;refGene_gn=
RBM4B;refGene_tx=XM_019812737.2
D1	111662997	111663409
refGene_gn=RBM4B;refGene_tx=XM_003993688.5;refGene_gn=RBM4B;refGene_tx

=XM_006937553.4;refGene_gn=RBM4B;refGene_tx=XM_019812736.2;refGene_gn=RBM4B;refGene_tx=XM_019812737.2
D1 111670431 111670665
refGene_gn=SPTBN2;refGene_tx=XM_011287043.3;refGene_gn=SPTBN2;refGene_tx=XM_011287044.3;refGene_gn=SPTBN2;refGene_tx=XM_019812716.2;refGene_gn=SPTBN2;refGene_tx=XM_023240087.1
D1 111670992 111671035
refGene_gn=SPTBN2;refGene_tx=XM_011287043.3;refGene_gn=SPTBN2;refGene_tx=XM_011287044.3;refGene_gn=SPTBN2;refGene_tx=XM_019812716.2;refGene_gn=SPTBN2;refGene_tx=XM_023240087.1
D1 111671535 111671709
refGene_gn=SPTBN2;refGene_tx=XM_011287043.3;refGene_gn=SPTBN2;refGene_tx=XM_011287044.3;refGene_gn=SPTBN2;refGene_tx=XM_019812716.2;refGene_gn=SPTBN2;refGene_tx=XM_023240087.1
D1 111672211 111672432
refGene_gn=SPTBN2;refGene_tx=XM_011287043.3;refGene_gn=SPTBN2;refGene_tx=XM_011287044.3;refGene_gn=SPTBN2;refGene_tx=XM_019812716.2;refGene_gn=SPTBN2;refGene_tx=XM_023240087.1
D1 111672653 111672704
refGene_gn=SPTBN2;refGene_tx=XM_011287043.3;refGene_gn=SPTBN2;refGene_tx=XM_011287044.3;refGene_gn=SPTBN2;refGene_tx=XM_019812716.2;refGene_gn=SPTBN2;refGene_tx=XM_023240087.1
D1 111672787 111672863
refGene_gn=SPTBN2;refGene_tx=XM_011287043.3;refGene_gn=SPTBN2;refGene_tx=XM_011287044.3;refGene_gn=SPTBN2;refGene_tx=XM_019812716.2;refGene_gn=SPTBN2;refGene_tx=XM_023240087.1
D1 111672953 111673096
refGene_gn=SPTBN2;refGene_tx=XM_011287043.3;refGene_gn=SPTBN2;refGene_tx=XM_011287044.3;refGene_gn=SPTBN2;refGene_tx=XM_019812716.2;refGene_gn=SPTBN2;refGene_tx=XM_023240087.1
D1 111673396 111673593
refGene_gn=SPTBN2;refGene_tx=XM_011287043.3;refGene_gn=SPTBN2;refGene_tx=XM_011287044.3;refGene_gn=SPTBN2;refGene_tx=XM_019812716.2;refGene_gn=SPTBN2;refGene_tx=XM_023240087.1
D1 111673867 111673952
refGene_gn=SPTBN2;refGene_tx=XM_011287043.3;refGene_gn=SPTBN2;refGene_tx=XM_011287044.3;refGene_gn=SPTBN2;refGene_tx=XM_019812716.2;refGene_gn=SPTBN2;refGene_tx=XM_023240087.1
D1 111674215 111674354
refGene_gn=SPTBN2;refGene_tx=XM_011287043.3;refGene_gn=SPTBN2;refGene_tx=XM_011287044.3;refGene_gn=SPTBN2;refGene_tx=XM_019812716.2;refGene_gn=SPTBN2;refGene_tx=XM_023240087.1
D1 111674448 111674693
refGene_gn=SPTBN2;refGene_tx=XM_011287043.3;refGene_gn=SPTBN2;refGene_tx=XM_011287044.3;refGene_gn=SPTBN2;refGene_tx=XM_019812716.2;refGene_gn=SPTBN2;refGene_tx=XM_023240087.1
D1 111675388 111675763
refGene_gn=SPTBN2;refGene_tx=XM_011287043.3;refGene_gn=SPTBN2;refGene_tx=XM_011287044.3;refGene_gn=SPTBN2;refGene_tx=XM_019812716.2;refGene_gn=SPTBN2;refGene_tx=XM_023240087.1

D1 111676320 111676525
refGene_gn=SPTBN2;refGene_tx=XM_011287043.3;refGene_gn=SPTBN2;refGene_tx=XM_011287044.3;refGene_gn=SPTBN2;refGene_tx=XM_019812716.2;refGene_gn=SPTBN2;refGene_tx=XM_023240087.1
D1 111676767 111676898
refGene_gn=SPTBN2;refGene_tx=XM_011287043.3;refGene_gn=SPTBN2;refGene_tx=XM_011287044.3;refGene_gn=SPTBN2;refGene_tx=XM_019812716.2;refGene_gn=SPTBN2;refGene_tx=XM_023240087.1
D1 111676983 111677244
refGene_gn=SPTBN2;refGene_tx=XM_011287043.3;refGene_gn=SPTBN2;refGene_tx=XM_011287044.3;refGene_gn=SPTBN2;refGene_tx=XM_019812716.2;refGene_gn=SPTBN2;refGene_tx=XM_023240087.1
D1 111677596 111677686
refGene_gn=SPTBN2;refGene_tx=XM_011287043.3;refGene_gn=SPTBN2;refGene_tx=XM_011287044.3;refGene_gn=SPTBN2;refGene_tx=XM_019812716.2;refGene_gn=SPTBN2;refGene_tx=XM_023240087.1
D1 111677999 111678224
refGene_gn=SPTBN2;refGene_tx=XM_011287043.3;refGene_gn=SPTBN2;refGene_tx=XM_011287044.3;refGene_gn=SPTBN2;refGene_tx=XM_019812716.2;refGene_gn=SPTBN2;refGene_tx=XM_023240087.1
D1 111679683 111679947
refGene_gn=SPTBN2;refGene_tx=XM_011287043.3;refGene_gn=SPTBN2;refGene_tx=XM_011287044.3;refGene_gn=SPTBN2;refGene_tx=XM_019812716.2;refGene_gn=SPTBN2;refGene_tx=XM_023240087.1
D1 111682371 111682518
refGene_gn=SPTBN2;refGene_tx=XM_011287043.3;refGene_gn=SPTBN2;refGene_tx=XM_011287044.3;refGene_gn=SPTBN2;refGene_tx=XM_019812716.2;refGene_gn=SPTBN2;refGene_tx=XM_023240087.1
D1 111682750 111682841
refGene_gn=SPTBN2;refGene_tx=XM_011287043.3;refGene_gn=SPTBN2;refGene_tx=XM_011287044.3;refGene_gn=SPTBN2;refGene_tx=XM_019812716.2;refGene_gn=SPTBN2;refGene_tx=XM_023240087.1
D1 111683224 111683427
refGene_gn=SPTBN2;refGene_tx=XM_011287043.3;refGene_gn=SPTBN2;refGene_tx=XM_011287044.3;refGene_gn=SPTBN2;refGene_tx=XM_019812716.2;refGene_gn=SPTBN2;refGene_tx=XM_023240087.1
D1 111684300 111685057
refGene_gn=SPTBN2;refGene_tx=XM_011287043.3;refGene_gn=SPTBN2;refGene_tx=XM_011287044.3;refGene_gn=SPTBN2;refGene_tx=XM_019812716.2;refGene_gn=SPTBN2;refGene_tx=XM_023240087.1
D1 111685422 111685560
refGene_gn=SPTBN2;refGene_tx=XM_011287043.3;refGene_gn=SPTBN2;refGene_tx=XM_011287044.3;refGene_gn=SPTBN2;refGene_tx=XM_019812716.2;refGene_gn=SPTBN2;refGene_tx=XM_023240087.1
D1 111687552 111688423
refGene_gn=SPTBN2;refGene_tx=XM_011287043.3;refGene_gn=SPTBN2;refGene_tx=XM_011287044.3;refGene_gn=SPTBN2;refGene_tx=XM_019812716.2;refGene_gn=SPTBN2;refGene_tx=XM_023240087.1
D1 111688668 111688822
refGene_gn=SPTBN2;refGene_tx=XM_011287043.3;refGene_gn=SPTBN2;refGene_

tx=XM_011287044.3;refGene_gn=SPTBN2;refGene_tx=XM_019812716.2;refGene_gn=SPTBN2;refGene_tx=XM_023240087.1
D1 111690111 111690414
refGene_gn=SPTBN2;refGene_tx=XM_011287043.3;refGene_gn=SPTBN2;refGene_tx=XM_011287044.3;refGene_gn=SPTBN2;refGene_tx=XM_019812716.2;refGene_gn=SPTBN2;refGene_tx=XM_023240087.1
D1 111690743 111690902
refGene_gn=SPTBN2;refGene_tx=XM_011287043.3;refGene_gn=SPTBN2;refGene_tx=XM_011287044.3;refGene_gn=SPTBN2;refGene_tx=XM_019812716.2;refGene_gn=SPTBN2;refGene_tx=XM_023240087.1
D1 111691478 111691596
refGene_gn=SPTBN2;refGene_tx=XM_011287043.3;refGene_gn=SPTBN2;refGene_tx=XM_011287044.3;refGene_gn=SPTBN2;refGene_tx=XM_019812716.2;refGene_gn=SPTBN2;refGene_tx=XM_023240087.1
D1 111692849 111693037
refGene_gn=SPTBN2;refGene_tx=XM_011287043.3;refGene_gn=SPTBN2;refGene_tx=XM_011287044.3;refGene_gn=SPTBN2;refGene_tx=XM_019812716.2;refGene_gn=SPTBN2;refGene_tx=XM_023240087.1
D1 111693171 111693284
refGene_gn=SPTBN2;refGene_tx=XM_011287043.3;refGene_gn=SPTBN2;refGene_tx=XM_011287044.3;refGene_gn=SPTBN2;refGene_tx=XM_019812716.2;refGene_gn=SPTBN2;refGene_tx=XM_023240087.1
D1 111695827 111695943
refGene_gn=SPTBN2;refGene_tx=XM_011287043.3;refGene_gn=SPTBN2;refGene_tx=XM_011287044.3;refGene_gn=SPTBN2;refGene_tx=XM_019812716.2;refGene_gn=SPTBN2;refGene_tx=XM_023240087.1
D1 111696268 111696349
refGene_gn=SPTBN2;refGene_tx=XM_011287043.3;refGene_gn=SPTBN2;refGene_tx=XM_011287044.3;refGene_gn=SPTBN2;refGene_tx=XM_019812716.2;refGene_gn=SPTBN2;refGene_tx=XM_023240087.1
D1 111696507 111696599
refGene_gn=SPTBN2;refGene_tx=XM_011287043.3;refGene_gn=SPTBN2;refGene_tx=XM_011287044.3;refGene_gn=SPTBN2;refGene_tx=XM_019812716.2;refGene_gn=SPTBN2;refGene_tx=XM_023240087.1
D1 111697421 111697595
refGene_gn=SPTBN2;refGene_tx=XM_011287043.3;refGene_gn=SPTBN2;refGene_tx=XM_011287044.3;refGene_gn=SPTBN2;refGene_tx=XM_019812716.2;refGene_gn=SPTBN2;refGene_tx=XM_023240087.1
D1 111698043 111698195
refGene_gn=SPTBN2;refGene_tx=XM_011287043.3;refGene_gn=SPTBN2;refGene_tx=XM_011287044.3;refGene_gn=SPTBN2;refGene_tx=XM_019812716.2;refGene_gn=SPTBN2;refGene_tx=XM_023240087.1
D1 111700176 111700354
refGene_gn=SPTBN2;refGene_tx=XM_011287043.3
D1 111702661 111702840
refGene_gn=SPTBN2;refGene_tx=XM_011287044.3;refGene_gn=SPTBN2;refGene_tx=XM_019812716.2;refGene_gn=SPTBN2;refGene_tx=XM_023240087.1
D1 111708493 111708510
refGene_gn=SPTBN2;refGene_tx=XM_019812716.2
D1 111723559 111723589

refGene_gn=CD1H11orf80;refGene_tx=XM_011287045.3;refGene_gn=CD1H11orf8
0;refGene_tx=XM_019812721.2;refGene_gn=CD1H11orf80;refGene_tx=XM_01981
2725.2
D1 111724542 111724545
refGene_gn=CD1H11orf80;refGene_tx=XM_011287046.3
D1 111725800 111725939
refGene_gn=CD1H11orf80;refGene_tx=XM_011287045.3;refGene_gn=CD1H11orf8
0;refGene_tx=XM_011287046.3;refGene_gn=CD1H11orf80;refGene_tx=XM_01981
2721.2;refGene_gn=CD1H11orf80;refGene_tx=XM_019812725.2;refGene_gn=CD1
H11orf80;refGene_tx=XM_019812727.2;refGene_gn=CD1H11orf80;refGene_tx=X
M_019812728.2;refGene_gn=CD1H11orf80;refGene_tx=XM_023240089.1;refGene
_gn=CD1H11orf80;refGene_tx=XM_023240090.1;refGene_gn=CD1H11orf80;refGe
ne_tx=XM_023240091.1;refGene_gn=CD1H11orf80;refGene_tx=XM_023240092.1;
refGene_gn=CD1H11orf80;refGene_tx=XM_023240093.1
D1 111732700 111732754
refGene_gn=CD1H11orf80;refGene_tx=XM_011287045.3;refGene_gn=CD1H11orf8
0;refGene_tx=XM_011287046.3;refGene_gn=CD1H11orf80;refGene_tx=XM_01981
2721.2;refGene_gn=CD1H11orf80;refGene_tx=XM_019812727.2;refGene_gn=CD1
H11orf80;refGene_tx=XM_019812728.2;refGene_gn=CD1H11orf80;refGene_tx=X
M_023240089.1;refGene_gn=CD1H11orf80;refGene_tx=XM_023240090.1;refGene
_gn=CD1H11orf80;refGene_tx=XM_023240091.1;refGene_gn=CD1H11orf80;refGe
ne_tx=XM_023240092.1;refGene_gn=CD1H11orf80;refGene_tx=XM_023240093.1
D1 111760310 111760332
refGene_gn=CD1H11orf80;refGene_tx=XM_019812731.2
D1 111777575 111777685
refGene_gn=CD1H11orf80;refGene_tx=XM_011287045.3;refGene_gn=CD1H11orf8
0;refGene_tx=XM_011287046.3;refGene_gn=CD1H11orf80;refGene_tx=XM_01981
2721.2;refGene_gn=CD1H11orf80;refGene_tx=XM_019812725.2;refGene_gn=CD1
H11orf80;refGene_tx=XM_019812727.2;refGene_gn=CD1H11orf80;refGene_tx=X
M_019812728.2;refGene_gn=CD1H11orf80;refGene_tx=XM_019812731.2;refGene
_gn=CD1H11orf80;refGene_tx=XM_019812732.2;refGene_gn=CD1H11orf80;refGe
ne_tx=XM_019812734.2;refGene_gn=CD1H11orf80;refGene_tx=XM_019812735.2;
refGene_gn=CD1H11orf80;refGene_tx=XM_023240089.1;refGene_gn=CD1H11orf8
0;refGene_tx=XM_023240090.1;refGene_gn=CD1H11orf80;refGene_tx=XM_02324
0091.1;refGene_gn=CD1H11orf80;refGene_tx=XM_023240092.1;refGene_gn=CD1
H11orf80;refGene_tx=XM_023240093.1;refGene_gn=CD1H11orf80;refGene_tx=X
M_023240094.1
D1 111783153 111783237
refGene_gn=CD1H11orf80;refGene_tx=XM_011287045.3;refGene_gn=CD1H11orf8
0;refGene_tx=XM_011287046.3;refGene_gn=CD1H11orf80;refGene_tx=XM_01981
2721.2;refGene_gn=CD1H11orf80;refGene_tx=XM_019812725.2;refGene_gn=CD1
H11orf80;refGene_tx=XM_019812727.2;refGene_gn=CD1H11orf80;refGene_tx=X
M_019812728.2;refGene_gn=CD1H11orf80;refGene_tx=XM_019812731.2;refGene
_gn=CD1H11orf80;refGene_tx=XM_019812732.2;refGene_gn=CD1H11orf80;refGe
ne_tx=XM_019812734.2;refGene_gn=CD1H11orf80;refGene_tx=XM_019812735.2;
refGene_gn=CD1H11orf80;refGene_tx=XM_023240089.1;refGene_gn=CD1H11orf8
0;refGene_tx=XM_023240090.1;refGene_gn=CD1H11orf80;refGene_tx=XM_02324
0091.1;refGene_gn=CD1H11orf80;refGene_tx=XM_023240092.1;refGene_gn=CD1
H11orf80;refGene_tx=XM_023240093.1;refGene_gn=CD1H11orf80;refGene_tx=X
M_023240094.1

D1 111784392 111784448
refGene_gn=CD1H11orf80;refGene_tx=XM_011287045.3;refGene_gn=CD1H11orf80;refGene_tx=XM_011287046.3;refGene_gn=CD1H11orf80;refGene_tx=XM_019812721.2;refGene_gn=CD1H11orf80;refGene_tx=XM_019812725.2;refGene_gn=CD1H11orf80;refGene_tx=XM_019812727.2;refGene_gn=CD1H11orf80;refGene_tx=XM_019812728.2;refGene_gn=CD1H11orf80;refGene_tx=XM_019812731.2;refGene_gn=CD1H11orf80;refGene_tx=XM_019812732.2;refGene_gn=CD1H11orf80;refGene_tx=XM_019812734.2;refGene_gn=CD1H11orf80;refGene_tx=XM_019812735.2;refGene_gn=CD1H11orf80;refGene_tx=XM_023240089.1;refGene_gn=CD1H11orf80;refGene_tx=XM_023240090.1;refGene_gn=CD1H11orf80;refGene_tx=XM_023240091.1;refGene_gn=CD1H11orf80;refGene_tx=XM_023240092.1;refGene_gn=CD1H11orf80;refGene_tx=XM_023240093.1;refGene_gn=CD1H11orf80;refGene_tx=XM_023240094.1
D1 111784822 111784894
refGene_gn=CD1H11orf80;refGene_tx=XM_011287045.3;refGene_gn=CD1H11orf80;refGene_tx=XM_011287046.3;refGene_gn=CD1H11orf80;refGene_tx=XM_019812721.2;refGene_gn=CD1H11orf80;refGene_tx=XM_019812725.2;refGene_gn=CD1H11orf80;refGene_tx=XM_019812727.2;refGene_gn=CD1H11orf80;refGene_tx=XM_019812728.2;refGene_gn=CD1H11orf80;refGene_tx=XM_019812731.2;refGene_gn=CD1H11orf80;refGene_tx=XM_019812732.2;refGene_gn=CD1H11orf80;refGene_tx=XM_019812734.2;refGene_gn=CD1H11orf80;refGene_tx=XM_019812735.2;refGene_gn=CD1H11orf80;refGene_tx=XM_023240089.1;refGene_gn=CD1H11orf80;refGene_tx=XM_023240090.1;refGene_gn=CD1H11orf80;refGene_tx=XM_023240091.1;refGene_gn=CD1H11orf80;refGene_tx=XM_023240092.1;refGene_gn=CD1H11orf80;refGene_tx=XM_023240093.1;refGene_gn=CD1H11orf80;refGene_tx=XM_023240094.1
D1 111786837 111787033
refGene_gn=CD1H11orf80;refGene_tx=XM_011287045.3;refGene_gn=CD1H11orf80;refGene_tx=XM_011287046.3;refGene_gn=CD1H11orf80;refGene_tx=XM_019812721.2;refGene_gn=CD1H11orf80;refGene_tx=XM_019812725.2;refGene_gn=CD1H11orf80;refGene_tx=XM_019812727.2;refGene_gn=CD1H11orf80;refGene_tx=XM_019812728.2;refGene_gn=CD1H11orf80;refGene_tx=XM_019812731.2;refGene_gn=CD1H11orf80;refGene_tx=XM_019812732.2;refGene_gn=CD1H11orf80;refGene_tx=XM_019812734.2;refGene_gn=CD1H11orf80;refGene_tx=XM_019812735.2;refGene_gn=CD1H11orf80;refGene_tx=XM_023240089.1;refGene_gn=CD1H11orf80;refGene_tx=XM_023240090.1;refGene_gn=CD1H11orf80;refGene_tx=XM_023240091.1;refGene_gn=CD1H11orf80;refGene_tx=XM_023240092.1;refGene_gn=CD1H11orf80;refGene_tx=XM_023240093.1;refGene_gn=CD1H11orf80;refGene_tx=XM_023240094.1
D1 111796574 111796725
refGene_gn=CD1H11orf80;refGene_tx=XM_011287045.3;refGene_gn=CD1H11orf80;refGene_tx=XM_011287046.3;refGene_gn=CD1H11orf80;refGene_tx=XM_019812721.2;refGene_gn=CD1H11orf80;refGene_tx=XM_019812725.2;refGene_gn=CD1H11orf80;refGene_tx=XM_019812727.2;refGene_gn=CD1H11orf80;refGene_tx=XM_019812728.2;refGene_gn=CD1H11orf80;refGene_tx=XM_019812731.2;refGene_gn=CD1H11orf80;refGene_tx=XM_019812732.2;refGene_gn=CD1H11orf80;refGene_tx=XM_019812734.2;refGene_gn=CD1H11orf80;refGene_tx=XM_019812735.2;refGene_gn=CD1H11orf80;refGene_tx=XM_023240089.1;refGene_gn=CD1H11orf80;refGene_tx=XM_023240090.1;refGene_gn=CD1H11orf80;refGene_tx=XM_023240091.1;refGene_gn=CD1H11orf80;refGene_tx=XM_023240092.1;refGene_gn=CD1

H11orf80;refGene_tx=XM_023240093.1;refGene_gn=CD1H11orf80;refGene_tx=XM_023240094.1
D1 111798606 111798753
refGene_gn=CD1H11orf80;refGene_tx=XM_011287045.3;refGene_gn=CD1H11orf80;refGene_tx=XM_011287046.3;refGene_gn=CD1H11orf80;refGene_tx=XM_019812721.2;refGene_gn=CD1H11orf80;refGene_tx=XM_019812725.2;refGene_gn=CD1H11orf80;refGene_tx=XM_019812727.2;refGene_gn=CD1H11orf80;refGene_tx=XM_019812728.2;refGene_gn=CD1H11orf80;refGene_tx=XM_019812731.2;refGene_gn=CD1H11orf80;refGene_tx=XM_019812732.2;refGene_gn=CD1H11orf80;refGene_tx=XM_019812734.2;refGene_gn=CD1H11orf80;refGene_tx=XM_019812735.2;refGene_gn=CD1H11orf80;refGene_tx=XM_023240089.1;refGene_gn=CD1H11orf80;refGene_tx=XM_023240090.1;refGene_gn=CD1H11orf80;refGene_tx=XM_023240091.1;refGene_gn=CD1H11orf80;refGene_tx=XM_023240092.1;refGene_gn=CD1H11orf80;refGene_tx=XM_023240093.1;refGene_gn=CD1H11orf80;refGene_tx=XM_023240094.1
D1 111802612 111802734
refGene_gn=CD1H11orf80;refGene_tx=XM_011287045.3;refGene_gn=CD1H11orf80;refGene_tx=XM_011287046.3;refGene_gn=CD1H11orf80;refGene_tx=XM_019812721.2;refGene_gn=CD1H11orf80;refGene_tx=XM_019812725.2;refGene_gn=CD1H11orf80;refGene_tx=XM_019812727.2;refGene_gn=CD1H11orf80;refGene_tx=XM_019812728.2;refGene_gn=CD1H11orf80;refGene_tx=XM_019812731.2;refGene_gn=CD1H11orf80;refGene_tx=XM_019812732.2;refGene_gn=CD1H11orf80;refGene_tx=XM_019812734.2;refGene_gn=CD1H11orf80;refGene_tx=XM_019812735.2;refGene_gn=CD1H11orf80;refGene_tx=XM_023240089.1;refGene_gn=CD1H11orf80;refGene_tx=XM_023240090.1;refGene_gn=CD1H11orf80;refGene_tx=XM_023240091.1;refGene_gn=CD1H11orf80;refGene_tx=XM_023240092.1;refGene_gn=CD1H11orf80;refGene_tx=XM_023240093.1;refGene_gn=CD1H11orf80;refGene_tx=XM_023240094.1
D1 111803880 111803991
refGene_gn=CD1H11orf80;refGene_tx=XM_011287045.3;refGene_gn=CD1H11orf80;refGene_tx=XM_011287046.3;refGene_gn=CD1H11orf80;refGene_tx=XM_019812721.2;refGene_gn=CD1H11orf80;refGene_tx=XM_019812725.2;refGene_gn=CD1H11orf80;refGene_tx=XM_019812727.2;refGene_gn=CD1H11orf80;refGene_tx=XM_019812728.2;refGene_gn=CD1H11orf80;refGene_tx=XM_019812731.2;refGene_gn=CD1H11orf80;refGene_tx=XM_019812732.2;refGene_gn=CD1H11orf80;refGene_tx=XM_019812734.2;refGene_gn=CD1H11orf80;refGene_tx=XM_019812735.2;refGene_gn=CD1H11orf80;refGene_tx=XM_023240089.1;refGene_gn=CD1H11orf80;refGene_tx=XM_023240090.1;refGene_gn=CD1H11orf80;refGene_tx=XM_023240091.1;refGene_gn=CD1H11orf80;refGene_tx=XM_023240092.1;refGene_gn=CD1H11orf80;refGene_tx=XM_023240093.1;refGene_gn=CD1H11orf80;refGene_tx=XM_023240094.1
D1 111805971 111806070
refGene_gn=CD1H11orf80;refGene_tx=XM_011287045.3;refGene_gn=CD1H11orf80;refGene_tx=XM_011287046.3;refGene_gn=CD1H11orf80;refGene_tx=XM_019812721.2;refGene_gn=CD1H11orf80;refGene_tx=XM_019812725.2;refGene_gn=CD1H11orf80;refGene_tx=XM_019812727.2;refGene_gn=CD1H11orf80;refGene_tx=XM_019812728.2;refGene_gn=CD1H11orf80;refGene_tx=XM_019812731.2;refGene_gn=CD1H11orf80;refGene_tx=XM_019812732.2;refGene_gn=CD1H11orf80;refGene_tx=XM_019812734.2;refGene_gn=CD1H11orf80;refGene_tx=XM_019812735.2;refGene_gn=CD1H11orf80;refGene_tx=XM_023240089.1;refGene_gn=CD1H11orf8

0;refGene_tx=XM_023240090.1;refGene_gn=CD1H11orf80;refGene_tx=XM_02324
0091.1;refGene_gn=CD1H11orf80;refGene_tx=XM_023240092.1;refGene_gn=CD1
H11orf80;refGene_tx=XM_023240093.1;refGene_gn=CD1H11orf80;refGene_tx=X
M_023240094.1
D1 111818065 111818141
refGene_gn=CD1H11orf80;refGene_tx=XM_011287045.3;refGene_gn=CD1H11orf8
0;refGene_tx=XM_011287046.3;refGene_gn=CD1H11orf80;refGene_tx=XM_01981
2721.2;refGene_gn=CD1H11orf80;refGene_tx=XM_019812725.2;refGene_gn=CD1
H11orf80;refGene_tx=XM_019812727.2;refGene_gn=CD1H11orf80;refGene_tx=X
M_019812728.2;refGene_gn=CD1H11orf80;refGene_tx=XM_019812731.2;refGene
_gn=CD1H11orf80;refGene_tx=XM_019812732.2;refGene_gn=CD1H11orf80;refGe
ne_tx=XM_019812734.2;refGene_gn=CD1H11orf80;refGene_tx=XM_019812735.2;
refGene_gn=CD1H11orf80;refGene_tx=XM_023240089.1;refGene_gn=CD1H11orf8
0;refGene_tx=XM_023240090.1;refGene_gn=CD1H11orf80;refGene_tx=XM_02324
0091.1;refGene_gn=CD1H11orf80;refGene_tx=XM_023240092.1;refGene_gn=CD1
H11orf80;refGene_tx=XM_023240093.1;refGene_gn=CD1H11orf80;refGene_tx=X
M_023240094.1
D1 111821442 111821630
refGene_gn=CD1H11orf80;refGene_tx=XM_011287045.3;refGene_gn=CD1H11orf8
0;refGene_tx=XM_011287046.3;refGene_gn=CD1H11orf80;refGene_tx=XM_01981
2721.2;refGene_gn=CD1H11orf80;refGene_tx=XM_019812725.2;refGene_gn=CD1
H11orf80;refGene_tx=XM_019812727.2;refGene_gn=CD1H11orf80;refGene_tx=X
M_019812728.2;refGene_gn=CD1H11orf80;refGene_tx=XM_019812731.2;refGene
_gn=CD1H11orf80;refGene_tx=XM_019812732.2;refGene_gn=CD1H11orf80;refGe
ne_tx=XM_019812734.2;refGene_gn=CD1H11orf80;refGene_tx=XM_019812735.2;
refGene_gn=CD1H11orf80;refGene_tx=XM_023240089.1;refGene_gn=CD1H11orf8
0;refGene_tx=XM_023240090.1;refGene_gn=CD1H11orf80;refGene_tx=XM_02324
0091.1;refGene_gn=CD1H11orf80;refGene_tx=XM_023240092.1;refGene_gn=CD1
H11orf80;refGene_tx=XM_023240093.1;refGene_gn=CD1H11orf80;refGene_tx=X
M_023240094.1
D1 111821727 111821793
refGene_gn=CD1H11orf80;refGene_tx=XM_011287045.3;refGene_gn=CD1H11orf8
0;refGene_tx=XM_011287046.3;refGene_gn=CD1H11orf80;refGene_tx=XM_01981
2721.2;refGene_gn=CD1H11orf80;refGene_tx=XM_019812725.2;refGene_gn=CD1
H11orf80;refGene_tx=XM_019812727.2;refGene_gn=CD1H11orf80;refGene_tx=X
M_019812728.2;refGene_gn=CD1H11orf80;refGene_tx=XM_019812731.2;refGene
_gn=CD1H11orf80;refGene_tx=XM_019812732.2;refGene_gn=CD1H11orf80;refGe
ne_tx=XM_019812734.2;refGene_gn=CD1H11orf80;refGene_tx=XM_019812735.2;
refGene_gn=CD1H11orf80;refGene_tx=XM_023240089.1;refGene_gn=CD1H11orf8
0;refGene_tx=XM_023240090.1;refGene_gn=CD1H11orf80;refGene_tx=XM_02324
0091.1;refGene_gn=CD1H11orf80;refGene_tx=XM_023240092.1;refGene_gn=CD1
H11orf80;refGene_tx=XM_023240093.1;refGene_gn=CD1H11orf80;refGene_tx=X
M_023240094.1
D1 111822022 111822207
refGene_gn=RCE1;refGene_tx=XM_023240095.1;refGene_gn=RCE1;refGene_tx=X
M_023240096.1;refGene_gn=RCE1;refGene_tx=XM_023240097.1;refGene_gn=RCE
1;refGene_tx=XM_023240098.1;refGene_gn=RCE1;refGene_tx=XM_023240099.1
D1 111822343 111822446
refGene_gn=RCE1;refGene_tx=XM_023240095.1;refGene_gn=RCE1;refGene_tx=X
M_023240096.1;refGene_gn=RCE1;refGene_tx=XM_023240097.1;refGene_gn=RCE

1;refGene_tx=XM_023240098.1;refGene_gn=RCE1;refGene_tx=XM_023240099.1
D1	111822539	111822623
refGene_gn=RCE1;refGene_tx=XM_023240095.1;refGene_gn=RCE1;refGene_tx=XM_023240096.1;refGene_gn=RCE1;refGene_tx=XM_023240097.1;refGene_gn=RCE1;refGene_tx=XM_023240098.1;refGene_gn=RCE1;refGene_tx=XM_023240099.1
D1	111822853	111822932
refGene_gn=RCE1;refGene_tx=XM_023240095.1;refGene_gn=RCE1;refGene_tx=XM_023240096.1;refGene_gn=RCE1;refGene_tx=XM_023240097.1;refGene_gn=RCE1;refGene_tx=XM_023240098.1;refGene_gn=RCE1;refGene_tx=XM_023240099.1
D1	111823460	111823628
refGene_gn=RCE1;refGene_tx=XM_023240095.1;refGene_gn=RCE1;refGene_tx=XM_023240096.1;refGene_gn=RCE1;refGene_tx=XM_023240097.1;refGene_gn=RCE1;refGene_tx=XM_023240098.1;refGene_gn=RCE1;refGene_tx=XM_023240099.1
D1	111823756	111823828
refGene_gn=RCE1;refGene_tx=XM_023240095.1;refGene_gn=RCE1;refGene_tx=XM_023240096.1;refGene_gn=RCE1;refGene_tx=XM_023240097.1;refGene_gn=RCE1;refGene_tx=XM_023240098.1;refGene_gn=RCE1;refGene_tx=XM_023240099.1
D1	111824099	111824162
refGene_gn=RCE1;refGene_tx=XM_023240095.1;refGene_gn=RCE1;refGene_tx=XM_023240097.1;refGene_gn=RCE1;refGene_tx=XM_023240099.1
D1	111824446	111824682
refGene_gn=RCE1;refGene_tx=XM_023240097.1;refGene_gn=RCE1;refGene_tx=XM_023240098.1
D1	111825623	111825682
refGene_gn=RCE1;refGene_tx=XM_023240095.1;refGene_gn=RCE1;refGene_tx=XM_023240096.1;refGene_gn=RCE1;refGene_tx=XM_023240099.1
D1	111827338	111827587
refGene_gn=PC;refGene_tx=XM_006937555.4;refGene_gn=PC;refGene_tx=XM_006937556.3;refGene_gn=PC;refGene_tx=XM_011287047.3;refGene_gn=PC;refGene_tx=XM_019812719.2;refGene_gn=PC;refGene_tx=XM_019812720.2
D1	111827669	111827810
refGene_gn=PC;refGene_tx=XM_006937555.4;refGene_gn=PC;refGene_tx=XM_006937556.3;refGene_gn=PC;refGene_tx=XM_011287047.3;refGene_gn=PC;refGene_tx=XM_019812719.2;refGene_gn=PC;refGene_tx=XM_019812720.2
D1	111828100	111828349
refGene_gn=PC;refGene_tx=XM_006937555.4;refGene_gn=PC;refGene_tx=XM_006937556.3;refGene_gn=PC;refGene_tx=XM_011287047.3;refGene_gn=PC;refGene_tx=XM_019812719.2;refGene_gn=PC;refGene_tx=XM_019812720.2
D1	111828423	111828603
refGene_gn=PC;refGene_tx=XM_006937555.4;refGene_gn=PC;refGene_tx=XM_006937556.3;refGene_gn=PC;refGene_tx=XM_011287047.3;refGene_gn=PC;refGene_tx=XM_019812719.2;refGene_gn=PC;refGene_tx=XM_019812720.2
D1	111828688	111828933
refGene_gn=PC;refGene_tx=XM_006937555.4;refGene_gn=PC;refGene_tx=XM_006937556.3;refGene_gn=PC;refGene_tx=XM_011287047.3;refGene_gn=PC;refGene_tx=XM_019812719.2;refGene_gn=PC;refGene_tx=XM_019812720.2
D1	111829136	111829386
refGene_gn=PC;refGene_tx=XM_006937555.4;refGene_gn=PC;refGene_tx=XM_006937556.3;refGene_gn=PC;refGene_tx=XM_011287047.3;refGene_gn=PC;refGene_tx=XM_019812719.2;refGene_gn=PC;refGene_tx=XM_019812720.2

D1 111829537 111829778
refGene_gn=PC;refGene_tx=XM_006937555.4;refGene_gn=PC;refGene_tx=XM_006937556.3;refGene_gn=PC;refGene_tx=XM_011287047.3;refGene_gn=PC;refGene_tx=XM_019812719.2;refGene_gn=PC;refGene_tx=XM_019812720.2
D1 111830290 111830447
refGene_gn=PC;refGene_tx=XM_006937555.4;refGene_gn=PC;refGene_tx=XM_006937556.3;refGene_gn=PC;refGene_tx=XM_011287047.3;refGene_gn=PC;refGene_tx=XM_019812719.2;refGene_gn=PC;refGene_tx=XM_019812720.2
D1 111830933 111831155
refGene_gn=PC;refGene_tx=XM_006937555.4;refGene_gn=PC;refGene_tx=XM_006937556.3;refGene_gn=PC;refGene_tx=XM_011287047.3;refGene_gn=PC;refGene_tx=XM_019812719.2;refGene_gn=PC;refGene_tx=XM_019812720.2
D1 111831244 111831334
refGene_gn=PC;refGene_tx=XM_006937555.4;refGene_gn=PC;refGene_tx=XM_006937556.3;refGene_gn=PC;refGene_tx=XM_011287047.3;refGene_gn=PC;refGene_tx=XM_019812719.2;refGene_gn=PC;refGene_tx=XM_019812720.2
D1 111831693 111831838
refGene_gn=PC;refGene_tx=XM_006937555.4;refGene_gn=PC;refGene_tx=XM_006937556.3;refGene_gn=PC;refGene_tx=XM_011287047.3;refGene_gn=PC;refGene_tx=XM_019812719.2;refGene_gn=PC;refGene_tx=XM_019812720.2
D1 111832467 111832731
refGene_gn=RCE1;refGene_tx=XM_023240095.1;refGene_gn=RCE1;refGene_tx=XM_023240096.1
D1 111833068 111833110
refGene_gn=RCE1;refGene_tx=XM_023240099.1
D1 111835931 111837280
refGene_gn=LRFN4;refGene_tx=XM_003993691.4;refGene_gn=LRFN4;refGene_tx=XM_023240088.1
D1 111837871 111838433
refGene_gn=LRFN4;refGene_tx=XM_003993691.4;refGene_gn=LRFN4;refGene_tx=XM_023240088.1
D1 111841667 111841850
refGene_gn=PC;refGene_tx=XM_006937555.4;refGene_gn=PC;refGene_tx=XM_006937556.3;refGene_gn=PC;refGene_tx=XM_011287047.3;refGene_gn=PC;refGene_tx=XM_019812719.2;refGene_gn=PC;refGene_tx=XM_019812720.2
D1 111844127 111844290
refGene_gn=PC;refGene_tx=XM_006937555.4;refGene_gn=PC;refGene_tx=XM_006937556.3;refGene_gn=PC;refGene_tx=XM_011287047.3;refGene_gn=PC;refGene_tx=XM_019812719.2;refGene_gn=PC;refGene_tx=XM_019812720.2
D1 111846902 111847021
refGene_gn=PC;refGene_tx=XM_006937555.4;refGene_gn=PC;refGene_tx=XM_006937556.3;refGene_gn=PC;refGene_tx=XM_011287047.3;refGene_gn=PC;refGene_tx=XM_019812719.2;refGene_gn=PC;refGene_tx=XM_019812720.2
D1 111848550 111848702
refGene_gn=PC;refGene_tx=XM_006937555.4;refGene_gn=PC;refGene_tx=XM_006937556.3;refGene_gn=PC;refGene_tx=XM_011287047.3;refGene_gn=PC;refGene_tx=XM_019812719.2;refGene_gn=PC;refGene_tx=XM_019812720.2
D1 111848976 111849094
refGene_gn=PC;refGene_tx=XM_006937555.4;refGene_gn=PC;refGene_tx=XM_006937556.3;refGene_gn=PC;refGene_tx=XM_011287047.3;refGene_gn=PC;refGen

e_tx=XM_019812719.2;refGene_gn=PC;refGene_tx=XM_019812720.2
D1 111849253 111849399
refGene_gn=PC;refGene_tx=XM_006937555.4;refGene_gn=PC;refGene_tx=XM_006937556.3;refGene_gn=PC;refGene_tx=XM_011287047.3;refGene_gn=PC;refGene_tx=XM_019812719.2;refGene_gn=PC;refGene_tx=XM_019812720.2
D1 111849511 111849677
refGene_gn=PC;refGene_tx=XM_006937555.4;refGene_gn=PC;refGene_tx=XM_006937556.3;refGene_gn=PC;refGene_tx=XM_011287047.3;refGene_gn=PC;refGene_tx=XM_019812719.2;refGene_gn=PC;refGene_tx=XM_019812720.2
D1 111849891 111850076
refGene_gn=PC;refGene_tx=XM_006937555.4;refGene_gn=PC;refGene_tx=XM_006937556.3;refGene_gn=PC;refGene_tx=XM_011287047.3;refGene_gn=PC;refGene_tx=XM_019812719.2;refGene_gn=PC;refGene_tx=XM_019812720.2
D1 111850225 111850361
refGene_gn=PC;refGene_tx=XM_006937555.4;refGene_gn=PC;refGene_tx=XM_006937556.3;refGene_gn=PC;refGene_tx=XM_011287047.3;refGene_gn=PC;refGene_tx=XM_019812719.2;refGene_gn=PC;refGene_tx=XM_019812720.2
D1 111877595 111877598
refGene_gn=PC;refGene_tx=XM_019812719.2
D1 111939425 111939695
refGene_gn=CD1H11orf86;refGene_tx=XM_003993735.3;refGene_gn=CD1H11orf86;refGene_tx=XM_023239030.1
D1 111940267 111940366
refGene_gn=CD1H11orf86;refGene_tx=XM_003993735.3;refGene_gn=CD1H11orf86;refGene_tx=XM_023239030.1
D1 111983182 111983216
refGene_gn=SYT12;refGene_tx=XM_011287048.3;refGene_gn=SYT12;refGene_tx=XM_019812743.2;refGene_gn=SYT12;refGene_tx=XM_023240100.1;refGene_gn=SYT12;refGene_tx=XM_023240101.1
D1 111987676 111987870
refGene_gn=SYT12;refGene_tx=XM_011287048.3;refGene_gn=SYT12;refGene_tx=XM_019812743.2;refGene_gn=SYT12;refGene_tx=XM_023240100.1;refGene_gn=SYT12;refGene_tx=XM_023240101.1
D1 111990056 111990449
refGene_gn=SYT12;refGene_tx=XM_011287048.3;refGene_gn=SYT12;refGene_tx=XM_019812743.2;refGene_gn=SYT12;refGene_tx=XM_023240100.1;refGene_gn=SYT12;refGene_tx=XM_023240101.1
D1 111993164 111993380
refGene_gn=SYT12;refGene_tx=XM_011287048.3;refGene_gn=SYT12;refGene_tx=XM_019812743.2;refGene_gn=SYT12;refGene_tx=XM_023240100.1;refGene_gn=SYT12;refGene_tx=XM_023240101.1
D1 111993914 111994035
refGene_gn=SYT12;refGene_tx=XM_011287048.3;refGene_gn=SYT12;refGene_tx=XM_019812743.2;refGene_gn=SYT12;refGene_tx=XM_023240100.1;refGene_gn=SYT12;refGene_tx=XM_023240101.1
D1 111994895 111995029
refGene_gn=SYT12;refGene_tx=XM_011287048.3;refGene_gn=SYT12;refGene_tx=XM_019812743.2;refGene_gn=SYT12;refGene_tx=XM_023240100.1;refGene_gn=SYT12;refGene_tx=XM_023240101.1
D1 111996336 111996342

refGene_gn=SYT12;refGene_tx=XM_023240101.1
D1 111996742 111996916
refGene_gn=SYT12;refGene_tx=XM_011287048.3;refGene_gn=SYT12;refGene_tx=XM_019812743.2;refGene_gn=SYT12;refGene_tx=XM_023240100.1
D1 112004371 112004503
refGene_gn=RHOD;refGene_tx=XM_006937609.2
D1 112008892 112008980
refGene_gn=RHOD;refGene_tx=XM_006937609.2
D1 112009751 112009861
refGene_gn=RHOD;refGene_tx=XM_006937609.2
D1 112012216 112012351
refGene_gn=RHOD;refGene_tx=XM_006937609.2
D1 112012903 112013071
refGene_gn=RHOD;refGene_tx=XM_006937609.2
D1 112046271 112046342
refGene_gn=KDM2A;refGene_tx=XM_023240102.1
D1 112046429 112046554
refGene_gn=KDM2A;refGene_tx=XM_019812744.2;refGene_gn=KDM2A;refGene_tx=XM_023240102.1
D1 112074965 112074999
refGene_gn=KDM2A;refGene_tx=XM_023240103.1
D1 112078202 112078351
refGene_gn=KDM2A;refGene_tx=XM_023240102.1;refGene_gn=KDM2A;refGene_tx=XM_023240103.1
D1 112083185 112083203
refGene_gn=KDM2A;refGene_tx=XM_019812745.1
D1 112109726 112109865
refGene_gn=KDM2A;refGene_tx=XM_019812744.2;refGene_gn=KDM2A;refGene_tx=XM_019812745.1;refGene_gn=KDM2A;refGene_tx=XM_019812746.2;refGene_gn=KDM2A;refGene_tx=XM_023240102.1;refGene_gn=KDM2A;refGene_tx=XM_023240103.1
D1 112111120 112111199
refGene_gn=KDM2A;refGene_tx=XM_019812744.2;refGene_gn=KDM2A;refGene_tx=XM_019812745.1;refGene_gn=KDM2A;refGene_tx=XM_019812746.2;refGene_gn=KDM2A;refGene_tx=XM_023240102.1;refGene_gn=KDM2A;refGene_tx=XM_023240103.1
D1 112111650 112111697
refGene_gn=KDM2A;refGene_tx=XM_019812744.2;refGene_gn=KDM2A;refGene_tx=XM_019812745.1;refGene_gn=KDM2A;refGene_tx=XM_019812746.2;refGene_gn=KDM2A;refGene_tx=XM_023240102.1;refGene_gn=KDM2A;refGene_tx=XM_023240103.1
D1 112116035 112116214
refGene_gn=KDM2A;refGene_tx=XM_019812744.2;refGene_gn=KDM2A;refGene_tx=XM_019812745.1;refGene_gn=KDM2A;refGene_tx=XM_019812746.2;refGene_gn=KDM2A;refGene_tx=XM_023240102.1;refGene_gn=KDM2A;refGene_tx=XM_023240103.1
D1 112122595 112122702
refGene_gn=KDM2A;refGene_tx=XM_019812744.2;refGene_gn=KDM2A;refGene_tx=XM_019812745.1;refGene_gn=KDM2A;refGene_tx=XM_019812746.2;refGene_gn=KDM2A;refGene_tx=XM_023240102.1;refGene_gn=KDM2A;refGene_tx=XM_0232401

03.1
D1 112123063 112123157
refGene_gn=KDM2A;refGene_tx=XM_019812744.2;refGene_gn=KDM2A;refGene_tx=XM_019812745.1;refGene_gn=KDM2A;refGene_tx=XM_019812746.2;refGene_gn=KDM2A;refGene_tx=XM_023240102.1;refGene_gn=KDM2A;refGene_tx=XM_023240103.1
D1 112124337 112124491
refGene_gn=KDM2A;refGene_tx=XM_019812744.2;refGene_gn=KDM2A;refGene_tx=XM_019812745.1;refGene_gn=KDM2A;refGene_tx=XM_019812746.2;refGene_gn=KDM2A;refGene_tx=XM_023240102.1;refGene_gn=KDM2A;refGene_tx=XM_023240103.1
D1 112125348 112125464
refGene_gn=KDM2A;refGene_tx=XM_019812744.2;refGene_gn=KDM2A;refGene_tx=XM_019812745.1;refGene_gn=KDM2A;refGene_tx=XM_019812746.2;refGene_gn=KDM2A;refGene_tx=XM_023240102.1;refGene_gn=KDM2A;refGene_tx=XM_023240103.1
D1 112131032 112131159
refGene_gn=KDM2A;refGene_tx=XM_019812744.2;refGene_gn=KDM2A;refGene_tx=XM_019812745.1;refGene_gn=KDM2A;refGene_tx=XM_019812746.2;refGene_gn=KDM2A;refGene_tx=XM_023240102.1;refGene_gn=KDM2A;refGene_tx=XM_023240103.1
D1 112134488 112134883
refGene_gn=KDM2A;refGene_tx=XM_019812744.2;refGene_gn=KDM2A;refGene_tx=XM_019812745.1;refGene_gn=KDM2A;refGene_tx=XM_019812746.2;refGene_gn=KDM2A;refGene_tx=XM_023240102.1;refGene_gn=KDM2A;refGene_tx=XM_023240103.1
D1 112145258 112145342
refGene_gn=KDM2A;refGene_tx=XM_019812744.2;refGene_gn=KDM2A;refGene_tx=XM_019812745.1;refGene_gn=KDM2A;refGene_tx=XM_019812746.2;refGene_gn=KDM2A;refGene_tx=XM_023240102.1;refGene_gn=KDM2A;refGene_tx=XM_023240103.1
D1 112146799 112147069
refGene_gn=KDM2A;refGene_tx=XM_019812744.2;refGene_gn=KDM2A;refGene_tx=XM_019812745.1;refGene_gn=KDM2A;refGene_tx=XM_019812746.2;refGene_gn=KDM2A;refGene_tx=XM_023240102.1;refGene_gn=KDM2A;refGene_tx=XM_023240103.1
D1 112147622 112147754
refGene_gn=KDM2A;refGene_tx=XM_019812744.2;refGene_gn=KDM2A;refGene_tx=XM_019812745.1;refGene_gn=KDM2A;refGene_tx=XM_019812746.2;refGene_gn=KDM2A;refGene_tx=XM_023240102.1;refGene_gn=KDM2A;refGene_tx=XM_023240103.1
D1 112149545 112149635
refGene_gn=KDM2A;refGene_tx=XM_019812744.2;refGene_gn=KDM2A;refGene_tx=XM_019812745.1;refGene_gn=KDM2A;refGene_tx=XM_019812746.2;refGene_gn=KDM2A;refGene_tx=XM_023240102.1;refGene_gn=KDM2A;refGene_tx=XM_023240103.1
D1 112151806 112152519
refGene_gn=KDM2A;refGene_tx=XM_019812744.2;refGene_gn=KDM2A;refGene_tx=XM_019812745.1;refGene_gn=KDM2A;refGene_tx=XM_019812746.2;refGene_gn=KDM2A;refGene_tx=XM_023240102.1;refGene_gn=KDM2A;refGene_tx=XM_0232401

03.1
D1 112154239 112154403
refGene_gn=KDM2A;refGene_tx=XM_019812744.2;refGene_gn=KDM2A;refGene_tx=XM_019812745.1;refGene_gn=KDM2A;refGene_tx=XM_019812746.2;refGene_gn=KDM2A;refGene_tx=XM_023240102.1;refGene_gn=KDM2A;refGene_tx=XM_023240103.1
D1 112154911 112155070
refGene_gn=KDM2A;refGene_tx=XM_019812744.2;refGene_gn=KDM2A;refGene_tx=XM_019812745.1;refGene_gn=KDM2A;refGene_tx=XM_019812746.2;refGene_gn=KDM2A;refGene_tx=XM_023240102.1;refGene_gn=KDM2A;refGene_tx=XM_023240103.1
D1 112155615 112155831
refGene_gn=KDM2A;refGene_tx=XM_019812744.2;refGene_gn=KDM2A;refGene_tx=XM_019812745.1;refGene_gn=KDM2A;refGene_tx=XM_019812746.2;refGene_gn=KDM2A;refGene_tx=XM_023240102.1;refGene_gn=KDM2A;refGene_tx=XM_023240103.1
D1 112156259 112156441
refGene_gn=KDM2A;refGene_tx=XM_019812744.2;refGene_gn=KDM2A;refGene_tx=XM_019812745.1;refGene_gn=KDM2A;refGene_tx=XM_019812746.2;refGene_gn=KDM2A;refGene_tx=XM_023240102.1;refGene_gn=KDM2A;refGene_tx=XM_023240103.1
D1 112170377 112170490
refGene_gn=GRK2;refGene_tx=XM_023240104.1;refGene_gn=GRK2;refGene_tx=XM_023240105.1
D1 112180707 112180784
refGene_gn=GRK2;refGene_tx=XM_023240104.1;refGene_gn=GRK2;refGene_tx=XM_023240105.1
D1 112182548 112182622
refGene_gn=GRK2;refGene_tx=XM_023240104.1;refGene_gn=GRK2;refGene_tx=XM_023240105.1
D1 112182734 112182836
refGene_gn=GRK2;refGene_tx=XM_023240104.1;refGene_gn=GRK2;refGene_tx=XM_023240105.1
D1 112182958 112183033
refGene_gn=GRK2;refGene_tx=XM_023240104.1;refGene_gn=GRK2;refGene_tx=XM_023240105.1
D1 112183158 112183220
refGene_gn=GRK2;refGene_tx=XM_023240104.1;refGene_gn=GRK2;refGene_tx=XM_023240105.1
D1 112183950 112184002
refGene_gn=GRK2;refGene_tx=XM_023240104.1;refGene_gn=GRK2;refGene_tx=XM_023240105.1
D1 112184277 112184369
refGene_gn=GRK2;refGene_tx=XM_023240104.1;refGene_gn=GRK2;refGene_tx=XM_023240105.1
D1 112184777 112184877
refGene_gn=GRK2;refGene_tx=XM_023240104.1;refGene_gn=GRK2;refGene_tx=XM_023240105.1
D1 112184971 112185050
refGene_gn=GRK2;refGene_tx=XM_023240104.1;refGene_gn=GRK2;refGene_tx=X

M_023240105.1
D1 112185139 112185270
refGene_gn=GRK2;refGene_tx=XM_023240104.1;refGene_gn=GRK2;refGene_tx=X
M_023240105.1
D1 112185556 112185651
refGene_gn=GRK2;refGene_tx=XM_023240104.1;refGene_gn=GRK2;refGene_tx=X
M_023240105.1
D1 112185725 112185833
refGene_gn=GRK2;refGene_tx=XM_023240104.1;refGene_gn=GRK2;refGene_tx=X
M_023240105.1
D1 112186032 112186099
refGene_gn=GRK2;refGene_tx=XM_023240104.1;refGene_gn=GRK2;refGene_tx=X
M_023240105.1
D1 112186380 112186481
refGene_gn=GRK2;refGene_tx=XM_023240104.1;refGene_gn=GRK2;refGene_tx=X
M_023240105.1
D1 112187023 112187090
refGene_gn=GRK2;refGene_tx=XM_023240104.1;refGene_gn=GRK2;refGene_tx=X
M_023240105.1
D1 112187169 112187265
refGene_gn=GRK2;refGene_tx=XM_023240104.1;refGene_gn=GRK2;refGene_tx=X
M_023240105.1
D1 112187474 112187637
refGene_gn=GRK2;refGene_tx=XM_023240104.1;refGene_gn=GRK2;refGene_tx=X
M_023240105.1
D1 112187793 112187930
refGene_gn=GRK2;refGene_tx=XM_023240104.1;refGene_gn=GRK2;refGene_tx=X
M_023240105.1
D1 112188040 112188154
refGene_gn=GRK2;refGene_tx=XM_023240104.1;refGene_gn=GRK2;refGene_tx=X
M_023240105.1
D1 112188260 112188425
refGene_gn=GRK2;refGene_tx=XM_023240104.1;refGene_gn=GRK2;refGene_tx=X
M_023240105.1
D1 112192463 112192553
refGene_gn=ANKRD13D;refGene_tx=XM_006937611.3;refGene_gn=ANKRD13D;refG
ene_tx=XM_019812751.2;refGene_gn=ANKRD13D;refGene_tx=XM_023240106.1
D1 112192976 112193112
refGene_gn=ANKRD13D;refGene_tx=XM_006937611.3;refGene_gn=ANKRD13D;refG
ene_tx=XM_019812751.2;refGene_gn=ANKRD13D;refGene_tx=XM_023240106.1
D1 112193228 112193353
refGene_gn=ANKRD13D;refGene_tx=XM_006937611.3;refGene_gn=ANKRD13D;refG
ene_tx=XM_019812751.2;refGene_gn=ANKRD13D;refGene_tx=XM_023240106.1;re
fGene_gn=ANKRD13D;refGene_tx=XM_023240107.1
D1 112194115 112194161
refGene_gn=ANKRD13D;refGene_tx=XM_006937611.3;refGene_gn=ANKRD13D;refG
ene_tx=XM_019812751.2;refGene_gn=ANKRD13D;refGene_tx=XM_023240106.1;re
fGene_gn=ANKRD13D;refGene_tx=XM_023240107.1
D1 112194244 112194388
refGene_gn=ANKRD13D;refGene_tx=XM_006937611.3;refGene_gn=ANKRD13D;refG

ene_tx=XM_019812751.2;refGene_gn=ANKRD13D;refGene_tx=XM_023240106.1;refGene_gn=ANKRD13D;refGene_tx=XM_023240107.1
D1 112194640 112194830
refGene_gn=ANKRD13D;refGene_tx=XM_006937611.3;refGene_gn=ANKRD13D;refGene_tx=XM_019812751.2;refGene_gn=ANKRD13D;refGene_tx=XM_023240106.1;refGene_gn=ANKRD13D;refGene_tx=XM_023240107.1
D1 112199360 112199436
refGene_gn=ANKRD13D;refGene_tx=XM_006937611.3;refGene_gn=ANKRD13D;refGene_tx=XM_019812751.2;refGene_gn=ANKRD13D;refGene_tx=XM_023240106.1;refGene_gn=ANKRD13D;refGene_tx=XM_023240107.1
D1 112199827 112199909
refGene_gn=ANKRD13D;refGene_tx=XM_006937611.3;refGene_gn=ANKRD13D;refGene_tx=XM_019812751.2;refGene_gn=ANKRD13D;refGene_tx=XM_023240106.1;refGene_gn=ANKRD13D;refGene_tx=XM_023240107.1
D1 112200126 112200188
refGene_gn=ANKRD13D;refGene_tx=XM_006937611.3;refGene_gn=ANKRD13D;refGene_tx=XM_019812751.2;refGene_gn=ANKRD13D;refGene_tx=XM_023240106.1;refGene_gn=ANKRD13D;refGene_tx=XM_023240107.1
D1 112200291 112200422
refGene_gn=ANKRD13D;refGene_tx=XM_006937611.3;refGene_gn=ANKRD13D;refGene_tx=XM_019812751.2;refGene_gn=ANKRD13D;refGene_tx=XM_023240106.1;refGene_gn=ANKRD13D;refGene_tx=XM_023240107.1
D1 112201367 112201525
refGene_gn=ANKRD13D;refGene_tx=XM_006937611.3;refGene_gn=ANKRD13D;refGene_tx=XM_019812751.2;refGene_gn=ANKRD13D;refGene_tx=XM_023240106.1;refGene_gn=ANKRD13D;refGene_tx=XM_023240107.1
D1 112201613 112201736
refGene_gn=ANKRD13D;refGene_tx=XM_006937611.3;refGene_gn=ANKRD13D;refGene_tx=XM_019812751.2;refGene_gn=ANKRD13D;refGene_tx=XM_023240107.1
D1 112201817 112201981
refGene_gn=ANKRD13D;refGene_tx=XM_006937611.3;refGene_gn=ANKRD13D;refGene_tx=XM_019812751.2;refGene_gn=ANKRD13D;refGene_tx=XM_023240106.1;refGene_gn=ANKRD13D;refGene_tx=XM_023240107.1
D1 112202060 112202152
refGene_gn=ANKRD13D;refGene_tx=XM_006937611.3;refGene_gn=ANKRD13D;refGene_tx=XM_019812751.2;refGene_gn=ANKRD13D;refGene_tx=XM_023240106.1;refGene_gn=ANKRD13D;refGene_tx=XM_023240107.1
D1 112202488 112202702
refGene_gn=ANKRD13D;refGene_tx=XM_006937611.3;refGene_gn=ANKRD13D;refGene_tx=XM_019812751.2;refGene_gn=ANKRD13D;refGene_tx=XM_023240106.1;refGene_gn=ANKRD13D;refGene_tx=XM_023240107.1
D1 112204072 112204138
refGene_gn=SSH3;refGene_tx=XM_003993695.5;refGene_gn=SSH3;refGene_tx=XM_006937562.4;refGene_gn=SSH3;refGene_tx=XM_019812752.2
D1 112204647 112204688
refGene_gn=SSH3;refGene_tx=XM_003993695.5;refGene_gn=SSH3;refGene_tx=XM_006937562.4;refGene_gn=SSH3;refGene_tx=XM_019812752.2
D1 112205278 112205510
refGene_gn=SSH3;refGene_tx=XM_003993695.5;refGene_gn=SSH3;refGene_tx=XM_006937562.4;refGene_gn=SSH3;refGene_tx=XM_019812752.2

D1 112206704 112206823
refGene_gn=SSH3;refGene_tx=XM_003993695.5;refGene_gn=SSH3;refGene_tx=XM_006937562.4;refGene_gn=SSH3;refGene_tx=XM_019812752.2
D1 112206903 112206975
refGene_gn=SSH3;refGene_tx=XM_003993695.5;refGene_gn=SSH3;refGene_tx=XM_006937562.4;refGene_gn=SSH3;refGene_tx=XM_019812752.2
D1 112207327 112207393
refGene_gn=SSH3;refGene_tx=XM_003993695.5;refGene_gn=SSH3;refGene_tx=XM_006937562.4;refGene_gn=SSH3;refGene_tx=XM_019812752.2
D1 112207508 112207697
refGene_gn=SSH3;refGene_tx=XM_003993695.5;refGene_gn=SSH3;refGene_tx=XM_006937562.4;refGene_gn=SSH3;refGene_tx=XM_019812752.2
D1 112207780 112207874
refGene_gn=SSH3;refGene_tx=XM_003993695.5;refGene_gn=SSH3;refGene_tx=XM_006937562.4;refGene_gn=SSH3;refGene_tx=XM_019812752.2
D1 112208265 112208394
refGene_gn=SSH3;refGene_tx=XM_003993695.5;refGene_gn=SSH3;refGene_tx=XM_006937562.4;refGene_gn=SSH3;refGene_tx=XM_019812752.2
D1 112208471 112208518
refGene_gn=SSH3;refGene_tx=XM_003993695.5;refGene_gn=SSH3;refGene_tx=XM_006937562.4;refGene_gn=SSH3;refGene_tx=XM_019812752.2
D1 112209237 112209384
refGene_gn=SSH3;refGene_tx=XM_003993695.5;refGene_gn=SSH3;refGene_tx=XM_006937562.4;refGene_gn=SSH3;refGene_tx=XM_019812752.2
D1 112209579 112209780
refGene_gn=SSH3;refGene_tx=XM_003993695.5;refGene_gn=SSH3;refGene_tx=XM_006937562.4;refGene_gn=SSH3;refGene_tx=XM_019812752.2
D1 112209864 112210195
refGene_gn=SSH3;refGene_tx=XM_003993695.5;refGene_gn=SSH3;refGene_tx=XM_006937562.4;refGene_gn=SSH3;refGene_tx=XM_019812752.2
D1 112211196 112211469
refGene_gn=SSH3;refGene_tx=XM_003993695.5;refGene_gn=SSH3;refGene_tx=XM_006937562.4
D1 112235175 112235200
refGene_gn=POLD4;refGene_tx=XM_003993696.4
D1 112235567 112235679
refGene_gn=POLD4;refGene_tx=XM_003993696.4
D1 112235858 112235948
refGene_gn=POLD4;refGene_tx=XM_003993696.4
D1 112236199 112236296
refGene_gn=POLD4;refGene_tx=XM_003993696.4
D1 112245017 112245512
refGene_gn=CLCF1;refGene_tx=XM_006937563.3;refGene_gn=CLCF1;refGene_tx=XM_011287135.3;refGene_gn=CLCF1;refGene_tx=XM_023240108.1
D1 112247350 112247517
refGene_gn=CLCF1;refGene_tx=XM_006937563.3;refGene_gn=CLCF1;refGene_tx=XM_011287135.3;refGene_gn=CLCF1;refGene_tx=XM_023240108.1
D1 112252685 112252716
refGene_gn=CLCF1;refGene_tx=XM_023240108.1
D1 112253385 112253401

refGene_gn=CLCF1;refGene_tx=XM_011287135.3
D1	112265727	112265761
refGene_gn=RAD9A;refGene_tx=XM_023240117.1
D1	112265847	112265918
refGene_gn=RAD9A;refGene_tx=XM_023240117.1
D1	112266322	112266451
refGene_gn=RAD9A;refGene_tx=XM_023240117.1
D1	112266878	112266993
refGene_gn=RAD9A;refGene_tx=XM_023240117.1
D1	112267073	112267173
refGene_gn=RAD9A;refGene_tx=XM_023240117.1
D1	112268246	112268868
refGene_gn=RAD9A;refGene_tx=XM_023240115.1;refGene_gn=RAD9A;refGene_tx=XM_023240116.1;refGene_gn=RAD9A;refGene_tx=XM_023240117.1
D1	112268986	112269096
refGene_gn=RAD9A;refGene_tx=XM_023240115.1;refGene_gn=RAD9A;refGene_tx=XM_023240116.1;refGene_gn=RAD9A;refGene_tx=XM_023240117.1
D1	112269175	112269240
refGene_gn=RAD9A;refGene_tx=XM_023240115.1;refGene_gn=RAD9A;refGene_tx=XM_023240116.1;refGene_gn=RAD9A;refGene_tx=XM_023240117.1
D1	112269313	112269454
refGene_gn=RAD9A;refGene_tx=XM_023240115.1;refGene_gn=RAD9A;refGene_tx=XM_023240116.1;refGene_gn=RAD9A;refGene_tx=XM_023240117.1
D1	112270255	112270460
refGene_gn=RAD9A;refGene_tx=XM_023240115.1;refGene_gn=RAD9A;refGene_tx=XM_023240116.1;refGene_gn=RAD9A;refGene_tx=XM_023240117.1
D1	112270545	112270641
refGene_gn=RAD9A;refGene_tx=XM_023240115.1;refGene_gn=RAD9A;refGene_tx=XM_023240116.1;refGene_gn=RAD9A;refGene_tx=XM_023240117.1
D1	112271621	112271732
refGene_gn=PPP1CA;refGene_tx=XM_023240118.1
D1	112271806	112271941
refGene_gn=PPP1CA;refGene_tx=XM_023240118.1
D1	112272026	112272250
refGene_gn=PPP1CA;refGene_tx=XM_023240118.1
D1	112272647	112272752
refGene_gn=PPP1CA;refGene_tx=XM_023240118.1
D1	112273338	112273569
refGene_gn=PPP1CA;refGene_tx=XM_023240118.1
D1	112273721	112273853
refGene_gn=PPP1CA;refGene_tx=XM_023240118.1
D1	112274391	112274446
refGene_gn=PPP1CA;refGene_tx=XM_023240118.1
D1	112275876	112276028
refGene_gn=TBC1D10C;refGene_tx=XM_006937566.3;refGene_gn=TBC1D10C;refGene_tx=XM_006937567.3;refGene_gn=TBC1D10C;refGene_tx=XM_019812755.1
D1	112276777	112276877
refGene_gn=TBC1D10C;refGene_tx=XM_006937566.3;refGene_gn=TBC1D10C;refGene_tx=XM_006937567.3;refGene_gn=TBC1D10C;refGene_tx=XM_019812755.1
D1	112277076	112277184

refGene_gn=TBC1D10C;refGene_tx=XM_006937566.3;refGene_gn=TBC1D10C;refGene_tx=XM_006937567.3;refGene_gn=TBC1D10C;refGene_tx=XM_019812755.1
D1 112277272 112277379
refGene_gn=TBC1D10C;refGene_tx=XM_006937566.3;refGene_gn=TBC1D10C;refGene_tx=XM_006937567.3;refGene_gn=TBC1D10C;refGene_tx=XM_019812755.1
D1 112277579 112277694
refGene_gn=TBC1D10C;refGene_tx=XM_006937566.3;refGene_gn=TBC1D10C;refGene_tx=XM_006937567.3;refGene_gn=TBC1D10C;refGene_tx=XM_019812755.1
D1 112278314 112278380
refGene_gn=TBC1D10C;refGene_tx=XM_006937566.3;refGene_gn=TBC1D10C;refGene_tx=XM_006937567.3;refGene_gn=TBC1D10C;refGene_tx=XM_019812755.1
D1 112278510 112278700
refGene_gn=TBC1D10C;refGene_tx=XM_006937566.3;refGene_gn=TBC1D10C;refGene_tx=XM_006937567.3;refGene_gn=TBC1D10C;refGene_tx=XM_019812755.1
D1 112280448 112280603
refGene_gn=TBC1D10C;refGene_tx=XM_006937566.3;refGene_gn=TBC1D10C;refGene_tx=XM_006937567.3;refGene_gn=TBC1D10C;refGene_tx=XM_019812755.1
D1 112280904 112281252
refGene_gn=TBC1D10C;refGene_tx=XM_006937566.3;refGene_gn=TBC1D10C;refGene_tx=XM_006937567.3;refGene_gn=TBC1D10C;refGene_tx=XM_019812755.1
D1 112287058 112287061
refGene_gn=CARNS1;refGene_tx=XM_023240109.1;refGene_gn=CARNS1;refGene_tx=XM_023240110.1;refGene_gn=CARNS1;refGene_tx=XM_023240113.1
D1 112288348 112288622
refGene_gn=CARNS1;refGene_tx=XM_023240109.1;refGene_gn=CARNS1;refGene_tx=XM_023240110.1;refGene_gn=CARNS1;refGene_tx=XM_023240113.1
D1 112289379 112289469
refGene_gn=CARNS1;refGene_tx=XM_023240109.1;refGene_gn=CARNS1;refGene_tx=XM_023240110.1;refGene_gn=CARNS1;refGene_tx=XM_023240113.1
D1 112289719 112290207
refGene_gn=CARNS1;refGene_tx=XM_023240109.1;refGene_gn=CARNS1;refGene_tx=XM_023240110.1;refGene_gn=CARNS1;refGene_tx=XM_023240111.1;refGene_gn=CARNS1;refGene_tx=XM_023240112.1;refGene_gn=CARNS1;refGene_tx=XM_023240113.1
D1 112290439 112290614
refGene_gn=CARNS1;refGene_tx=XM_023240109.1;refGene_gn=CARNS1;refGene_tx=XM_023240110.1;refGene_gn=CARNS1;refGene_tx=XM_023240111.1;refGene_gn=CARNS1;refGene_tx=XM_023240112.1;refGene_gn=CARNS1;refGene_tx=XM_023240113.1
D1 112290703 112290789
refGene_gn=CARNS1;refGene_tx=XM_023240109.1;refGene_gn=CARNS1;refGene_tx=XM_023240110.1;refGene_gn=CARNS1;refGene_tx=XM_023240111.1;refGene_gn=CARNS1;refGene_tx=XM_023240112.1;refGene_gn=CARNS1;refGene_tx=XM_023240113.1
D1 112291499 112291731
refGene_gn=CARNS1;refGene_tx=XM_023240109.1;refGene_gn=CARNS1;refGene_tx=XM_023240110.1;refGene_gn=CARNS1;refGene_tx=XM_023240111.1;refGene_gn=CARNS1;refGene_tx=XM_023240112.1
D1 112291817 112292098
refGene_gn=CARNS1;refGene_tx=XM_023240109.1;refGene_gn=CARNS1;refGene_

tx=XM_023240110.1;refGene_gn=CARNS1;refGene_tx=XM_023240111.1;refGene_gn=CARNS1;refGene_tx=XM_023240112.1
D1 112293412 112294639
refGene_gn=CARNS1;refGene_tx=XM_023240109.1;refGene_gn=CARNS1;refGene_tx=XM_023240110.1;refGene_gn=CARNS1;refGene_tx=XM_023240111.1;refGene_gn=CARNS1;refGene_tx=XM_023240112.1;refGene_gn=CARNS1;refGene_tx=XM_023240113.1
D1 112296796 112296868
refGene_gn=RPS6KB2;refGene_tx=XM_003993698.5;refGene_gn=RPS6KB2;refGene_tx=XM_011287134.2;refGene_gn=RPS6KB2;refGene_tx=XM_019812754.1
D1 112297194 112297235
refGene_gn=RPS6KB2;refGene_tx=XM_003993698.5;refGene_gn=RPS6KB2;refGene_tx=XM_011287134.2
D1 112297334 112297455
refGene_gn=RPS6KB2;refGene_tx=XM_003993698.5;refGene_gn=RPS6KB2;refGene_tx=XM_011287134.2
D1 112297737 112297806
refGene_gn=RPS6KB2;refGene_tx=XM_003993698.5;refGene_gn=RPS6KB2;refGene_tx=XM_011287134.2;refGene_gn=RPS6KB2;refGene_tx=XM_019812754.1
D1 112298325 112298473
refGene_gn=RPS6KB2;refGene_tx=XM_003993698.5;refGene_gn=RPS6KB2;refGene_tx=XM_011287134.2;refGene_gn=RPS6KB2;refGene_tx=XM_019812754.1
D1 112299547 112299605
refGene_gn=RPS6KB2;refGene_tx=XM_003993698.5;refGene_gn=RPS6KB2;refGene_tx=XM_011287134.2;refGene_gn=RPS6KB2;refGene_tx=XM_019812754.1
D1 112299707 112299808
refGene_gn=RPS6KB2;refGene_tx=XM_003993698.5;refGene_gn=RPS6KB2;refGene_tx=XM_011287134.2;refGene_gn=RPS6KB2;refGene_tx=XM_019812754.1
D1 112299932 112300023
refGene_gn=RPS6KB2;refGene_tx=XM_003993698.5;refGene_gn=RPS6KB2;refGene_tx=XM_011287134.2;refGene_gn=RPS6KB2;refGene_tx=XM_019812754.1
D1 112300108 112300199
refGene_gn=RPS6KB2;refGene_tx=XM_003993698.5;refGene_gn=RPS6KB2;refGene_tx=XM_011287134.2;refGene_gn=RPS6KB2;refGene_tx=XM_019812754.1
D1 112300322 112300430
refGene_gn=RPS6KB2;refGene_tx=XM_003993698.5;refGene_gn=RPS6KB2;refGene_tx=XM_011287134.2;refGene_gn=RPS6KB2;refGene_tx=XM_019812754.1
D1 112301130 112301193
refGene_gn=RPS6KB2;refGene_tx=XM_003993698.5;refGene_gn=RPS6KB2;refGene_tx=XM_011287134.2;refGene_gn=RPS6KB2;refGene_tx=XM_019812754.1
D1 112301330 112301408
refGene_gn=RPS6KB2;refGene_tx=XM_003993698.5;refGene_gn=RPS6KB2;refGene_tx=XM_011287134.2;refGene_gn=RPS6KB2;refGene_tx=XM_019812754.1
D1 112301497 112301605
refGene_gn=RPS6KB2;refGene_tx=XM_003993698.5;refGene_gn=RPS6KB2;refGene_tx=XM_011287134.2;refGene_gn=RPS6KB2;refGene_tx=XM_019812754.1
D1 112301711 112301824
refGene_gn=RPS6KB2;refGene_tx=XM_003993698.5;refGene_gn=RPS6KB2;refGene_tx=XM_011287134.2;refGene_gn=RPS6KB2;refGene_tx=XM_019812754.1
D1 112302085 112302269

refGene_gn=RPS6KB2;refGene_tx=XM_003993698.5;refGene_gn=RPS6KB2;refGene_tx=XM_011287134.2;refGene_gn=RPS6KB2;refGene_tx=XM_019812754.1
D1 112302854 112303478
refGene_gn=PTPRCAP;refGene_tx=XM_006937614.2
D1 112304763 112304766
refGene_gn=PTPRCAP;refGene_tx=XM_006937614.2
D1 112305592 112305718
refGene_gn=CORO1B;refGene_tx=XM_023240114.1
D1 112305902 112306181
refGene_gn=CORO1B;refGene_tx=XM_023240114.1
D1 112306928 112306986
refGene_gn=CORO1B;refGene_tx=XM_023240114.1
D1 112307230 112307376
refGene_gn=CORO1B;refGene_tx=XM_023240114.1
D1 112307458 112307560
refGene_gn=CORO1B;refGene_tx=XM_023240114.1
D1 112308231 112308351
refGene_gn=CORO1B;refGene_tx=XM_023240114.1
D1 112308438 112308620
refGene_gn=CORO1B;refGene_tx=XM_023240114.1
D1 112308802 112308932
refGene_gn=CORO1B;refGene_tx=XM_023240114.1
D1 112309038 112309161
refGene_gn=CORO1B;refGene_tx=XM_023240114.1
D1 112309412 112309613
refGene_gn=CORO1B;refGene_tx=XM_023240114.1
D1 112314851 112316465
refGene_gn=GPR152;refGene_tx=XM_023240119.1
D1 112319056 112319419
refGene_gn=CABP4;refGene_tx=XM_003993744.3
D1 112319796 112319827
refGene_gn=CABP4;refGene_tx=XM_003993744.3
D1 112319910 112320054
refGene_gn=CABP4;refGene_tx=XM_003993744.3
D1 112320872 112320982
refGene_gn=CABP4;refGene_tx=XM_003993744.3
D1 112321339 112321487
refGene_gn=CABP4;refGene_tx=XM_003993744.3
D1 112321599 112321628
refGene_gn=CABP4;refGene_tx=XM_003993744.3
D1 112327606 112327689
refGene_gn=TMEM134;refGene_tx=XM_023240120.1;refGene_gn=TMEM134;refGene_tx=XM_023240121.1;refGene_gn=TMEM134;refGene_tx=XM_023240122.1;refGene_gn=TMEM134;refGene_tx=XM_023240123.1;refGene_gn=TMEM134;refGene_tx=XM_023240124.1;refGene_gn=TMEM134;refGene_tx=XM_023240125.1
D1 112327775 112327829
refGene_gn=TMEM134;refGene_tx=XM_023240120.1;refGene_gn=TMEM134;refGene_tx=XM_023240121.1;refGene_gn=TMEM134;refGene_tx=XM_023240122.1;refGene_gn=TMEM134;refGene_tx=XM_023240123.1;refGene_gn=TMEM134;refGene_tx=XM_023240124.1;refGene_gn=TMEM134;refGene_tx=XM_023240125.1

D1 112328049 112328094
refGene_gn=TMEM134;refGene_tx=XM_023240120.1;refGene_gn=TMEM134;refGene_tx=XM_023240121.1;refGene_gn=TMEM134;refGene_tx=XM_023240123.1;refGene_gn=TMEM134;refGene_tx=XM_023240124.1
D1 112328259 112328343
refGene_gn=TMEM134;refGene_tx=XM_023240120.1;refGene_gn=TMEM134;refGene_tx=XM_023240121.1;refGene_gn=TMEM134;refGene_tx=XM_023240122.1
D1 112330042 112330164
refGene_gn=TMEM134;refGene_tx=XM_023240126.1
D1 112330298 112330375
refGene_gn=TMEM134;refGene_tx=XM_023240120.1;refGene_gn=TMEM134;refGene_tx=XM_023240121.1;refGene_gn=TMEM134;refGene_tx=XM_023240122.1;refGene_gn=TMEM134;refGene_tx=XM_023240123.1;refGene_gn=TMEM134;refGene_tx=XM_023240124.1;refGene_gn=TMEM134;refGene_tx=XM_023240125.1;refGene_gn=TMEM134;refGene_tx=XM_023240126.1
D1 112330527 112330617
refGene_gn=TMEM134;refGene_tx=XM_023240120.1;refGene_gn=TMEM134;refGene_tx=XM_023240121.1;refGene_gn=TMEM134;refGene_tx=XM_023240122.1;refGene_gn=TMEM134;refGene_tx=XM_023240123.1;refGene_gn=TMEM134;refGene_tx=XM_023240124.1;refGene_gn=TMEM134;refGene_tx=XM_023240125.1;refGene_gn=TMEM134;refGene_tx=XM_023240126.1
D1 112331110 112331175
refGene_gn=TMEM134;refGene_tx=XM_023240120.1;refGene_gn=TMEM134;refGene_tx=XM_023240121.1;refGene_gn=TMEM134;refGene_tx=XM_023240122.1;refGene_gn=TMEM134;refGene_tx=XM_023240123.1;refGene_gn=TMEM134;refGene_tx=XM_023240124.1;refGene_gn=TMEM134;refGene_tx=XM_023240125.1;refGene_gn=TMEM134;refGene_tx=XM_023240126.1
D1 112332069 112332243
refGene_gn=TMEM134;refGene_tx=XM_023240120.1;refGene_gn=TMEM134;refGene_tx=XM_023240121.1;refGene_gn=TMEM134;refGene_tx=XM_023240122.1;refGene_gn=TMEM134;refGene_tx=XM_023240123.1;refGene_gn=TMEM134;refGene_tx=XM_023240124.1;refGene_gn=TMEM134;refGene_tx=XM_023240125.1;refGene_gn=TMEM134;refGene_tx=XM_023240126.1
D1 112342397 112342496
refGene_gn=AIP;refGene_tx=XM_003993700.4;refGene_gn=AIP;refGene_tx=XM_011287132.3
D1 112345992 112346172
refGene_gn=AIP;refGene_tx=XM_003993700.4;refGene_gn=AIP;refGene_tx=XM_011287132.3
D1 112347385 112347574
refGene_gn=AIP;refGene_tx=XM_003993700.4;refGene_gn=AIP;refGene_tx=XM_011287132.3
D1 112348035 112348212
refGene_gn=AIP;refGene_tx=XM_003993700.4;refGene_gn=AIP;refGene_tx=XM_011287132.3
D1 112348298 112348440
refGene_gn=AIP;refGene_tx=XM_003993700.4;refGene_gn=AIP;refGene_tx=XM_011287132.3
D1 112348739 112348945
refGene_gn=AIP;refGene_tx=XM_003993700.4;refGene_gn=AIP;refGene_tx=XM_

011287132.3
D1 112349897 112350161
refGene_gn=PITPNM1;refGene_tx=XM_023240127.1;refGene_gn=PITPNM1;refGene_tx=XM_023240128.1;refGene_gn=PITPNM1;refGene_tx=XM_023240129.1
D1 112350796 112350925
refGene_gn=PITPNM1;refGene_tx=XM_023240127.1;refGene_gn=PITPNM1;refGene_tx=XM_023240128.1;refGene_gn=PITPNM1;refGene_tx=XM_023240129.1
D1 112351240 112351424
refGene_gn=PITPNM1;refGene_tx=XM_023240127.1;refGene_gn=PITPNM1;refGene_tx=XM_023240128.1;refGene_gn=PITPNM1;refGene_tx=XM_023240129.1
D1 112351504 112351654
refGene_gn=PITPNM1;refGene_tx=XM_023240127.1;refGene_gn=PITPNM1;refGene_tx=XM_023240128.1;refGene_gn=PITPNM1;refGene_tx=XM_023240129.1
D1 112351739 112351888
refGene_gn=PITPNM1;refGene_tx=XM_023240127.1;refGene_gn=PITPNM1;refGene_tx=XM_023240128.1;refGene_gn=PITPNM1;refGene_tx=XM_023240129.1
D1 112352051 112352168
refGene_gn=PITPNM1;refGene_tx=XM_023240127.1;refGene_gn=PITPNM1;refGene_tx=XM_023240128.1;refGene_gn=PITPNM1;refGene_tx=XM_023240129.1
D1 112352621 112352732
refGene_gn=PITPNM1;refGene_tx=XM_023240127.1;refGene_gn=PITPNM1;refGene_tx=XM_023240128.1;refGene_gn=PITPNM1;refGene_tx=XM_023240129.1
D1 112352849 112352998
refGene_gn=PITPNM1;refGene_tx=XM_023240127.1;refGene_gn=PITPNM1;refGene_tx=XM_023240128.1;refGene_gn=PITPNM1;refGene_tx=XM_023240129.1
D1 112353241 112353406
refGene_gn=PITPNM1;refGene_tx=XM_023240127.1;refGene_gn=PITPNM1;refGene_tx=XM_023240128.1;refGene_gn=PITPNM1;refGene_tx=XM_023240129.1
D1 112354051 112354222
refGene_gn=PITPNM1;refGene_tx=XM_023240127.1;refGene_gn=PITPNM1;refGene_tx=XM_023240128.1;refGene_gn=PITPNM1;refGene_tx=XM_023240129.1
D1 112354677 112354883
refGene_gn=PITPNM1;refGene_tx=XM_023240127.1;refGene_gn=PITPNM1;refGene_tx=XM_023240128.1;refGene_gn=PITPNM1;refGene_tx=XM_023240129.1
D1 112354970 112355128
refGene_gn=PITPNM1;refGene_tx=XM_023240127.1;refGene_gn=PITPNM1;refGene_tx=XM_023240128.1;refGene_gn=PITPNM1;refGene_tx=XM_023240129.1
D1 112355407 112355515
refGene_gn=PITPNM1;refGene_tx=XM_023240127.1;refGene_gn=PITPNM1;refGene_tx=XM_023240128.1;refGene_gn=PITPNM1;refGene_tx=XM_023240129.1
D1 112355597 112355787
refGene_gn=PITPNM1;refGene_tx=XM_023240127.1;refGene_gn=PITPNM1;refGene_tx=XM_023240128.1;refGene_gn=PITPNM1;refGene_tx=XM_023240129.1
D1 112356109 112356360
refGene_gn=PITPNM1;refGene_tx=XM_023240127.1;refGene_gn=PITPNM1;refGene_tx=XM_023240128.1;refGene_gn=PITPNM1;refGene_tx=XM_023240129.1
D1 112356452 112356514
refGene_gn=PITPNM1;refGene_tx=XM_023240127.1;refGene_gn=PITPNM1;refGene_tx=XM_023240128.1;refGene_gn=PITPNM1;refGene_tx=XM_023240129.1
D1 112357184 112357292

refGene_gn=PITPNM1;refGene_tx=XM_023240127.1;refGene_gn=PITPNM1;refGen
e_tx=XM_023240128.1;refGene_gn=PITPNM1;refGene_tx=XM_023240129.1
D1	112357374	112357470
refGene_gn=PITPNM1;refGene_tx=XM_023240127.1;refGene_gn=PITPNM1;refGen
e_tx=XM_023240128.1;refGene_gn=PITPNM1;refGene_tx=XM_023240129.1
D1	112357545	112357872
refGene_gn=PITPNM1;refGene_tx=XM_023240127.1;refGene_gn=PITPNM1;refGen
e_tx=XM_023240128.1;refGene_gn=PITPNM1;refGene_tx=XM_023240129.1
D1	112359335	112359560
refGene_gn=PITPNM1;refGene_tx=XM_023240127.1;refGene_gn=PITPNM1;refGen
e_tx=XM_023240128.1;refGene_gn=PITPNM1;refGene_tx=XM_023240129.1
D1	112359736	112359858
refGene_gn=PITPNM1;refGene_tx=XM_023240127.1;refGene_gn=PITPNM1;refGen
e_tx=XM_023240128.1;refGene_gn=PITPNM1;refGene_tx=XM_023240129.1
D1	112359942	112360157
refGene_gn=PITPNM1;refGene_tx=XM_023240127.1;refGene_gn=PITPNM1;refGen
e_tx=XM_023240128.1;refGene_gn=PITPNM1;refGene_tx=XM_023240129.1
D1	112361862	112361981
refGene_gn=PITPNM1;refGene_tx=XM_023240127.1;refGene_gn=PITPNM1;refGen
e_tx=XM_023240128.1;refGene_gn=PITPNM1;refGene_tx=XM_023240129.1
D1	112363746	112363750
refGene_gn=PITPNM1;refGene_tx=XM_023240127.1;refGene_gn=PITPNM1;refGen
e_tx=XM_023240128.1
D1	112364742	112364810
refGene_gn=CDK2AP2;refGene_tx=XM_003993701.4;refGene_gn=CDK2AP2;refGen
e_tx=XM_019812764.1
D1	112365184	112365317
refGene_gn=CDK2AP2;refGene_tx=XM_003993701.4;refGene_gn=CDK2AP2;refGen
e_tx=XM_019812764.1
D1	112365404	112365505
refGene_gn=CDK2AP2;refGene_tx=XM_003993701.4;refGene_gn=CDK2AP2;refGen
e_tx=XM_019812764.1
D1	112365802	112365884
refGene_gn=CDK2AP2;refGene_tx=XM_003993701.4
D1	112372900	112372926
refGene_gn=CABP2;refGene_tx=XM_003993704.5;refGene_gn=CABP2;refGene_tx
=XM_019812759.2;refGene_gn=CABP2;refGene_tx=XM_019812760.2;refGene_gn=
CABP2;refGene_tx=XM_019812761.2;refGene_gn=CABP2;refGene_tx=XM_0198127
62.1;refGene_gn=CABP2;refGene_tx=XM_019812763.2
D1	112373523	112373671
refGene_gn=CABP2;refGene_tx=XM_003993704.5;refGene_gn=CABP2;refGene_tx
=XM_019812759.2;refGene_gn=CABP2;refGene_tx=XM_019812761.2;refGene_gn=
CABP2;refGene_tx=XM_019812762.1;refGene_gn=CABP2;refGene_tx=XM_0198127
63.2
D1	112373785	112373895
refGene_gn=CABP2;refGene_tx=XM_003993704.5;refGene_gn=CABP2;refGene_tx
=XM_019812759.2;refGene_gn=CABP2;refGene_tx=XM_019812760.2;refGene_gn=
CABP2;refGene_tx=XM_019812761.2;refGene_gn=CABP2;refGene_tx=XM_0198127
62.1;refGene_gn=CABP2;refGene_tx=XM_019812763.2
D1	112374173	112374404

refGene_gn=CABP2;refGene_tx=XM_003993704.5;refGene_gn=CABP2;refGene_tx=XM_019812759.2;refGene_gn=CABP2;refGene_tx=XM_019812760.2;refGene_gn=CABP2;refGene_tx=XM_019812761.2;refGene_gn=CABP2;refGene_tx=XM_019812762.1;refGene_gn=CABP2;refGene_tx=XM_019812763.2
D1 112378127 112378158
refGene_gn=CABP2;refGene_tx=XM_003993704.5;refGene_gn=CABP2;refGene_tx=XM_019812759.2;refGene_gn=CABP2;refGene_tx=XM_019812760.2;refGene_gn=CABP2;refGene_tx=XM_019812761.2;refGene_gn=CABP2;refGene_tx=XM_019812762.1;refGene_gn=CABP2;refGene_tx=XM_019812763.2
D1 112378741 112378918
refGene_gn=CABP2;refGene_tx=XM_003993704.5;refGene_gn=CABP2;refGene_tx=XM_019812759.2;refGene_gn=CABP2;refGene_tx=XM_019812760.2
D1 112379479 112379536
refGene_gn=CABP2;refGene_tx=XM_003993704.5;refGene_gn=CABP2;refGene_tx=XM_019812759.2;refGene_gn=CABP2;refGene_tx=XM_019812760.2;refGene_gn=CABP2;refGene_tx=XM_019812761.2;refGene_gn=CABP2;refGene_tx=XM_019812763.2
D1 112380399 112380564
refGene_gn=CABP2;refGene_tx=XM_019812762.1
D1 112418831 112418832
refGene_gn=GSTP1;refGene_tx=XM_023240130.1;refGene_gn=GSTP1;refGene_tx=XM_023240131.1
D1 112419212 112419248
refGene_gn=GSTP1;refGene_tx=XM_023240130.1;refGene_gn=GSTP1;refGene_tx=XM_023240131.1
D1 112419557 112419664
refGene_gn=GSTP1;refGene_tx=XM_023240130.1;refGene_gn=GSTP1;refGene_tx=XM_023240131.1
D1 112419775 112419863
refGene_gn=GSTP1;refGene_tx=XM_023240130.1;refGene_gn=GSTP1;refGene_tx=XM_023240131.1
D1 112420198 112420302
refGene_gn=GSTP1;refGene_tx=XM_023240130.1;refGene_gn=GSTP1;refGene_tx=XM_023240131.1
D1 112420953 112421061
refGene_gn=GSTP1;refGene_tx=XM_023240130.1;refGene_gn=GSTP1;refGene_tx=XM_023240131.1
D1 112421239 112421428
refGene_gn=GSTP1;refGene_tx=XM_023240130.1;refGene_gn=GSTP1;refGene_tx=XM_023240131.1
D1 112433134 112433206
refGene_gn=NDUFV1;refGene_tx=XM_003993705.3
D1 112434291 112434374
refGene_gn=NDUFV1;refGene_tx=XM_003993705.3
D1 112434446 112434617
refGene_gn=NDUFV1;refGene_tx=XM_003993705.3
D1 112435343 112435527
refGene_gn=NDUFV1;refGene_tx=XM_003993705.3
D1 112436360 112436550
refGene_gn=NDUFV1;refGene_tx=XM_003993705.3

D1 112436949 112437162
refGene_gn=NDUFV1;refGene_tx=XM_003993705.3
D1 112437333 112437500
refGene_gn=NDUFV1;refGene_tx=XM_003993705.3
D1 112437737 112437819
refGene_gn=NDUFV1;refGene_tx=XM_003993705.3
D1 112438047 112438193
refGene_gn=NDUFV1;refGene_tx=XM_003993705.3
D1 112438296 112438383
refGene_gn=NDUFV1;refGene_tx=XM_003993705.3
D1 112439018 112439149
refGene_gn=LOC101082437;refGene_tx=XM_011287127.3;refGene_gn=LOC101082
437;refGene_tx=XM_019812769.2
D1 112439249 112439346
refGene_gn=LOC101082437;refGene_tx=XM_011287127.3;refGene_gn=LOC101082
437;refGene_tx=XM_019812769.2
D1 112439520 112439602
refGene_gn=LOC101082437;refGene_tx=XM_011287127.3;refGene_gn=LOC101082
437;refGene_tx=XM_019812769.2
D1 112439696 112439848
refGene_gn=LOC101082437;refGene_tx=XM_011287127.3;refGene_gn=LOC101082
437;refGene_tx=XM_019812769.2
D1 112441038 112441092
refGene_gn=LOC101082437;refGene_tx=XM_011287127.3;refGene_gn=LOC101082
437;refGene_tx=XM_019812769.2
D1 112441218 112441354
refGene_gn=LOC101082437;refGene_tx=XM_011287127.3;refGene_gn=LOC101082
437;refGene_tx=XM_019812769.2
D1 112441449 112441559
refGene_gn=LOC101082437;refGene_tx=XM_011287127.3;refGene_gn=LOC101082
437;refGene_tx=XM_019812769.2
D1 112441774 112441846
refGene_gn=LOC101082437;refGene_tx=XM_011287127.3;refGene_gn=LOC101082
437;refGene_tx=XM_019812769.2
D1 112441922 112442002
refGene_gn=LOC101082437;refGene_tx=XM_011287127.3;refGene_gn=LOC101082
437;refGene_tx=XM_019812769.2
D1 112442174 112442560
refGene_gn=LOC101082437;refGene_tx=XM_011287127.3;refGene_gn=LOC101082
437;refGene_tx=XM_019812769.2
D1 112442677 112442748
refGene_gn=LOC101082437;refGene_tx=XM_011287127.3
D1 112443517 112443706
refGene_gn=LOC101082437;refGene_tx=XM_011287127.3
D1 112444685 112444913
refGene_gn=NUDT8;refGene_tx=XM_023240132.1;refGene_gn=NUDT8;refGene_tx
=XM_023240133.1
D1 112445010 112445088
refGene_gn=NUDT8;refGene_tx=XM_023240132.1;refGene_gn=NUDT8;refGene_tx
=XM_023240133.1

D1	112445322	112445455
refGene_gn=NUDT8;refGene_tx=XM_023240132.1;refGene_gn=NUDT8;refGene_tx=XM_023240133.1
D1	112446091	112446285
refGene_gn=NUDT8;refGene_tx=XM_023240132.1;refGene_gn=NUDT8;refGene_tx=XM_023240133.1
D1	112447839	112448129
refGene_gn=TBX10;refGene_tx=XM_023239752.1
D1	112448571	112448666
refGene_gn=TBX10;refGene_tx=XM_023239752.1
D1	112448863	112448931
refGene_gn=TBX10;refGene_tx=XM_023239752.1
D1	112449232	112449388
refGene_gn=TBX10;refGene_tx=XM_023239752.1
D1	112450221	112450393
refGene_gn=TBX10;refGene_tx=XM_023239752.1
D1	112450776	112450878
refGene_gn=TBX10;refGene_tx=XM_023239752.1
D1	112450962	112451215
refGene_gn=TBX10;refGene_tx=XM_023239752.1
D1	112454824	112454878
refGene_gn=TBX10;refGene_tx=XM_023239752.1
D1	112455197	112455357
refGene_gn=TBX10;refGene_tx=XM_023239752.1
D1	112456359	112456406
refGene_gn=ACY3;refGene_tx=XM_023239753.1
D1	112457314	112457546
refGene_gn=ACY3;refGene_tx=XM_023239753.1
D1	112457547	112457610
refGene_gn=ACY3;refGene_tx=XM_023239753.1
D1	112459169	112459261
refGene_gn=ACY3;refGene_tx=XM_023239753.1
D1	112459696	112459762
refGene_gn=ACY3;refGene_tx=XM_023239753.1
D1	112459961	112460018
refGene_gn=ACY3;refGene_tx=XM_023239753.1
D1	112460020	112460056
refGene_gn=ACY3;refGene_tx=XM_023239753.1
D1	112460196	112460281
refGene_gn=ACY3;refGene_tx=XM_023239753.1
D1	112460554	112460587
refGene_gn=ACY3;refGene_tx=XM_023239753.1
D1	112460588	112460748
refGene_gn=ACY3;refGene_tx=XM_023239753.1
D1	112461445	112461705
refGene_gn=ACY3;refGene_tx=XM_023239753.1
D1	112464502	112464593
refGene_gn=LOC105259663;refGene_tx=XM_023239754.1
D1	112466748	112466961
refGene_gn=LOC105259663;refGene_tx=XM_023239754.1

D1 112471754 112472262
refGene_gn=LOC105259663;refGene_tx=XM_023239754.1
D1 112473975 112474057
refGene_gn=LOC105259663;refGene_tx=XM_023239754.1
D1 112474553 112474639
refGene_gn=LOC105259663;refGene_tx=XM_023239754.1
D1 112474899 112475020
refGene_gn=LOC105259663;refGene_tx=XM_023239754.1
D1 112475131 112475242
refGene_gn=LOC105259663;refGene_tx=XM_023239754.1
D1 112475604 112475682
refGene_gn=LOC105259663;refGene_tx=XM_023239754.1
D1 112477711 112477874
refGene_gn=LOC105259663;refGene_tx=XM_023239754.1
D1 112481252 112481311
refGene_gn=LOC105259663;refGene_tx=XM_023239754.1
D1 112511981 112512323
refGene_gn=UNC93B1;refGene_tx=XM_023240134.1
D1 112513068 112513187
refGene_gn=UNC93B1;refGene_tx=XM_023240134.1
D1 112515084 112515358
refGene_gn=UNC93B1;refGene_tx=XM_023240134.1
D1 112515844 112516027
refGene_gn=UNC93B1;refGene_tx=XM_023240134.1
D1 112516874 112516999
refGene_gn=UNC93B1;refGene_tx=XM_023240134.1
D1 112517553 112517647
refGene_gn=UNC93B1;refGene_tx=XM_023240134.1
D1 112517978 112518111
refGene_gn=UNC93B1;refGene_tx=XM_023240134.1
D1 112518315 112518477
refGene_gn=UNC93B1;refGene_tx=XM_023240134.1
D1 112520754 112520908
refGene_gn=UNC93B1;refGene_tx=XM_023240134.1
D1 112521417 112521559
refGene_gn=UNC93B1;refGene_tx=XM_023240134.1
D1 112521764 112521860
refGene_gn=UNC93B1;refGene_tx=XM_023240134.1
D1 112536061 112536223
refGene_gn=ALDH3B1;refGene_tx=XM_019812775.2;refGene_gn=ALDH3B1;refGene_tx=XM_023240135.1;refGene_gn=ALDH3B1;refGene_tx=XM_023240136.1;refGene_gn=ALDH3B1;refGene_tx=XM_023240137.1;refGene_gn=ALDH3B1;refGene_tx=XM_023240138.1;refGene_gn=ALDH3B1;refGene_tx=XM_023240139.1;refGene_gn=ALDH3B1;refGene_tx=XM_023240140.1
D1 112538911 112538989
refGene_gn=ALDH3B1;refGene_tx=XM_019812775.2;refGene_gn=ALDH3B1;refGene_tx=XM_023240135.1;refGene_gn=ALDH3B1;refGene_tx=XM_023240136.1;refGene_gn=ALDH3B1;refGene_tx=XM_023240137.1;refGene_gn=ALDH3B1;refGene_tx=XM_023240139.1;refGene_gn=ALDH3B1;refGene_tx=XM_023240140.1
D1 112539136 112539220

refGene_gn=ALDH3B1;refGene_tx=XM_011287120.3;refGene_gn=ALDH3B1;refGene_tx=XM_019812779.2
D1 112539351 112539462
refGene_gn=ALDH3B1;refGene_tx=XM_011287120.3;refGene_gn=ALDH3B1;refGene_tx=XM_019812775.2;refGene_gn=ALDH3B1;refGene_tx=XM_019812779.2;refGene_gn=ALDH3B1;refGene_tx=XM_023240135.1;refGene_gn=ALDH3B1;refGene_tx=XM_023240136.1;refGene_gn=ALDH3B1;refGene_tx=XM_023240137.1;refGene_gn=ALDH3B1;refGene_tx=XM_023240138.1;refGene_gn=ALDH3B1;refGene_tx=XM_023240139.1;refGene_gn=ALDH3B1;refGene_tx=XM_023240140.1
D1 112539572 112539693
refGene_gn=ALDH3B1;refGene_tx=XM_011287120.3;refGene_gn=ALDH3B1;refGene_tx=XM_019812775.2;refGene_gn=ALDH3B1;refGene_tx=XM_019812779.2;refGene_gn=ALDH3B1;refGene_tx=XM_023240135.1;refGene_gn=ALDH3B1;refGene_tx=XM_023240136.1;refGene_gn=ALDH3B1;refGene_tx=XM_023240137.1;refGene_gn=ALDH3B1;refGene_tx=XM_023240138.1;refGene_gn=ALDH3B1;refGene_tx=XM_023240139.1;refGene_gn=ALDH3B1;refGene_tx=XM_023240140.1
D1 112539958 112540044
refGene_gn=ALDH3B1;refGene_tx=XM_011287120.3;refGene_gn=ALDH3B1;refGene_tx=XM_019812775.2;refGene_gn=ALDH3B1;refGene_tx=XM_019812779.2;refGene_gn=ALDH3B1;refGene_tx=XM_023240135.1;refGene_gn=ALDH3B1;refGene_tx=XM_023240136.1;refGene_gn=ALDH3B1;refGene_tx=XM_023240137.1;refGene_gn=ALDH3B1;refGene_tx=XM_023240138.1;refGene_gn=ALDH3B1;refGene_tx=XM_023240139.1;refGene_gn=ALDH3B1;refGene_tx=XM_023240140.1
D1 112540540 112540622
refGene_gn=ALDH3B1;refGene_tx=XM_011287120.3;refGene_gn=ALDH3B1;refGene_tx=XM_019812775.2;refGene_gn=ALDH3B1;refGene_tx=XM_019812779.2;refGene_gn=ALDH3B1;refGene_tx=XM_023240135.1;refGene_gn=ALDH3B1;refGene_tx=XM_023240136.1;refGene_gn=ALDH3B1;refGene_tx=XM_023240137.1;refGene_gn=ALDH3B1;refGene_tx=XM_023240138.1;refGene_gn=ALDH3B1;refGene_tx=XM_023240139.1;refGene_gn=ALDH3B1;refGene_tx=XM_023240140.1
D1 112542328 112542715
refGene_gn=ALDH3B1;refGene_tx=XM_011287120.3;refGene_gn=ALDH3B1;refGene_tx=XM_011287121.3;refGene_gn=ALDH3B1;refGene_tx=XM_019812775.2;refGene_gn=ALDH3B1;refGene_tx=XM_019812779.2;refGene_gn=ALDH3B1;refGene_tx=XM_023240135.1;refGene_gn=ALDH3B1;refGene_tx=XM_023240136.1;refGene_gn=ALDH3B1;refGene_tx=XM_023240137.1;refGene_gn=ALDH3B1;refGene_tx=XM_023240138.1;refGene_gn=ALDH3B1;refGene_tx=XM_023240139.1;refGene_gn=ALDH3B1;refGene_tx=XM_023240140.1
D1 112543845 112544012
refGene_gn=ALDH3B1;refGene_tx=XM_011287120.3;refGene_gn=ALDH3B1;refGene_tx=XM_011287121.3;refGene_gn=ALDH3B1;refGene_tx=XM_019812775.2;refGene_gn=ALDH3B1;refGene_tx=XM_019812779.2;refGene_gn=ALDH3B1;refGene_tx=XM_023240135.1;refGene_gn=ALDH3B1;refGene_tx=XM_023240136.1;refGene_gn=ALDH3B1;refGene_tx=XM_023240137.1;refGene_gn=ALDH3B1;refGene_tx=XM_023240138.1;refGene_gn=ALDH3B1;refGene_tx=XM_023240139.1;refGene_gn=ALDH3B1;refGene_tx=XM_023240140.1
D1 112544733 112544763
refGene_gn=ALDH3B1;refGene_tx=XM_023240139.1;refGene_gn=ALDH3B1;refGene_tx=XM_023240140.1
D1 112546728 112546828

refGene_gn=ALDH3B1;refGene_tx=XM_011287120.3;refGene_gn=ALDH3B1;refGene_tx=XM_011287121.3;refGene_gn=ALDH3B1;refGene_tx=XM_019812775.2;refGene_gn=ALDH3B1;refGene_tx=XM_019812779.2;refGene_gn=ALDH3B1;refGene_tx=XM_023240135.1;refGene_gn=ALDH3B1;refGene_tx=XM_023240136.1;refGene_gn=ALDH3B1;refGene_tx=XM_023240137.1;refGene_gn=ALDH3B1;refGene_tx=XM_023240138.1
D1 112548192 112548383
refGene_gn=ALDH3B1;refGene_tx=XM_011287120.3;refGene_gn=ALDH3B1;refGene_tx=XM_011287121.3;refGene_gn=ALDH3B1;refGene_tx=XM_019812775.2;refGene_gn=ALDH3B1;refGene_tx=XM_019812779.2;refGene_gn=ALDH3B1;refGene_tx=XM_023240135.1;refGene_gn=ALDH3B1;refGene_tx=XM_023240136.1;refGene_gn=ALDH3B1;refGene_tx=XM_023240137.1;refGene_gn=ALDH3B1;refGene_tx=XM_023240138.1
D1 112550473 112550542
refGene_gn=NDUFS8;refGene_tx=XM_019812780.2
D1 112551281 112551339
refGene_gn=NDUFS8;refGene_tx=XM_003993710.4;refGene_gn=NDUFS8;refGene_tx=XM_006937584.4;refGene_gn=NDUFS8;refGene_tx=XM_019812780.2
D1 112551429 112551480
refGene_gn=NDUFS8;refGene_tx=XM_003993710.4;refGene_gn=NDUFS8;refGene_tx=XM_006937584.4;refGene_gn=NDUFS8;refGene_tx=XM_019812780.2
D1 112552310 112552400
refGene_gn=NDUFS8;refGene_tx=XM_003993710.4;refGene_gn=NDUFS8;refGene_tx=XM_006937584.4;refGene_gn=NDUFS8;refGene_tx=XM_019812780.2
D1 112552499 112552672
refGene_gn=NDUFS8;refGene_tx=XM_003993710.4;refGene_gn=NDUFS8;refGene_tx=XM_006937584.4;refGene_gn=NDUFS8;refGene_tx=XM_019812780.2
D1 112554106 112554235
refGene_gn=NDUFS8;refGene_tx=XM_003993710.4;refGene_gn=NDUFS8;refGene_tx=XM_006937584.4;refGene_gn=NDUFS8;refGene_tx=XM_019812780.2
D1 112554317 112554449
refGene_gn=NDUFS8;refGene_tx=XM_003993710.4;refGene_gn=NDUFS8;refGene_tx=XM_006937584.4;refGene_gn=NDUFS8;refGene_tx=XM_019812780.2
D1 112559579 112559696
refGene_gn=TCIRG1;refGene_tx=XM_023240141.1;refGene_gn=TCIRG1;refGene_tx=XM_023240142.1
D1 112560070 112560149
refGene_gn=TCIRG1;refGene_tx=XM_023240141.1;refGene_gn=TCIRG1;refGene_tx=XM_023240142.1
D1 112560804 112561025
refGene_gn=TCIRG1;refGene_tx=XM_023240141.1;refGene_gn=TCIRG1;refGene_tx=XM_023240142.1
D1 112561122 112561208
refGene_gn=TCIRG1;refGene_tx=XM_023240141.1;refGene_gn=TCIRG1;refGene_tx=XM_023240142.1
D1 112561728 112561855
refGene_gn=TCIRG1;refGene_tx=XM_023240141.1;refGene_gn=TCIRG1;refGene_tx=XM_023240142.1
D1 112561936 112562019
refGene_gn=TCIRG1;refGene_tx=XM_023240141.1;refGene_gn=TCIRG1;refGene_

tx=XM_023240142.1
D1 112562172 112562266
refGene_gn=TCIRG1;refGene_tx=XM_023240141.1;refGene_gn=TCIRG1;refGene_tx=XM_023240142.1
D1 112562346 112562559
refGene_gn=TCIRG1;refGene_tx=XM_023240141.1;refGene_gn=TCIRG1;refGene_tx=XM_023240142.1
D1 112563080 112563225
refGene_gn=TCIRG1;refGene_tx=XM_023240141.1;refGene_gn=TCIRG1;refGene_tx=XM_023240142.1
D1 112565409 112565549
refGene_gn=TCIRG1;refGene_tx=XM_023240141.1;refGene_gn=TCIRG1;refGene_tx=XM_023240142.1
D1 112565625 112565783
refGene_gn=TCIRG1;refGene_tx=XM_023240141.1;refGene_gn=TCIRG1;refGene_tx=XM_023240142.1
D1 112565864 112565955
refGene_gn=TCIRG1;refGene_tx=XM_023240141.1;refGene_gn=TCIRG1;refGene_tx=XM_023240142.1
D1 112566522 112566641
refGene_gn=TCIRG1;refGene_tx=XM_023240141.1;refGene_gn=TCIRG1;refGene_tx=XM_023240142.1
D1 112566732 112566946
refGene_gn=TCIRG1;refGene_tx=XM_023240141.1;refGene_gn=TCIRG1;refGene_tx=XM_023240142.1
D1 112567221 112567350
refGene_gn=TCIRG1;refGene_tx=XM_023240141.1;refGene_gn=TCIRG1;refGene_tx=XM_023240142.1
D1 112567494 112567599
refGene_gn=TCIRG1;refGene_tx=XM_023240141.1;refGene_gn=TCIRG1;refGene_tx=XM_023240142.1
D1 112567678 112567796
refGene_gn=TCIRG1;refGene_tx=XM_023240141.1;refGene_gn=TCIRG1;refGene_tx=XM_023240142.1
D1 112568042 112568220
refGene_gn=TCIRG1;refGene_tx=XM_023240141.1;refGene_gn=TCIRG1;refGene_tx=XM_023240142.1
D1 112568318 112568397
refGene_gn=TCIRG1;refGene_tx=XM_023240141.1;refGene_gn=TCIRG1;refGene_tx=XM_023240142.1
D1 112570785 112571041
refGene_gn=CHKA;refGene_tx=XM_023240143.1;refGene_gn=CHKA;refGene_tx=XM_023240144.1;refGene_gn=CHKA;refGene_tx=XM_023240146.1;refGene_gn=CHKA;refGene_tx=XM_023240147.1;refGene_gn=CHKA;refGene_tx=XM_023240148.1;refGene_gn=CHKA;refGene_tx=XM_023240149.1;refGene_gn=CHKA;refGene_tx=XM_023240150.1;refGene_gn=CHKA;refGene_tx=XM_023240151.1;refGene_gn=CHKA;refGene_tx=XM_023240152.1;refGene_gn=CHKA;refGene_tx=XM_023240154.1
D1 112577354 112577465
refGene_gn=CHKA;refGene_tx=XM_023240145.1
D1 112577774 112577856

refGene_gn=CHKA;refGene_tx=XM_023240145.1;refGene_gn=CHKA;refGene_tx=XM_023240147.1;refGene_gn=CHKA;refGene_tx=XM_023240149.1;refGene_gn=CHKA;refGene_tx=XM_023240151.1
D1 112579250 112579357
refGene_gn=CHKA;refGene_tx=XM_023240143.1;refGene_gn=CHKA;refGene_tx=XM_023240144.1;refGene_gn=CHKA;refGene_tx=XM_023240145.1;refGene_gn=CHKA;refGene_tx=XM_023240146.1;refGene_gn=CHKA;refGene_tx=XM_023240147.1;refGene_gn=CHKA;refGene_tx=XM_023240148.1;refGene_gn=CHKA;refGene_tx=XM_023240149.1;refGene_gn=CHKA;refGene_tx=XM_023240150.1;refGene_gn=CHKA;refGene_tx=XM_023240151.1;refGene_gn=CHKA;refGene_tx=XM_023240152.1
D1 112581366 112581459
refGene_gn=CHKA;refGene_tx=XM_023240153.1
D1 112581718 112581745
refGene_gn=CHKA;refGene_tx=XM_023240155.1
D1 112582075 112582184
refGene_gn=CHKA;refGene_tx=XM_023240143.1;refGene_gn=CHKA;refGene_tx=XM_023240144.1;refGene_gn=CHKA;refGene_tx=XM_023240145.1;refGene_gn=CHKA;refGene_tx=XM_023240146.1;refGene_gn=CHKA;refGene_tx=XM_023240147.1;refGene_gn=CHKA;refGene_tx=XM_023240148.1;refGene_gn=CHKA;refGene_tx=XM_023240149.1;refGene_gn=CHKA;refGene_tx=XM_023240150.1;refGene_gn=CHKA;refGene_tx=XM_023240151.1;refGene_gn=CHKA;refGene_tx=XM_023240152.1;refGene_gn=CHKA;refGene_tx=XM_023240153.1;refGene_gn=CHKA;refGene_tx=XM_023240154.1;refGene_gn=CHKA;refGene_tx=XM_023240155.1
D1 112582946 112583034
refGene_gn=CHKA;refGene_tx=XM_023240143.1;refGene_gn=CHKA;refGene_tx=XM_023240144.1;refGene_gn=CHKA;refGene_tx=XM_023240145.1;refGene_gn=CHKA;refGene_tx=XM_023240146.1;refGene_gn=CHKA;refGene_tx=XM_023240147.1;refGene_gn=CHKA;refGene_tx=XM_023240148.1;refGene_gn=CHKA;refGene_tx=XM_023240149.1;refGene_gn=CHKA;refGene_tx=XM_023240150.1;refGene_gn=CHKA;refGene_tx=XM_023240151.1;refGene_gn=CHKA;refGene_tx=XM_023240152.1;refGene_gn=CHKA;refGene_tx=XM_023240153.1;refGene_gn=CHKA;refGene_tx=XM_023240154.1;refGene_gn=CHKA;refGene_tx=XM_023240155.1
D1 112584879 112584938
refGene_gn=CHKA;refGene_tx=XM_023240143.1;refGene_gn=CHKA;refGene_tx=XM_023240144.1;refGene_gn=CHKA;refGene_tx=XM_023240145.1;refGene_gn=CHKA;refGene_tx=XM_023240146.1;refGene_gn=CHKA;refGene_tx=XM_023240147.1;refGene_gn=CHKA;refGene_tx=XM_023240148.1;refGene_gn=CHKA;refGene_tx=XM_023240149.1;refGene_gn=CHKA;refGene_tx=XM_023240150.1;refGene_gn=CHKA;refGene_tx=XM_023240151.1;refGene_gn=CHKA;refGene_tx=XM_023240152.1;refGene_gn=CHKA;refGene_tx=XM_023240153.1;refGene_gn=CHKA;refGene_tx=XM_023240154.1;refGene_gn=CHKA;refGene_tx=XM_023240155.1
D1 112585625 112585730
refGene_gn=CHKA;refGene_tx=XM_023240143.1;refGene_gn=CHKA;refGene_tx=XM_023240144.1;refGene_gn=CHKA;refGene_tx=XM_023240145.1;refGene_gn=CHKA;refGene_tx=XM_023240146.1;refGene_gn=CHKA;refGene_tx=XM_023240147.1;refGene_gn=CHKA;refGene_tx=XM_023240148.1;refGene_gn=CHKA;refGene_tx=XM_023240149.1;refGene_gn=CHKA;refGene_tx=XM_023240150.1;refGene_gn=CHKA;refGene_tx=XM_023240151.1;refGene_gn=CHKA;refGene_tx=XM_023240152.1;refGene_gn=CHKA;refGene_tx=XM_023240153.1;refGene_gn=CHKA;refGene_tx=XM_023240154.1;refGene_gn=CHKA;refGene_tx=XM_023240155.1

D1	112585982	112586116
refGene_gn=CHKA;refGene_tx=XM_023240143.1;refGene_gn=CHKA;refGene_tx=XM_023240144.1;refGene_gn=CHKA;refGene_tx=XM_023240145.1;refGene_gn=CHKA;refGene_tx=XM_023240146.1;refGene_gn=CHKA;refGene_tx=XM_023240147.1;refGene_gn=CHKA;refGene_tx=XM_023240148.1;refGene_gn=CHKA;refGene_tx=XM_023240149.1;refGene_gn=CHKA;refGene_tx=XM_023240150.1;refGene_gn=CHKA;refGene_tx=XM_023240151.1;refGene_gn=CHKA;refGene_tx=XM_023240152.1;refGene_gn=CHKA;refGene_tx=XM_023240153.1;refGene_gn=CHKA;refGene_tx=XM_023240154.1;refGene_gn=CHKA;refGene_tx=XM_023240155.1
D1	112589053	112589167
refGene_gn=CHKA;refGene_tx=XM_023240143.1;refGene_gn=CHKA;refGene_tx=XM_023240144.1;refGene_gn=CHKA;refGene_tx=XM_023240145.1;refGene_gn=CHKA;refGene_tx=XM_023240146.1;refGene_gn=CHKA;refGene_tx=XM_023240147.1;refGene_gn=CHKA;refGene_tx=XM_023240149.1;refGene_gn=CHKA;refGene_tx=XM_023240151.1;refGene_gn=CHKA;refGene_tx=XM_023240153.1;refGene_gn=CHKA;refGene_tx=XM_023240154.1;refGene_gn=CHKA;refGene_tx=XM_023240155.1
D1	112597057	112597111
refGene_gn=CHKA;refGene_tx=XM_023240143.1;refGene_gn=CHKA;refGene_tx=XM_023240144.1;refGene_gn=CHKA;refGene_tx=XM_023240145.1;refGene_gn=CHKA;refGene_tx=XM_023240147.1;refGene_gn=CHKA;refGene_tx=XM_023240148.1;refGene_gn=CHKA;refGene_tx=XM_023240149.1;refGene_gn=CHKA;refGene_tx=XM_023240150.1;refGene_gn=CHKA;refGene_tx=XM_023240153.1;refGene_gn=CHKA;refGene_tx=XM_023240154.1;refGene_gn=CHKA;refGene_tx=XM_023240155.1
D1	112598133	112598163
refGene_gn=CHKA;refGene_tx=XM_023240143.1;refGene_gn=CHKA;refGene_tx=XM_023240145.1;refGene_gn=CHKA;refGene_tx=XM_023240147.1;refGene_gn=CHKA;refGene_tx=XM_023240148.1;refGene_gn=CHKA;refGene_tx=XM_023240153.1;refGene_gn=CHKA;refGene_tx=XM_023240154.1;refGene_gn=CHKA;refGene_tx=XM_023240155.1
D1	112610589	112610701
refGene_gn=CHKA;refGene_tx=XM_023240143.1;refGene_gn=CHKA;refGene_tx=XM_023240144.1;refGene_gn=CHKA;refGene_tx=XM_023240145.1;refGene_gn=CHKA;refGene_tx=XM_023240146.1;refGene_gn=CHKA;refGene_tx=XM_023240147.1;refGene_gn=CHKA;refGene_tx=XM_023240148.1;refGene_gn=CHKA;refGene_tx=XM_023240149.1;refGene_gn=CHKA;refGene_tx=XM_023240150.1;refGene_gn=CHKA;refGene_tx=XM_023240151.1;refGene_gn=CHKA;refGene_tx=XM_023240152.1;refGene_gn=CHKA;refGene_tx=XM_023240153.1;refGene_gn=CHKA;refGene_tx=XM_023240154.1;refGene_gn=CHKA;refGene_tx=XM_023240155.1
D1	112635021	112635377
refGene_gn=CHKA;refGene_tx=XM_023240143.1;refGene_gn=CHKA;refGene_tx=XM_023240144.1;refGene_gn=CHKA;refGene_tx=XM_023240145.1;refGene_gn=CHKA;refGene_tx=XM_023240146.1;refGene_gn=CHKA;refGene_tx=XM_023240147.1;refGene_gn=CHKA;refGene_tx=XM_023240148.1;refGene_gn=CHKA;refGene_tx=XM_023240149.1;refGene_gn=CHKA;refGene_tx=XM_023240150.1;refGene_gn=CHKA;refGene_tx=XM_023240151.1;refGene_gn=CHKA;refGene_tx=XM_023240152.1;refGene_gn=CHKA;refGene_tx=XM_023240153.1;refGene_gn=CHKA;refGene_tx=XM_023240154.1;refGene_gn=CHKA;refGene_tx=XM_023240155.1
D1	112659443	112660912
refGene_gn=KMT5B;refGene_tx=XM_019812787.1;refGene_gn=KMT5B;refGene_tx=XM_023240156.1;refGene_gn=KMT5B;refGene_tx=XM_023240157.1;refGene_gn=

KMT5B;refGene_tx=XM_023240158.1
D1 112667693 112667898
refGene_gn=KMT5B;refGene_tx=XM_019812787.1;refGene_gn=KMT5B;refGene_tx=XM_023240156.1;refGene_gn=KMT5B;refGene_tx=XM_023240157.1;refGene_gn=KMT5B;refGene_tx=XM_023240158.1;refGene_gn=KMT5B;refGene_tx=XM_023240159.1
D1 112674407 112674544
refGene_gn=KMT5B;refGene_tx=XM_019812787.1;refGene_gn=KMT5B;refGene_tx=XM_023240156.1;refGene_gn=KMT5B;refGene_tx=XM_023240157.1;refGene_gn=KMT5B;refGene_tx=XM_023240158.1;refGene_gn=KMT5B;refGene_tx=XM_023240159.1
D1 112674625 112674645
refGene_gn=KMT5B;refGene_tx=XM_019812787.1;refGene_gn=KMT5B;refGene_tx=XM_023240156.1;refGene_gn=KMT5B;refGene_tx=XM_023240157.1;refGene_gn=KMT5B;refGene_tx=XM_023240158.1;refGene_gn=KMT5B;refGene_tx=XM_023240159.1
D1 112674939 112675106
refGene_gn=KMT5B;refGene_tx=XM_019812787.1;refGene_gn=KMT5B;refGene_tx=XM_023240156.1;refGene_gn=KMT5B;refGene_tx=XM_023240157.1;refGene_gn=KMT5B;refGene_tx=XM_023240158.1;refGene_gn=KMT5B;refGene_tx=XM_023240159.1
D1 112676619 112676729
refGene_gn=KMT5B;refGene_tx=XM_023240156.1;refGene_gn=KMT5B;refGene_tx=XM_023240157.1;refGene_gn=KMT5B;refGene_tx=XM_023240158.1;refGene_gn=KMT5B;refGene_tx=XM_023240159.1
D1 112677113 112677279
refGene_gn=KMT5B;refGene_tx=XM_023240156.1;refGene_gn=KMT5B;refGene_tx=XM_023240157.1;refGene_gn=KMT5B;refGene_tx=XM_023240158.1;refGene_gn=KMT5B;refGene_tx=XM_023240159.1
D1 112681504 112681573
refGene_gn=KMT5B;refGene_tx=XM_023240156.1;refGene_gn=KMT5B;refGene_tx=XM_023240157.1;refGene_gn=KMT5B;refGene_tx=XM_023240159.1
D1 112686533 112686681
refGene_gn=KMT5B;refGene_tx=XM_023240156.1;refGene_gn=KMT5B;refGene_tx=XM_023240157.1;refGene_gn=KMT5B;refGene_tx=XM_023240158.1;refGene_gn=KMT5B;refGene_tx=XM_023240159.1
D1 112692421 112692581
refGene_gn=KMT5B;refGene_tx=XM_023240156.1;refGene_gn=KMT5B;refGene_tx=XM_023240157.1;refGene_gn=KMT5B;refGene_tx=XM_023240158.1;refGene_gn=KMT5B;refGene_tx=XM_023240159.1
D1 112788719 112788783
refGene_gn=LRP5;refGene_tx=XM_023240160.1
D1 112818561 112818958
refGene_gn=LRP5;refGene_tx=XM_023240160.1
D1 112825471 112825669
refGene_gn=LRP5;refGene_tx=XM_023240160.1
D1 112832010 112832207
refGene_gn=LRP5;refGene_tx=XM_023240160.1
D1 112833334 112833466
refGene_gn=LRP5;refGene_tx=XM_023240160.1

D1 112849668 112850065
refGene_gn=LRP5;refGene_tx=XM_023240160.1
D1 112851515 112851687
refGene_gn=LRP5;refGene_tx=XM_023240160.1
D1 112861227 112861444
refGene_gn=LRP5;refGene_tx=XM_023240160.1
D1 112863505 112863795
refGene_gn=LRP5;refGene_tx=XM_023240160.1
D1 112864535 112864762
refGene_gn=LRP5;refGene_tx=XM_023240160.1
D1 112866108 112866293
refGene_gn=LRP5;refGene_tx=XM_023240160.1
D1 112868582 112868906
refGene_gn=LRP5;refGene_tx=XM_023240160.1
D1 112870128 112870328
refGene_gn=LRP5;refGene_tx=XM_023240160.1
D1 112874785 112874994
refGene_gn=LRP5;refGene_tx=XM_023240160.1
D1 112876160 112876351
refGene_gn=LRP5;refGene_tx=XM_023240160.1
D1 112876961 112877171
refGene_gn=LRP5;refGene_tx=XM_023240160.1
D1 112880042 112880168
refGene_gn=LRP5;refGene_tx=XM_023240160.1
D1 112883170 112883407
refGene_gn=LRP5;refGene_tx=XM_023240160.1
D1 112885503 112885614
refGene_gn=LRP5;refGene_tx=XM_023240160.1
D1 112887311 112887548
refGene_gn=LRP5;refGene_tx=XM_023240160.1
D1 112887731 112887871
refGene_gn=LRP5;refGene_tx=XM_023240160.1
D1 112892357 112892455
refGene_gn=LRP5;refGene_tx=XM_023240160.1
D1 112894386 112894648
refGene_gn=LRP5;refGene_tx=XM_023240160.1
D1 112971508 112971735
refGene_gn=PPP6R3;refGene_tx=XM_003993712.5;refGene_gn=PPP6R3;refGene_tx=XM_003993713.5;refGene_gn=PPP6R3;refGene_tx=XM_003993714.5;refGene_gn=PPP6R3;refGene_tx=XM_006937588.3;refGene_gn=PPP6R3;refGene_tx=XM_006937593.3;refGene_gn=PPP6R3;refGene_tx=XM_006937597.4;refGene_gn=PPP6R3;refGene_tx=XM_006937598.4;refGene_gn=PPP6R3;refGene_tx=XM_019812791.2;refGene_gn=PPP6R3;refGene_tx=XM_019812792.2;refGene_gn=PPP6R3;refGene_tx=XM_019812793.1;refGene_gn=PPP6R3;refGene_tx=XM_019812794.2;refGene_gn=PPP6R3;refGene_tx=XM_019812795.2;refGene_gn=PPP6R3;refGene_tx=XM_019812796.2;refGene_gn=PPP6R3;refGene_tx=XM_019812797.2;refGene_gn=PPP6R3;refGene_tx=XM_019812798.2;refGene_gn=PPP6R3;refGene_tx=XM_019812799.2;refGene_gn=PPP6R3;refGene_tx=XM_019812800.2;refGene_gn=PPP6R3;refGene_tx=XM_023240161.1;refGene_gn=PPP6R3;refGene_tx=XM_023240162.1;refGene_gn=PPP6R3;refGene_tx=XM_023240163.1;refGene_gn=PPP6R3;refGene_tx=X

M_023240164.1;refGene_gn=PPP6R3;refGene_tx=XM_023240165.1;refGene_gn=PPP6R3;refGene_tx=XM_023240166.1
D1 112976309 112976496
refGene_gn=PPP6R3;refGene_tx=XM_003993712.5;refGene_gn=PPP6R3;refGene_tx=XM_003993713.5;refGene_gn=PPP6R3;refGene_tx=XM_003993714.5;refGene_gn=PPP6R3;refGene_tx=XM_006937588.3;refGene_gn=PPP6R3;refGene_tx=XM_006937593.3;refGene_gn=PPP6R3;refGene_tx=XM_006937597.4;refGene_gn=PPP6R3;refGene_tx=XM_006937598.4;refGene_gn=PPP6R3;refGene_tx=XM_019812791.2;refGene_gn=PPP6R3;refGene_tx=XM_019812792.2;refGene_gn=PPP6R3;refGene_tx=XM_019812793.1;refGene_gn=PPP6R3;refGene_tx=XM_019812794.2;refGene_gn=PPP6R3;refGene_tx=XM_019812795.2;refGene_gn=PPP6R3;refGene_tx=XM_019812796.2;refGene_gn=PPP6R3;refGene_tx=XM_019812797.2;refGene_gn=PPP6R3;refGene_tx=XM_019812798.2;refGene_gn=PPP6R3;refGene_tx=XM_019812799.2;refGene_gn=PPP6R3;refGene_tx=XM_019812800.2;refGene_gn=PPP6R3;refGene_tx=XM_019812801.2;refGene_gn=PPP6R3;refGene_tx=XM_023240161.1;refGene_gn=PPP6R3;refGene_tx=XM_023240162.1;refGene_gn=PPP6R3;refGene_tx=XM_023240163.1;refGene_gn=PPP6R3;refGene_tx=XM_023240164.1;refGene_gn=PPP6R3;refGene_tx=XM_023240165.1;refGene_gn=PPP6R3;refGene_tx=XM_023240166.1
D1 112979598 112979736
refGene_gn=PPP6R3;refGene_tx=XM_003993712.5;refGene_gn=PPP6R3;refGene_tx=XM_003993713.5;refGene_gn=PPP6R3;refGene_tx=XM_003993714.5;refGene_gn=PPP6R3;refGene_tx=XM_006937588.3;refGene_gn=PPP6R3;refGene_tx=XM_006937593.3;refGene_gn=PPP6R3;refGene_tx=XM_006937597.4;refGene_gn=PPP6R3;refGene_tx=XM_006937598.4;refGene_gn=PPP6R3;refGene_tx=XM_019812791.2;refGene_gn=PPP6R3;refGene_tx=XM_019812792.2;refGene_gn=PPP6R3;refGene_tx=XM_019812793.1;refGene_gn=PPP6R3;refGene_tx=XM_019812794.2;refGene_gn=PPP6R3;refGene_tx=XM_019812795.2;refGene_gn=PPP6R3;refGene_tx=XM_019812796.2;refGene_gn=PPP6R3;refGene_tx=XM_019812797.2;refGene_gn=PPP6R3;refGene_tx=XM_019812798.2;refGene_gn=PPP6R3;refGene_tx=XM_019812799.2;refGene_gn=PPP6R3;refGene_tx=XM_019812800.2;refGene_gn=PPP6R3;refGene_tx=XM_019812801.2;refGene_gn=PPP6R3;refGene_tx=XM_019812802.2;refGene_gn=PPP6R3;refGene_tx=XM_023240161.1;refGene_gn=PPP6R3;refGene_tx=XM_023240162.1;refGene_gn=PPP6R3;refGene_tx=XM_023240163.1;refGene_gn=PPP6R3;refGene_tx=XM_023240164.1;refGene_gn=PPP6R3;refGene_tx=XM_023240165.1;refGene_gn=PPP6R3;refGene_tx=XM_023240166.1
D1 112981848 112981914
refGene_gn=PPP6R3;refGene_tx=XM_003993712.5;refGene_gn=PPP6R3;refGene_tx=XM_003993713.5;refGene_gn=PPP6R3;refGene_tx=XM_003993714.5;refGene_gn=PPP6R3;refGene_tx=XM_006937588.3;refGene_gn=PPP6R3;refGene_tx=XM_006937593.3;refGene_gn=PPP6R3;refGene_tx=XM_006937597.4;refGene_gn=PPP6R3;refGene_tx=XM_006937598.4;refGene_gn=PPP6R3;refGene_tx=XM_019812791.2;refGene_gn=PPP6R3;refGene_tx=XM_019812792.2;refGene_gn=PPP6R3;refGene_tx=XM_019812793.1;refGene_gn=PPP6R3;refGene_tx=XM_019812794.2;refGene_gn=PPP6R3;refGene_tx=XM_019812795.2;refGene_gn=PPP6R3;refGene_tx=XM_019812796.2;refGene_gn=PPP6R3;refGene_tx=XM_019812797.2;refGene_gn=PPP6R3;refGene_tx=XM_019812798.2;refGene_gn=PPP6R3;refGene_tx=XM_019812799.2;refGene_gn=PPP6R3;refGene_tx=XM_019812800.2;refGene_gn=PPP6R3;refGene_tx=XM_019812801.2;refGene_gn=PPP6R3;refGene_tx=XM_019812802.2;refGene_gn=PPP6R3;refGene_tx=XM_023240161.1;refGene_gn=PPP6R3;refGene_tx=X

M_023240162.1;refGene_gn=PPP6R3;refGene_tx=XM_023240163.1;refGene_gn=PPP6R3;refGene_tx=XM_023240164.1;refGene_gn=PPP6R3;refGene_tx=XM_023240165.1;refGene_gn=PPP6R3;refGene_tx=XM_023240166.1
D1 112984671 112984784
refGene_gn=PPP6R3;refGene_tx=XM_003993712.5;refGene_gn=PPP6R3;refGene_tx=XM_003993713.5;refGene_gn=PPP6R3;refGene_tx=XM_003993714.5;refGene_gn=PPP6R3;refGene_tx=XM_006937588.3;refGene_gn=PPP6R3;refGene_tx=XM_006937593.3;refGene_gn=PPP6R3;refGene_tx=XM_006937597.4;refGene_gn=PPP6R3;refGene_tx=XM_006937598.4;refGene_gn=PPP6R3;refGene_tx=XM_019812791.2;refGene_gn=PPP6R3;refGene_tx=XM_019812792.2;refGene_gn=PPP6R3;refGene_tx=XM_019812793.1;refGene_gn=PPP6R3;refGene_tx=XM_019812794.2;refGene_gn=PPP6R3;refGene_tx=XM_019812795.2;refGene_gn=PPP6R3;refGene_tx=XM_019812796.2;refGene_gn=PPP6R3;refGene_tx=XM_019812797.2;refGene_gn=PPP6R3;refGene_tx=XM_019812798.2;refGene_gn=PPP6R3;refGene_tx=XM_019812799.2;refGene_gn=PPP6R3;refGene_tx=XM_019812800.2;refGene_gn=PPP6R3;refGene_tx=XM_019812801.2;refGene_gn=PPP6R3;refGene_tx=XM_019812802.2;refGene_gn=PPP6R3;refGene_tx=XM_023240161.1;refGene_gn=PPP6R3;refGene_tx=XM_023240162.1;refGene_gn=PPP6R3;refGene_tx=XM_023240163.1;refGene_gn=PPP6R3;refGene_tx=XM_023240164.1;refGene_gn=PPP6R3;refGene_tx=XM_023240165.1;refGene_gn=PPP6R3;refGene_tx=XM_023240166.1
D1 112988295 112988409
refGene_gn=PPP6R3;refGene_tx=XM_003993712.5;refGene_gn=PPP6R3;refGene_tx=XM_003993713.5;refGene_gn=PPP6R3;refGene_tx=XM_003993714.5;refGene_gn=PPP6R3;refGene_tx=XM_006937588.3;refGene_gn=PPP6R3;refGene_tx=XM_006937593.3;refGene_gn=PPP6R3;refGene_tx=XM_006937597.4;refGene_gn=PPP6R3;refGene_tx=XM_006937598.4;refGene_gn=PPP6R3;refGene_tx=XM_019812791.2;refGene_gn=PPP6R3;refGene_tx=XM_019812792.2;refGene_gn=PPP6R3;refGene_tx=XM_019812793.1;refGene_gn=PPP6R3;refGene_tx=XM_019812794.2;refGene_gn=PPP6R3;refGene_tx=XM_019812795.2;refGene_gn=PPP6R3;refGene_tx=XM_019812796.2;refGene_gn=PPP6R3;refGene_tx=XM_019812797.2;refGene_gn=PPP6R3;refGene_tx=XM_019812798.2;refGene_gn=PPP6R3;refGene_tx=XM_019812799.2;refGene_gn=PPP6R3;refGene_tx=XM_019812800.2;refGene_gn=PPP6R3;refGene_tx=XM_019812801.2;refGene_gn=PPP6R3;refGene_tx=XM_019812802.2;refGene_gn=PPP6R3;refGene_tx=XM_023240161.1;refGene_gn=PPP6R3;refGene_tx=XM_023240162.1;refGene_gn=PPP6R3;refGene_tx=XM_023240163.1;refGene_gn=PPP6R3;refGene_tx=XM_023240164.1;refGene_gn=PPP6R3;refGene_tx=XM_023240165.1;refGene_gn=PPP6R3;refGene_tx=XM_023240166.1
D1 112998443 112998573
refGene_gn=PPP6R3;refGene_tx=XM_003993712.5;refGene_gn=PPP6R3;refGene_tx=XM_003993713.5;refGene_gn=PPP6R3;refGene_tx=XM_003993714.5;refGene_gn=PPP6R3;refGene_tx=XM_006937588.3;refGene_gn=PPP6R3;refGene_tx=XM_006937593.3;refGene_gn=PPP6R3;refGene_tx=XM_006937597.4;refGene_gn=PPP6R3;refGene_tx=XM_006937598.4;refGene_gn=PPP6R3;refGene_tx=XM_019812791.2;refGene_gn=PPP6R3;refGene_tx=XM_019812792.2;refGene_gn=PPP6R3;refGene_tx=XM_019812793.1;refGene_gn=PPP6R3;refGene_tx=XM_019812794.2;refGene_gn=PPP6R3;refGene_tx=XM_019812795.2;refGene_gn=PPP6R3;refGene_tx=XM_019812796.2;refGene_gn=PPP6R3;refGene_tx=XM_019812797.2;refGene_gn=PPP6R3;refGene_tx=XM_019812798.2;refGene_gn=PPP6R3;refGene_tx=XM_019812799.2;refGene_gn=PPP6R3;refGene_tx=XM_019812800.2;refGene_gn=PPP6R3;refGene_tx=XM_019812801.2;refGene_gn=PPP6R3;refGene_tx=XM_019812802.2;refG

ene_gn=PPP6R3;refGene_tx=XM_023240161.1;refGene_gn=PPP6R3;refGene_tx=X
M_023240162.1;refGene_gn=PPP6R3;refGene_tx=XM_023240163.1;refGene_gn=P
PP6R3;refGene_tx=XM_023240164.1;refGene_gn=PPP6R3;refGene_tx=XM_023240
165.1;refGene_gn=PPP6R3;refGene_tx=XM_023240166.1
D1 113000668 113000821
refGene_gn=PPP6R3;refGene_tx=XM_003993712.5;refGene_gn=PPP6R3;refGene_
tx=XM_003993713.5;refGene_gn=PPP6R3;refGene_tx=XM_003993714.5;refGene_
gn=PPP6R3;refGene_tx=XM_006937588.3;refGene_gn=PPP6R3;refGene_tx=XM_00
6937593.3;refGene_gn=PPP6R3;refGene_tx=XM_006937597.4;refGene_gn=PPP6R
3;refGene_tx=XM_006937598.4;refGene_gn=PPP6R3;refGene_tx=XM_019812791.
2;refGene_gn=PPP6R3;refGene_tx=XM_019812792.2;refGene_gn=PPP6R3;refGen
e_tx=XM_019812793.1;refGene_gn=PPP6R3;refGene_tx=XM_019812794.2;refGen
e_gn=PPP6R3;refGene_tx=XM_019812795.2;refGene_gn=PPP6R3;refGene_tx=XM_
019812796.2;refGene_gn=PPP6R3;refGene_tx=XM_019812797.2;refGene_gn=PPP
6R3;refGene_tx=XM_019812798.2;refGene_gn=PPP6R3;refGene_tx=XM_01981279
9.2;refGene_gn=PPP6R3;refGene_tx=XM_019812800.2;refGene_gn=PPP6R3;refG
ene_tx=XM_019812801.2;refGene_gn=PPP6R3;refGene_tx=XM_019812802.2;refG
ene_gn=PPP6R3;refGene_tx=XM_023240161.1;refGene_gn=PPP6R3;refGene_tx=X
M_023240162.1;refGene_gn=PPP6R3;refGene_tx=XM_023240163.1;refGene_gn=P
PP6R3;refGene_tx=XM_023240164.1;refGene_gn=PPP6R3;refGene_tx=XM_023240
165.1;refGene_gn=PPP6R3;refGene_tx=XM_023240166.1
D1 113002704 113002854
refGene_gn=PPP6R3;refGene_tx=XM_003993712.5;refGene_gn=PPP6R3;refGene_
tx=XM_003993713.5;refGene_gn=PPP6R3;refGene_tx=XM_003993714.5;refGene_
gn=PPP6R3;refGene_tx=XM_006937588.3;refGene_gn=PPP6R3;refGene_tx=XM_00
6937593.3;refGene_gn=PPP6R3;refGene_tx=XM_006937597.4;refGene_gn=PPP6R
3;refGene_tx=XM_006937598.4;refGene_gn=PPP6R3;refGene_tx=XM_019812791.
2;refGene_gn=PPP6R3;refGene_tx=XM_019812792.2;refGene_gn=PPP6R3;refGen
e_tx=XM_019812793.1;refGene_gn=PPP6R3;refGene_tx=XM_019812794.2;refGen
e_gn=PPP6R3;refGene_tx=XM_019812795.2;refGene_gn=PPP6R3;refGene_tx=XM_
019812796.2;refGene_gn=PPP6R3;refGene_tx=XM_019812797.2;refGene_gn=PPP
6R3;refGene_tx=XM_019812798.2;refGene_gn=PPP6R3;refGene_tx=XM_01981279
9.2;refGene_gn=PPP6R3;refGene_tx=XM_019812800.2;refGene_gn=PPP6R3;refG
ene_tx=XM_019812801.2;refGene_gn=PPP6R3;refGene_tx=XM_019812802.2;refG
ene_gn=PPP6R3;refGene_tx=XM_023240161.1;refGene_gn=PPP6R3;refGene_tx=X
M_023240162.1;refGene_gn=PPP6R3;refGene_tx=XM_023240163.1;refGene_gn=P
PP6R3;refGene_tx=XM_023240164.1;refGene_gn=PPP6R3;refGene_tx=XM_023240
165.1;refGene_gn=PPP6R3;refGene_tx=XM_023240166.1
D1 113003921 113003986
refGene_gn=PPP6R3;refGene_tx=XM_003993712.5;refGene_gn=PPP6R3;refGene_
tx=XM_003993713.5;refGene_gn=PPP6R3;refGene_tx=XM_003993714.5;refGene_
gn=PPP6R3;refGene_tx=XM_006937588.3;refGene_gn=PPP6R3;refGene_tx=XM_00
6937593.3;refGene_gn=PPP6R3;refGene_tx=XM_006937597.4;refGene_gn=PPP6R
3;refGene_tx=XM_006937598.4;refGene_gn=PPP6R3;refGene_tx=XM_019812791.
2;refGene_gn=PPP6R3;refGene_tx=XM_019812792.2;refGene_gn=PPP6R3;refGen
e_tx=XM_019812793.1;refGene_gn=PPP6R3;refGene_tx=XM_019812794.2;refGen
e_gn=PPP6R3;refGene_tx=XM_019812795.2;refGene_gn=PPP6R3;refGene_tx=XM_
019812796.2;refGene_gn=PPP6R3;refGene_tx=XM_019812797.2;refGene_gn=PPP
6R3;refGene_tx=XM_019812798.2;refGene_gn=PPP6R3;refGene_tx=XM_01981279
9.2;refGene_gn=PPP6R3;refGene_tx=XM_019812800.2;refGene_gn=PPP6R3;refG

ene_tx=XM_019812801.2;refGene_gn=PPP6R3;refGene_tx=XM_019812802.2;refGene_gn=PPP6R3;refGene_tx=XM_023240161.1;refGene_gn=PPP6R3;refGene_tx=XM_023240162.1;refGene_gn=PPP6R3;refGene_tx=XM_023240163.1;refGene_gn=PPP6R3;refGene_tx=XM_023240164.1;refGene_gn=PPP6R3;refGene_tx=XM_023240165.1;refGene_gn=PPP6R3;refGene_tx=XM_023240166.1
D1 113006166 113006282
refGene_gn=PPP6R3;refGene_tx=XM_003993712.5;refGene_gn=PPP6R3;refGene_tx=XM_003993713.5;refGene_gn=PPP6R3;refGene_tx=XM_003993714.5;refGene_gn=PPP6R3;refGene_tx=XM_006937588.3;refGene_gn=PPP6R3;refGene_tx=XM_006937593.3;refGene_gn=PPP6R3;refGene_tx=XM_006937597.4;refGene_gn=PPP6R3;refGene_tx=XM_006937598.4;refGene_gn=PPP6R3;refGene_tx=XM_019812791.2;refGene_gn=PPP6R3;refGene_tx=XM_019812792.2;refGene_gn=PPP6R3;refGene_tx=XM_019812793.1;refGene_gn=PPP6R3;refGene_tx=XM_019812794.2;refGene_gn=PPP6R3;refGene_tx=XM_019812795.2;refGene_gn=PPP6R3;refGene_tx=XM_019812796.2;refGene_gn=PPP6R3;refGene_tx=XM_019812797.2;refGene_gn=PPP6R3;refGene_tx=XM_019812798.2;refGene_gn=PPP6R3;refGene_tx=XM_019812799.2;refGene_gn=PPP6R3;refGene_tx=XM_019812800.2;refGene_gn=PPP6R3;refGene_tx=XM_019812801.2;refGene_gn=PPP6R3;refGene_tx=XM_019812802.2;refGene_gn=PPP6R3;refGene_tx=XM_023240161.1;refGene_gn=PPP6R3;refGene_tx=XM_023240162.1;refGene_gn=PPP6R3;refGene_tx=XM_023240163.1;refGene_gn=PPP6R3;refGene_tx=XM_023240164.1;refGene_gn=PPP6R3;refGene_tx=XM_023240165.1;refGene_gn=PPP6R3;refGene_tx=XM_023240166.1
D1 113007902 113007988
refGene_gn=PPP6R3;refGene_tx=XM_003993712.5;refGene_gn=PPP6R3;refGene_tx=XM_003993713.5;refGene_gn=PPP6R3;refGene_tx=XM_003993714.5;refGene_gn=PPP6R3;refGene_tx=XM_006937588.3;refGene_gn=PPP6R3;refGene_tx=XM_006937593.3;refGene_gn=PPP6R3;refGene_tx=XM_006937597.4;refGene_gn=PPP6R3;refGene_tx=XM_006937598.4;refGene_gn=PPP6R3;refGene_tx=XM_019812791.2;refGene_gn=PPP6R3;refGene_tx=XM_019812792.2;refGene_gn=PPP6R3;refGene_tx=XM_019812793.1;refGene_gn=PPP6R3;refGene_tx=XM_019812794.2;refGene_gn=PPP6R3;refGene_tx=XM_019812795.2;refGene_gn=PPP6R3;refGene_tx=XM_019812796.2;refGene_gn=PPP6R3;refGene_tx=XM_019812797.2;refGene_gn=PPP6R3;refGene_tx=XM_019812798.2;refGene_gn=PPP6R3;refGene_tx=XM_019812799.2;refGene_gn=PPP6R3;refGene_tx=XM_019812800.2;refGene_gn=PPP6R3;refGene_tx=XM_019812801.2;refGene_gn=PPP6R3;refGene_tx=XM_019812802.2;refGene_gn=PPP6R3;refGene_tx=XM_023240161.1;refGene_gn=PPP6R3;refGene_tx=XM_023240162.1;refGene_gn=PPP6R3;refGene_tx=XM_023240163.1;refGene_gn=PPP6R3;refGene_tx=XM_023240164.1;refGene_gn=PPP6R3;refGene_tx=XM_023240165.1;refGene_gn=PPP6R3;refGene_tx=XM_023240166.1
D1 113014090 113014177
refGene_gn=PPP6R3;refGene_tx=XM_003993712.5;refGene_gn=PPP6R3;refGene_tx=XM_003993714.5;refGene_gn=PPP6R3;refGene_tx=XM_006937588.3;refGene_gn=PPP6R3;refGene_tx=XM_019812791.2;refGene_gn=PPP6R3;refGene_tx=XM_019812792.2;refGene_gn=PPP6R3;refGene_tx=XM_019812793.1;refGene_gn=PPP6R3;refGene_tx=XM_019812794.2;refGene_gn=PPP6R3;refGene_tx=XM_019812795.2;refGene_gn=PPP6R3;refGene_tx=XM_019812796.2;refGene_gn=PPP6R3;refGene_tx=XM_019812800.2;refGene_gn=PPP6R3;refGene_tx=XM_019812801.2;refGene_gn=PPP6R3;refGene_tx=XM_019812802.2;refGene_gn=PPP6R3;refGene_tx=XM_023240161.1;refGene_gn=PPP6R3;refGene_tx=XM_023240162.1;refGene_gn=PPP6R3;refGene_tx=XM_023240163.1;refGene_gn=PPP6R3;refGene_tx=XM_02324016

4.1
D1 113018807 113018905
refGene_gn=PPP6R3;refGene_tx=XM_003993712.5;refGene_gn=PPP6R3;refGene_tx=XM_003993713.5;refGene_gn=PPP6R3;refGene_tx=XM_003993714.5;refGene_gn=PPP6R3;refGene_tx=XM_006937588.3;refGene_gn=PPP6R3;refGene_tx=XM_006937593.3;refGene_gn=PPP6R3;refGene_tx=XM_006937597.4;refGene_gn=PPP6R3;refGene_tx=XM_006937598.4;refGene_gn=PPP6R3;refGene_tx=XM_019812791.2;refGene_gn=PPP6R3;refGene_tx=XM_019812792.2;refGene_gn=PPP6R3;refGene_tx=XM_019812793.1;refGene_gn=PPP6R3;refGene_tx=XM_019812794.2;refGene_gn=PPP6R3;refGene_tx=XM_019812795.2;refGene_gn=PPP6R3;refGene_tx=XM_019812796.2;refGene_gn=PPP6R3;refGene_tx=XM_019812797.2;refGene_gn=PPP6R3;refGene_tx=XM_019812798.2;refGene_gn=PPP6R3;refGene_tx=XM_019812799.2;refGene_gn=PPP6R3;refGene_tx=XM_019812800.2;refGene_gn=PPP6R3;refGene_tx=XM_019812801.2;refGene_gn=PPP6R3;refGene_tx=XM_019812802.2;refGene_gn=PPP6R3;refGene_tx=XM_023240161.1;refGene_gn=PPP6R3;refGene_tx=XM_023240162.1;refGene_gn=PPP6R3;refGene_tx=XM_023240163.1;refGene_gn=PPP6R3;refGene_tx=XM_023240164.1;refGene_gn=PPP6R3;refGene_tx=XM_023240165.1;refGene_gn=PPP6R3;refGene_tx=XM_023240166.1
D1 113020736 113020791
refGene_gn=PPP6R3;refGene_tx=XM_003993712.5;refGene_gn=PPP6R3;refGene_tx=XM_003993713.5;refGene_gn=PPP6R3;refGene_tx=XM_003993714.5;refGene_gn=PPP6R3;refGene_tx=XM_006937588.3;refGene_gn=PPP6R3;refGene_tx=XM_006937593.3;refGene_gn=PPP6R3;refGene_tx=XM_006937597.4;refGene_gn=PPP6R3;refGene_tx=XM_006937598.4;refGene_gn=PPP6R3;refGene_tx=XM_019812791.2;refGene_gn=PPP6R3;refGene_tx=XM_019812792.2;refGene_gn=PPP6R3;refGene_tx=XM_019812793.1;refGene_gn=PPP6R3;refGene_tx=XM_019812794.2;refGene_gn=PPP6R3;refGene_tx=XM_019812795.2;refGene_gn=PPP6R3;refGene_tx=XM_019812796.2;refGene_gn=PPP6R3;refGene_tx=XM_019812797.2;refGene_gn=PPP6R3;refGene_tx=XM_019812798.2;refGene_gn=PPP6R3;refGene_tx=XM_019812799.2;refGene_gn=PPP6R3;refGene_tx=XM_019812800.2;refGene_gn=PPP6R3;refGene_tx=XM_019812801.2;refGene_gn=PPP6R3;refGene_tx=XM_019812802.2;refGene_gn=PPP6R3;refGene_tx=XM_023240161.1;refGene_gn=PPP6R3;refGene_tx=XM_023240162.1;refGene_gn=PPP6R3;refGene_tx=XM_023240163.1;refGene_gn=PPP6R3;refGene_tx=XM_023240164.1;refGene_gn=PPP6R3;refGene_tx=XM_023240165.1;refGene_gn=PPP6R3;refGene_tx=XM_023240166.1
D1 113021916 113022047
refGene_gn=PPP6R3;refGene_tx=XM_003993712.5;refGene_gn=PPP6R3;refGene_tx=XM_003993713.5;refGene_gn=PPP6R3;refGene_tx=XM_003993714.5;refGene_gn=PPP6R3;refGene_tx=XM_006937588.3;refGene_gn=PPP6R3;refGene_tx=XM_006937593.3;refGene_gn=PPP6R3;refGene_tx=XM_006937597.4;refGene_gn=PPP6R3;refGene_tx=XM_006937598.4;refGene_gn=PPP6R3;refGene_tx=XM_019812791.2;refGene_gn=PPP6R3;refGene_tx=XM_019812792.2;refGene_gn=PPP6R3;refGene_tx=XM_019812793.1;refGene_gn=PPP6R3;refGene_tx=XM_019812794.2;refGene_gn=PPP6R3;refGene_tx=XM_019812795.2;refGene_gn=PPP6R3;refGene_tx=XM_019812796.2;refGene_gn=PPP6R3;refGene_tx=XM_019812797.2;refGene_gn=PPP6R3;refGene_tx=XM_019812798.2;refGene_gn=PPP6R3;refGene_tx=XM_019812799.2;refGene_gn=PPP6R3;refGene_tx=XM_019812800.2;refGene_gn=PPP6R3;refGene_tx=XM_019812801.2;refGene_gn=PPP6R3;refGene_tx=XM_019812802.2;refGene_gn=PPP6R3;refGene_tx=XM_023240161.1;refGene_gn=PPP6R3;refGene_tx=XM_023240162.1;refGene_gn=PPP6R3;refGene_tx=XM_023240163.1;refGene_gn=P

PP6R3;refGene_tx=XM_023240164.1;refGene_gn=PPP6R3;refGene_tx=XM_023240165.1;refGene_gn=PPP6R3;refGene_tx=XM_023240166.1
D1	113027181	113027303
refGene_gn=PPP6R3;refGene_tx=XM_003993712.5;refGene_gn=PPP6R3;refGene_tx=XM_003993713.5;refGene_gn=PPP6R3;refGene_tx=XM_003993714.5;refGene_gn=PPP6R3;refGene_tx=XM_006937588.3;refGene_gn=PPP6R3;refGene_tx=XM_006937593.3;refGene_gn=PPP6R3;refGene_tx=XM_006937597.4;refGene_gn=PPP6R3;refGene_tx=XM_006937598.4;refGene_gn=PPP6R3;refGene_tx=XM_019812791.2;refGene_gn=PPP6R3;refGene_tx=XM_019812792.2;refGene_gn=PPP6R3;refGene_tx=XM_019812793.1;refGene_gn=PPP6R3;refGene_tx=XM_019812794.2;refGene_gn=PPP6R3;refGene_tx=XM_019812795.2;refGene_gn=PPP6R3;refGene_tx=XM_019812796.2;refGene_gn=PPP6R3;refGene_tx=XM_019812797.2;refGene_gn=PPP6R3;refGene_tx=XM_019812798.2;refGene_gn=PPP6R3;refGene_tx=XM_019812799.2;refGene_gn=PPP6R3;refGene_tx=XM_019812800.2;refGene_gn=PPP6R3;refGene_tx=XM_019812801.2;refGene_gn=PPP6R3;refGene_tx=XM_019812802.2;refGene_gn=PPP6R3;refGene_tx=XM_023240161.1;refGene_gn=PPP6R3;refGene_tx=XM_023240162.1;refGene_gn=PPP6R3;refGene_tx=XM_023240163.1;refGene_gn=PPP6R3;refGene_tx=XM_023240164.1;refGene_gn=PPP6R3;refGene_tx=XM_023240165.1;refGene_gn=PPP6R3;refGene_tx=XM_023240166.1
D1	113029758	113029912
refGene_gn=PPP6R3;refGene_tx=XM_003993712.5;refGene_gn=PPP6R3;refGene_tx=XM_003993713.5;refGene_gn=PPP6R3;refGene_tx=XM_003993714.5;refGene_gn=PPP6R3;refGene_tx=XM_006937588.3;refGene_gn=PPP6R3;refGene_tx=XM_006937593.3;refGene_gn=PPP6R3;refGene_tx=XM_006937597.4;refGene_gn=PPP6R3;refGene_tx=XM_006937598.4;refGene_gn=PPP6R3;refGene_tx=XM_019812791.2;refGene_gn=PPP6R3;refGene_tx=XM_019812792.2;refGene_gn=PPP6R3;refGene_tx=XM_019812793.1;refGene_gn=PPP6R3;refGene_tx=XM_019812794.2;refGene_gn=PPP6R3;refGene_tx=XM_019812795.2;refGene_gn=PPP6R3;refGene_tx=XM_019812796.2;refGene_gn=PPP6R3;refGene_tx=XM_019812797.2;refGene_gn=PPP6R3;refGene_tx=XM_019812798.2;refGene_gn=PPP6R3;refGene_tx=XM_019812799.2;refGene_gn=PPP6R3;refGene_tx=XM_019812800.2;refGene_gn=PPP6R3;refGene_tx=XM_019812801.2;refGene_gn=PPP6R3;refGene_tx=XM_019812802.2;refGene_gn=PPP6R3;refGene_tx=XM_023240161.1;refGene_gn=PPP6R3;refGene_tx=XM_023240162.1;refGene_gn=PPP6R3;refGene_tx=XM_023240163.1;refGene_gn=PPP6R3;refGene_tx=XM_023240164.1;refGene_gn=PPP6R3;refGene_tx=XM_023240165.1;refGene_gn=PPP6R3;refGene_tx=XM_023240166.1
D1	113031324	113031431
refGene_gn=PPP6R3;refGene_tx=XM_003993712.5;refGene_gn=PPP6R3;refGene_tx=XM_003993713.5;refGene_gn=PPP6R3;refGene_tx=XM_003993714.5;refGene_gn=PPP6R3;refGene_tx=XM_006937588.3;refGene_gn=PPP6R3;refGene_tx=XM_006937593.3;refGene_gn=PPP6R3;refGene_tx=XM_006937597.4;refGene_gn=PPP6R3;refGene_tx=XM_006937598.4;refGene_gn=PPP6R3;refGene_tx=XM_019812791.2;refGene_gn=PPP6R3;refGene_tx=XM_019812792.2;refGene_gn=PPP6R3;refGene_tx=XM_019812793.1;refGene_gn=PPP6R3;refGene_tx=XM_019812794.2;refGene_gn=PPP6R3;refGene_tx=XM_019812795.2;refGene_gn=PPP6R3;refGene_tx=XM_019812796.2;refGene_gn=PPP6R3;refGene_tx=XM_019812797.2;refGene_gn=PPP6R3;refGene_tx=XM_019812798.2;refGene_gn=PPP6R3;refGene_tx=XM_019812799.2;refGene_gn=PPP6R3;refGene_tx=XM_019812800.2;refGene_gn=PPP6R3;refGene_tx=XM_019812801.2;refGene_gn=PPP6R3;refGene_tx=XM_019812802.2;refGene_gn=PPP6R3;refGene_tx=XM_023240161.1;refGene_gn=PPP6R3;refGene_tx=X

M_023240162.1;refGene_gn=PPP6R3;refGene_tx=XM_023240163.1;refGene_gn=P
PP6R3;refGene_tx=XM_023240164.1;refGene_gn=PPP6R3;refGene_tx=XM_023240
165.1;refGene_gn=PPP6R3;refGene_tx=XM_023240166.1
D1 113032874 113033025
refGene_gn=PPP6R3;refGene_tx=XM_003993712.5;refGene_gn=PPP6R3;refGene_
tx=XM_003993713.5;refGene_gn=PPP6R3;refGene_tx=XM_003993714.5;refGene_
gn=PPP6R3;refGene_tx=XM_006937588.3;refGene_gn=PPP6R3;refGene_tx=XM_00
6937593.3;refGene_gn=PPP6R3;refGene_tx=XM_006937597.4;refGene_gn=PPP6R
3;refGene_tx=XM_006937598.4;refGene_gn=PPP6R3;refGene_tx=XM_019812791.
2;refGene_gn=PPP6R3;refGene_tx=XM_019812792.2;refGene_gn=PPP6R3;refGen
e_tx=XM_019812793.1;refGene_gn=PPP6R3;refGene_tx=XM_019812794.2;refGen
e_gn=PPP6R3;refGene_tx=XM_019812795.2;refGene_gn=PPP6R3;refGene_tx=XM_
019812796.2;refGene_gn=PPP6R3;refGene_tx=XM_019812797.2;refGene_gn=PPP
6R3;refGene_tx=XM_019812798.2;refGene_gn=PPP6R3;refGene_tx=XM_01981279
9.2;refGene_gn=PPP6R3;refGene_tx=XM_019812800.2;refGene_gn=PPP6R3;refG
ene_tx=XM_019812801.2;refGene_gn=PPP6R3;refGene_tx=XM_019812802.2;refG
ene_gn=PPP6R3;refGene_tx=XM_023240161.1;refGene_gn=PPP6R3;refGene_tx=X
M_023240162.1;refGene_gn=PPP6R3;refGene_tx=XM_023240163.1;refGene_gn=P
PP6R3;refGene_tx=XM_023240164.1;refGene_gn=PPP6R3;refGene_tx=XM_023240
165.1;refGene_gn=PPP6R3;refGene_tx=XM_023240166.1
D1 113034976 113034986
refGene_gn=PPP6R3;refGene_tx=XM_019812800.2
D1 113035115 113035133
refGene_gn=PPP6R3;refGene_tx=XM_019812791.2;refGene_gn=PPP6R3;refGene_
tx=XM_019812792.2;refGene_gn=PPP6R3;refGene_tx=XM_019812793.1;refGene_
gn=PPP6R3;refGene_tx=XM_019812794.2;refGene_gn=PPP6R3;refGene_tx=XM_01
9812795.2;refGene_gn=PPP6R3;refGene_tx=XM_019812796.2;refGene_gn=PPP6R
3;refGene_tx=XM_019812797.2;refGene_gn=PPP6R3;refGene_tx=XM_019812798.
2;refGene_gn=PPP6R3;refGene_tx=XM_019812799.2;refGene_gn=PPP6R3;refGen
e_tx=XM_019812801.2;refGene_gn=PPP6R3;refGene_tx=XM_019812802.2;refGen
e_gn=PPP6R3;refGene_tx=XM_023240162.1;refGene_gn=PPP6R3;refGene_tx=XM_
023240163.1;refGene_gn=PPP6R3;refGene_tx=XM_023240165.1;refGene_gn=PPP
6R3;refGene_tx=XM_023240166.1
D1 113035400 113035520
refGene_gn=PPP6R3;refGene_tx=XM_003993712.5;refGene_gn=PPP6R3;refGene_
tx=XM_003993713.5;refGene_gn=PPP6R3;refGene_tx=XM_003993714.5;refGene_
gn=PPP6R3;refGene_tx=XM_006937588.3;refGene_gn=PPP6R3;refGene_tx=XM_00
6937593.3;refGene_gn=PPP6R3;refGene_tx=XM_006937597.4;refGene_gn=PPP6R
3;refGene_tx=XM_006937598.4;refGene_gn=PPP6R3;refGene_tx=XM_019812791.
2;refGene_gn=PPP6R3;refGene_tx=XM_019812792.2;refGene_gn=PPP6R3;refGen
e_tx=XM_019812793.1;refGene_gn=PPP6R3;refGene_tx=XM_019812794.2;refGen
e_gn=PPP6R3;refGene_tx=XM_019812795.2;refGene_gn=PPP6R3;refGene_tx=XM_
019812796.2;refGene_gn=PPP6R3;refGene_tx=XM_019812797.2;refGene_gn=PPP
6R3;refGene_tx=XM_019812798.2;refGene_gn=PPP6R3;refGene_tx=XM_01981279
9.2;refGene_gn=PPP6R3;refGene_tx=XM_019812801.2;refGene_gn=PPP6R3;refG
ene_tx=XM_019812802.2;refGene_gn=PPP6R3;refGene_tx=XM_023240161.1;refG
ene_gn=PPP6R3;refGene_tx=XM_023240162.1;refGene_gn=PPP6R3;refGene_tx=X
M_023240163.1;refGene_gn=PPP6R3;refGene_tx=XM_023240164.1;refGene_gn=P
PP6R3;refGene_tx=XM_023240165.1;refGene_gn=PPP6R3;refGene_tx=XM_023240
166.1

D1	113036152	113036258
refGene_gn=PPP6R3;refGene_tx=XM_006937588.3;refGene_gn=PPP6R3;refGene_tx=XM_006937593.3;refGene_gn=PPP6R3;refGene_tx=XM_019812791.2;refGene_gn=PPP6R3;refGene_tx=XM_019812792.2;refGene_gn=PPP6R3;refGene_tx=XM_019812793.1;refGene_gn=PPP6R3;refGene_tx=XM_019812794.2;refGene_gn=PPP6R3;refGene_tx=XM_019812795.2;refGene_gn=PPP6R3;refGene_tx=XM_019812797.2;refGene_gn=PPP6R3;refGene_tx=XM_019812798.2;refGene_gn=PPP6R3;refGene_tx=XM_019812799.2;refGene_gn=PPP6R3;refGene_tx=XM_019812801.2;refGene_gn=PPP6R3;refGene_tx=XM_019812802.2;refGene_gn=PPP6R3;refGene_tx=XM_023240161.1
D1	113038589	113038641
refGene_gn=PPP6R3;refGene_tx=XM_003993712.5;refGene_gn=PPP6R3;refGene_tx=XM_003993713.5;refGene_gn=PPP6R3;refGene_tx=XM_003993714.5;refGene_gn=PPP6R3;refGene_tx=XM_006937597.4;refGene_gn=PPP6R3;refGene_tx=XM_006937598.4;refGene_gn=PPP6R3;refGene_tx=XM_019812796.2;refGene_gn=PPP6R3;refGene_tx=XM_023240162.1;refGene_gn=PPP6R3;refGene_tx=XM_023240163.1;refGene_gn=PPP6R3;refGene_tx=XM_023240164.1;refGene_gn=PPP6R3;refGene_tx=XM_023240165.1;refGene_gn=PPP6R3;refGene_tx=XM_023240166.1
D1	113076097	113076178
refGene_gn=GAL;refGene_tx=XM_023240167.1
D1	113076845	113076900
refGene_gn=GAL;refGene_tx=XM_023240167.1
D1	113080214	113080301
refGene_gn=GAL;refGene_tx=XM_023240167.1
D1	113080550	113080628
refGene_gn=GAL;refGene_tx=XM_023240167.1
D1	113082132	113082203
refGene_gn=GAL;refGene_tx=XM_023240167.1
D1	113090879	113090901
refGene_gn=TESMIN;refGene_tx=XM_019812806.2
D1	113091014	113091153
refGene_gn=TESMIN;refGene_tx=XM_019812806.2;refGene_gn=TESMIN;refGene_tx=XM_023240170.1
D1	113095707	113095897
refGene_gn=TESMIN;refGene_tx=XM_011287105.3;refGene_gn=TESMIN;refGene_tx=XM_011287108.3;refGene_gn=TESMIN;refGene_tx=XM_019812804.2;refGene_gn=TESMIN;refGene_tx=XM_019812805.2;refGene_gn=TESMIN;refGene_tx=XM_019812807.1;refGene_gn=TESMIN;refGene_tx=XM_023240168.1;refGene_gn=TESMIN;refGene_tx=XM_023240169.1
D1	113096289	113096465
refGene_gn=TESMIN;refGene_tx=XM_011287105.3;refGene_gn=TESMIN;refGene_tx=XM_011287108.3;refGene_gn=TESMIN;refGene_tx=XM_019812804.2;refGene_gn=TESMIN;refGene_tx=XM_019812805.2;refGene_gn=TESMIN;refGene_tx=XM_019812806.2;refGene_gn=TESMIN;refGene_tx=XM_019812807.1;refGene_gn=TESMIN;refGene_tx=XM_023240168.1;refGene_gn=TESMIN;refGene_tx=XM_023240169.1;refGene_gn=TESMIN;refGene_tx=XM_023240170.1
D1	113098769	113098907
refGene_gn=TESMIN;refGene_tx=XM_011287105.3;refGene_gn=TESMIN;refGene_tx=XM_019812804.2;refGene_gn=TESMIN;refGene_tx=XM_019812805.2;refGene_gn=TESMIN;refGene_tx=XM_019812806.2;refGene_gn=TESMIN;refGene_tx=XM_01

9812807.1;refGene_gn=TESMIN;refGene_tx=XM_023240168.1;refGene_gn=TESMI
N;refGene_tx=XM_023240169.1;refGene_gn=TESMIN;refGene_tx=XM_023240170.
1
D1 113101491 113101594
refGene_gn=TESMIN;refGene_tx=XM_011287105.3;refGene_gn=TESMIN;refGene_
tx=XM_011287108.3;refGene_gn=TESMIN;refGene_tx=XM_019812804.2;refGene_
gn=TESMIN;refGene_tx=XM_019812805.2;refGene_gn=TESMIN;refGene_tx=XM_01
9812806.2;refGene_gn=TESMIN;refGene_tx=XM_019812807.1;refGene_gn=TESMI
N;refGene_tx=XM_023240168.1;refGene_gn=TESMIN;refGene_tx=XM_023240169.
1;refGene_gn=TESMIN;refGene_tx=XM_023240170.1
D1 113115885 113115957
refGene_gn=TESMIN;refGene_tx=XM_011287105.3;refGene_gn=TESMIN;refGene_
tx=XM_011287108.3;refGene_gn=TESMIN;refGene_tx=XM_019812804.2;refGene_
gn=TESMIN;refGene_tx=XM_019812805.2;refGene_gn=TESMIN;refGene_tx=XM_01
9812806.2;refGene_gn=TESMIN;refGene_tx=XM_023240168.1;refGene_gn=TESMI
N;refGene_tx=XM_023240169.1;refGene_gn=TESMIN;refGene_tx=XM_023240170.
1
D1 113120442 113120538
refGene_gn=TESMIN;refGene_tx=XM_011287105.3;refGene_gn=TESMIN;refGene_
tx=XM_011287108.3;refGene_gn=TESMIN;refGene_tx=XM_019812804.2;refGene_
gn=TESMIN;refGene_tx=XM_019812805.2;refGene_gn=TESMIN;refGene_tx=XM_01
9812806.2;refGene_gn=TESMIN;refGene_tx=XM_019812807.1;refGene_gn=TESMI
N;refGene_tx=XM_019812808.2;refGene_gn=TESMIN;refGene_tx=XM_023240168.
1;refGene_gn=TESMIN;refGene_tx=XM_023240169.1;refGene_gn=TESMIN;refGen
e_tx=XM_023240170.1
D1 113123446 113123520
refGene_gn=TESMIN;refGene_tx=XM_011287105.3;refGene_gn=TESMIN;refGene_
tx=XM_011287108.3;refGene_gn=TESMIN;refGene_tx=XM_019812804.2;refGene_
gn=TESMIN;refGene_tx=XM_019812805.2;refGene_gn=TESMIN;refGene_tx=XM_01
9812806.2;refGene_gn=TESMIN;refGene_tx=XM_019812807.1;refGene_gn=TESMI
N;refGene_tx=XM_019812808.2;refGene_gn=TESMIN;refGene_tx=XM_023240168.
1;refGene_gn=TESMIN;refGene_tx=XM_023240169.1;refGene_gn=TESMIN;refGen
e_tx=XM_023240170.1
D1 113125579 113125700
refGene_gn=TESMIN;refGene_tx=XM_011287105.3;refGene_gn=TESMIN;refGene_
tx=XM_011287108.3;refGene_gn=TESMIN;refGene_tx=XM_019812804.2;refGene_
gn=TESMIN;refGene_tx=XM_019812805.2;refGene_gn=TESMIN;refGene_tx=XM_01
9812806.2;refGene_gn=TESMIN;refGene_tx=XM_019812807.1;refGene_gn=TESMI
N;refGene_tx=XM_019812808.2;refGene_gn=TESMIN;refGene_tx=XM_023240168.
1;refGene_gn=TESMIN;refGene_tx=XM_023240169.1;refGene_gn=TESMIN;refGen
e_tx=XM_023240170.1
D1 113128005 113128149
refGene_gn=TESMIN;refGene_tx=XM_011287105.3;refGene_gn=TESMIN;refGene_
tx=XM_011287108.3;refGene_gn=TESMIN;refGene_tx=XM_019812804.2;refGene_
gn=TESMIN;refGene_tx=XM_019812805.2;refGene_gn=TESMIN;refGene_tx=XM_01
9812806.2;refGene_gn=TESMIN;refGene_tx=XM_019812807.1;refGene_gn=TESMI
N;refGene_tx=XM_019812808.2;refGene_gn=TESMIN;refGene_tx=XM_023240168.
1;refGene_gn=TESMIN;refGene_tx=XM_023240169.1;refGene_gn=TESMIN;refGen
e_tx=XM_023240170.1
D1 113129846 113130317

refGene_gn=TESMIN;refGene_tx=XM_011287105.3;refGene_gn=TESMIN;refGene_tx=XM_011287108.3;refGene_gn=TESMIN;refGene_tx=XM_019812804.2;refGene_gn=TESMIN;refGene_tx=XM_019812805.2;refGene_gn=TESMIN;refGene_tx=XM_019812806.2;refGene_gn=TESMIN;refGene_tx=XM_019812807.1;refGene_gn=TESMIN;refGene_tx=XM_019812808.2;refGene_gn=TESMIN;refGene_tx=XM_023240168.1;refGene_gn=TESMIN;refGene_tx=XM_023240169.1;refGene_gn=TESMIN;refGene_tx=XM_023240170.1
D1 113136242 113136329
refGene_gn=CPT1A;refGene_tx=XM_023240171.1
D1 113137270 113137363
refGene_gn=CPT1A;refGene_tx=XM_023240171.1
D1 113137611 113137725
refGene_gn=CPT1A;refGene_tx=XM_023240171.1
D1 113138600 113138753
refGene_gn=CPT1A;refGene_tx=XM_023240171.1
D1 113139992 113140028
refGene_gn=CPT1A;refGene_tx=XM_023240171.1
D1 113140029 113140128
refGene_gn=CPT1A;refGene_tx=XM_023240171.1
D1 113147432 113147597
refGene_gn=CPT1A;refGene_tx=XM_023240171.1
D1 113149062 113149179
refGene_gn=CPT1A;refGene_tx=XM_023240171.1
D1 113152347 113152453
refGene_gn=CPT1A;refGene_tx=XM_023240171.1
D1 113153338 113153527
refGene_gn=CPT1A;refGene_tx=XM_023240171.1
D1 113155595 113155791
refGene_gn=CPT1A;refGene_tx=XM_023240171.1
D1 113158344 113158432
refGene_gn=CPT1A;refGene_tx=XM_023240171.1
D1 113160699 113160807
refGene_gn=CPT1A;refGene_tx=XM_023240171.1
D1 113161495 113161573
refGene_gn=CPT1A;refGene_tx=XM_023240171.1
D1 113162031 113162079
refGene_gn=CPT1A;refGene_tx=XM_023240171.1
D1 113162080 113162170
refGene_gn=CPT1A;refGene_tx=XM_023240171.1
D1 113164731 113164833
refGene_gn=CPT1A;refGene_tx=XM_023240171.1
D1 113166608 113166780
refGene_gn=CPT1A;refGene_tx=XM_023240171.1
D1 113170556 113170696
refGene_gn=CPT1A;refGene_tx=XM_023240171.1
D1 113173844 113173985
refGene_gn=CPT1A;refGene_tx=XM_023240171.1
D1 113224827 113224892
refGene_gn=MRPL21;refGene_tx=XM_011287104.3;refGene_gn=MRPL21;refGene_tx=XM_023240172.1

D1	113226280	113226384
refGene_gn=MRPL21;refGene_tx=XM_011287104.3;refGene_gn=MRPL21;refGene_tx=XM_023240172.1
D1	113227362	113227415
refGene_gn=MRPL21;refGene_tx=XM_011287104.3;refGene_gn=MRPL21;refGene_tx=XM_023240172.1
D1	113230020	113230184
refGene_gn=MRPL21;refGene_tx=XM_011287104.3;refGene_gn=MRPL21;refGene_tx=XM_023240172.1
D1	113231381	113231467
refGene_gn=MRPL21;refGene_tx=XM_011287104.3;refGene_gn=MRPL21;refGene_tx=XM_023240172.1
D1	113233516	113233574
refGene_gn=MRPL21;refGene_tx=XM_011287104.3;refGene_gn=MRPL21;refGene_tx=XM_023240172.1
D1	113237239	113237290
refGene_gn=MRPL21;refGene_tx=XM_023240172.1
D1	113237484	113237584
refGene_gn=MRPL21;refGene_tx=XM_011287104.3;refGene_gn=MRPL21;refGene_tx=XM_023240172.1
D1	113237720	113237806
refGene_gn=IGHMBP2;refGene_tx=XM_023240175.1
D1	113240066	113240236
refGene_gn=IGHMBP2;refGene_tx=XM_023240175.1
D1	113243832	113244016
refGene_gn=IGHMBP2;refGene_tx=XM_023240175.1
D1	113244217	113244315
refGene_gn=IGHMBP2;refGene_tx=XM_023240175.1
D1	113245625	113245789
refGene_gn=IGHMBP2;refGene_tx=XM_023240175.1
D1	113247992	113248205
refGene_gn=IGHMBP2;refGene_tx=XM_023240175.1
D1	113249783	113249919
refGene_gn=IGHMBP2;refGene_tx=XM_023240175.1
D1	113251343	113251518
refGene_gn=IGHMBP2;refGene_tx=XM_023240175.1
D1	113254115	113254298
refGene_gn=IGHMBP2;refGene_tx=XM_023240175.1
D1	113254611	113254730
refGene_gn=IGHMBP2;refGene_tx=XM_023240175.1
D1	113255150	113255245
refGene_gn=IGHMBP2;refGene_tx=XM_023240175.1
D1	113256185	113256309
refGene_gn=IGHMBP2;refGene_tx=XM_023240175.1
D1	113257142	113257988
refGene_gn=IGHMBP2;refGene_tx=XM_023240175.1
D1	113260188	113260361
refGene_gn=IGHMBP2;refGene_tx=XM_023240175.1
D1	113263053	113263251
refGene_gn=IGHMBP2;refGene_tx=XM_023240175.1

D1 113294860 113295692
refGene_gn=MRGPRD;refGene_tx=XM_023240174.1
D1 113295693 113296154
refGene_gn=MRGPRD;refGene_tx=XM_023240174.1
D1 113310757 113311741
refGene_gn=MRGPRF;refGene_tx=XM_003993757.5;refGene_gn=MRGPRF;refGene_tx=XM_006937623.4;refGene_gn=MRGPRF;refGene_tx=XM_011287103.3;refGene_gn=MRGPRF;refGene_tx=XM_019812811.2;refGene_gn=MRGPRF;refGene_tx=XM_023240173.1
D1 113314791 113314905
refGene_gn=MRGPRF;refGene_tx=XM_003993757.5;refGene_gn=MRGPRF;refGene_tx=XM_006937623.4;refGene_gn=MRGPRF;refGene_tx=XM_011287103.3;refGene_gn=MRGPRF;refGene_tx=XM_019812811.2;refGene_gn=MRGPRF;refGene_tx=XM_023240173.1
D1 113315015 113315064
refGene_gn=MRGPRF;refGene_tx=XM_023240173.1
D1 113316608 113316701
refGene_gn=MRGPRF;refGene_tx=XM_023240173.1
D1 113317117 113317239
refGene_gn=MRGPRF;refGene_tx=XM_023240173.1
D1 113343349 113343455
refGene_gn=TPCN2;refGene_tx=XM_023240176.1;refGene_gn=TPCN2;refGene_tx=XM_023240177.1;refGene_gn=TPCN2;refGene_tx=XM_023240178.1
D1 113347347 113347412
refGene_gn=TPCN2;refGene_tx=XM_023240176.1;refGene_gn=TPCN2;refGene_tx=XM_023240177.1;refGene_gn=TPCN2;refGene_tx=XM_023240178.1
D1 113348124 113348201
refGene_gn=TPCN2;refGene_tx=XM_023240176.1;refGene_gn=TPCN2;refGene_tx=XM_023240177.1;refGene_gn=TPCN2;refGene_tx=XM_023240178.1
D1 113348545 113348723
refGene_gn=TPCN2;refGene_tx=XM_023240176.1;refGene_gn=TPCN2;refGene_tx=XM_023240177.1;refGene_gn=TPCN2;refGene_tx=XM_023240178.1
D1 113350028 113350145
refGene_gn=TPCN2;refGene_tx=XM_023240176.1;refGene_gn=TPCN2;refGene_tx=XM_023240177.1;refGene_gn=TPCN2;refGene_tx=XM_023240178.1
D1 113353590 113353697
refGene_gn=TPCN2;refGene_tx=XM_023240176.1;refGene_gn=TPCN2;refGene_tx=XM_023240177.1;refGene_gn=TPCN2;refGene_tx=XM_023240178.1
D1 113354247 113354320
refGene_gn=TPCN2;refGene_tx=XM_023240176.1;refGene_gn=TPCN2;refGene_tx=XM_023240177.1;refGene_gn=TPCN2;refGene_tx=XM_023240178.1
D1 113355411 113355514
refGene_gn=TPCN2;refGene_tx=XM_023240176.1;refGene_gn=TPCN2;refGene_tx=XM_023240177.1;refGene_gn=TPCN2;refGene_tx=XM_023240178.1
D1 113356443 113356509
refGene_gn=TPCN2;refGene_tx=XM_023240176.1;refGene_gn=TPCN2;refGene_tx=XM_023240177.1;refGene_gn=TPCN2;refGene_tx=XM_023240178.1
D1 113357365 113357430
refGene_gn=TPCN2;refGene_tx=XM_023240176.1;refGene_gn=TPCN2;refGene_tx=XM_023240177.1;refGene_gn=TPCN2;refGene_tx=XM_023240178.1

D1	113357907	113358008
refGene_gn=TPCN2;refGene_tx=XM_023240176.1;refGene_gn=TPCN2;refGene_tx=XM_023240177.1;refGene_gn=TPCN2;refGene_tx=XM_023240178.1
D1	113358601	113358683
refGene_gn=TPCN2;refGene_tx=XM_023240176.1;refGene_gn=TPCN2;refGene_tx=XM_023240177.1;refGene_gn=TPCN2;refGene_tx=XM_023240178.1
D1	113358903	113358990
refGene_gn=TPCN2;refGene_tx=XM_023240176.1;refGene_gn=TPCN2;refGene_tx=XM_023240177.1;refGene_gn=TPCN2;refGene_tx=XM_023240178.1
D1	113359764	113359884
refGene_gn=TPCN2;refGene_tx=XM_023240176.1;refGene_gn=TPCN2;refGene_tx=XM_023240177.1;refGene_gn=TPCN2;refGene_tx=XM_023240178.1
D1	113360040	113360100
refGene_gn=TPCN2;refGene_tx=XM_023240176.1;refGene_gn=TPCN2;refGene_tx=XM_023240177.1;refGene_gn=TPCN2;refGene_tx=XM_023240178.1
D1	113361967	113362096
refGene_gn=TPCN2;refGene_tx=XM_023240176.1;refGene_gn=TPCN2;refGene_tx=XM_023240177.1;refGene_gn=TPCN2;refGene_tx=XM_023240178.1
D1	113363001	113363051
refGene_gn=TPCN2;refGene_tx=XM_023240176.1;refGene_gn=TPCN2;refGene_tx=XM_023240177.1
D1	113364372	113364472
refGene_gn=TPCN2;refGene_tx=XM_023240176.1;refGene_gn=TPCN2;refGene_tx=XM_023240177.1;refGene_gn=TPCN2;refGene_tx=XM_023240178.1
D1	113365016	113365091
refGene_gn=TPCN2;refGene_tx=XM_023240178.1
D1	113366350	113366422
refGene_gn=TPCN2;refGene_tx=XM_023240176.1;refGene_gn=TPCN2;refGene_tx=XM_023240177.1;refGene_gn=TPCN2;refGene_tx=XM_023240178.1
D1	113367727	113367801
refGene_gn=TPCN2;refGene_tx=XM_023240176.1;refGene_gn=TPCN2;refGene_tx=XM_023240177.1;refGene_gn=TPCN2;refGene_tx=XM_023240178.1
D1	113368297	113368379
refGene_gn=TPCN2;refGene_tx=XM_023240176.1;refGene_gn=TPCN2;refGene_tx=XM_023240177.1;refGene_gn=TPCN2;refGene_tx=XM_023240178.1
D1	113368468	113368551
refGene_gn=TPCN2;refGene_tx=XM_023240176.1;refGene_gn=TPCN2;refGene_tx=XM_023240177.1
D1	113368949	113369031
refGene_gn=TPCN2;refGene_tx=XM_023240176.1;refGene_gn=TPCN2;refGene_tx=XM_023240177.1
D1	113369331	113369426
refGene_gn=TPCN2;refGene_tx=XM_023240176.1;refGene_gn=TPCN2;refGene_tx=XM_023240177.1
D1	113370006	113370085
refGene_gn=TPCN2;refGene_tx=XM_023240176.1;refGene_gn=TPCN2;refGene_tx=XM_023240177.1
D1	113416500	113416935
refGene_gn=LOC109492308;refGene_tx=XM_023240179.1
D1	113417567	113417672

refGene_gn=LOC109492308;refGene_tx=XM_023240179.1
D1 113418113 113418557
refGene_gn=LOC109492308;refGene_tx=XM_023240179.1
D1 113810057 113810255
refGene_gn=CCND1;refGene_tx=XM_011287102.3
D1 113811808 113812024
refGene_gn=CCND1;refGene_tx=XM_011287102.3
D1 113812559 113812719
refGene_gn=CCND1;refGene_tx=XM_011287102.3
D1 113816766 113816915
refGene_gn=CCND1;refGene_tx=XM_011287102.3
D1 113819498 113819663
refGene_gn=CCND1;refGene_tx=XM_011287102.3
D1 113832174 113832243
refGene_gn=ORAOV1;refGene_tx=XM_023240180.1;refGene_gn=ORAOV1;refGene_tx=XM_023240181.1;refGene_gn=ORAOV1;refGene_tx=XM_023240182.1
D1 113832533 113832651
refGene_gn=ORAOV1;refGene_tx=XM_023240180.1;refGene_gn=ORAOV1;refGene_tx=XM_023240181.1;refGene_gn=ORAOV1;refGene_tx=XM_023240182.1
D1 113835479 113835550
refGene_gn=ORAOV1;refGene_tx=XM_023240180.1;refGene_gn=ORAOV1;refGene_tx=XM_023240181.1;refGene_gn=ORAOV1;refGene_tx=XM_023240182.1
D1 113837528 113837634
refGene_gn=ORAOV1;refGene_tx=XM_023240180.1;refGene_gn=ORAOV1;refGene_tx=XM_023240181.1;refGene_gn=ORAOV1;refGene_tx=XM_023240182.1
D1 113839564 113839614
refGene_gn=ORAOV1;refGene_tx=XM_023240180.1;refGene_gn=ORAOV1;refGene_tx=XM_023240181.1;refGene_gn=ORAOV1;refGene_tx=XM_023240182.1
D1 113852485 113852800
refGene_gn=FGF19;refGene_tx=XM_003993764.2
D1 113855576 113855680
refGene_gn=FGF19;refGene_tx=XM_003993764.2
D1 113856033 113856274
refGene_gn=FGF19;refGene_tx=XM_003993764.2
D1 113900598 113900775
refGene_gn=FGF4;refGene_tx=XM_023239032.1
D1 113901287 113901391
refGene_gn=FGF4;refGene_tx=XM_023239032.1
D1 113901910 113902250
refGene_gn=FGF4;refGene_tx=XM_023239032.1
D1 113924091 113924457
refGene_gn=FGF3;refGene_tx=XM_023239755.1
D1 113930360 113930464
refGene_gn=FGF3;refGene_tx=XM_023239755.1
D1 113932491 113932711
refGene_gn=FGF3;refGene_tx=XM_023239755.1
D1 114110435 114110543
refGene_gn=ANO1;refGene_tx=XM_023238951.1;refGene_gn=ANO1;refGene_tx=XM_023238952.1;refGene_gn=ANO1;refGene_tx=XM_023238953.1;refGene_gn=ANO1;refGene_tx=XM_023238954.1;refGene_gn=ANO1;refGene_tx=XM_023238956.1;

refGene_gn=ANO1;refGene_tx=XM_023238957.1;refGene_gn=ANO1;refGene_tx=XM_023238958.1;refGene_gn=ANO1;refGene_tx=XM_023238959.1;refGene_gn=ANO1;refGene_tx=XM_023238960.1;refGene_gn=ANO1;refGene_tx=XM_023238961.1;refGene_gn=ANO1;refGene_tx=XM_023238962.1
D1 114116293 114116359
refGene_gn=ANO1;refGene_tx=XM_023238955.1
D1 114117700 114118033
refGene_gn=ANO1;refGene_tx=XM_023238951.1;refGene_gn=ANO1;refGene_tx=XM_023238952.1;refGene_gn=ANO1;refGene_tx=XM_023238953.1;refGene_gn=ANO1;refGene_tx=XM_023238954.1;refGene_gn=ANO1;refGene_tx=XM_023238955.1;refGene_gn=ANO1;refGene_tx=XM_023238956.1;refGene_gn=ANO1;refGene_tx=XM_023238957.1;refGene_gn=ANO1;refGene_tx=XM_023238958.1;refGene_gn=ANO1;refGene_tx=XM_023238959.1;refGene_gn=ANO1;refGene_tx=XM_023238960.1;refGene_gn=ANO1;refGene_tx=XM_023238961.1
D1 114129533 114129632
refGene_gn=ANO1;refGene_tx=XM_023238951.1;refGene_gn=ANO1;refGene_tx=XM_023238952.1;refGene_gn=ANO1;refGene_tx=XM_023238953.1;refGene_gn=ANO1;refGene_tx=XM_023238954.1;refGene_gn=ANO1;refGene_tx=XM_023238955.1;refGene_gn=ANO1;refGene_tx=XM_023238956.1;refGene_gn=ANO1;refGene_tx=XM_023238957.1;refGene_gn=ANO1;refGene_tx=XM_023238958.1;refGene_gn=ANO1;refGene_tx=XM_023238959.1;refGene_gn=ANO1;refGene_tx=XM_023238960.1;refGene_gn=ANO1;refGene_tx=XM_023238961.1;refGene_gn=ANO1;refGene_tx=XM_023238962.1
D1 114130420 114130572
refGene_gn=ANO1;refGene_tx=XM_023238951.1;refGene_gn=ANO1;refGene_tx=XM_023238952.1;refGene_gn=ANO1;refGene_tx=XM_023238953.1;refGene_gn=ANO1;refGene_tx=XM_023238954.1;refGene_gn=ANO1;refGene_tx=XM_023238955.1;refGene_gn=ANO1;refGene_tx=XM_023238956.1;refGene_gn=ANO1;refGene_tx=XM_023238957.1;refGene_gn=ANO1;refGene_tx=XM_023238958.1;refGene_gn=ANO1;refGene_tx=XM_023238959.1;refGene_gn=ANO1;refGene_tx=XM_023238960.1;refGene_gn=ANO1;refGene_tx=XM_023238961.1;refGene_gn=ANO1;refGene_tx=XM_023238962.1
D1 114132093 114132148
refGene_gn=ANO1;refGene_tx=XM_023238951.1;refGene_gn=ANO1;refGene_tx=XM_023238952.1;refGene_gn=ANO1;refGene_tx=XM_023238953.1;refGene_gn=ANO1;refGene_tx=XM_023238954.1;refGene_gn=ANO1;refGene_tx=XM_023238955.1;refGene_gn=ANO1;refGene_tx=XM_023238956.1;refGene_gn=ANO1;refGene_tx=XM_023238957.1;refGene_gn=ANO1;refGene_tx=XM_023238958.1;refGene_gn=ANO1;refGene_tx=XM_023238959.1;refGene_gn=ANO1;refGene_tx=XM_023238960.1;refGene_gn=ANO1;refGene_tx=XM_023238961.1;refGene_gn=ANO1;refGene_tx=XM_023238962.1
D1 114135023 114135075
refGene_gn=ANO1;refGene_tx=XM_023238951.1;refGene_gn=ANO1;refGene_tx=XM_023238952.1;refGene_gn=ANO1;refGene_tx=XM_023238953.1;refGene_gn=ANO1;refGene_tx=XM_023238954.1;refGene_gn=ANO1;refGene_tx=XM_023238955.1;refGene_gn=ANO1;refGene_tx=XM_023238956.1;refGene_gn=ANO1;refGene_tx=XM_023238957.1;refGene_gn=ANO1;refGene_tx=XM_023238958.1;refGene_gn=ANO1;refGene_tx=XM_023238959.1;refGene_gn=ANO1;refGene_tx=XM_023238960.1;refGene_gn=ANO1;refGene_tx=XM_023238961.1;refGene_gn=ANO1;refGene_tx=XM_023238962.1

D1 114137472 114137538
refGene_gn=ANO1;refGene_tx=XM_023238951.1;refGene_gn=ANO1;refGene_tx=XM_023238952.1;refGene_gn=ANO1;refGene_tx=XM_023238953.1;refGene_gn=ANO1;refGene_tx=XM_023238954.1;refGene_gn=ANO1;refGene_tx=XM_023238955.1;refGene_gn=ANO1;refGene_tx=XM_023238957.1;refGene_gn=ANO1;refGene_tx=XM_023238958.1;refGene_gn=ANO1;refGene_tx=XM_023238960.1;refGene_gn=ANO1;refGene_tx=XM_023238962.1
D1 114138034 114138090
refGene_gn=ANO1;refGene_tx=XM_023238951.1;refGene_gn=ANO1;refGene_tx=XM_023238952.1;refGene_gn=ANO1;refGene_tx=XM_023238953.1;refGene_gn=ANO1;refGene_tx=XM_023238954.1;refGene_gn=ANO1;refGene_tx=XM_023238955.1;refGene_gn=ANO1;refGene_tx=XM_023238956.1;refGene_gn=ANO1;refGene_tx=XM_023238957.1;refGene_gn=ANO1;refGene_tx=XM_023238958.1;refGene_gn=ANO1;refGene_tx=XM_023238959.1;refGene_gn=ANO1;refGene_tx=XM_023238960.1;refGene_gn=ANO1;refGene_tx=XM_023238961.1;refGene_gn=ANO1;refGene_tx=XM_023238962.1
D1 114141305 114141347
refGene_gn=ANO1;refGene_tx=XM_023238951.1;refGene_gn=ANO1;refGene_tx=XM_023238952.1;refGene_gn=ANO1;refGene_tx=XM_023238953.1;refGene_gn=ANO1;refGene_tx=XM_023238954.1;refGene_gn=ANO1;refGene_tx=XM_023238955.1;refGene_gn=ANO1;refGene_tx=XM_023238956.1;refGene_gn=ANO1;refGene_tx=XM_023238957.1;refGene_gn=ANO1;refGene_tx=XM_023238958.1;refGene_gn=ANO1;refGene_tx=XM_023238959.1;refGene_gn=ANO1;refGene_tx=XM_023238960.1;refGene_gn=ANO1;refGene_tx=XM_023238961.1;refGene_gn=ANO1;refGene_tx=XM_023238962.1
D1 114147643 114147708
refGene_gn=ANO1;refGene_tx=XM_023238951.1;refGene_gn=ANO1;refGene_tx=XM_023238952.1;refGene_gn=ANO1;refGene_tx=XM_023238953.1;refGene_gn=ANO1;refGene_tx=XM_023238954.1;refGene_gn=ANO1;refGene_tx=XM_023238955.1;refGene_gn=ANO1;refGene_tx=XM_023238956.1;refGene_gn=ANO1;refGene_tx=XM_023238957.1;refGene_gn=ANO1;refGene_tx=XM_023238958.1;refGene_gn=ANO1;refGene_tx=XM_023238959.1;refGene_gn=ANO1;refGene_tx=XM_023238960.1;refGene_gn=ANO1;refGene_tx=XM_023238961.1;refGene_gn=ANO1;refGene_tx=XM_023238962.1
D1 114148354 114148489
refGene_gn=ANO1;refGene_tx=XM_023238951.1;refGene_gn=ANO1;refGene_tx=XM_023238952.1;refGene_gn=ANO1;refGene_tx=XM_023238953.1;refGene_gn=ANO1;refGene_tx=XM_023238954.1;refGene_gn=ANO1;refGene_tx=XM_023238955.1;refGene_gn=ANO1;refGene_tx=XM_023238956.1;refGene_gn=ANO1;refGene_tx=XM_023238957.1;refGene_gn=ANO1;refGene_tx=XM_023238958.1;refGene_gn=ANO1;refGene_tx=XM_023238959.1;refGene_gn=ANO1;refGene_tx=XM_023238960.1;refGene_gn=ANO1;refGene_tx=XM_023238961.1;refGene_gn=ANO1;refGene_tx=XM_023238962.1
D1 114151348 114151509
refGene_gn=ANO1;refGene_tx=XM_023238951.1;refGene_gn=ANO1;refGene_tx=XM_023238952.1;refGene_gn=ANO1;refGene_tx=XM_023238953.1;refGene_gn=ANO1;refGene_tx=XM_023238954.1;refGene_gn=ANO1;refGene_tx=XM_023238955.1;refGene_gn=ANO1;refGene_tx=XM_023238956.1;refGene_gn=ANO1;refGene_tx=XM_023238957.1;refGene_gn=ANO1;refGene_tx=XM_023238958.1;refGene_gn=ANO1;refGene_tx=XM_023238959.1;refGene_gn=ANO1;refGene_tx=XM_023238960.1;

refGene_gn=ANO1;refGene_tx=XM_023238961.1;refGene_gn=ANO1;refGene_tx=X
M_023238962.1
D1	114162484	114162567
refGene_gn=ANO1;refGene_tx=XM_023238951.1;refGene_gn=ANO1;refGene_tx=X
M_023238952.1;refGene_gn=ANO1;refGene_tx=XM_023238953.1;refGene_gn=ANO
1;refGene_tx=XM_023238954.1;refGene_gn=ANO1;refGene_tx=XM_023238955.1;
refGene_gn=ANO1;refGene_tx=XM_023238956.1;refGene_gn=ANO1;refGene_tx=X
M_023238957.1;refGene_gn=ANO1;refGene_tx=XM_023238958.1;refGene_gn=ANO
1;refGene_tx=XM_023238959.1;refGene_gn=ANO1;refGene_tx=XM_023238960.1;
refGene_gn=ANO1;refGene_tx=XM_023238961.1;refGene_gn=ANO1;refGene_tx=X
M_023238962.1
D1	114164783	114164795
refGene_gn=ANO1;refGene_tx=XM_023238951.1;refGene_gn=ANO1;refGene_tx=X
M_023238952.1;refGene_gn=ANO1;refGene_tx=XM_023238955.1;refGene_gn=ANO
1;refGene_tx=XM_023238956.1;refGene_gn=ANO1;refGene_tx=XM_023238957.1;
refGene_gn=ANO1;refGene_tx=XM_023238959.1;refGene_gn=ANO1;refGene_tx=X
M_023238960.1;refGene_gn=ANO1;refGene_tx=XM_023238962.1
D1	114165376	114165448
refGene_gn=ANO1;refGene_tx=XM_023238951.1;refGene_gn=ANO1;refGene_tx=X
M_023238952.1;refGene_gn=ANO1;refGene_tx=XM_023238953.1;refGene_gn=ANO
1;refGene_tx=XM_023238954.1;refGene_gn=ANO1;refGene_tx=XM_023238955.1;
refGene_gn=ANO1;refGene_tx=XM_023238956.1;refGene_gn=ANO1;refGene_tx=X
M_023238957.1;refGene_gn=ANO1;refGene_tx=XM_023238958.1;refGene_gn=ANO
1;refGene_tx=XM_023238959.1;refGene_gn=ANO1;refGene_tx=XM_023238960.1;
refGene_gn=ANO1;refGene_tx=XM_023238961.1;refGene_gn=ANO1;refGene_tx=X
M_023238962.1
D1	114167373	114167451
refGene_gn=ANO1;refGene_tx=XM_023238951.1;refGene_gn=ANO1;refGene_tx=X
M_023238952.1;refGene_gn=ANO1;refGene_tx=XM_023238953.1;refGene_gn=ANO
1;refGene_tx=XM_023238954.1;refGene_gn=ANO1;refGene_tx=XM_023238955.1;
refGene_gn=ANO1;refGene_tx=XM_023238956.1;refGene_gn=ANO1;refGene_tx=X
M_023238960.1;refGene_gn=ANO1;refGene_tx=XM_023238962.1
D1	114168440	114168515
refGene_gn=ANO1;refGene_tx=XM_023238951.1;refGene_gn=ANO1;refGene_tx=X
M_023238952.1;refGene_gn=ANO1;refGene_tx=XM_023238953.1;refGene_gn=ANO
1;refGene_tx=XM_023238954.1;refGene_gn=ANO1;refGene_tx=XM_023238955.1;
refGene_gn=ANO1;refGene_tx=XM_023238956.1;refGene_gn=ANO1;refGene_tx=X
M_023238957.1;refGene_gn=ANO1;refGene_tx=XM_023238958.1;refGene_gn=ANO
1;refGene_tx=XM_023238959.1;refGene_gn=ANO1;refGene_tx=XM_023238960.1;
refGene_gn=ANO1;refGene_tx=XM_023238961.1;refGene_gn=ANO1;refGene_tx=X
M_023238962.1
D1	114170923	114171125
refGene_gn=ANO1;refGene_tx=XM_023238951.1;refGene_gn=ANO1;refGene_tx=X
M_023238952.1;refGene_gn=ANO1;refGene_tx=XM_023238953.1;refGene_gn=ANO
1;refGene_tx=XM_023238954.1;refGene_gn=ANO1;refGene_tx=XM_023238955.1;
refGene_gn=ANO1;refGene_tx=XM_023238956.1;refGene_gn=ANO1;refGene_tx=X
M_023238957.1;refGene_gn=ANO1;refGene_tx=XM_023238958.1;refGene_gn=ANO
1;refGene_tx=XM_023238959.1;refGene_gn=ANO1;refGene_tx=XM_023238960.1;
refGene_gn=ANO1;refGene_tx=XM_023238961.1;refGene_gn=ANO1;refGene_tx=X
M_023238962.1

D1	114171253	114171365
refGene_gn=ANO1;refGene_tx=XM_023238951.1;refGene_gn=ANO1;refGene_tx=XM_023238952.1;refGene_gn=ANO1;refGene_tx=XM_023238953.1;refGene_gn=ANO1;refGene_tx=XM_023238954.1;refGene_gn=ANO1;refGene_tx=XM_023238955.1;refGene_gn=ANO1;refGene_tx=XM_023238956.1;refGene_gn=ANO1;refGene_tx=XM_023238957.1;refGene_gn=ANO1;refGene_tx=XM_023238958.1;refGene_gn=ANO1;refGene_tx=XM_023238959.1;refGene_gn=ANO1;refGene_tx=XM_023238960.1;refGene_gn=ANO1;refGene_tx=XM_023238961.1;refGene_gn=ANO1;refGene_tx=XM_023238962.1
D1	114173785	114173843
refGene_gn=ANO1;refGene_tx=XM_023238951.1;refGene_gn=ANO1;refGene_tx=XM_023238952.1;refGene_gn=ANO1;refGene_tx=XM_023238953.1;refGene_gn=ANO1;refGene_tx=XM_023238954.1;refGene_gn=ANO1;refGene_tx=XM_023238955.1;refGene_gn=ANO1;refGene_tx=XM_023238956.1;refGene_gn=ANO1;refGene_tx=XM_023238957.1;refGene_gn=ANO1;refGene_tx=XM_023238958.1;refGene_gn=ANO1;refGene_tx=XM_023238959.1;refGene_gn=ANO1;refGene_tx=XM_023238960.1;refGene_gn=ANO1;refGene_tx=XM_023238961.1;refGene_gn=ANO1;refGene_tx=XM_023238962.1
D1	114175857	114175958
refGene_gn=ANO1;refGene_tx=XM_023238951.1;refGene_gn=ANO1;refGene_tx=XM_023238952.1;refGene_gn=ANO1;refGene_tx=XM_023238953.1;refGene_gn=ANO1;refGene_tx=XM_023238954.1;refGene_gn=ANO1;refGene_tx=XM_023238955.1;refGene_gn=ANO1;refGene_tx=XM_023238956.1;refGene_gn=ANO1;refGene_tx=XM_023238957.1;refGene_gn=ANO1;refGene_tx=XM_023238958.1;refGene_gn=ANO1;refGene_tx=XM_023238959.1;refGene_gn=ANO1;refGene_tx=XM_023238960.1;refGene_gn=ANO1;refGene_tx=XM_023238961.1;refGene_gn=ANO1;refGene_tx=XM_023238962.1
D1	114176666	114176812
refGene_gn=ANO1;refGene_tx=XM_023238951.1;refGene_gn=ANO1;refGene_tx=XM_023238952.1;refGene_gn=ANO1;refGene_tx=XM_023238953.1;refGene_gn=ANO1;refGene_tx=XM_023238954.1;refGene_gn=ANO1;refGene_tx=XM_023238955.1;refGene_gn=ANO1;refGene_tx=XM_023238956.1;refGene_gn=ANO1;refGene_tx=XM_023238957.1;refGene_gn=ANO1;refGene_tx=XM_023238958.1;refGene_gn=ANO1;refGene_tx=XM_023238959.1;refGene_gn=ANO1;refGene_tx=XM_023238960.1;refGene_gn=ANO1;refGene_tx=XM_023238961.1;refGene_gn=ANO1;refGene_tx=XM_023238962.1
D1	114179503	114179656
refGene_gn=ANO1;refGene_tx=XM_023238951.1;refGene_gn=ANO1;refGene_tx=XM_023238952.1;refGene_gn=ANO1;refGene_tx=XM_023238953.1;refGene_gn=ANO1;refGene_tx=XM_023238954.1;refGene_gn=ANO1;refGene_tx=XM_023238955.1;refGene_gn=ANO1;refGene_tx=XM_023238956.1;refGene_gn=ANO1;refGene_tx=XM_023238957.1;refGene_gn=ANO1;refGene_tx=XM_023238958.1;refGene_gn=ANO1;refGene_tx=XM_023238959.1;refGene_gn=ANO1;refGene_tx=XM_023238960.1;refGene_gn=ANO1;refGene_tx=XM_023238961.1;refGene_gn=ANO1;refGene_tx=XM_023238962.1
D1	114183927	114183980
refGene_gn=ANO1;refGene_tx=XM_023238951.1;refGene_gn=ANO1;refGene_tx=XM_023238952.1;refGene_gn=ANO1;refGene_tx=XM_023238953.1;refGene_gn=ANO1;refGene_tx=XM_023238954.1;refGene_gn=ANO1;refGene_tx=XM_023238955.1;refGene_gn=ANO1;refGene_tx=XM_023238956.1;refGene_gn=ANO1;refGene_tx=X

M_023238957.1;refGene_gn=ANO1;refGene_tx=XM_023238958.1;refGene_gn=ANO1;refGene_tx=XM_023238959.1;refGene_gn=ANO1;refGene_tx=XM_023238960.1;refGene_gn=ANO1;refGene_tx=XM_023238961.1;refGene_gn=ANO1;refGene_tx=XM_023238962.1
D1 114185916 114186101
refGene_gn=ANO1;refGene_tx=XM_023238951.1;refGene_gn=ANO1;refGene_tx=XM_023238952.1;refGene_gn=ANO1;refGene_tx=XM_023238953.1;refGene_gn=ANO1;refGene_tx=XM_023238954.1;refGene_gn=ANO1;refGene_tx=XM_023238955.1;refGene_gn=ANO1;refGene_tx=XM_023238956.1;refGene_gn=ANO1;refGene_tx=XM_023238957.1;refGene_gn=ANO1;refGene_tx=XM_023238958.1;refGene_gn=ANO1;refGene_tx=XM_023238959.1;refGene_gn=ANO1;refGene_tx=XM_023238960.1;refGene_gn=ANO1;refGene_tx=XM_023238961.1;refGene_gn=ANO1;refGene_tx=XM_023238962.1
D1 114190326 114190432
refGene_gn=ANO1;refGene_tx=XM_023238951.1;refGene_gn=ANO1;refGene_tx=XM_023238952.1;refGene_gn=ANO1;refGene_tx=XM_023238953.1;refGene_gn=ANO1;refGene_tx=XM_023238954.1;refGene_gn=ANO1;refGene_tx=XM_023238955.1;refGene_gn=ANO1;refGene_tx=XM_023238956.1;refGene_gn=ANO1;refGene_tx=XM_023238957.1;refGene_gn=ANO1;refGene_tx=XM_023238958.1;refGene_gn=ANO1;refGene_tx=XM_023238959.1;refGene_gn=ANO1;refGene_tx=XM_023238960.1;refGene_gn=ANO1;refGene_tx=XM_023238961.1;refGene_gn=ANO1;refGene_tx=XM_023238962.1
D1 114192011 114192254
refGene_gn=ANO1;refGene_tx=XM_023238951.1;refGene_gn=ANO1;refGene_tx=XM_023238952.1;refGene_gn=ANO1;refGene_tx=XM_023238953.1;refGene_gn=ANO1;refGene_tx=XM_023238954.1;refGene_gn=ANO1;refGene_tx=XM_023238955.1;refGene_gn=ANO1;refGene_tx=XM_023238956.1;refGene_gn=ANO1;refGene_tx=XM_023238957.1;refGene_gn=ANO1;refGene_tx=XM_023238958.1;refGene_gn=ANO1;refGene_tx=XM_023238959.1;refGene_gn=ANO1;refGene_tx=XM_023238961.1;refGene_gn=ANO1;refGene_tx=XM_023238962.1
D1 114193249 114193342
refGene_gn=ANO1;refGene_tx=XM_023238960.1
D1 114206785 114207071
refGene_gn=FADD;refGene_tx=XM_011287100.3
D1 114208971 114209333
refGene_gn=FADD;refGene_tx=XM_011287100.3
D1 114283674 114283842
refGene_gn=LOC101088110;refGene_tx=XM_023239756.1
D1 114289894 114289996
refGene_gn=LOC101088110;refGene_tx=XM_023239756.1
D1 114290267 114290432
refGene_gn=LOC101088110;refGene_tx=XM_023239756.1
D1 114291020 114291158
refGene_gn=LOC101088110;refGene_tx=XM_023239756.1
D1 114292890 114293037
refGene_gn=LOC101088110;refGene_tx=XM_023239756.1
D1 114295832 114295904
refGene_gn=LOC101088110;refGene_tx=XM_023239756.1
D1 114299739 114299823
refGene_gn=LOC101088110;refGene_tx=XM_023239756.1

D1 114301315 114301470
refGene_gn=LOC101088110;refGene_tx=XM_023239756.1
D1 114303216 114303366
refGene_gn=LOC101088110;refGene_tx=XM_023239756.1
D1 114303894 114304011
refGene_gn=LOC101088110;refGene_tx=XM_023239756.1
D1 114310180 114310322
refGene_gn=LOC101088110;refGene_tx=XM_023239756.1
D1 114313228 114313512
refGene_gn=LOC101088110;refGene_tx=XM_023239756.1
D1 114315269 114315645
refGene_gn=LOC101088110;refGene_tx=XM_023239756.1
D1 114320180 114320257
refGene_gn=LOC101088110;refGene_tx=XM_023239756.1
D1 114328078 114328121
refGene_gn=LOC101088110;refGene_tx=XM_023239756.1
D1 114328389 114328590
refGene_gn=LOC101088110;refGene_tx=XM_023239756.1
D1 114328696 114328897
refGene_gn=LOC101088110;refGene_tx=XM_023239756.1
D1 114330451 114330514
refGene_gn=LOC101088110;refGene_tx=XM_023239756.1
D1 114331017 114331044
refGene_gn=LOC101088110;refGene_tx=XM_023239756.1
D1 114336087 114336263
refGene_gn=LOC101088110;refGene_tx=XM_023239756.1
D1 114336367 114336470
refGene_gn=LOC101088110;refGene_tx=XM_023239756.1
D1 114339956 114339983
refGene_gn=LOC111556516;refGene_tx=XM_023238978.1
D1 114341611 114341771
refGene_gn=LOC111556516;refGene_tx=XM_023238978.1
D1 114344005 114344074
refGene_gn=LOC111556516;refGene_tx=XM_023238978.1
D1 114345114 114345140
refGene_gn=LOC111556516;refGene_tx=XM_023238978.1
D1 114363340 114363427
refGene_gn=CTTN;refGene_tx=XM_023238972.1
D1 114363554 114363628
refGene_gn=CTTN;refGene_tx=XM_023238972.1
D1 114365550 114365680
refGene_gn=CTTN;refGene_tx=XM_023238972.1
D1 114368969 114369080
refGene_gn=CTTN;refGene_tx=XM_023238972.1
D1 114370033 114370088
refGene_gn=CTTN;refGene_tx=XM_023238972.1
D1 114370805 114370916
refGene_gn=CTTN;refGene_tx=XM_023238972.1
D1 114372979 114373090
refGene_gn=CTTN;refGene_tx=XM_023238972.1

D1 114374055 114374166
refGene_gn=CTTN;refGene_tx=XM_023238972.1
D1 114376482 114376538
refGene_gn=CTTN;refGene_tx=XM_023238972.1
D1 114378735 114378763
refGene_gn=CTTN;refGene_tx=XM_023238972.1
D1 114378765 114378804
refGene_gn=CTTN;refGene_tx=XM_023238972.1
D1 114380785 114380934
refGene_gn=CTTN;refGene_tx=XM_023238972.1
D1 114383318 114383408
refGene_gn=CTTN;refGene_tx=XM_023238972.1
D1 114384462 114384607
refGene_gn=CTTN;refGene_tx=XM_023238972.1
D1 114385402 114385474
refGene_gn=CTTN;refGene_tx=XM_023238972.1
D1 114386257 114386394
refGene_gn=CTTN;refGene_tx=XM_023238972.1
D1 114409024 114409595
refGene_gn=SHANK2;refGene_tx=XM_023238933.1;refGene_gn=SHANK2;refGene_tx=XM_023238934.1;refGene_gn=SHANK2;refGene_tx=XM_023238935.1;refGene_gn=SHANK2;refGene_tx=XM_023238936.1;refGene_gn=SHANK2;refGene_tx=XM_023238937.1;refGene_gn=SHANK2;refGene_tx=XM_023238939.1;refGene_gn=SHANK2;refGene_tx=XM_023238940.1;refGene_gn=SHANK2;refGene_tx=XM_023238941.1;refGene_gn=SHANK2;refGene_tx=XM_023238942.1;refGene_gn=SHANK2;refGene_tx=XM_023238943.1;refGene_gn=SHANK2;refGene_tx=XM_023238944.1;refGene_gn=SHANK2;refGene_tx=XM_023238948.1;refGene_gn=SHANK2;refGene_tx=XM_023238950.1
D1 114412246 114412322
refGene_gn=SHANK2;refGene_tx=XM_023238946.1;refGene_gn=SHANK2;refGene_tx=XM_023238947.1
D1 114412503 114412582
refGene_gn=SHANK2;refGene_tx=XM_023238945.1
D1 114418868 114421278
refGene_gn=SHANK2;refGene_tx=XM_023238933.1;refGene_gn=SHANK2;refGene_tx=XM_023238934.1;refGene_gn=SHANK2;refGene_tx=XM_023238935.1;refGene_gn=SHANK2;refGene_tx=XM_023238936.1;refGene_gn=SHANK2;refGene_tx=XM_023238937.1;refGene_gn=SHANK2;refGene_tx=XM_023238939.1;refGene_gn=SHANK2;refGene_tx=XM_023238940.1;refGene_gn=SHANK2;refGene_tx=XM_023238941.1;refGene_gn=SHANK2;refGene_tx=XM_023238942.1;refGene_gn=SHANK2;refGene_tx=XM_023238943.1;refGene_gn=SHANK2;refGene_tx=XM_023238944.1;refGene_gn=SHANK2;refGene_tx=XM_023238945.1;refGene_gn=SHANK2;refGene_tx=XM_023238946.1;refGene_gn=SHANK2;refGene_tx=XM_023238947.1;refGene_gn=SHANK2;refGene_tx=XM_023238948.1;refGene_gn=SHANK2;refGene_tx=XM_023238950.1
D1 114422649 114422670
refGene_gn=SHANK2;refGene_tx=XM_023238933.1;refGene_gn=SHANK2;refGene_tx=XM_023238934.1;refGene_gn=SHANK2;refGene_tx=XM_023238935.1;refGene_gn=SHANK2;refGene_tx=XM_023238937.1;refGene_gn=SHANK2;refGene_tx=XM_023238939.1;refGene_gn=SHANK2;refGene_tx=XM_023238940.1;refGene_gn=SHANK

2;refGene_tx=XM_023238942.1;refGene_gn=SHANK2;refGene_tx=XM_023238943.
1;refGene_gn=SHANK2;refGene_tx=XM_023238944.1;refGene_gn=SHANK2;refGen
e_tx=XM_023238945.1;refGene_gn=SHANK2;refGene_tx=XM_023238946.1;refGen
e_gn=SHANK2;refGene_tx=XM_023238947.1;refGene_gn=SHANK2;refGene_tx=XM_
023238948.1;refGene_gn=SHANK2;refGene_tx=XM_023238950.1
D1 114423475 114423587
refGene_gn=SHANK2;refGene_tx=XM_023238933.1;refGene_gn=SHANK2;refGene_
tx=XM_023238934.1;refGene_gn=SHANK2;refGene_tx=XM_023238935.1;refGene_
gn=SHANK2;refGene_tx=XM_023238936.1;refGene_gn=SHANK2;refGene_tx=XM_02
3238937.1;refGene_gn=SHANK2;refGene_tx=XM_023238939.1;refGene_gn=SHANK
2;refGene_tx=XM_023238940.1;refGene_gn=SHANK2;refGene_tx=XM_023238941.
1;refGene_gn=SHANK2;refGene_tx=XM_023238942.1;refGene_gn=SHANK2;refGen
e_tx=XM_023238943.1;refGene_gn=SHANK2;refGene_tx=XM_023238944.1;refGen
e_gn=SHANK2;refGene_tx=XM_023238945.1;refGene_gn=SHANK2;refGene_tx=XM_
023238946.1;refGene_gn=SHANK2;refGene_tx=XM_023238947.1;refGene_gn=SHA
NK2;refGene_tx=XM_023238948.1;refGene_gn=SHANK2;refGene_tx=XM_02323895
0.1
D1 114424634 114424765
refGene_gn=SHANK2;refGene_tx=XM_023238933.1;refGene_gn=SHANK2;refGene_
tx=XM_023238934.1;refGene_gn=SHANK2;refGene_tx=XM_023238935.1;refGene_
gn=SHANK2;refGene_tx=XM_023238936.1;refGene_gn=SHANK2;refGene_tx=XM_02
3238937.1;refGene_gn=SHANK2;refGene_tx=XM_023238939.1;refGene_gn=SHANK
2;refGene_tx=XM_023238940.1;refGene_gn=SHANK2;refGene_tx=XM_023238941.
1;refGene_gn=SHANK2;refGene_tx=XM_023238942.1;refGene_gn=SHANK2;refGen
e_tx=XM_023238943.1;refGene_gn=SHANK2;refGene_tx=XM_023238944.1;refGen
e_gn=SHANK2;refGene_tx=XM_023238945.1;refGene_gn=SHANK2;refGene_tx=XM_
023238946.1;refGene_gn=SHANK2;refGene_tx=XM_023238947.1;refGene_gn=SHA
NK2;refGene_tx=XM_023238948.1;refGene_gn=SHANK2;refGene_tx=XM_02323895
0.1
D1 114431491 114431512
refGene_gn=SHANK2;refGene_tx=XM_023238933.1;refGene_gn=SHANK2;refGene_
tx=XM_023238934.1;refGene_gn=SHANK2;refGene_tx=XM_023238936.1;refGene_
gn=SHANK2;refGene_tx=XM_023238937.1;refGene_gn=SHANK2;refGene_tx=XM_02
3238941.1;refGene_gn=SHANK2;refGene_tx=XM_023238942.1;refGene_gn=SHANK
2;refGene_tx=XM_023238943.1;refGene_gn=SHANK2;refGene_tx=XM_023238945.
1;refGene_gn=SHANK2;refGene_tx=XM_023238946.1;refGene_gn=SHANK2;refGen
e_tx=XM_023238947.1;refGene_gn=SHANK2;refGene_tx=XM_023238948.1;refGen
e_gn=SHANK2;refGene_tx=XM_023238950.1
D1 114432658 114432679
refGene_gn=SHANK2;refGene_tx=XM_023238933.1;refGene_gn=SHANK2;refGene_
tx=XM_023238934.1;refGene_gn=SHANK2;refGene_tx=XM_023238935.1;refGene_
gn=SHANK2;refGene_tx=XM_023238936.1;refGene_gn=SHANK2;refGene_tx=XM_02
3238937.1;refGene_gn=SHANK2;refGene_tx=XM_023238939.1;refGene_gn=SHANK
2;refGene_tx=XM_023238941.1;refGene_gn=SHANK2;refGene_tx=XM_023238942.
1;refGene_gn=SHANK2;refGene_tx=XM_023238943.1;refGene_gn=SHANK2;refGen
e_tx=XM_023238945.1;refGene_gn=SHANK2;refGene_tx=XM_023238946.1;refGen
e_gn=SHANK2;refGene_tx=XM_023238947.1;refGene_gn=SHANK2;refGene_tx=XM_
023238948.1;refGene_gn=SHANK2;refGene_tx=XM_023238950.1
D1 114432947 114433028
refGene_gn=SHANK2;refGene_tx=XM_023238933.1;refGene_gn=SHANK2;refGene_

tx=XM_023238934.1;refGene_gn=SHANK2;refGene_tx=XM_023238935.1;refGene_gn=SHANK2;refGene_tx=XM_023238936.1;refGene_gn=SHANK2;refGene_tx=XM_023238937.1;refGene_gn=SHANK2;refGene_tx=XM_023238939.1;refGene_gn=SHANK2;refGene_tx=XM_023238940.1;refGene_gn=SHANK2;refGene_tx=XM_023238941.1;refGene_gn=SHANK2;refGene_tx=XM_023238942.1;refGene_gn=SHANK2;refGene_tx=XM_023238943.1;refGene_gn=SHANK2;refGene_tx=XM_023238944.1;refGene_gn=SHANK2;refGene_tx=XM_023238945.1;refGene_gn=SHANK2;refGene_tx=XM_023238946.1;refGene_gn=SHANK2;refGene_tx=XM_023238947.1;refGene_gn=SHANK2;refGene_tx=XM_023238948.1;refGene_gn=SHANK2;refGene_tx=XM_023238950.1

D1 114433613 114433749
refGene_gn=SHANK2;refGene_tx=XM_023238933.1;refGene_gn=SHANK2;refGene_tx=XM_023238934.1;refGene_gn=SHANK2;refGene_tx=XM_023238935.1;refGene_gn=SHANK2;refGene_tx=XM_023238936.1;refGene_gn=SHANK2;refGene_tx=XM_023238937.1;refGene_gn=SHANK2;refGene_tx=XM_023238939.1;refGene_gn=SHANK2;refGene_tx=XM_023238940.1;refGene_gn=SHANK2;refGene_tx=XM_023238941.1;refGene_gn=SHANK2;refGene_tx=XM_023238942.1;refGene_gn=SHANK2;refGene_tx=XM_023238943.1;refGene_gn=SHANK2;refGene_tx=XM_023238944.1;refGene_gn=SHANK2;refGene_tx=XM_023238945.1;refGene_gn=SHANK2;refGene_tx=XM_023238946.1;refGene_gn=SHANK2;refGene_tx=XM_023238947.1;refGene_gn=SHANK2;refGene_tx=XM_023238948.1;refGene_gn=SHANK2;refGene_tx=XM_023238950.1

D1 114553475 114553600
refGene_gn=SHANK2;refGene_tx=XM_023238933.1;refGene_gn=SHANK2;refGene_tx=XM_023238934.1;refGene_gn=SHANK2;refGene_tx=XM_023238935.1;refGene_gn=SHANK2;refGene_tx=XM_023238936.1;refGene_gn=SHANK2;refGene_tx=XM_023238937.1;refGene_gn=SHANK2;refGene_tx=XM_023238939.1;refGene_gn=SHANK2;refGene_tx=XM_023238940.1;refGene_gn=SHANK2;refGene_tx=XM_023238941.1;refGene_gn=SHANK2;refGene_tx=XM_023238942.1;refGene_gn=SHANK2;refGene_tx=XM_023238943.1;refGene_gn=SHANK2;refGene_tx=XM_023238944.1;refGene_gn=SHANK2;refGene_tx=XM_023238945.1;refGene_gn=SHANK2;refGene_tx=XM_023238946.1;refGene_gn=SHANK2;refGene_tx=XM_023238947.1;refGene_gn=SHANK2;refGene_tx=XM_023238948.1;refGene_gn=SHANK2;refGene_tx=XM_023238950.1

D1 114555200 114555283
refGene_gn=SHANK2;refGene_tx=XM_023238933.1;refGene_gn=SHANK2;refGene_tx=XM_023238934.1;refGene_gn=SHANK2;refGene_tx=XM_023238935.1;refGene_gn=SHANK2;refGene_tx=XM_023238936.1;refGene_gn=SHANK2;refGene_tx=XM_023238937.1;refGene_gn=SHANK2;refGene_tx=XM_023238939.1;refGene_gn=SHANK2;refGene_tx=XM_023238940.1;refGene_gn=SHANK2;refGene_tx=XM_023238941.1;refGene_gn=SHANK2;refGene_tx=XM_023238942.1;refGene_gn=SHANK2;refGene_tx=XM_023238943.1;refGene_gn=SHANK2;refGene_tx=XM_023238944.1;refGene_gn=SHANK2;refGene_tx=XM_023238945.1;refGene_gn=SHANK2;refGene_tx=XM_023238946.1;refGene_gn=SHANK2;refGene_tx=XM_023238947.1;refGene_gn=SHANK2;refGene_tx=XM_023238948.1;refGene_gn=SHANK2;refGene_tx=XM_023238950.1

D1 114586836 114586912
refGene_gn=SHANK2;refGene_tx=XM_023238933.1;refGene_gn=SHANK2;refGene_tx=XM_023238934.1;refGene_gn=SHANK2;refGene_tx=XM_023238935.1;refGene_gn=SHANK2;refGene_tx=XM_023238936.1;refGene_gn=SHANK2;refGene_tx=XM_02

3238937.1;refGene_gn=SHANK2;refGene_tx=XM_023238939.1;refGene_gn=SHANK2;refGene_tx=XM_023238940.1;refGene_gn=SHANK2;refGene_tx=XM_023238941.1;refGene_gn=SHANK2;refGene_tx=XM_023238942.1;refGene_gn=SHANK2;refGene_tx=XM_023238943.1;refGene_gn=SHANK2;refGene_tx=XM_023238944.1;refGene_gn=SHANK2;refGene_tx=XM_023238945.1;refGene_gn=SHANK2;refGene_tx=XM_023238946.1;refGene_gn=SHANK2;refGene_tx=XM_023238947.1;refGene_gn=SHANK2;refGene_tx=XM_023238948.1;refGene_gn=SHANK2;refGene_tx=XM_023238950.1
D1 114663721 114663835
refGene_gn=SHANK2;refGene_tx=XM_023238933.1;refGene_gn=SHANK2;refGene_tx=XM_023238934.1;refGene_gn=SHANK2;refGene_tx=XM_023238935.1;refGene_gn=SHANK2;refGene_tx=XM_023238936.1;refGene_gn=SHANK2;refGene_tx=XM_023238937.1;refGene_gn=SHANK2;refGene_tx=XM_023238939.1;refGene_gn=SHANK2;refGene_tx=XM_023238940.1;refGene_gn=SHANK2;refGene_tx=XM_023238941.1;refGene_gn=SHANK2;refGene_tx=XM_023238942.1;refGene_gn=SHANK2;refGene_tx=XM_023238943.1;refGene_gn=SHANK2;refGene_tx=XM_023238944.1;refGene_gn=SHANK2;refGene_tx=XM_023238945.1;refGene_gn=SHANK2;refGene_tx=XM_023238946.1;refGene_gn=SHANK2;refGene_tx=XM_023238947.1;refGene_gn=SHANK2;refGene_tx=XM_023238948.1;refGene_gn=SHANK2;refGene_tx=XM_023238950.1
D1 114669762 114669932
refGene_gn=SHANK2;refGene_tx=XM_023238933.1;refGene_gn=SHANK2;refGene_tx=XM_023238934.1;refGene_gn=SHANK2;refGene_tx=XM_023238935.1;refGene_gn=SHANK2;refGene_tx=XM_023238936.1;refGene_gn=SHANK2;refGene_tx=XM_023238937.1;refGene_gn=SHANK2;refGene_tx=XM_023238939.1;refGene_gn=SHANK2;refGene_tx=XM_023238940.1;refGene_gn=SHANK2;refGene_tx=XM_023238941.1;refGene_gn=SHANK2;refGene_tx=XM_023238942.1;refGene_gn=SHANK2;refGene_tx=XM_023238943.1;refGene_gn=SHANK2;refGene_tx=XM_023238944.1;refGene_gn=SHANK2;refGene_tx=XM_023238945.1;refGene_gn=SHANK2;refGene_tx=XM_023238946.1;refGene_gn=SHANK2;refGene_tx=XM_023238947.1;refGene_gn=SHANK2;refGene_tx=XM_023238948.1;refGene_gn=SHANK2;refGene_tx=XM_023238950.1
D1 114681900 114682216
refGene_gn=SHANK2;refGene_tx=XM_023238933.1;refGene_gn=SHANK2;refGene_tx=XM_023238934.1;refGene_gn=SHANK2;refGene_tx=XM_023238935.1;refGene_gn=SHANK2;refGene_tx=XM_023238936.1;refGene_gn=SHANK2;refGene_tx=XM_023238937.1;refGene_gn=SHANK2;refGene_tx=XM_023238939.1;refGene_gn=SHANK2;refGene_tx=XM_023238940.1;refGene_gn=SHANK2;refGene_tx=XM_023238941.1;refGene_gn=SHANK2;refGene_tx=XM_023238942.1;refGene_gn=SHANK2;refGene_tx=XM_023238943.1;refGene_gn=SHANK2;refGene_tx=XM_023238944.1;refGene_gn=SHANK2;refGene_tx=XM_023238945.1;refGene_gn=SHANK2;refGene_tx=XM_023238946.1;refGene_gn=SHANK2;refGene_tx=XM_023238947.1;refGene_gn=SHANK2;refGene_tx=XM_023238948.1;refGene_gn=SHANK2;refGene_tx=XM_023238950.1
D1 114685937 114686133
refGene_gn=SHANK2;refGene_tx=XM_023238948.1;refGene_gn=SHANK2;refGene_tx=XM_023238950.1
D1 114737332 114737503
refGene_gn=SHANK2;refGene_tx=XM_023238933.1;refGene_gn=SHANK2;refGene_tx=XM_023238934.1;refGene_gn=SHANK2;refGene_tx=XM_023238935.1;refGene_

gn=SHANK2;refGene_tx=XM_023238936.1;refGene_gn=SHANK2;refGene_tx=XM_02
3238937.1;refGene_gn=SHANK2;refGene_tx=XM_023238939.1;refGene_gn=SHANK
2;refGene_tx=XM_023238940.1;refGene_gn=SHANK2;refGene_tx=XM_023238941.
1;refGene_gn=SHANK2;refGene_tx=XM_023238945.1;refGene_gn=SHANK2;refGen
e_tx=XM_023238946.1;refGene_gn=SHANK2;refGene_tx=XM_023238947.1
D1 114737875 114737942
refGene_gn=SHANK2;refGene_tx=XM_023238933.1;refGene_gn=SHANK2;refGene_
tx=XM_023238934.1;refGene_gn=SHANK2;refGene_tx=XM_023238935.1;refGene_
gn=SHANK2;refGene_tx=XM_023238936.1;refGene_gn=SHANK2;refGene_tx=XM_02
3238937.1;refGene_gn=SHANK2;refGene_tx=XM_023238939.1;refGene_gn=SHANK
2;refGene_tx=XM_023238940.1;refGene_gn=SHANK2;refGene_tx=XM_023238941.
1;refGene_gn=SHANK2;refGene_tx=XM_023238942.1;refGene_gn=SHANK2;refGen
e_tx=XM_023238943.1;refGene_gn=SHANK2;refGene_tx=XM_023238944.1;refGen
e_gn=SHANK2;refGene_tx=XM_023238945.1;refGene_gn=SHANK2;refGene_tx=XM_
023238946.1;refGene_gn=SHANK2;refGene_tx=XM_023238947.1
D1 114774165 114774243
refGene_gn=SHANK2;refGene_tx=XM_023238933.1;refGene_gn=SHANK2;refGene_
tx=XM_023238934.1;refGene_gn=SHANK2;refGene_tx=XM_023238935.1;refGene_
gn=SHANK2;refGene_tx=XM_023238936.1;refGene_gn=SHANK2;refGene_tx=XM_02
3238937.1;refGene_gn=SHANK2;refGene_tx=XM_023238939.1;refGene_gn=SHANK
2;refGene_tx=XM_023238940.1;refGene_gn=SHANK2;refGene_tx=XM_023238941.
1;refGene_gn=SHANK2;refGene_tx=XM_023238942.1;refGene_gn=SHANK2;refGen
e_tx=XM_023238943.1;refGene_gn=SHANK2;refGene_tx=XM_023238944.1;refGen
e_gn=SHANK2;refGene_tx=XM_023238945.1;refGene_gn=SHANK2;refGene_tx=XM_
023238946.1;refGene_gn=SHANK2;refGene_tx=XM_023238947.1
D1 114785985 114786102
refGene_gn=SHANK2;refGene_tx=XM_023238933.1;refGene_gn=SHANK2;refGene_
tx=XM_023238934.1;refGene_gn=SHANK2;refGene_tx=XM_023238935.1;refGene_
gn=SHANK2;refGene_tx=XM_023238936.1;refGene_gn=SHANK2;refGene_tx=XM_02
3238937.1;refGene_gn=SHANK2;refGene_tx=XM_023238939.1;refGene_gn=SHANK
2;refGene_tx=XM_023238940.1;refGene_gn=SHANK2;refGene_tx=XM_023238941.
1;refGene_gn=SHANK2;refGene_tx=XM_023238942.1;refGene_gn=SHANK2;refGen
e_tx=XM_023238943.1;refGene_gn=SHANK2;refGene_tx=XM_023238944.1;refGen
e_gn=SHANK2;refGene_tx=XM_023238945.1;refGene_gn=SHANK2;refGene_tx=XM_
023238946.1;refGene_gn=SHANK2;refGene_tx=XM_023238947.1
D1 114794812 114794980
refGene_gn=SHANK2;refGene_tx=XM_023238933.1;refGene_gn=SHANK2;refGene_
tx=XM_023238934.1;refGene_gn=SHANK2;refGene_tx=XM_023238935.1;refGene_
gn=SHANK2;refGene_tx=XM_023238936.1;refGene_gn=SHANK2;refGene_tx=XM_02
3238937.1;refGene_gn=SHANK2;refGene_tx=XM_023238939.1;refGene_gn=SHANK
2;refGene_tx=XM_023238940.1;refGene_gn=SHANK2;refGene_tx=XM_023238941.
1;refGene_gn=SHANK2;refGene_tx=XM_023238942.1;refGene_gn=SHANK2;refGen
e_tx=XM_023238943.1;refGene_gn=SHANK2;refGene_tx=XM_023238944.1;refGen
e_gn=SHANK2;refGene_tx=XM_023238945.1;refGene_gn=SHANK2;refGene_tx=XM_
023238946.1;refGene_gn=SHANK2;refGene_tx=XM_023238947.1
D1 114795920 114796072
refGene_gn=SHANK2;refGene_tx=XM_023238933.1;refGene_gn=SHANK2;refGene_
tx=XM_023238934.1;refGene_gn=SHANK2;refGene_tx=XM_023238935.1;refGene_
gn=SHANK2;refGene_tx=XM_023238936.1;refGene_gn=SHANK2;refGene_tx=XM_02
3238937.1;refGene_gn=SHANK2;refGene_tx=XM_023238939.1;refGene_gn=SHANK

2;refGene_tx=XM_023238940.1;refGene_gn=SHANK2;refGene_tx=XM_023238941.1;refGene_gn=SHANK2;refGene_tx=XM_023238942.1;refGene_gn=SHANK2;refGene_tx=XM_023238943.1;refGene_gn=SHANK2;refGene_tx=XM_023238944.1;refGene_gn=SHANK2;refGene_tx=XM_023238945.1;refGene_gn=SHANK2;refGene_tx=XM_023238946.1;refGene_gn=SHANK2;refGene_tx=XM_023238947.1
D1 114806566 114806675
refGene_gn=SHANK2;refGene_tx=XM_023238933.1;refGene_gn=SHANK2;refGene_tx=XM_023238934.1;refGene_gn=SHANK2;refGene_tx=XM_023238935.1;refGene_gn=SHANK2;refGene_tx=XM_023238936.1;refGene_gn=SHANK2;refGene_tx=XM_023238937.1;refGene_gn=SHANK2;refGene_tx=XM_023238939.1;refGene_gn=SHANK2;refGene_tx=XM_023238940.1;refGene_gn=SHANK2;refGene_tx=XM_023238941.1;refGene_gn=SHANK2;refGene_tx=XM_023238942.1;refGene_gn=SHANK2;refGene_tx=XM_023238943.1;refGene_gn=SHANK2;refGene_tx=XM_023238944.1;refGene_gn=SHANK2;refGene_tx=XM_023238945.1;refGene_gn=SHANK2;refGene_tx=XM_023238946.1;refGene_gn=SHANK2;refGene_tx=XM_023238947.1
D1 114808686 114808758
refGene_gn=SHANK2;refGene_tx=XM_023238933.1;refGene_gn=SHANK2;refGene_tx=XM_023238934.1;refGene_gn=SHANK2;refGene_tx=XM_023238935.1;refGene_gn=SHANK2;refGene_tx=XM_023238936.1;refGene_gn=SHANK2;refGene_tx=XM_023238937.1;refGene_gn=SHANK2;refGene_tx=XM_023238939.1;refGene_gn=SHANK2;refGene_tx=XM_023238940.1;refGene_gn=SHANK2;refGene_tx=XM_023238941.1;refGene_gn=SHANK2;refGene_tx=XM_023238942.1;refGene_gn=SHANK2;refGene_tx=XM_023238943.1;refGene_gn=SHANK2;refGene_tx=XM_023238944.1;refGene_gn=SHANK2;refGene_tx=XM_023238945.1;refGene_gn=SHANK2;refGene_tx=XM_023238946.1;refGene_gn=SHANK2;refGene_tx=XM_023238947.1
D1 114811897 114812101
refGene_gn=SHANK2;refGene_tx=XM_023238933.1;refGene_gn=SHANK2;refGene_tx=XM_023238934.1;refGene_gn=SHANK2;refGene_tx=XM_023238935.1;refGene_gn=SHANK2;refGene_tx=XM_023238936.1;refGene_gn=SHANK2;refGene_tx=XM_023238937.1;refGene_gn=SHANK2;refGene_tx=XM_023238939.1;refGene_gn=SHANK2;refGene_tx=XM_023238940.1;refGene_gn=SHANK2;refGene_tx=XM_023238941.1;refGene_gn=SHANK2;refGene_tx=XM_023238942.1;refGene_gn=SHANK2;refGene_tx=XM_023238943.1;refGene_gn=SHANK2;refGene_tx=XM_023238944.1;refGene_gn=SHANK2;refGene_tx=XM_023238945.1;refGene_gn=SHANK2;refGene_tx=XM_023238946.1;refGene_gn=SHANK2;refGene_tx=XM_023238947.1
D1 114834842 114835049
refGene_gn=SHANK2;refGene_tx=XM_023238933.1;refGene_gn=SHANK2;refGene_tx=XM_023238934.1;refGene_gn=SHANK2;refGene_tx=XM_023238935.1;refGene_gn=SHANK2;refGene_tx=XM_023238936.1;refGene_gn=SHANK2;refGene_tx=XM_023238937.1;refGene_gn=SHANK2;refGene_tx=XM_023238939.1;refGene_gn=SHANK2;refGene_tx=XM_023238940.1;refGene_gn=SHANK2;refGene_tx=XM_023238941.1;refGene_gn=SHANK2;refGene_tx=XM_023238942.1;refGene_gn=SHANK2;refGene_tx=XM_023238943.1;refGene_gn=SHANK2;refGene_tx=XM_023238944.1;refGene_gn=SHANK2;refGene_tx=XM_023238945.1;refGene_gn=SHANK2;refGene_tx=XM_023238946.1;refGene_gn=SHANK2;refGene_tx=XM_023238947.1
D1 115000622 115000766
refGene_gn=LOC109492241;refGene_tx=XM_019812835.1
D1 115006814 115006996
refGene_gn=LOC109492241;refGene_tx=XM_019812835.1
D1 115008836 115009025

refGene_gn=LOC109492241;refGene_tx=XM_019812835.1
D1 115009812 115009974
refGene_gn=LOC109492241;refGene_tx=XM_019812835.1
D1 115013374 115013459
refGene_gn=LOC109492241;refGene_tx=XM_019812835.1
D1 115017874 115017985
refGene_gn=LOC109492241;refGene_tx=XM_019812835.1
D1 115021166 115021295
refGene_gn=LOC109492241;refGene_tx=XM_019812835.1
D1 115025419 115025533
refGene_gn=LOC109492241;refGene_tx=XM_019812835.1
D1 115043833 115044298
refGene_gn=DHCR7;refGene_tx=XM_003993771.3;refGene_gn=DHCR7;refGene_tx
=XM_019812836.1;refGene_gn=DHCR7;refGene_tx=XM_019812837.2
D1 115047323 115047455
refGene_gn=DHCR7;refGene_tx=XM_003993771.3;refGene_gn=DHCR7;refGene_tx
=XM_019812836.1;refGene_gn=DHCR7;refGene_tx=XM_019812837.2
D1 115047874 115048079
refGene_gn=DHCR7;refGene_tx=XM_003993771.3;refGene_gn=DHCR7;refGene_tx
=XM_019812836.1;refGene_gn=DHCR7;refGene_tx=XM_019812837.2
D1 115050577 115050791
refGene_gn=DHCR7;refGene_tx=XM_003993771.3;refGene_gn=DHCR7;refGene_tx
=XM_019812836.1;refGene_gn=DHCR7;refGene_tx=XM_019812837.2
D1 115051591 115051682
refGene_gn=DHCR7;refGene_tx=XM_003993771.3;refGene_gn=DHCR7;refGene_tx
=XM_019812836.1;refGene_gn=DHCR7;refGene_tx=XM_019812837.2
D1 115053046 115053269
refGene_gn=DHCR7;refGene_tx=XM_003993771.3;refGene_gn=DHCR7;refGene_tx
=XM_019812836.1;refGene_gn=DHCR7;refGene_tx=XM_019812837.2
D1 115053832 115053930
refGene_gn=DHCR7;refGene_tx=XM_003993771.3;refGene_gn=DHCR7;refGene_tx
=XM_019812836.1;refGene_gn=DHCR7;refGene_tx=XM_019812837.2
D1 115064292 115064377
refGene_gn=NADSYN1;refGene_tx=XM_023238965.1;refGene_gn=NADSYN1;refGen
e_tx=XM_023238967.1;refGene_gn=NADSYN1;refGene_tx=XM_023238968.1;refGe
ne_gn=NADSYN1;refGene_tx=XM_023238969.1;refGene_gn=NADSYN1;refGene_tx=
XM_023238970.1
D1 115066103 115066164
refGene_gn=NADSYN1;refGene_tx=XM_023238965.1;refGene_gn=NADSYN1;refGen
e_tx=XM_023238967.1;refGene_gn=NADSYN1;refGene_tx=XM_023238968.1;refGe
ne_gn=NADSYN1;refGene_tx=XM_023238969.1;refGene_gn=NADSYN1;refGene_tx=
XM_023238970.1
D1 115069113 115069230
refGene_gn=NADSYN1;refGene_tx=XM_023238965.1;refGene_gn=NADSYN1;refGen
e_tx=XM_023238967.1;refGene_gn=NADSYN1;refGene_tx=XM_023238968.1;refGe
ne_gn=NADSYN1;refGene_tx=XM_023238969.1;refGene_gn=NADSYN1;refGene_tx=
XM_023238970.1
D1 115070918 115070972
refGene_gn=NADSYN1;refGene_tx=XM_023238965.1;refGene_gn=NADSYN1;refGen
e_tx=XM_023238967.1;refGene_gn=NADSYN1;refGene_tx=XM_023238968.1;refGe

ne_gn=NADSYN1;refGene_tx=XM_023238969.1;refGene_gn=NADSYN1;refGene_tx=XM_023238970.1
D1	115071386	115071476
refGene_gn=NADSYN1;refGene_tx=XM_023238965.1;refGene_gn=NADSYN1;refGene_tx=XM_023238968.1;refGene_gn=NADSYN1;refGene_tx=XM_023238969.1;refGene_gn=NADSYN1;refGene_tx=XM_023238970.1;refGene_gn=NADSYN1;refGene_tx=XM_023238971.1
D1	115076029	115076081
refGene_gn=NADSYN1;refGene_tx=XM_023238965.1;refGene_gn=NADSYN1;refGene_tx=XM_023238967.1;refGene_gn=NADSYN1;refGene_tx=XM_023238968.1;refGene_gn=NADSYN1;refGene_tx=XM_023238969.1;refGene_gn=NADSYN1;refGene_tx=XM_023238970.1;refGene_gn=NADSYN1;refGene_tx=XM_023238971.1
D1	115079243	115079332
refGene_gn=NADSYN1;refGene_tx=XM_023238965.1;refGene_gn=NADSYN1;refGene_tx=XM_023238967.1;refGene_gn=NADSYN1;refGene_tx=XM_023238968.1;refGene_gn=NADSYN1;refGene_tx=XM_023238969.1;refGene_gn=NADSYN1;refGene_tx=XM_023238970.1;refGene_gn=NADSYN1;refGene_tx=XM_023238971.1
D1	115079436	115079554
refGene_gn=NADSYN1;refGene_tx=XM_023238965.1;refGene_gn=NADSYN1;refGene_tx=XM_023238967.1;refGene_gn=NADSYN1;refGene_tx=XM_023238968.1;refGene_gn=NADSYN1;refGene_tx=XM_023238969.1;refGene_gn=NADSYN1;refGene_tx=XM_023238970.1;refGene_gn=NADSYN1;refGene_tx=XM_023238971.1
D1	115080234	115080366
refGene_gn=NADSYN1;refGene_tx=XM_023238965.1;refGene_gn=NADSYN1;refGene_tx=XM_023238967.1;refGene_gn=NADSYN1;refGene_tx=XM_023238968.1;refGene_gn=NADSYN1;refGene_tx=XM_023238969.1;refGene_gn=NADSYN1;refGene_tx=XM_023238970.1;refGene_gn=NADSYN1;refGene_tx=XM_023238971.1
D1	115082748	115082823
refGene_gn=NADSYN1;refGene_tx=XM_023238965.1;refGene_gn=NADSYN1;refGene_tx=XM_023238967.1;refGene_gn=NADSYN1;refGene_tx=XM_023238968.1;refGene_gn=NADSYN1;refGene_tx=XM_023238969.1;refGene_gn=NADSYN1;refGene_tx=XM_023238970.1;refGene_gn=NADSYN1;refGene_tx=XM_023238971.1
D1	115083567	115083692
refGene_gn=NADSYN1;refGene_tx=XM_023238965.1;refGene_gn=NADSYN1;refGene_tx=XM_023238967.1;refGene_gn=NADSYN1;refGene_tx=XM_023238968.1;refGene_gn=NADSYN1;refGene_tx=XM_023238969.1;refGene_gn=NADSYN1;refGene_tx=XM_023238970.1;refGene_gn=NADSYN1;refGene_tx=XM_023238971.1
D1	115084164	115084213
refGene_gn=NADSYN1;refGene_tx=XM_023238965.1;refGene_gn=NADSYN1;refGene_tx=XM_023238967.1;refGene_gn=NADSYN1;refGene_tx=XM_023238968.1;refGene_gn=NADSYN1;refGene_tx=XM_023238969.1;refGene_gn=NADSYN1;refGene_tx=XM_023238970.1;refGene_gn=NADSYN1;refGene_tx=XM_023238971.1
D1	115084630	115084733
refGene_gn=NADSYN1;refGene_tx=XM_023238965.1;refGene_gn=NADSYN1;refGene_tx=XM_023238967.1;refGene_gn=NADSYN1;refGene_tx=XM_023238968.1;refGene_gn=NADSYN1;refGene_tx=XM_023238969.1;refGene_gn=NADSYN1;refGene_tx=XM_023238970.1;refGene_gn=NADSYN1;refGene_tx=XM_023238971.1
D1	115085131	115085300
refGene_gn=NADSYN1;refGene_tx=XM_023238965.1;refGene_gn=NADSYN1;refGene_tx=XM_023238967.1;refGene_gn=NADSYN1;refGene_tx=XM_023238968.1;refGe

ne_gn=NADSYN1;refGene_tx=XM_023238969.1;refGene_gn=NADSYN1;refGene_tx=XM_023238970.1;refGene_gn=NADSYN1;refGene_tx=XM_023238971.1
D1 115086035 115086171
refGene_gn=NADSYN1;refGene_tx=XM_023238965.1;refGene_gn=NADSYN1;refGene_tx=XM_023238967.1;refGene_gn=NADSYN1;refGene_tx=XM_023238968.1;refGene_gn=NADSYN1;refGene_tx=XM_023238969.1;refGene_gn=NADSYN1;refGene_tx=XM_023238970.1;refGene_gn=NADSYN1;refGene_tx=XM_023238971.1
D1 115086975 115087082
refGene_gn=NADSYN1;refGene_tx=XM_023238965.1;refGene_gn=NADSYN1;refGene_tx=XM_023238967.1;refGene_gn=NADSYN1;refGene_tx=XM_023238968.1;refGene_gn=NADSYN1;refGene_tx=XM_023238969.1;refGene_gn=NADSYN1;refGene_tx=XM_023238970.1;refGene_gn=NADSYN1;refGene_tx=XM_023238971.1
D1 115096093 115096225
refGene_gn=NADSYN1;refGene_tx=XM_023238965.1;refGene_gn=NADSYN1;refGene_tx=XM_023238967.1;refGene_gn=NADSYN1;refGene_tx=XM_023238968.1;refGene_gn=NADSYN1;refGene_tx=XM_023238969.1;refGene_gn=NADSYN1;refGene_tx=XM_023238970.1;refGene_gn=NADSYN1;refGene_tx=XM_023238971.1
D1 115096770 115096840
refGene_gn=NADSYN1;refGene_tx=XM_023238965.1;refGene_gn=NADSYN1;refGene_tx=XM_023238967.1;refGene_gn=NADSYN1;refGene_tx=XM_023238968.1;refGene_gn=NADSYN1;refGene_tx=XM_023238969.1;refGene_gn=NADSYN1;refGene_tx=XM_023238970.1;refGene_gn=NADSYN1;refGene_tx=XM_023238971.1
D1 115097229 115097448
refGene_gn=NADSYN1;refGene_tx=XM_023238969.1;refGene_gn=NADSYN1;refGene_tx=XM_023238970.1
D1 115099915 115100044
refGene_gn=NADSYN1;refGene_tx=XM_023238965.1;refGene_gn=NADSYN1;refGene_tx=XM_023238967.1;refGene_gn=NADSYN1;refGene_tx=XM_023238968.1;refGene_gn=NADSYN1;refGene_tx=XM_023238971.1
D1 115100536 115100713
refGene_gn=NADSYN1;refGene_tx=XM_023238965.1;refGene_gn=NADSYN1;refGene_tx=XM_023238967.1;refGene_gn=NADSYN1;refGene_tx=XM_023238968.1;refGene_gn=NADSYN1;refGene_tx=XM_023238971.1
D1 115101614 115101785
refGene_gn=NADSYN1;refGene_tx=XM_023238965.1;refGene_gn=NADSYN1;refGene_tx=XM_023238967.1;refGene_gn=NADSYN1;refGene_tx=XM_023238971.1
D1 115101824 115101875
refGene_gn=NADSYN1;refGene_tx=XM_023238968.1
D1 115145341 115145387
refGene_gn=MRGPRG;refGene_tx=XM_019812998.2
D1 115145389 115145625
refGene_gn=MRGPRG;refGene_tx=XM_019812998.2
D1 115145627 115145772
refGene_gn=MRGPRG;refGene_tx=XM_019812998.2
D1 115145774 115145862
refGene_gn=MRGPRG;refGene_tx=XM_019812998.2
D1 115145863 115146028
refGene_gn=MRGPRG;refGene_tx=XM_019812998.2
D1 115146029 115146127
refGene_gn=MRGPRG;refGene_tx=XM_019812998.2

D1 115146129 115146179
refGene_gn=MRGPRG;refGene_tx=XM_019812998.2
D1 115162470 115162485
refGene_gn=OSBPL5;refGene_tx=XM_019812839.2
D1 115206928 115207000
refGene_gn=OSBPL5;refGene_tx=XM_019812838.2
D1 115216906 115217063
refGene_gn=OSBPL5;refGene_tx=XM_006937631.4;refGene_gn=OSBPL5;refGene_tx=XM_019812838.2;refGene_gn=OSBPL5;refGene_tx=XM_019812839.2;refGene_gn=OSBPL5;refGene_tx=XM_019812840.2;refGene_gn=OSBPL5;refGene_tx=XM_019812841.2;refGene_gn=OSBPL5;refGene_tx=XM_023238964.1
D1 115218994 115219077
refGene_gn=OSBPL5;refGene_tx=XM_006937631.4;refGene_gn=OSBPL5;refGene_tx=XM_019812838.2;refGene_gn=OSBPL5;refGene_tx=XM_019812839.2;refGene_gn=OSBPL5;refGene_tx=XM_019812840.2;refGene_gn=OSBPL5;refGene_tx=XM_019812841.2;refGene_gn=OSBPL5;refGene_tx=XM_023238964.1
D1 115224491 115224572
refGene_gn=OSBPL5;refGene_tx=XM_006937631.4;refGene_gn=OSBPL5;refGene_tx=XM_019812838.2;refGene_gn=OSBPL5;refGene_tx=XM_019812839.2;refGene_gn=OSBPL5;refGene_tx=XM_019812840.2;refGene_gn=OSBPL5;refGene_tx=XM_019812841.2;refGene_gn=OSBPL5;refGene_tx=XM_023238964.1
D1 115224828 115224930
refGene_gn=OSBPL5;refGene_tx=XM_006937631.4;refGene_gn=OSBPL5;refGene_tx=XM_019812838.2;refGene_gn=OSBPL5;refGene_tx=XM_019812839.2;refGene_gn=OSBPL5;refGene_tx=XM_019812840.2;refGene_gn=OSBPL5;refGene_tx=XM_019812841.2;refGene_gn=OSBPL5;refGene_tx=XM_023238964.1
D1 115226003 115226207
refGene_gn=OSBPL5;refGene_tx=XM_006937631.4;refGene_gn=OSBPL5;refGene_tx=XM_019812838.2;refGene_gn=OSBPL5;refGene_tx=XM_019812839.2;refGene_gn=OSBPL5;refGene_tx=XM_019812840.2;refGene_gn=OSBPL5;refGene_tx=XM_019812841.2;refGene_gn=OSBPL5;refGene_tx=XM_023238964.1
D1 115226908 115226993
refGene_gn=OSBPL5;refGene_tx=XM_006937631.4;refGene_gn=OSBPL5;refGene_tx=XM_019812838.2;refGene_gn=OSBPL5;refGene_tx=XM_019812839.2;refGene_gn=OSBPL5;refGene_tx=XM_019812840.2;refGene_gn=OSBPL5;refGene_tx=XM_019812841.2;refGene_gn=OSBPL5;refGene_tx=XM_023238964.1
D1 115228841 115229013
refGene_gn=OSBPL5;refGene_tx=XM_006937631.4;refGene_gn=OSBPL5;refGene_tx=XM_019812838.2;refGene_gn=OSBPL5;refGene_tx=XM_019812839.2;refGene_gn=OSBPL5;refGene_tx=XM_019812840.2;refGene_gn=OSBPL5;refGene_tx=XM_019812841.2;refGene_gn=OSBPL5;refGene_tx=XM_023238964.1
D1 115229339 115229535
refGene_gn=OSBPL5;refGene_tx=XM_006937631.4;refGene_gn=OSBPL5;refGene_tx=XM_019812838.2;refGene_gn=OSBPL5;refGene_tx=XM_019812839.2;refGene_gn=OSBPL5;refGene_tx=XM_019812840.2;refGene_gn=OSBPL5;refGene_tx=XM_019812841.2;refGene_gn=OSBPL5;refGene_tx=XM_023238964.1
D1 115231518 115231703
refGene_gn=OSBPL5;refGene_tx=XM_006937631.4;refGene_gn=OSBPL5;refGene_tx=XM_019812838.2;refGene_gn=OSBPL5;refGene_tx=XM_019812839.2;refGene_gn=OSBPL5;refGene_tx=XM_019812840.2;refGene_gn=OSBPL5;refGene_tx=XM_01

9812841.2;refGene_gn=OSBPL5;refGene_tx=XM_023238964.1
D1 115232330 115232412
refGene_gn=OSBPL5;refGene_tx=XM_006937631.4;refGene_gn=OSBPL5;refGene_tx=XM_019812838.2;refGene_gn=OSBPL5;refGene_tx=XM_019812839.2;refGene_gn=OSBPL5;refGene_tx=XM_019812840.2;refGene_gn=OSBPL5;refGene_tx=XM_019812841.2;refGene_gn=OSBPL5;refGene_tx=XM_023238964.1
D1 115234080 115234179
refGene_gn=OSBPL5;refGene_tx=XM_006937631.4;refGene_gn=OSBPL5;refGene_tx=XM_019812838.2;refGene_gn=OSBPL5;refGene_tx=XM_019812839.2;refGene_gn=OSBPL5;refGene_tx=XM_019812840.2;refGene_gn=OSBPL5;refGene_tx=XM_019812841.2;refGene_gn=OSBPL5;refGene_tx=XM_023238964.1
D1 115234523 115234620
refGene_gn=OSBPL5;refGene_tx=XM_006937631.4;refGene_gn=OSBPL5;refGene_tx=XM_019812838.2;refGene_gn=OSBPL5;refGene_tx=XM_019812839.2;refGene_gn=OSBPL5;refGene_tx=XM_019812840.2;refGene_gn=OSBPL5;refGene_tx=XM_019812841.2;refGene_gn=OSBPL5;refGene_tx=XM_023238964.1
D1 115235532 115235631
refGene_gn=OSBPL5;refGene_tx=XM_006937631.4;refGene_gn=OSBPL5;refGene_tx=XM_019812838.2;refGene_gn=OSBPL5;refGene_tx=XM_019812839.2;refGene_gn=OSBPL5;refGene_tx=XM_019812840.2;refGene_gn=OSBPL5;refGene_tx=XM_019812841.2;refGene_gn=OSBPL5;refGene_tx=XM_023238964.1
D1 115237196 115237294
refGene_gn=OSBPL5;refGene_tx=XM_006937631.4;refGene_gn=OSBPL5;refGene_tx=XM_019812838.2;refGene_gn=OSBPL5;refGene_tx=XM_019812839.2;refGene_gn=OSBPL5;refGene_tx=XM_019812840.2;refGene_gn=OSBPL5;refGene_tx=XM_019812841.2;refGene_gn=OSBPL5;refGene_tx=XM_023238964.1
D1 115237716 115237806
refGene_gn=OSBPL5;refGene_tx=XM_006937631.4;refGene_gn=OSBPL5;refGene_tx=XM_019812838.2;refGene_gn=OSBPL5;refGene_tx=XM_019812839.2;refGene_gn=OSBPL5;refGene_tx=XM_019812840.2;refGene_gn=OSBPL5;refGene_tx=XM_019812841.2;refGene_gn=OSBPL5;refGene_tx=XM_023238964.1
D1 115237893 115238030
refGene_gn=OSBPL5;refGene_tx=XM_006937631.4;refGene_gn=OSBPL5;refGene_tx=XM_019812838.2;refGene_gn=OSBPL5;refGene_tx=XM_019812839.2;refGene_gn=OSBPL5;refGene_tx=XM_019812840.2;refGene_gn=OSBPL5;refGene_tx=XM_019812841.2;refGene_gn=OSBPL5;refGene_tx=XM_023238964.1
D1 115238474 115238660
refGene_gn=OSBPL5;refGene_tx=XM_006937631.4;refGene_gn=OSBPL5;refGene_tx=XM_019812838.2;refGene_gn=OSBPL5;refGene_tx=XM_019812839.2;refGene_gn=OSBPL5;refGene_tx=XM_019812840.2;refGene_gn=OSBPL5;refGene_tx=XM_019812841.2;refGene_gn=OSBPL5;refGene_tx=XM_023238964.1
D1 115238904 115239031
refGene_gn=OSBPL5;refGene_tx=XM_006937631.4;refGene_gn=OSBPL5;refGene_tx=XM_019812838.2;refGene_gn=OSBPL5;refGene_tx=XM_019812839.2;refGene_gn=OSBPL5;refGene_tx=XM_019812840.2;refGene_gn=OSBPL5;refGene_tx=XM_019812841.2;refGene_gn=OSBPL5;refGene_tx=XM_023238964.1
D1 115240100 115240239
refGene_gn=OSBPL5;refGene_tx=XM_006937631.4;refGene_gn=OSBPL5;refGene_tx=XM_019812838.2;refGene_gn=OSBPL5;refGene_tx=XM_019812839.2;refGene_gn=OSBPL5;refGene_tx=XM_019812840.2;refGene_gn=OSBPL5;refGene_tx=XM_01

9812841.2;refGene_gn=OSBPL5;refGene_tx=XM_023238964.1
D1 115240786 115240889
refGene_gn=OSBPL5;refGene_tx=XM_006937631.4;refGene_gn=OSBPL5;refGene_tx=XM_019812838.2;refGene_gn=OSBPL5;refGene_tx=XM_019812839.2;refGene_gn=OSBPL5;refGene_tx=XM_019812840.2;refGene_gn=OSBPL5;refGene_tx=XM_019812841.2;refGene_gn=OSBPL5;refGene_tx=XM_023238964.1
D1 115241897 115242036
refGene_gn=OSBPL5;refGene_tx=XM_006937631.4;refGene_gn=OSBPL5;refGene_tx=XM_019812838.2;refGene_gn=OSBPL5;refGene_tx=XM_019812839.2;refGene_gn=OSBPL5;refGene_tx=XM_019812840.2;refGene_gn=OSBPL5;refGene_tx=XM_019812841.2;refGene_gn=OSBPL5;refGene_tx=XM_023238964.1
D1 115260433 115260493
refGene_gn=LOC105261012;refGene_tx=XM_011287091.3
D1 115265732 115265820
refGene_gn=LOC105261012;refGene_tx=XM_011287091.3
D1 115267068 115267197
refGene_gn=LOC105261012;refGene_tx=XM_011287091.3
D1 115269227 115269345
refGene_gn=LOC105261012;refGene_tx=XM_011287091.3
D1 115270547 115270627
refGene_gn=LOC105261012;refGene_tx=XM_011287091.3
D1 115272721 115272820
refGene_gn=LOC105261012;refGene_tx=XM_011287091.3
D1 115273419 115273461
refGene_gn=LOC105261012;refGene_tx=XM_011287091.3
D1 115273559 115273585
refGene_gn=LOC105261012;refGene_tx=XM_011287091.3
D1 115275082 115275376
refGene_gn=LOC105261012;refGene_tx=XM_011287091.3
D1 115276991 115277034
refGene_gn=LOC105261012;refGene_tx=XM_011287091.3
D1 115277902 115278207
refGene_gn=LOC105261012;refGene_tx=XM_011287091.3
D1 115280565 115280590
refGene_gn=CARS;refGene_tx=XM_011287089.2;refGene_gn=CARS;refGene_tx=XM_011287090.3
D1 115288185 115288285
refGene_gn=CARS;refGene_tx=XM_019812842.1
D1 115288632 115288881
refGene_gn=CARS;refGene_tx=XM_011287089.2;refGene_gn=CARS;refGene_tx=XM_019812842.1
D1 115292877 115292969
refGene_gn=CARS;refGene_tx=XM_011287089.2;refGene_gn=CARS;refGene_tx=XM_011287090.3;refGene_gn=CARS;refGene_tx=XM_019812842.1
D1 115293648 115293737
refGene_gn=CARS;refGene_tx=XM_011287089.2;refGene_gn=CARS;refGene_tx=XM_011287090.3;refGene_gn=CARS;refGene_tx=XM_019812842.1
D1 115294194 115294291
refGene_gn=CARS;refGene_tx=XM_011287089.2;refGene_gn=CARS;refGene_tx=XM_011287090.3;refGene_gn=CARS;refGene_tx=XM_019812842.1

D1 115294702 115294801
refGene_gn=CARS;refGene_tx=XM_011287089.2;refGene_gn=CARS;refGene_tx=XM_011287090.3;refGene_gn=CARS;refGene_tx=XM_019812842.1
D1 115295651 115295801
refGene_gn=CARS;refGene_tx=XM_011287089.2;refGene_gn=CARS;refGene_tx=XM_011287090.3;refGene_gn=CARS;refGene_tx=XM_019812842.1
D1 115303846 115303987
refGene_gn=CARS;refGene_tx=XM_011287089.2;refGene_gn=CARS;refGene_tx=XM_011287090.3;refGene_gn=CARS;refGene_tx=XM_019812842.1
D1 115304189 115304278
refGene_gn=CARS;refGene_tx=XM_011287089.2;refGene_gn=CARS;refGene_tx=XM_011287090.3;refGene_gn=CARS;refGene_tx=XM_019812842.1
D1 115305815 115305937
refGene_gn=CARS;refGene_tx=XM_011287089.2;refGene_gn=CARS;refGene_tx=XM_011287090.3;refGene_gn=CARS;refGene_tx=XM_019812842.1
D1 115309639 115309752
refGene_gn=CARS;refGene_tx=XM_011287089.2;refGene_gn=CARS;refGene_tx=XM_011287090.3;refGene_gn=CARS;refGene_tx=XM_019812842.1
D1 115310231 115310360
refGene_gn=CARS;refGene_tx=XM_011287089.2;refGene_gn=CARS;refGene_tx=XM_011287090.3;refGene_gn=CARS;refGene_tx=XM_019812842.1
D1 115310765 115310895
refGene_gn=CARS;refGene_tx=XM_011287089.2;refGene_gn=CARS;refGene_tx=XM_011287090.3;refGene_gn=CARS;refGene_tx=XM_019812842.1
D1 115311003 115311107
refGene_gn=CARS;refGene_tx=XM_011287089.2;refGene_gn=CARS;refGene_tx=XM_011287090.3;refGene_gn=CARS;refGene_tx=XM_019812842.1
D1 115311545 115311643
refGene_gn=CARS;refGene_tx=XM_011287089.2;refGene_gn=CARS;refGene_tx=XM_011287090.3;refGene_gn=CARS;refGene_tx=XM_019812842.1
D1 115311780 115311970
refGene_gn=CARS;refGene_tx=XM_011287089.2;refGene_gn=CARS;refGene_tx=XM_011287090.3;refGene_gn=CARS;refGene_tx=XM_019812842.1
D1 115312394 115312463
refGene_gn=CARS;refGene_tx=XM_011287089.2;refGene_gn=CARS;refGene_tx=XM_011287090.3;refGene_gn=CARS;refGene_tx=XM_019812842.1
D1 115314899 115314981
refGene_gn=CARS;refGene_tx=XM_011287089.2;refGene_gn=CARS;refGene_tx=XM_011287090.3;refGene_gn=CARS;refGene_tx=XM_019812842.1
D1 115321547 115321628
refGene_gn=CARS;refGene_tx=XM_011287089.2;refGene_gn=CARS;refGene_tx=XM_011287090.3;refGene_gn=CARS;refGene_tx=XM_019812842.1
D1 115322185 115322253
refGene_gn=CARS;refGene_tx=XM_011287089.2;refGene_gn=CARS;refGene_tx=XM_011287090.3;refGene_gn=CARS;refGene_tx=XM_019812842.1
D1 115323611 115323671
refGene_gn=CARS;refGene_tx=XM_011287089.2;refGene_gn=CARS;refGene_tx=XM_011287090.3;refGene_gn=CARS;refGene_tx=XM_019812842.1
D1 115324017 115324101
refGene_gn=CARS;refGene_tx=XM_011287089.2;refGene_gn=CARS;refGene_tx=X

M_011287090.3;refGene_gn=CARS;refGene_tx=XM_019812842.1
D1 115324644 115324779
refGene_gn=CARS;refGene_tx=XM_011287089.2;refGene_gn=CARS;refGene_tx=XM_011287090.3;refGene_gn=CARS;refGene_tx=XM_019812842.1
D1 115349692 115349706
refGene_gn=NAP1L4;refGene_tx=XM_023238975.1;refGene_gn=NAP1L4;refGene_tx=XM_023238976.1
D1 115350507 115350566
refGene_gn=NAP1L4;refGene_tx=XM_023238975.1;refGene_gn=NAP1L4;refGene_tx=XM_023238976.1
D1 115354275 115354375
refGene_gn=NAP1L4;refGene_tx=XM_023238975.1;refGene_gn=NAP1L4;refGene_tx=XM_023238976.1
D1 115356407 115356549
refGene_gn=NAP1L4;refGene_tx=XM_023238975.1;refGene_gn=NAP1L4;refGene_tx=XM_023238976.1
D1 115357025 115357112
refGene_gn=NAP1L4;refGene_tx=XM_023238975.1;refGene_gn=NAP1L4;refGene_tx=XM_023238976.1
D1 115358457 115358589
refGene_gn=NAP1L4;refGene_tx=XM_023238975.1;refGene_gn=NAP1L4;refGene_tx=XM_023238976.1
D1 115359328 115359400
refGene_gn=NAP1L4;refGene_tx=XM_023238975.1;refGene_gn=NAP1L4;refGene_tx=XM_023238976.1
D1 115363765 115363905
refGene_gn=NAP1L4;refGene_tx=XM_023238975.1;refGene_gn=NAP1L4;refGene_tx=XM_023238976.1
D1 115365109 115365255
refGene_gn=NAP1L4;refGene_tx=XM_023238975.1;refGene_gn=NAP1L4;refGene_tx=XM_023238976.1
D1 115367038 115367061
refGene_gn=NAP1L4;refGene_tx=XM_023238975.1;refGene_gn=NAP1L4;refGene_tx=XM_023238976.1
D1 115367314 115367434
refGene_gn=NAP1L4;refGene_tx=XM_023238975.1;refGene_gn=NAP1L4;refGene_tx=XM_023238976.1
D1 115369918 115369948
refGene_gn=NAP1L4;refGene_tx=XM_023238975.1;refGene_gn=NAP1L4;refGene_tx=XM_023238976.1
D1 115370342 115370399
refGene_gn=NAP1L4;refGene_tx=XM_023238975.1;refGene_gn=NAP1L4;refGene_tx=XM_023238976.1
D1 115376791 115376830
refGene_gn=NAP1L4;refGene_tx=XM_023238975.1;refGene_gn=NAP1L4;refGene_tx=XM_023238976.1
D1 115388880 115389276
refGene_gn=PHLDA2;refGene_tx=XM_023238977.1
D1 115391682 115391861
refGene_gn=SLC22A18;refGene_tx=XM_023238973.1

D1 115393736 115393857
refGene_gn=SLC22A18;refGene_tx=XM_023238973.1
D1 115394058 115394160
refGene_gn=SLC22A18;refGene_tx=XM_023238973.1
D1 115396303 115396404
refGene_gn=SLC22A18;refGene_tx=XM_023238973.1
D1 115397194 115397302
refGene_gn=SLC22A18;refGene_tx=XM_023238973.1
D1 115397992 115398111
refGene_gn=SLC22A18;refGene_tx=XM_023238973.1
D1 115405338 115405471
refGene_gn=SLC22A18;refGene_tx=XM_023238973.1
D1 115405821 115405982
refGene_gn=SLC22A18;refGene_tx=XM_023238973.1
D1 115406310 115406526
refGene_gn=SLC22A18;refGene_tx=XM_023238973.1
D1 115406733 115406831
refGene_gn=SLC22A18;refGene_tx=XM_023238973.1
D1 115409166 115409310
refGene_gn=SLC22A18;refGene_tx=XM_023238973.1
D1 115423789 115424450
refGene_gn=CDKN1C;refGene_tx=XM_023240183.1
D1 115425012 115425146
refGene_gn=CDKN1C;refGene_tx=XM_023240183.1
D1 115460142 115460364
refGene_gn=KCNQ1;refGene_tx=XM_011287088.3;refGene_gn=KCNQ1;refGene_tx=XM_023240184.1;refGene_gn=KCNQ1;refGene_tx=XM_023240185.1;refGene_gn=KCNQ1;refGene_tx=XM_023240186.1;refGene_gn=KCNQ1;refGene_tx=XM_023240187.1;refGene_gn=KCNQ1;refGene_tx=XM_023240188.1
D1 115461720 115461789
refGene_gn=KCNQ1;refGene_tx=XM_011287088.3;refGene_gn=KCNQ1;refGene_tx=XM_023240185.1;refGene_gn=KCNQ1;refGene_tx=XM_023240186.1
D1 115532194 115532256
refGene_gn=KCNQ1;refGene_tx=XM_011287088.3;refGene_gn=KCNQ1;refGene_tx=XM_023240184.1;refGene_gn=KCNQ1;refGene_tx=XM_023240185.1;refGene_gn=KCNQ1;refGene_tx=XM_023240186.1;refGene_gn=KCNQ1;refGene_tx=XM_023240187.1;refGene_gn=KCNQ1;refGene_tx=XM_023240188.1;refGene_gn=KCNQ1;refGene_tx=XM_023240189.1
D1 115533293 115533340
refGene_gn=KCNQ1;refGene_tx=XM_011287088.3;refGene_gn=KCNQ1;refGene_tx=XM_023240184.1;refGene_gn=KCNQ1;refGene_tx=XM_023240189.1
D1 115534276 115534371
refGene_gn=KCNQ1;refGene_tx=XM_011287088.3;refGene_gn=KCNQ1;refGene_tx=XM_023240184.1;refGene_gn=KCNQ1;refGene_tx=XM_023240185.1;refGene_gn=KCNQ1;refGene_tx=XM_023240186.1;refGene_gn=KCNQ1;refGene_tx=XM_023240187.1;refGene_gn=KCNQ1;refGene_tx=XM_023240188.1;refGene_gn=KCNQ1;refGene_tx=XM_023240189.1
D1 115539027 115539103
refGene_gn=KCNQ1;refGene_tx=XM_011287088.3;refGene_gn=KCNQ1;refGene_tx=XM_023240184.1;refGene_gn=KCNQ1;refGene_tx=XM_023240185.1;refGene_gn=

KCNQ1;refGene_tx=XM_023240186.1;refGene_gn=KCNQ1;refGene_tx=XM_0232401
87.1;refGene_gn=KCNQ1;refGene_tx=XM_023240188.1;refGene_gn=KCNQ1;refGe
ne_tx=XM_023240189.1
D1 115640037 115640155
refGene_gn=KCNQ1;refGene_tx=XM_011287088.3;refGene_gn=KCNQ1;refGene_tx
=XM_023240184.1;refGene_gn=KCNQ1;refGene_tx=XM_023240185.1;refGene_gn=
KCNQ1;refGene_tx=XM_023240187.1
D1 115685616 115685761
refGene_gn=KCNQ1;refGene_tx=XM_011287088.3;refGene_gn=KCNQ1;refGene_tx
=XM_023240184.1;refGene_gn=KCNQ1;refGene_tx=XM_023240185.1;refGene_gn=
KCNQ1;refGene_tx=XM_023240186.1;refGene_gn=KCNQ1;refGene_tx=XM_0232401
87.1;refGene_gn=KCNQ1;refGene_tx=XM_023240188.1;refGene_gn=KCNQ1;refGe
ne_tx=XM_023240189.1
D1 115686619 115686742
refGene_gn=KCNQ1;refGene_tx=XM_011287088.3;refGene_gn=KCNQ1;refGene_tx
=XM_023240184.1;refGene_gn=KCNQ1;refGene_tx=XM_023240185.1;refGene_gn=
KCNQ1;refGene_tx=XM_023240186.1;refGene_gn=KCNQ1;refGene_tx=XM_0232401
87.1;refGene_gn=KCNQ1;refGene_tx=XM_023240188.1;refGene_gn=KCNQ1;refGe
ne_tx=XM_023240189.1
D1 115688828 115688924
refGene_gn=KCNQ1;refGene_tx=XM_011287088.3;refGene_gn=KCNQ1;refGene_tx
=XM_023240184.1;refGene_gn=KCNQ1;refGene_tx=XM_023240185.1;refGene_gn=
KCNQ1;refGene_tx=XM_023240186.1;refGene_gn=KCNQ1;refGene_tx=XM_0232401
87.1;refGene_gn=KCNQ1;refGene_tx=XM_023240188.1;refGene_gn=KCNQ1;refGe
ne_tx=XM_023240189.1
D1 115689614 115689725
refGene_gn=KCNQ1;refGene_tx=XM_011287088.3;refGene_gn=KCNQ1;refGene_tx
=XM_023240184.1;refGene_gn=KCNQ1;refGene_tx=XM_023240185.1;refGene_gn=
KCNQ1;refGene_tx=XM_023240186.1;refGene_gn=KCNQ1;refGene_tx=XM_0232401
87.1;refGene_gn=KCNQ1;refGene_tx=XM_023240188.1;refGene_gn=KCNQ1;refGe
ne_tx=XM_023240189.1
D1 115698074 115698215
refGene_gn=KCNQ1;refGene_tx=XM_011287088.3;refGene_gn=KCNQ1;refGene_tx
=XM_023240184.1;refGene_gn=KCNQ1;refGene_tx=XM_023240185.1;refGene_gn=
KCNQ1;refGene_tx=XM_023240186.1;refGene_gn=KCNQ1;refGene_tx=XM_0232401
87.1;refGene_gn=KCNQ1;refGene_tx=XM_023240188.1;refGene_gn=KCNQ1;refGe
ne_tx=XM_023240189.1
D1 115698693 115698790
refGene_gn=KCNQ1;refGene_tx=XM_011287088.3;refGene_gn=KCNQ1;refGene_tx
=XM_023240184.1;refGene_gn=KCNQ1;refGene_tx=XM_023240185.1;refGene_gn=
KCNQ1;refGene_tx=XM_023240186.1;refGene_gn=KCNQ1;refGene_tx=XM_0232401
87.1;refGene_gn=KCNQ1;refGene_tx=XM_023240188.1;refGene_gn=KCNQ1;refGe
ne_tx=XM_023240189.1
D1 115699241 115699320
refGene_gn=KCNQ1;refGene_tx=XM_011287088.3;refGene_gn=KCNQ1;refGene_tx
=XM_023240184.1;refGene_gn=KCNQ1;refGene_tx=XM_023240185.1;refGene_gn=
KCNQ1;refGene_tx=XM_023240186.1;refGene_gn=KCNQ1;refGene_tx=XM_0232401
87.1;refGene_gn=KCNQ1;refGene_tx=XM_023240188.1;refGene_gn=KCNQ1;refGe
ne_tx=XM_023240189.1
D1 115699558 115699685

refGene_gn=KCNQ1;refGene_tx=XM_011287088.3;refGene_gn=KCNQ1;refGene_tx=XM_023240184.1;refGene_gn=KCNQ1;refGene_tx=XM_023240185.1;refGene_gn=KCNQ1;refGene_tx=XM_023240186.1;refGene_gn=KCNQ1;refGene_tx=XM_023240187.1;refGene_gn=KCNQ1;refGene_tx=XM_023240188.1;refGene_gn=KCNQ1;refGene_tx=XM_023240189.1
D1 115735949 115736040
refGene_gn=KCNQ1;refGene_tx=XM_011287088.3;refGene_gn=KCNQ1;refGene_tx=XM_023240184.1;refGene_gn=KCNQ1;refGene_tx=XM_023240185.1;refGene_gn=KCNQ1;refGene_tx=XM_023240186.1;refGene_gn=KCNQ1;refGene_tx=XM_023240187.1;refGene_gn=KCNQ1;refGene_tx=XM_023240188.1;refGene_gn=KCNQ1;refGene_tx=XM_023240189.1
D1 115780943 115781323
refGene_gn=KCNQ1;refGene_tx=XM_011287088.3;refGene_gn=KCNQ1;refGene_tx=XM_023240184.1;refGene_gn=KCNQ1;refGene_tx=XM_023240185.1;refGene_gn=KCNQ1;refGene_tx=XM_023240186.1;refGene_gn=KCNQ1;refGene_tx=XM_023240187.1;refGene_gn=KCNQ1;refGene_tx=XM_023240188.1;refGene_gn=KCNQ1;refGene_tx=XM_023240189.1
D1 115789346 115789604
refGene_gn=LOC111556744;refGene_tx=XM_023240190.1
D1 115789933 115790875
refGene_gn=LOC111556744;refGene_tx=XM_023240190.1
D1 115798446 115798554
refGene_gn=TRPM5;refGene_tx=XM_023239757.1
D1 115800991 115801181
refGene_gn=TRPM5;refGene_tx=XM_023239757.1
D1 115802312 115802377
refGene_gn=TRPM5;refGene_tx=XM_023239757.1
D1 115802538 115802730
refGene_gn=TRPM5;refGene_tx=XM_023239757.1
D1 115802939 115803042
refGene_gn=TRPM5;refGene_tx=XM_023239757.1
D1 115804177 115804296
refGene_gn=TRPM5;refGene_tx=XM_023239757.1
D1 115804517 115804841
refGene_gn=TRPM5;refGene_tx=XM_023239757.1
D1 115804921 115805062
refGene_gn=TRPM5;refGene_tx=XM_023239757.1
D1 115805156 115805196
refGene_gn=TRPM5;refGene_tx=XM_023239757.1
D1 115805198 115805279
refGene_gn=TRPM5;refGene_tx=XM_023239757.1
D1 115805506 115805652
refGene_gn=TRPM5;refGene_tx=XM_023239757.1
D1 115806090 115806203
refGene_gn=TRPM5;refGene_tx=XM_023239757.1
D1 115806358 115806451
refGene_gn=TRPM5;refGene_tx=XM_023239757.1
D1 115806550 115806809
refGene_gn=TRPM5;refGene_tx=XM_023239757.1
D1 115806947 115807066

refGene_gn=TRPM5;refGene_tx=XM_023239757.1
D1 115807255 115807388
refGene_gn=TRPM5;refGene_tx=XM_023239757.1
D1 115807497 115807672
refGene_gn=TRPM5;refGene_tx=XM_023239757.1
D1 115810309 115810463
refGene_gn=TRPM5;refGene_tx=XM_023239757.1
D1 115810906 115811088
refGene_gn=TRPM5;refGene_tx=XM_023239757.1
D1 115811387 115811520
refGene_gn=TRPM5;refGene_tx=XM_023239757.1
D1 115812088 115812161
refGene_gn=TRPM5;refGene_tx=XM_023239757.1
D1 115812469 115812524
refGene_gn=TRPM5;refGene_tx=XM_023239757.1
D1 115812854 115813168
refGene_gn=TRPM5;refGene_tx=XM_023239757.1
D1 115814261 115815224
refGene_gn=TSSC4;refGene_tx=XM_003993776.5;refGene_gn=TSSC4;refGene_tx=XM_011287083.3;refGene_gn=TSSC4;refGene_tx=XM_011287084.3;refGene_gn=TSSC4;refGene_tx=XM_011287087.3
D1 115819065 115819128
refGene_gn=CD81;refGene_tx=XM_023240191.1
D1 115819209 115819296
refGene_gn=CD81;refGene_tx=XM_023240191.1
D1 115820044 115820146
refGene_gn=CD81;refGene_tx=XM_023240191.1
D1 115820577 115820682
refGene_gn=CD81;refGene_tx=XM_023240191.1
D1 115821031 115821106
refGene_gn=CD81;refGene_tx=XM_023240191.1
D1 115821727 115821825
refGene_gn=CD81;refGene_tx=XM_023240191.1
D1 115825165 115825280
refGene_gn=CD81;refGene_tx=XM_023240191.1
D1 115835577 115835643
refGene_gn=CD81;refGene_tx=XM_023240191.1
D1 115849017 115849168
refGene_gn=TSPAN32;refGene_tx=XM_003993786.5
D1 115849704 115849796
refGene_gn=TSPAN32;refGene_tx=XM_003993786.5
D1 115850088 115850172
refGene_gn=TSPAN32;refGene_tx=XM_003993786.5
D1 115851749 115851836
refGene_gn=TSPAN32;refGene_tx=XM_003993786.5
D1 115852283 115852385
refGene_gn=TSPAN32;refGene_tx=XM_003993786.5
D1 115855843 115855918
refGene_gn=TSPAN32;refGene_tx=XM_003993786.5
D1 115860052 115860150

refGene_gn=TSPAN32;refGene_tx=XM_003993786.5
D1 115860930 115861045
refGene_gn=TSPAN32;refGene_tx=XM_003993786.5
D1 115861563 115861629
refGene_gn=TSPAN32;refGene_tx=XM_003993786.5
D1 115883174 115883768
refGene_gn=ASCL2;refGene_tx=XM_019812844.2;refGene_gn=ASCL2;refGene_tx
=XM_019812845.2
D1 115932476 115932566
refGene_gn=TH;refGene_tx=XM_003993778.3
D1 115933854 115934067
refGene_gn=TH;refGene_tx=XM_003993778.3
D1 115935119 115935294
refGene_gn=TH;refGene_tx=XM_003993778.3
D1 115935507 115935596
refGene_gn=TH;refGene_tx=XM_003993778.3
D1 115936007 115936075
refGene_gn=TH;refGene_tx=XM_003993778.3
D1 115936373 115936424
refGene_gn=TH;refGene_tx=XM_003993778.3
D1 115936829 115936975
refGene_gn=TH;refGene_tx=XM_003993778.3
D1 115937055 115937191
refGene_gn=TH;refGene_tx=XM_003993778.3
D1 115937330 115937400
refGene_gn=TH;refGene_tx=XM_003993778.3
D1 115938007 115938064
refGene_gn=TH;refGene_tx=XM_003993778.3
D1 115938304 115938400
refGene_gn=TH;refGene_tx=XM_003993778.3
D1 115938524 115938658
refGene_gn=TH;refGene_tx=XM_003993778.3
D1 115939478 115939638
refGene_gn=TH;refGene_tx=XM_003993778.3
D1 115942353 115942540
refGene_gn=INS;refGene_tx=NM_001009272.1;refGene_gn=INS;refGene_tx=XM_
019811180.2
D1 115942838 115942984
refGene_gn=INS;refGene_tx=NM_001009272.1;refGene_gn=INS;refGene_tx=XM_
019811180.2
D1 115961689 115961851
refGene_gn=IGF2;refGene_tx=XM_023240192.1;refGene_gn=IGF2;refGene_tx=X
M_023240193.1
D1 115966709 115966872
refGene_gn=IGF2;refGene_tx=XM_019812847.1;refGene_gn=IGF2;refGene_tx=X
M_019812848.1;refGene_gn=IGF2;refGene_tx=XM_023240192.1;refGene_gn=IGF
2;refGene_tx=XM_023240193.1
D1 115968702 115968863
refGene_gn=IGF2;refGene_tx=XM_019812847.1;refGene_gn=IGF2;refGene_tx=X
M_019812848.1;refGene_gn=IGF2;refGene_tx=XM_023240192.1;refGene_gn=IGF

2;refGene_tx=XM_023240193.1
D1 115969151 115969391
refGene_gn=IGF2;refGene_tx=XM_019812847.1;refGene_gn=IGF2;refGene_tx=XM_019812848.1;refGene_gn=IGF2;refGene_tx=XM_023240192.1;refGene_gn=IGF2;refGene_tx=XM_023240193.1
D1 116088244 116088412
refGene_gn=MRPL23;refGene_tx=XM_023240194.1
D1 116091694 116091768
refGene_gn=MRPL23;refGene_tx=XM_023240194.1
D1 116092351 116092434
refGene_gn=MRPL23;refGene_tx=XM_023240194.1
D1 116093281 116093404
refGene_gn=MRPL23;refGene_tx=XM_023240194.1
D1 116095342 116095359
refGene_gn=MRPL23;refGene_tx=XM_023240194.1
D1 116112827 116112882
refGene_gn=TNNT3;refGene_tx=XM_006937639.4;refGene_gn=TNNT3;refGene_tx=XM_006937640.4;refGene_gn=TNNT3;refGene_tx=XM_006937641.4;refGene_gn=TNNT3;refGene_tx=XM_006937642.4;refGene_gn=TNNT3;refGene_tx=XM_006937644.4;refGene_gn=TNNT3;refGene_tx=XM_006937645.4;refGene_gn=TNNT3;refGene_tx=XM_011287075.3;refGene_gn=TNNT3;refGene_tx=XM_011287077.3;refGene_gn=TNNT3;refGene_tx=XM_011287078.3;refGene_gn=TNNT3;refGene_tx=XM_023240198.1
D1 116114287 116114328
refGene_gn=TNNT3;refGene_tx=XM_006937639.4;refGene_gn=TNNT3;refGene_tx=XM_006937640.4;refGene_gn=TNNT3;refGene_tx=XM_006937641.4;refGene_gn=TNNT3;refGene_tx=XM_006937642.4;refGene_gn=TNNT3;refGene_tx=XM_006937644.4;refGene_gn=TNNT3;refGene_tx=XM_006937645.4;refGene_gn=TNNT3;refGene_tx=XM_023240198.1
D1 116114885 116114926
refGene_gn=TNNT3;refGene_tx=XM_011287075.3;refGene_gn=TNNT3;refGene_tx=XM_011287077.3;refGene_gn=TNNT3;refGene_tx=XM_011287078.3
D1 116116013 116116104
refGene_gn=TNNT3;refGene_tx=XM_006937639.4;refGene_gn=TNNT3;refGene_tx=XM_006937640.4;refGene_gn=TNNT3;refGene_tx=XM_006937641.4;refGene_gn=TNNT3;refGene_tx=XM_006937642.4;refGene_gn=TNNT3;refGene_tx=XM_006937644.4;refGene_gn=TNNT3;refGene_tx=XM_006937645.4;refGene_gn=TNNT3;refGene_tx=XM_011287075.3;refGene_gn=TNNT3;refGene_tx=XM_011287077.3;refGene_gn=TNNT3;refGene_tx=XM_011287078.3;refGene_gn=TNNT3;refGene_tx=XM_023240198.1
D1 116116223 116116333
refGene_gn=TNNT3;refGene_tx=XM_006937639.4;refGene_gn=TNNT3;refGene_tx=XM_006937640.4;refGene_gn=TNNT3;refGene_tx=XM_006937641.4;refGene_gn=TNNT3;refGene_tx=XM_006937642.4;refGene_gn=TNNT3;refGene_tx=XM_006937644.4;refGene_gn=TNNT3;refGene_tx=XM_006937645.4;refGene_gn=TNNT3;refGene_tx=XM_011287075.3;refGene_gn=TNNT3;refGene_tx=XM_011287077.3;refGene_gn=TNNT3;refGene_tx=XM_011287078.3;refGene_gn=TNNT3;refGene_tx=XM_023240198.1
D1 116116430 116116544
refGene_gn=TNNT3;refGene_tx=XM_006937639.4;refGene_gn=TNNT3;refGene_tx

=XM_006937640.4;refGene_gn=TNNT3;refGene_tx=XM_006937641.4;refGene_gn=TNNT3;refGene_tx=XM_006937642.4;refGene_gn=TNNT3;refGene_tx=XM_006937644.4;refGene_gn=TNNT3;refGene_tx=XM_006937645.4;refGene_gn=TNNT3;refGene_tx=XM_011287075.3;refGene_gn=TNNT3;refGene_tx=XM_011287077.3;refGene_gn=TNNT3;refGene_tx=XM_011287078.3;refGene_gn=TNNT3;refGene_tx=XM_023240198.1
D1 116116679 116116757
refGene_gn=TNNT3;refGene_tx=XM_006937639.4;refGene_gn=TNNT3;refGene_tx=XM_006937640.4;refGene_gn=TNNT3;refGene_tx=XM_006937641.4;refGene_gn=TNNT3;refGene_tx=XM_006937642.4;refGene_gn=TNNT3;refGene_tx=XM_006937644.4;refGene_gn=TNNT3;refGene_tx=XM_006937645.4;refGene_gn=TNNT3;refGene_tx=XM_011287075.3;refGene_gn=TNNT3;refGene_tx=XM_011287077.3;refGene_gn=TNNT3;refGene_tx=XM_011287078.3;refGene_gn=TNNT3;refGene_tx=XM_023240198.1
D1 116116859 116116976
refGene_gn=TNNT3;refGene_tx=XM_006937639.4;refGene_gn=TNNT3;refGene_tx=XM_006937640.4;refGene_gn=TNNT3;refGene_tx=XM_006937641.4;refGene_gn=TNNT3;refGene_tx=XM_006937642.4;refGene_gn=TNNT3;refGene_tx=XM_006937644.4;refGene_gn=TNNT3;refGene_tx=XM_006937645.4;refGene_gn=TNNT3;refGene_tx=XM_011287075.3;refGene_gn=TNNT3;refGene_tx=XM_011287077.3;refGene_gn=TNNT3;refGene_tx=XM_011287078.3;refGene_gn=TNNT3;refGene_tx=XM_023240198.1
D1 116117734 116117780
refGene_gn=TNNT3;refGene_tx=XM_006937639.4;refGene_gn=TNNT3;refGene_tx=XM_006937640.4;refGene_gn=TNNT3;refGene_tx=XM_006937641.4;refGene_gn=TNNT3;refGene_tx=XM_006937642.4;refGene_gn=TNNT3;refGene_tx=XM_006937644.4;refGene_gn=TNNT3;refGene_tx=XM_006937645.4;refGene_gn=TNNT3;refGene_tx=XM_011287075.3;refGene_gn=TNNT3;refGene_tx=XM_011287077.3;refGene_gn=TNNT3;refGene_tx=XM_011287078.3;refGene_gn=TNNT3;refGene_tx=XM_023240198.1
D1 116120152 116120171
refGene_gn=TNNT3;refGene_tx=XM_006937639.4;refGene_gn=TNNT3;refGene_tx=XM_006937640.4;refGene_gn=TNNT3;refGene_tx=XM_006937641.4;refGene_gn=TNNT3;refGene_tx=XM_006937642.4;refGene_gn=TNNT3;refGene_tx=XM_006937644.4;refGene_gn=TNNT3;refGene_tx=XM_006937645.4;refGene_gn=TNNT3;refGene_tx=XM_011287075.3;refGene_gn=TNNT3;refGene_tx=XM_011287077.3;refGene_gn=TNNT3;refGene_tx=XM_011287078.3;refGene_gn=TNNT3;refGene_tx=XM_023240198.1
D1 116122589 116122604
refGene_gn=TNNT3;refGene_tx=XM_006937639.4;refGene_gn=TNNT3;refGene_tx=XM_006937640.4;refGene_gn=TNNT3;refGene_tx=XM_006937641.4;refGene_gn=TNNT3;refGene_tx=XM_006937642.4;refGene_gn=TNNT3;refGene_tx=XM_006937644.4;refGene_gn=TNNT3;refGene_tx=XM_006937645.4;refGene_gn=TNNT3;refGene_tx=XM_011287075.3;refGene_gn=TNNT3;refGene_tx=XM_011287077.3;refGene_gn=TNNT3;refGene_tx=XM_011287078.3
D1 116122819 116122834
refGene_gn=TNNT3;refGene_tx=XM_006937639.4;refGene_gn=TNNT3;refGene_tx=XM_006937640.4;refGene_gn=TNNT3;refGene_tx=XM_006937641.4;refGene_gn=TNNT3;refGene_tx=XM_006937644.4;refGene_gn=TNNT3;refGene_tx=XM_011287075.3

D1 116123819 116123834
refGene_gn=TNNT3;refGene_tx=XM_006937639.4;refGene_gn=TNNT3;refGene_tx=XM_006937640.4;refGene_gn=TNNT3;refGene_tx=XM_006937642.4;refGene_gn=TNNT3;refGene_tx=XM_011287075.3;refGene_gn=TNNT3;refGene_tx=XM_011287077.3;refGene_gn=TNNT3;refGene_tx=XM_023240198.1
D1 116123986 116124004
refGene_gn=TNNT3;refGene_tx=XM_006937639.4;refGene_gn=TNNT3;refGene_tx=XM_006937640.4;refGene_gn=TNNT3;refGene_tx=XM_006937641.4;refGene_gn=TNNT3;refGene_tx=XM_006937642.4;refGene_gn=TNNT3;refGene_tx=XM_006937644.4;refGene_gn=TNNT3;refGene_tx=XM_006937645.4;refGene_gn=TNNT3;refGene_tx=XM_011287075.3;refGene_gn=TNNT3;refGene_tx=XM_011287077.3;refGene_gn=TNNT3;refGene_tx=XM_011287078.3;refGene_gn=TNNT3;refGene_tx=XM_023240198.1
D1 116125070 116125088
refGene_gn=TNNT3;refGene_tx=XM_006937639.4;refGene_gn=TNNT3;refGene_tx=XM_006937640.4;refGene_gn=TNNT3;refGene_tx=XM_006937641.4;refGene_gn=TNNT3;refGene_tx=XM_006937642.4;refGene_gn=TNNT3;refGene_tx=XM_006937644.4;refGene_gn=TNNT3;refGene_tx=XM_006937645.4;refGene_gn=TNNT3;refGene_tx=XM_011287075.3;refGene_gn=TNNT3;refGene_tx=XM_011287077.3;refGene_gn=TNNT3;refGene_tx=XM_011287078.3;refGene_gn=TNNT3;refGene_tx=XM_023240198.1
D1 116125610 116125624
refGene_gn=TNNT3;refGene_tx=XM_006937639.4;refGene_gn=TNNT3;refGene_tx=XM_006937640.4;refGene_gn=TNNT3;refGene_tx=XM_006937641.4;refGene_gn=TNNT3;refGene_tx=XM_006937642.4;refGene_gn=TNNT3;refGene_tx=XM_006937644.4;refGene_gn=TNNT3;refGene_tx=XM_006937645.4;refGene_gn=TNNT3;refGene_tx=XM_011287075.3;refGene_gn=TNNT3;refGene_tx=XM_011287077.3;refGene_gn=TNNT3;refGene_tx=XM_011287078.3;refGene_gn=TNNT3;refGene_tx=XM_023240198.1
D1 116125806 116125823
refGene_gn=TNNT3;refGene_tx=XM_006937639.4;refGene_gn=TNNT3;refGene_tx=XM_006937640.4;refGene_gn=TNNT3;refGene_tx=XM_006937641.4;refGene_gn=TNNT3;refGene_tx=XM_006937642.4;refGene_gn=TNNT3;refGene_tx=XM_006937644.4;refGene_gn=TNNT3;refGene_tx=XM_006937645.4;refGene_gn=TNNT3;refGene_tx=XM_011287075.3;refGene_gn=TNNT3;refGene_tx=XM_011287077.3;refGene_gn=TNNT3;refGene_tx=XM_011287078.3;refGene_gn=TNNT3;refGene_tx=XM_023240198.1
D1 116132363 116132366
refGene_gn=LOC111556748;refGene_tx=XM_023240195.1
D1 116134987 116136316
refGene_gn=LOC111556748;refGene_tx=XM_023240195.1;refGene_gn=LOC111556748;refGene_tx=XM_023240196.1;refGene_gn=LOC111556748;refGene_tx=XM_023240197.1
D1 116137576 116137669
refGene_gn=LSP1;refGene_tx=XM_006937648.3;refGene_gn=LSP1;refGene_tx=XM_011287070.2;refGene_gn=LSP1;refGene_tx=XM_011287073.2;refGene_gn=LSP1;refGene_tx=XM_019812850.2;refGene_gn=LSP1;refGene_tx=XM_019812852.2;refGene_gn=LSP1;refGene_tx=XM_019812853.2
D1 116137819 116137897
refGene_gn=LSP1;refGene_tx=XM_006937648.3;refGene_gn=LSP1;refGene_tx=X

M_011287070.2;refGene_gn=LSP1;refGene_tx=XM_011287073.2;refGene_gn=LSP1;refGene_tx=XM_019812850.2;refGene_gn=LSP1;refGene_tx=XM_019812852.2;refGene_gn=LSP1;refGene_tx=XM_019812853.2
D1 116138233 116138368
refGene_gn=LSP1;refGene_tx=XM_006937648.3;refGene_gn=LSP1;refGene_tx=XM_011287070.2;refGene_gn=LSP1;refGene_tx=XM_011287073.2;refGene_gn=LSP1;refGene_tx=XM_019812850.2;refGene_gn=LSP1;refGene_tx=XM_019812852.2;refGene_gn=LSP1;refGene_tx=XM_019812853.2
D1 116139392 116139474
refGene_gn=LSP1;refGene_tx=XM_006937648.3;refGene_gn=LSP1;refGene_tx=XM_011287070.2;refGene_gn=LSP1;refGene_tx=XM_011287073.2;refGene_gn=LSP1;refGene_tx=XM_019812850.2;refGene_gn=LSP1;refGene_tx=XM_019812852.2;refGene_gn=LSP1;refGene_tx=XM_019812853.2
D1 116139643 116139687
refGene_gn=LSP1;refGene_tx=XM_006937648.3;refGene_gn=LSP1;refGene_tx=XM_011287070.2;refGene_gn=LSP1;refGene_tx=XM_011287073.2;refGene_gn=LSP1;refGene_tx=XM_019812850.2;refGene_gn=LSP1;refGene_tx=XM_019812852.2;refGene_gn=LSP1;refGene_tx=XM_019812853.2
D1 116139955 116140048
refGene_gn=LSP1;refGene_tx=XM_006937648.3;refGene_gn=LSP1;refGene_tx=XM_011287070.2;refGene_gn=LSP1;refGene_tx=XM_011287073.2;refGene_gn=LSP1;refGene_tx=XM_019812850.2;refGene_gn=LSP1;refGene_tx=XM_019812852.2;refGene_gn=LSP1;refGene_tx=XM_019812853.2
D1 116140400 116140596
refGene_gn=LSP1;refGene_tx=XM_006937648.3;refGene_gn=LSP1;refGene_tx=XM_011287070.2;refGene_gn=LSP1;refGene_tx=XM_011287073.2;refGene_gn=LSP1;refGene_tx=XM_019812850.2;refGene_gn=LSP1;refGene_tx=XM_019812852.2;refGene_gn=LSP1;refGene_tx=XM_019812853.2
D1 116142323 116142491
refGene_gn=LSP1;refGene_tx=XM_006937648.3;refGene_gn=LSP1;refGene_tx=XM_011287070.2;refGene_gn=LSP1;refGene_tx=XM_011287073.2;refGene_gn=LSP1;refGene_tx=XM_019812850.2;refGene_gn=LSP1;refGene_tx=XM_019812852.2;refGene_gn=LSP1;refGene_tx=XM_019812853.2
D1 116144801 116144936
refGene_gn=LSP1;refGene_tx=XM_006937648.3;refGene_gn=LSP1;refGene_tx=XM_011287070.2;refGene_gn=LSP1;refGene_tx=XM_011287073.2;refGene_gn=LSP1;refGene_tx=XM_019812850.2;refGene_gn=LSP1;refGene_tx=XM_019812852.2;refGene_gn=LSP1;refGene_tx=XM_019812853.2
D1 116153530 116153661
refGene_gn=LSP1;refGene_tx=XM_011287070.2
D1 116155853 116155966
refGene_gn=LSP1;refGene_tx=XM_019812850.2;refGene_gn=LSP1;refGene_tx=XM_019812852.2;refGene_gn=LSP1;refGene_tx=XM_019812853.2
D1 116159133 116159195
refGene_gn=LSP1;refGene_tx=XM_011287073.2
D1 116171475 116171543
refGene_gn=LSP1;refGene_tx=XM_006937648.3
D1 116183164 116183260
refGene_gn=TNNI2;refGene_tx=XM_011287069.2
D1 116183410 116183587

refGene_gn=TNNI2;refGene_tx=XM_011287069.2
D1 116183696 116183786
refGene_gn=TNNI2;refGene_tx=XM_011287069.2
D1 116183911 116184040
refGene_gn=TNNI2;refGene_tx=XM_011287069.2
D1 116184136 116184178
refGene_gn=TNNI2;refGene_tx=XM_011287069.2
D1 116184684 116184691
refGene_gn=TNNI2;refGene_tx=XM_011287069.2
D1 116184876 116184884
refGene_gn=TNNI2;refGene_tx=XM_011287069.2
D1 116187049 116187283
refGene_gn=SYT8;refGene_tx=XM_023240199.1;refGene_gn=SYT8;refGene_tx=XM_023240200.1;refGene_gn=SYT8;refGene_tx=XM_023240201.1;refGene_gn=SYT8;refGene_tx=XM_023240202.1
D1 116187387 116187521
refGene_gn=SYT8;refGene_tx=XM_023240199.1;refGene_gn=SYT8;refGene_tx=XM_023240200.1;refGene_gn=SYT8;refGene_tx=XM_023240201.1;refGene_gn=SYT8;refGene_tx=XM_023240202.1
D1 116187618 116187724
refGene_gn=SYT8;refGene_tx=XM_023240199.1;refGene_gn=SYT8;refGene_tx=XM_023240200.1;refGene_gn=SYT8;refGene_tx=XM_023240201.1;refGene_gn=SYT8;refGene_tx=XM_023240202.1
D1 116187824 116187992
refGene_gn=SYT8;refGene_tx=XM_023240199.1;refGene_gn=SYT8;refGene_tx=XM_023240200.1;refGene_gn=SYT8;refGene_tx=XM_023240201.1;refGene_gn=SYT8;refGene_tx=XM_023240202.1
D1 116188070 116188238
refGene_gn=SYT8;refGene_tx=XM_023240199.1;refGene_gn=SYT8;refGene_tx=XM_023240200.1;refGene_gn=SYT8;refGene_tx=XM_023240201.1;refGene_gn=SYT8;refGene_tx=XM_023240202.1
D1 116188364 116188463
refGene_gn=SYT8;refGene_tx=XM_023240199.1;refGene_gn=SYT8;refGene_tx=XM_023240200.1;refGene_gn=SYT8;refGene_tx=XM_023240201.1;refGene_gn=SYT8;refGene_tx=XM_023240202.1
D1 116188824 116189003
refGene_gn=SYT8;refGene_tx=XM_023240199.1;refGene_gn=SYT8;refGene_tx=XM_023240200.1;refGene_gn=SYT8;refGene_tx=XM_023240201.1;refGene_gn=SYT8;refGene_tx=XM_023240202.1
D1 116189106 116189200
refGene_gn=SYT8;refGene_tx=XM_023240199.1;refGene_gn=SYT8;refGene_tx=XM_023240200.1;refGene_gn=SYT8;refGene_tx=XM_023240201.1;refGene_gn=SYT8;refGene_tx=XM_023240202.1
D1 116232586 116232654
refGene_gn=CTSD;refGene_tx=XM_011287067.2
D1 116234620 116234780
refGene_gn=CTSD;refGene_tx=XM_011287067.2
D1 116237012 116237136
refGene_gn=CTSD;refGene_tx=XM_011287067.2
D1 116237611 116237730

refGene_gn=CTSD;refGene_tx=XM_011287067.2
D1 116238697 116238918
refGene_gn=CTSD;refGene_tx=XM_011287067.2
D1 116240727 116240850
refGene_gn=CTSD;refGene_tx=XM_011287067.2
D1 116241220 116241365
refGene_gn=CTSD;refGene_tx=XM_011287067.2
D1 116241450 116241549
refGene_gn=CTSD;refGene_tx=XM_011287067.2
D1 116241702 116241870
refGene_gn=CTSD;refGene_tx=XM_011287067.2
D1 116243468 116243577
refGene_gn=IFITM10;refGene_tx=XM_019813001.2
D1 116244226 116244355
refGene_gn=IFITM10;refGene_tx=XM_019813001.2
D1 116246211 116246262
refGene_gn=IFITM10;refGene_tx=XM_019813001.2
D1 116246296 116246508
refGene_gn=IFITM10;refGene_tx=XM_019813001.2
D1 116247369 116247822
refGene_gn=IFITM10;refGene_tx=XM_019813001.2
D1 116253189 116253339
refGene_gn=IFITM10;refGene_tx=XM_019813001.2
D1 116432737 116433122
refGene_gn=LOC111556667;refGene_tx=XM_023239758.1
D1 116437003 116437057
refGene_gn=LOC111556667;refGene_tx=XM_023239758.1
D1 116437238 116437300
refGene_gn=LOC111556667;refGene_tx=XM_023239758.1
D1 116437814 116438179
refGene_gn=LOC111556667;refGene_tx=XM_023239758.1
D1 116438756 116438790
refGene_gn=LOC111556667;refGene_tx=XM_023239758.1
D1 116438865 116439096
refGene_gn=DUSP8;refGene_tx=XM_023240203.1
D1 116440881 116441020
refGene_gn=DUSP8;refGene_tx=XM_023240203.1
D1 116443332 116443447
refGene_gn=DUSP8;refGene_tx=XM_023240204.1
D1 116445756 116445923
refGene_gn=DUSP8;refGene_tx=XM_023240203.1;refGene_gn=DUSP8;refGene_tx
=XM_023240204.1
D1 116446370 116446530
refGene_gn=DUSP8;refGene_tx=XM_023240203.1;refGene_gn=DUSP8;refGene_tx
=XM_023240204.1
D1 116447597 116447721
refGene_gn=DUSP8;refGene_tx=XM_023240203.1;refGene_gn=DUSP8;refGene_tx
=XM_023240204.1
D1 116447959 116449016
refGene_gn=DUSP8;refGene_tx=XM_023240203.1;refGene_gn=DUSP8;refGene_tx

=XM_023240204.1
D1	116472585	116472602
refGene_gn=MOB2;refGene_tx=XM_023240206.1
D1	116530624	116530731
refGene_gn=MOB2;refGene_tx=XM_023240205.1
D1	116531255	116531365
refGene_gn=MOB2;refGene_tx=XM_019812861.2
D1	116535814	116535927
refGene_gn=MOB2;refGene_tx=XM_019812859.2
D1	116538440	116538601
refGene_gn=MOB2;refGene_tx=XM_019812859.2;refGene_gn=MOB2;refGene_tx=XM_019812861.2;refGene_gn=MOB2;refGene_tx=XM_023240205.1;refGene_gn=MOB2;refGene_tx=XM_023240206.1
D1	116539152	116539246
refGene_gn=MOB2;refGene_tx=XM_019812859.2;refGene_gn=MOB2;refGene_tx=XM_019812861.2;refGene_gn=MOB2;refGene_tx=XM_023240205.1;refGene_gn=MOB2;refGene_tx=XM_023240206.1
D1	116545685	116545810
refGene_gn=MOB2;refGene_tx=XM_019812859.2;refGene_gn=MOB2;refGene_tx=XM_019812861.2;refGene_gn=MOB2;refGene_tx=XM_023240205.1;refGene_gn=MOB2;refGene_tx=XM_023240206.1
D1	116546525	116546866
refGene_gn=MOB2;refGene_tx=XM_019812859.2;refGene_gn=MOB2;refGene_tx=XM_019812861.2;refGene_gn=MOB2;refGene_tx=XM_023240205.1;refGene_gn=MOB2;refGene_tx=XM_023240206.1
D1	116555313	116555333
refGene_gn=BRSK2;refGene_tx=XM_023240215.1;refGene_gn=BRSK2;refGene_tx=XM_023240217.1;refGene_gn=BRSK2;refGene_tx=XM_023240218.1;refGene_gn=BRSK2;refGene_tx=XM_023240219.1
D1	116555594	116555818
refGene_gn=BRSK2;refGene_tx=XM_023240207.1;refGene_gn=BRSK2;refGene_tx=XM_023240208.1;refGene_gn=BRSK2;refGene_tx=XM_023240209.1;refGene_gn=BRSK2;refGene_tx=XM_023240210.1;refGene_gn=BRSK2;refGene_tx=XM_023240211.1;refGene_gn=BRSK2;refGene_tx=XM_023240216.1
D1	116556991	116557039
refGene_gn=BRSK2;refGene_tx=XM_023240207.1;refGene_gn=BRSK2;refGene_tx=XM_023240208.1;refGene_gn=BRSK2;refGene_tx=XM_023240209.1;refGene_gn=BRSK2;refGene_tx=XM_023240211.1;refGene_gn=BRSK2;refGene_tx=XM_023240215.1;refGene_gn=BRSK2;refGene_tx=XM_023240216.1;refGene_gn=BRSK2;refGene_tx=XM_023240218.1
D1	116559333	116559419
refGene_gn=BRSK2;refGene_tx=XM_023240212.1;refGene_gn=BRSK2;refGene_tx=XM_023240213.1;refGene_gn=BRSK2;refGene_tx=XM_023240214.1
D1	116559609	116559699
refGene_gn=BRSK2;refGene_tx=XM_023240207.1;refGene_gn=BRSK2;refGene_tx=XM_023240208.1;refGene_gn=BRSK2;refGene_tx=XM_023240209.1;refGene_gn=BRSK2;refGene_tx=XM_023240210.1;refGene_gn=BRSK2;refGene_tx=XM_023240211.1;refGene_gn=BRSK2;refGene_tx=XM_023240212.1;refGene_gn=BRSK2;refGene_tx=XM_023240213.1;refGene_gn=BRSK2;refGene_tx=XM_023240214.1;refGene_gn=BRSK2;refGene_tx=XM_023240215.1;refGene_gn=BRSK2;refGene_tx=XM_02

3240216.1;refGene_gn=BRSK2;refGene_tx=XM_023240217.1;refGene_gn=BRSK2;
refGene_tx=XM_023240218.1;refGene_gn=BRSK2;refGene_tx=XM_023240219.1
D1 116559770 116559951
refGene_gn=BRSK2;refGene_tx=XM_023240207.1;refGene_gn=BRSK2;refGene_tx
=XM_023240208.1;refGene_gn=BRSK2;refGene_tx=XM_023240209.1;refGene_gn=
BRSK2;refGene_tx=XM_023240210.1;refGene_gn=BRSK2;refGene_tx=XM_0232402
11.1;refGene_gn=BRSK2;refGene_tx=XM_023240212.1;refGene_gn=BRSK2;refGe
ne_tx=XM_023240213.1;refGene_gn=BRSK2;refGene_tx=XM_023240214.1;refGen
e_gn=BRSK2;refGene_tx=XM_023240215.1;refGene_gn=BRSK2;refGene_tx=XM_02
3240216.1;refGene_gn=BRSK2;refGene_tx=XM_023240217.1;refGene_gn=BRSK2;
refGene_tx=XM_023240218.1;refGene_gn=BRSK2;refGene_tx=XM_023240219.1
D1 116561285 116561409
refGene_gn=BRSK2;refGene_tx=XM_023240207.1;refGene_gn=BRSK2;refGene_tx
=XM_023240208.1;refGene_gn=BRSK2;refGene_tx=XM_023240209.1;refGene_gn=
BRSK2;refGene_tx=XM_023240210.1;refGene_gn=BRSK2;refGene_tx=XM_0232402
11.1;refGene_gn=BRSK2;refGene_tx=XM_023240212.1;refGene_gn=BRSK2;refGe
ne_tx=XM_023240213.1;refGene_gn=BRSK2;refGene_tx=XM_023240214.1;refGen
e_gn=BRSK2;refGene_tx=XM_023240215.1;refGene_gn=BRSK2;refGene_tx=XM_02
3240216.1;refGene_gn=BRSK2;refGene_tx=XM_023240217.1;refGene_gn=BRSK2;
refGene_tx=XM_023240218.1;refGene_gn=BRSK2;refGene_tx=XM_023240219.1
D1 116564749 116564798
refGene_gn=BRSK2;refGene_tx=XM_023240207.1;refGene_gn=BRSK2;refGene_tx
=XM_023240208.1;refGene_gn=BRSK2;refGene_tx=XM_023240209.1;refGene_gn=
BRSK2;refGene_tx=XM_023240210.1;refGene_gn=BRSK2;refGene_tx=XM_0232402
11.1;refGene_gn=BRSK2;refGene_tx=XM_023240212.1;refGene_gn=BRSK2;refGe
ne_tx=XM_023240213.1;refGene_gn=BRSK2;refGene_tx=XM_023240214.1;refGen
e_gn=BRSK2;refGene_tx=XM_023240215.1;refGene_gn=BRSK2;refGene_tx=XM_02
3240216.1;refGene_gn=BRSK2;refGene_tx=XM_023240217.1;refGene_gn=BRSK2;
refGene_tx=XM_023240218.1;refGene_gn=BRSK2;refGene_tx=XM_023240219.1
D1 116565410 116565618
refGene_gn=BRSK2;refGene_tx=XM_023240207.1;refGene_gn=BRSK2;refGene_tx
=XM_023240208.1;refGene_gn=BRSK2;refGene_tx=XM_023240209.1;refGene_gn=
BRSK2;refGene_tx=XM_023240210.1;refGene_gn=BRSK2;refGene_tx=XM_0232402
11.1;refGene_gn=BRSK2;refGene_tx=XM_023240212.1;refGene_gn=BRSK2;refGe
ne_tx=XM_023240213.1;refGene_gn=BRSK2;refGene_tx=XM_023240214.1;refGen
e_gn=BRSK2;refGene_tx=XM_023240215.1;refGene_gn=BRSK2;refGene_tx=XM_02
3240216.1;refGene_gn=BRSK2;refGene_tx=XM_023240217.1;refGene_gn=BRSK2;
refGene_tx=XM_023240218.1;refGene_gn=BRSK2;refGene_tx=XM_023240219.1
D1 116566357 116566418
refGene_gn=BRSK2;refGene_tx=XM_023240207.1;refGene_gn=BRSK2;refGene_tx
=XM_023240208.1;refGene_gn=BRSK2;refGene_tx=XM_023240209.1;refGene_gn=
BRSK2;refGene_tx=XM_023240210.1;refGene_gn=BRSK2;refGene_tx=XM_0232402
11.1;refGene_gn=BRSK2;refGene_tx=XM_023240212.1;refGene_gn=BRSK2;refGe
ne_tx=XM_023240213.1;refGene_gn=BRSK2;refGene_tx=XM_023240214.1;refGen
e_gn=BRSK2;refGene_tx=XM_023240215.1;refGene_gn=BRSK2;refGene_tx=XM_02
3240216.1;refGene_gn=BRSK2;refGene_tx=XM_023240217.1;refGene_gn=BRSK2;
refGene_tx=XM_023240218.1;refGene_gn=BRSK2;refGene_tx=XM_023240219.1
D1 116568470 116568536
refGene_gn=BRSK2;refGene_tx=XM_023240207.1;refGene_gn=BRSK2;refGene_tx
=XM_023240208.1;refGene_gn=BRSK2;refGene_tx=XM_023240210.1;refGene_gn=

BRSK2;refGene_tx=XM_023240212.1;refGene_gn=BRSK2;refGene_tx=XM_0232402
13.1;refGene_gn=BRSK2;refGene_tx=XM_023240214.1;refGene_gn=BRSK2;refGe
ne_tx=XM_023240215.1;refGene_gn=BRSK2;refGene_tx=XM_023240216.1;refGen
e_gn=BRSK2;refGene_tx=XM_023240217.1
D1 116570540 116570691
refGene_gn=BRSK2;refGene_tx=XM_023240207.1;refGene_gn=BRSK2;refGene_tx
=XM_023240208.1;refGene_gn=BRSK2;refGene_tx=XM_023240209.1;refGene_gn=
BRSK2;refGene_tx=XM_023240210.1;refGene_gn=BRSK2;refGene_tx=XM_0232402
11.1;refGene_gn=BRSK2;refGene_tx=XM_023240212.1;refGene_gn=BRSK2;refGe
ne_tx=XM_023240213.1;refGene_gn=BRSK2;refGene_tx=XM_023240214.1;refGen
e_gn=BRSK2;refGene_tx=XM_023240215.1;refGene_gn=BRSK2;refGene_tx=XM_02
3240216.1;refGene_gn=BRSK2;refGene_tx=XM_023240217.1;refGene_gn=BRSK2;
refGene_tx=XM_023240218.1;refGene_gn=BRSK2;refGene_tx=XM_023240219.1
D1 116570776 116570874
refGene_gn=BRSK2;refGene_tx=XM_023240207.1;refGene_gn=BRSK2;refGene_tx
=XM_023240208.1;refGene_gn=BRSK2;refGene_tx=XM_023240209.1;refGene_gn=
BRSK2;refGene_tx=XM_023240210.1;refGene_gn=BRSK2;refGene_tx=XM_0232402
11.1;refGene_gn=BRSK2;refGene_tx=XM_023240212.1;refGene_gn=BRSK2;refGe
ne_tx=XM_023240213.1;refGene_gn=BRSK2;refGene_tx=XM_023240214.1;refGen
e_gn=BRSK2;refGene_tx=XM_023240215.1;refGene_gn=BRSK2;refGene_tx=XM_02
3240216.1;refGene_gn=BRSK2;refGene_tx=XM_023240217.1;refGene_gn=BRSK2;
refGene_tx=XM_023240218.1;refGene_gn=BRSK2;refGene_tx=XM_023240219.1
D1 116570966 116571131
refGene_gn=BRSK2;refGene_tx=XM_023240207.1;refGene_gn=BRSK2;refGene_tx
=XM_023240208.1;refGene_gn=BRSK2;refGene_tx=XM_023240209.1;refGene_gn=
BRSK2;refGene_tx=XM_023240210.1;refGene_gn=BRSK2;refGene_tx=XM_0232402
11.1;refGene_gn=BRSK2;refGene_tx=XM_023240212.1;refGene_gn=BRSK2;refGe
ne_tx=XM_023240213.1;refGene_gn=BRSK2;refGene_tx=XM_023240214.1;refGen
e_gn=BRSK2;refGene_tx=XM_023240215.1;refGene_gn=BRSK2;refGene_tx=XM_02
3240216.1;refGene_gn=BRSK2;refGene_tx=XM_023240217.1;refGene_gn=BRSK2;
refGene_tx=XM_023240218.1;refGene_gn=BRSK2;refGene_tx=XM_023240219.1
D1 116571463 116571495
refGene_gn=BRSK2;refGene_tx=XM_023240207.1;refGene_gn=BRSK2;refGene_tx
=XM_023240208.1;refGene_gn=BRSK2;refGene_tx=XM_023240209.1;refGene_gn=
BRSK2;refGene_tx=XM_023240210.1;refGene_gn=BRSK2;refGene_tx=XM_0232402
11.1;refGene_gn=BRSK2;refGene_tx=XM_023240212.1;refGene_gn=BRSK2;refGe
ne_tx=XM_023240213.1;refGene_gn=BRSK2;refGene_tx=XM_023240214.1;refGen
e_gn=BRSK2;refGene_tx=XM_023240215.1;refGene_gn=BRSK2;refGene_tx=XM_02
3240216.1;refGene_gn=BRSK2;refGene_tx=XM_023240217.1;refGene_gn=BRSK2;
refGene_tx=XM_023240218.1;refGene_gn=BRSK2;refGene_tx=XM_023240219.1
D1 116572592 116572739
refGene_gn=BRSK2;refGene_tx=XM_023240207.1;refGene_gn=BRSK2;refGene_tx
=XM_023240208.1;refGene_gn=BRSK2;refGene_tx=XM_023240209.1;refGene_gn=
BRSK2;refGene_tx=XM_023240210.1;refGene_gn=BRSK2;refGene_tx=XM_0232402
11.1;refGene_gn=BRSK2;refGene_tx=XM_023240212.1;refGene_gn=BRSK2;refGe
ne_tx=XM_023240213.1;refGene_gn=BRSK2;refGene_tx=XM_023240214.1;refGen
e_gn=BRSK2;refGene_tx=XM_023240215.1;refGene_gn=BRSK2;refGene_tx=XM_02
3240216.1;refGene_gn=BRSK2;refGene_tx=XM_023240217.1;refGene_gn=BRSK2;
refGene_tx=XM_023240218.1;refGene_gn=BRSK2;refGene_tx=XM_023240219.1
D1 116572838 116572907

refGene_gn=BRSK2;refGene_tx=XM_023240207.1;refGene_gn=BRSK2;refGene_tx=XM_023240208.1;refGene_gn=BRSK2;refGene_tx=XM_023240209.1;refGene_gn=BRSK2;refGene_tx=XM_023240210.1;refGene_gn=BRSK2;refGene_tx=XM_023240211.1;refGene_gn=BRSK2;refGene_tx=XM_023240212.1;refGene_gn=BRSK2;refGene_tx=XM_023240213.1;refGene_gn=BRSK2;refGene_tx=XM_023240214.1;refGene_gn=BRSK2;refGene_tx=XM_023240215.1;refGene_gn=BRSK2;refGene_tx=XM_023240216.1;refGene_gn=BRSK2;refGene_tx=XM_023240217.1;refGene_gn=BRSK2;refGene_tx=XM_023240218.1;refGene_gn=BRSK2;refGene_tx=XM_023240219.1
D1 116573078 116573112
refGene_gn=BRSK2;refGene_tx=XM_023240207.1;refGene_gn=BRSK2;refGene_tx=XM_023240208.1;refGene_gn=BRSK2;refGene_tx=XM_023240209.1;refGene_gn=BRSK2;refGene_tx=XM_023240210.1;refGene_gn=BRSK2;refGene_tx=XM_023240211.1;refGene_gn=BRSK2;refGene_tx=XM_023240212.1;refGene_gn=BRSK2;refGene_tx=XM_023240213.1;refGene_gn=BRSK2;refGene_tx=XM_023240214.1;refGene_gn=BRSK2;refGene_tx=XM_023240215.1;refGene_gn=BRSK2;refGene_tx=XM_023240216.1;refGene_gn=BRSK2;refGene_tx=XM_023240217.1;refGene_gn=BRSK2;refGene_tx=XM_023240218.1;refGene_gn=BRSK2;refGene_tx=XM_023240219.1
D1 116573789 116573906
refGene_gn=BRSK2;refGene_tx=XM_023240207.1;refGene_gn=BRSK2;refGene_tx=XM_023240208.1;refGene_gn=BRSK2;refGene_tx=XM_023240209.1;refGene_gn=BRSK2;refGene_tx=XM_023240210.1;refGene_gn=BRSK2;refGene_tx=XM_023240211.1;refGene_gn=BRSK2;refGene_tx=XM_023240212.1;refGene_gn=BRSK2;refGene_tx=XM_023240213.1;refGene_gn=BRSK2;refGene_tx=XM_023240214.1;refGene_gn=BRSK2;refGene_tx=XM_023240215.1;refGene_gn=BRSK2;refGene_tx=XM_023240216.1;refGene_gn=BRSK2;refGene_tx=XM_023240217.1;refGene_gn=BRSK2;refGene_tx=XM_023240218.1;refGene_gn=BRSK2;refGene_tx=XM_023240219.1
D1 116575829 116575970
refGene_gn=BRSK2;refGene_tx=XM_023240207.1;refGene_gn=BRSK2;refGene_tx=XM_023240208.1;refGene_gn=BRSK2;refGene_tx=XM_023240209.1;refGene_gn=BRSK2;refGene_tx=XM_023240210.1;refGene_gn=BRSK2;refGene_tx=XM_023240211.1;refGene_gn=BRSK2;refGene_tx=XM_023240212.1;refGene_gn=BRSK2;refGene_tx=XM_023240213.1;refGene_gn=BRSK2;refGene_tx=XM_023240214.1;refGene_gn=BRSK2;refGene_tx=XM_023240215.1;refGene_gn=BRSK2;refGene_tx=XM_023240216.1;refGene_gn=BRSK2;refGene_tx=XM_023240217.1;refGene_gn=BRSK2;refGene_tx=XM_023240218.1;refGene_gn=BRSK2;refGene_tx=XM_023240219.1
D1 116578544 116578634
refGene_gn=BRSK2;refGene_tx=XM_023240207.1;refGene_gn=BRSK2;refGene_tx=XM_023240208.1;refGene_gn=BRSK2;refGene_tx=XM_023240209.1;refGene_gn=BRSK2;refGene_tx=XM_023240210.1;refGene_gn=BRSK2;refGene_tx=XM_023240211.1;refGene_gn=BRSK2;refGene_tx=XM_023240212.1;refGene_gn=BRSK2;refGene_tx=XM_023240213.1;refGene_gn=BRSK2;refGene_tx=XM_023240214.1;refGene_gn=BRSK2;refGene_tx=XM_023240215.1;refGene_gn=BRSK2;refGene_tx=XM_023240216.1;refGene_gn=BRSK2;refGene_tx=XM_023240217.1;refGene_gn=BRSK2;refGene_tx=XM_023240218.1;refGene_gn=BRSK2;refGene_tx=XM_023240219.1
D1 116579886 116579981
refGene_gn=BRSK2;refGene_tx=XM_023240207.1;refGene_gn=BRSK2;refGene_tx=XM_023240208.1;refGene_gn=BRSK2;refGene_tx=XM_023240209.1;refGene_gn=BRSK2;refGene_tx=XM_023240210.1;refGene_gn=BRSK2;refGene_tx=XM_023240211.1;refGene_gn=BRSK2;refGene_tx=XM_023240212.1;refGene_gn=BRSK2;refGene_tx=XM_023240213.1;refGene_gn=BRSK2;refGene_tx=XM_023240214.1;refGen

e_gn=BRSK2;refGene_tx=XM_023240215.1;refGene_gn=BRSK2;refGene_tx=XM_023240217.1;refGene_gn=BRSK2;refGene_tx=XM_023240218.1;refGene_gn=BRSK2;refGene_tx=XM_023240219.1
D1 116615294 116615388
refGene_gn=BRSK2;refGene_tx=XM_023240207.1;refGene_gn=BRSK2;refGene_tx=XM_023240208.1;refGene_gn=BRSK2;refGene_tx=XM_023240209.1;refGene_gn=BRSK2;refGene_tx=XM_023240210.1;refGene_gn=BRSK2;refGene_tx=XM_023240211.1;refGene_gn=BRSK2;refGene_tx=XM_023240212.1;refGene_gn=BRSK2;refGene_tx=XM_023240213.1;refGene_gn=BRSK2;refGene_tx=XM_023240214.1;refGene_gn=BRSK2;refGene_tx=XM_023240215.1;refGene_gn=BRSK2;refGene_tx=XM_023240217.1;refGene_gn=BRSK2;refGene_tx=XM_023240218.1;refGene_gn=BRSK2;refGene_tx=XM_023240219.1
D1 116665732 116665765
refGene_gn=TOLLIP;refGene_tx=XM_003993793.5
D1 116675827 116675977
refGene_gn=TOLLIP;refGene_tx=XM_003993793.5
D1 116677148 116677331
refGene_gn=TOLLIP;refGene_tx=XM_003993793.5
D1 116678469 116678622
refGene_gn=TOLLIP;refGene_tx=XM_003993793.5
D1 116679437 116679528
refGene_gn=TOLLIP;refGene_tx=XM_003993793.5
D1 116681922 116682137
refGene_gn=TOLLIP;refGene_tx=XM_003993793.5
D1 116697581 116697786
refGene_gn=MUC5B;refGene_tx=XM_023239759.1
D1 116698196 116698305
refGene_gn=MUC5B;refGene_tx=XM_023239759.1
D1 116698500 116698546
refGene_gn=MUC5B;refGene_tx=XM_023239759.1
D1 116698798 116698921
refGene_gn=MUC5B;refGene_tx=XM_023239759.1
D1 116699051 116699138
refGene_gn=MUC5B;refGene_tx=XM_023239759.1
D1 116699292 116699409
refGene_gn=MUC5B;refGene_tx=XM_023239759.1
D1 116699931 116699969
refGene_gn=MUC5B;refGene_tx=XM_023239759.1
D1 116700071 116700176
refGene_gn=MUC5B;refGene_tx=XM_023239759.1
D1 116700449 116700630
refGene_gn=MUC5B;refGene_tx=XM_023239759.1
D1 116700899 116700931
refGene_gn=MUC5B;refGene_tx=XM_023239759.1
D1 116701225 116701326
refGene_gn=MUC5B;refGene_tx=XM_023239759.1
D1 116701522 116701592
refGene_gn=MUC5B;refGene_tx=XM_023239759.1
D1 116701659 116701835
refGene_gn=MUC5B;refGene_tx=XM_023239759.1

D1 116701915 116702123
refGene_gn=MUC5B;refGene_tx=XM_023239759.1
D1 116702206 116702393
refGene_gn=MUC5B;refGene_tx=XM_023239759.1
D1 116702634 116702718
refGene_gn=MUC5B;refGene_tx=XM_023239759.1
D1 116702719 116702892
refGene_gn=MUC5B;refGene_tx=XM_023239759.1
D1 116703742 116703914
refGene_gn=MUC5B;refGene_tx=XM_023239759.1
D1 116704177 116704359
refGene_gn=MUC5B;refGene_tx=XM_023239759.1
D1 116704528 116706290
refGene_gn=MUC5B;refGene_tx=XM_023239759.1
D1 116706291 116706363
refGene_gn=MUC5B;refGene_tx=XM_023239759.1
D1 116706500 116706767
refGene_gn=MUC5B;refGene_tx=XM_023239759.1
D1 116706885 116708486
refGene_gn=MUC5B;refGene_tx=XM_023239759.1
D1 116708488 116708789
refGene_gn=MUC5B;refGene_tx=XM_023239759.1
D1 116709160 116709986
refGene_gn=MUC5B;refGene_tx=XM_023239759.1
D1 116709987 116711435
refGene_gn=MUC5B;refGene_tx=XM_023239759.1
D1 116712080 116714710
refGene_gn=MUC5B;refGene_tx=XM_023239759.1
D1 116714897 116715119
refGene_gn=MUC5B;refGene_tx=XM_023239759.1
D1 116715201 116715245
refGene_gn=MUC5B;refGene_tx=XM_023239759.1
D1 116715325 116715470
refGene_gn=MUC5B;refGene_tx=XM_023239759.1
D1 116715572 116715701
refGene_gn=MUC5B;refGene_tx=XM_023239759.1
D1 116716241 116716398
refGene_gn=MUC5B;refGene_tx=XM_023239759.1
D1 116717190 116717430
refGene_gn=MUC5B;refGene_tx=XM_023239759.1
D1 116717857 116718037
refGene_gn=MUC5B;refGene_tx=XM_023239759.1
D1 116718778 116718898
refGene_gn=MUC5B;refGene_tx=XM_023239759.1
D1 116721031 116721170
refGene_gn=MUC5B;refGene_tx=XM_023239759.1
D1 116721523 116721675
refGene_gn=MUC5B;refGene_tx=XM_023239759.1
D1 116721765 116721866
refGene_gn=MUC5B;refGene_tx=XM_023239759.1

D1 116721960 116722016
refGene_gn=MUC5B;refGene_tx=XM_023239759.1
D1 116722394 116722650
refGene_gn=MUC5B;refGene_tx=XM_023239759.1
D1 116722895 116723022
refGene_gn=MUC5B;refGene_tx=XM_023239759.1
D1 116723992 116724087
refGene_gn=MUC5B;refGene_tx=XM_023239759.1
D1 116724422 116724587
refGene_gn=MUC5B;refGene_tx=XM_023239759.1
D1 116724998 116725136
refGene_gn=MUC5B;refGene_tx=XM_023239759.1
D1 116725539 116725609
refGene_gn=MUC5B;refGene_tx=XM_023239759.1
D1 116725781 116725892
refGene_gn=MUC5B;refGene_tx=XM_023239759.1
D1 116726233 116726372
refGene_gn=MUC5B;refGene_tx=XM_023239759.1
D1 116726470 116726588
refGene_gn=MUC5B;refGene_tx=XM_023239759.1
D1 116726731 116726857
refGene_gn=MUC5B;refGene_tx=XM_023239759.1
D1 116727261 116727463
refGene_gn=MUC5B;refGene_tx=XM_023239759.1
D1 116728218 116728325
refGene_gn=MUC5B;refGene_tx=XM_023239759.1
D1 116728573 116728664
refGene_gn=MUC5B;refGene_tx=XM_023239759.1
D1 116728748 116728863
refGene_gn=MUC5B;refGene_tx=XM_023239759.1
D1 116728988 116729401
refGene_gn=MUC5B;refGene_tx=XM_023239759.1
D1 116729492 116729572
refGene_gn=MUC5B;refGene_tx=XM_023239759.1
D1 116730064 116730115
refGene_gn=MUC5B;refGene_tx=XM_023239759.1
D1 116730227 116730339
refGene_gn=MUC5B;refGene_tx=XM_023239759.1
D1 116730609 116731281
refGene_gn=LOC109492311;refGene_tx=XM_023239760.1
D1 116731797 116731893
refGene_gn=LOC109492311;refGene_tx=XM_023239760.1
D1 116741988 116742177
refGene_gn=LOC109492311;refGene_tx=XM_023239760.1
D1 116748568 116748875
refGene_gn=MUC5AC;refGene_tx=XM_023240220.1
D1 116749263 116749378
refGene_gn=MUC5AC;refGene_tx=XM_023240220.1
D1 116749496 116749560
refGene_gn=MUC5AC;refGene_tx=XM_023240220.1

D1	116749776	116749896
refGene_gn=MUC5AC;refGene_tx=XM_023240220.1
D1	116750183	116750282
refGene_gn=MUC5AC;refGene_tx=XM_023240220.1
D1	116750366	116750489
refGene_gn=MUC5AC;refGene_tx=XM_023240220.1
D1	116750803	116750841
refGene_gn=MUC5AC;refGene_tx=XM_023240220.1
D1	116751053	116751155
refGene_gn=MUC5AC;refGene_tx=XM_023240220.1
D1	116751399	116751571
refGene_gn=MUC5AC;refGene_tx=XM_023240220.1
D1	116751958	116751990
refGene_gn=MUC5AC;refGene_tx=XM_023240220.1
D1	116752205	116752309
refGene_gn=MUC5AC;refGene_tx=XM_023240220.1
D1	116752416	116752504
refGene_gn=MUC5AC;refGene_tx=XM_023240220.1
D1	116752869	116753048
refGene_gn=MUC5AC;refGene_tx=XM_023240220.1
D1	116753289	116753497
refGene_gn=MUC5AC;refGene_tx=XM_023240220.1
D1	116753967	116754151
refGene_gn=MUC5AC;refGene_tx=XM_023240220.1
D1	116754246	116754497
refGene_gn=MUC5AC;refGene_tx=XM_023240220.1
D1	116754672	116754847
refGene_gn=MUC5AC;refGene_tx=XM_023240220.1
D1	116755275	116755481
refGene_gn=MUC5AC;refGene_tx=XM_023240220.1
D1	116755731	116755813
refGene_gn=MUC5AC;refGene_tx=XM_023240220.1
D1	116758103	116758989
refGene_gn=MUC5AC;refGene_tx=XM_023240220.1
D1	116761709	116761920
refGene_gn=MUC5AC;refGene_tx=XM_023240220.1
D1	116762588	116763092
refGene_gn=MUC5AC;refGene_tx=XM_023240220.1
D1	116763698	116763887
refGene_gn=MUC5AC;refGene_tx=XM_023240220.1
D1	116763981	116764025
refGene_gn=MUC5AC;refGene_tx=XM_023240220.1
D1	116764372	116764535
refGene_gn=MUC5AC;refGene_tx=XM_023240220.1
D1	116764732	116764861
refGene_gn=MUC5AC;refGene_tx=XM_023240220.1
D1	116765388	116765545
refGene_gn=MUC5AC;refGene_tx=XM_023240220.1
D1	116765765	116766005
refGene_gn=MUC5AC;refGene_tx=XM_023240220.1

D1 116766666 116766846
refGene_gn=MUC5AC;refGene_tx=XM_023240220.1
D1 116766959 116767073
refGene_gn=MUC5AC;refGene_tx=XM_023240220.1
D1 116767185 116767324
refGene_gn=MUC5AC;refGene_tx=XM_023240220.1
D1 116767401 116767553
refGene_gn=MUC5AC;refGene_tx=XM_023240220.1
D1 116767743 116767844
refGene_gn=MUC5AC;refGene_tx=XM_023240220.1
D1 116768111 116768164
refGene_gn=MUC5AC;refGene_tx=XM_023240220.1
D1 116768246 116768502
refGene_gn=MUC5AC;refGene_tx=XM_023240220.1
D1 116768693 116768820
refGene_gn=MUC5AC;refGene_tx=XM_023240220.1
D1 116769036 116769131
refGene_gn=MUC5AC;refGene_tx=XM_023240220.1
D1 116769445 116769610
refGene_gn=MUC5AC;refGene_tx=XM_023240220.1
D1 116769695 116769833
refGene_gn=MUC5AC;refGene_tx=XM_023240220.1
D1 116769909 116769979
refGene_gn=MUC5AC;refGene_tx=XM_023240220.1
D1 116770483 116770594
refGene_gn=MUC5AC;refGene_tx=XM_023240220.1
D1 116770800 116770939
refGene_gn=MUC5AC;refGene_tx=XM_023240220.1
D1 116771205 116771323
refGene_gn=MUC5AC;refGene_tx=XM_023240220.1
D1 116771473 116771599
refGene_gn=MUC5AC;refGene_tx=XM_023240220.1
D1 116771710 116771924
refGene_gn=MUC5AC;refGene_tx=XM_023240220.1
D1 116772032 116772142
refGene_gn=MUC5AC;refGene_tx=XM_023240220.1
D1 116772866 116772957
refGene_gn=MUC5AC;refGene_tx=XM_023240220.1
D1 116773323 116773438
refGene_gn=MUC5AC;refGene_tx=XM_023240220.1
D1 116773710 116773972
refGene_gn=MUC5AC;refGene_tx=XM_023240220.1
D1 116774185 116774262
refGene_gn=MUC5AC;refGene_tx=XM_023240220.1
D1 116774705 116774747
refGene_gn=MUC5AC;refGene_tx=XM_023240220.1
D1 116774981 116775000
refGene_gn=MUC5AC;refGene_tx=XM_023240220.1
D1 116776262 116776335
refGene_gn=MUC5AC;refGene_tx=XM_023240220.1

D1 116808803 116809041
refGene_gn=MUC2;refGene_tx=XM_023239761.1
D1 116809130 116809257
refGene_gn=MUC2;refGene_tx=XM_023239761.1
D1 116809564 116809604
refGene_gn=MUC2;refGene_tx=XM_023239761.1
D1 116809717 116809846
refGene_gn=MUC2;refGene_tx=XM_023239761.1
D1 116810497 116810596
refGene_gn=MUC2;refGene_tx=XM_023239761.1
D1 116810697 116810820
refGene_gn=MUC2;refGene_tx=XM_023239761.1
D1 116810914 116810955
refGene_gn=MUC2;refGene_tx=XM_023239761.1
D1 116811157 116811262
refGene_gn=MUC2;refGene_tx=XM_023239761.1
D1 116811461 116811636
refGene_gn=MUC2;refGene_tx=XM_023239761.1
D1 116813738 116813770
refGene_gn=MUC2;refGene_tx=XM_023239761.1
D1 116813864 116813965
refGene_gn=MUC2;refGene_tx=XM_023239761.1
D1 116814046 116814116
refGene_gn=MUC2;refGene_tx=XM_023239761.1
D1 116814489 116814668
refGene_gn=MUC2;refGene_tx=XM_023239761.1
D1 116815402 116815622
refGene_gn=MUC2;refGene_tx=XM_023239761.1
D1 116815936 116816120
refGene_gn=MUC2;refGene_tx=XM_023239761.1
D1 116816623 116816871
refGene_gn=MUC2;refGene_tx=XM_023239761.1
D1 116817298 116817473
refGene_gn=MUC2;refGene_tx=XM_023239761.1
D1 116817793 116818023
refGene_gn=MUC2;refGene_tx=XM_023239761.1
D1 116818288 116818370
refGene_gn=MUC2;refGene_tx=XM_023239761.1
D1 116818551 116818975
refGene_gn=MUC2;refGene_tx=XM_023239761.1
D1 116819728 116819855
refGene_gn=MUC2;refGene_tx=XM_023239761.1
D1 116820687 116820834
refGene_gn=MUC2;refGene_tx=XM_023239761.1
D1 116820835 116820866
refGene_gn=MUC2;refGene_tx=XM_023239761.1
D1 116821971 116822627
refGene_gn=MUC2;refGene_tx=XM_023239761.1
D1 116824906 116824950
refGene_gn=MUC2;refGene_tx=XM_023239761.1

D1 116825533 116825575
refGene_gn=MUC2;refGene_tx=XM_023239761.1
D1 116825950 116826136
refGene_gn=MUC2;refGene_tx=XM_023239761.1
D1 116826496 116826540
refGene_gn=MUC2;refGene_tx=XM_023239761.1
D1 116826912 116827060
refGene_gn=MUC2;refGene_tx=XM_023239761.1
D1 116827536 116827668
refGene_gn=MUC2;refGene_tx=XM_023239761.1
D1 116827858 116828015
refGene_gn=MUC2;refGene_tx=XM_023239761.1
D1 116828469 116828709
refGene_gn=MUC2;refGene_tx=XM_023239761.1
D1 116828820 116829000
refGene_gn=MUC2;refGene_tx=XM_023239761.1
D1 116829084 116829195
refGene_gn=MUC2;refGene_tx=XM_023239761.1
D1 116829419 116829558
refGene_gn=MUC2;refGene_tx=XM_023239761.1
D1 116829742 116829894
refGene_gn=MUC2;refGene_tx=XM_023239761.1
D1 116830265 116830366
refGene_gn=MUC2;refGene_tx=XM_023239761.1
D1 116830484 116830537
refGene_gn=MUC2;refGene_tx=XM_023239761.1
D1 116830628 116830884
refGene_gn=MUC2;refGene_tx=XM_023239761.1
D1 116831140 116831267
refGene_gn=MUC2;refGene_tx=XM_023239761.1
D1 116831520 116831615
refGene_gn=MUC2;refGene_tx=XM_023239761.1
D1 116832019 116832184
refGene_gn=MUC2;refGene_tx=XM_023239761.1
D1 116832282 116832420
refGene_gn=MUC2;refGene_tx=XM_023239761.1
D1 116832558 116832628
refGene_gn=MUC2;refGene_tx=XM_023239761.1
D1 116832714 116832822
refGene_gn=MUC2;refGene_tx=XM_023239761.1
D1 116832913 116833052
refGene_gn=MUC2;refGene_tx=XM_023239761.1
D1 116833368 116833486
refGene_gn=MUC2;refGene_tx=XM_023239761.1
D1 116834021 116834147
refGene_gn=MUC2;refGene_tx=XM_023239761.1
D1 116834967 116835178
refGene_gn=MUC2;refGene_tx=XM_023239761.1
D1 116835299 116835409
refGene_gn=MUC2;refGene_tx=XM_023239761.1

D1 116835492 116835583
refGene_gn=MUC2;refGene_tx=XM_023239761.1
D1 116835880 116835995
refGene_gn=MUC2;refGene_tx=XM_023239761.1
D1 116836567 116836838
refGene_gn=MUC2;refGene_tx=XM_023239761.1
D1 116837681 116837757
refGene_gn=MUC2;refGene_tx=XM_023239761.1
D1 116881876 116882890
refGene_gn=LOC102899254;refGene_tx=XM_006937664.2
D1 116882945 116883083
refGene_gn=LOC109492256;refGene_tx=XM_023239762.1
D1 116883142 116883706
refGene_gn=LOC109492256;refGene_tx=XM_023239762.1
D1 116913070 116913125
refGene_gn=SCGB1C1;refGene_tx=XM_023240221.1
D1 116913731 116913976
refGene_gn=SCGB1C1;refGene_tx=XM_023240221.1
D1 116916083 116916192
refGene_gn=ODF3;refGene_tx=XM_019812909.2
D1 116916590 116916791
refGene_gn=ODF3;refGene_tx=XM_019812909.2
D1 116917136 116917238
refGene_gn=ODF3;refGene_tx=XM_019812909.2
D1 116917416 116917538
refGene_gn=ODF3;refGene_tx=XM_019812909.2
D1 116918323 116918464
refGene_gn=ODF3;refGene_tx=XM_019812909.2
D1 116919065 116919122
refGene_gn=ODF3;refGene_tx=XM_019812909.2
D1 116922468 116922491
refGene_gn=BET1L;refGene_tx=XM_006937698.4
D1 116922876 116923044
refGene_gn=BET1L;refGene_tx=XM_003993823.5;refGene_gn=BET1L;refGene_tx=XM_006937698.4
D1 116923187 116923244
refGene_gn=BET1L;refGene_tx=XM_003993823.5;refGene_gn=BET1L;refGene_tx=XM_006937698.4
D1 116923523 116923615
refGene_gn=BET1L;refGene_tx=XM_003993823.5;refGene_gn=BET1L;refGene_tx=XM_006937698.4
D1 116924739 116924758
refGene_gn=BET1L;refGene_tx=XM_003993823.5;refGene_gn=BET1L;refGene_tx=XM_006937698.4
D1 116926308 116926626
refGene_gn=RIC8A;refGene_tx=XM_003993821.5
D1 116926851 116926899
refGene_gn=RIC8A;refGene_tx=XM_003993821.5
D1 116926986 116927574
refGene_gn=RIC8A;refGene_tx=XM_003993821.5

D1 116928029 116928121
refGene_gn=RIC8A;refGene_tx=XM_003993821.5
D1 116928882 116929033
refGene_gn=RIC8A;refGene_tx=XM_003993821.5
D1 116930104 116930200
refGene_gn=RIC8A;refGene_tx=XM_003993821.5
D1 116930285 116930430
refGene_gn=RIC8A;refGene_tx=XM_003993821.5
D1 116930504 116930649
refGene_gn=RIC8A;refGene_tx=XM_003993821.5
D1 116930996 116931116
refGene_gn=RIC8A;refGene_tx=XM_003993821.5
D1 116931467 116931588
refGene_gn=RIC8A;refGene_tx=XM_003993821.5
D1 116933341 116933362
refGene_gn=SIRT3;refGene_tx=NM_001290245.1
D1 116934802 116935012
refGene_gn=SIRT3;refGene_tx=NM_001290245.1
D1 116938717 116938879
refGene_gn=SIRT3;refGene_tx=NM_001290245.1
D1 116944404 116944505
refGene_gn=SIRT3;refGene_tx=NM_001290245.1
D1 116946070 116946303
refGene_gn=SIRT3;refGene_tx=NM_001290245.1
D1 116946398 116946587
refGene_gn=SIRT3;refGene_tx=NM_001290245.1
D1 116947585 116947733
refGene_gn=SIRT3;refGene_tx=NM_001290245.1
D1 116951730 116951785
refGene_gn=SIRT3;refGene_tx=NM_001290245.1
D1 116952398 116952493
refGene_gn=PSMD13;refGene_tx=XM_003993819.5
D1 116954063 116954142
refGene_gn=PSMD13;refGene_tx=XM_003993819.5
D1 116957416 116957451
refGene_gn=PSMD13;refGene_tx=XM_003993819.5
D1 116957535 116957591
refGene_gn=PSMD13;refGene_tx=XM_003993819.5
D1 116957828 116957872
refGene_gn=PSMD13;refGene_tx=XM_003993819.5
D1 116958061 116958148
refGene_gn=PSMD13;refGene_tx=XM_003993819.5
D1 116960322 116960494
refGene_gn=PSMD13;refGene_tx=XM_003993819.5
D1 116961468 116961548
refGene_gn=PSMD13;refGene_tx=XM_003993819.5
D1 116961626 116961752
refGene_gn=PSMD13;refGene_tx=XM_003993819.5
D1 116963292 116963355
refGene_gn=PSMD13;refGene_tx=XM_003993819.5

D1	116963567	116963648
refGene_gn=PSMD13;refGene_tx=XM_003993819.5
D1	116963835	116963952
refGene_gn=PSMD13;refGene_tx=XM_003993819.5
D1	116964519	116964615
refGene_gn=PSMD13;refGene_tx=XM_003993819.5
D1	116993187	116993482
refGene_gn=NLRP6;refGene_tx=XM_023239763.1
D1	116993756	116993795
refGene_gn=NLRP6;refGene_tx=XM_023239763.1
D1	116994046	116995793
refGene_gn=NLRP6;refGene_tx=XM_023239763.1
D1	116996598	116996691
refGene_gn=NLRP6;refGene_tx=XM_023239763.1
D1	116997813	116997984
refGene_gn=NLRP6;refGene_tx=XM_023239763.1
D1	116998084	116998252
refGene_gn=NLRP6;refGene_tx=XM_023239763.1
D1	116998723	116998832
refGene_gn=NLRP6;refGene_tx=XM_023239763.1
D1	117002015	117002274
refGene_gn=PGGHG;refGene_tx=XM_006937694.4;refGene_gn=PGGHG;refGene_tx
=XM_006937695.4;refGene_gn=PGGHG;refGene_tx=XM_019812907.2
D1	117002579	117002790
refGene_gn=PGGHG;refGene_tx=XM_006937694.4;refGene_gn=PGGHG;refGene_tx
=XM_006937695.4;refGene_gn=PGGHG;refGene_tx=XM_019812907.2
D1	117002865	117003301
refGene_gn=PGGHG;refGene_tx=XM_006937694.4;refGene_gn=PGGHG;refGene_tx
=XM_006937695.4;refGene_gn=PGGHG;refGene_tx=XM_019812907.2
D1	117004212	117004332
refGene_gn=PGGHG;refGene_tx=XM_006937694.4;refGene_gn=PGGHG;refGene_tx
=XM_006937695.4;refGene_gn=PGGHG;refGene_tx=XM_019812907.2
D1	117004765	117004897
refGene_gn=PGGHG;refGene_tx=XM_006937694.4;refGene_gn=PGGHG;refGene_tx
=XM_006937695.4;refGene_gn=PGGHG;refGene_tx=XM_019812907.2
D1	117005072	117005184
refGene_gn=PGGHG;refGene_tx=XM_006937694.4;refGene_gn=PGGHG;refGene_tx
=XM_006937695.4;refGene_gn=PGGHG;refGene_tx=XM_019812907.2
D1	117005426	117005499
refGene_gn=PGGHG;refGene_tx=XM_006937694.4;refGene_gn=PGGHG;refGene_tx
=XM_006937695.4;refGene_gn=PGGHG;refGene_tx=XM_019812907.2
D1	117005592	117005729
refGene_gn=PGGHG;refGene_tx=XM_006937694.4;refGene_gn=PGGHG;refGene_tx
=XM_006937695.4;refGene_gn=PGGHG;refGene_tx=XM_019812907.2
D1	117005854	117005988
refGene_gn=PGGHG;refGene_tx=XM_006937694.4;refGene_gn=PGGHG;refGene_tx
=XM_006937695.4;refGene_gn=PGGHG;refGene_tx=XM_019812907.2
D1	117006095	117006191
refGene_gn=PGGHG;refGene_tx=XM_006937694.4;refGene_gn=PGGHG;refGene_tx
=XM_006937695.4;refGene_gn=PGGHG;refGene_tx=XM_019812907.2

D1 117006368 117006466
refGene_gn=PGGHG;refGene_tx=XM_006937694.4;refGene_gn=PGGHG;refGene_tx
=XM_006937695.4;refGene_gn=PGGHG;refGene_tx=XM_019812907.2
D1 117006541 117006753
refGene_gn=PGGHG;refGene_tx=XM_006937694.4;refGene_gn=PGGHG;refGene_tx
=XM_006937695.4;refGene_gn=PGGHG;refGene_tx=XM_019812907.2
D1 117006846 117006902
refGene_gn=PGGHG;refGene_tx=XM_006937694.4;refGene_gn=PGGHG;refGene_tx
=XM_006937695.4;refGene_gn=PGGHG;refGene_tx=XM_019812907.2
D1 117010326 117010539
refGene_gn=IFITM5;refGene_tx=XM_003993817.4
D1 117011039 117011228
refGene_gn=IFITM5;refGene_tx=XM_003993817.4
D1 117035628 117035811
refGene_gn=LOC101086846;refGene_tx=XM_003993844.5
D1 117036316 117036511
refGene_gn=LOC101086846;refGene_tx=XM_003993844.5
D1 117045909 117046158
refGene_gn=LOC101100838;refGene_tx=XM_003993816.5
D1 117046659 117046851
refGene_gn=LOC101100838;refGene_tx=XM_003993816.5
D1 117051961 117052156
refGene_gn=LOC101086599;refGene_tx=XM_006937693.4
D1 117052730 117052916
refGene_gn=LOC101086599;refGene_tx=XM_006937693.4
D1 117105156 117105307
refGene_gn=B4GALNT4;refGene_tx=XM_023240223.1;refGene_gn=B4GALNT4;refG
ene_tx=XM_023240224.1
D1 117108348 117108452
refGene_gn=B4GALNT4;refGene_tx=XM_023240223.1;refGene_gn=B4GALNT4;refG
ene_tx=XM_023240224.1
D1 117108829 117108925
refGene_gn=B4GALNT4;refGene_tx=XM_023240223.1;refGene_gn=B4GALNT4;refG
ene_tx=XM_023240224.1;refGene_gn=B4GALNT4;refGene_tx=XM_023240225.1;re
fGene_gn=B4GALNT4;refGene_tx=XM_023240226.1
D1 117109013 117109109
refGene_gn=B4GALNT4;refGene_tx=XM_023240223.1;refGene_gn=B4GALNT4;refG
ene_tx=XM_023240224.1;refGene_gn=B4GALNT4;refGene_tx=XM_023240225.1;re
fGene_gn=B4GALNT4;refGene_tx=XM_023240226.1
D1 117109198 117109289
refGene_gn=B4GALNT4;refGene_tx=XM_023240223.1;refGene_gn=B4GALNT4;refG
ene_tx=XM_023240224.1;refGene_gn=B4GALNT4;refGene_tx=XM_023240225.1;re
fGene_gn=B4GALNT4;refGene_tx=XM_023240226.1
D1 117109371 117109472
refGene_gn=B4GALNT4;refGene_tx=XM_023240223.1;refGene_gn=B4GALNT4;refG
ene_tx=XM_023240224.1;refGene_gn=B4GALNT4;refGene_tx=XM_023240225.1;re
fGene_gn=B4GALNT4;refGene_tx=XM_023240226.1
D1 117109652 117109720
refGene_gn=B4GALNT4;refGene_tx=XM_023240223.1;refGene_gn=B4GALNT4;refG
ene_tx=XM_023240224.1;refGene_gn=B4GALNT4;refGene_tx=XM_023240225.1;re

fGene_gn=B4GALNT4;refGene_tx=XM_023240226.1
D1 117109932 117110011
refGene_gn=B4GALNT4;refGene_tx=XM_023240223.1;refGene_gn=B4GALNT4;refGene_tx=XM_023240224.1;refGene_gn=B4GALNT4;refGene_tx=XM_023240225.1;refGene_gn=B4GALNT4;refGene_tx=XM_023240226.1
D1 117111094 117111161
refGene_gn=B4GALNT4;refGene_tx=XM_023240223.1;refGene_gn=B4GALNT4;refGene_tx=XM_023240224.1;refGene_gn=B4GALNT4;refGene_tx=XM_023240225.1;refGene_gn=B4GALNT4;refGene_tx=XM_023240226.1
D1 117111308 117111443
refGene_gn=B4GALNT4;refGene_tx=XM_023240223.1;refGene_gn=B4GALNT4;refGene_tx=XM_023240224.1;refGene_gn=B4GALNT4;refGene_tx=XM_023240225.1;refGene_gn=B4GALNT4;refGene_tx=XM_023240226.1
D1 117111513 117111623
refGene_gn=B4GALNT4;refGene_tx=XM_023240223.1;refGene_gn=B4GALNT4;refGene_tx=XM_023240224.1;refGene_gn=B4GALNT4;refGene_tx=XM_023240225.1;refGene_gn=B4GALNT4;refGene_tx=XM_023240226.1
D1 117111703 117111804
refGene_gn=B4GALNT4;refGene_tx=XM_023240223.1;refGene_gn=B4GALNT4;refGene_tx=XM_023240224.1;refGene_gn=B4GALNT4;refGene_tx=XM_023240225.1;refGene_gn=B4GALNT4;refGene_tx=XM_023240226.1
D1 117111897 117111998
refGene_gn=B4GALNT4;refGene_tx=XM_023240223.1;refGene_gn=B4GALNT4;refGene_tx=XM_023240224.1;refGene_gn=B4GALNT4;refGene_tx=XM_023240225.1;refGene_gn=B4GALNT4;refGene_tx=XM_023240226.1
D1 117112179 117113074
refGene_gn=B4GALNT4;refGene_tx=XM_023240223.1;refGene_gn=B4GALNT4;refGene_tx=XM_023240224.1;refGene_gn=B4GALNT4;refGene_tx=XM_023240225.1;refGene_gn=B4GALNT4;refGene_tx=XM_023240226.1
D1 117115120 117115410
refGene_gn=B4GALNT4;refGene_tx=XM_023240223.1;refGene_gn=B4GALNT4;refGene_tx=XM_023240224.1;refGene_gn=B4GALNT4;refGene_tx=XM_023240225.1;refGene_gn=B4GALNT4;refGene_tx=XM_023240226.1
D1 117115554 117115708
refGene_gn=B4GALNT4;refGene_tx=XM_023240223.1;refGene_gn=B4GALNT4;refGene_tx=XM_023240224.1;refGene_gn=B4GALNT4;refGene_tx=XM_023240225.1;refGene_gn=B4GALNT4;refGene_tx=XM_023240226.1
D1 117115840 117115913
refGene_gn=B4GALNT4;refGene_tx=XM_023240223.1;refGene_gn=B4GALNT4;refGene_tx=XM_023240224.1;refGene_gn=B4GALNT4;refGene_tx=XM_023240225.1;refGene_gn=B4GALNT4;refGene_tx=XM_023240226.1
D1 117116024 117116178
refGene_gn=B4GALNT4;refGene_tx=XM_023240223.1;refGene_gn=B4GALNT4;refGene_tx=XM_023240224.1;refGene_gn=B4GALNT4;refGene_tx=XM_023240225.1;refGene_gn=B4GALNT4;refGene_tx=XM_023240226.1
D1 117116513 117116640
refGene_gn=B4GALNT4;refGene_tx=XM_023240223.1;refGene_gn=B4GALNT4;refGene_tx=XM_023240224.1;refGene_gn=B4GALNT4;refGene_tx=XM_023240225.1;refGene_gn=B4GALNT4;refGene_tx=XM_023240226.1
D1 117117049 117117173

refGene_gn=B4GALNT4;refGene_tx=XM_023240223.1;refGene_gn=B4GALNT4;refGene_tx=XM_023240224.1;refGene_gn=B4GALNT4;refGene_tx=XM_023240225.1;refGene_gn=B4GALNT4;refGene_tx=XM_023240226.1
D1	117125266	117125347
refGene_gn=PKP3;refGene_tx=XM_023240227.1
D1	117126945	117127141
refGene_gn=PKP3;refGene_tx=XM_023240227.1
D1	117128722	117129002
refGene_gn=PKP3;refGene_tx=XM_019812900.2
D1	117129264	117129344
refGene_gn=PKP3;refGene_tx=XM_019812900.2;refGene_gn=PKP3;refGene_tx=XM_023240227.1
D1	117129483	117130109
refGene_gn=PKP3;refGene_tx=XM_019812900.2;refGene_gn=PKP3;refGene_tx=XM_023240227.1
D1	117130202	117130326
refGene_gn=PKP3;refGene_tx=XM_019812900.2;refGene_gn=PKP3;refGene_tx=XM_023240227.1
D1	117130740	117130945
refGene_gn=PKP3;refGene_tx=XM_019812900.2;refGene_gn=PKP3;refGene_tx=XM_023240227.1
D1	117131284	117131459
refGene_gn=PKP3;refGene_tx=XM_019812900.2;refGene_gn=PKP3;refGene_tx=XM_023240227.1
D1	117131895	117132013
refGene_gn=PKP3;refGene_tx=XM_019812900.2;refGene_gn=PKP3;refGene_tx=XM_023240227.1
D1	117132084	117132255
refGene_gn=PKP3;refGene_tx=XM_019812900.2;refGene_gn=PKP3;refGene_tx=XM_023240227.1
D1	117132331	117132517
refGene_gn=PKP3;refGene_tx=XM_019812900.2;refGene_gn=PKP3;refGene_tx=XM_023240227.1
D1	117136064	117136218
refGene_gn=PKP3;refGene_tx=XM_019812900.2;refGene_gn=PKP3;refGene_tx=XM_023240227.1
D1	117136431	117136624
refGene_gn=PKP3;refGene_tx=XM_019812900.2;refGene_gn=PKP3;refGene_tx=XM_023240227.1
D1	117136701	117136789
refGene_gn=PKP3;refGene_tx=XM_019812900.2;refGene_gn=PKP3;refGene_tx=XM_023240227.1
D1	117136999	117137035
refGene_gn=PKP3;refGene_tx=XM_019812900.2;refGene_gn=PKP3;refGene_tx=XM_023240227.1
D1	117138282	117138446
refGene_gn=SIGIRR;refGene_tx=XM_023240228.1
D1	117138762	117138952
refGene_gn=SIGIRR;refGene_tx=XM_023240228.1
D1	117139223	117139374

refGene_gn=SIGIRR;refGene_tx=XM_023240228.1
D1	117139440	117139543
refGene_gn=SIGIRR;refGene_tx=XM_023240228.1
D1	117139913	117140058
refGene_gn=SIGIRR;refGene_tx=XM_023240228.1
D1	117140433	117140576
refGene_gn=SIGIRR;refGene_tx=XM_023240228.1
D1	117140696	117140830
refGene_gn=SIGIRR;refGene_tx=XM_023240228.1
D1	117141234	117141433
refGene_gn=SIGIRR;refGene_tx=XM_023240228.1
D1	117142342	117142349
refGene_gn=SIGIRR;refGene_tx=XM_023240228.1
D1	117147658	117147686
refGene_gn=ANO9;refGene_tx=XM_019812902.2;refGene_gn=ANO9;refGene_tx=XM_019812903.2
D1	117147792	117147905
refGene_gn=ANO9;refGene_tx=XM_019812902.2;refGene_gn=ANO9;refGene_tx=XM_019812903.2
D1	117148021	117148115
refGene_gn=ANO9;refGene_tx=XM_019812902.2;refGene_gn=ANO9;refGene_tx=XM_019812903.2
D1	117148196	117148298
refGene_gn=ANO9;refGene_tx=XM_019812902.2;refGene_gn=ANO9;refGene_tx=XM_019812903.2
D1	117148867	117149015
refGene_gn=ANO9;refGene_tx=XM_019812902.2;refGene_gn=ANO9;refGene_tx=XM_019812903.2
D1	117149761	117149914
refGene_gn=ANO9;refGene_tx=XM_019812902.2;refGene_gn=ANO9;refGene_tx=XM_019812903.2
D1	117150042	117150182
refGene_gn=ANO9;refGene_tx=XM_019812902.2;refGene_gn=ANO9;refGene_tx=XM_019812903.2
D1	117150334	117150432
refGene_gn=ANO9;refGene_tx=XM_019812902.2;refGene_gn=ANO9;refGene_tx=XM_019812903.2
D1	117150573	117150631
refGene_gn=ANO9;refGene_tx=XM_019812902.2;refGene_gn=ANO9;refGene_tx=XM_019812903.2
D1	117153120	117153232
refGene_gn=ANO9;refGene_tx=XM_019812902.2;refGene_gn=ANO9;refGene_tx=XM_019812903.2
D1	117153449	117153486
refGene_gn=ANO9;refGene_tx=XM_019812902.2;refGene_gn=ANO9;refGene_tx=XM_019812903.2
D1	117153574	117153736
refGene_gn=ANO9;refGene_tx=XM_019812902.2;refGene_gn=ANO9;refGene_tx=XM_019812903.2
D1	117153809	117153914

refGene_gn=ANO9;refGene_tx=XM_019812902.2;refGene_gn=ANO9;refGene_tx=X
M_019812903.2
D1 117154034 117154117
refGene_gn=ANO9;refGene_tx=XM_019812902.2;refGene_gn=ANO9;refGene_tx=X
M_019812903.2
D1 117154231 117154292
refGene_gn=ANO9;refGene_tx=XM_019812902.2;refGene_gn=ANO9;refGene_tx=X
M_019812903.2
D1 117154449 117154546
refGene_gn=ANO9;refGene_tx=XM_019812902.2;refGene_gn=ANO9;refGene_tx=X
M_019812903.2
D1 117154654 117154789
refGene_gn=ANO9;refGene_tx=XM_019812902.2;refGene_gn=ANO9;refGene_tx=X
M_019812903.2
D1 117155039 117155113
refGene_gn=ANO9;refGene_tx=XM_019812902.2;refGene_gn=ANO9;refGene_tx=X
M_019812903.2
D1 117155196 117155255
refGene_gn=ANO9;refGene_tx=XM_019812902.2;refGene_gn=ANO9;refGene_tx=X
M_019812903.2
D1 117155381 117155428
refGene_gn=ANO9;refGene_tx=XM_019812902.2;refGene_gn=ANO9;refGene_tx=X
M_019812903.2
D1 117156282 117156428
refGene_gn=ANO9;refGene_tx=XM_019812902.2;refGene_gn=ANO9;refGene_tx=X
M_019812903.2
D1 117156608 117156734
refGene_gn=ANO9;refGene_tx=XM_019812902.2;refGene_gn=ANO9;refGene_tx=X
M_019812903.2
D1 117156815 117156890
refGene_gn=ANO9;refGene_tx=XM_019812902.2;refGene_gn=ANO9;refGene_tx=X
M_019812903.2
D1 117161856 117161913
refGene_gn=ANO9;refGene_tx=XM_019812902.2
D1 117162502 117162508
refGene_gn=ANO9;refGene_tx=XM_019812903.2
D1 117176011 117176193
refGene_gn=PTDSS2;refGene_tx=XM_023240234.1
D1 117182969 117183071
refGene_gn=PTDSS2;refGene_tx=XM_023240234.1
D1 117187750 117187833
refGene_gn=PTDSS2;refGene_tx=XM_023240234.1
D1 117191669 117191737
refGene_gn=PTDSS2;refGene_tx=XM_023240234.1
D1 117193580 117193715
refGene_gn=PTDSS2;refGene_tx=XM_023240234.1
D1 117194121 117194172
refGene_gn=PTDSS2;refGene_tx=XM_023240234.1
D1 117194472 117194586
refGene_gn=PTDSS2;refGene_tx=XM_023240234.1;refGene_gn=PTDSS2;refGene_

tx=XM_023240235.1;refGene_gn=PTDSS2;refGene_tx=XM_023240236.1
D1 117194675 117194794
refGene_gn=PTDSS2;refGene_tx=XM_023240234.1;refGene_gn=PTDSS2;refGene_tx=XM_023240235.1;refGene_gn=PTDSS2;refGene_tx=XM_023240236.1
D1 117195031 117195146
refGene_gn=PTDSS2;refGene_tx=XM_023240234.1;refGene_gn=PTDSS2;refGene_tx=XM_023240235.1;refGene_gn=PTDSS2;refGene_tx=XM_023240236.1
D1 117195219 117195365
refGene_gn=PTDSS2;refGene_tx=XM_023240234.1;refGene_gn=PTDSS2;refGene_tx=XM_023240235.1;refGene_gn=PTDSS2;refGene_tx=XM_023240236.1
D1 117195456 117195642
refGene_gn=PTDSS2;refGene_tx=XM_023240234.1;refGene_gn=PTDSS2;refGene_tx=XM_023240235.1;refGene_gn=PTDSS2;refGene_tx=XM_023240236.1
D1 117195819 117195979
refGene_gn=PTDSS2;refGene_tx=XM_023240234.1;refGene_gn=PTDSS2;refGene_tx=XM_023240235.1;refGene_gn=PTDSS2;refGene_tx=XM_023240236.1
D1 117197005 117197093
refGene_gn=RNH1;refGene_tx=XM_003993813.4;refGene_gn=RNH1;refGene_tx=XM_006937689.3;refGene_gn=RNH1;refGene_tx=XM_006937690.3;refGene_gn=RNH1;refGene_tx=XM_011287166.2;refGene_gn=RNH1;refGene_tx=XM_011287167.3;refGene_gn=RNH1;refGene_tx=XM_023240229.1;refGene_gn=RNH1;refGene_tx=XM_023240230.1;refGene_gn=RNH1;refGene_tx=XM_023240231.1;refGene_gn=RNH1;refGene_tx=XM_023240232.1;refGene_gn=RNH1;refGene_tx=XM_023240233.1
D1 117197188 117197359
refGene_gn=RNH1;refGene_tx=XM_003993813.4;refGene_gn=RNH1;refGene_tx=XM_006937689.3;refGene_gn=RNH1;refGene_tx=XM_006937690.3;refGene_gn=RNH1;refGene_tx=XM_011287166.2;refGene_gn=RNH1;refGene_tx=XM_011287167.3;refGene_gn=RNH1;refGene_tx=XM_023240229.1;refGene_gn=RNH1;refGene_tx=XM_023240230.1;refGene_gn=RNH1;refGene_tx=XM_023240231.1;refGene_gn=RNH1;refGene_tx=XM_023240232.1;refGene_gn=RNH1;refGene_tx=XM_023240233.1
D1 117198553 117198724
refGene_gn=RNH1;refGene_tx=XM_003993813.4;refGene_gn=RNH1;refGene_tx=XM_006937689.3;refGene_gn=RNH1;refGene_tx=XM_006937690.3;refGene_gn=RNH1;refGene_tx=XM_011287166.2;refGene_gn=RNH1;refGene_tx=XM_011287167.3;refGene_gn=RNH1;refGene_tx=XM_023240229.1;refGene_gn=RNH1;refGene_tx=XM_023240230.1;refGene_gn=RNH1;refGene_tx=XM_023240231.1;refGene_gn=RNH1;refGene_tx=XM_023240232.1;refGene_gn=RNH1;refGene_tx=XM_023240233.1
D1 117198960 117199131
refGene_gn=RNH1;refGene_tx=XM_003993813.4;refGene_gn=RNH1;refGene_tx=XM_006937689.3;refGene_gn=RNH1;refGene_tx=XM_006937690.3;refGene_gn=RNH1;refGene_tx=XM_011287166.2;refGene_gn=RNH1;refGene_tx=XM_011287167.3;refGene_gn=RNH1;refGene_tx=XM_023240229.1;refGene_gn=RNH1;refGene_tx=XM_023240230.1;refGene_gn=RNH1;refGene_tx=XM_023240231.1;refGene_gn=RNH1;refGene_tx=XM_023240232.1;refGene_gn=RNH1;refGene_tx=XM_023240233.1
D1 117199255 117199426
refGene_gn=RNH1;refGene_tx=XM_003993813.4;refGene_gn=RNH1;refGene_tx=XM_006937689.3;refGene_gn=RNH1;refGene_tx=XM_006937690.3;refGene_gn=RNH1;refGene_tx=XM_011287166.2;refGene_gn=RNH1;refGene_tx=XM_011287167.3;refGene_gn=RNH1;refGene_tx=XM_023240229.1;refGene_gn=RNH1;refGene_tx=XM_023240230.1;refGene_gn=RNH1;refGene_tx=XM_023240231.1;refGene_gn=RNH

1;refGene_tx=XM_023240232.1;refGene_gn=RNH1;refGene_tx=XM_023240233.1
D1 117199512 117199683
refGene_gn=RNH1;refGene_tx=XM_003993813.4;refGene_gn=RNH1;refGene_tx=XM_006937689.3;refGene_gn=RNH1;refGene_tx=XM_006937690.3;refGene_gn=RNH1;refGene_tx=XM_011287166.2;refGene_gn=RNH1;refGene_tx=XM_011287167.3;refGene_gn=RNH1;refGene_tx=XM_023240229.1;refGene_gn=RNH1;refGene_tx=XM_023240230.1;refGene_gn=RNH1;refGene_tx=XM_023240231.1;refGene_gn=RNH1;refGene_tx=XM_023240232.1;refGene_gn=RNH1;refGene_tx=XM_023240233.1
D1 117200205 117200376
refGene_gn=RNH1;refGene_tx=XM_003993813.4;refGene_gn=RNH1;refGene_tx=XM_006937689.3;refGene_gn=RNH1;refGene_tx=XM_006937690.3;refGene_gn=RNH1;refGene_tx=XM_011287166.2;refGene_gn=RNH1;refGene_tx=XM_011287167.3;refGene_gn=RNH1;refGene_tx=XM_023240229.1;refGene_gn=RNH1;refGene_tx=XM_023240230.1;refGene_gn=RNH1;refGene_tx=XM_023240231.1;refGene_gn=RNH1;refGene_tx=XM_023240232.1;refGene_gn=RNH1;refGene_tx=XM_023240233.1
D1 117200830 117201001
refGene_gn=RNH1;refGene_tx=XM_003993813.4;refGene_gn=RNH1;refGene_tx=XM_006937689.3;refGene_gn=RNH1;refGene_tx=XM_006937690.3;refGene_gn=RNH1;refGene_tx=XM_011287166.2;refGene_gn=RNH1;refGene_tx=XM_011287167.3;refGene_gn=RNH1;refGene_tx=XM_023240229.1;refGene_gn=RNH1;refGene_tx=XM_023240230.1;refGene_gn=RNH1;refGene_tx=XM_023240231.1;refGene_gn=RNH1;refGene_tx=XM_023240232.1;refGene_gn=RNH1;refGene_tx=XM_023240233.1
D1 117201858 117202012
refGene_gn=RNH1;refGene_tx=XM_003993813.4;refGene_gn=RNH1;refGene_tx=XM_006937689.3;refGene_gn=RNH1;refGene_tx=XM_006937690.3;refGene_gn=RNH1;refGene_tx=XM_011287166.2;refGene_gn=RNH1;refGene_tx=XM_011287167.3;refGene_gn=RNH1;refGene_tx=XM_023240229.1;refGene_gn=RNH1;refGene_tx=XM_023240230.1;refGene_gn=RNH1;refGene_tx=XM_023240231.1;refGene_gn=RNH1;refGene_tx=XM_023240232.1;refGene_gn=RNH1;refGene_tx=XM_023240233.1
D1 117203890 117203990
refGene_gn=RNH1;refGene_tx=XM_011287166.2;refGene_gn=RNH1;refGene_tx=XM_023240229.1;refGene_gn=RNH1;refGene_tx=XM_023240230.1
D1 117243918 117244038
refGene_gn=HRAS;refGene_tx=XM_003993811.5;refGene_gn=HRAS;refGene_tx=XM_019812897.1;refGene_gn=HRAS;refGene_tx=XM_023240249.1
D1 117244490 117244572
refGene_gn=HRAS;refGene_tx=XM_019812897.1;refGene_gn=HRAS;refGene_tx=XM_023240250.1
D1 117244678 117244838
refGene_gn=HRAS;refGene_tx=XM_003993811.5;refGene_gn=HRAS;refGene_tx=XM_019812897.1;refGene_gn=HRAS;refGene_tx=XM_023240249.1;refGene_gn=HRAS;refGene_tx=XM_023240250.1
D1 117244992 117245171
refGene_gn=HRAS;refGene_tx=XM_003993811.5;refGene_gn=HRAS;refGene_tx=XM_023240249.1;refGene_gn=HRAS;refGene_tx=XM_023240250.1
D1 117245353 117245464
refGene_gn=HRAS;refGene_tx=XM_003993811.5;refGene_gn=HRAS;refGene_tx=XM_023240249.1;refGene_gn=HRAS;refGene_tx=XM_023240250.1
D1 117245641 117245712
refGene_gn=LOC111556668;refGene_tx=XM_023239764.1

D1	117246756	117247771
refGene_gn=LOC111556668;refGene_tx=XM_023239764.1
D1	117247929	117247938
refGene_gn=LOC111556668;refGene_tx=XM_023239764.1
D1	117251142	117251307
refGene_gn=LRRC56;refGene_tx=XM_006937683.4;refGene_gn=LRRC56;refGene_tx=XM_011287170.3;refGene_gn=LRRC56;refGene_tx=XM_019812884.2;refGene_gn=LRRC56;refGene_tx=XM_023240237.1;refGene_gn=LRRC56;refGene_tx=XM_023240238.1;refGene_gn=LRRC56;refGene_tx=XM_023240239.1;refGene_gn=LRRC56;refGene_tx=XM_023240240.1;refGene_gn=LRRC56;refGene_tx=XM_023240241.1;refGene_gn=LRRC56;refGene_tx=XM_023240242.1;refGene_gn=LRRC56;refGene_tx=XM_023240243.1;refGene_gn=LRRC56;refGene_tx=XM_023240244.1;refGene_gn=LRRC56;refGene_tx=XM_023240245.1;refGene_gn=LRRC56;refGene_tx=XM_023240246.1;refGene_gn=LRRC56;refGene_tx=XM_023240247.1;refGene_gn=LRRC56;refGene_tx=XM_023240248.1
D1	117254882	117254970
refGene_gn=LRRC56;refGene_tx=XM_006937683.4;refGene_gn=LRRC56;refGene_tx=XM_011287170.3;refGene_gn=LRRC56;refGene_tx=XM_019812884.2;refGene_gn=LRRC56;refGene_tx=XM_023240237.1;refGene_gn=LRRC56;refGene_tx=XM_023240238.1;refGene_gn=LRRC56;refGene_tx=XM_023240239.1;refGene_gn=LRRC56;refGene_tx=XM_023240240.1;refGene_gn=LRRC56;refGene_tx=XM_023240241.1;refGene_gn=LRRC56;refGene_tx=XM_023240242.1;refGene_gn=LRRC56;refGene_tx=XM_023240243.1;refGene_gn=LRRC56;refGene_tx=XM_023240244.1;refGene_gn=LRRC56;refGene_tx=XM_023240245.1;refGene_gn=LRRC56;refGene_tx=XM_023240246.1;refGene_gn=LRRC56;refGene_tx=XM_023240247.1;refGene_gn=LRRC56;refGene_tx=XM_023240248.1
D1	117255568	117255629
refGene_gn=LRRC56;refGene_tx=XM_006937683.4;refGene_gn=LRRC56;refGene_tx=XM_011287170.3;refGene_gn=LRRC56;refGene_tx=XM_019812884.2;refGene_gn=LRRC56;refGene_tx=XM_023240237.1;refGene_gn=LRRC56;refGene_tx=XM_023240238.1;refGene_gn=LRRC56;refGene_tx=XM_023240239.1;refGene_gn=LRRC56;refGene_tx=XM_023240240.1;refGene_gn=LRRC56;refGene_tx=XM_023240241.1;refGene_gn=LRRC56;refGene_tx=XM_023240242.1;refGene_gn=LRRC56;refGene_tx=XM_023240243.1;refGene_gn=LRRC56;refGene_tx=XM_023240244.1;refGene_gn=LRRC56;refGene_tx=XM_023240245.1;refGene_gn=LRRC56;refGene_tx=XM_023240246.1;refGene_gn=LRRC56;refGene_tx=XM_023240247.1;refGene_gn=LRRC56;refGene_tx=XM_023240248.1
D1	117262856	117262953
refGene_gn=LRRC56;refGene_tx=XM_006937683.4;refGene_gn=LRRC56;refGene_tx=XM_011287170.3;refGene_gn=LRRC56;refGene_tx=XM_019812884.2;refGene_gn=LRRC56;refGene_tx=XM_023240237.1;refGene_gn=LRRC56;refGene_tx=XM_023240238.1;refGene_gn=LRRC56;refGene_tx=XM_023240239.1;refGene_gn=LRRC56;refGene_tx=XM_023240240.1;refGene_gn=LRRC56;refGene_tx=XM_023240241.1;refGene_gn=LRRC56;refGene_tx=XM_023240242.1;refGene_gn=LRRC56;refGene_tx=XM_023240243.1;refGene_gn=LRRC56;refGene_tx=XM_023240244.1;refGene_gn=LRRC56;refGene_tx=XM_023240245.1;refGene_gn=LRRC56;refGene_tx=XM_023240246.1;refGene_gn=LRRC56;refGene_tx=XM_023240247.1;refGene_gn=LRRC56;refGene_tx=XM_023240248.1
D1	117263025	117263226
refGene_gn=LRRC56;refGene_tx=XM_006937683.4;refGene_gn=LRRC56;refGene_

tx=XM_011287170.3;refGene_gn=LRRC56;refGene_tx=XM_019812884.2;refGene_gn=LRRC56;refGene_tx=XM_023240237.1;refGene_gn=LRRC56;refGene_tx=XM_023240238.1;refGene_gn=LRRC56;refGene_tx=XM_023240239.1;refGene_gn=LRRC56;refGene_tx=XM_023240240.1;refGene_gn=LRRC56;refGene_tx=XM_023240241.1;refGene_gn=LRRC56;refGene_tx=XM_023240242.1;refGene_gn=LRRC56;refGene_tx=XM_023240243.1;refGene_gn=LRRC56;refGene_tx=XM_023240244.1;refGene_gn=LRRC56;refGene_tx=XM_023240245.1;refGene_gn=LRRC56;refGene_tx=XM_023240246.1;refGene_gn=LRRC56;refGene_tx=XM_023240247.1;refGene_gn=LRRC56;refGene_tx=XM_023240248.1
D1 117263707 117263807
refGene_gn=LRRC56;refGene_tx=XM_019812884.2;refGene_gn=LRRC56;refGene_tx=XM_023240237.1;refGene_gn=LRRC56;refGene_tx=XM_023240238.1;refGene_gn=LRRC56;refGene_tx=XM_023240239.1;refGene_gn=LRRC56;refGene_tx=XM_023240240.1;refGene_gn=LRRC56;refGene_tx=XM_023240241.1;refGene_gn=LRRC56;refGene_tx=XM_023240242.1;refGene_gn=LRRC56;refGene_tx=XM_023240243.1;refGene_gn=LRRC56;refGene_tx=XM_023240244.1;refGene_gn=LRRC56;refGene_tx=XM_023240245.1;refGene_gn=LRRC56;refGene_tx=XM_023240246.1;refGene_gn=LRRC56;refGene_tx=XM_023240247.1;refGene_gn=LRRC56;refGene_tx=XM_023240248.1
D1 117263885 117264140
refGene_gn=LRRC56;refGene_tx=XM_006937683.4;refGene_gn=LRRC56;refGene_tx=XM_011287170.3;refGene_gn=LRRC56;refGene_tx=XM_019812884.2;refGene_gn=LRRC56;refGene_tx=XM_023240237.1;refGene_gn=LRRC56;refGene_tx=XM_023240238.1;refGene_gn=LRRC56;refGene_tx=XM_023240239.1;refGene_gn=LRRC56;refGene_tx=XM_023240240.1;refGene_gn=LRRC56;refGene_tx=XM_023240241.1;refGene_gn=LRRC56;refGene_tx=XM_023240242.1;refGene_gn=LRRC56;refGene_tx=XM_023240243.1;refGene_gn=LRRC56;refGene_tx=XM_023240244.1;refGene_gn=LRRC56;refGene_tx=XM_023240245.1;refGene_gn=LRRC56;refGene_tx=XM_023240246.1;refGene_gn=LRRC56;refGene_tx=XM_023240247.1;refGene_gn=LRRC56;refGene_tx=XM_023240248.1
D1 117264524 117264701
refGene_gn=LRRC56;refGene_tx=XM_006937683.4;refGene_gn=LRRC56;refGene_tx=XM_011287170.3;refGene_gn=LRRC56;refGene_tx=XM_019812884.2;refGene_gn=LRRC56;refGene_tx=XM_023240237.1;refGene_gn=LRRC56;refGene_tx=XM_023240238.1;refGene_gn=LRRC56;refGene_tx=XM_023240239.1;refGene_gn=LRRC56;refGene_tx=XM_023240240.1;refGene_gn=LRRC56;refGene_tx=XM_023240241.1;refGene_gn=LRRC56;refGene_tx=XM_023240242.1;refGene_gn=LRRC56;refGene_tx=XM_023240243.1;refGene_gn=LRRC56;refGene_tx=XM_023240244.1;refGene_gn=LRRC56;refGene_tx=XM_023240246.1;refGene_gn=LRRC56;refGene_tx=XM_023240247.1;refGene_gn=LRRC56;refGene_tx=XM_023240248.1
D1 117264779 117264844
refGene_gn=LRRC56;refGene_tx=XM_006937683.4;refGene_gn=LRRC56;refGene_tx=XM_011287170.3;refGene_gn=LRRC56;refGene_tx=XM_019812884.2;refGene_gn=LRRC56;refGene_tx=XM_023240237.1;refGene_gn=LRRC56;refGene_tx=XM_023240238.1;refGene_gn=LRRC56;refGene_tx=XM_023240239.1;refGene_gn=LRRC56;refGene_tx=XM_023240240.1;refGene_gn=LRRC56;refGene_tx=XM_023240241.1;refGene_gn=LRRC56;refGene_tx=XM_023240242.1;refGene_gn=LRRC56;refGene_tx=XM_023240243.1;refGene_gn=LRRC56;refGene_tx=XM_023240244.1;refGene_gn=LRRC56;refGene_tx=XM_023240245.1;refGene_gn=LRRC56;refGene_tx=XM_023240246.1;refGene_gn=LRRC56;refGene_tx=XM_023240247.1;refGene_gn=LRR

C56;refGene_tx=XM_023240248.1
D1 117265057 117265194
refGene_gn=LRRC56;refGene_tx=XM_006937683.4;refGene_gn=LRRC56;refGene_tx=XM_011287170.3;refGene_gn=LRRC56;refGene_tx=XM_019812884.2;refGene_gn=LRRC56;refGene_tx=XM_023240237.1;refGene_gn=LRRC56;refGene_tx=XM_023240238.1;refGene_gn=LRRC56;refGene_tx=XM_023240239.1;refGene_gn=LRRC56;refGene_tx=XM_023240240.1;refGene_gn=LRRC56;refGene_tx=XM_023240241.1;refGene_gn=LRRC56;refGene_tx=XM_023240242.1;refGene_gn=LRRC56;refGene_tx=XM_023240243.1;refGene_gn=LRRC56;refGene_tx=XM_023240244.1;refGene_gn=LRRC56;refGene_tx=XM_023240245.1;refGene_gn=LRRC56;refGene_tx=XM_023240246.1;refGene_gn=LRRC56;refGene_tx=XM_023240247.1;refGene_gn=LRRC56;refGene_tx=XM_023240248.1
D1 117265413 117265535
refGene_gn=LRRC56;refGene_tx=XM_006937683.4;refGene_gn=LRRC56;refGene_tx=XM_011287170.3;refGene_gn=LRRC56;refGene_tx=XM_019812884.2;refGene_gn=LRRC56;refGene_tx=XM_023240237.1;refGene_gn=LRRC56;refGene_tx=XM_023240238.1;refGene_gn=LRRC56;refGene_tx=XM_023240239.1;refGene_gn=LRRC56;refGene_tx=XM_023240240.1;refGene_gn=LRRC56;refGene_tx=XM_023240241.1;refGene_gn=LRRC56;refGene_tx=XM_023240242.1;refGene_gn=LRRC56;refGene_tx=XM_023240243.1;refGene_gn=LRRC56;refGene_tx=XM_023240244.1;refGene_gn=LRRC56;refGene_tx=XM_023240245.1;refGene_gn=LRRC56;refGene_tx=XM_023240246.1;refGene_gn=LRRC56;refGene_tx=XM_023240247.1;refGene_gn=LRRC56;refGene_tx=XM_023240248.1
D1 117266917 117267249
refGene_gn=LRRC56;refGene_tx=XM_006937683.4;refGene_gn=LRRC56;refGene_tx=XM_011287170.3;refGene_gn=LRRC56;refGene_tx=XM_019812884.2;refGene_gn=LRRC56;refGene_tx=XM_023240237.1;refGene_gn=LRRC56;refGene_tx=XM_023240238.1;refGene_gn=LRRC56;refGene_tx=XM_023240239.1;refGene_gn=LRRC56;refGene_tx=XM_023240240.1;refGene_gn=LRRC56;refGene_tx=XM_023240241.1;refGene_gn=LRRC56;refGene_tx=XM_023240242.1;refGene_gn=LRRC56;refGene_tx=XM_023240243.1;refGene_gn=LRRC56;refGene_tx=XM_023240244.1;refGene_gn=LRRC56;refGene_tx=XM_023240245.1;refGene_gn=LRRC56;refGene_tx=XM_023240246.1;refGene_gn=LRRC56;refGene_tx=XM_023240247.1;refGene_gn=LRRC56;refGene_tx=XM_023240248.1
D1 117267868 117268000
refGene_gn=LMNTD2;refGene_tx=XM_019812882.2;refGene_gn=LMNTD2;refGene_tx=XM_019812883.2
D1 117268151 117268404
refGene_gn=LMNTD2;refGene_tx=XM_019812882.2;refGene_gn=LMNTD2;refGene_tx=XM_019812883.2
D1 117268754 117268951
refGene_gn=LMNTD2;refGene_tx=XM_019812882.2;refGene_gn=LMNTD2;refGene_tx=XM_019812883.2
D1 117269039 117269159
refGene_gn=LMNTD2;refGene_tx=XM_019812882.2;refGene_gn=LMNTD2;refGene_tx=XM_019812883.2
D1 117269242 117269432
refGene_gn=LMNTD2;refGene_tx=XM_019812882.2;refGene_gn=LMNTD2;refGene_tx=XM_019812883.2
D1 117269751 117269830

refGene_gn=LMNTD2;refGene_tx=XM_019812882.2;refGene_gn=LMNTD2;refGene_tx=XM_019812883.2
D1 117270085 117270378
refGene_gn=LMNTD2;refGene_tx=XM_019812882.2;refGene_gn=LMNTD2;refGene_tx=XM_019812883.2
D1 117270721 117270810
refGene_gn=LMNTD2;refGene_tx=XM_019812882.2;refGene_gn=LMNTD2;refGene_tx=XM_019812883.2
D1 117270892 117270964
refGene_gn=LMNTD2;refGene_tx=XM_019812882.2;refGene_gn=LMNTD2;refGene_tx=XM_019812883.2
D1 117271251 117271365
refGene_gn=LMNTD2;refGene_tx=XM_019812882.2;refGene_gn=LMNTD2;refGene_tx=XM_019812883.2
D1 117271493 117271581
refGene_gn=LMNTD2;refGene_tx=XM_019812882.2;refGene_gn=LMNTD2;refGene_tx=XM_019812883.2
D1 117271986 117272139
refGene_gn=LMNTD2;refGene_tx=XM_019812882.2;refGene_gn=LMNTD2;refGene_tx=XM_019812883.2
D1 117272212 117272457
refGene_gn=LMNTD2;refGene_tx=XM_019812882.2;refGene_gn=LMNTD2;refGene_tx=XM_019812883.2
D1 117275164 117275288
refGene_gn=RASSF7;refGene_tx=XM_006937681.4;refGene_gn=RASSF7;refGene_tx=XM_006937682.4
D1 117275470 117276183
refGene_gn=RASSF7;refGene_tx=XM_006937681.4;refGene_gn=RASSF7;refGene_tx=XM_006937682.4
D1 117276315 117276444
refGene_gn=RASSF7;refGene_tx=XM_006937681.4;refGene_gn=RASSF7;refGene_tx=XM_006937682.4
D1 117276521 117276716
refGene_gn=RASSF7;refGene_tx=XM_006937681.4;refGene_gn=RASSF7;refGene_tx=XM_006937682.4
D1 117290640 117290734
refGene_gn=PHRF1;refGene_tx=XM_023240251.1;refGene_gn=PHRF1;refGene_tx=XM_023240252.1;refGene_gn=PHRF1;refGene_tx=XM_023240253.1;refGene_gn=PHRF1;refGene_tx=XM_023240254.1;refGene_gn=PHRF1;refGene_tx=XM_023240255.1;refGene_gn=PHRF1;refGene_tx=XM_023240256.1;refGene_gn=PHRF1;refGene_tx=XM_023240257.1
D1 117291120 117291240
refGene_gn=PHRF1;refGene_tx=XM_023240251.1;refGene_gn=PHRF1;refGene_tx=XM_023240252.1;refGene_gn=PHRF1;refGene_tx=XM_023240253.1;refGene_gn=PHRF1;refGene_tx=XM_023240254.1;refGene_gn=PHRF1;refGene_tx=XM_023240255.1;refGene_gn=PHRF1;refGene_tx=XM_023240256.1;refGene_gn=PHRF1;refGene_tx=XM_023240257.1
D1 117295160 117295384
refGene_gn=PHRF1;refGene_tx=XM_023240251.1;refGene_gn=PHRF1;refGene_tx=XM_023240252.1;refGene_gn=PHRF1;refGene_tx=XM_023240253.1;refGene_gn=

PHRF1;refGene_tx=XM_023240254.1;refGene_gn=PHRF1;refGene_tx=XM_0232402
55.1;refGene_gn=PHRF1;refGene_tx=XM_023240256.1;refGene_gn=PHRF1;refGe
ne_tx=XM_023240257.1
D1 117298167 117298251
refGene_gn=PHRF1;refGene_tx=XM_023240251.1;refGene_gn=PHRF1;refGene_tx
=XM_023240252.1;refGene_gn=PHRF1;refGene_tx=XM_023240253.1;refGene_gn=
PHRF1;refGene_tx=XM_023240254.1;refGene_gn=PHRF1;refGene_tx=XM_0232402
55.1;refGene_gn=PHRF1;refGene_tx=XM_023240256.1;refGene_gn=PHRF1;refGe
ne_tx=XM_023240257.1
D1 117299760 117299879
refGene_gn=PHRF1;refGene_tx=XM_023240251.1;refGene_gn=PHRF1;refGene_tx
=XM_023240252.1;refGene_gn=PHRF1;refGene_tx=XM_023240253.1;refGene_gn=
PHRF1;refGene_tx=XM_023240254.1;refGene_gn=PHRF1;refGene_tx=XM_0232402
55.1;refGene_gn=PHRF1;refGene_tx=XM_023240256.1;refGene_gn=PHRF1;refGe
ne_tx=XM_023240257.1
D1 117303215 117303313
refGene_gn=PHRF1;refGene_tx=XM_023240251.1;refGene_gn=PHRF1;refGene_tx
=XM_023240252.1;refGene_gn=PHRF1;refGene_tx=XM_023240253.1;refGene_gn=
PHRF1;refGene_tx=XM_023240254.1;refGene_gn=PHRF1;refGene_tx=XM_0232402
55.1;refGene_gn=PHRF1;refGene_tx=XM_023240256.1;refGene_gn=PHRF1;refGe
ne_tx=XM_023240257.1
D1 117303504 117303680
refGene_gn=PHRF1;refGene_tx=XM_023240251.1;refGene_gn=PHRF1;refGene_tx
=XM_023240252.1;refGene_gn=PHRF1;refGene_tx=XM_023240253.1;refGene_gn=
PHRF1;refGene_tx=XM_023240254.1;refGene_gn=PHRF1;refGene_tx=XM_0232402
55.1;refGene_gn=PHRF1;refGene_tx=XM_023240256.1;refGene_gn=PHRF1;refGe
ne_tx=XM_023240257.1
D1 117304543 117304673
refGene_gn=PHRF1;refGene_tx=XM_023240251.1;refGene_gn=PHRF1;refGene_tx
=XM_023240252.1;refGene_gn=PHRF1;refGene_tx=XM_023240253.1;refGene_gn=
PHRF1;refGene_tx=XM_023240254.1;refGene_gn=PHRF1;refGene_tx=XM_0232402
55.1;refGene_gn=PHRF1;refGene_tx=XM_023240256.1;refGene_gn=PHRF1;refGe
ne_tx=XM_023240257.1
D1 117308086 117308214
refGene_gn=PHRF1;refGene_tx=XM_023240251.1;refGene_gn=PHRF1;refGene_tx
=XM_023240252.1;refGene_gn=PHRF1;refGene_tx=XM_023240253.1;refGene_gn=
PHRF1;refGene_tx=XM_023240254.1;refGene_gn=PHRF1;refGene_tx=XM_0232402
55.1;refGene_gn=PHRF1;refGene_tx=XM_023240256.1;refGene_gn=PHRF1;refGe
ne_tx=XM_023240257.1
D1 117309246 117309428
refGene_gn=PHRF1;refGene_tx=XM_023240251.1;refGene_gn=PHRF1;refGene_tx
=XM_023240252.1;refGene_gn=PHRF1;refGene_tx=XM_023240253.1;refGene_gn=
PHRF1;refGene_tx=XM_023240254.1;refGene_gn=PHRF1;refGene_tx=XM_0232402
55.1;refGene_gn=PHRF1;refGene_tx=XM_023240256.1;refGene_gn=PHRF1;refGe
ne_tx=XM_023240257.1
D1 117309781 117309901
refGene_gn=PHRF1;refGene_tx=XM_023240251.1;refGene_gn=PHRF1;refGene_tx
=XM_023240252.1;refGene_gn=PHRF1;refGene_tx=XM_023240253.1;refGene_gn=
PHRF1;refGene_tx=XM_023240254.1;refGene_gn=PHRF1;refGene_tx=XM_0232402
55.1;refGene_gn=PHRF1;refGene_tx=XM_023240256.1;refGene_gn=PHRF1;refGe

ne_tx=XM_023240257.1
D1	117310381	117310536
refGene_gn=PHRF1;refGene_tx=XM_023240251.1;refGene_gn=PHRF1;refGene_tx=XM_023240252.1;refGene_gn=PHRF1;refGene_tx=XM_023240253.1;refGene_gn=PHRF1;refGene_tx=XM_023240254.1;refGene_gn=PHRF1;refGene_tx=XM_023240255.1;refGene_gn=PHRF1;refGene_tx=XM_023240256.1;refGene_gn=PHRF1;refGene_tx=XM_023240257.1
D1	117311194	117313771
refGene_gn=PHRF1;refGene_tx=XM_023240251.1;refGene_gn=PHRF1;refGene_tx=XM_023240252.1;refGene_gn=PHRF1;refGene_tx=XM_023240253.1;refGene_gn=PHRF1;refGene_tx=XM_023240254.1;refGene_gn=PHRF1;refGene_tx=XM_023240255.1;refGene_gn=PHRF1;refGene_tx=XM_023240256.1;refGene_gn=PHRF1;refGene_tx=XM_023240257.1
D1	117314039	117314197
refGene_gn=PHRF1;refGene_tx=XM_023240251.1;refGene_gn=PHRF1;refGene_tx=XM_023240252.1;refGene_gn=PHRF1;refGene_tx=XM_023240253.1;refGene_gn=PHRF1;refGene_tx=XM_023240254.1;refGene_gn=PHRF1;refGene_tx=XM_023240255.1;refGene_gn=PHRF1;refGene_tx=XM_023240256.1;refGene_gn=PHRF1;refGene_tx=XM_023240257.1
D1	117314342	117314603
refGene_gn=PHRF1;refGene_tx=XM_023240251.1;refGene_gn=PHRF1;refGene_tx=XM_023240252.1;refGene_gn=PHRF1;refGene_tx=XM_023240253.1;refGene_gn=PHRF1;refGene_tx=XM_023240254.1;refGene_gn=PHRF1;refGene_tx=XM_023240255.1;refGene_gn=PHRF1;refGene_tx=XM_023240256.1;refGene_gn=PHRF1;refGene_tx=XM_023240257.1
D1	117314693	117314822
refGene_gn=PHRF1;refGene_tx=XM_023240251.1;refGene_gn=PHRF1;refGene_tx=XM_023240252.1;refGene_gn=PHRF1;refGene_tx=XM_023240253.1;refGene_gn=PHRF1;refGene_tx=XM_023240254.1;refGene_gn=PHRF1;refGene_tx=XM_023240255.1;refGene_gn=PHRF1;refGene_tx=XM_023240256.1;refGene_gn=PHRF1;refGene_tx=XM_023240257.1
D1	117315334	117315478
refGene_gn=PHRF1;refGene_tx=XM_023240251.1;refGene_gn=PHRF1;refGene_tx=XM_023240252.1;refGene_gn=PHRF1;refGene_tx=XM_023240253.1;refGene_gn=PHRF1;refGene_tx=XM_023240254.1;refGene_gn=PHRF1;refGene_tx=XM_023240255.1;refGene_gn=PHRF1;refGene_tx=XM_023240256.1;refGene_gn=PHRF1;refGene_tx=XM_023240257.1
D1	117316266	117316422
refGene_gn=IRF7;refGene_tx=XM_011287173.2;refGene_gn=IRF7;refGene_tx=XM_011287174.2
D1	117316495	117316614
refGene_gn=IRF7;refGene_tx=XM_011287173.2;refGene_gn=IRF7;refGene_tx=XM_011287174.2
D1	117316704	117317214
refGene_gn=IRF7;refGene_tx=XM_011287173.2;refGene_gn=IRF7;refGene_tx=XM_011287174.2
D1	117317291	117317387
refGene_gn=IRF7;refGene_tx=XM_011287173.2;refGene_gn=IRF7;refGene_tx=XM_011287174.2
D1	117317565	117317773

refGene_gn=IRF7;refGene_tx=XM_011287173.2;refGene_gn=IRF7;refGene_tx=XM_011287174.2
D1 117317863 117317922
refGene_gn=IRF7;refGene_tx=XM_011287173.2;refGene_gn=IRF7;refGene_tx=XM_011287174.2
D1 117318153 117318361
refGene_gn=IRF7;refGene_tx=XM_011287173.2;refGene_gn=IRF7;refGene_tx=XM_011287174.2
D1 117318443 117318606
refGene_gn=IRF7;refGene_tx=XM_011287173.2;refGene_gn=IRF7;refGene_tx=XM_011287174.2
D1 117318689 117318709
refGene_gn=IRF7;refGene_tx=XM_011287173.2;refGene_gn=IRF7;refGene_tx=XM_011287174.2
D1 117320536 117320980
refGene_gn=CDHR5;refGene_tx=XM_019812879.2
D1 117321317 117321523
refGene_gn=CDHR5;refGene_tx=XM_019812879.2
D1 117322060 117322372
refGene_gn=CDHR5;refGene_tx=XM_019812879.2
D1 117322506 117322594
refGene_gn=CDHR5;refGene_tx=XM_019812879.2
D1 117322671 117322785
refGene_gn=CDHR5;refGene_tx=XM_019812879.2
D1 117322865 117323066
refGene_gn=CDHR5;refGene_tx=XM_019812879.2
D1 117323167 117323265
refGene_gn=CDHR5;refGene_tx=XM_019812879.2
D1 117323438 117323529
refGene_gn=CDHR5;refGene_tx=XM_019812879.2
D1 117324111 117324282
refGene_gn=CDHR5;refGene_tx=XM_019812879.2
D1 117324371 117324482
refGene_gn=CDHR5;refGene_tx=XM_019812879.2
D1 117324593 117324695
refGene_gn=CDHR5;refGene_tx=XM_019812879.2
D1 117324820 117324913
refGene_gn=CDHR5;refGene_tx=XM_019812879.2
D1 117326236 117326287
refGene_gn=CDHR5;refGene_tx=XM_019812879.2
D1 117326683 117326859
refGene_gn=CDHR5;refGene_tx=XM_019812879.2
D1 117326940 117326995
refGene_gn=CDHR5;refGene_tx=XM_019812879.2
D1 117328562 117328686
refGene_gn=SCT;refGene_tx=XM_023240258.1;refGene_gn=SCT;refGene_tx=XM_023240259.1;refGene_gn=SCT;refGene_tx=XM_023240260.1;refGene_gn=SCT;refGene_tx=XM_023240261.1
D1 117328965 117329058
refGene_gn=SCT;refGene_tx=XM_023240258.1;refGene_gn=SCT;refGene_tx=XM_

023240259.1;refGene_gn=SCT;refGene_tx=XM_023240260.1;refGene_gn=SCT;refGene_tx=XM_023240261.1
D1 117329293 117329395
refGene_gn=SCT;refGene_tx=XM_023240258.1;refGene_gn=SCT;refGene_tx=XM_023240259.1;refGene_gn=SCT;refGene_tx=XM_023240260.1;refGene_gn=SCT;refGene_tx=XM_023240261.1
D1 117329482 117329562
refGene_gn=SCT;refGene_tx=XM_023240261.1
D1 117332440 117332442
refGene_gn=SCT;refGene_tx=XM_023240258.1;refGene_gn=SCT;refGene_tx=XM_023240259.1;refGene_gn=SCT;refGene_tx=XM_023240260.1
D1 117343147 117343417
refGene_gn=DRD4;refGene_tx=XM_023240262.1
D1 117346164 117346277
refGene_gn=DRD4;refGene_tx=XM_023240262.1
D1 117346352 117346831
refGene_gn=DRD4;refGene_tx=XM_023240262.1
D1 117346959 117347276
refGene_gn=DRD4;refGene_tx=XM_023240262.1
D1 117351112 117351217
refGene_gn=DEAF1;refGene_tx=XM_023240263.1
D1 117354991 117355081
refGene_gn=DEAF1;refGene_tx=XM_023240263.1
D1 117361671 117361689
refGene_gn=DEAF1;refGene_tx=XM_023240264.1
D1 117362242 117362490
refGene_gn=DEAF1;refGene_tx=XM_023240263.1;refGene_gn=DEAF1;refGene_tx=XM_023240264.1
D1 117363976 117364105
refGene_gn=DEAF1;refGene_tx=XM_023240263.1;refGene_gn=DEAF1;refGene_tx=XM_023240264.1
D1 117364562 117364691
refGene_gn=DEAF1;refGene_tx=XM_023240263.1;refGene_gn=DEAF1;refGene_tx=XM_023240264.1
D1 117365675 117365802
refGene_gn=DEAF1;refGene_tx=XM_023240263.1;refGene_gn=DEAF1;refGene_tx=XM_023240264.1
D1 117368575 117368641
refGene_gn=DEAF1;refGene_tx=XM_023240263.1;refGene_gn=DEAF1;refGene_tx=XM_023240264.1
D1 117369154 117369294
refGene_gn=DEAF1;refGene_tx=XM_023240263.1;refGene_gn=DEAF1;refGene_tx=XM_023240264.1
D1 117369820 117369967
refGene_gn=DEAF1;refGene_tx=XM_023240263.1;refGene_gn=DEAF1;refGene_tx=XM_023240264.1
D1 117370234 117370364
refGene_gn=DEAF1;refGene_tx=XM_023240263.1;refGene_gn=DEAF1;refGene_tx=XM_023240264.1
D1 117372417 117372515

refGene_gn=DEAF1;refGene_tx=XM_023240263.1;refGene_gn=DEAF1;refGene_tx=XM_023240264.1
D1 117374733 117375022
refGene_gn=DEAF1;refGene_tx=XM_023240263.1;refGene_gn=DEAF1;refGene_tx=XM_023240264.1
D1 117375379 117375668
refGene_gn=TMEM80;refGene_tx=XM_023239765.1
D1 117375670 117375735
refGene_gn=TMEM80;refGene_tx=XM_023239765.1
D1 117375737 117375770
refGene_gn=TMEM80;refGene_tx=XM_023239765.1
D1 117375772 117375824
refGene_gn=TMEM80;refGene_tx=XM_023239765.1
D1 117378496 117378516
refGene_gn=TMEM80;refGene_tx=XM_023239765.1
D1 117379285 117379379
refGene_gn=TMEM80;refGene_tx=XM_023239765.1
D1 117379663 117379756
refGene_gn=TMEM80;refGene_tx=XM_023239765.1
D1 117381243 117381449
refGene_gn=TMEM80;refGene_tx=XM_023239765.1
D1 117386996 117387040
refGene_gn=EPS8L2;refGene_tx=XM_023240265.1;refGene_gn=EPS8L2;refGene_tx=XM_023240266.1;refGene_gn=EPS8L2;refGene_tx=XM_023240267.1
D1 117387124 117387180
refGene_gn=EPS8L2;refGene_tx=XM_023240265.1;refGene_gn=EPS8L2;refGene_tx=XM_023240266.1;refGene_gn=EPS8L2;refGene_tx=XM_023240267.1
D1 117387943 117388008
refGene_gn=EPS8L2;refGene_tx=XM_023240265.1;refGene_gn=EPS8L2;refGene_tx=XM_023240266.1;refGene_gn=EPS8L2;refGene_tx=XM_023240267.1;refGene_gn=EPS8L2;refGene_tx=XM_023240268.1;refGene_gn=EPS8L2;refGene_tx=XM_023240269.1
D1 117394875 117395037
refGene_gn=EPS8L2;refGene_tx=XM_023240265.1;refGene_gn=EPS8L2;refGene_tx=XM_023240266.1;refGene_gn=EPS8L2;refGene_tx=XM_023240267.1;refGene_gn=EPS8L2;refGene_tx=XM_023240268.1;refGene_gn=EPS8L2;refGene_tx=XM_023240269.1;refGene_gn=EPS8L2;refGene_tx=XM_023240270.1
D1 117395620 117395770
refGene_gn=EPS8L2;refGene_tx=XM_023240265.1;refGene_gn=EPS8L2;refGene_tx=XM_023240266.1;refGene_gn=EPS8L2;refGene_tx=XM_023240267.1;refGene_gn=EPS8L2;refGene_tx=XM_023240268.1;refGene_gn=EPS8L2;refGene_tx=XM_023240269.1;refGene_gn=EPS8L2;refGene_tx=XM_023240270.1
D1 117395837 117395917
refGene_gn=EPS8L2;refGene_tx=XM_023240265.1;refGene_gn=EPS8L2;refGene_tx=XM_023240266.1;refGene_gn=EPS8L2;refGene_tx=XM_023240267.1;refGene_gn=EPS8L2;refGene_tx=XM_023240268.1;refGene_gn=EPS8L2;refGene_tx=XM_023240269.1;refGene_gn=EPS8L2;refGene_tx=XM_023240270.1
D1 117395997 117396140
refGene_gn=EPS8L2;refGene_tx=XM_023240265.1;refGene_gn=EPS8L2;refGene_tx=XM_023240266.1;refGene_gn=EPS8L2;refGene_tx=XM_023240267.1;refGene_

gn=EPS8L2;refGene_tx=XM_023240268.1;refGene_gn=EPS8L2;refGene_tx=XM_02
3240269.1;refGene_gn=EPS8L2;refGene_tx=XM_023240270.1
D1	117396216	117396287
refGene_gn=EPS8L2;refGene_tx=XM_023240265.1;refGene_gn=EPS8L2;refGene_
tx=XM_023240266.1;refGene_gn=EPS8L2;refGene_tx=XM_023240267.1;refGene_
gn=EPS8L2;refGene_tx=XM_023240268.1;refGene_gn=EPS8L2;refGene_tx=XM_02
3240269.1;refGene_gn=EPS8L2;refGene_tx=XM_023240270.1
D1	117396453	117396580
refGene_gn=EPS8L2;refGene_tx=XM_023240265.1;refGene_gn=EPS8L2;refGene_
tx=XM_023240266.1;refGene_gn=EPS8L2;refGene_tx=XM_023240267.1;refGene_
gn=EPS8L2;refGene_tx=XM_023240268.1;refGene_gn=EPS8L2;refGene_tx=XM_02
3240269.1;refGene_gn=EPS8L2;refGene_tx=XM_023240270.1
D1	117396804	117396893
refGene_gn=EPS8L2;refGene_tx=XM_023240265.1;refGene_gn=EPS8L2;refGene_
tx=XM_023240266.1;refGene_gn=EPS8L2;refGene_tx=XM_023240267.1;refGene_
gn=EPS8L2;refGene_tx=XM_023240268.1;refGene_gn=EPS8L2;refGene_tx=XM_02
3240269.1;refGene_gn=EPS8L2;refGene_tx=XM_023240270.1
D1	117397671	117397746
refGene_gn=EPS8L2;refGene_tx=XM_023240265.1;refGene_gn=EPS8L2;refGene_
tx=XM_023240266.1;refGene_gn=EPS8L2;refGene_tx=XM_023240267.1;refGene_
gn=EPS8L2;refGene_tx=XM_023240268.1;refGene_gn=EPS8L2;refGene_tx=XM_02
3240269.1;refGene_gn=EPS8L2;refGene_tx=XM_023240270.1
D1	117397951	117398100
refGene_gn=EPS8L2;refGene_tx=XM_023240265.1;refGene_gn=EPS8L2;refGene_
tx=XM_023240266.1;refGene_gn=EPS8L2;refGene_tx=XM_023240267.1;refGene_
gn=EPS8L2;refGene_tx=XM_023240268.1;refGene_gn=EPS8L2;refGene_tx=XM_02
3240269.1;refGene_gn=EPS8L2;refGene_tx=XM_023240270.1
D1	117398219	117398352
refGene_gn=EPS8L2;refGene_tx=XM_023240265.1;refGene_gn=EPS8L2;refGene_
tx=XM_023240266.1;refGene_gn=EPS8L2;refGene_tx=XM_023240267.1;refGene_
gn=EPS8L2;refGene_tx=XM_023240268.1;refGene_gn=EPS8L2;refGene_tx=XM_02
3240269.1;refGene_gn=EPS8L2;refGene_tx=XM_023240270.1
D1	117398509	117398622
refGene_gn=EPS8L2;refGene_tx=XM_023240265.1;refGene_gn=EPS8L2;refGene_
tx=XM_023240266.1;refGene_gn=EPS8L2;refGene_tx=XM_023240267.1;refGene_
gn=EPS8L2;refGene_tx=XM_023240268.1;refGene_gn=EPS8L2;refGene_tx=XM_02
3240269.1;refGene_gn=EPS8L2;refGene_tx=XM_023240270.1
D1	117399848	117399954
refGene_gn=EPS8L2;refGene_tx=XM_023240265.1;refGene_gn=EPS8L2;refGene_
tx=XM_023240266.1;refGene_gn=EPS8L2;refGene_tx=XM_023240267.1;refGene_
gn=EPS8L2;refGene_tx=XM_023240268.1;refGene_gn=EPS8L2;refGene_tx=XM_02
3240269.1;refGene_gn=EPS8L2;refGene_tx=XM_023240270.1
D1	117400615	117400729
refGene_gn=EPS8L2;refGene_tx=XM_023240265.1;refGene_gn=EPS8L2;refGene_
tx=XM_023240266.1;refGene_gn=EPS8L2;refGene_tx=XM_023240267.1;refGene_
gn=EPS8L2;refGene_tx=XM_023240268.1;refGene_gn=EPS8L2;refGene_tx=XM_02
3240269.1;refGene_gn=EPS8L2;refGene_tx=XM_023240270.1
D1	117401108	117401181
refGene_gn=EPS8L2;refGene_tx=XM_023240265.1;refGene_gn=EPS8L2;refGene_
tx=XM_023240266.1;refGene_gn=EPS8L2;refGene_tx=XM_023240267.1;refGene_

gn=EPS8L2;refGene_tx=XM_023240268.1;refGene_gn=EPS8L2;refGene_tx=XM_023240269.1;refGene_gn=EPS8L2;refGene_tx=XM_023240270.1
D1 117401337 117401518
refGene_gn=EPS8L2;refGene_tx=XM_023240265.1;refGene_gn=EPS8L2;refGene_tx=XM_023240266.1;refGene_gn=EPS8L2;refGene_tx=XM_023240267.1;refGene_gn=EPS8L2;refGene_tx=XM_023240268.1;refGene_gn=EPS8L2;refGene_tx=XM_023240269.1;refGene_gn=EPS8L2;refGene_tx=XM_023240270.1
D1 117401620 117401753
refGene_gn=EPS8L2;refGene_tx=XM_023240265.1;refGene_gn=EPS8L2;refGene_tx=XM_023240266.1;refGene_gn=EPS8L2;refGene_tx=XM_023240267.1;refGene_gn=EPS8L2;refGene_tx=XM_023240268.1;refGene_gn=EPS8L2;refGene_tx=XM_023240269.1;refGene_gn=EPS8L2;refGene_tx=XM_023240270.1
D1 117401899 117401983
refGene_gn=EPS8L2;refGene_tx=XM_023240265.1;refGene_gn=EPS8L2;refGene_tx=XM_023240266.1;refGene_gn=EPS8L2;refGene_tx=XM_023240267.1;refGene_gn=EPS8L2;refGene_tx=XM_023240268.1;refGene_gn=EPS8L2;refGene_tx=XM_023240269.1;refGene_gn=EPS8L2;refGene_tx=XM_023240270.1
D1 117411843 117411940
refGene_gn=TALDO1;refGene_tx=XM_023240271.1
D1 117414898 117415022
refGene_gn=TALDO1;refGene_tx=XM_023240271.1
D1 117416382 117416490
refGene_gn=TALDO1;refGene_tx=XM_023240271.1
D1 117416704 117416836
refGene_gn=TALDO1;refGene_tx=XM_023240271.1
D1 117419286 117419462
refGene_gn=TALDO1;refGene_tx=XM_023240271.1
D1 117419559 117419757
refGene_gn=TALDO1;refGene_tx=XM_023240271.1
D1 117419910 117420056
refGene_gn=TALDO1;refGene_tx=XM_023240271.1
D1 117420236 117420269
refGene_gn=TALDO1;refGene_tx=XM_023240271.1
D1 117423742 117423749
refGene_gn=GATD1;refGene_tx=XM_023240272.1
D1 117423845 117423961
refGene_gn=GATD1;refGene_tx=XM_023240272.1;refGene_gn=GATD1;refGene_tx=XM_023240273.1
D1 117424103 117424197
refGene_gn=GATD1;refGene_tx=XM_023240272.1;refGene_gn=GATD1;refGene_tx=XM_023240273.1
D1 117425035 117425130
refGene_gn=GATD1;refGene_tx=XM_023240272.1;refGene_gn=GATD1;refGene_tx=XM_023240273.1
D1 117425805 117425913
refGene_gn=GATD1;refGene_tx=XM_023240272.1;refGene_gn=GATD1;refGene_tx=XM_023240273.1
D1 117426073 117426179
refGene_gn=GATD1;refGene_tx=XM_023240272.1;refGene_gn=GATD1;refGene_tx=XM_023240273.1

D1 117426685 117426762
refGene_gn=GATD1;refGene_tx=XM_023240272.1;refGene_gn=GATD1;refGene_tx=XM_023240273.1
D1 117428449 117428513
refGene_gn=GATD1;refGene_tx=XM_023240272.1;refGene_gn=GATD1;refGene_tx=XM_023240273.1
D1 117437958 117438408
refGene_gn=CEND1;refGene_tx=XM_003993809.4;refGene_gn=CEND1;refGene_tx=XM_019812869.1;refGene_gn=CEND1;refGene_tx=XM_019812870.2;refGene_gn=CEND1;refGene_tx=XM_019812871.1
D1 117441799 117441953
refGene_gn=SLC25A22;refGene_tx=XM_003993808.5;refGene_gn=SLC25A22;refGene_tx=XM_019812867.2
D1 117442023 117442099
refGene_gn=SLC25A22;refGene_tx=XM_003993808.5;refGene_gn=SLC25A22;refGene_tx=XM_019812867.2
D1 117442208 117442363
refGene_gn=SLC25A22;refGene_tx=XM_003993808.5;refGene_gn=SLC25A22;refGene_tx=XM_019812867.2
D1 117442468 117442643
refGene_gn=SLC25A22;refGene_tx=XM_003993808.5;refGene_gn=SLC25A22;refGene_tx=XM_019812867.2
D1 117442751 117442870
refGene_gn=SLC25A22;refGene_tx=XM_003993808.5;refGene_gn=SLC25A22;refGene_tx=XM_019812867.2
D1 117443360 117443451
refGene_gn=SLC25A22;refGene_tx=XM_003993808.5;refGene_gn=SLC25A22;refGene_tx=XM_019812867.2
D1 117444078 117444134
refGene_gn=SLC25A22;refGene_tx=XM_003993808.5;refGene_gn=SLC25A22;refGene_tx=XM_019812867.2
D1 117444335 117444461
refGene_gn=SLC25A22;refGene_tx=XM_003993808.5;refGene_gn=SLC25A22;refGene_tx=XM_019812867.2
D1 117444562 117444582
refGene_gn=SLC25A22;refGene_tx=XM_003993808.5
D1 117447416 117447497
refGene_gn=PANO1;refGene_tx=XM_023240277.1
D1 117447498 117447615
refGene_gn=PANO1;refGene_tx=XM_023240277.1
D1 117447616 117447706
refGene_gn=PANO1;refGene_tx=XM_023240277.1
D1 117447708 117448074
refGene_gn=PANO1;refGene_tx=XM_023240277.1
D1 117448536 117448780
refGene_gn=PIDD1;refGene_tx=XM_023240274.1;refGene_gn=PIDD1;refGene_tx=XM_023240275.1;refGene_gn=PIDD1;refGene_tx=XM_023240276.1
D1 117449033 117449239
refGene_gn=PIDD1;refGene_tx=XM_023240274.1;refGene_gn=PIDD1;refGene_tx=XM_023240275.1;refGene_gn=PIDD1;refGene_tx=XM_023240276.1

D1	117449360	117449474
refGene_gn=PIDD1;refGene_tx=XM_023240274.1;refGene_gn=PIDD1;refGene_tx=XM_023240275.1;refGene_gn=PIDD1;refGene_tx=XM_023240276.1
D1	117449552	117449671
refGene_gn=PIDD1;refGene_tx=XM_023240274.1;refGene_gn=PIDD1;refGene_tx=XM_023240275.1;refGene_gn=PIDD1;refGene_tx=XM_023240276.1
D1	117449754	117449878
refGene_gn=PIDD1;refGene_tx=XM_023240274.1;refGene_gn=PIDD1;refGene_tx=XM_023240275.1;refGene_gn=PIDD1;refGene_tx=XM_023240276.1
D1	117449964	117450115
refGene_gn=PIDD1;refGene_tx=XM_023240274.1;refGene_gn=PIDD1;refGene_tx=XM_023240275.1;refGene_gn=PIDD1;refGene_tx=XM_023240276.1
D1	117450196	117450332
refGene_gn=PIDD1;refGene_tx=XM_023240274.1;refGene_gn=PIDD1;refGene_tx=XM_023240275.1;refGene_gn=PIDD1;refGene_tx=XM_023240276.1
D1	117450477	117450625
refGene_gn=PIDD1;refGene_tx=XM_023240274.1;refGene_gn=PIDD1;refGene_tx=XM_023240275.1;refGene_gn=PIDD1;refGene_tx=XM_023240276.1
D1	117450702	117450882
refGene_gn=PIDD1;refGene_tx=XM_023240274.1;refGene_gn=PIDD1;refGene_tx=XM_023240275.1;refGene_gn=PIDD1;refGene_tx=XM_023240276.1
D1	117450986	117451112
refGene_gn=PIDD1;refGene_tx=XM_023240274.1;refGene_gn=PIDD1;refGene_tx=XM_023240275.1;refGene_gn=PIDD1;refGene_tx=XM_023240276.1
D1	117451197	117451399
refGene_gn=PIDD1;refGene_tx=XM_023240274.1;refGene_gn=PIDD1;refGene_tx=XM_023240275.1;refGene_gn=PIDD1;refGene_tx=XM_023240276.1
D1	117451570	117451625
refGene_gn=PIDD1;refGene_tx=XM_023240274.1;refGene_gn=PIDD1;refGene_tx=XM_023240275.1;refGene_gn=PIDD1;refGene_tx=XM_023240276.1
D1	117451715	117451925
refGene_gn=PIDD1;refGene_tx=XM_023240274.1;refGene_gn=PIDD1;refGene_tx=XM_023240275.1;refGene_gn=PIDD1;refGene_tx=XM_023240276.1
D1	117452243	117452657
refGene_gn=PIDD1;refGene_tx=XM_023240274.1;refGene_gn=PIDD1;refGene_tx=XM_023240275.1;refGene_gn=PIDD1;refGene_tx=XM_023240276.1
D1	117452934	117453247
refGene_gn=PIDD1;refGene_tx=XM_023240274.1;refGene_gn=PIDD1;refGene_tx=XM_023240275.1;refGene_gn=PIDD1;refGene_tx=XM_023240276.1
D1	117456489	117456612
refGene_gn=RPLP2;refGene_tx=XM_003993807.4;refGene_gn=RPLP2;refGene_tx=XM_019812865.1
D1	117457181	117457230
refGene_gn=RPLP2;refGene_tx=XM_003993807.4;refGene_gn=RPLP2;refGene_tx=XM_019812865.1
D1	117458214	117458313
refGene_gn=RPLP2;refGene_tx=XM_003993807.4;refGene_gn=RPLP2;refGene_tx=XM_019812865.1
D1	117458396	117458473
refGene_gn=RPLP2;refGene_tx=XM_003993807.4;refGene_gn=RPLP2;refGene_tx

=XM_019812865.1
D1 117462031 117462218
refGene_gn=PNPLA2;refGene_tx=XM_023240283.1
D1 117463937 117464170
refGene_gn=PNPLA2;refGene_tx=XM_023240283.1
D1 117464258 117464324
refGene_gn=PNPLA2;refGene_tx=XM_023240283.1
D1 117464673 117464883
refGene_gn=PNPLA2;refGene_tx=XM_023240283.1
D1 117465140 117465201
refGene_gn=PNPLA2;refGene_tx=XM_023240283.1
D1 117465311 117465488
refGene_gn=PNPLA2;refGene_tx=XM_023240283.1
D1 117465601 117465734
refGene_gn=PNPLA2;refGene_tx=XM_023240283.1
D1 117465951 117466074
refGene_gn=PNPLA2;refGene_tx=XM_023240283.1
D1 117466167 117466438
refGene_gn=PNPLA2;refGene_tx=XM_023240283.1
D1 117468973 117469087
refGene_gn=CRACR2B;refGene_tx=XM_023240279.1;refGene_gn=CRACR2B;refGene_tx=XM_023240280.1
D1 117469912 117470104
refGene_gn=CRACR2B;refGene_tx=XM_023240279.1;refGene_gn=CRACR2B;refGene_tx=XM_023240280.1;refGene_gn=CRACR2B;refGene_tx=XM_023240281.1;refGene_gn=CRACR2B;refGene_tx=XM_023240282.1
D1 117470181 117470293
refGene_gn=CRACR2B;refGene_tx=XM_023240279.1;refGene_gn=CRACR2B;refGene_tx=XM_023240280.1;refGene_gn=CRACR2B;refGene_tx=XM_023240281.1;refGene_gn=CRACR2B;refGene_tx=XM_023240282.1
D1 117470774 117470949
refGene_gn=CRACR2B;refGene_tx=XM_023240279.1;refGene_gn=CRACR2B;refGene_tx=XM_023240280.1;refGene_gn=CRACR2B;refGene_tx=XM_023240281.1;refGene_gn=CRACR2B;refGene_tx=XM_023240282.1
D1 117471250 117471397
refGene_gn=CRACR2B;refGene_tx=XM_023240279.1;refGene_gn=CRACR2B;refGene_tx=XM_023240280.1;refGene_gn=CRACR2B;refGene_tx=XM_023240281.1;refGene_gn=CRACR2B;refGene_tx=XM_023240282.1
D1 117471507 117471598
refGene_gn=CRACR2B;refGene_tx=XM_023240279.1;refGene_gn=CRACR2B;refGene_tx=XM_023240280.1;refGene_gn=CRACR2B;refGene_tx=XM_023240281.1;refGene_gn=CRACR2B;refGene_tx=XM_023240282.1
D1 117471749 117471842
refGene_gn=CRACR2B;refGene_tx=XM_023240279.1;refGene_gn=CRACR2B;refGene_tx=XM_023240280.1;refGene_gn=CRACR2B;refGene_tx=XM_023240281.1;refGene_gn=CRACR2B;refGene_tx=XM_023240282.1
D1 117472073 117472240
refGene_gn=CRACR2B;refGene_tx=XM_023240279.1;refGene_gn=CRACR2B;refGene_tx=XM_023240280.1;refGene_gn=CRACR2B;refGene_tx=XM_023240281.1;refGene_gn=CRACR2B;refGene_tx=XM_023240282.1

D1	117472481	117472553
refGene_gn=CRACR2B;refGene_tx=XM_023240279.1;refGene_gn=CRACR2B;refGene_tx=XM_023240280.1;refGene_gn=CRACR2B;refGene_tx=XM_023240281.1;refGene_gn=CRACR2B;refGene_tx=XM_023240282.1
D1	117472788	117472969
refGene_gn=CRACR2B;refGene_tx=XM_023240279.1;refGene_gn=CRACR2B;refGene_tx=XM_023240281.1;refGene_gn=CRACR2B;refGene_tx=XM_023240282.1
D1	117473888	117473991
refGene_gn=CRACR2B;refGene_tx=XM_023240280.1
D1	117476354	117476435
refGene_gn=CD151;refGene_tx=XM_011287176.3
D1	117476544	117476736
refGene_gn=CD151;refGene_tx=XM_011287176.3
D1	117477075	117477150
refGene_gn=CD151;refGene_tx=XM_011287176.3
D1	117477597	117477702
refGene_gn=CD151;refGene_tx=XM_011287176.3
D1	117477809	117477968
refGene_gn=CD151;refGene_tx=XM_011287176.3
D1	117478084	117478171
refGene_gn=CD151;refGene_tx=XM_011287176.3
D1	117478260	117478320
refGene_gn=CD151;refGene_tx=XM_011287176.3
D1	117480521	117480630
refGene_gn=POLR2L;refGene_tx=XM_023240286.1
D1	117481799	117481894
refGene_gn=POLR2L;refGene_tx=XM_023240286.1
D1	117491015	117491078
refGene_gn=TSPAN4;refGene_tx=XM_011287177.3;refGene_gn=TSPAN4;refGene_tx=XM_011287178.3;refGene_gn=TSPAN4;refGene_tx=XM_011287180.3;refGene_gn=TSPAN4;refGene_tx=XM_023240284.1;refGene_gn=TSPAN4;refGene_tx=XM_023240285.1
D1	117500278	117500470
refGene_gn=TSPAN4;refGene_tx=XM_011287177.3;refGene_gn=TSPAN4;refGene_tx=XM_011287178.3;refGene_gn=TSPAN4;refGene_tx=XM_011287180.3;refGene_gn=TSPAN4;refGene_tx=XM_023240284.1;refGene_gn=TSPAN4;refGene_tx=XM_023240285.1
D1	117501668	117501743
refGene_gn=TSPAN4;refGene_tx=XM_011287177.3;refGene_gn=TSPAN4;refGene_tx=XM_011287178.3;refGene_gn=TSPAN4;refGene_tx=XM_023240284.1;refGene_gn=TSPAN4;refGene_tx=XM_023240285.1
D1	117502663	117502765
refGene_gn=TSPAN4;refGene_tx=XM_011287177.3;refGene_gn=TSPAN4;refGene_tx=XM_011287178.3;refGene_gn=TSPAN4;refGene_tx=XM_011287180.3;refGene_gn=TSPAN4;refGene_tx=XM_023240284.1;refGene_gn=TSPAN4;refGene_tx=XM_023240285.1
D1	117502860	117502992
refGene_gn=TSPAN4;refGene_tx=XM_011287177.3;refGene_gn=TSPAN4;refGene_tx=XM_011287178.3;refGene_gn=TSPAN4;refGene_tx=XM_011287180.3;refGene_gn=TSPAN4;refGene_tx=XM_023240284.1;refGene_gn=TSPAN4;refGene_tx=XM_02

3240285.1
D1	117503089	117503173
refGene_gn=TSPAN4;refGene_tx=XM_011287177.3;refGene_gn=TSPAN4;refGene_tx=XM_011287178.3;refGene_gn=TSPAN4;refGene_tx=XM_011287180.3;refGene_gn=TSPAN4;refGene_tx=XM_023240284.1;refGene_gn=TSPAN4;refGene_tx=XM_023240285.1
D1	117503706	117503775
refGene_gn=TSPAN4;refGene_tx=XM_011287177.3;refGene_gn=TSPAN4;refGene_tx=XM_011287178.3;refGene_gn=TSPAN4;refGene_tx=XM_011287180.3;refGene_gn=TSPAN4;refGene_tx=XM_023240284.1;refGene_gn=TSPAN4;refGene_tx=XM_023240285.1
D1	117506621	117506720
refGene_gn=CHID1;refGene_tx=XM_011287181.3;refGene_gn=CHID1;refGene_tx=XM_011287182.3;refGene_gn=CHID1;refGene_tx=XM_023240278.1
D1	117506904	117506947
refGene_gn=CHID1;refGene_tx=XM_011287181.3;refGene_gn=CHID1;refGene_tx=XM_011287182.3;refGene_gn=CHID1;refGene_tx=XM_023240278.1
D1	117507190	117507271
refGene_gn=CHID1;refGene_tx=XM_011287181.3;refGene_gn=CHID1;refGene_tx=XM_011287182.3;refGene_gn=CHID1;refGene_tx=XM_023240278.1
D1	117515000	117515156
refGene_gn=CHID1;refGene_tx=XM_011287181.3;refGene_gn=CHID1;refGene_tx=XM_011287182.3;refGene_gn=CHID1;refGene_tx=XM_023240278.1
D1	117515713	117515815
refGene_gn=CHID1;refGene_tx=XM_011287181.3;refGene_gn=CHID1;refGene_tx=XM_011287182.3;refGene_gn=CHID1;refGene_tx=XM_023240278.1
D1	117520001	117520094
refGene_gn=CHID1;refGene_tx=XM_011287181.3;refGene_gn=CHID1;refGene_tx=XM_011287182.3;refGene_gn=CHID1;refGene_tx=XM_023240278.1
D1	117522852	117522914
refGene_gn=CHID1;refGene_tx=XM_011287181.3;refGene_gn=CHID1;refGene_tx=XM_011287182.3;refGene_gn=CHID1;refGene_tx=XM_023240278.1
D1	117523292	117523399
refGene_gn=CHID1;refGene_tx=XM_011287181.3;refGene_gn=CHID1;refGene_tx=XM_011287182.3;refGene_gn=CHID1;refGene_tx=XM_023240278.1
D1	117523852	117523897
refGene_gn=CHID1;refGene_tx=XM_011287181.3;refGene_gn=CHID1;refGene_tx=XM_011287182.3;refGene_gn=CHID1;refGene_tx=XM_023240278.1
D1	117524563	117524745
refGene_gn=CHID1;refGene_tx=XM_011287181.3;refGene_gn=CHID1;refGene_tx=XM_011287182.3;refGene_gn=CHID1;refGene_tx=XM_023240278.1
D1	117525004	117525154
refGene_gn=CHID1;refGene_tx=XM_011287181.3;refGene_gn=CHID1;refGene_tx=XM_011287182.3;refGene_gn=CHID1;refGene_tx=XM_023240278.1
D1	117525683	117525794
refGene_gn=CHID1;refGene_tx=XM_011287181.3;refGene_gn=CHID1;refGene_tx=XM_023240278.1
D1	117539912	117539979
refGene_gn=AP2A2;refGene_tx=XM_006937669.3;refGene_gn=AP2A2;refGene_tx=XM_006937670.3

D1 117585366 117585435
refGene_gn=AP2A2;refGene_tx=XM_006937669.3;refGene_gn=AP2A2;refGene_tx
=XM_006937670.3
D1 117597436 117597579
refGene_gn=AP2A2;refGene_tx=XM_006937669.3;refGene_gn=AP2A2;refGene_tx
=XM_006937670.3
D1 117599200 117599394
refGene_gn=AP2A2;refGene_tx=XM_006937669.3;refGene_gn=AP2A2;refGene_tx
=XM_006937670.3
D1 117602532 117602662
refGene_gn=AP2A2;refGene_tx=XM_006937669.3;refGene_gn=AP2A2;refGene_tx
=XM_006937670.3
D1 117605923 117606025
refGene_gn=AP2A2;refGene_tx=XM_006937669.3;refGene_gn=AP2A2;refGene_tx
=XM_006937670.3
D1 117609419 117609528
refGene_gn=AP2A2;refGene_tx=XM_006937669.3;refGene_gn=AP2A2;refGene_tx
=XM_006937670.3
D1 117611090 117611241
refGene_gn=AP2A2;refGene_tx=XM_006937669.3;refGene_gn=AP2A2;refGene_tx
=XM_006937670.3
D1 117612544 117612713
refGene_gn=AP2A2;refGene_tx=XM_006937669.3;refGene_gn=AP2A2;refGene_tx
=XM_006937670.3
D1 117613675 117613813
refGene_gn=AP2A2;refGene_tx=XM_006937669.3;refGene_gn=AP2A2;refGene_tx
=XM_006937670.3
D1 117616533 117616716
refGene_gn=AP2A2;refGene_tx=XM_006937669.3;refGene_gn=AP2A2;refGene_tx
=XM_006937670.3
D1 117617210 117617308
refGene_gn=AP2A2;refGene_tx=XM_006937669.3;refGene_gn=AP2A2;refGene_tx
=XM_006937670.3
D1 117617735 117617967
refGene_gn=AP2A2;refGene_tx=XM_006937669.3;refGene_gn=AP2A2;refGene_tx
=XM_006937670.3
D1 117618070 117618238
refGene_gn=AP2A2;refGene_tx=XM_006937669.3;refGene_gn=AP2A2;refGene_tx
=XM_006937670.3
D1 117620902 117621063
refGene_gn=AP2A2;refGene_tx=XM_006937669.3;refGene_gn=AP2A2;refGene_tx
=XM_006937670.3
D1 117623671 117623754
refGene_gn=AP2A2;refGene_tx=XM_006937669.3;refGene_gn=AP2A2;refGene_tx
=XM_006937670.3
D1 117625660 117625750
refGene_gn=AP2A2;refGene_tx=XM_006937669.3;refGene_gn=AP2A2;refGene_tx
=XM_006937670.3
D1 117626558 117626682
refGene_gn=AP2A2;refGene_tx=XM_006937669.3;refGene_gn=AP2A2;refGene_tx

=XM_006937670.3
D1 117627722 117627839
refGene_gn=AP2A2;refGene_tx=XM_006937669.3;refGene_gn=AP2A2;refGene_tx
=XM_006937670.3
D1 117627918 117627988
refGene_gn=AP2A2;refGene_tx=XM_006937669.3;refGene_gn=AP2A2;refGene_tx
=XM_006937670.3
D1 117628135 117628270
refGene_gn=AP2A2;refGene_tx=XM_006937669.3;refGene_gn=AP2A2;refGene_tx
=XM_006937670.3
D1 117628961 117629039
refGene_gn=AP2A2;refGene_tx=XM_006937669.3;refGene_gn=AP2A2;refGene_tx
=XM_006937670.3
D1 117632542 117633946
refGene_gn=LOC111556669;refGene_tx=XM_023239767.1
D1 117634601 117635051
refGene_gn=LOC111556669;refGene_tx=XM_023239767.1
D1 117635982 117636162
refGene_gn=LOC111556669;refGene_tx=XM_023239767.1
D1 117636297 117636443
refGene_gn=LOC111556669;refGene_tx=XM_023239767.1
D1 117636868 117637063
refGene_gn=LOC111556669;refGene_tx=XM_023239767.1
D1 117637154 117637276
refGene_gn=LOC111556669;refGene_tx=XM_023239767.1
D1 117639021 117639251
refGene_gn=LOC111556669;refGene_tx=XM_023239767.1
D2 280954 281589
refGene_gn=LOC109492842;refGene_tx=XM_023240316.1
D2 281591 282270
refGene_gn=LOC109492842;refGene_tx=XM_023240316.1
D2 282271 282386
refGene_gn=LOC109492842;refGene_tx=XM_023240316.1
D2 282387 282595
refGene_gn=LOC109492842;refGene_tx=XM_023240316.1
D2 301713 301809
refGene_gn=LOC109492842;refGene_tx=XM_023240316.1
D2 302086 302231
refGene_gn=LOC109492842;refGene_tx=XM_023240316.1
D2 309738 309753
refGene_gn=LOC109492842;refGene_tx=XM_023240316.1
D2 1697237 1697278
refGene_gn=ZWINT;refGene_tx=XM_006937712.3;refGene_gn=ZWINT;refGene_tx
=XM_019814188.2;refGene_gn=ZWINT;refGene_tx=XM_019814189.2;refGene_gn=
ZWINT;refGene_tx=XM_019814191.2
D2 1698161 1698252
refGene_gn=ZWINT;refGene_tx=XM_006937712.3;refGene_gn=ZWINT;refGene_tx
=XM_019814188.2;refGene_gn=ZWINT;refGene_tx=XM_019814189.2;refGene_gn=
ZWINT;refGene_tx=XM_019814191.2
D2 1698342 1698466

refGene_gn=ZWINT;refGene_tx=XM_006937712.3;refGene_gn=ZWINT;refGene_tx=XM_019814188.2;refGene_gn=ZWINT;refGene_tx=XM_019814189.2;refGene_gn=ZWINT;refGene_tx=XM_019814191.2
D2	1698601	1698768	refGene_gn=ZWINT;refGene_tx=XM_006937712.3;refGene_gn=ZWINT;refGene_tx=XM_019814188.2;refGene_gn=ZWINT;refGene_tx=XM_019814189.2;refGene_gn=ZWINT;refGene_tx=XM_019814191.2
D2	1698878	1698935	refGene_gn=ZWINT;refGene_tx=XM_006937712.3;refGene_gn=ZWINT;refGene_tx=XM_019814188.2;refGene_gn=ZWINT;refGene_tx=XM_019814189.2;refGene_gn=ZWINT;refGene_tx=XM_019814191.2
D2	1699493	1699636	refGene_gn=ZWINT;refGene_tx=XM_006937712.3;refGene_gn=ZWINT;refGene_tx=XM_019814188.2;refGene_gn=ZWINT;refGene_tx=XM_019814189.2;refGene_gn=ZWINT;refGene_tx=XM_019814191.2
D2	1699712	1699872	refGene_gn=ZWINT;refGene_tx=XM_006937712.3;refGene_gn=ZWINT;refGene_tx=XM_019814188.2;refGene_gn=ZWINT;refGene_tx=XM_019814189.2;refGene_gn=ZWINT;refGene_tx=XM_019814191.2
D2	1699960	1700038	refGene_gn=ZWINT;refGene_tx=XM_006937712.3;refGene_gn=ZWINT;refGene_tx=XM_019814188.2;refGene_gn=ZWINT;refGene_tx=XM_019814189.2;refGene_gn=ZWINT;refGene_tx=XM_019814191.2
D2	3228765	3228856	refGene_gn=PCDH15;refGene_tx=XM_019814204.2
D2	3351147	3351162	refGene_gn=PCDH15;refGene_tx=XM_019814204.2
D2	3351671	3351737	refGene_gn=PCDH15;refGene_tx=XM_019814204.2
D2	3498649	3498810	refGene_gn=PCDH15;refGene_tx=XM_019814204.2
D2	3518143	3518299	refGene_gn=PCDH15;refGene_tx=XM_019814204.2
D2	3547042	3547162	refGene_gn=PCDH15;refGene_tx=XM_019814204.2
D2	3553078	3553189	refGene_gn=PCDH15;refGene_tx=XM_019814204.2
D2	3566114	3566285	refGene_gn=PCDH15;refGene_tx=XM_019814204.2
D2	3631877	3631986	refGene_gn=PCDH15;refGene_tx=XM_019814204.2
D2	3656068	3656181	refGene_gn=PCDH15;refGene_tx=XM_019814204.2
D2	3671811	3672018	refGene_gn=PCDH15;refGene_tx=XM_019814204.2
D2	3681361	3681496	refGene_gn=PCDH15;refGene_tx=XM_019814204.2
D2	3683301	3683388	refGene_gn=PCDH15;refGene_tx=XM_019814204.2
D2	3683389	3683452	refGene_gn=PCDH15;refGene_tx=XM_019814204.2
D2	3709471	3709665	refGene_gn=PCDH15;refGene_tx=XM_019814204.2
D2	3723389	3723522	refGene_gn=PCDH15;refGene_tx=XM_019814204.2
D2	3751973	3752053	refGene_gn=PCDH15;refGene_tx=XM_019814204.2
D2	3761800	3761894	refGene_gn=PCDH15;refGene_tx=XM_019814204.2
D2	3773084	3773213	refGene_gn=PCDH15;refGene_tx=XM_019814204.2
D2	3817543	3817849	refGene_gn=PCDH15;refGene_tx=XM_019814204.2
D2	3820745	3820970	refGene_gn=PCDH15;refGene_tx=XM_019814204.2
D2	3831136	3831253	refGene_gn=PCDH15;refGene_tx=XM_019814204.2
D2	3852939	3853080	refGene_gn=PCDH15;refGene_tx=XM_019814204.2
D2	3855007	3855120	refGene_gn=PCDH15;refGene_tx=XM_019814204.2
D2	3878605	3878715	refGene_gn=PCDH15;refGene_tx=XM_019814204.2
D2	3880065	3880206	refGene_gn=PCDH15;refGene_tx=XM_019814204.2
D2	3923352	3923480	refGene_gn=PCDH15;refGene_tx=XM_019814204.2

D2	3957583	3957799	refGene_gn=PCDH15;refGene_tx=XM_019814204.2
D2	3965316	3965405	refGene_gn=PCDH15;refGene_tx=XM_019814204.2
D2	3987142	3987319	refGene_gn=PCDH15;refGene_tx=XM_019814204.2
D2	3996238	3996457	refGene_gn=PCDH15;refGene_tx=XM_019814204.2
D2	3999095	3999104	refGene_gn=PCDH15;refGene_tx=XM_019814204.2
D2	3999916	4000069	refGene_gn=PCDH15;refGene_tx=XM_019814204.2
D2	4009574	4009593	refGene_gn=PCDH15;refGene_tx=XM_019814204.2
D2	4011207	4011246	refGene_gn=PCDH15;refGene_tx=XM_019814204.2
D2	4015478	4015549	refGene_gn=PCDH15;refGene_tx=XM_019814204.2
D2	4016408	4016517	refGene_gn=PCDH15;refGene_tx=XM_019814204.2
D2	4017288	4018305
refGene_gn=LOC111556972;refGene_tx=XM_023240317.1
D2	4019946	4020492	refGene_gn=PCDH15;refGene_tx=XM_019814204.2
D2	5202949	5203203	refGene_gn=DKK1;refGene_tx=XM_023240318.1
D2	5203323	5203467	refGene_gn=DKK1;refGene_tx=XM_023240318.1
D2	5204583	5204746	refGene_gn=DKK1;refGene_tx=XM_023240318.1
D2	5204950	5205196	refGene_gn=DKK1;refGene_tx=XM_023240318.1
D2	5263173	5263272
refGene_gn=PRKG1;refGene_tx=XM_023240287.1;refGene_gn=PRKG1;refGene_tx
=XM_023240288.1;refGene_gn=PRKG1;refGene_tx=XM_023240289.1;refGene_gn=
PRKG1;refGene_tx=XM_023240290.1
D2	5264658	5264725
refGene_gn=PRKG1;refGene_tx=XM_023240287.1;refGene_gn=PRKG1;refGene_tx
=XM_023240288.1;refGene_gn=PRKG1;refGene_tx=XM_023240289.1;refGene_gn=
PRKG1;refGene_tx=XM_023240290.1
D2	5266032	5266095
refGene_gn=PRKG1;refGene_tx=XM_023240287.1;refGene_gn=PRKG1;refGene_tx
=XM_023240288.1;refGene_gn=PRKG1;refGene_tx=XM_023240289.1;refGene_gn=
PRKG1;refGene_tx=XM_023240290.1
D2	5266174	5266297
refGene_gn=PRKG1;refGene_tx=XM_023240287.1;refGene_gn=PRKG1;refGene_tx
=XM_023240288.1;refGene_gn=PRKG1;refGene_tx=XM_023240289.1;refGene_gn=
PRKG1;refGene_tx=XM_023240290.1
D2	5272445	5272609
refGene_gn=PRKG1;refGene_tx=XM_023240287.1;refGene_gn=PRKG1;refGene_tx
=XM_023240288.1;refGene_gn=PRKG1;refGene_tx=XM_023240289.1;refGene_gn=
PRKG1;refGene_tx=XM_023240290.1
D2	5273811	5273953
refGene_gn=PRKG1;refGene_tx=XM_023240287.1;refGene_gn=PRKG1;refGene_tx
=XM_023240288.1;refGene_gn=PRKG1;refGene_tx=XM_023240289.1;refGene_gn=
PRKG1;refGene_tx=XM_023240290.1
D2	5287205	5287295
refGene_gn=PRKG1;refGene_tx=XM_023240287.1;refGene_gn=PRKG1;refGene_tx
=XM_023240288.1;refGene_gn=PRKG1;refGene_tx=XM_023240289.1;refGene_gn=
PRKG1;refGene_tx=XM_023240290.1
D2	5288437	5288577
refGene_gn=PRKG1;refGene_tx=XM_023240287.1;refGene_gn=PRKG1;refGene_tx
=XM_023240288.1;refGene_gn=PRKG1;refGene_tx=XM_023240289.1;refGene_gn=
PRKG1;refGene_tx=XM_023240290.1
D2	5308969	5309066

refGene_gn=PRKG1;refGene_tx=XM_023240287.1;refGene_gn=PRKG1;refGene_tx=XM_023240288.1;refGene_gn=PRKG1;refGene_tx=XM_023240289.1;refGene_gn=PRKG1;refGene_tx=XM_023240290.1
D2 5373094 5373169
refGene_gn=PRKG1;refGene_tx=XM_023240287.1;refGene_gn=PRKG1;refGene_tx=XM_023240288.1;refGene_gn=PRKG1;refGene_tx=XM_023240289.1;refGene_gn=PRKG1;refGene_tx=XM_023240290.1
D2 5404970 5405036
refGene_gn=PRKG1;refGene_tx=XM_023240287.1;refGene_gn=PRKG1;refGene_tx=XM_023240288.1;refGene_gn=PRKG1;refGene_tx=XM_023240289.1;refGene_gn=PRKG1;refGene_tx=XM_023240290.1
D2 5472882 5472977
refGene_gn=PRKG1;refGene_tx=XM_023240287.1;refGene_gn=PRKG1;refGene_tx=XM_023240288.1;refGene_gn=PRKG1;refGene_tx=XM_023240289.1;refGene_gn=PRKG1;refGene_tx=XM_023240290.1
D2 5482667 5482745
refGene_gn=PRKG1;refGene_tx=XM_023240287.1;refGene_gn=PRKG1;refGene_tx=XM_023240288.1;refGene_gn=PRKG1;refGene_tx=XM_023240289.1;refGene_gn=PRKG1;refGene_tx=XM_023240290.1
D2 5619973 5620037
refGene_gn=PRKG1;refGene_tx=XM_023240287.1;refGene_gn=PRKG1;refGene_tx=XM_023240288.1;refGene_gn=PRKG1;refGene_tx=XM_023240289.1;refGene_gn=PRKG1;refGene_tx=XM_023240290.1
D2 5722241 5722347
refGene_gn=PRKG1;refGene_tx=XM_023240287.1;refGene_gn=PRKG1;refGene_tx=XM_023240288.1;refGene_gn=PRKG1;refGene_tx=XM_023240289.1;refGene_gn=PRKG1;refGene_tx=XM_023240290.1
D2 5817272 5819264 refGene_gn=CSTF2T;refGene_tx=XM_006937713.4
D2 6056507 6056621
refGene_gn=PRKG1;refGene_tx=XM_023240287.1;refGene_gn=PRKG1;refGene_tx=XM_023240288.1;refGene_gn=PRKG1;refGene_tx=XM_023240289.1;refGene_gn=PRKG1;refGene_tx=XM_023240290.1
D2 6343655 6343822
refGene_gn=PRKG1;refGene_tx=XM_023240287.1;refGene_gn=PRKG1;refGene_tx=XM_023240288.1;refGene_gn=PRKG1;refGene_tx=XM_023240289.1;refGene_gn=PRKG1;refGene_tx=XM_023240290.1
D2 6345407 6345445 refGene_gn=PRKG1;refGene_tx=XM_023240289.1
D2 6419303 6419614 refGene_gn=PRKG1;refGene_tx=XM_023240287.1
D2 6513176 6513442 refGene_gn=PRKG1;refGene_tx=XM_023240288.1
D2 6653948 6654047
refGene_gn=A1CF;refGene_tx=XM_006937714.2;refGene_gn=A1CF;refGene_tx=XM_019814212.2;refGene_gn=A1CF;refGene_tx=XM_023240319.1
D2 6669989 6670124
refGene_gn=A1CF;refGene_tx=XM_006937714.2;refGene_gn=A1CF;refGene_tx=XM_019814212.2;refGene_gn=A1CF;refGene_tx=XM_023240319.1
D2 6672246 6672377
refGene_gn=A1CF;refGene_tx=XM_006937714.2;refGene_gn=A1CF;refGene_tx=XM_019814212.2;refGene_gn=A1CF;refGene_tx=XM_023240319.1
D2 6677811 6678050
refGene_gn=A1CF;refGene_tx=XM_006937714.2;refGene_gn=A1CF;refGene_tx=X

M_019814212.2;refGene_gn=A1CF;refGene_tx=XM_023240319.1
D2	6683160	6683325
refGene_gn=A1CF;refGene_tx=XM_006937714.2;refGene_gn=A1CF;refGene_tx=XM_019814212.2;refGene_gn=A1CF;refGene_tx=XM_023240319.1
D2	6690372	6690470
refGene_gn=A1CF;refGene_tx=XM_006937714.2;refGene_gn=A1CF;refGene_tx=XM_019814212.2;refGene_gn=A1CF;refGene_tx=XM_023240319.1
D2	6696107	6696381
refGene_gn=A1CF;refGene_tx=XM_006937714.2;refGene_gn=A1CF;refGene_tx=XM_019814212.2;refGene_gn=A1CF;refGene_tx=XM_023240319.1
D2	6698319	6698525
refGene_gn=A1CF;refGene_tx=XM_006937714.2;refGene_gn=A1CF;refGene_tx=XM_019814212.2;refGene_gn=A1CF;refGene_tx=XM_023240319.1
D2	6700895	6701032
refGene_gn=A1CF;refGene_tx=XM_006937714.2;refGene_gn=A1CF;refGene_tx=XM_019814212.2;refGene_gn=A1CF;refGene_tx=XM_023240319.1
D2	6702014	6702157
refGene_gn=A1CF;refGene_tx=XM_006937714.2;refGene_gn=A1CF;refGene_tx=XM_019814212.2;refGene_gn=A1CF;refGene_tx=XM_023240319.1
D2	6705340	6705492
refGene_gn=A1CF;refGene_tx=XM_006937714.2;refGene_gn=A1CF;refGene_tx=XM_019814212.2;refGene_gn=A1CF;refGene_tx=XM_023240319.1
D2	6744091	6744281
refGene_gn=ASAH2;refGene_tx=XM_023240320.1;refGene_gn=ASAH2;refGene_tx=XM_023240321.1;refGene_gn=ASAH2;refGene_tx=XM_023240322.1;refGene_gn=ASAH2;refGene_tx=XM_023240323.1;refGene_gn=ASAH2;refGene_tx=XM_023240324.1;refGene_gn=ASAH2;refGene_tx=XM_023240325.1
D2	6746263	6746352
refGene_gn=ASAH2;refGene_tx=XM_023240320.1;refGene_gn=ASAH2;refGene_tx=XM_023240321.1;refGene_gn=ASAH2;refGene_tx=XM_023240322.1;refGene_gn=ASAH2;refGene_tx=XM_023240323.1;refGene_gn=ASAH2;refGene_tx=XM_023240324.1;refGene_gn=ASAH2;refGene_tx=XM_023240325.1
D2	6749707	6749767
refGene_gn=ASAH2;refGene_tx=XM_023240320.1;refGene_gn=ASAH2;refGene_tx=XM_023240321.1;refGene_gn=ASAH2;refGene_tx=XM_023240322.1;refGene_gn=ASAH2;refGene_tx=XM_023240323.1;refGene_gn=ASAH2;refGene_tx=XM_023240324.1;refGene_gn=ASAH2;refGene_tx=XM_023240325.1
D2	6754791	6754938
refGene_gn=ASAH2;refGene_tx=XM_023240320.1;refGene_gn=ASAH2;refGene_tx=XM_023240321.1;refGene_gn=ASAH2;refGene_tx=XM_023240322.1;refGene_gn=ASAH2;refGene_tx=XM_023240323.1
D2	6756982	6757078
refGene_gn=ASAH2;refGene_tx=XM_023240320.1;refGene_gn=ASAH2;refGene_tx=XM_023240321.1;refGene_gn=ASAH2;refGene_tx=XM_023240322.1;refGene_gn=ASAH2;refGene_tx=XM_023240323.1;refGene_gn=ASAH2;refGene_tx=XM_023240324.1
D2	6760451	6760547
refGene_gn=ASAH2;refGene_tx=XM_023240320.1;refGene_gn=ASAH2;refGene_tx=XM_023240321.1;refGene_gn=ASAH2;refGene_tx=XM_023240322.1;refGene_gn=ASAH2;refGene_tx=XM_023240323.1;refGene_gn=ASAH2;refGene_tx=XM_0232403

24.1;refGene_gn=ASAH2;refGene_tx=XM_023240325.1
D2 6761574 6761614
refGene_gn=ASAH2;refGene_tx=XM_023240320.1;refGene_gn=ASAH2;refGene_tx=XM_023240321.1;refGene_gn=ASAH2;refGene_tx=XM_023240322.1;refGene_gn=ASAH2;refGene_tx=XM_023240323.1;refGene_gn=ASAH2;refGene_tx=XM_023240324.1;refGene_gn=ASAH2;refGene_tx=XM_023240325.1
D2 6762785 6762880
refGene_gn=ASAH2;refGene_tx=XM_023240320.1;refGene_gn=ASAH2;refGene_tx=XM_023240321.1;refGene_gn=ASAH2;refGene_tx=XM_023240322.1;refGene_gn=ASAH2;refGene_tx=XM_023240323.1;refGene_gn=ASAH2;refGene_tx=XM_023240324.1;refGene_gn=ASAH2;refGene_tx=XM_023240325.1
D2 6763929 6764045
refGene_gn=ASAH2;refGene_tx=XM_023240320.1;refGene_gn=ASAH2;refGene_tx=XM_023240321.1;refGene_gn=ASAH2;refGene_tx=XM_023240322.1;refGene_gn=ASAH2;refGene_tx=XM_023240323.1;refGene_gn=ASAH2;refGene_tx=XM_023240324.1;refGene_gn=ASAH2;refGene_tx=XM_023240325.1
D2 6766103 6766184
refGene_gn=ASAH2;refGene_tx=XM_023240320.1;refGene_gn=ASAH2;refGene_tx=XM_023240321.1;refGene_gn=ASAH2;refGene_tx=XM_023240322.1;refGene_gn=ASAH2;refGene_tx=XM_023240323.1;refGene_gn=ASAH2;refGene_tx=XM_023240324.1;refGene_gn=ASAH2;refGene_tx=XM_023240325.1
D2 6766323 6766429
refGene_gn=ASAH2;refGene_tx=XM_023240320.1;refGene_gn=ASAH2;refGene_tx=XM_023240321.1;refGene_gn=ASAH2;refGene_tx=XM_023240322.1;refGene_gn=ASAH2;refGene_tx=XM_023240323.1;refGene_gn=ASAH2;refGene_tx=XM_023240324.1;refGene_gn=ASAH2;refGene_tx=XM_023240325.1
D2 6768581 6768668
refGene_gn=ASAH2;refGene_tx=XM_023240320.1;refGene_gn=ASAH2;refGene_tx=XM_023240321.1;refGene_gn=ASAH2;refGene_tx=XM_023240322.1;refGene_gn=ASAH2;refGene_tx=XM_023240323.1;refGene_gn=ASAH2;refGene_tx=XM_023240324.1;refGene_gn=ASAH2;refGene_tx=XM_023240325.1
D2 6772633 6772759
refGene_gn=ASAH2;refGene_tx=XM_023240320.1;refGene_gn=ASAH2;refGene_tx=XM_023240321.1;refGene_gn=ASAH2;refGene_tx=XM_023240322.1;refGene_gn=ASAH2;refGene_tx=XM_023240323.1;refGene_gn=ASAH2;refGene_tx=XM_023240324.1;refGene_gn=ASAH2;refGene_tx=XM_023240325.1
D2 6774299 6774420
refGene_gn=ASAH2;refGene_tx=XM_023240320.1;refGene_gn=ASAH2;refGene_tx=XM_023240321.1;refGene_gn=ASAH2;refGene_tx=XM_023240322.1;refGene_gn=ASAH2;refGene_tx=XM_023240323.1;refGene_gn=ASAH2;refGene_tx=XM_023240324.1;refGene_gn=ASAH2;refGene_tx=XM_023240325.1
D2 6784816 6784894
refGene_gn=ASAH2;refGene_tx=XM_023240320.1;refGene_gn=ASAH2;refGene_tx=XM_023240321.1;refGene_gn=ASAH2;refGene_tx=XM_023240322.1;refGene_gn=ASAH2;refGene_tx=XM_023240323.1;refGene_gn=ASAH2;refGene_tx=XM_023240324.1;refGene_gn=ASAH2;refGene_tx=XM_023240325.1
D2 6786039 6786167
refGene_gn=ASAH2;refGene_tx=XM_023240320.1;refGene_gn=ASAH2;refGene_tx=XM_023240321.1;refGene_gn=ASAH2;refGene_tx=XM_023240322.1;refGene_gn=ASAH2;refGene_tx=XM_023240323.1;refGene_gn=ASAH2;refGene_tx=XM_0232403

24.1;refGene_gn=ASAH2;refGene_tx=XM_023240325.1
D2 6787435 6787612
refGene_gn=ASAH2;refGene_tx=XM_023240320.1;refGene_gn=ASAH2;refGene_tx=XM_023240321.1;refGene_gn=ASAH2;refGene_tx=XM_023240322.1;refGene_gn=ASAH2;refGene_tx=XM_023240323.1;refGene_gn=ASAH2;refGene_tx=XM_023240324.1;refGene_gn=ASAH2;refGene_tx=XM_023240325.1
D2 6795239 6795389
refGene_gn=ASAH2;refGene_tx=XM_023240320.1;refGene_gn=ASAH2;refGene_tx=XM_023240321.1;refGene_gn=ASAH2;refGene_tx=XM_023240322.1;refGene_gn=ASAH2;refGene_tx=XM_023240323.1;refGene_gn=ASAH2;refGene_tx=XM_023240324.1;refGene_gn=ASAH2;refGene_tx=XM_023240325.1
D2 6797029 6797235
refGene_gn=ASAH2;refGene_tx=XM_023240320.1;refGene_gn=ASAH2;refGene_tx=XM_023240321.1;refGene_gn=ASAH2;refGene_tx=XM_023240322.1;refGene_gn=ASAH2;refGene_tx=XM_023240323.1;refGene_gn=ASAH2;refGene_tx=XM_023240324.1;refGene_gn=ASAH2;refGene_tx=XM_023240325.1
D2 6799899 6800026
refGene_gn=ASAH2;refGene_tx=XM_023240320.1;refGene_gn=ASAH2;refGene_tx=XM_023240321.1;refGene_gn=ASAH2;refGene_tx=XM_023240322.1;refGene_gn=ASAH2;refGene_tx=XM_023240323.1;refGene_gn=ASAH2;refGene_tx=XM_023240324.1;refGene_gn=ASAH2;refGene_tx=XM_023240325.1
D2 6844394 6844574
refGene_gn=SGMS1;refGene_tx=XM_019814218.1;refGene_gn=SGMS1;refGene_tx=XM_019814219.1;refGene_gn=SGMS1;refGene_tx=XM_023240326.1
D2 6845252 6845419
refGene_gn=SGMS1;refGene_tx=XM_019814218.1;refGene_gn=SGMS1;refGene_tx=XM_019814219.1;refGene_gn=SGMS1;refGene_tx=XM_023240326.1
D2 6848502 6848656
refGene_gn=SGMS1;refGene_tx=XM_019814218.1;refGene_gn=SGMS1;refGene_tx=XM_019814219.1;refGene_gn=SGMS1;refGene_tx=XM_023240326.1
D2 6864676 6864794
refGene_gn=SGMS1;refGene_tx=XM_019814218.1;refGene_gn=SGMS1;refGene_tx=XM_019814219.1;refGene_gn=SGMS1;refGene_tx=XM_023240326.1
D2 6883037 6883642
refGene_gn=SGMS1;refGene_tx=XM_019814218.1;refGene_gn=SGMS1;refGene_tx=XM_019814219.1;refGene_gn=SGMS1;refGene_tx=XM_023240326.1
D2 7259545 7260182
refGene_gn=MINPP1;refGene_tx=XM_023240327.1;refGene_gn=MINPP1;refGene_tx=XM_023240328.1
D2 7263668 7263866
refGene_gn=MINPP1;refGene_tx=XM_023240327.1;refGene_gn=MINPP1;refGene_tx=XM_023240328.1
D2 7269403 7269501 refGene_gn=MINPP1;refGene_tx=XM_023240327.1
D2 7276578 7276712 refGene_gn=MINPP1;refGene_tx=XM_023240327.1
D2 7305693 7306090
refGene_gn=MINPP1;refGene_tx=XM_023240327.1;refGene_gn=MINPP1;refGene_tx=XM_023240328.1
D2 7390124 7390151
refGene_gn=PAPSS2;refGene_tx=XM_023240329.1;refGene_gn=PAPSS2;refGene_tx=XM_023240331.1

D2 7412240 7412264 refGene_gn=PAPSS2;refGene_tx=XM_023240330.1
D2 7430710 7430828
refGene_gn=PAPSS2;refGene_tx=XM_023240329.1;refGene_gn=PAPSS2;refGene_tx=XM_023240330.1;refGene_gn=PAPSS2;refGene_tx=XM_023240331.1
D2 7434317 7434553
refGene_gn=PAPSS2;refGene_tx=XM_023240329.1;refGene_gn=PAPSS2;refGene_tx=XM_023240330.1;refGene_gn=PAPSS2;refGene_tx=XM_023240331.1
D2 7435270 7435409
refGene_gn=PAPSS2;refGene_tx=XM_023240329.1;refGene_gn=PAPSS2;refGene_tx=XM_023240330.1;refGene_gn=PAPSS2;refGene_tx=XM_023240331.1
D2 7436258 7436377
refGene_gn=PAPSS2;refGene_tx=XM_023240329.1;refGene_gn=PAPSS2;refGene_tx=XM_023240330.1;refGene_gn=PAPSS2;refGene_tx=XM_023240331.1
D2 7436512 7436626
refGene_gn=PAPSS2;refGene_tx=XM_023240329.1;refGene_gn=PAPSS2;refGene_tx=XM_023240330.1;refGene_gn=PAPSS2;refGene_tx=XM_023240331.1
D2 7438265 7438377
refGene_gn=PAPSS2;refGene_tx=XM_023240329.1;refGene_gn=PAPSS2;refGene_tx=XM_023240330.1;refGene_gn=PAPSS2;refGene_tx=XM_023240331.1
D2 7443217 7443232
refGene_gn=PAPSS2;refGene_tx=XM_023240329.1;refGene_gn=PAPSS2;refGene_tx=XM_023240330.1
D2 7447375 7447581
refGene_gn=PAPSS2;refGene_tx=XM_023240329.1;refGene_gn=PAPSS2;refGene_tx=XM_023240330.1;refGene_gn=PAPSS2;refGene_tx=XM_023240331.1
D2 7456145 7456281
refGene_gn=PAPSS2;refGene_tx=XM_023240329.1;refGene_gn=PAPSS2;refGene_tx=XM_023240330.1;refGene_gn=PAPSS2;refGene_tx=XM_023240331.1
D2 7458091 7458360
refGene_gn=PAPSS2;refGene_tx=XM_023240329.1;refGene_gn=PAPSS2;refGene_tx=XM_023240330.1;refGene_gn=PAPSS2;refGene_tx=XM_023240331.1
D2 7459722 7459952
refGene_gn=PAPSS2;refGene_tx=XM_023240329.1;refGene_gn=PAPSS2;refGene_tx=XM_023240330.1;refGene_gn=PAPSS2;refGene_tx=XM_023240331.1
D2 7460440 7460582
refGene_gn=PAPSS2;refGene_tx=XM_023240329.1;refGene_gn=PAPSS2;refGene_tx=XM_023240330.1;refGene_gn=PAPSS2;refGene_tx=XM_023240331.1
D2 7471942 7472063 refGene_gn=ATAD1;refGene_tx=XM_003993862.5
D2 7473898 7474032 refGene_gn=ATAD1;refGene_tx=XM_003993862.5
D2 7483188 7483239 refGene_gn=ATAD1;refGene_tx=XM_003993862.5
D2 7486034 7486124 refGene_gn=ATAD1;refGene_tx=XM_003993862.5
D2 7494516 7494623 refGene_gn=ATAD1;refGene_tx=XM_003993862.5
D2 7502300 7502501 refGene_gn=ATAD1;refGene_tx=XM_003993862.5
D2 7507587 7507620 refGene_gn=ATAD1;refGene_tx=XM_003993862.5
D2 7507622 7507707 refGene_gn=ATAD1;refGene_tx=XM_003993862.5
D2 7509866 7509965 refGene_gn=ATAD1;refGene_tx=XM_003993862.5
D2 7523562 7523734 refGene_gn=ATAD1;refGene_tx=XM_003993862.5
D2 7524570 7524665 refGene_gn=ATAD1;refGene_tx=XM_003993862.5
D2 7526840 7527168
refGene_gn=LOC109492818;refGene_tx=XM_023240917.1

D2 7527276 7527647
refGene_gn=LOC109492818;refGene_tx=XM_023240917.1
D2 7528100 7528224
refGene_gn=LOC109492818;refGene_tx=XM_023240917.1
D2 7532115 7532120
refGene_gn=LOC109492818;refGene_tx=XM_023240917.1
D2 7563542 7563711 refGene_gn=KLLN;refGene_tx=XM_023240918.1
D2 7563712 7563933 refGene_gn=KLLN;refGene_tx=XM_023240918.1
D2 7563935 7564298 refGene_gn=KLLN;refGene_tx=XM_023240918.1
D2 7566036 7566115 refGene_gn=PTEN;refGene_tx=XM_019814225.2
D2 7592130 7592215 refGene_gn=PTEN;refGene_tx=XM_019814225.2
D2 7619623 7619625 refGene_gn=PTEN;refGene_tx=XM_019814226.2
D2 7622187 7622232 refGene_gn=PTEN;refGene_tx=XM_019814226.2
D2 7628018 7628062
refGene_gn=PTEN;refGene_tx=XM_019814225.2;refGene_gn=PTEN;refGene_tx=XM_019814226.2
D2 7630130 7630369
refGene_gn=PTEN;refGene_tx=XM_019814225.2;refGene_gn=PTEN;refGene_tx=XM_019814226.2
D2 7645191 7645333
refGene_gn=PTEN;refGene_tx=XM_019814225.2;refGene_gn=PTEN;refGene_tx=XM_019814226.2;refGene_gn=PTEN;refGene_tx=XM_023240332.1
D2 7650126 7650293
refGene_gn=PTEN;refGene_tx=XM_019814225.2;refGene_gn=PTEN;refGene_tx=XM_019814226.2;refGene_gn=PTEN;refGene_tx=XM_023240332.1
D2 7653295 7653520
refGene_gn=PTEN;refGene_tx=XM_019814225.2;refGene_gn=PTEN;refGene_tx=XM_019814226.2;refGene_gn=PTEN;refGene_tx=XM_023240332.1
D2 7656200 7656386
refGene_gn=PTEN;refGene_tx=XM_019814225.2;refGene_gn=PTEN;refGene_tx=XM_019814226.2;refGene_gn=PTEN;refGene_tx=XM_023240332.1
D2 7955073 7955226 refGene_gn=RNLS;refGene_tx=XM_023240333.1
D2 7979640 7979816 refGene_gn=RNLS;refGene_tx=XM_023240333.1
D2 8013411 8013517 refGene_gn=RNLS;refGene_tx=XM_023240333.1
D2 8013518 8013586 refGene_gn=RNLS;refGene_tx=XM_023240333.1
D2 8208834 8208993 refGene_gn=RNLS;refGene_tx=XM_023240333.1
D2 8217176 8217319 refGene_gn=RNLS;refGene_tx=XM_023240333.1
D2 8218045 8218151 refGene_gn=RNLS;refGene_tx=XM_023240333.1
D2 8219140 8219258 refGene_gn=RNLS;refGene_tx=XM_023240333.1
D2 8237345 8237450 refGene_gn=LIPJ;refGene_tx=XM_023240310.1
D2 8238033 8238151 refGene_gn=LIPJ;refGene_tx=XM_023240310.1
D2 8243111 8243310 refGene_gn=LIPJ;refGene_tx=XM_023240310.1
D2 8243976 8244086 refGene_gn=LIPJ;refGene_tx=XM_023240310.1
D2 8247390 8247527 refGene_gn=LIPJ;refGene_tx=XM_023240310.1
D2 8247888 8248035 refGene_gn=LIPJ;refGene_tx=XM_023240310.1
D2 8252439 8252511 refGene_gn=LIPJ;refGene_tx=XM_023240310.1
D2 8256517 8256589 refGene_gn=LIPJ;refGene_tx=XM_023240310.1
D2 8258120 8258354 refGene_gn=LIPJ;refGene_tx=XM_023240310.1
D2 8298981 8299086
refGene_gn=LIPF;refGene_tx=XM_003993865.3;refGene_gn=LIPF;refGene_tx=X

M_019814146.1
D2	8299263	8299381
refGene_gn=LIPF;refGene_tx=XM_003993865.3;refGene_gn=LIPF;refGene_tx=XM_019814146.1
D2	8300703	8300902
refGene_gn=LIPF;refGene_tx=XM_003993865.3;refGene_gn=LIPF;refGene_tx=XM_019814146.1
D2	8302769	8302879
refGene_gn=LIPF;refGene_tx=XM_003993865.3;refGene_gn=LIPF;refGene_tx=XM_019814146.1
D2	8307380	8307517
refGene_gn=LIPF;refGene_tx=XM_003993865.3;refGene_gn=LIPF;refGene_tx=XM_019814146.1
D2	8309131	8309278
refGene_gn=LIPF;refGene_tx=XM_003993865.3;refGene_gn=LIPF;refGene_tx=XM_019814146.1
D2	8311376	8311448
refGene_gn=LIPF;refGene_tx=XM_003993865.3;refGene_gn=LIPF;refGene_tx=XM_019814146.1
D2	8312104	8312176
refGene_gn=LIPF;refGene_tx=XM_003993865.3;refGene_gn=LIPF;refGene_tx=XM_019814146.1
D2	8314904	8315141
refGene_gn=LIPF;refGene_tx=XM_003993865.3;refGene_gn=LIPF;refGene_tx=XM_019814146.1
D2	8370294	8370399
refGene_gn=LIPK;refGene_tx=XM_003993866.2;refGene_gn=LIPK;refGene_tx=XM_019814247.1;refGene_gn=LIPK;refGene_tx=XM_019814248.2;refGene_gn=LIPK;refGene_tx=XM_019814249.1;refGene_gn=LIPK;refGene_tx=XM_019814250.1;refGene_gn=LIPK;refGene_tx=XM_019814251.1
D2	8372639	8372754
refGene_gn=LIPK;refGene_tx=XM_003993866.2;refGene_gn=LIPK;refGene_tx=XM_019814247.1;refGene_gn=LIPK;refGene_tx=XM_019814248.2;refGene_gn=LIPK;refGene_tx=XM_019814249.1;refGene_gn=LIPK;refGene_tx=XM_019814250.1;refGene_gn=LIPK;refGene_tx=XM_019814251.1
D2	8375530	8375729
refGene_gn=LIPK;refGene_tx=XM_003993866.2;refGene_gn=LIPK;refGene_tx=XM_019814247.1;refGene_gn=LIPK;refGene_tx=XM_019814248.2;refGene_gn=LIPK;refGene_tx=XM_019814249.1;refGene_gn=LIPK;refGene_tx=XM_019814250.1;refGene_gn=LIPK;refGene_tx=XM_019814251.1
D2	8377384	8377494
refGene_gn=LIPK;refGene_tx=XM_003993866.2;refGene_gn=LIPK;refGene_tx=XM_019814247.1;refGene_gn=LIPK;refGene_tx=XM_019814248.2;refGene_gn=LIPK;refGene_tx=XM_019814249.1;refGene_gn=LIPK;refGene_tx=XM_019814250.1;refGene_gn=LIPK;refGene_tx=XM_019814251.1
D2	8378777	8378914
refGene_gn=LIPK;refGene_tx=XM_003993866.2;refGene_gn=LIPK;refGene_tx=XM_019814247.1;refGene_gn=LIPK;refGene_tx=XM_019814248.2;refGene_gn=LIPK;refGene_tx=XM_019814249.1;refGene_gn=LIPK;refGene_tx=XM_019814250.1;refGene_gn=LIPK;refGene_tx=XM_019814251.1

D2	8384921	8385068
refGene_gn=LIPK;refGene_tx=XM_003993866.2;refGene_gn=LIPK;refGene_tx=XM_019814247.1;refGene_gn=LIPK;refGene_tx=XM_019814248.2;refGene_gn=LIPK;refGene_tx=XM_019814249.1;refGene_gn=LIPK;refGene_tx=XM_019814250.1;refGene_gn=LIPK;refGene_tx=XM_019814251.1
D2	8386899	8386971
refGene_gn=LIPK;refGene_tx=XM_003993866.2;refGene_gn=LIPK;refGene_tx=XM_019814247.1;refGene_gn=LIPK;refGene_tx=XM_019814248.2;refGene_gn=LIPK;refGene_tx=XM_019814249.1;refGene_gn=LIPK;refGene_tx=XM_019814251.1
D2	8389766	8389838
refGene_gn=LIPK;refGene_tx=XM_003993866.2;refGene_gn=LIPK;refGene_tx=XM_019814247.1;refGene_gn=LIPK;refGene_tx=XM_019814248.2;refGene_gn=LIPK;refGene_tx=XM_019814249.1;refGene_gn=LIPK;refGene_tx=XM_019814250.1
D2	8394380	8394620
refGene_gn=LIPK;refGene_tx=XM_003993866.2;refGene_gn=LIPK;refGene_tx=XM_019814247.1;refGene_gn=LIPK;refGene_tx=XM_019814248.2;refGene_gn=LIPK;refGene_tx=XM_019814249.1;refGene_gn=LIPK;refGene_tx=XM_019814250.1;refGene_gn=LIPK;refGene_tx=XM_019814251.1
D2	8410319	8410367
refGene_gn=LIPN;refGene_tx=XM_019814245.2;refGene_gn=LIPN;refGene_tx=XM_023240334.1
D2	8410832	8410937
refGene_gn=LIPN;refGene_tx=XM_003993867.4;refGene_gn=LIPN;refGene_tx=XM_019814244.2
D2	8411653	8411771
refGene_gn=LIPN;refGene_tx=XM_003993867.4;refGene_gn=LIPN;refGene_tx=XM_019814244.2;refGene_gn=LIPN;refGene_tx=XM_019814245.2;refGene_gn=LIPN;refGene_tx=XM_023240334.1
D2	8413803	8414002
refGene_gn=LIPN;refGene_tx=XM_003993867.4;refGene_gn=LIPN;refGene_tx=XM_019814244.2;refGene_gn=LIPN;refGene_tx=XM_019814245.2;refGene_gn=LIPN;refGene_tx=XM_023240334.1
D2	8415319	8415429
refGene_gn=LIPN;refGene_tx=XM_003993867.4;refGene_gn=LIPN;refGene_tx=XM_019814244.2;refGene_gn=LIPN;refGene_tx=XM_019814245.2;refGene_gn=LIPN;refGene_tx=XM_023240334.1
D2	8417695	8417832
refGene_gn=LIPN;refGene_tx=XM_003993867.4;refGene_gn=LIPN;refGene_tx=XM_019814244.2;refGene_gn=LIPN;refGene_tx=XM_019814245.2;refGene_gn=LIPN;refGene_tx=XM_023240334.1
D2	8418628	8418775
refGene_gn=LIPN;refGene_tx=XM_003993867.4;refGene_gn=LIPN;refGene_tx=XM_019814244.2;refGene_gn=LIPN;refGene_tx=XM_019814245.2;refGene_gn=LIPN;refGene_tx=XM_023240334.1
D2	8420399	8420471
refGene_gn=LIPN;refGene_tx=XM_003993867.4;refGene_gn=LIPN;refGene_tx=XM_019814244.2;refGene_gn=LIPN;refGene_tx=XM_019814245.2;refGene_gn=LIPN;refGene_tx=XM_023240334.1
D2	8421461	8421533
refGene_gn=LIPN;refGene_tx=XM_003993867.4;refGene_gn=LIPN;refGene_tx=X

M_019814244.2;refGene_gn=LIPN;refGene_tx=XM_019814245.2;refGene_gn=LIPN;refGene_tx=XM_023240334.1
D2 8425241 8425475
refGene_gn=LIPN;refGene_tx=XM_003993867.4;refGene_gn=LIPN;refGene_tx=XM_019814244.2;refGene_gn=LIPN;refGene_tx=XM_019814245.2;refGene_gn=LIPN;refGene_tx=XM_023240334.1
D2 8453855 8454002
refGene_gn=LIPM;refGene_tx=XM_003993868.4;refGene_gn=LIPM;refGene_tx=XM_019814253.2
D2 8456898 8457016
refGene_gn=LIPM;refGene_tx=XM_003993868.4;refGene_gn=LIPM;refGene_tx=XM_019814253.2
D2 8462149 8462348
refGene_gn=LIPM;refGene_tx=XM_003993868.4;refGene_gn=LIPM;refGene_tx=XM_019814253.2
D2 8463438 8463548
refGene_gn=LIPM;refGene_tx=XM_003993868.4;refGene_gn=LIPM;refGene_tx=XM_019814253.2
D2 8464443 8464580
refGene_gn=LIPM;refGene_tx=XM_003993868.4;refGene_gn=LIPM;refGene_tx=XM_019814253.2
D2 8465450 8465597
refGene_gn=LIPM;refGene_tx=XM_003993868.4;refGene_gn=LIPM;refGene_tx=XM_019814253.2
D2 8466857 8466929
refGene_gn=LIPM;refGene_tx=XM_003993868.4;refGene_gn=LIPM;refGene_tx=XM_019814253.2
D2 8468421 8468493 refGene_gn=LIPM;refGene_tx=XM_003993868.4
D2 8472351 8472621
refGene_gn=LIPM;refGene_tx=XM_003993868.4;refGene_gn=LIPM;refGene_tx=XM_019814253.2
D2 8475239 8475317
refGene_gn=ANKRD22;refGene_tx=XM_003993869.5;refGene_gn=ANKRD22;refGene_tx=XM_019814254.2;refGene_gn=ANKRD22;refGene_tx=XM_019814255.2;refGene_gn=ANKRD22;refGene_tx=XM_023240335.1
D2 8475594 8475693
refGene_gn=ANKRD22;refGene_tx=XM_003993869.5;refGene_gn=ANKRD22;refGene_tx=XM_019814254.2;refGene_gn=ANKRD22;refGene_tx=XM_019814255.2;refGene_gn=ANKRD22;refGene_tx=XM_023240335.1
D2 8478242 8478320
refGene_gn=ANKRD22;refGene_tx=XM_003993869.5;refGene_gn=ANKRD22;refGene_tx=XM_019814254.2;refGene_gn=ANKRD22;refGene_tx=XM_019814255.2;refGene_gn=ANKRD22;refGene_tx=XM_023240335.1
D2 8480565 8480673
refGene_gn=ANKRD22;refGene_tx=XM_003993869.5;refGene_gn=ANKRD22;refGene_tx=XM_019814254.2;refGene_gn=ANKRD22;refGene_tx=XM_019814255.2;refGene_gn=ANKRD22;refGene_tx=XM_023240335.1
D2 8482865 8483057
refGene_gn=ANKRD22;refGene_tx=XM_003993869.5;refGene_gn=ANKRD22;refGene_tx=XM_019814254.2;refGene_gn=ANKRD22;refGene_tx=XM_019814255.2;refGe

ne_gn=ANKRD22;refGene_tx=XM_023240335.1
D2 8496414 8496435
refGene_gn=ANKRD22;refGene_tx=XM_003993869.5;refGene_gn=ANKRD22;refGen
e_tx=XM_019814254.2;refGene_gn=ANKRD22;refGene_tx=XM_019814255.2;refGe
ne_gn=ANKRD22;refGene_tx=XM_023240335.1
D2 8548935 8548965
refGene_gn=STAMBPL1;refGene_tx=XM_003993870.5;refGene_gn=STAMBPL1;refG
ene_tx=XM_023240336.1;refGene_gn=STAMBPL1;refGene_tx=XM_023240337.1
D2 8553055 8553273
refGene_gn=STAMBPL1;refGene_tx=XM_003993870.5;refGene_gn=STAMBPL1;refG
ene_tx=XM_023240336.1;refGene_gn=STAMBPL1;refGene_tx=XM_023240337.1
D2 8556235 8556311
refGene_gn=STAMBPL1;refGene_tx=XM_003993870.5;refGene_gn=STAMBPL1;refG
ene_tx=XM_023240336.1;refGene_gn=STAMBPL1;refGene_tx=XM_023240337.1
D2 8558083 8558179
refGene_gn=STAMBPL1;refGene_tx=XM_003993870.5;refGene_gn=STAMBPL1;refG
ene_tx=XM_023240336.1;refGene_gn=STAMBPL1;refGene_tx=XM_023240337.1
D2 8560666 8561024
refGene_gn=STAMBPL1;refGene_tx=XM_003993870.5;refGene_gn=STAMBPL1;refG
ene_tx=XM_023240336.1;refGene_gn=STAMBPL1;refGene_tx=XM_023240337.1
D2 8563426 8563551
refGene_gn=STAMBPL1;refGene_tx=XM_003993870.5;refGene_gn=STAMBPL1;refG
ene_tx=XM_023240336.1;refGene_gn=STAMBPL1;refGene_tx=XM_023240337.1
D2 8565772 8565910
refGene_gn=STAMBPL1;refGene_tx=XM_003993870.5;refGene_gn=STAMBPL1;refG
ene_tx=XM_023240336.1;refGene_gn=STAMBPL1;refGene_tx=XM_023240337.1
D2 8570612 8570725
refGene_gn=STAMBPL1;refGene_tx=XM_003993870.5;refGene_gn=STAMBPL1;refG
ene_tx=XM_023240336.1;refGene_gn=STAMBPL1;refGene_tx=XM_023240337.1
D2 8571611 8571711
refGene_gn=STAMBPL1;refGene_tx=XM_003993870.5;refGene_gn=STAMBPL1;refG
ene_tx=XM_023240336.1;refGene_gn=STAMBPL1;refGene_tx=XM_023240337.1
D2 8572887 8572944
refGene_gn=STAMBPL1;refGene_tx=XM_003993870.5;refGene_gn=STAMBPL1;refG
ene_tx=XM_023240336.1;refGene_gn=STAMBPL1;refGene_tx=XM_023240337.1
D2 8582758 8582902 refGene_gn=ACTA2;refGene_tx=XM_003993871.5
D2 8585769 8585951 refGene_gn=ACTA2;refGene_tx=XM_003993871.5
D2 8587354 8587546 refGene_gn=ACTA2;refGene_tx=XM_003993871.5
D2 8588980 8589142 refGene_gn=ACTA2;refGene_tx=XM_003993871.5
D2 8589943 8590028 refGene_gn=ACTA2;refGene_tx=XM_003993871.5
D2 8592059 8592170 refGene_gn=ACTA2;refGene_tx=XM_003993871.5
D2 8594692 8594821 refGene_gn=ACTA2;refGene_tx=XM_003993871.5
D2 8596531 8596660 refGene_gn=ACTA2;refGene_tx=XM_003993871.5
D2 8634382 8634412 refGene_gn=FAS;refGene_tx=NM_001009314.1
D2 8648551 8648717 refGene_gn=FAS;refGene_tx=NM_001009314.1
D2 8653076 8653214 refGene_gn=FAS;refGene_tx=NM_001009314.1
D2 8654247 8654356 refGene_gn=FAS;refGene_tx=NM_001009314.1
D2 8657026 8657085 refGene_gn=FAS;refGene_tx=NM_001009314.1
D2 8657243 8657312 refGene_gn=FAS;refGene_tx=NM_001009314.1
D2 8658741 8658803 refGene_gn=FAS;refGene_tx=NM_001009314.1

D2 8659901 8659926 refGene_gn=FAS;refGene_tx=NM_001009314.1
D2 8660636 8660923 refGene_gn=FAS;refGene_tx=NM_001009314.1
D2 8836843 8837659 refGene_gn=CH25H;refGene_tx=XM_003993886.4
D2 8849205 8849439 refGene_gn=LIPA;refGene_tx=XM_003993874.4
D2 8850173 8850245 refGene_gn=LIPA;refGene_tx=XM_003993874.4
D2 8854430 8854502 refGene_gn=LIPA;refGene_tx=XM_003993874.4
D2 8855317 8855464 refGene_gn=LIPA;refGene_tx=XM_003993874.4
D2 8861210 8861347 refGene_gn=LIPA;refGene_tx=XM_003993874.4
D2 8862951 8863061 refGene_gn=LIPA;refGene_tx=XM_003993874.4
D2 8864415 8864614 refGene_gn=LIPA;refGene_tx=XM_003993874.4
D2 8878478 8878596 refGene_gn=LIPA;refGene_tx=XM_003993874.4
D2 8880570 8880681 refGene_gn=LIPA;refGene_tx=XM_003993874.4
D2 8947177 8947182 refGene_gn=IFIT2;refGene_tx=XM_006937718.4
D2 8951271 8952676 refGene_gn=IFIT2;refGene_tx=XM_006937718.4
D2 8967732 8967737
refGene_gn=IFIT1B;refGene_tx=XM_006937727.4;refGene_gn=IFIT3;refGene_t
x=XM_011287196.3
D2 8971266 8971271 refGene_gn=IFIT3;refGene_tx=XM_019814133.2
D2 8978029 8979494
refGene_gn=IFIT3;refGene_tx=XM_011287196.3;refGene_gn=IFIT3;refGene_tx
=XM_019814133.2
D2 8988034 8989454 refGene_gn=IFIT1B;refGene_tx=XM_006937727.4
D2 8999797 8999802
refGene_gn=LOC101094376;refGene_tx=XM_003993875.5
D2 9005234 9005895
refGene_gn=LOC101094376;refGene_tx=XM_003993875.5
D2 9005896 9006652
refGene_gn=LOC101094376;refGene_tx=XM_003993875.5
D2 9012535 9012630
refGene_gn=LOC101100156;refGene_tx=XM_003993890.5
D2 9021868 9023309
refGene_gn=LOC101100156;refGene_tx=XM_003993890.5;refGene_gn=LOC101100
156;refGene_tx=XM_023240357.1
D2 9023928 9024020
refGene_gn=LOC111556919;refGene_tx=XM_023240291.1
D2 9025641 9026482
refGene_gn=LOC111556919;refGene_tx=XM_023240291.1
D2 9035171 9036670
refGene_gn=ZNF248;refGene_tx=XM_006937723.4;refGene_gn=ZNF248;refGene_
tx=XM_006937724.4;refGene_gn=ZNF248;refGene_tx=XM_011287199.3;refGene_
gn=ZNF248;refGene_tx=XM_019814131.2;refGene_gn=ZNF248;refGene_tx=XM_01
9814132.2
D2 9042711 9042807
refGene_gn=ZNF248;refGene_tx=XM_006937723.4;refGene_gn=ZNF248;refGene_
tx=XM_006937724.4;refGene_gn=ZNF248;refGene_tx=XM_011287199.3;refGene_
gn=ZNF248;refGene_tx=XM_019814131.2;refGene_gn=ZNF248;refGene_tx=XM_01
9814132.2
D2 9043073 9043200
refGene_gn=ZNF248;refGene_tx=XM_006937723.4;refGene_gn=ZNF248;refGene_
tx=XM_006937724.4;refGene_gn=ZNF248;refGene_tx=XM_011287199.3;refGene_

gn=ZNF248;refGene_tx=XM_019814131.2;refGene_gn=ZNF248;refGene_tx=XM_01
9814132.2
D2 9050988 9051003
refGene_gn=ZNF248;refGene_tx=XM_006937723.4;refGene_gn=ZNF248;refGene_
tx=XM_006937724.4;refGene_gn=ZNF248;refGene_tx=XM_011287199.3;refGene_
gn=ZNF248;refGene_tx=XM_019814131.2;refGene_gn=ZNF248;refGene_tx=XM_01
9814132.2
D2 9070152 9070263
refGene_gn=LOC101100417;refGene_tx=XM_023240338.1;refGene_gn=LOC101100
417;refGene_tx=XM_023240339.1;refGene_gn=LOC101100417;refGene_tx=XM_02
3240340.1
D2 9072437 9072500
refGene_gn=LOC101100417;refGene_tx=XM_006937730.4;refGene_gn=LOC101100
417;refGene_tx=XM_023240338.1;refGene_gn=LOC101100417;refGene_tx=XM_02
3240339.1
D2 9084204 9084331
refGene_gn=LOC101100417;refGene_tx=XM_006937730.4;refGene_gn=LOC101100
417;refGene_tx=XM_019814116.2;refGene_gn=LOC101100417;refGene_tx=XM_01
9814118.2;refGene_gn=LOC101100417;refGene_tx=XM_023240338.1;refGene_gn
=LOC101100417;refGene_tx=XM_023240339.1;refGene_gn=LOC101100417;refGen
e_tx=XM_023240340.1;refGene_gn=LOC101100417;refGene_tx=XM_023240341.1
D2 9084766 9084862
refGene_gn=LOC101100417;refGene_tx=XM_006937730.4;refGene_gn=LOC101100
417;refGene_tx=XM_019814116.2;refGene_gn=LOC101100417;refGene_tx=XM_01
9814118.2;refGene_gn=LOC101100417;refGene_tx=XM_019814119.2;refGene_gn
=LOC101100417;refGene_tx=XM_023240338.1;refGene_gn=LOC101100417;refGen
e_tx=XM_023240339.1;refGene_gn=LOC101100417;refGene_tx=XM_023240340.1;
refGene_gn=LOC101100417;refGene_tx=XM_023240341.1
D2 9089312 9089368
refGene_gn=LOC101100417;refGene_tx=XM_019814119.2
D2 9096619 9098865
refGene_gn=LOC101100417;refGene_tx=XM_006937730.4;refGene_gn=LOC101100
417;refGene_tx=XM_019814116.2;refGene_gn=LOC101100417;refGene_tx=XM_01
9814118.2;refGene_gn=LOC101100417;refGene_tx=XM_019814119.2;refGene_gn
=LOC101100417;refGene_tx=XM_023240338.1;refGene_gn=LOC101100417;refGen
e_tx=XM_023240339.1;refGene_gn=LOC101100417;refGene_tx=XM_023240340.1;
refGene_gn=LOC101100417;refGene_tx=XM_023240341.1
D2 9121437 9122491
refGene_gn=ZNF25;refGene_tx=XM_011287209.3;refGene_gn=ZNF25;refGene_tx
=XM_019814136.2;refGene_gn=ZNF25;refGene_tx=XM_019814137.2
D2 9122952 9123016
refGene_gn=ZNF25;refGene_tx=XM_011287209.3;refGene_gn=ZNF25;refGene_tx
=XM_019814136.2;refGene_gn=ZNF25;refGene_tx=XM_019814137.2
D2 9125166 9125262
refGene_gn=ZNF25;refGene_tx=XM_011287209.3;refGene_gn=ZNF25;refGene_tx
=XM_019814136.2;refGene_gn=ZNF25;refGene_tx=XM_019814137.2
D2 9125548 9125675
refGene_gn=ZNF25;refGene_tx=XM_011287209.3;refGene_gn=ZNF25;refGene_tx
=XM_019814136.2;refGene_gn=ZNF25;refGene_tx=XM_019814137.2
D2 9143804 9143819

refGene_gn=ZNF25;refGene_tx=XM_011287209.3;refGene_gn=ZNF25;refGene_tx=XM_019814136.2;refGene_gn=ZNF25;refGene_tx=XM_019814137.2
D2 9159707 9159722
refGene_gn=LOC101083968;refGene_tx=XM_019814138.2;refGene_gn=LOC101083968;refGene_tx=XM_023240342.1;refGene_gn=LOC101083968;refGene_tx=XM_023240343.1;refGene_gn=LOC101083968;refGene_tx=XM_023240344.1;refGene_gn=LOC101083968;refGene_tx=XM_023240345.1;refGene_gn=LOC101083968;refGene_tx=XM_023240346.1;refGene_gn=LOC101083968;refGene_tx=XM_023240347.1;refGene_gn=LOC101083968;refGene_tx=XM_023240348.1;refGene_gn=LOC101083968;refGene_tx=XM_023240349.1;refGene_gn=LOC101083968;refGene_tx=XM_023240350.1;refGene_gn=LOC101083968;refGene_tx=XM_023240351.1;refGene_gn=LOC101083968;refGene_tx=XM_023240352.1;refGene_gn=LOC101083968;refGene_tx=XM_023240353.1;refGene_gn=LOC101083968;refGene_tx=XM_023240354.1;refGene_gn=LOC101083968;refGene_tx=XM_023240356.1
D2 9172117 9172244
refGene_gn=LOC101083968;refGene_tx=XM_019814138.2;refGene_gn=LOC101083968;refGene_tx=XM_023240342.1;refGene_gn=LOC101083968;refGene_tx=XM_023240343.1;refGene_gn=LOC101083968;refGene_tx=XM_023240344.1;refGene_gn=LOC101083968;refGene_tx=XM_023240345.1;refGene_gn=LOC101083968;refGene_tx=XM_023240346.1;refGene_gn=LOC101083968;refGene_tx=XM_023240347.1;refGene_gn=LOC101083968;refGene_tx=XM_023240348.1;refGene_gn=LOC101083968;refGene_tx=XM_023240349.1;refGene_gn=LOC101083968;refGene_tx=XM_023240350.1;refGene_gn=LOC101083968;refGene_tx=XM_023240351.1;refGene_gn=LOC101083968;refGene_tx=XM_023240352.1;refGene_gn=LOC101083968;refGene_tx=XM_023240353.1;refGene_gn=LOC101083968;refGene_tx=XM_023240354.1;refGene_gn=LOC101083968;refGene_tx=XM_023240356.1
D2 9172526 9172622
refGene_gn=LOC101083968;refGene_tx=XM_019814138.2;refGene_gn=LOC101083968;refGene_tx=XM_023240342.1;refGene_gn=LOC101083968;refGene_tx=XM_023240343.1;refGene_gn=LOC101083968;refGene_tx=XM_023240344.1;refGene_gn=LOC101083968;refGene_tx=XM_023240345.1;refGene_gn=LOC101083968;refGene_tx=XM_023240346.1;refGene_gn=LOC101083968;refGene_tx=XM_023240347.1;refGene_gn=LOC101083968;refGene_tx=XM_023240348.1;refGene_gn=LOC101083968;refGene_tx=XM_023240349.1;refGene_gn=LOC101083968;refGene_tx=XM_023240350.1;refGene_gn=LOC101083968;refGene_tx=XM_023240351.1;refGene_gn=LOC101083968;refGene_tx=XM_023240352.1;refGene_gn=LOC101083968;refGene_tx=XM_023240353.1;refGene_gn=LOC101083968;refGene_tx=XM_023240354.1;refGene_gn=LOC101083968;refGene_tx=XM_023240356.1
D2 9176614 9176799
refGene_gn=LOC101083968;refGene_tx=XM_019814138.2;refGene_gn=LOC101083968;refGene_tx=XM_023240342.1;refGene_gn=LOC101083968;refGene_tx=XM_023240343.1;refGene_gn=LOC101083968;refGene_tx=XM_023240344.1;refGene_gn=LOC101083968;refGene_tx=XM_023240345.1;refGene_gn=LOC101083968;refGene_tx=XM_023240346.1;refGene_gn=LOC101083968;refGene_tx=XM_023240347.1;refGene_gn=LOC101083968;refGene_tx=XM_023240348.1;refGene_gn=LOC101083968;refGene_tx=XM_023240349.1;refGene_gn=LOC101083968;refGene_tx=XM_023240350.1;refGene_gn=LOC101083968;refGene_tx=XM_023240351.1;refGene_gn=LOC101083968;refGene_tx=XM_023240352.1;refGene_gn=LOC101083968;refGene_tx=XM_023240353.1;refGene_gn=LOC101083968;refGene_tx=XM_023240354.1;refGene_gn=LOC101083968;refGene_tx=XM_023240356.1

D2 9178044 9178755
refGene_gn=LOC101083968;refGene_tx=XM_019814138.2;refGene_gn=LOC101083
968;refGene_tx=XM_023240342.1;refGene_gn=LOC101083968;refGene_tx=XM_02
3240343.1;refGene_gn=LOC101083968;refGene_tx=XM_023240344.1;refGene_gn
=LOC101083968;refGene_tx=XM_023240345.1;refGene_gn=LOC101083968;refGen
e_tx=XM_023240346.1;refGene_gn=LOC101083968;refGene_tx=XM_023240347.1;
refGene_gn=LOC101083968;refGene_tx=XM_023240348.1;refGene_gn=LOC101083
968;refGene_tx=XM_023240349.1;refGene_gn=LOC101083968;refGene_tx=XM_02
3240350.1;refGene_gn=LOC101083968;refGene_tx=XM_023240351.1;refGene_gn
=LOC101083968;refGene_tx=XM_023240352.1;refGene_gn=LOC101083968;refGen
e_tx=XM_023240353.1;refGene_gn=LOC101083968;refGene_tx=XM_023240354.1;
refGene_gn=LOC101083968;refGene_tx=XM_023240356.1
D2 9382356 9384010
refGene_gn=LOC111556921;refGene_tx=XM_023240292.1
D2 9384971 9385097
refGene_gn=LOC111556921;refGene_tx=XM_023240292.1
D2 9435506 9435586
refGene_gn=LOC111556921;refGene_tx=XM_023240292.1
D2 9438938 9440711
refGene_gn=CHRM3;refGene_tx=XM_011287200.3;refGene_gn=CHRM3;refGene_tx
=XM_019814130.2
D2 11287657 11287829 refGene_gn=ZP4;refGene_tx=NM_001009260.1
D2 11288091 11288379 refGene_gn=ZP4;refGene_tx=NM_001009260.1
D2 11289534 11289732 refGene_gn=ZP4;refGene_tx=NM_001009260.1
D2 11290428 11290616 refGene_gn=ZP4;refGene_tx=NM_001009260.1
D2 11291148 11291246 refGene_gn=ZP4;refGene_tx=NM_001009260.1
D2 11292114 11292245 refGene_gn=ZP4;refGene_tx=NM_001009260.1
D2 11292645 11292835 refGene_gn=ZP4;refGene_tx=NM_001009260.1
D2 11292911 11293059 refGene_gn=ZP4;refGene_tx=NM_001009260.1
D2 11293390 11293469 refGene_gn=ZP4;refGene_tx=NM_001009260.1
D2 11294681 11294798 refGene_gn=ZP4;refGene_tx=NM_001009260.1
D2 11295015 11295119 refGene_gn=ZP4;refGene_tx=NM_001009260.1
D2 11347982 11348078
refGene_gn=RYR2;refGene_tx=XM_023240358.1;refGene_gn=RYR2;refGene_tx=X
M_023240359.1;refGene_gn=RYR2;refGene_tx=XM_023240360.1;refGene_gn=RYR
2;refGene_tx=XM_023240361.1;refGene_gn=RYR2;refGene_tx=XM_023240362.1;
refGene_gn=RYR2;refGene_tx=XM_023240363.1;refGene_gn=RYR2;refGene_tx=X
M_023240364.1;refGene_gn=RYR2;refGene_tx=XM_023240365.1;refGene_gn=RYR
2;refGene_tx=XM_023240366.1;refGene_gn=RYR2;refGene_tx=XM_023240367.1;
refGene_gn=RYR2;refGene_tx=XM_023240368.1
D2 11348808 11348860
refGene_gn=RYR2;refGene_tx=XM_023240358.1;refGene_gn=RYR2;refGene_tx=X
M_023240359.1;refGene_gn=RYR2;refGene_tx=XM_023240360.1;refGene_gn=RYR
2;refGene_tx=XM_023240361.1;refGene_gn=RYR2;refGene_tx=XM_023240362.1;
refGene_gn=RYR2;refGene_tx=XM_023240363.1;refGene_gn=RYR2;refGene_tx=X
M_023240364.1;refGene_gn=RYR2;refGene_tx=XM_023240365.1;refGene_gn=RYR
2;refGene_tx=XM_023240366.1;refGene_gn=RYR2;refGene_tx=XM_023240367.1;
refGene_gn=RYR2;refGene_tx=XM_023240368.1
D2 11349850 11349951
refGene_gn=RYR2;refGene_tx=XM_023240358.1;refGene_gn=RYR2;refGene_tx=X

M_023240359.1;refGene_gn=RYR2;refGene_tx=XM_023240360.1;refGene_gn=RYR2;refGene_tx=XM_023240361.1;refGene_gn=RYR2;refGene_tx=XM_023240362.1;refGene_gn=RYR2;refGene_tx=XM_023240363.1;refGene_gn=RYR2;refGene_tx=XM_023240364.1;refGene_gn=RYR2;refGene_tx=XM_023240365.1;refGene_gn=RYR2;refGene_tx=XM_023240366.1;refGene_gn=RYR2;refGene_tx=XM_023240367.1;refGene_gn=RYR2;refGene_tx=XM_023240368.1
D2 11351587 11351652
refGene_gn=RYR2;refGene_tx=XM_023240358.1;refGene_gn=RYR2;refGene_tx=XM_023240359.1;refGene_gn=RYR2;refGene_tx=XM_023240360.1;refGene_gn=RYR2;refGene_tx=XM_023240361.1;refGene_gn=RYR2;refGene_tx=XM_023240362.1;refGene_gn=RYR2;refGene_tx=XM_023240363.1;refGene_gn=RYR2;refGene_tx=XM_023240364.1;refGene_gn=RYR2;refGene_tx=XM_023240365.1;refGene_gn=RYR2;refGene_tx=XM_023240366.1;refGene_gn=RYR2;refGene_tx=XM_023240367.1;refGene_gn=RYR2;refGene_tx=XM_023240368.1
D2 11353762 11353919
refGene_gn=RYR2;refGene_tx=XM_023240358.1;refGene_gn=RYR2;refGene_tx=XM_023240359.1;refGene_gn=RYR2;refGene_tx=XM_023240360.1;refGene_gn=RYR2;refGene_tx=XM_023240361.1;refGene_gn=RYR2;refGene_tx=XM_023240362.1;refGene_gn=RYR2;refGene_tx=XM_023240363.1;refGene_gn=RYR2;refGene_tx=XM_023240364.1;refGene_gn=RYR2;refGene_tx=XM_023240365.1;refGene_gn=RYR2;refGene_tx=XM_023240366.1;refGene_gn=RYR2;refGene_tx=XM_023240367.1;refGene_gn=RYR2;refGene_tx=XM_023240368.1
D2 11361387 11361522
refGene_gn=RYR2;refGene_tx=XM_023240358.1;refGene_gn=RYR2;refGene_tx=XM_023240359.1;refGene_gn=RYR2;refGene_tx=XM_023240360.1;refGene_gn=RYR2;refGene_tx=XM_023240361.1;refGene_gn=RYR2;refGene_tx=XM_023240362.1;refGene_gn=RYR2;refGene_tx=XM_023240363.1;refGene_gn=RYR2;refGene_tx=XM_023240364.1;refGene_gn=RYR2;refGene_tx=XM_023240365.1;refGene_gn=RYR2;refGene_tx=XM_023240366.1;refGene_gn=RYR2;refGene_tx=XM_023240367.1;refGene_gn=RYR2;refGene_tx=XM_023240368.1
D2 11363243 11363390
refGene_gn=RYR2;refGene_tx=XM_023240358.1;refGene_gn=RYR2;refGene_tx=XM_023240359.1;refGene_gn=RYR2;refGene_tx=XM_023240360.1;refGene_gn=RYR2;refGene_tx=XM_023240361.1;refGene_gn=RYR2;refGene_tx=XM_023240362.1;refGene_gn=RYR2;refGene_tx=XM_023240363.1;refGene_gn=RYR2;refGene_tx=XM_023240364.1;refGene_gn=RYR2;refGene_tx=XM_023240365.1;refGene_gn=RYR2;refGene_tx=XM_023240366.1;refGene_gn=RYR2;refGene_tx=XM_023240367.1;refGene_gn=RYR2;refGene_tx=XM_023240368.1
D2 11367022 11367083
refGene_gn=RYR2;refGene_tx=XM_023240358.1;refGene_gn=RYR2;refGene_tx=XM_023240359.1;refGene_gn=RYR2;refGene_tx=XM_023240360.1;refGene_gn=RYR2;refGene_tx=XM_023240361.1;refGene_gn=RYR2;refGene_tx=XM_023240362.1;refGene_gn=RYR2;refGene_tx=XM_023240363.1;refGene_gn=RYR2;refGene_tx=XM_023240364.1;refGene_gn=RYR2;refGene_tx=XM_023240365.1;refGene_gn=RYR2;refGene_tx=XM_023240366.1;refGene_gn=RYR2;refGene_tx=XM_023240367.1;refGene_gn=RYR2;refGene_tx=XM_023240368.1
D2 11371399 11371533
refGene_gn=RYR2;refGene_tx=XM_023240358.1;refGene_gn=RYR2;refGene_tx=XM_023240359.1;refGene_gn=RYR2;refGene_tx=XM_023240360.1;refGene_gn=RYR2;refGene_tx=XM_023240361.1;refGene_gn=RYR2;refGene_tx=XM_023240362.1;

refGene_gn=RYR2;refGene_tx=XM_023240363.1;refGene_gn=RYR2;refGene_tx=XM_023240364.1;refGene_gn=RYR2;refGene_tx=XM_023240365.1;refGene_gn=RYR2;refGene_tx=XM_023240366.1;refGene_gn=RYR2;refGene_tx=XM_023240367.1;refGene_gn=RYR2;refGene_tx=XM_023240368.1
D2 11374605 11374648
refGene_gn=RYR2;refGene_tx=XM_023240358.1;refGene_gn=RYR2;refGene_tx=XM_023240359.1;refGene_gn=RYR2;refGene_tx=XM_023240360.1;refGene_gn=RYR2;refGene_tx=XM_023240361.1;refGene_gn=RYR2;refGene_tx=XM_023240362.1;refGene_gn=RYR2;refGene_tx=XM_023240363.1;refGene_gn=RYR2;refGene_tx=XM_023240364.1;refGene_gn=RYR2;refGene_tx=XM_023240365.1;refGene_gn=RYR2;refGene_tx=XM_023240366.1;refGene_gn=RYR2;refGene_tx=XM_023240367.1;refGene_gn=RYR2;refGene_tx=XM_023240368.1
D2 11376000 11376131
refGene_gn=RYR2;refGene_tx=XM_023240358.1;refGene_gn=RYR2;refGene_tx=XM_023240359.1;refGene_gn=RYR2;refGene_tx=XM_023240360.1;refGene_gn=RYR2;refGene_tx=XM_023240361.1;refGene_gn=RYR2;refGene_tx=XM_023240362.1;refGene_gn=RYR2;refGene_tx=XM_023240363.1;refGene_gn=RYR2;refGene_tx=XM_023240364.1;refGene_gn=RYR2;refGene_tx=XM_023240365.1;refGene_gn=RYR2;refGene_tx=XM_023240366.1;refGene_gn=RYR2;refGene_tx=XM_023240367.1;refGene_gn=RYR2;refGene_tx=XM_023240368.1
D2 11377631 11377850
refGene_gn=RYR2;refGene_tx=XM_023240358.1;refGene_gn=RYR2;refGene_tx=XM_023240359.1;refGene_gn=RYR2;refGene_tx=XM_023240360.1;refGene_gn=RYR2;refGene_tx=XM_023240361.1;refGene_gn=RYR2;refGene_tx=XM_023240362.1;refGene_gn=RYR2;refGene_tx=XM_023240363.1;refGene_gn=RYR2;refGene_tx=XM_023240364.1;refGene_gn=RYR2;refGene_tx=XM_023240365.1;refGene_gn=RYR2;refGene_tx=XM_023240366.1;refGene_gn=RYR2;refGene_tx=XM_023240367.1;refGene_gn=RYR2;refGene_tx=XM_023240368.1
D2 11378453 11378540
refGene_gn=RYR2;refGene_tx=XM_023240358.1;refGene_gn=RYR2;refGene_tx=XM_023240359.1;refGene_gn=RYR2;refGene_tx=XM_023240360.1;refGene_gn=RYR2;refGene_tx=XM_023240361.1;refGene_gn=RYR2;refGene_tx=XM_023240362.1;refGene_gn=RYR2;refGene_tx=XM_023240363.1;refGene_gn=RYR2;refGene_tx=XM_023240364.1;refGene_gn=RYR2;refGene_tx=XM_023240365.1;refGene_gn=RYR2;refGene_tx=XM_023240366.1;refGene_gn=RYR2;refGene_tx=XM_023240367.1;refGene_gn=RYR2;refGene_tx=XM_023240368.1
D2 11381513 11381661
refGene_gn=RYR2;refGene_tx=XM_023240358.1;refGene_gn=RYR2;refGene_tx=XM_023240359.1;refGene_gn=RYR2;refGene_tx=XM_023240360.1;refGene_gn=RYR2;refGene_tx=XM_023240361.1;refGene_gn=RYR2;refGene_tx=XM_023240362.1;refGene_gn=RYR2;refGene_tx=XM_023240363.1;refGene_gn=RYR2;refGene_tx=XM_023240364.1;refGene_gn=RYR2;refGene_tx=XM_023240365.1;refGene_gn=RYR2;refGene_tx=XM_023240366.1;refGene_gn=RYR2;refGene_tx=XM_023240367.1;refGene_gn=RYR2;refGene_tx=XM_023240368.1
D2 11384248 11384316
refGene_gn=RYR2;refGene_tx=XM_023240358.1;refGene_gn=RYR2;refGene_tx=XM_023240359.1;refGene_gn=RYR2;refGene_tx=XM_023240360.1;refGene_gn=RYR2;refGene_tx=XM_023240361.1;refGene_gn=RYR2;refGene_tx=XM_023240362.1;refGene_gn=RYR2;refGene_tx=XM_023240363.1;refGene_gn=RYR2;refGene_tx=XM_023240364.1;refGene_gn=RYR2;refGene_tx=XM_023240365.1;refGene_gn=RYR

2;refGene_tx=XM_023240366.1;refGene_gn=RYR2;refGene_tx=XM_023240367.1;
refGene_gn=RYR2;refGene_tx=XM_023240368.1
D2 11385049 11386347
refGene_gn=RYR2;refGene_tx=XM_023240358.1;refGene_gn=RYR2;refGene_tx=X
M_023240359.1;refGene_gn=RYR2;refGene_tx=XM_023240360.1;refGene_gn=RYR
2;refGene_tx=XM_023240361.1;refGene_gn=RYR2;refGene_tx=XM_023240362.1;
refGene_gn=RYR2;refGene_tx=XM_023240363.1;refGene_gn=RYR2;refGene_tx=X
M_023240364.1;refGene_gn=RYR2;refGene_tx=XM_023240365.1;refGene_gn=RYR
2;refGene_tx=XM_023240366.1;refGene_gn=RYR2;refGene_tx=XM_023240367.1;
refGene_gn=RYR2;refGene_tx=XM_023240368.1
D2 11388663 11388745
refGene_gn=RYR2;refGene_tx=XM_023240358.1;refGene_gn=RYR2;refGene_tx=X
M_023240359.1;refGene_gn=RYR2;refGene_tx=XM_023240360.1;refGene_gn=RYR
2;refGene_tx=XM_023240361.1;refGene_gn=RYR2;refGene_tx=XM_023240362.1;
refGene_gn=RYR2;refGene_tx=XM_023240363.1;refGene_gn=RYR2;refGene_tx=X
M_023240364.1;refGene_gn=RYR2;refGene_tx=XM_023240365.1;refGene_gn=RYR
2;refGene_tx=XM_023240366.1;refGene_gn=RYR2;refGene_tx=XM_023240367.1;
refGene_gn=RYR2;refGene_tx=XM_023240368.1
D2 11391425 11391530
refGene_gn=RYR2;refGene_tx=XM_023240358.1;refGene_gn=RYR2;refGene_tx=X
M_023240359.1;refGene_gn=RYR2;refGene_tx=XM_023240360.1;refGene_gn=RYR
2;refGene_tx=XM_023240361.1;refGene_gn=RYR2;refGene_tx=XM_023240362.1;
refGene_gn=RYR2;refGene_tx=XM_023240363.1;refGene_gn=RYR2;refGene_tx=X
M_023240364.1;refGene_gn=RYR2;refGene_tx=XM_023240365.1;refGene_gn=RYR
2;refGene_tx=XM_023240366.1;refGene_gn=RYR2;refGene_tx=XM_023240367.1;
refGene_gn=RYR2;refGene_tx=XM_023240368.1
D2 11407394 11407523
refGene_gn=RYR2;refGene_tx=XM_023240358.1;refGene_gn=RYR2;refGene_tx=X
M_023240359.1;refGene_gn=RYR2;refGene_tx=XM_023240360.1;refGene_gn=RYR
2;refGene_tx=XM_023240361.1;refGene_gn=RYR2;refGene_tx=XM_023240362.1;
refGene_gn=RYR2;refGene_tx=XM_023240363.1;refGene_gn=RYR2;refGene_tx=X
M_023240364.1;refGene_gn=RYR2;refGene_tx=XM_023240365.1;refGene_gn=RYR
2;refGene_tx=XM_023240366.1;refGene_gn=RYR2;refGene_tx=XM_023240367.1;
refGene_gn=RYR2;refGene_tx=XM_023240368.1
D2 11409599 11409688
refGene_gn=RYR2;refGene_tx=XM_023240358.1;refGene_gn=RYR2;refGene_tx=X
M_023240359.1;refGene_gn=RYR2;refGene_tx=XM_023240360.1;refGene_gn=RYR
2;refGene_tx=XM_023240361.1;refGene_gn=RYR2;refGene_tx=XM_023240362.1;
refGene_gn=RYR2;refGene_tx=XM_023240363.1;refGene_gn=RYR2;refGene_tx=X
M_023240364.1;refGene_gn=RYR2;refGene_tx=XM_023240365.1;refGene_gn=RYR
2;refGene_tx=XM_023240366.1;refGene_gn=RYR2;refGene_tx=XM_023240367.1;
refGene_gn=RYR2;refGene_tx=XM_023240368.1
D2 11410424 11410505
refGene_gn=RYR2;refGene_tx=XM_023240358.1;refGene_gn=RYR2;refGene_tx=X
M_023240359.1;refGene_gn=RYR2;refGene_tx=XM_023240360.1;refGene_gn=RYR
2;refGene_tx=XM_023240361.1;refGene_gn=RYR2;refGene_tx=XM_023240362.1;
refGene_gn=RYR2;refGene_tx=XM_023240363.1;refGene_gn=RYR2;refGene_tx=X
M_023240364.1;refGene_gn=RYR2;refGene_tx=XM_023240365.1;refGene_gn=RYR
2;refGene_tx=XM_023240366.1;refGene_gn=RYR2;refGene_tx=XM_023240367.1;
refGene_gn=RYR2;refGene_tx=XM_023240368.1

D2 11411690 11411723
refGene_gn=RYR2;refGene_tx=XM_023240358.1;refGene_gn=RYR2;refGene_tx=XM_023240360.1;refGene_gn=RYR2;refGene_tx=XM_023240361.1;refGene_gn=RYR2;refGene_tx=XM_023240363.1;refGene_gn=RYR2;refGene_tx=XM_023240365.1
D2 11418211 11418229
refGene_gn=RYR2;refGene_tx=XM_023240359.1;refGene_gn=RYR2;refGene_tx=XM_023240366.1
D2 11420349 11420423
refGene_gn=RYR2;refGene_tx=XM_023240358.1;refGene_gn=RYR2;refGene_tx=XM_023240359.1;refGene_gn=RYR2;refGene_tx=XM_023240360.1;refGene_gn=RYR2;refGene_tx=XM_023240361.1;refGene_gn=RYR2;refGene_tx=XM_023240362.1;refGene_gn=RYR2;refGene_tx=XM_023240363.1;refGene_gn=RYR2;refGene_tx=XM_023240364.1;refGene_gn=RYR2;refGene_tx=XM_023240365.1;refGene_gn=RYR2;refGene_tx=XM_023240366.1;refGene_gn=RYR2;refGene_tx=XM_023240367.1;refGene_gn=RYR2;refGene_tx=XM_023240368.1
D2 11421213 11421290
refGene_gn=RYR2;refGene_tx=XM_023240358.1;refGene_gn=RYR2;refGene_tx=XM_023240359.1;refGene_gn=RYR2;refGene_tx=XM_023240360.1;refGene_gn=RYR2;refGene_tx=XM_023240361.1;refGene_gn=RYR2;refGene_tx=XM_023240362.1;refGene_gn=RYR2;refGene_tx=XM_023240363.1;refGene_gn=RYR2;refGene_tx=XM_023240364.1;refGene_gn=RYR2;refGene_tx=XM_023240365.1;refGene_gn=RYR2;refGene_tx=XM_023240366.1;refGene_gn=RYR2;refGene_tx=XM_023240367.1;refGene_gn=RYR2;refGene_tx=XM_023240368.1
D2 11424803 11424883
refGene_gn=RYR2;refGene_tx=XM_023240358.1;refGene_gn=RYR2;refGene_tx=XM_023240359.1;refGene_gn=RYR2;refGene_tx=XM_023240360.1;refGene_gn=RYR2;refGene_tx=XM_023240361.1;refGene_gn=RYR2;refGene_tx=XM_023240362.1;refGene_gn=RYR2;refGene_tx=XM_023240363.1;refGene_gn=RYR2;refGene_tx=XM_023240364.1;refGene_gn=RYR2;refGene_tx=XM_023240365.1;refGene_gn=RYR2;refGene_tx=XM_023240366.1;refGene_gn=RYR2;refGene_tx=XM_023240367.1;refGene_gn=RYR2;refGene_tx=XM_023240368.1
D2 11426350 11426450
refGene_gn=RYR2;refGene_tx=XM_023240358.1;refGene_gn=RYR2;refGene_tx=XM_023240359.1;refGene_gn=RYR2;refGene_tx=XM_023240360.1;refGene_gn=RYR2;refGene_tx=XM_023240361.1;refGene_gn=RYR2;refGene_tx=XM_023240362.1;refGene_gn=RYR2;refGene_tx=XM_023240363.1;refGene_gn=RYR2;refGene_tx=XM_023240364.1;refGene_gn=RYR2;refGene_tx=XM_023240365.1;refGene_gn=RYR2;refGene_tx=XM_023240366.1;refGene_gn=RYR2;refGene_tx=XM_023240367.1;refGene_gn=RYR2;refGene_tx=XM_023240368.1
D2 11437391 11437415
refGene_gn=RYR2;refGene_tx=XM_023240358.1;refGene_gn=RYR2;refGene_tx=XM_023240359.1;refGene_gn=RYR2;refGene_tx=XM_023240360.1;refGene_gn=RYR2;refGene_tx=XM_023240362.1;refGene_gn=RYR2;refGene_tx=XM_023240363.1;refGene_gn=RYR2;refGene_tx=XM_023240367.1
D2 11438665 11438719
refGene_gn=RYR2;refGene_tx=XM_023240358.1;refGene_gn=RYR2;refGene_tx=XM_023240359.1;refGene_gn=RYR2;refGene_tx=XM_023240360.1;refGene_gn=RYR2;refGene_tx=XM_023240361.1;refGene_gn=RYR2;refGene_tx=XM_023240362.1;refGene_gn=RYR2;refGene_tx=XM_023240363.1;refGene_gn=RYR2;refGene_tx=XM_023240364.1;refGene_gn=RYR2;refGene_tx=XM_023240365.1;refGene_gn=RYR

2;refGene_tx=XM_023240366.1;refGene_gn=RYR2;refGene_tx=XM_023240367.1;
refGene_gn=RYR2;refGene_tx=XM_023240368.1
D2 11449117 11449169
refGene_gn=RYR2;refGene_tx=XM_023240358.1;refGene_gn=RYR2;refGene_tx=X
M_023240359.1;refGene_gn=RYR2;refGene_tx=XM_023240360.1;refGene_gn=RYR
2;refGene_tx=XM_023240361.1;refGene_gn=RYR2;refGene_tx=XM_023240362.1;
refGene_gn=RYR2;refGene_tx=XM_023240363.1;refGene_gn=RYR2;refGene_tx=X
M_023240364.1;refGene_gn=RYR2;refGene_tx=XM_023240365.1;refGene_gn=RYR
2;refGene_tx=XM_023240366.1;refGene_gn=RYR2;refGene_tx=XM_023240367.1;
refGene_gn=RYR2;refGene_tx=XM_023240368.1
D2 11450697 11450801
refGene_gn=RYR2;refGene_tx=XM_023240358.1;refGene_gn=RYR2;refGene_tx=X
M_023240359.1;refGene_gn=RYR2;refGene_tx=XM_023240360.1;refGene_gn=RYR
2;refGene_tx=XM_023240361.1;refGene_gn=RYR2;refGene_tx=XM_023240362.1;
refGene_gn=RYR2;refGene_tx=XM_023240363.1;refGene_gn=RYR2;refGene_tx=X
M_023240364.1;refGene_gn=RYR2;refGene_tx=XM_023240365.1;refGene_gn=RYR
2;refGene_tx=XM_023240366.1;refGene_gn=RYR2;refGene_tx=XM_023240367.1;
refGene_gn=RYR2;refGene_tx=XM_023240368.1
D2 11452937 11453034
refGene_gn=RYR2;refGene_tx=XM_023240358.1;refGene_gn=RYR2;refGene_tx=X
M_023240359.1;refGene_gn=RYR2;refGene_tx=XM_023240360.1;refGene_gn=RYR
2;refGene_tx=XM_023240361.1;refGene_gn=RYR2;refGene_tx=XM_023240362.1;
refGene_gn=RYR2;refGene_tx=XM_023240363.1;refGene_gn=RYR2;refGene_tx=X
M_023240364.1;refGene_gn=RYR2;refGene_tx=XM_023240365.1;refGene_gn=RYR
2;refGene_tx=XM_023240366.1;refGene_gn=RYR2;refGene_tx=XM_023240367.1;
refGene_gn=RYR2;refGene_tx=XM_023240368.1
D2 11456069 11456182
refGene_gn=RYR2;refGene_tx=XM_023240358.1;refGene_gn=RYR2;refGene_tx=X
M_023240359.1;refGene_gn=RYR2;refGene_tx=XM_023240360.1;refGene_gn=RYR
2;refGene_tx=XM_023240361.1;refGene_gn=RYR2;refGene_tx=XM_023240362.1;
refGene_gn=RYR2;refGene_tx=XM_023240363.1;refGene_gn=RYR2;refGene_tx=X
M_023240364.1;refGene_gn=RYR2;refGene_tx=XM_023240365.1;refGene_gn=RYR
2;refGene_tx=XM_023240366.1;refGene_gn=RYR2;refGene_tx=XM_023240367.1;
refGene_gn=RYR2;refGene_tx=XM_023240368.1
D2 11456952 11456988
refGene_gn=RYR2;refGene_tx=XM_023240358.1;refGene_gn=RYR2;refGene_tx=X
M_023240359.1;refGene_gn=RYR2;refGene_tx=XM_023240360.1;refGene_gn=RYR
2;refGene_tx=XM_023240361.1;refGene_gn=RYR2;refGene_tx=XM_023240362.1;
refGene_gn=RYR2;refGene_tx=XM_023240364.1
D2 11460287 11460422
refGene_gn=RYR2;refGene_tx=XM_023240358.1;refGene_gn=RYR2;refGene_tx=X
M_023240359.1;refGene_gn=RYR2;refGene_tx=XM_023240360.1;refGene_gn=RYR
2;refGene_tx=XM_023240361.1;refGene_gn=RYR2;refGene_tx=XM_023240362.1;
refGene_gn=RYR2;refGene_tx=XM_023240363.1;refGene_gn=RYR2;refGene_tx=X
M_023240364.1;refGene_gn=RYR2;refGene_tx=XM_023240365.1;refGene_gn=RYR
2;refGene_tx=XM_023240366.1;refGene_gn=RYR2;refGene_tx=XM_023240367.1;
refGene_gn=RYR2;refGene_tx=XM_023240368.1
D2 11463015 11463075
refGene_gn=RYR2;refGene_tx=XM_023240358.1;refGene_gn=RYR2;refGene_tx=X
M_023240359.1;refGene_gn=RYR2;refGene_tx=XM_023240360.1;refGene_gn=RYR

2;refGene_tx=XM_023240361.1;refGene_gn=RYR2;refGene_tx=XM_023240362.1;
refGene_gn=RYR2;refGene_tx=XM_023240363.1;refGene_gn=RYR2;refGene_tx=X
M_023240364.1;refGene_gn=RYR2;refGene_tx=XM_023240365.1;refGene_gn=RYR
2;refGene_tx=XM_023240366.1;refGene_gn=RYR2;refGene_tx=XM_023240367.1;
refGene_gn=RYR2;refGene_tx=XM_023240368.1
D2 11464172 11464343
refGene_gn=RYR2;refGene_tx=XM_023240358.1;refGene_gn=RYR2;refGene_tx=X
M_023240359.1;refGene_gn=RYR2;refGene_tx=XM_023240360.1;refGene_gn=RYR
2;refGene_tx=XM_023240361.1;refGene_gn=RYR2;refGene_tx=XM_023240362.1;
refGene_gn=RYR2;refGene_tx=XM_023240363.1;refGene_gn=RYR2;refGene_tx=X
M_023240364.1;refGene_gn=RYR2;refGene_tx=XM_023240365.1;refGene_gn=RYR
2;refGene_tx=XM_023240366.1;refGene_gn=RYR2;refGene_tx=XM_023240367.1;
refGene_gn=RYR2;refGene_tx=XM_023240368.1
D2 11475174 11475267
refGene_gn=RYR2;refGene_tx=XM_023240358.1;refGene_gn=RYR2;refGene_tx=X
M_023240359.1;refGene_gn=RYR2;refGene_tx=XM_023240360.1;refGene_gn=RYR
2;refGene_tx=XM_023240361.1;refGene_gn=RYR2;refGene_tx=XM_023240362.1;
refGene_gn=RYR2;refGene_tx=XM_023240363.1;refGene_gn=RYR2;refGene_tx=X
M_023240364.1;refGene_gn=RYR2;refGene_tx=XM_023240365.1;refGene_gn=RYR
2;refGene_tx=XM_023240366.1;refGene_gn=RYR2;refGene_tx=XM_023240367.1;
refGene_gn=RYR2;refGene_tx=XM_023240368.1
D2 11478292 11478380
refGene_gn=RYR2;refGene_tx=XM_023240358.1;refGene_gn=RYR2;refGene_tx=X
M_023240359.1;refGene_gn=RYR2;refGene_tx=XM_023240360.1;refGene_gn=RYR
2;refGene_tx=XM_023240361.1;refGene_gn=RYR2;refGene_tx=XM_023240362.1;
refGene_gn=RYR2;refGene_tx=XM_023240363.1;refGene_gn=RYR2;refGene_tx=X
M_023240364.1;refGene_gn=RYR2;refGene_tx=XM_023240365.1;refGene_gn=RYR
2;refGene_tx=XM_023240366.1;refGene_gn=RYR2;refGene_tx=XM_023240367.1;
refGene_gn=RYR2;refGene_tx=XM_023240368.1
D2 11478974 11479215
refGene_gn=RYR2;refGene_tx=XM_023240358.1;refGene_gn=RYR2;refGene_tx=X
M_023240359.1;refGene_gn=RYR2;refGene_tx=XM_023240360.1;refGene_gn=RYR
2;refGene_tx=XM_023240361.1;refGene_gn=RYR2;refGene_tx=XM_023240362.1;
refGene_gn=RYR2;refGene_tx=XM_023240363.1;refGene_gn=RYR2;refGene_tx=X
M_023240364.1;refGene_gn=RYR2;refGene_tx=XM_023240365.1;refGene_gn=RYR
2;refGene_tx=XM_023240366.1;refGene_gn=RYR2;refGene_tx=XM_023240367.1;
refGene_gn=RYR2;refGene_tx=XM_023240368.1
D2 11480309 11480630
refGene_gn=RYR2;refGene_tx=XM_023240358.1;refGene_gn=RYR2;refGene_tx=X
M_023240359.1;refGene_gn=RYR2;refGene_tx=XM_023240360.1;refGene_gn=RYR
2;refGene_tx=XM_023240361.1;refGene_gn=RYR2;refGene_tx=XM_023240362.1;
refGene_gn=RYR2;refGene_tx=XM_023240363.1;refGene_gn=RYR2;refGene_tx=X
M_023240364.1;refGene_gn=RYR2;refGene_tx=XM_023240365.1;refGene_gn=RYR
2;refGene_tx=XM_023240366.1;refGene_gn=RYR2;refGene_tx=XM_023240367.1;
refGene_gn=RYR2;refGene_tx=XM_023240368.1
D2 11483573 11483704
refGene_gn=RYR2;refGene_tx=XM_023240358.1;refGene_gn=RYR2;refGene_tx=X
M_023240359.1;refGene_gn=RYR2;refGene_tx=XM_023240360.1;refGene_gn=RYR
2;refGene_tx=XM_023240361.1;refGene_gn=RYR2;refGene_tx=XM_023240362.1;
refGene_gn=RYR2;refGene_tx=XM_023240363.1;refGene_gn=RYR2;refGene_tx=X

M_023240364.1;refGene_gn=RYR2;refGene_tx=XM_023240365.1;refGene_gn=RYR2;refGene_tx=XM_023240366.1;refGene_gn=RYR2;refGene_tx=XM_023240367.1;refGene_gn=RYR2;refGene_tx=XM_023240368.1
D2 11490356 11490438
refGene_gn=RYR2;refGene_tx=XM_023240358.1;refGene_gn=RYR2;refGene_tx=XM_023240359.1;refGene_gn=RYR2;refGene_tx=XM_023240360.1;refGene_gn=RYR2;refGene_tx=XM_023240361.1;refGene_gn=RYR2;refGene_tx=XM_023240362.1;refGene_gn=RYR2;refGene_tx=XM_023240363.1;refGene_gn=RYR2;refGene_tx=XM_023240364.1;refGene_gn=RYR2;refGene_tx=XM_023240365.1;refGene_gn=RYR2;refGene_tx=XM_023240366.1;refGene_gn=RYR2;refGene_tx=XM_023240367.1;refGene_gn=RYR2;refGene_tx=XM_023240368.1
D2 11492103 11492342
refGene_gn=RYR2;refGene_tx=XM_023240358.1;refGene_gn=RYR2;refGene_tx=XM_023240359.1;refGene_gn=RYR2;refGene_tx=XM_023240360.1;refGene_gn=RYR2;refGene_tx=XM_023240361.1;refGene_gn=RYR2;refGene_tx=XM_023240362.1;refGene_gn=RYR2;refGene_tx=XM_023240363.1;refGene_gn=RYR2;refGene_tx=XM_023240364.1;refGene_gn=RYR2;refGene_tx=XM_023240365.1;refGene_gn=RYR2;refGene_tx=XM_023240366.1;refGene_gn=RYR2;refGene_tx=XM_023240367.1;refGene_gn=RYR2;refGene_tx=XM_023240368.1
D2 11493561 11493622
refGene_gn=RYR2;refGene_tx=XM_023240358.1;refGene_gn=RYR2;refGene_tx=XM_023240359.1;refGene_gn=RYR2;refGene_tx=XM_023240360.1;refGene_gn=RYR2;refGene_tx=XM_023240361.1;refGene_gn=RYR2;refGene_tx=XM_023240362.1;refGene_gn=RYR2;refGene_tx=XM_023240363.1;refGene_gn=RYR2;refGene_tx=XM_023240364.1;refGene_gn=RYR2;refGene_tx=XM_023240365.1;refGene_gn=RYR2;refGene_tx=XM_023240366.1;refGene_gn=RYR2;refGene_tx=XM_023240367.1;refGene_gn=RYR2;refGene_tx=XM_023240368.1
D2 11504282 11504332
refGene_gn=RYR2;refGene_tx=XM_023240358.1;refGene_gn=RYR2;refGene_tx=XM_023240359.1;refGene_gn=RYR2;refGene_tx=XM_023240360.1;refGene_gn=RYR2;refGene_tx=XM_023240361.1;refGene_gn=RYR2;refGene_tx=XM_023240362.1;refGene_gn=RYR2;refGene_tx=XM_023240363.1;refGene_gn=RYR2;refGene_tx=XM_023240364.1;refGene_gn=RYR2;refGene_tx=XM_023240365.1;refGene_gn=RYR2;refGene_tx=XM_023240366.1;refGene_gn=RYR2;refGene_tx=XM_023240367.1;refGene_gn=RYR2;refGene_tx=XM_023240368.1
D2 11510926 11511048
refGene_gn=RYR2;refGene_tx=XM_023240358.1;refGene_gn=RYR2;refGene_tx=XM_023240359.1;refGene_gn=RYR2;refGene_tx=XM_023240360.1;refGene_gn=RYR2;refGene_tx=XM_023240361.1;refGene_gn=RYR2;refGene_tx=XM_023240362.1;refGene_gn=RYR2;refGene_tx=XM_023240363.1;refGene_gn=RYR2;refGene_tx=XM_023240364.1;refGene_gn=RYR2;refGene_tx=XM_023240365.1;refGene_gn=RYR2;refGene_tx=XM_023240366.1;refGene_gn=RYR2;refGene_tx=XM_023240367.1;refGene_gn=RYR2;refGene_tx=XM_023240368.1
D2 11513314 11513379
refGene_gn=RYR2;refGene_tx=XM_023240358.1;refGene_gn=RYR2;refGene_tx=XM_023240359.1;refGene_gn=RYR2;refGene_tx=XM_023240360.1;refGene_gn=RYR2;refGene_tx=XM_023240361.1;refGene_gn=RYR2;refGene_tx=XM_023240362.1;refGene_gn=RYR2;refGene_tx=XM_023240363.1;refGene_gn=RYR2;refGene_tx=XM_023240364.1;refGene_gn=RYR2;refGene_tx=XM_023240365.1;refGene_gn=RYR2;refGene_tx=XM_023240366.1;refGene_gn=RYR2;refGene_tx=XM_023240367.1;

refGene_gn=RYR2;refGene_tx=XM_023240368.1
D2 11516521 11516637
refGene_gn=RYR2;refGene_tx=XM_023240358.1;refGene_gn=RYR2;refGene_tx=XM_023240359.1;refGene_gn=RYR2;refGene_tx=XM_023240360.1;refGene_gn=RYR2;refGene_tx=XM_023240361.1;refGene_gn=RYR2;refGene_tx=XM_023240362.1;refGene_gn=RYR2;refGene_tx=XM_023240363.1;refGene_gn=RYR2;refGene_tx=XM_023240364.1;refGene_gn=RYR2;refGene_tx=XM_023240365.1;refGene_gn=RYR2;refGene_tx=XM_023240366.1;refGene_gn=RYR2;refGene_tx=XM_023240367.1;refGene_gn=RYR2;refGene_tx=XM_023240368.1
D2 11517119 11517243
refGene_gn=RYR2;refGene_tx=XM_023240358.1;refGene_gn=RYR2;refGene_tx=XM_023240359.1;refGene_gn=RYR2;refGene_tx=XM_023240360.1;refGene_gn=RYR2;refGene_tx=XM_023240361.1;refGene_gn=RYR2;refGene_tx=XM_023240362.1;refGene_gn=RYR2;refGene_tx=XM_023240363.1;refGene_gn=RYR2;refGene_tx=XM_023240364.1;refGene_gn=RYR2;refGene_tx=XM_023240365.1;refGene_gn=RYR2;refGene_tx=XM_023240366.1;refGene_gn=RYR2;refGene_tx=XM_023240367.1;refGene_gn=RYR2;refGene_tx=XM_023240368.1
D2 11521173 11521249
refGene_gn=RYR2;refGene_tx=XM_023240358.1;refGene_gn=RYR2;refGene_tx=XM_023240359.1;refGene_gn=RYR2;refGene_tx=XM_023240360.1;refGene_gn=RYR2;refGene_tx=XM_023240361.1;refGene_gn=RYR2;refGene_tx=XM_023240362.1;refGene_gn=RYR2;refGene_tx=XM_023240363.1;refGene_gn=RYR2;refGene_tx=XM_023240364.1;refGene_gn=RYR2;refGene_tx=XM_023240365.1;refGene_gn=RYR2;refGene_tx=XM_023240366.1;refGene_gn=RYR2;refGene_tx=XM_023240367.1;refGene_gn=RYR2;refGene_tx=XM_023240368.1
D2 11522486 11522564
refGene_gn=RYR2;refGene_tx=XM_023240358.1;refGene_gn=RYR2;refGene_tx=XM_023240359.1;refGene_gn=RYR2;refGene_tx=XM_023240360.1;refGene_gn=RYR2;refGene_tx=XM_023240361.1;refGene_gn=RYR2;refGene_tx=XM_023240362.1;refGene_gn=RYR2;refGene_tx=XM_023240363.1;refGene_gn=RYR2;refGene_tx=XM_023240364.1;refGene_gn=RYR2;refGene_tx=XM_023240365.1;refGene_gn=RYR2;refGene_tx=XM_023240366.1;refGene_gn=RYR2;refGene_tx=XM_023240367.1;refGene_gn=RYR2;refGene_tx=XM_023240368.1
D2 11529338 11529476
refGene_gn=RYR2;refGene_tx=XM_023240358.1;refGene_gn=RYR2;refGene_tx=XM_023240359.1;refGene_gn=RYR2;refGene_tx=XM_023240360.1;refGene_gn=RYR2;refGene_tx=XM_023240361.1;refGene_gn=RYR2;refGene_tx=XM_023240362.1;refGene_gn=RYR2;refGene_tx=XM_023240363.1;refGene_gn=RYR2;refGene_tx=XM_023240364.1;refGene_gn=RYR2;refGene_tx=XM_023240365.1;refGene_gn=RYR2;refGene_tx=XM_023240366.1;refGene_gn=RYR2;refGene_tx=XM_023240367.1;refGene_gn=RYR2;refGene_tx=XM_023240368.1
D2 11530260 11530350
refGene_gn=RYR2;refGene_tx=XM_023240358.1;refGene_gn=RYR2;refGene_tx=XM_023240359.1;refGene_gn=RYR2;refGene_tx=XM_023240360.1;refGene_gn=RYR2;refGene_tx=XM_023240361.1;refGene_gn=RYR2;refGene_tx=XM_023240362.1;refGene_gn=RYR2;refGene_tx=XM_023240363.1;refGene_gn=RYR2;refGene_tx=XM_023240364.1;refGene_gn=RYR2;refGene_tx=XM_023240365.1;refGene_gn=RYR2;refGene_tx=XM_023240366.1;refGene_gn=RYR2;refGene_tx=XM_023240367.1;refGene_gn=RYR2;refGene_tx=XM_023240368.1
D2 11531888 11531967

refGene_gn=RYR2;refGene_tx=XM_023240358.1;refGene_gn=RYR2;refGene_tx=XM_023240359.1;refGene_gn=RYR2;refGene_tx=XM_023240360.1;refGene_gn=RYR2;refGene_tx=XM_023240361.1;refGene_gn=RYR2;refGene_tx=XM_023240362.1;refGene_gn=RYR2;refGene_tx=XM_023240363.1;refGene_gn=RYR2;refGene_tx=XM_023240364.1;refGene_gn=RYR2;refGene_tx=XM_023240365.1;refGene_gn=RYR2;refGene_tx=XM_023240366.1;refGene_gn=RYR2;refGene_tx=XM_023240367.1;refGene_gn=RYR2;refGene_tx=XM_023240368.1
D2 11534161 11534325
refGene_gn=RYR2;refGene_tx=XM_023240358.1;refGene_gn=RYR2;refGene_tx=XM_023240359.1;refGene_gn=RYR2;refGene_tx=XM_023240360.1;refGene_gn=RYR2;refGene_tx=XM_023240361.1;refGene_gn=RYR2;refGene_tx=XM_023240362.1;refGene_gn=RYR2;refGene_tx=XM_023240363.1;refGene_gn=RYR2;refGene_tx=XM_023240364.1;refGene_gn=RYR2;refGene_tx=XM_023240365.1;refGene_gn=RYR2;refGene_tx=XM_023240366.1;refGene_gn=RYR2;refGene_tx=XM_023240367.1;refGene_gn=RYR2;refGene_tx=XM_023240368.1
D2 11536183 11536324
refGene_gn=RYR2;refGene_tx=XM_023240358.1;refGene_gn=RYR2;refGene_tx=XM_023240359.1;refGene_gn=RYR2;refGene_tx=XM_023240360.1;refGene_gn=RYR2;refGene_tx=XM_023240361.1;refGene_gn=RYR2;refGene_tx=XM_023240362.1;refGene_gn=RYR2;refGene_tx=XM_023240363.1;refGene_gn=RYR2;refGene_tx=XM_023240364.1;refGene_gn=RYR2;refGene_tx=XM_023240365.1;refGene_gn=RYR2;refGene_tx=XM_023240366.1;refGene_gn=RYR2;refGene_tx=XM_023240367.1;refGene_gn=RYR2;refGene_tx=XM_023240368.1
D2 11538754 11538845
refGene_gn=RYR2;refGene_tx=XM_023240358.1;refGene_gn=RYR2;refGene_tx=XM_023240359.1;refGene_gn=RYR2;refGene_tx=XM_023240360.1;refGene_gn=RYR2;refGene_tx=XM_023240361.1;refGene_gn=RYR2;refGene_tx=XM_023240362.1;refGene_gn=RYR2;refGene_tx=XM_023240363.1;refGene_gn=RYR2;refGene_tx=XM_023240364.1;refGene_gn=RYR2;refGene_tx=XM_023240365.1;refGene_gn=RYR2;refGene_tx=XM_023240366.1;refGene_gn=RYR2;refGene_tx=XM_023240367.1;refGene_gn=RYR2;refGene_tx=XM_023240368.1
D2 11540155 11540376
refGene_gn=RYR2;refGene_tx=XM_023240358.1;refGene_gn=RYR2;refGene_tx=XM_023240359.1;refGene_gn=RYR2;refGene_tx=XM_023240360.1;refGene_gn=RYR2;refGene_tx=XM_023240361.1;refGene_gn=RYR2;refGene_tx=XM_023240362.1;refGene_gn=RYR2;refGene_tx=XM_023240363.1;refGene_gn=RYR2;refGene_tx=XM_023240364.1;refGene_gn=RYR2;refGene_tx=XM_023240365.1;refGene_gn=RYR2;refGene_tx=XM_023240366.1;refGene_gn=RYR2;refGene_tx=XM_023240367.1;refGene_gn=RYR2;refGene_tx=XM_023240368.1
D2 11541982 11542152
refGene_gn=RYR2;refGene_tx=XM_023240358.1;refGene_gn=RYR2;refGene_tx=XM_023240359.1;refGene_gn=RYR2;refGene_tx=XM_023240360.1;refGene_gn=RYR2;refGene_tx=XM_023240361.1;refGene_gn=RYR2;refGene_tx=XM_023240362.1;refGene_gn=RYR2;refGene_tx=XM_023240363.1;refGene_gn=RYR2;refGene_tx=XM_023240364.1;refGene_gn=RYR2;refGene_tx=XM_023240365.1;refGene_gn=RYR2;refGene_tx=XM_023240366.1;refGene_gn=RYR2;refGene_tx=XM_023240367.1;refGene_gn=RYR2;refGene_tx=XM_023240368.1
D2 11547323 11547444
refGene_gn=RYR2;refGene_tx=XM_023240358.1;refGene_gn=RYR2;refGene_tx=XM_023240359.1;refGene_gn=RYR2;refGene_tx=XM_023240360.1;refGene_gn=RYR

2;refGene_tx=XM_023240361.1;refGene_gn=RYR2;refGene_tx=XM_023240362.1;
refGene_gn=RYR2;refGene_tx=XM_023240363.1;refGene_gn=RYR2;refGene_tx=X
M_023240364.1;refGene_gn=RYR2;refGene_tx=XM_023240365.1;refGene_gn=RYR
2;refGene_tx=XM_023240366.1;refGene_gn=RYR2;refGene_tx=XM_023240367.1;
refGene_gn=RYR2;refGene_tx=XM_023240368.1
D2 11549875 11549981
refGene_gn=RYR2;refGene_tx=XM_023240358.1;refGene_gn=RYR2;refGene_tx=X
M_023240359.1;refGene_gn=RYR2;refGene_tx=XM_023240360.1;refGene_gn=RYR
2;refGene_tx=XM_023240361.1;refGene_gn=RYR2;refGene_tx=XM_023240362.1;
refGene_gn=RYR2;refGene_tx=XM_023240363.1;refGene_gn=RYR2;refGene_tx=X
M_023240364.1;refGene_gn=RYR2;refGene_tx=XM_023240365.1;refGene_gn=RYR
2;refGene_tx=XM_023240366.1;refGene_gn=RYR2;refGene_tx=XM_023240367.1;
refGene_gn=RYR2;refGene_tx=XM_023240368.1
D2 11552134 11552321
refGene_gn=RYR2;refGene_tx=XM_023240358.1;refGene_gn=RYR2;refGene_tx=X
M_023240359.1;refGene_gn=RYR2;refGene_tx=XM_023240360.1;refGene_gn=RYR
2;refGene_tx=XM_023240361.1;refGene_gn=RYR2;refGene_tx=XM_023240362.1;
refGene_gn=RYR2;refGene_tx=XM_023240363.1;refGene_gn=RYR2;refGene_tx=X
M_023240364.1;refGene_gn=RYR2;refGene_tx=XM_023240365.1;refGene_gn=RYR
2;refGene_tx=XM_023240366.1;refGene_gn=RYR2;refGene_tx=XM_023240367.1;
refGene_gn=RYR2;refGene_tx=XM_023240368.1
D2 11552866 11553002
refGene_gn=RYR2;refGene_tx=XM_023240358.1;refGene_gn=RYR2;refGene_tx=X
M_023240359.1;refGene_gn=RYR2;refGene_tx=XM_023240360.1;refGene_gn=RYR
2;refGene_tx=XM_023240361.1;refGene_gn=RYR2;refGene_tx=XM_023240362.1;
refGene_gn=RYR2;refGene_tx=XM_023240363.1;refGene_gn=RYR2;refGene_tx=X
M_023240364.1;refGene_gn=RYR2;refGene_tx=XM_023240365.1;refGene_gn=RYR
2;refGene_tx=XM_023240366.1;refGene_gn=RYR2;refGene_tx=XM_023240367.1;
refGene_gn=RYR2;refGene_tx=XM_023240368.1
D2 11556730 11556834
refGene_gn=RYR2;refGene_tx=XM_023240358.1;refGene_gn=RYR2;refGene_tx=X
M_023240359.1;refGene_gn=RYR2;refGene_tx=XM_023240360.1;refGene_gn=RYR
2;refGene_tx=XM_023240361.1;refGene_gn=RYR2;refGene_tx=XM_023240362.1;
refGene_gn=RYR2;refGene_tx=XM_023240363.1;refGene_gn=RYR2;refGene_tx=X
M_023240364.1;refGene_gn=RYR2;refGene_tx=XM_023240365.1;refGene_gn=RYR
2;refGene_tx=XM_023240366.1;refGene_gn=RYR2;refGene_tx=XM_023240367.1;
refGene_gn=RYR2;refGene_tx=XM_023240368.1
D2 11557038 11557171
refGene_gn=RYR2;refGene_tx=XM_023240358.1;refGene_gn=RYR2;refGene_tx=X
M_023240359.1;refGene_gn=RYR2;refGene_tx=XM_023240360.1;refGene_gn=RYR
2;refGene_tx=XM_023240361.1;refGene_gn=RYR2;refGene_tx=XM_023240362.1;
refGene_gn=RYR2;refGene_tx=XM_023240363.1;refGene_gn=RYR2;refGene_tx=X
M_023240364.1;refGene_gn=RYR2;refGene_tx=XM_023240365.1;refGene_gn=RYR
2;refGene_tx=XM_023240366.1;refGene_gn=RYR2;refGene_tx=XM_023240367.1;
refGene_gn=RYR2;refGene_tx=XM_023240368.1
D2 11558519 11558634
refGene_gn=RYR2;refGene_tx=XM_023240358.1;refGene_gn=RYR2;refGene_tx=X
M_023240359.1;refGene_gn=RYR2;refGene_tx=XM_023240360.1;refGene_gn=RYR
2;refGene_tx=XM_023240361.1;refGene_gn=RYR2;refGene_tx=XM_023240362.1;
refGene_gn=RYR2;refGene_tx=XM_023240363.1;refGene_gn=RYR2;refGene_tx=X

M_023240364.1;refGene_gn=RYR2;refGene_tx=XM_023240365.1;refGene_gn=RYR2;refGene_tx=XM_023240366.1;refGene_gn=RYR2;refGene_tx=XM_023240367.1;refGene_gn=RYR2;refGene_tx=XM_023240368.1
D2 11563835 11564109
refGene_gn=RYR2;refGene_tx=XM_023240358.1;refGene_gn=RYR2;refGene_tx=XM_023240359.1;refGene_gn=RYR2;refGene_tx=XM_023240360.1;refGene_gn=RYR2;refGene_tx=XM_023240361.1;refGene_gn=RYR2;refGene_tx=XM_023240362.1;refGene_gn=RYR2;refGene_tx=XM_023240363.1;refGene_gn=RYR2;refGene_tx=XM_023240364.1;refGene_gn=RYR2;refGene_tx=XM_023240365.1;refGene_gn=RYR2;refGene_tx=XM_023240366.1;refGene_gn=RYR2;refGene_tx=XM_023240367.1;refGene_gn=RYR2;refGene_tx=XM_023240368.1
D2 11566112 11566256
refGene_gn=RYR2;refGene_tx=XM_023240358.1;refGene_gn=RYR2;refGene_tx=XM_023240359.1;refGene_gn=RYR2;refGene_tx=XM_023240360.1;refGene_gn=RYR2;refGene_tx=XM_023240361.1;refGene_gn=RYR2;refGene_tx=XM_023240362.1;refGene_gn=RYR2;refGene_tx=XM_023240363.1;refGene_gn=RYR2;refGene_tx=XM_023240364.1;refGene_gn=RYR2;refGene_tx=XM_023240365.1;refGene_gn=RYR2;refGene_tx=XM_023240366.1;refGene_gn=RYR2;refGene_tx=XM_023240367.1;refGene_gn=RYR2;refGene_tx=XM_023240368.1
D2 11572961 11573067
refGene_gn=RYR2;refGene_tx=XM_023240358.1;refGene_gn=RYR2;refGene_tx=XM_023240359.1;refGene_gn=RYR2;refGene_tx=XM_023240360.1;refGene_gn=RYR2;refGene_tx=XM_023240361.1;refGene_gn=RYR2;refGene_tx=XM_023240362.1;refGene_gn=RYR2;refGene_tx=XM_023240363.1;refGene_gn=RYR2;refGene_tx=XM_023240364.1;refGene_gn=RYR2;refGene_tx=XM_023240365.1;refGene_gn=RYR2;refGene_tx=XM_023240366.1;refGene_gn=RYR2;refGene_tx=XM_023240367.1;refGene_gn=RYR2;refGene_tx=XM_023240368.1
D2 11579495 11579696
refGene_gn=RYR2;refGene_tx=XM_023240358.1;refGene_gn=RYR2;refGene_tx=XM_023240359.1;refGene_gn=RYR2;refGene_tx=XM_023240360.1;refGene_gn=RYR2;refGene_tx=XM_023240361.1;refGene_gn=RYR2;refGene_tx=XM_023240362.1;refGene_gn=RYR2;refGene_tx=XM_023240363.1;refGene_gn=RYR2;refGene_tx=XM_023240364.1;refGene_gn=RYR2;refGene_tx=XM_023240365.1;refGene_gn=RYR2;refGene_tx=XM_023240366.1;refGene_gn=RYR2;refGene_tx=XM_023240367.1;refGene_gn=RYR2;refGene_tx=XM_023240368.1
D2 11582672 11583477
refGene_gn=RYR2;refGene_tx=XM_023240358.1;refGene_gn=RYR2;refGene_tx=XM_023240359.1;refGene_gn=RYR2;refGene_tx=XM_023240360.1;refGene_gn=RYR2;refGene_tx=XM_023240361.1;refGene_gn=RYR2;refGene_tx=XM_023240362.1;refGene_gn=RYR2;refGene_tx=XM_023240363.1;refGene_gn=RYR2;refGene_tx=XM_023240364.1;refGene_gn=RYR2;refGene_tx=XM_023240365.1;refGene_gn=RYR2;refGene_tx=XM_023240366.1;refGene_gn=RYR2;refGene_tx=XM_023240367.1;refGene_gn=RYR2;refGene_tx=XM_023240368.1
D2 11586815 11587042
refGene_gn=RYR2;refGene_tx=XM_023240358.1;refGene_gn=RYR2;refGene_tx=XM_023240359.1;refGene_gn=RYR2;refGene_tx=XM_023240360.1;refGene_gn=RYR2;refGene_tx=XM_023240361.1;refGene_gn=RYR2;refGene_tx=XM_023240362.1;refGene_gn=RYR2;refGene_tx=XM_023240363.1;refGene_gn=RYR2;refGene_tx=XM_023240364.1;refGene_gn=RYR2;refGene_tx=XM_023240365.1;refGene_gn=RYR2;refGene_tx=XM_023240366.1;refGene_gn=RYR2;refGene_tx=XM_023240367.1;

refGene_gn=RYR2;refGene_tx=XM_023240368.1
D2 11589765 11589852
refGene_gn=RYR2;refGene_tx=XM_023240358.1;refGene_gn=RYR2;refGene_tx=XM_023240359.1;refGene_gn=RYR2;refGene_tx=XM_023240360.1;refGene_gn=RYR2;refGene_tx=XM_023240361.1;refGene_gn=RYR2;refGene_tx=XM_023240362.1;refGene_gn=RYR2;refGene_tx=XM_023240363.1;refGene_gn=RYR2;refGene_tx=XM_023240364.1;refGene_gn=RYR2;refGene_tx=XM_023240365.1;refGene_gn=RYR2;refGene_tx=XM_023240366.1;refGene_gn=RYR2;refGene_tx=XM_023240367.1;refGene_gn=RYR2;refGene_tx=XM_023240368.1
D2 11590232 11590392
refGene_gn=RYR2;refGene_tx=XM_023240358.1;refGene_gn=RYR2;refGene_tx=XM_023240359.1;refGene_gn=RYR2;refGene_tx=XM_023240360.1;refGene_gn=RYR2;refGene_tx=XM_023240361.1;refGene_gn=RYR2;refGene_tx=XM_023240362.1;refGene_gn=RYR2;refGene_tx=XM_023240363.1;refGene_gn=RYR2;refGene_tx=XM_023240364.1;refGene_gn=RYR2;refGene_tx=XM_023240365.1;refGene_gn=RYR2;refGene_tx=XM_023240366.1;refGene_gn=RYR2;refGene_tx=XM_023240367.1;refGene_gn=RYR2;refGene_tx=XM_023240368.1
D2 11592521 11592682
refGene_gn=RYR2;refGene_tx=XM_023240358.1;refGene_gn=RYR2;refGene_tx=XM_023240359.1;refGene_gn=RYR2;refGene_tx=XM_023240360.1;refGene_gn=RYR2;refGene_tx=XM_023240361.1;refGene_gn=RYR2;refGene_tx=XM_023240362.1;refGene_gn=RYR2;refGene_tx=XM_023240363.1;refGene_gn=RYR2;refGene_tx=XM_023240364.1;refGene_gn=RYR2;refGene_tx=XM_023240365.1;refGene_gn=RYR2;refGene_tx=XM_023240366.1;refGene_gn=RYR2;refGene_tx=XM_023240367.1;refGene_gn=RYR2;refGene_tx=XM_023240368.1
D2 11594220 11594335
refGene_gn=RYR2;refGene_tx=XM_023240358.1;refGene_gn=RYR2;refGene_tx=XM_023240359.1;refGene_gn=RYR2;refGene_tx=XM_023240360.1;refGene_gn=RYR2;refGene_tx=XM_023240361.1;refGene_gn=RYR2;refGene_tx=XM_023240362.1;refGene_gn=RYR2;refGene_tx=XM_023240363.1;refGene_gn=RYR2;refGene_tx=XM_023240364.1;refGene_gn=RYR2;refGene_tx=XM_023240365.1;refGene_gn=RYR2;refGene_tx=XM_023240366.1;refGene_gn=RYR2;refGene_tx=XM_023240367.1;refGene_gn=RYR2;refGene_tx=XM_023240368.1
D2 11595902 11596255
refGene_gn=RYR2;refGene_tx=XM_023240358.1;refGene_gn=RYR2;refGene_tx=XM_023240359.1;refGene_gn=RYR2;refGene_tx=XM_023240360.1;refGene_gn=RYR2;refGene_tx=XM_023240361.1;refGene_gn=RYR2;refGene_tx=XM_023240362.1;refGene_gn=RYR2;refGene_tx=XM_023240363.1;refGene_gn=RYR2;refGene_tx=XM_023240364.1;refGene_gn=RYR2;refGene_tx=XM_023240365.1;refGene_gn=RYR2;refGene_tx=XM_023240366.1;refGene_gn=RYR2;refGene_tx=XM_023240367.1;refGene_gn=RYR2;refGene_tx=XM_023240368.1
D2 11597247 11597456
refGene_gn=RYR2;refGene_tx=XM_023240358.1;refGene_gn=RYR2;refGene_tx=XM_023240359.1;refGene_gn=RYR2;refGene_tx=XM_023240360.1;refGene_gn=RYR2;refGene_tx=XM_023240361.1;refGene_gn=RYR2;refGene_tx=XM_023240362.1;refGene_gn=RYR2;refGene_tx=XM_023240363.1;refGene_gn=RYR2;refGene_tx=XM_023240364.1;refGene_gn=RYR2;refGene_tx=XM_023240365.1;refGene_gn=RYR2;refGene_tx=XM_023240366.1;refGene_gn=RYR2;refGene_tx=XM_023240367.1;refGene_gn=RYR2;refGene_tx=XM_023240368.1
D2 11618196 11618371

refGene_gn=RYR2;refGene_tx=XM_023240358.1;refGene_gn=RYR2;refGene_tx=XM_023240359.1;refGene_gn=RYR2;refGene_tx=XM_023240360.1;refGene_gn=RYR2;refGene_tx=XM_023240361.1;refGene_gn=RYR2;refGene_tx=XM_023240362.1;refGene_gn=RYR2;refGene_tx=XM_023240363.1;refGene_gn=RYR2;refGene_tx=XM_023240364.1;refGene_gn=RYR2;refGene_tx=XM_023240365.1;refGene_gn=RYR2;refGene_tx=XM_023240366.1;refGene_gn=RYR2;refGene_tx=XM_023240367.1;refGene_gn=RYR2;refGene_tx=XM_023240368.1
D2 11620489 11620698
refGene_gn=RYR2;refGene_tx=XM_023240358.1;refGene_gn=RYR2;refGene_tx=XM_023240359.1;refGene_gn=RYR2;refGene_tx=XM_023240360.1;refGene_gn=RYR2;refGene_tx=XM_023240361.1;refGene_gn=RYR2;refGene_tx=XM_023240362.1;refGene_gn=RYR2;refGene_tx=XM_023240363.1;refGene_gn=RYR2;refGene_tx=XM_023240364.1;refGene_gn=RYR2;refGene_tx=XM_023240365.1;refGene_gn=RYR2;refGene_tx=XM_023240366.1;refGene_gn=RYR2;refGene_tx=XM_023240367.1;refGene_gn=RYR2;refGene_tx=XM_023240368.1
D2 11637535 11637683
refGene_gn=RYR2;refGene_tx=XM_023240358.1;refGene_gn=RYR2;refGene_tx=XM_023240359.1;refGene_gn=RYR2;refGene_tx=XM_023240360.1;refGene_gn=RYR2;refGene_tx=XM_023240361.1;refGene_gn=RYR2;refGene_tx=XM_023240362.1;refGene_gn=RYR2;refGene_tx=XM_023240363.1;refGene_gn=RYR2;refGene_tx=XM_023240364.1;refGene_gn=RYR2;refGene_tx=XM_023240365.1;refGene_gn=RYR2;refGene_tx=XM_023240366.1;refGene_gn=RYR2;refGene_tx=XM_023240367.1;refGene_gn=RYR2;refGene_tx=XM_023240368.1
D2 11640807 11640967
refGene_gn=RYR2;refGene_tx=XM_023240358.1;refGene_gn=RYR2;refGene_tx=XM_023240359.1;refGene_gn=RYR2;refGene_tx=XM_023240360.1;refGene_gn=RYR2;refGene_tx=XM_023240361.1;refGene_gn=RYR2;refGene_tx=XM_023240362.1;refGene_gn=RYR2;refGene_tx=XM_023240363.1;refGene_gn=RYR2;refGene_tx=XM_023240364.1;refGene_gn=RYR2;refGene_tx=XM_023240365.1;refGene_gn=RYR2;refGene_tx=XM_023240366.1;refGene_gn=RYR2;refGene_tx=XM_023240367.1;refGene_gn=RYR2;refGene_tx=XM_023240368.1
D2 11656085 11656169
refGene_gn=RYR2;refGene_tx=XM_023240358.1;refGene_gn=RYR2;refGene_tx=XM_023240359.1;refGene_gn=RYR2;refGene_tx=XM_023240360.1;refGene_gn=RYR2;refGene_tx=XM_023240361.1;refGene_gn=RYR2;refGene_tx=XM_023240362.1;refGene_gn=RYR2;refGene_tx=XM_023240363.1;refGene_gn=RYR2;refGene_tx=XM_023240364.1;refGene_gn=RYR2;refGene_tx=XM_023240365.1;refGene_gn=RYR2;refGene_tx=XM_023240366.1;refGene_gn=RYR2;refGene_tx=XM_023240367.1;refGene_gn=RYR2;refGene_tx=XM_023240368.1
D2 11668080 11668184
refGene_gn=RYR2;refGene_tx=XM_023240358.1;refGene_gn=RYR2;refGene_tx=XM_023240359.1;refGene_gn=RYR2;refGene_tx=XM_023240360.1;refGene_gn=RYR2;refGene_tx=XM_023240361.1;refGene_gn=RYR2;refGene_tx=XM_023240362.1;refGene_gn=RYR2;refGene_tx=XM_023240363.1;refGene_gn=RYR2;refGene_tx=XM_023240364.1;refGene_gn=RYR2;refGene_tx=XM_023240365.1;refGene_gn=RYR2;refGene_tx=XM_023240366.1;refGene_gn=RYR2;refGene_tx=XM_023240367.1;refGene_gn=RYR2;refGene_tx=XM_023240368.1
D2 11671147 11671252
refGene_gn=RYR2;refGene_tx=XM_023240358.1;refGene_gn=RYR2;refGene_tx=XM_023240359.1;refGene_gn=RYR2;refGene_tx=XM_023240360.1;refGene_gn=RYR

2;refGene_tx=XM_023240361.1;refGene_gn=RYR2;refGene_tx=XM_023240362.1;
refGene_gn=RYR2;refGene_tx=XM_023240363.1;refGene_gn=RYR2;refGene_tx=X
M_023240364.1;refGene_gn=RYR2;refGene_tx=XM_023240365.1;refGene_gn=RYR
2;refGene_tx=XM_023240366.1;refGene_gn=RYR2;refGene_tx=XM_023240367.1;
refGene_gn=RYR2;refGene_tx=XM_023240368.1
D2 11673332 11673549
refGene_gn=RYR2;refGene_tx=XM_023240358.1;refGene_gn=RYR2;refGene_tx=X
M_023240359.1;refGene_gn=RYR2;refGene_tx=XM_023240360.1;refGene_gn=RYR
2;refGene_tx=XM_023240361.1;refGene_gn=RYR2;refGene_tx=XM_023240362.1;
refGene_gn=RYR2;refGene_tx=XM_023240363.1;refGene_gn=RYR2;refGene_tx=X
M_023240364.1;refGene_gn=RYR2;refGene_tx=XM_023240365.1;refGene_gn=RYR
2;refGene_tx=XM_023240366.1;refGene_gn=RYR2;refGene_tx=XM_023240367.1;
refGene_gn=RYR2;refGene_tx=XM_023240368.1
D2 11675483 11675676
refGene_gn=RYR2;refGene_tx=XM_023240358.1;refGene_gn=RYR2;refGene_tx=X
M_023240359.1;refGene_gn=RYR2;refGene_tx=XM_023240360.1;refGene_gn=RYR
2;refGene_tx=XM_023240361.1;refGene_gn=RYR2;refGene_tx=XM_023240362.1;
refGene_gn=RYR2;refGene_tx=XM_023240363.1;refGene_gn=RYR2;refGene_tx=X
M_023240364.1;refGene_gn=RYR2;refGene_tx=XM_023240365.1;refGene_gn=RYR
2;refGene_tx=XM_023240366.1;refGene_gn=RYR2;refGene_tx=XM_023240367.1;
refGene_gn=RYR2;refGene_tx=XM_023240368.1
D2 11681152 11681394
refGene_gn=RYR2;refGene_tx=XM_023240358.1;refGene_gn=RYR2;refGene_tx=X
M_023240359.1;refGene_gn=RYR2;refGene_tx=XM_023240360.1;refGene_gn=RYR
2;refGene_tx=XM_023240361.1;refGene_gn=RYR2;refGene_tx=XM_023240362.1;
refGene_gn=RYR2;refGene_tx=XM_023240363.1;refGene_gn=RYR2;refGene_tx=X
M_023240364.1;refGene_gn=RYR2;refGene_tx=XM_023240365.1;refGene_gn=RYR
2;refGene_tx=XM_023240366.1;refGene_gn=RYR2;refGene_tx=XM_023240367.1;
refGene_gn=RYR2;refGene_tx=XM_023240368.1
D2 11682860 11682994
refGene_gn=RYR2;refGene_tx=XM_023240358.1;refGene_gn=RYR2;refGene_tx=X
M_023240359.1;refGene_gn=RYR2;refGene_tx=XM_023240360.1;refGene_gn=RYR
2;refGene_tx=XM_023240361.1;refGene_gn=RYR2;refGene_tx=XM_023240362.1;
refGene_gn=RYR2;refGene_tx=XM_023240363.1;refGene_gn=RYR2;refGene_tx=X
M_023240364.1;refGene_gn=RYR2;refGene_tx=XM_023240365.1;refGene_gn=RYR
2;refGene_tx=XM_023240366.1;refGene_gn=RYR2;refGene_tx=XM_023240367.1;
refGene_gn=RYR2;refGene_tx=XM_023240368.1
D2 11683337 11683456
refGene_gn=RYR2;refGene_tx=XM_023240358.1;refGene_gn=RYR2;refGene_tx=X
M_023240359.1;refGene_gn=RYR2;refGene_tx=XM_023240360.1;refGene_gn=RYR
2;refGene_tx=XM_023240361.1;refGene_gn=RYR2;refGene_tx=XM_023240362.1;
refGene_gn=RYR2;refGene_tx=XM_023240363.1;refGene_gn=RYR2;refGene_tx=X
M_023240364.1;refGene_gn=RYR2;refGene_tx=XM_023240365.1;refGene_gn=RYR
2;refGene_tx=XM_023240366.1;refGene_gn=RYR2;refGene_tx=XM_023240367.1;
refGene_gn=RYR2;refGene_tx=XM_023240368.1
D2 11707787 11707883
refGene_gn=RYR2;refGene_tx=XM_023240358.1;refGene_gn=RYR2;refGene_tx=X
M_023240359.1;refGene_gn=RYR2;refGene_tx=XM_023240360.1;refGene_gn=RYR
2;refGene_tx=XM_023240361.1;refGene_gn=RYR2;refGene_tx=XM_023240362.1;
refGene_gn=RYR2;refGene_tx=XM_023240363.1;refGene_gn=RYR2;refGene_tx=X

M_023240364.1;refGene_gn=RYR2;refGene_tx=XM_023240365.1;refGene_gn=RYR2;refGene_tx=XM_023240366.1;refGene_gn=RYR2;refGene_tx=XM_023240367.1;refGene_gn=RYR2;refGene_tx=XM_023240368.1
D2 11724218 11724354
refGene_gn=RYR2;refGene_tx=XM_023240358.1;refGene_gn=RYR2;refGene_tx=XM_023240359.1;refGene_gn=RYR2;refGene_tx=XM_023240360.1;refGene_gn=RYR2;refGene_tx=XM_023240361.1;refGene_gn=RYR2;refGene_tx=XM_023240362.1;refGene_gn=RYR2;refGene_tx=XM_023240363.1;refGene_gn=RYR2;refGene_tx=XM_023240364.1;refGene_gn=RYR2;refGene_tx=XM_023240365.1;refGene_gn=RYR2;refGene_tx=XM_023240366.1;refGene_gn=RYR2;refGene_tx=XM_023240367.1;refGene_gn=RYR2;refGene_tx=XM_023240368.1
D2 11727548 11727732
refGene_gn=RYR2;refGene_tx=XM_023240358.1;refGene_gn=RYR2;refGene_tx=XM_023240359.1;refGene_gn=RYR2;refGene_tx=XM_023240360.1;refGene_gn=RYR2;refGene_tx=XM_023240361.1;refGene_gn=RYR2;refGene_tx=XM_023240362.1;refGene_gn=RYR2;refGene_tx=XM_023240363.1;refGene_gn=RYR2;refGene_tx=XM_023240364.1;refGene_gn=RYR2;refGene_tx=XM_023240365.1;refGene_gn=RYR2;refGene_tx=XM_023240366.1;refGene_gn=RYR2;refGene_tx=XM_023240367.1;refGene_gn=RYR2;refGene_tx=XM_023240368.1
D2 11732723 11732845
refGene_gn=RYR2;refGene_tx=XM_023240358.1;refGene_gn=RYR2;refGene_tx=XM_023240359.1;refGene_gn=RYR2;refGene_tx=XM_023240360.1;refGene_gn=RYR2;refGene_tx=XM_023240361.1;refGene_gn=RYR2;refGene_tx=XM_023240362.1;refGene_gn=RYR2;refGene_tx=XM_023240363.1;refGene_gn=RYR2;refGene_tx=XM_023240364.1;refGene_gn=RYR2;refGene_tx=XM_023240365.1;refGene_gn=RYR2;refGene_tx=XM_023240366.1;refGene_gn=RYR2;refGene_tx=XM_023240367.1;refGene_gn=RYR2;refGene_tx=XM_023240368.1
D2 11736482 11736647
refGene_gn=RYR2;refGene_tx=XM_023240358.1;refGene_gn=RYR2;refGene_tx=XM_023240359.1;refGene_gn=RYR2;refGene_tx=XM_023240360.1;refGene_gn=RYR2;refGene_tx=XM_023240361.1;refGene_gn=RYR2;refGene_tx=XM_023240362.1;refGene_gn=RYR2;refGene_tx=XM_023240363.1;refGene_gn=RYR2;refGene_tx=XM_023240364.1;refGene_gn=RYR2;refGene_tx=XM_023240365.1;refGene_gn=RYR2;refGene_tx=XM_023240366.1;refGene_gn=RYR2;refGene_tx=XM_023240367.1;refGene_gn=RYR2;refGene_tx=XM_023240368.1
D2 11746388 11746545
refGene_gn=RYR2;refGene_tx=XM_023240358.1;refGene_gn=RYR2;refGene_tx=XM_023240359.1;refGene_gn=RYR2;refGene_tx=XM_023240360.1;refGene_gn=RYR2;refGene_tx=XM_023240361.1;refGene_gn=RYR2;refGene_tx=XM_023240362.1;refGene_gn=RYR2;refGene_tx=XM_023240363.1;refGene_gn=RYR2;refGene_tx=XM_023240364.1;refGene_gn=RYR2;refGene_tx=XM_023240365.1;refGene_gn=RYR2;refGene_tx=XM_023240366.1;refGene_gn=RYR2;refGene_tx=XM_023240367.1;refGene_gn=RYR2;refGene_tx=XM_023240368.1
D2 11752861 11752936
refGene_gn=RYR2;refGene_tx=XM_023240358.1;refGene_gn=RYR2;refGene_tx=XM_023240359.1;refGene_gn=RYR2;refGene_tx=XM_023240360.1;refGene_gn=RYR2;refGene_tx=XM_023240361.1;refGene_gn=RYR2;refGene_tx=XM_023240362.1;refGene_gn=RYR2;refGene_tx=XM_023240363.1;refGene_gn=RYR2;refGene_tx=XM_023240364.1;refGene_gn=RYR2;refGene_tx=XM_023240365.1;refGene_gn=RYR2;refGene_tx=XM_023240366.1;refGene_gn=RYR2;refGene_tx=XM_023240367.1;

refGene_gn=RYR2;refGene_tx=XM_023240368.1
D2	11784203 11784300
refGene_gn=RYR2;refGene_tx=XM_023240358.1;refGene_gn=RYR2;refGene_tx=XM_023240359.1;refGene_gn=RYR2;refGene_tx=XM_023240360.1;refGene_gn=RYR2;refGene_tx=XM_023240361.1;refGene_gn=RYR2;refGene_tx=XM_023240362.1;refGene_gn=RYR2;refGene_tx=XM_023240363.1;refGene_gn=RYR2;refGene_tx=XM_023240364.1;refGene_gn=RYR2;refGene_tx=XM_023240365.1;refGene_gn=RYR2;refGene_tx=XM_023240366.1;refGene_gn=RYR2;refGene_tx=XM_023240367.1;refGene_gn=RYR2;refGene_tx=XM_023240368.1
D2	11785034 11785134
refGene_gn=RYR2;refGene_tx=XM_023240358.1;refGene_gn=RYR2;refGene_tx=XM_023240359.1;refGene_gn=RYR2;refGene_tx=XM_023240360.1;refGene_gn=RYR2;refGene_tx=XM_023240361.1;refGene_gn=RYR2;refGene_tx=XM_023240362.1;refGene_gn=RYR2;refGene_tx=XM_023240363.1;refGene_gn=RYR2;refGene_tx=XM_023240364.1;refGene_gn=RYR2;refGene_tx=XM_023240365.1;refGene_gn=RYR2;refGene_tx=XM_023240366.1;refGene_gn=RYR2;refGene_tx=XM_023240367.1;refGene_gn=RYR2;refGene_tx=XM_023240368.1
D2	11793001 11793114
refGene_gn=RYR2;refGene_tx=XM_023240358.1;refGene_gn=RYR2;refGene_tx=XM_023240359.1;refGene_gn=RYR2;refGene_tx=XM_023240360.1;refGene_gn=RYR2;refGene_tx=XM_023240361.1;refGene_gn=RYR2;refGene_tx=XM_023240362.1;refGene_gn=RYR2;refGene_tx=XM_023240363.1;refGene_gn=RYR2;refGene_tx=XM_023240364.1;refGene_gn=RYR2;refGene_tx=XM_023240365.1;refGene_gn=RYR2;refGene_tx=XM_023240366.1;refGene_gn=RYR2;refGene_tx=XM_023240367.1;refGene_gn=RYR2;refGene_tx=XM_023240368.1
D2	11796363 11796442
refGene_gn=RYR2;refGene_tx=XM_023240358.1;refGene_gn=RYR2;refGene_tx=XM_023240359.1;refGene_gn=RYR2;refGene_tx=XM_023240360.1;refGene_gn=RYR2;refGene_tx=XM_023240361.1;refGene_gn=RYR2;refGene_tx=XM_023240362.1;refGene_gn=RYR2;refGene_tx=XM_023240363.1;refGene_gn=RYR2;refGene_tx=XM_023240364.1;refGene_gn=RYR2;refGene_tx=XM_023240365.1;refGene_gn=RYR2;refGene_tx=XM_023240366.1;refGene_gn=RYR2;refGene_tx=XM_023240367.1;refGene_gn=RYR2;refGene_tx=XM_023240368.1
D2	11799752 11799827
refGene_gn=RYR2;refGene_tx=XM_023240358.1;refGene_gn=RYR2;refGene_tx=XM_023240359.1;refGene_gn=RYR2;refGene_tx=XM_023240360.1;refGene_gn=RYR2;refGene_tx=XM_023240361.1;refGene_gn=RYR2;refGene_tx=XM_023240362.1;refGene_gn=RYR2;refGene_tx=XM_023240363.1;refGene_gn=RYR2;refGene_tx=XM_023240364.1;refGene_gn=RYR2;refGene_tx=XM_023240365.1;refGene_gn=RYR2;refGene_tx=XM_023240366.1;refGene_gn=RYR2;refGene_tx=XM_023240367.1;refGene_gn=RYR2;refGene_tx=XM_023240368.1
D2	11807094 11807109
refGene_gn=RYR2;refGene_tx=XM_023240358.1;refGene_gn=RYR2;refGene_tx=XM_023240359.1;refGene_gn=RYR2;refGene_tx=XM_023240360.1;refGene_gn=RYR2;refGene_tx=XM_023240361.1;refGene_gn=RYR2;refGene_tx=XM_023240362.1;refGene_gn=RYR2;refGene_tx=XM_023240363.1;refGene_gn=RYR2;refGene_tx=XM_023240364.1;refGene_gn=RYR2;refGene_tx=XM_023240365.1;refGene_gn=RYR2;refGene_tx=XM_023240366.1;refGene_gn=RYR2;refGene_tx=XM_023240367.1;refGene_gn=RYR2;refGene_tx=XM_023240368.1
D2	11816557 11816578

refGene_gn=RYR2;refGene_tx=XM_023240358.1;refGene_gn=RYR2;refGene_tx=XM_023240359.1;refGene_gn=RYR2;refGene_tx=XM_023240361.1;refGene_gn=RYR2;refGene_tx=XM_023240362.1;refGene_gn=RYR2;refGene_tx=XM_023240363.1;refGene_gn=RYR2;refGene_tx=XM_023240364.1;refGene_gn=RYR2;refGene_tx=XM_023240365.1;refGene_gn=RYR2;refGene_tx=XM_023240366.1
D2 11840973 11841078
refGene_gn=RYR2;refGene_tx=XM_023240358.1;refGene_gn=RYR2;refGene_tx=XM_023240359.1;refGene_gn=RYR2;refGene_tx=XM_023240360.1;refGene_gn=RYR2;refGene_tx=XM_023240361.1;refGene_gn=RYR2;refGene_tx=XM_023240362.1;refGene_gn=RYR2;refGene_tx=XM_023240363.1;refGene_gn=RYR2;refGene_tx=XM_023240364.1;refGene_gn=RYR2;refGene_tx=XM_023240365.1;refGene_gn=RYR2;refGene_tx=XM_023240366.1;refGene_gn=RYR2;refGene_tx=XM_023240367.1;refGene_gn=RYR2;refGene_tx=XM_023240368.1
D2 11902846 11902966
refGene_gn=RYR2;refGene_tx=XM_023240358.1;refGene_gn=RYR2;refGene_tx=XM_023240359.1;refGene_gn=RYR2;refGene_tx=XM_023240360.1;refGene_gn=RYR2;refGene_tx=XM_023240361.1;refGene_gn=RYR2;refGene_tx=XM_023240362.1;refGene_gn=RYR2;refGene_tx=XM_023240363.1;refGene_gn=RYR2;refGene_tx=XM_023240364.1;refGene_gn=RYR2;refGene_tx=XM_023240365.1;refGene_gn=RYR2;refGene_tx=XM_023240366.1;refGene_gn=RYR2;refGene_tx=XM_023240367.1;refGene_gn=RYR2;refGene_tx=XM_023240368.1
D2 12128059 12128107
refGene_gn=RYR2;refGene_tx=XM_023240358.1;refGene_gn=RYR2;refGene_tx=XM_023240359.1;refGene_gn=RYR2;refGene_tx=XM_023240360.1;refGene_gn=RYR2;refGene_tx=XM_023240361.1;refGene_gn=RYR2;refGene_tx=XM_023240362.1;refGene_gn=RYR2;refGene_tx=XM_023240363.1;refGene_gn=RYR2;refGene_tx=XM_023240364.1;refGene_gn=RYR2;refGene_tx=XM_023240365.1;refGene_gn=RYR2;refGene_tx=XM_023240366.1;refGene_gn=RYR2;refGene_tx=XM_023240367.1;refGene_gn=RYR2;refGene_tx=XM_023240368.1
D2 12244854 12244941 refGene_gn=MTR;refGene_tx=XM_003993893.5
D2 12245387 12245500 refGene_gn=MTR;refGene_tx=XM_003993893.5
D2 12246781 12246974 refGene_gn=MTR;refGene_tx=XM_003993893.5
D2 12247535 12247736 refGene_gn=MTR;refGene_tx=XM_003993893.5
D2 12250491 12250688 refGene_gn=MTR;refGene_tx=XM_003993893.5
D2 12251949 12252105 refGene_gn=MTR;refGene_tx=XM_003993893.5
D2 12254829 12254905 refGene_gn=MTR;refGene_tx=XM_003993893.5
D2 12255969 12256068 refGene_gn=MTR;refGene_tx=XM_003993893.5
D2 12261151 12261233 refGene_gn=MTR;refGene_tx=XM_003993893.5
D2 12268638 12268759 refGene_gn=MTR;refGene_tx=XM_003993893.5
D2 12269344 12269412 refGene_gn=MTR;refGene_tx=XM_003993893.5
D2 12277481 12277582 refGene_gn=MTR;refGene_tx=XM_003993893.5
D2 12278662 12278770 refGene_gn=MTR;refGene_tx=XM_003993893.5
D2 12280747 12280846 refGene_gn=MTR;refGene_tx=XM_003993893.5
D2 12280847 12280901 refGene_gn=MTR;refGene_tx=XM_003993893.5
D2 12281975 12282065 refGene_gn=MTR;refGene_tx=XM_003993893.5
D2 12287773 12287914 refGene_gn=MTR;refGene_tx=XM_003993893.5
D2 12288395 12288512 refGene_gn=MTR;refGene_tx=XM_003993893.5
D2 12290859 12291039 refGene_gn=MTR;refGene_tx=XM_003993893.5
D2 12303174 12303360 refGene_gn=MTR;refGene_tx=XM_003993893.5
D2 12305862 12306003 refGene_gn=MTR;refGene_tx=XM_003993893.5

D2 12309436 12309549 refGene_gn=MTR;refGene_tx=XM_003993893.5
D2 12312257 12312337 refGene_gn=MTR;refGene_tx=XM_003993893.5
D2 12315450 12315518 refGene_gn=MTR;refGene_tx=XM_003993893.5
D2 12317233 12317295 refGene_gn=MTR;refGene_tx=XM_003993893.5
D2 12318384 12318485 refGene_gn=MTR;refGene_tx=XM_003993893.5
D2 12330673 12330768 refGene_gn=MTR;refGene_tx=XM_003993893.5
D2 12331515 12331575 refGene_gn=MTR;refGene_tx=XM_003993893.5
D2 12333540 12333647 refGene_gn=MTR;refGene_tx=XM_003993893.5
D2 12335090 12335183 refGene_gn=MTR;refGene_tx=XM_003993893.5
D2 12336937 12337007 refGene_gn=MTR;refGene_tx=XM_003993893.5
D2 12339449 12339539 refGene_gn=MTR;refGene_tx=XM_003993893.5
D2 12343395 12343610 refGene_gn=MTR;refGene_tx=XM_003993893.5
D2 12346747 12346781 refGene_gn=MTR;refGene_tx=XM_003993893.5
D2 12369478 12369637
refGene_gn=ACTN2;refGene_tx=XM_006937742.4;refGene_gn=ACTN2;refGene_tx
=XM_023240369.1
D2 12371293 12371452
refGene_gn=ACTN2;refGene_tx=XM_006937742.4;refGene_gn=ACTN2;refGene_tx
=XM_023240369.1
D2 12372551 12372617
refGene_gn=ACTN2;refGene_tx=XM_006937742.4;refGene_gn=ACTN2;refGene_tx
=XM_023240369.1
D2 12376330 12376477
refGene_gn=ACTN2;refGene_tx=XM_006937742.4;refGene_gn=ACTN2;refGene_tx
=XM_023240369.1
D2 12377118 12377298
refGene_gn=ACTN2;refGene_tx=XM_006937742.4;refGene_gn=ACTN2;refGene_tx
=XM_023240369.1
D2 12378425 12378560
refGene_gn=ACTN2;refGene_tx=XM_006937742.4;refGene_gn=ACTN2;refGene_tx
=XM_023240369.1
D2 12380365 12380548
refGene_gn=ACTN2;refGene_tx=XM_006937742.4;refGene_gn=ACTN2;refGene_tx
=XM_023240369.1
D2 12382720 12382861
refGene_gn=ACTN2;refGene_tx=XM_006937742.4;refGene_gn=ACTN2;refGene_tx
=XM_023240369.1
D2 12384536 12384645
refGene_gn=ACTN2;refGene_tx=XM_006937742.4;refGene_gn=ACTN2;refGene_tx
=XM_023240369.1
D2 12388461 12388612
refGene_gn=ACTN2;refGene_tx=XM_006937742.4;refGene_gn=ACTN2;refGene_tx
=XM_023240369.1
D2 12390566 12390714
refGene_gn=ACTN2;refGene_tx=XM_006937742.4;refGene_gn=ACTN2;refGene_tx
=XM_023240369.1
D2 12393422 12393653
refGene_gn=ACTN2;refGene_tx=XM_006937742.4;refGene_gn=ACTN2;refGene_tx
=XM_023240369.1
D2 12395412 12395505

refGene_gn=ACTN2;refGene_tx=XM_006937742.4;refGene_gn=ACTN2;refGene_tx=XM_023240369.1
D2	12397161	12397247	refGene_gn=ACTN2;refGene_tx=XM_006937742.4
D2	12398380	12398466	refGene_gn=ACTN2;refGene_tx=XM_023240369.1
D2	12401531	12401613	refGene_gn=ACTN2;refGene_tx=XM_006937742.4;refGene_gn=ACTN2;refGene_tx=XM_023240369.1
D2	12405409	12405488	refGene_gn=ACTN2;refGene_tx=XM_006937742.4;refGene_gn=ACTN2;refGene_tx=XM_023240369.1
D2	12407245	12407333	refGene_gn=ACTN2;refGene_tx=XM_006937742.4;refGene_gn=ACTN2;refGene_tx=XM_023240369.1
D2	12411053	12411140	refGene_gn=ACTN2;refGene_tx=XM_006937742.4;refGene_gn=ACTN2;refGene_tx=XM_023240369.1
D2	12411935	12412055	refGene_gn=ACTN2;refGene_tx=XM_006937742.4;refGene_gn=ACTN2;refGene_tx=XM_023240369.1
D2	12412981	12413096	refGene_gn=ACTN2;refGene_tx=XM_006937742.4;refGene_gn=ACTN2;refGene_tx=XM_023240369.1
D2	12442355	12442481	refGene_gn=ACTN2;refGene_tx=XM_006937742.4;refGene_gn=ACTN2;refGene_tx=XM_023240369.1
D2	12523717	12523859	refGene_gn=HEATR1;refGene_tx=XM_011287210.3
D2	12524383	12524600	refGene_gn=HEATR1;refGene_tx=XM_011287210.3
D2	12526621	12526763	refGene_gn=HEATR1;refGene_tx=XM_011287210.3
D2	12528244	12528346	refGene_gn=HEATR1;refGene_tx=XM_011287210.3
D2	12528773	12528914	refGene_gn=HEATR1;refGene_tx=XM_011287210.3
D2	12530143	12530355	refGene_gn=HEATR1;refGene_tx=XM_011287210.3
D2	12530510	12530644	refGene_gn=HEATR1;refGene_tx=XM_011287210.3
D2	12532502	12532605	refGene_gn=HEATR1;refGene_tx=XM_011287210.3
D2	12533332	12533443	refGene_gn=HEATR1;refGene_tx=XM_011287210.3
D2	12533752	12533870	refGene_gn=HEATR1;refGene_tx=XM_011287210.3
D2	12534925	12535033	refGene_gn=HEATR1;refGene_tx=XM_011287210.3
D2	12537670	12537766	refGene_gn=HEATR1;refGene_tx=XM_011287210.3
D2	12538200	12538289	refGene_gn=HEATR1;refGene_tx=XM_011287210.3
D2	12539580	12539792	refGene_gn=HEATR1;refGene_tx=XM_011287210.3
D2	12540442	12540564	refGene_gn=HEATR1;refGene_tx=XM_011287210.3
D2	12541357	12541549	refGene_gn=HEATR1;refGene_tx=XM_011287210.3
D2	12543087	12543271	refGene_gn=HEATR1;refGene_tx=XM_011287210.3
D2	12543396	12543533	refGene_gn=HEATR1;refGene_tx=XM_011287210.3
D2	12544254	12544447	refGene_gn=HEATR1;refGene_tx=XM_011287210.3
D2	12549258	12549428	refGene_gn=HEATR1;refGene_tx=XM_011287210.3
D2	12549824	12549983	refGene_gn=HEATR1;refGene_tx=XM_011287210.3
D2	12552185	12552428	refGene_gn=HEATR1;refGene_tx=XM_011287210.3
D2	12552704	12552836	refGene_gn=HEATR1;refGene_tx=XM_011287210.3
D2	12553816	12553920	refGene_gn=HEATR1;refGene_tx=XM_011287210.3
D2	12554090	12554234	refGene_gn=HEATR1;refGene_tx=XM_011287210.3

D2 12554860 12554979 refGene_gn=HEATR1;refGene_tx=XM_011287210.3
D2 12555076 12555198 refGene_gn=HEATR1;refGene_tx=XM_011287210.3
D2 12557153 12557282 refGene_gn=HEATR1;refGene_tx=XM_011287210.3
D2 12558704 12558935 refGene_gn=HEATR1;refGene_tx=XM_011287210.3
D2 12559680 12559807 refGene_gn=HEATR1;refGene_tx=XM_011287210.3
D2 12561567 12561728 refGene_gn=HEATR1;refGene_tx=XM_011287210.3
D2 12565140 12565187 refGene_gn=HEATR1;refGene_tx=XM_011287210.3
D2 12565303 12565427 refGene_gn=HEATR1;refGene_tx=XM_011287210.3
D2 12566026 12566167 refGene_gn=HEATR1;refGene_tx=XM_011287210.3
D2 12566661 12566719 refGene_gn=HEATR1;refGene_tx=XM_011287210.3
D2 12566721 12566947 refGene_gn=HEATR1;refGene_tx=XM_011287210.3
D2 12567898 12568049 refGene_gn=HEATR1;refGene_tx=XM_011287210.3
D2 12568887 12569046 refGene_gn=HEATR1;refGene_tx=XM_011287210.3
D2 12569203 12569338 refGene_gn=HEATR1;refGene_tx=XM_011287210.3
D2 12569481 12569586 refGene_gn=HEATR1;refGene_tx=XM_011287210.3
D2 12569675 12569756 refGene_gn=HEATR1;refGene_tx=XM_011287210.3
D2 12569757 12569845 refGene_gn=HEATR1;refGene_tx=XM_011287210.3
D2 12570350 12570505 refGene_gn=HEATR1;refGene_tx=XM_011287210.3
D2 12571415 12571574 refGene_gn=HEATR1;refGene_tx=XM_011287210.3
D2 12572978 12573087 refGene_gn=HEATR1;refGene_tx=XM_011287210.3
D2 12573890 12573979 refGene_gn=HEATR1;refGene_tx=XM_011287210.3
D2 12576958 12577108
refGene_gn=LGALS8;refGene_tx=XM_003993895.4;refGene_gn=LGALS8;refGene_tx=XM_006937747.3;refGene_gn=LGALS8;refGene_tx=XM_019814271.2;refGene_gn=LGALS8;refGene_tx=XM_019814273.2;refGene_gn=LGALS8;refGene_tx=XM_019814275.2;refGene_gn=LGALS8;refGene_tx=XM_019814276.2;refGene_gn=LGALS8;refGene_tx=XM_019814278.2;refGene_gn=LGALS8;refGene_tx=XM_019814279.2;refGene_gn=LGALS8;refGene_tx=XM_019814280.2;refGene_gn=LGALS8;refGene_tx=XM_019814281.2;refGene_gn=LGALS8;refGene_tx=XM_023240370.1;refGene_gn=LGALS8;refGene_tx=XM_023240371.1
D2 12579929 12580095
refGene_gn=LGALS8;refGene_tx=XM_003993895.4;refGene_gn=LGALS8;refGene_tx=XM_006937747.3;refGene_gn=LGALS8;refGene_tx=XM_019814271.2;refGene_gn=LGALS8;refGene_tx=XM_019814273.2;refGene_gn=LGALS8;refGene_tx=XM_019814275.2;refGene_gn=LGALS8;refGene_tx=XM_019814276.2;refGene_gn=LGALS8;refGene_tx=XM_019814278.2;refGene_gn=LGALS8;refGene_tx=XM_019814279.2;refGene_gn=LGALS8;refGene_tx=XM_019814280.2;refGene_gn=LGALS8;refGene_tx=XM_019814281.2;refGene_gn=LGALS8;refGene_tx=XM_023240370.1;refGene_gn=LGALS8;refGene_tx=XM_023240371.1
D2 12580821 12580910
refGene_gn=LGALS8;refGene_tx=XM_003993895.4;refGene_gn=LGALS8;refGene_tx=XM_006937747.3;refGene_gn=LGALS8;refGene_tx=XM_019814271.2;refGene_gn=LGALS8;refGene_tx=XM_019814273.2;refGene_gn=LGALS8;refGene_tx=XM_019814275.2;refGene_gn=LGALS8;refGene_tx=XM_019814276.2;refGene_gn=LGALS8;refGene_tx=XM_019814278.2;refGene_gn=LGALS8;refGene_tx=XM_019814279.2;refGene_gn=LGALS8;refGene_tx=XM_019814280.2;refGene_gn=LGALS8;refGene_tx=XM_019814281.2;refGene_gn=LGALS8;refGene_tx=XM_023240370.1;refGene_gn=LGALS8;refGene_tx=XM_023240371.1
D2 12581331 12581454
refGene_gn=LGALS8;refGene_tx=XM_003993895.4;refGene_gn=LGALS8;refGene_

tx=XM_006937747.3;refGene_gn=LGALS8;refGene_tx=XM_019814271.2;refGene_gn=LGALS8;refGene_tx=XM_019814273.2;refGene_gn=LGALS8;refGene_tx=XM_019814275.2;refGene_gn=LGALS8;refGene_tx=XM_019814276.2;refGene_gn=LGALS8;refGene_tx=XM_019814278.2;refGene_gn=LGALS8;refGene_tx=XM_019814279.2;refGene_gn=LGALS8;refGene_tx=XM_023240370.1;refGene_gn=LGALS8;refGene_tx=XM_023240371.1
D2 12581582 12581609
refGene_gn=LGALS8;refGene_tx=XM_003993895.4;refGene_gn=LGALS8;refGene_tx=XM_006937747.3;refGene_gn=LGALS8;refGene_tx=XM_019814271.2;refGene_gn=LGALS8;refGene_tx=XM_019814273.2;refGene_gn=LGALS8;refGene_tx=XM_019814275.2;refGene_gn=LGALS8;refGene_tx=XM_019814276.2;refGene_gn=LGALS8;refGene_tx=XM_019814278.2;refGene_gn=LGALS8;refGene_tx=XM_019814279.2;refGene_gn=LGALS8;refGene_tx=XM_019814280.2;refGene_gn=LGALS8;refGene_tx=XM_019814281.2;refGene_gn=LGALS8;refGene_tx=XM_023240370.1;refGene_gn=LGALS8;refGene_tx=XM_023240371.1
D2 12582560 12582617
refGene_gn=LGALS8;refGene_tx=XM_003993895.4;refGene_gn=LGALS8;refGene_tx=XM_006937747.3;refGene_gn=LGALS8;refGene_tx=XM_019814271.2;refGene_gn=LGALS8;refGene_tx=XM_019814273.2;refGene_gn=LGALS8;refGene_tx=XM_019814275.2;refGene_gn=LGALS8;refGene_tx=XM_019814276.2;refGene_gn=LGALS8;refGene_tx=XM_019814278.2;refGene_gn=LGALS8;refGene_tx=XM_019814280.2;refGene_gn=LGALS8;refGene_tx=XM_023240370.1;refGene_gn=LGALS8;refGene_tx=XM_023240371.1
D2 12583175 12583295
refGene_gn=LGALS8;refGene_tx=XM_003993895.4;refGene_gn=LGALS8;refGene_tx=XM_006937747.3;refGene_gn=LGALS8;refGene_tx=XM_019814271.2;refGene_gn=LGALS8;refGene_tx=XM_019814273.2;refGene_gn=LGALS8;refGene_tx=XM_019814275.2;refGene_gn=LGALS8;refGene_tx=XM_019814276.2;refGene_gn=LGALS8;refGene_tx=XM_019814278.2;refGene_gn=LGALS8;refGene_tx=XM_019814279.2;refGene_gn=LGALS8;refGene_tx=XM_019814280.2;refGene_gn=LGALS8;refGene_tx=XM_019814281.2;refGene_gn=LGALS8;refGene_tx=XM_023240370.1;refGene_gn=LGALS8;refGene_tx=XM_023240371.1
D2 12584672 12584883
refGene_gn=LGALS8;refGene_tx=XM_003993895.4;refGene_gn=LGALS8;refGene_tx=XM_006937747.3;refGene_gn=LGALS8;refGene_tx=XM_019814271.2;refGene_gn=LGALS8;refGene_tx=XM_019814273.2;refGene_gn=LGALS8;refGene_tx=XM_019814275.2;refGene_gn=LGALS8;refGene_tx=XM_019814276.2;refGene_gn=LGALS8;refGene_tx=XM_019814278.2;refGene_gn=LGALS8;refGene_tx=XM_019814279.2;refGene_gn=LGALS8;refGene_tx=XM_019814280.2;refGene_gn=LGALS8;refGene_tx=XM_019814281.2;refGene_gn=LGALS8;refGene_tx=XM_023240370.1;refGene_gn=LGALS8;refGene_tx=XM_023240371.1
D2 12585543 12585632
refGene_gn=LGALS8;refGene_tx=XM_003993895.4;refGene_gn=LGALS8;refGene_tx=XM_006937747.3;refGene_gn=LGALS8;refGene_tx=XM_019814271.2;refGene_gn=LGALS8;refGene_tx=XM_019814273.2;refGene_gn=LGALS8;refGene_tx=XM_019814275.2;refGene_gn=LGALS8;refGene_tx=XM_019814276.2;refGene_gn=LGALS8;refGene_tx=XM_019814278.2;refGene_gn=LGALS8;refGene_tx=XM_019814279.2;refGene_gn=LGALS8;refGene_tx=XM_019814280.2;refGene_gn=LGALS8;refGene_tx=XM_019814281.2;refGene_gn=LGALS8;refGene_tx=XM_023240370.1;refGene_gn=LGALS8;refGene_tx=XM_023240371.1

D2 12593543 12593588
refGene_gn=LGALS8;refGene_tx=XM_003993895.4;refGene_gn=LGALS8;refGene_tx=XM_006937747.3;refGene_gn=LGALS8;refGene_tx=XM_019814271.2;refGene_gn=LGALS8;refGene_tx=XM_019814273.2;refGene_gn=LGALS8;refGene_tx=XM_019814275.2;refGene_gn=LGALS8;refGene_tx=XM_019814276.2;refGene_gn=LGALS8;refGene_tx=XM_019814278.2;refGene_gn=LGALS8;refGene_tx=XM_019814279.2;refGene_gn=LGALS8;refGene_tx=XM_019814280.2;refGene_gn=LGALS8;refGene_tx=XM_019814281.2;refGene_gn=LGALS8;refGene_tx=XM_023240370.1;refGene_gn=LGALS8;refGene_tx=XM_023240371.1
D2 12642600 12642983
refGene_gn=EDARADD;refGene_tx=XM_006937751.4;refGene_gn=EDARADD;refGene_tx=XM_019814282.2;refGene_gn=EDARADD;refGene_tx=XM_019814283.2;refGene_gn=EDARADD;refGene_tx=XM_019814284.2;refGene_gn=EDARADD;refGene_tx=XM_019814285.2;refGene_gn=EDARADD;refGene_tx=XM_023240372.1
D2 12654295 12654313
refGene_gn=EDARADD;refGene_tx=XM_019814282.2;refGene_gn=EDARADD;refGene_tx=XM_019814283.2;refGene_gn=EDARADD;refGene_tx=XM_019814284.2;refGene_gn=EDARADD;refGene_tx=XM_019814285.2;refGene_gn=EDARADD;refGene_tx=XM_023240372.1
D2 12657369 12657415
refGene_gn=EDARADD;refGene_tx=XM_006937751.4;refGene_gn=EDARADD;refGene_tx=XM_019814282.2;refGene_gn=EDARADD;refGene_tx=XM_019814283.2;refGene_gn=EDARADD;refGene_tx=XM_019814284.2;refGene_gn=EDARADD;refGene_tx=XM_019814285.2;refGene_gn=EDARADD;refGene_tx=XM_023240372.1
D2 12683226 12683285
refGene_gn=EDARADD;refGene_tx=XM_006937751.4;refGene_gn=EDARADD;refGene_tx=XM_019814282.2;refGene_gn=EDARADD;refGene_tx=XM_019814283.2;refGene_gn=EDARADD;refGene_tx=XM_019814284.2;refGene_gn=EDARADD;refGene_tx=XM_019814285.2;refGene_gn=EDARADD;refGene_tx=XM_023240372.1
D2 12696916 12696956
refGene_gn=EDARADD;refGene_tx=XM_006937751.4;refGene_gn=EDARADD;refGene_tx=XM_019814282.2;refGene_gn=EDARADD;refGene_tx=XM_019814283.2;refGene_gn=EDARADD;refGene_tx=XM_019814284.2;refGene_gn=EDARADD;refGene_tx=XM_019814285.2;refGene_gn=EDARADD;refGene_tx=XM_023240372.1
D2 12700942 12701001
refGene_gn=EDARADD;refGene_tx=XM_006937751.4;refGene_gn=EDARADD;refGene_tx=XM_019814282.2;refGene_gn=EDARADD;refGene_tx=XM_019814283.2;refGene_gn=EDARADD;refGene_tx=XM_019814284.2;refGene_gn=EDARADD;refGene_tx=XM_019814285.2;refGene_gn=EDARADD;refGene_tx=XM_023240372.1
D2 12712129 12712160 refGene_gn=EDARADD;refGene_tx=XM_023240372.1
D2 12713473 12713537
refGene_gn=EDARADD;refGene_tx=XM_006937751.4;refGene_gn=EDARADD;refGene_tx=XM_019814282.2
D2 12814127 12814229
refGene_gn=ERO1B;refGene_tx=XM_006937755.4;refGene_gn=ERO1B;refGene_tx=XM_006937756.4
D2 12823786 12823906
refGene_gn=ERO1B;refGene_tx=XM_006937755.4;refGene_gn=ERO1B;refGene_tx=XM_006937756.4
D2 12829856 12829940

refGene_gn=ERO1B;refGene_tx=XM_006937755.4;refGene_gn=ERO1B;refGene_tx=XM_006937756.4
D2 12840142 12840184 refGene_gn=ERO1B;refGene_tx=XM_006937755.4
D2 12842431 12842514
refGene_gn=ERO1B;refGene_tx=XM_006937755.4;refGene_gn=ERO1B;refGene_tx=XM_006937756.4
D2 12847601 12847675
refGene_gn=ERO1B;refGene_tx=XM_006937755.4;refGene_gn=ERO1B;refGene_tx=XM_006937756.4
D2 12852591 12852712
refGene_gn=ERO1B;refGene_tx=XM_006937755.4;refGene_gn=ERO1B;refGene_tx=XM_006937756.4
D2 12852832 12852879
refGene_gn=ERO1B;refGene_tx=XM_006937755.4;refGene_gn=ERO1B;refGene_tx=XM_006937756.4
D2 12856454 12856466
refGene_gn=ERO1B;refGene_tx=XM_006937755.4;refGene_gn=ERO1B;refGene_tx=XM_006937756.4
D2 12858326 12858353
refGene_gn=ERO1B;refGene_tx=XM_006937755.4;refGene_gn=ERO1B;refGene_tx=XM_006937756.4
D2 12862170 12862263
refGene_gn=ERO1B;refGene_tx=XM_006937755.4;refGene_gn=ERO1B;refGene_tx=XM_006937756.4
D2 12862386 12862633
refGene_gn=ERO1B;refGene_tx=XM_006937755.4;refGene_gn=ERO1B;refGene_tx=XM_006937756.4
D2 12863935 12864005
refGene_gn=ERO1B;refGene_tx=XM_006937755.4;refGene_gn=ERO1B;refGene_tx=XM_006937756.4
D2 12866218 12866305
refGene_gn=ERO1B;refGene_tx=XM_006937755.4;refGene_gn=ERO1B;refGene_tx=XM_006937756.4
D2 12867229 12867363
refGene_gn=ERO1B;refGene_tx=XM_006937755.4;refGene_gn=ERO1B;refGene_tx=XM_006937756.4
D2 12867557 12867612 refGene_gn=GPR137B;refGene_tx=XM_019814287.2
D2 12872957 12873014
refGene_gn=ERO1B;refGene_tx=XM_006937755.4;refGene_gn=ERO1B;refGene_tx=XM_006937756.4
D2 12875606 12875778
refGene_gn=ERO1B;refGene_tx=XM_006937755.4;refGene_gn=ERO1B;refGene_tx=XM_006937756.4
D2 12882999 12883120
refGene_gn=GPR137B;refGene_tx=XM_003993898.4;refGene_gn=GPR137B;refGene_tx=XM_019814289.2
D2 12883999 12884009 refGene_gn=GPR137B;refGene_tx=XM_019814288.2
D2 12884599 12884724
refGene_gn=GPR137B;refGene_tx=XM_003993898.4;refGene_gn=GPR137B;refGene_tx=XM_019814287.2;refGene_gn=GPR137B;refGene_tx=XM_019814288.2;refGe

ne_gn=GPR137B;refGene_tx=XM_019814289.2
D2 12893297 12893333 refGene_gn=GPR137B;refGene_tx=XM_019814290.2
D2 12897169 12897298
refGene_gn=GPR137B;refGene_tx=XM_003993898.4;refGene_gn=GPR137B;refGene_tx=XM_019814287.2;refGene_gn=GPR137B;refGene_tx=XM_019814288.2;refGene_gn=GPR137B;refGene_tx=XM_019814290.2
D2 12900008 12900158
refGene_gn=GPR137B;refGene_tx=XM_003993898.4;refGene_gn=GPR137B;refGene_tx=XM_019814287.2;refGene_gn=GPR137B;refGene_tx=XM_019814288.2;refGene_gn=GPR137B;refGene_tx=XM_019814289.2;refGene_gn=GPR137B;refGene_tx=XM_019814290.2
D2 12902947 12903170
refGene_gn=GPR137B;refGene_tx=XM_003993898.4;refGene_gn=GPR137B;refGene_tx=XM_019814287.2;refGene_gn=GPR137B;refGene_tx=XM_019814288.2;refGene_gn=GPR137B;refGene_tx=XM_019814289.2;refGene_gn=GPR137B;refGene_tx=XM_019814290.2
D2 12909051 12909101
refGene_gn=GPR137B;refGene_tx=XM_003993898.4;refGene_gn=GPR137B;refGene_tx=XM_019814287.2;refGene_gn=GPR137B;refGene_tx=XM_019814288.2;refGene_gn=GPR137B;refGene_tx=XM_019814289.2;refGene_gn=GPR137B;refGene_tx=XM_019814290.2
D2 12931044 12931458
refGene_gn=GPR137B;refGene_tx=XM_003993898.4;refGene_gn=GPR137B;refGene_tx=XM_019814287.2;refGene_gn=GPR137B;refGene_tx=XM_019814288.2;refGene_gn=GPR137B;refGene_tx=XM_019814289.2;refGene_gn=GPR137B;refGene_tx=XM_019814290.2
D2 12999343 12999568 refGene_gn=NID1;refGene_tx=XM_003993899.5
D2 13028204 13028504 refGene_gn=NID1;refGene_tx=XM_003993899.5
D2 13030768 13030995 refGene_gn=NID1;refGene_tx=XM_003993899.5
D2 13033485 13033868 refGene_gn=NID1;refGene_tx=XM_003993899.5
D2 13036688 13036838 refGene_gn=NID1;refGene_tx=XM_003993899.5
D2 13038971 13039223 refGene_gn=NID1;refGene_tx=XM_003993899.5
D2 13041634 13041835 refGene_gn=NID1;refGene_tx=XM_003993899.5
D2 13044266 13044512 refGene_gn=NID1;refGene_tx=XM_003993899.5
D2 13045569 13045713 refGene_gn=NID1;refGene_tx=XM_003993899.5
D2 13050209 13050335 refGene_gn=NID1;refGene_tx=XM_003993899.5
D2 13056559 13056709 refGene_gn=NID1;refGene_tx=XM_003993899.5
D2 13058390 13058513 refGene_gn=NID1;refGene_tx=XM_003993899.5
D2 13067871 13068090 refGene_gn=NID1;refGene_tx=XM_003993899.5
D2 13070038 13070211 refGene_gn=NID1;refGene_tx=XM_003993899.5
D2 13073631 13073758 refGene_gn=NID1;refGene_tx=XM_003993899.5
D2 13077213 13077385 refGene_gn=NID1;refGene_tx=XM_003993899.5
D2 13078346 13078504 refGene_gn=NID1;refGene_tx=XM_003993899.5
D2 13079906 13080030 refGene_gn=NID1;refGene_tx=XM_003993899.5
D2 13080760 13080873 refGene_gn=NID1;refGene_tx=XM_003993899.5
D2 13081825 13081947 refGene_gn=NID1;refGene_tx=XM_003993899.5
D2 13199205 13199397 refGene_gn=LYST;refGene_tx=NM_001290242.1
D2 13213559 13213650 refGene_gn=LYST;refGene_tx=NM_001290242.1
D2 13215394 13217474 refGene_gn=LYST;refGene_tx=NM_001290242.1
D2 13218893 13219923 refGene_gn=LYST;refGene_tx=NM_001290242.1

| | | | |
|---|---|---|---|
| D2 | 13221824 | 13221986 | refGene_gn=LYST;refGene_tx=NM_001290242.1 |
| D2 | 13223431 | 13223588 | refGene_gn=LYST;refGene_tx=NM_001290242.1 |
| D2 | 13225804 | 13226031 | refGene_gn=LYST;refGene_tx=NM_001290242.1 |
| D2 | 13226558 | 13226625 | refGene_gn=LYST;refGene_tx=NM_001290242.1 |
| D2 | 13231519 | 13231629 | refGene_gn=LYST;refGene_tx=NM_001290242.1 |
| D2 | 13233722 | 13234149 | refGene_gn=LYST;refGene_tx=NM_001290242.1 |
| D2 | 13236808 | 13236953 | refGene_gn=LYST;refGene_tx=NM_001290242.1 |
| D2 | 13238591 | 13238765 | refGene_gn=LYST;refGene_tx=NM_001290242.1 |
| D2 | 13242299 | 13242460 | refGene_gn=LYST;refGene_tx=NM_001290242.1 |
| D2 | 13243574 | 13243765 | refGene_gn=LYST;refGene_tx=NM_001290242.1 |
| D2 | 13246489 | 13246735 | refGene_gn=LYST;refGene_tx=NM_001290242.1 |
| D2 | 13249137 | 13249311 | refGene_gn=LYST;refGene_tx=NM_001290242.1 |
| D2 | 13250236 | 13250386 | refGene_gn=LYST;refGene_tx=NM_001290242.1 |
| D2 | 13254641 | 13254779 | refGene_gn=LYST;refGene_tx=NM_001290242.1 |
| D2 | 13258674 | 13258873 | refGene_gn=LYST;refGene_tx=NM_001290242.1 |
| D2 | 13261476 | 13261608 | refGene_gn=LYST;refGene_tx=NM_001290242.1 |
| D2 | 13265877 | 13266493 | refGene_gn=LYST;refGene_tx=NM_001290242.1 |
| D2 | 13267208 | 13267386 | refGene_gn=LYST;refGene_tx=NM_001290242.1 |
| D2 | 13269499 | 13269669 | refGene_gn=LYST;refGene_tx=NM_001290242.1 |
| D2 | 13271184 | 13271415 | refGene_gn=LYST;refGene_tx=NM_001290242.1 |
| D2 | 13273740 | 13273907 | refGene_gn=LYST;refGene_tx=NM_001290242.1 |
| D2 | 13274556 | 13274709 | refGene_gn=LYST;refGene_tx=NM_001290242.1 |
| D2 | 13280257 | 13280449 | refGene_gn=LYST;refGene_tx=NM_001290242.1 |
| D2 | 13282627 | 13282806 | refGene_gn=LYST;refGene_tx=NM_001290242.1 |
| D2 | 13284428 | 13284635 | refGene_gn=LYST;refGene_tx=NM_001290242.1 |
| D2 | 13291921 | 13292098 | refGene_gn=LYST;refGene_tx=NM_001290242.1 |
| D2 | 13292858 | 13292935 | refGene_gn=LYST;refGene_tx=NM_001290242.1 |
| D2 | 13293080 | 13293269 | refGene_gn=LYST;refGene_tx=NM_001290242.1 |
| D2 | 13295929 | 13296075 | refGene_gn=LYST;refGene_tx=NM_001290242.1 |
| D2 | 13296161 | 13296258 | refGene_gn=LYST;refGene_tx=NM_001290242.1 |
| D2 | 13297463 | 13297525 | refGene_gn=LYST;refGene_tx=NM_001290242.1 |
| D2 | 13299425 | 13299481 | refGene_gn=LYST;refGene_tx=NM_001290242.1 |
| D2 | 13303318 | 13303471 | refGene_gn=LYST;refGene_tx=NM_001290242.1 |
| D2 | 13306867 | 13307112 | refGene_gn=LYST;refGene_tx=NM_001290242.1 |
| D2 | 13311802 | 13311869 | refGene_gn=LYST;refGene_tx=NM_001290242.1 |
| D2 | 13313549 | 13313706 | refGene_gn=LYST;refGene_tx=NM_001290242.1 |
| D2 | 13316979 | 13317120 | refGene_gn=LYST;refGene_tx=NM_001290242.1 |
| D2 | 13320200 | 13320418 | refGene_gn=LYST;refGene_tx=NM_001290242.1 |
| D2 | 13325127 | 13325358 | refGene_gn=LYST;refGene_tx=NM_001290242.1 |
| D2 | 13327288 | 13327478 | refGene_gn=LYST;refGene_tx=NM_001290242.1 |
| D2 | 13330807 | 13330944 | refGene_gn=LYST;refGene_tx=NM_001290242.1 |
| D2 | 13333855 | 13333954 | refGene_gn=LYST;refGene_tx=NM_001290242.1 |
| D2 | 13339442 | 13339582 | refGene_gn=LYST;refGene_tx=NM_001290242.1 |
| D2 | 13340287 | 13340304 | refGene_gn=LYST;refGene_tx=NM_001290242.1 |
| D2 | 13340305 | 13340386 | refGene_gn=LYST;refGene_tx=NM_001290242.1 |
| D2 | 13345177 | 13345334 | refGene_gn=LYST;refGene_tx=NM_001290242.1 |
| D2 | 13346123 | 13346195 | refGene_gn=LYST;refGene_tx=NM_001290242.1 |
| D2 | 13347079 | 13347218 | refGene_gn=LYST;refGene_tx=NM_001290242.1 |
| D2 | 13380393 | 13380466 | refGene_gn=GNG4;refGene_tx=XM_019814292.2 |
| D2 | 13399314 | 13399411 | refGene_gn=GNG4;refGene_tx=XM_019814292.2 |

D2	13402425	13402543
refGene_gn=GNG4;refGene_tx=XM_019814292.2;refGene_gn=GNG4;refGene_tx=XM_023240374.1
D2	13404325	13404433
refGene_gn=GNG4;refGene_tx=XM_019814292.2;refGene_gn=GNG4;refGene_tx=XM_019814293.2;refGene_gn=GNG4;refGene_tx=XM_023240373.1;refGene_gn=GNG4;refGene_tx=XM_023240374.1
D2	13428179	13428308
refGene_gn=GNG4;refGene_tx=XM_019814292.2;refGene_gn=GNG4;refGene_tx=XM_019814293.2;refGene_gn=GNG4;refGene_tx=XM_023240373.1;refGene_gn=GNG4;refGene_tx=XM_023240374.1
D2	13475938	13476017	refGene_gn=B3GALNT2;refGene_tx=XM_023240380.1
D2	13476699	13476811
refGene_gn=B3GALNT2;refGene_tx=XM_023240378.1;refGene_gn=B3GALNT2;refGene_tx=XM_023240379.1;refGene_gn=B3GALNT2;refGene_tx=XM_023240381.1
D2	13491675	13491823
refGene_gn=B3GALNT2;refGene_tx=XM_023240378.1;refGene_gn=B3GALNT2;refGene_tx=XM_023240379.1;refGene_gn=B3GALNT2;refGene_tx=XM_023240380.1;refGene_gn=B3GALNT2;refGene_tx=XM_023240381.1
D2	13499798	13499899
refGene_gn=B3GALNT2;refGene_tx=XM_023240378.1;refGene_gn=B3GALNT2;refGene_tx=XM_023240379.1;refGene_gn=B3GALNT2;refGene_tx=XM_023240380.1;refGene_gn=B3GALNT2;refGene_tx=XM_023240381.1
D2	13503533	13503727
refGene_gn=B3GALNT2;refGene_tx=XM_023240378.1;refGene_gn=B3GALNT2;refGene_tx=XM_023240379.1;refGene_gn=B3GALNT2;refGene_tx=XM_023240380.1;refGene_gn=B3GALNT2;refGene_tx=XM_023240381.1
D2	13508266	13508362
refGene_gn=B3GALNT2;refGene_tx=XM_023240378.1;refGene_gn=B3GALNT2;refGene_tx=XM_023240379.1;refGene_gn=B3GALNT2;refGene_tx=XM_023240380.1
D2	13518663	13518774
refGene_gn=B3GALNT2;refGene_tx=XM_023240378.1;refGene_gn=B3GALNT2;refGene_tx=XM_023240379.1;refGene_gn=B3GALNT2;refGene_tx=XM_023240380.1;refGene_gn=B3GALNT2;refGene_tx=XM_023240381.1
D2	13522055	13522134
refGene_gn=B3GALNT2;refGene_tx=XM_023240378.1;refGene_gn=B3GALNT2;refGene_tx=XM_023240379.1;refGene_gn=B3GALNT2;refGene_tx=XM_023240380.1;refGene_gn=B3GALNT2;refGene_tx=XM_023240381.1
D2	13527836	13528020
refGene_gn=B3GALNT2;refGene_tx=XM_023240378.1;refGene_gn=B3GALNT2;refGene_tx=XM_023240379.1;refGene_gn=B3GALNT2;refGene_tx=XM_023240380.1;refGene_gn=B3GALNT2;refGene_tx=XM_023240381.1
D2	13531223	13531349
refGene_gn=B3GALNT2;refGene_tx=XM_023240378.1;refGene_gn=B3GALNT2;refGene_tx=XM_023240379.1;refGene_gn=B3GALNT2;refGene_tx=XM_023240380.1;refGene_gn=B3GALNT2;refGene_tx=XM_023240381.1
D2	13531928	13532088
refGene_gn=B3GALNT2;refGene_tx=XM_023240378.1;refGene_gn=B3GALNT2;refGene_tx=XM_023240379.1;refGene_gn=B3GALNT2;refGene_tx=XM_023240380.1;refGene_gn=B3GALNT2;refGene_tx=XM_023240381.1

D2 13532702 13532759
refGene_gn=B3GALNT2;refGene_tx=XM_023240378.1;refGene_gn=B3GALNT2;refGene_tx=XM_023240379.1;refGene_gn=B3GALNT2;refGene_tx=XM_023240380.1;refGene_gn=B3GALNT2;refGene_tx=XM_023240381.1
D2 13534492 13534639
refGene_gn=B3GALNT2;refGene_tx=XM_023240378.1;refGene_gn=B3GALNT2;refGene_tx=XM_023240379.1;refGene_gn=B3GALNT2;refGene_tx=XM_023240380.1;refGene_gn=B3GALNT2;refGene_tx=XM_023240381.1
D2 13535963 13536056
refGene_gn=TBCE;refGene_tx=XM_003993902.5;refGene_gn=TBCE;refGene_tx=XM_019814294.2;refGene_gn=TBCE;refGene_tx=XM_023240375.1;refGene_gn=TBCE;refGene_tx=XM_023240377.1
D2 13536202 13536294
refGene_gn=TBCE;refGene_tx=XM_003993902.5;refGene_gn=TBCE;refGene_tx=XM_019814294.2;refGene_gn=TBCE;refGene_tx=XM_023240375.1;refGene_gn=TBCE;refGene_tx=XM_023240377.1
D2 13538285 13538323 refGene_gn=TBCE;refGene_tx=XM_023240376.1
D2 13539794 13539854
refGene_gn=TBCE;refGene_tx=XM_003993902.5;refGene_gn=TBCE;refGene_tx=XM_019814294.2;refGene_gn=TBCE;refGene_tx=XM_023240375.1;refGene_gn=TBCE;refGene_tx=XM_023240376.1;refGene_gn=TBCE;refGene_tx=XM_023240377.1
D2 13540652 13540721
refGene_gn=TBCE;refGene_tx=XM_003993902.5;refGene_gn=TBCE;refGene_tx=XM_019814294.2;refGene_gn=TBCE;refGene_tx=XM_023240375.1;refGene_gn=TBCE;refGene_tx=XM_023240376.1;refGene_gn=TBCE;refGene_tx=XM_023240377.1
D2 13542960 13543114
refGene_gn=TBCE;refGene_tx=XM_003993902.5;refGene_gn=TBCE;refGene_tx=XM_019814294.2;refGene_gn=TBCE;refGene_tx=XM_023240375.1;refGene_gn=TBCE;refGene_tx=XM_023240376.1;refGene_gn=TBCE;refGene_tx=XM_023240377.1
D2 13544117 13544270
refGene_gn=TBCE;refGene_tx=XM_003993902.5;refGene_gn=TBCE;refGene_tx=XM_019814294.2;refGene_gn=TBCE;refGene_tx=XM_023240375.1;refGene_gn=TBCE;refGene_tx=XM_023240376.1;refGene_gn=TBCE;refGene_tx=XM_023240377.1
D2 13546036 13546101
refGene_gn=TBCE;refGene_tx=XM_003993902.5;refGene_gn=TBCE;refGene_tx=XM_019814294.2;refGene_gn=TBCE;refGene_tx=XM_023240375.1;refGene_gn=TBCE;refGene_tx=XM_023240376.1;refGene_gn=TBCE;refGene_tx=XM_023240377.1
D2 13546196 13546261
refGene_gn=TBCE;refGene_tx=XM_003993902.5;refGene_gn=TBCE;refGene_tx=XM_019814294.2;refGene_gn=TBCE;refGene_tx=XM_023240375.1;refGene_gn=TBCE;refGene_tx=XM_023240376.1;refGene_gn=TBCE;refGene_tx=XM_023240377.1
D2 13546777 13546873
refGene_gn=TBCE;refGene_tx=XM_003993902.5;refGene_gn=TBCE;refGene_tx=XM_019814294.2;refGene_gn=TBCE;refGene_tx=XM_023240375.1;refGene_gn=TBCE;refGene_tx=XM_023240376.1;refGene_gn=TBCE;refGene_tx=XM_023240377.1
D2 13547891 13547968
refGene_gn=TBCE;refGene_tx=XM_003993902.5;refGene_gn=TBCE;refGene_tx=XM_019814294.2;refGene_gn=TBCE;refGene_tx=XM_023240375.1;refGene_gn=TBCE;refGene_tx=XM_023240376.1;refGene_gn=TBCE;refGene_tx=XM_023240377.1
D2 13549136 13549239

refGene_gn=TBCE;refGene_tx=XM_003993902.5;refGene_gn=TBCE;refGene_tx=XM_019814294.2;refGene_gn=TBCE;refGene_tx=XM_023240375.1;refGene_gn=TBCE;refGene_tx=XM_023240376.1;refGene_gn=TBCE;refGene_tx=XM_023240377.1
D2 13551447 13551547
refGene_gn=TBCE;refGene_tx=XM_003993902.5;refGene_gn=TBCE;refGene_tx=XM_023240375.1;refGene_gn=TBCE;refGene_tx=XM_023240376.1;refGene_gn=TBCE;refGene_tx=XM_023240377.1
D2 13555804 13555931
refGene_gn=TBCE;refGene_tx=XM_003993902.5;refGene_gn=TBCE;refGene_tx=XM_023240375.1;refGene_gn=TBCE;refGene_tx=XM_023240376.1;refGene_gn=TBCE;refGene_tx=XM_023240377.1
D2 13560985 13561171
refGene_gn=TBCE;refGene_tx=XM_003993902.5;refGene_gn=TBCE;refGene_tx=XM_019814294.2;refGene_gn=TBCE;refGene_tx=XM_023240375.1;refGene_gn=TBCE;refGene_tx=XM_023240376.1
D2 13574786 13574871
refGene_gn=TBCE;refGene_tx=XM_003993902.5;refGene_gn=TBCE;refGene_tx=XM_019814294.2;refGene_gn=TBCE;refGene_tx=XM_023240375.1;refGene_gn=TBCE;refGene_tx=XM_023240376.1
D2 13603728 13603828
refGene_gn=TBCE;refGene_tx=XM_003993902.5;refGene_gn=TBCE;refGene_tx=XM_019814294.2;refGene_gn=TBCE;refGene_tx=XM_023240375.1;refGene_gn=TBCE;refGene_tx=XM_023240376.1
D2 13646705 13647467
refGene_gn=GGPS1;refGene_tx=XM_003993903.5;refGene_gn=GGPS1;refGene_tx=XM_019814301.2;refGene_gn=GGPS1;refGene_tx=XM_019814302.2;refGene_gn=GGPS1;refGene_tx=XM_023240386.1
D2 13647713 13647784
refGene_gn=GGPS1;refGene_tx=XM_003993903.5;refGene_gn=GGPS1;refGene_tx=XM_019814301.2;refGene_gn=GGPS1;refGene_tx=XM_019814302.2
D2 13652879 13652949
refGene_gn=GGPS1;refGene_tx=XM_003993903.5;refGene_gn=GGPS1;refGene_tx=XM_019814301.2;refGene_gn=GGPS1;refGene_tx=XM_019814302.2
D2 13658346 13658352
refGene_gn=ARID4B;refGene_tx=XM_011287214.3;refGene_gn=ARID4B;refGene_tx=XM_019814296.2;refGene_gn=ARID4B;refGene_tx=XM_019814297.2;refGene_gn=ARID4B;refGene_tx=XM_019814298.2;refGene_gn=ARID4B;refGene_tx=XM_019814299.2;refGene_gn=ARID4B;refGene_tx=XM_019814300.2;refGene_gn=ARID4B;refGene_tx=XM_023240382.1
D2 13724122 13724233
refGene_gn=ARID4B;refGene_tx=XM_011287214.3;refGene_gn=ARID4B;refGene_tx=XM_019814296.2;refGene_gn=ARID4B;refGene_tx=XM_019814297.2;refGene_gn=ARID4B;refGene_tx=XM_019814298.2;refGene_gn=ARID4B;refGene_tx=XM_019814299.2;refGene_gn=ARID4B;refGene_tx=XM_019814300.2;refGene_gn=ARID4B;refGene_tx=XM_023240382.1
D2 13727309 13727375
refGene_gn=ARID4B;refGene_tx=XM_011287214.3;refGene_gn=ARID4B;refGene_tx=XM_019814296.2;refGene_gn=ARID4B;refGene_tx=XM_019814297.2;refGene_gn=ARID4B;refGene_tx=XM_019814298.2;refGene_gn=ARID4B;refGene_tx=XM_019814299.2;refGene_gn=ARID4B;refGene_tx=XM_019814300.2;refGene_gn=ARID4

B;refGene_tx=XM_023240382.1
D2 13730982 13731073
refGene_gn=ARID4B;refGene_tx=XM_011287214.3;refGene_gn=ARID4B;refGene_tx=XM_019814296.2;refGene_gn=ARID4B;refGene_tx=XM_019814297.2;refGene_gn=ARID4B;refGene_tx=XM_019814298.2;refGene_gn=ARID4B;refGene_tx=XM_019814299.2;refGene_gn=ARID4B;refGene_tx=XM_019814300.2;refGene_gn=ARID4B;refGene_tx=XM_023240382.1
D2 13733726 13733806
refGene_gn=ARID4B;refGene_tx=XM_011287214.3;refGene_gn=ARID4B;refGene_tx=XM_019814296.2;refGene_gn=ARID4B;refGene_tx=XM_019814297.2;refGene_gn=ARID4B;refGene_tx=XM_019814298.2;refGene_gn=ARID4B;refGene_tx=XM_019814299.2;refGene_gn=ARID4B;refGene_tx=XM_019814300.2;refGene_gn=ARID4B;refGene_tx=XM_023240382.1
D2 13739890 13739982
refGene_gn=ARID4B;refGene_tx=XM_011287214.3;refGene_gn=ARID4B;refGene_tx=XM_019814296.2;refGene_gn=ARID4B;refGene_tx=XM_019814297.2;refGene_gn=ARID4B;refGene_tx=XM_019814298.2;refGene_gn=ARID4B;refGene_tx=XM_019814299.2;refGene_gn=ARID4B;refGene_tx=XM_019814300.2;refGene_gn=ARID4B;refGene_tx=XM_023240382.1
D2 13748603 13748742
refGene_gn=ARID4B;refGene_tx=XM_011287214.3;refGene_gn=ARID4B;refGene_tx=XM_019814296.2;refGene_gn=ARID4B;refGene_tx=XM_019814297.2;refGene_gn=ARID4B;refGene_tx=XM_019814298.2;refGene_gn=ARID4B;refGene_tx=XM_019814299.2;refGene_gn=ARID4B;refGene_tx=XM_019814300.2;refGene_gn=ARID4B;refGene_tx=XM_023240382.1
D2 13754207 13754287
refGene_gn=ARID4B;refGene_tx=XM_011287214.3;refGene_gn=ARID4B;refGene_tx=XM_019814296.2;refGene_gn=ARID4B;refGene_tx=XM_019814297.2;refGene_gn=ARID4B;refGene_tx=XM_019814298.2;refGene_gn=ARID4B;refGene_tx=XM_019814299.2;refGene_gn=ARID4B;refGene_tx=XM_019814300.2;refGene_gn=ARID4B;refGene_tx=XM_023240382.1
D2 13756231 13756308
refGene_gn=ARID4B;refGene_tx=XM_011287214.3;refGene_gn=ARID4B;refGene_tx=XM_019814296.2;refGene_gn=ARID4B;refGene_tx=XM_019814297.2;refGene_gn=ARID4B;refGene_tx=XM_019814298.2;refGene_gn=ARID4B;refGene_tx=XM_019814299.2;refGene_gn=ARID4B;refGene_tx=XM_019814300.2;refGene_gn=ARID4B;refGene_tx=XM_023240382.1
D2 13757646 13757801
refGene_gn=ARID4B;refGene_tx=XM_011287214.3;refGene_gn=ARID4B;refGene_tx=XM_019814296.2;refGene_gn=ARID4B;refGene_tx=XM_019814297.2;refGene_gn=ARID4B;refGene_tx=XM_019814298.2;refGene_gn=ARID4B;refGene_tx=XM_019814299.2;refGene_gn=ARID4B;refGene_tx=XM_019814300.2;refGene_gn=ARID4B;refGene_tx=XM_023240382.1
D2 13760735 13760808
refGene_gn=ARID4B;refGene_tx=XM_011287214.3;refGene_gn=ARID4B;refGene_tx=XM_019814296.2;refGene_gn=ARID4B;refGene_tx=XM_019814297.2;refGene_gn=ARID4B;refGene_tx=XM_019814298.2;refGene_gn=ARID4B;refGene_tx=XM_019814299.2;refGene_gn=ARID4B;refGene_tx=XM_019814300.2;refGene_gn=ARID4B;refGene_tx=XM_023240382.1
D2 13762356 13762451

refGene_gn=ARID4B;refGene_tx=XM_011287214.3;refGene_gn=ARID4B;refGene_tx=XM_019814296.2;refGene_gn=ARID4B;refGene_tx=XM_019814297.2;refGene_gn=ARID4B;refGene_tx=XM_019814298.2;refGene_gn=ARID4B;refGene_tx=XM_019814299.2;refGene_gn=ARID4B;refGene_tx=XM_019814300.2;refGene_gn=ARID4B;refGene_tx=XM_023240382.1
D2 13763638 13763736
refGene_gn=ARID4B;refGene_tx=XM_011287214.3;refGene_gn=ARID4B;refGene_tx=XM_019814296.2;refGene_gn=ARID4B;refGene_tx=XM_019814297.2;refGene_gn=ARID4B;refGene_tx=XM_019814298.2;refGene_gn=ARID4B;refGene_tx=XM_019814299.2;refGene_gn=ARID4B;refGene_tx=XM_019814300.2;refGene_gn=ARID4B;refGene_tx=XM_023240382.1
D2 13764874 13765118
refGene_gn=ARID4B;refGene_tx=XM_011287214.3;refGene_gn=ARID4B;refGene_tx=XM_019814296.2;refGene_gn=ARID4B;refGene_tx=XM_019814297.2;refGene_gn=ARID4B;refGene_tx=XM_019814298.2;refGene_gn=ARID4B;refGene_tx=XM_019814299.2;refGene_gn=ARID4B;refGene_tx=XM_019814300.2;refGene_gn=ARID4B;refGene_tx=XM_023240382.1
D2 13765438 13765614
refGene_gn=ARID4B;refGene_tx=XM_011287214.3;refGene_gn=ARID4B;refGene_tx=XM_019814296.2;refGene_gn=ARID4B;refGene_tx=XM_019814297.2;refGene_gn=ARID4B;refGene_tx=XM_019814298.2;refGene_gn=ARID4B;refGene_tx=XM_019814299.2;refGene_gn=ARID4B;refGene_tx=XM_019814300.2;refGene_gn=ARID4B;refGene_tx=XM_023240382.1
D2 13769623 13769875
refGene_gn=ARID4B;refGene_tx=XM_011287214.3;refGene_gn=ARID4B;refGene_tx=XM_019814296.2;refGene_gn=ARID4B;refGene_tx=XM_019814297.2;refGene_gn=ARID4B;refGene_tx=XM_019814298.2;refGene_gn=ARID4B;refGene_tx=XM_023240382.1
D2 13784435 13784520
refGene_gn=ARID4B;refGene_tx=XM_011287214.3;refGene_gn=ARID4B;refGene_tx=XM_019814296.2;refGene_gn=ARID4B;refGene_tx=XM_019814297.2;refGene_gn=ARID4B;refGene_tx=XM_019814298.2;refGene_gn=ARID4B;refGene_tx=XM_019814299.2;refGene_gn=ARID4B;refGene_tx=XM_019814300.2;refGene_gn=ARID4B;refGene_tx=XM_023240382.1
D2 13786209 13786408
refGene_gn=ARID4B;refGene_tx=XM_011287214.3;refGene_gn=ARID4B;refGene_tx=XM_019814296.2;refGene_gn=ARID4B;refGene_tx=XM_019814297.2;refGene_gn=ARID4B;refGene_tx=XM_019814298.2;refGene_gn=ARID4B;refGene_tx=XM_019814299.2;refGene_gn=ARID4B;refGene_tx=XM_019814300.2;refGene_gn=ARID4B;refGene_tx=XM_023240382.1
D2 13794129 13795344
refGene_gn=ARID4B;refGene_tx=XM_011287214.3;refGene_gn=ARID4B;refGene_tx=XM_019814296.2;refGene_gn=ARID4B;refGene_tx=XM_019814297.2;refGene_gn=ARID4B;refGene_tx=XM_019814298.2;refGene_gn=ARID4B;refGene_tx=XM_019814299.2;refGene_gn=ARID4B;refGene_tx=XM_019814300.2;refGene_gn=ARID4B;refGene_tx=XM_023240382.1
D2 13795741 13795833 refGene_gn=ARID4B;refGene_tx=XM_023240382.1
D2 13798694 13798808
refGene_gn=ARID4B;refGene_tx=XM_011287214.3;refGene_gn=ARID4B;refGene_tx=XM_019814296.2;refGene_gn=ARID4B;refGene_tx=XM_019814297.2;refGene_

gn=ARID4B;refGene_tx=XM_019814298.2;refGene_gn=ARID4B;refGene_tx=XM_01
9814299.2;refGene_gn=ARID4B;refGene_tx=XM_019814300.2
D2 13801067 13801283
refGene_gn=ARID4B;refGene_tx=XM_011287214.3;refGene_gn=ARID4B;refGene_
tx=XM_019814296.2;refGene_gn=ARID4B;refGene_tx=XM_019814297.2;refGene_
gn=ARID4B;refGene_tx=XM_019814298.2;refGene_gn=ARID4B;refGene_tx=XM_01
9814299.2;refGene_gn=ARID4B;refGene_tx=XM_019814300.2
D2 13802873 13803020
refGene_gn=ARID4B;refGene_tx=XM_011287214.3;refGene_gn=ARID4B;refGene_
tx=XM_019814296.2;refGene_gn=ARID4B;refGene_tx=XM_019814297.2;refGene_
gn=ARID4B;refGene_tx=XM_019814298.2;refGene_gn=ARID4B;refGene_tx=XM_01
9814299.2;refGene_gn=ARID4B;refGene_tx=XM_019814300.2
D2 13808026 13808154
refGene_gn=ARID4B;refGene_tx=XM_011287214.3;refGene_gn=ARID4B;refGene_
tx=XM_019814300.2
D2 13812691 13812852
refGene_gn=ARID4B;refGene_tx=XM_019814296.2;refGene_gn=ARID4B;refGene_
tx=XM_019814297.2;refGene_gn=ARID4B;refGene_tx=XM_019814298.2;refGene_
gn=ARID4B;refGene_tx=XM_019814299.2
D2 13812906 13812965
refGene_gn=RBM34;refGene_tx=XM_023240383.1;refGene_gn=RBM34;refGene_tx
=XM_023240384.1
D2 13813074 13813249
refGene_gn=RBM34;refGene_tx=XM_023240383.1;refGene_gn=RBM34;refGene_tx
=XM_023240384.1;refGene_gn=RBM34;refGene_tx=XM_023240385.1
D2 13813464 13813601
refGene_gn=RBM34;refGene_tx=XM_023240383.1;refGene_gn=RBM34;refGene_tx
=XM_023240384.1
D2 13817643 13817875
refGene_gn=RBM34;refGene_tx=XM_023240383.1;refGene_gn=RBM34;refGene_tx
=XM_023240384.1;refGene_gn=RBM34;refGene_tx=XM_023240385.1
D2 13818581 13818641
refGene_gn=RBM34;refGene_tx=XM_023240383.1;refGene_gn=RBM34;refGene_tx
=XM_023240385.1
D2 13820937 13820981
refGene_gn=RBM34;refGene_tx=XM_023240383.1;refGene_gn=RBM34;refGene_tx
=XM_023240384.1;refGene_gn=RBM34;refGene_tx=XM_023240385.1
D2 13827352 13827436
refGene_gn=RBM34;refGene_tx=XM_023240383.1;refGene_gn=RBM34;refGene_tx
=XM_023240384.1;refGene_gn=RBM34;refGene_tx=XM_023240385.1
D2 13827592 13827656
refGene_gn=RBM34;refGene_tx=XM_023240383.1;refGene_gn=RBM34;refGene_tx
=XM_023240384.1;refGene_gn=RBM34;refGene_tx=XM_023240385.1
D2 13829204 13829244
refGene_gn=RBM34;refGene_tx=XM_023240383.1;refGene_gn=RBM34;refGene_tx
=XM_023240384.1;refGene_gn=RBM34;refGene_tx=XM_023240385.1
D2 13829472 13829591
refGene_gn=RBM34;refGene_tx=XM_023240383.1;refGene_gn=RBM34;refGene_tx
=XM_023240384.1;refGene_gn=RBM34;refGene_tx=XM_023240385.1
D2 13831418 13831697

refGene_gn=RBM34;refGene_tx=XM_023240383.1;refGene_gn=RBM34;refGene_tx=XM_023240384.1;refGene_gn=RBM34;refGene_tx=XM_023240385.1
D2 13837421 13837542
refGene_gn=TOMM20;refGene_tx=XM_003993904.5;refGene_gn=TOMM20;refGene_tx=XM_006937765.4
D2 13842801 13842848
refGene_gn=TOMM20;refGene_tx=XM_003993904.5;refGene_gn=TOMM20;refGene_tx=XM_006937765.4
D2 13845272 13845354
refGene_gn=TOMM20;refGene_tx=XM_003993904.5;refGene_gn=TOMM20;refGene_tx=XM_006937765.4
D2 13848811 13848954
refGene_gn=TOMM20;refGene_tx=XM_003993904.5;refGene_gn=TOMM20;refGene_tx=XM_006937765.4
D2 13852146 13852191 refGene_gn=TOMM20;refGene_tx=XM_003993904.5
D2 13870597 13870630 refGene_gn=TOMM20;refGene_tx=XM_006937765.4
D2 14290777 14291837
refGene_gn=IRF2BP2;refGene_tx=XM_023240387.1;refGene_gn=IRF2BP2;refGene_tx=XM_023240388.1
D2 14292422 14293138
refGene_gn=IRF2BP2;refGene_tx=XM_023240387.1;refGene_gn=IRF2BP2;refGene_tx=XM_023240388.1
D2 14345327 14345381
refGene_gn=LOC102902472;refGene_tx=XM_023240389.1
D2 14345770 14345861
refGene_gn=LOC102902472;refGene_tx=XM_023240389.1
D2 14346423 14346671
refGene_gn=LOC102902472;refGene_tx=XM_023240389.1
D2 14346903 14347245
refGene_gn=LOC102902472;refGene_tx=XM_023240389.1
D2 14348737 14348909
refGene_gn=LOC102902472;refGene_tx=XM_023240390.1;refGene_gn=LOC102902472;refGene_tx=XM_023240391.1;refGene_gn=LOC102902472;refGene_tx=XM_023240392.1;refGene_gn=LOC102902472;refGene_tx=XM_023240393.1;refGene_gn=LOC102902472;refGene_tx=XM_023240394.1
D2 14349108 14349216
refGene_gn=LOC102902472;refGene_tx=XM_023240390.1;refGene_gn=LOC102902472;refGene_tx=XM_023240391.1;refGene_gn=LOC102902472;refGene_tx=XM_023240392.1;refGene_gn=LOC102902472;refGene_tx=XM_023240393.1;refGene_gn=LOC102902472;refGene_tx=XM_023240394.1
D2 14360231 14360357
refGene_gn=LOC102902472;refGene_tx=XM_023240390.1;refGene_gn=LOC102902472;refGene_tx=XM_023240391.1;refGene_gn=LOC102902472;refGene_tx=XM_023240392.1;refGene_gn=LOC102902472;refGene_tx=XM_023240393.1;refGene_gn=LOC102902472;refGene_tx=XM_023240394.1
D2 14361153 14361245
refGene_gn=LOC102902472;refGene_tx=XM_023240394.1
D2 14364900 14365130
refGene_gn=LOC102902472;refGene_tx=XM_023240390.1;refGene_gn=LOC102902472;refGene_tx=XM_023240391.1;refGene_gn=LOC102902472;refGene_tx=XM_02

3240392.1;refGene_gn=LOC102902472;refGene_tx=XM_023240393.1
D2 14374285 14374334
refGene_gn=LOC102902472;refGene_tx=XM_023240392.1;refGene_gn=LOC102902472;refGene_tx=XM_023240393.1
D2 14408515 14409455 refGene_gn=TARBP1;refGene_tx=XM_023240395.1
D2 14415006 14415104 refGene_gn=TARBP1;refGene_tx=XM_023240395.1
D2 14416593 14416663 refGene_gn=TARBP1;refGene_tx=XM_023240395.1
D2 14419513 14419662 refGene_gn=TARBP1;refGene_tx=XM_023240395.1
D2 14420554 14420607 refGene_gn=TARBP1;refGene_tx=XM_023240395.1
D2 14422113 14422211 refGene_gn=TARBP1;refGene_tx=XM_023240395.1
D2 14425356 14425492 refGene_gn=TARBP1;refGene_tx=XM_023240395.1
D2 14426091 14426188 refGene_gn=TARBP1;refGene_tx=XM_023240395.1
D2 14427335 14427425 refGene_gn=TARBP1;refGene_tx=XM_023240395.1
D2 14433143 14433285 refGene_gn=TARBP1;refGene_tx=XM_023240395.1
D2 14438828 14438928 refGene_gn=TARBP1;refGene_tx=XM_023240395.1
D2 14439905 14440078 refGene_gn=TARBP1;refGene_tx=XM_023240395.1
D2 14445791 14445889 refGene_gn=TARBP1;refGene_tx=XM_023240395.1
D2 14449626 14449788 refGene_gn=TARBP1;refGene_tx=XM_023240395.1
D2 14455656 14455844 refGene_gn=TARBP1;refGene_tx=XM_023240395.1
D2 14456274 14456536 refGene_gn=TARBP1;refGene_tx=XM_023240395.1
D2 14456658 14456718 refGene_gn=TARBP1;refGene_tx=XM_023240395.1
D2 14456720 14456861 refGene_gn=TARBP1;refGene_tx=XM_023240395.1
D2 14459223 14459414 refGene_gn=TARBP1;refGene_tx=XM_023240395.1
D2 14459839 14459911 refGene_gn=TARBP1;refGene_tx=XM_023240395.1
D2 14461595 14461716 refGene_gn=TARBP1;refGene_tx=XM_023240395.1
D2 14468379 14468490 refGene_gn=TARBP1;refGene_tx=XM_023240395.1
D2 14471773 14471923 refGene_gn=TARBP1;refGene_tx=XM_023240395.1
D2 14479205 14479292 refGene_gn=TARBP1;refGene_tx=XM_023240395.1
D2 14487469 14487666 refGene_gn=TARBP1;refGene_tx=XM_023240395.1
D2 14492180 14492262 refGene_gn=TARBP1;refGene_tx=XM_023240395.1
D2 14494137 14494309 refGene_gn=TARBP1;refGene_tx=XM_023240395.1
D2 14497219 14497411 refGene_gn=TARBP1;refGene_tx=XM_023240395.1
D2 14497589 14497714 refGene_gn=TARBP1;refGene_tx=XM_023240395.1
D2 14498142 14498279 refGene_gn=TARBP1;refGene_tx=XM_023240395.1
D2 14498590 14498759 refGene_gn=TARBP1;refGene_tx=XM_023240395.1
D2 14506622 14506718
refGene_gn=LOC101082948;refGene_tx=XM_003993905.5;refGene_gn=LOC101082948;refGene_tx=XM_006937773.2;refGene_gn=LOC101082948;refGene_tx=XM_019813063.2
D2 14512364 14512580
refGene_gn=LOC101082948;refGene_tx=XM_003993905.5;refGene_gn=LOC101082948;refGene_tx=XM_006937773.2;refGene_gn=LOC101082948;refGene_tx=XM_019813063.2
D2 14513081 14513146
refGene_gn=LOC101082948;refGene_tx=XM_006937773.2;refGene_gn=LOC101082948;refGene_tx=XM_019813063.2
D2 14590427 14590663
refGene_gn=SLC35F3;refGene_tx=XM_011287215.3;refGene_gn=SLC35F3;refGene_tx=XM_023240396.1;refGene_gn=SLC35F3;refGene_tx=XM_023240397.1
D2 14593411 14593501

refGene_gn=SLC35F3;refGene_tx=XM_011287215.3;refGene_gn=SLC35F3;refGene_tx=XM_023240396.1;refGene_gn=SLC35F3;refGene_tx=XM_023240397.1
D2 14594306 14594499
refGene_gn=SLC35F3;refGene_tx=XM_011287215.3;refGene_gn=SLC35F3;refGene_tx=XM_023240396.1;refGene_gn=SLC35F3;refGene_tx=XM_023240397.1
D2 14597226 14597352
refGene_gn=SLC35F3;refGene_tx=XM_011287215.3;refGene_gn=SLC35F3;refGene_tx=XM_023240396.1;refGene_gn=SLC35F3;refGene_tx=XM_023240397.1
D2 14603894 14604114
refGene_gn=SLC35F3;refGene_tx=XM_011287215.3;refGene_gn=SLC35F3;refGene_tx=XM_023240396.1;refGene_gn=SLC35F3;refGene_tx=XM_023240397.1
D2 14692091 14692416
refGene_gn=SLC35F3;refGene_tx=XM_011287215.3;refGene_gn=SLC35F3;refGene_tx=XM_023240396.1;refGene_gn=SLC35F3;refGene_tx=XM_023240397.1
D2 14708827 14708903 refGene_gn=SLC35F3;refGene_tx=XM_011287215.3
D2 14988821 14989051
refGene_gn=SLC35F3;refGene_tx=XM_023240396.1;refGene_gn=SLC35F3;refGene_tx=XM_023240397.1
D2 14989402 14989455 refGene_gn=SLC35F3;refGene_tx=XM_023240396.1
D2 15209158 15209418 refGene_gn=KCNK1;refGene_tx=XM_023240398.1
D2 15212491 15212887 refGene_gn=KCNK1;refGene_tx=XM_023240398.1
D2 15261016 15261371 refGene_gn=KCNK1;refGene_tx=XM_023240398.1
D2 15470213 15470614
refGene_gn=MAP3K21;refGene_tx=XM_011287216.2;refGene_gn=MAP3K21;refGene_tx=XM_011287217.2
D2 15471923 15472709
refGene_gn=MAP3K21;refGene_tx=XM_011287216.2;refGene_gn=MAP3K21;refGene_tx=XM_011287217.2
D2 15476450 15476548
refGene_gn=MAP3K21;refGene_tx=XM_011287216.2;refGene_gn=MAP3K21;refGene_tx=XM_011287217.2
D2 15476912 15477063
refGene_gn=MAP3K21;refGene_tx=XM_011287216.2;refGene_gn=MAP3K21;refGene_tx=XM_011287217.2
D2 15489624 15489747
refGene_gn=MAP3K21;refGene_tx=XM_011287216.2;refGene_gn=MAP3K21;refGene_tx=XM_011287217.2
D2 15497534 15497775
refGene_gn=MAP3K21;refGene_tx=XM_011287216.2;refGene_gn=MAP3K21;refGene_tx=XM_011287217.2
D2 15507519 15507695
refGene_gn=MAP3K21;refGene_tx=XM_011287216.2;refGene_gn=MAP3K21;refGene_tx=XM_011287217.2
D2 15508544 15508693
refGene_gn=MAP3K21;refGene_tx=XM_011287216.2;refGene_gn=MAP3K21;refGene_tx=XM_011287217.2
D2 15514754 15514935
refGene_gn=MAP3K21;refGene_tx=XM_011287216.2;refGene_gn=MAP3K21;refGene_tx=XM_011287217.2
D2 15533539 15534362

refGene_gn=MAP3K21;refGene_tx=XM_011287216.2;refGene_gn=MAP3K21;refGene_tx=XM_011287217.2
D2 15562745 15562898
refGene_gn=PCNX2;refGene_tx=XM_023240399.1;refGene_gn=PCNX2;refGene_tx=XM_023240400.1;refGene_gn=PCNX2;refGene_tx=XM_023240401.1;refGene_gn=PCNX2;refGene_tx=XM_023240402.1
D2 15590228 15590434
refGene_gn=PCNX2;refGene_tx=XM_023240399.1;refGene_gn=PCNX2;refGene_tx=XM_023240400.1;refGene_gn=PCNX2;refGene_tx=XM_023240401.1;refGene_gn=PCNX2;refGene_tx=XM_023240402.1
D2 15591265 15591386
refGene_gn=PCNX2;refGene_tx=XM_023240399.1;refGene_gn=PCNX2;refGene_tx=XM_023240400.1;refGene_gn=PCNX2;refGene_tx=XM_023240401.1;refGene_gn=PCNX2;refGene_tx=XM_023240402.1
D2 15592374 15592411
refGene_gn=PCNX2;refGene_tx=XM_023240399.1;refGene_gn=PCNX2;refGene_tx=XM_023240400.1;refGene_gn=PCNX2;refGene_tx=XM_023240401.1;refGene_gn=PCNX2;refGene_tx=XM_023240402.1
D2 15601498 15602788
refGene_gn=PCNX2;refGene_tx=XM_023240399.1;refGene_gn=PCNX2;refGene_tx=XM_023240400.1;refGene_gn=PCNX2;refGene_tx=XM_023240401.1;refGene_gn=PCNX2;refGene_tx=XM_023240402.1
D2 15607044 15607192
refGene_gn=PCNX2;refGene_tx=XM_023240399.1;refGene_gn=PCNX2;refGene_tx=XM_023240400.1;refGene_gn=PCNX2;refGene_tx=XM_023240401.1;refGene_gn=PCNX2;refGene_tx=XM_023240402.1
D2 15607329 15607475
refGene_gn=PCNX2;refGene_tx=XM_023240399.1;refGene_gn=PCNX2;refGene_tx=XM_023240400.1;refGene_gn=PCNX2;refGene_tx=XM_023240401.1;refGene_gn=PCNX2;refGene_tx=XM_023240402.1
D2 15609747 15609841
refGene_gn=PCNX2;refGene_tx=XM_023240399.1;refGene_gn=PCNX2;refGene_tx=XM_023240400.1;refGene_gn=PCNX2;refGene_tx=XM_023240401.1;refGene_gn=PCNX2;refGene_tx=XM_023240402.1
D2 15623205 15623341
refGene_gn=PCNX2;refGene_tx=XM_023240399.1;refGene_gn=PCNX2;refGene_tx=XM_023240400.1;refGene_gn=PCNX2;refGene_tx=XM_023240401.1;refGene_gn=PCNX2;refGene_tx=XM_023240402.1
D2 15635975 15636121
refGene_gn=PCNX2;refGene_tx=XM_023240399.1;refGene_gn=PCNX2;refGene_tx=XM_023240400.1;refGene_gn=PCNX2;refGene_tx=XM_023240401.1;refGene_gn=PCNX2;refGene_tx=XM_023240402.1
D2 15644232 15644386
refGene_gn=PCNX2;refGene_tx=XM_023240399.1;refGene_gn=PCNX2;refGene_tx=XM_023240400.1;refGene_gn=PCNX2;refGene_tx=XM_023240401.1;refGene_gn=PCNX2;refGene_tx=XM_023240402.1
D2 15644472 15644505
refGene_gn=PCNX2;refGene_tx=XM_023240399.1;refGene_gn=PCNX2;refGene_tx=XM_023240400.1;refGene_gn=PCNX2;refGene_tx=XM_023240401.1;refGene_gn=PCNX2;refGene_tx=XM_023240402.1

D2 15652611 15652783
refGene_gn=PCNX2;refGene_tx=XM_023240399.1;refGene_gn=PCNX2;refGene_tx=XM_023240400.1;refGene_gn=PCNX2;refGene_tx=XM_023240401.1;refGene_gn=PCNX2;refGene_tx=XM_023240402.1
D2 15660641 15660752
refGene_gn=PCNX2;refGene_tx=XM_023240399.1;refGene_gn=PCNX2;refGene_tx=XM_023240400.1;refGene_gn=PCNX2;refGene_tx=XM_023240401.1;refGene_gn=PCNX2;refGene_tx=XM_023240402.1
D2 15661559 15661651
refGene_gn=PCNX2;refGene_tx=XM_023240399.1;refGene_gn=PCNX2;refGene_tx=XM_023240400.1;refGene_gn=PCNX2;refGene_tx=XM_023240401.1;refGene_gn=PCNX2;refGene_tx=XM_023240402.1
D2 15696255 15696365
refGene_gn=PCNX2;refGene_tx=XM_023240399.1;refGene_gn=PCNX2;refGene_tx=XM_023240400.1;refGene_gn=PCNX2;refGene_tx=XM_023240401.1;refGene_gn=PCNX2;refGene_tx=XM_023240402.1
D2 15697699 15697796
refGene_gn=PCNX2;refGene_tx=XM_023240399.1;refGene_gn=PCNX2;refGene_tx=XM_023240400.1;refGene_gn=PCNX2;refGene_tx=XM_023240401.1;refGene_gn=PCNX2;refGene_tx=XM_023240402.1
D2 15712150 15712243
refGene_gn=PCNX2;refGene_tx=XM_023240399.1;refGene_gn=PCNX2;refGene_tx=XM_023240400.1;refGene_gn=PCNX2;refGene_tx=XM_023240401.1;refGene_gn=PCNX2;refGene_tx=XM_023240402.1
D2 15713057 15713208
refGene_gn=PCNX2;refGene_tx=XM_023240399.1;refGene_gn=PCNX2;refGene_tx=XM_023240400.1;refGene_gn=PCNX2;refGene_tx=XM_023240401.1;refGene_gn=PCNX2;refGene_tx=XM_023240402.1
D2 15735463 15735605
refGene_gn=PCNX2;refGene_tx=XM_023240399.1;refGene_gn=PCNX2;refGene_tx=XM_023240400.1;refGene_gn=PCNX2;refGene_tx=XM_023240401.1;refGene_gn=PCNX2;refGene_tx=XM_023240402.1
D2 15739808 15739986
refGene_gn=PCNX2;refGene_tx=XM_023240399.1;refGene_gn=PCNX2;refGene_tx=XM_023240400.1;refGene_gn=PCNX2;refGene_tx=XM_023240401.1;refGene_gn=PCNX2;refGene_tx=XM_023240402.1
D2 15770952 15771061
refGene_gn=PCNX2;refGene_tx=XM_023240399.1;refGene_gn=PCNX2;refGene_tx=XM_023240400.1;refGene_gn=PCNX2;refGene_tx=XM_023240401.1;refGene_gn=PCNX2;refGene_tx=XM_023240402.1
D2 15776634 15776764
refGene_gn=PCNX2;refGene_tx=XM_023240399.1;refGene_gn=PCNX2;refGene_tx=XM_023240400.1;refGene_gn=PCNX2;refGene_tx=XM_023240401.1;refGene_gn=PCNX2;refGene_tx=XM_023240402.1
D2 15800386 15800445
refGene_gn=PCNX2;refGene_tx=XM_023240399.1;refGene_gn=PCNX2;refGene_tx=XM_023240400.1;refGene_gn=PCNX2;refGene_tx=XM_023240401.1;refGene_gn=PCNX2;refGene_tx=XM_023240402.1
D2 15804654 15804870
refGene_gn=PCNX2;refGene_tx=XM_023240399.1;refGene_gn=PCNX2;refGene_tx

=XM_023240400.1;refGene_gn=PCNX2;refGene_tx=XM_023240401.1;refGene_gn=PCNX2;refGene_tx=XM_023240402.1
D2 15826202 15826456
refGene_gn=PCNX2;refGene_tx=XM_023240399.1;refGene_gn=PCNX2;refGene_tx=XM_023240400.1;refGene_gn=PCNX2;refGene_tx=XM_023240401.1;refGene_gn=PCNX2;refGene_tx=XM_023240402.1
D2 15833038 15833272
refGene_gn=PCNX2;refGene_tx=XM_023240399.1;refGene_gn=PCNX2;refGene_tx=XM_023240400.1;refGene_gn=PCNX2;refGene_tx=XM_023240401.1;refGene_gn=PCNX2;refGene_tx=XM_023240402.1
D2 15834717 15834830
refGene_gn=PCNX2;refGene_tx=XM_023240399.1;refGene_gn=PCNX2;refGene_tx=XM_023240400.1;refGene_gn=PCNX2;refGene_tx=XM_023240401.1;refGene_gn=PCNX2;refGene_tx=XM_023240402.1
D2 15848646 15848791
refGene_gn=PCNX2;refGene_tx=XM_023240399.1;refGene_gn=PCNX2;refGene_tx=XM_023240400.1;refGene_gn=PCNX2;refGene_tx=XM_023240401.1;refGene_gn=PCNX2;refGene_tx=XM_023240402.1
D2 15849299 15849530
refGene_gn=PCNX2;refGene_tx=XM_023240399.1;refGene_gn=PCNX2;refGene_tx=XM_023240400.1;refGene_gn=PCNX2;refGene_tx=XM_023240401.1;refGene_gn=PCNX2;refGene_tx=XM_023240402.1
D2 15850061 15850336
refGene_gn=PCNX2;refGene_tx=XM_023240399.1;refGene_gn=PCNX2;refGene_tx=XM_023240400.1;refGene_gn=PCNX2;refGene_tx=XM_023240401.1;refGene_gn=PCNX2;refGene_tx=XM_023240402.1
D2 15851197 15851382
refGene_gn=PCNX2;refGene_tx=XM_023240399.1;refGene_gn=PCNX2;refGene_tx=XM_023240400.1;refGene_gn=PCNX2;refGene_tx=XM_023240401.1;refGene_gn=PCNX2;refGene_tx=XM_023240402.1
D2 15853977 15854889
refGene_gn=LOC111557016;refGene_tx=XM_023240403.1
D2 15860561 15861010
refGene_gn=PCNX2;refGene_tx=XM_023240399.1;refGene_gn=PCNX2;refGene_tx=XM_023240400.1;refGene_gn=PCNX2;refGene_tx=XM_023240401.1;refGene_gn=PCNX2;refGene_tx=XM_023240402.1
D2 15862461 15862635
refGene_gn=PCNX2;refGene_tx=XM_023240399.1;refGene_gn=PCNX2;refGene_tx=XM_023240400.1;refGene_gn=PCNX2;refGene_tx=XM_023240401.1;refGene_gn=PCNX2;refGene_tx=XM_023240402.1
D2 15868496 15868565
refGene_gn=NTPCR;refGene_tx=XM_023240404.1;refGene_gn=NTPCR;refGene_tx=XM_023240405.1;refGene_gn=NTPCR;refGene_tx=XM_023240406.1;refGene_gn=NTPCR;refGene_tx=XM_023240407.1;refGene_gn=NTPCR;refGene_tx=XM_023240408.1
D2 15876662 15876725
refGene_gn=NTPCR;refGene_tx=XM_023240404.1;refGene_gn=NTPCR;refGene_tx=XM_023240405.1;refGene_gn=NTPCR;refGene_tx=XM_023240406.1;refGene_gn=NTPCR;refGene_tx=XM_023240407.1
D2 15877259 15877469

refGene_gn=NTPCR;refGene_tx=XM_023240404.1;refGene_gn=NTPCR;refGene_tx=XM_023240405.1;refGene_gn=NTPCR;refGene_tx=XM_023240406.1;refGene_gn=NTPCR;refGene_tx=XM_023240407.1;refGene_gn=NTPCR;refGene_tx=XM_023240408.1
D2 15893253 15893350
refGene_gn=NTPCR;refGene_tx=XM_023240404.1;refGene_gn=NTPCR;refGene_tx=XM_023240405.1;refGene_gn=NTPCR;refGene_tx=XM_023240406.1;refGene_gn=NTPCR;refGene_tx=XM_023240407.1;refGene_gn=NTPCR;refGene_tx=XM_023240408.1
D2 15894112 15894275
refGene_gn=NTPCR;refGene_tx=XM_023240404.1;refGene_gn=NTPCR;refGene_tx=XM_023240405.1;refGene_gn=NTPCR;refGene_tx=XM_023240406.1;refGene_gn=NTPCR;refGene_tx=XM_023240407.1;refGene_gn=NTPCR;refGene_tx=XM_023240408.1
D2 15898464 15898498
refGene_gn=NTPCR;refGene_tx=XM_023240404.1;refGene_gn=NTPCR;refGene_tx=XM_023240405.1;refGene_gn=NTPCR;refGene_tx=XM_023240406.1;refGene_gn=NTPCR;refGene_tx=XM_023240407.1;refGene_gn=NTPCR;refGene_tx=XM_023240408.1
D2 16038592 16041301 refGene_gn=MAP10;refGene_tx=XM_006937779.3
D2 16306220 16307697
refGene_gn=SIPA1L2;refGene_tx=XM_023240409.1;refGene_gn=SIPA1L2;refGene_tx=XM_023240410.1;refGene_gn=SIPA1L2;refGene_tx=XM_023240411.1;refGene_gn=SIPA1L2;refGene_tx=XM_023240412.1;refGene_gn=SIPA1L2;refGene_tx=XM_023240413.1
D2 16324807 16324941
refGene_gn=SIPA1L2;refGene_tx=XM_023240409.1;refGene_gn=SIPA1L2;refGene_tx=XM_023240410.1;refGene_gn=SIPA1L2;refGene_tx=XM_023240411.1;refGene_gn=SIPA1L2;refGene_tx=XM_023240412.1;refGene_gn=SIPA1L2;refGene_tx=XM_023240413.1
D2 16326630 16326819
refGene_gn=SIPA1L2;refGene_tx=XM_023240409.1;refGene_gn=SIPA1L2;refGene_tx=XM_023240410.1;refGene_gn=SIPA1L2;refGene_tx=XM_023240411.1;refGene_gn=SIPA1L2;refGene_tx=XM_023240412.1;refGene_gn=SIPA1L2;refGene_tx=XM_023240413.1
D2 16332739 16332914
refGene_gn=SIPA1L2;refGene_tx=XM_023240409.1;refGene_gn=SIPA1L2;refGene_tx=XM_023240410.1;refGene_gn=SIPA1L2;refGene_tx=XM_023240411.1;refGene_gn=SIPA1L2;refGene_tx=XM_023240412.1;refGene_gn=SIPA1L2;refGene_tx=XM_023240413.1
D2 16337159 16337263
refGene_gn=SIPA1L2;refGene_tx=XM_023240409.1;refGene_gn=SIPA1L2;refGene_tx=XM_023240410.1;refGene_gn=SIPA1L2;refGene_tx=XM_023240411.1;refGene_gn=SIPA1L2;refGene_tx=XM_023240412.1;refGene_gn=SIPA1L2;refGene_tx=XM_023240413.1
D2 16347797 16347955
refGene_gn=SIPA1L2;refGene_tx=XM_023240409.1;refGene_gn=SIPA1L2;refGene_tx=XM_023240410.1;refGene_gn=SIPA1L2;refGene_tx=XM_023240411.1;refGene_gn=SIPA1L2;refGene_tx=XM_023240412.1;refGene_gn=SIPA1L2;refGene_tx=XM_023240413.1

D2 16354003 16354580
refGene_gn=SIPA1L2;refGene_tx=XM_023240409.1;refGene_gn=SIPA1L2;refGene_tx=XM_023240410.1;refGene_gn=SIPA1L2;refGene_tx=XM_023240411.1;refGene_gn=SIPA1L2;refGene_tx=XM_023240412.1;refGene_gn=SIPA1L2;refGene_tx=XM_023240413.1
D2 16357757 16358032
refGene_gn=SIPA1L2;refGene_tx=XM_023240409.1;refGene_gn=SIPA1L2;refGene_tx=XM_023240410.1;refGene_gn=SIPA1L2;refGene_tx=XM_023240411.1;refGene_gn=SIPA1L2;refGene_tx=XM_023240412.1;refGene_gn=SIPA1L2;refGene_tx=XM_023240413.1
D2 16371497 16371755
refGene_gn=SIPA1L2;refGene_tx=XM_023240409.1;refGene_gn=SIPA1L2;refGene_tx=XM_023240410.1;refGene_gn=SIPA1L2;refGene_tx=XM_023240411.1;refGene_gn=SIPA1L2;refGene_tx=XM_023240412.1;refGene_gn=SIPA1L2;refGene_tx=XM_023240413.1
D2 16373490 16373574
refGene_gn=SIPA1L2;refGene_tx=XM_023240409.1;refGene_gn=SIPA1L2;refGene_tx=XM_023240410.1;refGene_gn=SIPA1L2;refGene_tx=XM_023240411.1;refGene_gn=SIPA1L2;refGene_tx=XM_023240412.1;refGene_gn=SIPA1L2;refGene_tx=XM_023240413.1
D2 16375174 16375275
refGene_gn=SIPA1L2;refGene_tx=XM_023240409.1;refGene_gn=SIPA1L2;refGene_tx=XM_023240410.1;refGene_gn=SIPA1L2;refGene_tx=XM_023240411.1;refGene_gn=SIPA1L2;refGene_tx=XM_023240412.1;refGene_gn=SIPA1L2;refGene_tx=XM_023240413.1
D2 16375675 16375779
refGene_gn=SIPA1L2;refGene_tx=XM_023240409.1;refGene_gn=SIPA1L2;refGene_tx=XM_023240410.1;refGene_gn=SIPA1L2;refGene_tx=XM_023240411.1;refGene_gn=SIPA1L2;refGene_tx=XM_023240412.1;refGene_gn=SIPA1L2;refGene_tx=XM_023240413.1
D2 16377783 16378163
refGene_gn=SIPA1L2;refGene_tx=XM_023240409.1;refGene_gn=SIPA1L2;refGene_tx=XM_023240410.1;refGene_gn=SIPA1L2;refGene_tx=XM_023240411.1;refGene_gn=SIPA1L2;refGene_tx=XM_023240412.1;refGene_gn=SIPA1L2;refGene_tx=XM_023240413.1
D2 16385474 16385699
refGene_gn=SIPA1L2;refGene_tx=XM_023240409.1;refGene_gn=SIPA1L2;refGene_tx=XM_023240410.1;refGene_gn=SIPA1L2;refGene_tx=XM_023240411.1;refGene_gn=SIPA1L2;refGene_tx=XM_023240412.1;refGene_gn=SIPA1L2;refGene_tx=XM_023240413.1
D2 16389040 16389194
refGene_gn=SIPA1L2;refGene_tx=XM_023240409.1;refGene_gn=SIPA1L2;refGene_tx=XM_023240410.1;refGene_gn=SIPA1L2;refGene_tx=XM_023240411.1;refGene_gn=SIPA1L2;refGene_tx=XM_023240412.1;refGene_gn=SIPA1L2;refGene_tx=XM_023240413.1
D2 16391431 16391651
refGene_gn=SIPA1L2;refGene_tx=XM_023240409.1;refGene_gn=SIPA1L2;refGene_tx=XM_023240410.1;refGene_gn=SIPA1L2;refGene_tx=XM_023240411.1;refGene_gn=SIPA1L2;refGene_tx=XM_023240412.1;refGene_gn=SIPA1L2;refGene_tx=XM_023240413.1

D2 16399701 16399833
refGene_gn=SIPA1L2;refGene_tx=XM_023240409.1;refGene_gn=SIPA1L2;refGen
e_tx=XM_023240410.1;refGene_gn=SIPA1L2;refGene_tx=XM_023240411.1
D2 16410370 16410424
refGene_gn=SIPA1L2;refGene_tx=XM_023240409.1;refGene_gn=SIPA1L2;refGen
e_tx=XM_023240410.1;refGene_gn=SIPA1L2;refGene_tx=XM_023240412.1
D2 16411011 16411135
refGene_gn=SIPA1L2;refGene_tx=XM_023240409.1;refGene_gn=SIPA1L2;refGen
e_tx=XM_023240410.1;refGene_gn=SIPA1L2;refGene_tx=XM_023240411.1;refGe
ne_gn=SIPA1L2;refGene_tx=XM_023240412.1;refGene_gn=SIPA1L2;refGene_tx=
XM_023240413.1
D2 16412205 16412287
refGene_gn=SIPA1L2;refGene_tx=XM_023240409.1;refGene_gn=SIPA1L2;refGen
e_tx=XM_023240410.1;refGene_gn=SIPA1L2;refGene_tx=XM_023240411.1;refGe
ne_gn=SIPA1L2;refGene_tx=XM_023240412.1;refGene_gn=SIPA1L2;refGene_tx=
XM_023240413.1
D2 16414044 16414191
refGene_gn=SIPA1L2;refGene_tx=XM_023240409.1;refGene_gn=SIPA1L2;refGen
e_tx=XM_023240410.1;refGene_gn=SIPA1L2;refGene_tx=XM_023240411.1;refGe
ne_gn=SIPA1L2;refGene_tx=XM_023240412.1;refGene_gn=SIPA1L2;refGene_tx=
XM_023240413.1
D2 16697672 16697708 refGene_gn=DISC1;refGene_tx=XM_019813074.2
D2 16707454 16707594
refGene_gn=DISC1;refGene_tx=XM_003993922.4;refGene_gn=DISC1;refGene_tx
=XM_019813072.2;refGene_gn=DISC1;refGene_tx=XM_019813074.2
D2 16716874 16716992
refGene_gn=DISC1;refGene_tx=XM_003993922.4;refGene_gn=DISC1;refGene_tx
=XM_019813072.2;refGene_gn=DISC1;refGene_tx=XM_019813074.2
D2 16735933 16736210
refGene_gn=DISC1;refGene_tx=XM_003993922.4;refGene_gn=DISC1;refGene_tx
=XM_019813072.2;refGene_gn=DISC1;refGene_tx=XM_019813074.2
D2 16786672 16786733
refGene_gn=DISC1;refGene_tx=XM_003993922.4;refGene_gn=DISC1;refGene_tx
=XM_019813072.2;refGene_gn=DISC1;refGene_tx=XM_019813074.2
D2 16911205 16911394
refGene_gn=DISC1;refGene_tx=XM_003993922.4;refGene_gn=DISC1;refGene_tx
=XM_019813072.2;refGene_gn=DISC1;refGene_tx=XM_019813074.2
D2 16924445 16924548
refGene_gn=DISC1;refGene_tx=XM_003993922.4;refGene_gn=DISC1;refGene_tx
=XM_019813072.2;refGene_gn=DISC1;refGene_tx=XM_019813074.2
D2 16929233 16929288
refGene_gn=DISC1;refGene_tx=XM_003993922.4;refGene_gn=DISC1;refGene_tx
=XM_019813072.2;refGene_gn=DISC1;refGene_tx=XM_019813074.2
D2 16950447 16950683
refGene_gn=DISC1;refGene_tx=XM_003993922.4;refGene_gn=DISC1;refGene_tx
=XM_019813072.2;refGene_gn=DISC1;refGene_tx=XM_019813074.2
D2 16955453 16955583
refGene_gn=DISC1;refGene_tx=XM_003993922.4;refGene_gn=DISC1;refGene_tx
=XM_019813072.2;refGene_gn=DISC1;refGene_tx=XM_019813074.2
D2 16967069 16967094 refGene_gn=DISC1;refGene_tx=XM_019813085.2

D2 16970177 16970328
refGene_gn=DISC1;refGene_tx=XM_003993922.4;refGene_gn=DISC1;refGene_tx=XM_019813072.2;refGene_gn=DISC1;refGene_tx=XM_019813074.2;refGene_gn=DISC1;refGene_tx=XM_019813085.2
D2 17009075 17009145
refGene_gn=DISC1;refGene_tx=XM_003993922.4;refGene_gn=DISC1;refGene_tx=XM_019813072.2;refGene_gn=DISC1;refGene_tx=XM_019813074.2;refGene_gn=DISC1;refGene_tx=XM_019813085.2
D2 17016009 17017001
refGene_gn=DISC1;refGene_tx=XM_003993922.4;refGene_gn=DISC1;refGene_tx=XM_019813072.2;refGene_gn=DISC1;refGene_tx=XM_019813074.2;refGene_gn=DISC1;refGene_tx=XM_019813085.2
D2 17022280 17022361
refGene_gn=DISC1;refGene_tx=XM_019813072.2;refGene_gn=DISC1;refGene_tx=XM_019813074.2;refGene_gn=DISC1;refGene_tx=XM_019813085.2
D2 17065348 17065406
refGene_gn=DISC1;refGene_tx=XM_003993922.4;refGene_gn=DISC1;refGene_tx=XM_019813072.2;refGene_gn=DISC1;refGene_tx=XM_019813074.2;refGene_gn=DISC1;refGene_tx=XM_019813085.2
D2 17132737 17133115 refGene_gn=TSNAX;refGene_tx=XM_003993916.5
D2 17136161 17136289 refGene_gn=TSNAX;refGene_tx=XM_003993916.5
D2 17155755 17155886 refGene_gn=TSNAX;refGene_tx=XM_003993916.5
D2 17159363 17159478 refGene_gn=TSNAX;refGene_tx=XM_003993916.5
D2 17162796 17162901 refGene_gn=TSNAX;refGene_tx=XM_003993916.5
D2 17163349 17163365 refGene_gn=TSNAX;refGene_tx=XM_003993916.5
D2 17271839 17272769
refGene_gn=EGLN1;refGene_tx=XM_011287220.2;refGene_gn=EGLN1;refGene_tx=XM_023240414.1;refGene_gn=EGLN1;refGene_tx=XM_023240415.1
D2 17318836 17318956
refGene_gn=EGLN1;refGene_tx=XM_011287220.2;refGene_gn=EGLN1;refGene_tx=XM_023240414.1
D2 17322042 17322179
refGene_gn=EGLN1;refGene_tx=XM_011287220.2;refGene_gn=EGLN1;refGene_tx=XM_023240414.1;refGene_gn=EGLN1;refGene_tx=XM_023240415.1
D2 17324821 17324889
refGene_gn=EGLN1;refGene_tx=XM_011287220.2;refGene_gn=EGLN1;refGene_tx=XM_023240415.1
D2 17326462 17326527
refGene_gn=EGLN1;refGene_tx=XM_011287220.2;refGene_gn=EGLN1;refGene_tx=XM_023240414.1;refGene_gn=EGLN1;refGene_tx=XM_023240415.1
D2 17338181 17338711 refGene_gn=SPRTN;refGene_tx=XM_003993924.5
D2 17338713 17338926 refGene_gn=SPRTN;refGene_tx=XM_003993924.5
D2 17340024 17340286 refGene_gn=SPRTN;refGene_tx=XM_003993924.5
D2 17343604 17343733 refGene_gn=SPRTN;refGene_tx=XM_003993924.5
D2 17348213 17348313 refGene_gn=SPRTN;refGene_tx=XM_003993924.5
D2 17349762 17349983 refGene_gn=SPRTN;refGene_tx=XM_003993924.5
D2 17350632 17352813 refGene_gn=EXOC8;refGene_tx=XM_006937781.3
D2 17374451 17374495
refGene_gn=GNPAT;refGene_tx=XM_011287222.3;refGene_gn=GNPAT;refGene_tx=XM_019813088.1;refGene_gn=GNPAT;refGene_tx=XM_019813089.2;refGene_gn=

GNPAT;refGene_tx=XM_023240416.1
D2 17376533 17376595
refGene_gn=GNPAT;refGene_tx=XM_011287222.3;refGene_gn=GNPAT;refGene_tx=XM_019813088.1;refGene_gn=GNPAT;refGene_tx=XM_019813089.2;refGene_gn=GNPAT;refGene_tx=XM_023240416.1
D2 17377236 17377330
refGene_gn=GNPAT;refGene_tx=XM_011287222.3;refGene_gn=GNPAT;refGene_tx=XM_019813088.1;refGene_gn=GNPAT;refGene_tx=XM_019813089.2;refGene_gn=GNPAT;refGene_tx=XM_023240416.1
D2 17377409 17377509
refGene_gn=GNPAT;refGene_tx=XM_011287222.3;refGene_gn=GNPAT;refGene_tx=XM_019813088.1;refGene_gn=GNPAT;refGene_tx=XM_019813089.2;refGene_gn=GNPAT;refGene_tx=XM_023240416.1
D2 17378598 17378739
refGene_gn=GNPAT;refGene_tx=XM_011287222.3;refGene_gn=GNPAT;refGene_tx=XM_019813088.1;refGene_gn=GNPAT;refGene_tx=XM_019813089.2;refGene_gn=GNPAT;refGene_tx=XM_023240416.1
D2 17379794 17379874
refGene_gn=GNPAT;refGene_tx=XM_011287222.3;refGene_gn=GNPAT;refGene_tx=XM_019813088.1;refGene_gn=GNPAT;refGene_tx=XM_019813089.2;refGene_gn=GNPAT;refGene_tx=XM_023240416.1
D2 17381211 17381454
refGene_gn=GNPAT;refGene_tx=XM_011287222.3;refGene_gn=GNPAT;refGene_tx=XM_019813088.1;refGene_gn=GNPAT;refGene_tx=XM_019813089.2;refGene_gn=GNPAT;refGene_tx=XM_023240416.1
D2 17385225 17385449
refGene_gn=GNPAT;refGene_tx=XM_011287222.3;refGene_gn=GNPAT;refGene_tx=XM_019813088.1;refGene_gn=GNPAT;refGene_tx=XM_019813089.2;refGene_gn=GNPAT;refGene_tx=XM_023240416.1
D2 17386984 17387115
refGene_gn=GNPAT;refGene_tx=XM_011287222.3;refGene_gn=GNPAT;refGene_tx=XM_019813088.1;refGene_gn=GNPAT;refGene_tx=XM_019813089.2;refGene_gn=GNPAT;refGene_tx=XM_023240416.1
D2 17387252 17387404
refGene_gn=GNPAT;refGene_tx=XM_011287222.3;refGene_gn=GNPAT;refGene_tx=XM_019813088.1;refGene_gn=GNPAT;refGene_tx=XM_019813089.2;refGene_gn=GNPAT;refGene_tx=XM_023240416.1
D2 17387671 17387747
refGene_gn=GNPAT;refGene_tx=XM_011287222.3;refGene_gn=GNPAT;refGene_tx=XM_019813088.1;refGene_gn=GNPAT;refGene_tx=XM_019813089.2;refGene_gn=GNPAT;refGene_tx=XM_023240416.1
D2 17388233 17388361
refGene_gn=GNPAT;refGene_tx=XM_011287222.3;refGene_gn=GNPAT;refGene_tx=XM_019813088.1;refGene_gn=GNPAT;refGene_tx=XM_019813089.2;refGene_gn=GNPAT;refGene_tx=XM_023240416.1
D2 17389491 17389621
refGene_gn=GNPAT;refGene_tx=XM_011287222.3;refGene_gn=GNPAT;refGene_tx=XM_019813088.1;refGene_gn=GNPAT;refGene_tx=XM_019813089.2;refGene_gn=GNPAT;refGene_tx=XM_023240416.1
D2 17392220 17392397

refGene_gn=GNPAT;refGene_tx=XM_011287222.3;refGene_gn=GNPAT;refGene_tx=XM_019813088.1;refGene_gn=GNPAT;refGene_tx=XM_019813089.2;refGene_gn=GNPAT;refGene_tx=XM_023240416.1
D2 17399202 17399385
refGene_gn=GNPAT;refGene_tx=XM_011287222.3;refGene_gn=GNPAT;refGene_tx=XM_019813088.1;refGene_gn=GNPAT;refGene_tx=XM_019813089.2;refGene_gn=GNPAT;refGene_tx=XM_023240416.1
D2 17399932 17399977 refGene_gn=GNPAT;refGene_tx=XM_023240416.1
D2 17406643 17406649 refGene_gn=GNPAT;refGene_tx=XM_019813089.2
D2 17407475 17407553 refGene_gn=GNPAT;refGene_tx=XM_011287222.3
D2 17407827 17407915
refGene_gn=CD2H1orf131;refGene_tx=XM_011287221.3
D2 17408336 17408357 refGene_gn=GNPAT;refGene_tx=XM_019813088.1
D2 17409555 17409902
refGene_gn=CD2H1orf131;refGene_tx=XM_011287221.3
D2 17419906 17419963
refGene_gn=CD2H1orf131;refGene_tx=XM_011287221.3
D2 17420652 17420748
refGene_gn=CD2H1orf131;refGene_tx=XM_011287221.3
D2 17420889 17420976
refGene_gn=CD2H1orf131;refGene_tx=XM_011287221.3
D2 17421537 17421590
refGene_gn=CD2H1orf131;refGene_tx=XM_011287221.3
D2 17422699 17422838
refGene_gn=CD2H1orf131;refGene_tx=XM_011287221.3
D2 17431932 17431998
refGene_gn=TRIM67;refGene_tx=XM_019813091.2;refGene_gn=TRIM67;refGene_tx=XM_019813092.2
D2 17433139 17433302
refGene_gn=TRIM67;refGene_tx=XM_019813091.2;refGene_gn=TRIM67;refGene_tx=XM_019813092.2
D2 17435964 17436268
refGene_gn=TRIM67;refGene_tx=XM_019813091.2;refGene_gn=TRIM67;refGene_tx=XM_019813092.2
D2 17437598 17437737
refGene_gn=TRIM67;refGene_tx=XM_019813091.2;refGene_gn=TRIM67;refGene_tx=XM_019813092.2
D2 17440271 17440417
refGene_gn=TRIM67;refGene_tx=XM_019813091.2;refGene_gn=TRIM67;refGene_tx=XM_019813092.2
D2 17442672 17442832
refGene_gn=TRIM67;refGene_tx=XM_019813091.2;refGene_gn=TRIM67;refGene_tx=XM_019813092.2
D2 17444177 17444288
refGene_gn=TRIM67;refGene_tx=XM_019813091.2;refGene_gn=TRIM67;refGene_tx=XM_019813092.2
D2 17445366 17445489
refGene_gn=TRIM67;refGene_tx=XM_019813091.2;refGene_gn=TRIM67;refGene_tx=XM_019813092.2
D2 17446560 17446656

refGene_gn=TRIM67;refGene_tx=XM_019813091.2;refGene_gn=TRIM67;refGene_tx=XM_019813092.2
D2 17482635 17483679
refGene_gn=TRIM67;refGene_tx=XM_019813091.2;refGene_gn=TRIM67;refGene_tx=XM_019813092.2
D2 17607337 17607622 refGene_gn=FAM89A;refGene_tx=XM_023240417.1
D2 17627063 17627327 refGene_gn=FAM89A;refGene_tx=XM_023240417.1
D2 17640543 17640686 refGene_gn=ARV1;refGene_tx=XM_023240424.1
D2 17641724 17641949
refGene_gn=ARV1;refGene_tx=XM_023240424.1;refGene_gn=ARV1;refGene_tx=XM_023240425.1
D2 17644556 17644710
refGene_gn=ARV1;refGene_tx=XM_023240424.1;refGene_gn=ARV1;refGene_tx=XM_023240425.1
D2 17647412 17647532
refGene_gn=ARV1;refGene_tx=XM_023240424.1;refGene_gn=ARV1;refGene_tx=XM_023240425.1
D2 17655931 17656141
refGene_gn=ARV1;refGene_tx=XM_023240424.1;refGene_gn=ARV1;refGene_tx=XM_023240425.1
D2 17656415 17656686
refGene_gn=TTC13;refGene_tx=XM_023240418.1;refGene_gn=TTC13;refGene_tx=XM_023240419.1;refGene_gn=TTC13;refGene_tx=XM_023240420.1;refGene_gn=TTC13;refGene_tx=XM_023240421.1;refGene_gn=TTC13;refGene_tx=XM_023240422.1
D2 17675996 17676091
refGene_gn=TTC13;refGene_tx=XM_023240418.1;refGene_gn=TTC13;refGene_tx=XM_023240419.1;refGene_gn=TTC13;refGene_tx=XM_023240420.1;refGene_gn=TTC13;refGene_tx=XM_023240421.1;refGene_gn=TTC13;refGene_tx=XM_023240422.1
D2 17679873 17679949
refGene_gn=TTC13;refGene_tx=XM_023240418.1;refGene_gn=TTC13;refGene_tx=XM_023240419.1;refGene_gn=TTC13;refGene_tx=XM_023240420.1;refGene_gn=TTC13;refGene_tx=XM_023240421.1;refGene_gn=TTC13;refGene_tx=XM_023240422.1
D2 17684215 17684286
refGene_gn=TTC13;refGene_tx=XM_023240418.1;refGene_gn=TTC13;refGene_tx=XM_023240419.1;refGene_gn=TTC13;refGene_tx=XM_023240420.1;refGene_gn=TTC13;refGene_tx=XM_023240421.1;refGene_gn=TTC13;refGene_tx=XM_023240422.1;refGene_gn=TTC13;refGene_tx=XM_023240423.1
D2 17693610 17693676
refGene_gn=TTC13;refGene_tx=XM_023240418.1;refGene_gn=TTC13;refGene_tx=XM_023240419.1;refGene_gn=TTC13;refGene_tx=XM_023240420.1;refGene_gn=TTC13;refGene_tx=XM_023240423.1
D2 17695180 17695273
refGene_gn=TTC13;refGene_tx=XM_023240418.1;refGene_gn=TTC13;refGene_tx=XM_023240419.1;refGene_gn=TTC13;refGene_tx=XM_023240423.1
D2 17697942 17698059
refGene_gn=TTC13;refGene_tx=XM_023240418.1;refGene_gn=TTC13;refGene_tx=XM_023240419.1;refGene_gn=TTC13;refGene_tx=XM_023240420.1;refGene_gn=

TTC13;refGene_tx=XM_023240421.1;refGene_gn=TTC13;refGene_tx=XM_0232404
22.1;refGene_gn=TTC13;refGene_tx=XM_023240423.1
D2 17699118 17699229
refGene_gn=TTC13;refGene_tx=XM_023240418.1;refGene_gn=TTC13;refGene_tx
=XM_023240419.1;refGene_gn=TTC13;refGene_tx=XM_023240420.1;refGene_gn=
TTC13;refGene_tx=XM_023240421.1;refGene_gn=TTC13;refGene_tx=XM_0232404
22.1;refGene_gn=TTC13;refGene_tx=XM_023240423.1
D2 17704429 17704512
refGene_gn=TTC13;refGene_tx=XM_023240418.1;refGene_gn=TTC13;refGene_tx
=XM_023240419.1;refGene_gn=TTC13;refGene_tx=XM_023240420.1;refGene_gn=
TTC13;refGene_tx=XM_023240421.1;refGene_gn=TTC13;refGene_tx=XM_0232404
22.1;refGene_gn=TTC13;refGene_tx=XM_023240423.1
D2 17708418 17708560
refGene_gn=TTC13;refGene_tx=XM_023240418.1;refGene_gn=TTC13;refGene_tx
=XM_023240419.1;refGene_gn=TTC13;refGene_tx=XM_023240420.1;refGene_gn=
TTC13;refGene_tx=XM_023240421.1;refGene_gn=TTC13;refGene_tx=XM_0232404
22.1;refGene_gn=TTC13;refGene_tx=XM_023240423.1
D2 17708892 17709067
refGene_gn=TTC13;refGene_tx=XM_023240418.1;refGene_gn=TTC13;refGene_tx
=XM_023240419.1;refGene_gn=TTC13;refGene_tx=XM_023240420.1;refGene_gn=
TTC13;refGene_tx=XM_023240421.1;refGene_gn=TTC13;refGene_tx=XM_0232404
22.1;refGene_gn=TTC13;refGene_tx=XM_023240423.1
D2 17711169 17711326
refGene_gn=TTC13;refGene_tx=XM_023240418.1;refGene_gn=TTC13;refGene_tx
=XM_023240419.1;refGene_gn=TTC13;refGene_tx=XM_023240420.1;refGene_gn=
TTC13;refGene_tx=XM_023240421.1;refGene_gn=TTC13;refGene_tx=XM_0232404
22.1;refGene_gn=TTC13;refGene_tx=XM_023240423.1
D2 17715683 17715814
refGene_gn=TTC13;refGene_tx=XM_023240418.1;refGene_gn=TTC13;refGene_tx
=XM_023240419.1;refGene_gn=TTC13;refGene_tx=XM_023240420.1;refGene_gn=
TTC13;refGene_tx=XM_023240421.1;refGene_gn=TTC13;refGene_tx=XM_0232404
22.1;refGene_gn=TTC13;refGene_tx=XM_023240423.1
D2 17716585 17716718
refGene_gn=TTC13;refGene_tx=XM_023240418.1;refGene_gn=TTC13;refGene_tx
=XM_023240419.1;refGene_gn=TTC13;refGene_tx=XM_023240420.1;refGene_gn=
TTC13;refGene_tx=XM_023240421.1;refGene_gn=TTC13;refGene_tx=XM_0232404
22.1;refGene_gn=TTC13;refGene_tx=XM_023240423.1
D2 17717553 17717646
refGene_gn=TTC13;refGene_tx=XM_023240418.1;refGene_gn=TTC13;refGene_tx
=XM_023240419.1;refGene_gn=TTC13;refGene_tx=XM_023240420.1;refGene_gn=
TTC13;refGene_tx=XM_023240421.1;refGene_gn=TTC13;refGene_tx=XM_0232404
22.1;refGene_gn=TTC13;refGene_tx=XM_023240423.1
D2 17718950 17719034
refGene_gn=TTC13;refGene_tx=XM_023240418.1;refGene_gn=TTC13;refGene_tx
=XM_023240419.1;refGene_gn=TTC13;refGene_tx=XM_023240420.1;refGene_gn=
TTC13;refGene_tx=XM_023240421.1;refGene_gn=TTC13;refGene_tx=XM_0232404
22.1;refGene_gn=TTC13;refGene_tx=XM_023240423.1
D2 17720098 17720183
refGene_gn=TTC13;refGene_tx=XM_023240418.1;refGene_gn=TTC13;refGene_tx
=XM_023240419.1;refGene_gn=TTC13;refGene_tx=XM_023240420.1;refGene_gn=

TTC13;refGene_tx=XM_023240421.1;refGene_gn=TTC13;refGene_tx=XM_023240422.1;refGene_gn=TTC13;refGene_tx=XM_023240423.1
D2 17730013 17730123
refGene_gn=TTC13;refGene_tx=XM_023240418.1;refGene_gn=TTC13;refGene_tx=XM_023240419.1;refGene_gn=TTC13;refGene_tx=XM_023240420.1;refGene_gn=TTC13;refGene_tx=XM_023240421.1;refGene_gn=TTC13;refGene_tx=XM_023240422.1;refGene_gn=TTC13;refGene_tx=XM_023240423.1
D2 17732043 17732179
refGene_gn=TTC13;refGene_tx=XM_023240418.1;refGene_gn=TTC13;refGene_tx=XM_023240419.1;refGene_gn=TTC13;refGene_tx=XM_023240420.1;refGene_gn=TTC13;refGene_tx=XM_023240421.1;refGene_gn=TTC13;refGene_tx=XM_023240422.1;refGene_gn=TTC13;refGene_tx=XM_023240423.1
D2 17733337 17733417
refGene_gn=TTC13;refGene_tx=XM_023240418.1;refGene_gn=TTC13;refGene_tx=XM_023240419.1;refGene_gn=TTC13;refGene_tx=XM_023240420.1;refGene_gn=TTC13;refGene_tx=XM_023240421.1;refGene_gn=TTC13;refGene_tx=XM_023240422.1;refGene_gn=TTC13;refGene_tx=XM_023240423.1
D2 17736087 17736166
refGene_gn=TTC13;refGene_tx=XM_023240418.1;refGene_gn=TTC13;refGene_tx=XM_023240419.1;refGene_gn=TTC13;refGene_tx=XM_023240420.1;refGene_gn=TTC13;refGene_tx=XM_023240421.1;refGene_gn=TTC13;refGene_tx=XM_023240422.1;refGene_gn=TTC13;refGene_tx=XM_023240423.1
D2 17736315 17736395
refGene_gn=TTC13;refGene_tx=XM_023240418.1;refGene_gn=TTC13;refGene_tx=XM_023240419.1;refGene_gn=TTC13;refGene_tx=XM_023240420.1;refGene_gn=TTC13;refGene_tx=XM_023240421.1;refGene_gn=TTC13;refGene_tx=XM_023240422.1;refGene_gn=TTC13;refGene_tx=XM_023240423.1
D2 17737858 17737973
refGene_gn=TTC13;refGene_tx=XM_023240418.1;refGene_gn=TTC13;refGene_tx=XM_023240419.1;refGene_gn=TTC13;refGene_tx=XM_023240420.1;refGene_gn=TTC13;refGene_tx=XM_023240421.1;refGene_gn=TTC13;refGene_tx=XM_023240422.1;refGene_gn=TTC13;refGene_tx=XM_023240423.1
D2 17765965 17766307
refGene_gn=CD2H1orf198;refGene_tx=XM_023240426.1
D2 17780358 17780409
refGene_gn=CD2H1orf198;refGene_tx=XM_023240426.1
D2 17789811 17790345
refGene_gn=CD2H1orf198;refGene_tx=XM_023240426.1
D2 17793888 17793945
refGene_gn=CD2H1orf198;refGene_tx=XM_023240426.1
D2 17840359 17840386
refGene_gn=CAPN9;refGene_tx=XM_003993929.4;refGene_gn=CAPN9;refGene_tx=XM_003993930.4
D2 17843239 17843298
refGene_gn=CAPN9;refGene_tx=XM_003993929.4;refGene_gn=CAPN9;refGene_tx=XM_003993930.4
D2 17845932 17846049
refGene_gn=CAPN9;refGene_tx=XM_003993929.4;refGene_gn=CAPN9;refGene_tx=XM_003993930.4
D2 17847159 17847238

refGene_gn=CAPN9;refGene_tx=XM_003993929.4;refGene_gn=CAPN9;refGene_tx=XM_003993930.4
D2 17847558 17847627 refGene_gn=CAPN9;refGene_tx=XM_003993929.4;refGene_gn=CAPN9;refGene_tx=XM_003993930.4
D2 17848101 17848166 refGene_gn=CAPN9;refGene_tx=XM_003993929.4;refGene_gn=CAPN9;refGene_tx=XM_003993930.4
D2 17849648 17849706 refGene_gn=CAPN9;refGene_tx=XM_003993929.4;refGene_gn=CAPN9;refGene_tx=XM_003993930.4
D2 17851434 17851515 refGene_gn=CAPN9;refGene_tx=XM_003993929.4;refGene_gn=CAPN9;refGene_tx=XM_003993930.4
D2 17852559 17852596 refGene_gn=CAPN9;refGene_tx=XM_003993929.4;refGene_gn=CAPN9;refGene_tx=XM_003993930.4
D2 17855744 17855953 refGene_gn=CAPN9;refGene_tx=XM_003993929.4;refGene_gn=CAPN9;refGene_tx=XM_003993930.4
D2 17856109 17856267 refGene_gn=CAPN9;refGene_tx=XM_003993929.4;refGene_gn=CAPN9;refGene_tx=XM_003993930.4
D2 17857281 17857442 refGene_gn=CAPN9;refGene_tx=XM_003993929.4;refGene_gn=CAPN9;refGene_tx=XM_003993930.4
D2 17859398 17859476 refGene_gn=CAPN9;refGene_tx=XM_003993929.4
D2 17861773 17861859 refGene_gn=CAPN9;refGene_tx=XM_003993929.4;refGene_gn=CAPN9;refGene_tx=XM_003993930.4
D2 17866539 17866623 refGene_gn=CAPN9;refGene_tx=XM_003993929.4;refGene_gn=CAPN9;refGene_tx=XM_003993930.4
D2 17867521 17867690 refGene_gn=CAPN9;refGene_tx=XM_003993929.4;refGene_gn=CAPN9;refGene_tx=XM_003993930.4
D2 17871747 17871881 refGene_gn=CAPN9;refGene_tx=XM_003993929.4;refGene_gn=CAPN9;refGene_tx=XM_003993930.4
D2 17873355 17873474 refGene_gn=CAPN9;refGene_tx=XM_003993929.4;refGene_gn=CAPN9;refGene_tx=XM_003993930.4
D2 17877871 17877941 refGene_gn=CAPN9;refGene_tx=XM_003993929.4;refGene_gn=CAPN9;refGene_tx=XM_003993930.4
D2 17884279 17884492 refGene_gn=CAPN9;refGene_tx=XM_003993929.4;refGene_gn=CAPN9;refGene_tx=XM_003993930.4
D2 17913295 17914127 refGene_gn=AGT;refGene_tx=XM_011287235.3
D2 17917710 17917978 refGene_gn=AGT;refGene_tx=XM_011287235.3

D2 17919566 17919711 refGene_gn=AGT;refGene_tx=XM_011287235.3
D2 17920458 17920650 refGene_gn=AGT;refGene_tx=XM_011287235.3
D2 17930142 17930244
refGene_gn=COG2;refGene_tx=XM_003993932.4;refGene_gn=COG2;refGene_tx=XM_006937789.3
D2 17931140 17931311
refGene_gn=COG2;refGene_tx=XM_023240427.1;refGene_gn=COG2;refGene_tx=XM_023240428.1;refGene_gn=COG2;refGene_tx=XM_023240429.1;refGene_gn=COG2;refGene_tx=XM_023240430.1
D2 17931703 17931884
refGene_gn=COG2;refGene_tx=XM_003993932.4;refGene_gn=COG2;refGene_tx=XM_006937789.3;refGene_gn=COG2;refGene_tx=XM_023240427.1;refGene_gn=COG2;refGene_tx=XM_023240428.1;refGene_gn=COG2;refGene_tx=XM_023240429.1;refGene_gn=COG2;refGene_tx=XM_023240430.1
D2 17933155 17933295
refGene_gn=COG2;refGene_tx=XM_003993932.4;refGene_gn=COG2;refGene_tx=XM_006937789.3;refGene_gn=COG2;refGene_tx=XM_023240427.1;refGene_gn=COG2;refGene_tx=XM_023240428.1;refGene_gn=COG2;refGene_tx=XM_023240429.1;refGene_gn=COG2;refGene_tx=XM_023240430.1
D2 17935043 17935186
refGene_gn=COG2;refGene_tx=XM_003993932.4;refGene_gn=COG2;refGene_tx=XM_006937789.3;refGene_gn=COG2;refGene_tx=XM_023240427.1;refGene_gn=COG2;refGene_tx=XM_023240428.1;refGene_gn=COG2;refGene_tx=XM_023240429.1;refGene_gn=COG2;refGene_tx=XM_023240430.1
D2 17935485 17935558
refGene_gn=COG2;refGene_tx=XM_003993932.4;refGene_gn=COG2;refGene_tx=XM_006937789.3;refGene_gn=COG2;refGene_tx=XM_023240427.1;refGene_gn=COG2;refGene_tx=XM_023240428.1;refGene_gn=COG2;refGene_tx=XM_023240429.1;refGene_gn=COG2;refGene_tx=XM_023240430.1
D2 17936377 17936575
refGene_gn=COG2;refGene_tx=XM_003993932.4;refGene_gn=COG2;refGene_tx=XM_006937789.3;refGene_gn=COG2;refGene_tx=XM_023240427.1;refGene_gn=COG2;refGene_tx=XM_023240428.1;refGene_gn=COG2;refGene_tx=XM_023240429.1;refGene_gn=COG2;refGene_tx=XM_023240430.1
D2 17938659 17938811
refGene_gn=COG2;refGene_tx=XM_003993932.4;refGene_gn=COG2;refGene_tx=XM_006937789.3;refGene_gn=COG2;refGene_tx=XM_023240427.1;refGene_gn=COG2;refGene_tx=XM_023240428.1;refGene_gn=COG2;refGene_tx=XM_023240429.1;refGene_gn=COG2;refGene_tx=XM_023240430.1
D2 17940255 17940317
refGene_gn=COG2;refGene_tx=XM_003993932.4;refGene_gn=COG2;refGene_tx=XM_006937789.3;refGene_gn=COG2;refGene_tx=XM_023240427.1;refGene_gn=COG2;refGene_tx=XM_023240428.1;refGene_gn=COG2;refGene_tx=XM_023240429.1;refGene_gn=COG2;refGene_tx=XM_023240430.1
D2 17944972 17945112
refGene_gn=COG2;refGene_tx=XM_003993932.4;refGene_gn=COG2;refGene_tx=XM_006937789.3;refGene_gn=COG2;refGene_tx=XM_023240427.1;refGene_gn=COG2;refGene_tx=XM_023240428.1;refGene_gn=COG2;refGene_tx=XM_023240429.1
D2 17946994 17947121
refGene_gn=COG2;refGene_tx=XM_003993932.4;refGene_gn=COG2;refGene_tx=X

M_006937789.3;refGene_gn=COG2;refGene_tx=XM_023240427.1;refGene_gn=COG2;refGene_tx=XM_023240428.1;refGene_gn=COG2;refGene_tx=XM_023240429.1
D2 17951674 17951799
refGene_gn=COG2;refGene_tx=XM_003993932.4;refGene_gn=COG2;refGene_tx=XM_006937789.3;refGene_gn=COG2;refGene_tx=XM_023240427.1;refGene_gn=COG2;refGene_tx=XM_023240428.1;refGene_gn=COG2;refGene_tx=XM_023240429.1;refGene_gn=COG2;refGene_tx=XM_023240430.1
D2 17953461 17953641
refGene_gn=COG2;refGene_tx=XM_003993932.4;refGene_gn=COG2;refGene_tx=XM_006937789.3;refGene_gn=COG2;refGene_tx=XM_023240427.1;refGene_gn=COG2;refGene_tx=XM_023240428.1;refGene_gn=COG2;refGene_tx=XM_023240429.1;refGene_gn=COG2;refGene_tx=XM_023240430.1
D2 17954193 17954302
refGene_gn=COG2;refGene_tx=XM_003993932.4;refGene_gn=COG2;refGene_tx=XM_006937789.3;refGene_gn=COG2;refGene_tx=XM_023240427.1;refGene_gn=COG2;refGene_tx=XM_023240428.1;refGene_gn=COG2;refGene_tx=XM_023240429.1;refGene_gn=COG2;refGene_tx=XM_023240430.1
D2 17957065 17957169
refGene_gn=COG2;refGene_tx=XM_003993932.4;refGene_gn=COG2;refGene_tx=XM_006937789.3;refGene_gn=COG2;refGene_tx=XM_023240427.1;refGene_gn=COG2;refGene_tx=XM_023240428.1;refGene_gn=COG2;refGene_tx=XM_023240429.1;refGene_gn=COG2;refGene_tx=XM_023240430.1
D2 17958836 17958917
refGene_gn=COG2;refGene_tx=XM_003993932.4;refGene_gn=COG2;refGene_tx=XM_006937789.3;refGene_gn=COG2;refGene_tx=XM_023240427.1;refGene_gn=COG2;refGene_tx=XM_023240428.1;refGene_gn=COG2;refGene_tx=XM_023240430.1
D2 17961133 17961199
refGene_gn=COG2;refGene_tx=XM_003993932.4;refGene_gn=COG2;refGene_tx=XM_006937789.3;refGene_gn=COG2;refGene_tx=XM_023240427.1;refGene_gn=COG2;refGene_tx=XM_023240428.1;refGene_gn=COG2;refGene_tx=XM_023240429.1;refGene_gn=COG2;refGene_tx=XM_023240430.1
D2 17962317 17962479
refGene_gn=COG2;refGene_tx=XM_003993932.4;refGene_gn=COG2;refGene_tx=XM_006937789.3;refGene_gn=COG2;refGene_tx=XM_023240427.1;refGene_gn=COG2;refGene_tx=XM_023240428.1;refGene_gn=COG2;refGene_tx=XM_023240429.1;refGene_gn=COG2;refGene_tx=XM_023240430.1
D2 17975754 17975826
refGene_gn=COG2;refGene_tx=XM_003993932.4;refGene_gn=COG2;refGene_tx=XM_006937789.3;refGene_gn=COG2;refGene_tx=XM_023240427.1;refGene_gn=COG2;refGene_tx=XM_023240428.1;refGene_gn=COG2;refGene_tx=XM_023240429.1;refGene_gn=COG2;refGene_tx=XM_023240430.1
D2 18133671 18133996
refGene_gn=PGBD5;refGene_tx=XM_019814059.2;refGene_gn=PGBD5;refGene_tx=XM_023240431.1;refGene_gn=PGBD5;refGene_tx=XM_023240432.1
D2 18208259 18208687
refGene_gn=PGBD5;refGene_tx=XM_019814059.2;refGene_gn=PGBD5;refGene_tx=XM_023240431.1;refGene_gn=PGBD5;refGene_tx=XM_023240432.1;refGene_gn=PGBD5;refGene_tx=XM_023240433.1
D2 18216049 18216184
refGene_gn=PGBD5;refGene_tx=XM_019814059.2;refGene_gn=PGBD5;refGene_tx

=XM_023240431.1;refGene_gn=PGBD5;refGene_tx=XM_023240432.1;refGene_gn=PGBD5;refGene_tx=XM_023240433.1
D2 18229873 18230054
refGene_gn=PGBD5;refGene_tx=XM_019814059.2;refGene_gn=PGBD5;refGene_tx=XM_023240431.1;refGene_gn=PGBD5;refGene_tx=XM_023240432.1;refGene_gn=PGBD5;refGene_tx=XM_023240433.1
D2 18233879 18234077
refGene_gn=PGBD5;refGene_tx=XM_019814059.2;refGene_gn=PGBD5;refGene_tx=XM_023240431.1;refGene_gn=PGBD5;refGene_tx=XM_023240432.1;refGene_gn=PGBD5;refGene_tx=XM_023240433.1
D2 18235896 18236002
refGene_gn=PGBD5;refGene_tx=XM_019814059.2;refGene_gn=PGBD5;refGene_tx=XM_023240431.1;refGene_gn=PGBD5;refGene_tx=XM_023240432.1;refGene_gn=PGBD5;refGene_tx=XM_023240433.1
D2 18237457 18237653
refGene_gn=PGBD5;refGene_tx=XM_019814059.2;refGene_gn=PGBD5;refGene_tx=XM_023240431.1;refGene_gn=PGBD5;refGene_tx=XM_023240432.1;refGene_gn=PGBD5;refGene_tx=XM_023240433.1
D2 18237889 18237995
refGene_gn=PGBD5;refGene_tx=XM_023240431.1;refGene_gn=PGBD5;refGene_tx=XM_023240433.1
D2 18239181 18239185 refGene_gn=PGBD5;refGene_tx=XM_023240432.1
D2 18275235 18275391
refGene_gn=GALNT2;refGene_tx=XM_006937791.4;refGene_gn=GALNT2;refGene_tx=XM_023240434.1;refGene_gn=GALNT2;refGene_tx=XM_023240435.1
D2 18279495 18279615
refGene_gn=GALNT2;refGene_tx=XM_006937791.4;refGene_gn=GALNT2;refGene_tx=XM_023240434.1;refGene_gn=GALNT2;refGene_tx=XM_023240435.1
D2 18286057 18286184
refGene_gn=GALNT2;refGene_tx=XM_006937791.4;refGene_gn=GALNT2;refGene_tx=XM_023240434.1;refGene_gn=GALNT2;refGene_tx=XM_023240435.1
D2 18288676 18288760
refGene_gn=GALNT2;refGene_tx=XM_006937791.4;refGene_gn=GALNT2;refGene_tx=XM_023240434.1;refGene_gn=GALNT2;refGene_tx=XM_023240435.1
D2 18289093 18289186
refGene_gn=GALNT2;refGene_tx=XM_006937791.4;refGene_gn=GALNT2;refGene_tx=XM_023240434.1;refGene_gn=GALNT2;refGene_tx=XM_023240435.1
D2 18294270 18294397
refGene_gn=GALNT2;refGene_tx=XM_006937791.4;refGene_gn=GALNT2;refGene_tx=XM_023240434.1;refGene_gn=GALNT2;refGene_tx=XM_023240435.1
D2 18299140 18299244
refGene_gn=GALNT2;refGene_tx=XM_006937791.4;refGene_gn=GALNT2;refGene_tx=XM_023240434.1;refGene_gn=GALNT2;refGene_tx=XM_023240435.1
D2 18299848 18299936
refGene_gn=GALNT2;refGene_tx=XM_006937791.4;refGene_gn=GALNT2;refGene_tx=XM_023240434.1;refGene_gn=GALNT2;refGene_tx=XM_023240435.1
D2 18303501 18303589
refGene_gn=GALNT2;refGene_tx=XM_006937791.4;refGene_gn=GALNT2;refGene_tx=XM_023240434.1;refGene_gn=GALNT2;refGene_tx=XM_023240435.1
D2 18306047 18306169

refGene_gn=GALNT2;refGene_tx=XM_006937791.4;refGene_gn=GALNT2;refGene_tx=XM_023240434.1;refGene_gn=GALNT2;refGene_tx=XM_023240435.1
D2 18309687 18309753
refGene_gn=GALNT2;refGene_tx=XM_006937791.4;refGene_gn=GALNT2;refGene_tx=XM_023240434.1;refGene_gn=GALNT2;refGene_tx=XM_023240435.1
D2 18309975 18310043
refGene_gn=GALNT2;refGene_tx=XM_006937791.4;refGene_gn=GALNT2;refGene_tx=XM_023240434.1;refGene_gn=GALNT2;refGene_tx=XM_023240435.1
D2 18310551 18310650
refGene_gn=GALNT2;refGene_tx=XM_006937791.4;refGene_gn=GALNT2;refGene_tx=XM_023240434.1;refGene_gn=GALNT2;refGene_tx=XM_023240435.1
D2 18344114 18344268
refGene_gn=GALNT2;refGene_tx=XM_006937791.4;refGene_gn=GALNT2;refGene_tx=XM_023240434.1;refGene_gn=GALNT2;refGene_tx=XM_023240435.1
D2 18368103 18368197
refGene_gn=GALNT2;refGene_tx=XM_006937791.4;refGene_gn=GALNT2;refGene_tx=XM_023240434.1;refGene_gn=GALNT2;refGene_tx=XM_023240435.1
D2 18377071 18377083 refGene_gn=GALNT2;refGene_tx=XM_006937791.4
D2 18468473 18468599 refGene_gn=GALNT2;refGene_tx=XM_023240434.1
D2 18836431 18836629
refGene_gn=URB2;refGene_tx=XM_019813108.2;refGene_gn=URB2;refGene_tx=XM_023240436.1;refGene_gn=URB2;refGene_tx=XM_023240437.1;refGene_gn=URB2;refGene_tx=XM_023240438.1
D2 18840617 18840757
refGene_gn=URB2;refGene_tx=XM_019813108.2;refGene_gn=URB2;refGene_tx=XM_023240436.1;refGene_gn=URB2;refGene_tx=XM_023240437.1;refGene_gn=URB2;refGene_tx=XM_023240438.1;refGene_gn=URB2;refGene_tx=XM_023240439.1
D2 18844402 18844490
refGene_gn=URB2;refGene_tx=XM_019813108.2;refGene_gn=URB2;refGene_tx=XM_023240436.1;refGene_gn=URB2;refGene_tx=XM_023240437.1;refGene_gn=URB2;refGene_tx=XM_023240438.1
D2 18849003 18849246
refGene_gn=URB2;refGene_tx=XM_019813108.2;refGene_gn=URB2;refGene_tx=XM_023240436.1;refGene_gn=URB2;refGene_tx=XM_023240437.1;refGene_gn=URB2;refGene_tx=XM_023240438.1;refGene_gn=URB2;refGene_tx=XM_023240439.1
D2 18850754 18850865
refGene_gn=URB2;refGene_tx=XM_019813108.2;refGene_gn=URB2;refGene_tx=XM_023240436.1;refGene_gn=URB2;refGene_tx=XM_023240437.1;refGene_gn=URB2;refGene_tx=XM_023240438.1;refGene_gn=URB2;refGene_tx=XM_023240439.1
D2 18852726 18852887
refGene_gn=URB2;refGene_tx=XM_019813108.2;refGene_gn=URB2;refGene_tx=XM_023240436.1;refGene_gn=URB2;refGene_tx=XM_023240437.1;refGene_gn=URB2;refGene_tx=XM_023240438.1;refGene_gn=URB2;refGene_tx=XM_023240439.1
D2 18857059 18860381
refGene_gn=URB2;refGene_tx=XM_019813108.2;refGene_gn=URB2;refGene_tx=XM_023240436.1;refGene_gn=URB2;refGene_tx=XM_023240437.1;refGene_gn=URB2;refGene_tx=XM_023240438.1;refGene_gn=URB2;refGene_tx=XM_023240439.1
D2 18861542 18861719
refGene_gn=URB2;refGene_tx=XM_019813108.2;refGene_gn=URB2;refGene_tx=XM_023240436.1;refGene_gn=URB2;refGene_tx=XM_023240437.1;refGene_gn=URB

2;refGene_tx=XM_023240438.1;refGene_gn=URB2;refGene_tx=XM_023240439.1
D2 18863693 18863819
refGene_gn=URB2;refGene_tx=XM_019813108.2;refGene_gn=URB2;refGene_tx=XM_023240436.1;refGene_gn=URB2;refGene_tx=XM_023240437.1;refGene_gn=URB2;refGene_tx=XM_023240438.1;refGene_gn=URB2;refGene_tx=XM_023240439.1
D2 18875976 18876118
refGene_gn=TAF5L;refGene_tx=XM_003993933.5;refGene_gn=TAF5L;refGene_tx=XM_023240440.1;refGene_gn=TAF5L;refGene_tx=XM_023240441.1
D2 18880479 18880584
refGene_gn=TAF5L;refGene_tx=XM_003993933.5;refGene_gn=TAF5L;refGene_tx=XM_023240440.1;refGene_gn=TAF5L;refGene_tx=XM_023240441.1
D2 18888655 18889386
refGene_gn=TAF5L;refGene_tx=XM_003993933.5;refGene_gn=TAF5L;refGene_tx=XM_023240440.1;refGene_gn=TAF5L;refGene_tx=XM_023240441.1
D2 18896776 18897574
refGene_gn=TAF5L;refGene_tx=XM_003993933.5;refGene_gn=TAF5L;refGene_tx=XM_023240440.1
D2 18929545 18930062
refGene_gn=ABCB10;refGene_tx=XM_003993943.4;refGene_gn=ABCB10;refGene_tx=XM_023240442.1
D2 18936308 18936319 refGene_gn=ABCB10;refGene_tx=XM_023240443.1
D2 18939826 18940029
refGene_gn=ABCB10;refGene_tx=XM_023240443.1;refGene_gn=ABCB10;refGene_tx=XM_023240444.1
D2 18940286 18940487
refGene_gn=ABCB10;refGene_tx=XM_003993943.4;refGene_gn=ABCB10;refGene_tx=XM_023240442.1;refGene_gn=ABCB10;refGene_tx=XM_023240443.1;refGene_gn=ABCB10;refGene_tx=XM_023240444.1
D2 18941338 18941541
refGene_gn=ABCB10;refGene_tx=XM_003993943.4;refGene_gn=ABCB10;refGene_tx=XM_023240442.1;refGene_gn=ABCB10;refGene_tx=XM_023240443.1;refGene_gn=ABCB10;refGene_tx=XM_023240444.1
D2 18945514 18945649
refGene_gn=ABCB10;refGene_tx=XM_003993943.4;refGene_gn=ABCB10;refGene_tx=XM_023240442.1;refGene_gn=ABCB10;refGene_tx=XM_023240443.1;refGene_gn=ABCB10;refGene_tx=XM_023240444.1
D2 18946923 18947070
refGene_gn=ABCB10;refGene_tx=XM_003993943.4;refGene_gn=ABCB10;refGene_tx=XM_023240442.1;refGene_gn=ABCB10;refGene_tx=XM_023240443.1;refGene_gn=ABCB10;refGene_tx=XM_023240444.1
D2 18948068 18948204
refGene_gn=ABCB10;refGene_tx=XM_003993943.4;refGene_gn=ABCB10;refGene_tx=XM_023240442.1;refGene_gn=ABCB10;refGene_tx=XM_023240443.1;refGene_gn=ABCB10;refGene_tx=XM_023240444.1
D2 18953023 18953119
refGene_gn=ABCB10;refGene_tx=XM_003993943.4;refGene_gn=ABCB10;refGene_tx=XM_023240442.1;refGene_gn=ABCB10;refGene_tx=XM_023240443.1;refGene_gn=ABCB10;refGene_tx=XM_023240444.1
D2 18954497 18954707
refGene_gn=ABCB10;refGene_tx=XM_003993943.4;refGene_gn=ABCB10;refGene_

tx=XM_023240442.1;refGene_gn=ABCB10;refGene_tx=XM_023240443.1;refGene_gn=ABCB10;refGene_tx=XM_023240444.1
D2 18957019 18957099
refGene_gn=ABCB10;refGene_tx=XM_003993943.4;refGene_gn=ABCB10;refGene_tx=XM_023240442.1;refGene_gn=ABCB10;refGene_tx=XM_023240443.1;refGene_gn=ABCB10;refGene_tx=XM_023240444.1
D2 18958824 18959005
refGene_gn=ABCB10;refGene_tx=XM_003993943.4;refGene_gn=ABCB10;refGene_tx=XM_023240442.1;refGene_gn=ABCB10;refGene_tx=XM_023240443.1;refGene_gn=ABCB10;refGene_tx=XM_023240444.1
D2 18960399 18960443
refGene_gn=ABCB10;refGene_tx=XM_003993943.4;refGene_gn=ABCB10;refGene_tx=XM_023240442.1;refGene_gn=ABCB10;refGene_tx=XM_023240443.1;refGene_gn=ABCB10;refGene_tx=XM_023240444.1
D2 18961868 18961903
refGene_gn=ABCB10;refGene_tx=XM_003993943.4;refGene_gn=ABCB10;refGene_tx=XM_023240442.1;refGene_gn=ABCB10;refGene_tx=XM_023240443.1;refGene_gn=ABCB10;refGene_tx=XM_023240444.1
D2 18962352 18962572
refGene_gn=ABCB10;refGene_tx=XM_003993943.4;refGene_gn=ABCB10;refGene_tx=XM_023240442.1;refGene_gn=ABCB10;refGene_tx=XM_023240443.1;refGene_gn=ABCB10;refGene_tx=XM_023240444.1
D2 18968850 18969032
refGene_gn=NUP133;refGene_tx=XM_003993934.5;refGene_gn=NUP133;refGene_tx=XM_019813112.2;refGene_gn=NUP133;refGene_tx=XM_019813113.2
D2 18971564 18971683
refGene_gn=NUP133;refGene_tx=XM_003993934.5;refGene_gn=NUP133;refGene_tx=XM_019813112.2;refGene_gn=NUP133;refGene_tx=XM_019813113.2
D2 18973690 18973794
refGene_gn=NUP133;refGene_tx=XM_003993934.5;refGene_gn=NUP133;refGene_tx=XM_019813112.2;refGene_gn=NUP133;refGene_tx=XM_019813113.2
D2 18975122 18975230
refGene_gn=NUP133;refGene_tx=XM_003993934.5;refGene_gn=NUP133;refGene_tx=XM_019813112.2;refGene_gn=NUP133;refGene_tx=XM_019813113.2
D2 18976896 18977031
refGene_gn=NUP133;refGene_tx=XM_003993934.5;refGene_gn=NUP133;refGene_tx=XM_019813112.2;refGene_gn=NUP133;refGene_tx=XM_019813113.2
D2 18977656 18977827
refGene_gn=NUP133;refGene_tx=XM_003993934.5;refGene_gn=NUP133;refGene_tx=XM_019813112.2;refGene_gn=NUP133;refGene_tx=XM_019813113.2
D2 18979428 18979584
refGene_gn=NUP133;refGene_tx=XM_003993934.5;refGene_gn=NUP133;refGene_tx=XM_019813112.2;refGene_gn=NUP133;refGene_tx=XM_019813113.2
D2 18979884 18979955
refGene_gn=NUP133;refGene_tx=XM_003993934.5;refGene_gn=NUP133;refGene_tx=XM_019813112.2;refGene_gn=NUP133;refGene_tx=XM_019813113.2
D2 18986023 18986171
refGene_gn=NUP133;refGene_tx=XM_003993934.5;refGene_gn=NUP133;refGene_tx=XM_019813112.2;refGene_gn=NUP133;refGene_tx=XM_019813113.2
D2 18987727 18987875

refGene_gn=NUP133;refGene_tx=XM_003993934.5;refGene_gn=NUP133;refGene_tx=XM_019813112.2;refGene_gn=NUP133;refGene_tx=XM_019813113.2
D2 18989022 18989180 refGene_gn=NUP133;refGene_tx=XM_003993934.5;refGene_gn=NUP133;refGene_tx=XM_019813112.2;refGene_gn=NUP133;refGene_tx=XM_019813113.2
D2 18992729 18992821 refGene_gn=NUP133;refGene_tx=XM_003993934.5;refGene_gn=NUP133;refGene_tx=XM_019813112.2;refGene_gn=NUP133;refGene_tx=XM_019813113.2
D2 18994423 18994587 refGene_gn=NUP133;refGene_tx=XM_003993934.5;refGene_gn=NUP133;refGene_tx=XM_019813112.2;refGene_gn=NUP133;refGene_tx=XM_019813113.2
D2 18995825 18995920 refGene_gn=NUP133;refGene_tx=XM_003993934.5;refGene_gn=NUP133;refGene_tx=XM_019813112.2;refGene_gn=NUP133;refGene_tx=XM_019813113.2
D2 19000921 19001146 refGene_gn=NUP133;refGene_tx=XM_003993934.5;refGene_gn=NUP133;refGene_tx=XM_019813112.2;refGene_gn=NUP133;refGene_tx=XM_019813113.2
D2 19004118 19004241 refGene_gn=NUP133;refGene_tx=XM_003993934.5;refGene_gn=NUP133;refGene_tx=XM_019813112.2;refGene_gn=NUP133;refGene_tx=XM_019813113.2
D2 19004821 19004918 refGene_gn=NUP133;refGene_tx=XM_003993934.5;refGene_gn=NUP133;refGene_tx=XM_019813112.2;refGene_gn=NUP133;refGene_tx=XM_019813113.2
D2 19005584 19005836 refGene_gn=NUP133;refGene_tx=XM_003993934.5;refGene_gn=NUP133;refGene_tx=XM_019813112.2;refGene_gn=NUP133;refGene_tx=XM_019813113.2
D2 19006487 19006621 refGene_gn=NUP133;refGene_tx=XM_003993934.5;refGene_gn=NUP133;refGene_tx=XM_019813112.2;refGene_gn=NUP133;refGene_tx=XM_019813113.2
D2 19009147 19009306 refGene_gn=NUP133;refGene_tx=XM_003993934.5;refGene_gn=NUP133;refGene_tx=XM_019813112.2;refGene_gn=NUP133;refGene_tx=XM_019813113.2
D2 19010150 19010286 refGene_gn=NUP133;refGene_tx=XM_003993934.5;refGene_gn=NUP133;refGene_tx=XM_019813112.2;refGene_gn=NUP133;refGene_tx=XM_019813113.2
D2 19014565 19014684 refGene_gn=NUP133;refGene_tx=XM_003993934.5;refGene_gn=NUP133;refGene_tx=XM_019813112.2;refGene_gn=NUP133;refGene_tx=XM_019813113.2
D2 19015671 19015752 refGene_gn=NUP133;refGene_tx=XM_003993934.5;refGene_gn=NUP133;refGene_tx=XM_019813112.2;refGene_gn=NUP133;refGene_tx=XM_019813113.2
D2 19017010 19017075 refGene_gn=NUP133;refGene_tx=XM_003993934.5;refGene_gn=NUP133;refGene_tx=XM_019813112.2;refGene_gn=NUP133;refGene_tx=XM_019813113.2
D2 19018339 19018428 refGene_gn=NUP133;refGene_tx=XM_003993934.5;refGene_gn=NUP133;refGene_tx=XM_019813112.2;refGene_gn=NUP133;refGene_tx=XM_019813113.2
D2 19021169 19021207 refGene_gn=NUP133;refGene_tx=XM_019813112.2
D2 19023162 19023299 refGene_gn=NUP133;refGene_tx=XM_003993934.5
D2 19030735 19030864 refGene_gn=ACTA1;refGene_tx=XM_023240445.1

D2 19030988 19031313 refGene_gn=ACTA1;refGene_tx=XM_023240445.1
D2 19031437 19031599 refGene_gn=ACTA1;refGene_tx=XM_023240445.1
D2 19031695 19031887 refGene_gn=ACTA1;refGene_tx=XM_023240445.1
D2 19031982 19032164 refGene_gn=ACTA1;refGene_tx=XM_023240445.1
D2 19032247 19032391 refGene_gn=ACTA1;refGene_tx=XM_023240445.1
D2 19099074 19099456
refGene_gn=CCSAP;refGene_tx=XM_023240446.1;refGene_gn=CCSAP;refGene_tx=XM_023240447.1
D2 19113511 19113780
refGene_gn=CCSAP;refGene_tx=XM_023240446.1;refGene_gn=CCSAP;refGene_tx=XM_023240447.1
D2 19115319 19115496
refGene_gn=CCSAP;refGene_tx=XM_023240446.1;refGene_gn=CCSAP;refGene_tx=XM_023240447.1
D2 19144667 19144783
refGene_gn=RAB4A;refGene_tx=XM_003993935.5;refGene_gn=RAB4A;refGene_tx=XM_019813117.2;refGene_gn=RAB4A;refGene_tx=XM_023240448.1
D2 19149156 19149252
refGene_gn=RAB4A;refGene_tx=XM_003993935.5;refGene_gn=RAB4A;refGene_tx=XM_019813117.2;refGene_gn=RAB4A;refGene_tx=XM_023240448.1
D2 19150702 19150857
refGene_gn=RAB4A;refGene_tx=XM_003993935.5;refGene_gn=RAB4A;refGene_tx=XM_019813117.2;refGene_gn=RAB4A;refGene_tx=XM_023240448.1
D2 19152071 19152084 refGene_gn=RAB4A;refGene_tx=XM_023240448.1
D2 19152408 19152471
refGene_gn=RAB4A;refGene_tx=XM_003993935.5;refGene_gn=RAB4A;refGene_tx=XM_019813117.2
D2 19157313 19157428
refGene_gn=RAB4A;refGene_tx=XM_003993935.5;refGene_gn=RAB4A;refGene_tx=XM_019813117.2
D2 19159661 19159742
refGene_gn=RAB4A;refGene_tx=XM_003993935.5;refGene_gn=RAB4A;refGene_tx=XM_019813117.2
D2 19185617 19185648
refGene_gn=RAB4A;refGene_tx=XM_003993935.5;refGene_gn=RAB4A;refGene_tx=XM_019813117.2
D2 19632750 19633206 refGene_gn=RHOU;refGene_tx=XM_023240449.1
D2 19639196 19639255 refGene_gn=RHOU;refGene_tx=XM_023240449.1
D2 19640928 19641133 refGene_gn=RHOU;refGene_tx=XM_023240449.1
D2 19641134 19641170 refGene_gn=RHOU;refGene_tx=XM_023240449.1
D2 19641171 19641192 refGene_gn=RHOU;refGene_tx=XM_023240449.1
D2 19753584 19753596
refGene_gn=LOC109492447;refGene_tx=XM_023240450.1
D2 19753824 19753917
refGene_gn=LOC109492447;refGene_tx=XM_023240450.1;refGene_gn=LOC109492447;refGene_tx=XM_023240451.1;refGene_gn=LOC109492447;refGene_tx=XM_023240452.1
D2 19772103 19772205
refGene_gn=LOC109492447;refGene_tx=XM_023240450.1;refGene_gn=LOC109492447;refGene_tx=XM_023240451.1;refGene_gn=LOC109492447;refGene_tx=XM_02

3240452.1
D2 19772476 19772692
refGene_gn=LOC109492447;refGene_tx=XM_023240450.1;refGene_gn=LOC109492447;refGene_tx=XM_023240451.1;refGene_gn=LOC109492447;refGene_tx=XM_023240452.1
D2 21853608 21854231
refGene_gn=IPMK;refGene_tx=XM_011287388.3;refGene_gn=IPMK;refGene_tx=XM_019813125.2;refGene_gn=IPMK;refGene_tx=XM_019813126.2;refGene_gn=IPMK;refGene_tx=XM_019813128.2
D2 21855711 21855793
refGene_gn=IPMK;refGene_tx=XM_011287388.3;refGene_gn=IPMK;refGene_tx=XM_019813125.2;refGene_gn=IPMK;refGene_tx=XM_019813126.2;refGene_gn=IPMK;refGene_tx=XM_019813128.2
D2 21879862 21880035
refGene_gn=IPMK;refGene_tx=XM_011287388.3;refGene_gn=IPMK;refGene_tx=XM_019813125.2;refGene_gn=IPMK;refGene_tx=XM_019813126.2;refGene_gn=IPMK;refGene_tx=XM_019813128.2
D2 21885637 21885734
refGene_gn=IPMK;refGene_tx=XM_011287388.3;refGene_gn=IPMK;refGene_tx=XM_019813125.2;refGene_gn=IPMK;refGene_tx=XM_019813126.2;refGene_gn=IPMK;refGene_tx=XM_019813128.2
D2 21900083 21900178
refGene_gn=IPMK;refGene_tx=XM_011287388.3;refGene_gn=IPMK;refGene_tx=XM_019813125.2;refGene_gn=IPMK;refGene_tx=XM_019813126.2;refGene_gn=IPMK;refGene_tx=XM_019813128.2
D2 21901557 21901627
refGene_gn=IPMK;refGene_tx=XM_019813126.2;refGene_gn=IPMK;refGene_tx=XM_019813128.2
D2 21913469 21913680
refGene_gn=IPMK;refGene_tx=XM_011287388.3;refGene_gn=IPMK;refGene_tx=XM_019813125.2
D2 21914281 21914287
refGene_gn=IPMK;refGene_tx=XM_011287388.3;refGene_gn=IPMK;refGene_tx=XM_019813125.2
D2 21915100 21915324
refGene_gn=CISD1;refGene_tx=XM_003993947.5;refGene_gn=CISD1;refGene_tx=XM_011287387.3
D2 21920468 21920563 refGene_gn=CISD1;refGene_tx=XM_011287387.3
D2 21921964 21922170
refGene_gn=CISD1;refGene_tx=XM_003993947.5;refGene_gn=CISD1;refGene_tx=XM_011287387.3
D2 21931033 21931123
refGene_gn=CISD1;refGene_tx=XM_003993947.5;refGene_gn=CISD1;refGene_tx=XM_011287387.3
D2 22028569 22028593 refGene_gn=UBE2D1;refGene_tx=XM_023240453.1
D2 22049435 22049499 refGene_gn=UBE2D1;refGene_tx=XM_023240453.1
D2 22049592 22049624
refGene_gn=UBE2D1;refGene_tx=XM_023240453.1;refGene_gn=UBE2D1;refGene_tx=XM_023240454.1
D2 22052193 22052271

refGene_gn=UBE2D1;refGene_tx=XM_023240453.1;refGene_gn=UBE2D1;refGene_tx=XM_023240454.1
D2 22053572 22053678 refGene_gn=UBE2D1;refGene_tx=XM_023240453.1;refGene_gn=UBE2D1;refGene_tx=XM_023240454.1
D2 22056641 22056735 refGene_gn=UBE2D1;refGene_tx=XM_023240453.1;refGene_gn=UBE2D1;refGene_tx=XM_023240454.1
D2 22057706 22057752 refGene_gn=UBE2D1;refGene_tx=XM_023240453.1;refGene_gn=UBE2D1;refGene_tx=XM_023240454.1
D2 22080790 22080891 refGene_gn=TFAM;refGene_tx=XM_003993948.5;refGene_gn=TFAM;refGene_tx=XM_023240455.1
D2 22081453 22081572 refGene_gn=TFAM;refGene_tx=XM_003993948.5;refGene_gn=TFAM;refGene_tx=XM_023240455.1
D2 22083047 22083118 refGene_gn=TFAM;refGene_tx=XM_003993948.5;refGene_gn=TFAM;refGene_tx=XM_023240455.1
D2 22083796 22083946 refGene_gn=TFAM;refGene_tx=XM_003993948.5
D2 22087619 22087715 refGene_gn=TFAM;refGene_tx=XM_003993948.5;refGene_gn=TFAM;refGene_tx=XM_023240455.1
D2 22093392 22093449 refGene_gn=TFAM;refGene_tx=XM_003993948.5;refGene_gn=TFAM;refGene_tx=XM_023240455.1
D2 22093913 22094060 refGene_gn=TFAM;refGene_tx=XM_003993948.5;refGene_gn=TFAM;refGene_tx=XM_023240455.1
D2 22253269 22253459 refGene_gn=BICC1;refGene_tx=XM_011287385.3;refGene_gn=BICC1;refGene_tx=XM_019813130.2;refGene_gn=BICC1;refGene_tx=XM_019813131.2
D2 22355101 22355148 refGene_gn=BICC1;refGene_tx=XM_011287385.3;refGene_gn=BICC1;refGene_tx=XM_019813130.2;refGene_gn=BICC1;refGene_tx=XM_019813131.2
D2 22422526 22422596 refGene_gn=BICC1;refGene_tx=XM_011287385.3;refGene_gn=BICC1;refGene_tx=XM_019813130.2;refGene_gn=BICC1;refGene_tx=XM_019813131.2
D2 22496034 22496114 refGene_gn=BICC1;refGene_tx=XM_011287385.3;refGene_gn=BICC1;refGene_tx=XM_019813130.2;refGene_gn=BICC1;refGene_tx=XM_019813131.2
D2 22496122 22496224 refGene_gn=BICC1;refGene_tx=XM_019813132.1
D2 22497745 22497904 refGene_gn=BICC1;refGene_tx=XM_011287385.3;refGene_gn=BICC1;refGene_tx=XM_019813130.2;refGene_gn=BICC1;refGene_tx=XM_019813131.2;refGene_gn=BICC1;refGene_tx=XM_019813132.1
D2 22499763 22499817 refGene_gn=BICC1;refGene_tx=XM_011287385.3;refGene_gn=BICC1;refGene_tx=XM_019813130.2;refGene_gn=BICC1;refGene_tx=XM_019813131.2;refGene_gn=

BICC1;refGene_tx=XM_019813132.1
D2	22500256	22500451	refGene_gn=BICC1;refGene_tx=XM_011287385.3;refGene_gn=BICC1;refGene_tx=XM_019813130.2;refGene_gn=BICC1;refGene_tx=XM_019813131.2;refGene_gn=BICC1;refGene_tx=XM_019813132.1
D2	22500672	22500924	refGene_gn=BICC1;refGene_tx=XM_011287385.3;refGene_gn=BICC1;refGene_tx=XM_019813130.2;refGene_gn=BICC1;refGene_tx=XM_019813131.2;refGene_gn=BICC1;refGene_tx=XM_019813132.1
D2	22502979	22503111	refGene_gn=BICC1;refGene_tx=XM_011287385.3;refGene_gn=BICC1;refGene_tx=XM_019813130.2;refGene_gn=BICC1;refGene_tx=XM_019813131.2;refGene_gn=BICC1;refGene_tx=XM_019813132.1
D2	22505141	22505328	refGene_gn=BICC1;refGene_tx=XM_011287385.3;refGene_gn=BICC1;refGene_tx=XM_019813130.2;refGene_gn=BICC1;refGene_tx=XM_019813131.2;refGene_gn=BICC1;refGene_tx=XM_019813132.1
D2	22507170	22507332	refGene_gn=BICC1;refGene_tx=XM_011287385.3;refGene_gn=BICC1;refGene_tx=XM_019813130.2;refGene_gn=BICC1;refGene_tx=XM_019813131.2;refGene_gn=BICC1;refGene_tx=XM_019813132.1
D2	22507933	22508130	refGene_gn=BICC1;refGene_tx=XM_011287385.3;refGene_gn=BICC1;refGene_tx=XM_019813130.2;refGene_gn=BICC1;refGene_tx=XM_019813131.2;refGene_gn=BICC1;refGene_tx=XM_019813132.1
D2	22508267	22508400	refGene_gn=BICC1;refGene_tx=XM_011287385.3;refGene_gn=BICC1;refGene_tx=XM_019813130.2;refGene_gn=BICC1;refGene_tx=XM_019813131.2;refGene_gn=BICC1;refGene_tx=XM_019813132.1
D2	22508934	22509091	refGene_gn=BICC1;refGene_tx=XM_011287385.3;refGene_gn=BICC1;refGene_tx=XM_019813130.2;refGene_gn=BICC1;refGene_tx=XM_019813131.2;refGene_gn=BICC1;refGene_tx=XM_019813132.1
D2	22511093	22511259	refGene_gn=BICC1;refGene_tx=XM_011287385.3;refGene_gn=BICC1;refGene_tx=XM_019813130.2;refGene_gn=BICC1;refGene_tx=XM_019813131.2;refGene_gn=BICC1;refGene_tx=XM_019813132.1
D2	22513898	22513938	refGene_gn=BICC1;refGene_tx=XM_011287385.3;refGene_gn=BICC1;refGene_tx=XM_019813130.2;refGene_gn=BICC1;refGene_tx=XM_019813131.2;refGene_gn=BICC1;refGene_tx=XM_019813132.1
D2	22514384	22514539	refGene_gn=BICC1;refGene_tx=XM_011287385.3;refGene_gn=BICC1;refGene_tx=XM_019813130.2;refGene_gn=BICC1;refGene_tx=XM_019813131.2;refGene_gn=BICC1;refGene_tx=XM_019813132.1
D2	22520244	22520401	refGene_gn=BICC1;refGene_tx=XM_011287385.3;refGene_gn=BICC1;refGene_tx=XM_019813130.2;refGene_gn=BICC1;refGene_tx=XM_019813131.2;refGene_gn=BICC1;refGene_tx=XM_019813132.1
D2	22524109	22524270

refGene_gn=BICC1;refGene_tx=XM_011287385.3;refGene_gn=BICC1;refGene_tx=XM_019813130.2;refGene_gn=BICC1;refGene_tx=XM_019813131.2;refGene_gn=BICC1;refGene_tx=XM_019813132.1
D2 22526883 22526983 refGene_gn=BICC1;refGene_tx=XM_011287385.3;refGene_gn=BICC1;refGene_tx=XM_019813130.2;refGene_gn=BICC1;refGene_tx=XM_019813131.2;refGene_gn=BICC1;refGene_tx=XM_019813132.1
D2 22529966 22530022 refGene_gn=BICC1;refGene_tx=XM_019813131.2
D2 22532924 22533055 refGene_gn=BICC1;refGene_tx=XM_011287385.3;refGene_gn=BICC1;refGene_tx=XM_019813130.2;refGene_gn=BICC1;refGene_tx=XM_019813132.1
D2 22551566 22551607 refGene_gn=LOC102902473;refGene_tx=XM_019814061.1
D2 22556615 22558532 refGene_gn=LOC102902473;refGene_tx=XM_019814061.1
D2 22952152 22952258 refGene_gn=PHYHIPL;refGene_tx=XM_003993949.4
D2 23011644 23011841 refGene_gn=PHYHIPL;refGene_tx=XM_003993949.4;refGene_gn=PHYHIPL;refGene_tx=XM_011287381.2;refGene_gn=PHYHIPL;refGene_tx=XM_019813134.2
D2 23013736 23013911 refGene_gn=PHYHIPL;refGene_tx=XM_003993949.4;refGene_gn=PHYHIPL;refGene_tx=XM_011287381.2;refGene_gn=PHYHIPL;refGene_tx=XM_019813134.2
D2 23017779 23017897 refGene_gn=PHYHIPL;refGene_tx=XM_003993949.4;refGene_gn=PHYHIPL;refGene_tx=XM_011287381.2;refGene_gn=PHYHIPL;refGene_tx=XM_019813134.2
D2 23033247 23033782 refGene_gn=PHYHIPL;refGene_tx=XM_003993949.4;refGene_gn=PHYHIPL;refGene_tx=XM_011287381.2;refGene_gn=PHYHIPL;refGene_tx=XM_019813134.2
D2 23035752 23035876 refGene_gn=FAM13C;refGene_tx=XM_011287382.2;refGene_gn=FAM13C;refGene_tx=XM_011287383.2;refGene_gn=FAM13C;refGene_tx=XM_011287384.3
D2 23039065 23039167 refGene_gn=FAM13C;refGene_tx=XM_011287382.2;refGene_gn=FAM13C;refGene_tx=XM_011287383.2;refGene_gn=FAM13C;refGene_tx=XM_011287384.3
D2 23040771 23040971 refGene_gn=FAM13C;refGene_tx=XM_011287382.2;refGene_gn=FAM13C;refGene_tx=XM_011287383.2;refGene_gn=FAM13C;refGene_tx=XM_011287384.3
D2 23042490 23042586 refGene_gn=FAM13C;refGene_tx=XM_011287382.2;refGene_gn=FAM13C;refGene_tx=XM_011287383.2;refGene_gn=FAM13C;refGene_tx=XM_011287384.3
D2 23048576 23048788 refGene_gn=FAM13C;refGene_tx=XM_011287382.2;refGene_gn=FAM13C;refGene_tx=XM_011287383.2;refGene_gn=FAM13C;refGene_tx=XM_011287384.3
D2 23050434 23050516 refGene_gn=FAM13C;refGene_tx=XM_011287382.2;refGene_gn=FAM13C;refGene_tx=XM_011287383.2;refGene_gn=FAM13C;refGene_tx=XM_011287384.3
D2 23075879 23076018 refGene_gn=FAM13C;refGene_tx=XM_011287382.2;refGene_gn=FAM13C;refGene_tx=XM_011287383.2;refGene_gn=FAM13C;refGene_tx=XM_011287384.3
D2 23077109 23077320

refGene_gn=FAM13C;refGene_tx=XM_011287382.2;refGene_gn=FAM13C;refGene_tx=XM_011287383.2;refGene_gn=FAM13C;refGene_tx=XM_011287384.3
D2 23093573 23093658
refGene_gn=FAM13C;refGene_tx=XM_011287382.2;refGene_gn=FAM13C;refGene_tx=XM_011287383.2;refGene_gn=FAM13C;refGene_tx=XM_011287384.3
D2 23124463 23124527
refGene_gn=FAM13C;refGene_tx=XM_011287382.2;refGene_gn=FAM13C;refGene_tx=XM_011287383.2;refGene_gn=FAM13C;refGene_tx=XM_011287384.3
D2 23143392 23143511
refGene_gn=FAM13C;refGene_tx=XM_011287382.2;refGene_gn=FAM13C;refGene_tx=XM_011287383.2;refGene_gn=FAM13C;refGene_tx=XM_011287384.3
D2 23191519 23191721
refGene_gn=FAM13C;refGene_tx=XM_011287382.2;refGene_gn=FAM13C;refGene_tx=XM_011287383.2;refGene_gn=FAM13C;refGene_tx=XM_011287384.3
D2 23195293 23195350
refGene_gn=FAM13C;refGene_tx=XM_011287382.2;refGene_gn=FAM13C;refGene_tx=XM_011287383.2
D2 23202251 23202313
refGene_gn=FAM13C;refGene_tx=XM_011287382.2;refGene_gn=FAM13C;refGene_tx=XM_011287383.2
D2 23505784 23505963
refGene_gn=SLC16A9;refGene_tx=XM_023240456.1;refGene_gn=SLC16A9;refGene_tx=XM_023240457.1
D2 23506362 23507280
refGene_gn=SLC16A9;refGene_tx=XM_023240456.1;refGene_gn=SLC16A9;refGene_tx=XM_023240457.1
D2 23509817 23509913
refGene_gn=SLC16A9;refGene_tx=XM_023240456.1;refGene_gn=SLC16A9;refGene_tx=XM_023240457.1
D2 23521768 23521912 refGene_gn=SLC16A9;refGene_tx=XM_023240456.1
D2 23533114 23533310
refGene_gn=SLC16A9;refGene_tx=XM_023240456.1;refGene_gn=SLC16A9;refGene_tx=XM_023240457.1
D2 23578874 23579035
refGene_gn=MRLN;refGene_tx=XM_019813137.1;refGene_gn=MRLN;refGene_tx=XM_019813138.1;refGene_gn=MRLN;refGene_tx=XM_019813139.1;refGene_gn=MRLN;refGene_tx=XM_023240458.1
D2 23580221 23580231 refGene_gn=MRLN;refGene_tx=XM_019813139.1
D2 23580581 23580660
refGene_gn=MRLN;refGene_tx=XM_019813137.1;refGene_gn=MRLN;refGene_tx=XM_019813138.1;refGene_gn=MRLN;refGene_tx=XM_023240458.1
D2 23623392 23623587 refGene_gn=CCDC6;refGene_tx=XM_003993958.5
D2 23624614 23624739 refGene_gn=CCDC6;refGene_tx=XM_003993958.5
D2 23633885 23633923 refGene_gn=CCDC6;refGene_tx=XM_023240459.1
D2 23635807 23635908
refGene_gn=CCDC6;refGene_tx=XM_003993958.5;refGene_gn=CCDC6;refGene_tx=XM_023240459.1
D2 23637232 23637389
refGene_gn=CCDC6;refGene_tx=XM_003993958.5;refGene_gn=CCDC6;refGene_tx=XM_023240459.1

D2 23643025 23643186
refGene_gn=CCDC6;refGene_tx=XM_003993958.5;refGene_gn=CCDC6;refGene_tx=XM_023240459.1
D2 23645019 23645123
refGene_gn=CCDC6;refGene_tx=XM_003993958.5;refGene_gn=CCDC6;refGene_tx=XM_023240459.1
D2 23659229 23659358
refGene_gn=CCDC6;refGene_tx=XM_003993958.5;refGene_gn=CCDC6;refGene_tx=XM_023240459.1
D2 23677682 23677832
refGene_gn=CCDC6;refGene_tx=XM_003993958.5;refGene_gn=CCDC6;refGene_tx=XM_023240459.1
D2 23728502 23728811
refGene_gn=CCDC6;refGene_tx=XM_003993958.5;refGene_gn=CCDC6;refGene_tx=XM_023240459.1
D2 23852894 23852966
refGene_gn=ANK3;refGene_tx=XM_003993953.5;refGene_gn=ANK3;refGene_tx=XM_019813142.2;refGene_gn=ANK3;refGene_tx=XM_023240460.1;refGene_gn=ANK3;refGene_tx=XM_023240461.1;refGene_gn=ANK3;refGene_tx=XM_023240462.1;refGene_gn=ANK3;refGene_tx=XM_023240463.1;refGene_gn=ANK3;refGene_tx=XM_023240464.1;refGene_gn=ANK3;refGene_tx=XM_023240465.1;refGene_gn=ANK3;refGene_tx=XM_023240466.1;refGene_gn=ANK3;refGene_tx=XM_023240467.1;refGene_gn=ANK3;refGene_tx=XM_023240468.1;refGene_gn=ANK3;refGene_tx=XM_023240469.1;refGene_gn=ANK3;refGene_tx=XM_023240470.1;refGene_gn=ANK3;refGene_tx=XM_023240471.1;refGene_gn=ANK3;refGene_tx=XM_023240472.1
D2 23854457 23854517
refGene_gn=ANK3;refGene_tx=XM_019813142.2;refGene_gn=ANK3;refGene_tx=XM_023240460.1;refGene_gn=ANK3;refGene_tx=XM_023240463.1;refGene_gn=ANK3;refGene_tx=XM_023240464.1;refGene_gn=ANK3;refGene_tx=XM_023240465.1;refGene_gn=ANK3;refGene_tx=XM_023240466.1;refGene_gn=ANK3;refGene_tx=XM_023240467.1;refGene_gn=ANK3;refGene_tx=XM_023240468.1;refGene_gn=ANK3;refGene_tx=XM_023240469.1;refGene_gn=ANK3;refGene_tx=XM_023240470.1
D2 23854629 23854713
refGene_gn=ANK3;refGene_tx=XM_019813142.2;refGene_gn=ANK3;refGene_tx=XM_023240460.1;refGene_gn=ANK3;refGene_tx=XM_023240463.1;refGene_gn=ANK3;refGene_tx=XM_023240464.1;refGene_gn=ANK3;refGene_tx=XM_023240465.1;refGene_gn=ANK3;refGene_tx=XM_023240466.1;refGene_gn=ANK3;refGene_tx=XM_023240467.1;refGene_gn=ANK3;refGene_tx=XM_023240468.1;refGene_gn=ANK3;refGene_tx=XM_023240469.1;refGene_gn=ANK3;refGene_tx=XM_023240470.1
D2 23861202 23861289
refGene_gn=ANK3;refGene_tx=XM_019813142.2;refGene_gn=ANK3;refGene_tx=XM_023240460.1;refGene_gn=ANK3;refGene_tx=XM_023240463.1;refGene_gn=ANK3;refGene_tx=XM_023240464.1;refGene_gn=ANK3;refGene_tx=XM_023240465.1;refGene_gn=ANK3;refGene_tx=XM_023240466.1;refGene_gn=ANK3;refGene_tx=XM_023240467.1;refGene_gn=ANK3;refGene_tx=XM_023240468.1;refGene_gn=ANK3;refGene_tx=XM_023240469.1;refGene_gn=ANK3;refGene_tx=XM_023240470.1
D2 23863335 23863386
refGene_gn=ANK3;refGene_tx=XM_019813142.2;refGene_gn=ANK3;refGene_tx=XM_023240461.1;refGene_gn=ANK3;refGene_tx=XM_023240463.1;refGene_gn=ANK3;refGene_tx=XM_023240464.1;refGene_gn=ANK3;refGene_tx=XM_023240465.1;

refGene_gn=ANK3;refGene_tx=XM_023240466.1;refGene_gn=ANK3;refGene_tx=XM_023240467.1;refGene_gn=ANK3;refGene_tx=XM_023240468.1;refGene_gn=ANK3;refGene_tx=XM_023240469.1;refGene_gn=ANK3;refGene_tx=XM_023240470.1
D2 23865095 23865474
refGene_gn=ANK3;refGene_tx=XM_003993953.5;refGene_gn=ANK3;refGene_tx=XM_019813142.2;refGene_gn=ANK3;refGene_tx=XM_023240460.1;refGene_gn=ANK3;refGene_tx=XM_023240461.1;refGene_gn=ANK3;refGene_tx=XM_023240462.1;refGene_gn=ANK3;refGene_tx=XM_023240463.1;refGene_gn=ANK3;refGene_tx=XM_023240464.1;refGene_gn=ANK3;refGene_tx=XM_023240465.1;refGene_gn=ANK3;refGene_tx=XM_023240466.1;refGene_gn=ANK3;refGene_tx=XM_023240467.1;refGene_gn=ANK3;refGene_tx=XM_023240468.1;refGene_gn=ANK3;refGene_tx=XM_023240469.1;refGene_gn=ANK3;refGene_tx=XM_023240470.1;refGene_gn=ANK3;refGene_tx=XM_023240471.1;refGene_gn=ANK3;refGene_tx=XM_023240472.1
D2 23867508 23867599
refGene_gn=ANK3;refGene_tx=XM_003993953.5;refGene_gn=ANK3;refGene_tx=XM_019813142.2;refGene_gn=ANK3;refGene_tx=XM_023240460.1;refGene_gn=ANK3;refGene_tx=XM_023240461.1;refGene_gn=ANK3;refGene_tx=XM_023240462.1;refGene_gn=ANK3;refGene_tx=XM_023240463.1;refGene_gn=ANK3;refGene_tx=XM_023240464.1;refGene_gn=ANK3;refGene_tx=XM_023240465.1;refGene_gn=ANK3;refGene_tx=XM_023240466.1;refGene_gn=ANK3;refGene_tx=XM_023240467.1;refGene_gn=ANK3;refGene_tx=XM_023240468.1;refGene_gn=ANK3;refGene_tx=XM_023240469.1;refGene_gn=ANK3;refGene_tx=XM_023240470.1;refGene_gn=ANK3;refGene_tx=XM_023240471.1;refGene_gn=ANK3;refGene_tx=XM_023240472.1
D2 23867878 23868190
refGene_gn=ANK3;refGene_tx=XM_003993953.5;refGene_gn=ANK3;refGene_tx=XM_023240463.1;refGene_gn=ANK3;refGene_tx=XM_023240464.1;refGene_gn=ANK3;refGene_tx=XM_023240465.1;refGene_gn=ANK3;refGene_tx=XM_023240466.1;refGene_gn=ANK3;refGene_tx=XM_023240467.1;refGene_gn=ANK3;refGene_tx=XM_023240468.1;refGene_gn=ANK3;refGene_tx=XM_023240469.1;refGene_gn=ANK3;refGene_tx=XM_023240470.1;refGene_gn=ANK3;refGene_tx=XM_023240471.1;refGene_gn=ANK3;refGene_tx=XM_023240472.1
D2 23871054 23871198
refGene_gn=ANK3;refGene_tx=XM_003993953.5;refGene_gn=ANK3;refGene_tx=XM_019813142.2;refGene_gn=ANK3;refGene_tx=XM_023240460.1;refGene_gn=ANK3;refGene_tx=XM_023240461.1;refGene_gn=ANK3;refGene_tx=XM_023240462.1;refGene_gn=ANK3;refGene_tx=XM_023240463.1;refGene_gn=ANK3;refGene_tx=XM_023240464.1;refGene_gn=ANK3;refGene_tx=XM_023240465.1;refGene_gn=ANK3;refGene_tx=XM_023240466.1;refGene_gn=ANK3;refGene_tx=XM_023240467.1;refGene_gn=ANK3;refGene_tx=XM_023240468.1;refGene_gn=ANK3;refGene_tx=XM_023240469.1;refGene_gn=ANK3;refGene_tx=XM_023240470.1;refGene_gn=ANK3;refGene_tx=XM_023240471.1;refGene_gn=ANK3;refGene_tx=XM_023240472.1
D2 23872117 23872249
refGene_gn=ANK3;refGene_tx=XM_003993953.5;refGene_gn=ANK3;refGene_tx=XM_019813142.2;refGene_gn=ANK3;refGene_tx=XM_023240460.1;refGene_gn=ANK3;refGene_tx=XM_023240461.1;refGene_gn=ANK3;refGene_tx=XM_023240462.1;refGene_gn=ANK3;refGene_tx=XM_023240463.1;refGene_gn=ANK3;refGene_tx=XM_023240464.1;refGene_gn=ANK3;refGene_tx=XM_023240465.1;refGene_gn=ANK3;refGene_tx=XM_023240466.1;refGene_gn=ANK3;refGene_tx=XM_023240467.1;refGene_gn=ANK3;refGene_tx=XM_023240468.1;refGene_gn=ANK3;refGene_tx=XM_023240469.1;refGene_gn=ANK3;refGene_tx=XM_023240470.1;refGene_gn=ANK

3;refGene_tx=XM_023240471.1;refGene_gn=ANK3;refGene_tx=XM_023240472.1
D2 23875581 23875656
refGene_gn=ANK3;refGene_tx=XM_003993953.5;refGene_gn=ANK3;refGene_tx=XM_019813142.2;refGene_gn=ANK3;refGene_tx=XM_023240460.1;refGene_gn=ANK3;refGene_tx=XM_023240461.1;refGene_gn=ANK3;refGene_tx=XM_023240462.1;refGene_gn=ANK3;refGene_tx=XM_023240463.1;refGene_gn=ANK3;refGene_tx=XM_023240464.1;refGene_gn=ANK3;refGene_tx=XM_023240465.1;refGene_gn=ANK3;refGene_tx=XM_023240466.1;refGene_gn=ANK3;refGene_tx=XM_023240467.1;refGene_gn=ANK3;refGene_tx=XM_023240468.1;refGene_gn=ANK3;refGene_tx=XM_023240469.1;refGene_gn=ANK3;refGene_tx=XM_023240470.1;refGene_gn=ANK3;refGene_tx=XM_023240471.1;refGene_gn=ANK3;refGene_tx=XM_023240472.1
D2 23876280 23884086
refGene_gn=ANK3;refGene_tx=XM_019813142.2;refGene_gn=ANK3;refGene_tx=XM_023240460.1;refGene_gn=ANK3;refGene_tx=XM_023240461.1;refGene_gn=ANK3;refGene_tx=XM_023240462.1;refGene_gn=ANK3;refGene_tx=XM_023240463.1;refGene_gn=ANK3;refGene_tx=XM_023240464.1;refGene_gn=ANK3;refGene_tx=XM_023240465.1;refGene_gn=ANK3;refGene_tx=XM_023240466.1;refGene_gn=ANK3;refGene_tx=XM_023240467.1;refGene_gn=ANK3;refGene_tx=XM_023240468.1;refGene_gn=ANK3;refGene_tx=XM_023240469.1;refGene_gn=ANK3;refGene_tx=XM_023240470.1
D2 23888152 23888234
refGene_gn=ANK3;refGene_tx=XM_003993953.5;refGene_gn=ANK3;refGene_tx=XM_019813142.2;refGene_gn=ANK3;refGene_tx=XM_023240460.1;refGene_gn=ANK3;refGene_tx=XM_023240461.1;refGene_gn=ANK3;refGene_tx=XM_023240462.1;refGene_gn=ANK3;refGene_tx=XM_023240463.1;refGene_gn=ANK3;refGene_tx=XM_023240464.1;refGene_gn=ANK3;refGene_tx=XM_023240465.1;refGene_gn=ANK3;refGene_tx=XM_023240466.1;refGene_gn=ANK3;refGene_tx=XM_023240467.1;refGene_gn=ANK3;refGene_tx=XM_023240468.1;refGene_gn=ANK3;refGene_tx=XM_023240469.1;refGene_gn=ANK3;refGene_tx=XM_023240470.1;refGene_gn=ANK3;refGene_tx=XM_023240471.1;refGene_gn=ANK3;refGene_tx=XM_023240472.1
D2 23889294 23889321
refGene_gn=ANK3;refGene_tx=XM_019813142.2;refGene_gn=ANK3;refGene_tx=XM_023240460.1;refGene_gn=ANK3;refGene_tx=XM_023240461.1;refGene_gn=ANK3;refGene_tx=XM_023240462.1;refGene_gn=ANK3;refGene_tx=XM_023240463.1;refGene_gn=ANK3;refGene_tx=XM_023240464.1;refGene_gn=ANK3;refGene_tx=XM_023240465.1;refGene_gn=ANK3;refGene_tx=XM_023240466.1;refGene_gn=ANK3;refGene_tx=XM_023240467.1;refGene_gn=ANK3;refGene_tx=XM_023240468.1;refGene_gn=ANK3;refGene_tx=XM_023240469.1;refGene_gn=ANK3;refGene_tx=XM_023240470.1
D2 23889757 23889880
refGene_gn=ANK3;refGene_tx=XM_003993953.5;refGene_gn=ANK3;refGene_tx=XM_019813142.2;refGene_gn=ANK3;refGene_tx=XM_023240460.1;refGene_gn=ANK3;refGene_tx=XM_023240461.1;refGene_gn=ANK3;refGene_tx=XM_023240462.1;refGene_gn=ANK3;refGene_tx=XM_023240463.1;refGene_gn=ANK3;refGene_tx=XM_023240464.1;refGene_gn=ANK3;refGene_tx=XM_023240465.1;refGene_gn=ANK3;refGene_tx=XM_023240466.1;refGene_gn=ANK3;refGene_tx=XM_023240467.1;refGene_gn=ANK3;refGene_tx=XM_023240468.1;refGene_gn=ANK3;refGene_tx=XM_023240469.1;refGene_gn=ANK3;refGene_tx=XM_023240470.1;refGene_gn=ANK3;refGene_tx=XM_023240471.1;refGene_gn=ANK3;refGene_tx=XM_023240472.1
D2 23890809 23890935

refGene_gn=ANK3;refGene_tx=XM_003993953.5;refGene_gn=ANK3;refGene_tx=XM_019813142.2;refGene_gn=ANK3;refGene_tx=XM_023240460.1;refGene_gn=ANK3;refGene_tx=XM_023240461.1;refGene_gn=ANK3;refGene_tx=XM_023240462.1;refGene_gn=ANK3;refGene_tx=XM_023240463.1;refGene_gn=ANK3;refGene_tx=XM_023240464.1;refGene_gn=ANK3;refGene_tx=XM_023240465.1;refGene_gn=ANK3;refGene_tx=XM_023240466.1;refGene_gn=ANK3;refGene_tx=XM_023240467.1;refGene_gn=ANK3;refGene_tx=XM_023240468.1;refGene_gn=ANK3;refGene_tx=XM_023240469.1;refGene_gn=ANK3;refGene_tx=XM_023240470.1;refGene_gn=ANK3;refGene_tx=XM_023240471.1;refGene_gn=ANK3;refGene_tx=XM_023240472.1
D2 23891578 23891807
refGene_gn=ANK3;refGene_tx=XM_003993953.5;refGene_gn=ANK3;refGene_tx=XM_019813142.2;refGene_gn=ANK3;refGene_tx=XM_023240460.1;refGene_gn=ANK3;refGene_tx=XM_023240461.1;refGene_gn=ANK3;refGene_tx=XM_023240462.1;refGene_gn=ANK3;refGene_tx=XM_023240463.1;refGene_gn=ANK3;refGene_tx=XM_023240464.1;refGene_gn=ANK3;refGene_tx=XM_023240465.1;refGene_gn=ANK3;refGene_tx=XM_023240466.1;refGene_gn=ANK3;refGene_tx=XM_023240467.1;refGene_gn=ANK3;refGene_tx=XM_023240468.1;refGene_gn=ANK3;refGene_tx=XM_023240469.1;refGene_gn=ANK3;refGene_tx=XM_023240470.1;refGene_gn=ANK3;refGene_tx=XM_023240471.1;refGene_gn=ANK3;refGene_tx=XM_023240472.1
D2 23892495 23892592
refGene_gn=ANK3;refGene_tx=XM_003993953.5;refGene_gn=ANK3;refGene_tx=XM_019813142.2;refGene_gn=ANK3;refGene_tx=XM_023240460.1;refGene_gn=ANK3;refGene_tx=XM_023240461.1;refGene_gn=ANK3;refGene_tx=XM_023240462.1;refGene_gn=ANK3;refGene_tx=XM_023240463.1;refGene_gn=ANK3;refGene_tx=XM_023240464.1;refGene_gn=ANK3;refGene_tx=XM_023240465.1;refGene_gn=ANK3;refGene_tx=XM_023240466.1;refGene_gn=ANK3;refGene_tx=XM_023240467.1;refGene_gn=ANK3;refGene_tx=XM_023240468.1;refGene_gn=ANK3;refGene_tx=XM_023240469.1;refGene_gn=ANK3;refGene_tx=XM_023240470.1;refGene_gn=ANK3;refGene_tx=XM_023240471.1;refGene_gn=ANK3;refGene_tx=XM_023240472.1
D2 23895386 23895594
refGene_gn=ANK3;refGene_tx=XM_003993953.5;refGene_gn=ANK3;refGene_tx=XM_019813142.2;refGene_gn=ANK3;refGene_tx=XM_023240460.1;refGene_gn=ANK3;refGene_tx=XM_023240461.1;refGene_gn=ANK3;refGene_tx=XM_023240462.1;refGene_gn=ANK3;refGene_tx=XM_023240463.1;refGene_gn=ANK3;refGene_tx=XM_023240464.1;refGene_gn=ANK3;refGene_tx=XM_023240465.1;refGene_gn=ANK3;refGene_tx=XM_023240466.1;refGene_gn=ANK3;refGene_tx=XM_023240467.1;refGene_gn=ANK3;refGene_tx=XM_023240468.1;refGene_gn=ANK3;refGene_tx=XM_023240469.1;refGene_gn=ANK3;refGene_tx=XM_023240470.1;refGene_gn=ANK3;refGene_tx=XM_023240471.1;refGene_gn=ANK3;refGene_tx=XM_023240472.1
D2 23896824 23897036
refGene_gn=ANK3;refGene_tx=XM_003993953.5;refGene_gn=ANK3;refGene_tx=XM_019813142.2;refGene_gn=ANK3;refGene_tx=XM_023240460.1;refGene_gn=ANK3;refGene_tx=XM_023240461.1;refGene_gn=ANK3;refGene_tx=XM_023240462.1;refGene_gn=ANK3;refGene_tx=XM_023240463.1;refGene_gn=ANK3;refGene_tx=XM_023240464.1;refGene_gn=ANK3;refGene_tx=XM_023240465.1;refGene_gn=ANK3;refGene_tx=XM_023240466.1;refGene_gn=ANK3;refGene_tx=XM_023240467.1;refGene_gn=ANK3;refGene_tx=XM_023240468.1;refGene_gn=ANK3;refGene_tx=XM_023240469.1;refGene_gn=ANK3;refGene_tx=XM_023240470.1;refGene_gn=ANK3;refGene_tx=XM_023240471.1;refGene_gn=ANK3;refGene_tx=XM_023240472.1
D2 23914117 23914272

refGene_gn=ANK3;refGene_tx=XM_003993953.5;refGene_gn=ANK3;refGene_tx=XM_019813142.2;refGene_gn=ANK3;refGene_tx=XM_023240460.1;refGene_gn=ANK3;refGene_tx=XM_023240461.1;refGene_gn=ANK3;refGene_tx=XM_023240462.1;refGene_gn=ANK3;refGene_tx=XM_023240463.1;refGene_gn=ANK3;refGene_tx=XM_023240464.1;refGene_gn=ANK3;refGene_tx=XM_023240465.1;refGene_gn=ANK3;refGene_tx=XM_023240466.1;refGene_gn=ANK3;refGene_tx=XM_023240467.1;refGene_gn=ANK3;refGene_tx=XM_023240468.1;refGene_gn=ANK3;refGene_tx=XM_023240469.1;refGene_gn=ANK3;refGene_tx=XM_023240470.1;refGene_gn=ANK3;refGene_tx=XM_023240471.1;refGene_gn=ANK3;refGene_tx=XM_023240472.1
D2 23916443 23916542 refGene_gn=ANK3;refGene_tx=XM_023240470.1
D2 23917077 23917302
refGene_gn=ANK3;refGene_tx=XM_003993953.5;refGene_gn=ANK3;refGene_tx=XM_019813142.2;refGene_gn=ANK3;refGene_tx=XM_023240460.1;refGene_gn=ANK3;refGene_tx=XM_023240461.1;refGene_gn=ANK3;refGene_tx=XM_023240462.1;refGene_gn=ANK3;refGene_tx=XM_023240463.1;refGene_gn=ANK3;refGene_tx=XM_023240464.1;refGene_gn=ANK3;refGene_tx=XM_023240465.1;refGene_gn=ANK3;refGene_tx=XM_023240466.1;refGene_gn=ANK3;refGene_tx=XM_023240467.1;refGene_gn=ANK3;refGene_tx=XM_023240468.1;refGene_gn=ANK3;refGene_tx=XM_023240469.1;refGene_gn=ANK3;refGene_tx=XM_023240470.1;refGene_gn=ANK3;refGene_tx=XM_023240471.1;refGene_gn=ANK3;refGene_tx=XM_023240472.1
D2 23919998 23920031
refGene_gn=ANK3;refGene_tx=XM_023240463.1;refGene_gn=ANK3;refGene_tx=XM_023240467.1;refGene_gn=ANK3;refGene_tx=XM_023240470.1
D2 23921982 23922089
refGene_gn=ANK3;refGene_tx=XM_003993953.5;refGene_gn=ANK3;refGene_tx=XM_019813142.2;refGene_gn=ANK3;refGene_tx=XM_023240460.1;refGene_gn=ANK3;refGene_tx=XM_023240461.1;refGene_gn=ANK3;refGene_tx=XM_023240462.1;refGene_gn=ANK3;refGene_tx=XM_023240463.1;refGene_gn=ANK3;refGene_tx=XM_023240464.1;refGene_gn=ANK3;refGene_tx=XM_023240465.1;refGene_gn=ANK3;refGene_tx=XM_023240466.1;refGene_gn=ANK3;refGene_tx=XM_023240467.1;refGene_gn=ANK3;refGene_tx=XM_023240468.1;refGene_gn=ANK3;refGene_tx=XM_023240469.1;refGene_gn=ANK3;refGene_tx=XM_023240470.1;refGene_gn=ANK3;refGene_tx=XM_023240471.1;refGene_gn=ANK3;refGene_tx=XM_023240472.1
D2 23937696 23937799
refGene_gn=ANK3;refGene_tx=XM_003993953.5;refGene_gn=ANK3;refGene_tx=XM_019813142.2;refGene_gn=ANK3;refGene_tx=XM_023240460.1;refGene_gn=ANK3;refGene_tx=XM_023240461.1;refGene_gn=ANK3;refGene_tx=XM_023240462.1;refGene_gn=ANK3;refGene_tx=XM_023240463.1;refGene_gn=ANK3;refGene_tx=XM_023240464.1;refGene_gn=ANK3;refGene_tx=XM_023240465.1;refGene_gn=ANK3;refGene_tx=XM_023240466.1;refGene_gn=ANK3;refGene_tx=XM_023240467.1;refGene_gn=ANK3;refGene_tx=XM_023240468.1;refGene_gn=ANK3;refGene_tx=XM_023240469.1;refGene_gn=ANK3;refGene_tx=XM_023240470.1;refGene_gn=ANK3;refGene_tx=XM_023240471.1;refGene_gn=ANK3;refGene_tx=XM_023240472.1
D2 23941053 23941065 refGene_gn=ANK3;refGene_tx=XM_023240470.1
D2 23942712 23942836
refGene_gn=ANK3;refGene_tx=XM_003993953.5;refGene_gn=ANK3;refGene_tx=XM_019813142.2;refGene_gn=ANK3;refGene_tx=XM_023240460.1;refGene_gn=ANK3;refGene_tx=XM_023240461.1;refGene_gn=ANK3;refGene_tx=XM_023240462.1;refGene_gn=ANK3;refGene_tx=XM_023240463.1;refGene_gn=ANK3;refGene_tx=XM_023240464.1;refGene_gn=ANK3;refGene_tx=XM_023240465.1;refGene_gn=ANK

3;refGene_tx=XM_023240466.1;refGene_gn=ANK3;refGene_tx=XM_023240467.1;
refGene_gn=ANK3;refGene_tx=XM_023240468.1;refGene_gn=ANK3;refGene_tx=X
M_023240469.1;refGene_gn=ANK3;refGene_tx=XM_023240470.1;refGene_gn=ANK
3;refGene_tx=XM_023240471.1;refGene_gn=ANK3;refGene_tx=XM_023240472.1
D2 23944130 23944146 refGene_gn=ANK3;refGene_tx=XM_003993953.5
D2 23950179 23950233
refGene_gn=ANK3;refGene_tx=XM_023240470.1;refGene_gn=ANK3;refGene_tx=X
M_023240471.1
D2 23964156 23964219
refGene_gn=ANK3;refGene_tx=XM_019813142.2;refGene_gn=ANK3;refGene_tx=X
M_023240460.1;refGene_gn=ANK3;refGene_tx=XM_023240461.1;refGene_gn=ANK
3;refGene_tx=XM_023240462.1;refGene_gn=ANK3;refGene_tx=XM_023240470.1;
refGene_gn=ANK3;refGene_tx=XM_023240471.1;refGene_gn=ANK3;refGene_tx=X
M_023240472.1
D2 23964380 23964453
refGene_gn=ANK3;refGene_tx=XM_019813142.2;refGene_gn=ANK3;refGene_tx=X
M_023240460.1;refGene_gn=ANK3;refGene_tx=XM_023240461.1;refGene_gn=ANK
3;refGene_tx=XM_023240462.1;refGene_gn=ANK3;refGene_tx=XM_023240463.1;
refGene_gn=ANK3;refGene_tx=XM_023240464.1;refGene_gn=ANK3;refGene_tx=X
M_023240465.1;refGene_gn=ANK3;refGene_tx=XM_023240466.1;refGene_gn=ANK
3;refGene_tx=XM_023240467.1;refGene_gn=ANK3;refGene_tx=XM_023240468.1;
refGene_gn=ANK3;refGene_tx=XM_023240469.1;refGene_gn=ANK3;refGene_tx=X
M_023240470.1;refGene_gn=ANK3;refGene_tx=XM_023240471.1;refGene_gn=ANK
3;refGene_tx=XM_023240472.1
D2 23968737 23968833
refGene_gn=ANK3;refGene_tx=XM_019813142.2;refGene_gn=ANK3;refGene_tx=X
M_023240460.1;refGene_gn=ANK3;refGene_tx=XM_023240461.1;refGene_gn=ANK
3;refGene_tx=XM_023240462.1;refGene_gn=ANK3;refGene_tx=XM_023240463.1;
refGene_gn=ANK3;refGene_tx=XM_023240464.1;refGene_gn=ANK3;refGene_tx=X
M_023240465.1;refGene_gn=ANK3;refGene_tx=XM_023240466.1;refGene_gn=ANK
3;refGene_tx=XM_023240467.1;refGene_gn=ANK3;refGene_tx=XM_023240468.1;
refGene_gn=ANK3;refGene_tx=XM_023240469.1;refGene_gn=ANK3;refGene_tx=X
M_023240470.1;refGene_gn=ANK3;refGene_tx=XM_023240471.1;refGene_gn=ANK
3;refGene_tx=XM_023240472.1
D2 23969371 23969470
refGene_gn=ANK3;refGene_tx=XM_019813142.2;refGene_gn=ANK3;refGene_tx=X
M_023240460.1;refGene_gn=ANK3;refGene_tx=XM_023240461.1;refGene_gn=ANK
3;refGene_tx=XM_023240462.1;refGene_gn=ANK3;refGene_tx=XM_023240463.1;
refGene_gn=ANK3;refGene_tx=XM_023240464.1;refGene_gn=ANK3;refGene_tx=X
M_023240465.1;refGene_gn=ANK3;refGene_tx=XM_023240466.1;refGene_gn=ANK
3;refGene_tx=XM_023240467.1;refGene_gn=ANK3;refGene_tx=XM_023240468.1;
refGene_gn=ANK3;refGene_tx=XM_023240469.1;refGene_gn=ANK3;refGene_tx=X
M_023240470.1;refGene_gn=ANK3;refGene_tx=XM_023240471.1;refGene_gn=ANK
3;refGene_tx=XM_023240472.1
D2 23969559 23969658
refGene_gn=ANK3;refGene_tx=XM_019813142.2;refGene_gn=ANK3;refGene_tx=X
M_023240460.1;refGene_gn=ANK3;refGene_tx=XM_023240461.1;refGene_gn=ANK
3;refGene_tx=XM_023240462.1;refGene_gn=ANK3;refGene_tx=XM_023240463.1;
refGene_gn=ANK3;refGene_tx=XM_023240464.1;refGene_gn=ANK3;refGene_tx=X
M_023240465.1;refGene_gn=ANK3;refGene_tx=XM_023240466.1;refGene_gn=ANK

3;refGene_tx=XM_023240467.1;refGene_gn=ANK3;refGene_tx=XM_023240468.1;
refGene_gn=ANK3;refGene_tx=XM_023240469.1;refGene_gn=ANK3;refGene_tx=X
M_023240470.1;refGene_gn=ANK3;refGene_tx=XM_023240471.1;refGene_gn=ANK
3;refGene_tx=XM_023240472.1
D2 23978244 23978343
refGene_gn=ANK3;refGene_tx=XM_019813142.2;refGene_gn=ANK3;refGene_tx=X
M_023240460.1;refGene_gn=ANK3;refGene_tx=XM_023240461.1;refGene_gn=ANK
3;refGene_tx=XM_023240462.1;refGene_gn=ANK3;refGene_tx=XM_023240463.1;
refGene_gn=ANK3;refGene_tx=XM_023240464.1;refGene_gn=ANK3;refGene_tx=X
M_023240465.1;refGene_gn=ANK3;refGene_tx=XM_023240466.1;refGene_gn=ANK
3;refGene_tx=XM_023240467.1;refGene_gn=ANK3;refGene_tx=XM_023240468.1;
refGene_gn=ANK3;refGene_tx=XM_023240469.1;refGene_gn=ANK3;refGene_tx=X
M_023240470.1;refGene_gn=ANK3;refGene_tx=XM_023240471.1;refGene_gn=ANK
3;refGene_tx=XM_023240472.1
D2 23984202 23984400
refGene_gn=ANK3;refGene_tx=XM_019813142.2;refGene_gn=ANK3;refGene_tx=X
M_023240460.1;refGene_gn=ANK3;refGene_tx=XM_023240461.1;refGene_gn=ANK
3;refGene_tx=XM_023240462.1;refGene_gn=ANK3;refGene_tx=XM_023240463.1;
refGene_gn=ANK3;refGene_tx=XM_023240464.1;refGene_gn=ANK3;refGene_tx=X
M_023240465.1;refGene_gn=ANK3;refGene_tx=XM_023240466.1;refGene_gn=ANK
3;refGene_tx=XM_023240467.1;refGene_gn=ANK3;refGene_tx=XM_023240468.1;
refGene_gn=ANK3;refGene_tx=XM_023240469.1;refGene_gn=ANK3;refGene_tx=X
M_023240470.1;refGene_gn=ANK3;refGene_tx=XM_023240471.1;refGene_gn=ANK
3;refGene_tx=XM_023240472.1
D2 23989989 23990088
refGene_gn=ANK3;refGene_tx=XM_019813142.2;refGene_gn=ANK3;refGene_tx=X
M_023240460.1;refGene_gn=ANK3;refGene_tx=XM_023240461.1;refGene_gn=ANK
3;refGene_tx=XM_023240462.1;refGene_gn=ANK3;refGene_tx=XM_023240463.1;
refGene_gn=ANK3;refGene_tx=XM_023240464.1;refGene_gn=ANK3;refGene_tx=X
M_023240465.1;refGene_gn=ANK3;refGene_tx=XM_023240466.1;refGene_gn=ANK
3;refGene_tx=XM_023240470.1;refGene_gn=ANK3;refGene_tx=XM_023240471.1;
refGene_gn=ANK3;refGene_tx=XM_023240472.1
D2 23990369 23990468
refGene_gn=ANK3;refGene_tx=XM_019813142.2;refGene_gn=ANK3;refGene_tx=X
M_023240460.1;refGene_gn=ANK3;refGene_tx=XM_023240461.1;refGene_gn=ANK
3;refGene_tx=XM_023240462.1;refGene_gn=ANK3;refGene_tx=XM_023240463.1;
refGene_gn=ANK3;refGene_tx=XM_023240464.1;refGene_gn=ANK3;refGene_tx=X
M_023240465.1;refGene_gn=ANK3;refGene_tx=XM_023240466.1;refGene_gn=ANK
3;refGene_tx=XM_023240467.1;refGene_gn=ANK3;refGene_tx=XM_023240468.1;
refGene_gn=ANK3;refGene_tx=XM_023240469.1;refGene_gn=ANK3;refGene_tx=X
M_023240470.1;refGene_gn=ANK3;refGene_tx=XM_023240471.1;refGene_gn=ANK
3;refGene_tx=XM_023240472.1
D2 23992496 23992694
refGene_gn=ANK3;refGene_tx=XM_019813142.2;refGene_gn=ANK3;refGene_tx=X
M_023240460.1;refGene_gn=ANK3;refGene_tx=XM_023240461.1;refGene_gn=ANK
3;refGene_tx=XM_023240462.1;refGene_gn=ANK3;refGene_tx=XM_023240463.1;
refGene_gn=ANK3;refGene_tx=XM_023240464.1;refGene_gn=ANK3;refGene_tx=X
M_023240465.1;refGene_gn=ANK3;refGene_tx=XM_023240466.1;refGene_gn=ANK
3;refGene_tx=XM_023240467.1;refGene_gn=ANK3;refGene_tx=XM_023240468.1;
refGene_gn=ANK3;refGene_tx=XM_023240469.1;refGene_gn=ANK3;refGene_tx=X

M_023240470.1;refGene_gn=ANK3;refGene_tx=XM_023240471.1;refGene_gn=ANK3;refGene_tx=XM_023240472.1
D2 23995473 23995572
refGene_gn=ANK3;refGene_tx=XM_019813142.2;refGene_gn=ANK3;refGene_tx=XM_023240460.1;refGene_gn=ANK3;refGene_tx=XM_023240461.1;refGene_gn=ANK3;refGene_tx=XM_023240462.1;refGene_gn=ANK3;refGene_tx=XM_023240463.1;refGene_gn=ANK3;refGene_tx=XM_023240464.1;refGene_gn=ANK3;refGene_tx=XM_023240465.1;refGene_gn=ANK3;refGene_tx=XM_023240466.1;refGene_gn=ANK3;refGene_tx=XM_023240467.1;refGene_gn=ANK3;refGene_tx=XM_023240468.1;refGene_gn=ANK3;refGene_tx=XM_023240469.1;refGene_gn=ANK3;refGene_tx=XM_023240470.1;refGene_gn=ANK3;refGene_tx=XM_023240471.1;refGene_gn=ANK3;refGene_tx=XM_023240472.1
D2 23997431 23997530
refGene_gn=ANK3;refGene_tx=XM_019813142.2;refGene_gn=ANK3;refGene_tx=XM_023240460.1;refGene_gn=ANK3;refGene_tx=XM_023240461.1;refGene_gn=ANK3;refGene_tx=XM_023240462.1;refGene_gn=ANK3;refGene_tx=XM_023240463.1;refGene_gn=ANK3;refGene_tx=XM_023240464.1;refGene_gn=ANK3;refGene_tx=XM_023240465.1;refGene_gn=ANK3;refGene_tx=XM_023240466.1;refGene_gn=ANK3;refGene_tx=XM_023240467.1;refGene_gn=ANK3;refGene_tx=XM_023240468.1;refGene_gn=ANK3;refGene_tx=XM_023240469.1;refGene_gn=ANK3;refGene_tx=XM_023240470.1;refGene_gn=ANK3;refGene_tx=XM_023240471.1;refGene_gn=ANK3;refGene_tx=XM_023240472.1
D2 24000224 24000323
refGene_gn=ANK3;refGene_tx=XM_019813142.2;refGene_gn=ANK3;refGene_tx=XM_023240460.1;refGene_gn=ANK3;refGene_tx=XM_023240461.1;refGene_gn=ANK3;refGene_tx=XM_023240462.1;refGene_gn=ANK3;refGene_tx=XM_023240463.1;refGene_gn=ANK3;refGene_tx=XM_023240464.1;refGene_gn=ANK3;refGene_tx=XM_023240465.1;refGene_gn=ANK3;refGene_tx=XM_023240466.1;refGene_gn=ANK3;refGene_tx=XM_023240467.1;refGene_gn=ANK3;refGene_tx=XM_023240468.1;refGene_gn=ANK3;refGene_tx=XM_023240469.1;refGene_gn=ANK3;refGene_tx=XM_023240470.1;refGene_gn=ANK3;refGene_tx=XM_023240471.1;refGene_gn=ANK3;refGene_tx=XM_023240472.1
D2 24002412 24002610
refGene_gn=ANK3;refGene_tx=XM_019813142.2;refGene_gn=ANK3;refGene_tx=XM_023240460.1;refGene_gn=ANK3;refGene_tx=XM_023240461.1;refGene_gn=ANK3;refGene_tx=XM_023240462.1;refGene_gn=ANK3;refGene_tx=XM_023240463.1;refGene_gn=ANK3;refGene_tx=XM_023240464.1;refGene_gn=ANK3;refGene_tx=XM_023240465.1;refGene_gn=ANK3;refGene_tx=XM_023240466.1;refGene_gn=ANK3;refGene_tx=XM_023240467.1;refGene_gn=ANK3;refGene_tx=XM_023240468.1;refGene_gn=ANK3;refGene_tx=XM_023240469.1;refGene_gn=ANK3;refGene_tx=XM_023240470.1;refGene_gn=ANK3;refGene_tx=XM_023240471.1;refGene_gn=ANK3;refGene_tx=XM_023240472.1
D2 24008251 24008350
refGene_gn=ANK3;refGene_tx=XM_019813142.2;refGene_gn=ANK3;refGene_tx=XM_023240460.1;refGene_gn=ANK3;refGene_tx=XM_023240461.1;refGene_gn=ANK3;refGene_tx=XM_023240462.1;refGene_gn=ANK3;refGene_tx=XM_023240463.1;refGene_gn=ANK3;refGene_tx=XM_023240464.1;refGene_gn=ANK3;refGene_tx=XM_023240465.1;refGene_gn=ANK3;refGene_tx=XM_023240466.1;refGene_gn=ANK3;refGene_tx=XM_023240467.1;refGene_gn=ANK3;refGene_tx=XM_023240468.1;refGene_gn=ANK3;refGene_tx=XM_023240469.1;refGene_gn=ANK3;refGene_tx=X

M_023240470.1;refGene_gn=ANK3;refGene_tx=XM_023240471.1;refGene_gn=ANK
3;refGene_tx=XM_023240472.1
D2 24028932 24029031
refGene_gn=ANK3;refGene_tx=XM_019813142.2;refGene_gn=ANK3;refGene_tx=X
M_023240460.1;refGene_gn=ANK3;refGene_tx=XM_023240461.1;refGene_gn=ANK
3;refGene_tx=XM_023240462.1;refGene_gn=ANK3;refGene_tx=XM_023240463.1;
refGene_gn=ANK3;refGene_tx=XM_023240464.1;refGene_gn=ANK3;refGene_tx=X
M_023240465.1;refGene_gn=ANK3;refGene_tx=XM_023240466.1;refGene_gn=ANK
3;refGene_tx=XM_023240467.1;refGene_gn=ANK3;refGene_tx=XM_023240468.1;
refGene_gn=ANK3;refGene_tx=XM_023240469.1;refGene_gn=ANK3;refGene_tx=X
M_023240470.1;refGene_gn=ANK3;refGene_tx=XM_023240471.1;refGene_gn=ANK
3;refGene_tx=XM_023240472.1
D2 24050475 24050574
refGene_gn=ANK3;refGene_tx=XM_019813142.2;refGene_gn=ANK3;refGene_tx=X
M_023240460.1;refGene_gn=ANK3;refGene_tx=XM_023240461.1;refGene_gn=ANK
3;refGene_tx=XM_023240462.1;refGene_gn=ANK3;refGene_tx=XM_023240463.1;
refGene_gn=ANK3;refGene_tx=XM_023240464.1;refGene_gn=ANK3;refGene_tx=X
M_023240465.1;refGene_gn=ANK3;refGene_tx=XM_023240466.1;refGene_gn=ANK
3;refGene_tx=XM_023240467.1;refGene_gn=ANK3;refGene_tx=XM_023240468.1;
refGene_gn=ANK3;refGene_tx=XM_023240469.1;refGene_gn=ANK3;refGene_tx=X
M_023240470.1;refGene_gn=ANK3;refGene_tx=XM_023240471.1;refGene_gn=ANK
3;refGene_tx=XM_023240472.1
D2 24051735 24051759
refGene_gn=ANK3;refGene_tx=XM_023240463.1;refGene_gn=ANK3;refGene_tx=X
M_023240464.1;refGene_gn=ANK3;refGene_tx=XM_023240467.1;refGene_gn=ANK
3;refGene_tx=XM_023240468.1;refGene_gn=ANK3;refGene_tx=XM_023240470.1
D2 24052415 24052601
refGene_gn=ANK3;refGene_tx=XM_019813142.2;refGene_gn=ANK3;refGene_tx=X
M_023240460.1;refGene_gn=ANK3;refGene_tx=XM_023240461.1;refGene_gn=ANK
3;refGene_tx=XM_023240462.1;refGene_gn=ANK3;refGene_tx=XM_023240463.1;
refGene_gn=ANK3;refGene_tx=XM_023240464.1;refGene_gn=ANK3;refGene_tx=X
M_023240465.1;refGene_gn=ANK3;refGene_tx=XM_023240466.1;refGene_gn=ANK
3;refGene_tx=XM_023240467.1;refGene_gn=ANK3;refGene_tx=XM_023240468.1;
refGene_gn=ANK3;refGene_tx=XM_023240469.1;refGene_gn=ANK3;refGene_tx=X
M_023240470.1;refGene_gn=ANK3;refGene_tx=XM_023240471.1;refGene_gn=ANK
3;refGene_tx=XM_023240472.1
D2 24058085 24058184
refGene_gn=ANK3;refGene_tx=XM_019813142.2;refGene_gn=ANK3;refGene_tx=X
M_023240460.1;refGene_gn=ANK3;refGene_tx=XM_023240461.1;refGene_gn=ANK
3;refGene_tx=XM_023240462.1;refGene_gn=ANK3;refGene_tx=XM_023240463.1;
refGene_gn=ANK3;refGene_tx=XM_023240464.1;refGene_gn=ANK3;refGene_tx=X
M_023240465.1;refGene_gn=ANK3;refGene_tx=XM_023240466.1;refGene_gn=ANK
3;refGene_tx=XM_023240467.1;refGene_gn=ANK3;refGene_tx=XM_023240468.1;
refGene_gn=ANK3;refGene_tx=XM_023240469.1;refGene_gn=ANK3;refGene_tx=X
M_023240470.1;refGene_gn=ANK3;refGene_tx=XM_023240471.1;refGene_gn=ANK
3;refGene_tx=XM_023240472.1
D2 24066385 24066484
refGene_gn=ANK3;refGene_tx=XM_019813142.2;refGene_gn=ANK3;refGene_tx=X
M_023240460.1;refGene_gn=ANK3;refGene_tx=XM_023240461.1;refGene_gn=ANK
3;refGene_tx=XM_023240462.1;refGene_gn=ANK3;refGene_tx=XM_023240463.1;

refGene_gn=ANK3;refGene_tx=XM_023240464.1;refGene_gn=ANK3;refGene_tx=XM_023240465.1;refGene_gn=ANK3;refGene_tx=XM_023240466.1;refGene_gn=ANK3;refGene_tx=XM_023240467.1;refGene_gn=ANK3;refGene_tx=XM_023240468.1;refGene_gn=ANK3;refGene_tx=XM_023240469.1;refGene_gn=ANK3;refGene_tx=XM_023240470.1;refGene_gn=ANK3;refGene_tx=XM_023240471.1;refGene_gn=ANK3;refGene_tx=XM_023240472.1
D2 24066697 24066796
refGene_gn=ANK3;refGene_tx=XM_019813142.2;refGene_gn=ANK3;refGene_tx=XM_023240460.1;refGene_gn=ANK3;refGene_tx=XM_023240461.1;refGene_gn=ANK3;refGene_tx=XM_023240462.1;refGene_gn=ANK3;refGene_tx=XM_023240463.1;refGene_gn=ANK3;refGene_tx=XM_023240464.1;refGene_gn=ANK3;refGene_tx=XM_023240465.1;refGene_gn=ANK3;refGene_tx=XM_023240466.1;refGene_gn=ANK3;refGene_tx=XM_023240467.1;refGene_gn=ANK3;refGene_tx=XM_023240468.1;refGene_gn=ANK3;refGene_tx=XM_023240469.1;refGene_gn=ANK3;refGene_tx=XM_023240470.1;refGene_gn=ANK3;refGene_tx=XM_023240471.1;refGene_gn=ANK3;refGene_tx=XM_023240472.1
D2 24067202 24067304
refGene_gn=ANK3;refGene_tx=XM_019813142.2;refGene_gn=ANK3;refGene_tx=XM_023240460.1;refGene_gn=ANK3;refGene_tx=XM_023240461.1;refGene_gn=ANK3;refGene_tx=XM_023240462.1;refGene_gn=ANK3;refGene_tx=XM_023240463.1;refGene_gn=ANK3;refGene_tx=XM_023240464.1;refGene_gn=ANK3;refGene_tx=XM_023240465.1;refGene_gn=ANK3;refGene_tx=XM_023240466.1;refGene_gn=ANK3;refGene_tx=XM_023240467.1;refGene_gn=ANK3;refGene_tx=XM_023240468.1;refGene_gn=ANK3;refGene_tx=XM_023240469.1;refGene_gn=ANK3;refGene_tx=XM_023240470.1;refGene_gn=ANK3;refGene_tx=XM_023240471.1;refGene_gn=ANK3;refGene_tx=XM_023240472.1
D2 24173297 24173411
refGene_gn=ANK3;refGene_tx=XM_019813142.2;refGene_gn=ANK3;refGene_tx=XM_023240460.1;refGene_gn=ANK3;refGene_tx=XM_023240461.1;refGene_gn=ANK3;refGene_tx=XM_023240462.1;refGene_gn=ANK3;refGene_tx=XM_023240463.1;refGene_gn=ANK3;refGene_tx=XM_023240464.1;refGene_gn=ANK3;refGene_tx=XM_023240465.1;refGene_gn=ANK3;refGene_tx=XM_023240466.1;refGene_gn=ANK3;refGene_tx=XM_023240467.1;refGene_gn=ANK3;refGene_tx=XM_023240468.1;refGene_gn=ANK3;refGene_tx=XM_023240469.1;refGene_gn=ANK3;refGene_tx=XM_023240471.1;refGene_gn=ANK3;refGene_tx=XM_023240472.1
D2 24194708 24195055
refGene_gn=LOC101101340;refGene_tx=XM_023240473.1
D2 24195311 24195513
refGene_gn=LOC101101340;refGene_tx=XM_023240473.1
D2 24402957 24402996 refGene_gn=ANK3;refGene_tx=XM_023240470.1
D2 24522788 24522845 refGene_gn=ANK3;refGene_tx=XM_023240470.1
D2 24581144 24581181
refGene_gn=CDK1;refGene_tx=XM_006937802.4;refGene_gn=CDK1;refGene_tx=XM_006937803.2;refGene_gn=CDK1;refGene_tx=XM_019813147.2;refGene_gn=CDK1;refGene_tx=XM_019813148.2
D2 24586288 24586445
refGene_gn=CDK1;refGene_tx=XM_006937802.4;refGene_gn=CDK1;refGene_tx=XM_006937803.2;refGene_gn=CDK1;refGene_tx=XM_019813147.2;refGene_gn=CDK1;refGene_tx=XM_019813148.2
D2 24587291 24587415

refGene_gn=CDK1;refGene_tx=XM_006937802.4;refGene_gn=CDK1;refGene_tx=XM_006937803.2;refGene_gn=CDK1;refGene_tx=XM_019813147.2;refGene_gn=CDK1;refGene_tx=XM_019813148.2
D2 24589423 24589594 refGene_gn=CDK1;refGene_tx=XM_006937802.4;refGene_gn=CDK1;refGene_tx=XM_006937803.2;refGene_gn=CDK1;refGene_tx=XM_019813147.2;refGene_gn=CDK1;refGene_tx=XM_019813148.2
D2 24591292 24591456 refGene_gn=CDK1;refGene_tx=XM_006937802.4;refGene_gn=CDK1;refGene_tx=XM_006937803.2;refGene_gn=CDK1;refGene_tx=XM_019813147.2;refGene_gn=CDK1;refGene_tx=XM_019813148.2
D2 24591580 24591722 refGene_gn=CDK1;refGene_tx=XM_006937802.4;refGene_gn=CDK1;refGene_tx=XM_006937803.2;refGene_gn=CDK1;refGene_tx=XM_019813147.2;refGene_gn=CDK1;refGene_tx=XM_019813148.2
D2 24595140 24595239 refGene_gn=CDK1;refGene_tx=XM_006937802.4;refGene_gn=CDK1;refGene_tx=XM_006937803.2;refGene_gn=CDK1;refGene_tx=XM_019813147.2
D2 24606220 24606241 refGene_gn=CDK1;refGene_tx=XM_019813148.2
D2 24674644 24674814 refGene_gn=RHOBTB1;refGene_tx=XM_006937805.4;refGene_gn=RHOBTB1;refGene_tx=XM_006937807.4;refGene_gn=RHOBTB1;refGene_tx=XM_011287376.3;refGene_gn=RHOBTB1;refGene_tx=XM_019813149.2;refGene_gn=RHOBTB1;refGene_tx=XM_019813151.2;refGene_gn=RHOBTB1;refGene_tx=XM_023240474.1;refGene_gn=RHOBTB1;refGene_tx=XM_023240475.1;refGene_gn=RHOBTB1;refGene_tx=XM_023240476.1;refGene_gn=RHOBTB1;refGene_tx=XM_023240477.1;refGene_gn=RHOBTB1;refGene_tx=XM_023240478.1
D2 24675334 24675440 refGene_gn=RHOBTB1;refGene_tx=XM_006937805.4;refGene_gn=RHOBTB1;refGene_tx=XM_006937807.4;refGene_gn=RHOBTB1;refGene_tx=XM_011287376.3;refGene_gn=RHOBTB1;refGene_tx=XM_019813149.2;refGene_gn=RHOBTB1;refGene_tx=XM_019813151.2;refGene_gn=RHOBTB1;refGene_tx=XM_023240474.1;refGene_gn=RHOBTB1;refGene_tx=XM_023240475.1;refGene_gn=RHOBTB1;refGene_tx=XM_023240476.1;refGene_gn=RHOBTB1;refGene_tx=XM_023240477.1;refGene_gn=RHOBTB1;refGene_tx=XM_023240478.1
D2 24678179 24678268 refGene_gn=RHOBTB1;refGene_tx=XM_006937805.4;refGene_gn=RHOBTB1;refGene_tx=XM_006937807.4;refGene_gn=RHOBTB1;refGene_tx=XM_011287376.3;refGene_gn=RHOBTB1;refGene_tx=XM_019813149.2;refGene_gn=RHOBTB1;refGene_tx=XM_019813151.2;refGene_gn=RHOBTB1;refGene_tx=XM_023240474.1;refGene_gn=RHOBTB1;refGene_tx=XM_023240475.1;refGene_gn=RHOBTB1;refGene_tx=XM_023240476.1;refGene_gn=RHOBTB1;refGene_tx=XM_023240477.1;refGene_gn=RHOBTB1;refGene_tx=XM_023240478.1
D2 24679635 24679643 refGene_gn=RHOBTB1;refGene_tx=XM_023240479.1
D2 24681082 24681233 refGene_gn=RHOBTB1;refGene_tx=XM_006937805.4;refGene_gn=RHOBTB1;refGene_tx=XM_006937807.4;refGene_gn=RHOBTB1;refGene_tx=XM_011287376.3;refGene_gn=RHOBTB1;refGene_tx=XM_019813149.2;refGene_gn=RHOBTB1;refGene_tx=XM_019813151.2;refGene_gn=RHOBTB1;refGene_tx=XM_023240474.1;refGene_gn=RHOBTB1;refGene_tx=XM_023240475.1;refGene_gn=RHOBTB1;refGene_tx=XM_02

3240476.1;refGene_gn=RHOBTB1;refGene_tx=XM_023240477.1;refGene_gn=RHOBTB1;refGene_tx=XM_023240478.1;refGene_gn=RHOBTB1;refGene_tx=XM_023240479.1
D2 24687058 24687177
refGene_gn=RHOBTB1;refGene_tx=XM_006937805.4;refGene_gn=RHOBTB1;refGene_tx=XM_006937807.4;refGene_gn=RHOBTB1;refGene_tx=XM_011287376.3;refGene_gn=RHOBTB1;refGene_tx=XM_019813149.2;refGene_gn=RHOBTB1;refGene_tx=XM_019813151.2;refGene_gn=RHOBTB1;refGene_tx=XM_023240474.1;refGene_gn=RHOBTB1;refGene_tx=XM_023240475.1;refGene_gn=RHOBTB1;refGene_tx=XM_023240476.1;refGene_gn=RHOBTB1;refGene_tx=XM_023240477.1;refGene_gn=RHOBTB1;refGene_tx=XM_023240478.1;refGene_gn=RHOBTB1;refGene_tx=XM_023240479.1
D2 24688980 24689960
refGene_gn=RHOBTB1;refGene_tx=XM_006937805.4;refGene_gn=RHOBTB1;refGene_tx=XM_006937807.4;refGene_gn=RHOBTB1;refGene_tx=XM_011287376.3;refGene_gn=RHOBTB1;refGene_tx=XM_019813149.2;refGene_gn=RHOBTB1;refGene_tx=XM_019813151.2;refGene_gn=RHOBTB1;refGene_tx=XM_023240474.1;refGene_gn=RHOBTB1;refGene_tx=XM_023240475.1;refGene_gn=RHOBTB1;refGene_tx=XM_023240476.1;refGene_gn=RHOBTB1;refGene_tx=XM_023240477.1;refGene_gn=RHOBTB1;refGene_tx=XM_023240478.1;refGene_gn=RHOBTB1;refGene_tx=XM_023240479.1
D2 24693150 24693336
refGene_gn=RHOBTB1;refGene_tx=XM_006937805.4;refGene_gn=RHOBTB1;refGene_tx=XM_006937807.4;refGene_gn=RHOBTB1;refGene_tx=XM_011287376.3;refGene_gn=RHOBTB1;refGene_tx=XM_019813149.2;refGene_gn=RHOBTB1;refGene_tx=XM_019813151.2;refGene_gn=RHOBTB1;refGene_tx=XM_023240474.1;refGene_gn=RHOBTB1;refGene_tx=XM_023240475.1;refGene_gn=RHOBTB1;refGene_tx=XM_023240476.1;refGene_gn=RHOBTB1;refGene_tx=XM_023240477.1;refGene_gn=RHOBTB1;refGene_tx=XM_023240478.1;refGene_gn=RHOBTB1;refGene_tx=XM_023240479.1
D2 24698321 24698379 refGene_gn=RHOBTB1;refGene_tx=XM_006937807.4
D2 24710633 24710737
refGene_gn=RHOBTB1;refGene_tx=XM_006937805.4;refGene_gn=RHOBTB1;refGene_tx=XM_006937807.4;refGene_gn=RHOBTB1;refGene_tx=XM_011287376.3;refGene_gn=RHOBTB1;refGene_tx=XM_019813149.2;refGene_gn=RHOBTB1;refGene_tx=XM_019813151.2;refGene_gn=RHOBTB1;refGene_tx=XM_023240474.1;refGene_gn=RHOBTB1;refGene_tx=XM_023240475.1;refGene_gn=RHOBTB1;refGene_tx=XM_023240476.1;refGene_gn=RHOBTB1;refGene_tx=XM_023240477.1;refGene_gn=RHOBTB1;refGene_tx=XM_023240478.1;refGene_gn=RHOBTB1;refGene_tx=XM_023240479.1
D2 24711116 24711308
refGene_gn=RHOBTB1;refGene_tx=XM_006937805.4;refGene_gn=RHOBTB1;refGene_tx=XM_011287376.3;refGene_gn=RHOBTB1;refGene_tx=XM_019813149.2;refGene_gn=RHOBTB1;refGene_tx=XM_019813151.2;refGene_gn=RHOBTB1;refGene_tx=XM_023240474.1;refGene_gn=RHOBTB1;refGene_tx=XM_023240475.1;refGene_gn=RHOBTB1;refGene_tx=XM_023240476.1;refGene_gn=RHOBTB1;refGene_tx=XM_023240477.1;refGene_gn=RHOBTB1;refGene_tx=XM_023240478.1;refGene_gn=RHOBTB1;refGene_tx=XM_023240479.1
D2 25069124 25069377
refGene_gn=LOC101084319;refGene_tx=XM_011287405.1

D2 25069379 25069458
refGene_gn=LOC101084319;refGene_tx=XM_011287405.1
D2 25069459 25070210
refGene_gn=LOC101084319;refGene_tx=XM_011287405.1
D2 25203430 25203855
refGene_gn=TMEM26;refGene_tx=XM_003993964.5;refGene_gn=TMEM26;refGene_tx=XM_023240480.1
D2 25206612 25206689
refGene_gn=TMEM26;refGene_tx=XM_003993964.5;refGene_gn=TMEM26;refGene_tx=XM_023240480.1
D2 25223764 25223985
refGene_gn=TMEM26;refGene_tx=XM_003993964.5;refGene_gn=TMEM26;refGene_tx=XM_023240480.1
D2 25225904 25226018
refGene_gn=TMEM26;refGene_tx=XM_003993964.5;refGene_gn=TMEM26;refGene_tx=XM_023240480.1
D2 25231837 25231916
refGene_gn=TMEM26;refGene_tx=XM_003993964.5;refGene_gn=TMEM26;refGene_tx=XM_023240480.1
D2 25247374 25247565
refGene_gn=TMEM26;refGene_tx=XM_003993964.5;refGene_gn=TMEM26;refGene_tx=XM_023240480.1
D2 25429123 25429420 refGene_gn=CABCOCO1;refGene_tx=XM_023240481.1
D2 25441881 25441985 refGene_gn=CABCOCO1;refGene_tx=XM_023240481.1
D2 25447356 25447526 refGene_gn=CABCOCO1;refGene_tx=XM_023240481.1
D2 25459109 25459254 refGene_gn=CABCOCO1;refGene_tx=XM_023240481.1
D2 25464639 25464712 refGene_gn=CABCOCO1;refGene_tx=XM_023240481.1
D2 25542529 25542652 refGene_gn=CABCOCO1;refGene_tx=XM_023240481.1
D2 25543077 25543218 refGene_gn=CABCOCO1;refGene_tx=XM_023240481.1
D2 25548400 25548475 refGene_gn=CABCOCO1;refGene_tx=XM_023240481.1
D2 25681571 25681592 refGene_gn=ARID5B;refGene_tx=XM_003993966.5
D2 25682038 25682293 refGene_gn=ARID5B;refGene_tx=XM_003993966.5
D2 25712865 25713091 refGene_gn=ARID5B;refGene_tx=XM_003993966.5
D2 25770177 25770408
refGene_gn=ARID5B;refGene_tx=XM_003993966.5;refGene_gn=ARID5B;refGene_tx=XM_023240482.1
D2 25816145 25816149 refGene_gn=ARID5B;refGene_tx=XM_006937808.4
D2 25817553 25817666
refGene_gn=ARID5B;refGene_tx=XM_003993966.5;refGene_gn=ARID5B;refGene_tx=XM_006937808.4;refGene_gn=ARID5B;refGene_tx=XM_023240482.1
D2 25823643 25823848
refGene_gn=ARID5B;refGene_tx=XM_003993966.5;refGene_gn=ARID5B;refGene_tx=XM_006937808.4;refGene_gn=ARID5B;refGene_tx=XM_023240482.1
D2 25825774 25825827
refGene_gn=ARID5B;refGene_tx=XM_003993966.5;refGene_gn=ARID5B;refGene_tx=XM_006937808.4;refGene_gn=ARID5B;refGene_tx=XM_023240482.1
D2 25835519 25835617
refGene_gn=ARID5B;refGene_tx=XM_003993966.5;refGene_gn=ARID5B;refGene_tx=XM_006937808.4;refGene_gn=ARID5B;refGene_tx=XM_023240482.1
D2 25851530 25851729

refGene_gn=ARID5B;refGene_tx=XM_003993966.5;refGene_gn=ARID5B;refGene_tx=XM_006937808.4;refGene_gn=ARID5B;refGene_tx=XM_023240482.1
D2 25855152 25857318
refGene_gn=ARID5B;refGene_tx=XM_003993966.5;refGene_gn=ARID5B;refGene_tx=XM_006937808.4;refGene_gn=ARID5B;refGene_tx=XM_023240482.1
D2 25934854 25935390
refGene_gn=RTKN2;refGene_tx=XM_011287373.2;refGene_gn=RTKN2;refGene_tx=XM_019813154.2;refGene_gn=RTKN2;refGene_tx=XM_019813155.1
D2 25936659 25936767
refGene_gn=RTKN2;refGene_tx=XM_011287373.2;refGene_gn=RTKN2;refGene_tx=XM_019813154.2;refGene_gn=RTKN2;refGene_tx=XM_019813155.1
D2 25941064 25941230
refGene_gn=RTKN2;refGene_tx=XM_011287373.2;refGene_gn=RTKN2;refGene_tx=XM_019813154.2;refGene_gn=RTKN2;refGene_tx=XM_019813155.1
D2 25955286 25955418
refGene_gn=RTKN2;refGene_tx=XM_011287373.2;refGene_gn=RTKN2;refGene_tx=XM_019813154.2;refGene_gn=RTKN2;refGene_tx=XM_019813155.1
D2 25956336 25956443
refGene_gn=RTKN2;refGene_tx=XM_011287373.2;refGene_gn=RTKN2;refGene_tx=XM_019813154.2;refGene_gn=RTKN2;refGene_tx=XM_019813155.1
D2 25966629 25966724
refGene_gn=RTKN2;refGene_tx=XM_011287373.2;refGene_gn=RTKN2;refGene_tx=XM_019813154.2;refGene_gn=RTKN2;refGene_tx=XM_019813155.1
D2 25978320 25978515
refGene_gn=RTKN2;refGene_tx=XM_011287373.2;refGene_gn=RTKN2;refGene_tx=XM_019813154.2;refGene_gn=RTKN2;refGene_tx=XM_019813155.1
D2 25980884 25981002
refGene_gn=RTKN2;refGene_tx=XM_011287373.2;refGene_gn=RTKN2;refGene_tx=XM_019813154.2;refGene_gn=RTKN2;refGene_tx=XM_019813155.1
D2 25982329 25982383
refGene_gn=RTKN2;refGene_tx=XM_011287373.2;refGene_gn=RTKN2;refGene_tx=XM_019813154.2;refGene_gn=RTKN2;refGene_tx=XM_019813155.1
D2 25987475 25987534
refGene_gn=RTKN2;refGene_tx=XM_011287373.2;refGene_gn=RTKN2;refGene_tx=XM_019813154.2
D2 26000172 26000369
refGene_gn=RTKN2;refGene_tx=XM_011287373.2;refGene_gn=RTKN2;refGene_tx=XM_019813154.2
D2 26007264 26007324 refGene_gn=RTKN2;refGene_tx=XM_011287373.2
D2 26090434 26091177 refGene_gn=ZNF365;refGene_tx=XM_023240483.1
D2 26104218 26104399 refGene_gn=ZNF365;refGene_tx=XM_023240483.1
D2 26111280 26111318 refGene_gn=ZNF365;refGene_tx=XM_023240483.1
D2 26112055 26112317 refGene_gn=ZNF365;refGene_tx=XM_023240483.1
D2 26485111 26485924 refGene_gn=ADO;refGene_tx=XM_003993968.5
D2 26492775 26494028
refGene_gn=EGR2;refGene_tx=XM_019813157.2;refGene_gn=EGR2;refGene_tx=XM_019813158.2;refGene_gn=EGR2;refGene_tx=XM_019813159.2
D2 26495350 26495459
refGene_gn=EGR2;refGene_tx=XM_019813157.2;refGene_gn=EGR2;refGene_tx=XM_019813158.2;refGene_gn=EGR2;refGene_tx=XM_019813159.2

D2 26498018 26498117 refGene_gn=EGR2;refGene_tx=XM_019813157.2
D2 26769429 26769678 refGene_gn=NRBF2;refGene_tx=XM_003993970.4
D2 26787625 26787710 refGene_gn=NRBF2;refGene_tx=XM_003993970.4
D2 26792652 26792693 refGene_gn=NRBF2;refGene_tx=XM_003993970.4
D2 26804794 26805394 refGene_gn=NRBF2;refGene_tx=XM_003993970.4
D2 26805395 26805506 refGene_gn=NRBF2;refGene_tx=XM_003993970.4
D2 26821168 26821258
refGene_gn=JMJD1C;refGene_tx=XM_023240484.1;refGene_gn=JMJD1C;refGene_tx=XM_023240485.1;refGene_gn=JMJD1C;refGene_tx=XM_023240486.1;refGene_gn=JMJD1C;refGene_tx=XM_023240487.1
D2 26821563 26821695
refGene_gn=JMJD1C;refGene_tx=XM_023240484.1;refGene_gn=JMJD1C;refGene_tx=XM_023240485.1;refGene_gn=JMJD1C;refGene_tx=XM_023240486.1;refGene_gn=JMJD1C;refGene_tx=XM_023240487.1
D2 26829571 26829748
refGene_gn=JMJD1C;refGene_tx=XM_023240484.1;refGene_gn=JMJD1C;refGene_tx=XM_023240485.1;refGene_gn=JMJD1C;refGene_tx=XM_023240486.1;refGene_gn=JMJD1C;refGene_tx=XM_023240487.1
D2 26831490 26831630
refGene_gn=JMJD1C;refGene_tx=XM_023240484.1;refGene_gn=JMJD1C;refGene_tx=XM_023240485.1;refGene_gn=JMJD1C;refGene_tx=XM_023240486.1;refGene_gn=JMJD1C;refGene_tx=XM_023240487.1
D2 26834578 26834701
refGene_gn=JMJD1C;refGene_tx=XM_023240484.1;refGene_gn=JMJD1C;refGene_tx=XM_023240485.1;refGene_gn=JMJD1C;refGene_tx=XM_023240486.1;refGene_gn=JMJD1C;refGene_tx=XM_023240487.1
D2 26835083 26835214
refGene_gn=JMJD1C;refGene_tx=XM_023240484.1;refGene_gn=JMJD1C;refGene_tx=XM_023240485.1;refGene_gn=JMJD1C;refGene_tx=XM_023240486.1;refGene_gn=JMJD1C;refGene_tx=XM_023240487.1
D2 26836205 26836296
refGene_gn=JMJD1C;refGene_tx=XM_023240484.1;refGene_gn=JMJD1C;refGene_tx=XM_023240485.1;refGene_gn=JMJD1C;refGene_tx=XM_023240486.1;refGene_gn=JMJD1C;refGene_tx=XM_023240487.1
D2 26836853 26837022
refGene_gn=JMJD1C;refGene_tx=XM_023240484.1;refGene_gn=JMJD1C;refGene_tx=XM_023240485.1;refGene_gn=JMJD1C;refGene_tx=XM_023240486.1;refGene_gn=JMJD1C;refGene_tx=XM_023240487.1
D2 26839946 26840225
refGene_gn=JMJD1C;refGene_tx=XM_023240484.1;refGene_gn=JMJD1C;refGene_tx=XM_023240485.1;refGene_gn=JMJD1C;refGene_tx=XM_023240486.1;refGene_gn=JMJD1C;refGene_tx=XM_023240487.1
D2 26840912 26841127
refGene_gn=JMJD1C;refGene_tx=XM_023240484.1;refGene_gn=JMJD1C;refGene_tx=XM_023240485.1;refGene_gn=JMJD1C;refGene_tx=XM_023240486.1;refGene_gn=JMJD1C;refGene_tx=XM_023240487.1
D2 26841651 26841865
refGene_gn=JMJD1C;refGene_tx=XM_023240484.1;refGene_gn=JMJD1C;refGene_tx=XM_023240485.1;refGene_gn=JMJD1C;refGene_tx=XM_023240486.1;refGene_gn=JMJD1C;refGene_tx=XM_023240487.1

D2 26842054 26842182
refGene_gn=JMJD1C;refGene_tx=XM_023240484.1;refGene_gn=JMJD1C;refGene_tx=XM_023240485.1;refGene_gn=JMJD1C;refGene_tx=XM_023240486.1;refGene_gn=JMJD1C;refGene_tx=XM_023240487.1
D2 26842905 26842995
refGene_gn=JMJD1C;refGene_tx=XM_023240484.1;refGene_gn=JMJD1C;refGene_tx=XM_023240485.1;refGene_gn=JMJD1C;refGene_tx=XM_023240486.1;refGene_gn=JMJD1C;refGene_tx=XM_023240487.1
D2 26844981 26845134
refGene_gn=JMJD1C;refGene_tx=XM_023240484.1;refGene_gn=JMJD1C;refGene_tx=XM_023240485.1;refGene_gn=JMJD1C;refGene_tx=XM_023240486.1;refGene_gn=JMJD1C;refGene_tx=XM_023240487.1
D2 26846034 26846249
refGene_gn=JMJD1C;refGene_tx=XM_023240484.1;refGene_gn=JMJD1C;refGene_tx=XM_023240485.1;refGene_gn=JMJD1C;refGene_tx=XM_023240486.1;refGene_gn=JMJD1C;refGene_tx=XM_023240487.1
D2 26848332 26848534
refGene_gn=JMJD1C;refGene_tx=XM_023240484.1;refGene_gn=JMJD1C;refGene_tx=XM_023240485.1;refGene_gn=JMJD1C;refGene_tx=XM_023240486.1;refGene_gn=JMJD1C;refGene_tx=XM_023240487.1
D2 26853295 26855505
refGene_gn=JMJD1C;refGene_tx=XM_023240484.1;refGene_gn=JMJD1C;refGene_tx=XM_023240485.1;refGene_gn=JMJD1C;refGene_tx=XM_023240486.1;refGene_gn=JMJD1C;refGene_tx=XM_023240487.1
D2 26855784 26855957
refGene_gn=JMJD1C;refGene_tx=XM_023240484.1;refGene_gn=JMJD1C;refGene_tx=XM_023240485.1;refGene_gn=JMJD1C;refGene_tx=XM_023240486.1;refGene_gn=JMJD1C;refGene_tx=XM_023240487.1
D2 26859386 26861074
refGene_gn=JMJD1C;refGene_tx=XM_023240484.1;refGene_gn=JMJD1C;refGene_tx=XM_023240485.1;refGene_gn=JMJD1C;refGene_tx=XM_023240486.1;refGene_gn=JMJD1C;refGene_tx=XM_023240487.1
D2 26861188 26861381
refGene_gn=JMJD1C;refGene_tx=XM_023240484.1;refGene_gn=JMJD1C;refGene_tx=XM_023240485.1;refGene_gn=JMJD1C;refGene_tx=XM_023240486.1;refGene_gn=JMJD1C;refGene_tx=XM_023240487.1
D2 26861476 26861624
refGene_gn=JMJD1C;refGene_tx=XM_023240484.1;refGene_gn=JMJD1C;refGene_tx=XM_023240485.1;refGene_gn=JMJD1C;refGene_tx=XM_023240486.1;refGene_gn=JMJD1C;refGene_tx=XM_023240487.1
D2 26863696 26863822
refGene_gn=JMJD1C;refGene_tx=XM_023240484.1;refGene_gn=JMJD1C;refGene_tx=XM_023240485.1;refGene_gn=JMJD1C;refGene_tx=XM_023240486.1;refGene_gn=JMJD1C;refGene_tx=XM_023240487.1
D2 26907104 26907218
refGene_gn=JMJD1C;refGene_tx=XM_023240484.1;refGene_gn=JMJD1C;refGene_tx=XM_023240485.1;refGene_gn=JMJD1C;refGene_tx=XM_023240486.1;refGene_gn=JMJD1C;refGene_tx=XM_023240487.1
D2 26975235 26975478
refGene_gn=LOC111557048;refGene_tx=XM_023240489.1

D2 27019903 27020068
refGene_gn=JMJD1C;refGene_tx=XM_023240484.1;refGene_gn=JMJD1C;refGene_tx=XM_023240485.1;refGene_gn=JMJD1C;refGene_tx=XM_023240486.1;refGene_gn=JMJD1C;refGene_tx=XM_023240487.1
D2 27040475 27040523
refGene_gn=JMJD1C;refGene_tx=XM_023240484.1;refGene_gn=JMJD1C;refGene_tx=XM_023240486.1;refGene_gn=JMJD1C;refGene_tx=XM_023240487.1
D2 27085489 27085663
refGene_gn=JMJD1C;refGene_tx=XM_023240484.1;refGene_gn=JMJD1C;refGene_tx=XM_023240485.1;refGene_gn=JMJD1C;refGene_tx=XM_023240486.1;refGene_gn=JMJD1C;refGene_tx=XM_023240487.1
D2 27146665 27146697 refGene_gn=REEP3;refGene_tx=XM_023240488.1
D2 27192307 27192380 refGene_gn=REEP3;refGene_tx=XM_023240488.1
D2 27221015 27221092 refGene_gn=REEP3;refGene_tx=XM_023240488.1
D2 27223546 27223667 refGene_gn=REEP3;refGene_tx=XM_023240488.1
D2 27223838 27223952 refGene_gn=REEP3;refGene_tx=XM_023240488.1
D2 27236982 27237130 refGene_gn=REEP3;refGene_tx=XM_023240488.1
D2 27245425 27245571 refGene_gn=REEP3;refGene_tx=XM_023240488.1
D2 27246669 27246726 refGene_gn=REEP3;refGene_tx=XM_023240488.1
D2 28639031 28639185
refGene_gn=LOC109492866;refGene_tx=XM_019814147.2
D2 28639219 28639413
refGene_gn=LOC109492866;refGene_tx=XM_019814147.2
D2 29219967 29220255
refGene_gn=CTNNA3;refGene_tx=XM_023240490.1;refGene_gn=CTNNA3;refGene_tx=XM_023240491.1
D2 29287423 29287558
refGene_gn=CTNNA3;refGene_tx=XM_023240490.1;refGene_gn=CTNNA3;refGene_tx=XM_023240491.1
D2 29302405 29302511
refGene_gn=CTNNA3;refGene_tx=XM_023240490.1;refGene_gn=CTNNA3;refGene_tx=XM_023240491.1
D2 29366638 29366820
refGene_gn=CTNNA3;refGene_tx=XM_023240490.1;refGene_gn=CTNNA3;refGene_tx=XM_023240491.1
D2 29406505 29406598
refGene_gn=CTNNA3;refGene_tx=XM_023240490.1;refGene_gn=CTNNA3;refGene_tx=XM_023240491.1
D2 29561338 29561490
refGene_gn=CTNNA3;refGene_tx=XM_023240490.1;refGene_gn=CTNNA3;refGene_tx=XM_023240491.1
D2 29677600 29677801
refGene_gn=CTNNA3;refGene_tx=XM_023240490.1;refGene_gn=CTNNA3;refGene_tx=XM_023240491.1
D2 29802328 29802485
refGene_gn=CTNNA3;refGene_tx=XM_023240490.1;refGene_gn=CTNNA3;refGene_tx=XM_023240491.1
D2 29881079 29881172
refGene_gn=CTNNA3;refGene_tx=XM_023240490.1;refGene_gn=CTNNA3;refGene_tx=XM_023240491.1

D2 30043641 30043794
refGene_gn=CTNNA3;refGene_tx=XM_023240490.1;refGene_gn=CTNNA3;refGene_tx=XM_023240491.1
D2 30052320 30052401
refGene_gn=CTNNA3;refGene_tx=XM_023240490.1;refGene_gn=CTNNA3;refGene_tx=XM_023240491.1
D2 30207333 30207337 refGene_gn=LRRTM3;refGene_tx=XM_006937815.4
D2 30207696 30209228 refGene_gn=LRRTM3;refGene_tx=XM_006937815.4
D2 30366095 30366305 refGene_gn=LRRTM3;refGene_tx=XM_006937815.4
D2 30453337 30453541
refGene_gn=CTNNA3;refGene_tx=XM_023240490.1;refGene_gn=CTNNA3;refGene_tx=XM_023240491.1
D2 30511652 30511916
refGene_gn=CTNNA3;refGene_tx=XM_023240490.1;refGene_gn=CTNNA3;refGene_tx=XM_023240491.1
D2 30764470 30764590
refGene_gn=CTNNA3;refGene_tx=XM_023240490.1;refGene_gn=CTNNA3;refGene_tx=XM_023240491.1
D2 30800868 30801035
refGene_gn=CTNNA3;refGene_tx=XM_023240490.1;refGene_gn=CTNNA3;refGene_tx=XM_023240491.1
D2 30870870 30871063
refGene_gn=CTNNA3;refGene_tx=XM_023240490.1;refGene_gn=CTNNA3;refGene_tx=XM_023240491.1
D2 30903036 30903135
refGene_gn=CTNNA3;refGene_tx=XM_023240490.1;refGene_gn=CTNNA3;refGene_tx=XM_023240491.1
D2 31032387 31032482 refGene_gn=DNAJC12;refGene_tx=XM_003993977.3
D2 31037616 31037663 refGene_gn=DNAJC12;refGene_tx=XM_019813168.2
D2 31040607 31040812
refGene_gn=DNAJC12;refGene_tx=XM_003993977.3;refGene_gn=DNAJC12;refGene_tx=XM_011287368.2;refGene_gn=DNAJC12;refGene_tx=XM_019813168.2
D2 31048420 31048560
refGene_gn=DNAJC12;refGene_tx=XM_003993977.3;refGene_gn=DNAJC12;refGene_tx=XM_011287368.2;refGene_gn=DNAJC12;refGene_tx=XM_019813168.2
D2 31060551 31060630
refGene_gn=DNAJC12;refGene_tx=XM_003993977.3;refGene_gn=DNAJC12;refGene_tx=XM_011287368.2;refGene_gn=DNAJC12;refGene_tx=XM_019813168.2
D2 31070744 31070822
refGene_gn=DNAJC12;refGene_tx=XM_003993977.3;refGene_gn=DNAJC12;refGene_tx=XM_011287368.2;refGene_gn=DNAJC12;refGene_tx=XM_019813168.2
D2 31117241 31117659 refGene_gn=SIRT1;refGene_tx=NM_001290246.1
D2 31119060 31119177 refGene_gn=SIRT1;refGene_tx=NM_001290246.1
D2 31122551 31122770 refGene_gn=SIRT1;refGene_tx=NM_001290246.1
D2 31122771 31122794 refGene_gn=SIRT1;refGene_tx=NM_001290246.1
D2 31124586 31124739 refGene_gn=SIRT1;refGene_tx=NM_001290246.1
D2 31131923 31132071 refGene_gn=SIRT1;refGene_tx=NM_001290246.1
D2 31133312 31133392 refGene_gn=SIRT1;refGene_tx=NM_001290246.1
D2 31134739 31134926 refGene_gn=SIRT1;refGene_tx=NM_001290246.1
D2 31136408 31136972 refGene_gn=SIRT1;refGene_tx=NM_001290246.1

D2 31142325 31142654 refGene_gn=SIRT1;refGene_tx=NM_001290246.1
D2 31148519 31148728
refGene_gn=HERC4;refGene_tx=XM_011287363.2;refGene_gn=HERC4;refGene_tx=XM_011287365.2;refGene_gn=HERC4;refGene_tx=XM_011287366.2;refGene_gn=HERC4;refGene_tx=XM_019813173.1;refGene_gn=HERC4;refGene_tx=XM_023240492.1;refGene_gn=HERC4;refGene_tx=XM_023240493.1;refGene_gn=HERC4;refGene_tx=XM_023240494.1;refGene_gn=HERC4;refGene_tx=XM_023240495.1
D2 31149921 31150024
refGene_gn=HERC4;refGene_tx=XM_011287363.2;refGene_gn=HERC4;refGene_tx=XM_011287365.2;refGene_gn=HERC4;refGene_tx=XM_011287366.2;refGene_gn=HERC4;refGene_tx=XM_019813173.1;refGene_gn=HERC4;refGene_tx=XM_023240492.1;refGene_gn=HERC4;refGene_tx=XM_023240493.1;refGene_gn=HERC4;refGene_tx=XM_023240494.1;refGene_gn=HERC4;refGene_tx=XM_023240495.1
D2 31154741 31154925
refGene_gn=HERC4;refGene_tx=XM_011287363.2;refGene_gn=HERC4;refGene_tx=XM_011287365.2;refGene_gn=HERC4;refGene_tx=XM_011287366.2;refGene_gn=HERC4;refGene_tx=XM_019813173.1;refGene_gn=HERC4;refGene_tx=XM_023240492.1;refGene_gn=HERC4;refGene_tx=XM_023240493.1;refGene_gn=HERC4;refGene_tx=XM_023240494.1;refGene_gn=HERC4;refGene_tx=XM_023240495.1
D2 31159317 31159400
refGene_gn=HERC4;refGene_tx=XM_011287363.2;refGene_gn=HERC4;refGene_tx=XM_011287365.2;refGene_gn=HERC4;refGene_tx=XM_011287366.2;refGene_gn=HERC4;refGene_tx=XM_019813173.1;refGene_gn=HERC4;refGene_tx=XM_023240492.1;refGene_gn=HERC4;refGene_tx=XM_023240493.1;refGene_gn=HERC4;refGene_tx=XM_023240494.1;refGene_gn=HERC4;refGene_tx=XM_023240495.1
D2 31161790 31161857
refGene_gn=HERC4;refGene_tx=XM_011287363.2;refGene_gn=HERC4;refGene_tx=XM_011287365.2;refGene_gn=HERC4;refGene_tx=XM_011287366.2;refGene_gn=HERC4;refGene_tx=XM_019813173.1;refGene_gn=HERC4;refGene_tx=XM_023240492.1;refGene_gn=HERC4;refGene_tx=XM_023240493.1;refGene_gn=HERC4;refGene_tx=XM_023240494.1;refGene_gn=HERC4;refGene_tx=XM_023240495.1
D2 31162495 31162662
refGene_gn=HERC4;refGene_tx=XM_011287363.2;refGene_gn=HERC4;refGene_tx=XM_011287365.2;refGene_gn=HERC4;refGene_tx=XM_011287366.2;refGene_gn=HERC4;refGene_tx=XM_019813173.1;refGene_gn=HERC4;refGene_tx=XM_023240492.1;refGene_gn=HERC4;refGene_tx=XM_023240493.1;refGene_gn=HERC4;refGene_tx=XM_023240494.1;refGene_gn=HERC4;refGene_tx=XM_023240495.1
D2 31174615 31174759
refGene_gn=HERC4;refGene_tx=XM_011287363.2;refGene_gn=HERC4;refGene_tx=XM_011287365.2;refGene_gn=HERC4;refGene_tx=XM_011287366.2;refGene_gn=HERC4;refGene_tx=XM_019813173.1;refGene_gn=HERC4;refGene_tx=XM_023240492.1;refGene_gn=HERC4;refGene_tx=XM_023240493.1;refGene_gn=HERC4;refGene_tx=XM_023240494.1;refGene_gn=HERC4;refGene_tx=XM_023240495.1
D2 31174989 31175157
refGene_gn=HERC4;refGene_tx=XM_011287363.2;refGene_gn=HERC4;refGene_tx=XM_011287365.2;refGene_gn=HERC4;refGene_tx=XM_011287366.2;refGene_gn=HERC4;refGene_tx=XM_019813173.1;refGene_gn=HERC4;refGene_tx=XM_023240492.1;refGene_gn=HERC4;refGene_tx=XM_023240493.1;refGene_gn=HERC4;refGene_tx=XM_023240494.1;refGene_gn=HERC4;refGene_tx=XM_023240495.1
D2 31178059 31178158

refGene_gn=HERC4;refGene_tx=XM_011287363.2;refGene_gn=HERC4;refGene_tx=XM_011287365.2;refGene_gn=HERC4;refGene_tx=XM_011287366.2;refGene_gn=HERC4;refGene_tx=XM_019813173.1;refGene_gn=HERC4;refGene_tx=XM_023240492.1;refGene_gn=HERC4;refGene_tx=XM_023240493.1;refGene_gn=HERC4;refGene_tx=XM_023240494.1;refGene_gn=HERC4;refGene_tx=XM_023240495.1
D2 31180614 31180638
refGene_gn=HERC4;refGene_tx=XM_011287363.2;refGene_gn=HERC4;refGene_tx=XM_011287365.2;refGene_gn=HERC4;refGene_tx=XM_019813173.1;refGene_gn=HERC4;refGene_tx=XM_023240492.1;refGene_gn=HERC4;refGene_tx=XM_023240493.1;refGene_gn=HERC4;refGene_tx=XM_023240495.1
D2 31189176 31189296
refGene_gn=HERC4;refGene_tx=XM_011287363.2;refGene_gn=HERC4;refGene_tx=XM_011287365.2;refGene_gn=HERC4;refGene_tx=XM_011287366.2;refGene_gn=HERC4;refGene_tx=XM_019813173.1;refGene_gn=HERC4;refGene_tx=XM_023240492.1;refGene_gn=HERC4;refGene_tx=XM_023240493.1;refGene_gn=HERC4;refGene_tx=XM_023240494.1;refGene_gn=HERC4;refGene_tx=XM_023240495.1
D2 31203475 31203648
refGene_gn=HERC4;refGene_tx=XM_011287363.2;refGene_gn=HERC4;refGene_tx=XM_011287365.2;refGene_gn=HERC4;refGene_tx=XM_011287366.2;refGene_gn=HERC4;refGene_tx=XM_019813173.1;refGene_gn=HERC4;refGene_tx=XM_023240492.1;refGene_gn=HERC4;refGene_tx=XM_023240493.1;refGene_gn=HERC4;refGene_tx=XM_023240494.1;refGene_gn=HERC4;refGene_tx=XM_023240495.1
D2 31205129 31205319
refGene_gn=HERC4;refGene_tx=XM_011287363.2;refGene_gn=HERC4;refGene_tx=XM_011287365.2;refGene_gn=HERC4;refGene_tx=XM_011287366.2;refGene_gn=HERC4;refGene_tx=XM_019813173.1;refGene_gn=HERC4;refGene_tx=XM_023240492.1;refGene_gn=HERC4;refGene_tx=XM_023240493.1;refGene_gn=HERC4;refGene_tx=XM_023240494.1;refGene_gn=HERC4;refGene_tx=XM_023240495.1
D2 31205796 31205908
refGene_gn=HERC4;refGene_tx=XM_011287363.2;refGene_gn=HERC4;refGene_tx=XM_011287365.2;refGene_gn=HERC4;refGene_tx=XM_011287366.2;refGene_gn=HERC4;refGene_tx=XM_019813173.1;refGene_gn=HERC4;refGene_tx=XM_023240492.1;refGene_gn=HERC4;refGene_tx=XM_023240493.1;refGene_gn=HERC4;refGene_tx=XM_023240494.1;refGene_gn=HERC4;refGene_tx=XM_023240495.1
D2 31206044 31206104
refGene_gn=HERC4;refGene_tx=XM_011287363.2;refGene_gn=HERC4;refGene_tx=XM_011287365.2;refGene_gn=HERC4;refGene_tx=XM_011287366.2;refGene_gn=HERC4;refGene_tx=XM_019813173.1;refGene_gn=HERC4;refGene_tx=XM_023240492.1;refGene_gn=HERC4;refGene_tx=XM_023240493.1;refGene_gn=HERC4;refGene_tx=XM_023240494.1;refGene_gn=HERC4;refGene_tx=XM_023240495.1
D2 31207537 31207662
refGene_gn=HERC4;refGene_tx=XM_011287363.2;refGene_gn=HERC4;refGene_tx=XM_011287365.2;refGene_gn=HERC4;refGene_tx=XM_011287366.2;refGene_gn=HERC4;refGene_tx=XM_019813173.1;refGene_gn=HERC4;refGene_tx=XM_023240492.1;refGene_gn=HERC4;refGene_tx=XM_023240493.1;refGene_gn=HERC4;refGene_tx=XM_023240494.1;refGene_gn=HERC4;refGene_tx=XM_023240495.1
D2 31207958 31208035
refGene_gn=HERC4;refGene_tx=XM_011287363.2;refGene_gn=HERC4;refGene_tx=XM_011287365.2;refGene_gn=HERC4;refGene_tx=XM_011287366.2;refGene_gn=HERC4;refGene_tx=XM_019813173.1;refGene_gn=HERC4;refGene_tx=XM_0232404

92.1;refGene_gn=HERC4;refGene_tx=XM_023240493.1;refGene_gn=HERC4;refGene_tx=XM_023240494.1;refGene_gn=HERC4;refGene_tx=XM_023240495.1
D2 31222748 31222909
refGene_gn=HERC4;refGene_tx=XM_011287363.2;refGene_gn=HERC4;refGene_tx=XM_011287365.2;refGene_gn=HERC4;refGene_tx=XM_011287366.2;refGene_gn=HERC4;refGene_tx=XM_019813173.1;refGene_gn=HERC4;refGene_tx=XM_023240492.1;refGene_gn=HERC4;refGene_tx=XM_023240493.1;refGene_gn=HERC4;refGene_tx=XM_023240494.1;refGene_gn=HERC4;refGene_tx=XM_023240495.1
D2 31236510 31236641
refGene_gn=HERC4;refGene_tx=XM_011287363.2;refGene_gn=HERC4;refGene_tx=XM_011287365.2;refGene_gn=HERC4;refGene_tx=XM_011287366.2;refGene_gn=HERC4;refGene_tx=XM_019813173.1;refGene_gn=HERC4;refGene_tx=XM_023240492.1;refGene_gn=HERC4;refGene_tx=XM_023240493.1;refGene_gn=HERC4;refGene_tx=XM_023240494.1;refGene_gn=HERC4;refGene_tx=XM_023240495.1
D2 31246306 31246398
refGene_gn=HERC4;refGene_tx=XM_011287363.2;refGene_gn=HERC4;refGene_tx=XM_011287365.2;refGene_gn=HERC4;refGene_tx=XM_011287366.2;refGene_gn=HERC4;refGene_tx=XM_019813173.1;refGene_gn=HERC4;refGene_tx=XM_023240492.1;refGene_gn=HERC4;refGene_tx=XM_023240493.1;refGene_gn=HERC4;refGene_tx=XM_023240494.1;refGene_gn=HERC4;refGene_tx=XM_023240495.1
D2 31247762 31247984
refGene_gn=HERC4;refGene_tx=XM_011287363.2;refGene_gn=HERC4;refGene_tx=XM_011287365.2;refGene_gn=HERC4;refGene_tx=XM_011287366.2;refGene_gn=HERC4;refGene_tx=XM_019813173.1;refGene_gn=HERC4;refGene_tx=XM_023240492.1;refGene_gn=HERC4;refGene_tx=XM_023240493.1;refGene_gn=HERC4;refGene_tx=XM_023240494.1;refGene_gn=HERC4;refGene_tx=XM_023240495.1
D2 31250938 31251015
refGene_gn=HERC4;refGene_tx=XM_011287363.2;refGene_gn=HERC4;refGene_tx=XM_011287365.2;refGene_gn=HERC4;refGene_tx=XM_011287366.2;refGene_gn=HERC4;refGene_tx=XM_019813173.1;refGene_gn=HERC4;refGene_tx=XM_023240492.1;refGene_gn=HERC4;refGene_tx=XM_023240493.1;refGene_gn=HERC4;refGene_tx=XM_023240494.1;refGene_gn=HERC4;refGene_tx=XM_023240495.1
D2 31251883 31251973
refGene_gn=HERC4;refGene_tx=XM_019813173.1;refGene_gn=HERC4;refGene_tx=XM_023240492.1;refGene_gn=HERC4;refGene_tx=XM_023240493.1;refGene_gn=HERC4;refGene_tx=XM_023240494.1;refGene_gn=HERC4;refGene_tx=XM_023240495.1
D2 31257157 31257199
refGene_gn=HERC4;refGene_tx=XM_011287363.2;refGene_gn=HERC4;refGene_tx=XM_019813173.1;refGene_gn=HERC4;refGene_tx=XM_023240492.1;refGene_gn=HERC4;refGene_tx=XM_023240493.1;refGene_gn=HERC4;refGene_tx=XM_023240494.1;refGene_gn=HERC4;refGene_tx=XM_023240495.1
D2 31257943 31258103
refGene_gn=HERC4;refGene_tx=XM_011287363.2;refGene_gn=HERC4;refGene_tx=XM_011287365.2;refGene_gn=HERC4;refGene_tx=XM_011287366.2;refGene_gn=HERC4;refGene_tx=XM_019813173.1;refGene_gn=HERC4;refGene_tx=XM_023240492.1;refGene_gn=HERC4;refGene_tx=XM_023240493.1;refGene_gn=HERC4;refGene_tx=XM_023240494.1;refGene_gn=HERC4;refGene_tx=XM_023240495.1
D2 31279258 31279274 refGene_gn=HERC4;refGene_tx=XM_023240495.1
D2 31283921 31284147

refGene_gn=HERC4;refGene_tx=XM_011287363.2;refGene_gn=HERC4;refGene_tx=XM_011287365.2;refGene_gn=HERC4;refGene_tx=XM_011287366.2;refGene_gn=HERC4;refGene_tx=XM_019813173.1;refGene_gn=HERC4;refGene_tx=XM_023240492.1;refGene_gn=HERC4;refGene_tx=XM_023240493.1;refGene_gn=HERC4;refGene_tx=XM_023240494.1
D2 31288046 31288087
refGene_gn=MYPN;refGene_tx=XM_019813170.2;refGene_gn=MYPN;refGene_tx=XM_019813171.2;refGene_gn=MYPN;refGene_tx=XM_019813172.2
D2 31328291 31329194
refGene_gn=MYPN;refGene_tx=XM_003993981.5;refGene_gn=MYPN;refGene_tx=XM_006937819.4;refGene_gn=MYPN;refGene_tx=XM_006937820.4;refGene_gn=MYPN;refGene_tx=XM_019813170.2;refGene_gn=MYPN;refGene_tx=XM_019813171.2;refGene_gn=MYPN;refGene_tx=XM_019813172.2
D2 31347569 31347745
refGene_gn=MYPN;refGene_tx=XM_003993981.5;refGene_gn=MYPN;refGene_tx=XM_006937819.4;refGene_gn=MYPN;refGene_tx=XM_006937820.4;refGene_gn=MYPN;refGene_tx=XM_019813170.2;refGene_gn=MYPN;refGene_tx=XM_019813171.2;refGene_gn=MYPN;refGene_tx=XM_019813172.2
D2 31350413 31350465
refGene_gn=MYPN;refGene_tx=XM_003993981.5;refGene_gn=MYPN;refGene_tx=XM_006937819.4;refGene_gn=MYPN;refGene_tx=XM_006937820.4;refGene_gn=MYPN;refGene_tx=XM_019813170.2;refGene_gn=MYPN;refGene_tx=XM_019813171.2;refGene_gn=MYPN;refGene_tx=XM_019813172.2
D2 31354051 31354166
refGene_gn=MYPN;refGene_tx=XM_003993981.5;refGene_gn=MYPN;refGene_tx=XM_006937819.4;refGene_gn=MYPN;refGene_tx=XM_006937820.4;refGene_gn=MYPN;refGene_tx=XM_019813170.2;refGene_gn=MYPN;refGene_tx=XM_019813171.2;refGene_gn=MYPN;refGene_tx=XM_019813172.2
D2 31355271 31355343
refGene_gn=MYPN;refGene_tx=XM_003993981.5;refGene_gn=MYPN;refGene_tx=XM_006937819.4;refGene_gn=MYPN;refGene_tx=XM_006937820.4;refGene_gn=MYPN;refGene_tx=XM_019813170.2;refGene_gn=MYPN;refGene_tx=XM_019813171.2;refGene_gn=MYPN;refGene_tx=XM_019813172.2
D2 31362538 31362680
refGene_gn=MYPN;refGene_tx=XM_003993981.5;refGene_gn=MYPN;refGene_tx=XM_006937819.4;refGene_gn=MYPN;refGene_tx=XM_006937820.4;refGene_gn=MYPN;refGene_tx=XM_019813170.2;refGene_gn=MYPN;refGene_tx=XM_019813171.2;refGene_gn=MYPN;refGene_tx=XM_019813172.2
D2 31364830 31364854
refGene_gn=MYPN;refGene_tx=XM_003993981.5;refGene_gn=MYPN;refGene_tx=XM_006937819.4;refGene_gn=MYPN;refGene_tx=XM_006937820.4;refGene_gn=MYPN;refGene_tx=XM_019813170.2;refGene_gn=MYPN;refGene_tx=XM_019813171.2;refGene_gn=MYPN;refGene_tx=XM_019813172.2
D2 31367651 31367768
refGene_gn=MYPN;refGene_tx=XM_003993981.5;refGene_gn=MYPN;refGene_tx=XM_006937819.4;refGene_gn=MYPN;refGene_tx=XM_006937820.4;refGene_gn=MYPN;refGene_tx=XM_019813170.2;refGene_gn=MYPN;refGene_tx=XM_019813171.2;refGene_gn=MYPN;refGene_tx=XM_019813172.2
D2 31368253 31368626
refGene_gn=MYPN;refGene_tx=XM_003993981.5;refGene_gn=MYPN;refGene_tx=X

M_006937819.4;refGene_gn=MYPN;refGene_tx=XM_006937820.4;refGene_gn=MYPN;refGene_tx=XM_019813170.2;refGene_gn=MYPN;refGene_tx=XM_019813171.2;refGene_gn=MYPN;refGene_tx=XM_019813172.2
D2 31375619 31376210
refGene_gn=MYPN;refGene_tx=XM_003993981.5;refGene_gn=MYPN;refGene_tx=XM_006937819.4;refGene_gn=MYPN;refGene_tx=XM_006937820.4;refGene_gn=MYPN;refGene_tx=XM_019813170.2;refGene_gn=MYPN;refGene_tx=XM_019813171.2;refGene_gn=MYPN;refGene_tx=XM_019813172.2
D2 31376586 31376725
refGene_gn=MYPN;refGene_tx=XM_003993981.5;refGene_gn=MYPN;refGene_tx=XM_006937819.4;refGene_gn=MYPN;refGene_tx=XM_006937820.4;refGene_gn=MYPN;refGene_tx=XM_019813170.2;refGene_gn=MYPN;refGene_tx=XM_019813171.2;refGene_gn=MYPN;refGene_tx=XM_019813172.2
D2 31386516 31386738
refGene_gn=MYPN;refGene_tx=XM_003993981.5;refGene_gn=MYPN;refGene_tx=XM_006937819.4;refGene_gn=MYPN;refGene_tx=XM_006937820.4;refGene_gn=MYPN;refGene_tx=XM_019813170.2;refGene_gn=MYPN;refGene_tx=XM_019813171.2;refGene_gn=MYPN;refGene_tx=XM_019813172.2
D2 31387957 31388107
refGene_gn=MYPN;refGene_tx=XM_003993981.5;refGene_gn=MYPN;refGene_tx=XM_006937819.4;refGene_gn=MYPN;refGene_tx=XM_006937820.4;refGene_gn=MYPN;refGene_tx=XM_019813170.2;refGene_gn=MYPN;refGene_tx=XM_019813171.2;refGene_gn=MYPN;refGene_tx=XM_019813172.2
D2 31388981 31389064
refGene_gn=MYPN;refGene_tx=XM_003993981.5;refGene_gn=MYPN;refGene_tx=XM_006937819.4;refGene_gn=MYPN;refGene_tx=XM_006937820.4;refGene_gn=MYPN;refGene_tx=XM_019813170.2;refGene_gn=MYPN;refGene_tx=XM_019813171.2;refGene_gn=MYPN;refGene_tx=XM_019813172.2
D2 31390275 31390402
refGene_gn=MYPN;refGene_tx=XM_003993981.5;refGene_gn=MYPN;refGene_tx=XM_006937819.4;refGene_gn=MYPN;refGene_tx=XM_006937820.4;refGene_gn=MYPN;refGene_tx=XM_019813170.2;refGene_gn=MYPN;refGene_tx=XM_019813171.2;refGene_gn=MYPN;refGene_tx=XM_019813172.2
D2 31391655 31391863
refGene_gn=MYPN;refGene_tx=XM_003993981.5;refGene_gn=MYPN;refGene_tx=XM_006937819.4;refGene_gn=MYPN;refGene_tx=XM_006937820.4;refGene_gn=MYPN;refGene_tx=XM_019813170.2;refGene_gn=MYPN;refGene_tx=XM_019813171.2;refGene_gn=MYPN;refGene_tx=XM_019813172.2
D2 31393872 31394038
refGene_gn=MYPN;refGene_tx=XM_003993981.5;refGene_gn=MYPN;refGene_tx=XM_006937819.4;refGene_gn=MYPN;refGene_tx=XM_006937820.4;refGene_gn=MYPN;refGene_tx=XM_019813170.2;refGene_gn=MYPN;refGene_tx=XM_019813171.2;refGene_gn=MYPN;refGene_tx=XM_019813172.2
D2 31399715 31399849
refGene_gn=MYPN;refGene_tx=XM_003993981.5;refGene_gn=MYPN;refGene_tx=XM_006937819.4;refGene_gn=MYPN;refGene_tx=XM_006937820.4;refGene_gn=MYPN;refGene_tx=XM_019813170.2;refGene_gn=MYPN;refGene_tx=XM_019813171.2;refGene_gn=MYPN;refGene_tx=XM_019813172.2
D2 31403313 31403483
refGene_gn=MYPN;refGene_tx=XM_003993981.5;refGene_gn=MYPN;refGene_tx=X

M_006937819.4;refGene_gn=MYPN;refGene_tx=XM_006937820.4;refGene_gn=MYPN;refGene_tx=XM_019813170.2;refGene_gn=MYPN;refGene_tx=XM_019813171.2;refGene_gn=MYPN;refGene_tx=XM_019813172.2
D2 31416053 31416512 refGene_gn=ATOH7;refGene_tx=XM_023240496.1
D2 31453158 31453271
refGene_gn=PBLD;refGene_tx=XM_003993985.2;refGene_gn=PBLD;refGene_tx=XM_019813178.1
D2 31454378 31454441 refGene_gn=PBLD;refGene_tx=XM_003993985.2
D2 31457119 31457298
refGene_gn=PBLD;refGene_tx=XM_003993985.2;refGene_gn=PBLD;refGene_tx=XM_019813178.1
D2 31457564 31457653
refGene_gn=PBLD;refGene_tx=XM_003993985.2;refGene_gn=PBLD;refGene_tx=XM_019813178.1
D2 31460112 31460142
refGene_gn=PBLD;refGene_tx=XM_003993985.2;refGene_gn=PBLD;refGene_tx=XM_019813178.1
D2 31460229 31460339
refGene_gn=PBLD;refGene_tx=XM_003993985.2;refGene_gn=PBLD;refGene_tx=XM_019813178.1
D2 31463055 31463154
refGene_gn=PBLD;refGene_tx=XM_003993985.2;refGene_gn=PBLD;refGene_tx=XM_019813178.1
D2 31463508 31463608
refGene_gn=PBLD;refGene_tx=XM_003993985.2;refGene_gn=PBLD;refGene_tx=XM_019813178.1
D2 31473581 31473665
refGene_gn=PBLD;refGene_tx=XM_003993985.2;refGene_gn=PBLD;refGene_tx=XM_019813178.1
D2 31504833 31504945
refGene_gn=HNRNPH3;refGene_tx=XM_003993986.5;refGene_gn=HNRNPH3;refGene_tx=XM_003993987.5;refGene_gn=HNRNPH3;refGene_tx=XM_006937821.4;refGene_gn=HNRNPH3;refGene_tx=XM_006937822.4
D2 31505463 31505602
refGene_gn=HNRNPH3;refGene_tx=XM_003993986.5;refGene_gn=HNRNPH3;refGene_tx=XM_003993987.5;refGene_gn=HNRNPH3;refGene_tx=XM_006937821.4;refGene_gn=HNRNPH3;refGene_tx=XM_006937822.4
D2 31506115 31506300
refGene_gn=HNRNPH3;refGene_tx=XM_003993986.5;refGene_gn=HNRNPH3;refGene_tx=XM_003993987.5;refGene_gn=HNRNPH3;refGene_tx=XM_006937821.4;refGene_gn=HNRNPH3;refGene_tx=XM_006937822.4;refGene_gn=HNRNPH3;refGene_tx=XM_019813186.2
D2 31506754 31506841
refGene_gn=HNRNPH3;refGene_tx=XM_003993986.5;refGene_gn=HNRNPH3;refGene_tx=XM_003993987.5;refGene_gn=HNRNPH3;refGene_tx=XM_006937821.4;refGene_gn=HNRNPH3;refGene_tx=XM_006937822.4;refGene_gn=HNRNPH3;refGene_tx=XM_019813186.2
D2 31507152 31507268
refGene_gn=HNRNPH3;refGene_tx=XM_003993986.5;refGene_gn=HNRNPH3;refGene_tx=XM_003993987.5;refGene_gn=HNRNPH3;refGene_tx=XM_006937821.4;refGe

ne_gn=HNRNPH3;refGene_tx=XM_006937822.4;refGene_gn=HNRNPH3;refGene_tx=XM_019813186.2
D2 31508895 31509031
refGene_gn=HNRNPH3;refGene_tx=XM_003993986.5;refGene_gn=HNRNPH3;refGene_tx=XM_003993987.5;refGene_gn=HNRNPH3;refGene_tx=XM_006937821.4;refGene_gn=HNRNPH3;refGene_tx=XM_006937822.4;refGene_gn=HNRNPH3;refGene_tx=XM_019813186.2
D2 31510040 31510136
refGene_gn=HNRNPH3;refGene_tx=XM_003993986.5;refGene_gn=HNRNPH3;refGene_tx=XM_003993987.5;refGene_gn=HNRNPH3;refGene_tx=XM_006937821.4;refGene_gn=HNRNPH3;refGene_tx=XM_006937822.4;refGene_gn=HNRNPH3;refGene_tx=XM_019813186.2
D2 31510210 31510303
refGene_gn=HNRNPH3;refGene_tx=XM_003993986.5;refGene_gn=HNRNPH3;refGene_tx=XM_003993987.5;refGene_gn=HNRNPH3;refGene_tx=XM_006937821.4;refGene_gn=HNRNPH3;refGene_tx=XM_006937822.4;refGene_gn=HNRNPH3;refGene_tx=XM_019813186.2
D2 31510440 31510517
refGene_gn=HNRNPH3;refGene_tx=XM_003993986.5;refGene_gn=HNRNPH3;refGene_tx=XM_003993987.5;refGene_gn=HNRNPH3;refGene_tx=XM_006937821.4;refGene_gn=HNRNPH3;refGene_tx=XM_006937822.4;refGene_gn=HNRNPH3;refGene_tx=XM_019813186.2
D2 31514097 31514241 refGene_gn=RUFY2;refGene_tx=XM_003993989.5
D2 31514327 31514405 refGene_gn=RUFY2;refGene_tx=XM_003993989.5
D2 31519787 31519836 refGene_gn=RUFY2;refGene_tx=XM_003993989.5
D2 31526462 31526557 refGene_gn=RUFY2;refGene_tx=XM_003993989.5
D2 31526979 31527109 refGene_gn=RUFY2;refGene_tx=XM_003993989.5
D2 31535984 31536104 refGene_gn=RUFY2;refGene_tx=XM_003993989.5
D2 31538495 31538593 refGene_gn=RUFY2;refGene_tx=XM_003993989.5
D2 31539443 31539611 refGene_gn=RUFY2;refGene_tx=XM_003993989.5
D2 31541185 31541302 refGene_gn=RUFY2;refGene_tx=XM_003993989.5
D2 31541442 31541544 refGene_gn=RUFY2;refGene_tx=XM_003993989.5
D2 31543982 31544052 refGene_gn=RUFY2;refGene_tx=XM_003993989.5
D2 31548057 31548123 refGene_gn=RUFY2;refGene_tx=XM_003993989.5
D2 31549698 31549760 refGene_gn=RUFY2;refGene_tx=XM_003993989.5
D2 31549954 31550078 refGene_gn=RUFY2;refGene_tx=XM_003993989.5
D2 31551897 31551999 refGene_gn=RUFY2;refGene_tx=XM_003993989.5
D2 31555122 31555240 refGene_gn=RUFY2;refGene_tx=XM_003993989.5
D2 31559244 31559377 refGene_gn=RUFY2;refGene_tx=XM_003993989.5
D2 31559378 31559419 refGene_gn=RUFY2;refGene_tx=XM_003993989.5
D2 31560897 31561102 refGene_gn=RUFY2;refGene_tx=XM_003993989.5
D2 31568292 31568361
refGene_gn=DNA2;refGene_tx=XM_006937829.3;refGene_gn=DNA2;refGene_tx=XM_019813187.2
D2 31569514 31569661
refGene_gn=DNA2;refGene_tx=XM_006937829.3;refGene_gn=DNA2;refGene_tx=XM_019813187.2
D2 31572288 31572471
refGene_gn=DNA2;refGene_tx=XM_006937829.3;refGene_gn=DNA2;refGene_tx=XM_019813187.2

D2 31573272 31573362
refGene_gn=DNA2;refGene_tx=XM_006937829.3;refGene_gn=DNA2;refGene_tx=XM_019813187.2
D2 31578160 31578365
refGene_gn=DNA2;refGene_tx=XM_006937829.3;refGene_gn=DNA2;refGene_tx=XM_019813187.2
D2 31578449 31578539
refGene_gn=DNA2;refGene_tx=XM_006937829.3;refGene_gn=DNA2;refGene_tx=XM_019813187.2
D2 31578631 31578825
refGene_gn=DNA2;refGene_tx=XM_006937829.3;refGene_gn=DNA2;refGene_tx=XM_019813187.2
D2 31584032 31584257
refGene_gn=DNA2;refGene_tx=XM_006937829.3;refGene_gn=DNA2;refGene_tx=XM_019813187.2
D2 31585368 31585478
refGene_gn=DNA2;refGene_tx=XM_006937829.3;refGene_gn=DNA2;refGene_tx=XM_019813187.2
D2 31586287 31586397
refGene_gn=DNA2;refGene_tx=XM_006937829.3;refGene_gn=DNA2;refGene_tx=XM_019813187.2
D2 31586476 31586593
refGene_gn=DNA2;refGene_tx=XM_006937829.3;refGene_gn=DNA2;refGene_tx=XM_019813187.2
D2 31588788 31589019
refGene_gn=DNA2;refGene_tx=XM_006937829.3;refGene_gn=DNA2;refGene_tx=XM_019813187.2
D2 31590299 31590494
refGene_gn=DNA2;refGene_tx=XM_006937829.3;refGene_gn=DNA2;refGene_tx=XM_019813187.2
D2 31592782 31592948
refGene_gn=DNA2;refGene_tx=XM_006937829.3;refGene_gn=DNA2;refGene_tx=XM_019813187.2
D2 31593498 31593616
refGene_gn=DNA2;refGene_tx=XM_006937829.3;refGene_gn=DNA2;refGene_tx=XM_019813187.2
D2 31600822 31601042
refGene_gn=DNA2;refGene_tx=XM_006937829.3;refGene_gn=DNA2;refGene_tx=XM_019813187.2
D2 31607846 31607978
refGene_gn=DNA2;refGene_tx=XM_006937829.3;refGene_gn=DNA2;refGene_tx=XM_019813187.2
D2 31615354 31615500
refGene_gn=DNA2;refGene_tx=XM_006937829.3;refGene_gn=DNA2;refGene_tx=XM_019813187.2
D2 31617428 31617612
refGene_gn=DNA2;refGene_tx=XM_006937829.3;refGene_gn=DNA2;refGene_tx=XM_019813187.2
D2 31619444 31619627
refGene_gn=DNA2;refGene_tx=XM_006937829.3;refGene_gn=DNA2;refGene_tx=X

M_019813187.2
D2 31621299 31621370 refGene_gn=DNA2;refGene_tx=XM_006937829.3
D2 31633019 31633176 refGene_gn=SLC25A16;refGene_tx=XM_023240293.1
D2 31635909 31635978 refGene_gn=SLC25A16;refGene_tx=XM_023240293.1
D2 31641283 31641446 refGene_gn=SLC25A16;refGene_tx=XM_023240293.1
D2 31646286 31646353 refGene_gn=SLC25A16;refGene_tx=XM_023240293.1
D2 31646711 31646833 refGene_gn=SLC25A16;refGene_tx=XM_023240293.1
D2 31667214 31667278 refGene_gn=SLC25A16;refGene_tx=XM_023240293.1
D2 31668604 31668738 refGene_gn=SLC25A16;refGene_tx=XM_023240293.1
D2 31678414 31678507 refGene_gn=SLC25A16;refGene_tx=XM_023240293.1
D2 31683049 31683335 refGene_gn=SLC25A16;refGene_tx=XM_023240293.1
D2 31706762 31707531 refGene_gn=SLC25A16;refGene_tx=XM_023240293.1
D2 31708144 31708194 refGene_gn=SLC25A16;refGene_tx=XM_023240293.1
D2 31714741 31716649
refGene_gn=TET1;refGene_tx=XM_003993991.5;refGene_gn=TET1;refGene_tx=XM_023240497.1;refGene_gn=TET1;refGene_tx=XM_023240498.1;refGene_gn=TET1;refGene_tx=XM_023240499.1;refGene_gn=TET1;refGene_tx=XM_023240500.1
D2 31747287 31747341
refGene_gn=TET1;refGene_tx=XM_003993991.5;refGene_gn=TET1;refGene_tx=XM_023240497.1;refGene_gn=TET1;refGene_tx=XM_023240498.1;refGene_gn=TET1;refGene_tx=XM_023240499.1;refGene_gn=TET1;refGene_tx=XM_023240500.1
D2 31787173 31789499
refGene_gn=TET1;refGene_tx=XM_003993991.5;refGene_gn=TET1;refGene_tx=XM_023240497.1;refGene_gn=TET1;refGene_tx=XM_023240498.1;refGene_gn=TET1;refGene_tx=XM_023240499.1;refGene_gn=TET1;refGene_tx=XM_023240500.1
D2 31793526 31793617
refGene_gn=TET1;refGene_tx=XM_003993991.5;refGene_gn=TET1;refGene_tx=XM_023240497.1;refGene_gn=TET1;refGene_tx=XM_023240498.1;refGene_gn=TET1;refGene_tx=XM_023240499.1;refGene_gn=TET1;refGene_tx=XM_023240500.1
D2 31794170 31794264
refGene_gn=TET1;refGene_tx=XM_003993991.5;refGene_gn=TET1;refGene_tx=XM_023240497.1;refGene_gn=TET1;refGene_tx=XM_023240498.1;refGene_gn=TET1;refGene_tx=XM_023240499.1;refGene_gn=TET1;refGene_tx=XM_023240500.1
D2 31816430 31816642
refGene_gn=TET1;refGene_tx=XM_003993991.5;refGene_gn=TET1;refGene_tx=XM_023240497.1;refGene_gn=TET1;refGene_tx=XM_023240498.1;refGene_gn=TET1;refGene_tx=XM_023240499.1;refGene_gn=TET1;refGene_tx=XM_023240500.1
D2 31820648 31820799
refGene_gn=TET1;refGene_tx=XM_003993991.5;refGene_gn=TET1;refGene_tx=XM_023240497.1;refGene_gn=TET1;refGene_tx=XM_023240498.1;refGene_gn=TET1;refGene_tx=XM_023240499.1;refGene_gn=TET1;refGene_tx=XM_023240500.1
D2 31821135 31821222
refGene_gn=TET1;refGene_tx=XM_023240497.1;refGene_gn=TET1;refGene_tx=XM_023240498.1;refGene_gn=TET1;refGene_tx=XM_023240499.1;refGene_gn=TET1;refGene_tx=XM_023240500.1
D2 31824910 31825000
refGene_gn=TET1;refGene_tx=XM_003993991.5;refGene_gn=TET1;refGene_tx=XM_023240497.1;refGene_gn=TET1;refGene_tx=XM_023240498.1;refGene_gn=TET1;refGene_tx=XM_023240499.1;refGene_gn=TET1;refGene_tx=XM_023240500.1
D2 31827429 31827567

refGene_gn=TET1;refGene_tx=XM_003993991.5;refGene_gn=TET1;refGene_tx=XM_023240497.1;refGene_gn=TET1;refGene_tx=XM_023240498.1;refGene_gn=TET1;refGene_tx=XM_023240499.1;refGene_gn=TET1;refGene_tx=XM_023240500.1
D2 31830203 31830552
refGene_gn=TET1;refGene_tx=XM_003993991.5;refGene_gn=TET1;refGene_tx=XM_023240497.1;refGene_gn=TET1;refGene_tx=XM_023240498.1;refGene_gn=TET1;refGene_tx=XM_023240499.1;refGene_gn=TET1;refGene_tx=XM_023240500.1
D2 31836023 31837039
refGene_gn=TET1;refGene_tx=XM_003993991.5;refGene_gn=TET1;refGene_tx=XM_023240497.1;refGene_gn=TET1;refGene_tx=XM_023240498.1;refGene_gn=TET1;refGene_tx=XM_023240499.1;refGene_gn=TET1;refGene_tx=XM_023240500.1
D2 31874390 31874463
refGene_gn=CCAR1;refGene_tx=XM_006937830.4;refGene_gn=CCAR1;refGene_tx=XM_019813190.2;refGene_gn=CCAR1;refGene_tx=XM_019813192.2;refGene_gn=CCAR1;refGene_tx=XM_019813193.2
D2 31889987 31890160
refGene_gn=CCAR1;refGene_tx=XM_006937830.4;refGene_gn=CCAR1;refGene_tx=XM_019813190.2;refGene_gn=CCAR1;refGene_tx=XM_019813192.2;refGene_gn=CCAR1;refGene_tx=XM_019813193.2
D2 31890955 31891000
refGene_gn=CCAR1;refGene_tx=XM_006937830.4;refGene_gn=CCAR1;refGene_tx=XM_019813190.2;refGene_gn=CCAR1;refGene_tx=XM_019813192.2
D2 31895625 31895658
refGene_gn=CCAR1;refGene_tx=XM_006937830.4;refGene_gn=CCAR1;refGene_tx=XM_019813190.2;refGene_gn=CCAR1;refGene_tx=XM_019813192.2;refGene_gn=CCAR1;refGene_tx=XM_019813193.2
D2 31898103 31898297
refGene_gn=CCAR1;refGene_tx=XM_006937830.4;refGene_gn=CCAR1;refGene_tx=XM_019813190.2;refGene_gn=CCAR1;refGene_tx=XM_019813192.2;refGene_gn=CCAR1;refGene_tx=XM_019813193.2
D2 31899445 31899560
refGene_gn=CCAR1;refGene_tx=XM_006937830.4;refGene_gn=CCAR1;refGene_tx=XM_019813190.2;refGene_gn=CCAR1;refGene_tx=XM_019813192.2;refGene_gn=CCAR1;refGene_tx=XM_019813193.2
D2 31899674 31899867
refGene_gn=CCAR1;refGene_tx=XM_006937830.4;refGene_gn=CCAR1;refGene_tx=XM_019813190.2;refGene_gn=CCAR1;refGene_tx=XM_019813192.2;refGene_gn=CCAR1;refGene_tx=XM_019813193.2
D2 31902627 31902757
refGene_gn=CCAR1;refGene_tx=XM_006937830.4;refGene_gn=CCAR1;refGene_tx=XM_019813190.2;refGene_gn=CCAR1;refGene_tx=XM_019813192.2;refGene_gn=CCAR1;refGene_tx=XM_019813193.2
D2 31902993 31903155
refGene_gn=CCAR1;refGene_tx=XM_006937830.4;refGene_gn=CCAR1;refGene_tx=XM_019813190.2;refGene_gn=CCAR1;refGene_tx=XM_019813192.2;refGene_gn=CCAR1;refGene_tx=XM_019813193.2
D2 31906280 31906506
refGene_gn=CCAR1;refGene_tx=XM_006937830.4;refGene_gn=CCAR1;refGene_tx=XM_019813190.2;refGene_gn=CCAR1;refGene_tx=XM_019813192.2;refGene_gn=CCAR1;refGene_tx=XM_019813193.2

D2 31907086 31907200
refGene_gn=CCAR1;refGene_tx=XM_006937830.4;refGene_gn=CCAR1;refGene_tx=XM_019813190.2;refGene_gn=CCAR1;refGene_tx=XM_019813192.2;refGene_gn=CCAR1;refGene_tx=XM_019813193.2
D2 31907774 31907941
refGene_gn=CCAR1;refGene_tx=XM_006937830.4;refGene_gn=CCAR1;refGene_tx=XM_019813190.2;refGene_gn=CCAR1;refGene_tx=XM_019813192.2;refGene_gn=CCAR1;refGene_tx=XM_019813193.2
D2 31908679 31908890
refGene_gn=CCAR1;refGene_tx=XM_006937830.4;refGene_gn=CCAR1;refGene_tx=XM_019813190.2;refGene_gn=CCAR1;refGene_tx=XM_019813192.2;refGene_gn=CCAR1;refGene_tx=XM_019813193.2
D2 31909706 31909790
refGene_gn=CCAR1;refGene_tx=XM_006937830.4;refGene_gn=CCAR1;refGene_tx=XM_019813190.2;refGene_gn=CCAR1;refGene_tx=XM_019813192.2;refGene_gn=CCAR1;refGene_tx=XM_019813193.2
D2 31912157 31912343
refGene_gn=CCAR1;refGene_tx=XM_006937830.4;refGene_gn=CCAR1;refGene_tx=XM_019813190.2;refGene_gn=CCAR1;refGene_tx=XM_019813192.2;refGene_gn=CCAR1;refGene_tx=XM_019813193.2
D2 31920503 31920695
refGene_gn=CCAR1;refGene_tx=XM_006937830.4;refGene_gn=CCAR1;refGene_tx=XM_019813190.2;refGene_gn=CCAR1;refGene_tx=XM_019813192.2;refGene_gn=CCAR1;refGene_tx=XM_019813193.2
D2 31924453 31924693
refGene_gn=CCAR1;refGene_tx=XM_006937830.4;refGene_gn=CCAR1;refGene_tx=XM_019813190.2;refGene_gn=CCAR1;refGene_tx=XM_019813192.2;refGene_gn=CCAR1;refGene_tx=XM_019813193.2
D2 31925086 31925198
refGene_gn=CCAR1;refGene_tx=XM_006937830.4;refGene_gn=CCAR1;refGene_tx=XM_019813190.2;refGene_gn=CCAR1;refGene_tx=XM_019813192.2;refGene_gn=CCAR1;refGene_tx=XM_019813193.2
D2 31933103 31933186
refGene_gn=CCAR1;refGene_tx=XM_006937830.4;refGene_gn=CCAR1;refGene_tx=XM_019813190.2;refGene_gn=CCAR1;refGene_tx=XM_019813192.2;refGene_gn=CCAR1;refGene_tx=XM_019813193.2
D2 31933507 31933654
refGene_gn=CCAR1;refGene_tx=XM_006937830.4;refGene_gn=CCAR1;refGene_tx=XM_019813190.2;refGene_gn=CCAR1;refGene_tx=XM_019813192.2;refGene_gn=CCAR1;refGene_tx=XM_019813193.2
D2 31934939 31935060
refGene_gn=CCAR1;refGene_tx=XM_006937830.4;refGene_gn=CCAR1;refGene_tx=XM_019813190.2;refGene_gn=CCAR1;refGene_tx=XM_019813192.2;refGene_gn=CCAR1;refGene_tx=XM_019813193.2
D2 31935144 31935330
refGene_gn=CCAR1;refGene_tx=XM_006937830.4;refGene_gn=CCAR1;refGene_tx=XM_019813190.2;refGene_gn=CCAR1;refGene_tx=XM_019813192.2;refGene_gn=CCAR1;refGene_tx=XM_019813193.2
D2 31937292 31937498
refGene_gn=CCAR1;refGene_tx=XM_006937830.4;refGene_gn=CCAR1;refGene_tx

=XM_019813190.2;refGene_gn=CCAR1;refGene_tx=XM_019813192.2;refGene_gn=CCAR1;refGene_tx=XM_019813193.2
D2 31938067 31938127 refGene_gn=CCAR1;refGene_tx=XM_006937830.4;refGene_gn=CCAR1;refGene_tx=XM_019813190.2;refGene_gn=CCAR1;refGene_tx=XM_019813192.2;refGene_gn=CCAR1;refGene_tx=XM_019813193.2
D2 31971237 31971550 refGene_gn=STOX1;refGene_tx=XM_023240501.1;refGene_gn=STOX1;refGene_tx=XM_023240502.1
D2 32013152 32013305 refGene_gn=STOX1;refGene_tx=XM_023240501.1;refGene_gn=STOX1;refGene_tx=XM_023240502.1;refGene_gn=STOX1;refGene_tx=XM_023240504.1
D2 32013703 32013836 refGene_gn=STOX1;refGene_tx=XM_023240503.1
D2 32016532 32018888 refGene_gn=STOX1;refGene_tx=XM_023240501.1;refGene_gn=STOX1;refGene_tx=XM_023240502.1;refGene_gn=STOX1;refGene_tx=XM_023240503.1;refGene_gn=STOX1;refGene_tx=XM_023240504.1
D2 32022837 32022985 refGene_gn=STOX1;refGene_tx=XM_023240501.1;refGene_gn=STOX1;refGene_tx=XM_023240503.1;refGene_gn=STOX1;refGene_tx=XM_023240504.1
D2 32029405 32029430 refGene_gn=STOX1;refGene_tx=XM_023240502.1
D2 32030603 32030693 refGene_gn=DDX50;refGene_tx=XM_003993993.5
D2 32034166 32034466 refGene_gn=DDX50;refGene_tx=XM_003993993.5;refGene_gn=DDX50;refGene_tx=XM_006937832.2;refGene_gn=DDX50;refGene_tx=XM_019813196.2;refGene_gn=DDX50;refGene_tx=XM_019813197.2
D2 32037222 32037298 refGene_gn=DDX50;refGene_tx=XM_003993993.5;refGene_gn=DDX50;refGene_tx=XM_006937832.2;refGene_gn=DDX50;refGene_tx=XM_019813196.2;refGene_gn=DDX50;refGene_tx=XM_019813197.2
D2 32037662 32037841 refGene_gn=DDX50;refGene_tx=XM_003993993.5;refGene_gn=DDX50;refGene_tx=XM_006937832.2;refGene_gn=DDX50;refGene_tx=XM_019813196.2;refGene_gn=DDX50;refGene_tx=XM_019813197.2
D2 32039329 32039447 refGene_gn=DDX50;refGene_tx=XM_003993993.5;refGene_gn=DDX50;refGene_tx=XM_006937832.2;refGene_gn=DDX50;refGene_tx=XM_019813196.2;refGene_gn=DDX50;refGene_tx=XM_019813197.2
D2 32039563 32039749 refGene_gn=DDX50;refGene_tx=XM_003993993.5;refGene_gn=DDX50;refGene_tx=XM_006937832.2;refGene_gn=DDX50;refGene_tx=XM_019813196.2;refGene_gn=DDX50;refGene_tx=XM_019813197.2
D2 32040233 32040379 refGene_gn=DDX50;refGene_tx=XM_003993993.5;refGene_gn=DDX50;refGene_tx=XM_006937832.2;refGene_gn=DDX50;refGene_tx=XM_019813196.2;refGene_gn=DDX50;refGene_tx=XM_019813197.2
D2 32045044 32045194 refGene_gn=DDX50;refGene_tx=XM_003993993.5;refGene_gn=DDX50;refGene_tx=XM_006937832.2;refGene_gn=DDX50;refGene_tx=XM_019813196.2;refGene_gn=DDX50;refGene_tx=XM_019813197.2

D2 32052188 32052350
refGene_gn=DDX50;refGene_tx=XM_003993993.5;refGene_gn=DDX50;refGene_tx=XM_006937832.2;refGene_gn=DDX50;refGene_tx=XM_019813196.2;refGene_gn=DDX50;refGene_tx=XM_019813197.2
D2 32052813 32052933
refGene_gn=DDX50;refGene_tx=XM_003993993.5;refGene_gn=DDX50;refGene_tx=XM_006937832.2;refGene_gn=DDX50;refGene_tx=XM_019813196.2;refGene_gn=DDX50;refGene_tx=XM_019813197.2
D2 32053938 32054012
refGene_gn=DDX50;refGene_tx=XM_003993993.5;refGene_gn=DDX50;refGene_tx=XM_006937832.2;refGene_gn=DDX50;refGene_tx=XM_019813196.2;refGene_gn=DDX50;refGene_tx=XM_019813197.2
D2 32054236 32054396
refGene_gn=DDX50;refGene_tx=XM_003993993.5;refGene_gn=DDX50;refGene_tx=XM_006937832.2;refGene_gn=DDX50;refGene_tx=XM_019813196.2;refGene_gn=DDX50;refGene_tx=XM_019813197.2
D2 32057459 32057594
refGene_gn=DDX50;refGene_tx=XM_003993993.5;refGene_gn=DDX50;refGene_tx=XM_006937832.2;refGene_gn=DDX50;refGene_tx=XM_019813196.2;refGene_gn=DDX50;refGene_tx=XM_019813197.2
D2 32058749 32058794
refGene_gn=DDX50;refGene_tx=XM_003993993.5;refGene_gn=DDX50;refGene_tx=XM_006937832.2;refGene_gn=DDX50;refGene_tx=XM_019813196.2;refGene_gn=DDX50;refGene_tx=XM_019813197.2
D2 32061979 32062258
refGene_gn=DDX50;refGene_tx=XM_003993993.5;refGene_gn=DDX50;refGene_tx=XM_006937832.2;refGene_gn=DDX50;refGene_tx=XM_019813196.2;refGene_gn=DDX50;refGene_tx=XM_019813197.2
D2 32069194 32069281 refGene_gn=DDX21;refGene_tx=XM_003993995.5
D2 32071188 32071626 refGene_gn=DDX21;refGene_tx=XM_003993995.5
D2 32073852 32073870 refGene_gn=DDX21;refGene_tx=XM_003993995.5
D2 32073871 32073929 refGene_gn=DDX21;refGene_tx=XM_003993995.5
D2 32074819 32074998 refGene_gn=DDX21;refGene_tx=XM_003993995.5
D2 32076446 32076564 refGene_gn=DDX21;refGene_tx=XM_003993995.5
D2 32080132 32080318 refGene_gn=DDX21;refGene_tx=XM_003993995.5
D2 32081418 32081564 refGene_gn=DDX21;refGene_tx=XM_003993995.5
D2 32082391 32082541 refGene_gn=DDX21;refGene_tx=XM_003993995.5
D2 32082923 32083085 refGene_gn=DDX21;refGene_tx=XM_003993995.5
D2 32084588 32084708 refGene_gn=DDX21;refGene_tx=XM_003993995.5
D2 32085771 32085845 refGene_gn=DDX21;refGene_tx=XM_003993995.5
D2 32089155 32089315 refGene_gn=DDX21;refGene_tx=XM_003993995.5
D2 32090143 32090278 refGene_gn=DDX21;refGene_tx=XM_003993995.5
D2 32094552 32094597 refGene_gn=DDX21;refGene_tx=XM_003993995.5
D2 32095647 32095917 refGene_gn=DDX21;refGene_tx=XM_003993995.5
D2 32103952 32104378 refGene_gn=KIF1BP;refGene_tx=XM_003993997.4
D2 32114340 32114439 refGene_gn=KIF1BP;refGene_tx=XM_003993997.4
D2 32121387 32121467 refGene_gn=KIF1BP;refGene_tx=XM_003993997.4
D2 32121788 32121972 refGene_gn=KIF1BP;refGene_tx=XM_003993997.4
D2 32124642 32124724 refGene_gn=KIF1BP;refGene_tx=XM_003993997.4
D2 32132103 32132219 refGene_gn=KIF1BP;refGene_tx=XM_003993997.4

D2 32137323 32138199 refGene_gn=KIF1BP;refGene_tx=XM_003993997.4
D2 32204782 32204858 refGene_gn=SRGN;refGene_tx=XM_003993998.5
D2 32214635 32214783 refGene_gn=SRGN;refGene_tx=XM_003993998.5
D2 32218969 32219237 refGene_gn=SRGN;refGene_tx=XM_003993998.5
D2 32241172 32241175 refGene_gn=VPS26A;refGene_tx=XM_003993999.5
D2 32241423 32241525 refGene_gn=VPS26A;refGene_tx=XM_003994000.5
D2 32248427 32248577
refGene_gn=VPS26A;refGene_tx=XM_003993999.5;refGene_gn=VPS26A;refGene_tx=XM_003994000.5
D2 32259934 32260010
refGene_gn=VPS26A;refGene_tx=XM_003993999.5;refGene_gn=VPS26A;refGene_tx=XM_003994000.5
D2 32261172 32261329
refGene_gn=VPS26A;refGene_tx=XM_003993999.5;refGene_gn=VPS26A;refGene_tx=XM_003994000.5
D2 32261922 32262087
refGene_gn=VPS26A;refGene_tx=XM_003993999.5;refGene_gn=VPS26A;refGene_tx=XM_003994000.5
D2 32264660 32264767
refGene_gn=VPS26A;refGene_tx=XM_003993999.5;refGene_gn=VPS26A;refGene_tx=XM_003994000.5
D2 32267072 32267141
refGene_gn=VPS26A;refGene_tx=XM_003993999.5;refGene_gn=VPS26A;refGene_tx=XM_003994000.5
D2 32269961 32270104
refGene_gn=VPS26A;refGene_tx=XM_003993999.5;refGene_gn=VPS26A;refGene_tx=XM_003994000.5
D2 32273015 32273129
refGene_gn=VPS26A;refGene_tx=XM_003993999.5;refGene_gn=VPS26A;refGene_tx=XM_003994000.5
D2 32285470 32285750 refGene_gn=SUPV3L1;refGene_tx=XM_003994001.5
D2 32292528 32292606 refGene_gn=SUPV3L1;refGene_tx=XM_003994001.5
D2 32293433 32293541 refGene_gn=SUPV3L1;refGene_tx=XM_003994001.5
D2 32294835 32294950 refGene_gn=SUPV3L1;refGene_tx=XM_003994001.5
D2 32296115 32296284 refGene_gn=SUPV3L1;refGene_tx=XM_003994001.5
D2 32298136 32298248 refGene_gn=SUPV3L1;refGene_tx=XM_003994001.5
D2 32300563 32300641 refGene_gn=SUPV3L1;refGene_tx=XM_003994001.5
D2 32301993 32302085 refGene_gn=SUPV3L1;refGene_tx=XM_003994001.5
D2 32303467 32303648 refGene_gn=SUPV3L1;refGene_tx=XM_003994001.5
D2 32304195 32304289 refGene_gn=SUPV3L1;refGene_tx=XM_003994001.5
D2 32305456 32305676 refGene_gn=SUPV3L1;refGene_tx=XM_003994001.5
D2 32306948 32307029 refGene_gn=SUPV3L1;refGene_tx=XM_003994001.5
D2 32307406 32307583 refGene_gn=SUPV3L1;refGene_tx=XM_003994001.5
D2 32310623 32310772 refGene_gn=SUPV3L1;refGene_tx=XM_003994001.5
D2 32311407 32311840 refGene_gn=SUPV3L1;refGene_tx=XM_003994001.5
D2 32326497 32326560 refGene_gn=HKDC1;refGene_tx=XM_003994002.4
D2 32331516 32331679 refGene_gn=HKDC1;refGene_tx=XM_003994002.4
D2 32338346 32338495 refGene_gn=HKDC1;refGene_tx=XM_003994002.4
D2 32338618 32338738 refGene_gn=HKDC1;refGene_tx=XM_003994002.4
D2 32343740 32343836 refGene_gn=HKDC1;refGene_tx=XM_003994002.4

D2 32344859 32344959 refGene_gn=HKDC1;refGene_tx=XM_003994002.4
D2 32347346 32347530 refGene_gn=HKDC1;refGene_tx=XM_003994002.4
D2 32351141 32351297 refGene_gn=HKDC1;refGene_tx=XM_003994002.4
D2 32352403 32352637 refGene_gn=HKDC1;refGene_tx=XM_003994002.4
D2 32353863 32354168 refGene_gn=HKDC1;refGene_tx=XM_003994002.4
D2 32355772 32355918 refGene_gn=HKDC1;refGene_tx=XM_003994002.4
D2 32356012 32356132 refGene_gn=HKDC1;refGene_tx=XM_003994002.4
D2 32360249 32360345 refGene_gn=HKDC1;refGene_tx=XM_003994002.4
D2 32360558 32360658 refGene_gn=HKDC1;refGene_tx=XM_003994002.4
D2 32362322 32362506 refGene_gn=HKDC1;refGene_tx=XM_003994002.4
D2 32364518 32364674 refGene_gn=HKDC1;refGene_tx=XM_003994002.4
D2 32371734 32371968 refGene_gn=HKDC1;refGene_tx=XM_003994002.4
D2 32376419 32376567 refGene_gn=HKDC1;refGene_tx=XM_003994002.4
D2 32415415 32415442 refGene_gn=HK1;refGene_tx=XM_006937834.4
D2 32419016 32419067 refGene_gn=HK1;refGene_tx=XM_006937834.4
D2 32434208 32434271 refGene_gn=HK1;refGene_tx=XM_023240505.1
D2 32466772 32466935
refGene_gn=HK1;refGene_tx=XM_006937834.4;refGene_gn=HK1;refGene_tx=XM_023240505.1
D2 32478158 32478307
refGene_gn=HK1;refGene_tx=XM_006937834.4;refGene_gn=HK1;refGene_tx=XM_023240505.1
D2 32483255 32483375
refGene_gn=HK1;refGene_tx=XM_006937834.4;refGene_gn=HK1;refGene_tx=XM_023240505.1
D2 32486297 32486393
refGene_gn=HK1;refGene_tx=XM_006937834.4;refGene_gn=HK1;refGene_tx=XM_023240505.1
D2 32486968 32487068
refGene_gn=HK1;refGene_tx=XM_006937834.4;refGene_gn=HK1;refGene_tx=XM_023240505.1
D2 32487179 32487363
refGene_gn=HK1;refGene_tx=XM_006937834.4;refGene_gn=HK1;refGene_tx=XM_023240505.1
D2 32494144 32494300
refGene_gn=HK1;refGene_tx=XM_006937834.4;refGene_gn=HK1;refGene_tx=XM_023240505.1
D2 32496756 32496990
refGene_gn=HK1;refGene_tx=XM_006937834.4;refGene_gn=HK1;refGene_tx=XM_023240505.1
D2 32499100 32499405
refGene_gn=HK1;refGene_tx=XM_006937834.4;refGene_gn=HK1;refGene_tx=XM_023240505.1
D2 32500858 32501007
refGene_gn=HK1;refGene_tx=XM_006937834.4;refGene_gn=HK1;refGene_tx=XM_023240505.1
D2 32501325 32501445
refGene_gn=HK1;refGene_tx=XM_006937834.4;refGene_gn=HK1;refGene_tx=XM_023240505.1
D2 32502812 32502908

refGene_gn=HK1;refGene_tx=XM_006937834.4;refGene_gn=HK1;refGene_tx=XM_023240505.1
D2 32504759 32504859
refGene_gn=HK1;refGene_tx=XM_006937834.4;refGene_gn=HK1;refGene_tx=XM_023240505.1
D2 32507860 32508044
refGene_gn=HK1;refGene_tx=XM_006937834.4;refGene_gn=HK1;refGene_tx=XM_023240505.1
D2 32510420 32510576
refGene_gn=HK1;refGene_tx=XM_006937834.4;refGene_gn=HK1;refGene_tx=XM_023240505.1
D2 32514964 32515198
refGene_gn=HK1;refGene_tx=XM_006937834.4;refGene_gn=HK1;refGene_tx=XM_023240505.1
D2 32516855 32517000
refGene_gn=HK1;refGene_tx=XM_006937834.4;refGene_gn=HK1;refGene_tx=XM_023240505.1
D2 32519589 32519806 refGene_gn=TACR2;refGene_tx=XM_003994003.4
D2 32522894 32523091 refGene_gn=TACR2;refGene_tx=XM_003994003.4
D2 32524448 32524602 refGene_gn=TACR2;refGene_tx=XM_003994003.4
D2 32527895 32528090 refGene_gn=TACR2;refGene_tx=XM_003994003.4
D2 32532776 32533168 refGene_gn=TACR2;refGene_tx=XM_003994003.4
D2 32559035 32559131 refGene_gn=TSPAN15;refGene_tx=XM_003994004.5
D2 32586459 32586645 refGene_gn=TSPAN15;refGene_tx=XM_003994004.5
D2 32587921 32587996 refGene_gn=TSPAN15;refGene_tx=XM_003994004.5
D2 32598606 32598702 refGene_gn=TSPAN15;refGene_tx=XM_003994004.5
D2 32600894 32601011 refGene_gn=TSPAN15;refGene_tx=XM_003994004.5
D2 32605773 32605821 refGene_gn=TSPAN15;refGene_tx=XM_003994004.5
D2 32607182 32607299 refGene_gn=TSPAN15;refGene_tx=XM_003994004.5
D2 32607822 32607972 refGene_gn=TSPAN15;refGene_tx=XM_003994004.5
D2 32653339 32653987 refGene_gn=NEUROG3;refGene_tx=XM_004001305.1
D2 32696019 32696112
refGene_gn=FAM241B;refGene_tx=XM_006937836.4;refGene_gn=FAM241B;refGene_tx=XM_011287360.2
D2 32696798 32697068
refGene_gn=FAM241B;refGene_tx=XM_006937836.4;refGene_gn=FAM241B;refGene_tx=XM_011287360.2
D2 32834605 32834899
refGene_gn=COL13A1;refGene_tx=XM_023240506.1;refGene_gn=COL13A1;refGene_tx=XM_023240507.1;refGene_gn=COL13A1;refGene_tx=XM_023240508.1
D2 32851454 32851527
refGene_gn=COL13A1;refGene_tx=XM_023240506.1;refGene_gn=COL13A1;refGene_tx=XM_023240507.1;refGene_gn=COL13A1;refGene_tx=XM_023240508.1
D2 32896523 32896531
refGene_gn=COL13A1;refGene_tx=XM_023240506.1;refGene_gn=COL13A1;refGene_tx=XM_023240507.1;refGene_gn=COL13A1;refGene_tx=XM_023240508.1
D2 32901351 32901378
refGene_gn=COL13A1;refGene_tx=XM_023240506.1;refGene_gn=COL13A1;refGene_tx=XM_023240507.1;refGene_gn=COL13A1;refGene_tx=XM_023240508.1
D2 32904141 32904177

refGene_gn=COL13A1;refGene_tx=XM_023240506.1;refGene_gn=COL13A1;refGene_tx=XM_023240507.1;refGene_gn=COL13A1;refGene_tx=XM_023240508.1
D2 32906721 32906748
refGene_gn=COL13A1;refGene_tx=XM_023240506.1;refGene_gn=COL13A1;refGene_tx=XM_023240507.1;refGene_gn=COL13A1;refGene_tx=XM_023240508.1
D2 32908311 32908362
refGene_gn=COL13A1;refGene_tx=XM_023240506.1;refGene_gn=COL13A1;refGene_tx=XM_023240507.1;refGene_gn=COL13A1;refGene_tx=XM_023240508.1
D2 32915134 32915170
refGene_gn=COL13A1;refGene_tx=XM_023240506.1;refGene_gn=COL13A1;refGene_tx=XM_023240507.1;refGene_gn=COL13A1;refGene_tx=XM_023240508.1
D2 32915977 32916004
refGene_gn=COL13A1;refGene_tx=XM_023240506.1;refGene_gn=COL13A1;refGene_tx=XM_023240507.1;refGene_gn=COL13A1;refGene_tx=XM_023240508.1
D2 32916972 32916999
refGene_gn=COL13A1;refGene_tx=XM_023240506.1;refGene_gn=COL13A1;refGene_tx=XM_023240507.1;refGene_gn=COL13A1;refGene_tx=XM_023240508.1
D2 32921522 32921549
refGene_gn=COL13A1;refGene_tx=XM_023240506.1;refGene_gn=COL13A1;refGene_tx=XM_023240507.1;refGene_gn=COL13A1;refGene_tx=XM_023240508.1
D2 32921645 32921672
refGene_gn=COL13A1;refGene_tx=XM_023240506.1;refGene_gn=COL13A1;refGene_tx=XM_023240507.1;refGene_gn=COL13A1;refGene_tx=XM_023240508.1
D2 32922470 32922497
refGene_gn=COL13A1;refGene_tx=XM_023240506.1;refGene_gn=COL13A1;refGene_tx=XM_023240507.1;refGene_gn=COL13A1;refGene_tx=XM_023240508.1
D2 32925528 32925594
refGene_gn=COL13A1;refGene_tx=XM_023240506.1;refGene_gn=COL13A1;refGene_tx=XM_023240507.1;refGene_gn=COL13A1;refGene_tx=XM_023240508.1
D2 32929103 32929211
refGene_gn=COL13A1;refGene_tx=XM_023240506.1;refGene_gn=COL13A1;refGene_tx=XM_023240507.1;refGene_gn=COL13A1;refGene_tx=XM_023240508.1
D2 32931545 32931572
refGene_gn=COL13A1;refGene_tx=XM_023240506.1;refGene_gn=COL13A1;refGene_tx=XM_023240507.1;refGene_gn=COL13A1;refGene_tx=XM_023240508.1
D2 32932203 32932239
refGene_gn=COL13A1;refGene_tx=XM_023240506.1;refGene_gn=COL13A1;refGene_tx=XM_023240507.1;refGene_gn=COL13A1;refGene_tx=XM_023240508.1
D2 32941624 32941669
refGene_gn=COL13A1;refGene_tx=XM_023240506.1;refGene_gn=COL13A1;refGene_tx=XM_023240507.1;refGene_gn=COL13A1;refGene_tx=XM_023240508.1
D2 32942535 32942568
refGene_gn=COL13A1;refGene_tx=XM_023240506.1;refGene_gn=COL13A1;refGene_tx=XM_023240507.1;refGene_gn=COL13A1;refGene_tx=XM_023240508.1
D2 32943301 32943328
refGene_gn=COL13A1;refGene_tx=XM_023240506.1;refGene_gn=COL13A1;refGene_tx=XM_023240507.1;refGene_gn=COL13A1;refGene_tx=XM_023240508.1
D2 32946059 32946113
refGene_gn=COL13A1;refGene_tx=XM_023240506.1;refGene_gn=COL13A1;refGene_tx=XM_023240507.1;refGene_gn=COL13A1;refGene_tx=XM_023240508.1

D2 32946937 32947024
refGene_gn=COL13A1;refGene_tx=XM_023240506.1;refGene_gn=COL13A1;refGene_tx=XM_023240507.1;refGene_gn=COL13A1;refGene_tx=XM_023240508.1
D2 32948641 32948695
refGene_gn=COL13A1;refGene_tx=XM_023240506.1;refGene_gn=COL13A1;refGene_tx=XM_023240507.1;refGene_gn=COL13A1;refGene_tx=XM_023240508.1
D2 32949852 32949897
refGene_gn=COL13A1;refGene_tx=XM_023240506.1;refGene_gn=COL13A1;refGene_tx=XM_023240507.1;refGene_gn=COL13A1;refGene_tx=XM_023240508.1
D2 32950261 32950327
refGene_gn=COL13A1;refGene_tx=XM_023240506.1;refGene_gn=COL13A1;refGene_tx=XM_023240507.1;refGene_gn=COL13A1;refGene_tx=XM_023240508.1
D2 32951494 32951518
refGene_gn=COL13A1;refGene_tx=XM_023240506.1;refGene_gn=COL13A1;refGene_tx=XM_023240507.1;refGene_gn=COL13A1;refGene_tx=XM_023240508.1
D2 32953255 32953318
refGene_gn=COL13A1;refGene_tx=XM_023240506.1;refGene_gn=COL13A1;refGene_tx=XM_023240507.1;refGene_gn=COL13A1;refGene_tx=XM_023240508.1
D2 32954373 32954418
refGene_gn=COL13A1;refGene_tx=XM_023240506.1;refGene_gn=COL13A1;refGene_tx=XM_023240507.1;refGene_gn=COL13A1;refGene_tx=XM_023240508.1
D2 32954783 32954936
refGene_gn=COL13A1;refGene_tx=XM_023240506.1;refGene_gn=COL13A1;refGene_tx=XM_023240507.1;refGene_gn=COL13A1;refGene_tx=XM_023240508.1
D2 32955547 32955646
refGene_gn=COL13A1;refGene_tx=XM_023240506.1;refGene_gn=COL13A1;refGene_tx=XM_023240507.1;refGene_gn=COL13A1;refGene_tx=XM_023240508.1
D2 32957154 32957199
refGene_gn=COL13A1;refGene_tx=XM_023240506.1;refGene_gn=COL13A1;refGene_tx=XM_023240507.1;refGene_gn=COL13A1;refGene_tx=XM_023240508.1
D2 32959772 32959814
refGene_gn=COL13A1;refGene_tx=XM_023240506.1;refGene_gn=COL13A1;refGene_tx=XM_023240507.1;refGene_gn=COL13A1;refGene_tx=XM_023240508.1
D2 32960732 32960759
refGene_gn=COL13A1;refGene_tx=XM_023240506.1;refGene_gn=COL13A1;refGene_tx=XM_023240507.1;refGene_gn=COL13A1;refGene_tx=XM_023240508.1
D2 32961689 32961770
refGene_gn=COL13A1;refGene_tx=XM_023240506.1;refGene_gn=COL13A1;refGene_tx=XM_023240507.1;refGene_gn=COL13A1;refGene_tx=XM_023240508.1
D2 32965353 32965389
refGene_gn=COL13A1;refGene_tx=XM_023240506.1;refGene_gn=COL13A1;refGene_tx=XM_023240507.1
D2 32968199 32968253
refGene_gn=COL13A1;refGene_tx=XM_023240506.1;refGene_gn=COL13A1;refGene_tx=XM_023240507.1;refGene_gn=COL13A1;refGene_tx=XM_023240508.1
D2 32969266 32969320
refGene_gn=COL13A1;refGene_tx=XM_023240506.1;refGene_gn=COL13A1;refGene_tx=XM_023240507.1;refGene_gn=COL13A1;refGene_tx=XM_023240508.1
D2 32970340 32970376
refGene_gn=COL13A1;refGene_tx=XM_023240506.1;refGene_gn=COL13A1;refGen

e_tx=XM_023240507.1;refGene_gn=COL13A1;refGene_tx=XM_023240508.1
D2	32975254	32975341
refGene_gn=COL13A1;refGene_tx=XM_023240506.1;refGene_gn=COL13A1;refGene_tx=XM_023240508.1
D2	32976206	32976284	refGene_gn=COL13A1;refGene_tx=XM_023240507.1
D2	32979180	32979219
refGene_gn=COL13A1;refGene_tx=XM_023240506.1;refGene_gn=COL13A1;refGene_tx=XM_023240507.1;refGene_gn=COL13A1;refGene_tx=XM_023240508.1
D2	32981120	32981123
refGene_gn=COL13A1;refGene_tx=XM_023240506.1;refGene_gn=COL13A1;refGene_tx=XM_023240507.1;refGene_gn=COL13A1;refGene_tx=XM_023240508.1
D2	33051784	33051956
refGene_gn=LOC101095520;refGene_tx=XM_023240513.1
D2	33072086	33072193
refGene_gn=LOC101095520;refGene_tx=XM_023240513.1
D2	33073387	33073585
refGene_gn=LOC101095520;refGene_tx=XM_023240513.1
D2	33074748	33074859
refGene_gn=LOC101095520;refGene_tx=XM_023240513.1
D2	33076369	33076469
refGene_gn=LOC101095520;refGene_tx=XM_023240513.1
D2	33081420	33081510
refGene_gn=LOC101095520;refGene_tx=XM_023240513.1
D2	33087916	33088091
refGene_gn=LOC101095520;refGene_tx=XM_023240513.1
D2	33090130	33090296
refGene_gn=LOC101095520;refGene_tx=XM_023240513.1
D2	33092514	33092666
refGene_gn=AIFM2;refGene_tx=XM_006937839.4;refGene_gn=AIFM2;refGene_tx=XM_011287359.3;refGene_gn=AIFM2;refGene_tx=XM_023240509.1;refGene_gn=AIFM2;refGene_tx=XM_023240510.1;refGene_gn=AIFM2;refGene_tx=XM_023240511.1;refGene_gn=AIFM2;refGene_tx=XM_023240512.1
D2	33092954	33093155
refGene_gn=AIFM2;refGene_tx=XM_006937839.4;refGene_gn=AIFM2;refGene_tx=XM_011287359.3;refGene_gn=AIFM2;refGene_tx=XM_023240509.1;refGene_gn=AIFM2;refGene_tx=XM_023240510.1;refGene_gn=AIFM2;refGene_tx=XM_023240511.1;refGene_gn=AIFM2;refGene_tx=XM_023240512.1
D2	33093862	33093955
refGene_gn=AIFM2;refGene_tx=XM_006937839.4;refGene_gn=AIFM2;refGene_tx=XM_011287359.3;refGene_gn=AIFM2;refGene_tx=XM_023240509.1;refGene_gn=AIFM2;refGene_tx=XM_023240510.1;refGene_gn=AIFM2;refGene_tx=XM_023240511.1
D2	33094535	33094688
refGene_gn=AIFM2;refGene_tx=XM_006937839.4;refGene_gn=AIFM2;refGene_tx=XM_011287359.3;refGene_gn=AIFM2;refGene_tx=XM_023240509.1;refGene_gn=AIFM2;refGene_tx=XM_023240510.1;refGene_gn=AIFM2;refGene_tx=XM_023240511.1;refGene_gn=AIFM2;refGene_tx=XM_023240512.1
D2	33095653	33095762
refGene_gn=AIFM2;refGene_tx=XM_006937839.4;refGene_gn=AIFM2;refGene_tx=XM_011287359.3;refGene_gn=AIFM2;refGene_tx=XM_023240509.1;refGene_gn=

AIFM2;refGene_tx=XM_023240510.1;refGene_gn=AIFM2;refGene_tx=XM_0232405
11.1;refGene_gn=AIFM2;refGene_tx=XM_023240512.1
D2 33097450 33097543
refGene_gn=AIFM2;refGene_tx=XM_006937839.4;refGene_gn=AIFM2;refGene_tx
=XM_011287359.3;refGene_gn=AIFM2;refGene_tx=XM_023240509.1;refGene_gn=
AIFM2;refGene_tx=XM_023240510.1;refGene_gn=AIFM2;refGene_tx=XM_0232405
11.1;refGene_gn=AIFM2;refGene_tx=XM_023240512.1
D2 33098241 33098361
refGene_gn=AIFM2;refGene_tx=XM_006937839.4;refGene_gn=AIFM2;refGene_tx
=XM_011287359.3;refGene_gn=AIFM2;refGene_tx=XM_023240509.1;refGene_gn=
AIFM2;refGene_tx=XM_023240510.1;refGene_gn=AIFM2;refGene_tx=XM_0232405
11.1;refGene_gn=AIFM2;refGene_tx=XM_023240512.1
D2 33099690 33099806
refGene_gn=AIFM2;refGene_tx=XM_006937839.4;refGene_gn=AIFM2;refGene_tx
=XM_011287359.3;refGene_gn=AIFM2;refGene_tx=XM_023240509.1;refGene_gn=
AIFM2;refGene_tx=XM_023240510.1;refGene_gn=AIFM2;refGene_tx=XM_0232405
11.1;refGene_gn=AIFM2;refGene_tx=XM_023240512.1
D2 33100181 33100359
refGene_gn=AIFM2;refGene_tx=XM_006937839.4;refGene_gn=AIFM2;refGene_tx
=XM_011287359.3;refGene_gn=AIFM2;refGene_tx=XM_023240509.1;refGene_gn=
AIFM2;refGene_tx=XM_023240510.1;refGene_gn=AIFM2;refGene_tx=XM_0232405
11.1;refGene_gn=AIFM2;refGene_tx=XM_023240512.1
D2 33115723 33116175 refGene_gn=TYSND1;refGene_tx=XM_019814066.2
D2 33118343 33118529 refGene_gn=TYSND1;refGene_tx=XM_019814066.2
D2 33119535 33119666 refGene_gn=TYSND1;refGene_tx=XM_019814066.2
D2 33121709 33122887 refGene_gn=TYSND1;refGene_tx=XM_019814066.2
D2 33127354 33127471
refGene_gn=SAR1A;refGene_tx=XM_003994009.5;refGene_gn=SAR1A;refGene_tx
=XM_023240514.1
D2 33128828 33128960
refGene_gn=SAR1A;refGene_tx=XM_003994009.5;refGene_gn=SAR1A;refGene_tx
=XM_023240514.1
D2 33131905 33132009
refGene_gn=SAR1A;refGene_tx=XM_003994009.5;refGene_gn=SAR1A;refGene_tx
=XM_023240514.1
D2 33135291 33135357
refGene_gn=SAR1A;refGene_tx=XM_003994009.5;refGene_gn=SAR1A;refGene_tx
=XM_023240514.1
D2 33136025 33136145
refGene_gn=SAR1A;refGene_tx=XM_003994009.5;refGene_gn=SAR1A;refGene_tx
=XM_023240514.1
D2 33136270 33136328
refGene_gn=SAR1A;refGene_tx=XM_003994009.5;refGene_gn=SAR1A;refGene_tx
=XM_023240514.1
D2 33157614 33157646 refGene_gn=PPA1;refGene_tx=XM_023240515.1
D2 33158484 33158527 refGene_gn=PPA1;refGene_tx=XM_023240515.1
D2 33159548 33159618 refGene_gn=PPA1;refGene_tx=XM_023240515.1
D2 33165434 33165520 refGene_gn=PPA1;refGene_tx=XM_023240515.1
D2 33165791 33165919 refGene_gn=PPA1;refGene_tx=XM_023240515.1
D2 33168057 33168184 refGene_gn=PPA1;refGene_tx=XM_023240515.1

D2 33169078 33169165 refGene_gn=PPA1;refGene_tx=XM_023240515.1
D2 33171789 33171909 refGene_gn=PPA1;refGene_tx=XM_023240515.1
D2 33172789 33172843 refGene_gn=PPA1;refGene_tx=XM_023240515.1
D2 33184458 33184517 refGene_gn=PPA1;refGene_tx=XM_023240515.1
D2 33187540 33187604 refGene_gn=PPA1;refGene_tx=XM_023240515.1
D2 33205457 33206343 refGene_gn=NPFFR1;refGene_tx=XM_023240516.1
D2 33207483 33207583 refGene_gn=NPFFR1;refGene_tx=XM_023240516.1
D2 33209142 33209457 refGene_gn=NPFFR1;refGene_tx=XM_023240516.1
D2 33223588 33223595 refGene_gn=NPFFR1;refGene_tx=XM_023240516.1
D2 33235998 33236153
refGene_gn=LRRC20;refGene_tx=XM_011287357.2;refGene_gn=LRRC20;refGene_tx=XM_011287358.2
D2 33253656 33253841
refGene_gn=LRRC20;refGene_tx=XM_023240517.1;refGene_gn=LRRC20;refGene_tx=XM_023240518.1;refGene_gn=LRRC20;refGene_tx=XM_023240519.1
D2 33259490 33259658
refGene_gn=LRRC20;refGene_tx=XM_011287357.2;refGene_gn=LRRC20;refGene_tx=XM_011287358.2;refGene_gn=LRRC20;refGene_tx=XM_023240517.1;refGene_gn=LRRC20;refGene_tx=XM_023240518.1;refGene_gn=LRRC20;refGene_tx=XM_023240519.1
D2 33279870 33280020
refGene_gn=LRRC20;refGene_tx=XM_011287357.2;refGene_gn=LRRC20;refGene_tx=XM_023240517.1;refGene_gn=LRRC20;refGene_tx=XM_023240518.1;refGene_gn=LRRC20;refGene_tx=XM_023240519.1
D2 33317460 33317542
refGene_gn=LRRC20;refGene_tx=XM_011287357.2;refGene_gn=LRRC20;refGene_tx=XM_011287358.2;refGene_gn=LRRC20;refGene_tx=XM_023240517.1;refGene_gn=LRRC20;refGene_tx=XM_023240518.1;refGene_gn=LRRC20;refGene_tx=XM_023240519.1
D2 33349489 33349634 refGene_gn=EIF4EBP2;refGene_tx=XM_003994023.4
D2 33366410 33366596 refGene_gn=EIF4EBP2;refGene_tx=XM_003994023.4
D2 33367514 33367546 refGene_gn=EIF4EBP2;refGene_tx=XM_003994023.4
D2 33377704 33377857 refGene_gn=NODAL;refGene_tx=XM_019813208.2
D2 33379286 33379984 refGene_gn=NODAL;refGene_tx=XM_019813208.2
D2 33384069 33384265 refGene_gn=NODAL;refGene_tx=XM_019813208.2
D2 33446548 33446733
refGene_gn=PALD1;refGene_tx=XM_011287355.3;refGene_gn=PALD1;refGene_tx=XM_011287356.3
D2 33449303 33449406
refGene_gn=PALD1;refGene_tx=XM_011287355.3;refGene_gn=PALD1;refGene_tx=XM_011287356.3
D2 33450649 33450829
refGene_gn=PALD1;refGene_tx=XM_011287355.3;refGene_gn=PALD1;refGene_tx=XM_011287356.3
D2 33452466 33452631
refGene_gn=PALD1;refGene_tx=XM_011287355.3;refGene_gn=PALD1;refGene_tx=XM_011287356.3
D2 33453455 33453616
refGene_gn=PALD1;refGene_tx=XM_011287355.3;refGene_gn=PALD1;refGene_tx=XM_011287356.3

D2 33453846 33453922
refGene_gn=PALD1;refGene_tx=XM_011287355.3;refGene_gn=PALD1;refGene_tx
=XM_011287356.3
D2 33454682 33454834
refGene_gn=PALD1;refGene_tx=XM_011287355.3;refGene_gn=PALD1;refGene_tx
=XM_011287356.3
D2 33455186 33455286
refGene_gn=PALD1;refGene_tx=XM_011287355.3;refGene_gn=PALD1;refGene_tx
=XM_011287356.3
D2 33455487 33455592
refGene_gn=PALD1;refGene_tx=XM_011287355.3;refGene_gn=PALD1;refGene_tx
=XM_011287356.3
D2 33457863 33457959
refGene_gn=PALD1;refGene_tx=XM_011287355.3;refGene_gn=PALD1;refGene_tx
=XM_011287356.3
D2 33458398 33458527
refGene_gn=PALD1;refGene_tx=XM_011287355.3;refGene_gn=PALD1;refGene_tx
=XM_011287356.3
D2 33459028 33459145
refGene_gn=PALD1;refGene_tx=XM_011287355.3;refGene_gn=PALD1;refGene_tx
=XM_011287356.3
D2 33459222 33459378
refGene_gn=PALD1;refGene_tx=XM_011287355.3;refGene_gn=PALD1;refGene_tx
=XM_011287356.3
D2 33459688 33459871
refGene_gn=PALD1;refGene_tx=XM_011287355.3;refGene_gn=PALD1;refGene_tx
=XM_011287356.3
D2 33461010 33461151
refGene_gn=PALD1;refGene_tx=XM_011287355.3;refGene_gn=PALD1;refGene_tx
=XM_011287356.3
D2 33461388 33461460
refGene_gn=PALD1;refGene_tx=XM_011287355.3;refGene_gn=PALD1;refGene_tx
=XM_011287356.3
D2 33463483 33463624
refGene_gn=PALD1;refGene_tx=XM_011287355.3;refGene_gn=PALD1;refGene_tx
=XM_011287356.3
D2 33481842 33481998
refGene_gn=PALD1;refGene_tx=XM_011287355.3;refGene_gn=PALD1;refGene_tx
=XM_011287356.3
D2 33483973 33484126
refGene_gn=PALD1;refGene_tx=XM_011287355.3;refGene_gn=PALD1;refGene_tx
=XM_011287356.3
D2 33504116 33505245
refGene_gn=PRF1;refGene_tx=NM_001101660.1;refGene_gn=PRF1;refGene_tx=X
M_006937817.3
D2 33506133 33506672
refGene_gn=PRF1;refGene_tx=NM_001101660.1;refGene_gn=PRF1;refGene_tx=X
M_006937817.3
D2 33539956 33540038 refGene_gn=ADAMTS14;refGene_tx=XM_023240520.1
D2 33541737 33542180 refGene_gn=ADAMTS14;refGene_tx=XM_023240520.1

D2 33563308 33563465 refGene_gn=ADAMTS14;refGene_tx=XM_023240520.1
D2 33570799 33570990 refGene_gn=ADAMTS14;refGene_tx=XM_023240520.1
D2 33585345 33585429 refGene_gn=ADAMTS14;refGene_tx=XM_023240520.1
D2 33586140 33586288 refGene_gn=ADAMTS14;refGene_tx=XM_023240520.1
D2 33588591 33588697 refGene_gn=ADAMTS14;refGene_tx=XM_023240520.1
D2 33590398 33590542 refGene_gn=ADAMTS14;refGene_tx=XM_023240520.1
D2 33591043 33591176 refGene_gn=ADAMTS14;refGene_tx=XM_023240520.1
D2 33592623 33592737 refGene_gn=ADAMTS14;refGene_tx=XM_023240520.1
D2 33595002 33595151 refGene_gn=ADAMTS14;refGene_tx=XM_023240520.1
D2 33597002 33597178 refGene_gn=ADAMTS14;refGene_tx=XM_023240520.1
D2 33599635 33599769 refGene_gn=ADAMTS14;refGene_tx=XM_023240520.1
D2 33600153 33600277 refGene_gn=ADAMTS14;refGene_tx=XM_023240520.1
D2 33601256 33601337 refGene_gn=ADAMTS14;refGene_tx=XM_023240520.1
D2 33606598 33606762 refGene_gn=ADAMTS14;refGene_tx=XM_023240520.1
D2 33608177 33608346 refGene_gn=ADAMTS14;refGene_tx=XM_023240520.1
D2 33608829 33608962 refGene_gn=ADAMTS14;refGene_tx=XM_023240520.1
D2 33610678 33610886 refGene_gn=ADAMTS14;refGene_tx=XM_023240520.1
D2 33615419 33615549 refGene_gn=ADAMTS14;refGene_tx=XM_023240520.1
D2 33615676 33615787 refGene_gn=ADAMTS14;refGene_tx=XM_023240520.1
D2 33618269 33618751 refGene_gn=ADAMTS14;refGene_tx=XM_023240520.1
D2 33641802 33641888 refGene_gn=TBATA;refGene_tx=XM_023240525.1
D2 33642925 33642978 refGene_gn=TBATA;refGene_tx=XM_023240525.1
D2 33644454 33644599 refGene_gn=TBATA;refGene_tx=XM_023240525.1
D2 33645367 33645449 refGene_gn=TBATA;refGene_tx=XM_023240525.1
D2 33647353 33647527 refGene_gn=TBATA;refGene_tx=XM_023240525.1
D2 33648997 33649054 refGene_gn=TBATA;refGene_tx=XM_023240525.1
D2 33649056 33649091 refGene_gn=TBATA;refGene_tx=XM_023240525.1
D2 33649950 33650106 refGene_gn=TBATA;refGene_tx=XM_023240525.1
D2 33651050 33651286 refGene_gn=TBATA;refGene_tx=XM_023240525.1
D2 33652652 33652699 refGene_gn=TBATA;refGene_tx=XM_023240525.1
D2 33654128 33654402 refGene_gn=SGPL1;refGene_tx=XM_019813215.1
D2 33670871 33670933
refGene_gn=SGPL1;refGene_tx=XM_003994012.4;refGene_gn=SGPL1;refGene_tx=XM_019813215.1;refGene_gn=SGPL1;refGene_tx=XM_023240521.1;refGene_gn=SGPL1;refGene_tx=XM_023240522.1;refGene_gn=SGPL1;refGene_tx=XM_023240523.1;refGene_gn=SGPL1;refGene_tx=XM_023240524.1
D2 33695670 33695836
refGene_gn=SGPL1;refGene_tx=XM_003994012.4;refGene_gn=SGPL1;refGene_tx=XM_019813215.1;refGene_gn=SGPL1;refGene_tx=XM_023240521.1;refGene_gn=SGPL1;refGene_tx=XM_023240522.1;refGene_gn=SGPL1;refGene_tx=XM_023240523.1;refGene_gn=SGPL1;refGene_tx=XM_023240524.1
D2 33702071 33702139
refGene_gn=SGPL1;refGene_tx=XM_003994012.4;refGene_gn=SGPL1;refGene_tx=XM_019813215.1;refGene_gn=SGPL1;refGene_tx=XM_023240521.1;refGene_gn=SGPL1;refGene_tx=XM_023240522.1;refGene_gn=SGPL1;refGene_tx=XM_023240523.1;refGene_gn=SGPL1;refGene_tx=XM_023240524.1
D2 33705693 33705841
refGene_gn=SGPL1;refGene_tx=XM_003994012.4;refGene_gn=SGPL1;refGene_tx=XM_019813215.1;refGene_gn=SGPL1;refGene_tx=XM_023240521.1;refGene_gn=SGPL1;refGene_tx=XM_023240522.1;refGene_gn=SGPL1;refGene_tx=XM_0232405

23.1;refGene_gn=SGPL1;refGene_tx=XM_023240524.1
D2	33709070	33709147
refGene_gn=SGPL1;refGene_tx=XM_003994012.4;refGene_gn=SGPL1;refGene_tx=XM_019813215.1;refGene_gn=SGPL1;refGene_tx=XM_023240521.1;refGene_gn=SGPL1;refGene_tx=XM_023240522.1;refGene_gn=SGPL1;refGene_tx=XM_023240523.1;refGene_gn=SGPL1;refGene_tx=XM_023240524.1
D2	33711506	33711635
refGene_gn=SGPL1;refGene_tx=XM_003994012.4;refGene_gn=SGPL1;refGene_tx=XM_019813215.1;refGene_gn=SGPL1;refGene_tx=XM_023240521.1;refGene_gn=SGPL1;refGene_tx=XM_023240522.1;refGene_gn=SGPL1;refGene_tx=XM_023240523.1;refGene_gn=SGPL1;refGene_tx=XM_023240524.1
D2	33718466	33718555
refGene_gn=SGPL1;refGene_tx=XM_003994012.4;refGene_gn=SGPL1;refGene_tx=XM_019813215.1;refGene_gn=SGPL1;refGene_tx=XM_023240521.1;refGene_gn=SGPL1;refGene_tx=XM_023240522.1;refGene_gn=SGPL1;refGene_tx=XM_023240523.1;refGene_gn=SGPL1;refGene_tx=XM_023240524.1
D2	33719966	33720072
refGene_gn=SGPL1;refGene_tx=XM_003994012.4;refGene_gn=SGPL1;refGene_tx=XM_019813215.1;refGene_gn=SGPL1;refGene_tx=XM_023240521.1;refGene_gn=SGPL1;refGene_tx=XM_023240522.1;refGene_gn=SGPL1;refGene_tx=XM_023240523.1;refGene_gn=SGPL1;refGene_tx=XM_023240524.1
D2	33721441	33721540
refGene_gn=SGPL1;refGene_tx=XM_003994012.4;refGene_gn=SGPL1;refGene_tx=XM_019813215.1;refGene_gn=SGPL1;refGene_tx=XM_023240521.1;refGene_gn=SGPL1;refGene_tx=XM_023240522.1;refGene_gn=SGPL1;refGene_tx=XM_023240523.1;refGene_gn=SGPL1;refGene_tx=XM_023240524.1
D2	33722167	33722317
refGene_gn=SGPL1;refGene_tx=XM_003994012.4;refGene_gn=SGPL1;refGene_tx=XM_019813215.1;refGene_gn=SGPL1;refGene_tx=XM_023240521.1;refGene_gn=SGPL1;refGene_tx=XM_023240522.1;refGene_gn=SGPL1;refGene_tx=XM_023240523.1;refGene_gn=SGPL1;refGene_tx=XM_023240524.1
D2	33723630	33723869
refGene_gn=SGPL1;refGene_tx=XM_003994012.4;refGene_gn=SGPL1;refGene_tx=XM_019813215.1;refGene_gn=SGPL1;refGene_tx=XM_023240521.1;refGene_gn=SGPL1;refGene_tx=XM_023240522.1;refGene_gn=SGPL1;refGene_tx=XM_023240523.1;refGene_gn=SGPL1;refGene_tx=XM_023240524.1
D2	33725179	33725326
refGene_gn=SGPL1;refGene_tx=XM_003994012.4;refGene_gn=SGPL1;refGene_tx=XM_019813215.1;refGene_gn=SGPL1;refGene_tx=XM_023240521.1;refGene_gn=SGPL1;refGene_tx=XM_023240522.1;refGene_gn=SGPL1;refGene_tx=XM_023240523.1;refGene_gn=SGPL1;refGene_tx=XM_023240524.1
D2	33726294	33726415
refGene_gn=SGPL1;refGene_tx=XM_003994012.4;refGene_gn=SGPL1;refGene_tx=XM_019813215.1;refGene_gn=SGPL1;refGene_tx=XM_023240521.1;refGene_gn=SGPL1;refGene_tx=XM_023240522.1;refGene_gn=SGPL1;refGene_tx=XM_023240523.1;refGene_gn=SGPL1;refGene_tx=XM_023240524.1
D2	33726920	33727061
refGene_gn=SGPL1;refGene_tx=XM_003994012.4;refGene_gn=SGPL1;refGene_tx=XM_019813215.1;refGene_gn=SGPL1;refGene_tx=XM_023240521.1;refGene_gn=SGPL1;refGene_tx=XM_023240522.1;refGene_gn=SGPL1;refGene_tx=XM_0232405

23.1;refGene_gn=SGPL1;refGene_tx=XM_023240524.1
D2 33732585 33732684
refGene_gn=PCBD1;refGene_tx=XM_019813222.2;refGene_gn=PCBD1;refGene_tx=XM_023240526.1
D2 33733300 33733381
refGene_gn=PCBD1;refGene_tx=XM_019813222.2;refGene_gn=PCBD1;refGene_tx=XM_023240526.1
D2 33733902 33734034 refGene_gn=PCBD1;refGene_tx=XM_023240526.1
D2 33736782 33736785 refGene_gn=PCBD1;refGene_tx=XM_023240526.1
D2 34039559 34039638
refGene_gn=UNC5B;refGene_tx=XM_006937850.4;refGene_gn=UNC5B;refGene_tx=XM_023240527.1
D2 34100541 34100766
refGene_gn=UNC5B;refGene_tx=XM_006937850.4;refGene_gn=UNC5B;refGene_tx=XM_023240527.1
D2 34104641 34104785
refGene_gn=UNC5B;refGene_tx=XM_006937850.4;refGene_gn=UNC5B;refGene_tx=XM_023240527.1
D2 34105243 34105347
refGene_gn=UNC5B;refGene_tx=XM_006937850.4;refGene_gn=UNC5B;refGene_tx=XM_023240527.1
D2 34106104 34106285
refGene_gn=UNC5B;refGene_tx=XM_006937850.4;refGene_gn=UNC5B;refGene_tx=XM_023240527.1
D2 34106931 34107099
refGene_gn=UNC5B;refGene_tx=XM_006937850.4;refGene_gn=UNC5B;refGene_tx=XM_023240527.1
D2 34107917 34108082
refGene_gn=UNC5B;refGene_tx=XM_006937850.4;refGene_gn=UNC5B;refGene_tx=XM_023240527.1
D2 34108329 34108362 refGene_gn=UNC5B;refGene_tx=XM_006937850.4
D2 34110115 34110310
refGene_gn=UNC5B;refGene_tx=XM_006937850.4;refGene_gn=UNC5B;refGene_tx=XM_023240527.1
D2 34110519 34110909
refGene_gn=UNC5B;refGene_tx=XM_006937850.4;refGene_gn=UNC5B;refGene_tx=XM_023240527.1
D2 34111564 34111652
refGene_gn=UNC5B;refGene_tx=XM_006937850.4;refGene_gn=UNC5B;refGene_tx=XM_023240527.1
D2 34112537 34112706
refGene_gn=UNC5B;refGene_tx=XM_006937850.4;refGene_gn=UNC5B;refGene_tx=XM_023240527.1
D2 34112880 34113114
refGene_gn=UNC5B;refGene_tx=XM_006937850.4;refGene_gn=UNC5B;refGene_tx=XM_023240527.1
D2 34114397 34114547
refGene_gn=UNC5B;refGene_tx=XM_006937850.4;refGene_gn=UNC5B;refGene_tx=XM_023240527.1
D2 34115243 34115408

refGene_gn=UNC5B;refGene_tx=XM_006937850.4;refGene_gn=UNC5B;refGene_tx=XM_023240527.1
D2 34116168 34116350
refGene_gn=UNC5B;refGene_tx=XM_006937850.4;refGene_gn=UNC5B;refGene_tx=XM_023240527.1
D2 34117184 34117350
refGene_gn=UNC5B;refGene_tx=XM_006937850.4;refGene_gn=UNC5B;refGene_tx=XM_023240527.1
D2 34131365 34131366 refGene_gn=SLC29A3;refGene_tx=XM_023240528.1
D2 34134379 34134678 refGene_gn=SLC29A3;refGene_tx=XM_023240528.1
D2 34156134 34156217 refGene_gn=SLC29A3;refGene_tx=XM_023240528.1
D2 34163498 34163725 refGene_gn=SLC29A3;refGene_tx=XM_023240528.1
D2 34167076 34167239 refGene_gn=SLC29A3;refGene_tx=XM_023240528.1
D2 34172541 34173196 refGene_gn=SLC29A3;refGene_tx=XM_023240528.1
D2 34253842 34253909
refGene_gn=CDH23;refGene_tx=XM_023240529.1;refGene_gn=CDH23;refGene_tx=XM_023240530.1;refGene_gn=CDH23;refGene_tx=XM_023240531.1;refGene_gn=CDH23;refGene_tx=XM_023240532.1;refGene_gn=CDH23;refGene_tx=XM_023240533.1;refGene_gn=CDH23;refGene_tx=XM_023240534.1;refGene_gn=CDH23;refGene_tx=XM_023240535.1;refGene_gn=CDH23;refGene_tx=XM_023240536.1
D2 34261164 34261242
refGene_gn=CDH23;refGene_tx=XM_023240529.1;refGene_gn=CDH23;refGene_tx=XM_023240530.1;refGene_gn=CDH23;refGene_tx=XM_023240531.1;refGene_gn=CDH23;refGene_tx=XM_023240532.1;refGene_gn=CDH23;refGene_tx=XM_023240533.1;refGene_gn=CDH23;refGene_tx=XM_023240534.1;refGene_gn=CDH23;refGene_tx=XM_023240535.1;refGene_gn=CDH23;refGene_tx=XM_023240536.1
D2 34329445 34329588
refGene_gn=CDH23;refGene_tx=XM_023240529.1;refGene_gn=CDH23;refGene_tx=XM_023240530.1;refGene_gn=CDH23;refGene_tx=XM_023240531.1;refGene_gn=CDH23;refGene_tx=XM_023240532.1;refGene_gn=CDH23;refGene_tx=XM_023240533.1;refGene_gn=CDH23;refGene_tx=XM_023240534.1;refGene_gn=CDH23;refGene_tx=XM_023240535.1;refGene_gn=CDH23;refGene_tx=XM_023240536.1
D2 34330127 34330175
refGene_gn=CDH23;refGene_tx=XM_023240529.1;refGene_gn=CDH23;refGene_tx=XM_023240530.1;refGene_gn=CDH23;refGene_tx=XM_023240531.1;refGene_gn=CDH23;refGene_tx=XM_023240532.1;refGene_gn=CDH23;refGene_tx=XM_023240533.1;refGene_gn=CDH23;refGene_tx=XM_023240534.1;refGene_gn=CDH23;refGene_tx=XM_023240535.1;refGene_gn=CDH23;refGene_tx=XM_023240536.1
D2 34330289 34330382
refGene_gn=CDH23;refGene_tx=XM_023240529.1;refGene_gn=CDH23;refGene_tx=XM_023240530.1;refGene_gn=CDH23;refGene_tx=XM_023240531.1;refGene_gn=CDH23;refGene_tx=XM_023240532.1;refGene_gn=CDH23;refGene_tx=XM_023240533.1;refGene_gn=CDH23;refGene_tx=XM_023240534.1;refGene_gn=CDH23;refGene_tx=XM_023240535.1;refGene_gn=CDH23;refGene_tx=XM_023240536.1
D2 34386401 34386596
refGene_gn=CDH23;refGene_tx=XM_023240529.1;refGene_gn=CDH23;refGene_tx=XM_023240530.1;refGene_gn=CDH23;refGene_tx=XM_023240531.1;refGene_gn=CDH23;refGene_tx=XM_023240532.1;refGene_gn=CDH23;refGene_tx=XM_023240533.1;refGene_gn=CDH23;refGene_tx=XM_023240534.1;refGene_gn=CDH23;refGene_tx=XM_023240535.1;refGene_gn=CDH23;refGene_tx=XM_023240536.1

D2 34390142 34390271
refGene_gn=CDH23;refGene_tx=XM_023240529.1;refGene_gn=CDH23;refGene_tx=XM_023240530.1;refGene_gn=CDH23;refGene_tx=XM_023240531.1;refGene_gn=CDH23;refGene_tx=XM_023240532.1;refGene_gn=CDH23;refGene_tx=XM_023240533.1;refGene_gn=CDH23;refGene_tx=XM_023240534.1;refGene_gn=CDH23;refGene_tx=XM_023240535.1;refGene_gn=CDH23;refGene_tx=XM_023240536.1
D2 34398035 34398114
refGene_gn=CDH23;refGene_tx=XM_023240529.1;refGene_gn=CDH23;refGene_tx=XM_023240530.1;refGene_gn=CDH23;refGene_tx=XM_023240531.1;refGene_gn=CDH23;refGene_tx=XM_023240532.1;refGene_gn=CDH23;refGene_tx=XM_023240533.1;refGene_gn=CDH23;refGene_tx=XM_023240534.1;refGene_gn=CDH23;refGene_tx=XM_023240535.1;refGene_gn=CDH23;refGene_tx=XM_023240536.1
D2 34435426 34435539
refGene_gn=CDH23;refGene_tx=XM_023240529.1;refGene_gn=CDH23;refGene_tx=XM_023240530.1;refGene_gn=CDH23;refGene_tx=XM_023240531.1;refGene_gn=CDH23;refGene_tx=XM_023240532.1;refGene_gn=CDH23;refGene_tx=XM_023240533.1;refGene_gn=CDH23;refGene_tx=XM_023240534.1;refGene_gn=CDH23;refGene_tx=XM_023240535.1;refGene_gn=CDH23;refGene_tx=XM_023240536.1
D2 34437178 34437367
refGene_gn=CDH23;refGene_tx=XM_023240529.1;refGene_gn=CDH23;refGene_tx=XM_023240530.1;refGene_gn=CDH23;refGene_tx=XM_023240531.1;refGene_gn=CDH23;refGene_tx=XM_023240532.1;refGene_gn=CDH23;refGene_tx=XM_023240533.1;refGene_gn=CDH23;refGene_tx=XM_023240534.1;refGene_gn=CDH23;refGene_tx=XM_023240535.1;refGene_gn=CDH23;refGene_tx=XM_023240536.1
D2 34461654 34461660
refGene_gn=CDH23;refGene_tx=XM_023240529.1;refGene_gn=CDH23;refGene_tx=XM_023240530.1;refGene_gn=CDH23;refGene_tx=XM_023240532.1;refGene_gn=CDH23;refGene_tx=XM_023240533.1;refGene_gn=CDH23;refGene_tx=XM_023240534.1;refGene_gn=CDH23;refGene_tx=XM_023240535.1;refGene_gn=CDH23;refGene_tx=XM_023240536.1
D2 34463342 34463492
refGene_gn=CDH23;refGene_tx=XM_023240529.1;refGene_gn=CDH23;refGene_tx=XM_023240530.1;refGene_gn=CDH23;refGene_tx=XM_023240531.1;refGene_gn=CDH23;refGene_tx=XM_023240532.1;refGene_gn=CDH23;refGene_tx=XM_023240533.1;refGene_gn=CDH23;refGene_tx=XM_023240534.1;refGene_gn=CDH23;refGene_tx=XM_023240535.1;refGene_gn=CDH23;refGene_tx=XM_023240536.1
D2 34463960 34464119
refGene_gn=CDH23;refGene_tx=XM_023240529.1;refGene_gn=CDH23;refGene_tx=XM_023240530.1;refGene_gn=CDH23;refGene_tx=XM_023240531.1;refGene_gn=CDH23;refGene_tx=XM_023240532.1;refGene_gn=CDH23;refGene_tx=XM_023240533.1;refGene_gn=CDH23;refGene_tx=XM_023240534.1;refGene_gn=CDH23;refGene_tx=XM_023240535.1;refGene_gn=CDH23;refGene_tx=XM_023240536.1
D2 34494083 34494148
refGene_gn=CDH23;refGene_tx=XM_023240529.1;refGene_gn=CDH23;refGene_tx=XM_023240530.1;refGene_gn=CDH23;refGene_tx=XM_023240531.1;refGene_gn=CDH23;refGene_tx=XM_023240532.1;refGene_gn=CDH23;refGene_tx=XM_023240533.1;refGene_gn=CDH23;refGene_tx=XM_023240534.1;refGene_gn=CDH23;refGene_tx=XM_023240535.1;refGene_gn=CDH23;refGene_tx=XM_023240536.1
D2 34495831 34496069
refGene_gn=CDH23;refGene_tx=XM_023240529.1;refGene_gn=CDH23;refGene_tx

=XM_023240530.1;refGene_gn=CDH23;refGene_tx=XM_023240531.1;refGene_gn=CDH23;refGene_tx=XM_023240532.1;refGene_gn=CDH23;refGene_tx=XM_023240533.1;refGene_gn=CDH23;refGene_tx=XM_023240534.1;refGene_gn=CDH23;refGene_tx=XM_023240535.1;refGene_gn=CDH23;refGene_tx=XM_023240536.1
D2 34497650 34497756
refGene_gn=CDH23;refGene_tx=XM_023240529.1;refGene_gn=CDH23;refGene_tx=XM_023240530.1;refGene_gn=CDH23;refGene_tx=XM_023240531.1;refGene_gn=CDH23;refGene_tx=XM_023240532.1;refGene_gn=CDH23;refGene_tx=XM_023240533.1;refGene_gn=CDH23;refGene_tx=XM_023240534.1;refGene_gn=CDH23;refGene_tx=XM_023240535.1;refGene_gn=CDH23;refGene_tx=XM_023240536.1
D2 34500061 34500189
refGene_gn=CDH23;refGene_tx=XM_023240529.1;refGene_gn=CDH23;refGene_tx=XM_023240530.1;refGene_gn=CDH23;refGene_tx=XM_023240531.1;refGene_gn=CDH23;refGene_tx=XM_023240532.1;refGene_gn=CDH23;refGene_tx=XM_023240533.1;refGene_gn=CDH23;refGene_tx=XM_023240534.1;refGene_gn=CDH23;refGene_tx=XM_023240535.1;refGene_gn=CDH23;refGene_tx=XM_023240536.1
D2 34504815 34504888
refGene_gn=CDH23;refGene_tx=XM_023240529.1;refGene_gn=CDH23;refGene_tx=XM_023240530.1;refGene_gn=CDH23;refGene_tx=XM_023240531.1;refGene_gn=CDH23;refGene_tx=XM_023240532.1;refGene_gn=CDH23;refGene_tx=XM_023240533.1;refGene_gn=CDH23;refGene_tx=XM_023240534.1;refGene_gn=CDH23;refGene_tx=XM_023240535.1;refGene_gn=CDH23;refGene_tx=XM_023240536.1
D2 34506587 34506704
refGene_gn=CDH23;refGene_tx=XM_023240529.1;refGene_gn=CDH23;refGene_tx=XM_023240530.1;refGene_gn=CDH23;refGene_tx=XM_023240531.1;refGene_gn=CDH23;refGene_tx=XM_023240532.1;refGene_gn=CDH23;refGene_tx=XM_023240533.1;refGene_gn=CDH23;refGene_tx=XM_023240534.1;refGene_gn=CDH23;refGene_tx=XM_023240535.1;refGene_gn=CDH23;refGene_tx=XM_023240536.1
D2 34509838 34509951
refGene_gn=CDH23;refGene_tx=XM_023240529.1;refGene_gn=CDH23;refGene_tx=XM_023240530.1;refGene_gn=CDH23;refGene_tx=XM_023240531.1;refGene_gn=CDH23;refGene_tx=XM_023240532.1;refGene_gn=CDH23;refGene_tx=XM_023240533.1;refGene_gn=CDH23;refGene_tx=XM_023240534.1;refGene_gn=CDH23;refGene_tx=XM_023240535.1;refGene_gn=CDH23;refGene_tx=XM_023240536.1
D2 34511080 34511188
refGene_gn=CDH23;refGene_tx=XM_023240529.1;refGene_gn=CDH23;refGene_tx=XM_023240530.1;refGene_gn=CDH23;refGene_tx=XM_023240531.1;refGene_gn=CDH23;refGene_tx=XM_023240532.1;refGene_gn=CDH23;refGene_tx=XM_023240533.1;refGene_gn=CDH23;refGene_tx=XM_023240534.1;refGene_gn=CDH23;refGene_tx=XM_023240535.1;refGene_gn=CDH23;refGene_tx=XM_023240536.1
D2 34518244 34518434
refGene_gn=CDH23;refGene_tx=XM_023240529.1;refGene_gn=CDH23;refGene_tx=XM_023240530.1;refGene_gn=CDH23;refGene_tx=XM_023240531.1;refGene_gn=CDH23;refGene_tx=XM_023240532.1;refGene_gn=CDH23;refGene_tx=XM_023240533.1;refGene_gn=CDH23;refGene_tx=XM_023240534.1;refGene_gn=CDH23;refGene_tx=XM_023240535.1;refGene_gn=CDH23;refGene_tx=XM_023240536.1
D2 34518781 34518927
refGene_gn=CDH23;refGene_tx=XM_023240529.1;refGene_gn=CDH23;refGene_tx=XM_023240530.1;refGene_gn=CDH23;refGene_tx=XM_023240531.1;refGene_gn=CDH23;refGene_tx=XM_023240532.1;refGene_gn=CDH23;refGene_tx=XM_0232405

33.1;refGene_gn=CDH23;refGene_tx=XM_023240534.1;refGene_gn=CDH23;refGene_tx=XM_023240535.1;refGene_gn=CDH23;refGene_tx=XM_023240536.1
D2 34521192 34521412
refGene_gn=CDH23;refGene_tx=XM_023240529.1;refGene_gn=CDH23;refGene_tx=XM_023240530.1;refGene_gn=CDH23;refGene_tx=XM_023240531.1;refGene_gn=CDH23;refGene_tx=XM_023240532.1;refGene_gn=CDH23;refGene_tx=XM_023240533.1;refGene_gn=CDH23;refGene_tx=XM_023240534.1;refGene_gn=CDH23;refGene_tx=XM_023240535.1;refGene_gn=CDH23;refGene_tx=XM_023240536.1
D2 34522895 34523048
refGene_gn=CDH23;refGene_tx=XM_023240529.1;refGene_gn=CDH23;refGene_tx=XM_023240530.1;refGene_gn=CDH23;refGene_tx=XM_023240531.1;refGene_gn=CDH23;refGene_tx=XM_023240532.1;refGene_gn=CDH23;refGene_tx=XM_023240533.1;refGene_gn=CDH23;refGene_tx=XM_023240534.1;refGene_gn=CDH23;refGene_tx=XM_023240535.1;refGene_gn=CDH23;refGene_tx=XM_023240536.1
D2 34525059 34525173
refGene_gn=CDH23;refGene_tx=XM_023240529.1;refGene_gn=CDH23;refGene_tx=XM_023240530.1;refGene_gn=CDH23;refGene_tx=XM_023240531.1;refGene_gn=CDH23;refGene_tx=XM_023240532.1;refGene_gn=CDH23;refGene_tx=XM_023240533.1;refGene_gn=CDH23;refGene_tx=XM_023240534.1;refGene_gn=CDH23;refGene_tx=XM_023240535.1;refGene_gn=CDH23;refGene_tx=XM_023240536.1
D2 34528031 34528180
refGene_gn=CDH23;refGene_tx=XM_023240529.1;refGene_gn=CDH23;refGene_tx=XM_023240530.1;refGene_gn=CDH23;refGene_tx=XM_023240531.1;refGene_gn=CDH23;refGene_tx=XM_023240532.1;refGene_gn=CDH23;refGene_tx=XM_023240533.1;refGene_gn=CDH23;refGene_tx=XM_023240534.1;refGene_gn=CDH23;refGene_tx=XM_023240535.1;refGene_gn=CDH23;refGene_tx=XM_023240536.1
D2 34531279 34531648
refGene_gn=CD2H10orf105;refGene_tx=XM_019813232.2;refGene_gn=CD2H10orf105;refGene_tx=XM_019813234.2;refGene_gn=CD2H10orf105;refGene_tx=XM_019813235.2;refGene_gn=CD2H10orf105;refGene_tx=XM_019813236.2;refGene_gn=CD2H10orf105;refGene_tx=XM_019813237.2;refGene_gn=CD2H10orf105;refGene_tx=XM_019813238.2;refGene_gn=CD2H10orf105;refGene_tx=XM_019813239.2;refGene_gn=CD2H10orf105;refGene_tx=XM_019813240.2
D2 34536814 34536875
refGene_gn=CDH23;refGene_tx=XM_023240529.1;refGene_gn=CDH23;refGene_tx=XM_023240530.1;refGene_gn=CDH23;refGene_tx=XM_023240531.1;refGene_gn=CDH23;refGene_tx=XM_023240532.1;refGene_gn=CDH23;refGene_tx=XM_023240533.1;refGene_gn=CDH23;refGene_tx=XM_023240534.1;refGene_gn=CDH23;refGene_tx=XM_023240535.1;refGene_gn=CDH23;refGene_tx=XM_023240536.1
D2 34537805 34537954
refGene_gn=CDH23;refGene_tx=XM_023240529.1;refGene_gn=CDH23;refGene_tx=XM_023240530.1;refGene_gn=CDH23;refGene_tx=XM_023240531.1;refGene_gn=CDH23;refGene_tx=XM_023240532.1;refGene_gn=CDH23;refGene_tx=XM_023240533.1;refGene_gn=CDH23;refGene_tx=XM_023240534.1;refGene_gn=CDH23;refGene_tx=XM_023240535.1;refGene_gn=CDH23;refGene_tx=XM_023240536.1
D2 34542721 34542857
refGene_gn=CDH23;refGene_tx=XM_023240529.1;refGene_gn=CDH23;refGene_tx=XM_023240530.1;refGene_gn=CDH23;refGene_tx=XM_023240531.1;refGene_gn=CDH23;refGene_tx=XM_023240532.1;refGene_gn=CDH23;refGene_tx=XM_023240533.1;refGene_gn=CDH23;refGene_tx=XM_023240534.1;refGene_gn=CDH23;refGe

ne_tx=XM_023240535.1;refGene_gn=CDH23;refGene_tx=XM_023240536.1
D2	34544404	34544793
refGene_gn=CDH23;refGene_tx=XM_023240529.1;refGene_gn=CDH23;refGene_tx=XM_023240530.1;refGene_gn=CDH23;refGene_tx=XM_023240531.1;refGene_gn=CDH23;refGene_tx=XM_023240532.1;refGene_gn=CDH23;refGene_tx=XM_023240533.1;refGene_gn=CDH23;refGene_tx=XM_023240534.1;refGene_gn=CDH23;refGene_tx=XM_023240535.1;refGene_gn=CDH23;refGene_tx=XM_023240536.1
D2	34545908	34546010
refGene_gn=CDH23;refGene_tx=XM_023240529.1;refGene_gn=CDH23;refGene_tx=XM_023240530.1;refGene_gn=CDH23;refGene_tx=XM_023240531.1;refGene_gn=CDH23;refGene_tx=XM_023240532.1;refGene_gn=CDH23;refGene_tx=XM_023240533.1;refGene_gn=CDH23;refGene_tx=XM_023240534.1;refGene_gn=CDH23;refGene_tx=XM_023240535.1;refGene_gn=CDH23;refGene_tx=XM_023240536.1
D2	34546309	34546312
refGene_gn=CDH23;refGene_tx=XM_023240529.1;refGene_gn=CDH23;refGene_tx=XM_023240531.1;refGene_gn=CDH23;refGene_tx=XM_023240532.1;refGene_gn=CDH23;refGene_tx=XM_023240533.1;refGene_gn=CDH23;refGene_tx=XM_023240534.1;refGene_gn=CDH23;refGene_tx=XM_023240535.1;refGene_gn=CDH23;refGene_tx=XM_023240536.1
D2	34550216	34550366
refGene_gn=CDH23;refGene_tx=XM_023240529.1;refGene_gn=CDH23;refGene_tx=XM_023240530.1;refGene_gn=CDH23;refGene_tx=XM_023240531.1;refGene_gn=CDH23;refGene_tx=XM_023240532.1;refGene_gn=CDH23;refGene_tx=XM_023240533.1;refGene_gn=CDH23;refGene_tx=XM_023240534.1;refGene_gn=CDH23;refGene_tx=XM_023240535.1;refGene_gn=CDH23;refGene_tx=XM_023240536.1
D2	34551337	34551466
refGene_gn=CDH23;refGene_tx=XM_023240529.1;refGene_gn=CDH23;refGene_tx=XM_023240530.1;refGene_gn=CDH23;refGene_tx=XM_023240531.1;refGene_gn=CDH23;refGene_tx=XM_023240532.1;refGene_gn=CDH23;refGene_tx=XM_023240533.1;refGene_gn=CDH23;refGene_tx=XM_023240534.1;refGene_gn=CDH23;refGene_tx=XM_023240535.1;refGene_gn=CDH23;refGene_tx=XM_023240536.1
D2	34552381	34552510
refGene_gn=CDH23;refGene_tx=XM_023240529.1;refGene_gn=CDH23;refGene_tx=XM_023240530.1;refGene_gn=CDH23;refGene_tx=XM_023240531.1;refGene_gn=CDH23;refGene_tx=XM_023240532.1;refGene_gn=CDH23;refGene_tx=XM_023240533.1;refGene_gn=CDH23;refGene_tx=XM_023240534.1;refGene_gn=CDH23;refGene_tx=XM_023240535.1;refGene_gn=CDH23;refGene_tx=XM_023240536.1
D2	34553228	34553456
refGene_gn=CDH23;refGene_tx=XM_023240529.1;refGene_gn=CDH23;refGene_tx=XM_023240530.1;refGene_gn=CDH23;refGene_tx=XM_023240531.1;refGene_gn=CDH23;refGene_tx=XM_023240532.1;refGene_gn=CDH23;refGene_tx=XM_023240533.1;refGene_gn=CDH23;refGene_tx=XM_023240534.1;refGene_gn=CDH23;refGene_tx=XM_023240535.1;refGene_gn=CDH23;refGene_tx=XM_023240536.1
D2	34561052	34561232
refGene_gn=VSIR;refGene_tx=XM_011287352.3;refGene_gn=VSIR;refGene_tx=XM_019813231.2
D2	34561392	34561603
refGene_gn=VSIR;refGene_tx=XM_011287352.3;refGene_gn=VSIR;refGene_tx=XM_019813231.2
D2	34562587	34562615

refGene_gn=VSIR;refGene_tx=XM_011287352.3;refGene_gn=VSIR;refGene_tx=XM_019813231.2
D2 34565484 34565592
refGene_gn=VSIR;refGene_tx=XM_011287352.3;refGene_gn=VSIR;refGene_tx=XM_019813231.2
D2 34570811 34570868
refGene_gn=VSIR;refGene_tx=XM_011287352.3;refGene_gn=VSIR;refGene_tx=XM_019813231.2
D2 34571375 34571801
refGene_gn=VSIR;refGene_tx=XM_011287352.3;refGene_gn=VSIR;refGene_tx=XM_019813231.2
D2 34581725 34581785 refGene_gn=CDH23;refGene_tx=XM_019813229.2
D2 34581898 34581980 refGene_gn=VSIR;refGene_tx=XM_011287352.3
D2 34591513 34591735
refGene_gn=CDH23;refGene_tx=XM_019813229.2;refGene_gn=CDH23;refGene_tx=XM_023240529.1;refGene_gn=CDH23;refGene_tx=XM_023240530.1;refGene_gn=CDH23;refGene_tx=XM_023240531.1;refGene_gn=CDH23;refGene_tx=XM_023240532.1;refGene_gn=CDH23;refGene_tx=XM_023240533.1;refGene_gn=CDH23;refGene_tx=XM_023240534.1;refGene_gn=CDH23;refGene_tx=XM_023240535.1;refGene_gn=CDH23;refGene_tx=XM_023240536.1
D2 34592048 34592168
refGene_gn=CDH23;refGene_tx=XM_019813229.2;refGene_gn=CDH23;refGene_tx=XM_023240529.1;refGene_gn=CDH23;refGene_tx=XM_023240530.1;refGene_gn=CDH23;refGene_tx=XM_023240531.1;refGene_gn=CDH23;refGene_tx=XM_023240532.1;refGene_gn=CDH23;refGene_tx=XM_023240533.1;refGene_gn=CDH23;refGene_tx=XM_023240534.1;refGene_gn=CDH23;refGene_tx=XM_023240535.1;refGene_gn=CDH23;refGene_tx=XM_023240536.1
D2 34592807 34592988
refGene_gn=CDH23;refGene_tx=XM_019813229.2;refGene_gn=CDH23;refGene_tx=XM_023240529.1;refGene_gn=CDH23;refGene_tx=XM_023240530.1;refGene_gn=CDH23;refGene_tx=XM_023240531.1;refGene_gn=CDH23;refGene_tx=XM_023240532.1;refGene_gn=CDH23;refGene_tx=XM_023240533.1;refGene_gn=CDH23;refGene_tx=XM_023240534.1;refGene_gn=CDH23;refGene_tx=XM_023240535.1;refGene_gn=CDH23;refGene_tx=XM_023240536.1
D2 34596156 34596290
refGene_gn=CDH23;refGene_tx=XM_019813229.2;refGene_gn=CDH23;refGene_tx=XM_023240529.1;refGene_gn=CDH23;refGene_tx=XM_023240530.1;refGene_gn=CDH23;refGene_tx=XM_023240531.1;refGene_gn=CDH23;refGene_tx=XM_023240532.1;refGene_gn=CDH23;refGene_tx=XM_023240533.1;refGene_gn=CDH23;refGene_tx=XM_023240534.1;refGene_gn=CDH23;refGene_tx=XM_023240535.1;refGene_gn=CDH23;refGene_tx=XM_023240536.1
D2 34596835 34597045
refGene_gn=CDH23;refGene_tx=XM_019813229.2;refGene_gn=CDH23;refGene_tx=XM_023240529.1;refGene_gn=CDH23;refGene_tx=XM_023240530.1;refGene_gn=CDH23;refGene_tx=XM_023240531.1;refGene_gn=CDH23;refGene_tx=XM_023240532.1;refGene_gn=CDH23;refGene_tx=XM_023240533.1;refGene_gn=CDH23;refGene_tx=XM_023240534.1;refGene_gn=CDH23;refGene_tx=XM_023240535.1;refGene_gn=CDH23;refGene_tx=XM_023240536.1
D2 34597583 34597691
refGene_gn=CDH23;refGene_tx=XM_019813229.2;refGene_gn=CDH23;refGene_tx

=XM_023240529.1;refGene_gn=CDH23;refGene_tx=XM_023240530.1;refGene_gn=CDH23;refGene_tx=XM_023240531.1;refGene_gn=CDH23;refGene_tx=XM_023240532.1;refGene_gn=CDH23;refGene_tx=XM_023240533.1;refGene_gn=CDH23;refGene_tx=XM_023240534.1;refGene_gn=CDH23;refGene_tx=XM_023240535.1;refGene_gn=CDH23;refGene_tx=XM_023240536.1
D2 34599132 34599235
refGene_gn=CDH23;refGene_tx=XM_019813229.2;refGene_gn=CDH23;refGene_tx=XM_023240529.1;refGene_gn=CDH23;refGene_tx=XM_023240530.1;refGene_gn=CDH23;refGene_tx=XM_023240531.1;refGene_gn=CDH23;refGene_tx=XM_023240532.1;refGene_gn=CDH23;refGene_tx=XM_023240533.1;refGene_gn=CDH23;refGene_tx=XM_023240534.1;refGene_gn=CDH23;refGene_tx=XM_023240535.1;refGene_gn=CDH23;refGene_tx=XM_023240536.1
D2 34600570 34600696
refGene_gn=CDH23;refGene_tx=XM_019813229.2;refGene_gn=CDH23;refGene_tx=XM_023240529.1;refGene_gn=CDH23;refGene_tx=XM_023240530.1;refGene_gn=CDH23;refGene_tx=XM_023240531.1;refGene_gn=CDH23;refGene_tx=XM_023240532.1;refGene_gn=CDH23;refGene_tx=XM_023240533.1;refGene_gn=CDH23;refGene_tx=XM_023240534.1;refGene_gn=CDH23;refGene_tx=XM_023240535.1;refGene_gn=CDH23;refGene_tx=XM_023240536.1
D2 34601021 34601225
refGene_gn=CDH23;refGene_tx=XM_019813229.2;refGene_gn=CDH23;refGene_tx=XM_023240529.1;refGene_gn=CDH23;refGene_tx=XM_023240530.1;refGene_gn=CDH23;refGene_tx=XM_023240531.1;refGene_gn=CDH23;refGene_tx=XM_023240532.1;refGene_gn=CDH23;refGene_tx=XM_023240533.1;refGene_gn=CDH23;refGene_tx=XM_023240534.1;refGene_gn=CDH23;refGene_tx=XM_023240535.1;refGene_gn=CDH23;refGene_tx=XM_023240536.1
D2 34603315 34603774
refGene_gn=CDH23;refGene_tx=XM_019813229.2;refGene_gn=CDH23;refGene_tx=XM_023240529.1;refGene_gn=CDH23;refGene_tx=XM_023240530.1;refGene_gn=CDH23;refGene_tx=XM_023240531.1;refGene_gn=CDH23;refGene_tx=XM_023240532.1;refGene_gn=CDH23;refGene_tx=XM_023240533.1;refGene_gn=CDH23;refGene_tx=XM_023240534.1;refGene_gn=CDH23;refGene_tx=XM_023240535.1;refGene_gn=CDH23;refGene_tx=XM_023240536.1
D2 34605379 34605627 refGene_gn=CDH23;refGene_tx=XM_023240534.1
D2 34605666 34605680 refGene_gn=CDH23;refGene_tx=XM_023240535.1
D2 34606856 34606973
refGene_gn=CDH23;refGene_tx=XM_019813229.2;refGene_gn=CDH23;refGene_tx=XM_019813230.1;refGene_gn=CDH23;refGene_tx=XM_023240529.1;refGene_gn=CDH23;refGene_tx=XM_023240530.1;refGene_gn=CDH23;refGene_tx=XM_023240531.1;refGene_gn=CDH23;refGene_tx=XM_023240532.1;refGene_gn=CDH23;refGene_tx=XM_023240533.1;refGene_gn=CDH23;refGene_tx=XM_023240536.1
D2 34608061 34608286
refGene_gn=CDH23;refGene_tx=XM_019813229.2;refGene_gn=CDH23;refGene_tx=XM_019813230.1;refGene_gn=CDH23;refGene_tx=XM_023240529.1;refGene_gn=CDH23;refGene_tx=XM_023240530.1;refGene_gn=CDH23;refGene_tx=XM_023240531.1;refGene_gn=CDH23;refGene_tx=XM_023240532.1;refGene_gn=CDH23;refGene_tx=XM_023240533.1;refGene_gn=CDH23;refGene_tx=XM_023240536.1
D2 34608850 34609020
refGene_gn=CDH23;refGene_tx=XM_019813229.2;refGene_gn=CDH23;refGene_tx=XM_019813230.1;refGene_gn=CDH23;refGene_tx=XM_023240529.1;refGene_gn=

CDH23;refGene_tx=XM_023240530.1;refGene_gn=CDH23;refGene_tx=XM_0232405
31.1;refGene_gn=CDH23;refGene_tx=XM_023240532.1;refGene_gn=CDH23;refGe
ne_tx=XM_023240533.1;refGene_gn=CDH23;refGene_tx=XM_023240536.1
D2 34609242 34609380
refGene_gn=CDH23;refGene_tx=XM_019813229.2;refGene_gn=CDH23;refGene_tx
=XM_019813230.1;refGene_gn=CDH23;refGene_tx=XM_023240529.1;refGene_gn=
CDH23;refGene_tx=XM_023240530.1;refGene_gn=CDH23;refGene_tx=XM_0232405
31.1;refGene_gn=CDH23;refGene_tx=XM_023240532.1;refGene_gn=CDH23;refGe
ne_tx=XM_023240533.1;refGene_gn=CDH23;refGene_tx=XM_023240536.1
D2 34610253 34610373
refGene_gn=CDH23;refGene_tx=XM_019813229.2;refGene_gn=CDH23;refGene_tx
=XM_019813230.1;refGene_gn=CDH23;refGene_tx=XM_023240529.1;refGene_gn=
CDH23;refGene_tx=XM_023240530.1;refGene_gn=CDH23;refGene_tx=XM_0232405
31.1;refGene_gn=CDH23;refGene_tx=XM_023240532.1;refGene_gn=CDH23;refGe
ne_tx=XM_023240533.1;refGene_gn=CDH23;refGene_tx=XM_023240536.1
D2 34611909 34612087
refGene_gn=CDH23;refGene_tx=XM_019813229.2;refGene_gn=CDH23;refGene_tx
=XM_019813230.1;refGene_gn=CDH23;refGene_tx=XM_023240529.1;refGene_gn=
CDH23;refGene_tx=XM_023240530.1;refGene_gn=CDH23;refGene_tx=XM_0232405
31.1;refGene_gn=CDH23;refGene_tx=XM_023240532.1;refGene_gn=CDH23;refGe
ne_tx=XM_023240533.1;refGene_gn=CDH23;refGene_tx=XM_023240536.1
D2 34612214 34612426
refGene_gn=CDH23;refGene_tx=XM_019813229.2;refGene_gn=CDH23;refGene_tx
=XM_019813230.1;refGene_gn=CDH23;refGene_tx=XM_023240529.1;refGene_gn=
CDH23;refGene_tx=XM_023240530.1;refGene_gn=CDH23;refGene_tx=XM_0232405
31.1;refGene_gn=CDH23;refGene_tx=XM_023240532.1;refGene_gn=CDH23;refGe
ne_tx=XM_023240533.1;refGene_gn=CDH23;refGene_tx=XM_023240536.1
D2 34614453 34614645
refGene_gn=CDH23;refGene_tx=XM_019813229.2;refGene_gn=CDH23;refGene_tx
=XM_019813230.1;refGene_gn=CDH23;refGene_tx=XM_023240529.1;refGene_gn=
CDH23;refGene_tx=XM_023240530.1;refGene_gn=CDH23;refGene_tx=XM_0232405
31.1;refGene_gn=CDH23;refGene_tx=XM_023240532.1;refGene_gn=CDH23;refGe
ne_tx=XM_023240533.1;refGene_gn=CDH23;refGene_tx=XM_023240536.1
D2 34614799 34614913
refGene_gn=CDH23;refGene_tx=XM_019813229.2;refGene_gn=CDH23;refGene_tx
=XM_019813230.1;refGene_gn=CDH23;refGene_tx=XM_023240529.1;refGene_gn=
CDH23;refGene_tx=XM_023240530.1;refGene_gn=CDH23;refGene_tx=XM_0232405
31.1;refGene_gn=CDH23;refGene_tx=XM_023240532.1;refGene_gn=CDH23;refGe
ne_tx=XM_023240533.1;refGene_gn=CDH23;refGene_tx=XM_023240536.1
D2 34615766 34615896
refGene_gn=CDH23;refGene_tx=XM_019813229.2;refGene_gn=CDH23;refGene_tx
=XM_019813230.1;refGene_gn=CDH23;refGene_tx=XM_023240529.1;refGene_gn=
CDH23;refGene_tx=XM_023240530.1;refGene_gn=CDH23;refGene_tx=XM_0232405
31.1;refGene_gn=CDH23;refGene_tx=XM_023240532.1;refGene_gn=CDH23;refGe
ne_tx=XM_023240533.1;refGene_gn=CDH23;refGene_tx=XM_023240536.1
D2 34616004 34616256
refGene_gn=CDH23;refGene_tx=XM_019813229.2;refGene_gn=CDH23;refGene_tx
=XM_019813230.1;refGene_gn=CDH23;refGene_tx=XM_023240529.1;refGene_gn=
CDH23;refGene_tx=XM_023240530.1;refGene_gn=CDH23;refGene_tx=XM_0232405
31.1;refGene_gn=CDH23;refGene_tx=XM_023240532.1;refGene_gn=CDH23;refGe

ne_tx=XM_023240533.1;refGene_gn=CDH23;refGene_tx=XM_023240536.1
D2 34616336 34616498
refGene_gn=CDH23;refGene_tx=XM_019813229.2;refGene_gn=CDH23;refGene_tx=XM_019813230.1;refGene_gn=CDH23;refGene_tx=XM_023240529.1;refGene_gn=CDH23;refGene_tx=XM_023240530.1;refGene_gn=CDH23;refGene_tx=XM_023240531.1;refGene_gn=CDH23;refGene_tx=XM_023240532.1;refGene_gn=CDH23;refGene_tx=XM_023240533.1;refGene_gn=CDH23;refGene_tx=XM_023240536.1
D2 34618016 34618273
refGene_gn=CDH23;refGene_tx=XM_019813229.2;refGene_gn=CDH23;refGene_tx=XM_019813230.1;refGene_gn=CDH23;refGene_tx=XM_023240529.1;refGene_gn=CDH23;refGene_tx=XM_023240530.1;refGene_gn=CDH23;refGene_tx=XM_023240531.1;refGene_gn=CDH23;refGene_tx=XM_023240532.1;refGene_gn=CDH23;refGene_tx=XM_023240533.1;refGene_gn=CDH23;refGene_tx=XM_023240536.1
D2 34618634 34618732
refGene_gn=CDH23;refGene_tx=XM_019813229.2;refGene_gn=CDH23;refGene_tx=XM_019813230.1;refGene_gn=CDH23;refGene_tx=XM_023240529.1;refGene_gn=CDH23;refGene_tx=XM_023240530.1;refGene_gn=CDH23;refGene_tx=XM_023240531.1;refGene_gn=CDH23;refGene_tx=XM_023240532.1;refGene_gn=CDH23;refGene_tx=XM_023240533.1;refGene_gn=CDH23;refGene_tx=XM_023240536.1
D2 34619142 34619263
refGene_gn=CDH23;refGene_tx=XM_019813229.2;refGene_gn=CDH23;refGene_tx=XM_019813230.1;refGene_gn=CDH23;refGene_tx=XM_023240529.1;refGene_gn=CDH23;refGene_tx=XM_023240530.1;refGene_gn=CDH23;refGene_tx=XM_023240531.1;refGene_gn=CDH23;refGene_tx=XM_023240532.1;refGene_gn=CDH23;refGene_tx=XM_023240533.1;refGene_gn=CDH23;refGene_tx=XM_023240536.1
D2 34619347 34619427
refGene_gn=CDH23;refGene_tx=XM_019813229.2;refGene_gn=CDH23;refGene_tx=XM_019813230.1;refGene_gn=CDH23;refGene_tx=XM_023240529.1;refGene_gn=CDH23;refGene_tx=XM_023240530.1;refGene_gn=CDH23;refGene_tx=XM_023240531.1;refGene_gn=CDH23;refGene_tx=XM_023240532.1;refGene_gn=CDH23;refGene_tx=XM_023240533.1;refGene_gn=CDH23;refGene_tx=XM_023240536.1
D2 34619566 34619607
refGene_gn=CDH23;refGene_tx=XM_019813229.2;refGene_gn=CDH23;refGene_tx=XM_019813230.1;refGene_gn=CDH23;refGene_tx=XM_023240529.1;refGene_gn=CDH23;refGene_tx=XM_023240530.1;refGene_gn=CDH23;refGene_tx=XM_023240531.1;refGene_gn=CDH23;refGene_tx=XM_023240532.1;refGene_gn=CDH23;refGene_tx=XM_023240533.1;refGene_gn=CDH23;refGene_tx=XM_023240536.1
D2 34619775 34619851
refGene_gn=CDH23;refGene_tx=XM_019813229.2;refGene_gn=CDH23;refGene_tx=XM_019813230.1;refGene_gn=CDH23;refGene_tx=XM_023240529.1;refGene_gn=CDH23;refGene_tx=XM_023240530.1;refGene_gn=CDH23;refGene_tx=XM_023240531.1;refGene_gn=CDH23;refGene_tx=XM_023240532.1;refGene_gn=CDH23;refGene_tx=XM_023240533.1;refGene_gn=CDH23;refGene_tx=XM_023240536.1
D2 34620318 34620448
refGene_gn=CDH23;refGene_tx=XM_019813229.2;refGene_gn=CDH23;refGene_tx=XM_019813230.1;refGene_gn=CDH23;refGene_tx=XM_023240529.1;refGene_gn=CDH23;refGene_tx=XM_023240530.1;refGene_gn=CDH23;refGene_tx=XM_023240531.1;refGene_gn=CDH23;refGene_tx=XM_023240532.1;refGene_gn=CDH23;refGene_tx=XM_023240533.1;refGene_gn=CDH23;refGene_tx=XM_023240536.1
D2 34620650 34620773

refGene_gn=CDH23;refGene_tx=XM_019813229.2;refGene_gn=CDH23;refGene_tx=XM_019813230.1;refGene_gn=CDH23;refGene_tx=XM_023240529.1;refGene_gn=CDH23;refGene_tx=XM_023240530.1;refGene_gn=CDH23;refGene_tx=XM_023240531.1;refGene_gn=CDH23;refGene_tx=XM_023240532.1;refGene_gn=CDH23;refGene_tx=XM_023240533.1;refGene_gn=CDH23;refGene_tx=XM_023240536.1
D2 34621139 34621244 refGene_gn=CDH23;refGene_tx=XM_023240536.1
D2 34621632 34621665 refGene_gn=CDH23;refGene_tx=XM_023240533.1
D2 34622122 34622449
refGene_gn=CDH23;refGene_tx=XM_019813229.2;refGene_gn=CDH23;refGene_tx=XM_019813230.1;refGene_gn=CDH23;refGene_tx=XM_023240529.1;refGene_gn=CDH23;refGene_tx=XM_023240530.1;refGene_gn=CDH23;refGene_tx=XM_023240531.1;refGene_gn=CDH23;refGene_tx=XM_023240532.1;refGene_gn=CDH23;refGene_tx=XM_023240536.1
D2 34624476 34624512
refGene_gn=PSAP;refGene_tx=XM_003994031.4;refGene_gn=PSAP;refGene_tx=XM_003994032.4;refGene_gn=PSAP;refGene_tx=XM_006937852.3
D2 34625596 34625704
refGene_gn=PSAP;refGene_tx=XM_003994031.4;refGene_gn=PSAP;refGene_tx=XM_003994032.4;refGene_gn=PSAP;refGene_tx=XM_006937852.3
D2 34625949 34626030
refGene_gn=PSAP;refGene_tx=XM_003994031.4;refGene_gn=PSAP;refGene_tx=XM_003994032.4;refGene_gn=PSAP;refGene_tx=XM_006937852.3
D2 34626367 34626525
refGene_gn=PSAP;refGene_tx=XM_003994031.4;refGene_gn=PSAP;refGene_tx=XM_003994032.4;refGene_gn=PSAP;refGene_tx=XM_006937852.3
D2 34626608 34626795
refGene_gn=PSAP;refGene_tx=XM_003994031.4;refGene_gn=PSAP;refGene_tx=XM_003994032.4;refGene_gn=PSAP;refGene_tx=XM_006937852.3
D2 34627094 34627190
refGene_gn=PSAP;refGene_tx=XM_003994031.4;refGene_gn=PSAP;refGene_tx=XM_003994032.4;refGene_gn=PSAP;refGene_tx=XM_006937852.3
D2 34628652 34628784
refGene_gn=PSAP;refGene_tx=XM_003994031.4;refGene_gn=PSAP;refGene_tx=XM_003994032.4;refGene_gn=PSAP;refGene_tx=XM_006937852.3
D2 34630551 34630560
refGene_gn=PSAP;refGene_tx=XM_003994032.4;refGene_gn=PSAP;refGene_tx=XM_006937852.3
D2 34632564 34632621
refGene_gn=PSAP;refGene_tx=XM_003994031.4;refGene_gn=PSAP;refGene_tx=XM_003994032.4;refGene_gn=PSAP;refGene_tx=XM_006937852.3
D2 34633947 34634091
refGene_gn=PSAP;refGene_tx=XM_003994031.4;refGene_gn=PSAP;refGene_tx=XM_003994032.4;refGene_gn=PSAP;refGene_tx=XM_006937852.3
D2 34634650 34634851
refGene_gn=PSAP;refGene_tx=XM_003994031.4;refGene_gn=PSAP;refGene_tx=XM_003994032.4;refGene_gn=PSAP;refGene_tx=XM_006937852.3
D2 34635959 34636085
refGene_gn=PSAP;refGene_tx=XM_003994031.4;refGene_gn=PSAP;refGene_tx=XM_003994032.4;refGene_gn=PSAP;refGene_tx=XM_006937852.3
D2 34636669 34636744

refGene_gn=PSAP;refGene_tx=XM_003994031.4;refGene_gn=PSAP;refGene_tx=XM_003994032.4;refGene_gn=PSAP;refGene_tx=XM_006937852.3
D2 34639538 34639672
refGene_gn=PSAP;refGene_tx=XM_003994031.4;refGene_gn=PSAP;refGene_tx=XM_003994032.4;refGene_gn=PSAP;refGene_tx=XM_006937852.3
D2 34657823 34657863
refGene_gn=PSAP;refGene_tx=XM_003994031.4;refGene_gn=PSAP;refGene_tx=XM_003994032.4;refGene_gn=PSAP;refGene_tx=XM_006937852.3
D2 34742603 34742743
refGene_gn=CHST3;refGene_tx=XM_003994033.4;refGene_gn=CHST3;refGene_tx=XM_019813242.2
D2 34744241 34745541
refGene_gn=CHST3;refGene_tx=XM_003994033.4;refGene_gn=CHST3;refGene_tx=XM_019813242.2
D2 34803468 34803614 refGene_gn=SPOCK2;refGene_tx=XM_023240537.1
D2 34803741 34803879 refGene_gn=SPOCK2;refGene_tx=XM_023240537.1
D2 34805277 34805340 refGene_gn=SPOCK2;refGene_tx=XM_023240537.1
D2 34806692 34806911 refGene_gn=SPOCK2;refGene_tx=XM_023240537.1
D2 34807382 34807499 refGene_gn=SPOCK2;refGene_tx=XM_023240537.1
D2 34807844 34807959 refGene_gn=SPOCK2;refGene_tx=XM_023240537.1
D2 34809769 34809884 refGene_gn=SPOCK2;refGene_tx=XM_023240537.1
D2 34811166 34811281 refGene_gn=SPOCK2;refGene_tx=XM_023240537.1
D2 34811523 34811569 refGene_gn=SPOCK2;refGene_tx=XM_023240537.1
D2 34811911 34811920 refGene_gn=SPOCK2;refGene_tx=XM_023240537.1
D2 34825772 34825961 refGene_gn=SPOCK2;refGene_tx=XM_023240537.1
D2 34835108 34835225
refGene_gn=ASCC1;refGene_tx=XM_006937855.4;refGene_gn=ASCC1;refGene_tx=XM_019813249.1;refGene_gn=ASCC1;refGene_tx=XM_023240540.1
D2 34841416 34841557
refGene_gn=ASCC1;refGene_tx=XM_019813244.2;refGene_gn=ASCC1;refGene_tx=XM_019813245.2;refGene_gn=ASCC1;refGene_tx=XM_023240538.1;refGene_gn=ASCC1;refGene_tx=XM_023240539.1
D2 34849772 34849853 refGene_gn=ASCC1;refGene_tx=XM_019813246.2
D2 34857624 34857710
refGene_gn=ASCC1;refGene_tx=XM_006937855.4;refGene_gn=ASCC1;refGene_tx=XM_019813244.2;refGene_gn=ASCC1;refGene_tx=XM_019813245.2;refGene_gn=ASCC1;refGene_tx=XM_019813246.2;refGene_gn=ASCC1;refGene_tx=XM_023240538.1;refGene_gn=ASCC1;refGene_tx=XM_023240539.1;refGene_gn=ASCC1;refGene_tx=XM_023240540.1
D2 34863327 34863452
refGene_gn=ASCC1;refGene_tx=XM_006937855.4;refGene_gn=ASCC1;refGene_tx=XM_019813244.2;refGene_gn=ASCC1;refGene_tx=XM_019813245.2;refGene_gn=ASCC1;refGene_tx=XM_019813246.2;refGene_gn=ASCC1;refGene_tx=XM_019813249.1;refGene_gn=ASCC1;refGene_tx=XM_023240538.1;refGene_gn=ASCC1;refGene_tx=XM_023240539.1;refGene_gn=ASCC1;refGene_tx=XM_023240540.1
D2 34884808 34884928
refGene_gn=ASCC1;refGene_tx=XM_006937855.4;refGene_gn=ASCC1;refGene_tx=XM_019813244.2;refGene_gn=ASCC1;refGene_tx=XM_019813245.2;refGene_gn=ASCC1;refGene_tx=XM_019813246.2;refGene_gn=ASCC1;refGene_tx=XM_019813249.1;refGene_gn=ASCC1;refGene_tx=XM_023240538.1;refGene_gn=ASCC1;refGe

ne_tx=XM_023240539.1;refGene_gn=ASCC1;refGene_tx=XM_023240540.1
D2 34892309 34892446
refGene_gn=ASCC1;refGene_tx=XM_006937855.4;refGene_gn=ASCC1;refGene_tx=XM_019813244.2;refGene_gn=ASCC1;refGene_tx=XM_019813245.2;refGene_gn=ASCC1;refGene_tx=XM_019813246.2;refGene_gn=ASCC1;refGene_tx=XM_019813249.1;refGene_gn=ASCC1;refGene_tx=XM_023240538.1;refGene_gn=ASCC1;refGene_tx=XM_023240539.1;refGene_gn=ASCC1;refGene_tx=XM_023240540.1
D2 34924821 34925000
refGene_gn=ASCC1;refGene_tx=XM_006937855.4;refGene_gn=ASCC1;refGene_tx=XM_019813244.2;refGene_gn=ASCC1;refGene_tx=XM_019813245.2;refGene_gn=ASCC1;refGene_tx=XM_019813246.2;refGene_gn=ASCC1;refGene_tx=XM_019813249.1;refGene_gn=ASCC1;refGene_tx=XM_023240538.1;refGene_gn=ASCC1;refGene_tx=XM_023240539.1;refGene_gn=ASCC1;refGene_tx=XM_023240540.1
D2 34934504 34934602
refGene_gn=ASCC1;refGene_tx=XM_006937855.4;refGene_gn=ASCC1;refGene_tx=XM_019813244.2;refGene_gn=ASCC1;refGene_tx=XM_019813245.2;refGene_gn=ASCC1;refGene_tx=XM_019813246.2;refGene_gn=ASCC1;refGene_tx=XM_019813249.1;refGene_gn=ASCC1;refGene_tx=XM_023240538.1;refGene_gn=ASCC1;refGene_tx=XM_023240539.1;refGene_gn=ASCC1;refGene_tx=XM_023240540.1
D2 34940276 34940376
refGene_gn=ASCC1;refGene_tx=XM_006937855.4;refGene_gn=ASCC1;refGene_tx=XM_019813244.2;refGene_gn=ASCC1;refGene_tx=XM_019813245.2;refGene_gn=ASCC1;refGene_tx=XM_019813246.2;refGene_gn=ASCC1;refGene_tx=XM_019813249.1;refGene_gn=ASCC1;refGene_tx=XM_023240538.1;refGene_gn=ASCC1;refGene_tx=XM_023240539.1;refGene_gn=ASCC1;refGene_tx=XM_023240540.1
D2 34941910 34942022
refGene_gn=ASCC1;refGene_tx=XM_006937855.4;refGene_gn=ASCC1;refGene_tx=XM_019813244.2;refGene_gn=ASCC1;refGene_tx=XM_019813245.2;refGene_gn=ASCC1;refGene_tx=XM_019813246.2;refGene_gn=ASCC1;refGene_tx=XM_019813249.1;refGene_gn=ASCC1;refGene_tx=XM_023240538.1;refGene_gn=ASCC1;refGene_tx=XM_023240539.1;refGene_gn=ASCC1;refGene_tx=XM_023240540.1
D2 34948359 34948501 refGene_gn=ANAPC16;refGene_tx=XM_011287350.2
D2 34948950 34949083 refGene_gn=ANAPC16;refGene_tx=XM_006937854.1
D2 34954360 34954435
refGene_gn=ANAPC16;refGene_tx=XM_006937854.1;refGene_gn=ANAPC16;refGene_tx=XM_011287350.2
D2 34955215 34955331
refGene_gn=ANAPC16;refGene_tx=XM_006937854.1;refGene_gn=ANAPC16;refGene_tx=XM_011287350.2
D2 34983410 34983612
refGene_gn=DDIT4;refGene_tx=XM_003994039.4;refGene_gn=DDIT4;refGene_tx=XM_006937856.3
D2 34983898 34984392
refGene_gn=DDIT4;refGene_tx=XM_003994039.4;refGene_gn=DDIT4;refGene_tx=XM_006937856.3
D2 35032379 35032409 refGene_gn=DNAJB12;refGene_tx=XM_006937857.2
D2 35033588 35033710
refGene_gn=DNAJB12;refGene_tx=XM_003994040.4;refGene_gn=DNAJB12;refGene_tx=XM_006937857.2;refGene_gn=DNAJB12;refGene_tx=XM_006937858.2
D2 35034266 35034439

refGene_gn=DNAJB12;refGene_tx=XM_003994040.4;refGene_gn=DNAJB12;refGene_tx=XM_006937857.2;refGene_gn=DNAJB12;refGene_tx=XM_006937858.2
D2 35036058 35036168
refGene_gn=DNAJB12;refGene_tx=XM_003994040.4;refGene_gn=DNAJB12;refGene_tx=XM_006937857.2;refGene_gn=DNAJB12;refGene_tx=XM_006937858.2
D2 35037259 35037339
refGene_gn=DNAJB12;refGene_tx=XM_003994040.4;refGene_gn=DNAJB12;refGene_tx=XM_006937857.2;refGene_gn=DNAJB12;refGene_tx=XM_006937858.2
D2 35037465 35037651
refGene_gn=DNAJB12;refGene_tx=XM_003994040.4;refGene_gn=DNAJB12;refGene_tx=XM_006937857.2;refGene_gn=DNAJB12;refGene_tx=XM_006937858.2
D2 35039029 35039175
refGene_gn=DNAJB12;refGene_tx=XM_003994040.4;refGene_gn=DNAJB12;refGene_tx=XM_006937857.2;refGene_gn=DNAJB12;refGene_tx=XM_006937858.2
D2 35039874 35040046
refGene_gn=DNAJB12;refGene_tx=XM_003994040.4;refGene_gn=DNAJB12;refGene_tx=XM_006937857.2;refGene_gn=DNAJB12;refGene_tx=XM_006937858.2
D2 35048467 35048600
refGene_gn=DNAJB12;refGene_tx=XM_003994040.4;refGene_gn=DNAJB12;refGene_tx=XM_006937857.2;refGene_gn=DNAJB12;refGene_tx=XM_006937858.2
D2 35058905 35059066
refGene_gn=MICU1;refGene_tx=XM_003994041.3;refGene_gn=MICU1;refGene_tx=XM_006937859.2;refGene_gn=MICU1;refGene_tx=XM_006937861.2;refGene_gn=MICU1;refGene_tx=XM_019813250.1;refGene_gn=MICU1;refGene_tx=XM_019813251.1;refGene_gn=MICU1;refGene_tx=XM_019813253.1;refGene_gn=MICU1;refGene_tx=XM_023240541.1;refGene_gn=MICU1;refGene_tx=XM_023240542.1
D2 35063667 35063757
refGene_gn=MICU1;refGene_tx=XM_003994041.3;refGene_gn=MICU1;refGene_tx=XM_006937859.2;refGene_gn=MICU1;refGene_tx=XM_006937861.2;refGene_gn=MICU1;refGene_tx=XM_019813250.1;refGene_gn=MICU1;refGene_tx=XM_019813251.1;refGene_gn=MICU1;refGene_tx=XM_019813253.1;refGene_gn=MICU1;refGene_tx=XM_023240541.1;refGene_gn=MICU1;refGene_tx=XM_023240542.1
D2 35083204 35083313
refGene_gn=MICU1;refGene_tx=XM_003994041.3;refGene_gn=MICU1;refGene_tx=XM_006937859.2;refGene_gn=MICU1;refGene_tx=XM_006937861.2;refGene_gn=MICU1;refGene_tx=XM_019813250.1;refGene_gn=MICU1;refGene_tx=XM_019813251.1;refGene_gn=MICU1;refGene_tx=XM_019813253.1;refGene_gn=MICU1;refGene_tx=XM_023240541.1;refGene_gn=MICU1;refGene_tx=XM_023240542.1
D2 35097037 35097175
refGene_gn=MICU1;refGene_tx=XM_003994041.3;refGene_gn=MICU1;refGene_tx=XM_006937859.2;refGene_gn=MICU1;refGene_tx=XM_006937861.2;refGene_gn=MICU1;refGene_tx=XM_019813250.1;refGene_gn=MICU1;refGene_tx=XM_019813251.1;refGene_gn=MICU1;refGene_tx=XM_019813253.1;refGene_gn=MICU1;refGene_tx=XM_023240541.1;refGene_gn=MICU1;refGene_tx=XM_023240542.1
D2 35132464 35132662
refGene_gn=MICU1;refGene_tx=XM_003994041.3;refGene_gn=MICU1;refGene_tx=XM_006937859.2;refGene_gn=MICU1;refGene_tx=XM_006937861.2;refGene_gn=MICU1;refGene_tx=XM_019813250.1;refGene_gn=MICU1;refGene_tx=XM_019813251.1;refGene_gn=MICU1;refGene_tx=XM_019813253.1;refGene_gn=MICU1;refGene_tx=XM_023240541.1;refGene_gn=MICU1;refGene_tx=XM_023240542.1

D2 35145570 35145653
refGene_gn=MICU1;refGene_tx=XM_003994041.3;refGene_gn=MICU1;refGene_tx=XM_006937859.2;refGene_gn=MICU1;refGene_tx=XM_006937861.2;refGene_gn=MICU1;refGene_tx=XM_019813250.1;refGene_gn=MICU1;refGene_tx=XM_019813251.1;refGene_gn=MICU1;refGene_tx=XM_019813253.1;refGene_gn=MICU1;refGene_tx=XM_023240541.1;refGene_gn=MICU1;refGene_tx=XM_023240542.1
D2 35174730 35174845
refGene_gn=MICU1;refGene_tx=XM_003994041.3;refGene_gn=MICU1;refGene_tx=XM_006937859.2;refGene_gn=MICU1;refGene_tx=XM_006937861.2;refGene_gn=MICU1;refGene_tx=XM_019813250.1;refGene_gn=MICU1;refGene_tx=XM_019813251.1;refGene_gn=MICU1;refGene_tx=XM_023240541.1;refGene_gn=MICU1;refGene_tx=XM_023240542.1
D2 35176116 35176122
refGene_gn=MICU1;refGene_tx=XM_006937859.2;refGene_gn=MICU1;refGene_tx=XM_019813251.1
D2 35179354 35179381 refGene_gn=MICU1;refGene_tx=XM_023240542.1
D2 35191227 35191239
refGene_gn=MICU1;refGene_tx=XM_006937859.2;refGene_gn=MICU1;refGene_tx=XM_006937861.2
D2 35200528 35200702
refGene_gn=MICU1;refGene_tx=XM_019813250.1;refGene_gn=MICU1;refGene_tx=XM_023240541.1
D2 35201306 35201350
refGene_gn=MICU1;refGene_tx=XM_003994041.3;refGene_gn=MICU1;refGene_tx=XM_006937859.2;refGene_gn=MICU1;refGene_tx=XM_006937861.2;refGene_gn=MICU1;refGene_tx=XM_019813250.1;refGene_gn=MICU1;refGene_tx=XM_019813251.1;refGene_gn=MICU1;refGene_tx=XM_023240541.1
D2 35223698 35223861
refGene_gn=MICU1;refGene_tx=XM_003994041.3;refGene_gn=MICU1;refGene_tx=XM_006937859.2;refGene_gn=MICU1;refGene_tx=XM_006937861.2;refGene_gn=MICU1;refGene_tx=XM_019813250.1;refGene_gn=MICU1;refGene_tx=XM_019813251.1;refGene_gn=MICU1;refGene_tx=XM_019813253.1;refGene_gn=MICU1;refGene_tx=XM_023240541.1
D2 35235210 35235385
refGene_gn=MICU1;refGene_tx=XM_003994041.3;refGene_gn=MICU1;refGene_tx=XM_006937859.2;refGene_gn=MICU1;refGene_tx=XM_006937861.2;refGene_gn=MICU1;refGene_tx=XM_019813250.1;refGene_gn=MICU1;refGene_tx=XM_019813251.1;refGene_gn=MICU1;refGene_tx=XM_019813253.1;refGene_gn=MICU1;refGene_tx=XM_023240541.1
D2 35263580 35263741
refGene_gn=MICU1;refGene_tx=XM_003994041.3;refGene_gn=MICU1;refGene_tx=XM_006937859.2;refGene_gn=MICU1;refGene_tx=XM_006937861.2;refGene_gn=MICU1;refGene_tx=XM_019813250.1;refGene_gn=MICU1;refGene_tx=XM_019813251.1;refGene_gn=MICU1;refGene_tx=XM_019813253.1;refGene_gn=MICU1;refGene_tx=XM_023240541.1
D2 35373418 35373475 refGene_gn=MCU;refGene_tx=XM_023240544.1
D2 35373476 35373566 refGene_gn=MCU;refGene_tx=XM_023240544.1
D2 35529881 35529951 refGene_gn=MCU;refGene_tx=XM_023240544.1
D2 35550982 35551153 refGene_gn=MCU;refGene_tx=XM_023240544.1
D2 35552381 35552486 refGene_gn=MCU;refGene_tx=XM_023240544.1

D2 35563597 35563758 refGene_gn=MCU;refGene_tx=XM_023240544.1
D2 35565188 35565392 refGene_gn=MCU;refGene_tx=XM_023240544.1
D2 35570902 35571019 refGene_gn=MCU;refGene_tx=XM_023240544.1
D2 35573184 35573262 refGene_gn=MCU;refGene_tx=XM_023240544.1
D2 35579513 35579574 refGene_gn=OIT3;refGene_tx=XM_023240543.1
D2 35583684 35584059
refGene_gn=OIT3;refGene_tx=XM_019813255.1;refGene_gn=OIT3;refGene_tx=XM_023240543.1
D2 35585285 35585393
refGene_gn=OIT3;refGene_tx=XM_019813255.1;refGene_gn=OIT3;refGene_tx=XM_023240543.1
D2 35591149 35591272
refGene_gn=OIT3;refGene_tx=XM_019813255.1;refGene_gn=OIT3;refGene_tx=XM_023240543.1
D2 35595507 35595630
refGene_gn=OIT3;refGene_tx=XM_019813255.1;refGene_gn=OIT3;refGene_tx=XM_023240543.1
D2 35597353 35597514
refGene_gn=OIT3;refGene_tx=XM_019813255.1;refGene_gn=OIT3;refGene_tx=XM_023240543.1
D2 35599399 35599815
refGene_gn=OIT3;refGene_tx=XM_019813255.1;refGene_gn=OIT3;refGene_tx=XM_023240543.1
D2 35604507 35604607
refGene_gn=OIT3;refGene_tx=XM_019813255.1;refGene_gn=OIT3;refGene_tx=XM_023240543.1
D2 35606275 35606449
refGene_gn=OIT3;refGene_tx=XM_019813255.1;refGene_gn=OIT3;refGene_tx=XM_023240543.1
D2 35608585 35608707
refGene_gn=PLA2G12B;refGene_tx=XM_003994046.4;refGene_gn=PLA2G12B;refGene_tx=XM_019813261.1;refGene_gn=PLA2G12B;refGene_tx=XM_019813262.1
D2 35614636 35614802
refGene_gn=PLA2G12B;refGene_tx=XM_003994046.4;refGene_gn=PLA2G12B;refGene_tx=XM_019813261.1;refGene_gn=PLA2G12B;refGene_tx=XM_019813262.1
D2 35626521 35626610
refGene_gn=PLA2G12B;refGene_tx=XM_003994046.4;refGene_gn=PLA2G12B;refGene_tx=XM_019813261.1;refGene_gn=PLA2G12B;refGene_tx=XM_019813262.1
D2 35638799 35639010
refGene_gn=PLA2G12B;refGene_tx=XM_003994046.4;refGene_gn=PLA2G12B;refGene_tx=XM_019813261.1;refGene_gn=PLA2G12B;refGene_tx=XM_019813262.1
D2 35699507 35699578
refGene_gn=P4HA1;refGene_tx=XM_011287347.3;refGene_gn=P4HA1;refGene_tx=XM_011287348.3
D2 35700996 35701093
refGene_gn=P4HA1;refGene_tx=XM_011287347.3;refGene_gn=P4HA1;refGene_tx=XM_011287348.3
D2 35701613 35701682
refGene_gn=P4HA1;refGene_tx=XM_011287347.3;refGene_gn=P4HA1;refGene_tx=XM_011287348.3

D2 35705539 35705605
refGene_gn=P4HA1;refGene_tx=XM_011287347.3;refGene_gn=P4HA1;refGene_tx
=XM_011287348.3
D2 35708600 35708654
refGene_gn=P4HA1;refGene_tx=XM_011287347.3;refGene_gn=P4HA1;refGene_tx
=XM_011287348.3
D2 35721937 35722037
refGene_gn=P4HA1;refGene_tx=XM_011287347.3;refGene_gn=P4HA1;refGene_tx
=XM_011287348.3
D2 35736696 35736767 refGene_gn=P4HA1;refGene_tx=XM_011287347.3
D2 35737789 35737860 refGene_gn=P4HA1;refGene_tx=XM_011287348.3
D2 35739917 35740094
refGene_gn=P4HA1;refGene_tx=XM_011287347.3;refGene_gn=P4HA1;refGene_tx
=XM_011287348.3
D2 35742633 35742830
refGene_gn=P4HA1;refGene_tx=XM_011287347.3;refGene_gn=P4HA1;refGene_tx
=XM_011287348.3
D2 35745364 35745604
refGene_gn=P4HA1;refGene_tx=XM_011287347.3;refGene_gn=P4HA1;refGene_tx
=XM_011287348.3
D2 35765303 35765441
refGene_gn=P4HA1;refGene_tx=XM_011287347.3;refGene_gn=P4HA1;refGene_tx
=XM_011287348.3
D2 35768347 35768499
refGene_gn=P4HA1;refGene_tx=XM_011287347.3;refGene_gn=P4HA1;refGene_tx
=XM_011287348.3
D2 35770091 35770188
refGene_gn=P4HA1;refGene_tx=XM_011287347.3;refGene_gn=P4HA1;refGene_tx
=XM_011287348.3
D2 35771698 35771774
refGene_gn=P4HA1;refGene_tx=XM_011287347.3;refGene_gn=P4HA1;refGene_tx
=XM_011287348.3
D2 35804526 35804576
refGene_gn=ECD;refGene_tx=XM_003994047.5;refGene_gn=ECD;refGene_tx=XM_
019813263.2;refGene_gn=ECD;refGene_tx=XM_019813264.2;refGene_gn=ECD;re
fGene_tx=XM_019813265.2
D2 35806272 35806277 refGene_gn=NUDT13;refGene_tx=XM_019813275.1
D2 35808386 35808469
refGene_gn=NUDT13;refGene_tx=XM_003994049.5;refGene_gn=NUDT13;refGene_
tx=XM_019813273.1;refGene_gn=NUDT13;refGene_tx=XM_019813274.2;refGene_
gn=NUDT13;refGene_tx=XM_019813275.1;refGene_gn=NUDT13;refGene_tx=XM_02
3240547.1
D2 35811722 35811862
refGene_gn=NUDT13;refGene_tx=XM_003994049.5;refGene_gn=NUDT13;refGene_
tx=XM_019813273.1;refGene_gn=NUDT13;refGene_tx=XM_019813274.2;refGene_
gn=NUDT13;refGene_tx=XM_019813275.1;refGene_gn=NUDT13;refGene_tx=XM_02
3240547.1
D2 35813609 35813744
refGene_gn=NUDT13;refGene_tx=XM_003994049.5;refGene_gn=NUDT13;refGene_
tx=XM_019813273.1;refGene_gn=NUDT13;refGene_tx=XM_019813274.2;refGene_

gn=NUDT13;refGene_tx=XM_019813275.1;refGene_gn=NUDT13;refGene_tx=XM_023240547.1
D2 35814561 35814668
refGene_gn=NUDT13;refGene_tx=XM_003994049.5;refGene_gn=NUDT13;refGene_tx=XM_019813273.1;refGene_gn=NUDT13;refGene_tx=XM_019813274.2;refGene_gn=NUDT13;refGene_tx=XM_019813275.1;refGene_gn=NUDT13;refGene_tx=XM_019813276.2;refGene_gn=NUDT13;refGene_tx=XM_023240547.1
D2 35816711 35816837
refGene_gn=NUDT13;refGene_tx=XM_003994049.5;refGene_gn=NUDT13;refGene_tx=XM_019813273.1;refGene_gn=NUDT13;refGene_tx=XM_019813274.2;refGene_gn=NUDT13;refGene_tx=XM_019813275.1;refGene_gn=NUDT13;refGene_tx=XM_019813276.2;refGene_gn=NUDT13;refGene_tx=XM_023240547.1
D2 35817001 35817113
refGene_gn=NUDT13;refGene_tx=XM_003994049.5;refGene_gn=NUDT13;refGene_tx=XM_019813273.1;refGene_gn=NUDT13;refGene_tx=XM_019813274.2;refGene_gn=NUDT13;refGene_tx=XM_019813275.1;refGene_gn=NUDT13;refGene_tx=XM_019813276.2;refGene_gn=NUDT13;refGene_tx=XM_023240547.1
D2 35818163 35818174 refGene_gn=NUDT13;refGene_tx=XM_023240547.1
D2 35818194 35818349
refGene_gn=NUDT13;refGene_tx=XM_003994049.5;refGene_gn=NUDT13;refGene_tx=XM_019813273.1;refGene_gn=NUDT13;refGene_tx=XM_019813274.2;refGene_gn=NUDT13;refGene_tx=XM_019813275.1;refGene_gn=NUDT13;refGene_tx=XM_019813276.2
D2 35819299 35819494
refGene_gn=ECD;refGene_tx=XM_023240545.1;refGene_gn=NUDT13;refGene_tx=XM_003994049.5;refGene_gn=NUDT13;refGene_tx=XM_019813273.1;refGene_gn=NUDT13;refGene_tx=XM_019813274.2;refGene_gn=NUDT13;refGene_tx=XM_019813275.1;refGene_gn=NUDT13;refGene_tx=XM_019813276.2
D2 35821681 35821912
refGene_gn=ECD;refGene_tx=XM_003994047.5;refGene_gn=ECD;refGene_tx=XM_019813263.2;refGene_gn=ECD;refGene_tx=XM_019813264.2;refGene_gn=ECD;refGene_tx=XM_019813265.2;refGene_gn=ECD;refGene_tx=XM_023240545.1;refGene_gn=ECD;refGene_tx=XM_023240546.1
D2 35824418 35824633
refGene_gn=ECD;refGene_tx=XM_003994047.5;refGene_gn=ECD;refGene_tx=XM_019813263.2;refGene_gn=ECD;refGene_tx=XM_019813264.2;refGene_gn=ECD;refGene_tx=XM_019813265.2;refGene_gn=ECD;refGene_tx=XM_023240545.1;refGene_gn=ECD;refGene_tx=XM_023240546.1
D2 35826127 35826195
refGene_gn=ECD;refGene_tx=XM_003994047.5;refGene_gn=ECD;refGene_tx=XM_019813263.2;refGene_gn=ECD;refGene_tx=XM_019813264.2;refGene_gn=ECD;refGene_tx=XM_019813265.2;refGene_gn=ECD;refGene_tx=XM_023240545.1;refGene_gn=ECD;refGene_tx=XM_023240546.1
D2 35827798 35827982
refGene_gn=ECD;refGene_tx=XM_003994047.5;refGene_gn=ECD;refGene_tx=XM_019813263.2;refGene_gn=ECD;refGene_tx=XM_019813264.2;refGene_gn=ECD;refGene_tx=XM_019813265.2;refGene_gn=ECD;refGene_tx=XM_023240545.1;refGene_gn=ECD;refGene_tx=XM_023240546.1
D2 35828116 35828223
refGene_gn=ECD;refGene_tx=XM_003994047.5;refGene_gn=ECD;refGene_tx=XM_

019813263.2;refGene_gn=ECD;refGene_tx=XM_019813264.2;refGene_gn=ECD;refGene_tx=XM_019813265.2;refGene_gn=ECD;refGene_tx=XM_023240545.1;refGene_gn=ECD;refGene_tx=XM_023240546.1
D2 35838139 35838225
refGene_gn=ECD;refGene_tx=XM_003994047.5;refGene_gn=ECD;refGene_tx=XM_019813263.2;refGene_gn=ECD;refGene_tx=XM_019813264.2;refGene_gn=ECD;refGene_tx=XM_019813265.2;refGene_gn=ECD;refGene_tx=XM_023240545.1;refGene_gn=ECD;refGene_tx=XM_023240546.1
D2 35840122 35840251
refGene_gn=ECD;refGene_tx=XM_003994047.5;refGene_gn=ECD;refGene_tx=XM_019813263.2;refGene_gn=ECD;refGene_tx=XM_019813264.2;refGene_gn=ECD;refGene_tx=XM_019813265.2;refGene_gn=ECD;refGene_tx=XM_023240545.1;refGene_gn=ECD;refGene_tx=XM_023240546.1
D2 35842327 35842456
refGene_gn=ECD;refGene_tx=XM_003994047.5;refGene_gn=ECD;refGene_tx=XM_019813263.2;refGene_gn=ECD;refGene_tx=XM_019813264.2;refGene_gn=ECD;refGene_tx=XM_019813265.2;refGene_gn=ECD;refGene_tx=XM_023240545.1;refGene_gn=ECD;refGene_tx=XM_023240546.1
D2 35843002 35843195
refGene_gn=ECD;refGene_tx=XM_003994047.5;refGene_gn=ECD;refGene_tx=XM_019813263.2;refGene_gn=ECD;refGene_tx=XM_019813264.2;refGene_gn=ECD;refGene_tx=XM_019813265.2;refGene_gn=ECD;refGene_tx=XM_023240545.1;refGene_gn=ECD;refGene_tx=XM_023240546.1
D2 35844154 35844333
refGene_gn=ECD;refGene_tx=XM_003994047.5;refGene_gn=ECD;refGene_tx=XM_019813263.2;refGene_gn=ECD;refGene_tx=XM_019813264.2;refGene_gn=ECD;refGene_tx=XM_019813265.2;refGene_gn=ECD;refGene_tx=XM_023240545.1;refGene_gn=ECD;refGene_tx=XM_023240546.1
D2 35844437 35844525
refGene_gn=ECD;refGene_tx=XM_003994047.5;refGene_gn=ECD;refGene_tx=XM_019813263.2;refGene_gn=ECD;refGene_tx=XM_019813264.2;refGene_gn=ECD;refGene_tx=XM_019813265.2;refGene_gn=ECD;refGene_tx=XM_023240545.1;refGene_gn=ECD;refGene_tx=XM_023240546.1
D2 35847142 35847260
refGene_gn=ECD;refGene_tx=XM_003994047.5;refGene_gn=ECD;refGene_tx=XM_019813263.2;refGene_gn=ECD;refGene_tx=XM_019813264.2;refGene_gn=ECD;refGene_tx=XM_019813265.2;refGene_gn=ECD;refGene_tx=XM_023240545.1;refGene_gn=ECD;refGene_tx=XM_023240546.1
D2 35850266 35850471
refGene_gn=ECD;refGene_tx=XM_003994047.5;refGene_gn=ECD;refGene_tx=XM_019813263.2;refGene_gn=ECD;refGene_tx=XM_019813264.2;refGene_gn=ECD;refGene_tx=XM_019813265.2;refGene_gn=ECD;refGene_tx=XM_023240545.1;refGene_gn=ECD;refGene_tx=XM_023240546.1
D2 35852334 35852381
refGene_gn=FAM149B1;refGene_tx=XM_011287343.3;refGene_gn=FAM149B1;refGene_tx=XM_019813266.2;refGene_gn=FAM149B1;refGene_tx=XM_019813268.2;refGene_gn=FAM149B1;refGene_tx=XM_019813269.2;refGene_gn=FAM149B1;refGene_tx=XM_019813270.2;refGene_gn=FAM149B1;refGene_tx=XM_019813272.2
D2 35858477 35858582
refGene_gn=FAM149B1;refGene_tx=XM_011287343.3;refGene_gn=FAM149B1;refG

ene_tx=XM_019813266.2;refGene_gn=FAM149B1;refGene_tx=XM_019813268.2;refGene_gn=FAM149B1;refGene_tx=XM_019813269.2;refGene_gn=FAM149B1;refGene_tx=XM_019813270.2;refGene_gn=FAM149B1;refGene_tx=XM_019813271.2;refGene_gn=FAM149B1;refGene_tx=XM_019813272.2
D2 35860519 35860649
refGene_gn=FAM149B1;refGene_tx=XM_011287343.3;refGene_gn=FAM149B1;refGene_tx=XM_019813266.2;refGene_gn=FAM149B1;refGene_tx=XM_019813268.2;refGene_gn=FAM149B1;refGene_tx=XM_019813269.2;refGene_gn=FAM149B1;refGene_tx=XM_019813270.2;refGene_gn=FAM149B1;refGene_tx=XM_019813271.2;refGene_gn=FAM149B1;refGene_tx=XM_019813272.2
D2 35887346 35887489
refGene_gn=FAM149B1;refGene_tx=XM_011287343.3;refGene_gn=FAM149B1;refGene_tx=XM_019813266.2;refGene_gn=FAM149B1;refGene_tx=XM_019813268.2;refGene_gn=FAM149B1;refGene_tx=XM_019813269.2;refGene_gn=FAM149B1;refGene_tx=XM_019813270.2;refGene_gn=FAM149B1;refGene_tx=XM_019813271.2;refGene_gn=FAM149B1;refGene_tx=XM_019813272.2
D2 35888559 35888676
refGene_gn=FAM149B1;refGene_tx=XM_011287343.3;refGene_gn=FAM149B1;refGene_tx=XM_019813266.2;refGene_gn=FAM149B1;refGene_tx=XM_019813268.2;refGene_gn=FAM149B1;refGene_tx=XM_019813269.2;refGene_gn=FAM149B1;refGene_tx=XM_019813270.2;refGene_gn=FAM149B1;refGene_tx=XM_019813271.2;refGene_gn=FAM149B1;refGene_tx=XM_019813272.2
D2 35891050 35891218
refGene_gn=FAM149B1;refGene_tx=XM_011287343.3;refGene_gn=FAM149B1;refGene_tx=XM_019813266.2;refGene_gn=FAM149B1;refGene_tx=XM_019813268.2;refGene_gn=FAM149B1;refGene_tx=XM_019813269.2;refGene_gn=FAM149B1;refGene_tx=XM_019813270.2;refGene_gn=FAM149B1;refGene_tx=XM_019813271.2;refGene_gn=FAM149B1;refGene_tx=XM_019813272.2
D2 35894159 35894347
refGene_gn=FAM149B1;refGene_tx=XM_011287343.3;refGene_gn=FAM149B1;refGene_tx=XM_019813266.2;refGene_gn=FAM149B1;refGene_tx=XM_019813268.2;refGene_gn=FAM149B1;refGene_tx=XM_019813269.2;refGene_gn=FAM149B1;refGene_tx=XM_019813270.2;refGene_gn=FAM149B1;refGene_tx=XM_019813271.2
D2 35927803 35927928
refGene_gn=FAM149B1;refGene_tx=XM_011287343.3;refGene_gn=FAM149B1;refGene_tx=XM_019813266.2;refGene_gn=FAM149B1;refGene_tx=XM_019813268.2;refGene_gn=FAM149B1;refGene_tx=XM_019813269.2;refGene_gn=FAM149B1;refGene_tx=XM_019813270.2;refGene_gn=FAM149B1;refGene_tx=XM_019813271.2
D2 35929500 35929607
refGene_gn=FAM149B1;refGene_tx=XM_011287343.3;refGene_gn=FAM149B1;refGene_tx=XM_019813266.2;refGene_gn=FAM149B1;refGene_tx=XM_019813268.2;refGene_gn=FAM149B1;refGene_tx=XM_019813269.2;refGene_gn=FAM149B1;refGene_tx=XM_019813270.2;refGene_gn=FAM149B1;refGene_tx=XM_019813271.2
D2 35933094 35933319
refGene_gn=FAM149B1;refGene_tx=XM_011287343.3;refGene_gn=FAM149B1;refGene_tx=XM_019813266.2;refGene_gn=FAM149B1;refGene_tx=XM_019813268.2;refGene_gn=FAM149B1;refGene_tx=XM_019813269.2;refGene_gn=FAM149B1;refGene_tx=XM_019813270.2;refGene_gn=FAM149B1;refGene_tx=XM_019813271.2;refGene_gn=FAM149B1;refGene_tx=XM_019813272.2
D2 35934049 35934112

refGene_gn=FAM149B1;refGene_tx=XM_019813266.2;refGene_gn=FAM149B1;refGene_tx=XM_019813268.2;refGene_gn=FAM149B1;refGene_tx=XM_019813269.2;refGene_gn=FAM149B1;refGene_tx=XM_019813271.2;refGene_gn=FAM149B1;refGene_tx=XM_019813272.2
D2	35934596	35934720	refGene_gn=FAM149B1;refGene_tx=XM_011287343.3;refGene_gn=FAM149B1;refGene_tx=XM_019813266.2;refGene_gn=FAM149B1;refGene_tx=XM_019813268.2;refGene_gn=FAM149B1;refGene_tx=XM_019813269.2;refGene_gn=FAM149B1;refGene_tx=XM_019813270.2;refGene_gn=FAM149B1;refGene_tx=XM_019813271.2;refGene_gn=FAM149B1;refGene_tx=XM_019813272.2
D2	35934966	35935092	refGene_gn=FAM149B1;refGene_tx=XM_011287343.3;refGene_gn=FAM149B1;refGene_tx=XM_019813266.2;refGene_gn=FAM149B1;refGene_tx=XM_019813268.2;refGene_gn=FAM149B1;refGene_tx=XM_019813269.2;refGene_gn=FAM149B1;refGene_tx=XM_019813270.2;refGene_gn=FAM149B1;refGene_tx=XM_019813271.2;refGene_gn=FAM149B1;refGene_tx=XM_019813272.2
D2	35937631	35937826	refGene_gn=FAM149B1;refGene_tx=XM_011287343.3;refGene_gn=FAM149B1;refGene_tx=XM_019813266.2;refGene_gn=FAM149B1;refGene_tx=XM_019813268.2;refGene_gn=FAM149B1;refGene_tx=XM_019813269.2;refGene_gn=FAM149B1;refGene_tx=XM_019813270.2;refGene_gn=FAM149B1;refGene_tx=XM_019813271.2;refGene_gn=FAM149B1;refGene_tx=XM_019813272.2
D2	35938800	35938874	refGene_gn=FAM149B1;refGene_tx=XM_011287343.3;refGene_gn=FAM149B1;refGene_tx=XM_019813269.2
D2	35941269	35941389	refGene_gn=DNAJC9;refGene_tx=XM_003994050.5
D2	35941750	35941837	refGene_gn=DNAJC9;refGene_tx=XM_003994050.5
D2	35943790	35944045	refGene_gn=DNAJC9;refGene_tx=XM_003994050.5
D2	35944578	35944719	refGene_gn=DNAJC9;refGene_tx=XM_003994050.5
D2	35944892	35945072	refGene_gn=DNAJC9;refGene_tx=XM_003994050.5
D2	35947620	35947754	refGene_gn=MRPS16;refGene_tx=XM_003994051.4;refGene_gn=MRPS16;refGene_tx=XM_019813277.1
D2	35947953	35948214	refGene_gn=MRPS16;refGene_tx=XM_003994051.4;refGene_gn=MRPS16;refGene_tx=XM_019813277.1
D2	35948658	35948932	refGene_gn=MRPS16;refGene_tx=XM_003994051.4;refGene_gn=MRPS16;refGene_tx=XM_019813277.1
D2	35950237	35950318	refGene_gn=CFAP70;refGene_tx=XM_003994052.4;refGene_gn=CFAP70;refGene_tx=XM_011287342.3;refGene_gn=CFAP70;refGene_tx=XM_019813278.2;refGene_gn=CFAP70;refGene_tx=XM_019813279.2
D2	35952016	35952064	refGene_gn=CFAP70;refGene_tx=XM_003994052.4;refGene_gn=CFAP70;refGene_tx=XM_011287342.3;refGene_gn=CFAP70;refGene_tx=XM_019813278.2;refGene_gn=CFAP70;refGene_tx=XM_019813279.2
D2	35964162	35964264	refGene_gn=CFAP70;refGene_tx=XM_003994052.4;refGene_gn=CFAP70;refGene_tx=XM_011287342.3;refGene_gn=CFAP70;refGene_tx=XM_019813278.2;refGene_

gn=CFAP70;refGene_tx=XM_019813279.2
D2 35968462 35968552
refGene_gn=CFAP70;refGene_tx=XM_003994052.4;refGene_gn=CFAP70;refGene_tx=XM_011287342.3;refGene_gn=CFAP70;refGene_tx=XM_019813278.2;refGene_gn=CFAP70;refGene_tx=XM_019813279.2
D2 35970207 35970369
refGene_gn=CFAP70;refGene_tx=XM_003994052.4;refGene_gn=CFAP70;refGene_tx=XM_011287342.3;refGene_gn=CFAP70;refGene_tx=XM_019813278.2;refGene_gn=CFAP70;refGene_tx=XM_019813279.2
D2 35970896 35971049
refGene_gn=CFAP70;refGene_tx=XM_003994052.4;refGene_gn=CFAP70;refGene_tx=XM_011287342.3;refGene_gn=CFAP70;refGene_tx=XM_019813278.2;refGene_gn=CFAP70;refGene_tx=XM_019813279.2
D2 35976888 35977010
refGene_gn=CFAP70;refGene_tx=XM_003994052.4;refGene_gn=CFAP70;refGene_tx=XM_011287342.3;refGene_gn=CFAP70;refGene_tx=XM_019813278.2;refGene_gn=CFAP70;refGene_tx=XM_019813279.2
D2 35977755 35977914
refGene_gn=CFAP70;refGene_tx=XM_003994052.4;refGene_gn=CFAP70;refGene_tx=XM_019813278.2;refGene_gn=CFAP70;refGene_tx=XM_019813279.2
D2 35985508 35985730
refGene_gn=CFAP70;refGene_tx=XM_003994052.4;refGene_gn=CFAP70;refGene_tx=XM_011287342.3;refGene_gn=CFAP70;refGene_tx=XM_019813278.2;refGene_gn=CFAP70;refGene_tx=XM_019813279.2
D2 35985928 35986044
refGene_gn=CFAP70;refGene_tx=XM_003994052.4;refGene_gn=CFAP70;refGene_tx=XM_011287342.3;refGene_gn=CFAP70;refGene_tx=XM_019813278.2;refGene_gn=CFAP70;refGene_tx=XM_019813279.2
D2 35986165 35986299
refGene_gn=CFAP70;refGene_tx=XM_003994052.4;refGene_gn=CFAP70;refGene_tx=XM_011287342.3;refGene_gn=CFAP70;refGene_tx=XM_019813278.2;refGene_gn=CFAP70;refGene_tx=XM_019813279.2
D2 35992047 35992173
refGene_gn=CFAP70;refGene_tx=XM_003994052.4;refGene_gn=CFAP70;refGene_tx=XM_011287342.3;refGene_gn=CFAP70;refGene_tx=XM_019813278.2;refGene_gn=CFAP70;refGene_tx=XM_019813279.2
D2 35997416 35997548
refGene_gn=CFAP70;refGene_tx=XM_003994052.4;refGene_gn=CFAP70;refGene_tx=XM_011287342.3;refGene_gn=CFAP70;refGene_tx=XM_019813278.2;refGene_gn=CFAP70;refGene_tx=XM_019813279.2
D2 35999031 35999226
refGene_gn=CFAP70;refGene_tx=XM_003994052.4;refGene_gn=CFAP70;refGene_tx=XM_011287342.3;refGene_gn=CFAP70;refGene_tx=XM_019813278.2;refGene_gn=CFAP70;refGene_tx=XM_019813279.2
D2 35999542 35999603
refGene_gn=CFAP70;refGene_tx=XM_003994052.4;refGene_gn=CFAP70;refGene_tx=XM_011287342.3;refGene_gn=CFAP70;refGene_tx=XM_019813278.2;refGene_gn=CFAP70;refGene_tx=XM_019813279.2
D2 36004917 36005009
refGene_gn=CFAP70;refGene_tx=XM_003994052.4;refGene_gn=CFAP70;refGene_

tx=XM_011287342.3;refGene_gn=CFAP70;refGene_tx=XM_019813278.2;refGene_gn=CFAP70;refGene_tx=XM_019813279.2
D2 36006587 36006668 refGene_gn=CFAP70;refGene_tx=XM_003994052.4;refGene_gn=CFAP70;refGene_tx=XM_011287342.3;refGene_gn=CFAP70;refGene_tx=XM_019813278.2;refGene_gn=CFAP70;refGene_tx=XM_019813279.2
D2 36006848 36007019 refGene_gn=CFAP70;refGene_tx=XM_003994052.4;refGene_gn=CFAP70;refGene_tx=XM_011287342.3;refGene_gn=CFAP70;refGene_tx=XM_019813278.2;refGene_gn=CFAP70;refGene_tx=XM_019813279.2
D2 36014443 36014578 refGene_gn=CFAP70;refGene_tx=XM_003994052.4;refGene_gn=CFAP70;refGene_tx=XM_011287342.3;refGene_gn=CFAP70;refGene_tx=XM_019813278.2;refGene_gn=CFAP70;refGene_tx=XM_019813279.2
D2 36017325 36017425 refGene_gn=CFAP70;refGene_tx=XM_003994052.4;refGene_gn=CFAP70;refGene_tx=XM_011287342.3;refGene_gn=CFAP70;refGene_tx=XM_019813278.2;refGene_gn=CFAP70;refGene_tx=XM_019813279.2
D2 36021766 36021861 refGene_gn=CFAP70;refGene_tx=XM_003994052.4;refGene_gn=CFAP70;refGene_tx=XM_011287342.3;refGene_gn=CFAP70;refGene_tx=XM_019813278.2;refGene_gn=CFAP70;refGene_tx=XM_019813279.2
D2 36024820 36025003 refGene_gn=CFAP70;refGene_tx=XM_003994052.4;refGene_gn=CFAP70;refGene_tx=XM_011287342.3;refGene_gn=CFAP70;refGene_tx=XM_019813278.2;refGene_gn=CFAP70;refGene_tx=XM_019813279.2
D2 36027392 36027572 refGene_gn=CFAP70;refGene_tx=XM_003994052.4;refGene_gn=CFAP70;refGene_tx=XM_011287342.3;refGene_gn=CFAP70;refGene_tx=XM_019813278.2;refGene_gn=CFAP70;refGene_tx=XM_019813279.2
D2 36031266 36031346 refGene_gn=CFAP70;refGene_tx=XM_003994052.4;refGene_gn=CFAP70;refGene_tx=XM_011287342.3;refGene_gn=CFAP70;refGene_tx=XM_019813278.2;refGene_gn=CFAP70;refGene_tx=XM_019813279.2
D2 36031506 36031605 refGene_gn=CFAP70;refGene_tx=XM_003994052.4;refGene_gn=CFAP70;refGene_tx=XM_011287342.3;refGene_gn=CFAP70;refGene_tx=XM_019813278.2;refGene_gn=CFAP70;refGene_tx=XM_019813279.2
D2 36051206 36051393 refGene_gn=CFAP70;refGene_tx=XM_003994052.4;refGene_gn=CFAP70;refGene_tx=XM_011287342.3;refGene_gn=CFAP70;refGene_tx=XM_019813278.2;refGene_gn=CFAP70;refGene_tx=XM_019813279.2
D2 36052384 36052447 refGene_gn=CFAP70;refGene_tx=XM_003994052.4;refGene_gn=CFAP70;refGene_tx=XM_011287342.3;refGene_gn=CFAP70;refGene_tx=XM_019813278.2;refGene_gn=CFAP70;refGene_tx=XM_019813279.2
D2 36060935 36061058 refGene_gn=ANXA7;refGene_tx=XM_006937868.4
D2 36062743 36062856 refGene_gn=ANXA7;refGene_tx=XM_006937868.4
D2 36063893 36063969 refGene_gn=ANXA7;refGene_tx=XM_006937868.4
D2 36064048 36064219 refGene_gn=ANXA7;refGene_tx=XM_006937868.4

D2 36066415 36066586 refGene_gn=ANXA7;refGene_tx=XM_006937868.4
D2 36066816 36066930 refGene_gn=ANXA7;refGene_tx=XM_006937868.4
D2 36070518 36070613 refGene_gn=ANXA7;refGene_tx=XM_006937868.4
D2 36071099 36071202 refGene_gn=ANXA7;refGene_tx=XM_006937868.4
D2 36076478 36076544 refGene_gn=ANXA7;refGene_tx=XM_006937868.4
D2 36076951 36077016 refGene_gn=ANXA7;refGene_tx=XM_006937868.4
D2 36078025 36078139 refGene_gn=ANXA7;refGene_tx=XM_006937868.4
D2 36078746 36078771 refGene_gn=ANXA7;refGene_tx=XM_006937868.4
D2 36078772 36078952 refGene_gn=ANXA7;refGene_tx=XM_006937868.4
D2 36080700 36080754 refGene_gn=ANXA7;refGene_tx=XM_006937868.4
D2 36098523 36098743 refGene_gn=MSS51;refGene_tx=XM_003994057.4
D2 36099334 36099428 refGene_gn=MSS51;refGene_tx=XM_003994057.4
D2 36099769 36100336 refGene_gn=MSS51;refGene_tx=XM_003994057.4
D2 36100535 36100660 refGene_gn=MSS51;refGene_tx=XM_003994057.4
D2 36101675 36101831 refGene_gn=MSS51;refGene_tx=XM_003994057.4
D2 36102107 36102328 refGene_gn=MSS51;refGene_tx=XM_003994057.4
D2 36111180 36111359
refGene_gn=PPP3CB;refGene_tx=XM_023240548.1;refGene_gn=PPP3CB;refGene_tx=XM_023240549.1;refGene_gn=PPP3CB;refGene_tx=XM_023240550.1;refGene_gn=PPP3CB;refGene_tx=XM_023240551.1;refGene_gn=PPP3CB;refGene_tx=XM_023240552.1;refGene_gn=PPP3CB;refGene_tx=XM_023240553.1;refGene_gn=PPP3CB;refGene_tx=XM_023240554.1
D2 36112845 36112875
refGene_gn=PPP3CB;refGene_tx=XM_023240548.1;refGene_gn=PPP3CB;refGene_tx=XM_023240549.1;refGene_gn=PPP3CB;refGene_tx=XM_023240550.1;refGene_gn=PPP3CB;refGene_tx=XM_023240551.1
D2 36117556 36117654
refGene_gn=PPP3CB;refGene_tx=XM_023240548.1;refGene_gn=PPP3CB;refGene_tx=XM_023240549.1;refGene_gn=PPP3CB;refGene_tx=XM_023240550.1;refGene_gn=PPP3CB;refGene_tx=XM_023240551.1;refGene_gn=PPP3CB;refGene_tx=XM_023240552.1;refGene_gn=PPP3CB;refGene_tx=XM_023240553.1;refGene_gn=PPP3CB;refGene_tx=XM_023240554.1
D2 36118626 36118711
refGene_gn=PPP3CB;refGene_tx=XM_023240548.1;refGene_gn=PPP3CB;refGene_tx=XM_023240549.1;refGene_gn=PPP3CB;refGene_tx=XM_023240550.1;refGene_gn=PPP3CB;refGene_tx=XM_023240551.1;refGene_gn=PPP3CB;refGene_tx=XM_023240552.1;refGene_gn=PPP3CB;refGene_tx=XM_023240553.1;refGene_gn=PPP3CB;refGene_tx=XM_023240554.1
D2 36120785 36120812
refGene_gn=PPP3CB;refGene_tx=XM_023240548.1;refGene_gn=PPP3CB;refGene_tx=XM_023240549.1;refGene_gn=PPP3CB;refGene_tx=XM_023240552.1
D2 36125804 36125882
refGene_gn=PPP3CB;refGene_tx=XM_023240548.1;refGene_gn=PPP3CB;refGene_tx=XM_023240549.1;refGene_gn=PPP3CB;refGene_tx=XM_023240550.1;refGene_gn=PPP3CB;refGene_tx=XM_023240551.1;refGene_gn=PPP3CB;refGene_tx=XM_023240552.1;refGene_gn=PPP3CB;refGene_tx=XM_023240553.1;refGene_gn=PPP3CB;refGene_tx=XM_023240554.1
D2 36143786 36143912
refGene_gn=PPP3CB;refGene_tx=XM_023240548.1;refGene_gn=PPP3CB;refGene_tx=XM_023240549.1;refGene_gn=PPP3CB;refGene_tx=XM_023240550.1;refGene_

gn=PPP3CB;refGene_tx=XM_023240551.1;refGene_gn=PPP3CB;refGene_tx=XM_023240552.1;refGene_gn=PPP3CB;refGene_tx=XM_023240553.1;refGene_gn=PPP3CB;refGene_tx=XM_023240554.1
D2 36146659 36146754
refGene_gn=PPP3CB;refGene_tx=XM_023240548.1;refGene_gn=PPP3CB;refGene_tx=XM_023240549.1;refGene_gn=PPP3CB;refGene_tx=XM_023240550.1;refGene_gn=PPP3CB;refGene_tx=XM_023240551.1;refGene_gn=PPP3CB;refGene_tx=XM_023240552.1;refGene_gn=PPP3CB;refGene_tx=XM_023240553.1;refGene_gn=PPP3CB;refGene_tx=XM_023240554.1
D2 36146860 36146938
refGene_gn=PPP3CB;refGene_tx=XM_023240548.1;refGene_gn=PPP3CB;refGene_tx=XM_023240549.1;refGene_gn=PPP3CB;refGene_tx=XM_023240550.1;refGene_gn=PPP3CB;refGene_tx=XM_023240551.1;refGene_gn=PPP3CB;refGene_tx=XM_023240552.1;refGene_gn=PPP3CB;refGene_tx=XM_023240553.1;refGene_gn=PPP3CB;refGene_tx=XM_023240554.1
D2 36147041 36147181
refGene_gn=PPP3CB;refGene_tx=XM_023240548.1;refGene_gn=PPP3CB;refGene_tx=XM_023240549.1;refGene_gn=PPP3CB;refGene_tx=XM_023240550.1;refGene_gn=PPP3CB;refGene_tx=XM_023240551.1;refGene_gn=PPP3CB;refGene_tx=XM_023240552.1;refGene_gn=PPP3CB;refGene_tx=XM_023240553.1;refGene_gn=PPP3CB;refGene_tx=XM_023240554.1
D2 36147448 36147594
refGene_gn=PPP3CB;refGene_tx=XM_023240548.1;refGene_gn=PPP3CB;refGene_tx=XM_023240549.1;refGene_gn=PPP3CB;refGene_tx=XM_023240550.1;refGene_gn=PPP3CB;refGene_tx=XM_023240551.1;refGene_gn=PPP3CB;refGene_tx=XM_023240552.1;refGene_gn=PPP3CB;refGene_tx=XM_023240553.1;refGene_gn=PPP3CB;refGene_tx=XM_023240554.1
D2 36149494 36149606
refGene_gn=PPP3CB;refGene_tx=XM_023240548.1;refGene_gn=PPP3CB;refGene_tx=XM_023240549.1;refGene_gn=PPP3CB;refGene_tx=XM_023240550.1;refGene_gn=PPP3CB;refGene_tx=XM_023240551.1;refGene_gn=PPP3CB;refGene_tx=XM_023240552.1;refGene_gn=PPP3CB;refGene_tx=XM_023240553.1;refGene_gn=PPP3CB;refGene_tx=XM_023240554.1
D2 36152289 36152414
refGene_gn=PPP3CB;refGene_tx=XM_023240548.1;refGene_gn=PPP3CB;refGene_tx=XM_023240549.1;refGene_gn=PPP3CB;refGene_tx=XM_023240550.1;refGene_gn=PPP3CB;refGene_tx=XM_023240551.1;refGene_gn=PPP3CB;refGene_tx=XM_023240552.1;refGene_gn=PPP3CB;refGene_tx=XM_023240553.1;refGene_gn=PPP3CB;refGene_tx=XM_023240554.1
D2 36153095 36153296
refGene_gn=PPP3CB;refGene_tx=XM_023240548.1;refGene_gn=PPP3CB;refGene_tx=XM_023240549.1;refGene_gn=PPP3CB;refGene_tx=XM_023240550.1;refGene_gn=PPP3CB;refGene_tx=XM_023240551.1;refGene_gn=PPP3CB;refGene_tx=XM_023240552.1;refGene_gn=PPP3CB;refGene_tx=XM_023240553.1;refGene_gn=PPP3CB;refGene_tx=XM_023240554.1
D2 36168032 36168117
refGene_gn=PPP3CB;refGene_tx=XM_023240548.1;refGene_gn=PPP3CB;refGene_tx=XM_023240549.1;refGene_gn=PPP3CB;refGene_tx=XM_023240550.1;refGene_gn=PPP3CB;refGene_tx=XM_023240551.1;refGene_gn=PPP3CB;refGene_tx=XM_023240552.1;refGene_gn=PPP3CB;refGene_tx=XM_023240553.1;refGene_gn=PPP3C

B;refGene_tx=XM_023240554.1
D2 36170713 36171258
refGene_gn=USP54;refGene_tx=XM_019813281.2;refGene_gn=USP54;refGene_tx=XM_019813282.2;refGene_gn=USP54;refGene_tx=XM_019813283.2;refGene_gn=USP54;refGene_tx=XM_019813284.2;refGene_gn=USP54;refGene_tx=XM_019813287.2;refGene_gn=USP54;refGene_tx=XM_019813288.2;refGene_gn=USP54;refGene_tx=XM_019813294.1;refGene_gn=USP54;refGene_tx=XM_023240555.1;refGene_gn=USP54;refGene_tx=XM_023240556.1;refGene_gn=USP54;refGene_tx=XM_023240557.1;refGene_gn=USP54;refGene_tx=XM_023240558.1;refGene_gn=USP54;refGene_tx=XM_023240559.1;refGene_gn=USP54;refGene_tx=XM_023240560.1;refGene_gn=USP54;refGene_tx=XM_023240561.1;refGene_gn=USP54;refGene_tx=XM_023240562.1;refGene_gn=USP54;refGene_tx=XM_023240563.1
D2 36172885 36173072
refGene_gn=USP54;refGene_tx=XM_019813281.2;refGene_gn=USP54;refGene_tx=XM_019813282.2;refGene_gn=USP54;refGene_tx=XM_019813283.2;refGene_gn=USP54;refGene_tx=XM_019813284.2;refGene_gn=USP54;refGene_tx=XM_019813287.2;refGene_gn=USP54;refGene_tx=XM_019813288.2;refGene_gn=USP54;refGene_tx=XM_019813294.1;refGene_gn=USP54;refGene_tx=XM_023240555.1;refGene_gn=USP54;refGene_tx=XM_023240556.1;refGene_gn=USP54;refGene_tx=XM_023240557.1;refGene_gn=USP54;refGene_tx=XM_023240558.1;refGene_gn=USP54;refGene_tx=XM_023240559.1;refGene_gn=USP54;refGene_tx=XM_023240560.1;refGene_gn=USP54;refGene_tx=XM_023240561.1;refGene_gn=USP54;refGene_tx=XM_023240562.1;refGene_gn=USP54;refGene_tx=XM_023240564.1
D2 36176356 36176497
refGene_gn=USP54;refGene_tx=XM_019813281.2;refGene_gn=USP54;refGene_tx=XM_019813282.2;refGene_gn=USP54;refGene_tx=XM_019813283.2;refGene_gn=USP54;refGene_tx=XM_019813284.2;refGene_gn=USP54;refGene_tx=XM_019813287.2;refGene_gn=USP54;refGene_tx=XM_019813288.2;refGene_gn=USP54;refGene_tx=XM_019813294.1;refGene_gn=USP54;refGene_tx=XM_023240555.1;refGene_gn=USP54;refGene_tx=XM_023240556.1;refGene_gn=USP54;refGene_tx=XM_023240557.1;refGene_gn=USP54;refGene_tx=XM_023240558.1;refGene_gn=USP54;refGene_tx=XM_023240560.1;refGene_gn=USP54;refGene_tx=XM_023240561.1;refGene_gn=USP54;refGene_tx=XM_023240562.1
D2 36176814 36176933
refGene_gn=USP54;refGene_tx=XM_019813281.2;refGene_gn=USP54;refGene_tx=XM_019813282.2;refGene_gn=USP54;refGene_tx=XM_019813283.2;refGene_gn=USP54;refGene_tx=XM_019813284.2;refGene_gn=USP54;refGene_tx=XM_019813287.2;refGene_gn=USP54;refGene_tx=XM_019813288.2;refGene_gn=USP54;refGene_tx=XM_019813294.1;refGene_gn=USP54;refGene_tx=XM_023240555.1;refGene_gn=USP54;refGene_tx=XM_023240556.1;refGene_gn=USP54;refGene_tx=XM_023240557.1;refGene_gn=USP54;refGene_tx=XM_023240558.1;refGene_gn=USP54;refGene_tx=XM_023240559.1;refGene_gn=USP54;refGene_tx=XM_023240560.1;refGene_gn=USP54;refGene_tx=XM_023240561.1;refGene_gn=USP54;refGene_tx=XM_023240562.1;refGene_gn=USP54;refGene_tx=XM_023240563.1
D2 36185210 36186604
refGene_gn=USP54;refGene_tx=XM_019813281.2;refGene_gn=USP54;refGene_tx=XM_019813282.2;refGene_gn=USP54;refGene_tx=XM_019813283.2;refGene_gn=USP54;refGene_tx=XM_019813284.2;refGene_gn=USP54;refGene_tx=XM_019813287.2;refGene_gn=USP54;refGene_tx=XM_019813288.2;refGene_gn=USP54;refGene_tx=XM_019813294.1;refGene_gn=USP54;refGene_tx=XM_023240555.1;refGen

e_gn=USP54;refGene_tx=XM_023240556.1;refGene_gn=USP54;refGene_tx=XM_02
3240557.1;refGene_gn=USP54;refGene_tx=XM_023240558.1;refGene_gn=USP54;
refGene_tx=XM_023240559.1;refGene_gn=USP54;refGene_tx=XM_023240560.1;r
efGene_gn=USP54;refGene_tx=XM_023240561.1;refGene_gn=USP54;refGene_tx=
XM_023240562.1;refGene_gn=USP54;refGene_tx=XM_023240563.1;refGene_gn=U
SP54;refGene_tx=XM_023240564.1;refGene_gn=USP54;refGene_tx=XM_02324056
5.1;refGene_gn=USP54;refGene_tx=XM_023240566.1
D2 36188392 36188591
refGene_gn=USP54;refGene_tx=XM_019813281.2;refGene_gn=USP54;refGene_tx
=XM_019813282.2;refGene_gn=USP54;refGene_tx=XM_019813283.2;refGene_gn=
USP54;refGene_tx=XM_019813284.2;refGene_gn=USP54;refGene_tx=XM_0198132
87.2;refGene_gn=USP54;refGene_tx=XM_019813288.2;refGene_gn=USP54;refGe
ne_tx=XM_019813294.1;refGene_gn=USP54;refGene_tx=XM_023240555.1;refGen
e_gn=USP54;refGene_tx=XM_023240556.1;refGene_gn=USP54;refGene_tx=XM_02
3240557.1;refGene_gn=USP54;refGene_tx=XM_023240558.1;refGene_gn=USP54;
refGene_tx=XM_023240559.1;refGene_gn=USP54;refGene_tx=XM_023240560.1;r
efGene_gn=USP54;refGene_tx=XM_023240561.1;refGene_gn=USP54;refGene_tx=
XM_023240562.1;refGene_gn=USP54;refGene_tx=XM_023240563.1;refGene_gn=U
SP54;refGene_tx=XM_023240564.1
D2 36189510 36189645
refGene_gn=USP54;refGene_tx=XM_019813281.2;refGene_gn=USP54;refGene_tx
=XM_019813282.2;refGene_gn=USP54;refGene_tx=XM_019813283.2;refGene_gn=
USP54;refGene_tx=XM_019813284.2;refGene_gn=USP54;refGene_tx=XM_0198132
87.2;refGene_gn=USP54;refGene_tx=XM_019813288.2;refGene_gn=USP54;refGe
ne_tx=XM_019813294.1;refGene_gn=USP54;refGene_tx=XM_023240555.1;refGen
e_gn=USP54;refGene_tx=XM_023240556.1;refGene_gn=USP54;refGene_tx=XM_02
3240557.1;refGene_gn=USP54;refGene_tx=XM_023240559.1;refGene_gn=USP54;
refGene_tx=XM_023240560.1;refGene_gn=USP54;refGene_tx=XM_023240561.1;r
efGene_gn=USP54;refGene_tx=XM_023240563.1;refGene_gn=USP54;refGene_tx=
XM_023240564.1;refGene_gn=USP54;refGene_tx=XM_023240565.1
D2 36192083 36192251
refGene_gn=USP54;refGene_tx=XM_019813281.2;refGene_gn=USP54;refGene_tx
=XM_019813282.2;refGene_gn=USP54;refGene_tx=XM_019813283.2;refGene_gn=
USP54;refGene_tx=XM_019813284.2;refGene_gn=USP54;refGene_tx=XM_0198132
87.2;refGene_gn=USP54;refGene_tx=XM_019813288.2;refGene_gn=USP54;refGe
ne_tx=XM_019813294.1;refGene_gn=USP54;refGene_tx=XM_023240555.1;refGen
e_gn=USP54;refGene_tx=XM_023240556.1;refGene_gn=USP54;refGene_tx=XM_02
3240557.1;refGene_gn=USP54;refGene_tx=XM_023240558.1;refGene_gn=USP54;
refGene_tx=XM_023240559.1;refGene_gn=USP54;refGene_tx=XM_023240560.1;r
efGene_gn=USP54;refGene_tx=XM_023240561.1;refGene_gn=USP54;refGene_tx=
XM_023240562.1;refGene_gn=USP54;refGene_tx=XM_023240563.1;refGene_gn=U
SP54;refGene_tx=XM_023240564.1;refGene_gn=USP54;refGene_tx=XM_02324056
5.1;refGene_gn=USP54;refGene_tx=XM_023240566.1
D2 36195103 36195237
refGene_gn=USP54;refGene_tx=XM_019813281.2;refGene_gn=USP54;refGene_tx
=XM_019813282.2;refGene_gn=USP54;refGene_tx=XM_019813283.2;refGene_gn=
USP54;refGene_tx=XM_019813284.2;refGene_gn=USP54;refGene_tx=XM_0198132
87.2;refGene_gn=USP54;refGene_tx=XM_019813288.2;refGene_gn=USP54;refGe
ne_tx=XM_019813294.1;refGene_gn=USP54;refGene_tx=XM_023240555.1;refGen
e_gn=USP54;refGene_tx=XM_023240556.1;refGene_gn=USP54;refGene_tx=XM_02

3240557.1;refGene_gn=USP54;refGene_tx=XM_023240558.1;refGene_gn=USP54;
refGene_tx=XM_023240559.1;refGene_gn=USP54;refGene_tx=XM_023240560.1;r
efGene_gn=USP54;refGene_tx=XM_023240561.1;refGene_gn=USP54;refGene_tx=
XM_023240562.1;refGene_gn=USP54;refGene_tx=XM_023240563.1;refGene_gn=U
SP54;refGene_tx=XM_023240564.1;refGene_gn=USP54;refGene_tx=XM_02324056
5.1;refGene_gn=USP54;refGene_tx=XM_023240566.1
D2 36197169 36197401
refGene_gn=USP54;refGene_tx=XM_019813281.2;refGene_gn=USP54;refGene_tx
=XM_019813282.2;refGene_gn=USP54;refGene_tx=XM_019813283.2;refGene_gn=
USP54;refGene_tx=XM_019813284.2;refGene_gn=USP54;refGene_tx=XM_0198132
87.2;refGene_gn=USP54;refGene_tx=XM_019813288.2;refGene_gn=USP54;refGe
ne_tx=XM_019813294.1;refGene_gn=USP54;refGene_tx=XM_023240555.1;refGen
e_gn=USP54;refGene_tx=XM_023240556.1;refGene_gn=USP54;refGene_tx=XM_02
3240557.1;refGene_gn=USP54;refGene_tx=XM_023240558.1;refGene_gn=USP54;
refGene_tx=XM_023240559.1;refGene_gn=USP54;refGene_tx=XM_023240560.1;r
efGene_gn=USP54;refGene_tx=XM_023240561.1;refGene_gn=USP54;refGene_tx=
XM_023240562.1;refGene_gn=USP54;refGene_tx=XM_023240563.1;refGene_gn=U
SP54;refGene_tx=XM_023240564.1;refGene_gn=USP54;refGene_tx=XM_02324056
5.1;refGene_gn=USP54;refGene_tx=XM_023240566.1
D2 36197651 36198032
refGene_gn=USP54;refGene_tx=XM_019813281.2;refGene_gn=USP54;refGene_tx
=XM_019813282.2;refGene_gn=USP54;refGene_tx=XM_019813283.2;refGene_gn=
USP54;refGene_tx=XM_019813284.2;refGene_gn=USP54;refGene_tx=XM_0198132
87.2;refGene_gn=USP54;refGene_tx=XM_019813288.2;refGene_gn=USP54;refGe
ne_tx=XM_019813294.1;refGene_gn=USP54;refGene_tx=XM_023240555.1;refGen
e_gn=USP54;refGene_tx=XM_023240556.1;refGene_gn=USP54;refGene_tx=XM_02
3240557.1;refGene_gn=USP54;refGene_tx=XM_023240558.1;refGene_gn=USP54;
refGene_tx=XM_023240559.1;refGene_gn=USP54;refGene_tx=XM_023240560.1;r
efGene_gn=USP54;refGene_tx=XM_023240561.1;refGene_gn=USP54;refGene_tx=
XM_023240562.1;refGene_gn=USP54;refGene_tx=XM_023240563.1;refGene_gn=U
SP54;refGene_tx=XM_023240564.1;refGene_gn=USP54;refGene_tx=XM_02324056
5.1;refGene_gn=USP54;refGene_tx=XM_023240566.1
D2 36198204 36198336
refGene_gn=USP54;refGene_tx=XM_019813281.2;refGene_gn=USP54;refGene_tx
=XM_019813282.2;refGene_gn=USP54;refGene_tx=XM_019813283.2;refGene_gn=
USP54;refGene_tx=XM_019813284.2;refGene_gn=USP54;refGene_tx=XM_0198132
87.2;refGene_gn=USP54;refGene_tx=XM_019813288.2;refGene_gn=USP54;refGe
ne_tx=XM_019813294.1;refGene_gn=USP54;refGene_tx=XM_023240555.1;refGen
e_gn=USP54;refGene_tx=XM_023240556.1;refGene_gn=USP54;refGene_tx=XM_02
3240557.1;refGene_gn=USP54;refGene_tx=XM_023240558.1;refGene_gn=USP54;
refGene_tx=XM_023240559.1;refGene_gn=USP54;refGene_tx=XM_023240560.1;r
efGene_gn=USP54;refGene_tx=XM_023240561.1;refGene_gn=USP54;refGene_tx=
XM_023240562.1;refGene_gn=USP54;refGene_tx=XM_023240563.1;refGene_gn=U
SP54;refGene_tx=XM_023240564.1;refGene_gn=USP54;refGene_tx=XM_02324056
5.1;refGene_gn=USP54;refGene_tx=XM_023240566.1
D2 36201699 36201870
refGene_gn=USP54;refGene_tx=XM_019813281.2;refGene_gn=USP54;refGene_tx
=XM_019813282.2;refGene_gn=USP54;refGene_tx=XM_019813283.2;refGene_gn=
USP54;refGene_tx=XM_019813284.2;refGene_gn=USP54;refGene_tx=XM_0198132
87.2;refGene_gn=USP54;refGene_tx=XM_019813288.2;refGene_gn=USP54;refGe

ne_tx=XM_019813294.1;refGene_gn=USP54;refGene_tx=XM_023240555.1;refGene_gn=USP54;refGene_tx=XM_023240556.1;refGene_gn=USP54;refGene_tx=XM_023240557.1;refGene_gn=USP54;refGene_tx=XM_023240558.1;refGene_gn=USP54;refGene_tx=XM_023240559.1;refGene_gn=USP54;refGene_tx=XM_023240563.1;refGene_gn=USP54;refGene_tx=XM_023240564.1;refGene_gn=USP54;refGene_tx=XM_023240565.1;refGene_gn=USP54;refGene_tx=XM_023240566.1
D2 36203226 36203395
refGene_gn=USP54;refGene_tx=XM_019813281.2;refGene_gn=USP54;refGene_tx=XM_019813282.2;refGene_gn=USP54;refGene_tx=XM_019813283.2;refGene_gn=USP54;refGene_tx=XM_019813284.2;refGene_gn=USP54;refGene_tx=XM_019813287.2;refGene_gn=USP54;refGene_tx=XM_019813288.2;refGene_gn=USP54;refGene_tx=XM_019813294.1;refGene_gn=USP54;refGene_tx=XM_023240555.1;refGene_gn=USP54;refGene_tx=XM_023240556.1;refGene_gn=USP54;refGene_tx=XM_023240557.1;refGene_gn=USP54;refGene_tx=XM_023240558.1;refGene_gn=USP54;refGene_tx=XM_023240559.1;refGene_gn=USP54;refGene_tx=XM_023240560.1;refGene_gn=USP54;refGene_tx=XM_023240561.1;refGene_gn=USP54;refGene_tx=XM_023240562.1;refGene_gn=USP54;refGene_tx=XM_023240563.1;refGene_gn=USP54;refGene_tx=XM_023240564.1;refGene_gn=USP54;refGene_tx=XM_023240565.1;refGene_gn=USP54;refGene_tx=XM_023240566.1
D2 36205205 36205355
refGene_gn=USP54;refGene_tx=XM_019813281.2;refGene_gn=USP54;refGene_tx=XM_019813282.2;refGene_gn=USP54;refGene_tx=XM_019813283.2;refGene_gn=USP54;refGene_tx=XM_019813284.2;refGene_gn=USP54;refGene_tx=XM_019813287.2;refGene_gn=USP54;refGene_tx=XM_019813288.2;refGene_gn=USP54;refGene_tx=XM_019813294.1;refGene_gn=USP54;refGene_tx=XM_023240555.1;refGene_gn=USP54;refGene_tx=XM_023240556.1;refGene_gn=USP54;refGene_tx=XM_023240557.1;refGene_gn=USP54;refGene_tx=XM_023240558.1;refGene_gn=USP54;refGene_tx=XM_023240559.1;refGene_gn=USP54;refGene_tx=XM_023240560.1;refGene_gn=USP54;refGene_tx=XM_023240561.1;refGene_gn=USP54;refGene_tx=XM_023240562.1;refGene_gn=USP54;refGene_tx=XM_023240563.1;refGene_gn=USP54;refGene_tx=XM_023240564.1;refGene_gn=USP54;refGene_tx=XM_023240565.1;refGene_gn=USP54;refGene_tx=XM_023240566.1
D2 36206674 36206821
refGene_gn=USP54;refGene_tx=XM_019813281.2;refGene_gn=USP54;refGene_tx=XM_019813282.2;refGene_gn=USP54;refGene_tx=XM_019813283.2;refGene_gn=USP54;refGene_tx=XM_019813284.2;refGene_gn=USP54;refGene_tx=XM_019813287.2;refGene_gn=USP54;refGene_tx=XM_019813288.2;refGene_gn=USP54;refGene_tx=XM_019813294.1;refGene_gn=USP54;refGene_tx=XM_023240555.1;refGene_gn=USP54;refGene_tx=XM_023240556.1;refGene_gn=USP54;refGene_tx=XM_023240557.1;refGene_gn=USP54;refGene_tx=XM_023240558.1;refGene_gn=USP54;refGene_tx=XM_023240559.1;refGene_gn=USP54;refGene_tx=XM_023240560.1;refGene_gn=USP54;refGene_tx=XM_023240561.1;refGene_gn=USP54;refGene_tx=XM_023240562.1;refGene_gn=USP54;refGene_tx=XM_023240563.1;refGene_gn=USP54;refGene_tx=XM_023240564.1;refGene_gn=USP54;refGene_tx=XM_023240565.1;refGene_gn=USP54;refGene_tx=XM_023240566.1
D2 36206932 36207038
refGene_gn=USP54;refGene_tx=XM_019813281.2;refGene_gn=USP54;refGene_tx=XM_019813282.2;refGene_gn=USP54;refGene_tx=XM_019813283.2;refGene_gn=USP54;refGene_tx=XM_019813284.2;refGene_gn=USP54;refGene_tx=XM_019813287.2;refGene_gn=USP54;refGene_tx=XM_019813288.2;refGene_gn=USP54;refGe

ne_tx=XM_019813294.1;refGene_gn=USP54;refGene_tx=XM_023240555.1;refGen
e_gn=USP54;refGene_tx=XM_023240556.1;refGene_gn=USP54;refGene_tx=XM_02
3240557.1;refGene_gn=USP54;refGene_tx=XM_023240558.1;refGene_gn=USP54;
refGene_tx=XM_023240559.1;refGene_gn=USP54;refGene_tx=XM_023240560.1;r
efGene_gn=USP54;refGene_tx=XM_023240561.1;refGene_gn=USP54;refGene_tx=
XM_023240562.1;refGene_gn=USP54;refGene_tx=XM_023240563.1;refGene_gn=U
SP54;refGene_tx=XM_023240564.1;refGene_gn=USP54;refGene_tx=XM_02324056
5.1;refGene_gn=USP54;refGene_tx=XM_023240566.1
D2 36207483 36207566
refGene_gn=USP54;refGene_tx=XM_019813281.2;refGene_gn=USP54;refGene_tx
=XM_019813282.2;refGene_gn=USP54;refGene_tx=XM_019813283.2;refGene_gn=
USP54;refGene_tx=XM_019813284.2;refGene_gn=USP54;refGene_tx=XM_0198132
87.2;refGene_gn=USP54;refGene_tx=XM_019813288.2;refGene_gn=USP54;refGe
ne_tx=XM_019813294.1;refGene_gn=USP54;refGene_tx=XM_023240555.1;refGen
e_gn=USP54;refGene_tx=XM_023240556.1;refGene_gn=USP54;refGene_tx=XM_02
3240557.1;refGene_gn=USP54;refGene_tx=XM_023240558.1;refGene_gn=USP54;
refGene_tx=XM_023240559.1;refGene_gn=USP54;refGene_tx=XM_023240560.1;r
efGene_gn=USP54;refGene_tx=XM_023240561.1;refGene_gn=USP54;refGene_tx=
XM_023240562.1;refGene_gn=USP54;refGene_tx=XM_023240563.1;refGene_gn=U
SP54;refGene_tx=XM_023240564.1;refGene_gn=USP54;refGene_tx=XM_02324056
5.1;refGene_gn=USP54;refGene_tx=XM_023240566.1
D2 36207701 36207815
refGene_gn=USP54;refGene_tx=XM_019813281.2;refGene_gn=USP54;refGene_tx
=XM_019813282.2;refGene_gn=USP54;refGene_tx=XM_019813283.2;refGene_gn=
USP54;refGene_tx=XM_019813284.2;refGene_gn=USP54;refGene_tx=XM_0198132
87.2;refGene_gn=USP54;refGene_tx=XM_019813288.2;refGene_gn=USP54;refGe
ne_tx=XM_019813294.1;refGene_gn=USP54;refGene_tx=XM_023240555.1;refGen
e_gn=USP54;refGene_tx=XM_023240556.1;refGene_gn=USP54;refGene_tx=XM_02
3240557.1;refGene_gn=USP54;refGene_tx=XM_023240558.1;refGene_gn=USP54;
refGene_tx=XM_023240559.1;refGene_gn=USP54;refGene_tx=XM_023240560.1;r
efGene_gn=USP54;refGene_tx=XM_023240561.1;refGene_gn=USP54;refGene_tx=
XM_023240562.1;refGene_gn=USP54;refGene_tx=XM_023240563.1;refGene_gn=U
SP54;refGene_tx=XM_023240564.1;refGene_gn=USP54;refGene_tx=XM_02324056
5.1;refGene_gn=USP54;refGene_tx=XM_023240566.1
D2 36209407 36209542
refGene_gn=USP54;refGene_tx=XM_019813281.2;refGene_gn=USP54;refGene_tx
=XM_019813282.2;refGene_gn=USP54;refGene_tx=XM_019813283.2;refGene_gn=
USP54;refGene_tx=XM_019813284.2;refGene_gn=USP54;refGene_tx=XM_0198132
87.2;refGene_gn=USP54;refGene_tx=XM_019813288.2;refGene_gn=USP54;refGe
ne_tx=XM_019813294.1;refGene_gn=USP54;refGene_tx=XM_023240555.1;refGen
e_gn=USP54;refGene_tx=XM_023240556.1;refGene_gn=USP54;refGene_tx=XM_02
3240557.1;refGene_gn=USP54;refGene_tx=XM_023240558.1;refGene_gn=USP54;
refGene_tx=XM_023240559.1;refGene_gn=USP54;refGene_tx=XM_023240560.1;r
efGene_gn=USP54;refGene_tx=XM_023240561.1;refGene_gn=USP54;refGene_tx=
XM_023240562.1;refGene_gn=USP54;refGene_tx=XM_023240563.1;refGene_gn=U
SP54;refGene_tx=XM_023240564.1;refGene_gn=USP54;refGene_tx=XM_02324056
5.1;refGene_gn=USP54;refGene_tx=XM_023240566.1
D2 36228521 36228614
refGene_gn=USP54;refGene_tx=XM_019813281.2;refGene_gn=USP54;refGene_tx
=XM_019813282.2;refGene_gn=USP54;refGene_tx=XM_019813283.2;refGene_gn=

USP54;refGene_tx=XM_019813284.2;refGene_gn=USP54;refGene_tx=XM_019813287.2;refGene_gn=USP54;refGene_tx=XM_019813288.2;refGene_gn=USP54;refGene_tx=XM_023240555.1;refGene_gn=USP54;refGene_tx=XM_023240556.1;refGene_gn=USP54;refGene_tx=XM_023240557.1;refGene_gn=USP54;refGene_tx=XM_023240558.1;refGene_gn=USP54;refGene_tx=XM_023240559.1;refGene_gn=USP54;refGene_tx=XM_023240560.1;refGene_gn=USP54;refGene_tx=XM_023240561.1;refGene_gn=USP54;refGene_tx=XM_023240562.1;refGene_gn=USP54;refGene_tx=XM_023240563.1;refGene_gn=USP54;refGene_tx=XM_023240564.1;refGene_gn=USP54;refGene_tx=XM_023240565.1;refGene_gn=USP54;refGene_tx=XM_023240566.1
D2 36232427 36232574
refGene_gn=USP54;refGene_tx=XM_019813281.2;refGene_gn=USP54;refGene_tx=XM_019813282.2;refGene_gn=USP54;refGene_tx=XM_019813283.2;refGene_gn=USP54;refGene_tx=XM_019813284.2;refGene_gn=USP54;refGene_tx=XM_019813287.2;refGene_gn=USP54;refGene_tx=XM_019813288.2;refGene_gn=USP54;refGene_tx=XM_023240555.1;refGene_gn=USP54;refGene_tx=XM_023240556.1;refGene_gn=USP54;refGene_tx=XM_023240557.1;refGene_gn=USP54;refGene_tx=XM_023240558.1;refGene_gn=USP54;refGene_tx=XM_023240559.1;refGene_gn=USP54;refGene_tx=XM_023240560.1;refGene_gn=USP54;refGene_tx=XM_023240561.1;refGene_gn=USP54;refGene_tx=XM_023240562.1;refGene_gn=USP54;refGene_tx=XM_023240563.1;refGene_gn=USP54;refGene_tx=XM_023240564.1;refGene_gn=USP54;refGene_tx=XM_023240565.1;refGene_gn=USP54;refGene_tx=XM_023240566.1
D2 36304218 36304450 refGene_gn=MYOZ1;refGene_tx=XM_003994060.3
D2 36305356 36305522 refGene_gn=MYOZ1;refGene_tx=XM_003994060.3
D2 36305880 36306124 refGene_gn=MYOZ1;refGene_tx=XM_003994060.3
D2 36307713 36307892 refGene_gn=MYOZ1;refGene_tx=XM_003994060.3
D2 36308817 36308890 refGene_gn=MYOZ1;refGene_tx=XM_003994060.3
D2 36313884 36316034 refGene_gn=SYNPO2L;refGene_tx=XM_003994061.5
D2 36319270 36319785 refGene_gn=SYNPO2L;refGene_tx=XM_003994061.5
D2 36320251 36320403 refGene_gn=SYNPO2L;refGene_tx=XM_003994061.5
D2 36321587 36321692 refGene_gn=SYNPO2L;refGene_tx=XM_003994061.5
D2 36330593 36330765
refGene_gn=SEC24C;refGene_tx=XM_003994063.5;refGene_gn=SEC24C;refGene_tx=XM_006937877.4;refGene_gn=SEC24C;refGene_tx=XM_006937878.4;refGene_gn=SEC24C;refGene_tx=XM_006937879.4;refGene_gn=SEC24C;refGene_tx=XM_019813299.2;refGene_gn=SEC24C;refGene_tx=XM_019813300.2
D2 36332849 36332985
refGene_gn=SEC24C;refGene_tx=XM_003994063.5;refGene_gn=SEC24C;refGene_tx=XM_006937877.4;refGene_gn=SEC24C;refGene_tx=XM_006937878.4;refGene_gn=SEC24C;refGene_tx=XM_006937879.4;refGene_gn=SEC24C;refGene_tx=XM_019813299.2;refGene_gn=SEC24C;refGene_tx=XM_019813300.2
D2 36341696 36341872
refGene_gn=SEC24C;refGene_tx=XM_003994063.5;refGene_gn=SEC24C;refGene_tx=XM_006937877.4;refGene_gn=SEC24C;refGene_tx=XM_006937878.4;refGene_gn=SEC24C;refGene_tx=XM_006937879.4;refGene_gn=SEC24C;refGene_tx=XM_019813299.2;refGene_gn=SEC24C;refGene_tx=XM_019813300.2
D2 36342301 36342673
refGene_gn=SEC24C;refGene_tx=XM_003994063.5;refGene_gn=SEC24C;refGene_tx=XM_006937877.4;refGene_gn=SEC24C;refGene_tx=XM_006937878.4;refGene_

gn=SEC24C;refGene_tx=XM_006937879.4;refGene_gn=SEC24C;refGene_tx=XM_019813299.2;refGene_gn=SEC24C;refGene_tx=XM_019813300.2
D2 36342957 36343094
refGene_gn=SEC24C;refGene_tx=XM_003994063.5;refGene_gn=SEC24C;refGene_tx=XM_006937877.4;refGene_gn=SEC24C;refGene_tx=XM_006937878.4;refGene_gn=SEC24C;refGene_tx=XM_006937879.4;refGene_gn=SEC24C;refGene_tx=XM_019813299.2;refGene_gn=SEC24C;refGene_tx=XM_019813300.2
D2 36344329 36344398
refGene_gn=SEC24C;refGene_tx=XM_006937877.4;refGene_gn=SEC24C;refGene_tx=XM_006937878.4;refGene_gn=SEC24C;refGene_tx=XM_006937879.4;refGene_gn=SEC24C;refGene_tx=XM_019813299.2;refGene_gn=SEC24C;refGene_tx=XM_019813300.2
D2 36345710 36345822
refGene_gn=SEC24C;refGene_tx=XM_003994063.5;refGene_gn=SEC24C;refGene_tx=XM_006937877.4;refGene_gn=SEC24C;refGene_tx=XM_006937878.4;refGene_gn=SEC24C;refGene_tx=XM_006937879.4;refGene_gn=SEC24C;refGene_tx=XM_019813299.2;refGene_gn=SEC24C;refGene_tx=XM_019813300.2
D2 36346042 36346170
refGene_gn=SEC24C;refGene_tx=XM_003994063.5;refGene_gn=SEC24C;refGene_tx=XM_006937877.4;refGene_gn=SEC24C;refGene_tx=XM_006937878.4;refGene_gn=SEC24C;refGene_tx=XM_006937879.4;refGene_gn=SEC24C;refGene_tx=XM_019813299.2;refGene_gn=SEC24C;refGene_tx=XM_019813300.2
D2 36347330 36347469
refGene_gn=SEC24C;refGene_tx=XM_003994063.5;refGene_gn=SEC24C;refGene_tx=XM_006937877.4;refGene_gn=SEC24C;refGene_tx=XM_006937878.4;refGene_gn=SEC24C;refGene_tx=XM_006937879.4;refGene_gn=SEC24C;refGene_tx=XM_019813299.2;refGene_gn=SEC24C;refGene_tx=XM_019813300.2
D2 36347684 36347800
refGene_gn=SEC24C;refGene_tx=XM_003994063.5;refGene_gn=SEC24C;refGene_tx=XM_006937877.4;refGene_gn=SEC24C;refGene_tx=XM_006937878.4;refGene_gn=SEC24C;refGene_tx=XM_006937879.4;refGene_gn=SEC24C;refGene_tx=XM_019813299.2;refGene_gn=SEC24C;refGene_tx=XM_019813300.2
D2 36347972 36348097
refGene_gn=SEC24C;refGene_tx=XM_003994063.5;refGene_gn=SEC24C;refGene_tx=XM_006937877.4;refGene_gn=SEC24C;refGene_tx=XM_006937878.4;refGene_gn=SEC24C;refGene_tx=XM_006937879.4;refGene_gn=SEC24C;refGene_tx=XM_019813299.2;refGene_gn=SEC24C;refGene_tx=XM_019813300.2
D2 36348216 36348408
refGene_gn=SEC24C;refGene_tx=XM_003994063.5;refGene_gn=SEC24C;refGene_tx=XM_006937877.4;refGene_gn=SEC24C;refGene_tx=XM_006937878.4;refGene_gn=SEC24C;refGene_tx=XM_006937879.4;refGene_gn=SEC24C;refGene_tx=XM_019813299.2;refGene_gn=SEC24C;refGene_tx=XM_019813300.2
D2 36348617 36348711
refGene_gn=SEC24C;refGene_tx=XM_003994063.5;refGene_gn=SEC24C;refGene_tx=XM_006937877.4;refGene_gn=SEC24C;refGene_tx=XM_006937878.4;refGene_gn=SEC24C;refGene_tx=XM_006937879.4;refGene_gn=SEC24C;refGene_tx=XM_019813299.2;refGene_gn=SEC24C;refGene_tx=XM_019813300.2
D2 36348913 36349030
refGene_gn=SEC24C;refGene_tx=XM_003994063.5;refGene_gn=SEC24C;refGene_tx=XM_006937877.4;refGene_gn=SEC24C;refGene_tx=XM_006937878.4;refGene_

gn=SEC24C;refGene_tx=XM_006937879.4;refGene_gn=SEC24C;refGene_tx=XM_019813300.2
D2 36350043 36350214
refGene_gn=SEC24C;refGene_tx=XM_003994063.5;refGene_gn=SEC24C;refGene_tx=XM_006937877.4;refGene_gn=SEC24C;refGene_tx=XM_006937878.4;refGene_gn=SEC24C;refGene_tx=XM_006937879.4;refGene_gn=SEC24C;refGene_tx=XM_019813299.2;refGene_gn=SEC24C;refGene_tx=XM_019813300.2
D2 36350895 36350992
refGene_gn=SEC24C;refGene_tx=XM_003994063.5;refGene_gn=SEC24C;refGene_tx=XM_006937877.4;refGene_gn=SEC24C;refGene_tx=XM_006937878.4;refGene_gn=SEC24C;refGene_tx=XM_006937879.4;refGene_gn=SEC24C;refGene_tx=XM_019813299.2;refGene_gn=SEC24C;refGene_tx=XM_019813300.2
D2 36351094 36351240
refGene_gn=SEC24C;refGene_tx=XM_003994063.5;refGene_gn=SEC24C;refGene_tx=XM_006937877.4;refGene_gn=SEC24C;refGene_tx=XM_006937878.4;refGene_gn=SEC24C;refGene_tx=XM_006937879.4;refGene_gn=SEC24C;refGene_tx=XM_019813299.2;refGene_gn=SEC24C;refGene_tx=XM_019813300.2
D2 36351465 36351604
refGene_gn=SEC24C;refGene_tx=XM_003994063.5;refGene_gn=SEC24C;refGene_tx=XM_006937877.4;refGene_gn=SEC24C;refGene_tx=XM_006937878.4;refGene_gn=SEC24C;refGene_tx=XM_006937879.4;refGene_gn=SEC24C;refGene_tx=XM_019813299.2;refGene_gn=SEC24C;refGene_tx=XM_019813300.2
D2 36351752 36351871
refGene_gn=SEC24C;refGene_tx=XM_003994063.5;refGene_gn=SEC24C;refGene_tx=XM_006937877.4;refGene_gn=SEC24C;refGene_tx=XM_006937878.4;refGene_gn=SEC24C;refGene_tx=XM_006937879.4;refGene_gn=SEC24C;refGene_tx=XM_019813299.2;refGene_gn=SEC24C;refGene_tx=XM_019813300.2
D2 36351966 36352146
refGene_gn=SEC24C;refGene_tx=XM_003994063.5;refGene_gn=SEC24C;refGene_tx=XM_006937877.4;refGene_gn=SEC24C;refGene_tx=XM_006937878.4;refGene_gn=SEC24C;refGene_tx=XM_006937879.4;refGene_gn=SEC24C;refGene_tx=XM_019813299.2;refGene_gn=SEC24C;refGene_tx=XM_019813300.2
D2 36352393 36352585
refGene_gn=SEC24C;refGene_tx=XM_003994063.5;refGene_gn=SEC24C;refGene_tx=XM_006937877.4;refGene_gn=SEC24C;refGene_tx=XM_006937878.4;refGene_gn=SEC24C;refGene_tx=XM_006937879.4;refGene_gn=SEC24C;refGene_tx=XM_019813299.2;refGene_gn=SEC24C;refGene_tx=XM_019813300.2
D2 36352818 36352908
refGene_gn=SEC24C;refGene_tx=XM_003994063.5;refGene_gn=SEC24C;refGene_tx=XM_006937877.4;refGene_gn=SEC24C;refGene_tx=XM_006937878.4;refGene_gn=SEC24C;refGene_tx=XM_006937879.4;refGene_gn=SEC24C;refGene_tx=XM_019813299.2;refGene_gn=SEC24C;refGene_tx=XM_019813300.2
D2 36353053 36353194
refGene_gn=SEC24C;refGene_tx=XM_003994063.5;refGene_gn=SEC24C;refGene_tx=XM_006937877.4;refGene_gn=SEC24C;refGene_tx=XM_006937878.4;refGene_gn=SEC24C;refGene_tx=XM_006937879.4;refGene_gn=SEC24C;refGene_tx=XM_019813299.2;refGene_gn=SEC24C;refGene_tx=XM_019813300.2
D2 36354515 36355223 refGene_gn=FUT11;refGene_tx=XM_011287333.3
D2 36355342 36355968 refGene_gn=FUT11;refGene_tx=XM_011287333.3
D2 36357579 36357727 refGene_gn=FUT11;refGene_tx=XM_011287333.3

D2 36362803 36362927 refGene_gn=CHCHD1;refGene_tx=XM_003994066.4
D2 36363050 36363169 refGene_gn=CHCHD1;refGene_tx=XM_003994066.4
D2 36363734 36363848 refGene_gn=CHCHD1;refGene_tx=XM_003994066.4
D2 36366150 36366358
refGene_gn=ZSWIM8;refGene_tx=XM_011287326.2;refGene_gn=ZSWIM8;refGene_tx=XM_011287329.2;refGene_gn=ZSWIM8;refGene_tx=XM_011287330.3;refGene_gn=ZSWIM8;refGene_tx=XM_011287331.3
D2 36368703 36368857
refGene_gn=ZSWIM8;refGene_tx=XM_011287326.2;refGene_gn=ZSWIM8;refGene_tx=XM_011287329.2;refGene_gn=ZSWIM8;refGene_tx=XM_011287330.3;refGene_gn=ZSWIM8;refGene_tx=XM_011287331.3
D2 36369090 36369185
refGene_gn=ZSWIM8;refGene_tx=XM_011287326.2;refGene_gn=ZSWIM8;refGene_tx=XM_011287329.2;refGene_gn=ZSWIM8;refGene_tx=XM_011287330.3;refGene_gn=ZSWIM8;refGene_tx=XM_011287331.3
D2 36369361 36369534
refGene_gn=ZSWIM8;refGene_tx=XM_011287326.2;refGene_gn=ZSWIM8;refGene_tx=XM_011287329.2;refGene_gn=ZSWIM8;refGene_tx=XM_011287330.3;refGene_gn=ZSWIM8;refGene_tx=XM_011287331.3
D2 36369709 36369817
refGene_gn=ZSWIM8;refGene_tx=XM_011287326.2;refGene_gn=ZSWIM8;refGene_tx=XM_011287329.2;refGene_gn=ZSWIM8;refGene_tx=XM_011287330.3;refGene_gn=ZSWIM8;refGene_tx=XM_011287331.3
D2 36369932 36370014
refGene_gn=ZSWIM8;refGene_tx=XM_011287326.2;refGene_gn=ZSWIM8;refGene_tx=XM_011287329.2;refGene_gn=ZSWIM8;refGene_tx=XM_011287330.3;refGene_gn=ZSWIM8;refGene_tx=XM_011287331.3
D2 36370148 36370269
refGene_gn=ZSWIM8;refGene_tx=XM_011287326.2;refGene_gn=ZSWIM8;refGene_tx=XM_011287329.2;refGene_gn=ZSWIM8;refGene_tx=XM_011287330.3;refGene_gn=ZSWIM8;refGene_tx=XM_011287331.3
D2 36370508 36370710
refGene_gn=ZSWIM8;refGene_tx=XM_011287326.2;refGene_gn=ZSWIM8;refGene_tx=XM_011287329.2;refGene_gn=ZSWIM8;refGene_tx=XM_011287330.3;refGene_gn=ZSWIM8;refGene_tx=XM_011287331.3
D2 36370856 36371032
refGene_gn=ZSWIM8;refGene_tx=XM_011287326.2;refGene_gn=ZSWIM8;refGene_tx=XM_011287329.2;refGene_gn=ZSWIM8;refGene_tx=XM_011287330.3;refGene_gn=ZSWIM8;refGene_tx=XM_011287331.3
D2 36371404 36372398
refGene_gn=ZSWIM8;refGene_tx=XM_011287326.2;refGene_gn=ZSWIM8;refGene_tx=XM_011287329.2;refGene_gn=ZSWIM8;refGene_tx=XM_011287330.3;refGene_gn=ZSWIM8;refGene_tx=XM_011287331.3
D2 36373158 36373290
refGene_gn=ZSWIM8;refGene_tx=XM_011287326.2;refGene_gn=ZSWIM8;refGene_tx=XM_011287329.2;refGene_gn=ZSWIM8;refGene_tx=XM_011287330.3;refGene_gn=ZSWIM8;refGene_tx=XM_011287331.3
D2 36373426 36373606
refGene_gn=ZSWIM8;refGene_tx=XM_011287326.2;refGene_gn=ZSWIM8;refGene_tx=XM_011287329.2;refGene_gn=ZSWIM8;refGene_tx=XM_011287330.3;refGene_

gn=ZSWIM8;refGene_tx=XM_011287331.3
D2 36373717 36373901
refGene_gn=ZSWIM8;refGene_tx=XM_011287326.2;refGene_gn=ZSWIM8;refGene_tx=XM_011287329.2;refGene_gn=ZSWIM8;refGene_tx=XM_011287330.3;refGene_gn=ZSWIM8;refGene_tx=XM_011287331.3
D2 36374110 36374209
refGene_gn=ZSWIM8;refGene_tx=XM_011287326.2;refGene_gn=ZSWIM8;refGene_tx=XM_011287329.2;refGene_gn=ZSWIM8;refGene_tx=XM_011287330.3;refGene_gn=ZSWIM8;refGene_tx=XM_011287331.3
D2 36374765 36374890
refGene_gn=ZSWIM8;refGene_tx=XM_011287326.2;refGene_gn=ZSWIM8;refGene_tx=XM_011287329.2;refGene_gn=ZSWIM8;refGene_tx=XM_011287330.3;refGene_gn=ZSWIM8;refGene_tx=XM_011287331.3
D2 36375316 36375557
refGene_gn=ZSWIM8;refGene_tx=XM_011287326.2;refGene_gn=ZSWIM8;refGene_tx=XM_011287329.2;refGene_gn=ZSWIM8;refGene_tx=XM_011287330.3;refGene_gn=ZSWIM8;refGene_tx=XM_011287331.3
D2 36375656 36375815
refGene_gn=ZSWIM8;refGene_tx=XM_011287326.2;refGene_gn=ZSWIM8;refGene_tx=XM_011287329.2;refGene_gn=ZSWIM8;refGene_tx=XM_011287330.3;refGene_gn=ZSWIM8;refGene_tx=XM_011287331.3
D2 36375912 36376141
refGene_gn=ZSWIM8;refGene_tx=XM_011287326.2;refGene_gn=ZSWIM8;refGene_tx=XM_011287329.2;refGene_gn=ZSWIM8;refGene_tx=XM_011287330.3;refGene_gn=ZSWIM8;refGene_tx=XM_011287331.3
D2 36376310 36376600
refGene_gn=ZSWIM8;refGene_tx=XM_011287326.2;refGene_gn=ZSWIM8;refGene_tx=XM_011287329.2;refGene_gn=ZSWIM8;refGene_tx=XM_011287330.3;refGene_gn=ZSWIM8;refGene_tx=XM_011287331.3
D2 36376748 36376972
refGene_gn=ZSWIM8;refGene_tx=XM_011287326.2;refGene_gn=ZSWIM8;refGene_tx=XM_011287329.2;refGene_gn=ZSWIM8;refGene_tx=XM_011287330.3;refGene_gn=ZSWIM8;refGene_tx=XM_011287331.3
D2 36377487 36377976
refGene_gn=ZSWIM8;refGene_tx=XM_011287326.2;refGene_gn=ZSWIM8;refGene_tx=XM_011287329.2;refGene_gn=ZSWIM8;refGene_tx=XM_011287330.3;refGene_gn=ZSWIM8;refGene_tx=XM_011287331.3
D2 36378206 36378390
refGene_gn=ZSWIM8;refGene_tx=XM_011287326.2;refGene_gn=ZSWIM8;refGene_tx=XM_011287329.2;refGene_gn=ZSWIM8;refGene_tx=XM_011287330.3;refGene_gn=ZSWIM8;refGene_tx=XM_011287331.3
D2 36378494 36378671
refGene_gn=ZSWIM8;refGene_tx=XM_011287326.2;refGene_gn=ZSWIM8;refGene_tx=XM_011287329.2;refGene_gn=ZSWIM8;refGene_tx=XM_011287330.3;refGene_gn=ZSWIM8;refGene_tx=XM_011287331.3
D2 36378839 36378959
refGene_gn=ZSWIM8;refGene_tx=XM_011287326.2;refGene_gn=ZSWIM8;refGene_tx=XM_011287329.2;refGene_gn=ZSWIM8;refGene_tx=XM_011287330.3;refGene_gn=ZSWIM8;refGene_tx=XM_011287331.3
D2 36379214 36379393

refGene_gn=ZSWIM8;refGene_tx=XM_011287326.2;refGene_gn=ZSWIM8;refGene_tx=XM_011287329.2;refGene_gn=ZSWIM8;refGene_tx=XM_011287330.3;refGene_gn=ZSWIM8;refGene_tx=XM_011287331.3
D2 36379513 36379974
refGene_gn=ZSWIM8;refGene_tx=XM_011287326.2;refGene_gn=ZSWIM8;refGene_tx=XM_011287329.2;refGene_gn=ZSWIM8;refGene_tx=XM_011287330.3;refGene_gn=ZSWIM8;refGene_tx=XM_011287331.3
D2 36380620 36380746
refGene_gn=NDST2;refGene_tx=XM_023240567.1;refGene_gn=NDST2;refGene_tx=XM_023240568.1;refGene_gn=NDST2;refGene_tx=XM_023240569.1;refGene_gn=NDST2;refGene_tx=XM_023240570.1
D2 36380843 36380946
refGene_gn=NDST2;refGene_tx=XM_023240567.1;refGene_gn=NDST2;refGene_tx=XM_023240568.1;refGene_gn=NDST2;refGene_tx=XM_023240569.1;refGene_gn=NDST2;refGene_tx=XM_023240570.1
D2 36381116 36381226
refGene_gn=NDST2;refGene_tx=XM_023240567.1;refGene_gn=NDST2;refGene_tx=XM_023240568.1;refGene_gn=NDST2;refGene_tx=XM_023240569.1;refGene_gn=NDST2;refGene_tx=XM_023240570.1
D2 36381333 36381504
refGene_gn=NDST2;refGene_tx=XM_023240567.1;refGene_gn=NDST2;refGene_tx=XM_023240568.1;refGene_gn=NDST2;refGene_tx=XM_023240569.1;refGene_gn=NDST2;refGene_tx=XM_023240570.1
D2 36381694 36381869
refGene_gn=NDST2;refGene_tx=XM_023240567.1;refGene_gn=NDST2;refGene_tx=XM_023240568.1;refGene_gn=NDST2;refGene_tx=XM_023240569.1;refGene_gn=NDST2;refGene_tx=XM_023240570.1
D2 36382007 36382131
refGene_gn=NDST2;refGene_tx=XM_023240567.1;refGene_gn=NDST2;refGene_tx=XM_023240568.1;refGene_gn=NDST2;refGene_tx=XM_023240569.1;refGene_gn=NDST2;refGene_tx=XM_023240570.1
D2 36382492 36382589
refGene_gn=NDST2;refGene_tx=XM_023240567.1;refGene_gn=NDST2;refGene_tx=XM_023240568.1;refGene_gn=NDST2;refGene_tx=XM_023240569.1;refGene_gn=NDST2;refGene_tx=XM_023240570.1
D2 36382681 36382864
refGene_gn=NDST2;refGene_tx=XM_023240567.1;refGene_gn=NDST2;refGene_tx=XM_023240568.1;refGene_gn=NDST2;refGene_tx=XM_023240569.1;refGene_gn=NDST2;refGene_tx=XM_023240570.1
D2 36382995 36383124
refGene_gn=NDST2;refGene_tx=XM_023240567.1;refGene_gn=NDST2;refGene_tx=XM_023240568.1;refGene_gn=NDST2;refGene_tx=XM_023240569.1;refGene_gn=NDST2;refGene_tx=XM_023240570.1
D2 36383395 36383581
refGene_gn=NDST2;refGene_tx=XM_023240567.1;refGene_gn=NDST2;refGene_tx=XM_023240568.1;refGene_gn=NDST2;refGene_tx=XM_023240569.1;refGene_gn=NDST2;refGene_tx=XM_023240570.1
D2 36383766 36383921
refGene_gn=NDST2;refGene_tx=XM_023240567.1;refGene_gn=NDST2;refGene_tx=XM_023240568.1;refGene_gn=NDST2;refGene_tx=XM_023240569.1;refGene_gn=

NDST2;refGene_tx=XM_023240570.1
D2 36384232 36384320
refGene_gn=NDST2;refGene_tx=XM_023240567.1;refGene_gn=NDST2;refGene_tx=XM_023240568.1;refGene_gn=NDST2;refGene_tx=XM_023240569.1;refGene_gn=NDST2;refGene_tx=XM_023240570.1
D2 36384511 36385516
refGene_gn=NDST2;refGene_tx=XM_023240567.1;refGene_gn=NDST2;refGene_tx=XM_023240568.1;refGene_gn=NDST2;refGene_tx=XM_023240569.1;refGene_gn=NDST2;refGene_tx=XM_023240570.1
D2 36391364 36391597
refGene_gn=CAMK2G;refGene_tx=XM_019813323.1;refGene_gn=CAMK2G;refGene_tx=XM_023240571.1;refGene_gn=CAMK2G;refGene_tx=XM_023240572.1;refGene_gn=CAMK2G;refGene_tx=XM_023240573.1;refGene_gn=CAMK2G;refGene_tx=XM_023240574.1;refGene_gn=CAMK2G;refGene_tx=XM_023240575.1;refGene_gn=CAMK2G;refGene_tx=XM_023240576.1;refGene_gn=CAMK2G;refGene_tx=XM_023240577.1;refGene_gn=CAMK2G;refGene_tx=XM_023240578.1;refGene_gn=CAMK2G;refGene_tx=XM_023240579.1;refGene_gn=CAMK2G;refGene_tx=XM_023240580.1;refGene_gn=CAMK2G;refGene_tx=XM_023240581.1;refGene_gn=CAMK2G;refGene_tx=XM_023240582.1;refGene_gn=CAMK2G;refGene_tx=XM_023240583.1;refGene_gn=CAMK2G;refGene_tx=XM_023240584.1;refGene_gn=CAMK2G;refGene_tx=XM_023240585.1;refGene_gn=CAMK2G;refGene_tx=XM_023240586.1;refGene_gn=CAMK2G;refGene_tx=XM_023240587.1
D2 36393136 36393231
refGene_gn=CAMK2G;refGene_tx=XM_019813323.1;refGene_gn=CAMK2G;refGene_tx=XM_023240571.1;refGene_gn=CAMK2G;refGene_tx=XM_023240572.1;refGene_gn=CAMK2G;refGene_tx=XM_023240573.1;refGene_gn=CAMK2G;refGene_tx=XM_023240574.1;refGene_gn=CAMK2G;refGene_tx=XM_023240575.1;refGene_gn=CAMK2G;refGene_tx=XM_023240576.1;refGene_gn=CAMK2G;refGene_tx=XM_023240577.1;refGene_gn=CAMK2G;refGene_tx=XM_023240578.1;refGene_gn=CAMK2G;refGene_tx=XM_023240579.1;refGene_gn=CAMK2G;refGene_tx=XM_023240580.1;refGene_gn=CAMK2G;refGene_tx=XM_023240581.1;refGene_gn=CAMK2G;refGene_tx=XM_023240582.1;refGene_gn=CAMK2G;refGene_tx=XM_023240583.1;refGene_gn=CAMK2G;refGene_tx=XM_023240584.1;refGene_gn=CAMK2G;refGene_tx=XM_023240585.1;refGene_gn=CAMK2G;refGene_tx=XM_023240586.1;refGene_gn=CAMK2G;refGene_tx=XM_023240587.1
D2 36393600 36393676
refGene_gn=CAMK2G;refGene_tx=XM_019813323.1;refGene_gn=CAMK2G;refGene_tx=XM_023240571.1;refGene_gn=CAMK2G;refGene_tx=XM_023240572.1;refGene_gn=CAMK2G;refGene_tx=XM_023240573.1;refGene_gn=CAMK2G;refGene_tx=XM_023240574.1;refGene_gn=CAMK2G;refGene_tx=XM_023240575.1;refGene_gn=CAMK2G;refGene_tx=XM_023240576.1;refGene_gn=CAMK2G;refGene_tx=XM_023240577.1;refGene_gn=CAMK2G;refGene_tx=XM_023240578.1;refGene_gn=CAMK2G;refGene_tx=XM_023240579.1;refGene_gn=CAMK2G;refGene_tx=XM_023240580.1;refGene_gn=CAMK2G;refGene_tx=XM_023240581.1;refGene_gn=CAMK2G;refGene_tx=XM_023240582.1;refGene_gn=CAMK2G;refGene_tx=XM_023240583.1;refGene_gn=CAMK2G;refGene_tx=XM_023240584.1;refGene_gn=CAMK2G;refGene_tx=XM_023240585.1;refGene_gn=CAMK2G;refGene_tx=XM_023240586.1;refGene_gn=CAMK2G;refGene_tx=XM_023240587.1
D2 36395716 36395830
refGene_gn=CAMK2G;refGene_tx=XM_019813323.1;refGene_gn=CAMK2G;refGene_

tx=XM_023240571.1;refGene_gn=CAMK2G;refGene_tx=XM_023240572.1;refGene_gn=CAMK2G;refGene_tx=XM_023240573.1;refGene_gn=CAMK2G;refGene_tx=XM_023240574.1;refGene_gn=CAMK2G;refGene_tx=XM_023240575.1;refGene_gn=CAMK2G;refGene_tx=XM_023240576.1;refGene_gn=CAMK2G;refGene_tx=XM_023240578.1;refGene_gn=CAMK2G;refGene_tx=XM_023240580.1;refGene_gn=CAMK2G;refGene_tx=XM_023240586.1
D2 36397046 36397095
refGene_gn=CAMK2G;refGene_tx=XM_019813323.1;refGene_gn=CAMK2G;refGene_tx=XM_023240571.1;refGene_gn=CAMK2G;refGene_tx=XM_023240572.1;refGene_gn=CAMK2G;refGene_tx=XM_023240573.1;refGene_gn=CAMK2G;refGene_tx=XM_023240574.1;refGene_gn=CAMK2G;refGene_tx=XM_023240575.1;refGene_gn=CAMK2G;refGene_tx=XM_023240576.1;refGene_gn=CAMK2G;refGene_tx=XM_023240577.1;refGene_gn=CAMK2G;refGene_tx=XM_023240578.1;refGene_gn=CAMK2G;refGene_tx=XM_023240579.1;refGene_gn=CAMK2G;refGene_tx=XM_023240580.1;refGene_gn=CAMK2G;refGene_tx=XM_023240581.1;refGene_gn=CAMK2G;refGene_tx=XM_023240582.1;refGene_gn=CAMK2G;refGene_tx=XM_023240583.1;refGene_gn=CAMK2G;refGene_tx=XM_023240584.1;refGene_gn=CAMK2G;refGene_tx=XM_023240585.1;refGene_gn=CAMK2G;refGene_tx=XM_023240586.1;refGene_gn=CAMK2G;refGene_tx=XM_023240587.1
D2 36398941 36398986
refGene_gn=CAMK2G;refGene_tx=XM_019813323.1;refGene_gn=CAMK2G;refGene_tx=XM_023240571.1;refGene_gn=CAMK2G;refGene_tx=XM_023240572.1;refGene_gn=CAMK2G;refGene_tx=XM_023240573.1;refGene_gn=CAMK2G;refGene_tx=XM_023240574.1;refGene_gn=CAMK2G;refGene_tx=XM_023240575.1;refGene_gn=CAMK2G;refGene_tx=XM_023240576.1;refGene_gn=CAMK2G;refGene_tx=XM_023240577.1;refGene_gn=CAMK2G;refGene_tx=XM_023240578.1;refGene_gn=CAMK2G;refGene_tx=XM_023240579.1;refGene_gn=CAMK2G;refGene_tx=XM_023240580.1;refGene_gn=CAMK2G;refGene_tx=XM_023240581.1;refGene_gn=CAMK2G;refGene_tx=XM_023240582.1;refGene_gn=CAMK2G;refGene_tx=XM_023240583.1;refGene_gn=CAMK2G;refGene_tx=XM_023240584.1;refGene_gn=CAMK2G;refGene_tx=XM_023240585.1;refGene_gn=CAMK2G;refGene_tx=XM_023240586.1;refGene_gn=CAMK2G;refGene_tx=XM_023240587.1
D2 36400126 36400195
refGene_gn=CAMK2G;refGene_tx=XM_019813323.1;refGene_gn=CAMK2G;refGene_tx=XM_023240571.1;refGene_gn=CAMK2G;refGene_tx=XM_023240572.1;refGene_gn=CAMK2G;refGene_tx=XM_023240573.1;refGene_gn=CAMK2G;refGene_tx=XM_023240575.1;refGene_gn=CAMK2G;refGene_tx=XM_023240577.1;refGene_gn=CAMK2G;refGene_tx=XM_023240579.1;refGene_gn=CAMK2G;refGene_tx=XM_023240581.1;refGene_gn=CAMK2G;refGene_tx=XM_023240583.1;refGene_gn=CAMK2G;refGene_tx=XM_023240586.1
D2 36402406 36402439
refGene_gn=CAMK2G;refGene_tx=XM_019813323.1;refGene_gn=CAMK2G;refGene_tx=XM_023240571.1;refGene_gn=CAMK2G;refGene_tx=XM_023240573.1;refGene_gn=CAMK2G;refGene_tx=XM_023240574.1;refGene_gn=CAMK2G;refGene_tx=XM_023240577.1;refGene_gn=CAMK2G;refGene_tx=XM_023240578.1;refGene_gn=CAMK2G;refGene_tx=XM_023240581.1;refGene_gn=CAMK2G;refGene_tx=XM_023240582.1;refGene_gn=CAMK2G;refGene_tx=XM_023240585.1;refGene_gn=CAMK2G;refGene_tx=XM_023240586.1
D2 36409820 36409864
refGene_gn=CAMK2G;refGene_tx=XM_019813323.1;refGene_gn=CAMK2G;refGene_

tx=XM_023240571.1;refGene_gn=CAMK2G;refGene_tx=XM_023240572.1;refGene_gn=CAMK2G;refGene_tx=XM_023240573.1;refGene_gn=CAMK2G;refGene_tx=XM_023240574.1;refGene_gn=CAMK2G;refGene_tx=XM_023240575.1;refGene_gn=CAMK2G;refGene_tx=XM_023240576.1;refGene_gn=CAMK2G;refGene_tx=XM_023240577.1;refGene_gn=CAMK2G;refGene_tx=XM_023240578.1;refGene_gn=CAMK2G;refGene_tx=XM_023240579.1;refGene_gn=CAMK2G;refGene_tx=XM_023240580.1;refGene_gn=CAMK2G;refGene_tx=XM_023240581.1;refGene_gn=CAMK2G;refGene_tx=XM_023240582.1;refGene_gn=CAMK2G;refGene_tx=XM_023240583.1;refGene_gn=CAMK2G;refGene_tx=XM_023240584.1;refGene_gn=CAMK2G;refGene_tx=XM_023240585.1;refGene_gn=CAMK2G;refGene_tx=XM_023240586.1;refGene_gn=CAMK2G;refGene_tx=XM_023240587.1
D2 36411874 36411937
refGene_gn=CAMK2G;refGene_tx=XM_019813323.1;refGene_gn=CAMK2G;refGene_tx=XM_023240571.1;refGene_gn=CAMK2G;refGene_tx=XM_023240572.1;refGene_gn=CAMK2G;refGene_tx=XM_023240574.1;refGene_gn=CAMK2G;refGene_tx=XM_023240576.1;refGene_gn=CAMK2G;refGene_tx=XM_023240577.1;refGene_gn=CAMK2G;refGene_tx=XM_023240579.1;refGene_gn=CAMK2G;refGene_tx=XM_023240582.1;refGene_gn=CAMK2G;refGene_tx=XM_023240584.1;refGene_gn=CAMK2G;refGene_tx=XM_023240586.1
D2 36414470 36414513
refGene_gn=CAMK2G;refGene_tx=XM_019813323.1;refGene_gn=CAMK2G;refGene_tx=XM_023240571.1;refGene_gn=CAMK2G;refGene_tx=XM_023240572.1;refGene_gn=CAMK2G;refGene_tx=XM_023240573.1;refGene_gn=CAMK2G;refGene_tx=XM_023240574.1;refGene_gn=CAMK2G;refGene_tx=XM_023240575.1;refGene_gn=CAMK2G;refGene_tx=XM_023240576.1;refGene_gn=CAMK2G;refGene_tx=XM_023240577.1;refGene_gn=CAMK2G;refGene_tx=XM_023240578.1;refGene_gn=CAMK2G;refGene_tx=XM_023240579.1;refGene_gn=CAMK2G;refGene_tx=XM_023240580.1;refGene_gn=CAMK2G;refGene_tx=XM_023240581.1;refGene_gn=CAMK2G;refGene_tx=XM_023240582.1;refGene_gn=CAMK2G;refGene_tx=XM_023240583.1;refGene_gn=CAMK2G;refGene_tx=XM_023240584.1;refGene_gn=CAMK2G;refGene_tx=XM_023240585.1;refGene_gn=CAMK2G;refGene_tx=XM_023240586.1;refGene_gn=CAMK2G;refGene_tx=XM_023240587.1
D2 36414761 36414845
refGene_gn=CAMK2G;refGene_tx=XM_019813323.1;refGene_gn=CAMK2G;refGene_tx=XM_023240571.1;refGene_gn=CAMK2G;refGene_tx=XM_023240572.1;refGene_gn=CAMK2G;refGene_tx=XM_023240573.1;refGene_gn=CAMK2G;refGene_tx=XM_023240574.1;refGene_gn=CAMK2G;refGene_tx=XM_023240575.1;refGene_gn=CAMK2G;refGene_tx=XM_023240576.1;refGene_gn=CAMK2G;refGene_tx=XM_023240577.1;refGene_gn=CAMK2G;refGene_tx=XM_023240578.1;refGene_gn=CAMK2G;refGene_tx=XM_023240579.1;refGene_gn=CAMK2G;refGene_tx=XM_023240580.1;refGene_gn=CAMK2G;refGene_tx=XM_023240581.1;refGene_gn=CAMK2G;refGene_tx=XM_023240582.1;refGene_gn=CAMK2G;refGene_tx=XM_023240583.1;refGene_gn=CAMK2G;refGene_tx=XM_023240584.1;refGene_gn=CAMK2G;refGene_tx=XM_023240585.1;refGene_gn=CAMK2G;refGene_tx=XM_023240586.1;refGene_gn=CAMK2G;refGene_tx=XM_023240587.1
D2 36418433 36418556
refGene_gn=CAMK2G;refGene_tx=XM_019813323.1;refGene_gn=CAMK2G;refGene_tx=XM_023240571.1;refGene_gn=CAMK2G;refGene_tx=XM_023240572.1;refGene_gn=CAMK2G;refGene_tx=XM_023240573.1;refGene_gn=CAMK2G;refGene_tx=XM_023240574.1;refGene_gn=CAMK2G;refGene_tx=XM_023240575.1;refGene_gn=CAMK2

G;refGene_tx=XM_023240576.1;refGene_gn=CAMK2G;refGene_tx=XM_023240577.
1;refGene_gn=CAMK2G;refGene_tx=XM_023240578.1;refGene_gn=CAMK2G;refGen
e_tx=XM_023240579.1;refGene_gn=CAMK2G;refGene_tx=XM_023240580.1;refGen
e_gn=CAMK2G;refGene_tx=XM_023240581.1;refGene_gn=CAMK2G;refGene_tx=XM_
023240582.1;refGene_gn=CAMK2G;refGene_tx=XM_023240583.1;refGene_gn=CAM
K2G;refGene_tx=XM_023240584.1;refGene_gn=CAMK2G;refGene_tx=XM_02324058
5.1;refGene_gn=CAMK2G;refGene_tx=XM_023240586.1;refGene_gn=CAMK2G;refG
ene_tx=XM_023240587.1
D2 36419176 36419271
refGene_gn=CAMK2G;refGene_tx=XM_019813323.1;refGene_gn=CAMK2G;refGene_
tx=XM_023240571.1;refGene_gn=CAMK2G;refGene_tx=XM_023240572.1;refGene_
gn=CAMK2G;refGene_tx=XM_023240573.1;refGene_gn=CAMK2G;refGene_tx=XM_02
3240574.1;refGene_gn=CAMK2G;refGene_tx=XM_023240575.1;refGene_gn=CAMK2
G;refGene_tx=XM_023240576.1;refGene_gn=CAMK2G;refGene_tx=XM_023240577.
1;refGene_gn=CAMK2G;refGene_tx=XM_023240578.1;refGene_gn=CAMK2G;refGen
e_tx=XM_023240579.1;refGene_gn=CAMK2G;refGene_tx=XM_023240580.1;refGen
e_gn=CAMK2G;refGene_tx=XM_023240581.1;refGene_gn=CAMK2G;refGene_tx=XM_
023240582.1;refGene_gn=CAMK2G;refGene_tx=XM_023240583.1;refGene_gn=CAM
K2G;refGene_tx=XM_023240584.1;refGene_gn=CAMK2G;refGene_tx=XM_02324058
5.1;refGene_gn=CAMK2G;refGene_tx=XM_023240586.1;refGene_gn=CAMK2G;refG
ene_tx=XM_023240587.1
D2 36419717 36419801
refGene_gn=CAMK2G;refGene_tx=XM_019813323.1;refGene_gn=CAMK2G;refGene_
tx=XM_023240571.1;refGene_gn=CAMK2G;refGene_tx=XM_023240572.1;refGene_
gn=CAMK2G;refGene_tx=XM_023240573.1;refGene_gn=CAMK2G;refGene_tx=XM_02
3240574.1;refGene_gn=CAMK2G;refGene_tx=XM_023240575.1;refGene_gn=CAMK2
G;refGene_tx=XM_023240576.1;refGene_gn=CAMK2G;refGene_tx=XM_023240577.
1;refGene_gn=CAMK2G;refGene_tx=XM_023240578.1;refGene_gn=CAMK2G;refGen
e_tx=XM_023240579.1;refGene_gn=CAMK2G;refGene_tx=XM_023240580.1;refGen
e_gn=CAMK2G;refGene_tx=XM_023240581.1;refGene_gn=CAMK2G;refGene_tx=XM_
023240582.1;refGene_gn=CAMK2G;refGene_tx=XM_023240583.1;refGene_gn=CAM
K2G;refGene_tx=XM_023240584.1;refGene_gn=CAMK2G;refGene_tx=XM_02324058
5.1;refGene_gn=CAMK2G;refGene_tx=XM_023240586.1;refGene_gn=CAMK2G;refG
ene_tx=XM_023240587.1
D2 36420232 36420335
refGene_gn=CAMK2G;refGene_tx=XM_019813323.1;refGene_gn=CAMK2G;refGene_
tx=XM_023240571.1;refGene_gn=CAMK2G;refGene_tx=XM_023240572.1;refGene_
gn=CAMK2G;refGene_tx=XM_023240573.1;refGene_gn=CAMK2G;refGene_tx=XM_02
3240574.1;refGene_gn=CAMK2G;refGene_tx=XM_023240575.1;refGene_gn=CAMK2
G;refGene_tx=XM_023240576.1;refGene_gn=CAMK2G;refGene_tx=XM_023240577.
1;refGene_gn=CAMK2G;refGene_tx=XM_023240578.1;refGene_gn=CAMK2G;refGen
e_tx=XM_023240579.1;refGene_gn=CAMK2G;refGene_tx=XM_023240580.1;refGen
e_gn=CAMK2G;refGene_tx=XM_023240581.1;refGene_gn=CAMK2G;refGene_tx=XM_
023240582.1;refGene_gn=CAMK2G;refGene_tx=XM_023240583.1;refGene_gn=CAM
K2G;refGene_tx=XM_023240584.1;refGene_gn=CAMK2G;refGene_tx=XM_02324058
5.1;refGene_gn=CAMK2G;refGene_tx=XM_023240586.1;refGene_gn=CAMK2G;refG
ene_tx=XM_023240587.1
D2 36420478 36420551
refGene_gn=CAMK2G;refGene_tx=XM_019813323.1;refGene_gn=CAMK2G;refGene_
tx=XM_023240571.1;refGene_gn=CAMK2G;refGene_tx=XM_023240572.1;refGene_

gn=CAMK2G;refGene_tx=XM_023240573.1;refGene_gn=CAMK2G;refGene_tx=XM_023240574.1;refGene_gn=CAMK2G;refGene_tx=XM_023240575.1;refGene_gn=CAMK2G;refGene_tx=XM_023240576.1;refGene_gn=CAMK2G;refGene_tx=XM_023240577.1;refGene_gn=CAMK2G;refGene_tx=XM_023240578.1;refGene_gn=CAMK2G;refGene_tx=XM_023240579.1;refGene_gn=CAMK2G;refGene_tx=XM_023240580.1;refGene_gn=CAMK2G;refGene_tx=XM_023240581.1;refGene_gn=CAMK2G;refGene_tx=XM_023240582.1;refGene_gn=CAMK2G;refGene_tx=XM_023240583.1;refGene_gn=CAMK2G;refGene_tx=XM_023240584.1;refGene_gn=CAMK2G;refGene_tx=XM_023240585.1;refGene_gn=CAMK2G;refGene_tx=XM_023240586.1;refGene_gn=CAMK2G;refGene_tx=XM_023240587.1

D2 36423531 36423597 refGene_gn=CAMK2G;refGene_tx=XM_019813323.1;refGene_gn=CAMK2G;refGene_tx=XM_023240571.1;refGene_gn=CAMK2G;refGene_tx=XM_023240572.1;refGene_gn=CAMK2G;refGene_tx=XM_023240573.1;refGene_gn=CAMK2G;refGene_tx=XM_023240574.1;refGene_gn=CAMK2G;refGene_tx=XM_023240575.1;refGene_gn=CAMK2G;refGene_tx=XM_023240576.1;refGene_gn=CAMK2G;refGene_tx=XM_023240577.1;refGene_gn=CAMK2G;refGene_tx=XM_023240578.1;refGene_gn=CAMK2G;refGene_tx=XM_023240579.1;refGene_gn=CAMK2G;refGene_tx=XM_023240580.1;refGene_gn=CAMK2G;refGene_tx=XM_023240581.1;refGene_gn=CAMK2G;refGene_tx=XM_023240582.1;refGene_gn=CAMK2G;refGene_tx=XM_023240583.1;refGene_gn=CAMK2G;refGene_tx=XM_023240584.1;refGene_gn=CAMK2G;refGene_tx=XM_023240585.1;refGene_gn=CAMK2G;refGene_tx=XM_023240586.1;refGene_gn=CAMK2G;refGene_tx=XM_023240587.1

D2 36424481 36424536 refGene_gn=CAMK2G;refGene_tx=XM_023240571.1;refGene_gn=CAMK2G;refGene_tx=XM_023240572.1;refGene_gn=CAMK2G;refGene_tx=XM_023240573.1;refGene_gn=CAMK2G;refGene_tx=XM_023240574.1;refGene_gn=CAMK2G;refGene_tx=XM_023240575.1;refGene_gn=CAMK2G;refGene_tx=XM_023240576.1;refGene_gn=CAMK2G;refGene_tx=XM_023240577.1;refGene_gn=CAMK2G;refGene_tx=XM_023240578.1;refGene_gn=CAMK2G;refGene_tx=XM_023240579.1;refGene_gn=CAMK2G;refGene_tx=XM_023240580.1;refGene_gn=CAMK2G;refGene_tx=XM_023240581.1;refGene_gn=CAMK2G;refGene_tx=XM_023240582.1;refGene_gn=CAMK2G;refGene_tx=XM_023240583.1;refGene_gn=CAMK2G;refGene_tx=XM_023240584.1;refGene_gn=CAMK2G;refGene_tx=XM_023240585.1;refGene_gn=CAMK2G;refGene_tx=XM_023240587.1

D2 36431728 36431788 refGene_gn=CAMK2G;refGene_tx=XM_019813323.1;refGene_gn=CAMK2G;refGene_tx=XM_023240571.1;refGene_gn=CAMK2G;refGene_tx=XM_023240572.1;refGene_gn=CAMK2G;refGene_tx=XM_023240573.1;refGene_gn=CAMK2G;refGene_tx=XM_023240574.1;refGene_gn=CAMK2G;refGene_tx=XM_023240575.1;refGene_gn=CAMK2G;refGene_tx=XM_023240576.1;refGene_gn=CAMK2G;refGene_tx=XM_023240577.1;refGene_gn=CAMK2G;refGene_tx=XM_023240578.1;refGene_gn=CAMK2G;refGene_tx=XM_023240579.1;refGene_gn=CAMK2G;refGene_tx=XM_023240580.1;refGene_gn=CAMK2G;refGene_tx=XM_023240581.1;refGene_gn=CAMK2G;refGene_tx=XM_023240582.1;refGene_gn=CAMK2G;refGene_tx=XM_023240583.1;refGene_gn=CAMK2G;refGene_tx=XM_023240584.1;refGene_gn=CAMK2G;refGene_tx=XM_023240585.1;refGene_gn=CAMK2G;refGene_tx=XM_023240586.1;refGene_gn=CAMK2G;refGene_tx=XM_023240587.1

D2 36443250 36443345 refGene_gn=CAMK2G;refGene_tx=XM_023240571.1;refGene_gn=CAMK2G;refGene_

tx=XM_023240572.1;refGene_gn=CAMK2G;refGene_tx=XM_023240573.1;refGene_gn=CAMK2G;refGene_tx=XM_023240574.1;refGene_gn=CAMK2G;refGene_tx=XM_023240575.1;refGene_gn=CAMK2G;refGene_tx=XM_023240576.1;refGene_gn=CAMK2G;refGene_tx=XM_023240577.1;refGene_gn=CAMK2G;refGene_tx=XM_023240578.1;refGene_gn=CAMK2G;refGene_tx=XM_023240579.1;refGene_gn=CAMK2G;refGene_tx=XM_023240580.1;refGene_gn=CAMK2G;refGene_tx=XM_023240581.1;refGene_gn=CAMK2G;refGene_tx=XM_023240582.1;refGene_gn=CAMK2G;refGene_tx=XM_023240583.1;refGene_gn=CAMK2G;refGene_tx=XM_023240584.1;refGene_gn=CAMK2G;refGene_tx=XM_023240585.1;refGene_gn=CAMK2G;refGene_tx=XM_023240587.1
D2 36444680 36444745
refGene_gn=CAMK2G;refGene_tx=XM_023240571.1;refGene_gn=CAMK2G;refGene_tx=XM_023240572.1;refGene_gn=CAMK2G;refGene_tx=XM_023240573.1;refGene_gn=CAMK2G;refGene_tx=XM_023240574.1;refGene_gn=CAMK2G;refGene_tx=XM_023240575.1;refGene_gn=CAMK2G;refGene_tx=XM_023240576.1;refGene_gn=CAMK2G;refGene_tx=XM_023240577.1;refGene_gn=CAMK2G;refGene_tx=XM_023240578.1;refGene_gn=CAMK2G;refGene_tx=XM_023240579.1;refGene_gn=CAMK2G;refGene_tx=XM_023240580.1;refGene_gn=CAMK2G;refGene_tx=XM_023240581.1;refGene_gn=CAMK2G;refGene_tx=XM_023240582.1;refGene_gn=CAMK2G;refGene_tx=XM_023240583.1;refGene_gn=CAMK2G;refGene_tx=XM_023240584.1;refGene_gn=CAMK2G;refGene_tx=XM_023240585.1;refGene_gn=CAMK2G;refGene_tx=XM_023240587.1
D2 36475059 36475116 refGene_gn=PLAU;refGene_tx=XM_006937896.2
D2 36475564 36475598 refGene_gn=PLAU;refGene_tx=XM_006937896.2
D2 36475749 36475857 refGene_gn=PLAU;refGene_tx=XM_006937896.2
D2 36476173 36476348 refGene_gn=PLAU;refGene_tx=XM_006937896.2
D2 36476540 36476632 refGene_gn=PLAU;refGene_tx=XM_006937896.2
D2 36476815 36477059 refGene_gn=PLAU;refGene_tx=XM_006937896.2
D2 36477275 36477424 refGene_gn=PLAU;refGene_tx=XM_006937896.2
D2 36478014 36478155 refGene_gn=PLAU;refGene_tx=XM_006937896.2
D2 36478506 36478655 refGene_gn=PLAU;refGene_tx=XM_006937896.2
D2 36479322 36479499 refGene_gn=PLAU;refGene_tx=XM_006937896.2
D2 36550096 36550264
refGene_gn=VCL;refGene_tx=XM_003994068.5;refGene_gn=VCL;refGene_tx=XM_003994069.5;refGene_gn=VCL;refGene_tx=XM_006937897.4;refGene_gn=VCL;refGene_tx=XM_019813304.2;refGene_gn=VCL;refGene_tx=XM_019813305.2
D2 36585466 36585537
refGene_gn=VCL;refGene_tx=XM_003994068.5;refGene_gn=VCL;refGene_tx=XM_003994069.5;refGene_gn=VCL;refGene_tx=XM_006937897.4;refGene_gn=VCL;refGene_tx=XM_019813304.2;refGene_gn=VCL;refGene_tx=XM_019813305.2
D2 36613958 36614109
refGene_gn=VCL;refGene_tx=XM_003994068.5;refGene_gn=VCL;refGene_tx=XM_003994069.5;refGene_gn=VCL;refGene_tx=XM_006937897.4;refGene_gn=VCL;refGene_tx=XM_019813304.2;refGene_gn=VCL;refGene_tx=XM_019813305.2
D2 36614292 36614401
refGene_gn=VCL;refGene_tx=XM_003994068.5;refGene_gn=VCL;refGene_tx=XM_003994069.5;refGene_gn=VCL;refGene_tx=XM_006937897.4;refGene_gn=VCL;refGene_tx=XM_019813304.2;refGene_gn=VCL;refGene_tx=XM_019813305.2
D2 36616666 36616789
refGene_gn=VCL;refGene_tx=XM_003994068.5;refGene_gn=VCL;refGene_tx=XM_

003994069.5;refGene_gn=VCL;refGene_tx=XM_006937897.4;refGene_gn=VCL;refGene_tx=XM_019813304.2;refGene_gn=VCL;refGene_tx=XM_019813305.2
D2 36618376 36618537
refGene_gn=VCL;refGene_tx=XM_003994068.5;refGene_gn=VCL;refGene_tx=XM_003994069.5;refGene_gn=VCL;refGene_tx=XM_006937897.4;refGene_gn=VCL;refGene_tx=XM_019813304.2;refGene_gn=VCL;refGene_tx=XM_019813305.2
D2 36627679 36627773
refGene_gn=VCL;refGene_tx=XM_003994068.5;refGene_gn=VCL;refGene_tx=XM_003994069.5;refGene_gn=VCL;refGene_tx=XM_006937897.4;refGene_gn=VCL;refGene_tx=XM_019813304.2;refGene_gn=VCL;refGene_tx=XM_019813305.2
D2 36629038 36629186
refGene_gn=VCL;refGene_tx=XM_003994068.5;refGene_gn=VCL;refGene_tx=XM_003994069.5;refGene_gn=VCL;refGene_tx=XM_006937897.4;refGene_gn=VCL;refGene_tx=XM_019813304.2;refGene_gn=VCL;refGene_tx=XM_019813305.2
D2 36633060 36633214
refGene_gn=VCL;refGene_tx=XM_003994068.5;refGene_gn=VCL;refGene_tx=XM_003994069.5;refGene_gn=VCL;refGene_tx=XM_006937897.4;refGene_gn=VCL;refGene_tx=XM_019813304.2;refGene_gn=VCL;refGene_tx=XM_019813305.2
D2 36634649 36634825
refGene_gn=VCL;refGene_tx=XM_003994068.5;refGene_gn=VCL;refGene_tx=XM_003994069.5;refGene_gn=VCL;refGene_tx=XM_006937897.4;refGene_gn=VCL;refGene_tx=XM_019813304.2;refGene_gn=VCL;refGene_tx=XM_019813305.2
D2 36637816 36638007
refGene_gn=VCL;refGene_tx=XM_003994068.5;refGene_gn=VCL;refGene_tx=XM_003994069.5;refGene_gn=VCL;refGene_tx=XM_006937897.4;refGene_gn=VCL;refGene_tx=XM_019813304.2;refGene_gn=VCL;refGene_tx=XM_019813305.2
D2 36639900 36640100
refGene_gn=VCL;refGene_tx=XM_003994068.5;refGene_gn=VCL;refGene_tx=XM_003994069.5;refGene_gn=VCL;refGene_tx=XM_006937897.4;refGene_gn=VCL;refGene_tx=XM_019813304.2;refGene_gn=VCL;refGene_tx=XM_019813305.2
D2 36641223 36641352
refGene_gn=VCL;refGene_tx=XM_003994068.5;refGene_gn=VCL;refGene_tx=XM_003994069.5;refGene_gn=VCL;refGene_tx=XM_006937897.4;refGene_gn=VCL;refGene_tx=XM_019813304.2;refGene_gn=VCL;refGene_tx=XM_019813305.2
D2 36645021 36645171
refGene_gn=VCL;refGene_tx=XM_003994068.5;refGene_gn=VCL;refGene_tx=XM_003994069.5;refGene_gn=VCL;refGene_tx=XM_006937897.4;refGene_gn=VCL;refGene_tx=XM_019813304.2;refGene_gn=VCL;refGene_tx=XM_019813305.2
D2 36646891 36647000
refGene_gn=VCL;refGene_tx=XM_003994068.5;refGene_gn=VCL;refGene_tx=XM_003994069.5;refGene_gn=VCL;refGene_tx=XM_006937897.4;refGene_gn=VCL;refGene_tx=XM_019813304.2;refGene_gn=VCL;refGene_tx=XM_019813305.2
D2 36648123 36648426
refGene_gn=VCL;refGene_tx=XM_003994068.5;refGene_gn=VCL;refGene_tx=XM_003994069.5;refGene_gn=VCL;refGene_tx=XM_006937897.4;refGene_gn=VCL;refGene_tx=XM_019813304.2;refGene_gn=VCL;refGene_tx=XM_019813305.2
D2 36649654 36649779
refGene_gn=VCL;refGene_tx=XM_003994068.5;refGene_gn=VCL;refGene_tx=XM_003994069.5;refGene_gn=VCL;refGene_tx=XM_006937897.4;refGene_gn=VCL;refGene_tx=XM_019813304.2;refGene_gn=VCL;refGene_tx=XM_019813305.2

D2	36651083	36651269
refGene_gn=VCL;refGene_tx=XM_003994068.5;refGene_gn=VCL;refGene_tx=XM_003994069.5;refGene_gn=VCL;refGene_tx=XM_006937897.4;refGene_gn=VCL;refGene_tx=XM_019813304.2;refGene_gn=VCL;refGene_tx=XM_019813305.2
D2	36654333	36654537
refGene_gn=VCL;refGene_tx=XM_003994069.5;refGene_gn=VCL;refGene_tx=XM_019813304.2;refGene_gn=VCL;refGene_tx=XM_019813305.2
D2	36656562	36656766
refGene_gn=VCL;refGene_tx=XM_003994068.5;refGene_gn=VCL;refGene_tx=XM_003994069.5;refGene_gn=VCL;refGene_tx=XM_006937897.4;refGene_gn=VCL;refGene_tx=XM_019813304.2;refGene_gn=VCL;refGene_tx=XM_019813305.2
D2	36657146	36657251
refGene_gn=VCL;refGene_tx=XM_003994068.5;refGene_gn=VCL;refGene_tx=XM_003994069.5;refGene_gn=VCL;refGene_tx=XM_006937897.4;refGene_gn=VCL;refGene_tx=XM_019813304.2;refGene_gn=VCL;refGene_tx=XM_019813305.2
D2	36659480	36659627
refGene_gn=VCL;refGene_tx=XM_003994068.5;refGene_gn=VCL;refGene_tx=XM_003994069.5;refGene_gn=VCL;refGene_tx=XM_006937897.4;refGene_gn=VCL;refGene_tx=XM_019813304.2;refGene_gn=VCL;refGene_tx=XM_019813305.2
D2	36664901	36665002
refGene_gn=AP3M1;refGene_tx=XM_003994070.3;refGene_gn=AP3M1;refGene_tx=XM_006937899.4;refGene_gn=AP3M1;refGene_tx=XM_023240588.1
D2	36665441	36665586
refGene_gn=AP3M1;refGene_tx=XM_003994070.3;refGene_gn=AP3M1;refGene_tx=XM_006937899.4;refGene_gn=AP3M1;refGene_tx=XM_023240588.1
D2	36667272	36667480
refGene_gn=AP3M1;refGene_tx=XM_003994070.3;refGene_gn=AP3M1;refGene_tx=XM_006937899.4;refGene_gn=AP3M1;refGene_tx=XM_023240588.1
D2	36668337	36668471
refGene_gn=AP3M1;refGene_tx=XM_003994070.3;refGene_gn=AP3M1;refGene_tx=XM_006937899.4;refGene_gn=AP3M1;refGene_tx=XM_023240588.1
D2	36669083	36669169
refGene_gn=AP3M1;refGene_tx=XM_003994070.3;refGene_gn=AP3M1;refGene_tx=XM_006937899.4;refGene_gn=AP3M1;refGene_tx=XM_023240588.1
D2	36670583	36670721
refGene_gn=AP3M1;refGene_tx=XM_003994070.3;refGene_gn=AP3M1;refGene_tx=XM_006937899.4;refGene_gn=AP3M1;refGene_tx=XM_023240588.1
D2	36673626	36673798
refGene_gn=AP3M1;refGene_tx=XM_003994070.3;refGene_gn=AP3M1;refGene_tx=XM_006937899.4;refGene_gn=AP3M1;refGene_tx=XM_023240588.1
D2	36674506	36674779
refGene_gn=AP3M1;refGene_tx=XM_003994070.3;refGene_gn=AP3M1;refGene_tx=XM_006937899.4;refGene_gn=AP3M1;refGene_tx=XM_023240588.1
D2	36685561	36685626
refGene_gn=ADK;refGene_tx=XM_019813326.2;refGene_gn=ADK;refGene_tx=XM_019813327.1;refGene_gn=ADK;refGene_tx=XM_023240589.1
D2	36724057	36724119	refGene_gn=ADK;refGene_tx=XM_023240589.1
D2	36730475	36730551	refGene_gn=ADK;refGene_tx=XM_023240589.1
D2	36756058	36756112
refGene_gn=ADK;refGene_tx=XM_019813326.2;refGene_gn=ADK;refGene_tx=XM_

023240589.1
D2	36855713	36855792
refGene_gn=ADK;refGene_tx=XM_019813326.2;refGene_gn=ADK;refGene_tx=XM_019813327.1;refGene_gn=ADK;refGene_tx=XM_023240589.1
D2	36864799	36865051
refGene_gn=LOC101090551;refGene_tx=XM_019813328.2
D2	36939438	36939611
refGene_gn=ADK;refGene_tx=XM_019813326.2;refGene_gn=ADK;refGene_tx=XM_019813327.1;refGene_gn=ADK;refGene_tx=XM_023240589.1
D2	36944029	36944138
refGene_gn=ADK;refGene_tx=XM_019813326.2;refGene_gn=ADK;refGene_tx=XM_019813327.1;refGene_gn=ADK;refGene_tx=XM_023240589.1
D2	37047283	37047454
refGene_gn=ADK;refGene_tx=XM_019813326.2;refGene_gn=ADK;refGene_tx=XM_019813327.1;refGene_gn=ADK;refGene_tx=XM_023240589.1
D2	37108368	37108404
refGene_gn=ADK;refGene_tx=XM_019813326.2;refGene_gn=ADK;refGene_tx=XM_019813327.1;refGene_gn=ADK;refGene_tx=XM_023240589.1
D2	37117184	37117299
refGene_gn=ADK;refGene_tx=XM_019813326.2;refGene_gn=ADK;refGene_tx=XM_019813327.1;refGene_gn=ADK;refGene_tx=XM_023240589.1
D2	37176527	37176614
refGene_gn=ADK;refGene_tx=XM_019813326.2;refGene_gn=ADK;refGene_tx=XM_019813327.1;refGene_gn=ADK;refGene_tx=XM_023240589.1
D2	37217148	37217273
refGene_gn=ADK;refGene_tx=XM_019813326.2;refGene_gn=ADK;refGene_tx=XM_019813327.1;refGene_gn=ADK;refGene_tx=XM_023240589.1
D2	37314200	37314821	refGene_gn=KAT6B;refGene_tx=XM_019813329.2
D2	37420036	37420145	refGene_gn=KAT6B;refGene_tx=XM_019813329.2
D2	37431744	37431860	refGene_gn=KAT6B;refGene_tx=XM_019813329.2
D2	37432108	37432190	refGene_gn=KAT6B;refGene_tx=XM_019813329.2
D2	37434653	37434786	refGene_gn=KAT6B;refGene_tx=XM_019813329.2
D2	37437542	37438029	refGene_gn=KAT6B;refGene_tx=XM_019813329.2
D2	37438030	37438475	refGene_gn=KAT6B;refGene_tx=XM_019813329.2
D2	37439491	37439613	refGene_gn=KAT6B;refGene_tx=XM_019813329.2
D2	37441210	37441326	refGene_gn=KAT6B;refGene_tx=XM_019813329.2
D2	37443115	37443257	refGene_gn=KAT6B;refGene_tx=XM_019813329.2
D2	37446283	37446445	refGene_gn=KAT6B;refGene_tx=XM_019813329.2
D2	37449965	37450059	refGene_gn=KAT6B;refGene_tx=XM_019813329.2
D2	37476847	37477079	refGene_gn=KAT6B;refGene_tx=XM_019813329.2
D2	37477383	37477543	refGene_gn=KAT6B;refGene_tx=XM_019813329.2
D2	37478150	37478546	refGene_gn=KAT6B;refGene_tx=XM_019813329.2
D2	37481247	37481539	refGene_gn=KAT6B;refGene_tx=XM_019813329.2
D2	37484090	37486618	refGene_gn=KAT6B;refGene_tx=XM_019813329.2
D2	37493434	37493679
refGene_gn=DUPD1;refGene_tx=XM_003994074.5;refGene_gn=DUPD1;refGene_tx=XM_006937901.4;refGene_gn=DUPD1;refGene_tx=XM_006937902.4
D2	37501165	37501386
refGene_gn=DUPD1;refGene_tx=XM_003994074.5;refGene_gn=DUPD1;refGene_tx=XM_006937901.4;refGene_gn=DUPD1;refGene_tx=XM_006937902.4

D2 37515681 37515909
refGene_gn=DUPD1;refGene_tx=XM_003994074.5;refGene_gn=DUPD1;refGene_tx=XM_006937901.4;refGene_gn=DUPD1;refGene_tx=XM_006937902.4
D2 37528505 37528669 refGene_gn=DUPD1;refGene_tx=XM_003994074.5
D2 37549486 37549689
refGene_gn=LOC101091309;refGene_tx=XM_003994077.5;refGene_gn=LOC101091309;refGene_tx=XM_006937904.4;refGene_gn=LOC101091309;refGene_tx=XM_011287308.3;refGene_gn=LOC101091309;refGene_tx=XM_011287309.3
D2 37550410 37550631
refGene_gn=LOC101091309;refGene_tx=XM_003994077.5;refGene_gn=LOC101091309;refGene_tx=XM_006937904.4;refGene_gn=LOC101091309;refGene_tx=XM_011287308.3;refGene_gn=LOC101091309;refGene_tx=XM_011287309.3
D2 37553055 37553229
refGene_gn=LOC101091309;refGene_tx=XM_003994077.5;refGene_gn=LOC101091309;refGene_tx=XM_006937904.4;refGene_gn=LOC101091309;refGene_tx=XM_011287308.3;refGene_gn=LOC101091309;refGene_tx=XM_011287309.3
D2 37554087 37554440
refGene_gn=LOC101091309;refGene_tx=XM_006937904.4
D2 37556832 37556927
refGene_gn=LOC101091309;refGene_tx=XM_011287308.3;refGene_gn=LOC101091309;refGene_tx=XM_011287309.3
D2 37557933 37558137
refGene_gn=LOC101091309;refGene_tx=XM_011287308.3;refGene_gn=LOC101091309;refGene_tx=XM_011287309.3
D2 37560388 37560475
refGene_gn=LOC101091309;refGene_tx=XM_011287308.3;refGene_gn=LOC101091309;refGene_tx=XM_011287309.3
D2 37562134 37562369
refGene_gn=LOC109491364;refGene_tx=XM_019814069.1
D2 37563137 37563286
refGene_gn=LOC109491364;refGene_tx=XM_019814069.1
D2 37565970 37566144
refGene_gn=SAMD8;refGene_tx=XM_003994082.5;refGene_gn=SAMD8;refGene_tx=XM_019813332.2
D2 37566752 37566782 refGene_gn=SAMD8;refGene_tx=XM_019813332.2
D2 37592597 37593190
refGene_gn=SAMD8;refGene_tx=XM_003994082.5;refGene_gn=SAMD8;refGene_tx=XM_006937907.3;refGene_gn=SAMD8;refGene_tx=XM_006937908.3;refGene_gn=SAMD8;refGene_tx=XM_019813331.2;refGene_gn=SAMD8;refGene_tx=XM_019813332.2
D2 37605217 37605313
refGene_gn=SAMD8;refGene_tx=XM_003994082.5;refGene_gn=SAMD8;refGene_tx=XM_006937907.3;refGene_gn=SAMD8;refGene_tx=XM_006937908.3;refGene_gn=SAMD8;refGene_tx=XM_019813331.2;refGene_gn=SAMD8;refGene_tx=XM_019813332.2
D2 37607525 37607643
refGene_gn=SAMD8;refGene_tx=XM_003994082.5;refGene_gn=SAMD8;refGene_tx=XM_006937907.3;refGene_gn=SAMD8;refGene_tx=XM_006937908.3;refGene_gn=SAMD8;refGene_tx=XM_019813331.2;refGene_gn=SAMD8;refGene_tx=XM_019813332.2

D2 37611662 37611851
refGene_gn=SAMD8;refGene_tx=XM_003994082.5;refGene_gn=SAMD8;refGene_tx=XM_006937907.3;refGene_gn=SAMD8;refGene_tx=XM_006937908.3;refGene_gn=SAMD8;refGene_tx=XM_019813331.2;refGene_gn=SAMD8;refGene_tx=XM_019813332.2
D2 37611988 37612293
refGene_gn=SAMD8;refGene_tx=XM_003994082.5;refGene_gn=SAMD8;refGene_tx=XM_006937907.3;refGene_gn=SAMD8;refGene_tx=XM_006937908.3;refGene_gn=SAMD8;refGene_tx=XM_019813331.2
D2 37646075 37646106
refGene_gn=VDAC2;refGene_tx=XM_011287307.3;refGene_gn=VDAC2;refGene_tx=XM_019813333.2;refGene_gn=VDAC2;refGene_tx=XM_023240590.1
D2 37647205 37647274
refGene_gn=VDAC2;refGene_tx=XM_011287307.3;refGene_gn=VDAC2;refGene_tx=XM_019813333.2;refGene_gn=VDAC2;refGene_tx=XM_023240590.1
D2 37648091 37648141
refGene_gn=VDAC2;refGene_tx=XM_011287307.3;refGene_gn=VDAC2;refGene_tx=XM_019813333.2;refGene_gn=VDAC2;refGene_tx=XM_023240590.1
D2 37651226 37651379
refGene_gn=VDAC2;refGene_tx=XM_011287307.3;refGene_gn=VDAC2;refGene_tx=XM_019813333.2;refGene_gn=VDAC2;refGene_tx=XM_023240590.1
D2 37651472 37651525
refGene_gn=VDAC2;refGene_tx=XM_011287307.3;refGene_gn=VDAC2;refGene_tx=XM_019813333.2;refGene_gn=VDAC2;refGene_tx=XM_023240590.1
D2 37652128 37652356
refGene_gn=VDAC2;refGene_tx=XM_011287307.3;refGene_gn=VDAC2;refGene_tx=XM_019813333.2;refGene_gn=VDAC2;refGene_tx=XM_023240590.1
D2 37653486 37653637
refGene_gn=VDAC2;refGene_tx=XM_011287307.3;refGene_gn=VDAC2;refGene_tx=XM_019813333.2;refGene_gn=VDAC2;refGene_tx=XM_023240590.1
D2 37656182 37656240
refGene_gn=VDAC2;refGene_tx=XM_011287307.3;refGene_gn=VDAC2;refGene_tx=XM_019813333.2;refGene_gn=VDAC2;refGene_tx=XM_023240590.1
D2 37657382 37657474
refGene_gn=VDAC2;refGene_tx=XM_011287307.3;refGene_gn=VDAC2;refGene_tx=XM_019813333.2;refGene_gn=VDAC2;refGene_tx=XM_023240590.1
D2 37660827 37660980
refGene_gn=COMTD1;refGene_tx=XM_003994084.4;refGene_gn=COMTD1;refGene_tx=XM_006937909.3;refGene_gn=COMTD1;refGene_tx=XM_006937910.3;refGene_gn=COMTD1;refGene_tx=XM_006937912.4;refGene_gn=COMTD1;refGene_tx=XM_011287304.2;refGene_gn=COMTD1;refGene_tx=XM_019813334.1
D2 37661183 37661317
refGene_gn=COMTD1;refGene_tx=XM_003994084.4;refGene_gn=COMTD1;refGene_tx=XM_006937909.3;refGene_gn=COMTD1;refGene_tx=XM_006937910.3;refGene_gn=COMTD1;refGene_tx=XM_006937912.4;refGene_gn=COMTD1;refGene_tx=XM_011287304.2;refGene_gn=COMTD1;refGene_tx=XM_019813334.1
D2 37661586 37661641
refGene_gn=COMTD1;refGene_tx=XM_003994084.4;refGene_gn=COMTD1;refGene_tx=XM_006937909.3;refGene_gn=COMTD1;refGene_tx=XM_006937910.3;refGene_gn=COMTD1;refGene_tx=XM_006937912.4;refGene_gn=COMTD1;refGene_tx=XM_01

1287304.2;refGene_gn=COMTD1;refGene_tx=XM_019813334.1
D2 37661745 37661864
refGene_gn=COMTD1;refGene_tx=XM_003994084.4;refGene_gn=COMTD1;refGene_tx=XM_006937909.3;refGene_gn=COMTD1;refGene_tx=XM_006937910.3;refGene_gn=COMTD1;refGene_tx=XM_006937912.4;refGene_gn=COMTD1;refGene_tx=XM_011287304.2;refGene_gn=COMTD1;refGene_tx=XM_019813334.1
D2 37661952 37662058
refGene_gn=COMTD1;refGene_tx=XM_003994084.4;refGene_gn=COMTD1;refGene_tx=XM_006937909.3;refGene_gn=COMTD1;refGene_tx=XM_006937910.3;refGene_gn=COMTD1;refGene_tx=XM_006937912.4;refGene_gn=COMTD1;refGene_tx=XM_011287304.2;refGene_gn=COMTD1;refGene_tx=XM_019813334.1
D2 37662315 37662443
refGene_gn=COMTD1;refGene_tx=XM_003994084.4;refGene_gn=COMTD1;refGene_tx=XM_006937909.3;refGene_gn=COMTD1;refGene_tx=XM_006937910.3;refGene_gn=COMTD1;refGene_tx=XM_006937912.4;refGene_gn=COMTD1;refGene_tx=XM_011287304.2;refGene_gn=COMTD1;refGene_tx=XM_019813334.1
D2 37662599 37662693
refGene_gn=COMTD1;refGene_tx=XM_003994084.4;refGene_gn=COMTD1;refGene_tx=XM_006937909.3;refGene_gn=COMTD1;refGene_tx=XM_006937910.3;refGene_gn=COMTD1;refGene_tx=XM_006937912.4;refGene_gn=COMTD1;refGene_tx=XM_011287304.2;refGene_gn=COMTD1;refGene_tx=XM_019813334.1
D2 37781430 37783065 refGene_gn=ZNF503;refGene_tx=XM_003994085.5
D2 37783892 37784213 refGene_gn=ZNF503;refGene_tx=XM_003994085.5
D2 37808558 37808588
refGene_gn=LRMDA;refGene_tx=XM_023240595.1;refGene_gn=LRMDA;refGene_tx=XM_023240596.1
D2 37814585 37814686
refGene_gn=LRMDA;refGene_tx=XM_023240595.1;refGene_gn=LRMDA;refGene_tx=XM_023240596.1
D2 38020874 38020945
refGene_gn=LRMDA;refGene_tx=XM_023240591.1;refGene_gn=LRMDA;refGene_tx=XM_023240592.1
D2 38213864 38213931
refGene_gn=LRMDA;refGene_tx=XM_023240591.1;refGene_gn=LRMDA;refGene_tx=XM_023240593.1;refGene_gn=LRMDA;refGene_tx=XM_023240594.1
D2 38383327 38383454
refGene_gn=LRMDA;refGene_tx=XM_023240591.1;refGene_gn=LRMDA;refGene_tx=XM_023240592.1;refGene_gn=LRMDA;refGene_tx=XM_023240593.1;refGene_gn=LRMDA;refGene_tx=XM_023240594.1;refGene_gn=LRMDA;refGene_tx=XM_023240595.1;refGene_gn=LRMDA;refGene_tx=XM_023240596.1
D2 38394914 38395054
refGene_gn=LRMDA;refGene_tx=XM_023240591.1;refGene_gn=LRMDA;refGene_tx=XM_023240592.1;refGene_gn=LRMDA;refGene_tx=XM_023240593.1;refGene_gn=LRMDA;refGene_tx=XM_023240594.1;refGene_gn=LRMDA;refGene_tx=XM_023240595.1;refGene_gn=LRMDA;refGene_tx=XM_023240596.1
D2 38405702 38405820
refGene_gn=LRMDA;refGene_tx=XM_023240591.1;refGene_gn=LRMDA;refGene_tx=XM_023240592.1;refGene_gn=LRMDA;refGene_tx=XM_023240593.1;refGene_gn=LRMDA;refGene_tx=XM_023240594.1;refGene_gn=LRMDA;refGene_tx=XM_023240595.1;refGene_gn=LRMDA;refGene_tx=XM_023240596.1

D2 38407498 38407575
refGene_gn=LRMDA;refGene_tx=XM_023240591.1;refGene_gn=LRMDA;refGene_tx=XM_023240592.1;refGene_gn=LRMDA;refGene_tx=XM_023240593.1;refGene_gn=LRMDA;refGene_tx=XM_023240594.1
D2 38633725 38633810
refGene_gn=LRMDA;refGene_tx=XM_023240591.1;refGene_gn=LRMDA;refGene_tx=XM_023240592.1;refGene_gn=LRMDA;refGene_tx=XM_023240593.1;refGene_gn=LRMDA;refGene_tx=XM_023240594.1;refGene_gn=LRMDA;refGene_tx=XM_023240595.1;refGene_gn=LRMDA;refGene_tx=XM_023240596.1
D2 38832853 38832935
refGene_gn=LRMDA;refGene_tx=XM_023240591.1;refGene_gn=LRMDA;refGene_tx=XM_023240592.1;refGene_gn=LRMDA;refGene_tx=XM_023240593.1;refGene_gn=LRMDA;refGene_tx=XM_023240594.1;refGene_gn=LRMDA;refGene_tx=XM_023240595.1;refGene_gn=LRMDA;refGene_tx=XM_023240596.1
D2 38835451 38835549
refGene_gn=LRMDA;refGene_tx=XM_023240591.1;refGene_gn=LRMDA;refGene_tx=XM_023240592.1;refGene_gn=LRMDA;refGene_tx=XM_023240593.1;refGene_gn=LRMDA;refGene_tx=XM_023240594.1
D2 39072559 39072714
refGene_gn=KCNMA1;refGene_tx=XM_011287300.3;refGene_gn=KCNMA1;refGene_tx=XM_011287302.3;refGene_gn=KCNMA1;refGene_tx=XM_011287303.3;refGene_gn=KCNMA1;refGene_tx=XM_019813335.2;refGene_gn=KCNMA1;refGene_tx=XM_019813336.2;refGene_gn=KCNMA1;refGene_tx=XM_019813338.2;refGene_gn=KCNMA1;refGene_tx=XM_019813339.2;refGene_gn=KCNMA1;refGene_tx=XM_019813340.2;refGene_gn=KCNMA1;refGene_tx=XM_019813341.2;refGene_gn=KCNMA1;refGene_tx=XM_019813342.2;refGene_gn=KCNMA1;refGene_tx=XM_019813344.2;refGene_gn=KCNMA1;refGene_tx=XM_019813345.2;refGene_gn=KCNMA1;refGene_tx=XM_019813346.2;refGene_gn=KCNMA1;refGene_tx=XM_019813347.2;refGene_gn=KCNMA1;refGene_tx=XM_019813349.2;refGene_gn=KCNMA1;refGene_tx=XM_019813350.2;refGene_gn=KCNMA1;refGene_tx=XM_019813351.2;refGene_gn=KCNMA1;refGene_tx=XM_019813352.2;refGene_gn=KCNMA1;refGene_tx=XM_019813353.2;refGene_gn=KCNMA1;refGene_tx=XM_019813354.2;refGene_gn=KCNMA1;refGene_tx=XM_019813355.2;refGene_gn=KCNMA1;refGene_tx=XM_019813356.2;refGene_gn=KCNMA1;refGene_tx=XM_019813358.2;refGene_gn=KCNMA1;refGene_tx=XM_019813360.2;refGene_gn=KCNMA1;refGene_tx=XM_019813362.2;refGene_gn=KCNMA1;refGene_tx=XM_019813363.2;refGene_gn=KCNMA1;refGene_tx=XM_019813364.2;refGene_gn=KCNMA1;refGene_tx=XM_019813366.2;refGene_gn=KCNMA1;refGene_tx=XM_019813368.2;refGene_gn=KCNMA1;refGene_tx=XM_019813369.2;refGene_gn=KCNMA1;refGene_tx=XM_019813371.2;refGene_gn=KCNMA1;refGene_tx=XM_019813373.1;refGene_gn=KCNMA1;refGene_tx=XM_019813374.2;refGene_gn=KCNMA1;refGene_tx=XM_023240597.1;refGene_gn=KCNMA1;refGene_tx=XM_023240601.1
D2 39079825 39079854
refGene_gn=KCNMA1;refGene_tx=XM_019813335.2;refGene_gn=KCNMA1;refGene_tx=XM_019813336.2;refGene_gn=KCNMA1;refGene_tx=XM_019813338.2;refGene_gn=KCNMA1;refGene_tx=XM_019813339.2;refGene_gn=KCNMA1;refGene_tx=XM_019813340.2;refGene_gn=KCNMA1;refGene_tx=XM_019813341.2;refGene_gn=KCNMA1;refGene_tx=XM_019813342.2;refGene_gn=KCNMA1;refGene_tx=XM_019813346.2;refGene_gn=KCNMA1;refGene_tx=XM_019813347.2;refGene_gn=KCNMA1;refGene_tx=XM_019813349.2;refGene_gn=KCNMA1;refGene_tx=XM_019813350.2;refGene_gn=KCNMA1;refGene_tx=XM_019813351.2;refGene_gn=KCNMA1;refGene_tx=XM_

019813352.2;refGene_gn=KCNMA1;refGene_tx=XM_019813353.2;refGene_gn=KCNMA1;refGene_tx=XM_019813360.2;refGene_gn=KCNMA1;refGene_tx=XM_019813362.2;refGene_gn=KCNMA1;refGene_tx=XM_019813363.2;refGene_gn=KCNMA1;refGene_tx=XM_019813373.1;refGene_gn=KCNMA1;refGene_tx=XM_019813374.2
D2 39082130 39082380
refGene_gn=KCNMA1;refGene_tx=XM_011287300.3;refGene_gn=KCNMA1;refGene_tx=XM_011287302.3;refGene_gn=KCNMA1;refGene_tx=XM_011287303.3;refGene_gn=KCNMA1;refGene_tx=XM_019813335.2;refGene_gn=KCNMA1;refGene_tx=XM_019813336.2;refGene_gn=KCNMA1;refGene_tx=XM_019813338.2;refGene_gn=KCNMA1;refGene_tx=XM_019813339.2;refGene_gn=KCNMA1;refGene_tx=XM_019813340.2;refGene_gn=KCNMA1;refGene_tx=XM_019813341.2;refGene_gn=KCNMA1;refGene_tx=XM_019813342.2;refGene_gn=KCNMA1;refGene_tx=XM_019813344.2;refGene_gn=KCNMA1;refGene_tx=XM_019813345.2;refGene_gn=KCNMA1;refGene_tx=XM_019813346.2;refGene_gn=KCNMA1;refGene_tx=XM_019813347.2;refGene_gn=KCNMA1;refGene_tx=XM_019813349.2;refGene_gn=KCNMA1;refGene_tx=XM_019813350.2;refGene_gn=KCNMA1;refGene_tx=XM_019813351.2;refGene_gn=KCNMA1;refGene_tx=XM_019813352.2;refGene_gn=KCNMA1;refGene_tx=XM_019813353.2;refGene_gn=KCNMA1;refGene_tx=XM_019813354.2;refGene_gn=KCNMA1;refGene_tx=XM_019813355.2;refGene_gn=KCNMA1;refGene_tx=XM_019813356.2;refGene_gn=KCNMA1;refGene_tx=XM_019813357.2;refGene_gn=KCNMA1;refGene_tx=XM_019813358.2;refGene_gn=KCNMA1;refGene_tx=XM_019813360.2;refGene_gn=KCNMA1;refGene_tx=XM_019813362.2;refGene_gn=KCNMA1;refGene_tx=XM_019813363.2;refGene_gn=KCNMA1;refGene_tx=XM_019813364.2;refGene_gn=KCNMA1;refGene_tx=XM_019813366.2;refGene_gn=KCNMA1;refGene_tx=XM_019813368.2;refGene_gn=KCNMA1;refGene_tx=XM_019813369.2;refGene_gn=KCNMA1;refGene_tx=XM_019813371.2;refGene_gn=KCNMA1;refGene_tx=XM_019813372.2;refGene_gn=KCNMA1;refGene_tx=XM_019813373.1;refGene_gn=KCNMA1;refGene_tx=XM_019813374.2;refGene_gn=KCNMA1;refGene_tx=XM_023240597.1;refGene_gn=KCNMA1;refGene_tx=XM_023240599.1;refGene_gn=KCNMA1;refGene_tx=XM_023240600.1;refGene_gn=KCNMA1;refGene_tx=XM_023240601.1
D2 39084744 39084863
refGene_gn=KCNMA1;refGene_tx=XM_011287300.3;refGene_gn=KCNMA1;refGene_tx=XM_011287302.3;refGene_gn=KCNMA1;refGene_tx=XM_011287303.3;refGene_gn=KCNMA1;refGene_tx=XM_019813335.2;refGene_gn=KCNMA1;refGene_tx=XM_019813336.2;refGene_gn=KCNMA1;refGene_tx=XM_019813338.2;refGene_gn=KCNMA1;refGene_tx=XM_019813339.2;refGene_gn=KCNMA1;refGene_tx=XM_019813340.2;refGene_gn=KCNMA1;refGene_tx=XM_019813341.2;refGene_gn=KCNMA1;refGene_tx=XM_019813342.2;refGene_gn=KCNMA1;refGene_tx=XM_019813344.2;refGene_gn=KCNMA1;refGene_tx=XM_019813345.2;refGene_gn=KCNMA1;refGene_tx=XM_019813346.2;refGene_gn=KCNMA1;refGene_tx=XM_019813347.2;refGene_gn=KCNMA1;refGene_tx=XM_019813349.2;refGene_gn=KCNMA1;refGene_tx=XM_019813350.2;refGene_gn=KCNMA1;refGene_tx=XM_019813351.2;refGene_gn=KCNMA1;refGene_tx=XM_019813352.2;refGene_gn=KCNMA1;refGene_tx=XM_019813353.2;refGene_gn=KCNMA1;refGene_tx=XM_019813354.2;refGene_gn=KCNMA1;refGene_tx=XM_019813355.2;refGene_gn=KCNMA1;refGene_tx=XM_019813356.2;refGene_gn=KCNMA1;refGene_tx=XM_019813357.2;refGene_gn=KCNMA1;refGene_tx=XM_019813358.2;refGene_gn=KCNMA1;refGene_tx=XM_019813360.2;refGene_gn=KCNMA1;refGene_tx=XM_019813362.2;refGene_gn=KCNMA1;refGene_tx=XM_019813363.2;refGene_gn=KCNMA1;refGene_tx=XM_019813364.2;refGene_gn=KCNMA1;refGene_tx=XM_019813366.2;refGene_gn=KCNMA1;refGene_tx=XM_019813368.2;refGene_gn

=KCNMA1;refGene_tx=XM_019813369.2;refGene_gn=KCNMA1;refGene_tx=XM_0198
13371.2;refGene_gn=KCNMA1;refGene_tx=XM_019813372.2;refGene_gn=KCNMA1;
refGene_tx=XM_019813373.1;refGene_gn=KCNMA1;refGene_tx=XM_019813374.2;
refGene_gn=KCNMA1;refGene_tx=XM_023240597.1;refGene_gn=KCNMA1;refGene_
tx=XM_023240599.1;refGene_gn=KCNMA1;refGene_tx=XM_023240600.1;refGene_
gn=KCNMA1;refGene_tx=XM_023240601.1
D2 39086751 39086946
refGene_gn=KCNMA1;refGene_tx=XM_011287300.3;refGene_gn=KCNMA1;refGene_
tx=XM_011287302.3;refGene_gn=KCNMA1;refGene_tx=XM_011287303.3;refGene_
gn=KCNMA1;refGene_tx=XM_019813335.2;refGene_gn=KCNMA1;refGene_tx=XM_01
9813336.2;refGene_gn=KCNMA1;refGene_tx=XM_019813338.2;refGene_gn=KCNMA
1;refGene_tx=XM_019813339.2;refGene_gn=KCNMA1;refGene_tx=XM_019813340.
2;refGene_gn=KCNMA1;refGene_tx=XM_019813341.2;refGene_gn=KCNMA1;refGen
e_tx=XM_019813342.2;refGene_gn=KCNMA1;refGene_tx=XM_019813344.2;refGen
e_gn=KCNMA1;refGene_tx=XM_019813345.2;refGene_gn=KCNMA1;refGene_tx=XM_
019813346.2;refGene_gn=KCNMA1;refGene_tx=XM_019813347.2;refGene_gn=KCN
MA1;refGene_tx=XM_019813349.2;refGene_gn=KCNMA1;refGene_tx=XM_01981335
0.2;refGene_gn=KCNMA1;refGene_tx=XM_019813351.2;refGene_gn=KCNMA1;refG
ene_tx=XM_019813352.2;refGene_gn=KCNMA1;refGene_tx=XM_019813353.2;refG
ene_gn=KCNMA1;refGene_tx=XM_019813354.2;refGene_gn=KCNMA1;refGene_tx=X
M_019813355.2;refGene_gn=KCNMA1;refGene_tx=XM_019813356.2;refGene_gn=K
CNMA1;refGene_tx=XM_019813357.2;refGene_gn=KCNMA1;refGene_tx=XM_019813
358.2;refGene_gn=KCNMA1;refGene_tx=XM_019813360.2;refGene_gn=KCNMA1;re
fGene_tx=XM_019813362.2;refGene_gn=KCNMA1;refGene_tx=XM_019813363.2;re
fGene_gn=KCNMA1;refGene_tx=XM_019813364.2;refGene_gn=KCNMA1;refGene_tx
=XM_019813366.2;refGene_gn=KCNMA1;refGene_tx=XM_019813368.2;refGene_gn
=KCNMA1;refGene_tx=XM_019813369.2;refGene_gn=KCNMA1;refGene_tx=XM_0198
13371.2;refGene_gn=KCNMA1;refGene_tx=XM_019813372.2;refGene_gn=KCNMA1;
refGene_tx=XM_019813373.1;refGene_gn=KCNMA1;refGene_tx=XM_019813374.2;
refGene_gn=KCNMA1;refGene_tx=XM_023240597.1;refGene_gn=KCNMA1;refGene_
tx=XM_023240599.1;refGene_gn=KCNMA1;refGene_tx=XM_023240600.1;refGene_
gn=KCNMA1;refGene_tx=XM_023240601.1
D2 39107100 39107231
refGene_gn=KCNMA1;refGene_tx=XM_011287300.3;refGene_gn=KCNMA1;refGene_
tx=XM_011287302.3;refGene_gn=KCNMA1;refGene_tx=XM_011287303.3;refGene_
gn=KCNMA1;refGene_tx=XM_019813335.2;refGene_gn=KCNMA1;refGene_tx=XM_01
9813336.2;refGene_gn=KCNMA1;refGene_tx=XM_019813338.2;refGene_gn=KCNMA
1;refGene_tx=XM_019813339.2;refGene_gn=KCNMA1;refGene_tx=XM_019813340.
2;refGene_gn=KCNMA1;refGene_tx=XM_019813341.2;refGene_gn=KCNMA1;refGen
e_tx=XM_019813342.2;refGene_gn=KCNMA1;refGene_tx=XM_019813344.2;refGen
e_gn=KCNMA1;refGene_tx=XM_019813345.2;refGene_gn=KCNMA1;refGene_tx=XM_
019813346.2;refGene_gn=KCNMA1;refGene_tx=XM_019813347.2;refGene_gn=KCN
MA1;refGene_tx=XM_019813349.2;refGene_gn=KCNMA1;refGene_tx=XM_01981335
0.2;refGene_gn=KCNMA1;refGene_tx=XM_019813351.2;refGene_gn=KCNMA1;refG
ene_tx=XM_019813352.2;refGene_gn=KCNMA1;refGene_tx=XM_019813353.2;refG
ene_gn=KCNMA1;refGene_tx=XM_019813354.2;refGene_gn=KCNMA1;refGene_tx=X
M_019813355.2;refGene_gn=KCNMA1;refGene_tx=XM_019813356.2;refGene_gn=K
CNMA1;refGene_tx=XM_019813357.2;refGene_gn=KCNMA1;refGene_tx=XM_019813
358.2;refGene_gn=KCNMA1;refGene_tx=XM_019813360.2;refGene_gn=KCNMA1;re
fGene_tx=XM_019813362.2;refGene_gn=KCNMA1;refGene_tx=XM_019813363.2;re

fGene_gn=KCNMA1;refGene_tx=XM_019813364.2;refGene_gn=KCNMA1;refGene_tx=XM_019813366.2;refGene_gn=KCNMA1;refGene_tx=XM_019813368.2;refGene_gn=KCNMA1;refGene_tx=XM_019813369.2;refGene_gn=KCNMA1;refGene_tx=XM_019813371.2;refGene_gn=KCNMA1;refGene_tx=XM_019813372.2;refGene_gn=KCNMA1;refGene_tx=XM_019813373.1;refGene_gn=KCNMA1;refGene_tx=XM_019813374.2;refGene_gn=KCNMA1;refGene_tx=XM_023240597.1;refGene_gn=KCNMA1;refGene_tx=XM_023240599.1;refGene_gn=KCNMA1;refGene_tx=XM_023240600.1;refGene_gn=KCNMA1;refGene_tx=XM_023240601.1
D2 39111155 39111236
refGene_gn=KCNMA1;refGene_tx=XM_019813335.2;refGene_gn=KCNMA1;refGene_tx=XM_019813336.2;refGene_gn=KCNMA1;refGene_tx=XM_019813338.2;refGene_gn=KCNMA1;refGene_tx=XM_019813339.2;refGene_gn=KCNMA1;refGene_tx=XM_019813340.2;refGene_gn=KCNMA1;refGene_tx=XM_019813344.2;refGene_gn=KCNMA1;refGene_tx=XM_019813345.2;refGene_gn=KCNMA1;refGene_tx=XM_019813346.2;refGene_gn=KCNMA1;refGene_tx=XM_019813347.2;refGene_gn=KCNMA1;refGene_tx=XM_019813349.2;refGene_gn=KCNMA1;refGene_tx=XM_019813350.2;refGene_gn=KCNMA1;refGene_tx=XM_019813351.2;refGene_gn=KCNMA1;refGene_tx=XM_019813352.2;refGene_gn=KCNMA1;refGene_tx=XM_019813353.2;refGene_gn=KCNMA1;refGene_tx=XM_019813364.2;refGene_gn=KCNMA1;refGene_tx=XM_019813366.2;refGene_gn=KCNMA1;refGene_tx=XM_019813368.2;refGene_gn=KCNMA1;refGene_tx=XM_019813369.2;refGene_gn=KCNMA1;refGene_tx=XM_019813373.1
D2 39112018 39112132
refGene_gn=KCNMA1;refGene_tx=XM_011287300.3;refGene_gn=KCNMA1;refGene_tx=XM_011287302.3;refGene_gn=KCNMA1;refGene_tx=XM_011287303.3;refGene_gn=KCNMA1;refGene_tx=XM_019813335.2;refGene_gn=KCNMA1;refGene_tx=XM_019813336.2;refGene_gn=KCNMA1;refGene_tx=XM_019813338.2;refGene_gn=KCNMA1;refGene_tx=XM_019813339.2;refGene_gn=KCNMA1;refGene_tx=XM_019813340.2;refGene_gn=KCNMA1;refGene_tx=XM_019813341.2;refGene_gn=KCNMA1;refGene_tx=XM_019813342.2;refGene_gn=KCNMA1;refGene_tx=XM_019813344.2;refGene_gn=KCNMA1;refGene_tx=XM_019813345.2;refGene_gn=KCNMA1;refGene_tx=XM_019813346.2;refGene_gn=KCNMA1;refGene_tx=XM_019813347.2;refGene_gn=KCNMA1;refGene_tx=XM_019813349.2;refGene_gn=KCNMA1;refGene_tx=XM_019813350.2;refGene_gn=KCNMA1;refGene_tx=XM_019813351.2;refGene_gn=KCNMA1;refGene_tx=XM_019813352.2;refGene_gn=KCNMA1;refGene_tx=XM_019813353.2;refGene_gn=KCNMA1;refGene_tx=XM_019813354.2;refGene_gn=KCNMA1;refGene_tx=XM_019813355.2;refGene_gn=KCNMA1;refGene_tx=XM_019813356.2;refGene_gn=KCNMA1;refGene_tx=XM_019813357.2;refGene_gn=KCNMA1;refGene_tx=XM_019813358.2;refGene_gn=KCNMA1;refGene_tx=XM_019813360.2;refGene_gn=KCNMA1;refGene_tx=XM_019813362.2;refGene_gn=KCNMA1;refGene_tx=XM_019813363.2;refGene_gn=KCNMA1;refGene_tx=XM_019813364.2;refGene_gn=KCNMA1;refGene_tx=XM_019813366.2;refGene_gn=KCNMA1;refGene_tx=XM_019813368.2;refGene_gn=KCNMA1;refGene_tx=XM_019813369.2;refGene_gn=KCNMA1;refGene_tx=XM_019813371.2;refGene_gn=KCNMA1;refGene_tx=XM_019813372.2;refGene_gn=KCNMA1;refGene_tx=XM_019813373.1;refGene_gn=KCNMA1;refGene_tx=XM_019813374.2;refGene_gn=KCNMA1;refGene_tx=XM_023240597.1;refGene_gn=KCNMA1;refGene_tx=XM_023240599.1;refGene_gn=KCNMA1;refGene_tx=XM_023240600.1;refGene_gn=KCNMA1;refGene_tx=XM_023240601.1
D2 39141544 39141737
refGene_gn=KCNMA1;refGene_tx=XM_011287300.3;refGene_gn=KCNMA1;refGene_tx=XM_011287302.3;refGene_gn=KCNMA1;refGene_tx=XM_011287303.3;refGene_

gn=KCNMA1;refGene_tx=XM_019813335.2;refGene_gn=KCNMA1;refGene_tx=XM_019813336.2;refGene_gn=KCNMA1;refGene_tx=XM_019813338.2;refGene_gn=KCNMA1;refGene_tx=XM_019813339.2;refGene_gn=KCNMA1;refGene_tx=XM_019813340.2;refGene_gn=KCNMA1;refGene_tx=XM_019813341.2;refGene_gn=KCNMA1;refGene_tx=XM_019813342.2;refGene_gn=KCNMA1;refGene_tx=XM_019813344.2;refGene_gn=KCNMA1;refGene_tx=XM_019813345.2;refGene_gn=KCNMA1;refGene_tx=XM_019813346.2;refGene_gn=KCNMA1;refGene_tx=XM_019813347.2;refGene_gn=KCNMA1;refGene_tx=XM_019813349.2;refGene_gn=KCNMA1;refGene_tx=XM_019813350.2;refGene_gn=KCNMA1;refGene_tx=XM_019813351.2;refGene_gn=KCNMA1;refGene_tx=XM_019813352.2;refGene_gn=KCNMA1;refGene_tx=XM_019813353.2;refGene_gn=KCNMA1;refGene_tx=XM_019813354.2;refGene_gn=KCNMA1;refGene_tx=XM_019813355.2;refGene_gn=KCNMA1;refGene_tx=XM_019813356.2;refGene_gn=KCNMA1;refGene_tx=XM_019813357.2;refGene_gn=KCNMA1;refGene_tx=XM_019813358.2;refGene_gn=KCNMA1;refGene_tx=XM_019813360.2;refGene_gn=KCNMA1;refGene_tx=XM_019813362.2;refGene_gn=KCNMA1;refGene_tx=XM_019813363.2;refGene_gn=KCNMA1;refGene_tx=XM_019813364.2;refGene_gn=KCNMA1;refGene_tx=XM_019813366.2;refGene_gn=KCNMA1;refGene_tx=XM_019813368.2;refGene_gn=KCNMA1;refGene_tx=XM_019813369.2;refGene_gn=KCNMA1;refGene_tx=XM_019813371.2;refGene_gn=KCNMA1;refGene_tx=XM_019813372.2;refGene_gn=KCNMA1;refGene_tx=XM_019813373.1;refGene_gn=KCNMA1;refGene_tx=XM_019813374.2;refGene_gn=KCNMA1;refGene_tx=XM_023240597.1;refGene_gn=KCNMA1;refGene_tx=XM_023240599.1;refGene_gn=KCNMA1;refGene_tx=XM_023240600.1;refGene_gn=KCNMA1;refGene_tx=XM_023240601.1
D2 39144997 39145222
refGene_gn=KCNMA1;refGene_tx=XM_011287300.3;refGene_gn=KCNMA1;refGene_tx=XM_011287302.3;refGene_gn=KCNMA1;refGene_tx=XM_011287303.3;refGene_gn=KCNMA1;refGene_tx=XM_019813335.2;refGene_gn=KCNMA1;refGene_tx=XM_019813336.2;refGene_gn=KCNMA1;refGene_tx=XM_019813338.2;refGene_gn=KCNMA1;refGene_tx=XM_019813339.2;refGene_gn=KCNMA1;refGene_tx=XM_019813340.2;refGene_gn=KCNMA1;refGene_tx=XM_019813341.2;refGene_gn=KCNMA1;refGene_tx=XM_019813342.2;refGene_gn=KCNMA1;refGene_tx=XM_019813344.2;refGene_gn=KCNMA1;refGene_tx=XM_019813345.2;refGene_gn=KCNMA1;refGene_tx=XM_019813346.2;refGene_gn=KCNMA1;refGene_tx=XM_019813347.2;refGene_gn=KCNMA1;refGene_tx=XM_019813349.2;refGene_gn=KCNMA1;refGene_tx=XM_019813350.2;refGene_gn=KCNMA1;refGene_tx=XM_019813351.2;refGene_gn=KCNMA1;refGene_tx=XM_019813352.2;refGene_gn=KCNMA1;refGene_tx=XM_019813353.2;refGene_gn=KCNMA1;refGene_tx=XM_019813354.2;refGene_gn=KCNMA1;refGene_tx=XM_019813355.2;refGene_gn=KCNMA1;refGene_tx=XM_019813356.2;refGene_gn=KCNMA1;refGene_tx=XM_019813357.2;refGene_gn=KCNMA1;refGene_tx=XM_019813358.2;refGene_gn=KCNMA1;refGene_tx=XM_019813360.2;refGene_gn=KCNMA1;refGene_tx=XM_019813362.2;refGene_gn=KCNMA1;refGene_tx=XM_019813363.2;refGene_gn=KCNMA1;refGene_tx=XM_019813364.2;refGene_gn=KCNMA1;refGene_tx=XM_019813366.2;refGene_gn=KCNMA1;refGene_tx=XM_019813368.2;refGene_gn=KCNMA1;refGene_tx=XM_019813369.2;refGene_gn=KCNMA1;refGene_tx=XM_019813371.2;refGene_gn=KCNMA1;refGene_tx=XM_019813372.2;refGene_gn=KCNMA1;refGene_tx=XM_019813373.1;refGene_gn=KCNMA1;refGene_tx=XM_019813374.2;refGene_gn=KCNMA1;refGene_tx=XM_023240597.1;refGene_gn=KCNMA1;refGene_tx=XM_023240599.1;refGene_gn=KCNMA1;refGene_tx=XM_023240600.1;refGene_gn=KCNMA1;refGene_tx=XM_023240601.1
D2 39149253 39149377

refGene_gn=KCNMA1;refGene_tx=XM_011287300.3;refGene_gn=KCNMA1;refGene_tx=XM_011287302.3;refGene_gn=KCNMA1;refGene_tx=XM_011287303.3;refGene_gn=KCNMA1;refGene_tx=XM_019813335.2;refGene_gn=KCNMA1;refGene_tx=XM_019813336.2;refGene_gn=KCNMA1;refGene_tx=XM_019813338.2;refGene_gn=KCNMA1;refGene_tx=XM_019813339.2;refGene_gn=KCNMA1;refGene_tx=XM_019813340.2;refGene_gn=KCNMA1;refGene_tx=XM_019813341.2;refGene_gn=KCNMA1;refGene_tx=XM_019813342.2;refGene_gn=KCNMA1;refGene_tx=XM_019813344.2;refGene_gn=KCNMA1;refGene_tx=XM_019813345.2;refGene_gn=KCNMA1;refGene_tx=XM_019813346.2;refGene_gn=KCNMA1;refGene_tx=XM_019813347.2;refGene_gn=KCNMA1;refGene_tx=XM_019813349.2;refGene_gn=KCNMA1;refGene_tx=XM_019813350.2;refGene_gn=KCNMA1;refGene_tx=XM_019813351.2;refGene_gn=KCNMA1;refGene_tx=XM_019813352.2;refGene_gn=KCNMA1;refGene_tx=XM_019813353.2;refGene_gn=KCNMA1;refGene_tx=XM_019813354.2;refGene_gn=KCNMA1;refGene_tx=XM_019813355.2;refGene_gn=KCNMA1;refGene_tx=XM_019813356.2;refGene_gn=KCNMA1;refGene_tx=XM_019813357.2;refGene_gn=KCNMA1;refGene_tx=XM_019813358.2;refGene_gn=KCNMA1;refGene_tx=XM_019813360.2;refGene_gn=KCNMA1;refGene_tx=XM_019813362.2;refGene_gn=KCNMA1;refGene_tx=XM_019813363.2;refGene_gn=KCNMA1;refGene_tx=XM_019813364.2;refGene_gn=KCNMA1;refGene_tx=XM_019813366.2;refGene_gn=KCNMA1;refGene_tx=XM_019813368.2;refGene_gn=KCNMA1;refGene_tx=XM_019813369.2;refGene_gn=KCNMA1;refGene_tx=XM_019813371.2;refGene_gn=KCNMA1;refGene_tx=XM_019813372.2;refGene_gn=KCNMA1;refGene_tx=XM_019813373.1;refGene_gn=KCNMA1;refGene_tx=XM_019813374.2;refGene_gn=KCNMA1;refGene_tx=XM_023240597.1;refGene_gn=KCNMA1;refGene_tx=XM_023240599.1;refGene_gn=KCNMA1;refGene_tx=XM_023240600.1;refGene_gn=KCNMA1;refGene_tx=XM_023240601.1

D2 39163912 39164006

refGene_gn=KCNMA1;refGene_tx=XM_011287300.3;refGene_gn=KCNMA1;refGene_tx=XM_011287302.3;refGene_gn=KCNMA1;refGene_tx=XM_011287303.3;refGene_gn=KCNMA1;refGene_tx=XM_019813335.2;refGene_gn=KCNMA1;refGene_tx=XM_019813336.2;refGene_gn=KCNMA1;refGene_tx=XM_019813338.2;refGene_gn=KCNMA1;refGene_tx=XM_019813339.2;refGene_gn=KCNMA1;refGene_tx=XM_019813340.2;refGene_gn=KCNMA1;refGene_tx=XM_019813341.2;refGene_gn=KCNMA1;refGene_tx=XM_019813342.2;refGene_gn=KCNMA1;refGene_tx=XM_019813344.2;refGene_gn=KCNMA1;refGene_tx=XM_019813345.2;refGene_gn=KCNMA1;refGene_tx=XM_019813346.2;refGene_gn=KCNMA1;refGene_tx=XM_019813347.2;refGene_gn=KCNMA1;refGene_tx=XM_019813349.2;refGene_gn=KCNMA1;refGene_tx=XM_019813350.2;refGene_gn=KCNMA1;refGene_tx=XM_019813351.2;refGene_gn=KCNMA1;refGene_tx=XM_019813352.2;refGene_gn=KCNMA1;refGene_tx=XM_019813353.2;refGene_gn=KCNMA1;refGene_tx=XM_019813354.2;refGene_gn=KCNMA1;refGene_tx=XM_019813355.2;refGene_gn=KCNMA1;refGene_tx=XM_019813356.2;refGene_gn=KCNMA1;refGene_tx=XM_019813357.2;refGene_gn=KCNMA1;refGene_tx=XM_019813358.2;refGene_gn=KCNMA1;refGene_tx=XM_019813360.2;refGene_gn=KCNMA1;refGene_tx=XM_019813362.2;refGene_gn=KCNMA1;refGene_tx=XM_019813363.2;refGene_gn=KCNMA1;refGene_tx=XM_019813364.2;refGene_gn=KCNMA1;refGene_tx=XM_019813366.2;refGene_gn=KCNMA1;refGene_tx=XM_019813368.2;refGene_gn=KCNMA1;refGene_tx=XM_019813369.2;refGene_gn=KCNMA1;refGene_tx=XM_019813371.2;refGene_gn=KCNMA1;refGene_tx=XM_019813372.2;refGene_gn=KCNMA1;refGene_tx=XM_019813373.1;refGene_gn=KCNMA1;refGene_tx=XM_019813374.2;refGene_gn=KCNMA1;refGene_tx=XM_023240597.1;refGene_gn=KCNMA1;refGene_tx=XM_023240599.1;refGene_gn=KCNMA1;refGene_tx=XM_023240600.1;refGene_

gn=KCNMA1;refGene_tx=XM_023240601.1
D2 39168470 39168557
refGene_gn=KCNMA1;refGene_tx=XM_023240599.1;refGene_gn=KCNMA1;refGene_tx=XM_023240600.1;refGene_gn=KCNMA1;refGene_tx=XM_023240601.1
D2 39193030 39193204
refGene_gn=KCNMA1;refGene_tx=XM_019813335.2;refGene_gn=KCNMA1;refGene_tx=XM_019813336.2;refGene_gn=KCNMA1;refGene_tx=XM_019813338.2;refGene_gn=KCNMA1;refGene_tx=XM_019813339.2;refGene_gn=KCNMA1;refGene_tx=XM_019813340.2;refGene_gn=KCNMA1;refGene_tx=XM_019813341.2;refGene_gn=KCNMA1;refGene_tx=XM_019813342.2;refGene_gn=KCNMA1;refGene_tx=XM_019813344.2;refGene_gn=KCNMA1;refGene_tx=XM_019813345.2;refGene_gn=KCNMA1;refGene_tx=XM_019813347.2;refGene_gn=KCNMA1;refGene_tx=XM_019813354.2;refGene_gn=KCNMA1;refGene_tx=XM_019813355.2;refGene_gn=KCNMA1;refGene_tx=XM_019813356.2;refGene_gn=KCNMA1;refGene_tx=XM_019813357.2;refGene_gn=KCNMA1;refGene_tx=XM_019813358.2;refGene_gn=KCNMA1;refGene_tx=XM_019813373.1;refGene_gn=KCNMA1;refGene_tx=XM_023240597.1
D2 39199797 39199806
refGene_gn=KCNMA1;refGene_tx=XM_019813335.2;refGene_gn=KCNMA1;refGene_tx=XM_019813336.2;refGene_gn=KCNMA1;refGene_tx=XM_019813338.2;refGene_gn=KCNMA1;refGene_tx=XM_019813340.2;refGene_gn=KCNMA1;refGene_tx=XM_019813341.2;refGene_gn=KCNMA1;refGene_tx=XM_019813342.2;refGene_gn=KCNMA1;refGene_tx=XM_019813344.2;refGene_gn=KCNMA1;refGene_tx=XM_019813345.2;refGene_gn=KCNMA1;refGene_tx=XM_019813346.2;refGene_gn=KCNMA1;refGene_tx=XM_019813347.2;refGene_gn=KCNMA1;refGene_tx=XM_019813350.2;refGene_gn=KCNMA1;refGene_tx=XM_019813351.2;refGene_gn=KCNMA1;refGene_tx=XM_019813354.2;refGene_gn=KCNMA1;refGene_tx=XM_019813355.2;refGene_gn=KCNMA1;refGene_tx=XM_019813360.2;refGene_gn=KCNMA1;refGene_tx=XM_019813362.2;refGene_gn=KCNMA1;refGene_tx=XM_019813364.2;refGene_gn=KCNMA1;refGene_tx=XM_019813369.2;refGene_gn=KCNMA1;refGene_tx=XM_019813371.2;refGene_gn=KCNMA1;refGene_tx=XM_019813373.1
D2 39203636 39203713
refGene_gn=KCNMA1;refGene_tx=XM_011287300.3;refGene_gn=KCNMA1;refGene_tx=XM_011287302.3;refGene_gn=KCNMA1;refGene_tx=XM_011287303.3;refGene_gn=KCNMA1;refGene_tx=XM_019813335.2;refGene_gn=KCNMA1;refGene_tx=XM_019813336.2;refGene_gn=KCNMA1;refGene_tx=XM_019813338.2;refGene_gn=KCNMA1;refGene_tx=XM_019813339.2;refGene_gn=KCNMA1;refGene_tx=XM_019813340.2;refGene_gn=KCNMA1;refGene_tx=XM_019813341.2;refGene_gn=KCNMA1;refGene_tx=XM_019813342.2;refGene_gn=KCNMA1;refGene_tx=XM_019813344.2;refGene_gn=KCNMA1;refGene_tx=XM_019813345.2;refGene_gn=KCNMA1;refGene_tx=XM_019813346.2;refGene_gn=KCNMA1;refGene_tx=XM_019813347.2;refGene_gn=KCNMA1;refGene_tx=XM_019813349.2;refGene_gn=KCNMA1;refGene_tx=XM_019813350.2;refGene_gn=KCNMA1;refGene_tx=XM_019813351.2;refGene_gn=KCNMA1;refGene_tx=XM_019813352.2;refGene_gn=KCNMA1;refGene_tx=XM_019813353.2;refGene_gn=KCNMA1;refGene_tx=XM_019813354.2;refGene_gn=KCNMA1;refGene_tx=XM_019813355.2;refGene_gn=KCNMA1;refGene_tx=XM_019813356.2;refGene_gn=KCNMA1;refGene_tx=XM_019813357.2;refGene_gn=KCNMA1;refGene_tx=XM_019813358.2;refGene_gn=KCNMA1;refGene_tx=XM_019813360.2;refGene_gn=KCNMA1;refGene_tx=XM_019813362.2;refGene_gn=KCNMA1;refGene_tx=XM_019813363.2;refGene_gn=KCNMA1;refGene_tx=XM_019813364.2;refGene_gn=KCNMA1;refGene_tx=XM_019813366.2;refGene_gn=KCNMA1;refGene_tx=XM_019813368.2;refGene_gn

=KCNMA1;refGene_tx=XM_019813369.2;refGene_gn=KCNMA1;refGene_tx=XM_0198
13371.2;refGene_gn=KCNMA1;refGene_tx=XM_019813372.2;refGene_gn=KCNMA1;
refGene_tx=XM_019813373.1;refGene_gn=KCNMA1;refGene_tx=XM_019813374.2;
refGene_gn=KCNMA1;refGene_tx=XM_023240597.1;refGene_gn=KCNMA1;refGene_
tx=XM_023240599.1;refGene_gn=KCNMA1;refGene_tx=XM_023240600.1;refGene_
gn=KCNMA1;refGene_tx=XM_023240601.1
D2 39209903 39209990
refGene_gn=KCNMA1;refGene_tx=XM_011287300.3;refGene_gn=KCNMA1;refGene_
tx=XM_011287302.3;refGene_gn=KCNMA1;refGene_tx=XM_011287303.3;refGene_
gn=KCNMA1;refGene_tx=XM_019813335.2;refGene_gn=KCNMA1;refGene_tx=XM_01
9813336.2;refGene_gn=KCNMA1;refGene_tx=XM_019813338.2;refGene_gn=KCNMA
1;refGene_tx=XM_019813339.2;refGene_gn=KCNMA1;refGene_tx=XM_019813340.
2;refGene_gn=KCNMA1;refGene_tx=XM_019813341.2;refGene_gn=KCNMA1;refGen
e_tx=XM_019813342.2;refGene_gn=KCNMA1;refGene_tx=XM_019813344.2;refGen
e_gn=KCNMA1;refGene_tx=XM_019813345.2;refGene_gn=KCNMA1;refGene_tx=XM_
019813346.2;refGene_gn=KCNMA1;refGene_tx=XM_019813347.2;refGene_gn=KCN
MA1;refGene_tx=XM_019813349.2;refGene_gn=KCNMA1;refGene_tx=XM_01981335
0.2;refGene_gn=KCNMA1;refGene_tx=XM_019813351.2;refGene_gn=KCNMA1;refG
ene_tx=XM_019813352.2;refGene_gn=KCNMA1;refGene_tx=XM_019813353.2;refG
ene_gn=KCNMA1;refGene_tx=XM_019813354.2;refGene_gn=KCNMA1;refGene_tx=X
M_019813355.2;refGene_gn=KCNMA1;refGene_tx=XM_019813356.2;refGene_gn=K
CNMA1;refGene_tx=XM_019813357.2;refGene_gn=KCNMA1;refGene_tx=XM_019813
358.2;refGene_gn=KCNMA1;refGene_tx=XM_019813360.2;refGene_gn=KCNMA1;re
fGene_tx=XM_019813362.2;refGene_gn=KCNMA1;refGene_tx=XM_019813363.2;re
fGene_gn=KCNMA1;refGene_tx=XM_019813364.2;refGene_gn=KCNMA1;refGene_tx
=XM_019813366.2;refGene_gn=KCNMA1;refGene_tx=XM_019813368.2;refGene_gn
=KCNMA1;refGene_tx=XM_019813369.2;refGene_gn=KCNMA1;refGene_tx=XM_0198
13371.2;refGene_gn=KCNMA1;refGene_tx=XM_019813372.2;refGene_gn=KCNMA1;
refGene_tx=XM_019813373.1;refGene_gn=KCNMA1;refGene_tx=XM_019813374.2;
refGene_gn=KCNMA1;refGene_tx=XM_023240597.1;refGene_gn=KCNMA1;refGene_
tx=XM_023240599.1;refGene_gn=KCNMA1;refGene_tx=XM_023240600.1;refGene_
gn=KCNMA1;refGene_tx=XM_023240601.1
D2 39217109 39217121
refGene_gn=KCNMA1;refGene_tx=XM_011287300.3;refGene_gn=KCNMA1;refGene_
tx=XM_019813335.2;refGene_gn=KCNMA1;refGene_tx=XM_019813336.2;refGene_
gn=KCNMA1;refGene_tx=XM_019813338.2;refGene_gn=KCNMA1;refGene_tx=XM_01
9813339.2;refGene_gn=KCNMA1;refGene_tx=XM_019813341.2;refGene_gn=KCNMA
1;refGene_tx=XM_019813342.2;refGene_gn=KCNMA1;refGene_tx=XM_019813344.
2;refGene_gn=KCNMA1;refGene_tx=XM_019813345.2;refGene_gn=KCNMA1;refGen
e_tx=XM_019813346.2;refGene_gn=KCNMA1;refGene_tx=XM_019813347.2;refGen
e_gn=KCNMA1;refGene_tx=XM_019813349.2;refGene_gn=KCNMA1;refGene_tx=XM_
019813354.2;refGene_gn=KCNMA1;refGene_tx=XM_019813355.2;refGene_gn=KCN
MA1;refGene_tx=XM_019813356.2;refGene_gn=KCNMA1;refGene_tx=XM_01981336
4.2;refGene_gn=KCNMA1;refGene_tx=XM_019813366.2;refGene_gn=KCNMA1;refG
ene_tx=XM_019813368.2;refGene_gn=KCNMA1;refGene_tx=XM_019813373.1;refG
ene_gn=KCNMA1;refGene_tx=XM_023240597.1;refGene_gn=KCNMA1;refGene_tx=X
M_023240599.1
D2 39219494 39219563
refGene_gn=KCNMA1;refGene_tx=XM_011287300.3;refGene_gn=KCNMA1;refGene_
tx=XM_011287302.3;refGene_gn=KCNMA1;refGene_tx=XM_011287303.3;refGene_

gn=KCNMA1;refGene_tx=XM_019813335.2;refGene_gn=KCNMA1;refGene_tx=XM_01
9813336.2;refGene_gn=KCNMA1;refGene_tx=XM_019813338.2;refGene_gn=KCNMA
1;refGene_tx=XM_019813339.2;refGene_gn=KCNMA1;refGene_tx=XM_019813340.
2;refGene_gn=KCNMA1;refGene_tx=XM_019813341.2;refGene_gn=KCNMA1;refGen
e_tx=XM_019813342.2;refGene_gn=KCNMA1;refGene_tx=XM_019813344.2;refGen
e_gn=KCNMA1;refGene_tx=XM_019813345.2;refGene_gn=KCNMA1;refGene_tx=XM_
019813346.2;refGene_gn=KCNMA1;refGene_tx=XM_019813347.2;refGene_gn=KCN
MA1;refGene_tx=XM_019813349.2;refGene_gn=KCNMA1;refGene_tx=XM_01981335
0.2;refGene_gn=KCNMA1;refGene_tx=XM_019813351.2;refGene_gn=KCNMA1;refG
ene_tx=XM_019813352.2;refGene_gn=KCNMA1;refGene_tx=XM_019813353.2;refG
ene_gn=KCNMA1;refGene_tx=XM_019813354.2;refGene_gn=KCNMA1;refGene_tx=X
M_019813355.2;refGene_gn=KCNMA1;refGene_tx=XM_019813356.2;refGene_gn=K
CNMA1;refGene_tx=XM_019813357.2;refGene_gn=KCNMA1;refGene_tx=XM_019813
358.2;refGene_gn=KCNMA1;refGene_tx=XM_019813360.2;refGene_gn=KCNMA1;re
fGene_tx=XM_019813362.2;refGene_gn=KCNMA1;refGene_tx=XM_019813363.2;re
fGene_gn=KCNMA1;refGene_tx=XM_019813364.2;refGene_gn=KCNMA1;refGene_tx
=XM_019813366.2;refGene_gn=KCNMA1;refGene_tx=XM_019813368.2;refGene_gn
=KCNMA1;refGene_tx=XM_019813369.2;refGene_gn=KCNMA1;refGene_tx=XM_0198
13371.2;refGene_gn=KCNMA1;refGene_tx=XM_019813372.2;refGene_gn=KCNMA1;
refGene_tx=XM_019813373.1;refGene_gn=KCNMA1;refGene_tx=XM_019813374.2;
refGene_gn=KCNMA1;refGene_tx=XM_023240597.1;refGene_gn=KCNMA1;refGene_
tx=XM_023240599.1;refGene_gn=KCNMA1;refGene_tx=XM_023240600.1;refGene_
gn=KCNMA1;refGene_tx=XM_023240601.1
D2 39232538 39232648
refGene_gn=KCNMA1;refGene_tx=XM_011287300.3;refGene_gn=KCNMA1;refGene_
tx=XM_011287302.3;refGene_gn=KCNMA1;refGene_tx=XM_011287303.3;refGene_
gn=KCNMA1;refGene_tx=XM_019813335.2;refGene_gn=KCNMA1;refGene_tx=XM_01
9813336.2;refGene_gn=KCNMA1;refGene_tx=XM_019813338.2;refGene_gn=KCNMA
1;refGene_tx=XM_019813339.2;refGene_gn=KCNMA1;refGene_tx=XM_019813340.
2;refGene_gn=KCNMA1;refGene_tx=XM_019813341.2;refGene_gn=KCNMA1;refGen
e_tx=XM_019813342.2;refGene_gn=KCNMA1;refGene_tx=XM_019813344.2;refGen
e_gn=KCNMA1;refGene_tx=XM_019813345.2;refGene_gn=KCNMA1;refGene_tx=XM_
019813346.2;refGene_gn=KCNMA1;refGene_tx=XM_019813347.2;refGene_gn=KCN
MA1;refGene_tx=XM_019813349.2;refGene_gn=KCNMA1;refGene_tx=XM_01981335
0.2;refGene_gn=KCNMA1;refGene_tx=XM_019813351.2;refGene_gn=KCNMA1;refG
ene_tx=XM_019813352.2;refGene_gn=KCNMA1;refGene_tx=XM_019813353.2;refG
ene_gn=KCNMA1;refGene_tx=XM_019813354.2;refGene_gn=KCNMA1;refGene_tx=X
M_019813355.2;refGene_gn=KCNMA1;refGene_tx=XM_019813356.2;refGene_gn=K
CNMA1;refGene_tx=XM_019813357.2;refGene_gn=KCNMA1;refGene_tx=XM_019813
358.2;refGene_gn=KCNMA1;refGene_tx=XM_019813360.2;refGene_gn=KCNMA1;re
fGene_tx=XM_019813362.2;refGene_gn=KCNMA1;refGene_tx=XM_019813363.2;re
fGene_gn=KCNMA1;refGene_tx=XM_019813364.2;refGene_gn=KCNMA1;refGene_tx
=XM_019813366.2;refGene_gn=KCNMA1;refGene_tx=XM_019813368.2;refGene_gn
=KCNMA1;refGene_tx=XM_019813369.2;refGene_gn=KCNMA1;refGene_tx=XM_0198
13371.2;refGene_gn=KCNMA1;refGene_tx=XM_019813372.2;refGene_gn=KCNMA1;
refGene_tx=XM_019813373.1;refGene_gn=KCNMA1;refGene_tx=XM_019813374.2;
refGene_gn=KCNMA1;refGene_tx=XM_023240597.1;refGene_gn=KCNMA1;refGene_
tx=XM_023240599.1;refGene_gn=KCNMA1;refGene_tx=XM_023240600.1;refGene_
gn=KCNMA1;refGene_tx=XM_023240601.1
D2 39263910 39263952 refGene_gn=KCNMA1;refGene_tx=XM_023240598.1

D2 39265848 39266004
refGene_gn=KCNMA1;refGene_tx=XM_011287300.3;refGene_gn=KCNMA1;refGene_tx=XM_011287302.3;refGene_gn=KCNMA1;refGene_tx=XM_011287303.3;refGene_gn=KCNMA1;refGene_tx=XM_019813335.2;refGene_gn=KCNMA1;refGene_tx=XM_019813336.2;refGene_gn=KCNMA1;refGene_tx=XM_019813338.2;refGene_gn=KCNMA1;refGene_tx=XM_019813339.2;refGene_gn=KCNMA1;refGene_tx=XM_019813340.2;refGene_gn=KCNMA1;refGene_tx=XM_019813341.2;refGene_gn=KCNMA1;refGene_tx=XM_019813342.2;refGene_gn=KCNMA1;refGene_tx=XM_019813344.2;refGene_gn=KCNMA1;refGene_tx=XM_019813345.2;refGene_gn=KCNMA1;refGene_tx=XM_019813346.2;refGene_gn=KCNMA1;refGene_tx=XM_019813347.2;refGene_gn=KCNMA1;refGene_tx=XM_019813349.2;refGene_gn=KCNMA1;refGene_tx=XM_019813350.2;refGene_gn=KCNMA1;refGene_tx=XM_019813351.2;refGene_gn=KCNMA1;refGene_tx=XM_019813352.2;refGene_gn=KCNMA1;refGene_tx=XM_019813353.2;refGene_gn=KCNMA1;refGene_tx=XM_019813354.2;refGene_gn=KCNMA1;refGene_tx=XM_019813355.2;refGene_gn=KCNMA1;refGene_tx=XM_019813356.2;refGene_gn=KCNMA1;refGene_tx=XM_019813357.2;refGene_gn=KCNMA1;refGene_tx=XM_019813358.2;refGene_gn=KCNMA1;refGene_tx=XM_019813360.2;refGene_gn=KCNMA1;refGene_tx=XM_019813362.2;refGene_gn=KCNMA1;refGene_tx=XM_019813363.2;refGene_gn=KCNMA1;refGene_tx=XM_019813364.2;refGene_gn=KCNMA1;refGene_tx=XM_019813366.2;refGene_gn=KCNMA1;refGene_tx=XM_019813368.2;refGene_gn=KCNMA1;refGene_tx=XM_019813369.2;refGene_gn=KCNMA1;refGene_tx=XM_019813371.2;refGene_gn=KCNMA1;refGene_tx=XM_019813372.2;refGene_gn=KCNMA1;refGene_tx=XM_019813373.1;refGene_gn=KCNMA1;refGene_tx=XM_019813374.2;refGene_gn=KCNMA1;refGene_tx=XM_023240597.1;refGene_gn=KCNMA1;refGene_tx=XM_023240598.1;refGene_gn=KCNMA1;refGene_tx=XM_023240599.1;refGene_gn=KCNMA1;refGene_tx=XM_023240600.1;refGene_gn=KCNMA1;refGene_tx=XM_023240601.1
D2 39271950 39272020
refGene_gn=KCNMA1;refGene_tx=XM_011287300.3;refGene_gn=KCNMA1;refGene_tx=XM_011287302.3;refGene_gn=KCNMA1;refGene_tx=XM_011287303.3;refGene_gn=KCNMA1;refGene_tx=XM_019813335.2;refGene_gn=KCNMA1;refGene_tx=XM_019813336.2;refGene_gn=KCNMA1;refGene_tx=XM_019813338.2;refGene_gn=KCNMA1;refGene_tx=XM_019813339.2;refGene_gn=KCNMA1;refGene_tx=XM_019813340.2;refGene_gn=KCNMA1;refGene_tx=XM_019813341.2;refGene_gn=KCNMA1;refGene_tx=XM_019813342.2;refGene_gn=KCNMA1;refGene_tx=XM_019813344.2;refGene_gn=KCNMA1;refGene_tx=XM_019813345.2;refGene_gn=KCNMA1;refGene_tx=XM_019813346.2;refGene_gn=KCNMA1;refGene_tx=XM_019813347.2;refGene_gn=KCNMA1;refGene_tx=XM_019813349.2;refGene_gn=KCNMA1;refGene_tx=XM_019813350.2;refGene_gn=KCNMA1;refGene_tx=XM_019813351.2;refGene_gn=KCNMA1;refGene_tx=XM_019813352.2;refGene_gn=KCNMA1;refGene_tx=XM_019813353.2;refGene_gn=KCNMA1;refGene_tx=XM_019813354.2;refGene_gn=KCNMA1;refGene_tx=XM_019813355.2;refGene_gn=KCNMA1;refGene_tx=XM_019813356.2;refGene_gn=KCNMA1;refGene_tx=XM_019813357.2;refGene_gn=KCNMA1;refGene_tx=XM_019813358.2;refGene_gn=KCNMA1;refGene_tx=XM_019813360.2;refGene_gn=KCNMA1;refGene_tx=XM_019813362.2;refGene_gn=KCNMA1;refGene_tx=XM_019813363.2;refGene_gn=KCNMA1;refGene_tx=XM_019813364.2;refGene_gn=KCNMA1;refGene_tx=XM_019813366.2;refGene_gn=KCNMA1;refGene_tx=XM_019813368.2;refGene_gn=KCNMA1;refGene_tx=XM_019813369.2;refGene_gn=KCNMA1;refGene_tx=XM_019813371.2;refGene_gn=KCNMA1;refGene_tx=XM_019813372.2;refGene_gn=KCNMA1;refGene_tx=XM_019813373.1;refGene_gn=KCNMA1;refGene_tx=XM_019813374.2;

refGene_gn=KCNMA1;refGene_tx=XM_023240597.1;refGene_gn=KCNMA1;refGene_tx=XM_023240598.1;refGene_gn=KCNMA1;refGene_tx=XM_023240599.1;refGene_gn=KCNMA1;refGene_tx=XM_023240600.1;refGene_gn=KCNMA1;refGene_tx=XM_023240601.1
D2 39275539 39275622
refGene_gn=KCNMA1;refGene_tx=XM_011287300.3;refGene_gn=KCNMA1;refGene_tx=XM_011287302.3;refGene_gn=KCNMA1;refGene_tx=XM_011287303.3;refGene_gn=KCNMA1;refGene_tx=XM_019813335.2;refGene_gn=KCNMA1;refGene_tx=XM_019813336.2;refGene_gn=KCNMA1;refGene_tx=XM_019813338.2;refGene_gn=KCNMA1;refGene_tx=XM_019813339.2;refGene_gn=KCNMA1;refGene_tx=XM_019813340.2;refGene_gn=KCNMA1;refGene_tx=XM_019813341.2;refGene_gn=KCNMA1;refGene_tx=XM_019813342.2;refGene_gn=KCNMA1;refGene_tx=XM_019813344.2;refGene_gn=KCNMA1;refGene_tx=XM_019813345.2;refGene_gn=KCNMA1;refGene_tx=XM_019813346.2;refGene_gn=KCNMA1;refGene_tx=XM_019813347.2;refGene_gn=KCNMA1;refGene_tx=XM_019813349.2;refGene_gn=KCNMA1;refGene_tx=XM_019813350.2;refGene_gn=KCNMA1;refGene_tx=XM_019813351.2;refGene_gn=KCNMA1;refGene_tx=XM_019813352.2;refGene_gn=KCNMA1;refGene_tx=XM_019813353.2;refGene_gn=KCNMA1;refGene_tx=XM_019813354.2;refGene_gn=KCNMA1;refGene_tx=XM_019813355.2;refGene_gn=KCNMA1;refGene_tx=XM_019813356.2;refGene_gn=KCNMA1;refGene_tx=XM_019813357.2;refGene_gn=KCNMA1;refGene_tx=XM_019813358.2;refGene_gn=KCNMA1;refGene_tx=XM_019813360.2;refGene_gn=KCNMA1;refGene_tx=XM_019813362.2;refGene_gn=KCNMA1;refGene_tx=XM_019813363.2;refGene_gn=KCNMA1;refGene_tx=XM_019813364.2;refGene_gn=KCNMA1;refGene_tx=XM_019813366.2;refGene_gn=KCNMA1;refGene_tx=XM_019813368.2;refGene_gn=KCNMA1;refGene_tx=XM_019813369.2;refGene_gn=KCNMA1;refGene_tx=XM_019813371.2;refGene_gn=KCNMA1;refGene_tx=XM_019813372.2;refGene_gn=KCNMA1;refGene_tx=XM_019813373.1;refGene_gn=KCNMA1;refGene_tx=XM_019813374.2;refGene_gn=KCNMA1;refGene_tx=XM_023240597.1;refGene_gn=KCNMA1;refGene_tx=XM_023240598.1;refGene_gn=KCNMA1;refGene_tx=XM_023240599.1;refGene_gn=KCNMA1;refGene_tx=XM_023240600.1;refGene_gn=KCNMA1;refGene_tx=XM_023240601.1
D2 39277358 39277464
refGene_gn=KCNMA1;refGene_tx=XM_011287300.3;refGene_gn=KCNMA1;refGene_tx=XM_011287302.3;refGene_gn=KCNMA1;refGene_tx=XM_011287303.3;refGene_gn=KCNMA1;refGene_tx=XM_019813335.2;refGene_gn=KCNMA1;refGene_tx=XM_019813336.2;refGene_gn=KCNMA1;refGene_tx=XM_019813338.2;refGene_gn=KCNMA1;refGene_tx=XM_019813339.2;refGene_gn=KCNMA1;refGene_tx=XM_019813340.2;refGene_gn=KCNMA1;refGene_tx=XM_019813341.2;refGene_gn=KCNMA1;refGene_tx=XM_019813342.2;refGene_gn=KCNMA1;refGene_tx=XM_019813344.2;refGene_gn=KCNMA1;refGene_tx=XM_019813345.2;refGene_gn=KCNMA1;refGene_tx=XM_019813346.2;refGene_gn=KCNMA1;refGene_tx=XM_019813347.2;refGene_gn=KCNMA1;refGene_tx=XM_019813349.2;refGene_gn=KCNMA1;refGene_tx=XM_019813350.2;refGene_gn=KCNMA1;refGene_tx=XM_019813351.2;refGene_gn=KCNMA1;refGene_tx=XM_019813352.2;refGene_gn=KCNMA1;refGene_tx=XM_019813353.2;refGene_gn=KCNMA1;refGene_tx=XM_019813354.2;refGene_gn=KCNMA1;refGene_tx=XM_019813355.2;refGene_gn=KCNMA1;refGene_tx=XM_019813356.2;refGene_gn=KCNMA1;refGene_tx=XM_019813357.2;refGene_gn=KCNMA1;refGene_tx=XM_019813358.2;refGene_gn=KCNMA1;refGene_tx=XM_019813360.2;refGene_gn=KCNMA1;refGene_tx=XM_019813362.2;refGene_gn=KCNMA1;refGene_tx=XM_019813363.2;refGene_gn=KCNMA1;refGene_tx=XM_019813364.2;refGene_gn=KCNMA1;refGene_tx

=XM_019813366.2;refGene_gn=KCNMA1;refGene_tx=XM_019813368.2;refGene_gn
=KCNMA1;refGene_tx=XM_019813369.2;refGene_gn=KCNMA1;refGene_tx=XM_0198
13371.2;refGene_gn=KCNMA1;refGene_tx=XM_019813372.2;refGene_gn=KCNMA1;
refGene_tx=XM_019813373.1;refGene_gn=KCNMA1;refGene_tx=XM_019813374.2;
refGene_gn=KCNMA1;refGene_tx=XM_023240597.1;refGene_gn=KCNMA1;refGene_
tx=XM_023240598.1;refGene_gn=KCNMA1;refGene_tx=XM_023240599.1;refGene_
gn=KCNMA1;refGene_tx=XM_023240600.1;refGene_gn=KCNMA1;refGene_tx=XM_02
3240601.1
D2 39280034 39280145
refGene_gn=KCNMA1;refGene_tx=XM_011287300.3;refGene_gn=KCNMA1;refGene_
tx=XM_011287302.3;refGene_gn=KCNMA1;refGene_tx=XM_011287303.3;refGene_
gn=KCNMA1;refGene_tx=XM_019813335.2;refGene_gn=KCNMA1;refGene_tx=XM_01
9813336.2;refGene_gn=KCNMA1;refGene_tx=XM_019813338.2;refGene_gn=KCNMA
1;refGene_tx=XM_019813339.2;refGene_gn=KCNMA1;refGene_tx=XM_019813340.
2;refGene_gn=KCNMA1;refGene_tx=XM_019813341.2;refGene_gn=KCNMA1;refGen
e_tx=XM_019813342.2;refGene_gn=KCNMA1;refGene_tx=XM_019813344.2;refGen
e_gn=KCNMA1;refGene_tx=XM_019813345.2;refGene_gn=KCNMA1;refGene_tx=XM_
019813346.2;refGene_gn=KCNMA1;refGene_tx=XM_019813347.2;refGene_gn=KCN
MA1;refGene_tx=XM_019813349.2;refGene_gn=KCNMA1;refGene_tx=XM_01981335
0.2;refGene_gn=KCNMA1;refGene_tx=XM_019813351.2;refGene_gn=KCNMA1;refG
ene_tx=XM_019813352.2;refGene_gn=KCNMA1;refGene_tx=XM_019813353.2;refG
ene_gn=KCNMA1;refGene_tx=XM_019813354.2;refGene_gn=KCNMA1;refGene_tx=X
M_019813355.2;refGene_gn=KCNMA1;refGene_tx=XM_019813356.2;refGene_gn=K
CNMA1;refGene_tx=XM_019813357.2;refGene_gn=KCNMA1;refGene_tx=XM_019813
358.2;refGene_gn=KCNMA1;refGene_tx=XM_019813360.2;refGene_gn=KCNMA1;re
fGene_tx=XM_019813362.2;refGene_gn=KCNMA1;refGene_tx=XM_019813363.2;re
fGene_gn=KCNMA1;refGene_tx=XM_019813364.2;refGene_gn=KCNMA1;refGene_tx
=XM_019813366.2;refGene_gn=KCNMA1;refGene_tx=XM_019813368.2;refGene_gn
=KCNMA1;refGene_tx=XM_019813369.2;refGene_gn=KCNMA1;refGene_tx=XM_0198
13371.2;refGene_gn=KCNMA1;refGene_tx=XM_019813372.2;refGene_gn=KCNMA1;
refGene_tx=XM_019813373.1;refGene_gn=KCNMA1;refGene_tx=XM_019813374.2;
refGene_gn=KCNMA1;refGene_tx=XM_023240597.1;refGene_gn=KCNMA1;refGene_
tx=XM_023240598.1;refGene_gn=KCNMA1;refGene_tx=XM_023240599.1;refGene_
gn=KCNMA1;refGene_tx=XM_023240600.1;refGene_gn=KCNMA1;refGene_tx=XM_02
3240601.1
D2 39298451 39298543
refGene_gn=KCNMA1;refGene_tx=XM_019813336.2;refGene_gn=KCNMA1;refGene_
tx=XM_019813372.2
D2 39298642 39298734
refGene_gn=KCNMA1;refGene_tx=XM_011287300.3;refGene_gn=KCNMA1;refGene_
tx=XM_011287302.3;refGene_gn=KCNMA1;refGene_tx=XM_011287303.3;refGene_
gn=KCNMA1;refGene_tx=XM_019813335.2;refGene_gn=KCNMA1;refGene_tx=XM_01
9813338.2;refGene_gn=KCNMA1;refGene_tx=XM_019813339.2;refGene_gn=KCNMA
1;refGene_tx=XM_019813340.2;refGene_gn=KCNMA1;refGene_tx=XM_019813341.
2;refGene_gn=KCNMA1;refGene_tx=XM_019813342.2;refGene_gn=KCNMA1;refGen
e_tx=XM_019813344.2;refGene_gn=KCNMA1;refGene_tx=XM_019813345.2;refGen
e_gn=KCNMA1;refGene_tx=XM_019813346.2;refGene_gn=KCNMA1;refGene_tx=XM_
019813347.2;refGene_gn=KCNMA1;refGene_tx=XM_019813349.2;refGene_gn=KCN
MA1;refGene_tx=XM_019813350.2;refGene_gn=KCNMA1;refGene_tx=XM_01981335
1.2;refGene_gn=KCNMA1;refGene_tx=XM_019813352.2;refGene_gn=KCNMA1;refG

ene_tx=XM_019813353.2;refGene_gn=KCNMA1;refGene_tx=XM_019813354.2;refGene_gn=KCNMA1;refGene_tx=XM_019813355.2;refGene_gn=KCNMA1;refGene_tx=XM_019813356.2;refGene_gn=KCNMA1;refGene_tx=XM_019813357.2;refGene_gn=KCNMA1;refGene_tx=XM_019813358.2;refGene_gn=KCNMA1;refGene_tx=XM_019813360.2;refGene_gn=KCNMA1;refGene_tx=XM_019813362.2;refGene_gn=KCNMA1;refGene_tx=XM_019813363.2;refGene_gn=KCNMA1;refGene_tx=XM_019813364.2;refGene_gn=KCNMA1;refGene_tx=XM_019813366.2;refGene_gn=KCNMA1;refGene_tx=XM_019813368.2;refGene_gn=KCNMA1;refGene_tx=XM_019813369.2;refGene_gn=KCNMA1;refGene_tx=XM_019813371.2;refGene_gn=KCNMA1;refGene_tx=XM_019813373.1;refGene_gn=KCNMA1;refGene_tx=XM_019813374.2;refGene_gn=KCNMA1;refGene_tx=XM_023240597.1;refGene_gn=KCNMA1;refGene_tx=XM_023240598.1;refGene_gn=KCNMA1;refGene_tx=XM_023240599.1;refGene_gn=KCNMA1;refGene_tx=XM_023240600.1;refGene_gn=KCNMA1;refGene_tx=XM_023240601.1

D2 39300396 39300567

refGene_gn=KCNMA1;refGene_tx=XM_011287300.3;refGene_gn=KCNMA1;refGene_tx=XM_011287302.3;refGene_gn=KCNMA1;refGene_tx=XM_011287303.3;refGene_gn=KCNMA1;refGene_tx=XM_019813335.2;refGene_gn=KCNMA1;refGene_tx=XM_019813336.2;refGene_gn=KCNMA1;refGene_tx=XM_019813338.2;refGene_gn=KCNMA1;refGene_tx=XM_019813339.2;refGene_gn=KCNMA1;refGene_tx=XM_019813340.2;refGene_gn=KCNMA1;refGene_tx=XM_019813341.2;refGene_gn=KCNMA1;refGene_tx=XM_019813342.2;refGene_gn=KCNMA1;refGene_tx=XM_019813344.2;refGene_gn=KCNMA1;refGene_tx=XM_019813345.2;refGene_gn=KCNMA1;refGene_tx=XM_019813346.2;refGene_gn=KCNMA1;refGene_tx=XM_019813347.2;refGene_gn=KCNMA1;refGene_tx=XM_019813349.2;refGene_gn=KCNMA1;refGene_tx=XM_019813350.2;refGene_gn=KCNMA1;refGene_tx=XM_019813351.2;refGene_gn=KCNMA1;refGene_tx=XM_019813352.2;refGene_gn=KCNMA1;refGene_tx=XM_019813353.2;refGene_gn=KCNMA1;refGene_tx=XM_019813354.2;refGene_gn=KCNMA1;refGene_tx=XM_019813355.2;refGene_gn=KCNMA1;refGene_tx=XM_019813356.2;refGene_gn=KCNMA1;refGene_tx=XM_019813357.2;refGene_gn=KCNMA1;refGene_tx=XM_019813358.2;refGene_gn=KCNMA1;refGene_tx=XM_019813360.2;refGene_gn=KCNMA1;refGene_tx=XM_019813362.2;refGene_gn=KCNMA1;refGene_tx=XM_019813363.2;refGene_gn=KCNMA1;refGene_tx=XM_019813364.2;refGene_gn=KCNMA1;refGene_tx=XM_019813366.2;refGene_gn=KCNMA1;refGene_tx=XM_019813368.2;refGene_gn=KCNMA1;refGene_tx=XM_019813369.2;refGene_gn=KCNMA1;refGene_tx=XM_019813371.2;refGene_gn=KCNMA1;refGene_tx=XM_019813372.2;refGene_gn=KCNMA1;refGene_tx=XM_019813373.1;refGene_gn=KCNMA1;refGene_tx=XM_019813374.2;refGene_gn=KCNMA1;refGene_tx=XM_023240597.1;refGene_gn=KCNMA1;refGene_tx=XM_023240598.1;refGene_gn=KCNMA1;refGene_tx=XM_023240599.1;refGene_gn=KCNMA1;refGene_tx=XM_023240600.1;refGene_gn=KCNMA1;refGene_tx=XM_023240601.1

D2 39302704 39302780

refGene_gn=KCNMA1;refGene_tx=XM_011287300.3;refGene_gn=KCNMA1;refGene_tx=XM_011287302.3;refGene_gn=KCNMA1;refGene_tx=XM_011287303.3;refGene_gn=KCNMA1;refGene_tx=XM_019813335.2;refGene_gn=KCNMA1;refGene_tx=XM_019813336.2;refGene_gn=KCNMA1;refGene_tx=XM_019813338.2;refGene_gn=KCNMA1;refGene_tx=XM_019813339.2;refGene_gn=KCNMA1;refGene_tx=XM_019813340.2;refGene_gn=KCNMA1;refGene_tx=XM_019813341.2;refGene_gn=KCNMA1;refGene_tx=XM_019813342.2;refGene_gn=KCNMA1;refGene_tx=XM_019813344.2;refGene_gn=KCNMA1;refGene_tx=XM_019813345.2;refGene_gn=KCNMA1;refGene_tx=XM_019813346.2;refGene_gn=KCNMA1;refGene_tx=XM_019813347.2;refGene_gn=KCN

MA1;refGene_tx=XM_019813349.2;refGene_gn=KCNMA1;refGene_tx=XM_01981335
0.2;refGene_gn=KCNMA1;refGene_tx=XM_019813351.2;refGene_gn=KCNMA1;refG
ene_tx=XM_019813352.2;refGene_gn=KCNMA1;refGene_tx=XM_019813353.2;refG
ene_gn=KCNMA1;refGene_tx=XM_019813354.2;refGene_gn=KCNMA1;refGene_tx=X
M_019813355.2;refGene_gn=KCNMA1;refGene_tx=XM_019813356.2;refGene_gn=K
CNMA1;refGene_tx=XM_019813357.2;refGene_gn=KCNMA1;refGene_tx=XM_019813
358.2;refGene_gn=KCNMA1;refGene_tx=XM_019813360.2;refGene_gn=KCNMA1;re
fGene_tx=XM_019813362.2;refGene_gn=KCNMA1;refGene_tx=XM_019813363.2;re
fGene_gn=KCNMA1;refGene_tx=XM_019813364.2;refGene_gn=KCNMA1;refGene_tx
=XM_019813366.2;refGene_gn=KCNMA1;refGene_tx=XM_019813368.2;refGene_gn
=KCNMA1;refGene_tx=XM_019813369.2;refGene_gn=KCNMA1;refGene_tx=XM_0198
13371.2;refGene_gn=KCNMA1;refGene_tx=XM_019813372.2;refGene_gn=KCNMA1;
refGene_tx=XM_019813373.1;refGene_gn=KCNMA1;refGene_tx=XM_019813374.2;
refGene_gn=KCNMA1;refGene_tx=XM_023240597.1;refGene_gn=KCNMA1;refGene_
tx=XM_023240598.1;refGene_gn=KCNMA1;refGene_tx=XM_023240599.1;refGene_
gn=KCNMA1;refGene_tx=XM_023240600.1;refGene_gn=KCNMA1;refGene_tx=XM_02
3240601.1
D2 39312802 39312878
refGene_gn=KCNMA1;refGene_tx=XM_011287300.3;refGene_gn=KCNMA1;refGene_
tx=XM_011287302.3;refGene_gn=KCNMA1;refGene_tx=XM_011287303.3;refGene_
gn=KCNMA1;refGene_tx=XM_019813335.2;refGene_gn=KCNMA1;refGene_tx=XM_01
9813336.2;refGene_gn=KCNMA1;refGene_tx=XM_019813338.2;refGene_gn=KCNMA
1;refGene_tx=XM_019813339.2;refGene_gn=KCNMA1;refGene_tx=XM_019813340.
2;refGene_gn=KCNMA1;refGene_tx=XM_019813341.2;refGene_gn=KCNMA1;refGen
e_tx=XM_019813342.2;refGene_gn=KCNMA1;refGene_tx=XM_019813344.2;refGen
e_gn=KCNMA1;refGene_tx=XM_019813345.2;refGene_gn=KCNMA1;refGene_tx=XM_
019813346.2;refGene_gn=KCNMA1;refGene_tx=XM_019813347.2;refGene_gn=KCN
MA1;refGene_tx=XM_019813349.2;refGene_gn=KCNMA1;refGene_tx=XM_01981335
0.2;refGene_gn=KCNMA1;refGene_tx=XM_019813351.2;refGene_gn=KCNMA1;refG
ene_tx=XM_019813352.2;refGene_gn=KCNMA1;refGene_tx=XM_019813353.2;refG
ene_gn=KCNMA1;refGene_tx=XM_019813354.2;refGene_gn=KCNMA1;refGene_tx=X
M_019813355.2;refGene_gn=KCNMA1;refGene_tx=XM_019813356.2;refGene_gn=K
CNMA1;refGene_tx=XM_019813357.2;refGene_gn=KCNMA1;refGene_tx=XM_019813
358.2;refGene_gn=KCNMA1;refGene_tx=XM_019813360.2;refGene_gn=KCNMA1;re
fGene_tx=XM_019813362.2;refGene_gn=KCNMA1;refGene_tx=XM_019813363.2;re
fGene_gn=KCNMA1;refGene_tx=XM_019813364.2;refGene_gn=KCNMA1;refGene_tx
=XM_019813366.2;refGene_gn=KCNMA1;refGene_tx=XM_019813368.2;refGene_gn
=KCNMA1;refGene_tx=XM_019813369.2;refGene_gn=KCNMA1;refGene_tx=XM_0198
13371.2;refGene_gn=KCNMA1;refGene_tx=XM_019813372.2;refGene_gn=KCNMA1;
refGene_tx=XM_019813373.1;refGene_gn=KCNMA1;refGene_tx=XM_019813374.2;
refGene_gn=KCNMA1;refGene_tx=XM_023240597.1;refGene_gn=KCNMA1;refGene_
tx=XM_023240598.1;refGene_gn=KCNMA1;refGene_tx=XM_023240599.1;refGene_
gn=KCNMA1;refGene_tx=XM_023240600.1;refGene_gn=KCNMA1;refGene_tx=XM_02
3240601.1
D2 39313011 39313030 refGene_gn=KCNMA1;refGene_tx=XM_019813373.1
D2 39332595 39332806 refGene_gn=KCNMA1;refGene_tx=XM_019813347.2
D2 39371065 39371177
refGene_gn=KCNMA1;refGene_tx=XM_011287300.3;refGene_gn=KCNMA1;refGene_
tx=XM_011287302.3;refGene_gn=KCNMA1;refGene_tx=XM_011287303.3;refGene_
gn=KCNMA1;refGene_tx=XM_019813335.2;refGene_gn=KCNMA1;refGene_tx=XM_01

9813336.2;refGene_gn=KCNMA1;refGene_tx=XM_019813338.2;refGene_gn=KCNMA1;refGene_tx=XM_019813339.2;refGene_gn=KCNMA1;refGene_tx=XM_019813340.2;refGene_gn=KCNMA1;refGene_tx=XM_019813341.2;refGene_gn=KCNMA1;refGene_tx=XM_019813342.2;refGene_gn=KCNMA1;refGene_tx=XM_019813344.2;refGene_gn=KCNMA1;refGene_tx=XM_019813345.2;refGene_gn=KCNMA1;refGene_tx=XM_019813346.2;refGene_gn=KCNMA1;refGene_tx=XM_019813349.2;refGene_gn=KCNMA1;refGene_tx=XM_019813350.2;refGene_gn=KCNMA1;refGene_tx=XM_019813351.2;refGene_gn=KCNMA1;refGene_tx=XM_019813352.2;refGene_gn=KCNMA1;refGene_tx=XM_019813353.2;refGene_gn=KCNMA1;refGene_tx=XM_019813354.2;refGene_gn=KCNMA1;refGene_tx=XM_019813355.2;refGene_gn=KCNMA1;refGene_tx=XM_019813356.2;refGene_gn=KCNMA1;refGene_tx=XM_019813357.2;refGene_gn=KCNMA1;refGene_tx=XM_019813358.2;refGene_gn=KCNMA1;refGene_tx=XM_019813360.2;refGene_gn=KCNMA1;refGene_tx=XM_019813362.2;refGene_gn=KCNMA1;refGene_tx=XM_019813363.2;refGene_gn=KCNMA1;refGene_tx=XM_019813364.2;refGene_gn=KCNMA1;refGene_tx=XM_019813366.2;refGene_gn=KCNMA1;refGene_tx=XM_019813368.2;refGene_gn=KCNMA1;refGene_tx=XM_019813369.2;refGene_gn=KCNMA1;refGene_tx=XM_019813371.2;refGene_gn=KCNMA1;refGene_tx=XM_019813372.2;refGene_gn=KCNMA1;refGene_tx=XM_019813374.2;refGene_gn=KCNMA1;refGene_tx=XM_023240597.1;refGene_gn=KCNMA1;refGene_tx=XM_023240598.1;refGene_gn=KCNMA1;refGene_tx=XM_023240599.1;refGene_gn=KCNMA1;refGene_tx=XM_023240600.1;refGene_gn=KCNMA1;refGene_tx=XM_023240601.1

D2 39372032 39372126

refGene_gn=KCNMA1;refGene_tx=XM_011287300.3;refGene_gn=KCNMA1;refGene_tx=XM_011287302.3;refGene_gn=KCNMA1;refGene_tx=XM_011287303.3;refGene_gn=KCNMA1;refGene_tx=XM_019813335.2;refGene_gn=KCNMA1;refGene_tx=XM_019813336.2;refGene_gn=KCNMA1;refGene_tx=XM_019813338.2;refGene_gn=KCNMA1;refGene_tx=XM_019813339.2;refGene_gn=KCNMA1;refGene_tx=XM_019813340.2;refGene_gn=KCNMA1;refGene_tx=XM_019813341.2;refGene_gn=KCNMA1;refGene_tx=XM_019813342.2;refGene_gn=KCNMA1;refGene_tx=XM_019813344.2;refGene_gn=KCNMA1;refGene_tx=XM_019813345.2;refGene_gn=KCNMA1;refGene_tx=XM_019813346.2;refGene_gn=KCNMA1;refGene_tx=XM_019813349.2;refGene_gn=KCNMA1;refGene_tx=XM_019813350.2;refGene_gn=KCNMA1;refGene_tx=XM_019813351.2;refGene_gn=KCNMA1;refGene_tx=XM_019813352.2;refGene_gn=KCNMA1;refGene_tx=XM_019813353.2;refGene_gn=KCNMA1;refGene_tx=XM_019813354.2;refGene_gn=KCNMA1;refGene_tx=XM_019813355.2;refGene_gn=KCNMA1;refGene_tx=XM_019813356.2;refGene_gn=KCNMA1;refGene_tx=XM_019813357.2;refGene_gn=KCNMA1;refGene_tx=XM_019813358.2;refGene_gn=KCNMA1;refGene_tx=XM_019813360.2;refGene_gn=KCNMA1;refGene_tx=XM_019813362.2;refGene_gn=KCNMA1;refGene_tx=XM_019813363.2;refGene_gn=KCNMA1;refGene_tx=XM_019813364.2;refGene_gn=KCNMA1;refGene_tx=XM_019813366.2;refGene_gn=KCNMA1;refGene_tx=XM_019813368.2;refGene_gn=KCNMA1;refGene_tx=XM_019813369.2;refGene_gn=KCNMA1;refGene_tx=XM_019813371.2;refGene_gn=KCNMA1;refGene_tx=XM_019813372.2;refGene_gn=KCNMA1;refGene_tx=XM_019813374.2;refGene_gn=KCNMA1;refGene_tx=XM_023240597.1;refGene_gn=KCNMA1;refGene_tx=XM_023240598.1;refGene_gn=KCNMA1;refGene_tx=XM_023240599.1;refGene_gn=KCNMA1;refGene_tx=XM_023240600.1;refGene_gn=KCNMA1;refGene_tx=XM_023240601.1

D2 39434284 39434346

refGene_gn=KCNMA1;refGene_tx=XM_011287300.3;refGene_gn=KCNMA1;refGene_tx=XM_011287302.3;refGene_gn=KCNMA1;refGene_tx=XM_011287303.3;refGene_gn=KCNMA1;refGene_tx=XM_019813335.2;refGene_gn=KCNMA1;refGene_tx=XM_01

9813336.2;refGene_gn=KCNMA1;refGene_tx=XM_019813338.2;refGene_gn=KCNMA
1;refGene_tx=XM_019813339.2;refGene_gn=KCNMA1;refGene_tx=XM_019813340.
2;refGene_gn=KCNMA1;refGene_tx=XM_019813341.2;refGene_gn=KCNMA1;refGen
e_tx=XM_019813342.2;refGene_gn=KCNMA1;refGene_tx=XM_019813344.2;refGen
e_gn=KCNMA1;refGene_tx=XM_019813345.2;refGene_gn=KCNMA1;refGene_tx=XM_
019813346.2;refGene_gn=KCNMA1;refGene_tx=XM_019813349.2;refGene_gn=KCN
MA1;refGene_tx=XM_019813350.2;refGene_gn=KCNMA1;refGene_tx=XM_01981335
1.2;refGene_gn=KCNMA1;refGene_tx=XM_019813352.2;refGene_gn=KCNMA1;refG
ene_tx=XM_019813353.2;refGene_gn=KCNMA1;refGene_tx=XM_019813354.2;refG
ene_gn=KCNMA1;refGene_tx=XM_019813355.2;refGene_gn=KCNMA1;refGene_tx=X
M_019813356.2;refGene_gn=KCNMA1;refGene_tx=XM_019813357.2;refGene_gn=K
CNMA1;refGene_tx=XM_019813358.2;refGene_gn=KCNMA1;refGene_tx=XM_019813
360.2;refGene_gn=KCNMA1;refGene_tx=XM_019813362.2;refGene_gn=KCNMA1;re
fGene_tx=XM_019813363.2;refGene_gn=KCNMA1;refGene_tx=XM_019813364.2;re
fGene_gn=KCNMA1;refGene_tx=XM_019813366.2;refGene_gn=KCNMA1;refGene_tx
=XM_019813368.2;refGene_gn=KCNMA1;refGene_tx=XM_019813369.2;refGene_gn
=KCNMA1;refGene_tx=XM_019813371.2;refGene_gn=KCNMA1;refGene_tx=XM_0198
13372.2;refGene_gn=KCNMA1;refGene_tx=XM_019813374.2;refGene_gn=KCNMA1;
refGene_tx=XM_023240597.1;refGene_gn=KCNMA1;refGene_tx=XM_023240598.1;
refGene_gn=KCNMA1;refGene_tx=XM_023240599.1;refGene_gn=KCNMA1;refGene_
tx=XM_023240600.1;refGene_gn=KCNMA1;refGene_tx=XM_023240601.1
D2 39581841 39582003
refGene_gn=KCNMA1;refGene_tx=XM_011287300.3;refGene_gn=KCNMA1;refGene_
tx=XM_011287302.3;refGene_gn=KCNMA1;refGene_tx=XM_011287303.3;refGene_
gn=KCNMA1;refGene_tx=XM_019813335.2;refGene_gn=KCNMA1;refGene_tx=XM_01
9813336.2;refGene_gn=KCNMA1;refGene_tx=XM_019813338.2;refGene_gn=KCNMA
1;refGene_tx=XM_019813339.2;refGene_gn=KCNMA1;refGene_tx=XM_019813340.
2;refGene_gn=KCNMA1;refGene_tx=XM_019813341.2;refGene_gn=KCNMA1;refGen
e_tx=XM_019813342.2;refGene_gn=KCNMA1;refGene_tx=XM_019813344.2;refGen
e_gn=KCNMA1;refGene_tx=XM_019813345.2;refGene_gn=KCNMA1;refGene_tx=XM_
019813346.2;refGene_gn=KCNMA1;refGene_tx=XM_019813349.2;refGene_gn=KCN
MA1;refGene_tx=XM_019813350.2;refGene_gn=KCNMA1;refGene_tx=XM_01981335
1.2;refGene_gn=KCNMA1;refGene_tx=XM_019813352.2;refGene_gn=KCNMA1;refG
ene_tx=XM_019813353.2;refGene_gn=KCNMA1;refGene_tx=XM_019813354.2;refG
ene_gn=KCNMA1;refGene_tx=XM_019813355.2;refGene_gn=KCNMA1;refGene_tx=X
M_019813356.2;refGene_gn=KCNMA1;refGene_tx=XM_019813357.2;refGene_gn=K
CNMA1;refGene_tx=XM_019813358.2;refGene_gn=KCNMA1;refGene_tx=XM_019813
360.2;refGene_gn=KCNMA1;refGene_tx=XM_019813362.2;refGene_gn=KCNMA1;re
fGene_tx=XM_019813363.2;refGene_gn=KCNMA1;refGene_tx=XM_019813364.2;re
fGene_gn=KCNMA1;refGene_tx=XM_019813366.2;refGene_gn=KCNMA1;refGene_tx
=XM_019813368.2;refGene_gn=KCNMA1;refGene_tx=XM_019813369.2;refGene_gn
=KCNMA1;refGene_tx=XM_019813371.2;refGene_gn=KCNMA1;refGene_tx=XM_0198
13372.2;refGene_gn=KCNMA1;refGene_tx=XM_019813374.2;refGene_gn=KCNMA1;
refGene_tx=XM_023240597.1;refGene_gn=KCNMA1;refGene_tx=XM_023240598.1;
refGene_gn=KCNMA1;refGene_tx=XM_023240599.1;refGene_gn=KCNMA1;refGene_
tx=XM_023240600.1;refGene_gn=KCNMA1;refGene_tx=XM_023240601.1
D2 39802049 39802427
refGene_gn=KCNMA1;refGene_tx=XM_011287300.3;refGene_gn=KCNMA1;refGene_
tx=XM_011287302.3;refGene_gn=KCNMA1;refGene_tx=XM_011287303.3;refGene_
gn=KCNMA1;refGene_tx=XM_019813335.2;refGene_gn=KCNMA1;refGene_tx=XM_01

9813336.2;refGene_gn=KCNMA1;refGene_tx=XM_019813338.2;refGene_gn=KCNMA
1;refGene_tx=XM_019813339.2;refGene_gn=KCNMA1;refGene_tx=XM_019813340.
2;refGene_gn=KCNMA1;refGene_tx=XM_019813341.2;refGene_gn=KCNMA1;refGen
e_tx=XM_019813342.2;refGene_gn=KCNMA1;refGene_tx=XM_019813344.2;refGen
e_gn=KCNMA1;refGene_tx=XM_019813345.2;refGene_gn=KCNMA1;refGene_tx=XM_
019813346.2;refGene_gn=KCNMA1;refGene_tx=XM_019813349.2;refGene_gn=KCN
MA1;refGene_tx=XM_019813350.2;refGene_gn=KCNMA1;refGene_tx=XM_01981335
1.2;refGene_gn=KCNMA1;refGene_tx=XM_019813352.2;refGene_gn=KCNMA1;refG
ene_tx=XM_019813353.2;refGene_gn=KCNMA1;refGene_tx=XM_019813354.2;refG
ene_gn=KCNMA1;refGene_tx=XM_019813355.2;refGene_gn=KCNMA1;refGene_tx=X
M_019813356.2;refGene_gn=KCNMA1;refGene_tx=XM_019813357.2;refGene_gn=K
CNMA1;refGene_tx=XM_019813358.2;refGene_gn=KCNMA1;refGene_tx=XM_019813
360.2;refGene_gn=KCNMA1;refGene_tx=XM_019813362.2;refGene_gn=KCNMA1;re
fGene_tx=XM_019813363.2;refGene_gn=KCNMA1;refGene_tx=XM_019813364.2;re
fGene_gn=KCNMA1;refGene_tx=XM_019813366.2;refGene_gn=KCNMA1;refGene_tx
=XM_019813368.2;refGene_gn=KCNMA1;refGene_tx=XM_019813369.2;refGene_gn
=KCNMA1;refGene_tx=XM_019813371.2;refGene_gn=KCNMA1;refGene_tx=XM_0198
13372.2;refGene_gn=KCNMA1;refGene_tx=XM_019813374.2;refGene_gn=KCNMA1;
refGene_tx=XM_023240597.1;refGene_gn=KCNMA1;refGene_tx=XM_023240598.1;
refGene_gn=KCNMA1;refGene_tx=XM_023240599.1;refGene_gn=KCNMA1;refGene_
tx=XM_023240600.1;refGene_gn=KCNMA1;refGene_tx=XM_023240601.1
D2 39947129 39947233
refGene_gn=DLG5;refGene_tx=XM_023240602.1;refGene_gn=DLG5;refGene_tx=X
M_023240603.1;refGene_gn=DLG5;refGene_tx=XM_023240604.1;refGene_gn=DLG
5;refGene_tx=XM_023240605.1;refGene_gn=DLG5;refGene_tx=XM_023240606.1
D2 39948694 39948804
refGene_gn=DLG5;refGene_tx=XM_023240602.1;refGene_gn=DLG5;refGene_tx=X
M_023240603.1;refGene_gn=DLG5;refGene_tx=XM_023240604.1;refGene_gn=DLG
5;refGene_tx=XM_023240605.1;refGene_gn=DLG5;refGene_tx=XM_023240606.1
D2 39949495 39949605
refGene_gn=DLG5;refGene_tx=XM_023240602.1;refGene_gn=DLG5;refGene_tx=X
M_023240603.1;refGene_gn=DLG5;refGene_tx=XM_023240604.1;refGene_gn=DLG
5;refGene_tx=XM_023240605.1;refGene_gn=DLG5;refGene_tx=XM_023240606.1
D2 39950809 39950937
refGene_gn=DLG5;refGene_tx=XM_023240602.1;refGene_gn=DLG5;refGene_tx=X
M_023240603.1;refGene_gn=DLG5;refGene_tx=XM_023240604.1;refGene_gn=DLG
5;refGene_tx=XM_023240605.1;refGene_gn=DLG5;refGene_tx=XM_023240606.1
D2 39951152 39951296
refGene_gn=DLG5;refGene_tx=XM_023240602.1;refGene_gn=DLG5;refGene_tx=X
M_023240603.1;refGene_gn=DLG5;refGene_tx=XM_023240604.1;refGene_gn=DLG
5;refGene_tx=XM_023240605.1;refGene_gn=DLG5;refGene_tx=XM_023240606.1
D2 39962765 39962962
refGene_gn=DLG5;refGene_tx=XM_023240602.1;refGene_gn=DLG5;refGene_tx=X
M_023240603.1;refGene_gn=DLG5;refGene_tx=XM_023240604.1;refGene_gn=DLG
5;refGene_tx=XM_023240605.1;refGene_gn=DLG5;refGene_tx=XM_023240606.1
D2 39963826 39963997
refGene_gn=DLG5;refGene_tx=XM_023240602.1;refGene_gn=DLG5;refGene_tx=X
M_023240603.1;refGene_gn=DLG5;refGene_tx=XM_023240604.1;refGene_gn=DLG
5;refGene_tx=XM_023240605.1;refGene_gn=DLG5;refGene_tx=XM_023240606.1
D2 39964865 39965014

refGene_gn=DLG5;refGene_tx=XM_023240602.1;refGene_gn=DLG5;refGene_tx=XM_023240603.1;refGene_gn=DLG5;refGene_tx=XM_023240604.1;refGene_gn=DLG5;refGene_tx=XM_023240605.1;refGene_gn=DLG5;refGene_tx=XM_023240606.1
D2 39965620 39965804
refGene_gn=DLG5;refGene_tx=XM_023240602.1;refGene_gn=DLG5;refGene_tx=XM_023240603.1;refGene_gn=DLG5;refGene_tx=XM_023240604.1;refGene_gn=DLG5;refGene_tx=XM_023240605.1;refGene_gn=DLG5;refGene_tx=XM_023240606.1
D2 39967101 39967248
refGene_gn=DLG5;refGene_tx=XM_023240602.1;refGene_gn=DLG5;refGene_tx=XM_023240603.1;refGene_gn=DLG5;refGene_tx=XM_023240604.1;refGene_gn=DLG5;refGene_tx=XM_023240605.1;refGene_gn=DLG5;refGene_tx=XM_023240606.1
D2 39967924 39968058
refGene_gn=DLG5;refGene_tx=XM_023240602.1;refGene_gn=DLG5;refGene_tx=XM_023240603.1;refGene_gn=DLG5;refGene_tx=XM_023240604.1;refGene_gn=DLG5;refGene_tx=XM_023240605.1;refGene_gn=DLG5;refGene_tx=XM_023240606.1
D2 39968161 39968324
refGene_gn=DLG5;refGene_tx=XM_023240602.1;refGene_gn=DLG5;refGene_tx=XM_023240603.1;refGene_gn=DLG5;refGene_tx=XM_023240604.1;refGene_gn=DLG5;refGene_tx=XM_023240605.1;refGene_gn=DLG5;refGene_tx=XM_023240606.1
D2 39971994 39972145
refGene_gn=DLG5;refGene_tx=XM_023240602.1;refGene_gn=DLG5;refGene_tx=XM_023240603.1;refGene_gn=DLG5;refGene_tx=XM_023240604.1;refGene_gn=DLG5;refGene_tx=XM_023240605.1;refGene_gn=DLG5;refGene_tx=XM_023240606.1
D2 39972442 39972532
refGene_gn=DLG5;refGene_tx=XM_023240602.1;refGene_gn=DLG5;refGene_tx=XM_023240603.1;refGene_gn=DLG5;refGene_tx=XM_023240604.1;refGene_gn=DLG5;refGene_tx=XM_023240605.1;refGene_gn=DLG5;refGene_tx=XM_023240606.1
D2 39973193 39973306
refGene_gn=DLG5;refGene_tx=XM_023240602.1;refGene_gn=DLG5;refGene_tx=XM_023240603.1;refGene_gn=DLG5;refGene_tx=XM_023240604.1;refGene_gn=DLG5;refGene_tx=XM_023240605.1;refGene_gn=DLG5;refGene_tx=XM_023240606.1
D2 39975963 39976108
refGene_gn=DLG5;refGene_tx=XM_023240602.1;refGene_gn=DLG5;refGene_tx=XM_023240603.1;refGene_gn=DLG5;refGene_tx=XM_023240604.1;refGene_gn=DLG5;refGene_tx=XM_023240605.1;refGene_gn=DLG5;refGene_tx=XM_023240606.1
D2 39976458 39976582
refGene_gn=DLG5;refGene_tx=XM_023240602.1;refGene_gn=DLG5;refGene_tx=XM_023240603.1;refGene_gn=DLG5;refGene_tx=XM_023240604.1;refGene_gn=DLG5;refGene_tx=XM_023240605.1;refGene_gn=DLG5;refGene_tx=XM_023240606.1
D2 39977346 39978366
refGene_gn=DLG5;refGene_tx=XM_023240602.1;refGene_gn=DLG5;refGene_tx=XM_023240603.1;refGene_gn=DLG5;refGene_tx=XM_023240604.1;refGene_gn=DLG5;refGene_tx=XM_023240605.1;refGene_gn=DLG5;refGene_tx=XM_023240606.1
D2 39980312 39980405
refGene_gn=DLG5;refGene_tx=XM_023240602.1;refGene_gn=DLG5;refGene_tx=XM_023240603.1;refGene_gn=DLG5;refGene_tx=XM_023240604.1;refGene_gn=DLG5;refGene_tx=XM_023240606.1
D2 39983443 39983547
refGene_gn=DLG5;refGene_tx=XM_023240602.1;refGene_gn=DLG5;refGene_tx=XM_023240603.1;refGene_gn=DLG5;refGene_tx=XM_023240604.1;refGene_gn=DLG

5;refGene_tx=XM_023240605.1;refGene_gn=DLG5;refGene_tx=XM_023240606.1
D2 39983909 39984085
refGene_gn=DLG5;refGene_tx=XM_023240602.1;refGene_gn=DLG5;refGene_tx=XM_023240603.1;refGene_gn=DLG5;refGene_tx=XM_023240604.1;refGene_gn=DLG5;refGene_tx=XM_023240605.1;refGene_gn=DLG5;refGene_tx=XM_023240606.1
D2 39984742 39984870
refGene_gn=DLG5;refGene_tx=XM_023240602.1;refGene_gn=DLG5;refGene_tx=XM_023240603.1;refGene_gn=DLG5;refGene_tx=XM_023240604.1;refGene_gn=DLG5;refGene_tx=XM_023240605.1;refGene_gn=DLG5;refGene_tx=XM_023240606.1
D2 39985297 39985430
refGene_gn=DLG5;refGene_tx=XM_023240602.1;refGene_gn=DLG5;refGene_tx=XM_023240603.1;refGene_gn=DLG5;refGene_tx=XM_023240604.1;refGene_gn=DLG5;refGene_tx=XM_023240605.1;refGene_gn=DLG5;refGene_tx=XM_023240606.1
D2 39988021 39988147
refGene_gn=DLG5;refGene_tx=XM_023240602.1;refGene_gn=DLG5;refGene_tx=XM_023240603.1;refGene_gn=DLG5;refGene_tx=XM_023240604.1;refGene_gn=DLG5;refGene_tx=XM_023240605.1;refGene_gn=DLG5;refGene_tx=XM_023240606.1
D2 39989694 39989879
refGene_gn=DLG5;refGene_tx=XM_023240602.1;refGene_gn=DLG5;refGene_tx=XM_023240603.1;refGene_gn=DLG5;refGene_tx=XM_023240604.1;refGene_gn=DLG5;refGene_tx=XM_023240605.1;refGene_gn=DLG5;refGene_tx=XM_023240606.1
D2 39995069 39995382
refGene_gn=DLG5;refGene_tx=XM_023240602.1;refGene_gn=DLG5;refGene_tx=XM_023240603.1;refGene_gn=DLG5;refGene_tx=XM_023240604.1;refGene_gn=DLG5;refGene_tx=XM_023240605.1;refGene_gn=DLG5;refGene_tx=XM_023240606.1
D2 39996836 39997096
refGene_gn=DLG5;refGene_tx=XM_023240602.1;refGene_gn=DLG5;refGene_tx=XM_023240603.1;refGene_gn=DLG5;refGene_tx=XM_023240604.1;refGene_gn=DLG5;refGene_tx=XM_023240605.1;refGene_gn=DLG5;refGene_tx=XM_023240606.1
D2 40010689 40010873
refGene_gn=DLG5;refGene_tx=XM_023240602.1;refGene_gn=DLG5;refGene_tx=XM_023240603.1;refGene_gn=DLG5;refGene_tx=XM_023240604.1;refGene_gn=DLG5;refGene_tx=XM_023240605.1;refGene_gn=DLG5;refGene_tx=XM_023240606.1
D2 40011498 40011642
refGene_gn=DLG5;refGene_tx=XM_023240602.1;refGene_gn=DLG5;refGene_tx=XM_023240603.1;refGene_gn=DLG5;refGene_tx=XM_023240604.1;refGene_gn=DLG5;refGene_tx=XM_023240605.1;refGene_gn=DLG5;refGene_tx=XM_023240606.1
D2 40013478 40013641
refGene_gn=DLG5;refGene_tx=XM_023240602.1;refGene_gn=DLG5;refGene_tx=XM_023240603.1;refGene_gn=DLG5;refGene_tx=XM_023240604.1;refGene_gn=DLG5;refGene_tx=XM_023240605.1;refGene_gn=DLG5;refGene_tx=XM_023240606.1
D2 40024856 40024925
refGene_gn=DLG5;refGene_tx=XM_023240602.1;refGene_gn=DLG5;refGene_tx=XM_023240603.1;refGene_gn=DLG5;refGene_tx=XM_023240605.1;refGene_gn=DLG5;refGene_tx=XM_023240606.1
D2 40048476 40048525 refGene_gn=DLG5;refGene_tx=XM_023240606.1
D2 40075266 40075570
refGene_gn=DLG5;refGene_tx=XM_023240602.1;refGene_gn=DLG5;refGene_tx=XM_023240603.1;refGene_gn=DLG5;refGene_tx=XM_023240604.1;refGene_gn=DLG5;refGene_tx=XM_023240605.1

D2	40116070 40116219 refGene_gn=POLR3A;refGene_tx=XM_003994087.4
D2	40117914 40118047 refGene_gn=POLR3A;refGene_tx=XM_003994087.4
D2	40119061 40119193 refGene_gn=POLR3A;refGene_tx=XM_003994087.4
D2	40119408 40119573 refGene_gn=POLR3A;refGene_tx=XM_003994087.4
D2	40120022 40120187 refGene_gn=POLR3A;refGene_tx=XM_003994087.4
D2	40121464 40121557 refGene_gn=POLR3A;refGene_tx=XM_003994087.4
D2	40121742 40121836 refGene_gn=POLR3A;refGene_tx=XM_003994087.4
D2	40122926 40123097 refGene_gn=POLR3A;refGene_tx=XM_003994087.4
D2	40123671 40123754 refGene_gn=POLR3A;refGene_tx=XM_003994087.4
D2	40123844 40123931 refGene_gn=POLR3A;refGene_tx=XM_003994087.4
D2	40128364 40128478 refGene_gn=POLR3A;refGene_tx=XM_003994087.4
D2	40129394 40129565 refGene_gn=POLR3A;refGene_tx=XM_003994087.4
D2	40131390 40131528 refGene_gn=POLR3A;refGene_tx=XM_003994087.4
D2	40132655 40132774 refGene_gn=POLR3A;refGene_tx=XM_003994087.4
D2	40134720 40134832 refGene_gn=POLR3A;refGene_tx=XM_003994087.4
D2	40138410 40138583 refGene_gn=POLR3A;refGene_tx=XM_003994087.4
D2	40142188 40142353 refGene_gn=POLR3A;refGene_tx=XM_003994087.4
D2	40143610 40143749 refGene_gn=POLR3A;refGene_tx=XM_003994087.4
D2	40144090 40144218 refGene_gn=POLR3A;refGene_tx=XM_003994087.4
D2	40144860 40144930 refGene_gn=POLR3A;refGene_tx=XM_003994087.4
D2	40148716 40148857 refGene_gn=POLR3A;refGene_tx=XM_003994087.4
D2	40151787 40151929 refGene_gn=POLR3A;refGene_tx=XM_003994087.4
D2	40152793 40152897 refGene_gn=POLR3A;refGene_tx=XM_003994087.4
D2	40154209 40154346 refGene_gn=POLR3A;refGene_tx=XM_003994087.4
D2	40154524 40154687 refGene_gn=POLR3A;refGene_tx=XM_003994087.4
D2	40154796 40155036 refGene_gn=POLR3A;refGene_tx=XM_003994087.4
D2	40162736 40162891 refGene_gn=POLR3A;refGene_tx=XM_003994087.4
D2	40163138 40163310 refGene_gn=POLR3A;refGene_tx=XM_003994087.4
D2	40164468 40164606 refGene_gn=POLR3A;refGene_tx=XM_003994087.4
D2	40164836 40164972 refGene_gn=POLR3A;refGene_tx=XM_003994087.4
D2	40167528 40167572 refGene_gn=POLR3A;refGene_tx=XM_003994087.4
D2	40169620 40169623
refGene_gn=RPS24;refGene_tx=XM_003994088.5;refGene_gn=RPS24;refGene_tx
=XM_003994089.5;refGene_gn=RPS24;refGene_tx=XM_003994091.5;refGene_gn=
RPS24;refGene_tx=XM_011287296.3;refGene_gn=RPS24;refGene_tx=XM_0198133
76.2;refGene_gn=RPS24;refGene_tx=XM_019813377.2
D2	40171098 40171164
refGene_gn=RPS24;refGene_tx=XM_003994088.5;refGene_gn=RPS24;refGene_tx
=XM_003994089.5;refGene_gn=RPS24;refGene_tx=XM_003994091.5;refGene_gn=
RPS24;refGene_tx=XM_011287296.3;refGene_gn=RPS24;refGene_tx=XM_0198133
76.2;refGene_gn=RPS24;refGene_tx=XM_019813377.2
D2	40171253 40171463
refGene_gn=RPS24;refGene_tx=XM_003994088.5;refGene_gn=RPS24;refGene_tx
=XM_003994089.5;refGene_gn=RPS24;refGene_tx=XM_003994091.5;refGene_gn=
RPS24;refGene_tx=XM_011287296.3;refGene_gn=RPS24;refGene_tx=XM_0198133
76.2;refGene_gn=RPS24;refGene_tx=XM_019813377.2
D2	40172383 40172494
refGene_gn=RPS24;refGene_tx=XM_003994088.5;refGene_gn=RPS24;refGene_tx
=XM_003994089.5;refGene_gn=RPS24;refGene_tx=XM_003994091.5;refGene_gn=
RPS24;refGene_tx=XM_011287296.3;refGene_gn=RPS24;refGene_tx=XM_0198133

76.2;refGene_gn=RPS24;refGene_tx=XM_019813377.2
D2 40172619 40172622
refGene_gn=RPS24;refGene_tx=XM_003994089.5;refGene_gn=RPS24;refGene_tx=XM_003994091.5;refGene_gn=RPS24;refGene_tx=XM_019813376.2
D2 40173141 40173147
refGene_gn=RPS24;refGene_tx=XM_003994089.5;refGene_gn=RPS24;refGene_tx=XM_011287296.3
D2 40175363 40175366
refGene_gn=RPS24;refGene_tx=XM_003994088.5;refGene_gn=RPS24;refGene_tx=XM_003994091.5
D2 40175772 40175781
refGene_gn=RPS24;refGene_tx=XM_019813376.2;refGene_gn=RPS24;refGene_tx=XM_019813377.2
D2 41096113 41096151
refGene_gn=LOC105261021;refGene_tx=XM_019813378.2
D2 41102579 41103717
refGene_gn=LOC105261021;refGene_tx=XM_019813378.2
D2 41228481 41228541 refGene_gn=ZMIZ1;refGene_tx=XM_011287295.3
D2 41234971 41235085 refGene_gn=ZMIZ1;refGene_tx=XM_011287295.3
D2 41242140 41242246 refGene_gn=ZMIZ1;refGene_tx=XM_011287295.3
D2 41301539 41301684 refGene_gn=ZMIZ1;refGene_tx=XM_011287295.3
D2 41312304 41312419 refGene_gn=ZMIZ1;refGene_tx=XM_011287295.3
D2 41313419 41313637 refGene_gn=ZMIZ1;refGene_tx=XM_011287295.3
D2 41314342 41314541 refGene_gn=ZMIZ1;refGene_tx=XM_011287295.3
D2 41315473 41315623 refGene_gn=ZMIZ1;refGene_tx=XM_011287295.3
D2 41315624 41315747 refGene_gn=ZMIZ1;refGene_tx=XM_011287295.3
D2 41318506 41318689 refGene_gn=ZMIZ1;refGene_tx=XM_011287295.3
D2 41319869 41319947 refGene_gn=ZMIZ1;refGene_tx=XM_011287295.3
D2 41320621 41320796 refGene_gn=ZMIZ1;refGene_tx=XM_011287295.3
D2 41321434 41321576 refGene_gn=ZMIZ1;refGene_tx=XM_011287295.3
D2 41322766 41322977 refGene_gn=ZMIZ1;refGene_tx=XM_011287295.3
D2 41323798 41323904 refGene_gn=ZMIZ1;refGene_tx=XM_011287295.3
D2 41324839 41325000 refGene_gn=ZMIZ1;refGene_tx=XM_011287295.3
D2 41326020 41326088 refGene_gn=ZMIZ1;refGene_tx=XM_011287295.3
D2 41326397 41326466 refGene_gn=ZMIZ1;refGene_tx=XM_011287295.3
D2 41326950 41327195 refGene_gn=ZMIZ1;refGene_tx=XM_011287295.3
D2 41328325 41328492 refGene_gn=ZMIZ1;refGene_tx=XM_011287295.3
D2 41331722 41331980 refGene_gn=ZMIZ1;refGene_tx=XM_011287295.3
D2 41333530 41333638 refGene_gn=ZMIZ1;refGene_tx=XM_011287295.3
D2 41375491 41375677
refGene_gn=PPIF;refGene_tx=XM_003994109.5;refGene_gn=PPIF;refGene_tx=XM_011287292.3
D2 41375719 41376007
refGene_gn=PPIF;refGene_tx=XM_011287290.3;refGene_gn=PPIF;refGene_tx=XM_011287293.2;refGene_gn=PPIF;refGene_tx=XM_011287294.2
D2 41376911 41376942
refGene_gn=PPIF;refGene_tx=XM_003994109.5;refGene_gn=PPIF;refGene_tx=XM_011287290.3;refGene_gn=PPIF;refGene_tx=XM_011287292.3;refGene_gn=PPIF;refGene_tx=XM_011287293.2;refGene_gn=PPIF;refGene_tx=XM_011287294.2
D2 41377472 41377561

refGene_gn=PPIF;refGene_tx=XM_003994109.5;refGene_gn=PPIF;refGene_tx=XM_011287290.3;refGene_gn=PPIF;refGene_tx=XM_011287292.3;refGene_gn=PPIF;refGene_tx=XM_011287293.2;refGene_gn=PPIF;refGene_tx=XM_011287294.2
D2 41379310 41379407
refGene_gn=PPIF;refGene_tx=XM_003994109.5;refGene_gn=PPIF;refGene_tx=XM_011287290.3;refGene_gn=PPIF;refGene_tx=XM_011287292.3;refGene_gn=PPIF;refGene_tx=XM_011287293.2;refGene_gn=PPIF;refGene_tx=XM_011287294.2
D2 41379807 41379965
refGene_gn=PPIF;refGene_tx=XM_003994109.5;refGene_gn=PPIF;refGene_tx=XM_011287290.3;refGene_gn=PPIF;refGene_tx=XM_011287292.3;refGene_gn=PPIF;refGene_tx=XM_011287293.2;refGene_gn=PPIF;refGene_tx=XM_011287294.2
D2 41381168 41381507
refGene_gn=PPIF;refGene_tx=XM_003994109.5;refGene_gn=PPIF;refGene_tx=XM_011287290.3;refGene_gn=PPIF;refGene_tx=XM_011287292.3;refGene_gn=PPIF;refGene_tx=XM_011287293.2;refGene_gn=PPIF;refGene_tx=XM_011287294.2
D2 41406495 41406609 refGene_gn=ZCCHC24;refGene_tx=XM_023240607.1
D2 41414246 41414411 refGene_gn=ZCCHC24;refGene_tx=XM_023240607.1
D2 41453499 41453700 refGene_gn=ZCCHC24;refGene_tx=XM_023240607.1
D2 41465483 41465729 refGene_gn=ZCCHC24;refGene_tx=XM_023240607.1
D2 41559149 41559204
refGene_gn=ANXA11;refGene_tx=XM_023240608.1;refGene_gn=ANXA11;refGene_tx=XM_023240609.1
D2 41561138 41561251
refGene_gn=ANXA11;refGene_tx=XM_023240608.1;refGene_gn=ANXA11;refGene_tx=XM_023240609.1
D2 41562165 41562558
refGene_gn=ANXA11;refGene_tx=XM_023240608.1;refGene_gn=ANXA11;refGene_tx=XM_023240609.1
D2 41563873 41563961
refGene_gn=ANXA11;refGene_tx=XM_023240608.1;refGene_gn=ANXA11;refGene_tx=XM_023240609.1
D2 41564171 41564266
refGene_gn=ANXA11;refGene_tx=XM_023240608.1;refGene_gn=ANXA11;refGene_tx=XM_023240609.1
D2 41564976 41565090
refGene_gn=ANXA11;refGene_tx=XM_023240608.1;refGene_gn=ANXA11;refGene_tx=XM_023240609.1
D2 41566853 41566944
refGene_gn=ANXA11;refGene_tx=XM_023240608.1;refGene_gn=ANXA11;refGene_tx=XM_023240609.1
D2 41567313 41567393
refGene_gn=ANXA11;refGene_tx=XM_023240608.1;refGene_gn=ANXA11;refGene_tx=XM_023240609.1
D2 41567529 41567586
refGene_gn=ANXA11;refGene_tx=XM_023240608.1;refGene_gn=ANXA11;refGene_tx=XM_023240609.1
D2 41569881 41569975
refGene_gn=ANXA11;refGene_tx=XM_023240608.1;refGene_gn=ANXA11;refGene_tx=XM_023240609.1
D2 41573134 41573230

refGene_gn=ANXA11;refGene_tx=XM_023240608.1;refGene_gn=ANXA11;refGene_tx=XM_023240609.1
D2 41574189 41574248
refGene_gn=ANXA11;refGene_tx=XM_023240608.1;refGene_gn=ANXA11;refGene_tx=XM_023240609.1
D2 41574413 41574536
refGene_gn=ANXA11;refGene_tx=XM_023240608.1;refGene_gn=ANXA11;refGene_tx=XM_023240609.1
D2 41576525 41576585
refGene_gn=ANXA11;refGene_tx=XM_023240608.1;refGene_gn=ANXA11;refGene_tx=XM_023240609.1
D2 41589125 41589172 refGene_gn=PLAC9;refGene_tx=XM_006937919.3
D2 41589664 41589791 refGene_gn=PLAC9;refGene_tx=XM_006937919.3
D2 41591205 41591303
refGene_gn=PLAC9;refGene_tx=XM_006937919.3;refGene_gn=PLAC9;refGene_tx=XM_023240610.1
D2 41603161 41603620
refGene_gn=PLAC9;refGene_tx=XM_006937919.3;refGene_gn=PLAC9;refGene_tx=XM_023240610.1
D2 41634762 41634883 refGene_gn=TMEM254;refGene_tx=XM_003994095.5
D2 41639200 41639260 refGene_gn=TMEM254;refGene_tx=XM_003994095.5
D2 41639449 41639553 refGene_gn=TMEM254;refGene_tx=XM_003994095.5
D2 41641903 41641993 refGene_gn=TMEM254;refGene_tx=XM_003994095.5
D2 41699097 41699296 refGene_gn=SFTPD;refGene_tx=XM_003994112.3
D2 41704157 41704274 refGene_gn=SFTPD;refGene_tx=XM_003994112.3
D2 41704536 41704653 refGene_gn=SFTPD;refGene_tx=XM_003994112.3
D2 41704948 41705065 refGene_gn=SFTPD;refGene_tx=XM_003994112.3
D2 41705336 41705453 refGene_gn=SFTPD;refGene_tx=XM_003994112.3
D2 41706066 41706150 refGene_gn=SFTPD;refGene_tx=XM_003994112.3
D2 41708322 41708699 refGene_gn=SFTPD;refGene_tx=XM_003994112.3
D2 41723803 41724177
refGene_gn=MBL2;refGene_tx=XM_019813387.1;refGene_gn=MBL2;refGene_tx=XM_019813388.1
D2 41725081 41725156
refGene_gn=MBL2;refGene_tx=XM_019813387.1;refGene_gn=MBL2;refGene_tx=XM_019813388.1
D2 41726510 41726627
refGene_gn=MBL2;refGene_tx=XM_019813387.1;refGene_gn=MBL2;refGene_tx=XM_019813388.1
D2 41727040 41727206
refGene_gn=MBL2;refGene_tx=XM_019813387.1;refGene_gn=MBL2;refGene_tx=XM_019813388.1
D2 41737266 41737643
refGene_gn=LOC101101085;refGene_tx=XM_003994114.3;refGene_gn=LOC101101085;refGene_tx=XM_019813386.2
D2 41738345 41738423
refGene_gn=LOC101101085;refGene_tx=XM_003994114.3;refGene_gn=LOC101101085;refGene_tx=XM_019813386.2
D2 41739140 41739260
refGene_gn=LOC101101085;refGene_tx=XM_003994114.3;refGene_gn=LOC101101

085;refGene_tx=XM_019813386.2
D2	41739543	41739738
refGene_gn=LOC101101085;refGene_tx=XM_003994114.3;refGene_gn=LOC101101
085;refGene_tx=XM_019813386.2
D2	41740522	41740526
refGene_gn=LOC101101085;refGene_tx=XM_003994114.3
D2	41760368	41760471
refGene_gn=MAT1A;refGene_tx=XM_003994116.5;refGene_gn=MAT1A;refGene_tx
=XM_019813389.2;refGene_gn=MAT1A;refGene_tx=XM_019813391.2
D2	41760501	41760526	refGene_gn=MAT1A;refGene_tx=XM_019813390.2
D2	41761143	41761277
refGene_gn=MAT1A;refGene_tx=XM_003994116.5;refGene_gn=MAT1A;refGene_tx
=XM_019813389.2;refGene_gn=MAT1A;refGene_tx=XM_019813390.2;refGene_gn=
MAT1A;refGene_tx=XM_019813391.2
D2	41761633	41761816
refGene_gn=MAT1A;refGene_tx=XM_003994116.5;refGene_gn=MAT1A;refGene_tx
=XM_019813389.2;refGene_gn=MAT1A;refGene_tx=XM_019813390.2;refGene_gn=
MAT1A;refGene_tx=XM_019813391.2
D2	41763362	41763581
refGene_gn=MAT1A;refGene_tx=XM_003994116.5;refGene_gn=MAT1A;refGene_tx
=XM_019813389.2;refGene_gn=MAT1A;refGene_tx=XM_019813390.2;refGene_gn=
MAT1A;refGene_tx=XM_019813391.2
D2	41767794	41767938
refGene_gn=MAT1A;refGene_tx=XM_003994116.5;refGene_gn=MAT1A;refGene_tx
=XM_019813389.2;refGene_gn=MAT1A;refGene_tx=XM_019813390.2;refGene_gn=
MAT1A;refGene_tx=XM_019813391.2
D2	41768361	41768474
refGene_gn=MAT1A;refGene_tx=XM_003994116.5;refGene_gn=MAT1A;refGene_tx
=XM_019813389.2;refGene_gn=MAT1A;refGene_tx=XM_019813390.2;refGene_gn=
MAT1A;refGene_tx=XM_019813391.2
D2	41771319	41771442
refGene_gn=MAT1A;refGene_tx=XM_003994116.5;refGene_gn=MAT1A;refGene_tx
=XM_019813389.2;refGene_gn=MAT1A;refGene_tx=XM_019813390.2;refGene_gn=
MAT1A;refGene_tx=XM_019813391.2
D2	41772241	41772319
refGene_gn=MAT1A;refGene_tx=XM_003994116.5;refGene_gn=MAT1A;refGene_tx
=XM_019813389.2;refGene_gn=MAT1A;refGene_tx=XM_019813390.2
D2	41776042	41776136
refGene_gn=MAT1A;refGene_tx=XM_003994116.5;refGene_gn=MAT1A;refGene_tx
=XM_019813389.2;refGene_gn=MAT1A;refGene_tx=XM_019813390.2
D2	41794364	41794409
refGene_gn=DYDC1;refGene_tx=XM_019813392.2;refGene_gn=DYDC1;refGene_tx
=XM_019813393.2;refGene_gn=DYDC1;refGene_tx=XM_019813394.2;refGene_gn=
DYDC1;refGene_tx=XM_019813395.2;refGene_gn=DYDC1;refGene_tx=XM_0198133
96.2;refGene_gn=DYDC1;refGene_tx=XM_023240611.1
D2	41801688	41801718	refGene_gn=DYDC1;refGene_tx=XM_019813397.2
D2	41804386	41804491
refGene_gn=DYDC1;refGene_tx=XM_019813392.2;refGene_gn=DYDC1;refGene_tx
=XM_019813393.2;refGene_gn=DYDC1;refGene_tx=XM_019813394.2;refGene_gn=
DYDC1;refGene_tx=XM_019813395.2;refGene_gn=DYDC1;refGene_tx=XM_0198133

96.2;refGene_gn=DYDC1;refGene_tx=XM_019813397.2;refGene_gn=DYDC1;refGene_tx=XM_023240611.1
D2 41805387 41805444 refGene_gn=DYDC1;refGene_tx=XM_023240611.1
D2 41806406 41806463
refGene_gn=DYDC1;refGene_tx=XM_019813392.2;refGene_gn=DYDC1;refGene_tx=XM_019813393.2;refGene_gn=DYDC1;refGene_tx=XM_019813394.2;refGene_gn=DYDC1;refGene_tx=XM_019813395.2;refGene_gn=DYDC1;refGene_tx=XM_019813396.2;refGene_gn=DYDC1;refGene_tx=XM_019813397.2
D2 41808355 41808448
refGene_gn=DYDC1;refGene_tx=XM_019813392.2;refGene_gn=DYDC1;refGene_tx=XM_019813393.2;refGene_gn=DYDC1;refGene_tx=XM_019813394.2;refGene_gn=DYDC1;refGene_tx=XM_019813395.2;refGene_gn=DYDC1;refGene_tx=XM_019813396.2;refGene_gn=DYDC1;refGene_tx=XM_019813397.2;refGene_gn=DYDC1;refGene_tx=XM_023240611.1
D2 41814612 41814714
refGene_gn=DYDC1;refGene_tx=XM_019813392.2;refGene_gn=DYDC1;refGene_tx=XM_019813393.2;refGene_gn=DYDC1;refGene_tx=XM_019813394.2;refGene_gn=DYDC1;refGene_tx=XM_019813395.2;refGene_gn=DYDC1;refGene_tx=XM_019813396.2;refGene_gn=DYDC1;refGene_tx=XM_019813397.2;refGene_gn=DYDC1;refGene_tx=XM_023240611.1
D2 41815373 41815520
refGene_gn=DYDC1;refGene_tx=XM_019813392.2;refGene_gn=DYDC1;refGene_tx=XM_019813393.2;refGene_gn=DYDC1;refGene_tx=XM_019813394.2;refGene_gn=DYDC1;refGene_tx=XM_019813395.2;refGene_gn=DYDC1;refGene_tx=XM_019813396.2;refGene_gn=DYDC1;refGene_tx=XM_019813397.2;refGene_gn=DYDC1;refGene_tx=XM_023240611.1
D2 41818420 41818687
refGene_gn=LOC101080577;refGene_tx=XM_023240294.1
D2 41823120 41823243
refGene_gn=LOC101080577;refGene_tx=XM_023240294.1
D2 41826355 41826490
refGene_gn=LOC101080577;refGene_tx=XM_023240294.1
D2 41860640 41860643 refGene_gn=FAM213A;refGene_tx=XM_006937936.4
D2 41865397 41865575
refGene_gn=FAM213A;refGene_tx=XM_003994119.5;refGene_gn=FAM213A;refGene_tx=XM_006937936.4;refGene_gn=FAM213A;refGene_tx=XM_023240612.1;refGene_gn=FAM213A;refGene_tx=XM_023240614.1
D2 41867315 41867407
refGene_gn=FAM213A;refGene_tx=XM_003994119.5;refGene_gn=FAM213A;refGene_tx=XM_006937936.4;refGene_gn=FAM213A;refGene_tx=XM_023240612.1;refGene_gn=FAM213A;refGene_tx=XM_023240614.1
D2 41870482 41870623
refGene_gn=FAM213A;refGene_tx=XM_003994119.5;refGene_gn=FAM213A;refGene_tx=XM_006937936.4;refGene_gn=FAM213A;refGene_tx=XM_023240612.1;refGene_gn=FAM213A;refGene_tx=XM_023240614.1
D2 41871269 41871434
refGene_gn=FAM213A;refGene_tx=XM_003994119.5;refGene_gn=FAM213A;refGene_tx=XM_006937936.4;refGene_gn=FAM213A;refGene_tx=XM_023240612.1;refGene_gn=FAM213A;refGene_tx=XM_023240614.1
D2 41875004 41875118

refGene_gn=FAM213A;refGene_tx=XM_003994119.5;refGene_gn=FAM213A;refGene_tx=XM_006937936.4;refGene_gn=FAM213A;refGene_tx=XM_023240612.1;refGene_gn=FAM213A;refGene_tx=XM_023240614.1
D2 41929765 41929846
refGene_gn=TSPAN14;refGene_tx=XM_019813399.2;refGene_gn=TSPAN14;refGene_tx=XM_019813400.2;refGene_gn=TSPAN14;refGene_tx=XM_019813401.2;refGene_gn=TSPAN14;refGene_tx=XM_023240615.1
D2 41945950 41946001
refGene_gn=TSPAN14;refGene_tx=XM_019813399.2;refGene_gn=TSPAN14;refGene_tx=XM_019813400.2;refGene_gn=TSPAN14;refGene_tx=XM_019813401.2;refGene_gn=TSPAN14;refGene_tx=XM_023240615.1
D2 41948039 41948186
refGene_gn=TSPAN14;refGene_tx=XM_019813399.2;refGene_gn=TSPAN14;refGene_tx=XM_019813400.2;refGene_gn=TSPAN14;refGene_tx=XM_019813401.2;refGene_gn=TSPAN14;refGene_tx=XM_023240615.1
D2 41949709 41949880
refGene_gn=TSPAN14;refGene_tx=XM_019813399.2;refGene_gn=TSPAN14;refGene_tx=XM_019813400.2;refGene_gn=TSPAN14;refGene_tx=XM_019813401.2;refGene_gn=TSPAN14;refGene_tx=XM_023240615.1
D2 41951791 41951917
refGene_gn=TSPAN14;refGene_tx=XM_019813399.2;refGene_gn=TSPAN14;refGene_tx=XM_019813400.2;refGene_gn=TSPAN14;refGene_tx=XM_019813401.2;refGene_gn=TSPAN14;refGene_tx=XM_023240615.1
D2 41953093 41953138
refGene_gn=TSPAN14;refGene_tx=XM_019813399.2;refGene_gn=TSPAN14;refGene_tx=XM_019813400.2;refGene_gn=TSPAN14;refGene_tx=XM_019813401.2;refGene_gn=TSPAN14;refGene_tx=XM_023240615.1
D2 41955204 41955324
refGene_gn=TSPAN14;refGene_tx=XM_019813399.2;refGene_gn=TSPAN14;refGene_tx=XM_019813400.2;refGene_gn=TSPAN14;refGene_tx=XM_019813401.2;refGene_gn=TSPAN14;refGene_tx=XM_023240615.1
D2 41956393 41956465
refGene_gn=TSPAN14;refGene_tx=XM_019813399.2;refGene_gn=TSPAN14;refGene_tx=XM_019813400.2;refGene_gn=TSPAN14;refGene_tx=XM_019813401.2;refGene_gn=TSPAN14;refGene_tx=XM_023240615.1
D2 42007308 42007409 refGene_gn=SH2D4B;refGene_tx=XM_023240616.1
D2 42008761 42008909 refGene_gn=SH2D4B;refGene_tx=XM_023240616.1
D2 42023360 42023508 refGene_gn=SH2D4B;refGene_tx=XM_023240616.1
D2 42035678 42035895 refGene_gn=SH2D4B;refGene_tx=XM_023240616.1
D2 42038240 42038365 refGene_gn=SH2D4B;refGene_tx=XM_023240616.1
D2 42062272 42062493 refGene_gn=SH2D4B;refGene_tx=XM_023240616.1
D2 42066420 42066513 refGene_gn=SH2D4B;refGene_tx=XM_023240616.1
D2 42897444 42898285
refGene_gn=NRG3;refGene_tx=XM_019814071.2;refGene_gn=NRG3;refGene_tx=XM_023240617.1;refGene_gn=NRG3;refGene_tx=XM_023240618.1;refGene_gn=NRG3;refGene_tx=XM_023240619.1;refGene_gn=NRG3;refGene_tx=XM_023240620.1;refGene_gn=NRG3;refGene_tx=XM_023240621.1;refGene_gn=NRG3;refGene_tx=XM_023240622.1;refGene_gn=NRG3;refGene_tx=XM_023240623.1;refGene_gn=NRG3;refGene_tx=XM_023240624.1
D2 43374699 43374829

refGene_gn=NRG3;refGene_tx=XM_019814071.2;refGene_gn=NRG3;refGene_tx=XM_023240617.1;refGene_gn=NRG3;refGene_tx=XM_023240618.1;refGene_gn=NRG3;refGene_tx=XM_023240619.1;refGene_gn=NRG3;refGene_tx=XM_023240620.1;refGene_gn=NRG3;refGene_tx=XM_023240621.1;refGene_gn=NRG3;refGene_tx=XM_023240622.1;refGene_gn=NRG3;refGene_tx=XM_023240623.1;refGene_gn=NRG3;refGene_tx=XM_023240624.1
D2 43719504 43719578
refGene_gn=NRG3;refGene_tx=XM_019814071.2;refGene_gn=NRG3;refGene_tx=XM_023240617.1;refGene_gn=NRG3;refGene_tx=XM_023240618.1;refGene_gn=NRG3;refGene_tx=XM_023240619.1;refGene_gn=NRG3;refGene_tx=XM_023240620.1;refGene_gn=NRG3;refGene_tx=XM_023240621.1;refGene_gn=NRG3;refGene_tx=XM_023240622.1;refGene_gn=NRG3;refGene_tx=XM_023240623.1;refGene_gn=NRG3;refGene_tx=XM_023240624.1
D2 43835756 43835783
refGene_gn=NRG3;refGene_tx=XM_019814071.2;refGene_gn=NRG3;refGene_tx=XM_023240617.1;refGene_gn=NRG3;refGene_tx=XM_023240618.1;refGene_gn=NRG3;refGene_tx=XM_023240619.1;refGene_gn=NRG3;refGene_tx=XM_023240620.1;refGene_gn=NRG3;refGene_tx=XM_023240621.1;refGene_gn=NRG3;refGene_tx=XM_023240622.1;refGene_gn=NRG3;refGene_tx=XM_023240623.1;refGene_gn=NRG3;refGene_tx=XM_023240624.1
D2 43925927 43926030
refGene_gn=NRG3;refGene_tx=XM_019814071.2;refGene_gn=NRG3;refGene_tx=XM_023240617.1;refGene_gn=NRG3;refGene_tx=XM_023240618.1;refGene_gn=NRG3;refGene_tx=XM_023240619.1;refGene_gn=NRG3;refGene_tx=XM_023240620.1;refGene_gn=NRG3;refGene_tx=XM_023240621.1;refGene_gn=NRG3;refGene_tx=XM_023240622.1;refGene_gn=NRG3;refGene_tx=XM_023240623.1;refGene_gn=NRG3;refGene_tx=XM_023240624.1
D2 43933114 43933241
refGene_gn=NRG3;refGene_tx=XM_019814071.2;refGene_gn=NRG3;refGene_tx=XM_023240617.1;refGene_gn=NRG3;refGene_tx=XM_023240618.1;refGene_gn=NRG3;refGene_tx=XM_023240619.1;refGene_gn=NRG3;refGene_tx=XM_023240620.1;refGene_gn=NRG3;refGene_tx=XM_023240621.1;refGene_gn=NRG3;refGene_tx=XM_023240622.1;refGene_gn=NRG3;refGene_tx=XM_023240623.1;refGene_gn=NRG3;refGene_tx=XM_023240624.1
D2 43949602 43949736
refGene_gn=NRG3;refGene_tx=XM_019814071.2;refGene_gn=NRG3;refGene_tx=XM_023240617.1;refGene_gn=NRG3;refGene_tx=XM_023240618.1;refGene_gn=NRG3;refGene_tx=XM_023240619.1;refGene_gn=NRG3;refGene_tx=XM_023240620.1;refGene_gn=NRG3;refGene_tx=XM_023240621.1;refGene_gn=NRG3;refGene_tx=XM_023240622.1;refGene_gn=NRG3;refGene_tx=XM_023240623.1;refGene_gn=NRG3;refGene_tx=XM_023240624.1
D2 43954248 43954422
refGene_gn=NRG3;refGene_tx=XM_019814071.2;refGene_gn=NRG3;refGene_tx=XM_023240617.1;refGene_gn=NRG3;refGene_tx=XM_023240618.1;refGene_gn=NRG3;refGene_tx=XM_023240619.1;refGene_gn=NRG3;refGene_tx=XM_023240620.1;refGene_gn=NRG3;refGene_tx=XM_023240621.1;refGene_gn=NRG3;refGene_tx=XM_023240622.1;refGene_gn=NRG3;refGene_tx=XM_023240623.1;refGene_gn=NRG3;refGene_tx=XM_023240624.1
D2 43959640 43959712
refGene_gn=NRG3;refGene_tx=XM_019814071.2;refGene_gn=NRG3;refGene_tx=X

M_023240617.1;refGene_gn=NRG3;refGene_tx=XM_023240618.1;refGene_gn=NRG3;refGene_tx=XM_023240619.1;refGene_gn=NRG3;refGene_tx=XM_023240620.1
D2 43959953 43960461
refGene_gn=NRG3;refGene_tx=XM_019814071.2;refGene_gn=NRG3;refGene_tx=XM_023240617.1;refGene_gn=NRG3;refGene_tx=XM_023240618.1;refGene_gn=NRG3;refGene_tx=XM_023240619.1;refGene_gn=NRG3;refGene_tx=XM_023240620.1;refGene_gn=NRG3;refGene_tx=XM_023240621.1;refGene_gn=NRG3;refGene_tx=XM_023240622.1;refGene_gn=NRG3;refGene_tx=XM_023240623.1;refGene_gn=NRG3;refGene_tx=XM_023240624.1
D2 44806078 44806207
refGene_gn=GHITM;refGene_tx=XM_003994124.5;refGene_gn=GHITM;refGene_tx=XM_019813403.2
D2 44807442 44807542
refGene_gn=GHITM;refGene_tx=XM_003994124.5;refGene_gn=GHITM;refGene_tx=XM_019813403.2
D2 44808796 44808908
refGene_gn=GHITM;refGene_tx=XM_003994124.5;refGene_gn=GHITM;refGene_tx=XM_019813403.2
D2 44809548 44809690
refGene_gn=GHITM;refGene_tx=XM_003994124.5;refGene_gn=GHITM;refGene_tx=XM_019813403.2
D2 44812890 44812999
refGene_gn=GHITM;refGene_tx=XM_003994124.5;refGene_gn=GHITM;refGene_tx=XM_019813403.2
D2 44819296 44819485
refGene_gn=GHITM;refGene_tx=XM_003994124.5;refGene_gn=GHITM;refGene_tx=XM_019813403.2
D2 44820122 44820294
refGene_gn=GHITM;refGene_tx=XM_003994124.5;refGene_gn=GHITM;refGene_tx=XM_019813403.2
D2 44821574 44821659
refGene_gn=GHITM;refGene_tx=XM_003994124.5;refGene_gn=GHITM;refGene_tx=XM_019813403.2
D2 44832117 44832187
refGene_gn=CD2H10orf99;refGene_tx=XM_003994129.3
D2 44834047 44834137
refGene_gn=CD2H10orf99;refGene_tx=XM_003994129.3
D2 44839387 44839473
refGene_gn=CD2H10orf99;refGene_tx=XM_003994129.3
D2 44848609 44848664 refGene_gn=CDHR1;refGene_tx=XM_023240625.1
D2 44849264 44849360 refGene_gn=CDHR1;refGene_tx=XM_023240625.1
D2 44850316 44850462 refGene_gn=CDHR1;refGene_tx=XM_023240625.1
D2 44851449 44851500 refGene_gn=CDHR1;refGene_tx=XM_023240625.1
D2 44852693 44852783 refGene_gn=CDHR1;refGene_tx=XM_023240625.1
D2 44853982 44854069 refGene_gn=CDHR1;refGene_tx=XM_023240625.1
D2 44855294 44855411 refGene_gn=CDHR1;refGene_tx=XM_023240625.1
D2 44856498 44856642 refGene_gn=CDHR1;refGene_tx=XM_023240625.1
D2 44858467 44858546 refGene_gn=CDHR1;refGene_tx=XM_023240625.1
D2 44859875 44859976 refGene_gn=CDHR1;refGene_tx=XM_023240625.1
D2 44862558 44862762 refGene_gn=CDHR1;refGene_tx=XM_023240625.1

D2 44863259 44863412 refGene_gn=CDHR1;refGene_tx=XM_023240625.1
D2 44864931 44865096 refGene_gn=CDHR1;refGene_tx=XM_023240625.1
D2 44865589 44865657 refGene_gn=CDHR1;refGene_tx=XM_023240625.1
D2 44866142 44866305 refGene_gn=CDHR1;refGene_tx=XM_023240625.1
D2 44866306 44866372 refGene_gn=CDHR1;refGene_tx=XM_023240625.1
D2 44867502 44867760 refGene_gn=CDHR1;refGene_tx=XM_023240625.1
D2 44868442 44868979 refGene_gn=CDHR1;refGene_tx=XM_023240625.1
D2 44875488 44876249
refGene_gn=LRIT2;refGene_tx=XM_003994131.3;refGene_gn=LRIT2;refGene_tx
=XM_019813406.2;refGene_gn=LRIT2;refGene_tx=XM_019813407.2;refGene_gn=
LRIT2;refGene_tx=XM_019813408.2
D2 44877975 44878757
refGene_gn=LRIT2;refGene_tx=XM_003994131.3;refGene_gn=LRIT2;refGene_tx
=XM_019813406.2;refGene_gn=LRIT2;refGene_tx=XM_019813407.2;refGene_gn=
LRIT2;refGene_tx=XM_019813408.2
D2 44879033 44879143 refGene_gn=LRIT2;refGene_tx=XM_003994131.3
D2 44882973 44882975 refGene_gn=LRIT2;refGene_tx=XM_019813407.2
D2 44883130 44883147 refGene_gn=LRIT2;refGene_tx=XM_019813406.2
D2 44885020 44885985
refGene_gn=LRIT1;refGene_tx=XM_003994125.2;refGene_gn=LRIT1;refGene_tx
=XM_019813404.1;refGene_gn=LRIT1;refGene_tx=XM_019813405.1
D2 44887201 44887507
refGene_gn=LRIT1;refGene_tx=XM_003994125.2;refGene_gn=LRIT1;refGene_tx
=XM_019813404.1
D2 44891854 44892321
refGene_gn=LRIT1;refGene_tx=XM_003994125.2;refGene_gn=LRIT1;refGene_tx
=XM_019813404.1;refGene_gn=LRIT1;refGene_tx=XM_019813405.1
D2 44895783 44895905
refGene_gn=LRIT1;refGene_tx=XM_003994125.2;refGene_gn=LRIT1;refGene_tx
=XM_019813405.1
D2 44898436 44898515
refGene_gn=RGR;refGene_tx=XM_003994126.3;refGene_gn=RGR;refGene_tx=XM_
019813410.1
D2 44900102 44900259
refGene_gn=RGR;refGene_tx=XM_003994126.3;refGene_gn=RGR;refGene_tx=XM_
019813410.1
D2 44901374 44901508
refGene_gn=RGR;refGene_tx=XM_003994126.3;refGene_gn=RGR;refGene_tx=XM_
019813410.1
D2 44904199 44904353
refGene_gn=RGR;refGene_tx=XM_003994126.3;refGene_gn=RGR;refGene_tx=XM_
019813410.1
D2 44905764 44905882
refGene_gn=RGR;refGene_tx=XM_003994126.3;refGene_gn=RGR;refGene_tx=XM_
019813410.1
D2 44907868 44907982
refGene_gn=RGR;refGene_tx=XM_003994126.3;refGene_gn=RGR;refGene_tx=XM_
019813410.1
D2 44908403 44908535
refGene_gn=RGR;refGene_tx=XM_003994126.3;refGene_gn=RGR;refGene_tx=XM_

019813410.1
D2	45009495	45010906	refGene_gn=CCSER2;refGene_tx=XM_003994128.5
D2	45012070	45012267	refGene_gn=CCSER2;refGene_tx=XM_003994128.5
D2	45052130	45052173	refGene_gn=CCSER2;refGene_tx=XM_003994128.5
D2	45052175	45052220	refGene_gn=CCSER2;refGene_tx=XM_003994128.5
D2	45060092	45060255	refGene_gn=CCSER2;refGene_tx=XM_003994128.5
D2	45076121	45076317	refGene_gn=CCSER2;refGene_tx=XM_003994128.5
D2	45098054	45098138	refGene_gn=CCSER2;refGene_tx=XM_003994128.5
D2	45103641	45103728	refGene_gn=CCSER2;refGene_tx=XM_003994128.5
D2	45115969	45116059	refGene_gn=CCSER2;refGene_tx=XM_003994128.5
D2	45144926	45145745	refGene_gn=CCSER2;refGene_tx=XM_003994128.5
D2	45989609	45990038	refGene_gn=GRID1;refGene_tx=XM_023240627.1
D2	46000064	46000305	refGene_gn=GRID1;refGene_tx=XM_023240627.1
D2	46005515	46005682	refGene_gn=GRID1;refGene_tx=XM_023240627.1
D2	46029819	46030015	refGene_gn=GRID1;refGene_tx=XM_023240627.1
D2	46101508	46101647	refGene_gn=GRID1;refGene_tx=XM_023240627.1
D2	46102847	46102920	refGene_gn=GRID1;refGene_tx=XM_023240627.1
D2	46102921	46103173	refGene_gn=GRID1;refGene_tx=XM_023240627.1
D2	46106664	46106862	refGene_gn=GRID1;refGene_tx=XM_023240627.1
D2	46108303	46108405	refGene_gn=GRID1;refGene_tx=XM_023240627.1
D2	46200078	46200198	refGene_gn=GRID1;refGene_tx=XM_023240627.1
D2	46201705	46201867	refGene_gn=GRID1;refGene_tx=XM_023240627.1
D2	46214876	46215047	refGene_gn=GRID1;refGene_tx=XM_023240627.1
D2	46247881	46247935	refGene_gn=GRID1;refGene_tx=XM_023240627.1
D2	46457267	46457473	refGene_gn=GRID1;refGene_tx=XM_023240627.1
D2	46528660	46528945	refGene_gn=GRID1;refGene_tx=XM_023240627.1
D2	46684293	46684449	refGene_gn=GRID1;refGene_tx=XM_023240627.1
D2	46686597	46686676	refGene_gn=GRID1;refGene_tx=XM_023240627.1
D2	46773129	46773195
refGene_gn=WAPL;refGene_tx=XM_003994132.5;refGene_gn=WAPL;refGene_tx=X
M_003994133.5
D2	46773516	46773612
refGene_gn=WAPL;refGene_tx=XM_003994132.5;refGene_gn=WAPL;refGene_tx=X
M_003994133.5
D2	46777977	46778066
refGene_gn=WAPL;refGene_tx=XM_003994132.5;refGene_gn=WAPL;refGene_tx=X
M_003994133.5
D2	46780790	46780998
refGene_gn=WAPL;refGene_tx=XM_003994132.5;refGene_gn=WAPL;refGene_tx=X
M_003994133.5
D2	46783626	46783791
refGene_gn=WAPL;refGene_tx=XM_003994132.5;refGene_gn=WAPL;refGene_tx=X
M_003994133.5
D2	46784204	46784320
refGene_gn=WAPL;refGene_tx=XM_003994132.5;refGene_gn=WAPL;refGene_tx=X
M_003994133.5
D2	46784853	46785029
refGene_gn=WAPL;refGene_tx=XM_003994132.5;refGene_gn=WAPL;refGene_tx=X
M_003994133.5
D2	46789269	46789346

refGene_gn=WAPL;refGene_tx=XM_003994132.5;refGene_gn=WAPL;refGene_tx=XM_003994133.5
D2 46790288 46790386
refGene_gn=WAPL;refGene_tx=XM_003994132.5;refGene_gn=WAPL;refGene_tx=XM_003994133.5
D2 46791398 46791510
refGene_gn=WAPL;refGene_tx=XM_003994132.5;refGene_gn=WAPL;refGene_tx=XM_003994133.5
D2 46795029 46795257
refGene_gn=WAPL;refGene_tx=XM_003994132.5;refGene_gn=WAPL;refGene_tx=XM_003994133.5
D2 46797368 46797480
refGene_gn=WAPL;refGene_tx=XM_003994132.5;refGene_gn=WAPL;refGene_tx=XM_003994133.5
D2 46798196 46798333
refGene_gn=WAPL;refGene_tx=XM_003994132.5;refGene_gn=WAPL;refGene_tx=XM_003994133.5
D2 46798581 46798734
refGene_gn=WAPL;refGene_tx=XM_003994132.5;refGene_gn=WAPL;refGene_tx=XM_003994133.5
D2 46800016 46800112
refGene_gn=WAPL;refGene_tx=XM_003994132.5;refGene_gn=WAPL;refGene_tx=XM_003994133.5
D2 46833033 46833152
refGene_gn=WAPL;refGene_tx=XM_003994132.5;refGene_gn=WAPL;refGene_tx=XM_003994133.5
D2 46835702 46836746
refGene_gn=WAPL;refGene_tx=XM_003994132.5;refGene_gn=WAPL;refGene_tx=XM_003994133.5
D2 46862370 46862869
refGene_gn=WAPL;refGene_tx=XM_003994132.5;refGene_gn=WAPL;refGene_tx=XM_003994133.5
D2 46864420 46864486
refGene_gn=LOC102902576;refGene_tx=XM_023240295.1
D2 46865819 46866452
refGene_gn=LOC102902576;refGene_tx=XM_023240295.1
D2 46943650 46943779 refGene_gn=OPN4;refGene_tx=NM_001009325.2
D2 46945092 46945238 refGene_gn=OPN4;refGene_tx=NM_001009325.2
D2 46946527 46946661 refGene_gn=OPN4;refGene_tx=NM_001009325.2
D2 46946985 46947189 refGene_gn=OPN4;refGene_tx=NM_001009325.2
D2 46947830 46948002 refGene_gn=OPN4;refGene_tx=NM_001009325.2
D2 46948455 46948620 refGene_gn=OPN4;refGene_tx=NM_001009325.2
D2 46949553 46949661 refGene_gn=OPN4;refGene_tx=NM_001009325.2
D2 46950575 46950756 refGene_gn=OPN4;refGene_tx=NM_001009325.2
D2 46951585 46951729 refGene_gn=OPN4;refGene_tx=NM_001009325.2
D2 46954048 46954129 refGene_gn=OPN4;refGene_tx=NM_001009325.2
D2 46956746 46956839
refGene_gn=LDB3;refGene_tx=XM_006937946.3;refGene_gn=LDB3;refGene_tx=XM_011287279.3;refGene_gn=LDB3;refGene_tx=XM_011287280.3;refGene_gn=LDB3;refGene_tx=XM_019813415.2;refGene_gn=LDB3;refGene_tx=XM_019813416.2;

refGene_gn=LDB3;refGene_tx=XM_019813417.2;refGene_gn=LDB3;refGene_tx=XM_019813418.2;refGene_gn=LDB3;refGene_tx=XM_019813419.2;refGene_gn=LDB3;refGene_tx=XM_019813420.2
D2 46967507 46967659
refGene_gn=LDB3;refGene_tx=XM_006937946.3;refGene_gn=LDB3;refGene_tx=XM_011287279.3;refGene_gn=LDB3;refGene_tx=XM_011287280.3;refGene_gn=LDB3;refGene_tx=XM_019813415.2;refGene_gn=LDB3;refGene_tx=XM_019813416.2;refGene_gn=LDB3;refGene_tx=XM_019813417.2;refGene_gn=LDB3;refGene_tx=XM_019813418.2;refGene_gn=LDB3;refGene_tx=XM_019813419.2;refGene_gn=LDB3;refGene_tx=XM_019813420.2
D2 46968180 46968256
refGene_gn=LDB3;refGene_tx=XM_006937946.3;refGene_gn=LDB3;refGene_tx=XM_011287279.3;refGene_gn=LDB3;refGene_tx=XM_011287280.3;refGene_gn=LDB3;refGene_tx=XM_019813415.2;refGene_gn=LDB3;refGene_tx=XM_019813416.2;refGene_gn=LDB3;refGene_tx=XM_019813417.2;refGene_gn=LDB3;refGene_tx=XM_019813418.2;refGene_gn=LDB3;refGene_tx=XM_019813419.2;refGene_gn=LDB3;refGene_tx=XM_019813420.2
D2 46969685 46970053
refGene_gn=LDB3;refGene_tx=XM_019813415.2;refGene_gn=LDB3;refGene_tx=XM_019813416.2;refGene_gn=LDB3;refGene_tx=XM_019813419.2;refGene_gn=LDB3;refGene_tx=XM_019813420.2
D2 46974096 46974119
refGene_gn=LDB3;refGene_tx=XM_006937946.3;refGene_gn=LDB3;refGene_tx=XM_011287279.3;refGene_gn=LDB3;refGene_tx=XM_011287280.3;refGene_gn=LDB3;refGene_tx=XM_019813417.2;refGene_gn=LDB3;refGene_tx=XM_019813418.2
D2 46975191 46975395
refGene_gn=LDB3;refGene_tx=XM_006937946.3;refGene_gn=LDB3;refGene_tx=XM_011287279.3;refGene_gn=LDB3;refGene_tx=XM_011287280.3;refGene_gn=LDB3;refGene_tx=XM_019813417.2;refGene_gn=LDB3;refGene_tx=XM_019813418.2
D2 46977666 46977681
refGene_gn=LDB3;refGene_tx=XM_011287279.3;refGene_gn=LDB3;refGene_tx=XM_019813417.2
D2 46980422 46980592
refGene_gn=LDB3;refGene_tx=XM_006937946.3;refGene_gn=LDB3;refGene_tx=XM_011287279.3;refGene_gn=LDB3;refGene_tx=XM_011287280.3;refGene_gn=LDB3;refGene_tx=XM_019813415.2;refGene_gn=LDB3;refGene_tx=XM_019813416.2;refGene_gn=LDB3;refGene_tx=XM_019813417.2;refGene_gn=LDB3;refGene_tx=XM_019813418.2;refGene_gn=LDB3;refGene_tx=XM_019813419.2;refGene_gn=LDB3;refGene_tx=XM_019813420.2
D2 46981087 46981124
refGene_gn=LDB3;refGene_tx=XM_006937946.3;refGene_gn=LDB3;refGene_tx=XM_011287279.3;refGene_gn=LDB3;refGene_tx=XM_011287280.3;refGene_gn=LDB3;refGene_tx=XM_019813415.2;refGene_gn=LDB3;refGene_tx=XM_019813416.2;refGene_gn=LDB3;refGene_tx=XM_019813417.2;refGene_gn=LDB3;refGene_tx=XM_019813418.2;refGene_gn=LDB3;refGene_tx=XM_019813419.2;refGene_gn=LDB3;refGene_tx=XM_019813420.2
D2 46987202 46987299
refGene_gn=LDB3;refGene_tx=XM_006937946.3;refGene_gn=LDB3;refGene_tx=XM_019813420.2
D2 46993334 46993517

refGene_gn=LDB3;refGene_tx=XM_019813415.2;refGene_gn=LDB3;refGene_tx=XM_019813416.2;refGene_gn=LDB3;refGene_tx=XM_019813417.2;refGene_gn=LDB3;refGene_tx=XM_019813418.2
D2 46996799 46996942
refGene_gn=LDB3;refGene_tx=XM_011287279.3;refGene_gn=LDB3;refGene_tx=XM_011287280.3;refGene_gn=LDB3;refGene_tx=XM_019813415.2;refGene_gn=LDB3;refGene_tx=XM_019813416.2;refGene_gn=LDB3;refGene_tx=XM_019813417.2;refGene_gn=LDB3;refGene_tx=XM_019813418.2;refGene_gn=LDB3;refGene_tx=XM_019813419.2
D2 47003305 47003726
refGene_gn=LDB3;refGene_tx=XM_011287279.3;refGene_gn=LDB3;refGene_tx=XM_011287280.3;refGene_gn=LDB3;refGene_tx=XM_019813415.2;refGene_gn=LDB3;refGene_tx=XM_019813416.2;refGene_gn=LDB3;refGene_tx=XM_019813417.2;refGene_gn=LDB3;refGene_tx=XM_019813418.2;refGene_gn=LDB3;refGene_tx=XM_019813419.2
D2 47005415 47005596
refGene_gn=LDB3;refGene_tx=XM_011287279.3;refGene_gn=LDB3;refGene_tx=XM_011287280.3;refGene_gn=LDB3;refGene_tx=XM_019813415.2;refGene_gn=LDB3;refGene_tx=XM_019813416.2;refGene_gn=LDB3;refGene_tx=XM_019813417.2;refGene_gn=LDB3;refGene_tx=XM_019813418.2;refGene_gn=LDB3;refGene_tx=XM_019813419.2
D2 47006229 47006350
refGene_gn=LDB3;refGene_tx=XM_011287279.3;refGene_gn=LDB3;refGene_tx=XM_011287280.3;refGene_gn=LDB3;refGene_tx=XM_019813415.2;refGene_gn=LDB3;refGene_tx=XM_019813416.2;refGene_gn=LDB3;refGene_tx=XM_019813417.2;refGene_gn=LDB3;refGene_tx=XM_019813418.2;refGene_gn=LDB3;refGene_tx=XM_019813419.2
D2 47011907 47012023
refGene_gn=LDB3;refGene_tx=XM_011287279.3;refGene_gn=LDB3;refGene_tx=XM_011287280.3;refGene_gn=LDB3;refGene_tx=XM_019813415.2;refGene_gn=LDB3;refGene_tx=XM_019813416.2;refGene_gn=LDB3;refGene_tx=XM_019813417.2;refGene_gn=LDB3;refGene_tx=XM_019813418.2;refGene_gn=LDB3;refGene_tx=XM_019813419.2
D2 47017578 47017668
refGene_gn=LDB3;refGene_tx=XM_011287279.3;refGene_gn=LDB3;refGene_tx=XM_011287280.3;refGene_gn=LDB3;refGene_tx=XM_019813415.2;refGene_gn=LDB3;refGene_tx=XM_019813416.2;refGene_gn=LDB3;refGene_tx=XM_019813417.2;refGene_gn=LDB3;refGene_tx=XM_019813418.2;refGene_gn=LDB3;refGene_tx=XM_019813419.2
D2 47146386 47146453 refGene_gn=BMPR1A;refGene_tx=XM_023240628.1
D2 47154506 47154669 refGene_gn=BMPR1A;refGene_tx=XM_023240628.1
D2 47155487 47155590 refGene_gn=BMPR1A;refGene_tx=XM_023240628.1
D2 47159946 47160043 refGene_gn=BMPR1A;refGene_tx=XM_023240628.1
D2 47160205 47160305 refGene_gn=BMPR1A;refGene_tx=XM_023240628.1
D2 47165725 47165870 refGene_gn=BMPR1A;refGene_tx=XM_023240628.1
D2 47169751 47169944 refGene_gn=BMPR1A;refGene_tx=XM_023240628.1
D2 47171447 47171745 refGene_gn=BMPR1A;refGene_tx=XM_023240628.1
D2 47173891 47174067 refGene_gn=BMPR1A;refGene_tx=XM_023240628.1
D2 47175069 47175200 refGene_gn=BMPR1A;refGene_tx=XM_023240628.1
D2 47175284 47175410 refGene_gn=BMPR1A;refGene_tx=XM_023240628.1

D2 47187580 47187966 refGene_gn=MMRN2;refGene_tx=XM_023240629.1
D2 47190894 47192640 refGene_gn=MMRN2;refGene_tx=XM_023240629.1
D2 47192806 47192959 refGene_gn=MMRN2;refGene_tx=XM_023240629.1
D2 47193707 47193788 refGene_gn=MMRN2;refGene_tx=XM_023240629.1
D2 47193887 47193994 refGene_gn=MMRN2;refGene_tx=XM_023240629.1
D2 47194086 47194215 refGene_gn=MMRN2;refGene_tx=XM_023240629.1
D2 47203029 47203193 refGene_gn=MMRN2;refGene_tx=XM_023240629.1
D2 47204362 47204483
refGene_gn=SNCG;refGene_tx=XM_003994137.4;refGene_gn=SNCG;refGene_tx=XM_019813421.1
D2 47205303 47205345 refGene_gn=SNCG;refGene_tx=XM_003994137.4
D2 47205659 47205787
refGene_gn=SNCG;refGene_tx=XM_003994137.4;refGene_gn=SNCG;refGene_tx=XM_019813421.1
D2 47208210 47208282
refGene_gn=SNCG;refGene_tx=XM_003994137.4;refGene_gn=SNCG;refGene_tx=XM_019813421.1
D2 47208524 47208545
refGene_gn=SNCG;refGene_tx=XM_003994137.4;refGene_gn=SNCG;refGene_tx=XM_019813421.1
D2 47212574 47212635 refGene_gn=ADIRF;refGene_tx=XM_003994138.4
D2 47214341 47214404 refGene_gn=ADIRF;refGene_tx=XM_003994138.4
D2 47214606 47214710 refGene_gn=ADIRF;refGene_tx=XM_003994138.4
D2 47223422 47223495 refGene_gn=FAM25A;refGene_tx=XM_006937947.4
D2 47225147 47225210 refGene_gn=FAM25A;refGene_tx=XM_006937947.4
D2 47226151 47226285 refGene_gn=FAM25A;refGene_tx=XM_006937947.4
D2 47237163 47237283 refGene_gn=GLUD1;refGene_tx=XM_023240636.1
D2 47238170 47238233 refGene_gn=GLUD1;refGene_tx=XM_023240636.1
D2 47242326 47242418 refGene_gn=GLUD1;refGene_tx=XM_023240636.1
D2 47243200 47243324 refGene_gn=GLUD1;refGene_tx=XM_023240636.1
D2 47244661 47244742 refGene_gn=GLUD1;refGene_tx=XM_023240636.1
D2 47245173 47245311 refGene_gn=GLUD1;refGene_tx=XM_023240636.1
D2 47245516 47245654 refGene_gn=GLUD1;refGene_tx=XM_023240636.1
D2 47246385 47246565 refGene_gn=GLUD1;refGene_tx=XM_023240636.1
D2 47250872 47250968 refGene_gn=GLUD1;refGene_tx=XM_023240636.1
D2 47255698 47255761 refGene_gn=GLUD1;refGene_tx=XM_023240636.1
D2 47256568 47256624 refGene_gn=GLUD1;refGene_tx=XM_023240636.1
D2 47257135 47257216 refGene_gn=GLUD1;refGene_tx=XM_023240636.1
D2 47272979 47273445 refGene_gn=GLUD1;refGene_tx=XM_023240636.1
D2 47288530 47288567
refGene_gn=FAM35A;refGene_tx=XM_023240630.1;refGene_gn=FAM35A;refGene_tx=XM_023240634.1;refGene_gn=FAM35A;refGene_tx=XM_023240635.1
D2 47322282 47323818
refGene_gn=FAM35A;refGene_tx=XM_023240630.1;refGene_gn=FAM35A;refGene_tx=XM_023240631.1;refGene_gn=FAM35A;refGene_tx=XM_023240632.1;refGene_gn=FAM35A;refGene_tx=XM_023240633.1;refGene_gn=FAM35A;refGene_tx=XM_023240634.1;refGene_gn=FAM35A;refGene_tx=XM_023240635.1
D2 47327888 47327996
refGene_gn=FAM35A;refGene_tx=XM_023240630.1;refGene_gn=FAM35A;refGene_tx=XM_023240631.1;refGene_gn=FAM35A;refGene_tx=XM_023240632.1;refGene_

gn=FAM35A;refGene_tx=XM_023240633.1;refGene_gn=FAM35A;refGene_tx=XM_023240634.1;refGene_gn=FAM35A;refGene_tx=XM_023240635.1
D2 47334275 47334470 refGene_gn=FAM35A;refGene_tx=XM_023240630.1;refGene_gn=FAM35A;refGene_tx=XM_023240631.1;refGene_gn=FAM35A;refGene_tx=XM_023240632.1;refGene_gn=FAM35A;refGene_tx=XM_023240633.1;refGene_gn=FAM35A;refGene_tx=XM_023240634.1;refGene_gn=FAM35A;refGene_tx=XM_023240635.1
D2 47334632 47334767 refGene_gn=FAM35A;refGene_tx=XM_023240630.1;refGene_gn=FAM35A;refGene_tx=XM_023240631.1;refGene_gn=FAM35A;refGene_tx=XM_023240632.1;refGene_gn=FAM35A;refGene_tx=XM_023240633.1;refGene_gn=FAM35A;refGene_tx=XM_023240634.1;refGene_gn=FAM35A;refGene_tx=XM_023240635.1
D2 47341020 47341227 refGene_gn=FAM35A;refGene_tx=XM_023240630.1;refGene_gn=FAM35A;refGene_tx=XM_023240631.1;refGene_gn=FAM35A;refGene_tx=XM_023240632.1;refGene_gn=FAM35A;refGene_tx=XM_023240633.1;refGene_gn=FAM35A;refGene_tx=XM_023240635.1
D2 47346919 47347148 refGene_gn=FAM35A;refGene_tx=XM_023240630.1;refGene_gn=FAM35A;refGene_tx=XM_023240631.1;refGene_gn=FAM35A;refGene_tx=XM_023240632.1;refGene_gn=FAM35A;refGene_tx=XM_023240633.1;refGene_gn=FAM35A;refGene_tx=XM_023240634.1;refGene_gn=FAM35A;refGene_tx=XM_023240635.1
D2 47354924 47355040 refGene_gn=FAM35A;refGene_tx=XM_023240630.1;refGene_gn=FAM35A;refGene_tx=XM_023240631.1;refGene_gn=FAM35A;refGene_tx=XM_023240632.1;refGene_gn=FAM35A;refGene_tx=XM_023240633.1;refGene_gn=FAM35A;refGene_tx=XM_023240634.1;refGene_gn=FAM35A;refGene_tx=XM_023240635.1
D2 47356450 47356650 refGene_gn=FAM35A;refGene_tx=XM_023240630.1;refGene_gn=FAM35A;refGene_tx=XM_023240631.1;refGene_gn=FAM35A;refGene_tx=XM_023240632.1;refGene_gn=FAM35A;refGene_tx=XM_023240633.1;refGene_gn=FAM35A;refGene_tx=XM_023240634.1
D2 47357252 47357500 refGene_gn=FAM35A;refGene_tx=XM_023240635.1
D2 47362238 47362291 refGene_gn=LOC101090552;refGene_tx=XM_023240638.1;refGene_gn=LOC101090552;refGene_tx=XM_023240639.1
D2 47373458 47373601 refGene_gn=LOC101090552;refGene_tx=XM_019813428.2;refGene_gn=LOC101090552;refGene_tx=XM_023240637.1
D2 47374008 47374188 refGene_gn=LOC101090552;refGene_tx=XM_019813428.2;refGene_gn=LOC101090552;refGene_tx=XM_023240637.1;refGene_gn=LOC101090552;refGene_tx=XM_023240638.1;refGene_gn=LOC101090552;refGene_tx=XM_023240639.1
D2 47375122 47375243 refGene_gn=LOC101090552;refGene_tx=XM_019813428.2;refGene_gn=LOC101090552;refGene_tx=XM_023240637.1;refGene_gn=LOC101090552;refGene_tx=XM_023240638.1;refGene_gn=LOC101090552;refGene_tx=XM_023240639.1
D2 47375868 47376039 refGene_gn=LOC101090552;refGene_tx=XM_019813428.2;refGene_gn=LOC101090552;refGene_tx=XM_023240637.1;refGene_gn=LOC101090552;refGene_tx=XM_02

3240638.1;refGene_gn=LOC101090552;refGene_tx=XM_023240639.1
D2 47377560 47377847
refGene_gn=LOC101090552;refGene_tx=XM_019813428.2;refGene_gn=LOC101090552;refGene_tx=XM_023240637.1;refGene_gn=LOC101090552;refGene_tx=XM_023240638.1;refGene_gn=LOC101090552;refGene_tx=XM_023240639.1
D2 47379273 47379498
refGene_gn=LOC101090552;refGene_tx=XM_019813428.2;refGene_gn=LOC101090552;refGene_tx=XM_023240637.1;refGene_gn=LOC101090552;refGene_tx=XM_023240638.1;refGene_gn=LOC101090552;refGene_tx=XM_023240639.1
D2 47379913 47380109
refGene_gn=LOC101090552;refGene_tx=XM_019813428.2;refGene_gn=LOC101090552;refGene_tx=XM_023240637.1;refGene_gn=LOC101090552;refGene_tx=XM_023240638.1;refGene_gn=LOC101090552;refGene_tx=XM_023240639.1
D2 47381027 47381034
refGene_gn=LOC101090552;refGene_tx=XM_023240637.1;refGene_gn=LOC101090552;refGene_tx=XM_023240638.1;refGene_gn=LOC101090552;refGene_tx=XM_023240639.1
D2 47407875 47409249
refGene_gn=GPRIN2;refGene_tx=XM_003994139.5;refGene_gn=GPRIN2;refGene_tx=XM_023240640.1
D2 47455780 47456908
refGene_gn=LOC101086754;refGene_tx=XM_019813436.2;refGene_gn=LOC101086754;refGene_tx=XM_019813437.2
D2 47498920 47498980
refGene_gn=LOC101087010;refGene_tx=XM_003994141.5
D2 47501633 47501756
refGene_gn=LOC101087010;refGene_tx=XM_003994141.5
D2 47502892 47502951
refGene_gn=LOC101087010;refGene_tx=XM_003994141.5
D2 47504648 47504744
refGene_gn=LOC101087010;refGene_tx=XM_003994141.5
D2 47504998 47505092
refGene_gn=LOC101087010;refGene_tx=XM_003994141.5
D2 47505618 47505678
refGene_gn=LOC101087010;refGene_tx=XM_003994141.5
D2 47506181 47506261
refGene_gn=LOC101087010;refGene_tx=XM_003994141.5
D2 47507015 47507106
refGene_gn=LOC101087010;refGene_tx=XM_003994141.5
D2 47507863 47507977
refGene_gn=LOC101087010;refGene_tx=XM_003994141.5
D2 47509364 47509459
refGene_gn=LOC101087010;refGene_tx=XM_003994141.5
D2 47510467 47510558
refGene_gn=LOC101087010;refGene_tx=XM_003994141.5
D2 47515324 47515345
refGene_gn=LOC101087010;refGene_tx=XM_003994141.5
D2 47526950 47527205 refGene_gn=ANTXRL;refGene_tx=XM_011287273.3
D2 47532007 47532085 refGene_gn=ANTXRL;refGene_tx=XM_011287273.3
D2 47533524 47533605 refGene_gn=ANTXRL;refGene_tx=XM_011287273.3

D2 47534691 47534727 refGene_gn=ANTXRL;refGene_tx=XM_011287273.3
D2 47535956 47536142 refGene_gn=ANTXRL;refGene_tx=XM_011287273.3
D2 47537715 47537806 refGene_gn=ANTXRL;refGene_tx=XM_011287273.3
D2 47538367 47538437 refGene_gn=ANTXRL;refGene_tx=XM_011287273.3
D2 47539645 47539744 refGene_gn=ANTXRL;refGene_tx=XM_011287273.3
D2 47546132 47546193 refGene_gn=ANTXRL;refGene_tx=XM_011287273.3
D2 47546286 47546367 refGene_gn=ANTXRL;refGene_tx=XM_011287273.3
D2 47546706 47546775 refGene_gn=ANTXRL;refGene_tx=XM_011287273.3
D2 47546851 47546931 refGene_gn=ANTXRL;refGene_tx=XM_011287273.3
D2 47547994 47548028 refGene_gn=ANTXRL;refGene_tx=XM_011287273.3
D2 47548128 47548210 refGene_gn=ANTXRL;refGene_tx=XM_011287273.3
D2 47551912 47551984 refGene_gn=ANTXRL;refGene_tx=XM_011287273.3
D2 47553281 47553353 refGene_gn=ANTXRL;refGene_tx=XM_011287273.3
D2 47563135 47563410 refGene_gn=ANTXRL;refGene_tx=XM_011287273.3
D2 47579890 47580118
refGene_gn=LOC102900063;refGene_tx=XM_023240641.1;refGene_gn=LOC102900063;refGene_tx=XM_023240642.1;refGene_gn=LOC102900063;refGene_tx=XM_023240643.1;refGene_gn=LOC102900063;refGene_tx=XM_023240644.1
D2 47589879 47589957
refGene_gn=LOC102900063;refGene_tx=XM_023240641.1;refGene_gn=LOC102900063;refGene_tx=XM_023240642.1;refGene_gn=LOC102900063;refGene_tx=XM_023240643.1;refGene_gn=LOC102900063;refGene_tx=XM_023240644.1
D2 47591427 47591511
refGene_gn=LOC102900063;refGene_tx=XM_023240641.1;refGene_gn=LOC102900063;refGene_tx=XM_023240642.1;refGene_gn=LOC102900063;refGene_tx=XM_023240643.1;refGene_gn=LOC102900063;refGene_tx=XM_023240644.1
D2 47658753 47658789
refGene_gn=LOC102900063;refGene_tx=XM_023240641.1;refGene_gn=LOC102900063;refGene_tx=XM_023240642.1;refGene_gn=LOC102900063;refGene_tx=XM_023240643.1;refGene_gn=LOC102900063;refGene_tx=XM_023240644.1
D2 47660061 47660277
refGene_gn=LOC102900063;refGene_tx=XM_023240641.1;refGene_gn=LOC102900063;refGene_tx=XM_023240642.1;refGene_gn=LOC102900063;refGene_tx=XM_023240643.1;refGene_gn=LOC102900063;refGene_tx=XM_023240644.1
D2 47661876 47661955
refGene_gn=LOC102900063;refGene_tx=XM_023240641.1;refGene_gn=LOC102900063;refGene_tx=XM_023240642.1;refGene_gn=LOC102900063;refGene_tx=XM_023240643.1;refGene_gn=LOC102900063;refGene_tx=XM_023240644.1
D2 47662525 47662595
refGene_gn=LOC102900063;refGene_tx=XM_023240641.1;refGene_gn=LOC102900063;refGene_tx=XM_023240642.1;refGene_gn=LOC102900063;refGene_tx=XM_023240643.1;refGene_gn=LOC102900063;refGene_tx=XM_023240644.1
D2 47663825 47663921
refGene_gn=LOC102900063;refGene_tx=XM_023240641.1;refGene_gn=LOC102900063;refGene_tx=XM_023240642.1;refGene_gn=LOC102900063;refGene_tx=XM_023240644.1
D2 47682433 47682494
refGene_gn=LOC102900063;refGene_tx=XM_023240641.1;refGene_gn=LOC102900063;refGene_tx=XM_023240642.1;refGene_gn=LOC102900063;refGene_tx=XM_023240643.1;refGene_gn=LOC102900063;refGene_tx=XM_023240644.1

D2 47682597 47682678
refGene_gn=LOC102900063;refGene_tx=XM_023240641.1;refGene_gn=LOC102900063;refGene_tx=XM_023240642.1;refGene_gn=LOC102900063;refGene_tx=XM_023240643.1;refGene_gn=LOC102900063;refGene_tx=XM_023240644.1
D2 47683254 47683323
refGene_gn=LOC102900063;refGene_tx=XM_023240641.1;refGene_gn=LOC102900063;refGene_tx=XM_023240643.1;refGene_gn=LOC102900063;refGene_tx=XM_023240644.1
D2 47683399 47683479
refGene_gn=LOC102900063;refGene_tx=XM_023240641.1;refGene_gn=LOC102900063;refGene_tx=XM_023240642.1;refGene_gn=LOC102900063;refGene_tx=XM_023240643.1;refGene_gn=LOC102900063;refGene_tx=XM_023240644.1
D2 47684148 47684182
refGene_gn=LOC102900063;refGene_tx=XM_023240641.1;refGene_gn=LOC102900063;refGene_tx=XM_023240642.1;refGene_gn=LOC102900063;refGene_tx=XM_023240643.1;refGene_gn=LOC102900063;refGene_tx=XM_023240644.1
D2 47684282 47684364
refGene_gn=LOC102900063;refGene_tx=XM_023240641.1;refGene_gn=LOC102900063;refGene_tx=XM_023240642.1;refGene_gn=LOC102900063;refGene_tx=XM_023240643.1;refGene_gn=LOC102900063;refGene_tx=XM_023240644.1
D2 47686663 47686735
refGene_gn=LOC102900063;refGene_tx=XM_023240641.1;refGene_gn=LOC102900063;refGene_tx=XM_023240642.1;refGene_gn=LOC102900063;refGene_tx=XM_023240643.1;refGene_gn=LOC102900063;refGene_tx=XM_023240644.1
D2 47688061 47688133
refGene_gn=LOC102900063;refGene_tx=XM_023240641.1;refGene_gn=LOC102900063;refGene_tx=XM_023240642.1;refGene_gn=LOC102900063;refGene_tx=XM_023240643.1
D2 47709662 47709925
refGene_gn=LOC102900063;refGene_tx=XM_023240641.1;refGene_gn=LOC102900063;refGene_tx=XM_023240642.1;refGene_gn=LOC102900063;refGene_tx=XM_023240643.1
D2 47722811 47723783
refGene_gn=ZNF488;refGene_tx=XM_023240645.1;refGene_gn=ZNF488;refGene_tx=XM_023240646.1;refGene_gn=ZNF488;refGene_tx=XM_023240647.1
D2 47733983 47734378 refGene_gn=RBP3;refGene_tx=XM_003994154.4
D2 47735680 47735727 refGene_gn=RBP3;refGene_tx=XM_019813453.2
D2 47735906 47736049
refGene_gn=RBP3;refGene_tx=XM_003994154.4;refGene_gn=RBP3;refGene_tx=XM_019813453.2
D2 47737565 47737756
refGene_gn=RBP3;refGene_tx=XM_003994154.4;refGene_gn=RBP3;refGene_tx=XM_019813453.2
D2 47739940 47742988
refGene_gn=RBP3;refGene_tx=XM_003994154.4;refGene_gn=RBP3;refGene_tx=XM_019813453.2
D2 47758210 47759151 refGene_gn=GDF2;refGene_tx=XM_003994142.3
D2 47761936 47762282 refGene_gn=GDF2;refGene_tx=XM_003994142.3
D2 47773098 47773290
refGene_gn=GDF10;refGene_tx=XM_003994143.5;refGene_gn=GDF10;refGene_tx

=XM_019813454.2
D2 47775262 47776188
refGene_gn=GDF10;refGene_tx=XM_003994143.5;refGene_gn=GDF10;refGene_tx
=XM_019813454.2
D2 47784633 47784949
refGene_gn=GDF10;refGene_tx=XM_003994143.5;refGene_gn=GDF10;refGene_tx
=XM_019813454.2
D2 48033899 48033941 refGene_gn=PTPN20;refGene_tx=XM_006937952.4
D2 48034030 48034036 refGene_gn=PTPN20;refGene_tx=XM_019813473.2
D2 48035608 48035674
refGene_gn=PTPN20;refGene_tx=XM_019813464.2;refGene_gn=PTPN20;refGene_
tx=XM_019813465.2;refGene_gn=PTPN20;refGene_tx=XM_019813467.2;refGene_
gn=PTPN20;refGene_tx=XM_019813468.2;refGene_gn=PTPN20;refGene_tx=XM_01
9813470.2;refGene_gn=PTPN20;refGene_tx=XM_019813474.2;refGene_gn=PTPN2
0;refGene_tx=XM_019813475.2;refGene_gn=PTPN20;refGene_tx=XM_019813476.
2;refGene_gn=PTPN20;refGene_tx=XM_019813477.2;refGene_gn=PTPN20;refGen
e_tx=XM_019813478.2;refGene_gn=PTPN20;refGene_tx=XM_023240648.1
D2 48036538 48036601
refGene_gn=PTPN20;refGene_tx=XM_006937952.4;refGene_gn=PTPN20;refGene_
tx=XM_019813464.2;refGene_gn=PTPN20;refGene_tx=XM_019813465.2;refGene_
gn=PTPN20;refGene_tx=XM_019813467.2;refGene_gn=PTPN20;refGene_tx=XM_01
9813468.2;refGene_gn=PTPN20;refGene_tx=XM_019813470.2;refGene_gn=PTPN2
0;refGene_tx=XM_019813473.2;refGene_gn=PTPN20;refGene_tx=XM_019813474.
2;refGene_gn=PTPN20;refGene_tx=XM_019813475.2;refGene_gn=PTPN20;refGen
e_tx=XM_019813476.2;refGene_gn=PTPN20;refGene_tx=XM_019813477.2;refGen
e_gn=PTPN20;refGene_tx=XM_019813478.2;refGene_gn=PTPN20;refGene_tx=XM_
023240648.1
D2 48043074 48043290
refGene_gn=PTPN20;refGene_tx=XM_006937952.4;refGene_gn=PTPN20;refGene_
tx=XM_019813464.2;refGene_gn=PTPN20;refGene_tx=XM_019813465.2;refGene_
gn=PTPN20;refGene_tx=XM_019813467.2;refGene_gn=PTPN20;refGene_tx=XM_01
9813468.2;refGene_gn=PTPN20;refGene_tx=XM_019813470.2;refGene_gn=PTPN2
0;refGene_tx=XM_019813473.2;refGene_gn=PTPN20;refGene_tx=XM_019813474.
2;refGene_gn=PTPN20;refGene_tx=XM_019813475.2;refGene_gn=PTPN20;refGen
e_tx=XM_019813476.2;refGene_gn=PTPN20;refGene_tx=XM_019813477.2;refGen
e_gn=PTPN20;refGene_tx=XM_019813478.2;refGene_gn=PTPN20;refGene_tx=XM_
023240648.1
D2 48047268 48047603
refGene_gn=PTPN20;refGene_tx=XM_006937952.4;refGene_gn=PTPN20;refGene_
tx=XM_019813464.2;refGene_gn=PTPN20;refGene_tx=XM_019813465.2;refGene_
gn=PTPN20;refGene_tx=XM_019813467.2;refGene_gn=PTPN20;refGene_tx=XM_01
9813468.2;refGene_gn=PTPN20;refGene_tx=XM_019813470.2;refGene_gn=PTPN2
0;refGene_tx=XM_019813473.2;refGene_gn=PTPN20;refGene_tx=XM_019813474.
2;refGene_gn=PTPN20;refGene_tx=XM_019813475.2;refGene_gn=PTPN20;refGen
e_tx=XM_019813476.2;refGene_gn=PTPN20;refGene_tx=XM_019813477.2;refGen
e_gn=PTPN20;refGene_tx=XM_019813478.2;refGene_gn=PTPN20;refGene_tx=XM_
023240648.1
D2 48059451 48059542
refGene_gn=PTPN20;refGene_tx=XM_006937952.4;refGene_gn=PTPN20;refGene_
tx=XM_019813464.2;refGene_gn=PTPN20;refGene_tx=XM_019813465.2;refGene_

gn=PTPN20;refGene_tx=XM_019813467.2;refGene_gn=PTPN20;refGene_tx=XM_01
9813468.2;refGene_gn=PTPN20;refGene_tx=XM_019813470.2;refGene_gn=PTPN2
0;refGene_tx=XM_019813473.2;refGene_gn=PTPN20;refGene_tx=XM_019813474.
2;refGene_gn=PTPN20;refGene_tx=XM_019813475.2;refGene_gn=PTPN20;refGen
e_tx=XM_019813476.2;refGene_gn=PTPN20;refGene_tx=XM_019813477.2;refGen
e_gn=PTPN20;refGene_tx=XM_019813478.2;refGene_gn=PTPN20;refGene_tx=XM_
023240648.1
D2 48062520 48062669
refGene_gn=PTPN20;refGene_tx=XM_006937952.4;refGene_gn=PTPN20;refGene_
tx=XM_019813464.2;refGene_gn=PTPN20;refGene_tx=XM_019813465.2;refGene_
gn=PTPN20;refGene_tx=XM_019813467.2;refGene_gn=PTPN20;refGene_tx=XM_01
9813468.2;refGene_gn=PTPN20;refGene_tx=XM_019813470.2;refGene_gn=PTPN2
0;refGene_tx=XM_019813473.2;refGene_gn=PTPN20;refGene_tx=XM_019813474.
2;refGene_gn=PTPN20;refGene_tx=XM_023240648.1
D2 48080670 48080780
refGene_gn=PTPN20;refGene_tx=XM_006937952.4;refGene_gn=PTPN20;refGene_
tx=XM_019813464.2;refGene_gn=PTPN20;refGene_tx=XM_019813465.2;refGene_
gn=PTPN20;refGene_tx=XM_019813467.2;refGene_gn=PTPN20;refGene_tx=XM_01
9813468.2;refGene_gn=PTPN20;refGene_tx=XM_019813470.2;refGene_gn=PTPN2
0;refGene_tx=XM_019813473.2;refGene_gn=PTPN20;refGene_tx=XM_019813474.
2;refGene_gn=PTPN20;refGene_tx=XM_019813475.2;refGene_gn=PTPN20;refGen
e_tx=XM_019813476.2;refGene_gn=PTPN20;refGene_tx=XM_019813477.2;refGen
e_gn=PTPN20;refGene_tx=XM_019813478.2;refGene_gn=PTPN20;refGene_tx=XM_
023240648.1
D2 48088237 48088332
refGene_gn=PTPN20;refGene_tx=XM_006937952.4;refGene_gn=PTPN20;refGene_
tx=XM_019813464.2;refGene_gn=PTPN20;refGene_tx=XM_019813465.2;refGene_
gn=PTPN20;refGene_tx=XM_019813467.2;refGene_gn=PTPN20;refGene_tx=XM_01
9813468.2;refGene_gn=PTPN20;refGene_tx=XM_019813470.2;refGene_gn=PTPN2
0;refGene_tx=XM_019813473.2;refGene_gn=PTPN20;refGene_tx=XM_019813474.
2;refGene_gn=PTPN20;refGene_tx=XM_023240648.1
D2 48094074 48094172
refGene_gn=PTPN20;refGene_tx=XM_006937952.4;refGene_gn=PTPN20;refGene_
tx=XM_019813464.2;refGene_gn=PTPN20;refGene_tx=XM_019813465.2;refGene_
gn=PTPN20;refGene_tx=XM_019813467.2;refGene_gn=PTPN20;refGene_tx=XM_01
9813468.2;refGene_gn=PTPN20;refGene_tx=XM_019813470.2;refGene_gn=PTPN2
0;refGene_tx=XM_019813473.2;refGene_gn=PTPN20;refGene_tx=XM_019813474.
2;refGene_gn=PTPN20;refGene_tx=XM_023240648.1
D2 48107474 48107511 refGene_gn=PTPN20;refGene_tx=XM_019813474.2
D2 48123384 48123432
refGene_gn=PTPN20;refGene_tx=XM_006937952.4;refGene_gn=PTPN20;refGene_
tx=XM_019813464.2;refGene_gn=PTPN20;refGene_tx=XM_019813465.2;refGene_
gn=PTPN20;refGene_tx=XM_019813467.2;refGene_gn=PTPN20;refGene_tx=XM_01
9813468.2;refGene_gn=PTPN20;refGene_tx=XM_019813470.2;refGene_gn=PTPN2
0;refGene_tx=XM_019813473.2
D2 48123753 48123787
refGene_gn=PTPN20;refGene_tx=XM_006937952.4;refGene_gn=PTPN20;refGene_
tx=XM_019813464.2;refGene_gn=PTPN20;refGene_tx=XM_019813465.2;refGene_
gn=PTPN20;refGene_tx=XM_019813467.2;refGene_gn=PTPN20;refGene_tx=XM_01
9813468.2;refGene_gn=PTPN20;refGene_tx=XM_019813470.2;refGene_gn=PTPN2

0;refGene_tx=XM_019813473.2;refGene_gn=PTPN20;refGene_tx=XM_023240648.1
D2 48156993 48156997 refGene_gn=FRMPD2;refGene_tx=XM_023240649.1;refGene_gn=FRMPD2;refGene_tx=XM_023240653.1
D2 48160059 48160341 refGene_gn=FRMPD2;refGene_tx=XM_023240649.1
D2 48161255 48161304 refGene_gn=FRMPD2;refGene_tx=XM_023240651.1;refGene_gn=FRMPD2;refGene_tx=XM_023240652.1;refGene_gn=FRMPD2;refGene_tx=XM_023240654.1
D2 48165773 48166117 refGene_gn=FRMPD2;refGene_tx=XM_023240649.1;refGene_gn=FRMPD2;refGene_tx=XM_023240651.1;refGene_gn=FRMPD2;refGene_tx=XM_023240652.1;refGene_gn=FRMPD2;refGene_tx=XM_023240653.1;refGene_gn=FRMPD2;refGene_tx=XM_023240654.1
D2 48171184 48171283 refGene_gn=FRMPD2;refGene_tx=XM_023240649.1;refGene_gn=FRMPD2;refGene_tx=XM_023240651.1;refGene_gn=FRMPD2;refGene_tx=XM_023240652.1;refGene_gn=FRMPD2;refGene_tx=XM_023240653.1;refGene_gn=FRMPD2;refGene_tx=XM_023240654.1
D2 48173597 48173812 refGene_gn=FRMPD2;refGene_tx=XM_023240649.1;refGene_gn=FRMPD2;refGene_tx=XM_023240651.1;refGene_gn=FRMPD2;refGene_tx=XM_023240652.1;refGene_gn=FRMPD2;refGene_tx=XM_023240653.1;refGene_gn=FRMPD2;refGene_tx=XM_023240654.1
D2 48175761 48175909 refGene_gn=FRMPD2;refGene_tx=XM_023240649.1;refGene_gn=FRMPD2;refGene_tx=XM_023240651.1;refGene_gn=FRMPD2;refGene_tx=XM_023240652.1;refGene_gn=FRMPD2;refGene_tx=XM_023240653.1;refGene_gn=FRMPD2;refGene_tx=XM_023240654.1
D2 48179297 48179383 refGene_gn=FRMPD2;refGene_tx=XM_023240649.1;refGene_gn=FRMPD2;refGene_tx=XM_023240651.1;refGene_gn=FRMPD2;refGene_tx=XM_023240652.1;refGene_gn=FRMPD2;refGene_tx=XM_023240653.1;refGene_gn=FRMPD2;refGene_tx=XM_023240654.1
D2 48180246 48180340 refGene_gn=FRMPD2;refGene_tx=XM_023240649.1;refGene_gn=FRMPD2;refGene_tx=XM_023240651.1;refGene_gn=FRMPD2;refGene_tx=XM_023240652.1;refGene_gn=FRMPD2;refGene_tx=XM_023240653.1;refGene_gn=FRMPD2;refGene_tx=XM_023240654.1
D2 48181394 48181505 refGene_gn=FRMPD2;refGene_tx=XM_023240649.1;refGene_gn=FRMPD2;refGene_tx=XM_023240651.1;refGene_gn=FRMPD2;refGene_tx=XM_023240652.1;refGene_gn=FRMPD2;refGene_tx=XM_023240653.1;refGene_gn=FRMPD2;refGene_tx=XM_023240654.1
D2 48185303 48185476 refGene_gn=FRMPD2;refGene_tx=XM_023240649.1;refGene_gn=FRMPD2;refGene_tx=XM_023240651.1;refGene_gn=FRMPD2;refGene_tx=XM_023240652.1;refGene_gn=FRMPD2;refGene_tx=XM_023240653.1;refGene_gn=FRMPD2;refGene_tx=XM_023240654.1
D2 48188831 48188948

refGene_gn=FRMPD2;refGene_tx=XM_023240649.1;refGene_gn=FRMPD2;refGene_tx=XM_023240651.1;refGene_gn=FRMPD2;refGene_tx=XM_023240652.1;refGene_gn=FRMPD2;refGene_tx=XM_023240653.1;refGene_gn=FRMPD2;refGene_tx=XM_023240654.1
D2 48189037 48189145
refGene_gn=FRMPD2;refGene_tx=XM_023240649.1;refGene_gn=FRMPD2;refGene_tx=XM_023240651.1;refGene_gn=FRMPD2;refGene_tx=XM_023240652.1;refGene_gn=FRMPD2;refGene_tx=XM_023240653.1;refGene_gn=FRMPD2;refGene_tx=XM_023240654.1
D2 48189735 48189828
refGene_gn=FRMPD2;refGene_tx=XM_023240649.1;refGene_gn=FRMPD2;refGene_tx=XM_023240651.1;refGene_gn=FRMPD2;refGene_tx=XM_023240652.1;refGene_gn=FRMPD2;refGene_tx=XM_023240653.1;refGene_gn=FRMPD2;refGene_tx=XM_023240654.1
D2 48191195 48191296
refGene_gn=FRMPD2;refGene_tx=XM_023240649.1;refGene_gn=FRMPD2;refGene_tx=XM_023240651.1;refGene_gn=FRMPD2;refGene_tx=XM_023240652.1;refGene_gn=FRMPD2;refGene_tx=XM_023240653.1;refGene_gn=FRMPD2;refGene_tx=XM_023240654.1
D2 48196688 48196911
refGene_gn=FRMPD2;refGene_tx=XM_023240649.1;refGene_gn=FRMPD2;refGene_tx=XM_023240651.1;refGene_gn=FRMPD2;refGene_tx=XM_023240652.1;refGene_gn=FRMPD2;refGene_tx=XM_023240653.1;refGene_gn=FRMPD2;refGene_tx=XM_023240654.1
D2 48202836 48202984
refGene_gn=FRMPD2;refGene_tx=XM_023240649.1;refGene_gn=FRMPD2;refGene_tx=XM_023240651.1;refGene_gn=FRMPD2;refGene_tx=XM_023240652.1;refGene_gn=FRMPD2;refGene_tx=XM_023240653.1;refGene_gn=FRMPD2;refGene_tx=XM_023240654.1
D2 48208458 48208644
refGene_gn=FRMPD2;refGene_tx=XM_023240649.1;refGene_gn=FRMPD2;refGene_tx=XM_023240651.1;refGene_gn=FRMPD2;refGene_tx=XM_023240652.1;refGene_gn=FRMPD2;refGene_tx=XM_023240653.1;refGene_gn=FRMPD2;refGene_tx=XM_023240654.1
D2 48213578 48213734
refGene_gn=FRMPD2;refGene_tx=XM_023240649.1;refGene_gn=FRMPD2;refGene_tx=XM_023240651.1;refGene_gn=FRMPD2;refGene_tx=XM_023240652.1;refGene_gn=FRMPD2;refGene_tx=XM_023240653.1;refGene_gn=FRMPD2;refGene_tx=XM_023240654.1
D2 48220222 48220361
refGene_gn=FRMPD2;refGene_tx=XM_023240649.1;refGene_gn=FRMPD2;refGene_tx=XM_023240651.1;refGene_gn=FRMPD2;refGene_tx=XM_023240652.1;refGene_gn=FRMPD2;refGene_tx=XM_023240653.1;refGene_gn=FRMPD2;refGene_tx=XM_023240654.1
D2 48221003 48221154
refGene_gn=FRMPD2;refGene_tx=XM_023240649.1;refGene_gn=FRMPD2;refGene_tx=XM_023240651.1;refGene_gn=FRMPD2;refGene_tx=XM_023240652.1;refGene_gn=FRMPD2;refGene_tx=XM_023240653.1;refGene_gn=FRMPD2;refGene_tx=XM_023240654.1
D2 48224516 48224691

refGene_gn=FRMPD2;refGene_tx=XM_023240649.1;refGene_gn=FRMPD2;refGene_tx=XM_023240651.1;refGene_gn=FRMPD2;refGene_tx=XM_023240652.1;refGene_gn=FRMPD2;refGene_tx=XM_023240653.1;refGene_gn=FRMPD2;refGene_tx=XM_023240654.1
D2 48226797 48226869
refGene_gn=FRMPD2;refGene_tx=XM_023240649.1;refGene_gn=FRMPD2;refGene_tx=XM_023240651.1;refGene_gn=FRMPD2;refGene_tx=XM_023240652.1;refGene_gn=FRMPD2;refGene_tx=XM_023240653.1;refGene_gn=FRMPD2;refGene_tx=XM_023240654.1
D2 48228771 48228907
refGene_gn=FRMPD2;refGene_tx=XM_023240649.1;refGene_gn=FRMPD2;refGene_tx=XM_023240651.1;refGene_gn=FRMPD2;refGene_tx=XM_023240652.1;refGene_gn=FRMPD2;refGene_tx=XM_023240653.1;refGene_gn=FRMPD2;refGene_tx=XM_023240654.1
D2 48230932 48231020
refGene_gn=FRMPD2;refGene_tx=XM_023240649.1;refGene_gn=FRMPD2;refGene_tx=XM_023240651.1;refGene_gn=FRMPD2;refGene_tx=XM_023240652.1;refGene_gn=FRMPD2;refGene_tx=XM_023240653.1;refGene_gn=FRMPD2;refGene_tx=XM_023240654.1
D2 48231686 48231819
refGene_gn=FRMPD2;refGene_tx=XM_023240649.1;refGene_gn=FRMPD2;refGene_tx=XM_023240651.1;refGene_gn=FRMPD2;refGene_tx=XM_023240652.1;refGene_gn=FRMPD2;refGene_tx=XM_023240653.1;refGene_gn=FRMPD2;refGene_tx=XM_023240654.1
D2 48233370 48233562
refGene_gn=FRMPD2;refGene_tx=XM_023240649.1;refGene_gn=FRMPD2;refGene_tx=XM_023240651.1;refGene_gn=FRMPD2;refGene_tx=XM_023240652.1;refGene_gn=FRMPD2;refGene_tx=XM_023240653.1;refGene_gn=FRMPD2;refGene_tx=XM_023240654.1
D2 48236066 48236132
refGene_gn=FRMPD2;refGene_tx=XM_023240649.1;refGene_gn=FRMPD2;refGene_tx=XM_023240651.1;refGene_gn=FRMPD2;refGene_tx=XM_023240652.1;refGene_gn=FRMPD2;refGene_tx=XM_023240653.1;refGene_gn=FRMPD2;refGene_tx=XM_023240654.1
D2 48240306 48240464
refGene_gn=FRMPD2;refGene_tx=XM_023240649.1;refGene_gn=FRMPD2;refGene_tx=XM_023240651.1;refGene_gn=FRMPD2;refGene_tx=XM_023240652.1;refGene_gn=FRMPD2;refGene_tx=XM_023240653.1;refGene_gn=FRMPD2;refGene_tx=XM_023240654.1
D2 48242014 48242140
refGene_gn=FRMPD2;refGene_tx=XM_023240649.1;refGene_gn=FRMPD2;refGene_tx=XM_023240651.1;refGene_gn=FRMPD2;refGene_tx=XM_023240652.1;refGene_gn=FRMPD2;refGene_tx=XM_023240653.1;refGene_gn=FRMPD2;refGene_tx=XM_023240654.1
D2 48257038 48257063
refGene_gn=FRMPD2;refGene_tx=XM_023240649.1;refGene_gn=FRMPD2;refGene_tx=XM_023240651.1;refGene_gn=FRMPD2;refGene_tx=XM_023240652.1;refGene_gn=FRMPD2;refGene_tx=XM_023240653.1;refGene_gn=FRMPD2;refGene_tx=XM_023240654.1
D2 48340971 48341093

refGene_gn=MAPK8;refGene_tx=XM_003994155.5;refGene_gn=MAPK8;refGene_tx=XM_023240655.1;refGene_gn=MAPK8;refGene_tx=XM_023240656.1;refGene_gn=MAPK8;refGene_tx=XM_023240657.1
D2 48345736 48345866
refGene_gn=MAPK8;refGene_tx=XM_003994155.5;refGene_gn=MAPK8;refGene_tx=XM_023240655.1;refGene_gn=MAPK8;refGene_tx=XM_023240656.1;refGene_gn=MAPK8;refGene_tx=XM_023240657.1
D2 48351953 48352012
refGene_gn=MAPK8;refGene_tx=XM_003994155.5;refGene_gn=MAPK8;refGene_tx=XM_023240655.1;refGene_gn=MAPK8;refGene_tx=XM_023240656.1;refGene_gn=MAPK8;refGene_tx=XM_023240657.1
D2 48352101 48352240
refGene_gn=MAPK8;refGene_tx=XM_003994155.5;refGene_gn=MAPK8;refGene_tx=XM_023240655.1;refGene_gn=MAPK8;refGene_tx=XM_023240656.1;refGene_gn=MAPK8;refGene_tx=XM_023240657.1
D2 48366362 48366528
refGene_gn=MAPK8;refGene_tx=XM_003994155.5;refGene_gn=MAPK8;refGene_tx=XM_023240655.1;refGene_gn=MAPK8;refGene_tx=XM_023240656.1;refGene_gn=MAPK8;refGene_tx=XM_023240657.1
D2 48370320 48370392
refGene_gn=MAPK8;refGene_tx=XM_003994155.5;refGene_gn=MAPK8;refGene_tx=XM_023240655.1;refGene_gn=MAPK8;refGene_tx=XM_023240656.1;refGene_gn=MAPK8;refGene_tx=XM_023240657.1
D2 48371752 48371935
refGene_gn=MAPK8;refGene_tx=XM_003994155.5;refGene_gn=MAPK8;refGene_tx=XM_023240655.1;refGene_gn=MAPK8;refGene_tx=XM_023240656.1;refGene_gn=MAPK8;refGene_tx=XM_023240657.1
D2 48372264 48372389
refGene_gn=MAPK8;refGene_tx=XM_003994155.5;refGene_gn=MAPK8;refGene_tx=XM_023240655.1;refGene_gn=MAPK8;refGene_tx=XM_023240656.1;refGene_gn=MAPK8;refGene_tx=XM_023240657.1
D2 48372924 48372988
refGene_gn=MAPK8;refGene_tx=XM_003994155.5;refGene_gn=MAPK8;refGene_tx=XM_023240655.1;refGene_gn=MAPK8;refGene_tx=XM_023240656.1;refGene_gn=MAPK8;refGene_tx=XM_023240657.1
D2 48375926 48376004
refGene_gn=MAPK8;refGene_tx=XM_003994155.5;refGene_gn=MAPK8;refGene_tx=XM_023240655.1;refGene_gn=MAPK8;refGene_tx=XM_023240656.1;refGene_gn=MAPK8;refGene_tx=XM_023240657.1
D2 48379879 48380030
refGene_gn=MAPK8;refGene_tx=XM_003994155.5;refGene_gn=MAPK8;refGene_tx=XM_023240655.1;refGene_gn=MAPK8;refGene_tx=XM_023240656.1;refGene_gn=MAPK8;refGene_tx=XM_023240657.1
D2 48393297 48393526 refGene_gn=ARHGAP22;refGene_tx=XM_019813482.2
D2 48397579 48398489 refGene_gn=ARHGAP22;refGene_tx=XM_019813482.2
D2 48400495 48400617 refGene_gn=ARHGAP22;refGene_tx=XM_019813482.2
D2 48401197 48401271 refGene_gn=ARHGAP22;refGene_tx=XM_019813482.2
D2 48402285 48402418 refGene_gn=ARHGAP22;refGene_tx=XM_019813482.2
D2 48405084 48405292 refGene_gn=ARHGAP22;refGene_tx=XM_019813482.2
D2 48422086 48422128 refGene_gn=ARHGAP22;refGene_tx=XM_019813482.2

D2 48422130 48422262 refGene_gn=ARHGAP22;refGene_tx=XM_019813482.2
D2 48486486 48486574 refGene_gn=ARHGAP22;refGene_tx=XM_019813482.2
D2 48510833 48511033 refGene_gn=ARHGAP22;refGene_tx=XM_019813482.2
D2 48566572 48566621 refGene_gn=ARHGAP22;refGene_tx=XM_019813482.2
D2 48617550 48617784
refGene_gn=WDFY4;refGene_tx=XM_011287265.3;refGene_gn=WDFY4;refGene_tx=XM_023240659.1;refGene_gn=WDFY4;refGene_tx=XM_023240660.1;refGene_gn=WDFY4;refGene_tx=XM_023240661.1;refGene_gn=WDFY4;refGene_tx=XM_023240662.1
D2 48627602 48627717
refGene_gn=WDFY4;refGene_tx=XM_011287265.3;refGene_gn=WDFY4;refGene_tx=XM_023240659.1;refGene_gn=WDFY4;refGene_tx=XM_023240660.1;refGene_gn=WDFY4;refGene_tx=XM_023240661.1;refGene_gn=WDFY4;refGene_tx=XM_023240662.1
D2 48628627 48628734
refGene_gn=WDFY4;refGene_tx=XM_011287265.3;refGene_gn=WDFY4;refGene_tx=XM_023240659.1;refGene_gn=WDFY4;refGene_tx=XM_023240660.1;refGene_gn=WDFY4;refGene_tx=XM_023240661.1;refGene_gn=WDFY4;refGene_tx=XM_023240662.1
D2 48630540 48630675
refGene_gn=WDFY4;refGene_tx=XM_011287265.3;refGene_gn=WDFY4;refGene_tx=XM_023240659.1;refGene_gn=WDFY4;refGene_tx=XM_023240660.1;refGene_gn=WDFY4;refGene_tx=XM_023240661.1;refGene_gn=WDFY4;refGene_tx=XM_023240662.1
D2 48633293 48633483
refGene_gn=WDFY4;refGene_tx=XM_011287265.3;refGene_gn=WDFY4;refGene_tx=XM_023240659.1;refGene_gn=WDFY4;refGene_tx=XM_023240660.1;refGene_gn=WDFY4;refGene_tx=XM_023240661.1;refGene_gn=WDFY4;refGene_tx=XM_023240662.1
D2 48635311 48635501
refGene_gn=WDFY4;refGene_tx=XM_011287265.3;refGene_gn=WDFY4;refGene_tx=XM_023240659.1;refGene_gn=WDFY4;refGene_tx=XM_023240660.1;refGene_gn=WDFY4;refGene_tx=XM_023240661.1;refGene_gn=WDFY4;refGene_tx=XM_023240662.1
D2 48637027 48637185
refGene_gn=WDFY4;refGene_tx=XM_011287265.3;refGene_gn=WDFY4;refGene_tx=XM_023240659.1;refGene_gn=WDFY4;refGene_tx=XM_023240660.1;refGene_gn=WDFY4;refGene_tx=XM_023240661.1;refGene_gn=WDFY4;refGene_tx=XM_023240662.1
D2 48639165 48639618
refGene_gn=WDFY4;refGene_tx=XM_011287265.3;refGene_gn=WDFY4;refGene_tx=XM_023240659.1;refGene_gn=WDFY4;refGene_tx=XM_023240660.1;refGene_gn=WDFY4;refGene_tx=XM_023240661.1;refGene_gn=WDFY4;refGene_tx=XM_023240662.1
D2 48641928 48642033
refGene_gn=WDFY4;refGene_tx=XM_011287265.3;refGene_gn=WDFY4;refGene_tx=XM_023240659.1;refGene_gn=WDFY4;refGene_tx=XM_023240660.1;refGene_gn=WDFY4;refGene_tx=XM_023240661.1;refGene_gn=WDFY4;refGene_tx=XM_023240662.1
D2 48643871 48644062

refGene_gn=WDFY4;refGene_tx=XM_011287265.3;refGene_gn=WDFY4;refGene_tx
=XM_023240659.1;refGene_gn=WDFY4;refGene_tx=XM_023240660.1;refGene_gn=
WDFY4;refGene_tx=XM_023240661.1;refGene_gn=WDFY4;refGene_tx=XM_0232406
62.1
D2 48651408 48651989
refGene_gn=WDFY4;refGene_tx=XM_011287265.3;refGene_gn=WDFY4;refGene_tx
=XM_023240659.1;refGene_gn=WDFY4;refGene_tx=XM_023240660.1;refGene_gn=
WDFY4;refGene_tx=XM_023240661.1;refGene_gn=WDFY4;refGene_tx=XM_0232406
62.1
D2 48675279 48675373
refGene_gn=WDFY4;refGene_tx=XM_011287265.3;refGene_gn=WDFY4;refGene_tx
=XM_023240659.1;refGene_gn=WDFY4;refGene_tx=XM_023240660.1;refGene_gn=
WDFY4;refGene_tx=XM_023240661.1;refGene_gn=WDFY4;refGene_tx=XM_0232406
62.1
D2 48690514 48690729
refGene_gn=WDFY4;refGene_tx=XM_011287265.3;refGene_gn=WDFY4;refGene_tx
=XM_023240659.1;refGene_gn=WDFY4;refGene_tx=XM_023240660.1;refGene_gn=
WDFY4;refGene_tx=XM_023240661.1;refGene_gn=WDFY4;refGene_tx=XM_0232406
62.1
D2 48691564 48691659
refGene_gn=WDFY4;refGene_tx=XM_011287265.3;refGene_gn=WDFY4;refGene_tx
=XM_023240659.1;refGene_gn=WDFY4;refGene_tx=XM_023240660.1;refGene_gn=
WDFY4;refGene_tx=XM_023240661.1;refGene_gn=WDFY4;refGene_tx=XM_0232406
62.1
D2 48692570 48692808
refGene_gn=WDFY4;refGene_tx=XM_011287265.3;refGene_gn=WDFY4;refGene_tx
=XM_023240659.1;refGene_gn=WDFY4;refGene_tx=XM_023240660.1;refGene_gn=
WDFY4;refGene_tx=XM_023240661.1;refGene_gn=WDFY4;refGene_tx=XM_0232406
62.1
D2 48692995 48693072
refGene_gn=WDFY4;refGene_tx=XM_011287265.3;refGene_gn=WDFY4;refGene_tx
=XM_023240659.1;refGene_gn=WDFY4;refGene_tx=XM_023240660.1;refGene_gn=
WDFY4;refGene_tx=XM_023240661.1;refGene_gn=WDFY4;refGene_tx=XM_0232406
62.1
D2 48694130 48694352
refGene_gn=WDFY4;refGene_tx=XM_011287265.3;refGene_gn=WDFY4;refGene_tx
=XM_023240659.1;refGene_gn=WDFY4;refGene_tx=XM_023240660.1;refGene_gn=
WDFY4;refGene_tx=XM_023240661.1;refGene_gn=WDFY4;refGene_tx=XM_0232406
62.1
D2 48695035 48695214
refGene_gn=WDFY4;refGene_tx=XM_011287265.3;refGene_gn=WDFY4;refGene_tx
=XM_023240659.1;refGene_gn=WDFY4;refGene_tx=XM_023240660.1;refGene_gn=
WDFY4;refGene_tx=XM_023240661.1;refGene_gn=WDFY4;refGene_tx=XM_0232406
62.1
D2 48701314 48701546
refGene_gn=WDFY4;refGene_tx=XM_011287265.3;refGene_gn=WDFY4;refGene_tx
=XM_023240659.1;refGene_gn=WDFY4;refGene_tx=XM_023240660.1;refGene_gn=
WDFY4;refGene_tx=XM_023240661.1;refGene_gn=WDFY4;refGene_tx=XM_0232406
62.1
D2 48702889 48703035

refGene_gn=WDFY4;refGene_tx=XM_011287265.3;refGene_gn=WDFY4;refGene_tx=XM_023240659.1;refGene_gn=WDFY4;refGene_tx=XM_023240660.1;refGene_gn=WDFY4;refGene_tx=XM_023240661.1;refGene_gn=WDFY4;refGene_tx=XM_023240662.1
D2 48704065 48704177
refGene_gn=WDFY4;refGene_tx=XM_011287265.3;refGene_gn=WDFY4;refGene_tx=XM_023240659.1;refGene_gn=WDFY4;refGene_tx=XM_023240660.1;refGene_gn=WDFY4;refGene_tx=XM_023240661.1;refGene_gn=WDFY4;refGene_tx=XM_023240662.1
D2 48704878 48705069
refGene_gn=WDFY4;refGene_tx=XM_011287265.3;refGene_gn=WDFY4;refGene_tx=XM_023240659.1;refGene_gn=WDFY4;refGene_tx=XM_023240660.1;refGene_gn=WDFY4;refGene_tx=XM_023240661.1;refGene_gn=WDFY4;refGene_tx=XM_023240662.1
D2 48709472 48709625
refGene_gn=WDFY4;refGene_tx=XM_011287265.3;refGene_gn=WDFY4;refGene_tx=XM_023240659.1;refGene_gn=WDFY4;refGene_tx=XM_023240660.1;refGene_gn=WDFY4;refGene_tx=XM_023240661.1;refGene_gn=WDFY4;refGene_tx=XM_023240662.1
D2 48715757 48715831
refGene_gn=WDFY4;refGene_tx=XM_011287265.3;refGene_gn=WDFY4;refGene_tx=XM_023240659.1;refGene_gn=WDFY4;refGene_tx=XM_023240660.1;refGene_gn=WDFY4;refGene_tx=XM_023240661.1;refGene_gn=WDFY4;refGene_tx=XM_023240662.1
D2 48717635 48717797
refGene_gn=WDFY4;refGene_tx=XM_011287265.3;refGene_gn=WDFY4;refGene_tx=XM_023240659.1;refGene_gn=WDFY4;refGene_tx=XM_023240660.1;refGene_gn=WDFY4;refGene_tx=XM_023240661.1;refGene_gn=WDFY4;refGene_tx=XM_023240662.1
D2 48718394 48718486
refGene_gn=WDFY4;refGene_tx=XM_011287265.3;refGene_gn=WDFY4;refGene_tx=XM_023240659.1;refGene_gn=WDFY4;refGene_tx=XM_023240660.1;refGene_gn=WDFY4;refGene_tx=XM_023240661.1;refGene_gn=WDFY4;refGene_tx=XM_023240662.1
D2 48720153 48720253
refGene_gn=WDFY4;refGene_tx=XM_011287265.3;refGene_gn=WDFY4;refGene_tx=XM_023240659.1;refGene_gn=WDFY4;refGene_tx=XM_023240660.1;refGene_gn=WDFY4;refGene_tx=XM_023240661.1;refGene_gn=WDFY4;refGene_tx=XM_023240662.1
D2 48722210 48722416
refGene_gn=WDFY4;refGene_tx=XM_011287265.3;refGene_gn=WDFY4;refGene_tx=XM_023240659.1;refGene_gn=WDFY4;refGene_tx=XM_023240660.1;refGene_gn=WDFY4;refGene_tx=XM_023240661.1;refGene_gn=WDFY4;refGene_tx=XM_023240662.1
D2 48723245 48723415
refGene_gn=WDFY4;refGene_tx=XM_011287265.3;refGene_gn=WDFY4;refGene_tx=XM_023240659.1;refGene_gn=WDFY4;refGene_tx=XM_023240660.1;refGene_gn=WDFY4;refGene_tx=XM_023240661.1;refGene_gn=WDFY4;refGene_tx=XM_023240662.1
D2 48725542 48725668

refGene_gn=WDFY4;refGene_tx=XM_011287265.3;refGene_gn=WDFY4;refGene_tx=XM_023240659.1;refGene_gn=WDFY4;refGene_tx=XM_023240660.1;refGene_gn=WDFY4;refGene_tx=XM_023240661.1;refGene_gn=WDFY4;refGene_tx=XM_023240662.1
D2 48730682 48730847
refGene_gn=WDFY4;refGene_tx=XM_011287265.3;refGene_gn=WDFY4;refGene_tx=XM_023240659.1;refGene_gn=WDFY4;refGene_tx=XM_023240660.1;refGene_gn=WDFY4;refGene_tx=XM_023240661.1;refGene_gn=WDFY4;refGene_tx=XM_023240662.1
D2 48733644 48733848
refGene_gn=WDFY4;refGene_tx=XM_011287265.3;refGene_gn=WDFY4;refGene_tx=XM_023240659.1;refGene_gn=WDFY4;refGene_tx=XM_023240660.1;refGene_gn=WDFY4;refGene_tx=XM_023240661.1;refGene_gn=WDFY4;refGene_tx=XM_023240662.1
D2 48734548 48734663
refGene_gn=WDFY4;refGene_tx=XM_011287265.3;refGene_gn=WDFY4;refGene_tx=XM_023240659.1;refGene_gn=WDFY4;refGene_tx=XM_023240660.1;refGene_gn=WDFY4;refGene_tx=XM_023240661.1;refGene_gn=WDFY4;refGene_tx=XM_023240662.1
D2 48735944 48736102
refGene_gn=WDFY4;refGene_tx=XM_011287265.3;refGene_gn=WDFY4;refGene_tx=XM_023240659.1;refGene_gn=WDFY4;refGene_tx=XM_023240660.1;refGene_gn=WDFY4;refGene_tx=XM_023240661.1;refGene_gn=WDFY4;refGene_tx=XM_023240662.1
D2 48738917 48739156
refGene_gn=WDFY4;refGene_tx=XM_011287265.3;refGene_gn=WDFY4;refGene_tx=XM_023240659.1;refGene_gn=WDFY4;refGene_tx=XM_023240660.1;refGene_gn=WDFY4;refGene_tx=XM_023240661.1;refGene_gn=WDFY4;refGene_tx=XM_023240662.1
D2 48741165 48741287
refGene_gn=WDFY4;refGene_tx=XM_011287265.3;refGene_gn=WDFY4;refGene_tx=XM_023240659.1;refGene_gn=WDFY4;refGene_tx=XM_023240660.1;refGene_gn=WDFY4;refGene_tx=XM_023240661.1;refGene_gn=WDFY4;refGene_tx=XM_023240662.1
D2 48743633 48743819
refGene_gn=WDFY4;refGene_tx=XM_011287265.3;refGene_gn=WDFY4;refGene_tx=XM_023240659.1;refGene_gn=WDFY4;refGene_tx=XM_023240660.1;refGene_gn=WDFY4;refGene_tx=XM_023240661.1;refGene_gn=WDFY4;refGene_tx=XM_023240662.1
D2 48745186 48745323
refGene_gn=WDFY4;refGene_tx=XM_011287265.3;refGene_gn=WDFY4;refGene_tx=XM_023240659.1;refGene_gn=WDFY4;refGene_tx=XM_023240660.1;refGene_gn=WDFY4;refGene_tx=XM_023240661.1;refGene_gn=WDFY4;refGene_tx=XM_023240662.1
D2 48774179 48774257
refGene_gn=WDFY4;refGene_tx=XM_011287265.3;refGene_gn=WDFY4;refGene_tx=XM_023240659.1;refGene_gn=WDFY4;refGene_tx=XM_023240660.1;refGene_gn=WDFY4;refGene_tx=XM_023240661.1;refGene_gn=WDFY4;refGene_tx=XM_023240662.1
D2 48780255 48780462

refGene_gn=WDFY4;refGene_tx=XM_011287265.3;refGene_gn=WDFY4;refGene_tx=XM_023240659.1;refGene_gn=WDFY4;refGene_tx=XM_023240660.1;refGene_gn=WDFY4;refGene_tx=XM_023240661.1;refGene_gn=WDFY4;refGene_tx=XM_023240662.1
D2 48782216 48782268
refGene_gn=WDFY4;refGene_tx=XM_011287265.3;refGene_gn=WDFY4;refGene_tx=XM_023240659.1;refGene_gn=WDFY4;refGene_tx=XM_023240660.1;refGene_gn=WDFY4;refGene_tx=XM_023240661.1;refGene_gn=WDFY4;refGene_tx=XM_023240662.1
D2 48784983 48785150
refGene_gn=WDFY4;refGene_tx=XM_011287265.3;refGene_gn=WDFY4;refGene_tx=XM_023240659.1;refGene_gn=WDFY4;refGene_tx=XM_023240660.1;refGene_gn=WDFY4;refGene_tx=XM_023240661.1;refGene_gn=WDFY4;refGene_tx=XM_023240662.1
D2 48798850 48798999
refGene_gn=WDFY4;refGene_tx=XM_011287265.3;refGene_gn=WDFY4;refGene_tx=XM_023240659.1;refGene_gn=WDFY4;refGene_tx=XM_023240660.1;refGene_gn=WDFY4;refGene_tx=XM_023240661.1;refGene_gn=WDFY4;refGene_tx=XM_023240662.1
D2 48804697 48804818
refGene_gn=WDFY4;refGene_tx=XM_011287265.3;refGene_gn=WDFY4;refGene_tx=XM_023240659.1;refGene_gn=WDFY4;refGene_tx=XM_023240660.1;refGene_gn=WDFY4;refGene_tx=XM_023240661.1;refGene_gn=WDFY4;refGene_tx=XM_023240662.1
D2 48807345 48807431
refGene_gn=WDFY4;refGene_tx=XM_011287265.3;refGene_gn=WDFY4;refGene_tx=XM_023240659.1;refGene_gn=WDFY4;refGene_tx=XM_023240660.1;refGene_gn=WDFY4;refGene_tx=XM_023240661.1;refGene_gn=WDFY4;refGene_tx=XM_023240662.1
D2 48808943 48809006
refGene_gn=WDFY4;refGene_tx=XM_011287265.3;refGene_gn=WDFY4;refGene_tx=XM_023240659.1;refGene_gn=WDFY4;refGene_tx=XM_023240660.1;refGene_gn=WDFY4;refGene_tx=XM_023240661.1;refGene_gn=WDFY4;refGene_tx=XM_023240662.1
D2 48817310 48817332
refGene_gn=LRRC18;refGene_tx=XM_006937960.4;refGene_gn=LRRC18;refGene_tx=XM_011287264.3
D2 48820476 48821240
refGene_gn=LRRC18;refGene_tx=XM_006937960.4;refGene_gn=LRRC18;refGene_tx=XM_011287264.3
D2 48845800 48845843
refGene_gn=WDFY4;refGene_tx=XM_011287265.3;refGene_gn=WDFY4;refGene_tx=XM_023240659.1;refGene_gn=WDFY4;refGene_tx=XM_023240660.1;refGene_gn=WDFY4;refGene_tx=XM_023240661.1;refGene_gn=WDFY4;refGene_tx=XM_023240662.1
D2 48846966 48847086
refGene_gn=WDFY4;refGene_tx=XM_011287265.3;refGene_gn=WDFY4;refGene_tx=XM_023240659.1;refGene_gn=WDFY4;refGene_tx=XM_023240660.1;refGene_gn=WDFY4;refGene_tx=XM_023240661.1;refGene_gn=WDFY4;refGene_tx=XM_023240662.1

D2 48849056 48849174
refGene_gn=WDFY4;refGene_tx=XM_011287265.3;refGene_gn=WDFY4;refGene_tx=XM_023240659.1;refGene_gn=WDFY4;refGene_tx=XM_023240660.1;refGene_gn=WDFY4;refGene_tx=XM_023240661.1;refGene_gn=WDFY4;refGene_tx=XM_023240662.1
D2 48851503 48851613
refGene_gn=WDFY4;refGene_tx=XM_011287265.3;refGene_gn=WDFY4;refGene_tx=XM_023240659.1;refGene_gn=WDFY4;refGene_tx=XM_023240660.1;refGene_gn=WDFY4;refGene_tx=XM_023240661.1;refGene_gn=WDFY4;refGene_tx=XM_023240662.1
D2 48860658 48860812
refGene_gn=WDFY4;refGene_tx=XM_011287265.3;refGene_gn=WDFY4;refGene_tx=XM_023240659.1;refGene_gn=WDFY4;refGene_tx=XM_023240660.1;refGene_gn=WDFY4;refGene_tx=XM_023240661.1;refGene_gn=WDFY4;refGene_tx=XM_023240662.1
D2 48863235 48863327
refGene_gn=WDFY4;refGene_tx=XM_011287265.3;refGene_gn=WDFY4;refGene_tx=XM_023240659.1;refGene_gn=WDFY4;refGene_tx=XM_023240660.1;refGene_gn=WDFY4;refGene_tx=XM_023240661.1;refGene_gn=WDFY4;refGene_tx=XM_023240662.1
D2 48869182 48869395
refGene_gn=WDFY4;refGene_tx=XM_011287265.3;refGene_gn=WDFY4;refGene_tx=XM_023240659.1;refGene_gn=WDFY4;refGene_tx=XM_023240660.1;refGene_gn=WDFY4;refGene_tx=XM_023240661.1;refGene_gn=WDFY4;refGene_tx=XM_023240662.1
D2 48871888 48872036
refGene_gn=WDFY4;refGene_tx=XM_011287265.3;refGene_gn=WDFY4;refGene_tx=XM_023240659.1;refGene_gn=WDFY4;refGene_tx=XM_023240660.1;refGene_gn=WDFY4;refGene_tx=XM_023240661.1;refGene_gn=WDFY4;refGene_tx=XM_023240662.1
D2 48873113 48873298
refGene_gn=WDFY4;refGene_tx=XM_011287265.3;refGene_gn=WDFY4;refGene_tx=XM_023240659.1;refGene_gn=WDFY4;refGene_tx=XM_023240660.1;refGene_gn=WDFY4;refGene_tx=XM_023240661.1;refGene_gn=WDFY4;refGene_tx=XM_023240662.1
D2 48874169 48874328
refGene_gn=WDFY4;refGene_tx=XM_011287265.3;refGene_gn=WDFY4;refGene_tx=XM_023240659.1;refGene_gn=WDFY4;refGene_tx=XM_023240660.1;refGene_gn=WDFY4;refGene_tx=XM_023240661.1;refGene_gn=WDFY4;refGene_tx=XM_023240662.1
D2 48878814 48878994
refGene_gn=WDFY4;refGene_tx=XM_011287265.3;refGene_gn=WDFY4;refGene_tx=XM_023240659.1;refGene_gn=WDFY4;refGene_tx=XM_023240660.1;refGene_gn=WDFY4;refGene_tx=XM_023240661.1;refGene_gn=WDFY4;refGene_tx=XM_023240662.1
D2 48880857 48881040
refGene_gn=WDFY4;refGene_tx=XM_011287265.3;refGene_gn=WDFY4;refGene_tx=XM_023240659.1;refGene_gn=WDFY4;refGene_tx=XM_023240660.1;refGene_gn=WDFY4;refGene_tx=XM_023240661.1;refGene_gn=WDFY4;refGene_tx=XM_023240662.1

D2 48883207 48883292
refGene_gn=WDFY4;refGene_tx=XM_011287265.3;refGene_gn=WDFY4;refGene_tx=XM_023240659.1;refGene_gn=WDFY4;refGene_tx=XM_023240660.1;refGene_gn=WDFY4;refGene_tx=XM_023240661.1;refGene_gn=WDFY4;refGene_tx=XM_023240662.1
D2 48886941 48887053
refGene_gn=WDFY4;refGene_tx=XM_011287265.3;refGene_gn=WDFY4;refGene_tx=XM_023240659.1;refGene_gn=WDFY4;refGene_tx=XM_023240660.1;refGene_gn=WDFY4;refGene_tx=XM_023240661.1;refGene_gn=WDFY4;refGene_tx=XM_023240662.1
D2 48888076 48888143
refGene_gn=WDFY4;refGene_tx=XM_011287265.3;refGene_gn=WDFY4;refGene_tx=XM_023240659.1;refGene_gn=WDFY4;refGene_tx=XM_023240660.1;refGene_gn=WDFY4;refGene_tx=XM_023240661.1;refGene_gn=WDFY4;refGene_tx=XM_023240662.1
D2 48913388 48913514 refGene_gn=VSTM4;refGene_tx=XM_023240663.1
D2 48946357 48946419 refGene_gn=VSTM4;refGene_tx=XM_023240663.1
D2 48947874 48947981 refGene_gn=VSTM4;refGene_tx=XM_023240663.1
D2 48963179 48963213 refGene_gn=VSTM4;refGene_tx=XM_023240663.1
D2 48969716 48969824 refGene_gn=VSTM4;refGene_tx=XM_023240663.1
D2 48978806 48978875 refGene_gn=VSTM4;refGene_tx=XM_023240663.1
D2 49001450 49001849 refGene_gn=VSTM4;refGene_tx=XM_023240663.1
D2 49010231 49010286 refGene_gn=VSTM4;refGene_tx=XM_023240663.1
D2 49026058 49026795
refGene_gn=FAM170B;refGene_tx=XM_003994175.5;refGene_gn=FAM170B;refGene_tx=XM_019813488.2
D2 49028531 49028793
refGene_gn=FAM170B;refGene_tx=XM_003994175.5;refGene_gn=FAM170B;refGene_tx=XM_019813488.2
D2 49050251 49050276
refGene_gn=CD2H10orf128;refGene_tx=XM_006937961.4;refGene_gn=CD2H10orf128;refGene_tx=XM_023240665.1
D2 49058538 49058573
refGene_gn=CD2H10orf128;refGene_tx=XM_006937961.4;refGene_gn=CD2H10orf128;refGene_tx=XM_023240665.1;refGene_gn=CD2H10orf128;refGene_tx=XM_023240666.1
D2 49059777 49059918
refGene_gn=CD2H10orf128;refGene_tx=XM_006937961.4;refGene_gn=CD2H10orf128;refGene_tx=XM_023240665.1;refGene_gn=CD2H10orf128;refGene_tx=XM_023240666.1
D2 49060778 49060823
refGene_gn=CD2H10orf128;refGene_tx=XM_006937961.4;refGene_gn=CD2H10orf128;refGene_tx=XM_023240665.1;refGene_gn=CD2H10orf128;refGene_tx=XM_023240666.1
D2 49061811 49061842
refGene_gn=CD2H10orf128;refGene_tx=XM_023240665.1
D2 49083450 49083496
refGene_gn=CD2H10orf128;refGene_tx=XM_006937961.4;refGene_gn=CD2H10orf128;refGene_tx=XM_023240666.1
D2 49204621 49208881

refGene_gn=CD2H10orf71;refGene_tx=XM_006937962.4;refGene_gn=CD2H10orf71;refGene_tx=XM_006937965.4;refGene_gn=CD2H10orf71;refGene_tx=XM_011287262.3;refGene_gn=CD2H10orf71;refGene_tx=XM_019813495.2;refGene_gn=CD2H10orf71;refGene_tx=XM_019813496.2;refGene_gn=CD2H10orf71;refGene_tx=XM_023240667.1
D2 49238285 49238551
refGene_gn=DRGX;refGene_tx=XM_023240668.1;refGene_gn=DRGX;refGene_tx=XM_023240669.1;refGene_gn=DRGX;refGene_tx=XM_023240670.1;refGene_gn=DRGX;refGene_tx=XM_023240671.1
D2 49260756 49260872
refGene_gn=DRGX;refGene_tx=XM_023240668.1;refGene_gn=DRGX;refGene_tx=XM_023240669.1;refGene_gn=DRGX;refGene_tx=XM_023240670.1;refGene_gn=DRGX;refGene_tx=XM_023240671.1
D2 49260952 49261131
refGene_gn=DRGX;refGene_tx=XM_023240668.1;refGene_gn=DRGX;refGene_tx=XM_023240669.1;refGene_gn=DRGX;refGene_tx=XM_023240670.1;refGene_gn=DRGX;refGene_tx=XM_023240671.1
D2 49264090 49264192
refGene_gn=DRGX;refGene_tx=XM_023240668.1;refGene_gn=DRGX;refGene_tx=XM_023240669.1;refGene_gn=DRGX;refGene_tx=XM_023240670.1;refGene_gn=DRGX;refGene_tx=XM_023240671.1
D2 49265104 49265202
refGene_gn=DRGX;refGene_tx=XM_023240668.1;refGene_gn=DRGX;refGene_tx=XM_023240669.1;refGene_gn=DRGX;refGene_tx=XM_023240670.1;refGene_gn=DRGX;refGene_tx=XM_023240671.1
D2 49269281 49269315
refGene_gn=DRGX;refGene_tx=XM_023240668.1;refGene_gn=DRGX;refGene_tx=XM_023240669.1;refGene_gn=DRGX;refGene_tx=XM_023240670.1;refGene_gn=DRGX;refGene_tx=XM_023240671.1
D2 49322010 49322430 refGene_gn=ERCC6;refGene_tx=XM_011287260.3
D2 49323799 49323878 refGene_gn=ERCC6;refGene_tx=XM_011287260.3
D2 49324462 49324667 refGene_gn=ERCC6;refGene_tx=XM_011287260.3
D2 49326443 49327151 refGene_gn=ERCC6;refGene_tx=XM_011287260.3
D2 49327236 49327382 refGene_gn=ERCC6;refGene_tx=XM_011287260.3
D2 49329065 49329160 refGene_gn=ERCC6;refGene_tx=XM_011287260.3
D2 49329587 49329707 refGene_gn=ERCC6;refGene_tx=XM_011287260.3
D2 49330113 49330224 refGene_gn=ERCC6;refGene_tx=XM_011287260.3
D2 49330918 49331134 refGene_gn=ERCC6;refGene_tx=XM_011287260.3
D2 49333607 49333703 refGene_gn=ERCC6;refGene_tx=XM_011287260.3
D2 49337710 49337827 refGene_gn=ERCC6;refGene_tx=XM_011287260.3
D2 49342436 49342613 refGene_gn=ERCC6;refGene_tx=XM_011287260.3
D2 49343061 49343232 refGene_gn=ERCC6;refGene_tx=XM_011287260.3
D2 49352314 49352450 refGene_gn=ERCC6;refGene_tx=XM_011287260.3
D2 49359086 49359245 refGene_gn=ERCC6;refGene_tx=XM_011287260.3
D2 49364963 49365089 refGene_gn=ERCC6;refGene_tx=XM_011287260.3
D2 49382947 49383665 refGene_gn=ERCC6;refGene_tx=XM_011287260.3
D2 49387701 49387813 refGene_gn=ERCC6;refGene_tx=XM_011287260.3
D2 49390383 49390504 refGene_gn=ERCC6;refGene_tx=XM_011287260.3
D2 49392342 49392764 refGene_gn=ERCC6;refGene_tx=XM_011287260.3
D2 49440465 49442010 refGene_gn=SLC18A3;refGene_tx=XM_003994177.3

D2	49446275 49446341 refGene_gn=CHAT;refGene_tx=XM_019813499.2
D2	49449220 49449412 refGene_gn=CHAT;refGene_tx=XM_019813499.2
D2	49450233 49450352 refGene_gn=CHAT;refGene_tx=XM_019813499.2
D2	49451765 49451819 refGene_gn=CHAT;refGene_tx=XM_019813499.2
D2	49453290 49453471 refGene_gn=CHAT;refGene_tx=XM_019813499.2
D2	49455454 49455632 refGene_gn=CHAT;refGene_tx=XM_019813499.2
D2	49473845 49474015 refGene_gn=CHAT;refGene_tx=XM_019813499.2
D2	49476055 49476156 refGene_gn=CHAT;refGene_tx=XM_019813499.2
D2	49477012 49477141 refGene_gn=CHAT;refGene_tx=XM_019813499.2
D2	49479733 49479856 refGene_gn=CHAT;refGene_tx=XM_019813499.2
D2	49482816 49482958 refGene_gn=CHAT;refGene_tx=XM_019813499.2
D2	49483100 49483163 refGene_gn=CHAT;refGene_tx=XM_019813499.2
D2	49486195 49486333 refGene_gn=CHAT;refGene_tx=XM_019813499.2
D2	49488094 49488364 refGene_gn=CHAT;refGene_tx=XM_019813499.2
D2	49506090 49506187
refGene_gn=CD2H10orf53;refGene_tx=XM_003994162.5
D2	49515131 49515251
refGene_gn=CD2H10orf53;refGene_tx=XM_003994162.5
D2	49515947 49516012
refGene_gn=CD2H10orf53;refGene_tx=XM_003994162.5
D2	49548656 49548780
refGene_gn=OGDHL;refGene_tx=XM_023240672.1;refGene_gn=OGDHL;refGene_tx
=XM_023240673.1
D2	49549234 49549389
refGene_gn=OGDHL;refGene_tx=XM_023240672.1;refGene_gn=OGDHL;refGene_tx
=XM_023240673.1
D2	49549562 49549726
refGene_gn=OGDHL;refGene_tx=XM_023240672.1;refGene_gn=OGDHL;refGene_tx
=XM_023240673.1
D2	49551088 49551161
refGene_gn=OGDHL;refGene_tx=XM_023240672.1;refGene_gn=OGDHL;refGene_tx
=XM_023240673.1
D2	49551264 49551390
refGene_gn=OGDHL;refGene_tx=XM_023240672.1;refGene_gn=OGDHL;refGene_tx
=XM_023240673.1
D2	49551478 49551550
refGene_gn=OGDHL;refGene_tx=XM_023240672.1;refGene_gn=OGDHL;refGene_tx
=XM_023240673.1
D2	49553296 49553475
refGene_gn=OGDHL;refGene_tx=XM_023240672.1;refGene_gn=OGDHL;refGene_tx
=XM_023240673.1
D2	49554100 49554228
refGene_gn=OGDHL;refGene_tx=XM_023240672.1;refGene_gn=OGDHL;refGene_tx
=XM_023240673.1
D2	49554708 49554859
refGene_gn=OGDHL;refGene_tx=XM_023240672.1;refGene_gn=OGDHL;refGene_tx
=XM_023240673.1
D2	49555819 49555948
refGene_gn=OGDHL;refGene_tx=XM_023240672.1;refGene_gn=OGDHL;refGene_tx
=XM_023240673.1

D2 49556465 49556568
refGene_gn=OGDHL;refGene_tx=XM_023240672.1;refGene_gn=OGDHL;refGene_tx=XM_023240673.1
D2 49557107 49557260
refGene_gn=OGDHL;refGene_tx=XM_023240672.1;refGene_gn=OGDHL;refGene_tx=XM_023240673.1
D2 49557561 49557741
refGene_gn=OGDHL;refGene_tx=XM_023240672.1;refGene_gn=OGDHL;refGene_tx=XM_023240673.1
D2 49559491 49559620
refGene_gn=OGDHL;refGene_tx=XM_023240672.1;refGene_gn=OGDHL;refGene_tx=XM_023240673.1
D2 49559829 49560009
refGene_gn=OGDHL;refGene_tx=XM_023240672.1;refGene_gn=OGDHL;refGene_tx=XM_023240673.1
D2 49561072 49561163
refGene_gn=OGDHL;refGene_tx=XM_023240672.1;refGene_gn=OGDHL;refGene_tx=XM_023240673.1
D2 49562976 49563123
refGene_gn=OGDHL;refGene_tx=XM_023240672.1;refGene_gn=OGDHL;refGene_tx=XM_023240673.1
D2 49563946 49564101
refGene_gn=OGDHL;refGene_tx=XM_023240672.1;refGene_gn=OGDHL;refGene_tx=XM_023240673.1
D2 49564248 49564364
refGene_gn=OGDHL;refGene_tx=XM_023240672.1;refGene_gn=OGDHL;refGene_tx=XM_023240673.1
D2 49564758 49564861
refGene_gn=OGDHL;refGene_tx=XM_023240672.1;refGene_gn=OGDHL;refGene_tx=XM_023240673.1
D2 49571581 49571740
refGene_gn=OGDHL;refGene_tx=XM_023240672.1;refGene_gn=OGDHL;refGene_tx=XM_023240673.1
D2 49574292 49574496
refGene_gn=OGDHL;refGene_tx=XM_023240672.1;refGene_gn=OGDHL;refGene_tx=XM_023240673.1
D2 49619379 49619534
refGene_gn=PARG;refGene_tx=XM_011287257.3;refGene_gn=PARG;refGene_tx=XM_019813506.2;refGene_gn=PARG;refGene_tx=XM_023240674.1;refGene_gn=PARG;refGene_tx=XM_023240675.1;refGene_gn=PARG;refGene_tx=XM_023240676.1;refGene_gn=PARG;refGene_tx=XM_023240677.1;refGene_gn=PARG;refGene_tx=XM_023240678.1
D2 49620218 49620347
refGene_gn=PARG;refGene_tx=XM_011287257.3;refGene_gn=PARG;refGene_tx=XM_019813506.2;refGene_gn=PARG;refGene_tx=XM_023240674.1;refGene_gn=PARG;refGene_tx=XM_023240675.1;refGene_gn=PARG;refGene_tx=XM_023240676.1;refGene_gn=PARG;refGene_tx=XM_023240677.1;refGene_gn=PARG;refGene_tx=XM_023240678.1
D2 49635171 49635277
refGene_gn=PARG;refGene_tx=XM_011287257.3;refGene_gn=PARG;refGene_tx=X

M_019813506.2;refGene_gn=PARG;refGene_tx=XM_023240674.1;refGene_gn=PARG;refGene_tx=XM_023240675.1;refGene_gn=PARG;refGene_tx=XM_023240676.1;refGene_gn=PARG;refGene_tx=XM_023240677.1;refGene_gn=PARG;refGene_tx=XM_023240678.1
D2 49645828 49645937 refGene_gn=PARG;refGene_tx=XM_011287257.3;refGene_gn=PARG;refGene_tx=XM_019813506.2;refGene_gn=PARG;refGene_tx=XM_023240674.1;refGene_gn=PARG;refGene_tx=XM_023240675.1;refGene_gn=PARG;refGene_tx=XM_023240676.1;refGene_gn=PARG;refGene_tx=XM_023240677.1;refGene_gn=PARG;refGene_tx=XM_023240678.1
D2 49648425 49648504 refGene_gn=PARG;refGene_tx=XM_011287257.3;refGene_gn=PARG;refGene_tx=XM_019813506.2;refGene_gn=PARG;refGene_tx=XM_023240674.1;refGene_gn=PARG;refGene_tx=XM_023240675.1;refGene_gn=PARG;refGene_tx=XM_023240676.1;refGene_gn=PARG;refGene_tx=XM_023240677.1;refGene_gn=PARG;refGene_tx=XM_023240678.1
D2 49663529 49663677 refGene_gn=PARG;refGene_tx=XM_011287257.3;refGene_gn=PARG;refGene_tx=XM_019813506.2;refGene_gn=PARG;refGene_tx=XM_023240674.1;refGene_gn=PARG;refGene_tx=XM_023240675.1;refGene_gn=PARG;refGene_tx=XM_023240676.1;refGene_gn=PARG;refGene_tx=XM_023240677.1;refGene_gn=PARG;refGene_tx=XM_023240678.1
D2 49669918 49669994 refGene_gn=PARG;refGene_tx=XM_011287257.3;refGene_gn=PARG;refGene_tx=XM_019813506.2;refGene_gn=PARG;refGene_tx=XM_023240674.1;refGene_gn=PARG;refGene_tx=XM_023240675.1;refGene_gn=PARG;refGene_tx=XM_023240676.1;refGene_gn=PARG;refGene_tx=XM_023240677.1;refGene_gn=PARG;refGene_tx=XM_023240678.1
D2 49674038 49674099 refGene_gn=PARG;refGene_tx=XM_011287257.3;refGene_gn=PARG;refGene_tx=XM_019813506.2;refGene_gn=PARG;refGene_tx=XM_023240674.1;refGene_gn=PARG;refGene_tx=XM_023240675.1;refGene_gn=PARG;refGene_tx=XM_023240676.1;refGene_gn=PARG;refGene_tx=XM_023240677.1;refGene_gn=PARG;refGene_tx=XM_023240678.1
D2 49678493 49678573 refGene_gn=PARG;refGene_tx=XM_011287257.3;refGene_gn=PARG;refGene_tx=XM_019813506.2;refGene_gn=PARG;refGene_tx=XM_023240674.1;refGene_gn=PARG;refGene_tx=XM_023240675.1;refGene_gn=PARG;refGene_tx=XM_023240676.1;refGene_gn=PARG;refGene_tx=XM_023240677.1;refGene_gn=PARG;refGene_tx=XM_023240678.1
D2 49687917 49688075 refGene_gn=PARG;refGene_tx=XM_011287257.3;refGene_gn=PARG;refGene_tx=XM_019813506.2;refGene_gn=PARG;refGene_tx=XM_023240674.1;refGene_gn=PARG;refGene_tx=XM_023240675.1;refGene_gn=PARG;refGene_tx=XM_023240676.1;refGene_gn=PARG;refGene_tx=XM_023240677.1;refGene_gn=PARG;refGene_tx=XM_023240678.1
D2 49691329 49691422 refGene_gn=PARG;refGene_tx=XM_011287257.3;refGene_gn=PARG;refGene_tx=XM_019813506.2;refGene_gn=PARG;refGene_tx=XM_023240674.1;refGene_gn=PARG;refGene_tx=XM_023240675.1;refGene_gn=PARG;refGene_tx=XM_023240676.1;

refGene_gn=PARG;refGene_tx=XM_023240677.1;refGene_gn=PARG;refGene_tx=XM_023240678.1
D2 49725324 49725399
refGene_gn=PARG;refGene_tx=XM_011287257.3;refGene_gn=PARG;refGene_tx=XM_019813506.2;refGene_gn=PARG;refGene_tx=XM_023240674.1;refGene_gn=PARG;refGene_tx=XM_023240675.1;refGene_gn=PARG;refGene_tx=XM_023240676.1;refGene_gn=PARG;refGene_tx=XM_023240677.1;refGene_gn=PARG;refGene_tx=XM_023240678.1
D2 49731295 49731379
refGene_gn=PARG;refGene_tx=XM_011287257.3;refGene_gn=PARG;refGene_tx=XM_019813506.2;refGene_gn=PARG;refGene_tx=XM_023240674.1;refGene_gn=PARG;refGene_tx=XM_023240675.1;refGene_gn=PARG;refGene_tx=XM_023240676.1;refGene_gn=PARG;refGene_tx=XM_023240677.1;refGene_gn=PARG;refGene_tx=XM_023240678.1
D2 49731495 49731618
refGene_gn=PARG;refGene_tx=XM_011287257.3;refGene_gn=PARG;refGene_tx=XM_019813506.2;refGene_gn=PARG;refGene_tx=XM_023240674.1;refGene_gn=PARG;refGene_tx=XM_023240675.1;refGene_gn=PARG;refGene_tx=XM_023240676.1;refGene_gn=PARG;refGene_tx=XM_023240677.1;refGene_gn=PARG;refGene_tx=XM_023240678.1
D2 49738023 49738207
refGene_gn=PARG;refGene_tx=XM_011287257.3;refGene_gn=PARG;refGene_tx=XM_019813506.2;refGene_gn=PARG;refGene_tx=XM_023240674.1;refGene_gn=PARG;refGene_tx=XM_023240675.1;refGene_gn=PARG;refGene_tx=XM_023240676.1;refGene_gn=PARG;refGene_tx=XM_023240677.1;refGene_gn=PARG;refGene_tx=XM_023240678.1
D2 49739520 49740504
refGene_gn=PARG;refGene_tx=XM_011287257.3;refGene_gn=PARG;refGene_tx=XM_019813506.2;refGene_gn=PARG;refGene_tx=XM_023240674.1;refGene_gn=PARG;refGene_tx=XM_023240675.1;refGene_gn=PARG;refGene_tx=XM_023240676.1;refGene_gn=PARG;refGene_tx=XM_023240677.1;refGene_gn=PARG;refGene_tx=XM_023240678.1
D2 49741119 49741186
refGene_gn=PARG;refGene_tx=XM_011287257.3;refGene_gn=PARG;refGene_tx=XM_019813506.2;refGene_gn=PARG;refGene_tx=XM_023240674.1;refGene_gn=PARG;refGene_tx=XM_023240675.1;refGene_gn=PARG;refGene_tx=XM_023240676.1
D2 49747865 49748085
refGene_gn=PARG;refGene_tx=XM_011287257.3;refGene_gn=PARG;refGene_tx=XM_019813506.2;refGene_gn=PARG;refGene_tx=XM_023240674.1
D2 49748562 49748668
refGene_gn=LOC101094135;refGene_tx=XM_003994168.5
D2 49750932 49750991
refGene_gn=LOC101094135;refGene_tx=XM_003994168.5;refGene_gn=LOC101094135;refGene_tx=XM_019813516.2
D2 49758304 49758398
refGene_gn=LOC101094135;refGene_tx=XM_003994168.5;refGene_gn=LOC101094135;refGene_tx=XM_019813516.2
D2 49758940 49759025
refGene_gn=LOC101094135;refGene_tx=XM_003994168.5;refGene_gn=LOC101094135;refGene_tx=XM_019813516.2

D2 49761543 49761602
refGene_gn=LOC101094135;refGene_tx=XM_003994168.5;refGene_gn=LOC101094135;refGene_tx=XM_019813516.2
D2 49763367 49763478
refGene_gn=LOC101094135;refGene_tx=XM_003994168.5;refGene_gn=LOC101094135;refGene_tx=XM_019813516.2
D2 49774200 49774316
refGene_gn=LOC101094135;refGene_tx=XM_003994168.5;refGene_gn=LOC101094135;refGene_tx=XM_019813516.2
D2 49777678 49777756
refGene_gn=NCOA4;refGene_tx=XM_011287251.3;refGene_gn=NCOA4;refGene_tx=XM_011287252.3;refGene_gn=NCOA4;refGene_tx=XM_011287255.3
D2 49778199 49779195
refGene_gn=NCOA4;refGene_tx=XM_011287251.3;refGene_gn=NCOA4;refGene_tx=XM_011287252.3;refGene_gn=NCOA4;refGene_tx=XM_011287255.3
D2 49780285 49780432
refGene_gn=NCOA4;refGene_tx=XM_011287251.3;refGene_gn=NCOA4;refGene_tx=XM_011287252.3
D2 49780945 49781038
refGene_gn=NCOA4;refGene_tx=XM_011287251.3;refGene_gn=NCOA4;refGene_tx=XM_011287252.3;refGene_gn=NCOA4;refGene_tx=XM_011287255.3
D2 49782842 49782951
refGene_gn=NCOA4;refGene_tx=XM_011287251.3;refGene_gn=NCOA4;refGene_tx=XM_011287252.3;refGene_gn=NCOA4;refGene_tx=XM_011287255.3
D2 49783419 49783508
refGene_gn=NCOA4;refGene_tx=XM_011287251.3;refGene_gn=NCOA4;refGene_tx=XM_011287252.3;refGene_gn=NCOA4;refGene_tx=XM_011287255.3
D2 49783684 49783825
refGene_gn=NCOA4;refGene_tx=XM_011287251.3;refGene_gn=NCOA4;refGene_tx=XM_011287252.3;refGene_gn=NCOA4;refGene_tx=XM_011287255.3
D2 49785076 49785231
refGene_gn=NCOA4;refGene_tx=XM_011287251.3;refGene_gn=NCOA4;refGene_tx=XM_011287252.3;refGene_gn=NCOA4;refGene_tx=XM_011287255.3
D2 49791656 49791666
refGene_gn=NCOA4;refGene_tx=XM_011287251.3;refGene_gn=NCOA4;refGene_tx=XM_011287252.3;refGene_gn=NCOA4;refGene_tx=XM_011287255.3
D2 49802985 49803170 refGene_gn=MSMB;refGene_tx=XM_011287389.2
D2 49807842 49807987 refGene_gn=MSMB;refGene_tx=XM_011287389.2
D2 49837097 49837100
refGene_gn=LOC101096765;refGene_tx=XM_019813511.2;refGene_gn=LOC101096765;refGene_tx=XM_019813512.2;refGene_gn=LOC101096765;refGene_tx=XM_019813513.2;refGene_gn=LOC101096765;refGene_tx=XM_019813514.2
D2 49837191 49837314
refGene_gn=LOC101096765;refGene_tx=XM_019813511.2;refGene_gn=LOC101096765;refGene_tx=XM_019813512.2;refGene_gn=LOC101096765;refGene_tx=XM_019813513.2;refGene_gn=LOC101096765;refGene_tx=XM_019813514.2
D2 49838104 49838269
refGene_gn=LOC101096765;refGene_tx=XM_019813511.2;refGene_gn=LOC101096765;refGene_tx=XM_019813512.2;refGene_gn=LOC101096765;refGene_tx=XM_019813513.2;refGene_gn=LOC101096765;refGene_tx=XM_019813514.2

D2 49846860 49846923
refGene_gn=LOC101096765;refGene_tx=XM_019813511.2;refGene_gn=LOC101096765;refGene_tx=XM_019813512.2;refGene_gn=LOC101096765;refGene_tx=XM_019813513.2;refGene_gn=LOC101096765;refGene_tx=XM_019813514.2
D2 49849302 49849476
refGene_gn=LOC101096765;refGene_tx=XM_019813511.2;refGene_gn=LOC101096765;refGene_tx=XM_019813512.2;refGene_gn=LOC101096765;refGene_tx=XM_019813513.2;refGene_gn=LOC101096765;refGene_tx=XM_019813514.2
D2 49851881 49851975
refGene_gn=LOC101096765;refGene_tx=XM_019813511.2;refGene_gn=LOC101096765;refGene_tx=XM_019813512.2;refGene_gn=LOC101096765;refGene_tx=XM_019813513.2;refGene_gn=LOC101096765;refGene_tx=XM_019813514.2
D2 49852754 49852813
refGene_gn=LOC101096765;refGene_tx=XM_019813511.2;refGene_gn=LOC101096765;refGene_tx=XM_019813512.2;refGene_gn=LOC101096765;refGene_tx=XM_019813513.2;refGene_gn=LOC101096765;refGene_tx=XM_019813514.2
D2 49854019 49854067
refGene_gn=LOC101096765;refGene_tx=XM_019813511.2;refGene_gn=LOC101096765;refGene_tx=XM_019813512.2;refGene_gn=LOC101096765;refGene_tx=XM_019813513.2;refGene_gn=LOC101096765;refGene_tx=XM_019813514.2
D2 49856730 49856841
refGene_gn=LOC101096765;refGene_tx=XM_019813511.2;refGene_gn=LOC101096765;refGene_tx=XM_019813512.2;refGene_gn=LOC101096765;refGene_tx=XM_019813513.2;refGene_gn=LOC101096765;refGene_tx=XM_019813514.2
D2 49857609 49857691
refGene_gn=LOC101096765;refGene_tx=XM_019813511.2;refGene_gn=LOC101096765;refGene_tx=XM_019813512.2;refGene_gn=LOC101096765;refGene_tx=XM_019813513.2;refGene_gn=LOC101096765;refGene_tx=XM_019813514.2
D2 49858237 49858309
refGene_gn=LOC101096765;refGene_tx=XM_019813511.2;refGene_gn=LOC101096765;refGene_tx=XM_019813512.2;refGene_gn=LOC101096765;refGene_tx=XM_019813513.2;refGene_gn=LOC101096765;refGene_tx=XM_019813514.2
D2 49859018 49859137
refGene_gn=LOC101096765;refGene_tx=XM_019813511.2;refGene_gn=LOC101096765;refGene_tx=XM_019813512.2;refGene_gn=LOC101096765;refGene_tx=XM_019813513.2;refGene_gn=LOC101096765;refGene_tx=XM_019813514.2
D2 49859588 49859646
refGene_gn=LOC101096765;refGene_tx=XM_019813511.2;refGene_gn=LOC101096765;refGene_tx=XM_019813512.2;refGene_gn=LOC101096765;refGene_tx=XM_019813513.2;refGene_gn=LOC101096765;refGene_tx=XM_019813514.2
D2 49861149 49861209
refGene_gn=LOC101096765;refGene_tx=XM_019813511.2;refGene_gn=LOC101096765;refGene_tx=XM_019813512.2;refGene_gn=LOC101096765;refGene_tx=XM_019813513.2;refGene_gn=LOC101096765;refGene_tx=XM_019813514.2
D2 49861592 49861766
refGene_gn=LOC101096765;refGene_tx=XM_019813511.2;refGene_gn=LOC101096765;refGene_tx=XM_019813512.2;refGene_gn=LOC101096765;refGene_tx=XM_019813513.2;refGene_gn=LOC101096765;refGene_tx=XM_019813514.2
D2 49864451 49864579
refGene_gn=LOC101096765;refGene_tx=XM_019813511.2;refGene_gn=LOC101096

765;refGene_tx=XM_019813512.2;refGene_gn=LOC101096765;refGene_tx=XM_019813513.2;refGene_gn=LOC101096765;refGene_tx=XM_019813514.2
D2 49866127 49866214
refGene_gn=LOC101096765;refGene_tx=XM_019813511.2;refGene_gn=LOC101096765;refGene_tx=XM_019813512.2;refGene_gn=LOC101096765;refGene_tx=XM_019813513.2;refGene_gn=LOC101096765;refGene_tx=XM_019813514.2
D2 49869412 49869514
refGene_gn=LOC101096765;refGene_tx=XM_019813511.2;refGene_gn=LOC101096765;refGene_tx=XM_019813512.2;refGene_gn=LOC101096765;refGene_tx=XM_019813513.2;refGene_gn=LOC101096765;refGene_tx=XM_019813514.2
D2 49870919 49871051
refGene_gn=LOC101096765;refGene_tx=XM_019813511.2;refGene_gn=LOC101096765;refGene_tx=XM_019813512.2;refGene_gn=LOC101096765;refGene_tx=XM_019813513.2;refGene_gn=LOC101096765;refGene_tx=XM_019813514.2
D2 49874277 49874447
refGene_gn=LOC101096765;refGene_tx=XM_019813511.2;refGene_gn=LOC101096765;refGene_tx=XM_019813512.2;refGene_gn=LOC101096765;refGene_tx=XM_019813513.2;refGene_gn=LOC101096765;refGene_tx=XM_019813514.2
D2 49878811 49878914
refGene_gn=LOC101096765;refGene_tx=XM_019813511.2;refGene_gn=LOC101096765;refGene_tx=XM_019813512.2;refGene_gn=LOC101096765;refGene_tx=XM_019813513.2;refGene_gn=LOC101096765;refGene_tx=XM_019813514.2
D2 49884127 49884280
refGene_gn=LOC101096765;refGene_tx=XM_019813511.2;refGene_gn=LOC101096765;refGene_tx=XM_019813512.2;refGene_gn=LOC101096765;refGene_tx=XM_019813513.2;refGene_gn=LOC101096765;refGene_tx=XM_019813514.2
D2 49887128 49887311
refGene_gn=LOC101096765;refGene_tx=XM_019813511.2;refGene_gn=LOC101096765;refGene_tx=XM_019813512.2;refGene_gn=LOC101096765;refGene_tx=XM_019813513.2;refGene_gn=LOC101096765;refGene_tx=XM_019813514.2
D2 49890345 49890474
refGene_gn=LOC101096765;refGene_tx=XM_019813511.2;refGene_gn=LOC101096765;refGene_tx=XM_019813512.2;refGene_gn=LOC101096765;refGene_tx=XM_019813513.2;refGene_gn=LOC101096765;refGene_tx=XM_019813514.2
D2 49890604 49890685
refGene_gn=LOC101096765;refGene_tx=XM_019813511.2;refGene_gn=LOC101096765;refGene_tx=XM_019813512.2;refGene_gn=LOC101096765;refGene_tx=XM_019813513.2;refGene_gn=LOC101096765;refGene_tx=XM_019813514.2
D2 49891294 49891420
refGene_gn=LOC101096765;refGene_tx=XM_019813511.2;refGene_gn=LOC101096765;refGene_tx=XM_019813512.2;refGene_gn=LOC101096765;refGene_tx=XM_019813513.2;refGene_gn=LOC101096765;refGene_tx=XM_019813514.2
D2 49892815 49892878
refGene_gn=LOC101096765;refGene_tx=XM_019813511.2;refGene_gn=LOC101096765;refGene_tx=XM_019813512.2
D2 49893234 49893447
refGene_gn=LOC101096765;refGene_tx=XM_019813511.2;refGene_gn=LOC101096765;refGene_tx=XM_019813512.2;refGene_gn=LOC101096765;refGene_tx=XM_019813513.2
D2 49895122 49895740

refGene_gn=LOC101096765;refGene_tx=XM_019813511.2;refGene_gn=LOC101096765;refGene_tx=XM_019813512.2;refGene_gn=LOC101096765;refGene_tx=XM_019813513.2;refGene_gn=LOC101096765;refGene_tx=XM_019813514.2
D2 49896537 49896715
refGene_gn=LOC101096765;refGene_tx=XM_019813511.2;refGene_gn=LOC101096765;refGene_tx=XM_019813512.2;refGene_gn=LOC101096765;refGene_tx=XM_019813513.2;refGene_gn=LOC101096765;refGene_tx=XM_019813514.2
D2 49898481 49898621
refGene_gn=LOC101096765;refGene_tx=XM_019813511.2;refGene_gn=LOC101096765;refGene_tx=XM_019813512.2;refGene_gn=LOC101096765;refGene_tx=XM_019813513.2;refGene_gn=LOC101096765;refGene_tx=XM_019813514.2
D2 49910993 49911255
refGene_gn=ZFAND4;refGene_tx=XM_011287241.2;refGene_gn=ZFAND4;refGene_tx=XM_011287243.2;refGene_gn=ZFAND4;refGene_tx=XM_011287244.3;refGene_gn=ZFAND4;refGene_tx=XM_011287245.3;refGene_gn=ZFAND4;refGene_tx=XM_019813517.2
D2 49954127 49954203
refGene_gn=ZFAND4;refGene_tx=XM_011287241.2;refGene_gn=ZFAND4;refGene_tx=XM_011287243.2;refGene_gn=ZFAND4;refGene_tx=XM_011287244.3;refGene_gn=ZFAND4;refGene_tx=XM_011287245.3;refGene_gn=ZFAND4;refGene_tx=XM_019813517.2
D2 49957449 49957517
refGene_gn=ZFAND4;refGene_tx=XM_011287241.2;refGene_gn=ZFAND4;refGene_tx=XM_011287243.2;refGene_gn=ZFAND4;refGene_tx=XM_011287244.3;refGene_gn=ZFAND4;refGene_tx=XM_011287245.3;refGene_gn=ZFAND4;refGene_tx=XM_019813517.2;refGene_gn=ZFAND4;refGene_tx=XM_019813518.2
D2 49960875 49961062
refGene_gn=ZFAND4;refGene_tx=XM_011287241.2;refGene_gn=ZFAND4;refGene_tx=XM_011287243.2;refGene_gn=ZFAND4;refGene_tx=XM_011287244.3;refGene_gn=ZFAND4;refGene_tx=XM_011287245.3;refGene_gn=ZFAND4;refGene_tx=XM_019813517.2;refGene_gn=ZFAND4;refGene_tx=XM_019813518.2
D2 49969694 49969842
refGene_gn=ZFAND4;refGene_tx=XM_011287241.2;refGene_gn=ZFAND4;refGene_tx=XM_011287243.2;refGene_gn=ZFAND4;refGene_tx=XM_011287244.3;refGene_gn=ZFAND4;refGene_tx=XM_011287245.3;refGene_gn=ZFAND4;refGene_tx=XM_019813517.2;refGene_gn=ZFAND4;refGene_tx=XM_019813518.2
D2 49980621 49981797
refGene_gn=ZFAND4;refGene_tx=XM_011287241.2;refGene_gn=ZFAND4;refGene_tx=XM_011287243.2;refGene_gn=ZFAND4;refGene_tx=XM_011287244.3;refGene_gn=ZFAND4;refGene_tx=XM_011287245.3;refGene_gn=ZFAND4;refGene_tx=XM_019813517.2;refGene_gn=ZFAND4;refGene_tx=XM_019813518.2
D2 49982518 49982595
refGene_gn=ZFAND4;refGene_tx=XM_011287241.2;refGene_gn=ZFAND4;refGene_tx=XM_011287243.2;refGene_gn=ZFAND4;refGene_tx=XM_011287244.3;refGene_gn=ZFAND4;refGene_tx=XM_011287245.3;refGene_gn=ZFAND4;refGene_tx=XM_019813517.2;refGene_gn=ZFAND4;refGene_tx=XM_019813518.2
D2 49987940 49988062
refGene_gn=ZFAND4;refGene_tx=XM_011287241.2;refGene_gn=ZFAND4;refGene_tx=XM_011287243.2;refGene_gn=ZFAND4;refGene_tx=XM_011287244.3;refGene_gn=ZFAND4;refGene_tx=XM_011287245.3;refGene_gn=ZFAND4;refGene_tx=XM_01

9813518.2
D2	49989030	49989166
refGene_gn=ZFAND4;refGene_tx=XM_011287241.2;refGene_gn=ZFAND4;refGene_tx=XM_011287243.2;refGene_gn=ZFAND4;refGene_tx=XM_011287244.3;refGene_gn=ZFAND4;refGene_tx=XM_011287245.3;refGene_gn=ZFAND4;refGene_tx=XM_019813518.2
D2	50045920	50046022
refGene_gn=MARCH8;refGene_tx=XM_006937981.3;refGene_gn=MARCH8;refGene_tx=XM_006937983.3;refGene_gn=MARCH8;refGene_tx=XM_011287239.2;refGene_gn=MARCH8;refGene_tx=XM_019813519.1;refGene_gn=MARCH8;refGene_tx=XM_019813520.2;refGene_gn=MARCH8;refGene_tx=XM_019813521.2;refGene_gn=MARCH8;refGene_tx=XM_019813523.1
D2	50075980	50075992	refGene_gn=MARCH8;refGene_tx=XM_019813524.2
D2	50098616	50098667
refGene_gn=MARCH8;refGene_tx=XM_006937981.3;refGene_gn=MARCH8;refGene_tx=XM_006937982.2;refGene_gn=MARCH8;refGene_tx=XM_006937983.3;refGene_gn=MARCH8;refGene_tx=XM_011287239.2;refGene_gn=MARCH8;refGene_tx=XM_019813519.1;refGene_gn=MARCH8;refGene_tx=XM_019813520.2;refGene_gn=MARCH8;refGene_tx=XM_019813521.2;refGene_gn=MARCH8;refGene_tx=XM_019813523.1;refGene_gn=MARCH8;refGene_tx=XM_019813524.2;refGene_gn=MARCH8;refGene_tx=XM_023240679.1
D2	50112978	50113061
refGene_gn=MARCH8;refGene_tx=XM_006937981.3;refGene_gn=MARCH8;refGene_tx=XM_006937982.2;refGene_gn=MARCH8;refGene_tx=XM_006937983.3;refGene_gn=MARCH8;refGene_tx=XM_011287239.2;refGene_gn=MARCH8;refGene_tx=XM_019813519.1;refGene_gn=MARCH8;refGene_tx=XM_019813520.2;refGene_gn=MARCH8;refGene_tx=XM_019813521.2;refGene_gn=MARCH8;refGene_tx=XM_019813523.1;refGene_gn=MARCH8;refGene_tx=XM_019813524.2;refGene_gn=MARCH8;refGene_tx=XM_023240679.1
D2	50113308	50114148
refGene_gn=MARCH8;refGene_tx=XM_006937981.3;refGene_gn=MARCH8;refGene_tx=XM_006937982.2;refGene_gn=MARCH8;refGene_tx=XM_011287239.2;refGene_gn=MARCH8;refGene_tx=XM_019813519.1;refGene_gn=MARCH8;refGene_tx=XM_019813520.2;refGene_gn=MARCH8;refGene_tx=XM_019813521.2;refGene_gn=MARCH8;refGene_tx=XM_019813523.1;refGene_gn=MARCH8;refGene_tx=XM_019813524.2;refGene_gn=MARCH8;refGene_tx=XM_023240679.1
D2	50115570	50115751
refGene_gn=MARCH8;refGene_tx=XM_006937981.3;refGene_gn=MARCH8;refGene_tx=XM_006937982.2;refGene_gn=MARCH8;refGene_tx=XM_006937983.3;refGene_gn=MARCH8;refGene_tx=XM_011287239.2;refGene_gn=MARCH8;refGene_tx=XM_019813519.1;refGene_gn=MARCH8;refGene_tx=XM_019813520.2;refGene_gn=MARCH8;refGene_tx=XM_019813521.2;refGene_gn=MARCH8;refGene_tx=XM_019813523.1;refGene_gn=MARCH8;refGene_tx=XM_019813524.2;refGene_gn=MARCH8;refGene_tx=XM_023240679.1
D2	50117724	50117872
refGene_gn=MARCH8;refGene_tx=XM_006937981.3;refGene_gn=MARCH8;refGene_tx=XM_006937982.2;refGene_gn=MARCH8;refGene_tx=XM_006937983.3;refGene_gn=MARCH8;refGene_tx=XM_011287239.2;refGene_gn=MARCH8;refGene_tx=XM_019813519.1;refGene_gn=MARCH8;refGene_tx=XM_019813520.2;refGene_gn=MARCH8;refGene_tx=XM_019813521.2;refGene_gn=MARCH8;refGene_tx=XM_019813523.

1;refGene_gn=MARCH8;refGene_tx=XM_019813524.2;refGene_gn=MARCH8;refGene_tx=XM_023240679.1
D2 50118287 50118592
refGene_gn=MARCH8;refGene_tx=XM_006937981.3;refGene_gn=MARCH8;refGene_tx=XM_006937982.2;refGene_gn=MARCH8;refGene_tx=XM_006937983.3;refGene_gn=MARCH8;refGene_tx=XM_011287239.2;refGene_gn=MARCH8;refGene_tx=XM_019813519.1;refGene_gn=MARCH8;refGene_tx=XM_019813520.2;refGene_gn=MARCH8;refGene_tx=XM_019813521.2;refGene_gn=MARCH8;refGene_tx=XM_019813523.1;refGene_gn=MARCH8;refGene_tx=XM_019813524.2;refGene_gn=MARCH8;refGene_tx=XM_023240679.1
D2 50130156 50130336
refGene_gn=ALOX5;refGene_tx=XM_003994181.5;refGene_gn=ALOX5;refGene_tx=XM_019813525.2
D2 50131458 50131629
refGene_gn=ALOX5;refGene_tx=XM_003994181.5;refGene_gn=ALOX5;refGene_tx=XM_019813525.2
D2 50131956 50132057
refGene_gn=ALOX5;refGene_tx=XM_003994181.5;refGene_gn=ALOX5;refGene_tx=XM_019813525.2
D2 50132413 50132535
refGene_gn=ALOX5;refGene_tx=XM_003994181.5;refGene_gn=ALOX5;refGene_tx=XM_019813525.2
D2 50132821 50133000
refGene_gn=ALOX5;refGene_tx=XM_003994181.5;refGene_gn=ALOX5;refGene_tx=XM_019813525.2
D2 50134702 50134789
refGene_gn=ALOX5;refGene_tx=XM_003994181.5;refGene_gn=ALOX5;refGene_tx=XM_019813525.2
D2 50135528 50135732
refGene_gn=ALOX5;refGene_tx=XM_003994181.5;refGene_gn=ALOX5;refGene_tx=XM_019813525.2
D2 50142014 50142161
refGene_gn=ALOX5;refGene_tx=XM_003994181.5;refGene_gn=ALOX5;refGene_tx=XM_019813525.2
D2 50145234 50145407
refGene_gn=ALOX5;refGene_tx=XM_003994181.5;refGene_gn=ALOX5;refGene_tx=XM_019813525.2
D2 50147184 50147291
refGene_gn=ALOX5;refGene_tx=XM_003994181.5;refGene_gn=ALOX5;refGene_tx=XM_019813525.2
D2 50156910 50157033
refGene_gn=ALOX5;refGene_tx=XM_003994181.5;refGene_gn=ALOX5;refGene_tx=XM_019813525.2
D2 50167215 50167297 refGene_gn=ALOX5;refGene_tx=XM_003994181.5
D2 50172080 50172279 refGene_gn=ALOX5;refGene_tx=XM_003994181.5
D2 50178982 50179132 refGene_gn=ALOX5;refGene_tx=XM_003994181.5
D2 50243548 50244508
refGene_gn=LOC109496957;refGene_tx=XM_023240680.1;refGene_gn=LOC109496957;refGene_tx=XM_023240681.1
D2 50245344 50246331

refGene_gn=LOC111557100;refGene_tx=XM_023240682.1
D2 50273629 50274667
refGene_gn=LOC111557101;refGene_tx=XM_023240683.1
D2 50284779 50285741
refGene_gn=LOC105260014;refGene_tx=XM_023240684.1;refGene_gn=LOC105260014;refGene_tx=XM_023240685.1;refGene_gn=LOC105260014;refGene_tx=XM_023240686.1
D2 50295053 50295120
refGene_gn=LOC105260014;refGene_tx=XM_023240684.1;refGene_gn=LOC105260014;refGene_tx=XM_023240685.1;refGene_gn=LOC105260014;refGene_tx=XM_023240686.1
D2 50296500 50297463
refGene_gn=LOC109496939;refGene_tx=XM_019824705.2
D2 50338849 50339812
refGene_gn=LOC109492874;refGene_tx=XM_023240311.1
D2 50365503 50366472
refGene_gn=LOC109492827;refGene_tx=XM_019814073.2
D2 50401802 50402480
refGene_gn=ZNF22;refGene_tx=XM_003994185.3;refGene_gn=ZNF22;refGene_tx=XM_023240694.1
D2 50410577 50410688
refGene_gn=RASSF4;refGene_tx=XM_003994186.5;refGene_gn=RASSF4;refGene_tx=XM_023240692.1
D2 50411978 50412076 refGene_gn=RASSF4;refGene_tx=XM_003994186.5
D2 50412925 50413047
refGene_gn=RASSF4;refGene_tx=XM_003994186.5;refGene_gn=RASSF4;refGene_tx=XM_023240692.1
D2 50413562 50413614
refGene_gn=RASSF4;refGene_tx=XM_003994186.5;refGene_gn=RASSF4;refGene_tx=XM_023240692.1
D2 50413899 50414001
refGene_gn=RASSF4;refGene_tx=XM_003994186.5;refGene_gn=RASSF4;refGene_tx=XM_023240692.1
D2 50417507 50417665
refGene_gn=RASSF4;refGene_tx=XM_003994186.5;refGene_gn=RASSF4;refGene_tx=XM_023240692.1
D2 50418354 50418446
refGene_gn=RASSF4;refGene_tx=XM_003994186.5;refGene_gn=RASSF4;refGene_tx=XM_023240692.1
D2 50419754 50419897
refGene_gn=RASSF4;refGene_tx=XM_003994186.5;refGene_gn=RASSF4;refGene_tx=XM_023240692.1
D2 50423117 50423744 refGene_gn=DEPP1;refGene_tx=XM_003994187.5
D2 50429217 50429293
refGene_gn=RASSF4;refGene_tx=XM_003994186.5;refGene_gn=RASSF4;refGene_tx=XM_023240692.1
D2 50431716 50431778
refGene_gn=RASSF4;refGene_tx=XM_003994186.5;refGene_gn=RASSF4;refGene_tx=XM_023240692.1
D2 50461820 50462299 refGene_gn=TMEM72;refGene_tx=XM_003994188.4

D2 50463675 50463815 refGene_gn=TMEM72;refGene_tx=XM_003994188.4
D2 50465364 50465436 refGene_gn=TMEM72;refGene_tx=XM_003994188.4
D2 50469346 50469413 refGene_gn=TMEM72;refGene_tx=XM_003994188.4
D2 50487046 50487116 refGene_gn=TMEM72;refGene_tx=XM_003994188.4
D2 50847843 50847904
refGene_gn=CXCL12;refGene_tx=NM_001009847.1;refGene_gn=CXCL12;refGene_tx=XM_006937984.4
D2 50851940 50852058
refGene_gn=CXCL12;refGene_tx=NM_001009847.1;refGene_gn=CXCL12;refGene_tx=XM_006937984.4
D2 50854171 50854258
refGene_gn=CXCL12;refGene_tx=NM_001009847.1;refGene_gn=CXCL12;refGene_tx=XM_006937984.4
D2 50856945 50857039 refGene_gn=CXCL12;refGene_tx=XM_006937984.4
D2 50859549 50859565 refGene_gn=CXCL12;refGene_tx=NM_001009847.1
D2 51451115 51451185
refGene_gn=ZNF32;refGene_tx=XM_003994189.5;refGene_gn=ZNF32;refGene_tx=XM_006937987.4;refGene_gn=ZNF32;refGene_tx=XM_023240693.1
D2 51451415 51451458 refGene_gn=ZNF32;refGene_tx=XM_019814177.1
D2 51452346 51453140
refGene_gn=ZNF32;refGene_tx=XM_003994189.5;refGene_gn=ZNF32;refGene_tx=XM_006937987.4;refGene_gn=ZNF32;refGene_tx=XM_006937988.4;refGene_gn=ZNF32;refGene_tx=XM_019814177.1;refGene_gn=ZNF32;refGene_tx=XM_019814178.2;refGene_gn=ZNF32;refGene_tx=XM_023240693.1
D2 51461152 51462234 refGene_gn=ZNF485;refGene_tx=XM_019814165.2
D2 51467691 51467787 refGene_gn=ZNF485;refGene_tx=XM_019814165.2
D2 51468251 51468378 refGene_gn=ZNF485;refGene_tx=XM_019814165.2
D2 51470378 51470411 refGene_gn=ZNF485;refGene_tx=XM_019814165.2
D2 51497731 51498988
refGene_gn=ZNF239;refGene_tx=XM_019814169.2;refGene_gn=ZNF239;refGene_tx=XM_019814170.2
D2 51607124 51607427
refGene_gn=LOC111557102;refGene_tx=XM_023240697.1
D2 51628982 51630227
refGene_gn=HNRNPF;refGene_tx=XM_003994191.5;refGene_gn=HNRNPF;refGene_tx=XM_003994192.5;refGene_gn=HNRNPF;refGene_tx=XM_006937989.4;refGene_gn=HNRNPF;refGene_tx=XM_006937990.4;refGene_gn=HNRNPF;refGene_tx=XM_006937991.4;refGene_gn=HNRNPF;refGene_tx=XM_019814172.2;refGene_gn=HNRNPF;refGene_tx=XM_019814173.2;refGene_gn=HNRNPF;refGene_tx=XM_023240687.1;refGene_gn=HNRNPF;refGene_tx=XM_023240688.1;refGene_gn=HNRNPF;refGene_tx=XM_023240689.1;refGene_gn=HNRNPF;refGene_tx=XM_023240690.1;refGene_gn=HNRNPF;refGene_tx=XM_023240691.1
D2 51642310 51642330 refGene_gn=FXYD4;refGene_tx=XM_023240696.1
D2 51642467 51642567
refGene_gn=FXYD4;refGene_tx=XM_023240695.1;refGene_gn=FXYD4;refGene_tx=XM_023240696.1
D2 51642987 51643024
refGene_gn=FXYD4;refGene_tx=XM_023240695.1;refGene_gn=FXYD4;refGene_tx=XM_023240696.1
D2 51643133 51643208

refGene_gn=FXYD4;refGene_tx=XM_023240695.1;refGene_gn=FXYD4;refGene_tx=XM_023240696.1
D2 51643889 51643913
refGene_gn=FXYD4;refGene_tx=XM_023240695.1;refGene_gn=FXYD4;refGene_tx=XM_023240696.1
D2 51644054 51644087
refGene_gn=FXYD4;refGene_tx=XM_023240695.1;refGene_gn=FXYD4;refGene_tx=XM_023240696.1
D2 51644913 51644953
refGene_gn=FXYD4;refGene_tx=XM_023240695.1;refGene_gn=FXYD4;refGene_tx=XM_023240696.1
D2 51790784 51790802 refGene_gn=RASGEF1A;refGene_tx=XM_003994193.5
D2 51813339 51813540
refGene_gn=RASGEF1A;refGene_tx=XM_003994193.5;refGene_gn=RASGEF1A;refGene_tx=XM_011287411.3
D2 51816026 51816149
refGene_gn=RASGEF1A;refGene_tx=XM_003994193.5;refGene_gn=RASGEF1A;refGene_tx=XM_011287411.3
D2 51818299 51818437
refGene_gn=RASGEF1A;refGene_tx=XM_003994193.5;refGene_gn=RASGEF1A;refGene_tx=XM_011287411.3
D2 51819307 51819529
refGene_gn=RASGEF1A;refGene_tx=XM_003994193.5;refGene_gn=RASGEF1A;refGene_tx=XM_011287411.3
D2 51820080 51820155
refGene_gn=RASGEF1A;refGene_tx=XM_003994193.5;refGene_gn=RASGEF1A;refGene_tx=XM_011287411.3
D2 51820499 51820592
refGene_gn=RASGEF1A;refGene_tx=XM_003994193.5;refGene_gn=RASGEF1A;refGene_tx=XM_011287411.3
D2 51821052 51821155
refGene_gn=RASGEF1A;refGene_tx=XM_003994193.5;refGene_gn=RASGEF1A;refGene_tx=XM_011287411.3
D2 51821242 51821322
refGene_gn=RASGEF1A;refGene_tx=XM_003994193.5;refGene_gn=RASGEF1A;refGene_tx=XM_011287411.3
D2 51822065 51822257
refGene_gn=RASGEF1A;refGene_tx=XM_003994193.5;refGene_gn=RASGEF1A;refGene_tx=XM_011287411.3
D2 51823067 51823191
refGene_gn=RASGEF1A;refGene_tx=XM_003994193.5;refGene_gn=RASGEF1A;refGene_tx=XM_011287411.3
D2 51823597 51823670
refGene_gn=RASGEF1A;refGene_tx=XM_003994193.5;refGene_gn=RASGEF1A;refGene_tx=XM_011287411.3
D2 51823975 51824000
refGene_gn=RASGEF1A;refGene_tx=XM_003994193.5;refGene_gn=RASGEF1A;refGene_tx=XM_011287411.3
D2 51833670 51833963
refGene_gn=CSGALNACT2;refGene_tx=XM_003994194.5;refGene_gn=CSGALNACT2;

refGene_tx=XM_019814155.2;refGene_gn=CSGALNACT2;refGene_tx=XM_019814156.2;refGene_gn=CSGALNACT2;refGene_tx=XM_019814159.2
D2 51835111 51835172
refGene_gn=CSGALNACT2;refGene_tx=XM_019814155.2;refGene_gn=CSGALNACT2;refGene_tx=XM_019814156.2;refGene_gn=CSGALNACT2;refGene_tx=XM_019814157.2;refGene_gn=CSGALNACT2;refGene_tx=XM_019814158.2;refGene_gn=CSGALNACT2;refGene_tx=XM_019814160.2
D2 51835327 51835397
refGene_gn=CSGALNACT2;refGene_tx=XM_019814155.2;refGene_gn=CSGALNACT2;refGene_tx=XM_019814156.2;refGene_gn=CSGALNACT2;refGene_tx=XM_019814157.2
D2 51839103 51839185
refGene_gn=CSGALNACT2;refGene_tx=XM_003994194.5;refGene_gn=CSGALNACT2;refGene_tx=XM_019814155.2;refGene_gn=CSGALNACT2;refGene_tx=XM_019814156.2;refGene_gn=CSGALNACT2;refGene_tx=XM_019814157.2;refGene_gn=CSGALNACT2;refGene_tx=XM_019814158.2;refGene_gn=CSGALNACT2;refGene_tx=XM_019814159.2;refGene_gn=CSGALNACT2;refGene_tx=XM_019814160.2
D2 51849613 51849708
refGene_gn=CSGALNACT2;refGene_tx=XM_003994194.5;refGene_gn=CSGALNACT2;refGene_tx=XM_019814157.2;refGene_gn=CSGALNACT2;refGene_tx=XM_019814158.2
D2 51851659 51851838
refGene_gn=CSGALNACT2;refGene_tx=XM_003994194.5;refGene_gn=CSGALNACT2;refGene_tx=XM_019814155.2;refGene_gn=CSGALNACT2;refGene_tx=XM_019814156.2;refGene_gn=CSGALNACT2;refGene_tx=XM_019814157.2;refGene_gn=CSGALNACT2;refGene_tx=XM_019814158.2;refGene_gn=CSGALNACT2;refGene_tx=XM_019814159.2;refGene_gn=CSGALNACT2;refGene_tx=XM_019814160.2;refGene_gn=CSGALNACT2;refGene_tx=XM_019814164.2
D2 51853354 51853460
refGene_gn=CSGALNACT2;refGene_tx=XM_019814164.2;refGene_gn=CSGALNACT2;refGene_tx=XM_023240698.1
D2 51854148 51854209
refGene_gn=CSGALNACT2;refGene_tx=XM_019814162.2;refGene_gn=CSGALNACT2;refGene_tx=XM_019814163.2
D2 51855104 51855206
refGene_gn=CSGALNACT2;refGene_tx=XM_003994194.5;refGene_gn=CSGALNACT2;refGene_tx=XM_019814155.2;refGene_gn=CSGALNACT2;refGene_tx=XM_019814157.2;refGene_gn=CSGALNACT2;refGene_tx=XM_019814158.2;refGene_gn=CSGALNACT2;refGene_tx=XM_019814159.2;refGene_gn=CSGALNACT2;refGene_tx=XM_019814160.2;refGene_gn=CSGALNACT2;refGene_tx=XM_019814162.2;refGene_gn=CSGALNACT2;refGene_tx=XM_019814163.2;refGene_gn=CSGALNACT2;refGene_tx=XM_019814164.2;refGene_gn=CSGALNACT2;refGene_tx=XM_023240698.1
D2 51856552 51856769
refGene_gn=CSGALNACT2;refGene_tx=XM_003994194.5;refGene_gn=CSGALNACT2;refGene_tx=XM_019814155.2;refGene_gn=CSGALNACT2;refGene_tx=XM_019814156.2;refGene_gn=CSGALNACT2;refGene_tx=XM_019814157.2;refGene_gn=CSGALNACT2;refGene_tx=XM_019814158.2;refGene_gn=CSGALNACT2;refGene_tx=XM_019814159.2;refGene_gn=CSGALNACT2;refGene_tx=XM_019814160.2;refGene_gn=CSGALNACT2;refGene_tx=XM_019814162.2;refGene_gn=CSGALNACT2;refGene_tx=XM_019814163.2;refGene_gn=CSGALNACT2;refGene_tx=XM_019814164.2;refGene_gn

=CSGALNACT2;refGene_tx=XM_023240698.1
D2 51859512 51860173
refGene_gn=CSGALNACT2;refGene_tx=XM_003994194.5;refGene_gn=CSGALNACT2;refGene_tx=XM_019814155.2;refGene_gn=CSGALNACT2;refGene_tx=XM_019814156.2;refGene_gn=CSGALNACT2;refGene_tx=XM_019814157.2;refGene_gn=CSGALNACT2;refGene_tx=XM_019814158.2;refGene_gn=CSGALNACT2;refGene_tx=XM_019814159.2;refGene_gn=CSGALNACT2;refGene_tx=XM_019814160.2;refGene_gn=CSGALNACT2;refGene_tx=XM_019814162.2;refGene_gn=CSGALNACT2;refGene_tx=XM_019814163.2;refGene_gn=CSGALNACT2;refGene_tx=XM_019814164.2;refGene_gn=CSGALNACT2;refGene_tx=XM_023240698.1
D2 51888960 51889118 refGene_gn=RET;refGene_tx=XM_023240699.1
D2 51890469 51890617 refGene_gn=RET;refGene_tx=XM_023240699.1
D2 51892573 51892673 refGene_gn=RET;refGene_tx=XM_023240699.1
D2 51893778 51893916 refGene_gn=RET;refGene_tx=XM_023240699.1
D2 51895218 51895289 refGene_gn=RET;refGene_tx=XM_023240699.1
D2 51897522 51897645 refGene_gn=RET;refGene_tx=XM_023240699.1
D2 51897727 51897942 refGene_gn=RET;refGene_tx=XM_023240699.1
D2 51898927 51899035 refGene_gn=RET;refGene_tx=XM_023240699.1
D2 51900477 51900625 refGene_gn=RET;refGene_tx=XM_023240699.1
D2 51903577 51903834 refGene_gn=RET;refGene_tx=XM_023240699.1
D2 51904712 51904835 refGene_gn=RET;refGene_tx=XM_023240699.1
D2 51905413 51905524 refGene_gn=RET;refGene_tx=XM_023240699.1
D2 51906012 51906138 refGene_gn=RET;refGene_tx=XM_023240699.1
D2 51906871 51907130 refGene_gn=RET;refGene_tx=XM_023240699.1
D2 51909160 51909360 refGene_gn=RET;refGene_tx=XM_023240699.1
D2 51910723 51910919 refGene_gn=RET;refGene_tx=XM_023240699.1
D2 51912135 51912380 refGene_gn=RET;refGene_tx=XM_023240699.1
D2 51914414 51914702 refGene_gn=RET;refGene_tx=XM_023240699.1
D2 51916459 51916726 refGene_gn=RET;refGene_tx=XM_023240699.1
D2 51939989 51940062 refGene_gn=RET;refGene_tx=XM_023240699.1
D2 51986758 51987207
refGene_gn=LOC109492829;refGene_tx=XM_019814075.1
D2 51987369 51987463
refGene_gn=LOC109492829;refGene_tx=XM_019814075.1
D2 51987819 51988060
refGene_gn=LOC109492829;refGene_tx=XM_019814075.1
D2 51988522 51988643
refGene_gn=LOC109492829;refGene_tx=XM_019814075.1
D2 51989781 51989794
refGene_gn=LOC109492829;refGene_tx=XM_019814075.1
D2 52132970 52133201
refGene_gn=BMS1;refGene_tx=XM_011287413.3;refGene_gn=BMS1;refGene_tx=XM_023240700.1;refGene_gn=BMS1;refGene_tx=XM_023240701.1;refGene_gn=BMS1;refGene_tx=XM_023240702.1
D2 52133722 52133884
refGene_gn=BMS1;refGene_tx=XM_011287413.3;refGene_gn=BMS1;refGene_tx=XM_023240700.1;refGene_gn=BMS1;refGene_tx=XM_023240701.1;refGene_gn=BMS1;refGene_tx=XM_023240702.1
D2 52136163 52136339
refGene_gn=BMS1;refGene_tx=XM_011287413.3;refGene_gn=BMS1;refGene_tx=X

M_023240700.1;refGene_gn=BMS1;refGene_tx=XM_023240701.1;refGene_gn=BMS1;refGene_tx=XM_023240702.1
D2 52136681 52136829 refGene_gn=BMS1;refGene_tx=XM_011287413.3;refGene_gn=BMS1;refGene_tx=XM_023240700.1;refGene_gn=BMS1;refGene_tx=XM_023240701.1;refGene_gn=BMS1;refGene_tx=XM_023240702.1
D2 52137759 52137882 refGene_gn=BMS1;refGene_tx=XM_011287413.3;refGene_gn=BMS1;refGene_tx=XM_023240700.1;refGene_gn=BMS1;refGene_tx=XM_023240701.1;refGene_gn=BMS1;refGene_tx=XM_023240702.1
D2 52138657 52138716 refGene_gn=BMS1;refGene_tx=XM_011287413.3;refGene_gn=BMS1;refGene_tx=XM_023240700.1;refGene_gn=BMS1;refGene_tx=XM_023240701.1;refGene_gn=BMS1;refGene_tx=XM_023240702.1
D2 52138908 52139089 refGene_gn=BMS1;refGene_tx=XM_011287413.3;refGene_gn=BMS1;refGene_tx=XM_023240700.1;refGene_gn=BMS1;refGene_tx=XM_023240701.1;refGene_gn=BMS1;refGene_tx=XM_023240702.1
D2 52139183 52139372 refGene_gn=BMS1;refGene_tx=XM_011287413.3;refGene_gn=BMS1;refGene_tx=XM_023240700.1;refGene_gn=BMS1;refGene_tx=XM_023240701.1;refGene_gn=BMS1;refGene_tx=XM_023240702.1
D2 52141904 52142078 refGene_gn=BMS1;refGene_tx=XM_011287413.3;refGene_gn=BMS1;refGene_tx=XM_023240700.1;refGene_gn=BMS1;refGene_tx=XM_023240701.1;refGene_gn=BMS1;refGene_tx=XM_023240702.1
D2 52143183 52143257 refGene_gn=BMS1;refGene_tx=XM_011287413.3;refGene_gn=BMS1;refGene_tx=XM_023240700.1;refGene_gn=BMS1;refGene_tx=XM_023240701.1;refGene_gn=BMS1;refGene_tx=XM_023240702.1
D2 52158593 52158675 refGene_gn=BMS1;refGene_tx=XM_011287413.3;refGene_gn=BMS1;refGene_tx=XM_023240700.1;refGene_gn=BMS1;refGene_tx=XM_023240701.1;refGene_gn=BMS1;refGene_tx=XM_023240702.1
D2 52164629 52164787 refGene_gn=BMS1;refGene_tx=XM_011287413.3;refGene_gn=BMS1;refGene_tx=XM_023240700.1;refGene_gn=BMS1;refGene_tx=XM_023240701.1;refGene_gn=BMS1;refGene_tx=XM_023240702.1
D2 52165492 52165594 refGene_gn=BMS1;refGene_tx=XM_011287413.3;refGene_gn=BMS1;refGene_tx=XM_023240700.1;refGene_gn=BMS1;refGene_tx=XM_023240701.1;refGene_gn=BMS1;refGene_tx=XM_023240702.1
D2 52166017 52166784 refGene_gn=BMS1;refGene_tx=XM_011287413.3;refGene_gn=BMS1;refGene_tx=XM_023240700.1;refGene_gn=BMS1;refGene_tx=XM_023240701.1;refGene_gn=BMS1;refGene_tx=XM_023240702.1
D2 52168324 52168464 refGene_gn=BMS1;refGene_tx=XM_011287413.3;refGene_gn=BMS1;refGene_tx=XM_023240700.1;refGene_gn=BMS1;refGene_tx=XM_023240701.1;refGene_gn=BMS1;refGene_tx=XM_023240702.1

D2 52170021 52170215
refGene_gn=BMS1;refGene_tx=XM_011287413.3;refGene_gn=BMS1;refGene_tx=XM_023240700.1;refGene_gn=BMS1;refGene_tx=XM_023240701.1;refGene_gn=BMS1;refGene_tx=XM_023240702.1
D2 52170532 52170654
refGene_gn=BMS1;refGene_tx=XM_011287413.3;refGene_gn=BMS1;refGene_tx=XM_023240700.1;refGene_gn=BMS1;refGene_tx=XM_023240701.1;refGene_gn=BMS1;refGene_tx=XM_023240702.1
D2 52171396 52171539
refGene_gn=BMS1;refGene_tx=XM_011287413.3;refGene_gn=BMS1;refGene_tx=XM_023240700.1;refGene_gn=BMS1;refGene_tx=XM_023240701.1;refGene_gn=BMS1;refGene_tx=XM_023240702.1
D2 52172177 52172366
refGene_gn=BMS1;refGene_tx=XM_011287413.3;refGene_gn=BMS1;refGene_tx=XM_023240700.1;refGene_gn=BMS1;refGene_tx=XM_023240701.1;refGene_gn=BMS1;refGene_tx=XM_023240702.1
D2 52176218 52176298
refGene_gn=BMS1;refGene_tx=XM_011287413.3;refGene_gn=BMS1;refGene_tx=XM_023240700.1;refGene_gn=BMS1;refGene_tx=XM_023240701.1;refGene_gn=BMS1;refGene_tx=XM_023240702.1
D2 52177945 52178136
refGene_gn=BMS1;refGene_tx=XM_011287413.3;refGene_gn=BMS1;refGene_tx=XM_023240700.1;refGene_gn=BMS1;refGene_tx=XM_023240701.1;refGene_gn=BMS1;refGene_tx=XM_023240702.1
D2 52179055 52179231
refGene_gn=BMS1;refGene_tx=XM_011287413.3;refGene_gn=BMS1;refGene_tx=XM_023240700.1;refGene_gn=BMS1;refGene_tx=XM_023240701.1;refGene_gn=BMS1;refGene_tx=XM_023240702.1
D2 52203521 52204315 refGene_gn=ZNF37A;refGene_tx=XM_023240703.1
D2 52204317 52204410 refGene_gn=ZNF37A;refGene_tx=XM_023240703.1
D2 52204412 52204974 refGene_gn=ZNF37A;refGene_tx=XM_023240703.1
D2 52265027 52265032 refGene_gn=IFIT5;refGene_tx=XM_006937998.4
D2 52275010 52276454 refGene_gn=IFIT5;refGene_tx=XM_006937998.4
D2 52296315 52296578 refGene_gn=SLC16A12;refGene_tx=XM_023240704.1
D2 52298922 52299182 refGene_gn=SLC16A12;refGene_tx=XM_023240704.1
D2 52301130 52301710 refGene_gn=SLC16A12;refGene_tx=XM_023240704.1
D2 52304088 52304232 refGene_gn=SLC16A12;refGene_tx=XM_023240704.1
D2 52306931 52307035 refGene_gn=SLC16A12;refGene_tx=XM_023240704.1
D2 52316472 52316588 refGene_gn=SLC16A12;refGene_tx=XM_023240704.1
D2 52316589 52316646 refGene_gn=SLC16A12;refGene_tx=XM_023240704.1
D2 52316647 52316677 refGene_gn=SLC16A12;refGene_tx=XM_023240704.1
D2 52429711 52429771
refGene_gn=PANK1;refGene_tx=XM_019813527.2;refGene_gn=PANK1;refGene_tx=XM_019813529.2;refGene_gn=PANK1;refGene_tx=XM_019813530.2;refGene_gn=PANK1;refGene_tx=XM_019813531.2;refGene_gn=PANK1;refGene_tx=XM_019813532.2;refGene_gn=PANK1;refGene_tx=XM_023240708.1;refGene_gn=PANK1;refGene_tx=XM_023240709.1;refGene_gn=PANK1;refGene_tx=XM_023240710.1
D2 52434036 52434162
refGene_gn=PANK1;refGene_tx=XM_019813527.2;refGene_gn=PANK1;refGene_tx=XM_019813529.2;refGene_gn=PANK1;refGene_tx=XM_019813530.2;refGene_gn=

PANK1;refGene_tx=XM_019813531.2;refGene_gn=PANK1;refGene_tx=XM_019813532.2;refGene_gn=PANK1;refGene_tx=XM_023240708.1;refGene_gn=PANK1;refGene_tx=XM_023240709.1;refGene_gn=PANK1;refGene_tx=XM_023240710.1
D2 52438191 52438315
refGene_gn=PANK1;refGene_tx=XM_019813527.2;refGene_gn=PANK1;refGene_tx=XM_019813529.2;refGene_gn=PANK1;refGene_tx=XM_019813530.2;refGene_gn=PANK1;refGene_tx=XM_019813531.2;refGene_gn=PANK1;refGene_tx=XM_019813532.2;refGene_gn=PANK1;refGene_tx=XM_023240708.1;refGene_gn=PANK1;refGene_tx=XM_023240709.1;refGene_gn=PANK1;refGene_tx=XM_023240710.1
D2 52438828 52439005
refGene_gn=PANK1;refGene_tx=XM_019813527.2;refGene_gn=PANK1;refGene_tx=XM_019813529.2;refGene_gn=PANK1;refGene_tx=XM_019813530.2;refGene_gn=PANK1;refGene_tx=XM_019813531.2;refGene_gn=PANK1;refGene_tx=XM_019813532.2;refGene_gn=PANK1;refGene_tx=XM_023240708.1;refGene_gn=PANK1;refGene_tx=XM_023240709.1
D2 52443934 52444188
refGene_gn=PANK1;refGene_tx=XM_019813527.2;refGene_gn=PANK1;refGene_tx=XM_019813529.2;refGene_gn=PANK1;refGene_tx=XM_019813530.2;refGene_gn=PANK1;refGene_tx=XM_019813531.2;refGene_gn=PANK1;refGene_tx=XM_019813532.2;refGene_gn=PANK1;refGene_tx=XM_023240708.1;refGene_gn=PANK1;refGene_tx=XM_023240709.1;refGene_gn=PANK1;refGene_tx=XM_023240710.1
D2 52454080 52454433
refGene_gn=PANK1;refGene_tx=XM_019813527.2;refGene_gn=PANK1;refGene_tx=XM_019813529.2;refGene_gn=PANK1;refGene_tx=XM_019813530.2;refGene_gn=PANK1;refGene_tx=XM_019813531.2;refGene_gn=PANK1;refGene_tx=XM_019813532.2;refGene_gn=PANK1;refGene_tx=XM_023240708.1;refGene_gn=PANK1;refGene_tx=XM_023240709.1;refGene_gn=PANK1;refGene_tx=XM_023240710.1
D2 52469742 52469830 refGene_gn=PANK1;refGene_tx=XM_019813527.2
D2 52480052 52480080 refGene_gn=PANK1;refGene_tx=XM_019813529.2
D2 52480967 52481490
refGene_gn=PANK1;refGene_tx=XM_023240708.1;refGene_gn=PANK1;refGene_tx=XM_023240709.1;refGene_gn=PANK1;refGene_tx=XM_023240710.1
D2 52533662 52533809
refGene_gn=KIF20B;refGene_tx=XM_023240705.1;refGene_gn=KIF20B;refGene_tx=XM_023240706.1;refGene_gn=KIF20B;refGene_tx=XM_023240707.1
D2 52538669 52538756
refGene_gn=KIF20B;refGene_tx=XM_023240705.1;refGene_gn=KIF20B;refGene_tx=XM_023240706.1;refGene_gn=KIF20B;refGene_tx=XM_023240707.1
D2 52538871 52538988
refGene_gn=KIF20B;refGene_tx=XM_023240705.1;refGene_gn=KIF20B;refGene_tx=XM_023240706.1;refGene_gn=KIF20B;refGene_tx=XM_023240707.1
D2 52539617 52539756
refGene_gn=KIF20B;refGene_tx=XM_023240705.1;refGene_gn=KIF20B;refGene_tx=XM_023240706.1;refGene_gn=KIF20B;refGene_tx=XM_023240707.1
D2 52540309 52540494
refGene_gn=KIF20B;refGene_tx=XM_023240705.1;refGene_gn=KIF20B;refGene_tx=XM_023240706.1;refGene_gn=KIF20B;refGene_tx=XM_023240707.1
D2 52542152 52542189
refGene_gn=KIF20B;refGene_tx=XM_023240705.1;refGene_gn=KIF20B;refGene_tx=XM_023240706.1;refGene_gn=KIF20B;refGene_tx=XM_023240707.1

D2 52543379 52543607
refGene_gn=KIF20B;refGene_tx=XM_023240705.1;refGene_gn=KIF20B;refGene_tx=XM_023240706.1;refGene_gn=KIF20B;refGene_tx=XM_023240707.1
D2 52544767 52544879
refGene_gn=KIF20B;refGene_tx=XM_023240705.1;refGene_gn=KIF20B;refGene_tx=XM_023240706.1;refGene_gn=KIF20B;refGene_tx=XM_023240707.1
D2 52545720 52545792
refGene_gn=KIF20B;refGene_tx=XM_023240705.1;refGene_gn=KIF20B;refGene_tx=XM_023240706.1;refGene_gn=KIF20B;refGene_tx=XM_023240707.1
D2 52545892 52546039
refGene_gn=KIF20B;refGene_tx=XM_023240705.1;refGene_gn=KIF20B;refGene_tx=XM_023240706.1;refGene_gn=KIF20B;refGene_tx=XM_023240707.1
D2 52546719 52546882
refGene_gn=KIF20B;refGene_tx=XM_023240705.1;refGene_gn=KIF20B;refGene_tx=XM_023240706.1;refGene_gn=KIF20B;refGene_tx=XM_023240707.1
D2 52547584 52547875
refGene_gn=KIF20B;refGene_tx=XM_023240705.1;refGene_gn=KIF20B;refGene_tx=XM_023240706.1;refGene_gn=KIF20B;refGene_tx=XM_023240707.1
D2 52551271 52551411
refGene_gn=KIF20B;refGene_tx=XM_023240705.1;refGene_gn=KIF20B;refGene_tx=XM_023240706.1;refGene_gn=KIF20B;refGene_tx=XM_023240707.1
D2 52552407 52552546
refGene_gn=KIF20B;refGene_tx=XM_023240705.1;refGene_gn=KIF20B;refGene_tx=XM_023240706.1;refGene_gn=KIF20B;refGene_tx=XM_023240707.1
D2 52553431 52553660
refGene_gn=KIF20B;refGene_tx=XM_023240705.1;refGene_gn=KIF20B;refGene_tx=XM_023240706.1;refGene_gn=KIF20B;refGene_tx=XM_023240707.1
D2 52554818 52554871
refGene_gn=KIF20B;refGene_tx=XM_023240705.1;refGene_gn=KIF20B;refGene_tx=XM_023240706.1;refGene_gn=KIF20B;refGene_tx=XM_023240707.1
D2 52556223 52556343
refGene_gn=KIF20B;refGene_tx=XM_023240705.1;refGene_gn=KIF20B;refGene_tx=XM_023240706.1;refGene_gn=KIF20B;refGene_tx=XM_023240707.1
D2 52560250 52560404
refGene_gn=KIF20B;refGene_tx=XM_023240705.1;refGene_gn=KIF20B;refGene_tx=XM_023240706.1;refGene_gn=KIF20B;refGene_tx=XM_023240707.1
D2 52563755 52564989
refGene_gn=KIF20B;refGene_tx=XM_023240705.1;refGene_gn=KIF20B;refGene_tx=XM_023240706.1;refGene_gn=KIF20B;refGene_tx=XM_023240707.1
D2 52565324 52565463
refGene_gn=KIF20B;refGene_tx=XM_023240705.1;refGene_gn=KIF20B;refGene_tx=XM_023240706.1;refGene_gn=KIF20B;refGene_tx=XM_023240707.1
D2 52572050 52572170
refGene_gn=KIF20B;refGene_tx=XM_023240705.1;refGene_gn=KIF20B;refGene_tx=XM_023240706.1;refGene_gn=KIF20B;refGene_tx=XM_023240707.1
D2 52574124 52574185
refGene_gn=KIF20B;refGene_tx=XM_023240705.1;refGene_gn=KIF20B;refGene_tx=XM_023240706.1;refGene_gn=KIF20B;refGene_tx=XM_023240707.1
D2 52583046 52583172
refGene_gn=KIF20B;refGene_tx=XM_023240705.1;refGene_gn=KIF20B;refGene_

tx=XM_023240706.1;refGene_gn=KIF20B;refGene_tx=XM_023240707.1
D2 52583831 52583959
refGene_gn=KIF20B;refGene_tx=XM_023240705.1;refGene_gn=KIF20B;refGene_
tx=XM_023240706.1;refGene_gn=KIF20B;refGene_tx=XM_023240707.1
D2 52585608 52585764
refGene_gn=KIF20B;refGene_tx=XM_023240705.1;refGene_gn=KIF20B;refGene_
tx=XM_023240706.1;refGene_gn=KIF20B;refGene_tx=XM_023240707.1
D2 52589563 52589740
refGene_gn=KIF20B;refGene_tx=XM_023240705.1;refGene_gn=KIF20B;refGene_
tx=XM_023240706.1;refGene_gn=KIF20B;refGene_tx=XM_023240707.1
D2 52591359 52591470
refGene_gn=KIF20B;refGene_tx=XM_023240705.1;refGene_gn=KIF20B;refGene_
tx=XM_023240706.1;refGene_gn=KIF20B;refGene_tx=XM_023240707.1
D2 52593973 52594171
refGene_gn=KIF20B;refGene_tx=XM_023240705.1;refGene_gn=KIF20B;refGene_
tx=XM_023240706.1;refGene_gn=KIF20B;refGene_tx=XM_023240707.1
D2 52596470 52596572
refGene_gn=KIF20B;refGene_tx=XM_023240705.1;refGene_gn=KIF20B;refGene_
tx=XM_023240706.1;refGene_gn=KIF20B;refGene_tx=XM_023240707.1
D2 52596969 52597120
refGene_gn=KIF20B;refGene_tx=XM_023240705.1;refGene_gn=KIF20B;refGene_
tx=XM_023240706.1;refGene_gn=KIF20B;refGene_tx=XM_023240707.1
D2 52601470 52601613
refGene_gn=KIF20B;refGene_tx=XM_023240705.1;refGene_gn=KIF20B;refGene_
tx=XM_023240706.1;refGene_gn=KIF20B;refGene_tx=XM_023240707.1
D2 52603193 52603271
refGene_gn=KIF20B;refGene_tx=XM_023240705.1;refGene_gn=KIF20B;refGene_
tx=XM_023240706.1;refGene_gn=KIF20B;refGene_tx=XM_023240707.1
D2 53301413 53301438 refGene_gn=HTR7;refGene_tx=XM_023240712.1
D2 53306425 53306468 refGene_gn=HTR7;refGene_tx=XM_023240711.1
D2 53312691 53313454
refGene_gn=HTR7;refGene_tx=XM_019814076.2;refGene_gn=HTR7;refGene_tx=X
M_023240711.1;refGene_gn=HTR7;refGene_tx=XM_023240712.1
D2 53388307 53388852
refGene_gn=HTR7;refGene_tx=XM_019814076.2;refGene_gn=HTR7;refGene_tx=X
M_023240711.1;refGene_gn=HTR7;refGene_tx=XM_023240712.1
D2 53400495 53400577 refGene_gn=RPP30;refGene_tx=XM_003994205.4
D2 53401999 53402055 refGene_gn=RPP30;refGene_tx=XM_003994205.4
D2 53402664 53402721 refGene_gn=RPP30;refGene_tx=XM_003994205.4
D2 53403368 53403443 refGene_gn=RPP30;refGene_tx=XM_003994205.4
D2 53406707 53406779 refGene_gn=RPP30;refGene_tx=XM_003994205.4
D2 53413144 53413234 refGene_gn=RPP30;refGene_tx=XM_003994205.4
D2 53422048 53422165 refGene_gn=RPP30;refGene_tx=XM_003994205.4
D2 53422740 53422770 refGene_gn=RPP30;refGene_tx=XM_003994205.4
D2 53423164 53423202 refGene_gn=RPP30;refGene_tx=XM_003994205.4
D2 53423584 53423664 refGene_gn=RPP30;refGene_tx=XM_003994205.4
D2 53426826 53426936 refGene_gn=RPP30;refGene_tx=XM_003994205.4
D2 53435696 53435807 refGene_gn=ANKRD1;refGene_tx=XM_003994206.3
D2 53438351 53438450 refGene_gn=ANKRD1;refGene_tx=XM_003994206.3
D2 53438588 53438687 refGene_gn=ANKRD1;refGene_tx=XM_003994206.3

D2 53438895 53438994 refGene_gn=ANKRD1;refGene_tx=XM_003994206.3
D2 53440591 53440690 refGene_gn=ANKRD1;refGene_tx=XM_003994206.3
D2 53442162 53442270 refGene_gn=ANKRD1;refGene_tx=XM_003994206.3
D2 53442399 53442537 refGene_gn=ANKRD1;refGene_tx=XM_003994206.3
D2 53443427 53443607 refGene_gn=ANKRD1;refGene_tx=XM_003994206.3
D2 53444287 53444314 refGene_gn=ANKRD1;refGene_tx=XM_003994206.3
D2 53699813 53699925
refGene_gn=PCGF5;refGene_tx=XM_019813534.2;refGene_gn=PCGF5;refGene_tx=XM_023240713.1;refGene_gn=PCGF5;refGene_tx=XM_023240714.1;refGene_gn=PCGF5;refGene_tx=XM_023240715.1;refGene_gn=PCGF5;refGene_tx=XM_023240716.1;refGene_gn=PCGF5;refGene_tx=XM_023240717.1
D2 53717957 53718054
refGene_gn=PCGF5;refGene_tx=XM_019813534.2;refGene_gn=PCGF5;refGene_tx=XM_023240713.1;refGene_gn=PCGF5;refGene_tx=XM_023240714.1;refGene_gn=PCGF5;refGene_tx=XM_023240715.1;refGene_gn=PCGF5;refGene_tx=XM_023240716.1;refGene_gn=PCGF5;refGene_tx=XM_023240717.1
D2 53726262 53726318
refGene_gn=PCGF5;refGene_tx=XM_019813534.2;refGene_gn=PCGF5;refGene_tx=XM_023240713.1;refGene_gn=PCGF5;refGene_tx=XM_023240714.1;refGene_gn=PCGF5;refGene_tx=XM_023240715.1;refGene_gn=PCGF5;refGene_tx=XM_023240716.1;refGene_gn=PCGF5;refGene_tx=XM_023240717.1
D2 53726432 53726492
refGene_gn=PCGF5;refGene_tx=XM_019813534.2;refGene_gn=PCGF5;refGene_tx=XM_023240713.1;refGene_gn=PCGF5;refGene_tx=XM_023240714.1;refGene_gn=PCGF5;refGene_tx=XM_023240715.1;refGene_gn=PCGF5;refGene_tx=XM_023240716.1;refGene_gn=PCGF5;refGene_tx=XM_023240717.1
D2 53728980 53729129
refGene_gn=PCGF5;refGene_tx=XM_019813534.2;refGene_gn=PCGF5;refGene_tx=XM_023240713.1;refGene_gn=PCGF5;refGene_tx=XM_023240714.1;refGene_gn=PCGF5;refGene_tx=XM_023240715.1;refGene_gn=PCGF5;refGene_tx=XM_023240716.1;refGene_gn=PCGF5;refGene_tx=XM_023240717.1
D2 53747610 53747709
refGene_gn=PCGF5;refGene_tx=XM_019813534.2;refGene_gn=PCGF5;refGene_tx=XM_023240713.1;refGene_gn=PCGF5;refGene_tx=XM_023240714.1;refGene_gn=PCGF5;refGene_tx=XM_023240715.1;refGene_gn=PCGF5;refGene_tx=XM_023240716.1;refGene_gn=PCGF5;refGene_tx=XM_023240717.1
D2 53750576 53750666
refGene_gn=PCGF5;refGene_tx=XM_019813534.2;refGene_gn=PCGF5;refGene_tx=XM_023240713.1;refGene_gn=PCGF5;refGene_tx=XM_023240714.1;refGene_gn=PCGF5;refGene_tx=XM_023240715.1;refGene_gn=PCGF5;refGene_tx=XM_023240716.1;refGene_gn=PCGF5;refGene_tx=XM_023240717.1
D2 53757959 53758019
refGene_gn=PCGF5;refGene_tx=XM_019813534.2;refGene_gn=PCGF5;refGene_tx=XM_023240713.1;refGene_gn=PCGF5;refGene_tx=XM_023240714.1;refGene_gn=PCGF5;refGene_tx=XM_023240716.1
D2 53758390 53758555
refGene_gn=PCGF5;refGene_tx=XM_019813534.2;refGene_gn=PCGF5;refGene_tx=XM_023240713.1;refGene_gn=PCGF5;refGene_tx=XM_023240714.1
D2 53763858 53763906
refGene_gn=PCGF5;refGene_tx=XM_023240716.1;refGene_gn=PCGF5;refGene_tx

=XM_023240717.1
D2	53766637	53766763	refGene_gn=PCGF5;refGene_tx=XM_023240715.1
D2	53883305	53883443	refGene_gn=HECTD2;refGene_tx=XM_023240718.1;refGene_gn=HECTD2;refGene_tx=XM_023240719.1;refGene_gn=HECTD2;refGene_tx=XM_023240720.1
D2	53902347	53902477	refGene_gn=HECTD2;refGene_tx=XM_023240718.1;refGene_gn=HECTD2;refGene_tx=XM_023240719.1;refGene_gn=HECTD2;refGene_tx=XM_023240720.1
D2	53919065	53919204	refGene_gn=HECTD2;refGene_tx=XM_023240718.1;refGene_gn=HECTD2;refGene_tx=XM_023240719.1;refGene_gn=HECTD2;refGene_tx=XM_023240720.1
D2	53919908	53920011	refGene_gn=HECTD2;refGene_tx=XM_023240718.1;refGene_gn=HECTD2;refGene_tx=XM_023240719.1;refGene_gn=HECTD2;refGene_tx=XM_023240720.1
D2	53920670	53920760	refGene_gn=HECTD2;refGene_tx=XM_023240718.1;refGene_gn=HECTD2;refGene_tx=XM_023240719.1;refGene_gn=HECTD2;refGene_tx=XM_023240720.1
D2	53922809	53922874	refGene_gn=HECTD2;refGene_tx=XM_023240718.1;refGene_gn=HECTD2;refGene_tx=XM_023240719.1;refGene_gn=HECTD2;refGene_tx=XM_023240720.1
D2	53926574	53926620	refGene_gn=HECTD2;refGene_tx=XM_023240718.1;refGene_gn=HECTD2;refGene_tx=XM_023240719.1;refGene_gn=HECTD2;refGene_tx=XM_023240720.1
D2	53928314	53928424	refGene_gn=HECTD2;refGene_tx=XM_023240718.1;refGene_gn=HECTD2;refGene_tx=XM_023240719.1;refGene_gn=HECTD2;refGene_tx=XM_023240720.1
D2	53929844	53929993	refGene_gn=HECTD2;refGene_tx=XM_023240718.1;refGene_gn=HECTD2;refGene_tx=XM_023240719.1;refGene_gn=HECTD2;refGene_tx=XM_023240720.1
D2	53930788	53930912	refGene_gn=HECTD2;refGene_tx=XM_023240718.1;refGene_gn=HECTD2;refGene_tx=XM_023240719.1;refGene_gn=HECTD2;refGene_tx=XM_023240720.1
D2	53933193	53933290	refGene_gn=HECTD2;refGene_tx=XM_023240718.1;refGene_gn=HECTD2;refGene_tx=XM_023240719.1;refGene_gn=HECTD2;refGene_tx=XM_023240720.1
D2	53937117	53937225	refGene_gn=HECTD2;refGene_tx=XM_023240718.1;refGene_gn=HECTD2;refGene_tx=XM_023240719.1;refGene_gn=HECTD2;refGene_tx=XM_023240720.1
D2	53938084	53938217	refGene_gn=HECTD2;refGene_tx=XM_023240718.1;refGene_gn=HECTD2;refGene_tx=XM_023240719.1;refGene_gn=HECTD2;refGene_tx=XM_023240720.1
D2	53939139	53939228	refGene_gn=HECTD2;refGene_tx=XM_023240718.1;refGene_gn=HECTD2;refGene_tx=XM_023240719.1;refGene_gn=HECTD2;refGene_tx=XM_023240720.1
D2	53942072	53942090	refGene_gn=HECTD2;refGene_tx=XM_023240719.1
D2	53942932	53943100	refGene_gn=HECTD2;refGene_tx=XM_023240718.1;refGene_gn=HECTD2;refGene_tx=XM_023240720.1
D2	53946214	53946289	refGene_gn=HECTD2;refGene_tx=XM_023240718.1
D2	53947375	53947463	refGene_gn=HECTD2;refGene_tx=XM_023240718.1

D2 53947549 53947656 refGene_gn=HECTD2;refGene_tx=XM_023240718.1
D2 53948932 53949048 refGene_gn=HECTD2;refGene_tx=XM_023240718.1
D2 53956864 53957008 refGene_gn=HECTD2;refGene_tx=XM_023240718.1
D2 53958782 53958903 refGene_gn=HECTD2;refGene_tx=XM_023240718.1
D2 54052377 54053317 refGene_gn=PPP1R3C;refGene_tx=XM_003994208.4
D2 54055429 54055443 refGene_gn=PPP1R3C;refGene_tx=XM_003994208.4
D2 54167854 54168053
refGene_gn=TNKS2;refGene_tx=XM_019814078.2;refGene_gn=TNKS2;refGene_tx
=XM_023240721.1;refGene_gn=TNKS2;refGene_tx=XM_023240722.1
D2 54184402 54184627
refGene_gn=TNKS2;refGene_tx=XM_019814078.2;refGene_gn=TNKS2;refGene_tx
=XM_023240721.1;refGene_gn=TNKS2;refGene_tx=XM_023240722.1
D2 54187107 54187203
refGene_gn=TNKS2;refGene_tx=XM_019814078.2;refGene_gn=TNKS2;refGene_tx
=XM_023240721.1;refGene_gn=TNKS2;refGene_tx=XM_023240722.1
D2 54190393 54190430
refGene_gn=TNKS2;refGene_tx=XM_019814078.2;refGene_gn=TNKS2;refGene_tx
=XM_023240721.1;refGene_gn=TNKS2;refGene_tx=XM_023240722.1
D2 54190598 54190674
refGene_gn=TNKS2;refGene_tx=XM_019814078.2;refGene_gn=TNKS2;refGene_tx
=XM_023240721.1;refGene_gn=TNKS2;refGene_tx=XM_023240722.1
D2 54191258 54191353
refGene_gn=TNKS2;refGene_tx=XM_019814078.2;refGene_gn=TNKS2;refGene_tx
=XM_023240721.1;refGene_gn=TNKS2;refGene_tx=XM_023240722.1
D2 54193491 54193558
refGene_gn=TNKS2;refGene_tx=XM_019814078.2;refGene_gn=TNKS2;refGene_tx
=XM_023240721.1;refGene_gn=TNKS2;refGene_tx=XM_023240722.1
D2 54195458 54195645
refGene_gn=TNKS2;refGene_tx=XM_019814078.2;refGene_gn=TNKS2;refGene_tx
=XM_023240721.1;refGene_gn=TNKS2;refGene_tx=XM_023240722.1
D2 54196638 54196760
refGene_gn=TNKS2;refGene_tx=XM_019814078.2;refGene_gn=TNKS2;refGene_tx
=XM_023240721.1;refGene_gn=TNKS2;refGene_tx=XM_023240722.1
D2 54199132 54199224
refGene_gn=TNKS2;refGene_tx=XM_019814078.2;refGene_gn=TNKS2;refGene_tx
=XM_023240721.1
D2 54199311 54199390
refGene_gn=TNKS2;refGene_tx=XM_019814078.2;refGene_gn=TNKS2;refGene_tx
=XM_023240721.1
D2 54202012 54202184
refGene_gn=TNKS2;refGene_tx=XM_019814078.2;refGene_gn=TNKS2;refGene_tx
=XM_023240721.1;refGene_gn=TNKS2;refGene_tx=XM_023240722.1
D2 54204403 54204483
refGene_gn=TNKS2;refGene_tx=XM_019814078.2;refGene_gn=TNKS2;refGene_tx
=XM_023240721.1;refGene_gn=TNKS2;refGene_tx=XM_023240722.1
D2 54208582 54208728
refGene_gn=TNKS2;refGene_tx=XM_019814078.2;refGene_gn=TNKS2;refGene_tx
=XM_023240721.1;refGene_gn=TNKS2;refGene_tx=XM_023240722.1
D2 54210867 54211033
refGene_gn=TNKS2;refGene_tx=XM_019814078.2;refGene_gn=TNKS2;refGene_tx

=XM_023240721.1;refGene_gn=TNKS2;refGene_tx=XM_023240722.1
D2 54211908 54212128
refGene_gn=TNKS2;refGene_tx=XM_019814078.2;refGene_gn=TNKS2;refGene_tx=XM_023240721.1;refGene_gn=TNKS2;refGene_tx=XM_023240722.1
D2 54214731 54214841
refGene_gn=TNKS2;refGene_tx=XM_019814078.2;refGene_gn=TNKS2;refGene_tx=XM_023240721.1;refGene_gn=TNKS2;refGene_tx=XM_023240722.1
D2 54215807 54215996
refGene_gn=TNKS2;refGene_tx=XM_019814078.2;refGene_gn=TNKS2;refGene_tx=XM_023240721.1;refGene_gn=TNKS2;refGene_tx=XM_023240722.1
D2 54218387 54218640
refGene_gn=TNKS2;refGene_tx=XM_019814078.2;refGene_gn=TNKS2;refGene_tx=XM_023240721.1;refGene_gn=TNKS2;refGene_tx=XM_023240722.1
D2 54219354 54219437
refGene_gn=TNKS2;refGene_tx=XM_019814078.2;refGene_gn=TNKS2;refGene_tx=XM_023240721.1;refGene_gn=TNKS2;refGene_tx=XM_023240722.1
D2 54220838 54220959
refGene_gn=TNKS2;refGene_tx=XM_019814078.2;refGene_gn=TNKS2;refGene_tx=XM_023240721.1;refGene_gn=TNKS2;refGene_tx=XM_023240722.1
D2 54224388 54224486
refGene_gn=TNKS2;refGene_tx=XM_019814078.2;refGene_gn=TNKS2;refGene_tx=XM_023240721.1;refGene_gn=TNKS2;refGene_tx=XM_023240722.1
D2 54226243 54226318
refGene_gn=TNKS2;refGene_tx=XM_019814078.2;refGene_gn=TNKS2;refGene_tx=XM_023240721.1;refGene_gn=TNKS2;refGene_tx=XM_023240722.1
D2 54228071 54228177
refGene_gn=TNKS2;refGene_tx=XM_019814078.2;refGene_gn=TNKS2;refGene_tx=XM_023240721.1;refGene_gn=TNKS2;refGene_tx=XM_023240722.1
D2 54229805 54229992
refGene_gn=TNKS2;refGene_tx=XM_019814078.2;refGene_gn=TNKS2;refGene_tx=XM_023240721.1;refGene_gn=TNKS2;refGene_tx=XM_023240722.1
D2 54232851 54233008
refGene_gn=TNKS2;refGene_tx=XM_019814078.2;refGene_gn=TNKS2;refGene_tx=XM_023240721.1;refGene_gn=TNKS2;refGene_tx=XM_023240722.1
D2 54233844 54233907
refGene_gn=TNKS2;refGene_tx=XM_019814078.2;refGene_gn=TNKS2;refGene_tx=XM_023240721.1;refGene_gn=TNKS2;refGene_tx=XM_023240722.1
D2 54289662 54290445 refGene_gn=FGFBP3;refGene_tx=XM_023240723.1
D2 54302785 54302799 refGene_gn=BTAF1;refGene_tx=XM_011287417.3
D2 54316158 54316282 refGene_gn=BTAF1;refGene_tx=XM_011287417.3
D2 54322593 54322708 refGene_gn=BTAF1;refGene_tx=XM_011287417.3
D2 54325637 54325784
refGene_gn=BTAF1;refGene_tx=XM_011287417.3;refGene_gn=BTAF1;refGene_tx=XM_019813536.2
D2 54333327 54333491
refGene_gn=BTAF1;refGene_tx=XM_011287417.3;refGene_gn=BTAF1;refGene_tx=XM_019813536.2
D2 54335980 54336117
refGene_gn=BTAF1;refGene_tx=XM_011287417.3;refGene_gn=BTAF1;refGene_tx=XM_019813536.2

D2 54339477 54339607
refGene_gn=BTAF1;refGene_tx=XM_011287417.3;refGene_gn=BTAF1;refGene_tx=XM_019813536.2
D2 54339863 54339932
refGene_gn=BTAF1;refGene_tx=XM_011287417.3;refGene_gn=BTAF1;refGene_tx=XM_019813536.2
D2 54341032 54341122
refGene_gn=BTAF1;refGene_tx=XM_011287417.3;refGene_gn=BTAF1;refGene_tx=XM_019813536.2
D2 54341769 54341865
refGene_gn=BTAF1;refGene_tx=XM_011287417.3;refGene_gn=BTAF1;refGene_tx=XM_019813536.2
D2 54341961 54342138
refGene_gn=BTAF1;refGene_tx=XM_011287417.3;refGene_gn=BTAF1;refGene_tx=XM_019813536.2
D2 54343259 54343400
refGene_gn=BTAF1;refGene_tx=XM_011287417.3;refGene_gn=BTAF1;refGene_tx=XM_019813536.2
D2 54344408 54344533
refGene_gn=BTAF1;refGene_tx=XM_011287417.3;refGene_gn=BTAF1;refGene_tx=XM_019813536.2
D2 54347014 54347135
refGene_gn=BTAF1;refGene_tx=XM_011287417.3;refGene_gn=BTAF1;refGene_tx=XM_019813536.2
D2 54366020 54366125
refGene_gn=BTAF1;refGene_tx=XM_011287417.3;refGene_gn=BTAF1;refGene_tx=XM_019813536.2
D2 54367297 54367447
refGene_gn=BTAF1;refGene_tx=XM_011287417.3;refGene_gn=BTAF1;refGene_tx=XM_019813536.2
D2 54367758 54367901
refGene_gn=BTAF1;refGene_tx=XM_011287417.3;refGene_gn=BTAF1;refGene_tx=XM_019813536.2
D2 54368168 54368343
refGene_gn=BTAF1;refGene_tx=XM_011287417.3;refGene_gn=BTAF1;refGene_tx=XM_019813536.2
D2 54369800 54370004
refGene_gn=BTAF1;refGene_tx=XM_011287417.3;refGene_gn=BTAF1;refGene_tx=XM_019813536.2
D2 54379543 54379970
refGene_gn=BTAF1;refGene_tx=XM_011287417.3;refGene_gn=BTAF1;refGene_tx=XM_019813536.2
D2 54381129 54381320
refGene_gn=BTAF1;refGene_tx=XM_011287417.3;refGene_gn=BTAF1;refGene_tx=XM_019813536.2
D2 54382842 54382996
refGene_gn=BTAF1;refGene_tx=XM_011287417.3;refGene_gn=BTAF1;refGene_tx=XM_019813536.2
D2 54383074 54383184
refGene_gn=BTAF1;refGene_tx=XM_011287417.3;refGene_gn=BTAF1;refGene_tx

=XM_019813536.2
D2 54385191 54385393
refGene_gn=BTAF1;refGene_tx=XM_011287417.3;refGene_gn=BTAF1;refGene_tx=XM_019813536.2
D2 54386386 54386535
refGene_gn=BTAF1;refGene_tx=XM_011287417.3;refGene_gn=BTAF1;refGene_tx=XM_019813536.2
D2 54393576 54393729
refGene_gn=BTAF1;refGene_tx=XM_011287417.3;refGene_gn=BTAF1;refGene_tx=XM_019813536.2
D2 54393957 54394079
refGene_gn=BTAF1;refGene_tx=XM_011287417.3;refGene_gn=BTAF1;refGene_tx=XM_019813536.2
D2 54394185 54394353
refGene_gn=BTAF1;refGene_tx=XM_011287417.3;refGene_gn=BTAF1;refGene_tx=XM_019813536.2
D2 54406052 54406130
refGene_gn=BTAF1;refGene_tx=XM_011287417.3;refGene_gn=BTAF1;refGene_tx=XM_019813536.2
D2 54406258 54406388
refGene_gn=BTAF1;refGene_tx=XM_011287417.3;refGene_gn=BTAF1;refGene_tx=XM_019813536.2
D2 54408720 54408862
refGene_gn=BTAF1;refGene_tx=XM_011287417.3;refGene_gn=BTAF1;refGene_tx=XM_019813536.2
D2 54408950 54409081
refGene_gn=BTAF1;refGene_tx=XM_011287417.3;refGene_gn=BTAF1;refGene_tx=XM_019813536.2
D2 54411233 54411359
refGene_gn=BTAF1;refGene_tx=XM_011287417.3;refGene_gn=BTAF1;refGene_tx=XM_019813536.2
D2 54412948 54413101
refGene_gn=BTAF1;refGene_tx=XM_011287417.3;refGene_gn=BTAF1;refGene_tx=XM_019813536.2
D2 54418904 54419116
refGene_gn=BTAF1;refGene_tx=XM_011287417.3;refGene_gn=BTAF1;refGene_tx=XM_019813536.2
D2 54420832 54420992
refGene_gn=BTAF1;refGene_tx=XM_011287417.3;refGene_gn=BTAF1;refGene_tx=XM_019813536.2
D2 54421371 54421542
refGene_gn=BTAF1;refGene_tx=XM_011287417.3;refGene_gn=BTAF1;refGene_tx=XM_019813536.2
D2 54423347 54423491
refGene_gn=BTAF1;refGene_tx=XM_011287417.3;refGene_gn=BTAF1;refGene_tx=XM_019813536.2
D2 54453651 54453879
refGene_gn=CPEB3;refGene_tx=XM_023240724.1;refGene_gn=CPEB3;refGene_tx=XM_023240725.1;refGene_gn=CPEB3;refGene_tx=XM_023240726.1;refGene_gn=CPEB3;refGene_tx=XM_023240727.1;refGene_gn=CPEB3;refGene_tx=XM_0232407

28.1;refGene_gn=CPEB3;refGene_tx=XM_023240729.1;refGene_gn=CPEB3;refGene_tx=XM_023240730.1;refGene_gn=CPEB3;refGene_tx=XM_023240731.1;refGene_gn=CPEB3;refGene_tx=XM_023240732.1;refGene_gn=CPEB3;refGene_tx=XM_023240733.1;refGene_gn=CPEB3;refGene_tx=XM_023240734.1;refGene_gn=CPEB3;refGene_tx=XM_023240735.1;refGene_gn=CPEB3;refGene_tx=XM_023240736.1;refGene_gn=CPEB3;refGene_tx=XM_023240737.1
D2 54478079 54478261
refGene_gn=CPEB3;refGene_tx=XM_023240724.1;refGene_gn=CPEB3;refGene_tx=XM_023240725.1;refGene_gn=CPEB3;refGene_tx=XM_023240726.1;refGene_gn=CPEB3;refGene_tx=XM_023240727.1;refGene_gn=CPEB3;refGene_tx=XM_023240728.1;refGene_gn=CPEB3;refGene_tx=XM_023240729.1;refGene_gn=CPEB3;refGene_tx=XM_023240730.1;refGene_gn=CPEB3;refGene_tx=XM_023240731.1;refGene_gn=CPEB3;refGene_tx=XM_023240732.1;refGene_gn=CPEB3;refGene_tx=XM_023240733.1;refGene_gn=CPEB3;refGene_tx=XM_023240734.1;refGene_gn=CPEB3;refGene_tx=XM_023240735.1;refGene_gn=CPEB3;refGene_tx=XM_023240736.1;refGene_gn=CPEB3;refGene_tx=XM_023240737.1;refGene_gn=CPEB3;refGene_tx=XM_023240738.1
D2 54487720 54487835
refGene_gn=CPEB3;refGene_tx=XM_023240724.1;refGene_gn=CPEB3;refGene_tx=XM_023240725.1;refGene_gn=CPEB3;refGene_tx=XM_023240726.1;refGene_gn=CPEB3;refGene_tx=XM_023240727.1;refGene_gn=CPEB3;refGene_tx=XM_023240728.1;refGene_gn=CPEB3;refGene_tx=XM_023240729.1;refGene_gn=CPEB3;refGene_tx=XM_023240730.1;refGene_gn=CPEB3;refGene_tx=XM_023240731.1;refGene_gn=CPEB3;refGene_tx=XM_023240732.1;refGene_gn=CPEB3;refGene_tx=XM_023240733.1;refGene_gn=CPEB3;refGene_tx=XM_023240734.1;refGene_gn=CPEB3;refGene_tx=XM_023240735.1;refGene_gn=CPEB3;refGene_tx=XM_023240736.1;refGene_gn=CPEB3;refGene_tx=XM_023240737.1
D2 54502679 54502798
refGene_gn=CPEB3;refGene_tx=XM_023240724.1;refGene_gn=CPEB3;refGene_tx=XM_023240725.1;refGene_gn=CPEB3;refGene_tx=XM_023240726.1;refGene_gn=CPEB3;refGene_tx=XM_023240727.1;refGene_gn=CPEB3;refGene_tx=XM_023240728.1;refGene_gn=CPEB3;refGene_tx=XM_023240729.1;refGene_gn=CPEB3;refGene_tx=XM_023240730.1;refGene_gn=CPEB3;refGene_tx=XM_023240731.1;refGene_gn=CPEB3;refGene_tx=XM_023240732.1;refGene_gn=CPEB3;refGene_tx=XM_023240733.1;refGene_gn=CPEB3;refGene_tx=XM_023240734.1;refGene_gn=CPEB3;refGene_tx=XM_023240735.1;refGene_gn=CPEB3;refGene_tx=XM_023240736.1;refGene_gn=CPEB3;refGene_tx=XM_023240737.1;refGene_gn=CPEB3;refGene_tx=XM_023240738.1
D2 54529658 54529748
refGene_gn=CPEB3;refGene_tx=XM_023240724.1;refGene_gn=CPEB3;refGene_tx=XM_023240725.1;refGene_gn=CPEB3;refGene_tx=XM_023240726.1;refGene_gn=CPEB3;refGene_tx=XM_023240727.1;refGene_gn=CPEB3;refGene_tx=XM_023240728.1;refGene_gn=CPEB3;refGene_tx=XM_023240729.1;refGene_gn=CPEB3;refGene_tx=XM_023240730.1;refGene_gn=CPEB3;refGene_tx=XM_023240731.1;refGene_gn=CPEB3;refGene_tx=XM_023240732.1;refGene_gn=CPEB3;refGene_tx=XM_023240733.1;refGene_gn=CPEB3;refGene_tx=XM_023240734.1;refGene_gn=CPEB3;refGene_tx=XM_023240735.1;refGene_gn=CPEB3;refGene_tx=XM_023240738.1
D2 54531857 54532025
refGene_gn=CPEB3;refGene_tx=XM_023240724.1;refGene_gn=CPEB3;refGene_tx=XM_023240725.1;refGene_gn=CPEB3;refGene_tx=XM_023240726.1;refGene_gn=

CPEB3;refGene_tx=XM_023240727.1;refGene_gn=CPEB3;refGene_tx=XM_0232407
28.1;refGene_gn=CPEB3;refGene_tx=XM_023240729.1;refGene_gn=CPEB3;refGe
ne_tx=XM_023240730.1;refGene_gn=CPEB3;refGene_tx=XM_023240731.1;refGen
e_gn=CPEB3;refGene_tx=XM_023240732.1;refGene_gn=CPEB3;refGene_tx=XM_02
3240733.1;refGene_gn=CPEB3;refGene_tx=XM_023240734.1;refGene_gn=CPEB3;
refGene_tx=XM_023240735.1;refGene_gn=CPEB3;refGene_tx=XM_023240738.1
D2 54545572 54545596 refGene_gn=CPEB3;refGene_tx=XM_023240725.1
D2 54546377 54546401
refGene_gn=CPEB3;refGene_tx=XM_023240724.1;refGene_gn=CPEB3;refGene_tx
=XM_023240727.1;refGene_gn=CPEB3;refGene_tx=XM_023240729.1;refGene_gn=
CPEB3;refGene_tx=XM_023240730.1;refGene_gn=CPEB3;refGene_tx=XM_0232407
31.1;refGene_gn=CPEB3;refGene_tx=XM_023240732.1;refGene_gn=CPEB3;refGe
ne_tx=XM_023240733.1;refGene_gn=CPEB3;refGene_tx=XM_023240734.1;refGen
e_gn=CPEB3;refGene_tx=XM_023240735.1;refGene_gn=CPEB3;refGene_tx=XM_02
3240738.1
D2 54563800 54563857
refGene_gn=CPEB3;refGene_tx=XM_023240724.1;refGene_gn=CPEB3;refGene_tx
=XM_023240725.1;refGene_gn=CPEB3;refGene_tx=XM_023240726.1;refGene_gn=
CPEB3;refGene_tx=XM_023240727.1;refGene_gn=CPEB3;refGene_tx=XM_0232407
28.1;refGene_gn=CPEB3;refGene_tx=XM_023240729.1;refGene_gn=CPEB3;refGe
ne_tx=XM_023240730.1;refGene_gn=CPEB3;refGene_tx=XM_023240731.1;refGen
e_gn=CPEB3;refGene_tx=XM_023240732.1;refGene_gn=CPEB3;refGene_tx=XM_02
3240733.1;refGene_gn=CPEB3;refGene_tx=XM_023240734.1;refGene_gn=CPEB3;
refGene_tx=XM_023240735.1;refGene_gn=CPEB3;refGene_tx=XM_023240736.1;r
efGene_gn=CPEB3;refGene_tx=XM_023240737.1;refGene_gn=CPEB3;refGene_tx=
XM_023240738.1
D2 54573811 54573971
refGene_gn=CPEB3;refGene_tx=XM_023240724.1;refGene_gn=CPEB3;refGene_tx
=XM_023240725.1;refGene_gn=CPEB3;refGene_tx=XM_023240726.1;refGene_gn=
CPEB3;refGene_tx=XM_023240727.1;refGene_gn=CPEB3;refGene_tx=XM_0232407
28.1;refGene_gn=CPEB3;refGene_tx=XM_023240729.1;refGene_gn=CPEB3;refGe
ne_tx=XM_023240730.1;refGene_gn=CPEB3;refGene_tx=XM_023240731.1;refGen
e_gn=CPEB3;refGene_tx=XM_023240732.1;refGene_gn=CPEB3;refGene_tx=XM_02
3240733.1;refGene_gn=CPEB3;refGene_tx=XM_023240734.1;refGene_gn=CPEB3;
refGene_tx=XM_023240735.1;refGene_gn=CPEB3;refGene_tx=XM_023240736.1;r
efGene_gn=CPEB3;refGene_tx=XM_023240737.1;refGene_gn=CPEB3;refGene_tx=
XM_023240738.1
D2 54607858 54608871
refGene_gn=CPEB3;refGene_tx=XM_023240724.1;refGene_gn=CPEB3;refGene_tx
=XM_023240725.1;refGene_gn=CPEB3;refGene_tx=XM_023240726.1;refGene_gn=
CPEB3;refGene_tx=XM_023240727.1;refGene_gn=CPEB3;refGene_tx=XM_0232407
28.1;refGene_gn=CPEB3;refGene_tx=XM_023240729.1;refGene_gn=CPEB3;refGe
ne_tx=XM_023240730.1;refGene_gn=CPEB3;refGene_tx=XM_023240731.1;refGen
e_gn=CPEB3;refGene_tx=XM_023240732.1;refGene_gn=CPEB3;refGene_tx=XM_02
3240733.1;refGene_gn=CPEB3;refGene_tx=XM_023240734.1;refGene_gn=CPEB3;
refGene_tx=XM_023240735.1;refGene_gn=CPEB3;refGene_tx=XM_023240736.1;r
efGene_gn=CPEB3;refGene_tx=XM_023240737.1;refGene_gn=CPEB3;refGene_tx=
XM_023240738.1
D2 54644105 54644193
refGene_gn=CPEB3;refGene_tx=XM_023240724.1;refGene_gn=CPEB3;refGene_tx

=XM_023240725.1;refGene_gn=CPEB3;refGene_tx=XM_023240726.1;refGene_gn=CPEB3;refGene_tx=XM_023240727.1;refGene_gn=CPEB3;refGene_tx=XM_023240728.1;refGene_gn=CPEB3;refGene_tx=XM_023240736.1;refGene_gn=CPEB3;refGene_tx=XM_023240737.1;refGene_gn=CPEB3;refGene_tx=XM_023240738.1
D2	54644731 54644766 refGene_gn=MARCH5;refGene_tx=XM_003994212.5
D2	54663973 54664176 refGene_gn=MARCH5;refGene_tx=XM_003994212.5
D2	54688842 54688973 refGene_gn=MARCH5;refGene_tx=XM_003994212.5
D2	54697455 54697639 refGene_gn=MARCH5;refGene_tx=XM_003994212.5
D2	54697770 54697937 refGene_gn=MARCH5;refGene_tx=XM_003994212.5
D2	54698902 54699019 refGene_gn=MARCH5;refGene_tx=XM_003994212.5
D2	54791226 54791322
refGene_gn=IDE;refGene_tx=XM_019813543.2;refGene_gn=IDE;refGene_tx=XM_023240739.1;refGene_gn=IDE;refGene_tx=XM_023240740.1;refGene_gn=IDE;refGene_tx=XM_023240741.1
D2	54792016 54792084
refGene_gn=IDE;refGene_tx=XM_019813543.2;refGene_gn=IDE;refGene_tx=XM_023240739.1;refGene_gn=IDE;refGene_tx=XM_023240740.1;refGene_gn=IDE;refGene_tx=XM_023240741.1
D2	54792793 54792866
refGene_gn=IDE;refGene_tx=XM_019813543.2;refGene_gn=IDE;refGene_tx=XM_023240739.1;refGene_gn=IDE;refGene_tx=XM_023240740.1;refGene_gn=IDE;refGene_tx=XM_023240741.1
D2	54795495 54795557
refGene_gn=IDE;refGene_tx=XM_019813543.2;refGene_gn=IDE;refGene_tx=XM_023240739.1;refGene_gn=IDE;refGene_tx=XM_023240740.1;refGene_gn=IDE;refGene_tx=XM_023240741.1
D2	54796610 54796883
refGene_gn=IDE;refGene_tx=XM_019813543.2;refGene_gn=IDE;refGene_tx=XM_023240739.1;refGene_gn=IDE;refGene_tx=XM_023240740.1;refGene_gn=IDE;refGene_tx=XM_023240741.1
D2	54798505 54798673
refGene_gn=IDE;refGene_tx=XM_019813543.2;refGene_gn=IDE;refGene_tx=XM_023240739.1;refGene_gn=IDE;refGene_tx=XM_023240740.1;refGene_gn=IDE;refGene_tx=XM_023240741.1
D2	54801546 54801658
refGene_gn=IDE;refGene_tx=XM_019813543.2;refGene_gn=IDE;refGene_tx=XM_023240739.1;refGene_gn=IDE;refGene_tx=XM_023240740.1;refGene_gn=IDE;refGene_tx=XM_023240741.1
D2	54802676 54802768
refGene_gn=IDE;refGene_tx=XM_019813543.2;refGene_gn=IDE;refGene_tx=XM_023240739.1;refGene_gn=IDE;refGene_tx=XM_023240740.1;refGene_gn=IDE;refGene_tx=XM_023240741.1
D2	54806435 54806556
refGene_gn=IDE;refGene_tx=XM_019813543.2;refGene_gn=IDE;refGene_tx=XM_023240739.1;refGene_gn=IDE;refGene_tx=XM_023240740.1;refGene_gn=IDE;refGene_tx=XM_023240741.1
D2	54807896 54808007
refGene_gn=IDE;refGene_tx=XM_019813543.2;refGene_gn=IDE;refGene_tx=XM_023240739.1;refGene_gn=IDE;refGene_tx=XM_023240740.1;refGene_gn=IDE;refGene_tx=XM_023240741.1

D2 54809861 54810006 refGene_gn=IDE;refGene_tx=XM_023240740.1
D2 54810512 54810657
refGene_gn=IDE;refGene_tx=XM_019813543.2;refGene_gn=IDE;refGene_tx=XM_023240739.1;refGene_gn=IDE;refGene_tx=XM_023240741.1
D2 54814595 54814678
refGene_gn=IDE;refGene_tx=XM_019813543.2;refGene_gn=IDE;refGene_tx=XM_023240739.1;refGene_gn=IDE;refGene_tx=XM_023240740.1;refGene_gn=IDE;refGene_tx=XM_023240741.1
D2 54816980 54817103
refGene_gn=IDE;refGene_tx=XM_019813543.2;refGene_gn=IDE;refGene_tx=XM_023240739.1;refGene_gn=IDE;refGene_tx=XM_023240740.1;refGene_gn=IDE;refGene_tx=XM_023240741.1
D2 54819635 54819738
refGene_gn=IDE;refGene_tx=XM_019813543.2;refGene_gn=IDE;refGene_tx=XM_023240739.1;refGene_gn=IDE;refGene_tx=XM_023240740.1;refGene_gn=IDE;refGene_tx=XM_023240741.1
D2 54828176 54828280
refGene_gn=IDE;refGene_tx=XM_019813543.2;refGene_gn=IDE;refGene_tx=XM_023240739.1;refGene_gn=IDE;refGene_tx=XM_023240740.1;refGene_gn=IDE;refGene_tx=XM_023240741.1
D2 54830079 54830160
refGene_gn=IDE;refGene_tx=XM_019813543.2;refGene_gn=IDE;refGene_tx=XM_023240739.1;refGene_gn=IDE;refGene_tx=XM_023240740.1;refGene_gn=IDE;refGene_tx=XM_023240741.1
D2 54831917 54832009
refGene_gn=IDE;refGene_tx=XM_019813543.2;refGene_gn=IDE;refGene_tx=XM_023240739.1;refGene_gn=IDE;refGene_tx=XM_023240740.1;refGene_gn=IDE;refGene_tx=XM_023240741.1
D2 54832439 54832532
refGene_gn=IDE;refGene_tx=XM_019813543.2;refGene_gn=IDE;refGene_tx=XM_023240739.1;refGene_gn=IDE;refGene_tx=XM_023240740.1;refGene_gn=IDE;refGene_tx=XM_023240741.1
D2 54833687 54833850
refGene_gn=IDE;refGene_tx=XM_019813543.2;refGene_gn=IDE;refGene_tx=XM_023240739.1;refGene_gn=IDE;refGene_tx=XM_023240740.1;refGene_gn=IDE;refGene_tx=XM_023240741.1
D2 54834788 54834901
refGene_gn=IDE;refGene_tx=XM_019813543.2;refGene_gn=IDE;refGene_tx=XM_023240739.1;refGene_gn=IDE;refGene_tx=XM_023240740.1;refGene_gn=IDE;refGene_tx=XM_023240741.1
D2 54840954 54841077
refGene_gn=IDE;refGene_tx=XM_019813543.2;refGene_gn=IDE;refGene_tx=XM_023240739.1;refGene_gn=IDE;refGene_tx=XM_023240740.1;refGene_gn=IDE;refGene_tx=XM_023240741.1
D2 54857310 54857480
refGene_gn=IDE;refGene_tx=XM_019813543.2;refGene_gn=IDE;refGene_tx=XM_023240739.1;refGene_gn=IDE;refGene_tx=XM_023240740.1;refGene_gn=IDE;refGene_tx=XM_023240741.1
D2 54861237 54861445
refGene_gn=IDE;refGene_tx=XM_019813543.2;refGene_gn=IDE;refGene_tx=XM_

023240739.1;refGene_gn=IDE;refGene_tx=XM_023240740.1;refGene_gn=IDE;refGene_tx=XM_023240741.1
D2 54862738 54862923
refGene_gn=IDE;refGene_tx=XM_019813543.2;refGene_gn=IDE;refGene_tx=XM_023240739.1;refGene_gn=IDE;refGene_tx=XM_023240740.1;refGene_gn=IDE;refGene_tx=XM_023240741.1
D2 54900489 54900539 refGene_gn=IDE;refGene_tx=XM_023240741.1
D2 54901038 54901136
refGene_gn=IDE;refGene_tx=XM_023240739.1;refGene_gn=IDE;refGene_tx=XM_023240740.1
D2 54918982 54919059
refGene_gn=KIF11;refGene_tx=XM_011287418.3;refGene_gn=KIF11;refGene_tx=XM_023240742.1
D2 54925813 54925946
refGene_gn=KIF11;refGene_tx=XM_011287418.3;refGene_gn=KIF11;refGene_tx=XM_023240742.1
D2 54926182 54926280
refGene_gn=KIF11;refGene_tx=XM_011287418.3;refGene_gn=KIF11;refGene_tx=XM_023240742.1
D2 54926375 54926454
refGene_gn=KIF11;refGene_tx=XM_011287418.3;refGene_gn=KIF11;refGene_tx=XM_023240742.1
D2 54928031 54928217
refGene_gn=KIF11;refGene_tx=XM_011287418.3;refGene_gn=KIF11;refGene_tx=XM_023240742.1
D2 54928332 54928457
refGene_gn=KIF11;refGene_tx=XM_011287418.3;refGene_gn=KIF11;refGene_tx=XM_023240742.1
D2 54931152 54931243
refGene_gn=KIF11;refGene_tx=XM_011287418.3;refGene_gn=KIF11;refGene_tx=XM_023240742.1
D2 54932422 54932665
refGene_gn=KIF11;refGene_tx=XM_011287418.3;refGene_gn=KIF11;refGene_tx=XM_023240742.1
D2 54934536 54934632
refGene_gn=KIF11;refGene_tx=XM_011287418.3;refGene_gn=KIF11;refGene_tx=XM_023240742.1
D2 54936238 54936327
refGene_gn=KIF11;refGene_tx=XM_011287418.3;refGene_gn=KIF11;refGene_tx=XM_023240742.1
D2 54940145 54940233
refGene_gn=KIF11;refGene_tx=XM_011287418.3;refGene_gn=KIF11;refGene_tx=XM_023240742.1
D2 54942807 54942996
refGene_gn=KIF11;refGene_tx=XM_011287418.3;refGene_gn=KIF11;refGene_tx=XM_023240742.1
D2 54943873 54944081
refGene_gn=KIF11;refGene_tx=XM_011287418.3;refGene_gn=KIF11;refGene_tx=XM_023240742.1
D2 54945058 54945231

refGene_gn=KIF11;refGene_tx=XM_011287418.3;refGene_gn=KIF11;refGene_tx=XM_023240742.1
D2 54947533 54947659
refGene_gn=KIF11;refGene_tx=XM_011287418.3;refGene_gn=KIF11;refGene_tx=XM_023240742.1
D2 54948167 54948326
refGene_gn=KIF11;refGene_tx=XM_011287418.3;refGene_gn=KIF11;refGene_tx=XM_023240742.1
D2 54950747 54950854
refGene_gn=KIF11;refGene_tx=XM_011287418.3;refGene_gn=KIF11;refGene_tx=XM_023240742.1
D2 54954844 54955124
refGene_gn=KIF11;refGene_tx=XM_011287418.3;refGene_gn=KIF11;refGene_tx=XM_023240742.1
D2 54958559 54958782
refGene_gn=KIF11;refGene_tx=XM_011287418.3;refGene_gn=KIF11;refGene_tx=XM_023240742.1
D2 54959621 54959773
refGene_gn=KIF11;refGene_tx=XM_011287418.3;refGene_gn=KIF11;refGene_tx=XM_023240742.1
D2 54960832 54960949
refGene_gn=KIF11;refGene_tx=XM_011287418.3;refGene_gn=KIF11;refGene_tx=XM_023240742.1
D2 54961247 54961249 refGene_gn=KIF11;refGene_tx=XM_023240742.1
D2 54962857 54962990
refGene_gn=KIF11;refGene_tx=XM_011287418.3;refGene_gn=KIF11;refGene_tx=XM_023240742.1
D2 54985328 54985692 refGene_gn=HHEX;refGene_tx=XM_003994214.5
D2 54987831 54988010 refGene_gn=HHEX;refGene_tx=XM_003994214.5
D2 54988209 54988260 refGene_gn=HHEX;refGene_tx=XM_003994214.5
D2 54990060 54990279 refGene_gn=HHEX;refGene_tx=XM_003994214.5
D2 55113652 55113774
refGene_gn=EXOC6;refGene_tx=XM_019813544.2;refGene_gn=EXOC6;refGene_tx=XM_019813545.2;refGene_gn=EXOC6;refGene_tx=XM_019813548.2
D2 55129968 55130069
refGene_gn=EXOC6;refGene_tx=XM_023240743.1;refGene_gn=EXOC6;refGene_tx=XM_023240744.1
D2 55163210 55163382
refGene_gn=EXOC6;refGene_tx=XM_019813544.2;refGene_gn=EXOC6;refGene_tx=XM_019813545.2;refGene_gn=EXOC6;refGene_tx=XM_019813548.2;refGene_gn=EXOC6;refGene_tx=XM_023240743.1;refGene_gn=EXOC6;refGene_tx=XM_023240744.1;refGene_gn=EXOC6;refGene_tx=XM_023240745.1;refGene_gn=EXOC6;refGene_tx=XM_023240746.1
D2 55164872 55164920
refGene_gn=EXOC6;refGene_tx=XM_019813544.2;refGene_gn=EXOC6;refGene_tx=XM_019813545.2;refGene_gn=EXOC6;refGene_tx=XM_019813548.2;refGene_gn=EXOC6;refGene_tx=XM_023240743.1;refGene_gn=EXOC6;refGene_tx=XM_023240744.1;refGene_gn=EXOC6;refGene_tx=XM_023240745.1;refGene_gn=EXOC6;refGene_tx=XM_023240746.1
D2 55165009 55165100

refGene_gn=EXOC6;refGene_tx=XM_019813544.2;refGene_gn=EXOC6;refGene_tx=XM_019813545.2;refGene_gn=EXOC6;refGene_tx=XM_019813548.2;refGene_gn=EXOC6;refGene_tx=XM_023240743.1;refGene_gn=EXOC6;refGene_tx=XM_023240744.1;refGene_gn=EXOC6;refGene_tx=XM_023240745.1;refGene_gn=EXOC6;refGene_tx=XM_023240746.1
D2 55168229 55168275
refGene_gn=EXOC6;refGene_tx=XM_019813544.2;refGene_gn=EXOC6;refGene_tx=XM_019813545.2;refGene_gn=EXOC6;refGene_tx=XM_019813548.2;refGene_gn=EXOC6;refGene_tx=XM_023240743.1;refGene_gn=EXOC6;refGene_tx=XM_023240744.1;refGene_gn=EXOC6;refGene_tx=XM_023240745.1;refGene_gn=EXOC6;refGene_tx=XM_023240746.1
D2 55185444 55185649
refGene_gn=EXOC6;refGene_tx=XM_019813544.2;refGene_gn=EXOC6;refGene_tx=XM_019813545.2;refGene_gn=EXOC6;refGene_tx=XM_019813548.2;refGene_gn=EXOC6;refGene_tx=XM_023240743.1;refGene_gn=EXOC6;refGene_tx=XM_023240744.1;refGene_gn=EXOC6;refGene_tx=XM_023240745.1;refGene_gn=EXOC6;refGene_tx=XM_023240746.1
D2 55196466 55196619
refGene_gn=EXOC6;refGene_tx=XM_019813544.2;refGene_gn=EXOC6;refGene_tx=XM_019813545.2;refGene_gn=EXOC6;refGene_tx=XM_019813548.2;refGene_gn=EXOC6;refGene_tx=XM_023240743.1;refGene_gn=EXOC6;refGene_tx=XM_023240744.1;refGene_gn=EXOC6;refGene_tx=XM_023240745.1;refGene_gn=EXOC6;refGene_tx=XM_023240746.1
D2 55197964 55198033
refGene_gn=EXOC6;refGene_tx=XM_019813544.2;refGene_gn=EXOC6;refGene_tx=XM_019813545.2;refGene_gn=EXOC6;refGene_tx=XM_019813548.2;refGene_gn=EXOC6;refGene_tx=XM_023240743.1;refGene_gn=EXOC6;refGene_tx=XM_023240744.1;refGene_gn=EXOC6;refGene_tx=XM_023240745.1;refGene_gn=EXOC6;refGene_tx=XM_023240746.1
D2 55209863 55209947
refGene_gn=EXOC6;refGene_tx=XM_019813544.2;refGene_gn=EXOC6;refGene_tx=XM_019813545.2;refGene_gn=EXOC6;refGene_tx=XM_019813548.2;refGene_gn=EXOC6;refGene_tx=XM_023240743.1;refGene_gn=EXOC6;refGene_tx=XM_023240744.1;refGene_gn=EXOC6;refGene_tx=XM_023240745.1;refGene_gn=EXOC6;refGene_tx=XM_023240746.1
D2 55215207 55215254
refGene_gn=EXOC6;refGene_tx=XM_019813544.2;refGene_gn=EXOC6;refGene_tx=XM_019813545.2;refGene_gn=EXOC6;refGene_tx=XM_019813548.2;refGene_gn=EXOC6;refGene_tx=XM_023240743.1;refGene_gn=EXOC6;refGene_tx=XM_023240744.1;refGene_gn=EXOC6;refGene_tx=XM_023240745.1;refGene_gn=EXOC6;refGene_tx=XM_023240746.1
D2 55215385 55215506
refGene_gn=EXOC6;refGene_tx=XM_019813544.2;refGene_gn=EXOC6;refGene_tx=XM_019813545.2;refGene_gn=EXOC6;refGene_tx=XM_019813548.2;refGene_gn=EXOC6;refGene_tx=XM_023240743.1;refGene_gn=EXOC6;refGene_tx=XM_023240744.1;refGene_gn=EXOC6;refGene_tx=XM_023240745.1;refGene_gn=EXOC6;refGene_tx=XM_023240746.1
D2 55216553 55216625
refGene_gn=EXOC6;refGene_tx=XM_019813544.2;refGene_gn=EXOC6;refGene_tx=XM_019813545.2;refGene_gn=EXOC6;refGene_tx=XM_019813548.2;refGene_gn=

EXOC6;refGene_tx=XM_023240743.1;refGene_gn=EXOC6;refGene_tx=XM_0232407
44.1;refGene_gn=EXOC6;refGene_tx=XM_023240745.1;refGene_gn=EXOC6;refGe
ne_tx=XM_023240746.1
D2 55221297 55221395
refGene_gn=EXOC6;refGene_tx=XM_019813544.2;refGene_gn=EXOC6;refGene_tx
=XM_019813545.2;refGene_gn=EXOC6;refGene_tx=XM_019813548.2;refGene_gn=
EXOC6;refGene_tx=XM_023240743.1;refGene_gn=EXOC6;refGene_tx=XM_0232407
44.1;refGene_gn=EXOC6;refGene_tx=XM_023240745.1;refGene_gn=EXOC6;refGe
ne_tx=XM_023240746.1
D2 55227830 55227936
refGene_gn=EXOC6;refGene_tx=XM_019813544.2;refGene_gn=EXOC6;refGene_tx
=XM_019813545.2;refGene_gn=EXOC6;refGene_tx=XM_019813548.2;refGene_gn=
EXOC6;refGene_tx=XM_023240743.1;refGene_gn=EXOC6;refGene_tx=XM_0232407
44.1;refGene_gn=EXOC6;refGene_tx=XM_023240745.1;refGene_gn=EXOC6;refGe
ne_tx=XM_023240746.1
D2 55231917 55232027
refGene_gn=EXOC6;refGene_tx=XM_019813544.2;refGene_gn=EXOC6;refGene_tx
=XM_019813545.2;refGene_gn=EXOC6;refGene_tx=XM_019813548.2;refGene_gn=
EXOC6;refGene_tx=XM_023240743.1;refGene_gn=EXOC6;refGene_tx=XM_0232407
44.1;refGene_gn=EXOC6;refGene_tx=XM_023240745.1;refGene_gn=EXOC6;refGe
ne_tx=XM_023240746.1
D2 55241974 55242086
refGene_gn=EXOC6;refGene_tx=XM_019813544.2;refGene_gn=EXOC6;refGene_tx
=XM_019813545.2;refGene_gn=EXOC6;refGene_tx=XM_019813548.2;refGene_gn=
EXOC6;refGene_tx=XM_023240743.1;refGene_gn=EXOC6;refGene_tx=XM_0232407
44.1;refGene_gn=EXOC6;refGene_tx=XM_023240745.1;refGene_gn=EXOC6;refGe
ne_tx=XM_023240746.1
D2 55242614 55242749
refGene_gn=EXOC6;refGene_tx=XM_019813544.2;refGene_gn=EXOC6;refGene_tx
=XM_019813545.2;refGene_gn=EXOC6;refGene_tx=XM_019813548.2;refGene_gn=
EXOC6;refGene_tx=XM_023240743.1;refGene_gn=EXOC6;refGene_tx=XM_0232407
44.1;refGene_gn=EXOC6;refGene_tx=XM_023240745.1;refGene_gn=EXOC6;refGe
ne_tx=XM_023240746.1
D2 55262619 55262799
refGene_gn=EXOC6;refGene_tx=XM_019813544.2;refGene_gn=EXOC6;refGene_tx
=XM_019813545.2;refGene_gn=EXOC6;refGene_tx=XM_023240743.1;refGene_gn=
EXOC6;refGene_tx=XM_023240744.1;refGene_gn=EXOC6;refGene_tx=XM_0232407
45.1;refGene_gn=EXOC6;refGene_tx=XM_023240746.1
D2 55286304 55286446
refGene_gn=EXOC6;refGene_tx=XM_019813544.2;refGene_gn=EXOC6;refGene_tx
=XM_019813545.2;refGene_gn=EXOC6;refGene_tx=XM_019813548.2;refGene_gn=
EXOC6;refGene_tx=XM_023240743.1;refGene_gn=EXOC6;refGene_tx=XM_0232407
44.1;refGene_gn=EXOC6;refGene_tx=XM_023240745.1;refGene_gn=EXOC6;refGe
ne_tx=XM_023240746.1
D2 55305613 55305687
refGene_gn=EXOC6;refGene_tx=XM_019813544.2;refGene_gn=EXOC6;refGene_tx
=XM_019813545.2;refGene_gn=EXOC6;refGene_tx=XM_019813548.2;refGene_gn=
EXOC6;refGene_tx=XM_023240743.1;refGene_gn=EXOC6;refGene_tx=XM_0232407
44.1;refGene_gn=EXOC6;refGene_tx=XM_023240745.1;refGene_gn=EXOC6;refGe
ne_tx=XM_023240746.1

D2 55342643 55342756
refGene_gn=EXOC6;refGene_tx=XM_019813544.2;refGene_gn=EXOC6;refGene_tx=XM_019813545.2;refGene_gn=EXOC6;refGene_tx=XM_019813548.2;refGene_gn=EXOC6;refGene_tx=XM_023240743.1;refGene_gn=EXOC6;refGene_tx=XM_023240744.1;refGene_gn=EXOC6;refGene_tx=XM_023240745.1;refGene_gn=EXOC6;refGene_tx=XM_023240746.1
D2 55343546 55343558
refGene_gn=EXOC6;refGene_tx=XM_019813544.2;refGene_gn=EXOC6;refGene_tx=XM_019813548.2;refGene_gn=EXOC6;refGene_tx=XM_023240743.1;refGene_gn=EXOC6;refGene_tx=XM_023240745.1;refGene_gn=EXOC6;refGene_tx=XM_023240746.1
D2 55343862 55343995
refGene_gn=EXOC6;refGene_tx=XM_019813544.2;refGene_gn=EXOC6;refGene_tx=XM_019813545.2;refGene_gn=EXOC6;refGene_tx=XM_019813548.2;refGene_gn=EXOC6;refGene_tx=XM_023240743.1;refGene_gn=EXOC6;refGene_tx=XM_023240744.1;refGene_gn=EXOC6;refGene_tx=XM_023240745.1;refGene_gn=EXOC6;refGene_tx=XM_023240746.1
D2 55346819 55347023
refGene_gn=LOC101093894;refGene_tx=XM_023240747.1
D2 55347577 55347802
refGene_gn=LOC101093894;refGene_tx=XM_023240747.1
D2 55348285 55348561
refGene_gn=LOC101093894;refGene_tx=XM_023240747.1
D2 55349830 55349986
refGene_gn=LOC101093894;refGene_tx=XM_023240747.1
D2 55351890 55352220
refGene_gn=LOC101093894;refGene_tx=XM_023240747.1
D2 55354181 55354559
refGene_gn=LOC101093894;refGene_tx=XM_023240747.1
D2 55359374 55359563
refGene_gn=LOC101086422;refGene_tx=XM_003994216.5
D2 55359745 55359970
refGene_gn=LOC101086422;refGene_tx=XM_003994216.5
D2 55360203 55360494
refGene_gn=LOC101086422;refGene_tx=XM_003994216.5
D2 55360629 55360800
refGene_gn=LOC101086422;refGene_tx=XM_003994216.5
D2 55361306 55361441
refGene_gn=LOC101086422;refGene_tx=XM_003994216.5
D2 55362023 55362176
refGene_gn=LOC101086422;refGene_tx=XM_003994216.5
D2 55362430 55362772
refGene_gn=LOC101086422;refGene_tx=XM_003994216.5
D2 55538088 55538127
refGene_gn=MYOF;refGene_tx=XM_003994218.5;refGene_gn=MYOF;refGene_tx=XM_011287427.3;refGene_gn=MYOF;refGene_tx=XM_011287428.3;refGene_gn=MYOF;refGene_tx=XM_019813549.2;refGene_gn=MYOF;refGene_tx=XM_019813550.2
D2 55539524 55539672
refGene_gn=MYOF;refGene_tx=XM_003994218.5;refGene_gn=MYOF;refGene_tx=XM_011287427.3;refGene_gn=MYOF;refGene_tx=XM_011287428.3;refGene_gn=MYO

F;refGene_tx=XM_019813549.2;refGene_gn=MYOF;refGene_tx=XM_019813550.2
D2 55540051 55540161
refGene_gn=MYOF;refGene_tx=XM_003994218.5;refGene_gn=MYOF;refGene_tx=XM_011287427.3;refGene_gn=MYOF;refGene_tx=XM_011287428.3;refGene_gn=MYOF;refGene_tx=XM_019813549.2;refGene_gn=MYOF;refGene_tx=XM_019813550.2
D2 55542017 55542208
refGene_gn=MYOF;refGene_tx=XM_003994218.5;refGene_gn=MYOF;refGene_tx=XM_011287427.3;refGene_gn=MYOF;refGene_tx=XM_011287428.3;refGene_gn=MYOF;refGene_tx=XM_019813549.2;refGene_gn=MYOF;refGene_tx=XM_019813550.2
D2 55543362 55543462
refGene_gn=MYOF;refGene_tx=XM_003994218.5;refGene_gn=MYOF;refGene_tx=XM_011287427.3;refGene_gn=MYOF;refGene_tx=XM_011287428.3;refGene_gn=MYOF;refGene_tx=XM_019813549.2;refGene_gn=MYOF;refGene_tx=XM_019813550.2
D2 55545943 55546085
refGene_gn=MYOF;refGene_tx=XM_003994218.5;refGene_gn=MYOF;refGene_tx=XM_011287427.3;refGene_gn=MYOF;refGene_tx=XM_011287428.3;refGene_gn=MYOF;refGene_tx=XM_019813549.2;refGene_gn=MYOF;refGene_tx=XM_019813550.2
D2 55549796 55549892
refGene_gn=MYOF;refGene_tx=XM_003994218.5;refGene_gn=MYOF;refGene_tx=XM_011287427.3;refGene_gn=MYOF;refGene_tx=XM_011287428.3;refGene_gn=MYOF;refGene_tx=XM_019813549.2;refGene_gn=MYOF;refGene_tx=XM_019813550.2
D2 55549968 55550057
refGene_gn=MYOF;refGene_tx=XM_003994218.5;refGene_gn=MYOF;refGene_tx=XM_011287427.3;refGene_gn=MYOF;refGene_tx=XM_011287428.3;refGene_gn=MYOF;refGene_tx=XM_019813549.2;refGene_gn=MYOF;refGene_tx=XM_019813550.2
D2 55552173 55552313
refGene_gn=MYOF;refGene_tx=XM_003994218.5;refGene_gn=MYOF;refGene_tx=XM_011287427.3;refGene_gn=MYOF;refGene_tx=XM_011287428.3;refGene_gn=MYOF;refGene_tx=XM_019813549.2;refGene_gn=MYOF;refGene_tx=XM_019813550.2
D2 55554304 55554453
refGene_gn=MYOF;refGene_tx=XM_003994218.5;refGene_gn=MYOF;refGene_tx=XM_011287427.3;refGene_gn=MYOF;refGene_tx=XM_011287428.3;refGene_gn=MYOF;refGene_tx=XM_019813549.2;refGene_gn=MYOF;refGene_tx=XM_019813550.2
D2 55555015 55555186
refGene_gn=MYOF;refGene_tx=XM_003994218.5;refGene_gn=MYOF;refGene_tx=XM_011287427.3;refGene_gn=MYOF;refGene_tx=XM_011287428.3;refGene_gn=MYOF;refGene_tx=XM_019813549.2;refGene_gn=MYOF;refGene_tx=XM_019813550.2
D2 55558183 55558275
refGene_gn=MYOF;refGene_tx=XM_003994218.5;refGene_gn=MYOF;refGene_tx=XM_011287427.3;refGene_gn=MYOF;refGene_tx=XM_011287428.3;refGene_gn=MYOF;refGene_tx=XM_019813549.2;refGene_gn=MYOF;refGene_tx=XM_019813550.2
D2 55558739 55558895
refGene_gn=MYOF;refGene_tx=XM_003994218.5;refGene_gn=MYOF;refGene_tx=XM_011287427.3;refGene_gn=MYOF;refGene_tx=XM_011287428.3;refGene_gn=MYOF;refGene_tx=XM_019813549.2;refGene_gn=MYOF;refGene_tx=XM_019813550.2
D2 55561202 55561328
refGene_gn=MYOF;refGene_tx=XM_003994218.5;refGene_gn=MYOF;refGene_tx=XM_011287427.3;refGene_gn=MYOF;refGene_tx=XM_011287428.3;refGene_gn=MYOF;refGene_tx=XM_019813549.2;refGene_gn=MYOF;refGene_tx=XM_019813550.2
D2 55562859 55562958

refGene_gn=MYOF;refGene_tx=XM_003994218.5;refGene_gn=MYOF;refGene_tx=XM_011287427.3;refGene_gn=MYOF;refGene_tx=XM_011287428.3;refGene_gn=MYOF;refGene_tx=XM_019813549.2;refGene_gn=MYOF;refGene_tx=XM_019813550.2
D2 55564468 55564480
refGene_gn=MYOF;refGene_tx=XM_003994218.5;refGene_gn=MYOF;refGene_tx=XM_011287427.3;refGene_gn=MYOF;refGene_tx=XM_019813549.2;refGene_gn=MYOF;refGene_tx=XM_019813550.2
D2 55565929 55565986
refGene_gn=MYOF;refGene_tx=XM_011287427.3;refGene_gn=MYOF;refGene_tx=XM_019813549.2
D2 55568489 55568566
refGene_gn=MYOF;refGene_tx=XM_003994218.5;refGene_gn=MYOF;refGene_tx=XM_011287427.3;refGene_gn=MYOF;refGene_tx=XM_011287428.3;refGene_gn=MYOF;refGene_tx=XM_019813549.2;refGene_gn=MYOF;refGene_tx=XM_019813550.2
D2 55571934 55572100
refGene_gn=MYOF;refGene_tx=XM_003994218.5;refGene_gn=MYOF;refGene_tx=XM_011287427.3;refGene_gn=MYOF;refGene_tx=XM_011287428.3;refGene_gn=MYOF;refGene_tx=XM_019813549.2;refGene_gn=MYOF;refGene_tx=XM_019813550.2
D2 55573992 55574154
refGene_gn=MYOF;refGene_tx=XM_003994218.5;refGene_gn=MYOF;refGene_tx=XM_011287427.3;refGene_gn=MYOF;refGene_tx=XM_011287428.3;refGene_gn=MYOF;refGene_tx=XM_019813549.2;refGene_gn=MYOF;refGene_tx=XM_019813550.2
D2 55575135 55575234
refGene_gn=MYOF;refGene_tx=XM_003994218.5;refGene_gn=MYOF;refGene_tx=XM_011287427.3;refGene_gn=MYOF;refGene_tx=XM_011287428.3;refGene_gn=MYOF;refGene_tx=XM_019813549.2;refGene_gn=MYOF;refGene_tx=XM_019813550.2
D2 55575358 55575517
refGene_gn=MYOF;refGene_tx=XM_003994218.5;refGene_gn=MYOF;refGene_tx=XM_011287427.3;refGene_gn=MYOF;refGene_tx=XM_011287428.3;refGene_gn=MYOF;refGene_tx=XM_019813549.2;refGene_gn=MYOF;refGene_tx=XM_019813550.2
D2 55575610 55575792
refGene_gn=MYOF;refGene_tx=XM_003994218.5;refGene_gn=MYOF;refGene_tx=XM_011287427.3;refGene_gn=MYOF;refGene_tx=XM_011287428.3;refGene_gn=MYOF;refGene_tx=XM_019813549.2;refGene_gn=MYOF;refGene_tx=XM_019813550.2
D2 55579886 55579964
refGene_gn=MYOF;refGene_tx=XM_003994218.5;refGene_gn=MYOF;refGene_tx=XM_011287427.3;refGene_gn=MYOF;refGene_tx=XM_011287428.3;refGene_gn=MYOF;refGene_tx=XM_019813549.2;refGene_gn=MYOF;refGene_tx=XM_019813550.2
D2 55581210 55581319
refGene_gn=MYOF;refGene_tx=XM_003994218.5;refGene_gn=MYOF;refGene_tx=XM_011287427.3;refGene_gn=MYOF;refGene_tx=XM_011287428.3;refGene_gn=MYOF;refGene_tx=XM_019813549.2;refGene_gn=MYOF;refGene_tx=XM_019813550.2
D2 55582217 55582391
refGene_gn=MYOF;refGene_tx=XM_003994218.5;refGene_gn=MYOF;refGene_tx=XM_011287427.3;refGene_gn=MYOF;refGene_tx=XM_011287428.3;refGene_gn=MYOF;refGene_tx=XM_019813549.2;refGene_gn=MYOF;refGene_tx=XM_019813550.2
D2 55584580 55584726
refGene_gn=MYOF;refGene_tx=XM_003994218.5;refGene_gn=MYOF;refGene_tx=XM_011287427.3;refGene_gn=MYOF;refGene_tx=XM_011287428.3;refGene_gn=MYOF;refGene_tx=XM_019813549.2;refGene_gn=MYOF;refGene_tx=XM_019813550.2

D2 55585910 55586016
refGene_gn=MYOF;refGene_tx=XM_003994218.5;refGene_gn=MYOF;refGene_tx=XM_011287427.3;refGene_gn=MYOF;refGene_tx=XM_011287428.3;refGene_gn=MYOF;refGene_tx=XM_019813549.2;refGene_gn=MYOF;refGene_tx=XM_019813550.2
D2 55587530 55587645
refGene_gn=MYOF;refGene_tx=XM_003994218.5;refGene_gn=MYOF;refGene_tx=XM_011287427.3;refGene_gn=MYOF;refGene_tx=XM_011287428.3;refGene_gn=MYOF;refGene_tx=XM_019813549.2;refGene_gn=MYOF;refGene_tx=XM_019813550.2
D2 55589916 55590080
refGene_gn=MYOF;refGene_tx=XM_003994218.5;refGene_gn=MYOF;refGene_tx=XM_011287427.3;refGene_gn=MYOF;refGene_tx=XM_011287428.3;refGene_gn=MYOF;refGene_tx=XM_019813549.2;refGene_gn=MYOF;refGene_tx=XM_019813550.2
D2 55592402 55592534
refGene_gn=MYOF;refGene_tx=XM_003994218.5;refGene_gn=MYOF;refGene_tx=XM_011287427.3;refGene_gn=MYOF;refGene_tx=XM_011287428.3;refGene_gn=MYOF;refGene_tx=XM_019813549.2;refGene_gn=MYOF;refGene_tx=XM_019813550.2
D2 55594745 55594901
refGene_gn=MYOF;refGene_tx=XM_003994218.5;refGene_gn=MYOF;refGene_tx=XM_011287427.3;refGene_gn=MYOF;refGene_tx=XM_011287428.3;refGene_gn=MYOF;refGene_tx=XM_019813549.2;refGene_gn=MYOF;refGene_tx=XM_019813550.2
D2 55596046 55596239
refGene_gn=MYOF;refGene_tx=XM_003994218.5;refGene_gn=MYOF;refGene_tx=XM_011287427.3;refGene_gn=MYOF;refGene_tx=XM_011287428.3;refGene_gn=MYOF;refGene_tx=XM_019813549.2;refGene_gn=MYOF;refGene_tx=XM_019813550.2
D2 55598540 55598647
refGene_gn=MYOF;refGene_tx=XM_003994218.5;refGene_gn=MYOF;refGene_tx=XM_011287427.3;refGene_gn=MYOF;refGene_tx=XM_011287428.3;refGene_gn=MYOF;refGene_tx=XM_019813549.2;refGene_gn=MYOF;refGene_tx=XM_019813550.2
D2 55601514 55601639
refGene_gn=MYOF;refGene_tx=XM_003994218.5;refGene_gn=MYOF;refGene_tx=XM_011287427.3;refGene_gn=MYOF;refGene_tx=XM_011287428.3;refGene_gn=MYOF;refGene_tx=XM_019813549.2;refGene_gn=MYOF;refGene_tx=XM_019813550.2
D2 55602870 55603048
refGene_gn=MYOF;refGene_tx=XM_003994218.5;refGene_gn=MYOF;refGene_tx=XM_011287427.3;refGene_gn=MYOF;refGene_tx=XM_011287428.3;refGene_gn=MYOF;refGene_tx=XM_019813549.2;refGene_gn=MYOF;refGene_tx=XM_019813550.2
D2 55613448 55613565
refGene_gn=MYOF;refGene_tx=XM_003994218.5;refGene_gn=MYOF;refGene_tx=XM_011287427.3;refGene_gn=MYOF;refGene_tx=XM_011287428.3;refGene_gn=MYOF;refGene_tx=XM_019813549.2;refGene_gn=MYOF;refGene_tx=XM_019813550.2
D2 55614730 55614855
refGene_gn=MYOF;refGene_tx=XM_003994218.5;refGene_gn=MYOF;refGene_tx=XM_011287427.3;refGene_gn=MYOF;refGene_tx=XM_011287428.3;refGene_gn=MYOF;refGene_tx=XM_019813549.2;refGene_gn=MYOF;refGene_tx=XM_019813550.2
D2 55618717 55618756
refGene_gn=MYOF;refGene_tx=XM_003994218.5;refGene_gn=MYOF;refGene_tx=XM_011287427.3;refGene_gn=MYOF;refGene_tx=XM_011287428.3
D2 55620865 55620948
refGene_gn=MYOF;refGene_tx=XM_003994218.5;refGene_gn=MYOF;refGene_tx=XM_011287427.3;refGene_gn=MYOF;refGene_tx=XM_011287428.3;refGene_gn=MYO

F;refGene_tx=XM_019813549.2;refGene_gn=MYOF;refGene_tx=XM_019813550.2
D2 55623898 55623942
refGene_gn=MYOF;refGene_tx=XM_003994218.5;refGene_gn=MYOF;refGene_tx=XM_011287427.3;refGene_gn=MYOF;refGene_tx=XM_011287428.3;refGene_gn=MYOF;refGene_tx=XM_019813549.2;refGene_gn=MYOF;refGene_tx=XM_019813550.2
D2 55624040 55624109
refGene_gn=MYOF;refGene_tx=XM_003994218.5;refGene_gn=MYOF;refGene_tx=XM_011287427.3;refGene_gn=MYOF;refGene_tx=XM_011287428.3;refGene_gn=MYOF;refGene_tx=XM_019813549.2;refGene_gn=MYOF;refGene_tx=XM_019813550.2
D2 55625949 55626053
refGene_gn=MYOF;refGene_tx=XM_003994218.5;refGene_gn=MYOF;refGene_tx=XM_011287427.3;refGene_gn=MYOF;refGene_tx=XM_011287428.3;refGene_gn=MYOF;refGene_tx=XM_019813549.2;refGene_gn=MYOF;refGene_tx=XM_019813550.2
D2 55627376 55627503
refGene_gn=MYOF;refGene_tx=XM_003994218.5;refGene_gn=MYOF;refGene_tx=XM_011287427.3;refGene_gn=MYOF;refGene_tx=XM_011287428.3;refGene_gn=MYOF;refGene_tx=XM_019813549.2;refGene_gn=MYOF;refGene_tx=XM_019813550.2
D2 55628155 55628271
refGene_gn=MYOF;refGene_tx=XM_003994218.5;refGene_gn=MYOF;refGene_tx=XM_011287427.3;refGene_gn=MYOF;refGene_tx=XM_011287428.3;refGene_gn=MYOF;refGene_tx=XM_019813549.2;refGene_gn=MYOF;refGene_tx=XM_019813550.2
D2 55628743 55628774
refGene_gn=MYOF;refGene_tx=XM_003994218.5;refGene_gn=MYOF;refGene_tx=XM_011287427.3;refGene_gn=MYOF;refGene_tx=XM_011287428.3;refGene_gn=MYOF;refGene_tx=XM_019813549.2;refGene_gn=MYOF;refGene_tx=XM_019813550.2
D2 55630163 55630214
refGene_gn=MYOF;refGene_tx=XM_003994218.5;refGene_gn=MYOF;refGene_tx=XM_011287427.3;refGene_gn=MYOF;refGene_tx=XM_011287428.3;refGene_gn=MYOF;refGene_tx=XM_019813549.2;refGene_gn=MYOF;refGene_tx=XM_019813550.2
D2 55630299 55630362
refGene_gn=MYOF;refGene_tx=XM_003994218.5;refGene_gn=MYOF;refGene_tx=XM_011287427.3;refGene_gn=MYOF;refGene_tx=XM_011287428.3;refGene_gn=MYOF;refGene_tx=XM_019813549.2;refGene_gn=MYOF;refGene_tx=XM_019813550.2
D2 55632547 55632676
refGene_gn=MYOF;refGene_tx=XM_003994218.5;refGene_gn=MYOF;refGene_tx=XM_011287427.3;refGene_gn=MYOF;refGene_tx=XM_011287428.3;refGene_gn=MYOF;refGene_tx=XM_019813549.2;refGene_gn=MYOF;refGene_tx=XM_019813550.2
D2 55633280 55633447
refGene_gn=MYOF;refGene_tx=XM_003994218.5;refGene_gn=MYOF;refGene_tx=XM_011287427.3;refGene_gn=MYOF;refGene_tx=XM_011287428.3;refGene_gn=MYOF;refGene_tx=XM_019813549.2;refGene_gn=MYOF;refGene_tx=XM_019813550.2
D2 55644934 55645022
refGene_gn=MYOF;refGene_tx=XM_003994218.5;refGene_gn=MYOF;refGene_tx=XM_011287427.3;refGene_gn=MYOF;refGene_tx=XM_011287428.3;refGene_gn=MYOF;refGene_tx=XM_019813549.2;refGene_gn=MYOF;refGene_tx=XM_019813550.2
D2 55648377 55648486
refGene_gn=MYOF;refGene_tx=XM_003994218.5;refGene_gn=MYOF;refGene_tx=XM_011287427.3;refGene_gn=MYOF;refGene_tx=XM_011287428.3;refGene_gn=MYOF;refGene_tx=XM_019813549.2;refGene_gn=MYOF;refGene_tx=XM_019813550.2
D2 55662269 55662361

refGene_gn=MYOF;refGene_tx=XM_003994218.5;refGene_gn=MYOF;refGene_tx=XM_011287427.3;refGene_gn=MYOF;refGene_tx=XM_011287428.3;refGene_gn=MYOF;refGene_tx=XM_019813549.2;refGene_gn=MYOF;refGene_tx=XM_019813550.2
D2 55664843 55664899 refGene_gn=MYOF;refGene_tx=XM_003994218.5;refGene_gn=MYOF;refGene_tx=XM_011287427.3;refGene_gn=MYOF;refGene_tx=XM_011287428.3;refGene_gn=MYOF;refGene_tx=XM_019813549.2;refGene_gn=MYOF;refGene_tx=XM_019813550.2
D2 55689340 55689428 refGene_gn=MYOF;refGene_tx=XM_003994218.5;refGene_gn=MYOF;refGene_tx=XM_011287427.3;refGene_gn=MYOF;refGene_tx=XM_011287428.3;refGene_gn=MYOF;refGene_tx=XM_019813549.2;refGene_gn=MYOF;refGene_tx=XM_019813550.2
D2 55700398 55700405 refGene_gn=CEP55;refGene_tx=XM_019813551.2;refGene_gn=CEP55;refGene_tx=XM_019813552.2
D2 55700939 55701130 refGene_gn=CEP55;refGene_tx=XM_003994244.5;refGene_gn=CEP55;refGene_tx=XM_011287426.3;refGene_gn=CEP55;refGene_tx=XM_019813551.2;refGene_gn=CEP55;refGene_tx=XM_019813552.2
D2 55704745 55705021 refGene_gn=CEP55;refGene_tx=XM_003994244.5;refGene_gn=CEP55;refGene_tx=XM_011287426.3;refGene_gn=CEP55;refGene_tx=XM_019813551.2;refGene_gn=CEP55;refGene_tx=XM_019813552.2
D2 55708417 55708486 refGene_gn=CEP55;refGene_tx=XM_003994244.5;refGene_gn=CEP55;refGene_tx=XM_011287426.3;refGene_gn=CEP55;refGene_tx=XM_019813551.2;refGene_gn=CEP55;refGene_tx=XM_019813552.2
D2 55716058 55716209 refGene_gn=CEP55;refGene_tx=XM_003994244.5;refGene_gn=CEP55;refGene_tx=XM_011287426.3;refGene_gn=CEP55;refGene_tx=XM_019813551.2;refGene_gn=CEP55;refGene_tx=XM_019813552.2
D2 55717663 55717977 refGene_gn=CEP55;refGene_tx=XM_003994244.5;refGene_gn=CEP55;refGene_tx=XM_011287426.3;refGene_gn=CEP55;refGene_tx=XM_019813551.2;refGene_gn=CEP55;refGene_tx=XM_019813552.2
D2 55719581 55719653 refGene_gn=CEP55;refGene_tx=XM_003994244.5;refGene_gn=CEP55;refGene_tx=XM_011287426.3;refGene_gn=CEP55;refGene_tx=XM_019813551.2;refGene_gn=CEP55;refGene_tx=XM_019813552.2
D2 55720363 55720489 refGene_gn=CEP55;refGene_tx=XM_003994244.5;refGene_gn=CEP55;refGene_tx=XM_011287426.3;refGene_gn=CEP55;refGene_tx=XM_019813551.2;refGene_gn=CEP55;refGene_tx=XM_019813552.2
D2 55722215 55722416 refGene_gn=CEP55;refGene_tx=XM_003994244.5;refGene_gn=CEP55;refGene_tx=XM_011287426.3;refGene_gn=CEP55;refGene_tx=XM_019813551.2;refGene_gn=CEP55;refGene_tx=XM_019813552.2
D2 55749767 55750334 refGene_gn=FFAR4;refGene_tx=NM_001317359.1
D2 55758324 55758453 refGene_gn=FFAR4;refGene_tx=NM_001317359.1
D2 55767447 55767837 refGene_gn=FFAR4;refGene_tx=NM_001317359.1
D2 55771911 55771949 refGene_gn=RBP4;refGene_tx=NM_001290249.1

D2 55773953 55774166 refGene_gn=RBP4;refGene_tx=NM_001290249.1
D2 55779990 55780097 refGene_gn=RBP4;refGene_tx=NM_001290249.1
D2 55780241 55780378 refGene_gn=RBP4;refGene_tx=NM_001290249.1
D2 55780479 55780590 refGene_gn=RBP4;refGene_tx=NM_001290249.1
D2 55790709 55791144
refGene_gn=PDE6C;refGene_tx=XM_003994246.5;refGene_gn=PDE6C;refGene_tx=XM_019813553.2
D2 55798392 55798545
refGene_gn=PDE6C;refGene_tx=XM_003994246.5;refGene_gn=PDE6C;refGene_tx=XM_019813553.2
D2 55798648 55798738
refGene_gn=PDE6C;refGene_tx=XM_003994246.5;refGene_gn=PDE6C;refGene_tx=XM_019813553.2
D2 55799667 55799808
refGene_gn=PDE6C;refGene_tx=XM_003994246.5;refGene_gn=PDE6C;refGene_tx=XM_019813553.2
D2 55801086 55801161
refGene_gn=PDE6C;refGene_tx=XM_003994246.5;refGene_gn=PDE6C;refGene_tx=XM_019813553.2
D2 55802106 55802171
refGene_gn=PDE6C;refGene_tx=XM_003994246.5;refGene_gn=PDE6C;refGene_tx=XM_019813553.2
D2 55802275 55802342
refGene_gn=PDE6C;refGene_tx=XM_003994246.5;refGene_gn=PDE6C;refGene_tx=XM_019813553.2
D2 55805074 55805122
refGene_gn=PDE6C;refGene_tx=XM_003994246.5;refGene_gn=PDE6C;refGene_tx=XM_019813553.2
D2 55807615 55807765
refGene_gn=PDE6C;refGene_tx=XM_003994246.5;refGene_gn=PDE6C;refGene_tx=XM_019813553.2
D2 55808347 55808491
refGene_gn=PDE6C;refGene_tx=XM_003994246.5;refGene_gn=PDE6C;refGene_tx=XM_019813553.2
D2 55809782 55809851
refGene_gn=PDE6C;refGene_tx=XM_003994246.5;refGene_gn=PDE6C;refGene_tx=XM_019813553.2
D2 55814789 55814936
refGene_gn=PDE6C;refGene_tx=XM_003994246.5;refGene_gn=PDE6C;refGene_tx=XM_019813553.2
D2 55815175 55815283
refGene_gn=PDE6C;refGene_tx=XM_003994246.5;refGene_gn=PDE6C;refGene_tx=XM_019813553.2
D2 55815660 55815770
refGene_gn=PDE6C;refGene_tx=XM_003994246.5;refGene_gn=PDE6C;refGene_tx=XM_019813553.2
D2 55821402 55821490
refGene_gn=PDE6C;refGene_tx=XM_003994246.5;refGene_gn=PDE6C;refGene_tx=XM_019813553.2
D2 55829094 55829195

refGene_gn=PDE6C;refGene_tx=XM_003994246.5;refGene_gn=PDE6C;refGene_tx=XM_019813553.2
D2 55830961 55831084
refGene_gn=PDE6C;refGene_tx=XM_003994246.5;refGene_gn=PDE6C;refGene_tx=XM_019813553.2
D2 55831229 55831293
refGene_gn=PDE6C;refGene_tx=XM_003994246.5;refGene_gn=PDE6C;refGene_tx=XM_019813553.2
D2 55833449 55833524
refGene_gn=PDE6C;refGene_tx=XM_003994246.5;refGene_gn=PDE6C;refGene_tx=XM_019813553.2
D2 55833635 55833719
refGene_gn=PDE6C;refGene_tx=XM_003994246.5;refGene_gn=PDE6C;refGene_tx=XM_019813553.2
D2 55834089 55834246
refGene_gn=PDE6C;refGene_tx=XM_003994246.5;refGene_gn=PDE6C;refGene_tx=XM_019813553.2
D2 55836399 55836461
refGene_gn=PDE6C;refGene_tx=XM_003994246.5;refGene_gn=PDE6C;refGene_tx=XM_019813553.2
D2 55840825 55840868
refGene_gn=LOC101087786;refGene_tx=XM_019813554.2;refGene_gn=LOC101087786;refGene_tx=XM_023240748.1;refGene_gn=LOC101087786;refGene_tx=XM_023240749.1
D2 55843052 55843131
refGene_gn=LOC101087786;refGene_tx=XM_019813554.2;refGene_gn=LOC101087786;refGene_tx=XM_023240748.1;refGene_gn=LOC101087786;refGene_tx=XM_023240749.1
D2 55848911 55848950
refGene_gn=LOC101087786;refGene_tx=XM_019813554.2;refGene_gn=LOC101087786;refGene_tx=XM_023240748.1;refGene_gn=LOC101087786;refGene_tx=XM_023240749.1
D2 55853503 55853622
refGene_gn=LOC101087786;refGene_tx=XM_019813554.2;refGene_gn=LOC101087786;refGene_tx=XM_023240748.1;refGene_gn=LOC101087786;refGene_tx=XM_023240749.1
D2 55855862 55855905
refGene_gn=LOC101087786;refGene_tx=XM_019813554.2;refGene_gn=LOC101087786;refGene_tx=XM_023240748.1;refGene_gn=LOC101087786;refGene_tx=XM_023240749.1
D2 55857678 55857792
refGene_gn=LOC101087786;refGene_tx=XM_019813554.2;refGene_gn=LOC101087786;refGene_tx=XM_023240748.1;refGene_gn=LOC101087786;refGene_tx=XM_023240749.1
D2 55860059 55860105
refGene_gn=LOC101087786;refGene_tx=XM_019813554.2;refGene_gn=LOC101087786;refGene_tx=XM_023240748.1;refGene_gn=LOC101087786;refGene_tx=XM_023240749.1
D2 55869315 55869400
refGene_gn=LOC101087786;refGene_tx=XM_019813554.2;refGene_gn=LOC101087

786;refGene_tx=XM_023240748.1;refGene_gn=LOC101087786;refGene_tx=XM_02
3240749.1
D2 55869987 55870071
refGene_gn=LOC101087786;refGene_tx=XM_019813554.2;refGene_gn=LOC101087
786;refGene_tx=XM_023240748.1;refGene_gn=LOC101087786;refGene_tx=XM_02
3240749.1
D2 55871344 55871421
refGene_gn=LOC101087786;refGene_tx=XM_019813554.2;refGene_gn=LOC101087
786;refGene_tx=XM_023240748.1;refGene_gn=LOC101087786;refGene_tx=XM_02
3240749.1
D2 55873770 55873816
refGene_gn=LOC101087786;refGene_tx=XM_019813554.2;refGene_gn=LOC101087
786;refGene_tx=XM_023240748.1;refGene_gn=LOC101087786;refGene_tx=XM_02
3240749.1
D2 55873942 55874038
refGene_gn=LOC101087786;refGene_tx=XM_019813554.2;refGene_gn=LOC101087
786;refGene_tx=XM_023240748.1;refGene_gn=LOC101087786;refGene_tx=XM_02
3240749.1
D2 55875371 55875448
refGene_gn=LOC101087786;refGene_tx=XM_019813554.2;refGene_gn=LOC101087
786;refGene_tx=XM_023240748.1;refGene_gn=LOC101087786;refGene_tx=XM_02
3240749.1
D2 55914976 55915191
refGene_gn=LGI1;refGene_tx=XM_003994222.5;refGene_gn=LGI1;refGene_tx=X
M_019813555.2;refGene_gn=LGI1;refGene_tx=XM_019813556.2;refGene_gn=LGI
1;refGene_tx=XM_019813557.2
D2 55915601 55915673
refGene_gn=LGI1;refGene_tx=XM_003994222.5;refGene_gn=LGI1;refGene_tx=X
M_019813556.2;refGene_gn=LGI1;refGene_tx=XM_019813557.2
D2 55934291 55934363
refGene_gn=LGI1;refGene_tx=XM_003994222.5;refGene_gn=LGI1;refGene_tx=X
M_019813555.2;refGene_gn=LGI1;refGene_tx=XM_019813557.2
D2 55934458 55934530
refGene_gn=LGI1;refGene_tx=XM_003994222.5;refGene_gn=LGI1;refGene_tx=X
M_019813555.2;refGene_gn=LGI1;refGene_tx=XM_019813557.2
D2 55945813 55945885
refGene_gn=LGI1;refGene_tx=XM_003994222.5;refGene_gn=LGI1;refGene_tx=X
M_019813555.2;refGene_gn=LGI1;refGene_tx=XM_019813556.2;refGene_gn=LGI
1;refGene_tx=XM_019813557.2
D2 55948962 55949132
refGene_gn=LGI1;refGene_tx=XM_003994222.5;refGene_gn=LGI1;refGene_tx=X
M_019813555.2;refGene_gn=LGI1;refGene_tx=XM_019813556.2;refGene_gn=LGI
1;refGene_tx=XM_019813557.2
D2 55949408 55949573
refGene_gn=LGI1;refGene_tx=XM_003994222.5;refGene_gn=LGI1;refGene_tx=X
M_019813555.2;refGene_gn=LGI1;refGene_tx=XM_019813556.2;refGene_gn=LGI
1;refGene_tx=XM_019813557.2
D2 55952120 55952956
refGene_gn=LGI1;refGene_tx=XM_003994222.5;refGene_gn=LGI1;refGene_tx=X
M_019813555.2;refGene_gn=LGI1;refGene_tx=XM_019813556.2;refGene_gn=LGI

1;refGene_tx=XM_019813557.2
D2 56030008 56030186
refGene_gn=SLC35G1;refGene_tx=XM_003994224.4;refGene_gn=SLC35G1;refGene_tx=XM_019813559.2
D2 56034921 56035102
refGene_gn=SLC35G1;refGene_tx=XM_003994224.4;refGene_gn=SLC35G1;refGene_tx=XM_019813559.2
D2 56037107 56037846
refGene_gn=SLC35G1;refGene_tx=XM_003994224.4;refGene_gn=SLC35G1;refGene_tx=XM_019813559.2
D2 56155138 56156341
refGene_gn=PLCE1;refGene_tx=XM_019813560.2;refGene_gn=PLCE1;refGene_tx=XM_023240750.1;refGene_gn=PLCE1;refGene_tx=XM_023240751.1;refGene_gn=PLCE1;refGene_tx=XM_023240752.1;refGene_gn=PLCE1;refGene_tx=XM_023240753.1;refGene_gn=PLCE1;refGene_tx=XM_023240754.1;refGene_gn=PLCE1;refGene_tx=XM_023240755.1
D2 56210364 56210415
refGene_gn=PLCE1;refGene_tx=XM_023240756.1;refGene_gn=PLCE1;refGene_tx=XM_023240757.1
D2 56253317 56253606
refGene_gn=PLCE1;refGene_tx=XM_019813560.2;refGene_gn=PLCE1;refGene_tx=XM_023240750.1;refGene_gn=PLCE1;refGene_tx=XM_023240751.1;refGene_gn=PLCE1;refGene_tx=XM_023240752.1;refGene_gn=PLCE1;refGene_tx=XM_023240753.1;refGene_gn=PLCE1;refGene_tx=XM_023240754.1;refGene_gn=PLCE1;refGene_tx=XM_023240755.1;refGene_gn=PLCE1;refGene_tx=XM_023240756.1;refGene_gn=PLCE1;refGene_tx=XM_023240757.1
D2 56291547 56291864
refGene_gn=PLCE1;refGene_tx=XM_019813560.2;refGene_gn=PLCE1;refGene_tx=XM_023240750.1;refGene_gn=PLCE1;refGene_tx=XM_023240751.1;refGene_gn=PLCE1;refGene_tx=XM_023240752.1;refGene_gn=PLCE1;refGene_tx=XM_023240753.1;refGene_gn=PLCE1;refGene_tx=XM_023240754.1;refGene_gn=PLCE1;refGene_tx=XM_023240755.1;refGene_gn=PLCE1;refGene_tx=XM_023240756.1;refGene_gn=PLCE1;refGene_tx=XM_023240757.1
D2 56339440 56339586
refGene_gn=PLCE1;refGene_tx=XM_019813560.2;refGene_gn=PLCE1;refGene_tx=XM_023240750.1;refGene_gn=PLCE1;refGene_tx=XM_023240751.1;refGene_gn=PLCE1;refGene_tx=XM_023240752.1;refGene_gn=PLCE1;refGene_tx=XM_023240753.1;refGene_gn=PLCE1;refGene_tx=XM_023240754.1;refGene_gn=PLCE1;refGene_tx=XM_023240755.1;refGene_gn=PLCE1;refGene_tx=XM_023240756.1;refGene_gn=PLCE1;refGene_tx=XM_023240757.1
D2 56343942 56344201
refGene_gn=PLCE1;refGene_tx=XM_019813560.2;refGene_gn=PLCE1;refGene_tx=XM_023240750.1;refGene_gn=PLCE1;refGene_tx=XM_023240751.1;refGene_gn=PLCE1;refGene_tx=XM_023240752.1;refGene_gn=PLCE1;refGene_tx=XM_023240753.1;refGene_gn=PLCE1;refGene_tx=XM_023240754.1;refGene_gn=PLCE1;refGene_tx=XM_023240755.1;refGene_gn=PLCE1;refGene_tx=XM_023240756.1;refGene_gn=PLCE1;refGene_tx=XM_023240757.1
D2 56345801 56346007
refGene_gn=PLCE1;refGene_tx=XM_019813560.2;refGene_gn=PLCE1;refGene_tx=XM_023240750.1;refGene_gn=PLCE1;refGene_tx=XM_023240751.1;refGene_gn=

PLCE1;refGene_tx=XM_023240752.1;refGene_gn=PLCE1;refGene_tx=XM_0232407
53.1;refGene_gn=PLCE1;refGene_tx=XM_023240754.1;refGene_gn=PLCE1;refGe
ne_tx=XM_023240755.1;refGene_gn=PLCE1;refGene_tx=XM_023240756.1;refGen
e_gn=PLCE1;refGene_tx=XM_023240757.1
D2 56358375 56359051
refGene_gn=PLCE1;refGene_tx=XM_019813560.2;refGene_gn=PLCE1;refGene_tx
=XM_023240750.1;refGene_gn=PLCE1;refGene_tx=XM_023240751.1;refGene_gn=
PLCE1;refGene_tx=XM_023240752.1;refGene_gn=PLCE1;refGene_tx=XM_0232407
53.1;refGene_gn=PLCE1;refGene_tx=XM_023240754.1;refGene_gn=PLCE1;refGe
ne_tx=XM_023240755.1;refGene_gn=PLCE1;refGene_tx=XM_023240756.1;refGen
e_gn=PLCE1;refGene_tx=XM_023240757.1
D2 56366206 56366389
refGene_gn=PLCE1;refGene_tx=XM_019813560.2;refGene_gn=PLCE1;refGene_tx
=XM_023240750.1;refGene_gn=PLCE1;refGene_tx=XM_023240751.1;refGene_gn=
PLCE1;refGene_tx=XM_023240752.1;refGene_gn=PLCE1;refGene_tx=XM_0232407
53.1;refGene_gn=PLCE1;refGene_tx=XM_023240754.1;refGene_gn=PLCE1;refGe
ne_tx=XM_023240755.1;refGene_gn=PLCE1;refGene_tx=XM_023240756.1;refGen
e_gn=PLCE1;refGene_tx=XM_023240757.1
D2 56368593 56368711
refGene_gn=PLCE1;refGene_tx=XM_019813560.2;refGene_gn=PLCE1;refGene_tx
=XM_023240750.1;refGene_gn=PLCE1;refGene_tx=XM_023240751.1;refGene_gn=
PLCE1;refGene_tx=XM_023240752.1;refGene_gn=PLCE1;refGene_tx=XM_0232407
53.1;refGene_gn=PLCE1;refGene_tx=XM_023240754.1;refGene_gn=PLCE1;refGe
ne_tx=XM_023240755.1;refGene_gn=PLCE1;refGene_tx=XM_023240756.1;refGen
e_gn=PLCE1;refGene_tx=XM_023240757.1
D2 56371440 56371597
refGene_gn=PLCE1;refGene_tx=XM_019813560.2;refGene_gn=PLCE1;refGene_tx
=XM_023240750.1;refGene_gn=PLCE1;refGene_tx=XM_023240751.1;refGene_gn=
PLCE1;refGene_tx=XM_023240752.1;refGene_gn=PLCE1;refGene_tx=XM_0232407
53.1;refGene_gn=PLCE1;refGene_tx=XM_023240754.1;refGene_gn=PLCE1;refGe
ne_tx=XM_023240755.1;refGene_gn=PLCE1;refGene_tx=XM_023240756.1;refGen
e_gn=PLCE1;refGene_tx=XM_023240757.1
D2 56374132 56374255
refGene_gn=PLCE1;refGene_tx=XM_019813560.2;refGene_gn=PLCE1;refGene_tx
=XM_023240750.1;refGene_gn=PLCE1;refGene_tx=XM_023240751.1;refGene_gn=
PLCE1;refGene_tx=XM_023240752.1;refGene_gn=PLCE1;refGene_tx=XM_0232407
53.1;refGene_gn=PLCE1;refGene_tx=XM_023240754.1;refGene_gn=PLCE1;refGe
ne_tx=XM_023240755.1;refGene_gn=PLCE1;refGene_tx=XM_023240756.1;refGen
e_gn=PLCE1;refGene_tx=XM_023240757.1
D2 56374340 56374477
refGene_gn=PLCE1;refGene_tx=XM_019813560.2;refGene_gn=PLCE1;refGene_tx
=XM_023240750.1;refGene_gn=PLCE1;refGene_tx=XM_023240751.1;refGene_gn=
PLCE1;refGene_tx=XM_023240752.1;refGene_gn=PLCE1;refGene_tx=XM_0232407
53.1;refGene_gn=PLCE1;refGene_tx=XM_023240754.1;refGene_gn=PLCE1;refGe
ne_tx=XM_023240755.1;refGene_gn=PLCE1;refGene_tx=XM_023240756.1;refGen
e_gn=PLCE1;refGene_tx=XM_023240757.1
D2 56377140 56377379
refGene_gn=PLCE1;refGene_tx=XM_019813560.2;refGene_gn=PLCE1;refGene_tx
=XM_023240750.1;refGene_gn=PLCE1;refGene_tx=XM_023240751.1;refGene_gn=
PLCE1;refGene_tx=XM_023240752.1;refGene_gn=PLCE1;refGene_tx=XM_0232407

53.1;refGene_gn=PLCE1;refGene_tx=XM_023240754.1;refGene_gn=PLCE1;refGene_tx=XM_023240755.1;refGene_gn=PLCE1;refGene_tx=XM_023240756.1;refGene_gn=PLCE1;refGene_tx=XM_023240757.1
D2 56379280 56379342
refGene_gn=PLCE1;refGene_tx=XM_019813560.2;refGene_gn=PLCE1;refGene_tx=XM_023240750.1;refGene_gn=PLCE1;refGene_tx=XM_023240751.1;refGene_gn=PLCE1;refGene_tx=XM_023240752.1;refGene_gn=PLCE1;refGene_tx=XM_023240753.1;refGene_gn=PLCE1;refGene_tx=XM_023240754.1;refGene_gn=PLCE1;refGene_tx=XM_023240755.1;refGene_gn=PLCE1;refGene_tx=XM_023240756.1;refGene_gn=PLCE1;refGene_tx=XM_023240757.1
D2 56379424 56379590
refGene_gn=PLCE1;refGene_tx=XM_019813560.2;refGene_gn=PLCE1;refGene_tx=XM_023240750.1;refGene_gn=PLCE1;refGene_tx=XM_023240751.1;refGene_gn=PLCE1;refGene_tx=XM_023240752.1;refGene_gn=PLCE1;refGene_tx=XM_023240753.1;refGene_gn=PLCE1;refGene_tx=XM_023240754.1;refGene_gn=PLCE1;refGene_tx=XM_023240755.1;refGene_gn=PLCE1;refGene_tx=XM_023240756.1;refGene_gn=PLCE1;refGene_tx=XM_023240757.1
D2 56383208 56383316
refGene_gn=PLCE1;refGene_tx=XM_019813560.2;refGene_gn=PLCE1;refGene_tx=XM_023240750.1;refGene_gn=PLCE1;refGene_tx=XM_023240751.1;refGene_gn=PLCE1;refGene_tx=XM_023240752.1;refGene_gn=PLCE1;refGene_tx=XM_023240753.1;refGene_gn=PLCE1;refGene_tx=XM_023240754.1;refGene_gn=PLCE1;refGene_tx=XM_023240755.1;refGene_gn=PLCE1;refGene_tx=XM_023240756.1;refGene_gn=PLCE1;refGene_tx=XM_023240757.1
D2 56384427 56384544
refGene_gn=PLCE1;refGene_tx=XM_019813560.2;refGene_gn=PLCE1;refGene_tx=XM_023240750.1;refGene_gn=PLCE1;refGene_tx=XM_023240751.1;refGene_gn=PLCE1;refGene_tx=XM_023240752.1;refGene_gn=PLCE1;refGene_tx=XM_023240753.1;refGene_gn=PLCE1;refGene_tx=XM_023240754.1;refGene_gn=PLCE1;refGene_tx=XM_023240755.1;refGene_gn=PLCE1;refGene_tx=XM_023240756.1;refGene_gn=PLCE1;refGene_tx=XM_023240757.1
D2 56386524 56386683
refGene_gn=PLCE1;refGene_tx=XM_019813560.2;refGene_gn=PLCE1;refGene_tx=XM_023240750.1;refGene_gn=PLCE1;refGene_tx=XM_023240751.1;refGene_gn=PLCE1;refGene_tx=XM_023240752.1;refGene_gn=PLCE1;refGene_tx=XM_023240753.1;refGene_gn=PLCE1;refGene_tx=XM_023240754.1;refGene_gn=PLCE1;refGene_tx=XM_023240755.1;refGene_gn=PLCE1;refGene_tx=XM_023240756.1;refGene_gn=PLCE1;refGene_tx=XM_023240757.1
D2 56395332 56395462
refGene_gn=PLCE1;refGene_tx=XM_019813560.2;refGene_gn=PLCE1;refGene_tx=XM_023240750.1;refGene_gn=PLCE1;refGene_tx=XM_023240751.1;refGene_gn=PLCE1;refGene_tx=XM_023240752.1;refGene_gn=PLCE1;refGene_tx=XM_023240753.1;refGene_gn=PLCE1;refGene_tx=XM_023240754.1;refGene_gn=PLCE1;refGene_tx=XM_023240755.1;refGene_gn=PLCE1;refGene_tx=XM_023240756.1;refGene_gn=PLCE1;refGene_tx=XM_023240757.1
D2 56399093 56399135
refGene_gn=PLCE1;refGene_tx=XM_019813560.2;refGene_gn=PLCE1;refGene_tx=XM_023240750.1;refGene_gn=PLCE1;refGene_tx=XM_023240751.1;refGene_gn=PLCE1;refGene_tx=XM_023240753.1;refGene_gn=PLCE1;refGene_tx=XM_023240755.1;refGene_gn=PLCE1;refGene_tx=XM_023240756.1

D2 56400279 56400401
refGene_gn=PLCE1;refGene_tx=XM_019813560.2;refGene_gn=PLCE1;refGene_tx=XM_023240750.1;refGene_gn=PLCE1;refGene_tx=XM_023240751.1;refGene_gn=PLCE1;refGene_tx=XM_023240752.1;refGene_gn=PLCE1;refGene_tx=XM_023240753.1;refGene_gn=PLCE1;refGene_tx=XM_023240754.1;refGene_gn=PLCE1;refGene_tx=XM_023240755.1;refGene_gn=PLCE1;refGene_tx=XM_023240756.1;refGene_gn=PLCE1;refGene_tx=XM_023240757.1
D2 56401313 56401431
refGene_gn=PLCE1;refGene_tx=XM_019813560.2;refGene_gn=PLCE1;refGene_tx=XM_023240750.1;refGene_gn=PLCE1;refGene_tx=XM_023240751.1;refGene_gn=PLCE1;refGene_tx=XM_023240752.1;refGene_gn=PLCE1;refGene_tx=XM_023240753.1;refGene_gn=PLCE1;refGene_tx=XM_023240754.1;refGene_gn=PLCE1;refGene_tx=XM_023240755.1;refGene_gn=PLCE1;refGene_tx=XM_023240756.1;refGene_gn=PLCE1;refGene_tx=XM_023240757.1
D2 56406196 56406328
refGene_gn=PLCE1;refGene_tx=XM_019813560.2;refGene_gn=PLCE1;refGene_tx=XM_023240750.1;refGene_gn=PLCE1;refGene_tx=XM_023240751.1;refGene_gn=PLCE1;refGene_tx=XM_023240752.1;refGene_gn=PLCE1;refGene_tx=XM_023240753.1;refGene_gn=PLCE1;refGene_tx=XM_023240754.1;refGene_gn=PLCE1;refGene_tx=XM_023240755.1;refGene_gn=PLCE1;refGene_tx=XM_023240756.1;refGene_gn=PLCE1;refGene_tx=XM_023240757.1
D2 56409162 56409453
refGene_gn=PLCE1;refGene_tx=XM_019813560.2;refGene_gn=PLCE1;refGene_tx=XM_023240750.1;refGene_gn=PLCE1;refGene_tx=XM_023240751.1;refGene_gn=PLCE1;refGene_tx=XM_023240752.1;refGene_gn=PLCE1;refGene_tx=XM_023240753.1;refGene_gn=PLCE1;refGene_tx=XM_023240754.1;refGene_gn=PLCE1;refGene_tx=XM_023240755.1;refGene_gn=PLCE1;refGene_tx=XM_023240756.1;refGene_gn=PLCE1;refGene_tx=XM_023240757.1
D2 56413994 56414158
refGene_gn=PLCE1;refGene_tx=XM_019813560.2;refGene_gn=PLCE1;refGene_tx=XM_023240750.1;refGene_gn=PLCE1;refGene_tx=XM_023240751.1;refGene_gn=PLCE1;refGene_tx=XM_023240752.1;refGene_gn=PLCE1;refGene_tx=XM_023240753.1;refGene_gn=PLCE1;refGene_tx=XM_023240754.1;refGene_gn=PLCE1;refGene_tx=XM_023240755.1;refGene_gn=PLCE1;refGene_tx=XM_023240756.1;refGene_gn=PLCE1;refGene_tx=XM_023240757.1
D2 56415429 56415691
refGene_gn=PLCE1;refGene_tx=XM_019813560.2;refGene_gn=PLCE1;refGene_tx=XM_023240750.1;refGene_gn=PLCE1;refGene_tx=XM_023240751.1;refGene_gn=PLCE1;refGene_tx=XM_023240752.1;refGene_gn=PLCE1;refGene_tx=XM_023240753.1;refGene_gn=PLCE1;refGene_tx=XM_023240754.1;refGene_gn=PLCE1;refGene_tx=XM_023240755.1;refGene_gn=PLCE1;refGene_tx=XM_023240756.1;refGene_gn=PLCE1;refGene_tx=XM_023240757.1
D2 56418648 56418767
refGene_gn=PLCE1;refGene_tx=XM_019813560.2;refGene_gn=PLCE1;refGene_tx=XM_023240750.1;refGene_gn=PLCE1;refGene_tx=XM_023240751.1;refGene_gn=PLCE1;refGene_tx=XM_023240752.1;refGene_gn=PLCE1;refGene_tx=XM_023240753.1;refGene_gn=PLCE1;refGene_tx=XM_023240754.1;refGene_gn=PLCE1;refGene_tx=XM_023240755.1;refGene_gn=PLCE1;refGene_tx=XM_023240756.1;refGene_gn=PLCE1;refGene_tx=XM_023240757.1
D2 56423292 56423421

refGene_gn=PLCE1;refGene_tx=XM_019813560.2;refGene_gn=PLCE1;refGene_tx=XM_023240750.1;refGene_gn=PLCE1;refGene_tx=XM_023240751.1;refGene_gn=PLCE1;refGene_tx=XM_023240752.1;refGene_gn=PLCE1;refGene_tx=XM_023240753.1;refGene_gn=PLCE1;refGene_tx=XM_023240754.1;refGene_gn=PLCE1;refGene_tx=XM_023240755.1;refGene_gn=PLCE1;refGene_tx=XM_023240756.1;refGene_gn=PLCE1;refGene_tx=XM_023240757.1
D2 56425691 56425901
refGene_gn=PLCE1;refGene_tx=XM_019813560.2;refGene_gn=PLCE1;refGene_tx=XM_023240750.1;refGene_gn=PLCE1;refGene_tx=XM_023240751.1;refGene_gn=PLCE1;refGene_tx=XM_023240752.1;refGene_gn=PLCE1;refGene_tx=XM_023240753.1;refGene_gn=PLCE1;refGene_tx=XM_023240754.1;refGene_gn=PLCE1;refGene_tx=XM_023240755.1;refGene_gn=PLCE1;refGene_tx=XM_023240756.1;refGene_gn=PLCE1;refGene_tx=XM_023240757.1
D2 56428262 56428331 refGene_gn=PLCE1;refGene_tx=XM_023240755.1
D2 56430289 56430448
refGene_gn=PLCE1;refGene_tx=XM_019813560.2;refGene_gn=PLCE1;refGene_tx=XM_023240750.1;refGene_gn=PLCE1;refGene_tx=XM_023240751.1;refGene_gn=PLCE1;refGene_tx=XM_023240752.1;refGene_gn=PLCE1;refGene_tx=XM_023240753.1;refGene_gn=PLCE1;refGene_tx=XM_023240754.1;refGene_gn=PLCE1;refGene_tx=XM_023240756.1;refGene_gn=PLCE1;refGene_tx=XM_023240757.1
D2 56432810 56433026
refGene_gn=PLCE1;refGene_tx=XM_019813560.2;refGene_gn=PLCE1;refGene_tx=XM_023240750.1;refGene_gn=PLCE1;refGene_tx=XM_023240751.1;refGene_gn=PLCE1;refGene_tx=XM_023240752.1;refGene_gn=PLCE1;refGene_tx=XM_023240753.1;refGene_gn=PLCE1;refGene_tx=XM_023240754.1;refGene_gn=PLCE1;refGene_tx=XM_023240756.1;refGene_gn=PLCE1;refGene_tx=XM_023240757.1
D2 56433363 56433552
refGene_gn=PLCE1;refGene_tx=XM_019813560.2;refGene_gn=PLCE1;refGene_tx=XM_023240750.1;refGene_gn=PLCE1;refGene_tx=XM_023240751.1;refGene_gn=PLCE1;refGene_tx=XM_023240752.1;refGene_gn=PLCE1;refGene_tx=XM_023240753.1;refGene_gn=PLCE1;refGene_tx=XM_023240754.1;refGene_gn=PLCE1;refGene_tx=XM_023240756.1;refGene_gn=PLCE1;refGene_tx=XM_023240757.1
D2 56442725 56442857 refGene_gn=NOC3L;refGene_tx=XM_019813567.2
D2 56443161 56443246 refGene_gn=NOC3L;refGene_tx=XM_019813567.2
D2 56445706 56445804 refGene_gn=NOC3L;refGene_tx=XM_019813567.2
D2 56446723 56446852 refGene_gn=NOC3L;refGene_tx=XM_019813567.2
D2 56447359 56447541 refGene_gn=NOC3L;refGene_tx=XM_019813567.2
D2 56448115 56448187 refGene_gn=NOC3L;refGene_tx=XM_019813567.2
D2 56448322 56448386 refGene_gn=NOC3L;refGene_tx=XM_019813567.2
D2 56448471 56448544 refGene_gn=NOC3L;refGene_tx=XM_019813567.2
D2 56451786 56451887 refGene_gn=NOC3L;refGene_tx=XM_019813567.2
D2 56452203 56452284 refGene_gn=NOC3L;refGene_tx=XM_019813567.2
D2 56453754 56453886 refGene_gn=NOC3L;refGene_tx=XM_019813567.2
D2 56457872 56458001 refGene_gn=NOC3L;refGene_tx=XM_019813567.2
D2 56458383 56458559 refGene_gn=NOC3L;refGene_tx=XM_019813567.2
D2 56462064 56462158 refGene_gn=NOC3L;refGene_tx=XM_019813567.2
D2 56462339 56462501 refGene_gn=NOC3L;refGene_tx=XM_019813567.2
D2 56464555 56464686 refGene_gn=NOC3L;refGene_tx=XM_019813567.2
D2 56466697 56466751 refGene_gn=NOC3L;refGene_tx=XM_019813567.2
D2 56467355 56467513 refGene_gn=NOC3L;refGene_tx=XM_019813567.2

D2 56468776 56468909 refGene_gn=NOC3L;refGene_tx=XM_019813567.2
D2 56472188 56472396 refGene_gn=NOC3L;refGene_tx=XM_019813567.2
D2 56473139 56473148 refGene_gn=NOC3L;refGene_tx=XM_019813567.2
D2 56496084 56497016 refGene_gn=TBC1D12;refGene_tx=XM_023240758.1
D2 56531623 56531747 refGene_gn=TBC1D12;refGene_tx=XM_023240758.1
D2 56557837 56557953 refGene_gn=TBC1D12;refGene_tx=XM_023240758.1
D2 56569505 56569588 refGene_gn=TBC1D12;refGene_tx=XM_023240758.1
D2 56573160 56573255 refGene_gn=TBC1D12;refGene_tx=XM_023240758.1
D2 56573256 56573279 refGene_gn=TBC1D12;refGene_tx=XM_023240758.1
D2 56574487 56574594 refGene_gn=TBC1D12;refGene_tx=XM_023240758.1
D2 56579646 56579727 refGene_gn=TBC1D12;refGene_tx=XM_023240758.1
D2 56582619 56582708 refGene_gn=TBC1D12;refGene_tx=XM_023240758.1
D2 56583741 56583813 refGene_gn=TBC1D12;refGene_tx=XM_023240758.1
D2 56590780 56590909 refGene_gn=TBC1D12;refGene_tx=XM_023240758.1
D2 56591167 56591277 refGene_gn=TBC1D12;refGene_tx=XM_023240758.1
D2 56593693 56593952 refGene_gn=TBC1D12;refGene_tx=XM_023240758.1
D2 56594743 56594812 refGene_gn=TBC1D12;refGene_tx=XM_023240758.1
D2 56613231 56613262 refGene_gn=HELLS;refGene_tx=XM_006938008.4
D2 56613689 56613811 refGene_gn=HELLS;refGene_tx=XM_006938008.4
D2 56619175 56619298 refGene_gn=HELLS;refGene_tx=XM_006938008.4
D2 56622133 56622190 refGene_gn=HELLS;refGene_tx=XM_006938008.4
D2 56625535 56625572 refGene_gn=HELLS;refGene_tx=XM_006938008.4
D2 56625657 56625722 refGene_gn=HELLS;refGene_tx=XM_006938008.4
D2 56628440 56628482 refGene_gn=HELLS;refGene_tx=XM_006938008.4
D2 56631805 56632030 refGene_gn=HELLS;refGene_tx=XM_006938008.4
D2 56632394 56632577 refGene_gn=HELLS;refGene_tx=XM_006938008.4
D2 56633375 56633519 refGene_gn=HELLS;refGene_tx=XM_006938008.4
D2 56636297 56636494 refGene_gn=HELLS;refGene_tx=XM_006938008.4
D2 56636686 56636783 refGene_gn=HELLS;refGene_tx=XM_006938008.4
D2 56638325 56638487 refGene_gn=HELLS;refGene_tx=XM_006938008.4
D2 56639464 56639604 refGene_gn=HELLS;refGene_tx=XM_006938008.4
D2 56639690 56639829 refGene_gn=HELLS;refGene_tx=XM_006938008.4
D2 56641162 56641246 refGene_gn=HELLS;refGene_tx=XM_006938008.4
D2 56641327 56641447 refGene_gn=HELLS;refGene_tx=XM_006938008.4
D2 56643138 56643255 refGene_gn=HELLS;refGene_tx=XM_006938008.4
D2 56643562 56643722 refGene_gn=HELLS;refGene_tx=XM_006938008.4
D2 56646214 56646311 refGene_gn=HELLS;refGene_tx=XM_006938008.4
D2 56646389 56646466 refGene_gn=HELLS;refGene_tx=XM_006938008.4
D2 56650081 56650176 refGene_gn=HELLS;refGene_tx=XM_006938008.4
D2 56666614 56666782
refGene_gn=LOC101089396;refGene_tx=XM_019813568.2
D2 56668086 56668249
refGene_gn=LOC101089396;refGene_tx=XM_019813568.2;refGene_gn=LOC101089
396;refGene_tx=XM_023240759.1
D2 56668406 56668556
refGene_gn=LOC101089396;refGene_tx=XM_019813568.2;refGene_gn=LOC101089
396;refGene_tx=XM_023240759.1
D2 56672999 56673160
refGene_gn=LOC101089396;refGene_tx=XM_019813568.2;refGene_gn=LOC101089
396;refGene_tx=XM_023240759.1

D2 56676922 56677099
refGene_gn=LOC101089396;refGene_tx=XM_019813568.2;refGene_gn=LOC101089
396;refGene_tx=XM_023240759.1
D2 56681061 56681203
refGene_gn=LOC101089396;refGene_tx=XM_019813568.2;refGene_gn=LOC101089
396;refGene_tx=XM_023240759.1
D2 56687694 56687882
refGene_gn=LOC101089396;refGene_tx=XM_019813568.2;refGene_gn=LOC101089
396;refGene_tx=XM_023240759.1
D2 56696717 56696859
refGene_gn=LOC101089396;refGene_tx=XM_019813568.2;refGene_gn=LOC101089
396;refGene_tx=XM_023240759.1
D2 56699147 56699329
refGene_gn=LOC101089396;refGene_tx=XM_019813568.2;refGene_gn=LOC101089
396;refGene_tx=XM_023240759.1
D2 56716673 56716855
refGene_gn=LOC101089804;refGene_tx=XM_019813570.2
D2 56724962 56725104
refGene_gn=LOC101089804;refGene_tx=XM_019813570.2
D2 56729505 56729693
refGene_gn=LOC101089804;refGene_tx=XM_019813570.2
D2 56733329 56733471
refGene_gn=LOC101089804;refGene_tx=XM_019813570.2
D2 56734328 56734502
refGene_gn=LOC101089804;refGene_tx=XM_019813570.2
D2 56736275 56736436
refGene_gn=LOC101089804;refGene_tx=XM_019813570.2
D2 56738473 56738623
refGene_gn=LOC101089804;refGene_tx=XM_019813570.2
D2 56740408 56740571
refGene_gn=LOC101089804;refGene_tx=XM_019813570.2
D2 56745197 56745317
refGene_gn=LOC101089804;refGene_tx=XM_019813570.2
D2 56745318 56745441
refGene_gn=LOC101089804;refGene_tx=XM_019813570.2
D2 56771697 56771884 refGene_gn=PDLIM1;refGene_tx=XM_023240760.1
D2 56772297 56772415 refGene_gn=PDLIM1;refGene_tx=XM_023240760.1
D2 56779673 56779825 refGene_gn=PDLIM1;refGene_tx=XM_023240760.1
D2 56796265 56796465 refGene_gn=PDLIM1;refGene_tx=XM_023240760.1
D2 56801211 56801296 refGene_gn=PDLIM1;refGene_tx=XM_023240760.1
D2 56802903 56803055 refGene_gn=PDLIM1;refGene_tx=XM_023240760.1
D2 56819825 56819921 refGene_gn=PDLIM1;refGene_tx=XM_023240760.1
D2 56842457 56842525
refGene_gn=SORBS1;refGene_tx=XM_019813577.2;refGene_gn=SORBS1;refGene_
tx=XM_019813579.2;refGene_gn=SORBS1;refGene_tx=XM_023240761.1;refGene_
gn=SORBS1;refGene_tx=XM_023240762.1;refGene_gn=SORBS1;refGene_tx=XM_02
3240763.1;refGene_gn=SORBS1;refGene_tx=XM_023240764.1;refGene_gn=SORBS
1;refGene_tx=XM_023240765.1;refGene_gn=SORBS1;refGene_tx=XM_023240766.
1;refGene_gn=SORBS1;refGene_tx=XM_023240767.1;refGene_gn=SORBS1;refGen
e_tx=XM_023240769.1;refGene_gn=SORBS1;refGene_tx=XM_023240770.1;refGen

e_gn=SORBS1;refGene_tx=XM_023240771.1;refGene_gn=SORBS1;refGene_tx=XM_023240772.1;refGene_gn=SORBS1;refGene_tx=XM_023240773.1;refGene_gn=SORBS1;refGene_tx=XM_023240774.1;refGene_gn=SORBS1;refGene_tx=XM_023240775.1;refGene_gn=SORBS1;refGene_tx=XM_023240776.1;refGene_gn=SORBS1;refGene_tx=XM_023240777.1;refGene_gn=SORBS1;refGene_tx=XM_023240778.1;refGene_gn=SORBS1;refGene_tx=XM_023240779.1;refGene_gn=SORBS1;refGene_tx=XM_023240780.1;refGene_gn=SORBS1;refGene_tx=XM_023240781.1;refGene_gn=SORBS1;refGene_tx=XM_023240782.1
D2 56845694 56845807
refGene_gn=SORBS1;refGene_tx=XM_019813577.2;refGene_gn=SORBS1;refGene_tx=XM_019813579.2;refGene_gn=SORBS1;refGene_tx=XM_023240761.1;refGene_gn=SORBS1;refGene_tx=XM_023240762.1;refGene_gn=SORBS1;refGene_tx=XM_023240763.1;refGene_gn=SORBS1;refGene_tx=XM_023240764.1;refGene_gn=SORBS1;refGene_tx=XM_023240765.1;refGene_gn=SORBS1;refGene_tx=XM_023240766.1;refGene_gn=SORBS1;refGene_tx=XM_023240767.1;refGene_gn=SORBS1;refGene_tx=XM_023240769.1;refGene_gn=SORBS1;refGene_tx=XM_023240770.1;refGene_gn=SORBS1;refGene_tx=XM_023240771.1;refGene_gn=SORBS1;refGene_tx=XM_023240772.1;refGene_gn=SORBS1;refGene_tx=XM_023240773.1;refGene_gn=SORBS1;refGene_tx=XM_023240774.1;refGene_gn=SORBS1;refGene_tx=XM_023240775.1;refGene_gn=SORBS1;refGene_tx=XM_023240776.1;refGene_gn=SORBS1;refGene_tx=XM_023240777.1;refGene_gn=SORBS1;refGene_tx=XM_023240778.1;refGene_gn=SORBS1;refGene_tx=XM_023240779.1;refGene_gn=SORBS1;refGene_tx=XM_023240780.1;refGene_gn=SORBS1;refGene_tx=XM_023240781.1;refGene_gn=SORBS1;refGene_tx=XM_023240782.1
D2 56848445 56848636
refGene_gn=SORBS1;refGene_tx=XM_019813577.2;refGene_gn=SORBS1;refGene_tx=XM_019813579.2;refGene_gn=SORBS1;refGene_tx=XM_023240761.1;refGene_gn=SORBS1;refGene_tx=XM_023240762.1;refGene_gn=SORBS1;refGene_tx=XM_023240763.1;refGene_gn=SORBS1;refGene_tx=XM_023240764.1;refGene_gn=SORBS1;refGene_tx=XM_023240765.1;refGene_gn=SORBS1;refGene_tx=XM_023240766.1;refGene_gn=SORBS1;refGene_tx=XM_023240767.1;refGene_gn=SORBS1;refGene_tx=XM_023240769.1;refGene_gn=SORBS1;refGene_tx=XM_023240770.1;refGene_gn=SORBS1;refGene_tx=XM_023240771.1;refGene_gn=SORBS1;refGene_tx=XM_023240772.1;refGene_gn=SORBS1;refGene_tx=XM_023240773.1;refGene_gn=SORBS1;refGene_tx=XM_023240774.1;refGene_gn=SORBS1;refGene_tx=XM_023240775.1;refGene_gn=SORBS1;refGene_tx=XM_023240776.1;refGene_gn=SORBS1;refGene_tx=XM_023240777.1;refGene_gn=SORBS1;refGene_tx=XM_023240778.1;refGene_gn=SORBS1;refGene_tx=XM_023240779.1;refGene_gn=SORBS1;refGene_tx=XM_023240780.1;refGene_gn=SORBS1;refGene_tx=XM_023240781.1;refGene_gn=SORBS1;refGene_tx=XM_023240782.1
D2 56849212 56849272
refGene_gn=SORBS1;refGene_tx=XM_019813577.2;refGene_gn=SORBS1;refGene_tx=XM_019813579.2;refGene_gn=SORBS1;refGene_tx=XM_023240762.1;refGene_gn=SORBS1;refGene_tx=XM_023240763.1;refGene_gn=SORBS1;refGene_tx=XM_023240764.1;refGene_gn=SORBS1;refGene_tx=XM_023240765.1;refGene_gn=SORBS1;refGene_tx=XM_023240769.1;refGene_gn=SORBS1;refGene_tx=XM_023240770.1
D2 56860147 56860921
refGene_gn=SORBS1;refGene_tx=XM_019813577.2;refGene_gn=SORBS1;refGene_tx=XM_019813579.2;refGene_gn=SORBS1;refGene_tx=XM_023240761.1;refGene_

gn=SORBS1;refGene_tx=XM_023240762.1;refGene_gn=SORBS1;refGene_tx=XM_02
3240763.1;refGene_gn=SORBS1;refGene_tx=XM_023240764.1;refGene_gn=SORBS
1;refGene_tx=XM_023240765.1;refGene_gn=SORBS1;refGene_tx=XM_023240766.
1;refGene_gn=SORBS1;refGene_tx=XM_023240769.1
D2 56862432 56862627
refGene_gn=SORBS1;refGene_tx=XM_019813577.2;refGene_gn=SORBS1;refGene_
tx=XM_019813579.2;refGene_gn=SORBS1;refGene_tx=XM_023240761.1;refGene_
gn=SORBS1;refGene_tx=XM_023240762.1;refGene_gn=SORBS1;refGene_tx=XM_02
3240763.1;refGene_gn=SORBS1;refGene_tx=XM_023240764.1;refGene_gn=SORBS
1;refGene_tx=XM_023240765.1;refGene_gn=SORBS1;refGene_tx=XM_023240766.
1;refGene_gn=SORBS1;refGene_tx=XM_023240767.1;refGene_gn=SORBS1;refGen
e_tx=XM_023240769.1;refGene_gn=SORBS1;refGene_tx=XM_023240770.1;refGen
e_gn=SORBS1;refGene_tx=XM_023240771.1;refGene_gn=SORBS1;refGene_tx=XM_
023240772.1;refGene_gn=SORBS1;refGene_tx=XM_023240773.1;refGene_gn=SOR
BS1;refGene_tx=XM_023240774.1;refGene_gn=SORBS1;refGene_tx=XM_02324077
5.1;refGene_gn=SORBS1;refGene_tx=XM_023240776.1;refGene_gn=SORBS1;refG
ene_tx=XM_023240777.1;refGene_gn=SORBS1;refGene_tx=XM_023240778.1;refG
ene_gn=SORBS1;refGene_tx=XM_023240779.1;refGene_gn=SORBS1;refGene_tx=X
M_023240780.1;refGene_gn=SORBS1;refGene_tx=XM_023240781.1;refGene_gn=S
ORBS1;refGene_tx=XM_023240782.1
D2 56864548 56864674
refGene_gn=SORBS1;refGene_tx=XM_019813577.2;refGene_gn=SORBS1;refGene_
tx=XM_019813579.2;refGene_gn=SORBS1;refGene_tx=XM_023240761.1;refGene_
gn=SORBS1;refGene_tx=XM_023240762.1;refGene_gn=SORBS1;refGene_tx=XM_02
3240763.1;refGene_gn=SORBS1;refGene_tx=XM_023240764.1;refGene_gn=SORBS
1;refGene_tx=XM_023240765.1;refGene_gn=SORBS1;refGene_tx=XM_023240766.
1;refGene_gn=SORBS1;refGene_tx=XM_023240767.1;refGene_gn=SORBS1;refGen
e_tx=XM_023240769.1;refGene_gn=SORBS1;refGene_tx=XM_023240770.1;refGen
e_gn=SORBS1;refGene_tx=XM_023240771.1;refGene_gn=SORBS1;refGene_tx=XM_
023240772.1;refGene_gn=SORBS1;refGene_tx=XM_023240773.1;refGene_gn=SOR
BS1;refGene_tx=XM_023240774.1;refGene_gn=SORBS1;refGene_tx=XM_02324077
5.1;refGene_gn=SORBS1;refGene_tx=XM_023240776.1;refGene_gn=SORBS1;refG
ene_tx=XM_023240777.1;refGene_gn=SORBS1;refGene_tx=XM_023240778.1;refG
ene_gn=SORBS1;refGene_tx=XM_023240779.1;refGene_gn=SORBS1;refGene_tx=X
M_023240780.1;refGene_gn=SORBS1;refGene_tx=XM_023240781.1;refGene_gn=S
ORBS1;refGene_tx=XM_023240782.1
D2 56864821 56864936
refGene_gn=SORBS1;refGene_tx=XM_019813577.2;refGene_gn=SORBS1;refGene_
tx=XM_019813579.2;refGene_gn=SORBS1;refGene_tx=XM_023240761.1;refGene_
gn=SORBS1;refGene_tx=XM_023240762.1;refGene_gn=SORBS1;refGene_tx=XM_02
3240763.1;refGene_gn=SORBS1;refGene_tx=XM_023240764.1;refGene_gn=SORBS
1;refGene_tx=XM_023240765.1;refGene_gn=SORBS1;refGene_tx=XM_023240766.
1;refGene_gn=SORBS1;refGene_tx=XM_023240767.1;refGene_gn=SORBS1;refGen
e_tx=XM_023240769.1;refGene_gn=SORBS1;refGene_tx=XM_023240770.1;refGen
e_gn=SORBS1;refGene_tx=XM_023240771.1;refGene_gn=SORBS1;refGene_tx=XM_
023240772.1;refGene_gn=SORBS1;refGene_tx=XM_023240773.1;refGene_gn=SOR
BS1;refGene_tx=XM_023240774.1;refGene_gn=SORBS1;refGene_tx=XM_02324077
5.1;refGene_gn=SORBS1;refGene_tx=XM_023240776.1;refGene_gn=SORBS1;refG
ene_tx=XM_023240777.1;refGene_gn=SORBS1;refGene_tx=XM_023240778.1;refG
ene_gn=SORBS1;refGene_tx=XM_023240779.1;refGene_gn=SORBS1;refGene_tx=X

M_023240780.1;refGene_gn=SORBS1;refGene_tx=XM_023240781.1;refGene_gn=SORBS1;refGene_tx=XM_023240782.1
D2 56868319 56868366
refGene_gn=SORBS1;refGene_tx=XM_019813577.2;refGene_gn=SORBS1;refGene_tx=XM_019813579.2;refGene_gn=SORBS1;refGene_tx=XM_023240761.1;refGene_gn=SORBS1;refGene_tx=XM_023240762.1;refGene_gn=SORBS1;refGene_tx=XM_023240763.1;refGene_gn=SORBS1;refGene_tx=XM_023240764.1;refGene_gn=SORBS1;refGene_tx=XM_023240765.1;refGene_gn=SORBS1;refGene_tx=XM_023240766.1;refGene_gn=SORBS1;refGene_tx=XM_023240767.1;refGene_gn=SORBS1;refGene_tx=XM_023240769.1;refGene_gn=SORBS1;refGene_tx=XM_023240770.1;refGene_gn=SORBS1;refGene_tx=XM_023240771.1;refGene_gn=SORBS1;refGene_tx=XM_023240772.1;refGene_gn=SORBS1;refGene_tx=XM_023240773.1;refGene_gn=SORBS1;refGene_tx=XM_023240774.1;refGene_gn=SORBS1;refGene_tx=XM_023240775.1;refGene_gn=SORBS1;refGene_tx=XM_023240776.1;refGene_gn=SORBS1;refGene_tx=XM_023240777.1;refGene_gn=SORBS1;refGene_tx=XM_023240778.1;refGene_gn=SORBS1;refGene_tx=XM_023240779.1;refGene_gn=SORBS1;refGene_tx=XM_023240780.1;refGene_gn=SORBS1;refGene_tx=XM_023240781.1;refGene_gn=SORBS1;refGene_tx=XM_023240782.1
D2 56872386 56872554
refGene_gn=SORBS1;refGene_tx=XM_019813577.2;refGene_gn=SORBS1;refGene_tx=XM_019813579.2;refGene_gn=SORBS1;refGene_tx=XM_023240761.1;refGene_gn=SORBS1;refGene_tx=XM_023240762.1;refGene_gn=SORBS1;refGene_tx=XM_023240763.1;refGene_gn=SORBS1;refGene_tx=XM_023240764.1;refGene_gn=SORBS1;refGene_tx=XM_023240766.1;refGene_gn=SORBS1;refGene_tx=XM_023240767.1;refGene_gn=SORBS1;refGene_tx=XM_023240769.1;refGene_gn=SORBS1;refGene_tx=XM_023240770.1;refGene_gn=SORBS1;refGene_tx=XM_023240771.1;refGene_gn=SORBS1;refGene_tx=XM_023240773.1;refGene_gn=SORBS1;refGene_tx=XM_023240774.1;refGene_gn=SORBS1;refGene_tx=XM_023240776.1;refGene_gn=SORBS1;refGene_tx=XM_023240777.1;refGene_gn=SORBS1;refGene_tx=XM_023240778.1;refGene_gn=SORBS1;refGene_tx=XM_023240779.1;refGene_gn=SORBS1;refGene_tx=XM_023240782.1
D2 56875646 56875732
refGene_gn=SORBS1;refGene_tx=XM_019813577.2;refGene_gn=SORBS1;refGene_tx=XM_019813579.2;refGene_gn=SORBS1;refGene_tx=XM_023240761.1;refGene_gn=SORBS1;refGene_tx=XM_023240762.1;refGene_gn=SORBS1;refGene_tx=XM_023240763.1;refGene_gn=SORBS1;refGene_tx=XM_023240764.1;refGene_gn=SORBS1;refGene_tx=XM_023240765.1;refGene_gn=SORBS1;refGene_tx=XM_023240766.1;refGene_gn=SORBS1;refGene_tx=XM_023240767.1;refGene_gn=SORBS1;refGene_tx=XM_023240769.1;refGene_gn=SORBS1;refGene_tx=XM_023240770.1;refGene_gn=SORBS1;refGene_tx=XM_023240771.1;refGene_gn=SORBS1;refGene_tx=XM_023240772.1;refGene_gn=SORBS1;refGene_tx=XM_023240773.1;refGene_gn=SORBS1;refGene_tx=XM_023240774.1;refGene_gn=SORBS1;refGene_tx=XM_023240775.1;refGene_gn=SORBS1;refGene_tx=XM_023240776.1;refGene_gn=SORBS1;refGene_tx=XM_023240777.1;refGene_gn=SORBS1;refGene_tx=XM_023240778.1;refGene_gn=SORBS1;refGene_tx=XM_023240779.1;refGene_gn=SORBS1;refGene_tx=XM_023240780.1;refGene_gn=SORBS1;refGene_tx=XM_023240781.1;refGene_gn=SORBS1;refGene_tx=XM_023240782.1
D2 56876402 56877878
refGene_gn=SORBS1;refGene_tx=XM_019813577.2;refGene_gn=SORBS1;refGene_tx=XM_019813579.2;refGene_gn=SORBS1;refGene_tx=XM_023240761.1;refGene_

gn=SORBS1;refGene_tx=XM_023240762.1;refGene_gn=SORBS1;refGene_tx=XM_02
3240763.1
D2 56878419 56878592
refGene_gn=SORBS1;refGene_tx=XM_019813577.2;refGene_gn=SORBS1;refGene_
tx=XM_019813579.2;refGene_gn=SORBS1;refGene_tx=XM_023240761.1;refGene_
gn=SORBS1;refGene_tx=XM_023240762.1;refGene_gn=SORBS1;refGene_tx=XM_02
3240763.1;refGene_gn=SORBS1;refGene_tx=XM_023240764.1;refGene_gn=SORBS
1;refGene_tx=XM_023240765.1;refGene_gn=SORBS1;refGene_tx=XM_023240766.
1;refGene_gn=SORBS1;refGene_tx=XM_023240767.1;refGene_gn=SORBS1;refGen
e_tx=XM_023240769.1;refGene_gn=SORBS1;refGene_tx=XM_023240770.1;refGen
e_gn=SORBS1;refGene_tx=XM_023240771.1;refGene_gn=SORBS1;refGene_tx=XM_
023240772.1;refGene_gn=SORBS1;refGene_tx=XM_023240773.1;refGene_gn=SOR
BS1;refGene_tx=XM_023240774.1;refGene_gn=SORBS1;refGene_tx=XM_02324077
5.1;refGene_gn=SORBS1;refGene_tx=XM_023240776.1;refGene_gn=SORBS1;refG
ene_tx=XM_023240777.1;refGene_gn=SORBS1;refGene_tx=XM_023240778.1;refG
ene_gn=SORBS1;refGene_tx=XM_023240779.1;refGene_gn=SORBS1;refGene_tx=X
M_023240780.1;refGene_gn=SORBS1;refGene_tx=XM_023240781.1;refGene_gn=S
ORBS1;refGene_tx=XM_023240782.1
D2 56887668 56887718
refGene_gn=SORBS1;refGene_tx=XM_019813577.2;refGene_gn=SORBS1;refGene_
tx=XM_019813579.2;refGene_gn=SORBS1;refGene_tx=XM_023240761.1;refGene_
gn=SORBS1;refGene_tx=XM_023240762.1;refGene_gn=SORBS1;refGene_tx=XM_02
3240763.1;refGene_gn=SORBS1;refGene_tx=XM_023240764.1;refGene_gn=SORBS
1;refGene_tx=XM_023240765.1;refGene_gn=SORBS1;refGene_tx=XM_023240766.
1;refGene_gn=SORBS1;refGene_tx=XM_023240767.1;refGene_gn=SORBS1;refGen
e_tx=XM_023240769.1;refGene_gn=SORBS1;refGene_tx=XM_023240770.1;refGen
e_gn=SORBS1;refGene_tx=XM_023240771.1;refGene_gn=SORBS1;refGene_tx=XM_
023240772.1;refGene_gn=SORBS1;refGene_tx=XM_023240773.1;refGene_gn=SOR
BS1;refGene_tx=XM_023240774.1;refGene_gn=SORBS1;refGene_tx=XM_02324077
5.1;refGene_gn=SORBS1;refGene_tx=XM_023240776.1;refGene_gn=SORBS1;refG
ene_tx=XM_023240777.1;refGene_gn=SORBS1;refGene_tx=XM_023240778.1;refG
ene_gn=SORBS1;refGene_tx=XM_023240779.1;refGene_gn=SORBS1;refGene_tx=X
M_023240780.1;refGene_gn=SORBS1;refGene_tx=XM_023240781.1;refGene_gn=S
ORBS1;refGene_tx=XM_023240782.1
D2 56891634 56891736
refGene_gn=SORBS1;refGene_tx=XM_019813577.2;refGene_gn=SORBS1;refGene_
tx=XM_019813579.2;refGene_gn=SORBS1;refGene_tx=XM_023240761.1;refGene_
gn=SORBS1;refGene_tx=XM_023240762.1;refGene_gn=SORBS1;refGene_tx=XM_02
3240763.1;refGene_gn=SORBS1;refGene_tx=XM_023240764.1;refGene_gn=SORBS
1;refGene_tx=XM_023240765.1;refGene_gn=SORBS1;refGene_tx=XM_023240766.
1;refGene_gn=SORBS1;refGene_tx=XM_023240767.1;refGene_gn=SORBS1;refGen
e_tx=XM_023240769.1;refGene_gn=SORBS1;refGene_tx=XM_023240773.1;refGen
e_gn=SORBS1;refGene_tx=XM_023240776.1;refGene_gn=SORBS1;refGene_tx=XM_
023240782.1
D2 56892316 56892382
refGene_gn=SORBS1;refGene_tx=XM_019813577.2;refGene_gn=SORBS1;refGene_
tx=XM_019813579.2;refGene_gn=SORBS1;refGene_tx=XM_023240761.1;refGene_
gn=SORBS1;refGene_tx=XM_023240762.1;refGene_gn=SORBS1;refGene_tx=XM_02
3240763.1;refGene_gn=SORBS1;refGene_tx=XM_023240764.1;refGene_gn=SORBS
1;refGene_tx=XM_023240765.1;refGene_gn=SORBS1;refGene_tx=XM_023240766.

1;refGene_gn=SORBS1;refGene_tx=XM_023240767.1;refGene_gn=SORBS1;refGene_tx=XM_023240769.1;refGene_gn=SORBS1;refGene_tx=XM_023240770.1;refGene_gn=SORBS1;refGene_tx=XM_023240773.1
D2 56896421 56896505
refGene_gn=SORBS1;refGene_tx=XM_019813577.2;refGene_gn=SORBS1;refGene_tx=XM_019813579.2;refGene_gn=SORBS1;refGene_tx=XM_023240761.1;refGene_gn=SORBS1;refGene_tx=XM_023240762.1;refGene_gn=SORBS1;refGene_tx=XM_023240763.1;refGene_gn=SORBS1;refGene_tx=XM_023240764.1;refGene_gn=SORBS1;refGene_tx=XM_023240765.1;refGene_gn=SORBS1;refGene_tx=XM_023240766.1;refGene_gn=SORBS1;refGene_tx=XM_023240767.1;refGene_gn=SORBS1;refGene_tx=XM_023240769.1;refGene_gn=SORBS1;refGene_tx=XM_023240770.1;refGene_gn=SORBS1;refGene_tx=XM_023240773.1;refGene_gn=SORBS1;refGene_tx=XM_023240774.1;refGene_gn=SORBS1;refGene_tx=XM_023240776.1;refGene_gn=SORBS1;refGene_tx=XM_023240782.1
D2 56901960 56902075
refGene_gn=SORBS1;refGene_tx=XM_019813577.2;refGene_gn=SORBS1;refGene_tx=XM_019813579.2;refGene_gn=SORBS1;refGene_tx=XM_023240761.1;refGene_gn=SORBS1;refGene_tx=XM_023240762.1;refGene_gn=SORBS1;refGene_tx=XM_023240763.1;refGene_gn=SORBS1;refGene_tx=XM_023240764.1;refGene_gn=SORBS1;refGene_tx=XM_023240765.1;refGene_gn=SORBS1;refGene_tx=XM_023240766.1;refGene_gn=SORBS1;refGene_tx=XM_023240767.1;refGene_gn=SORBS1;refGene_tx=XM_023240769.1;refGene_gn=SORBS1;refGene_tx=XM_023240770.1;refGene_gn=SORBS1;refGene_tx=XM_023240771.1;refGene_gn=SORBS1;refGene_tx=XM_023240772.1;refGene_gn=SORBS1;refGene_tx=XM_023240773.1;refGene_gn=SORBS1;refGene_tx=XM_023240774.1;refGene_gn=SORBS1;refGene_tx=XM_023240775.1;refGene_gn=SORBS1;refGene_tx=XM_023240776.1;refGene_gn=SORBS1;refGene_tx=XM_023240777.1;refGene_gn=SORBS1;refGene_tx=XM_023240778.1;refGene_gn=SORBS1;refGene_tx=XM_023240779.1;refGene_gn=SORBS1;refGene_tx=XM_023240780.1;refGene_gn=SORBS1;refGene_tx=XM_023240781.1;refGene_gn=SORBS1;refGene_tx=XM_023240782.1
D2 56903579 56903709
refGene_gn=SORBS1;refGene_tx=XM_019813577.2;refGene_gn=SORBS1;refGene_tx=XM_019813579.2;refGene_gn=SORBS1;refGene_tx=XM_023240761.1;refGene_gn=SORBS1;refGene_tx=XM_023240762.1;refGene_gn=SORBS1;refGene_tx=XM_023240763.1;refGene_gn=SORBS1;refGene_tx=XM_023240764.1;refGene_gn=SORBS1;refGene_tx=XM_023240765.1;refGene_gn=SORBS1;refGene_tx=XM_023240766.1;refGene_gn=SORBS1;refGene_tx=XM_023240767.1;refGene_gn=SORBS1;refGene_tx=XM_023240769.1;refGene_gn=SORBS1;refGene_tx=XM_023240770.1;refGene_gn=SORBS1;refGene_tx=XM_023240771.1;refGene_gn=SORBS1;refGene_tx=XM_023240772.1;refGene_gn=SORBS1;refGene_tx=XM_023240773.1;refGene_gn=SORBS1;refGene_tx=XM_023240774.1;refGene_gn=SORBS1;refGene_tx=XM_023240775.1;refGene_gn=SORBS1;refGene_tx=XM_023240776.1;refGene_gn=SORBS1;refGene_tx=XM_023240777.1;refGene_gn=SORBS1;refGene_tx=XM_023240778.1;refGene_gn=SORBS1;refGene_tx=XM_023240779.1;refGene_gn=SORBS1;refGene_tx=XM_023240780.1;refGene_gn=SORBS1;refGene_tx=XM_023240781.1;refGene_gn=SORBS1;refGene_tx=XM_023240782.1
D2 56903833 56903881
refGene_gn=SORBS1;refGene_tx=XM_019813577.2;refGene_gn=SORBS1;refGene_tx=XM_023240761.1;refGene_gn=SORBS1;refGene_tx=XM_023240762.1;refGene_gn=SORBS1;refGene_tx=XM_023240763.1;refGene_gn=SORBS1;refGene_tx=XM_02

3240764.1;refGene_gn=SORBS1;refGene_tx=XM_023240765.1;refGene_gn=SORBS1;refGene_tx=XM_023240766.1;refGene_gn=SORBS1;refGene_tx=XM_023240767.1;refGene_gn=SORBS1;refGene_tx=XM_023240769.1;refGene_gn=SORBS1;refGene_tx=XM_023240770.1;refGene_gn=SORBS1;refGene_tx=XM_023240771.1;refGene_gn=SORBS1;refGene_tx=XM_023240772.1;refGene_gn=SORBS1;refGene_tx=XM_023240773.1;refGene_gn=SORBS1;refGene_tx=XM_023240774.1;refGene_gn=SORBS1;refGene_tx=XM_023240775.1;refGene_gn=SORBS1;refGene_tx=XM_023240776.1;refGene_gn=SORBS1;refGene_tx=XM_023240777.1;refGene_gn=SORBS1;refGene_tx=XM_023240778.1;refGene_gn=SORBS1;refGene_tx=XM_023240779.1;refGene_gn=SORBS1;refGene_tx=XM_023240780.1;refGene_gn=SORBS1;refGene_tx=XM_023240781.1;refGene_gn=SORBS1;refGene_tx=XM_023240782.1
D2 56913903 56913966
refGene_gn=SORBS1;refGene_tx=XM_019813577.2;refGene_gn=SORBS1;refGene_tx=XM_023240761.1;refGene_gn=SORBS1;refGene_tx=XM_023240762.1;refGene_gn=SORBS1;refGene_tx=XM_023240763.1;refGene_gn=SORBS1;refGene_tx=XM_023240764.1;refGene_gn=SORBS1;refGene_tx=XM_023240765.1;refGene_gn=SORBS1;refGene_tx=XM_023240766.1;refGene_gn=SORBS1;refGene_tx=XM_023240767.1;refGene_gn=SORBS1;refGene_tx=XM_023240774.1
D2 56914280 56914355
refGene_gn=SORBS1;refGene_tx=XM_019813577.2;refGene_gn=SORBS1;refGene_tx=XM_023240761.1;refGene_gn=SORBS1;refGene_tx=XM_023240762.1;refGene_gn=SORBS1;refGene_tx=XM_023240763.1;refGene_gn=SORBS1;refGene_tx=XM_023240764.1;refGene_gn=SORBS1;refGene_tx=XM_023240765.1;refGene_gn=SORBS1;refGene_tx=XM_023240766.1;refGene_gn=SORBS1;refGene_tx=XM_023240767.1;refGene_gn=SORBS1;refGene_tx=XM_023240773.1;refGene_gn=SORBS1;refGene_tx=XM_023240774.1
D2 56916062 56916126
refGene_gn=SORBS1;refGene_tx=XM_019813577.2;refGene_gn=SORBS1;refGene_tx=XM_023240761.1;refGene_gn=SORBS1;refGene_tx=XM_023240762.1;refGene_gn=SORBS1;refGene_tx=XM_023240763.1;refGene_gn=SORBS1;refGene_tx=XM_023240764.1;refGene_gn=SORBS1;refGene_tx=XM_023240765.1;refGene_gn=SORBS1;refGene_tx=XM_023240766.1;refGene_gn=SORBS1;refGene_tx=XM_023240767.1;refGene_gn=SORBS1;refGene_tx=XM_023240769.1;refGene_gn=SORBS1;refGene_tx=XM_023240770.1;refGene_gn=SORBS1;refGene_tx=XM_023240771.1;refGene_gn=SORBS1;refGene_tx=XM_023240772.1;refGene_gn=SORBS1;refGene_tx=XM_023240773.1;refGene_gn=SORBS1;refGene_tx=XM_023240774.1;refGene_gn=SORBS1;refGene_tx=XM_023240775.1;refGene_gn=SORBS1;refGene_tx=XM_023240776.1;refGene_gn=SORBS1;refGene_tx=XM_023240777.1;refGene_gn=SORBS1;refGene_tx=XM_023240778.1;refGene_gn=SORBS1;refGene_tx=XM_023240779.1;refGene_gn=SORBS1;refGene_tx=XM_023240780.1;refGene_gn=SORBS1;refGene_tx=XM_023240781.1;refGene_gn=SORBS1;refGene_tx=XM_023240782.1
D2 56917888 56918061
refGene_gn=SORBS1;refGene_tx=XM_019813577.2;refGene_gn=SORBS1;refGene_tx=XM_023240761.1;refGene_gn=SORBS1;refGene_tx=XM_023240762.1;refGene_gn=SORBS1;refGene_tx=XM_023240763.1;refGene_gn=SORBS1;refGene_tx=XM_023240764.1;refGene_gn=SORBS1;refGene_tx=XM_023240765.1;refGene_gn=SORBS1;refGene_tx=XM_023240766.1;refGene_gn=SORBS1;refGene_tx=XM_023240767.1;refGene_gn=SORBS1;refGene_tx=XM_023240769.1;refGene_gn=SORBS1;refGene_tx=XM_023240770.1;refGene_gn=SORBS1;refGene_tx=XM_023240771.1;refGene_gn=SORBS1;refGene_tx=XM_023240772.1;refGene_gn=SORBS1;refGene_tx=XM_

023240773.1;refGene_gn=SORBS1;refGene_tx=XM_023240774.1;refGene_gn=SORBS1;refGene_tx=XM_023240775.1;refGene_gn=SORBS1;refGene_tx=XM_023240776.1;refGene_gn=SORBS1;refGene_tx=XM_023240777.1;refGene_gn=SORBS1;refGene_tx=XM_023240778.1;refGene_gn=SORBS1;refGene_tx=XM_023240779.1;refGene_gn=SORBS1;refGene_tx=XM_023240780.1;refGene_gn=SORBS1;refGene_tx=XM_023240781.1;refGene_gn=SORBS1;refGene_tx=XM_023240782.1
D2 56923577 56923607
refGene_gn=SORBS1;refGene_tx=XM_019813577.2;refGene_gn=SORBS1;refGene_tx=XM_023240761.1;refGene_gn=SORBS1;refGene_tx=XM_023240762.1;refGene_gn=SORBS1;refGene_tx=XM_023240763.1;refGene_gn=SORBS1;refGene_tx=XM_023240764.1;refGene_gn=SORBS1;refGene_tx=XM_023240765.1;refGene_gn=SORBS1;refGene_tx=XM_023240766.1;refGene_gn=SORBS1;refGene_tx=XM_023240767.1;refGene_gn=SORBS1;refGene_tx=XM_023240769.1;refGene_gn=SORBS1;refGene_tx=XM_023240770.1;refGene_gn=SORBS1;refGene_tx=XM_023240771.1;refGene_gn=SORBS1;refGene_tx=XM_023240772.1;refGene_gn=SORBS1;refGene_tx=XM_023240773.1;refGene_gn=SORBS1;refGene_tx=XM_023240774.1;refGene_gn=SORBS1;refGene_tx=XM_023240775.1;refGene_gn=SORBS1;refGene_tx=XM_023240776.1;refGene_gn=SORBS1;refGene_tx=XM_023240777.1;refGene_gn=SORBS1;refGene_tx=XM_023240778.1;refGene_gn=SORBS1;refGene_tx=XM_023240779.1;refGene_gn=SORBS1;refGene_tx=XM_023240780.1;refGene_gn=SORBS1;refGene_tx=XM_023240781.1;refGene_gn=SORBS1;refGene_tx=XM_023240782.1
D2 56927478 56927623
refGene_gn=SORBS1;refGene_tx=XM_019813577.2;refGene_gn=SORBS1;refGene_tx=XM_023240761.1;refGene_gn=SORBS1;refGene_tx=XM_023240762.1;refGene_gn=SORBS1;refGene_tx=XM_023240763.1;refGene_gn=SORBS1;refGene_tx=XM_023240764.1;refGene_gn=SORBS1;refGene_tx=XM_023240765.1;refGene_gn=SORBS1;refGene_tx=XM_023240766.1;refGene_gn=SORBS1;refGene_tx=XM_023240767.1;refGene_gn=SORBS1;refGene_tx=XM_023240769.1;refGene_gn=SORBS1;refGene_tx=XM_023240770.1;refGene_gn=SORBS1;refGene_tx=XM_023240771.1;refGene_gn=SORBS1;refGene_tx=XM_023240772.1;refGene_gn=SORBS1;refGene_tx=XM_023240773.1;refGene_gn=SORBS1;refGene_tx=XM_023240774.1;refGene_gn=SORBS1;refGene_tx=XM_023240775.1;refGene_gn=SORBS1;refGene_tx=XM_023240776.1;refGene_gn=SORBS1;refGene_tx=XM_023240777.1;refGene_gn=SORBS1;refGene_tx=XM_023240778.1;refGene_gn=SORBS1;refGene_tx=XM_023240779.1;refGene_gn=SORBS1;refGene_tx=XM_023240780.1;refGene_gn=SORBS1;refGene_tx=XM_023240781.1;refGene_gn=SORBS1;refGene_tx=XM_023240782.1
D2 56931286 56931649
refGene_gn=SORBS1;refGene_tx=XM_023240761.1;refGene_gn=SORBS1;refGene_tx=XM_023240762.1;refGene_gn=SORBS1;refGene_tx=XM_023240764.1;refGene_gn=SORBS1;refGene_tx=XM_023240765.1;refGene_gn=SORBS1;refGene_tx=XM_023240766.1;refGene_gn=SORBS1;refGene_tx=XM_023240767.1;refGene_gn=SORBS1;refGene_tx=XM_023240769.1;refGene_gn=SORBS1;refGene_tx=XM_023240770.1;refGene_gn=SORBS1;refGene_tx=XM_023240771.1;refGene_gn=SORBS1;refGene_tx=XM_023240772.1;refGene_gn=SORBS1;refGene_tx=XM_023240773.1;refGene_gn=SORBS1;refGene_tx=XM_023240775.1;refGene_gn=SORBS1;refGene_tx=XM_023240781.1
D2 56932277 56932994
refGene_gn=SORBS1;refGene_tx=XM_023240761.1;refGene_gn=SORBS1;refGene_tx=XM_023240764.1;refGene_gn=SORBS1;refGene_tx=XM_023240765.1;refGene_gn=SORBS1;refGene_tx=XM_023240767.1;refGene_gn=SORBS1;refGene_tx=XM_02

3240770.1;refGene_gn=SORBS1;refGene_tx=XM_023240771.1;refGene_gn=SORBS1;refGene_tx=XM_023240772.1
D2 56937589 56937702
refGene_gn=SORBS1;refGene_tx=XM_023240761.1;refGene_gn=SORBS1;refGene_tx=XM_023240762.1;refGene_gn=SORBS1;refGene_tx=XM_023240763.1;refGene_gn=SORBS1;refGene_tx=XM_023240764.1;refGene_gn=SORBS1;refGene_tx=XM_023240765.1;refGene_gn=SORBS1;refGene_tx=XM_023240766.1;refGene_gn=SORBS1;refGene_tx=XM_023240767.1;refGene_gn=SORBS1;refGene_tx=XM_023240769.1;refGene_gn=SORBS1;refGene_tx=XM_023240770.1;refGene_gn=SORBS1;refGene_tx=XM_023240771.1;refGene_gn=SORBS1;refGene_tx=XM_023240772.1;refGene_gn=SORBS1;refGene_tx=XM_023240773.1;refGene_gn=SORBS1;refGene_tx=XM_023240774.1;refGene_gn=SORBS1;refGene_tx=XM_023240775.1;refGene_gn=SORBS1;refGene_tx=XM_023240776.1;refGene_gn=SORBS1;refGene_tx=XM_023240777.1;refGene_gn=SORBS1;refGene_tx=XM_023240778.1;refGene_gn=SORBS1;refGene_tx=XM_023240779.1;refGene_gn=SORBS1;refGene_tx=XM_023240780.1;refGene_gn=SORBS1;refGene_tx=XM_023240781.1;refGene_gn=SORBS1;refGene_tx=XM_023240782.1
D2 56937822 56937849
refGene_gn=SORBS1;refGene_tx=XM_023240761.1;refGene_gn=SORBS1;refGene_tx=XM_023240762.1;refGene_gn=SORBS1;refGene_tx=XM_023240763.1;refGene_gn=SORBS1;refGene_tx=XM_023240764.1;refGene_gn=SORBS1;refGene_tx=XM_023240765.1;refGene_gn=SORBS1;refGene_tx=XM_023240766.1;refGene_gn=SORBS1;refGene_tx=XM_023240767.1;refGene_gn=SORBS1;refGene_tx=XM_023240769.1;refGene_gn=SORBS1;refGene_tx=XM_023240770.1;refGene_gn=SORBS1;refGene_tx=XM_023240771.1;refGene_gn=SORBS1;refGene_tx=XM_023240772.1;refGene_gn=SORBS1;refGene_tx=XM_023240775.1;refGene_gn=SORBS1;refGene_tx=XM_023240776.1;refGene_gn=SORBS1;refGene_tx=XM_023240777.1;refGene_gn=SORBS1;refGene_tx=XM_023240778.1;refGene_gn=SORBS1;refGene_tx=XM_023240781.1;refGene_gn=SORBS1;refGene_tx=XM_023240782.1
D2 56947162 56947291
refGene_gn=SORBS1;refGene_tx=XM_019813577.2;refGene_gn=SORBS1;refGene_tx=XM_023240761.1;refGene_gn=SORBS1;refGene_tx=XM_023240762.1;refGene_gn=SORBS1;refGene_tx=XM_023240763.1;refGene_gn=SORBS1;refGene_tx=XM_023240764.1;refGene_gn=SORBS1;refGene_tx=XM_023240765.1;refGene_gn=SORBS1;refGene_tx=XM_023240766.1;refGene_gn=SORBS1;refGene_tx=XM_023240767.1;refGene_gn=SORBS1;refGene_tx=XM_023240769.1;refGene_gn=SORBS1;refGene_tx=XM_023240770.1;refGene_gn=SORBS1;refGene_tx=XM_023240771.1;refGene_gn=SORBS1;refGene_tx=XM_023240772.1;refGene_gn=SORBS1;refGene_tx=XM_023240773.1;refGene_gn=SORBS1;refGene_tx=XM_023240774.1;refGene_gn=SORBS1;refGene_tx=XM_023240775.1;refGene_gn=SORBS1;refGene_tx=XM_023240776.1;refGene_gn=SORBS1;refGene_tx=XM_023240777.1;refGene_gn=SORBS1;refGene_tx=XM_023240778.1;refGene_gn=SORBS1;refGene_tx=XM_023240779.1;refGene_gn=SORBS1;refGene_tx=XM_023240780.1;refGene_gn=SORBS1;refGene_tx=XM_023240781.1;refGene_gn=SORBS1;refGene_tx=XM_023240782.1
D2 56949435 56949531
refGene_gn=SORBS1;refGene_tx=XM_023240761.1;refGene_gn=SORBS1;refGene_tx=XM_023240762.1;refGene_gn=SORBS1;refGene_tx=XM_023240763.1;refGene_gn=SORBS1;refGene_tx=XM_023240764.1;refGene_gn=SORBS1;refGene_tx=XM_023240765.1;refGene_gn=SORBS1;refGene_tx=XM_023240766.1;refGene_gn=SORBS1;refGene_tx=XM_023240767.1;refGene_gn=SORBS1;refGene_tx=XM_023240769.

1;refGene_gn=SORBS1;refGene_tx=XM_023240770.1;refGene_gn=SORBS1;refGene_tx=XM_023240771.1;refGene_gn=SORBS1;refGene_tx=XM_023240772.1;refGene_gn=SORBS1;refGene_tx=XM_023240773.1;refGene_gn=SORBS1;refGene_tx=XM_023240774.1;refGene_gn=SORBS1;refGene_tx=XM_023240775.1;refGene_gn=SORBS1;refGene_tx=XM_023240776.1;refGene_gn=SORBS1;refGene_tx=XM_023240777.1

D2 56952293 56952362
refGene_gn=SORBS1;refGene_tx=XM_023240761.1;refGene_gn=SORBS1;refGene_tx=XM_023240762.1;refGene_gn=SORBS1;refGene_tx=XM_023240763.1;refGene_gn=SORBS1;refGene_tx=XM_023240764.1;refGene_gn=SORBS1;refGene_tx=XM_023240765.1;refGene_gn=SORBS1;refGene_tx=XM_023240766.1;refGene_gn=SORBS1;refGene_tx=XM_023240767.1;refGene_gn=SORBS1;refGene_tx=XM_023240769.1;refGene_gn=SORBS1;refGene_tx=XM_023240770.1;refGene_gn=SORBS1;refGene_tx=XM_023240771.1;refGene_gn=SORBS1;refGene_tx=XM_023240772.1;refGene_gn=SORBS1;refGene_tx=XM_023240773.1;refGene_gn=SORBS1;refGene_tx=XM_023240774.1;refGene_gn=SORBS1;refGene_tx=XM_023240775.1;refGene_gn=SORBS1;refGene_tx=XM_023240776.1;refGene_gn=SORBS1;refGene_tx=XM_023240777.1;refGene_gn=SORBS1;refGene_tx=XM_023240778.1;refGene_gn=SORBS1;refGene_tx=XM_023240779.1;refGene_gn=SORBS1;refGene_tx=XM_023240780.1;refGene_gn=SORBS1;refGene_tx=XM_023240781.1;refGene_gn=SORBS1;refGene_tx=XM_023240782.1

D2 56959118 56959128
refGene_gn=SORBS1;refGene_tx=XM_023240761.1;refGene_gn=SORBS1;refGene_tx=XM_023240762.1;refGene_gn=SORBS1;refGene_tx=XM_023240763.1;refGene_gn=SORBS1;refGene_tx=XM_023240764.1;refGene_gn=SORBS1;refGene_tx=XM_023240765.1;refGene_gn=SORBS1;refGene_tx=XM_023240766.1;refGene_gn=SORBS1;refGene_tx=XM_023240767.1;refGene_gn=SORBS1;refGene_tx=XM_023240769.1;refGene_gn=SORBS1;refGene_tx=XM_023240770.1;refGene_gn=SORBS1;refGene_tx=XM_023240771.1;refGene_gn=SORBS1;refGene_tx=XM_023240772.1;refGene_gn=SORBS1;refGene_tx=XM_023240773.1;refGene_gn=SORBS1;refGene_tx=XM_023240774.1;refGene_gn=SORBS1;refGene_tx=XM_023240775.1;refGene_gn=SORBS1;refGene_tx=XM_023240776.1;refGene_gn=SORBS1;refGene_tx=XM_023240777.1;refGene_gn=SORBS1;refGene_tx=XM_023240778.1;refGene_gn=SORBS1;refGene_tx=XM_023240779.1;refGene_gn=SORBS1;refGene_tx=XM_023240780.1;refGene_gn=SORBS1;refGene_tx=XM_023240781.1;refGene_gn=SORBS1;refGene_tx=XM_023240782.1

D2 57108552 57108734
refGene_gn=ALDH18A1;refGene_tx=XM_006938031.4;refGene_gn=ALDH18A1;refGene_tx=XM_006938032.4;refGene_gn=ALDH18A1;refGene_tx=XM_006938034.4;refGene_gn=ALDH18A1;refGene_tx=XM_019813599.2;refGene_gn=ALDH18A1;refGene_tx=XM_019813600.2

D2 57111144 57111240
refGene_gn=ALDH18A1;refGene_tx=XM_006938031.4;refGene_gn=ALDH18A1;refGene_tx=XM_006938032.4;refGene_gn=ALDH18A1;refGene_tx=XM_006938034.4;refGene_gn=ALDH18A1;refGene_tx=XM_019813599.2;refGene_gn=ALDH18A1;refGene_tx=XM_019813600.2

D2 57112265 57112452
refGene_gn=ALDH18A1;refGene_tx=XM_006938031.4;refGene_gn=ALDH18A1;refGene_tx=XM_006938032.4;refGene_gn=ALDH18A1;refGene_tx=XM_006938034.4;refGene_gn=ALDH18A1;refGene_tx=XM_019813599.2;refGene_gn=ALDH18A1;refGen

e_tx=XM_019813600.2
D2 57114467 57114589
refGene_gn=ALDH18A1;refGene_tx=XM_006938031.4;refGene_gn=ALDH18A1;refGene_tx=XM_006938032.4;refGene_gn=ALDH18A1;refGene_tx=XM_006938034.4;refGene_gn=ALDH18A1;refGene_tx=XM_019813599.2;refGene_gn=ALDH18A1;refGene_tx=XM_019813600.2
D2 57114694 57114890
refGene_gn=ALDH18A1;refGene_tx=XM_006938031.4;refGene_gn=ALDH18A1;refGene_tx=XM_006938032.4;refGene_gn=ALDH18A1;refGene_tx=XM_006938034.4;refGene_gn=ALDH18A1;refGene_tx=XM_019813599.2;refGene_gn=ALDH18A1;refGene_tx=XM_019813600.2
D2 57116741 57116879
refGene_gn=ALDH18A1;refGene_tx=XM_006938031.4;refGene_gn=ALDH18A1;refGene_tx=XM_006938032.4;refGene_gn=ALDH18A1;refGene_tx=XM_006938034.4;refGene_gn=ALDH18A1;refGene_tx=XM_019813599.2;refGene_gn=ALDH18A1;refGene_tx=XM_019813600.2
D2 57119695 57119916
refGene_gn=ALDH18A1;refGene_tx=XM_006938031.4;refGene_gn=ALDH18A1;refGene_tx=XM_006938032.4;refGene_gn=ALDH18A1;refGene_tx=XM_006938034.4;refGene_gn=ALDH18A1;refGene_tx=XM_019813599.2;refGene_gn=ALDH18A1;refGene_tx=XM_019813600.2
D2 57122327 57122421
refGene_gn=ALDH18A1;refGene_tx=XM_006938031.4;refGene_gn=ALDH18A1;refGene_tx=XM_006938032.4;refGene_gn=ALDH18A1;refGene_tx=XM_006938034.4;refGene_gn=ALDH18A1;refGene_tx=XM_019813599.2;refGene_gn=ALDH18A1;refGene_tx=XM_019813600.2
D2 57123547 57123621
refGene_gn=ALDH18A1;refGene_tx=XM_006938031.4;refGene_gn=ALDH18A1;refGene_tx=XM_006938032.4;refGene_gn=ALDH18A1;refGene_tx=XM_006938034.4;refGene_gn=ALDH18A1;refGene_tx=XM_019813599.2;refGene_gn=ALDH18A1;refGene_tx=XM_019813600.2
D2 57124241 57124386
refGene_gn=ALDH18A1;refGene_tx=XM_006938031.4;refGene_gn=ALDH18A1;refGene_tx=XM_006938032.4;refGene_gn=ALDH18A1;refGene_tx=XM_006938034.4;refGene_gn=ALDH18A1;refGene_tx=XM_019813599.2;refGene_gn=ALDH18A1;refGene_tx=XM_019813600.2
D2 57125791 57125916
refGene_gn=ALDH18A1;refGene_tx=XM_006938031.4;refGene_gn=ALDH18A1;refGene_tx=XM_006938032.4;refGene_gn=ALDH18A1;refGene_tx=XM_006938034.4;refGene_gn=ALDH18A1;refGene_tx=XM_019813599.2;refGene_gn=ALDH18A1;refGene_tx=XM_019813600.2
D2 57129653 57129744
refGene_gn=ALDH18A1;refGene_tx=XM_006938031.4;refGene_gn=ALDH18A1;refGene_tx=XM_006938032.4;refGene_gn=ALDH18A1;refGene_tx=XM_006938034.4;refGene_gn=ALDH18A1;refGene_tx=XM_019813599.2;refGene_gn=ALDH18A1;refGene_tx=XM_019813600.2
D2 57130157 57130316
refGene_gn=ALDH18A1;refGene_tx=XM_006938031.4;refGene_gn=ALDH18A1;refGene_tx=XM_006938032.4;refGene_gn=ALDH18A1;refGene_tx=XM_006938034.4;refGene_gn=ALDH18A1;refGene_tx=XM_019813599.2;refGene_gn=ALDH18A1;refGen

e_tx=XM_019813600.2
D2 57133937 57134042
refGene_gn=ALDH18A1;refGene_tx=XM_006938031.4;refGene_gn=ALDH18A1;refGene_tx=XM_006938032.4;refGene_gn=ALDH18A1;refGene_tx=XM_006938034.4;refGene_gn=ALDH18A1;refGene_tx=XM_019813599.2;refGene_gn=ALDH18A1;refGene_tx=XM_019813600.2
D2 57134132 57134282
refGene_gn=ALDH18A1;refGene_tx=XM_006938031.4;refGene_gn=ALDH18A1;refGene_tx=XM_006938032.4;refGene_gn=ALDH18A1;refGene_tx=XM_006938034.4;refGene_gn=ALDH18A1;refGene_tx=XM_019813599.2;refGene_gn=ALDH18A1;refGene_tx=XM_019813600.2
D2 57137763 57137987
refGene_gn=ALDH18A1;refGene_tx=XM_006938031.4;refGene_gn=ALDH18A1;refGene_tx=XM_006938032.4;refGene_gn=ALDH18A1;refGene_tx=XM_006938034.4;refGene_gn=ALDH18A1;refGene_tx=XM_019813599.2;refGene_gn=ALDH18A1;refGene_tx=XM_019813600.2
D2 57146933 57147027
refGene_gn=ALDH18A1;refGene_tx=XM_006938031.4;refGene_gn=ALDH18A1;refGene_tx=XM_006938032.4;refGene_gn=ALDH18A1;refGene_tx=XM_006938034.4;refGene_gn=ALDH18A1;refGene_tx=XM_019813599.2;refGene_gn=ALDH18A1;refGene_tx=XM_019813600.2
D2 57153927 57154158
refGene_gn=TCTN3;refGene_tx=XM_011287501.3;refGene_gn=TCTN3;refGene_tx=XM_011287502.3;refGene_gn=TCTN3;refGene_tx=XM_011287503.3
D2 57154231 57154240 refGene_gn=TCTN3;refGene_tx=XM_011287504.3
D2 57166956 57167094
refGene_gn=TCTN3;refGene_tx=XM_011287501.3;refGene_gn=TCTN3;refGene_tx=XM_011287502.3;refGene_gn=TCTN3;refGene_tx=XM_011287503.3;refGene_gn=TCTN3;refGene_tx=XM_011287504.3
D2 57170642 57170796
refGene_gn=TCTN3;refGene_tx=XM_011287501.3;refGene_gn=TCTN3;refGene_tx=XM_011287502.3;refGene_gn=TCTN3;refGene_tx=XM_011287503.3;refGene_gn=TCTN3;refGene_tx=XM_011287504.3
D2 57171290 57171385
refGene_gn=TCTN3;refGene_tx=XM_011287501.3;refGene_gn=TCTN3;refGene_tx=XM_011287502.3;refGene_gn=TCTN3;refGene_tx=XM_011287503.3;refGene_gn=TCTN3;refGene_tx=XM_011287504.3
D2 57171783 57171891
refGene_gn=TCTN3;refGene_tx=XM_011287501.3;refGene_gn=TCTN3;refGene_tx=XM_011287502.3;refGene_gn=TCTN3;refGene_tx=XM_011287504.3
D2 57172944 57173070
refGene_gn=TCTN3;refGene_tx=XM_011287501.3;refGene_gn=TCTN3;refGene_tx=XM_011287502.3;refGene_gn=TCTN3;refGene_tx=XM_011287503.3;refGene_gn=TCTN3;refGene_tx=XM_011287504.3
D2 57174469 57174550
refGene_gn=TCTN3;refGene_tx=XM_011287501.3;refGene_gn=TCTN3;refGene_tx=XM_011287502.3;refGene_gn=TCTN3;refGene_tx=XM_011287503.3;refGene_gn=TCTN3;refGene_tx=XM_011287504.3
D2 57175116 57175152
refGene_gn=TCTN3;refGene_tx=XM_011287501.3;refGene_gn=TCTN3;refGene_tx

=XM_011287502.3;refGene_gn=TCTN3;refGene_tx=XM_011287503.3;refGene_gn=TCTN3;refGene_tx=XM_011287504.3
D2 57176424 57176540
refGene_gn=TCTN3;refGene_tx=XM_011287501.3;refGene_gn=TCTN3;refGene_tx=XM_011287502.3;refGene_gn=TCTN3;refGene_tx=XM_011287503.3;refGene_gn=TCTN3;refGene_tx=XM_011287504.3
D2 57176953 57177062
refGene_gn=TCTN3;refGene_tx=XM_011287501.3;refGene_gn=TCTN3;refGene_tx=XM_011287502.3;refGene_gn=TCTN3;refGene_tx=XM_011287503.3;refGene_gn=TCTN3;refGene_tx=XM_011287504.3
D2 57177298 57177426
refGene_gn=TCTN3;refGene_tx=XM_011287501.3;refGene_gn=TCTN3;refGene_tx=XM_011287502.3;refGene_gn=TCTN3;refGene_tx=XM_011287503.3;refGene_gn=TCTN3;refGene_tx=XM_011287504.3
D2 57178624 57178743
refGene_gn=TCTN3;refGene_tx=XM_011287501.3;refGene_gn=TCTN3;refGene_tx=XM_011287502.3;refGene_gn=TCTN3;refGene_tx=XM_011287503.3;refGene_gn=TCTN3;refGene_tx=XM_011287504.3
D2 57179078 57179202
refGene_gn=TCTN3;refGene_tx=XM_011287501.3;refGene_gn=TCTN3;refGene_tx=XM_011287502.3;refGene_gn=TCTN3;refGene_tx=XM_011287503.3;refGene_gn=TCTN3;refGene_tx=XM_011287504.3
D2 57179365 57179618
refGene_gn=TCTN3;refGene_tx=XM_011287501.3;refGene_gn=TCTN3;refGene_tx=XM_011287502.3;refGene_gn=TCTN3;refGene_tx=XM_011287503.3;refGene_gn=TCTN3;refGene_tx=XM_011287504.3
D2 57219273 57219289 refGene_gn=ENTPD1;refGene_tx=XM_006938035.4
D2 57258162 57258307
refGene_gn=ENTPD1;refGene_tx=XM_019813614.2;refGene_gn=ENTPD1;refGene_tx=XM_023240784.1;refGene_gn=ENTPD1;refGene_tx=XM_023240785.1;refGene_gn=ENTPD1;refGene_tx=XM_023240786.1
D2 57265213 57265341
refGene_gn=ENTPD1;refGene_tx=XM_006938035.4;refGene_gn=ENTPD1;refGene_tx=XM_019813614.2;refGene_gn=ENTPD1;refGene_tx=XM_023240784.1;refGene_gn=ENTPD1;refGene_tx=XM_023240785.1;refGene_gn=ENTPD1;refGene_tx=XM_023240786.1
D2 57284431 57284549
refGene_gn=ENTPD1;refGene_tx=XM_006938035.4;refGene_gn=ENTPD1;refGene_tx=XM_019813614.2;refGene_gn=ENTPD1;refGene_tx=XM_023240784.1;refGene_gn=ENTPD1;refGene_tx=XM_023240785.1;refGene_gn=ENTPD1;refGene_tx=XM_023240786.1
D2 57287177 57287328
refGene_gn=ENTPD1;refGene_tx=XM_006938035.4;refGene_gn=ENTPD1;refGene_tx=XM_019813614.2;refGene_gn=ENTPD1;refGene_tx=XM_019813615.2;refGene_gn=ENTPD1;refGene_tx=XM_023240784.1;refGene_gn=ENTPD1;refGene_tx=XM_023240785.1;refGene_gn=ENTPD1;refGene_tx=XM_023240786.1
D2 57289314 57289474
refGene_gn=ENTPD1;refGene_tx=XM_006938035.4;refGene_gn=ENTPD1;refGene_tx=XM_019813614.2;refGene_gn=ENTPD1;refGene_tx=XM_019813615.2;refGene_gn=ENTPD1;refGene_tx=XM_023240784.1;refGene_gn=ENTPD1;refGene_tx=XM_02

3240785.1;refGene_gn=ENTPD1;refGene_tx=XM_023240786.1
D2	57290169	57290409
refGene_gn=ENTPD1;refGene_tx=XM_006938035.4;refGene_gn=ENTPD1;refGene_tx=XM_019813614.2;refGene_gn=ENTPD1;refGene_tx=XM_019813615.2;refGene_gn=ENTPD1;refGene_tx=XM_023240784.1;refGene_gn=ENTPD1;refGene_tx=XM_023240785.1;refGene_gn=ENTPD1;refGene_tx=XM_023240786.1
D2	57291612	57291870
refGene_gn=ENTPD1;refGene_tx=XM_006938035.4;refGene_gn=ENTPD1;refGene_tx=XM_019813614.2;refGene_gn=ENTPD1;refGene_tx=XM_019813615.2;refGene_gn=ENTPD1;refGene_tx=XM_023240784.1;refGene_gn=ENTPD1;refGene_tx=XM_023240786.1
D2	57292841	57292973
refGene_gn=ENTPD1;refGene_tx=XM_019813614.2;refGene_gn=ENTPD1;refGene_tx=XM_023240786.1
D2	57295482	57295483	refGene_gn=ENTPD1;refGene_tx=XM_019813614.2
D2	57302828	57302945
refGene_gn=ENTPD1;refGene_tx=XM_006938035.4;refGene_gn=ENTPD1;refGene_tx=XM_019813615.2;refGene_gn=ENTPD1;refGene_tx=XM_023240784.1;refGene_gn=ENTPD1;refGene_tx=XM_023240785.1
D2	57304724	57304862
refGene_gn=ENTPD1;refGene_tx=XM_006938035.4;refGene_gn=ENTPD1;refGene_tx=XM_019813615.2;refGene_gn=ENTPD1;refGene_tx=XM_023240784.1;refGene_gn=ENTPD1;refGene_tx=XM_023240785.1
D2	57306870	57307077
refGene_gn=ENTPD1;refGene_tx=XM_006938035.4;refGene_gn=ENTPD1;refGene_tx=XM_019813615.2;refGene_gn=ENTPD1;refGene_tx=XM_023240784.1;refGene_gn=ENTPD1;refGene_tx=XM_023240785.1
D2	57326369	57326411
refGene_gn=CC2D2B;refGene_tx=XM_019813601.2;refGene_gn=CC2D2B;refGene_tx=XM_019813602.2;refGene_gn=CC2D2B;refGene_tx=XM_019813605.2;refGene_gn=CC2D2B;refGene_tx=XM_019813607.2;refGene_gn=CC2D2B;refGene_tx=XM_019813608.2
D2	57330622	57330688
refGene_gn=CC2D2B;refGene_tx=XM_019813601.2;refGene_gn=CC2D2B;refGene_tx=XM_019813602.2;refGene_gn=CC2D2B;refGene_tx=XM_019813605.2;refGene_gn=CC2D2B;refGene_tx=XM_019813607.2;refGene_gn=CC2D2B;refGene_tx=XM_019813608.2
D2	57335519	57335603
refGene_gn=CC2D2B;refGene_tx=XM_019813601.2;refGene_gn=CC2D2B;refGene_tx=XM_019813602.2;refGene_gn=CC2D2B;refGene_tx=XM_019813607.2;refGene_gn=CC2D2B;refGene_tx=XM_019813608.2
D2	57338882	57339081
refGene_gn=CC2D2B;refGene_tx=XM_019813601.2;refGene_gn=CC2D2B;refGene_tx=XM_019813602.2;refGene_gn=CC2D2B;refGene_tx=XM_019813605.2;refGene_gn=CC2D2B;refGene_tx=XM_019813607.2;refGene_gn=CC2D2B;refGene_tx=XM_019813608.2
D2	57339387	57339524
refGene_gn=CC2D2B;refGene_tx=XM_019813601.2;refGene_gn=CC2D2B;refGene_tx=XM_019813602.2;refGene_gn=CC2D2B;refGene_tx=XM_019813605.2;refGene_gn=CC2D2B;refGene_tx=XM_019813607.2;refGene_gn=CC2D2B;refGene_tx=XM_01

9813608.2;refGene_gn=CC2D2B;refGene_tx=XM_019813609.2
D2 57339625 57339754
refGene_gn=CC2D2B;refGene_tx=XM_019813601.2;refGene_gn=CC2D2B;refGene_tx=XM_019813602.2;refGene_gn=CC2D2B;refGene_tx=XM_019813605.2;refGene_gn=CC2D2B;refGene_tx=XM_019813607.2;refGene_gn=CC2D2B;refGene_tx=XM_019813608.2;refGene_gn=CC2D2B;refGene_tx=XM_019813609.2
D2 57345542 57345752
refGene_gn=CC2D2B;refGene_tx=XM_019813601.2;refGene_gn=CC2D2B;refGene_tx=XM_019813602.2;refGene_gn=CC2D2B;refGene_tx=XM_019813605.2;refGene_gn=CC2D2B;refGene_tx=XM_019813607.2;refGene_gn=CC2D2B;refGene_tx=XM_019813608.2;refGene_gn=CC2D2B;refGene_tx=XM_019813609.2
D2 57347021 57347119
refGene_gn=CC2D2B;refGene_tx=XM_019813601.2;refGene_gn=CC2D2B;refGene_tx=XM_019813602.2;refGene_gn=CC2D2B;refGene_tx=XM_019813605.2;refGene_gn=CC2D2B;refGene_tx=XM_019813607.2;refGene_gn=CC2D2B;refGene_tx=XM_019813608.2;refGene_gn=CC2D2B;refGene_tx=XM_019813609.2
D2 57352531 57352672
refGene_gn=CC2D2B;refGene_tx=XM_019813601.2;refGene_gn=CC2D2B;refGene_tx=XM_019813602.2;refGene_gn=CC2D2B;refGene_tx=XM_019813605.2;refGene_gn=CC2D2B;refGene_tx=XM_019813607.2;refGene_gn=CC2D2B;refGene_tx=XM_019813608.2;refGene_gn=CC2D2B;refGene_tx=XM_019813609.2
D2 57354310 57354419
refGene_gn=CC2D2B;refGene_tx=XM_019813601.2;refGene_gn=CC2D2B;refGene_tx=XM_019813602.2;refGene_gn=CC2D2B;refGene_tx=XM_019813605.2;refGene_gn=CC2D2B;refGene_tx=XM_019813607.2;refGene_gn=CC2D2B;refGene_tx=XM_019813608.2;refGene_gn=CC2D2B;refGene_tx=XM_019813609.2
D2 57354619 57354738
refGene_gn=CC2D2B;refGene_tx=XM_019813601.2;refGene_gn=CC2D2B;refGene_tx=XM_019813602.2;refGene_gn=CC2D2B;refGene_tx=XM_019813605.2;refGene_gn=CC2D2B;refGene_tx=XM_019813607.2;refGene_gn=CC2D2B;refGene_tx=XM_019813608.2;refGene_gn=CC2D2B;refGene_tx=XM_019813609.2
D2 57356997 57357175
refGene_gn=CC2D2B;refGene_tx=XM_019813601.2;refGene_gn=CC2D2B;refGene_tx=XM_019813602.2;refGene_gn=CC2D2B;refGene_tx=XM_019813605.2;refGene_gn=CC2D2B;refGene_tx=XM_019813607.2;refGene_gn=CC2D2B;refGene_tx=XM_019813608.2;refGene_gn=CC2D2B;refGene_tx=XM_019813609.2
D2 57359785 57359936
refGene_gn=CC2D2B;refGene_tx=XM_019813601.2;refGene_gn=CC2D2B;refGene_tx=XM_019813602.2;refGene_gn=CC2D2B;refGene_tx=XM_019813605.2;refGene_gn=CC2D2B;refGene_tx=XM_019813607.2;refGene_gn=CC2D2B;refGene_tx=XM_019813608.2;refGene_gn=CC2D2B;refGene_tx=XM_019813609.2
D2 57360765 57360913
refGene_gn=CC2D2B;refGene_tx=XM_019813601.2;refGene_gn=CC2D2B;refGene_tx=XM_019813602.2;refGene_gn=CC2D2B;refGene_tx=XM_019813605.2;refGene_gn=CC2D2B;refGene_tx=XM_019813607.2;refGene_gn=CC2D2B;refGene_tx=XM_019813608.2;refGene_gn=CC2D2B;refGene_tx=XM_019813609.2
D2 57367494 57367633
refGene_gn=CC2D2B;refGene_tx=XM_019813601.2;refGene_gn=CC2D2B;refGene_tx=XM_019813602.2;refGene_gn=CC2D2B;refGene_tx=XM_019813605.2;refGene_gn=CC2D2B;refGene_tx=XM_019813607.2;refGene_gn=CC2D2B;refGene_tx=XM_01

9813608.2;refGene_gn=CC2D2B;refGene_tx=XM_019813609.2
D2 57368789 57368993
refGene_gn=CC2D2B;refGene_tx=XM_019813601.2;refGene_gn=CC2D2B;refGene_tx=XM_019813602.2;refGene_gn=CC2D2B;refGene_tx=XM_019813605.2;refGene_gn=CC2D2B;refGene_tx=XM_019813607.2;refGene_gn=CC2D2B;refGene_tx=XM_019813608.2;refGene_gn=CC2D2B;refGene_tx=XM_019813609.2
D2 57379129 57379222
refGene_gn=CC2D2B;refGene_tx=XM_019813601.2;refGene_gn=CC2D2B;refGene_tx=XM_019813602.2;refGene_gn=CC2D2B;refGene_tx=XM_019813605.2;refGene_gn=CC2D2B;refGene_tx=XM_019813607.2;refGene_gn=CC2D2B;refGene_tx=XM_019813608.2;refGene_gn=CC2D2B;refGene_tx=XM_019813609.2
D2 57379407 57379499
refGene_gn=CC2D2B;refGene_tx=XM_019813601.2;refGene_gn=CC2D2B;refGene_tx=XM_019813602.2;refGene_gn=CC2D2B;refGene_tx=XM_019813605.2;refGene_gn=CC2D2B;refGene_tx=XM_019813607.2;refGene_gn=CC2D2B;refGene_tx=XM_019813608.2;refGene_gn=CC2D2B;refGene_tx=XM_019813609.2
D2 57381296 57381467
refGene_gn=CC2D2B;refGene_tx=XM_019813601.2;refGene_gn=CC2D2B;refGene_tx=XM_019813602.2;refGene_gn=CC2D2B;refGene_tx=XM_019813605.2;refGene_gn=CC2D2B;refGene_tx=XM_019813608.2;refGene_gn=CC2D2B;refGene_tx=XM_019813609.2
D2 57382686 57382783
refGene_gn=CC2D2B;refGene_tx=XM_019813601.2;refGene_gn=CC2D2B;refGene_tx=XM_019813602.2;refGene_gn=CC2D2B;refGene_tx=XM_019813605.2;refGene_gn=CC2D2B;refGene_tx=XM_019813607.2;refGene_gn=CC2D2B;refGene_tx=XM_019813608.2;refGene_gn=CC2D2B;refGene_tx=XM_019813609.2
D2 57383668 57383778
refGene_gn=CC2D2B;refGene_tx=XM_019813601.2;refGene_gn=CC2D2B;refGene_tx=XM_019813602.2;refGene_gn=CC2D2B;refGene_tx=XM_019813605.2;refGene_gn=CC2D2B;refGene_tx=XM_019813607.2;refGene_gn=CC2D2B;refGene_tx=XM_019813608.2;refGene_gn=CC2D2B;refGene_tx=XM_019813609.2
D2 57388888 57388982
refGene_gn=CC2D2B;refGene_tx=XM_019813601.2;refGene_gn=CC2D2B;refGene_tx=XM_019813602.2;refGene_gn=CC2D2B;refGene_tx=XM_019813605.2;refGene_gn=CC2D2B;refGene_tx=XM_019813607.2;refGene_gn=CC2D2B;refGene_tx=XM_019813608.2;refGene_gn=CC2D2B;refGene_tx=XM_019813609.2
D2 57394867 57394966
refGene_gn=CC2D2B;refGene_tx=XM_019813601.2;refGene_gn=CC2D2B;refGene_tx=XM_019813602.2;refGene_gn=CC2D2B;refGene_tx=XM_019813605.2;refGene_gn=CC2D2B;refGene_tx=XM_019813607.2;refGene_gn=CC2D2B;refGene_tx=XM_019813608.2;refGene_gn=CC2D2B;refGene_tx=XM_019813609.2
D2 57397537 57397720
refGene_gn=CC2D2B;refGene_tx=XM_019813601.2;refGene_gn=CC2D2B;refGene_tx=XM_019813602.2;refGene_gn=CC2D2B;refGene_tx=XM_019813605.2;refGene_gn=CC2D2B;refGene_tx=XM_019813607.2;refGene_gn=CC2D2B;refGene_tx=XM_019813608.2;refGene_gn=CC2D2B;refGene_tx=XM_019813609.2
D2 57397882 57398080
refGene_gn=CC2D2B;refGene_tx=XM_019813601.2;refGene_gn=CC2D2B;refGene_tx=XM_019813602.2;refGene_gn=CC2D2B;refGene_tx=XM_019813605.2;refGene_gn=CC2D2B;refGene_tx=XM_019813607.2;refGene_gn=CC2D2B;refGene_tx=XM_01

9813608.2;refGene_gn=CC2D2B;refGene_tx=XM_019813609.2
D2 57400299 57400389
refGene_gn=CC2D2B;refGene_tx=XM_019813601.2;refGene_gn=CC2D2B;refGene_tx=XM_019813602.2;refGene_gn=CC2D2B;refGene_tx=XM_019813605.2;refGene_gn=CC2D2B;refGene_tx=XM_019813607.2;refGene_gn=CC2D2B;refGene_tx=XM_019813608.2;refGene_gn=CC2D2B;refGene_tx=XM_019813609.2
D2 57403208 57403322
refGene_gn=CC2D2B;refGene_tx=XM_019813601.2;refGene_gn=CC2D2B;refGene_tx=XM_019813602.2;refGene_gn=CC2D2B;refGene_tx=XM_019813605.2;refGene_gn=CC2D2B;refGene_tx=XM_019813607.2;refGene_gn=CC2D2B;refGene_tx=XM_019813608.2;refGene_gn=CC2D2B;refGene_tx=XM_019813609.2
D2 57405432 57405567
refGene_gn=CC2D2B;refGene_tx=XM_019813601.2;refGene_gn=CC2D2B;refGene_tx=XM_019813602.2;refGene_gn=CC2D2B;refGene_tx=XM_019813605.2;refGene_gn=CC2D2B;refGene_tx=XM_019813607.2;refGene_gn=CC2D2B;refGene_tx=XM_019813608.2;refGene_gn=CC2D2B;refGene_tx=XM_019813609.2
D2 57405945 57406068
refGene_gn=CC2D2B;refGene_tx=XM_019813601.2;refGene_gn=CC2D2B;refGene_tx=XM_019813602.2;refGene_gn=CC2D2B;refGene_tx=XM_019813605.2;refGene_gn=CC2D2B;refGene_tx=XM_019813607.2;refGene_gn=CC2D2B;refGene_tx=XM_019813608.2;refGene_gn=CC2D2B;refGene_tx=XM_019813609.2
D2 57410880 57410939
refGene_gn=CC2D2B;refGene_tx=XM_019813601.2;refGene_gn=CC2D2B;refGene_tx=XM_019813602.2;refGene_gn=CC2D2B;refGene_tx=XM_019813605.2;refGene_gn=CC2D2B;refGene_tx=XM_019813607.2;refGene_gn=CC2D2B;refGene_tx=XM_019813609.2
D2 57411002 57411035 refGene_gn=CC2D2B;refGene_tx=XM_019813608.2
D2 57412613 57412791
refGene_gn=CC2D2B;refGene_tx=XM_019813601.2;refGene_gn=CC2D2B;refGene_tx=XM_019813602.2;refGene_gn=CC2D2B;refGene_tx=XM_019813605.2;refGene_gn=CC2D2B;refGene_tx=XM_019813607.2;refGene_gn=CC2D2B;refGene_tx=XM_019813609.2
D2 57415100 57415289
refGene_gn=CC2D2B;refGene_tx=XM_019813601.2;refGene_gn=CC2D2B;refGene_tx=XM_019813602.2;refGene_gn=CC2D2B;refGene_tx=XM_019813605.2;refGene_gn=CC2D2B;refGene_tx=XM_019813607.2;refGene_gn=CC2D2B;refGene_tx=XM_019813609.2
D2 57426896 57426965
refGene_gn=CCNJ;refGene_tx=XM_023240787.1;refGene_gn=CCNJ;refGene_tx=XM_023240788.1;refGene_gn=CCNJ;refGene_tx=XM_023240789.1;refGene_gn=CCNJ;refGene_tx=XM_023240790.1
D2 57432233 57432444
refGene_gn=CCNJ;refGene_tx=XM_023240787.1;refGene_gn=CCNJ;refGene_tx=XM_023240788.1;refGene_gn=CCNJ;refGene_tx=XM_023240789.1;refGene_gn=CCNJ;refGene_tx=XM_023240790.1
D2 57440073 57440373
refGene_gn=CCNJ;refGene_tx=XM_023240787.1;refGene_gn=CCNJ;refGene_tx=XM_023240788.1;refGene_gn=CCNJ;refGene_tx=XM_023240789.1;refGene_gn=CCNJ;refGene_tx=XM_023240790.1
D2 57440460 57440620

refGene_gn=CCNJ;refGene_tx=XM_023240787.1;refGene_gn=CCNJ;refGene_tx=XM_023240788.1;refGene_gn=CCNJ;refGene_tx=XM_023240789.1;refGene_gn=CCNJ;refGene_tx=XM_023240790.1
D2 57441653 57442065
refGene_gn=CCNJ;refGene_tx=XM_023240787.1;refGene_gn=CCNJ;refGene_tx=XM_023240788.1;refGene_gn=CCNJ;refGene_tx=XM_023240789.1;refGene_gn=CCNJ;refGene_tx=XM_023240790.1
D2 57529037 57533498
refGene_gn=ZNF518A;refGene_tx=XM_003994239.4;refGene_gn=ZNF518A;refGene_tx=XM_019813616.2;refGene_gn=ZNF518A;refGene_tx=XM_019813617.2;refGene_gn=ZNF518A;refGene_tx=XM_019813619.2;refGene_gn=ZNF518A;refGene_tx=XM_023240791.1
D2 57561506 57561626
refGene_gn=BLNK;refGene_tx=XM_006938046.4;refGene_gn=BLNK;refGene_tx=XM_006938047.4
D2 57567074 57567230
refGene_gn=BLNK;refGene_tx=XM_006938046.4;refGene_gn=BLNK;refGene_tx=XM_006938047.4
D2 57569803 57569887
refGene_gn=BLNK;refGene_tx=XM_006938046.4;refGene_gn=BLNK;refGene_tx=XM_006938047.4
D2 57570707 57570775
refGene_gn=BLNK;refGene_tx=XM_006938046.4;refGene_gn=BLNK;refGene_tx=XM_006938047.4
D2 57574305 57574337
refGene_gn=BLNK;refGene_tx=XM_006938046.4;refGene_gn=BLNK;refGene_tx=XM_006938047.4
D2 57574765 57574850
refGene_gn=BLNK;refGene_tx=XM_006938046.4;refGene_gn=BLNK;refGene_tx=XM_006938047.4
D2 57577093 57577136
refGene_gn=BLNK;refGene_tx=XM_006938046.4;refGene_gn=BLNK;refGene_tx=XM_006938047.4
D2 57577982 57578010
refGene_gn=BLNK;refGene_tx=XM_006938046.4;refGene_gn=BLNK;refGene_tx=XM_006938047.4
D2 57579756 57579826
refGene_gn=BLNK;refGene_tx=XM_006938046.4;refGene_gn=BLNK;refGene_tx=XM_006938047.4
D2 57587027 57587099
refGene_gn=BLNK;refGene_tx=XM_006938046.4;refGene_gn=BLNK;refGene_tx=XM_006938047.4
D2 57588081 57588163
refGene_gn=BLNK;refGene_tx=XM_006938046.4;refGene_gn=BLNK;refGene_tx=XM_006938047.4
D2 57592428 57592589
refGene_gn=BLNK;refGene_tx=XM_006938046.4;refGene_gn=BLNK;refGene_tx=XM_006938047.4
D2 57595858 57596015
refGene_gn=BLNK;refGene_tx=XM_006938046.4;refGene_gn=BLNK;refGene_tx=X

M_006938047.4
D2	57596214	57596280	refGene_gn=BLNK;refGene_tx=XM_006938046.4
D2	57599974	57600015
refGene_gn=BLNK;refGene_tx=XM_006938046.4;refGene_gn=BLNK;refGene_tx=X
M_006938047.4
D2	57607709	57607759
refGene_gn=BLNK;refGene_tx=XM_006938046.4;refGene_gn=BLNK;refGene_tx=X
M_006938047.4
D2	57612363	57612429
refGene_gn=BLNK;refGene_tx=XM_006938046.4;refGene_gn=BLNK;refGene_tx=X
M_006938047.4
D2	57637277	57637324
refGene_gn=BLNK;refGene_tx=XM_006938046.4;refGene_gn=BLNK;refGene_tx=X
M_006938047.4
D2	57664973	57665272
refGene_gn=DNTT;refGene_tx=XM_019813620.2;refGene_gn=DNTT;refGene_tx=X
M_019813621.2;refGene_gn=DNTT;refGene_tx=XM_019813622.2;refGene_gn=DNT
T;refGene_tx=XM_019813623.2
D2	57676413	57676585
refGene_gn=DNTT;refGene_tx=XM_019813620.2;refGene_gn=DNTT;refGene_tx=X
M_019813621.2;refGene_gn=DNTT;refGene_tx=XM_019813622.2;refGene_gn=DNT
T;refGene_tx=XM_019813623.2
D2	57677525	57677654
refGene_gn=DNTT;refGene_tx=XM_019813620.2;refGene_gn=DNTT;refGene_tx=X
M_019813621.2;refGene_gn=DNTT;refGene_tx=XM_019813622.2;refGene_gn=DNT
T;refGene_tx=XM_019813623.2
D2	57679150	57679321
refGene_gn=DNTT;refGene_tx=XM_019813620.2;refGene_gn=DNTT;refGene_tx=X
M_019813621.2;refGene_gn=DNTT;refGene_tx=XM_019813622.2;refGene_gn=DNT
T;refGene_tx=XM_019813623.2
D2	57679629	57679701
refGene_gn=DNTT;refGene_tx=XM_019813620.2;refGene_gn=DNTT;refGene_tx=X
M_019813621.2;refGene_gn=DNTT;refGene_tx=XM_019813622.2;refGene_gn=DNT
T;refGene_tx=XM_019813623.2
D2	57680890	57681014
refGene_gn=DNTT;refGene_tx=XM_019813620.2;refGene_gn=DNTT;refGene_tx=X
M_019813621.2;refGene_gn=DNTT;refGene_tx=XM_019813622.2;refGene_gn=DNT
T;refGene_tx=XM_019813623.2
D2	57682867	57683000
refGene_gn=DNTT;refGene_tx=XM_019813620.2;refGene_gn=DNTT;refGene_tx=X
M_019813621.2;refGene_gn=DNTT;refGene_tx=XM_019813622.2;refGene_gn=DNT
T;refGene_tx=XM_019813623.2
D2	57684288	57684397
refGene_gn=DNTT;refGene_tx=XM_019813620.2;refGene_gn=DNTT;refGene_tx=X
M_019813621.2;refGene_gn=DNTT;refGene_tx=XM_019813622.2;refGene_gn=DNT
T;refGene_tx=XM_019813623.2
D2	57688757	57689003
refGene_gn=DNTT;refGene_tx=XM_019813620.2;refGene_gn=DNTT;refGene_tx=X
M_019813621.2;refGene_gn=DNTT;refGene_tx=XM_019813622.2;refGene_gn=DNT
T;refGene_tx=XM_019813623.2

D2 57693430 57693514
refGene_gn=DNTT;refGene_tx=XM_019813620.2;refGene_gn=DNTT;refGene_tx=XM_019813621.2;refGene_gn=DNTT;refGene_tx=XM_019813622.2;refGene_gn=DNTT;refGene_tx=XM_019813623.2
D2 57693621 57693756
refGene_gn=DNTT;refGene_tx=XM_019813620.2;refGene_gn=DNTT;refGene_tx=XM_019813621.2
D2 57695421 57695508
refGene_gn=DNTT;refGene_tx=XM_019813622.2;refGene_gn=DNTT;refGene_tx=XM_019813623.2
D2 57697395 57697572
refGene_gn=OPALIN;refGene_tx=XM_003994254.5;refGene_gn=OPALIN;refGene_tx=XM_019813624.2;refGene_gn=OPALIN;refGene_tx=XM_019813625.2
D2 57700209 57700266
refGene_gn=OPALIN;refGene_tx=XM_003994254.5;refGene_gn=OPALIN;refGene_tx=XM_019813624.2;refGene_gn=OPALIN;refGene_tx=XM_019813625.2
D2 57701639 57701759
refGene_gn=OPALIN;refGene_tx=XM_003994254.5;refGene_gn=OPALIN;refGene_tx=XM_019813624.2;refGene_gn=OPALIN;refGene_tx=XM_019813625.2
D2 57702761 57702797
refGene_gn=OPALIN;refGene_tx=XM_003994254.5;refGene_gn=OPALIN;refGene_tx=XM_019813624.2
D2 57706677 57706713 refGene_gn=OPALIN;refGene_tx=XM_003994254.5
D2 57709390 57709393
refGene_gn=OPALIN;refGene_tx=XM_003994254.5;refGene_gn=OPALIN;refGene_tx=XM_019813624.2;refGene_gn=OPALIN;refGene_tx=XM_019813625.2
D2 57714947 57715082 refGene_gn=TLL2;refGene_tx=XM_023240792.1
D2 57716661 57716912 refGene_gn=TLL2;refGene_tx=XM_023240792.1
D2 57719497 57719711 refGene_gn=TLL2;refGene_tx=XM_023240792.1
D2 57722675 57722803 refGene_gn=TLL2;refGene_tx=XM_023240792.1
D2 57725092 57725218 refGene_gn=TLL2;refGene_tx=XM_023240792.1
D2 57726058 57726239 refGene_gn=TLL2;refGene_tx=XM_023240792.1
D2 57726871 57727032 refGene_gn=TLL2;refGene_tx=XM_023240792.1
D2 57727775 57727901 refGene_gn=TLL2;refGene_tx=XM_023240792.1
D2 57735232 57735428 refGene_gn=TLL2;refGene_tx=XM_023240792.1
D2 57735846 57735992 refGene_gn=TLL2;refGene_tx=XM_023240792.1
D2 57737008 57737125 refGene_gn=TLL2;refGene_tx=XM_023240792.1
D2 57743987 57744090 refGene_gn=TLL2;refGene_tx=XM_023240792.1
D2 57749368 57749484 refGene_gn=TLL2;refGene_tx=XM_023240792.1
D2 57751968 57752093 refGene_gn=TLL2;refGene_tx=XM_023240792.1
D2 57759358 57759464 refGene_gn=TLL2;refGene_tx=XM_023240792.1
D2 57760464 57760643 refGene_gn=TLL2;refGene_tx=XM_023240792.1
D2 57764976 57765094 refGene_gn=TLL2;refGene_tx=XM_023240792.1
D2 57767845 57768001 refGene_gn=TLL2;refGene_tx=XM_023240792.1
D2 57783784 57783862 refGene_gn=TLL2;refGene_tx=XM_023240792.1
D2 57810491 57810602 refGene_gn=TLL2;refGene_tx=XM_023240792.1
D2 57835326 57835528 refGene_gn=TLL2;refGene_tx=XM_023240792.1
D2 57844111 57844179 refGene_gn=TM9SF3;refGene_tx=XM_023240793.1
D2 57848870 57848947 refGene_gn=TM9SF3;refGene_tx=XM_023240793.1
D2 57849275 57849359 refGene_gn=TM9SF3;refGene_tx=XM_023240793.1

D2 57849870 57850017 refGene_gn=TM9SF3;refGene_tx=XM_023240793.1
D2 57853015 57853084 refGene_gn=TM9SF3;refGene_tx=XM_023240793.1
D2 57855662 57855802 refGene_gn=TM9SF3;refGene_tx=XM_023240793.1
D2 57866315 57866446 refGene_gn=TM9SF3;refGene_tx=XM_023240793.1
D2 57869617 57869712 refGene_gn=TM9SF3;refGene_tx=XM_023240793.1
D2 57874716 57874883 refGene_gn=TM9SF3;refGene_tx=XM_023240793.1
D2 57876434 57876566 refGene_gn=TM9SF3;refGene_tx=XM_023240793.1
D2 57882766 57882844 refGene_gn=TM9SF3;refGene_tx=XM_023240793.1
D2 57884077 57884238 refGene_gn=TM9SF3;refGene_tx=XM_023240793.1
D2 57887345 57887468 refGene_gn=TM9SF3;refGene_tx=XM_023240793.1
D2 57897407 57897603 refGene_gn=TM9SF3;refGene_tx=XM_023240793.1
D2 57908416 57908515 refGene_gn=TM9SF3;refGene_tx=XM_023240793.1
D2 57915415 57915473 refGene_gn=PIK3AP1;refGene_tx=XM_023240794.1
D2 57922215 57922334 refGene_gn=PIK3AP1;refGene_tx=XM_023240794.1
D2 57924230 57924301 refGene_gn=PIK3AP1;refGene_tx=XM_023240794.1
D2 57926259 57926415 refGene_gn=PIK3AP1;refGene_tx=XM_023240794.1
D2 57933717 57933790 refGene_gn=PIK3AP1;refGene_tx=XM_023240794.1
D2 57937451 57937657 refGene_gn=PIK3AP1;refGene_tx=XM_023240794.1
D2 57939696 57939762 refGene_gn=PIK3AP1;refGene_tx=XM_023240794.1
D2 57942058 57942256 refGene_gn=PIK3AP1;refGene_tx=XM_023240794.1
D2 57944919 57945015 refGene_gn=PIK3AP1;refGene_tx=XM_023240794.1
D2 57961590 57961780 refGene_gn=PIK3AP1;refGene_tx=XM_023240794.1
D2 57964391 57964588 refGene_gn=PIK3AP1;refGene_tx=XM_023240794.1
D2 57966525 57966658 refGene_gn=PIK3AP1;refGene_tx=XM_023240794.1
D2 57967040 57967183 refGene_gn=PIK3AP1;refGene_tx=XM_023240794.1
D2 57969151 57969296 refGene_gn=PIK3AP1;refGene_tx=XM_023240794.1
D2 57971984 57972121 refGene_gn=PIK3AP1;refGene_tx=XM_023240794.1
D2 58022305 58022722 refGene_gn=PIK3AP1;refGene_tx=XM_023240794.1
D2 58030798 58030811 refGene_gn=PIK3AP1;refGene_tx=XM_023240794.1
D2 58126319 58126405
refGene_gn=LOC101098011;refGene_tx=XM_023240795.1;refGene_gn=LOC101098
011;refGene_tx=XM_023240798.1
D2 58126691 58126765
refGene_gn=LOC101098011;refGene_tx=XM_023240795.1;refGene_gn=LOC101098
011;refGene_tx=XM_023240798.1
D2 58189008 58189074
refGene_gn=LOC101098011;refGene_tx=XM_023240795.1;refGene_gn=LOC101098
011;refGene_tx=XM_023240798.1
D2 58189439 58189519
refGene_gn=LOC101098011;refGene_tx=XM_023240795.1;refGene_gn=LOC101098
011;refGene_tx=XM_023240798.1
D2 58214233 58214366
refGene_gn=LOC101098011;refGene_tx=XM_023240795.1;refGene_gn=LOC101098
011;refGene_tx=XM_023240798.1
D2 58218898 58219186
refGene_gn=LOC101098011;refGene_tx=XM_019813627.2;refGene_gn=LOC101098
011;refGene_tx=XM_019813628.2;refGene_gn=LOC101098011;refGene_tx=XM_02
3240795.1;refGene_gn=LOC101098011;refGene_tx=XM_023240798.1
D2 58222326 58222420
refGene_gn=LOC101098011;refGene_tx=XM_019813627.2;refGene_gn=LOC101098

011;refGene_tx=XM_019813628.2;refGene_gn=LOC101098011;refGene_tx=XM_02
3240795.1;refGene_gn=LOC101098011;refGene_tx=XM_023240798.1
D2	58224965	58225935
refGene_gn=LOC101098011;refGene_tx=XM_023240798.1
D2	58250071	58254401
refGene_gn=LOC101098011;refGene_tx=XM_019813627.2;refGene_gn=LOC101098
011;refGene_tx=XM_019813628.2;refGene_gn=LOC101098011;refGene_tx=XM_02
3240795.1;refGene_gn=LOC101098011;refGene_tx=XM_023240796.1;refGene_gn
=LOC101098011;refGene_tx=XM_023240797.1
D2	58270358	58270597	refGene_gn=SLIT1;refGene_tx=XM_019813631.2
D2	58271405	58271547	refGene_gn=SLIT1;refGene_tx=XM_019813631.2
D2	58271548	58271618	refGene_gn=SLIT1;refGene_tx=XM_019813631.2
D2	58271943	58272232	refGene_gn=SLIT1;refGene_tx=XM_019813631.2
D2	58273310	58273465	refGene_gn=SLIT1;refGene_tx=XM_019813631.2
D2	58273914	58274045	refGene_gn=SLIT1;refGene_tx=XM_019813631.2
D2	58275461	58275699	refGene_gn=SLIT1;refGene_tx=XM_019813631.2
D2	58276994	58277132	refGene_gn=SLIT1;refGene_tx=XM_019813631.2
D2	58279482	58279576	refGene_gn=SLIT1;refGene_tx=XM_019813631.2
D2	58279761	58279901	refGene_gn=SLIT1;refGene_tx=XM_019813631.2
D2	58282557	58282655	refGene_gn=SLIT1;refGene_tx=XM_019813631.2
D2	58282948	58283073	refGene_gn=SLIT1;refGene_tx=XM_019813631.2
D2	58285014	58285178	refGene_gn=SLIT1;refGene_tx=XM_019813631.2
D2	58292718	58292790	refGene_gn=SLIT1;refGene_tx=XM_019813631.2
D2	58293697	58293769	refGene_gn=SLIT1;refGene_tx=XM_019813631.2
D2	58296212	58296284	refGene_gn=SLIT1;refGene_tx=XM_019813631.2
D2	58298804	58298873	refGene_gn=SLIT1;refGene_tx=XM_019813631.2
D2	58300993	58301126	refGene_gn=SLIT1;refGene_tx=XM_019813631.2
D2	58303949	58304116	refGene_gn=SLIT1;refGene_tx=XM_019813631.2
D2	58304378	58304522	refGene_gn=SLIT1;refGene_tx=XM_019813631.2
D2	58307314	58307458	refGene_gn=SLIT1;refGene_tx=XM_019813631.2
D2	58307631	58307706	refGene_gn=SLIT1;refGene_tx=XM_019813631.2
D2	58308321	58308466	refGene_gn=SLIT1;refGene_tx=XM_019813631.2
D2	58308594	58308618	refGene_gn=SLIT1;refGene_tx=XM_019813631.2
D2	58309345	58309509	refGene_gn=SLIT1;refGene_tx=XM_019813631.2
D2	58316561	58316705	refGene_gn=SLIT1;refGene_tx=XM_019813631.2
D2	58317453	58317525	refGene_gn=SLIT1;refGene_tx=XM_019813631.2
D2	58319679	58319751	refGene_gn=SLIT1;refGene_tx=XM_019813631.2
D2	58320263	58320335	refGene_gn=SLIT1;refGene_tx=XM_019813631.2
D2	58320810	58320958	refGene_gn=SLIT1;refGene_tx=XM_019813631.2
D2	58322261	58322425	refGene_gn=SLIT1;refGene_tx=XM_019813631.2
D2	58322906	58322978	refGene_gn=SLIT1;refGene_tx=XM_019813631.2
D2	58323581	58323653	refGene_gn=SLIT1;refGene_tx=XM_019813631.2
D2	58325160	58325232	refGene_gn=SLIT1;refGene_tx=XM_019813631.2
D2	58414876	58414948	refGene_gn=SLIT1;refGene_tx=XM_019813631.2
D2	58418317	58418389	refGene_gn=SLIT1;refGene_tx=XM_019813631.2
D2	58420491	58420563	refGene_gn=SLIT1;refGene_tx=XM_019813631.2
D2	58442612	58442809	refGene_gn=SLIT1;refGene_tx=XM_019813631.2
D2	58477882	58477887	refGene_gn=ARHGAP19;refGene_tx=XM_011287496.3
D2	58485425	58485504	refGene_gn=ARHGAP19;refGene_tx=XM_011287496.3
D2	58486022	58486133	refGene_gn=ARHGAP19;refGene_tx=XM_011287496.3

D2 58491257 58491356 refGene_gn=ARHGAP19;refGene_tx=XM_011287496.3
D2 58496940 58497132 refGene_gn=ARHGAP19;refGene_tx=XM_011287496.3
D2 58499650 58499716 refGene_gn=ARHGAP19;refGene_tx=XM_011287496.3
D2 58509321 58509408 refGene_gn=ARHGAP19;refGene_tx=XM_011287496.3
D2 58515612 58515839 refGene_gn=ARHGAP19;refGene_tx=XM_011287496.3
D2 58518885 58518987 refGene_gn=ARHGAP19;refGene_tx=XM_011287496.3
D2 58518989 58519094 refGene_gn=ARHGAP19;refGene_tx=XM_011287496.3
D2 58520472 58520553 refGene_gn=ARHGAP19;refGene_tx=XM_011287496.3
D2 58521643 58521909 refGene_gn=ARHGAP19;refGene_tx=XM_011287496.3
D2 58547713 58547769 refGene_gn=ARHGAP19;refGene_tx=XM_011287496.3
D2 58562055 58562901 refGene_gn=FRAT1;refGene_tx=XM_004001583.3
D2 58575027 58575732 refGene_gn=FRAT2;refGene_tx=XM_023240799.1
D2 58597898 58598001
refGene_gn=RRP12;refGene_tx=XM_003994259.4;refGene_gn=RRP12;refGene_tx
=XM_023240800.1
D2 58598359 58598442
refGene_gn=RRP12;refGene_tx=XM_003994259.4;refGene_gn=RRP12;refGene_tx
=XM_023240800.1
D2 58598747 58598815
refGene_gn=RRP12;refGene_tx=XM_003994259.4;refGene_gn=RRP12;refGene_tx
=XM_023240800.1
D2 58600502 58600575
refGene_gn=RRP12;refGene_tx=XM_003994259.4;refGene_gn=RRP12;refGene_tx
=XM_023240800.1
D2 58602036 58602086
refGene_gn=RRP12;refGene_tx=XM_003994259.4;refGene_gn=RRP12;refGene_tx
=XM_023240800.1
D2 58604760 58604895
refGene_gn=RRP12;refGene_tx=XM_003994259.4;refGene_gn=RRP12;refGene_tx
=XM_023240800.1
D2 58605127 58605303
refGene_gn=RRP12;refGene_tx=XM_003994259.4;refGene_gn=RRP12;refGene_tx
=XM_023240800.1
D2 58605424 58605586
refGene_gn=RRP12;refGene_tx=XM_003994259.4;refGene_gn=RRP12;refGene_tx
=XM_023240800.1
D2 58605774 58605866
refGene_gn=RRP12;refGene_tx=XM_003994259.4;refGene_gn=RRP12;refGene_tx
=XM_023240800.1
D2 58607553 58607711
refGene_gn=RRP12;refGene_tx=XM_003994259.4;refGene_gn=RRP12;refGene_tx
=XM_023240800.1
D2 58608213 58608321
refGene_gn=RRP12;refGene_tx=XM_003994259.4;refGene_gn=RRP12;refGene_tx
=XM_023240800.1
D2 58608512 58608618
refGene_gn=RRP12;refGene_tx=XM_003994259.4;refGene_gn=RRP12;refGene_tx
=XM_023240800.1
D2 58608835 58608916
refGene_gn=RRP12;refGene_tx=XM_003994259.4;refGene_gn=RRP12;refGene_tx

=XM_023240800.1
D2 58609049 58609208
refGene_gn=RRP12;refGene_tx=XM_003994259.4;refGene_gn=RRP12;refGene_tx
=XM_023240800.1
D2 58609793 58609887
refGene_gn=RRP12;refGene_tx=XM_003994259.4;refGene_gn=RRP12;refGene_tx
=XM_023240800.1
D2 58610446 58610514
refGene_gn=RRP12;refGene_tx=XM_003994259.4;refGene_gn=RRP12;refGene_tx
=XM_023240800.1
D2 58610777 58610932
refGene_gn=RRP12;refGene_tx=XM_003994259.4;refGene_gn=RRP12;refGene_tx
=XM_023240800.1
D2 58611378 58611541
refGene_gn=RRP12;refGene_tx=XM_003994259.4;refGene_gn=RRP12;refGene_tx
=XM_023240800.1
D2 58611630 58611695
refGene_gn=RRP12;refGene_tx=XM_003994259.4;refGene_gn=RRP12;refGene_tx
=XM_023240800.1
D2 58612814 58612936
refGene_gn=RRP12;refGene_tx=XM_003994259.4;refGene_gn=RRP12;refGene_tx
=XM_023240800.1
D2 58613172 58613315
refGene_gn=RRP12;refGene_tx=XM_003994259.4;refGene_gn=RRP12;refGene_tx
=XM_023240800.1
D2 58615260 58615375
refGene_gn=RRP12;refGene_tx=XM_003994259.4;refGene_gn=RRP12;refGene_tx
=XM_023240800.1
D2 58616834 58616932
refGene_gn=RRP12;refGene_tx=XM_003994259.4;refGene_gn=RRP12;refGene_tx
=XM_023240800.1
D2 58617169 58617281
refGene_gn=RRP12;refGene_tx=XM_003994259.4;refGene_gn=RRP12;refGene_tx
=XM_023240800.1
D2 58618946 58619038
refGene_gn=RRP12;refGene_tx=XM_003994259.4;refGene_gn=RRP12;refGene_tx
=XM_023240800.1
D2 58619689 58619788
refGene_gn=RRP12;refGene_tx=XM_003994259.4;refGene_gn=RRP12;refGene_tx
=XM_023240800.1
D2 58620663 58620791
refGene_gn=RRP12;refGene_tx=XM_003994259.4;refGene_gn=RRP12;refGene_tx
=XM_023240800.1
D2 58620885 58621024
refGene_gn=RRP12;refGene_tx=XM_003994259.4;refGene_gn=RRP12;refGene_tx
=XM_023240800.1
D2 58622175 58622292
refGene_gn=RRP12;refGene_tx=XM_003994259.4;refGene_gn=RRP12;refGene_tx
=XM_023240800.1
D2 58622504 58622610

refGene_gn=RRP12;refGene_tx=XM_003994259.4;refGene_gn=RRP12;refGene_tx=XM_023240800.1
D2 58623366 58623443
refGene_gn=RRP12;refGene_tx=XM_003994259.4;refGene_gn=RRP12;refGene_tx=XM_023240800.1
D2 58624630 58624714 refGene_gn=RRP12;refGene_tx=XM_003994259.4
D2 58626702 58626932 refGene_gn=RRP12;refGene_tx=XM_003994259.4
D2 58627495 58627634 refGene_gn=RRP12;refGene_tx=XM_003994259.4
D2 58645758 58645897 refGene_gn=PGAM1;refGene_tx=XM_003994262.5
D2 58647062 58647193 refGene_gn=EXOSC1;refGene_tx=XM_019813635.2
D2 58649288 58649563 refGene_gn=PGAM1;refGene_tx=XM_003994262.5
D2 58650343 58650524 refGene_gn=PGAM1;refGene_tx=XM_003994262.5
D2 58651007 58651177 refGene_gn=PGAM1;refGene_tx=XM_003994262.5
D2 58656768 58656875
refGene_gn=EXOSC1;refGene_tx=XM_003994263.4;refGene_gn=EXOSC1;refGene_tx=XM_019813636.1
D2 58657206 58657291
refGene_gn=EXOSC1;refGene_tx=XM_003994263.4;refGene_gn=EXOSC1;refGene_tx=XM_019813635.2;refGene_gn=EXOSC1;refGene_tx=XM_019813636.1
D2 58657420 58657471
refGene_gn=EXOSC1;refGene_tx=XM_003994263.4;refGene_gn=EXOSC1;refGene_tx=XM_019813635.2;refGene_gn=EXOSC1;refGene_tx=XM_019813636.1
D2 58657995 58658036
refGene_gn=EXOSC1;refGene_tx=XM_003994263.4;refGene_gn=EXOSC1;refGene_tx=XM_019813635.2;refGene_gn=EXOSC1;refGene_tx=XM_019813636.1
D2 58659608 58659697
refGene_gn=EXOSC1;refGene_tx=XM_003994263.4;refGene_gn=EXOSC1;refGene_tx=XM_019813635.2;refGene_gn=EXOSC1;refGene_tx=XM_019813636.1
D2 58660922 58660997
refGene_gn=EXOSC1;refGene_tx=XM_003994263.4;refGene_gn=EXOSC1;refGene_tx=XM_019813635.2
D2 58663440 58663556
refGene_gn=EXOSC1;refGene_tx=XM_003994263.4;refGene_gn=EXOSC1;refGene_tx=XM_019813635.2
D2 58663661 58663728
refGene_gn=EXOSC1;refGene_tx=XM_003994263.4;refGene_gn=EXOSC1;refGene_tx=XM_019813635.2
D2 58667973 58668216
refGene_gn=ZDHHC16;refGene_tx=XM_003994264.4;refGene_gn=ZDHHC16;refGene_tx=XM_003994265.4;refGene_gn=ZDHHC16;refGene_tx=XM_019813640.2;refGene_gn=ZDHHC16;refGene_tx=XM_019813641.2
D2 58668394 58668589
refGene_gn=ZDHHC16;refGene_tx=XM_003994264.4;refGene_gn=ZDHHC16;refGene_tx=XM_003994265.4;refGene_gn=ZDHHC16;refGene_tx=XM_019813640.2;refGene_gn=ZDHHC16;refGene_tx=XM_019813641.2
D2 58668738 58668827
refGene_gn=ZDHHC16;refGene_tx=XM_003994264.4;refGene_gn=ZDHHC16;refGene_tx=XM_003994265.4;refGene_gn=ZDHHC16;refGene_tx=XM_019813640.2;refGene_gn=ZDHHC16;refGene_tx=XM_019813641.2
D2 58669210 58669239

refGene_gn=ZDHHC16;refGene_tx=XM_003994264.4;refGene_gn=ZDHHC16;refGene_tx=XM_003994265.4
D2 58669811 58669945
refGene_gn=ZDHHC16;refGene_tx=XM_003994264.4;refGene_gn=ZDHHC16;refGene_tx=XM_003994265.4;refGene_gn=ZDHHC16;refGene_tx=XM_019813640.2;refGene_gn=ZDHHC16;refGene_tx=XM_019813641.2
D2 58670079 58670127
refGene_gn=ZDHHC16;refGene_tx=XM_003994265.4;refGene_gn=ZDHHC16;refGene_tx=XM_019813640.2
D2 58670982 58671068
refGene_gn=ZDHHC16;refGene_tx=XM_003994264.4;refGene_gn=ZDHHC16;refGene_tx=XM_003994265.4;refGene_gn=ZDHHC16;refGene_tx=XM_019813640.2;refGene_gn=ZDHHC16;refGene_tx=XM_019813641.2
D2 58671710 58671834
refGene_gn=ZDHHC16;refGene_tx=XM_003994264.4;refGene_gn=ZDHHC16;refGene_tx=XM_003994265.4;refGene_gn=ZDHHC16;refGene_tx=XM_019813640.2;refGene_gn=ZDHHC16;refGene_tx=XM_019813641.2
D2 58672035 58672106
refGene_gn=ZDHHC16;refGene_tx=XM_003994264.4;refGene_gn=ZDHHC16;refGene_tx=XM_003994265.4;refGene_gn=ZDHHC16;refGene_tx=XM_019813640.2;refGene_gn=ZDHHC16;refGene_tx=XM_019813641.2
D2 58672914 58673029
refGene_gn=ZDHHC16;refGene_tx=XM_003994264.4;refGene_gn=ZDHHC16;refGene_tx=XM_003994265.4;refGene_gn=ZDHHC16;refGene_tx=XM_019813640.2;refGene_gn=ZDHHC16;refGene_tx=XM_019813641.2
D2 58674092 58674120
refGene_gn=MMS19;refGene_tx=XM_003994268.5;refGene_gn=MMS19;refGene_tx=XM_011287495.3;refGene_gn=MMS19;refGene_tx=XM_019813637.2;refGene_gn=MMS19;refGene_tx=XM_019813638.2
D2 58674199 58674300
refGene_gn=MMS19;refGene_tx=XM_003994268.5;refGene_gn=MMS19;refGene_tx=XM_011287495.3;refGene_gn=MMS19;refGene_tx=XM_019813637.2;refGene_gn=MMS19;refGene_tx=XM_019813638.2
D2 58674587 58674647
refGene_gn=MMS19;refGene_tx=XM_003994268.5;refGene_gn=MMS19;refGene_tx=XM_011287495.3;refGene_gn=MMS19;refGene_tx=XM_019813637.2;refGene_gn=MMS19;refGene_tx=XM_019813638.2
D2 58674734 58674899
refGene_gn=MMS19;refGene_tx=XM_003994268.5;refGene_gn=MMS19;refGene_tx=XM_011287495.3;refGene_gn=MMS19;refGene_tx=XM_019813637.2;refGene_gn=MMS19;refGene_tx=XM_019813638.2
D2 58675019 58675102
refGene_gn=MMS19;refGene_tx=XM_003994268.5;refGene_gn=MMS19;refGene_tx=XM_011287495.3;refGene_gn=MMS19;refGene_tx=XM_019813637.2;refGene_gn=MMS19;refGene_tx=XM_019813638.2
D2 58675412 58675599
refGene_gn=MMS19;refGene_tx=XM_003994268.5;refGene_gn=MMS19;refGene_tx=XM_011287495.3;refGene_gn=MMS19;refGene_tx=XM_019813637.2;refGene_gn=MMS19;refGene_tx=XM_019813638.2
D2 58675741 58675798

refGene_gn=MMS19;refGene_tx=XM_003994268.5;refGene_gn=MMS19;refGene_tx=XM_011287495.3;refGene_gn=MMS19;refGene_tx=XM_019813637.2;refGene_gn=MMS19;refGene_tx=XM_019813638.2
D2 58675944 58676045
refGene_gn=MMS19;refGene_tx=XM_003994268.5;refGene_gn=MMS19;refGene_tx=XM_011287495.3;refGene_gn=MMS19;refGene_tx=XM_019813637.2;refGene_gn=MMS19;refGene_tx=XM_019813638.2
D2 58676555 58676682
refGene_gn=MMS19;refGene_tx=XM_003994268.5;refGene_gn=MMS19;refGene_tx=XM_011287495.3;refGene_gn=MMS19;refGene_tx=XM_019813637.2;refGene_gn=MMS19;refGene_tx=XM_019813638.2
D2 58677120 58677189
refGene_gn=MMS19;refGene_tx=XM_003994268.5;refGene_gn=MMS19;refGene_tx=XM_011287495.3;refGene_gn=MMS19;refGene_tx=XM_019813637.2;refGene_gn=MMS19;refGene_tx=XM_019813638.2
D2 58677326 58677429
refGene_gn=MMS19;refGene_tx=XM_003994268.5;refGene_gn=MMS19;refGene_tx=XM_011287495.3;refGene_gn=MMS19;refGene_tx=XM_019813637.2;refGene_gn=MMS19;refGene_tx=XM_019813638.2
D2 58677904 58678004
refGene_gn=MMS19;refGene_tx=XM_003994268.5;refGene_gn=MMS19;refGene_tx=XM_011287495.3;refGene_gn=MMS19;refGene_tx=XM_019813637.2;refGene_gn=MMS19;refGene_tx=XM_019813638.2
D2 58678875 58679031
refGene_gn=MMS19;refGene_tx=XM_003994268.5;refGene_gn=MMS19;refGene_tx=XM_011287495.3;refGene_gn=MMS19;refGene_tx=XM_019813637.2;refGene_gn=MMS19;refGene_tx=XM_019813638.2
D2 58680147 58680297
refGene_gn=MMS19;refGene_tx=XM_003994268.5;refGene_gn=MMS19;refGene_tx=XM_011287495.3;refGene_gn=MMS19;refGene_tx=XM_019813637.2;refGene_gn=MMS19;refGene_tx=XM_019813638.2
D2 58680401 58680502
refGene_gn=MMS19;refGene_tx=XM_003994268.5;refGene_gn=MMS19;refGene_tx=XM_011287495.3;refGene_gn=MMS19;refGene_tx=XM_019813637.2;refGene_gn=MMS19;refGene_tx=XM_019813638.2
D2 58680784 58680866
refGene_gn=MMS19;refGene_tx=XM_003994268.5;refGene_gn=MMS19;refGene_tx=XM_011287495.3;refGene_gn=MMS19;refGene_tx=XM_019813637.2;refGene_gn=MMS19;refGene_tx=XM_019813638.2
D2 58681060 58681186
refGene_gn=MMS19;refGene_tx=XM_003994268.5;refGene_gn=MMS19;refGene_tx=XM_011287495.3;refGene_gn=MMS19;refGene_tx=XM_019813637.2;refGene_gn=MMS19;refGene_tx=XM_019813638.2
D2 58681471 58681550
refGene_gn=MMS19;refGene_tx=XM_003994268.5;refGene_gn=MMS19;refGene_tx=XM_011287495.3;refGene_gn=MMS19;refGene_tx=XM_019813637.2;refGene_gn=MMS19;refGene_tx=XM_019813638.2
D2 58681800 58681955
refGene_gn=MMS19;refGene_tx=XM_003994268.5;refGene_gn=MMS19;refGene_tx=XM_011287495.3;refGene_gn=MMS19;refGene_tx=XM_019813637.2;refGene_gn=

MMS19;refGene_tx=XM_019813638.2
D2 58682207 58682346
refGene_gn=MMS19;refGene_tx=XM_003994268.5;refGene_gn=MMS19;refGene_tx
=XM_011287495.3;refGene_gn=MMS19;refGene_tx=XM_019813637.2;refGene_gn=
MMS19;refGene_tx=XM_019813638.2
D2 58682878 58682956
refGene_gn=MMS19;refGene_tx=XM_003994268.5;refGene_gn=MMS19;refGene_tx
=XM_011287495.3;refGene_gn=MMS19;refGene_tx=XM_019813637.2;refGene_gn=
MMS19;refGene_tx=XM_019813638.2
D2 58683356 58683431
refGene_gn=MMS19;refGene_tx=XM_003994268.5;refGene_gn=MMS19;refGene_tx
=XM_011287495.3;refGene_gn=MMS19;refGene_tx=XM_019813637.2;refGene_gn=
MMS19;refGene_tx=XM_019813638.2
D2 58683873 58683960
refGene_gn=MMS19;refGene_tx=XM_003994268.5;refGene_gn=MMS19;refGene_tx
=XM_011287495.3;refGene_gn=MMS19;refGene_tx=XM_019813637.2;refGene_gn=
MMS19;refGene_tx=XM_019813638.2
D2 58688769 58688831
refGene_gn=MMS19;refGene_tx=XM_003994268.5;refGene_gn=MMS19;refGene_tx
=XM_011287495.3;refGene_gn=MMS19;refGene_tx=XM_019813637.2;refGene_gn=
MMS19;refGene_tx=XM_019813638.2
D2 58688921 58689050
refGene_gn=MMS19;refGene_tx=XM_003994268.5;refGene_gn=MMS19;refGene_tx
=XM_011287495.3;refGene_gn=MMS19;refGene_tx=XM_019813637.2;refGene_gn=
MMS19;refGene_tx=XM_019813638.2
D2 58689623 58689693
refGene_gn=MMS19;refGene_tx=XM_003994268.5;refGene_gn=MMS19;refGene_tx
=XM_011287495.3;refGene_gn=MMS19;refGene_tx=XM_019813637.2;refGene_gn=
MMS19;refGene_tx=XM_019813638.2
D2 58690094 58690169
refGene_gn=MMS19;refGene_tx=XM_003994268.5;refGene_gn=MMS19;refGene_tx
=XM_019813637.2;refGene_gn=MMS19;refGene_tx=XM_019813638.2
D2 58690551 58690637
refGene_gn=MMS19;refGene_tx=XM_003994268.5;refGene_gn=MMS19;refGene_tx
=XM_019813637.2;refGene_gn=MMS19;refGene_tx=XM_019813638.2
D2 58691555 58691656
refGene_gn=MMS19;refGene_tx=XM_003994268.5;refGene_gn=MMS19;refGene_tx
=XM_019813637.2;refGene_gn=MMS19;refGene_tx=XM_019813638.2
D2 58693472 58693521
refGene_gn=MMS19;refGene_tx=XM_003994268.5;refGene_gn=MMS19;refGene_tx
=XM_019813637.2;refGene_gn=MMS19;refGene_tx=XM_019813638.2
D2 58707195 58707274 refGene_gn=UBTD1;refGene_tx=XM_019813646.1
D2 58707602 58707714
refGene_gn=MMS19;refGene_tx=XM_003994268.5;refGene_gn=MMS19;refGene_tx
=XM_019813637.2;refGene_gn=MMS19;refGene_tx=XM_019813638.2
D2 58708574 58708644 refGene_gn=UBTD1;refGene_tx=XM_003994271.4
D2 58757946 58758174
refGene_gn=UBTD1;refGene_tx=XM_003994271.4;refGene_gn=UBTD1;refGene_tx
=XM_019813646.1
D2 58759521 58759907

refGene_gn=UBTD1;refGene_tx=XM_003994271.4;refGene_gn=UBTD1;refGene_tx=XM_019813646.1
D2 58762248 58762335 refGene_gn=ANKRD2;refGene_tx=XM_019813645.2
D2 58765826 58765928 refGene_gn=ANKRD2;refGene_tx=XM_019813645.2
D2 58766735 58766894 refGene_gn=ANKRD2;refGene_tx=XM_019813645.2
D2 58767009 58767117 refGene_gn=ANKRD2;refGene_tx=XM_019813645.2
D2 58768851 58768950 refGene_gn=ANKRD2;refGene_tx=XM_019813645.2
D2 58769280 58769379 refGene_gn=ANKRD2;refGene_tx=XM_019813645.2
D2 58770560 58770659 refGene_gn=ANKRD2;refGene_tx=XM_019813645.2
D2 58770765 58770864 refGene_gn=ANKRD2;refGene_tx=XM_019813645.2
D2 58771676 58771826 refGene_gn=ANKRD2;refGene_tx=XM_019813645.2
D2 58772703 58772914
refGene_gn=HOGA1;refGene_tx=XM_003994273.4;refGene_gn=HOGA1;refGene_tx=XM_006938055.4;refGene_gn=HOGA1;refGene_tx=XM_019813642.2;refGene_gn=HOGA1;refGene_tx=XM_019813643.2;refGene_gn=HOGA1;refGene_tx=XM_019813644.2
D2 58776992 58777679
refGene_gn=CD2H10orf62;refGene_tx=XM_006938054.3
D2 58784171 58784300
refGene_gn=HOGA1;refGene_tx=XM_003994273.4;refGene_gn=HOGA1;refGene_tx=XM_006938055.4;refGene_gn=HOGA1;refGene_tx=XM_019813642.2;refGene_gn=HOGA1;refGene_tx=XM_019813643.2
D2 58784503 58784631
refGene_gn=HOGA1;refGene_tx=XM_003994273.4;refGene_gn=HOGA1;refGene_tx=XM_006938055.4;refGene_gn=HOGA1;refGene_tx=XM_019813642.2;refGene_gn=HOGA1;refGene_tx=XM_019813643.2;refGene_gn=HOGA1;refGene_tx=XM_019813644.2
D2 58785142 58785277
refGene_gn=HOGA1;refGene_tx=XM_003994273.4;refGene_gn=HOGA1;refGene_tx=XM_006938055.4;refGene_gn=HOGA1;refGene_tx=XM_019813642.2;refGene_gn=HOGA1;refGene_tx=XM_019813643.2;refGene_gn=HOGA1;refGene_tx=XM_019813644.2
D2 58785528 58785625
refGene_gn=HOGA1;refGene_tx=XM_003994273.4;refGene_gn=HOGA1;refGene_tx=XM_006938055.4;refGene_gn=HOGA1;refGene_tx=XM_019813643.2;refGene_gn=HOGA1;refGene_tx=XM_019813644.2
D2 58787283 58787417
refGene_gn=HOGA1;refGene_tx=XM_003994273.4;refGene_gn=HOGA1;refGene_tx=XM_006938055.4;refGene_gn=HOGA1;refGene_tx=XM_019813642.2;refGene_gn=HOGA1;refGene_tx=XM_019813643.2;refGene_gn=HOGA1;refGene_tx=XM_019813644.2
D2 58788157 58788279
refGene_gn=HOGA1;refGene_tx=XM_019813642.2;refGene_gn=HOGA1;refGene_tx=XM_019813643.2
D2 58793168 58793378
refGene_gn=HOGA1;refGene_tx=XM_006938055.4;refGene_gn=HOGA1;refGene_tx=XM_019813644.2
D2 58794865 58795015
refGene_gn=HOGA1;refGene_tx=XM_003994273.4;refGene_gn=HOGA1;refGene_tx=XM_019813642.2;refGene_gn=HOGA1;refGene_tx=XM_019813643.2

D2 58798393 58798542
refGene_gn=MORN4;refGene_tx=XM_003994275.4;refGene_gn=MORN4;refGene_tx=XM_006938056.3;refGene_gn=MORN4;refGene_tx=XM_006938057.3;refGene_gn=MORN4;refGene_tx=XM_006938058.3;refGene_gn=MORN4;refGene_tx=XM_011287492.3;refGene_gn=MORN4;refGene_tx=XM_011287493.3;refGene_gn=MORN4;refGene_tx=XM_019813647.2;refGene_gn=MORN4;refGene_tx=XM_019813648.2;refGene_gn=MORN4;refGene_tx=XM_019813649.1;refGene_gn=MORN4;refGene_tx=XM_019813650.2;refGene_gn=MORN4;refGene_tx=XM_023240801.1
D2 58798790 58798900
refGene_gn=MORN4;refGene_tx=XM_003994275.4;refGene_gn=MORN4;refGene_tx=XM_006938056.3;refGene_gn=MORN4;refGene_tx=XM_006938057.3;refGene_gn=MORN4;refGene_tx=XM_006938058.3;refGene_gn=MORN4;refGene_tx=XM_011287492.3;refGene_gn=MORN4;refGene_tx=XM_011287493.3;refGene_gn=MORN4;refGene_tx=XM_019813647.2;refGene_gn=MORN4;refGene_tx=XM_019813648.2;refGene_gn=MORN4;refGene_tx=XM_019813649.1;refGene_gn=MORN4;refGene_tx=XM_019813650.2;refGene_gn=MORN4;refGene_tx=XM_023240801.1
D2 58799516 58799631
refGene_gn=MORN4;refGene_tx=XM_003994275.4;refGene_gn=MORN4;refGene_tx=XM_006938056.3;refGene_gn=MORN4;refGene_tx=XM_006938057.3;refGene_gn=MORN4;refGene_tx=XM_006938058.3;refGene_gn=MORN4;refGene_tx=XM_011287492.3;refGene_gn=MORN4;refGene_tx=XM_011287493.3;refGene_gn=MORN4;refGene_tx=XM_019813647.2;refGene_gn=MORN4;refGene_tx=XM_019813648.2;refGene_gn=MORN4;refGene_tx=XM_019813649.1;refGene_gn=MORN4;refGene_tx=XM_019813650.2;refGene_gn=MORN4;refGene_tx=XM_023240801.1
D2 58801115 58801182
refGene_gn=MORN4;refGene_tx=XM_003994275.4;refGene_gn=MORN4;refGene_tx=XM_006938056.3;refGene_gn=MORN4;refGene_tx=XM_006938057.3;refGene_gn=MORN4;refGene_tx=XM_006938058.3;refGene_gn=MORN4;refGene_tx=XM_011287493.3;refGene_gn=MORN4;refGene_tx=XM_019813649.1;refGene_gn=MORN4;refGene_tx=XM_019813650.2
D2 58804880 58805061
refGene_gn=MORN4;refGene_tx=XM_011287492.3;refGene_gn=MORN4;refGene_tx=XM_019813647.2;refGene_gn=MORN4;refGene_tx=XM_019813648.2;refGene_gn=MORN4;refGene_tx=XM_023240801.1
D2 58813996 58814431 refGene_gn=PI4K2A;refGene_tx=XM_023240802.1
D2 58823913 58824114 refGene_gn=PI4K2A;refGene_tx=XM_023240802.1
D2 58828594 58828726 refGene_gn=PI4K2A;refGene_tx=XM_023240802.1
D2 58829076 58829230 refGene_gn=PI4K2A;refGene_tx=XM_023240802.1
D2 58836624 58836686 refGene_gn=PI4K2A;refGene_tx=XM_023240802.1
D2 58838656 58838756 refGene_gn=PI4K2A;refGene_tx=XM_023240802.1
D2 58839482 58839616 refGene_gn=PI4K2A;refGene_tx=XM_023240802.1
D2 58840116 58840176 refGene_gn=PI4K2A;refGene_tx=XM_023240802.1
D2 58846167 58846329 refGene_gn=PI4K2A;refGene_tx=XM_023240802.1
D2 58850126 58850286
refGene_gn=AVPI1;refGene_tx=XM_003994276.3;refGene_gn=AVPI1;refGene_tx=XM_003994277.5
D2 58851286 58851573
refGene_gn=AVPI1;refGene_tx=XM_003994276.3;refGene_gn=AVPI1;refGene_tx=XM_003994277.5
D2 58874747 58875269

refGene_gn=MARVELD1;refGene_tx=XM_023240803.1;refGene_gn=MARVELD1;refGene_tx=XM_023240804.1;refGene_gn=MARVELD1;refGene_tx=XM_023240805.1;refGene_gn=MARVELD1;refGene_tx=XM_023240806.1
D2 58891501 58891698
refGene_gn=ZFYVE27;refGene_tx=XM_003994280.4;refGene_gn=ZFYVE27;refGene_tx=XM_019813652.1;refGene_gn=ZFYVE27;refGene_tx=XM_019813653.1
D2 58895945 58896016
refGene_gn=ZFYVE27;refGene_tx=XM_003994280.4;refGene_gn=ZFYVE27;refGene_tx=XM_019813652.1;refGene_gn=ZFYVE27;refGene_tx=XM_019813653.1
D2 58897612 58897799
refGene_gn=ZFYVE27;refGene_tx=XM_003994280.4;refGene_gn=ZFYVE27;refGene_tx=XM_019813652.1;refGene_gn=ZFYVE27;refGene_tx=XM_019813653.1
D2 58900655 58900751
refGene_gn=ZFYVE27;refGene_tx=XM_003994280.4;refGene_gn=ZFYVE27;refGene_tx=XM_019813652.1;refGene_gn=ZFYVE27;refGene_tx=XM_019813653.1
D2 58901096 58901209
refGene_gn=ZFYVE27;refGene_tx=XM_003994280.4;refGene_gn=ZFYVE27;refGene_tx=XM_019813652.1;refGene_gn=ZFYVE27;refGene_tx=XM_019813653.1
D2 58901943 58902083
refGene_gn=ZFYVE27;refGene_tx=XM_003994280.4;refGene_gn=ZFYVE27;refGene_tx=XM_019813652.1;refGene_gn=ZFYVE27;refGene_tx=XM_019813653.1
D2 58902667 58902739
refGene_gn=ZFYVE27;refGene_tx=XM_003994280.4;refGene_gn=ZFYVE27;refGene_tx=XM_019813652.1;refGene_gn=ZFYVE27;refGene_tx=XM_019813653.1
D2 58904324 58904345
refGene_gn=ZFYVE27;refGene_tx=XM_019813652.1;refGene_gn=ZFYVE27;refGene_tx=XM_019813653.1
D2 58904509 58904654
refGene_gn=ZFYVE27;refGene_tx=XM_003994280.4;refGene_gn=ZFYVE27;refGene_tx=XM_019813652.1;refGene_gn=ZFYVE27;refGene_tx=XM_019813653.1
D2 58908703 58908750
refGene_gn=ZFYVE27;refGene_tx=XM_003994280.4;refGene_gn=ZFYVE27;refGene_tx=XM_019813652.1;refGene_gn=ZFYVE27;refGene_tx=XM_019813653.1
D2 58909057 58909139
refGene_gn=ZFYVE27;refGene_tx=XM_003994280.4;refGene_gn=ZFYVE27;refGene_tx=XM_019813652.1;refGene_gn=ZFYVE27;refGene_tx=XM_019813653.1
D2 58909746 58909829 refGene_gn=ZFYVE27;refGene_tx=XM_019813652.1
D2 58910139 58910204
refGene_gn=ZFYVE27;refGene_tx=XM_003994280.4;refGene_gn=ZFYVE27;refGene_tx=XM_019813653.1
D2 58918875 58919222 refGene_gn=SFRP5;refGene_tx=XM_023240807.1
D2 58920929 58921007 refGene_gn=SFRP5;refGene_tx=XM_023240807.1
D2 58923114 58923637 refGene_gn=SFRP5;refGene_tx=XM_023240807.1
D2 58991773 58991860 refGene_gn=GOLGA7B;refGene_tx=XM_019813659.2
D2 58994831 58994854 refGene_gn=CRTAC1;refGene_tx=XM_019813657.2
D2 58997618 58997744 refGene_gn=GOLGA7B;refGene_tx=XM_019813659.2
D2 59000473 59000626 refGene_gn=GOLGA7B;refGene_tx=XM_019813659.2
D2 59000711 59000813 refGene_gn=GOLGA7B;refGene_tx=XM_019813659.2
D2 59001322 59001373 refGene_gn=GOLGA7B;refGene_tx=XM_019813659.2
D2 59001907 59002158

refGene_gn=CRTAC1;refGene_tx=XM_006938060.3;refGene_gn=CRTAC1;refGene_tx=XM_019813654.2
D2 59007484 59007513
refGene_gn=CRTAC1;refGene_tx=XM_006938062.4;refGene_gn=CRTAC1;refGene_tx=XM_019813655.2
D2 59009411 59009449 refGene_gn=CRTAC1;refGene_tx=XM_019813656.2
D2 59015015 59015134
refGene_gn=CRTAC1;refGene_tx=XM_003994283.5;refGene_gn=CRTAC1;refGene_tx=XM_019813654.2;refGene_gn=CRTAC1;refGene_tx=XM_019813655.2
D2 59015601 59015745
refGene_gn=CRTAC1;refGene_tx=XM_003994283.5;refGene_gn=CRTAC1;refGene_tx=XM_006938060.3;refGene_gn=CRTAC1;refGene_tx=XM_006938062.4;refGene_gn=CRTAC1;refGene_tx=XM_019813654.2;refGene_gn=CRTAC1;refGene_tx=XM_019813655.2;refGene_gn=CRTAC1;refGene_tx=XM_019813656.2;refGene_gn=CRTAC1;refGene_tx=XM_019813657.2
D2 59018657 59018700
refGene_gn=CRTAC1;refGene_tx=XM_003994283.5;refGene_gn=CRTAC1;refGene_tx=XM_006938060.3;refGene_gn=CRTAC1;refGene_tx=XM_006938062.4;refGene_gn=CRTAC1;refGene_tx=XM_019813654.2;refGene_gn=CRTAC1;refGene_tx=XM_019813655.2;refGene_gn=CRTAC1;refGene_tx=XM_019813656.2;refGene_gn=CRTAC1;refGene_tx=XM_019813657.2
D2 59019154 59019300
refGene_gn=CRTAC1;refGene_tx=XM_003994283.5;refGene_gn=CRTAC1;refGene_tx=XM_006938060.3;refGene_gn=CRTAC1;refGene_tx=XM_006938062.4;refGene_gn=CRTAC1;refGene_tx=XM_019813654.2;refGene_gn=CRTAC1;refGene_tx=XM_019813655.2;refGene_gn=CRTAC1;refGene_tx=XM_019813656.2;refGene_gn=CRTAC1;refGene_tx=XM_019813657.2
D2 59028654 59028823
refGene_gn=CRTAC1;refGene_tx=XM_003994283.5;refGene_gn=CRTAC1;refGene_tx=XM_006938060.3;refGene_gn=CRTAC1;refGene_tx=XM_006938062.4;refGene_gn=CRTAC1;refGene_tx=XM_019813654.2;refGene_gn=CRTAC1;refGene_tx=XM_019813655.2;refGene_gn=CRTAC1;refGene_tx=XM_019813656.2;refGene_gn=CRTAC1;refGene_tx=XM_019813657.2
D2 59029245 59029346
refGene_gn=CRTAC1;refGene_tx=XM_003994283.5;refGene_gn=CRTAC1;refGene_tx=XM_006938060.3;refGene_gn=CRTAC1;refGene_tx=XM_006938062.4;refGene_gn=CRTAC1;refGene_tx=XM_019813654.2;refGene_gn=CRTAC1;refGene_tx=XM_019813655.2;refGene_gn=CRTAC1;refGene_tx=XM_019813656.2;refGene_gn=CRTAC1;refGene_tx=XM_019813657.2
D2 59030226 59030309
refGene_gn=CRTAC1;refGene_tx=XM_003994283.5;refGene_gn=CRTAC1;refGene_tx=XM_006938060.3;refGene_gn=CRTAC1;refGene_tx=XM_006938062.4;refGene_gn=CRTAC1;refGene_tx=XM_019813654.2;refGene_gn=CRTAC1;refGene_tx=XM_019813655.2;refGene_gn=CRTAC1;refGene_tx=XM_019813656.2;refGene_gn=CRTAC1;refGene_tx=XM_019813657.2
D2 59033998 59034135
refGene_gn=CRTAC1;refGene_tx=XM_003994283.5;refGene_gn=CRTAC1;refGene_tx=XM_006938060.3;refGene_gn=CRTAC1;refGene_tx=XM_006938062.4;refGene_gn=CRTAC1;refGene_tx=XM_019813654.2;refGene_gn=CRTAC1;refGene_tx=XM_019813655.2;refGene_gn=CRTAC1;refGene_tx=XM_019813656.2;refGene_gn=CRTAC

1;refGene_tx=XM_019813657.2
D2 59036778 59036924
refGene_gn=CRTAC1;refGene_tx=XM_003994283.5;refGene_gn=CRTAC1;refGene_tx=XM_006938060.3;refGene_gn=CRTAC1;refGene_tx=XM_006938062.4;refGene_gn=CRTAC1;refGene_tx=XM_019813654.2;refGene_gn=CRTAC1;refGene_tx=XM_019813655.2;refGene_gn=CRTAC1;refGene_tx=XM_019813656.2;refGene_gn=CRTAC1;refGene_tx=XM_019813657.2
D2 59040249 59040384
refGene_gn=CRTAC1;refGene_tx=XM_003994283.5;refGene_gn=CRTAC1;refGene_tx=XM_006938060.3;refGene_gn=CRTAC1;refGene_tx=XM_006938062.4;refGene_gn=CRTAC1;refGene_tx=XM_019813654.2;refGene_gn=CRTAC1;refGene_tx=XM_019813655.2;refGene_gn=CRTAC1;refGene_tx=XM_019813656.2;refGene_gn=CRTAC1;refGene_tx=XM_019813657.2
D2 59048963 59049120
refGene_gn=CRTAC1;refGene_tx=XM_003994283.5;refGene_gn=CRTAC1;refGene_tx=XM_006938060.3;refGene_gn=CRTAC1;refGene_tx=XM_006938062.4;refGene_gn=CRTAC1;refGene_tx=XM_019813654.2;refGene_gn=CRTAC1;refGene_tx=XM_019813655.2;refGene_gn=CRTAC1;refGene_tx=XM_019813656.2;refGene_gn=CRTAC1;refGene_tx=XM_019813657.2
D2 59054589 59054726
refGene_gn=CRTAC1;refGene_tx=XM_003994283.5;refGene_gn=CRTAC1;refGene_tx=XM_006938060.3;refGene_gn=CRTAC1;refGene_tx=XM_006938062.4;refGene_gn=CRTAC1;refGene_tx=XM_019813654.2;refGene_gn=CRTAC1;refGene_tx=XM_019813655.2;refGene_gn=CRTAC1;refGene_tx=XM_019813656.2;refGene_gn=CRTAC1;refGene_tx=XM_019813657.2
D2 59067805 59068002
refGene_gn=CRTAC1;refGene_tx=XM_003994283.5;refGene_gn=CRTAC1;refGene_tx=XM_006938060.3;refGene_gn=CRTAC1;refGene_tx=XM_006938062.4;refGene_gn=CRTAC1;refGene_tx=XM_019813654.2;refGene_gn=CRTAC1;refGene_tx=XM_019813655.2;refGene_gn=CRTAC1;refGene_tx=XM_019813656.2;refGene_gn=CRTAC1;refGene_tx=XM_019813657.2
D2 59133536 59133736
refGene_gn=CRTAC1;refGene_tx=XM_003994283.5;refGene_gn=CRTAC1;refGene_tx=XM_006938060.3;refGene_gn=CRTAC1;refGene_tx=XM_006938062.4;refGene_gn=CRTAC1;refGene_tx=XM_019813654.2;refGene_gn=CRTAC1;refGene_tx=XM_019813655.2;refGene_gn=CRTAC1;refGene_tx=XM_019813656.2;refGene_gn=CRTAC1;refGene_tx=XM_019813657.2
D2 59152047 59152071
refGene_gn=CRTAC1;refGene_tx=XM_003994283.5;refGene_gn=CRTAC1;refGene_tx=XM_006938060.3;refGene_gn=CRTAC1;refGene_tx=XM_006938062.4;refGene_gn=CRTAC1;refGene_tx=XM_019813654.2;refGene_gn=CRTAC1;refGene_tx=XM_019813655.2;refGene_gn=CRTAC1;refGene_tx=XM_019813656.2;refGene_gn=CRTAC1;refGene_tx=XM_019813657.2
D2 59305300 59307109
refGene_gn=R3HCC1L;refGene_tx=XM_003994285.5;refGene_gn=R3HCC1L;refGene_tx=XM_006938064.3;refGene_gn=R3HCC1L;refGene_tx=XM_006938066.4;refGene_gn=R3HCC1L;refGene_tx=XM_006938067.4;refGene_gn=R3HCC1L;refGene_tx=XM_019813660.2;refGene_gn=R3HCC1L;refGene_tx=XM_019813662.2;refGene_gn=R3HCC1L;refGene_tx=XM_023240809.1;refGene_gn=R3HCC1L;refGene_tx=XM_023240810.1;refGene_gn=R3HCC1L;refGene_tx=XM_023240811.1

D2 59321896 59322072
refGene_gn=R3HCC1L;refGene_tx=XM_003994285.5;refGene_gn=R3HCC1L;refGene_tx=XM_006938064.3;refGene_gn=R3HCC1L;refGene_tx=XM_006938066.4;refGene_gn=R3HCC1L;refGene_tx=XM_006938067.4;refGene_gn=R3HCC1L;refGene_tx=XM_019813660.2;refGene_gn=R3HCC1L;refGene_tx=XM_019813662.2;refGene_gn=R3HCC1L;refGene_tx=XM_023240809.1;refGene_gn=R3HCC1L;refGene_tx=XM_023240810.1;refGene_gn=R3HCC1L;refGene_tx=XM_023240811.1
D2 59327026 59327097
refGene_gn=R3HCC1L;refGene_tx=XM_003994285.5;refGene_gn=R3HCC1L;refGene_tx=XM_006938064.3;refGene_gn=R3HCC1L;refGene_tx=XM_006938066.4;refGene_gn=R3HCC1L;refGene_tx=XM_006938067.4;refGene_gn=R3HCC1L;refGene_tx=XM_019813660.2;refGene_gn=R3HCC1L;refGene_tx=XM_019813662.2;refGene_gn=R3HCC1L;refGene_tx=XM_023240809.1;refGene_gn=R3HCC1L;refGene_tx=XM_023240810.1;refGene_gn=R3HCC1L;refGene_tx=XM_023240811.1
D2 59328017 59328113
refGene_gn=R3HCC1L;refGene_tx=XM_003994285.5;refGene_gn=R3HCC1L;refGene_tx=XM_006938064.3;refGene_gn=R3HCC1L;refGene_tx=XM_006938066.4;refGene_gn=R3HCC1L;refGene_tx=XM_006938067.4;refGene_gn=R3HCC1L;refGene_tx=XM_019813660.2;refGene_gn=R3HCC1L;refGene_tx=XM_019813662.2;refGene_gn=R3HCC1L;refGene_tx=XM_023240809.1;refGene_gn=R3HCC1L;refGene_tx=XM_023240810.1;refGene_gn=R3HCC1L;refGene_tx=XM_023240811.1
D2 59328608 59328749
refGene_gn=R3HCC1L;refGene_tx=XM_003994285.5;refGene_gn=R3HCC1L;refGene_tx=XM_006938064.3;refGene_gn=R3HCC1L;refGene_tx=XM_006938066.4;refGene_gn=R3HCC1L;refGene_tx=XM_006938067.4;refGene_gn=R3HCC1L;refGene_tx=XM_019813660.2;refGene_gn=R3HCC1L;refGene_tx=XM_019813662.2;refGene_gn=R3HCC1L;refGene_tx=XM_023240809.1;refGene_gn=R3HCC1L;refGene_tx=XM_023240810.1;refGene_gn=R3HCC1L;refGene_tx=XM_023240811.1
D2 59337399 59337467
refGene_gn=R3HCC1L;refGene_tx=XM_003994285.5;refGene_gn=R3HCC1L;refGene_tx=XM_006938064.3;refGene_gn=R3HCC1L;refGene_tx=XM_006938066.4;refGene_gn=R3HCC1L;refGene_tx=XM_006938067.4;refGene_gn=R3HCC1L;refGene_tx=XM_019813660.2;refGene_gn=R3HCC1L;refGene_tx=XM_019813662.2;refGene_gn=R3HCC1L;refGene_tx=XM_023240809.1;refGene_gn=R3HCC1L;refGene_tx=XM_023240810.1;refGene_gn=R3HCC1L;refGene_tx=XM_023240811.1
D2 59342151 59342222
refGene_gn=LOXL4;refGene_tx=XM_019813663.1;refGene_gn=LOXL4;refGene_tx=XM_023240813.1;refGene_gn=LOXL4;refGene_tx=XM_023240814.1;refGene_gn=LOXL4;refGene_tx=XM_023240815.1
D2 59344565 59344677
refGene_gn=LOXL4;refGene_tx=XM_019813663.1;refGene_gn=LOXL4;refGene_tx=XM_023240813.1;refGene_gn=LOXL4;refGene_tx=XM_023240814.1;refGene_gn=LOXL4;refGene_tx=XM_023240815.1
D2 59345039 59345176
refGene_gn=LOXL4;refGene_tx=XM_019813663.1;refGene_gn=LOXL4;refGene_tx=XM_023240813.1;refGene_gn=LOXL4;refGene_tx=XM_023240814.1;refGene_gn=LOXL4;refGene_tx=XM_023240815.1
D2 59345863 59345979
refGene_gn=LOXL4;refGene_tx=XM_019813663.1;refGene_gn=LOXL4;refGene_tx=XM_023240813.1;refGene_gn=LOXL4;refGene_tx=XM_023240814.1;refGene_gn=

LOXL4;refGene_tx=XM_023240815.1
D2 59347065 59347309
refGene_gn=LOXL4;refGene_tx=XM_019813663.1;refGene_gn=LOXL4;refGene_tx=XM_023240813.1;refGene_gn=LOXL4;refGene_tx=XM_023240814.1;refGene_gn=LOXL4;refGene_tx=XM_023240815.1
D2 59349542 59349705
refGene_gn=LOXL4;refGene_tx=XM_019813663.1;refGene_gn=LOXL4;refGene_tx=XM_023240813.1;refGene_gn=LOXL4;refGene_tx=XM_023240814.1;refGene_gn=LOXL4;refGene_tx=XM_023240815.1
D2 59350717 59350885
refGene_gn=LOXL4;refGene_tx=XM_019813663.1;refGene_gn=LOXL4;refGene_tx=XM_023240813.1;refGene_gn=LOXL4;refGene_tx=XM_023240814.1;refGene_gn=LOXL4;refGene_tx=XM_023240815.1
D2 59351597 59351752
refGene_gn=LOXL4;refGene_tx=XM_019813663.1;refGene_gn=LOXL4;refGene_tx=XM_023240813.1;refGene_gn=LOXL4;refGene_tx=XM_023240814.1;refGene_gn=LOXL4;refGene_tx=XM_023240815.1
D2 59351921 59352108
refGene_gn=LOXL4;refGene_tx=XM_019813663.1;refGene_gn=LOXL4;refGene_tx=XM_023240813.1;refGene_gn=LOXL4;refGene_tx=XM_023240814.1;refGene_gn=LOXL4;refGene_tx=XM_023240815.1
D2 59352915 59353135
refGene_gn=LOXL4;refGene_tx=XM_019813663.1;refGene_gn=LOXL4;refGene_tx=XM_023240813.1;refGene_gn=LOXL4;refGene_tx=XM_023240814.1;refGene_gn=LOXL4;refGene_tx=XM_023240815.1
D2 59353279 59353318
refGene_gn=LOXL4;refGene_tx=XM_019813663.1;refGene_gn=LOXL4;refGene_tx=XM_023240813.1;refGene_gn=LOXL4;refGene_tx=XM_023240814.1;refGene_gn=LOXL4;refGene_tx=XM_023240815.1
D2 59354697 59354903
refGene_gn=LOXL4;refGene_tx=XM_019813663.1;refGene_gn=LOXL4;refGene_tx=XM_023240813.1;refGene_gn=LOXL4;refGene_tx=XM_023240814.1;refGene_gn=LOXL4;refGene_tx=XM_023240815.1
D2 59355782 59355961
refGene_gn=LOXL4;refGene_tx=XM_019813663.1;refGene_gn=LOXL4;refGene_tx=XM_023240813.1;refGene_gn=LOXL4;refGene_tx=XM_023240814.1;refGene_gn=LOXL4;refGene_tx=XM_023240815.1
D2 59356506 59356786
refGene_gn=LOXL4;refGene_tx=XM_019813663.1;refGene_gn=LOXL4;refGene_tx=XM_023240813.1;refGene_gn=LOXL4;refGene_tx=XM_023240814.1
D2 59466132 59466220
refGene_gn=PYROXD2;refGene_tx=XM_003994289.3;refGene_gn=PYROXD2;refGene_tx=XM_019813672.1;refGene_gn=PYROXD2;refGene_tx=XM_019813673.1;refGene_gn=PYROXD2;refGene_tx=XM_019813674.1;refGene_gn=PYROXD2;refGene_tx=XM_019813675.1;refGene_gn=PYROXD2;refGene_tx=XM_019813676.1;refGene_gn=PYROXD2;refGene_tx=XM_019813679.1;refGene_gn=PYROXD2;refGene_tx=XM_019813680.1
D2 59466992 59467228
refGene_gn=PYROXD2;refGene_tx=XM_003994289.3;refGene_gn=PYROXD2;refGene_tx=XM_019813672.1;refGene_gn=PYROXD2;refGene_tx=XM_019813673.1;refGe

ne_gn=PYROXD2;refGene_tx=XM_019813674.1;refGene_gn=PYROXD2;refGene_tx=XM_019813675.1;refGene_gn=PYROXD2;refGene_tx=XM_019813676.1;refGene_gn=PYROXD2;refGene_tx=XM_019813679.1;refGene_gn=PYROXD2;refGene_tx=XM_019813680.1
D2 59467302 59467329 refGene_gn=PYROXD2;refGene_tx=XM_019813677.2
D2 59468602 59468709
refGene_gn=PYROXD2;refGene_tx=XM_003994289.3;refGene_gn=PYROXD2;refGene_tx=XM_019813672.1;refGene_gn=PYROXD2;refGene_tx=XM_019813673.1;refGene_gn=PYROXD2;refGene_tx=XM_019813674.1;refGene_gn=PYROXD2;refGene_tx=XM_019813675.1;refGene_gn=PYROXD2;refGene_tx=XM_019813676.1;refGene_gn=PYROXD2;refGene_tx=XM_019813677.2;refGene_gn=PYROXD2;refGene_tx=XM_019813679.1;refGene_gn=PYROXD2;refGene_tx=XM_019813680.1
D2 59469227 59469334 refGene_gn=PYROXD2;refGene_tx=XM_019813678.1
D2 59469421 59469576
refGene_gn=PYROXD2;refGene_tx=XM_003994289.3;refGene_gn=PYROXD2;refGene_tx=XM_019813672.1;refGene_gn=PYROXD2;refGene_tx=XM_019813673.1;refGene_gn=PYROXD2;refGene_tx=XM_019813674.1;refGene_gn=PYROXD2;refGene_tx=XM_019813675.1;refGene_gn=PYROXD2;refGene_tx=XM_019813676.1;refGene_gn=PYROXD2;refGene_tx=XM_019813677.2;refGene_gn=PYROXD2;refGene_tx=XM_019813678.1;refGene_gn=PYROXD2;refGene_tx=XM_019813679.1;refGene_gn=PYROXD2;refGene_tx=XM_019813680.1
D2 59472024 59472181
refGene_gn=PYROXD2;refGene_tx=XM_003994289.3;refGene_gn=PYROXD2;refGene_tx=XM_019813672.1;refGene_gn=PYROXD2;refGene_tx=XM_019813673.1;refGene_gn=PYROXD2;refGene_tx=XM_019813674.1;refGene_gn=PYROXD2;refGene_tx=XM_019813675.1;refGene_gn=PYROXD2;refGene_tx=XM_019813676.1;refGene_gn=PYROXD2;refGene_tx=XM_019813677.2;refGene_gn=PYROXD2;refGene_tx=XM_019813678.1;refGene_gn=PYROXD2;refGene_tx=XM_019813679.1;refGene_gn=PYROXD2;refGene_tx=XM_019813680.1
D2 59472268 59472280
refGene_gn=PYROXD2;refGene_tx=XM_019813672.1;refGene_gn=PYROXD2;refGene_tx=XM_019813674.1;refGene_gn=PYROXD2;refGene_tx=XM_019813675.1;refGene_gn=PYROXD2;refGene_tx=XM_019813676.1;refGene_gn=PYROXD2;refGene_tx=XM_019813677.2;refGene_gn=PYROXD2;refGene_tx=XM_019813678.1;refGene_gn=PYROXD2;refGene_tx=XM_019813679.1;refGene_gn=PYROXD2;refGene_tx=XM_019813680.1
D2 59472443 59472516
refGene_gn=PYROXD2;refGene_tx=XM_003994289.3;refGene_gn=PYROXD2;refGene_tx=XM_019813672.1;refGene_gn=PYROXD2;refGene_tx=XM_019813673.1;refGene_gn=PYROXD2;refGene_tx=XM_019813674.1;refGene_gn=PYROXD2;refGene_tx=XM_019813675.1;refGene_gn=PYROXD2;refGene_tx=XM_019813676.1;refGene_gn=PYROXD2;refGene_tx=XM_019813677.2;refGene_gn=PYROXD2;refGene_tx=XM_019813678.1;refGene_gn=PYROXD2;refGene_tx=XM_019813679.1;refGene_gn=PYROXD2;refGene_tx=XM_019813680.1
D2 59473788 59473923
refGene_gn=PYROXD2;refGene_tx=XM_003994289.3;refGene_gn=PYROXD2;refGene_tx=XM_019813672.1;refGene_gn=PYROXD2;refGene_tx=XM_019813673.1;refGene_gn=PYROXD2;refGene_tx=XM_019813674.1;refGene_gn=PYROXD2;refGene_tx=XM_019813675.1;refGene_gn=PYROXD2;refGene_tx=XM_019813676.1;refGene_gn=PYROXD2;refGene_tx=XM_019813677.2;refGene_gn=PYROXD2;refGene_tx=XM_01

9813678.1;refGene_gn=PYROXD2;refGene_tx=XM_019813679.1;refGene_gn=PYROXD2;refGene_tx=XM_019813680.1
D2 59474327 59474469
refGene_gn=PYROXD2;refGene_tx=XM_003994289.3;refGene_gn=PYROXD2;refGene_tx=XM_019813672.1;refGene_gn=PYROXD2;refGene_tx=XM_019813673.1;refGene_gn=PYROXD2;refGene_tx=XM_019813674.1;refGene_gn=PYROXD2;refGene_tx=XM_019813675.1;refGene_gn=PYROXD2;refGene_tx=XM_019813676.1;refGene_gn=PYROXD2;refGene_tx=XM_019813677.2;refGene_gn=PYROXD2;refGene_tx=XM_019813678.1;refGene_gn=PYROXD2;refGene_tx=XM_019813679.1;refGene_gn=PYROXD2;refGene_tx=XM_019813680.1
D2 59476307 59476405
refGene_gn=PYROXD2;refGene_tx=XM_003994289.3;refGene_gn=PYROXD2;refGene_tx=XM_019813672.1;refGene_gn=PYROXD2;refGene_tx=XM_019813673.1;refGene_gn=PYROXD2;refGene_tx=XM_019813674.1;refGene_gn=PYROXD2;refGene_tx=XM_019813675.1;refGene_gn=PYROXD2;refGene_tx=XM_019813676.1;refGene_gn=PYROXD2;refGene_tx=XM_019813677.2;refGene_gn=PYROXD2;refGene_tx=XM_019813678.1;refGene_gn=PYROXD2;refGene_tx=XM_019813679.1;refGene_gn=PYROXD2;refGene_tx=XM_019813680.1
D2 59476501 59476563
refGene_gn=PYROXD2;refGene_tx=XM_003994289.3;refGene_gn=PYROXD2;refGene_tx=XM_019813672.1;refGene_gn=PYROXD2;refGene_tx=XM_019813673.1;refGene_gn=PYROXD2;refGene_tx=XM_019813674.1;refGene_gn=PYROXD2;refGene_tx=XM_019813675.1;refGene_gn=PYROXD2;refGene_tx=XM_019813676.1;refGene_gn=PYROXD2;refGene_tx=XM_019813677.2;refGene_gn=PYROXD2;refGene_tx=XM_019813678.1;refGene_gn=PYROXD2;refGene_tx=XM_019813679.1;refGene_gn=PYROXD2;refGene_tx=XM_019813680.1
D2 59477395 59477549
refGene_gn=PYROXD2;refGene_tx=XM_003994289.3;refGene_gn=PYROXD2;refGene_tx=XM_019813672.1;refGene_gn=PYROXD2;refGene_tx=XM_019813673.1;refGene_gn=PYROXD2;refGene_tx=XM_019813674.1;refGene_gn=PYROXD2;refGene_tx=XM_019813675.1;refGene_gn=PYROXD2;refGene_tx=XM_019813676.1;refGene_gn=PYROXD2;refGene_tx=XM_019813677.2;refGene_gn=PYROXD2;refGene_tx=XM_019813678.1;refGene_gn=PYROXD2;refGene_tx=XM_019813679.1;refGene_gn=PYROXD2;refGene_tx=XM_019813680.1
D2 59479700 59479856
refGene_gn=PYROXD2;refGene_tx=XM_003994289.3;refGene_gn=PYROXD2;refGene_tx=XM_019813672.1;refGene_gn=PYROXD2;refGene_tx=XM_019813673.1;refGene_gn=PYROXD2;refGene_tx=XM_019813674.1;refGene_gn=PYROXD2;refGene_tx=XM_019813675.1;refGene_gn=PYROXD2;refGene_tx=XM_019813676.1;refGene_gn=PYROXD2;refGene_tx=XM_019813677.2;refGene_gn=PYROXD2;refGene_tx=XM_019813678.1;refGene_gn=PYROXD2;refGene_tx=XM_019813679.1;refGene_gn=PYROXD2;refGene_tx=XM_019813680.1
D2 59484895 59484969
refGene_gn=PYROXD2;refGene_tx=XM_003994289.3;refGene_gn=PYROXD2;refGene_tx=XM_019813672.1;refGene_gn=PYROXD2;refGene_tx=XM_019813673.1;refGene_gn=PYROXD2;refGene_tx=XM_019813674.1;refGene_gn=PYROXD2;refGene_tx=XM_019813675.1;refGene_gn=PYROXD2;refGene_tx=XM_019813676.1;refGene_gn=PYROXD2;refGene_tx=XM_019813677.2;refGene_gn=PYROXD2;refGene_tx=XM_019813678.1
D2 59485053 59485147

refGene_gn=PYROXD2;refGene_tx=XM_003994289.3;refGene_gn=PYROXD2;refGene_tx=XM_019813672.1;refGene_gn=PYROXD2;refGene_tx=XM_019813673.1;refGene_gn=PYROXD2;refGene_tx=XM_019813674.1;refGene_gn=PYROXD2;refGene_tx=XM_019813675.1;refGene_gn=PYROXD2;refGene_tx=XM_019813676.1;refGene_gn=PYROXD2;refGene_tx=XM_019813677.2;refGene_gn=PYROXD2;refGene_tx=XM_019813678.1
D2 59487678 59487698
refGene_gn=PYROXD2;refGene_tx=XM_003994289.3;refGene_gn=PYROXD2;refGene_tx=XM_019813672.1;refGene_gn=PYROXD2;refGene_tx=XM_019813673.1;refGene_gn=PYROXD2;refGene_tx=XM_019813675.1;refGene_gn=PYROXD2;refGene_tx=XM_019813676.1;refGene_gn=PYROXD2;refGene_tx=XM_019813677.2;refGene_gn=PYROXD2;refGene_tx=XM_019813678.1
D2 59487761 59487845 refGene_gn=PYROXD2;refGene_tx=XM_019813674.1
D2 59491999 59492126
refGene_gn=PYROXD2;refGene_tx=XM_003994289.3;refGene_gn=PYROXD2;refGene_tx=XM_019813672.1;refGene_gn=PYROXD2;refGene_tx=XM_019813673.1;refGene_gn=PYROXD2;refGene_tx=XM_019813676.1;refGene_gn=PYROXD2;refGene_tx=XM_019813677.2;refGene_gn=PYROXD2;refGene_tx=XM_019813678.1
D2 59492834 59492889 refGene_gn=PYROXD2;refGene_tx=XM_019813675.1
D2 59494540 59494703
refGene_gn=HPS1;refGene_tx=XM_011287485.2;refGene_gn=HPS1;refGene_tx=XM_019813664.2;refGene_gn=HPS1;refGene_tx=XM_019813665.1;refGene_gn=HPS1;refGene_tx=XM_019813666.2;refGene_gn=HPS1;refGene_tx=XM_019813667.2;refGene_gn=HPS1;refGene_tx=XM_019813668.1;refGene_gn=HPS1;refGene_tx=XM_019813669.2;refGene_gn=HPS1;refGene_tx=XM_019813670.1;refGene_gn=HPS1;refGene_tx=XM_019813671.1
D2 59495187 59495270
refGene_gn=HPS1;refGene_tx=XM_011287485.2;refGene_gn=HPS1;refGene_tx=XM_019813664.2;refGene_gn=HPS1;refGene_tx=XM_019813665.1;refGene_gn=HPS1;refGene_tx=XM_019813666.2;refGene_gn=HPS1;refGene_tx=XM_019813667.2;refGene_gn=HPS1;refGene_tx=XM_019813668.1;refGene_gn=HPS1;refGene_tx=XM_019813669.2;refGene_gn=HPS1;refGene_tx=XM_019813670.1;refGene_gn=HPS1;refGene_tx=XM_019813671.1
D2 59496767 59496881
refGene_gn=HPS1;refGene_tx=XM_011287485.2;refGene_gn=HPS1;refGene_tx=XM_019813664.2;refGene_gn=HPS1;refGene_tx=XM_019813665.1;refGene_gn=HPS1;refGene_tx=XM_019813666.2;refGene_gn=HPS1;refGene_tx=XM_019813667.2;refGene_gn=HPS1;refGene_tx=XM_019813668.1;refGene_gn=HPS1;refGene_tx=XM_019813669.2;refGene_gn=HPS1;refGene_tx=XM_019813671.1
D2 59498970 59499115
refGene_gn=HPS1;refGene_tx=XM_011287485.2;refGene_gn=HPS1;refGene_tx=XM_019813664.2;refGene_gn=HPS1;refGene_tx=XM_019813665.1;refGene_gn=HPS1;refGene_tx=XM_019813666.2;refGene_gn=HPS1;refGene_tx=XM_019813667.2;refGene_gn=HPS1;refGene_tx=XM_019813668.1;refGene_gn=HPS1;refGene_tx=XM_019813669.2;refGene_gn=HPS1;refGene_tx=XM_019813670.1;refGene_gn=HPS1;refGene_tx=XM_019813671.1
D2 59500209 59500275
refGene_gn=HPS1;refGene_tx=XM_011287485.2;refGene_gn=HPS1;refGene_tx=XM_019813664.2;refGene_gn=HPS1;refGene_tx=XM_019813665.1;refGene_gn=HPS1;refGene_tx=XM_019813666.2;refGene_gn=HPS1;refGene_tx=XM_019813667.2;

refGene_gn=HPS1;refGene_tx=XM_019813668.1;refGene_gn=HPS1;refGene_tx=XM_019813669.2;refGene_gn=HPS1;refGene_tx=XM_019813670.1;refGene_gn=HPS1;refGene_tx=XM_019813671.1
D2 59500358 59500493
refGene_gn=HPS1;refGene_tx=XM_011287485.2;refGene_gn=HPS1;refGene_tx=XM_019813664.2;refGene_gn=HPS1;refGene_tx=XM_019813665.1;refGene_gn=HPS1;refGene_tx=XM_019813666.2;refGene_gn=HPS1;refGene_tx=XM_019813667.2;refGene_gn=HPS1;refGene_tx=XM_019813668.1;refGene_gn=HPS1;refGene_tx=XM_019813669.2;refGene_gn=HPS1;refGene_tx=XM_019813670.1;refGene_gn=HPS1;refGene_tx=XM_019813671.1
D2 59501020 59501082
refGene_gn=HPS1;refGene_tx=XM_011287485.2;refGene_gn=HPS1;refGene_tx=XM_019813664.2;refGene_gn=HPS1;refGene_tx=XM_019813665.1;refGene_gn=HPS1;refGene_tx=XM_019813666.2;refGene_gn=HPS1;refGene_tx=XM_019813667.2;refGene_gn=HPS1;refGene_tx=XM_019813668.1;refGene_gn=HPS1;refGene_tx=XM_019813669.2;refGene_gn=HPS1;refGene_tx=XM_019813670.1;refGene_gn=HPS1;refGene_tx=XM_019813671.1
D2 59502166 59502346
refGene_gn=HPS1;refGene_tx=XM_011287485.2;refGene_gn=HPS1;refGene_tx=XM_019813664.2;refGene_gn=HPS1;refGene_tx=XM_019813665.1;refGene_gn=HPS1;refGene_tx=XM_019813666.2;refGene_gn=HPS1;refGene_tx=XM_019813667.2;refGene_gn=HPS1;refGene_tx=XM_019813668.1;refGene_gn=HPS1;refGene_tx=XM_019813669.2;refGene_gn=HPS1;refGene_tx=XM_019813670.1;refGene_gn=HPS1;refGene_tx=XM_019813671.1
D2 59502441 59502609
refGene_gn=HPS1;refGene_tx=XM_011287485.2;refGene_gn=HPS1;refGene_tx=XM_019813664.2;refGene_gn=HPS1;refGene_tx=XM_019813665.1;refGene_gn=HPS1;refGene_tx=XM_019813666.2;refGene_gn=HPS1;refGene_tx=XM_019813667.2;refGene_gn=HPS1;refGene_tx=XM_019813668.1;refGene_gn=HPS1;refGene_tx=XM_019813669.2;refGene_gn=HPS1;refGene_tx=XM_019813670.1;refGene_gn=HPS1;refGene_tx=XM_019813671.1
D2 59503919 59503975
refGene_gn=HPS1;refGene_tx=XM_011287485.2;refGene_gn=HPS1;refGene_tx=XM_019813664.2;refGene_gn=HPS1;refGene_tx=XM_019813665.1;refGene_gn=HPS1;refGene_tx=XM_019813666.2;refGene_gn=HPS1;refGene_tx=XM_019813667.2;refGene_gn=HPS1;refGene_tx=XM_019813668.1;refGene_gn=HPS1;refGene_tx=XM_019813669.2;refGene_gn=HPS1;refGene_tx=XM_019813670.1;refGene_gn=HPS1;refGene_tx=XM_019813671.1
D2 59505938 59506008
refGene_gn=HPS1;refGene_tx=XM_011287485.2;refGene_gn=HPS1;refGene_tx=XM_019813664.2;refGene_gn=HPS1;refGene_tx=XM_019813665.1;refGene_gn=HPS1;refGene_tx=XM_019813666.2;refGene_gn=HPS1;refGene_tx=XM_019813667.2;refGene_gn=HPS1;refGene_tx=XM_019813668.1;refGene_gn=HPS1;refGene_tx=XM_019813669.2;refGene_gn=HPS1;refGene_tx=XM_019813670.1;refGene_gn=HPS1;refGene_tx=XM_019813671.1
D2 59506133 59506232
refGene_gn=HPS1;refGene_tx=XM_011287485.2;refGene_gn=HPS1;refGene_tx=XM_019813664.2;refGene_gn=HPS1;refGene_tx=XM_019813665.1;refGene_gn=HPS1;refGene_tx=XM_019813666.2;refGene_gn=HPS1;refGene_tx=XM_019813667.2;refGene_gn=HPS1;refGene_tx=XM_019813668.1;refGene_gn=HPS1;refGene_tx=X

M_019813669.2;refGene_gn=HPS1;refGene_tx=XM_019813670.1;refGene_gn=HPS1;refGene_tx=XM_019813671.1
D2 59506793 59506896
refGene_gn=HPS1;refGene_tx=XM_011287485.2;refGene_gn=HPS1;refGene_tx=XM_019813664.2;refGene_gn=HPS1;refGene_tx=XM_019813665.1;refGene_gn=HPS1;refGene_tx=XM_019813666.2;refGene_gn=HPS1;refGene_tx=XM_019813667.2;refGene_gn=HPS1;refGene_tx=XM_019813668.1;refGene_gn=HPS1;refGene_tx=XM_019813669.2;refGene_gn=HPS1;refGene_tx=XM_019813670.1;refGene_gn=HPS1;refGene_tx=XM_019813671.1
D2 59507379 59507540
refGene_gn=HPS1;refGene_tx=XM_011287485.2;refGene_gn=HPS1;refGene_tx=XM_019813664.2;refGene_gn=HPS1;refGene_tx=XM_019813665.1;refGene_gn=HPS1;refGene_tx=XM_019813666.2;refGene_gn=HPS1;refGene_tx=XM_019813667.2;refGene_gn=HPS1;refGene_tx=XM_019813668.1;refGene_gn=HPS1;refGene_tx=XM_019813669.2;refGene_gn=HPS1;refGene_tx=XM_019813670.1;refGene_gn=HPS1;refGene_tx=XM_019813671.1
D2 59509745 59509854
refGene_gn=HPS1;refGene_tx=XM_011287485.2;refGene_gn=HPS1;refGene_tx=XM_019813664.2;refGene_gn=HPS1;refGene_tx=XM_019813665.1;refGene_gn=HPS1;refGene_tx=XM_019813666.2;refGene_gn=HPS1;refGene_tx=XM_019813667.2;refGene_gn=HPS1;refGene_tx=XM_019813668.1;refGene_gn=HPS1;refGene_tx=XM_019813669.2;refGene_gn=HPS1;refGene_tx=XM_019813670.1
D2 59510261 59510404
refGene_gn=HPS1;refGene_tx=XM_011287485.2;refGene_gn=HPS1;refGene_tx=XM_019813664.2;refGene_gn=HPS1;refGene_tx=XM_019813665.1;refGene_gn=HPS1;refGene_tx=XM_019813666.2;refGene_gn=HPS1;refGene_tx=XM_019813667.2;refGene_gn=HPS1;refGene_tx=XM_019813668.1;refGene_gn=HPS1;refGene_tx=XM_019813669.2;refGene_gn=HPS1;refGene_tx=XM_019813670.1
D2 59510620 59510758
refGene_gn=HPS1;refGene_tx=XM_011287485.2;refGene_gn=HPS1;refGene_tx=XM_019813664.2;refGene_gn=HPS1;refGene_tx=XM_019813665.1;refGene_gn=HPS1;refGene_tx=XM_019813666.2;refGene_gn=HPS1;refGene_tx=XM_019813667.2;refGene_gn=HPS1;refGene_tx=XM_019813668.1;refGene_gn=HPS1;refGene_tx=XM_019813669.2;refGene_gn=HPS1;refGene_tx=XM_019813670.1
D2 59515029 59515191
refGene_gn=HPS1;refGene_tx=XM_011287485.2;refGene_gn=HPS1;refGene_tx=XM_019813664.2;refGene_gn=HPS1;refGene_tx=XM_019813665.1;refGene_gn=HPS1;refGene_tx=XM_019813666.2;refGene_gn=HPS1;refGene_tx=XM_019813667.2;refGene_gn=HPS1;refGene_tx=XM_019813668.1;refGene_gn=HPS1;refGene_tx=XM_019813669.2;refGene_gn=HPS1;refGene_tx=XM_019813670.1
D2 59530257 59530423
refGene_gn=HPSE2;refGene_tx=XM_011287482.3;refGene_gn=HPSE2;refGene_tx=XM_019813681.2;refGene_gn=HPSE2;refGene_tx=XM_019813682.2;refGene_gn=HPSE2;refGene_tx=XM_023240819.1
D2 59549832 59549979
refGene_gn=HPSE2;refGene_tx=XM_011287482.3;refGene_gn=HPSE2;refGene_tx=XM_019813681.2;refGene_gn=HPSE2;refGene_tx=XM_019813682.2;refGene_gn=HPSE2;refGene_tx=XM_023240819.1
D2 59557637 59557783
refGene_gn=HPSE2;refGene_tx=XM_011287482.3;refGene_gn=HPSE2;refGene_tx

=XM_019813682.2;refGene_gn=HPSE2;refGene_tx=XM_023240819.1
D2 59657978 59658093
refGene_gn=HPSE2;refGene_tx=XM_011287482.3;refGene_gn=HPSE2;refGene_tx=XM_019813681.2;refGene_gn=HPSE2;refGene_tx=XM_019813682.2;refGene_gn=HPSE2;refGene_tx=XM_023240819.1
D2 59663239 59663346
refGene_gn=HPSE2;refGene_tx=XM_011287482.3;refGene_gn=HPSE2;refGene_tx=XM_019813681.2;refGene_gn=HPSE2;refGene_tx=XM_019813682.2;refGene_gn=HPSE2;refGene_tx=XM_023240819.1
D2 59682537 59682631
refGene_gn=HPSE2;refGene_tx=XM_011287482.3;refGene_gn=HPSE2;refGene_tx=XM_019813681.2;refGene_gn=HPSE2;refGene_tx=XM_019813682.2;refGene_gn=HPSE2;refGene_tx=XM_023240819.1
D2 59743964 59744012
refGene_gn=HPSE2;refGene_tx=XM_011287482.3;refGene_gn=HPSE2;refGene_tx=XM_019813681.2;refGene_gn=HPSE2;refGene_tx=XM_019813682.2;refGene_gn=HPSE2;refGene_tx=XM_023240819.1
D2 59764018 59764190
refGene_gn=HPSE2;refGene_tx=XM_011287482.3;refGene_gn=HPSE2;refGene_tx=XM_019813681.2;refGene_gn=HPSE2;refGene_tx=XM_019813682.2;refGene_gn=HPSE2;refGene_tx=XM_023240819.1
D2 59775908 59776082
refGene_gn=HPSE2;refGene_tx=XM_011287482.3;refGene_gn=HPSE2;refGene_tx=XM_019813681.2;refGene_gn=HPSE2;refGene_tx=XM_019813682.2;refGene_gn=HPSE2;refGene_tx=XM_023240819.1
D2 60125020 60125182
refGene_gn=HPSE2;refGene_tx=XM_011287482.3;refGene_gn=HPSE2;refGene_tx=XM_019813681.2;refGene_gn=HPSE2;refGene_tx=XM_019813682.2;refGene_gn=HPSE2;refGene_tx=XM_023240819.1
D2 60182336 60182494
refGene_gn=HPSE2;refGene_tx=XM_011287482.3;refGene_gn=HPSE2;refGene_tx=XM_019813681.2;refGene_gn=HPSE2;refGene_tx=XM_019813682.2;refGene_gn=HPSE2;refGene_tx=XM_023240819.1
D2 60185390 60185680
refGene_gn=HPSE2;refGene_tx=XM_011287482.3;refGene_gn=HPSE2;refGene_tx=XM_019813681.2
D2 60190428 60190433
refGene_gn=HPSE2;refGene_tx=XM_019813682.2;refGene_gn=HPSE2;refGene_tx=XM_023240819.1
D2 60251046 60252610
refGene_gn=CNNM1;refGene_tx=XM_023240821.1;refGene_gn=CNNM1;refGene_tx=XM_023240822.1;refGene_gn=CNNM1;refGene_tx=XM_023240823.1
D2 60276265 60276409
refGene_gn=CNNM1;refGene_tx=XM_023240821.1;refGene_gn=CNNM1;refGene_tx=XM_023240822.1;refGene_gn=CNNM1;refGene_tx=XM_023240823.1
D2 60279316 60279457
refGene_gn=CNNM1;refGene_tx=XM_023240821.1;refGene_gn=CNNM1;refGene_tx=XM_023240822.1;refGene_gn=CNNM1;refGene_tx=XM_023240823.1
D2 60281572 60281742
refGene_gn=CNNM1;refGene_tx=XM_023240821.1;refGene_gn=CNNM1;refGene_tx

=XM_023240822.1;refGene_gn=CNNM1;refGene_tx=XM_023240823.1
D2 60283445 60283545
refGene_gn=CNNM1;refGene_tx=XM_023240821.1;refGene_gn=CNNM1;refGene_tx
=XM_023240822.1
D2 60283951 60283999
refGene_gn=CNNM1;refGene_tx=XM_023240821.1;refGene_gn=CNNM1;refGene_tx
=XM_023240822.1;refGene_gn=CNNM1;refGene_tx=XM_023240823.1
D2 60287111 60287174 refGene_gn=CNNM1;refGene_tx=XM_023240821.1
D2 60295620 60295784
refGene_gn=CNNM1;refGene_tx=XM_023240821.1;refGene_gn=CNNM1;refGene_tx
=XM_023240822.1
D2 60305937 60306121
refGene_gn=CNNM1;refGene_tx=XM_023240821.1;refGene_gn=CNNM1;refGene_tx
=XM_023240822.1;refGene_gn=CNNM1;refGene_tx=XM_023240823.1
D2 60306272 60306422
refGene_gn=CNNM1;refGene_tx=XM_023240821.1;refGene_gn=CNNM1;refGene_tx
=XM_023240822.1;refGene_gn=CNNM1;refGene_tx=XM_023240823.1
D2 60308054 60308156
refGene_gn=CNNM1;refGene_tx=XM_023240821.1;refGene_gn=CNNM1;refGene_tx
=XM_023240822.1;refGene_gn=CNNM1;refGene_tx=XM_023240823.1
D2 60309139 60309219
refGene_gn=CNNM1;refGene_tx=XM_023240821.1;refGene_gn=CNNM1;refGene_tx
=XM_023240822.1;refGene_gn=CNNM1;refGene_tx=XM_023240823.1
D2 60314780 60314920 refGene_gn=GOT1;refGene_tx=XM_011287481.3
D2 60319144 60319287 refGene_gn=GOT1;refGene_tx=XM_011287481.3
D2 60320038 60320204 refGene_gn=GOT1;refGene_tx=XM_011287481.3
D2 60320290 60320441 refGene_gn=GOT1;refGene_tx=XM_011287481.3
D2 60322240 60322345 refGene_gn=GOT1;refGene_tx=XM_011287481.3
D2 60322584 60322697 refGene_gn=GOT1;refGene_tx=XM_011287481.3
D2 60323390 60323514 refGene_gn=GOT1;refGene_tx=XM_011287481.3
D2 60330111 60330293 refGene_gn=GOT1;refGene_tx=XM_011287481.3
D2 60339869 60339987 refGene_gn=GOT1;refGene_tx=XM_011287481.3
D2 60438369 60438730 refGene_gn=NKX2-3;refGene_tx=XM_023240825.1
D2 60440235 60440981 refGene_gn=NKX2-3;refGene_tx=XM_023240825.1
D2 60511871 60512389
refGene_gn=SLC25A28;refGene_tx=XM_019813687.2;refGene_gn=SLC25A28;refG
ene_tx=XM_019813688.1;refGene_gn=SLC25A28;refGene_tx=XM_019813689.1
D2 60513562 60513619
refGene_gn=SLC25A28;refGene_tx=XM_019813687.2;refGene_gn=SLC25A28;refG
ene_tx=XM_019813688.1;refGene_gn=SLC25A28;refGene_tx=XM_019813689.1
D2 60514693 60514922
refGene_gn=SLC25A28;refGene_tx=XM_019813687.2;refGene_gn=SLC25A28;refG
ene_tx=XM_019813688.1;refGene_gn=SLC25A28;refGene_tx=XM_019813689.1
D2 60520571 60520580 refGene_gn=SLC25A28;refGene_tx=XM_019813688.1
D2 60520879 60521170 refGene_gn=SLC25A28;refGene_tx=XM_019813687.2
D2 60521566 60521936
refGene_gn=LOC109492831;refGene_tx=XM_019814083.1
D2 60522065 60522545
refGene_gn=LOC109492831;refGene_tx=XM_019814083.1
D2 60524281 60524349

refGene_gn=LOC109492831;refGene_tx=XM_019814083.1
D2 60541719 60541727 refGene_gn=ENTPD7;refGene_tx=XM_011287480.2
D2 60542877 60543060 refGene_gn=ENTPD7;refGene_tx=XM_011287480.2
D2 60561229 60561435 refGene_gn=ENTPD7;refGene_tx=XM_011287480.2
D2 60561686 60561837 refGene_gn=ENTPD7;refGene_tx=XM_011287480.2
D2 60566568 60566672 refGene_gn=ENTPD7;refGene_tx=XM_011287480.2
D2 60569535 60569592 refGene_gn=ENTPD7;refGene_tx=XM_011287480.2
D2 60573453 60573587 refGene_gn=ENTPD7;refGene_tx=XM_011287480.2
D2 60576823 60576990 refGene_gn=ENTPD7;refGene_tx=XM_011287480.2
D2 60579446 60579771 refGene_gn=ENTPD7;refGene_tx=XM_011287480.2
D2 60581603 60581689 refGene_gn=ENTPD7;refGene_tx=XM_011287480.2
D2 60582750 60582912 refGene_gn=ENTPD7;refGene_tx=XM_011287480.2
D2 60584321 60584553 refGene_gn=ENTPD7;refGene_tx=XM_011287480.2
D2 60595238 60595370
refGene_gn=LOC101085408;refGene_tx=XM_003994297.4
D2 60596339 60596453
refGene_gn=LOC101085408;refGene_tx=XM_003994297.4
D2 60598193 60598265
refGene_gn=LOC101085408;refGene_tx=XM_003994297.4
D2 60598266 60598349
refGene_gn=LOC101085408;refGene_tx=XM_003994297.4
D2 60599689 60599771
refGene_gn=LOC101085408;refGene_tx=XM_003994297.4
D2 60601277 60601445
refGene_gn=LOC101085408;refGene_tx=XM_003994297.4
D2 60603283 60603470
refGene_gn=LOC101085408;refGene_tx=XM_003994297.4
D2 60603760 60603883
refGene_gn=LOC101085408;refGene_tx=XM_003994297.4
D2 60605617 60605808
refGene_gn=LOC101085408;refGene_tx=XM_003994297.4
D2 60607048 60607138
refGene_gn=LOC101085408;refGene_tx=XM_003994297.4
D2 60607438 60607499
refGene_gn=CUTC;refGene_tx=XM_003994298.5;refGene_gn=CUTC;refGene_tx=X
M_006938080.4
D2 60610343 60610415
refGene_gn=CUTC;refGene_tx=XM_003994298.5;refGene_gn=CUTC;refGene_tx=X
M_023240826.1
D2 60613262 60613322
refGene_gn=CUTC;refGene_tx=XM_003994298.5;refGene_gn=CUTC;refGene_tx=X
M_006938080.4;refGene_gn=CUTC;refGene_tx=XM_023240826.1
D2 60617089 60617299
refGene_gn=CUTC;refGene_tx=XM_003994298.5;refGene_gn=CUTC;refGene_tx=X
M_006938080.4;refGene_gn=CUTC;refGene_tx=XM_023240826.1
D2 60618015 60618051
refGene_gn=CUTC;refGene_tx=XM_003994298.5;refGene_gn=CUTC;refGene_tx=X
M_006938080.4;refGene_gn=CUTC;refGene_tx=XM_023240826.1
D2 60620723 60620857
refGene_gn=CUTC;refGene_tx=XM_003994298.5;refGene_gn=CUTC;refGene_tx=X

M_006938080.4;refGene_gn=CUTC;refGene_tx=XM_023240826.1
D2 60629479 60629507
refGene_gn=CUTC;refGene_tx=XM_003994298.5;refGene_gn=CUTC;refGene_tx=XM_006938080.4;refGene_gn=CUTC;refGene_tx=XM_023240826.1
D2 60631121 60631227
refGene_gn=CUTC;refGene_tx=XM_003994298.5;refGene_gn=CUTC;refGene_tx=XM_006938080.4;refGene_gn=CUTC;refGene_tx=XM_023240826.1
D2 60632218 60632333
refGene_gn=CUTC;refGene_tx=XM_003994298.5;refGene_gn=CUTC;refGene_tx=XM_006938080.4;refGene_gn=CUTC;refGene_tx=XM_023240826.1
D2 60649940 60649973
refGene_gn=ABCC2;refGene_tx=XM_019813691.2;refGene_gn=ABCC2;refGene_tx=XM_019813692.2;refGene_gn=ABCC2;refGene_tx=XM_019813693.1;refGene_gn=ABCC2;refGene_tx=XM_019813695.2
D2 60652095 60652269
refGene_gn=ABCC2;refGene_tx=XM_019813691.2;refGene_gn=ABCC2;refGene_tx=XM_019813692.2;refGene_gn=ABCC2;refGene_tx=XM_019813693.1;refGene_gn=ABCC2;refGene_tx=XM_019813695.2
D2 60665405 60665531
refGene_gn=ABCC2;refGene_tx=XM_019813691.2;refGene_gn=ABCC2;refGene_tx=XM_019813692.2;refGene_gn=ABCC2;refGene_tx=XM_019813693.1;refGene_gn=ABCC2;refGene_tx=XM_019813695.2
D2 60667710 60667845
refGene_gn=ABCC2;refGene_tx=XM_019813691.2;refGene_gn=ABCC2;refGene_tx=XM_019813692.2;refGene_gn=ABCC2;refGene_tx=XM_019813693.1;refGene_gn=ABCC2;refGene_tx=XM_019813695.2
D2 60668071 60668179
refGene_gn=ABCC2;refGene_tx=XM_019813691.2;refGene_gn=ABCC2;refGene_tx=XM_019813692.2;refGene_gn=ABCC2;refGene_tx=XM_019813693.1;refGene_gn=ABCC2;refGene_tx=XM_019813695.2
D2 60668639 60668695
refGene_gn=ABCC2;refGene_tx=XM_019813691.2;refGene_gn=ABCC2;refGene_tx=XM_019813692.2;refGene_gn=ABCC2;refGene_tx=XM_019813693.1;refGene_gn=ABCC2;refGene_tx=XM_019813695.2
D2 60669855 60670090
refGene_gn=ABCC2;refGene_tx=XM_019813691.2;refGene_gn=ABCC2;refGene_tx=XM_019813692.2;refGene_gn=ABCC2;refGene_tx=XM_019813693.1;refGene_gn=ABCC2;refGene_tx=XM_019813695.2
D2 60672151 60672312
refGene_gn=ABCC2;refGene_tx=XM_019813691.2;refGene_gn=ABCC2;refGene_tx=XM_019813692.2;refGene_gn=ABCC2;refGene_tx=XM_019813693.1;refGene_gn=ABCC2;refGene_tx=XM_019813695.2
D2 60673116 60673294
refGene_gn=ABCC2;refGene_tx=XM_019813691.2;refGene_gn=ABCC2;refGene_tx=XM_019813692.2;refGene_gn=ABCC2;refGene_tx=XM_019813693.1;refGene_gn=ABCC2;refGene_tx=XM_019813695.2
D2 60676942 60677197
refGene_gn=ABCC2;refGene_tx=XM_019813691.2;refGene_gn=ABCC2;refGene_tx=XM_019813692.2;refGene_gn=ABCC2;refGene_tx=XM_019813693.1
D2 60678274 60678340

refGene_gn=ABCC2;refGene_tx=XM_019813691.2;refGene_gn=ABCC2;refGene_tx=XM_019813693.1
D2 60680661 60680799 refGene_gn=ABCC2;refGene_tx=XM_019813691.2;refGene_gn=ABCC2;refGene_tx=XM_019813692.2;refGene_gn=ABCC2;refGene_tx=XM_019813693.1;refGene_gn=ABCC2;refGene_tx=XM_019813695.2
D2 60681320 60681467 refGene_gn=ABCC2;refGene_tx=XM_019813691.2;refGene_gn=ABCC2;refGene_tx=XM_019813692.2;refGene_gn=ABCC2;refGene_tx=XM_019813693.1;refGene_gn=ABCC2;refGene_tx=XM_019813695.2
D2 60683124 60683209 refGene_gn=ABCC2;refGene_tx=XM_019813691.2;refGene_gn=ABCC2;refGene_tx=XM_019813692.2;refGene_gn=ABCC2;refGene_tx=XM_019813693.1;refGene_gn=ABCC2;refGene_tx=XM_019813695.2
D2 60683920 60683987 refGene_gn=ABCC2;refGene_tx=XM_019813691.2;refGene_gn=ABCC2;refGene_tx=XM_019813692.2;refGene_gn=ABCC2;refGene_tx=XM_019813693.1;refGene_gn=ABCC2;refGene_tx=XM_019813695.2
D2 60686041 60686168 refGene_gn=ABCC2;refGene_tx=XM_019813691.2;refGene_gn=ABCC2;refGene_tx=XM_019813692.2;refGene_gn=ABCC2;refGene_tx=XM_019813693.1;refGene_gn=ABCC2;refGene_tx=XM_019813695.2
D2 60688589 60688766 refGene_gn=ABCC2;refGene_tx=XM_019813691.2;refGene_gn=ABCC2;refGene_tx=XM_019813692.2;refGene_gn=ABCC2;refGene_tx=XM_019813693.1;refGene_gn=ABCC2;refGene_tx=XM_019813695.2
D2 60690147 60690315 refGene_gn=ABCC2;refGene_tx=XM_019813691.2;refGene_gn=ABCC2;refGene_tx=XM_019813692.2;refGene_gn=ABCC2;refGene_tx=XM_019813693.1;refGene_gn=ABCC2;refGene_tx=XM_019813695.2
D2 60692732 60692913 refGene_gn=ABCC2;refGene_tx=XM_019813691.2;refGene_gn=ABCC2;refGene_tx=XM_019813692.2;refGene_gn=ABCC2;refGene_tx=XM_019813693.1;refGene_gn=ABCC2;refGene_tx=XM_019813695.2
D2 60698216 60698337 refGene_gn=ABCC2;refGene_tx=XM_019813691.2;refGene_gn=ABCC2;refGene_tx=XM_019813692.2;refGene_gn=ABCC2;refGene_tx=XM_019813693.1;refGene_gn=ABCC2;refGene_tx=XM_019813695.2
D2 60698628 60698764 refGene_gn=ABCC2;refGene_tx=XM_019813691.2;refGene_gn=ABCC2;refGene_tx=XM_019813692.2;refGene_gn=ABCC2;refGene_tx=XM_019813693.1;refGene_gn=ABCC2;refGene_tx=XM_019813695.2
D2 60699480 60699700 refGene_gn=ABCC2;refGene_tx=XM_019813691.2;refGene_gn=ABCC2;refGene_tx=XM_019813692.2;refGene_gn=ABCC2;refGene_tx=XM_019813693.1;refGene_gn=ABCC2;refGene_tx=XM_019813695.2
D2 60699852 60700007 refGene_gn=ABCC2;refGene_tx=XM_019813691.2;refGene_gn=ABCC2;refGene_tx=XM_019813692.2;refGene_gn=ABCC2;refGene_tx=XM_019813693.1;refGene_gn=ABCC2;refGene_tx=XM_019813695.2

D2 60703551 60703707
refGene_gn=ABCC2;refGene_tx=XM_019813691.2;refGene_gn=ABCC2;refGene_tx=XM_019813692.2;refGene_gn=ABCC2;refGene_tx=XM_019813693.1;refGene_gn=ABCC2;refGene_tx=XM_019813695.2
D2 60705167 60705367
refGene_gn=ABCC2;refGene_tx=XM_019813691.2;refGene_gn=ABCC2;refGene_tx=XM_019813692.2;refGene_gn=ABCC2;refGene_tx=XM_019813693.1;refGene_gn=ABCC2;refGene_tx=XM_019813695.2
D2 60707461 60707588
refGene_gn=ABCC2;refGene_tx=XM_019813691.2;refGene_gn=ABCC2;refGene_tx=XM_019813692.2;refGene_gn=ABCC2;refGene_tx=XM_019813693.1;refGene_gn=ABCC2;refGene_tx=XM_019813695.2
D2 60708365 60708467
refGene_gn=ABCC2;refGene_tx=XM_019813691.2;refGene_gn=ABCC2;refGene_tx=XM_019813692.2;refGene_gn=ABCC2;refGene_tx=XM_019813693.1;refGene_gn=ABCC2;refGene_tx=XM_019813695.2
D2 60708883 60709027
refGene_gn=ABCC2;refGene_tx=XM_019813691.2;refGene_gn=ABCC2;refGene_tx=XM_019813692.2;refGene_gn=ABCC2;refGene_tx=XM_019813693.1;refGene_gn=ABCC2;refGene_tx=XM_019813695.2
D2 60709834 60709993
refGene_gn=ABCC2;refGene_tx=XM_019813691.2;refGene_gn=ABCC2;refGene_tx=XM_019813692.2;refGene_gn=ABCC2;refGene_tx=XM_019813693.1;refGene_gn=ABCC2;refGene_tx=XM_019813695.2
D2 60710930 60711097
refGene_gn=ABCC2;refGene_tx=XM_019813691.2;refGene_gn=ABCC2;refGene_tx=XM_019813692.2;refGene_gn=ABCC2;refGene_tx=XM_019813693.1;refGene_gn=ABCC2;refGene_tx=XM_019813695.2
D2 60712917 60713116
refGene_gn=ABCC2;refGene_tx=XM_019813691.2;refGene_gn=ABCC2;refGene_tx=XM_019813692.2;refGene_gn=ABCC2;refGene_tx=XM_019813693.1;refGene_gn=ABCC2;refGene_tx=XM_019813695.2
D2 60713758 60713888
refGene_gn=ABCC2;refGene_tx=XM_019813691.2;refGene_gn=ABCC2;refGene_tx=XM_019813692.2;refGene_gn=ABCC2;refGene_tx=XM_019813695.2
D2 60726852 60727038
refGene_gn=DNMBP;refGene_tx=XM_019813697.2;refGene_gn=DNMBP;refGene_tx=XM_023240827.1
D2 60728672 60729214
refGene_gn=DNMBP;refGene_tx=XM_019813697.2;refGene_gn=DNMBP;refGene_tx=XM_023240827.1
D2 60732624 60732823
refGene_gn=DNMBP;refGene_tx=XM_019813697.2;refGene_gn=DNMBP;refGene_tx=XM_023240827.1
D2 60734396 60734576
refGene_gn=DNMBP;refGene_tx=XM_019813697.2;refGene_gn=DNMBP;refGene_tx=XM_023240827.1
D2 60734970 60735303
refGene_gn=DNMBP;refGene_tx=XM_019813697.2;refGene_gn=DNMBP;refGene_tx=XM_023240827.1

D2 60736834 60736963
refGene_gn=DNMBP;refGene_tx=XM_019813697.2;refGene_gn=DNMBP;refGene_tx=XM_023240827.1
D2 60742334 60742439
refGene_gn=DNMBP;refGene_tx=XM_019813697.2;refGene_gn=DNMBP;refGene_tx=XM_023240827.1
D2 60743385 60743516
refGene_gn=DNMBP;refGene_tx=XM_019813697.2;refGene_gn=DNMBP;refGene_tx=XM_023240827.1
D2 60745550 60745750
refGene_gn=DNMBP;refGene_tx=XM_019813697.2;refGene_gn=DNMBP;refGene_tx=XM_023240827.1
D2 60746459 60746477
refGene_gn=DNMBP;refGene_tx=XM_019813697.2;refGene_gn=DNMBP;refGene_tx=XM_023240827.1
D2 60747289 60747437
refGene_gn=DNMBP;refGene_tx=XM_019813697.2;refGene_gn=DNMBP;refGene_tx=XM_023240827.1
D2 60748599 60748699
refGene_gn=DNMBP;refGene_tx=XM_019813697.2;refGene_gn=DNMBP;refGene_tx=XM_023240827.1
D2 60749545 60749739
refGene_gn=DNMBP;refGene_tx=XM_019813697.2;refGene_gn=DNMBP;refGene_tx=XM_023240827.1
D2 60769264 60770426 refGene_gn=DNMBP;refGene_tx=XM_019813697.2
D2 60785377 60787354 refGene_gn=DNMBP;refGene_tx=XM_023240827.1
D2 60798271 60798394 refGene_gn=DNMBP;refGene_tx=XM_023240827.1
D2 60800121 60800266 refGene_gn=DNMBP;refGene_tx=XM_023240827.1
D2 60851691 60851805 refGene_gn=CPN1;refGene_tx=XM_003994312.5
D2 60854818 60854937 refGene_gn=CPN1;refGene_tx=XM_003994312.5
D2 60857290 60857390 refGene_gn=CPN1;refGene_tx=XM_003994312.5
D2 60858646 60858786 refGene_gn=CPN1;refGene_tx=XM_003994312.5
D2 60861395 60861507 refGene_gn=CPN1;refGene_tx=XM_003994312.5
D2 60862779 60862962 refGene_gn=CPN1;refGene_tx=XM_003994312.5
D2 60866138 60866294 refGene_gn=CPN1;refGene_tx=XM_003994312.5
D2 60871278 60871475 refGene_gn=CPN1;refGene_tx=XM_003994312.5
D2 60874041 60874264 refGene_gn=CPN1;refGene_tx=XM_003994312.5
D2 60876832 60876903
refGene_gn=ERLIN1;refGene_tx=XM_019813699.1;refGene_gn=ERLIN1;refGene_tx=XM_019813700.1
D2 60891544 60891576 refGene_gn=ERLIN1;refGene_tx=XM_019813701.1
D2 60892637 60892746
refGene_gn=ERLIN1;refGene_tx=XM_019813699.1;refGene_gn=ERLIN1;refGene_tx=XM_019813700.1;refGene_gn=ERLIN1;refGene_tx=XM_019813701.1
D2 60899538 60899760
refGene_gn=ERLIN1;refGene_tx=XM_011287475.3;refGene_gn=ERLIN1;refGene_tx=XM_023240828.1
D2 60902217 60902297
refGene_gn=ERLIN1;refGene_tx=XM_011287475.3;refGene_gn=ERLIN1;refGene_tx=XM_019813699.1;refGene_gn=ERLIN1;refGene_tx=XM_019813700.1;refGene_

gn=ERLIN1;refGene_tx=XM_019813701.1;refGene_gn=ERLIN1;refGene_tx=XM_02
3240828.1
D2 60903376 60903466
refGene_gn=ERLIN1;refGene_tx=XM_011287475.3;refGene_gn=ERLIN1;refGene_
tx=XM_019813699.1;refGene_gn=ERLIN1;refGene_tx=XM_019813700.1;refGene_
gn=ERLIN1;refGene_tx=XM_019813701.1;refGene_gn=ERLIN1;refGene_tx=XM_02
3240828.1
D2 60910578 60910670
refGene_gn=ERLIN1;refGene_tx=XM_011287475.3;refGene_gn=ERLIN1;refGene_
tx=XM_019813699.1;refGene_gn=ERLIN1;refGene_tx=XM_019813700.1;refGene_
gn=ERLIN1;refGene_tx=XM_019813701.1;refGene_gn=ERLIN1;refGene_tx=XM_02
3240828.1
D2 60912547 60912606
refGene_gn=ERLIN1;refGene_tx=XM_011287475.3;refGene_gn=ERLIN1;refGene_
tx=XM_019813699.1;refGene_gn=ERLIN1;refGene_tx=XM_019813700.1;refGene_
gn=ERLIN1;refGene_tx=XM_019813701.1;refGene_gn=ERLIN1;refGene_tx=XM_02
3240828.1
D2 60920245 60920319
refGene_gn=ERLIN1;refGene_tx=XM_011287475.3;refGene_gn=ERLIN1;refGene_
tx=XM_019813699.1;refGene_gn=ERLIN1;refGene_tx=XM_019813700.1;refGene_
gn=ERLIN1;refGene_tx=XM_019813701.1;refGene_gn=ERLIN1;refGene_tx=XM_02
3240828.1
D2 60922286 60922412
refGene_gn=ERLIN1;refGene_tx=XM_011287475.3;refGene_gn=ERLIN1;refGene_
tx=XM_019813699.1;refGene_gn=ERLIN1;refGene_tx=XM_019813700.1;refGene_
gn=ERLIN1;refGene_tx=XM_019813701.1;refGene_gn=ERLIN1;refGene_tx=XM_02
3240828.1
D2 60925035 60925097
refGene_gn=ERLIN1;refGene_tx=XM_011287475.3;refGene_gn=ERLIN1;refGene_
tx=XM_019813699.1;refGene_gn=ERLIN1;refGene_tx=XM_019813700.1;refGene_
gn=ERLIN1;refGene_tx=XM_019813701.1;refGene_gn=ERLIN1;refGene_tx=XM_02
3240828.1
D2 60926852 60926899
refGene_gn=ERLIN1;refGene_tx=XM_011287475.3;refGene_gn=ERLIN1;refGene_
tx=XM_019813699.1;refGene_gn=ERLIN1;refGene_tx=XM_019813700.1;refGene_
gn=ERLIN1;refGene_tx=XM_019813701.1
D2 60931124 60931206
refGene_gn=ERLIN1;refGene_tx=XM_011287475.3;refGene_gn=ERLIN1;refGene_
tx=XM_019813699.1;refGene_gn=ERLIN1;refGene_tx=XM_019813700.1;refGene_
gn=ERLIN1;refGene_tx=XM_019813701.1
D2 60932913 60933026
refGene_gn=ERLIN1;refGene_tx=XM_011287475.3;refGene_gn=ERLIN1;refGene_
tx=XM_019813699.1;refGene_gn=ERLIN1;refGene_tx=XM_019813700.1;refGene_
gn=ERLIN1;refGene_tx=XM_019813701.1
D2 60937428 60937458 refGene_gn=CHUK;refGene_tx=XM_023240831.1
D2 60937485 60937548
refGene_gn=CHUK;refGene_tx=XM_023240829.1;refGene_gn=CHUK;refGene_tx=X
M_023240830.1;refGene_gn=CHUK;refGene_tx=XM_023240834.1
D2 60938207 60938307
refGene_gn=CHUK;refGene_tx=XM_023240829.1;refGene_gn=CHUK;refGene_tx=X

M_023240830.1;refGene_gn=CHUK;refGene_tx=XM_023240831.1;refGene_gn=CHUK;refGene_tx=XM_023240832.1;refGene_gn=CHUK;refGene_tx=XM_023240833.1;refGene_gn=CHUK;refGene_tx=XM_023240834.1
D2 60940532 60940666
refGene_gn=CHUK;refGene_tx=XM_023240829.1;refGene_gn=CHUK;refGene_tx=XM_023240830.1;refGene_gn=CHUK;refGene_tx=XM_023240831.1;refGene_gn=CHUK;refGene_tx=XM_023240832.1;refGene_gn=CHUK;refGene_tx=XM_023240833.1;refGene_gn=CHUK;refGene_tx=XM_023240834.1
D2 60941216 60941364
refGene_gn=CHUK;refGene_tx=XM_023240829.1;refGene_gn=CHUK;refGene_tx=XM_023240830.1;refGene_gn=CHUK;refGene_tx=XM_023240831.1;refGene_gn=CHUK;refGene_tx=XM_023240832.1;refGene_gn=CHUK;refGene_tx=XM_023240833.1;refGene_gn=CHUK;refGene_tx=XM_023240834.1
D2 60941659 60941756
refGene_gn=CHUK;refGene_tx=XM_023240829.1;refGene_gn=CHUK;refGene_tx=XM_023240830.1;refGene_gn=CHUK;refGene_tx=XM_023240831.1;refGene_gn=CHUK;refGene_tx=XM_023240832.1;refGene_gn=CHUK;refGene_tx=XM_023240833.1;refGene_gn=CHUK;refGene_tx=XM_023240834.1
D2 60947896 60947946
refGene_gn=CHUK;refGene_tx=XM_023240829.1;refGene_gn=CHUK;refGene_tx=XM_023240830.1;refGene_gn=CHUK;refGene_tx=XM_023240831.1;refGene_gn=CHUK;refGene_tx=XM_023240832.1;refGene_gn=CHUK;refGene_tx=XM_023240833.1;refGene_gn=CHUK;refGene_tx=XM_023240834.1
D2 60952932 60953042
refGene_gn=CHUK;refGene_tx=XM_023240829.1;refGene_gn=CHUK;refGene_tx=XM_023240830.1;refGene_gn=CHUK;refGene_tx=XM_023240831.1;refGene_gn=CHUK;refGene_tx=XM_023240832.1;refGene_gn=CHUK;refGene_tx=XM_023240833.1;refGene_gn=CHUK;refGene_tx=XM_023240834.1
D2 60954001 60954063
refGene_gn=CHUK;refGene_tx=XM_023240829.1;refGene_gn=CHUK;refGene_tx=XM_023240830.1;refGene_gn=CHUK;refGene_tx=XM_023240831.1;refGene_gn=CHUK;refGene_tx=XM_023240832.1;refGene_gn=CHUK;refGene_tx=XM_023240833.1;refGene_gn=CHUK;refGene_tx=XM_023240834.1
D2 60955868 60956020
refGene_gn=CHUK;refGene_tx=XM_023240829.1;refGene_gn=CHUK;refGene_tx=XM_023240830.1;refGene_gn=CHUK;refGene_tx=XM_023240831.1;refGene_gn=CHUK;refGene_tx=XM_023240832.1;refGene_gn=CHUK;refGene_tx=XM_023240833.1;refGene_gn=CHUK;refGene_tx=XM_023240834.1
D2 60956423 60956547
refGene_gn=CHUK;refGene_tx=XM_023240829.1;refGene_gn=CHUK;refGene_tx=XM_023240830.1;refGene_gn=CHUK;refGene_tx=XM_023240831.1;refGene_gn=CHUK;refGene_tx=XM_023240832.1;refGene_gn=CHUK;refGene_tx=XM_023240833.1;refGene_gn=CHUK;refGene_tx=XM_023240834.1
D2 60958745 60958848
refGene_gn=CHUK;refGene_tx=XM_023240829.1;refGene_gn=CHUK;refGene_tx=XM_023240830.1;refGene_gn=CHUK;refGene_tx=XM_023240831.1;refGene_gn=CHUK;refGene_tx=XM_023240832.1;refGene_gn=CHUK;refGene_tx=XM_023240833.1;refGene_gn=CHUK;refGene_tx=XM_023240834.1
D2 60959815 60960010
refGene_gn=CHUK;refGene_tx=XM_023240829.1;refGene_gn=CHUK;refGene_tx=X

M_023240830.1;refGene_gn=CHUK;refGene_tx=XM_023240831.1;refGene_gn=CHUK;refGene_tx=XM_023240832.1;refGene_gn=CHUK;refGene_tx=XM_023240833.1;refGene_gn=CHUK;refGene_tx=XM_023240834.1
D2 60964372 60964508
refGene_gn=CHUK;refGene_tx=XM_023240829.1;refGene_gn=CHUK;refGene_tx=XM_023240830.1;refGene_gn=CHUK;refGene_tx=XM_023240831.1;refGene_gn=CHUK;refGene_tx=XM_023240832.1;refGene_gn=CHUK;refGene_tx=XM_023240833.1;refGene_gn=CHUK;refGene_tx=XM_023240834.1
D2 60965085 60965193
refGene_gn=CHUK;refGene_tx=XM_023240829.1;refGene_gn=CHUK;refGene_tx=XM_023240830.1;refGene_gn=CHUK;refGene_tx=XM_023240831.1;refGene_gn=CHUK;refGene_tx=XM_023240832.1;refGene_gn=CHUK;refGene_tx=XM_023240833.1;refGene_gn=CHUK;refGene_tx=XM_023240834.1
D2 60965383 60965508
refGene_gn=CHUK;refGene_tx=XM_023240829.1;refGene_gn=CHUK;refGene_tx=XM_023240830.1;refGene_gn=CHUK;refGene_tx=XM_023240831.1;refGene_gn=CHUK;refGene_tx=XM_023240832.1;refGene_gn=CHUK;refGene_tx=XM_023240833.1;refGene_gn=CHUK;refGene_tx=XM_023240834.1
D2 60965670 60965760
refGene_gn=CHUK;refGene_tx=XM_023240829.1;refGene_gn=CHUK;refGene_tx=XM_023240830.1;refGene_gn=CHUK;refGene_tx=XM_023240831.1;refGene_gn=CHUK;refGene_tx=XM_023240832.1;refGene_gn=CHUK;refGene_tx=XM_023240833.1;refGene_gn=CHUK;refGene_tx=XM_023240834.1
D2 60966944 60967033
refGene_gn=CHUK;refGene_tx=XM_023240829.1;refGene_gn=CHUK;refGene_tx=XM_023240830.1;refGene_gn=CHUK;refGene_tx=XM_023240831.1;refGene_gn=CHUK;refGene_tx=XM_023240832.1;refGene_gn=CHUK;refGene_tx=XM_023240833.1;refGene_gn=CHUK;refGene_tx=XM_023240834.1
D2 60968456 60968526
refGene_gn=CHUK;refGene_tx=XM_023240829.1;refGene_gn=CHUK;refGene_tx=XM_023240830.1;refGene_gn=CHUK;refGene_tx=XM_023240831.1;refGene_gn=CHUK;refGene_tx=XM_023240832.1;refGene_gn=CHUK;refGene_tx=XM_023240833.1;refGene_gn=CHUK;refGene_tx=XM_023240834.1
D2 60969181 60969296
refGene_gn=CHUK;refGene_tx=XM_023240829.1;refGene_gn=CHUK;refGene_tx=XM_023240830.1;refGene_gn=CHUK;refGene_tx=XM_023240831.1;refGene_gn=CHUK;refGene_tx=XM_023240832.1;refGene_gn=CHUK;refGene_tx=XM_023240833.1;refGene_gn=CHUK;refGene_tx=XM_023240834.1
D2 60971319 60971414
refGene_gn=CHUK;refGene_tx=XM_023240829.1;refGene_gn=CHUK;refGene_tx=XM_023240830.1;refGene_gn=CHUK;refGene_tx=XM_023240831.1;refGene_gn=CHUK;refGene_tx=XM_023240832.1;refGene_gn=CHUK;refGene_tx=XM_023240833.1
D2 60978560 60978665
refGene_gn=CHUK;refGene_tx=XM_023240829.1;refGene_gn=CHUK;refGene_tx=XM_023240830.1;refGene_gn=CHUK;refGene_tx=XM_023240831.1;refGene_gn=CHUK;refGene_tx=XM_023240832.1;refGene_gn=CHUK;refGene_tx=XM_023240833.1;refGene_gn=CHUK;refGene_tx=XM_023240834.1
D2 60981407 60981552
refGene_gn=CWF19L1;refGene_tx=XM_003994314.4;refGene_gn=CWF19L1;refGene_tx=XM_011287474.2

D2 60984436 60984534
refGene_gn=CWF19L1;refGene_tx=XM_003994314.4;refGene_gn=CWF19L1;refGene_tx=XM_011287474.2
D2 60984961 60985081
refGene_gn=CWF19L1;refGene_tx=XM_003994314.4;refGene_gn=CWF19L1;refGene_tx=XM_011287474.2
D2 60985922 60986132
refGene_gn=CWF19L1;refGene_tx=XM_003994314.4;refGene_gn=CWF19L1;refGene_tx=XM_011287474.2
D2 60989815 60989895
refGene_gn=CWF19L1;refGene_tx=XM_003994314.4;refGene_gn=CWF19L1;refGene_tx=XM_011287474.2
D2 60990653 60990768
refGene_gn=CWF19L1;refGene_tx=XM_003994314.4;refGene_gn=CWF19L1;refGene_tx=XM_011287474.2
D2 60993641 60993782
refGene_gn=CWF19L1;refGene_tx=XM_003994314.4;refGene_gn=CWF19L1;refGene_tx=XM_011287474.2
D2 60995377 60995462
refGene_gn=CWF19L1;refGene_tx=XM_003994314.4;refGene_gn=CWF19L1;refGene_tx=XM_011287474.2
D2 61003392 61003511
refGene_gn=CWF19L1;refGene_tx=XM_003994314.4;refGene_gn=CWF19L1;refGene_tx=XM_011287474.2
D2 61006008 61006223
refGene_gn=CWF19L1;refGene_tx=XM_003994314.4;refGene_gn=CWF19L1;refGene_tx=XM_011287474.2
D2 61009038 61009140 refGene_gn=CWF19L1;refGene_tx=XM_003994314.4
D2 61010628 61010707 refGene_gn=CWF19L1;refGene_tx=XM_003994314.4
D2 61011589 61011674 refGene_gn=CWF19L1;refGene_tx=XM_003994314.4
D2 61014190 61014213 refGene_gn=CWF19L1;refGene_tx=XM_003994314.4
D2 61021901 61021933 refGene_gn=BLOC1S2;refGene_tx=XM_003994315.5
D2 61023795 61023900 refGene_gn=BLOC1S2;refGene_tx=XM_003994315.5
D2 61024603 61024723 refGene_gn=BLOC1S2;refGene_tx=XM_003994315.5
D2 61027825 61027942 refGene_gn=BLOC1S2;refGene_tx=XM_003994315.5
D2 61028268 61028329 refGene_gn=BLOC1S2;refGene_tx=XM_003994315.5
D2 61030609 61030669 refGene_gn=PKD2L1;refGene_tx=XM_019813702.2
D2 61030986 61031110 refGene_gn=PKD2L1;refGene_tx=XM_019813702.2
D2 61031361 61031480 refGene_gn=PKD2L1;refGene_tx=XM_019813702.2
D2 61032268 61032395 refGene_gn=PKD2L1;refGene_tx=XM_019813702.2
D2 61033655 61033777 refGene_gn=PKD2L1;refGene_tx=XM_019813702.2
D2 61033984 61034083 refGene_gn=PKD2L1;refGene_tx=XM_019813702.2
D2 61034915 61035036 refGene_gn=PKD2L1;refGene_tx=XM_019813702.2
D2 61035346 61035528 refGene_gn=PKD2L1;refGene_tx=XM_019813702.2
D2 61035709 61035880 refGene_gn=PKD2L1;refGene_tx=XM_019813702.2
D2 61036530 61036759 refGene_gn=PKD2L1;refGene_tx=XM_019813702.2
D2 61036922 61037147 refGene_gn=PKD2L1;refGene_tx=XM_019813702.2
D2 61038117 61038371 refGene_gn=PKD2L1;refGene_tx=XM_019813702.2
D2 61038836 61038964 refGene_gn=PKD2L1;refGene_tx=XM_019813702.2
D2 61059705 61059819 refGene_gn=PKD2L1;refGene_tx=XM_019813702.2

D2 61060361 61060596 refGene_gn=PKD2L1;refGene_tx=XM_019813702.2
D2 61098717 61098744 refGene_gn=SCD;refGene_tx=XM_003994316.5
D2 61099264 61099547 refGene_gn=SCD;refGene_tx=XM_003994316.5
D2 61103482 61103613 refGene_gn=SCD;refGene_tx=XM_003994316.5
D2 61105065 61105271 refGene_gn=SCD;refGene_tx=XM_003994316.5
D2 61106748 61106981 refGene_gn=SCD;refGene_tx=XM_003994316.5
D2 61109971 61110171 refGene_gn=SCD;refGene_tx=XM_003994316.5
D2 61177739 61177807 refGene_gn=WNT8B;refGene_tx=XM_023240297.1
D2 61197281 61197315 refGene_gn=WNT8B;refGene_tx=XM_023240297.1
D2 61198002 61198246 refGene_gn=WNT8B;refGene_tx=XM_023240297.1
D2 61199098 61199224 refGene_gn=WNT8B;refGene_tx=XM_023240297.1
D2 61199970 61200113 refGene_gn=WNT8B;refGene_tx=XM_023240297.1
D2 61200347 61200893 refGene_gn=WNT8B;refGene_tx=XM_023240297.1
D2 61205792 61205972
refGene_gn=SEC31B;refGene_tx=XM_011287465.3;refGene_gn=SEC31B;refGene_tx=XM_011287466.3;refGene_gn=SEC31B;refGene_tx=XM_011287472.3;refGene_gn=SEC31B;refGene_tx=XM_019813704.2;refGene_gn=SEC31B;refGene_tx=XM_019813705.2;refGene_gn=SEC31B;refGene_tx=XM_019813706.2;refGene_gn=SEC31B;refGene_tx=XM_019813708.2;refGene_gn=SEC31B;refGene_tx=XM_019813709.2;refGene_gn=SEC31B;refGene_tx=XM_019813710.2;refGene_gn=SEC31B;refGene_tx=XM_019813711.2;refGene_gn=SEC31B;refGene_tx=XM_019813712.2;refGene_gn=SEC31B;refGene_tx=XM_019813713.2;refGene_gn=SEC31B;refGene_tx=XM_019813714.2;refGene_gn=SEC31B;refGene_tx=XM_019813715.2;refGene_gn=SEC31B;refGene_tx=XM_019813717.2;refGene_gn=SEC31B;refGene_tx=XM_019813718.2;refGene_gn=SEC31B;refGene_tx=XM_019813719.2;refGene_gn=SEC31B;refGene_tx=XM_019813720.2
D2 61206195 61206267
refGene_gn=SEC31B;refGene_tx=XM_011287465.3;refGene_gn=SEC31B;refGene_tx=XM_011287466.3;refGene_gn=SEC31B;refGene_tx=XM_011287472.3;refGene_gn=SEC31B;refGene_tx=XM_019813704.2;refGene_gn=SEC31B;refGene_tx=XM_019813705.2;refGene_gn=SEC31B;refGene_tx=XM_019813706.2;refGene_gn=SEC31B;refGene_tx=XM_019813708.2;refGene_gn=SEC31B;refGene_tx=XM_019813709.2;refGene_gn=SEC31B;refGene_tx=XM_019813710.2;refGene_gn=SEC31B;refGene_tx=XM_019813711.2;refGene_gn=SEC31B;refGene_tx=XM_019813712.2;refGene_gn=SEC31B;refGene_tx=XM_019813713.2;refGene_gn=SEC31B;refGene_tx=XM_019813714.2;refGene_gn=SEC31B;refGene_tx=XM_019813715.2;refGene_gn=SEC31B;refGene_tx=XM_019813717.2;refGene_gn=SEC31B;refGene_tx=XM_019813718.2;refGene_gn=SEC31B;refGene_tx=XM_019813719.2;refGene_gn=SEC31B;refGene_tx=XM_019813720.2
D2 61206722 61206842
refGene_gn=SEC31B;refGene_tx=XM_011287465.3;refGene_gn=SEC31B;refGene_tx=XM_011287466.3;refGene_gn=SEC31B;refGene_tx=XM_011287472.3;refGene_gn=SEC31B;refGene_tx=XM_019813704.2;refGene_gn=SEC31B;refGene_tx=XM_019813705.2;refGene_gn=SEC31B;refGene_tx=XM_019813706.2;refGene_gn=SEC31B;refGene_tx=XM_019813708.2;refGene_gn=SEC31B;refGene_tx=XM_019813709.2;refGene_gn=SEC31B;refGene_tx=XM_019813710.2;refGene_gn=SEC31B;refGene_tx=XM_019813711.2;refGene_gn=SEC31B;refGene_tx=XM_019813712.2;refGene_gn=SEC31B;refGene_tx=XM_019813713.2;refGene_gn=SEC31B;refGene_tx=XM_019813714.2;refGene_gn=SEC31B;refGene_tx=XM_019813715.2;refGene_gn=SEC31B;refGene_tx=XM_019813717.2;refGene_gn=SEC31B;refGene_tx=XM_01981371

8.2;refGene_gn=SEC31B;refGene_tx=XM_019813719.2;refGene_gn=SEC31B;refGene_tx=XM_019813720.2
D2 61207134 61207281
refGene_gn=SEC31B;refGene_tx=XM_011287465.3;refGene_gn=SEC31B;refGene_tx=XM_011287466.3;refGene_gn=SEC31B;refGene_tx=XM_011287472.3;refGene_gn=SEC31B;refGene_tx=XM_019813704.2;refGene_gn=SEC31B;refGene_tx=XM_019813705.2;refGene_gn=SEC31B;refGene_tx=XM_019813706.2;refGene_gn=SEC31B;refGene_tx=XM_019813708.2;refGene_gn=SEC31B;refGene_tx=XM_019813709.2;refGene_gn=SEC31B;refGene_tx=XM_019813710.2;refGene_gn=SEC31B;refGene_tx=XM_019813711.2;refGene_gn=SEC31B;refGene_tx=XM_019813712.2;refGene_gn=SEC31B;refGene_tx=XM_019813713.2;refGene_gn=SEC31B;refGene_tx=XM_019813714.2;refGene_gn=SEC31B;refGene_tx=XM_019813715.2;refGene_gn=SEC31B;refGene_tx=XM_019813717.2;refGene_gn=SEC31B;refGene_tx=XM_019813718.2;refGene_gn=SEC31B;refGene_tx=XM_019813719.2;refGene_gn=SEC31B;refGene_tx=XM_019813720.2
D2 61207579 61207638
refGene_gn=SEC31B;refGene_tx=XM_011287465.3;refGene_gn=SEC31B;refGene_tx=XM_011287466.3;refGene_gn=SEC31B;refGene_tx=XM_011287472.3;refGene_gn=SEC31B;refGene_tx=XM_019813704.2;refGene_gn=SEC31B;refGene_tx=XM_019813705.2;refGene_gn=SEC31B;refGene_tx=XM_019813706.2;refGene_gn=SEC31B;refGene_tx=XM_019813708.2;refGene_gn=SEC31B;refGene_tx=XM_019813709.2;refGene_gn=SEC31B;refGene_tx=XM_019813710.2;refGene_gn=SEC31B;refGene_tx=XM_019813711.2;refGene_gn=SEC31B;refGene_tx=XM_019813712.2;refGene_gn=SEC31B;refGene_tx=XM_019813713.2;refGene_gn=SEC31B;refGene_tx=XM_019813714.2;refGene_gn=SEC31B;refGene_tx=XM_019813715.2;refGene_gn=SEC31B;refGene_tx=XM_019813717.2;refGene_gn=SEC31B;refGene_tx=XM_019813718.2;refGene_gn=SEC31B;refGene_tx=XM_019813719.2;refGene_gn=SEC31B;refGene_tx=XM_019813720.2
D2 61207898 61208213
refGene_gn=SEC31B;refGene_tx=XM_011287465.3;refGene_gn=SEC31B;refGene_tx=XM_011287466.3;refGene_gn=SEC31B;refGene_tx=XM_011287472.3;refGene_gn=SEC31B;refGene_tx=XM_019813704.2;refGene_gn=SEC31B;refGene_tx=XM_019813705.2;refGene_gn=SEC31B;refGene_tx=XM_019813706.2;refGene_gn=SEC31B;refGene_tx=XM_019813708.2;refGene_gn=SEC31B;refGene_tx=XM_019813709.2;refGene_gn=SEC31B;refGene_tx=XM_019813710.2;refGene_gn=SEC31B;refGene_tx=XM_019813711.2;refGene_gn=SEC31B;refGene_tx=XM_019813712.2;refGene_gn=SEC31B;refGene_tx=XM_019813713.2;refGene_gn=SEC31B;refGene_tx=XM_019813714.2;refGene_gn=SEC31B;refGene_tx=XM_019813715.2;refGene_gn=SEC31B;refGene_tx=XM_019813718.2;refGene_gn=SEC31B;refGene_tx=XM_019813719.2;refGene_gn=SEC31B;refGene_tx=XM_019813720.2
D2 61208677 61208846
refGene_gn=SEC31B;refGene_tx=XM_011287465.3;refGene_gn=SEC31B;refGene_tx=XM_011287466.3;refGene_gn=SEC31B;refGene_tx=XM_011287472.3;refGene_gn=SEC31B;refGene_tx=XM_019813704.2;refGene_gn=SEC31B;refGene_tx=XM_019813705.2;refGene_gn=SEC31B;refGene_tx=XM_019813706.2;refGene_gn=SEC31B;refGene_tx=XM_019813708.2;refGene_gn=SEC31B;refGene_tx=XM_019813709.2;refGene_gn=SEC31B;refGene_tx=XM_019813710.2;refGene_gn=SEC31B;refGene_tx=XM_019813711.2;refGene_gn=SEC31B;refGene_tx=XM_019813712.2;refGene_gn=SEC31B;refGene_tx=XM_019813713.2;refGene_gn=SEC31B;refGene_tx=XM_019813714.2;refGene_gn=SEC31B;refGene_tx=XM_019813715.2;refGene_gn=SEC

31B;refGene_tx=XM_019813717.2;refGene_gn=SEC31B;refGene_tx=XM_019813718.2;refGene_gn=SEC31B;refGene_tx=XM_019813719.2;refGene_gn=SEC31B;refGene_tx=XM_019813720.2
D2	61213354	61213516
refGene_gn=SEC31B;refGene_tx=XM_011287465.3;refGene_gn=SEC31B;refGene_tx=XM_011287466.3;refGene_gn=SEC31B;refGene_tx=XM_011287472.3;refGene_gn=SEC31B;refGene_tx=XM_019813704.2;refGene_gn=SEC31B;refGene_tx=XM_019813705.2;refGene_gn=SEC31B;refGene_tx=XM_019813706.2;refGene_gn=SEC31B;refGene_tx=XM_019813708.2;refGene_gn=SEC31B;refGene_tx=XM_019813709.2;refGene_gn=SEC31B;refGene_tx=XM_019813710.2;refGene_gn=SEC31B;refGene_tx=XM_019813711.2;refGene_gn=SEC31B;refGene_tx=XM_019813713.2;refGene_gn=SEC31B;refGene_tx=XM_019813714.2;refGene_gn=SEC31B;refGene_tx=XM_019813715.2;refGene_gn=SEC31B;refGene_tx=XM_019813717.2;refGene_gn=SEC31B;refGene_tx=XM_019813718.2;refGene_gn=SEC31B;refGene_tx=XM_019813719.2;refGene_gn=SEC31B;refGene_tx=XM_019813720.2
D2	61214018	61214192
refGene_gn=SEC31B;refGene_tx=XM_011287465.3;refGene_gn=SEC31B;refGene_tx=XM_011287466.3;refGene_gn=SEC31B;refGene_tx=XM_011287472.3;refGene_gn=SEC31B;refGene_tx=XM_019813704.2;refGene_gn=SEC31B;refGene_tx=XM_019813705.2;refGene_gn=SEC31B;refGene_tx=XM_019813706.2;refGene_gn=SEC31B;refGene_tx=XM_019813708.2;refGene_gn=SEC31B;refGene_tx=XM_019813709.2;refGene_gn=SEC31B;refGene_tx=XM_019813710.2;refGene_gn=SEC31B;refGene_tx=XM_019813711.2;refGene_gn=SEC31B;refGene_tx=XM_019813712.2;refGene_gn=SEC31B;refGene_tx=XM_019813713.2;refGene_gn=SEC31B;refGene_tx=XM_019813714.2;refGene_gn=SEC31B;refGene_tx=XM_019813715.2;refGene_gn=SEC31B;refGene_tx=XM_019813717.2;refGene_gn=SEC31B;refGene_tx=XM_019813718.2;refGene_gn=SEC31B;refGene_tx=XM_019813719.2;refGene_gn=SEC31B;refGene_tx=XM_019813720.2
D2	61214893	61215039
refGene_gn=SEC31B;refGene_tx=XM_011287465.3;refGene_gn=SEC31B;refGene_tx=XM_011287466.3;refGene_gn=SEC31B;refGene_tx=XM_011287472.3;refGene_gn=SEC31B;refGene_tx=XM_019813704.2;refGene_gn=SEC31B;refGene_tx=XM_019813705.2;refGene_gn=SEC31B;refGene_tx=XM_019813706.2;refGene_gn=SEC31B;refGene_tx=XM_019813708.2;refGene_gn=SEC31B;refGene_tx=XM_019813709.2;refGene_gn=SEC31B;refGene_tx=XM_019813710.2;refGene_gn=SEC31B;refGene_tx=XM_019813711.2;refGene_gn=SEC31B;refGene_tx=XM_019813712.2;refGene_gn=SEC31B;refGene_tx=XM_019813713.2;refGene_gn=SEC31B;refGene_tx=XM_019813714.2;refGene_gn=SEC31B;refGene_tx=XM_019813715.2;refGene_gn=SEC31B;refGene_tx=XM_019813717.2;refGene_gn=SEC31B;refGene_tx=XM_019813718.2;refGene_gn=SEC31B;refGene_tx=XM_019813719.2;refGene_gn=SEC31B;refGene_tx=XM_019813720.2
D2	61215660	61215787
refGene_gn=SEC31B;refGene_tx=XM_011287465.3;refGene_gn=SEC31B;refGene_tx=XM_011287466.3;refGene_gn=SEC31B;refGene_tx=XM_011287472.3;refGene_gn=SEC31B;refGene_tx=XM_019813704.2;refGene_gn=SEC31B;refGene_tx=XM_019813705.2;refGene_gn=SEC31B;refGene_tx=XM_019813706.2;refGene_gn=SEC31B;refGene_tx=XM_019813708.2;refGene_gn=SEC31B;refGene_tx=XM_019813709.2;refGene_gn=SEC31B;refGene_tx=XM_019813710.2;refGene_gn=SEC31B;refGene_tx=XM_019813711.2;refGene_gn=SEC31B;refGene_tx=XM_019813712.2;refGene_gn=SEC31B;refGene_tx=XM_019813713.2;refGene_gn=SEC31B;refGene_tx=XM_

019813714.2;refGene_gn=SEC31B;refGene_tx=XM_019813715.2;refGene_gn=SEC31B;refGene_tx=XM_019813717.2;refGene_gn=SEC31B;refGene_tx=XM_019813718.2;refGene_gn=SEC31B;refGene_tx=XM_019813719.2;refGene_gn=SEC31B;refGene_tx=XM_019813720.2
D2	61216050	61216229
refGene_gn=SEC31B;refGene_tx=XM_011287465.3;refGene_gn=SEC31B;refGene_tx=XM_011287466.3;refGene_gn=SEC31B;refGene_tx=XM_011287472.3;refGene_gn=SEC31B;refGene_tx=XM_019813704.2;refGene_gn=SEC31B;refGene_tx=XM_019813705.2;refGene_gn=SEC31B;refGene_tx=XM_019813706.2;refGene_gn=SEC31B;refGene_tx=XM_019813708.2;refGene_gn=SEC31B;refGene_tx=XM_019813709.2;refGene_gn=SEC31B;refGene_tx=XM_019813710.2;refGene_gn=SEC31B;refGene_tx=XM_019813711.2;refGene_gn=SEC31B;refGene_tx=XM_019813712.2;refGene_gn=SEC31B;refGene_tx=XM_019813713.2;refGene_gn=SEC31B;refGene_tx=XM_019813714.2;refGene_gn=SEC31B;refGene_tx=XM_019813715.2;refGene_gn=SEC31B;refGene_tx=XM_019813717.2;refGene_gn=SEC31B;refGene_tx=XM_019813718.2;refGene_gn=SEC31B;refGene_tx=XM_019813719.2;refGene_gn=SEC31B;refGene_tx=XM_019813720.2
D2	61216757	61216857
refGene_gn=SEC31B;refGene_tx=XM_011287465.3;refGene_gn=SEC31B;refGene_tx=XM_011287466.3;refGene_gn=SEC31B;refGene_tx=XM_011287472.3;refGene_gn=SEC31B;refGene_tx=XM_019813704.2;refGene_gn=SEC31B;refGene_tx=XM_019813705.2;refGene_gn=SEC31B;refGene_tx=XM_019813706.2;refGene_gn=SEC31B;refGene_tx=XM_019813708.2;refGene_gn=SEC31B;refGene_tx=XM_019813709.2;refGene_gn=SEC31B;refGene_tx=XM_019813710.2;refGene_gn=SEC31B;refGene_tx=XM_019813711.2;refGene_gn=SEC31B;refGene_tx=XM_019813712.2;refGene_gn=SEC31B;refGene_tx=XM_019813713.2;refGene_gn=SEC31B;refGene_tx=XM_019813714.2;refGene_gn=SEC31B;refGene_tx=XM_019813715.2;refGene_gn=SEC31B;refGene_tx=XM_019813717.2;refGene_gn=SEC31B;refGene_tx=XM_019813718.2;refGene_gn=SEC31B;refGene_tx=XM_019813719.2;refGene_gn=SEC31B;refGene_tx=XM_019813720.2
D2	61217217	61217313
refGene_gn=SEC31B;refGene_tx=XM_011287465.3;refGene_gn=SEC31B;refGene_tx=XM_011287466.3;refGene_gn=SEC31B;refGene_tx=XM_011287472.3;refGene_gn=SEC31B;refGene_tx=XM_019813704.2;refGene_gn=SEC31B;refGene_tx=XM_019813705.2;refGene_gn=SEC31B;refGene_tx=XM_019813706.2;refGene_gn=SEC31B;refGene_tx=XM_019813708.2;refGene_gn=SEC31B;refGene_tx=XM_019813709.2;refGene_gn=SEC31B;refGene_tx=XM_019813710.2;refGene_gn=SEC31B;refGene_tx=XM_019813712.2;refGene_gn=SEC31B;refGene_tx=XM_019813713.2;refGene_gn=SEC31B;refGene_tx=XM_019813714.2;refGene_gn=SEC31B;refGene_tx=XM_019813715.2;refGene_gn=SEC31B;refGene_tx=XM_019813717.2;refGene_gn=SEC31B;refGene_tx=XM_019813718.2;refGene_gn=SEC31B;refGene_tx=XM_019813719.2;refGene_gn=SEC31B;refGene_tx=XM_019813720.2
D2	61217575	61217650
refGene_gn=SEC31B;refGene_tx=XM_011287465.3;refGene_gn=SEC31B;refGene_tx=XM_011287466.3;refGene_gn=SEC31B;refGene_tx=XM_011287472.3;refGene_gn=SEC31B;refGene_tx=XM_019813704.2;refGene_gn=SEC31B;refGene_tx=XM_019813705.2;refGene_gn=SEC31B;refGene_tx=XM_019813706.2;refGene_gn=SEC31B;refGene_tx=XM_019813708.2;refGene_gn=SEC31B;refGene_tx=XM_019813709.2;refGene_gn=SEC31B;refGene_tx=XM_019813710.2;refGene_gn=SEC31B;refGene_tx=XM_019813711.2;refGene_gn=SEC31B;refGene_tx=XM_019813712.2;refGen

e_gn=SEC31B;refGene_tx=XM_019813713.2;refGene_gn=SEC31B;refGene_tx=XM_019813714.2;refGene_gn=SEC31B;refGene_tx=XM_019813715.2;refGene_gn=SEC31B;refGene_tx=XM_019813717.2;refGene_gn=SEC31B;refGene_tx=XM_019813718.2;refGene_gn=SEC31B;refGene_tx=XM_019813719.2;refGene_gn=SEC31B;refGene_tx=XM_019813720.2
D2 61219576 61219807
refGene_gn=SEC31B;refGene_tx=XM_011287465.3;refGene_gn=SEC31B;refGene_tx=XM_011287466.3;refGene_gn=SEC31B;refGene_tx=XM_011287472.3;refGene_gn=SEC31B;refGene_tx=XM_019813704.2;refGene_gn=SEC31B;refGene_tx=XM_019813705.2;refGene_gn=SEC31B;refGene_tx=XM_019813706.2;refGene_gn=SEC31B;refGene_tx=XM_019813708.2;refGene_gn=SEC31B;refGene_tx=XM_019813709.2;refGene_gn=SEC31B;refGene_tx=XM_019813710.2;refGene_gn=SEC31B;refGene_tx=XM_019813711.2;refGene_gn=SEC31B;refGene_tx=XM_019813712.2;refGene_gn=SEC31B;refGene_tx=XM_019813713.2;refGene_gn=SEC31B;refGene_tx=XM_019813714.2;refGene_gn=SEC31B;refGene_tx=XM_019813715.2;refGene_gn=SEC31B;refGene_tx=XM_019813717.2;refGene_gn=SEC31B;refGene_tx=XM_019813718.2;refGene_gn=SEC31B;refGene_tx=XM_019813719.2;refGene_gn=SEC31B;refGene_tx=XM_019813720.2
D2 61227141 61227276
refGene_gn=SEC31B;refGene_tx=XM_011287465.3;refGene_gn=SEC31B;refGene_tx=XM_011287466.3;refGene_gn=SEC31B;refGene_tx=XM_011287472.3;refGene_gn=SEC31B;refGene_tx=XM_019813704.2;refGene_gn=SEC31B;refGene_tx=XM_019813705.2;refGene_gn=SEC31B;refGene_tx=XM_019813706.2;refGene_gn=SEC31B;refGene_tx=XM_019813708.2;refGene_gn=SEC31B;refGene_tx=XM_019813709.2;refGene_gn=SEC31B;refGene_tx=XM_019813710.2;refGene_gn=SEC31B;refGene_tx=XM_019813711.2;refGene_gn=SEC31B;refGene_tx=XM_019813712.2;refGene_gn=SEC31B;refGene_tx=XM_019813713.2;refGene_gn=SEC31B;refGene_tx=XM_019813714.2;refGene_gn=SEC31B;refGene_tx=XM_019813715.2;refGene_gn=SEC31B;refGene_tx=XM_019813717.2;refGene_gn=SEC31B;refGene_tx=XM_019813718.2;refGene_gn=SEC31B;refGene_tx=XM_019813719.2;refGene_gn=SEC31B;refGene_tx=XM_019813720.2
D2 61227783 61227945
refGene_gn=SEC31B;refGene_tx=XM_011287465.3;refGene_gn=SEC31B;refGene_tx=XM_011287466.3;refGene_gn=SEC31B;refGene_tx=XM_011287472.3;refGene_gn=SEC31B;refGene_tx=XM_019813704.2;refGene_gn=SEC31B;refGene_tx=XM_019813705.2;refGene_gn=SEC31B;refGene_tx=XM_019813706.2;refGene_gn=SEC31B;refGene_tx=XM_019813708.2;refGene_gn=SEC31B;refGene_tx=XM_019813709.2;refGene_gn=SEC31B;refGene_tx=XM_019813710.2;refGene_gn=SEC31B;refGene_tx=XM_019813711.2;refGene_gn=SEC31B;refGene_tx=XM_019813712.2;refGene_gn=SEC31B;refGene_tx=XM_019813713.2;refGene_gn=SEC31B;refGene_tx=XM_019813714.2;refGene_gn=SEC31B;refGene_tx=XM_019813715.2;refGene_gn=SEC31B;refGene_tx=XM_019813717.2;refGene_gn=SEC31B;refGene_tx=XM_019813718.2;refGene_gn=SEC31B;refGene_tx=XM_019813719.2;refGene_gn=SEC31B;refGene_tx=XM_019813720.2
D2 61228049 61228149
refGene_gn=SEC31B;refGene_tx=XM_011287465.3;refGene_gn=SEC31B;refGene_tx=XM_011287466.3;refGene_gn=SEC31B;refGene_tx=XM_011287472.3;refGene_gn=SEC31B;refGene_tx=XM_019813704.2;refGene_gn=SEC31B;refGene_tx=XM_019813705.2;refGene_gn=SEC31B;refGene_tx=XM_019813706.2;refGene_gn=SEC31B;refGene_tx=XM_019813708.2;refGene_gn=SEC31B;refGene_tx=XM_019813709.

2;refGene_gn=SEC31B;refGene_tx=XM_019813710.2;refGene_gn=SEC31B;refGene_tx=XM_019813711.2;refGene_gn=SEC31B;refGene_tx=XM_019813712.2;refGene_gn=SEC31B;refGene_tx=XM_019813713.2;refGene_gn=SEC31B;refGene_tx=XM_019813714.2;refGene_gn=SEC31B;refGene_tx=XM_019813715.2;refGene_gn=SEC31B;refGene_tx=XM_019813717.2;refGene_gn=SEC31B;refGene_tx=XM_019813718.2;refGene_gn=SEC31B;refGene_tx=XM_019813719.2;refGene_gn=SEC31B;refGene_tx=XM_019813720.2
D2 61229206 61229349
refGene_gn=SEC31B;refGene_tx=XM_011287465.3;refGene_gn=SEC31B;refGene_tx=XM_011287466.3;refGene_gn=SEC31B;refGene_tx=XM_011287472.3;refGene_gn=SEC31B;refGene_tx=XM_019813704.2;refGene_gn=SEC31B;refGene_tx=XM_019813705.2;refGene_gn=SEC31B;refGene_tx=XM_019813706.2;refGene_gn=SEC31B;refGene_tx=XM_019813708.2;refGene_gn=SEC31B;refGene_tx=XM_019813709.2;refGene_gn=SEC31B;refGene_tx=XM_019813710.2;refGene_gn=SEC31B;refGene_tx=XM_019813711.2;refGene_gn=SEC31B;refGene_tx=XM_019813712.2;refGene_gn=SEC31B;refGene_tx=XM_019813713.2;refGene_gn=SEC31B;refGene_tx=XM_019813714.2;refGene_gn=SEC31B;refGene_tx=XM_019813715.2;refGene_gn=SEC31B;refGene_tx=XM_019813717.2;refGene_gn=SEC31B;refGene_tx=XM_019813718.2;refGene_gn=SEC31B;refGene_tx=XM_019813719.2;refGene_gn=SEC31B;refGene_tx=XM_019813720.2
D2 61229666 61229810
refGene_gn=SEC31B;refGene_tx=XM_011287465.3;refGene_gn=SEC31B;refGene_tx=XM_011287466.3;refGene_gn=SEC31B;refGene_tx=XM_011287472.3;refGene_gn=SEC31B;refGene_tx=XM_019813704.2;refGene_gn=SEC31B;refGene_tx=XM_019813705.2;refGene_gn=SEC31B;refGene_tx=XM_019813706.2;refGene_gn=SEC31B;refGene_tx=XM_019813708.2;refGene_gn=SEC31B;refGene_tx=XM_019813709.2;refGene_gn=SEC31B;refGene_tx=XM_019813710.2;refGene_gn=SEC31B;refGene_tx=XM_019813711.2;refGene_gn=SEC31B;refGene_tx=XM_019813712.2;refGene_gn=SEC31B;refGene_tx=XM_019813713.2;refGene_gn=SEC31B;refGene_tx=XM_019813714.2;refGene_gn=SEC31B;refGene_tx=XM_019813715.2;refGene_gn=SEC31B;refGene_tx=XM_019813717.2;refGene_gn=SEC31B;refGene_tx=XM_019813718.2;refGene_gn=SEC31B;refGene_tx=XM_019813719.2
D2 61230228 61230237
refGene_gn=SEC31B;refGene_tx=XM_019813704.2;refGene_gn=SEC31B;refGene_tx=XM_019813705.2;refGene_gn=SEC31B;refGene_tx=XM_019813706.2;refGene_gn=SEC31B;refGene_tx=XM_019813708.2;refGene_gn=SEC31B;refGene_tx=XM_019813709.2;refGene_gn=SEC31B;refGene_tx=XM_019813710.2;refGene_gn=SEC31B;refGene_tx=XM_019813711.2;refGene_gn=SEC31B;refGene_tx=XM_019813712.2;refGene_gn=SEC31B;refGene_tx=XM_019813713.2;refGene_gn=SEC31B;refGene_tx=XM_019813714.2;refGene_gn=SEC31B;refGene_tx=XM_019813715.2;refGene_gn=SEC31B;refGene_tx=XM_019813717.2;refGene_gn=SEC31B;refGene_tx=XM_019813718.2
D2 61230742 61230838
refGene_gn=SEC31B;refGene_tx=XM_011287465.3;refGene_gn=SEC31B;refGene_tx=XM_011287466.3;refGene_gn=SEC31B;refGene_tx=XM_011287472.3;refGene_gn=SEC31B;refGene_tx=XM_019813704.2;refGene_gn=SEC31B;refGene_tx=XM_019813705.2;refGene_gn=SEC31B;refGene_tx=XM_019813706.2;refGene_gn=SEC31B;refGene_tx=XM_019813708.2;refGene_gn=SEC31B;refGene_tx=XM_019813709.2;refGene_gn=SEC31B;refGene_tx=XM_019813710.2;refGene_gn=SEC31B;refGene_tx=XM_019813711.2;refGene_gn=SEC31B;refGene_tx=XM_019813712.2;refGen

e_gn=SEC31B;refGene_tx=XM_019813713.2;refGene_gn=SEC31B;refGene_tx=XM_019813714.2;refGene_gn=SEC31B;refGene_tx=XM_019813715.2;refGene_gn=SEC31B;refGene_tx=XM_019813717.2;refGene_gn=SEC31B;refGene_tx=XM_019813718.2;refGene_gn=SEC31B;refGene_tx=XM_019813719.2
D2 61231051 61231247
refGene_gn=SEC31B;refGene_tx=XM_011287465.3;refGene_gn=SEC31B;refGene_tx=XM_011287466.3;refGene_gn=SEC31B;refGene_tx=XM_011287472.3;refGene_gn=SEC31B;refGene_tx=XM_019813704.2;refGene_gn=SEC31B;refGene_tx=XM_019813705.2;refGene_gn=SEC31B;refGene_tx=XM_019813706.2;refGene_gn=SEC31B;refGene_tx=XM_019813708.2;refGene_gn=SEC31B;refGene_tx=XM_019813709.2;refGene_gn=SEC31B;refGene_tx=XM_019813710.2;refGene_gn=SEC31B;refGene_tx=XM_019813711.2;refGene_gn=SEC31B;refGene_tx=XM_019813712.2;refGene_gn=SEC31B;refGene_tx=XM_019813713.2;refGene_gn=SEC31B;refGene_tx=XM_019813714.2;refGene_gn=SEC31B;refGene_tx=XM_019813715.2;refGene_gn=SEC31B;refGene_tx=XM_019813717.2
D2 61231882 61231992
refGene_gn=SEC31B;refGene_tx=XM_019813708.2;refGene_gn=SEC31B;refGene_tx=XM_019813709.2;refGene_gn=SEC31B;refGene_tx=XM_019813710.2
D2 61238072 61238196
refGene_gn=SEC31B;refGene_tx=XM_011287465.3;refGene_gn=SEC31B;refGene_tx=XM_011287466.3;refGene_gn=SEC31B;refGene_tx=XM_011287472.3;refGene_gn=SEC31B;refGene_tx=XM_019813704.2;refGene_gn=SEC31B;refGene_tx=XM_019813705.2;refGene_gn=SEC31B;refGene_tx=XM_019813706.2;refGene_gn=SEC31B;refGene_tx=XM_019813711.2;refGene_gn=SEC31B;refGene_tx=XM_019813712.2;refGene_gn=SEC31B;refGene_tx=XM_019813717.2
D2 61238548 61238627
refGene_gn=SEC31B;refGene_tx=XM_011287465.3;refGene_gn=SEC31B;refGene_tx=XM_011287466.3;refGene_gn=SEC31B;refGene_tx=XM_011287472.3;refGene_gn=SEC31B;refGene_tx=XM_019813704.2;refGene_gn=SEC31B;refGene_tx=XM_019813705.2;refGene_gn=SEC31B;refGene_tx=XM_019813706.2;refGene_gn=SEC31B;refGene_tx=XM_019813711.2;refGene_gn=SEC31B;refGene_tx=XM_019813712.2;refGene_gn=SEC31B;refGene_tx=XM_019813717.2
D2 61244301 61244394
refGene_gn=NDUFB8;refGene_tx=XM_003994317.4;refGene_gn=NDUFB8;refGene_tx=XM_019813722.1
D2 61246648 61246804
refGene_gn=NDUFB8;refGene_tx=XM_003994317.4;refGene_gn=NDUFB8;refGene_tx=XM_019813722.1
D2 61247239 61247339
refGene_gn=NDUFB8;refGene_tx=XM_003994317.4;refGene_gn=NDUFB8;refGene_tx=XM_019813722.1
D2 61248911 61249038
refGene_gn=NDUFB8;refGene_tx=XM_003994317.4;refGene_gn=NDUFB8;refGene_tx=XM_019813722.1
D2 61249298 61249383 refGene_gn=NDUFB8;refGene_tx=XM_003994317.4
D2 61249389 61249414 refGene_gn=NDUFB8;refGene_tx=XM_019813722.1
D2 61254113 61254290
refGene_gn=HIF1AN;refGene_tx=XM_003994318.5;refGene_gn=HIF1AN;refGene_tx=XM_019813721.2
D2 61254572 61254823

refGene_gn=HIF1AN;refGene_tx=XM_003994318.5;refGene_gn=HIF1AN;refGene_tx=XM_019813721.2
D2 61258267 61258416
refGene_gn=HIF1AN;refGene_tx=XM_003994318.5;refGene_gn=HIF1AN;refGene_tx=XM_019813721.2
D2 61260820 61260966
refGene_gn=HIF1AN;refGene_tx=XM_003994318.5;refGene_gn=HIF1AN;refGene_tx=XM_019813721.2
D2 61261808 61261915
refGene_gn=HIF1AN;refGene_tx=XM_003994318.5;refGene_gn=HIF1AN;refGene_tx=XM_019813721.2
D2 61262028 61262095 refGene_gn=HIF1AN;refGene_tx=XM_019813721.2
D2 61262335 61262399 refGene_gn=HIF1AN;refGene_tx=XM_003994318.5
D2 61262632 61262743 refGene_gn=HIF1AN;refGene_tx=XM_003994318.5
D2 61263551 61263596 refGene_gn=HIF1AN;refGene_tx=XM_003994318.5
D2 61420460 61420485
refGene_gn=PAX2;refGene_tx=XM_023240835.1;refGene_gn=PAX2;refGene_tx=XM_023240836.1;refGene_gn=PAX2;refGene_tx=XM_023240837.1;refGene_gn=PAX2;refGene_tx=XM_023240838.1;refGene_gn=PAX2;refGene_tx=XM_023240839.1;refGene_gn=PAX2;refGene_tx=XM_023240840.1;refGene_gn=PAX2;refGene_tx=XM_023240841.1
D2 61431203 61431314
refGene_gn=PAX2;refGene_tx=XM_023240835.1;refGene_gn=PAX2;refGene_tx=XM_023240836.1;refGene_gn=PAX2;refGene_tx=XM_023240837.1;refGene_gn=PAX2;refGene_tx=XM_023240838.1;refGene_gn=PAX2;refGene_tx=XM_023240839.1;refGene_gn=PAX2;refGene_tx=XM_023240840.1;refGene_gn=PAX2;refGene_tx=XM_023240841.1
D2 61434848 61435017
refGene_gn=PAX2;refGene_tx=XM_023240835.1;refGene_gn=PAX2;refGene_tx=XM_023240836.1;refGene_gn=PAX2;refGene_tx=XM_023240837.1;refGene_gn=PAX2;refGene_tx=XM_023240838.1;refGene_gn=PAX2;refGene_tx=XM_023240839.1;refGene_gn=PAX2;refGene_tx=XM_023240840.1;refGene_gn=PAX2;refGene_tx=XM_023240841.1
D2 61435816 61436014
refGene_gn=PAX2;refGene_tx=XM_023240835.1;refGene_gn=PAX2;refGene_tx=XM_023240836.1;refGene_gn=PAX2;refGene_tx=XM_023240837.1;refGene_gn=PAX2;refGene_tx=XM_023240838.1;refGene_gn=PAX2;refGene_tx=XM_023240839.1;refGene_gn=PAX2;refGene_tx=XM_023240840.1;refGene_gn=PAX2;refGene_tx=XM_023240841.1
D2 61462032 61462118
refGene_gn=PAX2;refGene_tx=XM_023240835.1;refGene_gn=PAX2;refGene_tx=XM_023240836.1;refGene_gn=PAX2;refGene_tx=XM_023240837.1;refGene_gn=PAX2;refGene_tx=XM_023240838.1;refGene_gn=PAX2;refGene_tx=XM_023240839.1;refGene_gn=PAX2;refGene_tx=XM_023240840.1;refGene_gn=PAX2;refGene_tx=XM_023240841.1
D2 61463372 61463492
refGene_gn=PAX2;refGene_tx=XM_023240835.1;refGene_gn=PAX2;refGene_tx=XM_023240836.1;refGene_gn=PAX2;refGene_tx=XM_023240837.1;refGene_gn=PAX2;refGene_tx=XM_023240838.1;refGene_gn=PAX2;refGene_tx=XM_023240839.1;refGene_gn=PAX2;refGene_tx=XM_023240840.1;refGene_gn=PAX2;refGene_tx=X

M_023240841.1
D2 61469067 61469136
refGene_gn=PAX2;refGene_tx=XM_023240835.1;refGene_gn=PAX2;refGene_tx=XM_023240836.1;refGene_gn=PAX2;refGene_tx=XM_023240837.1;refGene_gn=PAX2;refGene_tx=XM_023240838.1
D2 61488360 61488536
refGene_gn=PAX2;refGene_tx=XM_023240835.1;refGene_gn=PAX2;refGene_tx=XM_023240836.1;refGene_gn=PAX2;refGene_tx=XM_023240837.1;refGene_gn=PAX2;refGene_tx=XM_023240838.1;refGene_gn=PAX2;refGene_tx=XM_023240839.1;refGene_gn=PAX2;refGene_tx=XM_023240840.1;refGene_gn=PAX2;refGene_tx=XM_023240841.1
D2 61490706 61490833
refGene_gn=PAX2;refGene_tx=XM_023240835.1;refGene_gn=PAX2;refGene_tx=XM_023240836.1;refGene_gn=PAX2;refGene_tx=XM_023240837.1;refGene_gn=PAX2;refGene_tx=XM_023240838.1;refGene_gn=PAX2;refGene_tx=XM_023240839.1;refGene_gn=PAX2;refGene_tx=XM_023240840.1;refGene_gn=PAX2;refGene_tx=XM_023240841.1
D2 61505263 61505369
refGene_gn=PAX2;refGene_tx=XM_023240835.1;refGene_gn=PAX2;refGene_tx=XM_023240836.1;refGene_gn=PAX2;refGene_tx=XM_023240837.1;refGene_gn=PAX2;refGene_tx=XM_023240838.1;refGene_gn=PAX2;refGene_tx=XM_023240839.1;refGene_gn=PAX2;refGene_tx=XM_023240840.1;refGene_gn=PAX2;refGene_tx=XM_023240841.1
D2 61507554 61507637
refGene_gn=PAX2;refGene_tx=XM_023240835.1;refGene_gn=PAX2;refGene_tx=XM_023240836.1;refGene_gn=PAX2;refGene_tx=XM_023240837.1;refGene_gn=PAX2;refGene_tx=XM_023240838.1;refGene_gn=PAX2;refGene_tx=XM_023240839.1;refGene_gn=PAX2;refGene_tx=XM_023240840.1;refGene_gn=PAX2;refGene_tx=XM_023240841.1
D2 61508112 61508253
refGene_gn=PAX2;refGene_tx=XM_023240835.1;refGene_gn=PAX2;refGene_tx=XM_023240836.1;refGene_gn=PAX2;refGene_tx=XM_023240837.1;refGene_gn=PAX2;refGene_tx=XM_023240838.1;refGene_gn=PAX2;refGene_tx=XM_023240839.1;refGene_gn=PAX2;refGene_tx=XM_023240840.1;refGene_gn=PAX2;refGene_tx=XM_023240841.1
D2 61590135 61590275
refGene_gn=SLF2;refGene_tx=XM_003994324.5;refGene_gn=SLF2;refGene_tx=XM_011287462.3;refGene_gn=SLF2;refGene_tx=XM_023240842.1;refGene_gn=SLF2;refGene_tx=XM_023240843.1
D2 61592873 61592917
refGene_gn=SLF2;refGene_tx=XM_003994324.5;refGene_gn=SLF2;refGene_tx=XM_011287462.3;refGene_gn=SLF2;refGene_tx=XM_023240842.1;refGene_gn=SLF2;refGene_tx=XM_023240843.1
D2 61593417 61594151
refGene_gn=SLF2;refGene_tx=XM_003994324.5;refGene_gn=SLF2;refGene_tx=XM_011287462.3;refGene_gn=SLF2;refGene_tx=XM_019813733.2;refGene_gn=SLF2;refGene_tx=XM_023240842.1;refGene_gn=SLF2;refGene_tx=XM_023240843.1
D2 61595537 61595595
refGene_gn=SLF2;refGene_tx=XM_003994324.5;refGene_gn=SLF2;refGene_tx=XM_011287462.3;refGene_gn=SLF2;refGene_tx=XM_019813733.2;refGene_gn=SLF

2;refGene_tx=XM_023240842.1;refGene_gn=SLF2;refGene_tx=XM_023240843.1
D2 61600197 61601186
refGene_gn=SLF2;refGene_tx=XM_003994324.5;refGene_gn=SLF2;refGene_tx=XM_011287462.3;refGene_gn=SLF2;refGene_tx=XM_019813733.2;refGene_gn=SLF2;refGene_tx=XM_023240842.1;refGene_gn=SLF2;refGene_tx=XM_023240843.1
D2 61601693 61601699 refGene_gn=SLF2;refGene_tx=XM_011287464.3
D2 61602162 61602233
refGene_gn=SLF2;refGene_tx=XM_003994324.5;refGene_gn=SLF2;refGene_tx=XM_011287462.3;refGene_gn=SLF2;refGene_tx=XM_011287464.3;refGene_gn=SLF2;refGene_tx=XM_019813733.2;refGene_gn=SLF2;refGene_tx=XM_023240842.1;refGene_gn=SLF2;refGene_tx=XM_023240843.1
D2 61604485 61604608
refGene_gn=SLF2;refGene_tx=XM_003994324.5;refGene_gn=SLF2;refGene_tx=XM_011287462.3;refGene_gn=SLF2;refGene_tx=XM_011287464.3;refGene_gn=SLF2;refGene_tx=XM_019813733.2;refGene_gn=SLF2;refGene_tx=XM_023240842.1;refGene_gn=SLF2;refGene_tx=XM_023240843.1
D2 61604990 61605158
refGene_gn=SLF2;refGene_tx=XM_003994324.5;refGene_gn=SLF2;refGene_tx=XM_011287462.3;refGene_gn=SLF2;refGene_tx=XM_011287464.3;refGene_gn=SLF2;refGene_tx=XM_019813733.2;refGene_gn=SLF2;refGene_tx=XM_023240842.1;refGene_gn=SLF2;refGene_tx=XM_023240843.1
D2 61606158 61606261
refGene_gn=SLF2;refGene_tx=XM_003994324.5;refGene_gn=SLF2;refGene_tx=XM_011287462.3;refGene_gn=SLF2;refGene_tx=XM_011287464.3;refGene_gn=SLF2;refGene_tx=XM_019813733.2;refGene_gn=SLF2;refGene_tx=XM_023240842.1;refGene_gn=SLF2;refGene_tx=XM_023240843.1
D2 61611519 61611595
refGene_gn=SLF2;refGene_tx=XM_003994324.5;refGene_gn=SLF2;refGene_tx=XM_011287462.3;refGene_gn=SLF2;refGene_tx=XM_011287464.3;refGene_gn=SLF2;refGene_tx=XM_019813733.2;refGene_gn=SLF2;refGene_tx=XM_023240842.1;refGene_gn=SLF2;refGene_tx=XM_023240843.1
D2 61612391 61612533
refGene_gn=SLF2;refGene_tx=XM_003994324.5;refGene_gn=SLF2;refGene_tx=XM_011287462.3;refGene_gn=SLF2;refGene_tx=XM_011287464.3;refGene_gn=SLF2;refGene_tx=XM_019813733.2;refGene_gn=SLF2;refGene_tx=XM_023240842.1;refGene_gn=SLF2;refGene_tx=XM_023240843.1
D2 61617975 61618078
refGene_gn=SLF2;refGene_tx=XM_003994324.5;refGene_gn=SLF2;refGene_tx=XM_011287462.3;refGene_gn=SLF2;refGene_tx=XM_011287464.3;refGene_gn=SLF2;refGene_tx=XM_019813733.2;refGene_gn=SLF2;refGene_tx=XM_023240842.1;refGene_gn=SLF2;refGene_tx=XM_023240843.1
D2 61619055 61619232
refGene_gn=SLF2;refGene_tx=XM_003994324.5;refGene_gn=SLF2;refGene_tx=XM_011287462.3;refGene_gn=SLF2;refGene_tx=XM_011287464.3;refGene_gn=SLF2;refGene_tx=XM_019813733.2;refGene_gn=SLF2;refGene_tx=XM_023240842.1;refGene_gn=SLF2;refGene_tx=XM_023240843.1
D2 61620442 61620540
refGene_gn=SLF2;refGene_tx=XM_003994324.5;refGene_gn=SLF2;refGene_tx=XM_011287462.3;refGene_gn=SLF2;refGene_tx=XM_011287464.3;refGene_gn=SLF2;refGene_tx=XM_019813733.2;refGene_gn=SLF2;refGene_tx=XM_023240842.1;

refGene_gn=SLF2;refGene_tx=XM_023240843.1
D2 61621265 61621353
refGene_gn=SLF2;refGene_tx=XM_003994324.5;refGene_gn=SLF2;refGene_tx=XM_011287462.3;refGene_gn=SLF2;refGene_tx=XM_011287464.3;refGene_gn=SLF2;refGene_tx=XM_019813733.2;refGene_gn=SLF2;refGene_tx=XM_023240842.1;refGene_gn=SLF2;refGene_tx=XM_023240843.1
D2 61622843 61622975
refGene_gn=SLF2;refGene_tx=XM_003994324.5;refGene_gn=SLF2;refGene_tx=XM_011287462.3;refGene_gn=SLF2;refGene_tx=XM_011287464.3;refGene_gn=SLF2;refGene_tx=XM_019813733.2;refGene_gn=SLF2;refGene_tx=XM_023240842.1;refGene_gn=SLF2;refGene_tx=XM_023240843.1
D2 61623416 61623494
refGene_gn=SLF2;refGene_tx=XM_003994324.5;refGene_gn=SLF2;refGene_tx=XM_011287462.3;refGene_gn=SLF2;refGene_tx=XM_011287464.3;refGene_gn=SLF2;refGene_tx=XM_019813733.2;refGene_gn=SLF2;refGene_tx=XM_023240842.1;refGene_gn=SLF2;refGene_tx=XM_023240843.1
D2 61626847 61626934
refGene_gn=SLF2;refGene_tx=XM_003994324.5;refGene_gn=SLF2;refGene_tx=XM_011287462.3;refGene_gn=SLF2;refGene_tx=XM_011287464.3;refGene_gn=SLF2;refGene_tx=XM_019813733.2;refGene_gn=SLF2;refGene_tx=XM_023240842.1;refGene_gn=SLF2;refGene_tx=XM_023240843.1
D2 61630380 61630449
refGene_gn=SLF2;refGene_tx=XM_003994324.5;refGene_gn=SLF2;refGene_tx=XM_011287462.3;refGene_gn=SLF2;refGene_tx=XM_011287464.3;refGene_gn=SLF2;refGene_tx=XM_019813733.2;refGene_gn=SLF2;refGene_tx=XM_023240842.1;refGene_gn=SLF2;refGene_tx=XM_023240843.1
D2 61633519 61633555
refGene_gn=SLF2;refGene_tx=XM_003994324.5;refGene_gn=SLF2;refGene_tx=XM_011287462.3;refGene_gn=SLF2;refGene_tx=XM_011287464.3;refGene_gn=SLF2;refGene_tx=XM_019813733.2;refGene_gn=SLF2;refGene_tx=XM_023240842.1;refGene_gn=SLF2;refGene_tx=XM_023240843.1
D2 61644681 61644805
refGene_gn=SEMA4G;refGene_tx=XM_023240845.1;refGene_gn=SEMA4G;refGene_tx=XM_023240846.1
D2 61644910 61645059
refGene_gn=SEMA4G;refGene_tx=XM_019813736.2;refGene_gn=SEMA4G;refGene_tx=XM_023240845.1;refGene_gn=SEMA4G;refGene_tx=XM_023240846.1
D2 61645341 61645404
refGene_gn=SEMA4G;refGene_tx=XM_019813736.2;refGene_gn=SEMA4G;refGene_tx=XM_023240845.1;refGene_gn=SEMA4G;refGene_tx=XM_023240846.1
D2 61648666 61648787
refGene_gn=SEMA4G;refGene_tx=XM_019813736.2;refGene_gn=SEMA4G;refGene_tx=XM_023240845.1;refGene_gn=SEMA4G;refGene_tx=XM_023240846.1
D2 61648909 61649003
refGene_gn=SEMA4G;refGene_tx=XM_019813736.2;refGene_gn=SEMA4G;refGene_tx=XM_023240845.1;refGene_gn=SEMA4G;refGene_tx=XM_023240846.1
D2 61649154 61649268
refGene_gn=SEMA4G;refGene_tx=XM_019813736.2;refGene_gn=SEMA4G;refGene_tx=XM_023240845.1;refGene_gn=SEMA4G;refGene_tx=XM_023240846.1
D2 61649453 61649623

refGene_gn=SEMA4G;refGene_tx=XM_019813736.2;refGene_gn=SEMA4G;refGene_tx=XM_023240845.1;refGene_gn=SEMA4G;refGene_tx=XM_023240846.1
D2 61649698 61649868
refGene_gn=SEMA4G;refGene_tx=XM_019813736.2;refGene_gn=SEMA4G;refGene_tx=XM_023240845.1;refGene_gn=SEMA4G;refGene_tx=XM_023240846.1
D2 61650164 61650309
refGene_gn=SEMA4G;refGene_tx=XM_019813736.2;refGene_gn=SEMA4G;refGene_tx=XM_023240845.1;refGene_gn=SEMA4G;refGene_tx=XM_023240846.1
D2 61650451 61650674
refGene_gn=SEMA4G;refGene_tx=XM_019813736.2;refGene_gn=SEMA4G;refGene_tx=XM_023240845.1;refGene_gn=SEMA4G;refGene_tx=XM_023240846.1
D2 61651091 61651207
refGene_gn=SEMA4G;refGene_tx=XM_019813736.2;refGene_gn=SEMA4G;refGene_tx=XM_023240845.1;refGene_gn=SEMA4G;refGene_tx=XM_023240846.1
D2 61651329 61651493
refGene_gn=SEMA4G;refGene_tx=XM_019813736.2;refGene_gn=SEMA4G;refGene_tx=XM_023240845.1;refGene_gn=SEMA4G;refGene_tx=XM_023240846.1
D2 61651660 61651722
refGene_gn=SEMA4G;refGene_tx=XM_019813736.2;refGene_gn=SEMA4G;refGene_tx=XM_023240845.1;refGene_gn=SEMA4G;refGene_tx=XM_023240846.1
D2 61653159 61653989
refGene_gn=SEMA4G;refGene_tx=XM_019813736.2;refGene_gn=SEMA4G;refGene_tx=XM_023240845.1;refGene_gn=SEMA4G;refGene_tx=XM_023240846.1
D2 61656824 61657066 refGene_gn=MRPL43;refGene_tx=XM_003994327.4
D2 61657193 61657300 refGene_gn=MRPL43;refGene_tx=XM_003994327.4
D2 61657389 61657520 refGene_gn=MRPL43;refGene_tx=XM_003994327.4
D2 61658272 61659515
refGene_gn=TWNK;refGene_tx=XM_003994328.5;refGene_gn=TWNK;refGene_tx=XM_019813737.2
D2 61659679 61659942
refGene_gn=TWNK;refGene_tx=XM_003994328.5;refGene_gn=TWNK;refGene_tx=XM_019813737.2
D2 61660093 61660201
refGene_gn=TWNK;refGene_tx=XM_003994328.5;refGene_gn=TWNK;refGene_tx=XM_019813737.2
D2 61660517 61660659
refGene_gn=TWNK;refGene_tx=XM_003994328.5;refGene_gn=TWNK;refGene_tx=XM_019813737.2
D2 61661812 61662133
refGene_gn=TWNK;refGene_tx=XM_003994328.5;refGene_gn=TWNK;refGene_tx=XM_019813737.2
D2 61670823 61671231
refGene_gn=LZTS2;refGene_tx=XM_003994330.5;refGene_gn=LZTS2;refGene_tx=XM_006938099.3;refGene_gn=LZTS2;refGene_tx=XM_006938101.3;refGene_gn=LZTS2;refGene_tx=XM_006938103.3;refGene_gn=LZTS2;refGene_tx=XM_019813738.2;refGene_gn=LZTS2;refGene_tx=XM_023240847.1
D2 61671856 61672519
refGene_gn=LZTS2;refGene_tx=XM_003994330.5;refGene_gn=LZTS2;refGene_tx=XM_006938099.3;refGene_gn=LZTS2;refGene_tx=XM_006938101.3;refGene_gn=LZTS2;refGene_tx=XM_006938103.3;refGene_gn=LZTS2;refGene_tx=XM_0232408

47.1
D2	61673040	61673298
refGene_gn=LZTS2;refGene_tx=XM_003994330.5;refGene_gn=LZTS2;refGene_tx
=XM_006938099.3;refGene_gn=LZTS2;refGene_tx=XM_006938101.3;refGene_gn=
LZTS2;refGene_tx=XM_006938103.3;refGene_gn=LZTS2;refGene_tx=XM_0198137
38.2;refGene_gn=LZTS2;refGene_tx=XM_023240847.1
D2	61674103	61674787
refGene_gn=LZTS2;refGene_tx=XM_003994330.5;refGene_gn=LZTS2;refGene_tx
=XM_006938099.3;refGene_gn=LZTS2;refGene_tx=XM_006938101.3;refGene_gn=
LZTS2;refGene_tx=XM_006938103.3;refGene_gn=LZTS2;refGene_tx=XM_0198137
38.2;refGene_gn=LZTS2;refGene_tx=XM_023240847.1
D2	61676287	61676503	refGene_gn=PDZD7;refGene_tx=XM_006938098.4
D2	61676909	61677010	refGene_gn=PDZD7;refGene_tx=XM_006938098.4
D2	61677304	61677904
refGene_gn=PDZD7;refGene_tx=XM_006938098.4;refGene_gn=PDZD7;refGene_tx
=XM_019813734.2
D2	61678403	61678475
refGene_gn=PDZD7;refGene_tx=XM_006938098.4;refGene_gn=PDZD7;refGene_tx
=XM_019813734.2
D2	61678630	61678722
refGene_gn=PDZD7;refGene_tx=XM_006938098.4;refGene_gn=PDZD7;refGene_tx
=XM_019813734.2
D2	61678832	61678924
refGene_gn=PDZD7;refGene_tx=XM_006938098.4;refGene_gn=PDZD7;refGene_tx
=XM_019813734.2
D2	61681824	61682000
refGene_gn=PDZD7;refGene_tx=XM_006938098.4;refGene_gn=PDZD7;refGene_tx
=XM_019813734.2;refGene_gn=PDZD7;refGene_tx=XM_023240844.1
D2	61682575	61682623
refGene_gn=PDZD7;refGene_tx=XM_006938098.4;refGene_gn=PDZD7;refGene_tx
=XM_019813734.2;refGene_gn=PDZD7;refGene_tx=XM_023240844.1
D2	61684181	61684407
refGene_gn=PDZD7;refGene_tx=XM_006938098.4;refGene_gn=PDZD7;refGene_tx
=XM_019813734.2;refGene_gn=PDZD7;refGene_tx=XM_023240844.1
D2	61684890	61685298
refGene_gn=PDZD7;refGene_tx=XM_006938098.4;refGene_gn=PDZD7;refGene_tx
=XM_019813734.2;refGene_gn=PDZD7;refGene_tx=XM_023240844.1
D2	61685670	61685731
refGene_gn=PDZD7;refGene_tx=XM_006938098.4;refGene_gn=PDZD7;refGene_tx
=XM_019813734.2;refGene_gn=PDZD7;refGene_tx=XM_023240844.1
D2	61687051	61687199
refGene_gn=PDZD7;refGene_tx=XM_006938098.4;refGene_gn=PDZD7;refGene_tx
=XM_019813734.2;refGene_gn=PDZD7;refGene_tx=XM_023240844.1
D2	61687431	61687608
refGene_gn=PDZD7;refGene_tx=XM_006938098.4;refGene_gn=PDZD7;refGene_tx
=XM_019813734.2;refGene_gn=PDZD7;refGene_tx=XM_023240844.1
D2	61688142	61688317
refGene_gn=PDZD7;refGene_tx=XM_006938098.4;refGene_gn=PDZD7;refGene_tx
=XM_019813734.2;refGene_gn=PDZD7;refGene_tx=XM_023240844.1
D2	61688622	61688763

refGene_gn=PDZD7;refGene_tx=XM_006938098.4;refGene_gn=PDZD7;refGene_tx=XM_019813734.2;refGene_gn=PDZD7;refGene_tx=XM_023240844.1
D2 61694421 61694647 refGene_gn=PDZD7;refGene_tx=XM_006938098.4;refGene_gn=PDZD7;refGene_tx=XM_019813734.2;refGene_gn=PDZD7;refGene_tx=XM_023240844.1
D2 61698204 61698365 refGene_gn=SFXN3;refGene_tx=XM_006938104.4;refGene_gn=SFXN3;refGene_tx=XM_019813739.2
D2 61699030 61699201 refGene_gn=SFXN3;refGene_tx=XM_006938104.4;refGene_gn=SFXN3;refGene_tx=XM_019813739.2
D2 61699549 61699648 refGene_gn=SFXN3;refGene_tx=XM_006938104.4;refGene_gn=SFXN3;refGene_tx=XM_019813739.2
D2 61700020 61700096 refGene_gn=SFXN3;refGene_tx=XM_006938104.4;refGene_gn=SFXN3;refGene_tx=XM_019813739.2
D2 61700263 61700349 refGene_gn=SFXN3;refGene_tx=XM_006938104.4;refGene_gn=SFXN3;refGene_tx=XM_019813739.2
D2 61700610 61700738 refGene_gn=SFXN3;refGene_tx=XM_006938104.4;refGene_gn=SFXN3;refGene_tx=XM_019813739.2
D2 61700917 61700967 refGene_gn=SFXN3;refGene_tx=XM_006938104.4;refGene_gn=SFXN3;refGene_tx=XM_019813739.2
D2 61701763 61701813 refGene_gn=SFXN3;refGene_tx=XM_006938104.4;refGene_gn=SFXN3;refGene_tx=XM_019813739.2
D2 61702293 61702341 refGene_gn=SFXN3;refGene_tx=XM_006938104.4;refGene_gn=SFXN3;refGene_tx=XM_019813739.2
D2 61702608 61702705 refGene_gn=SFXN3;refGene_tx=XM_006938104.4;refGene_gn=SFXN3;refGene_tx=XM_019813739.2
D2 61724382 61724887 refGene_gn=KAZALD1;refGene_tx=XM_011287459.2;refGene_gn=KAZALD1;refGene_tx=XM_011287460.3
D2 61726071 61726232 refGene_gn=KAZALD1;refGene_tx=XM_011287459.2;refGene_gn=KAZALD1;refGene_tx=XM_011287460.3
D2 61726313 61726461 refGene_gn=KAZALD1;refGene_tx=XM_011287459.2;refGene_gn=KAZALD1;refGene_tx=XM_011287460.3
D2 61726652 61726747 refGene_gn=KAZALD1;refGene_tx=XM_011287459.2;refGene_gn=KAZALD1;refGene_tx=XM_011287460.3
D2 61786717 61787303 refGene_gn=TLX1;refGene_tx=XM_019814085.2
D2 61789308 61789510 refGene_gn=TLX1;refGene_tx=XM_019814085.2
D2 61791738 61791961 refGene_gn=TLX1;refGene_tx=XM_019814085.2

D2 61852985 61853506 refGene_gn=LBX1;refGene_tx=XM_023240848.1
D2 61854219 61854544 refGene_gn=LBX1;refGene_tx=XM_023240848.1
D2 61861009 61861051
refGene_gn=LOC109492650;refGene_tx=XM_023240298.1
D2 61861692 61862649
refGene_gn=LOC109492650;refGene_tx=XM_023240298.1
D2 61958486 61958534
refGene_gn=BTRC;refGene_tx=XM_003994340.5;refGene_gn=BTRC;refGene_tx=XM_003994341.5;refGene_gn=BTRC;refGene_tx=XM_006938110.4;refGene_gn=BTRC;refGene_tx=XM_006938111.4;refGene_gn=BTRC;refGene_tx=XM_019813740.2;refGene_gn=BTRC;refGene_tx=XM_019813741.2
D2 62021123 62021231
refGene_gn=BTRC;refGene_tx=XM_003994340.5;refGene_gn=BTRC;refGene_tx=XM_006938110.4;refGene_gn=BTRC;refGene_tx=XM_019813740.2;refGene_gn=BTRC;refGene_tx=XM_019813741.2;refGene_gn=BTRC;refGene_tx=XM_023240849.1
D2 62056870 62056948
refGene_gn=BTRC;refGene_tx=XM_003994340.5;refGene_gn=BTRC;refGene_tx=XM_003994341.5;refGene_gn=BTRC;refGene_tx=XM_019813740.2;refGene_gn=BTRC;refGene_tx=XM_019813741.2;refGene_gn=BTRC;refGene_tx=XM_023240849.1
D2 62072444 62072534
refGene_gn=BTRC;refGene_tx=XM_003994340.5;refGene_gn=BTRC;refGene_tx=XM_003994341.5;refGene_gn=BTRC;refGene_tx=XM_006938110.4;refGene_gn=BTRC;refGene_tx=XM_006938111.4;refGene_gn=BTRC;refGene_tx=XM_019813740.2;refGene_gn=BTRC;refGene_tx=XM_019813741.2;refGene_gn=BTRC;refGene_tx=XM_023240849.1
D2 62107938 62108170
refGene_gn=BTRC;refGene_tx=XM_003994340.5;refGene_gn=BTRC;refGene_tx=XM_003994341.5;refGene_gn=BTRC;refGene_tx=XM_006938110.4;refGene_gn=BTRC;refGene_tx=XM_006938111.4;refGene_gn=BTRC;refGene_tx=XM_019813740.2;refGene_gn=BTRC;refGene_tx=XM_019813741.2;refGene_gn=BTRC;refGene_tx=XM_023240849.1
D2 62111672 62111859
refGene_gn=BTRC;refGene_tx=XM_003994340.5;refGene_gn=BTRC;refGene_tx=XM_003994341.5;refGene_gn=BTRC;refGene_tx=XM_006938110.4;refGene_gn=BTRC;refGene_tx=XM_006938111.4;refGene_gn=BTRC;refGene_tx=XM_019813740.2;refGene_gn=BTRC;refGene_tx=XM_019813741.2;refGene_gn=BTRC;refGene_tx=XM_023240849.1
D2 62118334 62118431
refGene_gn=BTRC;refGene_tx=XM_003994340.5;refGene_gn=BTRC;refGene_tx=XM_003994341.5;refGene_gn=BTRC;refGene_tx=XM_006938110.4;refGene_gn=BTRC;refGene_tx=XM_006938111.4;refGene_gn=BTRC;refGene_tx=XM_019813740.2;refGene_gn=BTRC;refGene_tx=XM_019813741.2;refGene_gn=BTRC;refGene_tx=XM_023240849.1
D2 62119078 62119216
refGene_gn=BTRC;refGene_tx=XM_003994340.5;refGene_gn=BTRC;refGene_tx=XM_003994341.5;refGene_gn=BTRC;refGene_tx=XM_006938110.4;refGene_gn=BTRC;refGene_tx=XM_006938111.4;refGene_gn=BTRC;refGene_tx=XM_019813740.2;refGene_gn=BTRC;refGene_tx=XM_019813741.2;refGene_gn=BTRC;refGene_tx=XM_023240849.1
D2 62119743 62119862

refGene_gn=BTRC;refGene_tx=XM_003994340.5;refGene_gn=BTRC;refGene_tx=XM_003994341.5;refGene_gn=BTRC;refGene_tx=XM_006938110.4;refGene_gn=BTRC;refGene_tx=XM_006938111.4;refGene_gn=BTRC;refGene_tx=XM_019813740.2;refGene_gn=BTRC;refGene_tx=XM_019813741.2;refGene_gn=BTRC;refGene_tx=XM_023240849.1
D2 62122003 62122253 refGene_gn=BTRC;refGene_tx=XM_003994340.5;refGene_gn=BTRC;refGene_tx=XM_003994341.5;refGene_gn=BTRC;refGene_tx=XM_006938110.4;refGene_gn=BTRC;refGene_tx=XM_006938111.4;refGene_gn=BTRC;refGene_tx=XM_019813740.2;refGene_gn=BTRC;refGene_tx=XM_019813741.2;refGene_gn=BTRC;refGene_tx=XM_023240849.1
D2 62122765 62122884 refGene_gn=BTRC;refGene_tx=XM_003994340.5;refGene_gn=BTRC;refGene_tx=XM_003994341.5;refGene_gn=BTRC;refGene_tx=XM_006938110.4;refGene_gn=BTRC;refGene_tx=XM_006938111.4;refGene_gn=BTRC;refGene_tx=XM_019813740.2;refGene_gn=BTRC;refGene_tx=XM_019813741.2;refGene_gn=BTRC;refGene_tx=XM_023240849.1
D2 62124469 62124580 refGene_gn=BTRC;refGene_tx=XM_003994340.5;refGene_gn=BTRC;refGene_tx=XM_003994341.5;refGene_gn=BTRC;refGene_tx=XM_006938110.4;refGene_gn=BTRC;refGene_tx=XM_006938111.4;refGene_gn=BTRC;refGene_tx=XM_019813740.2;refGene_gn=BTRC;refGene_tx=XM_019813741.2;refGene_gn=BTRC;refGene_tx=XM_023240849.1
D2 62126486 62126565 refGene_gn=BTRC;refGene_tx=XM_003994340.5;refGene_gn=BTRC;refGene_tx=XM_003994341.5;refGene_gn=BTRC;refGene_tx=XM_006938110.4;refGene_gn=BTRC;refGene_tx=XM_006938111.4;refGene_gn=BTRC;refGene_tx=XM_019813740.2;refGene_gn=BTRC;refGene_tx=XM_019813741.2;refGene_gn=BTRC;refGene_tx=XM_023240849.1
D2 62127075 62127117 refGene_gn=BTRC;refGene_tx=XM_019813740.2
D2 62130468 62130477 refGene_gn=BTRC;refGene_tx=XM_019813741.2
D2 62133312 62133474 refGene_gn=BTRC;refGene_tx=XM_003994340.5;refGene_gn=BTRC;refGene_tx=XM_003994341.5;refGene_gn=BTRC;refGene_tx=XM_006938110.4;refGene_gn=BTRC;refGene_tx=XM_006938111.4;refGene_gn=BTRC;refGene_tx=XM_023240849.1
D2 62160729 62161094 refGene_gn=POLL;refGene_tx=XM_011287453.3;refGene_gn=POLL;refGene_tx=XM_011287454.3
D2 62161543 62161712 refGene_gn=POLL;refGene_tx=XM_011287453.3;refGene_gn=POLL;refGene_tx=XM_011287454.3
D2 62163777 62163906 refGene_gn=POLL;refGene_tx=XM_011287453.3;refGene_gn=POLL;refGene_tx=XM_011287454.3
D2 62164460 62164634 refGene_gn=POLL;refGene_tx=XM_011287453.3;refGene_gn=POLL;refGene_tx=XM_011287454.3
D2 62165398 62165716 refGene_gn=POLL;refGene_tx=XM_011287453.3;refGene_gn=POLL;refGene_tx=XM_011287454.3

D2 62166076 62166239
refGene_gn=POLL;refGene_tx=XM_011287453.3;refGene_gn=POLL;refGene_tx=XM_011287454.3
D2 62166426 62166721
refGene_gn=POLL;refGene_tx=XM_011287453.3;refGene_gn=POLL;refGene_tx=XM_011287454.3
D2 62167224 62167336
refGene_gn=POLL;refGene_tx=XM_011287453.3;refGene_gn=POLL;refGene_tx=XM_011287454.3
D2 62168359 62168423
refGene_gn=DPCD;refGene_tx=XM_003994342.5;refGene_gn=DPCD;refGene_tx=XM_019813743.2
D2 62173126 62173207
refGene_gn=DPCD;refGene_tx=XM_003994342.5;refGene_gn=DPCD;refGene_tx=XM_019813743.2
D2 62177825 62177950
refGene_gn=DPCD;refGene_tx=XM_003994342.5;refGene_gn=DPCD;refGene_tx=XM_019813743.2
D2 62178283 62178417
refGene_gn=DPCD;refGene_tx=XM_003994342.5;refGene_gn=DPCD;refGene_tx=XM_019813743.2
D2 62184985 62185088
refGene_gn=DPCD;refGene_tx=XM_003994342.5;refGene_gn=DPCD;refGene_tx=XM_019813743.2
D2 62185478 62185583
refGene_gn=DPCD;refGene_tx=XM_003994342.5;refGene_gn=DPCD;refGene_tx=XM_019813743.2
D2 62187004 62187124
refGene_gn=FBXW4;refGene_tx=XM_023240850.1;refGene_gn=FBXW4;refGene_tx=XM_023240851.1;refGene_gn=FBXW4;refGene_tx=XM_023240852.1
D2 62187261 62187403
refGene_gn=FBXW4;refGene_tx=XM_023240850.1;refGene_gn=FBXW4;refGene_tx=XM_023240851.1;refGene_gn=FBXW4;refGene_tx=XM_023240852.1
D2 62187817 62187958
refGene_gn=FBXW4;refGene_tx=XM_023240850.1;refGene_gn=FBXW4;refGene_tx=XM_023240851.1;refGene_gn=FBXW4;refGene_tx=XM_023240852.1
D2 62195475 62195540 refGene_gn=DPCD;refGene_tx=XM_019813743.2
D2 62198840 62198906
refGene_gn=FBXW4;refGene_tx=XM_023240850.1;refGene_gn=FBXW4;refGene_tx=XM_023240852.1
D2 62239570 62239665
refGene_gn=FBXW4;refGene_tx=XM_023240850.1;refGene_gn=FBXW4;refGene_tx=XM_023240851.1;refGene_gn=FBXW4;refGene_tx=XM_023240852.1
D2 62244557 62244690
refGene_gn=FBXW4;refGene_tx=XM_023240850.1;refGene_gn=FBXW4;refGene_tx=XM_023240851.1;refGene_gn=FBXW4;refGene_tx=XM_023240852.1
D2 62245152 62245338
refGene_gn=FBXW4;refGene_tx=XM_023240850.1;refGene_gn=FBXW4;refGene_tx=XM_023240851.1;refGene_gn=FBXW4;refGene_tx=XM_023240852.1
D2 62248204 62248300

refGene_gn=FBXW4;refGene_tx=XM_023240850.1;refGene_gn=FBXW4;refGene_tx=XM_023240851.1;refGene_gn=FBXW4;refGene_tx=XM_023240852.1
D2 62266595 62266843 refGene_gn=FBXW4;refGene_tx=XM_023240850.1;refGene_gn=FBXW4;refGene_tx=XM_023240851.1
D2 62324682 62324973 refGene_gn=FGF8;refGene_tx=XM_023240312.1;refGene_gn=FGF8;refGene_tx=XM_023240313.1
D2 62325821 62325928 refGene_gn=FGF8;refGene_tx=XM_023240312.1;refGene_gn=FGF8;refGene_tx=XM_023240313.1
D2 62329142 62329323 refGene_gn=FGF8;refGene_tx=XM_023240312.1;refGene_gn=FGF8;refGene_tx=XM_023240313.1
D2 62330179 62330216 refGene_gn=FGF8;refGene_tx=XM_023240312.1;refGene_gn=FGF8;refGene_tx=XM_023240313.1
D2 62330321 62330353 refGene_gn=FGF8;refGene_tx=XM_023240312.1;refGene_gn=FGF8;refGene_tx=XM_023240313.1
D2 62336202 62336315 refGene_gn=NPM3;refGene_tx=XM_003994343.4
D2 62336693 62336787 refGene_gn=NPM3;refGene_tx=XM_003994343.4
D2 62336923 62337043 refGene_gn=NPM3;refGene_tx=XM_003994343.4
D2 62337297 62337383 refGene_gn=NPM3;refGene_tx=XM_003994343.4
D2 62337682 62337800 refGene_gn=NPM3;refGene_tx=XM_003994343.4
D2 62340866 62341003 refGene_gn=MGEA5;refGene_tx=XM_003994344.5
D2 62343540 62343700 refGene_gn=MGEA5;refGene_tx=XM_003994344.5
D2 62345933 62346126 refGene_gn=MGEA5;refGene_tx=XM_003994344.5
D2 62346405 62346491 refGene_gn=MGEA5;refGene_tx=XM_003994344.5
D2 62347178 62347283 refGene_gn=MGEA5;refGene_tx=XM_003994344.5
D2 62348254 62348340 refGene_gn=MGEA5;refGene_tx=XM_003994344.5
D2 62351740 62351890 refGene_gn=MGEA5;refGene_tx=XM_003994344.5
D2 62351892 62351914 refGene_gn=MGEA5;refGene_tx=XM_003994344.5
D2 62352638 62353252 refGene_gn=MGEA5;refGene_tx=XM_003994344.5
D2 62353706 62353865 refGene_gn=MGEA5;refGene_tx=XM_003994344.5
D2 62356075 62356360 refGene_gn=MGEA5;refGene_tx=XM_003994344.5
D2 62357202 62357301 refGene_gn=MGEA5;refGene_tx=XM_003994344.5
D2 62359844 62360016 refGene_gn=MGEA5;refGene_tx=XM_003994344.5
D2 62361507 62361638 refGene_gn=MGEA5;refGene_tx=XM_003994344.5
D2 62363236 62363334 refGene_gn=MGEA5;refGene_tx=XM_003994344.5
D2 62363764 62363816 refGene_gn=MGEA5;refGene_tx=XM_003994344.5
D2 62367770 62367969 refGene_gn=MGEA5;refGene_tx=XM_003994344.5
D2 62373875 62373911 refGene_gn=KCNIP2;refGene_tx=XM_006938113.4
D2 62374799 62374847 refGene_gn=KCNIP2;refGene_tx=XM_003994346.5;refGene_gn=KCNIP2;refGene_tx=XM_003994347.5;refGene_gn=KCNIP2;refGene_tx=XM_003994348.5;refGene_gn=KCNIP2;refGene_tx=XM_019813747.2
D2 62375147 62375210 refGene_gn=KCNIP2;refGene_tx=XM_003994346.5;refGene_gn=KCNIP2;refGene_tx=XM_003994347.5;refGene_gn=KCNIP2;refGene_tx=XM_003994348.5;refGene_

gn=KCNIP2;refGene_tx=XM_006938113.4;refGene_gn=KCNIP2;refGene_tx=XM_01
9813744.2;refGene_gn=KCNIP2;refGene_tx=XM_019813745.2;refGene_gn=KCNIP
2;refGene_tx=XM_019813747.2
D2 62375329 62375448
refGene_gn=KCNIP2;refGene_tx=XM_003994346.5;refGene_gn=KCNIP2;refGene_
tx=XM_003994347.5;refGene_gn=KCNIP2;refGene_tx=XM_003994348.5;refGene_
gn=KCNIP2;refGene_tx=XM_006938113.4;refGene_gn=KCNIP2;refGene_tx=XM_01
9813744.2;refGene_gn=KCNIP2;refGene_tx=XM_019813745.2;refGene_gn=KCNIP
2;refGene_tx=XM_019813747.2
D2 62375632 62375740
refGene_gn=KCNIP2;refGene_tx=XM_003994346.5;refGene_gn=KCNIP2;refGene_
tx=XM_003994347.5;refGene_gn=KCNIP2;refGene_tx=XM_003994348.5;refGene_
gn=KCNIP2;refGene_tx=XM_006938113.4;refGene_gn=KCNIP2;refGene_tx=XM_01
9813744.2;refGene_gn=KCNIP2;refGene_tx=XM_019813745.2;refGene_gn=KCNIP
2;refGene_tx=XM_019813747.2
D2 62375871 62375942
refGene_gn=KCNIP2;refGene_tx=XM_003994346.5;refGene_gn=KCNIP2;refGene_
tx=XM_003994347.5;refGene_gn=KCNIP2;refGene_tx=XM_003994348.5;refGene_
gn=KCNIP2;refGene_tx=XM_006938113.4;refGene_gn=KCNIP2;refGene_tx=XM_01
9813744.2;refGene_gn=KCNIP2;refGene_tx=XM_019813745.2;refGene_gn=KCNIP
2;refGene_tx=XM_019813747.2
D2 62376100 62376170
refGene_gn=KCNIP2;refGene_tx=XM_003994346.5;refGene_gn=KCNIP2;refGene_
tx=XM_003994347.5;refGene_gn=KCNIP2;refGene_tx=XM_003994348.5;refGene_
gn=KCNIP2;refGene_tx=XM_006938113.4;refGene_gn=KCNIP2;refGene_tx=XM_01
9813744.2;refGene_gn=KCNIP2;refGene_tx=XM_019813745.2;refGene_gn=KCNIP
2;refGene_tx=XM_019813747.2
D2 62376821 62376946
refGene_gn=KCNIP2;refGene_tx=XM_003994346.5;refGene_gn=KCNIP2;refGene_
tx=XM_003994347.5;refGene_gn=KCNIP2;refGene_tx=XM_003994348.5;refGene_
gn=KCNIP2;refGene_tx=XM_006938113.4;refGene_gn=KCNIP2;refGene_tx=XM_01
9813744.2;refGene_gn=KCNIP2;refGene_tx=XM_019813745.2;refGene_gn=KCNIP
2;refGene_tx=XM_019813747.2
D2 62377600 62377654
refGene_gn=KCNIP2;refGene_tx=XM_003994346.5;refGene_gn=KCNIP2;refGene_
tx=XM_006938113.4;refGene_gn=KCNIP2;refGene_tx=XM_019813744.2;refGene_
gn=KCNIP2;refGene_tx=XM_019813745.2
D2 62378564 62378660
refGene_gn=KCNIP2;refGene_tx=XM_003994346.5;refGene_gn=KCNIP2;refGene_
tx=XM_003994348.5;refGene_gn=KCNIP2;refGene_tx=XM_006938113.4;refGene_
gn=KCNIP2;refGene_tx=XM_019813744.2;refGene_gn=KCNIP2;refGene_tx=XM_01
9813745.2
D2 62388167 62388255 refGene_gn=KCNIP2;refGene_tx=XM_019813747.2
D2 62391778 62391851
refGene_gn=KCNIP2;refGene_tx=XM_003994346.5;refGene_gn=KCNIP2;refGene_
tx=XM_003994347.5;refGene_gn=KCNIP2;refGene_tx=XM_003994348.5;refGene_
gn=KCNIP2;refGene_tx=XM_006938113.4;refGene_gn=KCNIP2;refGene_tx=XM_01
9813744.2;refGene_gn=KCNIP2;refGene_tx=XM_019813745.2
D2 62395367 62395460
refGene_gn=CD2H10orf76;refGene_tx=XM_003994351.5

D2 62397704 62397821
refGene_gn=CD2H10orf76;refGene_tx=XM_003994351.5
D2 62429607 62429686
refGene_gn=CD2H10orf76;refGene_tx=XM_003994351.5
D2 62483322 62483398
refGene_gn=CD2H10orf76;refGene_tx=XM_003994351.5
D2 62495540 62495667
refGene_gn=CD2H10orf76;refGene_tx=XM_003994351.5
D2 62496843 62496926
refGene_gn=CD2H10orf76;refGene_tx=XM_003994351.5
D2 62512277 62512366
refGene_gn=CD2H10orf76;refGene_tx=XM_003994351.5
D2 62517726 62517787
refGene_gn=CD2H10orf76;refGene_tx=XM_003994351.5
D2 62518861 62518931
refGene_gn=CD2H10orf76;refGene_tx=XM_003994351.5
D2 62519823 62519889
refGene_gn=CD2H10orf76;refGene_tx=XM_003994351.5
D2 62522552 62522611
refGene_gn=CD2H10orf76;refGene_tx=XM_003994351.5
D2 62526305 62526407
refGene_gn=CD2H10orf76;refGene_tx=XM_003994351.5
D2 62529785 62529879
refGene_gn=CD2H10orf76;refGene_tx=XM_003994351.5
D2 62531467 62531543
refGene_gn=CD2H10orf76;refGene_tx=XM_003994351.5
D2 62532078 62532125
refGene_gn=CD2H10orf76;refGene_tx=XM_003994351.5
D2 62534638 62534699
refGene_gn=CD2H10orf76;refGene_tx=XM_003994351.5
D2 62536513 62536557
refGene_gn=CD2H10orf76;refGene_tx=XM_003994351.5
D2 62537637 62537694
refGene_gn=CD2H10orf76;refGene_tx=XM_003994351.5
D2 62542369 62542456
refGene_gn=CD2H10orf76;refGene_tx=XM_003994351.5
D2 62542575 62542650
refGene_gn=CD2H10orf76;refGene_tx=XM_003994351.5
D2 62543642 62543735
refGene_gn=CD2H10orf76;refGene_tx=XM_003994351.5
D2 62547512 62547620
refGene_gn=CD2H10orf76;refGene_tx=XM_003994351.5
D2 62550738 62550885
refGene_gn=CD2H10orf76;refGene_tx=XM_003994351.5
D2 62551029 62551086
refGene_gn=CD2H10orf76;refGene_tx=XM_003994351.5
D2 62553496 62553598
refGene_gn=CD2H10orf76;refGene_tx=XM_003994351.5
D2 62571569 62573978 refGene_gn=HPS6;refGene_tx=XM_003994352.5
D2 62601610 62601841

refGene_gn=LDB1;refGene_tx=XM_003994353.5;refGene_gn=LDB1;refGene_tx=XM_023240853.1;refGene_gn=LDB1;refGene_tx=XM_023240854.1;refGene_gn=LDB1;refGene_tx=XM_023240855.1
D2 62602571 62602720
refGene_gn=LDB1;refGene_tx=XM_003994353.5;refGene_gn=LDB1;refGene_tx=XM_023240853.1;refGene_gn=LDB1;refGene_tx=XM_023240854.1;refGene_gn=LDB1;refGene_tx=XM_023240855.1
D2 62602924 62603048
refGene_gn=LDB1;refGene_tx=XM_003994353.5;refGene_gn=LDB1;refGene_tx=XM_023240853.1;refGene_gn=LDB1;refGene_tx=XM_023240854.1;refGene_gn=LDB1;refGene_tx=XM_023240855.1
D2 62603140 62603224
refGene_gn=LDB1;refGene_tx=XM_003994353.5;refGene_gn=LDB1;refGene_tx=XM_023240853.1;refGene_gn=LDB1;refGene_tx=XM_023240854.1;refGene_gn=LDB1;refGene_tx=XM_023240855.1
D2 62603456 62603579
refGene_gn=LDB1;refGene_tx=XM_003994353.5;refGene_gn=LDB1;refGene_tx=XM_023240853.1;refGene_gn=LDB1;refGene_tx=XM_023240854.1;refGene_gn=LDB1;refGene_tx=XM_023240855.1
D2 62604064 62604237
refGene_gn=LDB1;refGene_tx=XM_003994353.5;refGene_gn=LDB1;refGene_tx=XM_023240853.1;refGene_gn=LDB1;refGene_tx=XM_023240854.1;refGene_gn=LDB1;refGene_tx=XM_023240855.1
D2 62604410 62604513
refGene_gn=LDB1;refGene_tx=XM_003994353.5;refGene_gn=LDB1;refGene_tx=XM_023240853.1;refGene_gn=LDB1;refGene_tx=XM_023240854.1;refGene_gn=LDB1;refGene_tx=XM_023240855.1
D2 62604615 62604691
refGene_gn=LDB1;refGene_tx=XM_003994353.5;refGene_gn=LDB1;refGene_tx=XM_023240853.1;refGene_gn=LDB1;refGene_tx=XM_023240854.1;refGene_gn=LDB1;refGene_tx=XM_023240855.1
D2 62604810 62604855
refGene_gn=LDB1;refGene_tx=XM_003994353.5;refGene_gn=LDB1;refGene_tx=XM_023240853.1;refGene_gn=LDB1;refGene_tx=XM_023240854.1;refGene_gn=LDB1;refGene_tx=XM_023240855.1
D2 62604977 62605080
refGene_gn=LDB1;refGene_tx=XM_003994353.5;refGene_gn=LDB1;refGene_tx=XM_023240853.1;refGene_gn=LDB1;refGene_tx=XM_023240855.1
D2 62613197 62613222
refGene_gn=LDB1;refGene_tx=XM_023240853.1;refGene_gn=LDB1;refGene_tx=XM_023240855.1
D2 62625439 62625592
refGene_gn=PPRC1;refGene_tx=XM_003994354.5;refGene_gn=PPRC1;refGene_tx=XM_006938118.4;refGene_gn=PPRC1;refGene_tx=XM_006938119.4;refGene_gn=PPRC1;refGene_tx=XM_019813748.2
D2 62629280 62629469
refGene_gn=PPRC1;refGene_tx=XM_003994354.5;refGene_gn=PPRC1;refGene_tx=XM_006938118.4;refGene_gn=PPRC1;refGene_tx=XM_006938119.4;refGene_gn=PPRC1;refGene_tx=XM_019813748.2
D2 62630013 62630160

refGene_gn=PPRC1;refGene_tx=XM_003994354.5;refGene_gn=PPRC1;refGene_tx=XM_006938118.4;refGene_gn=PPRC1;refGene_tx=XM_006938119.4;refGene_gn=PPRC1;refGene_tx=XM_019813748.2
D2 62630274 62630376
refGene_gn=PPRC1;refGene_tx=XM_003994354.5;refGene_gn=PPRC1;refGene_tx=XM_006938118.4;refGene_gn=PPRC1;refGene_tx=XM_006938119.4;refGene_gn=PPRC1;refGene_tx=XM_019813748.2
D2 62630490 62633392
refGene_gn=PPRC1;refGene_tx=XM_003994354.5;refGene_gn=PPRC1;refGene_tx=XM_006938118.4;refGene_gn=PPRC1;refGene_tx=XM_006938119.4;refGene_gn=PPRC1;refGene_tx=XM_019813748.2
D2 62633716 62633770
refGene_gn=PPRC1;refGene_tx=XM_003994354.5;refGene_gn=PPRC1;refGene_tx=XM_006938118.4;refGene_gn=PPRC1;refGene_tx=XM_006938119.4;refGene_gn=PPRC1;refGene_tx=XM_019813748.2
D2 62634601 62634659
refGene_gn=PPRC1;refGene_tx=XM_003994354.5;refGene_gn=PPRC1;refGene_tx=XM_006938118.4;refGene_gn=PPRC1;refGene_tx=XM_006938119.4;refGene_gn=PPRC1;refGene_tx=XM_019813748.2
D2 62635436 62635507
refGene_gn=PPRC1;refGene_tx=XM_003994354.5;refGene_gn=PPRC1;refGene_tx=XM_006938118.4;refGene_gn=PPRC1;refGene_tx=XM_019813748.2
D2 62636473 62637197
refGene_gn=PPRC1;refGene_tx=XM_003994354.5;refGene_gn=PPRC1;refGene_tx=XM_006938118.4;refGene_gn=PPRC1;refGene_tx=XM_019813748.2
D2 62638626 62638773
refGene_gn=PPRC1;refGene_tx=XM_003994354.5;refGene_gn=PPRC1;refGene_tx=XM_006938118.4;refGene_gn=PPRC1;refGene_tx=XM_006938119.4;refGene_gn=PPRC1;refGene_tx=XM_019813748.2
D2 62638888 62638955
refGene_gn=PPRC1;refGene_tx=XM_003994354.5;refGene_gn=PPRC1;refGene_tx=XM_006938118.4;refGene_gn=PPRC1;refGene_tx=XM_006938119.4;refGene_gn=PPRC1;refGene_tx=XM_019813748.2
D2 62639072 62639194
refGene_gn=PPRC1;refGene_tx=XM_003994354.5;refGene_gn=PPRC1;refGene_tx=XM_006938118.4;refGene_gn=PPRC1;refGene_tx=XM_006938119.4;refGene_gn=PPRC1;refGene_tx=XM_019813748.2
D2 62639444 62639596
refGene_gn=PPRC1;refGene_tx=XM_003994354.5;refGene_gn=PPRC1;refGene_tx=XM_006938118.4;refGene_gn=PPRC1;refGene_tx=XM_006938119.4;refGene_gn=PPRC1;refGene_tx=XM_019813748.2
D2 62639882 62639986
refGene_gn=PPRC1;refGene_tx=XM_003994354.5;refGene_gn=PPRC1;refGene_tx=XM_006938118.4;refGene_gn=PPRC1;refGene_tx=XM_006938119.4;refGene_gn=PPRC1;refGene_tx=XM_019813748.2
D2 62641802 62641922
refGene_gn=NOLC1;refGene_tx=XM_006938120.4;refGene_gn=NOLC1;refGene_tx=XM_006938121.4
D2 62644089 62644145
refGene_gn=NOLC1;refGene_tx=XM_006938120.4;refGene_gn=NOLC1;refGene_tx

=XM_006938121.4
D2	62644254	62644397
refGene_gn=NOLC1;refGene_tx=XM_006938120.4;refGene_gn=NOLC1;refGene_tx
=XM_006938121.4
D2	62644499	62644627
refGene_gn=NOLC1;refGene_tx=XM_006938120.4;refGene_gn=NOLC1;refGene_tx
=XM_006938121.4
D2	62645275	62645441
refGene_gn=NOLC1;refGene_tx=XM_006938120.4;refGene_gn=NOLC1;refGene_tx
=XM_006938121.4
D2	62646696	62646821
refGene_gn=NOLC1;refGene_tx=XM_006938120.4;refGene_gn=NOLC1;refGene_tx
=XM_006938121.4
D2	62646940	62647076
refGene_gn=NOLC1;refGene_tx=XM_006938120.4;refGene_gn=NOLC1;refGene_tx
=XM_006938121.4
D2	62647397	62647526
refGene_gn=NOLC1;refGene_tx=XM_006938120.4;refGene_gn=NOLC1;refGene_tx
=XM_006938121.4
D2	62647718	62647829
refGene_gn=NOLC1;refGene_tx=XM_006938120.4;refGene_gn=NOLC1;refGene_tx
=XM_006938121.4
D2	62647911	62648553
refGene_gn=NOLC1;refGene_tx=XM_006938120.4;refGene_gn=NOLC1;refGene_tx
=XM_006938121.4
D2	62648705	62648812
refGene_gn=NOLC1;refGene_tx=XM_006938120.4;refGene_gn=NOLC1;refGene_tx
=XM_006938121.4
D2	62648988	62649081
refGene_gn=NOLC1;refGene_tx=XM_006938120.4;refGene_gn=NOLC1;refGene_tx
=XM_006938121.4
D2	62649258	62649417
refGene_gn=NOLC1;refGene_tx=XM_006938120.4;refGene_gn=NOLC1;refGene_tx
=XM_006938121.4
D2	62699330	62699431	refGene_gn=ELOVL3;refGene_tx=XM_003994357.4
D2	62700415	62700547	refGene_gn=ELOVL3;refGene_tx=XM_003994357.4
D2	62701424	62701576	refGene_gn=ELOVL3;refGene_tx=XM_003994357.4
D2	62701830	62702258	refGene_gn=ELOVL3;refGene_tx=XM_003994357.4
D2	62703532	62704120
refGene_gn=PITX3;refGene_tx=XM_011287448.3;refGene_gn=PITX3;refGene_tx
=XM_011287449.3
D2	62704680	62704883
refGene_gn=PITX3;refGene_tx=XM_011287448.3;refGene_gn=PITX3;refGene_tx
=XM_011287449.3
D2	62705060	62705178
refGene_gn=PITX3;refGene_tx=XM_011287448.3;refGene_gn=PITX3;refGene_tx
=XM_011287449.3
D2	62730176	62730272
refGene_gn=GBF1;refGene_tx=XM_003994358.5;refGene_gn=GBF1;refGene_tx=X
M_006938122.4;refGene_gn=GBF1;refGene_tx=XM_006938123.4;refGene_gn=GBF

1;refGene_tx=XM_006938124.4;refGene_gn=GBF1;refGene_tx=XM_006938125.4;
refGene_gn=GBF1;refGene_tx=XM_006938126.4;refGene_gn=GBF1;refGene_tx=X
M_006938128.4;refGene_gn=GBF1;refGene_tx=XM_006938129.4;refGene_gn=GBF
1;refGene_tx=XM_006938130.4;refGene_gn=GBF1;refGene_tx=XM_019813751.2;
refGene_gn=GBF1;refGene_tx=XM_019813752.2;refGene_gn=GBF1;refGene_tx=X
M_023240856.1
D2 62732503 62732570
refGene_gn=GBF1;refGene_tx=XM_003994358.5;refGene_gn=GBF1;refGene_tx=X
M_006938122.4;refGene_gn=GBF1;refGene_tx=XM_006938123.4;refGene_gn=GBF
1;refGene_tx=XM_006938124.4;refGene_gn=GBF1;refGene_tx=XM_006938125.4;
refGene_gn=GBF1;refGene_tx=XM_006938126.4;refGene_gn=GBF1;refGene_tx=X
M_006938128.4;refGene_gn=GBF1;refGene_tx=XM_006938129.4;refGene_gn=GBF
1;refGene_tx=XM_006938130.4;refGene_gn=GBF1;refGene_tx=XM_019813751.2;
refGene_gn=GBF1;refGene_tx=XM_019813752.2;refGene_gn=GBF1;refGene_tx=X
M_023240856.1
D2 62805014 62805146
refGene_gn=GBF1;refGene_tx=XM_003994358.5;refGene_gn=GBF1;refGene_tx=X
M_006938122.4;refGene_gn=GBF1;refGene_tx=XM_006938123.4;refGene_gn=GBF
1;refGene_tx=XM_006938124.4;refGene_gn=GBF1;refGene_tx=XM_006938125.4;
refGene_gn=GBF1;refGene_tx=XM_006938126.4;refGene_gn=GBF1;refGene_tx=X
M_006938128.4;refGene_gn=GBF1;refGene_tx=XM_006938129.4;refGene_gn=GBF
1;refGene_tx=XM_006938130.4;refGene_gn=GBF1;refGene_tx=XM_019813751.2;
refGene_gn=GBF1;refGene_tx=XM_019813752.2;refGene_gn=GBF1;refGene_tx=X
M_023240856.1
D2 62810054 62810173
refGene_gn=GBF1;refGene_tx=XM_003994358.5;refGene_gn=GBF1;refGene_tx=X
M_006938122.4;refGene_gn=GBF1;refGene_tx=XM_006938123.4;refGene_gn=GBF
1;refGene_tx=XM_006938124.4;refGene_gn=GBF1;refGene_tx=XM_006938125.4;
refGene_gn=GBF1;refGene_tx=XM_006938126.4;refGene_gn=GBF1;refGene_tx=X
M_006938128.4;refGene_gn=GBF1;refGene_tx=XM_006938129.4;refGene_gn=GBF
1;refGene_tx=XM_006938130.4;refGene_gn=GBF1;refGene_tx=XM_019813751.2;
refGene_gn=GBF1;refGene_tx=XM_019813752.2;refGene_gn=GBF1;refGene_tx=X
M_023240856.1
D2 62810706 62810815
refGene_gn=GBF1;refGene_tx=XM_003994358.5;refGene_gn=GBF1;refGene_tx=X
M_006938122.4;refGene_gn=GBF1;refGene_tx=XM_006938123.4;refGene_gn=GBF
1;refGene_tx=XM_006938124.4;refGene_gn=GBF1;refGene_tx=XM_006938125.4;
refGene_gn=GBF1;refGene_tx=XM_006938126.4;refGene_gn=GBF1;refGene_tx=X
M_006938128.4;refGene_gn=GBF1;refGene_tx=XM_006938129.4;refGene_gn=GBF
1;refGene_tx=XM_006938130.4;refGene_gn=GBF1;refGene_tx=XM_019813751.2;
refGene_gn=GBF1;refGene_tx=XM_019813752.2;refGene_gn=GBF1;refGene_tx=X
M_023240856.1
D2 62811343 62811404
refGene_gn=GBF1;refGene_tx=XM_003994358.5;refGene_gn=GBF1;refGene_tx=X
M_006938122.4;refGene_gn=GBF1;refGene_tx=XM_006938123.4;refGene_gn=GBF
1;refGene_tx=XM_006938124.4;refGene_gn=GBF1;refGene_tx=XM_006938125.4;
refGene_gn=GBF1;refGene_tx=XM_006938126.4;refGene_gn=GBF1;refGene_tx=X
M_006938128.4;refGene_gn=GBF1;refGene_tx=XM_006938129.4;refGene_gn=GBF
1;refGene_tx=XM_006938130.4;refGene_gn=GBF1;refGene_tx=XM_019813751.2;
refGene_gn=GBF1;refGene_tx=XM_019813752.2;refGene_gn=GBF1;refGene_tx=X

M_023240856.1
D2	62812683 62812738
refGene_gn=GBF1;refGene_tx=XM_003994358.5;refGene_gn=GBF1;refGene_tx=XM_006938122.4;refGene_gn=GBF1;refGene_tx=XM_006938123.4;refGene_gn=GBF1;refGene_tx=XM_006938124.4;refGene_gn=GBF1;refGene_tx=XM_006938125.4;refGene_gn=GBF1;refGene_tx=XM_006938126.4;refGene_gn=GBF1;refGene_tx=XM_006938128.4;refGene_gn=GBF1;refGene_tx=XM_006938129.4;refGene_gn=GBF1;refGene_tx=XM_006938130.4;refGene_gn=GBF1;refGene_tx=XM_019813751.2;refGene_gn=GBF1;refGene_tx=XM_019813752.2;refGene_gn=GBF1;refGene_tx=XM_023240856.1
D2	62813467 62813566
refGene_gn=GBF1;refGene_tx=XM_006938122.4;refGene_gn=GBF1;refGene_tx=XM_006938123.4;refGene_gn=GBF1;refGene_tx=XM_006938124.4;refGene_gn=GBF1;refGene_tx=XM_006938125.4;refGene_gn=GBF1;refGene_tx=XM_006938126.4;refGene_gn=GBF1;refGene_tx=XM_019813751.2;refGene_gn=GBF1;refGene_tx=XM_019813752.2;refGene_gn=GBF1;refGene_tx=XM_023240856.1
D2	62816448 62816596
refGene_gn=GBF1;refGene_tx=XM_003994358.5;refGene_gn=GBF1;refGene_tx=XM_006938122.4;refGene_gn=GBF1;refGene_tx=XM_006938123.4;refGene_gn=GBF1;refGene_tx=XM_006938124.4;refGene_gn=GBF1;refGene_tx=XM_006938125.4;refGene_gn=GBF1;refGene_tx=XM_006938126.4;refGene_gn=GBF1;refGene_tx=XM_006938128.4;refGene_gn=GBF1;refGene_tx=XM_006938129.4;refGene_gn=GBF1;refGene_tx=XM_006938130.4;refGene_gn=GBF1;refGene_tx=XM_019813751.2;refGene_gn=GBF1;refGene_tx=XM_019813752.2;refGene_gn=GBF1;refGene_tx=XM_019813753.2;refGene_gn=GBF1;refGene_tx=XM_023240856.1
D2	62816929 62817153
refGene_gn=GBF1;refGene_tx=XM_003994358.5;refGene_gn=GBF1;refGene_tx=XM_006938122.4;refGene_gn=GBF1;refGene_tx=XM_006938123.4;refGene_gn=GBF1;refGene_tx=XM_006938124.4;refGene_gn=GBF1;refGene_tx=XM_006938125.4;refGene_gn=GBF1;refGene_tx=XM_006938126.4;refGene_gn=GBF1;refGene_tx=XM_006938128.4;refGene_gn=GBF1;refGene_tx=XM_006938129.4;refGene_gn=GBF1;refGene_tx=XM_006938130.4;refGene_gn=GBF1;refGene_tx=XM_019813751.2;refGene_gn=GBF1;refGene_tx=XM_019813752.2;refGene_gn=GBF1;refGene_tx=XM_019813753.2;refGene_gn=GBF1;refGene_tx=XM_023240856.1
D2	62817682 62817851
refGene_gn=GBF1;refGene_tx=XM_003994358.5;refGene_gn=GBF1;refGene_tx=XM_006938122.4;refGene_gn=GBF1;refGene_tx=XM_006938123.4;refGene_gn=GBF1;refGene_tx=XM_006938124.4;refGene_gn=GBF1;refGene_tx=XM_006938125.4;refGene_gn=GBF1;refGene_tx=XM_006938126.4;refGene_gn=GBF1;refGene_tx=XM_006938128.4;refGene_gn=GBF1;refGene_tx=XM_006938129.4;refGene_gn=GBF1;refGene_tx=XM_006938130.4;refGene_gn=GBF1;refGene_tx=XM_019813751.2;refGene_gn=GBF1;refGene_tx=XM_019813752.2;refGene_gn=GBF1;refGene_tx=XM_019813753.2;refGene_gn=GBF1;refGene_tx=XM_023240856.1
D2	62818285 62818497
refGene_gn=GBF1;refGene_tx=XM_003994358.5;refGene_gn=GBF1;refGene_tx=XM_006938122.4;refGene_gn=GBF1;refGene_tx=XM_006938123.4;refGene_gn=GBF1;refGene_tx=XM_006938124.4;refGene_gn=GBF1;refGene_tx=XM_006938125.4;refGene_gn=GBF1;refGene_tx=XM_006938126.4;refGene_gn=GBF1;refGene_tx=XM_006938128.4;refGene_gn=GBF1;refGene_tx=XM_006938129.4;refGene_gn=GBF1;refGene_tx=XM_006938130.4;refGene_gn=GBF1;refGene_tx=XM_019813751.2;

refGene_gn=GBF1;refGene_tx=XM_019813752.2;refGene_gn=GBF1;refGene_tx=XM_019813753.2;refGene_gn=GBF1;refGene_tx=XM_023240856.1
D2 62818834 62818933
refGene_gn=GBF1;refGene_tx=XM_003994358.5;refGene_gn=GBF1;refGene_tx=XM_006938122.4;refGene_gn=GBF1;refGene_tx=XM_006938123.4;refGene_gn=GBF1;refGene_tx=XM_006938124.4;refGene_gn=GBF1;refGene_tx=XM_006938125.4;refGene_gn=GBF1;refGene_tx=XM_006938126.4;refGene_gn=GBF1;refGene_tx=XM_006938128.4;refGene_gn=GBF1;refGene_tx=XM_006938129.4;refGene_gn=GBF1;refGene_tx=XM_006938130.4;refGene_gn=GBF1;refGene_tx=XM_019813751.2;refGene_gn=GBF1;refGene_tx=XM_019813752.2;refGene_gn=GBF1;refGene_tx=XM_019813753.2;refGene_gn=GBF1;refGene_tx=XM_023240856.1
D2 62819520 62819715
refGene_gn=GBF1;refGene_tx=XM_003994358.5;refGene_gn=GBF1;refGene_tx=XM_006938122.4;refGene_gn=GBF1;refGene_tx=XM_006938123.4;refGene_gn=GBF1;refGene_tx=XM_006938124.4;refGene_gn=GBF1;refGene_tx=XM_006938125.4;refGene_gn=GBF1;refGene_tx=XM_006938126.4;refGene_gn=GBF1;refGene_tx=XM_006938128.4;refGene_gn=GBF1;refGene_tx=XM_006938129.4;refGene_gn=GBF1;refGene_tx=XM_006938130.4;refGene_gn=GBF1;refGene_tx=XM_019813751.2;refGene_gn=GBF1;refGene_tx=XM_019813752.2;refGene_gn=GBF1;refGene_tx=XM_019813753.2;refGene_gn=GBF1;refGene_tx=XM_023240856.1
D2 62819959 62820149
refGene_gn=GBF1;refGene_tx=XM_003994358.5;refGene_gn=GBF1;refGene_tx=XM_006938122.4;refGene_gn=GBF1;refGene_tx=XM_006938123.4;refGene_gn=GBF1;refGene_tx=XM_006938124.4;refGene_gn=GBF1;refGene_tx=XM_006938125.4;refGene_gn=GBF1;refGene_tx=XM_006938126.4;refGene_gn=GBF1;refGene_tx=XM_006938128.4;refGene_gn=GBF1;refGene_tx=XM_006938129.4;refGene_gn=GBF1;refGene_tx=XM_006938130.4;refGene_gn=GBF1;refGene_tx=XM_019813751.2;refGene_gn=GBF1;refGene_tx=XM_019813752.2;refGene_gn=GBF1;refGene_tx=XM_019813753.2;refGene_gn=GBF1;refGene_tx=XM_023240856.1
D2 62821562 62821703
refGene_gn=GBF1;refGene_tx=XM_003994358.5;refGene_gn=GBF1;refGene_tx=XM_006938122.4;refGene_gn=GBF1;refGene_tx=XM_006938123.4;refGene_gn=GBF1;refGene_tx=XM_006938124.4;refGene_gn=GBF1;refGene_tx=XM_006938125.4;refGene_gn=GBF1;refGene_tx=XM_006938126.4;refGene_gn=GBF1;refGene_tx=XM_006938128.4;refGene_gn=GBF1;refGene_tx=XM_006938129.4;refGene_gn=GBF1;refGene_tx=XM_006938130.4;refGene_gn=GBF1;refGene_tx=XM_019813751.2;refGene_gn=GBF1;refGene_tx=XM_019813752.2;refGene_gn=GBF1;refGene_tx=XM_019813753.2;refGene_gn=GBF1;refGene_tx=XM_023240856.1
D2 62822018 62822107
refGene_gn=GBF1;refGene_tx=XM_003994358.5;refGene_gn=GBF1;refGene_tx=XM_006938122.4;refGene_gn=GBF1;refGene_tx=XM_006938123.4;refGene_gn=GBF1;refGene_tx=XM_006938124.4;refGene_gn=GBF1;refGene_tx=XM_006938125.4;refGene_gn=GBF1;refGene_tx=XM_006938126.4;refGene_gn=GBF1;refGene_tx=XM_006938128.4;refGene_gn=GBF1;refGene_tx=XM_006938129.4;refGene_gn=GBF1;refGene_tx=XM_006938130.4;refGene_gn=GBF1;refGene_tx=XM_019813751.2;refGene_gn=GBF1;refGene_tx=XM_019813752.2;refGene_gn=GBF1;refGene_tx=XM_019813753.2;refGene_gn=GBF1;refGene_tx=XM_023240856.1
D2 62823292 62823495
refGene_gn=GBF1;refGene_tx=XM_003994358.5;refGene_gn=GBF1;refGene_tx=XM_006938122.4;refGene_gn=GBF1;refGene_tx=XM_006938123.4;refGene_gn=GBF

1;refGene_tx=XM_006938124.4;refGene_gn=GBF1;refGene_tx=XM_006938125.4;
refGene_gn=GBF1;refGene_tx=XM_006938126.4;refGene_gn=GBF1;refGene_tx=X
M_006938128.4;refGene_gn=GBF1;refGene_tx=XM_006938129.4;refGene_gn=GBF
1;refGene_tx=XM_006938130.4;refGene_gn=GBF1;refGene_tx=XM_019813751.2;
refGene_gn=GBF1;refGene_tx=XM_019813752.2;refGene_gn=GBF1;refGene_tx=X
M_019813753.2;refGene_gn=GBF1;refGene_tx=XM_023240856.1
D2 62823910 62824034
refGene_gn=GBF1;refGene_tx=XM_003994358.5;refGene_gn=GBF1;refGene_tx=X
M_006938122.4;refGene_gn=GBF1;refGene_tx=XM_006938123.4;refGene_gn=GBF
1;refGene_tx=XM_006938124.4;refGene_gn=GBF1;refGene_tx=XM_006938125.4;
refGene_gn=GBF1;refGene_tx=XM_006938126.4;refGene_gn=GBF1;refGene_tx=X
M_006938128.4;refGene_gn=GBF1;refGene_tx=XM_006938129.4;refGene_gn=GBF
1;refGene_tx=XM_006938130.4;refGene_gn=GBF1;refGene_tx=XM_019813751.2;
refGene_gn=GBF1;refGene_tx=XM_019813752.2;refGene_gn=GBF1;refGene_tx=X
M_019813753.2;refGene_gn=GBF1;refGene_tx=XM_023240856.1
D2 62824299 62824425
refGene_gn=GBF1;refGene_tx=XM_003994358.5;refGene_gn=GBF1;refGene_tx=X
M_006938122.4;refGene_gn=GBF1;refGene_tx=XM_006938123.4;refGene_gn=GBF
1;refGene_tx=XM_006938124.4;refGene_gn=GBF1;refGene_tx=XM_006938125.4;
refGene_gn=GBF1;refGene_tx=XM_006938126.4;refGene_gn=GBF1;refGene_tx=X
M_006938128.4;refGene_gn=GBF1;refGene_tx=XM_006938129.4;refGene_gn=GBF
1;refGene_tx=XM_006938130.4;refGene_gn=GBF1;refGene_tx=XM_019813751.2;
refGene_gn=GBF1;refGene_tx=XM_019813752.2;refGene_gn=GBF1;refGene_tx=X
M_019813753.2;refGene_gn=GBF1;refGene_tx=XM_023240856.1
D2 62824695 62824778
refGene_gn=GBF1;refGene_tx=XM_003994358.5;refGene_gn=GBF1;refGene_tx=X
M_006938122.4;refGene_gn=GBF1;refGene_tx=XM_006938123.4;refGene_gn=GBF
1;refGene_tx=XM_006938124.4;refGene_gn=GBF1;refGene_tx=XM_006938125.4;
refGene_gn=GBF1;refGene_tx=XM_006938126.4;refGene_gn=GBF1;refGene_tx=X
M_006938128.4;refGene_gn=GBF1;refGene_tx=XM_006938129.4;refGene_gn=GBF
1;refGene_tx=XM_006938130.4;refGene_gn=GBF1;refGene_tx=XM_019813751.2;
refGene_gn=GBF1;refGene_tx=XM_019813752.2;refGene_gn=GBF1;refGene_tx=X
M_019813753.2;refGene_gn=GBF1;refGene_tx=XM_023240856.1
D2 62825447 62825684
refGene_gn=GBF1;refGene_tx=XM_003994358.5;refGene_gn=GBF1;refGene_tx=X
M_006938122.4;refGene_gn=GBF1;refGene_tx=XM_006938123.4;refGene_gn=GBF
1;refGene_tx=XM_006938124.4;refGene_gn=GBF1;refGene_tx=XM_006938125.4;
refGene_gn=GBF1;refGene_tx=XM_006938126.4;refGene_gn=GBF1;refGene_tx=X
M_006938128.4;refGene_gn=GBF1;refGene_tx=XM_006938129.4;refGene_gn=GBF
1;refGene_tx=XM_006938130.4;refGene_gn=GBF1;refGene_tx=XM_019813751.2;
refGene_gn=GBF1;refGene_tx=XM_019813752.2;refGene_gn=GBF1;refGene_tx=X
M_019813753.2;refGene_gn=GBF1;refGene_tx=XM_023240856.1
D2 62825972 62826066
refGene_gn=GBF1;refGene_tx=XM_003994358.5;refGene_gn=GBF1;refGene_tx=X
M_006938122.4;refGene_gn=GBF1;refGene_tx=XM_006938123.4;refGene_gn=GBF
1;refGene_tx=XM_006938124.4;refGene_gn=GBF1;refGene_tx=XM_006938125.4;
refGene_gn=GBF1;refGene_tx=XM_006938126.4;refGene_gn=GBF1;refGene_tx=X
M_006938128.4;refGene_gn=GBF1;refGene_tx=XM_006938129.4;refGene_gn=GBF
1;refGene_tx=XM_006938130.4;refGene_gn=GBF1;refGene_tx=XM_019813751.2;
refGene_gn=GBF1;refGene_tx=XM_019813752.2;refGene_gn=GBF1;refGene_tx=X

M_019813753.2;refGene_gn=GBF1;refGene_tx=XM_023240856.1
D2 62826477 62826654
refGene_gn=GBF1;refGene_tx=XM_003994358.5;refGene_gn=GBF1;refGene_tx=XM_006938122.4;refGene_gn=GBF1;refGene_tx=XM_006938123.4;refGene_gn=GBF1;refGene_tx=XM_006938124.4;refGene_gn=GBF1;refGene_tx=XM_006938125.4;refGene_gn=GBF1;refGene_tx=XM_006938126.4;refGene_gn=GBF1;refGene_tx=XM_006938128.4;refGene_gn=GBF1;refGene_tx=XM_006938129.4;refGene_gn=GBF1;refGene_tx=XM_006938130.4;refGene_gn=GBF1;refGene_tx=XM_019813751.2;refGene_gn=GBF1;refGene_tx=XM_019813752.2;refGene_gn=GBF1;refGene_tx=XM_019813753.2;refGene_gn=GBF1;refGene_tx=XM_023240856.1
D2 62826976 62827041
refGene_gn=GBF1;refGene_tx=XM_003994358.5;refGene_gn=GBF1;refGene_tx=XM_006938122.4;refGene_gn=GBF1;refGene_tx=XM_006938123.4;refGene_gn=GBF1;refGene_tx=XM_006938124.4;refGene_gn=GBF1;refGene_tx=XM_006938125.4;refGene_gn=GBF1;refGene_tx=XM_006938126.4;refGene_gn=GBF1;refGene_tx=XM_006938128.4;refGene_gn=GBF1;refGene_tx=XM_006938129.4;refGene_gn=GBF1;refGene_tx=XM_006938130.4;refGene_gn=GBF1;refGene_tx=XM_019813751.2;refGene_gn=GBF1;refGene_tx=XM_019813752.2;refGene_gn=GBF1;refGene_tx=XM_019813753.2;refGene_gn=GBF1;refGene_tx=XM_023240856.1
D2 62827124 62827248
refGene_gn=GBF1;refGene_tx=XM_003994358.5;refGene_gn=GBF1;refGene_tx=XM_006938122.4;refGene_gn=GBF1;refGene_tx=XM_006938123.4;refGene_gn=GBF1;refGene_tx=XM_006938124.4;refGene_gn=GBF1;refGene_tx=XM_006938125.4;refGene_gn=GBF1;refGene_tx=XM_006938126.4;refGene_gn=GBF1;refGene_tx=XM_006938128.4;refGene_gn=GBF1;refGene_tx=XM_006938129.4;refGene_gn=GBF1;refGene_tx=XM_006938130.4;refGene_gn=GBF1;refGene_tx=XM_019813751.2;refGene_gn=GBF1;refGene_tx=XM_019813752.2;refGene_gn=GBF1;refGene_tx=XM_019813753.2;refGene_gn=GBF1;refGene_tx=XM_023240856.1
D2 62827447 62827519
refGene_gn=GBF1;refGene_tx=XM_003994358.5;refGene_gn=GBF1;refGene_tx=XM_006938122.4;refGene_gn=GBF1;refGene_tx=XM_006938123.4;refGene_gn=GBF1;refGene_tx=XM_006938124.4;refGene_gn=GBF1;refGene_tx=XM_006938125.4;refGene_gn=GBF1;refGene_tx=XM_006938126.4;refGene_gn=GBF1;refGene_tx=XM_006938128.4;refGene_gn=GBF1;refGene_tx=XM_006938129.4;refGene_gn=GBF1;refGene_tx=XM_006938130.4;refGene_gn=GBF1;refGene_tx=XM_019813751.2;refGene_gn=GBF1;refGene_tx=XM_019813752.2;refGene_gn=GBF1;refGene_tx=XM_019813753.2;refGene_gn=GBF1;refGene_tx=XM_023240856.1
D2 62827664 62827759
refGene_gn=GBF1;refGene_tx=XM_003994358.5;refGene_gn=GBF1;refGene_tx=XM_006938122.4;refGene_gn=GBF1;refGene_tx=XM_006938123.4;refGene_gn=GBF1;refGene_tx=XM_006938124.4;refGene_gn=GBF1;refGene_tx=XM_006938125.4;refGene_gn=GBF1;refGene_tx=XM_006938126.4;refGene_gn=GBF1;refGene_tx=XM_006938128.4;refGene_gn=GBF1;refGene_tx=XM_006938129.4;refGene_gn=GBF1;refGene_tx=XM_006938130.4;refGene_gn=GBF1;refGene_tx=XM_019813751.2;refGene_gn=GBF1;refGene_tx=XM_019813752.2;refGene_gn=GBF1;refGene_tx=XM_019813753.2;refGene_gn=GBF1;refGene_tx=XM_023240856.1
D2 62828018 62828172
refGene_gn=GBF1;refGene_tx=XM_003994358.5;refGene_gn=GBF1;refGene_tx=XM_006938122.4;refGene_gn=GBF1;refGene_tx=XM_006938123.4;refGene_gn=GBF1;refGene_tx=XM_006938124.4;refGene_gn=GBF1;refGene_tx=XM_006938125.4;

refGene_gn=GBF1;refGene_tx=XM_006938126.4;refGene_gn=GBF1;refGene_tx=XM_006938128.4;refGene_gn=GBF1;refGene_tx=XM_006938129.4;refGene_gn=GBF1;refGene_tx=XM_006938130.4;refGene_gn=GBF1;refGene_tx=XM_019813751.2;refGene_gn=GBF1;refGene_tx=XM_019813752.2;refGene_gn=GBF1;refGene_tx=XM_019813753.2;refGene_gn=GBF1;refGene_tx=XM_023240856.1
D2 62831374 62831600
refGene_gn=GBF1;refGene_tx=XM_003994358.5;refGene_gn=GBF1;refGene_tx=XM_006938122.4;refGene_gn=GBF1;refGene_tx=XM_006938123.4;refGene_gn=GBF1;refGene_tx=XM_006938124.4;refGene_gn=GBF1;refGene_tx=XM_006938125.4;refGene_gn=GBF1;refGene_tx=XM_006938126.4;refGene_gn=GBF1;refGene_tx=XM_006938128.4;refGene_gn=GBF1;refGene_tx=XM_006938129.4;refGene_gn=GBF1;refGene_tx=XM_006938130.4;refGene_gn=GBF1;refGene_tx=XM_019813751.2;refGene_gn=GBF1;refGene_tx=XM_019813752.2;refGene_gn=GBF1;refGene_tx=XM_019813753.2;refGene_gn=GBF1;refGene_tx=XM_023240856.1
D2 62832364 62832525
refGene_gn=GBF1;refGene_tx=XM_003994358.5;refGene_gn=GBF1;refGene_tx=XM_006938122.4;refGene_gn=GBF1;refGene_tx=XM_006938123.4;refGene_gn=GBF1;refGene_tx=XM_006938124.4;refGene_gn=GBF1;refGene_tx=XM_006938125.4;refGene_gn=GBF1;refGene_tx=XM_006938126.4;refGene_gn=GBF1;refGene_tx=XM_006938128.4;refGene_gn=GBF1;refGene_tx=XM_006938129.4;refGene_gn=GBF1;refGene_tx=XM_006938130.4;refGene_gn=GBF1;refGene_tx=XM_019813751.2;refGene_gn=GBF1;refGene_tx=XM_019813752.2;refGene_gn=GBF1;refGene_tx=XM_019813753.2;refGene_gn=GBF1;refGene_tx=XM_023240856.1
D2 62832659 62832906
refGene_gn=GBF1;refGene_tx=XM_003994358.5;refGene_gn=GBF1;refGene_tx=XM_006938122.4;refGene_gn=GBF1;refGene_tx=XM_006938123.4;refGene_gn=GBF1;refGene_tx=XM_006938124.4;refGene_gn=GBF1;refGene_tx=XM_006938125.4;refGene_gn=GBF1;refGene_tx=XM_006938126.4;refGene_gn=GBF1;refGene_tx=XM_006938128.4;refGene_gn=GBF1;refGene_tx=XM_006938129.4;refGene_gn=GBF1;refGene_tx=XM_006938130.4;refGene_gn=GBF1;refGene_tx=XM_019813751.2;refGene_gn=GBF1;refGene_tx=XM_019813752.2;refGene_gn=GBF1;refGene_tx=XM_019813753.2;refGene_gn=GBF1;refGene_tx=XM_023240856.1
D2 62833030 62833236
refGene_gn=GBF1;refGene_tx=XM_003994358.5;refGene_gn=GBF1;refGene_tx=XM_006938122.4;refGene_gn=GBF1;refGene_tx=XM_006938123.4;refGene_gn=GBF1;refGene_tx=XM_006938124.4;refGene_gn=GBF1;refGene_tx=XM_006938125.4;refGene_gn=GBF1;refGene_tx=XM_006938126.4;refGene_gn=GBF1;refGene_tx=XM_006938128.4;refGene_gn=GBF1;refGene_tx=XM_006938129.4;refGene_gn=GBF1;refGene_tx=XM_006938130.4;refGene_gn=GBF1;refGene_tx=XM_019813751.2;refGene_gn=GBF1;refGene_tx=XM_019813752.2;refGene_gn=GBF1;refGene_tx=XM_019813753.2;refGene_gn=GBF1;refGene_tx=XM_023240856.1
D2 62834358 62834509
refGene_gn=GBF1;refGene_tx=XM_003994358.5;refGene_gn=GBF1;refGene_tx=XM_006938122.4;refGene_gn=GBF1;refGene_tx=XM_006938123.4;refGene_gn=GBF1;refGene_tx=XM_006938124.4;refGene_gn=GBF1;refGene_tx=XM_006938125.4;refGene_gn=GBF1;refGene_tx=XM_006938126.4;refGene_gn=GBF1;refGene_tx=XM_006938128.4;refGene_gn=GBF1;refGene_tx=XM_006938129.4;refGene_gn=GBF1;refGene_tx=XM_006938130.4;refGene_gn=GBF1;refGene_tx=XM_019813751.2;refGene_gn=GBF1;refGene_tx=XM_019813752.2;refGene_gn=GBF1;refGene_tx=XM_019813753.2;refGene_gn=GBF1;refGene_tx=XM_023240856.1

D2	62834615 62834746
refGene_gn=GBF1;refGene_tx=XM_003994358.5;refGene_gn=GBF1;refGene_tx=XM_006938122.4;refGene_gn=GBF1;refGene_tx=XM_006938123.4;refGene_gn=GBF1;refGene_tx=XM_006938124.4;refGene_gn=GBF1;refGene_tx=XM_006938125.4;refGene_gn=GBF1;refGene_tx=XM_006938126.4;refGene_gn=GBF1;refGene_tx=XM_006938128.4;refGene_gn=GBF1;refGene_tx=XM_006938129.4;refGene_gn=GBF1;refGene_tx=XM_006938130.4;refGene_gn=GBF1;refGene_tx=XM_019813751.2;refGene_gn=GBF1;refGene_tx=XM_019813752.2;refGene_gn=GBF1;refGene_tx=XM_019813753.2;refGene_gn=GBF1;refGene_tx=XM_023240856.1
D2	62834952 62835054
refGene_gn=GBF1;refGene_tx=XM_003994358.5;refGene_gn=GBF1;refGene_tx=XM_006938122.4;refGene_gn=GBF1;refGene_tx=XM_006938123.4;refGene_gn=GBF1;refGene_tx=XM_006938124.4;refGene_gn=GBF1;refGene_tx=XM_006938125.4;refGene_gn=GBF1;refGene_tx=XM_006938126.4;refGene_gn=GBF1;refGene_tx=XM_006938128.4;refGene_gn=GBF1;refGene_tx=XM_006938129.4;refGene_gn=GBF1;refGene_tx=XM_006938130.4;refGene_gn=GBF1;refGene_tx=XM_019813751.2;refGene_gn=GBF1;refGene_tx=XM_019813752.2;refGene_gn=GBF1;refGene_tx=XM_019813753.2;refGene_gn=GBF1;refGene_tx=XM_023240856.1
D2	62835331 62835445
refGene_gn=GBF1;refGene_tx=XM_003994358.5;refGene_gn=GBF1;refGene_tx=XM_006938122.4;refGene_gn=GBF1;refGene_tx=XM_006938123.4;refGene_gn=GBF1;refGene_tx=XM_006938124.4;refGene_gn=GBF1;refGene_tx=XM_006938125.4;refGene_gn=GBF1;refGene_tx=XM_006938126.4;refGene_gn=GBF1;refGene_tx=XM_006938128.4;refGene_gn=GBF1;refGene_tx=XM_006938129.4;refGene_gn=GBF1;refGene_tx=XM_006938130.4;refGene_gn=GBF1;refGene_tx=XM_019813751.2;refGene_gn=GBF1;refGene_tx=XM_019813752.2;refGene_gn=GBF1;refGene_tx=XM_019813753.2;refGene_gn=GBF1;refGene_tx=XM_023240856.1
D2	62835576 62835757
refGene_gn=GBF1;refGene_tx=XM_003994358.5;refGene_gn=GBF1;refGene_tx=XM_006938122.4;refGene_gn=GBF1;refGene_tx=XM_006938123.4;refGene_gn=GBF1;refGene_tx=XM_006938124.4;refGene_gn=GBF1;refGene_tx=XM_006938125.4;refGene_gn=GBF1;refGene_tx=XM_006938126.4;refGene_gn=GBF1;refGene_tx=XM_006938128.4;refGene_gn=GBF1;refGene_tx=XM_006938129.4;refGene_gn=GBF1;refGene_tx=XM_006938130.4;refGene_gn=GBF1;refGene_tx=XM_019813751.2;refGene_gn=GBF1;refGene_tx=XM_019813752.2;refGene_gn=GBF1;refGene_tx=XM_019813753.2;refGene_gn=GBF1;refGene_tx=XM_023240856.1
D2	62835922 62836051
refGene_gn=GBF1;refGene_tx=XM_003994358.5;refGene_gn=GBF1;refGene_tx=XM_006938122.4;refGene_gn=GBF1;refGene_tx=XM_006938123.4;refGene_gn=GBF1;refGene_tx=XM_006938124.4;refGene_gn=GBF1;refGene_tx=XM_006938125.4;refGene_gn=GBF1;refGene_tx=XM_006938126.4;refGene_gn=GBF1;refGene_tx=XM_006938128.4;refGene_gn=GBF1;refGene_tx=XM_006938129.4;refGene_gn=GBF1;refGene_tx=XM_006938130.4;refGene_gn=GBF1;refGene_tx=XM_019813751.2;refGene_gn=GBF1;refGene_tx=XM_019813752.2;refGene_gn=GBF1;refGene_tx=XM_019813753.2;refGene_gn=GBF1;refGene_tx=XM_023240856.1
D2	62836543 62836824
refGene_gn=GBF1;refGene_tx=XM_003994358.5;refGene_gn=GBF1;refGene_tx=XM_006938122.4;refGene_gn=GBF1;refGene_tx=XM_006938123.4;refGene_gn=GBF1;refGene_tx=XM_006938124.4;refGene_gn=GBF1;refGene_tx=XM_006938125.4;refGene_gn=GBF1;refGene_tx=XM_006938126.4;refGene_gn=GBF1;refGene_tx=X

M_006938128.4;refGene_gn=GBF1;refGene_tx=XM_006938129.4;refGene_gn=GBF1;refGene_tx=XM_006938130.4;refGene_gn=GBF1;refGene_tx=XM_019813751.2;refGene_gn=GBF1;refGene_tx=XM_019813752.2;refGene_gn=GBF1;refGene_tx=XM_019813753.2;refGene_gn=GBF1;refGene_tx=XM_023240856.1
D2 62844320 62844341
refGene_gn=NFKB2;refGene_tx=XM_006938131.4;refGene_gn=NFKB2;refGene_tx=XM_006938132.4
D2 62844608 62844690
refGene_gn=NFKB2;refGene_tx=XM_006938131.4;refGene_gn=NFKB2;refGene_tx=XM_006938132.4
D2 62844806 62844847
refGene_gn=NFKB2;refGene_tx=XM_006938131.4;refGene_gn=NFKB2;refGene_tx=XM_006938132.4
D2 62845109 62845208
refGene_gn=NFKB2;refGene_tx=XM_006938131.4;refGene_gn=NFKB2;refGene_tx=XM_006938132.4
D2 62845287 62845439
refGene_gn=NFKB2;refGene_tx=XM_006938131.4;refGene_gn=NFKB2;refGene_tx=XM_006938132.4
D2 62845667 62845774
refGene_gn=NFKB2;refGene_tx=XM_006938131.4;refGene_gn=NFKB2;refGene_tx=XM_006938132.4;refGene_gn=NFKB2;refGene_tx=XM_019813757.2
D2 62845882 62846041
refGene_gn=NFKB2;refGene_tx=XM_006938131.4;refGene_gn=NFKB2;refGene_tx=XM_006938132.4;refGene_gn=NFKB2;refGene_tx=XM_019813757.2
D2 62846331 62846436
refGene_gn=NFKB2;refGene_tx=XM_006938131.4;refGene_gn=NFKB2;refGene_tx=XM_006938132.4;refGene_gn=NFKB2;refGene_tx=XM_019813757.2
D2 62846564 62846650
refGene_gn=NFKB2;refGene_tx=XM_006938131.4;refGene_gn=NFKB2;refGene_tx=XM_006938132.4;refGene_gn=NFKB2;refGene_tx=XM_019813757.2
D2 62846734 62846873
refGene_gn=NFKB2;refGene_tx=XM_006938131.4;refGene_gn=NFKB2;refGene_tx=XM_006938132.4;refGene_gn=NFKB2;refGene_tx=XM_019813757.2
D2 62847093 62847222
refGene_gn=NFKB2;refGene_tx=XM_006938131.4;refGene_gn=NFKB2;refGene_tx=XM_006938132.4;refGene_gn=NFKB2;refGene_tx=XM_019813757.2
D2 62847356 62847551
refGene_gn=NFKB2;refGene_tx=XM_006938131.4;refGene_gn=NFKB2;refGene_tx=XM_006938132.4;refGene_gn=NFKB2;refGene_tx=XM_019813757.2
D2 62847629 62847771
refGene_gn=NFKB2;refGene_tx=XM_006938131.4;refGene_gn=NFKB2;refGene_tx=XM_006938132.4;refGene_gn=NFKB2;refGene_tx=XM_019813757.2
D2 62848196 62848311
refGene_gn=NFKB2;refGene_tx=XM_006938131.4;refGene_gn=NFKB2;refGene_tx=XM_006938132.4;refGene_gn=NFKB2;refGene_tx=XM_019813757.2
D2 62848402 62848616
refGene_gn=NFKB2;refGene_tx=XM_006938131.4;refGene_gn=NFKB2;refGene_tx=XM_006938132.4;refGene_gn=NFKB2;refGene_tx=XM_019813757.2
D2 62848755 62848925

refGene_gn=NFKB2;refGene_tx=XM_006938131.4;refGene_gn=NFKB2;refGene_tx=XM_006938132.4;refGene_gn=NFKB2;refGene_tx=XM_019813757.2
D2 62849047 62849150
refGene_gn=NFKB2;refGene_tx=XM_006938131.4;refGene_gn=NFKB2;refGene_tx=XM_006938132.4;refGene_gn=NFKB2;refGene_tx=XM_019813757.2
D2 62849278 62849430
refGene_gn=NFKB2;refGene_tx=XM_006938131.4;refGene_gn=NFKB2;refGene_tx=XM_006938132.4;refGene_gn=NFKB2;refGene_tx=XM_019813757.2
D2 62849558 62849628
refGene_gn=NFKB2;refGene_tx=XM_006938131.4;refGene_gn=NFKB2;refGene_tx=XM_006938132.4;refGene_gn=NFKB2;refGene_tx=XM_019813757.2
D2 62849857 62850030
refGene_gn=NFKB2;refGene_tx=XM_006938131.4;refGene_gn=NFKB2;refGene_tx=XM_006938132.4;refGene_gn=NFKB2;refGene_tx=XM_019813757.2
D2 62850155 62850267
refGene_gn=NFKB2;refGene_tx=XM_006938131.4;refGene_gn=NFKB2;refGene_tx=XM_006938132.4;refGene_gn=NFKB2;refGene_tx=XM_019813757.2
D2 62850369 62850491
refGene_gn=NFKB2;refGene_tx=XM_006938131.4;refGene_gn=NFKB2;refGene_tx=XM_006938132.4;refGene_gn=NFKB2;refGene_tx=XM_019813757.2
D2 62851005 62851017 refGene_gn=PSD;refGene_tx=XM_019813756.2
D2 62851320 62851551
refGene_gn=PSD;refGene_tx=XM_006938134.3;refGene_gn=PSD;refGene_tx=XM_011287444.2;refGene_gn=PSD;refGene_tx=XM_019813754.1;refGene_gn=PSD;refGene_tx=XM_019813755.1;refGene_gn=PSD;refGene_tx=XM_023240857.1;refGene_gn=PSD;refGene_tx=XM_023240858.1
D2 62852040 62852184
refGene_gn=PSD;refGene_tx=XM_006938134.3;refGene_gn=PSD;refGene_tx=XM_011287444.2;refGene_gn=PSD;refGene_tx=XM_019813754.1;refGene_gn=PSD;refGene_tx=XM_019813755.1;refGene_gn=PSD;refGene_tx=XM_019813756.2;refGene_gn=PSD;refGene_tx=XM_023240857.1;refGene_gn=PSD;refGene_tx=XM_023240858.1
D2 62852470 62852615
refGene_gn=PSD;refGene_tx=XM_006938134.3;refGene_gn=PSD;refGene_tx=XM_011287444.2;refGene_gn=PSD;refGene_tx=XM_019813754.1;refGene_gn=PSD;refGene_tx=XM_019813755.1;refGene_gn=PSD;refGene_tx=XM_019813756.2;refGene_gn=PSD;refGene_tx=XM_023240857.1;refGene_gn=PSD;refGene_tx=XM_023240858.1
D2 62852787 62852945
refGene_gn=PSD;refGene_tx=XM_006938134.3;refGene_gn=PSD;refGene_tx=XM_011287444.2;refGene_gn=PSD;refGene_tx=XM_019813754.1;refGene_gn=PSD;refGene_tx=XM_019813755.1;refGene_gn=PSD;refGene_tx=XM_019813756.2;refGene_gn=PSD;refGene_tx=XM_023240857.1;refGene_gn=PSD;refGene_tx=XM_023240858.1
D2 62853054 62853125
refGene_gn=PSD;refGene_tx=XM_006938134.3;refGene_gn=PSD;refGene_tx=XM_011287444.2;refGene_gn=PSD;refGene_tx=XM_019813754.1;refGene_gn=PSD;refGene_tx=XM_019813755.1;refGene_gn=PSD;refGene_tx=XM_019813756.2;refGene_gn=PSD;refGene_tx=XM_023240857.1;refGene_gn=PSD;refGene_tx=XM_023240858.1

D2 62853209 62853400
refGene_gn=PSD;refGene_tx=XM_006938134.3;refGene_gn=PSD;refGene_tx=XM_011287444.2;refGene_gn=PSD;refGene_tx=XM_019813754.1;refGene_gn=PSD;refGene_tx=XM_019813755.1;refGene_gn=PSD;refGene_tx=XM_019813756.2;refGene_gn=PSD;refGene_tx=XM_023240857.1;refGene_gn=PSD;refGene_tx=XM_023240858.1
D2 62854787 62854831
refGene_gn=PSD;refGene_tx=XM_006938134.3;refGene_gn=PSD;refGene_tx=XM_011287444.2;refGene_gn=PSD;refGene_tx=XM_019813754.1;refGene_gn=PSD;refGene_tx=XM_019813755.1;refGene_gn=PSD;refGene_tx=XM_019813756.2;refGene_gn=PSD;refGene_tx=XM_023240857.1;refGene_gn=PSD;refGene_tx=XM_023240858.1
D2 62858527 62858617
refGene_gn=PSD;refGene_tx=XM_006938134.3;refGene_gn=PSD;refGene_tx=XM_011287444.2;refGene_gn=PSD;refGene_tx=XM_019813754.1;refGene_gn=PSD;refGene_tx=XM_019813755.1;refGene_gn=PSD;refGene_tx=XM_019813756.2;refGene_gn=PSD;refGene_tx=XM_023240857.1;refGene_gn=PSD;refGene_tx=XM_023240858.1
D2 62858724 62858783
refGene_gn=PSD;refGene_tx=XM_006938134.3;refGene_gn=PSD;refGene_tx=XM_011287444.2;refGene_gn=PSD;refGene_tx=XM_019813754.1;refGene_gn=PSD;refGene_tx=XM_019813755.1;refGene_gn=PSD;refGene_tx=XM_019813756.2;refGene_gn=PSD;refGene_tx=XM_023240857.1;refGene_gn=PSD;refGene_tx=XM_023240858.1
D2 62859349 62859462
refGene_gn=PSD;refGene_tx=XM_006938134.3;refGene_gn=PSD;refGene_tx=XM_011287444.2;refGene_gn=PSD;refGene_tx=XM_019813754.1;refGene_gn=PSD;refGene_tx=XM_019813755.1;refGene_gn=PSD;refGene_tx=XM_019813756.2;refGene_gn=PSD;refGene_tx=XM_023240857.1;refGene_gn=PSD;refGene_tx=XM_023240858.1
D2 62859668 62859749
refGene_gn=PSD;refGene_tx=XM_006938134.3;refGene_gn=PSD;refGene_tx=XM_011287444.2;refGene_gn=PSD;refGene_tx=XM_019813754.1;refGene_gn=PSD;refGene_tx=XM_019813755.1;refGene_gn=PSD;refGene_tx=XM_019813756.2;refGene_gn=PSD;refGene_tx=XM_023240857.1;refGene_gn=PSD;refGene_tx=XM_023240858.1
D2 62859913 62860108
refGene_gn=PSD;refGene_tx=XM_006938134.3;refGene_gn=PSD;refGene_tx=XM_011287444.2;refGene_gn=PSD;refGene_tx=XM_019813754.1;refGene_gn=PSD;refGene_tx=XM_019813755.1;refGene_gn=PSD;refGene_tx=XM_019813756.2;refGene_gn=PSD;refGene_tx=XM_023240857.1
D2 62861328 62861769
refGene_gn=PSD;refGene_tx=XM_006938134.3;refGene_gn=PSD;refGene_tx=XM_011287444.2;refGene_gn=PSD;refGene_tx=XM_019813754.1;refGene_gn=PSD;refGene_tx=XM_019813755.1;refGene_gn=PSD;refGene_tx=XM_019813756.2;refGene_gn=PSD;refGene_tx=XM_023240857.1;refGene_gn=PSD;refGene_tx=XM_023240858.1
D2 62862497 62862858
refGene_gn=PSD;refGene_tx=XM_006938134.3;refGene_gn=PSD;refGene_tx=XM_011287444.2;refGene_gn=PSD;refGene_tx=XM_019813754.1;refGene_gn=PSD;re

fGene_tx=XM_019813755.1;refGene_gn=PSD;refGene_tx=XM_019813756.2;refGene_gn=PSD;refGene_tx=XM_023240857.1;refGene_gn=PSD;refGene_tx=XM_023240858.1
D2	62863347	62863450	refGene_gn=PSD;refGene_tx=XM_006938134.3;refGene_gn=PSD;refGene_tx=XM_011287444.2;refGene_gn=PSD;refGene_tx=XM_019813754.1;refGene_gn=PSD;refGene_tx=XM_019813755.1;refGene_gn=PSD;refGene_tx=XM_019813756.2;refGene_gn=PSD;refGene_tx=XM_023240857.1;refGene_gn=PSD;refGene_tx=XM_023240858.1
D2	62863732	62864386	refGene_gn=PSD;refGene_tx=XM_006938134.3;refGene_gn=PSD;refGene_tx=XM_011287444.2;refGene_gn=PSD;refGene_tx=XM_019813754.1;refGene_gn=PSD;refGene_tx=XM_019813755.1;refGene_gn=PSD;refGene_tx=XM_019813756.2;refGene_gn=PSD;refGene_tx=XM_023240857.1;refGene_gn=PSD;refGene_tx=XM_023240858.1
D2	62868200	62868253	refGene_gn=FBXL15;refGene_tx=XM_003994361.4;refGene_gn=FBXL15;refGene_tx=XM_023240860.1
D2	62868424	62868578	refGene_gn=FBXL15;refGene_tx=XM_003994361.4;refGene_gn=FBXL15;refGene_tx=XM_023240860.1
D2	62868898	62869404	refGene_gn=FBXL15;refGene_tx=XM_003994361.4;refGene_gn=FBXL15;refGene_tx=XM_023240860.1
D2	62869914	62870104	refGene_gn=FBXL15;refGene_tx=XM_003994361.4;refGene_gn=FBXL15;refGene_tx=XM_023240860.1
D2	62870554	62870701	refGene_gn=CUEDC2;refGene_tx=XM_003994362.5;refGene_gn=CUEDC2;refGene_tx=XM_006938138.4;refGene_gn=CUEDC2;refGene_tx=XM_006938139.4;refGene_gn=CUEDC2;refGene_tx=XM_023240859.1
D2	62870786	62870847	refGene_gn=CUEDC2;refGene_tx=XM_003994362.5;refGene_gn=CUEDC2;refGene_tx=XM_006938138.4;refGene_gn=CUEDC2;refGene_tx=XM_006938139.4;refGene_gn=CUEDC2;refGene_tx=XM_023240859.1
D2	62870928	62870990	refGene_gn=CUEDC2;refGene_tx=XM_003994362.5;refGene_gn=CUEDC2;refGene_tx=XM_006938138.4;refGene_gn=CUEDC2;refGene_tx=XM_006938139.4;refGene_gn=CUEDC2;refGene_tx=XM_023240859.1
D2	62871129	62871312	refGene_gn=CUEDC2;refGene_tx=XM_003994362.5;refGene_gn=CUEDC2;refGene_tx=XM_006938138.4;refGene_gn=CUEDC2;refGene_tx=XM_006938139.4;refGene_gn=CUEDC2;refGene_tx=XM_023240859.1
D2	62871397	62871528	refGene_gn=CUEDC2;refGene_tx=XM_003994362.5;refGene_gn=CUEDC2;refGene_tx=XM_006938138.4;refGene_gn=CUEDC2;refGene_tx=XM_006938139.4;refGene_gn=CUEDC2;refGene_tx=XM_023240859.1
D2	62871612	62871674	refGene_gn=CUEDC2;refGene_tx=XM_003994362.5;refGene_gn=CUEDC2;refGene_tx=XM_006938138.4;refGene_gn=CUEDC2;refGene_tx=XM_006938139.4;refGene_

gn=CUEDC2;refGene_tx=XM_023240859.1
D2 62871767 62871936
refGene_gn=CUEDC2;refGene_tx=XM_003994362.5;refGene_gn=CUEDC2;refGene_tx=XM_006938138.4;refGene_gn=CUEDC2;refGene_tx=XM_006938139.4;refGene_gn=CUEDC2;refGene_tx=XM_023240859.1
D2 62872267 62872354
refGene_gn=CUEDC2;refGene_tx=XM_003994362.5;refGene_gn=CUEDC2;refGene_tx=XM_006938138.4;refGene_gn=CUEDC2;refGene_tx=XM_006938139.4;refGene_gn=CUEDC2;refGene_tx=XM_023240859.1
D2 62877891 62878003 refGene_gn=CUEDC2;refGene_tx=XM_023240859.1
D2 62886150 62886272
refGene_gn=CD2H10orf95;refGene_tx=XM_023240299.1
D2 62887210 62887947
refGene_gn=CD2H10orf95;refGene_tx=XM_023240299.1
D2 62890507 62890591
refGene_gn=CD2H10orf95;refGene_tx=XM_023240299.1
D2 62892494 62892505
refGene_gn=CD2H10orf95;refGene_tx=XM_023240299.1
D2 62902225 62902393
refGene_gn=MFSD13A;refGene_tx=XM_003994363.5;refGene_gn=MFSD13A;refGene_tx=XM_023240861.1
D2 62902740 62902823
refGene_gn=MFSD13A;refGene_tx=XM_003994363.5;refGene_gn=MFSD13A;refGene_tx=XM_023240861.1
D2 62903398 62903851
refGene_gn=MFSD13A;refGene_tx=XM_003994363.5;refGene_gn=MFSD13A;refGene_tx=XM_023240861.1
D2 62904002 62904126
refGene_gn=MFSD13A;refGene_tx=XM_003994363.5;refGene_gn=MFSD13A;refGene_tx=XM_023240861.1
D2 62904721 62904818
refGene_gn=MFSD13A;refGene_tx=XM_003994363.5;refGene_gn=MFSD13A;refGene_tx=XM_023240861.1
D2 62905602 62905774
refGene_gn=MFSD13A;refGene_tx=XM_003994363.5;refGene_gn=MFSD13A;refGene_tx=XM_023240861.1
D2 62905847 62906041
refGene_gn=MFSD13A;refGene_tx=XM_003994363.5;refGene_gn=MFSD13A;refGene_tx=XM_023240861.1
D2 62907129 62907392
refGene_gn=MFSD13A;refGene_tx=XM_003994363.5;refGene_gn=MFSD13A;refGene_tx=XM_023240861.1
D2 62910974 62911077 refGene_gn=ACTR1A;refGene_tx=XM_003994364.4
D2 62911250 62911291 refGene_gn=ACTR1A;refGene_tx=XM_003994364.4
D2 62911902 62911964 refGene_gn=ACTR1A;refGene_tx=XM_003994364.4
D2 62912065 62912240 refGene_gn=ACTR1A;refGene_tx=XM_003994364.4
D2 62912599 62912692 refGene_gn=ACTR1A;refGene_tx=XM_003994364.4
D2 62913437 62913654 refGene_gn=ACTR1A;refGene_tx=XM_003994364.4
D2 62915355 62915480 refGene_gn=ACTR1A;refGene_tx=XM_003994364.4
D2 62916334 62916460 refGene_gn=ACTR1A;refGene_tx=XM_003994364.4

D2 62917203 62917279 refGene_gn=ACTR1A;refGene_tx=XM_003994364.4
D2 62917753 62917818 refGene_gn=ACTR1A;refGene_tx=XM_003994364.4
D2 62925741 62925789 refGene_gn=ACTR1A;refGene_tx=XM_003994364.4
D2 62928036 62928218
refGene_gn=SUFU;refGene_tx=XM_003994375.5;refGene_gn=SUFU;refGene_tx=XM_023240862.1;refGene_gn=SUFU;refGene_tx=XM_023240863.1;refGene_gn=SUFU;refGene_tx=XM_023240864.1;refGene_gn=SUFU;refGene_tx=XM_023240865.1;refGene_gn=SUFU;refGene_tx=XM_023240866.1
D2 62931580 62931715
refGene_gn=SUFU;refGene_tx=XM_003994375.5;refGene_gn=SUFU;refGene_tx=XM_023240862.1;refGene_gn=SUFU;refGene_tx=XM_023240863.1;refGene_gn=SUFU;refGene_tx=XM_023240864.1;refGene_gn=SUFU;refGene_tx=XM_023240865.1;refGene_gn=SUFU;refGene_tx=XM_023240866.1
D2 62964469 62964606
refGene_gn=SUFU;refGene_tx=XM_003994375.5;refGene_gn=SUFU;refGene_tx=XM_023240862.1;refGene_gn=SUFU;refGene_tx=XM_023240863.1;refGene_gn=SUFU;refGene_tx=XM_023240864.1;refGene_gn=SUFU;refGene_tx=XM_023240865.1;refGene_gn=SUFU;refGene_tx=XM_023240866.1
D2 63007283 63007426
refGene_gn=SUFU;refGene_tx=XM_003994375.5;refGene_gn=SUFU;refGene_tx=XM_023240862.1;refGene_gn=SUFU;refGene_tx=XM_023240863.1;refGene_gn=SUFU;refGene_tx=XM_023240864.1;refGene_gn=SUFU;refGene_tx=XM_023240865.1;refGene_gn=SUFU;refGene_tx=XM_023240866.1
D2 63008362 63008448
refGene_gn=SUFU;refGene_tx=XM_003994375.5;refGene_gn=SUFU;refGene_tx=XM_023240862.1;refGene_gn=SUFU;refGene_tx=XM_023240863.1;refGene_gn=SUFU;refGene_tx=XM_023240864.1;refGene_gn=SUFU;refGene_tx=XM_023240865.1;refGene_gn=SUFU;refGene_tx=XM_023240866.1
D2 63008724 63008797
refGene_gn=SUFU;refGene_tx=XM_003994375.5;refGene_gn=SUFU;refGene_tx=XM_023240862.1;refGene_gn=SUFU;refGene_tx=XM_023240863.1;refGene_gn=SUFU;refGene_tx=XM_023240864.1;refGene_gn=SUFU;refGene_tx=XM_023240865.1;refGene_gn=SUFU;refGene_tx=XM_023240866.1
D2 63012379 63012536
refGene_gn=SUFU;refGene_tx=XM_003994375.5;refGene_gn=SUFU;refGene_tx=XM_023240862.1;refGene_gn=SUFU;refGene_tx=XM_023240863.1;refGene_gn=SUFU;refGene_tx=XM_023240864.1;refGene_gn=SUFU;refGene_tx=XM_023240865.1;refGene_gn=SUFU;refGene_tx=XM_023240866.1
D2 63014369 63014481
refGene_gn=SUFU;refGene_tx=XM_003994375.5;refGene_gn=SUFU;refGene_tx=XM_023240862.1;refGene_gn=SUFU;refGene_tx=XM_023240863.1;refGene_gn=SUFU;refGene_tx=XM_023240864.1;refGene_gn=SUFU;refGene_tx=XM_023240865.1;refGene_gn=SUFU;refGene_tx=XM_023240866.1
D2 63027874 63028009
refGene_gn=SUFU;refGene_tx=XM_003994375.5;refGene_gn=SUFU;refGene_tx=XM_023240862.1;refGene_gn=SUFU;refGene_tx=XM_023240863.1;refGene_gn=SUFU;refGene_tx=XM_023240864.1;refGene_gn=SUFU;refGene_tx=XM_023240865.1;refGene_gn=SUFU;refGene_tx=XM_023240866.1
D2 63029836 63029975
refGene_gn=SUFU;refGene_tx=XM_003994375.5;refGene_gn=SUFU;refGene_tx=X

M_023240862.1;refGene_gn=SUFU;refGene_tx=XM_023240863.1;refGene_gn=SUFU;refGene_tx=XM_023240864.1;refGene_gn=SUFU;refGene_tx=XM_023240865.1;refGene_gn=SUFU;refGene_tx=XM_023240866.1
D2 63038714 63038783 refGene_gn=SUFU;refGene_tx=XM_003994375.5;refGene_gn=SUFU;refGene_tx=XM_023240862.1;refGene_gn=SUFU;refGene_tx=XM_023240863.1;refGene_gn=SUFU;refGene_tx=XM_023240864.1;refGene_gn=SUFU;refGene_tx=XM_023240866.1
D2 63041202 63041292 refGene_gn=SUFU;refGene_tx=XM_003994375.5;refGene_gn=SUFU;refGene_tx=XM_023240864.1
D2 63041854 63041887 refGene_gn=SUFU;refGene_tx=XM_023240866.1
D2 63050877 63050985 refGene_gn=SUFU;refGene_tx=XM_023240862.1;refGene_gn=SUFU;refGene_tx=XM_023240863.1;refGene_gn=SUFU;refGene_tx=XM_023240865.1
D2 63053589 63053843 refGene_gn=TRIM8;refGene_tx=NM_001290250.1
D2 63053844 63054160 refGene_gn=TRIM8;refGene_tx=NM_001290250.1
D2 63063723 63063819 refGene_gn=TRIM8;refGene_tx=NM_001290250.1
D2 63064171 63064361 refGene_gn=TRIM8;refGene_tx=NM_001290250.1
D2 63064362 63064406 refGene_gn=TRIM8;refGene_tx=NM_001290250.1
D2 63065227 63065259 refGene_gn=TRIM8;refGene_tx=NM_001290250.1
D2 63065392 63065508 refGene_gn=TRIM8;refGene_tx=NM_001290250.1
D2 63065860 63066468 refGene_gn=TRIM8;refGene_tx=NM_001290250.1
D2 63081058 63081106 refGene_gn=ARL3;refGene_tx=XM_023240867.1
D2 63089959 63090145 refGene_gn=ARL3;refGene_tx=XM_023240867.1
D2 63091692 63091743 refGene_gn=ARL3;refGene_tx=XM_023240867.1
D2 63109037 63109058 refGene_gn=ARL3;refGene_tx=XM_023240867.1
D2 63109059 63109155 refGene_gn=ARL3;refGene_tx=XM_023240867.1
D2 63113794 63113938 refGene_gn=ARL3;refGene_tx=XM_023240867.1
D2 63116537 63116573 refGene_gn=ARL3;refGene_tx=XM_023240867.1
D2 63122214 63122235 refGene_gn=SFXN2;refGene_tx=XM_019813759.2;refGene_gn=SFXN2;refGene_tx=XM_019813760.2;refGene_gn=SFXN2;refGene_tx=XM_019813764.2
D2 63134944 63135129 refGene_gn=SFXN2;refGene_tx=XM_019813758.2;refGene_gn=SFXN2;refGene_tx=XM_019813759.2;refGene_gn=SFXN2;refGene_tx=XM_019813760.2;refGene_gn=SFXN2;refGene_tx=XM_019813761.2;refGene_gn=SFXN2;refGene_tx=XM_019813762.2;refGene_gn=SFXN2;refGene_tx=XM_019813763.2;refGene_gn=SFXN2;refGene_tx=XM_019813764.2
D2 63135303 63135492 refGene_gn=SFXN2;refGene_tx=XM_019813758.2;refGene_gn=SFXN2;refGene_tx=XM_019813759.2;refGene_gn=SFXN2;refGene_tx=XM_019813760.2;refGene_gn=SFXN2;refGene_tx=XM_019813761.2;refGene_gn=SFXN2;refGene_tx=XM_019813762.2;refGene_gn=SFXN2;refGene_tx=XM_019813763.2;refGene_gn=SFXN2;refGene_tx=XM_019813764.2
D2 63136458 63136557 refGene_gn=SFXN2;refGene_tx=XM_019813758.2;refGene_gn=SFXN2;refGene_tx=XM_019813759.2;refGene_gn=SFXN2;refGene_tx=XM_019813760.2;refGene_gn=SFXN2;refGene_tx=XM_019813761.2;refGene_gn=SFXN2;refGene_tx=XM_019813762.2;refGene_gn=SFXN2;refGene_tx=XM_019813763.2;refGene_gn=SFXN2;refGene_tx=XM_019813764.2

D2 63137301 63137377
refGene_gn=SFXN2;refGene_tx=XM_019813758.2;refGene_gn=SFXN2;refGene_tx=XM_019813759.2;refGene_gn=SFXN2;refGene_tx=XM_019813760.2;refGene_gn=SFXN2;refGene_tx=XM_019813761.2;refGene_gn=SFXN2;refGene_tx=XM_019813762.2;refGene_gn=SFXN2;refGene_tx=XM_019813763.2;refGene_gn=SFXN2;refGene_tx=XM_019813764.2
D2 63137652 63137738
refGene_gn=SFXN2;refGene_tx=XM_019813758.2;refGene_gn=SFXN2;refGene_tx=XM_019813759.2;refGene_gn=SFXN2;refGene_tx=XM_019813760.2;refGene_gn=SFXN2;refGene_tx=XM_019813761.2;refGene_gn=SFXN2;refGene_tx=XM_019813762.2;refGene_gn=SFXN2;refGene_tx=XM_019813763.2;refGene_gn=SFXN2;refGene_tx=XM_019813764.2
D2 63139018 63139079
refGene_gn=SFXN2;refGene_tx=XM_019813758.2;refGene_gn=SFXN2;refGene_tx=XM_019813759.2;refGene_gn=SFXN2;refGene_tx=XM_019813760.2;refGene_gn=SFXN2;refGene_tx=XM_019813761.2;refGene_gn=SFXN2;refGene_tx=XM_019813762.2;refGene_gn=SFXN2;refGene_tx=XM_019813763.2;refGene_gn=SFXN2;refGene_tx=XM_019813764.2
D2 63139445 63139512
refGene_gn=SFXN2;refGene_tx=XM_019813758.2;refGene_gn=SFXN2;refGene_tx=XM_019813759.2;refGene_gn=SFXN2;refGene_tx=XM_019813760.2;refGene_gn=SFXN2;refGene_tx=XM_019813761.2;refGene_gn=SFXN2;refGene_tx=XM_019813762.2;refGene_gn=SFXN2;refGene_tx=XM_019813763.2;refGene_gn=SFXN2;refGene_tx=XM_019813764.2
D2 63140235 63140285
refGene_gn=SFXN2;refGene_tx=XM_019813758.2;refGene_gn=SFXN2;refGene_tx=XM_019813759.2;refGene_gn=SFXN2;refGene_tx=XM_019813760.2;refGene_gn=SFXN2;refGene_tx=XM_019813761.2;refGene_gn=SFXN2;refGene_tx=XM_019813762.2;refGene_gn=SFXN2;refGene_tx=XM_019813763.2;refGene_gn=SFXN2;refGene_tx=XM_019813764.2
D2 63140619 63140669
refGene_gn=SFXN2;refGene_tx=XM_019813758.2;refGene_gn=SFXN2;refGene_tx=XM_019813759.2;refGene_gn=SFXN2;refGene_tx=XM_019813760.2;refGene_gn=SFXN2;refGene_tx=XM_019813761.2;refGene_gn=SFXN2;refGene_tx=XM_019813762.2;refGene_gn=SFXN2;refGene_tx=XM_019813763.2;refGene_gn=SFXN2;refGene_tx=XM_019813764.2
D2 63142699 63142747
refGene_gn=SFXN2;refGene_tx=XM_019813758.2;refGene_gn=SFXN2;refGene_tx=XM_019813759.2;refGene_gn=SFXN2;refGene_tx=XM_019813760.2;refGene_gn=SFXN2;refGene_tx=XM_019813761.2;refGene_gn=SFXN2;refGene_tx=XM_019813762.2;refGene_gn=SFXN2;refGene_tx=XM_019813763.2;refGene_gn=SFXN2;refGene_tx=XM_019813764.2
D2 63143582 63143682
refGene_gn=SFXN2;refGene_tx=XM_019813758.2;refGene_gn=SFXN2;refGene_tx=XM_019813759.2;refGene_gn=SFXN2;refGene_tx=XM_019813760.2;refGene_gn=SFXN2;refGene_tx=XM_019813761.2;refGene_gn=SFXN2;refGene_tx=XM_019813762.2;refGene_gn=SFXN2;refGene_tx=XM_019813763.2;refGene_gn=SFXN2;refGene_tx=XM_019813764.2
D2 63148750 63148840 refGene_gn=WBP1L;refGene_tx=XM_003994368.5
D2 63164946 63165015 refGene_gn=WBP1L;refGene_tx=XM_019813766.2

D2 63171591 63171618 refGene_gn=WBP1L;refGene_tx=XM_006938145.4
D2 63193524 63193627
refGene_gn=WBP1L;refGene_tx=XM_003994368.5;refGene_gn=WBP1L;refGene_tx=XM_006938145.4;refGene_gn=WBP1L;refGene_tx=XM_019813766.2
D2 63204230 63204392
refGene_gn=WBP1L;refGene_tx=XM_003994368.5;refGene_gn=WBP1L;refGene_tx=XM_006938145.4;refGene_gn=WBP1L;refGene_tx=XM_019813766.2
D2 63205731 63206474
refGene_gn=WBP1L;refGene_tx=XM_003994368.5;refGene_gn=WBP1L;refGene_tx=XM_006938145.4;refGene_gn=WBP1L;refGene_tx=XM_019813766.2
D2 63216852 63217136
refGene_gn=CYP17A1;refGene_tx=NM_001009371.2;refGene_gn=CYP17A1;refGene_tx=XM_011287431.2
D2 63217649 63217753
refGene_gn=CYP17A1;refGene_tx=NM_001009371.2;refGene_gn=CYP17A1;refGene_tx=XM_011287431.2
D2 63218579 63218749
refGene_gn=CYP17A1;refGene_tx=NM_001009371.2;refGene_gn=CYP17A1;refGene_tx=XM_011287431.2
D2 63219069 63219285
refGene_gn=CYP17A1;refGene_tx=NM_001009371.2;refGene_gn=CYP17A1;refGene_tx=XM_011287431.2
D2 63219672 63219759
refGene_gn=CYP17A1;refGene_tx=NM_001009371.2;refGene_gn=CYP17A1;refGene_tx=XM_011287431.2
D2 63220534 63220764
refGene_gn=CYP17A1;refGene_tx=NM_001009371.2;refGene_gn=CYP17A1;refGene_tx=XM_011287431.2
D2 63220974 63221113
refGene_gn=CYP17A1;refGene_tx=NM_001009371.2;refGene_gn=CYP17A1;refGene_tx=XM_011287431.2
D2 63222244 63222541
refGene_gn=CYP17A1;refGene_tx=NM_001009371.2;refGene_gn=CYP17A1;refGene_tx=XM_011287431.2
D2 63247426 63247564
refGene_gn=BORCS7;refGene_tx=XM_023240868.1;refGene_gn=BORCS7;refGene_tx=XM_023240869.1;refGene_gn=BORCS7;refGene_tx=XM_023240870.1;refGene_gn=BORCS7;refGene_tx=XM_023240871.1;refGene_gn=BORCS7;refGene_tx=XM_023240872.1
D2 63254631 63254771
refGene_gn=BORCS7;refGene_tx=XM_023240868.1;refGene_gn=BORCS7;refGene_tx=XM_023240869.1
D2 63255938 63256001
refGene_gn=BORCS7;refGene_tx=XM_023240868.1;refGene_gn=BORCS7;refGene_tx=XM_023240869.1;refGene_gn=BORCS7;refGene_tx=XM_023240870.1;refGene_gn=BORCS7;refGene_tx=XM_023240871.1;refGene_gn=BORCS7;refGene_tx=XM_023240872.1
D2 63256113 63256157
refGene_gn=BORCS7;refGene_tx=XM_023240870.1;refGene_gn=BORCS7;refGene_tx=XM_023240872.1

D2	63257115	63257152
refGene_gn=BORCS7;refGene_tx=XM_023240868.1;refGene_gn=BORCS7;refGene_tx=XM_023240869.1;refGene_gn=BORCS7;refGene_tx=XM_023240870.1;refGene_gn=BORCS7;refGene_tx=XM_023240871.1;refGene_gn=BORCS7;refGene_tx=XM_023240872.1
D2	63258988	63259040
refGene_gn=BORCS7;refGene_tx=XM_023240868.1;refGene_gn=BORCS7;refGene_tx=XM_023240870.1;refGene_gn=BORCS7;refGene_tx=XM_023240871.1
D2	63267086	63267087	refGene_gn=AS3MT;refGene_tx=XM_023240874.1
D2	63267280	63267321	refGene_gn=AS3MT;refGene_tx=XM_023240874.1
D2	63267530	63267751
refGene_gn=AS3MT;refGene_tx=XM_023240873.1;refGene_gn=AS3MT;refGene_tx=XM_023240874.1
D2	63268684	63268835
refGene_gn=AS3MT;refGene_tx=XM_023240873.1;refGene_gn=AS3MT;refGene_tx=XM_023240874.1
D2	63269451	63269588
refGene_gn=AS3MT;refGene_tx=XM_023240873.1;refGene_gn=AS3MT;refGene_tx=XM_023240874.1
D2	63270317	63270387
refGene_gn=AS3MT;refGene_tx=XM_023240873.1;refGene_gn=AS3MT;refGene_tx=XM_023240874.1
D2	63272226	63272308
refGene_gn=AS3MT;refGene_tx=XM_023240873.1;refGene_gn=AS3MT;refGene_tx=XM_023240874.1
D2	63274240	63274372
refGene_gn=AS3MT;refGene_tx=XM_023240873.1;refGene_gn=AS3MT;refGene_tx=XM_023240874.1
D2	63274710	63274853
refGene_gn=AS3MT;refGene_tx=XM_023240873.1;refGene_gn=AS3MT;refGene_tx=XM_023240874.1
D2	63277626	63277761
refGene_gn=AS3MT;refGene_tx=XM_023240873.1;refGene_gn=AS3MT;refGene_tx=XM_023240874.1
D2	63286135	63286240
refGene_gn=AS3MT;refGene_tx=XM_023240873.1;refGene_gn=AS3MT;refGene_tx=XM_023240874.1
D2	63301670	63302550	refGene_gn=CNNM2;refGene_tx=XM_023240875.1
D2	63302552	63303290	refGene_gn=CNNM2;refGene_tx=XM_023240875.1
D2	63420058	63420202	refGene_gn=CNNM2;refGene_tx=XM_023240875.1
D2	63423185	63423323	refGene_gn=CNNM2;refGene_tx=XM_023240875.1
D2	63425385	63425555	refGene_gn=CNNM2;refGene_tx=XM_023240875.1
D2	63437262	63437356	refGene_gn=CNNM2;refGene_tx=XM_023240875.1
D2	63444731	63444916	refGene_gn=CNNM2;refGene_tx=XM_023240875.1
D2	63445633	63445843	refGene_gn=CNNM2;refGene_tx=XM_023240875.1
D2	63458066	63458303
refGene_gn=NT5C2;refGene_tx=XM_011287440.3;refGene_gn=NT5C2;refGene_tx=XM_019813770.2;refGene_gn=NT5C2;refGene_tx=XM_023240876.1;refGene_gn=NT5C2;refGene_tx=XM_023240877.1;refGene_gn=NT5C2;refGene_tx=XM_023240878.1;refGene_gn=NT5C2;refGene_tx=XM_023240879.1;refGene_gn=NT5C2;refGe

ne_tx=XM_023240880.1
D2 63459054 63459231
refGene_gn=NT5C2;refGene_tx=XM_011287440.3;refGene_gn=NT5C2;refGene_tx=XM_019813770.2;refGene_gn=NT5C2;refGene_tx=XM_023240876.1;refGene_gn=NT5C2;refGene_tx=XM_023240877.1;refGene_gn=NT5C2;refGene_tx=XM_023240878.1;refGene_gn=NT5C2;refGene_tx=XM_023240879.1;refGene_gn=NT5C2;refGene_tx=XM_023240880.1
D2 63459361 63459422
refGene_gn=NT5C2;refGene_tx=XM_011287440.3;refGene_gn=NT5C2;refGene_tx=XM_019813770.2;refGene_gn=NT5C2;refGene_tx=XM_023240876.1;refGene_gn=NT5C2;refGene_tx=XM_023240877.1;refGene_gn=NT5C2;refGene_tx=XM_023240878.1;refGene_gn=NT5C2;refGene_tx=XM_023240879.1;refGene_gn=NT5C2;refGene_tx=XM_023240880.1
D2 63459548 63459600
refGene_gn=NT5C2;refGene_tx=XM_011287440.3;refGene_gn=NT5C2;refGene_tx=XM_019813770.2;refGene_gn=NT5C2;refGene_tx=XM_023240876.1;refGene_gn=NT5C2;refGene_tx=XM_023240877.1;refGene_gn=NT5C2;refGene_tx=XM_023240878.1;refGene_gn=NT5C2;refGene_tx=XM_023240879.1;refGene_gn=NT5C2;refGene_tx=XM_023240880.1
D2 63460872 63461043
refGene_gn=NT5C2;refGene_tx=XM_011287440.3;refGene_gn=NT5C2;refGene_tx=XM_019813770.2;refGene_gn=NT5C2;refGene_tx=XM_023240876.1;refGene_gn=NT5C2;refGene_tx=XM_023240877.1;refGene_gn=NT5C2;refGene_tx=XM_023240878.1;refGene_gn=NT5C2;refGene_tx=XM_023240879.1;refGene_gn=NT5C2;refGene_tx=XM_023240880.1
D2 63461748 63461815
refGene_gn=NT5C2;refGene_tx=XM_011287440.3;refGene_gn=NT5C2;refGene_tx=XM_019813770.2;refGene_gn=NT5C2;refGene_tx=XM_023240876.1;refGene_gn=NT5C2;refGene_tx=XM_023240877.1;refGene_gn=NT5C2;refGene_tx=XM_023240878.1;refGene_gn=NT5C2;refGene_tx=XM_023240879.1;refGene_gn=NT5C2;refGene_tx=XM_023240880.1
D2 63462086 63462194
refGene_gn=NT5C2;refGene_tx=XM_011287440.3;refGene_gn=NT5C2;refGene_tx=XM_019813770.2;refGene_gn=NT5C2;refGene_tx=XM_023240876.1;refGene_gn=NT5C2;refGene_tx=XM_023240877.1;refGene_gn=NT5C2;refGene_tx=XM_023240878.1;refGene_gn=NT5C2;refGene_tx=XM_023240879.1;refGene_gn=NT5C2;refGene_tx=XM_023240880.1
D2 63463295 63463337
refGene_gn=NT5C2;refGene_tx=XM_011287440.3;refGene_gn=NT5C2;refGene_tx=XM_019813770.2;refGene_gn=NT5C2;refGene_tx=XM_023240876.1;refGene_gn=NT5C2;refGene_tx=XM_023240877.1;refGene_gn=NT5C2;refGene_tx=XM_023240878.1;refGene_gn=NT5C2;refGene_tx=XM_023240879.1;refGene_gn=NT5C2;refGene_tx=XM_023240880.1
D2 63464389 63464473
refGene_gn=NT5C2;refGene_tx=XM_011287440.3;refGene_gn=NT5C2;refGene_tx=XM_019813770.2;refGene_gn=NT5C2;refGene_tx=XM_023240876.1;refGene_gn=NT5C2;refGene_tx=XM_023240877.1;refGene_gn=NT5C2;refGene_tx=XM_023240878.1;refGene_gn=NT5C2;refGene_tx=XM_023240879.1;refGene_gn=NT5C2;refGene_tx=XM_023240880.1
D2 63467007 63467061

refGene_gn=NT5C2;refGene_tx=XM_011287440.3;refGene_gn=NT5C2;refGene_tx=XM_019813770.2;refGene_gn=NT5C2;refGene_tx=XM_023240876.1;refGene_gn=NT5C2;refGene_tx=XM_023240877.1;refGene_gn=NT5C2;refGene_tx=XM_023240878.1;refGene_gn=NT5C2;refGene_tx=XM_023240879.1;refGene_gn=NT5C2;refGene_tx=XM_023240880.1
D2 63467682 63467776
refGene_gn=NT5C2;refGene_tx=XM_011287440.3;refGene_gn=NT5C2;refGene_tx=XM_019813770.2;refGene_gn=NT5C2;refGene_tx=XM_023240876.1;refGene_gn=NT5C2;refGene_tx=XM_023240877.1;refGene_gn=NT5C2;refGene_tx=XM_023240878.1;refGene_gn=NT5C2;refGene_tx=XM_023240879.1;refGene_gn=NT5C2;refGene_tx=XM_023240880.1
D2 63468814 63468872
refGene_gn=NT5C2;refGene_tx=XM_011287440.3;refGene_gn=NT5C2;refGene_tx=XM_019813770.2;refGene_gn=NT5C2;refGene_tx=XM_023240876.1;refGene_gn=NT5C2;refGene_tx=XM_023240877.1;refGene_gn=NT5C2;refGene_tx=XM_023240878.1;refGene_gn=NT5C2;refGene_tx=XM_023240879.1
D2 63469004 63469096
refGene_gn=NT5C2;refGene_tx=XM_011287440.3;refGene_gn=NT5C2;refGene_tx=XM_019813770.2;refGene_gn=NT5C2;refGene_tx=XM_023240876.1;refGene_gn=NT5C2;refGene_tx=XM_023240877.1;refGene_gn=NT5C2;refGene_tx=XM_023240878.1;refGene_gn=NT5C2;refGene_tx=XM_023240879.1
D2 63473465 63473489
refGene_gn=NT5C2;refGene_tx=XM_011287440.3;refGene_gn=NT5C2;refGene_tx=XM_019813770.2;refGene_gn=NT5C2;refGene_tx=XM_023240877.1;refGene_gn=NT5C2;refGene_tx=XM_023240878.1
D2 63473819 63473915
refGene_gn=NT5C2;refGene_tx=XM_011287440.3;refGene_gn=NT5C2;refGene_tx=XM_019813770.2;refGene_gn=NT5C2;refGene_tx=XM_023240876.1;refGene_gn=NT5C2;refGene_tx=XM_023240877.1;refGene_gn=NT5C2;refGene_tx=XM_023240878.1;refGene_gn=NT5C2;refGene_tx=XM_023240879.1
D2 63475182 63475300
refGene_gn=NT5C2;refGene_tx=XM_011287440.3;refGene_gn=NT5C2;refGene_tx=XM_019813770.2;refGene_gn=NT5C2;refGene_tx=XM_023240876.1;refGene_gn=NT5C2;refGene_tx=XM_023240877.1;refGene_gn=NT5C2;refGene_tx=XM_023240878.1;refGene_gn=NT5C2;refGene_tx=XM_023240879.1
D2 63503779 63503853
refGene_gn=NT5C2;refGene_tx=XM_011287440.3;refGene_gn=NT5C2;refGene_tx=XM_019813770.2;refGene_gn=NT5C2;refGene_tx=XM_023240876.1;refGene_gn=NT5C2;refGene_tx=XM_023240877.1;refGene_gn=NT5C2;refGene_tx=XM_023240878.1;refGene_gn=NT5C2;refGene_tx=XM_023240879.1
D2 63508736 63508741 refGene_gn=NT5C2;refGene_tx=XM_011287440.3
D2 63528404 63528466 refGene_gn=NT5C2;refGene_tx=XM_023240876.1
D2 63531005 63531106
refGene_gn=NT5C2;refGene_tx=XM_023240877.1;refGene_gn=NT5C2;refGene_tx=XM_023240878.1;refGene_gn=NT5C2;refGene_tx=XM_023240879.1
D2 63607074 63608136 refGene_gn=INA;refGene_tx=XM_003994377.5
D2 63616619 63616744 refGene_gn=INA;refGene_tx=XM_003994377.5
D2 63618515 63618594 refGene_gn=INA;refGene_tx=XM_003994377.5
D2 63618595 63618829 refGene_gn=INA;refGene_tx=XM_003994377.5
D2 63630217 63630274 refGene_gn=PCGF6;refGene_tx=XM_023240883.1

D2 63639410 63639497 refGene_gn=PCGF6;refGene_tx=XM_023240883.1
D2 63646273 63646372 refGene_gn=PCGF6;refGene_tx=XM_023240883.1
D2 63654536 63654564 refGene_gn=PCGF6;refGene_tx=XM_023240883.1
D2 63659544 63659653 refGene_gn=PCGF6;refGene_tx=XM_023240883.1
D2 63660533 63660593 refGene_gn=PCGF6;refGene_tx=XM_023240883.1
D2 63660693 63660749 refGene_gn=PCGF6;refGene_tx=XM_023240883.1
D2 63661564 63661661 refGene_gn=PCGF6;refGene_tx=XM_023240883.1
D2 63661751 63661851 refGene_gn=PCGF6;refGene_tx=XM_023240883.1
D2 63662651 63663008 refGene_gn=PCGF6;refGene_tx=XM_023240883.1
D2 63676516 63677078 refGene_gn=TAF5;refGene_tx=XM_003994378.5
D2 63679391 63679629 refGene_gn=TAF5;refGene_tx=XM_003994378.5
D2 63683380 63683696 refGene_gn=TAF5;refGene_tx=XM_003994378.5
D2 63684463 63684532 refGene_gn=TAF5;refGene_tx=XM_003994378.5
D2 63684533 63684628 refGene_gn=TAF5;refGene_tx=XM_003994378.5
D2 63684731 63684867 refGene_gn=TAF5;refGene_tx=XM_003994378.5
D2 63685508 63685629 refGene_gn=TAF5;refGene_tx=XM_003994378.5
D2 63686493 63686623 refGene_gn=TAF5;refGene_tx=XM_003994378.5
D2 63688152 63688317 refGene_gn=TAF5;refGene_tx=XM_003994378.5
D2 63688930 63689108 refGene_gn=TAF5;refGene_tx=XM_003994378.5
D2 63689265 63689443 refGene_gn=TAF5;refGene_tx=XM_003994378.5
D2 63689727 63689945 refGene_gn=TAF5;refGene_tx=XM_003994378.5
D2 63692615 63692705
refGene_gn=ATP5MD;refGene_tx=XM_003994380.3;refGene_gn=ATP5MD;refGene_
tx=XM_006938150.4;refGene_gn=ATP5MD;refGene_tx=XM_011287438.2
D2 63692794 63692881
refGene_gn=ATP5MD;refGene_tx=XM_003994380.3;refGene_gn=ATP5MD;refGene_
tx=XM_006938150.4;refGene_gn=ATP5MD;refGene_tx=XM_011287438.2
D2 63697867 63697960
refGene_gn=PDCD11;refGene_tx=XM_023240881.1;refGene_gn=PDCD11;refGene_
tx=XM_023240882.1
D2 63699013 63699145
refGene_gn=PDCD11;refGene_tx=XM_023240881.1;refGene_gn=PDCD11;refGene_
tx=XM_023240882.1
D2 63700988 63701156
refGene_gn=PDCD11;refGene_tx=XM_023240881.1;refGene_gn=PDCD11;refGene_
tx=XM_023240882.1
D2 63702618 63702780
refGene_gn=PDCD11;refGene_tx=XM_023240881.1;refGene_gn=PDCD11;refGene_
tx=XM_023240882.1
D2 63704088 63704212
refGene_gn=PDCD11;refGene_tx=XM_023240881.1;refGene_gn=PDCD11;refGene_
tx=XM_023240882.1
D2 63704707 63704889
refGene_gn=PDCD11;refGene_tx=XM_023240881.1;refGene_gn=PDCD11;refGene_
tx=XM_023240882.1
D2 63709065 63709173
refGene_gn=PDCD11;refGene_tx=XM_023240881.1;refGene_gn=PDCD11;refGene_
tx=XM_023240882.1
D2 63710704 63710911
refGene_gn=PDCD11;refGene_tx=XM_023240881.1;refGene_gn=PDCD11;refGene_

tx=XM_023240882.1
D2 63711557 63711682
refGene_gn=PDCD11;refGene_tx=XM_023240881.1;refGene_gn=PDCD11;refGene_tx=XM_023240882.1
D2 63711865 63711926
refGene_gn=PDCD11;refGene_tx=XM_023240881.1;refGene_gn=PDCD11;refGene_tx=XM_023240882.1
D2 63712575 63712722
refGene_gn=PDCD11;refGene_tx=XM_023240881.1;refGene_gn=PDCD11;refGene_tx=XM_023240882.1
D2 63713925 63714177
refGene_gn=PDCD11;refGene_tx=XM_023240881.1;refGene_gn=PDCD11;refGene_tx=XM_023240882.1
D2 63714746 63714881
refGene_gn=PDCD11;refGene_tx=XM_023240881.1;refGene_gn=PDCD11;refGene_tx=XM_023240882.1
D2 63715630 63715825
refGene_gn=PDCD11;refGene_tx=XM_023240881.1;refGene_gn=PDCD11;refGene_tx=XM_023240882.1
D2 63716672 63716843
refGene_gn=PDCD11;refGene_tx=XM_023240881.1;refGene_gn=PDCD11;refGene_tx=XM_023240882.1
D2 63717768 63717988
refGene_gn=PDCD11;refGene_tx=XM_023240881.1;refGene_gn=PDCD11;refGene_tx=XM_023240882.1
D2 63718770 63718920
refGene_gn=PDCD11;refGene_tx=XM_023240881.1;refGene_gn=PDCD11;refGene_tx=XM_023240882.1
D2 63719349 63719465
refGene_gn=PDCD11;refGene_tx=XM_023240881.1;refGene_gn=PDCD11;refGene_tx=XM_023240882.1
D2 63720716 63721258
refGene_gn=PDCD11;refGene_tx=XM_023240881.1;refGene_gn=PDCD11;refGene_tx=XM_023240882.1
D2 63722113 63722176
refGene_gn=PDCD11;refGene_tx=XM_023240881.1;refGene_gn=PDCD11;refGene_tx=XM_023240882.1
D2 63726137 63726246
refGene_gn=PDCD11;refGene_tx=XM_023240881.1;refGene_gn=PDCD11;refGene_tx=XM_023240882.1
D2 63727835 63727925
refGene_gn=PDCD11;refGene_tx=XM_023240881.1;refGene_gn=PDCD11;refGene_tx=XM_023240882.1
D2 63728153 63728256
refGene_gn=PDCD11;refGene_tx=XM_023240881.1;refGene_gn=PDCD11;refGene_tx=XM_023240882.1
D2 63728652 63728830
refGene_gn=PDCD11;refGene_tx=XM_023240881.1;refGene_gn=PDCD11;refGene_tx=XM_023240882.1
D2 63733092 63733149

refGene_gn=PDCD11;refGene_tx=XM_023240881.1;refGene_gn=PDCD11;refGene_tx=XM_023240882.1
D2 63733661 63733784
refGene_gn=PDCD11;refGene_tx=XM_023240881.1;refGene_gn=PDCD11;refGene_tx=XM_023240882.1
D2 63734631 63734754
refGene_gn=PDCD11;refGene_tx=XM_023240881.1;refGene_gn=PDCD11;refGene_tx=XM_023240882.1
D2 63735168 63735472
refGene_gn=PDCD11;refGene_tx=XM_023240881.1;refGene_gn=PDCD11;refGene_tx=XM_023240882.1
D2 63735694 63735811
refGene_gn=PDCD11;refGene_tx=XM_023240881.1;refGene_gn=PDCD11;refGene_tx=XM_023240882.1
D2 63736514 63736664
refGene_gn=PDCD11;refGene_tx=XM_023240881.1;refGene_gn=PDCD11;refGene_tx=XM_023240882.1
D2 63736870 63737118
refGene_gn=PDCD11;refGene_tx=XM_023240881.1;refGene_gn=PDCD11;refGene_tx=XM_023240882.1
D2 63737842 63738011
refGene_gn=PDCD11;refGene_tx=XM_023240881.1;refGene_gn=PDCD11;refGene_tx=XM_023240882.1
D2 63738584 63738738
refGene_gn=PDCD11;refGene_tx=XM_023240881.1;refGene_gn=PDCD11;refGene_tx=XM_023240882.1
D2 63739192 63739358
refGene_gn=PDCD11;refGene_tx=XM_023240881.1;refGene_gn=PDCD11;refGene_tx=XM_023240882.1
D2 63739949 63740121
refGene_gn=PDCD11;refGene_tx=XM_023240881.1;refGene_gn=PDCD11;refGene_tx=XM_023240882.1
D2 63742217 63742634
refGene_gn=CALHM2;refGene_tx=XM_006938151.4;refGene_gn=CALHM2;refGene_tx=XM_023240884.1;refGene_gn=CALHM2;refGene_tx=XM_023240885.1
D2 63743724 63744279
refGene_gn=CALHM2;refGene_tx=XM_006938151.4;refGene_gn=CALHM2;refGene_tx=XM_023240884.1;refGene_gn=CALHM2;refGene_tx=XM_023240885.1
D2 63749402 63749888 refGene_gn=CALHM1;refGene_tx=XM_003994382.4
D2 63752249 63752801 refGene_gn=CALHM1;refGene_tx=XM_003994382.4
D2 63763667 63764168
refGene_gn=CALHM3;refGene_tx=XM_003994383.4;refGene_gn=CALHM3;refGene_tx=XM_019814150.2
D2 63766987 63767243
refGene_gn=CALHM3;refGene_tx=XM_003994383.4;refGene_gn=CALHM3;refGene_tx=XM_019814150.2
D2 63768794 63769081
refGene_gn=CALHM3;refGene_tx=XM_003994383.4;refGene_gn=CALHM3;refGene_tx=XM_019814150.2
D2 63789033 63789118 refGene_gn=NEURL1;refGene_tx=XM_023240887.1

D2	63833590	63833624	refGene_gn=NEURL1;refGene_tx=XM_023240889.1
D2	63833639	63833706	refGene_gn=NEURL1;refGene_tx=XM_023240888.1
D2	63846959	63847201
refGene_gn=NEURL1;refGene_tx=XM_023240887.1;refGene_gn=NEURL1;refGene_tx=XM_023240888.1;refGene_gn=NEURL1;refGene_tx=XM_023240889.1;refGene_gn=NEURL1;refGene_tx=XM_023240890.1
D2	63847603	63847925
refGene_gn=NEURL1;refGene_tx=XM_023240887.1;refGene_gn=NEURL1;refGene_tx=XM_023240888.1;refGene_gn=NEURL1;refGene_tx=XM_023240889.1;refGene_gn=NEURL1;refGene_tx=XM_023240890.1
D2	63860228	63860918
refGene_gn=NEURL1;refGene_tx=XM_023240887.1;refGene_gn=NEURL1;refGene_tx=XM_023240888.1;refGene_gn=NEURL1;refGene_tx=XM_023240889.1;refGene_gn=NEURL1;refGene_tx=XM_023240890.1
D2	63864591	63864738
refGene_gn=NEURL1;refGene_tx=XM_023240887.1;refGene_gn=NEURL1;refGene_tx=XM_023240888.1;refGene_gn=NEURL1;refGene_tx=XM_023240889.1;refGene_gn=NEURL1;refGene_tx=XM_023240890.1
D2	63865214	63865453
refGene_gn=NEURL1;refGene_tx=XM_023240887.1;refGene_gn=NEURL1;refGene_tx=XM_023240888.1;refGene_gn=NEURL1;refGene_tx=XM_023240889.1;refGene_gn=NEURL1;refGene_tx=XM_023240890.1
D2	63876448	63878416
refGene_gn=SH3PXD2A;refGene_tx=XM_011287436.3;refGene_gn=SH3PXD2A;refGene_tx=XM_019813771.2;refGene_gn=SH3PXD2A;refGene_tx=XM_019813772.2;refGene_gn=SH3PXD2A;refGene_tx=XM_019813773.2;refGene_gn=SH3PXD2A;refGene_tx=XM_019813774.2;refGene_gn=SH3PXD2A;refGene_tx=XM_019813775.2;refGene_gn=SH3PXD2A;refGene_tx=XM_019813778.2;refGene_gn=SH3PXD2A;refGene_tx=XM_023240886.1
D2	63880382	63880502
refGene_gn=SH3PXD2A;refGene_tx=XM_011287436.3;refGene_gn=SH3PXD2A;refGene_tx=XM_019813771.2;refGene_gn=SH3PXD2A;refGene_tx=XM_019813772.2;refGene_gn=SH3PXD2A;refGene_tx=XM_019813773.2;refGene_gn=SH3PXD2A;refGene_tx=XM_019813774.2;refGene_gn=SH3PXD2A;refGene_tx=XM_019813775.2;refGene_gn=SH3PXD2A;refGene_tx=XM_019813778.2;refGene_gn=SH3PXD2A;refGene_tx=XM_023240886.1
D2	63883160	63883210
refGene_gn=SH3PXD2A;refGene_tx=XM_011287436.3;refGene_gn=SH3PXD2A;refGene_tx=XM_019813771.2;refGene_gn=SH3PXD2A;refGene_tx=XM_019813772.2;refGene_gn=SH3PXD2A;refGene_tx=XM_019813773.2;refGene_gn=SH3PXD2A;refGene_tx=XM_019813774.2;refGene_gn=SH3PXD2A;refGene_tx=XM_019813775.2;refGene_gn=SH3PXD2A;refGene_tx=XM_019813778.2;refGene_gn=SH3PXD2A;refGene_tx=XM_023240886.1
D2	63884158	63884496
refGene_gn=SH3PXD2A;refGene_tx=XM_011287436.3;refGene_gn=SH3PXD2A;refGene_tx=XM_019813771.2;refGene_gn=SH3PXD2A;refGene_tx=XM_019813772.2;refGene_gn=SH3PXD2A;refGene_tx=XM_019813773.2;refGene_gn=SH3PXD2A;refGene_tx=XM_019813774.2;refGene_gn=SH3PXD2A;refGene_tx=XM_019813775.2;refGene_gn=SH3PXD2A;refGene_tx=XM_019813778.2;refGene_gn=SH3PXD2A;refGene_tx=XM_023240886.1

D2 63888306 63888424
refGene_gn=SH3PXD2A;refGene_tx=XM_011287436.3;refGene_gn=SH3PXD2A;refGene_tx=XM_019813771.2;refGene_gn=SH3PXD2A;refGene_tx=XM_019813772.2;refGene_gn=SH3PXD2A;refGene_tx=XM_019813773.2;refGene_gn=SH3PXD2A;refGene_tx=XM_019813774.2;refGene_gn=SH3PXD2A;refGene_tx=XM_019813775.2;refGene_gn=SH3PXD2A;refGene_tx=XM_019813778.2;refGene_gn=SH3PXD2A;refGene_tx=XM_023240886.1
D2 63893499 63893583
refGene_gn=SH3PXD2A;refGene_tx=XM_011287436.3;refGene_gn=SH3PXD2A;refGene_tx=XM_019813771.2;refGene_gn=SH3PXD2A;refGene_tx=XM_019813773.2;refGene_gn=SH3PXD2A;refGene_tx=XM_019813774.2;refGene_gn=SH3PXD2A;refGene_tx=XM_019813775.2;refGene_gn=SH3PXD2A;refGene_tx=XM_019813778.2;refGene_gn=SH3PXD2A;refGene_tx=XM_023240886.1
D2 63897735 63897849
refGene_gn=SH3PXD2A;refGene_tx=XM_011287436.3;refGene_gn=SH3PXD2A;refGene_tx=XM_019813771.2;refGene_gn=SH3PXD2A;refGene_tx=XM_019813772.2;refGene_gn=SH3PXD2A;refGene_tx=XM_019813773.2;refGene_gn=SH3PXD2A;refGene_tx=XM_019813774.2;refGene_gn=SH3PXD2A;refGene_tx=XM_019813775.2;refGene_gn=SH3PXD2A;refGene_tx=XM_019813778.2;refGene_gn=SH3PXD2A;refGene_tx=XM_023240886.1
D2 63926775 63926907
refGene_gn=SH3PXD2A;refGene_tx=XM_011287436.3;refGene_gn=SH3PXD2A;refGene_tx=XM_019813771.2;refGene_gn=SH3PXD2A;refGene_tx=XM_019813772.2;refGene_gn=SH3PXD2A;refGene_tx=XM_019813773.2;refGene_gn=SH3PXD2A;refGene_tx=XM_019813774.2;refGene_gn=SH3PXD2A;refGene_tx=XM_019813775.2;refGene_gn=SH3PXD2A;refGene_tx=XM_019813778.2;refGene_gn=SH3PXD2A;refGene_tx=XM_023240886.1
D2 63933434 63933479
refGene_gn=SH3PXD2A;refGene_tx=XM_019813771.2;refGene_gn=SH3PXD2A;refGene_tx=XM_019813772.2;refGene_gn=SH3PXD2A;refGene_tx=XM_019813773.2;refGene_gn=SH3PXD2A;refGene_tx=XM_019813774.2;refGene_gn=SH3PXD2A;refGene_tx=XM_019813775.2
D2 63954615 63954644
refGene_gn=SH3PXD2A;refGene_tx=XM_011287436.3;refGene_gn=SH3PXD2A;refGene_tx=XM_019813771.2;refGene_gn=SH3PXD2A;refGene_tx=XM_019813772.2;refGene_gn=SH3PXD2A;refGene_tx=XM_019813773.2;refGene_gn=SH3PXD2A;refGene_tx=XM_019813774.2;refGene_gn=SH3PXD2A;refGene_tx=XM_019813775.2
D2 63980792 63980884
refGene_gn=SH3PXD2A;refGene_tx=XM_011287436.3;refGene_gn=SH3PXD2A;refGene_tx=XM_019813771.2;refGene_gn=SH3PXD2A;refGene_tx=XM_019813772.2;refGene_gn=SH3PXD2A;refGene_tx=XM_019813773.2;refGene_gn=SH3PXD2A;refGene_tx=XM_019813774.2;refGene_gn=SH3PXD2A;refGene_tx=XM_019813775.2
D2 63991028 63991105
refGene_gn=SH3PXD2A;refGene_tx=XM_011287436.3;refGene_gn=SH3PXD2A;refGene_tx=XM_019813771.2;refGene_gn=SH3PXD2A;refGene_tx=XM_019813772.2;refGene_gn=SH3PXD2A;refGene_tx=XM_019813773.2;refGene_gn=SH3PXD2A;refGene_tx=XM_019813774.2;refGene_gn=SH3PXD2A;refGene_tx=XM_019813775.2
D2 64021443 64021519
refGene_gn=SH3PXD2A;refGene_tx=XM_011287436.3;refGene_gn=SH3PXD2A;refGene_tx=XM_019813771.2;refGene_gn=SH3PXD2A;refGene_tx=XM_019813772.2;re

fGene_gn=SH3PXD2A;refGene_tx=XM_019813773.2;refGene_gn=SH3PXD2A;refGene_tx=XM_019813774.2;refGene_gn=SH3PXD2A;refGene_tx=XM_019813775.2
D2 64047359 64047392 refGene_gn=SH3PXD2A;refGene_tx=XM_019813774.2
D2 64056346 64056412 refGene_gn=SH3PXD2A;refGene_tx=XM_019813773.2
D2 64057694 64057775
refGene_gn=SH3PXD2A;refGene_tx=XM_011287436.3;refGene_gn=SH3PXD2A;refGene_tx=XM_019813771.2;refGene_gn=SH3PXD2A;refGene_tx=XM_019813772.2
D2 64111160 64111232
refGene_gn=SH3PXD2A;refGene_tx=XM_011287436.3;refGene_gn=SH3PXD2A;refGene_tx=XM_019813771.2;refGene_gn=SH3PXD2A;refGene_tx=XM_019813772.2
D2 64124073 64124126 refGene_gn=STN1;refGene_tx=XM_019813781.2
D2 64145868 64146026
refGene_gn=STN1;refGene_tx=XM_006938152.3;refGene_gn=STN1;refGene_tx=XM_011287432.3;refGene_gn=STN1;refGene_tx=XM_011287433.3;refGene_gn=STN1;refGene_tx=XM_019813779.2;refGene_gn=STN1;refGene_tx=XM_019813780.2;refGene_gn=STN1;refGene_tx=XM_019813782.2;refGene_gn=STN1;refGene_tx=XM_019813783.2
D2 64158042 64158115
refGene_gn=STN1;refGene_tx=XM_006938152.3;refGene_gn=STN1;refGene_tx=XM_011287432.3;refGene_gn=STN1;refGene_tx=XM_011287433.3;refGene_gn=STN1;refGene_tx=XM_019813779.2;refGene_gn=STN1;refGene_tx=XM_019813780.2;refGene_gn=STN1;refGene_tx=XM_019813781.2;refGene_gn=STN1;refGene_tx=XM_019813782.2;refGene_gn=STN1;refGene_tx=XM_019813783.2
D2 64161413 64161536
refGene_gn=STN1;refGene_tx=XM_006938152.3;refGene_gn=STN1;refGene_tx=XM_011287432.3;refGene_gn=STN1;refGene_tx=XM_011287433.3;refGene_gn=STN1;refGene_tx=XM_019813779.2;refGene_gn=STN1;refGene_tx=XM_019813780.2;refGene_gn=STN1;refGene_tx=XM_019813781.2;refGene_gn=STN1;refGene_tx=XM_019813783.2
D2 64167541 64167716
refGene_gn=STN1;refGene_tx=XM_006938152.3;refGene_gn=STN1;refGene_tx=XM_011287432.3;refGene_gn=STN1;refGene_tx=XM_011287433.3;refGene_gn=STN1;refGene_tx=XM_019813779.2;refGene_gn=STN1;refGene_tx=XM_019813780.2;refGene_gn=STN1;refGene_tx=XM_019813781.2;refGene_gn=STN1;refGene_tx=XM_019813782.2;refGene_gn=STN1;refGene_tx=XM_019813783.2
D2 64171441 64171556
refGene_gn=STN1;refGene_tx=XM_006938152.3;refGene_gn=STN1;refGene_tx=XM_011287432.3;refGene_gn=STN1;refGene_tx=XM_011287433.3;refGene_gn=STN1;refGene_tx=XM_019813779.2;refGene_gn=STN1;refGene_tx=XM_019813780.2;refGene_gn=STN1;refGene_tx=XM_019813781.2;refGene_gn=STN1;refGene_tx=XM_019813782.2;refGene_gn=STN1;refGene_tx=XM_019813783.2
D2 64172297 64172471
refGene_gn=STN1;refGene_tx=XM_006938152.3;refGene_gn=STN1;refGene_tx=XM_011287432.3;refGene_gn=STN1;refGene_tx=XM_011287433.3;refGene_gn=STN1;refGene_tx=XM_019813779.2;refGene_gn=STN1;refGene_tx=XM_019813780.2;refGene_gn=STN1;refGene_tx=XM_019813781.2;refGene_gn=STN1;refGene_tx=XM_019813782.2
D2 64177708 64177777
refGene_gn=STN1;refGene_tx=XM_006938152.3;refGene_gn=STN1;refGene_tx=XM_011287432.3;refGene_gn=STN1;refGene_tx=XM_011287433.3;refGene_gn=STN

1;refGene_tx=XM_019813779.2;refGene_gn=STN1;refGene_tx=XM_019813781.2;refGene_gn=STN1;refGene_tx=XM_019813782.2;refGene_gn=STN1;refGene_tx=XM_019813783.2
D2 64184092 64184188 refGene_gn=STN1;refGene_tx=XM_006938152.3;refGene_gn=STN1;refGene_tx=XM_011287432.3;refGene_gn=STN1;refGene_tx=XM_011287433.3;refGene_gn=STN1;refGene_tx=XM_019813779.2;refGene_gn=STN1;refGene_tx=XM_019813780.2;refGene_gn=STN1;refGene_tx=XM_019813781.2;refGene_gn=STN1;refGene_tx=XM_019813782.2;refGene_gn=STN1;refGene_tx=XM_019813783.2
D2 64190791 64190924 refGene_gn=STN1;refGene_tx=XM_006938152.3;refGene_gn=STN1;refGene_tx=XM_011287432.3;refGene_gn=STN1;refGene_tx=XM_011287433.3;refGene_gn=STN1;refGene_tx=XM_019813779.2;refGene_gn=STN1;refGene_tx=XM_019813780.2;refGene_gn=STN1;refGene_tx=XM_019813781.2;refGene_gn=STN1;refGene_tx=XM_019813782.2;refGene_gn=STN1;refGene_tx=XM_019813783.2
D2 64234708 64234858 refGene_gn=SLK;refGene_tx=XM_011287641.3;refGene_gn=SLK;refGene_tx=XM_019813784.2
D2 64255177 64255342 refGene_gn=SLK;refGene_tx=XM_011287641.3;refGene_gn=SLK;refGene_tx=XM_019813784.2
D2 64257274 64257323 refGene_gn=SLK;refGene_tx=XM_011287641.3;refGene_gn=SLK;refGene_tx=XM_019813784.2
D2 64257673 64257823 refGene_gn=SLK;refGene_tx=XM_011287641.3;refGene_gn=SLK;refGene_tx=XM_019813784.2
D2 64263180 64263253 refGene_gn=SLK;refGene_tx=XM_011287641.3;refGene_gn=SLK;refGene_tx=XM_019813784.2
D2 64263417 64263612 refGene_gn=SLK;refGene_tx=XM_011287641.3;refGene_gn=SLK;refGene_tx=XM_019813784.2
D2 64264382 64264464 refGene_gn=SLK;refGene_tx=XM_011287641.3;refGene_gn=SLK;refGene_tx=XM_019813784.2
D2 64265963 64266092 refGene_gn=SLK;refGene_tx=XM_011287641.3;refGene_gn=SLK;refGene_tx=XM_019813784.2
D2 64266426 64267800 refGene_gn=SLK;refGene_tx=XM_011287641.3;refGene_gn=SLK;refGene_tx=XM_019813784.2
D2 64269746 64269877 refGene_gn=SLK;refGene_tx=XM_011287641.3;refGene_gn=SLK;refGene_tx=XM_019813784.2
D2 64270085 64270209 refGene_gn=SLK;refGene_tx=XM_011287641.3;refGene_gn=SLK;refGene_tx=XM_019813784.2
D2 64271720 64271900 refGene_gn=SLK;refGene_tx=XM_011287641.3;refGene_gn=SLK;refGene_tx=XM_

019813784.2
D2	64274242 64274335 refGene_gn=SLK;refGene_tx=XM_019813784.2
D2	64280876 64281006
refGene_gn=SLK;refGene_tx=XM_011287641.3;refGene_gn=SLK;refGene_tx=XM_
019813784.2
D2	64281935 64282060
refGene_gn=SLK;refGene_tx=XM_011287641.3;refGene_gn=SLK;refGene_tx=XM_
019813784.2
D2	64282819 64283008
refGene_gn=SLK;refGene_tx=XM_011287641.3;refGene_gn=SLK;refGene_tx=XM_
019813784.2
D2	64283544 64283670
refGene_gn=SLK;refGene_tx=XM_011287641.3;refGene_gn=SLK;refGene_tx=XM_
019813784.2
D2	64284582 64284696
refGene_gn=SLK;refGene_tx=XM_011287641.3;refGene_gn=SLK;refGene_tx=XM_
019813784.2
D2	64288696 64288843
refGene_gn=SLK;refGene_tx=XM_011287641.3;refGene_gn=SLK;refGene_tx=XM_
019813784.2
D2	64295603 64295656
refGene_gn=COL17A1;refGene_tx=XM_006938156.4;refGene_gn=COL17A1;refGen
e_tx=XM_019813785.2;refGene_gn=COL17A1;refGene_tx=XM_019813786.2
D2	64296012 64296093
refGene_gn=COL17A1;refGene_tx=XM_006938156.4;refGene_gn=COL17A1;refGen
e_tx=XM_019813785.2;refGene_gn=COL17A1;refGene_tx=XM_019813786.2
D2	64296248 64296311
refGene_gn=COL17A1;refGene_tx=XM_006938156.4;refGene_gn=COL17A1;refGen
e_tx=XM_019813785.2;refGene_gn=COL17A1;refGene_tx=XM_019813786.2
D2	64296557 64296695
refGene_gn=COL17A1;refGene_tx=XM_006938156.4;refGene_gn=COL17A1;refGen
e_tx=XM_019813785.2;refGene_gn=COL17A1;refGene_tx=XM_019813786.2
D2	64297273 64297627
refGene_gn=COL17A1;refGene_tx=XM_006938156.4;refGene_gn=COL17A1;refGen
e_tx=XM_019813785.2;refGene_gn=COL17A1;refGene_tx=XM_019813786.2
D2	64297879 64298026
refGene_gn=COL17A1;refGene_tx=XM_006938156.4;refGene_gn=COL17A1;refGen
e_tx=XM_019813785.2;refGene_gn=COL17A1;refGene_tx=XM_019813786.2
D2	64298468 64298579
refGene_gn=COL17A1;refGene_tx=XM_006938156.4;refGene_gn=COL17A1;refGen
e_tx=XM_019813785.2;refGene_gn=COL17A1;refGene_tx=XM_019813786.2
D2	64298680 64298770
refGene_gn=COL17A1;refGene_tx=XM_006938156.4;refGene_gn=COL17A1;refGen
e_tx=XM_019813785.2;refGene_gn=COL17A1;refGene_tx=XM_019813786.2
D2	64299239 64299341
refGene_gn=COL17A1;refGene_tx=XM_006938156.4;refGene_gn=COL17A1;refGen
e_tx=XM_019813785.2;refGene_gn=COL17A1;refGene_tx=XM_019813786.2
D2	64299652 64299718
refGene_gn=COL17A1;refGene_tx=XM_006938156.4;refGene_gn=COL17A1;refGen
e_tx=XM_019813785.2;refGene_gn=COL17A1;refGene_tx=XM_019813786.2

D2 64300200 64300338
refGene_gn=COL17A1;refGene_tx=XM_006938156.4;refGene_gn=COL17A1;refGene_tx=XM_019813785.2;refGene_gn=COL17A1;refGene_tx=XM_019813786.2
D2 64301078 64301204
refGene_gn=COL17A1;refGene_tx=XM_006938156.4;refGene_gn=COL17A1;refGene_tx=XM_019813785.2;refGene_gn=COL17A1;refGene_tx=XM_019813786.2
D2 64301687 64301738
refGene_gn=COL17A1;refGene_tx=XM_006938156.4;refGene_gn=COL17A1;refGene_tx=XM_019813785.2;refGene_gn=COL17A1;refGene_tx=XM_019813786.2
D2 64302045 64302111
refGene_gn=COL17A1;refGene_tx=XM_006938156.4;refGene_gn=COL17A1;refGene_tx=XM_019813785.2;refGene_gn=COL17A1;refGene_tx=XM_019813786.2
D2 64302196 64302229
refGene_gn=COL17A1;refGene_tx=XM_006938156.4;refGene_gn=COL17A1;refGene_tx=XM_019813785.2;refGene_gn=COL17A1;refGene_tx=XM_019813786.2
D2 64302478 64302505
refGene_gn=COL17A1;refGene_tx=XM_006938156.4;refGene_gn=COL17A1;refGene_tx=XM_019813785.2;refGene_gn=COL17A1;refGene_tx=XM_019813786.2
D2 64302839 64302899
refGene_gn=COL17A1;refGene_tx=XM_006938156.4;refGene_gn=COL17A1;refGene_tx=XM_019813785.2;refGene_gn=COL17A1;refGene_tx=XM_019813786.2
D2 64303281 64303335
refGene_gn=COL17A1;refGene_tx=XM_006938156.4;refGene_gn=COL17A1;refGene_tx=XM_019813785.2;refGene_gn=COL17A1;refGene_tx=XM_019813786.2
D2 64303492 64303534
refGene_gn=COL17A1;refGene_tx=XM_006938156.4;refGene_gn=COL17A1;refGene_tx=XM_019813785.2;refGene_gn=COL17A1;refGene_tx=XM_019813786.2
D2 64303671 64303725
refGene_gn=COL17A1;refGene_tx=XM_006938156.4;refGene_gn=COL17A1;refGene_tx=XM_019813785.2;refGene_gn=COL17A1;refGene_tx=XM_019813786.2
D2 64304619 64304697
refGene_gn=COL17A1;refGene_tx=XM_006938156.4;refGene_gn=COL17A1;refGene_tx=XM_019813785.2;refGene_gn=COL17A1;refGene_tx=XM_019813786.2
D2 64305230 64305266
refGene_gn=COL17A1;refGene_tx=XM_006938156.4;refGene_gn=COL17A1;refGene_tx=XM_019813785.2;refGene_gn=COL17A1;refGene_tx=XM_019813786.2
D2 64306574 64306655
refGene_gn=COL17A1;refGene_tx=XM_006938156.4;refGene_gn=COL17A1;refGene_tx=XM_019813785.2;refGene_gn=COL17A1;refGene_tx=XM_019813786.2
D2 64306905 64306941
refGene_gn=COL17A1;refGene_tx=XM_006938156.4;refGene_gn=COL17A1;refGene_tx=XM_019813785.2;refGene_gn=COL17A1;refGene_tx=XM_019813786.2
D2 64308938 64308974
refGene_gn=COL17A1;refGene_tx=XM_006938156.4;refGene_gn=COL17A1;refGene_tx=XM_019813785.2;refGene_gn=COL17A1;refGene_tx=XM_019813786.2
D2 64309857 64309884
refGene_gn=COL17A1;refGene_tx=XM_006938156.4;refGene_gn=COL17A1;refGene_tx=XM_019813785.2;refGene_gn=COL17A1;refGene_tx=XM_019813786.2
D2 64310693 64310765
refGene_gn=COL17A1;refGene_tx=XM_006938156.4;refGene_gn=COL17A1;refGen

e_tx=XM_019813785.2;refGene_gn=COL17A1;refGene_tx=XM_019813786.2
D2 64311008 64311044
refGene_gn=COL17A1;refGene_tx=XM_006938156.4;refGene_gn=COL17A1;refGen
e_tx=XM_019813785.2;refGene_gn=COL17A1;refGene_tx=XM_019813786.2
D2 64312166 64312229
refGene_gn=COL17A1;refGene_tx=XM_006938156.4;refGene_gn=COL17A1;refGen
e_tx=XM_019813785.2;refGene_gn=COL17A1;refGene_tx=XM_019813786.2
D2 64312871 64312907
refGene_gn=COL17A1;refGene_tx=XM_006938156.4;refGene_gn=COL17A1;refGen
e_tx=XM_019813785.2;refGene_gn=COL17A1;refGene_tx=XM_019813786.2
D2 64313397 64313433
refGene_gn=COL17A1;refGene_tx=XM_006938156.4;refGene_gn=COL17A1;refGen
e_tx=XM_019813785.2;refGene_gn=COL17A1;refGene_tx=XM_019813786.2
D2 64313664 64313718
refGene_gn=COL17A1;refGene_tx=XM_006938156.4;refGene_gn=COL17A1;refGen
e_tx=XM_019813785.2;refGene_gn=COL17A1;refGene_tx=XM_019813786.2
D2 64314323 64314359
refGene_gn=COL17A1;refGene_tx=XM_006938156.4;refGene_gn=COL17A1;refGen
e_tx=XM_019813785.2;refGene_gn=COL17A1;refGene_tx=XM_019813786.2
D2 64314982 64315045
refGene_gn=COL17A1;refGene_tx=XM_006938156.4;refGene_gn=COL17A1;refGen
e_tx=XM_019813785.2;refGene_gn=COL17A1;refGene_tx=XM_019813786.2
D2 64315810 64315915
refGene_gn=COL17A1;refGene_tx=XM_006938156.4;refGene_gn=COL17A1;refGen
e_tx=XM_019813785.2;refGene_gn=COL17A1;refGene_tx=XM_019813786.2
D2 64316464 64316527
refGene_gn=COL17A1;refGene_tx=XM_006938156.4;refGene_gn=COL17A1;refGen
e_tx=XM_019813785.2;refGene_gn=COL17A1;refGene_tx=XM_019813786.2
D2 64316627 64316654
refGene_gn=COL17A1;refGene_tx=XM_006938156.4;refGene_gn=COL17A1;refGen
e_tx=XM_019813785.2;refGene_gn=COL17A1;refGene_tx=XM_019813786.2
D2 64318035 64318062
refGene_gn=COL17A1;refGene_tx=XM_006938156.4;refGene_gn=COL17A1;refGen
e_tx=XM_019813785.2;refGene_gn=COL17A1;refGene_tx=XM_019813786.2
D2 64318412 64318442
refGene_gn=COL17A1;refGene_tx=XM_006938156.4;refGene_gn=COL17A1;refGen
e_tx=XM_019813785.2;refGene_gn=COL17A1;refGene_tx=XM_019813786.2
D2 64318755 64318977
refGene_gn=COL17A1;refGene_tx=XM_006938156.4;refGene_gn=COL17A1;refGen
e_tx=XM_019813785.2;refGene_gn=COL17A1;refGene_tx=XM_019813786.2
D2 64319616 64319814
refGene_gn=COL17A1;refGene_tx=XM_006938156.4;refGene_gn=COL17A1;refGen
e_tx=XM_019813785.2;refGene_gn=COL17A1;refGene_tx=XM_019813786.2
D2 64320763 64320808
refGene_gn=COL17A1;refGene_tx=XM_006938156.4;refGene_gn=COL17A1;refGen
e_tx=XM_019813785.2;refGene_gn=COL17A1;refGene_tx=XM_019813786.2
D2 64322130 64322211
refGene_gn=COL17A1;refGene_tx=XM_006938156.4;refGene_gn=COL17A1;refGen
e_tx=XM_019813785.2;refGene_gn=COL17A1;refGene_tx=XM_019813786.2
D2 64322586 64322748

refGene_gn=COL17A1;refGene_tx=XM_006938156.4;refGene_gn=COL17A1;refGene_tx=XM_019813785.2;refGene_gn=COL17A1;refGene_tx=XM_019813786.2
D2 64324178 64324247
refGene_gn=COL17A1;refGene_tx=XM_006938156.4;refGene_gn=COL17A1;refGene_tx=XM_019813785.2;refGene_gn=COL17A1;refGene_tx=XM_019813786.2
D2 64325020 64325092
refGene_gn=COL17A1;refGene_tx=XM_006938156.4;refGene_gn=COL17A1;refGene_tx=XM_019813785.2;refGene_gn=COL17A1;refGene_tx=XM_019813786.2
D2 64326601 64326673
refGene_gn=COL17A1;refGene_tx=XM_006938156.4;refGene_gn=COL17A1;refGene_tx=XM_019813785.2;refGene_gn=COL17A1;refGene_tx=XM_019813786.2
D2 64327280 64327439
refGene_gn=COL17A1;refGene_tx=XM_006938156.4;refGene_gn=COL17A1;refGene_tx=XM_019813785.2;refGene_gn=COL17A1;refGene_tx=XM_019813786.2
D2 64329543 64329687
refGene_gn=COL17A1;refGene_tx=XM_006938156.4;refGene_gn=COL17A1;refGene_tx=XM_019813785.2;refGene_gn=COL17A1;refGene_tx=XM_019813786.2
D2 64331353 64331401
refGene_gn=COL17A1;refGene_tx=XM_006938156.4;refGene_gn=COL17A1;refGene_tx=XM_019813785.2;refGene_gn=COL17A1;refGene_tx=XM_019813786.2
D2 64332516 64332552
refGene_gn=COL17A1;refGene_tx=XM_006938156.4;refGene_gn=COL17A1;refGene_tx=XM_019813785.2;refGene_gn=COL17A1;refGene_tx=XM_019813786.2
D2 64333678 64333726
refGene_gn=COL17A1;refGene_tx=XM_006938156.4;refGene_gn=COL17A1;refGene_tx=XM_019813785.2;refGene_gn=COL17A1;refGene_tx=XM_019813786.2
D2 64336192 64336324
refGene_gn=COL17A1;refGene_tx=XM_006938156.4;refGene_gn=COL17A1;refGene_tx=XM_019813785.2;refGene_gn=COL17A1;refGene_tx=XM_019813786.2
D2 64337344 64337449
refGene_gn=COL17A1;refGene_tx=XM_006938156.4;refGene_gn=COL17A1;refGene_tx=XM_019813785.2;refGene_gn=COL17A1;refGene_tx=XM_019813786.2
D2 64338841 64338886
refGene_gn=COL17A1;refGene_tx=XM_006938156.4;refGene_gn=COL17A1;refGene_tx=XM_019813785.2;refGene_gn=COL17A1;refGene_tx=XM_019813786.2
D2 64341012 64341064
refGene_gn=COL17A1;refGene_tx=XM_006938156.4;refGene_gn=COL17A1;refGene_tx=XM_019813785.2;refGene_gn=COL17A1;refGene_tx=XM_019813786.2
D2 64395077 64395173 refGene_gn=SFR1;refGene_tx=XM_003994389.4
D2 64395785 64396196
refGene_gn=SFR1;refGene_tx=XM_003994389.4;refGene_gn=SFR1;refGene_tx=XM_011287543.3
D2 64397247 64397439
refGene_gn=SFR1;refGene_tx=XM_003994389.4;refGene_gn=SFR1;refGene_tx=XM_011287543.3
D2 64402724 64402891
refGene_gn=CFAP43;refGene_tx=XM_006938157.4;refGene_gn=CFAP43;refGene_tx=XM_023240891.1;refGene_gn=CFAP43;refGene_tx=XM_023240892.1;refGene_gn=CFAP43;refGene_tx=XM_023240893.1
D2 64403080 64403234

refGene_gn=CFAP43;refGene_tx=XM_006938157.4;refGene_gn=CFAP43;refGene_tx=XM_023240891.1;refGene_gn=CFAP43;refGene_tx=XM_023240892.1;refGene_gn=CFAP43;refGene_tx=XM_023240893.1
D2 64403864 64403945
refGene_gn=CFAP43;refGene_tx=XM_006938157.4;refGene_gn=CFAP43;refGene_tx=XM_023240891.1;refGene_gn=CFAP43;refGene_tx=XM_023240892.1;refGene_gn=CFAP43;refGene_tx=XM_023240893.1
D2 64405937 64406102
refGene_gn=CFAP43;refGene_tx=XM_006938157.4;refGene_gn=CFAP43;refGene_tx=XM_023240891.1;refGene_gn=CFAP43;refGene_tx=XM_023240892.1;refGene_gn=CFAP43;refGene_tx=XM_023240893.1
D2 64412356 64412516
refGene_gn=CFAP43;refGene_tx=XM_006938157.4;refGene_gn=CFAP43;refGene_tx=XM_023240891.1;refGene_gn=CFAP43;refGene_tx=XM_023240892.1;refGene_gn=CFAP43;refGene_tx=XM_023240893.1
D2 64413501 64413614
refGene_gn=CFAP43;refGene_tx=XM_006938157.4;refGene_gn=CFAP43;refGene_tx=XM_023240891.1;refGene_gn=CFAP43;refGene_tx=XM_023240892.1;refGene_gn=CFAP43;refGene_tx=XM_023240893.1
D2 64414748 64414962
refGene_gn=CFAP43;refGene_tx=XM_006938157.4;refGene_gn=CFAP43;refGene_tx=XM_023240891.1;refGene_gn=CFAP43;refGene_tx=XM_023240892.1;refGene_gn=CFAP43;refGene_tx=XM_023240893.1
D2 64416561 64416650
refGene_gn=CFAP43;refGene_tx=XM_006938157.4;refGene_gn=CFAP43;refGene_tx=XM_023240891.1;refGene_gn=CFAP43;refGene_tx=XM_023240892.1;refGene_gn=CFAP43;refGene_tx=XM_023240893.1
D2 64418408 64418495
refGene_gn=CFAP43;refGene_tx=XM_006938157.4;refGene_gn=CFAP43;refGene_tx=XM_023240891.1;refGene_gn=CFAP43;refGene_tx=XM_023240892.1;refGene_gn=CFAP43;refGene_tx=XM_023240893.1
D2 64420007 64420115
refGene_gn=CFAP43;refGene_tx=XM_006938157.4;refGene_gn=CFAP43;refGene_tx=XM_023240891.1;refGene_gn=CFAP43;refGene_tx=XM_023240892.1;refGene_gn=CFAP43;refGene_tx=XM_023240893.1
D2 64422840 64422960
refGene_gn=CFAP43;refGene_tx=XM_006938157.4;refGene_gn=CFAP43;refGene_tx=XM_023240891.1;refGene_gn=CFAP43;refGene_tx=XM_023240892.1;refGene_gn=CFAP43;refGene_tx=XM_023240893.1
D2 64430183 64430309
refGene_gn=CFAP43;refGene_tx=XM_006938157.4;refGene_gn=CFAP43;refGene_tx=XM_023240891.1;refGene_gn=CFAP43;refGene_tx=XM_023240892.1
D2 64431624 64431705
refGene_gn=CFAP43;refGene_tx=XM_006938157.4;refGene_gn=CFAP43;refGene_tx=XM_023240891.1;refGene_gn=CFAP43;refGene_tx=XM_023240892.1;refGene_gn=CFAP43;refGene_tx=XM_023240893.1
D2 64432001 64432091
refGene_gn=CFAP43;refGene_tx=XM_006938157.4;refGene_gn=CFAP43;refGene_tx=XM_023240891.1;refGene_gn=CFAP43;refGene_tx=XM_023240892.1;refGene_gn=CFAP43;refGene_tx=XM_023240893.1

D2 64433857 64434064
refGene_gn=CFAP43;refGene_tx=XM_006938157.4;refGene_gn=CFAP43;refGene_tx=XM_023240891.1;refGene_gn=CFAP43;refGene_tx=XM_023240892.1;refGene_gn=CFAP43;refGene_tx=XM_023240893.1
D2 64436170 64436398
refGene_gn=CFAP43;refGene_tx=XM_006938157.4;refGene_gn=CFAP43;refGene_tx=XM_023240891.1;refGene_gn=CFAP43;refGene_tx=XM_023240892.1;refGene_gn=CFAP43;refGene_tx=XM_023240893.1
D2 64437304 64437421
refGene_gn=CFAP43;refGene_tx=XM_006938157.4;refGene_gn=CFAP43;refGene_tx=XM_023240891.1;refGene_gn=CFAP43;refGene_tx=XM_023240892.1;refGene_gn=CFAP43;refGene_tx=XM_023240893.1
D2 64438648 64438753
refGene_gn=CFAP43;refGene_tx=XM_006938157.4;refGene_gn=CFAP43;refGene_tx=XM_023240892.1;refGene_gn=CFAP43;refGene_tx=XM_023240893.1
D2 64442852 64442978
refGene_gn=CFAP43;refGene_tx=XM_006938157.4;refGene_gn=CFAP43;refGene_tx=XM_023240891.1;refGene_gn=CFAP43;refGene_tx=XM_023240892.1;refGene_gn=CFAP43;refGene_tx=XM_023240893.1
D2 64461197 64461275
refGene_gn=CFAP43;refGene_tx=XM_006938157.4;refGene_gn=CFAP43;refGene_tx=XM_023240891.1;refGene_gn=CFAP43;refGene_tx=XM_023240892.1;refGene_gn=CFAP43;refGene_tx=XM_023240893.1
D2 64462042 64462144
refGene_gn=CFAP43;refGene_tx=XM_006938157.4;refGene_gn=CFAP43;refGene_tx=XM_023240891.1;refGene_gn=CFAP43;refGene_tx=XM_023240892.1;refGene_gn=CFAP43;refGene_tx=XM_023240893.1
D2 64464559 64464707
refGene_gn=CFAP43;refGene_tx=XM_006938157.4;refGene_gn=CFAP43;refGene_tx=XM_023240891.1;refGene_gn=CFAP43;refGene_tx=XM_023240892.1;refGene_gn=CFAP43;refGene_tx=XM_023240893.1
D2 64466638 64466769
refGene_gn=CFAP43;refGene_tx=XM_006938157.4;refGene_gn=CFAP43;refGene_tx=XM_023240891.1;refGene_gn=CFAP43;refGene_tx=XM_023240892.1;refGene_gn=CFAP43;refGene_tx=XM_023240893.1
D2 64469404 64469554
refGene_gn=CFAP43;refGene_tx=XM_006938157.4;refGene_gn=CFAP43;refGene_tx=XM_023240891.1;refGene_gn=CFAP43;refGene_tx=XM_023240892.1;refGene_gn=CFAP43;refGene_tx=XM_023240893.1
D2 64470579 64470752
refGene_gn=CFAP43;refGene_tx=XM_006938157.4;refGene_gn=CFAP43;refGene_tx=XM_023240891.1;refGene_gn=CFAP43;refGene_tx=XM_023240892.1;refGene_gn=CFAP43;refGene_tx=XM_023240893.1
D2 64471735 64471876
refGene_gn=CFAP43;refGene_tx=XM_006938157.4;refGene_gn=CFAP43;refGene_tx=XM_023240891.1;refGene_gn=CFAP43;refGene_tx=XM_023240892.1;refGene_gn=CFAP43;refGene_tx=XM_023240893.1
D2 64475855 64475959
refGene_gn=CFAP43;refGene_tx=XM_006938157.4;refGene_gn=CFAP43;refGene_tx=XM_023240891.1;refGene_gn=CFAP43;refGene_tx=XM_023240892.1;refGene_

gn=CFAP43;refGene_tx=XM_023240893.1
D2 64478356 64478505
refGene_gn=CFAP43;refGene_tx=XM_006938157.4;refGene_gn=CFAP43;refGene_tx=XM_023240891.1;refGene_gn=CFAP43;refGene_tx=XM_023240892.1;refGene_gn=CFAP43;refGene_tx=XM_023240893.1
D2 64481914 64481995
refGene_gn=CFAP43;refGene_tx=XM_006938157.4;refGene_gn=CFAP43;refGene_tx=XM_023240891.1;refGene_gn=CFAP43;refGene_tx=XM_023240892.1;refGene_gn=CFAP43;refGene_tx=XM_023240893.1
D2 64484919 64485036
refGene_gn=CFAP43;refGene_tx=XM_006938157.4;refGene_gn=CFAP43;refGene_tx=XM_023240891.1;refGene_gn=CFAP43;refGene_tx=XM_023240893.1
D2 64489982 64490114
refGene_gn=CFAP43;refGene_tx=XM_006938157.4;refGene_gn=CFAP43;refGene_tx=XM_023240891.1;refGene_gn=CFAP43;refGene_tx=XM_023240892.1;refGene_gn=CFAP43;refGene_tx=XM_023240893.1
D2 64492077 64492145
refGene_gn=CFAP43;refGene_tx=XM_006938157.4;refGene_gn=CFAP43;refGene_tx=XM_023240891.1;refGene_gn=CFAP43;refGene_tx=XM_023240892.1;refGene_gn=CFAP43;refGene_tx=XM_023240893.1
D2 64503265 64503425
refGene_gn=CFAP43;refGene_tx=XM_006938157.4;refGene_gn=CFAP43;refGene_tx=XM_023240891.1;refGene_gn=CFAP43;refGene_tx=XM_023240892.1;refGene_gn=CFAP43;refGene_tx=XM_023240893.1
D2 64509902 64510053
refGene_gn=CFAP43;refGene_tx=XM_006938157.4;refGene_gn=CFAP43;refGene_tx=XM_023240891.1;refGene_gn=CFAP43;refGene_tx=XM_023240892.1;refGene_gn=CFAP43;refGene_tx=XM_023240893.1
D2 64513006 64513174
refGene_gn=CFAP43;refGene_tx=XM_006938157.4;refGene_gn=CFAP43;refGene_tx=XM_023240891.1;refGene_gn=CFAP43;refGene_tx=XM_023240892.1;refGene_gn=CFAP43;refGene_tx=XM_023240893.1
D2 64518516 64518613
refGene_gn=CFAP43;refGene_tx=XM_006938157.4;refGene_gn=CFAP43;refGene_tx=XM_023240891.1;refGene_gn=CFAP43;refGene_tx=XM_023240892.1;refGene_gn=CFAP43;refGene_tx=XM_023240893.1
D2 64523385 64523639
refGene_gn=CFAP43;refGene_tx=XM_006938157.4;refGene_gn=CFAP43;refGene_tx=XM_023240891.1;refGene_gn=CFAP43;refGene_tx=XM_023240892.1;refGene_gn=CFAP43;refGene_tx=XM_023240893.1
D2 64524656 64524721
refGene_gn=CFAP43;refGene_tx=XM_006938157.4;refGene_gn=CFAP43;refGene_tx=XM_023240891.1;refGene_gn=CFAP43;refGene_tx=XM_023240892.1;refGene_gn=CFAP43;refGene_tx=XM_023240893.1
D2 64539352 64539386 refGene_gn=GSTO1;refGene_tx=XM_019813795.2
D2 64539606 64539715 refGene_gn=GSTO1;refGene_tx=XM_019813795.2
D2 64543268 64543491 refGene_gn=GSTO1;refGene_tx=XM_019813795.2
D2 64546455 64546554 refGene_gn=GSTO1;refGene_tx=XM_019813795.2
D2 64549880 64549987 refGene_gn=GSTO1;refGene_tx=XM_019813795.2
D2 64550744 64550898 refGene_gn=GSTO1;refGene_tx=XM_019813795.2

D2 64559028 64559059
refGene_gn=GSTO2;refGene_tx=XM_003994392.5;refGene_gn=GSTO2;refGene_tx=XM_006938158.4;refGene_gn=GSTO2;refGene_tx=XM_019813792.2;refGene_gn=GSTO2;refGene_tx=XM_019813793.2;refGene_gn=GSTO2;refGene_tx=XM_023240894.1
D2 64559329 64559438
refGene_gn=GSTO2;refGene_tx=XM_003994392.5;refGene_gn=GSTO2;refGene_tx=XM_006938158.4;refGene_gn=GSTO2;refGene_tx=XM_019813792.2;refGene_gn=GSTO2;refGene_tx=XM_019813793.2;refGene_gn=GSTO2;refGene_tx=XM_023240894.1
D2 64561786 64562009
refGene_gn=GSTO2;refGene_tx=XM_003994392.5;refGene_gn=GSTO2;refGene_tx=XM_006938158.4;refGene_gn=GSTO2;refGene_tx=XM_019813792.2;refGene_gn=GSTO2;refGene_tx=XM_019813793.2;refGene_gn=GSTO2;refGene_tx=XM_023240894.1
D2 64562943 64563045
refGene_gn=GSTO2;refGene_tx=XM_006938158.4;refGene_gn=GSTO2;refGene_tx=XM_019813792.2;refGene_gn=GSTO2;refGene_tx=XM_019813793.2;refGene_gn=GSTO2;refGene_tx=XM_023240894.1
D2 64569959 64570067
refGene_gn=GSTO2;refGene_tx=XM_019813792.2;refGene_gn=GSTO2;refGene_tx=XM_019813793.2;refGene_gn=GSTO2;refGene_tx=XM_023240894.1
D2 64576003 64576110
refGene_gn=GSTO2;refGene_tx=XM_003994392.5;refGene_gn=GSTO2;refGene_tx=XM_006938158.4;refGene_gn=GSTO2;refGene_tx=XM_019813792.2;refGene_gn=GSTO2;refGene_tx=XM_019813793.2;refGene_gn=GSTO2;refGene_tx=XM_023240894.1
D2 64577124 64577284
refGene_gn=GSTO2;refGene_tx=XM_003994392.5;refGene_gn=GSTO2;refGene_tx=XM_006938158.4;refGene_gn=GSTO2;refGene_tx=XM_019813792.2;refGene_gn=GSTO2;refGene_tx=XM_019813793.2;refGene_gn=GSTO2;refGene_tx=XM_023240894.1
D2 64588077 64589736
refGene_gn=ITPRIP;refGene_tx=XM_006938159.4;refGene_gn=ITPRIP;refGene_tx=XM_006938160.4;refGene_gn=ITPRIP;refGene_tx=XM_019813797.2
D2 64625881 64625890
refGene_gn=CFAP58;refGene_tx=XM_019813796.2;refGene_gn=CFAP58;refGene_tx=XM_023240896.1
D2 64631590 64631872
refGene_gn=CFAP58;refGene_tx=XM_019813796.2;refGene_gn=CFAP58;refGene_tx=XM_023240896.1
D2 64633935 64634084
refGene_gn=CFAP58;refGene_tx=XM_019813796.2;refGene_gn=CFAP58;refGene_tx=XM_023240896.1
D2 64636773 64636930
refGene_gn=CFAP58;refGene_tx=XM_019813796.2;refGene_gn=CFAP58;refGene_tx=XM_023240896.1
D2 64637779 64637974
refGene_gn=CFAP58;refGene_tx=XM_019813796.2;refGene_gn=CFAP58;refGene_tx=XM_023240896.1

D2 64640061 64640199
refGene_gn=CFAP58;refGene_tx=XM_019813796.2;refGene_gn=CFAP58;refGene_tx=XM_023240896.1
D2 64642639 64642799
refGene_gn=CFAP58;refGene_tx=XM_019813796.2;refGene_gn=CFAP58;refGene_tx=XM_023240896.1
D2 64646980 64647063
refGene_gn=CFAP58;refGene_tx=XM_019813796.2;refGene_gn=CFAP58;refGene_tx=XM_023240896.1
D2 64651929 64652121
refGene_gn=CFAP58;refGene_tx=XM_019813796.2;refGene_gn=CFAP58;refGene_tx=XM_023240896.1
D2 64662494 64662656
refGene_gn=CFAP58;refGene_tx=XM_019813796.2;refGene_gn=CFAP58;refGene_tx=XM_023240896.1
D2 64663372 64663519
refGene_gn=CFAP58;refGene_tx=XM_019813796.2;refGene_gn=CFAP58;refGene_tx=XM_023240896.1
D2 64673170 64673311
refGene_gn=CFAP58;refGene_tx=XM_019813796.2;refGene_gn=CFAP58;refGene_tx=XM_023240896.1
D2 64674563 64674787
refGene_gn=CFAP58;refGene_tx=XM_019813796.2;refGene_gn=CFAP58;refGene_tx=XM_023240896.1
D2 64677483 64677595
refGene_gn=CFAP58;refGene_tx=XM_019813796.2;refGene_gn=CFAP58;refGene_tx=XM_023240896.1
D2 64679709 64679814
refGene_gn=CFAP58;refGene_tx=XM_019813796.2;refGene_gn=CFAP58;refGene_tx=XM_023240896.1
D2 64712716 64712806 refGene_gn=CFAP58;refGene_tx=XM_023240896.1
D2 64723948 64724068 refGene_gn=CFAP58;refGene_tx=XM_019813796.2
D2 64726401 64726535 refGene_gn=CFAP58;refGene_tx=XM_019813796.2
D2 64731403 64731515 refGene_gn=CFAP58;refGene_tx=XM_019813796.2
D2 64899431 64900058
refGene_gn=SORCS3;refGene_tx=XM_023240898.1;refGene_gn=SORCS3;refGene_tx=XM_023240899.1
D2 65083506 65083574
refGene_gn=SORCS3;refGene_tx=XM_023240898.1;refGene_gn=SORCS3;refGene_tx=XM_023240899.1
D2 65159671 65159771
refGene_gn=SORCS3;refGene_tx=XM_023240898.1;refGene_gn=SORCS3;refGene_tx=XM_023240899.1
D2 65216966 65217125
refGene_gn=SORCS3;refGene_tx=XM_023240898.1;refGene_gn=SORCS3;refGene_tx=XM_023240899.1
D2 65283308 65283382
refGene_gn=SORCS3;refGene_tx=XM_023240898.1;refGene_gn=SORCS3;refGene_tx=XM_023240899.1
D2 65331152 65331217

refGene_gn=SORCS3;refGene_tx=XM_023240898.1;refGene_gn=SORCS3;refGene_tx=XM_023240899.1
D2 65351283 65351402
refGene_gn=SORCS3;refGene_tx=XM_023240898.1;refGene_gn=SORCS3;refGene_tx=XM_023240899.1
D2 65382692 65382782
refGene_gn=SORCS3;refGene_tx=XM_023240898.1;refGene_gn=SORCS3;refGene_tx=XM_023240899.1
D2 65391050 65391230
refGene_gn=SORCS3;refGene_tx=XM_023240898.1;refGene_gn=SORCS3;refGene_tx=XM_023240899.1
D2 65400605 65400752
refGene_gn=SORCS3;refGene_tx=XM_023240898.1;refGene_gn=SORCS3;refGene_tx=XM_023240899.1
D2 65402216 65402319
refGene_gn=SORCS3;refGene_tx=XM_023240898.1;refGene_gn=SORCS3;refGene_tx=XM_023240899.1
D2 65407895 65407972
refGene_gn=SORCS3;refGene_tx=XM_023240898.1;refGene_gn=SORCS3;refGene_tx=XM_023240899.1
D2 65411082 65411174
refGene_gn=SORCS3;refGene_tx=XM_023240898.1;refGene_gn=SORCS3;refGene_tx=XM_023240899.1
D2 65421070 65421178
refGene_gn=SORCS3;refGene_tx=XM_023240898.1;refGene_gn=SORCS3;refGene_tx=XM_023240899.1
D2 65447673 65447791
refGene_gn=SORCS3;refGene_tx=XM_023240898.1;refGene_gn=SORCS3;refGene_tx=XM_023240899.1
D2 65449040 65449174
refGene_gn=SORCS3;refGene_tx=XM_023240898.1;refGene_gn=SORCS3;refGene_tx=XM_023240899.1
D2 65459562 65459676
refGene_gn=SORCS3;refGene_tx=XM_023240898.1;refGene_gn=SORCS3;refGene_tx=XM_023240899.1
D2 65462823 65462995
refGene_gn=SORCS3;refGene_tx=XM_023240898.1;refGene_gn=SORCS3;refGene_tx=XM_023240899.1
D2 65465307 65465494
refGene_gn=SORCS3;refGene_tx=XM_023240898.1;refGene_gn=SORCS3;refGene_tx=XM_023240899.1
D2 65471082 65471216
refGene_gn=SORCS3;refGene_tx=XM_023240898.1;refGene_gn=SORCS3;refGene_tx=XM_023240899.1
D2 65480119 65480243
refGene_gn=SORCS3;refGene_tx=XM_023240898.1;refGene_gn=SORCS3;refGene_tx=XM_023240899.1
D2 65481853 65481966
refGene_gn=SORCS3;refGene_tx=XM_023240898.1;refGene_gn=SORCS3;refGene_tx=XM_023240899.1

D2 65486056 65486188
refGene_gn=SORCS3;refGene_tx=XM_023240898.1;refGene_gn=SORCS3;refGene_tx=XM_023240899.1
D2 65486532 65486625 refGene_gn=SORCS3;refGene_tx=XM_023240899.1
D2 65488052 65488152 refGene_gn=SORCS3;refGene_tx=XM_023240898.1
D2 65489138 65489244 refGene_gn=SORCS3;refGene_tx=XM_023240898.1
D2 65494964 65495125 refGene_gn=SORCS3;refGene_tx=XM_023240898.1
D2 65495917 65495982 refGene_gn=SORCS3;refGene_tx=XM_023240898.1
D2 66708583 66708648
refGene_gn=SORCS1;refGene_tx=XM_019813798.2;refGene_gn=SORCS1;refGene_tx=XM_023240902.1
D2 66708784 66708920
refGene_gn=SORCS1;refGene_tx=XM_023240900.1;refGene_gn=SORCS1;refGene_tx=XM_023240901.1
D2 66710460 66710621
refGene_gn=SORCS1;refGene_tx=XM_019813798.2;refGene_gn=SORCS1;refGene_tx=XM_023240900.1
D2 66710743 66710849
refGene_gn=SORCS1;refGene_tx=XM_019813798.2;refGene_gn=SORCS1;refGene_tx=XM_023240900.1;refGene_gn=SORCS1;refGene_tx=XM_023240901.1;refGene_gn=SORCS1;refGene_tx=XM_023240902.1
D2 66726796 66726896
refGene_gn=SORCS1;refGene_tx=XM_019813798.2;refGene_gn=SORCS1;refGene_tx=XM_023240900.1;refGene_gn=SORCS1;refGene_tx=XM_023240901.1;refGene_gn=SORCS1;refGene_tx=XM_023240902.1
D2 66738345 66738477
refGene_gn=SORCS1;refGene_tx=XM_019813798.2;refGene_gn=SORCS1;refGene_tx=XM_023240900.1;refGene_gn=SORCS1;refGene_tx=XM_023240901.1;refGene_gn=SORCS1;refGene_tx=XM_023240902.1
D2 66741188 66741301
refGene_gn=SORCS1;refGene_tx=XM_019813798.2;refGene_gn=SORCS1;refGene_tx=XM_023240900.1;refGene_gn=SORCS1;refGene_tx=XM_023240901.1;refGene_gn=SORCS1;refGene_tx=XM_023240902.1
D2 66749362 66749486
refGene_gn=SORCS1;refGene_tx=XM_019813798.2;refGene_gn=SORCS1;refGene_tx=XM_023240900.1;refGene_gn=SORCS1;refGene_tx=XM_023240901.1;refGene_gn=SORCS1;refGene_tx=XM_023240902.1
D2 66751692 66751826
refGene_gn=SORCS1;refGene_tx=XM_019813798.2;refGene_gn=SORCS1;refGene_tx=XM_023240900.1;refGene_gn=SORCS1;refGene_tx=XM_023240901.1;refGene_gn=SORCS1;refGene_tx=XM_023240902.1
D2 66759694 66759881
refGene_gn=SORCS1;refGene_tx=XM_019813798.2;refGene_gn=SORCS1;refGene_tx=XM_023240900.1;refGene_gn=SORCS1;refGene_tx=XM_023240901.1;refGene_gn=SORCS1;refGene_tx=XM_023240902.1
D2 66781790 66781962
refGene_gn=SORCS1;refGene_tx=XM_019813798.2;refGene_gn=SORCS1;refGene_tx=XM_023240900.1;refGene_gn=SORCS1;refGene_tx=XM_023240901.1;refGene_gn=SORCS1;refGene_tx=XM_023240902.1
D2 66794757 66794871

refGene_gn=SORCS1;refGene_tx=XM_019813798.2;refGene_gn=SORCS1;refGene_tx=XM_023240900.1;refGene_gn=SORCS1;refGene_tx=XM_023240901.1;refGene_gn=SORCS1;refGene_tx=XM_023240902.1
D2 66797982 66798113
refGene_gn=SORCS1;refGene_tx=XM_019813798.2;refGene_gn=SORCS1;refGene_tx=XM_023240900.1;refGene_gn=SORCS1;refGene_tx=XM_023240901.1;refGene_gn=SORCS1;refGene_tx=XM_023240902.1
D2 66799959 66800077
refGene_gn=SORCS1;refGene_tx=XM_019813798.2;refGene_gn=SORCS1;refGene_tx=XM_023240900.1;refGene_gn=SORCS1;refGene_tx=XM_023240901.1;refGene_gn=SORCS1;refGene_tx=XM_023240902.1
D2 66801555 66801663
refGene_gn=SORCS1;refGene_tx=XM_019813798.2;refGene_gn=SORCS1;refGene_tx=XM_023240900.1;refGene_gn=SORCS1;refGene_tx=XM_023240901.1;refGene_gn=SORCS1;refGene_tx=XM_023240902.1
D2 66802945 66803037
refGene_gn=SORCS1;refGene_tx=XM_019813798.2;refGene_gn=SORCS1;refGene_tx=XM_023240900.1;refGene_gn=SORCS1;refGene_tx=XM_023240901.1;refGene_gn=SORCS1;refGene_tx=XM_023240902.1
D2 66805028 66805105
refGene_gn=SORCS1;refGene_tx=XM_019813798.2;refGene_gn=SORCS1;refGene_tx=XM_023240900.1;refGene_gn=SORCS1;refGene_tx=XM_023240901.1;refGene_gn=SORCS1;refGene_tx=XM_023240902.1
D2 66805373 66805476
refGene_gn=SORCS1;refGene_tx=XM_019813798.2;refGene_gn=SORCS1;refGene_tx=XM_023240900.1;refGene_gn=SORCS1;refGene_tx=XM_023240901.1;refGene_gn=SORCS1;refGene_tx=XM_023240902.1
D2 66813495 66813642
refGene_gn=SORCS1;refGene_tx=XM_019813798.2;refGene_gn=SORCS1;refGene_tx=XM_023240900.1;refGene_gn=SORCS1;refGene_tx=XM_023240901.1;refGene_gn=SORCS1;refGene_tx=XM_023240902.1
D2 66823223 66823403
refGene_gn=SORCS1;refGene_tx=XM_019813798.2;refGene_gn=SORCS1;refGene_tx=XM_023240900.1;refGene_gn=SORCS1;refGene_tx=XM_023240901.1;refGene_gn=SORCS1;refGene_tx=XM_023240902.1
D2 66829989 66830079
refGene_gn=SORCS1;refGene_tx=XM_019813798.2;refGene_gn=SORCS1;refGene_tx=XM_023240900.1;refGene_gn=SORCS1;refGene_tx=XM_023240901.1;refGene_gn=SORCS1;refGene_tx=XM_023240902.1
D2 66832631 66832750
refGene_gn=SORCS1;refGene_tx=XM_019813798.2;refGene_gn=SORCS1;refGene_tx=XM_023240900.1;refGene_gn=SORCS1;refGene_tx=XM_023240901.1;refGene_gn=SORCS1;refGene_tx=XM_023240902.1
D2 66848109 66848174
refGene_gn=SORCS1;refGene_tx=XM_019813798.2;refGene_gn=SORCS1;refGene_tx=XM_023240900.1;refGene_gn=SORCS1;refGene_tx=XM_023240901.1;refGene_gn=SORCS1;refGene_tx=XM_023240902.1
D2 66879477 66879551
refGene_gn=SORCS1;refGene_tx=XM_019813798.2;refGene_gn=SORCS1;refGene_tx=XM_023240900.1;refGene_gn=SORCS1;refGene_tx=XM_023240901.1;refGene_

gn=SORCS1;refGene_tx=XM_023240902.1
D2 66895836 66895995
refGene_gn=SORCS1;refGene_tx=XM_019813798.2;refGene_gn=SORCS1;refGene_tx=XM_023240900.1;refGene_gn=SORCS1;refGene_tx=XM_023240901.1;refGene_gn=SORCS1;refGene_tx=XM_023240902.1
D2 66943495 66943595
refGene_gn=SORCS1;refGene_tx=XM_019813798.2;refGene_gn=SORCS1;refGene_tx=XM_023240900.1;refGene_gn=SORCS1;refGene_tx=XM_023240901.1;refGene_gn=SORCS1;refGene_tx=XM_023240902.1
D2 67028163 67028231
refGene_gn=SORCS1;refGene_tx=XM_019813798.2;refGene_gn=SORCS1;refGene_tx=XM_023240900.1;refGene_gn=SORCS1;refGene_tx=XM_023240901.1;refGene_gn=SORCS1;refGene_tx=XM_023240902.1
D2 67214366 67214924
refGene_gn=SORCS1;refGene_tx=XM_019813798.2;refGene_gn=SORCS1;refGene_tx=XM_023240900.1;refGene_gn=SORCS1;refGene_tx=XM_023240901.1;refGene_gn=SORCS1;refGene_tx=XM_023240902.1
D2 67762144 67762842
refGene_gn=LOC102900465;refGene_tx=XM_023240300.1
D2 67765669 67765811
refGene_gn=LOC102900465;refGene_tx=XM_023240300.1
D2 67842611 67842724
refGene_gn=LOC102900465;refGene_tx=XM_023240300.1
D2 67844901 67845093
refGene_gn=LOC102900465;refGene_tx=XM_023240300.1
D2 67849068 67849162
refGene_gn=LOC102900465;refGene_tx=XM_023240300.1
D2 69345850 69345979
refGene_gn=XPNPEP1;refGene_tx=XM_023240903.1;refGene_gn=XPNPEP1;refGene_tx=XM_023240904.1;refGene_gn=XPNPEP1;refGene_tx=XM_023240905.1;refGene_gn=XPNPEP1;refGene_tx=XM_023240906.1;refGene_gn=XPNPEP1;refGene_tx=XM_023240907.1
D2 69349632 69349731
refGene_gn=XPNPEP1;refGene_tx=XM_023240903.1;refGene_gn=XPNPEP1;refGene_tx=XM_023240904.1;refGene_gn=XPNPEP1;refGene_tx=XM_023240905.1;refGene_gn=XPNPEP1;refGene_tx=XM_023240906.1;refGene_gn=XPNPEP1;refGene_tx=XM_023240907.1
D2 69350919 69350996
refGene_gn=XPNPEP1;refGene_tx=XM_023240903.1;refGene_gn=XPNPEP1;refGene_tx=XM_023240904.1;refGene_gn=XPNPEP1;refGene_tx=XM_023240905.1;refGene_gn=XPNPEP1;refGene_tx=XM_023240906.1;refGene_gn=XPNPEP1;refGene_tx=XM_023240907.1
D2 69351910 69352084
refGene_gn=XPNPEP1;refGene_tx=XM_023240903.1;refGene_gn=XPNPEP1;refGene_tx=XM_023240904.1;refGene_gn=XPNPEP1;refGene_tx=XM_023240905.1;refGene_gn=XPNPEP1;refGene_tx=XM_023240906.1;refGene_gn=XPNPEP1;refGene_tx=XM_023240907.1
D2 69352969 69353039
refGene_gn=XPNPEP1;refGene_tx=XM_023240903.1;refGene_gn=XPNPEP1;refGene_tx=XM_023240904.1;refGene_gn=XPNPEP1;refGene_tx=XM_023240905.1;refGe

ne_gn=XPNPEP1;refGene_tx=XM_023240906.1;refGene_gn=XPNPEP1;refGene_tx=XM_023240907.1
D2 69354525 69354586
refGene_gn=XPNPEP1;refGene_tx=XM_023240903.1;refGene_gn=XPNPEP1;refGene_tx=XM_023240904.1;refGene_gn=XPNPEP1;refGene_tx=XM_023240905.1;refGene_gn=XPNPEP1;refGene_tx=XM_023240906.1;refGene_gn=XPNPEP1;refGene_tx=XM_023240907.1
D2 69357036 69357108
refGene_gn=XPNPEP1;refGene_tx=XM_023240903.1;refGene_gn=XPNPEP1;refGene_tx=XM_023240904.1;refGene_gn=XPNPEP1;refGene_tx=XM_023240905.1;refGene_gn=XPNPEP1;refGene_tx=XM_023240906.1;refGene_gn=XPNPEP1;refGene_tx=XM_023240907.1
D2 69359012 69359090
refGene_gn=XPNPEP1;refGene_tx=XM_023240903.1;refGene_gn=XPNPEP1;refGene_tx=XM_023240904.1;refGene_gn=XPNPEP1;refGene_tx=XM_023240905.1;refGene_gn=XPNPEP1;refGene_tx=XM_023240906.1;refGene_gn=XPNPEP1;refGene_tx=XM_023240907.1
D2 69359219 69359278
refGene_gn=XPNPEP1;refGene_tx=XM_023240903.1;refGene_gn=XPNPEP1;refGene_tx=XM_023240904.1;refGene_gn=XPNPEP1;refGene_tx=XM_023240905.1;refGene_gn=XPNPEP1;refGene_tx=XM_023240906.1;refGene_gn=XPNPEP1;refGene_tx=XM_023240907.1
D2 69360842 69360893
refGene_gn=XPNPEP1;refGene_tx=XM_023240903.1;refGene_gn=XPNPEP1;refGene_tx=XM_023240904.1;refGene_gn=XPNPEP1;refGene_tx=XM_023240905.1;refGene_gn=XPNPEP1;refGene_tx=XM_023240906.1;refGene_gn=XPNPEP1;refGene_tx=XM_023240907.1
D2 69361409 69361499
refGene_gn=XPNPEP1;refGene_tx=XM_023240903.1;refGene_gn=XPNPEP1;refGene_tx=XM_023240904.1;refGene_gn=XPNPEP1;refGene_tx=XM_023240905.1;refGene_gn=XPNPEP1;refGene_tx=XM_023240906.1;refGene_gn=XPNPEP1;refGene_tx=XM_023240907.1
D2 69363528 69363739
refGene_gn=XPNPEP1;refGene_tx=XM_023240903.1;refGene_gn=XPNPEP1;refGene_tx=XM_023240904.1;refGene_gn=XPNPEP1;refGene_tx=XM_023240905.1;refGene_gn=XPNPEP1;refGene_tx=XM_023240906.1;refGene_gn=XPNPEP1;refGene_tx=XM_023240907.1
D2 69365219 69365301
refGene_gn=XPNPEP1;refGene_tx=XM_023240903.1;refGene_gn=XPNPEP1;refGene_tx=XM_023240904.1;refGene_gn=XPNPEP1;refGene_tx=XM_023240905.1;refGene_gn=XPNPEP1;refGene_tx=XM_023240906.1;refGene_gn=XPNPEP1;refGene_tx=XM_023240907.1
D2 69368303 69368399
refGene_gn=XPNPEP1;refGene_tx=XM_023240903.1;refGene_gn=XPNPEP1;refGene_tx=XM_023240904.1;refGene_gn=XPNPEP1;refGene_tx=XM_023240905.1;refGene_gn=XPNPEP1;refGene_tx=XM_023240906.1;refGene_gn=XPNPEP1;refGene_tx=XM_023240907.1
D2 69370134 69370278
refGene_gn=XPNPEP1;refGene_tx=XM_023240903.1;refGene_gn=XPNPEP1;refGene_tx=XM_023240904.1;refGene_gn=XPNPEP1;refGene_tx=XM_023240905.1;refGe

ne_gn=XPNPEP1;refGene_tx=XM_023240906.1;refGene_gn=XPNPEP1;refGene_tx=XM_023240907.1
D2 69370614 69370707
refGene_gn=XPNPEP1;refGene_tx=XM_023240903.1;refGene_gn=XPNPEP1;refGene_tx=XM_023240904.1;refGene_gn=XPNPEP1;refGene_tx=XM_023240905.1;refGene_gn=XPNPEP1;refGene_tx=XM_023240906.1;refGene_gn=XPNPEP1;refGene_tx=XM_023240907.1
D2 69373129 69373234
refGene_gn=XPNPEP1;refGene_tx=XM_023240903.1;refGene_gn=XPNPEP1;refGene_tx=XM_023240904.1;refGene_gn=XPNPEP1;refGene_tx=XM_023240905.1;refGene_gn=XPNPEP1;refGene_tx=XM_023240906.1;refGene_gn=XPNPEP1;refGene_tx=XM_023240907.1
D2 69374633 69374697
refGene_gn=XPNPEP1;refGene_tx=XM_023240903.1;refGene_gn=XPNPEP1;refGene_tx=XM_023240904.1;refGene_gn=XPNPEP1;refGene_tx=XM_023240905.1;refGene_gn=XPNPEP1;refGene_tx=XM_023240906.1
D2 69385936 69386061
refGene_gn=XPNPEP1;refGene_tx=XM_023240903.1;refGene_gn=XPNPEP1;refGene_tx=XM_023240904.1;refGene_gn=XPNPEP1;refGene_tx=XM_023240905.1;refGene_gn=XPNPEP1;refGene_tx=XM_023240906.1
D2 69392960 69393049 refGene_gn=XPNPEP1;refGene_tx=XM_023240903.1
D2 69404047 69404079 refGene_gn=XPNPEP1;refGene_tx=XM_023240903.1
D2 69574571 69574766
refGene_gn=ADD3;refGene_tx=XM_003994400.5;refGene_gn=ADD3;refGene_tx=XM_006938168.4;refGene_gn=ADD3;refGene_tx=XM_011287553.3;refGene_gn=ADD3;refGene_tx=XM_011287554.3;refGene_gn=ADD3;refGene_tx=XM_011287555.3;refGene_gn=ADD3;refGene_tx=XM_019813801.2
D2 69586026 69586165
refGene_gn=ADD3;refGene_tx=XM_003994400.5;refGene_gn=ADD3;refGene_tx=XM_006938168.4;refGene_gn=ADD3;refGene_tx=XM_011287553.3;refGene_gn=ADD3;refGene_tx=XM_011287554.3;refGene_gn=ADD3;refGene_tx=XM_011287555.3;refGene_gn=ADD3;refGene_tx=XM_019813801.2;refGene_gn=ADD3;refGene_tx=XM_019813802.2
D2 69590100 69590252
refGene_gn=ADD3;refGene_tx=XM_003994400.5;refGene_gn=ADD3;refGene_tx=XM_006938168.4;refGene_gn=ADD3;refGene_tx=XM_011287553.3;refGene_gn=ADD3;refGene_tx=XM_011287554.3;refGene_gn=ADD3;refGene_tx=XM_011287555.3;refGene_gn=ADD3;refGene_tx=XM_019813801.2;refGene_gn=ADD3;refGene_tx=XM_019813802.2
D2 69591232 69591313
refGene_gn=ADD3;refGene_tx=XM_003994400.5;refGene_gn=ADD3;refGene_tx=XM_006938168.4;refGene_gn=ADD3;refGene_tx=XM_011287553.3;refGene_gn=ADD3;refGene_tx=XM_011287554.3;refGene_gn=ADD3;refGene_tx=XM_011287555.3;refGene_gn=ADD3;refGene_tx=XM_019813801.2;refGene_gn=ADD3;refGene_tx=XM_019813802.2
D2 69592378 69592528
refGene_gn=ADD3;refGene_tx=XM_003994400.5;refGene_gn=ADD3;refGene_tx=XM_006938168.4;refGene_gn=ADD3;refGene_tx=XM_011287553.3;refGene_gn=ADD3;refGene_tx=XM_011287554.3;refGene_gn=ADD3;refGene_tx=XM_011287555.3;refGene_gn=ADD3;refGene_tx=XM_019813801.2;refGene_gn=ADD3;refGene_tx=X

M_019813802.2
D2 69592958 69593102
refGene_gn=ADD3;refGene_tx=XM_003994400.5;refGene_gn=ADD3;refGene_tx=XM_006938168.4;refGene_gn=ADD3;refGene_tx=XM_011287553.3;refGene_gn=ADD3;refGene_tx=XM_011287554.3;refGene_gn=ADD3;refGene_tx=XM_011287555.3;refGene_gn=ADD3;refGene_tx=XM_019813801.2;refGene_gn=ADD3;refGene_tx=XM_019813802.2
D2 69593216 69593315
refGene_gn=ADD3;refGene_tx=XM_003994400.5;refGene_gn=ADD3;refGene_tx=XM_006938168.4;refGene_gn=ADD3;refGene_tx=XM_011287553.3;refGene_gn=ADD3;refGene_tx=XM_011287554.3;refGene_gn=ADD3;refGene_tx=XM_011287555.3;refGene_gn=ADD3;refGene_tx=XM_019813801.2;refGene_gn=ADD3;refGene_tx=XM_019813802.2
D2 69594416 69594599
refGene_gn=ADD3;refGene_tx=XM_003994400.5;refGene_gn=ADD3;refGene_tx=XM_006938168.4;refGene_gn=ADD3;refGene_tx=XM_011287553.3;refGene_gn=ADD3;refGene_tx=XM_011287554.3;refGene_gn=ADD3;refGene_tx=XM_011287555.3;refGene_gn=ADD3;refGene_tx=XM_019813801.2;refGene_gn=ADD3;refGene_tx=XM_019813802.2
D2 69595699 69595957
refGene_gn=ADD3;refGene_tx=XM_003994400.5;refGene_gn=ADD3;refGene_tx=XM_006938168.4;refGene_gn=ADD3;refGene_tx=XM_011287553.3;refGene_gn=ADD3;refGene_tx=XM_011287554.3;refGene_gn=ADD3;refGene_tx=XM_011287555.3;refGene_gn=ADD3;refGene_tx=XM_019813801.2;refGene_gn=ADD3;refGene_tx=XM_019813802.2
D2 69597243 69597363
refGene_gn=ADD3;refGene_tx=XM_003994400.5;refGene_gn=ADD3;refGene_tx=XM_006938168.4;refGene_gn=ADD3;refGene_tx=XM_011287553.3;refGene_gn=ADD3;refGene_tx=XM_011287554.3;refGene_gn=ADD3;refGene_tx=XM_011287555.3;refGene_gn=ADD3;refGene_tx=XM_019813801.2;refGene_gn=ADD3;refGene_tx=XM_019813802.2
D2 69598051 69598138
refGene_gn=ADD3;refGene_tx=XM_003994400.5;refGene_gn=ADD3;refGene_tx=XM_006938168.4;refGene_gn=ADD3;refGene_tx=XM_011287553.3;refGene_gn=ADD3;refGene_tx=XM_011287554.3;refGene_gn=ADD3;refGene_tx=XM_011287555.3;refGene_gn=ADD3;refGene_tx=XM_019813801.2;refGene_gn=ADD3;refGene_tx=XM_019813802.2
D2 69600425 69600549
refGene_gn=ADD3;refGene_tx=XM_003994400.5;refGene_gn=ADD3;refGene_tx=XM_006938168.4;refGene_gn=ADD3;refGene_tx=XM_011287553.3;refGene_gn=ADD3;refGene_tx=XM_011287554.3;refGene_gn=ADD3;refGene_tx=XM_011287555.3;refGene_gn=ADD3;refGene_tx=XM_019813801.2;refGene_gn=ADD3;refGene_tx=XM_019813802.2
D2 69602131 69602227
refGene_gn=ADD3;refGene_tx=XM_006938168.4;refGene_gn=ADD3;refGene_tx=XM_011287553.3;refGene_gn=ADD3;refGene_tx=XM_011287554.3;refGene_gn=ADD3;refGene_tx=XM_011287555.3;refGene_gn=ADD3;refGene_tx=XM_019813801.2;refGene_gn=ADD3;refGene_tx=XM_019813802.2
D2 69602858 69603151
refGene_gn=ADD3;refGene_tx=XM_003994400.5;refGene_gn=ADD3;refGene_tx=X

M_006938168.4;refGene_gn=ADD3;refGene_tx=XM_011287553.3;refGene_gn=ADD3;refGene_tx=XM_011287554.3;refGene_gn=ADD3;refGene_tx=XM_011287555.3;refGene_gn=ADD3;refGene_tx=XM_019813801.2;refGene_gn=ADD3;refGene_tx=XM_019813802.2
D2 69661598 69661860 refGene_gn=MXI1;refGene_tx=XM_003994402.5
D2 69668707 69668810 refGene_gn=MXI1;refGene_tx=XM_019813803.2
D2 69678420 69678493 refGene_gn=MXI1;refGene_tx=XM_023240909.1
D2 69679909 69680042 refGene_gn=MXI1;refGene_tx=XM_003994402.5;refGene_gn=MXI1;refGene_tx=XM_019813803.2;refGene_gn=MXI1;refGene_tx=XM_023240909.1
D2 69705082 69705112 refGene_gn=MXI1;refGene_tx=XM_003994402.5;refGene_gn=MXI1;refGene_tx=XM_019813803.2;refGene_gn=MXI1;refGene_tx=XM_023240909.1
D2 69734889 69735004 refGene_gn=MXI1;refGene_tx=XM_003994402.5;refGene_gn=MXI1;refGene_tx=XM_019813803.2;refGene_gn=MXI1;refGene_tx=XM_023240909.1
D2 69735622 69735794 refGene_gn=MXI1;refGene_tx=XM_003994402.5;refGene_gn=MXI1;refGene_tx=XM_019813803.2;refGene_gn=MXI1;refGene_tx=XM_023240909.1
D2 69740218 69740382 refGene_gn=MXI1;refGene_tx=XM_003994402.5;refGene_gn=MXI1;refGene_tx=XM_019813803.2;refGene_gn=MXI1;refGene_tx=XM_023240909.1
D2 69749482 69749620 refGene_gn=SMNDC1;refGene_tx=XM_006938169.4;refGene_gn=SMNDC1;refGene_tx=XM_011287559.3
D2 69750345 69750499 refGene_gn=SMNDC1;refGene_tx=XM_006938169.4;refGene_gn=SMNDC1;refGene_tx=XM_011287559.3
D2 69752691 69752853 refGene_gn=SMNDC1;refGene_tx=XM_006938169.4;refGene_gn=SMNDC1;refGene_tx=XM_011287559.3
D2 69753471 69753614 refGene_gn=SMNDC1;refGene_tx=XM_006938169.4;refGene_gn=SMNDC1;refGene_tx=XM_011287559.3
D2 69755545 69755548 refGene_gn=SMNDC1;refGene_tx=XM_011287559.3
D2 69758330 69758450 refGene_gn=SMNDC1;refGene_tx=XM_006938169.4
D2 69923906 69924285 refGene_gn=DUSP5;refGene_tx=XM_023240910.1
D2 69929259 69929408 refGene_gn=DUSP5;refGene_tx=XM_023240910.1
D2 69932267 69932487 refGene_gn=DUSP5;refGene_tx=XM_023240910.1
D2 69935265 69935666 refGene_gn=DUSP5;refGene_tx=XM_023240910.1
D2 69982847 69982862 refGene_gn=SMC3;refGene_tx=XM_003994404.5
D2 69983861 69983937 refGene_gn=SMC3;refGene_tx=XM_003994404.5
D2 69990205 69990244 refGene_gn=SMC3;refGene_tx=XM_003994404.5
D2 69991864 69991932 refGene_gn=SMC3;refGene_tx=XM_003994404.5
D2 69993798 69993870 refGene_gn=SMC3;refGene_tx=XM_003994404.5
D2 69994243 69994323 refGene_gn=SMC3;refGene_tx=XM_003994404.5
D2 69994591 69994670 refGene_gn=SMC3;refGene_tx=XM_003994404.5
D2 69997774 69997892 refGene_gn=SMC3;refGene_tx=XM_003994404.5;refGene_gn=SMC3;refGene_tx=XM_019813804.2

D2 69999432 69999608
refGene_gn=SMC3;refGene_tx=XM_003994404.5;refGene_gn=SMC3;refGene_tx=XM_019813804.2
D2 70000251 70000332
refGene_gn=SMC3;refGene_tx=XM_003994404.5;refGene_gn=SMC3;refGene_tx=XM_019813804.2
D2 70000807 70000972
refGene_gn=SMC3;refGene_tx=XM_003994404.5;refGene_gn=SMC3;refGene_tx=XM_019813804.2
D2 70001884 70002006
refGene_gn=SMC3;refGene_tx=XM_003994404.5;refGene_gn=SMC3;refGene_tx=XM_019813804.2
D2 70003020 70003234
refGene_gn=SMC3;refGene_tx=XM_003994404.5;refGene_gn=SMC3;refGene_tx=XM_019813804.2
D2 70006151 70006255
refGene_gn=SMC3;refGene_tx=XM_003994404.5;refGene_gn=SMC3;refGene_tx=XM_019813804.2
D2 70006562 70006662
refGene_gn=SMC3;refGene_tx=XM_003994404.5;refGene_gn=SMC3;refGene_tx=XM_019813804.2
D2 70007085 70007246
refGene_gn=SMC3;refGene_tx=XM_003994404.5;refGene_gn=SMC3;refGene_tx=XM_019813804.2
D2 70007568 70007710
refGene_gn=SMC3;refGene_tx=XM_003994404.5;refGene_gn=SMC3;refGene_tx=XM_019813804.2
D2 70009051 70009202
refGene_gn=SMC3;refGene_tx=XM_003994404.5;refGene_gn=SMC3;refGene_tx=XM_019813804.2
D2 70011541 70011694
refGene_gn=SMC3;refGene_tx=XM_003994404.5;refGene_gn=SMC3;refGene_tx=XM_019813804.2
D2 70013290 70013442
refGene_gn=SMC3;refGene_tx=XM_003994404.5;refGene_gn=SMC3;refGene_tx=XM_019813804.2
D2 70014030 70014189
refGene_gn=SMC3;refGene_tx=XM_003994404.5;refGene_gn=SMC3;refGene_tx=XM_019813804.2
D2 70014917 70015025
refGene_gn=SMC3;refGene_tx=XM_003994404.5;refGene_gn=SMC3;refGene_tx=XM_019813804.2
D2 70015532 70015641
refGene_gn=SMC3;refGene_tx=XM_003994404.5;refGene_gn=SMC3;refGene_tx=XM_019813804.2
D2 70016421 70016669
refGene_gn=SMC3;refGene_tx=XM_003994404.5;refGene_gn=SMC3;refGene_tx=XM_019813804.2
D2 70016753 70016966
refGene_gn=SMC3;refGene_tx=XM_003994404.5;refGene_gn=SMC3;refGene_tx=X

M_019813804.2
D2 70017306 70017498
refGene_gn=SMC3;refGene_tx=XM_003994404.5;refGene_gn=SMC3;refGene_tx=X
M_019813804.2
D2 70017654 70017832
refGene_gn=SMC3;refGene_tx=XM_003994404.5;refGene_gn=SMC3;refGene_tx=X
M_019813804.2
D2 70018923 70019030
refGene_gn=SMC3;refGene_tx=XM_003994404.5;refGene_gn=SMC3;refGene_tx=X
M_019813804.2
D2 70019627 70019699
refGene_gn=SMC3;refGene_tx=XM_003994404.5;refGene_gn=SMC3;refGene_tx=X
M_019813804.2
D2 70062138 70062329
refGene_gn=RBM20;refGene_tx=XM_023240911.1;refGene_gn=RBM20;refGene_tx
=XM_023240912.1;refGene_gn=RBM20;refGene_tx=XM_023240913.1
D2 70085093 70085122 refGene_gn=RBM20;refGene_tx=XM_019813807.2
D2 70187877 70187960 refGene_gn=RBM20;refGene_tx=XM_019813805.2
D2 70205257 70206335
refGene_gn=RBM20;refGene_tx=XM_019813805.2;refGene_gn=RBM20;refGene_tx
=XM_019813807.2;refGene_gn=RBM20;refGene_tx=XM_019813808.2;refGene_gn=
RBM20;refGene_tx=XM_019813809.2;refGene_gn=RBM20;refGene_tx=XM_0232409
11.1;refGene_gn=RBM20;refGene_tx=XM_023240912.1;refGene_gn=RBM20;refGe
ne_tx=XM_023240913.1
D2 70207750 70207812
refGene_gn=RBM20;refGene_tx=XM_019813805.2;refGene_gn=RBM20;refGene_tx
=XM_019813807.2;refGene_gn=RBM20;refGene_tx=XM_019813808.2;refGene_gn=
RBM20;refGene_tx=XM_019813809.2;refGene_gn=RBM20;refGene_tx=XM_0232409
11.1;refGene_gn=RBM20;refGene_tx=XM_023240912.1;refGene_gn=RBM20;refGe
ne_tx=XM_023240913.1
D2 70210078 70210173
refGene_gn=RBM20;refGene_tx=XM_019813805.2;refGene_gn=RBM20;refGene_tx
=XM_019813807.2;refGene_gn=RBM20;refGene_tx=XM_019813808.2;refGene_gn=
RBM20;refGene_tx=XM_019813809.2;refGene_gn=RBM20;refGene_tx=XM_0232409
11.1;refGene_gn=RBM20;refGene_tx=XM_023240912.1;refGene_gn=RBM20;refGe
ne_tx=XM_023240913.1
D2 70210756 70210854
refGene_gn=RBM20;refGene_tx=XM_019813805.2;refGene_gn=RBM20;refGene_tx
=XM_019813807.2;refGene_gn=RBM20;refGene_tx=XM_019813808.2;refGene_gn=
RBM20;refGene_tx=XM_019813809.2;refGene_gn=RBM20;refGene_tx=XM_0232409
11.1;refGene_gn=RBM20;refGene_tx=XM_023240912.1;refGene_gn=RBM20;refGe
ne_tx=XM_023240913.1
D2 70222849 70222993
refGene_gn=RBM20;refGene_tx=XM_019813805.2;refGene_gn=RBM20;refGene_tx
=XM_019813807.2;refGene_gn=RBM20;refGene_tx=XM_019813808.2;refGene_gn=
RBM20;refGene_tx=XM_019813809.2;refGene_gn=RBM20;refGene_tx=XM_0232409
11.1;refGene_gn=RBM20;refGene_tx=XM_023240912.1;refGene_gn=RBM20;refGe
ne_tx=XM_023240913.1
D2 70225111 70225243
refGene_gn=RBM20;refGene_tx=XM_019813805.2;refGene_gn=RBM20;refGene_tx

=XM_019813807.2;refGene_gn=RBM20;refGene_tx=XM_019813808.2;refGene_gn=RBM20;refGene_tx=XM_019813809.2;refGene_gn=RBM20;refGene_tx=XM_023240911.1;refGene_gn=RBM20;refGene_tx=XM_023240912.1;refGene_gn=RBM20;refGene_tx=XM_023240913.1
D2 70231100 70231180
refGene_gn=RBM20;refGene_tx=XM_019813805.2;refGene_gn=RBM20;refGene_tx=XM_019813807.2;refGene_gn=RBM20;refGene_tx=XM_019813808.2;refGene_gn=RBM20;refGene_tx=XM_019813809.2;refGene_gn=RBM20;refGene_tx=XM_023240911.1;refGene_gn=RBM20;refGene_tx=XM_023240912.1;refGene_gn=RBM20;refGene_tx=XM_023240913.1
D2 70232931 70233625
refGene_gn=RBM20;refGene_tx=XM_019813805.2;refGene_gn=RBM20;refGene_tx=XM_019813807.2;refGene_gn=RBM20;refGene_tx=XM_019813808.2;refGene_gn=RBM20;refGene_tx=XM_019813809.2;refGene_gn=RBM20;refGene_tx=XM_023240911.1;refGene_gn=RBM20;refGene_tx=XM_023240912.1;refGene_gn=RBM20;refGene_tx=XM_023240913.1
D2 70239574 70239676
refGene_gn=RBM20;refGene_tx=XM_019813805.2;refGene_gn=RBM20;refGene_tx=XM_019813807.2;refGene_gn=RBM20;refGene_tx=XM_019813808.2;refGene_gn=RBM20;refGene_tx=XM_019813809.2;refGene_gn=RBM20;refGene_tx=XM_023240911.1;refGene_gn=RBM20;refGene_tx=XM_023240912.1;refGene_gn=RBM20;refGene_tx=XM_023240913.1
D2 70240783 70241444
refGene_gn=RBM20;refGene_tx=XM_019813805.2;refGene_gn=RBM20;refGene_tx=XM_019813807.2;refGene_gn=RBM20;refGene_tx=XM_019813808.2;refGene_gn=RBM20;refGene_tx=XM_019813809.2;refGene_gn=RBM20;refGene_tx=XM_023240911.1;refGene_gn=RBM20;refGene_tx=XM_023240912.1;refGene_gn=RBM20;refGene_tx=XM_023240913.1
D2 70243009 70243144
refGene_gn=RBM20;refGene_tx=XM_019813805.2;refGene_gn=RBM20;refGene_tx=XM_019813807.2;refGene_gn=RBM20;refGene_tx=XM_019813808.2;refGene_gn=RBM20;refGene_tx=XM_019813809.2;refGene_gn=RBM20;refGene_tx=XM_023240911.1;refGene_gn=RBM20;refGene_tx=XM_023240912.1;refGene_gn=RBM20;refGene_tx=XM_023240913.1
D2 70250955 70251077
refGene_gn=RBM20;refGene_tx=XM_019813805.2;refGene_gn=RBM20;refGene_tx=XM_019813807.2;refGene_gn=RBM20;refGene_tx=XM_019813808.2;refGene_gn=RBM20;refGene_tx=XM_019813809.2;refGene_gn=RBM20;refGene_tx=XM_023240911.1;refGene_gn=RBM20;refGene_tx=XM_023240912.1;refGene_gn=RBM20;refGene_tx=XM_023240913.1
D2 70255764 70255875
refGene_gn=RBM20;refGene_tx=XM_019813805.2;refGene_gn=RBM20;refGene_tx=XM_019813807.2;refGene_gn=RBM20;refGene_tx=XM_019813808.2;refGene_gn=RBM20;refGene_tx=XM_019813809.2;refGene_gn=RBM20;refGene_tx=XM_023240911.1;refGene_gn=RBM20;refGene_tx=XM_023240912.1;refGene_gn=RBM20;refGene_tx=XM_023240913.1
D2 70299552 70299595 refGene_gn=PDCD4;refGene_tx=XM_011287560.3
D2 70303084 70303387 refGene_gn=PDCD4;refGene_tx=XM_011287560.3
D2 70306010 70306105 refGene_gn=PDCD4;refGene_tx=XM_011287560.3
D2 70308469 70308583 refGene_gn=PDCD4;refGene_tx=XM_011287560.3

D2 70311495 70311717 refGene_gn=PDCD4;refGene_tx=XM_011287560.3
D2 70313654 70313752 refGene_gn=PDCD4;refGene_tx=XM_011287560.3
D2 70314554 70314669 refGene_gn=PDCD4;refGene_tx=XM_011287560.3
D2 70317970 70318078 refGene_gn=PDCD4;refGene_tx=XM_011287560.3
D2 70318340 70318451 refGene_gn=PDCD4;refGene_tx=XM_011287560.3
D2 70319078 70319218 refGene_gn=PDCD4;refGene_tx=XM_011287560.3
D2 70321588 70321649 refGene_gn=PDCD4;refGene_tx=XM_011287560.3
D2 70386109 70386812
refGene_gn=SHOC2;refGene_tx=XM_006938171.4;refGene_gn=SHOC2;refGene_tx=XM_019813811.2;refGene_gn=SHOC2;refGene_tx=XM_023240914.1;refGene_gn=SHOC2;refGene_tx=XM_023240915.1
D2 70415238 70415376
refGene_gn=SHOC2;refGene_tx=XM_006938171.4;refGene_gn=SHOC2;refGene_tx=XM_019813811.2;refGene_gn=SHOC2;refGene_tx=XM_023240914.1;refGene_gn=SHOC2;refGene_tx=XM_023240915.1
D2 70428243 70428374
refGene_gn=SHOC2;refGene_tx=XM_006938171.4;refGene_gn=SHOC2;refGene_tx=XM_019813811.2;refGene_gn=SHOC2;refGene_tx=XM_023240914.1;refGene_gn=SHOC2;refGene_tx=XM_023240915.1
D2 70432916 70433105
refGene_gn=SHOC2;refGene_tx=XM_006938171.4;refGene_gn=SHOC2;refGene_tx=XM_019813811.2;refGene_gn=SHOC2;refGene_tx=XM_023240914.1;refGene_gn=SHOC2;refGene_tx=XM_023240915.1
D2 70436964 70437087
refGene_gn=SHOC2;refGene_tx=XM_006938171.4;refGene_gn=SHOC2;refGene_tx=XM_019813811.2;refGene_gn=SHOC2;refGene_tx=XM_023240914.1;refGene_gn=SHOC2;refGene_tx=XM_023240915.1
D2 70440336 70440474
refGene_gn=SHOC2;refGene_tx=XM_006938171.4;refGene_gn=SHOC2;refGene_tx=XM_019813811.2;refGene_gn=SHOC2;refGene_tx=XM_023240914.1;refGene_gn=SHOC2;refGene_tx=XM_023240915.1
D2 70440802 70440920
refGene_gn=SHOC2;refGene_tx=XM_006938171.4;refGene_gn=SHOC2;refGene_tx=XM_019813811.2;refGene_gn=SHOC2;refGene_tx=XM_023240914.1;refGene_gn=SHOC2;refGene_tx=XM_023240915.1
D2 70443213 70443422
refGene_gn=SHOC2;refGene_tx=XM_006938171.4;refGene_gn=SHOC2;refGene_tx=XM_019813811.2;refGene_gn=SHOC2;refGene_tx=XM_023240914.1;refGene_gn=SHOC2;refGene_tx=XM_023240915.1
D2 70478210 70478941 refGene_gn=ADRA2A;refGene_tx=NM_001039955.1
D2 70478943 70479562 refGene_gn=ADRA2A;refGene_tx=NM_001039955.1
D2 71405788 71405905
refGene_gn=GPAM;refGene_tx=XM_011287561.3;refGene_gn=GPAM;refGene_tx=XM_023240919.1;refGene_gn=GPAM;refGene_tx=XM_023240920.1;refGene_gn=GPAM;refGene_tx=XM_023240921.1;refGene_gn=GPAM;refGene_tx=XM_023240922.1
D2 71406574 71406633
refGene_gn=GPAM;refGene_tx=XM_011287561.3;refGene_gn=GPAM;refGene_tx=XM_023240919.1;refGene_gn=GPAM;refGene_tx=XM_023240920.1;refGene_gn=GPAM;refGene_tx=XM_023240921.1;refGene_gn=GPAM;refGene_tx=XM_023240922.1
D2 71407793 71407983

refGene_gn=GPAM;refGene_tx=XM_011287561.3;refGene_gn=GPAM;refGene_tx=XM_023240919.1;refGene_gn=GPAM;refGene_tx=XM_023240920.1;refGene_gn=GPAM;refGene_tx=XM_023240921.1;refGene_gn=GPAM;refGene_tx=XM_023240922.1
D2 71409576 71409717
refGene_gn=GPAM;refGene_tx=XM_011287561.3;refGene_gn=GPAM;refGene_tx=XM_023240919.1;refGene_gn=GPAM;refGene_tx=XM_023240920.1;refGene_gn=GPAM;refGene_tx=XM_023240921.1;refGene_gn=GPAM;refGene_tx=XM_023240922.1
D2 71410353 71410431
refGene_gn=GPAM;refGene_tx=XM_011287561.3;refGene_gn=GPAM;refGene_tx=XM_023240919.1;refGene_gn=GPAM;refGene_tx=XM_023240920.1;refGene_gn=GPAM;refGene_tx=XM_023240921.1;refGene_gn=GPAM;refGene_tx=XM_023240922.1
D2 71411609 71411752
refGene_gn=GPAM;refGene_tx=XM_011287561.3;refGene_gn=GPAM;refGene_tx=XM_023240919.1;refGene_gn=GPAM;refGene_tx=XM_023240920.1;refGene_gn=GPAM;refGene_tx=XM_023240921.1;refGene_gn=GPAM;refGene_tx=XM_023240922.1
D2 71412228 71412493
refGene_gn=GPAM;refGene_tx=XM_011287561.3;refGene_gn=GPAM;refGene_tx=XM_023240919.1;refGene_gn=GPAM;refGene_tx=XM_023240920.1;refGene_gn=GPAM;refGene_tx=XM_023240921.1;refGene_gn=GPAM;refGene_tx=XM_023240922.1
D2 71413305 71413376
refGene_gn=GPAM;refGene_tx=XM_011287561.3;refGene_gn=GPAM;refGene_tx=XM_023240919.1;refGene_gn=GPAM;refGene_tx=XM_023240920.1;refGene_gn=GPAM;refGene_tx=XM_023240921.1;refGene_gn=GPAM;refGene_tx=XM_023240922.1
D2 71415461 71415577
refGene_gn=GPAM;refGene_tx=XM_011287561.3;refGene_gn=GPAM;refGene_tx=XM_023240919.1;refGene_gn=GPAM;refGene_tx=XM_023240920.1;refGene_gn=GPAM;refGene_tx=XM_023240921.1;refGene_gn=GPAM;refGene_tx=XM_023240922.1
D2 71416760 71416846
refGene_gn=GPAM;refGene_tx=XM_011287561.3;refGene_gn=GPAM;refGene_tx=XM_023240919.1;refGene_gn=GPAM;refGene_tx=XM_023240920.1;refGene_gn=GPAM;refGene_tx=XM_023240921.1;refGene_gn=GPAM;refGene_tx=XM_023240922.1
D2 71418332 71418446
refGene_gn=GPAM;refGene_tx=XM_011287561.3;refGene_gn=GPAM;refGene_tx=XM_023240919.1;refGene_gn=GPAM;refGene_tx=XM_023240920.1;refGene_gn=GPAM;refGene_tx=XM_023240921.1;refGene_gn=GPAM;refGene_tx=XM_023240922.1
D2 71421641 71421854
refGene_gn=GPAM;refGene_tx=XM_011287561.3;refGene_gn=GPAM;refGene_tx=XM_023240919.1;refGene_gn=GPAM;refGene_tx=XM_023240920.1;refGene_gn=GPAM;refGene_tx=XM_023240921.1;refGene_gn=GPAM;refGene_tx=XM_023240922.1
D2 71422184 71422284
refGene_gn=GPAM;refGene_tx=XM_011287561.3;refGene_gn=GPAM;refGene_tx=XM_023240919.1;refGene_gn=GPAM;refGene_tx=XM_023240920.1;refGene_gn=GPAM;refGene_tx=XM_023240921.1;refGene_gn=GPAM;refGene_tx=XM_023240922.1
D2 71425093 71425230
refGene_gn=GPAM;refGene_tx=XM_011287561.3;refGene_gn=GPAM;refGene_tx=XM_023240919.1;refGene_gn=GPAM;refGene_tx=XM_023240920.1;refGene_gn=GPAM;refGene_tx=XM_023240921.1;refGene_gn=GPAM;refGene_tx=XM_023240922.1
D2 71425891 71425988
refGene_gn=GPAM;refGene_tx=XM_011287561.3;refGene_gn=GPAM;refGene_tx=XM_023240919.1;refGene_gn=GPAM;refGene_tx=XM_023240920.1;refGene_gn=GPA

M;refGene_tx=XM_023240921.1;refGene_gn=GPAM;refGene_tx=XM_023240922.1
D2 71426591 71426738
refGene_gn=GPAM;refGene_tx=XM_011287561.3;refGene_gn=GPAM;refGene_tx=XM_023240919.1;refGene_gn=GPAM;refGene_tx=XM_023240920.1;refGene_gn=GPAM;refGene_tx=XM_023240921.1;refGene_gn=GPAM;refGene_tx=XM_023240922.1
D2 71429135 71429249
refGene_gn=GPAM;refGene_tx=XM_011287561.3;refGene_gn=GPAM;refGene_tx=XM_023240919.1;refGene_gn=GPAM;refGene_tx=XM_023240920.1;refGene_gn=GPAM;refGene_tx=XM_023240921.1;refGene_gn=GPAM;refGene_tx=XM_023240922.1
D2 71432506 71432580
refGene_gn=GPAM;refGene_tx=XM_011287561.3;refGene_gn=GPAM;refGene_tx=XM_023240919.1;refGene_gn=GPAM;refGene_tx=XM_023240920.1;refGene_gn=GPAM;refGene_tx=XM_023240921.1;refGene_gn=GPAM;refGene_tx=XM_023240922.1
D2 71435122 71435245
refGene_gn=GPAM;refGene_tx=XM_011287561.3;refGene_gn=GPAM;refGene_tx=XM_023240919.1;refGene_gn=GPAM;refGene_tx=XM_023240920.1;refGene_gn=GPAM;refGene_tx=XM_023240921.1;refGene_gn=GPAM;refGene_tx=XM_023240922.1
D2 71436352 71436454
refGene_gn=GPAM;refGene_tx=XM_011287561.3;refGene_gn=GPAM;refGene_tx=XM_023240919.1;refGene_gn=GPAM;refGene_tx=XM_023240920.1;refGene_gn=GPAM;refGene_tx=XM_023240921.1;refGene_gn=GPAM;refGene_tx=XM_023240922.1
D2 71512248 71512324 refGene_gn=TECTB;refGene_tx=XM_006938175.3
D2 71513039 71513230 refGene_gn=TECTB;refGene_tx=XM_006938175.3
D2 71514421 71514564 refGene_gn=TECTB;refGene_tx=XM_006938175.3
D2 71514669 71514742 refGene_gn=TECTB;refGene_tx=XM_006938175.3
D2 71519898 71520002 refGene_gn=TECTB;refGene_tx=XM_006938175.3
D2 71520122 71520206 refGene_gn=TECTB;refGene_tx=XM_006938175.3
D2 71524328 71524491 refGene_gn=TECTB;refGene_tx=XM_006938175.3
D2 71526001 71526074 refGene_gn=TECTB;refGene_tx=XM_006938175.3
D2 71527264 71527297 refGene_gn=TECTB;refGene_tx=XM_006938175.3
D2 71528443 71528493 refGene_gn=TECTB;refGene_tx=XM_006938175.3
D2 71533037 71533142
refGene_gn=LOC101101087;refGene_tx=XM_011287645.2
D2 71534467 71534559
refGene_gn=LOC101101087;refGene_tx=XM_011287645.2
D2 71535805 71535904
refGene_gn=LOC101101087;refGene_tx=XM_011287645.2
D2 71537309 71537484
refGene_gn=LOC101101087;refGene_tx=XM_011287645.2
D2 71539443 71539637
refGene_gn=LOC101101087;refGene_tx=XM_011287645.2
D2 71540533 71540673
refGene_gn=LOC101101087;refGene_tx=XM_011287645.2
D2 71542268 71542446
refGene_gn=LOC101101087;refGene_tx=XM_011287645.2
D2 71548143 71548299
refGene_gn=LOC101101087;refGene_tx=XM_011287645.2
D2 71550381 71550405
refGene_gn=LOC101101087;refGene_tx=XM_011287645.2
D2 71550407 71550546

refGene_gn=LOC101101087;refGene_tx=XM_011287645.2
D2 71557256 71557328
refGene_gn=LOC101101087;refGene_tx=XM_011287645.2
D2 71557850 71557955
refGene_gn=LOC101101087;refGene_tx=XM_011287645.2
D2 71558674 71558758
refGene_gn=LOC101101087;refGene_tx=XM_011287645.2
D2 71561192 71561330
refGene_gn=LOC101101087;refGene_tx=XM_011287645.2
D2 71563687 71563851
refGene_gn=LOC101101087;refGene_tx=XM_011287645.2
D2 71564493 71564615
refGene_gn=LOC101101087;refGene_tx=XM_011287645.2
D2 71565736 71565890
refGene_gn=LOC101101087;refGene_tx=XM_011287645.2
D2 71567534 71567661
refGene_gn=LOC101101087;refGene_tx=XM_011287645.2
D2 71569088 71569274
refGene_gn=LOC101101087;refGene_tx=XM_011287645.2
D2 71572487 71572597
refGene_gn=LOC101101087;refGene_tx=XM_011287645.2
D2 71574148 71574494
refGene_gn=LOC101101087;refGene_tx=XM_011287645.2
D2 71575919 71576300
refGene_gn=LOC101101087;refGene_tx=XM_011287645.2
D2 71606626 71606782 refGene_gn=ACSL5;refGene_tx=XM_023240924.1
D2 71609473 71609579 refGene_gn=ACSL5;refGene_tx=XM_023240924.1
D2 71612362 71612427 refGene_gn=ACSL5;refGene_tx=XM_023240924.1
D2 71612552 71612654 refGene_gn=ACSL5;refGene_tx=XM_023240924.1
D2 71614283 71614383 refGene_gn=ACSL5;refGene_tx=XM_023240924.1
D2 71615377 71615556 refGene_gn=ACSL5;refGene_tx=XM_023240924.1
D2 71616477 71616510 refGene_gn=ACSL5;refGene_tx=XM_023240924.1
D2 71616598 71616650 refGene_gn=ACSL5;refGene_tx=XM_023240924.1
D2 71617402 71617483 refGene_gn=ACSL5;refGene_tx=XM_023240924.1
D2 71619493 71619585 refGene_gn=ACSL5;refGene_tx=XM_023240924.1
D2 71622056 71622191 refGene_gn=ACSL5;refGene_tx=XM_023240924.1
D2 71622716 71622812 refGene_gn=ACSL5;refGene_tx=XM_023240924.1
D2 71624592 71624665 refGene_gn=ACSL5;refGene_tx=XM_023240924.1
D2 71624943 71625032 refGene_gn=ACSL5;refGene_tx=XM_023240924.1
D2 71625333 71625450 refGene_gn=ACSL5;refGene_tx=XM_023240924.1
D2 71626733 71626877 refGene_gn=ACSL5;refGene_tx=XM_023240924.1
D2 71627531 71627633 refGene_gn=ACSL5;refGene_tx=XM_023240924.1
D2 71628112 71628184 refGene_gn=ACSL5;refGene_tx=XM_023240924.1
D2 71628546 71628687 refGene_gn=ACSL5;refGene_tx=XM_023240924.1
D2 71631465 71631569
refGene_gn=ZDHHC6;refGene_tx=XM_003994413.5;refGene_gn=ZDHHC6;refGene_
tx=XM_006938181.4;refGene_gn=ZDHHC6;refGene_tx=XM_006938182.4;refGene_
gn=ZDHHC6;refGene_tx=XM_023240923.1
D2 71632812 71632859
refGene_gn=ZDHHC6;refGene_tx=XM_003994413.5;refGene_gn=ZDHHC6;refGene_

tx=XM_006938181.4;refGene_gn=ZDHHC6;refGene_tx=XM_006938182.4;refGene_gn=ZDHHC6;refGene_tx=XM_023240923.1
D2 71632948 71633094
refGene_gn=ZDHHC6;refGene_tx=XM_003994413.5;refGene_gn=ZDHHC6;refGene_tx=XM_006938181.4;refGene_gn=ZDHHC6;refGene_tx=XM_006938182.4;refGene_gn=ZDHHC6;refGene_tx=XM_023240923.1
D2 71633769 71633811
refGene_gn=ZDHHC6;refGene_tx=XM_003994413.5;refGene_gn=ZDHHC6;refGene_tx=XM_006938181.4;refGene_gn=ZDHHC6;refGene_tx=XM_006938182.4;refGene_gn=ZDHHC6;refGene_tx=XM_023240923.1
D2 71634819 71634987
refGene_gn=ZDHHC6;refGene_tx=XM_003994413.5;refGene_gn=ZDHHC6;refGene_tx=XM_006938181.4;refGene_gn=ZDHHC6;refGene_tx=XM_006938182.4;refGene_gn=ZDHHC6;refGene_tx=XM_023240923.1
D2 71639455 71639509
refGene_gn=ZDHHC6;refGene_tx=XM_003994413.5;refGene_gn=ZDHHC6;refGene_tx=XM_006938181.4;refGene_gn=ZDHHC6;refGene_tx=XM_006938182.4;refGene_gn=ZDHHC6;refGene_tx=XM_023240923.1
D2 71641142 71641304
refGene_gn=ZDHHC6;refGene_tx=XM_003994413.5;refGene_gn=ZDHHC6;refGene_tx=XM_006938181.4;refGene_gn=ZDHHC6;refGene_tx=XM_006938182.4;refGene_gn=ZDHHC6;refGene_tx=XM_023240923.1
D2 71642769 71642929
refGene_gn=ZDHHC6;refGene_tx=XM_003994413.5;refGene_gn=ZDHHC6;refGene_tx=XM_006938181.4;refGene_gn=ZDHHC6;refGene_tx=XM_006938182.4;refGene_gn=ZDHHC6;refGene_tx=XM_023240923.1
D2 71644477 71644569
refGene_gn=ZDHHC6;refGene_tx=XM_003994413.5;refGene_gn=ZDHHC6;refGene_tx=XM_006938181.4;refGene_gn=ZDHHC6;refGene_tx=XM_006938182.4;refGene_gn=ZDHHC6;refGene_tx=XM_023240923.1
D2 71646295 71646562
refGene_gn=ZDHHC6;refGene_tx=XM_003994413.5;refGene_gn=ZDHHC6;refGene_tx=XM_006938181.4;refGene_gn=ZDHHC6;refGene_tx=XM_006938182.4;refGene_gn=ZDHHC6;refGene_tx=XM_023240923.1
D2 71648663 71648757
refGene_gn=VTI1A;refGene_tx=XM_003994414.5;refGene_gn=VTI1A;refGene_tx=XM_006938177.4;refGene_gn=VTI1A;refGene_tx=XM_006938178.4;refGene_gn=VTI1A;refGene_tx=XM_019813824.2;refGene_gn=VTI1A;refGene_tx=XM_019813825.2
D2 71659209 71659268
refGene_gn=VTI1A;refGene_tx=XM_003994414.5;refGene_gn=VTI1A;refGene_tx=XM_006938177.4;refGene_gn=VTI1A;refGene_tx=XM_006938178.4;refGene_gn=VTI1A;refGene_tx=XM_019813824.2;refGene_gn=VTI1A;refGene_tx=XM_019813825.2
D2 71663469 71663580
refGene_gn=VTI1A;refGene_tx=XM_003994414.5;refGene_gn=VTI1A;refGene_tx=XM_006938177.4;refGene_gn=VTI1A;refGene_tx=XM_006938178.4;refGene_gn=VTI1A;refGene_tx=XM_019813824.2;refGene_gn=VTI1A;refGene_tx=XM_019813825.2
D2 71726534 71726612

refGene_gn=VTI1A;refGene_tx=XM_003994414.5;refGene_gn=VTI1A;refGene_tx=XM_006938177.4;refGene_gn=VTI1A;refGene_tx=XM_006938178.4;refGene_gn=VTI1A;refGene_tx=XM_019813824.2;refGene_gn=VTI1A;refGene_tx=XM_019813825.2
D2 71733038 71733059 refGene_gn=VTI1A;refGene_tx=XM_006938177.4;refGene_gn=VTI1A;refGene_tx=XM_006938178.4;refGene_gn=VTI1A;refGene_tx=XM_019813824.2
D2 71737837 71737922 refGene_gn=VTI1A;refGene_tx=XM_003994414.5;refGene_gn=VTI1A;refGene_tx=XM_006938177.4;refGene_gn=VTI1A;refGene_tx=XM_006938178.4;refGene_gn=VTI1A;refGene_tx=XM_019813824.2;refGene_gn=VTI1A;refGene_tx=XM_019813825.2
D2 71857280 71857351 refGene_gn=VTI1A;refGene_tx=XM_003994414.5;refGene_gn=VTI1A;refGene_tx=XM_006938177.4;refGene_gn=VTI1A;refGene_tx=XM_006938178.4;refGene_gn=VTI1A;refGene_tx=XM_019813824.2;refGene_gn=VTI1A;refGene_tx=XM_019813825.2
D2 71858020 71858086 refGene_gn=VTI1A;refGene_tx=XM_003994414.5;refGene_gn=VTI1A;refGene_tx=XM_006938177.4;refGene_gn=VTI1A;refGene_tx=XM_006938178.4;refGene_gn=VTI1A;refGene_tx=XM_019813824.2;refGene_gn=VTI1A;refGene_tx=XM_019813825.2
D2 71978337 71978356 refGene_gn=VTI1A;refGene_tx=XM_019813824.2
D2 71997643 71997737 refGene_gn=VTI1A;refGene_tx=XM_003994414.5;refGene_gn=VTI1A;refGene_tx=XM_006938177.4
D2 72046165 72046887 refGene_gn=LOC101101594;refGene_tx=XM_011287646.3
D2 72046888 72047279 refGene_gn=LOC101101594;refGene_tx=XM_011287646.3
D2 72117134 72117323 refGene_gn=TCF7L2;refGene_tx=XM_023240925.1;refGene_gn=TCF7L2;refGene_tx=XM_023240927.1;refGene_gn=TCF7L2;refGene_tx=XM_023240928.1;refGene_gn=TCF7L2;refGene_tx=XM_023240929.1;refGene_gn=TCF7L2;refGene_tx=XM_023240930.1;refGene_gn=TCF7L2;refGene_tx=XM_023240931.1;refGene_gn=TCF7L2;refGene_tx=XM_023240932.1;refGene_gn=TCF7L2;refGene_tx=XM_023240933.1;refGene_gn=TCF7L2;refGene_tx=XM_023240934.1;refGene_gn=TCF7L2;refGene_tx=XM_023240935.1;refGene_gn=TCF7L2;refGene_tx=XM_023240936.1;refGene_gn=TCF7L2;refGene_tx=XM_023240937.1;refGene_gn=TCF7L2;refGene_tx=XM_023240938.1;refGene_gn=TCF7L2;refGene_tx=XM_023240939.1;refGene_gn=TCF7L2;refGene_tx=XM_023240940.1;refGene_gn=TCF7L2;refGene_tx=XM_023240941.1;refGene_gn=TCF7L2;refGene_tx=XM_023240942.1;refGene_gn=TCF7L2;refGene_tx=XM_023240943.1;refGene_gn=TCF7L2;refGene_tx=XM_023240945.1;refGene_gn=TCF7L2;refGene_tx=XM_023240946.1;refGene_gn=TCF7L2;refGene_tx=XM_023240947.1;refGene_gn=TCF7L2;refGene_tx=XM_023240949.1;refGene_gn=TCF7L2;refGene_tx=XM_023240950.1;refGene_gn=TCF7L2;refGene_tx=XM_023240951.1;refGene_gn=TCF7L2;refGene_tx=XM_023240952.1;refGene_gn=TCF7L2;refGene_tx=XM_023240953.1
D2 72117588 72117655 refGene_gn=TCF7L2;refGene_tx=XM_023240925.1;refGene_gn=TCF7L2;refGene_

tx=XM_023240927.1;refGene_gn=TCF7L2;refGene_tx=XM_023240928.1;refGene_gn=TCF7L2;refGene_tx=XM_023240929.1;refGene_gn=TCF7L2;refGene_tx=XM_023240930.1;refGene_gn=TCF7L2;refGene_tx=XM_023240931.1;refGene_gn=TCF7L2;refGene_tx=XM_023240932.1;refGene_gn=TCF7L2;refGene_tx=XM_023240933.1;refGene_gn=TCF7L2;refGene_tx=XM_023240934.1;refGene_gn=TCF7L2;refGene_tx=XM_023240935.1;refGene_gn=TCF7L2;refGene_tx=XM_023240936.1;refGene_gn=TCF7L2;refGene_tx=XM_023240937.1;refGene_gn=TCF7L2;refGene_tx=XM_023240938.1;refGene_gn=TCF7L2;refGene_tx=XM_023240939.1;refGene_gn=TCF7L2;refGene_tx=XM_023240940.1;refGene_gn=TCF7L2;refGene_tx=XM_023240941.1;refGene_gn=TCF7L2;refGene_tx=XM_023240942.1;refGene_gn=TCF7L2;refGene_tx=XM_023240943.1;refGene_gn=TCF7L2;refGene_tx=XM_023240945.1;refGene_gn=TCF7L2;refGene_tx=XM_023240946.1;refGene_gn=TCF7L2;refGene_tx=XM_023240947.1;refGene_gn=TCF7L2;refGene_tx=XM_023240949.1;refGene_gn=TCF7L2;refGene_tx=XM_023240950.1;refGene_gn=TCF7L2;refGene_tx=XM_023240951.1;refGene_gn=TCF7L2;refGene_tx=XM_023240952.1;refGene_gn=TCF7L2;refGene_tx=XM_023240953.1

D2 72117864 72117989 refGene_gn=TCF7L2;refGene_tx=XM_023240925.1;refGene_gn=TCF7L2;refGene_tx=XM_023240927.1;refGene_gn=TCF7L2;refGene_tx=XM_023240928.1;refGene_gn=TCF7L2;refGene_tx=XM_023240929.1;refGene_gn=TCF7L2;refGene_tx=XM_023240930.1;refGene_gn=TCF7L2;refGene_tx=XM_023240931.1;refGene_gn=TCF7L2;refGene_tx=XM_023240932.1;refGene_gn=TCF7L2;refGene_tx=XM_023240933.1;refGene_gn=TCF7L2;refGene_tx=XM_023240934.1;refGene_gn=TCF7L2;refGene_tx=XM_023240935.1;refGene_gn=TCF7L2;refGene_tx=XM_023240936.1;refGene_gn=TCF7L2;refGene_tx=XM_023240937.1;refGene_gn=TCF7L2;refGene_tx=XM_023240938.1;refGene_gn=TCF7L2;refGene_tx=XM_023240939.1;refGene_gn=TCF7L2;refGene_tx=XM_023240940.1;refGene_gn=TCF7L2;refGene_tx=XM_023240941.1;refGene_gn=TCF7L2;refGene_tx=XM_023240942.1;refGene_gn=TCF7L2;refGene_tx=XM_023240943.1;refGene_gn=TCF7L2;refGene_tx=XM_023240945.1;refGene_gn=TCF7L2;refGene_tx=XM_023240946.1;refGene_gn=TCF7L2;refGene_tx=XM_023240947.1;refGene_gn=TCF7L2;refGene_tx=XM_023240949.1;refGene_gn=TCF7L2;refGene_tx=XM_023240950.1;refGene_gn=TCF7L2;refGene_tx=XM_023240951.1;refGene_gn=TCF7L2;refGene_tx=XM_023240952.1;refGene_gn=TCF7L2;refGene_tx=XM_023240953.1

D2 72130400 72130469 refGene_gn=TCF7L2;refGene_tx=XM_023240925.1;refGene_gn=TCF7L2;refGene_tx=XM_023240927.1;refGene_gn=TCF7L2;refGene_tx=XM_023240928.1;refGene_gn=TCF7L2;refGene_tx=XM_023240929.1;refGene_gn=TCF7L2;refGene_tx=XM_023240930.1;refGene_gn=TCF7L2;refGene_tx=XM_023240931.1;refGene_gn=TCF7L2;refGene_tx=XM_023240932.1;refGene_gn=TCF7L2;refGene_tx=XM_023240934.1;refGene_gn=TCF7L2;refGene_tx=XM_023240935.1;refGene_gn=TCF7L2;refGene_tx=XM_023240936.1;refGene_gn=TCF7L2;refGene_tx=XM_023240937.1;refGene_gn=TCF7L2;refGene_tx=XM_023240940.1;refGene_gn=TCF7L2;refGene_tx=XM_023240945.1;refGene_gn=TCF7L2;refGene_tx=XM_023240946.1;refGene_gn=TCF7L2;refGene_tx=XM_023240950.1

D2 72187897 72187963 refGene_gn=TCF7L2;refGene_tx=XM_023240944.1

D2 72194860 72194962 refGene_gn=TCF7L2;refGene_tx=XM_023240925.1;refGene_gn=TCF7L2;refGene_tx=XM_023240927.1;refGene_gn=TCF7L2;refGene_tx=XM_023240928.1;refGene_gn=TCF7L2;refGene_tx=XM_023240929.1;refGene_gn=TCF7L2;refGene_tx=XM_02

3240930.1;refGene_gn=TCF7L2;refGene_tx=XM_023240931.1;refGene_gn=TCF7L2;refGene_tx=XM_023240932.1;refGene_gn=TCF7L2;refGene_tx=XM_023240933.1;refGene_gn=TCF7L2;refGene_tx=XM_023240934.1;refGene_gn=TCF7L2;refGene_tx=XM_023240935.1;refGene_gn=TCF7L2;refGene_tx=XM_023240936.1;refGene_gn=TCF7L2;refGene_tx=XM_023240937.1;refGene_gn=TCF7L2;refGene_tx=XM_023240938.1;refGene_gn=TCF7L2;refGene_tx=XM_023240939.1;refGene_gn=TCF7L2;refGene_tx=XM_023240941.1;refGene_gn=TCF7L2;refGene_tx=XM_023240942.1;refGene_gn=TCF7L2;refGene_tx=XM_023240943.1;refGene_gn=TCF7L2;refGene_tx=XM_023240944.1;refGene_gn=TCF7L2;refGene_tx=XM_023240945.1;refGene_gn=TCF7L2;refGene_tx=XM_023240946.1;refGene_gn=TCF7L2;refGene_tx=XM_023240947.1;refGene_gn=TCF7L2;refGene_tx=XM_023240949.1;refGene_gn=TCF7L2;refGene_tx=XM_023240950.1;refGene_gn=TCF7L2;refGene_tx=XM_023240951.1;refGene_gn=TCF7L2;refGene_tx=XM_023240952.1;refGene_gn=TCF7L2;refGene_tx=XM_023240953.1

D2 72243834 72243978 refGene_gn=TCF7L2;refGene_tx=XM_023240925.1;refGene_gn=TCF7L2;refGene_tx=XM_023240927.1;refGene_gn=TCF7L2;refGene_tx=XM_023240928.1;refGene_gn=TCF7L2;refGene_tx=XM_023240929.1;refGene_gn=TCF7L2;refGene_tx=XM_023240930.1;refGene_gn=TCF7L2;refGene_tx=XM_023240931.1;refGene_gn=TCF7L2;refGene_tx=XM_023240932.1;refGene_gn=TCF7L2;refGene_tx=XM_023240933.1;refGene_gn=TCF7L2;refGene_tx=XM_023240944.1;refGene_gn=TCF7L2;refGene_tx=XM_023240945.1;refGene_gn=TCF7L2;refGene_tx=XM_023240946.1;refGene_gn=TCF7L2;refGene_tx=XM_023240947.1

D2 72281251 72281374 refGene_gn=TCF7L2;refGene_tx=XM_023240948.1

D2 72292825 72292958 refGene_gn=TCF7L2;refGene_tx=XM_023240925.1;refGene_gn=TCF7L2;refGene_tx=XM_023240927.1;refGene_gn=TCF7L2;refGene_tx=XM_023240928.1;refGene_gn=TCF7L2;refGene_tx=XM_023240929.1;refGene_gn=TCF7L2;refGene_tx=XM_023240930.1;refGene_gn=TCF7L2;refGene_tx=XM_023240931.1;refGene_gn=TCF7L2;refGene_tx=XM_023240932.1;refGene_gn=TCF7L2;refGene_tx=XM_023240933.1;refGene_gn=TCF7L2;refGene_tx=XM_023240934.1;refGene_gn=TCF7L2;refGene_tx=XM_023240935.1;refGene_gn=TCF7L2;refGene_tx=XM_023240936.1;refGene_gn=TCF7L2;refGene_tx=XM_023240937.1;refGene_gn=TCF7L2;refGene_tx=XM_023240938.1;refGene_gn=TCF7L2;refGene_tx=XM_023240939.1;refGene_gn=TCF7L2;refGene_tx=XM_023240940.1;refGene_gn=TCF7L2;refGene_tx=XM_023240941.1;refGene_gn=TCF7L2;refGene_tx=XM_023240942.1;refGene_gn=TCF7L2;refGene_tx=XM_023240943.1;refGene_gn=TCF7L2;refGene_tx=XM_023240944.1;refGene_gn=TCF7L2;refGene_tx=XM_023240945.1;refGene_gn=TCF7L2;refGene_tx=XM_023240946.1;refGene_gn=TCF7L2;refGene_tx=XM_023240947.1;refGene_gn=TCF7L2;refGene_tx=XM_023240948.1;refGene_gn=TCF7L2;refGene_tx=XM_023240949.1;refGene_gn=TCF7L2;refGene_tx=XM_023240950.1;refGene_gn=TCF7L2;refGene_tx=XM_023240951.1;refGene_gn=TCF7L2;refGene_tx=XM_023240952.1;refGene_gn=TCF7L2;refGene_tx=XM_023240953.1

D2 72295492 72295595 refGene_gn=TCF7L2;refGene_tx=XM_023240925.1;refGene_gn=TCF7L2;refGene_tx=XM_023240927.1;refGene_gn=TCF7L2;refGene_tx=XM_023240928.1;refGene_gn=TCF7L2;refGene_tx=XM_023240929.1;refGene_gn=TCF7L2;refGene_tx=XM_023240930.1;refGene_gn=TCF7L2;refGene_tx=XM_023240931.1;refGene_gn=TCF7L2;refGene_tx=XM_023240932.1;refGene_gn=TCF7L2;refGene_tx=XM_023240933.1;refGene_gn=TCF7L2;refGene_tx=XM_023240934.1;refGene_gn=TCF7L2;refGen e_tx=XM_023240935.1;refGene_gn=TCF7L2;refGene_tx=XM_023240936.1;refGene_gn=TCF7L2;refGene_tx=XM_023240937.1;refGene_gn=TCF7L2;refGene_tx=XM_023240938.1;refGene_gn=TCF7L2;refGene_tx=XM_023240939.1;refGene_gn=TCF7L2;refGene_tx=XM_023240940.1;refGene_gn=TCF7L2;refGene_tx=XM_023240941.1;refGene_gn=TCF7L2;refGene_tx=XM_023240942.1;refGene_gn=TCF7L2;refGene_tx=XM_023240943.1;refGene_gn=TCF7L2;refGene_tx=XM_023240944.1;refGene_gn=TCF7L2;refGene_tx=XM_023240945.1;refGene_gn=TCF7L2;refGene_tx=XM_023240946.1;refGene_gn=TCF7L2;refGene_tx=XM_023240947.1;refGene_gn=TCF7L2;refGene_tx=XM_023240948.1;refGene_gn=TCF7L2;refGene_tx=XM_023240949.1;refGene_gn=TCF7L2;refGene_tx=XM_023240950.1;refGene_gn=TCF7L2;refGene_tx=XM_023240951.1;refGene_gn=TCF7L2;refGene_tx=XM_023240952.1;refGene_gn=TCF7L2;refGene_tx=XM_023240953.1

D2 72297856 72297943

refGene_gn=TCF7L2;refGene_tx=XM_023240925.1;refGene_gn=TCF7L2;refGene_tx=XM_023240927.1;refGene_gn=TCF7L2;refGene_tx=XM_023240928.1;refGene_gn=TCF7L2;refGene_tx=XM_023240929.1;refGene_gn=TCF7L2;refGene_tx=XM_023240930.1;refGene_gn=TCF7L2;refGene_tx=XM_023240931.1;refGene_gn=TCF7L2;refGene_tx=XM_023240932.1;refGene_gn=TCF7L2;refGene_tx=XM_023240933.1;refGene_gn=TCF7L2;refGene_tx=XM_023240934.1;refGene_gn=TCF7L2;refGene_tx=XM_023240935.1;refGene_gn=TCF7L2;refGene_tx=XM_023240936.1;refGene_gn=TCF7L2;refGene_tx=XM_023240937.1;refGene_gn=TCF7L2;refGene_tx=XM_023240938.1;refGene_gn=TCF7L2;refGene_tx=XM_023240939.1;refGene_gn=TCF7L2;refGene_tx=XM_023240940.1;refGene_gn=TCF7L2;refGene_tx=XM_023240941.1;refGene_gn=TCF7L2;refGene_tx=XM_023240942.1;refGene_gn=TCF7L2;refGene_tx=XM_023240943.1;refGene_gn=TCF7L2;refGene_tx=XM_023240944.1;refGene_gn=TCF7L2;refGene_tx=XM_023240945.1;refGene_gn=TCF7L2;refGene_tx=XM_023240946.1;refGene_gn=TCF7L2;refGene_tx=XM_023240947.1;refGene_gn=TCF7L2;refGene_tx=XM_023240948.1;refGene_gn=TCF7L2;refGene_tx=XM_023240949.1;refGene_gn=TCF7L2;refGene_tx=XM_023240950.1;refGene_gn=TCF7L2;refGene_tx=XM_023240951.1;refGene_gn=TCF7L2;refGene_tx=XM_023240952.1;refGene_gn=TCF7L2;refGene_tx=XM_023240953.1

D2 72303536 72303677

refGene_gn=TCF7L2;refGene_tx=XM_023240925.1;refGene_gn=TCF7L2;refGene_tx=XM_023240927.1;refGene_gn=TCF7L2;refGene_tx=XM_023240928.1;refGene_gn=TCF7L2;refGene_tx=XM_023240929.1;refGene_gn=TCF7L2;refGene_tx=XM_023240930.1;refGene_gn=TCF7L2;refGene_tx=XM_023240931.1;refGene_gn=TCF7L2;refGene_tx=XM_023240932.1;refGene_gn=TCF7L2;refGene_tx=XM_023240933.1;refGene_gn=TCF7L2;refGene_tx=XM_023240934.1;refGene_gn=TCF7L2;refGene_tx=XM_023240935.1;refGene_gn=TCF7L2;refGene_tx=XM_023240936.1;refGene_gn=TCF7L2;refGene_tx=XM_023240937.1;refGene_gn=TCF7L2;refGene_tx=XM_023240938.1;refGene_gn=TCF7L2;refGene_tx=XM_023240939.1;refGene_gn=TCF7L2;refGene_tx=XM_023240940.1;refGene_gn=TCF7L2;refGene_tx=XM_023240941.1;refGene_gn=TCF7L2;refGene_tx=XM_023240942.1;refGene_gn=TCF7L2;refGene_tx=XM_023240943.1;refGene_gn=TCF7L2;refGene_tx=XM_023240944.1;refGene_gn=TCF7L2;refGene_tx=XM_023240945.1;refGene_gn=TCF7L2;refGene_tx=XM_023240946.1;refGene_gn=TCF7L2;refGene_tx=XM_023240947.1;refGene_gn=TCF7L2;refGene_tx=XM_023240948.1;refGene_gn=TCF7L2;refGene_tx=XM_023240949.1;refGene_gn=TCF7L2;refGene_tx=XM_023240950.1;refGene_gn=TCF7L2;refGene_tx=XM_023240951.1;refGene_gn=TCF7L2;refGene_tx=XM_023240952.1;refGene_gn=TCF7L2;refGene_tx=XM_023240953.1

D2 72304260 72304420
refGene_gn=TCF7L2;refGene_tx=XM_023240925.1;refGene_gn=TCF7L2;refGene_tx=XM_023240927.1;refGene_gn=TCF7L2;refGene_tx=XM_023240928.1;refGene_gn=TCF7L2;refGene_tx=XM_023240929.1;refGene_gn=TCF7L2;refGene_tx=XM_023240930.1;refGene_gn=TCF7L2;refGene_tx=XM_023240931.1;refGene_gn=TCF7L2;refGene_tx=XM_023240932.1;refGene_gn=TCF7L2;refGene_tx=XM_023240933.1;refGene_gn=TCF7L2;refGene_tx=XM_023240934.1;refGene_gn=TCF7L2;refGene_tx=XM_023240935.1;refGene_gn=TCF7L2;refGene_tx=XM_023240936.1;refGene_gn=TCF7L2;refGene_tx=XM_023240937.1;refGene_gn=TCF7L2;refGene_tx=XM_023240938.1;refGene_gn=TCF7L2;refGene_tx=XM_023240939.1;refGene_gn=TCF7L2;refGene_tx=XM_023240940.1;refGene_gn=TCF7L2;refGene_tx=XM_023240941.1;refGene_gn=TCF7L2;refGene_tx=XM_023240942.1;refGene_gn=TCF7L2;refGene_tx=XM_023240943.1;refGene_gn=TCF7L2;refGene_tx=XM_023240944.1;refGene_gn=TCF7L2;refGene_tx=XM_023240945.1;refGene_gn=TCF7L2;refGene_tx=XM_023240946.1;refGene_gn=TCF7L2;refGene_tx=XM_023240947.1;refGene_gn=TCF7L2;refGene_tx=XM_023240948.1;refGene_gn=TCF7L2;refGene_tx=XM_023240949.1;refGene_gn=TCF7L2;refGene_tx=XM_023240950.1;refGene_gn=TCF7L2;refGene_tx=XM_023240951.1;refGene_gn=TCF7L2;refGene_tx=XM_023240952.1;refGene_gn=TCF7L2;refGene_tx=XM_023240953.1
D2 72304768 72304876
refGene_gn=TCF7L2;refGene_tx=XM_023240925.1;refGene_gn=TCF7L2;refGene_tx=XM_023240927.1;refGene_gn=TCF7L2;refGene_tx=XM_023240928.1;refGene_gn=TCF7L2;refGene_tx=XM_023240929.1;refGene_gn=TCF7L2;refGene_tx=XM_023240930.1;refGene_gn=TCF7L2;refGene_tx=XM_023240931.1;refGene_gn=TCF7L2;refGene_tx=XM_023240932.1;refGene_gn=TCF7L2;refGene_tx=XM_023240933.1;refGene_gn=TCF7L2;refGene_tx=XM_023240934.1;refGene_gn=TCF7L2;refGene_tx=XM_023240935.1;refGene_gn=TCF7L2;refGene_tx=XM_023240936.1;refGene_gn=TCF7L2;refGene_tx=XM_023240937.1;refGene_gn=TCF7L2;refGene_tx=XM_023240938.1;refGene_gn=TCF7L2;refGene_tx=XM_023240939.1;refGene_gn=TCF7L2;refGene_tx=XM_023240940.1;refGene_gn=TCF7L2;refGene_tx=XM_023240941.1;refGene_gn=TCF7L2;refGene_tx=XM_023240942.1;refGene_gn=TCF7L2;refGene_tx=XM_023240943.1;refGene_gn=TCF7L2;refGene_tx=XM_023240944.1;refGene_gn=TCF7L2;refGene_tx=XM_023240945.1;refGene_gn=TCF7L2;refGene_tx=XM_023240946.1;refGene_gn=TCF7L2;refGene_tx=XM_023240947.1;refGene_gn=TCF7L2;refGene_tx=XM_023240948.1;refGene_gn=TCF7L2;refGene_tx=XM_023240949.1;refGene_gn=TCF7L2;refGene_tx=XM_023240950.1;refGene_gn=TCF7L2;refGene_tx=XM_023240951.1;refGene_gn=TCF7L2;refGene_tx=XM_023240952.1;refGene_gn=TCF7L2;refGene_tx=XM_023240953.1
D2 72310077 72310126
refGene_gn=TCF7L2;refGene_tx=XM_023240925.1;refGene_gn=TCF7L2;refGene_tx=XM_023240927.1;refGene_gn=TCF7L2;refGene_tx=XM_023240928.1;refGene_gn=TCF7L2;refGene_tx=XM_023240929.1;refGene_gn=TCF7L2;refGene_tx=XM_023240930.1;refGene_gn=TCF7L2;refGene_tx=XM_023240931.1;refGene_gn=TCF7L2;refGene_tx=XM_023240932.1;refGene_gn=TCF7L2;refGene_tx=XM_023240933.1;refGene_gn=TCF7L2;refGene_tx=XM_023240934.1;refGene_gn=TCF7L2;refGene_tx=XM_023240935.1;refGene_gn=TCF7L2;refGene_tx=XM_023240936.1;refGene_gn=TCF7L2;refGene_tx=XM_023240937.1;refGene_gn=TCF7L2;refGene_tx=XM_023240938.1;refGene_gn=TCF7L2;refGene_tx=XM_023240939.1;refGene_gn=TCF7L2;refGene_tx=XM_023240940.1;refGene_gn=TCF7L2;refGene_tx=XM_023240941.1;refGene_gn=TCF7L2;refGene_tx=XM_023240942.1;refGene_gn=TCF7L2;refG

ene_tx=XM_023240943.1;refGene_gn=TCF7L2;refGene_tx=XM_023240944.1;refGene_gn=TCF7L2;refGene_tx=XM_023240945.1;refGene_gn=TCF7L2;refGene_tx=XM_023240946.1;refGene_gn=TCF7L2;refGene_tx=XM_023240947.1;refGene_gn=TCF7L2;refGene_tx=XM_023240948.1;refGene_gn=TCF7L2;refGene_tx=XM_023240949.1;refGene_gn=TCF7L2;refGene_tx=XM_023240950.1;refGene_gn=TCF7L2;refGene_tx=XM_023240951.1;refGene_gn=TCF7L2;refGene_tx=XM_023240952.1;refGene_gn=TCF7L2;refGene_tx=XM_023240953.1
D2 72310715 72310766
refGene_gn=TCF7L2;refGene_tx=XM_023240925.1;refGene_gn=TCF7L2;refGene_tx=XM_023240927.1;refGene_gn=TCF7L2;refGene_tx=XM_023240928.1;refGene_gn=TCF7L2;refGene_tx=XM_023240929.1;refGene_gn=TCF7L2;refGene_tx=XM_023240930.1;refGene_gn=TCF7L2;refGene_tx=XM_023240933.1;refGene_gn=TCF7L2;refGene_tx=XM_023240934.1;refGene_gn=TCF7L2;refGene_tx=XM_023240935.1;refGene_gn=TCF7L2;refGene_tx=XM_023240938.1;refGene_gn=TCF7L2;refGene_tx=XM_023240939.1;refGene_gn=TCF7L2;refGene_tx=XM_023240940.1;refGene_gn=TCF7L2;refGene_tx=XM_023240944.1;refGene_gn=TCF7L2;refGene_tx=XM_023240945.1;refGene_gn=TCF7L2;refGene_tx=XM_023240946.1;refGene_gn=TCF7L2;refGene_tx=XM_023240948.1;refGene_gn=TCF7L2;refGene_tx=XM_023240949.1;refGene_gn=TCF7L2;refGene_tx=XM_023240951.1;refGene_gn=TCF7L2;refGene_tx=XM_023240952.1
D2 72312090 72312163
refGene_gn=TCF7L2;refGene_tx=XM_023240925.1;refGene_gn=TCF7L2;refGene_tx=XM_023240927.1;refGene_gn=TCF7L2;refGene_tx=XM_023240928.1;refGene_gn=TCF7L2;refGene_tx=XM_023240929.1;refGene_gn=TCF7L2;refGene_tx=XM_023240930.1;refGene_gn=TCF7L2;refGene_tx=XM_023240931.1;refGene_gn=TCF7L2;refGene_tx=XM_023240933.1;refGene_gn=TCF7L2;refGene_tx=XM_023240934.1;refGene_gn=TCF7L2;refGene_tx=XM_023240935.1;refGene_gn=TCF7L2;refGene_tx=XM_023240938.1;refGene_gn=TCF7L2;refGene_tx=XM_023240939.1;refGene_gn=TCF7L2;refGene_tx=XM_023240940.1;refGene_gn=TCF7L2;refGene_tx=XM_023240942.1;refGene_gn=TCF7L2;refGene_tx=XM_023240944.1;refGene_gn=TCF7L2;refGene_tx=XM_023240945.1;refGene_gn=TCF7L2;refGene_tx=XM_023240946.1;refGene_gn=TCF7L2;refGene_tx=XM_023240948.1;refGene_gn=TCF7L2;refGene_tx=XM_023240949.1
D2 72312809 72312882
refGene_gn=TCF7L2;refGene_tx=XM_023240932.1;refGene_gn=TCF7L2;refGene_tx=XM_023240936.1;refGene_gn=TCF7L2;refGene_tx=XM_023240937.1;refGene_gn=TCF7L2;refGene_tx=XM_023240941.1;refGene_gn=TCF7L2;refGene_tx=XM_023240943.1;refGene_gn=TCF7L2;refGene_tx=XM_023240945.1;refGene_gn=TCF7L2;refGene_tx=XM_023240946.1;refGene_gn=TCF7L2;refGene_tx=XM_023240949.1
D2 72317798 72318234
refGene_gn=TCF7L2;refGene_tx=XM_023240925.1;refGene_gn=TCF7L2;refGene_tx=XM_023240927.1;refGene_gn=TCF7L2;refGene_tx=XM_023240928.1;refGene_gn=TCF7L2;refGene_tx=XM_023240929.1;refGene_gn=TCF7L2;refGene_tx=XM_023240930.1;refGene_gn=TCF7L2;refGene_tx=XM_023240931.1;refGene_gn=TCF7L2;refGene_tx=XM_023240932.1;refGene_gn=TCF7L2;refGene_tx=XM_023240933.1;refGene_gn=TCF7L2;refGene_tx=XM_023240934.1;refGene_gn=TCF7L2;refGene_tx=XM_023240935.1;refGene_gn=TCF7L2;refGene_tx=XM_023240936.1;refGene_gn=TCF7L2;refGene_tx=XM_023240937.1;refGene_gn=TCF7L2;refGene_tx=XM_023240938.1;refGene_gn=TCF7L2;refGene_tx=XM_023240939.1;refGene_gn=TCF

7L2;refGene_tx=XM_023240940.1;refGene_gn=TCF7L2;refGene_tx=XM_02324094
1.1;refGene_gn=TCF7L2;refGene_tx=XM_023240942.1;refGene_gn=TCF7L2;refG
ene_tx=XM_023240943.1;refGene_gn=TCF7L2;refGene_tx=XM_023240944.1;refG
ene_gn=TCF7L2;refGene_tx=XM_023240947.1;refGene_gn=TCF7L2;refGene_tx=X
M_023240948.1;refGene_gn=TCF7L2;refGene_tx=XM_023240950.1;refGene_gn=T
CF7L2;refGene_tx=XM_023240951.1;refGene_gn=TCF7L2;refGene_tx=XM_023240
952.1;refGene_gn=TCF7L2;refGene_tx=XM_023240953.1
D2 72659004 72659073 refGene_gn=HABP2;refGene_tx=XM_006938184.4
D2 72671359 72671396 refGene_gn=HABP2;refGene_tx=XM_006938184.4
D2 72678526 72678637 refGene_gn=HABP2;refGene_tx=XM_006938184.4
D2 72680066 72680174 refGene_gn=HABP2;refGene_tx=XM_006938184.4
D2 72681379 72681496 refGene_gn=HABP2;refGene_tx=XM_006938184.4
D2 72682705 72682825 refGene_gn=HABP2;refGene_tx=XM_006938184.4
D2 72683302 72683474 refGene_gn=HABP2;refGene_tx=XM_006938184.4
D2 72684656 72684754 refGene_gn=HABP2;refGene_tx=XM_006938184.4
D2 72685867 72686117 refGene_gn=HABP2;refGene_tx=XM_006938184.4
D2 72687151 72687294 refGene_gn=HABP2;refGene_tx=XM_006938184.4
D2 72688048 72688183 refGene_gn=HABP2;refGene_tx=XM_006938184.4
D2 72689810 72689956 refGene_gn=HABP2;refGene_tx=XM_006938184.4
D2 72691563 72691728 refGene_gn=HABP2;refGene_tx=XM_006938184.4
D2 72692299 72692404
refGene_gn=NRAP;refGene_tx=XM_006938185.3;refGene_gn=NRAP;refGene_tx=X
M_006938188.3;refGene_gn=NRAP;refGene_tx=XM_006938191.3;refGene_gn=NRA
P;refGene_tx=XM_011287564.2;refGene_gn=NRAP;refGene_tx=XM_011287565.2;
refGene_gn=NRAP;refGene_tx=XM_011287566.2;refGene_gn=NRAP;refGene_tx=X
M_011287567.2;refGene_gn=NRAP;refGene_tx=XM_011287568.2;refGene_gn=NRA
P;refGene_tx=XM_019813827.1;refGene_gn=NRAP;refGene_tx=XM_019813828.1;
refGene_gn=NRAP;refGene_tx=XM_023240955.1
D2 72692981 72693113
refGene_gn=NRAP;refGene_tx=XM_006938185.3;refGene_gn=NRAP;refGene_tx=X
M_006938188.3;refGene_gn=NRAP;refGene_tx=XM_006938191.3;refGene_gn=NRA
P;refGene_tx=XM_011287564.2;refGene_gn=NRAP;refGene_tx=XM_011287565.2;
refGene_gn=NRAP;refGene_tx=XM_011287566.2;refGene_gn=NRAP;refGene_tx=X
M_011287567.2;refGene_gn=NRAP;refGene_tx=XM_011287568.2;refGene_gn=NRA
P;refGene_tx=XM_019813827.1;refGene_gn=NRAP;refGene_tx=XM_019813828.1;
refGene_gn=NRAP;refGene_tx=XM_023240955.1
D2 72693261 72693573
refGene_gn=NRAP;refGene_tx=XM_006938185.3;refGene_gn=NRAP;refGene_tx=X
M_006938188.3;refGene_gn=NRAP;refGene_tx=XM_006938191.3;refGene_gn=NRA
P;refGene_tx=XM_011287564.2;refGene_gn=NRAP;refGene_tx=XM_011287565.2;
refGene_gn=NRAP;refGene_tx=XM_011287566.2;refGene_gn=NRAP;refGene_tx=X
M_011287567.2;refGene_gn=NRAP;refGene_tx=XM_011287568.2;refGene_gn=NRA
P;refGene_tx=XM_019813827.1;refGene_gn=NRAP;refGene_tx=XM_019813828.1;
refGene_gn=NRAP;refGene_tx=XM_023240955.1
D2 72695494 72695602
refGene_gn=NRAP;refGene_tx=XM_006938185.3;refGene_gn=NRAP;refGene_tx=X
M_006938188.3;refGene_gn=NRAP;refGene_tx=XM_006938191.3;refGene_gn=NRA
P;refGene_tx=XM_011287564.2;refGene_gn=NRAP;refGene_tx=XM_011287565.2;
refGene_gn=NRAP;refGene_tx=XM_011287566.2;refGene_gn=NRAP;refGene_tx=X
M_011287567.2;refGene_gn=NRAP;refGene_tx=XM_011287568.2;refGene_gn=NRA

P;refGene_tx=XM_019813827.1;refGene_gn=NRAP;refGene_tx=XM_019813828.1;
refGene_gn=NRAP;refGene_tx=XM_023240955.1
D2 72698918 72699023
refGene_gn=NRAP;refGene_tx=XM_006938185.3;refGene_gn=NRAP;refGene_tx=X
M_006938188.3;refGene_gn=NRAP;refGene_tx=XM_006938191.3;refGene_gn=NRA
P;refGene_tx=XM_011287564.2;refGene_gn=NRAP;refGene_tx=XM_011287565.2;
refGene_gn=NRAP;refGene_tx=XM_011287566.2;refGene_gn=NRAP;refGene_tx=X
M_011287567.2;refGene_gn=NRAP;refGene_tx=XM_011287568.2;refGene_gn=NRA
P;refGene_tx=XM_019813827.1;refGene_gn=NRAP;refGene_tx=XM_019813828.1;
refGene_gn=NRAP;refGene_tx=XM_023240955.1
D2 72700345 72700444
refGene_gn=NRAP;refGene_tx=XM_006938185.3;refGene_gn=NRAP;refGene_tx=X
M_006938188.3;refGene_gn=NRAP;refGene_tx=XM_006938191.3;refGene_gn=NRA
P;refGene_tx=XM_011287564.2;refGene_gn=NRAP;refGene_tx=XM_011287565.2;
refGene_gn=NRAP;refGene_tx=XM_011287566.2;refGene_gn=NRAP;refGene_tx=X
M_011287567.2;refGene_gn=NRAP;refGene_tx=XM_011287568.2;refGene_gn=NRA
P;refGene_tx=XM_019813827.1;refGene_gn=NRAP;refGene_tx=XM_019813828.1;
refGene_gn=NRAP;refGene_tx=XM_023240955.1
D2 72701279 72701384
refGene_gn=NRAP;refGene_tx=XM_006938185.3;refGene_gn=NRAP;refGene_tx=X
M_006938188.3;refGene_gn=NRAP;refGene_tx=XM_006938191.3;refGene_gn=NRA
P;refGene_tx=XM_011287564.2;refGene_gn=NRAP;refGene_tx=XM_011287565.2;
refGene_gn=NRAP;refGene_tx=XM_011287566.2;refGene_gn=NRAP;refGene_tx=X
M_011287567.2;refGene_gn=NRAP;refGene_tx=XM_011287568.2;refGene_gn=NRA
P;refGene_tx=XM_019813827.1;refGene_gn=NRAP;refGene_tx=XM_019813828.1;
refGene_gn=NRAP;refGene_tx=XM_023240955.1
D2 72707068 72707380
refGene_gn=NRAP;refGene_tx=XM_006938185.3;refGene_gn=NRAP;refGene_tx=X
M_006938188.3;refGene_gn=NRAP;refGene_tx=XM_006938191.3;refGene_gn=NRA
P;refGene_tx=XM_011287564.2;refGene_gn=NRAP;refGene_tx=XM_011287565.2;
refGene_gn=NRAP;refGene_tx=XM_011287566.2;refGene_gn=NRAP;refGene_tx=X
M_011287567.2;refGene_gn=NRAP;refGene_tx=XM_019813827.1;refGene_gn=NRA
P;refGene_tx=XM_019813828.1;refGene_gn=NRAP;refGene_tx=XM_023240955.1
D2 72708248 72708356
refGene_gn=NRAP;refGene_tx=XM_006938185.3;refGene_gn=NRAP;refGene_tx=X
M_006938188.3;refGene_gn=NRAP;refGene_tx=XM_006938191.3;refGene_gn=NRA
P;refGene_tx=XM_011287564.2;refGene_gn=NRAP;refGene_tx=XM_011287565.2;
refGene_gn=NRAP;refGene_tx=XM_011287566.2;refGene_gn=NRAP;refGene_tx=X
M_011287567.2;refGene_gn=NRAP;refGene_tx=XM_011287568.2;refGene_gn=NRA
P;refGene_tx=XM_019813827.1;refGene_gn=NRAP;refGene_tx=XM_019813828.1;
refGene_gn=NRAP;refGene_tx=XM_023240955.1
D2 72708709 72708814
refGene_gn=NRAP;refGene_tx=XM_006938185.3;refGene_gn=NRAP;refGene_tx=X
M_006938188.3;refGene_gn=NRAP;refGene_tx=XM_006938191.3;refGene_gn=NRA
P;refGene_tx=XM_011287564.2;refGene_gn=NRAP;refGene_tx=XM_011287565.2;
refGene_gn=NRAP;refGene_tx=XM_011287566.2;refGene_gn=NRAP;refGene_tx=X
M_011287567.2;refGene_gn=NRAP;refGene_tx=XM_011287568.2;refGene_gn=NRA
P;refGene_tx=XM_019813827.1;refGene_gn=NRAP;refGene_tx=XM_019813828.1;
refGene_gn=NRAP;refGene_tx=XM_023240955.1
D2 72710775 72710874

refGene_gn=NRAP;refGene_tx=XM_006938185.3;refGene_gn=NRAP;refGene_tx=X
M_006938188.3;refGene_gn=NRAP;refGene_tx=XM_006938191.3;refGene_gn=NRA
P;refGene_tx=XM_011287564.2;refGene_gn=NRAP;refGene_tx=XM_011287565.2;
refGene_gn=NRAP;refGene_tx=XM_011287566.2;refGene_gn=NRAP;refGene_tx=X
M_011287567.2;refGene_gn=NRAP;refGene_tx=XM_011287568.2;refGene_gn=NRA
P;refGene_tx=XM_019813827.1;refGene_gn=NRAP;refGene_tx=XM_019813828.1;
refGene_gn=NRAP;refGene_tx=XM_023240955.1
D2 72712144 72712249
refGene_gn=NRAP;refGene_tx=XM_006938185.3;refGene_gn=NRAP;refGene_tx=X
M_006938188.3;refGene_gn=NRAP;refGene_tx=XM_006938191.3;refGene_gn=NRA
P;refGene_tx=XM_011287564.2;refGene_gn=NRAP;refGene_tx=XM_011287565.2;
refGene_gn=NRAP;refGene_tx=XM_011287566.2;refGene_gn=NRAP;refGene_tx=X
M_011287567.2;refGene_gn=NRAP;refGene_tx=XM_011287568.2;refGene_gn=NRA
P;refGene_tx=XM_019813827.1;refGene_gn=NRAP;refGene_tx=XM_019813828.1;
refGene_gn=NRAP;refGene_tx=XM_023240955.1
D2 72713914 72714112
refGene_gn=NRAP;refGene_tx=XM_006938185.3;refGene_gn=NRAP;refGene_tx=X
M_006938188.3;refGene_gn=NRAP;refGene_tx=XM_006938191.3;refGene_gn=NRA
P;refGene_tx=XM_011287564.2;refGene_gn=NRAP;refGene_tx=XM_011287565.2;
refGene_gn=NRAP;refGene_tx=XM_011287566.2;refGene_gn=NRAP;refGene_tx=X
M_011287567.2;refGene_gn=NRAP;refGene_tx=XM_011287568.2;refGene_gn=NRA
P;refGene_tx=XM_019813827.1;refGene_gn=NRAP;refGene_tx=XM_019813828.1;
refGene_gn=NRAP;refGene_tx=XM_023240955.1
D2 72716060 72716174
refGene_gn=NRAP;refGene_tx=XM_006938185.3;refGene_gn=NRAP;refGene_tx=X
M_006938188.3;refGene_gn=NRAP;refGene_tx=XM_006938191.3;refGene_gn=NRA
P;refGene_tx=XM_011287564.2;refGene_gn=NRAP;refGene_tx=XM_011287565.2;
refGene_gn=NRAP;refGene_tx=XM_011287566.2;refGene_gn=NRAP;refGene_tx=X
M_011287567.2;refGene_gn=NRAP;refGene_tx=XM_011287568.2;refGene_gn=NRA
P;refGene_tx=XM_019813827.1;refGene_gn=NRAP;refGene_tx=XM_019813828.1;
refGene_gn=NRAP;refGene_tx=XM_023240955.1
D2 72716717 72716825
refGene_gn=NRAP;refGene_tx=XM_006938185.3;refGene_gn=NRAP;refGene_tx=X
M_006938188.3;refGene_gn=NRAP;refGene_tx=XM_006938191.3;refGene_gn=NRA
P;refGene_tx=XM_011287564.2;refGene_gn=NRAP;refGene_tx=XM_011287565.2;
refGene_gn=NRAP;refGene_tx=XM_011287566.2;refGene_gn=NRAP;refGene_tx=X
M_011287567.2;refGene_gn=NRAP;refGene_tx=XM_011287568.2;refGene_gn=NRA
P;refGene_tx=XM_019813827.1;refGene_gn=NRAP;refGene_tx=XM_019813828.1;
refGene_gn=NRAP;refGene_tx=XM_023240955.1
D2 72717559 72717664
refGene_gn=NRAP;refGene_tx=XM_006938185.3;refGene_gn=NRAP;refGene_tx=X
M_006938188.3;refGene_gn=NRAP;refGene_tx=XM_006938191.3;refGene_gn=NRA
P;refGene_tx=XM_011287564.2;refGene_gn=NRAP;refGene_tx=XM_011287565.2;
refGene_gn=NRAP;refGene_tx=XM_011287566.2;refGene_gn=NRAP;refGene_tx=X
M_011287567.2;refGene_gn=NRAP;refGene_tx=XM_011287568.2;refGene_gn=NRA
P;refGene_tx=XM_019813827.1;refGene_gn=NRAP;refGene_tx=XM_019813828.1;
refGene_gn=NRAP;refGene_tx=XM_023240955.1
D2 72719275 72719374
refGene_gn=NRAP;refGene_tx=XM_006938185.3;refGene_gn=NRAP;refGene_tx=X
M_006938188.3;refGene_gn=NRAP;refGene_tx=XM_006938191.3;refGene_gn=NRA

P;refGene_tx=XM_011287564.2;refGene_gn=NRAP;refGene_tx=XM_011287565.2;
refGene_gn=NRAP;refGene_tx=XM_011287566.2;refGene_gn=NRAP;refGene_tx=X
M_011287567.2;refGene_gn=NRAP;refGene_tx=XM_011287568.2;refGene_gn=NRA
P;refGene_tx=XM_019813827.1;refGene_gn=NRAP;refGene_tx=XM_019813828.1;
refGene_gn=NRAP;refGene_tx=XM_023240955.1
D2 72722187 72722292
refGene_gn=NRAP;refGene_tx=XM_006938185.3;refGene_gn=NRAP;refGene_tx=X
M_006938188.3;refGene_gn=NRAP;refGene_tx=XM_006938191.3;refGene_gn=NRA
P;refGene_tx=XM_011287564.2;refGene_gn=NRAP;refGene_tx=XM_011287565.2;
refGene_gn=NRAP;refGene_tx=XM_011287566.2;refGene_gn=NRAP;refGene_tx=X
M_011287567.2;refGene_gn=NRAP;refGene_tx=XM_011287568.2;refGene_gn=NRA
P;refGene_tx=XM_019813827.1;refGene_gn=NRAP;refGene_tx=XM_019813828.1;
refGene_gn=NRAP;refGene_tx=XM_023240955.1
D2 72723385 72723697
refGene_gn=NRAP;refGene_tx=XM_006938185.3;refGene_gn=NRAP;refGene_tx=X
M_006938188.3;refGene_gn=NRAP;refGene_tx=XM_006938191.3;refGene_gn=NRA
P;refGene_tx=XM_011287564.2;refGene_gn=NRAP;refGene_tx=XM_011287565.2;
refGene_gn=NRAP;refGene_tx=XM_011287566.2;refGene_gn=NRAP;refGene_tx=X
M_011287567.2;refGene_gn=NRAP;refGene_tx=XM_011287568.2;refGene_gn=NRA
P;refGene_tx=XM_019813827.1;refGene_gn=NRAP;refGene_tx=XM_019813828.1;
refGene_gn=NRAP;refGene_tx=XM_023240955.1
D2 72725055 72725163
refGene_gn=NRAP;refGene_tx=XM_006938185.3;refGene_gn=NRAP;refGene_tx=X
M_006938188.3;refGene_gn=NRAP;refGene_tx=XM_006938191.3;refGene_gn=NRA
P;refGene_tx=XM_011287564.2;refGene_gn=NRAP;refGene_tx=XM_011287565.2;
refGene_gn=NRAP;refGene_tx=XM_011287566.2;refGene_gn=NRAP;refGene_tx=X
M_011287567.2;refGene_gn=NRAP;refGene_tx=XM_011287568.2;refGene_gn=NRA
P;refGene_tx=XM_023240955.1
D2 72726311 72726416
refGene_gn=NRAP;refGene_tx=XM_006938185.3;refGene_gn=NRAP;refGene_tx=X
M_006938188.3;refGene_gn=NRAP;refGene_tx=XM_006938191.3;refGene_gn=NRA
P;refGene_tx=XM_011287564.2;refGene_gn=NRAP;refGene_tx=XM_011287565.2;
refGene_gn=NRAP;refGene_tx=XM_011287566.2;refGene_gn=NRAP;refGene_tx=X
M_011287567.2;refGene_gn=NRAP;refGene_tx=XM_011287568.2;refGene_gn=NRA
P;refGene_tx=XM_019813827.1;refGene_gn=NRAP;refGene_tx=XM_019813828.1;
refGene_gn=NRAP;refGene_tx=XM_023240955.1
D2 72727543 72727642
refGene_gn=NRAP;refGene_tx=XM_006938185.3;refGene_gn=NRAP;refGene_tx=X
M_006938188.3;refGene_gn=NRAP;refGene_tx=XM_006938191.3;refGene_gn=NRA
P;refGene_tx=XM_011287564.2;refGene_gn=NRAP;refGene_tx=XM_011287565.2;
refGene_gn=NRAP;refGene_tx=XM_011287566.2;refGene_gn=NRAP;refGene_tx=X
M_011287567.2;refGene_gn=NRAP;refGene_tx=XM_011287568.2;refGene_gn=NRA
P;refGene_tx=XM_019813827.1;refGene_gn=NRAP;refGene_tx=XM_019813828.1;
refGene_gn=NRAP;refGene_tx=XM_023240955.1
D2 72729943 72730048
refGene_gn=NRAP;refGene_tx=XM_006938185.3;refGene_gn=NRAP;refGene_tx=X
M_006938188.3;refGene_gn=NRAP;refGene_tx=XM_006938191.3;refGene_gn=NRA
P;refGene_tx=XM_011287564.2;refGene_gn=NRAP;refGene_tx=XM_011287565.2;
refGene_gn=NRAP;refGene_tx=XM_011287566.2;refGene_gn=NRAP;refGene_tx=X
M_011287567.2;refGene_gn=NRAP;refGene_tx=XM_011287568.2;refGene_gn=NRA

P;refGene_tx=XM_019813827.1;refGene_gn=NRAP;refGene_tx=XM_019813828.1;
refGene_gn=NRAP;refGene_tx=XM_023240955.1
D2 72730615 72730813
refGene_gn=NRAP;refGene_tx=XM_006938185.3;refGene_gn=NRAP;refGene_tx=X
M_006938188.3;refGene_gn=NRAP;refGene_tx=XM_006938191.3;refGene_gn=NRA
P;refGene_tx=XM_011287564.2;refGene_gn=NRAP;refGene_tx=XM_011287565.2;
refGene_gn=NRAP;refGene_tx=XM_011287566.2;refGene_gn=NRAP;refGene_tx=X
M_011287567.2;refGene_gn=NRAP;refGene_tx=XM_011287568.2;refGene_gn=NRA
P;refGene_tx=XM_019813827.1;refGene_gn=NRAP;refGene_tx=XM_019813828.1
D2 72732610 72732712
refGene_gn=NRAP;refGene_tx=XM_006938185.3;refGene_gn=NRAP;refGene_tx=X
M_006938188.3;refGene_gn=NRAP;refGene_tx=XM_006938191.3;refGene_gn=NRA
P;refGene_tx=XM_011287564.2;refGene_gn=NRAP;refGene_tx=XM_011287565.2;
refGene_gn=NRAP;refGene_tx=XM_011287566.2;refGene_gn=NRAP;refGene_tx=X
M_011287567.2;refGene_gn=NRAP;refGene_tx=XM_011287568.2;refGene_gn=NRA
P;refGene_tx=XM_019813827.1;refGene_gn=NRAP;refGene_tx=XM_019813828.1;
refGene_gn=NRAP;refGene_tx=XM_023240955.1
D2 72732958 72733066
refGene_gn=NRAP;refGene_tx=XM_006938185.3;refGene_gn=NRAP;refGene_tx=X
M_006938188.3;refGene_gn=NRAP;refGene_tx=XM_006938191.3;refGene_gn=NRA
P;refGene_tx=XM_011287564.2;refGene_gn=NRAP;refGene_tx=XM_011287565.2;
refGene_gn=NRAP;refGene_tx=XM_011287566.2;refGene_gn=NRAP;refGene_tx=X
M_011287567.2;refGene_gn=NRAP;refGene_tx=XM_011287568.2;refGene_gn=NRA
P;refGene_tx=XM_019813827.1;refGene_gn=NRAP;refGene_tx=XM_019813828.1;
refGene_gn=NRAP;refGene_tx=XM_023240955.1
D2 72734215 72734320
refGene_gn=NRAP;refGene_tx=XM_006938185.3;refGene_gn=NRAP;refGene_tx=X
M_006938188.3;refGene_gn=NRAP;refGene_tx=XM_006938191.3;refGene_gn=NRA
P;refGene_tx=XM_011287564.2;refGene_gn=NRAP;refGene_tx=XM_011287565.2;
refGene_gn=NRAP;refGene_tx=XM_011287566.2;refGene_gn=NRAP;refGene_tx=X
M_011287567.2;refGene_gn=NRAP;refGene_tx=XM_011287568.2;refGene_gn=NRA
P;refGene_tx=XM_019813827.1;refGene_gn=NRAP;refGene_tx=XM_019813828.1;
refGene_gn=NRAP;refGene_tx=XM_023240955.1
D2 72735600 72735699
refGene_gn=NRAP;refGene_tx=XM_006938185.3;refGene_gn=NRAP;refGene_tx=X
M_006938188.3;refGene_gn=NRAP;refGene_tx=XM_006938191.3;refGene_gn=NRA
P;refGene_tx=XM_011287564.2;refGene_gn=NRAP;refGene_tx=XM_011287565.2;
refGene_gn=NRAP;refGene_tx=XM_011287566.2;refGene_gn=NRAP;refGene_tx=X
M_011287567.2;refGene_gn=NRAP;refGene_tx=XM_011287568.2;refGene_gn=NRA
P;refGene_tx=XM_019813827.1;refGene_gn=NRAP;refGene_tx=XM_019813828.1;
refGene_gn=NRAP;refGene_tx=XM_023240955.1
D2 72740762 72740867
refGene_gn=NRAP;refGene_tx=XM_006938185.3;refGene_gn=NRAP;refGene_tx=X
M_006938188.3;refGene_gn=NRAP;refGene_tx=XM_006938191.3;refGene_gn=NRA
P;refGene_tx=XM_011287564.2;refGene_gn=NRAP;refGene_tx=XM_011287565.2;
refGene_gn=NRAP;refGene_tx=XM_011287566.2;refGene_gn=NRAP;refGene_tx=X
M_011287567.2;refGene_gn=NRAP;refGene_tx=XM_011287568.2;refGene_gn=NRA
P;refGene_tx=XM_019813827.1;refGene_gn=NRAP;refGene_tx=XM_019813828.1;
refGene_gn=NRAP;refGene_tx=XM_023240955.1
D2 72741840 72741948

refGene_gn=NRAP;refGene_tx=XM_006938185.3;refGene_gn=NRAP;refGene_tx=X
M_006938188.3;refGene_gn=NRAP;refGene_tx=XM_006938191.3;refGene_gn=NRA
P;refGene_tx=XM_011287564.2;refGene_gn=NRAP;refGene_tx=XM_011287565.2;
refGene_gn=NRAP;refGene_tx=XM_011287566.2;refGene_gn=NRAP;refGene_tx=X
M_011287567.2;refGene_gn=NRAP;refGene_tx=XM_011287568.2;refGene_gn=NRA
P;refGene_tx=XM_019813827.1;refGene_gn=NRAP;refGene_tx=XM_019813828.1;
refGene_gn=NRAP;refGene_tx=XM_023240955.1
D2 72743642 72743747
refGene_gn=NRAP;refGene_tx=XM_006938185.3;refGene_gn=NRAP;refGene_tx=X
M_006938188.3;refGene_gn=NRAP;refGene_tx=XM_011287564.2;refGene_gn=NRA
P;refGene_tx=XM_011287565.2;refGene_gn=NRAP;refGene_tx=XM_011287566.2;
refGene_gn=NRAP;refGene_tx=XM_011287568.2;refGene_gn=NRAP;refGene_tx=X
M_019813827.1;refGene_gn=NRAP;refGene_tx=XM_019813828.1;refGene_gn=NRA
P;refGene_tx=XM_023240955.1
D2 72746204 72746321
refGene_gn=NRAP;refGene_tx=XM_006938185.3;refGene_gn=NRAP;refGene_tx=X
M_006938188.3;refGene_gn=NRAP;refGene_tx=XM_006938191.3;refGene_gn=NRA
P;refGene_tx=XM_011287564.2;refGene_gn=NRAP;refGene_tx=XM_011287565.2;
refGene_gn=NRAP;refGene_tx=XM_011287566.2;refGene_gn=NRAP;refGene_tx=X
M_011287567.2;refGene_gn=NRAP;refGene_tx=XM_011287568.2;refGene_gn=NRA
P;refGene_tx=XM_019813827.1;refGene_gn=NRAP;refGene_tx=XM_019813828.1;
refGene_gn=NRAP;refGene_tx=XM_023240955.1
D2 72747281 72747389
refGene_gn=NRAP;refGene_tx=XM_006938185.3;refGene_gn=NRAP;refGene_tx=X
M_006938188.3;refGene_gn=NRAP;refGene_tx=XM_006938191.3;refGene_gn=NRA
P;refGene_tx=XM_011287564.2;refGene_gn=NRAP;refGene_tx=XM_011287565.2;
refGene_gn=NRAP;refGene_tx=XM_011287566.2;refGene_gn=NRAP;refGene_tx=X
M_011287567.2;refGene_gn=NRAP;refGene_tx=XM_011287568.2;refGene_gn=NRA
P;refGene_tx=XM_019813827.1;refGene_gn=NRAP;refGene_tx=XM_019813828.1;
refGene_gn=NRAP;refGene_tx=XM_023240955.1
D2 72750751 72750856
refGene_gn=NRAP;refGene_tx=XM_006938185.3;refGene_gn=NRAP;refGene_tx=X
M_006938188.3;refGene_gn=NRAP;refGene_tx=XM_006938191.3;refGene_gn=NRA
P;refGene_tx=XM_011287564.2;refGene_gn=NRAP;refGene_tx=XM_011287565.2;
refGene_gn=NRAP;refGene_tx=XM_011287566.2;refGene_gn=NRAP;refGene_tx=X
M_011287567.2;refGene_gn=NRAP;refGene_tx=XM_011287568.2;refGene_gn=NRA
P;refGene_tx=XM_019813827.1;refGene_gn=NRAP;refGene_tx=XM_019813828.1;
refGene_gn=NRAP;refGene_tx=XM_023240955.1
D2 72751166 72751274
refGene_gn=NRAP;refGene_tx=XM_006938185.3;refGene_gn=NRAP;refGene_tx=X
M_006938188.3;refGene_gn=NRAP;refGene_tx=XM_006938191.3;refGene_gn=NRA
P;refGene_tx=XM_011287564.2;refGene_gn=NRAP;refGene_tx=XM_011287565.2;
refGene_gn=NRAP;refGene_tx=XM_011287566.2;refGene_gn=NRAP;refGene_tx=X
M_011287567.2;refGene_gn=NRAP;refGene_tx=XM_011287568.2;refGene_gn=NRA
P;refGene_tx=XM_019813827.1;refGene_gn=NRAP;refGene_tx=XM_019813828.1;
refGene_gn=NRAP;refGene_tx=XM_023240955.1
D2 72752034 72752139
refGene_gn=NRAP;refGene_tx=XM_006938185.3;refGene_gn=NRAP;refGene_tx=X
M_006938188.3;refGene_gn=NRAP;refGene_tx=XM_006938191.3;refGene_gn=NRA
P;refGene_tx=XM_011287564.2;refGene_gn=NRAP;refGene_tx=XM_011287565.2;

refGene_gn=NRAP;refGene_tx=XM_011287566.2;refGene_gn=NRAP;refGene_tx=XM_011287567.2;refGene_gn=NRAP;refGene_tx=XM_011287568.2;refGene_gn=NRAP;refGene_tx=XM_019813827.1;refGene_gn=NRAP;refGene_tx=XM_019813828.1;refGene_gn=NRAP;refGene_tx=XM_023240955.1
D2 72752854 72752962
refGene_gn=NRAP;refGene_tx=XM_006938185.3;refGene_gn=NRAP;refGene_tx=XM_006938188.3;refGene_gn=NRAP;refGene_tx=XM_006938191.3;refGene_gn=NRAP;refGene_tx=XM_011287564.2;refGene_gn=NRAP;refGene_tx=XM_011287565.2;refGene_gn=NRAP;refGene_tx=XM_011287566.2;refGene_gn=NRAP;refGene_tx=XM_011287567.2;refGene_gn=NRAP;refGene_tx=XM_011287568.2;refGene_gn=NRAP;refGene_tx=XM_019813827.1;refGene_gn=NRAP;refGene_tx=XM_019813828.1;refGene_gn=NRAP;refGene_tx=XM_023240955.1
D2 72754059 72754164
refGene_gn=NRAP;refGene_tx=XM_006938185.3;refGene_gn=NRAP;refGene_tx=XM_006938188.3;refGene_gn=NRAP;refGene_tx=XM_006938191.3;refGene_gn=NRAP;refGene_tx=XM_011287564.2;refGene_gn=NRAP;refGene_tx=XM_011287565.2;refGene_gn=NRAP;refGene_tx=XM_011287566.2;refGene_gn=NRAP;refGene_tx=XM_011287567.2;refGene_gn=NRAP;refGene_tx=XM_011287568.2;refGene_gn=NRAP;refGene_tx=XM_019813827.1;refGene_gn=NRAP;refGene_tx=XM_019813828.1;refGene_gn=NRAP;refGene_tx=XM_023240955.1
D2 72756330 72756447
refGene_gn=NRAP;refGene_tx=XM_006938185.3;refGene_gn=NRAP;refGene_tx=XM_006938188.3;refGene_gn=NRAP;refGene_tx=XM_006938191.3;refGene_gn=NRAP;refGene_tx=XM_011287564.2;refGene_gn=NRAP;refGene_tx=XM_011287565.2;refGene_gn=NRAP;refGene_tx=XM_011287566.2;refGene_gn=NRAP;refGene_tx=XM_011287567.2;refGene_gn=NRAP;refGene_tx=XM_011287568.2;refGene_gn=NRAP;refGene_tx=XM_019813827.1;refGene_gn=NRAP;refGene_tx=XM_019813828.1;refGene_gn=NRAP;refGene_tx=XM_023240955.1
D2 72761401 72761489
refGene_gn=NRAP;refGene_tx=XM_006938185.3;refGene_gn=NRAP;refGene_tx=XM_006938188.3;refGene_gn=NRAP;refGene_tx=XM_006938191.3;refGene_gn=NRAP;refGene_tx=XM_011287564.2;refGene_gn=NRAP;refGene_tx=XM_011287565.2;refGene_gn=NRAP;refGene_tx=XM_011287566.2;refGene_gn=NRAP;refGene_tx=XM_011287567.2;refGene_gn=NRAP;refGene_tx=XM_011287568.2;refGene_gn=NRAP;refGene_tx=XM_019813827.1;refGene_gn=NRAP;refGene_tx=XM_019813828.1;refGene_gn=NRAP;refGene_tx=XM_023240955.1
D2 72762295 72762390
refGene_gn=NRAP;refGene_tx=XM_006938185.3;refGene_gn=NRAP;refGene_tx=XM_006938188.3;refGene_gn=NRAP;refGene_tx=XM_006938191.3;refGene_gn=NRAP;refGene_tx=XM_011287564.2;refGene_gn=NRAP;refGene_tx=XM_011287565.2;refGene_gn=NRAP;refGene_tx=XM_011287566.2;refGene_gn=NRAP;refGene_tx=XM_011287567.2;refGene_gn=NRAP;refGene_tx=XM_011287568.2;refGene_gn=NRAP;refGene_tx=XM_019813827.1;refGene_gn=NRAP;refGene_tx=XM_019813828.1;refGene_gn=NRAP;refGene_tx=XM_023240955.1
D2 72762711 72762783
refGene_gn=NRAP;refGene_tx=XM_006938185.3;refGene_gn=NRAP;refGene_tx=XM_006938188.3;refGene_gn=NRAP;refGene_tx=XM_006938191.3;refGene_gn=NRAP;refGene_tx=XM_011287564.2;refGene_gn=NRAP;refGene_tx=XM_011287565.2;refGene_gn=NRAP;refGene_tx=XM_011287566.2;refGene_gn=NRAP;refGene_tx=XM_011287567.2;refGene_gn=NRAP;refGene_tx=XM_011287568.2;refGene_gn=NRA

P;refGene_tx=XM_019813827.1;refGene_gn=NRAP;refGene_tx=XM_019813828.1;refGene_gn=NRAP;refGene_tx=XM_023240955.1
D2 72774764 72774874 refGene_gn=CASP7;refGene_tx=XM_006938193.4
D2 72784478 72784513 refGene_gn=CASP7;refGene_tx=XM_019813834.2
D2 72792604 72792741 refGene_gn=CASP7;refGene_tx=XM_006938193.4;refGene_gn=CASP7;refGene_tx=XM_019813834.2
D2 72793421 72793550 refGene_gn=CASP7;refGene_tx=XM_006938193.4;refGene_gn=CASP7;refGene_tx=XM_019813834.2
D2 72794545 72794721 refGene_gn=CASP7;refGene_tx=XM_006938193.4;refGene_gn=CASP7;refGene_tx=XM_019813834.2
D2 72795078 72795208 refGene_gn=CASP7;refGene_tx=XM_006938193.4;refGene_gn=CASP7;refGene_tx=XM_019813834.2
D2 72798701 72798925 refGene_gn=CASP7;refGene_tx=XM_006938193.4;refGene_gn=CASP7;refGene_tx=XM_019813834.2
D2 72815243 72815271 refGene_gn=PLEKHS1;refGene_tx=XM_023240956.1;refGene_gn=PLEKHS1;refGene_tx=XM_023240957.1;refGene_gn=PLEKHS1;refGene_tx=XM_023240958.1;refGene_gn=PLEKHS1;refGene_tx=XM_023240959.1;refGene_gn=PLEKHS1;refGene_tx=XM_023240960.1
D2 72823878 72823967 refGene_gn=PLEKHS1;refGene_tx=XM_023240956.1;refGene_gn=PLEKHS1;refGene_tx=XM_023240957.1;refGene_gn=PLEKHS1;refGene_tx=XM_023240958.1;refGene_gn=PLEKHS1;refGene_tx=XM_023240959.1;refGene_gn=PLEKHS1;refGene_tx=XM_023240960.1
D2 72824078 72824185 refGene_gn=PLEKHS1;refGene_tx=XM_023240956.1;refGene_gn=PLEKHS1;refGene_tx=XM_023240957.1;refGene_gn=PLEKHS1;refGene_tx=XM_023240958.1;refGene_gn=PLEKHS1;refGene_tx=XM_023240959.1;refGene_gn=PLEKHS1;refGene_tx=XM_023240960.1
D2 72824801 72824936 refGene_gn=PLEKHS1;refGene_tx=XM_023240956.1;refGene_gn=PLEKHS1;refGene_tx=XM_023240957.1;refGene_gn=PLEKHS1;refGene_tx=XM_023240958.1;refGene_gn=PLEKHS1;refGene_tx=XM_023240959.1;refGene_gn=PLEKHS1;refGene_tx=XM_023240960.1
D2 72826290 72826366 refGene_gn=PLEKHS1;refGene_tx=XM_023240956.1;refGene_gn=PLEKHS1;refGene_tx=XM_023240957.1;refGene_gn=PLEKHS1;refGene_tx=XM_023240958.1;refGene_gn=PLEKHS1;refGene_tx=XM_023240959.1;refGene_gn=PLEKHS1;refGene_tx=XM_023240960.1
D2 72827228 72827348 refGene_gn=PLEKHS1;refGene_tx=XM_023240956.1;refGene_gn=PLEKHS1;refGene_tx=XM_023240957.1;refGene_gn=PLEKHS1;refGene_tx=XM_023240958.1;refGene_gn=PLEKHS1;refGene_tx=XM_023240959.1;refGene_gn=PLEKHS1;refGene_tx=XM_023240960.1
D2 72829121 72829247

refGene_gn=PLEKHS1;refGene_tx=XM_023240956.1;refGene_gn=PLEKHS1;refGene_tx=XM_023240957.1;refGene_gn=PLEKHS1;refGene_tx=XM_023240958.1;refGene_gn=PLEKHS1;refGene_tx=XM_023240959.1;refGene_gn=PLEKHS1;refGene_tx=XM_023240960.1
D2 72830279 72830386
refGene_gn=PLEKHS1;refGene_tx=XM_023240956.1;refGene_gn=PLEKHS1;refGene_tx=XM_023240957.1;refGene_gn=PLEKHS1;refGene_tx=XM_023240958.1;refGene_gn=PLEKHS1;refGene_tx=XM_023240959.1;refGene_gn=PLEKHS1;refGene_tx=XM_023240960.1
D2 72830877 72831084
refGene_gn=PLEKHS1;refGene_tx=XM_023240956.1;refGene_gn=PLEKHS1;refGene_tx=XM_023240957.1;refGene_gn=PLEKHS1;refGene_tx=XM_023240958.1;refGene_gn=PLEKHS1;refGene_tx=XM_023240959.1;refGene_gn=PLEKHS1;refGene_tx=XM_023240960.1
D2 72831783 72831885
refGene_gn=PLEKHS1;refGene_tx=XM_023240956.1;refGene_gn=PLEKHS1;refGene_tx=XM_023240957.1;refGene_gn=PLEKHS1;refGene_tx=XM_023240958.1;refGene_gn=PLEKHS1;refGene_tx=XM_023240959.1;refGene_gn=PLEKHS1;refGene_tx=XM_023240960.1
D2 72832998 72833134
refGene_gn=PLEKHS1;refGene_tx=XM_023240956.1;refGene_gn=PLEKHS1;refGene_tx=XM_023240957.1;refGene_gn=PLEKHS1;refGene_tx=XM_023240958.1;refGene_gn=PLEKHS1;refGene_tx=XM_023240959.1;refGene_gn=PLEKHS1;refGene_tx=XM_023240960.1
D2 72833264 72833267 refGene_gn=PLEKHS1;refGene_tx=XM_023240960.1
D2 72835784 72835985
refGene_gn=PLEKHS1;refGene_tx=XM_023240956.1;refGene_gn=PLEKHS1;refGene_tx=XM_023240957.1;refGene_gn=PLEKHS1;refGene_tx=XM_023240958.1;refGene_gn=PLEKHS1;refGene_tx=XM_023240959.1
D2 72876130 72876291
refGene_gn=DCLRE1A;refGene_tx=XM_003994417.5;refGene_gn=DCLRE1A;refGene_tx=XM_019813835.2;refGene_gn=DCLRE1A;refGene_tx=XM_019813836.1
D2 72877729 72877871
refGene_gn=DCLRE1A;refGene_tx=XM_003994417.5;refGene_gn=DCLRE1A;refGene_tx=XM_019813835.2;refGene_gn=DCLRE1A;refGene_tx=XM_019813836.1
D2 72883472 72883627
refGene_gn=DCLRE1A;refGene_tx=XM_003994417.5;refGene_gn=DCLRE1A;refGene_tx=XM_019813835.2;refGene_gn=DCLRE1A;refGene_tx=XM_019813836.1
D2 72884412 72884558
refGene_gn=DCLRE1A;refGene_tx=XM_003994417.5;refGene_gn=DCLRE1A;refGene_tx=XM_019813835.2;refGene_gn=DCLRE1A;refGene_tx=XM_019813836.1
D2 72886365 72886506
refGene_gn=DCLRE1A;refGene_tx=XM_003994417.5;refGene_gn=DCLRE1A;refGene_tx=XM_019813835.2;refGene_gn=DCLRE1A;refGene_tx=XM_019813836.1
D2 72889622 72889741
refGene_gn=DCLRE1A;refGene_tx=XM_003994417.5;refGene_gn=DCLRE1A;refGene_tx=XM_019813835.2;refGene_gn=DCLRE1A;refGene_tx=XM_019813836.1
D2 72891468 72891602
refGene_gn=DCLRE1A;refGene_tx=XM_003994417.5;refGene_gn=DCLRE1A;refGene_tx=XM_019813835.2;refGene_gn=DCLRE1A;refGene_tx=XM_019813836.1

D2 72893129 72894815
refGene_gn=DCLRE1A;refGene_tx=XM_003994417.5;refGene_gn=DCLRE1A;refGene_tx=XM_019813835.2;refGene_gn=DCLRE1A;refGene_tx=XM_019813836.1
D2 72897061 72897527
refGene_gn=DCLRE1A;refGene_tx=XM_003994417.5;refGene_gn=DCLRE1A;refGene_tx=XM_019813835.2;refGene_gn=DCLRE1A;refGene_tx=XM_019813836.1
D2 72899291 72899469 refGene_gn=NHLRC2;refGene_tx=XM_011287569.3
D2 72902487 72902640 refGene_gn=NHLRC2;refGene_tx=XM_011287569.3
D2 72918432 72918888 refGene_gn=NHLRC2;refGene_tx=XM_011287569.3
D2 72920664 72920786 refGene_gn=NHLRC2;refGene_tx=XM_011287569.3
D2 72929034 72929164 refGene_gn=NHLRC2;refGene_tx=XM_011287569.3
D2 72941743 72941843 refGene_gn=NHLRC2;refGene_tx=XM_011287569.3
D2 72945229 72945461 refGene_gn=NHLRC2;refGene_tx=XM_011287569.3
D2 72946260 72946383 refGene_gn=NHLRC2;refGene_tx=XM_011287569.3
D2 72947295 72947505 refGene_gn=NHLRC2;refGene_tx=XM_011287569.3
D2 72948516 72948736 refGene_gn=NHLRC2;refGene_tx=XM_011287569.3
D2 72951772 72952029 refGene_gn=NHLRC2;refGene_tx=XM_011287569.3
D2 73093098 73094040
refGene_gn=LOC111556931;refGene_tx=XM_023240301.1
D2 73094178 73095603 refGene_gn=ADRB1;refGene_tx=NM_001009375.1
D2 73164330 73164414
refGene_gn=CCDC186;refGene_tx=XM_003994418.5;refGene_gn=CCDC186;refGene_tx=XM_019813837.2
D2 73165251 73165471
refGene_gn=CCDC186;refGene_tx=XM_003994418.5;refGene_gn=CCDC186;refGene_tx=XM_019813837.2
D2 73166756 73166967
refGene_gn=CCDC186;refGene_tx=XM_003994418.5;refGene_gn=CCDC186;refGene_tx=XM_019813837.2
D2 73168843 73168924
refGene_gn=CCDC186;refGene_tx=XM_003994418.5;refGene_gn=CCDC186;refGene_tx=XM_019813837.2
D2 73170054 73170244
refGene_gn=CCDC186;refGene_tx=XM_003994418.5;refGene_gn=CCDC186;refGene_tx=XM_019813837.2
D2 73171394 73171650
refGene_gn=CCDC186;refGene_tx=XM_003994418.5;refGene_gn=CCDC186;refGene_tx=XM_019813837.2
D2 73174803 73174946
refGene_gn=CCDC186;refGene_tx=XM_003994418.5;refGene_gn=CCDC186;refGene_tx=XM_019813837.2
D2 73175784 73175871
refGene_gn=CCDC186;refGene_tx=XM_003994418.5;refGene_gn=CCDC186;refGene_tx=XM_019813837.2
D2 73176054 73176153
refGene_gn=CCDC186;refGene_tx=XM_003994418.5;refGene_gn=CCDC186;refGene_tx=XM_019813837.2
D2 73176593 73176698
refGene_gn=CCDC186;refGene_tx=XM_003994418.5;refGene_gn=CCDC186;refGene_tx=XM_019813837.2

D2 73184012 73184132
refGene_gn=CCDC186;refGene_tx=XM_003994418.5;refGene_gn=CCDC186;refGene_tx=XM_019813837.2
D2 73185059 73185272
refGene_gn=CCDC186;refGene_tx=XM_003994418.5;refGene_gn=CCDC186;refGene_tx=XM_019813837.2
D2 73190307 73190436
refGene_gn=CCDC186;refGene_tx=XM_003994418.5;refGene_gn=CCDC186;refGene_tx=XM_019813837.2
D2 73197174 73197301
refGene_gn=CCDC186;refGene_tx=XM_003994418.5;refGene_gn=CCDC186;refGene_tx=XM_019813837.2
D2 73203452 73204145
refGene_gn=CCDC186;refGene_tx=XM_003994418.5;refGene_gn=CCDC186;refGene_tx=XM_019813837.2
D2 73213671 73213820 refGene_gn=CCDC186;refGene_tx=XM_019813837.2
D2 73218638 73218767
refGene_gn=TDRD1;refGene_tx=XM_023240961.1;refGene_gn=TDRD1;refGene_tx=XM_023240964.1;refGene_gn=TDRD1;refGene_tx=XM_023240965.1
D2 73226924 73227255
refGene_gn=TDRD1;refGene_tx=XM_023240961.1;refGene_gn=TDRD1;refGene_tx=XM_023240962.1;refGene_gn=TDRD1;refGene_tx=XM_023240963.1;refGene_gn=TDRD1;refGene_tx=XM_023240964.1;refGene_gn=TDRD1;refGene_tx=XM_023240965.1
D2 73229740 73229790
refGene_gn=TDRD1;refGene_tx=XM_023240961.1;refGene_gn=TDRD1;refGene_tx=XM_023240962.1;refGene_gn=TDRD1;refGene_tx=XM_023240963.1;refGene_gn=TDRD1;refGene_tx=XM_023240964.1;refGene_gn=TDRD1;refGene_tx=XM_023240965.1
D2 73236928 73237073
refGene_gn=TDRD1;refGene_tx=XM_023240961.1;refGene_gn=TDRD1;refGene_tx=XM_023240962.1;refGene_gn=TDRD1;refGene_tx=XM_023240963.1;refGene_gn=TDRD1;refGene_tx=XM_023240964.1;refGene_gn=TDRD1;refGene_tx=XM_023240965.1
D2 73239660 73239766
refGene_gn=TDRD1;refGene_tx=XM_023240961.1;refGene_gn=TDRD1;refGene_tx=XM_023240962.1;refGene_gn=TDRD1;refGene_tx=XM_023240963.1;refGene_gn=TDRD1;refGene_tx=XM_023240964.1;refGene_gn=TDRD1;refGene_tx=XM_023240965.1
D2 73240487 73240551
refGene_gn=TDRD1;refGene_tx=XM_023240961.1;refGene_gn=TDRD1;refGene_tx=XM_023240962.1;refGene_gn=TDRD1;refGene_tx=XM_023240963.1;refGene_gn=TDRD1;refGene_tx=XM_023240964.1;refGene_gn=TDRD1;refGene_tx=XM_023240965.1
D2 73241512 73241617
refGene_gn=TDRD1;refGene_tx=XM_023240961.1;refGene_gn=TDRD1;refGene_tx=XM_023240962.1;refGene_gn=TDRD1;refGene_tx=XM_023240963.1;refGene_gn=TDRD1;refGene_tx=XM_023240964.1;refGene_gn=TDRD1;refGene_tx=XM_023240965.1
D2 73241843 73242026

refGene_gn=TDRD1;refGene_tx=XM_023240961.1;refGene_gn=TDRD1;refGene_tx=XM_023240962.1;refGene_gn=TDRD1;refGene_tx=XM_023240963.1;refGene_gn=TDRD1;refGene_tx=XM_023240964.1;refGene_gn=TDRD1;refGene_tx=XM_023240965.1
D2	73242538	73242682	refGene_gn=TDRD1;refGene_tx=XM_023240961.1;refGene_gn=TDRD1;refGene_tx=XM_023240962.1;refGene_gn=TDRD1;refGene_tx=XM_023240963.1;refGene_gn=TDRD1;refGene_tx=XM_023240964.1;refGene_gn=TDRD1;refGene_tx=XM_023240965.1
D2	73243460	73243632	refGene_gn=TDRD1;refGene_tx=XM_023240961.1;refGene_gn=TDRD1;refGene_tx=XM_023240962.1;refGene_gn=TDRD1;refGene_tx=XM_023240963.1;refGene_gn=TDRD1;refGene_tx=XM_023240964.1;refGene_gn=TDRD1;refGene_tx=XM_023240965.1
D2	73244708	73244795	refGene_gn=TDRD1;refGene_tx=XM_023240961.1;refGene_gn=TDRD1;refGene_tx=XM_023240962.1;refGene_gn=TDRD1;refGene_tx=XM_023240963.1;refGene_gn=TDRD1;refGene_tx=XM_023240964.1;refGene_gn=TDRD1;refGene_tx=XM_023240965.1
D2	73246633	73246801	refGene_gn=TDRD1;refGene_tx=XM_023240961.1;refGene_gn=TDRD1;refGene_tx=XM_023240962.1;refGene_gn=TDRD1;refGene_tx=XM_023240963.1;refGene_gn=TDRD1;refGene_tx=XM_023240964.1;refGene_gn=TDRD1;refGene_tx=XM_023240965.1
D2	73246922	73247030	refGene_gn=TDRD1;refGene_tx=XM_023240961.1;refGene_gn=TDRD1;refGene_tx=XM_023240962.1;refGene_gn=TDRD1;refGene_tx=XM_023240963.1;refGene_gn=TDRD1;refGene_tx=XM_023240964.1;refGene_gn=TDRD1;refGene_tx=XM_023240965.1
D2	73247903	73248074	refGene_gn=TDRD1;refGene_tx=XM_023240961.1;refGene_gn=TDRD1;refGene_tx=XM_023240962.1;refGene_gn=TDRD1;refGene_tx=XM_023240963.1;refGene_gn=TDRD1;refGene_tx=XM_023240964.1;refGene_gn=TDRD1;refGene_tx=XM_023240965.1
D2	73249671	73249914	refGene_gn=TDRD1;refGene_tx=XM_023240961.1;refGene_gn=TDRD1;refGene_tx=XM_023240962.1;refGene_gn=TDRD1;refGene_tx=XM_023240963.1;refGene_gn=TDRD1;refGene_tx=XM_023240965.1
D2	73250753	73250891	refGene_gn=TDRD1;refGene_tx=XM_023240961.1;refGene_gn=TDRD1;refGene_tx=XM_023240962.1;refGene_gn=TDRD1;refGene_tx=XM_023240963.1;refGene_gn=TDRD1;refGene_tx=XM_023240964.1;refGene_gn=TDRD1;refGene_tx=XM_023240965.1
D2	73253737	73253848	refGene_gn=TDRD1;refGene_tx=XM_023240961.1;refGene_gn=TDRD1;refGene_tx=XM_023240962.1;refGene_gn=TDRD1;refGene_tx=XM_023240963.1;refGene_gn=TDRD1;refGene_tx=XM_023240964.1;refGene_gn=TDRD1;refGene_tx=XM_023240965.1
D2	73254823	73254994	refGene_gn=TDRD1;refGene_tx=XM_023240961.1;refGene_gn=TDRD1;refGene_tx

=XM_023240962.1;refGene_gn=TDRD1;refGene_tx=XM_023240963.1;refGene_gn=
TDRD1;refGene_tx=XM_023240964.1;refGene_gn=TDRD1;refGene_tx=XM_0232409
65.1
D2 73256894 73257170
refGene_gn=TDRD1;refGene_tx=XM_023240961.1;refGene_gn=TDRD1;refGene_tx
=XM_023240962.1;refGene_gn=TDRD1;refGene_tx=XM_023240963.1;refGene_gn=
TDRD1;refGene_tx=XM_023240964.1;refGene_gn=TDRD1;refGene_tx=XM_0232409
65.1
D2 73257889 73258009
refGene_gn=TDRD1;refGene_tx=XM_023240961.1;refGene_gn=TDRD1;refGene_tx
=XM_023240962.1;refGene_gn=TDRD1;refGene_tx=XM_023240963.1;refGene_gn=
TDRD1;refGene_tx=XM_023240964.1;refGene_gn=TDRD1;refGene_tx=XM_0232409
65.1
D2 73260166 73260297
refGene_gn=TDRD1;refGene_tx=XM_023240961.1;refGene_gn=TDRD1;refGene_tx
=XM_023240962.1;refGene_gn=TDRD1;refGene_tx=XM_023240963.1;refGene_gn=
TDRD1;refGene_tx=XM_023240964.1;refGene_gn=TDRD1;refGene_tx=XM_0232409
65.1
D2 73263428 73263596
refGene_gn=TDRD1;refGene_tx=XM_023240961.1;refGene_gn=TDRD1;refGene_tx
=XM_023240962.1;refGene_gn=TDRD1;refGene_tx=XM_023240963.1;refGene_gn=
TDRD1;refGene_tx=XM_023240964.1
D2 73264628 73264856
refGene_gn=TDRD1;refGene_tx=XM_023240961.1;refGene_gn=TDRD1;refGene_tx
=XM_023240962.1;refGene_gn=TDRD1;refGene_tx=XM_023240963.1;refGene_gn=
TDRD1;refGene_tx=XM_023240964.1
D2 73265472 73265519
refGene_gn=TDRD1;refGene_tx=XM_023240961.1;refGene_gn=TDRD1;refGene_tx
=XM_023240962.1;refGene_gn=TDRD1;refGene_tx=XM_023240963.1;refGene_gn=
TDRD1;refGene_tx=XM_023240964.1
D2 73265600 73265693
refGene_gn=TDRD1;refGene_tx=XM_023240961.1;refGene_gn=TDRD1;refGene_tx
=XM_023240962.1;refGene_gn=TDRD1;refGene_tx=XM_023240963.1;refGene_gn=
TDRD1;refGene_tx=XM_023240964.1
D2 73283530 73283582 refGene_gn=VWA2;refGene_tx=XM_023240966.1
D2 73287491 73287566 refGene_gn=VWA2;refGene_tx=XM_023240966.1
D2 73288898 73289032 refGene_gn=VWA2;refGene_tx=XM_023240966.1
D2 73294473 73294583
refGene_gn=VWA2;refGene_tx=XM_023240966.1;refGene_gn=VWA2;refGene_tx=X
M_023240967.1
D2 73303688 73303883
refGene_gn=VWA2;refGene_tx=XM_023240966.1;refGene_gn=VWA2;refGene_tx=X
M_023240967.1
D2 73308154 73308288
refGene_gn=VWA2;refGene_tx=XM_023240966.1;refGene_gn=VWA2;refGene_tx=X
M_023240967.1
D2 73308902 73309035
refGene_gn=VWA2;refGene_tx=XM_023240966.1;refGene_gn=VWA2;refGene_tx=X
M_023240967.1
D2 73311411 73311467

refGene_gn=VWA2;refGene_tx=XM_023240966.1;refGene_gn=VWA2;refGene_tx=XM_023240967.1
D2 73314244 73314352
refGene_gn=VWA2;refGene_tx=XM_023240966.1;refGene_gn=VWA2;refGene_tx=XM_023240967.1
D2 73315638 73316211
refGene_gn=VWA2;refGene_tx=XM_023240966.1;refGene_gn=VWA2;refGene_tx=XM_023240967.1
D2 73317403 73317955
refGene_gn=VWA2;refGene_tx=XM_023240966.1;refGene_gn=VWA2;refGene_tx=XM_023240967.1
D2 73318645 73318771
refGene_gn=VWA2;refGene_tx=XM_023240966.1;refGene_gn=VWA2;refGene_tx=XM_023240967.1
D2 73319637 73319726
refGene_gn=VWA2;refGene_tx=XM_023240966.1;refGene_gn=VWA2;refGene_tx=XM_023240967.1
D2 73324027 73324054 refGene_gn=AFAP1L2;refGene_tx=XM_023240968.1
D2 73324929 73325052 refGene_gn=AFAP1L2;refGene_tx=XM_023240968.1
D2 73325166 73325360 refGene_gn=AFAP1L2;refGene_tx=XM_023240968.1
D2 73327632 73327788 refGene_gn=AFAP1L2;refGene_tx=XM_023240968.1
D2 73328046 73328221 refGene_gn=AFAP1L2;refGene_tx=XM_023240968.1
D2 73328303 73328540 refGene_gn=AFAP1L2;refGene_tx=XM_023240968.1
D2 73329154 73329266 refGene_gn=AFAP1L2;refGene_tx=XM_023240968.1
D2 73329858 73330004 refGene_gn=AFAP1L2;refGene_tx=XM_023240968.1
D2 73331591 73331803 refGene_gn=AFAP1L2;refGene_tx=XM_023240968.1
D2 73334036 73334141 refGene_gn=AFAP1L2;refGene_tx=XM_023240968.1
D2 73334677 73334762 refGene_gn=AFAP1L2;refGene_tx=XM_023240968.1
D2 73336431 73336521 refGene_gn=AFAP1L2;refGene_tx=XM_023240968.1
D2 73339841 73340021 refGene_gn=AFAP1L2;refGene_tx=XM_023240968.1
D2 73341134 73341194 refGene_gn=AFAP1L2;refGene_tx=XM_023240968.1
D2 73341195 73341341 refGene_gn=AFAP1L2;refGene_tx=XM_023240968.1
D2 73344136 73344220 refGene_gn=AFAP1L2;refGene_tx=XM_023240968.1
D2 73347085 73347176 refGene_gn=AFAP1L2;refGene_tx=XM_023240968.1
D2 73353407 73353502 refGene_gn=AFAP1L2;refGene_tx=XM_023240968.1
D2 73355073 73355148 refGene_gn=AFAP1L2;refGene_tx=XM_023240968.1
D2 73361462 73361591 refGene_gn=AFAP1L2;refGene_tx=XM_023240968.1
D2 73414788 73414842 refGene_gn=AFAP1L2;refGene_tx=XM_023240968.1
D2 73420785 73420801 refGene_gn=AFAP1L2;refGene_tx=XM_023240968.1
D2 73449452 73449566
refGene_gn=ABLIM1;refGene_tx=XM_019813857.2;refGene_gn=ABLIM1;refGene_tx=XM_019813858.2;refGene_gn=ABLIM1;refGene_tx=XM_019813859.2;refGene_gn=ABLIM1;refGene_tx=XM_019813860.1;refGene_gn=ABLIM1;refGene_tx=XM_019813861.2;refGene_gn=ABLIM1;refGene_tx=XM_019813862.2;refGene_gn=ABLIM1;refGene_tx=XM_019813863.2;refGene_gn=ABLIM1;refGene_tx=XM_019813864.2;refGene_gn=ABLIM1;refGene_tx=XM_019813865.1;refGene_gn=ABLIM1;refGene_tx=XM_019813866.2;refGene_gn=ABLIM1;refGene_tx=XM_019813867.2;refGene_gn=ABLIM1;refGene_tx=XM_019813868.2;refGene_gn=ABLIM1;refGene_tx=XM_019813869.1;refGene_gn=ABLIM1;refGene_tx=XM_019813873.2;refGene_gn=ABLIM1;refGene_tx=XM_019813874.2;refGene_gn=ABLIM1;refGene_tx=XM_01981387

5.2;refGene_gn=ABLIM1;refGene_tx=XM_019813876.2;refGene_gn=ABLIM1;refGene_tx=XM_019813877.2;refGene_gn=ABLIM1;refGene_tx=XM_019813878.2;refGene_gn=ABLIM1;refGene_tx=XM_019813879.2;refGene_gn=ABLIM1;refGene_tx=XM_019813880.2;refGene_gn=ABLIM1;refGene_tx=XM_019813881.2;refGene_gn=ABLIM1;refGene_tx=XM_019813882.2;refGene_gn=ABLIM1;refGene_tx=XM_023240969.1;refGene_gn=ABLIM1;refGene_tx=XM_023240970.1
D2 73450684 73450765
refGene_gn=ABLIM1;refGene_tx=XM_019813857.2;refGene_gn=ABLIM1;refGene_tx=XM_019813858.2;refGene_gn=ABLIM1;refGene_tx=XM_019813859.2;refGene_gn=ABLIM1;refGene_tx=XM_019813860.1;refGene_gn=ABLIM1;refGene_tx=XM_019813861.2;refGene_gn=ABLIM1;refGene_tx=XM_019813862.2;refGene_gn=ABLIM1;refGene_tx=XM_019813863.2;refGene_gn=ABLIM1;refGene_tx=XM_019813864.2;refGene_gn=ABLIM1;refGene_tx=XM_019813865.1;refGene_gn=ABLIM1;refGene_tx=XM_019813866.2;refGene_gn=ABLIM1;refGene_tx=XM_019813867.2;refGene_gn=ABLIM1;refGene_tx=XM_019813868.2;refGene_gn=ABLIM1;refGene_tx=XM_019813869.1;refGene_gn=ABLIM1;refGene_tx=XM_019813873.2;refGene_gn=ABLIM1;refGene_tx=XM_019813874.2;refGene_gn=ABLIM1;refGene_tx=XM_019813875.2;refGene_gn=ABLIM1;refGene_tx=XM_019813876.2;refGene_gn=ABLIM1;refGene_tx=XM_019813877.2;refGene_gn=ABLIM1;refGene_tx=XM_019813878.2;refGene_gn=ABLIM1;refGene_tx=XM_019813879.2;refGene_gn=ABLIM1;refGene_tx=XM_019813880.2;refGene_gn=ABLIM1;refGene_tx=XM_019813881.2;refGene_gn=ABLIM1;refGene_tx=XM_019813882.2;refGene_gn=ABLIM1;refGene_tx=XM_023240969.1;refGene_gn=ABLIM1;refGene_tx=XM_023240970.1
D2 73451686 73451761
refGene_gn=ABLIM1;refGene_tx=XM_019813857.2;refGene_gn=ABLIM1;refGene_tx=XM_019813858.2;refGene_gn=ABLIM1;refGene_tx=XM_019813859.2;refGene_gn=ABLIM1;refGene_tx=XM_019813860.1;refGene_gn=ABLIM1;refGene_tx=XM_019813861.2;refGene_gn=ABLIM1;refGene_tx=XM_019813862.2;refGene_gn=ABLIM1;refGene_tx=XM_019813863.2;refGene_gn=ABLIM1;refGene_tx=XM_019813864.2;refGene_gn=ABLIM1;refGene_tx=XM_019813865.1;refGene_gn=ABLIM1;refGene_tx=XM_019813866.2;refGene_gn=ABLIM1;refGene_tx=XM_019813867.2;refGene_gn=ABLIM1;refGene_tx=XM_019813868.2;refGene_gn=ABLIM1;refGene_tx=XM_019813869.1;refGene_gn=ABLIM1;refGene_tx=XM_019813873.2;refGene_gn=ABLIM1;refGene_tx=XM_019813874.2;refGene_gn=ABLIM1;refGene_tx=XM_019813875.2;refGene_gn=ABLIM1;refGene_tx=XM_019813876.2;refGene_gn=ABLIM1;refGene_tx=XM_019813877.2;refGene_gn=ABLIM1;refGene_tx=XM_019813878.2;refGene_gn=ABLIM1;refGene_tx=XM_019813879.2;refGene_gn=ABLIM1;refGene_tx=XM_019813880.2;refGene_gn=ABLIM1;refGene_tx=XM_019813881.2;refGene_gn=ABLIM1;refGene_tx=XM_019813882.2;refGene_gn=ABLIM1;refGene_tx=XM_023240969.1;refGene_gn=ABLIM1;refGene_tx=XM_023240970.1
D2 73452781 73452789
refGene_gn=ABLIM1;refGene_tx=XM_019813857.2;refGene_gn=ABLIM1;refGene_tx=XM_019813858.2;refGene_gn=ABLIM1;refGene_tx=XM_019813859.2;refGene_gn=ABLIM1;refGene_tx=XM_019813860.1;refGene_gn=ABLIM1;refGene_tx=XM_019813861.2;refGene_gn=ABLIM1;refGene_tx=XM_019813862.2;refGene_gn=ABLIM1;refGene_tx=XM_019813863.2;refGene_gn=ABLIM1;refGene_tx=XM_019813864.2;refGene_gn=ABLIM1;refGene_tx=XM_019813865.1;refGene_gn=ABLIM1;refGene_tx=XM_019813866.2;refGene_gn=ABLIM1;refGene_tx=XM_019813867.2;refGene_gn=ABLIM1;refGene_tx=XM_019813868.2;refGene_gn=ABLIM1;refGene_tx=XM_019813869.1;refGene_gn=ABLIM1;refGene_tx=XM_019813873.2;refGene_gn=ABL

IM1;refGene_tx=XM_019813874.2;refGene_gn=ABLIM1;refGene_tx=XM_01981387
5.2;refGene_gn=ABLIM1;refGene_tx=XM_019813876.2;refGene_gn=ABLIM1;refG
ene_tx=XM_019813877.2;refGene_gn=ABLIM1;refGene_tx=XM_019813878.2;refG
ene_gn=ABLIM1;refGene_tx=XM_019813879.2;refGene_gn=ABLIM1;refGene_tx=X
M_019813880.2;refGene_gn=ABLIM1;refGene_tx=XM_019813881.2;refGene_gn=A
BLIM1;refGene_tx=XM_019813882.2;refGene_gn=ABLIM1;refGene_tx=XM_023240
969.1;refGene_gn=ABLIM1;refGene_tx=XM_023240970.1
D2 73453860 73453921
refGene_gn=ABLIM1;refGene_tx=XM_019813857.2;refGene_gn=ABLIM1;refGene_
tx=XM_019813858.2;refGene_gn=ABLIM1;refGene_tx=XM_019813859.2;refGene_
gn=ABLIM1;refGene_tx=XM_019813860.1;refGene_gn=ABLIM1;refGene_tx=XM_01
9813861.2;refGene_gn=ABLIM1;refGene_tx=XM_019813862.2;refGene_gn=ABLIM
1;refGene_tx=XM_019813863.2;refGene_gn=ABLIM1;refGene_tx=XM_019813864.
2;refGene_gn=ABLIM1;refGene_tx=XM_019813865.1;refGene_gn=ABLIM1;refGen
e_tx=XM_019813866.2;refGene_gn=ABLIM1;refGene_tx=XM_019813867.2;refGen
e_gn=ABLIM1;refGene_tx=XM_019813868.2;refGene_gn=ABLIM1;refGene_tx=XM_
019813869.1;refGene_gn=ABLIM1;refGene_tx=XM_019813873.2;refGene_gn=ABL
IM1;refGene_tx=XM_019813874.2;refGene_gn=ABLIM1;refGene_tx=XM_01981387
5.2;refGene_gn=ABLIM1;refGene_tx=XM_019813876.2;refGene_gn=ABLIM1;refG
ene_tx=XM_019813877.2;refGene_gn=ABLIM1;refGene_tx=XM_019813878.2;refG
ene_gn=ABLIM1;refGene_tx=XM_019813879.2;refGene_gn=ABLIM1;refGene_tx=X
M_019813880.2;refGene_gn=ABLIM1;refGene_tx=XM_019813881.2;refGene_gn=A
BLIM1;refGene_tx=XM_019813882.2;refGene_gn=ABLIM1;refGene_tx=XM_023240
969.1;refGene_gn=ABLIM1;refGene_tx=XM_023240970.1
D2 73454542 73454607
refGene_gn=ABLIM1;refGene_tx=XM_019813857.2;refGene_gn=ABLIM1;refGene_
tx=XM_019813858.2;refGene_gn=ABLIM1;refGene_tx=XM_019813859.2;refGene_
gn=ABLIM1;refGene_tx=XM_019813860.1;refGene_gn=ABLIM1;refGene_tx=XM_01
9813861.2;refGene_gn=ABLIM1;refGene_tx=XM_019813862.2;refGene_gn=ABLIM
1;refGene_tx=XM_019813863.2;refGene_gn=ABLIM1;refGene_tx=XM_019813864.
2;refGene_gn=ABLIM1;refGene_tx=XM_019813865.1;refGene_gn=ABLIM1;refGen
e_tx=XM_019813866.2;refGene_gn=ABLIM1;refGene_tx=XM_019813867.2;refGen
e_gn=ABLIM1;refGene_tx=XM_019813868.2;refGene_gn=ABLIM1;refGene_tx=XM_
019813869.1;refGene_gn=ABLIM1;refGene_tx=XM_019813873.2;refGene_gn=ABL
IM1;refGene_tx=XM_019813874.2;refGene_gn=ABLIM1;refGene_tx=XM_01981387
5.2;refGene_gn=ABLIM1;refGene_tx=XM_019813876.2;refGene_gn=ABLIM1;refG
ene_tx=XM_019813877.2;refGene_gn=ABLIM1;refGene_tx=XM_019813878.2;refG
ene_gn=ABLIM1;refGene_tx=XM_019813879.2;refGene_gn=ABLIM1;refGene_tx=X
M_019813880.2;refGene_gn=ABLIM1;refGene_tx=XM_019813881.2;refGene_gn=A
BLIM1;refGene_tx=XM_019813882.2;refGene_gn=ABLIM1;refGene_tx=XM_023240
969.1;refGene_gn=ABLIM1;refGene_tx=XM_023240970.1
D2 73456970 73457076
refGene_gn=ABLIM1;refGene_tx=XM_019813857.2;refGene_gn=ABLIM1;refGene_
tx=XM_019813858.2;refGene_gn=ABLIM1;refGene_tx=XM_019813859.2;refGene_
gn=ABLIM1;refGene_tx=XM_019813860.1;refGene_gn=ABLIM1;refGene_tx=XM_01
9813861.2;refGene_gn=ABLIM1;refGene_tx=XM_019813862.2;refGene_gn=ABLIM
1;refGene_tx=XM_019813863.2;refGene_gn=ABLIM1;refGene_tx=XM_019813864.
2;refGene_gn=ABLIM1;refGene_tx=XM_019813865.1;refGene_gn=ABLIM1;refGen
e_tx=XM_019813866.2;refGene_gn=ABLIM1;refGene_tx=XM_019813867.2;refGen
e_gn=ABLIM1;refGene_tx=XM_019813868.2;refGene_gn=ABLIM1;refGene_tx=XM_

019813869.1;refGene_gn=ABLIM1;refGene_tx=XM_019813873.2;refGene_gn=ABLIM1;refGene_tx=XM_019813874.2;refGene_gn=ABLIM1;refGene_tx=XM_019813875.2;refGene_gn=ABLIM1;refGene_tx=XM_019813876.2;refGene_gn=ABLIM1;refGene_tx=XM_019813877.2;refGene_gn=ABLIM1;refGene_tx=XM_019813878.2;refGene_gn=ABLIM1;refGene_tx=XM_019813879.2;refGene_gn=ABLIM1;refGene_tx=XM_019813880.2;refGene_gn=ABLIM1;refGene_tx=XM_019813881.2;refGene_gn=ABLIM1;refGene_tx=XM_019813882.2;refGene_gn=ABLIM1;refGene_tx=XM_023240969.1;refGene_gn=ABLIM1;refGene_tx=XM_023240970.1
D2 73458301 73458393
refGene_gn=ABLIM1;refGene_tx=XM_019813857.2;refGene_gn=ABLIM1;refGene_tx=XM_019813858.2;refGene_gn=ABLIM1;refGene_tx=XM_019813859.2;refGene_gn=ABLIM1;refGene_tx=XM_019813860.1;refGene_gn=ABLIM1;refGene_tx=XM_019813861.2;refGene_gn=ABLIM1;refGene_tx=XM_019813862.2;refGene_gn=ABLIM1;refGene_tx=XM_019813863.2;refGene_gn=ABLIM1;refGene_tx=XM_019813864.2;refGene_gn=ABLIM1;refGene_tx=XM_019813865.1;refGene_gn=ABLIM1;refGene_tx=XM_019813866.2;refGene_gn=ABLIM1;refGene_tx=XM_019813867.2;refGene_gn=ABLIM1;refGene_tx=XM_019813868.2;refGene_gn=ABLIM1;refGene_tx=XM_019813869.1;refGene_gn=ABLIM1;refGene_tx=XM_019813873.2;refGene_gn=ABLIM1;refGene_tx=XM_019813874.2;refGene_gn=ABLIM1;refGene_tx=XM_019813875.2;refGene_gn=ABLIM1;refGene_tx=XM_019813876.2;refGene_gn=ABLIM1;refGene_tx=XM_019813877.2;refGene_gn=ABLIM1;refGene_tx=XM_019813878.2;refGene_gn=ABLIM1;refGene_tx=XM_019813879.2;refGene_gn=ABLIM1;refGene_tx=XM_019813880.2;refGene_gn=ABLIM1;refGene_tx=XM_019813881.2;refGene_gn=ABLIM1;refGene_tx=XM_019813882.2;refGene_gn=ABLIM1;refGene_tx=XM_023240969.1;refGene_gn=ABLIM1;refGene_tx=XM_023240970.1
D2 73460525 73460672
refGene_gn=ABLIM1;refGene_tx=XM_019813857.2;refGene_gn=ABLIM1;refGene_tx=XM_019813858.2;refGene_gn=ABLIM1;refGene_tx=XM_019813859.2;refGene_gn=ABLIM1;refGene_tx=XM_019813860.1;refGene_gn=ABLIM1;refGene_tx=XM_019813861.2;refGene_gn=ABLIM1;refGene_tx=XM_019813862.2;refGene_gn=ABLIM1;refGene_tx=XM_019813864.2;refGene_gn=ABLIM1;refGene_tx=XM_019813865.1;refGene_gn=ABLIM1;refGene_tx=XM_019813866.2;refGene_gn=ABLIM1;refGene_tx=XM_019813867.2;refGene_gn=ABLIM1;refGene_tx=XM_019813868.2;refGene_gn=ABLIM1;refGene_tx=XM_019813869.1;refGene_gn=ABLIM1;refGene_tx=XM_019813873.2;refGene_gn=ABLIM1;refGene_tx=XM_019813875.2;refGene_gn=ABLIM1;refGene_tx=XM_019813878.2;refGene_gn=ABLIM1;refGene_tx=XM_019813879.2;refGene_gn=ABLIM1;refGene_tx=XM_023240969.1;refGene_gn=ABLIM1;refGene_tx=XM_023240970.1
D2 73463569 73463617
refGene_gn=ABLIM1;refGene_tx=XM_019813857.2;refGene_gn=ABLIM1;refGene_tx=XM_019813858.2;refGene_gn=ABLIM1;refGene_tx=XM_019813859.2;refGene_gn=ABLIM1;refGene_tx=XM_019813860.1;refGene_gn=ABLIM1;refGene_tx=XM_019813861.2;refGene_gn=ABLIM1;refGene_tx=XM_019813862.2;refGene_gn=ABLIM1;refGene_tx=XM_019813863.2;refGene_gn=ABLIM1;refGene_tx=XM_019813864.2;refGene_gn=ABLIM1;refGene_tx=XM_019813865.1;refGene_gn=ABLIM1;refGene_tx=XM_019813866.2;refGene_gn=ABLIM1;refGene_tx=XM_019813867.2;refGene_gn=ABLIM1;refGene_tx=XM_019813868.2;refGene_gn=ABLIM1;refGene_tx=XM_019813869.1;refGene_gn=ABLIM1;refGene_tx=XM_019813873.2;refGene_gn=ABLIM1;refGene_tx=XM_019813874.2;refGene_gn=ABLIM1;refGene_tx=XM_019813875.2;refGene_gn=ABLIM1;refGene_tx=XM_019813876.2;refGene_gn=ABLIM1;refG

ene_tx=XM_019813877.2;refGene_gn=ABLIM1;refGene_tx=XM_019813878.2;refGene_gn=ABLIM1;refGene_tx=XM_019813879.2;refGene_gn=ABLIM1;refGene_tx=XM_019813880.2;refGene_gn=ABLIM1;refGene_tx=XM_019813881.2;refGene_gn=ABLIM1;refGene_tx=XM_019813882.2;refGene_gn=ABLIM1;refGene_tx=XM_023240969.1;refGene_gn=ABLIM1;refGene_tx=XM_023240970.1
D2 73465377 73465482
refGene_gn=ABLIM1;refGene_tx=XM_019813857.2;refGene_gn=ABLIM1;refGene_tx=XM_019813858.2;refGene_gn=ABLIM1;refGene_tx=XM_019813859.2;refGene_gn=ABLIM1;refGene_tx=XM_019813860.1;refGene_gn=ABLIM1;refGene_tx=XM_019813861.2;refGene_gn=ABLIM1;refGene_tx=XM_019813862.2;refGene_gn=ABLIM1;refGene_tx=XM_019813863.2;refGene_gn=ABLIM1;refGene_tx=XM_019813864.2;refGene_gn=ABLIM1;refGene_tx=XM_019813865.1;refGene_gn=ABLIM1;refGene_tx=XM_019813866.2;refGene_gn=ABLIM1;refGene_tx=XM_019813869.1;refGene_gn=ABLIM1;refGene_tx=XM_019813873.2;refGene_gn=ABLIM1;refGene_tx=XM_019813874.2;refGene_gn=ABLIM1;refGene_tx=XM_019813876.2;refGene_gn=ABLIM1;refGene_tx=XM_019813877.2;refGene_gn=ABLIM1;refGene_tx=XM_019813881.2;refGene_gn=ABLIM1;refGene_tx=XM_023240969.1;refGene_gn=ABLIM1;refGene_tx=XM_023240970.1
D2 73473773 73473881
refGene_gn=ABLIM1;refGene_tx=XM_019813857.2;refGene_gn=ABLIM1;refGene_tx=XM_019813858.2;refGene_gn=ABLIM1;refGene_tx=XM_019813859.2;refGene_gn=ABLIM1;refGene_tx=XM_019813860.1;refGene_gn=ABLIM1;refGene_tx=XM_019813862.2;refGene_gn=ABLIM1;refGene_tx=XM_019813863.2;refGene_gn=ABLIM1;refGene_tx=XM_019813866.2;refGene_gn=ABLIM1;refGene_tx=XM_019813869.1;refGene_gn=ABLIM1;refGene_tx=XM_019813876.2;refGene_gn=ABLIM1;refGene_tx=XM_023240969.1
D2 73476122 73476252
refGene_gn=ABLIM1;refGene_tx=XM_019813857.2;refGene_gn=ABLIM1;refGene_tx=XM_019813858.2;refGene_gn=ABLIM1;refGene_tx=XM_019813859.2;refGene_gn=ABLIM1;refGene_tx=XM_019813860.1;refGene_gn=ABLIM1;refGene_tx=XM_019813861.2;refGene_gn=ABLIM1;refGene_tx=XM_019813862.2;refGene_gn=ABLIM1;refGene_tx=XM_019813863.2;refGene_gn=ABLIM1;refGene_tx=XM_019813864.2;refGene_gn=ABLIM1;refGene_tx=XM_019813865.1;refGene_gn=ABLIM1;refGene_tx=XM_019813866.2;refGene_gn=ABLIM1;refGene_tx=XM_019813867.2;refGene_gn=ABLIM1;refGene_tx=XM_019813868.2;refGene_gn=ABLIM1;refGene_tx=XM_019813869.1;refGene_gn=ABLIM1;refGene_tx=XM_019813873.2;refGene_gn=ABLIM1;refGene_tx=XM_019813874.2;refGene_gn=ABLIM1;refGene_tx=XM_019813875.2;refGene_gn=ABLIM1;refGene_tx=XM_019813876.2;refGene_gn=ABLIM1;refGene_tx=XM_019813877.2;refGene_gn=ABLIM1;refGene_tx=XM_019813878.2;refGene_gn=ABLIM1;refGene_tx=XM_019813879.2;refGene_gn=ABLIM1;refGene_tx=XM_019813880.2;refGene_gn=ABLIM1;refGene_tx=XM_019813881.2;refGene_gn=ABLIM1;refGene_tx=XM_019813882.2;refGene_gn=ABLIM1;refGene_tx=XM_023240969.1;refGene_gn=ABLIM1;refGene_tx=XM_023240970.1
D2 73477157 73477193
refGene_gn=ABLIM1;refGene_tx=XM_019813857.2;refGene_gn=ABLIM1;refGene_tx=XM_019813858.2;refGene_gn=ABLIM1;refGene_tx=XM_019813859.2;refGene_gn=ABLIM1;refGene_tx=XM_019813860.1;refGene_gn=ABLIM1;refGene_tx=XM_019813861.2;refGene_gn=ABLIM1;refGene_tx=XM_019813862.2;refGene_gn=ABLIM1;refGene_tx=XM_019813863.2;refGene_gn=ABLIM1;refGene_tx=XM_019813864.2;refGene_gn=ABLIM1;refGene_tx=XM_019813865.1;refGene_gn=ABLIM1;refGen

e_tx=XM_019813866.2;refGene_gn=ABLIM1;refGene_tx=XM_019813867.2;refGen
e_gn=ABLIM1;refGene_tx=XM_019813868.2;refGene_gn=ABLIM1;refGene_tx=XM_
019813869.1;refGene_gn=ABLIM1;refGene_tx=XM_019813873.2;refGene_gn=ABL
IM1;refGene_tx=XM_019813874.2;refGene_gn=ABLIM1;refGene_tx=XM_01981387
5.2;refGene_gn=ABLIM1;refGene_tx=XM_019813876.2;refGene_gn=ABLIM1;refG
ene_tx=XM_019813877.2;refGene_gn=ABLIM1;refGene_tx=XM_019813878.2;refG
ene_gn=ABLIM1;refGene_tx=XM_019813879.2;refGene_gn=ABLIM1;refGene_tx=X
M_019813880.2;refGene_gn=ABLIM1;refGene_tx=XM_019813881.2;refGene_gn=A
BLIM1;refGene_tx=XM_019813882.2;refGene_gn=ABLIM1;refGene_tx=XM_023240
969.1;refGene_gn=ABLIM1;refGene_tx=XM_023240970.1
D2 73480647 73480803
refGene_gn=ABLIM1;refGene_tx=XM_019813857.2;refGene_gn=ABLIM1;refGene_
tx=XM_019813858.2;refGene_gn=ABLIM1;refGene_tx=XM_019813859.2;refGene_
gn=ABLIM1;refGene_tx=XM_019813860.1;refGene_gn=ABLIM1;refGene_tx=XM_01
9813861.2;refGene_gn=ABLIM1;refGene_tx=XM_019813862.2;refGene_gn=ABLIM
1;refGene_tx=XM_019813863.2;refGene_gn=ABLIM1;refGene_tx=XM_019813864.
2;refGene_gn=ABLIM1;refGene_tx=XM_019813865.1;refGene_gn=ABLIM1;refGen
e_tx=XM_019813866.2;refGene_gn=ABLIM1;refGene_tx=XM_019813867.2;refGen
e_gn=ABLIM1;refGene_tx=XM_019813868.2;refGene_gn=ABLIM1;refGene_tx=XM_
019813869.1;refGene_gn=ABLIM1;refGene_tx=XM_019813873.2;refGene_gn=ABL
IM1;refGene_tx=XM_019813874.2;refGene_gn=ABLIM1;refGene_tx=XM_01981387
5.2;refGene_gn=ABLIM1;refGene_tx=XM_019813876.2;refGene_gn=ABLIM1;refG
ene_tx=XM_019813877.2;refGene_gn=ABLIM1;refGene_tx=XM_019813878.2;refG
ene_gn=ABLIM1;refGene_tx=XM_019813879.2;refGene_gn=ABLIM1;refGene_tx=X
M_019813880.2;refGene_gn=ABLIM1;refGene_tx=XM_019813881.2;refGene_gn=A
BLIM1;refGene_tx=XM_019813882.2;refGene_gn=ABLIM1;refGene_tx=XM_023240
969.1;refGene_gn=ABLIM1;refGene_tx=XM_023240970.1
D2 73481528 73481606
refGene_gn=ABLIM1;refGene_tx=XM_019813857.2;refGene_gn=ABLIM1;refGene_
tx=XM_019813858.2;refGene_gn=ABLIM1;refGene_tx=XM_019813859.2;refGene_
gn=ABLIM1;refGene_tx=XM_019813860.1;refGene_gn=ABLIM1;refGene_tx=XM_01
9813861.2;refGene_gn=ABLIM1;refGene_tx=XM_019813862.2;refGene_gn=ABLIM
1;refGene_tx=XM_019813863.2;refGene_gn=ABLIM1;refGene_tx=XM_019813864.
2;refGene_gn=ABLIM1;refGene_tx=XM_019813865.1;refGene_gn=ABLIM1;refGen
e_tx=XM_019813866.2;refGene_gn=ABLIM1;refGene_tx=XM_019813867.2;refGen
e_gn=ABLIM1;refGene_tx=XM_019813868.2;refGene_gn=ABLIM1;refGene_tx=XM_
019813869.1;refGene_gn=ABLIM1;refGene_tx=XM_019813873.2;refGene_gn=ABL
IM1;refGene_tx=XM_019813874.2;refGene_gn=ABLIM1;refGene_tx=XM_01981387
5.2;refGene_gn=ABLIM1;refGene_tx=XM_019813876.2;refGene_gn=ABLIM1;refG
ene_tx=XM_019813877.2;refGene_gn=ABLIM1;refGene_tx=XM_019813878.2;refG
ene_gn=ABLIM1;refGene_tx=XM_019813879.2;refGene_gn=ABLIM1;refGene_tx=X
M_019813880.2;refGene_gn=ABLIM1;refGene_tx=XM_019813881.2;refGene_gn=A
BLIM1;refGene_tx=XM_019813882.2;refGene_gn=ABLIM1;refGene_tx=XM_023240
969.1;refGene_gn=ABLIM1;refGene_tx=XM_023240970.1
D2 73487969 73488017
refGene_gn=ABLIM1;refGene_tx=XM_019813857.2;refGene_gn=ABLIM1;refGene_
tx=XM_019813858.2;refGene_gn=ABLIM1;refGene_tx=XM_019813860.1;refGene_
gn=ABLIM1;refGene_tx=XM_019813861.2;refGene_gn=ABLIM1;refGene_tx=XM_01
9813863.2;refGene_gn=ABLIM1;refGene_tx=XM_019813866.2;refGene_gn=ABLIM
1;refGene_tx=XM_019813867.2;refGene_gn=ABLIM1;refGene_tx=XM_019813868.

2;refGene_gn=ABLIM1;refGene_tx=XM_019813869.1;refGene_gn=ABLIM1;refGene_tx=XM_019813874.2;refGene_gn=ABLIM1;refGene_tx=XM_019813880.2;refGene_gn=ABLIM1;refGene_tx=XM_023240969.1
D2 73491025 73491109
refGene_gn=ABLIM1;refGene_tx=XM_019813857.2;refGene_gn=ABLIM1;refGene_tx=XM_019813858.2;refGene_gn=ABLIM1;refGene_tx=XM_019813859.2;refGene_gn=ABLIM1;refGene_tx=XM_019813861.2;refGene_gn=ABLIM1;refGene_tx=XM_019813863.2;refGene_gn=ABLIM1;refGene_tx=XM_019813864.2;refGene_gn=ABLIM1;refGene_tx=XM_019813865.1;refGene_gn=ABLIM1;refGene_tx=XM_019813866.2;refGene_gn=ABLIM1;refGene_tx=XM_019813867.2;refGene_gn=ABLIM1;refGene_tx=XM_019813868.2;refGene_gn=ABLIM1;refGene_tx=XM_019813869.1;refGene_gn=ABLIM1;refGene_tx=XM_019813874.2;refGene_gn=ABLIM1;refGene_tx=XM_019813875.2;refGene_gn=ABLIM1;refGene_tx=XM_019813877.2;refGene_gn=ABLIM1;refGene_tx=XM_019813880.2;refGene_gn=ABLIM1;refGene_tx=XM_023240969.1
D2 73493580 73493639
refGene_gn=ABLIM1;refGene_tx=XM_019813857.2;refGene_gn=ABLIM1;refGene_tx=XM_019813858.2;refGene_gn=ABLIM1;refGene_tx=XM_019813859.2;refGene_gn=ABLIM1;refGene_tx=XM_019813860.1;refGene_gn=ABLIM1;refGene_tx=XM_019813861.2;refGene_gn=ABLIM1;refGene_tx=XM_019813862.2;refGene_gn=ABLIM1;refGene_tx=XM_019813863.2;refGene_gn=ABLIM1;refGene_tx=XM_019813864.2;refGene_gn=ABLIM1;refGene_tx=XM_019813865.1;refGene_gn=ABLIM1;refGene_tx=XM_019813866.2;refGene_gn=ABLIM1;refGene_tx=XM_019813867.2;refGene_gn=ABLIM1;refGene_tx=XM_019813868.2;refGene_gn=ABLIM1;refGene_tx=XM_019813869.1;refGene_gn=ABLIM1;refGene_tx=XM_019813873.2;refGene_gn=ABLIM1;refGene_tx=XM_019813874.2;refGene_gn=ABLIM1;refGene_tx=XM_019813875.2;refGene_gn=ABLIM1;refGene_tx=XM_019813876.2;refGene_gn=ABLIM1;refGene_tx=XM_019813877.2;refGene_gn=ABLIM1;refGene_tx=XM_019813878.2;refGene_gn=ABLIM1;refGene_tx=XM_019813879.2;refGene_gn=ABLIM1;refGene_tx=XM_019813880.2;refGene_gn=ABLIM1;refGene_tx=XM_019813881.2;refGene_gn=ABLIM1;refGene_tx=XM_019813882.2;refGene_gn=ABLIM1;refGene_tx=XM_023240969.1;refGene_gn=ABLIM1;refGene_tx=XM_023240970.1
D2 73497046 73497134
refGene_gn=ABLIM1;refGene_tx=XM_019813857.2;refGene_gn=ABLIM1;refGene_tx=XM_019813858.2;refGene_gn=ABLIM1;refGene_tx=XM_019813859.2;refGene_gn=ABLIM1;refGene_tx=XM_019813860.1;refGene_gn=ABLIM1;refGene_tx=XM_019813861.2;refGene_gn=ABLIM1;refGene_tx=XM_019813862.2;refGene_gn=ABLIM1;refGene_tx=XM_019813863.2;refGene_gn=ABLIM1;refGene_tx=XM_019813864.2;refGene_gn=ABLIM1;refGene_tx=XM_019813865.1;refGene_gn=ABLIM1;refGene_tx=XM_019813866.2;refGene_gn=ABLIM1;refGene_tx=XM_019813867.2;refGene_gn=ABLIM1;refGene_tx=XM_019813868.2;refGene_gn=ABLIM1;refGene_tx=XM_019813869.1;refGene_gn=ABLIM1;refGene_tx=XM_019813873.2;refGene_gn=ABLIM1;refGene_tx=XM_019813874.2;refGene_gn=ABLIM1;refGene_tx=XM_019813875.2;refGene_gn=ABLIM1;refGene_tx=XM_019813876.2;refGene_gn=ABLIM1;refGene_tx=XM_019813877.2;refGene_gn=ABLIM1;refGene_tx=XM_019813878.2;refGene_gn=ABLIM1;refGene_tx=XM_019813879.2;refGene_gn=ABLIM1;refGene_tx=XM_019813880.2;refGene_gn=ABLIM1;refGene_tx=XM_019813881.2;refGene_gn=ABLIM1;refGene_tx=XM_019813882.2;refGene_gn=ABLIM1;refGene_tx=XM_023240969.1;refGene_gn=ABLIM1;refGene_tx=XM_023240970.1
D2 73541835 73541929

refGene_gn=ABLIM1;refGene_tx=XM_019813857.2;refGene_gn=ABLIM1;refGene_tx=XM_019813858.2;refGene_gn=ABLIM1;refGene_tx=XM_019813859.2;refGene_gn=ABLIM1;refGene_tx=XM_019813860.1;refGene_gn=ABLIM1;refGene_tx=XM_019813861.2;refGene_gn=ABLIM1;refGene_tx=XM_019813862.2;refGene_gn=ABLIM1;refGene_tx=XM_019813863.2;refGene_gn=ABLIM1;refGene_tx=XM_019813864.2;refGene_gn=ABLIM1;refGene_tx=XM_019813865.1;refGene_gn=ABLIM1;refGene_tx=XM_019813866.2;refGene_gn=ABLIM1;refGene_tx=XM_019813867.2;refGene_gn=ABLIM1;refGene_tx=XM_019813868.2;refGene_gn=ABLIM1;refGene_tx=XM_019813869.1;refGene_gn=ABLIM1;refGene_tx=XM_019813873.2;refGene_gn=ABLIM1;refGene_tx=XM_019813874.2;refGene_gn=ABLIM1;refGene_tx=XM_019813875.2;refGene_gn=ABLIM1;refGene_tx=XM_019813876.2;refGene_gn=ABLIM1;refGene_tx=XM_019813877.2;refGene_gn=ABLIM1;refGene_tx=XM_019813878.2;refGene_gn=ABLIM1;refGene_tx=XM_019813879.2;refGene_gn=ABLIM1;refGene_tx=XM_019813880.2;refGene_gn=ABLIM1;refGene_tx=XM_019813881.2;refGene_gn=ABLIM1;refGene_tx=XM_019813882.2;refGene_gn=ABLIM1;refGene_tx=XM_023240969.1;refGene_gn=ABLIM1;refGene_tx=XM_023240970.1
D2 73543823 73543950
refGene_gn=ABLIM1;refGene_tx=XM_019813857.2;refGene_gn=ABLIM1;refGene_tx=XM_019813858.2;refGene_gn=ABLIM1;refGene_tx=XM_019813859.2;refGene_gn=ABLIM1;refGene_tx=XM_019813860.1;refGene_gn=ABLIM1;refGene_tx=XM_019813861.2;refGene_gn=ABLIM1;refGene_tx=XM_019813862.2;refGene_gn=ABLIM1;refGene_tx=XM_019813863.2;refGene_gn=ABLIM1;refGene_tx=XM_019813864.2;refGene_gn=ABLIM1;refGene_tx=XM_019813865.1;refGene_gn=ABLIM1;refGene_tx=XM_019813866.2;refGene_gn=ABLIM1;refGene_tx=XM_019813867.2;refGene_gn=ABLIM1;refGene_tx=XM_019813868.2;refGene_gn=ABLIM1;refGene_tx=XM_019813869.1;refGene_gn=ABLIM1;refGene_tx=XM_019813873.2;refGene_gn=ABLIM1;refGene_tx=XM_019813874.2;refGene_gn=ABLIM1;refGene_tx=XM_019813875.2;refGene_gn=ABLIM1;refGene_tx=XM_019813876.2;refGene_gn=ABLIM1;refGene_tx=XM_019813877.2;refGene_gn=ABLIM1;refGene_tx=XM_019813878.2;refGene_gn=ABLIM1;refGene_tx=XM_019813879.2;refGene_gn=ABLIM1;refGene_tx=XM_019813880.2;refGene_gn=ABLIM1;refGene_tx=XM_019813881.2;refGene_gn=ABLIM1;refGene_tx=XM_019813882.2;refGene_gn=ABLIM1;refGene_tx=XM_023240969.1;refGene_gn=ABLIM1;refGene_tx=XM_023240970.1
D2 73563532 73563642
refGene_gn=ABLIM1;refGene_tx=XM_019813857.2;refGene_gn=ABLIM1;refGene_tx=XM_019813858.2;refGene_gn=ABLIM1;refGene_tx=XM_019813859.2;refGene_gn=ABLIM1;refGene_tx=XM_019813860.1;refGene_gn=ABLIM1;refGene_tx=XM_019813861.2;refGene_gn=ABLIM1;refGene_tx=XM_019813862.2;refGene_gn=ABLIM1;refGene_tx=XM_019813863.2;refGene_gn=ABLIM1;refGene_tx=XM_019813864.2;refGene_gn=ABLIM1;refGene_tx=XM_019813865.1;refGene_gn=ABLIM1;refGene_tx=XM_019813866.2;refGene_gn=ABLIM1;refGene_tx=XM_019813867.2;refGene_gn=ABLIM1;refGene_tx=XM_019813868.2;refGene_gn=ABLIM1;refGene_tx=XM_019813869.1;refGene_gn=ABLIM1;refGene_tx=XM_019813873.2;refGene_gn=ABLIM1;refGene_tx=XM_019813874.2;refGene_gn=ABLIM1;refGene_tx=XM_019813875.2;refGene_gn=ABLIM1;refGene_tx=XM_019813876.2;refGene_gn=ABLIM1;refGene_tx=XM_019813877.2;refGene_gn=ABLIM1;refGene_tx=XM_019813878.2;refGene_gn=ABLIM1;refGene_tx=XM_019813879.2;refGene_gn=ABLIM1;refGene_tx=XM_019813880.2;refGene_gn=ABLIM1;refGene_tx=XM_019813881.2;refGene_gn=ABLIM1;refGene_tx=XM_019813882.2;refGene_gn=ABLIM1;refGene_tx=XM_023240969.1;refGene_gn=ABLIM1;refGene_tx=XM_023240970.1

D2	73567796	73567980
refGene_gn=ABLIM1;refGene_tx=XM_019813857.2;refGene_gn=ABLIM1;refGene_tx=XM_019813858.2;refGene_gn=ABLIM1;refGene_tx=XM_019813859.2;refGene_gn=ABLIM1;refGene_tx=XM_019813860.1;refGene_gn=ABLIM1;refGene_tx=XM_019813861.2;refGene_gn=ABLIM1;refGene_tx=XM_019813862.2;refGene_gn=ABLIM1;refGene_tx=XM_019813863.2;refGene_gn=ABLIM1;refGene_tx=XM_019813864.2;refGene_gn=ABLIM1;refGene_tx=XM_019813865.1;refGene_gn=ABLIM1;refGene_tx=XM_019813866.2;refGene_gn=ABLIM1;refGene_tx=XM_019813867.2;refGene_gn=ABLIM1;refGene_tx=XM_019813868.2;refGene_gn=ABLIM1;refGene_tx=XM_019813869.1;refGene_gn=ABLIM1;refGene_tx=XM_019813873.2;refGene_gn=ABLIM1;refGene_tx=XM_019813874.2;refGene_gn=ABLIM1;refGene_tx=XM_019813875.2;refGene_gn=ABLIM1;refGene_tx=XM_019813876.2;refGene_gn=ABLIM1;refGene_tx=XM_019813877.2;refGene_gn=ABLIM1;refGene_tx=XM_019813878.2;refGene_gn=ABLIM1;refGene_tx=XM_019813879.2;refGene_gn=ABLIM1;refGene_tx=XM_019813880.2;refGene_gn=ABLIM1;refGene_tx=XM_019813881.2;refGene_gn=ABLIM1;refGene_tx=XM_019813882.2;refGene_gn=ABLIM1;refGene_tx=XM_023240969.1;refGene_gn=ABLIM1;refGene_tx=XM_023240970.1
D2	73586624	73586759
refGene_gn=ABLIM1;refGene_tx=XM_019813857.2;refGene_gn=ABLIM1;refGene_tx=XM_019813858.2;refGene_gn=ABLIM1;refGene_tx=XM_019813859.2;refGene_gn=ABLIM1;refGene_tx=XM_019813860.1;refGene_gn=ABLIM1;refGene_tx=XM_019813861.2;refGene_gn=ABLIM1;refGene_tx=XM_019813862.2;refGene_gn=ABLIM1;refGene_tx=XM_019813863.2;refGene_gn=ABLIM1;refGene_tx=XM_019813864.2;refGene_gn=ABLIM1;refGene_tx=XM_019813865.1;refGene_gn=ABLIM1;refGene_tx=XM_019813866.2;refGene_gn=ABLIM1;refGene_tx=XM_019813867.2;refGene_gn=ABLIM1;refGene_tx=XM_019813868.2;refGene_gn=ABLIM1;refGene_tx=XM_019813869.1;refGene_gn=ABLIM1;refGene_tx=XM_019813873.2;refGene_gn=ABLIM1;refGene_tx=XM_019813874.2;refGene_gn=ABLIM1;refGene_tx=XM_019813875.2;refGene_gn=ABLIM1;refGene_tx=XM_019813876.2;refGene_gn=ABLIM1;refGene_tx=XM_019813877.2;refGene_gn=ABLIM1;refGene_tx=XM_019813878.2;refGene_gn=ABLIM1;refGene_tx=XM_019813879.2;refGene_gn=ABLIM1;refGene_tx=XM_019813880.2;refGene_gn=ABLIM1;refGene_tx=XM_019813881.2;refGene_gn=ABLIM1;refGene_tx=XM_019813882.2;refGene_gn=ABLIM1;refGene_tx=XM_023240969.1;refGene_gn=ABLIM1;refGene_tx=XM_023240970.1
D2	73613565	73613581	refGene_gn=ABLIM1;refGene_tx=XM_019813869.1
D2	73638547	73638791
refGene_gn=ABLIM1;refGene_tx=XM_019813857.2;refGene_gn=ABLIM1;refGene_tx=XM_019813858.2;refGene_gn=ABLIM1;refGene_tx=XM_019813859.2;refGene_gn=ABLIM1;refGene_tx=XM_019813860.1;refGene_gn=ABLIM1;refGene_tx=XM_019813861.2;refGene_gn=ABLIM1;refGene_tx=XM_019813862.2;refGene_gn=ABLIM1;refGene_tx=XM_019813863.2;refGene_gn=ABLIM1;refGene_tx=XM_019813864.2;refGene_gn=ABLIM1;refGene_tx=XM_019813865.1;refGene_gn=ABLIM1;refGene_tx=XM_019813867.2;refGene_gn=ABLIM1;refGene_tx=XM_019813868.2;refGene_gn=ABLIM1;refGene_tx=XM_019813873.2;refGene_gn=ABLIM1;refGene_tx=XM_019813874.2;refGene_gn=ABLIM1;refGene_tx=XM_019813875.2;refGene_gn=ABLIM1;refGene_tx=XM_019813876.2;refGene_gn=ABLIM1;refGene_tx=XM_019813877.2;refGene_gn=ABLIM1;refGene_tx=XM_019813878.2;refGene_gn=ABLIM1;refGene_tx=XM_019813879.2;refGene_gn=ABLIM1;refGene_tx=XM_019813880.2;refGene_gn=ABLIM1;refGene_tx=XM_019813881.2;refGene_gn=ABLIM1;refGene_tx=XM_019813882.2;refGene_gn=ABLIM1;refGene_tx=XM_023240970.1

D2 73665214 73665272 refGene_gn=ABLIM1;refGene_tx=XM_019813866.2
D2 73751047 73751086 refGene_gn=FAM160B1;refGene_tx=XM_023240972.1
D2 73751606 73751619 refGene_gn=ABLIM1;refGene_tx=XM_023240969.1
D2 73806047 73806092 refGene_gn=FAM160B1;refGene_tx=XM_023240971.1
D2 73812684 73812763
refGene_gn=FAM160B1;refGene_tx=XM_023240971.1;refGene_gn=FAM160B1;refGene_tx=XM_023240972.1
D2 73818783 73818953
refGene_gn=FAM160B1;refGene_tx=XM_023240971.1;refGene_gn=FAM160B1;refGene_tx=XM_023240972.1
D2 73820682 73820787
refGene_gn=FAM160B1;refGene_tx=XM_023240971.1;refGene_gn=FAM160B1;refGene_tx=XM_023240972.1
D2 73821229 73821352
refGene_gn=FAM160B1;refGene_tx=XM_023240971.1;refGene_gn=FAM160B1;refGene_tx=XM_023240972.1
D2 73826894 73827194
refGene_gn=FAM160B1;refGene_tx=XM_023240971.1;refGene_gn=FAM160B1;refGene_tx=XM_023240972.1
D2 73827409 73827606
refGene_gn=FAM160B1;refGene_tx=XM_023240971.1;refGene_gn=FAM160B1;refGene_tx=XM_023240972.1
D2 73828871 73828986
refGene_gn=FAM160B1;refGene_tx=XM_023240971.1;refGene_gn=FAM160B1;refGene_tx=XM_023240972.1
D2 73829534 73829611
refGene_gn=FAM160B1;refGene_tx=XM_023240971.1;refGene_gn=FAM160B1;refGene_tx=XM_023240972.1
D2 73829697 73829890
refGene_gn=FAM160B1;refGene_tx=XM_023240971.1;refGene_gn=FAM160B1;refGene_tx=XM_023240972.1
D2 73830342 73830512
refGene_gn=FAM160B1;refGene_tx=XM_023240971.1;refGene_gn=FAM160B1;refGene_tx=XM_023240972.1
D2 73830890 73831034
refGene_gn=FAM160B1;refGene_tx=XM_023240971.1;refGene_gn=FAM160B1;refGene_tx=XM_023240972.1
D2 73831904 73831995
refGene_gn=FAM160B1;refGene_tx=XM_023240971.1;refGene_gn=FAM160B1;refGene_tx=XM_023240972.1
D2 73835261 73835405
refGene_gn=FAM160B1;refGene_tx=XM_023240971.1;refGene_gn=FAM160B1;refGene_tx=XM_023240972.1
D2 73840407 73840548
refGene_gn=FAM160B1;refGene_tx=XM_023240971.1;refGene_gn=FAM160B1;refGene_tx=XM_023240972.1
D2 73841182 73841286
refGene_gn=FAM160B1;refGene_tx=XM_023240971.1;refGene_gn=FAM160B1;refGene_tx=XM_023240972.1
D2 73841383 73841489

refGene_gn=FAM160B1;refGene_tx=XM_023240971.1;refGene_gn=FAM160B1;refGene_tx=XM_023240972.1
D2 73904619 73904902 refGene_gn=TRUB1;refGene_tx=XM_003994437.5
D2 73908035 73908134 refGene_gn=TRUB1;refGene_tx=XM_003994437.5
D2 73915744 73915800 refGene_gn=TRUB1;refGene_tx=XM_003994437.5
D2 73924495 73924577 refGene_gn=TRUB1;refGene_tx=XM_003994437.5
D2 73935313 73935386 refGene_gn=TRUB1;refGene_tx=XM_003994437.5
D2 73936744 73936884 refGene_gn=TRUB1;refGene_tx=XM_003994437.5
D2 73938951 73939008 refGene_gn=TRUB1;refGene_tx=XM_003994437.5
D2 73939766 73940026 refGene_gn=TRUB1;refGene_tx=XM_003994437.5
D2 74035453 74035770 refGene_gn=ATRNL1;refGene_tx=XM_023240973.1;refGene_gn=ATRNL1;refGene_tx=XM_023240974.1
D2 74066303 74066387 refGene_gn=ATRNL1;refGene_tx=XM_023240973.1;refGene_gn=ATRNL1;refGene_tx=XM_023240974.1
D2 74067803 74067917 refGene_gn=ATRNL1;refGene_tx=XM_023240973.1;refGene_gn=ATRNL1;refGene_tx=XM_023240974.1
D2 74073768 74073897 refGene_gn=ATRNL1;refGene_tx=XM_023240973.1;refGene_gn=ATRNL1;refGene_tx=XM_023240974.1
D2 74075543 74075752 refGene_gn=ATRNL1;refGene_tx=XM_023240973.1;refGene_gn=ATRNL1;refGene_tx=XM_023240974.1
D2 74093670 74093845 refGene_gn=ATRNL1;refGene_tx=XM_023240973.1;refGene_gn=ATRNL1;refGene_tx=XM_023240974.1
D2 74098219 74098307 refGene_gn=ATRNL1;refGene_tx=XM_023240973.1;refGene_gn=ATRNL1;refGene_tx=XM_023240974.1
D2 74104161 74104417 refGene_gn=ATRNL1;refGene_tx=XM_023240973.1;refGene_gn=ATRNL1;refGene_tx=XM_023240974.1
D2 74135720 74135904 refGene_gn=ATRNL1;refGene_tx=XM_023240973.1;refGene_gn=ATRNL1;refGene_tx=XM_023240974.1
D2 74163025 74163180 refGene_gn=ATRNL1;refGene_tx=XM_023240973.1;refGene_gn=ATRNL1;refGene_tx=XM_023240974.1
D2 74185734 74185819 refGene_gn=ATRNL1;refGene_tx=XM_023240973.1;refGene_gn=ATRNL1;refGene_tx=XM_023240974.1
D2 74187141 74187350 refGene_gn=ATRNL1;refGene_tx=XM_023240973.1;refGene_gn=ATRNL1;refGene_tx=XM_023240974.1
D2 74189435 74189554 refGene_gn=ATRNL1;refGene_tx=XM_023240973.1;refGene_gn=ATRNL1;refGene_tx=XM_023240974.1
D2 74193949 74194082

refGene_gn=ATRNL1;refGene_tx=XM_023240973.1;refGene_gn=ATRNL1;refGene_tx=XM_023240974.1
D2	74198478	74198660	refGene_gn=ATRNL1;refGene_tx=XM_023240973.1;refGene_gn=ATRNL1;refGene_tx=XM_023240974.1
D2	74203860	74204074	refGene_gn=ATRNL1;refGene_tx=XM_023240973.1;refGene_gn=ATRNL1;refGene_tx=XM_023240974.1
D2	74206338	74206527	refGene_gn=ATRNL1;refGene_tx=XM_023240973.1;refGene_gn=ATRNL1;refGene_tx=XM_023240974.1
D2	74217921	74218140	refGene_gn=ATRNL1;refGene_tx=XM_023240973.1;refGene_gn=ATRNL1;refGene_tx=XM_023240974.1
D2	74247831	74247969	refGene_gn=ATRNL1;refGene_tx=XM_023240973.1;refGene_gn=ATRNL1;refGene_tx=XM_023240974.1
D2	74276450	74276544	refGene_gn=ATRNL1;refGene_tx=XM_023240973.1;refGene_gn=ATRNL1;refGene_tx=XM_023240974.1
D2	74299601	74299654	refGene_gn=ATRNL1;refGene_tx=XM_023240973.1;refGene_gn=ATRNL1;refGene_tx=XM_023240974.1
D2	74326141	74326236	refGene_gn=ATRNL1;refGene_tx=XM_023240973.1;refGene_gn=ATRNL1;refGene_tx=XM_023240974.1
D2	74341411	74341490	refGene_gn=ATRNL1;refGene_tx=XM_023240973.1;refGene_gn=ATRNL1;refGene_tx=XM_023240974.1
D2	74343499	74343657	refGene_gn=ATRNL1;refGene_tx=XM_023240973.1;refGene_gn=ATRNL1;refGene_tx=XM_023240974.1
D2	74388222	74388284	refGene_gn=ATRNL1;refGene_tx=XM_023240973.1;refGene_gn=ATRNL1;refGene_tx=XM_023240974.1
D2	74410115	74410194	refGene_gn=ATRNL1;refGene_tx=XM_023240973.1;refGene_gn=ATRNL1;refGene_tx=XM_023240974.1
D2	74559427	74559535	refGene_gn=ATRNL1;refGene_tx=XM_023240973.1;refGene_gn=ATRNL1;refGene_tx=XM_023240974.1
D2	74696501	74696616	refGene_gn=ATRNL1;refGene_tx=XM_023240973.1
D2	74804413	74804538	refGene_gn=ATRNL1;refGene_tx=XM_023240973.1;refGene_gn=ATRNL1;refGene_tx=XM_023240974.1
D2	74915955	74916111	refGene_gn=GFRA1;refGene_tx=XM_023240975.1;refGene_gn=GFRA1;refGene_tx=XM_023240977.1;refGene_gn=GFRA1;refGene_tx=XM_023240978.1
D2	74917241	74917295	refGene_gn=GFRA1;refGene_tx=XM_023240975.1;refGene_gn=GFRA1;refGene_tx

=XM_023240977.1;refGene_gn=GFRA1;refGene_tx=XM_023240978.1
D2 74941953 74942135
refGene_gn=GFRA1;refGene_tx=XM_023240975.1;refGene_gn=GFRA1;refGene_tx=XM_023240977.1;refGene_gn=GFRA1;refGene_tx=XM_023240978.1
D2 74946080 74946215
refGene_gn=GFRA1;refGene_tx=XM_023240975.1;refGene_gn=GFRA1;refGene_tx=XM_023240977.1;refGene_gn=GFRA1;refGene_tx=XM_023240978.1
D2 74948893 74949003
refGene_gn=GFRA1;refGene_tx=XM_023240975.1;refGene_gn=GFRA1;refGene_tx=XM_023240977.1;refGene_gn=GFRA1;refGene_tx=XM_023240978.1
D2 74976377 74976714
refGene_gn=GFRA1;refGene_tx=XM_023240975.1;refGene_gn=GFRA1;refGene_tx=XM_023240977.1;refGene_gn=GFRA1;refGene_tx=XM_023240978.1
D2 75061889 75061904
refGene_gn=GFRA1;refGene_tx=XM_023240975.1;refGene_gn=GFRA1;refGene_tx=XM_023240977.1
D2 75120362 75120446
refGene_gn=GFRA1;refGene_tx=XM_023240975.1;refGene_gn=GFRA1;refGene_tx=XM_023240977.1;refGene_gn=GFRA1;refGene_tx=XM_023240978.1
D2 75121636 75121930
refGene_gn=GFRA1;refGene_tx=XM_023240975.1;refGene_gn=GFRA1;refGene_tx=XM_023240977.1;refGene_gn=GFRA1;refGene_tx=XM_023240978.1
D2 75122746 75122786
refGene_gn=GFRA1;refGene_tx=XM_023240975.1;refGene_gn=GFRA1;refGene_tx=XM_023240977.1;refGene_gn=GFRA1;refGene_tx=XM_023240978.1
D2 75168409 75168412 refGene_gn=CCDC172;refGene_tx=XM_003994438.5
D2 75168819 75168958 refGene_gn=CCDC172;refGene_tx=XM_003994438.5
D2 75169139 75169225 refGene_gn=CCDC172;refGene_tx=XM_003994438.5
D2 75193378 75193495 refGene_gn=CCDC172;refGene_tx=XM_003994438.5
D2 75194660 75194826 refGene_gn=CCDC172;refGene_tx=XM_003994438.5
D2 75211872 75211974 refGene_gn=CCDC172;refGene_tx=XM_003994438.5
D2 75212332 75212435 refGene_gn=CCDC172;refGene_tx=XM_003994438.5
D2 75226080 75226168 refGene_gn=CCDC172;refGene_tx=XM_003994438.5
D2 75227002 75227038 refGene_gn=CCDC172;refGene_tx=XM_003994438.5
D2 75270451 75270500
refGene_gn=PNLIPRP3;refGene_tx=XM_003994448.3;refGene_gn=PNLIPRP3;refGene_tx=XM_019813886.2
D2 75280657 75280812
refGene_gn=PNLIPRP3;refGene_tx=XM_003994448.3;refGene_gn=PNLIPRP3;refGene_tx=XM_019813886.2
D2 75291203 75291323
refGene_gn=PNLIPRP3;refGene_tx=XM_003994448.3;refGene_gn=PNLIPRP3;refGene_tx=XM_019813886.2
D2 75291976 75292108
refGene_gn=PNLIPRP3;refGene_tx=XM_003994448.3;refGene_gn=PNLIPRP3;refGene_tx=XM_019813886.2
D2 75294382 75294491
refGene_gn=PNLIPRP3;refGene_tx=XM_003994448.3;refGene_gn=PNLIPRP3;refGene_tx=XM_019813886.2
D2 75298641 75298761

refGene_gn=PNLIPRP3;refGene_tx=XM_003994448.3;refGene_gn=PNLIPRP3;refGene_tx=XM_019813886.2
D2	75298844	75298967	refGene_gn=PNLIPRP3;refGene_tx=XM_003994448.3
D2	75302061	75302180	refGene_gn=PNLIPRP3;refGene_tx=XM_003994448.3;refGene_gn=PNLIPRP3;refGene_tx=XM_019813886.2
D2	75305193	75305326	refGene_gn=PNLIPRP3;refGene_tx=XM_003994448.3;refGene_gn=PNLIPRP3;refGene_tx=XM_019813886.2
D2	75307130	75307242	refGene_gn=PNLIPRP3;refGene_tx=XM_003994448.3;refGene_gn=PNLIPRP3;refGene_tx=XM_019813886.2
D2	75309824	75309992	refGene_gn=PNLIPRP3;refGene_tx=XM_003994448.3;refGene_gn=PNLIPRP3;refGene_tx=XM_019813886.2
D2	75310248	75310330	refGene_gn=PNLIPRP3;refGene_tx=XM_003994448.3;refGene_gn=PNLIPRP3;refGene_tx=XM_019813886.2
D2	75361774	75361777	refGene_gn=PNLIP;refGene_tx=XM_006938207.2
D2	75361919	75361965	refGene_gn=PNLIP;refGene_tx=XM_006938207.2
D2	75363128	75363283	refGene_gn=PNLIP;refGene_tx=XM_006938207.2
D2	75368215	75368338	refGene_gn=PNLIP;refGene_tx=XM_006938207.2
D2	75370896	75371031	refGene_gn=PNLIP;refGene_tx=XM_006938207.2
D2	75372291	75372403	refGene_gn=PNLIP;refGene_tx=XM_006938207.2
D2	75373902	75374022	refGene_gn=PNLIP;refGene_tx=XM_006938207.2
D2	75374138	75374258	refGene_gn=PNLIP;refGene_tx=XM_006938207.2
D2	75374753	75374872	refGene_gn=PNLIP;refGene_tx=XM_006938207.2
D2	75377709	75377839	refGene_gn=PNLIP;refGene_tx=XM_006938207.2
D2	75379115	75379224	refGene_gn=PNLIP;refGene_tx=XM_006938207.2
D2	75379708	75379873	refGene_gn=PNLIP;refGene_tx=XM_006938207.2
D2	75381065	75381129	refGene_gn=PNLIP;refGene_tx=XM_006938207.2
D2	75392517	75392566	refGene_gn=PNLIPRP1;refGene_tx=XM_006938208.2
D2	75393173	75393328	refGene_gn=PNLIPRP1;refGene_tx=XM_006938208.2
D2	75393844	75393970	refGene_gn=PNLIPRP1;refGene_tx=XM_006938208.2
D2	75395563	75395698	refGene_gn=PNLIPRP1;refGene_tx=XM_006938208.2
D2	75397093	75397202	refGene_gn=PNLIPRP1;refGene_tx=XM_006938208.2
D2	75398714	75398834	refGene_gn=PNLIPRP1;refGene_tx=XM_006938208.2
D2	75398937	75399057	refGene_gn=PNLIPRP1;refGene_tx=XM_006938208.2
D2	75400507	75400626	refGene_gn=PNLIPRP1;refGene_tx=XM_006938208.2
D2	75401366	75401496	refGene_gn=PNLIPRP1;refGene_tx=XM_006938208.2
D2	75403695	75403804	refGene_gn=PNLIPRP1;refGene_tx=XM_006938208.2
D2	75404425	75404593	refGene_gn=PNLIPRP1;refGene_tx=XM_006938208.2
D2	75407294	75407358	refGene_gn=PNLIPRP1;refGene_tx=XM_006938208.2
D2	75416144	75416193	refGene_gn=LOC101083714;refGene_tx=XM_003994441.3
D2	75418612	75418767	refGene_gn=LOC101083714;refGene_tx=XM_003994441.3
D2	75420452	75420578	refGene_gn=LOC101083714;refGene_tx=XM_003994441.3
D2	75423303	75423438

refGene_gn=LOC101083714;refGene_tx=XM_003994441.3
D2 75424089 75424201
refGene_gn=LOC101083714;refGene_tx=XM_003994441.3
D2 75425662 75425782
refGene_gn=LOC101083714;refGene_tx=XM_003994441.3
D2 75426988 75427108
refGene_gn=LOC101083714;refGene_tx=XM_003994441.3
D2 75429826 75429945
refGene_gn=LOC101083714;refGene_tx=XM_003994441.3
D2 75431845 75431975
refGene_gn=LOC101083714;refGene_tx=XM_003994441.3
D2 75434399 75434511
refGene_gn=LOC101083714;refGene_tx=XM_003994441.3
D2 75435398 75435566
refGene_gn=LOC101083714;refGene_tx=XM_003994441.3
D2 75438817 75438881
refGene_gn=LOC101083714;refGene_tx=XM_003994441.3
D2 75450772 75450918
refGene_gn=CD2H10orf82;refGene_tx=XM_003994442.4;refGene_gn=CD2H10orf82;refGene_tx=XM_019813888.2;refGene_gn=CD2H10orf82;refGene_tx=XM_019813889.2
D2 75451042 75451143
refGene_gn=CD2H10orf82;refGene_tx=XM_003994442.4;refGene_gn=CD2H10orf82;refGene_tx=XM_019813888.2;refGene_gn=CD2H10orf82;refGene_tx=XM_019813889.2
D2 75451649 75451836
refGene_gn=CD2H10orf82;refGene_tx=XM_003994442.4;refGene_gn=CD2H10orf82;refGene_tx=XM_019813888.2;refGene_gn=CD2H10orf82;refGene_tx=XM_019813889.2
D2 75453271 75453490
refGene_gn=CD2H10orf82;refGene_tx=XM_003994442.4;refGene_gn=CD2H10orf82;refGene_tx=XM_019813888.2;refGene_gn=CD2H10orf82;refGene_tx=XM_019813889.2
D2 75456387 75456445
refGene_gn=CD2H10orf82;refGene_tx=XM_003994442.4;refGene_gn=CD2H10orf82;refGene_tx=XM_019813888.2;refGene_gn=CD2H10orf82;refGene_tx=XM_019813889.2
D2 75462663 75463301
refGene_gn=HSPA12A;refGene_tx=XM_023240980.1;refGene_gn=HSPA12A;refGene_tx=XM_023240981.1
D2 75464444 75464548
refGene_gn=HSPA12A;refGene_tx=XM_023240980.1;refGene_gn=HSPA12A;refGene_tx=XM_023240981.1
D2 75467355 75467614
refGene_gn=HSPA12A;refGene_tx=XM_023240980.1;refGene_gn=HSPA12A;refGene_tx=XM_023240981.1
D2 75468616 75468721
refGene_gn=HSPA12A;refGene_tx=XM_023240980.1;refGene_gn=HSPA12A;refGene_tx=XM_023240981.1
D2 75469265 75469352

refGene_gn=HSPA12A;refGene_tx=XM_023240980.1;refGene_gn=HSPA12A;refGene_tx=XM_023240981.1
D2 75473216 75473388 refGene_gn=HSPA12A;refGene_tx=XM_023240980.1;refGene_gn=HSPA12A;refGene_tx=XM_023240981.1
D2 75481558 75481675 refGene_gn=HSPA12A;refGene_tx=XM_023240980.1;refGene_gn=HSPA12A;refGene_tx=XM_023240981.1
D2 75487026 75487131 refGene_gn=HSPA12A;refGene_tx=XM_023240980.1;refGene_gn=HSPA12A;refGene_tx=XM_023240981.1
D2 75489710 75489897 refGene_gn=HSPA12A;refGene_tx=XM_023240980.1;refGene_gn=HSPA12A;refGene_tx=XM_023240981.1
D2 75493527 75493655 refGene_gn=HSPA12A;refGene_tx=XM_023240980.1;refGene_gn=HSPA12A;refGene_tx=XM_023240981.1
D2 75495378 75495464 refGene_gn=HSPA12A;refGene_tx=XM_023240980.1;refGene_gn=HSPA12A;refGene_tx=XM_023240981.1
D2 75529230 75529273 refGene_gn=HSPA12A;refGene_tx=XM_023240981.1
D2 75614247 75614335 refGene_gn=HSPA12A;refGene_tx=XM_023240980.1
D2 75627785 75627951 refGene_gn=ENO4;refGene_tx=XM_023240985.1;refGene_gn=ENO4;refGene_tx=XM_023240986.1;refGene_gn=ENO4;refGene_tx=XM_023240987.1;refGene_gn=ENO4;refGene_tx=XM_023240988.1;refGene_gn=HSPA12A;refGene_tx=XM_023240980.1
D2 75634819 75634948 refGene_gn=ENO4;refGene_tx=XM_023240985.1;refGene_gn=ENO4;refGene_tx=XM_023240986.1;refGene_gn=ENO4;refGene_tx=XM_023240987.1;refGene_gn=ENO4;refGene_tx=XM_023240988.1
D2 75635714 75635905 refGene_gn=ENO4;refGene_tx=XM_023240985.1;refGene_gn=ENO4;refGene_tx=XM_023240986.1;refGene_gn=ENO4;refGene_tx=XM_023240987.1;refGene_gn=ENO4;refGene_tx=XM_023240988.1
D2 75637801 75637935 refGene_gn=ENO4;refGene_tx=XM_023240985.1;refGene_gn=ENO4;refGene_tx=XM_023240986.1;refGene_gn=ENO4;refGene_tx=XM_023240987.1;refGene_gn=ENO4;refGene_tx=XM_023240988.1
D2 75639584 75639754 refGene_gn=ENO4;refGene_tx=XM_023240985.1;refGene_gn=ENO4;refGene_tx=XM_023240986.1;refGene_gn=ENO4;refGene_tx=XM_023240987.1;refGene_gn=ENO4;refGene_tx=XM_023240988.1
D2 75639843 75639975 refGene_gn=ENO4;refGene_tx=XM_023240985.1;refGene_gn=ENO4;refGene_tx=XM_023240986.1;refGene_gn=ENO4;refGene_tx=XM_023240987.1
D2 75641319 75641373 refGene_gn=ENO4;refGene_tx=XM_023240985.1;refGene_gn=ENO4;refGene_tx=XM_023240986.1;refGene_gn=ENO4;refGene_tx=XM_023240987.1;refGene_gn=ENO4;refGene_tx=XM_023240988.1

D2 75642584 75642641
refGene_gn=ENO4;refGene_tx=XM_023240985.1;refGene_gn=ENO4;refGene_tx=XM_023240986.1;refGene_gn=ENO4;refGene_tx=XM_023240987.1;refGene_gn=ENO4;refGene_tx=XM_023240988.1
D2 75644730 75644898
refGene_gn=ENO4;refGene_tx=XM_023240985.1;refGene_gn=ENO4;refGene_tx=XM_023240986.1;refGene_gn=ENO4;refGene_tx=XM_023240987.1;refGene_gn=ENO4;refGene_tx=XM_023240988.1
D2 75647998 75648124
refGene_gn=ENO4;refGene_tx=XM_023240985.1;refGene_gn=ENO4;refGene_tx=XM_023240986.1;refGene_gn=ENO4;refGene_tx=XM_023240988.1
D2 75649640 75649836
refGene_gn=ENO4;refGene_tx=XM_023240985.1;refGene_gn=ENO4;refGene_tx=XM_023240986.1;refGene_gn=ENO4;refGene_tx=XM_023240987.1;refGene_gn=ENO4;refGene_tx=XM_023240988.1
D2 75652183 75652251
refGene_gn=ENO4;refGene_tx=XM_023240985.1;refGene_gn=ENO4;refGene_tx=XM_023240986.1;refGene_gn=ENO4;refGene_tx=XM_023240987.1;refGene_gn=ENO4;refGene_tx=XM_023240988.1
D2 75652774 75652892
refGene_gn=ENO4;refGene_tx=XM_023240985.1;refGene_gn=ENO4;refGene_tx=XM_023240986.1;refGene_gn=ENO4;refGene_tx=XM_023240987.1;refGene_gn=ENO4;refGene_tx=XM_023240988.1
D2 75655379 75655510
refGene_gn=ENO4;refGene_tx=XM_023240985.1;refGene_gn=ENO4;refGene_tx=XM_023240986.1;refGene_gn=ENO4;refGene_tx=XM_023240987.1;refGene_gn=ENO4;refGene_tx=XM_023240988.1
D2 75660015 75660238
refGene_gn=SHTN1;refGene_tx=XM_023240982.1;refGene_gn=SHTN1;refGene_tx=XM_023240983.1;refGene_gn=SHTN1;refGene_tx=XM_023240984.1
D2 75675636 75675829
refGene_gn=SHTN1;refGene_tx=XM_023240982.1;refGene_gn=SHTN1;refGene_tx=XM_023240983.1
D2 75679878 75679999
refGene_gn=SHTN1;refGene_tx=XM_023240982.1;refGene_gn=SHTN1;refGene_tx=XM_023240983.1
D2 75685743 75685797
refGene_gn=SHTN1;refGene_tx=XM_023240982.1;refGene_gn=SHTN1;refGene_tx=XM_023240983.1;refGene_gn=SHTN1;refGene_tx=XM_023240984.1
D2 75688987 75689097
refGene_gn=SHTN1;refGene_tx=XM_023240982.1;refGene_gn=SHTN1;refGene_tx=XM_023240983.1;refGene_gn=SHTN1;refGene_tx=XM_023240984.1
D2 75695778 75695861
refGene_gn=SHTN1;refGene_tx=XM_023240982.1;refGene_gn=SHTN1;refGene_tx=XM_023240983.1;refGene_gn=SHTN1;refGene_tx=XM_023240984.1
D2 75700937 75701037
refGene_gn=SHTN1;refGene_tx=XM_023240982.1;refGene_gn=SHTN1;refGene_tx=XM_023240983.1;refGene_gn=SHTN1;refGene_tx=XM_023240984.1
D2 75702731 75702885
refGene_gn=SHTN1;refGene_tx=XM_023240982.1;refGene_gn=SHTN1;refGene_tx

=XM_023240983.1;refGene_gn=SHTN1;refGene_tx=XM_023240984.1
D2	75705778	75705925
refGene_gn=SHTN1;refGene_tx=XM_023240982.1;refGene_gn=SHTN1;refGene_tx=XM_023240983.1;refGene_gn=SHTN1;refGene_tx=XM_023240984.1
D2	75712619	75712714
refGene_gn=SHTN1;refGene_tx=XM_023240982.1;refGene_gn=SHTN1;refGene_tx=XM_023240983.1;refGene_gn=SHTN1;refGene_tx=XM_023240984.1
D2	75716312	75716394
refGene_gn=SHTN1;refGene_tx=XM_023240982.1;refGene_gn=SHTN1;refGene_tx=XM_023240983.1;refGene_gn=SHTN1;refGene_tx=XM_023240984.1
D2	75718356	75718454
refGene_gn=SHTN1;refGene_tx=XM_023240982.1;refGene_gn=SHTN1;refGene_tx=XM_023240983.1;refGene_gn=SHTN1;refGene_tx=XM_023240984.1
D2	75720482	75720651
refGene_gn=SHTN1;refGene_tx=XM_023240982.1;refGene_gn=SHTN1;refGene_tx=XM_023240983.1;refGene_gn=SHTN1;refGene_tx=XM_023240984.1
D2	75726480	75726575
refGene_gn=SHTN1;refGene_tx=XM_023240982.1;refGene_gn=SHTN1;refGene_tx=XM_023240984.1
D2	75734044	75734105
refGene_gn=SHTN1;refGene_tx=XM_023240982.1;refGene_gn=SHTN1;refGene_tx=XM_023240983.1;refGene_gn=SHTN1;refGene_tx=XM_023240984.1
D2	75745544	75745597
refGene_gn=SHTN1;refGene_tx=XM_023240982.1;refGene_gn=SHTN1;refGene_tx=XM_023240983.1;refGene_gn=SHTN1;refGene_tx=XM_023240984.1
D2	75760034	75760092
refGene_gn=SHTN1;refGene_tx=XM_023240982.1;refGene_gn=SHTN1;refGene_tx=XM_023240984.1
D2	75856798	75857386	refGene_gn=VAX1;refGene_tx=XM_011287584.3
D2	75859226	75859414	refGene_gn=VAX1;refGene_tx=XM_011287584.3
D2	75860583	75860824	refGene_gn=VAX1;refGene_tx=XM_011287584.3
D2	75919726	75919946	refGene_gn=KCNK18;refGene_tx=XM_003994444.4
D2	75922490	75922619	refGene_gn=KCNK18;refGene_tx=XM_003994444.4
D2	75931466	75932281	refGene_gn=KCNK18;refGene_tx=XM_003994444.4
D2	75958205	75958326	refGene_gn=SLC18A2;refGene_tx=XM_023240989.1
D2	75959976	75960328	refGene_gn=SLC18A2;refGene_tx=XM_023240989.1
D2	75969616	75969675	refGene_gn=SLC18A2;refGene_tx=XM_023240989.1
D2	75969959	75970043	refGene_gn=SLC18A2;refGene_tx=XM_023240989.1
D2	75970373	75970466	refGene_gn=SLC18A2;refGene_tx=XM_023240989.1
D2	75971250	75971340	refGene_gn=SLC18A2;refGene_tx=XM_023240989.1
D2	75971457	75971501	refGene_gn=SLC18A2;refGene_tx=XM_023240989.1
D2	75971574	75971635	refGene_gn=SLC18A2;refGene_tx=XM_023240989.1
D2	75974214	75974310	refGene_gn=SLC18A2;refGene_tx=XM_023240989.1
D2	75984950	75985029	refGene_gn=SLC18A2;refGene_tx=XM_023240989.1
D2	75985267	75985319	refGene_gn=SLC18A2;refGene_tx=XM_023240989.1
D2	75986231	75986295	refGene_gn=SLC18A2;refGene_tx=XM_023240989.1
D2	75988010	75988130	refGene_gn=SLC18A2;refGene_tx=XM_023240989.1
D2	75988271	75988405	refGene_gn=SLC18A2;refGene_tx=XM_023240989.1
D2	75994302	75994407	refGene_gn=SLC18A2;refGene_tx=XM_023240989.1
D2	75999551	76001755	refGene_gn=PDZD8;refGene_tx=XM_004001358.5

D2 76005589 76005752 refGene_gn=PDZD8;refGene_tx=XM_004001358.5
D2 76029703 76029806 refGene_gn=PDZD8;refGene_tx=XM_004001358.5
D2 76044116 76044242 refGene_gn=PDZD8;refGene_tx=XM_004001358.5
D2 76068245 76069114 refGene_gn=PDZD8;refGene_tx=XM_004001358.5
D2 76219433 76219563 refGene_gn=EMX2;refGene_tx=XM_023240990.1
D2 76219565 76219785 refGene_gn=EMX2;refGene_tx=XM_023240990.1
D2 76219786 76219839 refGene_gn=EMX2;refGene_tx=XM_023240990.1
D2 76221726 76221911 refGene_gn=EMX2;refGene_tx=XM_023240990.1
D2 76224150 76224318 refGene_gn=EMX2;refGene_tx=XM_023240990.1
D2 76642331 76642559
refGene_gn=RAB11FIP2;refGene_tx=XM_003994445.5
D2 76648372 76648418
refGene_gn=RAB11FIP2;refGene_tx=XM_003994445.5
D2 76669164 76669303
refGene_gn=RAB11FIP2;refGene_tx=XM_019813894.2
D2 76672043 76672077
refGene_gn=RAB11FIP2;refGene_tx=XM_006938211.4
D2 76672785 76673254
refGene_gn=RAB11FIP2;refGene_tx=XM_003994445.5;refGene_gn=RAB11FIP2;re
fGene_tx=XM_006938211.4;refGene_gn=RAB11FIP2;refGene_tx=XM_019813894.2
D2 76673981 76674424
refGene_gn=RAB11FIP2;refGene_tx=XM_003994445.5;refGene_gn=RAB11FIP2;re
fGene_tx=XM_006938211.4;refGene_gn=RAB11FIP2;refGene_tx=XM_019813894.2
D2 76680132 76680485
refGene_gn=RAB11FIP2;refGene_tx=XM_003994445.5;refGene_gn=RAB11FIP2;re
fGene_tx=XM_006938211.4;refGene_gn=RAB11FIP2;refGene_tx=XM_019813894.2
D2 76929160 76929212
refGene_gn=FAM204A;refGene_tx=XM_003994454.4;refGene_gn=FAM204A;refGen
e_tx=XM_019813895.2
D2 76929507 76929614
refGene_gn=FAM204A;refGene_tx=XM_003994454.4;refGene_gn=FAM204A;refGen
e_tx=XM_019813895.2
D2 76937542 76937632 refGene_gn=FAM204A;refGene_tx=XM_006938216.4
D2 76943970 76944060
refGene_gn=FAM204A;refGene_tx=XM_003994454.4;refGene_gn=FAM204A;refGen
e_tx=XM_006938216.4;refGene_gn=FAM204A;refGene_tx=XM_019813895.2
D2 76955103 76955203
refGene_gn=FAM204A;refGene_tx=XM_003994454.4;refGene_gn=FAM204A;refGen
e_tx=XM_006938216.4;refGene_gn=FAM204A;refGene_tx=XM_019813895.2
D2 76955364 76955395
refGene_gn=FAM204A;refGene_tx=XM_003994454.4;refGene_gn=FAM204A;refGen
e_tx=XM_006938216.4;refGene_gn=FAM204A;refGene_tx=XM_019813895.2
D2 76955534 76955622
refGene_gn=FAM204A;refGene_tx=XM_003994454.4;refGene_gn=FAM204A;refGen
e_tx=XM_006938216.4;refGene_gn=FAM204A;refGene_tx=XM_019813895.2
D2 76956167 76956409
refGene_gn=FAM204A;refGene_tx=XM_003994454.4;refGene_gn=FAM204A;refGen
e_tx=XM_006938216.4;refGene_gn=FAM204A;refGene_tx=XM_019813895.2
D2 76961168 76961241
refGene_gn=FAM204A;refGene_tx=XM_003994454.4;refGene_gn=FAM204A;refGen

e_tx=XM_006938216.4
D2 77192215 77193328 refGene_gn=PRLHR;refGene_tx=XM_023240991.1
D2 77265282 77265323 refGene_gn=CACUL1;refGene_tx=XM_003994457.5
D2 77265755 77265799 refGene_gn=CACUL1;refGene_tx=XM_003994457.5
D2 77272093 77272232 refGene_gn=CACUL1;refGene_tx=XM_003994457.5
D2 77276491 77276581 refGene_gn=CACUL1;refGene_tx=XM_003994457.5
D2 77282206 77282309 refGene_gn=CACUL1;refGene_tx=XM_003994457.5
D2 77284545 77284641 refGene_gn=CACUL1;refGene_tx=XM_003994457.5
D2 77304532 77304635 refGene_gn=CACUL1;refGene_tx=XM_003994457.5
D2 77305550 77305677 refGene_gn=CACUL1;refGene_tx=XM_003994457.5
D2 77332057 77332424 refGene_gn=CACUL1;refGene_tx=XM_003994457.5
D2 77536658 77537483 refGene_gn=NANOS1;refGene_tx=XM_023240992.1
D2 77537484 77537550 refGene_gn=NANOS1;refGene_tx=XM_023240992.1
D2 77543459 77543689 refGene_gn=EIF3A;refGene_tx=XM_011287585.3
D2 77544559 77544741 refGene_gn=EIF3A;refGene_tx=XM_011287585.3
D2 77545476 77545690 refGene_gn=EIF3A;refGene_tx=XM_011287585.3
D2 77549038 77549503 refGene_gn=EIF3A;refGene_tx=XM_011287585.3
D2 77549505 77549699 refGene_gn=EIF3A;refGene_tx=XM_011287585.3
D2 77550530 77550619 refGene_gn=EIF3A;refGene_tx=XM_011287585.3
D2 77554164 77554349 refGene_gn=EIF3A;refGene_tx=XM_011287585.3
D2 77554909 77555063 refGene_gn=EIF3A;refGene_tx=XM_011287585.3
D2 77555571 77555694 refGene_gn=EIF3A;refGene_tx=XM_011287585.3
D2 77556945 77557059 refGene_gn=EIF3A;refGene_tx=XM_011287585.3
D2 77557147 77557252 refGene_gn=EIF3A;refGene_tx=XM_011287585.3
D2 77557682 77558030 refGene_gn=EIF3A;refGene_tx=XM_011287585.3
D2 77558544 77558730 refGene_gn=EIF3A;refGene_tx=XM_011287585.3
D2 77559232 77559349 refGene_gn=EIF3A;refGene_tx=XM_011287585.3
D2 77560481 77560580 refGene_gn=EIF3A;refGene_tx=XM_011287585.3
D2 77560964 77561069 refGene_gn=EIF3A;refGene_tx=XM_011287585.3
D2 77564670 77564842 refGene_gn=EIF3A;refGene_tx=XM_011287585.3
D2 77566557 77566766 refGene_gn=EIF3A;refGene_tx=XM_011287585.3
D2 77568621 77568821 refGene_gn=EIF3A;refGene_tx=XM_011287585.3
D2 77570865 77571029 refGene_gn=EIF3A;refGene_tx=XM_011287585.3
D2 77571448 77571585 refGene_gn=EIF3A;refGene_tx=XM_011287585.3
D2 77571757 77571948 refGene_gn=EIF3A;refGene_tx=XM_011287585.3
D2 77577936 77577985 refGene_gn=EIF3A;refGene_tx=XM_011287585.3
D2 77596533 77596588
refGene_gn=FAM45A;refGene_tx=XM_003994458.4;refGene_gn=FAM45A;refGene_
tx=XM_023240993.1
D2 77599106 77599303
refGene_gn=FAM45A;refGene_tx=XM_003994458.4;refGene_gn=FAM45A;refGene_
tx=XM_023240993.1
D2 77601671 77601751
refGene_gn=FAM45A;refGene_tx=XM_003994458.4;refGene_gn=FAM45A;refGene_
tx=XM_023240993.1
D2 77605606 77605755
refGene_gn=FAM45A;refGene_tx=XM_003994458.4;refGene_gn=FAM45A;refGene_
tx=XM_023240993.1;refGene_gn=FAM45A;refGene_tx=XM_023240994.1
D2 77607729 77607841
refGene_gn=FAM45A;refGene_tx=XM_003994458.4;refGene_gn=FAM45A;refGene_

tx=XM_023240993.1;refGene_gn=FAM45A;refGene_tx=XM_023240994.1
D2 77609554 77609655
refGene_gn=FAM45A;refGene_tx=XM_003994458.4;refGene_gn=FAM45A;refGene_tx=XM_023240993.1;refGene_gn=FAM45A;refGene_tx=XM_023240994.1
D2 77611630 77611738
refGene_gn=FAM45A;refGene_tx=XM_003994458.4;refGene_gn=FAM45A;refGene_tx=XM_023240993.1;refGene_gn=FAM45A;refGene_tx=XM_023240994.1
D2 77614875 77614970
refGene_gn=FAM45A;refGene_tx=XM_003994458.4;refGene_gn=FAM45A;refGene_tx=XM_023240993.1;refGene_gn=FAM45A;refGene_tx=XM_023240994.1
D2 77616293 77616413
refGene_gn=FAM45A;refGene_tx=XM_023240993.1;refGene_gn=FAM45A;refGene_tx=XM_023240994.1
D2 77617442 77617619
refGene_gn=FAM45A;refGene_tx=XM_003994458.4;refGene_gn=FAM45A;refGene_tx=XM_023240993.1;refGene_gn=FAM45A;refGene_tx=XM_023240994.1
D2 77621175 77621253
refGene_gn=SFXN4;refGene_tx=XM_003994482.3;refGene_gn=SFXN4;refGene_tx=XM_006938217.2
D2 77626606 77626724
refGene_gn=SFXN4;refGene_tx=XM_003994482.3;refGene_gn=SFXN4;refGene_tx=XM_006938217.2
D2 77628345 77628431
refGene_gn=SFXN4;refGene_tx=XM_003994482.3;refGene_gn=SFXN4;refGene_tx=XM_006938217.2
D2 77634851 77634967
refGene_gn=SFXN4;refGene_tx=XM_003994482.3;refGene_gn=SFXN4;refGene_tx=XM_006938217.2
D2 77635843 77635916
refGene_gn=SFXN4;refGene_tx=XM_003994482.3;refGene_gn=SFXN4;refGene_tx=XM_006938217.2
D2 77636735 77636801 refGene_gn=SFXN4;refGene_tx=XM_006938217.2
D2 77636933 77636990
refGene_gn=SFXN4;refGene_tx=XM_003994482.3;refGene_gn=SFXN4;refGene_tx=XM_006938217.2
D2 77637075 77637129
refGene_gn=SFXN4;refGene_tx=XM_003994482.3;refGene_gn=SFXN4;refGene_tx=XM_006938217.2
D2 77637846 77637872
refGene_gn=SFXN4;refGene_tx=XM_003994482.3;refGene_gn=SFXN4;refGene_tx=XM_006938217.2
D2 77640145 77640200
refGene_gn=SFXN4;refGene_tx=XM_003994482.3;refGene_gn=SFXN4;refGene_tx=XM_006938217.2
D2 77640286 77640313
refGene_gn=SFXN4;refGene_tx=XM_003994482.3;refGene_gn=SFXN4;refGene_tx=XM_006938217.2
D2 77641702 77641777
refGene_gn=SFXN4;refGene_tx=XM_003994482.3;refGene_gn=SFXN4;refGene_tx=XM_006938217.2

D2 77643607 77643673
refGene_gn=SFXN4;refGene_tx=XM_003994482.3;refGene_gn=SFXN4;refGene_tx=XM_006938217.2
D2 77645067 77645106
refGene_gn=SFXN4;refGene_tx=XM_003994482.3;refGene_gn=SFXN4;refGene_tx=XM_006938217.2
D2 77648310 77648364 refGene_gn=PRDX3;refGene_tx=XM_003994460.3
D2 77648698 77648864 refGene_gn=PRDX3;refGene_tx=XM_003994460.3
D2 77651102 77651206 refGene_gn=PRDX3;refGene_tx=XM_003994460.3
D2 77652018 77652154 refGene_gn=PRDX3;refGene_tx=XM_003994460.3
D2 77653177 77653319 refGene_gn=PRDX3;refGene_tx=XM_003994460.3
D2 77655567 77655703 refGene_gn=PRDX3;refGene_tx=XM_003994460.3
D2 77657331 77657367 refGene_gn=PRDX3;refGene_tx=XM_003994460.3
D2 77681258 77681310
refGene_gn=GRK5;refGene_tx=XM_003994462.5;refGene_gn=GRK5;refGene_tx=XM_023240995.1
D2 77779560 77779656
refGene_gn=GRK5;refGene_tx=XM_003994462.5;refGene_gn=GRK5;refGene_tx=XM_023240995.1
D2 77827694 77827807
refGene_gn=GRK5;refGene_tx=XM_003994462.5;refGene_gn=GRK5;refGene_tx=XM_023240995.1
D2 77839331 77839409 refGene_gn=GRK5;refGene_tx=XM_003994462.5
D2 77865971 77866072
refGene_gn=GRK5;refGene_tx=XM_003994462.5;refGene_gn=GRK5;refGene_tx=XM_023240995.1
D2 77868074 77868167
refGene_gn=GRK5;refGene_tx=XM_003994462.5;refGene_gn=GRK5;refGene_tx=XM_023240995.1
D2 77872801 77872865
refGene_gn=GRK5;refGene_tx=XM_003994462.5;refGene_gn=GRK5;refGene_tx=XM_023240995.1
D2 77874105 77874246
refGene_gn=GRK5;refGene_tx=XM_003994462.5;refGene_gn=GRK5;refGene_tx=XM_023240995.1
D2 77876790 77876981
refGene_gn=GRK5;refGene_tx=XM_003994462.5;refGene_gn=GRK5;refGene_tx=XM_023240995.1
D2 77881057 77881095
refGene_gn=GRK5;refGene_tx=XM_003994462.5;refGene_gn=GRK5;refGene_tx=XM_023240995.1
D2 77882790 77882880
refGene_gn=GRK5;refGene_tx=XM_003994462.5;refGene_gn=GRK5;refGene_tx=XM_023240995.1
D2 77884814 77885023
refGene_gn=GRK5;refGene_tx=XM_003994462.5;refGene_gn=GRK5;refGene_tx=XM_023240995.1
D2 77889426 77889564
refGene_gn=GRK5;refGene_tx=XM_003994462.5;refGene_gn=GRK5;refGene_tx=XM_023240995.1

D2 77890371 77890509
refGene_gn=GRK5;refGene_tx=XM_003994462.5;refGene_gn=GRK5;refGene_tx=XM_023240995.1
D2 77891072 77891204
refGene_gn=GRK5;refGene_tx=XM_003994462.5;refGene_gn=GRK5;refGene_tx=XM_023240995.1
D2 77893416 77893515
refGene_gn=GRK5;refGene_tx=XM_003994462.5;refGene_gn=GRK5;refGene_tx=XM_023240995.1
D2 77940542 77940689
refGene_gn=RGS10;refGene_tx=XM_011287586.3;refGene_gn=RGS10;refGene_tx=XM_023240996.1
D2 77941096 77941156 refGene_gn=RGS10;refGene_tx=XM_023240998.1
D2 77951082 77951154 refGene_gn=RGS10;refGene_tx=XM_023240997.1
D2 77955592 77955736
refGene_gn=RGS10;refGene_tx=XM_011287586.3;refGene_gn=RGS10;refGene_tx=XM_023240996.1;refGene_gn=RGS10;refGene_tx=XM_023240997.1;refGene_gn=RGS10;refGene_tx=XM_023240998.1
D2 77967457 77967544
refGene_gn=RGS10;refGene_tx=XM_011287586.3;refGene_gn=RGS10;refGene_tx=XM_023240996.1;refGene_gn=RGS10;refGene_tx=XM_023240997.1;refGene_gn=RGS10;refGene_tx=XM_023240998.1
D2 77968784 77968903
refGene_gn=RGS10;refGene_tx=XM_011287586.3;refGene_gn=RGS10;refGene_tx=XM_023240996.1;refGene_gn=RGS10;refGene_tx=XM_023240997.1;refGene_gn=RGS10;refGene_tx=XM_023240998.1
D2 77975398 77975405 refGene_gn=RGS10;refGene_tx=XM_011287586.3
D2 77981410 77981459
refGene_gn=RGS10;refGene_tx=XM_023240996.1;refGene_gn=RGS10;refGene_tx=XM_023240997.1;refGene_gn=RGS10;refGene_tx=XM_023240998.1
D2 78002710 78002837
refGene_gn=TIAL1;refGene_tx=XM_011287587.2;refGene_gn=TIAL1;refGene_tx=XM_011287588.3;refGene_gn=TIAL1;refGene_tx=XM_019813902.2;refGene_gn=TIAL1;refGene_tx=XM_019813904.2;refGene_gn=TIAL1;refGene_tx=XM_019813908.2;refGene_gn=TIAL1;refGene_tx=XM_019813909.2;refGene_gn=TIAL1;refGene_tx=XM_023240999.1;refGene_gn=TIAL1;refGene_tx=XM_023241000.1;refGene_gn=TIAL1;refGene_tx=XM_023241001.1
D2 78003644 78003784
refGene_gn=TIAL1;refGene_tx=XM_011287587.2;refGene_gn=TIAL1;refGene_tx=XM_011287588.3;refGene_gn=TIAL1;refGene_tx=XM_019813902.2;refGene_gn=TIAL1;refGene_tx=XM_019813904.2;refGene_gn=TIAL1;refGene_tx=XM_019813908.2;refGene_gn=TIAL1;refGene_tx=XM_019813909.2;refGene_gn=TIAL1;refGene_tx=XM_023240999.1;refGene_gn=TIAL1;refGene_tx=XM_023241000.1;refGene_gn=TIAL1;refGene_tx=XM_023241001.1
D2 78003919 78003937
refGene_gn=TIAL1;refGene_tx=XM_019813906.2;refGene_gn=TIAL1;refGene_tx=XM_019813907.2
D2 78004126 78004250
refGene_gn=TIAL1;refGene_tx=XM_011287587.2;refGene_gn=TIAL1;refGene_tx=XM_011287588.3;refGene_gn=TIAL1;refGene_tx=XM_019813902.2;refGene_gn=

TIAL1;refGene_tx=XM_019813904.2;refGene_gn=TIAL1;refGene_tx=XM_0198139
06.2;refGene_gn=TIAL1;refGene_tx=XM_019813907.2;refGene_gn=TIAL1;refGe
ne_tx=XM_019813908.2;refGene_gn=TIAL1;refGene_tx=XM_019813909.2;refGen
e_gn=TIAL1;refGene_tx=XM_023240999.1;refGene_gn=TIAL1;refGene_tx=XM_02
3241000.1;refGene_gn=TIAL1;refGene_tx=XM_023241001.1
D2 78004484 78004569
refGene_gn=TIAL1;refGene_tx=XM_011287587.2;refGene_gn=TIAL1;refGene_tx
=XM_011287588.3;refGene_gn=TIAL1;refGene_tx=XM_019813902.2;refGene_gn=
TIAL1;refGene_tx=XM_019813904.2;refGene_gn=TIAL1;refGene_tx=XM_0198139
06.2;refGene_gn=TIAL1;refGene_tx=XM_019813907.2;refGene_gn=TIAL1;refGe
ne_tx=XM_019813908.2;refGene_gn=TIAL1;refGene_tx=XM_019813909.2;refGen
e_gn=TIAL1;refGene_tx=XM_023240999.1;refGene_gn=TIAL1;refGene_tx=XM_02
3241000.1;refGene_gn=TIAL1;refGene_tx=XM_023241001.1
D2 78004669 78004765
refGene_gn=TIAL1;refGene_tx=XM_011287587.2;refGene_gn=TIAL1;refGene_tx
=XM_011287588.3;refGene_gn=TIAL1;refGene_tx=XM_019813902.2;refGene_gn=
TIAL1;refGene_tx=XM_019813904.2;refGene_gn=TIAL1;refGene_tx=XM_0198139
06.2;refGene_gn=TIAL1;refGene_tx=XM_019813907.2;refGene_gn=TIAL1;refGe
ne_tx=XM_019813908.2;refGene_gn=TIAL1;refGene_tx=XM_019813909.2;refGen
e_gn=TIAL1;refGene_tx=XM_023240999.1;refGene_gn=TIAL1;refGene_tx=XM_02
3241000.1;refGene_gn=TIAL1;refGene_tx=XM_023241001.1
D2 78005078 78005187
refGene_gn=TIAL1;refGene_tx=XM_011287587.2;refGene_gn=TIAL1;refGene_tx
=XM_011287588.3;refGene_gn=TIAL1;refGene_tx=XM_019813902.2;refGene_gn=
TIAL1;refGene_tx=XM_019813904.2;refGene_gn=TIAL1;refGene_tx=XM_0198139
06.2;refGene_gn=TIAL1;refGene_tx=XM_019813907.2;refGene_gn=TIAL1;refGe
ne_tx=XM_019813908.2;refGene_gn=TIAL1;refGene_tx=XM_019813909.2;refGen
e_gn=TIAL1;refGene_tx=XM_023240999.1;refGene_gn=TIAL1;refGene_tx=XM_02
3241000.1;refGene_gn=TIAL1;refGene_tx=XM_023241001.1
D2 78006353 78006429
refGene_gn=TIAL1;refGene_tx=XM_011287587.2;refGene_gn=TIAL1;refGene_tx
=XM_011287588.3;refGene_gn=TIAL1;refGene_tx=XM_019813902.2;refGene_gn=
TIAL1;refGene_tx=XM_019813904.2;refGene_gn=TIAL1;refGene_tx=XM_0198139
06.2;refGene_gn=TIAL1;refGene_tx=XM_019813907.2;refGene_gn=TIAL1;refGe
ne_tx=XM_019813908.2;refGene_gn=TIAL1;refGene_tx=XM_019813909.2;refGen
e_gn=TIAL1;refGene_tx=XM_023240999.1;refGene_gn=TIAL1;refGene_tx=XM_02
3241000.1;refGene_gn=TIAL1;refGene_tx=XM_023241001.1
D2 78006890 78006892
refGene_gn=TIAL1;refGene_tx=XM_019813908.2;refGene_gn=TIAL1;refGene_tx
=XM_019813909.2
D2 78008324 78008412
refGene_gn=TIAL1;refGene_tx=XM_011287587.2;refGene_gn=TIAL1;refGene_tx
=XM_011287588.3;refGene_gn=TIAL1;refGene_tx=XM_019813902.2;refGene_gn=
TIAL1;refGene_tx=XM_019813904.2;refGene_gn=TIAL1;refGene_tx=XM_0198139
06.2;refGene_gn=TIAL1;refGene_tx=XM_019813907.2;refGene_gn=TIAL1;refGe
ne_tx=XM_023240999.1;refGene_gn=TIAL1;refGene_tx=XM_023241000.1;refGen
e_gn=TIAL1;refGene_tx=XM_023241001.1
D2 78008564 78008619
refGene_gn=TIAL1;refGene_tx=XM_011287587.2;refGene_gn=TIAL1;refGene_tx
=XM_011287588.3;refGene_gn=TIAL1;refGene_tx=XM_019813902.2;refGene_gn=

TIAL1;refGene_tx=XM_019813904.2;refGene_gn=TIAL1;refGene_tx=XM_019813906.2;refGene_gn=TIAL1;refGene_tx=XM_019813907.2;refGene_gn=TIAL1;refGene_tx=XM_023240999.1;refGene_gn=TIAL1;refGene_tx=XM_023241000.1;refGene_gn=TIAL1;refGene_tx=XM_023241001.1
D2 78008849 78008999
refGene_gn=TIAL1;refGene_tx=XM_011287587.2;refGene_gn=TIAL1;refGene_tx=XM_011287588.3;refGene_gn=TIAL1;refGene_tx=XM_019813902.2;refGene_gn=TIAL1;refGene_tx=XM_019813904.2;refGene_gn=TIAL1;refGene_tx=XM_019813906.2;refGene_gn=TIAL1;refGene_tx=XM_019813907.2;refGene_gn=TIAL1;refGene_tx=XM_023240999.1;refGene_gn=TIAL1;refGene_tx=XM_023241000.1;refGene_gn=TIAL1;refGene_tx=XM_023241001.1
D2 78013941 78014038
refGene_gn=TIAL1;refGene_tx=XM_011287587.2;refGene_gn=TIAL1;refGene_tx=XM_011287588.3;refGene_gn=TIAL1;refGene_tx=XM_019813902.2;refGene_gn=TIAL1;refGene_tx=XM_019813904.2;refGene_gn=TIAL1;refGene_tx=XM_019813906.2;refGene_gn=TIAL1;refGene_tx=XM_019813907.2;refGene_gn=TIAL1;refGene_tx=XM_023240999.1;refGene_gn=TIAL1;refGene_tx=XM_023241000.1;refGene_gn=TIAL1;refGene_tx=XM_023241001.1
D2 78020889 78020921
refGene_gn=TIAL1;refGene_tx=XM_011287587.2;refGene_gn=TIAL1;refGene_tx=XM_019813906.2;refGene_gn=TIAL1;refGene_tx=XM_019813907.2;refGene_gn=TIAL1;refGene_tx=XM_023240999.1
D2 78068922 78069108
refGene_gn=BAG3;refGene_tx=XM_023241002.1;refGene_gn=BAG3;refGene_tx=XM_023241004.1
D2 78085440 78085770
refGene_gn=BAG3;refGene_tx=XM_023241002.1;refGene_gn=BAG3;refGene_tx=XM_023241004.1
D2 78087113 78087518
refGene_gn=BAG3;refGene_tx=XM_023241002.1;refGene_gn=BAG3;refGene_tx=XM_023241004.1
D2 78091470 78092289
refGene_gn=BAG3;refGene_tx=XM_023241002.1;refGene_gn=BAG3;refGene_tx=XM_023241004.1
D2 78141974 78142071
refGene_gn=INPP5F;refGene_tx=XM_023241005.1;refGene_gn=INPP5F;refGene_tx=XM_023241006.1
D2 78163101 78163182
refGene_gn=INPP5F;refGene_tx=XM_023241005.1;refGene_gn=INPP5F;refGene_tx=XM_023241006.1
D2 78181464 78181601
refGene_gn=INPP5F;refGene_tx=XM_023241005.1;refGene_gn=INPP5F;refGene_tx=XM_023241006.1
D2 78190980 78191109
refGene_gn=INPP5F;refGene_tx=XM_023241005.1;refGene_gn=INPP5F;refGene_tx=XM_023241006.1
D2 78191318 78191486
refGene_gn=INPP5F;refGene_tx=XM_023241005.1;refGene_gn=INPP5F;refGene_tx=XM_023241006.1
D2 78191602 78191659

refGene_gn=INPP5F;refGene_tx=XM_023241005.1;refGene_gn=INPP5F;refGene_tx=XM_023241006.1
D2 78194226 78194425
refGene_gn=INPP5F;refGene_tx=XM_023241005.1;refGene_gn=INPP5F;refGene_tx=XM_023241006.1
D2 78195192 78195372 refGene_gn=INPP5F;refGene_tx=XM_023241005.1
D2 78196277 78196345 refGene_gn=INPP5F;refGene_tx=XM_023241005.1
D2 78203353 78203478 refGene_gn=INPP5F;refGene_tx=XM_023241005.1
D2 78204250 78204328 refGene_gn=INPP5F;refGene_tx=XM_023241005.1
D2 78205281 78205402 refGene_gn=INPP5F;refGene_tx=XM_023241005.1
D2 78206639 78206768 refGene_gn=INPP5F;refGene_tx=XM_023241005.1
D2 78213165 78213283 refGene_gn=INPP5F;refGene_tx=XM_023241005.1
D2 78215443 78215642 refGene_gn=INPP5F;refGene_tx=XM_023241005.1
D2 78223410 78223482 refGene_gn=INPP5F;refGene_tx=XM_023241005.1
D2 78224522 78224596 refGene_gn=INPP5F;refGene_tx=XM_023241005.1
D2 78225032 78225161 refGene_gn=INPP5F;refGene_tx=XM_023241005.1
D2 78225771 78225859 refGene_gn=INPP5F;refGene_tx=XM_023241005.1
D2 78228362 78229515 refGene_gn=INPP5F;refGene_tx=XM_023241005.1
D2 78233204 78233331
refGene_gn=MCMBP;refGene_tx=XM_023241007.1;refGene_gn=MCMBP;refGene_tx=XM_023241008.1
D2 78233706 78233795
refGene_gn=MCMBP;refGene_tx=XM_023241007.1;refGene_gn=MCMBP;refGene_tx=XM_023241008.1
D2 78236513 78236678
refGene_gn=MCMBP;refGene_tx=XM_023241007.1;refGene_gn=MCMBP;refGene_tx=XM_023241008.1
D2 78237887 78238021
refGene_gn=MCMBP;refGene_tx=XM_023241007.1;refGene_gn=MCMBP;refGene_tx=XM_023241008.1
D2 78238925 78239091
refGene_gn=MCMBP;refGene_tx=XM_023241007.1;refGene_gn=MCMBP;refGene_tx=XM_023241008.1
D2 78241804 78241922
refGene_gn=MCMBP;refGene_tx=XM_023241007.1;refGene_gn=MCMBP;refGene_tx=XM_023241008.1
D2 78243720 78243844
refGene_gn=MCMBP;refGene_tx=XM_023241007.1;refGene_gn=MCMBP;refGene_tx=XM_023241008.1
D2 78244211 78244390
refGene_gn=MCMBP;refGene_tx=XM_023241007.1;refGene_gn=MCMBP;refGene_tx=XM_023241008.1
D2 78247635 78247736
refGene_gn=MCMBP;refGene_tx=XM_023241007.1;refGene_gn=MCMBP;refGene_tx=XM_023241008.1
D2 78249115 78249267
refGene_gn=MCMBP;refGene_tx=XM_023241007.1;refGene_gn=MCMBP;refGene_tx=XM_023241008.1
D2 78251891 78252036
refGene_gn=MCMBP;refGene_tx=XM_023241007.1;refGene_gn=MCMBP;refGene_tx

=XM_023241008.1
D2 78256575 78256677
refGene_gn=MCMBP;refGene_tx=XM_023241007.1;refGene_gn=MCMBP;refGene_tx=XM_023241008.1
D2 78258903 78258945
refGene_gn=MCMBP;refGene_tx=XM_023241007.1;refGene_gn=MCMBP;refGene_tx=XM_023241008.1
D2 78259074 78259215
refGene_gn=MCMBP;refGene_tx=XM_023241007.1;refGene_gn=MCMBP;refGene_tx=XM_023241008.1
D2 78260049 78260135
refGene_gn=MCMBP;refGene_tx=XM_023241007.1;refGene_gn=MCMBP;refGene_tx=XM_023241008.1
D2 78280708 78280766
refGene_gn=MCMBP;refGene_tx=XM_023241007.1;refGene_gn=MCMBP;refGene_tx=XM_023241008.1
D2 78293879 78294042
refGene_gn=SEC23IP;refGene_tx=XM_011287592.3;refGene_gn=SEC23IP;refGene_tx=XM_023241009.1;refGene_gn=SEC23IP;refGene_tx=XM_023241010.1;refGene_gn=SEC23IP;refGene_tx=XM_023241011.1
D2 78297362 78297895
refGene_gn=SEC23IP;refGene_tx=XM_011287592.3;refGene_gn=SEC23IP;refGene_tx=XM_023241009.1;refGene_gn=SEC23IP;refGene_tx=XM_023241010.1;refGene_gn=SEC23IP;refGene_tx=XM_023241011.1
D2 78301194 78301417
refGene_gn=SEC23IP;refGene_tx=XM_011287592.3;refGene_gn=SEC23IP;refGene_tx=XM_023241009.1;refGene_gn=SEC23IP;refGene_tx=XM_023241010.1;refGene_gn=SEC23IP;refGene_tx=XM_023241011.1
D2 78302592 78302786
refGene_gn=SEC23IP;refGene_tx=XM_011287592.3;refGene_gn=SEC23IP;refGene_tx=XM_023241009.1;refGene_gn=SEC23IP;refGene_tx=XM_023241010.1;refGene_gn=SEC23IP;refGene_tx=XM_023241011.1
D2 78304739 78304829
refGene_gn=SEC23IP;refGene_tx=XM_011287592.3;refGene_gn=SEC23IP;refGene_tx=XM_023241009.1;refGene_gn=SEC23IP;refGene_tx=XM_023241010.1;refGene_gn=SEC23IP;refGene_tx=XM_023241011.1
D2 78306915 78307036
refGene_gn=SEC23IP;refGene_tx=XM_011287592.3;refGene_gn=SEC23IP;refGene_tx=XM_023241009.1;refGene_gn=SEC23IP;refGene_tx=XM_023241010.1;refGene_gn=SEC23IP;refGene_tx=XM_023241011.1
D2 78308549 78308639
refGene_gn=SEC23IP;refGene_tx=XM_011287592.3;refGene_gn=SEC23IP;refGene_tx=XM_023241009.1;refGene_gn=SEC23IP;refGene_tx=XM_023241010.1;refGene_gn=SEC23IP;refGene_tx=XM_023241011.1
D2 78309892 78310034
refGene_gn=SEC23IP;refGene_tx=XM_011287592.3;refGene_gn=SEC23IP;refGene_tx=XM_023241009.1;refGene_gn=SEC23IP;refGene_tx=XM_023241010.1;refGene_gn=SEC23IP;refGene_tx=XM_023241011.1
D2 78311424 78311633
refGene_gn=SEC23IP;refGene_tx=XM_011287592.3;refGene_gn=SEC23IP;refGen

e_tx=XM_023241009.1;refGene_gn=SEC23IP;refGene_tx=XM_023241010.1;refGene_gn=SEC23IP;refGene_tx=XM_023241011.1
D2	78312214	78312333
refGene_gn=SEC23IP;refGene_tx=XM_011287592.3;refGene_gn=SEC23IP;refGene_tx=XM_023241009.1;refGene_gn=SEC23IP;refGene_tx=XM_023241010.1;refGene_gn=SEC23IP;refGene_tx=XM_023241011.1
D2	78312860	78313013
refGene_gn=SEC23IP;refGene_tx=XM_011287592.3;refGene_gn=SEC23IP;refGene_tx=XM_023241009.1;refGene_gn=SEC23IP;refGene_tx=XM_023241010.1;refGene_gn=SEC23IP;refGene_tx=XM_023241011.1
D2	78314284	78314380
refGene_gn=SEC23IP;refGene_tx=XM_011287592.3;refGene_gn=SEC23IP;refGene_tx=XM_023241009.1;refGene_gn=SEC23IP;refGene_tx=XM_023241010.1;refGene_gn=SEC23IP;refGene_tx=XM_023241011.1
D2	78319246	78319444
refGene_gn=SEC23IP;refGene_tx=XM_011287592.3;refGene_gn=SEC23IP;refGene_tx=XM_023241009.1;refGene_gn=SEC23IP;refGene_tx=XM_023241010.1;refGene_gn=SEC23IP;refGene_tx=XM_023241011.1
D2	78324605	78324761
refGene_gn=SEC23IP;refGene_tx=XM_011287592.3;refGene_gn=SEC23IP;refGene_tx=XM_023241009.1;refGene_gn=SEC23IP;refGene_tx=XM_023241010.1;refGene_gn=SEC23IP;refGene_tx=XM_023241011.1
D2	78325185	78325288
refGene_gn=SEC23IP;refGene_tx=XM_011287592.3;refGene_gn=SEC23IP;refGene_tx=XM_023241009.1;refGene_gn=SEC23IP;refGene_tx=XM_023241010.1;refGene_gn=SEC23IP;refGene_tx=XM_023241011.1
D2	78326999	78327185
refGene_gn=SEC23IP;refGene_tx=XM_011287592.3;refGene_gn=SEC23IP;refGene_tx=XM_023241009.1;refGene_gn=SEC23IP;refGene_tx=XM_023241010.1;refGene_gn=SEC23IP;refGene_tx=XM_023241011.1
D2	78327853	78328016
refGene_gn=SEC23IP;refGene_tx=XM_011287592.3;refGene_gn=SEC23IP;refGene_tx=XM_023241009.1;refGene_gn=SEC23IP;refGene_tx=XM_023241010.1;refGene_gn=SEC23IP;refGene_tx=XM_023241011.1
D2	78328494	78328576
refGene_gn=SEC23IP;refGene_tx=XM_011287592.3;refGene_gn=SEC23IP;refGene_tx=XM_023241009.1;refGene_gn=SEC23IP;refGene_tx=XM_023241010.1;refGene_gn=SEC23IP;refGene_tx=XM_023241011.1
D2	78685361	78685390
refGene_gn=PLPP4;refGene_tx=XM_019813914.1;refGene_gn=PLPP4;refGene_tx=XM_019813916.2
D2	78685849	78685942
refGene_gn=PLPP4;refGene_tx=XM_019813914.1;refGene_gn=PLPP4;refGene_tx=XM_019813916.2;refGene_gn=PLPP4;refGene_tx=XM_023241014.1
D2	78787545	78787601
refGene_gn=PLPP4;refGene_tx=XM_023241012.1;refGene_gn=PLPP4;refGene_tx=XM_023241015.1
D2	78834025	78834134
refGene_gn=PLPP4;refGene_tx=XM_019813914.1;refGene_gn=PLPP4;refGene_tx=XM_019813916.2;refGene_gn=PLPP4;refGene_tx=XM_023241012.1;refGene_gn=

PLPP4;refGene_tx=XM_023241014.1;refGene_gn=PLPP4;refGene_tx=XM_0232410
15.1
D2 78843833 78843924
refGene_gn=PLPP4;refGene_tx=XM_019813914.1;refGene_gn=PLPP4;refGene_tx
=XM_019813916.2;refGene_gn=PLPP4;refGene_tx=XM_023241012.1
D2 78848690 78848754
refGene_gn=PLPP4;refGene_tx=XM_019813914.1;refGene_gn=PLPP4;refGene_tx
=XM_019813916.2;refGene_gn=PLPP4;refGene_tx=XM_023241012.1;refGene_gn=
PLPP4;refGene_tx=XM_023241014.1
D2 78850341 78850466
refGene_gn=PLPP4;refGene_tx=XM_019813914.1;refGene_gn=PLPP4;refGene_tx
=XM_019813916.2;refGene_gn=PLPP4;refGene_tx=XM_023241012.1;refGene_gn=
PLPP4;refGene_tx=XM_023241014.1
D2 78904174 78904345
refGene_gn=PLPP4;refGene_tx=XM_019813914.1;refGene_gn=PLPP4;refGene_tx
=XM_023241012.1;refGene_gn=PLPP4;refGene_tx=XM_023241014.1;refGene_gn=
PLPP4;refGene_tx=XM_023241015.1
D2 78915034 78915234
refGene_gn=PLPP4;refGene_tx=XM_019813914.1;refGene_gn=PLPP4;refGene_tx
=XM_019813916.2;refGene_gn=PLPP4;refGene_tx=XM_023241012.1;refGene_gn=
PLPP4;refGene_tx=XM_023241014.1
D2 79113492 79113578 refGene_gn=WDR11;refGene_tx=XM_011287593.2
D2 79114411 79114523 refGene_gn=WDR11;refGene_tx=XM_011287593.2
D2 79119792 79119946 refGene_gn=WDR11;refGene_tx=XM_011287593.2
D2 79120857 79121031 refGene_gn=WDR11;refGene_tx=XM_011287593.2
D2 79123058 79123245 refGene_gn=WDR11;refGene_tx=XM_011287593.2
D2 79125720 79125886 refGene_gn=WDR11;refGene_tx=XM_011287593.2
D2 79126312 79126427 refGene_gn=WDR11;refGene_tx=XM_011287593.2
D2 79127454 79127650 refGene_gn=WDR11;refGene_tx=XM_011287593.2
D2 79127946 79128050 refGene_gn=WDR11;refGene_tx=XM_011287593.2
D2 79131469 79131646 refGene_gn=WDR11;refGene_tx=XM_011287593.2
D2 79134842 79134927 refGene_gn=WDR11;refGene_tx=XM_011287593.2
D2 79149049 79149156 refGene_gn=WDR11;refGene_tx=XM_011287593.2
D2 79151340 79151416 refGene_gn=WDR11;refGene_tx=XM_011287593.2
D2 79156163 79156272 refGene_gn=WDR11;refGene_tx=XM_011287593.2
D2 79158075 79158200 refGene_gn=WDR11;refGene_tx=XM_011287593.2
D2 79159158 79159306 refGene_gn=WDR11;refGene_tx=XM_011287593.2
D2 79163458 79163565 refGene_gn=WDR11;refGene_tx=XM_011287593.2
D2 79163978 79164093 refGene_gn=WDR11;refGene_tx=XM_011287593.2
D2 79164394 79164566 refGene_gn=WDR11;refGene_tx=XM_011287593.2
D2 79172542 79172651 refGene_gn=WDR11;refGene_tx=XM_011287593.2
D2 79173534 79173597 refGene_gn=WDR11;refGene_tx=XM_011287593.2
D2 79174800 79174866 refGene_gn=WDR11;refGene_tx=XM_011287593.2
D2 79175892 79176058 refGene_gn=WDR11;refGene_tx=XM_011287593.2
D2 79176716 79176812 refGene_gn=WDR11;refGene_tx=XM_011287593.2
D2 79177340 79177506 refGene_gn=WDR11;refGene_tx=XM_011287593.2
D2 79177979 79178077 refGene_gn=WDR11;refGene_tx=XM_011287593.2
D2 79178522 79178668 refGene_gn=WDR11;refGene_tx=XM_011287593.2
D2 79179624 79179704 refGene_gn=WDR11;refGene_tx=XM_011287593.2
D2 79180873 79181031 refGene_gn=WDR11;refGene_tx=XM_011287593.2

D2	79667234	79667399
refGene_gn=FGFR2;refGene_tx=XM_019813918.2;refGene_gn=FGFR2;refGene_tx=XM_019813921.2;refGene_gn=FGFR2;refGene_tx=XM_019813922.2;refGene_gn=FGFR2;refGene_tx=XM_019813923.2;refGene_gn=FGFR2;refGene_tx=XM_023241016.1;refGene_gn=FGFR2;refGene_tx=XM_023241017.1;refGene_gn=FGFR2;refGene_tx=XM_023241018.1;refGene_gn=FGFR2;refGene_tx=XM_023241019.1;refGene_gn=FGFR2;refGene_tx=XM_023241020.1
D2	79670455	79670561
refGene_gn=FGFR2;refGene_tx=XM_019813918.2;refGene_gn=FGFR2;refGene_tx=XM_019813921.2;refGene_gn=FGFR2;refGene_tx=XM_019813922.2;refGene_gn=FGFR2;refGene_tx=XM_019813923.2;refGene_gn=FGFR2;refGene_tx=XM_023241016.1;refGene_gn=FGFR2;refGene_tx=XM_023241017.1;refGene_gn=FGFR2;refGene_tx=XM_023241018.1;refGene_gn=FGFR2;refGene_tx=XM_023241019.1;refGene_gn=FGFR2;refGene_tx=XM_023241020.1
D2	79672206	79672344
refGene_gn=FGFR2;refGene_tx=XM_019813918.2;refGene_gn=FGFR2;refGene_tx=XM_019813921.2;refGene_gn=FGFR2;refGene_tx=XM_019813922.2;refGene_gn=FGFR2;refGene_tx=XM_019813923.2;refGene_gn=FGFR2;refGene_tx=XM_023241016.1;refGene_gn=FGFR2;refGene_tx=XM_023241017.1;refGene_gn=FGFR2;refGene_tx=XM_023241018.1;refGene_gn=FGFR2;refGene_tx=XM_023241019.1;refGene_gn=FGFR2;refGene_tx=XM_023241020.1
D2	79674576	79674647
refGene_gn=FGFR2;refGene_tx=XM_019813918.2;refGene_gn=FGFR2;refGene_tx=XM_019813921.2;refGene_gn=FGFR2;refGene_tx=XM_019813922.2;refGene_gn=FGFR2;refGene_tx=XM_019813923.2;refGene_gn=FGFR2;refGene_tx=XM_023241016.1;refGene_gn=FGFR2;refGene_tx=XM_023241017.1;refGene_gn=FGFR2;refGene_tx=XM_023241018.1;refGene_gn=FGFR2;refGene_tx=XM_023241019.1;refGene_gn=FGFR2;refGene_tx=XM_023241020.1
D2	79675397	79675520
refGene_gn=FGFR2;refGene_tx=XM_019813918.2;refGene_gn=FGFR2;refGene_tx=XM_019813921.2;refGene_gn=FGFR2;refGene_tx=XM_019813922.2;refGene_gn=FGFR2;refGene_tx=XM_019813923.2;refGene_gn=FGFR2;refGene_tx=XM_023241016.1;refGene_gn=FGFR2;refGene_tx=XM_023241017.1;refGene_gn=FGFR2;refGene_tx=XM_023241018.1;refGene_gn=FGFR2;refGene_tx=XM_023241019.1;refGene_gn=FGFR2;refGene_tx=XM_023241020.1
D2	79683183	79683374
refGene_gn=FGFR2;refGene_tx=XM_019813918.2;refGene_gn=FGFR2;refGene_tx=XM_019813921.2;refGene_gn=FGFR2;refGene_tx=XM_019813922.2;refGene_gn=FGFR2;refGene_tx=XM_019813923.2;refGene_gn=FGFR2;refGene_tx=XM_023241016.1;refGene_gn=FGFR2;refGene_tx=XM_023241017.1;refGene_gn=FGFR2;refGene_tx=XM_023241018.1;refGene_gn=FGFR2;refGene_tx=XM_023241019.1;refGene_gn=FGFR2;refGene_tx=XM_023241020.1
D2	79684852	79684963
refGene_gn=FGFR2;refGene_tx=XM_019813918.2;refGene_gn=FGFR2;refGene_tx=XM_019813921.2;refGene_gn=FGFR2;refGene_tx=XM_019813922.2;refGene_gn=FGFR2;refGene_tx=XM_019813923.2;refGene_gn=FGFR2;refGene_tx=XM_023241016.1;refGene_gn=FGFR2;refGene_tx=XM_023241017.1;refGene_gn=FGFR2;refGene_tx=XM_023241018.1;refGene_gn=FGFR2;refGene_tx=XM_023241019.1;refGene_gn=FGFR2;refGene_tx=XM_023241020.1
D2	79687269	79687391

refGene_gn=FGFR2;refGene_tx=XM_019813918.2;refGene_gn=FGFR2;refGene_tx=XM_019813921.2;refGene_gn=FGFR2;refGene_tx=XM_019813922.2;refGene_gn=FGFR2;refGene_tx=XM_019813923.2;refGene_gn=FGFR2;refGene_tx=XM_023241016.1;refGene_gn=FGFR2;refGene_tx=XM_023241017.1;refGene_gn=FGFR2;refGene_tx=XM_023241018.1;refGene_gn=FGFR2;refGene_tx=XM_023241019.1;refGene_gn=FGFR2;refGene_tx=XM_023241020.1
D2 79690266 79690418
refGene_gn=FGFR2;refGene_tx=XM_019813918.2;refGene_gn=FGFR2;refGene_tx=XM_019813921.2;refGene_gn=FGFR2;refGene_tx=XM_019813922.2;refGene_gn=FGFR2;refGene_tx=XM_019813923.2;refGene_gn=FGFR2;refGene_tx=XM_023241016.1;refGene_gn=FGFR2;refGene_tx=XM_023241017.1;refGene_gn=FGFR2;refGene_tx=XM_023241018.1;refGene_gn=FGFR2;refGene_tx=XM_023241019.1;refGene_gn=FGFR2;refGene_tx=XM_023241020.1
D2 79701221 79701424
refGene_gn=FGFR2;refGene_tx=XM_019813918.2;refGene_gn=FGFR2;refGene_tx=XM_019813921.2;refGene_gn=FGFR2;refGene_tx=XM_019813922.2;refGene_gn=FGFR2;refGene_tx=XM_019813923.2;refGene_gn=FGFR2;refGene_tx=XM_023241016.1;refGene_gn=FGFR2;refGene_tx=XM_023241017.1;refGene_gn=FGFR2;refGene_tx=XM_023241018.1;refGene_gn=FGFR2;refGene_tx=XM_023241019.1;refGene_gn=FGFR2;refGene_tx=XM_023241020.1
D2 79703470 79703615
refGene_gn=FGFR2;refGene_tx=XM_019813921.2;refGene_gn=FGFR2;refGene_tx=XM_019813923.2;refGene_gn=FGFR2;refGene_tx=XM_023241016.1;refGene_gn=FGFR2;refGene_tx=XM_023241018.1;refGene_gn=FGFR2;refGene_tx=XM_023241020.1
D2 79704870 79705018
refGene_gn=FGFR2;refGene_tx=XM_019813918.2;refGene_gn=FGFR2;refGene_tx=XM_019813922.2;refGene_gn=FGFR2;refGene_tx=XM_023241017.1;refGene_gn=FGFR2;refGene_tx=XM_023241019.1
D2 79706122 79706313
refGene_gn=FGFR2;refGene_tx=XM_019813918.2;refGene_gn=FGFR2;refGene_tx=XM_019813921.2;refGene_gn=FGFR2;refGene_tx=XM_019813922.2;refGene_gn=FGFR2;refGene_tx=XM_019813923.2;refGene_gn=FGFR2;refGene_tx=XM_023241016.1;refGene_gn=FGFR2;refGene_tx=XM_023241017.1;refGene_gn=FGFR2;refGene_tx=XM_023241018.1;refGene_gn=FGFR2;refGene_tx=XM_023241019.1;refGene_gn=FGFR2;refGene_tx=XM_023241020.1
D2 79721975 79722099
refGene_gn=FGFR2;refGene_tx=XM_019813918.2;refGene_gn=FGFR2;refGene_tx=XM_019813921.2;refGene_gn=FGFR2;refGene_tx=XM_019813922.2;refGene_gn=FGFR2;refGene_tx=XM_019813923.2;refGene_gn=FGFR2;refGene_tx=XM_023241016.1;refGene_gn=FGFR2;refGene_tx=XM_023241017.1;refGene_gn=FGFR2;refGene_tx=XM_023241018.1;refGene_gn=FGFR2;refGene_tx=XM_023241019.1;refGene_gn=FGFR2;refGene_tx=XM_023241020.1
D2 79736904 79737074
refGene_gn=FGFR2;refGene_tx=XM_019813918.2;refGene_gn=FGFR2;refGene_tx=XM_019813921.2;refGene_gn=FGFR2;refGene_tx=XM_019813922.2;refGene_gn=FGFR2;refGene_tx=XM_019813923.2;refGene_gn=FGFR2;refGene_tx=XM_023241016.1;refGene_gn=FGFR2;refGene_tx=XM_023241017.1;refGene_gn=FGFR2;refGene_tx=XM_023241018.1;refGene_gn=FGFR2;refGene_tx=XM_023241019.1;refGene_gn=FGFR2;refGene_tx=XM_023241020.1

D2 79747711 79747786
refGene_gn=FGFR2;refGene_tx=XM_019813918.2;refGene_gn=FGFR2;refGene_tx
=XM_019813921.2;refGene_gn=FGFR2;refGene_tx=XM_023241016.1;refGene_gn=
FGFR2;refGene_tx=XM_023241017.1;refGene_gn=FGFR2;refGene_tx=XM_0232410
18.1;refGene_gn=FGFR2;refGene_tx=XM_023241019.1
D2 79748641 79748908
refGene_gn=FGFR2;refGene_tx=XM_019813918.2;refGene_gn=FGFR2;refGene_tx
=XM_023241016.1;refGene_gn=FGFR2;refGene_tx=XM_023241017.1;refGene_gn=
FGFR2;refGene_tx=XM_023241018.1
D2 79773137 79773306
refGene_gn=FGFR2;refGene_tx=XM_019813918.2;refGene_gn=FGFR2;refGene_tx
=XM_019813921.2;refGene_gn=FGFR2;refGene_tx=XM_019813922.2;refGene_gn=
FGFR2;refGene_tx=XM_019813923.2;refGene_gn=FGFR2;refGene_tx=XM_0232410
16.1;refGene_gn=FGFR2;refGene_tx=XM_023241017.1;refGene_gn=FGFR2;refGe
ne_tx=XM_023241018.1;refGene_gn=FGFR2;refGene_tx=XM_023241019.1;refGen
e_gn=FGFR2;refGene_tx=XM_023241020.1
D2 79907647 79907826
refGene_gn=ATE1;refGene_tx=XM_019813925.2;refGene_gn=ATE1;refGene_tx=X
M_023241022.1;refGene_gn=ATE1;refGene_tx=XM_023241023.1;refGene_gn=ATE
1;refGene_tx=XM_023241024.1;refGene_gn=ATE1;refGene_tx=XM_023241025.1;
refGene_gn=ATE1;refGene_tx=XM_023241026.1;refGene_gn=ATE1;refGene_tx=X
M_023241027.1;refGene_gn=ATE1;refGene_tx=XM_023241028.1
D2 79944276 79944397
refGene_gn=ATE1;refGene_tx=XM_019813925.2;refGene_gn=ATE1;refGene_tx=X
M_023241022.1;refGene_gn=ATE1;refGene_tx=XM_023241023.1;refGene_gn=ATE
1;refGene_tx=XM_023241024.1;refGene_gn=ATE1;refGene_tx=XM_023241025.1;
refGene_gn=ATE1;refGene_tx=XM_023241027.1;refGene_gn=ATE1;refGene_tx=X
M_023241028.1
D2 79985314 79985414
refGene_gn=ATE1;refGene_tx=XM_019813925.2;refGene_gn=ATE1;refGene_tx=X
M_023241022.1;refGene_gn=ATE1;refGene_tx=XM_023241023.1;refGene_gn=ATE
1;refGene_tx=XM_023241024.1;refGene_gn=ATE1;refGene_tx=XM_023241025.1;
refGene_gn=ATE1;refGene_tx=XM_023241026.1;refGene_gn=ATE1;refGene_tx=X
M_023241027.1;refGene_gn=ATE1;refGene_tx=XM_023241028.1
D2 79989011 79989193
refGene_gn=ATE1;refGene_tx=XM_019813925.2;refGene_gn=ATE1;refGene_tx=X
M_023241022.1;refGene_gn=ATE1;refGene_tx=XM_023241023.1;refGene_gn=ATE
1;refGene_tx=XM_023241024.1;refGene_gn=ATE1;refGene_tx=XM_023241025.1;
refGene_gn=ATE1;refGene_tx=XM_023241026.1;refGene_gn=ATE1;refGene_tx=X
M_023241027.1;refGene_gn=ATE1;refGene_tx=XM_023241028.1
D2 80013644 80013677
refGene_gn=ATE1;refGene_tx=XM_019813925.2;refGene_gn=ATE1;refGene_tx=X
M_023241022.1;refGene_gn=ATE1;refGene_tx=XM_023241023.1;refGene_gn=ATE
1;refGene_tx=XM_023241024.1;refGene_gn=ATE1;refGene_tx=XM_023241025.1;
refGene_gn=ATE1;refGene_tx=XM_023241026.1;refGene_gn=ATE1;refGene_tx=X
M_023241027.1;refGene_gn=ATE1;refGene_tx=XM_023241028.1
D2 80039359 80039488
refGene_gn=ATE1;refGene_tx=XM_019813925.2;refGene_gn=ATE1;refGene_tx=X
M_023241022.1;refGene_gn=ATE1;refGene_tx=XM_023241023.1;refGene_gn=ATE
1;refGene_tx=XM_023241025.1;refGene_gn=ATE1;refGene_tx=XM_023241026.1;

refGene_gn=ATE1;refGene_tx=XM_023241027.1
D2 80040402 80040531
refGene_gn=ATE1;refGene_tx=XM_019813925.2;refGene_gn=ATE1;refGene_tx=XM_023241022.1;refGene_gn=ATE1;refGene_tx=XM_023241024.1;refGene_gn=ATE1;refGene_tx=XM_023241025.1;refGene_gn=ATE1;refGene_tx=XM_023241026.1;refGene_gn=ATE1;refGene_tx=XM_023241028.1
D2 80042314 80042547
refGene_gn=ATE1;refGene_tx=XM_019813925.2;refGene_gn=ATE1;refGene_tx=XM_023241022.1;refGene_gn=ATE1;refGene_tx=XM_023241023.1;refGene_gn=ATE1;refGene_tx=XM_023241024.1;refGene_gn=ATE1;refGene_tx=XM_023241025.1;refGene_gn=ATE1;refGene_tx=XM_023241026.1;refGene_gn=ATE1;refGene_tx=XM_023241027.1;refGene_gn=ATE1;refGene_tx=XM_023241028.1
D2 80048278 80048533
refGene_gn=ATE1;refGene_tx=XM_019813925.2;refGene_gn=ATE1;refGene_tx=XM_023241022.1;refGene_gn=ATE1;refGene_tx=XM_023241023.1;refGene_gn=ATE1;refGene_tx=XM_023241024.1;refGene_gn=ATE1;refGene_tx=XM_023241025.1;refGene_gn=ATE1;refGene_tx=XM_023241026.1;refGene_gn=ATE1;refGene_tx=XM_023241027.1;refGene_gn=ATE1;refGene_tx=XM_023241028.1
D2 80050289 80050393
refGene_gn=ATE1;refGene_tx=XM_019813925.2;refGene_gn=ATE1;refGene_tx=XM_023241022.1;refGene_gn=ATE1;refGene_tx=XM_023241023.1;refGene_gn=ATE1;refGene_tx=XM_023241024.1;refGene_gn=ATE1;refGene_tx=XM_023241025.1;refGene_gn=ATE1;refGene_tx=XM_023241026.1;refGene_gn=ATE1;refGene_tx=XM_023241027.1;refGene_gn=ATE1;refGene_tx=XM_023241028.1
D2 80054475 80054538
refGene_gn=ATE1;refGene_tx=XM_019813925.2;refGene_gn=ATE1;refGene_tx=XM_023241022.1;refGene_gn=ATE1;refGene_tx=XM_023241023.1;refGene_gn=ATE1;refGene_tx=XM_023241024.1;refGene_gn=ATE1;refGene_tx=XM_023241025.1;refGene_gn=ATE1;refGene_tx=XM_023241026.1;refGene_gn=ATE1;refGene_tx=XM_023241027.1;refGene_gn=ATE1;refGene_tx=XM_023241028.1
D2 80057574 80057638
refGene_gn=ATE1;refGene_tx=XM_019813925.2;refGene_gn=ATE1;refGene_tx=XM_023241022.1;refGene_gn=ATE1;refGene_tx=XM_023241023.1;refGene_gn=ATE1;refGene_tx=XM_023241024.1;refGene_gn=ATE1;refGene_tx=XM_023241025.1;refGene_gn=ATE1;refGene_tx=XM_023241026.1;refGene_gn=ATE1;refGene_tx=XM_023241027.1;refGene_gn=ATE1;refGene_tx=XM_023241028.1
D2 80060104 80060372
refGene_gn=ATE1;refGene_tx=XM_023241022.1;refGene_gn=ATE1;refGene_tx=XM_023241023.1;refGene_gn=ATE1;refGene_tx=XM_023241024.1;refGene_gn=ATE1;refGene_tx=XM_023241026.1
D2 80060620 80060705
refGene_gn=ATE1;refGene_tx=XM_023241025.1;refGene_gn=ATE1;refGene_tx=XM_023241027.1;refGene_gn=ATE1;refGene_tx=XM_023241028.1
D2 80082391 80082466 refGene_gn=NSMCE4A;refGene_tx=XM_023241029.1
D2 80082566 80082661 refGene_gn=NSMCE4A;refGene_tx=XM_023241029.1
D2 80083450 80083503 refGene_gn=NSMCE4A;refGene_tx=XM_023241030.1
D2 80083602 80083651
refGene_gn=NSMCE4A;refGene_tx=XM_023241029.1;refGene_gn=NSMCE4A;refGene_tx=XM_023241030.1
D2 80085774 80085869

refGene_gn=NSMCE4A;refGene_tx=XM_023241029.1;refGene_gn=NSMCE4A;refGene_tx=XM_023241030.1
D2	80086745	80086836
refGene_gn=NSMCE4A;refGene_tx=XM_023241029.1;refGene_gn=NSMCE4A;refGene_tx=XM_023241030.1
D2	80087504	80087604
refGene_gn=NSMCE4A;refGene_tx=XM_023241029.1;refGene_gn=NSMCE4A;refGene_tx=XM_023241030.1
D2	80089507	80089659
refGene_gn=NSMCE4A;refGene_tx=XM_023241029.1;refGene_gn=NSMCE4A;refGene_tx=XM_023241030.1
D2	80091753	80091884
refGene_gn=NSMCE4A;refGene_tx=XM_023241029.1;refGene_gn=NSMCE4A;refGene_tx=XM_023241030.1
D2	80094601	80094679
refGene_gn=NSMCE4A;refGene_tx=XM_023241029.1;refGene_gn=NSMCE4A;refGene_tx=XM_023241030.1
D2	80095383	80095675
refGene_gn=NSMCE4A;refGene_tx=XM_023241029.1;refGene_gn=NSMCE4A;refGene_tx=XM_023241030.1
D2	80110160	80110178	refGene_gn=TACC2;refGene_tx=XM_019813932.1
D2	80125845	80125878
refGene_gn=TACC2;refGene_tx=XM_006938226.2;refGene_gn=TACC2;refGene_tx=XM_019813927.1;refGene_gn=TACC2;refGene_tx=XM_019813928.1;refGene_gn=TACC2;refGene_tx=XM_019813929.1;refGene_gn=TACC2;refGene_tx=XM_019813930.1;refGene_gn=TACC2;refGene_tx=XM_019813931.1;refGene_gn=TACC2;refGene_tx=XM_019813933.1;refGene_gn=TACC2;refGene_tx=XM_019813934.1;refGene_gn=TACC2;refGene_tx=XM_019813935.1;refGene_gn=TACC2;refGene_tx=XM_019813936.1;refGene_gn=TACC2;refGene_tx=XM_019813937.1;refGene_gn=TACC2;refGene_tx=XM_019813938.1
D2	80132766	80132826
refGene_gn=TACC2;refGene_tx=XM_006938226.2;refGene_gn=TACC2;refGene_tx=XM_019813927.1;refGene_gn=TACC2;refGene_tx=XM_019813928.1;refGene_gn=TACC2;refGene_tx=XM_019813929.1;refGene_gn=TACC2;refGene_tx=XM_019813930.1;refGene_gn=TACC2;refGene_tx=XM_019813933.1;refGene_gn=TACC2;refGene_tx=XM_019813934.1;refGene_gn=TACC2;refGene_tx=XM_019813935.1;refGene_gn=TACC2;refGene_tx=XM_019813936.1;refGene_gn=TACC2;refGene_tx=XM_019813937.1;refGene_gn=TACC2;refGene_tx=XM_019813938.1
D2	80147394	80147507
refGene_gn=TACC2;refGene_tx=XM_006938226.2;refGene_gn=TACC2;refGene_tx=XM_019813927.1;refGene_gn=TACC2;refGene_tx=XM_019813928.1;refGene_gn=TACC2;refGene_tx=XM_019813929.1;refGene_gn=TACC2;refGene_tx=XM_019813930.1;refGene_gn=TACC2;refGene_tx=XM_019813931.1;refGene_gn=TACC2;refGene_tx=XM_019813932.1;refGene_gn=TACC2;refGene_tx=XM_019813933.1;refGene_gn=TACC2;refGene_tx=XM_019813934.1;refGene_gn=TACC2;refGene_tx=XM_019813935.1;refGene_gn=TACC2;refGene_tx=XM_019813936.1;refGene_gn=TACC2;refGene_tx=XM_019813937.1;refGene_gn=TACC2;refGene_tx=XM_019813938.1
D2	80170595	80175842
refGene_gn=TACC2;refGene_tx=XM_006938226.2;refGene_gn=TACC2;refGene_tx=XM_019813927.1;refGene_gn=TACC2;refGene_tx=XM_019813928.1;refGene_gn=

TACC2;refGene_tx=XM_019813929.1;refGene_gn=TACC2;refGene_tx=XM_0198139
30.1;refGene_gn=TACC2;refGene_tx=XM_019813931.1;refGene_gn=TACC2;refGe
ne_tx=XM_019813932.1;refGene_gn=TACC2;refGene_tx=XM_019813933.1;refGen
e_gn=TACC2;refGene_tx=XM_019813934.1;refGene_gn=TACC2;refGene_tx=XM_01
9813935.1;refGene_gn=TACC2;refGene_tx=XM_019813936.1;refGene_gn=TACC2;
refGene_tx=XM_019813937.1;refGene_gn=TACC2;refGene_tx=XM_019813938.1
D2 80176403 80176529
refGene_gn=TACC2;refGene_tx=XM_006938226.2;refGene_gn=TACC2;refGene_tx
=XM_019813927.1;refGene_gn=TACC2;refGene_tx=XM_019813928.1;refGene_gn=
TACC2;refGene_tx=XM_019813929.1;refGene_gn=TACC2;refGene_tx=XM_0198139
30.1;refGene_gn=TACC2;refGene_tx=XM_019813931.1;refGene_gn=TACC2;refGe
ne_tx=XM_019813932.1;refGene_gn=TACC2;refGene_tx=XM_019813933.1;refGen
e_gn=TACC2;refGene_tx=XM_019813934.1;refGene_gn=TACC2;refGene_tx=XM_01
9813935.1;refGene_gn=TACC2;refGene_tx=XM_019813936.1;refGene_gn=TACC2;
refGene_tx=XM_019813937.1;refGene_gn=TACC2;refGene_tx=XM_019813938.1
D2 80214026 80214152
refGene_gn=TACC2;refGene_tx=XM_006938226.2;refGene_gn=TACC2;refGene_tx
=XM_019813927.1;refGene_gn=TACC2;refGene_tx=XM_019813928.1;refGene_gn=
TACC2;refGene_tx=XM_019813929.1;refGene_gn=TACC2;refGene_tx=XM_0198139
30.1;refGene_gn=TACC2;refGene_tx=XM_019813931.1;refGene_gn=TACC2;refGe
ne_tx=XM_019813932.1;refGene_gn=TACC2;refGene_tx=XM_019813933.1;refGen
e_gn=TACC2;refGene_tx=XM_019813934.1;refGene_gn=TACC2;refGene_tx=XM_01
9813935.1;refGene_gn=TACC2;refGene_tx=XM_019813936.1;refGene_gn=TACC2;
refGene_tx=XM_019813937.1;refGene_gn=TACC2;refGene_tx=XM_019813938.1
D2 80225664 80225799
refGene_gn=TACC2;refGene_tx=XM_006938226.2;refGene_gn=TACC2;refGene_tx
=XM_019813927.1;refGene_gn=TACC2;refGene_tx=XM_019813928.1;refGene_gn=
TACC2;refGene_tx=XM_019813929.1;refGene_gn=TACC2;refGene_tx=XM_0198139
30.1;refGene_gn=TACC2;refGene_tx=XM_019813931.1;refGene_gn=TACC2;refGe
ne_tx=XM_019813932.1;refGene_gn=TACC2;refGene_tx=XM_019813933.1;refGen
e_gn=TACC2;refGene_tx=XM_019813934.1;refGene_gn=TACC2;refGene_tx=XM_01
9813936.1
D2 80241645 80241713
refGene_gn=TACC2;refGene_tx=XM_019813940.2;refGene_gn=TACC2;refGene_tx
=XM_019813941.2
D2 80274002 80274148
refGene_gn=TACC2;refGene_tx=XM_006938226.2;refGene_gn=TACC2;refGene_tx
=XM_019813927.1;refGene_gn=TACC2;refGene_tx=XM_019813928.1;refGene_gn=
TACC2;refGene_tx=XM_019813929.1;refGene_gn=TACC2;refGene_tx=XM_0198139
30.1;refGene_gn=TACC2;refGene_tx=XM_019813931.1;refGene_gn=TACC2;refGe
ne_tx=XM_019813932.1;refGene_gn=TACC2;refGene_tx=XM_019813933.1;refGen
e_gn=TACC2;refGene_tx=XM_019813934.1;refGene_gn=TACC2;refGene_tx=XM_01
9813935.1;refGene_gn=TACC2;refGene_tx=XM_019813936.1;refGene_gn=TACC2;
refGene_tx=XM_019813937.1;refGene_gn=TACC2;refGene_tx=XM_019813938.1;r
efGene_gn=TACC2;refGene_tx=XM_019813940.2;refGene_gn=TACC2;refGene_tx=
XM_019813941.2
D2 80284543 80285840
refGene_gn=TACC2;refGene_tx=XM_006938226.2;refGene_gn=TACC2;refGene_tx
=XM_019813927.1;refGene_gn=TACC2;refGene_tx=XM_019813928.1;refGene_gn=
TACC2;refGene_tx=XM_019813929.1;refGene_gn=TACC2;refGene_tx=XM_0198139

30.1;refGene_gn=TACC2;refGene_tx=XM_019813931.1;refGene_gn=TACC2;refGene_tx=XM_019813932.1;refGene_gn=TACC2;refGene_tx=XM_019813933.1;refGene_gn=TACC2;refGene_tx=XM_019813934.1;refGene_gn=TACC2;refGene_tx=XM_019813935.1;refGene_gn=TACC2;refGene_tx=XM_019813936.1;refGene_gn=TACC2;refGene_tx=XM_019813937.1;refGene_gn=TACC2;refGene_tx=XM_019813938.1;refGene_gn=TACC2;refGene_tx=XM_019813940.2;refGene_gn=TACC2;refGene_tx=XM_019813941.2
D2 80289786 80289847
refGene_gn=TACC2;refGene_tx=XM_006938226.2;refGene_gn=TACC2;refGene_tx=XM_019813927.1;refGene_gn=TACC2;refGene_tx=XM_019813928.1;refGene_gn=TACC2;refGene_tx=XM_019813929.1;refGene_gn=TACC2;refGene_tx=XM_019813930.1;refGene_gn=TACC2;refGene_tx=XM_019813931.1;refGene_gn=TACC2;refGene_tx=XM_019813932.1;refGene_gn=TACC2;refGene_tx=XM_019813933.1;refGene_gn=TACC2;refGene_tx=XM_019813934.1;refGene_gn=TACC2;refGene_tx=XM_019813935.1;refGene_gn=TACC2;refGene_tx=XM_019813936.1;refGene_gn=TACC2;refGene_tx=XM_019813937.1;refGene_gn=TACC2;refGene_tx=XM_019813938.1;refGene_gn=TACC2;refGene_tx=XM_019813940.2;refGene_gn=TACC2;refGene_tx=XM_019813941.2
D2 80291004 80291203
refGene_gn=TACC2;refGene_tx=XM_006938226.2;refGene_gn=TACC2;refGene_tx=XM_019813927.1;refGene_gn=TACC2;refGene_tx=XM_019813928.1;refGene_gn=TACC2;refGene_tx=XM_019813929.1;refGene_gn=TACC2;refGene_tx=XM_019813930.1;refGene_gn=TACC2;refGene_tx=XM_019813931.1;refGene_gn=TACC2;refGene_tx=XM_019813932.1;refGene_gn=TACC2;refGene_tx=XM_019813933.1;refGene_gn=TACC2;refGene_tx=XM_019813934.1;refGene_gn=TACC2;refGene_tx=XM_019813935.1;refGene_gn=TACC2;refGene_tx=XM_019813936.1;refGene_gn=TACC2;refGene_tx=XM_019813937.1;refGene_gn=TACC2;refGene_tx=XM_019813938.1;refGene_gn=TACC2;refGene_tx=XM_019813940.2;refGene_gn=TACC2;refGene_tx=XM_019813941.2
D2 80298241 80298303
refGene_gn=TACC2;refGene_tx=XM_006938226.2;refGene_gn=TACC2;refGene_tx=XM_019813927.1;refGene_gn=TACC2;refGene_tx=XM_019813928.1;refGene_gn=TACC2;refGene_tx=XM_019813929.1;refGene_gn=TACC2;refGene_tx=XM_019813930.1;refGene_gn=TACC2;refGene_tx=XM_019813931.1;refGene_gn=TACC2;refGene_tx=XM_019813932.1;refGene_gn=TACC2;refGene_tx=XM_019813933.1;refGene_gn=TACC2;refGene_tx=XM_019813934.1;refGene_gn=TACC2;refGene_tx=XM_019813935.1;refGene_gn=TACC2;refGene_tx=XM_019813936.1;refGene_gn=TACC2;refGene_tx=XM_019813937.1;refGene_gn=TACC2;refGene_tx=XM_019813938.1;refGene_gn=TACC2;refGene_tx=XM_019813940.2;refGene_gn=TACC2;refGene_tx=XM_019813941.2
D2 80299722 80299838
refGene_gn=TACC2;refGene_tx=XM_006938226.2;refGene_gn=TACC2;refGene_tx=XM_019813927.1;refGene_gn=TACC2;refGene_tx=XM_019813928.1;refGene_gn=TACC2;refGene_tx=XM_019813929.1;refGene_gn=TACC2;refGene_tx=XM_019813930.1;refGene_gn=TACC2;refGene_tx=XM_019813931.1;refGene_gn=TACC2;refGene_tx=XM_019813932.1;refGene_gn=TACC2;refGene_tx=XM_019813933.1;refGene_gn=TACC2;refGene_tx=XM_019813934.1;refGene_gn=TACC2;refGene_tx=XM_019813935.1;refGene_gn=TACC2;refGene_tx=XM_019813936.1;refGene_gn=TACC2;refGene_tx=XM_019813937.1;refGene_gn=TACC2;refGene_tx=XM_019813938.1;refGene_gn=TACC2;refGene_tx=XM_019813940.2;refGene_gn=TACC2;refGene_tx=

XM_019813941.2
D2 80300974 80301146
refGene_gn=TACC2;refGene_tx=XM_006938226.2;refGene_gn=TACC2;refGene_tx=XM_019813927.1;refGene_gn=TACC2;refGene_tx=XM_019813928.1;refGene_gn=TACC2;refGene_tx=XM_019813929.1;refGene_gn=TACC2;refGene_tx=XM_019813930.1;refGene_gn=TACC2;refGene_tx=XM_019813931.1;refGene_gn=TACC2;refGene_tx=XM_019813932.1;refGene_gn=TACC2;refGene_tx=XM_019813933.1;refGene_gn=TACC2;refGene_tx=XM_019813934.1;refGene_gn=TACC2;refGene_tx=XM_019813935.1;refGene_gn=TACC2;refGene_tx=XM_019813936.1;refGene_gn=TACC2;refGene_tx=XM_019813937.1;refGene_gn=TACC2;refGene_tx=XM_019813938.1;refGene_gn=TACC2;refGene_tx=XM_019813940.2;refGene_gn=TACC2;refGene_tx=XM_019813941.2
D2 80302431 80302572
refGene_gn=TACC2;refGene_tx=XM_006938226.2;refGene_gn=TACC2;refGene_tx=XM_019813927.1;refGene_gn=TACC2;refGene_tx=XM_019813928.1;refGene_gn=TACC2;refGene_tx=XM_019813929.1;refGene_gn=TACC2;refGene_tx=XM_019813930.1;refGene_gn=TACC2;refGene_tx=XM_019813931.1;refGene_gn=TACC2;refGene_tx=XM_019813932.1;refGene_gn=TACC2;refGene_tx=XM_019813933.1;refGene_gn=TACC2;refGene_tx=XM_019813934.1;refGene_gn=TACC2;refGene_tx=XM_019813935.1;refGene_gn=TACC2;refGene_tx=XM_019813940.2;refGene_gn=TACC2;refGene_tx=XM_019813941.2
D2 80303466 80303556
refGene_gn=TACC2;refGene_tx=XM_006938226.2;refGene_gn=TACC2;refGene_tx=XM_019813927.1;refGene_gn=TACC2;refGene_tx=XM_019813928.1;refGene_gn=TACC2;refGene_tx=XM_019813929.1;refGene_gn=TACC2;refGene_tx=XM_019813930.1;refGene_gn=TACC2;refGene_tx=XM_019813931.1;refGene_gn=TACC2;refGene_tx=XM_019813932.1;refGene_gn=TACC2;refGene_tx=XM_019813935.1;refGene_gn=TACC2;refGene_tx=XM_019813940.2
D2 80309977 80310121
refGene_gn=TACC2;refGene_tx=XM_006938226.2;refGene_gn=TACC2;refGene_tx=XM_019813927.1;refGene_gn=TACC2;refGene_tx=XM_019813928.1;refGene_gn=TACC2;refGene_tx=XM_019813929.1;refGene_gn=TACC2;refGene_tx=XM_019813930.1;refGene_gn=TACC2;refGene_tx=XM_019813931.1;refGene_gn=TACC2;refGene_tx=XM_019813932.1;refGene_gn=TACC2;refGene_tx=XM_019813933.1;refGene_gn=TACC2;refGene_tx=XM_019813934.1;refGene_gn=TACC2;refGene_tx=XM_019813935.1;refGene_gn=TACC2;refGene_tx=XM_019813936.1;refGene_gn=TACC2;refGene_tx=XM_019813937.1;refGene_gn=TACC2;refGene_tx=XM_019813938.1;refGene_gn=TACC2;refGene_tx=XM_019813940.2;refGene_gn=TACC2;refGene_tx=XM_019813941.2
D2 80310582 80310659
refGene_gn=TACC2;refGene_tx=XM_006938226.2;refGene_gn=TACC2;refGene_tx=XM_019813927.1;refGene_gn=TACC2;refGene_tx=XM_019813928.1;refGene_gn=TACC2;refGene_tx=XM_019813929.1;refGene_gn=TACC2;refGene_tx=XM_019813930.1;refGene_gn=TACC2;refGene_tx=XM_019813931.1;refGene_gn=TACC2;refGene_tx=XM_019813932.1;refGene_gn=TACC2;refGene_tx=XM_019813933.1;refGene_gn=TACC2;refGene_tx=XM_019813934.1;refGene_gn=TACC2;refGene_tx=XM_019813935.1;refGene_gn=TACC2;refGene_tx=XM_019813936.1;refGene_gn=TACC2;refGene_tx=XM_019813937.1;refGene_gn=TACC2;refGene_tx=XM_019813938.1;refGene_gn=TACC2;refGene_tx=XM_019813940.2;refGene_gn=TACC2;refGene_tx=XM_019813941.2

D2 80315382 80315426
refGene_gn=TACC2;refGene_tx=XM_006938226.2;refGene_gn=TACC2;refGene_tx=XM_019813927.1;refGene_gn=TACC2;refGene_tx=XM_019813928.1;refGene_gn=TACC2;refGene_tx=XM_019813929.1;refGene_gn=TACC2;refGene_tx=XM_019813930.1;refGene_gn=TACC2;refGene_tx=XM_019813931.1;refGene_gn=TACC2;refGene_tx=XM_019813932.1;refGene_gn=TACC2;refGene_tx=XM_019813933.1;refGene_gn=TACC2;refGene_tx=XM_019813934.1;refGene_gn=TACC2;refGene_tx=XM_019813935.1;refGene_gn=TACC2;refGene_tx=XM_019813936.1;refGene_gn=TACC2;refGene_tx=XM_019813937.1;refGene_gn=TACC2;refGene_tx=XM_019813938.1;refGene_gn=TACC2;refGene_tx=XM_019813940.2;refGene_gn=TACC2;refGene_tx=XM_019813941.2
D2 80318045 80318054
refGene_gn=TACC2;refGene_tx=XM_019813927.1;refGene_gn=TACC2;refGene_tx=XM_019813928.1;refGene_gn=TACC2;refGene_tx=XM_019813931.1;refGene_gn=TACC2;refGene_tx=XM_019813932.1
D2 80320633 80320794
refGene_gn=TACC2;refGene_tx=XM_006938226.2;refGene_gn=TACC2;refGene_tx=XM_019813927.1;refGene_gn=TACC2;refGene_tx=XM_019813928.1;refGene_gn=TACC2;refGene_tx=XM_019813929.1;refGene_gn=TACC2;refGene_tx=XM_019813930.1;refGene_gn=TACC2;refGene_tx=XM_019813931.1;refGene_gn=TACC2;refGene_tx=XM_019813932.1;refGene_gn=TACC2;refGene_tx=XM_019813933.1;refGene_gn=TACC2;refGene_tx=XM_019813934.1;refGene_gn=TACC2;refGene_tx=XM_019813935.1;refGene_gn=TACC2;refGene_tx=XM_019813936.1;refGene_gn=TACC2;refGene_tx=XM_019813937.1;refGene_gn=TACC2;refGene_tx=XM_019813938.1;refGene_gn=TACC2;refGene_tx=XM_019813940.2;refGene_gn=TACC2;refGene_tx=XM_019813941.2
D2 80320982 80321089
refGene_gn=TACC2;refGene_tx=XM_006938226.2;refGene_gn=TACC2;refGene_tx=XM_019813927.1;refGene_gn=TACC2;refGene_tx=XM_019813928.1;refGene_gn=TACC2;refGene_tx=XM_019813929.1;refGene_gn=TACC2;refGene_tx=XM_019813930.1;refGene_gn=TACC2;refGene_tx=XM_019813931.1;refGene_gn=TACC2;refGene_tx=XM_019813932.1;refGene_gn=TACC2;refGene_tx=XM_019813933.1;refGene_gn=TACC2;refGene_tx=XM_019813934.1;refGene_gn=TACC2;refGene_tx=XM_019813935.1;refGene_gn=TACC2;refGene_tx=XM_019813936.1;refGene_gn=TACC2;refGene_tx=XM_019813937.1;refGene_gn=TACC2;refGene_tx=XM_019813938.1;refGene_gn=TACC2;refGene_tx=XM_019813940.2;refGene_gn=TACC2;refGene_tx=XM_019813941.2
D2 80321490 80321611
refGene_gn=TACC2;refGene_tx=XM_006938226.2;refGene_gn=TACC2;refGene_tx=XM_019813927.1;refGene_gn=TACC2;refGene_tx=XM_019813928.1;refGene_gn=TACC2;refGene_tx=XM_019813929.1;refGene_gn=TACC2;refGene_tx=XM_019813930.1;refGene_gn=TACC2;refGene_tx=XM_019813931.1;refGene_gn=TACC2;refGene_tx=XM_019813932.1;refGene_gn=TACC2;refGene_tx=XM_019813933.1;refGene_gn=TACC2;refGene_tx=XM_019813934.1;refGene_gn=TACC2;refGene_tx=XM_019813935.1;refGene_gn=TACC2;refGene_tx=XM_019813936.1;refGene_gn=TACC2;refGene_tx=XM_019813937.1;refGene_gn=TACC2;refGene_tx=XM_019813938.1;refGene_gn=TACC2;refGene_tx=XM_019813940.2;refGene_gn=TACC2;refGene_tx=XM_019813941.2
D2 80325126 80325192
refGene_gn=TACC2;refGene_tx=XM_006938226.2;refGene_gn=TACC2;refGene_tx

=XM_019813927.1;refGene_gn=TACC2;refGene_tx=XM_019813928.1;refGene_gn=TACC2;refGene_tx=XM_019813929.1;refGene_gn=TACC2;refGene_tx=XM_019813930.1;refGene_gn=TACC2;refGene_tx=XM_019813931.1;refGene_gn=TACC2;refGene_tx=XM_019813932.1;refGene_gn=TACC2;refGene_tx=XM_019813933.1;refGene_gn=TACC2;refGene_tx=XM_019813934.1;refGene_gn=TACC2;refGene_tx=XM_019813935.1;refGene_gn=TACC2;refGene_tx=XM_019813936.1;refGene_gn=TACC2;refGene_tx=XM_019813937.1;refGene_gn=TACC2;refGene_tx=XM_019813938.1;refGene_gn=TACC2;refGene_tx=XM_019813940.2;refGene_gn=TACC2;refGene_tx=XM_019813941.2
D2 80335711 80335729
refGene_gn=BTBD16;refGene_tx=XM_006938230.3;refGene_gn=BTBD16;refGene_tx=XM_019813942.2;refGene_gn=BTBD16;refGene_tx=XM_019813943.2
D2 80337511 80337660
refGene_gn=BTBD16;refGene_tx=XM_006938230.3;refGene_gn=BTBD16;refGene_tx=XM_019813942.2;refGene_gn=BTBD16;refGene_tx=XM_019813943.2
D2 80341122 80341196
refGene_gn=BTBD16;refGene_tx=XM_006938230.3;refGene_gn=BTBD16;refGene_tx=XM_019813942.2;refGene_gn=BTBD16;refGene_tx=XM_019813943.2
D2 80343386 80343530
refGene_gn=BTBD16;refGene_tx=XM_006938230.3;refGene_gn=BTBD16;refGene_tx=XM_019813942.2;refGene_gn=BTBD16;refGene_tx=XM_019813943.2;refGene_gn=BTBD16;refGene_tx=XM_023241033.1
D2 80347193 80347283
refGene_gn=BTBD16;refGene_tx=XM_006938230.3;refGene_gn=BTBD16;refGene_tx=XM_019813942.2;refGene_gn=BTBD16;refGene_tx=XM_019813943.2;refGene_gn=BTBD16;refGene_tx=XM_023241033.1
D2 80349382 80349497
refGene_gn=BTBD16;refGene_tx=XM_006938230.3;refGene_gn=BTBD16;refGene_tx=XM_019813942.2;refGene_gn=BTBD16;refGene_tx=XM_019813943.2;refGene_gn=BTBD16;refGene_tx=XM_023241033.1
D2 80353202 80353272
refGene_gn=BTBD16;refGene_tx=XM_006938230.3;refGene_gn=BTBD16;refGene_tx=XM_019813942.2;refGene_gn=BTBD16;refGene_tx=XM_019813943.2;refGene_gn=BTBD16;refGene_tx=XM_023241033.1
D2 80354451 80354582
refGene_gn=BTBD16;refGene_tx=XM_006938230.3;refGene_gn=BTBD16;refGene_tx=XM_019813942.2;refGene_gn=BTBD16;refGene_tx=XM_019813943.2;refGene_gn=BTBD16;refGene_tx=XM_023241033.1
D2 80361794 80361914
refGene_gn=BTBD16;refGene_tx=XM_006938230.3;refGene_gn=BTBD16;refGene_tx=XM_019813942.2;refGene_gn=BTBD16;refGene_tx=XM_019813943.2;refGene_gn=BTBD16;refGene_tx=XM_023241033.1
D2 80371672 80371795
refGene_gn=BTBD16;refGene_tx=XM_019813942.2;refGene_gn=BTBD16;refGene_tx=XM_019813943.2;refGene_gn=BTBD16;refGene_tx=XM_023241033.1
D2 80373805 80373897
refGene_gn=BTBD16;refGene_tx=XM_006938230.3;refGene_gn=BTBD16;refGene_tx=XM_019813942.2;refGene_gn=BTBD16;refGene_tx=XM_019813943.2;refGene_gn=BTBD16;refGene_tx=XM_023241033.1
D2 80374775 80374858

refGene_gn=BTBD16;refGene_tx=XM_006938230.3;refGene_gn=BTBD16;refGene_tx=XM_019813942.2;refGene_gn=BTBD16;refGene_tx=XM_019813943.2;refGene_gn=BTBD16;refGene_tx=XM_023241033.1
D2 80377085 80377163
refGene_gn=BTBD16;refGene_tx=XM_006938230.3;refGene_gn=BTBD16;refGene_tx=XM_019813942.2;refGene_gn=BTBD16;refGene_tx=XM_023241033.1
D2 80378305 80378404
refGene_gn=BTBD16;refGene_tx=XM_006938230.3;refGene_gn=BTBD16;refGene_tx=XM_019813942.2;refGene_gn=BTBD16;refGene_tx=XM_019813943.2;refGene_gn=BTBD16;refGene_tx=XM_023241033.1
D2 80379722 80379896
refGene_gn=BTBD16;refGene_tx=XM_006938230.3;refGene_gn=BTBD16;refGene_tx=XM_019813942.2;refGene_gn=BTBD16;refGene_tx=XM_019813943.2;refGene_gn=BTBD16;refGene_tx=XM_023241033.1
D2 80380935 80381004
refGene_gn=BTBD16;refGene_tx=XM_006938230.3;refGene_gn=BTBD16;refGene_tx=XM_019813942.2;refGene_gn=BTBD16;refGene_tx=XM_019813943.2;refGene_gn=BTBD16;refGene_tx=XM_023241033.1
D2 80432891 80433032
refGene_gn=PLEKHA1;refGene_tx=XM_023241034.1;refGene_gn=PLEKHA1;refGene_tx=XM_023241035.1;refGene_gn=PLEKHA1;refGene_tx=XM_023241036.1
D2 80438646 80438703
refGene_gn=PLEKHA1;refGene_tx=XM_023241034.1;refGene_gn=PLEKHA1;refGene_tx=XM_023241035.1;refGene_gn=PLEKHA1;refGene_tx=XM_023241036.1
D2 80441128 80441174
refGene_gn=PLEKHA1;refGene_tx=XM_023241034.1;refGene_gn=PLEKHA1;refGene_tx=XM_023241035.1;refGene_gn=PLEKHA1;refGene_tx=XM_023241036.1
D2 80448208 80448306
refGene_gn=PLEKHA1;refGene_tx=XM_023241034.1;refGene_gn=PLEKHA1;refGene_tx=XM_023241035.1;refGene_gn=PLEKHA1;refGene_tx=XM_023241036.1
D2 80454271 80454397
refGene_gn=PLEKHA1;refGene_tx=XM_023241034.1;refGene_gn=PLEKHA1;refGene_tx=XM_023241035.1;refGene_gn=PLEKHA1;refGene_tx=XM_023241036.1
D2 80458216 80458360
refGene_gn=PLEKHA1;refGene_tx=XM_023241034.1;refGene_gn=PLEKHA1;refGene_tx=XM_023241035.1;refGene_gn=PLEKHA1;refGene_tx=XM_023241036.1
D2 80460157 80460226
refGene_gn=PLEKHA1;refGene_tx=XM_023241034.1;refGene_gn=PLEKHA1;refGene_tx=XM_023241035.1;refGene_gn=PLEKHA1;refGene_tx=XM_023241036.1
D2 80466550 80466615
refGene_gn=PLEKHA1;refGene_tx=XM_023241034.1;refGene_gn=PLEKHA1;refGene_tx=XM_023241035.1;refGene_gn=PLEKHA1;refGene_tx=XM_023241036.1
D2 80467229 80467293
refGene_gn=PLEKHA1;refGene_tx=XM_023241034.1;refGene_gn=PLEKHA1;refGene_tx=XM_023241035.1;refGene_gn=PLEKHA1;refGene_tx=XM_023241036.1
D2 80468916 80469006
refGene_gn=PLEKHA1;refGene_tx=XM_023241034.1;refGene_gn=PLEKHA1;refGene_tx=XM_023241035.1;refGene_gn=PLEKHA1;refGene_tx=XM_023241036.1
D2 80470257 80470328
refGene_gn=PLEKHA1;refGene_tx=XM_023241035.1;refGene_gn=PLEKHA1;refGen

e_tx=XM_023241036.1
D2 80471614 80471929
refGene_gn=PLEKHA1;refGene_tx=XM_023241034.1;refGene_gn=PLEKHA1;refGen
e_tx=XM_023241035.1;refGene_gn=PLEKHA1;refGene_tx=XM_023241036.1
D2 80498206 80498690 refGene_gn=HTRA1;refGene_tx=XM_023241037.1
D2 80525927 80526027 refGene_gn=HTRA1;refGene_tx=XM_023241037.1
D2 80526391 80526596 refGene_gn=HTRA1;refGene_tx=XM_023241037.1
D2 80543053 80543248 refGene_gn=HTRA1;refGene_tx=XM_023241037.1
D2 80544434 80544467 refGene_gn=HTRA1;refGene_tx=XM_023241037.1
D2 80545803 80545918 refGene_gn=HTRA1;refGene_tx=XM_023241037.1
D2 80547068 80547126 refGene_gn=HTRA1;refGene_tx=XM_023241037.1
D2 80548609 80548705 refGene_gn=HTRA1;refGene_tx=XM_023241037.1
D2 80550071 80550240 refGene_gn=HTRA1;refGene_tx=XM_023241037.1
D2 80595910 80595971
refGene_gn=LOC111556932;refGene_tx=XM_023240302.1
D2 80604599 80604656
refGene_gn=LOC111556932;refGene_tx=XM_023240302.1
D2 80606593 80606623
refGene_gn=LOC111556932;refGene_tx=XM_023240302.1
D2 80611783 80611813
refGene_gn=LOC111556932;refGene_tx=XM_023240302.1
D2 80613943 80614255
refGene_gn=LOC111556932;refGene_tx=XM_023240302.1
D2 80615505 80615553
refGene_gn=LOC111556932;refGene_tx=XM_023240302.1
D2 80617290 80617329
refGene_gn=LOC111556932;refGene_tx=XM_023240302.1
D2 80619379 80619409
refGene_gn=LOC111556932;refGene_tx=XM_023240302.1
D2 80620248 80620278
refGene_gn=LOC111556932;refGene_tx=XM_023240302.1
D2 80622640 80622654
refGene_gn=LOC111556932;refGene_tx=XM_023240302.1
D2 80622799 80623415
refGene_gn=LOC111556934;refGene_tx=XM_023240303.1
D2 80624226 80624256
refGene_gn=LOC111556934;refGene_tx=XM_023240303.1
D2 80624871 80624898
refGene_gn=LOC111556934;refGene_tx=XM_023240303.1
D2 80627518 80627874
refGene_gn=LOC111556934;refGene_tx=XM_023240303.1
D2 80672380 80672456
refGene_gn=LOC101096171;refGene_tx=XM_023241038.1
D2 80672860 80673181
refGene_gn=LOC101096171;refGene_tx=XM_023241038.1
D2 80673831 80673852
refGene_gn=LOC101096171;refGene_tx=XM_023241038.1
D2 80674943 80674973
refGene_gn=LOC101096171;refGene_tx=XM_023241038.1
D2 80675185 80675336

refGene_gn=LOC101096171;refGene_tx=XM_023241038.1
D2	80676608	80676817
refGene_gn=LOC101096171;refGene_tx=XM_023241038.1
D2	80677645	80677830
refGene_gn=LOC101096171;refGene_tx=XM_023241038.1
D2	80682268	80682657
refGene_gn=LOC101096171;refGene_tx=XM_023241038.1
D2	80685259	80685512
refGene_gn=LOC101096171;refGene_tx=XM_023241038.1
D2	80706280	80707024
refGene_gn=CD2H10orf120;refGene_tx=XM_006938233.3
D2	80707413	80707489
refGene_gn=CD2H10orf120;refGene_tx=XM_006938233.3
D2	80707683	80707868
refGene_gn=CD2H10orf120;refGene_tx=XM_006938233.3
D2	80756897	80756904
refGene_gn=LOC111556935;refGene_tx=XM_023240304.1
D2	80756999	80757317
refGene_gn=LOC111556935;refGene_tx=XM_023240304.1
D2	80757436	80757466
refGene_gn=LOC111556935;refGene_tx=XM_023240304.1
D2	80767280	80767726
refGene_gn=LOC111556935;refGene_tx=XM_023240304.1
D2	80775442	80775541
refGene_gn=LOC101096674;refGene_tx=XM_023240305.1
D2	80778819	80779326
refGene_gn=LOC101096674;refGene_tx=XM_023240305.1
D2	80816206	80816584
refGene_gn=LOC111557129;refGene_tx=XM_023241045.1
D2	80827751	80827854
refGene_gn=LOC109492837;refGene_tx=XM_023241039.1;refGene_gn=LOC109492837;refGene_tx=XM_023241040.1
D2	80830129	80830171
refGene_gn=LOC109492837;refGene_tx=XM_023241039.1;refGene_gn=LOC109492837;refGene_tx=XM_023241040.1
D2	80831036	80831075
refGene_gn=LOC109492837;refGene_tx=XM_023241039.1;refGene_gn=LOC109492837;refGene_tx=XM_023241040.1
D2	80831586	80831619
refGene_gn=LOC109492837;refGene_tx=XM_023241039.1;refGene_gn=LOC109492837;refGene_tx=XM_023241040.1
D2	80833003	80833036
refGene_gn=LOC109492837;refGene_tx=XM_023241039.1;refGene_gn=LOC109492837;refGene_tx=XM_023241040.1
D2	80833152	80833191
refGene_gn=LOC109492837;refGene_tx=XM_023241039.1;refGene_gn=LOC109492837;refGene_tx=XM_023241040.1
D2	80833374	80833413
refGene_gn=LOC109492837;refGene_tx=XM_023241039.1;refGene_gn=LOC109492837;refGene_tx=XM_023241040.1

D2 80833612 80833859
refGene_gn=LOC109492837;refGene_tx=XM_023241039.1;refGene_gn=LOC109492837;refGene_tx=XM_023241040.1;refGene_gn=LOC109492837;refGene_tx=XM_023241041.1
D2 80835775 80835814
refGene_gn=LOC109492837;refGene_tx=XM_023241039.1;refGene_gn=LOC109492837;refGene_tx=XM_023241040.1;refGene_gn=LOC109492837;refGene_tx=XM_023241041.1
D2 80836565 80836604
refGene_gn=LOC109492837;refGene_tx=XM_023241039.1;refGene_gn=LOC109492837;refGene_tx=XM_023241040.1;refGene_gn=LOC109492837;refGene_tx=XM_023241041.1;refGene_gn=LOC109492837;refGene_tx=XM_023241042.1
D2 80838129 80838441
refGene_gn=LOC109492837;refGene_tx=XM_023241039.1;refGene_gn=LOC109492837;refGene_tx=XM_023241040.1;refGene_gn=LOC109492837;refGene_tx=XM_023241041.1;refGene_gn=LOC109492837;refGene_tx=XM_023241042.1
D2 80839056 80839098
refGene_gn=LOC109492837;refGene_tx=XM_023241039.1;refGene_gn=LOC109492837;refGene_tx=XM_023241040.1;refGene_gn=LOC109492837;refGene_tx=XM_023241041.1;refGene_gn=LOC109492837;refGene_tx=XM_023241042.1
D2 80840216 80840392
refGene_gn=LOC109492837;refGene_tx=XM_023241039.1;refGene_gn=LOC109492837;refGene_tx=XM_023241040.1;refGene_gn=LOC109492837;refGene_tx=XM_023241041.1;refGene_gn=LOC109492837;refGene_tx=XM_023241042.1
D2 80843047 80843415
refGene_gn=LOC109492837;refGene_tx=XM_023241039.1;refGene_gn=LOC109492837;refGene_tx=XM_023241040.1;refGene_gn=LOC109492837;refGene_tx=XM_023241041.1;refGene_gn=LOC109492837;refGene_tx=XM_023241042.1
D2 80846121 80846413
refGene_gn=LOC109492837;refGene_tx=XM_023241039.1;refGene_gn=LOC109492837;refGene_tx=XM_023241040.1;refGene_gn=LOC109492837;refGene_tx=XM_023241041.1;refGene_gn=LOC109492837;refGene_tx=XM_023241042.1
D2 80850909 80851085 refGene_gn=CUZD1;refGene_tx=XM_011287600.3
D2 80852823 80853092 refGene_gn=CUZD1;refGene_tx=XM_011287600.3
D2 80853872 80854264 refGene_gn=CUZD1;refGene_tx=XM_011287600.3
D2 80855314 80855487 refGene_gn=CUZD1;refGene_tx=XM_011287600.3
D2 80856427 80856645 refGene_gn=CUZD1;refGene_tx=XM_011287600.3
D2 80857174 80857325 refGene_gn=CUZD1;refGene_tx=XM_011287600.3
D2 80859182 80859397 refGene_gn=CUZD1;refGene_tx=XM_011287600.3
D2 80861291 80861442 refGene_gn=CUZD1;refGene_tx=XM_011287600.3
D2 80863481 80863520 refGene_gn=CUZD1;refGene_tx=XM_011287600.3
D2 80863522 80863562 refGene_gn=CUZD1;refGene_tx=XM_011287600.3
D2 80867810 80867985
refGene_gn=LOC101091815;refGene_tx=XM_011287598.3
D2 80868541 80868662
refGene_gn=LOC101091815;refGene_tx=XM_011287598.3
D2 80883736 80884307
refGene_gn=LOC101091815;refGene_tx=XM_011287598.3;refGene_gn=LOC111557128;refGene_tx=XM_023241043.1;refGene_gn=LOC111557128;refGene_tx=XM_023241044.1

D2 80884898 80885105
refGene_gn=LOC111557128;refGene_tx=XM_023241043.1;refGene_gn=LOC111557128;refGene_tx=XM_023241044.1
D2 80886125 80886198
refGene_gn=LOC111557128;refGene_tx=XM_023241043.1;refGene_gn=LOC111557128;refGene_tx=XM_023241044.1
D2 80888748 80888955
refGene_gn=LOC111557128;refGene_tx=XM_023241043.1;refGene_gn=LOC111557128;refGene_tx=XM_023241044.1
D2 80893236 80893691
refGene_gn=LOC111557128;refGene_tx=XM_023241043.1;refGene_gn=LOC111557128;refGene_tx=XM_023241044.1
D2 80899137 80899372
refGene_gn=LOC111557128;refGene_tx=XM_023241043.1
D2 80923114 80923349
refGene_gn=LOC101092296;refGene_tx=XM_003994475.5
D2 80925113 80925565
refGene_gn=LOC101092296;refGene_tx=XM_003994475.5
D2 80932883 80933090
refGene_gn=LOC101092296;refGene_tx=XM_003994475.5
D2 80936872 80936942
refGene_gn=LOC101092296;refGene_tx=XM_003994475.5
D2 80938607 80938814
refGene_gn=LOC101092296;refGene_tx=XM_003994475.5
D2 80939406 80939570
refGene_gn=LOC101092296;refGene_tx=XM_003994475.5
D2 80950654 80950870
refGene_gn=PSTK;refGene_tx=XM_011287601.2;refGene_gn=PSTK;refGene_tx=XM_019813959.2
D2 80951023 80951098 refGene_gn=PSTK;refGene_tx=XM_011287602.3
D2 80952942 80953234
refGene_gn=PSTK;refGene_tx=XM_011287601.2;refGene_gn=PSTK;refGene_tx=XM_011287602.3;refGene_gn=PSTK;refGene_tx=XM_019813959.2
D2 80953467 80953666
refGene_gn=PSTK;refGene_tx=XM_011287601.2;refGene_gn=PSTK;refGene_tx=XM_011287602.3;refGene_gn=PSTK;refGene_tx=XM_019813959.2
D2 80955236 80955312
refGene_gn=PSTK;refGene_tx=XM_011287601.2;refGene_gn=PSTK;refGene_tx=XM_011287602.3;refGene_gn=PSTK;refGene_tx=XM_019813959.2
D2 80955807 80955901
refGene_gn=PSTK;refGene_tx=XM_011287601.2;refGene_gn=PSTK;refGene_tx=XM_011287602.3;refGene_gn=PSTK;refGene_tx=XM_019813959.2
D2 80959079 80959276
refGene_gn=PSTK;refGene_tx=XM_011287601.2;refGene_gn=PSTK;refGene_tx=XM_011287602.3
D2 80963011 80963880
refGene_gn=IKZF5;refGene_tx=XM_003994476.5;refGene_gn=IKZF5;refGene_tx=XM_019813954.2;refGene_gn=IKZF5;refGene_tx=XM_019813955.2;refGene_gn=IKZF5;refGene_tx=XM_019813957.2
D2 80965102 80965285

refGene_gn=IKZF5;refGene_tx=XM_003994476.5;refGene_gn=IKZF5;refGene_tx=XM_019813954.2;refGene_gn=IKZF5;refGene_tx=XM_019813955.2;refGene_gn=IKZF5;refGene_tx=XM_019813957.2
D2 80967307 80967440 refGene_gn=IKZF5;refGene_tx=XM_003994476.5;refGene_gn=IKZF5;refGene_tx=XM_019813954.2;refGene_gn=IKZF5;refGene_tx=XM_019813955.2
D2 80984834 80984876 refGene_gn=ACADSB;refGene_tx=XM_003994477.5;refGene_gn=ACADSB;refGene_tx=XM_023241046.1
D2 81003209 81003242 refGene_gn=ACADSB;refGene_tx=XM_019813952.1
D2 81010099 81010259 refGene_gn=ACADSB;refGene_tx=XM_003994477.5;refGene_gn=ACADSB;refGene_tx=XM_019813952.1;refGene_gn=ACADSB;refGene_tx=XM_019813953.2;refGene_gn=ACADSB;refGene_tx=XM_023241046.1
D2 81010473 81010574 refGene_gn=ACADSB;refGene_tx=XM_003994477.5;refGene_gn=ACADSB;refGene_tx=XM_019813952.1;refGene_gn=ACADSB;refGene_tx=XM_019813953.2;refGene_gn=ACADSB;refGene_tx=XM_023241046.1
D2 81013804 81014011 refGene_gn=ACADSB;refGene_tx=XM_003994477.5;refGene_gn=ACADSB;refGene_tx=XM_019813952.1;refGene_gn=ACADSB;refGene_tx=XM_019813953.2;refGene_gn=ACADSB;refGene_tx=XM_023241046.1
D2 81014777 81014948 refGene_gn=ACADSB;refGene_tx=XM_003994477.5;refGene_gn=ACADSB;refGene_tx=XM_019813952.1;refGene_gn=ACADSB;refGene_tx=XM_019813953.2;refGene_gn=ACADSB;refGene_tx=XM_023241046.1
D2 81015935 81016061 refGene_gn=ACADSB;refGene_tx=XM_003994477.5;refGene_gn=ACADSB;refGene_tx=XM_019813952.1;refGene_gn=ACADSB;refGene_tx=XM_019813953.2;refGene_gn=ACADSB;refGene_tx=XM_023241046.1
D2 81017746 81017839 refGene_gn=ACADSB;refGene_tx=XM_003994477.5;refGene_gn=ACADSB;refGene_tx=XM_019813952.1;refGene_gn=ACADSB;refGene_tx=XM_019813953.2;refGene_gn=ACADSB;refGene_tx=XM_023241046.1
D2 81021332 81021422 refGene_gn=ACADSB;refGene_tx=XM_003994477.5;refGene_gn=ACADSB;refGene_tx=XM_019813952.1;refGene_gn=ACADSB;refGene_tx=XM_019813953.2;refGene_gn=ACADSB;refGene_tx=XM_023241046.1
D2 81025288 81025426 refGene_gn=ACADSB;refGene_tx=XM_003994477.5;refGene_gn=ACADSB;refGene_tx=XM_019813952.1;refGene_gn=ACADSB;refGene_tx=XM_019813953.2
D2 81027428 81027528 refGene_gn=ACADSB;refGene_tx=XM_003994477.5;refGene_gn=ACADSB;refGene_tx=XM_019813952.1;refGene_gn=ACADSB;refGene_tx=XM_019813953.2
D2 81028262 81028333 refGene_gn=ACADSB;refGene_tx=XM_003994477.5;refGene_gn=ACADSB;refGene_tx=XM_019813952.1;refGene_gn=ACADSB;refGene_tx=XM_019813953.2
D2 81031427 81031430 refGene_gn=ACADSB;refGene_tx=XM_023241046.1
D2 81037423 81037528 refGene_gn=ACADSB;refGene_tx=XM_023241046.1
D2 81104712 81105112 refGene_gn=HMX3;refGene_tx=XM_019814102.2

D2 81105739 81106413 refGene_gn=HMX3;refGene_tx=XM_019814102.2
D2 81116899 81117167
refGene_gn=HMX2;refGene_tx=XM_023241047.1;refGene_gn=HMX2;refGene_tx=XM_023241048.1
D2 81118109 81118663
refGene_gn=HMX2;refGene_tx=XM_023241047.1;refGene_gn=HMX2;refGene_tx=XM_023241048.1
D2 81123458 81123563 refGene_gn=BUB3;refGene_tx=XM_011287604.3
D2 81123564 81123654 refGene_gn=BUB3;refGene_tx=XM_011287604.3
D2 81124249 81124319 refGene_gn=BUB3;refGene_tx=XM_011287604.3
D2 81126161 81126313 refGene_gn=BUB3;refGene_tx=XM_011287604.3
D2 81128931 81129090 refGene_gn=BUB3;refGene_tx=XM_011287604.3
D2 81130750 81130928 refGene_gn=BUB3;refGene_tx=XM_011287604.3
D2 81131210 81131427 refGene_gn=BUB3;refGene_tx=XM_011287604.3
D2 81133656 81133666 refGene_gn=BUB3;refGene_tx=XM_011287604.3
D2 81597624 81598292
refGene_gn=GPR26;refGene_tx=XM_003994496.5;refGene_gn=GPR26;refGene_tx=XM_023241049.1;refGene_gn=GPR26;refGene_tx=XM_023241050.1
D2 81607933 81608047
refGene_gn=GPR26;refGene_tx=XM_003994496.5;refGene_gn=GPR26;refGene_tx=XM_023241049.1;refGene_gn=GPR26;refGene_tx=XM_023241050.1
D2 81623192 81623424
refGene_gn=GPR26;refGene_tx=XM_003994496.5;refGene_gn=GPR26;refGene_tx=XM_023241049.1
D2 81645171 81645268 refGene_gn=GPR26;refGene_tx=XM_023241050.1
D2 81663412 81663681
refGene_gn=CPXM2;refGene_tx=XM_006938252.3;refGene_gn=CPXM2;refGene_tx=XM_023241051.1;refGene_gn=CPXM2;refGene_tx=XM_023241052.1;refGene_gn=CPXM2;refGene_tx=XM_023241053.1
D2 81671094 81671194
refGene_gn=CPXM2;refGene_tx=XM_006938252.3;refGene_gn=CPXM2;refGene_tx=XM_023241051.1;refGene_gn=CPXM2;refGene_tx=XM_023241052.1;refGene_gn=CPXM2;refGene_tx=XM_023241053.1
D2 81673069 81673209
refGene_gn=CPXM2;refGene_tx=XM_006938252.3;refGene_gn=CPXM2;refGene_tx=XM_023241051.1;refGene_gn=CPXM2;refGene_tx=XM_023241052.1;refGene_gn=CPXM2;refGene_tx=XM_023241053.1
D2 81678187 81678485
refGene_gn=CPXM2;refGene_tx=XM_006938252.3;refGene_gn=CPXM2;refGene_tx=XM_023241051.1;refGene_gn=CPXM2;refGene_tx=XM_023241052.1;refGene_gn=CPXM2;refGene_tx=XM_023241053.1
D2 81684213 81684393
refGene_gn=CPXM2;refGene_tx=XM_006938252.3;refGene_gn=CPXM2;refGene_tx=XM_023241051.1;refGene_gn=CPXM2;refGene_tx=XM_023241052.1;refGene_gn=CPXM2;refGene_tx=XM_023241053.1
D2 81685396 81685593
refGene_gn=CPXM2;refGene_tx=XM_006938252.3;refGene_gn=CPXM2;refGene_tx=XM_023241051.1;refGene_gn=CPXM2;refGene_tx=XM_023241052.1;refGene_gn=CPXM2;refGene_tx=XM_023241053.1
D2 81688110 81688239

refGene_gn=CPXM2;refGene_tx=XM_006938252.3;refGene_gn=CPXM2;refGene_tx=XM_023241051.1;refGene_gn=CPXM2;refGene_tx=XM_023241052.1;refGene_gn=CPXM2;refGene_tx=XM_023241053.1
D2 81695068 81695157
refGene_gn=CPXM2;refGene_tx=XM_006938252.3;refGene_gn=CPXM2;refGene_tx=XM_023241051.1;refGene_gn=CPXM2;refGene_tx=XM_023241052.1;refGene_gn=CPXM2;refGene_tx=XM_023241053.1
D2 81707655 81707806
refGene_gn=CPXM2;refGene_tx=XM_006938252.3;refGene_gn=CPXM2;refGene_tx=XM_023241051.1;refGene_gn=CPXM2;refGene_tx=XM_023241052.1;refGene_gn=CPXM2;refGene_tx=XM_023241053.1
D2 81708760 81708845
refGene_gn=CPXM2;refGene_tx=XM_006938252.3;refGene_gn=CPXM2;refGene_tx=XM_023241051.1;refGene_gn=CPXM2;refGene_tx=XM_023241052.1;refGene_gn=CPXM2;refGene_tx=XM_023241053.1
D2 81744395 81744577 refGene_gn=CPXM2;refGene_tx=XM_006938252.3
D2 81750259 81750399
refGene_gn=CPXM2;refGene_tx=XM_023241051.1;refGene_gn=CPXM2;refGene_tx=XM_023241052.1;refGene_gn=CPXM2;refGene_tx=XM_023241053.1
D2 81773053 81773163
refGene_gn=CPXM2;refGene_tx=XM_023241051.1;refGene_gn=CPXM2;refGene_tx=XM_023241052.1;refGene_gn=CPXM2;refGene_tx=XM_023241053.1
D2 81797566 81797665
refGene_gn=CPXM2;refGene_tx=XM_023241051.1;refGene_gn=CPXM2;refGene_tx=XM_023241052.1
D2 81807394 81807701
refGene_gn=CPXM2;refGene_tx=XM_023241051.1;refGene_gn=CPXM2;refGene_tx=XM_023241052.1;refGene_gn=CPXM2;refGene_tx=XM_023241053.1
D2 81895254 81895445
refGene_gn=CHST15;refGene_tx=XM_011287605.3;refGene_gn=CHST15;refGene_tx=XM_019813964.2;refGene_gn=CHST15;refGene_tx=XM_023241054.1;refGene_gn=CHST15;refGene_tx=XM_023241055.1;refGene_gn=CHST15;refGene_tx=XM_023241056.1
D2 81897029 81897177
refGene_gn=CHST15;refGene_tx=XM_011287605.3;refGene_gn=CHST15;refGene_tx=XM_019813964.2;refGene_gn=CHST15;refGene_tx=XM_023241054.1;refGene_gn=CHST15;refGene_tx=XM_023241055.1;refGene_gn=CHST15;refGene_tx=XM_023241056.1
D2 81905077 81905234
refGene_gn=CHST15;refGene_tx=XM_011287605.3;refGene_gn=CHST15;refGene_tx=XM_019813964.2;refGene_gn=CHST15;refGene_tx=XM_023241054.1;refGene_gn=CHST15;refGene_tx=XM_023241055.1;refGene_gn=CHST15;refGene_tx=XM_023241056.1
D2 81921409 81921566
refGene_gn=CHST15;refGene_tx=XM_011287605.3;refGene_gn=CHST15;refGene_tx=XM_019813964.2;refGene_gn=CHST15;refGene_tx=XM_023241054.1;refGene_gn=CHST15;refGene_tx=XM_023241055.1;refGene_gn=CHST15;refGene_tx=XM_023241056.1
D2 81925123 81925270
refGene_gn=CHST15;refGene_tx=XM_011287605.3;refGene_gn=CHST15;refGene_

tx=XM_019813964.2;refGene_gn=CHST15;refGene_tx=XM_023241054.1;refGene_gn=CHST15;refGene_tx=XM_023241055.1;refGene_gn=CHST15;refGene_tx=XM_023241056.1
D2 81927007 81927347
refGene_gn=CHST15;refGene_tx=XM_011287605.3;refGene_gn=CHST15;refGene_tx=XM_019813964.2;refGene_gn=CHST15;refGene_tx=XM_023241054.1;refGene_gn=CHST15;refGene_tx=XM_023241055.1;refGene_gn=CHST15;refGene_tx=XM_023241056.1
D2 81928058 81928604
refGene_gn=CHST15;refGene_tx=XM_011287605.3;refGene_gn=CHST15;refGene_tx=XM_019813964.2;refGene_gn=CHST15;refGene_tx=XM_023241054.1;refGene_gn=CHST15;refGene_tx=XM_023241055.1;refGene_gn=CHST15;refGene_tx=XM_023241056.1
D2 82255595 82255756
refGene_gn=OAT;refGene_tx=XM_023241057.1;refGene_gn=OAT;refGene_tx=XM_023241058.1
D2 82257083 82257228
refGene_gn=OAT;refGene_tx=XM_023241057.1;refGene_gn=OAT;refGene_tx=XM_023241058.1
D2 82257963 82258077
refGene_gn=OAT;refGene_tx=XM_023241057.1;refGene_gn=OAT;refGene_tx=XM_023241058.1
D2 82258861 82258990
refGene_gn=OAT;refGene_tx=XM_023241057.1;refGene_gn=OAT;refGene_tx=XM_023241058.1
D2 82259631 82259754
refGene_gn=OAT;refGene_tx=XM_023241057.1;refGene_gn=OAT;refGene_tx=XM_023241058.1
D2 82260771 82260899
refGene_gn=OAT;refGene_tx=XM_023241057.1;refGene_gn=OAT;refGene_tx=XM_023241058.1
D2 82264183 82264279
refGene_gn=OAT;refGene_tx=XM_023241057.1;refGene_gn=OAT;refGene_tx=XM_023241058.1
D2 82264382 82264607
refGene_gn=OAT;refGene_tx=XM_023241057.1;refGene_gn=OAT;refGene_tx=XM_023241058.1
D2 82267014 82267213
refGene_gn=OAT;refGene_tx=XM_023241057.1;refGene_gn=OAT;refGene_tx=XM_023241058.1
D2 82294928 82295650 refGene_gn=NKX1-2;refGene_tx=XM_023241059.1
D2 82297316 82297530 refGene_gn=NKX1-2;refGene_tx=XM_023241059.1
D2 82311507 82311632
refGene_gn=LHPP;refGene_tx=XM_023241062.1;refGene_gn=LHPP;refGene_tx=XM_023241063.1;refGene_gn=LHPP;refGene_tx=XM_023241064.1;refGene_gn=LHPP;refGene_tx=XM_023241065.1
D2 82330505 82330693
refGene_gn=LHPP;refGene_tx=XM_023241062.1;refGene_gn=LHPP;refGene_tx=XM_023241063.1;refGene_gn=LHPP;refGene_tx=XM_023241064.1;refGene_gn=LHPP;refGene_tx=XM_023241065.1

D2 82334016 82334170
refGene_gn=LHPP;refGene_tx=XM_023241062.1;refGene_gn=LHPP;refGene_tx=XM_023241063.1;refGene_gn=LHPP;refGene_tx=XM_023241064.1;refGene_gn=LHPP;refGene_tx=XM_023241065.1
D2 82340280 82340344
refGene_gn=LHPP;refGene_tx=XM_023241062.1;refGene_gn=LHPP;refGene_tx=XM_023241063.1;refGene_gn=LHPP;refGene_tx=XM_023241064.1
D2 82341192 82341285
refGene_gn=LHPP;refGene_tx=XM_023241062.1;refGene_gn=LHPP;refGene_tx=XM_023241063.1;refGene_gn=LHPP;refGene_tx=XM_023241064.1
D2 82356334 82356426
refGene_gn=LHPP;refGene_tx=XM_023241062.1;refGene_gn=LHPP;refGene_tx=XM_023241063.1
D2 82365940 82366103 refGene_gn=LHPP;refGene_tx=XM_023241062.1
D2 82428973 82429070
refGene_gn=LHPP;refGene_tx=XM_023241063.1;refGene_gn=LHPP;refGene_tx=XM_023241064.1;refGene_gn=LHPP;refGene_tx=XM_023241065.1
D2 82437667 82438036
refGene_gn=FAM53B;refGene_tx=XM_023241060.1;refGene_gn=FAM53B;refGene_tx=XM_023241061.1
D2 82487628 82488407
refGene_gn=FAM53B;refGene_tx=XM_023241060.1;refGene_gn=FAM53B;refGene_tx=XM_023241061.1
D2 82501624 82501679
refGene_gn=FAM53B;refGene_tx=XM_023241060.1;refGene_gn=FAM53B;refGene_tx=XM_023241061.1
D2 82509170 82509248
refGene_gn=FAM53B;refGene_tx=XM_023241060.1;refGene_gn=FAM53B;refGene_tx=XM_023241061.1
D2 82541849 82541893
refGene_gn=EEF1AKMT2;refGene_tx=XM_006938255.4
D2 82553215 82553277
refGene_gn=EEF1AKMT2;refGene_tx=XM_023241066.1
D2 82560388 82560483
refGene_gn=EEF1AKMT2;refGene_tx=XM_006938253.1
D2 82561694 82561741
refGene_gn=EEF1AKMT2;refGene_tx=XM_019813966.2
D2 82564423 82564640
refGene_gn=EEF1AKMT2;refGene_tx=XM_006938253.1;refGene_gn=EEF1AKMT2;refGene_tx=XM_006938255.4;refGene_gn=EEF1AKMT2;refGene_tx=XM_019813966.2;refGene_gn=EEF1AKMT2;refGene_tx=XM_023241066.1
D2 82567077 82567185
refGene_gn=EEF1AKMT2;refGene_tx=XM_006938253.1;refGene_gn=EEF1AKMT2;refGene_tx=XM_006938255.4;refGene_gn=EEF1AKMT2;refGene_tx=XM_019813966.2;refGene_gn=EEF1AKMT2;refGene_tx=XM_023241066.1
D2 82576817 82576932
refGene_gn=EEF1AKMT2;refGene_tx=XM_006938253.1;refGene_gn=EEF1AKMT2;refGene_tx=XM_006938255.4;refGene_gn=EEF1AKMT2;refGene_tx=XM_019813966.2;refGene_gn=EEF1AKMT2;refGene_tx=XM_023241066.1
D2 82578190 82578256

refGene_gn=EEF1AKMT2;refGene_tx=XM_006938253.1;refGene_gn=EEF1AKMT2;refGene_tx=XM_006938255.4;refGene_gn=EEF1AKMT2;refGene_tx=XM_019813966.2;refGene_gn=EEF1AKMT2;refGene_tx=XM_023241066.1
D2 82579324 82579443
refGene_gn=EEF1AKMT2;refGene_tx=XM_006938253.1;refGene_gn=EEF1AKMT2;refGene_tx=XM_006938255.4;refGene_gn=EEF1AKMT2;refGene_tx=XM_019813966.2;refGene_gn=EEF1AKMT2;refGene_tx=XM_023241066.1
D2 82588080 82588179 refGene_gn=ABRAXAS2;refGene_tx=XM_003994500.4
D2 82593071 82593162 refGene_gn=ABRAXAS2;refGene_tx=XM_003994500.4
D2 82598366 82598403 refGene_gn=ABRAXAS2;refGene_tx=XM_003994500.4
D2 82601103 82601170 refGene_gn=ABRAXAS2;refGene_tx=XM_003994500.4
D2 82610323 82610514 refGene_gn=ABRAXAS2;refGene_tx=XM_003994500.4
D2 82611988 82612108 refGene_gn=ABRAXAS2;refGene_tx=XM_003994500.4
D2 82612623 82612708 refGene_gn=ABRAXAS2;refGene_tx=XM_003994500.4
D2 82614256 82614371 refGene_gn=ABRAXAS2;refGene_tx=XM_003994500.4
D2 82617801 82618271 refGene_gn=ABRAXAS2;refGene_tx=XM_003994500.4
D2 82712834 82712875 refGene_gn=ZRANB1;refGene_tx=XM_003994501.5
D2 82714168 82715019
refGene_gn=ZRANB1;refGene_tx=XM_003994501.5;refGene_gn=ZRANB1;refGene_tx=XM_023241070.1;refGene_gn=ZRANB1;refGene_tx=XM_023241071.1
D2 82735215 82735403
refGene_gn=ZRANB1;refGene_tx=XM_003994501.5;refGene_gn=ZRANB1;refGene_tx=XM_019813968.2;refGene_gn=ZRANB1;refGene_tx=XM_023241070.1;refGene_gn=ZRANB1;refGene_tx=XM_023241071.1
D2 82739639 82739793
refGene_gn=ZRANB1;refGene_tx=XM_003994501.5;refGene_gn=ZRANB1;refGene_tx=XM_019813968.2;refGene_gn=ZRANB1;refGene_tx=XM_023241070.1;refGene_gn=ZRANB1;refGene_tx=XM_023241071.1
D2 82742166 82742238
refGene_gn=ZRANB1;refGene_tx=XM_003994501.5;refGene_gn=ZRANB1;refGene_tx=XM_019813968.2;refGene_gn=ZRANB1;refGene_tx=XM_023241070.1;refGene_gn=ZRANB1;refGene_tx=XM_023241071.1
D2 82743234 82743433
refGene_gn=ZRANB1;refGene_tx=XM_003994501.5;refGene_gn=ZRANB1;refGene_tx=XM_019813968.2;refGene_gn=ZRANB1;refGene_tx=XM_023241070.1;refGene_gn=ZRANB1;refGene_tx=XM_023241071.1
D2 82747175 82747296
refGene_gn=ZRANB1;refGene_tx=XM_003994501.5;refGene_gn=ZRANB1;refGene_tx=XM_019813968.2;refGene_gn=ZRANB1;refGene_tx=XM_023241070.1;refGene_gn=ZRANB1;refGene_tx=XM_023241071.1
D2 82748488 82748618
refGene_gn=ZRANB1;refGene_tx=XM_003994501.5;refGene_gn=ZRANB1;refGene_tx=XM_019813968.2;refGene_gn=ZRANB1;refGene_tx=XM_023241070.1;refGene_gn=ZRANB1;refGene_tx=XM_023241071.1
D2 82748726 82748956
refGene_gn=ZRANB1;refGene_tx=XM_003994501.5;refGene_gn=ZRANB1;refGene_tx=XM_019813968.2;refGene_gn=ZRANB1;refGene_tx=XM_023241070.1;refGene_gn=ZRANB1;refGene_tx=XM_023241071.1
D2 82750561 82750780
refGene_gn=ZRANB1;refGene_tx=XM_003994501.5;refGene_gn=ZRANB1;refGene_

tx=XM_019813968.2;refGene_gn=ZRANB1;refGene_tx=XM_023241070.1;refGene_gn=ZRANB1;refGene_tx=XM_023241071.1
D2 82755247 82755428
refGene_gn=CTBP2;refGene_tx=XM_011287608.3;refGene_gn=CTBP2;refGene_tx=XM_019813967.2;refGene_gn=CTBP2;refGene_tx=XM_023241068.1;refGene_gn=CTBP2;refGene_tx=XM_023241069.1
D2 82757136 82757254
refGene_gn=CTBP2;refGene_tx=XM_011287608.3;refGene_gn=CTBP2;refGene_tx=XM_019813967.2;refGene_gn=CTBP2;refGene_tx=XM_023241068.1;refGene_gn=CTBP2;refGene_tx=XM_023241069.1
D2 82757984 82758112
refGene_gn=CTBP2;refGene_tx=XM_011287608.3;refGene_gn=CTBP2;refGene_tx=XM_019813967.2;refGene_gn=CTBP2;refGene_tx=XM_023241068.1;refGene_gn=CTBP2;refGene_tx=XM_023241069.1
D2 82758675 82758806
refGene_gn=CTBP2;refGene_tx=XM_011287608.3;refGene_gn=CTBP2;refGene_tx=XM_019813967.2;refGene_gn=CTBP2;refGene_tx=XM_023241068.1;refGene_gn=CTBP2;refGene_tx=XM_023241069.1
D2 82759274 82759489
refGene_gn=CTBP2;refGene_tx=XM_011287608.3;refGene_gn=CTBP2;refGene_tx=XM_019813967.2;refGene_gn=CTBP2;refGene_tx=XM_023241068.1;refGene_gn=CTBP2;refGene_tx=XM_023241069.1
D2 82762681 82762888
refGene_gn=CTBP2;refGene_tx=XM_011287608.3;refGene_gn=CTBP2;refGene_tx=XM_019813967.2;refGene_gn=CTBP2;refGene_tx=XM_023241068.1;refGene_gn=CTBP2;refGene_tx=XM_023241069.1
D2 82766180 82766325
refGene_gn=CTBP2;refGene_tx=XM_011287608.3;refGene_gn=CTBP2;refGene_tx=XM_019813967.2;refGene_gn=CTBP2;refGene_tx=XM_023241068.1;refGene_gn=CTBP2;refGene_tx=XM_023241069.1
D2 82766536 82766691
refGene_gn=CTBP2;refGene_tx=XM_011287608.3;refGene_gn=CTBP2;refGene_tx=XM_019813967.2;refGene_gn=CTBP2;refGene_tx=XM_023241068.1;refGene_gn=CTBP2;refGene_tx=XM_023241069.1
D2 82790052 82791706 refGene_gn=CTBP2;refGene_tx=XM_011287608.3
D2 82804105 82804163
refGene_gn=CTBP2;refGene_tx=XM_019813967.2;refGene_gn=CTBP2;refGene_tx=XM_023241068.1;refGene_gn=CTBP2;refGene_tx=XM_023241069.1
D2 83354488 83354524
refGene_gn=TEX36;refGene_tx=XM_003994511.4;refGene_gn=TEX36;refGene_tx=XM_019813969.2;refGene_gn=TEX36;refGene_tx=XM_019813971.2
D2 83355030 83355279
refGene_gn=TEX36;refGene_tx=XM_003994511.4;refGene_gn=TEX36;refGene_tx=XM_019813969.2
D2 83360925 83361006
refGene_gn=TEX36;refGene_tx=XM_003994511.4;refGene_gn=TEX36;refGene_tx=XM_019813969.2;refGene_gn=TEX36;refGene_tx=XM_019813971.2
D2 83361722 83361854
refGene_gn=TEX36;refGene_tx=XM_003994511.4;refGene_gn=TEX36;refGene_tx=XM_019813969.2;refGene_gn=TEX36;refGene_tx=XM_019813971.2

D2	83366838	83366889
refGene_gn=TEX36;refGene_tx=XM_003994511.4;refGene_gn=TEX36;refGene_tx=XM_019813969.2;refGene_gn=TEX36;refGene_tx=XM_019813971.2
D2	83400782	83400890
refGene_gn=EDRF1;refGene_tx=XM_003994512.4;refGene_gn=EDRF1;refGene_tx=XM_006938262.3;refGene_gn=EDRF1;refGene_tx=XM_019813972.2;refGene_gn=EDRF1;refGene_tx=XM_019813973.2
D2	83401944	83402153
refGene_gn=EDRF1;refGene_tx=XM_003994512.4;refGene_gn=EDRF1;refGene_tx=XM_006938262.3;refGene_gn=EDRF1;refGene_tx=XM_019813972.2;refGene_gn=EDRF1;refGene_tx=XM_019813973.2
D2	83404030	83404097
refGene_gn=EDRF1;refGene_tx=XM_003994512.4;refGene_gn=EDRF1;refGene_tx=XM_006938262.3;refGene_gn=EDRF1;refGene_tx=XM_019813972.2;refGene_gn=EDRF1;refGene_tx=XM_019813973.2
D2	83404767	83404893
refGene_gn=EDRF1;refGene_tx=XM_003994512.4;refGene_gn=EDRF1;refGene_tx=XM_006938262.3;refGene_gn=EDRF1;refGene_tx=XM_019813972.2;refGene_gn=EDRF1;refGene_tx=XM_019813973.2
D2	83405816	83405941
refGene_gn=EDRF1;refGene_tx=XM_003994512.4;refGene_gn=EDRF1;refGene_tx=XM_006938262.3;refGene_gn=EDRF1;refGene_tx=XM_019813972.2;refGene_gn=EDRF1;refGene_tx=XM_019813973.2
D2	83406180	83406337
refGene_gn=EDRF1;refGene_tx=XM_003994512.4;refGene_gn=EDRF1;refGene_tx=XM_006938262.3;refGene_gn=EDRF1;refGene_tx=XM_019813972.2;refGene_gn=EDRF1;refGene_tx=XM_019813973.2
D2	83409609	83409711
refGene_gn=EDRF1;refGene_tx=XM_003994512.4;refGene_gn=EDRF1;refGene_tx=XM_019813972.2;refGene_gn=EDRF1;refGene_tx=XM_019813973.2
D2	83409980	83410102
refGene_gn=EDRF1;refGene_tx=XM_003994512.4;refGene_gn=EDRF1;refGene_tx=XM_006938262.3;refGene_gn=EDRF1;refGene_tx=XM_019813972.2;refGene_gn=EDRF1;refGene_tx=XM_019813973.2
D2	83410776	83410888
refGene_gn=EDRF1;refGene_tx=XM_003994512.4;refGene_gn=EDRF1;refGene_tx=XM_006938262.3;refGene_gn=EDRF1;refGene_tx=XM_019813972.2;refGene_gn=EDRF1;refGene_tx=XM_019813973.2
D2	83413633	83413781
refGene_gn=EDRF1;refGene_tx=XM_003994512.4;refGene_gn=EDRF1;refGene_tx=XM_006938262.3;refGene_gn=EDRF1;refGene_tx=XM_019813972.2;refGene_gn=EDRF1;refGene_tx=XM_019813973.2
D2	83413852	83413961
refGene_gn=EDRF1;refGene_tx=XM_003994512.4;refGene_gn=EDRF1;refGene_tx=XM_006938262.3;refGene_gn=EDRF1;refGene_tx=XM_019813972.2;refGene_gn=EDRF1;refGene_tx=XM_019813973.2
D2	83414247	83414359
refGene_gn=EDRF1;refGene_tx=XM_003994512.4;refGene_gn=EDRF1;refGene_tx=XM_006938262.3;refGene_gn=EDRF1;refGene_tx=XM_019813972.2;refGene_gn=EDRF1;refGene_tx=XM_019813973.2

D2 83415800 83416061
refGene_gn=EDRF1;refGene_tx=XM_003994512.4;refGene_gn=EDRF1;refGene_tx=XM_006938262.3;refGene_gn=EDRF1;refGene_tx=XM_019813972.2;refGene_gn=EDRF1;refGene_tx=XM_019813973.2
D2 83418205 83418277
refGene_gn=EDRF1;refGene_tx=XM_003994512.4;refGene_gn=EDRF1;refGene_tx=XM_006938262.3;refGene_gn=EDRF1;refGene_tx=XM_019813972.2;refGene_gn=EDRF1;refGene_tx=XM_019813973.2
D2 83418462 83418613
refGene_gn=EDRF1;refGene_tx=XM_003994512.4;refGene_gn=EDRF1;refGene_tx=XM_006938262.3;refGene_gn=EDRF1;refGene_tx=XM_019813972.2;refGene_gn=EDRF1;refGene_tx=XM_019813973.2
D2 83420625 83420814
refGene_gn=EDRF1;refGene_tx=XM_003994512.4;refGene_gn=EDRF1;refGene_tx=XM_006938262.3;refGene_gn=EDRF1;refGene_tx=XM_019813972.2;refGene_gn=EDRF1;refGene_tx=XM_019813973.2
D2 83421142 83421343
refGene_gn=EDRF1;refGene_tx=XM_003994512.4;refGene_gn=EDRF1;refGene_tx=XM_006938262.3;refGene_gn=EDRF1;refGene_tx=XM_019813972.2;refGene_gn=EDRF1;refGene_tx=XM_019813973.2
D2 83422818 83423037
refGene_gn=EDRF1;refGene_tx=XM_003994512.4;refGene_gn=EDRF1;refGene_tx=XM_006938262.3;refGene_gn=EDRF1;refGene_tx=XM_019813972.2;refGene_gn=EDRF1;refGene_tx=XM_019813973.2
D2 83425103 83425327
refGene_gn=EDRF1;refGene_tx=XM_003994512.4;refGene_gn=EDRF1;refGene_tx=XM_006938262.3;refGene_gn=EDRF1;refGene_tx=XM_019813972.2;refGene_gn=EDRF1;refGene_tx=XM_019813973.2
D2 83427484 83427643
refGene_gn=EDRF1;refGene_tx=XM_003994512.4;refGene_gn=EDRF1;refGene_tx=XM_006938262.3;refGene_gn=EDRF1;refGene_tx=XM_019813972.2;refGene_gn=EDRF1;refGene_tx=XM_019813973.2
D2 83427790 83427940
refGene_gn=EDRF1;refGene_tx=XM_003994512.4;refGene_gn=EDRF1;refGene_tx=XM_006938262.3;refGene_gn=EDRF1;refGene_tx=XM_019813972.2;refGene_gn=EDRF1;refGene_tx=XM_019813973.2
D2 83429251 83429405
refGene_gn=EDRF1;refGene_tx=XM_003994512.4;refGene_gn=EDRF1;refGene_tx=XM_006938262.3;refGene_gn=EDRF1;refGene_tx=XM_019813972.2;refGene_gn=EDRF1;refGene_tx=XM_019813973.2
D2 83430848 83430964
refGene_gn=EDRF1;refGene_tx=XM_003994512.4;refGene_gn=EDRF1;refGene_tx=XM_006938262.3;refGene_gn=EDRF1;refGene_tx=XM_019813972.2;refGene_gn=EDRF1;refGene_tx=XM_019813973.2
D2 83432052 83432204
refGene_gn=EDRF1;refGene_tx=XM_003994512.4;refGene_gn=EDRF1;refGene_tx=XM_006938262.3;refGene_gn=EDRF1;refGene_tx=XM_019813972.2;refGene_gn=EDRF1;refGene_tx=XM_019813973.2
D2 83437898 83437911 refGene_gn=EDRF1;refGene_tx=XM_019813973.2
D2 83439819 83439976

refGene_gn=EDRF1;refGene_tx=XM_003994512.4;refGene_gn=EDRF1;refGene_tx=XM_006938262.3;refGene_gn=EDRF1;refGene_tx=XM_019813972.2
D2 83442376 83442676 refGene_gn=MMP21;refGene_tx=XM_023241072.1
D2 83442931 83443104 refGene_gn=MMP21;refGene_tx=XM_023241072.1
D2 83445552 83445810 refGene_gn=MMP21;refGene_tx=XM_023241072.1
D2 83447000 83447142 refGene_gn=MMP21;refGene_tx=XM_023241072.1
D2 83447342 83447482 refGene_gn=MMP21;refGene_tx=XM_023241072.1
D2 83448235 83448746 refGene_gn=MMP21;refGene_tx=XM_023241072.1
D2 83449167 83449329 refGene_gn=MMP21;refGene_tx=XM_023241072.1
D2 83457961 83458096
refGene_gn=UROS;refGene_tx=XM_003994514.4;refGene_gn=UROS;refGene_tx=XM_006938265.3;refGene_gn=UROS;refGene_tx=XM_006938267.3;refGene_gn=UROS;refGene_tx=XM_019813980.2;refGene_gn=UROS;refGene_tx=XM_019813981.2;refGene_gn=UROS;refGene_tx=XM_019813982.2;refGene_gn=UROS;refGene_tx=XM_023241082.1
D2 83460436 83460535
refGene_gn=UROS;refGene_tx=XM_003994514.4;refGene_gn=UROS;refGene_tx=XM_006938265.3;refGene_gn=UROS;refGene_tx=XM_006938267.3;refGene_gn=UROS;refGene_tx=XM_019813980.2;refGene_gn=UROS;refGene_tx=XM_019813981.2;refGene_gn=UROS;refGene_tx=XM_019813982.2;refGene_gn=UROS;refGene_tx=XM_023241082.1
D2 83461396 83461482
refGene_gn=UROS;refGene_tx=XM_003994514.4;refGene_gn=UROS;refGene_tx=XM_006938265.3;refGene_gn=UROS;refGene_tx=XM_006938267.3;refGene_gn=UROS;refGene_tx=XM_019813980.2;refGene_gn=UROS;refGene_tx=XM_019813981.2;refGene_gn=UROS;refGene_tx=XM_019813982.2;refGene_gn=UROS;refGene_tx=XM_023241082.1
D2 83462410 83462491
refGene_gn=UROS;refGene_tx=XM_003994514.4;refGene_gn=UROS;refGene_tx=XM_006938265.3;refGene_gn=UROS;refGene_tx=XM_006938267.3;refGene_gn=UROS;refGene_tx=XM_019813980.2;refGene_gn=UROS;refGene_tx=XM_019813981.2;refGene_gn=UROS;refGene_tx=XM_019813982.2;refGene_gn=UROS;refGene_tx=XM_023241082.1
D2 83467325 83467414 refGene_gn=UROS;refGene_tx=XM_019813983.2
D2 83467781 83467856
refGene_gn=UROS;refGene_tx=XM_003994514.4;refGene_gn=UROS;refGene_tx=XM_006938265.3;refGene_gn=UROS;refGene_tx=XM_006938267.3;refGene_gn=UROS;refGene_tx=XM_019813980.2;refGene_gn=UROS;refGene_tx=XM_019813981.2;refGene_gn=UROS;refGene_tx=XM_019813982.2;refGene_gn=UROS;refGene_tx=XM_019813983.2;refGene_gn=UROS;refGene_tx=XM_023241082.1
D2 83477466 83477541
refGene_gn=UROS;refGene_tx=XM_003994514.4;refGene_gn=UROS;refGene_tx=XM_006938265.3;refGene_gn=UROS;refGene_tx=XM_006938267.3;refGene_gn=UROS;refGene_tx=XM_019813980.2;refGene_gn=UROS;refGene_tx=XM_019813981.2;refGene_gn=UROS;refGene_tx=XM_019813982.2;refGene_gn=UROS;refGene_tx=XM_019813983.2;refGene_gn=UROS;refGene_tx=XM_023241082.1
D2 83480649 83480746
refGene_gn=UROS;refGene_tx=XM_003994514.4;refGene_gn=UROS;refGene_tx=XM_006938265.3;refGene_gn=UROS;refGene_tx=XM_006938267.3;refGene_gn=UROS;refGene_tx=XM_019813980.2;refGene_gn=UROS;refGene_tx=XM_019813981.2;

refGene_gn=UROS;refGene_tx=XM_019813982.2;refGene_gn=UROS;refGene_tx=XM_019813983.2;refGene_gn=UROS;refGene_tx=XM_023241082.1
D2 83482386 83482470
refGene_gn=UROS;refGene_tx=XM_003994514.4;refGene_gn=UROS;refGene_tx=XM_006938265.3;refGene_gn=UROS;refGene_tx=XM_006938267.3;refGene_gn=UROS;refGene_tx=XM_019813980.2;refGene_gn=UROS;refGene_tx=XM_019813981.2;refGene_gn=UROS;refGene_tx=XM_019813982.2;refGene_gn=UROS;refGene_tx=XM_019813983.2;refGene_gn=UROS;refGene_tx=XM_023241082.1
D2 83482652 83482741
refGene_gn=UROS;refGene_tx=XM_003994514.4;refGene_gn=UROS;refGene_tx=XM_006938265.3;refGene_gn=UROS;refGene_tx=XM_006938267.3;refGene_gn=UROS;refGene_tx=XM_019813980.2;refGene_gn=UROS;refGene_tx=XM_019813981.2;refGene_gn=UROS;refGene_tx=XM_019813982.2;refGene_gn=UROS;refGene_tx=XM_019813983.2;refGene_gn=UROS;refGene_tx=XM_023241082.1
D2 83486782 83486813
refGene_gn=UROS;refGene_tx=XM_006938265.3;refGene_gn=UROS;refGene_tx=XM_019813982.2
D2 83487480 83487556
refGene_gn=UROS;refGene_tx=XM_019813980.2;refGene_gn=UROS;refGene_tx=XM_019813981.2;refGene_gn=UROS;refGene_tx=XM_019813983.2;refGene_gn=UROS;refGene_tx=XM_023241082.1
D2 83493191 83493359
refGene_gn=BCCIP;refGene_tx=XM_003994509.5;refGene_gn=BCCIP;refGene_tx=XM_019813979.2
D2 83498065 83498140
refGene_gn=BCCIP;refGene_tx=XM_003994509.5;refGene_gn=BCCIP;refGene_tx=XM_019813979.2
D2 83499032 83499113
refGene_gn=BCCIP;refGene_tx=XM_003994509.5;refGene_gn=BCCIP;refGene_tx=XM_019813979.2
D2 83503139 83503229
refGene_gn=BCCIP;refGene_tx=XM_003994509.5;refGene_gn=BCCIP;refGene_tx=XM_019813979.2
D2 83503882 83504070
refGene_gn=BCCIP;refGene_tx=XM_003994509.5;refGene_gn=BCCIP;refGene_tx=XM_019813979.2
D2 83505371 83505546
refGene_gn=BCCIP;refGene_tx=XM_003994509.5;refGene_gn=BCCIP;refGene_tx=XM_019813979.2
D2 83506779 83506950
refGene_gn=BCCIP;refGene_tx=XM_003994509.5;refGene_gn=BCCIP;refGene_tx=XM_019813979.2
D2 83507390 83507562
refGene_gn=DHX32;refGene_tx=XM_011287615.3;refGene_gn=DHX32;refGene_tx=XM_011287616.3;refGene_gn=DHX32;refGene_tx=XM_011287617.3;refGene_gn=DHX32;refGene_tx=XM_011287618.3;refGene_gn=DHX32;refGene_tx=XM_023241075.1;refGene_gn=DHX32;refGene_tx=XM_023241076.1;refGene_gn=DHX32;refGene_tx=XM_023241077.1;refGene_gn=DHX32;refGene_tx=XM_023241078.1
D2 83508544 83508548 refGene_gn=DHX32;refGene_tx=XM_023241079.1
D2 83508629 83508811

refGene_gn=DHX32;refGene_tx=XM_011287615.3;refGene_gn=DHX32;refGene_tx
=XM_011287616.3;refGene_gn=DHX32;refGene_tx=XM_011287617.3;refGene_gn=
DHX32;refGene_tx=XM_011287618.3;refGene_gn=DHX32;refGene_tx=XM_0232410
75.1;refGene_gn=DHX32;refGene_tx=XM_023241076.1;refGene_gn=DHX32;refGe
ne_tx=XM_023241077.1;refGene_gn=DHX32;refGene_tx=XM_023241078.1;refGen
e_gn=DHX32;refGene_tx=XM_023241079.1
D2 83510654 83510842
refGene_gn=DHX32;refGene_tx=XM_011287615.3;refGene_gn=DHX32;refGene_tx
=XM_011287616.3;refGene_gn=DHX32;refGene_tx=XM_011287617.3;refGene_gn=
DHX32;refGene_tx=XM_011287618.3;refGene_gn=DHX32;refGene_tx=XM_0232410
75.1;refGene_gn=DHX32;refGene_tx=XM_023241076.1;refGene_gn=DHX32;refGe
ne_tx=XM_023241077.1;refGene_gn=DHX32;refGene_tx=XM_023241078.1;refGen
e_gn=DHX32;refGene_tx=XM_023241079.1
D2 83512342 83512492
refGene_gn=DHX32;refGene_tx=XM_011287615.3;refGene_gn=DHX32;refGene_tx
=XM_011287616.3;refGene_gn=DHX32;refGene_tx=XM_011287617.3;refGene_gn=
DHX32;refGene_tx=XM_011287618.3;refGene_gn=DHX32;refGene_tx=XM_0232410
75.1;refGene_gn=DHX32;refGene_tx=XM_023241076.1;refGene_gn=DHX32;refGe
ne_tx=XM_023241077.1;refGene_gn=DHX32;refGene_tx=XM_023241078.1;refGen
e_gn=DHX32;refGene_tx=XM_023241079.1
D2 83513168 83513360
refGene_gn=DHX32;refGene_tx=XM_011287615.3;refGene_gn=DHX32;refGene_tx
=XM_011287616.3;refGene_gn=DHX32;refGene_tx=XM_011287617.3;refGene_gn=
DHX32;refGene_tx=XM_011287618.3;refGene_gn=DHX32;refGene_tx=XM_0232410
75.1;refGene_gn=DHX32;refGene_tx=XM_023241076.1;refGene_gn=DHX32;refGe
ne_tx=XM_023241077.1;refGene_gn=DHX32;refGene_tx=XM_023241078.1;refGen
e_gn=DHX32;refGene_tx=XM_023241079.1
D2 83522178 83522337
refGene_gn=DHX32;refGene_tx=XM_011287615.3;refGene_gn=DHX32;refGene_tx
=XM_011287616.3;refGene_gn=DHX32;refGene_tx=XM_011287617.3;refGene_gn=
DHX32;refGene_tx=XM_011287618.3;refGene_gn=DHX32;refGene_tx=XM_0232410
75.1;refGene_gn=DHX32;refGene_tx=XM_023241076.1;refGene_gn=DHX32;refGe
ne_tx=XM_023241077.1;refGene_gn=DHX32;refGene_tx=XM_023241078.1;refGen
e_gn=DHX32;refGene_tx=XM_023241079.1
D2 83522424 83522524
refGene_gn=DHX32;refGene_tx=XM_011287615.3;refGene_gn=DHX32;refGene_tx
=XM_011287616.3;refGene_gn=DHX32;refGene_tx=XM_011287617.3;refGene_gn=
DHX32;refGene_tx=XM_011287618.3;refGene_gn=DHX32;refGene_tx=XM_0232410
75.1;refGene_gn=DHX32;refGene_tx=XM_023241076.1;refGene_gn=DHX32;refGe
ne_tx=XM_023241077.1;refGene_gn=DHX32;refGene_tx=XM_023241078.1;refGen
e_gn=DHX32;refGene_tx=XM_023241079.1
D2 83523835 83524081
refGene_gn=DHX32;refGene_tx=XM_011287615.3;refGene_gn=DHX32;refGene_tx
=XM_011287616.3;refGene_gn=DHX32;refGene_tx=XM_011287617.3;refGene_gn=
DHX32;refGene_tx=XM_011287618.3;refGene_gn=DHX32;refGene_tx=XM_0232410
75.1;refGene_gn=DHX32;refGene_tx=XM_023241076.1;refGene_gn=DHX32;refGe
ne_tx=XM_023241077.1;refGene_gn=DHX32;refGene_tx=XM_023241078.1;refGen
e_gn=DHX32;refGene_tx=XM_023241079.1
D2 83529993 83530366
refGene_gn=DHX32;refGene_tx=XM_011287615.3;refGene_gn=DHX32;refGene_tx

=XM_011287616.3;refGene_gn=DHX32;refGene_tx=XM_011287617.3;refGene_gn=DHX32;refGene_tx=XM_011287618.3;refGene_gn=DHX32;refGene_tx=XM_023241075.1;refGene_gn=DHX32;refGene_tx=XM_023241076.1;refGene_gn=DHX32;refGene_tx=XM_023241077.1;refGene_gn=DHX32;refGene_tx=XM_023241078.1;refGene_gn=DHX32;refGene_tx=XM_023241079.1
D2 83533832 83534026
refGene_gn=DHX32;refGene_tx=XM_011287615.3;refGene_gn=DHX32;refGene_tx=XM_011287616.3;refGene_gn=DHX32;refGene_tx=XM_011287617.3;refGene_gn=DHX32;refGene_tx=XM_011287618.3;refGene_gn=DHX32;refGene_tx=XM_023241075.1;refGene_gn=DHX32;refGene_tx=XM_023241076.1;refGene_gn=DHX32;refGene_tx=XM_023241077.1;refGene_gn=DHX32;refGene_tx=XM_023241078.1;refGene_gn=DHX32;refGene_tx=XM_023241079.1
D2 83545361 83545643
refGene_gn=DHX32;refGene_tx=XM_011287615.3;refGene_gn=DHX32;refGene_tx=XM_011287616.3;refGene_gn=DHX32;refGene_tx=XM_011287617.3;refGene_gn=DHX32;refGene_tx=XM_011287618.3;refGene_gn=DHX32;refGene_tx=XM_023241075.1;refGene_gn=DHX32;refGene_tx=XM_023241076.1;refGene_gn=DHX32;refGene_tx=XM_023241077.1;refGene_gn=DHX32;refGene_tx=XM_023241078.1;refGene_gn=DHX32;refGene_tx=XM_023241079.1
D2 83561063 83561076
refGene_gn=FANK1;refGene_tx=XM_023241080.1;refGene_gn=FANK1;refGene_tx=XM_023241081.1
D2 83642314 83642492
refGene_gn=FANK1;refGene_tx=XM_023241080.1;refGene_gn=FANK1;refGene_tx=XM_023241081.1
D2 83648854 83648979
refGene_gn=FANK1;refGene_tx=XM_023241080.1;refGene_gn=FANK1;refGene_tx=XM_023241081.1
D2 83656045 83656127
refGene_gn=FANK1;refGene_tx=XM_023241080.1;refGene_gn=FANK1;refGene_tx=XM_023241081.1
D2 83657146 83657221
refGene_gn=FANK1;refGene_tx=XM_023241080.1;refGene_gn=FANK1;refGene_tx=XM_023241081.1
D2 83658369 83658435
refGene_gn=FANK1;refGene_tx=XM_023241080.1;refGene_gn=FANK1;refGene_tx=XM_023241081.1
D2 83664679 83664845
refGene_gn=FANK1;refGene_tx=XM_023241080.1;refGene_gn=FANK1;refGene_tx=XM_023241081.1
D2 83667450 83667594
refGene_gn=FANK1;refGene_tx=XM_023241080.1;refGene_gn=FANK1;refGene_tx=XM_023241081.1
D2 83668118 83668196
refGene_gn=FANK1;refGene_tx=XM_023241080.1;refGene_gn=FANK1;refGene_tx=XM_023241081.1
D2 83668392 83668452
refGene_gn=FANK1;refGene_tx=XM_023241080.1;refGene_gn=FANK1;refGene_tx=XM_023241081.1
D2 83668539 83668605

refGene_gn=FANK1;refGene_tx=XM_023241080.1;refGene_gn=FANK1;refGene_tx=XM_023241081.1
D2 83677230 83677291 refGene_gn=ADAM12;refGene_tx=XM_023241073.1
D2 83679445 83679576 refGene_gn=ADAM12;refGene_tx=XM_023241073.1
D2 83689182 83689380 refGene_gn=ADAM12;refGene_tx=XM_023241074.1
D2 83698608 83698788 refGene_gn=ADAM12;refGene_tx=XM_023241073.1;refGene_gn=ADAM12;refGene_tx=XM_023241074.1
D2 83700805 83700914 refGene_gn=ADAM12;refGene_tx=XM_023241073.1;refGene_gn=ADAM12;refGene_tx=XM_023241074.1
D2 83701542 83701678 refGene_gn=ADAM12;refGene_tx=XM_023241073.1;refGene_gn=ADAM12;refGene_tx=XM_023241074.1
D2 83704551 83704660 refGene_gn=ADAM12;refGene_tx=XM_023241073.1;refGene_gn=ADAM12;refGene_tx=XM_023241074.1
D2 83707339 83707417 refGene_gn=ADAM12;refGene_tx=XM_023241073.1;refGene_gn=ADAM12;refGene_tx=XM_023241074.1
D2 83709767 83709966 refGene_gn=ADAM12;refGene_tx=XM_023241073.1;refGene_gn=ADAM12;refGene_tx=XM_023241074.1
D2 83710072 83710181 refGene_gn=ADAM12;refGene_tx=XM_023241073.1;refGene_gn=ADAM12;refGene_tx=XM_023241074.1
D2 83725317 83725513 refGene_gn=ADAM12;refGene_tx=XM_023241073.1;refGene_gn=ADAM12;refGene_tx=XM_023241074.1
D2 83727473 83727563 refGene_gn=ADAM12;refGene_tx=XM_023241073.1;refGene_gn=ADAM12;refGene_tx=XM_023241074.1
D2 83731938 83732116 refGene_gn=ADAM12;refGene_tx=XM_023241073.1;refGene_gn=ADAM12;refGene_tx=XM_023241074.1
D2 83753235 83753384 refGene_gn=ADAM12;refGene_tx=XM_023241073.1;refGene_gn=ADAM12;refGene_tx=XM_023241074.1
D2 83757557 83757642 refGene_gn=ADAM12;refGene_tx=XM_023241073.1;refGene_gn=ADAM12;refGene_tx=XM_023241074.1
D2 83760414 83760584 refGene_gn=ADAM12;refGene_tx=XM_023241073.1;refGene_gn=ADAM12;refGene_tx=XM_023241074.1
D2 83765321 83765393 refGene_gn=ADAM12;refGene_tx=XM_023241073.1;refGene_gn=ADAM12;refGene_tx=XM_023241074.1
D2 83766440 83766506 refGene_gn=ADAM12;refGene_tx=XM_023241073.1;refGene_gn=ADAM12;refGene_tx=XM_023241074.1

D2 83774129 83774316
refGene_gn=ADAM12;refGene_tx=XM_023241073.1;refGene_gn=ADAM12;refGene_tx=XM_023241074.1
D2 83789886 83789963
refGene_gn=ADAM12;refGene_tx=XM_023241073.1;refGene_gn=ADAM12;refGene_tx=XM_023241074.1
D2 83807201 83807280
refGene_gn=ADAM12;refGene_tx=XM_023241073.1;refGene_gn=ADAM12;refGene_tx=XM_023241074.1
D2 83929479 83929553
refGene_gn=ADAM12;refGene_tx=XM_023241073.1;refGene_gn=ADAM12;refGene_tx=XM_023241074.1
D2 83974071 83974208
refGene_gn=ADAM12;refGene_tx=XM_023241073.1;refGene_gn=ADAM12;refGene_tx=XM_023241074.1
D2 84020542 84020630
refGene_gn=ADAM12;refGene_tx=XM_023241073.1;refGene_gn=ADAM12;refGene_tx=XM_023241074.1
D2 84463924 84463970
refGene_gn=DOCK1;refGene_tx=XM_011287620.3;refGene_gn=DOCK1;refGene_tx=XM_011287621.3;refGene_gn=DOCK1;refGene_tx=XM_019813984.2
D2 84526745 84526829
refGene_gn=DOCK1;refGene_tx=XM_011287620.3;refGene_gn=DOCK1;refGene_tx=XM_011287621.3;refGene_gn=DOCK1;refGene_tx=XM_019813984.2
D2 84533647 84533688
refGene_gn=DOCK1;refGene_tx=XM_011287620.3;refGene_gn=DOCK1;refGene_tx=XM_011287621.3;refGene_gn=DOCK1;refGene_tx=XM_019813984.2
D2 84536728 84536784
refGene_gn=DOCK1;refGene_tx=XM_011287620.3;refGene_gn=DOCK1;refGene_tx=XM_011287621.3;refGene_gn=DOCK1;refGene_tx=XM_019813984.2
D2 84540366 84540463
refGene_gn=DOCK1;refGene_tx=XM_011287620.3;refGene_gn=DOCK1;refGene_tx=XM_011287621.3;refGene_gn=DOCK1;refGene_tx=XM_019813984.2
D2 84542815 84542964
refGene_gn=DOCK1;refGene_tx=XM_011287620.3;refGene_gn=DOCK1;refGene_tx=XM_011287621.3;refGene_gn=DOCK1;refGene_tx=XM_019813984.2
D2 84547392 84547528
refGene_gn=DOCK1;refGene_tx=XM_011287620.3;refGene_gn=DOCK1;refGene_tx=XM_011287621.3;refGene_gn=DOCK1;refGene_tx=XM_019813984.2
D2 84548692 84548850
refGene_gn=DOCK1;refGene_tx=XM_011287620.3;refGene_gn=DOCK1;refGene_tx=XM_011287621.3;refGene_gn=DOCK1;refGene_tx=XM_019813984.2
D2 84550071 84550153
refGene_gn=DOCK1;refGene_tx=XM_011287620.3;refGene_gn=DOCK1;refGene_tx=XM_011287621.3;refGene_gn=DOCK1;refGene_tx=XM_019813984.2
D2 84550566 84550702
refGene_gn=DOCK1;refGene_tx=XM_011287620.3;refGene_gn=DOCK1;refGene_tx=XM_011287621.3;refGene_gn=DOCK1;refGene_tx=XM_019813984.2
D2 84553849 84553922
refGene_gn=DOCK1;refGene_tx=XM_011287620.3;refGene_gn=DOCK1;refGene_tx

=XM_011287621.3;refGene_gn=DOCK1;refGene_tx=XM_019813984.2
D2	84557019	84557162
refGene_gn=DOCK1;refGene_tx=XM_011287620.3;refGene_gn=DOCK1;refGene_tx=XM_011287621.3;refGene_gn=DOCK1;refGene_tx=XM_019813984.2
D2	84563909	84564035
refGene_gn=DOCK1;refGene_tx=XM_011287620.3;refGene_gn=DOCK1;refGene_tx=XM_011287621.3;refGene_gn=DOCK1;refGene_tx=XM_019813984.2
D2	84568767	84568892
refGene_gn=DOCK1;refGene_tx=XM_011287620.3;refGene_gn=DOCK1;refGene_tx=XM_011287621.3;refGene_gn=DOCK1;refGene_tx=XM_019813984.2
D2	84570059	84570158
refGene_gn=DOCK1;refGene_tx=XM_011287620.3;refGene_gn=DOCK1;refGene_tx=XM_011287621.3;refGene_gn=DOCK1;refGene_tx=XM_019813984.2
D2	84571123	84571196
refGene_gn=DOCK1;refGene_tx=XM_011287620.3;refGene_gn=DOCK1;refGene_tx=XM_011287621.3;refGene_gn=DOCK1;refGene_tx=XM_019813984.2
D2	84576543	84576647
refGene_gn=DOCK1;refGene_tx=XM_011287620.3;refGene_gn=DOCK1;refGene_tx=XM_011287621.3;refGene_gn=DOCK1;refGene_tx=XM_019813984.2
D2	84577043	84577227
refGene_gn=DOCK1;refGene_tx=XM_011287620.3;refGene_gn=DOCK1;refGene_tx=XM_011287621.3;refGene_gn=DOCK1;refGene_tx=XM_019813984.2
D2	84582029	84582127
refGene_gn=DOCK1;refGene_tx=XM_011287620.3;refGene_gn=DOCK1;refGene_tx=XM_011287621.3;refGene_gn=DOCK1;refGene_tx=XM_019813984.2
D2	84586283	84586373
refGene_gn=DOCK1;refGene_tx=XM_011287620.3;refGene_gn=DOCK1;refGene_tx=XM_011287621.3;refGene_gn=DOCK1;refGene_tx=XM_019813984.2
D2	84586685	84586786
refGene_gn=DOCK1;refGene_tx=XM_011287620.3;refGene_gn=DOCK1;refGene_tx=XM_011287621.3;refGene_gn=DOCK1;refGene_tx=XM_019813984.2
D2	84591344	84591479
refGene_gn=DOCK1;refGene_tx=XM_011287620.3;refGene_gn=DOCK1;refGene_tx=XM_011287621.3;refGene_gn=DOCK1;refGene_tx=XM_019813984.2
D2	84600391	84600500
refGene_gn=DOCK1;refGene_tx=XM_011287620.3;refGene_gn=DOCK1;refGene_tx=XM_011287621.3;refGene_gn=DOCK1;refGene_tx=XM_019813984.2
D2	84651515	84651586
refGene_gn=DOCK1;refGene_tx=XM_011287620.3;refGene_gn=DOCK1;refGene_tx=XM_011287621.3;refGene_gn=DOCK1;refGene_tx=XM_019813984.2
D2	84655388	84655495
refGene_gn=DOCK1;refGene_tx=XM_011287620.3;refGene_gn=DOCK1;refGene_tx=XM_011287621.3;refGene_gn=DOCK1;refGene_tx=XM_019813984.2
D2	84672857	84672985
refGene_gn=DOCK1;refGene_tx=XM_011287620.3;refGene_gn=DOCK1;refGene_tx=XM_011287621.3;refGene_gn=DOCK1;refGene_tx=XM_019813984.2
D2	84675044	84675140
refGene_gn=DOCK1;refGene_tx=XM_011287620.3;refGene_gn=DOCK1;refGene_tx=XM_011287621.3;refGene_gn=DOCK1;refGene_tx=XM_019813984.2
D2	84685023	84685207

refGene_gn=FAM196A;refGene_tx=XM_023241083.1;refGene_gn=FAM196A;refGene_tx=XM_023241084.1;refGene_gn=FAM196A;refGene_tx=XM_023241085.1;refGene_gn=FAM196A;refGene_tx=XM_023241086.1
D2 84700614 84700686
refGene_gn=FAM196A;refGene_tx=XM_023241083.1;refGene_gn=FAM196A;refGene_tx=XM_023241084.1;refGene_gn=FAM196A;refGene_tx=XM_023241085.1
D2 84720625 84721794
refGene_gn=FAM196A;refGene_tx=XM_023241083.1;refGene_gn=FAM196A;refGene_tx=XM_023241084.1;refGene_gn=FAM196A;refGene_tx=XM_023241085.1;refGene_gn=FAM196A;refGene_tx=XM_023241086.1
D2 84795045 84795147
refGene_gn=DOCK1;refGene_tx=XM_011287620.3;refGene_gn=DOCK1;refGene_tx=XM_011287621.3
D2 84802533 84802628
refGene_gn=DOCK1;refGene_tx=XM_011287620.3;refGene_gn=DOCK1;refGene_tx=XM_011287621.3;refGene_gn=DOCK1;refGene_tx=XM_019813984.2
D2 84890384 84890463
refGene_gn=DOCK1;refGene_tx=XM_011287620.3;refGene_gn=DOCK1;refGene_tx=XM_011287621.3;refGene_gn=DOCK1;refGene_tx=XM_019813984.2
D2 84895456 84895557
refGene_gn=DOCK1;refGene_tx=XM_011287620.3;refGene_gn=DOCK1;refGene_tx=XM_011287621.3;refGene_gn=DOCK1;refGene_tx=XM_019813984.2
D2 84905197 84905256
refGene_gn=DOCK1;refGene_tx=XM_011287620.3;refGene_gn=DOCK1;refGene_tx=XM_011287621.3;refGene_gn=DOCK1;refGene_tx=XM_019813984.2
D2 84916769 84916918
refGene_gn=DOCK1;refGene_tx=XM_011287620.3;refGene_gn=DOCK1;refGene_tx=XM_011287621.3;refGene_gn=DOCK1;refGene_tx=XM_019813984.2
D2 84927445 84927531
refGene_gn=DOCK1;refGene_tx=XM_011287620.3;refGene_gn=DOCK1;refGene_tx=XM_011287621.3;refGene_gn=DOCK1;refGene_tx=XM_019813984.2
D2 84927715 84927872
refGene_gn=DOCK1;refGene_tx=XM_011287620.3;refGene_gn=DOCK1;refGene_tx=XM_011287621.3;refGene_gn=DOCK1;refGene_tx=XM_019813984.2
D2 84934246 84934287
refGene_gn=DOCK1;refGene_tx=XM_011287620.3;refGene_gn=DOCK1;refGene_tx=XM_011287621.3;refGene_gn=DOCK1;refGene_tx=XM_019813984.2
D2 84935745 84935836
refGene_gn=DOCK1;refGene_tx=XM_011287620.3;refGene_gn=DOCK1;refGene_tx=XM_011287621.3;refGene_gn=DOCK1;refGene_tx=XM_019813984.2
D2 84940070 84940190
refGene_gn=DOCK1;refGene_tx=XM_011287620.3;refGene_gn=DOCK1;refGene_tx=XM_011287621.3;refGene_gn=DOCK1;refGene_tx=XM_019813984.2
D2 84956338 84956428
refGene_gn=DOCK1;refGene_tx=XM_011287620.3;refGene_gn=DOCK1;refGene_tx=XM_011287621.3;refGene_gn=DOCK1;refGene_tx=XM_019813984.2
D2 84957246 84957351
refGene_gn=DOCK1;refGene_tx=XM_011287620.3;refGene_gn=DOCK1;refGene_tx=XM_011287621.3;refGene_gn=DOCK1;refGene_tx=XM_019813984.2
D2 84961832 84961974

refGene_gn=DOCK1;refGene_tx=XM_011287620.3;refGene_gn=DOCK1;refGene_tx=XM_011287621.3;refGene_gn=DOCK1;refGene_tx=XM_019813984.2
D2 84962108 84962187
refGene_gn=DOCK1;refGene_tx=XM_011287620.3;refGene_gn=DOCK1;refGene_tx=XM_011287621.3;refGene_gn=DOCK1;refGene_tx=XM_019813984.2
D2 84963664 84963749
refGene_gn=DOCK1;refGene_tx=XM_011287620.3;refGene_gn=DOCK1;refGene_tx=XM_011287621.3;refGene_gn=DOCK1;refGene_tx=XM_019813984.2
D2 84966718 84966805
refGene_gn=DOCK1;refGene_tx=XM_011287620.3;refGene_gn=DOCK1;refGene_tx=XM_011287621.3;refGene_gn=DOCK1;refGene_tx=XM_019813984.2
D2 84969563 84969740
refGene_gn=DOCK1;refGene_tx=XM_011287620.3;refGene_gn=DOCK1;refGene_tx=XM_011287621.3;refGene_gn=DOCK1;refGene_tx=XM_019813984.2
D2 84970918 84971002
refGene_gn=DOCK1;refGene_tx=XM_011287620.3;refGene_gn=DOCK1;refGene_tx=XM_011287621.3;refGene_gn=DOCK1;refGene_tx=XM_019813984.2
D2 84975997 84976135
refGene_gn=DOCK1;refGene_tx=XM_011287620.3;refGene_gn=DOCK1;refGene_tx=XM_011287621.3;refGene_gn=DOCK1;refGene_tx=XM_019813984.2
D2 84983277 84983423
refGene_gn=DOCK1;refGene_tx=XM_011287620.3;refGene_gn=DOCK1;refGene_tx=XM_011287621.3;refGene_gn=DOCK1;refGene_tx=XM_019813984.2
D2 84986472 84986671
refGene_gn=DOCK1;refGene_tx=XM_011287620.3;refGene_gn=DOCK1;refGene_tx=XM_011287621.3;refGene_gn=DOCK1;refGene_tx=XM_019813984.2
D2 84989555 84989709
refGene_gn=DOCK1;refGene_tx=XM_011287620.3;refGene_gn=DOCK1;refGene_tx=XM_011287621.3;refGene_gn=DOCK1;refGene_tx=XM_019813984.2
D2 84992675 84992836
refGene_gn=DOCK1;refGene_tx=XM_011287620.3;refGene_gn=DOCK1;refGene_tx=XM_011287621.3;refGene_gn=DOCK1;refGene_tx=XM_019813984.2
D2 84996790 84996886
refGene_gn=DOCK1;refGene_tx=XM_011287620.3;refGene_gn=DOCK1;refGene_tx=XM_011287621.3;refGene_gn=DOCK1;refGene_tx=XM_019813984.2
D2 85086213 85086221 refGene_gn=NPS;refGene_tx=XM_019814154.2
D2 85086334 85086416 refGene_gn=NPS;refGene_tx=XM_019814154.2
D2 85089574 85089754 refGene_gn=NPS;refGene_tx=XM_019814154.2
D2 85237206 85237750 refGene_gn=FOXI2;refGene_tx=XM_023240306.1
D2 85238439 85238882 refGene_gn=FOXI2;refGene_tx=XM_023240306.1
D2 85319704 85319985 refGene_gn=CLRN3;refGene_tx=XM_003994520.3
D2 85325207 85325387 refGene_gn=CLRN3;refGene_tx=XM_003994520.3
D2 85331755 85331984 refGene_gn=CLRN3;refGene_tx=XM_003994520.3
D2 85411352 85411472 refGene_gn=PTPRE;refGene_tx=XM_023241087.1
D2 85411737 85411760 refGene_gn=PTPRE;refGene_tx=XM_023241087.1
D2 85465841 85465960
refGene_gn=PTPRE;refGene_tx=XM_023241087.1;refGene_gn=PTPRE;refGene_tx=XM_023241088.1;refGene_gn=PTPRE;refGene_tx=XM_023241090.1;refGene_gn=PTPRE;refGene_tx=XM_023241091.1;refGene_gn=PTPRE;refGene_tx=XM_023241092.1;refGene_gn=PTPRE;refGene_tx=XM_023241094.1

D2 85466909 85467021 refGene_gn=PTPRE;refGene_tx=XM_023241089.1
D2 85473419 85473519
refGene_gn=PTPRE;refGene_tx=XM_023241087.1;refGene_gn=PTPRE;refGene_tx=XM_023241088.1;refGene_gn=PTPRE;refGene_tx=XM_023241089.1;refGene_gn=PTPRE;refGene_tx=XM_023241090.1;refGene_gn=PTPRE;refGene_tx=XM_023241091.1;refGene_gn=PTPRE;refGene_tx=XM_023241092.1;refGene_gn=PTPRE;refGene_tx=XM_023241094.1
D2 85473695 85473730 refGene_gn=PTPRE;refGene_tx=XM_023241093.1
D2 85473807 85473881
refGene_gn=PTPRE;refGene_tx=XM_023241087.1;refGene_gn=PTPRE;refGene_tx=XM_023241088.1;refGene_gn=PTPRE;refGene_tx=XM_023241089.1;refGene_gn=PTPRE;refGene_tx=XM_023241090.1;refGene_gn=PTPRE;refGene_tx=XM_023241091.1;refGene_gn=PTPRE;refGene_tx=XM_023241092.1;refGene_gn=PTPRE;refGene_tx=XM_023241093.1;refGene_gn=PTPRE;refGene_tx=XM_023241094.1
D2 85475778 85475915
refGene_gn=PTPRE;refGene_tx=XM_023241087.1;refGene_gn=PTPRE;refGene_tx=XM_023241088.1;refGene_gn=PTPRE;refGene_tx=XM_023241089.1;refGene_gn=PTPRE;refGene_tx=XM_023241090.1;refGene_gn=PTPRE;refGene_tx=XM_023241091.1;refGene_gn=PTPRE;refGene_tx=XM_023241092.1;refGene_gn=PTPRE;refGene_tx=XM_023241093.1;refGene_gn=PTPRE;refGene_tx=XM_023241094.1
D2 85481829 85481920
refGene_gn=PTPRE;refGene_tx=XM_023241087.1;refGene_gn=PTPRE;refGene_tx=XM_023241088.1;refGene_gn=PTPRE;refGene_tx=XM_023241089.1;refGene_gn=PTPRE;refGene_tx=XM_023241090.1;refGene_gn=PTPRE;refGene_tx=XM_023241091.1;refGene_gn=PTPRE;refGene_tx=XM_023241092.1;refGene_gn=PTPRE;refGene_tx=XM_023241093.1;refGene_gn=PTPRE;refGene_tx=XM_023241094.1
D2 85485818 85485895
refGene_gn=PTPRE;refGene_tx=XM_023241087.1;refGene_gn=PTPRE;refGene_tx=XM_023241088.1;refGene_gn=PTPRE;refGene_tx=XM_023241089.1;refGene_gn=PTPRE;refGene_tx=XM_023241090.1;refGene_gn=PTPRE;refGene_tx=XM_023241091.1;refGene_gn=PTPRE;refGene_tx=XM_023241092.1;refGene_gn=PTPRE;refGene_tx=XM_023241093.1;refGene_gn=PTPRE;refGene_tx=XM_023241094.1
D2 85486795 85486832
refGene_gn=PTPRE;refGene_tx=XM_023241087.1;refGene_gn=PTPRE;refGene_tx=XM_023241088.1;refGene_gn=PTPRE;refGene_tx=XM_023241089.1;refGene_gn=PTPRE;refGene_tx=XM_023241090.1;refGene_gn=PTPRE;refGene_tx=XM_023241091.1;refGene_gn=PTPRE;refGene_tx=XM_023241092.1;refGene_gn=PTPRE;refGene_tx=XM_023241093.1;refGene_gn=PTPRE;refGene_tx=XM_023241094.1
D2 85488607 85488705
refGene_gn=PTPRE;refGene_tx=XM_023241087.1;refGene_gn=PTPRE;refGene_tx=XM_023241088.1;refGene_gn=PTPRE;refGene_tx=XM_023241089.1;refGene_gn=PTPRE;refGene_tx=XM_023241090.1;refGene_gn=PTPRE;refGene_tx=XM_023241091.1;refGene_gn=PTPRE;refGene_tx=XM_023241092.1;refGene_gn=PTPRE;refGene_tx=XM_023241093.1;refGene_gn=PTPRE;refGene_tx=XM_023241094.1
D2 85492630 85492750
refGene_gn=PTPRE;refGene_tx=XM_023241087.1;refGene_gn=PTPRE;refGene_tx=XM_023241088.1;refGene_gn=PTPRE;refGene_tx=XM_023241089.1;refGene_gn=PTPRE;refGene_tx=XM_023241090.1;refGene_gn=PTPRE;refGene_tx=XM_023241091.1;refGene_gn=PTPRE;refGene_tx=XM_023241092.1;refGene_gn=PTPRE;refGene_tx=XM_023241093.1;refGene_gn=PTPRE;refGene_tx=XM_023241094.1

D2	85493963	85494127
refGene_gn=PTPRE;refGene_tx=XM_023241087.1;refGene_gn=PTPRE;refGene_tx=XM_023241088.1;refGene_gn=PTPRE;refGene_tx=XM_023241089.1;refGene_gn=PTPRE;refGene_tx=XM_023241090.1;refGene_gn=PTPRE;refGene_tx=XM_023241091.1;refGene_gn=PTPRE;refGene_tx=XM_023241092.1;refGene_gn=PTPRE;refGene_tx=XM_023241093.1;refGene_gn=PTPRE;refGene_tx=XM_023241094.1
D2	85498039	85498175
refGene_gn=PTPRE;refGene_tx=XM_023241087.1;refGene_gn=PTPRE;refGene_tx=XM_023241088.1;refGene_gn=PTPRE;refGene_tx=XM_023241089.1;refGene_gn=PTPRE;refGene_tx=XM_023241090.1;refGene_gn=PTPRE;refGene_tx=XM_023241091.1;refGene_gn=PTPRE;refGene_tx=XM_023241092.1;refGene_gn=PTPRE;refGene_tx=XM_023241093.1;refGene_gn=PTPRE;refGene_tx=XM_023241094.1
D2	85498497	85498647
refGene_gn=PTPRE;refGene_tx=XM_023241087.1;refGene_gn=PTPRE;refGene_tx=XM_023241088.1;refGene_gn=PTPRE;refGene_tx=XM_023241089.1;refGene_gn=PTPRE;refGene_tx=XM_023241090.1;refGene_gn=PTPRE;refGene_tx=XM_023241091.1;refGene_gn=PTPRE;refGene_tx=XM_023241092.1;refGene_gn=PTPRE;refGene_tx=XM_023241093.1;refGene_gn=PTPRE;refGene_tx=XM_023241094.1
D2	85498978	85499072
refGene_gn=PTPRE;refGene_tx=XM_023241087.1;refGene_gn=PTPRE;refGene_tx=XM_023241088.1;refGene_gn=PTPRE;refGene_tx=XM_023241089.1;refGene_gn=PTPRE;refGene_tx=XM_023241090.1;refGene_gn=PTPRE;refGene_tx=XM_023241091.1;refGene_gn=PTPRE;refGene_tx=XM_023241092.1;refGene_gn=PTPRE;refGene_tx=XM_023241093.1;refGene_gn=PTPRE;refGene_tx=XM_023241094.1
D2	85500344	85500421
refGene_gn=PTPRE;refGene_tx=XM_023241087.1;refGene_gn=PTPRE;refGene_tx=XM_023241088.1;refGene_gn=PTPRE;refGene_tx=XM_023241089.1;refGene_gn=PTPRE;refGene_tx=XM_023241090.1;refGene_gn=PTPRE;refGene_tx=XM_023241091.1;refGene_gn=PTPRE;refGene_tx=XM_023241092.1;refGene_gn=PTPRE;refGene_tx=XM_023241093.1;refGene_gn=PTPRE;refGene_tx=XM_023241094.1
D2	85501280	85501415
refGene_gn=PTPRE;refGene_tx=XM_023241087.1;refGene_gn=PTPRE;refGene_tx=XM_023241088.1;refGene_gn=PTPRE;refGene_tx=XM_023241089.1;refGene_gn=PTPRE;refGene_tx=XM_023241090.1;refGene_gn=PTPRE;refGene_tx=XM_023241091.1;refGene_gn=PTPRE;refGene_tx=XM_023241092.1;refGene_gn=PTPRE;refGene_tx=XM_023241093.1;refGene_gn=PTPRE;refGene_tx=XM_023241094.1
D2	85504206	85504332
refGene_gn=PTPRE;refGene_tx=XM_023241087.1;refGene_gn=PTPRE;refGene_tx=XM_023241088.1;refGene_gn=PTPRE;refGene_tx=XM_023241089.1;refGene_gn=PTPRE;refGene_tx=XM_023241090.1;refGene_gn=PTPRE;refGene_tx=XM_023241091.1;refGene_gn=PTPRE;refGene_tx=XM_023241092.1;refGene_gn=PTPRE;refGene_tx=XM_023241093.1;refGene_gn=PTPRE;refGene_tx=XM_023241094.1
D2	85505064	85505231
refGene_gn=PTPRE;refGene_tx=XM_023241087.1;refGene_gn=PTPRE;refGene_tx=XM_023241088.1;refGene_gn=PTPRE;refGene_tx=XM_023241089.1;refGene_gn=PTPRE;refGene_tx=XM_023241090.1;refGene_gn=PTPRE;refGene_tx=XM_023241091.1;refGene_gn=PTPRE;refGene_tx=XM_023241092.1;refGene_gn=PTPRE;refGene_tx=XM_023241093.1;refGene_gn=PTPRE;refGene_tx=XM_023241094.1
D2	85507011	85507147
refGene_gn=PTPRE;refGene_tx=XM_023241087.1;refGene_gn=PTPRE;refGene_tx

=XM_023241088.1;refGene_gn=PTPRE;refGene_tx=XM_023241089.1;refGene_gn=PTPRE;refGene_tx=XM_023241090.1;refGene_gn=PTPRE;refGene_tx=XM_023241091.1;refGene_gn=PTPRE;refGene_tx=XM_023241092.1;refGene_gn=PTPRE;refGene_tx=XM_023241093.1;refGene_gn=PTPRE;refGene_tx=XM_023241094.1
D2 85509795 85509870
refGene_gn=PTPRE;refGene_tx=XM_023241087.1;refGene_gn=PTPRE;refGene_tx=XM_023241088.1;refGene_gn=PTPRE;refGene_tx=XM_023241089.1;refGene_gn=PTPRE;refGene_tx=XM_023241090.1;refGene_gn=PTPRE;refGene_tx=XM_023241091.1;refGene_gn=PTPRE;refGene_tx=XM_023241092.1;refGene_gn=PTPRE;refGene_tx=XM_023241093.1;refGene_gn=PTPRE;refGene_tx=XM_023241094.1
D2 85518790 85518856 refGene_gn=MKI67;refGene_tx=XM_023241095.1
D2 85521161 85521605 refGene_gn=MKI67;refGene_tx=XM_023241095.1
D2 85522446 85525865 refGene_gn=MKI67;refGene_tx=XM_023241095.1
D2 85525867 85528123 refGene_gn=MKI67;refGene_tx=XM_023241095.1
D2 85529101 85529260 refGene_gn=MKI67;refGene_tx=XM_023241095.1
D2 85532246 85532418 refGene_gn=MKI67;refGene_tx=XM_023241095.1
D2 85532521 85532640 refGene_gn=MKI67;refGene_tx=XM_023241095.1
D2 85532728 85533047 refGene_gn=MKI67;refGene_tx=XM_023241095.1
D2 85534221 85534391 refGene_gn=MKI67;refGene_tx=XM_023241095.1
D2 85535823 85536532 refGene_gn=MKI67;refGene_tx=XM_023241095.1
D2 85536533 85536895 refGene_gn=MKI67;refGene_tx=XM_023241095.1
D2 85537389 85537435 refGene_gn=MKI67;refGene_tx=XM_023241095.1
D2 85540063 85540130 refGene_gn=MKI67;refGene_tx=XM_023241095.1
D2 85542597 85542713 refGene_gn=MKI67;refGene_tx=XM_023241095.1
D2 85542824 85542903 refGene_gn=MKI67;refGene_tx=XM_023241095.1
D2 85545515 85545607 refGene_gn=MKI67;refGene_tx=XM_023241095.1
D2 86764485 86764662 refGene_gn=MGMT;refGene_tx=XM_023241096.1
D2 86824142 86824279
refGene_gn=MGMT;refGene_tx=XM_023241096.1;refGene_gn=MGMT;refGene_tx=XM_023241098.1;refGene_gn=MGMT;refGene_tx=XM_023241099.1;refGene_gn=MGMT;refGene_tx=XM_023241100.1;refGene_gn=MGMT;refGene_tx=XM_023241101.1
D2 87009846 87010007
refGene_gn=MGMT;refGene_tx=XM_023241096.1;refGene_gn=MGMT;refGene_tx=XM_023241098.1;refGene_gn=MGMT;refGene_tx=XM_023241099.1;refGene_gn=MGMT;refGene_tx=XM_023241100.1;refGene_gn=MGMT;refGene_tx=XM_023241101.1
D2 87048068 87048208
refGene_gn=MGMT;refGene_tx=XM_023241096.1;refGene_gn=MGMT;refGene_tx=XM_023241098.1;refGene_gn=MGMT;refGene_tx=XM_023241099.1;refGene_gn=MGMT;refGene_tx=XM_023241100.1;refGene_gn=MGMT;refGene_tx=XM_023241101.1
D2 87055524 87055725
refGene_gn=MGMT;refGene_tx=XM_023241096.1;refGene_gn=MGMT;refGene_tx=XM_023241098.1;refGene_gn=MGMT;refGene_tx=XM_023241099.1;refGene_gn=MGMT;refGene_tx=XM_023241100.1;refGene_gn=MGMT;refGene_tx=XM_023241101.1
D2 87125091 87125123
refGene_gn=EBF3;refGene_tx=XM_023241102.1;refGene_gn=EBF3;refGene_tx=XM_023241103.1;refGene_gn=EBF3;refGene_tx=XM_023241104.1;refGene_gn=EBF3;refGene_tx=XM_023241105.1;refGene_gn=EBF3;refGene_tx=XM_023241107.1;refGene_gn=EBF3;refGene_tx=XM_023241109.1;refGene_gn=EBF3;refGene_tx=XM_023241110.1;refGene_gn=EBF3;refGene_tx=XM_023241112.1;refGene_gn=EBF3;refGene_tx=XM_023241113.1;refGene_gn=EBF3;refGene_tx=XM_023241114.1

D2 87126174 87126287
refGene_gn=EBF3;refGene_tx=XM_023241102.1;refGene_gn=EBF3;refGene_tx=XM_023241103.1;refGene_gn=EBF3;refGene_tx=XM_023241104.1;refGene_gn=EBF3;refGene_tx=XM_023241107.1;refGene_gn=EBF3;refGene_tx=XM_023241113.1
D2 87126362 87126394
refGene_gn=EBF3;refGene_tx=XM_023241106.1;refGene_gn=EBF3;refGene_tx=XM_023241111.1
D2 87127279 87127477
refGene_gn=EBF3;refGene_tx=XM_023241102.1;refGene_gn=EBF3;refGene_tx=XM_023241103.1;refGene_gn=EBF3;refGene_tx=XM_023241104.1;refGene_gn=EBF3;refGene_tx=XM_023241105.1;refGene_gn=EBF3;refGene_tx=XM_023241106.1;refGene_gn=EBF3;refGene_tx=XM_023241107.1;refGene_gn=EBF3;refGene_tx=XM_023241109.1;refGene_gn=EBF3;refGene_tx=XM_023241110.1;refGene_gn=EBF3;refGene_tx=XM_023241111.1;refGene_gn=EBF3;refGene_tx=XM_023241112.1;refGene_gn=EBF3;refGene_tx=XM_023241113.1;refGene_gn=EBF3;refGene_tx=XM_023241114.1
D2 87127916 87128105
refGene_gn=EBF3;refGene_tx=XM_023241102.1;refGene_gn=EBF3;refGene_tx=XM_023241103.1;refGene_gn=EBF3;refGene_tx=XM_023241104.1;refGene_gn=EBF3;refGene_tx=XM_023241105.1;refGene_gn=EBF3;refGene_tx=XM_023241106.1;refGene_gn=EBF3;refGene_tx=XM_023241107.1;refGene_gn=EBF3;refGene_tx=XM_023241109.1;refGene_gn=EBF3;refGene_tx=XM_023241110.1;refGene_gn=EBF3;refGene_tx=XM_023241111.1;refGene_gn=EBF3;refGene_tx=XM_023241112.1
D2 87129083 87129261
refGene_gn=EBF3;refGene_tx=XM_023241102.1;refGene_gn=EBF3;refGene_tx=XM_023241103.1;refGene_gn=EBF3;refGene_tx=XM_023241104.1;refGene_gn=EBF3;refGene_tx=XM_023241105.1;refGene_gn=EBF3;refGene_tx=XM_023241106.1;refGene_gn=EBF3;refGene_tx=XM_023241107.1;refGene_gn=EBF3;refGene_tx=XM_023241109.1;refGene_gn=EBF3;refGene_tx=XM_023241110.1;refGene_gn=EBF3;refGene_tx=XM_023241111.1;refGene_gn=EBF3;refGene_tx=XM_023241112.1;refGene_gn=EBF3;refGene_tx=XM_023241113.1;refGene_gn=EBF3;refGene_tx=XM_023241114.1
D2 87129921 87129987
refGene_gn=EBF3;refGene_tx=XM_023241102.1;refGene_gn=EBF3;refGene_tx=XM_023241103.1;refGene_gn=EBF3;refGene_tx=XM_023241104.1;refGene_gn=EBF3;refGene_tx=XM_023241105.1;refGene_gn=EBF3;refGene_tx=XM_023241106.1;refGene_gn=EBF3;refGene_tx=XM_023241107.1;refGene_gn=EBF3;refGene_tx=XM_023241109.1;refGene_gn=EBF3;refGene_tx=XM_023241110.1;refGene_gn=EBF3;refGene_tx=XM_023241111.1;refGene_gn=EBF3;refGene_tx=XM_023241112.1;refGene_gn=EBF3;refGene_tx=XM_023241113.1;refGene_gn=EBF3;refGene_tx=XM_023241114.1
D2 87135199 87135288
refGene_gn=EBF3;refGene_tx=XM_023241102.1;refGene_gn=EBF3;refGene_tx=XM_023241103.1;refGene_gn=EBF3;refGene_tx=XM_023241104.1;refGene_gn=EBF3;refGene_tx=XM_023241105.1;refGene_gn=EBF3;refGene_tx=XM_023241106.1;refGene_gn=EBF3;refGene_tx=XM_023241107.1;refGene_gn=EBF3;refGene_tx=XM_023241109.1;refGene_gn=EBF3;refGene_tx=XM_023241110.1;refGene_gn=EBF3;refGene_tx=XM_023241111.1;refGene_gn=EBF3;refGene_tx=XM_023241112.1;refGene_gn=EBF3;refGene_tx=XM_023241113.1;refGene_gn=EBF3;refGene_tx=XM_023241114.1

D2 87152997 87153124
refGene_gn=EBF3;refGene_tx=XM_023241102.1;refGene_gn=EBF3;refGene_tx=XM_023241103.1;refGene_gn=EBF3;refGene_tx=XM_023241104.1;refGene_gn=EBF3;refGene_tx=XM_023241105.1;refGene_gn=EBF3;refGene_tx=XM_023241106.1;refGene_gn=EBF3;refGene_tx=XM_023241107.1;refGene_gn=EBF3;refGene_tx=XM_023241109.1;refGene_gn=EBF3;refGene_tx=XM_023241110.1;refGene_gn=EBF3;refGene_tx=XM_023241111.1;refGene_gn=EBF3;refGene_tx=XM_023241112.1;refGene_gn=EBF3;refGene_tx=XM_023241113.1;refGene_gn=EBF3;refGene_tx=XM_023241114.1
D2 87153586 87153717
refGene_gn=EBF3;refGene_tx=XM_023241102.1;refGene_gn=EBF3;refGene_tx=XM_023241103.1;refGene_gn=EBF3;refGene_tx=XM_023241104.1;refGene_gn=EBF3;refGene_tx=XM_023241105.1;refGene_gn=EBF3;refGene_tx=XM_023241106.1;refGene_gn=EBF3;refGene_tx=XM_023241107.1;refGene_gn=EBF3;refGene_tx=XM_023241109.1;refGene_gn=EBF3;refGene_tx=XM_023241110.1;refGene_gn=EBF3;refGene_tx=XM_023241111.1;refGene_gn=EBF3;refGene_tx=XM_023241112.1;refGene_gn=EBF3;refGene_tx=XM_023241113.1;refGene_gn=EBF3;refGene_tx=XM_023241114.1
D2 87159104 87159249
refGene_gn=EBF3;refGene_tx=XM_023241102.1;refGene_gn=EBF3;refGene_tx=XM_023241103.1;refGene_gn=EBF3;refGene_tx=XM_023241104.1;refGene_gn=EBF3;refGene_tx=XM_023241105.1;refGene_gn=EBF3;refGene_tx=XM_023241106.1;refGene_gn=EBF3;refGene_tx=XM_023241107.1;refGene_gn=EBF3;refGene_tx=XM_023241109.1;refGene_gn=EBF3;refGene_tx=XM_023241110.1;refGene_gn=EBF3;refGene_tx=XM_023241111.1;refGene_gn=EBF3;refGene_tx=XM_023241112.1;refGene_gn=EBF3;refGene_tx=XM_023241113.1;refGene_gn=EBF3;refGene_tx=XM_023241114.1
D2 87162745 87162827
refGene_gn=EBF3;refGene_tx=XM_023241102.1;refGene_gn=EBF3;refGene_tx=XM_023241103.1;refGene_gn=EBF3;refGene_tx=XM_023241104.1;refGene_gn=EBF3;refGene_tx=XM_023241105.1;refGene_gn=EBF3;refGene_tx=XM_023241106.1;refGene_gn=EBF3;refGene_tx=XM_023241107.1;refGene_gn=EBF3;refGene_tx=XM_023241109.1;refGene_gn=EBF3;refGene_tx=XM_023241110.1;refGene_gn=EBF3;refGene_tx=XM_023241111.1;refGene_gn=EBF3;refGene_tx=XM_023241112.1;refGene_gn=EBF3;refGene_tx=XM_023241113.1;refGene_gn=EBF3;refGene_tx=XM_023241114.1
D2 87235585 87235654
refGene_gn=EBF3;refGene_tx=XM_023241102.1;refGene_gn=EBF3;refGene_tx=XM_023241103.1;refGene_gn=EBF3;refGene_tx=XM_023241104.1;refGene_gn=EBF3;refGene_tx=XM_023241105.1;refGene_gn=EBF3;refGene_tx=XM_023241106.1;refGene_gn=EBF3;refGene_tx=XM_023241107.1;refGene_gn=EBF3;refGene_tx=XM_023241109.1;refGene_gn=EBF3;refGene_tx=XM_023241110.1;refGene_gn=EBF3;refGene_tx=XM_023241111.1;refGene_gn=EBF3;refGene_tx=XM_023241112.1;refGene_gn=EBF3;refGene_tx=XM_023241113.1;refGene_gn=EBF3;refGene_tx=XM_023241114.1
D2 87237265 87237339
refGene_gn=EBF3;refGene_tx=XM_023241102.1;refGene_gn=EBF3;refGene_tx=XM_023241103.1;refGene_gn=EBF3;refGene_tx=XM_023241104.1;refGene_gn=EBF3;refGene_tx=XM_023241105.1;refGene_gn=EBF3;refGene_tx=XM_023241106.1;refGene_gn=EBF3;refGene_tx=XM_023241107.1;refGene_gn=EBF3;refGene_tx=X

M_023241109.1;refGene_gn=EBF3;refGene_tx=XM_023241110.1;refGene_gn=EBF3;refGene_tx=XM_023241111.1;refGene_gn=EBF3;refGene_tx=XM_023241112.1;refGene_gn=EBF3;refGene_tx=XM_023241113.1;refGene_gn=EBF3;refGene_tx=XM_023241114.1
D2 87240241 87240297
refGene_gn=EBF3;refGene_tx=XM_023241102.1;refGene_gn=EBF3;refGene_tx=XM_023241103.1;refGene_gn=EBF3;refGene_tx=XM_023241104.1;refGene_gn=EBF3;refGene_tx=XM_023241105.1;refGene_gn=EBF3;refGene_tx=XM_023241106.1;refGene_gn=EBF3;refGene_tx=XM_023241107.1;refGene_gn=EBF3;refGene_tx=XM_023241109.1;refGene_gn=EBF3;refGene_tx=XM_023241110.1;refGene_gn=EBF3;refGene_tx=XM_023241111.1;refGene_gn=EBF3;refGene_tx=XM_023241112.1;refGene_gn=EBF3;refGene_tx=XM_023241113.1;refGene_gn=EBF3;refGene_tx=XM_023241114.1
D2 87241005 87241069
refGene_gn=EBF3;refGene_tx=XM_023241102.1;refGene_gn=EBF3;refGene_tx=XM_023241103.1;refGene_gn=EBF3;refGene_tx=XM_023241104.1;refGene_gn=EBF3;refGene_tx=XM_023241105.1;refGene_gn=EBF3;refGene_tx=XM_023241106.1;refGene_gn=EBF3;refGene_tx=XM_023241107.1;refGene_gn=EBF3;refGene_tx=XM_023241109.1;refGene_gn=EBF3;refGene_tx=XM_023241110.1;refGene_gn=EBF3;refGene_tx=XM_023241111.1;refGene_gn=EBF3;refGene_tx=XM_023241112.1;refGene_gn=EBF3;refGene_tx=XM_023241113.1;refGene_gn=EBF3;refGene_tx=XM_023241114.1
D2 87241435 87241592
refGene_gn=EBF3;refGene_tx=XM_023241102.1;refGene_gn=EBF3;refGene_tx=XM_023241103.1;refGene_gn=EBF3;refGene_tx=XM_023241104.1;refGene_gn=EBF3;refGene_tx=XM_023241105.1;refGene_gn=EBF3;refGene_tx=XM_023241106.1;refGene_gn=EBF3;refGene_tx=XM_023241107.1;refGene_gn=EBF3;refGene_tx=XM_023241109.1;refGene_gn=EBF3;refGene_tx=XM_023241110.1;refGene_gn=EBF3;refGene_tx=XM_023241111.1;refGene_gn=EBF3;refGene_tx=XM_023241112.1;refGene_gn=EBF3;refGene_tx=XM_023241113.1;refGene_gn=EBF3;refGene_tx=XM_023241114.1
D2 87241677 87241811
refGene_gn=EBF3;refGene_tx=XM_023241102.1;refGene_gn=EBF3;refGene_tx=XM_023241103.1;refGene_gn=EBF3;refGene_tx=XM_023241104.1;refGene_gn=EBF3;refGene_tx=XM_023241105.1;refGene_gn=EBF3;refGene_tx=XM_023241106.1;refGene_gn=EBF3;refGene_tx=XM_023241107.1;refGene_gn=EBF3;refGene_tx=XM_023241109.1;refGene_gn=EBF3;refGene_tx=XM_023241110.1;refGene_gn=EBF3;refGene_tx=XM_023241111.1;refGene_gn=EBF3;refGene_tx=XM_023241112.1;refGene_gn=EBF3;refGene_tx=XM_023241113.1;refGene_gn=EBF3;refGene_tx=XM_023241114.1
D2 87383917 87384006 refGene_gn=GLRX3;refGene_tx=XM_023241115.1
D2 87390049 87390158 refGene_gn=GLRX3;refGene_tx=XM_023241115.1
D2 87398812 87398887 refGene_gn=GLRX3;refGene_tx=XM_023241115.1
D2 87399631 87399833 refGene_gn=GLRX3;refGene_tx=XM_023241115.1
D2 87404854 87405027 refGene_gn=GLRX3;refGene_tx=XM_023241115.1
D2 87405265 87405327 refGene_gn=GLRX3;refGene_tx=XM_023241115.1
D2 87407382 87407440 refGene_gn=GLRX3;refGene_tx=XM_023241115.1
D2 87409172 87409225 refGene_gn=GLRX3;refGene_tx=XM_023241115.1
D2 87411828 87411868 refGene_gn=GLRX3;refGene_tx=XM_023241115.1
D2 87411962 87412055 refGene_gn=GLRX3;refGene_tx=XM_023241115.1

D2 87415311 87415362 refGene_gn=GLRX3;refGene_tx=XM_023241115.1
D2 88120101 88120258
refGene_gn=LOC101085835;refGene_tx=XM_023241116.1
D2 88124236 88124355
refGene_gn=LOC101085835;refGene_tx=XM_023241116.1
D2 88128148 88128238
refGene_gn=LOC101085835;refGene_tx=XM_023241116.1
D2 88140615 88140751
refGene_gn=LOC101085835;refGene_tx=XM_023241116.1
D2 88160482 88160552
refGene_gn=LOC101085835;refGene_tx=XM_023241116.1
D2 88171467 88171622
refGene_gn=LOC101085835;refGene_tx=XM_023241116.1
D2 88184901 88184990
refGene_gn=LOC101085835;refGene_tx=XM_023241116.1
D2 88188020 88188068
refGene_gn=LOC101085835;refGene_tx=XM_023241116.1
D2 88216600 88216746 refGene_gn=TCERG1L;refGene_tx=XM_023240307.1
D2 88272952 88273132 refGene_gn=TCERG1L;refGene_tx=XM_023240307.1
D2 88310042 88310223 refGene_gn=TCERG1L;refGene_tx=XM_023240307.1
D2 88310980 88311127 refGene_gn=TCERG1L;refGene_tx=XM_023240307.1
D2 88313180 88313450 refGene_gn=TCERG1L;refGene_tx=XM_023240307.1
D2 88707327 88707334 refGene_gn=PPP2R2D;refGene_tx=XM_023241118.1
D2 88707439 88707532 refGene_gn=PPP2R2D;refGene_tx=XM_023241118.1
D2 88738444 88738542 refGene_gn=PPP2R2D;refGene_tx=XM_023241118.1
D2 88742146 88742312 refGene_gn=PPP2R2D;refGene_tx=XM_023241118.1
D2 88742726 88742839 refGene_gn=PPP2R2D;refGene_tx=XM_023241118.1
D2 88745935 88746113 refGene_gn=PPP2R2D;refGene_tx=XM_023241118.1
D2 88748982 88749147 refGene_gn=PPP2R2D;refGene_tx=XM_023241118.1
D2 88751081 88751251 refGene_gn=PPP2R2D;refGene_tx=XM_023241118.1
D2 88751323 88751415 refGene_gn=PPP2R2D;refGene_tx=XM_023241118.1
D2 88756633 88756913 refGene_gn=PPP2R2D;refGene_tx=XM_023241118.1
D2 88759035 88759233
refGene_gn=LOC109492797;refGene_tx=XM_023241119.1
D2 88759581 88759904
refGene_gn=LOC109492797;refGene_tx=XM_023241119.1
D2 88760016 88760156
refGene_gn=LOC109492797;refGene_tx=XM_023241119.1
D2 88762876 88762962
refGene_gn=LOC109492797;refGene_tx=XM_023241119.1
D2 88763082 88763181
refGene_gn=LOC109492797;refGene_tx=XM_023241119.1
D2 88763843 88763889 refGene_gn=BNIP3;refGene_tx=XM_011287628.3
D2 88765203 88765353 refGene_gn=BNIP3;refGene_tx=XM_011287628.3
D2 88765425 88765532 refGene_gn=BNIP3;refGene_tx=XM_011287628.3
D2 88767531 88767616 refGene_gn=BNIP3;refGene_tx=XM_011287628.3
D2 88768218 88768369 refGene_gn=BNIP3;refGene_tx=XM_011287628.3
D2 88775999 88776045 refGene_gn=BNIP3;refGene_tx=XM_011287628.3
D2 88853948 88854083
refGene_gn=JAKMIP3;refGene_tx=XM_023241120.1;refGene_gn=JAKMIP3;refGen

e_tx=XM_023241121.1;refGene_gn=JAKMIP3;refGene_tx=XM_023241122.1;refGene_gn=JAKMIP3;refGene_tx=XM_023241123.1;refGene_gn=JAKMIP3;refGene_tx=XM_023241124.1;refGene_gn=JAKMIP3;refGene_tx=XM_023241125.1;refGene_gn=JAKMIP3;refGene_tx=XM_023241126.1;refGene_gn=JAKMIP3;refGene_tx=XM_023241127.1;refGene_gn=JAKMIP3;refGene_tx=XM_023241128.1;refGene_gn=JAKMIP3;refGene_tx=XM_023241129.1;refGene_gn=JAKMIP3;refGene_tx=XM_023241130.1;refGene_gn=JAKMIP3;refGene_tx=XM_023241131.1;refGene_gn=JAKMIP3;refGene_tx=XM_023241132.1;refGene_gn=JAKMIP3;refGene_tx=XM_023241133.1;refGene_gn=JAKMIP3;refGene_tx=XM_023241134.1;refGene_gn=JAKMIP3;refGene_tx=XM_023241135.1

D2 88862457 88862955

refGene_gn=JAKMIP3;refGene_tx=XM_023241120.1;refGene_gn=JAKMIP3;refGene_tx=XM_023241121.1;refGene_gn=JAKMIP3;refGene_tx=XM_023241122.1;refGene_gn=JAKMIP3;refGene_tx=XM_023241123.1;refGene_gn=JAKMIP3;refGene_tx=XM_023241124.1;refGene_gn=JAKMIP3;refGene_tx=XM_023241125.1;refGene_gn=JAKMIP3;refGene_tx=XM_023241126.1;refGene_gn=JAKMIP3;refGene_tx=XM_023241127.1;refGene_gn=JAKMIP3;refGene_tx=XM_023241128.1;refGene_gn=JAKMIP3;refGene_tx=XM_023241129.1;refGene_gn=JAKMIP3;refGene_tx=XM_023241130.1;refGene_gn=JAKMIP3;refGene_tx=XM_023241131.1;refGene_gn=JAKMIP3;refGene_tx=XM_023241132.1;refGene_gn=JAKMIP3;refGene_tx=XM_023241133.1;refGene_gn=JAKMIP3;refGene_tx=XM_023241134.1;refGene_gn=JAKMIP3;refGene_tx=XM_023241135.1

D2 88874513 88874729

refGene_gn=JAKMIP3;refGene_tx=XM_023241120.1;refGene_gn=JAKMIP3;refGene_tx=XM_023241121.1;refGene_gn=JAKMIP3;refGene_tx=XM_023241122.1;refGene_gn=JAKMIP3;refGene_tx=XM_023241123.1;refGene_gn=JAKMIP3;refGene_tx=XM_023241124.1;refGene_gn=JAKMIP3;refGene_tx=XM_023241125.1;refGene_gn=JAKMIP3;refGene_tx=XM_023241126.1;refGene_gn=JAKMIP3;refGene_tx=XM_023241127.1;refGene_gn=JAKMIP3;refGene_tx=XM_023241128.1;refGene_gn=JAKMIP3;refGene_tx=XM_023241129.1;refGene_gn=JAKMIP3;refGene_tx=XM_023241130.1

D2 88876144 88876264

refGene_gn=JAKMIP3;refGene_tx=XM_023241120.1;refGene_gn=JAKMIP3;refGene_tx=XM_023241121.1;refGene_gn=JAKMIP3;refGene_tx=XM_023241122.1;refGene_gn=JAKMIP3;refGene_tx=XM_023241123.1;refGene_gn=JAKMIP3;refGene_tx=XM_023241124.1;refGene_gn=JAKMIP3;refGene_tx=XM_023241125.1;refGene_gn=JAKMIP3;refGene_tx=XM_023241126.1;refGene_gn=JAKMIP3;refGene_tx=XM_023241127.1;refGene_gn=JAKMIP3;refGene_tx=XM_023241128.1;refGene_gn=JAKMIP3;refGene_tx=XM_023241129.1;refGene_gn=JAKMIP3;refGene_tx=XM_023241130.1;refGene_gn=JAKMIP3;refGene_tx=XM_023241131.1;refGene_gn=JAKMIP3;refGene_tx=XM_023241132.1;refGene_gn=JAKMIP3;refGene_tx=XM_023241133.1;refGene_gn=JAKMIP3;refGene_tx=XM_023241134.1;refGene_gn=JAKMIP3;refGene_tx=XM_023241135.1

D2 88876812 88876959

refGene_gn=JAKMIP3;refGene_tx=XM_023241120.1;refGene_gn=JAKMIP3;refGene_tx=XM_023241121.1;refGene_gn=JAKMIP3;refGene_tx=XM_023241122.1;refGene_gn=JAKMIP3;refGene_tx=XM_023241123.1;refGene_gn=JAKMIP3;refGene_tx=XM_023241124.1;refGene_gn=JAKMIP3;refGene_tx=XM_023241125.1;refGene_gn=JAKMIP3;refGene_tx=XM_023241126.1;refGene_gn=JAKMIP3;refGene_tx=XM_023241127.1;refGene_gn=JAKMIP3;refGene_tx=XM_023241128.1;refGene_gn=JAKM

IP3;refGene_tx=XM_023241129.1;refGene_gn=JAKMIP3;refGene_tx=XM_0232411
30.1;refGene_gn=JAKMIP3;refGene_tx=XM_023241131.1;refGene_gn=JAKMIP3;r
efGene_tx=XM_023241132.1;refGene_gn=JAKMIP3;refGene_tx=XM_023241133.1;
refGene_gn=JAKMIP3;refGene_tx=XM_023241134.1;refGene_gn=JAKMIP3;refGen
e_tx=XM_023241135.1
D2 88877745 88877877
refGene_gn=JAKMIP3;refGene_tx=XM_023241120.1;refGene_gn=JAKMIP3;refGen
e_tx=XM_023241121.1;refGene_gn=JAKMIP3;refGene_tx=XM_023241122.1;refGe
ne_gn=JAKMIP3;refGene_tx=XM_023241123.1;refGene_gn=JAKMIP3;refGene_tx=
XM_023241124.1;refGene_gn=JAKMIP3;refGene_tx=XM_023241125.1;refGene_gn
=JAKMIP3;refGene_tx=XM_023241126.1;refGene_gn=JAKMIP3;refGene_tx=XM_02
3241127.1;refGene_gn=JAKMIP3;refGene_tx=XM_023241128.1;refGene_gn=JAKM
IP3;refGene_tx=XM_023241129.1;refGene_gn=JAKMIP3;refGene_tx=XM_0232411
30.1;refGene_gn=JAKMIP3;refGene_tx=XM_023241131.1;refGene_gn=JAKMIP3;r
efGene_tx=XM_023241132.1;refGene_gn=JAKMIP3;refGene_tx=XM_023241133.1;
refGene_gn=JAKMIP3;refGene_tx=XM_023241134.1;refGene_gn=JAKMIP3;refGen
e_tx=XM_023241135.1
D2 88880334 88880370
refGene_gn=JAKMIP3;refGene_tx=XM_023241120.1;refGene_gn=JAKMIP3;refGen
e_tx=XM_023241121.1;refGene_gn=JAKMIP3;refGene_tx=XM_023241122.1;refGe
ne_gn=JAKMIP3;refGene_tx=XM_023241123.1;refGene_gn=JAKMIP3;refGene_tx=
XM_023241124.1;refGene_gn=JAKMIP3;refGene_tx=XM_023241125.1;refGene_gn
=JAKMIP3;refGene_tx=XM_023241126.1;refGene_gn=JAKMIP3;refGene_tx=XM_02
3241127.1;refGene_gn=JAKMIP3;refGene_tx=XM_023241128.1;refGene_gn=JAKM
IP3;refGene_tx=XM_023241129.1;refGene_gn=JAKMIP3;refGene_tx=XM_0232411
30.1;refGene_gn=JAKMIP3;refGene_tx=XM_023241131.1;refGene_gn=JAKMIP3;r
efGene_tx=XM_023241132.1;refGene_gn=JAKMIP3;refGene_tx=XM_023241133.1;
refGene_gn=JAKMIP3;refGene_tx=XM_023241134.1;refGene_gn=JAKMIP3;refGen
e_tx=XM_023241135.1
D2 88881135 88881195
refGene_gn=JAKMIP3;refGene_tx=XM_023241120.1;refGene_gn=JAKMIP3;refGen
e_tx=XM_023241121.1;refGene_gn=JAKMIP3;refGene_tx=XM_023241122.1;refGe
ne_gn=JAKMIP3;refGene_tx=XM_023241123.1;refGene_gn=JAKMIP3;refGene_tx=
XM_023241124.1;refGene_gn=JAKMIP3;refGene_tx=XM_023241125.1;refGene_gn
=JAKMIP3;refGene_tx=XM_023241126.1;refGene_gn=JAKMIP3;refGene_tx=XM_02
3241127.1;refGene_gn=JAKMIP3;refGene_tx=XM_023241128.1;refGene_gn=JAKM
IP3;refGene_tx=XM_023241129.1;refGene_gn=JAKMIP3;refGene_tx=XM_0232411
30.1;refGene_gn=JAKMIP3;refGene_tx=XM_023241131.1;refGene_gn=JAKMIP3;r
efGene_tx=XM_023241132.1;refGene_gn=JAKMIP3;refGene_tx=XM_023241133.1;
refGene_gn=JAKMIP3;refGene_tx=XM_023241134.1;refGene_gn=JAKMIP3;refGen
e_tx=XM_023241135.1
D2 88882939 88883071
refGene_gn=JAKMIP3;refGene_tx=XM_023241120.1;refGene_gn=JAKMIP3;refGen
e_tx=XM_023241121.1;refGene_gn=JAKMIP3;refGene_tx=XM_023241122.1;refGe
ne_gn=JAKMIP3;refGene_tx=XM_023241123.1;refGene_gn=JAKMIP3;refGene_tx=
XM_023241124.1;refGene_gn=JAKMIP3;refGene_tx=XM_023241125.1;refGene_gn
=JAKMIP3;refGene_tx=XM_023241126.1;refGene_gn=JAKMIP3;refGene_tx=XM_02
3241127.1;refGene_gn=JAKMIP3;refGene_tx=XM_023241128.1;refGene_gn=JAKM
IP3;refGene_tx=XM_023241129.1;refGene_gn=JAKMIP3;refGene_tx=XM_0232411
30.1;refGene_gn=JAKMIP3;refGene_tx=XM_023241131.1;refGene_gn=JAKMIP3;r

efGene_tx=XM_023241132.1;refGene_gn=JAKMIP3;refGene_tx=XM_023241133.1;refGene_gn=JAKMIP3;refGene_tx=XM_023241134.1;refGene_gn=JAKMIP3;refGene_tx=XM_023241135.1
D2 88883967 88884096
refGene_gn=JAKMIP3;refGene_tx=XM_023241120.1;refGene_gn=JAKMIP3;refGene_tx=XM_023241121.1;refGene_gn=JAKMIP3;refGene_tx=XM_023241122.1;refGene_gn=JAKMIP3;refGene_tx=XM_023241123.1;refGene_gn=JAKMIP3;refGene_tx=XM_023241124.1;refGene_gn=JAKMIP3;refGene_tx=XM_023241125.1;refGene_gn=JAKMIP3;refGene_tx=XM_023241126.1;refGene_gn=JAKMIP3;refGene_tx=XM_023241127.1;refGene_gn=JAKMIP3;refGene_tx=XM_023241128.1;refGene_gn=JAKMIP3;refGene_tx=XM_023241129.1;refGene_gn=JAKMIP3;refGene_tx=XM_023241130.1;refGene_gn=JAKMIP3;refGene_tx=XM_023241131.1;refGene_gn=JAKMIP3;refGene_tx=XM_023241132.1;refGene_gn=JAKMIP3;refGene_tx=XM_023241133.1;refGene_gn=JAKMIP3;refGene_tx=XM_023241134.1;refGene_gn=JAKMIP3;refGene_tx=XM_023241135.1
D2 88885858 88885942
refGene_gn=JAKMIP3;refGene_tx=XM_023241120.1;refGene_gn=JAKMIP3;refGene_tx=XM_023241121.1;refGene_gn=JAKMIP3;refGene_tx=XM_023241122.1;refGene_gn=JAKMIP3;refGene_tx=XM_023241123.1;refGene_gn=JAKMIP3;refGene_tx=XM_023241124.1;refGene_gn=JAKMIP3;refGene_tx=XM_023241125.1;refGene_gn=JAKMIP3;refGene_tx=XM_023241126.1;refGene_gn=JAKMIP3;refGene_tx=XM_023241127.1;refGene_gn=JAKMIP3;refGene_tx=XM_023241128.1;refGene_gn=JAKMIP3;refGene_tx=XM_023241129.1;refGene_gn=JAKMIP3;refGene_tx=XM_023241130.1;refGene_gn=JAKMIP3;refGene_tx=XM_023241131.1;refGene_gn=JAKMIP3;refGene_tx=XM_023241132.1;refGene_gn=JAKMIP3;refGene_tx=XM_023241133.1;refGene_gn=JAKMIP3;refGene_tx=XM_023241134.1;refGene_gn=JAKMIP3;refGene_tx=XM_023241135.1
D2 88887522 88887585
refGene_gn=JAKMIP3;refGene_tx=XM_023241120.1;refGene_gn=JAKMIP3;refGene_tx=XM_023241121.1;refGene_gn=JAKMIP3;refGene_tx=XM_023241122.1;refGene_gn=JAKMIP3;refGene_tx=XM_023241123.1;refGene_gn=JAKMIP3;refGene_tx=XM_023241124.1;refGene_gn=JAKMIP3;refGene_tx=XM_023241125.1;refGene_gn=JAKMIP3;refGene_tx=XM_023241126.1;refGene_gn=JAKMIP3;refGene_tx=XM_023241127.1;refGene_gn=JAKMIP3;refGene_tx=XM_023241128.1;refGene_gn=JAKMIP3;refGene_tx=XM_023241129.1;refGene_gn=JAKMIP3;refGene_tx=XM_023241130.1;refGene_gn=JAKMIP3;refGene_tx=XM_023241131.1;refGene_gn=JAKMIP3;refGene_tx=XM_023241132.1;refGene_gn=JAKMIP3;refGene_tx=XM_023241133.1;refGene_gn=JAKMIP3;refGene_tx=XM_023241134.1;refGene_gn=JAKMIP3;refGene_tx=XM_023241135.1
D2 88890126 88890225
refGene_gn=JAKMIP3;refGene_tx=XM_023241120.1;refGene_gn=JAKMIP3;refGene_tx=XM_023241121.1;refGene_gn=JAKMIP3;refGene_tx=XM_023241122.1;refGene_gn=JAKMIP3;refGene_tx=XM_023241123.1;refGene_gn=JAKMIP3;refGene_tx=XM_023241124.1;refGene_gn=JAKMIP3;refGene_tx=XM_023241125.1;refGene_gn=JAKMIP3;refGene_tx=XM_023241126.1;refGene_gn=JAKMIP3;refGene_tx=XM_023241127.1;refGene_gn=JAKMIP3;refGene_tx=XM_023241128.1;refGene_gn=JAKMIP3;refGene_tx=XM_023241129.1;refGene_gn=JAKMIP3;refGene_tx=XM_023241130.1;refGene_gn=JAKMIP3;refGene_tx=XM_023241131.1;refGene_gn=JAKMIP3;refGene_tx=XM_023241132.1;refGene_gn=JAKMIP3;refGene_tx=XM_023241133.1;refGene_gn=JAKMIP3;refGene_tx=XM_023241134.1;refGene_gn=JAKMIP3;refGen

e_tx=XM_023241135.1
D2 88891261 88891360
refGene_gn=JAKMIP3;refGene_tx=XM_023241120.1;refGene_gn=JAKMIP3;refGene_tx=XM_023241121.1;refGene_gn=JAKMIP3;refGene_tx=XM_023241122.1;refGene_gn=JAKMIP3;refGene_tx=XM_023241123.1;refGene_gn=JAKMIP3;refGene_tx=XM_023241124.1;refGene_gn=JAKMIP3;refGene_tx=XM_023241125.1;refGene_gn=JAKMIP3;refGene_tx=XM_023241126.1;refGene_gn=JAKMIP3;refGene_tx=XM_023241127.1;refGene_gn=JAKMIP3;refGene_tx=XM_023241128.1;refGene_gn=JAKMIP3;refGene_tx=XM_023241129.1;refGene_gn=JAKMIP3;refGene_tx=XM_023241130.1;refGene_gn=JAKMIP3;refGene_tx=XM_023241131.1;refGene_gn=JAKMIP3;refGene_tx=XM_023241132.1;refGene_gn=JAKMIP3;refGene_tx=XM_023241133.1;refGene_gn=JAKMIP3;refGene_tx=XM_023241134.1;refGene_gn=JAKMIP3;refGene_tx=XM_023241135.1
D2 88891451 88891511
refGene_gn=JAKMIP3;refGene_tx=XM_023241120.1;refGene_gn=JAKMIP3;refGene_tx=XM_023241121.1;refGene_gn=JAKMIP3;refGene_tx=XM_023241122.1;refGene_gn=JAKMIP3;refGene_tx=XM_023241123.1;refGene_gn=JAKMIP3;refGene_tx=XM_023241124.1;refGene_gn=JAKMIP3;refGene_tx=XM_023241125.1;refGene_gn=JAKMIP3;refGene_tx=XM_023241126.1;refGene_gn=JAKMIP3;refGene_tx=XM_023241127.1;refGene_gn=JAKMIP3;refGene_tx=XM_023241128.1;refGene_gn=JAKMIP3;refGene_tx=XM_023241129.1;refGene_gn=JAKMIP3;refGene_tx=XM_023241130.1;refGene_gn=JAKMIP3;refGene_tx=XM_023241131.1;refGene_gn=JAKMIP3;refGene_tx=XM_023241132.1;refGene_gn=JAKMIP3;refGene_tx=XM_023241133.1;refGene_gn=JAKMIP3;refGene_tx=XM_023241134.1;refGene_gn=JAKMIP3;refGene_tx=XM_023241135.1
D2 88895515 88895581
refGene_gn=JAKMIP3;refGene_tx=XM_023241120.1;refGene_gn=JAKMIP3;refGene_tx=XM_023241121.1;refGene_gn=JAKMIP3;refGene_tx=XM_023241122.1;refGene_gn=JAKMIP3;refGene_tx=XM_023241123.1;refGene_gn=JAKMIP3;refGene_tx=XM_023241124.1;refGene_gn=JAKMIP3;refGene_tx=XM_023241125.1;refGene_gn=JAKMIP3;refGene_tx=XM_023241126.1;refGene_gn=JAKMIP3;refGene_tx=XM_023241127.1;refGene_gn=JAKMIP3;refGene_tx=XM_023241128.1;refGene_gn=JAKMIP3;refGene_tx=XM_023241129.1;refGene_gn=JAKMIP3;refGene_tx=XM_023241130.1;refGene_gn=JAKMIP3;refGene_tx=XM_023241131.1;refGene_gn=JAKMIP3;refGene_tx=XM_023241132.1;refGene_gn=JAKMIP3;refGene_tx=XM_023241133.1;refGene_gn=JAKMIP3;refGene_tx=XM_023241134.1;refGene_gn=JAKMIP3;refGene_tx=XM_023241135.1
D2 88896310 88896379
refGene_gn=JAKMIP3;refGene_tx=XM_023241120.1;refGene_gn=JAKMIP3;refGene_tx=XM_023241121.1;refGene_gn=JAKMIP3;refGene_tx=XM_023241122.1;refGene_gn=JAKMIP3;refGene_tx=XM_023241123.1;refGene_gn=JAKMIP3;refGene_tx=XM_023241124.1;refGene_gn=JAKMIP3;refGene_tx=XM_023241127.1;refGene_gn=JAKMIP3;refGene_tx=XM_023241128.1;refGene_gn=JAKMIP3;refGene_tx=XM_023241129.1;refGene_gn=JAKMIP3;refGene_tx=XM_023241130.1;refGene_gn=JAKMIP3;refGene_tx=XM_023241131.1;refGene_gn=JAKMIP3;refGene_tx=XM_023241132.1;refGene_gn=JAKMIP3;refGene_tx=XM_023241133.1;refGene_gn=JAKMIP3;refGene_tx=XM_023241134.1;refGene_gn=JAKMIP3;refGene_tx=XM_023241135.1
D2 88896465 88896543
refGene_gn=JAKMIP3;refGene_tx=XM_023241120.1;refGene_gn=JAKMIP3;refGene_tx=XM_023241121.1;refGene_gn=JAKMIP3;refGene_tx=XM_023241122.1;refGe

ne_gn=JAKMIP3;refGene_tx=XM_023241123.1;refGene_gn=JAKMIP3;refGene_tx=
XM_023241124.1;refGene_gn=JAKMIP3;refGene_tx=XM_023241125.1;refGene_gn
=JAKMIP3;refGene_tx=XM_023241126.1;refGene_gn=JAKMIP3;refGene_tx=XM_02
3241127.1;refGene_gn=JAKMIP3;refGene_tx=XM_023241128.1;refGene_gn=JAKM
IP3;refGene_tx=XM_023241129.1;refGene_gn=JAKMIP3;refGene_tx=XM_0232411
30.1;refGene_gn=JAKMIP3;refGene_tx=XM_023241131.1;refGene_gn=JAKMIP3;r
efGene_tx=XM_023241132.1;refGene_gn=JAKMIP3;refGene_tx=XM_023241133.1;
refGene_gn=JAKMIP3;refGene_tx=XM_023241134.1;refGene_gn=JAKMIP3;refGen
e_tx=XM_023241135.1
D2 88902319 88902523
refGene_gn=JAKMIP3;refGene_tx=XM_023241120.1;refGene_gn=JAKMIP3;refGen
e_tx=XM_023241121.1;refGene_gn=JAKMIP3;refGene_tx=XM_023241122.1;refGe
ne_gn=JAKMIP3;refGene_tx=XM_023241123.1;refGene_gn=JAKMIP3;refGene_tx=
XM_023241124.1;refGene_gn=JAKMIP3;refGene_tx=XM_023241125.1;refGene_gn
=JAKMIP3;refGene_tx=XM_023241126.1;refGene_gn=JAKMIP3;refGene_tx=XM_02
3241127.1;refGene_gn=JAKMIP3;refGene_tx=XM_023241128.1;refGene_gn=JAKM
IP3;refGene_tx=XM_023241129.1;refGene_gn=JAKMIP3;refGene_tx=XM_0232411
30.1;refGene_gn=JAKMIP3;refGene_tx=XM_023241131.1;refGene_gn=JAKMIP3;r
efGene_tx=XM_023241132.1;refGene_gn=JAKMIP3;refGene_tx=XM_023241133.1;
refGene_gn=JAKMIP3;refGene_tx=XM_023241134.1;refGene_gn=JAKMIP3;refGen
e_tx=XM_023241135.1
D2 88904073 88904139
refGene_gn=JAKMIP3;refGene_tx=XM_023241120.1;refGene_gn=JAKMIP3;refGen
e_tx=XM_023241121.1;refGene_gn=JAKMIP3;refGene_tx=XM_023241122.1;refGe
ne_gn=JAKMIP3;refGene_tx=XM_023241123.1;refGene_gn=JAKMIP3;refGene_tx=
XM_023241124.1;refGene_gn=JAKMIP3;refGene_tx=XM_023241125.1;refGene_gn
=JAKMIP3;refGene_tx=XM_023241126.1;refGene_gn=JAKMIP3;refGene_tx=XM_02
3241127.1;refGene_gn=JAKMIP3;refGene_tx=XM_023241128.1;refGene_gn=JAKM
IP3;refGene_tx=XM_023241129.1;refGene_gn=JAKMIP3;refGene_tx=XM_0232411
30.1;refGene_gn=JAKMIP3;refGene_tx=XM_023241131.1;refGene_gn=JAKMIP3;r
efGene_tx=XM_023241132.1;refGene_gn=JAKMIP3;refGene_tx=XM_023241133.1;
refGene_gn=JAKMIP3;refGene_tx=XM_023241134.1;refGene_gn=JAKMIP3;refGen
e_tx=XM_023241135.1
D2 88905950 88906023
refGene_gn=JAKMIP3;refGene_tx=XM_023241120.1;refGene_gn=JAKMIP3;refGen
e_tx=XM_023241121.1;refGene_gn=JAKMIP3;refGene_tx=XM_023241122.1;refGe
ne_gn=JAKMIP3;refGene_tx=XM_023241123.1;refGene_gn=JAKMIP3;refGene_tx=
XM_023241125.1;refGene_gn=JAKMIP3;refGene_tx=XM_023241126.1;refGene_gn
=JAKMIP3;refGene_tx=XM_023241127.1;refGene_gn=JAKMIP3;refGene_tx=XM_02
3241128.1;refGene_gn=JAKMIP3;refGene_tx=XM_023241129.1;refGene_gn=JAKM
IP3;refGene_tx=XM_023241130.1;refGene_gn=JAKMIP3;refGene_tx=XM_0232411
31.1;refGene_gn=JAKMIP3;refGene_tx=XM_023241132.1;refGene_gn=JAKMIP3;r
efGene_tx=XM_023241133.1;refGene_gn=JAKMIP3;refGene_tx=XM_023241134.1;
refGene_gn=JAKMIP3;refGene_tx=XM_023241135.1
D2 88906710 88906869
refGene_gn=JAKMIP3;refGene_tx=XM_023241120.1;refGene_gn=JAKMIP3;refGen
e_tx=XM_023241121.1;refGene_gn=JAKMIP3;refGene_tx=XM_023241122.1;refGe
ne_gn=JAKMIP3;refGene_tx=XM_023241123.1;refGene_gn=JAKMIP3;refGene_tx=
XM_023241124.1;refGene_gn=JAKMIP3;refGene_tx=XM_023241125.1;refGene_gn
=JAKMIP3;refGene_tx=XM_023241126.1;refGene_gn=JAKMIP3;refGene_tx=XM_02

3241131.1;refGene_gn=JAKMIP3;refGene_tx=XM_023241132.1;refGene_gn=JAKMIP3;refGene_tx=XM_023241133.1;refGene_gn=JAKMIP3;refGene_tx=XM_023241134.1
D2 88923674 88923932 refGene_gn=DPYSL4;refGene_tx=XM_023241136.1
D2 88925034 88925073 refGene_gn=DPYSL4;refGene_tx=XM_023241137.1
D2 88927349 88927438 refGene_gn=DPYSL4;refGene_tx=XM_023241136.1;refGene_gn=DPYSL4;refGene_tx=XM_023241137.1
D2 88927927 88928112 refGene_gn=DPYSL4;refGene_tx=XM_023241136.1;refGene_gn=DPYSL4;refGene_tx=XM_023241137.1
D2 88930107 88930272 refGene_gn=DPYSL4;refGene_tx=XM_023241136.1;refGene_gn=DPYSL4;refGene_tx=XM_023241137.1
D2 88930847 88930909 refGene_gn=DPYSL4;refGene_tx=XM_023241136.1;refGene_gn=DPYSL4;refGene_tx=XM_023241137.1
D2 88931003 88931084 refGene_gn=DPYSL4;refGene_tx=XM_023241136.1;refGene_gn=DPYSL4;refGene_tx=XM_023241137.1
D2 88932512 88932581 refGene_gn=DPYSL4;refGene_tx=XM_023241136.1;refGene_gn=DPYSL4;refGene_tx=XM_023241137.1
D2 88932936 88933057 refGene_gn=DPYSL4;refGene_tx=XM_023241136.1;refGene_gn=DPYSL4;refGene_tx=XM_023241137.1
D2 88933898 88934055 refGene_gn=DPYSL4;refGene_tx=XM_023241136.1;refGene_gn=DPYSL4;refGene_tx=XM_023241137.1
D2 88934345 88934487 refGene_gn=DPYSL4;refGene_tx=XM_023241136.1;refGene_gn=DPYSL4;refGene_tx=XM_023241137.1
D2 88935148 88935319 refGene_gn=DPYSL4;refGene_tx=XM_023241136.1;refGene_gn=DPYSL4;refGene_tx=XM_023241137.1
D2 88935613 88935793 refGene_gn=DPYSL4;refGene_tx=XM_023241136.1;refGene_gn=DPYSL4;refGene_tx=XM_023241137.1
D2 88937449 88937615 refGene_gn=DPYSL4;refGene_tx=XM_023241136.1;refGene_gn=DPYSL4;refGene_tx=XM_023241137.1
D2 88938446 88938538 refGene_gn=DPYSL4;refGene_tx=XM_023241136.1;refGene_gn=DPYSL4;refGene_tx=XM_023241137.1
D2 88941298 88941437 refGene_gn=STK32C;refGene_tx=XM_019814031.2;refGene_gn=STK32C;refGene_tx=XM_023241145.1;refGene_gn=STK32C;refGene_tx=XM_023241146.1
D2 88942359 88942427 refGene_gn=STK32C;refGene_tx=XM_019814031.2;refGene_gn=STK32C;refGene_tx=XM_023241145.1;refGene_gn=STK32C;refGene_tx=XM_023241146.1

D2 88945897 88946029
refGene_gn=STK32C;refGene_tx=XM_019814031.2;refGene_gn=STK32C;refGene_tx=XM_023241145.1;refGene_gn=STK32C;refGene_tx=XM_023241146.1
D2 88946113 88946239
refGene_gn=STK32C;refGene_tx=XM_019814031.2;refGene_gn=STK32C;refGene_tx=XM_023241145.1;refGene_gn=STK32C;refGene_tx=XM_023241146.1
D2 88947445 88947562
refGene_gn=STK32C;refGene_tx=XM_019814031.2;refGene_gn=STK32C;refGene_tx=XM_023241145.1;refGene_gn=STK32C;refGene_tx=XM_023241146.1
D2 88948218 88948322
refGene_gn=STK32C;refGene_tx=XM_019814031.2;refGene_gn=STK32C;refGene_tx=XM_023241145.1;refGene_gn=STK32C;refGene_tx=XM_023241146.1
D2 88948459 88948624
refGene_gn=STK32C;refGene_tx=XM_019814031.2;refGene_gn=STK32C;refGene_tx=XM_023241145.1;refGene_gn=STK32C;refGene_tx=XM_023241146.1
D2 88948725 88948763
refGene_gn=STK32C;refGene_tx=XM_019814031.2;refGene_gn=STK32C;refGene_tx=XM_023241145.1;refGene_gn=STK32C;refGene_tx=XM_023241146.1
D2 88949639 88949813
refGene_gn=STK32C;refGene_tx=XM_019814031.2;refGene_gn=STK32C;refGene_tx=XM_023241145.1;refGene_gn=STK32C;refGene_tx=XM_023241146.1
D2 88952470 88952622
refGene_gn=STK32C;refGene_tx=XM_019814031.2;refGene_gn=STK32C;refGene_tx=XM_023241145.1;refGene_gn=STK32C;refGene_tx=XM_023241146.1
D2 88972773 88972829
refGene_gn=STK32C;refGene_tx=XM_023241145.1;refGene_gn=STK32C;refGene_tx=XM_023241146.1
D2 89018688 89018950
refGene_gn=STK32C;refGene_tx=XM_023241145.1;refGene_gn=STK32C;refGene_tx=XM_023241146.1
D2 89040515 89040716
refGene_gn=LRRC27;refGene_tx=XM_006938284.4;refGene_gn=LRRC27;refGene_tx=XM_019814024.2;refGene_gn=LRRC27;refGene_tx=XM_019814025.2;refGene_gn=LRRC27;refGene_tx=XM_023241138.1;refGene_gn=LRRC27;refGene_tx=XM_023241140.1;refGene_gn=LRRC27;refGene_tx=XM_023241141.1;refGene_gn=LRRC27;refGene_tx=XM_023241142.1;refGene_gn=LRRC27;refGene_tx=XM_023241143.1;refGene_gn=LRRC27;refGene_tx=XM_023241144.1
D2 89043211 89043342
refGene_gn=LRRC27;refGene_tx=XM_006938284.4;refGene_gn=LRRC27;refGene_tx=XM_019814024.2;refGene_gn=LRRC27;refGene_tx=XM_019814025.2;refGene_gn=LRRC27;refGene_tx=XM_023241138.1;refGene_gn=LRRC27;refGene_tx=XM_023241140.1;refGene_gn=LRRC27;refGene_tx=XM_023241141.1;refGene_gn=LRRC27;refGene_tx=XM_023241142.1;refGene_gn=LRRC27;refGene_tx=XM_023241143.1;refGene_gn=LRRC27;refGene_tx=XM_023241144.1
D2 89046604 89046663
refGene_gn=LRRC27;refGene_tx=XM_006938284.4;refGene_gn=LRRC27;refGene_tx=XM_019814024.2;refGene_gn=LRRC27;refGene_tx=XM_019814025.2;refGene_gn=LRRC27;refGene_tx=XM_019814029.2;refGene_gn=LRRC27;refGene_tx=XM_023241138.1;refGene_gn=LRRC27;refGene_tx=XM_023241140.1;refGene_gn=LRRC27;refGene_tx=XM_023241141.1;refGene_gn=LRRC27;refGene_tx=XM_023241142.

1;refGene_gn=LRRC27;refGene_tx=XM_023241143.1;refGene_gn=LRRC27;refGene_tx=XM_023241144.1
D2 89052181 89052334
refGene_gn=LRRC27;refGene_tx=XM_006938284.4;refGene_gn=LRRC27;refGene_tx=XM_019814024.2;refGene_gn=LRRC27;refGene_tx=XM_019814025.2;refGene_gn=LRRC27;refGene_tx=XM_019814029.2;refGene_gn=LRRC27;refGene_tx=XM_023241138.1;refGene_gn=LRRC27;refGene_tx=XM_023241140.1;refGene_gn=LRRC27;refGene_tx=XM_023241141.1;refGene_gn=LRRC27;refGene_tx=XM_023241142.1;refGene_gn=LRRC27;refGene_tx=XM_023241143.1
D2 89054262 89054569
refGene_gn=LRRC27;refGene_tx=XM_006938284.4;refGene_gn=LRRC27;refGene_tx=XM_019814024.2;refGene_gn=LRRC27;refGene_tx=XM_019814025.2;refGene_gn=LRRC27;refGene_tx=XM_019814029.2;refGene_gn=LRRC27;refGene_tx=XM_023241138.1;refGene_gn=LRRC27;refGene_tx=XM_023241140.1;refGene_gn=LRRC27;refGene_tx=XM_023241141.1;refGene_gn=LRRC27;refGene_tx=XM_023241142.1;refGene_gn=LRRC27;refGene_tx=XM_023241143.1;refGene_gn=LRRC27;refGene_tx=XM_023241144.1
D2 89055465 89055633
refGene_gn=LRRC27;refGene_tx=XM_006938284.4;refGene_gn=LRRC27;refGene_tx=XM_019814024.2;refGene_gn=LRRC27;refGene_tx=XM_019814025.2;refGene_gn=LRRC27;refGene_tx=XM_019814029.2;refGene_gn=LRRC27;refGene_tx=XM_023241138.1;refGene_gn=LRRC27;refGene_tx=XM_023241140.1;refGene_gn=LRRC27;refGene_tx=XM_023241141.1;refGene_gn=LRRC27;refGene_tx=XM_023241142.1;refGene_gn=LRRC27;refGene_tx=XM_023241143.1;refGene_gn=LRRC27;refGene_tx=XM_023241144.1
D2 89056971 89057714 refGene_gn=LRRC27;refGene_tx=XM_019814027.2
D2 89058698 89058897
refGene_gn=LRRC27;refGene_tx=XM_006938284.4;refGene_gn=LRRC27;refGene_tx=XM_019814024.2;refGene_gn=LRRC27;refGene_tx=XM_019814025.2;refGene_gn=LRRC27;refGene_tx=XM_019814027.2;refGene_gn=LRRC27;refGene_tx=XM_019814029.2;refGene_gn=LRRC27;refGene_tx=XM_023241138.1;refGene_gn=LRRC27;refGene_tx=XM_023241140.1;refGene_gn=LRRC27;refGene_tx=XM_023241141.1;refGene_gn=LRRC27;refGene_tx=XM_023241142.1;refGene_gn=LRRC27;refGene_tx=XM_023241143.1;refGene_gn=LRRC27;refGene_tx=XM_023241144.1
D2 89066230 89066349
refGene_gn=LRRC27;refGene_tx=XM_006938284.4;refGene_gn=LRRC27;refGene_tx=XM_019814024.2;refGene_gn=LRRC27;refGene_tx=XM_019814025.2;refGene_gn=LRRC27;refGene_tx=XM_019814027.2;refGene_gn=LRRC27;refGene_tx=XM_019814029.2;refGene_gn=LRRC27;refGene_tx=XM_023241138.1;refGene_gn=LRRC27;refGene_tx=XM_023241140.1;refGene_gn=LRRC27;refGene_tx=XM_023241141.1;refGene_gn=LRRC27;refGene_tx=XM_023241142.1;refGene_gn=LRRC27;refGene_tx=XM_023241143.1;refGene_gn=LRRC27;refGene_tx=XM_023241144.1
D2 89068093 89068223
refGene_gn=LRRC27;refGene_tx=XM_006938284.4;refGene_gn=LRRC27;refGene_tx=XM_019814024.2;refGene_gn=LRRC27;refGene_tx=XM_019814025.2;refGene_gn=LRRC27;refGene_tx=XM_019814027.2;refGene_gn=LRRC27;refGene_tx=XM_019814029.2;refGene_gn=LRRC27;refGene_tx=XM_023241138.1;refGene_gn=LRRC27;refGene_tx=XM_023241140.1;refGene_gn=LRRC27;refGene_tx=XM_023241141.1;refGene_gn=LRRC27;refGene_tx=XM_023241142.1;refGene_gn=LRRC27;refGene_tx=XM_023241143.1;refGene_gn=LRRC27;refGene_tx=XM_023241144.1

D2 89069417 89069501 refGene_gn=LRRC27;refGene_tx=XM_023241143.1
D2 89073852 89073999
refGene_gn=LRRC27;refGene_tx=XM_006938284.4;refGene_gn=LRRC27;refGene_tx=XM_019814024.2;refGene_gn=LRRC27;refGene_tx=XM_019814025.2;refGene_gn=LRRC27;refGene_tx=XM_019814027.2;refGene_gn=LRRC27;refGene_tx=XM_019814029.2;refGene_gn=LRRC27;refGene_tx=XM_023241138.1;refGene_gn=LRRC27;refGene_tx=XM_023241140.1;refGene_gn=LRRC27;refGene_tx=XM_023241141.1;refGene_gn=LRRC27;refGene_tx=XM_023241142.1;refGene_gn=LRRC27;refGene_tx=XM_023241144.1
D2 89082040 89082165
refGene_gn=PWWP2B;refGene_tx=XM_023241147.1;refGene_gn=PWWP2B;refGene_tx=XM_023241151.1
D2 89091110 89092692
refGene_gn=PWWP2B;refGene_tx=XM_023241147.1;refGene_gn=PWWP2B;refGene_tx=XM_023241148.1;refGene_gn=PWWP2B;refGene_tx=XM_023241149.1;refGene_gn=PWWP2B;refGene_tx=XM_023241150.1;refGene_gn=PWWP2B;refGene_tx=XM_023241151.1
D2 89101688 89102837
refGene_gn=LOC111557137;refGene_tx=XM_023241152.1;refGene_gn=LOC111557137;refGene_tx=XM_023241153.1
D2 89106120 89106134 refGene_gn=PWWP2B;refGene_tx=XM_023241151.1
D2 89191408 89191483
refGene_gn=INPP5A;refGene_tx=XM_003994524.5;refGene_gn=INPP5A;refGene_tx=XM_019814032.2;refGene_gn=INPP5A;refGene_tx=XM_019814033.2;refGene_gn=INPP5A;refGene_tx=XM_019814035.2
D2 89235839 89235881
refGene_gn=INPP5A;refGene_tx=XM_003994524.5;refGene_gn=INPP5A;refGene_tx=XM_019814032.2;refGene_gn=INPP5A;refGene_tx=XM_019814033.2;refGene_gn=INPP5A;refGene_tx=XM_019814035.2
D2 89256304 89256405
refGene_gn=INPP5A;refGene_tx=XM_003994524.5;refGene_gn=INPP5A;refGene_tx=XM_019814032.2;refGene_gn=INPP5A;refGene_tx=XM_019814033.2;refGene_gn=INPP5A;refGene_tx=XM_019814034.2
D2 89261548 89261636
refGene_gn=INPP5A;refGene_tx=XM_003994524.5;refGene_gn=INPP5A;refGene_tx=XM_019814032.2;refGene_gn=INPP5A;refGene_tx=XM_019814033.2;refGene_gn=INPP5A;refGene_tx=XM_019814034.2
D2 89296704 89296768
refGene_gn=INPP5A;refGene_tx=XM_003994524.5;refGene_gn=INPP5A;refGene_tx=XM_019814032.2;refGene_gn=INPP5A;refGene_tx=XM_019814033.2;refGene_gn=INPP5A;refGene_tx=XM_019814034.2;refGene_gn=INPP5A;refGene_tx=XM_019814035.2
D2 89306391 89306495
refGene_gn=INPP5A;refGene_tx=XM_003994524.5;refGene_gn=INPP5A;refGene_tx=XM_019814032.2;refGene_gn=INPP5A;refGene_tx=XM_019814033.2;refGene_gn=INPP5A;refGene_tx=XM_019814034.2;refGene_gn=INPP5A;refGene_tx=XM_019814035.2
D2 89313622 89313675
refGene_gn=INPP5A;refGene_tx=XM_003994524.5;refGene_gn=INPP5A;refGene_tx=XM_019814032.2;refGene_gn=INPP5A;refGene_tx=XM_019814033.2;refGene_

gn=INPP5A;refGene_tx=XM_019814034.2;refGene_gn=INPP5A;refGene_tx=XM_019814035.2
D2 89315476 89315596 refGene_gn=INPP5A;refGene_tx=XM_003994524.5;refGene_gn=INPP5A;refGene_tx=XM_019814032.2;refGene_gn=INPP5A;refGene_tx=XM_019814033.2;refGene_gn=INPP5A;refGene_tx=XM_019814034.2;refGene_gn=INPP5A;refGene_tx=XM_019814035.2
D2 89326404 89326489 refGene_gn=INPP5A;refGene_tx=XM_003994524.5;refGene_gn=INPP5A;refGene_tx=XM_019814032.2;refGene_gn=INPP5A;refGene_tx=XM_019814033.2;refGene_gn=INPP5A;refGene_tx=XM_019814034.2;refGene_gn=INPP5A;refGene_tx=XM_019814035.2
D2 89341454 89341550 refGene_gn=INPP5A;refGene_tx=XM_003994524.5;refGene_gn=INPP5A;refGene_tx=XM_019814032.2;refGene_gn=INPP5A;refGene_tx=XM_019814033.2;refGene_gn=INPP5A;refGene_tx=XM_019814034.2;refGene_gn=INPP5A;refGene_tx=XM_019814035.2
D2 89341653 89341728 refGene_gn=INPP5A;refGene_tx=XM_003994524.5;refGene_gn=INPP5A;refGene_tx=XM_019814032.2;refGene_gn=INPP5A;refGene_tx=XM_019814033.2;refGene_gn=INPP5A;refGene_tx=XM_019814034.2;refGene_gn=INPP5A;refGene_tx=XM_019814035.2
D2 89349056 89349130 refGene_gn=INPP5A;refGene_tx=XM_003994524.5;refGene_gn=INPP5A;refGene_tx=XM_019814032.2;refGene_gn=INPP5A;refGene_tx=XM_019814033.2;refGene_gn=INPP5A;refGene_tx=XM_019814034.2;refGene_gn=INPP5A;refGene_tx=XM_019814035.2
D2 89359435 89359547 refGene_gn=INPP5A;refGene_tx=XM_003994524.5;refGene_gn=INPP5A;refGene_tx=XM_019814032.2;refGene_gn=INPP5A;refGene_tx=XM_019814033.2;refGene_gn=INPP5A;refGene_tx=XM_019814034.2;refGene_gn=INPP5A;refGene_tx=XM_019814035.2
D2 89359842 89359923 refGene_gn=INPP5A;refGene_tx=XM_019814033.2
D2 89361989 89362058 refGene_gn=INPP5A;refGene_tx=XM_003994524.5;refGene_gn=INPP5A;refGene_tx=XM_019814032.2;refGene_gn=INPP5A;refGene_tx=XM_019814034.2;refGene_gn=INPP5A;refGene_tx=XM_019814035.2
D2 89362842 89362923 refGene_gn=INPP5A;refGene_tx=XM_003994524.5;refGene_gn=INPP5A;refGene_tx=XM_019814032.2;refGene_gn=INPP5A;refGene_tx=XM_019814034.2;refGene_gn=INPP5A;refGene_tx=XM_019814035.2
D2 89363017 89363067 refGene_gn=INPP5A;refGene_tx=XM_019814032.2;refGene_gn=INPP5A;refGene_tx=XM_019814034.2;refGene_gn=INPP5A;refGene_tx=XM_019814035.2
D2 89365997 89366252 refGene_gn=NKX6-2;refGene_tx=XM_023241155.1
D2 89366556 89366725 refGene_gn=NKX6-2;refGene_tx=XM_023241155.1
D2 89366827 89367066 refGene_gn=NKX6-2;refGene_tx=XM_023241155.1
D2 89367067 89367235 refGene_gn=NKX6-2;refGene_tx=XM_023241155.1
D2 89385727 89385852 refGene_gn=CFAP46;refGene_tx=XM_023241156.1
D2 89386878 89386959 refGene_gn=CFAP46;refGene_tx=XM_023241156.1

| | | | |
|---|---|---|---|
| D2 | 89387037 | 89387174 | refGene_gn=CFAP46;refGene_tx=XM_023241156.1 |
| D2 | 89387733 | 89387815 | refGene_gn=CFAP46;refGene_tx=XM_023241156.1 |
| D2 | 89389116 | 89389229 | refGene_gn=CFAP46;refGene_tx=XM_023241156.1 |
| D2 | 89390353 | 89390456 | refGene_gn=CFAP46;refGene_tx=XM_023241156.1 |
| D2 | 89391057 | 89391093 | refGene_gn=CFAP46;refGene_tx=XM_023241156.1 |
| D2 | 89391163 | 89391224 | refGene_gn=CFAP46;refGene_tx=XM_023241156.1 |
| D2 | 89391325 | 89391399 | refGene_gn=CFAP46;refGene_tx=XM_023241156.1 |
| D2 | 89394514 | 89394673 | refGene_gn=CFAP46;refGene_tx=XM_023241156.1 |
| D2 | 89396101 | 89396181 | refGene_gn=CFAP46;refGene_tx=XM_023241156.1 |
| D2 | 89397029 | 89397151 | refGene_gn=CFAP46;refGene_tx=XM_023241156.1 |
| D2 | 89397648 | 89397779 | refGene_gn=CFAP46;refGene_tx=XM_023241156.1 |
| D2 | 89398261 | 89398338 | refGene_gn=CFAP46;refGene_tx=XM_023241156.1 |
| D2 | 89398776 | 89398874 | refGene_gn=CFAP46;refGene_tx=XM_023241156.1 |
| D2 | 89405269 | 89405440 | refGene_gn=CFAP46;refGene_tx=XM_023241156.1 |
| D2 | 89405861 | 89406041 | refGene_gn=CFAP46;refGene_tx=XM_023241156.1 |
| D2 | 89406109 | 89406244 | refGene_gn=CFAP46;refGene_tx=XM_023241156.1 |
| D2 | 89407229 | 89407418 | refGene_gn=CFAP46;refGene_tx=XM_023241156.1 |
| D2 | 89408093 | 89408282 | refGene_gn=CFAP46;refGene_tx=XM_023241156.1 |
| D2 | 89415961 | 89416097 | refGene_gn=CFAP46;refGene_tx=XM_023241156.1 |
| D2 | 89417281 | 89417458 | refGene_gn=CFAP46;refGene_tx=XM_023241156.1 |
| D2 | 89418701 | 89418808 | refGene_gn=CFAP46;refGene_tx=XM_023241156.1 |
| D2 | 89418895 | 89419096 | refGene_gn=CFAP46;refGene_tx=XM_023241156.1 |
| D2 | 89420989 | 89421136 | refGene_gn=CFAP46;refGene_tx=XM_023241156.1 |
| D2 | 89421880 | 89422013 | refGene_gn=CFAP46;refGene_tx=XM_023241156.1 |
| D2 | 89422932 | 89423031 | refGene_gn=CFAP46;refGene_tx=XM_023241156.1 |
| D2 | 89423819 | 89423968 | refGene_gn=CFAP46;refGene_tx=XM_023241156.1 |
| D2 | 89427048 | 89427123 | refGene_gn=CFAP46;refGene_tx=XM_023241156.1 |
| D2 | 89427124 | 89427199 | refGene_gn=CFAP46;refGene_tx=XM_023241156.1 |
| D2 | 89427842 | 89427989 | refGene_gn=CFAP46;refGene_tx=XM_023241156.1 |
| D2 | 89429065 | 89429271 | refGene_gn=CFAP46;refGene_tx=XM_023241156.1 |
| D2 | 89429570 | 89429730 | refGene_gn=CFAP46;refGene_tx=XM_023241156.1 |
| D2 | 89433228 | 89433412 | refGene_gn=CFAP46;refGene_tx=XM_023241156.1 |
| D2 | 89433748 | 89433887 | refGene_gn=CFAP46;refGene_tx=XM_023241156.1 |
| D2 | 89436082 | 89436167 | refGene_gn=CFAP46;refGene_tx=XM_023241156.1 |
| D2 | 89441687 | 89441850 | refGene_gn=CFAP46;refGene_tx=XM_023241156.1 |
| D2 | 89442242 | 89442374 | refGene_gn=CFAP46;refGene_tx=XM_023241156.1 |
| D2 | 89452724 | 89452891 | refGene_gn=CFAP46;refGene_tx=XM_023241156.1 |
| D2 | 89453177 | 89453285 | refGene_gn=CFAP46;refGene_tx=XM_023241156.1 |
| D2 | 89453662 | 89453812 | refGene_gn=CFAP46;refGene_tx=XM_023241156.1 |
| D2 | 89454210 | 89454376 | refGene_gn=CFAP46;refGene_tx=XM_023241156.1 |
| D2 | 89455830 | 89456043 | refGene_gn=CFAP46;refGene_tx=XM_023241156.1 |
| D2 | 89456787 | 89456921 | refGene_gn=CFAP46;refGene_tx=XM_023241156.1 |
| D2 | 89457753 | 89457881 | refGene_gn=CFAP46;refGene_tx=XM_023241156.1 |
| D2 | 89458724 | 89458855 | refGene_gn=CFAP46;refGene_tx=XM_023241156.1 |
| D2 | 89459032 | 89459156 | refGene_gn=CFAP46;refGene_tx=XM_023241156.1 |
| D2 | 89460157 | 89460278 | refGene_gn=CFAP46;refGene_tx=XM_023241156.1 |
| D2 | 89460412 | 89460641 | refGene_gn=CFAP46;refGene_tx=XM_023241156.1 |
| D2 | 89465407 | 89465598 | refGene_gn=CFAP46;refGene_tx=XM_023241156.1 |
| D2 | 89467329 | 89467428 | refGene_gn=CFAP46;refGene_tx=XM_023241156.1 |
| D2 | 89473482 | 89473582 | refGene_gn=CFAP46;refGene_tx=XM_023241156.1 |

D2 89476109 89476220 refGene_gn=CFAP46;refGene_tx=XM_023241156.1
D2 89476986 89477081 refGene_gn=CFAP46;refGene_tx=XM_023241156.1
D2 89477550 89477674 refGene_gn=CFAP46;refGene_tx=XM_023241156.1
D2 89478534 89478699 refGene_gn=CFAP46;refGene_tx=XM_023241156.1
D2 89480351 89480416 refGene_gn=CFAP46;refGene_tx=XM_023241156.1
D2 89480992 89481124 refGene_gn=CFAP46;refGene_tx=XM_023241156.1
D2 89481338 89481463 refGene_gn=CFAP46;refGene_tx=XM_023241156.1
D2 89481918 89481967 refGene_gn=CFAP46;refGene_tx=XM_023241156.1
D2 89606450 89606650 refGene_gn=ADGRA1;refGene_tx=XM_006938288.4
D2 89607260 89607462 refGene_gn=ADGRA1;refGene_tx=XM_006938288.4
D2 89614576 89614704 refGene_gn=ADGRA1;refGene_tx=XM_006938288.4
D2 89616844 89616968 refGene_gn=ADGRA1;refGene_tx=XM_006938288.4
D2 89624214 89624336 refGene_gn=ADGRA1;refGene_tx=XM_006938288.4
D2 89638367 89638466 refGene_gn=ADGRA1;refGene_tx=XM_006938288.4
D2 89638987 89640191 refGene_gn=ADGRA1;refGene_tx=XM_006938288.4
D2 89656087 89656189
refGene_gn=KNDC1;refGene_tx=XM_019814037.2;refGene_gn=KNDC1;refGene_tx=XM_019814038.2;refGene_gn=KNDC1;refGene_tx=XM_023241157.1
D2 89663063 89663262
refGene_gn=KNDC1;refGene_tx=XM_019814037.2;refGene_gn=KNDC1;refGene_tx=XM_019814038.2;refGene_gn=KNDC1;refGene_tx=XM_023241157.1
D2 89663818 89663877
refGene_gn=KNDC1;refGene_tx=XM_019814037.2;refGene_gn=KNDC1;refGene_tx=XM_019814038.2;refGene_gn=KNDC1;refGene_tx=XM_023241157.1
D2 89676900 89677053
refGene_gn=KNDC1;refGene_tx=XM_019814037.2;refGene_gn=KNDC1;refGene_tx=XM_019814038.2;refGene_gn=KNDC1;refGene_tx=XM_023241157.1
D2 89677436 89677554
refGene_gn=KNDC1;refGene_tx=XM_019814037.2;refGene_gn=KNDC1;refGene_tx=XM_019814038.2;refGene_gn=KNDC1;refGene_tx=XM_023241157.1
D2 89679028 89679678
refGene_gn=KNDC1;refGene_tx=XM_019814037.2;refGene_gn=KNDC1;refGene_tx=XM_019814038.2;refGene_gn=KNDC1;refGene_tx=XM_023241157.1
D2 89682877 89682992
refGene_gn=KNDC1;refGene_tx=XM_019814037.2;refGene_gn=KNDC1;refGene_tx=XM_019814038.2;refGene_gn=KNDC1;refGene_tx=XM_023241157.1
D2 89683721 89683793
refGene_gn=KNDC1;refGene_tx=XM_019814037.2;refGene_gn=KNDC1;refGene_tx=XM_019814038.2;refGene_gn=KNDC1;refGene_tx=XM_023241157.1
D2 89683885 89683947
refGene_gn=KNDC1;refGene_tx=XM_019814037.2;refGene_gn=KNDC1;refGene_tx=XM_019814038.2;refGene_gn=KNDC1;refGene_tx=XM_023241157.1
D2 89690915 89691074
refGene_gn=KNDC1;refGene_tx=XM_019814037.2;refGene_gn=KNDC1;refGene_tx=XM_019814038.2;refGene_gn=KNDC1;refGene_tx=XM_023241157.1
D2 89692971 89693049
refGene_gn=KNDC1;refGene_tx=XM_019814037.2;refGene_gn=KNDC1;refGene_tx=XM_019814038.2;refGene_gn=KNDC1;refGene_tx=XM_023241157.1
D2 89693633 89693727
refGene_gn=KNDC1;refGene_tx=XM_019814037.2;refGene_gn=KNDC1;refGene_tx

=XM_019814038.2;refGene_gn=KNDC1;refGene_tx=XM_023241157.1
D2 89694223 89694386
refGene_gn=KNDC1;refGene_tx=XM_019814037.2;refGene_gn=KNDC1;refGene_tx=XM_019814038.2;refGene_gn=KNDC1;refGene_tx=XM_023241157.1
D2 89694463 89695152
refGene_gn=KNDC1;refGene_tx=XM_019814037.2;refGene_gn=KNDC1;refGene_tx=XM_019814038.2;refGene_gn=KNDC1;refGene_tx=XM_023241157.1
D2 89695487 89695632
refGene_gn=KNDC1;refGene_tx=XM_019814037.2;refGene_gn=KNDC1;refGene_tx=XM_019814038.2;refGene_gn=KNDC1;refGene_tx=XM_023241157.1
D2 89696649 89696747
refGene_gn=KNDC1;refGene_tx=XM_019814037.2;refGene_gn=KNDC1;refGene_tx=XM_019814038.2;refGene_gn=KNDC1;refGene_tx=XM_023241157.1
D2 89697596 89697985
refGene_gn=KNDC1;refGene_tx=XM_019814037.2;refGene_gn=KNDC1;refGene_tx=XM_019814038.2;refGene_gn=KNDC1;refGene_tx=XM_023241157.1
D2 89698816 89698882 refGene_gn=KNDC1;refGene_tx=XM_019814038.2
D2 89702229 89702259 refGene_gn=KNDC1;refGene_tx=XM_019814037.2
D2 89706578 89706672
refGene_gn=KNDC1;refGene_tx=XM_019814037.2;refGene_gn=KNDC1;refGene_tx=XM_023241157.1
D2 89706773 89706871
refGene_gn=KNDC1;refGene_tx=XM_019814037.2;refGene_gn=KNDC1;refGene_tx=XM_023241157.1
D2 89707359 89707580
refGene_gn=KNDC1;refGene_tx=XM_019814037.2;refGene_gn=KNDC1;refGene_tx=XM_023241157.1
D2 89709647 89709761
refGene_gn=KNDC1;refGene_tx=XM_019814037.2;refGene_gn=KNDC1;refGene_tx=XM_023241157.1
D2 89710866 89711014
refGene_gn=KNDC1;refGene_tx=XM_019814037.2;refGene_gn=KNDC1;refGene_tx=XM_023241157.1
D2 89711130 89711310
refGene_gn=KNDC1;refGene_tx=XM_019814037.2;refGene_gn=KNDC1;refGene_tx=XM_023241157.1
D2 89711834 89712041
refGene_gn=KNDC1;refGene_tx=XM_019814037.2;refGene_gn=KNDC1;refGene_tx=XM_023241157.1
D2 89712478 89712561
refGene_gn=KNDC1;refGene_tx=XM_019814037.2;refGene_gn=KNDC1;refGene_tx=XM_023241157.1
D2 89712771 89712922
refGene_gn=KNDC1;refGene_tx=XM_019814037.2;refGene_gn=KNDC1;refGene_tx=XM_023241157.1
D2 89717008 89717131
refGene_gn=KNDC1;refGene_tx=XM_019814037.2;refGene_gn=KNDC1;refGene_tx=XM_023241157.1
D2 89717215 89717275
refGene_gn=KNDC1;refGene_tx=XM_019814037.2;refGene_gn=KNDC1;refGene_tx

=XM_023241157.1
D2 89718053 89718211
refGene_gn=KNDC1;refGene_tx=XM_019814037.2;refGene_gn=KNDC1;refGene_tx
=XM_023241157.1
D2 89719496 89719728
refGene_gn=KNDC1;refGene_tx=XM_019814037.2;refGene_gn=KNDC1;refGene_tx
=XM_023241157.1
D2 89725928 89726502 refGene_gn=UTF1;refGene_tx=XM_023240314.1
D2 89726617 89727102 refGene_gn=UTF1;refGene_tx=XM_023240314.1
D2 89735292 89735596
refGene_gn=VENTX;refGene_tx=XM_023241158.1;refGene_gn=VENTX;refGene_tx
=XM_023241159.1
D2 89737366 89737527
refGene_gn=VENTX;refGene_tx=XM_023241158.1;refGene_gn=VENTX;refGene_tx
=XM_023241159.1
D2 89737625 89738006
refGene_gn=VENTX;refGene_tx=XM_023241158.1;refGene_gn=VENTX;refGene_tx
=XM_023241159.1
D2 89754614 89754710
refGene_gn=LOC102901922;refGene_tx=XM_019814040.1
D2 89756703 89756806
refGene_gn=LOC102901922;refGene_tx=XM_019814040.1
D2 89756893 89756984
refGene_gn=LOC102901922;refGene_tx=XM_019814040.1
D2 89757330 89757397
refGene_gn=LOC102901922;refGene_tx=XM_019814040.1
D2 89757745 89757861
refGene_gn=LOC102901922;refGene_tx=XM_019814040.1
D2 89758849 89759041
refGene_gn=LOC102901922;refGene_tx=XM_019814040.1
D2 89759506 89759648
refGene_gn=LOC102901922;refGene_tx=XM_019814040.1
D2 89760322 89760373
refGene_gn=ADAM8;refGene_tx=XM_011287631.3;refGene_gn=ADAM8;refGene_tx
=XM_011287633.3
D2 89760564 89760573 refGene_gn=ADAM8;refGene_tx=XM_011287632.3
D2 89760777 89760858
refGene_gn=ADAM8;refGene_tx=XM_011287631.3;refGene_gn=ADAM8;refGene_tx
=XM_011287632.3;refGene_gn=ADAM8;refGene_tx=XM_011287633.3
D2 89761605 89761656
refGene_gn=ADAM8;refGene_tx=XM_011287631.3;refGene_gn=ADAM8;refGene_tx
=XM_011287632.3;refGene_gn=ADAM8;refGene_tx=XM_011287633.3
D2 89761826 89761892
refGene_gn=ADAM8;refGene_tx=XM_011287631.3;refGene_gn=ADAM8;refGene_tx
=XM_011287632.3;refGene_gn=ADAM8;refGene_tx=XM_011287633.3
D2 89762330 89762490
refGene_gn=ADAM8;refGene_tx=XM_011287631.3;refGene_gn=ADAM8;refGene_tx
=XM_011287632.3;refGene_gn=ADAM8;refGene_tx=XM_011287633.3
D2 89762894 89763006
refGene_gn=ADAM8;refGene_tx=XM_011287631.3;refGene_gn=ADAM8;refGene_tx

=XM_011287632.3;refGene_gn=ADAM8;refGene_tx=XM_011287633.3
D2 89763140 89763275
refGene_gn=ADAM8;refGene_tx=XM_011287631.3;refGene_gn=ADAM8;refGene_tx=XM_011287632.3
D2 89763580 89763665
refGene_gn=ADAM8;refGene_tx=XM_011287631.3;refGene_gn=ADAM8;refGene_tx=XM_011287632.3;refGene_gn=ADAM8;refGene_tx=XM_011287633.3
D2 89764230 89764308
refGene_gn=ADAM8;refGene_tx=XM_011287631.3;refGene_gn=ADAM8;refGene_tx=XM_011287632.3;refGene_gn=ADAM8;refGene_tx=XM_011287633.3
D2 89764757 89764920
refGene_gn=ADAM8;refGene_tx=XM_011287631.3;refGene_gn=ADAM8;refGene_tx=XM_011287632.3;refGene_gn=ADAM8;refGene_tx=XM_011287633.3
D2 89765180 89765262
refGene_gn=ADAM8;refGene_tx=XM_011287631.3;refGene_gn=ADAM8;refGene_tx=XM_011287632.3;refGene_gn=ADAM8;refGene_tx=XM_011287633.3
D2 89765338 89765528
refGene_gn=ADAM8;refGene_tx=XM_011287631.3;refGene_gn=ADAM8;refGene_tx=XM_011287632.3;refGene_gn=ADAM8;refGene_tx=XM_011287633.3
D2 89765673 89765763
refGene_gn=ADAM8;refGene_tx=XM_011287631.3;refGene_gn=ADAM8;refGene_tx=XM_011287632.3;refGene_gn=ADAM8;refGene_tx=XM_011287633.3
D2 89765964 89766142
refGene_gn=ADAM8;refGene_tx=XM_011287631.3;refGene_gn=ADAM8;refGene_tx=XM_011287632.3;refGene_gn=ADAM8;refGene_tx=XM_011287633.3
D2 89766224 89766373
refGene_gn=ADAM8;refGene_tx=XM_011287631.3;refGene_gn=ADAM8;refGene_tx=XM_011287632.3;refGene_gn=ADAM8;refGene_tx=XM_011287633.3
D2 89766675 89766757
refGene_gn=ADAM8;refGene_tx=XM_011287631.3;refGene_gn=ADAM8;refGene_tx=XM_011287632.3;refGene_gn=ADAM8;refGene_tx=XM_011287633.3
D2 89766883 89767053
refGene_gn=ADAM8;refGene_tx=XM_011287631.3;refGene_gn=ADAM8;refGene_tx=XM_011287632.3;refGene_gn=ADAM8;refGene_tx=XM_011287633.3
D2 89767214 89767283
refGene_gn=ADAM8;refGene_tx=XM_011287631.3;refGene_gn=ADAM8;refGene_tx=XM_011287632.3;refGene_gn=ADAM8;refGene_tx=XM_011287633.3
D2 89767375 89767438
refGene_gn=ADAM8;refGene_tx=XM_011287631.3;refGene_gn=ADAM8;refGene_tx=XM_011287632.3;refGene_gn=ADAM8;refGene_tx=XM_011287633.3
D2 89767787 89767971
refGene_gn=ADAM8;refGene_tx=XM_011287631.3;refGene_gn=ADAM8;refGene_tx=XM_011287632.3;refGene_gn=ADAM8;refGene_tx=XM_011287633.3
D2 89768318 89768395
refGene_gn=ADAM8;refGene_tx=XM_011287631.3;refGene_gn=ADAM8;refGene_tx=XM_011287632.3;refGene_gn=ADAM8;refGene_tx=XM_011287633.3
D2 89768484 89768563
refGene_gn=ADAM8;refGene_tx=XM_011287631.3;refGene_gn=ADAM8;refGene_tx=XM_011287632.3;refGene_gn=ADAM8;refGene_tx=XM_011287633.3
D2 89768647 89768724

refGene_gn=ADAM8;refGene_tx=XM_011287631.3;refGene_gn=ADAM8;refGene_tx=XM_011287632.3;refGene_gn=ADAM8;refGene_tx=XM_011287633.3
D2 89769608 89769712
refGene_gn=ADAM8;refGene_tx=XM_011287631.3;refGene_gn=ADAM8;refGene_tx=XM_011287632.3;refGene_gn=ADAM8;refGene_tx=XM_011287633.3
D2 89770809 89770852
refGene_gn=ADAM8;refGene_tx=XM_011287631.3;refGene_gn=ADAM8;refGene_tx=XM_011287632.3;refGene_gn=ADAM8;refGene_tx=XM_011287633.3
D2 89773359 89773495
refGene_gn=TUBGCP2;refGene_tx=XM_011287637.2;refGene_gn=TUBGCP2;refGene_tx=XM_019814039.1;refGene_gn=TUBGCP2;refGene_tx=XM_023241160.1;refGene_gn=TUBGCP2;refGene_tx=XM_023241161.1
D2 89774452 89774616
refGene_gn=TUBGCP2;refGene_tx=XM_011287637.2;refGene_gn=TUBGCP2;refGene_tx=XM_019814039.1;refGene_gn=TUBGCP2;refGene_tx=XM_023241160.1;refGene_gn=TUBGCP2;refGene_tx=XM_023241161.1
D2 89775280 89775409
refGene_gn=TUBGCP2;refGene_tx=XM_011287637.2;refGene_gn=TUBGCP2;refGene_tx=XM_019814039.1;refGene_gn=TUBGCP2;refGene_tx=XM_023241160.1;refGene_gn=TUBGCP2;refGene_tx=XM_023241161.1
D2 89775924 89776068
refGene_gn=TUBGCP2;refGene_tx=XM_011287637.2;refGene_gn=TUBGCP2;refGene_tx=XM_019814039.1;refGene_gn=TUBGCP2;refGene_tx=XM_023241160.1;refGene_gn=TUBGCP2;refGene_tx=XM_023241161.1
D2 89776418 89776539
refGene_gn=TUBGCP2;refGene_tx=XM_011287637.2;refGene_gn=TUBGCP2;refGene_tx=XM_019814039.1;refGene_gn=TUBGCP2;refGene_tx=XM_023241160.1;refGene_gn=TUBGCP2;refGene_tx=XM_023241161.1
D2 89776948 89777077
refGene_gn=TUBGCP2;refGene_tx=XM_011287637.2;refGene_gn=TUBGCP2;refGene_tx=XM_019814039.1;refGene_gn=TUBGCP2;refGene_tx=XM_023241160.1;refGene_gn=TUBGCP2;refGene_tx=XM_023241161.1
D2 89777366 89777539
refGene_gn=TUBGCP2;refGene_tx=XM_011287637.2;refGene_gn=TUBGCP2;refGene_tx=XM_019814039.1;refGene_gn=TUBGCP2;refGene_tx=XM_023241160.1;refGene_gn=TUBGCP2;refGene_tx=XM_023241161.1
D2 89778270 89778451
refGene_gn=TUBGCP2;refGene_tx=XM_011287637.2;refGene_gn=TUBGCP2;refGene_tx=XM_019814039.1;refGene_gn=TUBGCP2;refGene_tx=XM_023241160.1;refGene_gn=TUBGCP2;refGene_tx=XM_023241161.1
D2 89779063 89779244
refGene_gn=TUBGCP2;refGene_tx=XM_011287637.2;refGene_gn=TUBGCP2;refGene_tx=XM_019814039.1;refGene_gn=TUBGCP2;refGene_tx=XM_023241160.1;refGene_gn=TUBGCP2;refGene_tx=XM_023241161.1
D2 89780628 89780774
refGene_gn=TUBGCP2;refGene_tx=XM_011287637.2;refGene_gn=TUBGCP2;refGene_tx=XM_019814039.1;refGene_gn=TUBGCP2;refGene_tx=XM_023241160.1;refGene_gn=TUBGCP2;refGene_tx=XM_023241161.1
D2 89781952 89782142
refGene_gn=TUBGCP2;refGene_tx=XM_011287637.2;refGene_gn=TUBGCP2;refGen

e_tx=XM_019814039.1;refGene_gn=TUBGCP2;refGene_tx=XM_023241160.1;refGene_gn=TUBGCP2;refGene_tx=XM_023241161.1
D2 89782451 89782651
refGene_gn=TUBGCP2;refGene_tx=XM_011287637.2;refGene_gn=TUBGCP2;refGene_tx=XM_019814039.1;refGene_gn=TUBGCP2;refGene_tx=XM_023241160.1;refGene_gn=TUBGCP2;refGene_tx=XM_023241161.1
D2 89782914 89783122
refGene_gn=TUBGCP2;refGene_tx=XM_011287637.2;refGene_gn=TUBGCP2;refGene_tx=XM_019814039.1;refGene_gn=TUBGCP2;refGene_tx=XM_023241160.1;refGene_gn=TUBGCP2;refGene_tx=XM_023241161.1
D2 89785107 89785267
refGene_gn=TUBGCP2;refGene_tx=XM_011287637.2;refGene_gn=TUBGCP2;refGene_tx=XM_019814039.1;refGene_gn=TUBGCP2;refGene_tx=XM_023241160.1;refGene_gn=TUBGCP2;refGene_tx=XM_023241161.1
D2 89786397 89786574
refGene_gn=TUBGCP2;refGene_tx=XM_011287637.2;refGene_gn=TUBGCP2;refGene_tx=XM_019814039.1;refGene_gn=TUBGCP2;refGene_tx=XM_023241160.1;refGene_gn=TUBGCP2;refGene_tx=XM_023241161.1
D2 89787375 89787504 refGene_gn=TUBGCP2;refGene_tx=XM_023241161.1
D2 89789468 89789618 refGene_gn=TUBGCP2;refGene_tx=XM_023241161.1
D2 89793031 89793184
refGene_gn=ZNF511;refGene_tx=XM_023241162.1;refGene_gn=ZNF511;refGene_tx=XM_023241163.1
D2 89793446 89793520
refGene_gn=ZNF511;refGene_tx=XM_023241162.1;refGene_gn=ZNF511;refGene_tx=XM_023241163.1
D2 89793815 89794017
refGene_gn=ZNF511;refGene_tx=XM_023241162.1;refGene_gn=ZNF511;refGene_tx=XM_023241163.1
D2 89794202 89794327
refGene_gn=ZNF511;refGene_tx=XM_023241162.1;refGene_gn=ZNF511;refGene_tx=XM_023241163.1
D2 89794790 89794875 refGene_gn=ZNF511;refGene_tx=XM_023241163.1
D2 89795331 89795481 refGene_gn=ZNF511;refGene_tx=XM_023241162.1
D2 89796278 89796357 refGene_gn=ZNF511;refGene_tx=XM_023241162.1
D2 89803612 89804013
refGene_gn=LOC101089397;refGene_tx=XM_023241164.1
D2 89804597 89804811
refGene_gn=LOC101089397;refGene_tx=XM_023241164.1
D2 89838055 89839285
refGene_gn=LOC102902043;refGene_tx=XM_006938294.2
D2 89848852 89849161 refGene_gn=CALY;refGene_tx=XM_023241166.1
D2 89849357 89849471 refGene_gn=CALY;refGene_tx=XM_023241166.1
D2 89850068 89850182 refGene_gn=CALY;refGene_tx=XM_023241166.1
D2 89851192 89851327 refGene_gn=CALY;refGene_tx=XM_023241166.1
D2 89864793 89864801 refGene_gn=PRAP1;refGene_tx=XM_019814041.2
D2 89867195 89867262 refGene_gn=PRAP1;refGene_tx=XM_019814041.2
D2 89867986 89868039 refGene_gn=PRAP1;refGene_tx=XM_019814041.2
D2 89868832 89869046 refGene_gn=PRAP1;refGene_tx=XM_019814041.2
D2 89871513 89871580 refGene_gn=FUOM;refGene_tx=XM_003994526.5

D2 89871844 89871918 refGene_gn=FUOM;refGene_tx=XM_003994526.5
D2 89872549 89872648 refGene_gn=FUOM;refGene_tx=XM_003994526.5
D2 89872837 89872908 refGene_gn=FUOM;refGene_tx=XM_003994526.5
D2 89874349 89874418 refGene_gn=FUOM;refGene_tx=XM_003994526.5
D2 89875148 89875233 refGene_gn=FUOM;refGene_tx=XM_003994526.5
D2 89881325 89881391 refGene_gn=ECHS1;refGene_tx=XM_023241167.1
D2 89882384 89882452 refGene_gn=ECHS1;refGene_tx=XM_023241167.1
D2 89883971 89884091 refGene_gn=ECHS1;refGene_tx=XM_023241167.1
D2 89884566 89884671 refGene_gn=ECHS1;refGene_tx=XM_023241167.1
D2 89885844 89885944 refGene_gn=ECHS1;refGene_tx=XM_023241167.1
D2 89886958 89887086 refGene_gn=ECHS1;refGene_tx=XM_023241167.1
D2 89887587 89887785 refGene_gn=ECHS1;refGene_tx=XM_023241167.1
D2 89888945 89889033 refGene_gn=ECHS1;refGene_tx=XM_023241167.1
D2 89898044 89898228 refGene_gn=PAOX;refGene_tx=XM_023241168.1
D2 89898816 89899303 refGene_gn=PAOX;refGene_tx=XM_023241168.1
D2 89901098 89901298 refGene_gn=PAOX;refGene_tx=XM_023241168.1
D2 89903209 89903462 refGene_gn=PAOX;refGene_tx=XM_023241168.1
D2 89904943 89905056 refGene_gn=PAOX;refGene_tx=XM_023241168.1
D2 89905663 89905821 refGene_gn=PAOX;refGene_tx=XM_023241168.1
D2 89908035 89908173 refGene_gn=PAOX;refGene_tx=XM_023241168.1
D2 89910934 89911046
refGene_gn=MTG1;refGene_tx=XM_003994527.5;refGene_gn=MTG1;refGene_tx=XM_006938300.2
D2 89912096 89912220
refGene_gn=MTG1;refGene_tx=XM_003994527.5;refGene_gn=MTG1;refGene_tx=XM_006938297.3;refGene_gn=MTG1;refGene_tx=XM_006938298.3;refGene_gn=MTG1;refGene_tx=XM_006938299.3;refGene_gn=MTG1;refGene_tx=XM_006938300.2;refGene_gn=MTG1;refGene_tx=XM_023241169.1
D2 89915101 89915206
refGene_gn=MTG1;refGene_tx=XM_003994527.5;refGene_gn=MTG1;refGene_tx=XM_006938297.3;refGene_gn=MTG1;refGene_tx=XM_006938298.3;refGene_gn=MTG1;refGene_tx=XM_006938299.3;refGene_gn=MTG1;refGene_tx=XM_006938300.2;refGene_gn=MTG1;refGene_tx=XM_023241169.1
D2 89917603 89917684
refGene_gn=MTG1;refGene_tx=XM_003994527.5;refGene_gn=MTG1;refGene_tx=XM_006938297.3;refGene_gn=MTG1;refGene_tx=XM_006938298.3;refGene_gn=MTG1;refGene_tx=XM_006938299.3;refGene_gn=MTG1;refGene_tx=XM_006938300.2;refGene_gn=MTG1;refGene_tx=XM_023241169.1
D2 89918226 89918283
refGene_gn=MTG1;refGene_tx=XM_003994527.5;refGene_gn=MTG1;refGene_tx=XM_006938297.3;refGene_gn=MTG1;refGene_tx=XM_006938298.3;refGene_gn=MTG1;refGene_tx=XM_006938299.3;refGene_gn=MTG1;refGene_tx=XM_006938300.2;refGene_gn=MTG1;refGene_tx=XM_023241169.1
D2 89918635 89918726
refGene_gn=MTG1;refGene_tx=XM_003994527.5;refGene_gn=MTG1;refGene_tx=XM_006938297.3;refGene_gn=MTG1;refGene_tx=XM_006938298.3;refGene_gn=MTG1;refGene_tx=XM_006938299.3;refGene_gn=MTG1;refGene_tx=XM_006938300.2;refGene_gn=MTG1;refGene_tx=XM_023241169.1
D2 89920938 89921000
refGene_gn=MTG1;refGene_tx=XM_003994527.5;refGene_gn=MTG1;refGene_tx=X

M_006938297.3;refGene_gn=MTG1;refGene_tx=XM_006938298.3;refGene_gn=MTG1;refGene_tx=XM_006938299.3;refGene_gn=MTG1;refGene_tx=XM_006938300.2;refGene_gn=MTG1;refGene_tx=XM_023241169.1
D2 89921438 89921535
refGene_gn=MTG1;refGene_tx=XM_003994527.5;refGene_gn=MTG1;refGene_tx=XM_006938297.3;refGene_gn=MTG1;refGene_tx=XM_006938298.3;refGene_gn=MTG1;refGene_tx=XM_006938299.3;refGene_gn=MTG1;refGene_tx=XM_006938300.2;refGene_gn=MTG1;refGene_tx=XM_023241169.1
D2 89921918 89922000
refGene_gn=MTG1;refGene_tx=XM_003994527.5;refGene_gn=MTG1;refGene_tx=XM_006938297.3;refGene_gn=MTG1;refGene_tx=XM_006938298.3;refGene_gn=MTG1;refGene_tx=XM_006938299.3;refGene_gn=MTG1;refGene_tx=XM_006938300.2;refGene_gn=MTG1;refGene_tx=XM_023241169.1
D2 89925804 89925917
refGene_gn=MTG1;refGene_tx=XM_003994527.5;refGene_gn=MTG1;refGene_tx=XM_006938297.3;refGene_gn=MTG1;refGene_tx=XM_006938298.3;refGene_gn=MTG1;refGene_tx=XM_006938299.3;refGene_gn=MTG1;refGene_tx=XM_006938300.2;refGene_gn=MTG1;refGene_tx=XM_023241169.1
D2 89926308 89926442
refGene_gn=MTG1;refGene_tx=XM_003994527.5;refGene_gn=MTG1;refGene_tx=XM_006938297.3;refGene_gn=MTG1;refGene_tx=XM_006938298.3;refGene_gn=MTG1;refGene_tx=XM_006938299.3;refGene_gn=MTG1;refGene_tx=XM_006938300.2;refGene_gn=MTG1;refGene_tx=XM_023241169.1
D2 89929376 89929814
refGene_gn=SPRN;refGene_tx=XM_023241171.1;refGene_gn=SPRN;refGene_tx=XM_023241172.1
D2 89948953 89949880
refGene_gn=LOC101090391;refGene_tx=XM_003994550.5
D2 89975553 89976750
refGene_gn=LOC101090648;refGene_tx=XM_003994551.5
D2 89988634 89988701
refGene_gn=LOC111556939;refGene_tx=XM_023240308.1
D2 89990910 89991260
refGene_gn=LOC111556939;refGene_tx=XM_023240308.1
D2 89992365 89992759
refGene_gn=LOC101090899;refGene_tx=XM_023241173.1;refGene_gn=LOC101090899;refGene_tx=XM_023241174.1;refGene_gn=LOC101090899;refGene_tx=XM_023241175.1
D2 89993650 89993959
refGene_gn=LOC101090899;refGene_tx=XM_023241173.1;refGene_gn=LOC101090899;refGene_tx=XM_023241174.1;refGene_gn=LOC101090899;refGene_tx=XM_023241175.1
D2 89994345 89994651
refGene_gn=LOC101090899;refGene_tx=XM_023241173.1;refGene_gn=LOC101090899;refGene_tx=XM_023241174.1;refGene_gn=LOC101090899;refGene_tx=XM_023241175.1
D2 89994766 89995084
refGene_gn=LOC101090899;refGene_tx=XM_023241173.1;refGene_gn=LOC101090899;refGene_tx=XM_023241174.1;refGene_gn=LOC101090899;refGene_tx=XM_023241175.1

D2 89995152 89995467
refGene_gn=LOC101090899;refGene_tx=XM_023241173.1;refGene_gn=LOC101090
899;refGene_tx=XM_023241174.1;refGene_gn=LOC101090899;refGene_tx=XM_02
3241175.1
D2 89998810 89999125
refGene_gn=LOC101090899;refGene_tx=XM_023241173.1;refGene_gn=LOC101090
899;refGene_tx=XM_023241174.1;refGene_gn=LOC101090899;refGene_tx=XM_02
3241175.1
D2 89999272 89999338
refGene_gn=LOC101090899;refGene_tx=XM_023241173.1;refGene_gn=LOC101090
899;refGene_tx=XM_023241174.1;refGene_gn=LOC101090899;refGene_tx=XM_02
3241175.1
D2 89999413 89999728
refGene_gn=LOC101090899;refGene_tx=XM_023241173.1;refGene_gn=LOC101090
899;refGene_tx=XM_023241174.1;refGene_gn=LOC101090899;refGene_tx=XM_02
3241175.1
D2 89999985 90000033
refGene_gn=LOC101090899;refGene_tx=XM_023241173.1;refGene_gn=LOC101090
899;refGene_tx=XM_023241174.1;refGene_gn=LOC101090899;refGene_tx=XM_02
3241175.1
D2 90000529 90000671
refGene_gn=LOC101090899;refGene_tx=XM_023241173.1;refGene_gn=LOC101090
899;refGene_tx=XM_023241174.1;refGene_gn=LOC101090899;refGene_tx=XM_02
3241175.1
D2 90001059 90001331
refGene_gn=LOC101090899;refGene_tx=XM_023241173.1;refGene_gn=LOC101090
899;refGene_tx=XM_023241174.1;refGene_gn=LOC101090899;refGene_tx=XM_02
3241175.1
D2 90001711 90001782
refGene_gn=LOC101090899;refGene_tx=XM_023241174.1
D2 90006690 90007212
refGene_gn=LOC101091153;refGene_tx=XM_023240309.1
D2 90007383 90007608
refGene_gn=LOC101091153;refGene_tx=XM_023240309.1
D2 90020207 90021116
refGene_gn=LOC101091405;refGene_tx=XM_003994554.4
D2 90088409 90088586 refGene_gn=CYP2E2;refGene_tx=NM_001048010.1
D2 90089526 90089686 refGene_gn=CYP2E2;refGene_tx=NM_001048010.1
D2 90092709 90092859 refGene_gn=CYP2E2;refGene_tx=NM_001048010.1
D2 90093247 90093408 refGene_gn=CYP2E2;refGene_tx=NM_001048010.1
D2 90093859 90094036 refGene_gn=CYP2E2;refGene_tx=NM_001048010.1
D2 90095050 90095192 refGene_gn=CYP2E2;refGene_tx=NM_001048010.1
D2 90098676 90098864 refGene_gn=CYP2E2;refGene_tx=NM_001048010.1
D2 90099297 90099439 refGene_gn=CYP2E2;refGene_tx=NM_001048010.1
D2 90100386 90100577 refGene_gn=CYP2E2;refGene_tx=NM_001048010.1
D2 90142011 90142104
refGene_gn=LOC101091903;refGene_tx=XM_023241176.1;refGene_gn=LOC101091
903;refGene_tx=XM_023241177.1;refGene_gn=LOC101091903;refGene_tx=XM_02
3241178.1;refGene_gn=LOC101091903;refGene_tx=XM_023241179.1;refGene_gn
=LOC101091903;refGene_tx=XM_023241180.1;refGene_gn=LOC101091903;refGen

e_tx=XM_023241181.1;refGene_gn=LOC101091903;refGene_tx=XM_023241182.1;refGene_gn=LOC101091903;refGene_tx=XM_023241183.1;refGene_gn=LOC101091903;refGene_tx=XM_023241184.1;refGene_gn=LOC101091903;refGene_tx=XM_023241185.1
D2 90148530 90148657
refGene_gn=LOC101091903;refGene_tx=XM_023241176.1;refGene_gn=LOC101091903;refGene_tx=XM_023241177.1;refGene_gn=LOC101091903;refGene_tx=XM_023241178.1;refGene_gn=LOC101091903;refGene_tx=XM_023241179.1;refGene_gn=LOC101091903;refGene_tx=XM_023241180.1;refGene_gn=LOC101091903;refGene_tx=XM_023241181.1;refGene_gn=LOC101091903;refGene_tx=XM_023241182.1;refGene_gn=LOC101091903;refGene_tx=XM_023241183.1;refGene_gn=LOC101091903;refGene_tx=XM_023241184.1;refGene_gn=LOC101091903;refGene_tx=XM_023241185.1
D2 90148891 90148984
refGene_gn=LOC101091903;refGene_tx=XM_023241176.1;refGene_gn=LOC101091903;refGene_tx=XM_023241177.1;refGene_gn=LOC101091903;refGene_tx=XM_023241178.1;refGene_gn=LOC101091903;refGene_tx=XM_023241179.1;refGene_gn=LOC101091903;refGene_tx=XM_023241180.1;refGene_gn=LOC101091903;refGene_tx=XM_023241181.1;refGene_gn=LOC101091903;refGene_tx=XM_023241182.1;refGene_gn=LOC101091903;refGene_tx=XM_023241183.1;refGene_gn=LOC101091903;refGene_tx=XM_023241184.1;refGene_gn=LOC101091903;refGene_tx=XM_023241185.1
D2 90150289 90150345
refGene_gn=LOC101091903;refGene_tx=XM_023241185.1
D2 90150856 90152274
refGene_gn=LOC101091903;refGene_tx=XM_023241176.1;refGene_gn=LOC101091903;refGene_tx=XM_023241177.1;refGene_gn=LOC101091903;refGene_tx=XM_023241178.1;refGene_gn=LOC101091903;refGene_tx=XM_023241179.1;refGene_gn=LOC101091903;refGene_tx=XM_023241180.1;refGene_gn=LOC101091903;refGene_tx=XM_023241181.1;refGene_gn=LOC101091903;refGene_tx=XM_023241182.1;refGene_gn=LOC101091903;refGene_tx=XM_023241183.1
D2 90156353 90156503
refGene_gn=LOC101091903;refGene_tx=XM_023241184.1
D2 90156659 90156691
refGene_gn=LOC101091903;refGene_tx=XM_023241184.1
D2 90157045 90157143
refGene_gn=SYCE1;refGene_tx=XM_023241186.1;refGene_gn=SYCE1;refGene_tx=XM_023241187.1;refGene_gn=SYCE1;refGene_tx=XM_023241188.1
D2 90157369 90157457
refGene_gn=SYCE1;refGene_tx=XM_023241186.1;refGene_gn=SYCE1;refGene_tx=XM_023241187.1;refGene_gn=SYCE1;refGene_tx=XM_023241188.1
D2 90157612 90157723
refGene_gn=SYCE1;refGene_tx=XM_023241186.1;refGene_gn=SYCE1;refGene_tx=XM_023241187.1;refGene_gn=SYCE1;refGene_tx=XM_023241188.1
D2 90157798 90157922
refGene_gn=SYCE1;refGene_tx=XM_023241186.1;refGene_gn=SYCE1;refGene_tx=XM_023241187.1;refGene_gn=SYCE1;refGene_tx=XM_023241188.1
D2 90157994 90158061
refGene_gn=SYCE1;refGene_tx=XM_023241186.1;refGene_gn=SYCE1;refGene_tx=XM_023241187.1;refGene_gn=SYCE1;refGene_tx=XM_023241188.1

D2 90158843 90158907
refGene_gn=SYCE1;refGene_tx=XM_023241186.1;refGene_gn=SYCE1;refGene_tx=XM_023241187.1;refGene_gn=SYCE1;refGene_tx=XM_023241188.1
D2 90159173 90159263
refGene_gn=SYCE1;refGene_tx=XM_023241186.1;refGene_gn=SYCE1;refGene_tx=XM_023241187.1;refGene_gn=SYCE1;refGene_tx=XM_023241188.1
D2 90159972 90160027
refGene_gn=SYCE1;refGene_tx=XM_023241186.1;refGene_gn=SYCE1;refGene_tx=XM_023241187.1;refGene_gn=SYCE1;refGene_tx=XM_023241188.1
D2 90160284 90160332
refGene_gn=SYCE1;refGene_tx=XM_023241186.1;refGene_gn=SYCE1;refGene_tx=XM_023241187.1;refGene_gn=SYCE1;refGene_tx=XM_023241188.1
D2 90160963 90161038
refGene_gn=SYCE1;refGene_tx=XM_023241186.1;refGene_gn=SYCE1;refGene_tx=XM_023241187.1;refGene_gn=SYCE1;refGene_tx=XM_023241188.1
D2 90161435 90161495
refGene_gn=SYCE1;refGene_tx=XM_023241186.1;refGene_gn=SYCE1;refGene_tx=XM_023241187.1;refGene_gn=SYCE1;refGene_tx=XM_023241188.1
D2 90162125 90162188
refGene_gn=SYCE1;refGene_tx=XM_023241186.1;refGene_gn=SYCE1;refGene_tx=XM_023241187.1;refGene_gn=SYCE1;refGene_tx=XM_023241188.1
D2 90165684 90165754
refGene_gn=SYCE1;refGene_tx=XM_023241186.1;refGene_gn=SYCE1;refGene_tx=XM_023241187.1;refGene_gn=SYCE1;refGene_tx=XM_023241188.1
D3 133983 134730
refGene_gn=LOC109493088;refGene_tx=XM_023242147.1
D3 136355 136412
refGene_gn=LOC109493088;refGene_tx=XM_023242147.1
D3 155939 156026
refGene_gn=ANHX;refGene_tx=XM_023241211.1;refGene_gn=ANHX;refGene_tx=XM_023241212.1
D3 156290 156372
refGene_gn=ANHX;refGene_tx=XM_023241211.1;refGene_gn=ANHX;refGene_tx=XM_023241212.1
D3 158113 158261
refGene_gn=ANHX;refGene_tx=XM_023241211.1;refGene_gn=ANHX;refGene_tx=XM_023241212.1;refGene_gn=ANHX;refGene_tx=XM_023241214.1
D3 159423 159554 refGene_gn=ANHX;refGene_tx=XM_023241213.1
D3 161145 161438
refGene_gn=ANHX;refGene_tx=XM_023241211.1;refGene_gn=ANHX;refGene_tx=XM_023241212.1;refGene_gn=ANHX;refGene_tx=XM_023241213.1;refGene_gn=ANHX;refGene_tx=XM_023241214.1
D3 162997 163118
refGene_gn=ANHX;refGene_tx=XM_023241211.1;refGene_gn=ANHX;refGene_tx=XM_023241212.1;refGene_gn=ANHX;refGene_tx=XM_023241213.1;refGene_gn=ANHX;refGene_tx=XM_023241214.1
D3 163576 163793
refGene_gn=ANHX;refGene_tx=XM_023241211.1;refGene_gn=ANHX;refGene_tx=XM_023241212.1;refGene_gn=ANHX;refGene_tx=XM_023241213.1;refGene_gn=ANHX;refGene_tx=XM_023241214.1

D3 164827 164951
refGene_gn=ANHX;refGene_tx=XM_023241211.1;refGene_gn=ANHX;refGene_tx=XM_023241212.1;refGene_gn=ANHX;refGene_tx=XM_023241213.1;refGene_gn=ANHX;refGene_tx=XM_023241214.1
D3 167051 167179
refGene_gn=ANHX;refGene_tx=XM_023241211.1;refGene_gn=ANHX;refGene_tx=XM_023241212.1;refGene_gn=ANHX;refGene_tx=XM_023241213.1;refGene_gn=ANHX;refGene_tx=XM_023241214.1
D3 169424 169693
refGene_gn=ANHX;refGene_tx=XM_023241211.1;refGene_gn=ANHX;refGene_tx=XM_023241212.1;refGene_gn=ANHX;refGene_tx=XM_023241213.1;refGene_gn=ANHX;refGene_tx=XM_023241214.1
D3 185657 185667
refGene_gn=ANHX;refGene_tx=XM_023241211.1;refGene_gn=ANHX;refGene_tx=XM_023241212.1;refGene_gn=ANHX;refGene_tx=XM_023241213.1;refGene_gn=ANHX;refGene_tx=XM_023241214.1
D3 189566 192004
refGene_gn=ZNF268;refGene_tx=XM_006938318.4;refGene_gn=ZNF268;refGene_tx=XM_023241210.1
D3 196080 196176
refGene_gn=ZNF268;refGene_tx=XM_006938318.4;refGene_gn=ZNF268;refGene_tx=XM_023241210.1
D3 196463 196590
refGene_gn=ZNF268;refGene_tx=XM_006938318.4;refGene_gn=ZNF268;refGene_tx=XM_023241210.1
D3 199594 199768
refGene_gn=ZNF268;refGene_tx=XM_006938318.4;refGene_gn=ZNF268;refGene_tx=XM_023241210.1
D3 201563 201566 refGene_gn=ZNF268;refGene_tx=XM_023241210.1
D3 218855 218888 refGene_gn=ZNF268;refGene_tx=XM_006938318.4
D3 227633 229099
refGene_gn=ZNF10;refGene_tx=XM_006938319.4;refGene_gn=ZNF10;refGene_tx=XM_019814374.2;refGene_gn=ZNF10;refGene_tx=XM_019814375.2;refGene_gn=ZNF10;refGene_tx=XM_019814376.2
D3 233177 233273
refGene_gn=ZNF10;refGene_tx=XM_006938319.4;refGene_gn=ZNF10;refGene_tx=XM_019814374.2;refGene_gn=ZNF10;refGene_tx=XM_019814375.2;refGene_gn=ZNF10;refGene_tx=XM_019814376.2
D3 233556 233683
refGene_gn=ZNF10;refGene_tx=XM_006938319.4;refGene_gn=ZNF10;refGene_tx=XM_019814374.2;refGene_gn=ZNF10;refGene_tx=XM_019814375.2;refGene_gn=ZNF10;refGene_tx=XM_019814376.2
D3 235430 235507
refGene_gn=ZNF10;refGene_tx=XM_006938319.4;refGene_gn=ZNF10;refGene_tx=XM_019814374.2;refGene_gn=ZNF10;refGene_tx=XM_019814375.2;refGene_gn=ZNF10;refGene_tx=XM_019814376.2
D3 236304 236371 refGene_gn=ZNF10;refGene_tx=XM_006938319.4
D3 261178 261290 refGene_gn=ZNF891;refGene_tx=XM_019814400.1
D3 261292 261379 refGene_gn=ZNF891;refGene_tx=XM_019814400.1
D3 261381 261604 refGene_gn=ZNF891;refGene_tx=XM_019814400.1

D3 261605 261718 refGene_gn=ZNF891;refGene_tx=XM_019814400.1
D3 261720 261947 refGene_gn=ZNF891;refGene_tx=XM_019814400.1
D3 276088 277099
refGene_gn=ZNF140;refGene_tx=XM_006938320.4;refGene_gn=ZNF140;refGene_tx=XM_006938321.4;refGene_gn=ZNF140;refGene_tx=XM_019814391.2;refGene_gn=ZNF140;refGene_tx=XM_019814392.2;refGene_gn=ZNF140;refGene_tx=XM_019814393.2;refGene_gn=ZNF140;refGene_tx=XM_019814394.2;refGene_gn=ZNF140;refGene_tx=XM_023241216.1;refGene_gn=ZNF140;refGene_tx=XM_023241217.1
D3 290046 290053 refGene_gn=ZNF140;refGene_tx=XM_023241218.1
D3 291517 291644 refGene_gn=ZNF140;refGene_tx=XM_023241218.1
D3 292444 292525 refGene_gn=ZNF140;refGene_tx=XM_023241218.1
D3 292794 292849 refGene_gn=ZNF140;refGene_tx=XM_023241218.1
D3 293349 293400 refGene_gn=ZNF140;refGene_tx=XM_023241218.1
D3 294447 294738 refGene_gn=ZNF140;refGene_tx=XM_023241218.1
D3 312688 314673
refGene_gn=LOC101095441;refGene_tx=XM_019814386.2;refGene_gn=LOC101095441;refGene_tx=XM_019814390.2
D3 319805 319901
refGene_gn=LOC101095441;refGene_tx=XM_019814386.2
D3 320521 320648
refGene_gn=LOC101095441;refGene_tx=XM_019814386.2
D3 331447 331462
refGene_gn=LOC101095441;refGene_tx=XM_019814386.2
D3 352852 354204
refGene_gn=LOC101095441;refGene_tx=XM_019814387.1;refGene_gn=LOC101095441;refGene_tx=XM_019814388.1;refGene_gn=LOC101095441;refGene_tx=XM_019814389.2;refGene_gn=LOC101095441;refGene_tx=XM_019814390.2
D3 359141 359237
refGene_gn=LOC101095441;refGene_tx=XM_019814387.1;refGene_gn=LOC101095441;refGene_tx=XM_019814388.1;refGene_gn=LOC101095441;refGene_tx=XM_019814389.2;refGene_gn=LOC101095441;refGene_tx=XM_019814390.2
D3 359502 359629
refGene_gn=LOC101095441;refGene_tx=XM_019814387.1;refGene_gn=LOC101095441;refGene_tx=XM_019814388.1;refGene_gn=LOC101095441;refGene_tx=XM_019814389.2;refGene_gn=LOC101095441;refGene_tx=XM_019814390.2
D3 369210 369243
refGene_gn=LOC101095441;refGene_tx=XM_019814387.1;refGene_gn=LOC101095441;refGene_tx=XM_019814388.1;refGene_gn=LOC101095441;refGene_tx=XM_019814390.2
D3 381574 381589 refGene_gn=ZNF605;refGene_tx=XM_011287675.3
D3 388320 388441 refGene_gn=ZNF605;refGene_tx=XM_011287675.3
D3 388668 388758 refGene_gn=ZNF605;refGene_tx=XM_011287675.3
D3 392189 393985 refGene_gn=ZNF605;refGene_tx=XM_011287675.3
D3 403989 404004
refGene_gn=LOC101095686;refGene_tx=XM_011287676.3
D3 412414 412541
refGene_gn=LOC101095686;refGene_tx=XM_011287676.3
D3 413032 413125
refGene_gn=LOC101095686;refGene_tx=XM_011287676.3

D3	414198	414632
refGene_gn=LOC101095686;refGene_tx=XM_011287676.3
D3	421471	422674
refGene_gn=LOC101095921;refGene_tx=XM_023242148.1
D3	432908	433048
refGene_gn=LOC101095921;refGene_tx=XM_023242148.1
D3	436123	436367
refGene_gn=LOC101095921;refGene_tx=XM_023242148.1
D3	436606	436840
refGene_gn=LOC101095921;refGene_tx=XM_023242148.1
D3	436901	437036
refGene_gn=LOC101095921;refGene_tx=XM_023242148.1
D3	442846	443451	refGene_gn=CHFR;refGene_tx=XM_023241219.1
D3	443644	443788
refGene_gn=CHFR;refGene_tx=XM_023241219.1;refGene_gn=CHFR;refGene_tx=XM_023241220.1
D3	447759	447859	refGene_gn=CHFR;refGene_tx=XM_023241220.1
D3	450289	450399
refGene_gn=CHFR;refGene_tx=XM_023241219.1;refGene_gn=CHFR;refGene_tx=XM_023241220.1
D3	451503	451563
refGene_gn=CHFR;refGene_tx=XM_023241219.1;refGene_gn=CHFR;refGene_tx=XM_023241220.1
D3	451717	451927
refGene_gn=CHFR;refGene_tx=XM_023241219.1;refGene_gn=CHFR;refGene_tx=XM_023241220.1
D3	455619	455787
refGene_gn=CHFR;refGene_tx=XM_023241219.1;refGene_gn=CHFR;refGene_tx=XM_023241220.1
D3	457664	457824
refGene_gn=CHFR;refGene_tx=XM_023241219.1;refGene_gn=CHFR;refGene_tx=XM_023241220.1
D3	458795	458950
refGene_gn=CHFR;refGene_tx=XM_023241219.1;refGene_gn=CHFR;refGene_tx=XM_023241220.1
D3	459040	459203
refGene_gn=CHFR;refGene_tx=XM_023241219.1;refGene_gn=CHFR;refGene_tx=XM_023241220.1
D3	461173	461316
refGene_gn=CHFR;refGene_tx=XM_023241219.1;refGene_gn=CHFR;refGene_tx=XM_023241220.1
D3	462730	462847
refGene_gn=CHFR;refGene_tx=XM_023241219.1;refGene_gn=CHFR;refGene_tx=XM_023241220.1
D3	463302	463386
refGene_gn=CHFR;refGene_tx=XM_023241219.1;refGene_gn=CHFR;refGene_tx=XM_023241220.1
D3	464270	464341
refGene_gn=CHFR;refGene_tx=XM_023241219.1;refGene_gn=CHFR;refGene_tx=XM_023241220.1

D3 465299 465387
refGene_gn=CHFR;refGene_tx=XM_023241219.1;refGene_gn=CHFR;refGene_tx=XM_023241220.1
D3 467779 467887
refGene_gn=CHFR;refGene_tx=XM_023241219.1;refGene_gn=CHFR;refGene_tx=XM_023241220.1
D3 468316 468389
refGene_gn=CHFR;refGene_tx=XM_023241219.1;refGene_gn=CHFR;refGene_tx=XM_023241220.1
D3 469625 469668
refGene_gn=CHFR;refGene_tx=XM_023241219.1;refGene_gn=CHFR;refGene_tx=XM_023241220.1
D3 483061 483170 refGene_gn=GOLGA3;refGene_tx=XM_006938338.4
D3 485998 486277 refGene_gn=GOLGA3;refGene_tx=XM_006938338.4
D3 488859 488972 refGene_gn=GOLGA3;refGene_tx=XM_006938338.4
D3 493284 493943 refGene_gn=GOLGA3;refGene_tx=XM_006938338.4
D3 494321 494433 refGene_gn=GOLGA3;refGene_tx=XM_006938338.4
D3 496066 496373 refGene_gn=GOLGA3;refGene_tx=XM_006938338.4
D3 498316 498519 refGene_gn=GOLGA3;refGene_tx=XM_006938338.4
D3 501364 501502 refGene_gn=GOLGA3;refGene_tx=XM_006938338.4
D3 502857 503019 refGene_gn=GOLGA3;refGene_tx=XM_006938338.4
D3 503279 503648 refGene_gn=GOLGA3;refGene_tx=XM_006938338.4
D3 504625 504703 refGene_gn=GOLGA3;refGene_tx=XM_006938338.4
D3 505453 505657 refGene_gn=GOLGA3;refGene_tx=XM_006938338.4
D3 505658 505718 refGene_gn=GOLGA3;refGene_tx=XM_006938338.4
D3 506785 506880 refGene_gn=GOLGA3;refGene_tx=XM_006938338.4
D3 507548 507765 refGene_gn=GOLGA3;refGene_tx=XM_006938338.4
D3 510265 510409 refGene_gn=GOLGA3;refGene_tx=XM_006938338.4
D3 511023 511221 refGene_gn=GOLGA3;refGene_tx=XM_006938338.4
D3 512058 512175 refGene_gn=GOLGA3;refGene_tx=XM_006938338.4
D3 514712 514852 refGene_gn=GOLGA3;refGene_tx=XM_006938338.4
D3 515400 515533 refGene_gn=GOLGA3;refGene_tx=XM_006938338.4
D3 515771 515894 refGene_gn=GOLGA3;refGene_tx=XM_006938338.4
D3 516908 517073 refGene_gn=GOLGA3;refGene_tx=XM_006938338.4
D3 517892 518056 refGene_gn=GOLGA3;refGene_tx=XM_006938338.4
D3 518966 519186 refGene_gn=GOLGA3;refGene_tx=XM_006938338.4
D3 526620 526807 refGene_gn=ANKLE2;refGene_tx=XM_023241222.1
D3 534829 535303
refGene_gn=ANKLE2;refGene_tx=XM_006938342.3;refGene_gn=ANKLE2;refGene_tx=XM_023241222.1
D3 537494 537701
refGene_gn=ANKLE2;refGene_tx=XM_006938342.3;refGene_gn=ANKLE2;refGene_tx=XM_023241222.1
D3 538824 539018
refGene_gn=ANKLE2;refGene_tx=XM_006938342.3;refGene_gn=ANKLE2;refGene_tx=XM_023241222.1
D3 539098 539287
refGene_gn=ANKLE2;refGene_tx=XM_006938342.3;refGene_gn=ANKLE2;refGene_tx=XM_023241222.1
D3 541788 541911

refGene_gn=ANKLE2;refGene_tx=XM_006938342.3;refGene_gn=ANKLE2;refGene_tx=XM_023241222.1
D3 544289 544356
refGene_gn=ANKLE2;refGene_tx=XM_006938342.3;refGene_gn=ANKLE2;refGene_tx=XM_023241222.1
D3 545898 546071
refGene_gn=ANKLE2;refGene_tx=XM_006938342.3;refGene_gn=ANKLE2;refGene_tx=XM_023241222.1
D3 547752 547859
refGene_gn=ANKLE2;refGene_tx=XM_006938342.3;refGene_gn=ANKLE2;refGene_tx=XM_023241222.1
D3 548707 548892
refGene_gn=ANKLE2;refGene_tx=XM_006938342.3;refGene_gn=ANKLE2;refGene_tx=XM_023241222.1
D3 549288 549934
refGene_gn=ANKLE2;refGene_tx=XM_006938342.3;refGene_gn=ANKLE2;refGene_tx=XM_023241222.1
D3 551260 551392
refGene_gn=ANKLE2;refGene_tx=XM_006938342.3;refGene_gn=ANKLE2;refGene_tx=XM_023241222.1
D3 551753 551958
refGene_gn=ANKLE2;refGene_tx=XM_006938342.3;refGene_gn=ANKLE2;refGene_tx=XM_023241222.1
D3 556455 556606
refGene_gn=PGAM5;refGene_tx=XM_023241223.1;refGene_gn=PGAM5;refGene_tx=XM_023241224.1
D3 557907 557950 refGene_gn=PGAM5;refGene_tx=XM_023241225.1
D3 558290 558424
refGene_gn=PGAM5;refGene_tx=XM_023241223.1;refGene_gn=PGAM5;refGene_tx=XM_023241224.1;refGene_gn=PGAM5;refGene_tx=XM_023241225.1
D3 558542 558631
refGene_gn=PGAM5;refGene_tx=XM_023241223.1;refGene_gn=PGAM5;refGene_tx=XM_023241224.1;refGene_gn=PGAM5;refGene_tx=XM_023241225.1
D3 558735 558861
refGene_gn=PGAM5;refGene_tx=XM_023241223.1;refGene_gn=PGAM5;refGene_tx=XM_023241224.1;refGene_gn=PGAM5;refGene_tx=XM_023241225.1
D3 559801 559980
refGene_gn=PGAM5;refGene_tx=XM_023241223.1;refGene_gn=PGAM5;refGene_tx=XM_023241224.1;refGene_gn=PGAM5;refGene_tx=XM_023241225.1
D3 562717 562908
refGene_gn=PGAM5;refGene_tx=XM_023241223.1;refGene_gn=PGAM5;refGene_tx=XM_023241224.1;refGene_gn=PGAM5;refGene_tx=XM_023241225.1
D3 566267 566336 refGene_gn=PXMP2;refGene_tx=XM_023241227.1
D3 566754 566769 refGene_gn=PXMP2;refGene_tx=XM_023241228.1
D3 568582 569047
refGene_gn=PXMP2;refGene_tx=XM_023241226.1;refGene_gn=PXMP2;refGene_tx=XM_023241227.1;refGene_gn=PXMP2;refGene_tx=XM_023241228.1
D3 572470 572633
refGene_gn=PXMP2;refGene_tx=XM_023241226.1;refGene_gn=PXMP2;refGene_tx=XM_023241227.1;refGene_gn=PXMP2;refGene_tx=XM_023241228.1

D3 574142 574256
refGene_gn=PXMP2;refGene_tx=XM_023241226.1;refGene_gn=PXMP2;refGene_tx=XM_023241227.1;refGene_gn=PXMP2;refGene_tx=XM_023241228.1
D3 575840 575962
refGene_gn=PXMP2;refGene_tx=XM_023241226.1;refGene_gn=PXMP2;refGene_tx=XM_023241227.1;refGene_gn=PXMP2;refGene_tx=XM_023241228.1
D3 576292 576354 refGene_gn=POLE;refGene_tx=XM_023241229.1
D3 578727 578869 refGene_gn=POLE;refGene_tx=XM_023241229.1
D3 579533 579614 refGene_gn=POLE;refGene_tx=XM_023241229.1
D3 580220 580265 refGene_gn=POLE;refGene_tx=XM_023241229.1
D3 580412 580505 refGene_gn=POLE;refGene_tx=XM_023241229.1
D3 580812 580967 refGene_gn=POLE;refGene_tx=XM_023241229.1
D3 585163 585305 refGene_gn=POLE;refGene_tx=XM_023241229.1
D3 585419 585500 refGene_gn=POLE;refGene_tx=XM_023241229.1
D3 586480 586588 refGene_gn=POLE;refGene_tx=XM_023241229.1
D3 586879 586990 refGene_gn=POLE;refGene_tx=XM_023241229.1
D3 587261 587347 refGene_gn=POLE;refGene_tx=XM_023241229.1
D3 587582 587702 refGene_gn=POLE;refGene_tx=XM_023241229.1
D3 588954 589087 refGene_gn=POLE;refGene_tx=XM_023241229.1
D3 589180 589294 refGene_gn=POLE;refGene_tx=XM_023241229.1
D3 589423 589636 refGene_gn=POLE;refGene_tx=XM_023241229.1
D3 589888 589996 refGene_gn=POLE;refGene_tx=XM_023241229.1
D3 594906 595035 refGene_gn=POLE;refGene_tx=XM_023241229.1
D3 595119 595222 refGene_gn=POLE;refGene_tx=XM_023241229.1
D3 595313 595460 refGene_gn=POLE;refGene_tx=XM_023241229.1
D3 595856 596002 refGene_gn=POLE;refGene_tx=XM_023241229.1
D3 598043 598192 refGene_gn=POLE;refGene_tx=XM_023241229.1
D3 599112 599205 refGene_gn=POLE;refGene_tx=XM_023241229.1
D3 599398 599543 refGene_gn=POLE;refGene_tx=XM_023241229.1
D3 602235 602393 refGene_gn=POLE;refGene_tx=XM_023241229.1
D3 602759 602955 refGene_gn=POLE;refGene_tx=XM_023241229.1
D3 604034 604249 refGene_gn=POLE;refGene_tx=XM_023241229.1
D3 606779 606882 refGene_gn=POLE;refGene_tx=XM_023241229.1
D3 607295 607376 refGene_gn=POLE;refGene_tx=XM_023241229.1
D3 607495 607618 refGene_gn=POLE;refGene_tx=XM_023241229.1
D3 608365 608578 refGene_gn=POLE;refGene_tx=XM_023241229.1
D3 608829 609039 refGene_gn=POLE;refGene_tx=XM_023241229.1
D3 609243 609387 refGene_gn=POLE;refGene_tx=XM_023241229.1
D3 610602 610743 refGene_gn=POLE;refGene_tx=XM_023241229.1
D3 610907 611061 refGene_gn=POLE;refGene_tx=XM_023241229.1
D3 611137 611244 refGene_gn=POLE;refGene_tx=XM_023241229.1
D3 611403 611580 refGene_gn=POLE;refGene_tx=XM_023241229.1
D3 611662 611886 refGene_gn=POLE;refGene_tx=XM_023241229.1
D3 611980 612201 refGene_gn=POLE;refGene_tx=XM_023241229.1
D3 612489 612694 refGene_gn=POLE;refGene_tx=XM_023241229.1
D3 613119 613293 refGene_gn=POLE;refGene_tx=XM_023241229.1
D3 614239 614365 refGene_gn=POLE;refGene_tx=XM_023241229.1
D3 615451 615584 refGene_gn=POLE;refGene_tx=XM_023241229.1
D3 616954 617141 refGene_gn=POLE;refGene_tx=XM_023241229.1
D3 618011 618143 refGene_gn=POLE;refGene_tx=XM_023241229.1

D3	618283	618477	refGene_gn=POLE;refGene_tx=XM_023241229.1
D3	621724	621925	refGene_gn=POLE;refGene_tx=XM_023241229.1
D3	622140	622266	refGene_gn=POLE;refGene_tx=XM_023241229.1
D3	622949	623039	refGene_gn=POLE;refGene_tx=XM_023241229.1
D3	623171	623285	refGene_gn=POLE;refGene_tx=XM_023241229.1
D3	626525	626867
refGene_gn=P2RX2;refGene_tx=XM_023241231.1;refGene_gn=P2RX2;refGene_tx=XM_023241232.1;refGene_gn=P2RX2;refGene_tx=XM_023241233.1;refGene_gn=P2RX2;refGene_tx=XM_023241234.1;refGene_gn=P2RX2;refGene_tx=XM_023241235.1
D3	626947	627013
refGene_gn=P2RX2;refGene_tx=XM_023241231.1;refGene_gn=P2RX2;refGene_tx=XM_023241233.1;refGene_gn=P2RX2;refGene_tx=XM_023241234.1;refGene_gn=P2RX2;refGene_tx=XM_023241235.1
D3	627129	627220
refGene_gn=P2RX2;refGene_tx=XM_023241231.1;refGene_gn=P2RX2;refGene_tx=XM_023241232.1;refGene_gn=P2RX2;refGene_tx=XM_023241233.1;refGene_gn=P2RX2;refGene_tx=XM_023241234.1;refGene_gn=P2RX2;refGene_tx=XM_023241235.1
D3	627373	627504
refGene_gn=P2RX2;refGene_tx=XM_023241231.1;refGene_gn=P2RX2;refGene_tx=XM_023241232.1;refGene_gn=P2RX2;refGene_tx=XM_023241233.1;refGene_gn=P2RX2;refGene_tx=XM_023241234.1;refGene_gn=P2RX2;refGene_tx=XM_023241235.1
D3	627900	628039
refGene_gn=P2RX2;refGene_tx=XM_023241231.1;refGene_gn=P2RX2;refGene_tx=XM_023241232.1;refGene_gn=P2RX2;refGene_tx=XM_023241233.1;refGene_gn=P2RX2;refGene_tx=XM_023241234.1;refGene_gn=P2RX2;refGene_tx=XM_023241235.1
D3	628164	628245
refGene_gn=P2RX2;refGene_tx=XM_023241231.1;refGene_gn=P2RX2;refGene_tx=XM_023241232.1;refGene_gn=P2RX2;refGene_tx=XM_023241233.1;refGene_gn=P2RX2;refGene_tx=XM_023241234.1;refGene_gn=P2RX2;refGene_tx=XM_023241235.1
D3	628403	628500
refGene_gn=P2RX2;refGene_tx=XM_023241231.1;refGene_gn=P2RX2;refGene_tx=XM_023241232.1;refGene_gn=P2RX2;refGene_tx=XM_023241233.1;refGene_gn=P2RX2;refGene_tx=XM_023241234.1;refGene_gn=P2RX2;refGene_tx=XM_023241235.1
D3	628570	628646
refGene_gn=P2RX2;refGene_tx=XM_023241231.1;refGene_gn=P2RX2;refGene_tx=XM_023241232.1;refGene_gn=P2RX2;refGene_tx=XM_023241233.1;refGene_gn=P2RX2;refGene_tx=XM_023241234.1
D3	628753	628825
refGene_gn=P2RX2;refGene_tx=XM_023241231.1;refGene_gn=P2RX2;refGene_tx=XM_023241232.1;refGene_gn=P2RX2;refGene_tx=XM_023241233.1;refGene_gn=P2RX2;refGene_tx=XM_023241234.1
D3	628925	629061
refGene_gn=P2RX2;refGene_tx=XM_023241231.1;refGene_gn=P2RX2;refGene_tx=XM_023241232.1;refGene_gn=P2RX2;refGene_tx=XM_023241233.1;refGene_gn=

P2RX2;refGene_tx=XM_023241234.1;refGene_gn=P2RX2;refGene_tx=XM_0232412
35.1
D3 629181 629357
refGene_gn=P2RX2;refGene_tx=XM_023241231.1;refGene_gn=P2RX2;refGene_tx
=XM_023241232.1;refGene_gn=P2RX2;refGene_tx=XM_023241233.1;refGene_gn=
P2RX2;refGene_tx=XM_023241234.1;refGene_gn=P2RX2;refGene_tx=XM_0232412
35.1
D3 639126 639216
refGene_gn=LRCOL1;refGene_tx=XM_023241236.1;refGene_gn=LRCOL1;refGene_
tx=XM_023241237.1;refGene_gn=LRCOL1;refGene_tx=XM_023241238.1
D3 640132 640700
refGene_gn=LRCOL1;refGene_tx=XM_023241236.1;refGene_gn=LRCOL1;refGene_
tx=XM_023241237.1;refGene_gn=LRCOL1;refGene_tx=XM_023241238.1
D3 641453 641614
refGene_gn=LRCOL1;refGene_tx=XM_023241236.1;refGene_gn=LRCOL1;refGene_
tx=XM_023241238.1
D3 686615 687576
refGene_gn=FBRSL1;refGene_tx=XM_023241239.1;refGene_gn=FBRSL1;refGene_
tx=XM_023241240.1;refGene_gn=FBRSL1;refGene_tx=XM_023241241.1;refGene_
gn=FBRSL1;refGene_tx=XM_023241242.1;refGene_gn=FBRSL1;refGene_tx=XM_02
3241243.1;refGene_gn=FBRSL1;refGene_tx=XM_023241244.1;refGene_gn=FBRSL
1;refGene_tx=XM_023241245.1;refGene_gn=FBRSL1;refGene_tx=XM_023241246.
1;refGene_gn=FBRSL1;refGene_tx=XM_023241247.1
D3 688467 688672
refGene_gn=FBRSL1;refGene_tx=XM_023241239.1;refGene_gn=FBRSL1;refGene_
tx=XM_023241240.1;refGene_gn=FBRSL1;refGene_tx=XM_023241241.1;refGene_
gn=FBRSL1;refGene_tx=XM_023241242.1;refGene_gn=FBRSL1;refGene_tx=XM_02
3241243.1;refGene_gn=FBRSL1;refGene_tx=XM_023241244.1;refGene_gn=FBRSL
1;refGene_tx=XM_023241245.1;refGene_gn=FBRSL1;refGene_tx=XM_023241246.
1;refGene_gn=FBRSL1;refGene_tx=XM_023241247.1
D3 688923 689007
refGene_gn=FBRSL1;refGene_tx=XM_023241239.1;refGene_gn=FBRSL1;refGene_
tx=XM_023241240.1;refGene_gn=FBRSL1;refGene_tx=XM_023241241.1;refGene_
gn=FBRSL1;refGene_tx=XM_023241242.1;refGene_gn=FBRSL1;refGene_tx=XM_02
3241243.1;refGene_gn=FBRSL1;refGene_tx=XM_023241244.1;refGene_gn=FBRSL
1;refGene_tx=XM_023241245.1;refGene_gn=FBRSL1;refGene_tx=XM_023241246.
1;refGene_gn=FBRSL1;refGene_tx=XM_023241247.1
D3 689210 689288
refGene_gn=FBRSL1;refGene_tx=XM_023241239.1;refGene_gn=FBRSL1;refGene_
tx=XM_023241240.1;refGene_gn=FBRSL1;refGene_tx=XM_023241241.1;refGene_
gn=FBRSL1;refGene_tx=XM_023241242.1;refGene_gn=FBRSL1;refGene_tx=XM_02
3241243.1;refGene_gn=FBRSL1;refGene_tx=XM_023241244.1;refGene_gn=FBRSL
1;refGene_tx=XM_023241245.1;refGene_gn=FBRSL1;refGene_tx=XM_023241246.
1;refGene_gn=FBRSL1;refGene_tx=XM_023241247.1
D3 691308 691441
refGene_gn=FBRSL1;refGene_tx=XM_023241239.1;refGene_gn=FBRSL1;refGene_
tx=XM_023241240.1;refGene_gn=FBRSL1;refGene_tx=XM_023241241.1;refGene_
gn=FBRSL1;refGene_tx=XM_023241242.1;refGene_gn=FBRSL1;refGene_tx=XM_02
3241243.1;refGene_gn=FBRSL1;refGene_tx=XM_023241244.1;refGene_gn=FBRSL
1;refGene_tx=XM_023241245.1;refGene_gn=FBRSL1;refGene_tx=XM_023241246.

1;refGene_gn=FBRSL1;refGene_tx=XM_023241247.1
D3 692323 692395
refGene_gn=FBRSL1;refGene_tx=XM_023241239.1;refGene_gn=FBRSL1;refGene_tx=XM_023241240.1;refGene_gn=FBRSL1;refGene_tx=XM_023241241.1;refGene_gn=FBRSL1;refGene_tx=XM_023241242.1;refGene_gn=FBRSL1;refGene_tx=XM_023241243.1;refGene_gn=FBRSL1;refGene_tx=XM_023241244.1;refGene_gn=FBRSL1;refGene_tx=XM_023241245.1;refGene_gn=FBRSL1;refGene_tx=XM_023241246.1;refGene_gn=FBRSL1;refGene_tx=XM_023241247.1
D3 692594 692624
refGene_gn=FBRSL1;refGene_tx=XM_023241239.1;refGene_gn=FBRSL1;refGene_tx=XM_023241240.1;refGene_gn=FBRSL1;refGene_tx=XM_023241241.1;refGene_gn=FBRSL1;refGene_tx=XM_023241242.1;refGene_gn=FBRSL1;refGene_tx=XM_023241243.1;refGene_gn=FBRSL1;refGene_tx=XM_023241244.1;refGene_gn=FBRSL1;refGene_tx=XM_023241245.1;refGene_gn=FBRSL1;refGene_tx=XM_023241246.1;refGene_gn=FBRSL1;refGene_tx=XM_023241247.1
D3 692792 692861
refGene_gn=FBRSL1;refGene_tx=XM_023241239.1;refGene_gn=FBRSL1;refGene_tx=XM_023241240.1;refGene_gn=FBRSL1;refGene_tx=XM_023241241.1;refGene_gn=FBRSL1;refGene_tx=XM_023241242.1;refGene_gn=FBRSL1;refGene_tx=XM_023241243.1;refGene_gn=FBRSL1;refGene_tx=XM_023241244.1;refGene_gn=FBRSL1;refGene_tx=XM_023241245.1;refGene_gn=FBRSL1;refGene_tx=XM_023241246.1;refGene_gn=FBRSL1;refGene_tx=XM_023241247.1
D3 693400 693496
refGene_gn=FBRSL1;refGene_tx=XM_023241239.1;refGene_gn=FBRSL1;refGene_tx=XM_023241240.1;refGene_gn=FBRSL1;refGene_tx=XM_023241241.1;refGene_gn=FBRSL1;refGene_tx=XM_023241242.1;refGene_gn=FBRSL1;refGene_tx=XM_023241243.1;refGene_gn=FBRSL1;refGene_tx=XM_023241246.1
D3 693687 693738
refGene_gn=FBRSL1;refGene_tx=XM_023241239.1;refGene_gn=FBRSL1;refGene_tx=XM_023241240.1;refGene_gn=FBRSL1;refGene_tx=XM_023241241.1;refGene_gn=FBRSL1;refGene_tx=XM_023241242.1;refGene_gn=FBRSL1;refGene_tx=XM_023241243.1;refGene_gn=FBRSL1;refGene_tx=XM_023241244.1;refGene_gn=FBRSL1;refGene_tx=XM_023241245.1;refGene_gn=FBRSL1;refGene_tx=XM_023241246.1;refGene_gn=FBRSL1;refGene_tx=XM_023241247.1
D3 694526 694741
refGene_gn=FBRSL1;refGene_tx=XM_023241239.1;refGene_gn=FBRSL1;refGene_tx=XM_023241240.1;refGene_gn=FBRSL1;refGene_tx=XM_023241241.1;refGene_gn=FBRSL1;refGene_tx=XM_023241242.1;refGene_gn=FBRSL1;refGene_tx=XM_023241243.1;refGene_gn=FBRSL1;refGene_tx=XM_023241244.1;refGene_gn=FBRSL1;refGene_tx=XM_023241245.1
D3 694816 694980
refGene_gn=FBRSL1;refGene_tx=XM_023241239.1;refGene_gn=FBRSL1;refGene_tx=XM_023241240.1;refGene_gn=FBRSL1;refGene_tx=XM_023241241.1;refGene_gn=FBRSL1;refGene_tx=XM_023241242.1;refGene_gn=FBRSL1;refGene_tx=XM_023241244.1;refGene_gn=FBRSL1;refGene_tx=XM_023241245.1;refGene_gn=FBRSL1;refGene_tx=XM_023241246.1;refGene_gn=FBRSL1;refGene_tx=XM_023241247.1
D3 695673 695894
refGene_gn=FBRSL1;refGene_tx=XM_023241239.1;refGene_gn=FBRSL1;refGene_tx=XM_023241240.1;refGene_gn=FBRSL1;refGene_tx=XM_023241241.1;refGene_

gn=FBRSL1;refGene_tx=XM_023241242.1;refGene_gn=FBRSL1;refGene_tx=XM_02
3241243.1;refGene_gn=FBRSL1;refGene_tx=XM_023241244.1;refGene_gn=FBRSL
1;refGene_tx=XM_023241245.1;refGene_gn=FBRSL1;refGene_tx=XM_023241246.
1;refGene_gn=FBRSL1;refGene_tx=XM_023241247.1
D3 695972 696294
refGene_gn=FBRSL1;refGene_tx=XM_023241239.1;refGene_gn=FBRSL1;refGene_
tx=XM_023241240.1;refGene_gn=FBRSL1;refGene_tx=XM_023241241.1;refGene_
gn=FBRSL1;refGene_tx=XM_023241242.1;refGene_gn=FBRSL1;refGene_tx=XM_02
3241243.1;refGene_gn=FBRSL1;refGene_tx=XM_023241244.1;refGene_gn=FBRSL
1;refGene_tx=XM_023241245.1;refGene_gn=FBRSL1;refGene_tx=XM_023241246.
1;refGene_gn=FBRSL1;refGene_tx=XM_023241247.1
D3 698810 698856
refGene_gn=FBRSL1;refGene_tx=XM_023241239.1;refGene_gn=FBRSL1;refGene_
tx=XM_023241240.1;refGene_gn=FBRSL1;refGene_tx=XM_023241241.1;refGene_
gn=FBRSL1;refGene_tx=XM_023241242.1;refGene_gn=FBRSL1;refGene_tx=XM_02
3241243.1;refGene_gn=FBRSL1;refGene_tx=XM_023241244.1;refGene_gn=FBRSL
1;refGene_tx=XM_023241245.1;refGene_gn=FBRSL1;refGene_tx=XM_023241246.
1;refGene_gn=FBRSL1;refGene_tx=XM_023241247.1
D3 716764 716794
refGene_gn=FBRSL1;refGene_tx=XM_023241239.1;refGene_gn=FBRSL1;refGene_
tx=XM_023241240.1;refGene_gn=FBRSL1;refGene_tx=XM_023241241.1;refGene_
gn=FBRSL1;refGene_tx=XM_023241243.1;refGene_gn=FBRSL1;refGene_tx=XM_02
3241244.1;refGene_gn=FBRSL1;refGene_tx=XM_023241245.1;refGene_gn=FBRSL
1;refGene_tx=XM_023241246.1;refGene_gn=FBRSL1;refGene_tx=XM_023241247.
1
D3 731538 731574
refGene_gn=FBRSL1;refGene_tx=XM_023241239.1;refGene_gn=FBRSL1;refGene_
tx=XM_023241240.1;refGene_gn=FBRSL1;refGene_tx=XM_023241241.1;refGene_
gn=FBRSL1;refGene_tx=XM_023241242.1;refGene_gn=FBRSL1;refGene_tx=XM_02
3241243.1;refGene_gn=FBRSL1;refGene_tx=XM_023241244.1;refGene_gn=FBRSL
1;refGene_tx=XM_023241245.1;refGene_gn=FBRSL1;refGene_tx=XM_023241246.
1;refGene_gn=FBRSL1;refGene_tx=XM_023241247.1
D3 733523 733628
refGene_gn=FBRSL1;refGene_tx=XM_023241239.1;refGene_gn=FBRSL1;refGene_
tx=XM_023241240.1;refGene_gn=FBRSL1;refGene_tx=XM_023241241.1;refGene_
gn=FBRSL1;refGene_tx=XM_023241242.1;refGene_gn=FBRSL1;refGene_tx=XM_02
3241243.1;refGene_gn=FBRSL1;refGene_tx=XM_023241244.1;refGene_gn=FBRSL
1;refGene_tx=XM_023241245.1;refGene_gn=FBRSL1;refGene_tx=XM_023241246.
1;refGene_gn=FBRSL1;refGene_tx=XM_023241247.1
D3 747892 748093
refGene_gn=FBRSL1;refGene_tx=XM_023241239.1;refGene_gn=FBRSL1;refGene_
tx=XM_023241240.1;refGene_gn=FBRSL1;refGene_tx=XM_023241241.1;refGene_
gn=FBRSL1;refGene_tx=XM_023241242.1;refGene_gn=FBRSL1;refGene_tx=XM_02
3241243.1;refGene_gn=FBRSL1;refGene_tx=XM_023241244.1;refGene_gn=FBRSL
1;refGene_tx=XM_023241245.1;refGene_gn=FBRSL1;refGene_tx=XM_023241246.
1;refGene_gn=FBRSL1;refGene_tx=XM_023241247.1
D3 764213 764492
refGene_gn=FBRSL1;refGene_tx=XM_023241239.1;refGene_gn=FBRSL1;refGene_
tx=XM_023241240.1;refGene_gn=FBRSL1;refGene_tx=XM_023241241.1;refGene_
gn=FBRSL1;refGene_tx=XM_023241242.1;refGene_gn=FBRSL1;refGene_tx=XM_02

3241243.1;refGene_gn=FBRSL1;refGene_tx=XM_023241244.1;refGene_gn=FBRSL1;refGene_tx=XM_023241245.1;refGene_gn=FBRSL1;refGene_tx=XM_023241246.1;refGene_gn=FBRSL1;refGene_tx=XM_023241247.1
D3 853899 854140 refGene_gn=GALNT9;refGene_tx=XM_023241248.1
D3 880025 880206 refGene_gn=GALNT9;refGene_tx=XM_023241248.1;refGene_gn=GALNT9;refGene_tx=XM_023241249.1
D3 896589 896756 refGene_gn=GALNT9;refGene_tx=XM_023241248.1;refGene_gn=GALNT9;refGene_tx=XM_023241249.1
D3 898326 898501 refGene_gn=GALNT9;refGene_tx=XM_023241248.1;refGene_gn=GALNT9;refGene_tx=XM_023241249.1
D3 902330 902528 refGene_gn=GALNT9;refGene_tx=XM_023241248.1;refGene_gn=GALNT9;refGene_tx=XM_023241249.1
D3 908439 908557 refGene_gn=GALNT9;refGene_tx=XM_023241248.1;refGene_gn=GALNT9;refGene_tx=XM_023241249.1
D3 926325 926511 refGene_gn=GALNT9;refGene_tx=XM_023241248.1;refGene_gn=GALNT9;refGene_tx=XM_023241249.1
D3 929317 929455 refGene_gn=GALNT9;refGene_tx=XM_023241248.1;refGene_gn=GALNT9;refGene_tx=XM_023241249.1
D3 930398 930494 refGene_gn=GALNT9;refGene_tx=XM_023241248.1;refGene_gn=GALNT9;refGene_tx=XM_023241249.1
D3 933109 933277 refGene_gn=GALNT9;refGene_tx=XM_023241248.1;refGene_gn=GALNT9;refGene_tx=XM_023241249.1
D3 935205 935352 refGene_gn=GALNT9;refGene_tx=XM_023241248.1;refGene_gn=GALNT9;refGene_tx=XM_023241249.1
D3 959999 960122 refGene_gn=NOC4L;refGene_tx=XM_003994565.4
D3 960201 960315 refGene_gn=NOC4L;refGene_tx=XM_003994565.4
D3 960454 960537 refGene_gn=NOC4L;refGene_tx=XM_003994565.4
D3 960765 960926 refGene_gn=NOC4L;refGene_tx=XM_003994565.4
D3 961023 961134 refGene_gn=NOC4L;refGene_tx=XM_003994565.4
D3 961351 961412 refGene_gn=NOC4L;refGene_tx=XM_003994565.4
D3 961614 961726 refGene_gn=NOC4L;refGene_tx=XM_003994565.4
D3 961829 961880 refGene_gn=NOC4L;refGene_tx=XM_003994565.4
D3 962282 962317 refGene_gn=NOC4L;refGene_tx=XM_003994565.4
D3 962402 962502 refGene_gn=NOC4L;refGene_tx=XM_003994565.4
D3 962574 962724 refGene_gn=NOC4L;refGene_tx=XM_003994565.4
D3 962916 963024 refGene_gn=NOC4L;refGene_tx=XM_003994565.4
D3 963302 963409 refGene_gn=NOC4L;refGene_tx=XM_003994565.4
D3 964087 964208 refGene_gn=NOC4L;refGene_tx=XM_003994565.4
D3 964413 964530 refGene_gn=NOC4L;refGene_tx=XM_003994565.4
D3 964695 964987

refGene_gn=DDX51;refGene_tx=XM_023241250.1;refGene_gn=DDX51;refGene_tx=XM_023241251.1
D3 965074 965226
refGene_gn=DDX51;refGene_tx=XM_023241250.1;refGene_gn=DDX51;refGene_tx=XM_023241251.1
D3 966148 966299
refGene_gn=DDX51;refGene_tx=XM_023241250.1;refGene_gn=DDX51;refGene_tx=XM_023241251.1
D3 966614 966760
refGene_gn=DDX51;refGene_tx=XM_023241250.1;refGene_gn=DDX51;refGene_tx=XM_023241251.1
D3 966853 966925
refGene_gn=DDX51;refGene_tx=XM_023241250.1;refGene_gn=DDX51;refGene_tx=XM_023241251.1
D3 967015 967122
refGene_gn=DDX51;refGene_tx=XM_023241250.1;refGene_gn=DDX51;refGene_tx=XM_023241251.1
D3 967278 967387
refGene_gn=DDX51;refGene_tx=XM_023241250.1;refGene_gn=DDX51;refGene_tx=XM_023241251.1
D3 967472 967618
refGene_gn=DDX51;refGene_tx=XM_023241250.1;refGene_gn=DDX51;refGene_tx=XM_023241251.1
D3 967792 967985
refGene_gn=DDX51;refGene_tx=XM_023241250.1;refGene_gn=DDX51;refGene_tx=XM_023241251.1
D3 968061 968177
refGene_gn=DDX51;refGene_tx=XM_023241250.1;refGene_gn=DDX51;refGene_tx=XM_023241251.1
D3 968272 968393
refGene_gn=DDX51;refGene_tx=XM_023241250.1;refGene_gn=DDX51;refGene_tx=XM_023241251.1
D3 968564 968666 refGene_gn=DDX51;refGene_tx=XM_023241250.1
D3 968769 968833 refGene_gn=DDX51;refGene_tx=XM_023241250.1
D3 968919 969054 refGene_gn=DDX51;refGene_tx=XM_023241250.1
D3 969268 969295 refGene_gn=DDX51;refGene_tx=XM_023241250.1
D3 991021 991294 refGene_gn=EP400;refGene_tx=XM_023241252.1
D3 992076 992154 refGene_gn=EP400;refGene_tx=XM_023241252.1
D3 996462 996609 refGene_gn=EP400;refGene_tx=XM_023241252.1
D3 998492 998617 refGene_gn=EP400;refGene_tx=XM_023241252.1
D3 999082 999278 refGene_gn=EP400;refGene_tx=XM_023241252.1
D3 1001044 1001263 refGene_gn=EP400;refGene_tx=XM_023241252.1
D3 1002842 1003039 refGene_gn=EP400;refGene_tx=XM_023241252.1
D3 1003168 1003376 refGene_gn=EP400;refGene_tx=XM_023241252.1
D3 1008004 1008114 refGene_gn=EP400;refGene_tx=XM_023241252.1
D3 1008188 1008234 refGene_gn=EP400;refGene_tx=XM_023241252.1
D3 1009205 1009457 refGene_gn=EP400;refGene_tx=XM_023241252.1
D3 1009884 1009963 refGene_gn=EP400;refGene_tx=XM_023241252.1
D3 1012402 1012459 refGene_gn=EP400;refGene_tx=XM_023241252.1
D3 1012575 1012670 refGene_gn=EP400;refGene_tx=XM_023241252.1

D3 1012671 1012713 refGene_gn=EP400;refGene_tx=XM_023241252.1
D3 1014560 1014734 refGene_gn=EP400;refGene_tx=XM_023241252.1
D3 1014843 1015085 refGene_gn=EP400;refGene_tx=XM_023241252.1
D3 1015300 1015454 refGene_gn=EP400;refGene_tx=XM_023241252.1
D3 1015533 1015578 refGene_gn=EP400;refGene_tx=XM_023241252.1
D3 1015796 1015931 refGene_gn=EP400;refGene_tx=XM_023241252.1
D3 1016252 1016336 refGene_gn=EP400;refGene_tx=XM_023241252.1
D3 1016545 1016686 refGene_gn=EP400;refGene_tx=XM_023241252.1
D3 1020181 1020325 refGene_gn=EP400;refGene_tx=XM_023241252.1
D3 1020498 1020610 refGene_gn=EP400;refGene_tx=XM_023241252.1
D3 1023634 1023831 refGene_gn=EP400;refGene_tx=XM_023241252.1
D3 1025044 1025214 refGene_gn=EP400;refGene_tx=XM_023241252.1
D3 1025525 1025728 refGene_gn=EP400;refGene_tx=XM_023241252.1
D3 1029282 1029554 refGene_gn=EP400;refGene_tx=XM_023241252.1
D3 1030012 1030107 refGene_gn=EP400;refGene_tx=XM_023241252.1
D3 1031554 1031710 refGene_gn=EP400;refGene_tx=XM_023241252.1
D3 1033423 1033588 refGene_gn=EP400;refGene_tx=XM_023241252.1
D3 1035402 1035612 refGene_gn=EP400;refGene_tx=XM_023241252.1
D3 1036375 1036545 refGene_gn=EP400;refGene_tx=XM_023241252.1
D3 1037912 1038079 refGene_gn=EP400;refGene_tx=XM_023241252.1
D3 1039562 1039749 refGene_gn=EP400;refGene_tx=XM_023241252.1
D3 1041808 1041945 refGene_gn=EP400;refGene_tx=XM_023241252.1
D3 1042042 1042217 refGene_gn=EP400;refGene_tx=XM_023241252.1
D3 1042501 1042671 refGene_gn=EP400;refGene_tx=XM_023241252.1
D3 1043516 1043653 refGene_gn=EP400;refGene_tx=XM_023241252.1
D3 1046029 1046207 refGene_gn=EP400;refGene_tx=XM_023241252.1
D3 1046452 1046637 refGene_gn=EP400;refGene_tx=XM_023241252.1
D3 1047671 1047779 refGene_gn=EP400;refGene_tx=XM_023241252.1
D3 1050088 1050178 refGene_gn=EP400;refGene_tx=XM_023241252.1
D3 1050896 1050954 refGene_gn=EP400;refGene_tx=XM_023241252.1
D3 1051323 1051373 refGene_gn=EP400;refGene_tx=XM_023241252.1
D3 1052079 1052158 refGene_gn=EP400;refGene_tx=XM_023241252.1
D3 1052610 1052751 refGene_gn=EP400;refGene_tx=XM_023241252.1
D3 1054676 1054862 refGene_gn=EP400;refGene_tx=XM_023241252.1
D3 1055793 1056084 refGene_gn=EP400;refGene_tx=XM_023241252.1
D3 1058023 1058406 refGene_gn=EP400;refGene_tx=XM_023241252.1
D3 1058890 1058998 refGene_gn=EP400;refGene_tx=XM_023241252.1
D3 1060546 1060637 refGene_gn=EP400;refGene_tx=XM_023241252.1
D3 1072336 1073662 refGene_gn=EP400;refGene_tx=XM_023241252.1
D3 1079084 1079134
refGene_gn=LOC111557362;refGene_tx=XM_023242149.1
D3 1082741 1082956
refGene_gn=LOC111557362;refGene_tx=XM_023242149.1
D3 1086756 1087574
refGene_gn=LOC111557362;refGene_tx=XM_023242149.1
D3 1092851 1092881
refGene_gn=PUS1;refGene_tx=XM_023241253.1;refGene_gn=PUS1;refGene_tx=X
M_023241254.1;refGene_gn=PUS1;refGene_tx=XM_023241255.1
D3 1094095 1094787
refGene_gn=PUS1;refGene_tx=XM_023241253.1;refGene_gn=PUS1;refGene_tx=X

M_023241254.1;refGene_gn=PUS1;refGene_tx=XM_023241255.1
D3 1096856 1096959
refGene_gn=PUS1;refGene_tx=XM_023241253.1;refGene_gn=PUS1;refGene_tx=XM_023241254.1;refGene_gn=PUS1;refGene_tx=XM_023241255.1
D3 1100495 1100633
refGene_gn=PUS1;refGene_tx=XM_023241253.1;refGene_gn=PUS1;refGene_tx=XM_023241254.1;refGene_gn=PUS1;refGene_tx=XM_023241255.1
D3 1101575 1101795
refGene_gn=PUS1;refGene_tx=XM_023241253.1;refGene_gn=PUS1;refGene_tx=XM_023241254.1;refGene_gn=PUS1;refGene_tx=XM_023241255.1
D3 1101884 1101973 refGene_gn=PUS1;refGene_tx=XM_023241253.1
D3 1106568 1106624 refGene_gn=ULK1;refGene_tx=XM_023241256.1
D3 1106704 1106840 refGene_gn=ULK1;refGene_tx=XM_023241256.1
D3 1107528 1107686 refGene_gn=ULK1;refGene_tx=XM_023241256.1
D3 1108052 1108171 refGene_gn=ULK1;refGene_tx=XM_023241256.1
D3 1108273 1108446 refGene_gn=ULK1;refGene_tx=XM_023241256.1
D3 1108528 1108722 refGene_gn=ULK1;refGene_tx=XM_023241256.1
D3 1109098 1109242 refGene_gn=ULK1;refGene_tx=XM_023241256.1
D3 1109325 1109435 refGene_gn=ULK1;refGene_tx=XM_023241256.1
D3 1109729 1109923 refGene_gn=ULK1;refGene_tx=XM_023241256.1
D3 1110436 1110705 refGene_gn=ULK1;refGene_tx=XM_023241256.1
D3 1111024 1111111 refGene_gn=ULK1;refGene_tx=XM_023241256.1
D3 1111234 1111383 refGene_gn=ULK1;refGene_tx=XM_023241256.1
D3 1112119 1112242 refGene_gn=ULK1;refGene_tx=XM_023241256.1
D3 1112794 1112884 refGene_gn=ULK1;refGene_tx=XM_023241256.1
D3 1113019 1113080 refGene_gn=ULK1;refGene_tx=XM_023241256.1
D3 1113866 1114020 refGene_gn=ULK1;refGene_tx=XM_023241256.1
D3 1114700 1114789 refGene_gn=ULK1;refGene_tx=XM_023241256.1
D3 1115039 1115090 refGene_gn=ULK1;refGene_tx=XM_023241256.1
D3 1115343 1115426 refGene_gn=ULK1;refGene_tx=XM_023241256.1
D3 1115507 1115566 refGene_gn=ULK1;refGene_tx=XM_023241256.1
D3 1116485 1116587 refGene_gn=ULK1;refGene_tx=XM_023241256.1
D3 1116799 1116873 refGene_gn=ULK1;refGene_tx=XM_023241256.1
D3 1116963 1117137 refGene_gn=ULK1;refGene_tx=XM_023241256.1
D3 1117853 1117890 refGene_gn=ULK1;refGene_tx=XM_023241256.1
D3 1118219 1118252 refGene_gn=ULK1;refGene_tx=XM_023241256.1
D3 1126229 1126271 refGene_gn=ULK1;refGene_tx=XM_023241256.1
D3 1126354 1126447 refGene_gn=ULK1;refGene_tx=XM_023241256.1
D3 1126822 1126933 refGene_gn=ULK1;refGene_tx=XM_023241256.1
D3 1148596 1148937
refGene_gn=MMP17;refGene_tx=XM_023241257.1;refGene_gn=MMP17;refGene_tx=XM_023241258.1;refGene_gn=MMP17;refGene_tx=XM_023241259.1;refGene_gn=MMP17;refGene_tx=XM_023241260.1;refGene_gn=MMP17;refGene_tx=XM_023241261.1
D3 1149560 1149818
refGene_gn=MMP17;refGene_tx=XM_023241257.1;refGene_gn=MMP17;refGene_tx=XM_023241258.1;refGene_gn=MMP17;refGene_tx=XM_023241259.1;refGene_gn=MMP17;refGene_tx=XM_023241260.1;refGene_gn=MMP17;refGene_tx=XM_023241261.1
D3 1154256 1154441

refGene_gn=MMP17;refGene_tx=XM_023241257.1;refGene_gn=MMP17;refGene_tx=XM_023241258.1;refGene_gn=MMP17;refGene_tx=XM_023241259.1;refGene_gn=MMP17;refGene_tx=XM_023241260.1;refGene_gn=MMP17;refGene_tx=XM_023241261.1;refGene_gn=MMP17;refGene_tx=XM_023241262.1
D3 1154540 1154623 refGene_gn=MMP17;refGene_tx=XM_023241257.1;refGene_gn=MMP17;refGene_tx=XM_023241258.1;refGene_gn=MMP17;refGene_tx=XM_023241259.1;refGene_gn=MMP17;refGene_tx=XM_023241260.1;refGene_gn=MMP17;refGene_tx=XM_023241261.1;refGene_gn=MMP17;refGene_tx=XM_023241262.1
D3 1155160 1155260 refGene_gn=MMP17;refGene_tx=XM_023241257.1;refGene_gn=MMP17;refGene_tx=XM_023241258.1;refGene_gn=MMP17;refGene_tx=XM_023241259.1;refGene_gn=MMP17;refGene_tx=XM_023241260.1;refGene_gn=MMP17;refGene_tx=XM_023241261.1;refGene_gn=MMP17;refGene_tx=XM_023241262.1
D3 1156853 1157030 refGene_gn=MMP17;refGene_tx=XM_023241257.1;refGene_gn=MMP17;refGene_tx=XM_023241258.1;refGene_gn=MMP17;refGene_tx=XM_023241259.1;refGene_gn=MMP17;refGene_tx=XM_023241260.1;refGene_gn=MMP17;refGene_tx=XM_023241261.1;refGene_gn=MMP17;refGene_tx=XM_023241262.1
D3 1157309 1157593 refGene_gn=MMP17;refGene_tx=XM_023241257.1;refGene_gn=MMP17;refGene_tx=XM_023241258.1;refGene_gn=MMP17;refGene_tx=XM_023241259.1;refGene_gn=MMP17;refGene_tx=XM_023241260.1;refGene_gn=MMP17;refGene_tx=XM_023241261.1;refGene_gn=MMP17;refGene_tx=XM_023241262.1
D3 1158019 1158149 refGene_gn=MMP17;refGene_tx=XM_023241257.1;refGene_gn=MMP17;refGene_tx=XM_023241258.1;refGene_gn=MMP17;refGene_tx=XM_023241259.1;refGene_gn=MMP17;refGene_tx=XM_023241260.1;refGene_gn=MMP17;refGene_tx=XM_023241261.1;refGene_gn=MMP17;refGene_tx=XM_023241262.1
D3 1158911 1159044 refGene_gn=MMP17;refGene_tx=XM_023241257.1;refGene_gn=MMP17;refGene_tx=XM_023241258.1;refGene_gn=MMP17;refGene_tx=XM_023241259.1;refGene_gn=MMP17;refGene_tx=XM_023241260.1;refGene_gn=MMP17;refGene_tx=XM_023241261.1;refGene_gn=MMP17;refGene_tx=XM_023241262.1
D3 1165293 1165725 refGene_gn=MMP17;refGene_tx=XM_023241257.1;refGene_gn=MMP17;refGene_tx=XM_023241258.1;refGene_gn=MMP17;refGene_tx=XM_023241262.1
D3 1168106 1168250 refGene_gn=MMP17;refGene_tx=XM_023241259.1
D3 1176366 1176432 refGene_gn=SFSWAP;refGene_tx=XM_003994566.4;refGene_gn=SFSWAP;refGene_tx=XM_006938344.3
D3 1176571 1176644 refGene_gn=SFSWAP;refGene_tx=XM_003994566.4;refGene_gn=SFSWAP;refGene_tx=XM_006938344.3
D3 1177594 1177759 refGene_gn=SFSWAP;refGene_tx=XM_003994566.4;refGene_gn=SFSWAP;refGene_tx=XM_006938344.3
D3 1187835 1187961 refGene_gn=SFSWAP;refGene_tx=XM_003994566.4;refGene_gn=SFSWAP;refGene_tx=XM_006938344.3

D3 1194302 1194568
refGene_gn=SFSWAP;refGene_tx=XM_003994566.4;refGene_gn=SFSWAP;refGene_tx=XM_006938344.3
D3 1203216 1203407
refGene_gn=SFSWAP;refGene_tx=XM_003994566.4;refGene_gn=SFSWAP;refGene_tx=XM_006938344.3
D3 1204440 1204671
refGene_gn=SFSWAP;refGene_tx=XM_003994566.4;refGene_gn=SFSWAP;refGene_tx=XM_006938344.3
D3 1210792 1210964
refGene_gn=SFSWAP;refGene_tx=XM_003994566.4;refGene_gn=SFSWAP;refGene_tx=XM_006938344.3
D3 1211881 1211975
refGene_gn=SFSWAP;refGene_tx=XM_003994566.4;refGene_gn=SFSWAP;refGene_tx=XM_006938344.3
D3 1212968 1213100
refGene_gn=SFSWAP;refGene_tx=XM_003994566.4;refGene_gn=SFSWAP;refGene_tx=XM_006938344.3
D3 1214259 1214497
refGene_gn=SFSWAP;refGene_tx=XM_003994566.4;refGene_gn=SFSWAP;refGene_tx=XM_006938344.3
D3 1229323 1229456
refGene_gn=SFSWAP;refGene_tx=XM_003994566.4;refGene_gn=SFSWAP;refGene_tx=XM_006938344.3
D3 1229896 1230009
refGene_gn=SFSWAP;refGene_tx=XM_003994566.4;refGene_gn=SFSWAP;refGene_tx=XM_006938344.3
D3 1231179 1231405
refGene_gn=SFSWAP;refGene_tx=XM_003994566.4;refGene_gn=SFSWAP;refGene_tx=XM_006938344.3
D3 1236600 1236686
refGene_gn=SFSWAP;refGene_tx=XM_003994566.4;refGene_gn=SFSWAP;refGene_tx=XM_006938344.3
D3 1240998 1241130
refGene_gn=SFSWAP;refGene_tx=XM_003994566.4;refGene_gn=SFSWAP;refGene_tx=XM_006938344.3
D3 1241709 1241879
refGene_gn=SFSWAP;refGene_tx=XM_003994566.4;refGene_gn=SFSWAP;refGene_tx=XM_006938344.3
D3 1244468 1244686 refGene_gn=SFSWAP;refGene_tx=XM_003994566.4
D3 1656087 1656183
refGene_gn=ADGRD1;refGene_tx=XM_011287690.3;refGene_gn=ADGRD1;refGene_tx=XM_019814407.2
D3 1657389 1657482
refGene_gn=ADGRD1;refGene_tx=XM_011287690.3;refGene_gn=ADGRD1;refGene_tx=XM_019814407.2
D3 1658607 1658649
refGene_gn=ADGRD1;refGene_tx=XM_011287690.3;refGene_gn=ADGRD1;refGene_tx=XM_019814407.2
D3 1659930 1660057

refGene_gn=ADGRD1;refGene_tx=XM_011287690.3;refGene_gn=ADGRD1;refGene_tx=XM_019814407.2
D3 1661746 1661838
refGene_gn=ADGRD1;refGene_tx=XM_011287690.3;refGene_gn=ADGRD1;refGene_tx=XM_019814407.2
D3 1671588 1671655
refGene_gn=ADGRD1;refGene_tx=XM_011287690.3;refGene_gn=ADGRD1;refGene_tx=XM_019814407.2
D3 1673071 1673138
refGene_gn=ADGRD1;refGene_tx=XM_011287690.3;refGene_gn=ADGRD1;refGene_tx=XM_019814407.2
D3 1677720 1677874
refGene_gn=ADGRD1;refGene_tx=XM_011287690.3;refGene_gn=ADGRD1;refGene_tx=XM_019814407.2
D3 1679068 1679180
refGene_gn=ADGRD1;refGene_tx=XM_011287690.3;refGene_gn=ADGRD1;refGene_tx=XM_019814407.2
D3 1680591 1680695
refGene_gn=ADGRD1;refGene_tx=XM_011287690.3;refGene_gn=ADGRD1;refGene_tx=XM_019814407.2
D3 1694003 1694127
refGene_gn=ADGRD1;refGene_tx=XM_011287690.3;refGene_gn=ADGRD1;refGene_tx=XM_019814407.2
D3 1698648 1698722
refGene_gn=ADGRD1;refGene_tx=XM_011287690.3;refGene_gn=ADGRD1;refGene_tx=XM_019814407.2
D3 1729420 1729562
refGene_gn=ADGRD1;refGene_tx=XM_011287690.3;refGene_gn=ADGRD1;refGene_tx=XM_019814407.2
D3 1735356 1735432
refGene_gn=ADGRD1;refGene_tx=XM_011287690.3;refGene_gn=ADGRD1;refGene_tx=XM_019814407.2
D3 1737267 1737378
refGene_gn=ADGRD1;refGene_tx=XM_011287690.3;refGene_gn=ADGRD1;refGene_tx=XM_019814407.2
D3 1738646 1738764
refGene_gn=ADGRD1;refGene_tx=XM_011287690.3;refGene_gn=ADGRD1;refGene_tx=XM_019814407.2
D3 1741179 1741239
refGene_gn=ADGRD1;refGene_tx=XM_011287690.3;refGene_gn=ADGRD1;refGene_tx=XM_019814407.2
D3 1743617 1743773
refGene_gn=ADGRD1;refGene_tx=XM_011287690.3;refGene_gn=ADGRD1;refGene_tx=XM_019814407.2
D3 1744498 1744563
refGene_gn=ADGRD1;refGene_tx=XM_011287690.3;refGene_gn=ADGRD1;refGene_tx=XM_019814407.2
D3 1748293 1748548
refGene_gn=ADGRD1;refGene_tx=XM_011287690.3;refGene_gn=ADGRD1;refGene_tx=XM_019814407.2

D3 1751473 1751653
refGene_gn=ADGRD1;refGene_tx=XM_011287690.3;refGene_gn=ADGRD1;refGene_
tx=XM_019814407.2
D3 1763160 1763283
refGene_gn=ADGRD1;refGene_tx=XM_011287690.3;refGene_gn=ADGRD1;refGene_
tx=XM_019814407.2
D3 1765672 1765768 refGene_gn=ADGRD1;refGene_tx=XM_019814407.2
D3 1767795 1767879
refGene_gn=ADGRD1;refGene_tx=XM_011287690.3;refGene_gn=ADGRD1;refGene_
tx=XM_019814407.2
D3 1777658 1777695
refGene_gn=ADGRD1;refGene_tx=XM_011287690.3;refGene_gn=ADGRD1;refGene_
tx=XM_019814407.2
D3 1777787 1777853
refGene_gn=ADGRD1;refGene_tx=XM_011287690.3;refGene_gn=ADGRD1;refGene_
tx=XM_019814407.2
D3 1835648 1835693 refGene_gn=RAN;refGene_tx=XM_023241263.1
D3 1835781 1835952 refGene_gn=RAN;refGene_tx=XM_023241263.1
D3 1836358 1836546 refGene_gn=RAN;refGene_tx=XM_023241263.1
D3 1837706 1837832 refGene_gn=RAN;refGene_tx=XM_023241263.1
D3 1837907 1837992 refGene_gn=RAN;refGene_tx=XM_023241263.1
D3 1838364 1838400 refGene_gn=RAN;refGene_tx=XM_023241263.1
D3 1850003 1850033
refGene_gn=STX2;refGene_tx=XM_023241264.1;refGene_gn=STX2;refGene_tx=X
M_023241265.1;refGene_gn=STX2;refGene_tx=XM_023241266.1;refGene_gn=STX
2;refGene_tx=XM_023241267.1
D3 1861531 1861606
refGene_gn=STX2;refGene_tx=XM_023241264.1;refGene_gn=STX2;refGene_tx=X
M_023241265.1;refGene_gn=STX2;refGene_tx=XM_023241266.1;refGene_gn=STX
2;refGene_tx=XM_023241267.1
D3 1865459 1865559
refGene_gn=STX2;refGene_tx=XM_023241264.1;refGene_gn=STX2;refGene_tx=X
M_023241265.1;refGene_gn=STX2;refGene_tx=XM_023241266.1;refGene_gn=STX
2;refGene_tx=XM_023241267.1
D3 1872910 1872985
refGene_gn=STX2;refGene_tx=XM_023241264.1;refGene_gn=STX2;refGene_tx=X
M_023241265.1;refGene_gn=STX2;refGene_tx=XM_023241266.1;refGene_gn=STX
2;refGene_tx=XM_023241267.1
D3 1874537 1874611
refGene_gn=STX2;refGene_tx=XM_023241264.1;refGene_gn=STX2;refGene_tx=X
M_023241265.1;refGene_gn=STX2;refGene_tx=XM_023241266.1;refGene_gn=STX
2;refGene_tx=XM_023241267.1
D3 1875103 1875212
refGene_gn=STX2;refGene_tx=XM_023241264.1;refGene_gn=STX2;refGene_tx=X
M_023241265.1;refGene_gn=STX2;refGene_tx=XM_023241266.1;refGene_gn=STX
2;refGene_tx=XM_023241267.1
D3 1877441 1877515
refGene_gn=STX2;refGene_tx=XM_023241264.1;refGene_gn=STX2;refGene_tx=X
M_023241265.1;refGene_gn=STX2;refGene_tx=XM_023241266.1;refGene_gn=STX
2;refGene_tx=XM_023241267.1

D3	1877617	1877755
refGene_gn=STX2;refGene_tx=XM_023241264.1;refGene_gn=STX2;refGene_tx=XM_023241265.1;refGene_gn=STX2;refGene_tx=XM_023241266.1;refGene_gn=STX2;refGene_tx=XM_023241267.1
D3	1879810	1879921
refGene_gn=STX2;refGene_tx=XM_023241264.1;refGene_gn=STX2;refGene_tx=XM_023241265.1;refGene_gn=STX2;refGene_tx=XM_023241266.1;refGene_gn=STX2;refGene_tx=XM_023241267.1
D3	1881660	1881741	refGene_gn=STX2;refGene_tx=XM_023241265.1
D3	1884994	1885072	refGene_gn=STX2;refGene_tx=XM_023241266.1
D3	1889449	1889602
refGene_gn=STX2;refGene_tx=XM_023241264.1;refGene_gn=STX2;refGene_tx=XM_023241267.1
D3	1920925	1920931	refGene_gn=STX2;refGene_tx=XM_023241264.1
D3	1928179	1928274	refGene_gn=STX2;refGene_tx=XM_023241267.1
D3	1929562	1929740	refGene_gn=STX2;refGene_tx=XM_023241267.1
D3	2054037	2054580
refGene_gn=RIMBP2;refGene_tx=XM_019814410.2;refGene_gn=RIMBP2;refGene_tx=XM_019814411.2;refGene_gn=RIMBP2;refGene_tx=XM_023241268.1;refGene_gn=RIMBP2;refGene_tx=XM_023241269.1
D3	2057525	2057615
refGene_gn=RIMBP2;refGene_tx=XM_019814410.2;refGene_gn=RIMBP2;refGene_tx=XM_019814411.2;refGene_gn=RIMBP2;refGene_tx=XM_023241268.1;refGene_gn=RIMBP2;refGene_tx=XM_023241269.1
D3	2066768	2066891
refGene_gn=RIMBP2;refGene_tx=XM_019814410.2;refGene_gn=RIMBP2;refGene_tx=XM_019814411.2;refGene_gn=RIMBP2;refGene_tx=XM_023241268.1;refGene_gn=RIMBP2;refGene_tx=XM_023241269.1
D3	2082705	2082810
refGene_gn=RIMBP2;refGene_tx=XM_019814410.2;refGene_gn=RIMBP2;refGene_tx=XM_019814411.2;refGene_gn=RIMBP2;refGene_tx=XM_019814413.2;refGene_gn=RIMBP2;refGene_tx=XM_023241268.1;refGene_gn=RIMBP2;refGene_tx=XM_023241269.1
D3	2090261	2090312
refGene_gn=RIMBP2;refGene_tx=XM_019814410.2;refGene_gn=RIMBP2;refGene_tx=XM_019814411.2;refGene_gn=RIMBP2;refGene_tx=XM_019814413.2;refGene_gn=RIMBP2;refGene_tx=XM_023241268.1;refGene_gn=RIMBP2;refGene_tx=XM_023241269.1
D3	2091751	2091772
refGene_gn=RIMBP2;refGene_tx=XM_019814412.2;refGene_gn=RIMBP2;refGene_tx=XM_019814413.2
D3	2101174	2101376
refGene_gn=RIMBP2;refGene_tx=XM_019814410.2;refGene_gn=RIMBP2;refGene_tx=XM_019814411.2;refGene_gn=RIMBP2;refGene_tx=XM_019814412.2;refGene_gn=RIMBP2;refGene_tx=XM_019814413.2;refGene_gn=RIMBP2;refGene_tx=XM_023241268.1;refGene_gn=RIMBP2;refGene_tx=XM_023241269.1
D3	2104410	2104556
refGene_gn=RIMBP2;refGene_tx=XM_019814410.2;refGene_gn=RIMBP2;refGene_tx=XM_019814411.2;refGene_gn=RIMBP2;refGene_tx=XM_019814412.2;refGene_gn=RIMBP2;refGene_tx=XM_019814413.2;refGene_gn=RIMBP2;refGene_tx=XM_02

3241268.1;refGene_gn=RIMBP2;refGene_tx=XM_023241269.1
D3 2105185 2105265
refGene_gn=RIMBP2;refGene_tx=XM_019814410.2;refGene_gn=RIMBP2;refGene_tx=XM_019814411.2;refGene_gn=RIMBP2;refGene_tx=XM_019814412.2;refGene_gn=RIMBP2;refGene_tx=XM_019814413.2;refGene_gn=RIMBP2;refGene_tx=XM_023241268.1;refGene_gn=RIMBP2;refGene_tx=XM_023241269.1
D3 2108809 2108919
refGene_gn=RIMBP2;refGene_tx=XM_019814410.2;refGene_gn=RIMBP2;refGene_tx=XM_019814411.2;refGene_gn=RIMBP2;refGene_tx=XM_019814412.2;refGene_gn=RIMBP2;refGene_tx=XM_019814413.2;refGene_gn=RIMBP2;refGene_tx=XM_023241268.1;refGene_gn=RIMBP2;refGene_tx=XM_023241269.1
D3 2111261 2112074
refGene_gn=RIMBP2;refGene_tx=XM_019814410.2;refGene_gn=RIMBP2;refGene_tx=XM_019814411.2;refGene_gn=RIMBP2;refGene_tx=XM_019814412.2;refGene_gn=RIMBP2;refGene_tx=XM_019814413.2;refGene_gn=RIMBP2;refGene_tx=XM_023241268.1;refGene_gn=RIMBP2;refGene_tx=XM_023241269.1
D3 2120370 2120522
refGene_gn=RIMBP2;refGene_tx=XM_019814410.2;refGene_gn=RIMBP2;refGene_tx=XM_019814411.2;refGene_gn=RIMBP2;refGene_tx=XM_019814412.2;refGene_gn=RIMBP2;refGene_tx=XM_019814413.2;refGene_gn=RIMBP2;refGene_tx=XM_023241268.1;refGene_gn=RIMBP2;refGene_tx=XM_023241269.1
D3 2122036 2122498
refGene_gn=RIMBP2;refGene_tx=XM_019814410.2;refGene_gn=RIMBP2;refGene_tx=XM_019814411.2;refGene_gn=RIMBP2;refGene_tx=XM_019814412.2;refGene_gn=RIMBP2;refGene_tx=XM_019814413.2;refGene_gn=RIMBP2;refGene_tx=XM_023241268.1;refGene_gn=RIMBP2;refGene_tx=XM_023241269.1
D3 2123678 2123822
refGene_gn=RIMBP2;refGene_tx=XM_019814410.2;refGene_gn=RIMBP2;refGene_tx=XM_019814411.2;refGene_gn=RIMBP2;refGene_tx=XM_019814412.2;refGene_gn=RIMBP2;refGene_tx=XM_019814413.2;refGene_gn=RIMBP2;refGene_tx=XM_023241268.1;refGene_gn=RIMBP2;refGene_tx=XM_023241269.1
D3 2130255 2130414
refGene_gn=RIMBP2;refGene_tx=XM_019814410.2;refGene_gn=RIMBP2;refGene_tx=XM_019814411.2;refGene_gn=RIMBP2;refGene_tx=XM_019814412.2;refGene_gn=RIMBP2;refGene_tx=XM_019814413.2;refGene_gn=RIMBP2;refGene_tx=XM_023241268.1;refGene_gn=RIMBP2;refGene_tx=XM_023241269.1
D3 2132925 2133612
refGene_gn=RIMBP2;refGene_tx=XM_019814410.2;refGene_gn=RIMBP2;refGene_tx=XM_019814411.2;refGene_gn=RIMBP2;refGene_tx=XM_019814412.2;refGene_gn=RIMBP2;refGene_tx=XM_019814413.2
D3 2135204 2135322
refGene_gn=RIMBP2;refGene_tx=XM_019814410.2;refGene_gn=RIMBP2;refGene_tx=XM_019814411.2;refGene_gn=RIMBP2;refGene_tx=XM_019814412.2;refGene_gn=RIMBP2;refGene_tx=XM_019814413.2;refGene_gn=RIMBP2;refGene_tx=XM_023241268.1;refGene_gn=RIMBP2;refGene_tx=XM_023241269.1
D3 2140275 2140460
refGene_gn=RIMBP2;refGene_tx=XM_019814410.2;refGene_gn=RIMBP2;refGene_tx=XM_019814411.2;refGene_gn=RIMBP2;refGene_tx=XM_019814412.2;refGene_gn=RIMBP2;refGene_tx=XM_019814413.2;refGene_gn=RIMBP2;refGene_tx=XM_023241268.1;refGene_gn=RIMBP2;refGene_tx=XM_023241269.1

D3 2141546 2141715
refGene_gn=RIMBP2;refGene_tx=XM_019814410.2;refGene_gn=RIMBP2;refGene_tx=XM_019814411.2;refGene_gn=RIMBP2;refGene_tx=XM_019814412.2;refGene_gn=RIMBP2;refGene_tx=XM_019814413.2;refGene_gn=RIMBP2;refGene_tx=XM_023241268.1;refGene_gn=RIMBP2;refGene_tx=XM_023241269.1
D3 2146181 2146282
refGene_gn=RIMBP2;refGene_tx=XM_019814410.2;refGene_gn=RIMBP2;refGene_tx=XM_019814411.2;refGene_gn=RIMBP2;refGene_tx=XM_019814412.2;refGene_gn=RIMBP2;refGene_tx=XM_019814413.2;refGene_gn=RIMBP2;refGene_tx=XM_023241268.1;refGene_gn=RIMBP2;refGene_tx=XM_023241269.1
D3 2147310 2147382
refGene_gn=RIMBP2;refGene_tx=XM_019814410.2;refGene_gn=RIMBP2;refGene_tx=XM_019814411.2;refGene_gn=RIMBP2;refGene_tx=XM_019814412.2;refGene_gn=RIMBP2;refGene_tx=XM_019814413.2;refGene_gn=RIMBP2;refGene_tx=XM_023241268.1;refGene_gn=RIMBP2;refGene_tx=XM_023241269.1
D3 2151365 2151500
refGene_gn=RIMBP2;refGene_tx=XM_019814410.2;refGene_gn=RIMBP2;refGene_tx=XM_019814411.2;refGene_gn=RIMBP2;refGene_tx=XM_019814412.2;refGene_gn=RIMBP2;refGene_tx=XM_019814413.2;refGene_gn=RIMBP2;refGene_tx=XM_023241268.1;refGene_gn=RIMBP2;refGene_tx=XM_023241269.1
D3 2151977 2152004
refGene_gn=RIMBP2;refGene_tx=XM_019814411.2;refGene_gn=RIMBP2;refGene_tx=XM_023241269.1
D3 2153543 2153732
refGene_gn=RIMBP2;refGene_tx=XM_019814410.2;refGene_gn=RIMBP2;refGene_tx=XM_019814412.2;refGene_gn=RIMBP2;refGene_tx=XM_019814413.2;refGene_gn=RIMBP2;refGene_tx=XM_023241268.1
D3 2166121 2166238 refGene_gn=PIWIL1;refGene_tx=XM_006938364.3
D3 2166373 2166521 refGene_gn=PIWIL1;refGene_tx=XM_006938364.3
D3 2169116 2169242 refGene_gn=PIWIL1;refGene_tx=XM_006938364.3
D3 2171523 2172022
refGene_gn=PIWIL1;refGene_tx=XM_006938362.4;refGene_gn=PIWIL1;refGene_tx=XM_006938363.4;refGene_gn=PIWIL1;refGene_tx=XM_019814415.2
D3 2172390 2172544
refGene_gn=PIWIL1;refGene_tx=XM_006938362.4;refGene_gn=PIWIL1;refGene_tx=XM_006938363.4;refGene_gn=PIWIL1;refGene_tx=XM_006938364.3;refGene_gn=PIWIL1;refGene_tx=XM_019814415.2
D3 2172638 2172709
refGene_gn=PIWIL1;refGene_tx=XM_006938362.4;refGene_gn=PIWIL1;refGene_tx=XM_006938363.4;refGene_gn=PIWIL1;refGene_tx=XM_006938364.3;refGene_gn=PIWIL1;refGene_tx=XM_019814415.2
D3 2173563 2173667
refGene_gn=PIWIL1;refGene_tx=XM_006938362.4;refGene_gn=PIWIL1;refGene_tx=XM_006938363.4;refGene_gn=PIWIL1;refGene_tx=XM_006938364.3;refGene_gn=PIWIL1;refGene_tx=XM_019814415.2
D3 2173776 2173977
refGene_gn=PIWIL1;refGene_tx=XM_006938362.4;refGene_gn=PIWIL1;refGene_tx=XM_006938363.4;refGene_gn=PIWIL1;refGene_tx=XM_006938364.3;refGene_gn=PIWIL1;refGene_tx=XM_019814415.2
D3 2177332 2177405

refGene_gn=PIWIL1;refGene_tx=XM_006938362.4;refGene_gn=PIWIL1;refGene_tx=XM_006938363.4;refGene_gn=PIWIL1;refGene_tx=XM_006938364.3;refGene_gn=PIWIL1;refGene_tx=XM_019814415.2
D3 2177777 2177965
refGene_gn=PIWIL1;refGene_tx=XM_006938362.4;refGene_gn=PIWIL1;refGene_tx=XM_006938363.4;refGene_gn=PIWIL1;refGene_tx=XM_006938364.3;refGene_gn=PIWIL1;refGene_tx=XM_019814415.2
D3 2179028 2179143
refGene_gn=PIWIL1;refGene_tx=XM_006938362.4;refGene_gn=PIWIL1;refGene_tx=XM_006938363.4;refGene_gn=PIWIL1;refGene_tx=XM_006938364.3;refGene_gn=PIWIL1;refGene_tx=XM_019814415.2
D3 2179648 2179766
refGene_gn=PIWIL1;refGene_tx=XM_006938362.4;refGene_gn=PIWIL1;refGene_tx=XM_006938363.4;refGene_gn=PIWIL1;refGene_tx=XM_006938364.3;refGene_gn=PIWIL1;refGene_tx=XM_019814415.2
D3 2179967 2180094
refGene_gn=PIWIL1;refGene_tx=XM_006938362.4;refGene_gn=PIWIL1;refGene_tx=XM_006938363.4;refGene_gn=PIWIL1;refGene_tx=XM_006938364.3;refGene_gn=PIWIL1;refGene_tx=XM_019814415.2
D3 2184113 2184225
refGene_gn=PIWIL1;refGene_tx=XM_006938362.4;refGene_gn=PIWIL1;refGene_tx=XM_006938363.4;refGene_gn=PIWIL1;refGene_tx=XM_006938364.3;refGene_gn=PIWIL1;refGene_tx=XM_019814415.2
D3 2184647 2184845
refGene_gn=PIWIL1;refGene_tx=XM_006938362.4;refGene_gn=PIWIL1;refGene_tx=XM_006938363.4;refGene_gn=PIWIL1;refGene_tx=XM_006938364.3;refGene_gn=PIWIL1;refGene_tx=XM_019814415.2
D3 2185485 2185566
refGene_gn=PIWIL1;refGene_tx=XM_006938362.4;refGene_gn=PIWIL1;refGene_tx=XM_006938363.4;refGene_gn=PIWIL1;refGene_tx=XM_006938364.3;refGene_gn=PIWIL1;refGene_tx=XM_019814415.2
D3 2185705 2185827
refGene_gn=PIWIL1;refGene_tx=XM_006938362.4;refGene_gn=PIWIL1;refGene_tx=XM_006938363.4;refGene_gn=PIWIL1;refGene_tx=XM_006938364.3;refGene_gn=PIWIL1;refGene_tx=XM_019814415.2
D3 2186403 2186618
refGene_gn=PIWIL1;refGene_tx=XM_006938362.4;refGene_gn=PIWIL1;refGene_tx=XM_006938363.4;refGene_gn=PIWIL1;refGene_tx=XM_006938364.3;refGene_gn=PIWIL1;refGene_tx=XM_019814415.2
D3 2187125 2187251
refGene_gn=PIWIL1;refGene_tx=XM_006938362.4;refGene_gn=PIWIL1;refGene_tx=XM_006938363.4;refGene_gn=PIWIL1;refGene_tx=XM_006938364.3;refGene_gn=PIWIL1;refGene_tx=XM_019814415.2
D3 2191924 2192036
refGene_gn=PIWIL1;refGene_tx=XM_006938362.4;refGene_gn=PIWIL1;refGene_tx=XM_006938363.4;refGene_gn=PIWIL1;refGene_tx=XM_006938364.3;refGene_gn=PIWIL1;refGene_tx=XM_019814415.2
D3 2192585 2192663
refGene_gn=PIWIL1;refGene_tx=XM_006938362.4;refGene_gn=PIWIL1;refGene_tx=XM_006938363.4;refGene_gn=PIWIL1;refGene_tx=XM_006938364.3;refGene_

gn=PIWIL1;refGene_tx=XM_019814415.2
D3	2302826	2304572	refGene_gn=FZD10;refGene_tx=XM_023241270.1
D3	2463239	2463318
refGene_gn=TMEM132D;refGene_tx=XM_023241271.1;refGene_gn=TMEM132D;refGene_tx=XM_023241272.1
D3	2601630	2601673	refGene_gn=TMEM132D;refGene_tx=XM_019814417.2
D3	2615862	2616754
refGene_gn=TMEM132D;refGene_tx=XM_019814417.2;refGene_gn=TMEM132D;refGene_tx=XM_023241271.1;refGene_gn=TMEM132D;refGene_tx=XM_023241272.1
D3	2728846	2728993
refGene_gn=TMEM132D;refGene_tx=XM_019814417.2;refGene_gn=TMEM132D;refGene_tx=XM_023241271.1
D3	2853960	2854144
refGene_gn=TMEM132D;refGene_tx=XM_019814417.2;refGene_gn=TMEM132D;refGene_tx=XM_023241271.1;refGene_gn=TMEM132D;refGene_tx=XM_023241272.1
D3	2946751	2946895
refGene_gn=TMEM132D;refGene_tx=XM_019814417.2;refGene_gn=TMEM132D;refGene_tx=XM_023241271.1;refGene_gn=TMEM132D;refGene_tx=XM_023241272.1
D3	3021221	3021427
refGene_gn=TMEM132D;refGene_tx=XM_019814417.2;refGene_gn=TMEM132D;refGene_tx=XM_023241271.1;refGene_gn=TMEM132D;refGene_tx=XM_023241272.1
D3	3023615	3023892
refGene_gn=TMEM132D;refGene_tx=XM_019814417.2;refGene_gn=TMEM132D;refGene_tx=XM_023241271.1;refGene_gn=TMEM132D;refGene_tx=XM_023241272.1
D3	3025977	3026169
refGene_gn=TMEM132D;refGene_tx=XM_019814417.2;refGene_gn=TMEM132D;refGene_tx=XM_023241271.1;refGene_gn=TMEM132D;refGene_tx=XM_023241272.1
D3	3029099	3030281
refGene_gn=TMEM132D;refGene_tx=XM_019814417.2;refGene_gn=TMEM132D;refGene_tx=XM_023241271.1;refGene_gn=TMEM132D;refGene_tx=XM_023241272.1
D3	3079647	3079809
refGene_gn=GLT1D1;refGene_tx=XM_006938365.3;refGene_gn=GLT1D1;refGene_tx=XM_023241275.1
D3	3088210	3088309
refGene_gn=GLT1D1;refGene_tx=XM_006938365.3;refGene_gn=GLT1D1;refGene_tx=XM_023241275.1
D3	3098429	3098550
refGene_gn=GLT1D1;refGene_tx=XM_006938365.3;refGene_gn=GLT1D1;refGene_tx=XM_023241275.1
D3	3099084	3099128
refGene_gn=GLT1D1;refGene_tx=XM_006938365.3;refGene_gn=GLT1D1;refGene_tx=XM_023241275.1
D3	3112171	3112221
refGene_gn=GLT1D1;refGene_tx=XM_006938365.3;refGene_gn=GLT1D1;refGene_tx=XM_023241275.1
D3	3113645	3113741
refGene_gn=GLT1D1;refGene_tx=XM_006938365.3;refGene_gn=GLT1D1;refGene_tx=XM_023241275.1
D3	3121610	3121667
refGene_gn=GLT1D1;refGene_tx=XM_006938365.3;refGene_gn=GLT1D1;refGene_

tx=XM_023241275.1
D3	3123639	3123676
refGene_gn=GLT1D1;refGene_tx=XM_006938365.3;refGene_gn=GLT1D1;refGene_tx=XM_023241275.1
D3	3135355	3135407
refGene_gn=GLT1D1;refGene_tx=XM_006938365.3;refGene_gn=GLT1D1;refGene_tx=XM_023241275.1
D3	3139348	3139454
refGene_gn=GLT1D1;refGene_tx=XM_006938365.3;refGene_gn=GLT1D1;refGene_tx=XM_023241275.1
D3	3150339	3150488
refGene_gn=GLT1D1;refGene_tx=XM_006938365.3;refGene_gn=GLT1D1;refGene_tx=XM_023241275.1
D3	3163148	3163216	refGene_gn=GLT1D1;refGene_tx=XM_023241275.1
D3	3172739	3172771	refGene_gn=GLT1D1;refGene_tx=XM_006938365.3
D3	3173837	3174362
refGene_gn=SLC15A4;refGene_tx=XM_023241273.1;refGene_gn=SLC15A4;refGene_tx=XM_023241274.1
D3	3180503	3180799
refGene_gn=SLC15A4;refGene_tx=XM_023241273.1;refGene_gn=SLC15A4;refGene_tx=XM_023241274.1
D3	3184786	3184955
refGene_gn=SLC15A4;refGene_tx=XM_023241273.1;refGene_gn=SLC15A4;refGene_tx=XM_023241274.1
D3	3185328	3185406
refGene_gn=SLC15A4;refGene_tx=XM_023241273.1;refGene_gn=SLC15A4;refGene_tx=XM_023241274.1
D3	3185809	3185978
refGene_gn=SLC15A4;refGene_tx=XM_023241273.1;refGene_gn=SLC15A4;refGene_tx=XM_023241274.1
D3	3193921	3194077
refGene_gn=SLC15A4;refGene_tx=XM_023241273.1;refGene_gn=SLC15A4;refGene_tx=XM_023241274.1
D3	3195148	3195307
refGene_gn=SLC15A4;refGene_tx=XM_023241273.1;refGene_gn=SLC15A4;refGene_tx=XM_023241274.1
D3	3199875	3200024	refGene_gn=SLC15A4;refGene_tx=XM_023241274.1
D3	3255254	3255475	refGene_gn=SLC15A4;refGene_tx=XM_023241273.1
D3	3261561	3262749
refGene_gn=TMEM132C;refGene_tx=XM_019814428.1;refGene_gn=TMEM132C;refGene_tx=XM_023241276.1
D3	3267989	3268181
refGene_gn=TMEM132C;refGene_tx=XM_019814428.1;refGene_gn=TMEM132C;refGene_tx=XM_023241276.1
D3	3269171	3269445
refGene_gn=TMEM132C;refGene_tx=XM_019814428.1;refGene_gn=TMEM132C;refGene_tx=XM_023241276.1
D3	3270829	3271035
refGene_gn=TMEM132C;refGene_tx=XM_019814428.1;refGene_gn=TMEM132C;refGene_tx=XM_023241276.1

D3 3291799 3291943
refGene_gn=TMEM132C;refGene_tx=XM_019814428.1;refGene_gn=TMEM132C;refGene_tx=XM_023241276.1
D3 3332916 3333100
refGene_gn=TMEM132C;refGene_tx=XM_019814428.1;refGene_gn=TMEM132C;refGene_tx=XM_023241276.1
D3 3378691 3378838 refGene_gn=TMEM132C;refGene_tx=XM_023241276.1
D3 3466854 3467746 refGene_gn=TMEM132C;refGene_tx=XM_023241276.1
D3 3574673 3574758 refGene_gn=TMEM132C;refGene_tx=XM_023241276.1
D3 4061008 4061906
refGene_gn=LOC102900410;refGene_tx=XM_023242150.1
D3 4062832 4063020
refGene_gn=LOC102900410;refGene_tx=XM_023242150.1
D3 4091663 4091875
refGene_gn=LOC102900410;refGene_tx=XM_023242150.1
D3 4092895 4092972
refGene_gn=LOC102900410;refGene_tx=XM_023242150.1
D3 4095710 4095782
refGene_gn=LOC102900410;refGene_tx=XM_023242150.1
D3 4095922 4095975
refGene_gn=LOC102900410;refGene_tx=XM_023242150.1
D3 4852847 4853996
refGene_gn=LOC101082698;refGene_tx=XM_023241277.1;refGene_gn=LOC101082698;refGene_tx=XM_023241278.1
D3 4854920 4855112
refGene_gn=LOC101082698;refGene_tx=XM_023241277.1;refGene_gn=LOC101082698;refGene_tx=XM_023241278.1
D3 4857143 4857414
refGene_gn=LOC101082698;refGene_tx=XM_023241277.1;refGene_gn=LOC101082698;refGene_tx=XM_023241278.1
D3 4860706 4860912
refGene_gn=LOC101082698;refGene_tx=XM_023241277.1;refGene_gn=LOC101082698;refGene_tx=XM_023241278.1
D3 4892071 4892215
refGene_gn=LOC101082698;refGene_tx=XM_023241277.1;refGene_gn=LOC101082698;refGene_tx=XM_023241278.1
D3 4912105 4912108
refGene_gn=LOC101082698;refGene_tx=XM_023241278.1
D3 4941997 4942184
refGene_gn=LOC101082698;refGene_tx=XM_023241277.1
D3 5027394 5027541
refGene_gn=LOC101082698;refGene_tx=XM_023241277.1
D3 5071987 5072879
refGene_gn=LOC101082698;refGene_tx=XM_023241277.1
D3 5222703 5222770
refGene_gn=LOC101082698;refGene_tx=XM_023241277.1
D3 5258370 5258555
refGene_gn=AACS;refGene_tx=XM_023241279.1;refGene_gn=AACS;refGene_tx=XM_023241280.1
D3 5260970 5261173

refGene_gn=AACS;refGene_tx=XM_023241279.1;refGene_gn=AACS;refGene_tx=XM_023241280.1
D3 5261923 5261982
refGene_gn=AACS;refGene_tx=XM_023241279.1;refGene_gn=AACS;refGene_tx=XM_023241280.1
D3 5262459 5262529
refGene_gn=AACS;refGene_tx=XM_023241279.1;refGene_gn=AACS;refGene_tx=XM_023241280.1
D3 5266229 5266355
refGene_gn=AACS;refGene_tx=XM_023241279.1;refGene_gn=AACS;refGene_tx=XM_023241280.1
D3 5268893 5269007
refGene_gn=AACS;refGene_tx=XM_023241279.1;refGene_gn=AACS;refGene_tx=XM_023241280.1
D3 5271615 5271738
refGene_gn=AACS;refGene_tx=XM_023241279.1;refGene_gn=AACS;refGene_tx=XM_023241280.1
D3 5271888 5271953
refGene_gn=AACS;refGene_tx=XM_023241279.1;refGene_gn=AACS;refGene_tx=XM_023241280.1
D3 5274809 5274934
refGene_gn=AACS;refGene_tx=XM_023241279.1;refGene_gn=AACS;refGene_tx=XM_023241280.1
D3 5277787 5277868
refGene_gn=AACS;refGene_tx=XM_023241279.1;refGene_gn=AACS;refGene_tx=XM_023241280.1
D3 5283520 5283668
refGene_gn=AACS;refGene_tx=XM_023241279.1;refGene_gn=AACS;refGene_tx=XM_023241280.1
D3 5287999 5288081
refGene_gn=AACS;refGene_tx=XM_023241279.1;refGene_gn=AACS;refGene_tx=XM_023241280.1
D3 5288247 5288362
refGene_gn=AACS;refGene_tx=XM_023241279.1;refGene_gn=AACS;refGene_tx=XM_023241280.1
D3 5292164 5292262
refGene_gn=AACS;refGene_tx=XM_023241279.1;refGene_gn=AACS;refGene_tx=XM_023241280.1
D3 5295259 5295373
refGene_gn=AACS;refGene_tx=XM_023241279.1;refGene_gn=AACS;refGene_tx=XM_023241280.1
D3 5301372 5301493
refGene_gn=AACS;refGene_tx=XM_023241279.1;refGene_gn=AACS;refGene_tx=XM_023241280.1
D3 5303118 5303222
refGene_gn=AACS;refGene_tx=XM_023241279.1;refGene_gn=AACS;refGene_tx=XM_023241280.1
D3 5309390 5309523
refGene_gn=AACS;refGene_tx=XM_023241279.1;refGene_gn=AACS;refGene_tx=XM_023241280.1

D3 5329900 5330322 refGene_gn=BRI3BP;refGene_tx=XM_023241281.1
D3 5342471 5342574 refGene_gn=BRI3BP;refGene_tx=XM_023241281.1
D3 5354966 5355176 refGene_gn=BRI3BP;refGene_tx=XM_023241281.1
D3 5358306 5358412 refGene_gn=DHX37;refGene_tx=XM_023241282.1
D3 5361925 5362095 refGene_gn=DHX37;refGene_tx=XM_023241282.1
D3 5364622 5364735 refGene_gn=DHX37;refGene_tx=XM_023241282.1
D3 5366188 5366519 refGene_gn=DHX37;refGene_tx=XM_023241282.1
D3 5368389 5368538 refGene_gn=DHX37;refGene_tx=XM_023241282.1
D3 5369624 5369717 refGene_gn=DHX37;refGene_tx=XM_023241282.1
D3 5371366 5371463 refGene_gn=DHX37;refGene_tx=XM_023241282.1
D3 5372336 5372450 refGene_gn=DHX37;refGene_tx=XM_023241282.1
D3 5372630 5372732 refGene_gn=DHX37;refGene_tx=XM_023241282.1
D3 5372976 5373091 refGene_gn=DHX37;refGene_tx=XM_023241282.1
D3 5374292 5374388 refGene_gn=DHX37;refGene_tx=XM_023241282.1
D3 5374571 5374654 refGene_gn=DHX37;refGene_tx=XM_023241282.1
D3 5375355 5375506 refGene_gn=DHX37;refGene_tx=XM_023241282.1
D3 5376120 5376197 refGene_gn=DHX37;refGene_tx=XM_023241282.1
D3 5376531 5376764 refGene_gn=DHX37;refGene_tx=XM_023241282.1
D3 5378259 5378371 refGene_gn=DHX37;refGene_tx=XM_023241282.1
D3 5379792 5379899 refGene_gn=DHX37;refGene_tx=XM_023241282.1
D3 5379989 5380178 refGene_gn=DHX37;refGene_tx=XM_023241282.1
D3 5381155 5381280 refGene_gn=DHX37;refGene_tx=XM_023241282.1
D3 5381434 5381551 refGene_gn=DHX37;refGene_tx=XM_023241282.1
D3 5382809 5382982 refGene_gn=DHX37;refGene_tx=XM_023241282.1
D3 5383660 5383775 refGene_gn=DHX37;refGene_tx=XM_023241282.1
D3 5383923 5384061 refGene_gn=DHX37;refGene_tx=XM_023241282.1
D3 5384251 5384346 refGene_gn=DHX37;refGene_tx=XM_023241282.1
D3 5384434 5384508 refGene_gn=DHX37;refGene_tx=XM_023241282.1
D3 5385719 5385817 refGene_gn=DHX37;refGene_tx=XM_023241282.1
D3 5385973 5386059 refGene_gn=DHX37;refGene_tx=XM_023241282.1
D3 5403116 5403493 refGene_gn=UBC;refGene_tx=XM_023241283.1
D3 5433400 5433526
refGene_gn=SCARB1;refGene_tx=XM_023241284.1;refGene_gn=SCARB1;refGene_tx=XM_023241285.1;refGene_gn=SCARB1;refGene_tx=XM_023241286.1;refGene_gn=SCARB1;refGene_tx=XM_023241287.1;refGene_gn=SCARB1;refGene_tx=XM_023241288.1;refGene_gn=SCARB1;refGene_tx=XM_023241289.1
D3 5467764 5467922
refGene_gn=SCARB1;refGene_tx=XM_023241284.1;refGene_gn=SCARB1;refGene_tx=XM_023241285.1;refGene_gn=SCARB1;refGene_tx=XM_023241286.1;refGene_gn=SCARB1;refGene_tx=XM_023241287.1;refGene_gn=SCARB1;refGene_tx=XM_023241288.1;refGene_gn=SCARB1;refGene_tx=XM_023241289.1
D3 5469579 5469721
refGene_gn=SCARB1;refGene_tx=XM_023241284.1;refGene_gn=SCARB1;refGene_tx=XM_023241285.1;refGene_gn=SCARB1;refGene_tx=XM_023241286.1;refGene_gn=SCARB1;refGene_tx=XM_023241287.1;refGene_gn=SCARB1;refGene_tx=XM_023241288.1;refGene_gn=SCARB1;refGene_tx=XM_023241289.1
D3 5471924 5472128
refGene_gn=SCARB1;refGene_tx=XM_023241284.1;refGene_gn=SCARB1;refGene_tx=XM_023241285.1;refGene_gn=SCARB1;refGene_tx=XM_023241286.1;refGene_gn=SCARB1;refGene_tx=XM_023241287.1;refGene_gn=SCARB1;refGene_tx=XM_02

3241288.1;refGene_gn=SCARB1;refGene_tx=XM_023241289.1
D3 5474027 5474123
refGene_gn=SCARB1;refGene_tx=XM_023241284.1;refGene_gn=SCARB1;refGene_tx=XM_023241285.1;refGene_gn=SCARB1;refGene_tx=XM_023241286.1;refGene_gn=SCARB1;refGene_tx=XM_023241287.1;refGene_gn=SCARB1;refGene_tx=XM_023241288.1;refGene_gn=SCARB1;refGene_tx=XM_023241289.1
D3 5475596 5475712
refGene_gn=SCARB1;refGene_tx=XM_023241284.1;refGene_gn=SCARB1;refGene_tx=XM_023241285.1;refGene_gn=SCARB1;refGene_tx=XM_023241286.1;refGene_gn=SCARB1;refGene_tx=XM_023241287.1;refGene_gn=SCARB1;refGene_tx=XM_023241288.1;refGene_gn=SCARB1;refGene_tx=XM_023241289.1
D3 5478047 5478214
refGene_gn=SCARB1;refGene_tx=XM_023241284.1;refGene_gn=SCARB1;refGene_tx=XM_023241285.1;refGene_gn=SCARB1;refGene_tx=XM_023241286.1;refGene_gn=SCARB1;refGene_tx=XM_023241287.1;refGene_gn=SCARB1;refGene_tx=XM_023241288.1;refGene_gn=SCARB1;refGene_tx=XM_023241289.1
D3 5483390 5483509
refGene_gn=SCARB1;refGene_tx=XM_023241284.1;refGene_gn=SCARB1;refGene_tx=XM_023241285.1;refGene_gn=SCARB1;refGene_tx=XM_023241286.1;refGene_gn=SCARB1;refGene_tx=XM_023241287.1;refGene_gn=SCARB1;refGene_tx=XM_023241288.1;refGene_gn=SCARB1;refGene_tx=XM_023241289.1
D3 5484427 5484538 refGene_gn=SCARB1;refGene_tx=XM_023241288.1
D3 5486519 5486593
refGene_gn=SCARB1;refGene_tx=XM_023241284.1;refGene_gn=SCARB1;refGene_tx=XM_023241285.1;refGene_gn=SCARB1;refGene_tx=XM_023241286.1;refGene_gn=SCARB1;refGene_tx=XM_023241287.1;refGene_gn=SCARB1;refGene_tx=XM_023241289.1
D3 5486771 5486838 refGene_gn=SCARB1;refGene_tx=XM_023241287.1
D3 5489674 5489726
refGene_gn=SCARB1;refGene_tx=XM_023241284.1;refGene_gn=SCARB1;refGene_tx=XM_023241285.1;refGene_gn=SCARB1;refGene_tx=XM_023241286.1;refGene_gn=SCARB1;refGene_tx=XM_023241289.1
D3 5490818 5490965
refGene_gn=SCARB1;refGene_tx=XM_023241284.1;refGene_gn=SCARB1;refGene_tx=XM_023241285.1;refGene_gn=SCARB1;refGene_tx=XM_023241286.1;refGene_gn=SCARB1;refGene_tx=XM_023241289.1
D3 5494143 5494272
refGene_gn=SCARB1;refGene_tx=XM_023241284.1;refGene_gn=SCARB1;refGene_tx=XM_023241289.1
D3 5498122 5498242 refGene_gn=SCARB1;refGene_tx=XM_023241285.1
D3 5517971 5518034 refGene_gn=SCARB1;refGene_tx=XM_023241286.1
D3 5573962 5574366
refGene_gn=LOC109493140;refGene_tx=XM_019814440.2;refGene_gn=LOC109493140;refGene_tx=XM_019814443.2
D3 5576380 5576481
refGene_gn=LOC109493140;refGene_tx=XM_019814443.2
D3 5579145 5579317
refGene_gn=LOC109493140;refGene_tx=XM_019814440.2;refGene_gn=LOC109493140;refGene_tx=XM_019814443.2
D3 5579841 5579973

refGene_gn=LOC109493140;refGene_tx=XM_019814440.2;refGene_gn=LOC109493140;refGene_tx=XM_019814443.2
D3 5716506 5716611
refGene_gn=NCOR2;refGene_tx=XM_023241290.1;refGene_gn=NCOR2;refGene_tx=XM_023241291.1;refGene_gn=NCOR2;refGene_tx=XM_023241292.1;refGene_gn=NCOR2;refGene_tx=XM_023241293.1;refGene_gn=NCOR2;refGene_tx=XM_023241294.1;refGene_gn=NCOR2;refGene_tx=XM_023241295.1;refGene_gn=NCOR2;refGene_tx=XM_023241296.1;refGene_gn=NCOR2;refGene_tx=XM_023241297.1;refGene_gn=NCOR2;refGene_tx=XM_023241298.1;refGene_gn=NCOR2;refGene_tx=XM_023241299.1;refGene_gn=NCOR2;refGene_tx=XM_023241300.1;refGene_gn=NCOR2;refGene_tx=XM_023241301.1;refGene_gn=NCOR2;refGene_tx=XM_023241302.1;refGene_gn=NCOR2;refGene_tx=XM_023241303.1;refGene_gn=NCOR2;refGene_tx=XM_023241304.1;refGene_gn=NCOR2;refGene_tx=XM_023241305.1;refGene_gn=NCOR2;refGene_tx=XM_023241306.1;refGene_gn=NCOR2;refGene_tx=XM_023241307.1;refGene_gn=NCOR2;refGene_tx=XM_023241308.1;refGene_gn=NCOR2;refGene_tx=XM_023241309.1;refGene_gn=NCOR2;refGene_tx=XM_023241310.1;refGene_gn=NCOR2;refGene_tx=XM_023241311.1;refGene_gn=NCOR2;refGene_tx=XM_023241312.1
D3 5724158 5724286
refGene_gn=NCOR2;refGene_tx=XM_023241290.1;refGene_gn=NCOR2;refGene_tx=XM_023241291.1;refGene_gn=NCOR2;refGene_tx=XM_023241292.1;refGene_gn=NCOR2;refGene_tx=XM_023241293.1;refGene_gn=NCOR2;refGene_tx=XM_023241294.1;refGene_gn=NCOR2;refGene_tx=XM_023241295.1;refGene_gn=NCOR2;refGene_tx=XM_023241296.1;refGene_gn=NCOR2;refGene_tx=XM_023241297.1;refGene_gn=NCOR2;refGene_tx=XM_023241298.1;refGene_gn=NCOR2;refGene_tx=XM_023241299.1;refGene_gn=NCOR2;refGene_tx=XM_023241300.1;refGene_gn=NCOR2;refGene_tx=XM_023241301.1;refGene_gn=NCOR2;refGene_tx=XM_023241302.1;refGene_gn=NCOR2;refGene_tx=XM_023241303.1;refGene_gn=NCOR2;refGene_tx=XM_023241304.1;refGene_gn=NCOR2;refGene_tx=XM_023241305.1;refGene_gn=NCOR2;refGene_tx=XM_023241306.1;refGene_gn=NCOR2;refGene_tx=XM_023241307.1;refGene_gn=NCOR2;refGene_tx=XM_023241308.1;refGene_gn=NCOR2;refGene_tx=XM_023241309.1;refGene_gn=NCOR2;refGene_tx=XM_023241310.1;refGene_gn=NCOR2;refGene_tx=XM_023241311.1;refGene_gn=NCOR2;refGene_tx=XM_023241312.1
D3 5725854 5726032
refGene_gn=NCOR2;refGene_tx=XM_023241290.1;refGene_gn=NCOR2;refGene_tx=XM_023241291.1;refGene_gn=NCOR2;refGene_tx=XM_023241292.1;refGene_gn=NCOR2;refGene_tx=XM_023241293.1;refGene_gn=NCOR2;refGene_tx=XM_023241294.1;refGene_gn=NCOR2;refGene_tx=XM_023241295.1;refGene_gn=NCOR2;refGene_tx=XM_023241296.1;refGene_gn=NCOR2;refGene_tx=XM_023241297.1;refGene_gn=NCOR2;refGene_tx=XM_023241298.1;refGene_gn=NCOR2;refGene_tx=XM_023241299.1;refGene_gn=NCOR2;refGene_tx=XM_023241300.1;refGene_gn=NCOR2;refGene_tx=XM_023241301.1;refGene_gn=NCOR2;refGene_tx=XM_023241302.1;refGene_gn=NCOR2;refGene_tx=XM_023241303.1;refGene_gn=NCOR2;refGene_tx=XM_023241304.1;refGene_gn=NCOR2;refGene_tx=XM_023241305.1;refGene_gn=NCOR2;refGene_tx=XM_023241306.1;refGene_gn=NCOR2;refGene_tx=XM_023241307.1;refGene_gn=NCOR2;refGene_tx=XM_023241308.1;refGene_gn=NCOR2;refGene_tx=XM_023241309.1;refGene_gn=NCOR2;refGene_tx=XM_023241310.1;refGene_gn=NCOR2;refGene_tx=XM_023241311.1;refGene_gn=NCOR2;refGene_tx=XM_023241312.1

D3	5732418	5732598
refGene_gn=NCOR2;refGene_tx=XM_023241290.1;refGene_gn=NCOR2;refGene_tx=XM_023241291.1;refGene_gn=NCOR2;refGene_tx=XM_023241292.1;refGene_gn=NCOR2;refGene_tx=XM_023241293.1;refGene_gn=NCOR2;refGene_tx=XM_023241294.1;refGene_gn=NCOR2;refGene_tx=XM_023241295.1;refGene_gn=NCOR2;refGene_tx=XM_023241296.1;refGene_gn=NCOR2;refGene_tx=XM_023241297.1;refGene_gn=NCOR2;refGene_tx=XM_023241298.1;refGene_gn=NCOR2;refGene_tx=XM_023241299.1;refGene_gn=NCOR2;refGene_tx=XM_023241300.1;refGene_gn=NCOR2;refGene_tx=XM_023241301.1;refGene_gn=NCOR2;refGene_tx=XM_023241302.1;refGene_gn=NCOR2;refGene_tx=XM_023241303.1;refGene_gn=NCOR2;refGene_tx=XM_023241304.1;refGene_gn=NCOR2;refGene_tx=XM_023241305.1;refGene_gn=NCOR2;refGene_tx=XM_023241306.1;refGene_gn=NCOR2;refGene_tx=XM_023241307.1;refGene_gn=NCOR2;refGene_tx=XM_023241308.1;refGene_gn=NCOR2;refGene_tx=XM_023241309.1;refGene_gn=NCOR2;refGene_tx=XM_023241310.1;refGene_gn=NCOR2;refGene_tx=XM_023241311.1;refGene_gn=NCOR2;refGene_tx=XM_023241312.1
D3	5736514	5736628
refGene_gn=NCOR2;refGene_tx=XM_023241290.1;refGene_gn=NCOR2;refGene_tx=XM_023241291.1;refGene_gn=NCOR2;refGene_tx=XM_023241292.1;refGene_gn=NCOR2;refGene_tx=XM_023241293.1;refGene_gn=NCOR2;refGene_tx=XM_023241294.1;refGene_gn=NCOR2;refGene_tx=XM_023241295.1;refGene_gn=NCOR2;refGene_tx=XM_023241296.1;refGene_gn=NCOR2;refGene_tx=XM_023241297.1;refGene_gn=NCOR2;refGene_tx=XM_023241298.1;refGene_gn=NCOR2;refGene_tx=XM_023241299.1;refGene_gn=NCOR2;refGene_tx=XM_023241300.1;refGene_gn=NCOR2;refGene_tx=XM_023241301.1;refGene_gn=NCOR2;refGene_tx=XM_023241302.1;refGene_gn=NCOR2;refGene_tx=XM_023241304.1;refGene_gn=NCOR2;refGene_tx=XM_023241305.1;refGene_gn=NCOR2;refGene_tx=XM_023241306.1;refGene_gn=NCOR2;refGene_tx=XM_023241307.1;refGene_gn=NCOR2;refGene_tx=XM_023241308.1;refGene_gn=NCOR2;refGene_tx=XM_023241309.1;refGene_gn=NCOR2;refGene_tx=XM_023241310.1;refGene_gn=NCOR2;refGene_tx=XM_023241311.1;refGene_gn=NCOR2;refGene_tx=XM_023241312.1
D3	5746526	5746583
refGene_gn=NCOR2;refGene_tx=XM_023241290.1;refGene_gn=NCOR2;refGene_tx=XM_023241291.1;refGene_gn=NCOR2;refGene_tx=XM_023241292.1;refGene_gn=NCOR2;refGene_tx=XM_023241293.1;refGene_gn=NCOR2;refGene_tx=XM_023241294.1;refGene_gn=NCOR2;refGene_tx=XM_023241295.1;refGene_gn=NCOR2;refGene_tx=XM_023241296.1;refGene_gn=NCOR2;refGene_tx=XM_023241297.1;refGene_gn=NCOR2;refGene_tx=XM_023241298.1;refGene_gn=NCOR2;refGene_tx=XM_023241299.1;refGene_gn=NCOR2;refGene_tx=XM_023241300.1;refGene_gn=NCOR2;refGene_tx=XM_023241301.1;refGene_gn=NCOR2;refGene_tx=XM_023241302.1;refGene_gn=NCOR2;refGene_tx=XM_023241303.1;refGene_gn=NCOR2;refGene_tx=XM_023241304.1;refGene_gn=NCOR2;refGene_tx=XM_023241305.1;refGene_gn=NCOR2;refGene_tx=XM_023241306.1;refGene_gn=NCOR2;refGene_tx=XM_023241307.1;refGene_gn=NCOR2;refGene_tx=XM_023241308.1;refGene_gn=NCOR2;refGene_tx=XM_023241309.1;refGene_gn=NCOR2;refGene_tx=XM_023241310.1;refGene_gn=NCOR2;refGene_tx=XM_023241311.1;refGene_gn=NCOR2;refGene_tx=XM_023241312.1
D3	5753769	5753822
refGene_gn=NCOR2;refGene_tx=XM_023241290.1;refGene_gn=NCOR2;refGene_tx=XM_023241291.1;refGene_gn=NCOR2;refGene_tx=XM_023241292.1;refGene_gn=

NCOR2;refGene_tx=XM_023241293.1;refGene_gn=NCOR2;refGene_tx=XM_0232412
94.1;refGene_gn=NCOR2;refGene_tx=XM_023241295.1;refGene_gn=NCOR2;refGe
ne_tx=XM_023241296.1;refGene_gn=NCOR2;refGene_tx=XM_023241297.1;refGen
e_gn=NCOR2;refGene_tx=XM_023241298.1;refGene_gn=NCOR2;refGene_tx=XM_02
3241299.1;refGene_gn=NCOR2;refGene_tx=XM_023241300.1;refGene_gn=NCOR2;
refGene_tx=XM_023241301.1;refGene_gn=NCOR2;refGene_tx=XM_023241302.1;r
efGene_gn=NCOR2;refGene_tx=XM_023241303.1;refGene_gn=NCOR2;refGene_tx=
XM_023241304.1;refGene_gn=NCOR2;refGene_tx=XM_023241305.1;refGene_gn=N
COR2;refGene_tx=XM_023241306.1;refGene_gn=NCOR2;refGene_tx=XM_02324130
7.1;refGene_gn=NCOR2;refGene_tx=XM_023241308.1;refGene_gn=NCOR2;refGen
e_tx=XM_023241309.1;refGene_gn=NCOR2;refGene_tx=XM_023241310.1;refGene
_gn=NCOR2;refGene_tx=XM_023241311.1;refGene_gn=NCOR2;refGene_tx=XM_023
241312.1
D3 5763248 5763315
refGene_gn=NCOR2;refGene_tx=XM_023241290.1;refGene_gn=NCOR2;refGene_tx
=XM_023241291.1;refGene_gn=NCOR2;refGene_tx=XM_023241292.1;refGene_gn=
NCOR2;refGene_tx=XM_023241293.1;refGene_gn=NCOR2;refGene_tx=XM_0232412
94.1;refGene_gn=NCOR2;refGene_tx=XM_023241295.1;refGene_gn=NCOR2;refGe
ne_tx=XM_023241296.1;refGene_gn=NCOR2;refGene_tx=XM_023241297.1;refGen
e_gn=NCOR2;refGene_tx=XM_023241298.1;refGene_gn=NCOR2;refGene_tx=XM_02
3241299.1;refGene_gn=NCOR2;refGene_tx=XM_023241300.1;refGene_gn=NCOR2;
refGene_tx=XM_023241301.1;refGene_gn=NCOR2;refGene_tx=XM_023241302.1;r
efGene_gn=NCOR2;refGene_tx=XM_023241303.1;refGene_gn=NCOR2;refGene_tx=
XM_023241304.1;refGene_gn=NCOR2;refGene_tx=XM_023241305.1;refGene_gn=N
COR2;refGene_tx=XM_023241306.1;refGene_gn=NCOR2;refGene_tx=XM_02324130
7.1;refGene_gn=NCOR2;refGene_tx=XM_023241308.1;refGene_gn=NCOR2;refGen
e_tx=XM_023241309.1;refGene_gn=NCOR2;refGene_tx=XM_023241310.1;refGene
_gn=NCOR2;refGene_tx=XM_023241311.1;refGene_gn=NCOR2;refGene_tx=XM_023
241312.1
D3 5769273 5769446
refGene_gn=NCOR2;refGene_tx=XM_023241290.1;refGene_gn=NCOR2;refGene_tx
=XM_023241291.1;refGene_gn=NCOR2;refGene_tx=XM_023241292.1;refGene_gn=
NCOR2;refGene_tx=XM_023241293.1;refGene_gn=NCOR2;refGene_tx=XM_0232412
94.1;refGene_gn=NCOR2;refGene_tx=XM_023241295.1;refGene_gn=NCOR2;refGe
ne_tx=XM_023241296.1;refGene_gn=NCOR2;refGene_tx=XM_023241297.1;refGen
e_gn=NCOR2;refGene_tx=XM_023241298.1;refGene_gn=NCOR2;refGene_tx=XM_02
3241299.1;refGene_gn=NCOR2;refGene_tx=XM_023241300.1;refGene_gn=NCOR2;
refGene_tx=XM_023241301.1;refGene_gn=NCOR2;refGene_tx=XM_023241302.1;r
efGene_gn=NCOR2;refGene_tx=XM_023241303.1;refGene_gn=NCOR2;refGene_tx=
XM_023241304.1;refGene_gn=NCOR2;refGene_tx=XM_023241305.1;refGene_gn=N
COR2;refGene_tx=XM_023241306.1;refGene_gn=NCOR2;refGene_tx=XM_02324130
7.1;refGene_gn=NCOR2;refGene_tx=XM_023241308.1;refGene_gn=NCOR2;refGen
e_tx=XM_023241309.1;refGene_gn=NCOR2;refGene_tx=XM_023241310.1;refGene
_gn=NCOR2;refGene_tx=XM_023241311.1;refGene_gn=NCOR2;refGene_tx=XM_023
241312.1
D3 5770579 5770673
refGene_gn=NCOR2;refGene_tx=XM_023241290.1;refGene_gn=NCOR2;refGene_tx
=XM_023241291.1;refGene_gn=NCOR2;refGene_tx=XM_023241292.1;refGene_gn=
NCOR2;refGene_tx=XM_023241293.1;refGene_gn=NCOR2;refGene_tx=XM_0232412
94.1;refGene_gn=NCOR2;refGene_tx=XM_023241295.1;refGene_gn=NCOR2;refGe

ne_tx=XM_023241296.1;refGene_gn=NCOR2;refGene_tx=XM_023241297.1;refGene_gn=NCOR2;refGene_tx=XM_023241298.1;refGene_gn=NCOR2;refGene_tx=XM_023241299.1;refGene_gn=NCOR2;refGene_tx=XM_023241300.1;refGene_gn=NCOR2;refGene_tx=XM_023241301.1;refGene_gn=NCOR2;refGene_tx=XM_023241302.1;refGene_gn=NCOR2;refGene_tx=XM_023241303.1;refGene_gn=NCOR2;refGene_tx=XM_023241304.1;refGene_gn=NCOR2;refGene_tx=XM_023241305.1;refGene_gn=NCOR2;refGene_tx=XM_023241306.1;refGene_gn=NCOR2;refGene_tx=XM_023241307.1;refGene_gn=NCOR2;refGene_tx=XM_023241308.1;refGene_gn=NCOR2;refGene_tx=XM_023241309.1;refGene_gn=NCOR2;refGene_tx=XM_023241310.1;refGene_gn=NCOR2;refGene_tx=XM_023241311.1;refGene_gn=NCOR2;refGene_tx=XM_023241312.1

D3 5773421 5773600 refGene_gn=NCOR2;refGene_tx=XM_023241290.1;refGene_gn=NCOR2;refGene_tx=XM_023241291.1;refGene_gn=NCOR2;refGene_tx=XM_023241292.1;refGene_gn=NCOR2;refGene_tx=XM_023241293.1;refGene_gn=NCOR2;refGene_tx=XM_023241294.1;refGene_gn=NCOR2;refGene_tx=XM_023241295.1;refGene_gn=NCOR2;refGene_tx=XM_023241296.1;refGene_gn=NCOR2;refGene_tx=XM_023241297.1;refGene_gn=NCOR2;refGene_tx=XM_023241298.1;refGene_gn=NCOR2;refGene_tx=XM_023241299.1;refGene_gn=NCOR2;refGene_tx=XM_023241300.1;refGene_gn=NCOR2;refGene_tx=XM_023241301.1;refGene_gn=NCOR2;refGene_tx=XM_023241302.1;refGene_gn=NCOR2;refGene_tx=XM_023241303.1;refGene_gn=NCOR2;refGene_tx=XM_023241304.1;refGene_gn=NCOR2;refGene_tx=XM_023241305.1;refGene_gn=NCOR2;refGene_tx=XM_023241306.1;refGene_gn=NCOR2;refGene_tx=XM_023241307.1;refGene_gn=NCOR2;refGene_tx=XM_023241308.1;refGene_gn=NCOR2;refGene_tx=XM_023241309.1;refGene_gn=NCOR2;refGene_tx=XM_023241310.1;refGene_gn=NCOR2;refGene_tx=XM_023241311.1;refGene_gn=NCOR2;refGene_tx=XM_023241312.1

D3 5776980 5777035 refGene_gn=NCOR2;refGene_tx=XM_023241290.1;refGene_gn=NCOR2;refGene_tx=XM_023241291.1;refGene_gn=NCOR2;refGene_tx=XM_023241292.1;refGene_gn=NCOR2;refGene_tx=XM_023241293.1;refGene_gn=NCOR2;refGene_tx=XM_023241294.1;refGene_gn=NCOR2;refGene_tx=XM_023241295.1;refGene_gn=NCOR2;refGene_tx=XM_023241296.1;refGene_gn=NCOR2;refGene_tx=XM_023241297.1;refGene_gn=NCOR2;refGene_tx=XM_023241298.1;refGene_gn=NCOR2;refGene_tx=XM_023241299.1;refGene_gn=NCOR2;refGene_tx=XM_023241300.1;refGene_gn=NCOR2;refGene_tx=XM_023241301.1;refGene_gn=NCOR2;refGene_tx=XM_023241302.1;refGene_gn=NCOR2;refGene_tx=XM_023241303.1;refGene_gn=NCOR2;refGene_tx=XM_023241304.1;refGene_gn=NCOR2;refGene_tx=XM_023241305.1;refGene_gn=NCOR2;refGene_tx=XM_023241306.1;refGene_gn=NCOR2;refGene_tx=XM_023241307.1;refGene_gn=NCOR2;refGene_tx=XM_023241308.1;refGene_gn=NCOR2;refGene_tx=XM_023241309.1;refGene_gn=NCOR2;refGene_tx=XM_023241310.1;refGene_gn=NCOR2;refGene_tx=XM_023241311.1;refGene_gn=NCOR2;refGene_tx=XM_023241312.1

D3 5779347 5779446 refGene_gn=NCOR2;refGene_tx=XM_023241290.1;refGene_gn=NCOR2;refGene_tx=XM_023241291.1;refGene_gn=NCOR2;refGene_tx=XM_023241292.1;refGene_gn=NCOR2;refGene_tx=XM_023241293.1;refGene_gn=NCOR2;refGene_tx=XM_023241294.1;refGene_gn=NCOR2;refGene_tx=XM_023241295.1;refGene_gn=NCOR2;refGene_tx=XM_023241296.1;refGene_gn=NCOR2;refGene_tx=XM_023241297.1;refGene_gn=NCOR2;refGene_tx=XM_023241298.1;refGene_gn=NCOR2;refGene_tx=XM_02

3241299.1;refGene_gn=NCOR2;refGene_tx=XM_023241300.1;refGene_gn=NCOR2;
refGene_tx=XM_023241301.1;refGene_gn=NCOR2;refGene_tx=XM_023241302.1;r
efGene_gn=NCOR2;refGene_tx=XM_023241303.1;refGene_gn=NCOR2;refGene_tx=
XM_023241304.1;refGene_gn=NCOR2;refGene_tx=XM_023241305.1;refGene_gn=N
COR2;refGene_tx=XM_023241306.1;refGene_gn=NCOR2;refGene_tx=XM_02324130
7.1;refGene_gn=NCOR2;refGene_tx=XM_023241308.1;refGene_gn=NCOR2;refGen
e_tx=XM_023241309.1;refGene_gn=NCOR2;refGene_tx=XM_023241310.1;refGene
_gn=NCOR2;refGene_tx=XM_023241311.1;refGene_gn=NCOR2;refGene_tx=XM_023
241312.1
D3 5794656 5794802
refGene_gn=NCOR2;refGene_tx=XM_023241290.1;refGene_gn=NCOR2;refGene_tx
=XM_023241291.1;refGene_gn=NCOR2;refGene_tx=XM_023241292.1;refGene_gn=
NCOR2;refGene_tx=XM_023241293.1;refGene_gn=NCOR2;refGene_tx=XM_0232412
94.1;refGene_gn=NCOR2;refGene_tx=XM_023241295.1;refGene_gn=NCOR2;refGe
ne_tx=XM_023241296.1;refGene_gn=NCOR2;refGene_tx=XM_023241297.1;refGen
e_gn=NCOR2;refGene_tx=XM_023241298.1;refGene_gn=NCOR2;refGene_tx=XM_02
3241299.1;refGene_gn=NCOR2;refGene_tx=XM_023241300.1;refGene_gn=NCOR2;
refGene_tx=XM_023241301.1;refGene_gn=NCOR2;refGene_tx=XM_023241302.1;r
efGene_gn=NCOR2;refGene_tx=XM_023241303.1;refGene_gn=NCOR2;refGene_tx=
XM_023241304.1;refGene_gn=NCOR2;refGene_tx=XM_023241305.1;refGene_gn=N
COR2;refGene_tx=XM_023241306.1;refGene_gn=NCOR2;refGene_tx=XM_02324130
7.1;refGene_gn=NCOR2;refGene_tx=XM_023241308.1;refGene_gn=NCOR2;refGen
e_tx=XM_023241309.1;refGene_gn=NCOR2;refGene_tx=XM_023241310.1;refGene
_gn=NCOR2;refGene_tx=XM_023241311.1;refGene_gn=NCOR2;refGene_tx=XM_023
241312.1
D3 5796150 5796323
refGene_gn=NCOR2;refGene_tx=XM_023241290.1;refGene_gn=NCOR2;refGene_tx
=XM_023241291.1;refGene_gn=NCOR2;refGene_tx=XM_023241292.1;refGene_gn=
NCOR2;refGene_tx=XM_023241293.1;refGene_gn=NCOR2;refGene_tx=XM_0232412
94.1;refGene_gn=NCOR2;refGene_tx=XM_023241295.1;refGene_gn=NCOR2;refGe
ne_tx=XM_023241296.1;refGene_gn=NCOR2;refGene_tx=XM_023241297.1;refGen
e_gn=NCOR2;refGene_tx=XM_023241298.1;refGene_gn=NCOR2;refGene_tx=XM_02
3241299.1;refGene_gn=NCOR2;refGene_tx=XM_023241300.1;refGene_gn=NCOR2;
refGene_tx=XM_023241301.1;refGene_gn=NCOR2;refGene_tx=XM_023241302.1;r
efGene_gn=NCOR2;refGene_tx=XM_023241303.1;refGene_gn=NCOR2;refGene_tx=
XM_023241304.1;refGene_gn=NCOR2;refGene_tx=XM_023241305.1;refGene_gn=N
COR2;refGene_tx=XM_023241306.1;refGene_gn=NCOR2;refGene_tx=XM_02324130
7.1;refGene_gn=NCOR2;refGene_tx=XM_023241308.1;refGene_gn=NCOR2;refGen
e_tx=XM_023241309.1;refGene_gn=NCOR2;refGene_tx=XM_023241310.1;refGene
_gn=NCOR2;refGene_tx=XM_023241311.1;refGene_gn=NCOR2;refGene_tx=XM_023
241312.1
D3 5798217 5798280
refGene_gn=NCOR2;refGene_tx=XM_023241290.1;refGene_gn=NCOR2;refGene_tx
=XM_023241291.1;refGene_gn=NCOR2;refGene_tx=XM_023241292.1;refGene_gn=
NCOR2;refGene_tx=XM_023241293.1;refGene_gn=NCOR2;refGene_tx=XM_0232412
94.1;refGene_gn=NCOR2;refGene_tx=XM_023241295.1;refGene_gn=NCOR2;refGe
ne_tx=XM_023241296.1;refGene_gn=NCOR2;refGene_tx=XM_023241297.1;refGen
e_gn=NCOR2;refGene_tx=XM_023241298.1;refGene_gn=NCOR2;refGene_tx=XM_02
3241299.1;refGene_gn=NCOR2;refGene_tx=XM_023241300.1;refGene_gn=NCOR2;
refGene_tx=XM_023241301.1;refGene_gn=NCOR2;refGene_tx=XM_023241302.1;r

efGene_gn=NCOR2;refGene_tx=XM_023241303.1;refGene_gn=NCOR2;refGene_tx=XM_023241304.1;refGene_gn=NCOR2;refGene_tx=XM_023241305.1;refGene_gn=NCOR2;refGene_tx=XM_023241306.1;refGene_gn=NCOR2;refGene_tx=XM_023241307.1;refGene_gn=NCOR2;refGene_tx=XM_023241308.1;refGene_gn=NCOR2;refGene_tx=XM_023241309.1;refGene_gn=NCOR2;refGene_tx=XM_023241310.1;refGene_gn=NCOR2;refGene_tx=XM_023241311.1;refGene_gn=NCOR2;refGene_tx=XM_023241312.1
D3 5810237 5810380
refGene_gn=NCOR2;refGene_tx=XM_023241290.1;refGene_gn=NCOR2;refGene_tx=XM_023241291.1;refGene_gn=NCOR2;refGene_tx=XM_023241292.1;refGene_gn=NCOR2;refGene_tx=XM_023241293.1;refGene_gn=NCOR2;refGene_tx=XM_023241294.1;refGene_gn=NCOR2;refGene_tx=XM_023241295.1;refGene_gn=NCOR2;refGene_tx=XM_023241296.1;refGene_gn=NCOR2;refGene_tx=XM_023241297.1;refGene_gn=NCOR2;refGene_tx=XM_023241298.1;refGene_gn=NCOR2;refGene_tx=XM_023241299.1;refGene_gn=NCOR2;refGene_tx=XM_023241300.1;refGene_gn=NCOR2;refGene_tx=XM_023241301.1;refGene_gn=NCOR2;refGene_tx=XM_023241302.1;refGene_gn=NCOR2;refGene_tx=XM_023241303.1;refGene_gn=NCOR2;refGene_tx=XM_023241304.1;refGene_gn=NCOR2;refGene_tx=XM_023241305.1;refGene_gn=NCOR2;refGene_tx=XM_023241306.1;refGene_gn=NCOR2;refGene_tx=XM_023241307.1;refGene_gn=NCOR2;refGene_tx=XM_023241308.1;refGene_gn=NCOR2;refGene_tx=XM_023241309.1;refGene_gn=NCOR2;refGene_tx=XM_023241310.1;refGene_gn=NCOR2;refGene_tx=XM_023241311.1;refGene_gn=NCOR2;refGene_tx=XM_023241312.1
D3 5817793 5817941
refGene_gn=NCOR2;refGene_tx=XM_023241290.1;refGene_gn=NCOR2;refGene_tx=XM_023241291.1;refGene_gn=NCOR2;refGene_tx=XM_023241292.1;refGene_gn=NCOR2;refGene_tx=XM_023241293.1;refGene_gn=NCOR2;refGene_tx=XM_023241294.1;refGene_gn=NCOR2;refGene_tx=XM_023241295.1;refGene_gn=NCOR2;refGene_tx=XM_023241296.1;refGene_gn=NCOR2;refGene_tx=XM_023241297.1;refGene_gn=NCOR2;refGene_tx=XM_023241298.1;refGene_gn=NCOR2;refGene_tx=XM_023241299.1;refGene_gn=NCOR2;refGene_tx=XM_023241300.1;refGene_gn=NCOR2;refGene_tx=XM_023241301.1;refGene_gn=NCOR2;refGene_tx=XM_023241302.1;refGene_gn=NCOR2;refGene_tx=XM_023241303.1;refGene_gn=NCOR2;refGene_tx=XM_023241304.1;refGene_gn=NCOR2;refGene_tx=XM_023241305.1;refGene_gn=NCOR2;refGene_tx=XM_023241306.1;refGene_gn=NCOR2;refGene_tx=XM_023241307.1;refGene_gn=NCOR2;refGene_tx=XM_023241308.1;refGene_gn=NCOR2;refGene_tx=XM_023241309.1;refGene_gn=NCOR2;refGene_tx=XM_023241310.1;refGene_gn=NCOR2;refGene_tx=XM_023241311.1;refGene_gn=NCOR2;refGene_tx=XM_023241312.1
D3 5820375 5820426
refGene_gn=NCOR2;refGene_tx=XM_023241290.1;refGene_gn=NCOR2;refGene_tx=XM_023241291.1;refGene_gn=NCOR2;refGene_tx=XM_023241292.1;refGene_gn=NCOR2;refGene_tx=XM_023241293.1;refGene_gn=NCOR2;refGene_tx=XM_023241294.1;refGene_gn=NCOR2;refGene_tx=XM_023241295.1;refGene_gn=NCOR2;refGene_tx=XM_023241296.1;refGene_gn=NCOR2;refGene_tx=XM_023241297.1;refGene_gn=NCOR2;refGene_tx=XM_023241298.1;refGene_gn=NCOR2;refGene_tx=XM_023241299.1;refGene_gn=NCOR2;refGene_tx=XM_023241301.1;refGene_gn=NCOR2;refGene_tx=XM_023241302.1;refGene_gn=NCOR2;refGene_tx=XM_023241303.1;refGene_gn=NCOR2;refGene_tx=XM_023241304.1;refGene_gn=NCOR2;refGene_tx=XM_023241305.1;refGene_gn=NCOR2;refGene_tx=XM_023241306.1;refGene_gn=N

COR2;refGene_tx=XM_023241307.1
D3 5821407 5822035
refGene_gn=NCOR2;refGene_tx=XM_023241290.1;refGene_gn=NCOR2;refGene_tx=XM_023241291.1;refGene_gn=NCOR2;refGene_tx=XM_023241292.1;refGene_gn=NCOR2;refGene_tx=XM_023241293.1;refGene_gn=NCOR2;refGene_tx=XM_023241294.1;refGene_gn=NCOR2;refGene_tx=XM_023241295.1;refGene_gn=NCOR2;refGene_tx=XM_023241296.1;refGene_gn=NCOR2;refGene_tx=XM_023241297.1;refGene_gn=NCOR2;refGene_tx=XM_023241298.1;refGene_gn=NCOR2;refGene_tx=XM_023241299.1;refGene_gn=NCOR2;refGene_tx=XM_023241300.1;refGene_gn=NCOR2;refGene_tx=XM_023241301.1;refGene_gn=NCOR2;refGene_tx=XM_023241302.1;refGene_gn=NCOR2;refGene_tx=XM_023241303.1;refGene_gn=NCOR2;refGene_tx=XM_023241304.1;refGene_gn=NCOR2;refGene_tx=XM_023241305.1;refGene_gn=NCOR2;refGene_tx=XM_023241306.1;refGene_gn=NCOR2;refGene_tx=XM_023241307.1;refGene_gn=NCOR2;refGene_tx=XM_023241308.1;refGene_gn=NCOR2;refGene_tx=XM_023241309.1;refGene_gn=NCOR2;refGene_tx=XM_023241310.1;refGene_gn=NCOR2;refGene_tx=XM_023241311.1;refGene_gn=NCOR2;refGene_tx=XM_023241312.1
D3 5828860 5828981
refGene_gn=NCOR2;refGene_tx=XM_023241290.1;refGene_gn=NCOR2;refGene_tx=XM_023241291.1;refGene_gn=NCOR2;refGene_tx=XM_023241292.1;refGene_gn=NCOR2;refGene_tx=XM_023241293.1;refGene_gn=NCOR2;refGene_tx=XM_023241294.1;refGene_gn=NCOR2;refGene_tx=XM_023241295.1;refGene_gn=NCOR2;refGene_tx=XM_023241296.1;refGene_gn=NCOR2;refGene_tx=XM_023241297.1;refGene_gn=NCOR2;refGene_tx=XM_023241298.1;refGene_gn=NCOR2;refGene_tx=XM_023241299.1;refGene_gn=NCOR2;refGene_tx=XM_023241300.1;refGene_gn=NCOR2;refGene_tx=XM_023241301.1;refGene_gn=NCOR2;refGene_tx=XM_023241302.1;refGene_gn=NCOR2;refGene_tx=XM_023241303.1;refGene_gn=NCOR2;refGene_tx=XM_023241304.1;refGene_gn=NCOR2;refGene_tx=XM_023241305.1;refGene_gn=NCOR2;refGene_tx=XM_023241306.1;refGene_gn=NCOR2;refGene_tx=XM_023241307.1;refGene_gn=NCOR2;refGene_tx=XM_023241308.1;refGene_gn=NCOR2;refGene_tx=XM_023241309.1;refGene_gn=NCOR2;refGene_tx=XM_023241310.1;refGene_gn=NCOR2;refGene_tx=XM_023241311.1;refGene_gn=NCOR2;refGene_tx=XM_023241312.1
D3 5829948 5830105
refGene_gn=NCOR2;refGene_tx=XM_023241290.1;refGene_gn=NCOR2;refGene_tx=XM_023241291.1;refGene_gn=NCOR2;refGene_tx=XM_023241292.1;refGene_gn=NCOR2;refGene_tx=XM_023241293.1;refGene_gn=NCOR2;refGene_tx=XM_023241294.1;refGene_gn=NCOR2;refGene_tx=XM_023241295.1;refGene_gn=NCOR2;refGene_tx=XM_023241296.1;refGene_gn=NCOR2;refGene_tx=XM_023241297.1;refGene_gn=NCOR2;refGene_tx=XM_023241298.1;refGene_gn=NCOR2;refGene_tx=XM_023241299.1;refGene_gn=NCOR2;refGene_tx=XM_023241300.1;refGene_gn=NCOR2;refGene_tx=XM_023241301.1;refGene_gn=NCOR2;refGene_tx=XM_023241302.1;refGene_gn=NCOR2;refGene_tx=XM_023241303.1;refGene_gn=NCOR2;refGene_tx=XM_023241304.1;refGene_gn=NCOR2;refGene_tx=XM_023241305.1;refGene_gn=NCOR2;refGene_tx=XM_023241306.1;refGene_gn=NCOR2;refGene_tx=XM_023241307.1;refGene_gn=NCOR2;refGene_tx=XM_023241308.1;refGene_gn=NCOR2;refGene_tx=XM_023241309.1;refGene_gn=NCOR2;refGene_tx=XM_023241310.1;refGene_gn=NCOR2;refGene_tx=XM_023241311.1;refGene_gn=NCOR2;refGene_tx=XM_023241312.1
D3 5834100 5834265

refGene_gn=NCOR2;refGene_tx=XM_023241290.1;refGene_gn=NCOR2;refGene_tx
=XM_023241291.1;refGene_gn=NCOR2;refGene_tx=XM_023241292.1;refGene_gn=
NCOR2;refGene_tx=XM_023241293.1;refGene_gn=NCOR2;refGene_tx=XM_0232412
94.1;refGene_gn=NCOR2;refGene_tx=XM_023241295.1;refGene_gn=NCOR2;refGe
ne_tx=XM_023241296.1;refGene_gn=NCOR2;refGene_tx=XM_023241297.1;refGen
e_gn=NCOR2;refGene_tx=XM_023241298.1;refGene_gn=NCOR2;refGene_tx=XM_02
3241299.1;refGene_gn=NCOR2;refGene_tx=XM_023241300.1;refGene_gn=NCOR2;
refGene_tx=XM_023241301.1;refGene_gn=NCOR2;refGene_tx=XM_023241302.1;r
efGene_gn=NCOR2;refGene_tx=XM_023241303.1;refGene_gn=NCOR2;refGene_tx=
XM_023241304.1;refGene_gn=NCOR2;refGene_tx=XM_023241305.1;refGene_gn=N
COR2;refGene_tx=XM_023241306.1;refGene_gn=NCOR2;refGene_tx=XM_02324130
7.1;refGene_gn=NCOR2;refGene_tx=XM_023241308.1;refGene_gn=NCOR2;refGen
e_tx=XM_023241309.1;refGene_gn=NCOR2;refGene_tx=XM_023241310.1;refGene
_gn=NCOR2;refGene_tx=XM_023241311.1;refGene_gn=NCOR2;refGene_tx=XM_023
241312.1
D3 5835270 5835410
refGene_gn=NCOR2;refGene_tx=XM_023241290.1;refGene_gn=NCOR2;refGene_tx
=XM_023241291.1;refGene_gn=NCOR2;refGene_tx=XM_023241292.1;refGene_gn=
NCOR2;refGene_tx=XM_023241293.1;refGene_gn=NCOR2;refGene_tx=XM_0232412
94.1;refGene_gn=NCOR2;refGene_tx=XM_023241295.1;refGene_gn=NCOR2;refGe
ne_tx=XM_023241296.1;refGene_gn=NCOR2;refGene_tx=XM_023241297.1;refGen
e_gn=NCOR2;refGene_tx=XM_023241298.1;refGene_gn=NCOR2;refGene_tx=XM_02
3241299.1;refGene_gn=NCOR2;refGene_tx=XM_023241300.1;refGene_gn=NCOR2;
refGene_tx=XM_023241301.1;refGene_gn=NCOR2;refGene_tx=XM_023241302.1;r
efGene_gn=NCOR2;refGene_tx=XM_023241303.1;refGene_gn=NCOR2;refGene_tx=
XM_023241304.1;refGene_gn=NCOR2;refGene_tx=XM_023241305.1;refGene_gn=N
COR2;refGene_tx=XM_023241306.1;refGene_gn=NCOR2;refGene_tx=XM_02324130
7.1;refGene_gn=NCOR2;refGene_tx=XM_023241308.1;refGene_gn=NCOR2;refGen
e_tx=XM_023241309.1;refGene_gn=NCOR2;refGene_tx=XM_023241310.1;refGene
_gn=NCOR2;refGene_tx=XM_023241311.1;refGene_gn=NCOR2;refGene_tx=XM_023
241312.1
D3 5836119 5836222
refGene_gn=NCOR2;refGene_tx=XM_023241290.1;refGene_gn=NCOR2;refGene_tx
=XM_023241291.1;refGene_gn=NCOR2;refGene_tx=XM_023241292.1;refGene_gn=
NCOR2;refGene_tx=XM_023241293.1;refGene_gn=NCOR2;refGene_tx=XM_0232412
94.1;refGene_gn=NCOR2;refGene_tx=XM_023241295.1;refGene_gn=NCOR2;refGe
ne_tx=XM_023241296.1;refGene_gn=NCOR2;refGene_tx=XM_023241297.1;refGen
e_gn=NCOR2;refGene_tx=XM_023241298.1;refGene_gn=NCOR2;refGene_tx=XM_02
3241299.1;refGene_gn=NCOR2;refGene_tx=XM_023241300.1;refGene_gn=NCOR2;
refGene_tx=XM_023241301.1;refGene_gn=NCOR2;refGene_tx=XM_023241302.1;r
efGene_gn=NCOR2;refGene_tx=XM_023241303.1;refGene_gn=NCOR2;refGene_tx=
XM_023241304.1;refGene_gn=NCOR2;refGene_tx=XM_023241305.1;refGene_gn=N
COR2;refGene_tx=XM_023241306.1;refGene_gn=NCOR2;refGene_tx=XM_02324130
7.1;refGene_gn=NCOR2;refGene_tx=XM_023241308.1;refGene_gn=NCOR2;refGen
e_tx=XM_023241309.1;refGene_gn=NCOR2;refGene_tx=XM_023241310.1;refGene
_gn=NCOR2;refGene_tx=XM_023241311.1;refGene_gn=NCOR2;refGene_tx=XM_023
241312.1
D3 5836393 5836498
refGene_gn=NCOR2;refGene_tx=XM_023241290.1;refGene_gn=NCOR2;refGene_tx
=XM_023241291.1;refGene_gn=NCOR2;refGene_tx=XM_023241292.1;refGene_gn=

NCOR2;refGene_tx=XM_023241293.1;refGene_gn=NCOR2;refGene_tx=XM_0232412
94.1;refGene_gn=NCOR2;refGene_tx=XM_023241295.1;refGene_gn=NCOR2;refGe
ne_tx=XM_023241296.1;refGene_gn=NCOR2;refGene_tx=XM_023241297.1;refGen
e_gn=NCOR2;refGene_tx=XM_023241298.1;refGene_gn=NCOR2;refGene_tx=XM_02
3241299.1;refGene_gn=NCOR2;refGene_tx=XM_023241300.1;refGene_gn=NCOR2;
refGene_tx=XM_023241301.1;refGene_gn=NCOR2;refGene_tx=XM_023241302.1;r
efGene_gn=NCOR2;refGene_tx=XM_023241303.1;refGene_gn=NCOR2;refGene_tx=
XM_023241304.1;refGene_gn=NCOR2;refGene_tx=XM_023241305.1;refGene_gn=N
COR2;refGene_tx=XM_023241306.1;refGene_gn=NCOR2;refGene_tx=XM_02324130
7.1;refGene_gn=NCOR2;refGene_tx=XM_023241308.1;refGene_gn=NCOR2;refGen
e_tx=XM_023241309.1;refGene_gn=NCOR2;refGene_tx=XM_023241310.1;refGene
_gn=NCOR2;refGene_tx=XM_023241311.1;refGene_gn=NCOR2;refGene_tx=XM_023
241312.1
D3 5836818 5836922
refGene_gn=NCOR2;refGene_tx=XM_023241290.1;refGene_gn=NCOR2;refGene_tx
=XM_023241291.1;refGene_gn=NCOR2;refGene_tx=XM_023241292.1;refGene_gn=
NCOR2;refGene_tx=XM_023241293.1;refGene_gn=NCOR2;refGene_tx=XM_0232412
94.1;refGene_gn=NCOR2;refGene_tx=XM_023241295.1;refGene_gn=NCOR2;refGe
ne_tx=XM_023241296.1;refGene_gn=NCOR2;refGene_tx=XM_023241297.1;refGen
e_gn=NCOR2;refGene_tx=XM_023241298.1;refGene_gn=NCOR2;refGene_tx=XM_02
3241299.1;refGene_gn=NCOR2;refGene_tx=XM_023241300.1;refGene_gn=NCOR2;
refGene_tx=XM_023241301.1;refGene_gn=NCOR2;refGene_tx=XM_023241302.1;r
efGene_gn=NCOR2;refGene_tx=XM_023241303.1;refGene_gn=NCOR2;refGene_tx=
XM_023241304.1;refGene_gn=NCOR2;refGene_tx=XM_023241305.1;refGene_gn=N
COR2;refGene_tx=XM_023241306.1;refGene_gn=NCOR2;refGene_tx=XM_02324130
7.1;refGene_gn=NCOR2;refGene_tx=XM_023241308.1;refGene_gn=NCOR2;refGen
e_tx=XM_023241309.1;refGene_gn=NCOR2;refGene_tx=XM_023241310.1;refGene
_gn=NCOR2;refGene_tx=XM_023241311.1;refGene_gn=NCOR2;refGene_tx=XM_023
241312.1
D3 5840415 5840566
refGene_gn=NCOR2;refGene_tx=XM_023241290.1;refGene_gn=NCOR2;refGene_tx
=XM_023241291.1;refGene_gn=NCOR2;refGene_tx=XM_023241292.1;refGene_gn=
NCOR2;refGene_tx=XM_023241293.1;refGene_gn=NCOR2;refGene_tx=XM_0232412
94.1;refGene_gn=NCOR2;refGene_tx=XM_023241295.1;refGene_gn=NCOR2;refGe
ne_tx=XM_023241296.1;refGene_gn=NCOR2;refGene_tx=XM_023241297.1;refGen
e_gn=NCOR2;refGene_tx=XM_023241298.1;refGene_gn=NCOR2;refGene_tx=XM_02
3241299.1;refGene_gn=NCOR2;refGene_tx=XM_023241300.1;refGene_gn=NCOR2;
refGene_tx=XM_023241301.1;refGene_gn=NCOR2;refGene_tx=XM_023241302.1;r
efGene_gn=NCOR2;refGene_tx=XM_023241303.1;refGene_gn=NCOR2;refGene_tx=
XM_023241304.1;refGene_gn=NCOR2;refGene_tx=XM_023241305.1;refGene_gn=N
COR2;refGene_tx=XM_023241306.1;refGene_gn=NCOR2;refGene_tx=XM_02324130
7.1;refGene_gn=NCOR2;refGene_tx=XM_023241308.1;refGene_gn=NCOR2;refGen
e_tx=XM_023241309.1;refGene_gn=NCOR2;refGene_tx=XM_023241310.1;refGene
_gn=NCOR2;refGene_tx=XM_023241311.1;refGene_gn=NCOR2;refGene_tx=XM_023
241312.1
D3 5842898 5843039
refGene_gn=NCOR2;refGene_tx=XM_023241290.1;refGene_gn=NCOR2;refGene_tx
=XM_023241291.1;refGene_gn=NCOR2;refGene_tx=XM_023241292.1;refGene_gn=
NCOR2;refGene_tx=XM_023241293.1;refGene_gn=NCOR2;refGene_tx=XM_0232412
94.1;refGene_gn=NCOR2;refGene_tx=XM_023241295.1;refGene_gn=NCOR2;refGe

ne_tx=XM_023241296.1;refGene_gn=NCOR2;refGene_tx=XM_023241297.1;refGen
e_gn=NCOR2;refGene_tx=XM_023241298.1;refGene_gn=NCOR2;refGene_tx=XM_02
3241299.1;refGene_gn=NCOR2;refGene_tx=XM_023241300.1;refGene_gn=NCOR2;
refGene_tx=XM_023241301.1;refGene_gn=NCOR2;refGene_tx=XM_023241302.1;r
efGene_gn=NCOR2;refGene_tx=XM_023241303.1;refGene_gn=NCOR2;refGene_tx=
XM_023241304.1;refGene_gn=NCOR2;refGene_tx=XM_023241305.1;refGene_gn=N
COR2;refGene_tx=XM_023241306.1;refGene_gn=NCOR2;refGene_tx=XM_02324130
7.1;refGene_gn=NCOR2;refGene_tx=XM_023241308.1;refGene_gn=NCOR2;refGen
e_tx=XM_023241309.1;refGene_gn=NCOR2;refGene_tx=XM_023241310.1;refGene
_gn=NCOR2;refGene_tx=XM_023241311.1;refGene_gn=NCOR2;refGene_tx=XM_023
241312.1
D3 5843317 5843404
refGene_gn=NCOR2;refGene_tx=XM_023241290.1;refGene_gn=NCOR2;refGene_tx
=XM_023241291.1;refGene_gn=NCOR2;refGene_tx=XM_023241292.1;refGene_gn=
NCOR2;refGene_tx=XM_023241293.1;refGene_gn=NCOR2;refGene_tx=XM_0232412
94.1;refGene_gn=NCOR2;refGene_tx=XM_023241295.1;refGene_gn=NCOR2;refGe
ne_tx=XM_023241296.1;refGene_gn=NCOR2;refGene_tx=XM_023241297.1;refGen
e_gn=NCOR2;refGene_tx=XM_023241298.1;refGene_gn=NCOR2;refGene_tx=XM_02
3241299.1;refGene_gn=NCOR2;refGene_tx=XM_023241300.1;refGene_gn=NCOR2;
refGene_tx=XM_023241301.1;refGene_gn=NCOR2;refGene_tx=XM_023241302.1;r
efGene_gn=NCOR2;refGene_tx=XM_023241303.1;refGene_gn=NCOR2;refGene_tx=
XM_023241304.1;refGene_gn=NCOR2;refGene_tx=XM_023241305.1;refGene_gn=N
COR2;refGene_tx=XM_023241306.1;refGene_gn=NCOR2;refGene_tx=XM_02324130
7.1;refGene_gn=NCOR2;refGene_tx=XM_023241308.1;refGene_gn=NCOR2;refGen
e_tx=XM_023241309.1;refGene_gn=NCOR2;refGene_tx=XM_023241310.1;refGene
_gn=NCOR2;refGene_tx=XM_023241311.1;refGene_gn=NCOR2;refGene_tx=XM_023
241312.1
D3 5844033 5844329
refGene_gn=NCOR2;refGene_tx=XM_023241290.1;refGene_gn=NCOR2;refGene_tx
=XM_023241291.1;refGene_gn=NCOR2;refGene_tx=XM_023241292.1;refGene_gn=
NCOR2;refGene_tx=XM_023241293.1;refGene_gn=NCOR2;refGene_tx=XM_0232412
94.1;refGene_gn=NCOR2;refGene_tx=XM_023241295.1;refGene_gn=NCOR2;refGe
ne_tx=XM_023241296.1;refGene_gn=NCOR2;refGene_tx=XM_023241297.1;refGen
e_gn=NCOR2;refGene_tx=XM_023241298.1;refGene_gn=NCOR2;refGene_tx=XM_02
3241299.1;refGene_gn=NCOR2;refGene_tx=XM_023241300.1;refGene_gn=NCOR2;
refGene_tx=XM_023241301.1;refGene_gn=NCOR2;refGene_tx=XM_023241302.1;r
efGene_gn=NCOR2;refGene_tx=XM_023241303.1;refGene_gn=NCOR2;refGene_tx=
XM_023241304.1;refGene_gn=NCOR2;refGene_tx=XM_023241305.1;refGene_gn=N
COR2;refGene_tx=XM_023241306.1;refGene_gn=NCOR2;refGene_tx=XM_02324130
7.1;refGene_gn=NCOR2;refGene_tx=XM_023241308.1;refGene_gn=NCOR2;refGen
e_tx=XM_023241309.1;refGene_gn=NCOR2;refGene_tx=XM_023241310.1;refGene
_gn=NCOR2;refGene_tx=XM_023241311.1;refGene_gn=NCOR2;refGene_tx=XM_023
241312.1
D3 5845780 5846135
refGene_gn=NCOR2;refGene_tx=XM_023241290.1;refGene_gn=NCOR2;refGene_tx
=XM_023241291.1;refGene_gn=NCOR2;refGene_tx=XM_023241292.1;refGene_gn=
NCOR2;refGene_tx=XM_023241293.1;refGene_gn=NCOR2;refGene_tx=XM_0232412
94.1;refGene_gn=NCOR2;refGene_tx=XM_023241295.1;refGene_gn=NCOR2;refGe
ne_tx=XM_023241296.1;refGene_gn=NCOR2;refGene_tx=XM_023241297.1;refGen
e_gn=NCOR2;refGene_tx=XM_023241298.1;refGene_gn=NCOR2;refGene_tx=XM_02

3241299.1;refGene_gn=NCOR2;refGene_tx=XM_023241300.1;refGene_gn=NCOR2;
refGene_tx=XM_023241301.1;refGene_gn=NCOR2;refGene_tx=XM_023241302.1;r
efGene_gn=NCOR2;refGene_tx=XM_023241303.1;refGene_gn=NCOR2;refGene_tx=
XM_023241304.1;refGene_gn=NCOR2;refGene_tx=XM_023241305.1;refGene_gn=N
COR2;refGene_tx=XM_023241306.1;refGene_gn=NCOR2;refGene_tx=XM_02324130
7.1;refGene_gn=NCOR2;refGene_tx=XM_023241308.1;refGene_gn=NCOR2;refGen
e_tx=XM_023241309.1;refGene_gn=NCOR2;refGene_tx=XM_023241310.1;refGene
_gn=NCOR2;refGene_tx=XM_023241311.1;refGene_gn=NCOR2;refGene_tx=XM_023
241312.1
D3 5847271 5847493
refGene_gn=NCOR2;refGene_tx=XM_023241290.1;refGene_gn=NCOR2;refGene_tx
=XM_023241291.1;refGene_gn=NCOR2;refGene_tx=XM_023241292.1;refGene_gn=
NCOR2;refGene_tx=XM_023241293.1;refGene_gn=NCOR2;refGene_tx=XM_0232412
94.1;refGene_gn=NCOR2;refGene_tx=XM_023241295.1;refGene_gn=NCOR2;refGe
ne_tx=XM_023241296.1;refGene_gn=NCOR2;refGene_tx=XM_023241297.1;refGen
e_gn=NCOR2;refGene_tx=XM_023241298.1;refGene_gn=NCOR2;refGene_tx=XM_02
3241299.1;refGene_gn=NCOR2;refGene_tx=XM_023241300.1;refGene_gn=NCOR2;
refGene_tx=XM_023241301.1;refGene_gn=NCOR2;refGene_tx=XM_023241302.1;r
efGene_gn=NCOR2;refGene_tx=XM_023241303.1;refGene_gn=NCOR2;refGene_tx=
XM_023241304.1;refGene_gn=NCOR2;refGene_tx=XM_023241305.1;refGene_gn=N
COR2;refGene_tx=XM_023241306.1;refGene_gn=NCOR2;refGene_tx=XM_02324130
7.1;refGene_gn=NCOR2;refGene_tx=XM_023241308.1;refGene_gn=NCOR2;refGen
e_tx=XM_023241309.1;refGene_gn=NCOR2;refGene_tx=XM_023241310.1;refGene
_gn=NCOR2;refGene_tx=XM_023241311.1;refGene_gn=NCOR2;refGene_tx=XM_023
241312.1
D3 5847906 5848158
refGene_gn=NCOR2;refGene_tx=XM_023241290.1;refGene_gn=NCOR2;refGene_tx
=XM_023241291.1;refGene_gn=NCOR2;refGene_tx=XM_023241292.1;refGene_gn=
NCOR2;refGene_tx=XM_023241293.1;refGene_gn=NCOR2;refGene_tx=XM_0232412
94.1;refGene_gn=NCOR2;refGene_tx=XM_023241295.1;refGene_gn=NCOR2;refGe
ne_tx=XM_023241296.1;refGene_gn=NCOR2;refGene_tx=XM_023241297.1;refGen
e_gn=NCOR2;refGene_tx=XM_023241298.1;refGene_gn=NCOR2;refGene_tx=XM_02
3241299.1;refGene_gn=NCOR2;refGene_tx=XM_023241300.1;refGene_gn=NCOR2;
refGene_tx=XM_023241301.1;refGene_gn=NCOR2;refGene_tx=XM_023241302.1;r
efGene_gn=NCOR2;refGene_tx=XM_023241303.1;refGene_gn=NCOR2;refGene_tx=
XM_023241304.1;refGene_gn=NCOR2;refGene_tx=XM_023241305.1;refGene_gn=N
COR2;refGene_tx=XM_023241306.1;refGene_gn=NCOR2;refGene_tx=XM_02324130
7.1;refGene_gn=NCOR2;refGene_tx=XM_023241308.1;refGene_gn=NCOR2;refGen
e_tx=XM_023241309.1;refGene_gn=NCOR2;refGene_tx=XM_023241310.1;refGene
_gn=NCOR2;refGene_tx=XM_023241311.1;refGene_gn=NCOR2;refGene_tx=XM_023
241312.1
D3 5848890 5849043
refGene_gn=NCOR2;refGene_tx=XM_023241290.1;refGene_gn=NCOR2;refGene_tx
=XM_023241291.1;refGene_gn=NCOR2;refGene_tx=XM_023241292.1;refGene_gn=
NCOR2;refGene_tx=XM_023241293.1;refGene_gn=NCOR2;refGene_tx=XM_0232412
94.1;refGene_gn=NCOR2;refGene_tx=XM_023241295.1;refGene_gn=NCOR2;refGe
ne_tx=XM_023241296.1;refGene_gn=NCOR2;refGene_tx=XM_023241297.1;refGen
e_gn=NCOR2;refGene_tx=XM_023241298.1;refGene_gn=NCOR2;refGene_tx=XM_02
3241299.1;refGene_gn=NCOR2;refGene_tx=XM_023241300.1;refGene_gn=NCOR2;
refGene_tx=XM_023241301.1;refGene_gn=NCOR2;refGene_tx=XM_023241302.1;r

efGene_gn=NCOR2;refGene_tx=XM_023241303.1;refGene_gn=NCOR2;refGene_tx=
XM_023241304.1;refGene_gn=NCOR2;refGene_tx=XM_023241305.1;refGene_gn=N
COR2;refGene_tx=XM_023241306.1;refGene_gn=NCOR2;refGene_tx=XM_02324130
7.1;refGene_gn=NCOR2;refGene_tx=XM_023241308.1;refGene_gn=NCOR2;refGen
e_tx=XM_023241309.1;refGene_gn=NCOR2;refGene_tx=XM_023241310.1;refGene
_gn=NCOR2;refGene_tx=XM_023241311.1;refGene_gn=NCOR2;refGene_tx=XM_023
241312.1
D3 5849168 5849324
refGene_gn=NCOR2;refGene_tx=XM_023241290.1;refGene_gn=NCOR2;refGene_tx
=XM_023241291.1;refGene_gn=NCOR2;refGene_tx=XM_023241292.1;refGene_gn=
NCOR2;refGene_tx=XM_023241293.1;refGene_gn=NCOR2;refGene_tx=XM_0232412
94.1;refGene_gn=NCOR2;refGene_tx=XM_023241295.1;refGene_gn=NCOR2;refGe
ne_tx=XM_023241296.1;refGene_gn=NCOR2;refGene_tx=XM_023241297.1;refGen
e_gn=NCOR2;refGene_tx=XM_023241298.1;refGene_gn=NCOR2;refGene_tx=XM_02
3241299.1;refGene_gn=NCOR2;refGene_tx=XM_023241300.1;refGene_gn=NCOR2;
refGene_tx=XM_023241301.1;refGene_gn=NCOR2;refGene_tx=XM_023241302.1;r
efGene_gn=NCOR2;refGene_tx=XM_023241303.1;refGene_gn=NCOR2;refGene_tx=
XM_023241304.1;refGene_gn=NCOR2;refGene_tx=XM_023241305.1;refGene_gn=N
COR2;refGene_tx=XM_023241306.1;refGene_gn=NCOR2;refGene_tx=XM_02324130
7.1;refGene_gn=NCOR2;refGene_tx=XM_023241308.1;refGene_gn=NCOR2;refGen
e_tx=XM_023241309.1;refGene_gn=NCOR2;refGene_tx=XM_023241310.1;refGene
_gn=NCOR2;refGene_tx=XM_023241311.1;refGene_gn=NCOR2;refGene_tx=XM_023
241312.1
D3 5849413 5849627
refGene_gn=NCOR2;refGene_tx=XM_023241290.1;refGene_gn=NCOR2;refGene_tx
=XM_023241291.1;refGene_gn=NCOR2;refGene_tx=XM_023241292.1;refGene_gn=
NCOR2;refGene_tx=XM_023241293.1;refGene_gn=NCOR2;refGene_tx=XM_0232412
94.1;refGene_gn=NCOR2;refGene_tx=XM_023241295.1;refGene_gn=NCOR2;refGe
ne_tx=XM_023241296.1;refGene_gn=NCOR2;refGene_tx=XM_023241297.1;refGen
e_gn=NCOR2;refGene_tx=XM_023241298.1;refGene_gn=NCOR2;refGene_tx=XM_02
3241299.1;refGene_gn=NCOR2;refGene_tx=XM_023241300.1;refGene_gn=NCOR2;
refGene_tx=XM_023241301.1;refGene_gn=NCOR2;refGene_tx=XM_023241302.1;r
efGene_gn=NCOR2;refGene_tx=XM_023241303.1;refGene_gn=NCOR2;refGene_tx=
XM_023241304.1;refGene_gn=NCOR2;refGene_tx=XM_023241305.1;refGene_gn=N
COR2;refGene_tx=XM_023241306.1;refGene_gn=NCOR2;refGene_tx=XM_02324130
7.1;refGene_gn=NCOR2;refGene_tx=XM_023241308.1;refGene_gn=NCOR2;refGen
e_tx=XM_023241309.1;refGene_gn=NCOR2;refGene_tx=XM_023241310.1;refGene
_gn=NCOR2;refGene_tx=XM_023241311.1;refGene_gn=NCOR2;refGene_tx=XM_023
241312.1
D3 5852254 5852796
refGene_gn=NCOR2;refGene_tx=XM_023241290.1;refGene_gn=NCOR2;refGene_tx
=XM_023241291.1;refGene_gn=NCOR2;refGene_tx=XM_023241292.1;refGene_gn=
NCOR2;refGene_tx=XM_023241293.1;refGene_gn=NCOR2;refGene_tx=XM_0232412
94.1;refGene_gn=NCOR2;refGene_tx=XM_023241295.1;refGene_gn=NCOR2;refGe
ne_tx=XM_023241296.1;refGene_gn=NCOR2;refGene_tx=XM_023241297.1;refGen
e_gn=NCOR2;refGene_tx=XM_023241298.1;refGene_gn=NCOR2;refGene_tx=XM_02
3241299.1;refGene_gn=NCOR2;refGene_tx=XM_023241300.1;refGene_gn=NCOR2;
refGene_tx=XM_023241301.1;refGene_gn=NCOR2;refGene_tx=XM_023241302.1;r
efGene_gn=NCOR2;refGene_tx=XM_023241303.1;refGene_gn=NCOR2;refGene_tx=
XM_023241304.1;refGene_gn=NCOR2;refGene_tx=XM_023241305.1;refGene_gn=N

COR2;refGene_tx=XM_023241306.1;refGene_gn=NCOR2;refGene_tx=XM_02324130
7.1;refGene_gn=NCOR2;refGene_tx=XM_023241308.1;refGene_gn=NCOR2;refGen
e_tx=XM_023241309.1;refGene_gn=NCOR2;refGene_tx=XM_023241310.1;refGene
_gn=NCOR2;refGene_tx=XM_023241311.1;refGene_gn=NCOR2;refGene_tx=XM_023
241312.1
D3 5853855 5854005
refGene_gn=NCOR2;refGene_tx=XM_023241290.1;refGene_gn=NCOR2;refGene_tx
=XM_023241291.1;refGene_gn=NCOR2;refGene_tx=XM_023241292.1;refGene_gn=
NCOR2;refGene_tx=XM_023241293.1;refGene_gn=NCOR2;refGene_tx=XM_0232412
94.1;refGene_gn=NCOR2;refGene_tx=XM_023241295.1;refGene_gn=NCOR2;refGe
ne_tx=XM_023241296.1;refGene_gn=NCOR2;refGene_tx=XM_023241297.1;refGen
e_gn=NCOR2;refGene_tx=XM_023241298.1;refGene_gn=NCOR2;refGene_tx=XM_02
3241299.1;refGene_gn=NCOR2;refGene_tx=XM_023241300.1;refGene_gn=NCOR2;
refGene_tx=XM_023241301.1;refGene_gn=NCOR2;refGene_tx=XM_023241302.1;r
efGene_gn=NCOR2;refGene_tx=XM_023241303.1;refGene_gn=NCOR2;refGene_tx=
XM_023241304.1;refGene_gn=NCOR2;refGene_tx=XM_023241305.1;refGene_gn=N
COR2;refGene_tx=XM_023241306.1;refGene_gn=NCOR2;refGene_tx=XM_02324130
7.1;refGene_gn=NCOR2;refGene_tx=XM_023241308.1;refGene_gn=NCOR2;refGen
e_tx=XM_023241309.1;refGene_gn=NCOR2;refGene_tx=XM_023241310.1;refGene
_gn=NCOR2;refGene_tx=XM_023241311.1;refGene_gn=NCOR2;refGene_tx=XM_023
241312.1
D3 5854242 5854388
refGene_gn=NCOR2;refGene_tx=XM_023241290.1;refGene_gn=NCOR2;refGene_tx
=XM_023241291.1;refGene_gn=NCOR2;refGene_tx=XM_023241292.1;refGene_gn=
NCOR2;refGene_tx=XM_023241293.1;refGene_gn=NCOR2;refGene_tx=XM_0232412
94.1;refGene_gn=NCOR2;refGene_tx=XM_023241295.1;refGene_gn=NCOR2;refGe
ne_tx=XM_023241296.1;refGene_gn=NCOR2;refGene_tx=XM_023241297.1;refGen
e_gn=NCOR2;refGene_tx=XM_023241298.1;refGene_gn=NCOR2;refGene_tx=XM_02
3241299.1;refGene_gn=NCOR2;refGene_tx=XM_023241300.1;refGene_gn=NCOR2;
refGene_tx=XM_023241301.1;refGene_gn=NCOR2;refGene_tx=XM_023241302.1;r
efGene_gn=NCOR2;refGene_tx=XM_023241303.1;refGene_gn=NCOR2;refGene_tx=
XM_023241304.1;refGene_gn=NCOR2;refGene_tx=XM_023241305.1;refGene_gn=N
COR2;refGene_tx=XM_023241306.1;refGene_gn=NCOR2;refGene_tx=XM_02324130
7.1;refGene_gn=NCOR2;refGene_tx=XM_023241308.1;refGene_gn=NCOR2;refGen
e_tx=XM_023241309.1;refGene_gn=NCOR2;refGene_tx=XM_023241310.1;refGene
_gn=NCOR2;refGene_tx=XM_023241311.1;refGene_gn=NCOR2;refGene_tx=XM_023
241312.1
D3 5854863 5855057
refGene_gn=NCOR2;refGene_tx=XM_023241290.1;refGene_gn=NCOR2;refGene_tx
=XM_023241291.1;refGene_gn=NCOR2;refGene_tx=XM_023241292.1;refGene_gn=
NCOR2;refGene_tx=XM_023241293.1;refGene_gn=NCOR2;refGene_tx=XM_0232412
94.1;refGene_gn=NCOR2;refGene_tx=XM_023241295.1;refGene_gn=NCOR2;refGe
ne_tx=XM_023241296.1;refGene_gn=NCOR2;refGene_tx=XM_023241297.1;refGen
e_gn=NCOR2;refGene_tx=XM_023241298.1;refGene_gn=NCOR2;refGene_tx=XM_02
3241299.1;refGene_gn=NCOR2;refGene_tx=XM_023241300.1;refGene_gn=NCOR2;
refGene_tx=XM_023241301.1;refGene_gn=NCOR2;refGene_tx=XM_023241302.1;r
efGene_gn=NCOR2;refGene_tx=XM_023241303.1;refGene_gn=NCOR2;refGene_tx=
XM_023241304.1;refGene_gn=NCOR2;refGene_tx=XM_023241305.1;refGene_gn=N
COR2;refGene_tx=XM_023241306.1;refGene_gn=NCOR2;refGene_tx=XM_02324130
7.1;refGene_gn=NCOR2;refGene_tx=XM_023241308.1;refGene_gn=NCOR2;refGen

e_tx=XM_023241309.1;refGene_gn=NCOR2;refGene_tx=XM_023241310.1;refGene_gn=NCOR2;refGene_tx=XM_023241311.1;refGene_gn=NCOR2;refGene_tx=XM_023241312.1
D3 5855274 5855358
refGene_gn=NCOR2;refGene_tx=XM_023241290.1;refGene_gn=NCOR2;refGene_tx=XM_023241291.1;refGene_gn=NCOR2;refGene_tx=XM_023241292.1;refGene_gn=NCOR2;refGene_tx=XM_023241293.1;refGene_gn=NCOR2;refGene_tx=XM_023241294.1;refGene_gn=NCOR2;refGene_tx=XM_023241295.1;refGene_gn=NCOR2;refGene_tx=XM_023241296.1;refGene_gn=NCOR2;refGene_tx=XM_023241297.1;refGene_gn=NCOR2;refGene_tx=XM_023241298.1;refGene_gn=NCOR2;refGene_tx=XM_023241299.1;refGene_gn=NCOR2;refGene_tx=XM_023241300.1;refGene_gn=NCOR2;refGene_tx=XM_023241302.1;refGene_gn=NCOR2;refGene_tx=XM_023241303.1;refGene_gn=NCOR2;refGene_tx=XM_023241304.1;refGene_gn=NCOR2;refGene_tx=XM_023241307.1
D3 5855897 5856047
refGene_gn=NCOR2;refGene_tx=XM_023241290.1;refGene_gn=NCOR2;refGene_tx=XM_023241291.1;refGene_gn=NCOR2;refGene_tx=XM_023241292.1;refGene_gn=NCOR2;refGene_tx=XM_023241293.1;refGene_gn=NCOR2;refGene_tx=XM_023241294.1;refGene_gn=NCOR2;refGene_tx=XM_023241295.1;refGene_gn=NCOR2;refGene_tx=XM_023241296.1;refGene_gn=NCOR2;refGene_tx=XM_023241297.1;refGene_gn=NCOR2;refGene_tx=XM_023241298.1;refGene_gn=NCOR2;refGene_tx=XM_023241299.1;refGene_gn=NCOR2;refGene_tx=XM_023241300.1;refGene_gn=NCOR2;refGene_tx=XM_023241301.1;refGene_gn=NCOR2;refGene_tx=XM_023241302.1;refGene_gn=NCOR2;refGene_tx=XM_023241303.1;refGene_gn=NCOR2;refGene_tx=XM_023241304.1;refGene_gn=NCOR2;refGene_tx=XM_023241305.1;refGene_gn=NCOR2;refGene_tx=XM_023241306.1;refGene_gn=NCOR2;refGene_tx=XM_023241307.1;refGene_gn=NCOR2;refGene_tx=XM_023241308.1;refGene_gn=NCOR2;refGene_tx=XM_023241309.1;refGene_gn=NCOR2;refGene_tx=XM_023241310.1;refGene_gn=NCOR2;refGene_tx=XM_023241311.1;refGene_gn=NCOR2;refGene_tx=XM_023241312.1
D3 5856660 5856809
refGene_gn=NCOR2;refGene_tx=XM_023241290.1;refGene_gn=NCOR2;refGene_tx=XM_023241291.1;refGene_gn=NCOR2;refGene_tx=XM_023241292.1;refGene_gn=NCOR2;refGene_tx=XM_023241293.1;refGene_gn=NCOR2;refGene_tx=XM_023241294.1;refGene_gn=NCOR2;refGene_tx=XM_023241295.1;refGene_gn=NCOR2;refGene_tx=XM_023241296.1;refGene_gn=NCOR2;refGene_tx=XM_023241297.1;refGene_gn=NCOR2;refGene_tx=XM_023241298.1;refGene_gn=NCOR2;refGene_tx=XM_023241299.1;refGene_gn=NCOR2;refGene_tx=XM_023241300.1;refGene_gn=NCOR2;refGene_tx=XM_023241301.1;refGene_gn=NCOR2;refGene_tx=XM_023241302.1;refGene_gn=NCOR2;refGene_tx=XM_023241303.1;refGene_gn=NCOR2;refGene_tx=XM_023241304.1;refGene_gn=NCOR2;refGene_tx=XM_023241305.1;refGene_gn=NCOR2;refGene_tx=XM_023241306.1;refGene_gn=NCOR2;refGene_tx=XM_023241307.1;refGene_gn=NCOR2;refGene_tx=XM_023241308.1;refGene_gn=NCOR2;refGene_tx=XM_023241309.1;refGene_gn=NCOR2;refGene_tx=XM_023241310.1;refGene_gn=NCOR2;refGene_tx=XM_023241311.1;refGene_gn=NCOR2;refGene_tx=XM_023241312.1
D3 5857357 5857411
refGene_gn=NCOR2;refGene_tx=XM_023241290.1;refGene_gn=NCOR2;refGene_tx=XM_023241291.1;refGene_gn=NCOR2;refGene_tx=XM_023241292.1;refGene_gn=NCOR2;refGene_tx=XM_023241293.1;refGene_gn=NCOR2;refGene_tx=XM_0232412

94.1;refGene_gn=NCOR2;refGene_tx=XM_023241295.1;refGene_gn=NCOR2;refGe
ne_tx=XM_023241296.1;refGene_gn=NCOR2;refGene_tx=XM_023241297.1;refGen
e_gn=NCOR2;refGene_tx=XM_023241298.1;refGene_gn=NCOR2;refGene_tx=XM_02
3241299.1;refGene_gn=NCOR2;refGene_tx=XM_023241300.1;refGene_gn=NCOR2;
refGene_tx=XM_023241301.1;refGene_gn=NCOR2;refGene_tx=XM_023241302.1;r
efGene_gn=NCOR2;refGene_tx=XM_023241303.1;refGene_gn=NCOR2;refGene_tx=
XM_023241304.1;refGene_gn=NCOR2;refGene_tx=XM_023241305.1;refGene_gn=N
COR2;refGene_tx=XM_023241306.1;refGene_gn=NCOR2;refGene_tx=XM_02324130
7.1;refGene_gn=NCOR2;refGene_tx=XM_023241308.1;refGene_gn=NCOR2;refGen
e_tx=XM_023241309.1;refGene_gn=NCOR2;refGene_tx=XM_023241310.1;refGene
_gn=NCOR2;refGene_tx=XM_023241311.1;refGene_gn=NCOR2;refGene_tx=XM_023
241312.1
D3 5860380 5860602
refGene_gn=NCOR2;refGene_tx=XM_023241290.1;refGene_gn=NCOR2;refGene_tx
=XM_023241291.1;refGene_gn=NCOR2;refGene_tx=XM_023241292.1;refGene_gn=
NCOR2;refGene_tx=XM_023241293.1;refGene_gn=NCOR2;refGene_tx=XM_0232412
94.1;refGene_gn=NCOR2;refGene_tx=XM_023241295.1;refGene_gn=NCOR2;refGe
ne_tx=XM_023241296.1;refGene_gn=NCOR2;refGene_tx=XM_023241297.1;refGen
e_gn=NCOR2;refGene_tx=XM_023241298.1;refGene_gn=NCOR2;refGene_tx=XM_02
3241299.1;refGene_gn=NCOR2;refGene_tx=XM_023241300.1;refGene_gn=NCOR2;
refGene_tx=XM_023241301.1;refGene_gn=NCOR2;refGene_tx=XM_023241302.1;r
efGene_gn=NCOR2;refGene_tx=XM_023241303.1;refGene_gn=NCOR2;refGene_tx=
XM_023241304.1;refGene_gn=NCOR2;refGene_tx=XM_023241305.1;refGene_gn=N
COR2;refGene_tx=XM_023241306.1;refGene_gn=NCOR2;refGene_tx=XM_02324130
8.1;refGene_gn=NCOR2;refGene_tx=XM_023241309.1;refGene_gn=NCOR2;refGen
e_tx=XM_023241310.1;refGene_gn=NCOR2;refGene_tx=XM_023241311.1;refGene
_gn=NCOR2;refGene_tx=XM_023241312.1
D3 5861675 5861855
refGene_gn=NCOR2;refGene_tx=XM_023241290.1;refGene_gn=NCOR2;refGene_tx
=XM_023241291.1;refGene_gn=NCOR2;refGene_tx=XM_023241292.1;refGene_gn=
NCOR2;refGene_tx=XM_023241293.1;refGene_gn=NCOR2;refGene_tx=XM_0232412
94.1;refGene_gn=NCOR2;refGene_tx=XM_023241295.1;refGene_gn=NCOR2;refGe
ne_tx=XM_023241296.1;refGene_gn=NCOR2;refGene_tx=XM_023241297.1;refGen
e_gn=NCOR2;refGene_tx=XM_023241298.1;refGene_gn=NCOR2;refGene_tx=XM_02
3241299.1;refGene_gn=NCOR2;refGene_tx=XM_023241300.1;refGene_gn=NCOR2;
refGene_tx=XM_023241301.1;refGene_gn=NCOR2;refGene_tx=XM_023241302.1;r
efGene_gn=NCOR2;refGene_tx=XM_023241303.1;refGene_gn=NCOR2;refGene_tx=
XM_023241304.1;refGene_gn=NCOR2;refGene_tx=XM_023241305.1;refGene_gn=N
COR2;refGene_tx=XM_023241306.1;refGene_gn=NCOR2;refGene_tx=XM_02324130
7.1;refGene_gn=NCOR2;refGene_tx=XM_023241308.1;refGene_gn=NCOR2;refGen
e_tx=XM_023241309.1;refGene_gn=NCOR2;refGene_tx=XM_023241310.1;refGene
_gn=NCOR2;refGene_tx=XM_023241311.1;refGene_gn=NCOR2;refGene_tx=XM_023
241312.1
D3 5862412 5862594
refGene_gn=NCOR2;refGene_tx=XM_023241290.1;refGene_gn=NCOR2;refGene_tx
=XM_023241291.1;refGene_gn=NCOR2;refGene_tx=XM_023241292.1;refGene_gn=
NCOR2;refGene_tx=XM_023241293.1;refGene_gn=NCOR2;refGene_tx=XM_0232412
94.1;refGene_gn=NCOR2;refGene_tx=XM_023241295.1;refGene_gn=NCOR2;refGe
ne_tx=XM_023241296.1;refGene_gn=NCOR2;refGene_tx=XM_023241297.1;refGen
e_gn=NCOR2;refGene_tx=XM_023241298.1;refGene_gn=NCOR2;refGene_tx=XM_02

3241299.1;refGene_gn=NCOR2;refGene_tx=XM_023241300.1;refGene_gn=NCOR2;
refGene_tx=XM_023241301.1;refGene_gn=NCOR2;refGene_tx=XM_023241302.1;r
efGene_gn=NCOR2;refGene_tx=XM_023241303.1;refGene_gn=NCOR2;refGene_tx=
XM_023241304.1;refGene_gn=NCOR2;refGene_tx=XM_023241305.1;refGene_gn=N
COR2;refGene_tx=XM_023241306.1;refGene_gn=NCOR2;refGene_tx=XM_02324130
7.1;refGene_gn=NCOR2;refGene_tx=XM_023241308.1;refGene_gn=NCOR2;refGen
e_tx=XM_023241309.1;refGene_gn=NCOR2;refGene_tx=XM_023241310.1;refGene
_gn=NCOR2;refGene_tx=XM_023241311.1;refGene_gn=NCOR2;refGene_tx=XM_023
241312.1
D3 5869950 5870263
refGene_gn=LOC101084234;refGene_tx=XM_023241313.1
D3 5872148 5872258
refGene_gn=LOC101084234;refGene_tx=XM_023241313.1
D3 5884736 5884943
refGene_gn=LOC101084234;refGene_tx=XM_023241313.1
D3 6073035 6073821
refGene_gn=ZNF664;refGene_tx=XM_019814447.2;refGene_gn=ZNF664;refGene_
tx=XM_023241316.1;refGene_gn=ZNF664;refGene_tx=XM_023241317.1;refGene_
gn=ZNF664;refGene_tx=XM_023241318.1
D3 6140316 6140446
refGene_gn=CCDC92;refGene_tx=XM_006938374.4;refGene_gn=CCDC92;refGene_
tx=XM_023241315.1
D3 6141002 6141044
refGene_gn=CCDC92;refGene_tx=XM_006938374.4;refGene_gn=CCDC92;refGene_
tx=XM_023241315.1
D3 6143721 6144497
refGene_gn=CCDC92;refGene_tx=XM_006938374.4;refGene_gn=CCDC92;refGene_
tx=XM_023241315.1
D3 6146059 6146206 refGene_gn=DNAH10;refGene_tx=XM_023241314.1
D3 6146791 6146937 refGene_gn=DNAH10;refGene_tx=XM_023241314.1
D3 6147429 6147610 refGene_gn=DNAH10;refGene_tx=XM_023241314.1
D3 6149061 6149229 refGene_gn=DNAH10;refGene_tx=XM_023241314.1
D3 6149320 6149532 refGene_gn=DNAH10;refGene_tx=XM_023241314.1
D3 6149714 6149846 refGene_gn=DNAH10;refGene_tx=XM_023241314.1
D3 6150316 6150488 refGene_gn=DNAH10;refGene_tx=XM_023241314.1
D3 6151158 6151254 refGene_gn=DNAH10;refGene_tx=XM_023241314.1
D3 6151436 6151646 refGene_gn=DNAH10;refGene_tx=XM_023241314.1
D3 6152336 6152537 refGene_gn=DNAH10;refGene_tx=XM_023241314.1
D3 6153099 6153283 refGene_gn=DNAH10;refGene_tx=XM_023241314.1
D3 6154351 6154506 refGene_gn=DNAH10;refGene_tx=XM_023241314.1
D3 6154986 6155141 refGene_gn=DNAH10;refGene_tx=XM_023241314.1
D3 6155509 6155614 refGene_gn=DNAH10;refGene_tx=XM_023241314.1
D3 6156930 6157204 refGene_gn=DNAH10;refGene_tx=XM_023241314.1
D3 6158667 6158897 refGene_gn=DNAH10;refGene_tx=XM_023241314.1
D3 6159569 6159849 refGene_gn=DNAH10;refGene_tx=XM_023241314.1
D3 6161872 6162020 refGene_gn=DNAH10;refGene_tx=XM_023241314.1
D3 6162269 6162491 refGene_gn=DNAH10;refGene_tx=XM_023241314.1
D3 6163520 6163738 refGene_gn=DNAH10;refGene_tx=XM_023241314.1
D3 6164887 6165024 refGene_gn=DNAH10;refGene_tx=XM_023241314.1
D3 6165987 6166169 refGene_gn=DNAH10;refGene_tx=XM_023241314.1

| | | | |
|---|---|---|---|
| D3 | 6171555 | 6171730 | refGene_gn=DNAH10;refGene_tx=XM_023241314.1 |
| D3 | 6173424 | 6173586 | refGene_gn=DNAH10;refGene_tx=XM_023241314.1 |
| D3 | 6174106 | 6174304 | refGene_gn=DNAH10;refGene_tx=XM_023241314.1 |
| D3 | 6177586 | 6177667 | refGene_gn=DNAH10;refGene_tx=XM_023241314.1 |
| D3 | 6178297 | 6178501 | refGene_gn=DNAH10;refGene_tx=XM_023241314.1 |
| D3 | 6182960 | 6183132 | refGene_gn=DNAH10;refGene_tx=XM_023241314.1 |
| D3 | 6187661 | 6187850 | refGene_gn=DNAH10;refGene_tx=XM_023241314.1 |
| D3 | 6188385 | 6188553 | refGene_gn=DNAH10;refGene_tx=XM_023241314.1 |
| D3 | 6188861 | 6188955 | refGene_gn=DNAH10;refGene_tx=XM_023241314.1 |
| D3 | 6189959 | 6190132 | refGene_gn=DNAH10;refGene_tx=XM_023241314.1 |
| D3 | 6192135 | 6192396 | refGene_gn=DNAH10;refGene_tx=XM_023241314.1 |
| D3 | 6193737 | 6193890 | refGene_gn=DNAH10;refGene_tx=XM_023241314.1 |
| D3 | 6196853 | 6196999 | refGene_gn=DNAH10;refGene_tx=XM_023241314.1 |
| D3 | 6198028 | 6198148 | refGene_gn=DNAH10;refGene_tx=XM_023241314.1 |
| D3 | 6200852 | 6201069 | refGene_gn=DNAH10;refGene_tx=XM_023241314.1 |
| D3 | 6201352 | 6201487 | refGene_gn=DNAH10;refGene_tx=XM_023241314.1 |
| D3 | 6202982 | 6203105 | refGene_gn=DNAH10;refGene_tx=XM_023241314.1 |
| D3 | 6204284 | 6204420 | refGene_gn=DNAH10;refGene_tx=XM_023241314.1 |
| D3 | 6205273 | 6205432 | refGene_gn=DNAH10;refGene_tx=XM_023241314.1 |
| D3 | 6207556 | 6207675 | refGene_gn=DNAH10;refGene_tx=XM_023241314.1 |
| D3 | 6209266 | 6209458 | refGene_gn=DNAH10;refGene_tx=XM_023241314.1 |
| D3 | 6210016 | 6210163 | refGene_gn=DNAH10;refGene_tx=XM_023241314.1 |
| D3 | 6210687 | 6210876 | refGene_gn=DNAH10;refGene_tx=XM_023241314.1 |
| D3 | 6212633 | 6212786 | refGene_gn=DNAH10;refGene_tx=XM_023241314.1 |
| D3 | 6213403 | 6213513 | refGene_gn=DNAH10;refGene_tx=XM_023241314.1 |
| D3 | 6213515 | 6213537 | refGene_gn=DNAH10;refGene_tx=XM_023241314.1 |
| D3 | 6215128 | 6215312 | refGene_gn=DNAH10;refGene_tx=XM_023241314.1 |
| D3 | 6215410 | 6215680 | refGene_gn=DNAH10;refGene_tx=XM_023241314.1 |
| D3 | 6218236 | 6218460 | refGene_gn=DNAH10;refGene_tx=XM_023241314.1 |
| D3 | 6221357 | 6221591 | refGene_gn=DNAH10;refGene_tx=XM_023241314.1 |
| D3 | 6223199 | 6223322 | refGene_gn=DNAH10;refGene_tx=XM_023241314.1 |
| D3 | 6225589 | 6225823 | refGene_gn=DNAH10;refGene_tx=XM_023241314.1 |
| D3 | 6227285 | 6227439 | refGene_gn=DNAH10;refGene_tx=XM_023241314.1 |
| D3 | 6229940 | 6230152 | refGene_gn=DNAH10;refGene_tx=XM_023241314.1 |
| D3 | 6233514 | 6233693 | refGene_gn=DNAH10;refGene_tx=XM_023241314.1 |
| D3 | 6234342 | 6234445 | refGene_gn=DNAH10;refGene_tx=XM_023241314.1 |
| D3 | 6234530 | 6234647 | refGene_gn=DNAH10;refGene_tx=XM_023241314.1 |
| D3 | 6236541 | 6236700 | refGene_gn=DNAH10;refGene_tx=XM_023241314.1 |
| D3 | 6236802 | 6237279 | refGene_gn=DNAH10;refGene_tx=XM_023241314.1 |
| D3 | 6239862 | 6240019 | refGene_gn=DNAH10;refGene_tx=XM_023241314.1 |
| D3 | 6241992 | 6242200 | refGene_gn=DNAH10;refGene_tx=XM_023241314.1 |
| D3 | 6242818 | 6242983 | refGene_gn=DNAH10;refGene_tx=XM_023241314.1 |
| D3 | 6247794 | 6247946 | refGene_gn=DNAH10;refGene_tx=XM_023241314.1 |
| D3 | 6248381 | 6248554 | refGene_gn=DNAH10;refGene_tx=XM_023241314.1 |
| D3 | 6249422 | 6249548 | refGene_gn=DNAH10;refGene_tx=XM_023241314.1 |
| D3 | 6250519 | 6250696 | refGene_gn=DNAH10;refGene_tx=XM_023241314.1 |
| D3 | 6253339 | 6253510 | refGene_gn=DNAH10;refGene_tx=XM_023241314.1 |
| D3 | 6256883 | 6257078 | refGene_gn=DNAH10;refGene_tx=XM_023241314.1 |
| D3 | 6267469 | 6267668 | refGene_gn=DNAH10;refGene_tx=XM_023241314.1 |
| D3 | 6269402 | 6269593 | refGene_gn=DNAH10;refGene_tx=XM_023241314.1 |

D3	6271834	6272065	refGene_gn=DNAH10;refGene_tx=XM_023241314.1
D3	6272811	6272969	refGene_gn=DNAH10;refGene_tx=XM_023241314.1
D3	6278713	6278924	refGene_gn=DNAH10;refGene_tx=XM_023241314.1
D3	6282432	6282548	refGene_gn=DNAH10;refGene_tx=XM_023241314.1
D3	6285833	6285942	refGene_gn=DNAH10;refGene_tx=XM_023241314.1
D3	6287005	6287106	refGene_gn=DNAH10;refGene_tx=XM_023241314.1
D3	6290233	6290305	refGene_gn=DNAH10;refGene_tx=XM_023241314.1
D3	6293108	6293322	refGene_gn=DNAH10;refGene_tx=XM_023241314.1
D3	6297511	6297611
refGene_gn=ATP6V0A2;refGene_tx=XM_003994607.5;refGene_gn=ATP6V0A2;refGene_tx=XM_011287700.3;refGene_gn=ATP6V0A2;refGene_tx=XM_019814450.2
D3	6298137	6298309
refGene_gn=ATP6V0A2;refGene_tx=XM_003994607.5;refGene_gn=ATP6V0A2;refGene_tx=XM_011287700.3;refGene_gn=ATP6V0A2;refGene_tx=XM_019814450.2
D3	6298734	6298852
refGene_gn=ATP6V0A2;refGene_tx=XM_003994607.5;refGene_gn=ATP6V0A2;refGene_tx=XM_011287700.3;refGene_gn=ATP6V0A2;refGene_tx=XM_019814450.2
D3	6300606	6300726
refGene_gn=ATP6V0A2;refGene_tx=XM_003994607.5;refGene_gn=ATP6V0A2;refGene_tx=XM_011287700.3;refGene_gn=ATP6V0A2;refGene_tx=XM_019814450.2
D3	6301964	6302084
refGene_gn=ATP6V0A2;refGene_tx=XM_003994607.5;refGene_gn=ATP6V0A2;refGene_tx=XM_011287700.3
D3	6304407	6304618
refGene_gn=ATP6V0A2;refGene_tx=XM_003994607.5;refGene_gn=ATP6V0A2;refGene_tx=XM_011287700.3;refGene_gn=ATP6V0A2;refGene_tx=XM_019814450.2
D3	6306020	6306139
refGene_gn=ATP6V0A2;refGene_tx=XM_003994607.5;refGene_gn=ATP6V0A2;refGene_tx=XM_011287700.3;refGene_gn=ATP6V0A2;refGene_tx=XM_019814450.2
D3	6308355	6308446
refGene_gn=ATP6V0A2;refGene_tx=XM_003994607.5;refGene_gn=ATP6V0A2;refGene_tx=XM_011287700.3;refGene_gn=ATP6V0A2;refGene_tx=XM_019814450.2
D3	6308548	6308736
refGene_gn=ATP6V0A2;refGene_tx=XM_003994607.5;refGene_gn=ATP6V0A2;refGene_tx=XM_011287700.3;refGene_gn=ATP6V0A2;refGene_tx=XM_019814450.2
D3	6308978	6309115
refGene_gn=ATP6V0A2;refGene_tx=XM_003994607.5;refGene_gn=ATP6V0A2;refGene_tx=XM_011287700.3;refGene_gn=ATP6V0A2;refGene_tx=XM_019814450.2
D3	6309355	6309506
refGene_gn=ATP6V0A2;refGene_tx=XM_003994607.5;refGene_gn=ATP6V0A2;refGene_tx=XM_011287700.3;refGene_gn=ATP6V0A2;refGene_tx=XM_019814450.2
D3	6316211	6316424
refGene_gn=ATP6V0A2;refGene_tx=XM_003994607.5;refGene_gn=ATP6V0A2;refGene_tx=XM_011287700.3;refGene_gn=ATP6V0A2;refGene_tx=XM_019814450.2
D3	6317106	6317200
refGene_gn=ATP6V0A2;refGene_tx=XM_003994607.5;refGene_gn=ATP6V0A2;refGene_tx=XM_011287700.3;refGene_gn=ATP6V0A2;refGene_tx=XM_019814450.2
D3	6318957	6319040
refGene_gn=ATP6V0A2;refGene_tx=XM_003994607.5;refGene_gn=ATP6V0A2;refGene_tx=XM_011287700.3;refGene_gn=ATP6V0A2;refGene_tx=XM_019814450.2

D3 6325714 6325841
refGene_gn=ATP6V0A2;refGene_tx=XM_003994607.5;refGene_gn=ATP6V0A2;refGene_tx=XM_011287700.3;refGene_gn=ATP6V0A2;refGene_tx=XM_019814450.2
D3 6326613 6326702
refGene_gn=ATP6V0A2;refGene_tx=XM_003994607.5;refGene_gn=ATP6V0A2;refGene_tx=XM_011287700.3;refGene_gn=ATP6V0A2;refGene_tx=XM_019814450.2
D3 6327403 6327541
refGene_gn=ATP6V0A2;refGene_tx=XM_003994607.5;refGene_gn=ATP6V0A2;refGene_tx=XM_011287700.3;refGene_gn=ATP6V0A2;refGene_tx=XM_019814450.2
D3 6329545 6329643
refGene_gn=ATP6V0A2;refGene_tx=XM_003994607.5;refGene_gn=ATP6V0A2;refGene_tx=XM_011287700.3;refGene_gn=ATP6V0A2;refGene_tx=XM_019814450.2
D3 6332389 6332468
refGene_gn=ATP6V0A2;refGene_tx=XM_003994607.5;refGene_gn=ATP6V0A2;refGene_tx=XM_011287700.3;refGene_gn=ATP6V0A2;refGene_tx=XM_019814450.2
D3 6338059 6338176
refGene_gn=ATP6V0A2;refGene_tx=XM_003994607.5;refGene_gn=ATP6V0A2;refGene_tx=XM_011287700.3;refGene_gn=ATP6V0A2;refGene_tx=XM_019814450.2
D3 6340990 6341094
refGene_gn=TCTN2;refGene_tx=XM_023241319.1;refGene_gn=TCTN2;refGene_tx=XM_023241320.1;refGene_gn=TCTN2;refGene_tx=XM_023241321.1;refGene_gn=TCTN2;refGene_tx=XM_023241322.1;refGene_gn=TCTN2;refGene_tx=XM_023241323.1;refGene_gn=TCTN2;refGene_tx=XM_023241324.1;refGene_gn=TCTN2;refGene_tx=XM_023241325.1;refGene_gn=TCTN2;refGene_tx=XM_023241326.1;refGene_gn=TCTN2;refGene_tx=XM_023241327.1
D3 6341497 6341586
refGene_gn=TCTN2;refGene_tx=XM_023241319.1;refGene_gn=TCTN2;refGene_tx=XM_023241320.1;refGene_gn=TCTN2;refGene_tx=XM_023241321.1;refGene_gn=TCTN2;refGene_tx=XM_023241322.1;refGene_gn=TCTN2;refGene_tx=XM_023241323.1;refGene_gn=TCTN2;refGene_tx=XM_023241324.1;refGene_gn=TCTN2;refGene_tx=XM_023241325.1;refGene_gn=TCTN2;refGene_tx=XM_023241326.1;refGene_gn=TCTN2;refGene_tx=XM_023241327.1
D3 6341735 6341861
refGene_gn=TCTN2;refGene_tx=XM_023241319.1;refGene_gn=TCTN2;refGene_tx=XM_023241320.1;refGene_gn=TCTN2;refGene_tx=XM_023241321.1;refGene_gn=TCTN2;refGene_tx=XM_023241322.1;refGene_gn=TCTN2;refGene_tx=XM_023241323.1;refGene_gn=TCTN2;refGene_tx=XM_023241324.1;refGene_gn=TCTN2;refGene_tx=XM_023241325.1;refGene_gn=TCTN2;refGene_tx=XM_023241326.1;refGene_gn=TCTN2;refGene_tx=XM_023241327.1
D3 6342631 6342791
refGene_gn=TCTN2;refGene_tx=XM_023241319.1;refGene_gn=TCTN2;refGene_tx=XM_023241320.1;refGene_gn=TCTN2;refGene_tx=XM_023241321.1;refGene_gn=TCTN2;refGene_tx=XM_023241322.1;refGene_gn=TCTN2;refGene_tx=XM_023241323.1;refGene_gn=TCTN2;refGene_tx=XM_023241324.1;refGene_gn=TCTN2;refGene_tx=XM_023241325.1;refGene_gn=TCTN2;refGene_tx=XM_023241326.1;refGene_gn=TCTN2;refGene_tx=XM_023241327.1
D3 6345635 6345742
refGene_gn=TCTN2;refGene_tx=XM_023241319.1;refGene_gn=TCTN2;refGene_tx=XM_023241320.1;refGene_gn=TCTN2;refGene_tx=XM_023241321.1;refGene_gn=TCTN2;refGene_tx=XM_023241322.1;refGene_gn=TCTN2;refGene_tx=XM_0232413

23.1;refGene_gn=TCTN2;refGene_tx=XM_023241324.1;refGene_gn=TCTN2;refGene_tx=XM_023241325.1;refGene_gn=TCTN2;refGene_tx=XM_023241326.1;refGene_gn=TCTN2;refGene_tx=XM_023241327.1
D3 6348553 6348665
refGene_gn=TCTN2;refGene_tx=XM_023241319.1;refGene_gn=TCTN2;refGene_tx=XM_023241320.1;refGene_gn=TCTN2;refGene_tx=XM_023241321.1;refGene_gn=TCTN2;refGene_tx=XM_023241322.1;refGene_gn=TCTN2;refGene_tx=XM_023241323.1;refGene_gn=TCTN2;refGene_tx=XM_023241324.1;refGene_gn=TCTN2;refGene_tx=XM_023241325.1;refGene_gn=TCTN2;refGene_tx=XM_023241326.1;refGene_gn=TCTN2;refGene_tx=XM_023241327.1
D3 6349210 6349291
refGene_gn=TCTN2;refGene_tx=XM_023241319.1;refGene_gn=TCTN2;refGene_tx=XM_023241320.1;refGene_gn=TCTN2;refGene_tx=XM_023241321.1;refGene_gn=TCTN2;refGene_tx=XM_023241322.1;refGene_gn=TCTN2;refGene_tx=XM_023241323.1;refGene_gn=TCTN2;refGene_tx=XM_023241324.1;refGene_gn=TCTN2;refGene_tx=XM_023241325.1;refGene_gn=TCTN2;refGene_tx=XM_023241326.1;refGene_gn=TCTN2;refGene_tx=XM_023241327.1
D3 6353415 6353493
refGene_gn=TCTN2;refGene_tx=XM_023241319.1;refGene_gn=TCTN2;refGene_tx=XM_023241320.1;refGene_gn=TCTN2;refGene_tx=XM_023241321.1;refGene_gn=TCTN2;refGene_tx=XM_023241322.1;refGene_gn=TCTN2;refGene_tx=XM_023241323.1;refGene_gn=TCTN2;refGene_tx=XM_023241324.1;refGene_gn=TCTN2;refGene_tx=XM_023241325.1;refGene_gn=TCTN2;refGene_tx=XM_023241326.1;refGene_gn=TCTN2;refGene_tx=XM_023241327.1
D3 6353734 6353869
refGene_gn=TCTN2;refGene_tx=XM_023241319.1;refGene_gn=TCTN2;refGene_tx=XM_023241320.1;refGene_gn=TCTN2;refGene_tx=XM_023241321.1;refGene_gn=TCTN2;refGene_tx=XM_023241322.1;refGene_gn=TCTN2;refGene_tx=XM_023241323.1;refGene_gn=TCTN2;refGene_tx=XM_023241324.1;refGene_gn=TCTN2;refGene_tx=XM_023241326.1;refGene_gn=TCTN2;refGene_tx=XM_023241327.1
D3 6355566 6355632
refGene_gn=TCTN2;refGene_tx=XM_023241319.1;refGene_gn=TCTN2;refGene_tx=XM_023241320.1;refGene_gn=TCTN2;refGene_tx=XM_023241321.1;refGene_gn=TCTN2;refGene_tx=XM_023241322.1;refGene_gn=TCTN2;refGene_tx=XM_023241323.1;refGene_gn=TCTN2;refGene_tx=XM_023241324.1;refGene_gn=TCTN2;refGene_tx=XM_023241325.1;refGene_gn=TCTN2;refGene_tx=XM_023241326.1;refGene_gn=TCTN2;refGene_tx=XM_023241327.1
D3 6357133 6357275
refGene_gn=TCTN2;refGene_tx=XM_023241319.1;refGene_gn=TCTN2;refGene_tx=XM_023241320.1;refGene_gn=TCTN2;refGene_tx=XM_023241321.1;refGene_gn=TCTN2;refGene_tx=XM_023241322.1;refGene_gn=TCTN2;refGene_tx=XM_023241323.1;refGene_gn=TCTN2;refGene_tx=XM_023241324.1;refGene_gn=TCTN2;refGene_tx=XM_023241325.1;refGene_gn=TCTN2;refGene_tx=XM_023241326.1;refGene_gn=TCTN2;refGene_tx=XM_023241327.1
D3 6358306 6358433
refGene_gn=TCTN2;refGene_tx=XM_023241319.1;refGene_gn=TCTN2;refGene_tx=XM_023241320.1;refGene_gn=TCTN2;refGene_tx=XM_023241321.1;refGene_gn=TCTN2;refGene_tx=XM_023241322.1;refGene_gn=TCTN2;refGene_tx=XM_023241323.1;refGene_gn=TCTN2;refGene_tx=XM_023241324.1;refGene_gn=TCTN2;refGene_tx=XM_023241325.1;refGene_gn=TCTN2;refGene_tx=XM_023241326.1;refGen

e_gn=TCTN2;refGene_tx=XM_023241327.1
D3 6358820 6359062
refGene_gn=TCTN2;refGene_tx=XM_023241319.1;refGene_gn=TCTN2;refGene_tx=XM_023241320.1;refGene_gn=TCTN2;refGene_tx=XM_023241321.1;refGene_gn=TCTN2;refGene_tx=XM_023241322.1;refGene_gn=TCTN2;refGene_tx=XM_023241323.1;refGene_gn=TCTN2;refGene_tx=XM_023241324.1;refGene_gn=TCTN2;refGene_tx=XM_023241325.1;refGene_gn=TCTN2;refGene_tx=XM_023241326.1;refGene_gn=TCTN2;refGene_tx=XM_023241327.1
D3 6363734 6363835
refGene_gn=TCTN2;refGene_tx=XM_023241319.1;refGene_gn=TCTN2;refGene_tx=XM_023241320.1;refGene_gn=TCTN2;refGene_tx=XM_023241321.1;refGene_gn=TCTN2;refGene_tx=XM_023241322.1;refGene_gn=TCTN2;refGene_tx=XM_023241323.1;refGene_gn=TCTN2;refGene_tx=XM_023241324.1;refGene_gn=TCTN2;refGene_tx=XM_023241325.1;refGene_gn=TCTN2;refGene_tx=XM_023241326.1;refGene_gn=TCTN2;refGene_tx=XM_023241327.1
D3 6364988 6365057
refGene_gn=TCTN2;refGene_tx=XM_023241319.1;refGene_gn=TCTN2;refGene_tx=XM_023241320.1;refGene_gn=TCTN2;refGene_tx=XM_023241321.1;refGene_gn=TCTN2;refGene_tx=XM_023241325.1;refGene_gn=TCTN2;refGene_tx=XM_023241326.1
D3 6365979 6366013
refGene_gn=TCTN2;refGene_tx=XM_023241326.1;refGene_gn=TCTN2;refGene_tx=XM_023241327.1
D3 6367122 6367318
refGene_gn=TCTN2;refGene_tx=XM_023241319.1;refGene_gn=TCTN2;refGene_tx=XM_023241320.1;refGene_gn=TCTN2;refGene_tx=XM_023241321.1;refGene_gn=TCTN2;refGene_tx=XM_023241322.1;refGene_gn=TCTN2;refGene_tx=XM_023241323.1;refGene_gn=TCTN2;refGene_tx=XM_023241324.1;refGene_gn=TCTN2;refGene_tx=XM_023241325.1
D3 6368144 6368233
refGene_gn=TCTN2;refGene_tx=XM_023241319.1;refGene_gn=TCTN2;refGene_tx=XM_023241320.1;refGene_gn=TCTN2;refGene_tx=XM_023241321.1;refGene_gn=TCTN2;refGene_tx=XM_023241322.1;refGene_gn=TCTN2;refGene_tx=XM_023241323.1;refGene_gn=TCTN2;refGene_tx=XM_023241324.1;refGene_gn=TCTN2;refGene_tx=XM_023241325.1
D3 6368708 6368816
refGene_gn=TCTN2;refGene_tx=XM_023241319.1;refGene_gn=TCTN2;refGene_tx=XM_023241320.1;refGene_gn=TCTN2;refGene_tx=XM_023241321.1;refGene_gn=TCTN2;refGene_tx=XM_023241322.1;refGene_gn=TCTN2;refGene_tx=XM_023241323.1;refGene_gn=TCTN2;refGene_tx=XM_023241324.1;refGene_gn=TCTN2;refGene_tx=XM_023241325.1
D3 6368946 6369028
refGene_gn=TCTN2;refGene_tx=XM_023241319.1;refGene_gn=TCTN2;refGene_tx=XM_023241320.1;refGene_gn=TCTN2;refGene_tx=XM_023241321.1;refGene_gn=TCTN2;refGene_tx=XM_023241322.1;refGene_gn=TCTN2;refGene_tx=XM_023241323.1;refGene_gn=TCTN2;refGene_tx=XM_023241324.1;refGene_gn=TCTN2;refGene_tx=XM_023241325.1
D3 6374589 6374659
refGene_gn=GTF2H3;refGene_tx=XM_011287707.3;refGene_gn=GTF2H3;refGene_tx=XM_011287708.3;refGene_gn=GTF2H3;refGene_tx=XM_019814458.2;refGene_

gn=GTF2H3;refGene_tx=XM_019814459.2
D3 6374745 6374782
refGene_gn=GTF2H3;refGene_tx=XM_011287707.3;refGene_gn=GTF2H3;refGene_tx=XM_011287708.3;refGene_gn=GTF2H3;refGene_tx=XM_019814458.2;refGene_gn=GTF2H3;refGene_tx=XM_019814459.2
D3 6374893 6375029
refGene_gn=GTF2H3;refGene_tx=XM_011287707.3;refGene_gn=GTF2H3;refGene_tx=XM_011287708.3;refGene_gn=GTF2H3;refGene_tx=XM_019814458.2;refGene_gn=GTF2H3;refGene_tx=XM_019814459.2
D3 6375425 6375494
refGene_gn=GTF2H3;refGene_tx=XM_011287707.3;refGene_gn=GTF2H3;refGene_tx=XM_011287708.3;refGene_gn=GTF2H3;refGene_tx=XM_019814459.2
D3 6378631 6378685
refGene_gn=GTF2H3;refGene_tx=XM_011287707.3;refGene_gn=GTF2H3;refGene_tx=XM_011287708.3;refGene_gn=GTF2H3;refGene_tx=XM_019814458.2;refGene_gn=GTF2H3;refGene_tx=XM_019814459.2
D3 6379755 6379830
refGene_gn=GTF2H3;refGene_tx=XM_011287707.3;refGene_gn=GTF2H3;refGene_tx=XM_011287708.3;refGene_gn=GTF2H3;refGene_tx=XM_019814458.2;refGene_gn=GTF2H3;refGene_tx=XM_019814459.2
D3 6382135 6382164
refGene_gn=GTF2H3;refGene_tx=XM_011287707.3;refGene_gn=GTF2H3;refGene_tx=XM_011287708.3;refGene_gn=GTF2H3;refGene_tx=XM_019814458.2;refGene_gn=GTF2H3;refGene_tx=XM_019814459.2
D3 6382311 6382341
refGene_gn=GTF2H3;refGene_tx=XM_011287707.3;refGene_gn=GTF2H3;refGene_tx=XM_011287708.3;refGene_gn=GTF2H3;refGene_tx=XM_019814458.2;refGene_gn=GTF2H3;refGene_tx=XM_019814459.2
D3 6383231 6383294
refGene_gn=GTF2H3;refGene_tx=XM_011287707.3;refGene_gn=GTF2H3;refGene_tx=XM_011287708.3;refGene_gn=GTF2H3;refGene_tx=XM_019814458.2;refGene_gn=GTF2H3;refGene_tx=XM_019814459.2
D3 6385484 6385651
refGene_gn=GTF2H3;refGene_tx=XM_011287707.3;refGene_gn=GTF2H3;refGene_tx=XM_011287708.3;refGene_gn=GTF2H3;refGene_tx=XM_019814458.2;refGene_gn=GTF2H3;refGene_tx=XM_019814459.2
D3 6387419 6387526
refGene_gn=GTF2H3;refGene_tx=XM_011287707.3;refGene_gn=GTF2H3;refGene_tx=XM_011287708.3;refGene_gn=GTF2H3;refGene_tx=XM_019814458.2;refGene_gn=GTF2H3;refGene_tx=XM_019814459.2
D3 6394225 6394305
refGene_gn=GTF2H3;refGene_tx=XM_011287707.3;refGene_gn=GTF2H3;refGene_tx=XM_019814458.2
D3 6397439 6397452
refGene_gn=GTF2H3;refGene_tx=XM_011287707.3;refGene_gn=GTF2H3;refGene_tx=XM_019814458.2
D3 6397730 6397743
refGene_gn=EIF2B1;refGene_tx=XM_006938380.4;refGene_gn=EIF2B1;refGene_tx=XM_006938381.4;refGene_gn=EIF2B1;refGene_tx=XM_006938382.4;refGene_gn=EIF2B1;refGene_tx=XM_006938383.4;refGene_gn=EIF2B1;refGene_tx=XM_00

6938384.4;refGene_gn=EIF2B1;refGene_tx=XM_006938385.4
D3 6398783 6398885
refGene_gn=EIF2B1;refGene_tx=XM_006938380.4;refGene_gn=EIF2B1;refGene_tx=XM_006938381.4;refGene_gn=EIF2B1;refGene_tx=XM_006938382.4;refGene_gn=EIF2B1;refGene_tx=XM_006938383.4;refGene_gn=EIF2B1;refGene_tx=XM_006938384.4;refGene_gn=EIF2B1;refGene_tx=XM_006938385.4
D3 6400425 6400562
refGene_gn=EIF2B1;refGene_tx=XM_006938380.4;refGene_gn=EIF2B1;refGene_tx=XM_006938381.4;refGene_gn=EIF2B1;refGene_tx=XM_006938382.4;refGene_gn=EIF2B1;refGene_tx=XM_006938383.4;refGene_gn=EIF2B1;refGene_tx=XM_006938384.4;refGene_gn=EIF2B1;refGene_tx=XM_006938385.4
D3 6400660 6400777
refGene_gn=EIF2B1;refGene_tx=XM_006938380.4;refGene_gn=EIF2B1;refGene_tx=XM_006938381.4;refGene_gn=EIF2B1;refGene_tx=XM_006938382.4;refGene_gn=EIF2B1;refGene_tx=XM_006938383.4;refGene_gn=EIF2B1;refGene_tx=XM_006938384.4;refGene_gn=EIF2B1;refGene_tx=XM_006938385.4
D3 6402447 6402560
refGene_gn=EIF2B1;refGene_tx=XM_006938380.4;refGene_gn=EIF2B1;refGene_tx=XM_006938381.4;refGene_gn=EIF2B1;refGene_tx=XM_006938382.4;refGene_gn=EIF2B1;refGene_tx=XM_006938383.4;refGene_gn=EIF2B1;refGene_tx=XM_006938384.4;refGene_gn=EIF2B1;refGene_tx=XM_006938385.4
D3 6403325 6403394
refGene_gn=EIF2B1;refGene_tx=XM_006938380.4;refGene_gn=EIF2B1;refGene_tx=XM_006938381.4;refGene_gn=EIF2B1;refGene_tx=XM_006938382.4;refGene_gn=EIF2B1;refGene_tx=XM_006938383.4;refGene_gn=EIF2B1;refGene_tx=XM_006938384.4;refGene_gn=EIF2B1;refGene_tx=XM_006938385.4
D3 6404600 6404676
refGene_gn=EIF2B1;refGene_tx=XM_006938380.4;refGene_gn=EIF2B1;refGene_tx=XM_006938381.4;refGene_gn=EIF2B1;refGene_tx=XM_006938382.4;refGene_gn=EIF2B1;refGene_tx=XM_006938383.4;refGene_gn=EIF2B1;refGene_tx=XM_006938384.4;refGene_gn=EIF2B1;refGene_tx=XM_006938385.4
D3 6405955 6406081
refGene_gn=EIF2B1;refGene_tx=XM_006938380.4;refGene_gn=EIF2B1;refGene_tx=XM_006938381.4;refGene_gn=EIF2B1;refGene_tx=XM_006938382.4;refGene_gn=EIF2B1;refGene_tx=XM_006938383.4;refGene_gn=EIF2B1;refGene_tx=XM_006938384.4;refGene_gn=EIF2B1;refGene_tx=XM_006938385.4
D3 6406819 6406984
refGene_gn=EIF2B1;refGene_tx=XM_006938380.4;refGene_gn=EIF2B1;refGene_tx=XM_006938381.4;refGene_gn=EIF2B1;refGene_tx=XM_006938382.4;refGene_gn=EIF2B1;refGene_tx=XM_006938383.4;refGene_gn=EIF2B1;refGene_tx=XM_006938384.4;refGene_gn=EIF2B1;refGene_tx=XM_006938385.4
D3 6408505 6408682 refGene_gn=DDX55;refGene_tx=XM_003994610.5
D3 6408874 6409167 refGene_gn=DDX55;refGene_tx=XM_003994610.5
D3 6409950 6410119 refGene_gn=DDX55;refGene_tx=XM_003994610.5
D3 6410592 6410707 refGene_gn=DDX55;refGene_tx=XM_003994610.5
D3 6411333 6411426 refGene_gn=DDX55;refGene_tx=XM_003994610.5
D3 6412190 6412322 refGene_gn=DDX55;refGene_tx=XM_003994610.5
D3 6413057 6413140 refGene_gn=DDX55;refGene_tx=XM_003994610.5
D3 6414718 6414908 refGene_gn=DDX55;refGene_tx=XM_003994610.5
D3 6415429 6415579 refGene_gn=DDX55;refGene_tx=XM_003994610.5

D3	6417781	6417844	refGene_gn=DDX55;refGene_tx=XM_003994610.5
D3	6417923	6418015	refGene_gn=DDX55;refGene_tx=XM_003994610.5
D3	6418848	6418935	refGene_gn=DDX55;refGene_tx=XM_003994610.5
D3	6419029	6419080	refGene_gn=DDX55;refGene_tx=XM_003994610.5
D3	6421825	6421933	refGene_gn=DDX55;refGene_tx=XM_003994610.5
D3	6425495	6425620
refGene_gn=TMED2;refGene_tx=XM_023241328.1;refGene_gn=TMED2;refGene_tx=XM_023241329.1
D3	6428457	6428565
refGene_gn=TMED2;refGene_tx=XM_023241328.1;refGene_gn=TMED2;refGene_tx=XM_023241329.1
D3	6431258	6431279	refGene_gn=TMED2;refGene_tx=XM_023241328.1
D3	6431687	6431880
refGene_gn=TMED2;refGene_tx=XM_023241328.1;refGene_gn=TMED2;refGene_tx=XM_023241329.1
D3	6433350	6433530
refGene_gn=TMED2;refGene_tx=XM_023241328.1;refGene_gn=TMED2;refGene_tx=XM_023241329.1
D3	6456872	6457181
refGene_gn=RILPL1;refGene_tx=XM_003994586.4;refGene_gn=RILPL1;refGene_tx=XM_003994587.4;refGene_gn=RILPL1;refGene_tx=XM_006938386.3;refGene_gn=RILPL1;refGene_tx=XM_006938387.3;refGene_gn=RILPL1;refGene_tx=XM_006938388.3;refGene_gn=RILPL1;refGene_tx=XM_006938389.3
D3	6464361	6464512
refGene_gn=RILPL1;refGene_tx=XM_003994586.4;refGene_gn=RILPL1;refGene_tx=XM_003994587.4;refGene_gn=RILPL1;refGene_tx=XM_006938386.3;refGene_gn=RILPL1;refGene_tx=XM_006938387.3;refGene_gn=RILPL1;refGene_tx=XM_006938388.3;refGene_gn=RILPL1;refGene_tx=XM_006938389.3
D3	6475414	6475533
refGene_gn=RILPL1;refGene_tx=XM_003994586.4;refGene_gn=RILPL1;refGene_tx=XM_003994587.4;refGene_gn=RILPL1;refGene_tx=XM_006938386.3;refGene_gn=RILPL1;refGene_tx=XM_006938387.3;refGene_gn=RILPL1;refGene_tx=XM_006938388.3;refGene_gn=RILPL1;refGene_tx=XM_006938389.3
D3	6476116	6476338
refGene_gn=RILPL1;refGene_tx=XM_003994586.4;refGene_gn=RILPL1;refGene_tx=XM_003994587.4;refGene_gn=RILPL1;refGene_tx=XM_006938386.3;refGene_gn=RILPL1;refGene_tx=XM_006938387.3;refGene_gn=RILPL1;refGene_tx=XM_006938388.3;refGene_gn=RILPL1;refGene_tx=XM_006938389.3
D3	6484645	6484818
refGene_gn=RILPL1;refGene_tx=XM_003994586.4;refGene_gn=RILPL1;refGene_tx=XM_003994587.4;refGene_gn=RILPL1;refGene_tx=XM_006938386.3;refGene_gn=RILPL1;refGene_tx=XM_006938387.3;refGene_gn=RILPL1;refGene_tx=XM_006938388.3;refGene_gn=RILPL1;refGene_tx=XM_006938389.3
D3	6486491	6486584
refGene_gn=RILPL1;refGene_tx=XM_003994586.4;refGene_gn=RILPL1;refGene_tx=XM_003994587.4;refGene_gn=RILPL1;refGene_tx=XM_006938386.3;refGene_gn=RILPL1;refGene_tx=XM_006938387.3;refGene_gn=RILPL1;refGene_tx=XM_006938388.3;refGene_gn=RILPL1;refGene_tx=XM_006938389.3
D3	6492557	6492645
refGene_gn=RILPL1;refGene_tx=XM_003994587.4;refGene_gn=RILPL1;refGene_

tx=XM_006938388.3
D3	6492649	6492704	refGene_gn=RILPL1;refGene_tx=XM_006938389.3
D3	6492911	6492935
refGene_gn=RILPL1;refGene_tx=XM_006938386.3;refGene_gn=RILPL1;refGene_tx=XM_006938387.3;refGene_gn=RILPL1;refGene_tx=XM_006938388.3
D3	6493123	6493205	refGene_gn=RILPL1;refGene_tx=XM_006938387.3
D3	6494780	6494925
refGene_gn=RILPL1;refGene_tx=XM_003994586.4;refGene_gn=RILPL1;refGene_tx=XM_003994587.4;refGene_gn=RILPL1;refGene_tx=XM_006938386.3
D3	6498612	6499353	refGene_gn=SNRNP35;refGene_tx=XM_003994588.5
D3	6525319	6525661
refGene_gn=RILPL2;refGene_tx=XM_006938391.4;refGene_gn=RILPL2;refGene_tx=XM_006938393.2;refGene_gn=RILPL2;refGene_tx=XM_023241330.1;refGene_gn=RILPL2;refGene_tx=XM_023241331.1
D3	6532833	6532985
refGene_gn=RILPL2;refGene_tx=XM_006938391.4;refGene_gn=RILPL2;refGene_tx=XM_006938393.2;refGene_gn=RILPL2;refGene_tx=XM_023241330.1;refGene_gn=RILPL2;refGene_tx=XM_023241331.1
D3	6538234	6538348
refGene_gn=RILPL2;refGene_tx=XM_006938391.4;refGene_gn=RILPL2;refGene_tx=XM_006938393.2;refGene_gn=RILPL2;refGene_tx=XM_023241330.1;refGene_gn=RILPL2;refGene_tx=XM_023241331.1
D3	6541283	6541308
refGene_gn=RILPL2;refGene_tx=XM_006938391.4;refGene_gn=RILPL2;refGene_tx=XM_006938393.2;refGene_gn=RILPL2;refGene_tx=XM_023241330.1;refGene_gn=RILPL2;refGene_tx=XM_023241331.1
D3	6547664	6547875
refGene_gn=LOC101086008;refGene_tx=XM_023241332.1
D3	6548639	6548830
refGene_gn=LOC101086008;refGene_tx=XM_023241332.1
D3	6549608	6549668
refGene_gn=LOC101086008;refGene_tx=XM_023241332.1
D3	6554316	6554404
refGene_gn=LOC101086008;refGene_tx=XM_023241332.1
D3	6554749	6554934
refGene_gn=LOC101086008;refGene_tx=XM_023241332.1
D3	6555427	6555620
refGene_gn=LOC111557363;refGene_tx=XM_023242151.1
D3	6555966	6556186
refGene_gn=LOC111557363;refGene_tx=XM_023242151.1
D3	6560087	6560244
refGene_gn=LOC111557363;refGene_tx=XM_023242151.1
D3	6561414	6561536
refGene_gn=LOC111557363;refGene_tx=XM_023242151.1
D3	6561804	6562225
refGene_gn=LOC111557363;refGene_tx=XM_023242151.1
D3	6567172	6567298
refGene_gn=LOC111557363;refGene_tx=XM_023242151.1
D3	6589745	6589877
refGene_gn=SBNO1;refGene_tx=XM_023241333.1;refGene_gn=SBNO1;refGene_tx

=XM_023241334.1;refGene_gn=SBNO1;refGene_tx=XM_023241335.1;refGene_gn=SBNO1;refGene_tx=XM_023241336.1
D3 6590597 6590702
refGene_gn=SBNO1;refGene_tx=XM_023241333.1;refGene_gn=SBNO1;refGene_tx=XM_023241334.1;refGene_gn=SBNO1;refGene_tx=XM_023241335.1
D3 6592627 6592940
refGene_gn=SBNO1;refGene_tx=XM_019814465.2;refGene_gn=SBNO1;refGene_tx=XM_023241333.1;refGene_gn=SBNO1;refGene_tx=XM_023241334.1;refGene_gn=SBNO1;refGene_tx=XM_023241335.1;refGene_gn=SBNO1;refGene_tx=XM_023241336.1
D3 6596529 6596630
refGene_gn=SBNO1;refGene_tx=XM_019814465.2;refGene_gn=SBNO1;refGene_tx=XM_023241333.1;refGene_gn=SBNO1;refGene_tx=XM_023241334.1;refGene_gn=SBNO1;refGene_tx=XM_023241335.1;refGene_gn=SBNO1;refGene_tx=XM_023241336.1
D3 6600800 6600897
refGene_gn=SBNO1;refGene_tx=XM_019814465.2;refGene_gn=SBNO1;refGene_tx=XM_023241333.1;refGene_gn=SBNO1;refGene_tx=XM_023241334.1;refGene_gn=SBNO1;refGene_tx=XM_023241335.1;refGene_gn=SBNO1;refGene_tx=XM_023241336.1
D3 6603449 6603610
refGene_gn=SBNO1;refGene_tx=XM_019814465.2;refGene_gn=SBNO1;refGene_tx=XM_023241333.1;refGene_gn=SBNO1;refGene_tx=XM_023241334.1;refGene_gn=SBNO1;refGene_tx=XM_023241335.1;refGene_gn=SBNO1;refGene_tx=XM_023241336.1
D3 6604793 6604927
refGene_gn=SBNO1;refGene_tx=XM_019814465.2;refGene_gn=SBNO1;refGene_tx=XM_023241333.1;refGene_gn=SBNO1;refGene_tx=XM_023241334.1;refGene_gn=SBNO1;refGene_tx=XM_023241335.1;refGene_gn=SBNO1;refGene_tx=XM_023241336.1
D3 6605128 6605219
refGene_gn=SBNO1;refGene_tx=XM_019814465.2;refGene_gn=SBNO1;refGene_tx=XM_023241333.1;refGene_gn=SBNO1;refGene_tx=XM_023241334.1;refGene_gn=SBNO1;refGene_tx=XM_023241335.1;refGene_gn=SBNO1;refGene_tx=XM_023241336.1
D3 6606686 6606848
refGene_gn=SBNO1;refGene_tx=XM_019814465.2;refGene_gn=SBNO1;refGene_tx=XM_023241333.1;refGene_gn=SBNO1;refGene_tx=XM_023241334.1;refGene_gn=SBNO1;refGene_tx=XM_023241335.1;refGene_gn=SBNO1;refGene_tx=XM_023241336.1
D3 6607726 6607862
refGene_gn=SBNO1;refGene_tx=XM_019814465.2;refGene_gn=SBNO1;refGene_tx=XM_023241333.1;refGene_gn=SBNO1;refGene_tx=XM_023241334.1;refGene_gn=SBNO1;refGene_tx=XM_023241335.1;refGene_gn=SBNO1;refGene_tx=XM_023241336.1
D3 6607942 6608048
refGene_gn=SBNO1;refGene_tx=XM_019814465.2;refGene_gn=SBNO1;refGene_tx=XM_023241333.1;refGene_gn=SBNO1;refGene_tx=XM_023241334.1;refGene_gn=SBNO1;refGene_tx=XM_023241335.1;refGene_gn=SBNO1;refGene_tx=XM_023241336.1

D3	6608183	6608337
refGene_gn=SBNO1;refGene_tx=XM_019814465.2;refGene_gn=SBNO1;refGene_tx=XM_023241333.1;refGene_gn=SBNO1;refGene_tx=XM_023241334.1;refGene_gn=SBNO1;refGene_tx=XM_023241335.1;refGene_gn=SBNO1;refGene_tx=XM_023241336.1
D3	6609567	6609750
refGene_gn=SBNO1;refGene_tx=XM_019814465.2;refGene_gn=SBNO1;refGene_tx=XM_023241333.1;refGene_gn=SBNO1;refGene_tx=XM_023241334.1;refGene_gn=SBNO1;refGene_tx=XM_023241335.1;refGene_gn=SBNO1;refGene_tx=XM_023241336.1
D3	6610267	6610365
refGene_gn=SBNO1;refGene_tx=XM_019814465.2;refGene_gn=SBNO1;refGene_tx=XM_023241333.1;refGene_gn=SBNO1;refGene_tx=XM_023241334.1;refGene_gn=SBNO1;refGene_tx=XM_023241335.1;refGene_gn=SBNO1;refGene_tx=XM_023241336.1
D3	6611462	6611614
refGene_gn=SBNO1;refGene_tx=XM_019814465.2;refGene_gn=SBNO1;refGene_tx=XM_023241333.1;refGene_gn=SBNO1;refGene_tx=XM_023241334.1;refGene_gn=SBNO1;refGene_tx=XM_023241335.1;refGene_gn=SBNO1;refGene_tx=XM_023241336.1
D3	6613581	6613779
refGene_gn=SBNO1;refGene_tx=XM_019814465.2;refGene_gn=SBNO1;refGene_tx=XM_023241333.1;refGene_gn=SBNO1;refGene_tx=XM_023241334.1;refGene_gn=SBNO1;refGene_tx=XM_023241335.1;refGene_gn=SBNO1;refGene_tx=XM_023241336.1
D3	6614168	6614336
refGene_gn=SBNO1;refGene_tx=XM_019814465.2;refGene_gn=SBNO1;refGene_tx=XM_023241333.1;refGene_gn=SBNO1;refGene_tx=XM_023241334.1;refGene_gn=SBNO1;refGene_tx=XM_023241335.1;refGene_gn=SBNO1;refGene_tx=XM_023241336.1
D3	6614446	6614622
refGene_gn=SBNO1;refGene_tx=XM_019814465.2;refGene_gn=SBNO1;refGene_tx=XM_023241333.1;refGene_gn=SBNO1;refGene_tx=XM_023241334.1;refGene_gn=SBNO1;refGene_tx=XM_023241335.1;refGene_gn=SBNO1;refGene_tx=XM_023241336.1
D3	6615090	6615222
refGene_gn=SBNO1;refGene_tx=XM_019814465.2;refGene_gn=SBNO1;refGene_tx=XM_023241333.1;refGene_gn=SBNO1;refGene_tx=XM_023241334.1;refGene_gn=SBNO1;refGene_tx=XM_023241335.1;refGene_gn=SBNO1;refGene_tx=XM_023241336.1
D3	6617067	6617203
refGene_gn=SBNO1;refGene_tx=XM_019814465.2;refGene_gn=SBNO1;refGene_tx=XM_023241333.1;refGene_gn=SBNO1;refGene_tx=XM_023241334.1;refGene_gn=SBNO1;refGene_tx=XM_023241335.1;refGene_gn=SBNO1;refGene_tx=XM_023241336.1
D3	6618140	6618253
refGene_gn=SBNO1;refGene_tx=XM_019814465.2;refGene_gn=SBNO1;refGene_tx=XM_023241333.1;refGene_gn=SBNO1;refGene_tx=XM_023241334.1;refGene_gn=SBNO1;refGene_tx=XM_023241335.1;refGene_gn=SBNO1;refGene_tx=XM_023241336.1

D3	6618344	6618416
refGene_gn=SBNO1;refGene_tx=XM_019814465.2;refGene_gn=SBNO1;refGene_tx=XM_023241333.1;refGene_gn=SBNO1;refGene_tx=XM_023241334.1;refGene_gn=SBNO1;refGene_tx=XM_023241335.1;refGene_gn=SBNO1;refGene_tx=XM_023241336.1
D3	6618936	6619036
refGene_gn=SBNO1;refGene_tx=XM_019814465.2;refGene_gn=SBNO1;refGene_tx=XM_023241333.1;refGene_gn=SBNO1;refGene_tx=XM_023241334.1;refGene_gn=SBNO1;refGene_tx=XM_023241335.1;refGene_gn=SBNO1;refGene_tx=XM_023241336.1
D3	6623376	6623451
refGene_gn=SBNO1;refGene_tx=XM_019814465.2;refGene_gn=SBNO1;refGene_tx=XM_023241333.1;refGene_gn=SBNO1;refGene_tx=XM_023241334.1;refGene_gn=SBNO1;refGene_tx=XM_023241335.1;refGene_gn=SBNO1;refGene_tx=XM_023241336.1
D3	6624359	6624506
refGene_gn=SBNO1;refGene_tx=XM_019814465.2;refGene_gn=SBNO1;refGene_tx=XM_023241333.1;refGene_gn=SBNO1;refGene_tx=XM_023241334.1;refGene_gn=SBNO1;refGene_tx=XM_023241335.1;refGene_gn=SBNO1;refGene_tx=XM_023241336.1
D3	6624591	6624686
refGene_gn=SBNO1;refGene_tx=XM_019814465.2;refGene_gn=SBNO1;refGene_tx=XM_023241333.1;refGene_gn=SBNO1;refGene_tx=XM_023241334.1;refGene_gn=SBNO1;refGene_tx=XM_023241335.1;refGene_gn=SBNO1;refGene_tx=XM_023241336.1
D3	6624783	6624876
refGene_gn=SBNO1;refGene_tx=XM_019814465.2;refGene_gn=SBNO1;refGene_tx=XM_023241333.1;refGene_gn=SBNO1;refGene_tx=XM_023241334.1;refGene_gn=SBNO1;refGene_tx=XM_023241335.1;refGene_gn=SBNO1;refGene_tx=XM_023241336.1
D3	6630095	6630233
refGene_gn=SBNO1;refGene_tx=XM_019814465.2;refGene_gn=SBNO1;refGene_tx=XM_023241333.1;refGene_gn=SBNO1;refGene_tx=XM_023241334.1;refGene_gn=SBNO1;refGene_tx=XM_023241335.1;refGene_gn=SBNO1;refGene_tx=XM_023241336.1
D3	6630844	6630921
refGene_gn=SBNO1;refGene_tx=XM_019814465.2;refGene_gn=SBNO1;refGene_tx=XM_023241333.1;refGene_gn=SBNO1;refGene_tx=XM_023241334.1;refGene_gn=SBNO1;refGene_tx=XM_023241335.1;refGene_gn=SBNO1;refGene_tx=XM_023241336.1
D3	6632551	6632745
refGene_gn=SBNO1;refGene_tx=XM_019814465.2;refGene_gn=SBNO1;refGene_tx=XM_023241333.1;refGene_gn=SBNO1;refGene_tx=XM_023241334.1;refGene_gn=SBNO1;refGene_tx=XM_023241335.1;refGene_gn=SBNO1;refGene_tx=XM_023241336.1
D3	6633971	6634114
refGene_gn=SBNO1;refGene_tx=XM_019814465.2;refGene_gn=SBNO1;refGene_tx=XM_023241333.1;refGene_gn=SBNO1;refGene_tx=XM_023241334.1;refGene_gn=SBNO1;refGene_tx=XM_023241335.1;refGene_gn=SBNO1;refGene_tx=XM_023241336.1

D3 6653327 6653382 refGene_gn=CDK2AP1;refGene_tx=XM_023241337.1
D3 6657964 6658062
refGene_gn=CDK2AP1;refGene_tx=XM_006938406.3;refGene_gn=CDK2AP1;refGene_tx=XM_023241337.1
D3 6659701 6659828
refGene_gn=CDK2AP1;refGene_tx=XM_006938406.3;refGene_gn=CDK2AP1;refGene_tx=XM_023241337.1
D3 6662911 6662979
refGene_gn=CDK2AP1;refGene_tx=XM_006938406.3;refGene_gn=CDK2AP1;refGene_tx=XM_023241337.1
D3 6666828 6667047
refGene_gn=CD3H12orf65;refGene_tx=XM_003994613.4;refGene_gn=CD3H12orf65;refGene_tx=XM_006938408.3;refGene_gn=CD3H12orf65;refGene_tx=XM_006938409.2;refGene_gn=CD3H12orf65;refGene_tx=XM_023241339.1
D3 6668738 6669049
refGene_gn=CD3H12orf65;refGene_tx=XM_003994613.4;refGene_gn=CD3H12orf65;refGene_tx=XM_006938408.3;refGene_gn=CD3H12orf65;refGene_tx=XM_006938409.2;refGene_gn=CD3H12orf65;refGene_tx=XM_023241339.1
D3 6680407 6680474
refGene_gn=CD3H12orf65;refGene_tx=XM_006938408.3
D3 6680561 6680864
refGene_gn=MPHOSPH9;refGene_tx=XM_019814466.2;refGene_gn=MPHOSPH9;refGene_tx=XM_019814467.2
D3 6683699 6683806
refGene_gn=MPHOSPH9;refGene_tx=XM_011287712.3;refGene_gn=MPHOSPH9;refGene_tx=XM_023241338.1
D3 6686274 6686428
refGene_gn=MPHOSPH9;refGene_tx=XM_011287712.3;refGene_gn=MPHOSPH9;refGene_tx=XM_019814468.2;refGene_gn=MPHOSPH9;refGene_tx=XM_023241338.1
D3 6692101 6692191
refGene_gn=MPHOSPH9;refGene_tx=XM_011287712.3;refGene_gn=MPHOSPH9;refGene_tx=XM_019814466.2;refGene_gn=MPHOSPH9;refGene_tx=XM_019814467.2;refGene_gn=MPHOSPH9;refGene_tx=XM_019814468.2;refGene_gn=MPHOSPH9;refGene_tx=XM_023241338.1
D3 6692395 6692916
refGene_gn=MPHOSPH9;refGene_tx=XM_011287712.3;refGene_gn=MPHOSPH9;refGene_tx=XM_019814466.2;refGene_gn=MPHOSPH9;refGene_tx=XM_019814467.2;refGene_gn=MPHOSPH9;refGene_tx=XM_019814468.2;refGene_gn=MPHOSPH9;refGene_tx=XM_023241338.1
D3 6693569 6693693
refGene_gn=MPHOSPH9;refGene_tx=XM_011287712.3;refGene_gn=MPHOSPH9;refGene_tx=XM_019814466.2;refGene_gn=MPHOSPH9;refGene_tx=XM_019814467.2;refGene_gn=MPHOSPH9;refGene_tx=XM_019814468.2;refGene_gn=MPHOSPH9;refGene_tx=XM_023241338.1
D3 6695778 6695869
refGene_gn=MPHOSPH9;refGene_tx=XM_011287712.3;refGene_gn=MPHOSPH9;refGene_tx=XM_019814466.2;refGene_gn=MPHOSPH9;refGene_tx=XM_019814467.2;refGene_gn=MPHOSPH9;refGene_tx=XM_019814468.2;refGene_gn=MPHOSPH9;refGene_tx=XM_023241338.1
D3 6697600 6697704

refGene_gn=MPHOSPH9;refGene_tx=XM_011287712.3;refGene_gn=MPHOSPH9;refGene_tx=XM_019814466.2;refGene_gn=MPHOSPH9;refGene_tx=XM_019814467.2;refGene_gn=MPHOSPH9;refGene_tx=XM_019814468.2;refGene_gn=MPHOSPH9;refGene_tx=XM_023241338.1
D3 6702530 6702656
refGene_gn=MPHOSPH9;refGene_tx=XM_011287712.3;refGene_gn=MPHOSPH9;refGene_tx=XM_019814466.2;refGene_gn=MPHOSPH9;refGene_tx=XM_019814468.2;refGene_gn=MPHOSPH9;refGene_tx=XM_023241338.1
D3 6702819 6703277
refGene_gn=MPHOSPH9;refGene_tx=XM_011287712.3;refGene_gn=MPHOSPH9;refGene_tx=XM_019814466.2;refGene_gn=MPHOSPH9;refGene_tx=XM_019814467.2;refGene_gn=MPHOSPH9;refGene_tx=XM_019814468.2;refGene_gn=MPHOSPH9;refGene_tx=XM_023241338.1
D3 6703608 6703764
refGene_gn=MPHOSPH9;refGene_tx=XM_011287712.3;refGene_gn=MPHOSPH9;refGene_tx=XM_019814466.2;refGene_gn=MPHOSPH9;refGene_tx=XM_019814467.2;refGene_gn=MPHOSPH9;refGene_tx=XM_019814468.2;refGene_gn=MPHOSPH9;refGene_tx=XM_023241338.1
D3 6708083 6708171
refGene_gn=MPHOSPH9;refGene_tx=XM_011287712.3;refGene_gn=MPHOSPH9;refGene_tx=XM_019814466.2;refGene_gn=MPHOSPH9;refGene_tx=XM_019814467.2;refGene_gn=MPHOSPH9;refGene_tx=XM_019814468.2;refGene_gn=MPHOSPH9;refGene_tx=XM_023241338.1
D3 6710676 6710892
refGene_gn=MPHOSPH9;refGene_tx=XM_011287712.3;refGene_gn=MPHOSPH9;refGene_tx=XM_019814466.2;refGene_gn=MPHOSPH9;refGene_tx=XM_019814467.2;refGene_gn=MPHOSPH9;refGene_tx=XM_019814468.2;refGene_gn=MPHOSPH9;refGene_tx=XM_023241338.1
D3 6721753 6721801
refGene_gn=MPHOSPH9;refGene_tx=XM_011287712.3;refGene_gn=MPHOSPH9;refGene_tx=XM_019814466.2;refGene_gn=MPHOSPH9;refGene_tx=XM_019814467.2;refGene_gn=MPHOSPH9;refGene_tx=XM_019814468.2;refGene_gn=MPHOSPH9;refGene_tx=XM_023241338.1
D3 6722750 6722815
refGene_gn=MPHOSPH9;refGene_tx=XM_011287712.3;refGene_gn=MPHOSPH9;refGene_tx=XM_019814466.2;refGene_gn=MPHOSPH9;refGene_tx=XM_019814467.2;refGene_gn=MPHOSPH9;refGene_tx=XM_019814468.2;refGene_gn=MPHOSPH9;refGene_tx=XM_023241338.1
D3 6724259 6724361
refGene_gn=MPHOSPH9;refGene_tx=XM_011287712.3;refGene_gn=MPHOSPH9;refGene_tx=XM_019814466.2;refGene_gn=MPHOSPH9;refGene_tx=XM_019814467.2;refGene_gn=MPHOSPH9;refGene_tx=XM_019814468.2;refGene_gn=MPHOSPH9;refGene_tx=XM_023241338.1
D3 6731543 6731678
refGene_gn=MPHOSPH9;refGene_tx=XM_011287712.3;refGene_gn=MPHOSPH9;refGene_tx=XM_019814466.2;refGene_gn=MPHOSPH9;refGene_tx=XM_019814467.2;refGene_gn=MPHOSPH9;refGene_tx=XM_019814468.2;refGene_gn=MPHOSPH9;refGene_tx=XM_023241338.1
D3 6732554 6732730
refGene_gn=MPHOSPH9;refGene_tx=XM_011287712.3;refGene_gn=MPHOSPH9;refG

ene_tx=XM_019814466.2;refGene_gn=MPHOSPH9;refGene_tx=XM_019814467.2;refGene_gn=MPHOSPH9;refGene_tx=XM_019814468.2;refGene_gn=MPHOSPH9;refGene_tx=XM_023241338.1
D3 6733471 6733618
refGene_gn=MPHOSPH9;refGene_tx=XM_011287712.3;refGene_gn=MPHOSPH9;refGene_tx=XM_019814466.2;refGene_gn=MPHOSPH9;refGene_tx=XM_019814467.2;refGene_gn=MPHOSPH9;refGene_tx=XM_019814468.2;refGene_gn=MPHOSPH9;refGene_tx=XM_023241338.1
D3 6734395 6734516
refGene_gn=MPHOSPH9;refGene_tx=XM_011287712.3;refGene_gn=MPHOSPH9;refGene_tx=XM_019814466.2;refGene_gn=MPHOSPH9;refGene_tx=XM_019814467.2;refGene_gn=MPHOSPH9;refGene_tx=XM_019814468.2;refGene_gn=MPHOSPH9;refGene_tx=XM_023241338.1
D3 6735278 6735382
refGene_gn=MPHOSPH9;refGene_tx=XM_011287712.3;refGene_gn=MPHOSPH9;refGene_tx=XM_019814466.2;refGene_gn=MPHOSPH9;refGene_tx=XM_019814467.2;refGene_gn=MPHOSPH9;refGene_tx=XM_019814468.2;refGene_gn=MPHOSPH9;refGene_tx=XM_023241338.1
D3 6736116 6736364
refGene_gn=MPHOSPH9;refGene_tx=XM_011287712.3;refGene_gn=MPHOSPH9;refGene_tx=XM_019814466.2;refGene_gn=MPHOSPH9;refGene_tx=XM_019814467.2;refGene_gn=MPHOSPH9;refGene_tx=XM_019814468.2;refGene_gn=MPHOSPH9;refGene_tx=XM_023241338.1
D3 6736678 6736747
refGene_gn=MPHOSPH9;refGene_tx=XM_011287712.3;refGene_gn=MPHOSPH9;refGene_tx=XM_019814466.2;refGene_gn=MPHOSPH9;refGene_tx=XM_019814467.2;refGene_gn=MPHOSPH9;refGene_tx=XM_019814468.2;refGene_gn=MPHOSPH9;refGene_tx=XM_023241338.1
D3 6739963 6740065
refGene_gn=MPHOSPH9;refGene_tx=XM_011287712.3;refGene_gn=MPHOSPH9;refGene_tx=XM_019814466.2;refGene_gn=MPHOSPH9;refGene_tx=XM_019814467.2;refGene_gn=MPHOSPH9;refGene_tx=XM_019814468.2;refGene_gn=MPHOSPH9;refGene_tx=XM_023241338.1
D3 6852803 6852881
refGene_gn=PITPNM2;refGene_tx=XM_023241340.1;refGene_gn=PITPNM2;refGene_tx=XM_023241341.1;refGene_gn=PITPNM2;refGene_tx=XM_023241342.1;refGene_gn=PITPNM2;refGene_tx=XM_023241343.1;refGene_gn=PITPNM2;refGene_tx=XM_023241344.1;refGene_gn=PITPNM2;refGene_tx=XM_023241345.1;refGene_gn=PITPNM2;refGene_tx=XM_023241346.1;refGene_gn=PITPNM2;refGene_tx=XM_023241347.1;refGene_gn=PITPNM2;refGene_tx=XM_023241348.1;refGene_gn=PITPNM2;refGene_tx=XM_023241349.1;refGene_gn=PITPNM2;refGene_tx=XM_023241350.1;refGene_gn=PITPNM2;refGene_tx=XM_023241351.1
D3 6861685 6861900
refGene_gn=PITPNM2;refGene_tx=XM_023241340.1;refGene_gn=PITPNM2;refGene_tx=XM_023241341.1;refGene_gn=PITPNM2;refGene_tx=XM_023241342.1;refGene_gn=PITPNM2;refGene_tx=XM_023241343.1;refGene_gn=PITPNM2;refGene_tx=XM_023241344.1;refGene_gn=PITPNM2;refGene_tx=XM_023241345.1;refGene_gn=PITPNM2;refGene_tx=XM_023241346.1;refGene_gn=PITPNM2;refGene_tx=XM_023241347.1;refGene_gn=PITPNM2;refGene_tx=XM_023241348.1;refGene_gn=PITPNM2;refGene_tx=XM_023241349.1;refGene_gn=PITPNM2;refGene_tx=XM_0232413

50.1;refGene_gn=PITPNM2;refGene_tx=XM_023241351.1
D3 6862962 6863084
refGene_gn=PITPNM2;refGene_tx=XM_023241340.1;refGene_gn=PITPNM2;refGene_tx=XM_023241341.1;refGene_gn=PITPNM2;refGene_tx=XM_023241342.1;refGene_gn=PITPNM2;refGene_tx=XM_023241343.1;refGene_gn=PITPNM2;refGene_tx=XM_023241344.1;refGene_gn=PITPNM2;refGene_tx=XM_023241345.1;refGene_gn=PITPNM2;refGene_tx=XM_023241346.1;refGene_gn=PITPNM2;refGene_tx=XM_023241347.1;refGene_gn=PITPNM2;refGene_tx=XM_023241348.1;refGene_gn=PITPNM2;refGene_tx=XM_023241349.1;refGene_gn=PITPNM2;refGene_tx=XM_023241350.1;refGene_gn=PITPNM2;refGene_tx=XM_023241351.1
D3 6865612 6865840
refGene_gn=PITPNM2;refGene_tx=XM_023241340.1;refGene_gn=PITPNM2;refGene_tx=XM_023241341.1;refGene_gn=PITPNM2;refGene_tx=XM_023241342.1;refGene_gn=PITPNM2;refGene_tx=XM_023241343.1;refGene_gn=PITPNM2;refGene_tx=XM_023241344.1;refGene_gn=PITPNM2;refGene_tx=XM_023241345.1;refGene_gn=PITPNM2;refGene_tx=XM_023241346.1;refGene_gn=PITPNM2;refGene_tx=XM_023241347.1;refGene_gn=PITPNM2;refGene_tx=XM_023241348.1;refGene_gn=PITPNM2;refGene_tx=XM_023241349.1;refGene_gn=PITPNM2;refGene_tx=XM_023241350.1;refGene_gn=PITPNM2;refGene_tx=XM_023241351.1
D3 6868094 6868412
refGene_gn=PITPNM2;refGene_tx=XM_023241340.1;refGene_gn=PITPNM2;refGene_tx=XM_023241341.1;refGene_gn=PITPNM2;refGene_tx=XM_023241342.1;refGene_gn=PITPNM2;refGene_tx=XM_023241343.1;refGene_gn=PITPNM2;refGene_tx=XM_023241344.1;refGene_gn=PITPNM2;refGene_tx=XM_023241345.1;refGene_gn=PITPNM2;refGene_tx=XM_023241346.1;refGene_gn=PITPNM2;refGene_tx=XM_023241347.1;refGene_gn=PITPNM2;refGene_tx=XM_023241348.1;refGene_gn=PITPNM2;refGene_tx=XM_023241349.1;refGene_gn=PITPNM2;refGene_tx=XM_023241350.1;refGene_gn=PITPNM2;refGene_tx=XM_023241351.1
D3 6869072 6869132
refGene_gn=PITPNM2;refGene_tx=XM_023241340.1;refGene_gn=PITPNM2;refGene_tx=XM_023241341.1;refGene_gn=PITPNM2;refGene_tx=XM_023241342.1;refGene_gn=PITPNM2;refGene_tx=XM_023241343.1;refGene_gn=PITPNM2;refGene_tx=XM_023241345.1;refGene_gn=PITPNM2;refGene_tx=XM_023241346.1;refGene_gn=PITPNM2;refGene_tx=XM_023241347.1;refGene_gn=PITPNM2;refGene_tx=XM_023241350.1
D3 6869401 6869497
refGene_gn=PITPNM2;refGene_tx=XM_023241340.1;refGene_gn=PITPNM2;refGene_tx=XM_023241341.1;refGene_gn=PITPNM2;refGene_tx=XM_023241342.1;refGene_gn=PITPNM2;refGene_tx=XM_023241343.1;refGene_gn=PITPNM2;refGene_tx=XM_023241344.1;refGene_gn=PITPNM2;refGene_tx=XM_023241345.1;refGene_gn=PITPNM2;refGene_tx=XM_023241346.1;refGene_gn=PITPNM2;refGene_tx=XM_023241347.1;refGene_gn=PITPNM2;refGene_tx=XM_023241348.1;refGene_gn=PITPNM2;refGene_tx=XM_023241349.1;refGene_gn=PITPNM2;refGene_tx=XM_023241350.1;refGene_gn=PITPNM2;refGene_tx=XM_023241351.1
D3 6871914 6872019
refGene_gn=PITPNM2;refGene_tx=XM_023241340.1;refGene_gn=PITPNM2;refGene_tx=XM_023241341.1;refGene_gn=PITPNM2;refGene_tx=XM_023241342.1;refGene_gn=PITPNM2;refGene_tx=XM_023241343.1;refGene_gn=PITPNM2;refGene_tx=XM_023241344.1;refGene_gn=PITPNM2;refGene_tx=XM_023241345.1;refGene_gn=PITPNM2;refGene_tx=XM_023241346.1;refGene_gn=PITPNM2;refGene_tx=XM_02

3241347.1;refGene_gn=PITPNM2;refGene_tx=XM_023241348.1;refGene_gn=PITPNM2;refGene_tx=XM_023241349.1;refGene_gn=PITPNM2;refGene_tx=XM_023241350.1;refGene_gn=PITPNM2;refGene_tx=XM_023241351.1
D3 6872243 6872314
refGene_gn=PITPNM2;refGene_tx=XM_023241340.1;refGene_gn=PITPNM2;refGene_tx=XM_023241341.1;refGene_gn=PITPNM2;refGene_tx=XM_023241342.1;refGene_gn=PITPNM2;refGene_tx=XM_023241343.1;refGene_gn=PITPNM2;refGene_tx=XM_023241344.1;refGene_gn=PITPNM2;refGene_tx=XM_023241345.1;refGene_gn=PITPNM2;refGene_tx=XM_023241346.1;refGene_gn=PITPNM2;refGene_tx=XM_023241347.1;refGene_gn=PITPNM2;refGene_tx=XM_023241348.1;refGene_gn=PITPNM2;refGene_tx=XM_023241349.1;refGene_gn=PITPNM2;refGene_tx=XM_023241350.1;refGene_gn=PITPNM2;refGene_tx=XM_023241351.1
D3 6875064 6875312
refGene_gn=PITPNM2;refGene_tx=XM_023241340.1;refGene_gn=PITPNM2;refGene_tx=XM_023241341.1;refGene_gn=PITPNM2;refGene_tx=XM_023241342.1;refGene_gn=PITPNM2;refGene_tx=XM_023241343.1;refGene_gn=PITPNM2;refGene_tx=XM_023241344.1;refGene_gn=PITPNM2;refGene_tx=XM_023241345.1;refGene_gn=PITPNM2;refGene_tx=XM_023241346.1;refGene_gn=PITPNM2;refGene_tx=XM_023241347.1;refGene_gn=PITPNM2;refGene_tx=XM_023241348.1;refGene_gn=PITPNM2;refGene_tx=XM_023241349.1;refGene_gn=PITPNM2;refGene_tx=XM_023241350.1;refGene_gn=PITPNM2;refGene_tx=XM_023241351.1
D3 6875709 6875899
refGene_gn=PITPNM2;refGene_tx=XM_023241340.1;refGene_gn=PITPNM2;refGene_tx=XM_023241341.1;refGene_gn=PITPNM2;refGene_tx=XM_023241342.1;refGene_gn=PITPNM2;refGene_tx=XM_023241343.1;refGene_gn=PITPNM2;refGene_tx=XM_023241344.1;refGene_gn=PITPNM2;refGene_tx=XM_023241345.1;refGene_gn=PITPNM2;refGene_tx=XM_023241346.1;refGene_gn=PITPNM2;refGene_tx=XM_023241347.1;refGene_gn=PITPNM2;refGene_tx=XM_023241348.1;refGene_gn=PITPNM2;refGene_tx=XM_023241349.1;refGene_gn=PITPNM2;refGene_tx=XM_023241350.1;refGene_gn=PITPNM2;refGene_tx=XM_023241351.1
D3 6876023 6876143
refGene_gn=PITPNM2;refGene_tx=XM_023241340.1;refGene_gn=PITPNM2;refGene_tx=XM_023241341.1;refGene_gn=PITPNM2;refGene_tx=XM_023241342.1;refGene_gn=PITPNM2;refGene_tx=XM_023241343.1;refGene_gn=PITPNM2;refGene_tx=XM_023241344.1;refGene_gn=PITPNM2;refGene_tx=XM_023241345.1;refGene_gn=PITPNM2;refGene_tx=XM_023241346.1;refGene_gn=PITPNM2;refGene_tx=XM_023241347.1;refGene_gn=PITPNM2;refGene_tx=XM_023241348.1;refGene_gn=PITPNM2;refGene_tx=XM_023241349.1;refGene_gn=PITPNM2;refGene_tx=XM_023241350.1;refGene_gn=PITPNM2;refGene_tx=XM_023241351.1
D3 6876740 6876958
refGene_gn=PITPNM2;refGene_tx=XM_023241340.1;refGene_gn=PITPNM2;refGene_tx=XM_023241341.1;refGene_gn=PITPNM2;refGene_tx=XM_023241342.1;refGene_gn=PITPNM2;refGene_tx=XM_023241343.1;refGene_gn=PITPNM2;refGene_tx=XM_023241344.1;refGene_gn=PITPNM2;refGene_tx=XM_023241345.1;refGene_gn=PITPNM2;refGene_tx=XM_023241346.1;refGene_gn=PITPNM2;refGene_tx=XM_023241347.1;refGene_gn=PITPNM2;refGene_tx=XM_023241348.1;refGene_gn=PITPNM2;refGene_tx=XM_023241349.1;refGene_gn=PITPNM2;refGene_tx=XM_023241350.1;refGene_gn=PITPNM2;refGene_tx=XM_023241351.1
D3 6877410 6877589
refGene_gn=PITPNM2;refGene_tx=XM_023241340.1;refGene_gn=PITPNM2;refGen

e_tx=XM_023241341.1;refGene_gn=PITPNM2;refGene_tx=XM_023241342.1;refGene_gn=PITPNM2;refGene_tx=XM_023241343.1;refGene_gn=PITPNM2;refGene_tx=XM_023241344.1;refGene_gn=PITPNM2;refGene_tx=XM_023241345.1;refGene_gn=PITPNM2;refGene_tx=XM_023241346.1;refGene_gn=PITPNM2;refGene_tx=XM_023241347.1;refGene_gn=PITPNM2;refGene_tx=XM_023241348.1;refGene_gn=PITPNM2;refGene_tx=XM_023241349.1;refGene_gn=PITPNM2;refGene_tx=XM_023241350.1;refGene_gn=PITPNM2;refGene_tx=XM_023241351.1
D3 6878674 6878845
refGene_gn=PITPNM2;refGene_tx=XM_023241340.1;refGene_gn=PITPNM2;refGene_tx=XM_023241341.1;refGene_gn=PITPNM2;refGene_tx=XM_023241342.1;refGene_gn=PITPNM2;refGene_tx=XM_023241343.1;refGene_gn=PITPNM2;refGene_tx=XM_023241344.1;refGene_gn=PITPNM2;refGene_tx=XM_023241345.1;refGene_gn=PITPNM2;refGene_tx=XM_023241346.1;refGene_gn=PITPNM2;refGene_tx=XM_023241347.1;refGene_gn=PITPNM2;refGene_tx=XM_023241348.1;refGene_gn=PITPNM2;refGene_tx=XM_023241349.1;refGene_gn=PITPNM2;refGene_tx=XM_023241350.1;refGene_gn=PITPNM2;refGene_tx=XM_023241351.1
D3 6879361 6879511
refGene_gn=PITPNM2;refGene_tx=XM_023241340.1;refGene_gn=PITPNM2;refGene_tx=XM_023241341.1;refGene_gn=PITPNM2;refGene_tx=XM_023241342.1;refGene_gn=PITPNM2;refGene_tx=XM_023241343.1;refGene_gn=PITPNM2;refGene_tx=XM_023241344.1;refGene_gn=PITPNM2;refGene_tx=XM_023241346.1;refGene_gn=PITPNM2;refGene_tx=XM_023241347.1;refGene_gn=PITPNM2;refGene_tx=XM_023241349.1
D3 6880566 6880731
refGene_gn=PITPNM2;refGene_tx=XM_023241340.1;refGene_gn=PITPNM2;refGene_tx=XM_023241341.1;refGene_gn=PITPNM2;refGene_tx=XM_023241342.1;refGene_gn=PITPNM2;refGene_tx=XM_023241343.1;refGene_gn=PITPNM2;refGene_tx=XM_023241344.1;refGene_gn=PITPNM2;refGene_tx=XM_023241345.1;refGene_gn=PITPNM2;refGene_tx=XM_023241346.1;refGene_gn=PITPNM2;refGene_tx=XM_023241347.1;refGene_gn=PITPNM2;refGene_tx=XM_023241348.1;refGene_gn=PITPNM2;refGene_tx=XM_023241349.1;refGene_gn=PITPNM2;refGene_tx=XM_023241350.1;refGene_gn=PITPNM2;refGene_tx=XM_023241351.1
D3 6881412 6881574
refGene_gn=PITPNM2;refGene_tx=XM_023241340.1;refGene_gn=PITPNM2;refGene_tx=XM_023241341.1;refGene_gn=PITPNM2;refGene_tx=XM_023241342.1;refGene_gn=PITPNM2;refGene_tx=XM_023241343.1;refGene_gn=PITPNM2;refGene_tx=XM_023241344.1;refGene_gn=PITPNM2;refGene_tx=XM_023241345.1;refGene_gn=PITPNM2;refGene_tx=XM_023241348.1
D3 6882391 6882540
refGene_gn=PITPNM2;refGene_tx=XM_023241340.1;refGene_gn=PITPNM2;refGene_tx=XM_023241341.1;refGene_gn=PITPNM2;refGene_tx=XM_023241342.1;refGene_gn=PITPNM2;refGene_tx=XM_023241343.1;refGene_gn=PITPNM2;refGene_tx=XM_023241344.1;refGene_gn=PITPNM2;refGene_tx=XM_023241345.1;refGene_gn=PITPNM2;refGene_tx=XM_023241346.1;refGene_gn=PITPNM2;refGene_tx=XM_023241347.1;refGene_gn=PITPNM2;refGene_tx=XM_023241348.1;refGene_gn=PITPNM2;refGene_tx=XM_023241349.1;refGene_gn=PITPNM2;refGene_tx=XM_023241350.1;refGene_gn=PITPNM2;refGene_tx=XM_023241351.1
D3 6882859 6882976
refGene_gn=PITPNM2;refGene_tx=XM_023241340.1;refGene_gn=PITPNM2;refGene_tx=XM_023241341.1;refGene_gn=PITPNM2;refGene_tx=XM_023241342.1;refGe

ne_gn=PITPNM2;refGene_tx=XM_023241343.1;refGene_gn=PITPNM2;refGene_tx=XM_023241344.1;refGene_gn=PITPNM2;refGene_tx=XM_023241345.1;refGene_gn=PITPNM2;refGene_tx=XM_023241346.1;refGene_gn=PITPNM2;refGene_tx=XM_023241347.1;refGene_gn=PITPNM2;refGene_tx=XM_023241348.1;refGene_gn=PITPNM2;refGene_tx=XM_023241349.1;refGene_gn=PITPNM2;refGene_tx=XM_023241350.1;refGene_gn=PITPNM2;refGene_tx=XM_023241351.1
D3 6883289 6883406
refGene_gn=PITPNM2;refGene_tx=XM_023241340.1;refGene_gn=PITPNM2;refGene_tx=XM_023241341.1;refGene_gn=PITPNM2;refGene_tx=XM_023241342.1;refGene_gn=PITPNM2;refGene_tx=XM_023241343.1;refGene_gn=PITPNM2;refGene_tx=XM_023241344.1;refGene_gn=PITPNM2;refGene_tx=XM_023241345.1;refGene_gn=PITPNM2;refGene_tx=XM_023241346.1;refGene_gn=PITPNM2;refGene_tx=XM_023241347.1;refGene_gn=PITPNM2;refGene_tx=XM_023241348.1;refGene_gn=PITPNM2;refGene_tx=XM_023241349.1;refGene_gn=PITPNM2;refGene_tx=XM_023241350.1;refGene_gn=PITPNM2;refGene_tx=XM_023241351.1
D3 6883508 6883657
refGene_gn=PITPNM2;refGene_tx=XM_023241340.1;refGene_gn=PITPNM2;refGene_tx=XM_023241341.1;refGene_gn=PITPNM2;refGene_tx=XM_023241342.1;refGene_gn=PITPNM2;refGene_tx=XM_023241343.1;refGene_gn=PITPNM2;refGene_tx=XM_023241344.1;refGene_gn=PITPNM2;refGene_tx=XM_023241345.1;refGene_gn=PITPNM2;refGene_tx=XM_023241346.1;refGene_gn=PITPNM2;refGene_tx=XM_023241347.1;refGene_gn=PITPNM2;refGene_tx=XM_023241348.1;refGene_gn=PITPNM2;refGene_tx=XM_023241349.1;refGene_gn=PITPNM2;refGene_tx=XM_023241350.1;refGene_gn=PITPNM2;refGene_tx=XM_023241351.1
D3 6883750 6883900
refGene_gn=PITPNM2;refGene_tx=XM_023241340.1;refGene_gn=PITPNM2;refGene_tx=XM_023241341.1;refGene_gn=PITPNM2;refGene_tx=XM_023241342.1;refGene_gn=PITPNM2;refGene_tx=XM_023241343.1;refGene_gn=PITPNM2;refGene_tx=XM_023241344.1;refGene_gn=PITPNM2;refGene_tx=XM_023241345.1;refGene_gn=PITPNM2;refGene_tx=XM_023241346.1;refGene_gn=PITPNM2;refGene_tx=XM_023241347.1;refGene_gn=PITPNM2;refGene_tx=XM_023241348.1;refGene_gn=PITPNM2;refGene_tx=XM_023241349.1;refGene_gn=PITPNM2;refGene_tx=XM_023241350.1;refGene_gn=PITPNM2;refGene_tx=XM_023241351.1
D3 6884261 6884445
refGene_gn=PITPNM2;refGene_tx=XM_023241340.1;refGene_gn=PITPNM2;refGene_tx=XM_023241341.1;refGene_gn=PITPNM2;refGene_tx=XM_023241342.1;refGene_gn=PITPNM2;refGene_tx=XM_023241343.1;refGene_gn=PITPNM2;refGene_tx=XM_023241344.1;refGene_gn=PITPNM2;refGene_tx=XM_023241345.1;refGene_gn=PITPNM2;refGene_tx=XM_023241346.1;refGene_gn=PITPNM2;refGene_tx=XM_023241347.1;refGene_gn=PITPNM2;refGene_tx=XM_023241348.1;refGene_gn=PITPNM2;refGene_tx=XM_023241349.1;refGene_gn=PITPNM2;refGene_tx=XM_023241350.1;refGene_gn=PITPNM2;refGene_tx=XM_023241351.1
D3 6884531 6884660
refGene_gn=PITPNM2;refGene_tx=XM_023241340.1;refGene_gn=PITPNM2;refGene_tx=XM_023241341.1;refGene_gn=PITPNM2;refGene_tx=XM_023241342.1;refGene_gn=PITPNM2;refGene_tx=XM_023241343.1;refGene_gn=PITPNM2;refGene_tx=XM_023241344.1;refGene_gn=PITPNM2;refGene_tx=XM_023241345.1;refGene_gn=PITPNM2;refGene_tx=XM_023241346.1;refGene_gn=PITPNM2;refGene_tx=XM_023241347.1;refGene_gn=PITPNM2;refGene_tx=XM_023241348.1;refGene_gn=PITPNM2;refGene_tx=XM_023241349.1;refGene_gn=PITPNM2;refGene_tx=XM_0232413

50.1;refGene_gn=PITPNM2;refGene_tx=XM_023241351.1
D3 6884761 6885085
refGene_gn=PITPNM2;refGene_tx=XM_023241340.1;refGene_gn=PITPNM2;refGene_tx=XM_023241341.1;refGene_gn=PITPNM2;refGene_tx=XM_023241342.1;refGene_gn=PITPNM2;refGene_tx=XM_023241343.1;refGene_gn=PITPNM2;refGene_tx=XM_023241344.1;refGene_gn=PITPNM2;refGene_tx=XM_023241345.1;refGene_gn=PITPNM2;refGene_tx=XM_023241346.1;refGene_gn=PITPNM2;refGene_tx=XM_023241347.1;refGene_gn=PITPNM2;refGene_tx=XM_023241348.1;refGene_gn=PITPNM2;refGene_tx=XM_023241349.1;refGene_gn=PITPNM2;refGene_tx=XM_023241350.1;refGene_gn=PITPNM2;refGene_tx=XM_023241351.1
D3 6888570 6888627
refGene_gn=ARL6IP4;refGene_tx=XM_006938410.3;refGene_gn=ARL6IP4;refGene_tx=XM_006938412.3
D3 6888706 6888776
refGene_gn=ARL6IP4;refGene_tx=XM_006938410.3;refGene_gn=ARL6IP4;refGene_tx=XM_006938412.3
D3 6889175 6889293
refGene_gn=ARL6IP4;refGene_tx=XM_006938410.3;refGene_gn=ARL6IP4;refGene_tx=XM_006938412.3
D3 6889376 6889691
refGene_gn=ARL6IP4;refGene_tx=XM_006938410.3;refGene_gn=ARL6IP4;refGene_tx=XM_006938412.3
D3 6890116 6890279
refGene_gn=ARL6IP4;refGene_tx=XM_006938410.3;refGene_gn=ARL6IP4;refGene_tx=XM_006938412.3
D3 6892164 6892428 refGene_gn=OGFOD2;refGene_tx=XM_003994615.5
D3 6892502 6892760 refGene_gn=OGFOD2;refGene_tx=XM_003994615.5
D3 6893014 6893142 refGene_gn=OGFOD2;refGene_tx=XM_003994615.5
D3 6894610 6894710 refGene_gn=OGFOD2;refGene_tx=XM_003994615.5
D3 6894799 6894913 refGene_gn=OGFOD2;refGene_tx=XM_003994615.5
D3 6895617 6895674 refGene_gn=OGFOD2;refGene_tx=XM_003994615.5
D3 6896108 6896240 refGene_gn=OGFOD2;refGene_tx=XM_003994615.5
D3 6909896 6910497
refGene_gn=ABCB9;refGene_tx=XM_023241352.1;refGene_gn=ABCB9;refGene_tx=XM_023241353.1;refGene_gn=ABCB9;refGene_tx=XM_023241354.1
D3 6915354 6915469
refGene_gn=ABCB9;refGene_tx=XM_023241352.1;refGene_gn=ABCB9;refGene_tx=XM_023241353.1;refGene_gn=ABCB9;refGene_tx=XM_023241354.1
D3 6915967 6916098
refGene_gn=ABCB9;refGene_tx=XM_023241352.1;refGene_gn=ABCB9;refGene_tx=XM_023241353.1;refGene_gn=ABCB9;refGene_tx=XM_023241354.1
D3 6916979 6917185
refGene_gn=ABCB9;refGene_tx=XM_023241352.1;refGene_gn=ABCB9;refGene_tx=XM_023241353.1;refGene_gn=ABCB9;refGene_tx=XM_023241354.1
D3 6919568 6919766
refGene_gn=ABCB9;refGene_tx=XM_023241352.1;refGene_gn=ABCB9;refGene_tx=XM_023241353.1;refGene_gn=ABCB9;refGene_tx=XM_023241354.1
D3 6920961 6921090
refGene_gn=ABCB9;refGene_tx=XM_023241352.1;refGene_gn=ABCB9;refGene_tx=XM_023241353.1;refGene_gn=ABCB9;refGene_tx=XM_023241354.1

D3 6925809 6925998
refGene_gn=ABCB9;refGene_tx=XM_023241352.1;refGene_gn=ABCB9;refGene_tx
=XM_023241353.1;refGene_gn=ABCB9;refGene_tx=XM_023241354.1
D3 6926484 6926658
refGene_gn=ABCB9;refGene_tx=XM_023241352.1;refGene_gn=ABCB9;refGene_tx
=XM_023241353.1;refGene_gn=ABCB9;refGene_tx=XM_023241354.1
D3 6928529 6928689
refGene_gn=ABCB9;refGene_tx=XM_023241352.1;refGene_gn=ABCB9;refGene_tx
=XM_023241353.1;refGene_gn=ABCB9;refGene_tx=XM_023241354.1
D3 6930689 6930826
refGene_gn=ABCB9;refGene_tx=XM_023241352.1;refGene_gn=ABCB9;refGene_tx
=XM_023241353.1;refGene_gn=ABCB9;refGene_tx=XM_023241354.1
D3 6933491 6933755
refGene_gn=ABCB9;refGene_tx=XM_023241352.1;refGene_gn=ABCB9;refGene_tx
=XM_023241353.1;refGene_gn=ABCB9;refGene_tx=XM_023241354.1
D3 6961961 6962072 refGene_gn=VPS37B;refGene_tx=XM_023241356.1
D3 6983875 6984047
refGene_gn=VPS37B;refGene_tx=XM_019814482.2;refGene_gn=VPS37B;refGene_
tx=XM_023241356.1
D3 6986017 6986100
refGene_gn=VPS37B;refGene_tx=XM_019814482.2;refGene_gn=VPS37B;refGene_
tx=XM_023241356.1
D3 6986909 6987407
refGene_gn=VPS37B;refGene_tx=XM_019814482.2;refGene_gn=VPS37B;refGene_
tx=XM_023241356.1
D3 6991903 6991951 refGene_gn=HIP1R;refGene_tx=XM_023241355.1
D3 6992159 6992366 refGene_gn=HIP1R;refGene_tx=XM_023241355.1
D3 6992478 6992540 refGene_gn=HIP1R;refGene_tx=XM_023241355.1
D3 6992615 6992739 refGene_gn=HIP1R;refGene_tx=XM_023241355.1
D3 6992836 6992942 refGene_gn=HIP1R;refGene_tx=XM_023241355.1
D3 6993063 6993164 refGene_gn=HIP1R;refGene_tx=XM_023241355.1
D3 6993472 6993535 refGene_gn=HIP1R;refGene_tx=XM_023241355.1
D3 6993619 6993650 refGene_gn=HIP1R;refGene_tx=XM_023241355.1
D3 6993890 6993949 refGene_gn=HIP1R;refGene_tx=XM_023241355.1
D3 6994159 6994270 refGene_gn=HIP1R;refGene_tx=XM_023241355.1
D3 6994515 6994652 refGene_gn=HIP1R;refGene_tx=XM_023241355.1
D3 6994791 6994899 refGene_gn=HIP1R;refGene_tx=XM_023241355.1
D3 6995258 6995345 refGene_gn=HIP1R;refGene_tx=XM_023241355.1
D3 6995461 6995609 refGene_gn=HIP1R;refGene_tx=XM_023241355.1
D3 6996811 6997006 refGene_gn=HIP1R;refGene_tx=XM_023241355.1
D3 6997280 6997382 refGene_gn=HIP1R;refGene_tx=XM_023241355.1
D3 6997478 6997595 refGene_gn=HIP1R;refGene_tx=XM_023241355.1
D3 6997686 6997775 refGene_gn=HIP1R;refGene_tx=XM_023241355.1
D3 6997869 6998053 refGene_gn=HIP1R;refGene_tx=XM_023241355.1
D3 6998147 6998220 refGene_gn=HIP1R;refGene_tx=XM_023241355.1
D3 6998304 6998366 refGene_gn=HIP1R;refGene_tx=XM_023241355.1
D3 6998466 6998607 refGene_gn=HIP1R;refGene_tx=XM_023241355.1
D3 6998773 6998849 refGene_gn=HIP1R;refGene_tx=XM_023241355.1
D3 6998946 6999004 refGene_gn=HIP1R;refGene_tx=XM_023241355.1
D3 7000020 7000161 refGene_gn=HIP1R;refGene_tx=XM_023241355.1

D3	7001905	7001967	refGene_gn=HIP1R;refGene_tx=XM_023241355.1
D3	7002292	7002369	refGene_gn=HIP1R;refGene_tx=XM_023241355.1
D3	7002770	7002851	refGene_gn=HIP1R;refGene_tx=XM_023241355.1
D3	7004070	7004127	refGene_gn=HIP1R;refGene_tx=XM_023241355.1
D3	7004365	7004508	refGene_gn=HIP1R;refGene_tx=XM_023241355.1
D3	7005046	7005110	refGene_gn=HIP1R;refGene_tx=XM_023241355.1
D3	7019691	7019784	refGene_gn=HIP1R;refGene_tx=XM_023241355.1
D3	7025425	7025479	refGene_gn=CCDC62;refGene_tx=XM_011287845.3
D3	7038725	7038851	refGene_gn=CCDC62;refGene_tx=XM_011287845.3
D3	7044766	7044923	refGene_gn=CCDC62;refGene_tx=XM_011287845.3
D3	7048326	7048519	refGene_gn=CCDC62;refGene_tx=XM_011287845.3
D3	7048521	7049054	refGene_gn=CCDC62;refGene_tx=XM_011287845.3
D3	7051150	7051266	refGene_gn=CCDC62;refGene_tx=XM_011287845.3
D3	7051607	7051696	refGene_gn=CCDC62;refGene_tx=XM_011287845.3
D3	7055884	7055986	refGene_gn=CCDC62;refGene_tx=XM_011287845.3
D3	7059117	7059289	refGene_gn=CCDC62;refGene_tx=XM_011287845.3
D3	7060984	7061086	refGene_gn=CCDC62;refGene_tx=XM_011287845.3
D3	7063103	7063270	refGene_gn=CCDC62;refGene_tx=XM_011287845.3
D3	7066962	7067155	refGene_gn=CCDC62;refGene_tx=XM_011287845.3
D3	7069217	7069253	refGene_gn=CCDC62;refGene_tx=XM_011287845.3
D3	7073257	7073302
refGene_gn=DENR;refGene_tx=XM_011287842.2;refGene_gn=DENR;refGene_tx=XM_011287843.2;refGene_gn=DENR;refGene_tx=XM_011287844.2
D3	7073409	7073549
refGene_gn=DENR;refGene_tx=XM_011287842.2;refGene_gn=DENR;refGene_tx=XM_011287843.2;refGene_gn=DENR;refGene_tx=XM_011287844.2
D3	7074448	7074565
refGene_gn=DENR;refGene_tx=XM_011287842.2;refGene_gn=DENR;refGene_tx=XM_011287843.2;refGene_gn=DENR;refGene_tx=XM_011287844.2
D3	7076828	7076912
refGene_gn=DENR;refGene_tx=XM_011287842.2;refGene_gn=DENR;refGene_tx=XM_011287843.2;refGene_gn=DENR;refGene_tx=XM_011287844.2
D3	7078650	7078735
refGene_gn=DENR;refGene_tx=XM_011287842.2;refGene_gn=DENR;refGene_tx=XM_011287843.2;refGene_gn=DENR;refGene_tx=XM_011287844.2
D3	7079436	7079456
refGene_gn=DENR;refGene_tx=XM_011287842.2;refGene_gn=DENR;refGene_tx=XM_011287843.2;refGene_gn=DENR;refGene_tx=XM_011287844.2
D3	7080531	7080571	refGene_gn=DENR;refGene_tx=XM_011287843.2
D3	7084493	7084599
refGene_gn=DENR;refGene_tx=XM_011287842.2;refGene_gn=DENR;refGene_tx=XM_011287844.2
D3	7105329	7105716	refGene_gn=HCAR1;refGene_tx=XM_006938419.3
D3	7105802	7106834
refGene_gn=HCAR1;refGene_tx=XM_006938419.3;refGene_gn=HCAR1;refGene_tx=XM_019814483.1
D3	7120106	7121198	refGene_gn=HCAR2;refGene_tx=NM_001309044.1
D3	7155864	7155885
refGene_gn=KNTC1;refGene_tx=XM_006938420.4;refGene_gn=KNTC1;refGene_tx=XM_019814484.1

D3	7157670	7157761
refGene_gn=KNTC1;refGene_tx=XM_006938420.4;refGene_gn=KNTC1;refGene_tx=XM_019814484.1
D3	7158506	7158652
refGene_gn=KNTC1;refGene_tx=XM_006938420.4;refGene_gn=KNTC1;refGene_tx=XM_019814484.1
D3	7159123	7159213
refGene_gn=KNTC1;refGene_tx=XM_006938420.4;refGene_gn=KNTC1;refGene_tx=XM_019814484.1
D3	7160267	7160397
refGene_gn=KNTC1;refGene_tx=XM_006938420.4;refGene_gn=KNTC1;refGene_tx=XM_019814484.1
D3	7161183	7161247
refGene_gn=KNTC1;refGene_tx=XM_006938420.4;refGene_gn=KNTC1;refGene_tx=XM_019814484.1
D3	7161364	7161419
refGene_gn=KNTC1;refGene_tx=XM_006938420.4;refGene_gn=KNTC1;refGene_tx=XM_019814484.1
D3	7164933	7164990
refGene_gn=KNTC1;refGene_tx=XM_006938420.4;refGene_gn=KNTC1;refGene_tx=XM_019814484.1
D3	7165356	7165452
refGene_gn=KNTC1;refGene_tx=XM_006938420.4;refGene_gn=KNTC1;refGene_tx=XM_019814484.1
D3	7166135	7166271
refGene_gn=KNTC1;refGene_tx=XM_006938420.4;refGene_gn=KNTC1;refGene_tx=XM_019814484.1
D3	7166644	7166763
refGene_gn=KNTC1;refGene_tx=XM_006938420.4;refGene_gn=KNTC1;refGene_tx=XM_019814484.1
D3	7168170	7168249
refGene_gn=KNTC1;refGene_tx=XM_006938420.4;refGene_gn=KNTC1;refGene_tx=XM_019814484.1
D3	7168288	7168439	refGene_gn=KNTC1;refGene_tx=XM_023241357.1
D3	7169637	7169684
refGene_gn=KNTC1;refGene_tx=XM_006938420.4;refGene_gn=KNTC1;refGene_tx=XM_019814484.1;refGene_gn=KNTC1;refGene_tx=XM_023241357.1
D3	7171992	7172102
refGene_gn=KNTC1;refGene_tx=XM_006938420.4;refGene_gn=KNTC1;refGene_tx=XM_019814484.1;refGene_gn=KNTC1;refGene_tx=XM_023241357.1
D3	7172222	7172433
refGene_gn=KNTC1;refGene_tx=XM_006938420.4;refGene_gn=KNTC1;refGene_tx=XM_019814484.1;refGene_gn=KNTC1;refGene_tx=XM_023241357.1
D3	7172709	7172783
refGene_gn=KNTC1;refGene_tx=XM_006938420.4;refGene_gn=KNTC1;refGene_tx=XM_019814484.1;refGene_gn=KNTC1;refGene_tx=XM_023241357.1
D3	7173810	7174027
refGene_gn=KNTC1;refGene_tx=XM_006938420.4;refGene_gn=KNTC1;refGene_tx=XM_019814484.1;refGene_gn=KNTC1;refGene_tx=XM_023241357.1
D3	7174375	7174434

refGene_gn=KNTC1;refGene_tx=XM_006938420.4;refGene_gn=KNTC1;refGene_tx=XM_019814484.1;refGene_gn=KNTC1;refGene_tx=XM_023241357.1
D3 7174521 7174693
refGene_gn=KNTC1;refGene_tx=XM_006938420.4;refGene_gn=KNTC1;refGene_tx=XM_019814484.1;refGene_gn=KNTC1;refGene_tx=XM_023241357.1
D3 7176013 7176103
refGene_gn=KNTC1;refGene_tx=XM_006938420.4;refGene_gn=KNTC1;refGene_tx=XM_019814484.1;refGene_gn=KNTC1;refGene_tx=XM_023241357.1
D3 7177447 7177655
refGene_gn=KNTC1;refGene_tx=XM_006938420.4;refGene_gn=KNTC1;refGene_tx=XM_019814484.1;refGene_gn=KNTC1;refGene_tx=XM_023241357.1
D3 7182771 7182881
refGene_gn=KNTC1;refGene_tx=XM_006938420.4;refGene_gn=KNTC1;refGene_tx=XM_019814484.1;refGene_gn=KNTC1;refGene_tx=XM_023241357.1
D3 7185045 7185162
refGene_gn=KNTC1;refGene_tx=XM_006938420.4;refGene_gn=KNTC1;refGene_tx=XM_019814484.1;refGene_gn=KNTC1;refGene_tx=XM_023241357.1
D3 7185781 7185910
refGene_gn=KNTC1;refGene_tx=XM_006938420.4;refGene_gn=KNTC1;refGene_tx=XM_019814484.1;refGene_gn=KNTC1;refGene_tx=XM_023241357.1
D3 7186308 7186413
refGene_gn=KNTC1;refGene_tx=XM_006938420.4;refGene_gn=KNTC1;refGene_tx=XM_019814484.1;refGene_gn=KNTC1;refGene_tx=XM_023241357.1
D3 7187444 7187608
refGene_gn=KNTC1;refGene_tx=XM_006938420.4;refGene_gn=KNTC1;refGene_tx=XM_019814484.1;refGene_gn=KNTC1;refGene_tx=XM_023241357.1
D3 7188104 7188161
refGene_gn=KNTC1;refGene_tx=XM_006938420.4;refGene_gn=KNTC1;refGene_tx=XM_019814484.1;refGene_gn=KNTC1;refGene_tx=XM_023241357.1
D3 7188682 7188821
refGene_gn=KNTC1;refGene_tx=XM_006938420.4;refGene_gn=KNTC1;refGene_tx=XM_019814484.1;refGene_gn=KNTC1;refGene_tx=XM_023241357.1
D3 7189487 7189585
refGene_gn=KNTC1;refGene_tx=XM_006938420.4;refGene_gn=KNTC1;refGene_tx=XM_019814484.1;refGene_gn=KNTC1;refGene_tx=XM_023241357.1
D3 7190027 7190200
refGene_gn=KNTC1;refGene_tx=XM_006938420.4;refGene_gn=KNTC1;refGene_tx=XM_019814484.1;refGene_gn=KNTC1;refGene_tx=XM_023241357.1
D3 7191040 7191321
refGene_gn=KNTC1;refGene_tx=XM_006938420.4;refGene_gn=KNTC1;refGene_tx=XM_019814484.1;refGene_gn=KNTC1;refGene_tx=XM_023241357.1
D3 7194680 7194748
refGene_gn=KNTC1;refGene_tx=XM_006938420.4;refGene_gn=KNTC1;refGene_tx=XM_019814484.1;refGene_gn=KNTC1;refGene_tx=XM_023241357.1
D3 7195465 7195538
refGene_gn=KNTC1;refGene_tx=XM_006938420.4;refGene_gn=KNTC1;refGene_tx=XM_019814484.1;refGene_gn=KNTC1;refGene_tx=XM_023241357.1
D3 7197464 7197584
refGene_gn=KNTC1;refGene_tx=XM_006938420.4;refGene_gn=KNTC1;refGene_tx=XM_019814484.1;refGene_gn=KNTC1;refGene_tx=XM_023241357.1

D3 7198130 7198265
refGene_gn=KNTC1;refGene_tx=XM_006938420.4;refGene_gn=KNTC1;refGene_tx=XM_019814484.1;refGene_gn=KNTC1;refGene_tx=XM_023241357.1
D3 7198808 7198908
refGene_gn=KNTC1;refGene_tx=XM_006938420.4;refGene_gn=KNTC1;refGene_tx=XM_019814484.1;refGene_gn=KNTC1;refGene_tx=XM_023241357.1
D3 7199061 7199165
refGene_gn=KNTC1;refGene_tx=XM_006938420.4;refGene_gn=KNTC1;refGene_tx=XM_019814484.1;refGene_gn=KNTC1;refGene_tx=XM_023241357.1
D3 7200282 7200381
refGene_gn=KNTC1;refGene_tx=XM_006938420.4;refGene_gn=KNTC1;refGene_tx=XM_019814484.1;refGene_gn=KNTC1;refGene_tx=XM_023241357.1
D3 7200895 7201039
refGene_gn=KNTC1;refGene_tx=XM_006938420.4;refGene_gn=KNTC1;refGene_tx=XM_019814484.1;refGene_gn=KNTC1;refGene_tx=XM_023241357.1
D3 7201134 7201254
refGene_gn=KNTC1;refGene_tx=XM_006938420.4;refGene_gn=KNTC1;refGene_tx=XM_019814484.1;refGene_gn=KNTC1;refGene_tx=XM_023241357.1
D3 7203029 7203131
refGene_gn=KNTC1;refGene_tx=XM_006938420.4;refGene_gn=KNTC1;refGene_tx=XM_019814484.1;refGene_gn=KNTC1;refGene_tx=XM_023241357.1
D3 7203223 7203280
refGene_gn=KNTC1;refGene_tx=XM_006938420.4;refGene_gn=KNTC1;refGene_tx=XM_019814484.1;refGene_gn=KNTC1;refGene_tx=XM_023241357.1
D3 7203467 7203611
refGene_gn=KNTC1;refGene_tx=XM_006938420.4;refGene_gn=KNTC1;refGene_tx=XM_019814484.1;refGene_gn=KNTC1;refGene_tx=XM_023241357.1
D3 7204412 7204524
refGene_gn=KNTC1;refGene_tx=XM_006938420.4;refGene_gn=KNTC1;refGene_tx=XM_019814484.1;refGene_gn=KNTC1;refGene_tx=XM_023241357.1
D3 7207329 7207391
refGene_gn=KNTC1;refGene_tx=XM_006938420.4;refGene_gn=KNTC1;refGene_tx=XM_019814484.1;refGene_gn=KNTC1;refGene_tx=XM_023241357.1
D3 7207965 7208019
refGene_gn=KNTC1;refGene_tx=XM_006938420.4;refGene_gn=KNTC1;refGene_tx=XM_019814484.1;refGene_gn=KNTC1;refGene_tx=XM_023241357.1
D3 7208616 7208706
refGene_gn=KNTC1;refGene_tx=XM_006938420.4;refGene_gn=KNTC1;refGene_tx=XM_019814484.1;refGene_gn=KNTC1;refGene_tx=XM_023241357.1
D3 7208796 7208922
refGene_gn=KNTC1;refGene_tx=XM_006938420.4;refGene_gn=KNTC1;refGene_tx=XM_019814484.1;refGene_gn=KNTC1;refGene_tx=XM_023241357.1
D3 7211899 7211975
refGene_gn=KNTC1;refGene_tx=XM_006938420.4;refGene_gn=KNTC1;refGene_tx=XM_019814484.1;refGene_gn=KNTC1;refGene_tx=XM_023241357.1
D3 7212073 7212139
refGene_gn=KNTC1;refGene_tx=XM_006938420.4;refGene_gn=KNTC1;refGene_tx=XM_019814484.1;refGene_gn=KNTC1;refGene_tx=XM_023241357.1
D3 7212241 7212285
refGene_gn=KNTC1;refGene_tx=XM_006938420.4;refGene_gn=KNTC1;refGene_tx

=XM_019814484.1;refGene_gn=KNTC1;refGene_tx=XM_023241357.1
D3 7213773 7213872
refGene_gn=KNTC1;refGene_tx=XM_006938420.4;refGene_gn=KNTC1;refGene_tx=XM_019814484.1;refGene_gn=KNTC1;refGene_tx=XM_023241357.1
D3 7214812 7214864
refGene_gn=KNTC1;refGene_tx=XM_006938420.4;refGene_gn=KNTC1;refGene_tx=XM_019814484.1;refGene_gn=KNTC1;refGene_tx=XM_023241357.1
D3 7215190 7215306
refGene_gn=KNTC1;refGene_tx=XM_006938420.4;refGene_gn=KNTC1;refGene_tx=XM_019814484.1;refGene_gn=KNTC1;refGene_tx=XM_023241357.1
D3 7215512 7215565
refGene_gn=KNTC1;refGene_tx=XM_006938420.4;refGene_gn=KNTC1;refGene_tx=XM_019814484.1;refGene_gn=KNTC1;refGene_tx=XM_023241357.1
D3 7216214 7216308
refGene_gn=KNTC1;refGene_tx=XM_006938420.4;refGene_gn=KNTC1;refGene_tx=XM_019814484.1;refGene_gn=KNTC1;refGene_tx=XM_023241357.1
D3 7217081 7217192
refGene_gn=KNTC1;refGene_tx=XM_006938420.4;refGene_gn=KNTC1;refGene_tx=XM_019814484.1;refGene_gn=KNTC1;refGene_tx=XM_023241357.1
D3 7218207 7218242
refGene_gn=KNTC1;refGene_tx=XM_006938420.4;refGene_gn=KNTC1;refGene_tx=XM_019814484.1;refGene_gn=KNTC1;refGene_tx=XM_023241357.1
D3 7219765 7219843
refGene_gn=KNTC1;refGene_tx=XM_006938420.4;refGene_gn=KNTC1;refGene_tx=XM_019814484.1;refGene_gn=KNTC1;refGene_tx=XM_023241357.1
D3 7220438 7220517
refGene_gn=KNTC1;refGene_tx=XM_006938420.4;refGene_gn=KNTC1;refGene_tx=XM_023241357.1
D3 7220879 7220995
refGene_gn=KNTC1;refGene_tx=XM_006938420.4;refGene_gn=KNTC1;refGene_tx=XM_023241357.1
D3 7222713 7222834
refGene_gn=KNTC1;refGene_tx=XM_006938420.4;refGene_gn=KNTC1;refGene_tx=XM_023241357.1
D3 7227028 7227157
refGene_gn=KNTC1;refGene_tx=XM_006938420.4;refGene_gn=KNTC1;refGene_tx=XM_023241357.1
D3 7229189 7229195
refGene_gn=RSRC2;refGene_tx=XM_003994618.4;refGene_gn=RSRC2;refGene_tx=XM_006938421.3
D3 7232954 7233111
refGene_gn=RSRC2;refGene_tx=XM_003994618.4;refGene_gn=RSRC2;refGene_tx=XM_006938421.3;refGene_gn=RSRC2;refGene_tx=XM_019814486.2;refGene_gn=RSRC2;refGene_tx=XM_019814487.2
D3 7234430 7234474
refGene_gn=RSRC2;refGene_tx=XM_003994618.4;refGene_gn=RSRC2;refGene_tx=XM_006938421.3;refGene_gn=RSRC2;refGene_tx=XM_019814486.2;refGene_gn=RSRC2;refGene_tx=XM_019814487.2
D3 7235366 7235374 refGene_gn=RSRC2;refGene_tx=XM_006938422.2
D3 7236934 7237147

refGene_gn=RSRC2;refGene_tx=XM_003994618.4;refGene_gn=RSRC2;refGene_tx
=XM_006938421.3;refGene_gn=RSRC2;refGene_tx=XM_006938422.2;refGene_gn=
RSRC2;refGene_tx=XM_019814486.2;refGene_gn=RSRC2;refGene_tx=XM_0198144
87.2
D3 7238176 7238380
refGene_gn=RSRC2;refGene_tx=XM_003994618.4;refGene_gn=RSRC2;refGene_tx
=XM_006938421.3;refGene_gn=RSRC2;refGene_tx=XM_006938422.2;refGene_gn=
RSRC2;refGene_tx=XM_019814486.2;refGene_gn=RSRC2;refGene_tx=XM_0198144
87.2
D3 7240321 7240444
refGene_gn=RSRC2;refGene_tx=XM_003994618.4;refGene_gn=RSRC2;refGene_tx
=XM_006938421.3;refGene_gn=RSRC2;refGene_tx=XM_006938422.2;refGene_gn=
RSRC2;refGene_tx=XM_019814486.2;refGene_gn=RSRC2;refGene_tx=XM_0198144
87.2
D3 7242441 7242521
refGene_gn=RSRC2;refGene_tx=XM_003994618.4;refGene_gn=RSRC2;refGene_tx
=XM_006938421.3;refGene_gn=RSRC2;refGene_tx=XM_006938422.2;refGene_gn=
RSRC2;refGene_tx=XM_019814486.2;refGene_gn=RSRC2;refGene_tx=XM_0198144
87.2
D3 7243741 7243974
refGene_gn=RSRC2;refGene_tx=XM_003994618.4;refGene_gn=RSRC2;refGene_tx
=XM_006938421.3;refGene_gn=RSRC2;refGene_tx=XM_006938422.2;refGene_gn=
RSRC2;refGene_tx=XM_019814486.2;refGene_gn=RSRC2;refGene_tx=XM_0198144
87.2
D3 7244419 7244509
refGene_gn=RSRC2;refGene_tx=XM_003994618.4;refGene_gn=RSRC2;refGene_tx
=XM_006938421.3;refGene_gn=RSRC2;refGene_tx=XM_006938422.2;refGene_gn=
RSRC2;refGene_tx=XM_019814486.2;refGene_gn=RSRC2;refGene_tx=XM_0198144
87.2
D3 7245562 7245742
refGene_gn=RSRC2;refGene_tx=XM_003994618.4;refGene_gn=RSRC2;refGene_tx
=XM_006938421.3;refGene_gn=RSRC2;refGene_tx=XM_006938422.2;refGene_gn=
RSRC2;refGene_tx=XM_019814486.2;refGene_gn=RSRC2;refGene_tx=XM_0198144
87.2
D3 7251270 7251469
refGene_gn=ZCCHC8;refGene_tx=XM_003994621.5;refGene_gn=ZCCHC8;refGene_
tx=XM_019814490.2
D3 7252957 7253000
refGene_gn=ZCCHC8;refGene_tx=XM_003994621.5;refGene_gn=ZCCHC8;refGene_
tx=XM_019814490.2
D3 7256871 7256946
refGene_gn=ZCCHC8;refGene_tx=XM_003994621.5;refGene_gn=ZCCHC8;refGene_
tx=XM_019814490.2;refGene_gn=ZCCHC8;refGene_tx=XM_023241358.1
D3 7258767 7258873
refGene_gn=ZCCHC8;refGene_tx=XM_003994621.5;refGene_gn=ZCCHC8;refGene_
tx=XM_019814490.2;refGene_gn=ZCCHC8;refGene_tx=XM_023241358.1
D3 7260115 7260193
refGene_gn=ZCCHC8;refGene_tx=XM_003994621.5;refGene_gn=ZCCHC8;refGene_
tx=XM_019814490.2;refGene_gn=ZCCHC8;refGene_tx=XM_023241358.1
D3 7263469 7263573

refGene_gn=ZCCHC8;refGene_tx=XM_003994621.5;refGene_gn=ZCCHC8;refGene_tx=XM_023241358.1
D3 7263690 7263756
refGene_gn=ZCCHC8;refGene_tx=XM_003994621.5;refGene_gn=ZCCHC8;refGene_tx=XM_023241358.1
D3 7264338 7264399
refGene_gn=ZCCHC8;refGene_tx=XM_003994621.5;refGene_gn=ZCCHC8;refGene_tx=XM_023241358.1;refGene_gn=ZCCHC8;refGene_tx=XM_023241359.1
D3 7264876 7265019
refGene_gn=ZCCHC8;refGene_tx=XM_003994621.5;refGene_gn=ZCCHC8;refGene_tx=XM_019814490.2;refGene_gn=ZCCHC8;refGene_tx=XM_023241358.1;refGene_gn=ZCCHC8;refGene_tx=XM_023241359.1
D3 7265333 7265476
refGene_gn=ZCCHC8;refGene_tx=XM_003994621.5;refGene_gn=ZCCHC8;refGene_tx=XM_019814490.2;refGene_gn=ZCCHC8;refGene_tx=XM_023241358.1;refGene_gn=ZCCHC8;refGene_tx=XM_023241359.1
D3 7268350 7268472
refGene_gn=ZCCHC8;refGene_tx=XM_003994621.5;refGene_gn=ZCCHC8;refGene_tx=XM_019814490.2;refGene_gn=ZCCHC8;refGene_tx=XM_023241358.1;refGene_gn=ZCCHC8;refGene_tx=XM_023241359.1
D3 7270578 7270665
refGene_gn=ZCCHC8;refGene_tx=XM_003994621.5;refGene_gn=ZCCHC8;refGene_tx=XM_019814490.2;refGene_gn=ZCCHC8;refGene_tx=XM_023241358.1;refGene_gn=ZCCHC8;refGene_tx=XM_023241359.1
D3 7270927 7271045
refGene_gn=ZCCHC8;refGene_tx=XM_003994621.5;refGene_gn=ZCCHC8;refGene_tx=XM_019814490.2;refGene_gn=ZCCHC8;refGene_tx=XM_023241358.1;refGene_gn=ZCCHC8;refGene_tx=XM_023241359.1
D3 7274105 7274890
refGene_gn=ZCCHC8;refGene_tx=XM_003994621.5;refGene_gn=ZCCHC8;refGene_tx=XM_019814490.2;refGene_gn=ZCCHC8;refGene_tx=XM_023241358.1;refGene_gn=ZCCHC8;refGene_tx=XM_023241359.1
D3 7345773 7345858
refGene_gn=CLIP1;refGene_tx=XM_023241360.1;refGene_gn=CLIP1;refGene_tx=XM_023241361.1;refGene_gn=CLIP1;refGene_tx=XM_023241362.1;refGene_gn=CLIP1;refGene_tx=XM_023241363.1;refGene_gn=CLIP1;refGene_tx=XM_023241364.1;refGene_gn=CLIP1;refGene_tx=XM_023241365.1;refGene_gn=CLIP1;refGene_tx=XM_023241366.1;refGene_gn=CLIP1;refGene_tx=XM_023241367.1;refGene_gn=CLIP1;refGene_tx=XM_023241368.1;refGene_gn=CLIP1;refGene_tx=XM_023241369.1;refGene_gn=CLIP1;refGene_tx=XM_023241370.1;refGene_gn=CLIP1;refGene_tx=XM_023241371.1;refGene_gn=CLIP1;refGene_tx=XM_023241373.1;refGene_gn=CLIP1;refGene_tx=XM_023241374.1;refGene_gn=CLIP1;refGene_tx=XM_023241375.1;refGene_gn=CLIP1;refGene_tx=XM_023241376.1;refGene_gn=CLIP1;refGene_tx=XM_023241377.1
D3 7350358 7350930
refGene_gn=CLIP1;refGene_tx=XM_023241360.1;refGene_gn=CLIP1;refGene_tx=XM_023241361.1;refGene_gn=CLIP1;refGene_tx=XM_023241362.1;refGene_gn=CLIP1;refGene_tx=XM_023241363.1;refGene_gn=CLIP1;refGene_tx=XM_023241364.1;refGene_gn=CLIP1;refGene_tx=XM_023241365.1;refGene_gn=CLIP1;refGene_tx=XM_023241366.1;refGene_gn=CLIP1;refGene_tx=XM_023241367.1;refGen

e_gn=CLIP1;refGene_tx=XM_023241368.1;refGene_gn=CLIP1;refGene_tx=XM_023241369.1;refGene_gn=CLIP1;refGene_tx=XM_023241370.1;refGene_gn=CLIP1;refGene_tx=XM_023241371.1;refGene_gn=CLIP1;refGene_tx=XM_023241373.1;refGene_gn=CLIP1;refGene_tx=XM_023241374.1;refGene_gn=CLIP1;refGene_tx=XM_023241375.1;refGene_gn=CLIP1;refGene_tx=XM_023241376.1;refGene_gn=CLIP1;refGene_tx=XM_023241377.1
D3	7369166	7369291
refGene_gn=CLIP1;refGene_tx=XM_023241360.1;refGene_gn=CLIP1;refGene_tx=XM_023241361.1;refGene_gn=CLIP1;refGene_tx=XM_023241362.1;refGene_gn=CLIP1;refGene_tx=XM_023241363.1;refGene_gn=CLIP1;refGene_tx=XM_023241364.1;refGene_gn=CLIP1;refGene_tx=XM_023241365.1;refGene_gn=CLIP1;refGene_tx=XM_023241366.1;refGene_gn=CLIP1;refGene_tx=XM_023241367.1;refGene_gn=CLIP1;refGene_tx=XM_023241368.1;refGene_gn=CLIP1;refGene_tx=XM_023241369.1;refGene_gn=CLIP1;refGene_tx=XM_023241370.1;refGene_gn=CLIP1;refGene_tx=XM_023241371.1;refGene_gn=CLIP1;refGene_tx=XM_023241373.1;refGene_gn=CLIP1;refGene_tx=XM_023241374.1;refGene_gn=CLIP1;refGene_tx=XM_023241375.1;refGene_gn=CLIP1;refGene_tx=XM_023241376.1;refGene_gn=CLIP1;refGene_tx=XM_023241377.1
D3	7369972	7370195
refGene_gn=CLIP1;refGene_tx=XM_023241360.1;refGene_gn=CLIP1;refGene_tx=XM_023241361.1;refGene_gn=CLIP1;refGene_tx=XM_023241362.1;refGene_gn=CLIP1;refGene_tx=XM_023241363.1;refGene_gn=CLIP1;refGene_tx=XM_023241364.1;refGene_gn=CLIP1;refGene_tx=XM_023241365.1;refGene_gn=CLIP1;refGene_tx=XM_023241366.1;refGene_gn=CLIP1;refGene_tx=XM_023241367.1;refGene_gn=CLIP1;refGene_tx=XM_023241368.1;refGene_gn=CLIP1;refGene_tx=XM_023241369.1;refGene_gn=CLIP1;refGene_tx=XM_023241370.1;refGene_gn=CLIP1;refGene_tx=XM_023241371.1;refGene_gn=CLIP1;refGene_tx=XM_023241373.1;refGene_gn=CLIP1;refGene_tx=XM_023241374.1;refGene_gn=CLIP1;refGene_tx=XM_023241375.1;refGene_gn=CLIP1;refGene_tx=XM_023241376.1;refGene_gn=CLIP1;refGene_tx=XM_023241377.1
D3	7374431	7374629
refGene_gn=CLIP1;refGene_tx=XM_023241360.1;refGene_gn=CLIP1;refGene_tx=XM_023241361.1;refGene_gn=CLIP1;refGene_tx=XM_023241362.1;refGene_gn=CLIP1;refGene_tx=XM_023241363.1;refGene_gn=CLIP1;refGene_tx=XM_023241364.1;refGene_gn=CLIP1;refGene_tx=XM_023241365.1;refGene_gn=CLIP1;refGene_tx=XM_023241366.1;refGene_gn=CLIP1;refGene_tx=XM_023241367.1;refGene_gn=CLIP1;refGene_tx=XM_023241369.1;refGene_gn=CLIP1;refGene_tx=XM_023241370.1;refGene_gn=CLIP1;refGene_tx=XM_023241371.1;refGene_gn=CLIP1;refGene_tx=XM_023241373.1;refGene_gn=CLIP1;refGene_tx=XM_023241374.1;refGene_gn=CLIP1;refGene_tx=XM_023241375.1;refGene_gn=CLIP1;refGene_tx=XM_023241376.1;refGene_gn=CLIP1;refGene_tx=XM_023241377.1
D3	7375246	7375350
refGene_gn=CLIP1;refGene_tx=XM_023241360.1;refGene_gn=CLIP1;refGene_tx=XM_023241361.1;refGene_gn=CLIP1;refGene_tx=XM_023241362.1;refGene_gn=CLIP1;refGene_tx=XM_023241363.1;refGene_gn=CLIP1;refGene_tx=XM_023241364.1;refGene_gn=CLIP1;refGene_tx=XM_023241365.1;refGene_gn=CLIP1;refGene_tx=XM_023241366.1;refGene_gn=CLIP1;refGene_tx=XM_023241367.1;refGene_gn=CLIP1;refGene_tx=XM_023241368.1;refGene_gn=CLIP1;refGene_tx=XM_023241369.1;refGene_gn=CLIP1;refGene_tx=XM_023241370.1;refGene_gn=CLIP1;refGene_tx=XM_023241371.1;refGene_gn=CLIP1;refGene_tx=XM_023241373.1;r

efGene_gn=CLIP1;refGene_tx=XM_023241374.1;refGene_gn=CLIP1;refGene_tx=XM_023241375.1;refGene_gn=CLIP1;refGene_tx=XM_023241376.1;refGene_gn=CLIP1;refGene_tx=XM_023241377.1
D3 7377883 7377944
refGene_gn=CLIP1;refGene_tx=XM_023241360.1;refGene_gn=CLIP1;refGene_tx=XM_023241361.1;refGene_gn=CLIP1;refGene_tx=XM_023241362.1;refGene_gn=CLIP1;refGene_tx=XM_023241363.1;refGene_gn=CLIP1;refGene_tx=XM_023241364.1;refGene_gn=CLIP1;refGene_tx=XM_023241365.1;refGene_gn=CLIP1;refGene_tx=XM_023241366.1;refGene_gn=CLIP1;refGene_tx=XM_023241367.1;refGene_gn=CLIP1;refGene_tx=XM_023241368.1;refGene_gn=CLIP1;refGene_tx=XM_023241369.1;refGene_gn=CLIP1;refGene_tx=XM_023241370.1;refGene_gn=CLIP1;refGene_tx=XM_023241371.1;refGene_gn=CLIP1;refGene_tx=XM_023241373.1;refGene_gn=CLIP1;refGene_tx=XM_023241374.1;refGene_gn=CLIP1;refGene_tx=XM_023241375.1;refGene_gn=CLIP1;refGene_tx=XM_023241376.1;refGene_gn=CLIP1;refGene_tx=XM_023241377.1
D3 7379218 7379251
refGene_gn=CLIP1;refGene_tx=XM_023241360.1;refGene_gn=CLIP1;refGene_tx=XM_023241361.1;refGene_gn=CLIP1;refGene_tx=XM_023241362.1;refGene_gn=CLIP1;refGene_tx=XM_023241363.1;refGene_gn=CLIP1;refGene_tx=XM_023241364.1;refGene_gn=CLIP1;refGene_tx=XM_023241366.1;refGene_gn=CLIP1;refGene_tx=XM_023241368.1;refGene_gn=CLIP1;refGene_tx=XM_023241369.1;refGene_gn=CLIP1;refGene_tx=XM_023241370.1;refGene_gn=CLIP1;refGene_tx=XM_023241373.1;refGene_gn=CLIP1;refGene_tx=XM_023241374.1
D3 7382311 7382416
refGene_gn=CLIP1;refGene_tx=XM_023241360.1;refGene_gn=CLIP1;refGene_tx=XM_023241361.1;refGene_gn=CLIP1;refGene_tx=XM_023241362.1;refGene_gn=CLIP1;refGene_tx=XM_023241363.1;refGene_gn=CLIP1;refGene_tx=XM_023241364.1;refGene_gn=CLIP1;refGene_tx=XM_023241365.1;refGene_gn=CLIP1;refGene_tx=XM_023241368.1;refGene_gn=CLIP1;refGene_tx=XM_023241369.1;refGene_gn=CLIP1;refGene_tx=XM_023241370.1;refGene_gn=CLIP1;refGene_tx=XM_023241371.1;refGene_gn=CLIP1;refGene_tx=XM_023241373.1;refGene_gn=CLIP1;refGene_tx=XM_023241374.1;refGene_gn=CLIP1;refGene_tx=XM_023241375.1;refGene_gn=CLIP1;refGene_tx=XM_023241377.1
D3 7386181 7387126
refGene_gn=CLIP1;refGene_tx=XM_023241360.1;refGene_gn=CLIP1;refGene_tx=XM_023241361.1;refGene_gn=CLIP1;refGene_tx=XM_023241362.1;refGene_gn=CLIP1;refGene_tx=XM_023241363.1;refGene_gn=CLIP1;refGene_tx=XM_023241364.1;refGene_gn=CLIP1;refGene_tx=XM_023241365.1;refGene_gn=CLIP1;refGene_tx=XM_023241366.1;refGene_gn=CLIP1;refGene_tx=XM_023241367.1;refGene_gn=CLIP1;refGene_tx=XM_023241368.1;refGene_gn=CLIP1;refGene_tx=XM_023241369.1;refGene_gn=CLIP1;refGene_tx=XM_023241370.1;refGene_gn=CLIP1;refGene_tx=XM_023241371.1;refGene_gn=CLIP1;refGene_tx=XM_023241372.1;refGene_gn=CLIP1;refGene_tx=XM_023241373.1;refGene_gn=CLIP1;refGene_tx=XM_023241374.1;refGene_gn=CLIP1;refGene_tx=XM_023241375.1;refGene_gn=CLIP1;refGene_tx=XM_023241376.1;refGene_gn=CLIP1;refGene_tx=XM_023241377.1
D3 7388913 7389030
refGene_gn=CLIP1;refGene_tx=XM_023241360.1;refGene_gn=CLIP1;refGene_tx=XM_023241361.1;refGene_gn=CLIP1;refGene_tx=XM_023241362.1;refGene_gn=CLIP1;refGene_tx=XM_023241363.1;refGene_gn=CLIP1;refGene_tx=XM_0232413

64.1;refGene_gn=CLIP1;refGene_tx=XM_023241365.1;refGene_gn=CLIP1;refGene_tx=XM_023241366.1;refGene_gn=CLIP1;refGene_tx=XM_023241367.1;refGene_gn=CLIP1;refGene_tx=XM_023241368.1;refGene_gn=CLIP1;refGene_tx=XM_023241369.1;refGene_gn=CLIP1;refGene_tx=XM_023241372.1;refGene_gn=CLIP1;refGene_tx=XM_023241373.1;refGene_gn=CLIP1;refGene_tx=XM_023241374.1;refGene_gn=CLIP1;refGene_tx=XM_023241375.1;refGene_gn=CLIP1;refGene_tx=XM_023241376.1
D3 7391365 7391423
refGene_gn=CLIP1;refGene_tx=XM_023241360.1;refGene_gn=CLIP1;refGene_tx=XM_023241361.1;refGene_gn=CLIP1;refGene_tx=XM_023241362.1;refGene_gn=CLIP1;refGene_tx=XM_023241363.1;refGene_gn=CLIP1;refGene_tx=XM_023241364.1;refGene_gn=CLIP1;refGene_tx=XM_023241365.1;refGene_gn=CLIP1;refGene_tx=XM_023241366.1;refGene_gn=CLIP1;refGene_tx=XM_023241367.1;refGene_gn=CLIP1;refGene_tx=XM_023241368.1;refGene_gn=CLIP1;refGene_tx=XM_023241369.1;refGene_gn=CLIP1;refGene_tx=XM_023241370.1;refGene_gn=CLIP1;refGene_tx=XM_023241371.1;refGene_gn=CLIP1;refGene_tx=XM_023241372.1;refGene_gn=CLIP1;refGene_tx=XM_023241373.1;refGene_gn=CLIP1;refGene_tx=XM_023241374.1;refGene_gn=CLIP1;refGene_tx=XM_023241375.1;refGene_gn=CLIP1;refGene_tx=XM_023241376.1;refGene_gn=CLIP1;refGene_tx=XM_023241377.1
D3 7392180 7392264
refGene_gn=CLIP1;refGene_tx=XM_023241360.1;refGene_gn=CLIP1;refGene_tx=XM_023241361.1;refGene_gn=CLIP1;refGene_tx=XM_023241362.1;refGene_gn=CLIP1;refGene_tx=XM_023241363.1;refGene_gn=CLIP1;refGene_tx=XM_023241364.1;refGene_gn=CLIP1;refGene_tx=XM_023241365.1;refGene_gn=CLIP1;refGene_tx=XM_023241366.1;refGene_gn=CLIP1;refGene_tx=XM_023241367.1;refGene_gn=CLIP1;refGene_tx=XM_023241368.1;refGene_gn=CLIP1;refGene_tx=XM_023241369.1;refGene_gn=CLIP1;refGene_tx=XM_023241370.1;refGene_gn=CLIP1;refGene_tx=XM_023241371.1;refGene_gn=CLIP1;refGene_tx=XM_023241372.1;refGene_gn=CLIP1;refGene_tx=XM_023241373.1;refGene_gn=CLIP1;refGene_tx=XM_023241374.1;refGene_gn=CLIP1;refGene_tx=XM_023241375.1;refGene_gn=CLIP1;refGene_tx=XM_023241376.1;refGene_gn=CLIP1;refGene_tx=XM_023241377.1
D3 7393217 7393374
refGene_gn=CLIP1;refGene_tx=XM_023241360.1;refGene_gn=CLIP1;refGene_tx=XM_023241361.1;refGene_gn=CLIP1;refGene_tx=XM_023241362.1;refGene_gn=CLIP1;refGene_tx=XM_023241363.1;refGene_gn=CLIP1;refGene_tx=XM_023241364.1;refGene_gn=CLIP1;refGene_tx=XM_023241365.1;refGene_gn=CLIP1;refGene_tx=XM_023241366.1;refGene_gn=CLIP1;refGene_tx=XM_023241367.1;refGene_gn=CLIP1;refGene_tx=XM_023241368.1;refGene_gn=CLIP1;refGene_tx=XM_023241369.1;refGene_gn=CLIP1;refGene_tx=XM_023241370.1;refGene_gn=CLIP1;refGene_tx=XM_023241371.1;refGene_gn=CLIP1;refGene_tx=XM_023241372.1;refGene_gn=CLIP1;refGene_tx=XM_023241373.1;refGene_gn=CLIP1;refGene_tx=XM_023241374.1;refGene_gn=CLIP1;refGene_tx=XM_023241375.1;refGene_gn=CLIP1;refGene_tx=XM_023241376.1;refGene_gn=CLIP1;refGene_tx=XM_023241377.1
D3 7394725 7394891
refGene_gn=CLIP1;refGene_tx=XM_023241360.1;refGene_gn=CLIP1;refGene_tx=XM_023241361.1;refGene_gn=CLIP1;refGene_tx=XM_023241362.1;refGene_gn=CLIP1;refGene_tx=XM_023241363.1;refGene_gn=CLIP1;refGene_tx=XM_0232413

64.1;refGene_gn=CLIP1;refGene_tx=XM_023241365.1;refGene_gn=CLIP1;refGene_tx=XM_023241366.1;refGene_gn=CLIP1;refGene_tx=XM_023241367.1;refGene_gn=CLIP1;refGene_tx=XM_023241368.1;refGene_gn=CLIP1;refGene_tx=XM_023241369.1;refGene_gn=CLIP1;refGene_tx=XM_023241370.1;refGene_gn=CLIP1;refGene_tx=XM_023241371.1;refGene_gn=CLIP1;refGene_tx=XM_023241372.1;refGene_gn=CLIP1;refGene_tx=XM_023241373.1;refGene_gn=CLIP1;refGene_tx=XM_023241374.1;refGene_gn=CLIP1;refGene_tx=XM_023241375.1;refGene_gn=CLIP1;refGene_tx=XM_023241376.1;refGene_gn=CLIP1;refGene_tx=XM_023241377.1

D3 7394988 7395204

refGene_gn=CLIP1;refGene_tx=XM_023241360.1;refGene_gn=CLIP1;refGene_tx=XM_023241361.1;refGene_gn=CLIP1;refGene_tx=XM_023241362.1;refGene_gn=CLIP1;refGene_tx=XM_023241363.1;refGene_gn=CLIP1;refGene_tx=XM_023241364.1;refGene_gn=CLIP1;refGene_tx=XM_023241365.1;refGene_gn=CLIP1;refGene_tx=XM_023241366.1;refGene_gn=CLIP1;refGene_tx=XM_023241367.1;refGene_gn=CLIP1;refGene_tx=XM_023241368.1;refGene_gn=CLIP1;refGene_tx=XM_023241369.1;refGene_gn=CLIP1;refGene_tx=XM_023241370.1;refGene_gn=CLIP1;refGene_tx=XM_023241371.1;refGene_gn=CLIP1;refGene_tx=XM_023241372.1;refGene_gn=CLIP1;refGene_tx=XM_023241373.1;refGene_gn=CLIP1;refGene_tx=XM_023241374.1;refGene_gn=CLIP1;refGene_tx=XM_023241375.1;refGene_gn=CLIP1;refGene_tx=XM_023241376.1;refGene_gn=CLIP1;refGene_tx=XM_023241377.1

D3 7397285 7399553

refGene_gn=CLIP1;refGene_tx=XM_023241360.1;refGene_gn=CLIP1;refGene_tx=XM_023241361.1;refGene_gn=CLIP1;refGene_tx=XM_023241362.1;refGene_gn=CLIP1;refGene_tx=XM_023241363.1;refGene_gn=CLIP1;refGene_tx=XM_023241364.1;refGene_gn=CLIP1;refGene_tx=XM_023241365.1;refGene_gn=CLIP1;refGene_tx=XM_023241366.1;refGene_gn=CLIP1;refGene_tx=XM_023241367.1;refGene_gn=CLIP1;refGene_tx=XM_023241368.1;refGene_gn=CLIP1;refGene_tx=XM_023241369.1;refGene_gn=CLIP1;refGene_tx=XM_023241370.1;refGene_gn=CLIP1;refGene_tx=XM_023241371.1;refGene_gn=CLIP1;refGene_tx=XM_023241372.1

D3 7400705 7400822

refGene_gn=CLIP1;refGene_tx=XM_023241360.1;refGene_gn=CLIP1;refGene_tx=XM_023241361.1;refGene_gn=CLIP1;refGene_tx=XM_023241362.1;refGene_gn=CLIP1;refGene_tx=XM_023241363.1;refGene_gn=CLIP1;refGene_tx=XM_023241364.1;refGene_gn=CLIP1;refGene_tx=XM_023241365.1;refGene_gn=CLIP1;refGene_tx=XM_023241366.1;refGene_gn=CLIP1;refGene_tx=XM_023241367.1;refGene_gn=CLIP1;refGene_tx=XM_023241368.1;refGene_gn=CLIP1;refGene_tx=XM_023241369.1;refGene_gn=CLIP1;refGene_tx=XM_023241370.1;refGene_gn=CLIP1;refGene_tx=XM_023241371.1;refGene_gn=CLIP1;refGene_tx=XM_023241372.1;refGene_gn=CLIP1;refGene_tx=XM_023241373.1;refGene_gn=CLIP1;refGene_tx=XM_023241374.1;refGene_gn=CLIP1;refGene_tx=XM_023241375.1;refGene_gn=CLIP1;refGene_tx=XM_023241376.1;refGene_gn=CLIP1;refGene_tx=XM_023241377.1

D3 7402875 7402982

refGene_gn=CLIP1;refGene_tx=XM_023241360.1;refGene_gn=CLIP1;refGene_tx=XM_023241361.1;refGene_gn=CLIP1;refGene_tx=XM_023241362.1;refGene_gn=CLIP1;refGene_tx=XM_023241363.1;refGene_gn=CLIP1;refGene_tx=XM_023241364.1;refGene_gn=CLIP1;refGene_tx=XM_023241365.1;refGene_gn=CLIP1;refGene_tx=XM_023241366.1;refGene_gn=CLIP1;refGene_tx=XM_023241367.1;refGen

e_gn=CLIP1;refGene_tx=XM_023241368.1;refGene_gn=CLIP1;refGene_tx=XM_02
3241369.1;refGene_gn=CLIP1;refGene_tx=XM_023241370.1;refGene_gn=CLIP1;
refGene_tx=XM_023241371.1;refGene_gn=CLIP1;refGene_tx=XM_023241372.1;r
efGene_gn=CLIP1;refGene_tx=XM_023241373.1;refGene_gn=CLIP1;refGene_tx=
XM_023241374.1;refGene_gn=CLIP1;refGene_tx=XM_023241375.1;refGene_gn=C
LIP1;refGene_tx=XM_023241376.1;refGene_gn=CLIP1;refGene_tx=XM_02324137
7.1
D3 7408124 7408245
refGene_gn=CLIP1;refGene_tx=XM_023241360.1;refGene_gn=CLIP1;refGene_tx
=XM_023241361.1;refGene_gn=CLIP1;refGene_tx=XM_023241362.1;refGene_gn=
CLIP1;refGene_tx=XM_023241363.1;refGene_gn=CLIP1;refGene_tx=XM_0232413
64.1;refGene_gn=CLIP1;refGene_tx=XM_023241365.1;refGene_gn=CLIP1;refGe
ne_tx=XM_023241366.1;refGene_gn=CLIP1;refGene_tx=XM_023241367.1;refGen
e_gn=CLIP1;refGene_tx=XM_023241368.1;refGene_gn=CLIP1;refGene_tx=XM_02
3241369.1;refGene_gn=CLIP1;refGene_tx=XM_023241370.1;refGene_gn=CLIP1;
refGene_tx=XM_023241371.1;refGene_gn=CLIP1;refGene_tx=XM_023241372.1;r
efGene_gn=CLIP1;refGene_tx=XM_023241373.1;refGene_gn=CLIP1;refGene_tx=
XM_023241374.1;refGene_gn=CLIP1;refGene_tx=XM_023241375.1;refGene_gn=C
LIP1;refGene_tx=XM_023241376.1;refGene_gn=CLIP1;refGene_tx=XM_02324137
7.1
D3 7422751 7422804
refGene_gn=CLIP1;refGene_tx=XM_023241360.1;refGene_gn=CLIP1;refGene_tx
=XM_023241361.1;refGene_gn=CLIP1;refGene_tx=XM_023241362.1;refGene_gn=
CLIP1;refGene_tx=XM_023241363.1;refGene_gn=CLIP1;refGene_tx=XM_0232413
64.1;refGene_gn=CLIP1;refGene_tx=XM_023241365.1;refGene_gn=CLIP1;refGe
ne_tx=XM_023241366.1;refGene_gn=CLIP1;refGene_tx=XM_023241367.1;refGen
e_gn=CLIP1;refGene_tx=XM_023241368.1;refGene_gn=CLIP1;refGene_tx=XM_02
3241369.1;refGene_gn=CLIP1;refGene_tx=XM_023241370.1;refGene_gn=CLIP1;
refGene_tx=XM_023241371.1;refGene_gn=CLIP1;refGene_tx=XM_023241372.1;r
efGene_gn=CLIP1;refGene_tx=XM_023241373.1;refGene_gn=CLIP1;refGene_tx=
XM_023241374.1;refGene_gn=CLIP1;refGene_tx=XM_023241375.1;refGene_gn=C
LIP1;refGene_tx=XM_023241376.1;refGene_gn=CLIP1;refGene_tx=XM_02324137
7.1
D3 7428901 7429019
refGene_gn=CLIP1;refGene_tx=XM_023241360.1;refGene_gn=CLIP1;refGene_tx
=XM_023241361.1;refGene_gn=CLIP1;refGene_tx=XM_023241362.1;refGene_gn=
CLIP1;refGene_tx=XM_023241363.1;refGene_gn=CLIP1;refGene_tx=XM_0232413
64.1;refGene_gn=CLIP1;refGene_tx=XM_023241365.1;refGene_gn=CLIP1;refGe
ne_tx=XM_023241366.1;refGene_gn=CLIP1;refGene_tx=XM_023241367.1;refGen
e_gn=CLIP1;refGene_tx=XM_023241368.1;refGene_gn=CLIP1;refGene_tx=XM_02
3241369.1;refGene_gn=CLIP1;refGene_tx=XM_023241370.1;refGene_gn=CLIP1;
refGene_tx=XM_023241371.1;refGene_gn=CLIP1;refGene_tx=XM_023241372.1;r
efGene_gn=CLIP1;refGene_tx=XM_023241373.1;refGene_gn=CLIP1;refGene_tx=
XM_023241374.1;refGene_gn=CLIP1;refGene_tx=XM_023241375.1;refGene_gn=C
LIP1;refGene_tx=XM_023241376.1;refGene_gn=CLIP1;refGene_tx=XM_02324137
7.1
D3 7429103 7429254
refGene_gn=CLIP1;refGene_tx=XM_023241360.1;refGene_gn=CLIP1;refGene_tx
=XM_023241361.1;refGene_gn=CLIP1;refGene_tx=XM_023241362.1;refGene_gn=
CLIP1;refGene_tx=XM_023241363.1;refGene_gn=CLIP1;refGene_tx=XM_0232413

64.1;refGene_gn=CLIP1;refGene_tx=XM_023241365.1;refGene_gn=CLIP1;refGene_tx=XM_023241366.1;refGene_gn=CLIP1;refGene_tx=XM_023241367.1;refGene_gn=CLIP1;refGene_tx=XM_023241368.1;refGene_gn=CLIP1;refGene_tx=XM_023241369.1;refGene_gn=CLIP1;refGene_tx=XM_023241370.1;refGene_gn=CLIP1;refGene_tx=XM_023241371.1;refGene_gn=CLIP1;refGene_tx=XM_023241372.1;refGene_gn=CLIP1;refGene_tx=XM_023241373.1;refGene_gn=CLIP1;refGene_tx=XM_023241374.1;refGene_gn=CLIP1;refGene_tx=XM_023241375.1;refGene_gn=CLIP1;refGene_tx=XM_023241376.1;refGene_gn=CLIP1;refGene_tx=XM_023241377.1
D3 7429826 7429876
refGene_gn=CLIP1;refGene_tx=XM_023241360.1;refGene_gn=CLIP1;refGene_tx=XM_023241361.1;refGene_gn=CLIP1;refGene_tx=XM_023241362.1;refGene_gn=CLIP1;refGene_tx=XM_023241363.1;refGene_gn=CLIP1;refGene_tx=XM_023241364.1;refGene_gn=CLIP1;refGene_tx=XM_023241365.1;refGene_gn=CLIP1;refGene_tx=XM_023241366.1;refGene_gn=CLIP1;refGene_tx=XM_023241367.1;refGene_gn=CLIP1;refGene_tx=XM_023241368.1;refGene_gn=CLIP1;refGene_tx=XM_023241369.1;refGene_gn=CLIP1;refGene_tx=XM_023241370.1;refGene_gn=CLIP1;refGene_tx=XM_023241371.1;refGene_gn=CLIP1;refGene_tx=XM_023241372.1;refGene_gn=CLIP1;refGene_tx=XM_023241373.1;refGene_gn=CLIP1;refGene_tx=XM_023241374.1;refGene_gn=CLIP1;refGene_tx=XM_023241375.1;refGene_gn=CLIP1;refGene_tx=XM_023241376.1;refGene_gn=CLIP1;refGene_tx=XM_023241377.1
D3 7431805 7431811
refGene_gn=CLIP1;refGene_tx=XM_023241360.1;refGene_gn=CLIP1;refGene_tx=XM_023241361.1;refGene_gn=CLIP1;refGene_tx=XM_023241362.1;refGene_gn=CLIP1;refGene_tx=XM_023241363.1;refGene_gn=CLIP1;refGene_tx=XM_023241365.1;refGene_gn=CLIP1;refGene_tx=XM_023241366.1;refGene_gn=CLIP1;refGene_tx=XM_023241367.1;refGene_gn=CLIP1;refGene_tx=XM_023241368.1;refGene_gn=CLIP1;refGene_tx=XM_023241369.1;refGene_gn=CLIP1;refGene_tx=XM_023241370.1;refGene_gn=CLIP1;refGene_tx=XM_023241371.1;refGene_gn=CLIP1;refGene_tx=XM_023241372.1;refGene_gn=CLIP1;refGene_tx=XM_023241373.1;refGene_gn=CLIP1;refGene_tx=XM_023241377.1
D3 7433018 7433143
refGene_gn=CLIP1;refGene_tx=XM_023241360.1;refGene_gn=CLIP1;refGene_tx=XM_023241361.1;refGene_gn=CLIP1;refGene_tx=XM_023241362.1;refGene_gn=CLIP1;refGene_tx=XM_023241363.1;refGene_gn=CLIP1;refGene_tx=XM_023241364.1;refGene_gn=CLIP1;refGene_tx=XM_023241365.1;refGene_gn=CLIP1;refGene_tx=XM_023241366.1;refGene_gn=CLIP1;refGene_tx=XM_023241367.1;refGene_gn=CLIP1;refGene_tx=XM_023241368.1;refGene_gn=CLIP1;refGene_tx=XM_023241369.1;refGene_gn=CLIP1;refGene_tx=XM_023241370.1;refGene_gn=CLIP1;refGene_tx=XM_023241371.1;refGene_gn=CLIP1;refGene_tx=XM_023241372.1;refGene_gn=CLIP1;refGene_tx=XM_023241373.1;refGene_gn=CLIP1;refGene_tx=XM_023241374.1;refGene_gn=CLIP1;refGene_tx=XM_023241375.1;refGene_gn=CLIP1;refGene_tx=XM_023241376.1;refGene_gn=CLIP1;refGene_tx=XM_023241377.1
D3 7433459 7433685
refGene_gn=CLIP1;refGene_tx=XM_023241360.1;refGene_gn=CLIP1;refGene_tx=XM_023241361.1;refGene_gn=CLIP1;refGene_tx=XM_023241362.1;refGene_gn=CLIP1;refGene_tx=XM_023241363.1;refGene_gn=CLIP1;refGene_tx=XM_023241364.1;refGene_gn=CLIP1;refGene_tx=XM_023241365.1;refGene_gn=CLIP1;refGe

ne_tx=XM_023241366.1;refGene_gn=CLIP1;refGene_tx=XM_023241367.1;refGene_gn=CLIP1;refGene_tx=XM_023241368.1;refGene_gn=CLIP1;refGene_tx=XM_023241369.1;refGene_gn=CLIP1;refGene_tx=XM_023241370.1;refGene_gn=CLIP1;refGene_tx=XM_023241371.1;refGene_gn=CLIP1;refGene_tx=XM_023241372.1;refGene_gn=CLIP1;refGene_tx=XM_023241373.1;refGene_gn=CLIP1;refGene_tx=XM_023241374.1;refGene_gn=CLIP1;refGene_tx=XM_023241375.1;refGene_gn=CLIP1;refGene_tx=XM_023241376.1;refGene_gn=CLIP1;refGene_tx=XM_023241377.1
D3 7437312 7437414 refGene_gn=VPS33A;refGene_tx=XM_011287839.3;refGene_gn=VPS33A;refGene_tx=XM_019814491.2
D3 7439190 7439256 refGene_gn=VPS33A;refGene_tx=XM_011287839.3;refGene_gn=VPS33A;refGene_tx=XM_019814491.2
D3 7439704 7439832 refGene_gn=VPS33A;refGene_tx=XM_011287839.3;refGene_gn=VPS33A;refGene_tx=XM_019814491.2
D3 7440713 7440900 refGene_gn=VPS33A;refGene_tx=XM_011287839.3;refGene_gn=VPS33A;refGene_tx=XM_019814491.2
D3 7447898 7448015 refGene_gn=VPS33A;refGene_tx=XM_011287839.3;refGene_gn=VPS33A;refGene_tx=XM_019814491.2
D3 7448854 7449029 refGene_gn=VPS33A;refGene_tx=XM_011287839.3;refGene_gn=VPS33A;refGene_tx=XM_019814491.2
D3 7450017 7450211 refGene_gn=VPS33A;refGene_tx=XM_011287839.3;refGene_gn=VPS33A;refGene_tx=XM_019814491.2
D3 7451815 7451942 refGene_gn=VPS33A;refGene_tx=XM_011287839.3;refGene_gn=VPS33A;refGene_tx=XM_019814491.2
D3 7453307 7453375 refGene_gn=VPS33A;refGene_tx=XM_011287839.3;refGene_gn=VPS33A;refGene_tx=XM_019814491.2
D3 7453954 7454092 refGene_gn=VPS33A;refGene_tx=XM_011287839.3;refGene_gn=VPS33A;refGene_tx=XM_019814491.2
D3 7457000 7457138 refGene_gn=VPS33A;refGene_tx=XM_011287839.3;refGene_gn=VPS33A;refGene_tx=XM_019814491.2
D3 7459151 7459166 refGene_gn=VPS33A;refGene_tx=XM_019814491.2
D3 7459177 7459346 refGene_gn=VPS33A;refGene_tx=XM_011287839.3
D3 7459720 7459902 refGene_gn=VPS33A;refGene_tx=XM_011287839.3
D3 7465755 7465805 refGene_gn=DIABLO;refGene_tx=XM_003994629.5
D3 7466646 7466773 refGene_gn=DIABLO;refGene_tx=XM_003994629.5
D3 7470744 7470876 refGene_gn=DIABLO;refGene_tx=XM_003994629.5
D3 7471627 7471738 refGene_gn=DIABLO;refGene_tx=XM_003994629.5
D3 7471910 7472007 refGene_gn=DIABLO;refGene_tx=XM_003994629.5
D3 7480885 7481082 refGene_gn=DIABLO;refGene_tx=XM_003994629.5

D3 7482050 7483112 refGene_gn=B3GNT4;refGene_tx=XM_003994631.5
D3 7486515 7486643 refGene_gn=LRRC43;refGene_tx=XM_023241378.1
D3 7493134 7493168 refGene_gn=LRRC43;refGene_tx=XM_023241378.1
D3 7493497 7493683 refGene_gn=LRRC43;refGene_tx=XM_023241378.1
D3 7493791 7493947 refGene_gn=LRRC43;refGene_tx=XM_023241378.1
D3 7494077 7494219 refGene_gn=LRRC43;refGene_tx=XM_023241378.1
D3 7496120 7496359 refGene_gn=LRRC43;refGene_tx=XM_023241378.1
D3 7496957 7497145 refGene_gn=LRRC43;refGene_tx=XM_023241378.1
D3 7497778 7498017 refGene_gn=LRRC43;refGene_tx=XM_023241378.1
D3 7500196 7500336 refGene_gn=LRRC43;refGene_tx=XM_023241378.1
D3 7503200 7503311 refGene_gn=LRRC43;refGene_tx=XM_023241378.1
D3 7504298 7504424 refGene_gn=LRRC43;refGene_tx=XM_023241378.1
D3 7504425 7504560 refGene_gn=LRRC43;refGene_tx=XM_023241378.1
D3 7505852 7505945 refGene_gn=LRRC43;refGene_tx=XM_023241378.1
D3 7512242 7512258 refGene_gn=IL31;refGene_tx=XM_011287838.2
D3 7512429 7512578 refGene_gn=IL31;refGene_tx=XM_011287838.2
D3 7513961 7514276 refGene_gn=IL31;refGene_tx=XM_011287838.2
D3 7532329 7532451 refGene_gn=MLXIP;refGene_tx=XM_023241379.1
D3 7533100 7533230 refGene_gn=MLXIP;refGene_tx=XM_023241379.1
D3 7534696 7534820 refGene_gn=MLXIP;refGene_tx=XM_023241379.1
D3 7535062 7535190 refGene_gn=MLXIP;refGene_tx=XM_023241379.1
D3 7535298 7535400 refGene_gn=MLXIP;refGene_tx=XM_023241379.1
D3 7535835 7535960 refGene_gn=MLXIP;refGene_tx=XM_023241379.1
D3 7537422 7537576 refGene_gn=MLXIP;refGene_tx=XM_023241379.1
D3 7537696 7537818 refGene_gn=MLXIP;refGene_tx=XM_023241379.1
D3 7538309 7538961 refGene_gn=MLXIP;refGene_tx=XM_023241379.1
D3 7539903 7539995 refGene_gn=MLXIP;refGene_tx=XM_023241379.1
D3 7540570 7540660 refGene_gn=MLXIP;refGene_tx=XM_023241379.1
D3 7541387 7541559 refGene_gn=MLXIP;refGene_tx=XM_023241379.1
D3 7541894 7541936 refGene_gn=MLXIP;refGene_tx=XM_023241379.1
D3 7542331 7542421 refGene_gn=MLXIP;refGene_tx=XM_023241379.1
D3 7546144 7546230 refGene_gn=MLXIP;refGene_tx=XM_023241379.1
D3 7546948 7547055 refGene_gn=MLXIP;refGene_tx=XM_023241379.1
D3 7588889 7589302 refGene_gn=MLXIP;refGene_tx=XM_023241379.1
D3 7605341 7605413
refGene_gn=BCL7A;refGene_tx=XM_023241380.1;refGene_gn=BCL7A;refGene_tx
=XM_023241381.1
D3 7609456 7609641
refGene_gn=BCL7A;refGene_tx=XM_023241380.1;refGene_gn=BCL7A;refGene_tx
=XM_023241381.1
D3 7616799 7616967
refGene_gn=BCL7A;refGene_tx=XM_023241380.1;refGene_gn=BCL7A;refGene_tx
=XM_023241381.1
D3 7623580 7623677
refGene_gn=BCL7A;refGene_tx=XM_023241380.1;refGene_gn=BCL7A;refGene_tx
=XM_023241381.1
D3 7626900 7626982
refGene_gn=BCL7A;refGene_tx=XM_023241380.1;refGene_gn=BCL7A;refGene_tx
=XM_023241381.1
D3 7634198 7634290

refGene_gn=BCL7A;refGene_tx=XM_023241380.1;refGene_gn=BCL7A;refGene_tx=XM_023241381.1
D3 7645667 7645804
refGene_gn=WDR66;refGene_tx=XM_023241382.1;refGene_gn=WDR66;refGene_tx=XM_023241383.1;refGene_gn=WDR66;refGene_tx=XM_023241384.1;refGene_gn=WDR66;refGene_tx=XM_023241388.1;refGene_gn=WDR66;refGene_tx=XM_023241389.1
D3 7647100 7647202
refGene_gn=WDR66;refGene_tx=XM_023241382.1;refGene_gn=WDR66;refGene_tx=XM_023241384.1;refGene_gn=WDR66;refGene_tx=XM_023241388.1;refGene_gn=WDR66;refGene_tx=XM_023241389.1
D3 7648619 7648848
refGene_gn=WDR66;refGene_tx=XM_023241382.1;refGene_gn=WDR66;refGene_tx=XM_023241383.1;refGene_gn=WDR66;refGene_tx=XM_023241384.1;refGene_gn=WDR66;refGene_tx=XM_023241388.1;refGene_gn=WDR66;refGene_tx=XM_023241389.1
D3 7664128 7664146 refGene_gn=WDR66;refGene_tx=XM_023241385.1
D3 7667444 7667588
refGene_gn=WDR66;refGene_tx=XM_023241382.1;refGene_gn=WDR66;refGene_tx=XM_023241383.1;refGene_gn=WDR66;refGene_tx=XM_023241384.1;refGene_gn=WDR66;refGene_tx=XM_023241385.1;refGene_gn=WDR66;refGene_tx=XM_023241388.1;refGene_gn=WDR66;refGene_tx=XM_023241389.1
D3 7667797 7667888
refGene_gn=WDR66;refGene_tx=XM_023241382.1;refGene_gn=WDR66;refGene_tx=XM_023241383.1;refGene_gn=WDR66;refGene_tx=XM_023241384.1;refGene_gn=WDR66;refGene_tx=XM_023241385.1;refGene_gn=WDR66;refGene_tx=XM_023241388.1;refGene_gn=WDR66;refGene_tx=XM_023241389.1
D3 7667983 7668035 refGene_gn=WDR66;refGene_tx=XM_023241387.1
D3 7672448 7672612
refGene_gn=WDR66;refGene_tx=XM_023241382.1;refGene_gn=WDR66;refGene_tx=XM_023241383.1;refGene_gn=WDR66;refGene_tx=XM_023241384.1;refGene_gn=WDR66;refGene_tx=XM_023241385.1;refGene_gn=WDR66;refGene_tx=XM_023241387.1;refGene_gn=WDR66;refGene_tx=XM_023241388.1;refGene_gn=WDR66;refGene_tx=XM_023241389.1
D3 7673471 7673586
refGene_gn=WDR66;refGene_tx=XM_023241382.1;refGene_gn=WDR66;refGene_tx=XM_023241383.1;refGene_gn=WDR66;refGene_tx=XM_023241384.1;refGene_gn=WDR66;refGene_tx=XM_023241385.1;refGene_gn=WDR66;refGene_tx=XM_023241387.1;refGene_gn=WDR66;refGene_tx=XM_023241388.1;refGene_gn=WDR66;refGene_tx=XM_023241389.1
D3 7679657 7679842
refGene_gn=WDR66;refGene_tx=XM_023241382.1;refGene_gn=WDR66;refGene_tx=XM_023241383.1;refGene_gn=WDR66;refGene_tx=XM_023241384.1;refGene_gn=WDR66;refGene_tx=XM_023241385.1;refGene_gn=WDR66;refGene_tx=XM_023241387.1;refGene_gn=WDR66;refGene_tx=XM_023241388.1;refGene_gn=WDR66;refGene_tx=XM_023241389.1
D3 7681047 7681221
refGene_gn=WDR66;refGene_tx=XM_023241382.1;refGene_gn=WDR66;refGene_tx=XM_023241383.1;refGene_gn=WDR66;refGene_tx=XM_023241384.1;refGene_gn=WDR66;refGene_tx=XM_023241385.1;refGene_gn=WDR66;refGene_tx=XM_0232413

87.1;refGene_gn=WDR66;refGene_tx=XM_023241388.1;refGene_gn=WDR66;refGene_tx=XM_023241389.1
D3 7682805 7682957
refGene_gn=WDR66;refGene_tx=XM_023241382.1;refGene_gn=WDR66;refGene_tx=XM_023241383.1;refGene_gn=WDR66;refGene_tx=XM_023241384.1;refGene_gn=WDR66;refGene_tx=XM_023241385.1;refGene_gn=WDR66;refGene_tx=XM_023241387.1;refGene_gn=WDR66;refGene_tx=XM_023241388.1;refGene_gn=WDR66;refGene_tx=XM_023241389.1
D3 7683302 7683553
refGene_gn=WDR66;refGene_tx=XM_023241382.1;refGene_gn=WDR66;refGene_tx=XM_023241383.1;refGene_gn=WDR66;refGene_tx=XM_023241384.1;refGene_gn=WDR66;refGene_tx=XM_023241385.1;refGene_gn=WDR66;refGene_tx=XM_023241387.1;refGene_gn=WDR66;refGene_tx=XM_023241388.1;refGene_gn=WDR66;refGene_tx=XM_023241389.1
D3 7684283 7684478
refGene_gn=WDR66;refGene_tx=XM_023241382.1;refGene_gn=WDR66;refGene_tx=XM_023241383.1;refGene_gn=WDR66;refGene_tx=XM_023241384.1;refGene_gn=WDR66;refGene_tx=XM_023241385.1;refGene_gn=WDR66;refGene_tx=XM_023241387.1;refGene_gn=WDR66;refGene_tx=XM_023241388.1;refGene_gn=WDR66;refGene_tx=XM_023241389.1
D3 7686648 7686863
refGene_gn=WDR66;refGene_tx=XM_023241382.1;refGene_gn=WDR66;refGene_tx=XM_023241383.1;refGene_gn=WDR66;refGene_tx=XM_023241384.1;refGene_gn=WDR66;refGene_tx=XM_023241385.1;refGene_gn=WDR66;refGene_tx=XM_023241387.1;refGene_gn=WDR66;refGene_tx=XM_023241388.1;refGene_gn=WDR66;refGene_tx=XM_023241389.1
D3 7689025 7689076
refGene_gn=WDR66;refGene_tx=XM_023241382.1;refGene_gn=WDR66;refGene_tx=XM_023241383.1;refGene_gn=WDR66;refGene_tx=XM_023241384.1;refGene_gn=WDR66;refGene_tx=XM_023241385.1;refGene_gn=WDR66;refGene_tx=XM_023241387.1;refGene_gn=WDR66;refGene_tx=XM_023241388.1;refGene_gn=WDR66;refGene_tx=XM_023241389.1
D3 7690591 7690669
refGene_gn=WDR66;refGene_tx=XM_023241382.1;refGene_gn=WDR66;refGene_tx=XM_023241383.1;refGene_gn=WDR66;refGene_tx=XM_023241384.1;refGene_gn=WDR66;refGene_tx=XM_023241385.1;refGene_gn=WDR66;refGene_tx=XM_023241387.1;refGene_gn=WDR66;refGene_tx=XM_023241388.1;refGene_gn=WDR66;refGene_tx=XM_023241389.1
D3 7695768 7695849
refGene_gn=WDR66;refGene_tx=XM_023241382.1;refGene_gn=WDR66;refGene_tx=XM_023241383.1;refGene_gn=WDR66;refGene_tx=XM_023241384.1;refGene_gn=WDR66;refGene_tx=XM_023241385.1;refGene_gn=WDR66;refGene_tx=XM_023241387.1;refGene_gn=WDR66;refGene_tx=XM_023241388.1;refGene_gn=WDR66;refGene_tx=XM_023241389.1
D3 7696106 7696218
refGene_gn=WDR66;refGene_tx=XM_023241382.1;refGene_gn=WDR66;refGene_tx=XM_023241383.1;refGene_gn=WDR66;refGene_tx=XM_023241384.1;refGene_gn=WDR66;refGene_tx=XM_023241385.1;refGene_gn=WDR66;refGene_tx=XM_023241387.1;refGene_gn=WDR66;refGene_tx=XM_023241388.1;refGene_gn=WDR66;refGene_tx=XM_023241389.1

D3	7703476	7703586
refGene_gn=WDR66;refGene_tx=XM_023241382.1;refGene_gn=WDR66;refGene_tx=XM_023241383.1;refGene_gn=WDR66;refGene_tx=XM_023241384.1;refGene_gn=WDR66;refGene_tx=XM_023241385.1;refGene_gn=WDR66;refGene_tx=XM_023241387.1
D3	7705851	7705992
refGene_gn=WDR66;refGene_tx=XM_023241382.1;refGene_gn=WDR66;refGene_tx=XM_023241383.1;refGene_gn=WDR66;refGene_tx=XM_023241384.1;refGene_gn=WDR66;refGene_tx=XM_023241385.1;refGene_gn=WDR66;refGene_tx=XM_023241387.1;refGene_gn=WDR66;refGene_tx=XM_023241388.1;refGene_gn=WDR66;refGene_tx=XM_023241389.1
D3	7710957	7711341
refGene_gn=WDR66;refGene_tx=XM_023241382.1;refGene_gn=WDR66;refGene_tx=XM_023241383.1;refGene_gn=WDR66;refGene_tx=XM_023241384.1;refGene_gn=WDR66;refGene_tx=XM_023241385.1;refGene_gn=WDR66;refGene_tx=XM_023241387.1
D3	7712025	7712226
refGene_gn=WDR66;refGene_tx=XM_023241382.1;refGene_gn=WDR66;refGene_tx=XM_023241383.1;refGene_gn=WDR66;refGene_tx=XM_023241385.1;refGene_gn=WDR66;refGene_tx=XM_023241387.1
D3	7716038	7716066	refGene_gn=PSMD9;refGene_tx=XM_003994632.5
D3	7716771	7716860	refGene_gn=PSMD9;refGene_tx=XM_003994632.5
D3	7723874	7723976	refGene_gn=PSMD9;refGene_tx=XM_003994632.5
D3	7726116	7726328	refGene_gn=PSMD9;refGene_tx=XM_003994632.5
D3	7730377	7730480	refGene_gn=PSMD9;refGene_tx=XM_003994632.5
D3	7735131	7735380	refGene_gn=PSMD9;refGene_tx=XM_003994632.5
D3	7765917	7765920	refGene_gn=HPD;refGene_tx=XM_003994634.3
D3	7766037	7766064	refGene_gn=HPD;refGene_tx=XM_003994634.3
D3	7766705	7766768	refGene_gn=HPD;refGene_tx=XM_003994634.3
D3	7767123	7767228	refGene_gn=HPD;refGene_tx=XM_003994634.3
D3	7767805	7767848	refGene_gn=HPD;refGene_tx=XM_003994634.3
D3	7768045	7768128	refGene_gn=HPD;refGene_tx=XM_003994634.3
D3	7769072	7769162	refGene_gn=HPD;refGene_tx=XM_003994634.3
D3	7770170	7770274	refGene_gn=HPD;refGene_tx=XM_003994634.3
D3	7770688	7770766	refGene_gn=HPD;refGene_tx=XM_003994634.3
D3	7771324	7771487	refGene_gn=HPD;refGene_tx=XM_003994634.3
D3	7771618	7771690	refGene_gn=HPD;refGene_tx=XM_003994634.3
D3	7773840	7773963	refGene_gn=HPD;refGene_tx=XM_003994634.3
D3	7776043	7776160	refGene_gn=HPD;refGene_tx=XM_003994634.3
D3	7776251	7776362	refGene_gn=HPD;refGene_tx=XM_003994634.3
D3	7784615	7784789	refGene_gn=SETD1B;refGene_tx=XM_023241390.1
D3	7788091	7788229	refGene_gn=SETD1B;refGene_tx=XM_023241390.1
D3	7788307	7788427	refGene_gn=SETD1B;refGene_tx=XM_023241390.1
D3	7788512	7788644	refGene_gn=SETD1B;refGene_tx=XM_023241390.1
D3	7790931	7791098	refGene_gn=SETD1B;refGene_tx=XM_023241390.1
D3	7791966	7793226	refGene_gn=SETD1B;refGene_tx=XM_023241390.1
D3	7794579	7795080	refGene_gn=SETD1B;refGene_tx=XM_023241390.1
D3	7796135	7796241	refGene_gn=SETD1B;refGene_tx=XM_023241390.1
D3	7796334	7796702	refGene_gn=SETD1B;refGene_tx=XM_023241390.1
D3	7796780	7797042	refGene_gn=SETD1B;refGene_tx=XM_023241390.1

D3 7797428 7797991 refGene_gn=SETD1B;refGene_tx=XM_023241390.1
D3 7797993 7798255 refGene_gn=SETD1B;refGene_tx=XM_023241390.1
D3 7801790 7803029 refGene_gn=SETD1B;refGene_tx=XM_023241390.1
D3 7803901 7804014 refGene_gn=SETD1B;refGene_tx=XM_023241390.1
D3 7806101 7806372 refGene_gn=SETD1B;refGene_tx=XM_023241390.1
D3 7807008 7807107 refGene_gn=SETD1B;refGene_tx=XM_023241390.1
D3 7807351 7807525 refGene_gn=SETD1B;refGene_tx=XM_023241390.1
D3 7826991 7827117 refGene_gn=RHOF;refGene_tx=XM_003994635.5
D3 7827401 7827489 refGene_gn=RHOF;refGene_tx=XM_003994635.5
D3 7836457 7836567 refGene_gn=RHOF;refGene_tx=XM_003994635.5
D3 7836654 7836789 refGene_gn=RHOF;refGene_tx=XM_003994635.5
D3 7837628 7837793 refGene_gn=RHOF;refGene_tx=XM_003994635.5
D3 7840729 7840843 refGene_gn=TMEM120B;refGene_tx=XM_003994636.4
D3 7841365 7841434 refGene_gn=TMEM120B;refGene_tx=XM_003994636.4
D3 7841781 7841846 refGene_gn=TMEM120B;refGene_tx=XM_003994636.4
D3 7842963 7843056 refGene_gn=TMEM120B;refGene_tx=XM_003994636.4
D3 7844231 7844293 refGene_gn=TMEM120B;refGene_tx=XM_003994636.4
D3 7844789 7844855 refGene_gn=TMEM120B;refGene_tx=XM_003994636.4
D3 7852734 7852824 refGene_gn=TMEM120B;refGene_tx=XM_003994636.4
D3 7858379 7858475 refGene_gn=TMEM120B;refGene_tx=XM_003994636.4
D3 7859136 7859196 refGene_gn=TMEM120B;refGene_tx=XM_003994636.4
D3 7862761 7862878 refGene_gn=TMEM120B;refGene_tx=XM_003994636.4
D3 7864561 7864680 refGene_gn=TMEM120B;refGene_tx=XM_003994636.4
D3 7882273 7882342 refGene_gn=TMEM120B;refGene_tx=XM_003994636.4
D3 7905369 7905514
refGene_gn=MORN3;refGene_tx=XM_003994637.4;refGene_gn=MORN3;refGene_tx=XM_019814501.2;refGene_gn=MORN3;refGene_tx=XM_019814502.2;refGene_gn=MORN3;refGene_tx=XM_023241391.1
D3 7910486 7910644
refGene_gn=MORN3;refGene_tx=XM_003994637.4;refGene_gn=MORN3;refGene_tx=XM_019814501.2;refGene_gn=MORN3;refGene_tx=XM_019814502.2;refGene_gn=MORN3;refGene_tx=XM_023241391.1
D3 7912178 7912338
refGene_gn=MORN3;refGene_tx=XM_003994637.4;refGene_gn=MORN3;refGene_tx=XM_019814501.2;refGene_gn=MORN3;refGene_tx=XM_019814502.2;refGene_gn=MORN3;refGene_tx=XM_023241391.1
D3 7915165 7915350
refGene_gn=MORN3;refGene_tx=XM_003994637.4;refGene_gn=MORN3;refGene_tx=XM_019814501.2;refGene_gn=MORN3;refGene_tx=XM_019814502.2;refGene_gn=MORN3;refGene_tx=XM_023241391.1
D3 7915628 7915706
refGene_gn=MORN3;refGene_tx=XM_003994637.4;refGene_gn=MORN3;refGene_tx=XM_019814501.2;refGene_gn=MORN3;refGene_tx=XM_019814502.2;refGene_gn=MORN3;refGene_tx=XM_023241391.1
D3 7920716 7921322 refGene_gn=ORAI1;refGene_tx=XM_003994647.5
D3 7933883 7934183 refGene_gn=ORAI1;refGene_tx=XM_003994647.5
D3 7963175 7963208 refGene_gn=KDM2B;refGene_tx=XM_023241392.1
D3 7963592 7963607 refGene_gn=KDM2B;refGene_tx=XM_023241393.1
D3 7964276 7964421
refGene_gn=KDM2B;refGene_tx=XM_023241392.1;refGene_gn=KDM2B;refGene_tx

=XM_023241393.1
D3	7967375	7967454
refGene_gn=KDM2B;refGene_tx=XM_023241392.1;refGene_gn=KDM2B;refGene_tx=XM_023241393.1
D3	7968741	7968788
refGene_gn=KDM2B;refGene_tx=XM_023241392.1;refGene_gn=KDM2B;refGene_tx=XM_023241393.1
D3	7993608	7993787
refGene_gn=KDM2B;refGene_tx=XM_023241392.1;refGene_gn=KDM2B;refGene_tx=XM_023241393.1
D3	7994000	7994107
refGene_gn=KDM2B;refGene_tx=XM_023241392.1;refGene_gn=KDM2B;refGene_tx=XM_023241393.1
D3	8004885	8004979
refGene_gn=KDM2B;refGene_tx=XM_023241392.1;refGene_gn=KDM2B;refGene_tx=XM_023241393.1
D3	8005952	8006106
refGene_gn=KDM2B;refGene_tx=XM_023241392.1;refGene_gn=KDM2B;refGene_tx=XM_023241393.1
D3	8014659	8014775
refGene_gn=KDM2B;refGene_tx=XM_023241392.1;refGene_gn=KDM2B;refGene_tx=XM_023241393.1
D3	8020415	8020546
refGene_gn=KDM2B;refGene_tx=XM_023241392.1;refGene_gn=KDM2B;refGene_tx=XM_023241393.1
D3	8023141	8023634
refGene_gn=KDM2B;refGene_tx=XM_023241392.1;refGene_gn=KDM2B;refGene_tx=XM_023241393.1
D3	8041205	8041292
refGene_gn=KDM2B;refGene_tx=XM_023241392.1;refGene_gn=KDM2B;refGene_tx=XM_023241393.1
D3	8070306	8070531
refGene_gn=KDM2B;refGene_tx=XM_023241392.1;refGene_gn=KDM2B;refGene_tx=XM_023241393.1
D3	8075920	8076064
refGene_gn=KDM2B;refGene_tx=XM_023241392.1;refGene_gn=KDM2B;refGene_tx=XM_023241393.1
D3	8077467	8077554
refGene_gn=KDM2B;refGene_tx=XM_023241392.1;refGene_gn=KDM2B;refGene_tx=XM_023241393.1
D3	8077726	8077987
refGene_gn=KDM2B;refGene_tx=XM_023241392.1;refGene_gn=KDM2B;refGene_tx=XM_023241393.1
D3	8078191	8078305
refGene_gn=KDM2B;refGene_tx=XM_023241392.1;refGene_gn=KDM2B;refGene_tx=XM_023241393.1
D3	8078966	8079005
refGene_gn=KDM2B;refGene_tx=XM_023241392.1;refGene_gn=KDM2B;refGene_tx=XM_023241393.1
D3	8079117	8079797

refGene_gn=KDM2B;refGene_tx=XM_023241392.1;refGene_gn=KDM2B;refGene_tx=XM_023241393.1
D3 8080503 8080667
refGene_gn=KDM2B;refGene_tx=XM_023241392.1;refGene_gn=KDM2B;refGene_tx=XM_023241393.1
D3 8080752 8080914
refGene_gn=KDM2B;refGene_tx=XM_023241392.1;refGene_gn=KDM2B;refGene_tx=XM_023241393.1
D3 8081587 8081806
refGene_gn=KDM2B;refGene_tx=XM_023241392.1;refGene_gn=KDM2B;refGene_tx=XM_023241393.1
D3 8088477 8088659
refGene_gn=KDM2B;refGene_tx=XM_023241392.1;refGene_gn=KDM2B;refGene_tx=XM_023241393.1
D3 8093702 8093893
refGene_gn=RNF34;refGene_tx=XM_003994639.5;refGene_gn=RNF34;refGene_tx=XM_006938444.4;refGene_gn=RNF34;refGene_tx=XM_019814503.2
D3 8095986 8096188
refGene_gn=RNF34;refGene_tx=XM_003994639.5;refGene_gn=RNF34;refGene_tx=XM_006938444.4;refGene_gn=RNF34;refGene_tx=XM_019814503.2
D3 8096469 8096592
refGene_gn=RNF34;refGene_tx=XM_003994639.5;refGene_gn=RNF34;refGene_tx=XM_006938444.4;refGene_gn=RNF34;refGene_tx=XM_019814503.2
D3 8098078 8098486
refGene_gn=RNF34;refGene_tx=XM_003994639.5;refGene_gn=RNF34;refGene_tx=XM_006938444.4;refGene_gn=RNF34;refGene_tx=XM_019814503.2
D3 8100275 8100494
refGene_gn=RNF34;refGene_tx=XM_003994639.5;refGene_gn=RNF34;refGene_tx=XM_006938444.4
D3 8111978 8111984
refGene_gn=RNF34;refGene_tx=XM_003994639.5;refGene_gn=RNF34;refGene_tx=XM_019814503.2
D3 8141073 8141280 refGene_gn=ANAPC5;refGene_tx=XM_011287834.3
D3 8144764 8144844 refGene_gn=ANAPC5;refGene_tx=XM_011287834.3
D3 8145730 8145840 refGene_gn=ANAPC5;refGene_tx=XM_011287834.3
D3 8146650 8146843 refGene_gn=ANAPC5;refGene_tx=XM_011287834.3
D3 8156440 8156507 refGene_gn=ANAPC5;refGene_tx=XM_011287834.3
D3 8158852 8158954 refGene_gn=ANAPC5;refGene_tx=XM_011287834.3
D3 8159728 8159919 refGene_gn=ANAPC5;refGene_tx=XM_011287834.3
D3 8163769 8163851 refGene_gn=ANAPC5;refGene_tx=XM_011287834.3
D3 8164500 8164590 refGene_gn=ANAPC5;refGene_tx=XM_011287834.3
D3 8166294 8166476 refGene_gn=ANAPC5;refGene_tx=XM_011287834.3
D3 8167541 8167677 refGene_gn=ANAPC5;refGene_tx=XM_011287834.3
D3 8171580 8171655 refGene_gn=ANAPC5;refGene_tx=XM_011287834.3
D3 8174383 8174505 refGene_gn=ANAPC5;refGene_tx=XM_011287834.3
D3 8174962 8175070 refGene_gn=ANAPC5;refGene_tx=XM_011287834.3
D3 8175153 8175301 refGene_gn=ANAPC5;refGene_tx=XM_011287834.3
D3 8183912 8184075 refGene_gn=ANAPC5;refGene_tx=XM_011287834.3
D3 8184475 8184687 refGene_gn=ANAPC5;refGene_tx=XM_011287834.3
D3 8206546 8206999

refGene_gn=CAMKK2;refGene_tx=XM_023241394.1;refGene_gn=CAMKK2;refGene_tx=XM_023241395.1;refGene_gn=CAMKK2;refGene_tx=XM_023241396.1;refGene_gn=CAMKK2;refGene_tx=XM_023241397.1;refGene_gn=CAMKK2;refGene_tx=XM_023241398.1
D3 8211056 8211104
refGene_gn=CAMKK2;refGene_tx=XM_023241394.1;refGene_gn=CAMKK2;refGene_tx=XM_023241395.1;refGene_gn=CAMKK2;refGene_tx=XM_023241396.1;refGene_gn=CAMKK2;refGene_tx=XM_023241397.1;refGene_gn=CAMKK2;refGene_tx=XM_023241398.1
D3 8211787 8211841
refGene_gn=CAMKK2;refGene_tx=XM_023241394.1;refGene_gn=CAMKK2;refGene_tx=XM_023241395.1;refGene_gn=CAMKK2;refGene_tx=XM_023241396.1;refGene_gn=CAMKK2;refGene_tx=XM_023241397.1;refGene_gn=CAMKK2;refGene_tx=XM_023241398.1
D3 8212567 8212619
refGene_gn=CAMKK2;refGene_tx=XM_023241394.1;refGene_gn=CAMKK2;refGene_tx=XM_023241395.1;refGene_gn=CAMKK2;refGene_tx=XM_023241396.1;refGene_gn=CAMKK2;refGene_tx=XM_023241397.1;refGene_gn=CAMKK2;refGene_tx=XM_023241398.1
D3 8212977 8213111
refGene_gn=CAMKK2;refGene_tx=XM_023241394.1;refGene_gn=CAMKK2;refGene_tx=XM_023241395.1;refGene_gn=CAMKK2;refGene_tx=XM_023241396.1;refGene_gn=CAMKK2;refGene_tx=XM_023241397.1;refGene_gn=CAMKK2;refGene_tx=XM_023241398.1
D3 8215684 8215721
refGene_gn=CAMKK2;refGene_tx=XM_023241394.1;refGene_gn=CAMKK2;refGene_tx=XM_023241395.1;refGene_gn=CAMKK2;refGene_tx=XM_023241396.1;refGene_gn=CAMKK2;refGene_tx=XM_023241397.1;refGene_gn=CAMKK2;refGene_tx=XM_023241398.1
D3 8221634 8221656
refGene_gn=CAMKK2;refGene_tx=XM_023241394.1;refGene_gn=CAMKK2;refGene_tx=XM_023241395.1;refGene_gn=CAMKK2;refGene_tx=XM_023241396.1;refGene_gn=CAMKK2;refGene_tx=XM_023241397.1;refGene_gn=CAMKK2;refGene_tx=XM_023241398.1
D3 8221805 8221894
refGene_gn=CAMKK2;refGene_tx=XM_023241394.1;refGene_gn=CAMKK2;refGene_tx=XM_023241395.1;refGene_gn=CAMKK2;refGene_tx=XM_023241396.1;refGene_gn=CAMKK2;refGene_tx=XM_023241397.1;refGene_gn=CAMKK2;refGene_tx=XM_023241398.1
D3 8223086 8223286
refGene_gn=CAMKK2;refGene_tx=XM_023241394.1;refGene_gn=CAMKK2;refGene_tx=XM_023241395.1;refGene_gn=CAMKK2;refGene_tx=XM_023241396.1;refGene_gn=CAMKK2;refGene_tx=XM_023241397.1;refGene_gn=CAMKK2;refGene_tx=XM_023241398.1
D3 8223902 8223956
refGene_gn=CAMKK2;refGene_tx=XM_023241394.1;refGene_gn=CAMKK2;refGene_tx=XM_023241395.1;refGene_gn=CAMKK2;refGene_tx=XM_023241396.1;refGene_gn=CAMKK2;refGene_tx=XM_023241397.1;refGene_gn=CAMKK2;refGene_tx=XM_023241398.1
D3 8226042 8226116

refGene_gn=CAMKK2;refGene_tx=XM_023241394.1;refGene_gn=CAMKK2;refGene_tx=XM_023241395.1;refGene_gn=CAMKK2;refGene_tx=XM_023241396.1;refGene_gn=CAMKK2;refGene_tx=XM_023241397.1;refGene_gn=CAMKK2;refGene_tx=XM_023241398.1
D3 8226211 8226299
refGene_gn=CAMKK2;refGene_tx=XM_023241394.1;refGene_gn=CAMKK2;refGene_tx=XM_023241395.1;refGene_gn=CAMKK2;refGene_tx=XM_023241396.1;refGene_gn=CAMKK2;refGene_tx=XM_023241397.1;refGene_gn=CAMKK2;refGene_tx=XM_023241398.1
D3 8227445 8227574
refGene_gn=CAMKK2;refGene_tx=XM_023241394.1;refGene_gn=CAMKK2;refGene_tx=XM_023241396.1;refGene_gn=CAMKK2;refGene_tx=XM_023241397.1
D3 8232455 8232556
refGene_gn=CAMKK2;refGene_tx=XM_023241394.1;refGene_gn=CAMKK2;refGene_tx=XM_023241395.1;refGene_gn=CAMKK2;refGene_tx=XM_023241396.1;refGene_gn=CAMKK2;refGene_tx=XM_023241397.1;refGene_gn=CAMKK2;refGene_tx=XM_023241398.1
D3 8233062 8233105
refGene_gn=CAMKK2;refGene_tx=XM_023241394.1;refGene_gn=CAMKK2;refGene_tx=XM_023241395.1;refGene_gn=CAMKK2;refGene_tx=XM_023241397.1
D3 8235941 8236106
refGene_gn=CAMKK2;refGene_tx=XM_023241394.1;refGene_gn=CAMKK2;refGene_tx=XM_023241395.1;refGene_gn=CAMKK2;refGene_tx=XM_023241396.1;refGene_gn=CAMKK2;refGene_tx=XM_023241398.1
D3 8236255 8236279 refGene_gn=CAMKK2;refGene_tx=XM_023241397.1
D3 8241567 8241594
refGene_gn=P2RX4;refGene_tx=XM_023241408.1;refGene_gn=P2RX4;refGene_tx=XM_023241409.1
D3 8242162 8242258
refGene_gn=P2RX4;refGene_tx=XM_023241408.1;refGene_gn=P2RX4;refGene_tx=XM_023241409.1
D3 8242592 8242658
refGene_gn=P2RX4;refGene_tx=XM_023241408.1;refGene_gn=P2RX4;refGene_tx=XM_023241409.1
D3 8242819 8242913
refGene_gn=P2RX4;refGene_tx=XM_023241408.1;refGene_gn=P2RX4;refGene_tx=XM_023241409.1
D3 8245476 8245613
refGene_gn=P2RX4;refGene_tx=XM_023241408.1;refGene_gn=P2RX4;refGene_tx=XM_023241409.1
D3 8245711 8245853
refGene_gn=P2RX4;refGene_tx=XM_023241408.1;refGene_gn=P2RX4;refGene_tx=XM_023241409.1
D3 8245955 8246036
refGene_gn=P2RX4;refGene_tx=XM_023241408.1;refGene_gn=P2RX4;refGene_tx=XM_023241409.1
D3 8249963 8250060
refGene_gn=P2RX4;refGene_tx=XM_023241408.1;refGene_gn=P2RX4;refGene_tx=XM_023241409.1
D3 8250473 8250546

refGene_gn=P2RX4;refGene_tx=XM_023241408.1;refGene_gn=P2RX4;refGene_tx=XM_023241409.1
D3 8250637 8250709
refGene_gn=P2RX4;refGene_tx=XM_023241408.1;refGene_gn=P2RX4;refGene_tx=XM_023241409.1
D3 8255261 8255409
refGene_gn=P2RX4;refGene_tx=XM_023241408.1;refGene_gn=P2RX4;refGene_tx=XM_023241409.1
D3 8259700 8259834 refGene_gn=P2RX4;refGene_tx=XM_023241408.1
D3 8280030 8280060 refGene_gn=P2RX7;refGene_tx=XM_019814513.2
D3 8285376 8285868
refGene_gn=P2RX7;refGene_tx=XM_003994650.5;refGene_gn=P2RX7;refGene_tx=XM_011287830.2;refGene_gn=P2RX7;refGene_tx=XM_019814510.2;refGene_gn=P2RX7;refGene_tx=XM_019814511.2;refGene_gn=P2RX7;refGene_tx=XM_019814512.1
D3 8289271 8289373
refGene_gn=P2RX7;refGene_tx=XM_003994650.5;refGene_gn=P2RX7;refGene_tx=XM_011287830.2;refGene_gn=P2RX7;refGene_tx=XM_019814510.2;refGene_gn=P2RX7;refGene_tx=XM_019814511.2;refGene_gn=P2RX7;refGene_tx=XM_019814512.1;refGene_gn=P2RX7;refGene_tx=XM_019814513.2
D3 8291156 8291306
refGene_gn=P2RX7;refGene_tx=XM_003994650.5;refGene_gn=P2RX7;refGene_tx=XM_011287830.2;refGene_gn=P2RX7;refGene_tx=XM_019814510.2;refGene_gn=P2RX7;refGene_tx=XM_019814511.2;refGene_gn=P2RX7;refGene_tx=XM_019814512.1;refGene_gn=P2RX7;refGene_tx=XM_019814513.2
D3 8291392 8291458
refGene_gn=P2RX7;refGene_tx=XM_003994650.5;refGene_gn=P2RX7;refGene_tx=XM_011287830.2;refGene_gn=P2RX7;refGene_tx=XM_019814510.2;refGene_gn=P2RX7;refGene_tx=XM_019814511.2;refGene_gn=P2RX7;refGene_tx=XM_019814512.1;refGene_gn=P2RX7;refGene_tx=XM_019814513.2
D3 8293003 8293094
refGene_gn=P2RX7;refGene_tx=XM_003994650.5;refGene_gn=P2RX7;refGene_tx=XM_011287830.2;refGene_gn=P2RX7;refGene_tx=XM_019814510.2;refGene_gn=P2RX7;refGene_tx=XM_019814511.2;refGene_gn=P2RX7;refGene_tx=XM_019814512.1;refGene_gn=P2RX7;refGene_tx=XM_019814513.2
D3 8302075 8302212
refGene_gn=P2RX7;refGene_tx=XM_003994650.5;refGene_gn=P2RX7;refGene_tx=XM_011287830.2;refGene_gn=P2RX7;refGene_tx=XM_019814510.2;refGene_gn=P2RX7;refGene_tx=XM_019814511.2;refGene_gn=P2RX7;refGene_tx=XM_019814512.1;refGene_gn=P2RX7;refGene_tx=XM_019814513.2
D3 8306614 8306744
refGene_gn=P2RX7;refGene_tx=XM_003994650.5;refGene_gn=P2RX7;refGene_tx=XM_011287830.2;refGene_gn=P2RX7;refGene_tx=XM_019814510.2;refGene_gn=P2RX7;refGene_tx=XM_019814511.2;refGene_gn=P2RX7;refGene_tx=XM_019814512.1;refGene_gn=P2RX7;refGene_tx=XM_019814513.2
D3 8307381 8307462
refGene_gn=P2RX7;refGene_tx=XM_003994650.5;refGene_gn=P2RX7;refGene_tx=XM_011287830.2;refGene_gn=P2RX7;refGene_tx=XM_019814510.2;refGene_gn=P2RX7;refGene_tx=XM_019814511.2;refGene_gn=P2RX7;refGene_tx=XM_019814512.1;refGene_gn=P2RX7;refGene_tx=XM_019814513.2

D3	8310133	8310230
refGene_gn=P2RX7;refGene_tx=XM_003994650.5;refGene_gn=P2RX7;refGene_tx=XM_011287830.2;refGene_gn=P2RX7;refGene_tx=XM_019814510.2;refGene_gn=P2RX7;refGene_tx=XM_019814511.2;refGene_gn=P2RX7;refGene_tx=XM_019814512.1;refGene_gn=P2RX7;refGene_tx=XM_019814513.2
D3	8311275	8311348
refGene_gn=P2RX7;refGene_tx=XM_003994650.5;refGene_gn=P2RX7;refGene_tx=XM_011287830.2;refGene_gn=P2RX7;refGene_tx=XM_019814510.2;refGene_gn=P2RX7;refGene_tx=XM_019814511.2;refGene_gn=P2RX7;refGene_tx=XM_019814512.1;refGene_gn=P2RX7;refGene_tx=XM_019814513.2
D3	8312340	8312409
refGene_gn=P2RX7;refGene_tx=XM_003994650.5;refGene_gn=P2RX7;refGene_tx=XM_011287830.2;refGene_gn=P2RX7;refGene_tx=XM_019814510.2;refGene_gn=P2RX7;refGene_tx=XM_019814511.2;refGene_gn=P2RX7;refGene_tx=XM_019814512.1;refGene_gn=P2RX7;refGene_tx=XM_019814513.2
D3	8313577	8313746
refGene_gn=P2RX7;refGene_tx=XM_003994650.5;refGene_gn=P2RX7;refGene_tx=XM_011287830.2;refGene_gn=P2RX7;refGene_tx=XM_019814510.2;refGene_gn=P2RX7;refGene_tx=XM_019814511.2;refGene_gn=P2RX7;refGene_tx=XM_019814513.2
D3	8318604	8318675
refGene_gn=P2RX7;refGene_tx=XM_011287830.2;refGene_gn=P2RX7;refGene_tx=XM_019814510.2
D3	8330523	8330648
refGene_gn=P2RX7;refGene_tx=XM_003994650.5;refGene_gn=P2RX7;refGene_tx=XM_019814513.2
D3	8369341	8369485
refGene_gn=IFT81;refGene_tx=XM_023241399.1;refGene_gn=IFT81;refGene_tx=XM_023241400.1;refGene_gn=IFT81;refGene_tx=XM_023241401.1;refGene_gn=IFT81;refGene_tx=XM_023241402.1;refGene_gn=IFT81;refGene_tx=XM_023241403.1;refGene_gn=IFT81;refGene_tx=XM_023241404.1;refGene_gn=IFT81;refGene_tx=XM_023241405.1;refGene_gn=IFT81;refGene_tx=XM_023241406.1;refGene_gn=IFT81;refGene_tx=XM_023241407.1
D3	8370004	8370108
refGene_gn=IFT81;refGene_tx=XM_023241399.1;refGene_gn=IFT81;refGene_tx=XM_023241400.1;refGene_gn=IFT81;refGene_tx=XM_023241401.1;refGene_gn=IFT81;refGene_tx=XM_023241402.1;refGene_gn=IFT81;refGene_tx=XM_023241403.1;refGene_gn=IFT81;refGene_tx=XM_023241404.1;refGene_gn=IFT81;refGene_tx=XM_023241405.1;refGene_gn=IFT81;refGene_tx=XM_023241406.1;refGene_gn=IFT81;refGene_tx=XM_023241407.1
D3	8370512	8370693
refGene_gn=IFT81;refGene_tx=XM_023241399.1;refGene_gn=IFT81;refGene_tx=XM_023241400.1;refGene_gn=IFT81;refGene_tx=XM_023241401.1;refGene_gn=IFT81;refGene_tx=XM_023241402.1;refGene_gn=IFT81;refGene_tx=XM_023241403.1;refGene_gn=IFT81;refGene_tx=XM_023241404.1;refGene_gn=IFT81;refGene_tx=XM_023241405.1;refGene_gn=IFT81;refGene_tx=XM_023241406.1;refGene_gn=IFT81;refGene_tx=XM_023241407.1
D3	8372823	8372913
refGene_gn=IFT81;refGene_tx=XM_023241399.1;refGene_gn=IFT81;refGene_tx=XM_023241400.1;refGene_gn=IFT81;refGene_tx=XM_023241401.1;refGene_gn=

IFT81;refGene_tx=XM_023241402.1;refGene_gn=IFT81;refGene_tx=XM_0232414
03.1;refGene_gn=IFT81;refGene_tx=XM_023241404.1;refGene_gn=IFT81;refGe
ne_tx=XM_023241405.1;refGene_gn=IFT81;refGene_tx=XM_023241406.1;refGen
e_gn=IFT81;refGene_tx=XM_023241407.1
D3 8374771 8374837
refGene_gn=IFT81;refGene_tx=XM_023241399.1;refGene_gn=IFT81;refGene_tx
=XM_023241400.1;refGene_gn=IFT81;refGene_tx=XM_023241401.1;refGene_gn=
IFT81;refGene_tx=XM_023241402.1;refGene_gn=IFT81;refGene_tx=XM_0232414
03.1;refGene_gn=IFT81;refGene_tx=XM_023241404.1;refGene_gn=IFT81;refGe
ne_tx=XM_023241405.1;refGene_gn=IFT81;refGene_tx=XM_023241406.1;refGen
e_gn=IFT81;refGene_tx=XM_023241407.1
D3 8375149 8375260
refGene_gn=IFT81;refGene_tx=XM_023241399.1;refGene_gn=IFT81;refGene_tx
=XM_023241400.1;refGene_gn=IFT81;refGene_tx=XM_023241401.1;refGene_gn=
IFT81;refGene_tx=XM_023241402.1;refGene_gn=IFT81;refGene_tx=XM_0232414
03.1;refGene_gn=IFT81;refGene_tx=XM_023241404.1;refGene_gn=IFT81;refGe
ne_tx=XM_023241405.1;refGene_gn=IFT81;refGene_tx=XM_023241406.1;refGen
e_gn=IFT81;refGene_tx=XM_023241407.1
D3 8377082 8377167
refGene_gn=IFT81;refGene_tx=XM_023241399.1;refGene_gn=IFT81;refGene_tx
=XM_023241400.1;refGene_gn=IFT81;refGene_tx=XM_023241401.1;refGene_gn=
IFT81;refGene_tx=XM_023241402.1;refGene_gn=IFT81;refGene_tx=XM_0232414
03.1;refGene_gn=IFT81;refGene_tx=XM_023241404.1;refGene_gn=IFT81;refGe
ne_tx=XM_023241405.1;refGene_gn=IFT81;refGene_tx=XM_023241406.1;refGen
e_gn=IFT81;refGene_tx=XM_023241407.1
D3 8383327 8383491
refGene_gn=IFT81;refGene_tx=XM_023241399.1;refGene_gn=IFT81;refGene_tx
=XM_023241400.1;refGene_gn=IFT81;refGene_tx=XM_023241401.1;refGene_gn=
IFT81;refGene_tx=XM_023241402.1;refGene_gn=IFT81;refGene_tx=XM_0232414
03.1;refGene_gn=IFT81;refGene_tx=XM_023241404.1;refGene_gn=IFT81;refGe
ne_tx=XM_023241405.1;refGene_gn=IFT81;refGene_tx=XM_023241406.1;refGen
e_gn=IFT81;refGene_tx=XM_023241407.1
D3 8385127 8385223
refGene_gn=IFT81;refGene_tx=XM_023241399.1;refGene_gn=IFT81;refGene_tx
=XM_023241400.1;refGene_gn=IFT81;refGene_tx=XM_023241401.1;refGene_gn=
IFT81;refGene_tx=XM_023241402.1;refGene_gn=IFT81;refGene_tx=XM_0232414
03.1;refGene_gn=IFT81;refGene_tx=XM_023241404.1;refGene_gn=IFT81;refGe
ne_tx=XM_023241405.1;refGene_gn=IFT81;refGene_tx=XM_023241406.1;refGen
e_gn=IFT81;refGene_tx=XM_023241407.1
D3 8392499 8392646
refGene_gn=IFT81;refGene_tx=XM_023241399.1;refGene_gn=IFT81;refGene_tx
=XM_023241400.1;refGene_gn=IFT81;refGene_tx=XM_023241401.1;refGene_gn=
IFT81;refGene_tx=XM_023241402.1;refGene_gn=IFT81;refGene_tx=XM_0232414
03.1;refGene_gn=IFT81;refGene_tx=XM_023241404.1;refGene_gn=IFT81;refGe
ne_tx=XM_023241405.1;refGene_gn=IFT81;refGene_tx=XM_023241406.1;refGen
e_gn=IFT81;refGene_tx=XM_023241407.1
D3 8414012 8414162
refGene_gn=IFT81;refGene_tx=XM_023241399.1;refGene_gn=IFT81;refGene_tx
=XM_023241400.1;refGene_gn=IFT81;refGene_tx=XM_023241401.1;refGene_gn=
IFT81;refGene_tx=XM_023241402.1;refGene_gn=IFT81;refGene_tx=XM_0232414

03.1;refGene_gn=IFT81;refGene_tx=XM_023241404.1;refGene_gn=IFT81;refGene_tx=XM_023241405.1;refGene_gn=IFT81;refGene_tx=XM_023241406.1
D3	8423585	8423714
refGene_gn=IFT81;refGene_tx=XM_023241399.1;refGene_gn=IFT81;refGene_tx=XM_023241400.1;refGene_gn=IFT81;refGene_tx=XM_023241401.1;refGene_gn=IFT81;refGene_tx=XM_023241402.1;refGene_gn=IFT81;refGene_tx=XM_023241403.1;refGene_gn=IFT81;refGene_tx=XM_023241404.1;refGene_gn=IFT81;refGene_tx=XM_023241405.1;refGene_gn=IFT81;refGene_tx=XM_023241406.1;refGene_gn=IFT81;refGene_tx=XM_023241407.1
D3	8425459	8425549
refGene_gn=IFT81;refGene_tx=XM_023241399.1;refGene_gn=IFT81;refGene_tx=XM_023241400.1;refGene_gn=IFT81;refGene_tx=XM_023241401.1;refGene_gn=IFT81;refGene_tx=XM_023241402.1;refGene_gn=IFT81;refGene_tx=XM_023241403.1;refGene_gn=IFT81;refGene_tx=XM_023241404.1;refGene_gn=IFT81;refGene_tx=XM_023241405.1;refGene_gn=IFT81;refGene_tx=XM_023241406.1;refGene_gn=IFT81;refGene_tx=XM_023241407.1
D3	8436682	8436769
refGene_gn=IFT81;refGene_tx=XM_023241399.1;refGene_gn=IFT81;refGene_tx=XM_023241400.1;refGene_gn=IFT81;refGene_tx=XM_023241401.1;refGene_gn=IFT81;refGene_tx=XM_023241402.1;refGene_gn=IFT81;refGene_tx=XM_023241403.1;refGene_gn=IFT81;refGene_tx=XM_023241404.1;refGene_gn=IFT81;refGene_tx=XM_023241405.1;refGene_gn=IFT81;refGene_tx=XM_023241406.1;refGene_gn=IFT81;refGene_tx=XM_023241407.1
D3	8437799	8437871
refGene_gn=IFT81;refGene_tx=XM_023241399.1;refGene_gn=IFT81;refGene_tx=XM_023241400.1;refGene_gn=IFT81;refGene_tx=XM_023241401.1;refGene_gn=IFT81;refGene_tx=XM_023241402.1;refGene_gn=IFT81;refGene_tx=XM_023241403.1;refGene_gn=IFT81;refGene_tx=XM_023241404.1;refGene_gn=IFT81;refGene_tx=XM_023241405.1;refGene_gn=IFT81;refGene_tx=XM_023241406.1;refGene_gn=IFT81;refGene_tx=XM_023241407.1
D3	8437955	8438041
refGene_gn=IFT81;refGene_tx=XM_023241399.1;refGene_gn=IFT81;refGene_tx=XM_023241400.1;refGene_gn=IFT81;refGene_tx=XM_023241401.1;refGene_gn=IFT81;refGene_tx=XM_023241402.1;refGene_gn=IFT81;refGene_tx=XM_023241403.1;refGene_gn=IFT81;refGene_tx=XM_023241404.1;refGene_gn=IFT81;refGene_tx=XM_023241405.1;refGene_gn=IFT81;refGene_tx=XM_023241406.1;refGene_gn=IFT81;refGene_tx=XM_023241407.1
D3	8446074	8446120
refGene_gn=IFT81;refGene_tx=XM_023241399.1;refGene_gn=IFT81;refGene_tx=XM_023241400.1;refGene_gn=IFT81;refGene_tx=XM_023241401.1;refGene_gn=IFT81;refGene_tx=XM_023241402.1;refGene_gn=IFT81;refGene_tx=XM_023241403.1;refGene_gn=IFT81;refGene_tx=XM_023241404.1;refGene_gn=IFT81;refGene_tx=XM_023241405.1;refGene_gn=IFT81;refGene_tx=XM_023241406.1;refGene_gn=IFT81;refGene_tx=XM_023241407.1
D3	8453686	8453830
refGene_gn=IFT81;refGene_tx=XM_023241399.1;refGene_gn=IFT81;refGene_tx=XM_023241400.1;refGene_gn=IFT81;refGene_tx=XM_023241401.1;refGene_gn=IFT81;refGene_tx=XM_023241402.1;refGene_gn=IFT81;refGene_tx=XM_023241403.1;refGene_gn=IFT81;refGene_tx=XM_023241404.1;refGene_gn=IFT81;refGene_tx=XM_023241405.1;refGene_gn=IFT81;refGene_tx=XM_023241406.1;refGen

e_gn=IFT81;refGene_tx=XM_023241407.1
D3	8454232	8454283	refGene_gn=IFT81;refGene_tx=XM_023241404.1
D3	8455648	8455693	refGene_gn=IFT81;refGene_tx=XM_023241405.1
D3	8456834	8456840	refGene_gn=IFT81;refGene_tx=XM_023241406.1
D3	8457174	8457276
refGene_gn=IFT81;refGene_tx=XM_023241399.1;refGene_gn=IFT81;refGene_tx=XM_023241400.1;refGene_gn=IFT81;refGene_tx=XM_023241401.1;refGene_gn=IFT81;refGene_tx=XM_023241402.1;refGene_gn=IFT81;refGene_tx=XM_023241403.1;refGene_gn=IFT81;refGene_tx=XM_023241407.1
D3	8502637	8502755	refGene_gn=ATP2A2;refGene_tx=NM_001009216.1
D3	8503464	8503482	refGene_gn=ATP2A2;refGene_tx=NM_001009216.1
D3	8503567	8503650	refGene_gn=ATP2A2;refGene_tx=NM_001009216.1
D3	8511685	8511790	refGene_gn=ATP2A2;refGene_tx=NM_001009216.1
D3	8516601	8516740	refGene_gn=ATP2A2;refGene_tx=NM_001009216.1
D3	8537208	8537289	refGene_gn=ATP2A2;refGene_tx=NM_001009216.1
D3	8539160	8539246	refGene_gn=ATP2A2;refGene_tx=NM_001009216.1
D3	8540232	8540697	refGene_gn=ATP2A2;refGene_tx=NM_001009216.1
D3	8545297	8545386	refGene_gn=ATP2A2;refGene_tx=NM_001009216.1
D3	8545770	8545873	refGene_gn=ATP2A2;refGene_tx=NM_001009216.1
D3	8546693	8546825	refGene_gn=ATP2A2;refGene_tx=NM_001009216.1
D3	8550445	8550568	refGene_gn=ATP2A2;refGene_tx=NM_001009216.1
D3	8550668	8550887	refGene_gn=ATP2A2;refGene_tx=NM_001009216.1
D3	8551664	8552000	refGene_gn=ATP2A2;refGene_tx=NM_001009216.1
D3	8553034	8553255	refGene_gn=ATP2A2;refGene_tx=NM_001009216.1
D3	8553662	8553865	refGene_gn=ATP2A2;refGene_tx=NM_001009216.1
D3	8555769	8555855	refGene_gn=ATP2A2;refGene_tx=NM_001009216.1
D3	8556120	8556254	refGene_gn=ATP2A2;refGene_tx=NM_001009216.1
D3	8556866	8556984	refGene_gn=ATP2A2;refGene_tx=NM_001009216.1
D3	8557072	8557193	refGene_gn=ATP2A2;refGene_tx=NM_001009216.1
D3	8560906	8560920	refGene_gn=ATP2A2;refGene_tx=NM_001009216.1
D3	8577001	8577191
refGene_gn=ANAPC7;refGene_tx=XM_003994652.4;refGene_gn=ANAPC7;refGene_tx=XM_023241410.1
D3	8578832	8578983
refGene_gn=ANAPC7;refGene_tx=XM_003994652.4;refGene_gn=ANAPC7;refGene_tx=XM_023241410.1
D3	8579642	8579867
refGene_gn=ANAPC7;refGene_tx=XM_003994652.4;refGene_gn=ANAPC7;refGene_tx=XM_023241410.1
D3	8582110	8582307
refGene_gn=ANAPC7;refGene_tx=XM_003994652.4;refGene_gn=ANAPC7;refGene_tx=XM_023241410.1
D3	8583020	8583138
refGene_gn=ANAPC7;refGene_tx=XM_003994652.4;refGene_gn=ANAPC7;refGene_tx=XM_023241410.1
D3	8584833	8584976
refGene_gn=ANAPC7;refGene_tx=XM_003994652.4;refGene_gn=ANAPC7;refGene_tx=XM_023241410.1
D3	8585723	8585877
refGene_gn=ANAPC7;refGene_tx=XM_003994652.4;refGene_gn=ANAPC7;refGene_

tx=XM_023241410.1
D3 8586434 8586546
refGene_gn=ANAPC7;refGene_tx=XM_003994652.4;refGene_gn=ANAPC7;refGene_
tx=XM_023241410.1
D3 8589713 8589833
refGene_gn=ANAPC7;refGene_tx=XM_003994652.4;refGene_gn=ANAPC7;refGene_
tx=XM_023241410.1
D3 8592175 8592362
refGene_gn=ANAPC7;refGene_tx=XM_003994652.4;refGene_gn=ANAPC7;refGene_
tx=XM_023241410.1
D3 8595911 8596111 refGene_gn=ANAPC7;refGene_tx=XM_003994652.4
D3 8614027 8614090 refGene_gn=ARPC3;refGene_tx=XM_003994653.3
D3 8614567 8614662 refGene_gn=ARPC3;refGene_tx=XM_003994653.3
D3 8615013 8615140 refGene_gn=ARPC3;refGene_tx=XM_003994653.3
D3 8615993 8616062 refGene_gn=ARPC3;refGene_tx=XM_003994653.3
D3 8617903 8617980 refGene_gn=ARPC3;refGene_tx=XM_003994653.3
D3 8620874 8620974 refGene_gn=ARPC3;refGene_tx=XM_003994653.3
D3 8623146 8623152 refGene_gn=ARPC3;refGene_tx=XM_003994653.3
D3 8627303 8627366
refGene_gn=GPN3;refGene_tx=XM_003994654.5;refGene_gn=GPN3;refGene_tx=X
M_006938458.4;refGene_gn=GPN3;refGene_tx=XM_006938459.4;refGene_gn=GPN
3;refGene_tx=XM_023241411.1
D3 8627926 8628055
refGene_gn=GPN3;refGene_tx=XM_003994654.5;refGene_gn=GPN3;refGene_tx=X
M_006938458.4;refGene_gn=GPN3;refGene_tx=XM_006938459.4;refGene_gn=GPN
3;refGene_tx=XM_023241411.1
D3 8628150 8628159 refGene_gn=GPN3;refGene_tx=XM_019814514.2
D3 8629007 8629104
refGene_gn=GPN3;refGene_tx=XM_003994654.5;refGene_gn=GPN3;refGene_tx=X
M_006938458.4;refGene_gn=GPN3;refGene_tx=XM_006938459.4;refGene_gn=GPN
3;refGene_tx=XM_019814514.2;refGene_gn=GPN3;refGene_tx=XM_023241411.1
D3 8629225 8629341
refGene_gn=GPN3;refGene_tx=XM_003994654.5;refGene_gn=GPN3;refGene_tx=X
M_006938458.4;refGene_gn=GPN3;refGene_tx=XM_006938459.4;refGene_gn=GPN
3;refGene_tx=XM_019814514.2;refGene_gn=GPN3;refGene_tx=XM_023241411.1
D3 8630041 8630166
refGene_gn=GPN3;refGene_tx=XM_003994654.5;refGene_gn=GPN3;refGene_tx=X
M_006938458.4;refGene_gn=GPN3;refGene_tx=XM_006938459.4;refGene_gn=GPN
3;refGene_tx=XM_019814514.2;refGene_gn=GPN3;refGene_tx=XM_023241411.1
D3 8631410 8631578
refGene_gn=GPN3;refGene_tx=XM_003994654.5;refGene_gn=GPN3;refGene_tx=X
M_006938458.4;refGene_gn=GPN3;refGene_tx=XM_006938459.4;refGene_gn=GPN
3;refGene_tx=XM_019814514.2;refGene_gn=GPN3;refGene_tx=XM_023241411.1
D3 8633283 8633355
refGene_gn=GPN3;refGene_tx=XM_006938458.4;refGene_gn=GPN3;refGene_tx=X
M_006938459.4;refGene_gn=GPN3;refGene_tx=XM_019814514.2;refGene_gn=GPN
3;refGene_tx=XM_023241411.1
D3 8636308 8636417
refGene_gn=GPN3;refGene_tx=XM_003994654.5;refGene_gn=GPN3;refGene_tx=X
M_006938458.4;refGene_gn=GPN3;refGene_tx=XM_006938459.4;refGene_gn=GPN

3;refGene_tx=XM_019814514.2;refGene_gn=GPN3;refGene_tx=XM_023241411.1
D3	8640113	8640161
refGene_gn=GPN3;refGene_tx=XM_003994654.5;refGene_gn=GPN3;refGene_tx=XM_006938458.4;refGene_gn=GPN3;refGene_tx=XM_019814514.2
D3	8640915	8641007
refGene_gn=FAM216A;refGene_tx=XM_003994656.5;refGene_gn=FAM216A;refGene_tx=XM_006938456.4;refGene_gn=FAM216A;refGene_tx=XM_006938457.4
D3	8642578	8642619
refGene_gn=FAM216A;refGene_tx=XM_003994656.5;refGene_gn=FAM216A;refGene_tx=XM_006938456.4;refGene_gn=FAM216A;refGene_tx=XM_006938457.4
D3	8646593	8646709
refGene_gn=FAM216A;refGene_tx=XM_003994656.5;refGene_gn=FAM216A;refGene_tx=XM_006938456.4;refGene_gn=FAM216A;refGene_tx=XM_006938457.4
D3	8648262	8648392
refGene_gn=FAM216A;refGene_tx=XM_003994656.5;refGene_gn=FAM216A;refGene_tx=XM_006938456.4;refGene_gn=FAM216A;refGene_tx=XM_006938457.4
D3	8648471	8648655
refGene_gn=FAM216A;refGene_tx=XM_003994656.5;refGene_gn=FAM216A;refGene_tx=XM_006938456.4;refGene_gn=FAM216A;refGene_tx=XM_006938457.4
D3	8649214	8649297
refGene_gn=FAM216A;refGene_tx=XM_003994656.5;refGene_gn=FAM216A;refGene_tx=XM_006938456.4;refGene_gn=FAM216A;refGene_tx=XM_006938457.4
D3	8650927	8651043
refGene_gn=FAM216A;refGene_tx=XM_003994656.5;refGene_gn=FAM216A;refGene_tx=XM_006938456.4;refGene_gn=FAM216A;refGene_tx=XM_006938457.4
D3	8652972	8653090
refGene_gn=VPS29;refGene_tx=XM_003994657.4;refGene_gn=VPS29;refGene_tx=XM_003994658.4
D3	8653325	8653561
refGene_gn=VPS29;refGene_tx=XM_003994657.4;refGene_gn=VPS29;refGene_tx=XM_003994658.4
D3	8656430	8656622
refGene_gn=VPS29;refGene_tx=XM_003994657.4;refGene_gn=VPS29;refGene_tx=XM_003994658.4
D3	8658868	8658880	refGene_gn=VPS29;refGene_tx=XM_003994658.4
D3	8661170	8661173
refGene_gn=VPS29;refGene_tx=XM_003994657.4;refGene_gn=VPS29;refGene_tx=XM_003994658.4
D3	8661463	8661500	refGene_gn=RAD9B;refGene_tx=XM_003994659.4
D3	8663349	8663420	refGene_gn=RAD9B;refGene_tx=XM_003994659.4
D3	8664216	8664369
refGene_gn=RAD9B;refGene_tx=XM_003994659.4;refGene_gn=RAD9B;refGene_tx=XM_006938460.3;refGene_gn=RAD9B;refGene_tx=XM_019814516.2;refGene_gn=RAD9B;refGene_tx=XM_023241412.1
D3	8664575	8664690
refGene_gn=RAD9B;refGene_tx=XM_003994659.4;refGene_gn=RAD9B;refGene_tx=XM_006938460.3;refGene_gn=RAD9B;refGene_tx=XM_019814516.2;refGene_gn=RAD9B;refGene_tx=XM_023241412.1
D3	8672410	8672510
refGene_gn=RAD9B;refGene_tx=XM_003994659.4;refGene_gn=RAD9B;refGene_tx

=XM_006938460.3;refGene_gn=RAD9B;refGene_tx=XM_019814516.2;refGene_gn=RAD9B;refGene_tx=XM_023241412.1
D3	8674189	8674296	refGene_gn=RAD9B;refGene_tx=XM_003994659.4;refGene_gn=RAD9B;refGene_tx=XM_006938460.3;refGene_gn=RAD9B;refGene_tx=XM_019814516.2;refGene_gn=RAD9B;refGene_tx=XM_023241412.1
D3	8680611	8680718	refGene_gn=RAD9B;refGene_tx=XM_003994659.4;refGene_gn=RAD9B;refGene_tx=XM_006938460.3;refGene_gn=RAD9B;refGene_tx=XM_019814516.2;refGene_gn=RAD9B;refGene_tx=XM_023241412.1
D3	8680800	8680865	refGene_gn=RAD9B;refGene_tx=XM_003994659.4;refGene_gn=RAD9B;refGene_tx=XM_006938460.3;refGene_gn=RAD9B;refGene_tx=XM_019814516.2;refGene_gn=RAD9B;refGene_tx=XM_023241412.1
D3	8681415	8681538	refGene_gn=RAD9B;refGene_tx=XM_003994659.4;refGene_gn=RAD9B;refGene_tx=XM_006938460.3;refGene_gn=RAD9B;refGene_tx=XM_019814516.2;refGene_gn=RAD9B;refGene_tx=XM_023241412.1
D3	8682903	8683144	refGene_gn=RAD9B;refGene_tx=XM_003994659.4;refGene_gn=RAD9B;refGene_tx=XM_006938460.3;refGene_gn=RAD9B;refGene_tx=XM_019814516.2;refGene_gn=RAD9B;refGene_tx=XM_023241412.1
D3	8690738	8690867	refGene_gn=RAD9B;refGene_tx=XM_003994659.4;refGene_gn=RAD9B;refGene_tx=XM_006938460.3;refGene_gn=RAD9B;refGene_tx=XM_019814516.2;refGene_gn=RAD9B;refGene_tx=XM_023241412.1
D3	8696649	8696708	refGene_gn=PPTC7;refGene_tx=XM_011287829.3
D3	8697822	8697952	refGene_gn=PPTC7;refGene_tx=XM_011287829.3
D3	8699618	8699742	refGene_gn=PPTC7;refGene_tx=XM_011287829.3
D3	8703432	8703631	refGene_gn=PPTC7;refGene_tx=XM_011287829.3
D3	8708964	8709144	refGene_gn=PPTC7;refGene_tx=XM_011287829.3
D3	8729616	8729839	refGene_gn=PPTC7;refGene_tx=XM_011287829.3
D3	8752062	8752282	refGene_gn=TCTN1;refGene_tx=XM_006938463.3;refGene_gn=TCTN1;refGene_tx=XM_019814520.2;refGene_gn=TCTN1;refGene_tx=XM_023241413.1
D3	8755803	8755924	refGene_gn=TCTN1;refGene_tx=XM_006938463.3;refGene_gn=TCTN1;refGene_tx=XM_019814520.2;refGene_gn=TCTN1;refGene_tx=XM_023241413.1
D3	8762671	8762802	refGene_gn=TCTN1;refGene_tx=XM_006938463.3;refGene_gn=TCTN1;refGene_tx=XM_019814520.2;refGene_gn=TCTN1;refGene_tx=XM_023241413.1
D3	8764158	8764310	refGene_gn=TCTN1;refGene_tx=XM_006938463.3;refGene_gn=TCTN1;refGene_tx=XM_019814520.2;refGene_gn=TCTN1;refGene_tx=XM_019814521.1;refGene_gn=TCTN1;refGene_tx=XM_019814522.1;refGene_gn=TCTN1;refGene_tx=XM_023241413.1
D3	8766013	8766101	refGene_gn=TCTN1;refGene_tx=XM_006938463.3;refGene_gn=TCTN1;refGene_tx=XM_019814520.2;refGene_gn=TCTN1;refGene_tx=XM_019814521.1;refGene_gn=TCTN1;refGene_tx=XM_019814522.1;refGene_gn=TCTN1;refGene_tx=XM_0232414

13.1
D3 8767628 8767738
refGene_gn=TCTN1;refGene_tx=XM_006938463.3;refGene_gn=TCTN1;refGene_tx
=XM_019814520.2;refGene_gn=TCTN1;refGene_tx=XM_019814521.1;refGene_gn=
TCTN1;refGene_tx=XM_019814522.1;refGene_gn=TCTN1;refGene_tx=XM_0232414
13.1
D3 8769515 8769542
refGene_gn=TCTN1;refGene_tx=XM_006938463.3;refGene_gn=TCTN1;refGene_tx
=XM_019814520.2;refGene_gn=TCTN1;refGene_tx=XM_019814521.1;refGene_gn=
TCTN1;refGene_tx=XM_019814522.1;refGene_gn=TCTN1;refGene_tx=XM_0232414
13.1
D3 8773847 8773982
refGene_gn=TCTN1;refGene_tx=XM_006938463.3;refGene_gn=TCTN1;refGene_tx
=XM_019814520.2;refGene_gn=TCTN1;refGene_tx=XM_019814521.1;refGene_gn=
TCTN1;refGene_tx=XM_019814522.1
D3 8774519 8774645
refGene_gn=TCTN1;refGene_tx=XM_006938463.3;refGene_gn=TCTN1;refGene_tx
=XM_019814520.2;refGene_gn=TCTN1;refGene_tx=XM_019814521.1;refGene_gn=
TCTN1;refGene_tx=XM_019814522.1;refGene_gn=TCTN1;refGene_tx=XM_0232414
13.1
D3 8775034 8775120
refGene_gn=TCTN1;refGene_tx=XM_006938463.3;refGene_gn=TCTN1;refGene_tx
=XM_019814520.2;refGene_gn=TCTN1;refGene_tx=XM_019814521.1;refGene_gn=
TCTN1;refGene_tx=XM_019814522.1;refGene_gn=TCTN1;refGene_tx=XM_0232414
13.1
D3 8775660 8775801
refGene_gn=TCTN1;refGene_tx=XM_006938463.3;refGene_gn=TCTN1;refGene_tx
=XM_019814520.2;refGene_gn=TCTN1;refGene_tx=XM_019814521.1;refGene_gn=
TCTN1;refGene_tx=XM_019814522.1;refGene_gn=TCTN1;refGene_tx=XM_0232414
13.1
D3 8776951 8777114
refGene_gn=TCTN1;refGene_tx=XM_006938463.3;refGene_gn=TCTN1;refGene_tx
=XM_019814520.2;refGene_gn=TCTN1;refGene_tx=XM_019814521.1;refGene_gn=
TCTN1;refGene_tx=XM_019814522.1;refGene_gn=TCTN1;refGene_tx=XM_0232414
13.1
D3 8779354 8779480
refGene_gn=TCTN1;refGene_tx=XM_006938463.3;refGene_gn=TCTN1;refGene_tx
=XM_019814520.2;refGene_gn=TCTN1;refGene_tx=XM_019814521.1;refGene_gn=
TCTN1;refGene_tx=XM_019814522.1;refGene_gn=TCTN1;refGene_tx=XM_0232414
13.1
D3 8780062 8780206
refGene_gn=TCTN1;refGene_tx=XM_006938463.3;refGene_gn=TCTN1;refGene_tx
=XM_019814521.1;refGene_gn=TCTN1;refGene_tx=XM_019814522.1;refGene_gn=
TCTN1;refGene_tx=XM_023241413.1
D3 8781067 8781139 refGene_gn=TCTN1;refGene_tx=XM_019814520.2
D3 8781473 8781539
refGene_gn=HVCN1;refGene_tx=XM_006938464.3;refGene_gn=HVCN1;refGene_tx
=XM_006938465.3;refGene_gn=HVCN1;refGene_tx=XM_006938466.3;refGene_gn=
HVCN1;refGene_tx=XM_006938467.2;refGene_gn=HVCN1;refGene_tx=XM_0069384
68.4;refGene_gn=HVCN1;refGene_tx=XM_011287827.2;refGene_gn=HVCN1;refGe

ne_tx=XM_019814523.1;refGene_gn=HVCN1;refGene_tx=XM_019814524.2;refGene_gn=HVCN1;refGene_tx=XM_019814525.1;refGene_gn=HVCN1;refGene_tx=XM_019814526.1
D3 8781891 8782004
refGene_gn=HVCN1;refGene_tx=XM_006938464.3;refGene_gn=HVCN1;refGene_tx=XM_006938465.3;refGene_gn=HVCN1;refGene_tx=XM_006938466.3;refGene_gn=HVCN1;refGene_tx=XM_006938467.2;refGene_gn=HVCN1;refGene_tx=XM_006938468.4;refGene_gn=HVCN1;refGene_tx=XM_011287827.2;refGene_gn=HVCN1;refGene_tx=XM_019814523.1;refGene_gn=HVCN1;refGene_tx=XM_019814524.2;refGene_gn=HVCN1;refGene_tx=XM_019814525.1;refGene_gn=HVCN1;refGene_tx=XM_019814526.1
D3 8784252 8784484
refGene_gn=HVCN1;refGene_tx=XM_006938464.3;refGene_gn=HVCN1;refGene_tx=XM_006938465.3;refGene_gn=HVCN1;refGene_tx=XM_006938466.3;refGene_gn=HVCN1;refGene_tx=XM_006938467.2;refGene_gn=HVCN1;refGene_tx=XM_006938468.4;refGene_gn=HVCN1;refGene_tx=XM_011287827.2;refGene_gn=HVCN1;refGene_tx=XM_019814523.1;refGene_gn=HVCN1;refGene_tx=XM_019814524.2;refGene_gn=HVCN1;refGene_tx=XM_019814525.1;refGene_gn=HVCN1;refGene_tx=XM_019814526.1
D3 8786866 8786965
refGene_gn=HVCN1;refGene_tx=XM_006938464.3;refGene_gn=HVCN1;refGene_tx=XM_006938465.3;refGene_gn=HVCN1;refGene_tx=XM_006938466.3;refGene_gn=HVCN1;refGene_tx=XM_006938467.2;refGene_gn=HVCN1;refGene_tx=XM_006938468.4;refGene_gn=HVCN1;refGene_tx=XM_011287827.2;refGene_gn=HVCN1;refGene_tx=XM_019814523.1;refGene_gn=HVCN1;refGene_tx=XM_019814524.2;refGene_gn=HVCN1;refGene_tx=XM_019814525.1;refGene_gn=HVCN1;refGene_tx=XM_019814526.1
D3 8791465 8791747
refGene_gn=HVCN1;refGene_tx=XM_006938464.3;refGene_gn=HVCN1;refGene_tx=XM_006938465.3;refGene_gn=HVCN1;refGene_tx=XM_006938466.3;refGene_gn=HVCN1;refGene_tx=XM_006938467.2;refGene_gn=HVCN1;refGene_tx=XM_006938468.4;refGene_gn=HVCN1;refGene_tx=XM_011287827.2;refGene_gn=HVCN1;refGene_tx=XM_019814523.1;refGene_gn=HVCN1;refGene_tx=XM_019814524.2;refGene_gn=HVCN1;refGene_tx=XM_019814525.1;refGene_gn=HVCN1;refGene_tx=XM_019814526.1
D3 8810371 8810392
refGene_gn=HVCN1;refGene_tx=XM_006938464.3;refGene_gn=HVCN1;refGene_tx=XM_006938465.3;refGene_gn=HVCN1;refGene_tx=XM_011287827.2;refGene_gn=HVCN1;refGene_tx=XM_019814524.2;refGene_gn=HVCN1;refGene_tx=XM_019814525.1
D3 8842231 8842302 refGene_gn=PPP1CC;refGene_tx=XM_003994662.5
D3 8843192 8843253 refGene_gn=PPP1CC;refGene_tx=XM_003994662.5
D3 8844211 8844346 refGene_gn=PPP1CC;refGene_tx=XM_003994662.5
D3 8844554 8844778 refGene_gn=PPP1CC;refGene_tx=XM_003994662.5
D3 8846183 8846288 refGene_gn=PPP1CC;refGene_tx=XM_003994662.5
D3 8850675 8850906 refGene_gn=PPP1CC;refGene_tx=XM_003994662.5
D3 8851872 8852004 refGene_gn=PPP1CC;refGene_tx=XM_003994662.5
D3 8862647 8862702 refGene_gn=PPP1CC;refGene_tx=XM_003994662.5
D3 8940631 8940640
refGene_gn=CCDC63;refGene_tx=XM_006938471.4;refGene_gn=CCDC63;refGene_

tx=XM_019814539.2;refGene_gn=CCDC63;refGene_tx=XM_023241414.1
D3 8941034 8941204
refGene_gn=CCDC63;refGene_tx=XM_006938471.4;refGene_gn=CCDC63;refGene_tx=XM_019814539.2;refGene_gn=CCDC63;refGene_tx=XM_023241414.1
D3 8944150 8944343
refGene_gn=CCDC63;refGene_tx=XM_006938471.4;refGene_gn=CCDC63;refGene_tx=XM_019814539.2;refGene_gn=CCDC63;refGene_tx=XM_023241414.1
D3 8949922 8950042
refGene_gn=CCDC63;refGene_tx=XM_006938471.4;refGene_gn=CCDC63;refGene_tx=XM_019814539.2;refGene_gn=CCDC63;refGene_tx=XM_023241414.1
D3 8953965 8954147
refGene_gn=CCDC63;refGene_tx=XM_006938471.4;refGene_gn=CCDC63;refGene_tx=XM_019814539.2;refGene_gn=CCDC63;refGene_tx=XM_023241414.1
D3 8954937 8955119
refGene_gn=CCDC63;refGene_tx=XM_006938471.4;refGene_gn=CCDC63;refGene_tx=XM_019814539.2;refGene_gn=CCDC63;refGene_tx=XM_023241414.1
D3 8957048 8957269
refGene_gn=CCDC63;refGene_tx=XM_006938471.4;refGene_gn=CCDC63;refGene_tx=XM_019814539.2;refGene_gn=CCDC63;refGene_tx=XM_023241414.1
D3 8963521 8963596
refGene_gn=CCDC63;refGene_tx=XM_006938471.4;refGene_gn=CCDC63;refGene_tx=XM_019814539.2;refGene_gn=CCDC63;refGene_tx=XM_023241414.1
D3 8967114 8967307
refGene_gn=CCDC63;refGene_tx=XM_006938471.4;refGene_gn=CCDC63;refGene_tx=XM_019814539.2
D3 8969880 8970084
refGene_gn=CCDC63;refGene_tx=XM_006938471.4;refGene_gn=CCDC63;refGene_tx=XM_019814539.2;refGene_gn=CCDC63;refGene_tx=XM_023241414.1
D3 8972328 8972448
refGene_gn=CCDC63;refGene_tx=XM_006938471.4;refGene_gn=CCDC63;refGene_tx=XM_019814539.2;refGene_gn=CCDC63;refGene_tx=XM_023241414.1
D3 8973333 8973432 refGene_gn=MYL2;refGene_tx=XM_003994663.5
D3 8975082 8975131 refGene_gn=MYL2;refGene_tx=XM_003994663.5
D3 8975221 8975300 refGene_gn=MYL2;refGene_tx=XM_003994663.5
D3 8975895 8976000 refGene_gn=MYL2;refGene_tx=XM_003994663.5
D3 8977134 8977210 refGene_gn=MYL2;refGene_tx=XM_003994663.5
D3 8979609 8979699 refGene_gn=MYL2;refGene_tx=XM_003994663.5
D3 8980953 8980956 refGene_gn=MYL2;refGene_tx=XM_003994663.5
D3 9076201 9076264 refGene_gn=CUX2;refGene_tx=XM_019814530.2
D3 9234295 9234406 refGene_gn=CUX2;refGene_tx=XM_019814530.2
D3 9238105 9238153 refGene_gn=CUX2;refGene_tx=XM_019814530.2
D3 9277961 9278040 refGene_gn=CUX2;refGene_tx=XM_019814530.2
D3 9298211 9298346 refGene_gn=CUX2;refGene_tx=XM_019814530.2
D3 9301363 9301487 refGene_gn=CUX2;refGene_tx=XM_019814530.2
D3 9303288 9303365 refGene_gn=CUX2;refGene_tx=XM_019814530.2
D3 9304233 9304300 refGene_gn=CUX2;refGene_tx=XM_019814530.2
D3 9306809 9306858 refGene_gn=CUX2;refGene_tx=XM_019814530.2
D3 9310369 9310474 refGene_gn=CUX2;refGene_tx=XM_019814530.2
D3 9313628 9313820 refGene_gn=CUX2;refGene_tx=XM_019814530.2
D3 9313921 9313980 refGene_gn=CUX2;refGene_tx=XM_019814530.2

| | | | |
|---|---|---|---|
| D3 | 9314470 | 9314519 | refGene_gn=CUX2;refGene_tx=XM_019814530.2 |
| D3 | 9314612 | 9314712 | refGene_gn=CUX2;refGene_tx=XM_019814530.2 |
| D3 | 9315895 | 9316570 | refGene_gn=CUX2;refGene_tx=XM_019814530.2 |
| D3 | 9317433 | 9317535 | refGene_gn=CUX2;refGene_tx=XM_019814530.2 |
| D3 | 9321982 | 9322776 | refGene_gn=CUX2;refGene_tx=XM_019814530.2 |
| D3 | 9323516 | 9323624 | refGene_gn=CUX2;refGene_tx=XM_019814530.2 |
| D3 | 9323625 | 9323677 | refGene_gn=CUX2;refGene_tx=XM_019814530.2 |
| D3 | 9330343 | 9330613 | refGene_gn=CUX2;refGene_tx=XM_019814530.2 |
| D3 | 9331810 | 9331999 | refGene_gn=CUX2;refGene_tx=XM_019814530.2 |
| D3 | 9333217 | 9333491 | refGene_gn=CUX2;refGene_tx=XM_019814530.2 |
| D3 | 9337184 | 9337986 | refGene_gn=CUX2;refGene_tx=XM_019814530.2 |
| D3 | 9349928 | 9350576 | refGene_gn=FAM109A;refGene_tx=XM_023241415.1;refGene_gn=FAM109A;refGene_tx=XM_023241416.1;refGene_gn=FAM109A;refGene_tx=XM_023241417.1;refGene_gn=FAM109A;refGene_tx=XM_023241418.1;refGene_gn=FAM109A;refGene_tx=XM_023241419.1 |
| D3 | 9384633 | 9385398 | refGene_gn=SH2B3;refGene_tx=XM_019814546.2 |
| D3 | 9408352 | 9408454 | refGene_gn=SH2B3;refGene_tx=XM_019814546.2 |
| D3 | 9408552 | 9408644 | refGene_gn=SH2B3;refGene_tx=XM_019814546.2 |
| D3 | 9408734 | 9408829 | refGene_gn=SH2B3;refGene_tx=XM_019814546.2 |
| D3 | 9408932 | 9409147 | refGene_gn=SH2B3;refGene_tx=XM_019814546.2 |
| D3 | 9409240 | 9409412 | refGene_gn=SH2B3;refGene_tx=XM_019814546.2 |
| D3 | 9409580 | 9409876 | refGene_gn=SH2B3;refGene_tx=XM_019814546.2 |
| D3 | 9414233 | 9414262 | refGene_gn=ATXN2;refGene_tx=XM_019814545.2 |
| D3 | 9415110 | 9415279 | refGene_gn=ATXN2;refGene_tx=XM_019814545.2 |
| D3 | 9417639 | 9417867 | refGene_gn=ATXN2;refGene_tx=XM_019814545.2 |
| D3 | 9418544 | 9418690 | refGene_gn=ATXN2;refGene_tx=XM_019814545.2 |
| D3 | 9424044 | 9424098 | refGene_gn=ATXN2;refGene_tx=XM_019814545.2 |
| D3 | 9427686 | 9427819 | refGene_gn=ATXN2;refGene_tx=XM_019814545.2 |
| D3 | 9428130 | 9428315 | refGene_gn=ATXN2;refGene_tx=XM_019814545.2 |
| D3 | 9441800 | 9441861 | refGene_gn=ATXN2;refGene_tx=XM_019814545.2 |
| D3 | 9442246 | 9442399 | refGene_gn=ATXN2;refGene_tx=XM_019814545.2 |
| D3 | 9443442 | 9443506 | refGene_gn=ATXN2;refGene_tx=XM_019814545.2 |
| D3 | 9444669 | 9444974 | refGene_gn=ATXN2;refGene_tx=XM_019814545.2 |
| D3 | 9452996 | 9453067 | refGene_gn=ATXN2;refGene_tx=XM_019814545.2 |
| D3 | 9453327 | 9453435 | refGene_gn=ATXN2;refGene_tx=XM_019814545.2 |
| D3 | 9453811 | 9454009 | refGene_gn=ATXN2;refGene_tx=XM_019814545.2 |
| D3 | 9456521 | 9456704 | refGene_gn=ATXN2;refGene_tx=XM_019814545.2 |
| D3 | 9459359 | 9459569 | refGene_gn=ATXN2;refGene_tx=XM_019814545.2 |
| D3 | 9461441 | 9461620 | refGene_gn=ATXN2;refGene_tx=XM_019814545.2 |
| D3 | 9463086 | 9463284 | refGene_gn=ATXN2;refGene_tx=XM_019814545.2 |
| D3 | 9463773 | 9463865 | refGene_gn=ATXN2;refGene_tx=XM_019814545.2 |
| D3 | 9465845 | 9465970 | refGene_gn=ATXN2;refGene_tx=XM_019814545.2 |
| D3 | 9497951 | 9498102 | refGene_gn=ATXN2;refGene_tx=XM_019814545.2 |
| D3 | 9498654 | 9498726 | refGene_gn=ATXN2;refGene_tx=XM_019814545.2 |
| D3 | 9499219 | 9499279 | refGene_gn=ATXN2;refGene_tx=XM_019814545.2 |
| D3 | 9500734 | 9500768 | refGene_gn=ATXN2;refGene_tx=XM_019814545.2 |
| D3 | 9533921 | 9534426 | refGene_gn=ATXN2;refGene_tx=XM_019814545.2 |
| D3 | 9534428 | 9534609 | refGene_gn=ATXN2;refGene_tx=XM_019814545.2 |
| D3 | 9569931 | 9570295 | refGene_gn=BRAP;refGene_tx=XM_003994672.3 |

D3	9572481	9572585	refGene_gn=BRAP;refGene_tx=XM_003994672.3
D3	9575875	9575965	refGene_gn=BRAP;refGene_tx=XM_003994672.3
D3	9578359	9578469	refGene_gn=BRAP;refGene_tx=XM_003994672.3
D3	9578842	9578981	refGene_gn=BRAP;refGene_tx=XM_003994672.3
D3	9579434	9579510	refGene_gn=BRAP;refGene_tx=XM_003994672.3
D3	9585474	9585623	refGene_gn=BRAP;refGene_tx=XM_003994672.3
D3	9587104	9587218	refGene_gn=BRAP;refGene_tx=XM_003994672.3
D3	9592235	9592425	refGene_gn=BRAP;refGene_tx=XM_003994672.3
D3	9593703	9593902	refGene_gn=BRAP;refGene_tx=XM_003994672.3
D3	9594596	9594758	refGene_gn=BRAP;refGene_tx=XM_003994672.3
D3	9596511	9596593	refGene_gn=BRAP;refGene_tx=XM_003994672.3
D3	9601779	9601966
refGene_gn=ACAD10;refGene_tx=XM_023241420.1;refGene_gn=ACAD10;refGene_tx=XM_023241421.1
D3	9606312	9606461
refGene_gn=ACAD10;refGene_tx=XM_023241420.1;refGene_gn=ACAD10;refGene_tx=XM_023241421.1
D3	9608557	9608752
refGene_gn=ACAD10;refGene_tx=XM_023241420.1;refGene_gn=ACAD10;refGene_tx=XM_023241421.1
D3	9612424	9612583
refGene_gn=ACAD10;refGene_tx=XM_023241420.1;refGene_gn=ACAD10;refGene_tx=XM_023241421.1
D3	9613303	9613463
refGene_gn=ACAD10;refGene_tx=XM_023241420.1;refGene_gn=ACAD10;refGene_tx=XM_023241421.1
D3	9614668	9614810
refGene_gn=ACAD10;refGene_tx=XM_023241420.1;refGene_gn=ACAD10;refGene_tx=XM_023241421.1
D3	9619988	9620057	refGene_gn=ACAD10;refGene_tx=XM_023241420.1
D3	9621833	9622015
refGene_gn=ACAD10;refGene_tx=XM_023241420.1;refGene_gn=ACAD10;refGene_tx=XM_023241421.1
D3	9628174	9628325
refGene_gn=ACAD10;refGene_tx=XM_023241420.1;refGene_gn=ACAD10;refGene_tx=XM_023241421.1
D3	9629400	9629552
refGene_gn=ACAD10;refGene_tx=XM_023241420.1;refGene_gn=ACAD10;refGene_tx=XM_023241421.1
D3	9631140	9631314
refGene_gn=ACAD10;refGene_tx=XM_023241420.1;refGene_gn=ACAD10;refGene_tx=XM_023241421.1
D3	9637809	9637994
refGene_gn=ACAD10;refGene_tx=XM_023241420.1;refGene_gn=ACAD10;refGene_tx=XM_023241421.1
D3	9662864	9662909	refGene_gn=ALDH2;refGene_tx=XM_003994673.5
D3	9662910	9662979	refGene_gn=ALDH2;refGene_tx=XM_003994673.5
D3	9674488	9674593	refGene_gn=ALDH2;refGene_tx=XM_003994673.5
D3	9675649	9675790	refGene_gn=ALDH2;refGene_tx=XM_003994673.5
D3	9676881	9676961	refGene_gn=ALDH2;refGene_tx=XM_003994673.5

```
D3	9679083	9679195	refGene_gn=ALDH2;refGene_tx=XM_003994673.5
D3	9679661	9679790	refGene_gn=ALDH2;refGene_tx=XM_003994673.5
D3	9680224	9680338	refGene_gn=ALDH2;refGene_tx=XM_003994673.5
D3	9682004	9682107	refGene_gn=ALDH2;refGene_tx=XM_003994673.5
D3	9682674	9682859	refGene_gn=ALDH2;refGene_tx=XM_003994673.5
D3	9685095	9685260	refGene_gn=ALDH2;refGene_tx=XM_003994673.5
D3	9686927	9687085	refGene_gn=ALDH2;refGene_tx=XM_003994673.5
D3	9688420	9688535	refGene_gn=ALDH2;refGene_tx=XM_003994673.5
D3	9690600	9690633	refGene_gn=ALDH2;refGene_tx=XM_003994673.5
D3	9717188	9717224
refGene_gn=MAPKAPK5;refGene_tx=XM_003994675.5;refGene_gn=MAPKAPK5;refG
ene_tx=XM_006938476.4;refGene_gn=MAPKAPK5;refGene_tx=XM_023241423.1
D3	9728811	9728885
refGene_gn=MAPKAPK5;refGene_tx=XM_003994675.5;refGene_gn=MAPKAPK5;refG
ene_tx=XM_006938476.4;refGene_gn=MAPKAPK5;refGene_tx=XM_023241423.1
D3	9729590	9729666
refGene_gn=MAPKAPK5;refGene_tx=XM_003994675.5;refGene_gn=MAPKAPK5;refG
ene_tx=XM_006938476.4;refGene_gn=MAPKAPK5;refGene_tx=XM_023241423.1
D3	9730229	9730327
refGene_gn=MAPKAPK5;refGene_tx=XM_003994675.5;refGene_gn=MAPKAPK5;refG
ene_tx=XM_006938476.4;refGene_gn=MAPKAPK5;refGene_tx=XM_023241422.1
D3	9731904	9732013
refGene_gn=MAPKAPK5;refGene_tx=XM_003994675.5;refGene_gn=MAPKAPK5;refG
ene_tx=XM_006938476.4;refGene_gn=MAPKAPK5;refGene_tx=XM_023241422.1
D3	9733646	9733736
refGene_gn=MAPKAPK5;refGene_tx=XM_003994675.5;refGene_gn=MAPKAPK5;refG
ene_tx=XM_006938476.4;refGene_gn=MAPKAPK5;refGene_tx=XM_023241422.1;re
fGene_gn=MAPKAPK5;refGene_tx=XM_023241423.1
D3	9734526	9734622
refGene_gn=MAPKAPK5;refGene_tx=XM_003994675.5;refGene_gn=MAPKAPK5;refG
ene_tx=XM_006938476.4;refGene_gn=MAPKAPK5;refGene_tx=XM_023241422.1;re
fGene_gn=MAPKAPK5;refGene_tx=XM_023241423.1
D3	9748393	9748474
refGene_gn=MAPKAPK5;refGene_tx=XM_003994675.5;refGene_gn=MAPKAPK5;refG
ene_tx=XM_006938476.4;refGene_gn=MAPKAPK5;refGene_tx=XM_023241422.1;re
fGene_gn=MAPKAPK5;refGene_tx=XM_023241423.1
D3	9751071	9751259
refGene_gn=MAPKAPK5;refGene_tx=XM_003994675.5;refGene_gn=MAPKAPK5;refG
ene_tx=XM_006938476.4;refGene_gn=MAPKAPK5;refGene_tx=XM_023241422.1;re
fGene_gn=MAPKAPK5;refGene_tx=XM_023241423.1
D3	9753464	9753585
refGene_gn=MAPKAPK5;refGene_tx=XM_003994675.5;refGene_gn=MAPKAPK5;refG
ene_tx=XM_006938476.4;refGene_gn=MAPKAPK5;refGene_tx=XM_023241422.1;re
fGene_gn=MAPKAPK5;refGene_tx=XM_023241423.1
D3	9756207	9756338
refGene_gn=MAPKAPK5;refGene_tx=XM_003994675.5;refGene_gn=MAPKAPK5;refG
ene_tx=XM_006938476.4;refGene_gn=MAPKAPK5;refGene_tx=XM_023241422.1;re
fGene_gn=MAPKAPK5;refGene_tx=XM_023241423.1
D3	9756599	9756721
refGene_gn=MAPKAPK5;refGene_tx=XM_003994675.5;refGene_gn=MAPKAPK5;refG
```

ene_tx=XM_006938476.4;refGene_gn=MAPKAPK5;refGene_tx=XM_023241422.1;refGene_gn=MAPKAPK5;refGene_tx=XM_023241423.1
D3 9757767 9757872 refGene_gn=MAPKAPK5;refGene_tx=XM_003994675.5;refGene_gn=MAPKAPK5;refGene_tx=XM_006938476.4;refGene_gn=MAPKAPK5;refGene_tx=XM_023241422.1;refGene_gn=MAPKAPK5;refGene_tx=XM_023241423.1
D3 9758204 9758299 refGene_gn=MAPKAPK5;refGene_tx=XM_003994675.5;refGene_gn=MAPKAPK5;refGene_tx=XM_006938476.4;refGene_gn=MAPKAPK5;refGene_tx=XM_023241422.1;refGene_gn=MAPKAPK5;refGene_tx=XM_023241423.1
D3 9773876 9774025 refGene_gn=TMEM116;refGene_tx=XM_023241424.1;refGene_gn=TMEM116;refGene_tx=XM_023241425.1;refGene_gn=TMEM116;refGene_tx=XM_023241426.1;refGene_gn=TMEM116;refGene_tx=XM_023241427.1
D3 9779100 9779224 refGene_gn=TMEM116;refGene_tx=XM_023241424.1;refGene_gn=TMEM116;refGene_tx=XM_023241425.1;refGene_gn=TMEM116;refGene_tx=XM_023241426.1;refGene_gn=TMEM116;refGene_tx=XM_023241427.1
D3 9785793 9785871 refGene_gn=TMEM116;refGene_tx=XM_023241428.1
D3 9810879 9811131 refGene_gn=TMEM116;refGene_tx=XM_019814547.2
D3 9811890 9811964 refGene_gn=TMEM116;refGene_tx=XM_019814547.2;refGene_gn=TMEM116;refGene_tx=XM_019814550.2;refGene_gn=TMEM116;refGene_tx=XM_023241424.1;refGene_gn=TMEM116;refGene_tx=XM_023241425.1;refGene_gn=TMEM116;refGene_tx=XM_023241426.1;refGene_gn=TMEM116;refGene_tx=XM_023241427.1;refGene_gn=TMEM116;refGene_tx=XM_023241428.1
D3 9812556 9812701 refGene_gn=TMEM116;refGene_tx=XM_019814547.2;refGene_gn=TMEM116;refGene_tx=XM_019814550.2;refGene_gn=TMEM116;refGene_tx=XM_023241424.1;refGene_gn=TMEM116;refGene_tx=XM_023241425.1;refGene_gn=TMEM116;refGene_tx=XM_023241426.1;refGene_gn=TMEM116;refGene_tx=XM_023241427.1;refGene_gn=TMEM116;refGene_tx=XM_023241428.1
D3 9814439 9814575 refGene_gn=TMEM116;refGene_tx=XM_019814547.2;refGene_gn=TMEM116;refGene_tx=XM_019814550.2;refGene_gn=TMEM116;refGene_tx=XM_023241424.1;refGene_gn=TMEM116;refGene_tx=XM_023241425.1;refGene_gn=TMEM116;refGene_tx=XM_023241426.1;refGene_gn=TMEM116;refGene_tx=XM_023241427.1;refGene_gn=TMEM116;refGene_tx=XM_023241428.1
D3 9814933 9815017 refGene_gn=TMEM116;refGene_tx=XM_019814547.2;refGene_gn=TMEM116;refGene_tx=XM_019814550.2;refGene_gn=TMEM116;refGene_tx=XM_023241424.1;refGene_gn=TMEM116;refGene_tx=XM_023241425.1;refGene_gn=TMEM116;refGene_tx=XM_023241426.1;refGene_gn=TMEM116;refGene_tx=XM_023241427.1;refGene_gn=TMEM116;refGene_tx=XM_023241428.1
D3 9815999 9816049 refGene_gn=TMEM116;refGene_tx=XM_019814547.2;refGene_gn=TMEM116;refGene_tx=XM_019814550.2;refGene_gn=TMEM116;refGene_tx=XM_023241424.1;refGene_gn=TMEM116;refGene_tx=XM_023241425.1;refGene_gn=TMEM116;refGene_tx=XM_023241426.1;refGene_gn=TMEM116;refGene_tx=XM_023241427.1;refGene_gn=TMEM116;refGene_tx=XM_023241428.1

D3 9820455 9820560
refGene_gn=TMEM116;refGene_tx=XM_019814547.2;refGene_gn=TMEM116;refGene_tx=XM_019814550.2;refGene_gn=TMEM116;refGene_tx=XM_023241424.1;refGene_gn=TMEM116;refGene_tx=XM_023241425.1;refGene_gn=TMEM116;refGene_tx=XM_023241426.1;refGene_gn=TMEM116;refGene_tx=XM_023241427.1;refGene_gn=TMEM116;refGene_tx=XM_023241428.1
D3 9865629 9865761
refGene_gn=TMEM116;refGene_tx=XM_019814547.2;refGene_gn=TMEM116;refGene_tx=XM_019814550.2;refGene_gn=TMEM116;refGene_tx=XM_023241424.1;refGene_gn=TMEM116;refGene_tx=XM_023241425.1;refGene_gn=TMEM116;refGene_tx=XM_023241426.1;refGene_gn=TMEM116;refGene_tx=XM_023241427.1;refGene_gn=TMEM116;refGene_tx=XM_023241428.1
D3 9873501 9873565
refGene_gn=TMEM116;refGene_tx=XM_019814547.2;refGene_gn=TMEM116;refGene_tx=XM_019814550.2;refGene_gn=TMEM116;refGene_tx=XM_023241424.1;refGene_gn=TMEM116;refGene_tx=XM_023241425.1;refGene_gn=TMEM116;refGene_tx=XM_023241428.1
D3 9874254 9874301
refGene_gn=TMEM116;refGene_tx=XM_019814547.2;refGene_gn=TMEM116;refGene_tx=XM_019814550.2;refGene_gn=TMEM116;refGene_tx=XM_023241424.1;refGene_gn=TMEM116;refGene_tx=XM_023241425.1;refGene_gn=TMEM116;refGene_tx=XM_023241428.1
D3 9874847 9874895
refGene_gn=TMEM116;refGene_tx=XM_019814547.2;refGene_gn=TMEM116;refGene_tx=XM_019814550.2;refGene_gn=TMEM116;refGene_tx=XM_023241424.1;refGene_gn=TMEM116;refGene_tx=XM_023241428.1
D3 9892483 9892627 refGene_gn=ERP29;refGene_tx=XM_003994679.5
D3 9897142 9897281 refGene_gn=ERP29;refGene_tx=XM_003994679.5
D3 9900482 9900985 refGene_gn=ERP29;refGene_tx=XM_003994679.5
D3 9908934 9909057
refGene_gn=NAA25;refGene_tx=XM_023241429.1;refGene_gn=NAA25;refGene_tx=XM_023241430.1
D3 9912233 9912380
refGene_gn=NAA25;refGene_tx=XM_023241429.1;refGene_gn=NAA25;refGene_tx=XM_023241430.1
D3 9916745 9916856
refGene_gn=NAA25;refGene_tx=XM_023241429.1;refGene_gn=NAA25;refGene_tx=XM_023241430.1
D3 9917935 9918033
refGene_gn=NAA25;refGene_tx=XM_023241429.1;refGene_gn=NAA25;refGene_tx=XM_023241430.1
D3 9925936 9926002
refGene_gn=NAA25;refGene_tx=XM_023241429.1;refGene_gn=NAA25;refGene_tx=XM_023241430.1
D3 9926685 9926809
refGene_gn=NAA25;refGene_tx=XM_023241429.1;refGene_gn=NAA25;refGene_tx=XM_023241430.1
D3 9927190 9927434
refGene_gn=NAA25;refGene_tx=XM_023241429.1;refGene_gn=NAA25;refGene_tx=XM_023241430.1

D3 9930714 9930840
refGene_gn=NAA25;refGene_tx=XM_023241429.1;refGene_gn=NAA25;refGene_tx
=XM_023241430.1
D3 9931399 9931551
refGene_gn=NAA25;refGene_tx=XM_023241429.1;refGene_gn=NAA25;refGene_tx
=XM_023241430.1
D3 9936762 9936862
refGene_gn=NAA25;refGene_tx=XM_023241429.1;refGene_gn=NAA25;refGene_tx
=XM_023241430.1
D3 9938706 9938887
refGene_gn=NAA25;refGene_tx=XM_023241429.1;refGene_gn=NAA25;refGene_tx
=XM_023241430.1
D3 9944189 9944279
refGene_gn=NAA25;refGene_tx=XM_023241429.1;refGene_gn=NAA25;refGene_tx
=XM_023241430.1
D3 9945072 9945280
refGene_gn=NAA25;refGene_tx=XM_023241429.1;refGene_gn=NAA25;refGene_tx
=XM_023241430.1
D3 9952746 9952859
refGene_gn=NAA25;refGene_tx=XM_023241429.1;refGene_gn=NAA25;refGene_tx
=XM_023241430.1
D3 9956426 9956596
refGene_gn=NAA25;refGene_tx=XM_023241429.1;refGene_gn=NAA25;refGene_tx
=XM_023241430.1
D3 9957789 9957879
refGene_gn=NAA25;refGene_tx=XM_023241429.1;refGene_gn=NAA25;refGene_tx
=XM_023241430.1
D3 9958469 9958581
refGene_gn=NAA25;refGene_tx=XM_023241429.1;refGene_gn=NAA25;refGene_tx
=XM_023241430.1
D3 9962623 9962702
refGene_gn=NAA25;refGene_tx=XM_023241429.1;refGene_gn=NAA25;refGene_tx
=XM_023241430.1
D3 9962994 9963102
refGene_gn=NAA25;refGene_tx=XM_023241429.1;refGene_gn=NAA25;refGene_tx
=XM_023241430.1
D3 9964344 9964419
refGene_gn=NAA25;refGene_tx=XM_023241429.1;refGene_gn=NAA25;refGene_tx
=XM_023241430.1
D3 9968841 9968960
refGene_gn=NAA25;refGene_tx=XM_023241429.1;refGene_gn=NAA25;refGene_tx
=XM_023241430.1
D3 9972212 9972351 refGene_gn=NAA25;refGene_tx=XM_023241429.1
D3 9973302 9973388 refGene_gn=NAA25;refGene_tx=XM_023241429.1
D3 9986457 9986515 refGene_gn=NAA25;refGene_tx=XM_023241429.1
D3 10012120 10012167 refGene_gn=TRAFD1;refGene_tx=XM_003994681.4
D3 10018546 10018682 refGene_gn=TRAFD1;refGene_tx=XM_003994681.4
D3 10018791 10018845 refGene_gn=TRAFD1;refGene_tx=XM_003994681.4
D3 10019478 10019884 refGene_gn=TRAFD1;refGene_tx=XM_003994681.4
D3 10020836 10021043 refGene_gn=TRAFD1;refGene_tx=XM_003994681.4

D3 10028611 10028688 refGene_gn=TRAFD1;refGene_tx=XM_003994681.4
D3 10030154 10030385 refGene_gn=TRAFD1;refGene_tx=XM_003994681.4
D3 10031686 10031807 refGene_gn=TRAFD1;refGene_tx=XM_003994681.4
D3 10032680 10033020 refGene_gn=TRAFD1;refGene_tx=XM_003994681.4
D3 10033276 10033349 refGene_gn=TRAFD1;refGene_tx=XM_003994681.4
D3 10033572 10033632 refGene_gn=TRAFD1;refGene_tx=XM_003994681.4
D3 10048405 10048542
refGene_gn=HECTD4;refGene_tx=XM_023241431.1;refGene_gn=HECTD4;refGene_tx=XM_023241432.1;refGene_gn=HECTD4;refGene_tx=XM_023241433.1;refGene_gn=HECTD4;refGene_tx=XM_023241434.1;refGene_gn=HECTD4;refGene_tx=XM_023241435.1
D3 10049320 10049543
refGene_gn=HECTD4;refGene_tx=XM_023241431.1;refGene_gn=HECTD4;refGene_tx=XM_023241432.1;refGene_gn=HECTD4;refGene_tx=XM_023241433.1;refGene_gn=HECTD4;refGene_tx=XM_023241434.1;refGene_gn=HECTD4;refGene_tx=XM_023241435.1
D3 10050128 10050324
refGene_gn=HECTD4;refGene_tx=XM_023241431.1;refGene_gn=HECTD4;refGene_tx=XM_023241432.1;refGene_gn=HECTD4;refGene_tx=XM_023241433.1;refGene_gn=HECTD4;refGene_tx=XM_023241434.1;refGene_gn=HECTD4;refGene_tx=XM_023241435.1
D3 10050571 10050735
refGene_gn=HECTD4;refGene_tx=XM_023241431.1;refGene_gn=HECTD4;refGene_tx=XM_023241432.1;refGene_gn=HECTD4;refGene_tx=XM_023241433.1;refGene_gn=HECTD4;refGene_tx=XM_023241434.1;refGene_gn=HECTD4;refGene_tx=XM_023241435.1
D3 10053052 10053274
refGene_gn=HECTD4;refGene_tx=XM_023241431.1;refGene_gn=HECTD4;refGene_tx=XM_023241432.1;refGene_gn=HECTD4;refGene_tx=XM_023241433.1;refGene_gn=HECTD4;refGene_tx=XM_023241434.1;refGene_gn=HECTD4;refGene_tx=XM_023241435.1
D3 10053381 10053485
refGene_gn=HECTD4;refGene_tx=XM_023241431.1;refGene_gn=HECTD4;refGene_tx=XM_023241432.1;refGene_gn=HECTD4;refGene_tx=XM_023241433.1;refGene_gn=HECTD4;refGene_tx=XM_023241434.1;refGene_gn=HECTD4;refGene_tx=XM_023241435.1
D3 10054865 10055021
refGene_gn=HECTD4;refGene_tx=XM_023241431.1;refGene_gn=HECTD4;refGene_tx=XM_023241432.1;refGene_gn=HECTD4;refGene_tx=XM_023241433.1;refGene_gn=HECTD4;refGene_tx=XM_023241434.1;refGene_gn=HECTD4;refGene_tx=XM_023241435.1
D3 10055979 10056099
refGene_gn=HECTD4;refGene_tx=XM_023241431.1;refGene_gn=HECTD4;refGene_tx=XM_023241432.1;refGene_gn=HECTD4;refGene_tx=XM_023241433.1;refGene_gn=HECTD4;refGene_tx=XM_023241434.1;refGene_gn=HECTD4;refGene_tx=XM_023241435.1
D3 10056669 10056816
refGene_gn=HECTD4;refGene_tx=XM_023241431.1;refGene_gn=HECTD4;refGene_tx=XM_023241432.1;refGene_gn=HECTD4;refGene_tx=XM_023241433.1;refGene_gn=HECTD4;refGene_tx=XM_023241434.1;refGene_gn=HECTD4;refGene_tx=XM_02

3241435.1
D3 10057858 10058049
refGene_gn=HECTD4;refGene_tx=XM_023241431.1;refGene_gn=HECTD4;refGene_tx=XM_023241432.1;refGene_gn=HECTD4;refGene_tx=XM_023241433.1;refGene_gn=HECTD4;refGene_tx=XM_023241434.1;refGene_gn=HECTD4;refGene_tx=XM_023241435.1
D3 10060307 10060431
refGene_gn=HECTD4;refGene_tx=XM_023241431.1;refGene_gn=HECTD4;refGene_tx=XM_023241432.1;refGene_gn=HECTD4;refGene_tx=XM_023241433.1;refGene_gn=HECTD4;refGene_tx=XM_023241434.1;refGene_gn=HECTD4;refGene_tx=XM_023241435.1
D3 10060904 10061011
refGene_gn=HECTD4;refGene_tx=XM_023241431.1;refGene_gn=HECTD4;refGene_tx=XM_023241432.1;refGene_gn=HECTD4;refGene_tx=XM_023241433.1;refGene_gn=HECTD4;refGene_tx=XM_023241434.1;refGene_gn=HECTD4;refGene_tx=XM_023241435.1
D3 10062391 10062543
refGene_gn=HECTD4;refGene_tx=XM_023241431.1;refGene_gn=HECTD4;refGene_tx=XM_023241432.1;refGene_gn=HECTD4;refGene_tx=XM_023241433.1;refGene_gn=HECTD4;refGene_tx=XM_023241434.1;refGene_gn=HECTD4;refGene_tx=XM_023241435.1
D3 10062635 10062859
refGene_gn=HECTD4;refGene_tx=XM_023241431.1;refGene_gn=HECTD4;refGene_tx=XM_023241432.1;refGene_gn=HECTD4;refGene_tx=XM_023241433.1;refGene_gn=HECTD4;refGene_tx=XM_023241434.1;refGene_gn=HECTD4;refGene_tx=XM_023241435.1
D3 10065061 10065269
refGene_gn=HECTD4;refGene_tx=XM_023241431.1;refGene_gn=HECTD4;refGene_tx=XM_023241432.1;refGene_gn=HECTD4;refGene_tx=XM_023241433.1;refGene_gn=HECTD4;refGene_tx=XM_023241434.1;refGene_gn=HECTD4;refGene_tx=XM_023241435.1
D3 10066265 10067572
refGene_gn=HECTD4;refGene_tx=XM_023241431.1;refGene_gn=HECTD4;refGene_tx=XM_023241432.1;refGene_gn=HECTD4;refGene_tx=XM_023241433.1;refGene_gn=HECTD4;refGene_tx=XM_023241434.1;refGene_gn=HECTD4;refGene_tx=XM_023241435.1
D3 10070166 10070346
refGene_gn=HECTD4;refGene_tx=XM_023241431.1;refGene_gn=HECTD4;refGene_tx=XM_023241432.1;refGene_gn=HECTD4;refGene_tx=XM_023241433.1;refGene_gn=HECTD4;refGene_tx=XM_023241434.1;refGene_gn=HECTD4;refGene_tx=XM_023241435.1
D3 10072173 10072379
refGene_gn=HECTD4;refGene_tx=XM_023241431.1;refGene_gn=HECTD4;refGene_tx=XM_023241432.1;refGene_gn=HECTD4;refGene_tx=XM_023241433.1;refGene_gn=HECTD4;refGene_tx=XM_023241434.1;refGene_gn=HECTD4;refGene_tx=XM_023241435.1
D3 10072891 10073022
refGene_gn=HECTD4;refGene_tx=XM_023241431.1;refGene_gn=HECTD4;refGene_tx=XM_023241432.1;refGene_gn=HECTD4;refGene_tx=XM_023241433.1;refGene_gn=HECTD4;refGene_tx=XM_023241434.1;refGene_gn=HECTD4;refGene_tx=XM_02

3241435.1
D3 10073326 10073532
refGene_gn=HECTD4;refGene_tx=XM_023241431.1;refGene_gn=HECTD4;refGene_tx=XM_023241432.1;refGene_gn=HECTD4;refGene_tx=XM_023241433.1;refGene_gn=HECTD4;refGene_tx=XM_023241434.1;refGene_gn=HECTD4;refGene_tx=XM_023241435.1
D3 10074695 10074877
refGene_gn=HECTD4;refGene_tx=XM_023241431.1;refGene_gn=HECTD4;refGene_tx=XM_023241432.1;refGene_gn=HECTD4;refGene_tx=XM_023241433.1;refGene_gn=HECTD4;refGene_tx=XM_023241434.1;refGene_gn=HECTD4;refGene_tx=XM_023241435.1
D3 10077300 10077461
refGene_gn=HECTD4;refGene_tx=XM_023241431.1;refGene_gn=HECTD4;refGene_tx=XM_023241432.1;refGene_gn=HECTD4;refGene_tx=XM_023241433.1;refGene_gn=HECTD4;refGene_tx=XM_023241434.1;refGene_gn=HECTD4;refGene_tx=XM_023241435.1
D3 10080008 10080145
refGene_gn=HECTD4;refGene_tx=XM_023241431.1;refGene_gn=HECTD4;refGene_tx=XM_023241432.1;refGene_gn=HECTD4;refGene_tx=XM_023241433.1;refGene_gn=HECTD4;refGene_tx=XM_023241434.1;refGene_gn=HECTD4;refGene_tx=XM_023241435.1
D3 10081605 10081743
refGene_gn=HECTD4;refGene_tx=XM_023241431.1;refGene_gn=HECTD4;refGene_tx=XM_023241432.1;refGene_gn=HECTD4;refGene_tx=XM_023241433.1;refGene_gn=HECTD4;refGene_tx=XM_023241434.1;refGene_gn=HECTD4;refGene_tx=XM_023241435.1
D3 10085115 10085242
refGene_gn=HECTD4;refGene_tx=XM_023241431.1;refGene_gn=HECTD4;refGene_tx=XM_023241432.1;refGene_gn=HECTD4;refGene_tx=XM_023241433.1;refGene_gn=HECTD4;refGene_tx=XM_023241434.1;refGene_gn=HECTD4;refGene_tx=XM_023241435.1
D3 10085658 10085795
refGene_gn=HECTD4;refGene_tx=XM_023241431.1;refGene_gn=HECTD4;refGene_tx=XM_023241432.1;refGene_gn=HECTD4;refGene_tx=XM_023241433.1;refGene_gn=HECTD4;refGene_tx=XM_023241434.1;refGene_gn=HECTD4;refGene_tx=XM_023241435.1
D3 10086780 10087018
refGene_gn=HECTD4;refGene_tx=XM_023241431.1;refGene_gn=HECTD4;refGene_tx=XM_023241432.1;refGene_gn=HECTD4;refGene_tx=XM_023241433.1;refGene_gn=HECTD4;refGene_tx=XM_023241434.1;refGene_gn=HECTD4;refGene_tx=XM_023241435.1
D3 10087665 10087829
refGene_gn=HECTD4;refGene_tx=XM_023241431.1;refGene_gn=HECTD4;refGene_tx=XM_023241432.1;refGene_gn=HECTD4;refGene_tx=XM_023241433.1;refGene_gn=HECTD4;refGene_tx=XM_023241434.1;refGene_gn=HECTD4;refGene_tx=XM_023241435.1
D3 10089626 10089706
refGene_gn=HECTD4;refGene_tx=XM_023241431.1;refGene_gn=HECTD4;refGene_tx=XM_023241432.1;refGene_gn=HECTD4;refGene_tx=XM_023241433.1;refGene_gn=HECTD4;refGene_tx=XM_023241434.1;refGene_gn=HECTD4;refGene_tx=XM_02

3241435.1
D3 10089985 10090134
refGene_gn=HECTD4;refGene_tx=XM_023241431.1;refGene_gn=HECTD4;refGene_tx=XM_023241432.1;refGene_gn=HECTD4;refGene_tx=XM_023241433.1;refGene_gn=HECTD4;refGene_tx=XM_023241434.1;refGene_gn=HECTD4;refGene_tx=XM_023241435.1
D3 10090247 10090409
refGene_gn=HECTD4;refGene_tx=XM_023241431.1;refGene_gn=HECTD4;refGene_tx=XM_023241432.1;refGene_gn=HECTD4;refGene_tx=XM_023241433.1;refGene_gn=HECTD4;refGene_tx=XM_023241434.1;refGene_gn=HECTD4;refGene_tx=XM_023241435.1
D3 10092203 10092307
refGene_gn=HECTD4;refGene_tx=XM_023241431.1;refGene_gn=HECTD4;refGene_tx=XM_023241432.1;refGene_gn=HECTD4;refGene_tx=XM_023241433.1;refGene_gn=HECTD4;refGene_tx=XM_023241434.1;refGene_gn=HECTD4;refGene_tx=XM_023241435.1
D3 10099862 10099978
refGene_gn=HECTD4;refGene_tx=XM_023241431.1;refGene_gn=HECTD4;refGene_tx=XM_023241432.1;refGene_gn=HECTD4;refGene_tx=XM_023241433.1;refGene_gn=HECTD4;refGene_tx=XM_023241434.1;refGene_gn=HECTD4;refGene_tx=XM_023241435.1
D3 10100904 10101074
refGene_gn=HECTD4;refGene_tx=XM_023241431.1;refGene_gn=HECTD4;refGene_tx=XM_023241432.1;refGene_gn=HECTD4;refGene_tx=XM_023241433.1;refGene_gn=HECTD4;refGene_tx=XM_023241434.1;refGene_gn=HECTD4;refGene_tx=XM_023241435.1
D3 10101468 10101633
refGene_gn=HECTD4;refGene_tx=XM_023241431.1;refGene_gn=HECTD4;refGene_tx=XM_023241432.1;refGene_gn=HECTD4;refGene_tx=XM_023241433.1;refGene_gn=HECTD4;refGene_tx=XM_023241434.1;refGene_gn=HECTD4;refGene_tx=XM_023241435.1
D3 10102118 10102301
refGene_gn=HECTD4;refGene_tx=XM_023241431.1;refGene_gn=HECTD4;refGene_tx=XM_023241432.1;refGene_gn=HECTD4;refGene_tx=XM_023241433.1;refGene_gn=HECTD4;refGene_tx=XM_023241434.1;refGene_gn=HECTD4;refGene_tx=XM_023241435.1
D3 10103124 10103260
refGene_gn=HECTD4;refGene_tx=XM_023241431.1;refGene_gn=HECTD4;refGene_tx=XM_023241432.1;refGene_gn=HECTD4;refGene_tx=XM_023241433.1;refGene_gn=HECTD4;refGene_tx=XM_023241434.1;refGene_gn=HECTD4;refGene_tx=XM_023241435.1
D3 10103913 10104116
refGene_gn=HECTD4;refGene_tx=XM_023241431.1;refGene_gn=HECTD4;refGene_tx=XM_023241432.1;refGene_gn=HECTD4;refGene_tx=XM_023241433.1;refGene_gn=HECTD4;refGene_tx=XM_023241434.1;refGene_gn=HECTD4;refGene_tx=XM_023241435.1
D3 10105796 10105878
refGene_gn=HECTD4;refGene_tx=XM_023241431.1;refGene_gn=HECTD4;refGene_tx=XM_023241432.1;refGene_gn=HECTD4;refGene_tx=XM_023241433.1;refGene_gn=HECTD4;refGene_tx=XM_023241434.1;refGene_gn=HECTD4;refGene_tx=XM_02

3241435.1
D3	10106907	10107097
refGene_gn=HECTD4;refGene_tx=XM_023241431.1;refGene_gn=HECTD4;refGene_tx=XM_023241432.1;refGene_gn=HECTD4;refGene_tx=XM_023241433.1;refGene_gn=HECTD4;refGene_tx=XM_023241434.1;refGene_gn=HECTD4;refGene_tx=XM_023241435.1
D3	10107318	10107599
refGene_gn=HECTD4;refGene_tx=XM_023241431.1;refGene_gn=HECTD4;refGene_tx=XM_023241432.1;refGene_gn=HECTD4;refGene_tx=XM_023241433.1;refGene_gn=HECTD4;refGene_tx=XM_023241434.1;refGene_gn=HECTD4;refGene_tx=XM_023241435.1
D3	10108534	10108688
refGene_gn=HECTD4;refGene_tx=XM_023241431.1;refGene_gn=HECTD4;refGene_tx=XM_023241432.1;refGene_gn=HECTD4;refGene_tx=XM_023241433.1;refGene_gn=HECTD4;refGene_tx=XM_023241434.1;refGene_gn=HECTD4;refGene_tx=XM_023241435.1
D3	10110921	10111106
refGene_gn=HECTD4;refGene_tx=XM_023241431.1;refGene_gn=HECTD4;refGene_tx=XM_023241432.1;refGene_gn=HECTD4;refGene_tx=XM_023241433.1;refGene_gn=HECTD4;refGene_tx=XM_023241434.1;refGene_gn=HECTD4;refGene_tx=XM_023241435.1
D3	10111715	10111862
refGene_gn=HECTD4;refGene_tx=XM_023241431.1;refGene_gn=HECTD4;refGene_tx=XM_023241432.1;refGene_gn=HECTD4;refGene_tx=XM_023241433.1;refGene_gn=HECTD4;refGene_tx=XM_023241434.1;refGene_gn=HECTD4;refGene_tx=XM_023241435.1
D3	10114807	10114974
refGene_gn=HECTD4;refGene_tx=XM_023241431.1;refGene_gn=HECTD4;refGene_tx=XM_023241432.1;refGene_gn=HECTD4;refGene_tx=XM_023241433.1;refGene_gn=HECTD4;refGene_tx=XM_023241434.1;refGene_gn=HECTD4;refGene_tx=XM_023241435.1
D3	10115074	10115216
refGene_gn=HECTD4;refGene_tx=XM_023241431.1;refGene_gn=HECTD4;refGene_tx=XM_023241432.1;refGene_gn=HECTD4;refGene_tx=XM_023241433.1;refGene_gn=HECTD4;refGene_tx=XM_023241434.1;refGene_gn=HECTD4;refGene_tx=XM_023241435.1
D3	10115327	10115463
refGene_gn=HECTD4;refGene_tx=XM_023241431.1;refGene_gn=HECTD4;refGene_tx=XM_023241432.1;refGene_gn=HECTD4;refGene_tx=XM_023241433.1;refGene_gn=HECTD4;refGene_tx=XM_023241434.1;refGene_gn=HECTD4;refGene_tx=XM_023241435.1
D3	10117567	10117743
refGene_gn=HECTD4;refGene_tx=XM_023241431.1;refGene_gn=HECTD4;refGene_tx=XM_023241432.1;refGene_gn=HECTD4;refGene_tx=XM_023241433.1;refGene_gn=HECTD4;refGene_tx=XM_023241434.1;refGene_gn=HECTD4;refGene_tx=XM_023241435.1
D3	10118122	10118211
refGene_gn=HECTD4;refGene_tx=XM_023241431.1;refGene_gn=HECTD4;refGene_tx=XM_023241432.1;refGene_gn=HECTD4;refGene_tx=XM_023241433.1;refGene_gn=HECTD4;refGene_tx=XM_023241434.1;refGene_gn=HECTD4;refGene_tx=XM_02

3241435.1
D3	10118709	10118814
refGene_gn=HECTD4;refGene_tx=XM_023241431.1;refGene_gn=HECTD4;refGene_tx=XM_023241432.1;refGene_gn=HECTD4;refGene_tx=XM_023241433.1;refGene_gn=HECTD4;refGene_tx=XM_023241434.1;refGene_gn=HECTD4;refGene_tx=XM_023241435.1
D3	10119578	10119771
refGene_gn=HECTD4;refGene_tx=XM_023241431.1;refGene_gn=HECTD4;refGene_tx=XM_023241432.1;refGene_gn=HECTD4;refGene_tx=XM_023241433.1;refGene_gn=HECTD4;refGene_tx=XM_023241434.1;refGene_gn=HECTD4;refGene_tx=XM_023241435.1
D3	10120752	10120986
refGene_gn=HECTD4;refGene_tx=XM_023241431.1;refGene_gn=HECTD4;refGene_tx=XM_023241432.1;refGene_gn=HECTD4;refGene_tx=XM_023241433.1;refGene_gn=HECTD4;refGene_tx=XM_023241434.1;refGene_gn=HECTD4;refGene_tx=XM_023241435.1
D3	10121560	10121724
refGene_gn=HECTD4;refGene_tx=XM_023241431.1;refGene_gn=HECTD4;refGene_tx=XM_023241432.1;refGene_gn=HECTD4;refGene_tx=XM_023241433.1;refGene_gn=HECTD4;refGene_tx=XM_023241434.1;refGene_gn=HECTD4;refGene_tx=XM_023241435.1
D3	10122831	10122936
refGene_gn=HECTD4;refGene_tx=XM_023241431.1;refGene_gn=HECTD4;refGene_tx=XM_023241432.1;refGene_gn=HECTD4;refGene_tx=XM_023241433.1;refGene_gn=HECTD4;refGene_tx=XM_023241434.1;refGene_gn=HECTD4;refGene_tx=XM_023241435.1
D3	10124063	10124210
refGene_gn=HECTD4;refGene_tx=XM_023241431.1;refGene_gn=HECTD4;refGene_tx=XM_023241432.1;refGene_gn=HECTD4;refGene_tx=XM_023241433.1;refGene_gn=HECTD4;refGene_tx=XM_023241434.1;refGene_gn=HECTD4;refGene_tx=XM_023241435.1
D3	10126290	10126489
refGene_gn=HECTD4;refGene_tx=XM_023241431.1;refGene_gn=HECTD4;refGene_tx=XM_023241432.1;refGene_gn=HECTD4;refGene_tx=XM_023241433.1;refGene_gn=HECTD4;refGene_tx=XM_023241434.1;refGene_gn=HECTD4;refGene_tx=XM_023241435.1
D3	10128142	10128243
refGene_gn=HECTD4;refGene_tx=XM_023241431.1;refGene_gn=HECTD4;refGene_tx=XM_023241432.1;refGene_gn=HECTD4;refGene_tx=XM_023241433.1;refGene_gn=HECTD4;refGene_tx=XM_023241434.1;refGene_gn=HECTD4;refGene_tx=XM_023241435.1
D3	10128759	10128913
refGene_gn=HECTD4;refGene_tx=XM_023241431.1;refGene_gn=HECTD4;refGene_tx=XM_023241432.1;refGene_gn=HECTD4;refGene_tx=XM_023241433.1;refGene_gn=HECTD4;refGene_tx=XM_023241434.1;refGene_gn=HECTD4;refGene_tx=XM_023241435.1
D3	10130386	10130511
refGene_gn=HECTD4;refGene_tx=XM_023241431.1;refGene_gn=HECTD4;refGene_tx=XM_023241432.1;refGene_gn=HECTD4;refGene_tx=XM_023241433.1;refGene_gn=HECTD4;refGene_tx=XM_023241434.1;refGene_gn=HECTD4;refGene_tx=XM_02

3241435.1
D3 10132435 10132564
refGene_gn=HECTD4;refGene_tx=XM_023241431.1;refGene_gn=HECTD4;refGene_tx=XM_023241432.1;refGene_gn=HECTD4;refGene_tx=XM_023241433.1;refGene_gn=HECTD4;refGene_tx=XM_023241434.1;refGene_gn=HECTD4;refGene_tx=XM_023241435.1
D3 10133276 10133397
refGene_gn=HECTD4;refGene_tx=XM_023241431.1;refGene_gn=HECTD4;refGene_tx=XM_023241432.1;refGene_gn=HECTD4;refGene_tx=XM_023241433.1;refGene_gn=HECTD4;refGene_tx=XM_023241434.1;refGene_gn=HECTD4;refGene_tx=XM_023241435.1
D3 10133972 10134078
refGene_gn=HECTD4;refGene_tx=XM_023241431.1;refGene_gn=HECTD4;refGene_tx=XM_023241432.1;refGene_gn=HECTD4;refGene_tx=XM_023241433.1;refGene_gn=HECTD4;refGene_tx=XM_023241434.1;refGene_gn=HECTD4;refGene_tx=XM_023241435.1
D3 10134945 10135016
refGene_gn=HECTD4;refGene_tx=XM_023241431.1;refGene_gn=HECTD4;refGene_tx=XM_023241432.1;refGene_gn=HECTD4;refGene_tx=XM_023241433.1;refGene_gn=HECTD4;refGene_tx=XM_023241434.1;refGene_gn=HECTD4;refGene_tx=XM_023241435.1
D3 10136587 10136733
refGene_gn=HECTD4;refGene_tx=XM_023241431.1;refGene_gn=HECTD4;refGene_tx=XM_023241432.1;refGene_gn=HECTD4;refGene_tx=XM_023241433.1;refGene_gn=HECTD4;refGene_tx=XM_023241434.1;refGene_gn=HECTD4;refGene_tx=XM_023241435.1
D3 10137110 10137343
refGene_gn=HECTD4;refGene_tx=XM_023241431.1;refGene_gn=HECTD4;refGene_tx=XM_023241432.1;refGene_gn=HECTD4;refGene_tx=XM_023241433.1;refGene_gn=HECTD4;refGene_tx=XM_023241434.1;refGene_gn=HECTD4;refGene_tx=XM_023241435.1
D3 10142661 10142802
refGene_gn=HECTD4;refGene_tx=XM_023241431.1;refGene_gn=HECTD4;refGene_tx=XM_023241432.1;refGene_gn=HECTD4;refGene_tx=XM_023241433.1;refGene_gn=HECTD4;refGene_tx=XM_023241434.1;refGene_gn=HECTD4;refGene_tx=XM_023241435.1
D3 10143516 10143630
refGene_gn=HECTD4;refGene_tx=XM_023241431.1;refGene_gn=HECTD4;refGene_tx=XM_023241432.1;refGene_gn=HECTD4;refGene_tx=XM_023241433.1;refGene_gn=HECTD4;refGene_tx=XM_023241434.1;refGene_gn=HECTD4;refGene_tx=XM_023241435.1
D3 10147734 10147893
refGene_gn=HECTD4;refGene_tx=XM_023241431.1;refGene_gn=HECTD4;refGene_tx=XM_023241432.1;refGene_gn=HECTD4;refGene_tx=XM_023241433.1;refGene_gn=HECTD4;refGene_tx=XM_023241434.1;refGene_gn=HECTD4;refGene_tx=XM_023241435.1
D3 10151598 10151791
refGene_gn=HECTD4;refGene_tx=XM_023241431.1;refGene_gn=HECTD4;refGene_tx=XM_023241432.1;refGene_gn=HECTD4;refGene_tx=XM_023241433.1;refGene_gn=HECTD4;refGene_tx=XM_023241434.1;refGene_gn=HECTD4;refGene_tx=XM_02

3241435.1
D3	10165761	10165932
refGene_gn=HECTD4;refGene_tx=XM_023241431.1;refGene_gn=HECTD4;refGene_tx=XM_023241432.1;refGene_gn=HECTD4;refGene_tx=XM_023241433.1;refGene_gn=HECTD4;refGene_tx=XM_023241434.1;refGene_gn=HECTD4;refGene_tx=XM_023241435.1
D3	10168956	10169095
refGene_gn=HECTD4;refGene_tx=XM_023241431.1;refGene_gn=HECTD4;refGene_tx=XM_023241432.1;refGene_gn=HECTD4;refGene_tx=XM_023241433.1;refGene_gn=HECTD4;refGene_tx=XM_023241434.1;refGene_gn=HECTD4;refGene_tx=XM_023241435.1
D3	10169974	10170083
refGene_gn=HECTD4;refGene_tx=XM_023241431.1;refGene_gn=HECTD4;refGene_tx=XM_023241432.1;refGene_gn=HECTD4;refGene_tx=XM_023241433.1;refGene_gn=HECTD4;refGene_tx=XM_023241434.1;refGene_gn=HECTD4;refGene_tx=XM_023241435.1
D3	10172564	10172695
refGene_gn=HECTD4;refGene_tx=XM_023241431.1;refGene_gn=HECTD4;refGene_tx=XM_023241432.1;refGene_gn=HECTD4;refGene_tx=XM_023241433.1;refGene_gn=HECTD4;refGene_tx=XM_023241434.1;refGene_gn=HECTD4;refGene_tx=XM_023241435.1
D3	10173715	10173805
refGene_gn=HECTD4;refGene_tx=XM_023241431.1;refGene_gn=HECTD4;refGene_tx=XM_023241432.1;refGene_gn=HECTD4;refGene_tx=XM_023241433.1;refGene_gn=HECTD4;refGene_tx=XM_023241434.1;refGene_gn=HECTD4;refGene_tx=XM_023241435.1
D3	10178114	10178632
refGene_gn=HECTD4;refGene_tx=XM_023241431.1;refGene_gn=HECTD4;refGene_tx=XM_023241432.1;refGene_gn=HECTD4;refGene_tx=XM_023241433.1;refGene_gn=HECTD4;refGene_tx=XM_023241434.1;refGene_gn=HECTD4;refGene_tx=XM_023241435.1
D3	10238723	10238900
refGene_gn=HECTD4;refGene_tx=XM_023241431.1;refGene_gn=HECTD4;refGene_tx=XM_023241432.1;refGene_gn=HECTD4;refGene_tx=XM_023241433.1;refGene_gn=HECTD4;refGene_tx=XM_023241434.1;refGene_gn=HECTD4;refGene_tx=XM_023241435.1
D3	10253883	10254036	refGene_gn=RPL6;refGene_tx=XM_006938486.4
D3	10254513	10254698	refGene_gn=RPL6;refGene_tx=XM_006938486.4
D3	10255329	10255378	refGene_gn=RPL6;refGene_tx=XM_006938486.4
D3	10255908	10256052	refGene_gn=RPL6;refGene_tx=XM_006938486.4
D3	10257197	10257296	refGene_gn=RPL6;refGene_tx=XM_006938486.4
D3	10257392	10257629	refGene_gn=RPL6;refGene_tx=XM_006938486.4
D3	10268069	10268083
refGene_gn=PTPN11;refGene_tx=XM_023241436.1;refGene_gn=PTPN11;refGene_tx=XM_023241437.1
D3	10288894	10289017
refGene_gn=PTPN11;refGene_tx=XM_023241436.1;refGene_gn=PTPN11;refGene_tx=XM_023241437.1
D3	10293815	10294010
refGene_gn=PTPN11;refGene_tx=XM_023241436.1;refGene_gn=PTPN11;refGene_

tx=XM_023241437.1
D3 10296277 10296470
refGene_gn=PTPN11;refGene_tx=XM_023241436.1;refGene_gn=PTPN11;refGene_tx=XM_023241437.1
D3 10300302 10300419
refGene_gn=PTPN11;refGene_tx=XM_023241436.1;refGene_gn=PTPN11;refGene_tx=XM_023241437.1
D3 10301103 10301217
refGene_gn=PTPN11;refGene_tx=XM_023241436.1;refGene_gn=PTPN11;refGene_tx=XM_023241437.1
D3 10317098 10317195
refGene_gn=PTPN11;refGene_tx=XM_023241436.1;refGene_gn=PTPN11;refGene_tx=XM_023241437.1
D3 10320273 10320353
refGene_gn=PTPN11;refGene_tx=XM_023241436.1;refGene_gn=PTPN11;refGene_tx=XM_023241437.1
D3 10320476 10320635
refGene_gn=PTPN11;refGene_tx=XM_023241436.1;refGene_gn=PTPN11;refGene_tx=XM_023241437.1
D3 10325132 10325264
refGene_gn=PTPN11;refGene_tx=XM_023241436.1;refGene_gn=PTPN11;refGene_tx=XM_023241437.1
D3 10329292 10329459
refGene_gn=PTPN11;refGene_tx=XM_023241436.1;refGene_gn=PTPN11;refGene_tx=XM_023241437.1
D3 10330907 10330975
refGene_gn=PTPN11;refGene_tx=XM_023241436.1;refGene_gn=PTPN11;refGene_tx=XM_023241437.1
D3 10331466 10331618
refGene_gn=PTPN11;refGene_tx=XM_023241436.1;refGene_gn=PTPN11;refGene_tx=XM_023241437.1
D3 10341301 10341414
refGene_gn=PTPN11;refGene_tx=XM_023241436.1;refGene_gn=PTPN11;refGene_tx=XM_023241437.1
D3 10343234 10343304
refGene_gn=PTPN11;refGene_tx=XM_023241436.1;refGene_gn=PTPN11;refGene_tx=XM_023241437.1
D3 10606788 10606859
refGene_gn=RPH3A;refGene_tx=XM_006938490.4;refGene_gn=RPH3A;refGene_tx=XM_019814557.2;refGene_gn=RPH3A;refGene_tx=XM_019814558.2;refGene_gn=RPH3A;refGene_tx=XM_019814559.2;refGene_gn=RPH3A;refGene_tx=XM_019814560.2;refGene_gn=RPH3A;refGene_tx=XM_023241438.1
D3 10611254 10611266
refGene_gn=RPH3A;refGene_tx=XM_006938490.4;refGene_gn=RPH3A;refGene_tx=XM_019814557.2;refGene_gn=RPH3A;refGene_tx=XM_019814558.2;refGene_gn=RPH3A;refGene_tx=XM_019814559.2;refGene_gn=RPH3A;refGene_tx=XM_023241438.1
D3 10611901 10611939 refGene_gn=RPH3A;refGene_tx=XM_019814556.2
D3 10619461 10619608
refGene_gn=RPH3A;refGene_tx=XM_006938490.4;refGene_gn=RPH3A;refGene_tx

=XM_019814556.2;refGene_gn=RPH3A;refGene_tx=XM_019814557.2;refGene_gn=RPH3A;refGene_tx=XM_019814558.2;refGene_gn=RPH3A;refGene_tx=XM_019814559.2;refGene_gn=RPH3A;refGene_tx=XM_019814560.2;refGene_gn=RPH3A;refGene_tx=XM_023241438.1
D3 10634409 10634539
refGene_gn=RPH3A;refGene_tx=XM_006938490.4;refGene_gn=RPH3A;refGene_tx=XM_019814556.2;refGene_gn=RPH3A;refGene_tx=XM_019814557.2;refGene_gn=RPH3A;refGene_tx=XM_019814558.2;refGene_gn=RPH3A;refGene_tx=XM_019814559.2;refGene_gn=RPH3A;refGene_tx=XM_019814560.2;refGene_gn=RPH3A;refGene_tx=XM_023241438.1
D3 10635886 10635973
refGene_gn=RPH3A;refGene_tx=XM_006938490.4;refGene_gn=RPH3A;refGene_tx=XM_019814556.2;refGene_gn=RPH3A;refGene_tx=XM_019814557.2;refGene_gn=RPH3A;refGene_tx=XM_019814558.2;refGene_gn=RPH3A;refGene_tx=XM_019814559.2;refGene_gn=RPH3A;refGene_tx=XM_019814560.2;refGene_gn=RPH3A;refGene_tx=XM_023241438.1
D3 10636480 10636646
refGene_gn=RPH3A;refGene_tx=XM_006938490.4;refGene_gn=RPH3A;refGene_tx=XM_019814556.2;refGene_gn=RPH3A;refGene_tx=XM_019814557.2;refGene_gn=RPH3A;refGene_tx=XM_019814558.2;refGene_gn=RPH3A;refGene_tx=XM_019814559.2;refGene_gn=RPH3A;refGene_tx=XM_019814560.2;refGene_gn=RPH3A;refGene_tx=XM_023241438.1
D3 10639994 10640033
refGene_gn=RPH3A;refGene_tx=XM_006938490.4;refGene_gn=RPH3A;refGene_tx=XM_019814556.2;refGene_gn=RPH3A;refGene_tx=XM_019814557.2;refGene_gn=RPH3A;refGene_tx=XM_019814558.2;refGene_gn=RPH3A;refGene_tx=XM_019814559.2;refGene_gn=RPH3A;refGene_tx=XM_019814560.2;refGene_gn=RPH3A;refGene_tx=XM_023241438.1
D3 10640115 10640268
refGene_gn=RPH3A;refGene_tx=XM_006938490.4;refGene_gn=RPH3A;refGene_tx=XM_019814556.2;refGene_gn=RPH3A;refGene_tx=XM_019814557.2;refGene_gn=RPH3A;refGene_tx=XM_019814558.2;refGene_gn=RPH3A;refGene_tx=XM_019814559.2;refGene_gn=RPH3A;refGene_tx=XM_019814560.2;refGene_gn=RPH3A;refGene_tx=XM_023241438.1
D3 10645258 10645345
refGene_gn=RPH3A;refGene_tx=XM_006938490.4;refGene_gn=RPH3A;refGene_tx=XM_019814556.2;refGene_gn=RPH3A;refGene_tx=XM_019814557.2;refGene_gn=RPH3A;refGene_tx=XM_019814558.2;refGene_gn=RPH3A;refGene_tx=XM_019814559.2;refGene_gn=RPH3A;refGene_tx=XM_019814560.2;refGene_gn=RPH3A;refGene_tx=XM_023241438.1
D3 10646661 10646721
refGene_gn=RPH3A;refGene_tx=XM_006938490.4;refGene_gn=RPH3A;refGene_tx=XM_019814556.2;refGene_gn=RPH3A;refGene_tx=XM_019814557.2;refGene_gn=RPH3A;refGene_tx=XM_019814558.2;refGene_gn=RPH3A;refGene_tx=XM_019814559.2;refGene_gn=RPH3A;refGene_tx=XM_019814560.2;refGene_gn=RPH3A;refGene_tx=XM_023241438.1
D3 10648629 10648869
refGene_gn=RPH3A;refGene_tx=XM_006938490.4;refGene_gn=RPH3A;refGene_tx=XM_019814556.2;refGene_gn=RPH3A;refGene_tx=XM_019814557.2;refGene_gn=RPH3A;refGene_tx=XM_019814558.2;refGene_gn=RPH3A;refGene_tx=XM_0198145

59.2;refGene_gn=RPH3A;refGene_tx=XM_019814560.2;refGene_gn=RPH3A;refGene_tx=XM_023241438.1
D3 10651416 10651496
refGene_gn=RPH3A;refGene_tx=XM_006938490.4;refGene_gn=RPH3A;refGene_tx=XM_019814556.2;refGene_gn=RPH3A;refGene_tx=XM_019814557.2;refGene_gn=RPH3A;refGene_tx=XM_019814558.2;refGene_gn=RPH3A;refGene_tx=XM_019814559.2;refGene_gn=RPH3A;refGene_tx=XM_019814560.2;refGene_gn=RPH3A;refGene_tx=XM_023241438.1
D3 10654602 10654677
refGene_gn=RPH3A;refGene_tx=XM_006938490.4;refGene_gn=RPH3A;refGene_tx=XM_019814556.2;refGene_gn=RPH3A;refGene_tx=XM_019814557.2;refGene_gn=RPH3A;refGene_tx=XM_019814558.2;refGene_gn=RPH3A;refGene_tx=XM_019814559.2;refGene_gn=RPH3A;refGene_tx=XM_019814560.2;refGene_gn=RPH3A;refGene_tx=XM_023241438.1
D3 10656088 10656198
refGene_gn=RPH3A;refGene_tx=XM_006938490.4;refGene_gn=RPH3A;refGene_tx=XM_019814556.2;refGene_gn=RPH3A;refGene_tx=XM_019814557.2;refGene_gn=RPH3A;refGene_tx=XM_019814558.2;refGene_gn=RPH3A;refGene_tx=XM_019814559.2;refGene_gn=RPH3A;refGene_tx=XM_019814560.2;refGene_gn=RPH3A;refGene_tx=XM_023241438.1
D3 10659293 10659420
refGene_gn=RPH3A;refGene_tx=XM_006938490.4;refGene_gn=RPH3A;refGene_tx=XM_019814556.2;refGene_gn=RPH3A;refGene_tx=XM_019814557.2;refGene_gn=RPH3A;refGene_tx=XM_019814558.2;refGene_gn=RPH3A;refGene_tx=XM_019814559.2;refGene_gn=RPH3A;refGene_tx=XM_019814560.2;refGene_gn=RPH3A;refGene_tx=XM_023241438.1
D3 10662834 10662891
refGene_gn=RPH3A;refGene_tx=XM_006938490.4;refGene_gn=RPH3A;refGene_tx=XM_019814556.2;refGene_gn=RPH3A;refGene_tx=XM_019814557.2;refGene_gn=RPH3A;refGene_tx=XM_019814558.2;refGene_gn=RPH3A;refGene_tx=XM_019814559.2;refGene_gn=RPH3A;refGene_tx=XM_019814560.2;refGene_gn=RPH3A;refGene_tx=XM_023241438.1
D3 10663594 10663749
refGene_gn=RPH3A;refGene_tx=XM_006938490.4;refGene_gn=RPH3A;refGene_tx=XM_019814556.2;refGene_gn=RPH3A;refGene_tx=XM_019814557.2;refGene_gn=RPH3A;refGene_tx=XM_019814558.2;refGene_gn=RPH3A;refGene_tx=XM_019814559.2;refGene_gn=RPH3A;refGene_tx=XM_019814560.2;refGene_gn=RPH3A;refGene_tx=XM_023241438.1
D3 10667357 10667439
refGene_gn=RPH3A;refGene_tx=XM_006938490.4;refGene_gn=RPH3A;refGene_tx=XM_019814556.2;refGene_gn=RPH3A;refGene_tx=XM_019814557.2;refGene_gn=RPH3A;refGene_tx=XM_019814558.2;refGene_gn=RPH3A;refGene_tx=XM_019814559.2;refGene_gn=RPH3A;refGene_tx=XM_019814560.2;refGene_gn=RPH3A;refGene_tx=XM_023241438.1
D3 10671069 10671166
refGene_gn=RPH3A;refGene_tx=XM_006938490.4;refGene_gn=RPH3A;refGene_tx=XM_019814556.2;refGene_gn=RPH3A;refGene_tx=XM_019814557.2;refGene_gn=RPH3A;refGene_tx=XM_019814558.2;refGene_gn=RPH3A;refGene_tx=XM_019814559.2;refGene_gn=RPH3A;refGene_tx=XM_019814560.2;refGene_gn=RPH3A;refGene_tx=XM_023241438.1

D3 10671362 10671427 refGene_gn=RPH3A;refGene_tx=XM_023241438.1
D3 10671956 10672087
refGene_gn=RPH3A;refGene_tx=XM_006938490.4;refGene_gn=RPH3A;refGene_tx=XM_019814556.2;refGene_gn=RPH3A;refGene_tx=XM_019814557.2;refGene_gn=RPH3A;refGene_tx=XM_019814558.2;refGene_gn=RPH3A;refGene_tx=XM_019814559.2;refGene_gn=RPH3A;refGene_tx=XM_019814560.2
D3 10679713 10679899 refGene_gn=OAS1;refGene_tx=XM_006938491.3
D3 10682444 10682733 refGene_gn=OAS1;refGene_tx=XM_006938491.3
D3 10685067 10685252 refGene_gn=OAS1;refGene_tx=XM_006938491.3
D3 10689433 10689666 refGene_gn=OAS1;refGene_tx=XM_006938491.3
D3 10690425 10690579 refGene_gn=OAS1;refGene_tx=XM_006938491.3
D3 10691984 10692143 refGene_gn=OAS1;refGene_tx=XM_006938491.3
D3 10698759 10699098 refGene_gn=OAS3;refGene_tx=XM_023241439.1
D3 10702032 10702315 refGene_gn=OAS3;refGene_tx=XM_023241439.1
D3 10703904 10704080 refGene_gn=OAS3;refGene_tx=XM_023241439.1
D3 10705370 10705609 refGene_gn=OAS3;refGene_tx=XM_023241439.1
D3 10706892 10707046 refGene_gn=OAS3;refGene_tx=XM_023241439.1
D3 10707468 10707821 refGene_gn=OAS3;refGene_tx=XM_023241439.1
D3 10708915 10709199 refGene_gn=OAS3;refGene_tx=XM_023241439.1
D3 10709535 10709711 refGene_gn=OAS3;refGene_tx=XM_023241439.1
D3 10710527 10710778 refGene_gn=OAS3;refGene_tx=XM_023241439.1
D3 10711183 10711328 refGene_gn=OAS3;refGene_tx=XM_023241439.1
D3 10712036 10712210 refGene_gn=OAS3;refGene_tx=XM_023241439.1
D3 10713535 10713821 refGene_gn=OAS3;refGene_tx=XM_023241439.1
D3 10714889 10715065 refGene_gn=OAS3;refGene_tx=XM_023241439.1
D3 10715383 10715622 refGene_gn=OAS3;refGene_tx=XM_023241439.1
D3 10716386 10716534 refGene_gn=OAS3;refGene_tx=XM_023241439.1
D3 10716709 10716721 refGene_gn=OAS3;refGene_tx=XM_023241439.1
D3 10723021 10723198 refGene_gn=OAS2;refGene_tx=XM_006938494.4
D3 10724880 10725151 refGene_gn=OAS2;refGene_tx=XM_006938494.4
D3 10728804 10728983 refGene_gn=OAS2;refGene_tx=XM_006938494.4
D3 10730000 10730236 refGene_gn=OAS2;refGene_tx=XM_006938494.4
D3 10730943 10731085 refGene_gn=OAS2;refGene_tx=XM_006938494.4
D3 10732596 10732770 refGene_gn=OAS2;refGene_tx=XM_006938494.4
D3 10735029 10735312 refGene_gn=OAS2;refGene_tx=XM_006938494.4
D3 10737719 10737907 refGene_gn=OAS2;refGene_tx=XM_006938494.4
D3 10738973 10739212 refGene_gn=OAS2;refGene_tx=XM_006938494.4
D3 10740366 10740520 refGene_gn=OAS2;refGene_tx=XM_006938494.4
D3 10741128 10741212 refGene_gn=OAS2;refGene_tx=XM_006938494.4
D3 10762519 10762778 refGene_gn=DTX1;refGene_tx=XM_023241443.1
D3 10778595 10779295 refGene_gn=DTX1;refGene_tx=XM_023241443.1
D3 10791668 10791730 refGene_gn=DTX1;refGene_tx=XM_023241443.1
D3 10792090 10792252 refGene_gn=DTX1;refGene_tx=XM_023241443.1
D3 10792523 10792585 refGene_gn=DTX1;refGene_tx=XM_023241443.1
D3 10793065 10793224 refGene_gn=DTX1;refGene_tx=XM_023241443.1
D3 10793300 10793462 refGene_gn=DTX1;refGene_tx=XM_023241443.1
D3 10793559 10793649 refGene_gn=DTX1;refGene_tx=XM_023241443.1
D3 10794290 10794515 refGene_gn=DTX1;refGene_tx=XM_023241443.1
D3 10797113 10797253
refGene_gn=RASAL1;refGene_tx=XM_023241440.1;refGene_gn=RASAL1;refGene_

tx=XM_023241441.1;refGene_gn=RASAL1;refGene_tx=XM_023241442.1
D3	10797513	10797590
refGene_gn=RASAL1;refGene_tx=XM_023241440.1;refGene_gn=RASAL1;refGene_tx=XM_023241441.1;refGene_gn=RASAL1;refGene_tx=XM_023241442.1
D3	10798448	10798569
refGene_gn=RASAL1;refGene_tx=XM_023241440.1;refGene_gn=RASAL1;refGene_tx=XM_023241441.1;refGene_gn=RASAL1;refGene_tx=XM_023241442.1
D3	10799354	10799490
refGene_gn=RASAL1;refGene_tx=XM_023241440.1;refGene_gn=RASAL1;refGene_tx=XM_023241441.1;refGene_gn=RASAL1;refGene_tx=XM_023241442.1
D3	10799580	10799718
refGene_gn=RASAL1;refGene_tx=XM_023241440.1;refGene_gn=RASAL1;refGene_tx=XM_023241441.1;refGene_gn=RASAL1;refGene_tx=XM_023241442.1
D3	10800676	10800849
refGene_gn=RASAL1;refGene_tx=XM_023241440.1;refGene_gn=RASAL1;refGene_tx=XM_023241441.1;refGene_gn=RASAL1;refGene_tx=XM_023241442.1
D3	10802013	10802155
refGene_gn=RASAL1;refGene_tx=XM_023241440.1;refGene_gn=RASAL1;refGene_tx=XM_023241441.1;refGene_gn=RASAL1;refGene_tx=XM_023241442.1
D3	10802951	10803089
refGene_gn=RASAL1;refGene_tx=XM_023241440.1;refGene_gn=RASAL1;refGene_tx=XM_023241441.1;refGene_gn=RASAL1;refGene_tx=XM_023241442.1
D3	10806091	10806284
refGene_gn=RASAL1;refGene_tx=XM_023241440.1;refGene_gn=RASAL1;refGene_tx=XM_023241441.1;refGene_gn=RASAL1;refGene_tx=XM_023241442.1
D3	10807267	10807380
refGene_gn=RASAL1;refGene_tx=XM_023241440.1;refGene_gn=RASAL1;refGene_tx=XM_023241441.1;refGene_gn=RASAL1;refGene_tx=XM_023241442.1
D3	10807663	10807728
refGene_gn=RASAL1;refGene_tx=XM_023241440.1;refGene_gn=RASAL1;refGene_tx=XM_023241441.1;refGene_gn=RASAL1;refGene_tx=XM_023241442.1
D3	10808101	10808255
refGene_gn=RASAL1;refGene_tx=XM_023241440.1;refGene_gn=RASAL1;refGene_tx=XM_023241441.1;refGene_gn=RASAL1;refGene_tx=XM_023241442.1
D3	10808377	10808495
refGene_gn=RASAL1;refGene_tx=XM_023241440.1;refGene_gn=RASAL1;refGene_tx=XM_023241441.1;refGene_gn=RASAL1;refGene_tx=XM_023241442.1
D3	10809188	10809277
refGene_gn=RASAL1;refGene_tx=XM_023241440.1;refGene_gn=RASAL1;refGene_tx=XM_023241441.1;refGene_gn=RASAL1;refGene_tx=XM_023241442.1
D3	10810942	10811074
refGene_gn=RASAL1;refGene_tx=XM_023241440.1;refGene_gn=RASAL1;refGene_tx=XM_023241441.1;refGene_gn=RASAL1;refGene_tx=XM_023241442.1
D3	10811154	10811604
refGene_gn=RASAL1;refGene_tx=XM_023241440.1;refGene_gn=RASAL1;refGene_tx=XM_023241441.1;refGene_gn=RASAL1;refGene_tx=XM_023241442.1
D3	10813293	10813423
refGene_gn=RASAL1;refGene_tx=XM_023241440.1;refGene_gn=RASAL1;refGene_tx=XM_023241441.1
D3	10818296	10818358

refGene_gn=RASAL1;refGene_tx=XM_023241440.1;refGene_gn=RASAL1;refGene_tx=XM_023241441.1
D3 10818496 10818792
refGene_gn=RASAL1;refGene_tx=XM_023241440.1;refGene_gn=RASAL1;refGene_tx=XM_023241441.1
D3 10820760 10820817 refGene_gn=RASAL1;refGene_tx=XM_023241441.1
D3 10825025 10825090 refGene_gn=RASAL1;refGene_tx=XM_023241441.1
D3 10835592 10835648
refGene_gn=CFAP73;refGene_tx=XM_019814575.2;refGene_gn=CFAP73;refGene_tx=XM_023241444.1;refGene_gn=CFAP73;refGene_tx=XM_023241445.1
D3 10837512 10837627
refGene_gn=CFAP73;refGene_tx=XM_019814575.2;refGene_gn=CFAP73;refGene_tx=XM_023241444.1;refGene_gn=CFAP73;refGene_tx=XM_023241445.1
D3 10838144 10838249
refGene_gn=CFAP73;refGene_tx=XM_019814575.2;refGene_gn=CFAP73;refGene_tx=XM_023241444.1;refGene_gn=CFAP73;refGene_tx=XM_023241445.1
D3 10838580 10838781
refGene_gn=CFAP73;refGene_tx=XM_019814575.2;refGene_gn=CFAP73;refGene_tx=XM_023241444.1;refGene_gn=CFAP73;refGene_tx=XM_023241445.1
D3 10839348 10839570
refGene_gn=CFAP73;refGene_tx=XM_019814575.2;refGene_gn=CFAP73;refGene_tx=XM_023241444.1;refGene_gn=CFAP73;refGene_tx=XM_023241445.1
D3 10839879 10840116
refGene_gn=CFAP73;refGene_tx=XM_019814575.2;refGene_gn=CFAP73;refGene_tx=XM_023241444.1;refGene_gn=CFAP73;refGene_tx=XM_023241445.1
D3 10842707 10842940 refGene_gn=DDX54;refGene_tx=XM_011287868.3
D3 10844873 10844986 refGene_gn=DDX54;refGene_tx=XM_011287868.3
D3 10845211 10845316 refGene_gn=DDX54;refGene_tx=XM_011287868.3
D3 10845982 10846096 refGene_gn=DDX54;refGene_tx=XM_011287868.3
D3 10846185 10846328 refGene_gn=DDX54;refGene_tx=XM_011287868.3
D3 10847413 10847620 refGene_gn=DDX54;refGene_tx=XM_011287868.3
D3 10848341 10848415 refGene_gn=DDX54;refGene_tx=XM_011287868.3
D3 10848506 10848737 refGene_gn=DDX54;refGene_tx=XM_011287868.3
D3 10850164 10850299 refGene_gn=DDX54;refGene_tx=XM_011287868.3
D3 10852441 10852652 refGene_gn=DDX54;refGene_tx=XM_011287868.3
D3 10853991 10854123 refGene_gn=DDX54;refGene_tx=XM_011287868.3
D3 10854203 10854265 refGene_gn=DDX54;refGene_tx=XM_011287868.3
D3 10854385 10854507 refGene_gn=DDX54;refGene_tx=XM_011287868.3
D3 10855169 10855265 refGene_gn=DDX54;refGene_tx=XM_011287868.3
D3 10855389 10855431 refGene_gn=DDX54;refGene_tx=XM_011287868.3
D3 10856198 10856248 refGene_gn=DDX54;refGene_tx=XM_011287868.3
D3 10856350 10856539 refGene_gn=DDX54;refGene_tx=XM_011287868.3
D3 10857111 10857182 refGene_gn=DDX54;refGene_tx=XM_011287868.3
D3 10858585 10858715 refGene_gn=DDX54;refGene_tx=XM_011287868.3
D3 10861530 10861704 refGene_gn=DDX54;refGene_tx=XM_011287868.3
D3 10862964 10863275 refGene_gn=RITA1;refGene_tx=XM_006938497.4
D3 10865658 10866166 refGene_gn=RITA1;refGene_tx=XM_006938497.4
D3 10867830 10868136
refGene_gn=IQCD;refGene_tx=XM_006938495.4;refGene_gn=IQCD;refGene_tx=XM_006938496.4

D3 10868920 10868992
refGene_gn=IQCD;refGene_tx=XM_006938495.4;refGene_gn=IQCD;refGene_tx=XM_006938496.4
D3 10870858 10871092
refGene_gn=IQCD;refGene_tx=XM_006938495.4;refGene_gn=IQCD;refGene_tx=XM_006938496.4
D3 10873243 10873972
refGene_gn=IQCD;refGene_tx=XM_006938495.4;refGene_gn=IQCD;refGene_tx=XM_006938496.4
D3 10894767 10894879 refGene_gn=TPCN1;refGene_tx=XM_023241446.1
D3 10922810 10922938 refGene_gn=TPCN1;refGene_tx=XM_023241446.1
D3 10927998 10928175 refGene_gn=TPCN1;refGene_tx=XM_023241446.1
D3 10928563 10928677 refGene_gn=TPCN1;refGene_tx=XM_023241446.1
D3 10929479 10929610 refGene_gn=TPCN1;refGene_tx=XM_023241446.1
D3 10930488 10930577 refGene_gn=TPCN1;refGene_tx=XM_023241446.1
D3 10932591 10932626 refGene_gn=TPCN1;refGene_tx=XM_023241446.1
D3 10933108 10933167 refGene_gn=TPCN1;refGene_tx=XM_023241446.1
D3 10933439 10933539 refGene_gn=TPCN1;refGene_tx=XM_023241446.1
D3 10935453 10935570 refGene_gn=TPCN1;refGene_tx=XM_023241446.1
D3 10935824 10935949 refGene_gn=TPCN1;refGene_tx=XM_023241446.1
D3 10936797 10936846 refGene_gn=TPCN1;refGene_tx=XM_023241446.1
D3 10937381 10937445 refGene_gn=TPCN1;refGene_tx=XM_023241446.1
D3 10938662 10938707 refGene_gn=TPCN1;refGene_tx=XM_023241446.1
D3 10941434 10941491 refGene_gn=TPCN1;refGene_tx=XM_023241446.1
D3 10941572 10941640 refGene_gn=TPCN1;refGene_tx=XM_023241446.1
D3 10942280 10942354 refGene_gn=TPCN1;refGene_tx=XM_023241446.1
D3 10943317 10943425 refGene_gn=TPCN1;refGene_tx=XM_023241446.1
D3 10944252 10944324 refGene_gn=TPCN1;refGene_tx=XM_023241446.1
D3 10944869 10944959 refGene_gn=TPCN1;refGene_tx=XM_023241446.1
D3 10946281 10946397 refGene_gn=TPCN1;refGene_tx=XM_023241446.1
D3 10947161 10947208 refGene_gn=TPCN1;refGene_tx=XM_023241446.1
D3 10947827 10947896 refGene_gn=TPCN1;refGene_tx=XM_023241446.1
D3 10948070 10948155 refGene_gn=TPCN1;refGene_tx=XM_023241446.1
D3 10948960 10949100 refGene_gn=TPCN1;refGene_tx=XM_023241446.1
D3 10949220 10949301 refGene_gn=TPCN1;refGene_tx=XM_023241446.1
D3 10951337 10951460 refGene_gn=TPCN1;refGene_tx=XM_023241446.1
D3 10954951 10955149 refGene_gn=SLC8B1;refGene_tx=XM_019814572.2
D3 10957143 10957194 refGene_gn=SLC8B1;refGene_tx=XM_019814573.2
D3 10957278 10957343
refGene_gn=SLC8B1;refGene_tx=XM_019814572.2;refGene_gn=SLC8B1;refGene_tx=XM_019814573.2
D3 10958768 10958849
refGene_gn=SLC8B1;refGene_tx=XM_019814572.2;refGene_gn=SLC8B1;refGene_tx=XM_019814573.2
D3 10959537 10959691
refGene_gn=SLC8B1;refGene_tx=XM_019814572.2;refGene_gn=SLC8B1;refGene_tx=XM_019814573.2
D3 10961190 10961312
refGene_gn=SLC8B1;refGene_tx=XM_019814572.2;refGene_gn=SLC8B1;refGene_tx=XM_019814573.2

D3 10963598 10963740
refGene_gn=SLC8B1;refGene_tx=XM_019814572.2;refGene_gn=SLC8B1;refGene_tx=XM_019814573.2
D3 10966564 10966695
refGene_gn=SLC8B1;refGene_tx=XM_019814572.2;refGene_gn=SLC8B1;refGene_tx=XM_019814573.2
D3 10966800 10966854
refGene_gn=SLC8B1;refGene_tx=XM_019814572.2;refGene_gn=SLC8B1;refGene_tx=XM_019814573.2
D3 10968057 10968165
refGene_gn=SLC8B1;refGene_tx=XM_019814572.2;refGene_gn=SLC8B1;refGene_tx=XM_019814573.2
D3 10969099 10969267
refGene_gn=SLC8B1;refGene_tx=XM_019814572.2;refGene_gn=SLC8B1;refGene_tx=XM_019814573.2
D3 10969345 10969451
refGene_gn=SLC8B1;refGene_tx=XM_019814572.2;refGene_gn=SLC8B1;refGene_tx=XM_019814573.2
D3 10969605 10969663
refGene_gn=SLC8B1;refGene_tx=XM_019814572.2;refGene_gn=SLC8B1;refGene_tx=XM_019814573.2
D3 10969813 10969866
refGene_gn=SLC8B1;refGene_tx=XM_019814572.2;refGene_gn=SLC8B1;refGene_tx=XM_019814573.2
D3 10969957 10970110
refGene_gn=SLC8B1;refGene_tx=XM_019814572.2;refGene_gn=SLC8B1;refGene_tx=XM_019814573.2
D3 10973555 10973660
refGene_gn=SLC8B1;refGene_tx=XM_019814572.2;refGene_gn=SLC8B1;refGene_tx=XM_019814573.2
D3 10975556 10975724
refGene_gn=SLC8B1;refGene_tx=XM_019814572.2;refGene_gn=SLC8B1;refGene_tx=XM_019814573.2
D3 10993326 10993616 refGene_gn=PLBD2;refGene_tx=XM_011287867.3
D3 11000530 11000624 refGene_gn=PLBD2;refGene_tx=XM_011287867.3
D3 11003405 11003564 refGene_gn=PLBD2;refGene_tx=XM_011287867.3
D3 11005464 11005565 refGene_gn=PLBD2;refGene_tx=XM_011287867.3
D3 11005780 11005995 refGene_gn=PLBD2;refGene_tx=XM_011287867.3
D3 11010474 11010572 refGene_gn=PLBD2;refGene_tx=XM_011287867.3
D3 11012357 11012518 refGene_gn=PLBD2;refGene_tx=XM_011287867.3
D3 11013138 11013234 refGene_gn=PLBD2;refGene_tx=XM_011287867.3
D3 11013481 11013553 refGene_gn=PLBD2;refGene_tx=XM_011287867.3
D3 11014197 11014350 refGene_gn=PLBD2;refGene_tx=XM_011287867.3
D3 11014927 11015090 refGene_gn=PLBD2;refGene_tx=XM_011287867.3
D3 11015310 11015478 refGene_gn=PLBD2;refGene_tx=XM_011287867.3
D3 11018072 11018281 refGene_gn=SDS;refGene_tx=XM_003994691.4
D3 11018914 11018975 refGene_gn=SDS;refGene_tx=XM_003994691.4
D3 11018976 11019040 refGene_gn=SDS;refGene_tx=XM_003994691.4
D3 11020815 11021043 refGene_gn=SDS;refGene_tx=XM_003994691.4
D3 11021281 11021373 refGene_gn=SDS;refGene_tx=XM_003994691.4

D3 11022205 11022345 refGene_gn=SDS;refGene_tx=XM_003994691.4
D3 11022589 11022629 refGene_gn=SDS;refGene_tx=XM_003994691.4
D3 11022924 11023077 refGene_gn=SDS;refGene_tx=XM_003994691.4
D3 11040183 11040357
refGene_gn=SDSL;refGene_tx=XM_003994699.4;refGene_gn=SDSL;refGene_tx=XM_019814576.1;refGene_gn=SDSL;refGene_tx=XM_019814577.1
D3 11040588 11040628
refGene_gn=SDSL;refGene_tx=XM_003994699.4;refGene_gn=SDSL;refGene_tx=XM_019814576.1;refGene_gn=SDSL;refGene_tx=XM_019814577.1
D3 11041164 11041203
refGene_gn=SDSL;refGene_tx=XM_019814576.1;refGene_gn=SDSL;refGene_tx=XM_019814577.1
D3 11041316 11041456
refGene_gn=SDSL;refGene_tx=XM_003994699.4;refGene_gn=SDSL;refGene_tx=XM_019814576.1;refGene_gn=SDSL;refGene_tx=XM_019814577.1
D3 11043607 11043696
refGene_gn=SDSL;refGene_tx=XM_003994699.4;refGene_gn=SDSL;refGene_tx=XM_019814576.1;refGene_gn=SDSL;refGene_tx=XM_019814577.1
D3 11044406 11044634
refGene_gn=SDSL;refGene_tx=XM_003994699.4;refGene_gn=SDSL;refGene_tx=XM_019814576.1;refGene_gn=SDSL;refGene_tx=XM_019814577.1
D3 11047813 11047938
refGene_gn=SDSL;refGene_tx=XM_003994699.4;refGene_gn=SDSL;refGene_tx=XM_019814576.1;refGene_gn=SDSL;refGene_tx=XM_019814577.1
D3 11049106 11049300
refGene_gn=SDSL;refGene_tx=XM_003994699.4;refGene_gn=SDSL;refGene_tx=XM_019814576.1;refGene_gn=SDSL;refGene_tx=XM_019814577.1
D3 11066412 11066780 refGene_gn=LHX5;refGene_tx=XM_023241192.1
D3 11070499 11070665 refGene_gn=LHX5;refGene_tx=XM_023241192.1
D3 11071355 11071633 refGene_gn=LHX5;refGene_tx=XM_023241192.1
D3 11072345 11072569 refGene_gn=LHX5;refGene_tx=XM_023241192.1
D3 11074614 11074787 refGene_gn=LHX5;refGene_tx=XM_023241192.1
D3 11305225 11305294 refGene_gn=RBM19;refGene_tx=XM_019814579.2
D3 11314886 11314898 refGene_gn=RBM19;refGene_tx=XM_006938504.4
D3 11329723 11329729 refGene_gn=RBM19;refGene_tx=XM_019814580.2
D3 11330340 11330453
refGene_gn=RBM19;refGene_tx=XM_006938504.4;refGene_gn=RBM19;refGene_tx=XM_019814579.2;refGene_gn=RBM19;refGene_tx=XM_019814580.2
D3 11349187 11349308
refGene_gn=RBM19;refGene_tx=XM_006938504.4;refGene_gn=RBM19;refGene_tx=XM_019814579.2;refGene_gn=RBM19;refGene_tx=XM_019814580.2
D3 11361791 11361897
refGene_gn=RBM19;refGene_tx=XM_006938504.4;refGene_gn=RBM19;refGene_tx=XM_019814579.2;refGene_gn=RBM19;refGene_tx=XM_019814580.2
D3 11374662 11375637
refGene_gn=LOC111557270;refGene_tx=XM_023241447.1
D3 11408096 11408213
refGene_gn=RBM19;refGene_tx=XM_006938504.4;refGene_gn=RBM19;refGene_tx=XM_019814579.2;refGene_gn=RBM19;refGene_tx=XM_019814580.2
D3 11411327 11411383

refGene_gn=RBM19;refGene_tx=XM_006938504.4;refGene_gn=RBM19;refGene_tx=XM_019814579.2;refGene_gn=RBM19;refGene_tx=XM_019814580.2
D3 11413921 11414001
refGene_gn=RBM19;refGene_tx=XM_006938504.4;refGene_gn=RBM19;refGene_tx=XM_019814579.2;refGene_gn=RBM19;refGene_tx=XM_019814580.2
D3 11417363 11417424
refGene_gn=RBM19;refGene_tx=XM_006938504.4;refGene_gn=RBM19;refGene_tx=XM_019814579.2;refGene_gn=RBM19;refGene_tx=XM_019814580.2
D3 11419254 11419427
refGene_gn=RBM19;refGene_tx=XM_006938504.4;refGene_gn=RBM19;refGene_tx=XM_019814579.2;refGene_gn=RBM19;refGene_tx=XM_019814580.2
D3 11427867 11427997
refGene_gn=RBM19;refGene_tx=XM_006938504.4;refGene_gn=RBM19;refGene_tx=XM_019814579.2;refGene_gn=RBM19;refGene_tx=XM_019814580.2
D3 11429255 11429456
refGene_gn=RBM19;refGene_tx=XM_006938504.4;refGene_gn=RBM19;refGene_tx=XM_019814579.2;refGene_gn=RBM19;refGene_tx=XM_019814580.2
D3 11431468 11431579
refGene_gn=RBM19;refGene_tx=XM_006938504.4;refGene_gn=RBM19;refGene_tx=XM_019814579.2;refGene_gn=RBM19;refGene_tx=XM_019814580.2
D3 11433384 11433481
refGene_gn=RBM19;refGene_tx=XM_006938504.4;refGene_gn=RBM19;refGene_tx=XM_019814579.2;refGene_gn=RBM19;refGene_tx=XM_019814580.2
D3 11433857 11433979
refGene_gn=RBM19;refGene_tx=XM_006938504.4;refGene_gn=RBM19;refGene_tx=XM_019814579.2;refGene_gn=RBM19;refGene_tx=XM_019814580.2
D3 11435010 11435141
refGene_gn=RBM19;refGene_tx=XM_006938504.4;refGene_gn=RBM19;refGene_tx=XM_019814579.2;refGene_gn=RBM19;refGene_tx=XM_019814580.2
D3 11436449 11436653
refGene_gn=RBM19;refGene_tx=XM_006938504.4;refGene_gn=RBM19;refGene_tx=XM_019814579.2;refGene_gn=RBM19;refGene_tx=XM_019814580.2
D3 11437589 11437661
refGene_gn=RBM19;refGene_tx=XM_006938504.4;refGene_gn=RBM19;refGene_tx=XM_019814579.2;refGene_gn=RBM19;refGene_tx=XM_019814580.2
D3 11439817 11439896
refGene_gn=RBM19;refGene_tx=XM_006938504.4;refGene_gn=RBM19;refGene_tx=XM_019814579.2;refGene_gn=RBM19;refGene_tx=XM_019814580.2
D3 11448114 11448195
refGene_gn=RBM19;refGene_tx=XM_006938504.4;refGene_gn=RBM19;refGene_tx=XM_019814579.2;refGene_gn=RBM19;refGene_tx=XM_019814580.2
D3 11449786 11450028
refGene_gn=RBM19;refGene_tx=XM_006938504.4;refGene_gn=RBM19;refGene_tx=XM_019814579.2;refGene_gn=RBM19;refGene_tx=XM_019814580.2
D3 11451101 11451294
refGene_gn=RBM19;refGene_tx=XM_006938504.4;refGene_gn=RBM19;refGene_tx=XM_019814579.2;refGene_gn=RBM19;refGene_tx=XM_019814580.2
D3 11451595 11451634
refGene_gn=RBM19;refGene_tx=XM_006938504.4;refGene_gn=RBM19;refGene_tx=XM_019814579.2;refGene_gn=RBM19;refGene_tx=XM_019814580.2

D3 11451832 11451958
refGene_gn=RBM19;refGene_tx=XM_006938504.4;refGene_gn=RBM19;refGene_tx=XM_019814579.2;refGene_gn=RBM19;refGene_tx=XM_019814580.2
D3 11453617 11453800
refGene_gn=RBM19;refGene_tx=XM_006938504.4;refGene_gn=RBM19;refGene_tx=XM_019814579.2;refGene_gn=RBM19;refGene_tx=XM_019814580.2
D3 11457647 11457683
refGene_gn=RBM19;refGene_tx=XM_006938504.4;refGene_gn=RBM19;refGene_tx=XM_019814579.2;refGene_gn=RBM19;refGene_tx=XM_019814580.2
D3 11766533 11767111
refGene_gn=TBX5;refGene_tx=XM_006938507.4;refGene_gn=TBX5;refGene_tx=XM_011287807.3;refGene_gn=TBX5;refGene_tx=XM_011287808.3;refGene_gn=TBX5;refGene_tx=XM_011287809.3;refGene_gn=TBX5;refGene_tx=XM_019814582.2;refGene_gn=TBX5;refGene_tx=XM_019814583.2;refGene_gn=TBX5;refGene_tx=XM_019814585.2
D3 11775931 11776158
refGene_gn=TBX5;refGene_tx=XM_006938507.4;refGene_gn=TBX5;refGene_tx=XM_011287807.3;refGene_gn=TBX5;refGene_tx=XM_011287808.3;refGene_gn=TBX5;refGene_tx=XM_011287809.3;refGene_gn=TBX5;refGene_tx=XM_019814582.2;refGene_gn=TBX5;refGene_tx=XM_019814583.2;refGene_gn=TBX5;refGene_tx=XM_019814585.2
D3 11792045 11792137
refGene_gn=TBX5;refGene_tx=XM_006938507.4;refGene_gn=TBX5;refGene_tx=XM_011287807.3;refGene_gn=TBX5;refGene_tx=XM_011287808.3;refGene_gn=TBX5;refGene_tx=XM_011287809.3;refGene_gn=TBX5;refGene_tx=XM_019814582.2;refGene_gn=TBX5;refGene_tx=XM_019814583.2;refGene_gn=TBX5;refGene_tx=XM_019814585.2
D3 11800318 11800471
refGene_gn=TBX5;refGene_tx=XM_006938507.4;refGene_gn=TBX5;refGene_tx=XM_011287807.3;refGene_gn=TBX5;refGene_tx=XM_011287808.3;refGene_gn=TBX5;refGene_tx=XM_011287809.3;refGene_gn=TBX5;refGene_tx=XM_019814582.2;refGene_gn=TBX5;refGene_tx=XM_019814583.2;refGene_gn=TBX5;refGene_tx=XM_019814585.2
D3 11803951 11804099
refGene_gn=TBX5;refGene_tx=XM_006938507.4;refGene_gn=TBX5;refGene_tx=XM_011287807.3;refGene_gn=TBX5;refGene_tx=XM_011287808.3;refGene_gn=TBX5;refGene_tx=XM_011287809.3;refGene_gn=TBX5;refGene_tx=XM_019814582.2;refGene_gn=TBX5;refGene_tx=XM_019814583.2;refGene_gn=TBX5;refGene_tx=XM_019814585.2
D3 11804892 11805012
refGene_gn=TBX5;refGene_tx=XM_006938507.4;refGene_gn=TBX5;refGene_tx=XM_011287807.3;refGene_gn=TBX5;refGene_tx=XM_011287808.3;refGene_gn=TBX5;refGene_tx=XM_011287809.3;refGene_gn=TBX5;refGene_tx=XM_019814582.2;refGene_gn=TBX5;refGene_tx=XM_019814583.2;refGene_gn=TBX5;refGene_tx=XM_019814585.2
D3 11807177 11807272
refGene_gn=TBX5;refGene_tx=XM_006938507.4;refGene_gn=TBX5;refGene_tx=XM_011287807.3;refGene_gn=TBX5;refGene_tx=XM_011287808.3;refGene_gn=TBX5;refGene_tx=XM_011287809.3;refGene_gn=TBX5;refGene_tx=XM_019814582.2;refGene_gn=TBX5;refGene_tx=XM_019814583.2;refGene_gn=TBX5;refGene_tx=X

M_019814585.2
D3 11808932 11809079
refGene_gn=TBX5;refGene_tx=XM_006938507.4;refGene_gn=TBX5;refGene_tx=XM_011287807.3;refGene_gn=TBX5;refGene_tx=XM_011287808.3;refGene_gn=TBX5;refGene_tx=XM_011287809.3;refGene_gn=TBX5;refGene_tx=XM_019814582.2;refGene_gn=TBX5;refGene_tx=XM_019814583.2;refGene_gn=TBX5;refGene_tx=XM_019814585.2
D3 11988509 11988971
refGene_gn=TBX3;refGene_tx=XM_023241448.1;refGene_gn=TBX3;refGene_tx=XM_023241449.1;refGene_gn=TBX3;refGene_tx=XM_023241450.1
D3 11990619 11991296
refGene_gn=TBX3;refGene_tx=XM_023241448.1;refGene_gn=TBX3;refGene_tx=XM_023241449.1;refGene_gn=TBX3;refGene_tx=XM_023241450.1
D3 11992764 11992922
refGene_gn=TBX3;refGene_tx=XM_023241448.1;refGene_gn=TBX3;refGene_tx=XM_023241449.1;refGene_gn=TBX3;refGene_tx=XM_023241450.1
D3 11994013 11994090
refGene_gn=TBX3;refGene_tx=XM_023241448.1;refGene_gn=TBX3;refGene_tx=XM_023241449.1;refGene_gn=TBX3;refGene_tx=XM_023241450.1
D3 11995980 11996127
refGene_gn=TBX3;refGene_tx=XM_023241448.1;refGene_gn=TBX3;refGene_tx=XM_023241449.1;refGene_gn=TBX3;refGene_tx=XM_023241450.1
D3 11996421 11996481
refGene_gn=TBX3;refGene_tx=XM_023241448.1;refGene_gn=TBX3;refGene_tx=XM_023241449.1
D3 11997390 11997658
refGene_gn=TBX3;refGene_tx=XM_023241448.1;refGene_gn=TBX3;refGene_tx=XM_023241449.1;refGene_gn=TBX3;refGene_tx=XM_023241450.1
D3 11999284 11999673
refGene_gn=TBX3;refGene_tx=XM_023241448.1;refGene_gn=TBX3;refGene_tx=XM_023241449.1;refGene_gn=TBX3;refGene_tx=XM_023241450.1
D3 13073769 13073902
refGene_gn=MED13L;refGene_tx=XM_023241451.1;refGene_gn=MED13L;refGene_tx=XM_023241452.1
D3 13074963 13075076
refGene_gn=MED13L;refGene_tx=XM_023241451.1;refGene_gn=MED13L;refGene_tx=XM_023241452.1
D3 13077164 13077326
refGene_gn=MED13L;refGene_tx=XM_023241451.1;refGene_gn=MED13L;refGene_tx=XM_023241452.1
D3 13079980 13080138
refGene_gn=MED13L;refGene_tx=XM_023241451.1;refGene_gn=MED13L;refGene_tx=XM_023241452.1
D3 13081354 13081531
refGene_gn=MED13L;refGene_tx=XM_023241451.1;refGene_gn=MED13L;refGene_tx=XM_023241452.1
D3 13083068 13083227
refGene_gn=MED13L;refGene_tx=XM_023241451.1;refGene_gn=MED13L;refGene_tx=XM_023241452.1
D3 13086357 13086500

refGene_gn=MED13L;refGene_tx=XM_023241451.1;refGene_gn=MED13L;refGene_tx=XM_023241452.1
D3 13086698 13086922
refGene_gn=MED13L;refGene_tx=XM_023241451.1;refGene_gn=MED13L;refGene_tx=XM_023241452.1
D3 13089975 13090164
refGene_gn=MED13L;refGene_tx=XM_023241451.1;refGene_gn=MED13L;refGene_tx=XM_023241452.1
D3 13091631 13091851
refGene_gn=MED13L;refGene_tx=XM_023241451.1;refGene_gn=MED13L;refGene_tx=XM_023241452.1
D3 13092404 13092837
refGene_gn=MED13L;refGene_tx=XM_023241451.1;refGene_gn=MED13L;refGene_tx=XM_023241452.1
D3 13093499 13093692
refGene_gn=MED13L;refGene_tx=XM_023241451.1;refGene_gn=MED13L;refGene_tx=XM_023241452.1
D3 13095577 13095801
refGene_gn=MED13L;refGene_tx=XM_023241451.1;refGene_gn=MED13L;refGene_tx=XM_023241452.1
D3 13096401 13096581
refGene_gn=MED13L;refGene_tx=XM_023241451.1;refGene_gn=MED13L;refGene_tx=XM_023241452.1
D3 13100510 13101448
refGene_gn=MED13L;refGene_tx=XM_023241451.1;refGene_gn=MED13L;refGene_tx=XM_023241452.1
D3 13104943 13105149
refGene_gn=MED13L;refGene_tx=XM_023241451.1;refGene_gn=MED13L;refGene_tx=XM_023241452.1
D3 13105454 13105675
refGene_gn=MED13L;refGene_tx=XM_023241451.1;refGene_gn=MED13L;refGene_tx=XM_023241452.1
D3 13111326 13111426
refGene_gn=MED13L;refGene_tx=XM_023241451.1;refGene_gn=MED13L;refGene_tx=XM_023241452.1
D3 13114804 13114929
refGene_gn=MED13L;refGene_tx=XM_023241451.1;refGene_gn=MED13L;refGene_tx=XM_023241452.1
D3 13115236 13115342
refGene_gn=MED13L;refGene_tx=XM_023241451.1;refGene_gn=MED13L;refGene_tx=XM_023241452.1
D3 13116343 13116569
refGene_gn=MED13L;refGene_tx=XM_023241451.1;refGene_gn=MED13L;refGene_tx=XM_023241452.1
D3 13117628 13118357
refGene_gn=MED13L;refGene_tx=XM_023241451.1;refGene_gn=MED13L;refGene_tx=XM_023241452.1
D3 13122115 13122220
refGene_gn=MED13L;refGene_tx=XM_023241451.1;refGene_gn=MED13L;refGene_tx=XM_023241452.1

D3 13124390 13124556
refGene_gn=MED13L;refGene_tx=XM_023241451.1;refGene_gn=MED13L;refGene_tx=XM_023241452.1
D3 13128243 13128432
refGene_gn=MED13L;refGene_tx=XM_023241451.1;refGene_gn=MED13L;refGene_tx=XM_023241452.1
D3 13128822 13129017
refGene_gn=MED13L;refGene_tx=XM_023241451.1;refGene_gn=MED13L;refGene_tx=XM_023241452.1
D3 13131593 13131739
refGene_gn=MED13L;refGene_tx=XM_023241451.1;refGene_gn=MED13L;refGene_tx=XM_023241452.1
D3 13204659 13204743
refGene_gn=MED13L;refGene_tx=XM_023241451.1;refGene_gn=MED13L;refGene_tx=XM_023241452.1
D3 13217877 13217962
refGene_gn=MED13L;refGene_tx=XM_023241451.1;refGene_gn=MED13L;refGene_tx=XM_023241452.1
D3 13342670 13342908
refGene_gn=MED13L;refGene_tx=XM_023241451.1;refGene_gn=MED13L;refGene_tx=XM_023241452.1
D3 13380256 13380328
refGene_gn=MED13L;refGene_tx=XM_023241451.1;refGene_gn=MED13L;refGene_tx=XM_023241452.1
D3 13616447 13616468
refGene_gn=CD3H12orf49;refGene_tx=XM_023241456.1
D3 13705768 13705774
refGene_gn=CD3H12orf49;refGene_tx=XM_023241457.1
D3 13721420 13721591
refGene_gn=CD3H12orf49;refGene_tx=XM_023241453.1;refGene_gn=CD3H12orf49;refGene_tx=XM_023241454.1
D3 13723577 13723661
refGene_gn=CD3H12orf49;refGene_tx=XM_011287804.2;refGene_gn=CD3H12orf49;refGene_tx=XM_023241455.1;refGene_gn=CD3H12orf49;refGene_tx=XM_023241458.1
D3 13725562 13725676
refGene_gn=CD3H12orf49;refGene_tx=XM_011287804.2;refGene_gn=CD3H12orf49;refGene_tx=XM_023241453.1;refGene_gn=CD3H12orf49;refGene_tx=XM_023241454.1;refGene_gn=CD3H12orf49;refGene_tx=XM_023241455.1;refGene_gn=CD3H12orf49;refGene_tx=XM_023241456.1;refGene_gn=CD3H12orf49;refGene_tx=XM_023241457.1;refGene_gn=CD3H12orf49;refGene_tx=XM_023241458.1
D3 13726266 13726418
refGene_gn=CD3H12orf49;refGene_tx=XM_011287804.2;refGene_gn=CD3H12orf49;refGene_tx=XM_023241453.1;refGene_gn=CD3H12orf49;refGene_tx=XM_023241454.1;refGene_gn=CD3H12orf49;refGene_tx=XM_023241455.1;refGene_gn=CD3H12orf49;refGene_tx=XM_023241456.1;refGene_gn=CD3H12orf49;refGene_tx=XM_023241457.1;refGene_gn=CD3H12orf49;refGene_tx=XM_023241458.1
D3 13729933 13730090
refGene_gn=CD3H12orf49;refGene_tx=XM_011287804.2;refGene_gn=CD3H12orf49;refGene_tx=XM_023241453.1;refGene_gn=CD3H12orf49;refGene_tx=XM_02324

1454.1;refGene_gn=CD3H12orf49;refGene_tx=XM_023241455.1;refGene_gn=CD3H12orf49;refGene_tx=XM_023241456.1;refGene_gn=CD3H12orf49;refGene_tx=XM_023241457.1
D3 13738361 13738539
refGene_gn=CD3H12orf49;refGene_tx=XM_011287804.2;refGene_gn=CD3H12orf49;refGene_tx=XM_023241453.1;refGene_gn=CD3H12orf49;refGene_tx=XM_023241454.1;refGene_gn=CD3H12orf49;refGene_tx=XM_023241455.1;refGene_gn=CD3H12orf49;refGene_tx=XM_023241456.1;refGene_gn=CD3H12orf49;refGene_tx=XM_023241457.1;refGene_gn=CD3H12orf49;refGene_tx=XM_023241458.1
D3 13740718 13740742 refGene_gn=RNFT2;refGene_tx=XM_003994702.4
D3 13741282 13741341 refGene_gn=RNFT2;refGene_tx=XM_003994702.4
D3 13747145 13747615 refGene_gn=RNFT2;refGene_tx=XM_003994702.4
D3 13748010 13748087 refGene_gn=RNFT2;refGene_tx=XM_003994702.4
D3 13751685 13751786 refGene_gn=RNFT2;refGene_tx=XM_003994702.4
D3 13755352 13755506 refGene_gn=RNFT2;refGene_tx=XM_003994702.4
D3 13794537 13794687 refGene_gn=RNFT2;refGene_tx=XM_003994702.4
D3 13796005 13796071 refGene_gn=RNFT2;refGene_tx=XM_003994702.4
D3 13796199 13796301 refGene_gn=RNFT2;refGene_tx=XM_003994702.4
D3 13807208 13807343 refGene_gn=RNFT2;refGene_tx=XM_003994702.4
D3 13833438 13833714 refGene_gn=HRK;refGene_tx=XM_023241459.1
D3 13857563 13857881
refGene_gn=FBXW8;refGene_tx=XM_023241460.1;refGene_gn=FBXW8;refGene_tx=XM_023241461.1
D3 13872287 13872392
refGene_gn=FBXW8;refGene_tx=XM_023241460.1;refGene_gn=FBXW8;refGene_tx=XM_023241461.1
D3 13890136 13890301
refGene_gn=FBXW8;refGene_tx=XM_023241460.1;refGene_gn=FBXW8;refGene_tx=XM_023241461.1
D3 13892888 13892977
refGene_gn=FBXW8;refGene_tx=XM_023241460.1;refGene_gn=FBXW8;refGene_tx=XM_023241461.1
D3 13907134 13907292
refGene_gn=FBXW8;refGene_tx=XM_023241460.1;refGene_gn=FBXW8;refGene_tx=XM_023241461.1
D3 13936954 13937151
refGene_gn=FBXW8;refGene_tx=XM_023241460.1;refGene_gn=FBXW8;refGene_tx=XM_023241461.1
D3 13940103 13940310
refGene_gn=FBXW8;refGene_tx=XM_023241460.1;refGene_gn=FBXW8;refGene_tx=XM_023241461.1
D3 13959147 13959275
refGene_gn=FBXW8;refGene_tx=XM_023241460.1;refGene_gn=FBXW8;refGene_tx=XM_023241461.1
D3 13974372 13974546
refGene_gn=FBXW8;refGene_tx=XM_023241460.1;refGene_gn=FBXW8;refGene_tx=XM_023241461.1
D3 13976908 13977016
refGene_gn=FBXW8;refGene_tx=XM_023241460.1;refGene_gn=FBXW8;refGene_tx=XM_023241461.1

D3 13977132 13977282 refGene_gn=FBXW8;refGene_tx=XM_023241460.1
D3 13977433 13977578
refGene_gn=FBXW8;refGene_tx=XM_023241460.1;refGene_gn=FBXW8;refGene_tx
=XM_023241461.1
D3 13985060 13985138 refGene_gn=TESC;refGene_tx=XM_019814591.1
D3 13987065 13987113 refGene_gn=TESC;refGene_tx=XM_019814591.1
D3 13990763 13990871 refGene_gn=TESC;refGene_tx=XM_019814591.1
D3 13990973 13991035 refGene_gn=TESC;refGene_tx=XM_019814591.1
D3 13992653 13992793 refGene_gn=TESC;refGene_tx=XM_019814591.1
D3 13997025 13997106 refGene_gn=TESC;refGene_tx=XM_019814591.1
D3 14012597 14012667 refGene_gn=TESC;refGene_tx=XM_019814591.1
D3 14046095 14046306 refGene_gn=TESC;refGene_tx=XM_019814591.1
D3 14056142 14056567
refGene_gn=FBXO21;refGene_tx=XM_023241462.1;refGene_gn=FBXO21;refGene_
tx=XM_023241463.1;refGene_gn=FBXO21;refGene_tx=XM_023241464.1
D3 14059579 14059857
refGene_gn=FBXO21;refGene_tx=XM_023241462.1;refGene_gn=FBXO21;refGene_
tx=XM_023241463.1;refGene_gn=FBXO21;refGene_tx=XM_023241464.1;refGene_
gn=FBXO21;refGene_tx=XM_023241465.1
D3 14059923 14059940 refGene_gn=FBXO21;refGene_tx=XM_023241467.1
D3 14060415 14060448
refGene_gn=FBXO21;refGene_tx=XM_023241462.1;refGene_gn=FBXO21;refGene_
tx=XM_023241463.1;refGene_gn=FBXO21;refGene_tx=XM_023241464.1;refGene_
gn=FBXO21;refGene_tx=XM_023241465.1;refGene_gn=FBXO21;refGene_tx=XM_02
3241467.1
D3 14069147 14069335
refGene_gn=FBXO21;refGene_tx=XM_006938515.4;refGene_gn=FBXO21;refGene_
tx=XM_023241466.1
D3 14079364 14079522
refGene_gn=FBXO21;refGene_tx=XM_006938515.4;refGene_gn=FBXO21;refGene_
tx=XM_023241462.1;refGene_gn=FBXO21;refGene_tx=XM_023241463.1;refGene_
gn=FBXO21;refGene_tx=XM_023241464.1;refGene_gn=FBXO21;refGene_tx=XM_02
3241465.1;refGene_gn=FBXO21;refGene_tx=XM_023241466.1;refGene_gn=FBXO2
1;refGene_tx=XM_023241467.1
D3 14081045 14081257
refGene_gn=FBXO21;refGene_tx=XM_006938515.4;refGene_gn=FBXO21;refGene_
tx=XM_023241462.1;refGene_gn=FBXO21;refGene_tx=XM_023241463.1;refGene_
gn=FBXO21;refGene_tx=XM_023241464.1;refGene_gn=FBXO21;refGene_tx=XM_02
3241465.1;refGene_gn=FBXO21;refGene_tx=XM_023241466.1;refGene_gn=FBXO2
1;refGene_tx=XM_023241467.1
D3 14089170 14089303
refGene_gn=FBXO21;refGene_tx=XM_006938515.4;refGene_gn=FBXO21;refGene_
tx=XM_023241462.1;refGene_gn=FBXO21;refGene_tx=XM_023241463.1;refGene_
gn=FBXO21;refGene_tx=XM_023241464.1;refGene_gn=FBXO21;refGene_tx=XM_02
3241465.1;refGene_gn=FBXO21;refGene_tx=XM_023241466.1;refGene_gn=FBXO2
1;refGene_tx=XM_023241467.1
D3 14090411 14090591
refGene_gn=FBXO21;refGene_tx=XM_006938515.4;refGene_gn=FBXO21;refGene_
tx=XM_023241462.1;refGene_gn=FBXO21;refGene_tx=XM_023241463.1;refGene_
gn=FBXO21;refGene_tx=XM_023241464.1;refGene_gn=FBXO21;refGene_tx=XM_02

3241465.1;refGene_gn=FBXO21;refGene_tx=XM_023241466.1;refGene_gn=FBXO2
1;refGene_tx=XM_023241467.1
D3 14099188 14099325
refGene_gn=FBXO21;refGene_tx=XM_006938515.4;refGene_gn=FBXO21;refGene_
tx=XM_023241462.1;refGene_gn=FBXO21;refGene_tx=XM_023241463.1;refGene_
gn=FBXO21;refGene_tx=XM_023241464.1;refGene_gn=FBXO21;refGene_tx=XM_02
3241465.1;refGene_gn=FBXO21;refGene_tx=XM_023241466.1;refGene_gn=FBXO2
1;refGene_tx=XM_023241467.1
D3 14100424 14100561
refGene_gn=FBXO21;refGene_tx=XM_006938515.4;refGene_gn=FBXO21;refGene_
tx=XM_023241462.1;refGene_gn=FBXO21;refGene_tx=XM_023241463.1;refGene_
gn=FBXO21;refGene_tx=XM_023241464.1;refGene_gn=FBXO21;refGene_tx=XM_02
3241465.1;refGene_gn=FBXO21;refGene_tx=XM_023241466.1;refGene_gn=FBXO2
1;refGene_tx=XM_023241467.1
D3 14100864 14101011
refGene_gn=FBXO21;refGene_tx=XM_006938515.4;refGene_gn=FBXO21;refGene_
tx=XM_023241462.1;refGene_gn=FBXO21;refGene_tx=XM_023241463.1;refGene_
gn=FBXO21;refGene_tx=XM_023241464.1;refGene_gn=FBXO21;refGene_tx=XM_02
3241465.1;refGene_gn=FBXO21;refGene_tx=XM_023241466.1;refGene_gn=FBXO2
1;refGene_tx=XM_023241467.1
D3 14103982 14104104
refGene_gn=FBXO21;refGene_tx=XM_006938515.4;refGene_gn=FBXO21;refGene_
tx=XM_023241462.1;refGene_gn=FBXO21;refGene_tx=XM_023241463.1;refGene_
gn=FBXO21;refGene_tx=XM_023241464.1;refGene_gn=FBXO21;refGene_tx=XM_02
3241465.1;refGene_gn=FBXO21;refGene_tx=XM_023241466.1;refGene_gn=FBXO2
1;refGene_tx=XM_023241467.1
D3 14110789 14110884
refGene_gn=FBXO21;refGene_tx=XM_006938515.4;refGene_gn=FBXO21;refGene_
tx=XM_023241462.1;refGene_gn=FBXO21;refGene_tx=XM_023241463.1;refGene_
gn=FBXO21;refGene_tx=XM_023241464.1;refGene_gn=FBXO21;refGene_tx=XM_02
3241465.1;refGene_gn=FBXO21;refGene_tx=XM_023241466.1;refGene_gn=FBXO2
1;refGene_tx=XM_023241467.1
D3 14113209 14113345
refGene_gn=FBXO21;refGene_tx=XM_006938515.4;refGene_gn=FBXO21;refGene_
tx=XM_023241462.1;refGene_gn=FBXO21;refGene_tx=XM_023241463.1;refGene_
gn=FBXO21;refGene_tx=XM_023241464.1;refGene_gn=FBXO21;refGene_tx=XM_02
3241465.1;refGene_gn=FBXO21;refGene_tx=XM_023241466.1;refGene_gn=FBXO2
1;refGene_tx=XM_023241467.1
D3 14113760 14113885 refGene_gn=FBXO21;refGene_tx=XM_023241464.1
D3 14114217 14114456
refGene_gn=FBXO21;refGene_tx=XM_006938515.4;refGene_gn=FBXO21;refGene_
tx=XM_023241462.1;refGene_gn=FBXO21;refGene_tx=XM_023241463.1;refGene_
gn=FBXO21;refGene_tx=XM_023241465.1;refGene_gn=FBXO21;refGene_tx=XM_02
3241466.1;refGene_gn=FBXO21;refGene_tx=XM_023241467.1
D3 14131498 14131514
refGene_gn=NOS1;refGene_tx=XM_011287802.3;refGene_gn=NOS1;refGene_tx=X
M_011287803.3;refGene_gn=NOS1;refGene_tx=XM_023241468.1
D3 14132688 14132807
refGene_gn=NOS1;refGene_tx=XM_011287802.3;refGene_gn=NOS1;refGene_tx=X
M_011287803.3;refGene_gn=NOS1;refGene_tx=XM_023241468.1

D3 14134490 14134685
refGene_gn=NOS1;refGene_tx=XM_011287802.3;refGene_gn=NOS1;refGene_tx=XM_011287803.3;refGene_gn=NOS1;refGene_tx=XM_023241468.1
D3 14135721 14135870
refGene_gn=NOS1;refGene_tx=XM_011287802.3;refGene_gn=NOS1;refGene_tx=XM_011287803.3;refGene_gn=NOS1;refGene_tx=XM_023241468.1
D3 14136929 14137051
refGene_gn=NOS1;refGene_tx=XM_011287802.3;refGene_gn=NOS1;refGene_tx=XM_011287803.3;refGene_gn=NOS1;refGene_tx=XM_023241468.1
D3 14138639 14138727
refGene_gn=NOS1;refGene_tx=XM_011287802.3;refGene_gn=NOS1;refGene_tx=XM_011287803.3;refGene_gn=NOS1;refGene_tx=XM_023241468.1
D3 14139427 14139638
refGene_gn=NOS1;refGene_tx=XM_011287802.3;refGene_gn=NOS1;refGene_tx=XM_011287803.3;refGene_gn=NOS1;refGene_tx=XM_023241468.1
D3 14142154 14142324
refGene_gn=NOS1;refGene_tx=XM_011287802.3;refGene_gn=NOS1;refGene_tx=XM_011287803.3;refGene_gn=NOS1;refGene_tx=XM_023241468.1
D3 14144259 14144453
refGene_gn=NOS1;refGene_tx=XM_011287802.3;refGene_gn=NOS1;refGene_tx=XM_011287803.3;refGene_gn=NOS1;refGene_tx=XM_023241468.1
D3 14150347 14150426
refGene_gn=NOS1;refGene_tx=XM_011287802.3;refGene_gn=NOS1;refGene_tx=XM_011287803.3;refGene_gn=NOS1;refGene_tx=XM_023241468.1
D3 14150849 14150988
refGene_gn=NOS1;refGene_tx=XM_011287802.3;refGene_gn=NOS1;refGene_tx=XM_011287803.3;refGene_gn=NOS1;refGene_tx=XM_023241468.1
D3 14155787 14155962
refGene_gn=NOS1;refGene_tx=XM_011287802.3;refGene_gn=NOS1;refGene_tx=XM_011287803.3;refGene_gn=NOS1;refGene_tx=XM_023241468.1
D3 14159666 14159783
refGene_gn=NOS1;refGene_tx=XM_011287802.3;refGene_gn=NOS1;refGene_tx=XM_011287803.3;refGene_gn=NOS1;refGene_tx=XM_023241468.1
D3 14162279 14162381
refGene_gn=NOS1;refGene_tx=XM_011287802.3;refGene_gn=NOS1;refGene_tx=XM_023241468.1
D3 14163340 14163399
refGene_gn=NOS1;refGene_tx=XM_011287802.3;refGene_gn=NOS1;refGene_tx=XM_011287803.3;refGene_gn=NOS1;refGene_tx=XM_023241468.1
D3 14164308 14164413
refGene_gn=NOS1;refGene_tx=XM_011287802.3;refGene_gn=NOS1;refGene_tx=XM_011287803.3;refGene_gn=NOS1;refGene_tx=XM_023241468.1
D3 14165690 14165835
refGene_gn=NOS1;refGene_tx=XM_011287802.3;refGene_gn=NOS1;refGene_tx=XM_011287803.3;refGene_gn=NOS1;refGene_tx=XM_023241468.1
D3 14168154 14168240
refGene_gn=NOS1;refGene_tx=XM_011287802.3;refGene_gn=NOS1;refGene_tx=XM_011287803.3;refGene_gn=NOS1;refGene_tx=XM_023241468.1
D3 14168675 14168870
refGene_gn=NOS1;refGene_tx=XM_011287802.3;refGene_gn=NOS1;refGene_tx=X

M_011287803.3;refGene_gn=NOS1;refGene_tx=XM_023241468.1
D3 14170833 14170935
refGene_gn=NOS1;refGene_tx=XM_011287802.3;refGene_gn=NOS1;refGene_tx=X
M_011287803.3;refGene_gn=NOS1;refGene_tx=XM_023241468.1
D3 14175120 14175295
refGene_gn=NOS1;refGene_tx=XM_011287802.3;refGene_gn=NOS1;refGene_tx=X
M_011287803.3;refGene_gn=NOS1;refGene_tx=XM_023241468.1
D3 14177395 14177535
refGene_gn=NOS1;refGene_tx=XM_011287802.3;refGene_gn=NOS1;refGene_tx=X
M_011287803.3;refGene_gn=NOS1;refGene_tx=XM_023241468.1
D3 14179515 14179657
refGene_gn=NOS1;refGene_tx=XM_011287802.3;refGene_gn=NOS1;refGene_tx=X
M_011287803.3;refGene_gn=NOS1;refGene_tx=XM_023241468.1
D3 14182785 14182877
refGene_gn=NOS1;refGene_tx=XM_011287802.3;refGene_gn=NOS1;refGene_tx=X
M_011287803.3;refGene_gn=NOS1;refGene_tx=XM_023241468.1
D3 14183868 14184031
refGene_gn=NOS1;refGene_tx=XM_011287802.3;refGene_gn=NOS1;refGene_tx=X
M_011287803.3;refGene_gn=NOS1;refGene_tx=XM_023241468.1
D3 14186045 14186191
refGene_gn=NOS1;refGene_tx=XM_011287802.3;refGene_gn=NOS1;refGene_tx=X
M_011287803.3;refGene_gn=NOS1;refGene_tx=XM_023241468.1
D3 14187572 14187701
refGene_gn=NOS1;refGene_tx=XM_011287802.3;refGene_gn=NOS1;refGene_tx=X
M_011287803.3;refGene_gn=NOS1;refGene_tx=XM_023241468.1
D3 14202882 14203009
refGene_gn=NOS1;refGene_tx=XM_011287802.3;refGene_gn=NOS1;refGene_tx=X
M_011287803.3;refGene_gn=NOS1;refGene_tx=XM_023241468.1
D3 14217319 14218029
refGene_gn=NOS1;refGene_tx=XM_011287802.3;refGene_gn=NOS1;refGene_tx=X
M_011287803.3;refGene_gn=NOS1;refGene_tx=XM_023241468.1
D3 14346634 14346772 refGene_gn=KSR2;refGene_tx=XM_023241469.1
D3 14347798 14347928 refGene_gn=KSR2;refGene_tx=XM_023241469.1
D3 14352738 14352870 refGene_gn=KSR2;refGene_tx=XM_023241469.1
D3 14358467 14358601 refGene_gn=KSR2;refGene_tx=XM_023241469.1
D3 14359647 14359744 refGene_gn=KSR2;refGene_tx=XM_023241469.1
D3 14405686 14406054 refGene_gn=KSR2;refGene_tx=XM_023241469.1
D3 14407994 14408043 refGene_gn=KSR2;refGene_tx=XM_023241469.1
D3 14411794 14411867 refGene_gn=KSR2;refGene_tx=XM_023241469.1
D3 14412529 14412571 refGene_gn=KSR2;refGene_tx=XM_023241469.1
D3 14420409 14420575 refGene_gn=KSR2;refGene_tx=XM_023241469.1
D3 14429878 14430003 refGene_gn=KSR2;refGene_tx=XM_023241469.1
D3 14432852 14432920 refGene_gn=KSR2;refGene_tx=XM_023241469.1
D3 14452201 14452285 refGene_gn=KSR2;refGene_tx=XM_023241469.1
D3 14455556 14455626 refGene_gn=KSR2;refGene_tx=XM_023241469.1
D3 14520103 14520288 refGene_gn=KSR2;refGene_tx=XM_023241469.1
D3 14600696 14601210 refGene_gn=KSR2;refGene_tx=XM_023241469.1
D3 14679025 14679029 refGene_gn=KSR2;refGene_tx=XM_023241469.1
D3 14681628 14681775 refGene_gn=KSR2;refGene_tx=XM_023241469.1
D3 14685692 14685833 refGene_gn=KSR2;refGene_tx=XM_023241469.1

D3 14774439 14774619 refGene_gn=KSR2;refGene_tx=XM_023241469.1
D3 14807122 14807187 refGene_gn=RFC5;refGene_tx=XM_003994707.5
D3 14808972 14809037 refGene_gn=RFC5;refGene_tx=XM_003994707.5
D3 14809306 14809443 refGene_gn=RFC5;refGene_tx=XM_003994707.5
D3 14810599 14810679 refGene_gn=RFC5;refGene_tx=XM_003994707.5
D3 14811780 14811854 refGene_gn=RFC5;refGene_tx=XM_003994707.5
D3 14813906 14814066 refGene_gn=RFC5;refGene_tx=XM_003994707.5
D3 14814806 14814888 refGene_gn=RFC5;refGene_tx=XM_003994707.5
D3 14815283 14815413 refGene_gn=RFC5;refGene_tx=XM_003994707.5
D3 14816190 14816268 refGene_gn=RFC5;refGene_tx=XM_003994707.5
D3 14817306 14817361 refGene_gn=RFC5;refGene_tx=XM_003994707.5
D3 14818422 14818519 refGene_gn=RFC5;refGene_tx=XM_003994707.5
D3 14821012 14821175
refGene_gn=WSB2;refGene_tx=XM_003994708.4;refGene_gn=WSB2;refGene_tx=XM_023241470.1
D3 14821852 14821960
refGene_gn=WSB2;refGene_tx=XM_003994708.4;refGene_gn=WSB2;refGene_tx=XM_023241470.1
D3 14822083 14822194
refGene_gn=WSB2;refGene_tx=XM_003994708.4;refGene_gn=WSB2;refGene_tx=XM_023241470.1
D3 14822939 14823112
refGene_gn=WSB2;refGene_tx=XM_003994708.4;refGene_gn=WSB2;refGene_tx=XM_023241470.1
D3 14824297 14824398
refGene_gn=WSB2;refGene_tx=XM_003994708.4;refGene_gn=WSB2;refGene_tx=XM_023241470.1
D3 14827651 14827783
refGene_gn=WSB2;refGene_tx=XM_003994708.4;refGene_gn=WSB2;refGene_tx=XM_023241470.1
D3 14827963 14828208
refGene_gn=WSB2;refGene_tx=XM_003994708.4;refGene_gn=WSB2;refGene_tx=XM_023241470.1
D3 14834863 14835051
refGene_gn=WSB2;refGene_tx=XM_003994708.4;refGene_gn=WSB2;refGene_tx=XM_023241470.1
D3 14840916 14840929 refGene_gn=WSB2;refGene_tx=XM_023241470.1
D3 14845884 14845937
refGene_gn=VSIG10;refGene_tx=XM_011287799.3;refGene_gn=VSIG10;refGene_tx=XM_011287800.3;refGene_gn=VSIG10;refGene_tx=XM_019814601.2
D3 14847358 14847576
refGene_gn=VSIG10;refGene_tx=XM_011287799.3;refGene_gn=VSIG10;refGene_tx=XM_011287800.3;refGene_gn=VSIG10;refGene_tx=XM_019814601.2
D3 14849002 14849018
refGene_gn=VSIG10;refGene_tx=XM_011287799.3;refGene_gn=VSIG10;refGene_tx=XM_011287800.3;refGene_gn=VSIG10;refGene_tx=XM_019814601.2
D3 14849848 14849959
refGene_gn=VSIG10;refGene_tx=XM_011287799.3;refGene_gn=VSIG10;refGene_tx=XM_011287800.3
D3 14852354 14852648

refGene_gn=VSIG10;refGene_tx=XM_011287799.3;refGene_gn=VSIG10;refGene_tx=XM_011287800.3;refGene_gn=VSIG10;refGene_tx=XM_019814601.2
D3 14859909 14860170
refGene_gn=VSIG10;refGene_tx=XM_011287799.3;refGene_gn=VSIG10;refGene_tx=XM_011287800.3;refGene_gn=VSIG10;refGene_tx=XM_019814601.2
D3 14861823 14862126
refGene_gn=VSIG10;refGene_tx=XM_011287799.3;refGene_gn=VSIG10;refGene_tx=XM_011287800.3;refGene_gn=VSIG10;refGene_tx=XM_019814601.2
D3 14873498 14873780
refGene_gn=VSIG10;refGene_tx=XM_011287799.3;refGene_gn=VSIG10;refGene_tx=XM_011287800.3;refGene_gn=VSIG10;refGene_tx=XM_019814601.2
D3 14879910 14879989
refGene_gn=VSIG10;refGene_tx=XM_011287799.3;refGene_gn=VSIG10;refGene_tx=XM_011287800.3;refGene_gn=VSIG10;refGene_tx=XM_019814601.2
D3 14906693 14906828 refGene_gn=PEBP1;refGene_tx=XM_003994709.5
D3 14908001 14908111 refGene_gn=PEBP1;refGene_tx=XM_003994709.5
D3 14908796 14908897 refGene_gn=PEBP1;refGene_tx=XM_003994709.5
D3 14910559 14910777 refGene_gn=PEBP1;refGene_tx=XM_003994709.5
D3 14924008 14924170
refGene_gn=TAOK3;refGene_tx=XM_011287798.3;refGene_gn=TAOK3;refGene_tx=XM_019814602.2;refGene_gn=TAOK3;refGene_tx=XM_019814603.2;refGene_gn=TAOK3;refGene_tx=XM_023241471.1
D3 14925462 14925645
refGene_gn=TAOK3;refGene_tx=XM_011287798.3;refGene_gn=TAOK3;refGene_tx=XM_019814602.2;refGene_gn=TAOK3;refGene_tx=XM_019814603.2;refGene_gn=TAOK3;refGene_tx=XM_023241471.1
D3 14931585 14931798
refGene_gn=TAOK3;refGene_tx=XM_011287798.3;refGene_gn=TAOK3;refGene_tx=XM_019814602.2;refGene_gn=TAOK3;refGene_tx=XM_019814603.2;refGene_gn=TAOK3;refGene_tx=XM_023241471.1
D3 14933262 14933502
refGene_gn=TAOK3;refGene_tx=XM_011287798.3;refGene_gn=TAOK3;refGene_tx=XM_019814602.2;refGene_gn=TAOK3;refGene_tx=XM_019814603.2;refGene_gn=TAOK3;refGene_tx=XM_023241471.1
D3 14940999 14941203
refGene_gn=TAOK3;refGene_tx=XM_011287798.3;refGene_gn=TAOK3;refGene_tx=XM_019814602.2;refGene_gn=TAOK3;refGene_tx=XM_019814603.2;refGene_gn=TAOK3;refGene_tx=XM_023241471.1
D3 14946749 14946878
refGene_gn=TAOK3;refGene_tx=XM_011287798.3;refGene_gn=TAOK3;refGene_tx=XM_019814602.2;refGene_gn=TAOK3;refGene_tx=XM_019814603.2;refGene_gn=TAOK3;refGene_tx=XM_023241471.1
D3 14948583 14948820
refGene_gn=TAOK3;refGene_tx=XM_011287798.3;refGene_gn=TAOK3;refGene_tx=XM_019814602.2;refGene_gn=TAOK3;refGene_tx=XM_019814603.2;refGene_gn=TAOK3;refGene_tx=XM_023241471.1
D3 14957074 14957209
refGene_gn=TAOK3;refGene_tx=XM_011287798.3;refGene_gn=TAOK3;refGene_tx=XM_019814602.2;refGene_gn=TAOK3;refGene_tx=XM_023241471.1
D3 14969794 14970001

refGene_gn=TAOK3;refGene_tx=XM_011287798.3;refGene_gn=TAOK3;refGene_tx=XM_019814602.2;refGene_gn=TAOK3;refGene_tx=XM_023241471.1
D3 14972684 14972852
refGene_gn=TAOK3;refGene_tx=XM_011287798.3;refGene_gn=TAOK3;refGene_tx=XM_019814602.2;refGene_gn=TAOK3;refGene_tx=XM_023241471.1
D3 14980130 14980212
refGene_gn=TAOK3;refGene_tx=XM_011287798.3;refGene_gn=TAOK3;refGene_tx=XM_019814602.2;refGene_gn=TAOK3;refGene_tx=XM_023241471.1
D3 14981240 14981334
refGene_gn=TAOK3;refGene_tx=XM_011287798.3;refGene_gn=TAOK3;refGene_tx=XM_019814602.2;refGene_gn=TAOK3;refGene_tx=XM_023241471.1
D3 14993982 14994074
refGene_gn=TAOK3;refGene_tx=XM_011287798.3;refGene_gn=TAOK3;refGene_tx=XM_019814602.2;refGene_gn=TAOK3;refGene_tx=XM_023241471.1
D3 14996155 14996269
refGene_gn=TAOK3;refGene_tx=XM_011287798.3;refGene_gn=TAOK3;refGene_tx=XM_019814602.2;refGene_gn=TAOK3;refGene_tx=XM_023241471.1
D3 14999135 14999232
refGene_gn=TAOK3;refGene_tx=XM_011287798.3;refGene_gn=TAOK3;refGene_tx=XM_019814602.2;refGene_gn=TAOK3;refGene_tx=XM_023241471.1
D3 15000453 15000499
refGene_gn=TAOK3;refGene_tx=XM_011287798.3;refGene_gn=TAOK3;refGene_tx=XM_019814602.2;refGene_gn=TAOK3;refGene_tx=XM_023241471.1
D3 15003791 15003893
refGene_gn=TAOK3;refGene_tx=XM_011287798.3;refGene_gn=TAOK3;refGene_tx=XM_019814602.2;refGene_gn=TAOK3;refGene_tx=XM_023241471.1
D3 15004280 15004352
refGene_gn=TAOK3;refGene_tx=XM_011287798.3;refGene_gn=TAOK3;refGene_tx=XM_019814602.2;refGene_gn=TAOK3;refGene_tx=XM_023241471.1
D3 15011098 15011218
refGene_gn=TAOK3;refGene_tx=XM_011287798.3;refGene_gn=TAOK3;refGene_tx=XM_019814602.2;refGene_gn=TAOK3;refGene_tx=XM_023241471.1
D3 15109128 15109270
refGene_gn=SUDS3;refGene_tx=XM_003994711.5;refGene_gn=SUDS3;refGene_tx=XM_006938518.3;refGene_gn=SUDS3;refGene_tx=XM_019814608.2
D3 15111154 15111224
refGene_gn=SUDS3;refGene_tx=XM_003994711.5;refGene_gn=SUDS3;refGene_tx=XM_006938518.3;refGene_gn=SUDS3;refGene_tx=XM_019814608.2;refGene_gn=SUDS3;refGene_tx=XM_023241472.1
D3 15113733 15113789
refGene_gn=SUDS3;refGene_tx=XM_003994711.5;refGene_gn=SUDS3;refGene_tx=XM_006938518.3;refGene_gn=SUDS3;refGene_tx=XM_019814608.2;refGene_gn=SUDS3;refGene_tx=XM_023241472.1
D3 15115126 15115198
refGene_gn=SUDS3;refGene_tx=XM_003994711.5;refGene_gn=SUDS3;refGene_tx=XM_006938518.3;refGene_gn=SUDS3;refGene_tx=XM_019814608.2;refGene_gn=SUDS3;refGene_tx=XM_023241472.1
D3 15118155 15118175
refGene_gn=SUDS3;refGene_tx=XM_003994711.5;refGene_gn=SUDS3;refGene_tx=XM_006938518.3;refGene_gn=SUDS3;refGene_tx=XM_019814608.2;refGene_gn=

SUDS3;refGene_tx=XM_023241472.1
D3 15119386 15119543
refGene_gn=SUDS3;refGene_tx=XM_003994711.5;refGene_gn=SUDS3;refGene_tx=XM_006938518.3;refGene_gn=SUDS3;refGene_tx=XM_019814608.2;refGene_gn=SUDS3;refGene_tx=XM_023241472.1
D3 15127681 15127777
refGene_gn=SUDS3;refGene_tx=XM_003994711.5;refGene_gn=SUDS3;refGene_tx=XM_006938518.3;refGene_gn=SUDS3;refGene_tx=XM_019814608.2;refGene_gn=SUDS3;refGene_tx=XM_023241472.1
D3 15128764 15128826
refGene_gn=SUDS3;refGene_tx=XM_003994711.5;refGene_gn=SUDS3;refGene_tx=XM_006938518.3;refGene_gn=SUDS3;refGene_tx=XM_019814608.2;refGene_gn=SUDS3;refGene_tx=XM_023241472.1
D3 15128987 15129009
refGene_gn=SUDS3;refGene_tx=XM_003994711.5;refGene_gn=SUDS3;refGene_tx=XM_006938518.3;refGene_gn=SUDS3;refGene_tx=XM_019814608.2;refGene_gn=SUDS3;refGene_tx=XM_023241472.1
D3 15130175 15130281
refGene_gn=SUDS3;refGene_tx=XM_003994711.5;refGene_gn=SUDS3;refGene_tx=XM_006938518.3;refGene_gn=SUDS3;refGene_tx=XM_019814608.2;refGene_gn=SUDS3;refGene_tx=XM_023241472.1
D3 15137166 15137251
refGene_gn=SUDS3;refGene_tx=XM_003994711.5;refGene_gn=SUDS3;refGene_tx=XM_006938518.3;refGene_gn=SUDS3;refGene_tx=XM_019814608.2;refGene_gn=SUDS3;refGene_tx=XM_023241472.1
D3 15140506 15140605
refGene_gn=SUDS3;refGene_tx=XM_003994711.5;refGene_gn=SUDS3;refGene_tx=XM_006938518.3;refGene_gn=SUDS3;refGene_tx=XM_019814608.2;refGene_gn=SUDS3;refGene_tx=XM_023241472.1
D3 15145853 15145896
refGene_gn=SUDS3;refGene_tx=XM_006938518.3;refGene_gn=SUDS3;refGene_tx=XM_023241472.1
D3 15149570 15149574 refGene_gn=SUDS3;refGene_tx=XM_019814608.2
D3 15580665 15580796
refGene_gn=SRRM4;refGene_tx=XM_023241473.1;refGene_gn=SRRM4;refGene_tx=XM_023241474.1
D3 15689828 15689975 refGene_gn=SRRM4;refGene_tx=XM_023241473.1
D3 15694030 15694117
refGene_gn=SRRM4;refGene_tx=XM_023241473.1;refGene_gn=SRRM4;refGene_tx=XM_023241474.1
D3 15696380 15696452
refGene_gn=SRRM4;refGene_tx=XM_023241473.1;refGene_gn=SRRM4;refGene_tx=XM_023241474.1
D3 15699024 15699051
refGene_gn=SRRM4;refGene_tx=XM_023241473.1;refGene_gn=SRRM4;refGene_tx=XM_023241474.1
D3 15700865 15700916
refGene_gn=SRRM4;refGene_tx=XM_023241473.1;refGene_gn=SRRM4;refGene_tx=XM_023241474.1
D3 15704118 15704217

refGene_gn=SRRM4;refGene_tx=XM_023241473.1;refGene_gn=SRRM4;refGene_tx=XM_023241474.1
D3 15707757 15707914
refGene_gn=SRRM4;refGene_tx=XM_023241473.1;refGene_gn=SRRM4;refGene_tx=XM_023241474.1
D3 15721436 15721741
refGene_gn=SRRM4;refGene_tx=XM_023241473.1;refGene_gn=SRRM4;refGene_tx=XM_023241474.1
D3 15726071 15726275
refGene_gn=SRRM4;refGene_tx=XM_023241473.1;refGene_gn=SRRM4;refGene_tx=XM_023241474.1
D3 15726989 15727100
refGene_gn=SRRM4;refGene_tx=XM_023241473.1;refGene_gn=SRRM4;refGene_tx=XM_023241474.1
D3 15727605 15727746
refGene_gn=SRRM4;refGene_tx=XM_023241473.1;refGene_gn=SRRM4;refGene_tx=XM_023241474.1
D3 15729413 15729759
refGene_gn=SRRM4;refGene_tx=XM_023241473.1;refGene_gn=SRRM4;refGene_tx=XM_023241474.1
D3 15750104 15750471 refGene_gn=HSPB8;refGene_tx=XM_003994712.5
D3 15756871 15756935 refGene_gn=HSPB8;refGene_tx=XM_003994712.5
D3 15762578 15762738 refGene_gn=HSPB8;refGene_tx=XM_003994712.5
D3 15866573 15866663 refGene_gn=CCDC60;refGene_tx=XM_003994742.5
D3 15936618 15936698 refGene_gn=CCDC60;refGene_tx=XM_003994742.5
D3 15974901 15975072 refGene_gn=CCDC60;refGene_tx=XM_003994742.5
D3 15984763 15984871 refGene_gn=CCDC60;refGene_tx=XM_003994742.5
D3 15992401 15992509 refGene_gn=CCDC60;refGene_tx=XM_003994742.5
D3 16003184 16003275 refGene_gn=CCDC60;refGene_tx=XM_003994742.5
D3 16008564 16008799 refGene_gn=CCDC60;refGene_tx=XM_003994742.5
D3 16016179 16016264 refGene_gn=CCDC60;refGene_tx=XM_003994742.5
D3 16019578 16019650 refGene_gn=CCDC60;refGene_tx=XM_003994742.5
D3 16022192 16022255 refGene_gn=CCDC60;refGene_tx=XM_003994742.5
D3 16022883 16023006 refGene_gn=CCDC60;refGene_tx=XM_003994742.5
D3 16024893 16025025 refGene_gn=CCDC60;refGene_tx=XM_003994742.5
D3 16027288 16027478 refGene_gn=CCDC60;refGene_tx=XM_003994742.5
D3 16029199 16029301 refGene_gn=CCDC60;refGene_tx=XM_003994742.5
D3 16075249 16075438 refGene_gn=TMEM233;refGene_tx=XM_003994743.4
D3 16101629 16101766 refGene_gn=TMEM233;refGene_tx=XM_003994743.4
D3 16110469 16110476 refGene_gn=TMEM233;refGene_tx=XM_003994743.4
D3 16130317 16130476 refGene_gn=PRKAB1;refGene_tx=XM_003994713.5
D3 16133350 16133514 refGene_gn=PRKAB1;refGene_tx=XM_003994713.5
D3 16134718 16134812 refGene_gn=PRKAB1;refGene_tx=XM_003994713.5
D3 16135096 16135211 refGene_gn=PRKAB1;refGene_tx=XM_003994713.5
D3 16137546 16137680 refGene_gn=PRKAB1;refGene_tx=XM_003994713.5
D3 16138878 16138947 refGene_gn=PRKAB1;refGene_tx=XM_003994713.5
D3 16139220 16139298 refGene_gn=PRKAB1;refGene_tx=XM_003994713.5
D3 16147734 16147758
refGene_gn=CIT;refGene_tx=XM_006938519.3;refGene_gn=CIT;refGene_tx=XM_019814610.1;refGene_gn=CIT;refGene_tx=XM_019814611.1;refGene_gn=CIT;re

fGene_tx=XM_019814612.1;refGene_gn=CIT;refGene_tx=XM_019814613.1;refGene_gn=CIT;refGene_tx=XM_019814614.1;refGene_gn=CIT;refGene_tx=XM_019814615.1;refGene_gn=CIT;refGene_tx=XM_019814616.1;refGene_gn=CIT;refGene_tx=XM_019814617.2;refGene_gn=CIT;refGene_tx=XM_023241475.1
D3 16148769 16149070
refGene_gn=CIT;refGene_tx=XM_006938519.3;refGene_gn=CIT;refGene_tx=XM_019814610.1;refGene_gn=CIT;refGene_tx=XM_019814611.1;refGene_gn=CIT;refGene_tx=XM_019814612.1;refGene_gn=CIT;refGene_tx=XM_019814613.1;refGene_gn=CIT;refGene_tx=XM_019814614.1;refGene_gn=CIT;refGene_tx=XM_019814615.1;refGene_gn=CIT;refGene_tx=XM_019814616.1;refGene_gn=CIT;refGene_tx=XM_019814617.2;refGene_gn=CIT;refGene_tx=XM_023241475.1
D3 16154254 16154434
refGene_gn=CIT;refGene_tx=XM_006938519.3;refGene_gn=CIT;refGene_tx=XM_019814610.1;refGene_gn=CIT;refGene_tx=XM_019814611.1;refGene_gn=CIT;refGene_tx=XM_019814612.1;refGene_gn=CIT;refGene_tx=XM_019814613.1;refGene_gn=CIT;refGene_tx=XM_019814614.1;refGene_gn=CIT;refGene_tx=XM_019814615.1;refGene_gn=CIT;refGene_tx=XM_019814616.1;refGene_gn=CIT;refGene_tx=XM_019814617.2;refGene_gn=CIT;refGene_tx=XM_023241475.1
D3 16154561 16154640
refGene_gn=CIT;refGene_tx=XM_006938519.3;refGene_gn=CIT;refGene_tx=XM_019814610.1;refGene_gn=CIT;refGene_tx=XM_019814611.1;refGene_gn=CIT;refGene_tx=XM_019814612.1;refGene_gn=CIT;refGene_tx=XM_019814613.1;refGene_gn=CIT;refGene_tx=XM_019814614.1;refGene_gn=CIT;refGene_tx=XM_019814615.1;refGene_gn=CIT;refGene_tx=XM_019814616.1;refGene_gn=CIT;refGene_tx=XM_019814617.2;refGene_gn=CIT;refGene_tx=XM_023241475.1
D3 16156226 16156307
refGene_gn=CIT;refGene_tx=XM_006938519.3;refGene_gn=CIT;refGene_tx=XM_019814610.1;refGene_gn=CIT;refGene_tx=XM_019814611.1;refGene_gn=CIT;refGene_tx=XM_019814612.1;refGene_gn=CIT;refGene_tx=XM_019814613.1;refGene_gn=CIT;refGene_tx=XM_019814614.1;refGene_gn=CIT;refGene_tx=XM_019814615.1;refGene_gn=CIT;refGene_tx=XM_019814616.1;refGene_gn=CIT;refGene_tx=XM_019814617.2;refGene_gn=CIT;refGene_tx=XM_023241475.1
D3 16156811 16156940
refGene_gn=CIT;refGene_tx=XM_006938519.3;refGene_gn=CIT;refGene_tx=XM_019814610.1;refGene_gn=CIT;refGene_tx=XM_019814611.1;refGene_gn=CIT;refGene_tx=XM_019814612.1;refGene_gn=CIT;refGene_tx=XM_019814613.1;refGene_gn=CIT;refGene_tx=XM_019814614.1;refGene_gn=CIT;refGene_tx=XM_019814615.1;refGene_gn=CIT;refGene_tx=XM_019814616.1;refGene_gn=CIT;refGene_tx=XM_019814617.2;refGene_gn=CIT;refGene_tx=XM_023241475.1
D3 16157037 16157146
refGene_gn=CIT;refGene_tx=XM_006938519.3;refGene_gn=CIT;refGene_tx=XM_019814610.1;refGene_gn=CIT;refGene_tx=XM_019814611.1;refGene_gn=CIT;refGene_tx=XM_019814612.1;refGene_gn=CIT;refGene_tx=XM_019814613.1;refGene_gn=CIT;refGene_tx=XM_019814614.1;refGene_gn=CIT;refGene_tx=XM_019814615.1;refGene_gn=CIT;refGene_tx=XM_019814616.1;refGene_gn=CIT;refGene_tx=XM_019814617.2;refGene_gn=CIT;refGene_tx=XM_023241475.1
D3 16158652 16158745
refGene_gn=CIT;refGene_tx=XM_006938519.3;refGene_gn=CIT;refGene_tx=XM_019814610.1;refGene_gn=CIT;refGene_tx=XM_019814611.1;refGene_gn=CIT;refGene_tx=XM_019814612.1;refGene_gn=CIT;refGene_tx=XM_019814613.1;refGe

ne_gn=CIT;refGene_tx=XM_019814614.1;refGene_gn=CIT;refGene_tx=XM_019814615.1;refGene_gn=CIT;refGene_tx=XM_019814616.1;refGene_gn=CIT;refGene_tx=XM_019814617.2;refGene_gn=CIT;refGene_tx=XM_023241475.1
D3 16160146 16160286
refGene_gn=CIT;refGene_tx=XM_006938519.3;refGene_gn=CIT;refGene_tx=XM_019814610.1;refGene_gn=CIT;refGene_tx=XM_019814611.1;refGene_gn=CIT;refGene_tx=XM_019814612.1;refGene_gn=CIT;refGene_tx=XM_019814613.1;refGene_gn=CIT;refGene_tx=XM_019814614.1;refGene_gn=CIT;refGene_tx=XM_019814615.1;refGene_gn=CIT;refGene_tx=XM_019814616.1;refGene_gn=CIT;refGene_tx=XM_019814617.2;refGene_gn=CIT;refGene_tx=XM_023241475.1
D3 16161927 16162063
refGene_gn=CIT;refGene_tx=XM_006938519.3;refGene_gn=CIT;refGene_tx=XM_019814610.1;refGene_gn=CIT;refGene_tx=XM_019814611.1;refGene_gn=CIT;refGene_tx=XM_019814612.1;refGene_gn=CIT;refGene_tx=XM_019814613.1;refGene_gn=CIT;refGene_tx=XM_019814614.1;refGene_gn=CIT;refGene_tx=XM_019814615.1;refGene_gn=CIT;refGene_tx=XM_019814616.1;refGene_gn=CIT;refGene_tx=XM_019814617.2;refGene_gn=CIT;refGene_tx=XM_023241475.1
D3 16162193 16162274
refGene_gn=CIT;refGene_tx=XM_006938519.3;refGene_gn=CIT;refGene_tx=XM_019814610.1;refGene_gn=CIT;refGene_tx=XM_019814611.1;refGene_gn=CIT;refGene_tx=XM_019814612.1;refGene_gn=CIT;refGene_tx=XM_019814613.1;refGene_gn=CIT;refGene_tx=XM_019814614.1;refGene_gn=CIT;refGene_tx=XM_019814615.1;refGene_gn=CIT;refGene_tx=XM_019814616.1;refGene_gn=CIT;refGene_tx=XM_019814617.2;refGene_gn=CIT;refGene_tx=XM_023241475.1
D3 16162737 16162782
refGene_gn=CIT;refGene_tx=XM_019814610.1;refGene_gn=CIT;refGene_tx=XM_019814611.1;refGene_gn=CIT;refGene_tx=XM_019814613.1;refGene_gn=CIT;refGene_tx=XM_019814615.1;refGene_gn=CIT;refGene_tx=XM_019814617.2
D3 16163507 16163677
refGene_gn=CIT;refGene_tx=XM_006938519.3;refGene_gn=CIT;refGene_tx=XM_019814610.1;refGene_gn=CIT;refGene_tx=XM_019814611.1;refGene_gn=CIT;refGene_tx=XM_019814612.1;refGene_gn=CIT;refGene_tx=XM_019814613.1;refGene_gn=CIT;refGene_tx=XM_019814614.1;refGene_gn=CIT;refGene_tx=XM_019814615.1;refGene_gn=CIT;refGene_tx=XM_019814616.1;refGene_gn=CIT;refGene_tx=XM_019814617.2;refGene_gn=CIT;refGene_tx=XM_023241475.1
D3 16163944 16164049
refGene_gn=CIT;refGene_tx=XM_006938519.3;refGene_gn=CIT;refGene_tx=XM_019814610.1;refGene_gn=CIT;refGene_tx=XM_019814611.1;refGene_gn=CIT;refGene_tx=XM_019814612.1;refGene_gn=CIT;refGene_tx=XM_019814613.1;refGene_gn=CIT;refGene_tx=XM_019814614.1;refGene_gn=CIT;refGene_tx=XM_019814615.1;refGene_gn=CIT;refGene_tx=XM_019814616.1;refGene_gn=CIT;refGene_tx=XM_019814617.2;refGene_gn=CIT;refGene_tx=XM_023241475.1
D3 16164593 16164685
refGene_gn=CIT;refGene_tx=XM_006938519.3;refGene_gn=CIT;refGene_tx=XM_019814610.1;refGene_gn=CIT;refGene_tx=XM_019814611.1;refGene_gn=CIT;refGene_tx=XM_019814612.1;refGene_gn=CIT;refGene_tx=XM_019814613.1;refGene_gn=CIT;refGene_tx=XM_019814614.1;refGene_gn=CIT;refGene_tx=XM_019814615.1;refGene_gn=CIT;refGene_tx=XM_019814616.1;refGene_gn=CIT;refGene_tx=XM_019814617.2;refGene_gn=CIT;refGene_tx=XM_023241475.1
D3 16165335 16165516

refGene_gn=CIT;refGene_tx=XM_006938519.3;refGene_gn=CIT;refGene_tx=XM_019814610.1;refGene_gn=CIT;refGene_tx=XM_019814611.1;refGene_gn=CIT;refGene_tx=XM_019814612.1;refGene_gn=CIT;refGene_tx=XM_019814613.1;refGene_gn=CIT;refGene_tx=XM_019814614.1;refGene_gn=CIT;refGene_tx=XM_019814615.1;refGene_gn=CIT;refGene_tx=XM_019814616.1;refGene_gn=CIT;refGene_tx=XM_019814617.2;refGene_gn=CIT;refGene_tx=XM_023241475.1
D3 16166067 16166205
refGene_gn=CIT;refGene_tx=XM_006938519.3;refGene_gn=CIT;refGene_tx=XM_019814610.1;refGene_gn=CIT;refGene_tx=XM_019814611.1;refGene_gn=CIT;refGene_tx=XM_019814612.1;refGene_gn=CIT;refGene_tx=XM_019814613.1;refGene_gn=CIT;refGene_tx=XM_019814614.1;refGene_gn=CIT;refGene_tx=XM_019814615.1;refGene_gn=CIT;refGene_tx=XM_019814616.1;refGene_gn=CIT;refGene_tx=XM_019814617.2;refGene_gn=CIT;refGene_tx=XM_023241475.1
D3 16168376 16168541
refGene_gn=CIT;refGene_tx=XM_006938519.3;refGene_gn=CIT;refGene_tx=XM_019814610.1;refGene_gn=CIT;refGene_tx=XM_019814611.1;refGene_gn=CIT;refGene_tx=XM_019814612.1;refGene_gn=CIT;refGene_tx=XM_019814613.1;refGene_gn=CIT;refGene_tx=XM_019814614.1;refGene_gn=CIT;refGene_tx=XM_019814615.1;refGene_gn=CIT;refGene_tx=XM_019814616.1;refGene_gn=CIT;refGene_tx=XM_019814617.2;refGene_gn=CIT;refGene_tx=XM_023241475.1
D3 16168818 16168981
refGene_gn=CIT;refGene_tx=XM_006938519.3;refGene_gn=CIT;refGene_tx=XM_019814610.1;refGene_gn=CIT;refGene_tx=XM_019814611.1;refGene_gn=CIT;refGene_tx=XM_019814612.1;refGene_gn=CIT;refGene_tx=XM_019814613.1;refGene_gn=CIT;refGene_tx=XM_019814614.1;refGene_gn=CIT;refGene_tx=XM_019814615.1;refGene_gn=CIT;refGene_tx=XM_019814616.1;refGene_gn=CIT;refGene_tx=XM_019814617.2;refGene_gn=CIT;refGene_tx=XM_023241475.1
D3 16170662 16170770
refGene_gn=CIT;refGene_tx=XM_006938519.3;refGene_gn=CIT;refGene_tx=XM_019814610.1;refGene_gn=CIT;refGene_tx=XM_019814611.1;refGene_gn=CIT;refGene_tx=XM_019814612.1;refGene_gn=CIT;refGene_tx=XM_019814613.1;refGene_gn=CIT;refGene_tx=XM_019814614.1;refGene_gn=CIT;refGene_tx=XM_019814615.1;refGene_gn=CIT;refGene_tx=XM_019814616.1;refGene_gn=CIT;refGene_tx=XM_019814617.2;refGene_gn=CIT;refGene_tx=XM_023241475.1
D3 16171206 16171347
refGene_gn=CIT;refGene_tx=XM_006938519.3;refGene_gn=CIT;refGene_tx=XM_019814610.1;refGene_gn=CIT;refGene_tx=XM_019814611.1;refGene_gn=CIT;refGene_tx=XM_019814612.1;refGene_gn=CIT;refGene_tx=XM_019814613.1;refGene_gn=CIT;refGene_tx=XM_019814614.1;refGene_gn=CIT;refGene_tx=XM_019814615.1;refGene_gn=CIT;refGene_tx=XM_019814616.1;refGene_gn=CIT;refGene_tx=XM_019814617.2;refGene_gn=CIT;refGene_tx=XM_023241475.1
D3 16178781 16178886
refGene_gn=CIT;refGene_tx=XM_006938519.3;refGene_gn=CIT;refGene_tx=XM_019814610.1;refGene_gn=CIT;refGene_tx=XM_019814611.1;refGene_gn=CIT;refGene_tx=XM_019814612.1;refGene_gn=CIT;refGene_tx=XM_019814613.1;refGene_gn=CIT;refGene_tx=XM_019814614.1;refGene_gn=CIT;refGene_tx=XM_019814615.1;refGene_gn=CIT;refGene_tx=XM_019814616.1;refGene_gn=CIT;refGene_tx=XM_019814617.2;refGene_gn=CIT;refGene_tx=XM_023241475.1
D3 16183113 16183249
refGene_gn=CIT;refGene_tx=XM_006938519.3;refGene_gn=CIT;refGene_tx=XM_

019814610.1;refGene_gn=CIT;refGene_tx=XM_019814611.1;refGene_gn=CIT;refGene_tx=XM_019814612.1;refGene_gn=CIT;refGene_tx=XM_019814613.1;refGene_gn=CIT;refGene_tx=XM_019814614.1;refGene_gn=CIT;refGene_tx=XM_019814615.1;refGene_gn=CIT;refGene_tx=XM_019814616.1;refGene_gn=CIT;refGene_tx=XM_019814617.2;refGene_gn=CIT;refGene_tx=XM_023241475.1
D3 16186033 16186254
refGene_gn=CIT;refGene_tx=XM_006938519.3;refGene_gn=CIT;refGene_tx=XM_019814610.1;refGene_gn=CIT;refGene_tx=XM_019814611.1;refGene_gn=CIT;refGene_tx=XM_019814612.1;refGene_gn=CIT;refGene_tx=XM_019814613.1;refGene_gn=CIT;refGene_tx=XM_019814614.1;refGene_gn=CIT;refGene_tx=XM_019814615.1;refGene_gn=CIT;refGene_tx=XM_019814616.1;refGene_gn=CIT;refGene_tx=XM_019814617.2;refGene_gn=CIT;refGene_tx=XM_023241475.1
D3 16187134 16187332
refGene_gn=CIT;refGene_tx=XM_006938519.3;refGene_gn=CIT;refGene_tx=XM_019814610.1;refGene_gn=CIT;refGene_tx=XM_019814611.1;refGene_gn=CIT;refGene_tx=XM_019814612.1;refGene_gn=CIT;refGene_tx=XM_019814613.1;refGene_gn=CIT;refGene_tx=XM_019814614.1;refGene_gn=CIT;refGene_tx=XM_019814615.1;refGene_gn=CIT;refGene_tx=XM_019814616.1;refGene_gn=CIT;refGene_tx=XM_019814617.2;refGene_gn=CIT;refGene_tx=XM_023241475.1
D3 16194935 16194989
refGene_gn=CIT;refGene_tx=XM_006938519.3;refGene_gn=CIT;refGene_tx=XM_019814610.1;refGene_gn=CIT;refGene_tx=XM_019814611.1;refGene_gn=CIT;refGene_tx=XM_019814612.1;refGene_gn=CIT;refGene_tx=XM_019814613.1;refGene_gn=CIT;refGene_tx=XM_019814614.1;refGene_gn=CIT;refGene_tx=XM_019814615.1;refGene_gn=CIT;refGene_tx=XM_019814616.1;refGene_gn=CIT;refGene_tx=XM_019814617.2;refGene_gn=CIT;refGene_tx=XM_023241475.1
D3 16203106 16203304
refGene_gn=CIT;refGene_tx=XM_006938519.3;refGene_gn=CIT;refGene_tx=XM_019814610.1;refGene_gn=CIT;refGene_tx=XM_019814611.1;refGene_gn=CIT;refGene_tx=XM_019814612.1;refGene_gn=CIT;refGene_tx=XM_019814613.1;refGene_gn=CIT;refGene_tx=XM_019814614.1;refGene_gn=CIT;refGene_tx=XM_019814615.1;refGene_gn=CIT;refGene_tx=XM_019814616.1;refGene_gn=CIT;refGene_tx=XM_019814617.2;refGene_gn=CIT;refGene_tx=XM_023241475.1
D3 16207222 16207397
refGene_gn=CIT;refGene_tx=XM_006938519.3;refGene_gn=CIT;refGene_tx=XM_019814610.1;refGene_gn=CIT;refGene_tx=XM_019814611.1;refGene_gn=CIT;refGene_tx=XM_019814612.1;refGene_gn=CIT;refGene_tx=XM_019814613.1;refGene_gn=CIT;refGene_tx=XM_019814614.1;refGene_gn=CIT;refGene_tx=XM_019814615.1;refGene_gn=CIT;refGene_tx=XM_019814616.1;refGene_gn=CIT;refGene_tx=XM_019814617.2;refGene_gn=CIT;refGene_tx=XM_023241475.1
D3 16208401 16208511
refGene_gn=CIT;refGene_tx=XM_006938519.3;refGene_gn=CIT;refGene_tx=XM_019814610.1;refGene_gn=CIT;refGene_tx=XM_019814611.1;refGene_gn=CIT;refGene_tx=XM_019814612.1;refGene_gn=CIT;refGene_tx=XM_019814613.1;refGene_gn=CIT;refGene_tx=XM_019814614.1;refGene_gn=CIT;refGene_tx=XM_019814615.1;refGene_gn=CIT;refGene_tx=XM_019814616.1;refGene_gn=CIT;refGene_tx=XM_019814617.2;refGene_gn=CIT;refGene_tx=XM_023241475.1
D3 16209499 16209616
refGene_gn=CIT;refGene_tx=XM_006938519.3;refGene_gn=CIT;refGene_tx=XM_019814610.1;refGene_gn=CIT;refGene_tx=XM_019814611.1;refGene_gn=CIT;re

fGene_tx=XM_019814612.1;refGene_gn=CIT;refGene_tx=XM_019814613.1;refGene_gn=CIT;refGene_tx=XM_019814614.1;refGene_gn=CIT;refGene_tx=XM_019814615.1;refGene_gn=CIT;refGene_tx=XM_019814616.1;refGene_gn=CIT;refGene_tx=XM_019814617.2;refGene_gn=CIT;refGene_tx=XM_023241475.1
D3 16214528 16214624
refGene_gn=CIT;refGene_tx=XM_006938519.3;refGene_gn=CIT;refGene_tx=XM_019814610.1;refGene_gn=CIT;refGene_tx=XM_019814611.1;refGene_gn=CIT;refGene_tx=XM_019814612.1;refGene_gn=CIT;refGene_tx=XM_019814613.1;refGene_gn=CIT;refGene_tx=XM_019814614.1;refGene_gn=CIT;refGene_tx=XM_019814615.1;refGene_gn=CIT;refGene_tx=XM_019814616.1;refGene_gn=CIT;refGene_tx=XM_019814617.2;refGene_gn=CIT;refGene_tx=XM_023241475.1
D3 16217984 16218110
refGene_gn=CIT;refGene_tx=XM_006938519.3;refGene_gn=CIT;refGene_tx=XM_019814610.1;refGene_gn=CIT;refGene_tx=XM_019814611.1;refGene_gn=CIT;refGene_tx=XM_019814612.1;refGene_gn=CIT;refGene_tx=XM_019814613.1;refGene_gn=CIT;refGene_tx=XM_019814614.1;refGene_gn=CIT;refGene_tx=XM_019814617.2
D3 16219913 16220054
refGene_gn=CIT;refGene_tx=XM_006938519.3;refGene_gn=CIT;refGene_tx=XM_019814610.1;refGene_gn=CIT;refGene_tx=XM_019814611.1;refGene_gn=CIT;refGene_tx=XM_019814612.1;refGene_gn=CIT;refGene_tx=XM_019814613.1;refGene_gn=CIT;refGene_tx=XM_019814614.1;refGene_gn=CIT;refGene_tx=XM_019814615.1;refGene_gn=CIT;refGene_tx=XM_019814616.1;refGene_gn=CIT;refGene_tx=XM_019814617.2;refGene_gn=CIT;refGene_tx=XM_023241475.1
D3 16222647 16222701
refGene_gn=CIT;refGene_tx=XM_006938519.3;refGene_gn=CIT;refGene_tx=XM_019814610.1;refGene_gn=CIT;refGene_tx=XM_019814611.1;refGene_gn=CIT;refGene_tx=XM_019814612.1;refGene_gn=CIT;refGene_tx=XM_019814613.1;refGene_gn=CIT;refGene_tx=XM_019814614.1;refGene_gn=CIT;refGene_tx=XM_019814615.1;refGene_gn=CIT;refGene_tx=XM_019814616.1;refGene_gn=CIT;refGene_tx=XM_019814617.2;refGene_gn=CIT;refGene_tx=XM_023241475.1
D3 16223221 16223272
refGene_gn=CIT;refGene_tx=XM_006938519.3;refGene_gn=CIT;refGene_tx=XM_019814610.1;refGene_gn=CIT;refGene_tx=XM_019814611.1;refGene_gn=CIT;refGene_tx=XM_019814612.1;refGene_gn=CIT;refGene_tx=XM_019814613.1;refGene_gn=CIT;refGene_tx=XM_019814614.1;refGene_gn=CIT;refGene_tx=XM_019814615.1;refGene_gn=CIT;refGene_tx=XM_019814616.1;refGene_gn=CIT;refGene_tx=XM_019814617.2;refGene_gn=CIT;refGene_tx=XM_023241475.1
D3 16223574 16223745
refGene_gn=CIT;refGene_tx=XM_006938519.3;refGene_gn=CIT;refGene_tx=XM_019814610.1;refGene_gn=CIT;refGene_tx=XM_019814611.1;refGene_gn=CIT;refGene_tx=XM_019814612.1;refGene_gn=CIT;refGene_tx=XM_019814613.1;refGene_gn=CIT;refGene_tx=XM_019814614.1;refGene_gn=CIT;refGene_tx=XM_019814615.1;refGene_gn=CIT;refGene_tx=XM_019814616.1;refGene_gn=CIT;refGene_tx=XM_019814617.2;refGene_gn=CIT;refGene_tx=XM_023241475.1
D3 16228365 16228485
refGene_gn=CIT;refGene_tx=XM_006938519.3;refGene_gn=CIT;refGene_tx=XM_019814610.1;refGene_gn=CIT;refGene_tx=XM_019814611.1;refGene_gn=CIT;refGene_tx=XM_019814612.1;refGene_gn=CIT;refGene_tx=XM_019814613.1;refGene_gn=CIT;refGene_tx=XM_019814614.1;refGene_gn=CIT;refGene_tx=XM_01981

4615.1;refGene_gn=CIT;refGene_tx=XM_019814616.1;refGene_gn=CIT;refGene_tx=XM_019814617.2;refGene_gn=CIT;refGene_tx=XM_023241475.1
D3 16229720 16229864
refGene_gn=CIT;refGene_tx=XM_006938519.3;refGene_gn=CIT;refGene_tx=XM_019814610.1;refGene_gn=CIT;refGene_tx=XM_019814611.1;refGene_gn=CIT;refGene_tx=XM_019814612.1;refGene_gn=CIT;refGene_tx=XM_019814613.1;refGene_gn=CIT;refGene_tx=XM_019814614.1;refGene_gn=CIT;refGene_tx=XM_019814615.1;refGene_gn=CIT;refGene_tx=XM_019814616.1;refGene_gn=CIT;refGene_tx=XM_019814617.2;refGene_gn=CIT;refGene_tx=XM_023241475.1
D3 16230774 16230880
refGene_gn=CIT;refGene_tx=XM_006938519.3;refGene_gn=CIT;refGene_tx=XM_019814610.1;refGene_gn=CIT;refGene_tx=XM_019814611.1;refGene_gn=CIT;refGene_tx=XM_019814612.1;refGene_gn=CIT;refGene_tx=XM_019814613.1;refGene_gn=CIT;refGene_tx=XM_019814614.1;refGene_gn=CIT;refGene_tx=XM_019814615.1;refGene_gn=CIT;refGene_tx=XM_019814616.1;refGene_gn=CIT;refGene_tx=XM_019814617.2;refGene_gn=CIT;refGene_tx=XM_023241475.1
D3 16239861 16239893 refGene_gn=CIT;refGene_tx=XM_019814617.2
D3 16246813 16246997
refGene_gn=CIT;refGene_tx=XM_006938519.3;refGene_gn=CIT;refGene_tx=XM_019814610.1;refGene_gn=CIT;refGene_tx=XM_019814611.1;refGene_gn=CIT;refGene_tx=XM_019814612.1;refGene_gn=CIT;refGene_tx=XM_019814613.1;refGene_gn=CIT;refGene_tx=XM_019814614.1;refGene_gn=CIT;refGene_tx=XM_019814615.1;refGene_gn=CIT;refGene_tx=XM_019814616.1;refGene_gn=CIT;refGene_tx=XM_023241475.1
D3 16267482 16267636
refGene_gn=CIT;refGene_tx=XM_006938519.3;refGene_gn=CIT;refGene_tx=XM_019814610.1;refGene_gn=CIT;refGene_tx=XM_019814611.1;refGene_gn=CIT;refGene_tx=XM_019814612.1;refGene_gn=CIT;refGene_tx=XM_019814613.1;refGene_gn=CIT;refGene_tx=XM_019814614.1;refGene_gn=CIT;refGene_tx=XM_019814615.1;refGene_gn=CIT;refGene_tx=XM_019814616.1;refGene_gn=CIT;refGene_tx=XM_023241475.1
D3 16269483 16269687
refGene_gn=CIT;refGene_tx=XM_006938519.3;refGene_gn=CIT;refGene_tx=XM_019814610.1;refGene_gn=CIT;refGene_tx=XM_019814611.1;refGene_gn=CIT;refGene_tx=XM_019814612.1;refGene_gn=CIT;refGene_tx=XM_019814613.1;refGene_gn=CIT;refGene_tx=XM_019814614.1;refGene_gn=CIT;refGene_tx=XM_019814615.1;refGene_gn=CIT;refGene_tx=XM_019814616.1;refGene_gn=CIT;refGene_tx=XM_023241475.1
D3 16276039 16276133
refGene_gn=CIT;refGene_tx=XM_006938519.3;refGene_gn=CIT;refGene_tx=XM_019814610.1;refGene_gn=CIT;refGene_tx=XM_019814611.1;refGene_gn=CIT;refGene_tx=XM_019814612.1;refGene_gn=CIT;refGene_tx=XM_019814613.1;refGene_gn=CIT;refGene_tx=XM_019814614.1;refGene_gn=CIT;refGene_tx=XM_019814615.1;refGene_gn=CIT;refGene_tx=XM_019814616.1;refGene_gn=CIT;refGene_tx=XM_023241475.1
D3 16277319 16277462
refGene_gn=CIT;refGene_tx=XM_006938519.3;refGene_gn=CIT;refGene_tx=XM_019814610.1;refGene_gn=CIT;refGene_tx=XM_019814611.1;refGene_gn=CIT;refGene_tx=XM_019814612.1;refGene_gn=CIT;refGene_tx=XM_019814613.1;refGene_gn=CIT;refGene_tx=XM_019814614.1;refGene_gn=CIT;refGene_tx=XM_01981

4615.1;refGene_gn=CIT;refGene_tx=XM_019814616.1;refGene_gn=CIT;refGene_tx=XM_023241475.1
D3 16287614 16287716
refGene_gn=CIT;refGene_tx=XM_006938519.3;refGene_gn=CIT;refGene_tx=XM_019814610.1;refGene_gn=CIT;refGene_tx=XM_019814611.1;refGene_gn=CIT;refGene_tx=XM_019814612.1;refGene_gn=CIT;refGene_tx=XM_019814613.1;refGene_gn=CIT;refGene_tx=XM_019814614.1;refGene_gn=CIT;refGene_tx=XM_019814615.1;refGene_gn=CIT;refGene_tx=XM_019814616.1;refGene_gn=CIT;refGene_tx=XM_023241475.1
D3 16293112 16293288
refGene_gn=CIT;refGene_tx=XM_006938519.3;refGene_gn=CIT;refGene_tx=XM_019814610.1;refGene_gn=CIT;refGene_tx=XM_019814611.1;refGene_gn=CIT;refGene_tx=XM_019814612.1;refGene_gn=CIT;refGene_tx=XM_019814613.1;refGene_gn=CIT;refGene_tx=XM_019814614.1;refGene_gn=CIT;refGene_tx=XM_019814615.1;refGene_gn=CIT;refGene_tx=XM_019814616.1;refGene_gn=CIT;refGene_tx=XM_023241475.1
D3 16300638 16300780
refGene_gn=CIT;refGene_tx=XM_006938519.3;refGene_gn=CIT;refGene_tx=XM_019814610.1;refGene_gn=CIT;refGene_tx=XM_019814611.1;refGene_gn=CIT;refGene_tx=XM_019814612.1;refGene_gn=CIT;refGene_tx=XM_019814613.1;refGene_gn=CIT;refGene_tx=XM_019814614.1;refGene_gn=CIT;refGene_tx=XM_019814615.1;refGene_gn=CIT;refGene_tx=XM_019814616.1;refGene_gn=CIT;refGene_tx=XM_023241475.1
D3 16307105 16307201
refGene_gn=CIT;refGene_tx=XM_006938519.3;refGene_gn=CIT;refGene_tx=XM_019814610.1;refGene_gn=CIT;refGene_tx=XM_019814611.1;refGene_gn=CIT;refGene_tx=XM_019814612.1;refGene_gn=CIT;refGene_tx=XM_019814613.1;refGene_gn=CIT;refGene_tx=XM_019814614.1;refGene_gn=CIT;refGene_tx=XM_019814615.1;refGene_gn=CIT;refGene_tx=XM_019814616.1;refGene_gn=CIT;refGene_tx=XM_023241475.1
D3 16385886 16386993
refGene_gn=LOC109493235;refGene_tx=XM_023242152.1
D3 16387039 16387508
refGene_gn=BICDL1;refGene_tx=XM_011287795.3;refGene_gn=BICDL1;refGene_tx=XM_023241476.1;refGene_gn=BICDL1;refGene_tx=XM_023241477.1;refGene_gn=BICDL1;refGene_tx=XM_023241478.1;refGene_gn=LOC109493235;refGene_tx=XM_023242152.1
D3 16396904 16397120
refGene_gn=BICDL1;refGene_tx=XM_011287795.3;refGene_gn=BICDL1;refGene_tx=XM_023241476.1;refGene_gn=BICDL1;refGene_tx=XM_023241477.1;refGene_gn=BICDL1;refGene_tx=XM_023241478.1
D3 16468546 16468663
refGene_gn=BICDL1;refGene_tx=XM_011287795.3;refGene_gn=BICDL1;refGene_tx=XM_023241476.1;refGene_gn=BICDL1;refGene_tx=XM_023241477.1;refGene_gn=BICDL1;refGene_tx=XM_023241478.1
D3 16469636 16469648
refGene_gn=BICDL1;refGene_tx=XM_011287795.3;refGene_gn=BICDL1;refGene_tx=XM_023241477.1
D3 16471544 16471691
refGene_gn=BICDL1;refGene_tx=XM_011287795.3;refGene_gn=BICDL1;refGene_

tx=XM_023241476.1;refGene_gn=BICDL1;refGene_tx=XM_023241477.1;refGene_gn=BICDL1;refGene_tx=XM_023241478.1
D3 16477753 16477933
refGene_gn=BICDL1;refGene_tx=XM_011287795.3;refGene_gn=BICDL1;refGene_tx=XM_023241476.1;refGene_gn=BICDL1;refGene_tx=XM_023241477.1;refGene_gn=BICDL1;refGene_tx=XM_023241478.1
D3 16478533 16478752
refGene_gn=BICDL1;refGene_tx=XM_011287795.3;refGene_gn=BICDL1;refGene_tx=XM_023241476.1;refGene_gn=BICDL1;refGene_tx=XM_023241477.1;refGene_gn=BICDL1;refGene_tx=XM_023241478.1
D3 16479804 16479948
refGene_gn=BICDL1;refGene_tx=XM_011287795.3;refGene_gn=BICDL1;refGene_tx=XM_023241476.1
D3 16484750 16484881
refGene_gn=BICDL1;refGene_tx=XM_011287795.3;refGene_gn=BICDL1;refGene_tx=XM_023241476.1;refGene_gn=BICDL1;refGene_tx=XM_023241477.1;refGene_gn=BICDL1;refGene_tx=XM_023241478.1
D3 16489649 16489770
refGene_gn=BICDL1;refGene_tx=XM_011287795.3;refGene_gn=BICDL1;refGene_tx=XM_023241476.1;refGene_gn=BICDL1;refGene_tx=XM_023241477.1;refGene_gn=BICDL1;refGene_tx=XM_023241478.1
D3 16492565 16492727
refGene_gn=BICDL1;refGene_tx=XM_011287795.3;refGene_gn=BICDL1;refGene_tx=XM_023241476.1;refGene_gn=BICDL1;refGene_tx=XM_023241477.1;refGene_gn=BICDL1;refGene_tx=XM_023241478.1
D3 16496850 16496979
refGene_gn=RAB35;refGene_tx=XM_023241479.1;refGene_gn=RAB35;refGene_tx=XM_023241480.1
D3 16498359 16498484 refGene_gn=RAB35;refGene_tx=XM_023241479.1
D3 16498573 16498698
refGene_gn=RAB35;refGene_tx=XM_023241479.1;refGene_gn=RAB35;refGene_tx=XM_023241480.1
D3 16502792 16502916
refGene_gn=RAB35;refGene_tx=XM_023241479.1;refGene_gn=RAB35;refGene_tx=XM_023241480.1
D3 16505594 16505645
refGene_gn=RAB35;refGene_tx=XM_023241479.1;refGene_gn=RAB35;refGene_tx=XM_023241480.1
D3 16511052 16511104
refGene_gn=RAB35;refGene_tx=XM_023241479.1;refGene_gn=RAB35;refGene_tx=XM_023241480.1
D3 16521330 16521456 refGene_gn=GCN1;refGene_tx=XM_023241481.1
D3 16521975 16522194 refGene_gn=GCN1;refGene_tx=XM_023241481.1
D3 16524290 16524398 refGene_gn=GCN1;refGene_tx=XM_023241481.1
D3 16524944 16525093 refGene_gn=GCN1;refGene_tx=XM_023241481.1
D3 16525435 16525532 refGene_gn=GCN1;refGene_tx=XM_023241481.1
D3 16527661 16527776 refGene_gn=GCN1;refGene_tx=XM_023241481.1
D3 16527918 16528112 refGene_gn=GCN1;refGene_tx=XM_023241481.1
D3 16530086 16530317 refGene_gn=GCN1;refGene_tx=XM_023241481.1
D3 16530757 16530871 refGene_gn=GCN1;refGene_tx=XM_023241481.1

| | | | |
|---|---|---|---|
| D3 | 16531112 | 16531382 | refGene_gn=GCN1;refGene_tx=XM_023241481.1 |
| D3 | 16531477 | 16531621 | refGene_gn=GCN1;refGene_tx=XM_023241481.1 |
| D3 | 16531849 | 16531942 | refGene_gn=GCN1;refGene_tx=XM_023241481.1 |
| D3 | 16532228 | 16532390 | refGene_gn=GCN1;refGene_tx=XM_023241481.1 |
| D3 | 16533565 | 16533730 | refGene_gn=GCN1;refGene_tx=XM_023241481.1 |
| D3 | 16535245 | 16535461 | refGene_gn=GCN1;refGene_tx=XM_023241481.1 |
| D3 | 16535573 | 16535691 | refGene_gn=GCN1;refGene_tx=XM_023241481.1 |
| D3 | 16538431 | 16538574 | refGene_gn=GCN1;refGene_tx=XM_023241481.1 |
| D3 | 16538771 | 16538968 | refGene_gn=GCN1;refGene_tx=XM_023241481.1 |
| D3 | 16539057 | 16539196 | refGene_gn=GCN1;refGene_tx=XM_023241481.1 |
| D3 | 16539399 | 16539468 | refGene_gn=GCN1;refGene_tx=XM_023241481.1 |
| D3 | 16540900 | 16541121 | refGene_gn=GCN1;refGene_tx=XM_023241481.1 |
| D3 | 16543515 | 16543695 | refGene_gn=GCN1;refGene_tx=XM_023241481.1 |
| D3 | 16544534 | 16544649 | refGene_gn=GCN1;refGene_tx=XM_023241481.1 |
| D3 | 16544981 | 16545103 | refGene_gn=GCN1;refGene_tx=XM_023241481.1 |
| D3 | 16545555 | 16545802 | refGene_gn=GCN1;refGene_tx=XM_023241481.1 |
| D3 | 16546752 | 16546947 | refGene_gn=GCN1;refGene_tx=XM_023241481.1 |
| D3 | 16547298 | 16547464 | refGene_gn=GCN1;refGene_tx=XM_023241481.1 |
| D3 | 16548103 | 16548174 | refGene_gn=GCN1;refGene_tx=XM_023241481.1 |
| D3 | 16548383 | 16548573 | refGene_gn=GCN1;refGene_tx=XM_023241481.1 |
| D3 | 16548756 | 16548884 | refGene_gn=GCN1;refGene_tx=XM_023241481.1 |
| D3 | 16549909 | 16550053 | refGene_gn=GCN1;refGene_tx=XM_023241481.1 |
| D3 | 16550417 | 16550498 | refGene_gn=GCN1;refGene_tx=XM_023241481.1 |
| D3 | 16551919 | 16552101 | refGene_gn=GCN1;refGene_tx=XM_023241481.1 |
| D3 | 16552587 | 16552743 | refGene_gn=GCN1;refGene_tx=XM_023241481.1 |
| D3 | 16553629 | 16553828 | refGene_gn=GCN1;refGene_tx=XM_023241481.1 |
| D3 | 16553920 | 16554034 | refGene_gn=GCN1;refGene_tx=XM_023241481.1 |
| D3 | 16555155 | 16555249 | refGene_gn=GCN1;refGene_tx=XM_023241481.1 |
| D3 | 16555554 | 16555733 | refGene_gn=GCN1;refGene_tx=XM_023241481.1 |
| D3 | 16556534 | 16556659 | refGene_gn=GCN1;refGene_tx=XM_023241481.1 |
| D3 | 16556757 | 16556947 | refGene_gn=GCN1;refGene_tx=XM_023241481.1 |
| D3 | 16557982 | 16558142 | refGene_gn=GCN1;refGene_tx=XM_023241481.1 |
| D3 | 16558287 | 16558363 | refGene_gn=GCN1;refGene_tx=XM_023241481.1 |
| D3 | 16559440 | 16559533 | refGene_gn=GCN1;refGene_tx=XM_023241481.1 |
| D3 | 16560679 | 16560832 | refGene_gn=GCN1;refGene_tx=XM_023241481.1 |
| D3 | 16562367 | 16562541 | refGene_gn=GCN1;refGene_tx=XM_023241481.1 |
| D3 | 16562760 | 16562859 | refGene_gn=GCN1;refGene_tx=XM_023241481.1 |
| D3 | 16563144 | 16563195 | refGene_gn=GCN1;refGene_tx=XM_023241481.1 |
| D3 | 16563714 | 16563843 | refGene_gn=GCN1;refGene_tx=XM_023241481.1 |
| D3 | 16564492 | 16564567 | refGene_gn=GCN1;refGene_tx=XM_023241481.1 |
| D3 | 16565038 | 16565147 | refGene_gn=GCN1;refGene_tx=XM_023241481.1 |
| D3 | 16565269 | 16565338 | refGene_gn=GCN1;refGene_tx=XM_023241481.1 |
| D3 | 16566211 | 16566346 | refGene_gn=GCN1;refGene_tx=XM_023241481.1 |
| D3 | 16566432 | 16566531 | refGene_gn=GCN1;refGene_tx=XM_023241481.1 |
| D3 | 16569874 | 16569983 | refGene_gn=GCN1;refGene_tx=XM_023241481.1 |
| D3 | 16570496 | 16570628 | refGene_gn=GCN1;refGene_tx=XM_023241481.1 |
| D3 | 16571139 | 16571203 | refGene_gn=GCN1;refGene_tx=XM_023241481.1 |
| D3 | 16575073 | 16575091 | refGene_gn=GCN1;refGene_tx=XM_023241481.1 |
| D3 | 16575093 | 16575175 | refGene_gn=GCN1;refGene_tx=XM_023241481.1 |
| D3 | 16580056 | 16580074 | refGene_gn=GCN1;refGene_tx=XM_023241481.1 |

D3 16582598 16582760 refGene_gn=RPLP0;refGene_tx=XM_003994717.5
D3 16583326 16583467 refGene_gn=RPLP0;refGene_tx=XM_003994717.5
D3 16584320 16584506 refGene_gn=RPLP0;refGene_tx=XM_003994717.5
D3 16584607 16584754 refGene_gn=RPLP0;refGene_tx=XM_003994717.5
D3 16584877 16584965 refGene_gn=RPLP0;refGene_tx=XM_003994717.5
D3 16585077 16585253 refGene_gn=RPLP0;refGene_tx=XM_003994717.5
D3 16586240 16586294 refGene_gn=RPLP0;refGene_tx=XM_003994717.5
D3 16591912 16592179
refGene_gn=PXN;refGene_tx=XM_011287790.2;refGene_gn=PXN;refGene_tx=XM_011287791.2;refGene_gn=PXN;refGene_tx=XM_023241482.1;refGene_gn=PXN;refGene_tx=XM_023241483.1;refGene_gn=PXN;refGene_tx=XM_023241484.1
D3 16592942 16593091
refGene_gn=PXN;refGene_tx=XM_011287790.2;refGene_gn=PXN;refGene_tx=XM_011287791.2;refGene_gn=PXN;refGene_tx=XM_023241482.1;refGene_gn=PXN;refGene_tx=XM_023241483.1;refGene_gn=PXN;refGene_tx=XM_023241484.1
D3 16593251 16593333
refGene_gn=PXN;refGene_tx=XM_011287790.2;refGene_gn=PXN;refGene_tx=XM_011287791.2;refGene_gn=PXN;refGene_tx=XM_023241482.1;refGene_gn=PXN;refGene_tx=XM_023241483.1;refGene_gn=PXN;refGene_tx=XM_023241484.1
D3 16593903 16594077
refGene_gn=PXN;refGene_tx=XM_011287790.2;refGene_gn=PXN;refGene_tx=XM_011287791.2;refGene_gn=PXN;refGene_tx=XM_023241482.1;refGene_gn=PXN;refGene_tx=XM_023241483.1;refGene_gn=PXN;refGene_tx=XM_023241484.1
D3 16594195 16594366
refGene_gn=PXN;refGene_tx=XM_011287790.2;refGene_gn=PXN;refGene_tx=XM_011287791.2;refGene_gn=PXN;refGene_tx=XM_023241482.1;refGene_gn=PXN;refGene_tx=XM_023241483.1;refGene_gn=PXN;refGene_tx=XM_023241484.1
D3 16594635 16594734
refGene_gn=PXN;refGene_tx=XM_011287790.2;refGene_gn=PXN;refGene_tx=XM_011287791.2;refGene_gn=PXN;refGene_tx=XM_023241482.1;refGene_gn=PXN;refGene_tx=XM_023241483.1
D3 16595363 16595675
refGene_gn=PXN;refGene_tx=XM_011287790.2;refGene_gn=PXN;refGene_tx=XM_011287791.2;refGene_gn=PXN;refGene_tx=XM_023241482.1
D3 16595939 16596215
refGene_gn=PXN;refGene_tx=XM_011287790.2;refGene_gn=PXN;refGene_tx=XM_023241482.1
D3 16597347 16598256
refGene_gn=PXN;refGene_tx=XM_011287790.2;refGene_gn=PXN;refGene_tx=XM_023241482.1
D3 16599736 16599872
refGene_gn=PXN;refGene_tx=XM_011287790.2;refGene_gn=PXN;refGene_tx=XM_011287791.2;refGene_gn=PXN;refGene_tx=XM_023241482.1;refGene_gn=PXN;refGene_tx=XM_023241483.1;refGene_gn=PXN;refGene_tx=XM_023241484.1
D3 16600729 16600931
refGene_gn=PXN;refGene_tx=XM_011287790.2;refGene_gn=PXN;refGene_tx=XM_011287791.2;refGene_gn=PXN;refGene_tx=XM_023241482.1;refGene_gn=PXN;refGene_tx=XM_023241483.1;refGene_gn=PXN;refGene_tx=XM_023241484.1
D3 16601040 16601177
refGene_gn=PXN;refGene_tx=XM_011287790.2;refGene_gn=PXN;refGene_tx=XM_

011287791.2;refGene_gn=PXN;refGene_tx=XM_023241482.1;refGene_gn=PXN;refGene_tx=XM_023241483.1;refGene_gn=PXN;refGene_tx=XM_023241484.1
D3	16601529	16601645	refGene_gn=PXN;refGene_tx=XM_023241482.1
D3	16601935	16602162	refGene_gn=PXN;refGene_tx=XM_023241482.1
D3	16637205	16637218	refGene_gn=PXN;refGene_tx=XM_023241482.1
D3	16666040	16666531	refGene_gn=SIRT4;refGene_tx=XM_003994719.5;refGene_gn=SIRT4;refGene_tx=XM_006938523.4;refGene_gn=SIRT4;refGene_tx=XM_019814619.2
D3	16668561	16668853	refGene_gn=SIRT4;refGene_tx=XM_003994719.5;refGene_gn=SIRT4;refGene_tx=XM_006938523.4;refGene_gn=SIRT4;refGene_tx=XM_019814619.2
D3	16668975	16669128	refGene_gn=SIRT4;refGene_tx=XM_003994719.5;refGene_gn=SIRT4;refGene_tx=XM_006938523.4;refGene_gn=SIRT4;refGene_tx=XM_019814619.2
D3	16685390	16685515	refGene_gn=PLA2G1B;refGene_tx=XM_019814620.1
D3	16689210	16689338	refGene_gn=PLA2G1B;refGene_tx=XM_019814620.1
D3	16689850	16690010	refGene_gn=PLA2G1B;refGene_tx=XM_019814620.1
D3	16694041	16694075	refGene_gn=PLA2G1B;refGene_tx=XM_019814620.1
D3	16708280	16708322	refGene_gn=MSI1;refGene_tx=XM_023241485.1;refGene_gn=MSI1;refGene_tx=XM_023241486.1;refGene_gn=MSI1;refGene_tx=XM_023241487.1;refGene_gn=MSI1;refGene_tx=XM_023241488.1
D3	16708717	16708905	refGene_gn=MSI1;refGene_tx=XM_023241485.1;refGene_gn=MSI1;refGene_tx=XM_023241486.1;refGene_gn=MSI1;refGene_tx=XM_023241487.1;refGene_gn=MSI1;refGene_tx=XM_023241488.1
D3	16709651	16709720	refGene_gn=MSI1;refGene_tx=XM_023241485.1;refGene_gn=MSI1;refGene_tx=XM_023241486.1;refGene_gn=MSI1;refGene_tx=XM_023241487.1;refGene_gn=MSI1;refGene_tx=XM_023241488.1
D3	16712803	16712860	refGene_gn=MSI1;refGene_tx=XM_023241485.1;refGene_gn=MSI1;refGene_tx=XM_023241486.1
D3	16714140	16714221	refGene_gn=MSI1;refGene_tx=XM_023241485.1;refGene_gn=MSI1;refGene_tx=XM_023241486.1;refGene_gn=MSI1;refGene_tx=XM_023241487.1;refGene_gn=MSI1;refGene_tx=XM_023241488.1
D3	16716972	16717090	refGene_gn=MSI1;refGene_tx=XM_023241485.1;refGene_gn=MSI1;refGene_tx=XM_023241486.1;refGene_gn=MSI1;refGene_tx=XM_023241487.1;refGene_gn=MSI1;refGene_tx=XM_023241488.1
D3	16717562	16717645	refGene_gn=MSI1;refGene_tx=XM_023241485.1;refGene_gn=MSI1;refGene_tx=XM_023241486.1;refGene_gn=MSI1;refGene_tx=XM_023241487.1;refGene_gn=MSI1;refGene_tx=XM_023241488.1
D3	16718835	16718884	refGene_gn=MSI1;refGene_tx=XM_023241485.1;refGene_gn=MSI1;refGene_tx=XM_023241486.1;refGene_gn=MSI1;refGene_tx=XM_023241487.1;refGene_gn=MSI1;refGene_tx=XM_023241488.1
D3	16722515	16722608

refGene_gn=MSI1;refGene_tx=XM_023241485.1;refGene_gn=MSI1;refGene_tx=XM_023241486.1;refGene_gn=MSI1;refGene_tx=XM_023241487.1;refGene_gn=MSI1;refGene_tx=XM_023241488.1
D3 16724152 16724194
refGene_gn=MSI1;refGene_tx=XM_023241485.1;refGene_gn=MSI1;refGene_tx=XM_023241486.1;refGene_gn=MSI1;refGene_tx=XM_023241487.1;refGene_gn=MSI1;refGene_tx=XM_023241488.1
D3 16727084 16727169
refGene_gn=MSI1;refGene_tx=XM_023241485.1;refGene_gn=MSI1;refGene_tx=XM_023241486.1;refGene_gn=MSI1;refGene_tx=XM_023241487.1;refGene_gn=MSI1;refGene_tx=XM_023241488.1
D3 16727267 16727349
refGene_gn=MSI1;refGene_tx=XM_023241485.1;refGene_gn=MSI1;refGene_tx=XM_023241486.1;refGene_gn=MSI1;refGene_tx=XM_023241487.1;refGene_gn=MSI1;refGene_tx=XM_023241488.1
D3 16727914 16727955
refGene_gn=MSI1;refGene_tx=XM_023241485.1;refGene_gn=MSI1;refGene_tx=XM_023241486.1;refGene_gn=MSI1;refGene_tx=XM_023241487.1;refGene_gn=MSI1;refGene_tx=XM_023241488.1
D3 16728124 16728183
refGene_gn=MSI1;refGene_tx=XM_023241485.1;refGene_gn=MSI1;refGene_tx=XM_023241486.1;refGene_gn=MSI1;refGene_tx=XM_023241487.1;refGene_gn=MSI1;refGene_tx=XM_023241488.1
D3 16782512 16782615
refGene_gn=LOC101095773;refGene_tx=XM_003994721.5
D3 16782765 16782908
refGene_gn=LOC101095773;refGene_tx=XM_003994721.5
D3 16784046 16784130
refGene_gn=LOC101095773;refGene_tx=XM_003994721.5
D3 16786710 16786794 refGene_gn=TRIAP1;refGene_tx=XM_003994722.5
D3 16787749 16787896 refGene_gn=TRIAP1;refGene_tx=XM_003994722.5
D3 16787995 16788076 refGene_gn=GATC;refGene_tx=XM_003994723.3
D3 16788170 16788343 refGene_gn=GATC;refGene_tx=XM_003994723.3
D3 16794468 16794572 refGene_gn=GATC;refGene_tx=XM_003994723.3
D3 16796521 16796574 refGene_gn=GATC;refGene_tx=XM_003994723.3
D3 16798763 16798907 refGene_gn=SRSF9;refGene_tx=XM_006938532.3
D3 16800684 16800857 refGene_gn=SRSF9;refGene_tx=XM_006938532.3
D3 16802617 16802778 refGene_gn=SRSF9;refGene_tx=XM_006938532.3
D3 16805489 16805677 refGene_gn=SRSF9;refGene_tx=XM_006938532.3
D3 16830596 16830728
refGene_gn=DYNLL1;refGene_tx=XM_003994724.3;refGene_gn=DYNLL1;refGene_tx=XM_019814621.2
D3 16832288 16832426
refGene_gn=DYNLL1;refGene_tx=XM_003994724.3;refGene_gn=DYNLL1;refGene_tx=XM_019814621.2
D3 16836108 16836210 refGene_gn=COQ5;refGene_tx=XM_003994726.3
D3 16836290 16836402 refGene_gn=COQ5;refGene_tx=XM_003994726.3
D3 16837167 16837256 refGene_gn=COQ5;refGene_tx=XM_003994726.3
D3 16839775 16839882 refGene_gn=COQ5;refGene_tx=XM_003994726.3
D3 16844394 16844616 refGene_gn=COQ5;refGene_tx=XM_003994726.3

D3 16848958 16849108 refGene_gn=COQ5;refGene_tx=XM_003994726.3
D3 16853866 16854113 refGene_gn=COQ5;refGene_tx=XM_003994726.3
D3 16857999 16858156 refGene_gn=RNF10;refGene_tx=XM_003994750.4;refGene_gn=RNF10;refGene_tx=XM_023241489.1;refGene_gn=RNF10;refGene_tx=XM_023241490.1
D3 16869730 16869927 refGene_gn=RNF10;refGene_tx=XM_003994750.4;refGene_gn=RNF10;refGene_tx=XM_023241489.1;refGene_gn=RNF10;refGene_tx=XM_023241490.1
D3 16875135 16875335 refGene_gn=RNF10;refGene_tx=XM_003994750.4;refGene_gn=RNF10;refGene_tx=XM_023241489.1;refGene_gn=RNF10;refGene_tx=XM_023241490.1
D3 16876282 16876373 refGene_gn=RNF10;refGene_tx=XM_003994750.4;refGene_gn=RNF10;refGene_tx=XM_023241489.1;refGene_gn=RNF10;refGene_tx=XM_023241490.1
D3 16877627 16877812 refGene_gn=RNF10;refGene_tx=XM_003994750.4;refGene_gn=RNF10;refGene_tx=XM_023241489.1;refGene_gn=RNF10;refGene_tx=XM_023241490.1
D3 16877891 16878028 refGene_gn=RNF10;refGene_tx=XM_003994750.4;refGene_gn=RNF10;refGene_tx=XM_023241489.1;refGene_gn=RNF10;refGene_tx=XM_023241490.1
D3 16880822 16880983 refGene_gn=RNF10;refGene_tx=XM_003994750.4;refGene_gn=RNF10;refGene_tx=XM_023241489.1;refGene_gn=RNF10;refGene_tx=XM_023241490.1
D3 16882226 16882352 refGene_gn=RNF10;refGene_tx=XM_003994750.4;refGene_gn=RNF10;refGene_tx=XM_023241489.1;refGene_gn=RNF10;refGene_tx=XM_023241490.1
D3 16882608 16882870 refGene_gn=RNF10;refGene_tx=XM_003994750.4;refGene_gn=RNF10;refGene_tx=XM_023241489.1;refGene_gn=RNF10;refGene_tx=XM_023241490.1
D3 16883084 16883218 refGene_gn=RNF10;refGene_tx=XM_003994750.4;refGene_gn=RNF10;refGene_tx=XM_023241489.1;refGene_gn=RNF10;refGene_tx=XM_023241490.1
D3 16884180 16884298 refGene_gn=RNF10;refGene_tx=XM_003994750.4;refGene_gn=RNF10;refGene_tx=XM_023241489.1;refGene_gn=RNF10;refGene_tx=XM_023241490.1
D3 16884525 16884627 refGene_gn=RNF10;refGene_tx=XM_003994750.4;refGene_gn=RNF10;refGene_tx=XM_023241489.1;refGene_gn=RNF10;refGene_tx=XM_023241490.1
D3 16886976 16887132 refGene_gn=RNF10;refGene_tx=XM_003994750.4;refGene_gn=RNF10;refGene_tx=XM_023241489.1;refGene_gn=RNF10;refGene_tx=XM_023241490.1
D3 16888644 16888745 refGene_gn=RNF10;refGene_tx=XM_003994750.4;refGene_gn=RNF10;refGene_tx=XM_023241489.1;refGene_gn=RNF10;refGene_tx=XM_023241490.1
D3 16891833 16891891 refGene_gn=RNF10;refGene_tx=XM_003994750.4;refGene_gn=RNF10;refGene_tx=XM_023241489.1;refGene_gn=RNF10;refGene_tx=XM_023241490.1
D3 16891963 16892181 refGene_gn=RNF10;refGene_tx=XM_003994750.4;refGene_gn=RNF10;refGene_tx=XM_023241489.1;refGene_gn=RNF10;refGene_tx=XM_023241490.1

D3 16892717 16893068
refGene_gn=RNF10;refGene_tx=XM_003994750.4;refGene_gn=RNF10;refGene_tx=XM_023241489.1;refGene_gn=RNF10;refGene_tx=XM_023241490.1
D3 16895000 16895116 refGene_gn=POP5;refGene_tx=XM_003994727.4
D3 16895213 16895297 refGene_gn=POP5;refGene_tx=XM_003994727.4
D3 16895477 16895627 refGene_gn=POP5;refGene_tx=XM_003994727.4
D3 16896781 16896924 refGene_gn=POP5;refGene_tx=XM_003994727.4
D3 16897009 16897029 refGene_gn=POP5;refGene_tx=XM_003994727.4
D3 16933671 16934319
refGene_gn=CABP1;refGene_tx=XM_023241491.1;refGene_gn=CABP1;refGene_tx=XM_023241492.1
D3 16934780 16934843 refGene_gn=CABP1;refGene_tx=XM_023241493.1
D3 16941772 16941817 refGene_gn=CABP1;refGene_tx=XM_011287785.3
D3 16946393 16946576
refGene_gn=CABP1;refGene_tx=XM_011287785.3;refGene_gn=CABP1;refGene_tx=XM_023241491.1
D3 16949604 16949635
refGene_gn=CABP1;refGene_tx=XM_011287785.3;refGene_gn=CABP1;refGene_tx=XM_023241491.1;refGene_gn=CABP1;refGene_tx=XM_023241492.1;refGene_gn=CABP1;refGene_tx=XM_023241493.1
D3 16949909 16950053
refGene_gn=CABP1;refGene_tx=XM_011287785.3;refGene_gn=CABP1;refGene_tx=XM_023241491.1;refGene_gn=CABP1;refGene_tx=XM_023241492.1;refGene_gn=CABP1;refGene_tx=XM_023241493.1
D3 16951087 16951197
refGene_gn=CABP1;refGene_tx=XM_011287785.3;refGene_gn=CABP1;refGene_tx=XM_023241491.1;refGene_gn=CABP1;refGene_tx=XM_023241492.1;refGene_gn=CABP1;refGene_tx=XM_023241493.1
D3 16951424 16951572
refGene_gn=CABP1;refGene_tx=XM_011287785.3;refGene_gn=CABP1;refGene_tx=XM_023241491.1;refGene_gn=CABP1;refGene_tx=XM_023241492.1;refGene_gn=CABP1;refGene_tx=XM_023241493.1
D3 16954085 16954111
refGene_gn=CABP1;refGene_tx=XM_011287785.3;refGene_gn=CABP1;refGene_tx=XM_023241491.1;refGene_gn=CABP1;refGene_tx=XM_023241492.1;refGene_gn=CABP1;refGene_tx=XM_023241493.1
D3 16967914 16967918 refGene_gn=MLEC;refGene_tx=XM_011287783.3
D3 16968154 16968383
refGene_gn=MLEC;refGene_tx=XM_003994728.4;refGene_gn=MLEC;refGene_tx=XM_011287784.3
D3 16974765 16974944
refGene_gn=MLEC;refGene_tx=XM_003994728.4;refGene_gn=MLEC;refGene_tx=XM_011287783.3;refGene_gn=MLEC;refGene_tx=XM_011287784.3
D3 16975533 16975710
refGene_gn=MLEC;refGene_tx=XM_003994728.4;refGene_gn=MLEC;refGene_tx=XM_011287783.3
D3 16975816 16975874
refGene_gn=MLEC;refGene_tx=XM_003994728.4;refGene_gn=MLEC;refGene_tx=XM_011287783.3
D3 16977284 16977599

refGene_gn=MLEC;refGene_tx=XM_003994728.4;refGene_gn=MLEC;refGene_tx=XM_011287783.3;refGene_gn=MLEC;refGene_tx=XM_011287784.3
D3 16989118 16989362 refGene_gn=UNC119B;refGene_tx=XM_003994729.3
D3 16992020 16992134 refGene_gn=UNC119B;refGene_tx=XM_003994729.3
D3 16994695 16994807 refGene_gn=UNC119B;refGene_tx=XM_003994729.3
D3 16994932 16995105 refGene_gn=UNC119B;refGene_tx=XM_003994729.3
D3 16997262 16997375 refGene_gn=UNC119B;refGene_tx=XM_003994729.3
D3 17003289 17003335 refGene_gn=ACADS;refGene_tx=XM_023241494.1
D3 17004874 17005038 refGene_gn=ACADS;refGene_tx=XM_023241494.1
D3 17012007 17012157 refGene_gn=ACADS;refGene_tx=XM_023241494.1
D3 17012395 17012507 refGene_gn=ACADS;refGene_tx=XM_023241494.1
D3 17012968 17013120 refGene_gn=ACADS;refGene_tx=XM_023241494.1
D3 17013420 17013591 refGene_gn=ACADS;refGene_tx=XM_023241494.1
D3 17013662 17013800 refGene_gn=ACADS;refGene_tx=XM_023241494.1
D3 17013976 17014072 refGene_gn=ACADS;refGene_tx=XM_023241494.1
D3 17014273 17014330 refGene_gn=ACADS;refGene_tx=XM_023241494.1
D3 17014489 17014642 refGene_gn=ACADS;refGene_tx=XM_023241494.1
D3 17060068 17060140
refGene_gn=SPPL3;refGene_tx=XM_019814624.2;refGene_gn=SPPL3;refGene_tx=XM_019814625.1;refGene_gn=SPPL3;refGene_tx=XM_023241495.1
D3 17060793 17060903
refGene_gn=SPPL3;refGene_tx=XM_019814624.2;refGene_gn=SPPL3;refGene_tx=XM_019814625.1;refGene_gn=SPPL3;refGene_tx=XM_023241495.1
D3 17061893 17062093
refGene_gn=SPPL3;refGene_tx=XM_019814624.2;refGene_gn=SPPL3;refGene_tx=XM_019814625.1;refGene_gn=SPPL3;refGene_tx=XM_023241495.1
D3 17062846 17063010
refGene_gn=SPPL3;refGene_tx=XM_019814624.2;refGene_gn=SPPL3;refGene_tx=XM_019814625.1;refGene_gn=SPPL3;refGene_tx=XM_023241495.1
D3 17063270 17063377
refGene_gn=SPPL3;refGene_tx=XM_019814624.2;refGene_gn=SPPL3;refGene_tx=XM_019814625.1;refGene_gn=SPPL3;refGene_tx=XM_023241495.1
D3 17072571 17072684
refGene_gn=SPPL3;refGene_tx=XM_019814624.2;refGene_gn=SPPL3;refGene_tx=XM_019814625.1;refGene_gn=SPPL3;refGene_tx=XM_023241495.1
D3 17073601 17073680
refGene_gn=SPPL3;refGene_tx=XM_019814624.2;refGene_gn=SPPL3;refGene_tx=XM_019814625.1;refGene_gn=SPPL3;refGene_tx=XM_023241495.1
D3 17074919 17075039
refGene_gn=SPPL3;refGene_tx=XM_019814624.2;refGene_gn=SPPL3;refGene_tx=XM_019814625.1;refGene_gn=SPPL3;refGene_tx=XM_023241495.1
D3 17078931 17078968 refGene_gn=SPPL3;refGene_tx=XM_019814625.1
D3 17082693 17082782
refGene_gn=SPPL3;refGene_tx=XM_019814624.2;refGene_gn=SPPL3;refGene_tx=XM_023241495.1
D3 17107608 17107686 refGene_gn=SPPL3;refGene_tx=XM_023241495.1
D3 17196401 17196424 refGene_gn=SPPL3;refGene_tx=XM_023241495.1
D3 17232727 17232781
refGene_gn=CD3H12orf43;refGene_tx=XM_006938527.3
D3 17232954 17232985

refGene_gn=CD3H12orf43;refGene_tx=XM_019814627.1
D3	17236661 17236990 refGene_gn=HNF1A;refGene_tx=XM_003994731.4
D3	17237502 17237545
refGene_gn=CD3H12orf43;refGene_tx=XM_006938527.3
D3	17238445 17238577
refGene_gn=CD3H12orf43;refGene_tx=XM_006938527.3;refGene_gn=CD3H12orf4
3;refGene_tx=XM_019814627.1
D3	17243995 17244195 refGene_gn=HNF1A;refGene_tx=XM_003994731.4
D3	17244319 17244409
refGene_gn=CD3H12orf43;refGene_tx=XM_006938527.3;refGene_gn=CD3H12orf4
3;refGene_tx=XM_019814627.1
D3	17249894 17250081 refGene_gn=HNF1A;refGene_tx=XM_003994731.4
D3	17250525 17250767 refGene_gn=HNF1A;refGene_tx=XM_003994731.4
D3	17251603 17251755 refGene_gn=HNF1A;refGene_tx=XM_003994731.4
D3	17251896 17252098 refGene_gn=HNF1A;refGene_tx=XM_003994731.4
D3	17253015 17253207 refGene_gn=HNF1A;refGene_tx=XM_003994731.4
D3	17254327 17254449 refGene_gn=HNF1A;refGene_tx=XM_003994731.4
D3	17254537 17254682 refGene_gn=HNF1A;refGene_tx=XM_003994731.4
D3	17256892 17257020 refGene_gn=HNF1A;refGene_tx=XM_003994731.4
D3	17259260 17259618
refGene_gn=CD3H12orf43;refGene_tx=XM_003994733.4;refGene_gn=CD3H12orf4
3;refGene_tx=XM_006938526.3
D3	17260089 17260180
refGene_gn=CD3H12orf43;refGene_tx=XM_003994733.4;refGene_gn=CD3H12orf4
3;refGene_tx=XM_006938526.3;refGene_gn=CD3H12orf43;refGene_tx=XM_00693
8527.3;refGene_gn=CD3H12orf43;refGene_tx=XM_019814627.1
D3	17260702 17260776
refGene_gn=CD3H12orf43;refGene_tx=XM_003994733.4;refGene_gn=CD3H12orf4
3;refGene_tx=XM_006938526.3;refGene_gn=CD3H12orf43;refGene_tx=XM_00693
8527.3;refGene_gn=CD3H12orf43;refGene_tx=XM_019814627.1
D3	17263323 17263422
refGene_gn=CD3H12orf43;refGene_tx=XM_003994733.4;refGene_gn=CD3H12orf4
3;refGene_tx=XM_006938526.3;refGene_gn=CD3H12orf43;refGene_tx=XM_00693
8527.3;refGene_gn=CD3H12orf43;refGene_tx=XM_019814627.1
D3	17263615 17263658
refGene_gn=CD3H12orf43;refGene_tx=XM_003994733.4;refGene_gn=CD3H12orf4
3;refGene_tx=XM_006938526.3;refGene_gn=CD3H12orf43;refGene_tx=XM_00693
8527.3;refGene_gn=CD3H12orf43;refGene_tx=XM_019814627.1
D3	17268500 17268645
refGene_gn=CD3H12orf43;refGene_tx=XM_003994733.4;refGene_gn=CD3H12orf4
3;refGene_tx=XM_006938526.3;refGene_gn=CD3H12orf43;refGene_tx=XM_00693
8527.3;refGene_gn=CD3H12orf43;refGene_tx=XM_019814627.1
D3	17270754 17271255 refGene_gn=OASL;refGene_tx=XM_003994734.4
D3	17272279 17272427 refGene_gn=OASL;refGene_tx=XM_003994734.4
D3	17275562 17275804 refGene_gn=OASL;refGene_tx=XM_003994734.4
D3	17282634 17282810 refGene_gn=OASL;refGene_tx=XM_003994734.4
D3	17283964 17284247 refGene_gn=OASL;refGene_tx=XM_003994734.4
D3	17288034 17288232 refGene_gn=OASL;refGene_tx=XM_003994734.4
D3	17291741 17292257
refGene_gn=LOC101099332;refGene_tx=XM_003994735.4

D3 17293741 17293883
refGene_gn=LOC101099332;refGene_tx=XM_003994735.4
D3 17294362 17294604
refGene_gn=LOC101099332;refGene_tx=XM_003994735.4
D3 17298865 17299041
refGene_gn=LOC101099332;refGene_tx=XM_003994735.4
D3 17300268 17300557
refGene_gn=LOC101099332;refGene_tx=XM_003994735.4
D3 17306498 17306696
refGene_gn=LOC101099332;refGene_tx=XM_003994735.4
D3 17557608 17557987
refGene_gn=WSCD2;refGene_tx=XM_003994869.5;refGene_gn=WSCD2;refGene_tx=XM_019814629.2;refGene_gn=WSCD2;refGene_tx=XM_019814630.2;refGene_gn=WSCD2;refGene_tx=XM_019814631.2;refGene_gn=WSCD2;refGene_tx=XM_019814632.2;refGene_gn=WSCD2;refGene_tx=XM_019814633.2;refGene_gn=WSCD2;refGene_tx=XM_019814634.2;refGene_gn=WSCD2;refGene_tx=XM_019814635.2;refGene_gn=WSCD2;refGene_tx=XM_023241496.1
D3 17569054 17569228
refGene_gn=WSCD2;refGene_tx=XM_019814629.2;refGene_gn=WSCD2;refGene_tx=XM_019814630.2;refGene_gn=WSCD2;refGene_tx=XM_019814631.2;refGene_gn=WSCD2;refGene_tx=XM_019814632.2;refGene_gn=WSCD2;refGene_tx=XM_019814633.2;refGene_gn=WSCD2;refGene_tx=XM_019814634.2;refGene_gn=WSCD2;refGene_tx=XM_023241496.1
D3 17569679 17569794
refGene_gn=WSCD2;refGene_tx=XM_003994869.5;refGene_gn=WSCD2;refGene_tx=XM_019814629.2;refGene_gn=WSCD2;refGene_tx=XM_019814630.2;refGene_gn=WSCD2;refGene_tx=XM_019814631.2;refGene_gn=WSCD2;refGene_tx=XM_019814632.2;refGene_gn=WSCD2;refGene_tx=XM_019814633.2;refGene_gn=WSCD2;refGene_tx=XM_019814634.2;refGene_gn=WSCD2;refGene_tx=XM_023241496.1
D3 17574000 17574185
refGene_gn=WSCD2;refGene_tx=XM_003994869.5;refGene_gn=WSCD2;refGene_tx=XM_019814629.2;refGene_gn=WSCD2;refGene_tx=XM_019814630.2;refGene_gn=WSCD2;refGene_tx=XM_019814631.2;refGene_gn=WSCD2;refGene_tx=XM_019814632.2;refGene_gn=WSCD2;refGene_tx=XM_019814633.2;refGene_gn=WSCD2;refGene_tx=XM_019814634.2;refGene_gn=WSCD2;refGene_tx=XM_023241496.1
D3 17588876 17588998
refGene_gn=WSCD2;refGene_tx=XM_003994869.5;refGene_gn=WSCD2;refGene_tx=XM_019814629.2;refGene_gn=WSCD2;refGene_tx=XM_019814630.2;refGene_gn=WSCD2;refGene_tx=XM_019814631.2;refGene_gn=WSCD2;refGene_tx=XM_019814632.2;refGene_gn=WSCD2;refGene_tx=XM_019814633.2;refGene_gn=WSCD2;refGene_tx=XM_019814634.2;refGene_gn=WSCD2;refGene_tx=XM_019814635.2;refGene_gn=WSCD2;refGene_tx=XM_023241496.1
D3 17591429 17591604
refGene_gn=WSCD2;refGene_tx=XM_003994869.5;refGene_gn=WSCD2;refGene_tx=XM_019814629.2;refGene_gn=WSCD2;refGene_tx=XM_019814630.2;refGene_gn=WSCD2;refGene_tx=XM_019814631.2;refGene_gn=WSCD2;refGene_tx=XM_019814632.2;refGene_gn=WSCD2;refGene_tx=XM_019814633.2;refGene_gn=WSCD2;refGene_tx=XM_019814634.2;refGene_gn=WSCD2;refGene_tx=XM_019814635.2;refGene_gn=WSCD2;refGene_tx=XM_023241496.1
D3 17597687 17597852

refGene_gn=WSCD2;refGene_tx=XM_003994869.5;refGene_gn=WSCD2;refGene_tx=XM_019814629.2;refGene_gn=WSCD2;refGene_tx=XM_019814630.2;refGene_gn=WSCD2;refGene_tx=XM_019814631.2;refGene_gn=WSCD2;refGene_tx=XM_019814632.2;refGene_gn=WSCD2;refGene_tx=XM_019814633.2;refGene_gn=WSCD2;refGene_tx=XM_019814634.2;refGene_gn=WSCD2;refGene_tx=XM_019814635.2;refGene_gn=WSCD2;refGene_tx=XM_023241496.1
D3 17604974 17605175
refGene_gn=WSCD2;refGene_tx=XM_003994869.5;refGene_gn=WSCD2;refGene_tx=XM_019814629.2;refGene_gn=WSCD2;refGene_tx=XM_019814630.2;refGene_gn=WSCD2;refGene_tx=XM_019814631.2;refGene_gn=WSCD2;refGene_tx=XM_019814632.2;refGene_gn=WSCD2;refGene_tx=XM_019814633.2;refGene_gn=WSCD2;refGene_tx=XM_019814634.2;refGene_gn=WSCD2;refGene_tx=XM_019814635.2;refGene_gn=WSCD2;refGene_tx=XM_023241496.1
D3 17611225 17611578
refGene_gn=WSCD2;refGene_tx=XM_003994869.5;refGene_gn=WSCD2;refGene_tx=XM_019814629.2;refGene_gn=WSCD2;refGene_tx=XM_019814630.2;refGene_gn=WSCD2;refGene_tx=XM_019814631.2;refGene_gn=WSCD2;refGene_tx=XM_019814632.2;refGene_gn=WSCD2;refGene_tx=XM_019814633.2;refGene_gn=WSCD2;refGene_tx=XM_019814634.2;refGene_gn=WSCD2;refGene_tx=XM_019814635.2;refGene_gn=WSCD2;refGene_tx=XM_023241496.1
D3 17654441 17655539
refGene_gn=CMKLR1;refGene_tx=XM_023241497.1;refGene_gn=CMKLR1;refGene_tx=XM_023241498.1;refGene_gn=CMKLR1;refGene_tx=XM_023241499.1;refGene_gn=CMKLR1;refGene_tx=XM_023241500.1;refGene_gn=CMKLR1;refGene_tx=XM_023241501.1;refGene_gn=CMKLR1;refGene_tx=XM_023241502.1
D3 17833718 17833840 refGene_gn=FICD;refGene_tx=XM_003994867.5
D3 17835323 17835676
refGene_gn=FICD;refGene_tx=XM_003994867.5;refGene_gn=FICD;refGene_tx=XM_023241503.1
D3 17836918 17837994
refGene_gn=FICD;refGene_tx=XM_003994867.5;refGene_gn=FICD;refGene_tx=XM_023241503.1
D3 17841975 17842153
refGene_gn=SART3;refGene_tx=XM_003994865.4;refGene_gn=SART3;refGene_tx=XM_006938660.3
D3 17842901 17843092
refGene_gn=SART3;refGene_tx=XM_003994865.4;refGene_gn=SART3;refGene_tx=XM_006938660.3
D3 17844003 17844156
refGene_gn=SART3;refGene_tx=XM_003994865.4;refGene_gn=SART3;refGene_tx=XM_006938660.3
D3 17844919 17845374
refGene_gn=SART3;refGene_tx=XM_003994865.4;refGene_gn=SART3;refGene_tx=XM_006938660.3
D3 17847609 17847778
refGene_gn=SART3;refGene_tx=XM_003994865.4;refGene_gn=SART3;refGene_tx=XM_006938660.3
D3 17848368 17848445
refGene_gn=SART3;refGene_tx=XM_003994865.4;refGene_gn=SART3;refGene_tx=XM_006938660.3

D3 17849388 17849501
refGene_gn=SART3;refGene_tx=XM_003994865.4;refGene_gn=SART3;refGene_tx=XM_006938660.3
D3 17853636 17853746
refGene_gn=SART3;refGene_tx=XM_003994865.4;refGene_gn=SART3;refGene_tx=XM_006938660.3
D3 17854914 17854973
refGene_gn=SART3;refGene_tx=XM_003994865.4;refGene_gn=SART3;refGene_tx=XM_006938660.3
D3 17855097 17855175
refGene_gn=SART3;refGene_tx=XM_003994865.4;refGene_gn=SART3;refGene_tx=XM_006938660.3
D3 17855840 17855948
refGene_gn=SART3;refGene_tx=XM_003994865.4;refGene_gn=SART3;refGene_tx=XM_006938660.3
D3 17856433 17856572
refGene_gn=SART3;refGene_tx=XM_003994865.4;refGene_gn=SART3;refGene_tx=XM_006938660.3
D3 17857279 17857435
refGene_gn=SART3;refGene_tx=XM_003994865.4;refGene_gn=SART3;refGene_tx=XM_006938660.3
D3 17861176 17861301
refGene_gn=SART3;refGene_tx=XM_003994865.4;refGene_gn=SART3;refGene_tx=XM_006938660.3
D3 17862554 17862660
refGene_gn=SART3;refGene_tx=XM_003994865.4;refGene_gn=SART3;refGene_tx=XM_006938660.3
D3 17864055 17864240
refGene_gn=SART3;refGene_tx=XM_003994865.4;refGene_gn=SART3;refGene_tx=XM_006938660.3
D3 17867656 17867761
refGene_gn=SART3;refGene_tx=XM_003994865.4;refGene_gn=SART3;refGene_tx=XM_006938660.3
D3 17869282 17869409
refGene_gn=SART3;refGene_tx=XM_003994865.4;refGene_gn=SART3;refGene_tx=XM_006938660.3
D3 17877755 17878070
refGene_gn=SART3;refGene_tx=XM_003994865.4;refGene_gn=SART3;refGene_tx=XM_006938660.3
D3 17883081 17883195 refGene_gn=ISCU;refGene_tx=XM_003994864.5
D3 17884773 17884887 refGene_gn=ISCU;refGene_tx=XM_003994864.5
D3 17885733 17885844 refGene_gn=ISCU;refGene_tx=XM_003994864.5
D3 17887629 17887708 refGene_gn=ISCU;refGene_tx=XM_003994864.5
D3 17889245 17889331 refGene_gn=ISCU;refGene_tx=XM_003994864.5
D3 17909573 17910413
refGene_gn=TMEM119;refGene_tx=XM_011287780.3;refGene_gn=TMEM119;refGene_tx=XM_023241504.1
D3 17932063 17933496 refGene_gn=SELPLG;refGene_tx=XM_006938659.4
D3 17943761 17943907 refGene_gn=SELPLG;refGene_tx=XM_006938659.4
D3 17944270 17944464 refGene_gn=SELPLG;refGene_tx=XM_006938659.4

D3 17958953 17959073
refGene_gn=CORO1C;refGene_tx=XM_023241509.1;refGene_gn=CORO1C;refGene_tx=XM_023241510.1
D3 17960251 17960497
refGene_gn=CORO1C;refGene_tx=XM_023241509.1;refGene_gn=CORO1C;refGene_tx=XM_023241510.1
D3 17960599 17960657
refGene_gn=CORO1C;refGene_tx=XM_023241509.1;refGene_gn=CORO1C;refGene_tx=XM_023241510.1
D3 17963100 17963246
refGene_gn=CORO1C;refGene_tx=XM_023241509.1;refGene_gn=CORO1C;refGene_tx=XM_023241510.1
D3 17965189 17965294
refGene_gn=CORO1C;refGene_tx=XM_023241509.1;refGene_gn=CORO1C;refGene_tx=XM_023241510.1
D3 17966350 17966470
refGene_gn=CORO1C;refGene_tx=XM_023241509.1;refGene_gn=CORO1C;refGene_tx=XM_023241510.1
D3 17967677 17967859
refGene_gn=CORO1C;refGene_tx=XM_023241509.1;refGene_gn=CORO1C;refGene_tx=XM_023241510.1
D3 17971076 17971206
refGene_gn=CORO1C;refGene_tx=XM_023241509.1;refGene_gn=CORO1C;refGene_tx=XM_023241510.1
D3 17985479 17985602
refGene_gn=CORO1C;refGene_tx=XM_023241509.1;refGene_gn=CORO1C;refGene_tx=XM_023241510.1
D3 18005366 18005561 refGene_gn=CORO1C;refGene_tx=XM_023241509.1
D3 18066005 18067244
refGene_gn=SSH1;refGene_tx=XM_019814643.2;refGene_gn=SSH1;refGene_tx=XM_019814644.2;refGene_gn=SSH1;refGene_tx=XM_019814645.2;refGene_gn=SSH1;refGene_tx=XM_023241505.1;refGene_gn=SSH1;refGene_tx=XM_023241506.1;refGene_gn=SSH1;refGene_tx=XM_023241507.1;refGene_gn=SSH1;refGene_tx=XM_023241508.1
D3 18070035 18070603
refGene_gn=SSH1;refGene_tx=XM_019814643.2;refGene_gn=SSH1;refGene_tx=XM_019814644.2;refGene_gn=SSH1;refGene_tx=XM_019814645.2;refGene_gn=SSH1;refGene_tx=XM_023241505.1;refGene_gn=SSH1;refGene_tx=XM_023241506.1;refGene_gn=SSH1;refGene_tx=XM_023241507.1;refGene_gn=SSH1;refGene_tx=XM_023241508.1
D3 18072909 18073110
refGene_gn=SSH1;refGene_tx=XM_019814643.2;refGene_gn=SSH1;refGene_tx=XM_019814644.2;refGene_gn=SSH1;refGene_tx=XM_019814645.2;refGene_gn=SSH1;refGene_tx=XM_023241505.1;refGene_gn=SSH1;refGene_tx=XM_023241506.1;refGene_gn=SSH1;refGene_tx=XM_023241507.1;refGene_gn=SSH1;refGene_tx=XM_023241508.1
D3 18074702 18074849
refGene_gn=SSH1;refGene_tx=XM_019814643.2;refGene_gn=SSH1;refGene_tx=XM_019814644.2;refGene_gn=SSH1;refGene_tx=XM_019814645.2;refGene_gn=SSH1;refGene_tx=XM_023241505.1;refGene_gn=SSH1;refGene_tx=XM_023241506.1;

refGene_gn=SSH1;refGene_tx=XM_023241507.1;refGene_gn=SSH1;refGene_tx=XM_023241508.1
D3 18076250 18076297
refGene_gn=SSH1;refGene_tx=XM_019814643.2;refGene_gn=SSH1;refGene_tx=XM_019814644.2;refGene_gn=SSH1;refGene_tx=XM_019814645.2;refGene_gn=SSH1;refGene_tx=XM_023241505.1;refGene_gn=SSH1;refGene_tx=XM_023241506.1;refGene_gn=SSH1;refGene_tx=XM_023241507.1;refGene_gn=SSH1;refGene_tx=XM_023241508.1
D3 18079012 18079141
refGene_gn=SSH1;refGene_tx=XM_019814643.2;refGene_gn=SSH1;refGene_tx=XM_019814644.2;refGene_gn=SSH1;refGene_tx=XM_019814645.2;refGene_gn=SSH1;refGene_tx=XM_023241505.1;refGene_gn=SSH1;refGene_tx=XM_023241506.1;refGene_gn=SSH1;refGene_tx=XM_023241507.1;refGene_gn=SSH1;refGene_tx=XM_023241508.1
D3 18080328 18080422
refGene_gn=SSH1;refGene_tx=XM_019814643.2;refGene_gn=SSH1;refGene_tx=XM_019814644.2;refGene_gn=SSH1;refGene_tx=XM_019814645.2;refGene_gn=SSH1;refGene_tx=XM_023241505.1;refGene_gn=SSH1;refGene_tx=XM_023241506.1;refGene_gn=SSH1;refGene_tx=XM_023241507.1;refGene_gn=SSH1;refGene_tx=XM_023241508.1
D3 18081641 18081836
refGene_gn=SSH1;refGene_tx=XM_019814643.2;refGene_gn=SSH1;refGene_tx=XM_019814644.2;refGene_gn=SSH1;refGene_tx=XM_019814645.2;refGene_gn=SSH1;refGene_tx=XM_023241505.1;refGene_gn=SSH1;refGene_tx=XM_023241506.1;refGene_gn=SSH1;refGene_tx=XM_023241507.1;refGene_gn=SSH1;refGene_tx=XM_023241508.1
D3 18084639 18084705
refGene_gn=SSH1;refGene_tx=XM_019814643.2;refGene_gn=SSH1;refGene_tx=XM_019814644.2;refGene_gn=SSH1;refGene_tx=XM_019814645.2;refGene_gn=SSH1;refGene_tx=XM_023241505.1;refGene_gn=SSH1;refGene_tx=XM_023241506.1;refGene_gn=SSH1;refGene_tx=XM_023241507.1;refGene_gn=SSH1;refGene_tx=XM_023241508.1
D3 18085804 18085873
refGene_gn=SSH1;refGene_tx=XM_019814643.2;refGene_gn=SSH1;refGene_tx=XM_019814644.2;refGene_gn=SSH1;refGene_tx=XM_019814645.2;refGene_gn=SSH1;refGene_tx=XM_023241505.1;refGene_gn=SSH1;refGene_tx=XM_023241506.1;refGene_gn=SSH1;refGene_tx=XM_023241507.1;refGene_gn=SSH1;refGene_tx=XM_023241508.1
D3 18087068 18087190
refGene_gn=SSH1;refGene_tx=XM_019814643.2;refGene_gn=SSH1;refGene_tx=XM_019814644.2;refGene_gn=SSH1;refGene_tx=XM_019814645.2;refGene_gn=SSH1;refGene_tx=XM_023241505.1;refGene_gn=SSH1;refGene_tx=XM_023241506.1;refGene_gn=SSH1;refGene_tx=XM_023241507.1;refGene_gn=SSH1;refGene_tx=XM_023241508.1
D3 18087644 18087709
refGene_gn=SSH1;refGene_tx=XM_019814643.2;refGene_gn=SSH1;refGene_tx=XM_023241505.1;refGene_gn=SSH1;refGene_tx=XM_023241506.1;refGene_gn=SSH1;refGene_tx=XM_023241507.1;refGene_gn=SSH1;refGene_tx=XM_023241508.1
D3 18092102 18092206
refGene_gn=SSH1;refGene_tx=XM_019814643.2;refGene_gn=SSH1;refGene_tx=X

M_023241505.1;refGene_gn=SSH1;refGene_tx=XM_023241506.1;refGene_gn=SSH1;refGene_tx=XM_023241507.1
D3 18093720 18093725 refGene_gn=SSH1;refGene_tx=XM_019814643.2
D3 18117738 18117779
refGene_gn=SSH1;refGene_tx=XM_023241505.1;refGene_gn=SSH1;refGene_tx=XM_023241507.1
D3 18122259 18122328
refGene_gn=SSH1;refGene_tx=XM_023241505.1;refGene_gn=SSH1;refGene_tx=XM_023241507.1
D3 18138401 18138595
refGene_gn=DAO;refGene_tx=XM_003994861.4;refGene_gn=DAO;refGene_tx=XM_006938655.3;refGene_gn=DAO;refGene_tx=XM_019814649.2;refGene_gn=DAO;refGene_tx=XM_019814650.2
D3 18141172 18141287
refGene_gn=DAO;refGene_tx=XM_003994861.4;refGene_gn=DAO;refGene_tx=XM_006938655.3;refGene_gn=DAO;refGene_tx=XM_019814649.2;refGene_gn=DAO;refGene_tx=XM_019814650.2
D3 18142554 18142631
refGene_gn=DAO;refGene_tx=XM_003994861.4;refGene_gn=DAO;refGene_tx=XM_006938655.3;refGene_gn=DAO;refGene_tx=XM_019814649.2;refGene_gn=DAO;refGene_tx=XM_019814650.2
D3 18144324 18144390
refGene_gn=DAO;refGene_tx=XM_003994861.4;refGene_gn=DAO;refGene_tx=XM_006938655.3;refGene_gn=DAO;refGene_tx=XM_019814649.2;refGene_gn=DAO;refGene_tx=XM_019814650.2
D3 18146270 18146325
refGene_gn=DAO;refGene_tx=XM_003994861.4;refGene_gn=DAO;refGene_tx=XM_006938655.3;refGene_gn=DAO;refGene_tx=XM_019814649.2;refGene_gn=DAO;refGene_tx=XM_019814650.2
D3 18147546 18147651
refGene_gn=DAO;refGene_tx=XM_003994861.4;refGene_gn=DAO;refGene_tx=XM_006938655.3;refGene_gn=DAO;refGene_tx=XM_019814649.2
D3 18147696 18147889
refGene_gn=DAO;refGene_tx=XM_006938655.3;refGene_gn=DAO;refGene_tx=XM_019814649.2;refGene_gn=DAO;refGene_tx=XM_019814650.2
D3 18148060 18148203
refGene_gn=DAO;refGene_tx=XM_006938655.3;refGene_gn=DAO;refGene_tx=XM_019814649.2;refGene_gn=DAO;refGene_tx=XM_019814650.2
D3 18148555 18148638
refGene_gn=DAO;refGene_tx=XM_003994861.4;refGene_gn=DAO;refGene_tx=XM_006938655.3;refGene_gn=DAO;refGene_tx=XM_019814649.2;refGene_gn=DAO;refGene_tx=XM_019814650.2
D3 18149553 18149671
refGene_gn=DAO;refGene_tx=XM_003994861.4;refGene_gn=DAO;refGene_tx=XM_006938655.3;refGene_gn=DAO;refGene_tx=XM_019814649.2;refGene_gn=DAO;refGene_tx=XM_019814650.2
D3 18150208 18150307
refGene_gn=DAO;refGene_tx=XM_003994861.4;refGene_gn=DAO;refGene_tx=XM_006938655.3;refGene_gn=DAO;refGene_tx=XM_019814649.2;refGene_gn=DAO;refGene_tx=XM_019814650.2

D3 18151540 18151672
refGene_gn=DAO;refGene_tx=XM_003994861.4;refGene_gn=DAO;refGene_tx=XM_006938655.3;refGene_gn=DAO;refGene_tx=XM_019814649.2;refGene_gn=DAO;refGene_tx=XM_019814650.2
D3 18157494 18157701 refGene_gn=SVOP;refGene_tx=XM_023241512.1
D3 18159104 18159344 refGene_gn=SVOP;refGene_tx=XM_023241511.1
D3 18159478 18159568
refGene_gn=SVOP;refGene_tx=XM_023241511.1;refGene_gn=SVOP;refGene_tx=XM_023241512.1
D3 18164342 18164424
refGene_gn=SVOP;refGene_tx=XM_023241511.1;refGene_gn=SVOP;refGene_tx=XM_023241512.1
D3 18165329 18165441
refGene_gn=SVOP;refGene_tx=XM_023241511.1;refGene_gn=SVOP;refGene_tx=XM_023241512.1
D3 18168982 18169090
refGene_gn=SVOP;refGene_tx=XM_023241511.1;refGene_gn=SVOP;refGene_tx=XM_023241512.1
D3 18174105 18174182
refGene_gn=SVOP;refGene_tx=XM_023241511.1;refGene_gn=SVOP;refGene_tx=XM_023241512.1
D3 18177501 18177575
refGene_gn=SVOP;refGene_tx=XM_023241511.1;refGene_gn=SVOP;refGene_tx=XM_023241512.1
D3 18179134 18179263
refGene_gn=SVOP;refGene_tx=XM_023241511.1;refGene_gn=SVOP;refGene_tx=XM_023241512.1
D3 18179951 18180077
refGene_gn=SVOP;refGene_tx=XM_023241511.1;refGene_gn=SVOP;refGene_tx=XM_023241512.1
D3 18181976 18182040
refGene_gn=SVOP;refGene_tx=XM_023241511.1;refGene_gn=SVOP;refGene_tx=XM_023241512.1
D3 18196053 18196178
refGene_gn=SVOP;refGene_tx=XM_023241511.1;refGene_gn=SVOP;refGene_tx=XM_023241512.1
D3 18205805 18205877
refGene_gn=SVOP;refGene_tx=XM_023241511.1;refGene_gn=SVOP;refGene_tx=XM_023241512.1
D3 18211265 18211364
refGene_gn=SVOP;refGene_tx=XM_023241511.1;refGene_gn=SVOP;refGene_tx=XM_023241512.1
D3 18212669 18212755
refGene_gn=SVOP;refGene_tx=XM_023241511.1;refGene_gn=SVOP;refGene_tx=XM_023241512.1
D3 18217615 18217776
refGene_gn=SVOP;refGene_tx=XM_023241511.1;refGene_gn=SVOP;refGene_tx=XM_023241512.1
D3 18237075 18237110
refGene_gn=SVOP;refGene_tx=XM_023241511.1;refGene_gn=SVOP;refGene_tx=X

M_023241512.1
D3 18250034 18250117 refGene_gn=USP30;refGene_tx=XM_003994860.5;refGene_gn=USP30;refGene_tx=XM_006938653.4;refGene_gn=USP30;refGene_tx=XM_006938654.4
D3 18254172 18254282 refGene_gn=USP30;refGene_tx=XM_003994860.5;refGene_gn=USP30;refGene_tx=XM_006938653.4;refGene_gn=USP30;refGene_tx=XM_006938654.4
D3 18255400 18255583 refGene_gn=USP30;refGene_tx=XM_003994860.5;refGene_gn=USP30;refGene_tx=XM_006938653.4;refGene_gn=USP30;refGene_tx=XM_006938654.4
D3 18260349 18260453 refGene_gn=USP30;refGene_tx=XM_003994860.5;refGene_gn=USP30;refGene_tx=XM_006938653.4;refGene_gn=USP30;refGene_tx=XM_006938654.4
D3 18264025 18264124 refGene_gn=USP30;refGene_tx=XM_003994860.5;refGene_gn=USP30;refGene_tx=XM_006938653.4;refGene_gn=USP30;refGene_tx=XM_006938654.4
D3 18264931 18264977 refGene_gn=USP30;refGene_tx=XM_003994860.5;refGene_gn=USP30;refGene_tx=XM_006938653.4;refGene_gn=USP30;refGene_tx=XM_006938654.4
D3 18266516 18266611 refGene_gn=USP30;refGene_tx=XM_003994860.5;refGene_gn=USP30;refGene_tx=XM_006938653.4;refGene_gn=USP30;refGene_tx=XM_006938654.4
D3 18272278 18272338 refGene_gn=USP30;refGene_tx=XM_003994860.5;refGene_gn=USP30;refGene_tx=XM_006938653.4;refGene_gn=USP30;refGene_tx=XM_006938654.4
D3 18272821 18272908 refGene_gn=USP30;refGene_tx=XM_003994860.5;refGene_gn=USP30;refGene_tx=XM_006938653.4;refGene_gn=USP30;refGene_tx=XM_006938654.4
D3 18273502 18273583 refGene_gn=USP30;refGene_tx=XM_003994860.5;refGene_gn=USP30;refGene_tx=XM_006938653.4;refGene_gn=USP30;refGene_tx=XM_006938654.4
D3 18273678 18273898 refGene_gn=USP30;refGene_tx=XM_003994860.5;refGene_gn=USP30;refGene_tx=XM_006938653.4;refGene_gn=USP30;refGene_tx=XM_006938654.4
D3 18276046 18276167 refGene_gn=USP30;refGene_tx=XM_003994860.5;refGene_gn=USP30;refGene_tx=XM_006938653.4;refGene_gn=USP30;refGene_tx=XM_006938654.4
D3 18276722 18276987 refGene_gn=USP30;refGene_tx=XM_003994860.5;refGene_gn=USP30;refGene_tx=XM_006938653.4;refGene_gn=USP30;refGene_tx=XM_006938654.4
D3 18279345 18279652 refGene_gn=ALKBH2;refGene_tx=XM_019814652.2;refGene_gn=ALKBH2;refGene_tx=XM_023241513.1
D3 18280696 18280895 refGene_gn=ALKBH2;refGene_tx=XM_019814652.2;refGene_gn=ALKBH2;refGene_tx=XM_023241513.1
D3 18281876 18282153 refGene_gn=ALKBH2;refGene_tx=XM_019814652.2;refGene_gn=ALKBH2;refGene_tx=XM_023241513.1
D3 18285023 18285155 refGene_gn=UNG;refGene_tx=XM_023241514.1

D3	18285795	18286002	refGene_gn=UNG;refGene_tx=XM_023241514.1
D3	18286796	18286892	refGene_gn=UNG;refGene_tx=XM_023241514.1
D3	18289162	18289260	refGene_gn=UNG;refGene_tx=XM_023241514.1
D3	18289931	18290020	refGene_gn=UNG;refGene_tx=XM_023241514.1
D3	18291547	18291726	refGene_gn=UNG;refGene_tx=XM_023241514.1
D3	18294322	18294463	refGene_gn=UNG;refGene_tx=XM_023241514.1
D3	18322856	18323503
refGene_gn=ACACB;refGene_tx=XM_003994858.5;refGene_gn=ACACB;refGene_tx
=XM_006938651.4;refGene_gn=ACACB;refGene_tx=XM_019814654.2
D3	18329183	18329254	refGene_gn=ACACB;refGene_tx=XM_006938649.4
D3	18334297	18334344	refGene_gn=ACACB;refGene_tx=XM_006938650.4
D3	18345372	18345505
refGene_gn=ACACB;refGene_tx=XM_003994858.5;refGene_gn=ACACB;refGene_tx
=XM_006938649.4;refGene_gn=ACACB;refGene_tx=XM_006938650.4;refGene_gn=
ACACB;refGene_tx=XM_006938651.4;refGene_gn=ACACB;refGene_tx=XM_0198146
54.2
D3	18349038	18349177
refGene_gn=ACACB;refGene_tx=XM_003994858.5;refGene_gn=ACACB;refGene_tx
=XM_006938649.4;refGene_gn=ACACB;refGene_tx=XM_006938650.4;refGene_gn=
ACACB;refGene_tx=XM_006938651.4;refGene_gn=ACACB;refGene_tx=XM_0198146
54.2
D3	18352250	18352360
refGene_gn=ACACB;refGene_tx=XM_003994858.5;refGene_gn=ACACB;refGene_tx
=XM_006938649.4;refGene_gn=ACACB;refGene_tx=XM_006938650.4;refGene_gn=
ACACB;refGene_tx=XM_006938651.4;refGene_gn=ACACB;refGene_tx=XM_0198146
54.2
D3	18352572	18352654
refGene_gn=ACACB;refGene_tx=XM_003994858.5;refGene_gn=ACACB;refGene_tx
=XM_006938649.4;refGene_gn=ACACB;refGene_tx=XM_006938650.4;refGene_gn=
ACACB;refGene_tx=XM_006938651.4;refGene_gn=ACACB;refGene_tx=XM_0198146
54.2
D3	18353911	18354010
refGene_gn=ACACB;refGene_tx=XM_003994858.5;refGene_gn=ACACB;refGene_tx
=XM_006938649.4;refGene_gn=ACACB;refGene_tx=XM_006938650.4;refGene_gn=
ACACB;refGene_tx=XM_006938651.4;refGene_gn=ACACB;refGene_tx=XM_0198146
54.2
D3	18355033	18355143
refGene_gn=ACACB;refGene_tx=XM_003994858.5;refGene_gn=ACACB;refGene_tx
=XM_006938649.4;refGene_gn=ACACB;refGene_tx=XM_006938650.4;refGene_gn=
ACACB;refGene_tx=XM_006938651.4;refGene_gn=ACACB;refGene_tx=XM_0198146
54.2
D3	18355239	18355350
refGene_gn=ACACB;refGene_tx=XM_003994858.5;refGene_gn=ACACB;refGene_tx
=XM_006938649.4;refGene_gn=ACACB;refGene_tx=XM_006938650.4;refGene_gn=
ACACB;refGene_tx=XM_006938651.4;refGene_gn=ACACB;refGene_tx=XM_0198146
54.2
D3	18358436	18358646
refGene_gn=ACACB;refGene_tx=XM_003994858.5;refGene_gn=ACACB;refGene_tx
=XM_006938649.4;refGene_gn=ACACB;refGene_tx=XM_006938650.4;refGene_gn=
ACACB;refGene_tx=XM_006938651.4;refGene_gn=ACACB;refGene_tx=XM_0198146

54.2
D3 18358950 18359121
refGene_gn=ACACB;refGene_tx=XM_003994858.5;refGene_gn=ACACB;refGene_tx
=XM_006938649.4;refGene_gn=ACACB;refGene_tx=XM_006938650.4;refGene_gn=
ACACB;refGene_tx=XM_006938651.4;refGene_gn=ACACB;refGene_tx=XM_0198146
54.2
D3 18361632 18361794
refGene_gn=ACACB;refGene_tx=XM_003994858.5;refGene_gn=ACACB;refGene_tx
=XM_006938649.4;refGene_gn=ACACB;refGene_tx=XM_006938650.4;refGene_gn=
ACACB;refGene_tx=XM_006938651.4;refGene_gn=ACACB;refGene_tx=XM_0198146
54.2
D3 18363169 18363333
refGene_gn=ACACB;refGene_tx=XM_003994858.5;refGene_gn=ACACB;refGene_tx
=XM_006938649.4;refGene_gn=ACACB;refGene_tx=XM_006938650.4;refGene_gn=
ACACB;refGene_tx=XM_006938651.4;refGene_gn=ACACB;refGene_tx=XM_0198146
54.2
D3 18365072 18365223
refGene_gn=ACACB;refGene_tx=XM_003994858.5;refGene_gn=ACACB;refGene_tx
=XM_006938649.4;refGene_gn=ACACB;refGene_tx=XM_006938650.4;refGene_gn=
ACACB;refGene_tx=XM_006938651.4;refGene_gn=ACACB;refGene_tx=XM_0198146
54.2
D3 18365320 18365424
refGene_gn=ACACB;refGene_tx=XM_003994858.5;refGene_gn=ACACB;refGene_tx
=XM_006938649.4;refGene_gn=ACACB;refGene_tx=XM_006938650.4;refGene_gn=
ACACB;refGene_tx=XM_006938651.4;refGene_gn=ACACB;refGene_tx=XM_0198146
54.2
D3 18366672 18366754
refGene_gn=ACACB;refGene_tx=XM_003994858.5;refGene_gn=ACACB;refGene_tx
=XM_006938649.4;refGene_gn=ACACB;refGene_tx=XM_006938650.4;refGene_gn=
ACACB;refGene_tx=XM_006938651.4;refGene_gn=ACACB;refGene_tx=XM_0198146
54.2
D3 18369109 18369255
refGene_gn=ACACB;refGene_tx=XM_003994858.5;refGene_gn=ACACB;refGene_tx
=XM_006938649.4;refGene_gn=ACACB;refGene_tx=XM_006938650.4;refGene_gn=
ACACB;refGene_tx=XM_006938651.4;refGene_gn=ACACB;refGene_tx=XM_0198146
54.2
D3 18370422 18370573
refGene_gn=ACACB;refGene_tx=XM_003994858.5;refGene_gn=ACACB;refGene_tx
=XM_006938649.4;refGene_gn=ACACB;refGene_tx=XM_006938650.4;refGene_gn=
ACACB;refGene_tx=XM_006938651.4;refGene_gn=ACACB;refGene_tx=XM_0198146
54.2
D3 18372595 18372730
refGene_gn=ACACB;refGene_tx=XM_003994858.5;refGene_gn=ACACB;refGene_tx
=XM_006938649.4;refGene_gn=ACACB;refGene_tx=XM_006938650.4;refGene_gn=
ACACB;refGene_tx=XM_006938651.4;refGene_gn=ACACB;refGene_tx=XM_0198146
54.2
D3 18375707 18375854
refGene_gn=ACACB;refGene_tx=XM_003994858.5;refGene_gn=ACACB;refGene_tx
=XM_006938649.4;refGene_gn=ACACB;refGene_tx=XM_006938650.4;refGene_gn=
ACACB;refGene_tx=XM_006938651.4;refGene_gn=ACACB;refGene_tx=XM_0198146

54.2
D3 18377176 18377365
refGene_gn=ACACB;refGene_tx=XM_003994858.5;refGene_gn=ACACB;refGene_tx=XM_006938649.4;refGene_gn=ACACB;refGene_tx=XM_006938650.4;refGene_gn=ACACB;refGene_tx=XM_006938651.4;refGene_gn=ACACB;refGene_tx=XM_019814654.2
D3 18379215 18379316
refGene_gn=ACACB;refGene_tx=XM_003994858.5;refGene_gn=ACACB;refGene_tx=XM_006938649.4;refGene_gn=ACACB;refGene_tx=XM_006938650.4;refGene_gn=ACACB;refGene_tx=XM_006938651.4;refGene_gn=ACACB;refGene_tx=XM_019814654.2
D3 18381928 18382017
refGene_gn=ACACB;refGene_tx=XM_003994858.5;refGene_gn=ACACB;refGene_tx=XM_006938649.4;refGene_gn=ACACB;refGene_tx=XM_006938650.4;refGene_gn=ACACB;refGene_tx=XM_006938651.4;refGene_gn=ACACB;refGene_tx=XM_019814654.2
D3 18382096 18382221
refGene_gn=ACACB;refGene_tx=XM_003994858.5;refGene_gn=ACACB;refGene_tx=XM_006938649.4;refGene_gn=ACACB;refGene_tx=XM_006938650.4;refGene_gn=ACACB;refGene_tx=XM_006938651.4;refGene_gn=ACACB;refGene_tx=XM_019814654.2
D3 18386262 18386376
refGene_gn=ACACB;refGene_tx=XM_003994858.5;refGene_gn=ACACB;refGene_tx=XM_006938649.4;refGene_gn=ACACB;refGene_tx=XM_006938650.4;refGene_gn=ACACB;refGene_tx=XM_006938651.4;refGene_gn=ACACB;refGene_tx=XM_019814654.2
D3 18386509 18386623
refGene_gn=ACACB;refGene_tx=XM_003994858.5;refGene_gn=ACACB;refGene_tx=XM_006938649.4;refGene_gn=ACACB;refGene_tx=XM_006938650.4;refGene_gn=ACACB;refGene_tx=XM_006938651.4;refGene_gn=ACACB;refGene_tx=XM_019814654.2
D3 18387487 18387577
refGene_gn=ACACB;refGene_tx=XM_003994858.5;refGene_gn=ACACB;refGene_tx=XM_006938649.4;refGene_gn=ACACB;refGene_tx=XM_006938650.4;refGene_gn=ACACB;refGene_tx=XM_006938651.4;refGene_gn=ACACB;refGene_tx=XM_019814654.2
D3 18391235 18391354
refGene_gn=ACACB;refGene_tx=XM_003994858.5;refGene_gn=ACACB;refGene_tx=XM_006938649.4;refGene_gn=ACACB;refGene_tx=XM_006938650.4;refGene_gn=ACACB;refGene_tx=XM_006938651.4;refGene_gn=ACACB;refGene_tx=XM_019814654.2
D3 18396064 18396202
refGene_gn=ACACB;refGene_tx=XM_003994858.5;refGene_gn=ACACB;refGene_tx=XM_006938649.4;refGene_gn=ACACB;refGene_tx=XM_006938650.4;refGene_gn=ACACB;refGene_tx=XM_006938651.4;refGene_gn=ACACB;refGene_tx=XM_019814654.2
D3 18396925 18397025
refGene_gn=ACACB;refGene_tx=XM_003994858.5;refGene_gn=ACACB;refGene_tx=XM_006938649.4;refGene_gn=ACACB;refGene_tx=XM_006938650.4;refGene_gn=ACACB;refGene_tx=XM_006938651.4;refGene_gn=ACACB;refGene_tx=XM_0198146

54.2
D3 18397114 18397222
refGene_gn=ACACB;refGene_tx=XM_003994858.5;refGene_gn=ACACB;refGene_tx=XM_006938649.4;refGene_gn=ACACB;refGene_tx=XM_006938650.4;refGene_gn=ACACB;refGene_tx=XM_006938651.4;refGene_gn=ACACB;refGene_tx=XM_019814654.2
D3 18401407 18401464
refGene_gn=ACACB;refGene_tx=XM_003994858.5;refGene_gn=ACACB;refGene_tx=XM_006938649.4;refGene_gn=ACACB;refGene_tx=XM_006938650.4;refGene_gn=ACACB;refGene_tx=XM_006938651.4;refGene_gn=ACACB;refGene_tx=XM_019814654.2
D3 18401704 18401746
refGene_gn=ACACB;refGene_tx=XM_003994858.5;refGene_gn=ACACB;refGene_tx=XM_006938649.4;refGene_gn=ACACB;refGene_tx=XM_006938650.4;refGene_gn=ACACB;refGene_tx=XM_006938651.4;refGene_gn=ACACB;refGene_tx=XM_019814654.2
D3 18402749 18402965
refGene_gn=ACACB;refGene_tx=XM_003994858.5;refGene_gn=ACACB;refGene_tx=XM_006938649.4;refGene_gn=ACACB;refGene_tx=XM_006938650.4;refGene_gn=ACACB;refGene_tx=XM_006938651.4;refGene_gn=ACACB;refGene_tx=XM_019814654.2
D3 18404561 18404717
refGene_gn=ACACB;refGene_tx=XM_003994858.5;refGene_gn=ACACB;refGene_tx=XM_006938649.4;refGene_gn=ACACB;refGene_tx=XM_006938650.4;refGene_gn=ACACB;refGene_tx=XM_006938651.4;refGene_gn=ACACB;refGene_tx=XM_019814654.2
D3 18405454 18405658
refGene_gn=ACACB;refGene_tx=XM_003994858.5;refGene_gn=ACACB;refGene_tx=XM_006938649.4;refGene_gn=ACACB;refGene_tx=XM_006938650.4;refGene_gn=ACACB;refGene_tx=XM_006938651.4;refGene_gn=ACACB;refGene_tx=XM_019814654.2
D3 18406497 18406653
refGene_gn=ACACB;refGene_tx=XM_003994858.5;refGene_gn=ACACB;refGene_tx=XM_006938649.4;refGene_gn=ACACB;refGene_tx=XM_006938650.4;refGene_gn=ACACB;refGene_tx=XM_006938651.4;refGene_gn=ACACB;refGene_tx=XM_019814654.2
D3 18409799 18409922
refGene_gn=ACACB;refGene_tx=XM_003994858.5;refGene_gn=ACACB;refGene_tx=XM_006938649.4;refGene_gn=ACACB;refGene_tx=XM_006938650.4;refGene_gn=ACACB;refGene_tx=XM_006938651.4;refGene_gn=ACACB;refGene_tx=XM_019814654.2
D3 18410025 18410295
refGene_gn=ACACB;refGene_tx=XM_003994858.5;refGene_gn=ACACB;refGene_tx=XM_006938649.4;refGene_gn=ACACB;refGene_tx=XM_006938650.4;refGene_gn=ACACB;refGene_tx=XM_006938651.4;refGene_gn=ACACB;refGene_tx=XM_019814654.2
D3 18411209 18411307
refGene_gn=ACACB;refGene_tx=XM_003994858.5;refGene_gn=ACACB;refGene_tx=XM_006938649.4;refGene_gn=ACACB;refGene_tx=XM_006938650.4;refGene_gn=ACACB;refGene_tx=XM_006938651.4;refGene_gn=ACACB;refGene_tx=XM_0198146

54.2
D3 18412412 18412533
refGene_gn=ACACB;refGene_tx=XM_003994858.5;refGene_gn=ACACB;refGene_tx
=XM_006938649.4;refGene_gn=ACACB;refGene_tx=XM_006938650.4;refGene_gn=
ACACB;refGene_tx=XM_006938651.4;refGene_gn=ACACB;refGene_tx=XM_0198146
54.2
D3 18414380 18414491
refGene_gn=ACACB;refGene_tx=XM_003994858.5;refGene_gn=ACACB;refGene_tx
=XM_006938649.4;refGene_gn=ACACB;refGene_tx=XM_006938650.4;refGene_gn=
ACACB;refGene_tx=XM_006938651.4;refGene_gn=ACACB;refGene_tx=XM_0198146
54.2
D3 18415354 18415498
refGene_gn=ACACB;refGene_tx=XM_003994858.5;refGene_gn=ACACB;refGene_tx
=XM_006938649.4;refGene_gn=ACACB;refGene_tx=XM_006938650.4;refGene_gn=
ACACB;refGene_tx=XM_006938651.4;refGene_gn=ACACB;refGene_tx=XM_0198146
54.2
D3 18416344 18416465
refGene_gn=ACACB;refGene_tx=XM_003994858.5;refGene_gn=ACACB;refGene_tx
=XM_006938649.4;refGene_gn=ACACB;refGene_tx=XM_006938650.4;refGene_gn=
ACACB;refGene_tx=XM_006938651.4;refGene_gn=ACACB;refGene_tx=XM_0198146
54.2
D3 18417856 18417953
refGene_gn=ACACB;refGene_tx=XM_003994858.5;refGene_gn=ACACB;refGene_tx
=XM_006938649.4;refGene_gn=ACACB;refGene_tx=XM_006938650.4;refGene_gn=
ACACB;refGene_tx=XM_006938651.4;refGene_gn=ACACB;refGene_tx=XM_0198146
54.2
D3 18419525 18419622
refGene_gn=ACACB;refGene_tx=XM_003994858.5;refGene_gn=ACACB;refGene_tx
=XM_006938649.4;refGene_gn=ACACB;refGene_tx=XM_006938650.4;refGene_gn=
ACACB;refGene_tx=XM_006938651.4;refGene_gn=ACACB;refGene_tx=XM_0198146
54.2
D3 18420172 18420308
refGene_gn=ACACB;refGene_tx=XM_003994858.5;refGene_gn=ACACB;refGene_tx
=XM_006938649.4;refGene_gn=ACACB;refGene_tx=XM_006938650.4;refGene_gn=
ACACB;refGene_tx=XM_006938651.4
D3 18420677 18420682 refGene_gn=ACACB;refGene_tx=XM_006938651.4
D3 18421144 18421322
refGene_gn=ACACB;refGene_tx=XM_003994858.5;refGene_gn=ACACB;refGene_tx
=XM_006938649.4;refGene_gn=ACACB;refGene_tx=XM_006938650.4;refGene_gn=
ACACB;refGene_tx=XM_019814654.2
D3 18422901 18423014
refGene_gn=ACACB;refGene_tx=XM_003994858.5;refGene_gn=ACACB;refGene_tx
=XM_006938649.4;refGene_gn=ACACB;refGene_tx=XM_006938650.4
D3 18423986 18424141
refGene_gn=ACACB;refGene_tx=XM_003994858.5;refGene_gn=ACACB;refGene_tx
=XM_006938649.4;refGene_gn=ACACB;refGene_tx=XM_006938650.4
D3 18424764 18424935
refGene_gn=ACACB;refGene_tx=XM_003994858.5;refGene_gn=ACACB;refGene_tx
=XM_006938649.4;refGene_gn=ACACB;refGene_tx=XM_006938650.4
D3 18425044 18425181

refGene_gn=ACACB;refGene_tx=XM_003994858.5;refGene_gn=ACACB;refGene_tx=XM_006938649.4;refGene_gn=ACACB;refGene_tx=XM_006938650.4
D3	18425820	18425947	refGene_gn=ACACB;refGene_tx=XM_003994858.5;refGene_gn=ACACB;refGene_tx=XM_006938649.4;refGene_gn=ACACB;refGene_tx=XM_006938650.4
D3	18435831	18436091	refGene_gn=FOXN4;refGene_tx=XM_023241515.1
D3	18437154	18437547	refGene_gn=FOXN4;refGene_tx=XM_023241515.1
D3	18440060	18440268	refGene_gn=FOXN4;refGene_tx=XM_023241515.1
D3	18441509	18441606	refGene_gn=FOXN4;refGene_tx=XM_023241515.1
D3	18442249	18442377	refGene_gn=FOXN4;refGene_tx=XM_023241515.1
D3	18442707	18442818	refGene_gn=FOXN4;refGene_tx=XM_023241515.1
D3	18442924	18442968	refGene_gn=FOXN4;refGene_tx=XM_023241515.1
D3	18442970	18443048	refGene_gn=FOXN4;refGene_tx=XM_023241515.1
D3	18444816	18444965	refGene_gn=FOXN4;refGene_tx=XM_023241515.1
D3	18458993	18459079	refGene_gn=FOXN4;refGene_tx=XM_023241515.1
D3	18532845	18532971	refGene_gn=MYO1H;refGene_tx=XM_023241516.1
D3	18535239	18535355	refGene_gn=MYO1H;refGene_tx=XM_023241516.1
D3	18536718	18536917	refGene_gn=MYO1H;refGene_tx=XM_023241516.1
D3	18537813	18537894	refGene_gn=MYO1H;refGene_tx=XM_023241516.1
D3	18540427	18540607	refGene_gn=MYO1H;refGene_tx=XM_023241516.1
D3	18542868	18542967	refGene_gn=MYO1H;refGene_tx=XM_023241516.1
D3	18544435	18544549	refGene_gn=MYO1H;refGene_tx=XM_023241516.1
D3	18544889	18544961	refGene_gn=MYO1H;refGene_tx=XM_023241516.1
D3	18545809	18545929	refGene_gn=MYO1H;refGene_tx=XM_023241516.1
D3	18547302	18547370	refGene_gn=MYO1H;refGene_tx=XM_023241516.1
D3	18548135	18548241	refGene_gn=MYO1H;refGene_tx=XM_023241516.1
D3	18548562	18548643	refGene_gn=MYO1H;refGene_tx=XM_023241516.1
D3	18550362	18550454	refGene_gn=MYO1H;refGene_tx=XM_023241516.1
D3	18552689	18552784	refGene_gn=MYO1H;refGene_tx=XM_023241516.1
D3	18556861	18556908	refGene_gn=MYO1H;refGene_tx=XM_023241516.1
D3	18560475	18560556	refGene_gn=MYO1H;refGene_tx=XM_023241516.1
D3	18561209	18561315	refGene_gn=MYO1H;refGene_tx=XM_023241516.1
D3	18562310	18562428	refGene_gn=MYO1H;refGene_tx=XM_023241516.1
D3	18565461	18565575	refGene_gn=MYO1H;refGene_tx=XM_023241516.1
D3	18568999	18569076	refGene_gn=MYO1H;refGene_tx=XM_023241516.1
D3	18570273	18570342	refGene_gn=MYO1H;refGene_tx=XM_023241516.1
D3	18571664	18571749	refGene_gn=MYO1H;refGene_tx=XM_023241516.1
D3	18572598	18572758	refGene_gn=MYO1H;refGene_tx=XM_023241516.1
D3	18573588	18573672	refGene_gn=MYO1H;refGene_tx=XM_023241516.1
D3	18574447	18574541	refGene_gn=MYO1H;refGene_tx=XM_023241516.1
D3	18574727	18574783	refGene_gn=MYO1H;refGene_tx=XM_023241516.1
D3	18575354	18575490	refGene_gn=MYO1H;refGene_tx=XM_023241516.1
D3	18575903	18575974	refGene_gn=MYO1H;refGene_tx=XM_023241516.1
D3	18576338	18576436	refGene_gn=MYO1H;refGene_tx=XM_023241516.1
D3	18577328	18577428	refGene_gn=MYO1H;refGene_tx=XM_023241516.1
D3	18578475	18578502	refGene_gn=MYO1H;refGene_tx=XM_023241516.1
D3	18581731	18581956	refGene_gn=KCTD10;refGene_tx=XM_003994856.5;refGene_gn=KCTD10;refGene_tx=XM_006938646.4
D3	18585885	18586081

refGene_gn=KCTD10;refGene_tx=XM_003994856.5;refGene_gn=KCTD10;refGene_tx=XM_006938646.4
D3 18587311 18587364
refGene_gn=KCTD10;refGene_tx=XM_003994856.5;refGene_gn=KCTD10;refGene_tx=XM_006938646.4
D3 18587671 18587761
refGene_gn=KCTD10;refGene_tx=XM_003994856.5;refGene_gn=KCTD10;refGene_tx=XM_006938646.4
D3 18590508 18590678
refGene_gn=KCTD10;refGene_tx=XM_003994856.5;refGene_gn=KCTD10;refGene_tx=XM_006938646.4
D3 18599655 18599869
refGene_gn=KCTD10;refGene_tx=XM_003994856.5;refGene_gn=KCTD10;refGene_tx=XM_006938646.4
D3 18605739 18605742
refGene_gn=KCTD10;refGene_tx=XM_003994856.5;refGene_gn=KCTD10;refGene_tx=XM_006938646.4
D3 18609627 18609681
refGene_gn=UBE3B;refGene_tx=XM_006938641.4;refGene_gn=UBE3B;refGene_tx=XM_019814658.2
D3 18610768 18610950
refGene_gn=UBE3B;refGene_tx=XM_003994855.5;refGene_gn=UBE3B;refGene_tx=XM_006938641.4;refGene_gn=UBE3B;refGene_tx=XM_006938642.4;refGene_gn=UBE3B;refGene_tx=XM_019814656.2;refGene_gn=UBE3B;refGene_tx=XM_019814657.2;refGene_gn=UBE3B;refGene_tx=XM_019814658.2
D3 18611098 18611219
refGene_gn=UBE3B;refGene_tx=XM_003994855.5;refGene_gn=UBE3B;refGene_tx=XM_006938641.4;refGene_gn=UBE3B;refGene_tx=XM_006938642.4;refGene_gn=UBE3B;refGene_tx=XM_019814656.2;refGene_gn=UBE3B;refGene_tx=XM_019814657.2;refGene_gn=UBE3B;refGene_tx=XM_019814658.2
D3 18613013 18613073
refGene_gn=UBE3B;refGene_tx=XM_003994855.5;refGene_gn=UBE3B;refGene_tx=XM_006938641.4;refGene_gn=UBE3B;refGene_tx=XM_006938642.4;refGene_gn=UBE3B;refGene_tx=XM_019814656.2;refGene_gn=UBE3B;refGene_tx=XM_019814657.2;refGene_gn=UBE3B;refGene_tx=XM_019814658.2
D3 18613493 18613598
refGene_gn=UBE3B;refGene_tx=XM_003994855.5;refGene_gn=UBE3B;refGene_tx=XM_006938641.4;refGene_gn=UBE3B;refGene_tx=XM_006938642.4;refGene_gn=UBE3B;refGene_tx=XM_019814656.2;refGene_gn=UBE3B;refGene_tx=XM_019814657.2
D3 18615534 18615631
refGene_gn=UBE3B;refGene_tx=XM_003994855.5;refGene_gn=UBE3B;refGene_tx=XM_006938641.4;refGene_gn=UBE3B;refGene_tx=XM_006938642.4;refGene_gn=UBE3B;refGene_tx=XM_019814656.2;refGene_gn=UBE3B;refGene_tx=XM_019814657.2;refGene_gn=UBE3B;refGene_tx=XM_019814658.2
D3 18616913 18616999
refGene_gn=UBE3B;refGene_tx=XM_003994855.5;refGene_gn=UBE3B;refGene_tx=XM_006938641.4;refGene_gn=UBE3B;refGene_tx=XM_006938642.4;refGene_gn=UBE3B;refGene_tx=XM_019814656.2;refGene_gn=UBE3B;refGene_tx=XM_019814657.2;refGene_gn=UBE3B;refGene_tx=XM_019814658.2

D3 18617904 18617987
refGene_gn=UBE3B;refGene_tx=XM_003994855.5;refGene_gn=UBE3B;refGene_tx=XM_006938641.4;refGene_gn=UBE3B;refGene_tx=XM_006938642.4;refGene_gn=UBE3B;refGene_tx=XM_019814656.2;refGene_gn=UBE3B;refGene_tx=XM_019814657.2;refGene_gn=UBE3B;refGene_tx=XM_019814658.2
D3 18622036 18622142
refGene_gn=UBE3B;refGene_tx=XM_003994855.5;refGene_gn=UBE3B;refGene_tx=XM_006938641.4;refGene_gn=UBE3B;refGene_tx=XM_006938642.4;refGene_gn=UBE3B;refGene_tx=XM_019814656.2;refGene_gn=UBE3B;refGene_tx=XM_019814657.2;refGene_gn=UBE3B;refGene_tx=XM_019814658.2
D3 18622445 18622566
refGene_gn=UBE3B;refGene_tx=XM_003994855.5;refGene_gn=UBE3B;refGene_tx=XM_006938641.4;refGene_gn=UBE3B;refGene_tx=XM_006938642.4;refGene_gn=UBE3B;refGene_tx=XM_019814656.2;refGene_gn=UBE3B;refGene_tx=XM_019814657.2;refGene_gn=UBE3B;refGene_tx=XM_019814658.2
D3 18623474 18623652
refGene_gn=UBE3B;refGene_tx=XM_003994855.5;refGene_gn=UBE3B;refGene_tx=XM_006938641.4;refGene_gn=UBE3B;refGene_tx=XM_006938642.4;refGene_gn=UBE3B;refGene_tx=XM_019814656.2;refGene_gn=UBE3B;refGene_tx=XM_019814657.2;refGene_gn=UBE3B;refGene_tx=XM_019814658.2
D3 18625438 18625602
refGene_gn=UBE3B;refGene_tx=XM_003994855.5;refGene_gn=UBE3B;refGene_tx=XM_006938641.4;refGene_gn=UBE3B;refGene_tx=XM_006938642.4;refGene_gn=UBE3B;refGene_tx=XM_019814656.2;refGene_gn=UBE3B;refGene_tx=XM_019814657.2;refGene_gn=UBE3B;refGene_tx=XM_019814658.2
D3 18627136 18627304
refGene_gn=UBE3B;refGene_tx=XM_003994855.5;refGene_gn=UBE3B;refGene_tx=XM_006938641.4;refGene_gn=UBE3B;refGene_tx=XM_006938642.4;refGene_gn=UBE3B;refGene_tx=XM_019814656.2;refGene_gn=UBE3B;refGene_tx=XM_019814657.2;refGene_gn=UBE3B;refGene_tx=XM_019814658.2
D3 18630560 18630732
refGene_gn=UBE3B;refGene_tx=XM_003994855.5;refGene_gn=UBE3B;refGene_tx=XM_006938641.4;refGene_gn=UBE3B;refGene_tx=XM_006938642.4;refGene_gn=UBE3B;refGene_tx=XM_019814656.2;refGene_gn=UBE3B;refGene_tx=XM_019814657.2;refGene_gn=UBE3B;refGene_tx=XM_019814658.2
D3 18632309 18632428
refGene_gn=UBE3B;refGene_tx=XM_003994855.5;refGene_gn=UBE3B;refGene_tx=XM_006938641.4;refGene_gn=UBE3B;refGene_tx=XM_006938642.4;refGene_gn=UBE3B;refGene_tx=XM_019814656.2;refGene_gn=UBE3B;refGene_tx=XM_019814657.2;refGene_gn=UBE3B;refGene_tx=XM_019814658.2
D3 18633048 18633163
refGene_gn=UBE3B;refGene_tx=XM_003994855.5;refGene_gn=UBE3B;refGene_tx=XM_006938641.4;refGene_gn=UBE3B;refGene_tx=XM_006938642.4;refGene_gn=UBE3B;refGene_tx=XM_019814656.2;refGene_gn=UBE3B;refGene_tx=XM_019814657.2;refGene_gn=UBE3B;refGene_tx=XM_019814658.2
D3 18634019 18634119
refGene_gn=UBE3B;refGene_tx=XM_003994855.5;refGene_gn=UBE3B;refGene_tx=XM_006938641.4;refGene_gn=UBE3B;refGene_tx=XM_006938642.4;refGene_gn=UBE3B;refGene_tx=XM_019814656.2;refGene_gn=UBE3B;refGene_tx=XM_019814657.2;refGene_gn=UBE3B;refGene_tx=XM_019814658.2

D3 18639318 18639438
refGene_gn=UBE3B;refGene_tx=XM_003994855.5;refGene_gn=UBE3B;refGene_tx=XM_006938641.4;refGene_gn=UBE3B;refGene_tx=XM_006938642.4;refGene_gn=UBE3B;refGene_tx=XM_019814656.2;refGene_gn=UBE3B;refGene_tx=XM_019814657.2;refGene_gn=UBE3B;refGene_tx=XM_019814658.2
D3 18644008 18644185
refGene_gn=UBE3B;refGene_tx=XM_003994855.5;refGene_gn=UBE3B;refGene_tx=XM_006938641.4;refGene_gn=UBE3B;refGene_tx=XM_006938642.4;refGene_gn=UBE3B;refGene_tx=XM_019814656.2;refGene_gn=UBE3B;refGene_tx=XM_019814657.2;refGene_gn=UBE3B;refGene_tx=XM_019814658.2
D3 18644314 18644425
refGene_gn=UBE3B;refGene_tx=XM_003994855.5;refGene_gn=UBE3B;refGene_tx=XM_006938641.4;refGene_gn=UBE3B;refGene_tx=XM_006938642.4;refGene_gn=UBE3B;refGene_tx=XM_019814656.2;refGene_gn=UBE3B;refGene_tx=XM_019814657.2;refGene_gn=UBE3B;refGene_tx=XM_019814658.2
D3 18646870 18647008
refGene_gn=UBE3B;refGene_tx=XM_003994855.5;refGene_gn=UBE3B;refGene_tx=XM_006938641.4;refGene_gn=UBE3B;refGene_tx=XM_006938642.4;refGene_gn=UBE3B;refGene_tx=XM_019814656.2;refGene_gn=UBE3B;refGene_tx=XM_019814657.2;refGene_gn=UBE3B;refGene_tx=XM_019814658.2
D3 18647283 18647349
refGene_gn=UBE3B;refGene_tx=XM_003994855.5;refGene_gn=UBE3B;refGene_tx=XM_006938641.4;refGene_gn=UBE3B;refGene_tx=XM_006938642.4;refGene_gn=UBE3B;refGene_tx=XM_019814656.2;refGene_gn=UBE3B;refGene_tx=XM_019814657.2;refGene_gn=UBE3B;refGene_tx=XM_019814658.2
D3 18649258 18649317
refGene_gn=UBE3B;refGene_tx=XM_003994855.5;refGene_gn=UBE3B;refGene_tx=XM_006938641.4;refGene_gn=UBE3B;refGene_tx=XM_006938642.4;refGene_gn=UBE3B;refGene_tx=XM_019814656.2;refGene_gn=UBE3B;refGene_tx=XM_019814657.2;refGene_gn=UBE3B;refGene_tx=XM_019814658.2
D3 18651695 18651878
refGene_gn=UBE3B;refGene_tx=XM_003994855.5;refGene_gn=UBE3B;refGene_tx=XM_006938641.4;refGene_gn=UBE3B;refGene_tx=XM_006938642.4;refGene_gn=UBE3B;refGene_tx=XM_019814656.2;refGene_gn=UBE3B;refGene_tx=XM_019814657.2;refGene_gn=UBE3B;refGene_tx=XM_019814658.2
D3 18652371 18652483
refGene_gn=UBE3B;refGene_tx=XM_003994855.5;refGene_gn=UBE3B;refGene_tx=XM_006938641.4;refGene_gn=UBE3B;refGene_tx=XM_006938642.4;refGene_gn=UBE3B;refGene_tx=XM_019814656.2;refGene_gn=UBE3B;refGene_tx=XM_019814657.2;refGene_gn=UBE3B;refGene_tx=XM_019814658.2
D3 18655334 18655427
refGene_gn=UBE3B;refGene_tx=XM_003994855.5;refGene_gn=UBE3B;refGene_tx=XM_006938641.4;refGene_gn=UBE3B;refGene_tx=XM_006938642.4;refGene_gn=UBE3B;refGene_tx=XM_019814656.2;refGene_gn=UBE3B;refGene_tx=XM_019814657.2;refGene_gn=UBE3B;refGene_tx=XM_019814658.2
D3 18656365 18656557
refGene_gn=UBE3B;refGene_tx=XM_003994855.5;refGene_gn=UBE3B;refGene_tx=XM_006938641.4;refGene_gn=UBE3B;refGene_tx=XM_006938642.4;refGene_gn=UBE3B;refGene_tx=XM_019814656.2;refGene_gn=UBE3B;refGene_tx=XM_019814657.2;refGene_gn=UBE3B;refGene_tx=XM_019814658.2

D3 18673975 18674075
refGene_gn=MMAB;refGene_tx=XM_003994918.5;refGene_gn=MMAB;refGene_tx=XM_011287770.3
D3 18675448 18675508
refGene_gn=MMAB;refGene_tx=XM_003994918.5;refGene_gn=MMAB;refGene_tx=XM_011287770.3;refGene_gn=MMAB;refGene_tx=XM_011287772.3
D3 18677291 18677356
refGene_gn=MMAB;refGene_tx=XM_003994918.5;refGene_gn=MMAB;refGene_tx=XM_011287770.3;refGene_gn=MMAB;refGene_tx=XM_011287772.3
D3 18677732 18677830
refGene_gn=MMAB;refGene_tx=XM_003994918.5;refGene_gn=MMAB;refGene_tx=XM_011287770.3;refGene_gn=MMAB;refGene_tx=XM_011287772.3
D3 18678022 18678132
refGene_gn=MMAB;refGene_tx=XM_003994918.5;refGene_gn=MMAB;refGene_tx=XM_011287770.3;refGene_gn=MMAB;refGene_tx=XM_011287772.3
D3 18680550 18680608
refGene_gn=MMAB;refGene_tx=XM_003994918.5;refGene_gn=MMAB;refGene_tx=XM_011287770.3;refGene_gn=MMAB;refGene_tx=XM_011287772.3
D3 18682453 18682547
refGene_gn=MMAB;refGene_tx=XM_003994918.5;refGene_gn=MMAB;refGene_tx=XM_011287770.3;refGene_gn=MMAB;refGene_tx=XM_011287772.3
D3 18687073 18687135 refGene_gn=MMAB;refGene_tx=XM_003994918.5
D3 18688274 18688470
refGene_gn=MMAB;refGene_tx=XM_003994918.5;refGene_gn=MMAB;refGene_tx=XM_011287770.3;refGene_gn=MMAB;refGene_tx=XM_011287772.3
D3 18688991 18688997 refGene_gn=MVK;refGene_tx=XM_023242153.1
D3 18689611 18689785 refGene_gn=MVK;refGene_tx=XM_023242153.1
D3 18692758 18693012 refGene_gn=MVK;refGene_tx=XM_023242153.1
D3 18695006 18695162 refGene_gn=MVK;refGene_tx=XM_023242153.1
D3 18698274 18698378 refGene_gn=MVK;refGene_tx=XM_023242153.1
D3 18698994 18699040 refGene_gn=MVK;refGene_tx=XM_023242153.1
D3 18703278 18703369 refGene_gn=MVK;refGene_tx=XM_023242153.1
D3 18703760 18703877 refGene_gn=MVK;refGene_tx=XM_023242153.1
D3 18707446 18707600 refGene_gn=MVK;refGene_tx=XM_023242153.1
D3 18708722 18708874 refGene_gn=MVK;refGene_tx=XM_023242153.1
D3 18872312 18872470 refGene_gn=TRPV4;refGene_tx=XM_023241517.1
D3 18872917 18873039 refGene_gn=TRPV4;refGene_tx=XM_023241517.1
D3 18874434 18874562 refGene_gn=TRPV4;refGene_tx=XM_023241517.1
D3 18875617 18875934 refGene_gn=TRPV4;refGene_tx=XM_023241517.1
D3 18878462 18878529 refGene_gn=TRPV4;refGene_tx=XM_023241517.1
D3 18878753 18878919 refGene_gn=TRPV4;refGene_tx=XM_023241517.1
D3 18879618 18879692 refGene_gn=TRPV4;refGene_tx=XM_023241517.1
D3 18879932 18880025 refGene_gn=TRPV4;refGene_tx=XM_023241517.1
D3 18880286 18880445 refGene_gn=TRPV4;refGene_tx=XM_023241517.1
D3 18881675 18881855 refGene_gn=TRPV4;refGene_tx=XM_023241517.1
D3 18884090 18884389 refGene_gn=TRPV4;refGene_tx=XM_023241517.1
D3 18885192 18885333 refGene_gn=TRPV4;refGene_tx=XM_023241517.1
D3 18886090 18886243 refGene_gn=TRPV4;refGene_tx=XM_023241517.1
D3 18890009 18890182 refGene_gn=TRPV4;refGene_tx=XM_023241517.1
D3 18892504 18892890 refGene_gn=TRPV4;refGene_tx=XM_023241517.1

D3	18918731 18918914 refGene_gn=GLTP;refGene_tx=XM_006938684.3
D3	18920750 18920901 refGene_gn=GLTP;refGene_tx=XM_006938684.3
D3	18923608 18923742 refGene_gn=GLTP;refGene_tx=XM_006938684.3
D3	18924784 18924843 refGene_gn=GLTP;refGene_tx=XM_006938684.3
D3	18940442 18940545 refGene_gn=GLTP;refGene_tx=XM_006938684.3
D3	18961891 18962085 refGene_gn=TCHP;refGene_tx=XM_003994848.5
D3	18963280 18963491 refGene_gn=TCHP;refGene_tx=XM_003994848.5
D3	18964083 18964140 refGene_gn=TCHP;refGene_tx=XM_003994848.5
D3	18965834 18965903 refGene_gn=TCHP;refGene_tx=XM_003994848.5
D3	18966572 18966746 refGene_gn=TCHP;refGene_tx=XM_003994848.5
D3	18967909 18968022 refGene_gn=TCHP;refGene_tx=XM_003994848.5
D3	18968168 18968235 refGene_gn=TCHP;refGene_tx=XM_003994848.5
D3	18969444 18969617 refGene_gn=TCHP;refGene_tx=XM_003994848.5
D3	18970473 18970555 refGene_gn=TCHP;refGene_tx=XM_003994848.5
D3	18972390 18972576 refGene_gn=TCHP;refGene_tx=XM_003994848.5
D3	18973154 18973298 refGene_gn=TCHP;refGene_tx=XM_003994848.5
D3	18974435 18974468 refGene_gn=TCHP;refGene_tx=XM_003994848.5
D3	18987747 18987960
refGene_gn=GIT2;refGene_tx=XM_023241518.1;refGene_gn=GIT2;refGene_tx=XM_023241519.1;refGene_gn=GIT2;refGene_tx=XM_023241520.1;refGene_gn=GIT2;refGene_tx=XM_023241521.1;refGene_gn=GIT2;refGene_tx=XM_023241522.1;refGene_gn=GIT2;refGene_tx=XM_023241523.1;refGene_gn=GIT2;refGene_tx=XM_023241524.1;refGene_gn=GIT2;refGene_tx=XM_023241525.1;refGene_gn=GIT2;refGene_tx=XM_023241526.1;refGene_gn=GIT2;refGene_tx=XM_023241527.1;refGene_gn=GIT2;refGene_tx=XM_023241528.1;refGene_gn=GIT2;refGene_tx=XM_023241529.1;refGene_gn=GIT2;refGene_tx=XM_023241530.1;refGene_gn=GIT2;refGene_tx=XM_023241531.1;refGene_gn=GIT2;refGene_tx=XM_023241532.1;refGene_gn=GIT2;refGene_tx=XM_023241533.1;refGene_gn=GIT2;refGene_tx=XM_023241534.1;refGene_gn=GIT2;refGene_tx=XM_023241535.1;refGene_gn=GIT2;refGene_tx=XM_023241536.1;refGene_gn=GIT2;refGene_tx=XM_023241537.1
D3	18988673 18988737
refGene_gn=GIT2;refGene_tx=XM_023241518.1;refGene_gn=GIT2;refGene_tx=XM_023241519.1;refGene_gn=GIT2;refGene_tx=XM_023241520.1;refGene_gn=GIT2;refGene_tx=XM_023241521.1;refGene_gn=GIT2;refGene_tx=XM_023241522.1;refGene_gn=GIT2;refGene_tx=XM_023241523.1;refGene_gn=GIT2;refGene_tx=XM_023241524.1;refGene_gn=GIT2;refGene_tx=XM_023241525.1;refGene_gn=GIT2;refGene_tx=XM_023241526.1;refGene_gn=GIT2;refGene_tx=XM_023241527.1;refGene_gn=GIT2;refGene_tx=XM_023241528.1;refGene_gn=GIT2;refGene_tx=XM_023241529.1;refGene_gn=GIT2;refGene_tx=XM_023241530.1;refGene_gn=GIT2;refGene_tx=XM_023241532.1;refGene_gn=GIT2;refGene_tx=XM_023241533.1;refGene_gn=GIT2;refGene_tx=XM_023241534.1;refGene_gn=GIT2;refGene_tx=XM_023241535.1;refGene_gn=GIT2;refGene_tx=XM_023241536.1;refGene_gn=GIT2;refGene_tx=XM_023241537.1
D3	18993458 18993647
refGene_gn=GIT2;refGene_tx=XM_023241518.1;refGene_gn=GIT2;refGene_tx=XM_023241519.1;refGene_gn=GIT2;refGene_tx=XM_023241520.1;refGene_gn=GIT2;refGene_tx=XM_023241521.1;refGene_gn=GIT2;refGene_tx=XM_023241522.1;refGene_gn=GIT2;refGene_tx=XM_023241523.1;refGene_gn=GIT2;refGene_tx=XM_023241524.1;refGene_gn=GIT2;refGene_tx=XM_023241525.1;refGene_gn=GIT2;refGene_tx=XM_023241526.1;refGene_gn=GIT2;refGene_tx=XM_023241527.1;

refGene_gn=GIT2;refGene_tx=XM_023241528.1;refGene_gn=GIT2;refGene_tx=XM_023241529.1;refGene_gn=GIT2;refGene_tx=XM_023241530.1;refGene_gn=GIT2;refGene_tx=XM_023241531.1;refGene_gn=GIT2;refGene_tx=XM_023241532.1;refGene_gn=GIT2;refGene_tx=XM_023241533.1;refGene_gn=GIT2;refGene_tx=XM_023241534.1;refGene_gn=GIT2;refGene_tx=XM_023241535.1;refGene_gn=GIT2;refGene_tx=XM_023241536.1;refGene_gn=GIT2;refGene_tx=XM_023241537.1
D3 18994232 18994315
refGene_gn=GIT2;refGene_tx=XM_023241518.1;refGene_gn=GIT2;refGene_tx=XM_023241519.1;refGene_gn=GIT2;refGene_tx=XM_023241520.1;refGene_gn=GIT2;refGene_tx=XM_023241521.1;refGene_gn=GIT2;refGene_tx=XM_023241522.1;refGene_gn=GIT2;refGene_tx=XM_023241523.1;refGene_gn=GIT2;refGene_tx=XM_023241524.1;refGene_gn=GIT2;refGene_tx=XM_023241525.1;refGene_gn=GIT2;refGene_tx=XM_023241526.1;refGene_gn=GIT2;refGene_tx=XM_023241527.1;refGene_gn=GIT2;refGene_tx=XM_023241528.1;refGene_gn=GIT2;refGene_tx=XM_023241529.1;refGene_gn=GIT2;refGene_tx=XM_023241530.1;refGene_gn=GIT2;refGene_tx=XM_023241531.1;refGene_gn=GIT2;refGene_tx=XM_023241532.1;refGene_gn=GIT2;refGene_tx=XM_023241533.1;refGene_gn=GIT2;refGene_tx=XM_023241534.1;refGene_gn=GIT2;refGene_tx=XM_023241535.1;refGene_gn=GIT2;refGene_tx=XM_023241536.1;refGene_gn=GIT2;refGene_tx=XM_023241537.1
D3 18998270 18998360
refGene_gn=GIT2;refGene_tx=XM_023241518.1;refGene_gn=GIT2;refGene_tx=XM_023241519.1;refGene_gn=GIT2;refGene_tx=XM_023241520.1;refGene_gn=GIT2;refGene_tx=XM_023241521.1;refGene_gn=GIT2;refGene_tx=XM_023241524.1;refGene_gn=GIT2;refGene_tx=XM_023241525.1;refGene_gn=GIT2;refGene_tx=XM_023241526.1;refGene_gn=GIT2;refGene_tx=XM_023241527.1;refGene_gn=GIT2;refGene_tx=XM_023241530.1;refGene_gn=GIT2;refGene_tx=XM_023241531.1;refGene_gn=GIT2;refGene_tx=XM_023241532.1;refGene_gn=GIT2;refGene_tx=XM_023241536.1
D3 18999955 19000204
refGene_gn=GIT2;refGene_tx=XM_023241518.1;refGene_gn=GIT2;refGene_tx=XM_023241519.1;refGene_gn=GIT2;refGene_tx=XM_023241520.1;refGene_gn=GIT2;refGene_tx=XM_023241521.1;refGene_gn=GIT2;refGene_tx=XM_023241522.1;refGene_gn=GIT2;refGene_tx=XM_023241523.1;refGene_gn=GIT2;refGene_tx=XM_023241524.1;refGene_gn=GIT2;refGene_tx=XM_023241525.1;refGene_gn=GIT2;refGene_tx=XM_023241526.1;refGene_gn=GIT2;refGene_tx=XM_023241527.1;refGene_gn=GIT2;refGene_tx=XM_023241528.1;refGene_gn=GIT2;refGene_tx=XM_023241529.1;refGene_gn=GIT2;refGene_tx=XM_023241531.1
D3 19001518 19001542 refGene_gn=GIT2;refGene_tx=XM_023241538.1
D3 19003897 19004047
refGene_gn=GIT2;refGene_tx=XM_023241518.1;refGene_gn=GIT2;refGene_tx=XM_023241519.1;refGene_gn=GIT2;refGene_tx=XM_023241520.1;refGene_gn=GIT2;refGene_tx=XM_023241521.1;refGene_gn=GIT2;refGene_tx=XM_023241522.1;refGene_gn=GIT2;refGene_tx=XM_023241523.1;refGene_gn=GIT2;refGene_tx=XM_023241526.1;refGene_gn=GIT2;refGene_tx=XM_023241527.1;refGene_gn=GIT2;refGene_tx=XM_023241530.1;refGene_gn=GIT2;refGene_tx=XM_023241531.1;refGene_gn=GIT2;refGene_tx=XM_023241532.1;refGene_gn=GIT2;refGene_tx=XM_023241533.1;refGene_gn=GIT2;refGene_tx=XM_023241534.1;refGene_gn=GIT2;refGene_tx=XM_023241535.1;refGene_gn=GIT2;refGene_tx=XM_023241538.1
D3 19006129 19006272
refGene_gn=GIT2;refGene_tx=XM_023241518.1;refGene_gn=GIT2;refGene_tx=X

M_023241519.1;refGene_gn=GIT2;refGene_tx=XM_023241520.1;refGene_gn=GIT
2;refGene_tx=XM_023241521.1;refGene_gn=GIT2;refGene_tx=XM_023241522.1;
refGene_gn=GIT2;refGene_tx=XM_023241523.1;refGene_gn=GIT2;refGene_tx=X
M_023241524.1;refGene_gn=GIT2;refGene_tx=XM_023241525.1;refGene_gn=GIT
2;refGene_tx=XM_023241526.1;refGene_gn=GIT2;refGene_tx=XM_023241527.1;
refGene_gn=GIT2;refGene_tx=XM_023241528.1;refGene_gn=GIT2;refGene_tx=X
M_023241529.1;refGene_gn=GIT2;refGene_tx=XM_023241530.1;refGene_gn=GIT
2;refGene_tx=XM_023241531.1;refGene_gn=GIT2;refGene_tx=XM_023241532.1;
refGene_gn=GIT2;refGene_tx=XM_023241533.1;refGene_gn=GIT2;refGene_tx=X
M_023241534.1;refGene_gn=GIT2;refGene_tx=XM_023241535.1;refGene_gn=GIT
2;refGene_tx=XM_023241536.1;refGene_gn=GIT2;refGene_tx=XM_023241537.1;
refGene_gn=GIT2;refGene_tx=XM_023241538.1
D3 19010597 19010709
refGene_gn=GIT2;refGene_tx=XM_023241518.1;refGene_gn=GIT2;refGene_tx=X
M_023241519.1;refGene_gn=GIT2;refGene_tx=XM_023241520.1;refGene_gn=GIT
2;refGene_tx=XM_023241521.1;refGene_gn=GIT2;refGene_tx=XM_023241522.1;
refGene_gn=GIT2;refGene_tx=XM_023241523.1;refGene_gn=GIT2;refGene_tx=X
M_023241524.1;refGene_gn=GIT2;refGene_tx=XM_023241525.1;refGene_gn=GIT
2;refGene_tx=XM_023241526.1;refGene_gn=GIT2;refGene_tx=XM_023241527.1;
refGene_gn=GIT2;refGene_tx=XM_023241528.1;refGene_gn=GIT2;refGene_tx=X
M_023241529.1;refGene_gn=GIT2;refGene_tx=XM_023241530.1;refGene_gn=GIT
2;refGene_tx=XM_023241531.1;refGene_gn=GIT2;refGene_tx=XM_023241532.1;
refGene_gn=GIT2;refGene_tx=XM_023241533.1;refGene_gn=GIT2;refGene_tx=X
M_023241534.1;refGene_gn=GIT2;refGene_tx=XM_023241535.1;refGene_gn=GIT
2;refGene_tx=XM_023241536.1;refGene_gn=GIT2;refGene_tx=XM_023241537.1;
refGene_gn=GIT2;refGene_tx=XM_023241538.1
D3 19011981 19012079
refGene_gn=GIT2;refGene_tx=XM_023241518.1;refGene_gn=GIT2;refGene_tx=X
M_023241519.1;refGene_gn=GIT2;refGene_tx=XM_023241520.1;refGene_gn=GIT
2;refGene_tx=XM_023241521.1;refGene_gn=GIT2;refGene_tx=XM_023241522.1;
refGene_gn=GIT2;refGene_tx=XM_023241523.1;refGene_gn=GIT2;refGene_tx=X
M_023241524.1;refGene_gn=GIT2;refGene_tx=XM_023241525.1;refGene_gn=GIT
2;refGene_tx=XM_023241528.1;refGene_gn=GIT2;refGene_tx=XM_023241529.1;
refGene_gn=GIT2;refGene_tx=XM_023241530.1;refGene_gn=GIT2;refGene_tx=X
M_023241531.1;refGene_gn=GIT2;refGene_tx=XM_023241532.1;refGene_gn=GIT
2;refGene_tx=XM_023241533.1;refGene_gn=GIT2;refGene_tx=XM_023241534.1;
refGene_gn=GIT2;refGene_tx=XM_023241535.1;refGene_gn=GIT2;refGene_tx=X
M_023241536.1;refGene_gn=GIT2;refGene_tx=XM_023241537.1;refGene_gn=GIT
2;refGene_tx=XM_023241538.1
D3 19012331 19012404
refGene_gn=GIT2;refGene_tx=XM_023241518.1;refGene_gn=GIT2;refGene_tx=X
M_023241519.1;refGene_gn=GIT2;refGene_tx=XM_023241520.1;refGene_gn=GIT
2;refGene_tx=XM_023241521.1;refGene_gn=GIT2;refGene_tx=XM_023241522.1;
refGene_gn=GIT2;refGene_tx=XM_023241523.1;refGene_gn=GIT2;refGene_tx=X
M_023241524.1;refGene_gn=GIT2;refGene_tx=XM_023241525.1;refGene_gn=GIT
2;refGene_tx=XM_023241528.1;refGene_gn=GIT2;refGene_tx=XM_023241529.1;
refGene_gn=GIT2;refGene_tx=XM_023241530.1;refGene_gn=GIT2;refGene_tx=X
M_023241531.1;refGene_gn=GIT2;refGene_tx=XM_023241532.1;refGene_gn=GIT
2;refGene_tx=XM_023241533.1;refGene_gn=GIT2;refGene_tx=XM_023241534.1;
refGene_gn=GIT2;refGene_tx=XM_023241535.1;refGene_gn=GIT2;refGene_tx=X

M_023241536.1;refGene_gn=GIT2;refGene_tx=XM_023241537.1;refGene_gn=GIT2;refGene_tx=XM_023241538.1
D3 19015716 19015768
refGene_gn=GIT2;refGene_tx=XM_023241518.1;refGene_gn=GIT2;refGene_tx=XM_023241519.1;refGene_gn=GIT2;refGene_tx=XM_023241520.1;refGene_gn=GIT2;refGene_tx=XM_023241521.1;refGene_gn=GIT2;refGene_tx=XM_023241522.1;refGene_gn=GIT2;refGene_tx=XM_023241523.1;refGene_gn=GIT2;refGene_tx=XM_023241524.1;refGene_gn=GIT2;refGene_tx=XM_023241525.1;refGene_gn=GIT2;refGene_tx=XM_023241526.1;refGene_gn=GIT2;refGene_tx=XM_023241527.1;refGene_gn=GIT2;refGene_tx=XM_023241528.1;refGene_gn=GIT2;refGene_tx=XM_023241529.1;refGene_gn=GIT2;refGene_tx=XM_023241530.1;refGene_gn=GIT2;refGene_tx=XM_023241531.1;refGene_gn=GIT2;refGene_tx=XM_023241532.1;refGene_gn=GIT2;refGene_tx=XM_023241533.1;refGene_gn=GIT2;refGene_tx=XM_023241534.1;refGene_gn=GIT2;refGene_tx=XM_023241535.1;refGene_gn=GIT2;refGene_tx=XM_023241536.1;refGene_gn=GIT2;refGene_tx=XM_023241537.1;refGene_gn=GIT2;refGene_tx=XM_023241538.1
D3 19016765 19016771
refGene_gn=GIT2;refGene_tx=XM_023241518.1;refGene_gn=GIT2;refGene_tx=XM_023241521.1;refGene_gn=GIT2;refGene_tx=XM_023241522.1;refGene_gn=GIT2;refGene_tx=XM_023241524.1;refGene_gn=GIT2;refGene_tx=XM_023241528.1;refGene_gn=GIT2;refGene_tx=XM_023241530.1;refGene_gn=GIT2;refGene_tx=XM_023241531.1;refGene_gn=GIT2;refGene_tx=XM_023241532.1;refGene_gn=GIT2;refGene_tx=XM_023241533.1;refGene_gn=GIT2;refGene_tx=XM_023241534.1;refGene_gn=GIT2;refGene_tx=XM_023241536.1;refGene_gn=GIT2;refGene_tx=XM_023241537.1
D3 19016909 19016955
refGene_gn=GIT2;refGene_tx=XM_023241518.1;refGene_gn=GIT2;refGene_tx=XM_023241519.1;refGene_gn=GIT2;refGene_tx=XM_023241520.1;refGene_gn=GIT2;refGene_tx=XM_023241521.1;refGene_gn=GIT2;refGene_tx=XM_023241522.1;refGene_gn=GIT2;refGene_tx=XM_023241523.1;refGene_gn=GIT2;refGene_tx=XM_023241524.1;refGene_gn=GIT2;refGene_tx=XM_023241525.1;refGene_gn=GIT2;refGene_tx=XM_023241526.1;refGene_gn=GIT2;refGene_tx=XM_023241527.1;refGene_gn=GIT2;refGene_tx=XM_023241528.1;refGene_gn=GIT2;refGene_tx=XM_023241529.1;refGene_gn=GIT2;refGene_tx=XM_023241530.1;refGene_gn=GIT2;refGene_tx=XM_023241531.1;refGene_gn=GIT2;refGene_tx=XM_023241532.1;refGene_gn=GIT2;refGene_tx=XM_023241533.1;refGene_gn=GIT2;refGene_tx=XM_023241534.1;refGene_gn=GIT2;refGene_tx=XM_023241535.1;refGene_gn=GIT2;refGene_tx=XM_023241536.1;refGene_gn=GIT2;refGene_tx=XM_023241537.1;refGene_gn=GIT2;refGene_tx=XM_023241538.1
D3 19024842 19024937
refGene_gn=GIT2;refGene_tx=XM_023241518.1;refGene_gn=GIT2;refGene_tx=XM_023241519.1;refGene_gn=GIT2;refGene_tx=XM_023241520.1;refGene_gn=GIT2;refGene_tx=XM_023241521.1;refGene_gn=GIT2;refGene_tx=XM_023241522.1;refGene_gn=GIT2;refGene_tx=XM_023241523.1;refGene_gn=GIT2;refGene_tx=XM_023241524.1;refGene_gn=GIT2;refGene_tx=XM_023241525.1;refGene_gn=GIT2;refGene_tx=XM_023241526.1;refGene_gn=GIT2;refGene_tx=XM_023241527.1;refGene_gn=GIT2;refGene_tx=XM_023241528.1;refGene_gn=GIT2;refGene_tx=XM_023241529.1;refGene_gn=GIT2;refGene_tx=XM_023241530.1;refGene_gn=GIT2;refGene_tx=XM_023241531.1;refGene_gn=GIT2;refGene_tx=XM_023241532.1;refGene_gn=GIT2;refGene_tx=XM_023241533.1;refGene_gn=GIT2;refGene_tx=X

M_023241534.1;refGene_gn=GIT2;refGene_tx=XM_023241535.1;refGene_gn=GIT2;refGene_tx=XM_023241536.1;refGene_gn=GIT2;refGene_tx=XM_023241537.1;refGene_gn=GIT2;refGene_tx=XM_023241538.1
D3 19027100 19027231
refGene_gn=GIT2;refGene_tx=XM_023241518.1;refGene_gn=GIT2;refGene_tx=XM_023241519.1;refGene_gn=GIT2;refGene_tx=XM_023241520.1;refGene_gn=GIT2;refGene_tx=XM_023241521.1;refGene_gn=GIT2;refGene_tx=XM_023241522.1;refGene_gn=GIT2;refGene_tx=XM_023241523.1;refGene_gn=GIT2;refGene_tx=XM_023241524.1;refGene_gn=GIT2;refGene_tx=XM_023241525.1;refGene_gn=GIT2;refGene_tx=XM_023241526.1;refGene_gn=GIT2;refGene_tx=XM_023241527.1;refGene_gn=GIT2;refGene_tx=XM_023241528.1;refGene_gn=GIT2;refGene_tx=XM_023241529.1;refGene_gn=GIT2;refGene_tx=XM_023241530.1;refGene_gn=GIT2;refGene_tx=XM_023241531.1;refGene_gn=GIT2;refGene_tx=XM_023241532.1;refGene_gn=GIT2;refGene_tx=XM_023241533.1;refGene_gn=GIT2;refGene_tx=XM_023241534.1;refGene_gn=GIT2;refGene_tx=XM_023241535.1;refGene_gn=GIT2;refGene_tx=XM_023241536.1;refGene_gn=GIT2;refGene_tx=XM_023241537.1;refGene_gn=GIT2;refGene_tx=XM_023241538.1
D3 19027344 19027431
refGene_gn=GIT2;refGene_tx=XM_023241518.1;refGene_gn=GIT2;refGene_tx=XM_023241519.1;refGene_gn=GIT2;refGene_tx=XM_023241520.1;refGene_gn=GIT2;refGene_tx=XM_023241521.1;refGene_gn=GIT2;refGene_tx=XM_023241522.1;refGene_gn=GIT2;refGene_tx=XM_023241523.1;refGene_gn=GIT2;refGene_tx=XM_023241524.1;refGene_gn=GIT2;refGene_tx=XM_023241525.1;refGene_gn=GIT2;refGene_tx=XM_023241526.1;refGene_gn=GIT2;refGene_tx=XM_023241527.1;refGene_gn=GIT2;refGene_tx=XM_023241528.1;refGene_gn=GIT2;refGene_tx=XM_023241529.1;refGene_gn=GIT2;refGene_tx=XM_023241530.1;refGene_gn=GIT2;refGene_tx=XM_023241531.1;refGene_gn=GIT2;refGene_tx=XM_023241532.1;refGene_gn=GIT2;refGene_tx=XM_023241533.1;refGene_gn=GIT2;refGene_tx=XM_023241534.1;refGene_gn=GIT2;refGene_tx=XM_023241535.1;refGene_gn=GIT2;refGene_tx=XM_023241536.1;refGene_gn=GIT2;refGene_tx=XM_023241537.1;refGene_gn=GIT2;refGene_tx=XM_023241538.1
D3 19029420 19029526
refGene_gn=GIT2;refGene_tx=XM_023241518.1;refGene_gn=GIT2;refGene_tx=XM_023241519.1;refGene_gn=GIT2;refGene_tx=XM_023241520.1;refGene_gn=GIT2;refGene_tx=XM_023241521.1;refGene_gn=GIT2;refGene_tx=XM_023241522.1;refGene_gn=GIT2;refGene_tx=XM_023241523.1;refGene_gn=GIT2;refGene_tx=XM_023241524.1;refGene_gn=GIT2;refGene_tx=XM_023241525.1;refGene_gn=GIT2;refGene_tx=XM_023241526.1;refGene_gn=GIT2;refGene_tx=XM_023241527.1;refGene_gn=GIT2;refGene_tx=XM_023241528.1;refGene_gn=GIT2;refGene_tx=XM_023241529.1;refGene_gn=GIT2;refGene_tx=XM_023241530.1;refGene_gn=GIT2;refGene_tx=XM_023241531.1;refGene_gn=GIT2;refGene_tx=XM_023241532.1;refGene_gn=GIT2;refGene_tx=XM_023241533.1;refGene_gn=GIT2;refGene_tx=XM_023241534.1;refGene_gn=GIT2;refGene_tx=XM_023241535.1;refGene_gn=GIT2;refGene_tx=XM_023241536.1;refGene_gn=GIT2;refGene_tx=XM_023241537.1;refGene_gn=GIT2;refGene_tx=XM_023241538.1
D3 19030392 19030505
refGene_gn=GIT2;refGene_tx=XM_023241518.1;refGene_gn=GIT2;refGene_tx=XM_023241519.1;refGene_gn=GIT2;refGene_tx=XM_023241520.1;refGene_gn=GIT2;refGene_tx=XM_023241521.1;refGene_gn=GIT2;refGene_tx=XM_023241522.1;refGene_gn=GIT2;refGene_tx=XM_023241523.1;refGene_gn=GIT2;refGene_tx=X

M_023241524.1;refGene_gn=GIT2;refGene_tx=XM_023241525.1;refGene_gn=GIT
2;refGene_tx=XM_023241526.1;refGene_gn=GIT2;refGene_tx=XM_023241527.1;
refGene_gn=GIT2;refGene_tx=XM_023241528.1;refGene_gn=GIT2;refGene_tx=X
M_023241529.1;refGene_gn=GIT2;refGene_tx=XM_023241530.1;refGene_gn=GIT
2;refGene_tx=XM_023241531.1;refGene_gn=GIT2;refGene_tx=XM_023241532.1;
refGene_gn=GIT2;refGene_tx=XM_023241533.1;refGene_gn=GIT2;refGene_tx=X
M_023241534.1;refGene_gn=GIT2;refGene_tx=XM_023241535.1;refGene_gn=GIT
2;refGene_tx=XM_023241536.1;refGene_gn=GIT2;refGene_tx=XM_023241537.1;
refGene_gn=GIT2;refGene_tx=XM_023241538.1
D3 19032022 19032156
refGene_gn=GIT2;refGene_tx=XM_023241518.1;refGene_gn=GIT2;refGene_tx=X
M_023241519.1;refGene_gn=GIT2;refGene_tx=XM_023241520.1;refGene_gn=GIT
2;refGene_tx=XM_023241521.1;refGene_gn=GIT2;refGene_tx=XM_023241522.1;
refGene_gn=GIT2;refGene_tx=XM_023241523.1;refGene_gn=GIT2;refGene_tx=X
M_023241524.1;refGene_gn=GIT2;refGene_tx=XM_023241525.1;refGene_gn=GIT
2;refGene_tx=XM_023241526.1;refGene_gn=GIT2;refGene_tx=XM_023241527.1;
refGene_gn=GIT2;refGene_tx=XM_023241528.1;refGene_gn=GIT2;refGene_tx=X
M_023241529.1;refGene_gn=GIT2;refGene_tx=XM_023241530.1;refGene_gn=GIT
2;refGene_tx=XM_023241531.1;refGene_gn=GIT2;refGene_tx=XM_023241532.1;
refGene_gn=GIT2;refGene_tx=XM_023241533.1;refGene_gn=GIT2;refGene_tx=X
M_023241534.1;refGene_gn=GIT2;refGene_tx=XM_023241535.1;refGene_gn=GIT
2;refGene_tx=XM_023241536.1;refGene_gn=GIT2;refGene_tx=XM_023241537.1;
refGene_gn=GIT2;refGene_tx=XM_023241538.1
D3 19036588 19036640
refGene_gn=GIT2;refGene_tx=XM_023241518.1;refGene_gn=GIT2;refGene_tx=X
M_023241519.1;refGene_gn=GIT2;refGene_tx=XM_023241520.1;refGene_gn=GIT
2;refGene_tx=XM_023241521.1;refGene_gn=GIT2;refGene_tx=XM_023241522.1;
refGene_gn=GIT2;refGene_tx=XM_023241523.1;refGene_gn=GIT2;refGene_tx=X
M_023241524.1;refGene_gn=GIT2;refGene_tx=XM_023241525.1;refGene_gn=GIT
2;refGene_tx=XM_023241526.1;refGene_gn=GIT2;refGene_tx=XM_023241527.1;
refGene_gn=GIT2;refGene_tx=XM_023241528.1;refGene_gn=GIT2;refGene_tx=X
M_023241529.1;refGene_gn=GIT2;refGene_tx=XM_023241530.1;refGene_gn=GIT
2;refGene_tx=XM_023241531.1;refGene_gn=GIT2;refGene_tx=XM_023241532.1;
refGene_gn=GIT2;refGene_tx=XM_023241533.1;refGene_gn=GIT2;refGene_tx=X
M_023241534.1;refGene_gn=GIT2;refGene_tx=XM_023241535.1;refGene_gn=GIT
2;refGene_tx=XM_023241536.1;refGene_gn=GIT2;refGene_tx=XM_023241537.1;
refGene_gn=GIT2;refGene_tx=XM_023241538.1
D3 19039628 19039724
refGene_gn=ANKRD13A;refGene_tx=XM_023241539.1;refGene_gn=ANKRD13A;refG
ene_tx=XM_023241540.1
D3 19049986 19050119
refGene_gn=ANKRD13A;refGene_tx=XM_023241539.1;refGene_gn=ANKRD13A;refG
ene_tx=XM_023241540.1
D3 19050946 19051071
refGene_gn=ANKRD13A;refGene_tx=XM_023241539.1;refGene_gn=ANKRD13A;refG
ene_tx=XM_023241540.1
D3 19054475 19054521
refGene_gn=ANKRD13A;refGene_tx=XM_023241539.1;refGene_gn=ANKRD13A;refG
ene_tx=XM_023241540.1
D3 19055673 19055817

refGene_gn=ANKRD13A;refGene_tx=XM_023241539.1;refGene_gn=ANKRD13A;refGene_tx=XM_023241540.1
D3 19056883 19057073
refGene_gn=ANKRD13A;refGene_tx=XM_023241539.1;refGene_gn=ANKRD13A;refGene_tx=XM_023241540.1
D3 19060911 19060978
refGene_gn=ANKRD13A;refGene_tx=XM_023241539.1;refGene_gn=ANKRD13A;refGene_tx=XM_023241540.1
D3 19062345 19062427
refGene_gn=ANKRD13A;refGene_tx=XM_023241539.1;refGene_gn=ANKRD13A;refGene_tx=XM_023241540.1
D3 19063267 19063329
refGene_gn=ANKRD13A;refGene_tx=XM_023241539.1;refGene_gn=ANKRD13A;refGene_tx=XM_023241540.1
D3 19064019 19064150
refGene_gn=ANKRD13A;refGene_tx=XM_023241539.1;refGene_gn=ANKRD13A;refGene_tx=XM_023241540.1
D3 19064659 19064817
refGene_gn=ANKRD13A;refGene_tx=XM_023241539.1;refGene_gn=ANKRD13A;refGene_tx=XM_023241540.1
D3 19065528 19065642
refGene_gn=ANKRD13A;refGene_tx=XM_023241539.1;refGene_gn=ANKRD13A;refGene_tx=XM_023241540.1
D3 19067539 19067700
refGene_gn=ANKRD13A;refGene_tx=XM_023241539.1;refGene_gn=ANKRD13A;refGene_tx=XM_023241540.1
D3 19068293 19068716 refGene_gn=ANKRD13A;refGene_tx=XM_023241539.1
D3 19069254 19069322 refGene_gn=ANKRD13A;refGene_tx=XM_023241540.1
D3 19070040 19070236 refGene_gn=ANKRD13A;refGene_tx=XM_023241540.1
D3 19076018 19076085
refGene_gn=CD3H12orf76;refGene_tx=XM_023241541.1
D3 19080818 19081213
refGene_gn=CD3H12orf76;refGene_tx=XM_023241541.1
D3 19102268 19102283
refGene_gn=LOC111557364;refGene_tx=XM_023242154.1
D3 19102541 19103058
refGene_gn=LOC111557364;refGene_tx=XM_023242154.1
D3 19117758 19117862
refGene_gn=LOC101092798;refGene_tx=XM_023242155.1
D3 19118959 19119080
refGene_gn=LOC101092798;refGene_tx=XM_023242155.1
D3 19119081 19119159
refGene_gn=LOC101092798;refGene_tx=XM_023242155.1
D3 19119237 19119411
refGene_gn=LOC101092798;refGene_tx=XM_023242155.1
D3 19119715 19119925
refGene_gn=LOC101092798;refGene_tx=XM_023242155.1
D3 19130439 19130551
refGene_gn=LOC101092550;refGene_tx=XM_006938638.4
D3 19130636 19130724

refGene_gn=LOC101092550;refGene_tx=XM_006938638.4
D3 19131674 19131825
refGene_gn=LOC101092550;refGene_tx=XM_006938638.4
D3 19132107 19132284
refGene_gn=LOC101092550;refGene_tx=XM_006938638.4
D3 19132368 19132569
refGene_gn=LOC101092550;refGene_tx=XM_006938638.4
D3 19133350 19133561
refGene_gn=LOC101083623;refGene_tx=XM_003994846.5
D3 19134922 19135010
refGene_gn=LOC101083623;refGene_tx=XM_003994846.5
D3 19137772 19137923
refGene_gn=LOC101083623;refGene_tx=XM_003994846.5
D3 19138483 19138660
refGene_gn=LOC101083623;refGene_tx=XM_003994846.5
D3 19139833 19140031
refGene_gn=LOC101083623;refGene_tx=XM_003994846.5
D3 19146833 19146996
refGene_gn=LOC101083370;refGene_tx=XM_003994845.5
D3 19148830 19148918
refGene_gn=LOC101083370;refGene_tx=XM_003994845.5
D3 19149508 19149659
refGene_gn=LOC101083370;refGene_tx=XM_003994845.5
D3 19157574 19157751
refGene_gn=LOC101083370;refGene_tx=XM_003994845.5;refGene_gn=LOC101083
370;refGene_tx=XM_019814694.2
D3 19157937 19158132
refGene_gn=LOC101083370;refGene_tx=XM_003994845.5;refGene_gn=LOC101083
370;refGene_tx=XM_019814694.2
D3 19160856 19160968
refGene_gn=LOC101083114;refGene_tx=XM_003994844.4
D3 19162332 19162420
refGene_gn=LOC101083114;refGene_tx=XM_003994844.4
D3 19164471 19164622
refGene_gn=LOC101083114;refGene_tx=XM_003994844.4
D3 19166669 19166846
refGene_gn=LOC101083114;refGene_tx=XM_003994844.4
D3 19167178 19167373
refGene_gn=LOC101083114;refGene_tx=XM_003994844.4
D3 19170142 19170250
refGene_gn=LOC101082852;refGene_tx=XM_003994843.4
D3 19170578 19170754
refGene_gn=LOC101082852;refGene_tx=XM_003994843.4
D3 19172317 19172390
refGene_gn=LOC101082852;refGene_tx=XM_003994843.4
D3 19188845 19188848
refGene_gn=CABIN1;refGene_tx=XM_006938634.4;refGene_gn=CABIN1;refGene_
tx=XM_006938635.4;refGene_gn=CABIN1;refGene_tx=XM_006938636.4;refGene_
gn=CABIN1;refGene_tx=XM_019814696.2;refGene_gn=CABIN1;refGene_tx=XM_01
9814700.2;refGene_gn=CABIN1;refGene_tx=XM_019814701.2;refGene_gn=CABIN

1;refGene_tx=XM_023241542.1
D3 19189366 19189459
refGene_gn=CABIN1;refGene_tx=XM_006938634.4;refGene_gn=CABIN1;refGene_tx=XM_006938635.4;refGene_gn=CABIN1;refGene_tx=XM_006938636.4;refGene_gn=CABIN1;refGene_tx=XM_019814696.2;refGene_gn=CABIN1;refGene_tx=XM_019814700.2;refGene_gn=CABIN1;refGene_tx=XM_019814701.2;refGene_gn=CABIN1;refGene_tx=XM_023241542.1
D3 19190740 19190854
refGene_gn=CABIN1;refGene_tx=XM_006938634.4;refGene_gn=CABIN1;refGene_tx=XM_006938635.4;refGene_gn=CABIN1;refGene_tx=XM_006938636.4;refGene_gn=CABIN1;refGene_tx=XM_019814696.2;refGene_gn=CABIN1;refGene_tx=XM_019814700.2;refGene_gn=CABIN1;refGene_tx=XM_019814701.2;refGene_gn=CABIN1;refGene_tx=XM_023241542.1
D3 19193628 19193763
refGene_gn=CABIN1;refGene_tx=XM_006938634.4;refGene_gn=CABIN1;refGene_tx=XM_006938635.4;refGene_gn=CABIN1;refGene_tx=XM_006938636.4;refGene_gn=CABIN1;refGene_tx=XM_019814696.2;refGene_gn=CABIN1;refGene_tx=XM_019814700.2;refGene_gn=CABIN1;refGene_tx=XM_019814701.2
D3 19194895 19195076
refGene_gn=CABIN1;refGene_tx=XM_006938634.4;refGene_gn=CABIN1;refGene_tx=XM_006938635.4;refGene_gn=CABIN1;refGene_tx=XM_006938636.4;refGene_gn=CABIN1;refGene_tx=XM_019814696.2;refGene_gn=CABIN1;refGene_tx=XM_019814700.2;refGene_gn=CABIN1;refGene_tx=XM_019814701.2;refGene_gn=CABIN1;refGene_tx=XM_023241542.1
D3 19198532 19198662
refGene_gn=CABIN1;refGene_tx=XM_006938634.4;refGene_gn=CABIN1;refGene_tx=XM_006938635.4;refGene_gn=CABIN1;refGene_tx=XM_006938636.4;refGene_gn=CABIN1;refGene_tx=XM_019814696.2;refGene_gn=CABIN1;refGene_tx=XM_019814700.2;refGene_gn=CABIN1;refGene_tx=XM_019814701.2;refGene_gn=CABIN1;refGene_tx=XM_023241542.1
D3 19200122 19200272
refGene_gn=CABIN1;refGene_tx=XM_006938634.4;refGene_gn=CABIN1;refGene_tx=XM_006938635.4;refGene_gn=CABIN1;refGene_tx=XM_006938636.4;refGene_gn=CABIN1;refGene_tx=XM_019814696.2;refGene_gn=CABIN1;refGene_tx=XM_019814700.2;refGene_gn=CABIN1;refGene_tx=XM_019814701.2;refGene_gn=CABIN1;refGene_tx=XM_023241542.1
D3 19203189 19203476
refGene_gn=CABIN1;refGene_tx=XM_006938634.4;refGene_gn=CABIN1;refGene_tx=XM_006938635.4;refGene_gn=CABIN1;refGene_tx=XM_006938636.4;refGene_gn=CABIN1;refGene_tx=XM_019814696.2;refGene_gn=CABIN1;refGene_tx=XM_019814700.2;refGene_gn=CABIN1;refGene_tx=XM_019814701.2;refGene_gn=CABIN1;refGene_tx=XM_023241542.1
D3 19204909 19205078
refGene_gn=CABIN1;refGene_tx=XM_006938634.4;refGene_gn=CABIN1;refGene_tx=XM_006938635.4;refGene_gn=CABIN1;refGene_tx=XM_006938636.4;refGene_gn=CABIN1;refGene_tx=XM_019814696.2;refGene_gn=CABIN1;refGene_tx=XM_019814700.2;refGene_gn=CABIN1;refGene_tx=XM_019814701.2;refGene_gn=CABIN1;refGene_tx=XM_023241542.1
D3 19208251 19208388
refGene_gn=CABIN1;refGene_tx=XM_006938634.4;refGene_gn=CABIN1;refGene_

tx=XM_006938635.4;refGene_gn=CABIN1;refGene_tx=XM_006938636.4;refGene_gn=CABIN1;refGene_tx=XM_019814696.2;refGene_gn=CABIN1;refGene_tx=XM_019814700.2;refGene_gn=CABIN1;refGene_tx=XM_019814701.2;refGene_gn=CABIN1;refGene_tx=XM_023241542.1
D3 19208923 19209141
refGene_gn=CABIN1;refGene_tx=XM_006938634.4;refGene_gn=CABIN1;refGene_tx=XM_006938635.4;refGene_gn=CABIN1;refGene_tx=XM_006938636.4;refGene_gn=CABIN1;refGene_tx=XM_019814696.2;refGene_gn=CABIN1;refGene_tx=XM_019814700.2;refGene_gn=CABIN1;refGene_tx=XM_019814701.2;refGene_gn=CABIN1;refGene_tx=XM_023241542.1
D3 19210878 19210957
refGene_gn=CABIN1;refGene_tx=XM_006938634.4;refGene_gn=CABIN1;refGene_tx=XM_006938635.4;refGene_gn=CABIN1;refGene_tx=XM_006938636.4;refGene_gn=CABIN1;refGene_tx=XM_019814696.2;refGene_gn=CABIN1;refGene_tx=XM_019814700.2;refGene_gn=CABIN1;refGene_tx=XM_019814701.2;refGene_gn=CABIN1;refGene_tx=XM_023241542.1
D3 19211540 19211728
refGene_gn=CABIN1;refGene_tx=XM_006938634.4;refGene_gn=CABIN1;refGene_tx=XM_006938635.4;refGene_gn=CABIN1;refGene_tx=XM_006938636.4;refGene_gn=CABIN1;refGene_tx=XM_019814696.2;refGene_gn=CABIN1;refGene_tx=XM_019814700.2;refGene_gn=CABIN1;refGene_tx=XM_019814701.2;refGene_gn=CABIN1;refGene_tx=XM_023241542.1
D3 19212552 19212705
refGene_gn=CABIN1;refGene_tx=XM_006938634.4;refGene_gn=CABIN1;refGene_tx=XM_006938635.4;refGene_gn=CABIN1;refGene_tx=XM_006938636.4;refGene_gn=CABIN1;refGene_tx=XM_019814696.2;refGene_gn=CABIN1;refGene_tx=XM_019814700.2;refGene_gn=CABIN1;refGene_tx=XM_019814701.2;refGene_gn=CABIN1;refGene_tx=XM_023241542.1
D3 19214480 19214675
refGene_gn=CABIN1;refGene_tx=XM_006938634.4;refGene_gn=CABIN1;refGene_tx=XM_006938635.4;refGene_gn=CABIN1;refGene_tx=XM_006938636.4;refGene_gn=CABIN1;refGene_tx=XM_019814696.2;refGene_gn=CABIN1;refGene_tx=XM_019814700.2;refGene_gn=CABIN1;refGene_tx=XM_019814701.2;refGene_gn=CABIN1;refGene_tx=XM_023241542.1
D3 19218722 19218965
refGene_gn=CABIN1;refGene_tx=XM_006938634.4;refGene_gn=CABIN1;refGene_tx=XM_006938635.4;refGene_gn=CABIN1;refGene_tx=XM_006938636.4;refGene_gn=CABIN1;refGene_tx=XM_019814696.2;refGene_gn=CABIN1;refGene_tx=XM_019814700.2;refGene_gn=CABIN1;refGene_tx=XM_019814701.2;refGene_gn=CABIN1;refGene_tx=XM_023241542.1
D3 19220270 19220427
refGene_gn=CABIN1;refGene_tx=XM_006938634.4;refGene_gn=CABIN1;refGene_tx=XM_006938635.4;refGene_gn=CABIN1;refGene_tx=XM_006938636.4;refGene_gn=CABIN1;refGene_tx=XM_019814696.2;refGene_gn=CABIN1;refGene_tx=XM_019814700.2;refGene_gn=CABIN1;refGene_tx=XM_019814701.2;refGene_gn=CABIN1;refGene_tx=XM_023241542.1
D3 19224747 19224863
refGene_gn=CABIN1;refGene_tx=XM_006938634.4;refGene_gn=CABIN1;refGene_tx=XM_006938635.4;refGene_gn=CABIN1;refGene_tx=XM_006938636.4;refGene_gn=CABIN1;refGene_tx=XM_019814696.2;refGene_gn=CABIN1;refGene_tx=XM_01

9814700.2;refGene_gn=CABIN1;refGene_tx=XM_019814701.2;refGene_gn=CABIN1;refGene_tx=XM_023241542.1
D3 19230319 19230481
refGene_gn=CABIN1;refGene_tx=XM_006938634.4;refGene_gn=CABIN1;refGene_tx=XM_006938635.4;refGene_gn=CABIN1;refGene_tx=XM_006938636.4;refGene_gn=CABIN1;refGene_tx=XM_019814696.2;refGene_gn=CABIN1;refGene_tx=XM_019814700.2;refGene_gn=CABIN1;refGene_tx=XM_019814701.2;refGene_gn=CABIN1;refGene_tx=XM_023241542.1
D3 19231245 19231452
refGene_gn=CABIN1;refGene_tx=XM_006938634.4;refGene_gn=CABIN1;refGene_tx=XM_006938635.4;refGene_gn=CABIN1;refGene_tx=XM_006938636.4;refGene_gn=CABIN1;refGene_tx=XM_019814696.2;refGene_gn=CABIN1;refGene_tx=XM_019814700.2;refGene_gn=CABIN1;refGene_tx=XM_019814701.2;refGene_gn=CABIN1;refGene_tx=XM_023241542.1
D3 19231672 19231818
refGene_gn=CABIN1;refGene_tx=XM_006938634.4;refGene_gn=CABIN1;refGene_tx=XM_006938635.4;refGene_gn=CABIN1;refGene_tx=XM_006938636.4;refGene_gn=CABIN1;refGene_tx=XM_019814696.2;refGene_gn=CABIN1;refGene_tx=XM_019814700.2;refGene_gn=CABIN1;refGene_tx=XM_019814701.2;refGene_gn=CABIN1;refGene_tx=XM_023241542.1
D3 19235122 19235384
refGene_gn=CABIN1;refGene_tx=XM_006938634.4;refGene_gn=CABIN1;refGene_tx=XM_006938635.4;refGene_gn=CABIN1;refGene_tx=XM_006938636.4;refGene_gn=CABIN1;refGene_tx=XM_019814696.2;refGene_gn=CABIN1;refGene_tx=XM_019814700.2;refGene_gn=CABIN1;refGene_tx=XM_019814701.2;refGene_gn=CABIN1;refGene_tx=XM_023241542.1
D3 19241654 19241915
refGene_gn=CABIN1;refGene_tx=XM_006938634.4;refGene_gn=CABIN1;refGene_tx=XM_006938635.4;refGene_gn=CABIN1;refGene_tx=XM_006938636.4;refGene_gn=CABIN1;refGene_tx=XM_019814696.2;refGene_gn=CABIN1;refGene_tx=XM_019814700.2;refGene_gn=CABIN1;refGene_tx=XM_019814701.2;refGene_gn=CABIN1;refGene_tx=XM_023241542.1
D3 19245296 19245448
refGene_gn=CABIN1;refGene_tx=XM_006938634.4;refGene_gn=CABIN1;refGene_tx=XM_006938635.4;refGene_gn=CABIN1;refGene_tx=XM_006938636.4;refGene_gn=CABIN1;refGene_tx=XM_019814696.2;refGene_gn=CABIN1;refGene_tx=XM_019814700.2;refGene_gn=CABIN1;refGene_tx=XM_019814701.2;refGene_gn=CABIN1;refGene_tx=XM_023241542.1
D3 19247155 19247334
refGene_gn=CABIN1;refGene_tx=XM_006938634.4;refGene_gn=CABIN1;refGene_tx=XM_006938635.4;refGene_gn=CABIN1;refGene_tx=XM_006938636.4;refGene_gn=CABIN1;refGene_tx=XM_019814696.2;refGene_gn=CABIN1;refGene_tx=XM_019814700.2;refGene_gn=CABIN1;refGene_tx=XM_019814701.2;refGene_gn=CABIN1;refGene_tx=XM_023241542.1
D3 19261539 19261722
refGene_gn=CABIN1;refGene_tx=XM_006938634.4;refGene_gn=CABIN1;refGene_tx=XM_006938635.4;refGene_gn=CABIN1;refGene_tx=XM_006938636.4;refGene_gn=CABIN1;refGene_tx=XM_019814696.2;refGene_gn=CABIN1;refGene_tx=XM_019814700.2;refGene_gn=CABIN1;refGene_tx=XM_023241542.1
D3 19266969 19267301

refGene_gn=CABIN1;refGene_tx=XM_006938634.4;refGene_gn=CABIN1;refGene_tx=XM_006938635.4;refGene_gn=CABIN1;refGene_tx=XM_006938636.4;refGene_gn=CABIN1;refGene_tx=XM_019814696.2;refGene_gn=CABIN1;refGene_tx=XM_019814700.2;refGene_gn=CABIN1;refGene_tx=XM_019814701.2;refGene_gn=CABIN1;refGene_tx=XM_023241542.1
D3 19282538 19282652
refGene_gn=CABIN1;refGene_tx=XM_006938634.4;refGene_gn=CABIN1;refGene_tx=XM_006938635.4;refGene_gn=CABIN1;refGene_tx=XM_006938636.4;refGene_gn=CABIN1;refGene_tx=XM_019814696.2;refGene_gn=CABIN1;refGene_tx=XM_019814700.2;refGene_gn=CABIN1;refGene_tx=XM_019814701.2;refGene_gn=CABIN1;refGene_tx=XM_023241542.1
D3 19314298 19314462
refGene_gn=CABIN1;refGene_tx=XM_006938634.4;refGene_gn=CABIN1;refGene_tx=XM_006938635.4;refGene_gn=CABIN1;refGene_tx=XM_006938636.4;refGene_gn=CABIN1;refGene_tx=XM_019814696.2;refGene_gn=CABIN1;refGene_tx=XM_019814700.2;refGene_gn=CABIN1;refGene_tx=XM_019814701.2;refGene_gn=CABIN1;refGene_tx=XM_023241542.1
D3 19315320 19315417
refGene_gn=CABIN1;refGene_tx=XM_006938634.4;refGene_gn=CABIN1;refGene_tx=XM_006938635.4;refGene_gn=CABIN1;refGene_tx=XM_006938636.4;refGene_gn=CABIN1;refGene_tx=XM_019814696.2;refGene_gn=CABIN1;refGene_tx=XM_019814700.2;refGene_gn=CABIN1;refGene_tx=XM_019814701.2;refGene_gn=CABIN1;refGene_tx=XM_023241542.1
D3 19316311 19316986
refGene_gn=CABIN1;refGene_tx=XM_006938634.4;refGene_gn=CABIN1;refGene_tx=XM_006938635.4;refGene_gn=CABIN1;refGene_tx=XM_006938636.4;refGene_gn=CABIN1;refGene_tx=XM_019814696.2;refGene_gn=CABIN1;refGene_tx=XM_019814700.2;refGene_gn=CABIN1;refGene_tx=XM_019814701.2;refGene_gn=CABIN1;refGene_tx=XM_023241542.1
D3 19318693 19318768
refGene_gn=CABIN1;refGene_tx=XM_006938634.4;refGene_gn=CABIN1;refGene_tx=XM_006938635.4;refGene_gn=CABIN1;refGene_tx=XM_006938636.4;refGene_gn=CABIN1;refGene_tx=XM_019814696.2;refGene_gn=CABIN1;refGene_tx=XM_019814700.2;refGene_gn=CABIN1;refGene_tx=XM_019814701.2;refGene_gn=CABIN1;refGene_tx=XM_023241542.1
D3 19321430 19321713
refGene_gn=CABIN1;refGene_tx=XM_006938634.4;refGene_gn=CABIN1;refGene_tx=XM_006938635.4;refGene_gn=CABIN1;refGene_tx=XM_006938636.4;refGene_gn=CABIN1;refGene_tx=XM_019814696.2;refGene_gn=CABIN1;refGene_tx=XM_019814700.2;refGene_gn=CABIN1;refGene_tx=XM_019814701.2;refGene_gn=CABIN1;refGene_tx=XM_023241542.1
D3 19326711 19326876
refGene_gn=CABIN1;refGene_tx=XM_006938634.4;refGene_gn=CABIN1;refGene_tx=XM_006938635.4;refGene_gn=CABIN1;refGene_tx=XM_006938636.4;refGene_gn=CABIN1;refGene_tx=XM_019814696.2;refGene_gn=CABIN1;refGene_tx=XM_019814700.2;refGene_gn=CABIN1;refGene_tx=XM_019814701.2;refGene_gn=CABIN1;refGene_tx=XM_023241542.1
D3 19327936 19328163
refGene_gn=CABIN1;refGene_tx=XM_006938634.4;refGene_gn=CABIN1;refGene_tx=XM_006938635.4;refGene_gn=CABIN1;refGene_tx=XM_006938636.4;refGene_

gn=CABIN1;refGene_tx=XM_019814696.2;refGene_gn=CABIN1;refGene_tx=XM_019814700.2;refGene_gn=CABIN1;refGene_tx=XM_019814701.2;refGene_gn=CABIN1;refGene_tx=XM_023241542.1
D3 19328271 19328415 refGene_gn=CABIN1;refGene_tx=XM_006938634.4;refGene_gn=CABIN1;refGene_tx=XM_006938635.4;refGene_gn=CABIN1;refGene_tx=XM_006938636.4;refGene_gn=CABIN1;refGene_tx=XM_019814696.2;refGene_gn=CABIN1;refGene_tx=XM_019814700.2;refGene_gn=CABIN1;refGene_tx=XM_019814701.2;refGene_gn=CABIN1;refGene_tx=XM_023241542.1
D3 19331251 19331333 refGene_gn=SUSD2;refGene_tx=XM_011287767.3
D3 19332854 19333062 refGene_gn=SUSD2;refGene_tx=XM_011287767.3
D3 19333210 19333362 refGene_gn=SUSD2;refGene_tx=XM_011287767.3
D3 19333787 19333955 refGene_gn=SUSD2;refGene_tx=XM_011287767.3
D3 19334478 19334653 refGene_gn=SUSD2;refGene_tx=XM_011287767.3
D3 19334786 19334988 refGene_gn=SUSD2;refGene_tx=XM_011287767.3
D3 19335208 19335294 refGene_gn=SUSD2;refGene_tx=XM_011287767.3
D3 19335387 19335656 refGene_gn=SUSD2;refGene_tx=XM_011287767.3
D3 19335730 19335874 refGene_gn=SUSD2;refGene_tx=XM_011287767.3
D3 19335968 19336127 refGene_gn=SUSD2;refGene_tx=XM_011287767.3
D3 19336987 19337236 refGene_gn=SUSD2;refGene_tx=XM_011287767.3
D3 19337351 19337624 refGene_gn=SUSD2;refGene_tx=XM_011287767.3
D3 19337758 19337935 refGene_gn=SUSD2;refGene_tx=XM_011287767.3
D3 19338024 19338127 refGene_gn=SUSD2;refGene_tx=XM_011287767.3
D3 19338240 19338265 refGene_gn=SUSD2;refGene_tx=XM_011287767.3
D3 19340817 19340877 refGene_gn=GGT5;refGene_tx=XM_019814703.2
D3 19351901 19352048 refGene_gn=GGT5;refGene_tx=XM_006938633.4;refGene_gn=GGT5;refGene_tx=XM_019814704.2
D3 19354437 19354548 refGene_gn=GGT5;refGene_tx=XM_006938633.4;refGene_gn=GGT5;refGene_tx=XM_019814703.2;refGene_gn=GGT5;refGene_tx=XM_019814704.2
D3 19354657 19354776 refGene_gn=GGT5;refGene_tx=XM_006938633.4;refGene_gn=GGT5;refGene_tx=XM_019814703.2;refGene_gn=GGT5;refGene_tx=XM_019814704.2
D3 19354969 19355076 refGene_gn=GGT5;refGene_tx=XM_006938633.4;refGene_gn=GGT5;refGene_tx=XM_019814703.2;refGene_gn=GGT5;refGene_tx=XM_019814704.2
D3 19355480 19355671 refGene_gn=GGT5;refGene_tx=XM_006938633.4;refGene_gn=GGT5;refGene_tx=XM_019814703.2;refGene_gn=GGT5;refGene_tx=XM_019814704.2
D3 19356051 19356188 refGene_gn=GGT5;refGene_tx=XM_006938633.4;refGene_gn=GGT5;refGene_tx=XM_019814703.2;refGene_gn=GGT5;refGene_tx=XM_019814704.2
D3 19357077 19357224 refGene_gn=GGT5;refGene_tx=XM_006938633.4;refGene_gn=GGT5;refGene_tx=XM_019814703.2;refGene_gn=GGT5;refGene_tx=XM_019814704.2
D3 19357762 19357920 refGene_gn=GGT5;refGene_tx=XM_006938633.4;refGene_gn=GGT5;refGene_tx=XM_019814703.2;refGene_gn=GGT5;refGene_tx=XM_019814704.2
D3 19358517 19358713

refGene_gn=GGT5;refGene_tx=XM_006938633.4;refGene_gn=GGT5;refGene_tx=XM_019814703.2;refGene_gn=GGT5;refGene_tx=XM_019814704.2
D3 19359227 19359323
refGene_gn=GGT5;refGene_tx=XM_006938633.4;refGene_gn=GGT5;refGene_tx=XM_019814703.2
D3 19360217 19360348
refGene_gn=GGT5;refGene_tx=XM_006938633.4;refGene_gn=GGT5;refGene_tx=XM_019814703.2;refGene_gn=GGT5;refGene_tx=XM_019814704.2
D3 19381220 19381393
refGene_gn=GGT5;refGene_tx=XM_006938633.4;refGene_gn=GGT5;refGene_tx=XM_019814703.2;refGene_gn=GGT5;refGene_tx=XM_019814704.2
D3 19384620 19384767 refGene_gn=GGT1;refGene_tx=XM_019814705.2
D3 19385098 19385212 refGene_gn=GGT1;refGene_tx=XM_019814705.2
D3 19385292 19385405 refGene_gn=GGT1;refGene_tx=XM_019814705.2
D3 19385501 19385629 refGene_gn=GGT1;refGene_tx=XM_019814705.2
D3 19385845 19386033 refGene_gn=GGT1;refGene_tx=XM_019814705.2
D3 19388863 19389000 refGene_gn=GGT1;refGene_tx=XM_019814705.2
D3 19389373 19389523 refGene_gn=GGT1;refGene_tx=XM_019814705.2
D3 19390532 19390690 refGene_gn=GGT1;refGene_tx=XM_019814705.2
D3 19391026 19391219 refGene_gn=GGT1;refGene_tx=XM_019814705.2
D3 19394035 19394122 refGene_gn=GGT1;refGene_tx=XM_019814705.2
D3 19394277 19394408 refGene_gn=GGT1;refGene_tx=XM_019814705.2
D3 19395009 19395170 refGene_gn=GGT1;refGene_tx=XM_019814705.2
D3 19409160 19409337 refGene_gn=LRRC75B;refGene_tx=XM_023241543.1
D3 19411926 19412055 refGene_gn=LRRC75B;refGene_tx=XM_023241543.1
D3 19413500 19413616 refGene_gn=LRRC75B;refGene_tx=XM_023241543.1
D3 19415126 19415652 refGene_gn=LRRC75B;refGene_tx=XM_023241543.1
D3 19424465 19424681
refGene_gn=LOC101081501;refGene_tx=XM_011287766.3
D3 19432757 19432819 refGene_gn=SNRPD3;refGene_tx=XM_003994836.5
D3 19434684 19434814 refGene_gn=SNRPD3;refGene_tx=XM_003994836.5
D3 19434816 19434879 refGene_gn=SNRPD3;refGene_tx=XM_003994836.5
D3 19446345 19446471 refGene_gn=SNRPD3;refGene_tx=XM_003994836.5
D3 19449040 19449083
refGene_gn=GUCD1;refGene_tx=XM_023241544.1;refGene_gn=GUCD1;refGene_tx=XM_023241545.1;refGene_gn=GUCD1;refGene_tx=XM_023241546.1
D3 19454062 19454147
refGene_gn=GUCD1;refGene_tx=XM_023241544.1;refGene_gn=GUCD1;refGene_tx=XM_023241545.1;refGene_gn=GUCD1;refGene_tx=XM_023241546.1
D3 19454921 19455087
refGene_gn=GUCD1;refGene_tx=XM_023241544.1;refGene_gn=GUCD1;refGene_tx=XM_023241545.1;refGene_gn=GUCD1;refGene_tx=XM_023241546.1
D3 19456262 19456354
refGene_gn=GUCD1;refGene_tx=XM_023241544.1;refGene_gn=GUCD1;refGene_tx=XM_023241545.1;refGene_gn=GUCD1;refGene_tx=XM_023241546.1
D3 19458453 19458695
refGene_gn=GUCD1;refGene_tx=XM_023241544.1;refGene_gn=GUCD1;refGene_tx=XM_023241545.1;refGene_gn=GUCD1;refGene_tx=XM_023241546.1
D3 19459214 19459307 refGene_gn=GUCD1;refGene_tx=XM_023241544.1
D3 19459396 19459491

refGene_gn=GUCD1;refGene_tx=XM_023241544.1;refGene_gn=GUCD1;refGene_tx=XM_023241545.1;refGene_gn=GUCD1;refGene_tx=XM_023241546.1
D3 19480264 19480348
refGene_gn=UPB1;refGene_tx=XM_003994835.5;refGene_gn=UPB1;refGene_tx=XM_019814709.2
D3 19481917 19482072
refGene_gn=UPB1;refGene_tx=XM_003994835.5;refGene_gn=UPB1;refGene_tx=XM_019814709.2
D3 19483326 19483369
refGene_gn=UPB1;refGene_tx=XM_003994835.5;refGene_gn=UPB1;refGene_tx=XM_019814709.2
D3 19484373 19484455
refGene_gn=UPB1;refGene_tx=XM_003994835.5;refGene_gn=UPB1;refGene_tx=XM_019814709.2
D3 19488266 19488436
refGene_gn=UPB1;refGene_tx=XM_003994835.5;refGene_gn=UPB1;refGene_tx=XM_019814709.2
D3 19490519 19490681
refGene_gn=UPB1;refGene_tx=XM_003994835.5;refGene_gn=UPB1;refGene_tx=XM_019814709.2
D3 19493024 19493119
refGene_gn=UPB1;refGene_tx=XM_003994835.5;refGene_gn=UPB1;refGene_tx=XM_019814709.2
D3 19499965 19500053
refGene_gn=UPB1;refGene_tx=XM_003994835.5;refGene_gn=UPB1;refGene_tx=XM_019814709.2
D3 19502305 19502477
refGene_gn=UPB1;refGene_tx=XM_003994835.5;refGene_gn=UPB1;refGene_tx=XM_019814709.2
D3 19508432 19508536 refGene_gn=UPB1;refGene_tx=XM_003994835.5
D3 19554582 19554636 refGene_gn=ADORA2A;refGene_tx=XM_019814711.2
D3 19554665 19555572
refGene_gn=ADORA2A;refGene_tx=XM_003994834.5;refGene_gn=ADORA2A;refGene_tx=XM_006938627.4;refGene_gn=ADORA2A;refGene_tx=XM_019814711.2
D3 19562214 19562546
refGene_gn=ADORA2A;refGene_tx=XM_003994834.5;refGene_gn=ADORA2A;refGene_tx=XM_006938627.4;refGene_gn=ADORA2A;refGene_tx=XM_019814711.2
D3 19580553 19580643
refGene_gn=SPECC1L;refGene_tx=XM_006938626.4;refGene_gn=SPECC1L;refGene_tx=XM_023241547.1
D3 19582338 19582398
refGene_gn=SPECC1L;refGene_tx=XM_006938626.4;refGene_gn=SPECC1L;refGene_tx=XM_023241547.1
D3 19583227 19583344
refGene_gn=SPECC1L;refGene_tx=XM_006938626.4;refGene_gn=SPECC1L;refGene_tx=XM_023241547.1
D3 19614864 19614967
refGene_gn=SPECC1L;refGene_tx=XM_006938626.4;refGene_gn=SPECC1L;refGene_tx=XM_023241547.1
D3 19618646 19618803

refGene_gn=SPECC1L;refGene_tx=XM_006938626.4;refGene_gn=SPECC1L;refGene_tx=XM_023241547.1
D3 19621078 19621162
refGene_gn=SPECC1L;refGene_tx=XM_006938626.4;refGene_gn=SPECC1L;refGene_tx=XM_023241547.1
D3 19634559 19634650
refGene_gn=SPECC1L;refGene_tx=XM_006938626.4;refGene_gn=SPECC1L;refGene_tx=XM_023241547.1
D3 19640699 19640791
refGene_gn=SPECC1L;refGene_tx=XM_006938626.4;refGene_gn=SPECC1L;refGene_tx=XM_023241547.1
D3 19645085 19645249
refGene_gn=SPECC1L;refGene_tx=XM_006938626.4;refGene_gn=SPECC1L;refGene_tx=XM_023241547.1
D3 19649705 19649881
refGene_gn=SPECC1L;refGene_tx=XM_006938626.4;refGene_gn=SPECC1L;refGene_tx=XM_023241547.1
D3 19650960 19651034
refGene_gn=SPECC1L;refGene_tx=XM_006938626.4;refGene_gn=SPECC1L;refGene_tx=XM_023241547.1
D3 19662528 19662736
refGene_gn=SPECC1L;refGene_tx=XM_006938626.4;refGene_gn=SPECC1L;refGene_tx=XM_023241547.1
D3 19663781 19665415
refGene_gn=SPECC1L;refGene_tx=XM_006938626.4;refGene_gn=SPECC1L;refGene_tx=XM_023241547.1
D3 19669674 19669828
refGene_gn=SPECC1L;refGene_tx=XM_006938626.4;refGene_gn=SPECC1L;refGene_tx=XM_023241547.1
D3 19685742 19685895
refGene_gn=SPECC1L;refGene_tx=XM_006938626.4;refGene_gn=SPECC1L;refGene_tx=XM_023241547.1
D3 19734182 19734272 refGene_gn=BCR;refGene_tx=XM_003994832.5
D3 19734619 19734782 refGene_gn=BCR;refGene_tx=XM_003994832.5
D3 19735271 19735377 refGene_gn=BCR;refGene_tx=XM_003994832.5
D3 19736140 19736275 refGene_gn=BCR;refGene_tx=XM_003994832.5
D3 19737148 19737288 refGene_gn=BCR;refGene_tx=XM_003994832.5
D3 19738579 19738689 refGene_gn=BCR;refGene_tx=XM_003994832.5
D3 19739377 19739437 refGene_gn=BCR;refGene_tx=XM_003994832.5
D3 19744931 19744982 refGene_gn=BCR;refGene_tx=XM_006938625.4
D3 19748696 19748783 refGene_gn=BCR;refGene_tx=XM_019814712.2
D3 19751276 19751408
refGene_gn=BCR;refGene_tx=XM_003994832.5;refGene_gn=BCR;refGene_tx=XM_006938625.4;refGene_gn=BCR;refGene_tx=XM_019814712.2
D3 19753241 19753339
refGene_gn=BCR;refGene_tx=XM_003994832.5;refGene_gn=BCR;refGene_tx=XM_006938625.4;refGene_gn=BCR;refGene_tx=XM_019814712.2
D3 19756765 19756840
refGene_gn=BCR;refGene_tx=XM_003994832.5;refGene_gn=BCR;refGene_tx=XM_006938625.4;refGene_gn=BCR;refGene_tx=XM_019814712.2

D3 19757487 19757592
refGene_gn=BCR;refGene_tx=XM_003994832.5;refGene_gn=BCR;refGene_tx=XM_006938625.4;refGene_gn=BCR;refGene_tx=XM_019814712.2
D3 19759014 19759090
refGene_gn=BCR;refGene_tx=XM_003994832.5;refGene_gn=BCR;refGene_tx=XM_006938625.4;refGene_gn=BCR;refGene_tx=XM_019814712.2
D3 19759969 19760089
refGene_gn=BCR;refGene_tx=XM_003994832.5;refGene_gn=BCR;refGene_tx=XM_006938625.4;refGene_gn=BCR;refGene_tx=XM_019814712.2
D3 19760649 19760818
refGene_gn=BCR;refGene_tx=XM_003994832.5;refGene_gn=BCR;refGene_tx=XM_006938625.4;refGene_gn=BCR;refGene_tx=XM_019814712.2
D3 19762038 19762160
refGene_gn=BCR;refGene_tx=XM_003994832.5;refGene_gn=BCR;refGene_tx=XM_006938625.4;refGene_gn=BCR;refGene_tx=XM_019814712.2
D3 19773426 19773567
refGene_gn=BCR;refGene_tx=XM_003994832.5;refGene_gn=BCR;refGene_tx=XM_006938625.4;refGene_gn=BCR;refGene_tx=XM_019814712.2
D3 19775489 19775542
refGene_gn=BCR;refGene_tx=XM_003994832.5;refGene_gn=BCR;refGene_tx=XM_006938625.4;refGene_gn=BCR;refGene_tx=XM_019814712.2
D3 19776826 19776887
refGene_gn=BCR;refGene_tx=XM_003994832.5;refGene_gn=BCR;refGene_tx=XM_006938625.4;refGene_gn=BCR;refGene_tx=XM_019814712.2
D3 19780806 19780914
refGene_gn=BCR;refGene_tx=XM_003994832.5;refGene_gn=BCR;refGene_tx=XM_006938625.4;refGene_gn=BCR;refGene_tx=XM_019814712.2
D3 19785854 19786040
refGene_gn=BCR;refGene_tx=XM_003994832.5;refGene_gn=BCR;refGene_tx=XM_006938625.4;refGene_gn=BCR;refGene_tx=XM_019814712.2
D3 19786394 19786499
refGene_gn=BCR;refGene_tx=XM_003994832.5;refGene_gn=BCR;refGene_tx=XM_006938625.4;refGene_gn=BCR;refGene_tx=XM_019814712.2
D3 19794578 19794760
refGene_gn=BCR;refGene_tx=XM_003994832.5;refGene_gn=BCR;refGene_tx=XM_006938625.4;refGene_gn=BCR;refGene_tx=XM_019814712.2
D3 19859468 19860780
refGene_gn=BCR;refGene_tx=XM_003994832.5;refGene_gn=BCR;refGene_tx=XM_006938625.4;refGene_gn=BCR;refGene_tx=XM_019814712.2
D3 19872970 19873108
refGene_gn=RAB36;refGene_tx=XM_019814716.2;refGene_gn=RAB36;refGene_tx=XM_019814717.2;refGene_gn=RAB36;refGene_tx=XM_019814718.2
D3 19873290 19873730
refGene_gn=RAB36;refGene_tx=XM_019814713.2;refGene_gn=RAB36;refGene_tx=XM_019814714.2;refGene_gn=RAB36;refGene_tx=XM_019814715.2;refGene_gn=RAB36;refGene_tx=XM_019814716.2;refGene_gn=RAB36;refGene_tx=XM_019814717.2;refGene_gn=RAB36;refGene_tx=XM_019814718.2
D3 19875281 19875372
refGene_gn=RAB36;refGene_tx=XM_019814713.2;refGene_gn=RAB36;refGene_tx=XM_019814714.2;refGene_gn=RAB36;refGene_tx=XM_019814715.2;refGene_gn=

RAB36;refGene_tx=XM_019814716.2;refGene_gn=RAB36;refGene_tx=XM_0198147
17.2;refGene_gn=RAB36;refGene_tx=XM_019814718.2
D3 19875699 19875781
refGene_gn=RAB36;refGene_tx=XM_019814713.2;refGene_gn=RAB36;refGene_tx
=XM_019814714.2;refGene_gn=RAB36;refGene_tx=XM_019814715.2;refGene_gn=
RAB36;refGene_tx=XM_019814716.2;refGene_gn=RAB36;refGene_tx=XM_0198147
17.2;refGene_gn=RAB36;refGene_tx=XM_019814718.2
D3 19876828 19876880
refGene_gn=RAB36;refGene_tx=XM_019814713.2;refGene_gn=RAB36;refGene_tx
=XM_019814714.2;refGene_gn=RAB36;refGene_tx=XM_019814716.2;refGene_gn=
RAB36;refGene_tx=XM_019814717.2;refGene_gn=RAB36;refGene_tx=XM_0198147
18.2
D3 19878483 19878548
refGene_gn=RAB36;refGene_tx=XM_019814713.2;refGene_gn=RAB36;refGene_tx
=XM_019814714.2;refGene_gn=RAB36;refGene_tx=XM_019814716.2;refGene_gn=
RAB36;refGene_tx=XM_019814717.2;refGene_gn=RAB36;refGene_tx=XM_0198147
18.2
D3 19880960 19881062
refGene_gn=RAB36;refGene_tx=XM_019814713.2;refGene_gn=RAB36;refGene_tx
=XM_019814714.2;refGene_gn=RAB36;refGene_tx=XM_019814715.2;refGene_gn=
RAB36;refGene_tx=XM_019814716.2;refGene_gn=RAB36;refGene_tx=XM_0198147
17.2;refGene_gn=RAB36;refGene_tx=XM_019814718.2
D3 19881491 19881557
refGene_gn=RAB36;refGene_tx=XM_019814713.2;refGene_gn=RAB36;refGene_tx
=XM_019814715.2;refGene_gn=RAB36;refGene_tx=XM_019814716.2;refGene_gn=
RAB36;refGene_tx=XM_019814717.2;refGene_gn=RAB36;refGene_tx=XM_0198147
18.2
D3 19883431 19883523
refGene_gn=RAB36;refGene_tx=XM_019814713.2;refGene_gn=RAB36;refGene_tx
=XM_019814714.2;refGene_gn=RAB36;refGene_tx=XM_019814715.2;refGene_gn=
RAB36;refGene_tx=XM_019814716.2;refGene_gn=RAB36;refGene_tx=XM_0198147
17.2;refGene_gn=RAB36;refGene_tx=XM_019814718.2
D3 19886548 19886617
refGene_gn=RAB36;refGene_tx=XM_019814713.2;refGene_gn=RAB36;refGene_tx
=XM_019814714.2;refGene_gn=RAB36;refGene_tx=XM_019814715.2;refGene_gn=
RAB36;refGene_tx=XM_019814716.2;refGene_gn=RAB36;refGene_tx=XM_0198147
17.2;refGene_gn=RAB36;refGene_tx=XM_019814718.2
D3 19894860 19895059
refGene_gn=RSPH14;refGene_tx=XM_003994830.4;refGene_gn=RSPH14;refGene_
tx=XM_023241549.1
D3 19896120 19896223
refGene_gn=RSPH14;refGene_tx=XM_003994830.4;refGene_gn=RSPH14;refGene_
tx=XM_023241549.1
D3 19899642 19899761
refGene_gn=RSPH14;refGene_tx=XM_003994830.4;refGene_gn=RSPH14;refGene_
tx=XM_023241549.1
D3 19909191 19909536 refGene_gn=GNAZ;refGene_tx=XM_023241548.1
D3 19936913 19937636 refGene_gn=GNAZ;refGene_tx=XM_023241548.1
D3 19967926 19968152
refGene_gn=RSPH14;refGene_tx=XM_003994830.4;refGene_gn=RSPH14;refGene_

tx=XM_023241549.1
D3 19970020 19970157
refGene_gn=RSPH14;refGene_tx=XM_003994830.4;refGene_gn=RSPH14;refGene_
tx=XM_023241549.1
D3 19972431 19972670
refGene_gn=RSPH14;refGene_tx=XM_003994830.4;refGene_gn=RSPH14;refGene_
tx=XM_023241549.1
D3 19977133 19977561
refGene_gn=LOC101091563;refGene_tx=XM_023241550.1
D3 19979871 19979971
refGene_gn=LOC101091563;refGene_tx=XM_023241550.1
D3 19982264 19982480
refGene_gn=LOC101091563;refGene_tx=XM_023241550.1
D3 19984877 19985046
refGene_gn=LOC101091563;refGene_tx=XM_023241550.1
D3 19995177 19995328
refGene_gn=LOC101091563;refGene_tx=XM_023241550.1
D3 19996799 19996907
refGene_gn=LOC101091563;refGene_tx=XM_023241550.1
D3 19999363 19999499
refGene_gn=LOC101091563;refGene_tx=XM_023241550.1
D3 20006207 20006440
refGene_gn=LOC101091563;refGene_tx=XM_023241550.1
D3 20011202 20011283
refGene_gn=LOC101091563;refGene_tx=XM_023241550.1
D3 20012545 20012651
refGene_gn=LOC101091563;refGene_tx=XM_023241550.1
D3 20014526 20014631
refGene_gn=LOC101091563;refGene_tx=XM_023241550.1
D3 20024203 20024263
refGene_gn=LOC101091563;refGene_tx=XM_023241550.1
D3 20025558 20025663
refGene_gn=LOC101091563;refGene_tx=XM_023241550.1
D3 20030345 20030417
refGene_gn=LOC101091563;refGene_tx=XM_023241550.1
D3 20034890 20035013
refGene_gn=LOC101091563;refGene_tx=XM_023241550.1
D3 20097014 20810341
refGene_gn=LOC101099449;refGene_gn=LOC105260055;refGene_gn=LOC10526028
9;refGene_gn=LOC109497182;refGene_tx=XM_023242171.1;refGene_gn=LOC1094
97182;refGene_tx=XM_023242172.1;refGene_gn=LOC109497182;refGene_tx=XM_
023242174.1;refGene_gn=LOC109497182;refGene_tx=XM_023242175.1;refGene_
gn=LOC109497182;refGene_tx=XM_023242176.1;refGene_gn=LOC109497182;refG
ene_tx=XM_023242177.1;refGene_gn=LOC109497182;refGene_tx=XM_023242178.
1;refGene_gn=LOC109497182;refGene_tx=XM_023242179.1;refGene_gn=LOC1094
97182;refGene_tx=XM_023242180.1;refGene_gn=LOC109497182;refGene_tx=XM_
023242181.1;refGene_gn=LOC109497182;refGene_tx=XM_023242182.1;refGene_
gn=LOC109497182;refGene_tx=XM_023242183.1;refGene_gn=LOC109497182;refG
ene_tx=XM_023242184.1;refGene_gn=LOC111557359;refGene_gn=LOC111557360;
refGene_gn=LOC111557369;refGene_tx=XM_023242185.1;refGene_gn=LOC111557

370;refGene_gn=LOC111557372;refGene_gn=LOC111557373;refGene_gn=PRAME;refGene_tx=XM_006938621.4;refGene_gn=PRAME;refGene_tx=XM_006938622.4;refGene_gn=PRAME;refGene_tx=XM_011287761.3;refGene_gn=PRAME;refGene_tx=XM_019814722.2;refGene_gn=PRAME;refGene_tx=XM_023242169.1;refGene_gn=ZNF280B;refGene_tx=XM_023242166.1;refGene_gn=ZNF280B;refGene_tx=XM_023242168.1
D3	20845634	20845941	refGene_gn=LOC111557369;refGene_tx=XM_023242185.1
D3	20846052	20846110	refGene_gn=LOC111557369;refGene_tx=XM_023242185.1
D3	20869882	20870212	refGene_gn=LOC109497182;refGene_tx=XM_023242171.1
D3	20870328	20870374	refGene_gn=LOC109497182;refGene_tx=XM_023242171.1
D3	20877952	20878566	refGene_gn=LOC105260295
D3	20881447	20884847	refGene_gn=LOC111557371
D3	20907171	20913326	refGene_gn=LOC109493262
D3	21010838	21010973	refGene_gn=LOC101100420;refGene_tx=XM_003994827.3
D3	21011760	21011832	refGene_gn=LOC101100420;refGene_tx=XM_003994827.3
D3	21013490	21013595	refGene_gn=LOC101100420;refGene_tx=XM_003994827.3
D3	21015590	21015650	refGene_gn=LOC101100420;refGene_tx=XM_003994827.3
D3	21017868	21017973	refGene_gn=LOC101100420;refGene_tx=XM_003994827.3
D3	21021931	21022037	refGene_gn=LOC101100420;refGene_tx=XM_003994827.3
D3	21030079	21030160	refGene_gn=LOC101100420;refGene_tx=XM_003994827.3
D3	21032601	21032822	refGene_gn=LOC101100420;refGene_tx=XM_003994827.3
D3	21035850	21035986	refGene_gn=LOC101100420;refGene_tx=XM_003994827.3
D3	21037012	21037120	refGene_gn=LOC101100420;refGene_tx=XM_003994827.3
D3	21039162	21039313	refGene_gn=LOC101100420;refGene_tx=XM_003994827.3
D3	21041860	21042029	refGene_gn=LOC101100420;refGene_tx=XM_003994827.3
D3	21043629	21043845	refGene_gn=LOC101100420;refGene_tx=XM_003994827.3
D3	21046244	21046347	refGene_gn=LOC101100420;refGene_tx=XM_003994827.3
D3	21049968	21050183	refGene_gn=LOC101100420;refGene_tx=XM_003994827.3
D3	21076620	21076838	refGene_gn=SLC5A1;refGene_tx=NM_001195405.1
D3	21090920	21091026	refGene_gn=SLC5A1;refGene_tx=NM_001195405.1
D3	21092714	21092930	refGene_gn=SLC5A1;refGene_tx=NM_001195405.1

D3 21099300 21099469 refGene_gn=SLC5A1;refGene_tx=NM_001195405.1
D3 21101710 21101861 refGene_gn=SLC5A1;refGene_tx=NM_001195405.1
D3 21106917 21107025 refGene_gn=SLC5A1;refGene_tx=NM_001195405.1
D3 21108415 21108551 refGene_gn=SLC5A1;refGene_tx=NM_001195405.1
D3 21108792 21109013 refGene_gn=SLC5A1;refGene_tx=NM_001195405.1
D3 21110894 21110975 refGene_gn=SLC5A1;refGene_tx=NM_001195405.1
D3 21114595 21114701 refGene_gn=SLC5A1;refGene_tx=NM_001195405.1
D3 21124275 21124380 refGene_gn=SLC5A1;refGene_tx=NM_001195405.1
D3 21124999 21125059 refGene_gn=SLC5A1;refGene_tx=NM_001195405.1
D3 21126053 21126158 refGene_gn=SLC5A1;refGene_tx=NM_001195405.1
D3 21144511 21144583 refGene_gn=SLC5A1;refGene_tx=NM_001195405.1
D3 21151772 21151904 refGene_gn=SLC5A1;refGene_tx=NM_001195405.1
D3 21235880 21236534 refGene_gn=YWHAH;refGene_tx=XM_023241551.1
D3 21246244 21246331 refGene_gn=YWHAH;refGene_tx=XM_023241551.1
D3 21277430 21277723
refGene_gn=DEPDC5;refGene_tx=XM_019814733.2;refGene_gn=DEPDC5;refGene_tx=XM_019814734.2;refGene_gn=DEPDC5;refGene_tx=XM_019814735.2;refGene_gn=DEPDC5;refGene_tx=XM_019814736.2;refGene_gn=DEPDC5;refGene_tx=XM_019814737.2;refGene_gn=DEPDC5;refGene_tx=XM_019814738.2;refGene_gn=DEPDC5;refGene_tx=XM_019814740.2;refGene_gn=DEPDC5;refGene_tx=XM_019814741.2;refGene_gn=DEPDC5;refGene_tx=XM_023241552.1
D3 21277884 21277967
refGene_gn=DEPDC5;refGene_tx=XM_019814733.2;refGene_gn=DEPDC5;refGene_tx=XM_019814734.2;refGene_gn=DEPDC5;refGene_tx=XM_019814735.2;refGene_gn=DEPDC5;refGene_tx=XM_019814736.2;refGene_gn=DEPDC5;refGene_tx=XM_019814737.2;refGene_gn=DEPDC5;refGene_tx=XM_019814738.2;refGene_gn=DEPDC5;refGene_tx=XM_019814740.2;refGene_gn=DEPDC5;refGene_tx=XM_019814741.2;refGene_gn=DEPDC5;refGene_tx=XM_023241552.1
D3 21280887 21280948
refGene_gn=DEPDC5;refGene_tx=XM_019814733.2;refGene_gn=DEPDC5;refGene_tx=XM_019814734.2;refGene_gn=DEPDC5;refGene_tx=XM_019814735.2;refGene_gn=DEPDC5;refGene_tx=XM_019814736.2;refGene_gn=DEPDC5;refGene_tx=XM_019814737.2;refGene_gn=DEPDC5;refGene_tx=XM_019814738.2;refGene_gn=DEPDC5;refGene_tx=XM_019814740.2;refGene_gn=DEPDC5;refGene_tx=XM_019814741.2;refGene_gn=DEPDC5;refGene_tx=XM_023241552.1
D3 21283458 21283630
refGene_gn=DEPDC5;refGene_tx=XM_019814733.2;refGene_gn=DEPDC5;refGene_tx=XM_019814734.2;refGene_gn=DEPDC5;refGene_tx=XM_019814735.2;refGene_gn=DEPDC5;refGene_tx=XM_019814736.2;refGene_gn=DEPDC5;refGene_tx=XM_019814737.2;refGene_gn=DEPDC5;refGene_tx=XM_019814738.2;refGene_gn=DEPDC5;refGene_tx=XM_019814740.2;refGene_gn=DEPDC5;refGene_tx=XM_019814741.2;refGene_gn=DEPDC5;refGene_tx=XM_023241552.1
D3 21287977 21288147
refGene_gn=DEPDC5;refGene_tx=XM_019814733.2;refGene_gn=DEPDC5;refGene_tx=XM_019814734.2;refGene_gn=DEPDC5;refGene_tx=XM_019814735.2;refGene_gn=DEPDC5;refGene_tx=XM_019814736.2;refGene_gn=DEPDC5;refGene_tx=XM_019814737.2;refGene_gn=DEPDC5;refGene_tx=XM_019814738.2;refGene_gn=DEPDC5;refGene_tx=XM_019814740.2;refGene_gn=DEPDC5;refGene_tx=XM_019814741.2;refGene_gn=DEPDC5;refGene_tx=XM_023241552.1
D3 21291936 21292067 refGene_gn=DEPDC5;refGene_tx=XM_019814742.2

D3	21292880 21293105
refGene_gn=DEPDC5;refGene_tx=XM_019814733.2;refGene_gn=DEPDC5;refGene_tx=XM_019814734.2;refGene_gn=DEPDC5;refGene_tx=XM_019814735.2;refGene_gn=DEPDC5;refGene_tx=XM_019814736.2;refGene_gn=DEPDC5;refGene_tx=XM_019814737.2;refGene_gn=DEPDC5;refGene_tx=XM_019814738.2;refGene_gn=DEPDC5;refGene_tx=XM_019814740.2;refGene_gn=DEPDC5;refGene_tx=XM_019814741.2;refGene_gn=DEPDC5;refGene_tx=XM_019814742.2;refGene_gn=DEPDC5;refGene_tx=XM_023241552.1
D3	21294576 21294685
refGene_gn=DEPDC5;refGene_tx=XM_019814733.2;refGene_gn=DEPDC5;refGene_tx=XM_019814734.2;refGene_gn=DEPDC5;refGene_tx=XM_019814735.2;refGene_gn=DEPDC5;refGene_tx=XM_019814736.2;refGene_gn=DEPDC5;refGene_tx=XM_019814737.2;refGene_gn=DEPDC5;refGene_tx=XM_019814738.2;refGene_gn=DEPDC5;refGene_tx=XM_019814740.2;refGene_gn=DEPDC5;refGene_tx=XM_019814741.2;refGene_gn=DEPDC5;refGene_tx=XM_019814742.2;refGene_gn=DEPDC5;refGene_tx=XM_023241552.1
D3	21296784 21296917
refGene_gn=DEPDC5;refGene_tx=XM_019814733.2;refGene_gn=DEPDC5;refGene_tx=XM_019814734.2;refGene_gn=DEPDC5;refGene_tx=XM_019814735.2;refGene_gn=DEPDC5;refGene_tx=XM_019814736.2;refGene_gn=DEPDC5;refGene_tx=XM_019814737.2;refGene_gn=DEPDC5;refGene_tx=XM_019814738.2;refGene_gn=DEPDC5;refGene_tx=XM_019814740.2;refGene_gn=DEPDC5;refGene_tx=XM_019814741.2;refGene_gn=DEPDC5;refGene_tx=XM_019814742.2;refGene_gn=DEPDC5;refGene_tx=XM_023241552.1
D3	21297424 21297502
refGene_gn=DEPDC5;refGene_tx=XM_019814733.2;refGene_gn=DEPDC5;refGene_tx=XM_019814734.2;refGene_gn=DEPDC5;refGene_tx=XM_019814735.2;refGene_gn=DEPDC5;refGene_tx=XM_019814736.2;refGene_gn=DEPDC5;refGene_tx=XM_019814737.2;refGene_gn=DEPDC5;refGene_tx=XM_019814738.2;refGene_gn=DEPDC5;refGene_tx=XM_019814740.2;refGene_gn=DEPDC5;refGene_tx=XM_019814741.2;refGene_gn=DEPDC5;refGene_tx=XM_019814742.2;refGene_gn=DEPDC5;refGene_tx=XM_023241552.1
D3	21301415 21301570
refGene_gn=DEPDC5;refGene_tx=XM_019814733.2;refGene_gn=DEPDC5;refGene_tx=XM_019814734.2;refGene_gn=DEPDC5;refGene_tx=XM_019814735.2;refGene_gn=DEPDC5;refGene_tx=XM_019814736.2;refGene_gn=DEPDC5;refGene_tx=XM_019814737.2;refGene_gn=DEPDC5;refGene_tx=XM_019814738.2;refGene_gn=DEPDC5;refGene_tx=XM_019814740.2;refGene_gn=DEPDC5;refGene_tx=XM_019814741.2;refGene_gn=DEPDC5;refGene_tx=XM_019814742.2;refGene_gn=DEPDC5;refGene_tx=XM_023241552.1
D3	21311933 21311999
refGene_gn=DEPDC5;refGene_tx=XM_019814733.2;refGene_gn=DEPDC5;refGene_tx=XM_019814734.2;refGene_gn=DEPDC5;refGene_tx=XM_019814735.2;refGene_gn=DEPDC5;refGene_tx=XM_019814738.2;refGene_gn=DEPDC5;refGene_tx=XM_019814740.2;refGene_gn=DEPDC5;refGene_tx=XM_019814742.2;refGene_gn=DEPDC5;refGene_tx=XM_023241552.1
D3	21314899 21315008
refGene_gn=DEPDC5;refGene_tx=XM_019814733.2;refGene_gn=DEPDC5;refGene_tx=XM_019814734.2;refGene_gn=DEPDC5;refGene_tx=XM_019814735.2;refGene_gn=DEPDC5;refGene_tx=XM_019814736.2;refGene_gn=DEPDC5;refGene_tx=XM_01

9814737.2;refGene_gn=DEPDC5;refGene_tx=XM_019814738.2;refGene_gn=DEPDC5;refGene_tx=XM_019814740.2;refGene_gn=DEPDC5;refGene_tx=XM_019814741.2;refGene_gn=DEPDC5;refGene_tx=XM_019814742.2;refGene_gn=DEPDC5;refGene_tx=XM_023241552.1
D3 21321608 21321742
refGene_gn=DEPDC5;refGene_tx=XM_019814733.2;refGene_gn=DEPDC5;refGene_tx=XM_019814734.2;refGene_gn=DEPDC5;refGene_tx=XM_019814735.2;refGene_gn=DEPDC5;refGene_tx=XM_019814736.2;refGene_gn=DEPDC5;refGene_tx=XM_019814737.2;refGene_gn=DEPDC5;refGene_tx=XM_019814738.2;refGene_gn=DEPDC5;refGene_tx=XM_019814740.2;refGene_gn=DEPDC5;refGene_tx=XM_019814741.2;refGene_gn=DEPDC5;refGene_tx=XM_019814742.2;refGene_gn=DEPDC5;refGene_tx=XM_023241552.1
D3 21323077 21323297
refGene_gn=DEPDC5;refGene_tx=XM_019814733.2;refGene_gn=DEPDC5;refGene_tx=XM_019814734.2;refGene_gn=DEPDC5;refGene_tx=XM_019814735.2;refGene_gn=DEPDC5;refGene_tx=XM_019814736.2;refGene_gn=DEPDC5;refGene_tx=XM_019814737.2;refGene_gn=DEPDC5;refGene_tx=XM_019814738.2;refGene_gn=DEPDC5;refGene_tx=XM_019814740.2;refGene_gn=DEPDC5;refGene_tx=XM_019814741.2;refGene_gn=DEPDC5;refGene_tx=XM_019814742.2;refGene_gn=DEPDC5;refGene_tx=XM_023241552.1
D3 21323968 21324136
refGene_gn=DEPDC5;refGene_tx=XM_019814733.2;refGene_gn=DEPDC5;refGene_tx=XM_019814734.2;refGene_gn=DEPDC5;refGene_tx=XM_019814735.2;refGene_gn=DEPDC5;refGene_tx=XM_019814736.2;refGene_gn=DEPDC5;refGene_tx=XM_019814737.2;refGene_gn=DEPDC5;refGene_tx=XM_019814738.2;refGene_gn=DEPDC5;refGene_tx=XM_019814740.2;refGene_gn=DEPDC5;refGene_tx=XM_019814741.2;refGene_gn=DEPDC5;refGene_tx=XM_019814742.2;refGene_gn=DEPDC5;refGene_tx=XM_023241552.1
D3 21324600 21324718
refGene_gn=DEPDC5;refGene_tx=XM_019814733.2;refGene_gn=DEPDC5;refGene_tx=XM_019814734.2;refGene_gn=DEPDC5;refGene_tx=XM_019814735.2;refGene_gn=DEPDC5;refGene_tx=XM_019814736.2;refGene_gn=DEPDC5;refGene_tx=XM_019814737.2;refGene_gn=DEPDC5;refGene_tx=XM_019814738.2;refGene_gn=DEPDC5;refGene_tx=XM_019814740.2;refGene_gn=DEPDC5;refGene_tx=XM_019814741.2;refGene_gn=DEPDC5;refGene_tx=XM_019814742.2;refGene_gn=DEPDC5;refGene_tx=XM_023241552.1
D3 21328680 21328841
refGene_gn=DEPDC5;refGene_tx=XM_019814733.2;refGene_gn=DEPDC5;refGene_tx=XM_019814734.2;refGene_gn=DEPDC5;refGene_tx=XM_019814735.2;refGene_gn=DEPDC5;refGene_tx=XM_019814736.2;refGene_gn=DEPDC5;refGene_tx=XM_019814737.2;refGene_gn=DEPDC5;refGene_tx=XM_019814738.2;refGene_gn=DEPDC5;refGene_tx=XM_019814740.2;refGene_gn=DEPDC5;refGene_tx=XM_019814741.2;refGene_gn=DEPDC5;refGene_tx=XM_019814742.2;refGene_gn=DEPDC5;refGene_tx=XM_023241552.1
D3 21329680 21329864
refGene_gn=DEPDC5;refGene_tx=XM_019814733.2;refGene_gn=DEPDC5;refGene_tx=XM_019814734.2;refGene_gn=DEPDC5;refGene_tx=XM_019814735.2;refGene_gn=DEPDC5;refGene_tx=XM_019814736.2;refGene_gn=DEPDC5;refGene_tx=XM_019814737.2;refGene_gn=DEPDC5;refGene_tx=XM_019814738.2;refGene_gn=DEPDC5;refGene_tx=XM_019814740.2;refGene_gn=DEPDC5;refGene_tx=XM_019814741.

2;refGene_gn=DEPDC5;refGene_tx=XM_019814742.2;refGene_gn=DEPDC5;refGene_tx=XM_023241552.1
D3 21333750 21333816
refGene_gn=DEPDC5;refGene_tx=XM_019814733.2;refGene_gn=DEPDC5;refGene_tx=XM_019814734.2;refGene_gn=DEPDC5;refGene_tx=XM_019814735.2;refGene_gn=DEPDC5;refGene_tx=XM_019814736.2;refGene_gn=DEPDC5;refGene_tx=XM_019814737.2;refGene_gn=DEPDC5;refGene_tx=XM_019814738.2;refGene_gn=DEPDC5;refGene_tx=XM_019814740.2;refGene_gn=DEPDC5;refGene_tx=XM_019814741.2;refGene_gn=DEPDC5;refGene_tx=XM_019814742.2;refGene_gn=DEPDC5;refGene_tx=XM_023241552.1
D3 21343781 21343879
refGene_gn=DEPDC5;refGene_tx=XM_019814733.2;refGene_gn=DEPDC5;refGene_tx=XM_019814734.2;refGene_gn=DEPDC5;refGene_tx=XM_019814735.2;refGene_gn=DEPDC5;refGene_tx=XM_019814736.2;refGene_gn=DEPDC5;refGene_tx=XM_019814737.2;refGene_gn=DEPDC5;refGene_tx=XM_019814742.2;refGene_gn=DEPDC5;refGene_tx=XM_023241552.1
D3 21344875 21345011
refGene_gn=DEPDC5;refGene_tx=XM_019814733.2;refGene_gn=DEPDC5;refGene_tx=XM_019814734.2;refGene_gn=DEPDC5;refGene_tx=XM_019814735.2;refGene_gn=DEPDC5;refGene_tx=XM_019814736.2;refGene_gn=DEPDC5;refGene_tx=XM_019814737.2;refGene_gn=DEPDC5;refGene_tx=XM_019814742.2;refGene_gn=DEPDC5;refGene_tx=XM_023241552.1
D3 21346919 21347123
refGene_gn=DEPDC5;refGene_tx=XM_019814733.2;refGene_gn=DEPDC5;refGene_tx=XM_019814734.2;refGene_gn=DEPDC5;refGene_tx=XM_019814735.2;refGene_gn=DEPDC5;refGene_tx=XM_019814736.2;refGene_gn=DEPDC5;refGene_tx=XM_019814737.2;refGene_gn=DEPDC5;refGene_tx=XM_019814738.2;refGene_gn=DEPDC5;refGene_tx=XM_019814740.2;refGene_gn=DEPDC5;refGene_tx=XM_019814741.2;refGene_gn=DEPDC5;refGene_tx=XM_019814742.2;refGene_gn=DEPDC5;refGene_tx=XM_023241552.1
D3 21352139 21352360
refGene_gn=DEPDC5;refGene_tx=XM_019814733.2;refGene_gn=DEPDC5;refGene_tx=XM_019814734.2;refGene_gn=DEPDC5;refGene_tx=XM_019814735.2;refGene_gn=DEPDC5;refGene_tx=XM_019814736.2;refGene_gn=DEPDC5;refGene_tx=XM_019814737.2;refGene_gn=DEPDC5;refGene_tx=XM_019814738.2;refGene_gn=DEPDC5;refGene_tx=XM_019814740.2;refGene_gn=DEPDC5;refGene_tx=XM_019814741.2;refGene_gn=DEPDC5;refGene_tx=XM_019814742.2;refGene_gn=DEPDC5;refGene_tx=XM_023241552.1
D3 21354719 21354840
refGene_gn=DEPDC5;refGene_tx=XM_019814733.2;refGene_gn=DEPDC5;refGene_tx=XM_019814734.2;refGene_gn=DEPDC5;refGene_tx=XM_019814735.2;refGene_gn=DEPDC5;refGene_tx=XM_019814736.2;refGene_gn=DEPDC5;refGene_tx=XM_019814737.2;refGene_gn=DEPDC5;refGene_tx=XM_019814738.2;refGene_gn=DEPDC5;refGene_tx=XM_019814740.2;refGene_gn=DEPDC5;refGene_tx=XM_019814741.2;refGene_gn=DEPDC5;refGene_tx=XM_019814742.2;refGene_gn=DEPDC5;refGene_tx=XM_023241552.1
D3 21355363 21355400
refGene_gn=DEPDC5;refGene_tx=XM_019814733.2;refGene_gn=DEPDC5;refGene_tx=XM_019814734.2;refGene_gn=DEPDC5;refGene_tx=XM_019814735.2;refGene_gn=DEPDC5;refGene_tx=XM_019814736.2;refGene_gn=DEPDC5;refGene_tx=XM_01

9814737.2;refGene_gn=DEPDC5;refGene_tx=XM_019814738.2;refGene_gn=DEPDC5;refGene_tx=XM_019814740.2;refGene_gn=DEPDC5;refGene_tx=XM_019814741.2;refGene_gn=DEPDC5;refGene_tx=XM_019814742.2;refGene_gn=DEPDC5;refGene_tx=XM_023241552.1
D3 21357400 21357470
refGene_gn=DEPDC5;refGene_tx=XM_019814733.2;refGene_gn=DEPDC5;refGene_tx=XM_019814734.2;refGene_gn=DEPDC5;refGene_tx=XM_019814735.2;refGene_gn=DEPDC5;refGene_tx=XM_019814736.2;refGene_gn=DEPDC5;refGene_tx=XM_019814737.2;refGene_gn=DEPDC5;refGene_tx=XM_019814738.2;refGene_gn=DEPDC5;refGene_tx=XM_019814740.2;refGene_gn=DEPDC5;refGene_tx=XM_019814741.2;refGene_gn=DEPDC5;refGene_tx=XM_019814742.2;refGene_gn=DEPDC5;refGene_tx=XM_023241552.1
D3 21363326 21363400
refGene_gn=DEPDC5;refGene_tx=XM_019814733.2;refGene_gn=DEPDC5;refGene_tx=XM_019814734.2;refGene_gn=DEPDC5;refGene_tx=XM_019814735.2;refGene_gn=DEPDC5;refGene_tx=XM_019814736.2;refGene_gn=DEPDC5;refGene_tx=XM_019814737.2;refGene_gn=DEPDC5;refGene_tx=XM_019814738.2;refGene_gn=DEPDC5;refGene_tx=XM_019814740.2;refGene_gn=DEPDC5;refGene_tx=XM_019814741.2;refGene_gn=DEPDC5;refGene_tx=XM_019814742.2;refGene_gn=DEPDC5;refGene_tx=XM_023241552.1
D3 21363543 21363605
refGene_gn=DEPDC5;refGene_tx=XM_019814733.2;refGene_gn=DEPDC5;refGene_tx=XM_019814734.2;refGene_gn=DEPDC5;refGene_tx=XM_019814735.2;refGene_gn=DEPDC5;refGene_tx=XM_019814736.2;refGene_gn=DEPDC5;refGene_tx=XM_019814737.2;refGene_gn=DEPDC5;refGene_tx=XM_019814738.2;refGene_gn=DEPDC5;refGene_tx=XM_019814740.2;refGene_gn=DEPDC5;refGene_tx=XM_019814741.2;refGene_gn=DEPDC5;refGene_tx=XM_019814742.2;refGene_gn=DEPDC5;refGene_tx=XM_023241552.1
D3 21364433 21364568
refGene_gn=DEPDC5;refGene_tx=XM_019814733.2;refGene_gn=DEPDC5;refGene_tx=XM_019814734.2;refGene_gn=DEPDC5;refGene_tx=XM_019814735.2;refGene_gn=DEPDC5;refGene_tx=XM_019814736.2;refGene_gn=DEPDC5;refGene_tx=XM_019814737.2;refGene_gn=DEPDC5;refGene_tx=XM_019814738.2;refGene_gn=DEPDC5;refGene_tx=XM_019814740.2;refGene_gn=DEPDC5;refGene_tx=XM_019814741.2;refGene_gn=DEPDC5;refGene_tx=XM_019814742.2;refGene_gn=DEPDC5;refGene_tx=XM_023241552.1
D3 21367247 21367322
refGene_gn=DEPDC5;refGene_tx=XM_019814733.2;refGene_gn=DEPDC5;refGene_tx=XM_019814734.2;refGene_gn=DEPDC5;refGene_tx=XM_019814735.2;refGene_gn=DEPDC5;refGene_tx=XM_019814736.2;refGene_gn=DEPDC5;refGene_tx=XM_019814737.2;refGene_gn=DEPDC5;refGene_tx=XM_019814738.2;refGene_gn=DEPDC5;refGene_tx=XM_019814740.2;refGene_gn=DEPDC5;refGene_tx=XM_019814741.2;refGene_gn=DEPDC5;refGene_tx=XM_019814742.2;refGene_gn=DEPDC5;refGene_tx=XM_023241552.1
D3 21367609 21367713
refGene_gn=DEPDC5;refGene_tx=XM_019814733.2;refGene_gn=DEPDC5;refGene_tx=XM_019814734.2;refGene_gn=DEPDC5;refGene_tx=XM_019814735.2;refGene_gn=DEPDC5;refGene_tx=XM_019814736.2;refGene_gn=DEPDC5;refGene_tx=XM_019814737.2;refGene_gn=DEPDC5;refGene_tx=XM_019814738.2;refGene_gn=DEPDC5;refGene_tx=XM_019814740.2;refGene_gn=DEPDC5;refGene_tx=XM_019814741.

2;refGene_gn=DEPDC5;refGene_tx=XM_019814742.2;refGene_gn=DEPDC5;refGene_tx=XM_023241552.1
D3 21377086 21377159
refGene_gn=DEPDC5;refGene_tx=XM_019814733.2;refGene_gn=DEPDC5;refGene_tx=XM_019814734.2;refGene_gn=DEPDC5;refGene_tx=XM_019814735.2;refGene_gn=DEPDC5;refGene_tx=XM_019814736.2;refGene_gn=DEPDC5;refGene_tx=XM_019814737.2;refGene_gn=DEPDC5;refGene_tx=XM_019814738.2;refGene_gn=DEPDC5;refGene_tx=XM_019814740.2;refGene_gn=DEPDC5;refGene_tx=XM_019814741.2;refGene_gn=DEPDC5;refGene_tx=XM_019814742.2;refGene_gn=DEPDC5;refGene_tx=XM_023241552.1
D3 21377970 21378040
refGene_gn=DEPDC5;refGene_tx=XM_019814733.2;refGene_gn=DEPDC5;refGene_tx=XM_019814734.2;refGene_gn=DEPDC5;refGene_tx=XM_019814735.2;refGene_gn=DEPDC5;refGene_tx=XM_019814736.2;refGene_gn=DEPDC5;refGene_tx=XM_019814737.2;refGene_gn=DEPDC5;refGene_tx=XM_019814738.2;refGene_gn=DEPDC5;refGene_tx=XM_019814740.2;refGene_gn=DEPDC5;refGene_tx=XM_019814741.2;refGene_gn=DEPDC5;refGene_tx=XM_019814742.2;refGene_gn=DEPDC5;refGene_tx=XM_023241552.1
D3 21382937 21382999
refGene_gn=DEPDC5;refGene_tx=XM_019814733.2;refGene_gn=DEPDC5;refGene_tx=XM_019814734.2;refGene_gn=DEPDC5;refGene_tx=XM_019814735.2;refGene_gn=DEPDC5;refGene_tx=XM_019814736.2;refGene_gn=DEPDC5;refGene_tx=XM_019814737.2;refGene_gn=DEPDC5;refGene_tx=XM_019814738.2;refGene_gn=DEPDC5;refGene_tx=XM_019814740.2;refGene_gn=DEPDC5;refGene_tx=XM_019814741.2;refGene_gn=DEPDC5;refGene_tx=XM_019814742.2;refGene_gn=DEPDC5;refGene_tx=XM_023241552.1
D3 21384381 21384460
refGene_gn=DEPDC5;refGene_tx=XM_019814733.2;refGene_gn=DEPDC5;refGene_tx=XM_019814734.2;refGene_gn=DEPDC5;refGene_tx=XM_019814735.2;refGene_gn=DEPDC5;refGene_tx=XM_019814736.2;refGene_gn=DEPDC5;refGene_tx=XM_019814737.2;refGene_gn=DEPDC5;refGene_tx=XM_019814738.2;refGene_gn=DEPDC5;refGene_tx=XM_019814740.2;refGene_gn=DEPDC5;refGene_tx=XM_019814741.2;refGene_gn=DEPDC5;refGene_tx=XM_019814742.2;refGene_gn=DEPDC5;refGene_tx=XM_023241552.1
D3 21389360 21389430
refGene_gn=DEPDC5;refGene_tx=XM_019814733.2;refGene_gn=DEPDC5;refGene_tx=XM_019814734.2;refGene_gn=DEPDC5;refGene_tx=XM_019814735.2;refGene_gn=DEPDC5;refGene_tx=XM_019814736.2;refGene_gn=DEPDC5;refGene_tx=XM_019814737.2;refGene_gn=DEPDC5;refGene_tx=XM_019814738.2;refGene_gn=DEPDC5;refGene_tx=XM_019814740.2;refGene_gn=DEPDC5;refGene_tx=XM_019814741.2;refGene_gn=DEPDC5;refGene_tx=XM_019814742.2;refGene_gn=DEPDC5;refGene_tx=XM_023241552.1
D3 21395816 21395866
refGene_gn=DEPDC5;refGene_tx=XM_019814733.2;refGene_gn=DEPDC5;refGene_tx=XM_019814734.2;refGene_gn=DEPDC5;refGene_tx=XM_019814735.2;refGene_gn=DEPDC5;refGene_tx=XM_019814736.2;refGene_gn=DEPDC5;refGene_tx=XM_019814737.2;refGene_gn=DEPDC5;refGene_tx=XM_019814738.2;refGene_gn=DEPDC5;refGene_tx=XM_019814740.2;refGene_gn=DEPDC5;refGene_tx=XM_019814741.2;refGene_gn=DEPDC5;refGene_tx=XM_019814742.2;refGene_gn=DEPDC5;refGene_tx=XM_023241552.1

D3 21396870 21396954
refGene_gn=DEPDC5;refGene_tx=XM_019814733.2;refGene_gn=DEPDC5;refGene_tx=XM_019814734.2;refGene_gn=DEPDC5;refGene_tx=XM_019814735.2;refGene_gn=DEPDC5;refGene_tx=XM_019814736.2;refGene_gn=DEPDC5;refGene_tx=XM_019814737.2;refGene_gn=DEPDC5;refGene_tx=XM_019814738.2;refGene_gn=DEPDC5;refGene_tx=XM_019814740.2;refGene_gn=DEPDC5;refGene_tx=XM_019814741.2;refGene_gn=DEPDC5;refGene_tx=XM_019814742.2;refGene_gn=DEPDC5;refGene_tx=XM_023241552.1
D3 21397851 21397937
refGene_gn=DEPDC5;refGene_tx=XM_019814733.2;refGene_gn=DEPDC5;refGene_tx=XM_019814734.2;refGene_gn=DEPDC5;refGene_tx=XM_019814735.2;refGene_gn=DEPDC5;refGene_tx=XM_019814736.2;refGene_gn=DEPDC5;refGene_tx=XM_019814737.2;refGene_gn=DEPDC5;refGene_tx=XM_019814738.2;refGene_gn=DEPDC5;refGene_tx=XM_019814740.2;refGene_gn=DEPDC5;refGene_tx=XM_019814741.2;refGene_gn=DEPDC5;refGene_tx=XM_019814742.2
D3 21399638 21399685
refGene_gn=DEPDC5;refGene_tx=XM_019814733.2;refGene_gn=DEPDC5;refGene_tx=XM_019814734.2;refGene_gn=DEPDC5;refGene_tx=XM_019814735.2;refGene_gn=DEPDC5;refGene_tx=XM_019814736.2;refGene_gn=DEPDC5;refGene_tx=XM_019814737.2;refGene_gn=DEPDC5;refGene_tx=XM_019814738.2;refGene_gn=DEPDC5;refGene_tx=XM_019814740.2;refGene_gn=DEPDC5;refGene_tx=XM_019814741.2;refGene_gn=DEPDC5;refGene_tx=XM_019814742.2
D3 21400074 21400162
refGene_gn=DEPDC5;refGene_tx=XM_019814733.2;refGene_gn=DEPDC5;refGene_tx=XM_019814734.2;refGene_gn=DEPDC5;refGene_tx=XM_019814735.2;refGene_gn=DEPDC5;refGene_tx=XM_019814736.2;refGene_gn=DEPDC5;refGene_tx=XM_019814737.2;refGene_gn=DEPDC5;refGene_tx=XM_019814738.2;refGene_gn=DEPDC5;refGene_tx=XM_019814740.2;refGene_gn=DEPDC5;refGene_tx=XM_019814741.2;refGene_gn=DEPDC5;refGene_tx=XM_019814742.2
D3 21402952 21403010
refGene_gn=DEPDC5;refGene_tx=XM_019814733.2;refGene_gn=DEPDC5;refGene_tx=XM_019814734.2;refGene_gn=DEPDC5;refGene_tx=XM_019814735.2;refGene_gn=DEPDC5;refGene_tx=XM_019814736.2;refGene_gn=DEPDC5;refGene_tx=XM_019814737.2;refGene_gn=DEPDC5;refGene_tx=XM_019814738.2;refGene_gn=DEPDC5;refGene_tx=XM_019814740.2;refGene_gn=DEPDC5;refGene_tx=XM_019814741.2;refGene_gn=DEPDC5;refGene_tx=XM_019814742.2
D3 21417432 21417906
refGene_gn=PRR14L;refGene_tx=XM_023241553.1;refGene_gn=PRR14L;refGene_tx=XM_023241554.1;refGene_gn=PRR14L;refGene_tx=XM_023241555.1
D3 21430609 21430682
refGene_gn=PRR14L;refGene_tx=XM_023241553.1;refGene_gn=PRR14L;refGene_tx=XM_023241554.1;refGene_gn=PRR14L;refGene_tx=XM_023241555.1;refGene_gn=PRR14L;refGene_tx=XM_023241556.1;refGene_gn=PRR14L;refGene_tx=XM_023241557.1;refGene_gn=PRR14L;refGene_tx=XM_023241558.1
D3 21432550 21432560 refGene_gn=PRR14L;refGene_tx=XM_023241559.1
D3 21441648 21446839
refGene_gn=PRR14L;refGene_tx=XM_023241553.1;refGene_gn=PRR14L;refGene_tx=XM_023241554.1;refGene_gn=PRR14L;refGene_tx=XM_023241555.1;refGene_gn=PRR14L;refGene_tx=XM_023241556.1;refGene_gn=PRR14L;refGene_tx=XM_023241557.1;refGene_gn=PRR14L;refGene_tx=XM_023241558.1;refGene_gn=PRR14

L;refGene_tx=XM_023241559.1
D3 21450936 21451008
refGene_gn=PRR14L;refGene_tx=XM_023241553.1;refGene_gn=PRR14L;refGene_tx=XM_023241554.1;refGene_gn=PRR14L;refGene_tx=XM_023241555.1;refGene_gn=PRR14L;refGene_tx=XM_023241556.1;refGene_gn=PRR14L;refGene_tx=XM_023241557.1;refGene_gn=PRR14L;refGene_tx=XM_023241558.1;refGene_gn=PRR14L;refGene_tx=XM_023241559.1
D3 21451717 21451889
refGene_gn=PRR14L;refGene_tx=XM_023241553.1;refGene_gn=PRR14L;refGene_tx=XM_023241554.1;refGene_gn=PRR14L;refGene_tx=XM_023241555.1;refGene_gn=PRR14L;refGene_tx=XM_023241556.1;refGene_gn=PRR14L;refGene_tx=XM_023241557.1;refGene_gn=PRR14L;refGene_tx=XM_023241558.1;refGene_gn=PRR14L;refGene_tx=XM_023241559.1
D3 21452464 21452571
refGene_gn=PRR14L;refGene_tx=XM_023241553.1;refGene_gn=PRR14L;refGene_tx=XM_023241554.1;refGene_gn=PRR14L;refGene_tx=XM_023241555.1;refGene_gn=PRR14L;refGene_tx=XM_023241556.1;refGene_gn=PRR14L;refGene_tx=XM_023241557.1;refGene_gn=PRR14L;refGene_tx=XM_023241558.1;refGene_gn=PRR14L;refGene_tx=XM_023241559.1
D3 21454602 21454671
refGene_gn=PRR14L;refGene_tx=XM_023241553.1;refGene_gn=PRR14L;refGene_tx=XM_023241554.1;refGene_gn=PRR14L;refGene_tx=XM_023241556.1;refGene_gn=PRR14L;refGene_tx=XM_023241557.1;refGene_gn=PRR14L;refGene_tx=XM_023241558.1;refGene_gn=PRR14L;refGene_tx=XM_023241559.1
D3 21463376 21463448
refGene_gn=PRR14L;refGene_tx=XM_023241553.1;refGene_gn=PRR14L;refGene_tx=XM_023241554.1;refGene_gn=PRR14L;refGene_tx=XM_023241555.1;refGene_gn=PRR14L;refGene_tx=XM_023241556.1;refGene_gn=PRR14L;refGene_tx=XM_023241557.1;refGene_gn=PRR14L;refGene_tx=XM_023241558.1;refGene_gn=PRR14L;refGene_tx=XM_023241559.1
D3 21464659 21464939
refGene_gn=PRR14L;refGene_tx=XM_023241553.1;refGene_gn=PRR14L;refGene_tx=XM_023241554.1;refGene_gn=PRR14L;refGene_tx=XM_023241555.1;refGene_gn=PRR14L;refGene_tx=XM_023241556.1;refGene_gn=PRR14L;refGene_tx=XM_023241557.1;refGene_gn=PRR14L;refGene_tx=XM_023241558.1;refGene_gn=PRR14L;refGene_tx=XM_023241559.1
D3 21499710 21499775
refGene_gn=PISD;refGene_tx=XM_011287756.2;refGene_gn=PISD;refGene_tx=XM_019814756.1
D3 21508477 21508557
refGene_gn=PISD;refGene_tx=XM_011287756.2;refGene_gn=PISD;refGene_tx=XM_019814756.1
D3 21512154 21512341
refGene_gn=PISD;refGene_tx=XM_011287756.2;refGene_gn=PISD;refGene_tx=XM_011287757.2;refGene_gn=PISD;refGene_tx=XM_019814756.1
D3 21529144 21529194 refGene_gn=PISD;refGene_tx=XM_006938605.4
D3 21530738 21530904
refGene_gn=PISD;refGene_tx=XM_006938605.4;refGene_gn=PISD;refGene_tx=XM_011287757.2;refGene_gn=PISD;refGene_tx=XM_019814756.1
D3 21532297 21532534

refGene_gn=PISD;refGene_tx=XM_006938605.4;refGene_gn=PISD;refGene_tx=XM_011287756.2;refGene_gn=PISD;refGene_tx=XM_011287757.2;refGene_gn=PISD;refGene_tx=XM_019814756.1
D3 21532700 21532839
refGene_gn=PISD;refGene_tx=XM_006938605.4;refGene_gn=PISD;refGene_tx=XM_011287756.2;refGene_gn=PISD;refGene_tx=XM_011287757.2;refGene_gn=PISD;refGene_tx=XM_019814756.1
D3 21533040 21533187
refGene_gn=PISD;refGene_tx=XM_006938605.4;refGene_gn=PISD;refGene_tx=XM_011287756.2;refGene_gn=PISD;refGene_tx=XM_011287757.2;refGene_gn=PISD;refGene_tx=XM_019814756.1
D3 21533407 21533568
refGene_gn=PISD;refGene_tx=XM_006938605.4;refGene_gn=PISD;refGene_tx=XM_011287756.2;refGene_gn=PISD;refGene_tx=XM_011287757.2;refGene_gn=PISD;refGene_tx=XM_019814756.1
D3 21534163 21534388
refGene_gn=PISD;refGene_tx=XM_006938605.4;refGene_gn=PISD;refGene_tx=XM_011287756.2;refGene_gn=PISD;refGene_tx=XM_011287757.2;refGene_gn=PISD;refGene_tx=XM_019814756.1
D3 21535505 21535794
refGene_gn=SFI1;refGene_tx=XM_006938606.4;refGene_gn=SFI1;refGene_tx=XM_006938607.4;refGene_gn=SFI1;refGene_tx=XM_006938608.4;refGene_gn=SFI1;refGene_tx=XM_006938609.4;refGene_gn=SFI1;refGene_tx=XM_006938610.4;refGene_gn=SFI1;refGene_tx=XM_006938612.4;refGene_gn=SFI1;refGene_tx=XM_006938613.4;refGene_gn=SFI1;refGene_tx=XM_019814747.2;refGene_gn=SFI1;refGene_tx=XM_019814748.2;refGene_gn=SFI1;refGene_tx=XM_019814749.2;refGene_gn=SFI1;refGene_tx=XM_019814750.2;refGene_gn=SFI1;refGene_tx=XM_019814751.2;refGene_gn=SFI1;refGene_tx=XM_019814752.2;refGene_gn=SFI1;refGene_tx=XM_019814754.2;refGene_gn=SFI1;refGene_tx=XM_023241560.1
D3 21536259 21536591
refGene_gn=SFI1;refGene_tx=XM_006938606.4;refGene_gn=SFI1;refGene_tx=XM_006938607.4;refGene_gn=SFI1;refGene_tx=XM_006938608.4;refGene_gn=SFI1;refGene_tx=XM_006938609.4;refGene_gn=SFI1;refGene_tx=XM_006938610.4;refGene_gn=SFI1;refGene_tx=XM_006938612.4;refGene_gn=SFI1;refGene_tx=XM_006938613.4;refGene_gn=SFI1;refGene_tx=XM_019814747.2;refGene_gn=SFI1;refGene_tx=XM_019814748.2;refGene_gn=SFI1;refGene_tx=XM_019814749.2;refGene_gn=SFI1;refGene_tx=XM_019814750.2;refGene_gn=SFI1;refGene_tx=XM_019814751.2;refGene_gn=SFI1;refGene_tx=XM_019814752.2;refGene_gn=SFI1;refGene_tx=XM_019814753.2;refGene_gn=SFI1;refGene_tx=XM_019814754.2;refGene_gn=SFI1;refGene_tx=XM_023241560.1;refGene_gn=SFI1;refGene_tx=XM_023241561.1
D3 21537814 21538046
refGene_gn=SFI1;refGene_tx=XM_006938606.4;refGene_gn=SFI1;refGene_tx=XM_006938607.4;refGene_gn=SFI1;refGene_tx=XM_006938608.4;refGene_gn=SFI1;refGene_tx=XM_006938609.4;refGene_gn=SFI1;refGene_tx=XM_006938610.4;refGene_gn=SFI1;refGene_tx=XM_006938612.4;refGene_gn=SFI1;refGene_tx=XM_006938613.4;refGene_gn=SFI1;refGene_tx=XM_019814747.2;refGene_gn=SFI1;refGene_tx=XM_019814748.2;refGene_gn=SFI1;refGene_tx=XM_019814749.2;refGene_gn=SFI1;refGene_tx=XM_019814750.2;refGene_gn=SFI1;refGene_tx=XM_019814751.2;refGene_gn=SFI1;refGene_tx=XM_019814752.2;refGene_gn=SFI

1;refGene_tx=XM_019814754.2;refGene_gn=SFI1;refGene_tx=XM_023241560.1
D3 21538143 21538248
refGene_gn=SFI1;refGene_tx=XM_006938606.4;refGene_gn=SFI1;refGene_tx=XM_006938607.4;refGene_gn=SFI1;refGene_tx=XM_006938608.4;refGene_gn=SFI1;refGene_tx=XM_006938609.4;refGene_gn=SFI1;refGene_tx=XM_006938610.4;refGene_gn=SFI1;refGene_tx=XM_006938612.4;refGene_gn=SFI1;refGene_tx=XM_006938613.4;refGene_gn=SFI1;refGene_tx=XM_019814747.2;refGene_gn=SFI1;refGene_tx=XM_019814748.2;refGene_gn=SFI1;refGene_tx=XM_019814749.2;refGene_gn=SFI1;refGene_tx=XM_019814750.2;refGene_gn=SFI1;refGene_tx=XM_019814751.2;refGene_gn=SFI1;refGene_tx=XM_019814752.2;refGene_gn=SFI1;refGene_tx=XM_019814753.2;refGene_gn=SFI1;refGene_tx=XM_019814754.2;refGene_gn=SFI1;refGene_tx=XM_023241560.1
D3 21538711 21538715 refGene_gn=SFI1;refGene_tx=XM_023241562.1
D3 21538998 21539258
refGene_gn=SFI1;refGene_tx=XM_006938606.4;refGene_gn=SFI1;refGene_tx=XM_006938607.4;refGene_gn=SFI1;refGene_tx=XM_006938608.4;refGene_gn=SFI1;refGene_tx=XM_006938609.4;refGene_gn=SFI1;refGene_tx=XM_006938610.4;refGene_gn=SFI1;refGene_tx=XM_006938612.4;refGene_gn=SFI1;refGene_tx=XM_006938613.4;refGene_gn=SFI1;refGene_tx=XM_019814747.2;refGene_gn=SFI1;refGene_tx=XM_019814748.2;refGene_gn=SFI1;refGene_tx=XM_019814749.2;refGene_gn=SFI1;refGene_tx=XM_019814750.2;refGene_gn=SFI1;refGene_tx=XM_019814751.2;refGene_gn=SFI1;refGene_tx=XM_019814752.2;refGene_gn=SFI1;refGene_tx=XM_019814753.2;refGene_gn=SFI1;refGene_tx=XM_019814754.2;refGene_gn=SFI1;refGene_tx=XM_023241560.1;refGene_gn=SFI1;refGene_tx=XM_023241561.1;refGene_gn=SFI1;refGene_tx=XM_023241562.1
D3 21539335 21539512
refGene_gn=SFI1;refGene_tx=XM_006938606.4;refGene_gn=SFI1;refGene_tx=XM_006938607.4;refGene_gn=SFI1;refGene_tx=XM_006938608.4;refGene_gn=SFI1;refGene_tx=XM_006938609.4;refGene_gn=SFI1;refGene_tx=XM_006938610.4;refGene_gn=SFI1;refGene_tx=XM_006938612.4;refGene_gn=SFI1;refGene_tx=XM_006938613.4;refGene_gn=SFI1;refGene_tx=XM_019814747.2;refGene_gn=SFI1;refGene_tx=XM_019814748.2;refGene_gn=SFI1;refGene_tx=XM_019814749.2;refGene_gn=SFI1;refGene_tx=XM_019814750.2;refGene_gn=SFI1;refGene_tx=XM_019814751.2;refGene_gn=SFI1;refGene_tx=XM_019814752.2;refGene_gn=SFI1;refGene_tx=XM_019814753.2;refGene_gn=SFI1;refGene_tx=XM_019814754.2;refGene_gn=SFI1;refGene_tx=XM_023241560.1;refGene_gn=SFI1;refGene_tx=XM_023241561.1;refGene_gn=SFI1;refGene_tx=XM_023241562.1
D3 21539635 21539710
refGene_gn=SFI1;refGene_tx=XM_006938606.4;refGene_gn=SFI1;refGene_tx=XM_006938607.4;refGene_gn=SFI1;refGene_tx=XM_006938608.4;refGene_gn=SFI1;refGene_tx=XM_006938609.4;refGene_gn=SFI1;refGene_tx=XM_006938610.4;refGene_gn=SFI1;refGene_tx=XM_006938612.4;refGene_gn=SFI1;refGene_tx=XM_006938613.4;refGene_gn=SFI1;refGene_tx=XM_019814747.2;refGene_gn=SFI1;refGene_tx=XM_019814748.2;refGene_gn=SFI1;refGene_tx=XM_019814749.2;refGene_gn=SFI1;refGene_tx=XM_019814750.2;refGene_gn=SFI1;refGene_tx=XM_019814751.2;refGene_gn=SFI1;refGene_tx=XM_019814752.2;refGene_gn=SFI1;refGene_tx=XM_019814753.2;refGene_gn=SFI1;refGene_tx=XM_019814754.2;refGene_gn=SFI1;refGene_tx=XM_023241560.1;refGene_gn=SFI1;refGene_tx=XM_023241561.1;refGene_gn=SFI1;refGene_tx=XM_023241562.1
D3 21540584 21540659

refGene_gn=SFI1;refGene_tx=XM_006938606.4;refGene_gn=SFI1;refGene_tx=XM_006938607.4;refGene_gn=SFI1;refGene_tx=XM_006938608.4;refGene_gn=SFI1;refGene_tx=XM_006938609.4;refGene_gn=SFI1;refGene_tx=XM_006938610.4;refGene_gn=SFI1;refGene_tx=XM_006938612.4;refGene_gn=SFI1;refGene_tx=XM_006938613.4;refGene_gn=SFI1;refGene_tx=XM_019814747.2;refGene_gn=SFI1;refGene_tx=XM_019814748.2;refGene_gn=SFI1;refGene_tx=XM_019814749.2;refGene_gn=SFI1;refGene_tx=XM_019814750.2;refGene_gn=SFI1;refGene_tx=XM_019814751.2;refGene_gn=SFI1;refGene_tx=XM_019814752.2;refGene_gn=SFI1;refGene_tx=XM_019814753.2;refGene_gn=SFI1;refGene_tx=XM_019814754.2;refGene_gn=SFI1;refGene_tx=XM_023241560.1;refGene_gn=SFI1;refGene_tx=XM_023241561.1;refGene_gn=SFI1;refGene_tx=XM_023241562.1
D3 21541108 21541269
refGene_gn=SFI1;refGene_tx=XM_006938606.4;refGene_gn=SFI1;refGene_tx=XM_006938607.4;refGene_gn=SFI1;refGene_tx=XM_006938608.4;refGene_gn=SFI1;refGene_tx=XM_006938609.4;refGene_gn=SFI1;refGene_tx=XM_006938610.4;refGene_gn=SFI1;refGene_tx=XM_006938612.4;refGene_gn=SFI1;refGene_tx=XM_006938613.4;refGene_gn=SFI1;refGene_tx=XM_019814747.2;refGene_gn=SFI1;refGene_tx=XM_019814748.2;refGene_gn=SFI1;refGene_tx=XM_019814749.2;refGene_gn=SFI1;refGene_tx=XM_019814750.2;refGene_gn=SFI1;refGene_tx=XM_019814751.2;refGene_gn=SFI1;refGene_tx=XM_019814752.2;refGene_gn=SFI1;refGene_tx=XM_019814753.2;refGene_gn=SFI1;refGene_tx=XM_019814754.2;refGene_gn=SFI1;refGene_tx=XM_023241560.1;refGene_gn=SFI1;refGene_tx=XM_023241561.1;refGene_gn=SFI1;refGene_tx=XM_023241562.1
D3 21544158 21544255
refGene_gn=SFI1;refGene_tx=XM_006938606.4;refGene_gn=SFI1;refGene_tx=XM_006938607.4;refGene_gn=SFI1;refGene_tx=XM_006938608.4;refGene_gn=SFI1;refGene_tx=XM_006938609.4;refGene_gn=SFI1;refGene_tx=XM_006938610.4;refGene_gn=SFI1;refGene_tx=XM_006938612.4;refGene_gn=SFI1;refGene_tx=XM_006938613.4;refGene_gn=SFI1;refGene_tx=XM_019814747.2;refGene_gn=SFI1;refGene_tx=XM_019814748.2;refGene_gn=SFI1;refGene_tx=XM_019814749.2;refGene_gn=SFI1;refGene_tx=XM_019814750.2;refGene_gn=SFI1;refGene_tx=XM_019814751.2;refGene_gn=SFI1;refGene_tx=XM_019814752.2;refGene_gn=SFI1;refGene_tx=XM_019814753.2;refGene_gn=SFI1;refGene_tx=XM_019814754.2;refGene_gn=SFI1;refGene_tx=XM_023241560.1;refGene_gn=SFI1;refGene_tx=XM_023241561.1;refGene_gn=SFI1;refGene_tx=XM_023241562.1
D3 21546100 21546203
refGene_gn=SFI1;refGene_tx=XM_006938606.4;refGene_gn=SFI1;refGene_tx=XM_006938607.4;refGene_gn=SFI1;refGene_tx=XM_006938608.4;refGene_gn=SFI1;refGene_tx=XM_006938609.4;refGene_gn=SFI1;refGene_tx=XM_006938610.4;refGene_gn=SFI1;refGene_tx=XM_006938612.4;refGene_gn=SFI1;refGene_tx=XM_006938613.4;refGene_gn=SFI1;refGene_tx=XM_019814747.2;refGene_gn=SFI1;refGene_tx=XM_019814748.2;refGene_gn=SFI1;refGene_tx=XM_019814749.2;refGene_gn=SFI1;refGene_tx=XM_019814750.2;refGene_gn=SFI1;refGene_tx=XM_019814751.2;refGene_gn=SFI1;refGene_tx=XM_019814752.2;refGene_gn=SFI1;refGene_tx=XM_019814753.2;refGene_gn=SFI1;refGene_tx=XM_019814754.2;refGene_gn=SFI1;refGene_tx=XM_023241560.1;refGene_gn=SFI1;refGene_tx=XM_023241561.1;refGene_gn=SFI1;refGene_tx=XM_023241562.1
D3 21547210 21547287
refGene_gn=SFI1;refGene_tx=XM_006938606.4;refGene_gn=SFI1;refGene_tx=XM_006938607.4;refGene_gn=SFI1;refGene_tx=XM_006938608.4;refGene_gn=SFI

1;refGene_tx=XM_006938609.4;refGene_gn=SFI1;refGene_tx=XM_006938610.4;
refGene_gn=SFI1;refGene_tx=XM_006938612.4;refGene_gn=SFI1;refGene_tx=X
M_006938613.4;refGene_gn=SFI1;refGene_tx=XM_019814747.2;refGene_gn=SFI
1;refGene_tx=XM_019814748.2;refGene_gn=SFI1;refGene_tx=XM_019814749.2;
refGene_gn=SFI1;refGene_tx=XM_019814750.2;refGene_gn=SFI1;refGene_tx=X
M_019814751.2;refGene_gn=SFI1;refGene_tx=XM_019814752.2;refGene_gn=SFI
1;refGene_tx=XM_019814753.2;refGene_gn=SFI1;refGene_tx=XM_019814754.2;
refGene_gn=SFI1;refGene_tx=XM_023241560.1;refGene_gn=SFI1;refGene_tx=X
M_023241561.1;refGene_gn=SFI1;refGene_tx=XM_023241562.1
D3 21547688 21547784
refGene_gn=SFI1;refGene_tx=XM_006938606.4;refGene_gn=SFI1;refGene_tx=X
M_006938607.4;refGene_gn=SFI1;refGene_tx=XM_006938608.4;refGene_gn=SFI
1;refGene_tx=XM_006938609.4;refGene_gn=SFI1;refGene_tx=XM_006938610.4;
refGene_gn=SFI1;refGene_tx=XM_006938612.4;refGene_gn=SFI1;refGene_tx=X
M_006938613.4;refGene_gn=SFI1;refGene_tx=XM_019814747.2;refGene_gn=SFI
1;refGene_tx=XM_019814748.2;refGene_gn=SFI1;refGene_tx=XM_019814749.2;
refGene_gn=SFI1;refGene_tx=XM_019814750.2;refGene_gn=SFI1;refGene_tx=X
M_019814751.2;refGene_gn=SFI1;refGene_tx=XM_019814752.2;refGene_gn=SFI
1;refGene_tx=XM_019814753.2;refGene_gn=SFI1;refGene_tx=XM_019814754.2;
refGene_gn=SFI1;refGene_tx=XM_023241560.1;refGene_gn=SFI1;refGene_tx=X
M_023241561.1;refGene_gn=SFI1;refGene_tx=XM_023241562.1
D3 21548250 21548326
refGene_gn=SFI1;refGene_tx=XM_006938606.4;refGene_gn=SFI1;refGene_tx=X
M_006938607.4;refGene_gn=SFI1;refGene_tx=XM_006938608.4;refGene_gn=SFI
1;refGene_tx=XM_006938609.4;refGene_gn=SFI1;refGene_tx=XM_006938610.4;
refGene_gn=SFI1;refGene_tx=XM_006938612.4;refGene_gn=SFI1;refGene_tx=X
M_006938613.4;refGene_gn=SFI1;refGene_tx=XM_019814747.2;refGene_gn=SFI
1;refGene_tx=XM_019814748.2;refGene_gn=SFI1;refGene_tx=XM_019814749.2;
refGene_gn=SFI1;refGene_tx=XM_019814750.2;refGene_gn=SFI1;refGene_tx=X
M_019814751.2;refGene_gn=SFI1;refGene_tx=XM_019814752.2;refGene_gn=SFI
1;refGene_tx=XM_019814753.2;refGene_gn=SFI1;refGene_tx=XM_019814754.2;
refGene_gn=SFI1;refGene_tx=XM_023241560.1;refGene_gn=SFI1;refGene_tx=X
M_023241561.1;refGene_gn=SFI1;refGene_tx=XM_023241562.1
D3 21549093 21549272
refGene_gn=SFI1;refGene_tx=XM_006938606.4;refGene_gn=SFI1;refGene_tx=X
M_006938607.4;refGene_gn=SFI1;refGene_tx=XM_006938608.4;refGene_gn=SFI
1;refGene_tx=XM_006938609.4;refGene_gn=SFI1;refGene_tx=XM_006938610.4;
refGene_gn=SFI1;refGene_tx=XM_006938612.4;refGene_gn=SFI1;refGene_tx=X
M_006938613.4;refGene_gn=SFI1;refGene_tx=XM_019814747.2;refGene_gn=SFI
1;refGene_tx=XM_019814748.2;refGene_gn=SFI1;refGene_tx=XM_019814749.2;
refGene_gn=SFI1;refGene_tx=XM_019814750.2;refGene_gn=SFI1;refGene_tx=X
M_019814751.2;refGene_gn=SFI1;refGene_tx=XM_019814752.2;refGene_gn=SFI
1;refGene_tx=XM_019814753.2;refGene_gn=SFI1;refGene_tx=XM_019814754.2;
refGene_gn=SFI1;refGene_tx=XM_023241560.1;refGene_gn=SFI1;refGene_tx=X
M_023241561.1;refGene_gn=SFI1;refGene_tx=XM_023241562.1
D3 21549588 21549670
refGene_gn=SFI1;refGene_tx=XM_006938606.4;refGene_gn=SFI1;refGene_tx=X
M_006938607.4;refGene_gn=SFI1;refGene_tx=XM_006938608.4;refGene_gn=SFI
1;refGene_tx=XM_006938609.4;refGene_gn=SFI1;refGene_tx=XM_006938610.4;
refGene_gn=SFI1;refGene_tx=XM_006938612.4;refGene_gn=SFI1;refGene_tx=X

M_006938613.4;refGene_gn=SFI1;refGene_tx=XM_019814747.2;refGene_gn=SFI1;refGene_tx=XM_019814748.2;refGene_gn=SFI1;refGene_tx=XM_019814749.2;refGene_gn=SFI1;refGene_tx=XM_019814750.2;refGene_gn=SFI1;refGene_tx=XM_019814751.2;refGene_gn=SFI1;refGene_tx=XM_019814752.2;refGene_gn=SFI1;refGene_tx=XM_019814753.2;refGene_gn=SFI1;refGene_tx=XM_019814754.2;refGene_gn=SFI1;refGene_tx=XM_023241560.1;refGene_gn=SFI1;refGene_tx=XM_023241561.1;refGene_gn=SFI1;refGene_tx=XM_023241562.1
D3 21562338 21562469
refGene_gn=SFI1;refGene_tx=XM_006938606.4;refGene_gn=SFI1;refGene_tx=XM_006938607.4;refGene_gn=SFI1;refGene_tx=XM_006938608.4;refGene_gn=SFI1;refGene_tx=XM_006938609.4;refGene_gn=SFI1;refGene_tx=XM_006938610.4;refGene_gn=SFI1;refGene_tx=XM_006938612.4;refGene_gn=SFI1;refGene_tx=XM_006938613.4;refGene_gn=SFI1;refGene_tx=XM_019814747.2;refGene_gn=SFI1;refGene_tx=XM_019814748.2;refGene_gn=SFI1;refGene_tx=XM_019814749.2;refGene_gn=SFI1;refGene_tx=XM_019814750.2;refGene_gn=SFI1;refGene_tx=XM_019814751.2;refGene_gn=SFI1;refGene_tx=XM_019814752.2;refGene_gn=SFI1;refGene_tx=XM_019814753.2;refGene_gn=SFI1;refGene_tx=XM_019814754.2;refGene_gn=SFI1;refGene_tx=XM_023241560.1;refGene_gn=SFI1;refGene_tx=XM_023241561.1;refGene_gn=SFI1;refGene_tx=XM_023241562.1
D3 21566527 21566594
refGene_gn=SFI1;refGene_tx=XM_006938606.4;refGene_gn=SFI1;refGene_tx=XM_006938607.4;refGene_gn=SFI1;refGene_tx=XM_006938608.4;refGene_gn=SFI1;refGene_tx=XM_006938609.4;refGene_gn=SFI1;refGene_tx=XM_006938610.4;refGene_gn=SFI1;refGene_tx=XM_006938612.4;refGene_gn=SFI1;refGene_tx=XM_006938613.4;refGene_gn=SFI1;refGene_tx=XM_019814747.2;refGene_gn=SFI1;refGene_tx=XM_019814748.2;refGene_gn=SFI1;refGene_tx=XM_019814749.2;refGene_gn=SFI1;refGene_tx=XM_019814750.2;refGene_gn=SFI1;refGene_tx=XM_019814751.2;refGene_gn=SFI1;refGene_tx=XM_019814752.2;refGene_gn=SFI1;refGene_tx=XM_019814753.2;refGene_gn=SFI1;refGene_tx=XM_019814754.2;refGene_gn=SFI1;refGene_tx=XM_023241560.1;refGene_gn=SFI1;refGene_tx=XM_023241561.1;refGene_gn=SFI1;refGene_tx=XM_023241562.1
D3 21568004 21568102
refGene_gn=SFI1;refGene_tx=XM_006938606.4;refGene_gn=SFI1;refGene_tx=XM_006938607.4;refGene_gn=SFI1;refGene_tx=XM_006938608.4;refGene_gn=SFI1;refGene_tx=XM_006938609.4;refGene_gn=SFI1;refGene_tx=XM_006938610.4;refGene_gn=SFI1;refGene_tx=XM_006938612.4;refGene_gn=SFI1;refGene_tx=XM_006938613.4;refGene_gn=SFI1;refGene_tx=XM_019814747.2;refGene_gn=SFI1;refGene_tx=XM_019814748.2;refGene_gn=SFI1;refGene_tx=XM_019814749.2;refGene_gn=SFI1;refGene_tx=XM_019814750.2;refGene_gn=SFI1;refGene_tx=XM_019814751.2;refGene_gn=SFI1;refGene_tx=XM_019814752.2;refGene_gn=SFI1;refGene_tx=XM_019814753.2;refGene_gn=SFI1;refGene_tx=XM_019814754.2;refGene_gn=SFI1;refGene_tx=XM_023241560.1;refGene_gn=SFI1;refGene_tx=XM_023241561.1;refGene_gn=SFI1;refGene_tx=XM_023241562.1
D3 21570391 21570484
refGene_gn=SFI1;refGene_tx=XM_006938606.4;refGene_gn=SFI1;refGene_tx=XM_006938607.4;refGene_gn=SFI1;refGene_tx=XM_006938608.4;refGene_gn=SFI1;refGene_tx=XM_006938609.4;refGene_gn=SFI1;refGene_tx=XM_006938610.4;refGene_gn=SFI1;refGene_tx=XM_006938612.4;refGene_gn=SFI1;refGene_tx=XM_006938613.4;refGene_gn=SFI1;refGene_tx=XM_019814747.2;refGene_gn=SFI1;refGene_tx=XM_019814748.2;refGene_gn=SFI1;refGene_tx=XM_019814749.2;

refGene_gn=SFI1;refGene_tx=XM_019814750.2;refGene_gn=SFI1;refGene_tx=XM_019814751.2;refGene_gn=SFI1;refGene_tx=XM_019814752.2;refGene_gn=SFI1;refGene_tx=XM_019814753.2;refGene_gn=SFI1;refGene_tx=XM_019814754.2;refGene_gn=SFI1;refGene_tx=XM_023241560.1;refGene_gn=SFI1;refGene_tx=XM_023241561.1;refGene_gn=SFI1;refGene_tx=XM_023241562.1
D3 21571798 21571869
refGene_gn=SFI1;refGene_tx=XM_006938606.4;refGene_gn=SFI1;refGene_tx=XM_006938607.4;refGene_gn=SFI1;refGene_tx=XM_006938608.4;refGene_gn=SFI1;refGene_tx=XM_006938609.4;refGene_gn=SFI1;refGene_tx=XM_006938610.4;refGene_gn=SFI1;refGene_tx=XM_006938612.4;refGene_gn=SFI1;refGene_tx=XM_006938613.4;refGene_gn=SFI1;refGene_tx=XM_019814747.2;refGene_gn=SFI1;refGene_tx=XM_019814748.2;refGene_gn=SFI1;refGene_tx=XM_019814749.2;refGene_gn=SFI1;refGene_tx=XM_019814750.2;refGene_gn=SFI1;refGene_tx=XM_019814751.2;refGene_gn=SFI1;refGene_tx=XM_019814752.2;refGene_gn=SFI1;refGene_tx=XM_019814753.2;refGene_gn=SFI1;refGene_tx=XM_019814754.2;refGene_gn=SFI1;refGene_tx=XM_023241560.1;refGene_gn=SFI1;refGene_tx=XM_023241561.1;refGene_gn=SFI1;refGene_tx=XM_023241562.1
D3 21573687 21573849
refGene_gn=SFI1;refGene_tx=XM_006938606.4;refGene_gn=SFI1;refGene_tx=XM_006938607.4;refGene_gn=SFI1;refGene_tx=XM_006938608.4;refGene_gn=SFI1;refGene_tx=XM_006938609.4;refGene_gn=SFI1;refGene_tx=XM_006938610.4;refGene_gn=SFI1;refGene_tx=XM_006938612.4;refGene_gn=SFI1;refGene_tx=XM_006938613.4;refGene_gn=SFI1;refGene_tx=XM_019814747.2;refGene_gn=SFI1;refGene_tx=XM_019814748.2;refGene_gn=SFI1;refGene_tx=XM_019814749.2;refGene_gn=SFI1;refGene_tx=XM_019814750.2;refGene_gn=SFI1;refGene_tx=XM_019814751.2;refGene_gn=SFI1;refGene_tx=XM_019814752.2;refGene_gn=SFI1;refGene_tx=XM_019814753.2;refGene_gn=SFI1;refGene_tx=XM_019814754.2;refGene_gn=SFI1;refGene_tx=XM_023241560.1;refGene_gn=SFI1;refGene_tx=XM_023241561.1;refGene_gn=SFI1;refGene_tx=XM_023241562.1
D3 21575207 21575364
refGene_gn=SFI1;refGene_tx=XM_006938606.4;refGene_gn=SFI1;refGene_tx=XM_006938607.4;refGene_gn=SFI1;refGene_tx=XM_006938608.4;refGene_gn=SFI1;refGene_tx=XM_006938609.4;refGene_gn=SFI1;refGene_tx=XM_006938610.4;refGene_gn=SFI1;refGene_tx=XM_006938612.4;refGene_gn=SFI1;refGene_tx=XM_006938613.4;refGene_gn=SFI1;refGene_tx=XM_019814747.2;refGene_gn=SFI1;refGene_tx=XM_019814748.2;refGene_gn=SFI1;refGene_tx=XM_019814749.2;refGene_gn=SFI1;refGene_tx=XM_019814750.2;refGene_gn=SFI1;refGene_tx=XM_019814751.2;refGene_gn=SFI1;refGene_tx=XM_019814752.2;refGene_gn=SFI1;refGene_tx=XM_019814753.2;refGene_gn=SFI1;refGene_tx=XM_019814754.2;refGene_gn=SFI1;refGene_tx=XM_023241560.1;refGene_gn=SFI1;refGene_tx=XM_023241561.1;refGene_gn=SFI1;refGene_tx=XM_023241562.1
D3 21585297 21585400
refGene_gn=SFI1;refGene_tx=XM_006938606.4;refGene_gn=SFI1;refGene_tx=XM_006938607.4;refGene_gn=SFI1;refGene_tx=XM_006938608.4;refGene_gn=SFI1;refGene_tx=XM_006938609.4;refGene_gn=SFI1;refGene_tx=XM_006938610.4;refGene_gn=SFI1;refGene_tx=XM_006938612.4;refGene_gn=SFI1;refGene_tx=XM_006938613.4;refGene_gn=SFI1;refGene_tx=XM_019814747.2;refGene_gn=SFI1;refGene_tx=XM_019814748.2;refGene_gn=SFI1;refGene_tx=XM_019814749.2;refGene_gn=SFI1;refGene_tx=XM_019814750.2;refGene_gn=SFI1;refGene_tx=XM_019814751.2;refGene_gn=SFI1;refGene_tx=XM_019814752.2;refGene_gn=SFI

1;refGene_tx=XM_019814753.2;refGene_gn=SFI1;refGene_tx=XM_019814754.2;
refGene_gn=SFI1;refGene_tx=XM_023241560.1;refGene_gn=SFI1;refGene_tx=X
M_023241561.1;refGene_gn=SFI1;refGene_tx=XM_023241562.1
D3 21587680 21587798
refGene_gn=SFI1;refGene_tx=XM_006938606.4;refGene_gn=SFI1;refGene_tx=X
M_006938607.4;refGene_gn=SFI1;refGene_tx=XM_006938608.4;refGene_gn=SFI
1;refGene_tx=XM_006938609.4;refGene_gn=SFI1;refGene_tx=XM_006938610.4;
refGene_gn=SFI1;refGene_tx=XM_006938612.4;refGene_gn=SFI1;refGene_tx=X
M_006938613.4;refGene_gn=SFI1;refGene_tx=XM_019814747.2;refGene_gn=SFI
1;refGene_tx=XM_019814748.2;refGene_gn=SFI1;refGene_tx=XM_019814749.2;
refGene_gn=SFI1;refGene_tx=XM_019814750.2;refGene_gn=SFI1;refGene_tx=X
M_019814751.2;refGene_gn=SFI1;refGene_tx=XM_019814752.2;refGene_gn=SFI
1;refGene_tx=XM_019814753.2;refGene_gn=SFI1;refGene_tx=XM_019814754.2;
refGene_gn=SFI1;refGene_tx=XM_023241560.1;refGene_gn=SFI1;refGene_tx=X
M_023241561.1;refGene_gn=SFI1;refGene_tx=XM_023241562.1
D3 21600335 21600430
refGene_gn=SFI1;refGene_tx=XM_006938606.4;refGene_gn=SFI1;refGene_tx=X
M_006938607.4;refGene_gn=SFI1;refGene_tx=XM_006938608.4;refGene_gn=SFI
1;refGene_tx=XM_006938609.4;refGene_gn=SFI1;refGene_tx=XM_006938610.4;
refGene_gn=SFI1;refGene_tx=XM_006938612.4;refGene_gn=SFI1;refGene_tx=X
M_006938613.4;refGene_gn=SFI1;refGene_tx=XM_019814750.2;refGene_gn=SFI
1;refGene_tx=XM_019814751.2;refGene_gn=SFI1;refGene_tx=XM_019814754.2;
refGene_gn=SFI1;refGene_tx=XM_023241560.1
D3 21605109 21605220
refGene_gn=SFI1;refGene_tx=XM_006938606.4;refGene_gn=SFI1;refGene_tx=X
M_006938607.4;refGene_gn=SFI1;refGene_tx=XM_006938608.4;refGene_gn=SFI
1;refGene_tx=XM_006938609.4;refGene_gn=SFI1;refGene_tx=XM_006938610.4;
refGene_gn=SFI1;refGene_tx=XM_006938612.4;refGene_gn=SFI1;refGene_tx=X
M_019814747.2;refGene_gn=SFI1;refGene_tx=XM_019814748.2;refGene_gn=SFI
1;refGene_tx=XM_019814749.2;refGene_gn=SFI1;refGene_tx=XM_019814750.2;
refGene_gn=SFI1;refGene_tx=XM_019814752.2;refGene_gn=SFI1;refGene_tx=X
M_019814753.2;refGene_gn=SFI1;refGene_tx=XM_019814754.2;refGene_gn=SFI
1;refGene_tx=XM_023241560.1;refGene_gn=SFI1;refGene_tx=XM_023241561.1;
refGene_gn=SFI1;refGene_tx=XM_023241562.1
D3 21613693 21613765
refGene_gn=SFI1;refGene_tx=XM_006938606.4;refGene_gn=SFI1;refGene_tx=X
M_006938607.4;refGene_gn=SFI1;refGene_tx=XM_006938608.4;refGene_gn=SFI
1;refGene_tx=XM_006938609.4;refGene_gn=SFI1;refGene_tx=XM_006938610.4;
refGene_gn=SFI1;refGene_tx=XM_006938612.4;refGene_gn=SFI1;refGene_tx=X
M_006938613.4
D3 21615659 21615833
refGene_gn=SFI1;refGene_tx=XM_006938606.4;refGene_gn=SFI1;refGene_tx=X
M_006938607.4;refGene_gn=SFI1;refGene_tx=XM_006938608.4;refGene_gn=SFI
1;refGene_tx=XM_006938609.4;refGene_gn=SFI1;refGene_tx=XM_006938610.4;
refGene_gn=SFI1;refGene_tx=XM_006938612.4;refGene_gn=SFI1;refGene_tx=X
M_006938613.4;refGene_gn=SFI1;refGene_tx=XM_019814747.2;refGene_gn=SFI
1;refGene_tx=XM_019814748.2;refGene_gn=SFI1;refGene_tx=XM_019814749.2;
refGene_gn=SFI1;refGene_tx=XM_019814750.2;refGene_gn=SFI1;refGene_tx=X
M_019814751.2;refGene_gn=SFI1;refGene_tx=XM_019814753.2;refGene_gn=SFI
1;refGene_tx=XM_019814754.2;refGene_gn=SFI1;refGene_tx=XM_023241561.1;

refGene_gn=SFI1;refGene_tx=XM_023241562.1
D3	21630463	21630587
refGene_gn=SFI1;refGene_tx=XM_006938606.4;refGene_gn=SFI1;refGene_tx=XM_006938607.4;refGene_gn=SFI1;refGene_tx=XM_006938608.4;refGene_gn=SFI1;refGene_tx=XM_006938609.4;refGene_gn=SFI1;refGene_tx=XM_006938610.4;refGene_gn=SFI1;refGene_tx=XM_006938612.4;refGene_gn=SFI1;refGene_tx=XM_006938613.4;refGene_gn=SFI1;refGene_tx=XM_019814747.2;refGene_gn=SFI1;refGene_tx=XM_019814748.2;refGene_gn=SFI1;refGene_tx=XM_019814749.2;refGene_gn=SFI1;refGene_tx=XM_019814750.2;refGene_gn=SFI1;refGene_tx=XM_019814751.2;refGene_gn=SFI1;refGene_tx=XM_019814753.2;refGene_gn=SFI1;refGene_tx=XM_019814754.2;refGene_gn=SFI1;refGene_tx=XM_023241560.1;refGene_gn=SFI1;refGene_tx=XM_023241561.1;refGene_gn=SFI1;refGene_tx=XM_023241562.1
D3	21639502	21639504
refGene_gn=SFI1;refGene_tx=XM_019814747.2;refGene_gn=SFI1;refGene_tx=XM_019814748.2;refGene_gn=SFI1;refGene_tx=XM_019814749.2;refGene_gn=SFI1;refGene_tx=XM_019814753.2;refGene_gn=SFI1;refGene_tx=XM_023241561.1;refGene_gn=SFI1;refGene_tx=XM_023241562.1
D3	21639635	21639825
refGene_gn=SFI1;refGene_tx=XM_019814747.2;refGene_gn=SFI1;refGene_tx=XM_019814748.2;refGene_gn=SFI1;refGene_tx=XM_019814749.2;refGene_gn=SFI1;refGene_tx=XM_019814753.2;refGene_gn=SFI1;refGene_tx=XM_023241561.1;refGene_gn=SFI1;refGene_tx=XM_023241562.1
D3	21645553	21645649
refGene_gn=EIF4ENIF1;refGene_tx=XM_023241563.1;refGene_gn=EIF4ENIF1;refGene_tx=XM_023241564.1;refGene_gn=EIF4ENIF1;refGene_tx=XM_023241565.1;refGene_gn=EIF4ENIF1;refGene_tx=XM_023241566.1;refGene_gn=EIF4ENIF1;refGene_tx=XM_023241567.1
D3	21657530	21657604
refGene_gn=EIF4ENIF1;refGene_tx=XM_023241563.1;refGene_gn=EIF4ENIF1;refGene_tx=XM_023241564.1;refGene_gn=EIF4ENIF1;refGene_tx=XM_023241565.1;refGene_gn=EIF4ENIF1;refGene_tx=XM_023241566.1;refGene_gn=EIF4ENIF1;refGene_tx=XM_023241567.1
D3	21660694	21660822
refGene_gn=EIF4ENIF1;refGene_tx=XM_023241563.1;refGene_gn=EIF4ENIF1;refGene_tx=XM_023241564.1;refGene_gn=EIF4ENIF1;refGene_tx=XM_023241565.1;refGene_gn=EIF4ENIF1;refGene_tx=XM_023241566.1;refGene_gn=EIF4ENIF1;refGene_tx=XM_023241567.1
D3	21663547	21663834
refGene_gn=EIF4ENIF1;refGene_tx=XM_023241563.1;refGene_gn=EIF4ENIF1;refGene_tx=XM_023241564.1;refGene_gn=EIF4ENIF1;refGene_tx=XM_023241565.1;refGene_gn=EIF4ENIF1;refGene_tx=XM_023241566.1;refGene_gn=EIF4ENIF1;refGene_tx=XM_023241567.1
D3	21664126	21664328
refGene_gn=EIF4ENIF1;refGene_tx=XM_023241563.1;refGene_gn=EIF4ENIF1;refGene_tx=XM_023241564.1;refGene_gn=EIF4ENIF1;refGene_tx=XM_023241565.1;refGene_gn=EIF4ENIF1;refGene_tx=XM_023241566.1;refGene_gn=EIF4ENIF1;refGene_tx=XM_023241567.1
D3	21669012	21669188
refGene_gn=EIF4ENIF1;refGene_tx=XM_023241563.1;refGene_gn=EIF4ENIF1;re

fGene_tx=XM_023241564.1;refGene_gn=EIF4ENIF1;refGene_tx=XM_023241565.1
;refGene_gn=EIF4ENIF1;refGene_tx=XM_023241566.1;refGene_gn=EIF4ENIF1;r
efGene_tx=XM_023241567.1
D3 21674379 21674515
refGene_gn=EIF4ENIF1;refGene_tx=XM_023241563.1;refGene_gn=EIF4ENIF1;re
fGene_tx=XM_023241564.1;refGene_gn=EIF4ENIF1;refGene_tx=XM_023241565.1
;refGene_gn=EIF4ENIF1;refGene_tx=XM_023241566.1;refGene_gn=EIF4ENIF1;r
efGene_tx=XM_023241567.1
D3 21674728 21674908
refGene_gn=EIF4ENIF1;refGene_tx=XM_023241563.1;refGene_gn=EIF4ENIF1;re
fGene_tx=XM_023241564.1;refGene_gn=EIF4ENIF1;refGene_tx=XM_023241565.1
;refGene_gn=EIF4ENIF1;refGene_tx=XM_023241566.1;refGene_gn=EIF4ENIF1;r
efGene_tx=XM_023241567.1
D3 21675643 21675876
refGene_gn=EIF4ENIF1;refGene_tx=XM_023241563.1;refGene_gn=EIF4ENIF1;re
fGene_tx=XM_023241564.1;refGene_gn=EIF4ENIF1;refGene_tx=XM_023241565.1
;refGene_gn=EIF4ENIF1;refGene_tx=XM_023241566.1;refGene_gn=EIF4ENIF1;r
efGene_tx=XM_023241567.1
D3 21678744 21678816
refGene_gn=EIF4ENIF1;refGene_tx=XM_023241563.1;refGene_gn=EIF4ENIF1;re
fGene_tx=XM_023241564.1;refGene_gn=EIF4ENIF1;refGene_tx=XM_023241565.1
;refGene_gn=EIF4ENIF1;refGene_tx=XM_023241566.1
D3 21679271 21679455
refGene_gn=EIF4ENIF1;refGene_tx=XM_023241563.1;refGene_gn=EIF4ENIF1;re
fGene_tx=XM_023241564.1;refGene_gn=EIF4ENIF1;refGene_tx=XM_023241565.1
;refGene_gn=EIF4ENIF1;refGene_tx=XM_023241566.1;refGene_gn=EIF4ENIF1;r
efGene_tx=XM_023241567.1
D3 21680425 21680505
refGene_gn=EIF4ENIF1;refGene_tx=XM_023241563.1;refGene_gn=EIF4ENIF1;re
fGene_tx=XM_023241564.1;refGene_gn=EIF4ENIF1;refGene_tx=XM_023241565.1
;refGene_gn=EIF4ENIF1;refGene_tx=XM_023241566.1;refGene_gn=EIF4ENIF1;r
efGene_tx=XM_023241567.1
D3 21680833 21680976
refGene_gn=EIF4ENIF1;refGene_tx=XM_023241563.1;refGene_gn=EIF4ENIF1;re
fGene_tx=XM_023241564.1;refGene_gn=EIF4ENIF1;refGene_tx=XM_023241565.1
;refGene_gn=EIF4ENIF1;refGene_tx=XM_023241566.1;refGene_gn=EIF4ENIF1;r
efGene_tx=XM_023241567.1
D3 21683659 21683744
refGene_gn=EIF4ENIF1;refGene_tx=XM_023241563.1;refGene_gn=EIF4ENIF1;re
fGene_tx=XM_023241564.1;refGene_gn=EIF4ENIF1;refGene_tx=XM_023241565.1
;refGene_gn=EIF4ENIF1;refGene_tx=XM_023241566.1;refGene_gn=EIF4ENIF1;r
efGene_tx=XM_023241567.1
D3 21685023 21685162
refGene_gn=EIF4ENIF1;refGene_tx=XM_023241563.1;refGene_gn=EIF4ENIF1;re
fGene_tx=XM_023241564.1;refGene_gn=EIF4ENIF1;refGene_tx=XM_023241565.1
;refGene_gn=EIF4ENIF1;refGene_tx=XM_023241566.1;refGene_gn=EIF4ENIF1;r
efGene_tx=XM_023241567.1
D3 21686205 21686550
refGene_gn=EIF4ENIF1;refGene_tx=XM_023241563.1;refGene_gn=EIF4ENIF1;re
fGene_tx=XM_023241564.1;refGene_gn=EIF4ENIF1;refGene_tx=XM_023241565.1

;refGene_gn=EIF4ENIF1;refGene_tx=XM_023241566.1;refGene_gn=EIF4ENIF1;refGene_tx=XM_023241567.1
D3 21686825 21686990
refGene_gn=EIF4ENIF1;refGene_tx=XM_023241563.1;refGene_gn=EIF4ENIF1;refGene_tx=XM_023241564.1;refGene_gn=EIF4ENIF1;refGene_tx=XM_023241565.1;refGene_gn=EIF4ENIF1;refGene_tx=XM_023241566.1;refGene_gn=EIF4ENIF1;refGene_tx=XM_023241567.1
D3 21687573 21687815
refGene_gn=EIF4ENIF1;refGene_tx=XM_023241563.1;refGene_gn=EIF4ENIF1;refGene_tx=XM_023241564.1;refGene_gn=EIF4ENIF1;refGene_tx=XM_023241565.1;refGene_gn=EIF4ENIF1;refGene_tx=XM_023241566.1;refGene_gn=EIF4ENIF1;refGene_tx=XM_023241567.1
D3 21694175 21694275 refGene_gn=DRG1;refGene_tx=XM_003994820.5
D3 21700216 21700339 refGene_gn=DRG1;refGene_tx=XM_003994820.5
D3 21700613 21700781 refGene_gn=DRG1;refGene_tx=XM_003994820.5
D3 21702901 21703032 refGene_gn=DRG1;refGene_tx=XM_003994820.5
D3 21704795 21704965 refGene_gn=DRG1;refGene_tx=XM_003994820.5
D3 21708204 21708274 refGene_gn=DRG1;refGene_tx=XM_003994820.5
D3 21714589 21714765 refGene_gn=DRG1;refGene_tx=XM_003994820.5
D3 21716668 21716792 refGene_gn=DRG1;refGene_tx=XM_003994820.5
D3 21717732 21717774 refGene_gn=DRG1;refGene_tx=XM_003994820.5
D3 21752038 21753309
refGene_gn=PATZ1;refGene_tx=XM_023241568.1;refGene_gn=PATZ1;refGene_tx=XM_023241569.1;refGene_gn=PATZ1;refGene_tx=XM_023241570.1;refGene_gn=PATZ1;refGene_tx=XM_023241571.1;refGene_gn=PATZ1;refGene_tx=XM_023241572.1;refGene_gn=PATZ1;refGene_tx=XM_023241573.1
D3 21754708 21754772
refGene_gn=PATZ1;refGene_tx=XM_023241568.1;refGene_gn=PATZ1;refGene_tx=XM_023241569.1;refGene_gn=PATZ1;refGene_tx=XM_023241570.1;refGene_gn=PATZ1;refGene_tx=XM_023241571.1;refGene_gn=PATZ1;refGene_tx=XM_023241572.1;refGene_gn=PATZ1;refGene_tx=XM_023241573.1
D3 21756022 21756301 refGene_gn=PATZ1;refGene_tx=XM_023241573.1
D3 21761114 21761286
refGene_gn=PATZ1;refGene_tx=XM_023241568.1;refGene_gn=PATZ1;refGene_tx=XM_023241569.1;refGene_gn=PATZ1;refGene_tx=XM_023241570.1;refGene_gn=PATZ1;refGene_tx=XM_023241571.1;refGene_gn=PATZ1;refGene_tx=XM_023241572.1
D3 21767098 21767236
refGene_gn=PATZ1;refGene_tx=XM_023241568.1;refGene_gn=PATZ1;refGene_tx=XM_023241569.1;refGene_gn=PATZ1;refGene_tx=XM_023241572.1
D3 21768710 21769129
refGene_gn=PATZ1;refGene_tx=XM_023241568.1;refGene_gn=PATZ1;refGene_tx=XM_023241569.1;refGene_gn=PATZ1;refGene_tx=XM_023241570.1;refGene_gn=PATZ1;refGene_tx=XM_023241571.1;refGene_gn=PATZ1;refGene_tx=XM_023241572.1
D3 21796008 21796078
refGene_gn=PIK3IP1;refGene_tx=XM_006938599.4;refGene_gn=PIK3IP1;refGene_tx=XM_006938600.4;refGene_gn=PIK3IP1;refGene_tx=XM_006938601.4
D3 21796994 21797255
refGene_gn=PIK3IP1;refGene_tx=XM_006938599.4;refGene_gn=PIK3IP1;refGen

e_tx=XM_006938600.4;refGene_gn=PIK3IP1;refGene_tx=XM_006938601.4
D3 21797341 21797461
refGene_gn=PIK3IP1;refGene_tx=XM_006938599.4;refGene_gn=PIK3IP1;refGene_tx=XM_006938600.4;refGene_gn=PIK3IP1;refGene_tx=XM_006938601.4
D3 21798414 21798615
refGene_gn=PIK3IP1;refGene_tx=XM_006938599.4;refGene_gn=PIK3IP1;refGene_tx=XM_006938600.4;refGene_gn=PIK3IP1;refGene_tx=XM_006938601.4
D3 21798712 21798791
refGene_gn=PIK3IP1;refGene_tx=XM_006938599.4;refGene_gn=PIK3IP1;refGene_tx=XM_006938600.4;refGene_gn=PIK3IP1;refGene_tx=XM_006938601.4
D3 21805669 21805874
refGene_gn=PIK3IP1;refGene_tx=XM_006938599.4;refGene_gn=PIK3IP1;refGene_tx=XM_006938600.4;refGene_gn=PIK3IP1;refGene_tx=XM_006938601.4
D3 21810198 21810343
refGene_gn=LIMK2;refGene_tx=XM_003994812.5;refGene_gn=LIMK2;refGene_tx=XM_003994813.5;refGene_gn=LIMK2;refGene_tx=XM_019814757.2
D3 21812990 21813148
refGene_gn=LIMK2;refGene_tx=XM_003994812.5;refGene_gn=LIMK2;refGene_tx=XM_003994813.5;refGene_gn=LIMK2;refGene_tx=XM_019814757.2
D3 21814211 21814267
refGene_gn=LIMK2;refGene_tx=XM_003994812.5;refGene_gn=LIMK2;refGene_tx=XM_003994813.5;refGene_gn=LIMK2;refGene_tx=XM_019814757.2
D3 21815218 21815393
refGene_gn=LIMK2;refGene_tx=XM_003994812.5;refGene_gn=LIMK2;refGene_tx=XM_003994813.5;refGene_gn=LIMK2;refGene_tx=XM_019814757.2
D3 21816347 21816413
refGene_gn=LIMK2;refGene_tx=XM_003994812.5;refGene_gn=LIMK2;refGene_tx=XM_003994813.5;refGene_gn=LIMK2;refGene_tx=XM_019814757.2
D3 21818298 21818355
refGene_gn=LIMK2;refGene_tx=XM_003994812.5;refGene_gn=LIMK2;refGene_tx=XM_003994813.5;refGene_gn=LIMK2;refGene_tx=XM_019814757.2
D3 21818517 21818649
refGene_gn=LIMK2;refGene_tx=XM_003994812.5;refGene_gn=LIMK2;refGene_tx=XM_003994813.5;refGene_gn=LIMK2;refGene_tx=XM_019814757.2
D3 21819808 21819895
refGene_gn=LIMK2;refGene_tx=XM_003994812.5;refGene_gn=LIMK2;refGene_tx=XM_003994813.5;refGene_gn=LIMK2;refGene_tx=XM_019814757.2
D3 21820730 21820917
refGene_gn=LIMK2;refGene_tx=XM_003994812.5;refGene_gn=LIMK2;refGene_tx=XM_003994813.5;refGene_gn=LIMK2;refGene_tx=XM_019814757.2
D3 21822634 21822831
refGene_gn=LIMK2;refGene_tx=XM_003994812.5;refGene_gn=LIMK2;refGene_tx=XM_003994813.5;refGene_gn=LIMK2;refGene_tx=XM_019814757.2
D3 21823190 21823296
refGene_gn=LIMK2;refGene_tx=XM_003994812.5;refGene_gn=LIMK2;refGene_tx=XM_003994813.5;refGene_gn=LIMK2;refGene_tx=XM_019814757.2
D3 21824764 21824953
refGene_gn=LIMK2;refGene_tx=XM_003994812.5;refGene_gn=LIMK2;refGene_tx=XM_003994813.5;refGene_gn=LIMK2;refGene_tx=XM_019814757.2
D3 21825548 21825658

refGene_gn=LIMK2;refGene_tx=XM_003994812.5;refGene_gn=LIMK2;refGene_tx=XM_003994813.5;refGene_gn=LIMK2;refGene_tx=XM_019814757.2
D3	21826609	21826745	refGene_gn=LIMK2;refGene_tx=XM_003994812.5;refGene_gn=LIMK2;refGene_tx=XM_003994813.5;refGene_gn=LIMK2;refGene_tx=XM_019814757.2
D3	21836055	21836108	refGene_gn=LIMK2;refGene_tx=XM_003994813.5
D3	21836242	21836319	refGene_gn=LIMK2;refGene_tx=XM_019814757.2
D3	21856039	21856139	refGene_gn=LIMK2;refGene_tx=XM_003994812.5
D3	21867468	21867484	refGene_gn=LIMK2;refGene_tx=XM_003994812.5
D3	21874286	21874384	refGene_gn=RNF185;refGene_tx=XM_011287752.3;refGene_gn=RNF185;refGene_tx=XM_019814761.2
D3	21876332	21876450	refGene_gn=RNF185;refGene_tx=XM_011287752.3;refGene_gn=RNF185;refGene_tx=XM_019814761.2
D3	21879532	21879587	refGene_gn=RNF185;refGene_tx=XM_011287752.3;refGene_gn=RNF185;refGene_tx=XM_019814761.2
D3	21880977	21881090	refGene_gn=RNF185;refGene_tx=XM_011287752.3;refGene_gn=RNF185;refGene_tx=XM_019814761.2
D3	21882721	21882740	refGene_gn=RNF185;refGene_tx=XM_011287752.3;refGene_gn=RNF185;refGene_tx=XM_019814761.2
D3	21885346	21885522	refGene_gn=RNF185;refGene_tx=XM_011287752.3;refGene_gn=RNF185;refGene_tx=XM_019814761.2
D3	21923616	21924157	refGene_gn=PLA2G3;refGene_tx=XM_019814758.2;refGene_gn=PLA2G3;refGene_tx=XM_023241574.1
D3	21925171	21925304	refGene_gn=PLA2G3;refGene_tx=XM_019814758.2;refGene_gn=PLA2G3;refGene_tx=XM_023241574.1
D3	21925552	21925687	refGene_gn=PLA2G3;refGene_tx=XM_019814758.2
D3	21925950	21926258	refGene_gn=PLA2G3;refGene_tx=XM_019814758.2;refGene_gn=PLA2G3;refGene_tx=XM_023241574.1
D3	21926949	21927079	refGene_gn=PLA2G3;refGene_tx=XM_019814758.2;refGene_gn=PLA2G3;refGene_tx=XM_023241574.1
D3	21927188	21927305	refGene_gn=PLA2G3;refGene_tx=XM_019814758.2;refGene_gn=PLA2G3;refGene_tx=XM_023241574.1
D3	21927954	21928159	refGene_gn=PLA2G3;refGene_tx=XM_019814758.2;refGene_gn=PLA2G3;refGene_tx=XM_023241574.1
D3	21929401	21929908	refGene_gn=INPP5J;refGene_tx=XM_003994807.5
D3	21930106	21930211	refGene_gn=INPP5J;refGene_tx=XM_003994807.5
D3	21930327	21930405	refGene_gn=INPP5J;refGene_tx=XM_003994807.5
D3	21930566	21930704	refGene_gn=INPP5J;refGene_tx=XM_003994807.5

D3 21932619 21932803 refGene_gn=INPP5J;refGene_tx=XM_003994807.5
D3 21932941 21933051 refGene_gn=INPP5J;refGene_tx=XM_003994807.5
D3 21933191 21933303 refGene_gn=INPP5J;refGene_tx=XM_003994807.5
D3 21933715 21933891 refGene_gn=INPP5J;refGene_tx=XM_003994807.5
D3 21934226 21934343 refGene_gn=INPP5J;refGene_tx=XM_003994807.5
D3 21934552 21934661 refGene_gn=INPP5J;refGene_tx=XM_003994807.5
D3 21934792 21934909 refGene_gn=INPP5J;refGene_tx=XM_003994807.5
D3 21935255 21936409 refGene_gn=INPP5J;refGene_tx=XM_003994807.5
D3 21938040 21938145 refGene_gn=INPP5J;refGene_tx=XM_003994807.5
D3 21945873 21945993 refGene_gn=SELENOM;refGene_tx=XM_006938597.4
D3 21947192 21947228 refGene_gn=SELENOM;refGene_tx=XM_006938597.4
D3 21947411 21947446 refGene_gn=SELENOM;refGene_tx=XM_006938597.4
D3 21947798 21947877 refGene_gn=SELENOM;refGene_tx=XM_006938597.4
D3 21947954 21948047 refGene_gn=SELENOM;refGene_tx=XM_006938597.4
D3 21948048 21948114 refGene_gn=SELENOM;refGene_tx=XM_006938597.4
D3 21948617 21948768
refGene_gn=SMTN;refGene_tx=XM_019814762.2;refGene_gn=SMTN;refGene_tx=XM_019814764.2;refGene_gn=SMTN;refGene_tx=XM_019814765.2;refGene_gn=SMTN;refGene_tx=XM_019814766.2;refGene_gn=SMTN;refGene_tx=XM_019814767.2
D3 21951789 21951934
refGene_gn=SMTN;refGene_tx=XM_011287750.3;refGene_gn=SMTN;refGene_tx=XM_019814763.2
D3 21952886 21953038
refGene_gn=SMTN;refGene_tx=XM_011287750.3;refGene_gn=SMTN;refGene_tx=XM_019814762.2;refGene_gn=SMTN;refGene_tx=XM_019814763.2;refGene_gn=SMTN;refGene_tx=XM_019814764.2;refGene_gn=SMTN;refGene_tx=XM_019814765.2;refGene_gn=SMTN;refGene_tx=XM_019814766.2;refGene_gn=SMTN;refGene_tx=XM_019814767.2
D3 21953629 21953747
refGene_gn=SMTN;refGene_tx=XM_011287750.3;refGene_gn=SMTN;refGene_tx=XM_019814762.2;refGene_gn=SMTN;refGene_tx=XM_019814763.2;refGene_gn=SMTN;refGene_tx=XM_019814764.2;refGene_gn=SMTN;refGene_tx=XM_019814765.2;refGene_gn=SMTN;refGene_tx=XM_019814766.2;refGene_gn=SMTN;refGene_tx=XM_019814767.2
D3 21953935 21954109
refGene_gn=SMTN;refGene_tx=XM_011287750.3;refGene_gn=SMTN;refGene_tx=XM_019814762.2;refGene_gn=SMTN;refGene_tx=XM_019814763.2;refGene_gn=SMTN;refGene_tx=XM_019814764.2;refGene_gn=SMTN;refGene_tx=XM_019814765.2;refGene_gn=SMTN;refGene_tx=XM_019814766.2;refGene_gn=SMTN;refGene_tx=XM_019814767.2
D3 21955063 21955130
refGene_gn=SMTN;refGene_tx=XM_011287750.3;refGene_gn=SMTN;refGene_tx=XM_019814762.2;refGene_gn=SMTN;refGene_tx=XM_019814763.2;refGene_gn=SMTN;refGene_tx=XM_019814764.2;refGene_gn=SMTN;refGene_tx=XM_019814765.2;refGene_gn=SMTN;refGene_tx=XM_019814766.2;refGene_gn=SMTN;refGene_tx=XM_019814767.2
D3 21955352 21955415
refGene_gn=SMTN;refGene_tx=XM_011287750.3;refGene_gn=SMTN;refGene_tx=XM_019814762.2;refGene_gn=SMTN;refGene_tx=XM_019814763.2;refGene_gn=SMTN;refGene_tx=XM_019814764.2;refGene_gn=SMTN;refGene_tx=XM_019814765.2;

refGene_gn=SMTN;refGene_tx=XM_019814767.2
D3 21955512 21955677
refGene_gn=SMTN;refGene_tx=XM_011287750.3;refGene_gn=SMTN;refGene_tx=XM_019814762.2;refGene_gn=SMTN;refGene_tx=XM_019814763.2;refGene_gn=SMTN;refGene_tx=XM_019814764.2;refGene_gn=SMTN;refGene_tx=XM_019814765.2;refGene_gn=SMTN;refGene_tx=XM_019814767.2
D3 21956764 21956840
refGene_gn=SMTN;refGene_tx=XM_011287750.3;refGene_gn=SMTN;refGene_tx=XM_019814762.2;refGene_gn=SMTN;refGene_tx=XM_019814763.2;refGene_gn=SMTN;refGene_tx=XM_019814764.2;refGene_gn=SMTN;refGene_tx=XM_019814765.2;refGene_gn=SMTN;refGene_tx=XM_019814766.2;refGene_gn=SMTN;refGene_tx=XM_019814767.2
D3 21956927 21957080
refGene_gn=SMTN;refGene_tx=XM_011287750.3;refGene_gn=SMTN;refGene_tx=XM_019814762.2;refGene_gn=SMTN;refGene_tx=XM_019814763.2;refGene_gn=SMTN;refGene_tx=XM_019814764.2;refGene_gn=SMTN;refGene_tx=XM_019814765.2;refGene_gn=SMTN;refGene_tx=XM_019814766.2;refGene_gn=SMTN;refGene_tx=XM_019814767.2
D3 21958190 21958283 refGene_gn=SMTN;refGene_tx=XM_019814767.2
D3 21960219 21960392
refGene_gn=SMTN;refGene_tx=XM_011287750.3;refGene_gn=SMTN;refGene_tx=XM_019814762.2;refGene_gn=SMTN;refGene_tx=XM_019814763.2;refGene_gn=SMTN;refGene_tx=XM_019814764.2;refGene_gn=SMTN;refGene_tx=XM_019814765.2;refGene_gn=SMTN;refGene_tx=XM_019814766.2;refGene_gn=SMTN;refGene_tx=XM_019814767.2
D3 21960581 21961103
refGene_gn=SMTN;refGene_tx=XM_011287750.3;refGene_gn=SMTN;refGene_tx=XM_019814762.2;refGene_gn=SMTN;refGene_tx=XM_019814763.2;refGene_gn=SMTN;refGene_tx=XM_019814764.2;refGene_gn=SMTN;refGene_tx=XM_019814765.2;refGene_gn=SMTN;refGene_tx=XM_019814766.2;refGene_gn=SMTN;refGene_tx=XM_019814767.2
D3 21961183 21961276
refGene_gn=SMTN;refGene_tx=XM_011287750.3;refGene_gn=SMTN;refGene_tx=XM_019814762.2;refGene_gn=SMTN;refGene_tx=XM_019814763.2;refGene_gn=SMTN;refGene_tx=XM_019814764.2;refGene_gn=SMTN;refGene_tx=XM_019814766.2;refGene_gn=SMTN;refGene_tx=XM_019814767.2
D3 21961849 21961922
refGene_gn=SMTN;refGene_tx=XM_011287750.3;refGene_gn=SMTN;refGene_tx=XM_019814762.2;refGene_gn=SMTN;refGene_tx=XM_019814763.2;refGene_gn=SMTN;refGene_tx=XM_019814764.2;refGene_gn=SMTN;refGene_tx=XM_019814765.2;refGene_gn=SMTN;refGene_tx=XM_019814766.2;refGene_gn=SMTN;refGene_tx=XM_019814767.2
D3 21962010 21962331
refGene_gn=SMTN;refGene_tx=XM_011287750.3;refGene_gn=SMTN;refGene_tx=XM_019814762.2;refGene_gn=SMTN;refGene_tx=XM_019814763.2;refGene_gn=SMTN;refGene_tx=XM_019814764.2;refGene_gn=SMTN;refGene_tx=XM_019814765.2;refGene_gn=SMTN;refGene_tx=XM_019814766.2;refGene_gn=SMTN;refGene_tx=XM_019814767.2
D3 21963255 21963353
refGene_gn=SMTN;refGene_tx=XM_011287750.3;refGene_gn=SMTN;refGene_tx=X

M_019814762.2;refGene_gn=SMTN;refGene_tx=XM_019814763.2;refGene_gn=SMTN;refGene_tx=XM_019814764.2;refGene_gn=SMTN;refGene_tx=XM_019814765.2;refGene_gn=SMTN;refGene_tx=XM_019814766.2;refGene_gn=SMTN;refGene_tx=XM_019814767.2
D3 21963448 21963527
refGene_gn=SMTN;refGene_tx=XM_011287750.3;refGene_gn=SMTN;refGene_tx=XM_019814762.2;refGene_gn=SMTN;refGene_tx=XM_019814763.2;refGene_gn=SMTN;refGene_tx=XM_019814764.2;refGene_gn=SMTN;refGene_tx=XM_019814765.2;refGene_gn=SMTN;refGene_tx=XM_019814766.2;refGene_gn=SMTN;refGene_tx=XM_019814767.2
D3 21963613 21963707
refGene_gn=SMTN;refGene_tx=XM_011287750.3;refGene_gn=SMTN;refGene_tx=XM_019814762.2;refGene_gn=SMTN;refGene_tx=XM_019814763.2;refGene_gn=SMTN;refGene_tx=XM_019814764.2;refGene_gn=SMTN;refGene_tx=XM_019814765.2;refGene_gn=SMTN;refGene_tx=XM_019814766.2;refGene_gn=SMTN;refGene_tx=XM_019814767.2
D3 21964096 21964245
refGene_gn=SMTN;refGene_tx=XM_011287750.3;refGene_gn=SMTN;refGene_tx=XM_019814762.2;refGene_gn=SMTN;refGene_tx=XM_019814763.2;refGene_gn=SMTN;refGene_tx=XM_019814764.2;refGene_gn=SMTN;refGene_tx=XM_019814765.2;refGene_gn=SMTN;refGene_tx=XM_019814766.2;refGene_gn=SMTN;refGene_tx=XM_019814767.2
D3 21968794 21968845
refGene_gn=SMTN;refGene_tx=XM_011287750.3;refGene_gn=SMTN;refGene_tx=XM_019814762.2;refGene_gn=SMTN;refGene_tx=XM_019814763.2;refGene_gn=SMTN;refGene_tx=XM_019814764.2;refGene_gn=SMTN;refGene_tx=XM_019814765.2;refGene_gn=SMTN;refGene_tx=XM_019814766.2;refGene_gn=SMTN;refGene_tx=XM_019814767.2
D3 22070171 22070239
refGene_gn=MORC2;refGene_tx=XM_003994803.5;refGene_gn=MORC2;refGene_tx=XM_006938594.4;refGene_gn=MORC2;refGene_tx=XM_006938595.4;refGene_gn=MORC2;refGene_tx=XM_006938596.4
D3 22077407 22077461
refGene_gn=MORC2;refGene_tx=XM_003994803.5;refGene_gn=MORC2;refGene_tx=XM_006938594.4;refGene_gn=MORC2;refGene_tx=XM_006938595.4;refGene_gn=MORC2;refGene_tx=XM_006938596.4
D3 22079818 22079853
refGene_gn=MORC2;refGene_tx=XM_003994803.5;refGene_gn=MORC2;refGene_tx=XM_006938594.4;refGene_gn=MORC2;refGene_tx=XM_006938595.4;refGene_gn=MORC2;refGene_tx=XM_006938596.4
D3 22085350 22085419
refGene_gn=MORC2;refGene_tx=XM_003994803.5;refGene_gn=MORC2;refGene_tx=XM_006938594.4;refGene_gn=MORC2;refGene_tx=XM_006938595.4;refGene_gn=MORC2;refGene_tx=XM_006938596.4
D3 22085891 22085982
refGene_gn=MORC2;refGene_tx=XM_003994803.5;refGene_gn=MORC2;refGene_tx=XM_006938594.4;refGene_gn=MORC2;refGene_tx=XM_006938595.4;refGene_gn=MORC2;refGene_tx=XM_006938596.4
D3 22089408 22089517
refGene_gn=MORC2;refGene_tx=XM_003994803.5;refGene_gn=MORC2;refGene_tx

=XM_006938594.4;refGene_gn=MORC2;refGene_tx=XM_006938595.4;refGene_gn=MORC2;refGene_tx=XM_006938596.4
D3 22093281 22093441
refGene_gn=MORC2;refGene_tx=XM_003994803.5;refGene_gn=MORC2;refGene_tx=XM_006938594.4;refGene_gn=MORC2;refGene_tx=XM_006938595.4;refGene_gn=MORC2;refGene_tx=XM_006938596.4
D3 22093549 22093661
refGene_gn=MORC2;refGene_tx=XM_003994803.5;refGene_gn=MORC2;refGene_tx=XM_006938594.4;refGene_gn=MORC2;refGene_tx=XM_006938595.4;refGene_gn=MORC2;refGene_tx=XM_006938596.4
D3 22094179 22094305
refGene_gn=MORC2;refGene_tx=XM_003994803.5;refGene_gn=MORC2;refGene_tx=XM_006938594.4;refGene_gn=MORC2;refGene_tx=XM_006938595.4;refGene_gn=MORC2;refGene_tx=XM_006938596.4
D3 22094910 22094990
refGene_gn=MORC2;refGene_tx=XM_003994803.5;refGene_gn=MORC2;refGene_tx=XM_006938594.4;refGene_gn=MORC2;refGene_tx=XM_006938595.4;refGene_gn=MORC2;refGene_tx=XM_006938596.4
D3 22095700 22095783
refGene_gn=MORC2;refGene_tx=XM_003994803.5;refGene_gn=MORC2;refGene_tx=XM_006938594.4;refGene_gn=MORC2;refGene_tx=XM_006938595.4;refGene_gn=MORC2;refGene_tx=XM_006938596.4
D3 22096324 22096410
refGene_gn=MORC2;refGene_tx=XM_003994803.5;refGene_gn=MORC2;refGene_tx=XM_006938594.4;refGene_gn=MORC2;refGene_tx=XM_006938595.4;refGene_gn=MORC2;refGene_tx=XM_006938596.4
D3 22097788 22097929
refGene_gn=MORC2;refGene_tx=XM_003994803.5;refGene_gn=MORC2;refGene_tx=XM_006938594.4;refGene_gn=MORC2;refGene_tx=XM_006938595.4;refGene_gn=MORC2;refGene_tx=XM_006938596.4
D3 22098027 22098182
refGene_gn=MORC2;refGene_tx=XM_003994803.5;refGene_gn=MORC2;refGene_tx=XM_006938594.4;refGene_gn=MORC2;refGene_tx=XM_006938595.4;refGene_gn=MORC2;refGene_tx=XM_006938596.4
D3 22098297 22098426
refGene_gn=MORC2;refGene_tx=XM_003994803.5;refGene_gn=MORC2;refGene_tx=XM_006938594.4;refGene_gn=MORC2;refGene_tx=XM_006938595.4;refGene_gn=MORC2;refGene_tx=XM_006938596.4
D3 22098940 22099046
refGene_gn=MORC2;refGene_tx=XM_003994803.5;refGene_gn=MORC2;refGene_tx=XM_006938594.4;refGene_gn=MORC2;refGene_tx=XM_006938595.4;refGene_gn=MORC2;refGene_tx=XM_006938596.4
D3 22099572 22099705
refGene_gn=MORC2;refGene_tx=XM_003994803.5;refGene_gn=MORC2;refGene_tx=XM_006938594.4;refGene_gn=MORC2;refGene_tx=XM_006938595.4;refGene_gn=MORC2;refGene_tx=XM_006938596.4
D3 22101051 22101126
refGene_gn=MORC2;refGene_tx=XM_003994803.5;refGene_gn=MORC2;refGene_tx=XM_006938594.4;refGene_gn=MORC2;refGene_tx=XM_006938595.4;refGene_gn=MORC2;refGene_tx=XM_006938596.4

D3 22101211 22101592
refGene_gn=MORC2;refGene_tx=XM_003994803.5;refGene_gn=MORC2;refGene_tx=XM_006938594.4;refGene_gn=MORC2;refGene_tx=XM_006938595.4;refGene_gn=MORC2;refGene_tx=XM_006938596.4
D3 22102099 22102231
refGene_gn=MORC2;refGene_tx=XM_003994803.5;refGene_gn=MORC2;refGene_tx=XM_006938594.4;refGene_gn=MORC2;refGene_tx=XM_006938595.4;refGene_gn=MORC2;refGene_tx=XM_006938596.4
D3 22102677 22102732
refGene_gn=MORC2;refGene_tx=XM_003994803.5;refGene_gn=MORC2;refGene_tx=XM_006938594.4;refGene_gn=MORC2;refGene_tx=XM_006938595.4;refGene_gn=MORC2;refGene_tx=XM_006938596.4
D3 22103152 22103294
refGene_gn=MORC2;refGene_tx=XM_003994803.5;refGene_gn=MORC2;refGene_tx=XM_006938594.4;refGene_gn=MORC2;refGene_tx=XM_006938595.4;refGene_gn=MORC2;refGene_tx=XM_006938596.4
D3 22103420 22103663
refGene_gn=MORC2;refGene_tx=XM_003994803.5;refGene_gn=MORC2;refGene_tx=XM_006938594.4;refGene_gn=MORC2;refGene_tx=XM_006938595.4;refGene_gn=MORC2;refGene_tx=XM_006938596.4
D3 22103750 22103844
refGene_gn=MORC2;refGene_tx=XM_003994803.5;refGene_gn=MORC2;refGene_tx=XM_006938594.4;refGene_gn=MORC2;refGene_tx=XM_006938595.4;refGene_gn=MORC2;refGene_tx=XM_006938596.4
D3 22107299 22107488
refGene_gn=MORC2;refGene_tx=XM_003994803.5;refGene_gn=MORC2;refGene_tx=XM_006938594.4;refGene_gn=MORC2;refGene_tx=XM_006938595.4;refGene_gn=MORC2;refGene_tx=XM_006938596.4
D3 22107581 22107674
refGene_gn=MORC2;refGene_tx=XM_006938594.4;refGene_gn=MORC2;refGene_tx=XM_006938595.4
D3 22108745 22108814
refGene_gn=MORC2;refGene_tx=XM_003994803.5;refGene_gn=MORC2;refGene_tx=XM_006938594.4;refGene_gn=MORC2;refGene_tx=XM_006938595.4;refGene_gn=MORC2;refGene_tx=XM_006938596.4
D3 22124985 22125131
refGene_gn=OSBP2;refGene_tx=XM_019814768.2;refGene_gn=OSBP2;refGene_tx=XM_019814769.2;refGene_gn=OSBP2;refGene_tx=XM_023241575.1
D3 22125351 22125578
refGene_gn=OSBP2;refGene_tx=XM_019814768.2;refGene_gn=OSBP2;refGene_tx=XM_019814769.2;refGene_gn=OSBP2;refGene_tx=XM_023241575.1
D3 22134903 22135088
refGene_gn=OSBP2;refGene_tx=XM_019814768.2;refGene_gn=OSBP2;refGene_tx=XM_019814769.2;refGene_gn=OSBP2;refGene_tx=XM_023241575.1
D3 22135170 22135266
refGene_gn=OSBP2;refGene_tx=XM_019814768.2;refGene_gn=OSBP2;refGene_tx=XM_019814769.2;refGene_gn=OSBP2;refGene_tx=XM_023241575.1
D3 22135338 22135442
refGene_gn=OSBP2;refGene_tx=XM_019814768.2;refGene_gn=OSBP2;refGene_tx=XM_019814769.2;refGene_gn=OSBP2;refGene_tx=XM_023241575.1

D3 22135727 22135848
refGene_gn=OSBP2;refGene_tx=XM_019814768.2;refGene_gn=OSBP2;refGene_tx=XM_019814769.2;refGene_gn=OSBP2;refGene_tx=XM_023241575.1
D3 22138581 22138827
refGene_gn=OSBP2;refGene_tx=XM_019814768.2;refGene_gn=OSBP2;refGene_tx=XM_019814769.2;refGene_gn=OSBP2;refGene_tx=XM_023241575.1
D3 22139930 22140077
refGene_gn=OSBP2;refGene_tx=XM_019814768.2;refGene_gn=OSBP2;refGene_tx=XM_019814769.2;refGene_gn=OSBP2;refGene_tx=XM_023241575.1
D3 22140329 22140381
refGene_gn=OSBP2;refGene_tx=XM_019814768.2;refGene_gn=OSBP2;refGene_tx=XM_019814769.2;refGene_gn=OSBP2;refGene_tx=XM_023241575.1
D3 22142628 22142746
refGene_gn=OSBP2;refGene_tx=XM_019814768.2;refGene_gn=OSBP2;refGene_tx=XM_019814769.2;refGene_gn=OSBP2;refGene_tx=XM_023241575.1
D3 22143374 22143567
refGene_gn=OSBP2;refGene_tx=XM_019814768.2;refGene_gn=OSBP2;refGene_tx=XM_019814769.2;refGene_gn=OSBP2;refGene_tx=XM_023241575.1
D3 22156445 22156702
refGene_gn=OSBP2;refGene_tx=XM_019814768.2;refGene_gn=OSBP2;refGene_tx=XM_019814769.2;refGene_gn=OSBP2;refGene_tx=XM_023241575.1
D3 22206861 22206922 refGene_gn=OSBP2;refGene_tx=XM_023241575.1
D3 22269498 22269707
refGene_gn=OSBP2;refGene_tx=XM_019814768.2;refGene_gn=OSBP2;refGene_tx=XM_019814769.2
D3 22302643 22303320
refGene_gn=OSBP2;refGene_tx=XM_019814768.2;refGene_gn=OSBP2;refGene_tx=XM_019814769.2
D3 22319848 22319947
refGene_gn=CD3H5orf52;refGene_tx=XM_006938593.4
D3 22323036 22323145
refGene_gn=CD3H5orf52;refGene_tx=XM_006938593.4
D3 22325254 22325520
refGene_gn=CD3H5orf52;refGene_tx=XM_006938593.4
D3 22328784 22329351 refGene_gn=DUSP18;refGene_tx=XM_003994799.5
D3 22344462 22344896 refGene_gn=SLC35E4;refGene_tx=XM_003994798.4
D3 22349106 22349725 refGene_gn=SLC35E4;refGene_tx=XM_003994798.4
D3 22355455 22355517
refGene_gn=TCN2;refGene_tx=XM_003994797.5;refGene_gn=TCN2;refGene_tx=XM_019814772.1;refGene_gn=TCN2;refGene_tx=XM_019814773.2
D3 22357998 22358114
refGene_gn=TCN2;refGene_tx=XM_003994797.5;refGene_gn=TCN2;refGene_tx=XM_019814772.1;refGene_gn=TCN2;refGene_tx=XM_019814773.2
D3 22359879 22360045
refGene_gn=TCN2;refGene_tx=XM_003994797.5;refGene_gn=TCN2;refGene_tx=XM_019814772.1;refGene_gn=TCN2;refGene_tx=XM_019814773.2
D3 22360893 22361080
refGene_gn=TCN2;refGene_tx=XM_003994797.5;refGene_gn=TCN2;refGene_tx=XM_019814772.1;refGene_gn=TCN2;refGene_tx=XM_019814773.2
D3 22361217 22361390

refGene_gn=TCN2;refGene_tx=XM_003994797.5;refGene_gn=TCN2;refGene_tx=XM_019814772.1;refGene_gn=TCN2;refGene_tx=XM_019814773.2
D3 22362461 22362623
refGene_gn=TCN2;refGene_tx=XM_003994797.5;refGene_gn=TCN2;refGene_tx=XM_019814772.1;refGene_gn=TCN2;refGene_tx=XM_019814773.2
D3 22363661 22363831
refGene_gn=TCN2;refGene_tx=XM_003994797.5;refGene_gn=TCN2;refGene_tx=XM_019814772.1;refGene_gn=TCN2;refGene_tx=XM_019814773.2
D3 22364835 22365028
refGene_gn=TCN2;refGene_tx=XM_003994797.5;refGene_gn=TCN2;refGene_tx=XM_019814772.1;refGene_gn=TCN2;refGene_tx=XM_019814773.2
D3 22368954 22369018
refGene_gn=TCN2;refGene_tx=XM_003994797.5;refGene_gn=TCN2;refGene_tx=XM_019814772.1;refGene_gn=TCN2;refGene_tx=XM_019814773.2
D3 22406777 22406801
refGene_gn=PES1;refGene_tx=XM_003994796.5;refGene_gn=PES1;refGene_tx=XM_019814771.2
D3 22409262 22409342
refGene_gn=PES1;refGene_tx=XM_003994796.5;refGene_gn=PES1;refGene_tx=XM_019814771.2
D3 22410106 22410260
refGene_gn=PES1;refGene_tx=XM_003994796.5;refGene_gn=PES1;refGene_tx=XM_019814771.2
D3 22411427 22411537
refGene_gn=PES1;refGene_tx=XM_003994796.5;refGene_gn=PES1;refGene_tx=XM_019814771.2
D3 22413861 22414033
refGene_gn=PES1;refGene_tx=XM_003994796.5;refGene_gn=PES1;refGene_tx=XM_019814771.2
D3 22414118 22414208
refGene_gn=PES1;refGene_tx=XM_003994796.5;refGene_gn=PES1;refGene_tx=XM_019814771.2
D3 22416778 22416895
refGene_gn=PES1;refGene_tx=XM_003994796.5;refGene_gn=PES1;refGene_tx=XM_019814771.2
D3 22417007 22417082
refGene_gn=PES1;refGene_tx=XM_003994796.5;refGene_gn=PES1;refGene_tx=XM_019814771.2
D3 22417303 22417390
refGene_gn=PES1;refGene_tx=XM_003994796.5;refGene_gn=PES1;refGene_tx=XM_019814771.2
D3 22417526 22417657
refGene_gn=PES1;refGene_tx=XM_003994796.5;refGene_gn=PES1;refGene_tx=XM_019814771.2
D3 22418032 22418158
refGene_gn=PES1;refGene_tx=XM_003994796.5;refGene_gn=PES1;refGene_tx=XM_019814771.2
D3 22418277 22418462
refGene_gn=PES1;refGene_tx=XM_003994796.5;refGene_gn=PES1;refGene_tx=XM_019814771.2

D3 22418917 22419084
refGene_gn=PES1;refGene_tx=XM_003994796.5;refGene_gn=PES1;refGene_tx=XM_019814771.2
D3 22419223 22419385
refGene_gn=PES1;refGene_tx=XM_003994796.5;refGene_gn=PES1;refGene_tx=XM_019814771.2
D3 22420011 22420095
refGene_gn=PES1;refGene_tx=XM_003994796.5;refGene_gn=PES1;refGene_tx=XM_019814771.2
D3 22435626 22435757
refGene_gn=GAL3ST1;refGene_tx=XM_011287748.3;refGene_gn=GAL3ST1;refGene_tx=XM_019814774.2;refGene_gn=GAL3ST1;refGene_tx=XM_023241576.1;refGene_gn=GAL3ST1;refGene_tx=XM_023241577.1;refGene_gn=GAL3ST1;refGene_tx=XM_023241578.1
D3 22436553 22437697
refGene_gn=GAL3ST1;refGene_tx=XM_011287748.3;refGene_gn=GAL3ST1;refGene_tx=XM_019814774.2;refGene_gn=GAL3ST1;refGene_tx=XM_023241576.1;refGene_gn=GAL3ST1;refGene_tx=XM_023241577.1;refGene_gn=GAL3ST1;refGene_tx=XM_023241578.1
D3 22446674 22446718
refGene_gn=LOC101095136;refGene_tx=XM_023242156.1
D3 22450129 22450235
refGene_gn=LOC101095136;refGene_tx=XM_023242156.1
D3 22450311 22450451
refGene_gn=LOC101095136;refGene_tx=XM_023242156.1
D3 22450542 22450562
refGene_gn=LOC101095136;refGene_tx=XM_023242156.1
D3 22450564 22450711
refGene_gn=LOC101095136;refGene_tx=XM_023242156.1
D3 22450951 22451064
refGene_gn=LOC101095136;refGene_tx=XM_023242156.1
D3 22455686 22455752
refGene_gn=LOC101095136;refGene_tx=XM_023242156.1
D3 22467425 22467521
refGene_gn=LOC101094885;refGene_tx=XM_019814777.2;refGene_gn=LOC101094885;refGene_tx=XM_019814778.2
D3 22467641 22467772
refGene_gn=LOC101094885;refGene_tx=XM_003994889.5;refGene_gn=LOC101094885;refGene_tx=XM_019814777.2
D3 22470436 22470512
refGene_gn=LOC101094885;refGene_tx=XM_003994889.5;refGene_gn=LOC101094885;refGene_tx=XM_019814777.2;refGene_gn=LOC101094885;refGene_tx=XM_019814778.2;refGene_gn=LOC101094885;refGene_tx=XM_019814779.2;refGene_gn=LOC101094885;refGene_tx=XM_019814780.2
D3 22475837 22475881
refGene_gn=LOC101094885;refGene_tx=XM_003994889.5;refGene_gn=LOC101094885;refGene_tx=XM_019814777.2;refGene_gn=LOC101094885;refGene_tx=XM_019814778.2;refGene_gn=LOC101094885;refGene_tx=XM_019814779.2;refGene_gn=LOC101094885;refGene_tx=XM_019814780.2
D3 22476138 22476253

refGene_gn=LOC101094885;refGene_tx=XM_003994889.5;refGene_gn=LOC101094885;refGene_tx=XM_019814777.2;refGene_gn=LOC101094885;refGene_tx=XM_019814778.2;refGene_gn=LOC101094885;refGene_tx=XM_019814779.2;refGene_gn=LOC101094885;refGene_tx=XM_019814780.2
D3 22476375 22476564
refGene_gn=LOC101094885;refGene_tx=XM_003994889.5;refGene_gn=LOC101094885;refGene_tx=XM_019814777.2;refGene_gn=LOC101094885;refGene_tx=XM_019814778.2;refGene_gn=LOC101094885;refGene_tx=XM_019814779.2;refGene_gn=LOC101094885;refGene_tx=XM_019814780.2
D3 22478003 22478099
refGene_gn=LOC101094885;refGene_tx=XM_003994889.5;refGene_gn=LOC101094885;refGene_tx=XM_019814777.2;refGene_gn=LOC101094885;refGene_tx=XM_019814778.2;refGene_gn=LOC101094885;refGene_tx=XM_019814779.2;refGene_gn=LOC101094885;refGene_tx=XM_019814780.2
D3 22478398 22478459
refGene_gn=LOC101094885;refGene_tx=XM_003994889.5;refGene_gn=LOC101094885;refGene_tx=XM_019814777.2;refGene_gn=LOC101094885;refGene_tx=XM_019814778.2;refGene_gn=LOC101094885;refGene_tx=XM_019814779.2;refGene_gn=LOC101094885;refGene_tx=XM_019814780.2
D3 22479425 22479509
refGene_gn=LOC101094885;refGene_tx=XM_003994889.5;refGene_gn=LOC101094885;refGene_tx=XM_019814777.2;refGene_gn=LOC101094885;refGene_tx=XM_019814778.2;refGene_gn=LOC101094885;refGene_tx=XM_019814779.2;refGene_gn=LOC101094885;refGene_tx=XM_019814780.2
D3 22479841 22479948
refGene_gn=LOC101094885;refGene_tx=XM_003994889.5;refGene_gn=LOC101094885;refGene_tx=XM_019814777.2;refGene_gn=LOC101094885;refGene_tx=XM_019814778.2;refGene_gn=LOC101094885;refGene_tx=XM_019814779.2;refGene_gn=LOC101094885;refGene_tx=XM_019814780.2
D3 22480024 22480164
refGene_gn=LOC101094885;refGene_tx=XM_003994889.5;refGene_gn=LOC101094885;refGene_tx=XM_019814777.2;refGene_gn=LOC101094885;refGene_tx=XM_019814778.2;refGene_gn=LOC101094885;refGene_tx=XM_019814779.2;refGene_gn=LOC101094885;refGene_tx=XM_019814780.2
D3 22480256 22480426
refGene_gn=LOC101094885;refGene_tx=XM_003994889.5;refGene_gn=LOC101094885;refGene_tx=XM_019814777.2;refGene_gn=LOC101094885;refGene_tx=XM_019814778.2;refGene_gn=LOC101094885;refGene_tx=XM_019814779.2;refGene_gn=LOC101094885;refGene_tx=XM_019814780.2
D3 22481553 22481693
refGene_gn=LOC101094885;refGene_tx=XM_003994889.5;refGene_gn=LOC101094885;refGene_tx=XM_019814777.2;refGene_gn=LOC101094885;refGene_tx=XM_019814778.2;refGene_gn=LOC101094885;refGene_tx=XM_019814779.2;refGene_gn=LOC101094885;refGene_tx=XM_019814780.2
D3 22492655 22492709
refGene_gn=LOC101092402;refGene_tx=XM_003994793.3
D3 22493691 22493767
refGene_gn=LOC101092402;refGene_tx=XM_003994793.3
D3 22494043 22494087
refGene_gn=LOC101092402;refGene_tx=XM_003994793.3

D3 22494208 22494268
refGene_gn=LOC101092402;refGene_tx=XM_003994793.3
D3 22495287 22495476
refGene_gn=LOC101092402;refGene_tx=XM_003994793.3
D3 22496417 22496513
refGene_gn=LOC101092402;refGene_tx=XM_003994793.3
D3 22497170 22497231
refGene_gn=LOC101092402;refGene_tx=XM_003994793.3
D3 22498400 22498484
refGene_gn=LOC101092402;refGene_tx=XM_003994793.3
D3 22499360 22499467
refGene_gn=LOC101092402;refGene_tx=XM_003994793.3
D3 22499857 22499997
refGene_gn=LOC101092402;refGene_tx=XM_003994793.3
D3 22500093 22500263
refGene_gn=LOC101092402;refGene_tx=XM_003994793.3
D3 22501099 22501221
refGene_gn=LOC101092402;refGene_tx=XM_003994793.3
D3 22518387 22518502
refGene_gn=MTFP1;refGene_tx=XM_003994792.5;refGene_gn=MTFP1;refGene_tx
=XM_006938590.4
D3 22519477 22519710
refGene_gn=MTFP1;refGene_tx=XM_003994792.5;refGene_gn=MTFP1;refGene_tx
=XM_006938590.4
D3 22520013 22520141
refGene_gn=MTFP1;refGene_tx=XM_003994792.5;refGene_gn=MTFP1;refGene_tx
=XM_006938590.4
D3 22520883 22520950
refGene_gn=MTFP1;refGene_tx=XM_003994792.5;refGene_gn=MTFP1;refGene_tx
=XM_006938590.4
D3 22524203 22524334
refGene_gn=SEC14L2;refGene_tx=XM_003994790.5;refGene_gn=SEC14L2;refGen
e_tx=XM_006938587.4;refGene_gn=SEC14L2;refGene_tx=XM_006938588.4;refGe
ne_gn=SEC14L2;refGene_tx=XM_006938589.4
D3 22529951 22530121
refGene_gn=SEC14L2;refGene_tx=XM_003994790.5;refGene_gn=SEC14L2;refGen
e_tx=XM_006938587.4;refGene_gn=SEC14L2;refGene_tx=XM_006938588.4;refGe
ne_gn=SEC14L2;refGene_tx=XM_006938589.4
D3 22530260 22530400
refGene_gn=SEC14L2;refGene_tx=XM_003994790.5;refGene_gn=SEC14L2;refGen
e_tx=XM_006938587.4;refGene_gn=SEC14L2;refGene_tx=XM_006938588.4;refGe
ne_gn=SEC14L2;refGene_tx=XM_006938589.4
D3 22530505 22530612
refGene_gn=SEC14L2;refGene_tx=XM_003994790.5;refGene_gn=SEC14L2;refGen
e_tx=XM_006938587.4;refGene_gn=SEC14L2;refGene_tx=XM_006938588.4;refGe
ne_gn=SEC14L2;refGene_tx=XM_006938589.4
D3 22535079 22535163
refGene_gn=SEC14L2;refGene_tx=XM_003994790.5;refGene_gn=SEC14L2;refGen
e_tx=XM_006938587.4;refGene_gn=SEC14L2;refGene_tx=XM_006938588.4;refGe
ne_gn=SEC14L2;refGene_tx=XM_006938589.4

D3 22535793 22535854
refGene_gn=SEC14L2;refGene_tx=XM_003994790.5;refGene_gn=SEC14L2;refGene_tx=XM_006938587.4;refGene_gn=SEC14L2;refGene_tx=XM_006938588.4;refGene_gn=SEC14L2;refGene_tx=XM_006938589.4
D3 22536040 22536136
refGene_gn=SEC14L2;refGene_tx=XM_003994790.5;refGene_gn=SEC14L2;refGene_tx=XM_006938587.4;refGene_gn=SEC14L2;refGene_tx=XM_006938588.4;refGene_gn=SEC14L2;refGene_tx=XM_006938589.4
D3 22537132 22537321
refGene_gn=SEC14L2;refGene_tx=XM_003994790.5;refGene_gn=SEC14L2;refGene_tx=XM_006938587.4;refGene_gn=SEC14L2;refGene_tx=XM_006938588.4;refGene_gn=SEC14L2;refGene_tx=XM_006938589.4
D3 22537588 22537648
refGene_gn=SEC14L2;refGene_tx=XM_003994790.5;refGene_gn=SEC14L2;refGene_tx=XM_006938587.4;refGene_gn=SEC14L2;refGene_tx=XM_006938588.4;refGene_gn=SEC14L2;refGene_tx=XM_006938589.4
D3 22538050 22538094
refGene_gn=SEC14L2;refGene_tx=XM_003994790.5;refGene_gn=SEC14L2;refGene_tx=XM_006938587.4;refGene_gn=SEC14L2;refGene_tx=XM_006938588.4;refGene_gn=SEC14L2;refGene_tx=XM_006938589.4
D3 22542091 22542140 refGene_gn=SEC14L2;refGene_tx=XM_006938589.4
D3 22542451 22542527 refGene_gn=SEC14L2;refGene_tx=XM_003994790.5
D3 22544653 22544707 refGene_gn=SEC14L2;refGene_tx=XM_003994790.5
D3 22559498 22559783 refGene_gn=RNF215;refGene_tx=XM_003994789.4
D3 22560017 22560161 refGene_gn=RNF215;refGene_tx=XM_003994789.4
D3 22560637 22560709 refGene_gn=RNF215;refGene_tx=XM_003994789.4
D3 22560789 22560875 refGene_gn=RNF215;refGene_tx=XM_003994789.4
D3 22561333 22561490 refGene_gn=RNF215;refGene_tx=XM_003994789.4
D3 22564907 22565027 refGene_gn=RNF215;refGene_tx=XM_003994789.4
D3 22565106 22565250 refGene_gn=RNF215;refGene_tx=XM_003994789.4
D3 22565497 22565600 refGene_gn=RNF215;refGene_tx=XM_003994789.4
D3 22565688 22565711 refGene_gn=RNF215;refGene_tx=XM_003994789.4
D3 22568180 22568483
refGene_gn=CCDC157;refGene_tx=XM_019814781.2;refGene_gn=CCDC157;refGene_tx=XM_019814782.2;refGene_gn=CCDC157;refGene_tx=XM_019814783.2;refGene_gn=CCDC157;refGene_tx=XM_019814784.2;refGene_gn=CCDC157;refGene_tx=XM_019814785.2;refGene_gn=CCDC157;refGene_tx=XM_019814786.2;refGene_gn=CCDC157;refGene_tx=XM_019814787.2;refGene_gn=CCDC157;refGene_tx=XM_019814788.2;refGene_gn=CCDC157;refGene_tx=XM_019814789.2;refGene_gn=CCDC157;refGene_tx=XM_019814790.2;refGene_gn=CCDC157;refGene_tx=XM_019814791.2;refGene_gn=CCDC157;refGene_tx=XM_019814792.2;refGene_gn=CCDC157;refGene_tx=XM_019814793.2;refGene_gn=CCDC157;refGene_tx=XM_019814794.2;refGene_gn=CCDC157;refGene_tx=XM_019814795.2
D3 22568570 22568659
refGene_gn=CCDC157;refGene_tx=XM_019814781.2;refGene_gn=CCDC157;refGene_tx=XM_019814782.2;refGene_gn=CCDC157;refGene_tx=XM_019814783.2;refGene_gn=CCDC157;refGene_tx=XM_019814784.2;refGene_gn=CCDC157;refGene_tx=XM_019814785.2;refGene_gn=CCDC157;refGene_tx=XM_019814786.2;refGene_gn=CCDC157;refGene_tx=XM_019814787.2;refGene_gn=CCDC157;refGene_tx=XM_019814788.2;refGene_gn=CCDC157;refGene_tx=XM_019814789.2;refGene_gn=CCDC

157;refGene_tx=XM_019814790.2;refGene_gn=CCDC157;refGene_tx=XM_019814791.2;refGene_gn=CCDC157;refGene_tx=XM_019814792.2;refGene_gn=CCDC157;refGene_tx=XM_019814793.2;refGene_gn=CCDC157;refGene_tx=XM_019814794.2;refGene_gn=CCDC157;refGene_tx=XM_019814795.2
D3 22569245 22569440
refGene_gn=CCDC157;refGene_tx=XM_019814781.2;refGene_gn=CCDC157;refGene_tx=XM_019814782.2;refGene_gn=CCDC157;refGene_tx=XM_019814783.2;refGene_gn=CCDC157;refGene_tx=XM_019814784.2;refGene_gn=CCDC157;refGene_tx=XM_019814785.2;refGene_gn=CCDC157;refGene_tx=XM_019814786.2;refGene_gn=CCDC157;refGene_tx=XM_019814787.2;refGene_gn=CCDC157;refGene_tx=XM_019814788.2;refGene_gn=CCDC157;refGene_tx=XM_019814789.2;refGene_gn=CCDC157;refGene_tx=XM_019814790.2;refGene_gn=CCDC157;refGene_tx=XM_019814791.2;refGene_gn=CCDC157;refGene_tx=XM_019814792.2;refGene_gn=CCDC157;refGene_tx=XM_019814793.2;refGene_gn=CCDC157;refGene_tx=XM_019814794.2;refGene_gn=CCDC157;refGene_tx=XM_019814795.2
D3 22570510 22570679
refGene_gn=CCDC157;refGene_tx=XM_019814781.2;refGene_gn=CCDC157;refGene_tx=XM_019814782.2;refGene_gn=CCDC157;refGene_tx=XM_019814783.2;refGene_gn=CCDC157;refGene_tx=XM_019814784.2;refGene_gn=CCDC157;refGene_tx=XM_019814785.2;refGene_gn=CCDC157;refGene_tx=XM_019814786.2;refGene_gn=CCDC157;refGene_tx=XM_019814787.2;refGene_gn=CCDC157;refGene_tx=XM_019814788.2;refGene_gn=CCDC157;refGene_tx=XM_019814789.2;refGene_gn=CCDC157;refGene_tx=XM_019814790.2;refGene_gn=CCDC157;refGene_tx=XM_019814791.2;refGene_gn=CCDC157;refGene_tx=XM_019814792.2;refGene_gn=CCDC157;refGene_tx=XM_019814793.2;refGene_gn=CCDC157;refGene_tx=XM_019814794.2;refGene_gn=CCDC157;refGene_tx=XM_019814795.2
D3 22570845 22571022
refGene_gn=CCDC157;refGene_tx=XM_019814781.2;refGene_gn=CCDC157;refGene_tx=XM_019814782.2;refGene_gn=CCDC157;refGene_tx=XM_019814783.2;refGene_gn=CCDC157;refGene_tx=XM_019814784.2;refGene_gn=CCDC157;refGene_tx=XM_019814785.2;refGene_gn=CCDC157;refGene_tx=XM_019814786.2;refGene_gn=CCDC157;refGene_tx=XM_019814787.2;refGene_gn=CCDC157;refGene_tx=XM_019814788.2;refGene_gn=CCDC157;refGene_tx=XM_019814789.2;refGene_gn=CCDC157;refGene_tx=XM_019814790.2;refGene_gn=CCDC157;refGene_tx=XM_019814791.2;refGene_gn=CCDC157;refGene_tx=XM_019814792.2;refGene_gn=CCDC157;refGene_tx=XM_019814793.2;refGene_gn=CCDC157;refGene_tx=XM_019814794.2;refGene_gn=CCDC157;refGene_tx=XM_019814795.2
D3 22572305 22572496
refGene_gn=CCDC157;refGene_tx=XM_019814781.2;refGene_gn=CCDC157;refGene_tx=XM_019814782.2;refGene_gn=CCDC157;refGene_tx=XM_019814783.2;refGene_gn=CCDC157;refGene_tx=XM_019814784.2;refGene_gn=CCDC157;refGene_tx=XM_019814785.2;refGene_gn=CCDC157;refGene_tx=XM_019814786.2;refGene_gn=CCDC157;refGene_tx=XM_019814787.2;refGene_gn=CCDC157;refGene_tx=XM_019814788.2;refGene_gn=CCDC157;refGene_tx=XM_019814789.2;refGene_gn=CCDC157;refGene_tx=XM_019814790.2;refGene_gn=CCDC157;refGene_tx=XM_019814791.2;refGene_gn=CCDC157;refGene_tx=XM_019814792.2;refGene_gn=CCDC157;refGene_tx=XM_019814793.2;refGene_gn=CCDC157;refGene_tx=XM_019814794.2;refGene_gn=CCDC157;refGene_tx=XM_019814795.2
D3 22572870 22572948
refGene_gn=CCDC157;refGene_tx=XM_019814781.2;refGene_gn=CCDC157;refGen

e_tx=XM_019814782.2;refGene_gn=CCDC157;refGene_tx=XM_019814783.2;refGene_gn=CCDC157;refGene_tx=XM_019814784.2;refGene_gn=CCDC157;refGene_tx=XM_019814785.2;refGene_gn=CCDC157;refGene_tx=XM_019814786.2;refGene_gn=CCDC157;refGene_tx=XM_019814787.2;refGene_gn=CCDC157;refGene_tx=XM_019814788.2;refGene_gn=CCDC157;refGene_tx=XM_019814789.2;refGene_gn=CCDC157;refGene_tx=XM_019814790.2;refGene_gn=CCDC157;refGene_tx=XM_019814791.2;refGene_gn=CCDC157;refGene_tx=XM_019814792.2;refGene_gn=CCDC157;refGene_tx=XM_019814793.2;refGene_gn=CCDC157;refGene_tx=XM_019814794.2;refGene_gn=CCDC157;refGene_tx=XM_019814795.2
D3 22573661 22574307
refGene_gn=CCDC157;refGene_tx=XM_019814781.2;refGene_gn=CCDC157;refGene_tx=XM_019814782.2;refGene_gn=CCDC157;refGene_tx=XM_019814783.2;refGene_gn=CCDC157;refGene_tx=XM_019814784.2;refGene_gn=CCDC157;refGene_tx=XM_019814785.2;refGene_gn=CCDC157;refGene_tx=XM_019814786.2;refGene_gn=CCDC157;refGene_tx=XM_019814787.2;refGene_gn=CCDC157;refGene_tx=XM_019814788.2;refGene_gn=CCDC157;refGene_tx=XM_019814789.2;refGene_gn=CCDC157;refGene_tx=XM_019814790.2;refGene_gn=CCDC157;refGene_tx=XM_019814791.2;refGene_gn=CCDC157;refGene_tx=XM_019814792.2;refGene_gn=CCDC157;refGene_tx=XM_019814793.2;refGene_gn=CCDC157;refGene_tx=XM_019814794.2;refGene_gn=CCDC157;refGene_tx=XM_019814795.2
D3 22575065 22575237
refGene_gn=CCDC157;refGene_tx=XM_019814781.2;refGene_gn=CCDC157;refGene_tx=XM_019814782.2;refGene_gn=CCDC157;refGene_tx=XM_019814783.2;refGene_gn=CCDC157;refGene_tx=XM_019814784.2;refGene_gn=CCDC157;refGene_tx=XM_019814785.2;refGene_gn=CCDC157;refGene_tx=XM_019814786.2;refGene_gn=CCDC157;refGene_tx=XM_019814787.2;refGene_gn=CCDC157;refGene_tx=XM_019814788.2;refGene_gn=CCDC157;refGene_tx=XM_019814789.2;refGene_gn=CCDC157;refGene_tx=XM_019814790.2;refGene_gn=CCDC157;refGene_tx=XM_019814791.2;refGene_gn=CCDC157;refGene_tx=XM_019814792.2;refGene_gn=CCDC157;refGene_tx=XM_019814793.2;refGene_gn=CCDC157;refGene_tx=XM_019814794.2;refGene_gn=CCDC157;refGene_tx=XM_019814795.2
D3 22576995 22577243
refGene_gn=CCDC157;refGene_tx=XM_019814781.2;refGene_gn=CCDC157;refGene_tx=XM_019814782.2;refGene_gn=CCDC157;refGene_tx=XM_019814783.2;refGene_gn=CCDC157;refGene_tx=XM_019814784.2;refGene_gn=CCDC157;refGene_tx=XM_019814785.2;refGene_gn=CCDC157;refGene_tx=XM_019814786.2;refGene_gn=CCDC157;refGene_tx=XM_019814787.2;refGene_gn=CCDC157;refGene_tx=XM_019814788.2;refGene_gn=CCDC157;refGene_tx=XM_019814789.2;refGene_gn=CCDC157;refGene_tx=XM_019814790.2;refGene_gn=CCDC157;refGene_tx=XM_019814791.2;refGene_gn=CCDC157;refGene_tx=XM_019814792.2;refGene_gn=CCDC157;refGene_tx=XM_019814793.2;refGene_gn=CCDC157;refGene_tx=XM_019814794.2;refGene_gn=CCDC157;refGene_tx=XM_019814795.2
D3 22582650 22582713 refGene_gn=SF3A1;refGene_tx=XM_003994786.5
D3 22585345 22585467 refGene_gn=SF3A1;refGene_tx=XM_003994786.5
D3 22589667 22589875 refGene_gn=SF3A1;refGene_tx=XM_003994786.5
D3 22591001 22591259 refGene_gn=SF3A1;refGene_tx=XM_003994786.5
D3 22593772 22593847 refGene_gn=SF3A1;refGene_tx=XM_003994786.5
D3 22594799 22594950 refGene_gn=SF3A1;refGene_tx=XM_003994786.5
D3 22595305 22595499 refGene_gn=SF3A1;refGene_tx=XM_003994786.5
D3 22596385 22596503 refGene_gn=SF3A1;refGene_tx=XM_003994786.5

D3 22596801 22596987 refGene_gn=SF3A1;refGene_tx=XM_003994786.5
D3 22597628 22597750 refGene_gn=SF3A1;refGene_tx=XM_003994786.5
D3 22597853 22598099 refGene_gn=SF3A1;refGene_tx=XM_003994786.5
D3 22598652 22598860 refGene_gn=SF3A1;refGene_tx=XM_003994786.5
D3 22599371 22599526 refGene_gn=SF3A1;refGene_tx=XM_003994786.5
D3 22600760 22600862 refGene_gn=SF3A1;refGene_tx=XM_003994786.5
D3 22600975 22601047 refGene_gn=SF3A1;refGene_tx=XM_003994786.5
D3 22601839 22601941 refGene_gn=SF3A1;refGene_tx=XM_003994786.5
D3 22609215 22609424 refGene_gn=TBC1D10A;refGene_tx=XM_023241579.1
D3 22628735 22628835 refGene_gn=TBC1D10A;refGene_tx=XM_023241579.1
D3 22632854 22632962 refGene_gn=TBC1D10A;refGene_tx=XM_023241579.1
D3 22636451 22636558 refGene_gn=TBC1D10A;refGene_tx=XM_023241579.1
D3 22637235 22637350 refGene_gn=TBC1D10A;refGene_tx=XM_023241579.1
D3 22637429 22637495 refGene_gn=TBC1D10A;refGene_tx=XM_023241579.1
D3 22638181 22638371 refGene_gn=TBC1D10A;refGene_tx=XM_023241579.1
D3 22638477 22638632 refGene_gn=TBC1D10A;refGene_tx=XM_023241579.1
D3 22639231 22639738 refGene_gn=TBC1D10A;refGene_tx=XM_023241579.1
D3 22642600 22642713
refGene_gn=CASTOR1;refGene_tx=XM_003994785.5;refGene_gn=CASTOR1;refGene_tx=XM_011287745.3
D3 22642986 22643126
refGene_gn=CASTOR1;refGene_tx=XM_011287743.3;refGene_gn=CASTOR1;refGene_tx=XM_011287744.3;refGene_gn=CASTOR1;refGene_tx=XM_011287746.2;refGene_gn=CASTOR1;refGene_tx=XM_019814798.2
D3 22643344 22643415
refGene_gn=CASTOR1;refGene_tx=XM_003994785.5;refGene_gn=CASTOR1;refGene_tx=XM_011287743.3;refGene_gn=CASTOR1;refGene_tx=XM_011287744.3;refGene_gn=CASTOR1;refGene_tx=XM_011287745.3;refGene_gn=CASTOR1;refGene_tx=XM_011287746.2;refGene_gn=CASTOR1;refGene_tx=XM_019814798.2;refGene_gn=CASTOR1;refGene_tx=XM_019814799.2
D3 22644597 22644785
refGene_gn=CASTOR1;refGene_tx=XM_003994785.5;refGene_gn=CASTOR1;refGene_tx=XM_011287743.3;refGene_gn=CASTOR1;refGene_tx=XM_011287744.3;refGene_gn=CASTOR1;refGene_tx=XM_011287745.3;refGene_gn=CASTOR1;refGene_tx=XM_011287746.2;refGene_gn=CASTOR1;refGene_tx=XM_019814798.2;refGene_gn=CASTOR1;refGene_tx=XM_019814799.2
D3 22644870 22645003
refGene_gn=CASTOR1;refGene_tx=XM_003994785.5;refGene_gn=CASTOR1;refGene_tx=XM_011287743.3;refGene_gn=CASTOR1;refGene_tx=XM_011287744.3;refGene_gn=CASTOR1;refGene_tx=XM_011287745.3;refGene_gn=CASTOR1;refGene_tx=XM_011287746.2;refGene_gn=CASTOR1;refGene_tx=XM_019814798.2;refGene_gn=CASTOR1;refGene_tx=XM_019814799.2
D3 22645260 22645387
refGene_gn=CASTOR1;refGene_tx=XM_003994785.5;refGene_gn=CASTOR1;refGene_tx=XM_011287743.3;refGene_gn=CASTOR1;refGene_tx=XM_011287744.3;refGene_gn=CASTOR1;refGene_tx=XM_011287745.3;refGene_gn=CASTOR1;refGene_tx=XM_011287746.2;refGene_gn=CASTOR1;refGene_tx=XM_019814798.2;refGene_gn=CASTOR1;refGene_tx=XM_019814799.2
D3 22645469 22645695 refGene_gn=CASTOR1;refGene_tx=XM_019814798.2
D3 22645808 22645922

refGene_gn=CASTOR1;refGene_tx=XM_003994785.5;refGene_gn=CASTOR1;refGene_tx=XM_011287743.3;refGene_gn=CASTOR1;refGene_tx=XM_011287744.3;refGene_gn=CASTOR1;refGene_tx=XM_011287745.3;refGene_gn=CASTOR1;refGene_tx=XM_019814799.2
D3 22646095 22646178
refGene_gn=CASTOR1;refGene_tx=XM_003994785.5;refGene_gn=CASTOR1;refGene_tx=XM_011287743.3;refGene_gn=CASTOR1;refGene_tx=XM_011287744.3;refGene_gn=CASTOR1;refGene_tx=XM_011287745.3;refGene_gn=CASTOR1;refGene_tx=XM_011287746.2;refGene_gn=CASTOR1;refGene_tx=XM_019814799.2
D3 22646276 22646371
refGene_gn=CASTOR1;refGene_tx=XM_003994785.5;refGene_gn=CASTOR1;refGene_tx=XM_011287743.3;refGene_gn=CASTOR1;refGene_tx=XM_011287744.3;refGene_gn=CASTOR1;refGene_tx=XM_011287745.3;refGene_gn=CASTOR1;refGene_tx=XM_011287746.2;refGene_gn=CASTOR1;refGene_tx=XM_019814799.2
D3 22646501 22646570
refGene_gn=CASTOR1;refGene_tx=XM_003994785.5;refGene_gn=CASTOR1;refGene_tx=XM_011287743.3;refGene_gn=CASTOR1;refGene_tx=XM_011287744.3;refGene_gn=CASTOR1;refGene_tx=XM_011287745.3;refGene_gn=CASTOR1;refGene_tx=XM_011287746.2;refGene_gn=CASTOR1;refGene_tx=XM_019814799.2
D3 22660402 22660436 refGene_gn=OSM;refGene_tx=XM_006938582.3
D3 22661461 22661483 refGene_gn=OSM;refGene_tx=XM_023241580.1
D3 22661966 22662100
refGene_gn=OSM;refGene_tx=XM_006938582.3;refGene_gn=OSM;refGene_tx=XM_023241580.1
D3 22662751 22663300
refGene_gn=OSM;refGene_tx=XM_006938582.3;refGene_gn=OSM;refGene_tx=XM_023241580.1
D3 22678828 22678847
refGene_gn=LIF;refGene_tx=XM_003994784.5;refGene_gn=LIF;refGene_tx=XM_006938581.4
D3 22680639 22680818
refGene_gn=LIF;refGene_tx=XM_003994784.5;refGene_gn=LIF;refGene_tx=XM_006938581.4
D3 22681551 22681962
refGene_gn=LIF;refGene_tx=XM_003994784.5;refGene_gn=LIF;refGene_tx=XM_006938581.4
D3 22745987 22746092
refGene_gn=HORMAD2;refGene_tx=XM_011287741.2;refGene_gn=HORMAD2;refGene_tx=XM_023241581.1;refGene_gn=HORMAD2;refGene_tx=XM_023241582.1;refGene_gn=HORMAD2;refGene_tx=XM_023241583.1;refGene_gn=HORMAD2;refGene_tx=XM_023241584.1
D3 22799144 22799398
refGene_gn=HORMAD2;refGene_tx=XM_011287741.2;refGene_gn=HORMAD2;refGene_tx=XM_023241581.1;refGene_gn=HORMAD2;refGene_tx=XM_023241582.1;refGene_gn=HORMAD2;refGene_tx=XM_023241583.1;refGene_gn=HORMAD2;refGene_tx=XM_023241584.1
D3 22799793 22799945
refGene_gn=HORMAD2;refGene_tx=XM_011287741.2;refGene_gn=HORMAD2;refGene_tx=XM_023241581.1;refGene_gn=HORMAD2;refGene_tx=XM_023241582.1;refGene_gn=HORMAD2;refGene_tx=XM_023241583.1;refGene_gn=HORMAD2;refGene_tx=

XM_023241584.1
D3 22803526 22803594
refGene_gn=HORMAD2;refGene_tx=XM_011287741.2;refGene_gn=HORMAD2;refGene_tx=XM_023241581.1;refGene_gn=HORMAD2;refGene_tx=XM_023241582.1;refGene_gn=HORMAD2;refGene_tx=XM_023241583.1;refGene_gn=HORMAD2;refGene_tx=XM_023241584.1
D3 22810595 22810622
refGene_gn=HORMAD2;refGene_tx=XM_011287741.2;refGene_gn=HORMAD2;refGene_tx=XM_023241581.1;refGene_gn=HORMAD2;refGene_tx=XM_023241582.1;refGene_gn=HORMAD2;refGene_tx=XM_023241583.1;refGene_gn=HORMAD2;refGene_tx=XM_023241584.1
D3 22811338 22811359
refGene_gn=HORMAD2;refGene_tx=XM_011287741.2;refGene_gn=HORMAD2;refGene_tx=XM_023241581.1;refGene_gn=HORMAD2;refGene_tx=XM_023241582.1;refGene_gn=HORMAD2;refGene_tx=XM_023241584.1
D3 22815906 22815943
refGene_gn=HORMAD2;refGene_tx=XM_011287741.2;refGene_gn=HORMAD2;refGene_tx=XM_023241581.1;refGene_gn=HORMAD2;refGene_tx=XM_023241582.1;refGene_gn=HORMAD2;refGene_tx=XM_023241583.1;refGene_gn=HORMAD2;refGene_tx=XM_023241584.1
D3 22816607 22816671
refGene_gn=HORMAD2;refGene_tx=XM_011287741.2;refGene_gn=HORMAD2;refGene_tx=XM_023241581.1;refGene_gn=HORMAD2;refGene_tx=XM_023241582.1;refGene_gn=HORMAD2;refGene_tx=XM_023241583.1;refGene_gn=HORMAD2;refGene_tx=XM_023241584.1
D3 22819871 22820013
refGene_gn=HORMAD2;refGene_tx=XM_011287741.2;refGene_gn=HORMAD2;refGene_tx=XM_023241581.1;refGene_gn=HORMAD2;refGene_tx=XM_023241582.1;refGene_gn=HORMAD2;refGene_tx=XM_023241583.1;refGene_gn=HORMAD2;refGene_tx=XM_023241584.1
D3 22825774 22825825
refGene_gn=HORMAD2;refGene_tx=XM_011287741.2;refGene_gn=HORMAD2;refGene_tx=XM_023241581.1;refGene_gn=HORMAD2;refGene_tx=XM_023241582.1;refGene_gn=HORMAD2;refGene_tx=XM_023241583.1;refGene_gn=HORMAD2;refGene_tx=XM_023241584.1
D3 22902143 22902315
refGene_gn=MTMR3;refGene_tx=XM_003994780.4;refGene_gn=MTMR3;refGene_tx=XM_003994781.5;refGene_gn=MTMR3;refGene_tx=XM_003994782.5;refGene_gn=MTMR3;refGene_tx=XM_006938574.3;refGene_gn=MTMR3;refGene_tx=XM_006938575.4;refGene_gn=MTMR3;refGene_tx=XM_006938576.3;refGene_gn=MTMR3;refGene_tx=XM_006938577.3;refGene_gn=MTMR3;refGene_tx=XM_006938578.3;refGene_gn=MTMR3;refGene_tx=XM_006938579.3;refGene_gn=MTMR3;refGene_tx=XM_019814803.2;refGene_gn=MTMR3;refGene_tx=XM_019814804.2
D3 22904207 22904234
refGene_gn=MTMR3;refGene_tx=XM_003994782.5;refGene_gn=MTMR3;refGene_tx=XM_006938574.3;refGene_gn=MTMR3;refGene_tx=XM_006938575.4;refGene_gn=MTMR3;refGene_tx=XM_006938576.3;refGene_gn=MTMR3;refGene_tx=XM_006938578.3;refGene_gn=MTMR3;refGene_tx=XM_006938579.3;refGene_gn=MTMR3;refGene_tx=XM_019814804.2
D3 22905012 22905101

refGene_gn=MTMR3;refGene_tx=XM_003994780.4;refGene_gn=MTMR3;refGene_tx=XM_003994781.5;refGene_gn=MTMR3;refGene_tx=XM_003994782.5;refGene_gn=MTMR3;refGene_tx=XM_006938574.3;refGene_gn=MTMR3;refGene_tx=XM_006938575.4;refGene_gn=MTMR3;refGene_tx=XM_006938576.3;refGene_gn=MTMR3;refGene_tx=XM_006938577.3;refGene_gn=MTMR3;refGene_tx=XM_006938578.3;refGene_gn=MTMR3;refGene_tx=XM_006938579.3;refGene_gn=MTMR3;refGene_tx=XM_019814803.2;refGene_gn=MTMR3;refGene_tx=XM_019814804.2
D3 22905573 22905684
refGene_gn=MTMR3;refGene_tx=XM_003994780.4;refGene_gn=MTMR3;refGene_tx=XM_006938574.3;refGene_gn=MTMR3;refGene_tx=XM_006938575.4;refGene_gn=MTMR3;refGene_tx=XM_006938576.3;refGene_gn=MTMR3;refGene_tx=XM_006938577.3;refGene_gn=MTMR3;refGene_tx=XM_006938578.3;refGene_gn=MTMR3;refGene_tx=XM_019814804.2
D3 22906846 22908263
refGene_gn=MTMR3;refGene_tx=XM_003994780.4;refGene_gn=MTMR3;refGene_tx=XM_003994781.5;refGene_gn=MTMR3;refGene_tx=XM_003994782.5;refGene_gn=MTMR3;refGene_tx=XM_006938574.3;refGene_gn=MTMR3;refGene_tx=XM_006938575.4;refGene_gn=MTMR3;refGene_tx=XM_006938576.3;refGene_gn=MTMR3;refGene_tx=XM_006938577.3;refGene_gn=MTMR3;refGene_tx=XM_006938578.3;refGene_gn=MTMR3;refGene_tx=XM_006938579.3;refGene_gn=MTMR3;refGene_tx=XM_019814803.2;refGene_gn=MTMR3;refGene_tx=XM_019814804.2
D3 22909275 22909421
refGene_gn=MTMR3;refGene_tx=XM_003994780.4;refGene_gn=MTMR3;refGene_tx=XM_003994781.5;refGene_gn=MTMR3;refGene_tx=XM_003994782.5;refGene_gn=MTMR3;refGene_tx=XM_006938574.3;refGene_gn=MTMR3;refGene_tx=XM_006938575.4;refGene_gn=MTMR3;refGene_tx=XM_006938576.3;refGene_gn=MTMR3;refGene_tx=XM_006938577.3;refGene_gn=MTMR3;refGene_tx=XM_006938578.3;refGene_gn=MTMR3;refGene_tx=XM_006938579.3;refGene_gn=MTMR3;refGene_tx=XM_019814803.2;refGene_gn=MTMR3;refGene_tx=XM_019814804.2
D3 22910528 22910699
refGene_gn=MTMR3;refGene_tx=XM_003994780.4;refGene_gn=MTMR3;refGene_tx=XM_003994781.5;refGene_gn=MTMR3;refGene_tx=XM_003994782.5;refGene_gn=MTMR3;refGene_tx=XM_006938574.3;refGene_gn=MTMR3;refGene_tx=XM_006938575.4;refGene_gn=MTMR3;refGene_tx=XM_006938576.3;refGene_gn=MTMR3;refGene_tx=XM_006938577.3;refGene_gn=MTMR3;refGene_tx=XM_006938578.3;refGene_gn=MTMR3;refGene_tx=XM_006938579.3;refGene_gn=MTMR3;refGene_tx=XM_019814803.2;refGene_gn=MTMR3;refGene_tx=XM_019814804.2
D3 22913748 22913934
refGene_gn=MTMR3;refGene_tx=XM_003994780.4;refGene_gn=MTMR3;refGene_tx=XM_003994781.5;refGene_gn=MTMR3;refGene_tx=XM_003994782.5;refGene_gn=MTMR3;refGene_tx=XM_006938574.3;refGene_gn=MTMR3;refGene_tx=XM_006938575.4;refGene_gn=MTMR3;refGene_tx=XM_006938576.3;refGene_gn=MTMR3;refGene_tx=XM_006938577.3;refGene_gn=MTMR3;refGene_tx=XM_006938578.3;refGene_gn=MTMR3;refGene_tx=XM_006938579.3;refGene_gn=MTMR3;refGene_tx=XM_019814803.2;refGene_gn=MTMR3;refGene_tx=XM_019814804.2
D3 22914728 22914924
refGene_gn=MTMR3;refGene_tx=XM_003994780.4;refGene_gn=MTMR3;refGene_tx=XM_003994781.5;refGene_gn=MTMR3;refGene_tx=XM_003994782.5;refGene_gn=MTMR3;refGene_tx=XM_006938574.3;refGene_gn=MTMR3;refGene_tx=XM_006938575.4;refGene_gn=MTMR3;refGene_tx=XM_006938576.3;refGene_gn=MTMR3;refGe

ne_tx=XM_006938577.3;refGene_gn=MTMR3;refGene_tx=XM_006938578.3;refGene_gn=MTMR3;refGene_tx=XM_006938579.3;refGene_gn=MTMR3;refGene_tx=XM_019814803.2;refGene_gn=MTMR3;refGene_tx=XM_019814804.2
D3 22917887 22917999 refGene_gn=MTMR3;refGene_tx=XM_003994780.4;refGene_gn=MTMR3;refGene_tx=XM_003994781.5;refGene_gn=MTMR3;refGene_tx=XM_003994782.5;refGene_gn=MTMR3;refGene_tx=XM_006938574.3;refGene_gn=MTMR3;refGene_tx=XM_006938575.4;refGene_gn=MTMR3;refGene_tx=XM_006938576.3;refGene_gn=MTMR3;refGene_tx=XM_006938577.3;refGene_gn=MTMR3;refGene_tx=XM_006938578.3;refGene_gn=MTMR3;refGene_tx=XM_006938579.3;refGene_gn=MTMR3;refGene_tx=XM_019814803.2;refGene_gn=MTMR3;refGene_tx=XM_019814804.2
D3 22919254 22919386 refGene_gn=MTMR3;refGene_tx=XM_003994780.4;refGene_gn=MTMR3;refGene_tx=XM_003994781.5;refGene_gn=MTMR3;refGene_tx=XM_003994782.5;refGene_gn=MTMR3;refGene_tx=XM_006938574.3;refGene_gn=MTMR3;refGene_tx=XM_006938575.4;refGene_gn=MTMR3;refGene_tx=XM_006938576.3;refGene_gn=MTMR3;refGene_tx=XM_006938577.3;refGene_gn=MTMR3;refGene_tx=XM_006938578.3;refGene_gn=MTMR3;refGene_tx=XM_006938579.3;refGene_gn=MTMR3;refGene_tx=XM_019814803.2;refGene_gn=MTMR3;refGene_tx=XM_019814804.2
D3 22919892 22920098 refGene_gn=MTMR3;refGene_tx=XM_003994780.4;refGene_gn=MTMR3;refGene_tx=XM_003994781.5;refGene_gn=MTMR3;refGene_tx=XM_003994782.5;refGene_gn=MTMR3;refGene_tx=XM_006938574.3;refGene_gn=MTMR3;refGene_tx=XM_006938575.4;refGene_gn=MTMR3;refGene_tx=XM_006938576.3;refGene_gn=MTMR3;refGene_tx=XM_006938577.3;refGene_gn=MTMR3;refGene_tx=XM_006938578.3;refGene_gn=MTMR3;refGene_tx=XM_006938579.3;refGene_gn=MTMR3;refGene_tx=XM_019814803.2;refGene_gn=MTMR3;refGene_tx=XM_019814804.2
D3 22924210 22924324 refGene_gn=MTMR3;refGene_tx=XM_003994780.4;refGene_gn=MTMR3;refGene_tx=XM_003994781.5;refGene_gn=MTMR3;refGene_tx=XM_003994782.5;refGene_gn=MTMR3;refGene_tx=XM_006938574.3;refGene_gn=MTMR3;refGene_tx=XM_006938575.4;refGene_gn=MTMR3;refGene_tx=XM_006938576.3;refGene_gn=MTMR3;refGene_tx=XM_006938577.3;refGene_gn=MTMR3;refGene_tx=XM_006938578.3;refGene_gn=MTMR3;refGene_tx=XM_006938579.3;refGene_gn=MTMR3;refGene_tx=XM_019814803.2;refGene_gn=MTMR3;refGene_tx=XM_019814804.2
D3 22927021 22927118 refGene_gn=MTMR3;refGene_tx=XM_003994780.4;refGene_gn=MTMR3;refGene_tx=XM_003994781.5;refGene_gn=MTMR3;refGene_tx=XM_003994782.5;refGene_gn=MTMR3;refGene_tx=XM_006938574.3;refGene_gn=MTMR3;refGene_tx=XM_006938575.4;refGene_gn=MTMR3;refGene_tx=XM_006938576.3;refGene_gn=MTMR3;refGene_tx=XM_006938577.3;refGene_gn=MTMR3;refGene_tx=XM_006938578.3;refGene_gn=MTMR3;refGene_tx=XM_006938579.3;refGene_gn=MTMR3;refGene_tx=XM_019814803.2;refGene_gn=MTMR3;refGene_tx=XM_019814804.2
D3 22935418 22935585 refGene_gn=MTMR3;refGene_tx=XM_003994780.4;refGene_gn=MTMR3;refGene_tx=XM_003994781.5;refGene_gn=MTMR3;refGene_tx=XM_003994782.5;refGene_gn=MTMR3;refGene_tx=XM_006938574.3;refGene_gn=MTMR3;refGene_tx=XM_006938575.4;refGene_gn=MTMR3;refGene_tx=XM_006938576.3;refGene_gn=MTMR3;refGene_tx=XM_006938577.3;refGene_gn=MTMR3;refGene_tx=XM_006938578.3;refGene_gn=MTMR3;refGene_tx=XM_006938579.3;refGene_gn=MTMR3;refGene_tx=XM_01

9814803.2;refGene_gn=MTMR3;refGene_tx=XM_019814804.2
D3 22937734 22937817
refGene_gn=MTMR3;refGene_tx=XM_003994780.4;refGene_gn=MTMR3;refGene_tx=XM_003994781.5;refGene_gn=MTMR3;refGene_tx=XM_003994782.5;refGene_gn=MTMR3;refGene_tx=XM_006938574.3;refGene_gn=MTMR3;refGene_tx=XM_006938575.4;refGene_gn=MTMR3;refGene_tx=XM_006938576.3;refGene_gn=MTMR3;refGene_tx=XM_006938577.3;refGene_gn=MTMR3;refGene_tx=XM_006938578.3;refGene_gn=MTMR3;refGene_tx=XM_006938579.3;refGene_gn=MTMR3;refGene_tx=XM_019814803.2
D3 22942967 22943000
refGene_gn=MTMR3;refGene_tx=XM_006938574.3;refGene_gn=MTMR3;refGene_tx=XM_006938575.4;refGene_gn=MTMR3;refGene_tx=XM_006938576.3;refGene_gn=MTMR3;refGene_tx=XM_006938577.3;refGene_gn=MTMR3;refGene_tx=XM_006938579.3;refGene_gn=MTMR3;refGene_tx=XM_019814803.2
D3 22945949 22946066
refGene_gn=MTMR3;refGene_tx=XM_003994780.4;refGene_gn=MTMR3;refGene_tx=XM_003994781.5;refGene_gn=MTMR3;refGene_tx=XM_003994782.5;refGene_gn=MTMR3;refGene_tx=XM_006938574.3;refGene_gn=MTMR3;refGene_tx=XM_006938575.4;refGene_gn=MTMR3;refGene_tx=XM_006938576.3;refGene_gn=MTMR3;refGene_tx=XM_006938577.3;refGene_gn=MTMR3;refGene_tx=XM_006938578.3;refGene_gn=MTMR3;refGene_tx=XM_006938579.3;refGene_gn=MTMR3;refGene_tx=XM_019814803.2;refGene_gn=MTMR3;refGene_tx=XM_019814804.2
D3 22946473 22946563
refGene_gn=MTMR3;refGene_tx=XM_003994780.4;refGene_gn=MTMR3;refGene_tx=XM_003994781.5;refGene_gn=MTMR3;refGene_tx=XM_003994782.5;refGene_gn=MTMR3;refGene_tx=XM_006938574.3;refGene_gn=MTMR3;refGene_tx=XM_006938575.4;refGene_gn=MTMR3;refGene_tx=XM_006938576.3;refGene_gn=MTMR3;refGene_tx=XM_006938577.3;refGene_gn=MTMR3;refGene_tx=XM_006938578.3;refGene_gn=MTMR3;refGene_tx=XM_006938579.3;refGene_gn=MTMR3;refGene_tx=XM_019814803.2;refGene_gn=MTMR3;refGene_tx=XM_019814804.2
D3 22956291 22956294
refGene_gn=MTMR3;refGene_tx=XM_003994780.4;refGene_gn=MTMR3;refGene_tx=XM_003994781.5;refGene_gn=MTMR3;refGene_tx=XM_003994782.5;refGene_gn=MTMR3;refGene_tx=XM_006938574.3;refGene_gn=MTMR3;refGene_tx=XM_006938575.4;refGene_gn=MTMR3;refGene_tx=XM_006938576.3;refGene_gn=MTMR3;refGene_tx=XM_006938577.3;refGene_gn=MTMR3;refGene_tx=XM_006938578.3;refGene_gn=MTMR3;refGene_tx=XM_006938579.3;refGene_gn=MTMR3;refGene_tx=XM_019814803.2
D3 23097301 23097382
refGene_gn=ASCC2;refGene_tx=XM_006938572.4;refGene_gn=ASCC2;refGene_tx=XM_006938573.4
D3 23103691 23103850 refGene_gn=ASCC2;refGene_tx=XM_006938572.4
D3 23104219 23104390
refGene_gn=ASCC2;refGene_tx=XM_006938572.4;refGene_gn=ASCC2;refGene_tx=XM_006938573.4
D3 23105720 23105850
refGene_gn=ASCC2;refGene_tx=XM_006938572.4;refGene_gn=ASCC2;refGene_tx=XM_006938573.4
D3 23111139 23111207
refGene_gn=ASCC2;refGene_tx=XM_006938572.4;refGene_gn=ASCC2;refGene_tx

=XM_006938573.4
D3 23112169 23112280
refGene_gn=ASCC2;refGene_tx=XM_006938572.4;refGene_gn=ASCC2;refGene_tx
=XM_006938573.4
D3 23113350 23113463
refGene_gn=ASCC2;refGene_tx=XM_006938572.4;refGene_gn=ASCC2;refGene_tx
=XM_006938573.4
D3 23118781 23118856
refGene_gn=ASCC2;refGene_tx=XM_006938572.4;refGene_gn=ASCC2;refGene_tx
=XM_006938573.4
D3 23119276 23119384
refGene_gn=ASCC2;refGene_tx=XM_006938572.4;refGene_gn=ASCC2;refGene_tx
=XM_006938573.4
D3 23119628 23119697
refGene_gn=ASCC2;refGene_tx=XM_006938572.4;refGene_gn=ASCC2;refGene_tx
=XM_006938573.4
D3 23119852 23119927
refGene_gn=ASCC2;refGene_tx=XM_006938572.4;refGene_gn=ASCC2;refGene_tx
=XM_006938573.4
D3 23121296 23121477
refGene_gn=ASCC2;refGene_tx=XM_006938572.4;refGene_gn=ASCC2;refGene_tx
=XM_006938573.4
D3 23122894 23123109
refGene_gn=ASCC2;refGene_tx=XM_006938572.4;refGene_gn=ASCC2;refGene_tx
=XM_006938573.4
D3 23124024 23124144
refGene_gn=ASCC2;refGene_tx=XM_006938572.4;refGene_gn=ASCC2;refGene_tx
=XM_006938573.4
D3 23129994 23130088
refGene_gn=ASCC2;refGene_tx=XM_006938572.4;refGene_gn=ASCC2;refGene_tx
=XM_006938573.4
D3 23130177 23130308
refGene_gn=ASCC2;refGene_tx=XM_006938572.4;refGene_gn=ASCC2;refGene_tx
=XM_006938573.4
D3 23130856 23130959
refGene_gn=ASCC2;refGene_tx=XM_006938572.4;refGene_gn=ASCC2;refGene_tx
=XM_006938573.4
D3 23132483 23132563
refGene_gn=ASCC2;refGene_tx=XM_006938572.4;refGene_gn=ASCC2;refGene_tx
=XM_006938573.4
D3 23133035 23133207
refGene_gn=ASCC2;refGene_tx=XM_006938572.4;refGene_gn=ASCC2;refGene_tx
=XM_006938573.4
D3 23149501 23149543
refGene_gn=LOC105261050;refGene_tx=XM_011287738.3
D3 23150915 23150921
refGene_gn=LOC105261050;refGene_tx=XM_011287739.3
D3 23151067 23151430
refGene_gn=LOC105261050;refGene_tx=XM_011287738.3;refGene_gn=LOC105261
050;refGene_tx=XM_011287739.3

D3 23164084 23164211
refGene_gn=ZMAT5;refGene_tx=XM_003994776.5;refGene_gn=ZMAT5;refGene_tx=XM_006938571.4;refGene_gn=ZMAT5;refGene_tx=XM_019814812.2
D3 23169663 23169726
refGene_gn=ZMAT5;refGene_tx=XM_003994776.5;refGene_gn=ZMAT5;refGene_tx=XM_006938571.4;refGene_gn=ZMAT5;refGene_tx=XM_019814812.2
D3 23171346 23171427
refGene_gn=ZMAT5;refGene_tx=XM_003994776.5;refGene_gn=ZMAT5;refGene_tx=XM_006938571.4;refGene_gn=ZMAT5;refGene_tx=XM_019814812.2
D3 23173178 23173290
refGene_gn=ZMAT5;refGene_tx=XM_003994776.5;refGene_gn=ZMAT5;refGene_tx=XM_006938571.4;refGene_gn=ZMAT5;refGene_tx=XM_019814812.2
D3 23178596 23178726
refGene_gn=ZMAT5;refGene_tx=XM_003994776.5;refGene_gn=ZMAT5;refGene_tx=XM_006938571.4;refGene_gn=ZMAT5;refGene_tx=XM_019814812.2
D3 23180332 23180460 refGene_gn=CABP7;refGene_tx=XM_011287735.3
D3 23180689 23180843 refGene_gn=CABP7;refGene_tx=XM_011287735.3
D3 23181156 23181269 refGene_gn=CABP7;refGene_tx=XM_011287735.3
D3 23182157 23182301 refGene_gn=CABP7;refGene_tx=XM_011287735.3
D3 23188602 23188711 refGene_gn=CABP7;refGene_tx=XM_011287735.3
D3 23213003 23213054
refGene_gn=NF2;refGene_tx=XM_003994773.4;refGene_gn=NF2;refGene_tx=XM_023241585.1;refGene_gn=NF2;refGene_tx=XM_023241588.1;refGene_gn=NF2;refGene_tx=XM_023241589.1;refGene_gn=NF2;refGene_tx=XM_023241590.1;refGene_gn=NF2;refGene_tx=XM_023241592.1;refGene_gn=NF2;refGene_tx=XM_023241593.1
D3 23223460 23223496
refGene_gn=NF2;refGene_tx=XM_023241586.1;refGene_gn=NF2;refGene_tx=XM_023241587.1;refGene_gn=NF2;refGene_tx=XM_023241591.1
D3 23224914 23225080
refGene_gn=NF2;refGene_tx=XM_003994773.4;refGene_gn=NF2;refGene_tx=XM_023241585.1;refGene_gn=NF2;refGene_tx=XM_023241586.1;refGene_gn=NF2;refGene_tx=XM_023241587.1;refGene_gn=NF2;refGene_tx=XM_023241588.1;refGene_gn=NF2;refGene_tx=XM_023241589.1;refGene_gn=NF2;refGene_tx=XM_023241590.1;refGene_gn=NF2;refGene_tx=XM_023241591.1;refGene_gn=NF2;refGene_tx=XM_023241592.1;refGene_gn=NF2;refGene_tx=XM_023241593.1
D3 23227276 23227404
refGene_gn=NF2;refGene_tx=XM_003994773.4;refGene_gn=NF2;refGene_tx=XM_023241585.1;refGene_gn=NF2;refGene_tx=XM_023241586.1;refGene_gn=NF2;refGene_tx=XM_023241587.1;refGene_gn=NF2;refGene_tx=XM_023241588.1;refGene_gn=NF2;refGene_tx=XM_023241589.1;refGene_gn=NF2;refGene_tx=XM_023241590.1;refGene_gn=NF2;refGene_tx=XM_023241591.1;refGene_gn=NF2;refGene_tx=XM_023241592.1;refGene_gn=NF2;refGene_tx=XM_023241593.1
D3 23231373 23231479
refGene_gn=NF2;refGene_tx=XM_003994773.4;refGene_gn=NF2;refGene_tx=XM_023241585.1;refGene_gn=NF2;refGene_tx=XM_023241586.1;refGene_gn=NF2;refGene_tx=XM_023241587.1;refGene_gn=NF2;refGene_tx=XM_023241588.1;refGene_gn=NF2;refGene_tx=XM_023241589.1;refGene_gn=NF2;refGene_tx=XM_023241590.1;refGene_gn=NF2;refGene_tx=XM_023241591.1;refGene_gn=NF2;refGene_tx=XM_023241592.1;refGene_gn=NF2;refGene_tx=XM_023241593.1

D3 23232856 23233074
refGene_gn=NF2;refGene_tx=XM_003994773.4;refGene_gn=NF2;refGene_tx=XM_023241585.1;refGene_gn=NF2;refGene_tx=XM_023241586.1;refGene_gn=NF2;refGene_tx=XM_023241587.1;refGene_gn=NF2;refGene_tx=XM_023241588.1;refGene_gn=NF2;refGene_tx=XM_023241589.1;refGene_gn=NF2;refGene_tx=XM_023241590.1;refGene_gn=NF2;refGene_tx=XM_023241591.1;refGene_gn=NF2;refGene_tx=XM_023241592.1;refGene_gn=NF2;refGene_tx=XM_023241593.1
D3 23233746 23233869
refGene_gn=NF2;refGene_tx=XM_003994773.4;refGene_gn=NF2;refGene_tx=XM_023241585.1;refGene_gn=NF2;refGene_tx=XM_023241586.1;refGene_gn=NF2;refGene_tx=XM_023241587.1;refGene_gn=NF2;refGene_tx=XM_023241588.1;refGene_gn=NF2;refGene_tx=XM_023241589.1;refGene_gn=NF2;refGene_tx=XM_023241590.1;refGene_gn=NF2;refGene_tx=XM_023241591.1;refGene_gn=NF2;refGene_tx=XM_023241592.1;refGene_gn=NF2;refGene_tx=XM_023241593.1
D3 23236398 23236512
refGene_gn=NF2;refGene_tx=XM_003994773.4;refGene_gn=NF2;refGene_tx=XM_023241585.1;refGene_gn=NF2;refGene_tx=XM_023241586.1;refGene_gn=NF2;refGene_tx=XM_023241587.1;refGene_gn=NF2;refGene_tx=XM_023241589.1;refGene_gn=NF2;refGene_tx=XM_023241590.1;refGene_gn=NF2;refGene_tx=XM_023241591.1;refGene_gn=NF2;refGene_tx=XM_023241592.1
D3 23239257 23239332
refGene_gn=NF2;refGene_tx=XM_003994773.4;refGene_gn=NF2;refGene_tx=XM_023241585.1;refGene_gn=NF2;refGene_tx=XM_023241586.1;refGene_gn=NF2;refGene_tx=XM_023241587.1;refGene_gn=NF2;refGene_tx=XM_023241588.1;refGene_gn=NF2;refGene_tx=XM_023241589.1;refGene_gn=NF2;refGene_tx=XM_023241590.1;refGene_gn=NF2;refGene_tx=XM_023241591.1;refGene_gn=NF2;refGene_tx=XM_023241593.1
D3 23242361 23242496
refGene_gn=NF2;refGene_tx=XM_003994773.4;refGene_gn=NF2;refGene_tx=XM_023241585.1;refGene_gn=NF2;refGene_tx=XM_023241586.1;refGene_gn=NF2;refGene_tx=XM_023241587.1;refGene_gn=NF2;refGene_tx=XM_023241588.1;refGene_gn=NF2;refGene_tx=XM_023241589.1;refGene_gn=NF2;refGene_tx=XM_023241591.1
D3 23247157 23247233
refGene_gn=NF2;refGene_tx=XM_003994773.4;refGene_gn=NF2;refGene_tx=XM_023241585.1;refGene_gn=NF2;refGene_tx=XM_023241586.1;refGene_gn=NF2;refGene_tx=XM_023241587.1;refGene_gn=NF2;refGene_tx=XM_023241588.1;refGene_gn=NF2;refGene_tx=XM_023241589.1;refGene_gn=NF2;refGene_tx=XM_023241590.1;refGene_gn=NF2;refGene_tx=XM_023241591.1;refGene_gn=NF2;refGene_tx=XM_023241592.1;refGene_gn=NF2;refGene_tx=XM_023241593.1
D3 23249092 23249175
refGene_gn=NF2;refGene_tx=XM_003994773.4;refGene_gn=NF2;refGene_tx=XM_023241585.1;refGene_gn=NF2;refGene_tx=XM_023241586.1;refGene_gn=NF2;refGene_tx=XM_023241587.1;refGene_gn=NF2;refGene_tx=XM_023241588.1;refGene_gn=NF2;refGene_tx=XM_023241589.1;refGene_gn=NF2;refGene_tx=XM_023241590.1;refGene_gn=NF2;refGene_tx=XM_023241591.1;refGene_gn=NF2;refGene_tx=XM_023241592.1;refGene_gn=NF2;refGene_tx=XM_023241593.1
D3 23250019 23250088
refGene_gn=NF2;refGene_tx=XM_003994773.4;refGene_gn=NF2;refGene_tx=XM_023241585.1;refGene_gn=NF2;refGene_tx=XM_023241586.1;refGene_gn=NF2;re

fGene_tx=XM_023241587.1;refGene_gn=NF2;refGene_tx=XM_023241588.1;refGene_gn=NF2;refGene_tx=XM_023241589.1;refGene_gn=NF2;refGene_tx=XM_023241590.1;refGene_gn=NF2;refGene_tx=XM_023241591.1;refGene_gn=NF2;refGene_tx=XM_023241592.1;refGene_gn=NF2;refGene_tx=XM_023241593.1
D3 23258225 23258309 refGene_gn=NF2;refGene_tx=XM_003994773.4;refGene_gn=NF2;refGene_tx=XM_023241585.1;refGene_gn=NF2;refGene_tx=XM_023241586.1;refGene_gn=NF2;refGene_tx=XM_023241587.1;refGene_gn=NF2;refGene_tx=XM_023241588.1;refGene_gn=NF2;refGene_tx=XM_023241589.1;refGene_gn=NF2;refGene_tx=XM_023241590.1;refGene_gn=NF2;refGene_tx=XM_023241591.1;refGene_gn=NF2;refGene_tx=XM_023241592.1;refGene_gn=NF2;refGene_tx=XM_023241593.1
D3 23260309 23260432 refGene_gn=NF2;refGene_tx=XM_003994773.4;refGene_gn=NF2;refGene_tx=XM_023241585.1;refGene_gn=NF2;refGene_tx=XM_023241586.1;refGene_gn=NF2;refGene_tx=XM_023241587.1;refGene_gn=NF2;refGene_tx=XM_023241588.1;refGene_gn=NF2;refGene_tx=XM_023241589.1;refGene_gn=NF2;refGene_tx=XM_023241590.1;refGene_gn=NF2;refGene_tx=XM_023241591.1;refGene_gn=NF2;refGene_tx=XM_023241592.1;refGene_gn=NF2;refGene_tx=XM_023241593.1
D3 23262378 23262504 refGene_gn=NF2;refGene_tx=XM_003994773.4;refGene_gn=NF2;refGene_tx=XM_023241585.1;refGene_gn=NF2;refGene_tx=XM_023241586.1;refGene_gn=NF2;refGene_tx=XM_023241587.1;refGene_gn=NF2;refGene_tx=XM_023241588.1;refGene_gn=NF2;refGene_tx=XM_023241590.1;refGene_gn=NF2;refGene_tx=XM_023241592.1;refGene_gn=NF2;refGene_tx=XM_023241593.1
D3 23291489 23291603 refGene_gn=NF2;refGene_tx=XM_023241585.1;refGene_gn=NF2;refGene_tx=XM_023241586.1;refGene_gn=NF2;refGene_tx=XM_023241587.1;refGene_gn=NF2;refGene_tx=XM_023241588.1;refGene_gn=NF2;refGene_tx=XM_023241589.1;refGene_gn=NF2;refGene_tx=XM_023241590.1;refGene_gn=NF2;refGene_tx=XM_023241591.1;refGene_gn=NF2;refGene_tx=XM_023241592.1;refGene_gn=NF2;refGene_tx=XM_023241593.1
D3 23311242 23311334 refGene_gn=NIPSNAP1;refGene_tx=XM_011287734.3
D3 23317966 23318094 refGene_gn=NIPSNAP1;refGene_tx=XM_011287734.3
D3 23318292 23318338 refGene_gn=NIPSNAP1;refGene_tx=XM_011287734.3
D3 23318598 23318693 refGene_gn=NIPSNAP1;refGene_tx=XM_011287734.3
D3 23322094 23322165 refGene_gn=NIPSNAP1;refGene_tx=XM_011287734.3
D3 23322312 23322453 refGene_gn=NIPSNAP1;refGene_tx=XM_011287734.3
D3 23322773 23322805 refGene_gn=NIPSNAP1;refGene_tx=XM_011287734.3
D3 23323145 23323240 refGene_gn=NIPSNAP1;refGene_tx=XM_011287734.3
D3 23324390 23324474 refGene_gn=NIPSNAP1;refGene_tx=XM_011287734.3
D3 23326183 23326248 refGene_gn=NIPSNAP1;refGene_tx=XM_011287734.3
D3 23330243 23330339 refGene_gn=THOC5;refGene_tx=XM_006938564.3;refGene_gn=THOC5;refGene_tx=XM_006938565.3;refGene_gn=THOC5;refGene_tx=XM_006938566.3;refGene_gn=THOC5;refGene_tx=XM_006938567.3
D3 23332275 23332419 refGene_gn=THOC5;refGene_tx=XM_006938564.3;refGene_gn=THOC5;refGene_tx=XM_006938565.3;refGene_gn=THOC5;refGene_tx=XM_006938566.3;refGene_gn=THOC5;refGene_tx=XM_006938567.3;refGene_gn=THOC5;refGene_tx=XM_019814818.1

D3 23333599 23333713
refGene_gn=THOC5;refGene_tx=XM_006938564.3;refGene_gn=THOC5;refGene_tx=XM_006938565.3;refGene_gn=THOC5;refGene_tx=XM_006938566.3;refGene_gn=THOC5;refGene_tx=XM_006938567.3;refGene_gn=THOC5;refGene_tx=XM_019814818.1
D3 23333851 23333949
refGene_gn=THOC5;refGene_tx=XM_006938564.3;refGene_gn=THOC5;refGene_tx=XM_006938565.3;refGene_gn=THOC5;refGene_tx=XM_006938566.3;refGene_gn=THOC5;refGene_tx=XM_006938567.3;refGene_gn=THOC5;refGene_tx=XM_019814818.1
D3 23335955 23336102
refGene_gn=THOC5;refGene_tx=XM_006938564.3;refGene_gn=THOC5;refGene_tx=XM_006938565.3;refGene_gn=THOC5;refGene_tx=XM_006938566.3;refGene_gn=THOC5;refGene_tx=XM_006938567.3;refGene_gn=THOC5;refGene_tx=XM_019814818.1
D3 23337499 23337614
refGene_gn=THOC5;refGene_tx=XM_006938564.3;refGene_gn=THOC5;refGene_tx=XM_006938565.3;refGene_gn=THOC5;refGene_tx=XM_006938566.3;refGene_gn=THOC5;refGene_tx=XM_006938567.3;refGene_gn=THOC5;refGene_tx=XM_019814818.1
D3 23341074 23341207
refGene_gn=THOC5;refGene_tx=XM_006938564.3;refGene_gn=THOC5;refGene_tx=XM_006938565.3;refGene_gn=THOC5;refGene_tx=XM_006938566.3;refGene_gn=THOC5;refGene_tx=XM_006938567.3;refGene_gn=THOC5;refGene_tx=XM_019814818.1
D3 23341928 23341961
refGene_gn=THOC5;refGene_tx=XM_006938564.3;refGene_gn=THOC5;refGene_tx=XM_006938565.3;refGene_gn=THOC5;refGene_tx=XM_006938566.3;refGene_gn=THOC5;refGene_tx=XM_019814818.1
D3 23343465 23343543
refGene_gn=THOC5;refGene_tx=XM_006938564.3;refGene_gn=THOC5;refGene_tx=XM_006938565.3;refGene_gn=THOC5;refGene_tx=XM_006938566.3;refGene_gn=THOC5;refGene_tx=XM_006938567.3;refGene_gn=THOC5;refGene_tx=XM_019814818.1
D3 23344455 23344496
refGene_gn=THOC5;refGene_tx=XM_006938564.3;refGene_gn=THOC5;refGene_tx=XM_006938565.3;refGene_gn=THOC5;refGene_tx=XM_006938566.3;refGene_gn=THOC5;refGene_tx=XM_006938567.3;refGene_gn=THOC5;refGene_tx=XM_019814818.1
D3 23344739 23344839
refGene_gn=THOC5;refGene_tx=XM_006938564.3;refGene_gn=THOC5;refGene_tx=XM_006938565.3;refGene_gn=THOC5;refGene_tx=XM_006938566.3;refGene_gn=THOC5;refGene_tx=XM_006938567.3;refGene_gn=THOC5;refGene_tx=XM_019814818.1
D3 23346234 23346343
refGene_gn=THOC5;refGene_tx=XM_006938564.3;refGene_gn=THOC5;refGene_tx=XM_006938565.3;refGene_gn=THOC5;refGene_tx=XM_006938566.3;refGene_gn=THOC5;refGene_tx=XM_006938567.3;refGene_gn=THOC5;refGene_tx=XM_019814818.1
D3 23349946 23350048

refGene_gn=THOC5;refGene_tx=XM_006938564.3;refGene_gn=THOC5;refGene_tx
=XM_006938565.3;refGene_gn=THOC5;refGene_tx=XM_006938566.3;refGene_gn=
THOC5;refGene_tx=XM_006938567.3;refGene_gn=THOC5;refGene_tx=XM_0198148
18.1
D3 23350617 23350714
refGene_gn=THOC5;refGene_tx=XM_006938564.3;refGene_gn=THOC5;refGene_tx
=XM_006938565.3;refGene_gn=THOC5;refGene_tx=XM_006938566.3;refGene_gn=
THOC5;refGene_tx=XM_006938567.3;refGene_gn=THOC5;refGene_tx=XM_0198148
18.1
D3 23351307 23351422
refGene_gn=THOC5;refGene_tx=XM_006938564.3;refGene_gn=THOC5;refGene_tx
=XM_006938565.3;refGene_gn=THOC5;refGene_tx=XM_006938566.3;refGene_gn=
THOC5;refGene_tx=XM_006938567.3;refGene_gn=THOC5;refGene_tx=XM_0198148
18.1
D3 23352707 23352811
refGene_gn=THOC5;refGene_tx=XM_006938564.3;refGene_gn=THOC5;refGene_tx
=XM_006938565.3;refGene_gn=THOC5;refGene_tx=XM_006938566.3;refGene_gn=
THOC5;refGene_tx=XM_006938567.3;refGene_gn=THOC5;refGene_tx=XM_0198148
18.1
D3 23352945 23353033
refGene_gn=THOC5;refGene_tx=XM_006938564.3;refGene_gn=THOC5;refGene_tx
=XM_006938565.3;refGene_gn=THOC5;refGene_tx=XM_006938566.3;refGene_gn=
THOC5;refGene_tx=XM_006938567.3;refGene_gn=THOC5;refGene_tx=XM_0198148
18.1
D3 23355616 23355732
refGene_gn=THOC5;refGene_tx=XM_006938564.3;refGene_gn=THOC5;refGene_tx
=XM_006938565.3;refGene_gn=THOC5;refGene_tx=XM_006938566.3;refGene_gn=
THOC5;refGene_tx=XM_006938567.3;refGene_gn=THOC5;refGene_tx=XM_0198148
18.1
D3 23356393 23356584
refGene_gn=THOC5;refGene_tx=XM_006938564.3;refGene_gn=THOC5;refGene_tx
=XM_006938565.3;refGene_gn=THOC5;refGene_tx=XM_006938566.3;refGene_gn=
THOC5;refGene_tx=XM_006938567.3;refGene_gn=THOC5;refGene_tx=XM_0198148
18.1
D3 23359493 23359557
refGene_gn=THOC5;refGene_tx=XM_006938564.3;refGene_gn=THOC5;refGene_tx
=XM_006938565.3;refGene_gn=THOC5;refGene_tx=XM_006938566.3;refGene_gn=
THOC5;refGene_tx=XM_006938567.3;refGene_gn=THOC5;refGene_tx=XM_0198148
18.1
D3 23373529 23374687
refGene_gn=NEFH;refGene_tx=XM_023241594.1;refGene_gn=NEFH;refGene_tx=X
M_023241595.1;refGene_gn=NEFH;refGene_tx=XM_023241596.1;refGene_gn=NEF
H;refGene_tx=XM_023241597.1;refGene_gn=NEFH;refGene_tx=XM_023241598.1
D3 23374752 23375004
refGene_gn=NEFH;refGene_tx=XM_023241594.1;refGene_gn=NEFH;refGene_tx=X
M_023241595.1;refGene_gn=NEFH;refGene_tx=XM_023241596.1;refGene_gn=NEF
H;refGene_tx=XM_023241597.1;refGene_gn=NEFH;refGene_tx=XM_023241598.1
D3 23375457 23376229
refGene_gn=NEFH;refGene_tx=XM_023241594.1;refGene_gn=NEFH;refGene_tx=X
M_023241595.1;refGene_gn=NEFH;refGene_tx=XM_023241596.1;refGene_gn=NEF

H;refGene_tx=XM_023241597.1;refGene_gn=NEFH;refGene_tx=XM_023241598.1
D3 23377864 23377989
refGene_gn=NEFH;refGene_tx=XM_023241594.1;refGene_gn=NEFH;refGene_tx=XM_023241595.1;refGene_gn=NEFH;refGene_tx=XM_023241596.1;refGene_gn=NEFH;refGene_tx=XM_023241597.1;refGene_gn=NEFH;refGene_tx=XM_023241598.1
D3 23378976 23379176
refGene_gn=NEFH;refGene_tx=XM_023241594.1;refGene_gn=NEFH;refGene_tx=XM_023241595.1;refGene_gn=NEFH;refGene_tx=XM_023241596.1;refGene_gn=NEFH;refGene_tx=XM_023241597.1;refGene_gn=NEFH;refGene_tx=XM_023241598.1
D3 23381015 23381889
refGene_gn=NEFH;refGene_tx=XM_023241594.1;refGene_gn=NEFH;refGene_tx=XM_023241595.1;refGene_gn=NEFH;refGene_tx=XM_023241596.1;refGene_gn=NEFH;refGene_tx=XM_023241597.1;refGene_gn=NEFH;refGene_tx=XM_023241598.1
D3 23391594 23391758 refGene_gn=AP1B1;refGene_tx=XM_023241599.1
D3 23441450 23441513
refGene_gn=AP1B1;refGene_tx=XM_003994768.5;refGene_gn=AP1B1;refGene_tx=XM_006938560.4;refGene_gn=AP1B1;refGene_tx=XM_006938561.4;refGene_gn=AP1B1;refGene_tx=XM_006938562.4;refGene_gn=AP1B1;refGene_tx=XM_006938563.4;refGene_gn=AP1B1;refGene_tx=XM_019814823.2;refGene_gn=AP1B1;refGene_tx=XM_023241599.1;refGene_gn=AP1B1;refGene_tx=XM_023241600.1
D3 23444454 23444560
refGene_gn=AP1B1;refGene_tx=XM_003994768.5;refGene_gn=AP1B1;refGene_tx=XM_006938560.4;refGene_gn=AP1B1;refGene_tx=XM_006938561.4;refGene_gn=AP1B1;refGene_tx=XM_006938562.4;refGene_gn=AP1B1;refGene_tx=XM_006938563.4;refGene_gn=AP1B1;refGene_tx=XM_019814823.2;refGene_gn=AP1B1;refGene_tx=XM_023241600.1
D3 23447406 23447542
refGene_gn=AP1B1;refGene_tx=XM_003994768.5;refGene_gn=AP1B1;refGene_tx=XM_006938560.4;refGene_gn=AP1B1;refGene_tx=XM_006938561.4;refGene_gn=AP1B1;refGene_tx=XM_006938562.4;refGene_gn=AP1B1;refGene_tx=XM_006938563.4;refGene_gn=AP1B1;refGene_tx=XM_019814823.2;refGene_gn=AP1B1;refGene_tx=XM_023241599.1;refGene_gn=AP1B1;refGene_tx=XM_023241600.1
D3 23448329 23448575
refGene_gn=AP1B1;refGene_tx=XM_003994768.5;refGene_gn=AP1B1;refGene_tx=XM_006938560.4;refGene_gn=AP1B1;refGene_tx=XM_006938561.4;refGene_gn=AP1B1;refGene_tx=XM_006938562.4;refGene_gn=AP1B1;refGene_tx=XM_006938563.4;refGene_gn=AP1B1;refGene_tx=XM_019814823.2;refGene_gn=AP1B1;refGene_tx=XM_023241599.1;refGene_gn=AP1B1;refGene_tx=XM_023241600.1
D3 23450378 23450569
refGene_gn=AP1B1;refGene_tx=XM_003994768.5;refGene_gn=AP1B1;refGene_tx=XM_006938560.4;refGene_gn=AP1B1;refGene_tx=XM_006938561.4;refGene_gn=AP1B1;refGene_tx=XM_006938562.4;refGene_gn=AP1B1;refGene_tx=XM_006938563.4;refGene_gn=AP1B1;refGene_tx=XM_019814823.2;refGene_gn=AP1B1;refGene_tx=XM_023241599.1;refGene_gn=AP1B1;refGene_tx=XM_023241600.1
D3 23451373 23451595
refGene_gn=AP1B1;refGene_tx=XM_003994768.5;refGene_gn=AP1B1;refGene_tx=XM_006938560.4;refGene_gn=AP1B1;refGene_tx=XM_006938561.4;refGene_gn=AP1B1;refGene_tx=XM_006938562.4;refGene_gn=AP1B1;refGene_tx=XM_006938563.4;refGene_gn=AP1B1;refGene_tx=XM_019814823.2;refGene_gn=AP1B1;refGene_tx=XM_023241599.1;refGene_gn=AP1B1;refGene_tx=XM_023241600.1

D3 23454114 23454235
refGene_gn=AP1B1;refGene_tx=XM_003994768.5;refGene_gn=AP1B1;refGene_tx=XM_006938560.4;refGene_gn=AP1B1;refGene_tx=XM_006938561.4;refGene_gn=AP1B1;refGene_tx=XM_006938562.4;refGene_gn=AP1B1;refGene_tx=XM_006938563.4;refGene_gn=AP1B1;refGene_tx=XM_019814823.2;refGene_gn=AP1B1;refGene_tx=XM_023241599.1;refGene_gn=AP1B1;refGene_tx=XM_023241600.1
D3 23454699 23454795
refGene_gn=AP1B1;refGene_tx=XM_003994768.5;refGene_gn=AP1B1;refGene_tx=XM_006938560.4;refGene_gn=AP1B1;refGene_tx=XM_006938561.4;refGene_gn=AP1B1;refGene_tx=XM_006938562.4;refGene_gn=AP1B1;refGene_tx=XM_006938563.4;refGene_gn=AP1B1;refGene_tx=XM_019814823.2;refGene_gn=AP1B1;refGene_tx=XM_023241599.1;refGene_gn=AP1B1;refGene_tx=XM_023241600.1
D3 23455784 23455900
refGene_gn=AP1B1;refGene_tx=XM_003994768.5;refGene_gn=AP1B1;refGene_tx=XM_006938560.4;refGene_gn=AP1B1;refGene_tx=XM_006938561.4;refGene_gn=AP1B1;refGene_tx=XM_006938562.4;refGene_gn=AP1B1;refGene_tx=XM_006938563.4;refGene_gn=AP1B1;refGene_tx=XM_019814823.2;refGene_gn=AP1B1;refGene_tx=XM_023241599.1;refGene_gn=AP1B1;refGene_tx=XM_023241600.1
D3 23456248 23456414
refGene_gn=AP1B1;refGene_tx=XM_003994768.5;refGene_gn=AP1B1;refGene_tx=XM_006938560.4;refGene_gn=AP1B1;refGene_tx=XM_006938561.4;refGene_gn=AP1B1;refGene_tx=XM_006938562.4;refGene_gn=AP1B1;refGene_tx=XM_006938563.4;refGene_gn=AP1B1;refGene_tx=XM_019814823.2;refGene_gn=AP1B1;refGene_tx=XM_023241599.1;refGene_gn=AP1B1;refGene_tx=XM_023241600.1
D3 23463121 23463220
refGene_gn=AP1B1;refGene_tx=XM_003994768.5;refGene_gn=AP1B1;refGene_tx=XM_006938560.4;refGene_gn=AP1B1;refGene_tx=XM_006938561.4;refGene_gn=AP1B1;refGene_tx=XM_006938562.4;refGene_gn=AP1B1;refGene_tx=XM_006938563.4;refGene_gn=AP1B1;refGene_tx=XM_019814823.2;refGene_gn=AP1B1;refGene_tx=XM_023241599.1;refGene_gn=AP1B1;refGene_tx=XM_023241600.1
D3 23463756 23464016
refGene_gn=AP1B1;refGene_tx=XM_003994768.5;refGene_gn=AP1B1;refGene_tx=XM_006938560.4;refGene_gn=AP1B1;refGene_tx=XM_006938561.4;refGene_gn=AP1B1;refGene_tx=XM_006938562.4;refGene_gn=AP1B1;refGene_tx=XM_006938563.4;refGene_gn=AP1B1;refGene_tx=XM_019814823.2;refGene_gn=AP1B1;refGene_tx=XM_023241599.1;refGene_gn=AP1B1;refGene_tx=XM_023241600.1
D3 23464668 23464870
refGene_gn=AP1B1;refGene_tx=XM_003994768.5;refGene_gn=AP1B1;refGene_tx=XM_006938560.4;refGene_gn=AP1B1;refGene_tx=XM_006938561.4;refGene_gn=AP1B1;refGene_tx=XM_006938562.4;refGene_gn=AP1B1;refGene_tx=XM_006938563.4;refGene_gn=AP1B1;refGene_tx=XM_019814823.2;refGene_gn=AP1B1;refGene_tx=XM_023241599.1;refGene_gn=AP1B1;refGene_tx=XM_023241600.1
D3 23465652 23465673
refGene_gn=AP1B1;refGene_tx=XM_003994768.5;refGene_gn=AP1B1;refGene_tx=XM_006938560.4;refGene_gn=AP1B1;refGene_tx=XM_006938561.4;refGene_gn=AP1B1;refGene_tx=XM_019814823.2;refGene_gn=AP1B1;refGene_tx=XM_023241599.1;refGene_gn=AP1B1;refGene_tx=XM_023241600.1
D3 23466223 23466367
refGene_gn=AP1B1;refGene_tx=XM_003994768.5;refGene_gn=AP1B1;refGene_tx=XM_006938560.4;refGene_gn=AP1B1;refGene_tx=XM_006938561.4;refGene_gn=

AP1B1;refGene_tx=XM_006938562.4;refGene_gn=AP1B1;refGene_tx=XM_0069385
63.4;refGene_gn=AP1B1;refGene_tx=XM_019814823.2;refGene_gn=AP1B1;refGe
ne_tx=XM_023241599.1;refGene_gn=AP1B1;refGene_tx=XM_023241600.1
D3 23471167 23471313
refGene_gn=AP1B1;refGene_tx=XM_003994768.5;refGene_gn=AP1B1;refGene_tx
=XM_006938560.4;refGene_gn=AP1B1;refGene_tx=XM_006938561.4;refGene_gn=
AP1B1;refGene_tx=XM_006938562.4;refGene_gn=AP1B1;refGene_tx=XM_0069385
63.4;refGene_gn=AP1B1;refGene_tx=XM_019814823.2;refGene_gn=AP1B1;refGe
ne_tx=XM_023241599.1;refGene_gn=AP1B1;refGene_tx=XM_023241600.1
D3 23473079 23473209
refGene_gn=AP1B1;refGene_tx=XM_003994768.5;refGene_gn=AP1B1;refGene_tx
=XM_006938560.4;refGene_gn=AP1B1;refGene_tx=XM_006938561.4;refGene_gn=
AP1B1;refGene_tx=XM_006938562.4;refGene_gn=AP1B1;refGene_tx=XM_0069385
63.4;refGene_gn=AP1B1;refGene_tx=XM_019814823.2;refGene_gn=AP1B1;refGe
ne_tx=XM_023241599.1;refGene_gn=AP1B1;refGene_tx=XM_023241600.1
D3 23473506 23473591
refGene_gn=AP1B1;refGene_tx=XM_003994768.5;refGene_gn=AP1B1;refGene_tx
=XM_006938560.4;refGene_gn=AP1B1;refGene_tx=XM_006938561.4;refGene_gn=
AP1B1;refGene_tx=XM_006938562.4;refGene_gn=AP1B1;refGene_tx=XM_0069385
63.4;refGene_gn=AP1B1;refGene_tx=XM_019814823.2;refGene_gn=AP1B1;refGe
ne_tx=XM_023241599.1;refGene_gn=AP1B1;refGene_tx=XM_023241600.1
D3 23474288 23474375
refGene_gn=AP1B1;refGene_tx=XM_003994768.5;refGene_gn=AP1B1;refGene_tx
=XM_006938560.4;refGene_gn=AP1B1;refGene_tx=XM_006938561.4;refGene_gn=
AP1B1;refGene_tx=XM_006938562.4;refGene_gn=AP1B1;refGene_tx=XM_0069385
63.4;refGene_gn=AP1B1;refGene_tx=XM_019814823.2;refGene_gn=AP1B1;refGe
ne_tx=XM_023241599.1;refGene_gn=AP1B1;refGene_tx=XM_023241600.1
D3 23474478 23474633
refGene_gn=AP1B1;refGene_tx=XM_003994768.5;refGene_gn=AP1B1;refGene_tx
=XM_006938560.4;refGene_gn=AP1B1;refGene_tx=XM_006938561.4;refGene_gn=
AP1B1;refGene_tx=XM_006938562.4;refGene_gn=AP1B1;refGene_tx=XM_0069385
63.4;refGene_gn=AP1B1;refGene_tx=XM_019814823.2;refGene_gn=AP1B1;refGe
ne_tx=XM_023241599.1;refGene_gn=AP1B1;refGene_tx=XM_023241600.1
D3 23475330 23475339
refGene_gn=AP1B1;refGene_tx=XM_003994768.5;refGene_gn=AP1B1;refGene_tx
=XM_006938560.4;refGene_gn=AP1B1;refGene_tx=XM_006938562.4;refGene_gn=
AP1B1;refGene_tx=XM_019814823.2;refGene_gn=AP1B1;refGene_tx=XM_0232416
00.1
D3 23476200 23476275
refGene_gn=AP1B1;refGene_tx=XM_003994768.5;refGene_gn=AP1B1;refGene_tx
=XM_006938560.4;refGene_gn=AP1B1;refGene_tx=XM_006938561.4;refGene_gn=
AP1B1;refGene_tx=XM_006938562.4;refGene_gn=AP1B1;refGene_tx=XM_0069385
63.4;refGene_gn=AP1B1;refGene_tx=XM_019814823.2;refGene_gn=AP1B1;refGe
ne_tx=XM_023241599.1;refGene_gn=AP1B1;refGene_tx=XM_023241600.1
D3 23487625 23487844
refGene_gn=RASL10A;refGene_tx=XM_019814333.2;refGene_gn=RASL10A;refGen
e_tx=XM_023241609.1
D3 23488017 23488053 refGene_gn=RASL10A;refGene_tx=XM_019814333.2
D3 23489103 23489228
refGene_gn=RASL10A;refGene_tx=XM_019814333.2;refGene_gn=RASL10A;refGen

e_tx=XM_023241609.1
D3 23489511 23489779
refGene_gn=RASL10A;refGene_tx=XM_019814333.2;refGene_gn=RASL10A;refGene_tx=XM_023241609.1
D3 23490594 23491630 refGene_gn=GAS2L1;refGene_tx=XM_023241601.1
D3 23492165 23492337 refGene_gn=GAS2L1;refGene_tx=XM_023241601.1
D3 23492440 23492537 refGene_gn=GAS2L1;refGene_tx=XM_023241601.1
D3 23492631 23492739 refGene_gn=GAS2L1;refGene_tx=XM_023241601.1
D3 23494027 23494660 refGene_gn=GAS2L1;refGene_tx=XM_023241601.1
D3 23503081 23503121
refGene_gn=EWSR1;refGene_tx=XM_011287726.3;refGene_gn=EWSR1;refGene_tx=XM_011287727.3;refGene_gn=EWSR1;refGene_tx=XM_019814825.2;refGene_gn=EWSR1;refGene_tx=XM_019814826.2
D3 23503275 23503528
refGene_gn=EWSR1;refGene_tx=XM_011287726.3;refGene_gn=EWSR1;refGene_tx=XM_011287727.3;refGene_gn=EWSR1;refGene_tx=XM_019814825.2;refGene_gn=EWSR1;refGene_tx=XM_019814826.2
D3 23503828 23503926
refGene_gn=EWSR1;refGene_tx=XM_011287726.3;refGene_gn=EWSR1;refGene_tx=XM_011287727.3;refGene_gn=EWSR1;refGene_tx=XM_019814825.2;refGene_gn=EWSR1;refGene_tx=XM_019814826.2
D3 23504335 23504498
refGene_gn=EWSR1;refGene_tx=XM_011287726.3;refGene_gn=EWSR1;refGene_tx=XM_011287727.3;refGene_gn=EWSR1;refGene_tx=XM_019814825.2;refGene_gn=EWSR1;refGene_tx=XM_019814826.2
D3 23504996 23505119
refGene_gn=EWSR1;refGene_tx=XM_011287726.3;refGene_gn=EWSR1;refGene_tx=XM_011287727.3;refGene_gn=EWSR1;refGene_tx=XM_019814825.2;refGene_gn=EWSR1;refGene_tx=XM_019814826.2
D3 23505971 23506101
refGene_gn=EWSR1;refGene_tx=XM_011287726.3;refGene_gn=EWSR1;refGene_tx=XM_011287727.3;refGene_gn=EWSR1;refGene_tx=XM_019814825.2;refGene_gn=EWSR1;refGene_tx=XM_019814826.2
D3 23510238 23510357
refGene_gn=EWSR1;refGene_tx=XM_011287726.3;refGene_gn=EWSR1;refGene_tx=XM_011287727.3;refGene_gn=EWSR1;refGene_tx=XM_019814825.2;refGene_gn=EWSR1;refGene_tx=XM_019814826.2
D3 23510673 23510706
refGene_gn=EWSR1;refGene_tx=XM_011287726.3;refGene_gn=EWSR1;refGene_tx=XM_011287727.3;refGene_gn=EWSR1;refGene_tx=XM_019814825.2;refGene_gn=EWSR1;refGene_tx=XM_019814826.2
D3 23511241 23511279
refGene_gn=EWSR1;refGene_tx=XM_011287726.3;refGene_gn=EWSR1;refGene_tx=XM_011287727.3;refGene_gn=EWSR1;refGene_tx=XM_019814825.2;refGene_gn=EWSR1;refGene_tx=XM_019814826.2
D3 23512318 23512409 refGene_gn=EWSR1;refGene_tx=XM_011287728.3
D3 23514060 23514241
refGene_gn=EWSR1;refGene_tx=XM_011287726.3;refGene_gn=EWSR1;refGene_tx=XM_011287727.3;refGene_gn=EWSR1;refGene_tx=XM_011287728.3;refGene_gn=EWSR1;refGene_tx=XM_019814825.2;refGene_gn=EWSR1;refGene_tx=XM_0198148

26.2
D3	23515251	23515463
refGene_gn=EWSR1;refGene_tx=XM_011287726.3;refGene_gn=EWSR1;refGene_tx=XM_011287727.3;refGene_gn=EWSR1;refGene_tx=XM_011287728.3;refGene_gn=EWSR1;refGene_tx=XM_019814825.2;refGene_gn=EWSR1;refGene_tx=XM_019814826.2
D3	23517199	23517367
refGene_gn=EWSR1;refGene_tx=XM_011287726.3;refGene_gn=EWSR1;refGene_tx=XM_011287727.3;refGene_gn=EWSR1;refGene_tx=XM_011287728.3;refGene_gn=EWSR1;refGene_tx=XM_019814825.2;refGene_gn=EWSR1;refGene_tx=XM_019814826.2
D3	23519312	23519499
refGene_gn=EWSR1;refGene_tx=XM_011287726.3;refGene_gn=EWSR1;refGene_tx=XM_011287727.3;refGene_gn=EWSR1;refGene_tx=XM_011287728.3;refGene_gn=EWSR1;refGene_tx=XM_019814825.2;refGene_gn=EWSR1;refGene_tx=XM_019814826.2
D3	23523400	23523418
refGene_gn=EWSR1;refGene_tx=XM_019814825.2;refGene_gn=EWSR1;refGene_tx=XM_019814826.2
D3	23523811	23523935
refGene_gn=EWSR1;refGene_tx=XM_011287726.3;refGene_gn=EWSR1;refGene_tx=XM_011287727.3;refGene_gn=EWSR1;refGene_tx=XM_011287728.3;refGene_gn=EWSR1;refGene_tx=XM_019814825.2;refGene_gn=EWSR1;refGene_tx=XM_019814826.2
D3	23525724	23525776
refGene_gn=EWSR1;refGene_tx=XM_011287726.3;refGene_gn=EWSR1;refGene_tx=XM_011287727.3;refGene_gn=EWSR1;refGene_tx=XM_011287728.3;refGene_gn=EWSR1;refGene_tx=XM_019814825.2;refGene_gn=EWSR1;refGene_tx=XM_019814826.2
D3	23525918	23525955
refGene_gn=EWSR1;refGene_tx=XM_011287726.3;refGene_gn=EWSR1;refGene_tx=XM_011287727.3;refGene_gn=EWSR1;refGene_tx=XM_011287728.3;refGene_gn=EWSR1;refGene_tx=XM_019814825.2;refGene_gn=EWSR1;refGene_tx=XM_019814826.2
D3	23529900	23529913
refGene_gn=EWSR1;refGene_tx=XM_011287726.3;refGene_gn=EWSR1;refGene_tx=XM_011287727.3;refGene_gn=EWSR1;refGene_tx=XM_011287728.3;refGene_gn=EWSR1;refGene_tx=XM_019814825.2;refGene_gn=EWSR1;refGene_tx=XM_019814826.2
D3	23532532	23532680
refGene_gn=RHBDD3;refGene_tx=XM_011287731.3;refGene_gn=RHBDD3;refGene_tx=XM_011287732.3;refGene_gn=RHBDD3;refGene_tx=XM_023241608.1
D3	23533307	23533691
refGene_gn=RHBDD3;refGene_tx=XM_011287731.3;refGene_gn=RHBDD3;refGene_tx=XM_011287732.3;refGene_gn=RHBDD3;refGene_tx=XM_023241608.1
D3	23534591	23534754
refGene_gn=RHBDD3;refGene_tx=XM_011287731.3;refGene_gn=RHBDD3;refGene_tx=XM_011287732.3;refGene_gn=RHBDD3;refGene_tx=XM_023241608.1
D3	23534842	23535130
refGene_gn=RHBDD3;refGene_tx=XM_011287731.3;refGene_gn=RHBDD3;refGene_

tx=XM_011287732.3;refGene_gn=RHBDD3;refGene_tx=XM_023241608.1
D3 23535216 23535415
refGene_gn=RHBDD3;refGene_tx=XM_011287731.3;refGene_gn=RHBDD3;refGene_
tx=XM_011287732.3;refGene_gn=RHBDD3;refGene_tx=XM_023241608.1
D3 23537214 23537436
refGene_gn=EMID1;refGene_tx=XM_023241602.1;refGene_gn=EMID1;refGene_tx
=XM_023241603.1;refGene_gn=EMID1;refGene_tx=XM_023241604.1;refGene_gn=
EMID1;refGene_tx=XM_023241605.1;refGene_gn=EMID1;refGene_tx=XM_0232416
06.1
D3 23540661 23540752 refGene_gn=EMID1;refGene_tx=XM_023241603.1
D3 23541731 23541816
refGene_gn=EMID1;refGene_tx=XM_023241604.1;refGene_gn=EMID1;refGene_tx
=XM_023241606.1
D3 23546824 23546869
refGene_gn=EMID1;refGene_tx=XM_023241602.1;refGene_gn=EMID1;refGene_tx
=XM_023241603.1;refGene_gn=EMID1;refGene_tx=XM_023241604.1;refGene_gn=
EMID1;refGene_tx=XM_023241605.1;refGene_gn=EMID1;refGene_tx=XM_0232416
06.1
D3 23560401 23560536 refGene_gn=EMID1;refGene_tx=XM_023241607.1
D3 23562154 23562199
refGene_gn=EMID1;refGene_tx=XM_023241602.1;refGene_gn=EMID1;refGene_tx
=XM_023241603.1;refGene_gn=EMID1;refGene_tx=XM_023241604.1;refGene_gn=
EMID1;refGene_tx=XM_023241605.1;refGene_gn=EMID1;refGene_tx=XM_0232416
06.1;refGene_gn=EMID1;refGene_tx=XM_023241607.1
D3 23562305 23562368
refGene_gn=EMID1;refGene_tx=XM_023241602.1;refGene_gn=EMID1;refGene_tx
=XM_023241603.1;refGene_gn=EMID1;refGene_tx=XM_023241604.1;refGene_gn=
EMID1;refGene_tx=XM_023241605.1;refGene_gn=EMID1;refGene_tx=XM_0232416
06.1;refGene_gn=EMID1;refGene_tx=XM_023241607.1
D3 23562851 23562904
refGene_gn=EMID1;refGene_tx=XM_023241602.1;refGene_gn=EMID1;refGene_tx
=XM_023241603.1;refGene_gn=EMID1;refGene_tx=XM_023241604.1;refGene_gn=
EMID1;refGene_tx=XM_023241605.1;refGene_gn=EMID1;refGene_tx=XM_0232416
06.1;refGene_gn=EMID1;refGene_tx=XM_023241607.1
D3 23563048 23563138
refGene_gn=EMID1;refGene_tx=XM_023241602.1;refGene_gn=EMID1;refGene_tx
=XM_023241603.1;refGene_gn=EMID1;refGene_tx=XM_023241604.1;refGene_gn=
EMID1;refGene_tx=XM_023241605.1;refGene_gn=EMID1;refGene_tx=XM_0232416
06.1;refGene_gn=EMID1;refGene_tx=XM_023241607.1
D3 23564118 23564274
refGene_gn=EMID1;refGene_tx=XM_023241602.1;refGene_gn=EMID1;refGene_tx
=XM_023241603.1;refGene_gn=EMID1;refGene_tx=XM_023241604.1;refGene_gn=
EMID1;refGene_tx=XM_023241605.1;refGene_gn=EMID1;refGene_tx=XM_0232416
06.1;refGene_gn=EMID1;refGene_tx=XM_023241607.1
D3 23565053 23565143
refGene_gn=EMID1;refGene_tx=XM_023241602.1;refGene_gn=EMID1;refGene_tx
=XM_023241603.1;refGene_gn=EMID1;refGene_tx=XM_023241604.1;refGene_gn=
EMID1;refGene_tx=XM_023241605.1;refGene_gn=EMID1;refGene_tx=XM_0232416
06.1;refGene_gn=EMID1;refGene_tx=XM_023241607.1
D3 23565226 23565277

refGene_gn=EMID1;refGene_tx=XM_023241602.1;refGene_gn=EMID1;refGene_tx=XM_023241603.1;refGene_gn=EMID1;refGene_tx=XM_023241604.1;refGene_gn=EMID1;refGene_tx=XM_023241607.1
D3 23565480 23565601
refGene_gn=EMID1;refGene_tx=XM_023241602.1;refGene_gn=EMID1;refGene_tx=XM_023241603.1;refGene_gn=EMID1;refGene_tx=XM_023241604.1;refGene_gn=EMID1;refGene_tx=XM_023241605.1;refGene_gn=EMID1;refGene_tx=XM_023241606.1;refGene_gn=EMID1;refGene_tx=XM_023241607.1
D3 23567578 23567640
refGene_gn=EMID1;refGene_tx=XM_023241602.1;refGene_gn=EMID1;refGene_tx=XM_023241603.1;refGene_gn=EMID1;refGene_tx=XM_023241604.1;refGene_gn=EMID1;refGene_tx=XM_023241605.1;refGene_gn=EMID1;refGene_tx=XM_023241606.1;refGene_gn=EMID1;refGene_tx=XM_023241607.1
D3 23568758 23568842
refGene_gn=EMID1;refGene_tx=XM_023241602.1;refGene_gn=EMID1;refGene_tx=XM_023241603.1;refGene_gn=EMID1;refGene_tx=XM_023241604.1;refGene_gn=EMID1;refGene_tx=XM_023241605.1;refGene_gn=EMID1;refGene_tx=XM_023241606.1;refGene_gn=EMID1;refGene_tx=XM_023241607.1
D3 23576406 23576510
refGene_gn=EMID1;refGene_tx=XM_023241602.1;refGene_gn=EMID1;refGene_tx=XM_023241603.1;refGene_gn=EMID1;refGene_tx=XM_023241604.1;refGene_gn=EMID1;refGene_tx=XM_023241605.1;refGene_gn=EMID1;refGene_tx=XM_023241606.1;refGene_gn=EMID1;refGene_tx=XM_023241607.1
D3 23577003 23577117
refGene_gn=EMID1;refGene_tx=XM_023241602.1;refGene_gn=EMID1;refGene_tx=XM_023241603.1;refGene_gn=EMID1;refGene_tx=XM_023241604.1;refGene_gn=EMID1;refGene_tx=XM_023241605.1;refGene_gn=EMID1;refGene_tx=XM_023241606.1;refGene_gn=EMID1;refGene_tx=XM_023241607.1
D3 23585307 23585408
refGene_gn=EMID1;refGene_tx=XM_023241602.1;refGene_gn=EMID1;refGene_tx=XM_023241603.1;refGene_gn=EMID1;refGene_tx=XM_023241604.1;refGene_gn=EMID1;refGene_tx=XM_023241605.1;refGene_gn=EMID1;refGene_tx=XM_023241606.1;refGene_gn=EMID1;refGene_tx=XM_023241607.1
D3 23633097 23633266
refGene_gn=KREMEN1;refGene_tx=XM_023241610.1;refGene_gn=KREMEN1;refGene_tx=XM_023241611.1
D3 23634976 23635061
refGene_gn=KREMEN1;refGene_tx=XM_023241610.1;refGene_gn=KREMEN1;refGene_tx=XM_023241611.1
D3 23636158 23636320
refGene_gn=KREMEN1;refGene_tx=XM_023241610.1;refGene_gn=KREMEN1;refGene_tx=XM_023241611.1
D3 23637388 23637721
refGene_gn=KREMEN1;refGene_tx=XM_023241610.1;refGene_gn=KREMEN1;refGene_tx=XM_023241611.1
D3 23648763 23648917
refGene_gn=KREMEN1;refGene_tx=XM_023241610.1;refGene_gn=KREMEN1;refGene_tx=XM_023241611.1
D3 23656724 23656849
refGene_gn=KREMEN1;refGene_tx=XM_023241610.1;refGene_gn=KREMEN1;refGen

e_tx=XM_023241611.1
D3 23676009 23676101
refGene_gn=KREMEN1;refGene_tx=XM_023241610.1;refGene_gn=KREMEN1;refGene_tx=XM_023241611.1
D3 23680628 23680791
refGene_gn=KREMEN1;refGene_tx=XM_023241610.1;refGene_gn=KREMEN1;refGene_tx=XM_023241611.1
D3 23702720 23702817
refGene_gn=KREMEN1;refGene_tx=XM_023241610.1;refGene_gn=KREMEN1;refGene_tx=XM_023241611.1
D3 23710663 23710666
refGene_gn=CD3H22orf31;refGene_tx=XM_019814368.1
D3 23711543 23711990
refGene_gn=CD3H22orf31;refGene_tx=XM_019814368.1;refGene_gn=CD3H22orf31;refGene_tx=XM_019814369.2
D3 23712890 23713328
refGene_gn=CD3H22orf31;refGene_tx=XM_019814368.1;refGene_gn=CD3H22orf31;refGene_tx=XM_019814369.2
D3 23717610 23717651
refGene_gn=ZNRF3;refGene_tx=XM_003994766.5;refGene_gn=ZNRF3;refGene_tx=XM_006938551.4;refGene_gn=ZNRF3;refGene_tx=XM_006938553.4;refGene_gn=ZNRF3;refGene_tx=XM_006938554.4;refGene_gn=ZNRF3;refGene_tx=XM_023241612.1;refGene_gn=ZNRF3;refGene_tx=XM_023241613.1
D3 23719049 23720819
refGene_gn=ZNRF3;refGene_tx=XM_003994766.5;refGene_gn=ZNRF3;refGene_tx=XM_006938551.4;refGene_gn=ZNRF3;refGene_tx=XM_006938553.4;refGene_gn=ZNRF3;refGene_tx=XM_006938554.4;refGene_gn=ZNRF3;refGene_tx=XM_023241612.1;refGene_gn=ZNRF3;refGene_tx=XM_023241613.1
D3 23721571 23721674
refGene_gn=ZNRF3;refGene_tx=XM_003994766.5;refGene_gn=ZNRF3;refGene_tx=XM_006938551.4;refGene_gn=ZNRF3;refGene_tx=XM_006938553.4;refGene_gn=ZNRF3;refGene_tx=XM_006938554.4;refGene_gn=ZNRF3;refGene_tx=XM_023241612.1;refGene_gn=ZNRF3;refGene_tx=XM_023241613.1
D3 23723553 23723721
refGene_gn=ZNRF3;refGene_tx=XM_003994766.5;refGene_gn=ZNRF3;refGene_tx=XM_006938551.4;refGene_gn=ZNRF3;refGene_tx=XM_006938553.4;refGene_gn=ZNRF3;refGene_tx=XM_006938554.4;refGene_gn=ZNRF3;refGene_tx=XM_023241612.1;refGene_gn=ZNRF3;refGene_tx=XM_023241613.1
D3 23724705 23724816
refGene_gn=ZNRF3;refGene_tx=XM_003994766.5;refGene_gn=ZNRF3;refGene_tx=XM_006938551.4;refGene_gn=ZNRF3;refGene_tx=XM_006938553.4;refGene_gn=ZNRF3;refGene_tx=XM_006938554.4;refGene_gn=ZNRF3;refGene_tx=XM_023241612.1;refGene_gn=ZNRF3;refGene_tx=XM_023241613.1
D3 23726431 23726563
refGene_gn=ZNRF3;refGene_tx=XM_003994766.5;refGene_gn=ZNRF3;refGene_tx=XM_006938551.4;refGene_gn=ZNRF3;refGene_tx=XM_006938553.4;refGene_gn=ZNRF3;refGene_tx=XM_006938554.4;refGene_gn=ZNRF3;refGene_tx=XM_023241612.1;refGene_gn=ZNRF3;refGene_tx=XM_023241613.1
D3 23726983 23727058
refGene_gn=ZNRF3;refGene_tx=XM_003994766.5;refGene_gn=ZNRF3;refGene_tx

=XM_006938551.4;refGene_gn=ZNRF3;refGene_tx=XM_006938553.4;refGene_gn=ZNRF3;refGene_tx=XM_006938554.4;refGene_gn=ZNRF3;refGene_tx=XM_023241612.1;refGene_gn=ZNRF3;refGene_tx=XM_023241613.1
D3 23730449 23730554 refGene_gn=ZNRF3;refGene_tx=XM_006938553.4
D3 23734525 23734561
refGene_gn=ZNRF3;refGene_tx=XM_006938554.4;refGene_gn=ZNRF3;refGene_tx=XM_023241613.1
D3 23769600 23769720
refGene_gn=ZNRF3;refGene_tx=XM_006938551.4;refGene_gn=ZNRF3;refGene_tx=XM_023241612.1
D3 23775473 23775599
refGene_gn=ZNRF3;refGene_tx=XM_003994766.5;refGene_gn=ZNRF3;refGene_tx=XM_006938551.4;refGene_gn=ZNRF3;refGene_tx=XM_023241612.1
D3 23870509 23870809 refGene_gn=ZNRF3;refGene_tx=XM_003994766.5
D3 23942482 23942724 refGene_gn=XBP1;refGene_tx=XM_023241614.1
D3 23943689 23943786 refGene_gn=XBP1;refGene_tx=XM_023241614.1
D3 23944594 23944729 refGene_gn=XBP1;refGene_tx=XM_023241614.1
D3 23945730 23945766 refGene_gn=XBP1;refGene_tx=XM_023241614.1
D3 23945768 23945876 refGene_gn=XBP1;refGene_tx=XM_023241614.1
D3 23946190 23946748 refGene_gn=XBP1;refGene_tx=XM_023241614.1
D3 23952699 23952937 refGene_gn=CCDC117;refGene_tx=XM_023241615.1
D3 23956244 23956382 refGene_gn=CCDC117;refGene_tx=XM_023241615.1
D3 23956820 23957045 refGene_gn=CCDC117;refGene_tx=XM_023241615.1
D3 23961716 23961767 refGene_gn=CCDC117;refGene_tx=XM_023241615.1
D3 23962439 23962621 refGene_gn=CCDC117;refGene_tx=XM_023241615.1
D3 23970922 23970960 refGene_gn=HSCB;refGene_tx=XM_019814842.2
D3 23980400 23980492
refGene_gn=HSCB;refGene_tx=XM_003994765.4;refGene_gn=HSCB;refGene_tx=XM_019814841.1;refGene_gn=HSCB;refGene_tx=XM_019814845.1;refGene_gn=HSCB;refGene_tx=XM_023241622.1
D3 23986453 23986485 refGene_gn=HSCB;refGene_tx=XM_019814843.2
D3 23988787 23988835
refGene_gn=HSCB;refGene_tx=XM_003994765.4;refGene_gn=HSCB;refGene_tx=XM_019814842.2;refGene_gn=HSCB;refGene_tx=XM_019814843.2;refGene_gn=HSCB;refGene_tx=XM_019814845.1;refGene_gn=HSCB;refGene_tx=XM_023241622.1
D3 23992731 23992876
refGene_gn=HSCB;refGene_tx=XM_003994765.4;refGene_gn=HSCB;refGene_tx=XM_019814841.1;refGene_gn=HSCB;refGene_tx=XM_019814842.2;refGene_gn=HSCB;refGene_tx=XM_019814843.2;refGene_gn=HSCB;refGene_tx=XM_019814844.1;refGene_gn=HSCB;refGene_tx=XM_019814845.1;refGene_gn=HSCB;refGene_tx=XM_023241622.1
D3 23993505 23993600
refGene_gn=HSCB;refGene_tx=XM_003994765.4;refGene_gn=HSCB;refGene_tx=XM_019814841.1;refGene_gn=HSCB;refGene_tx=XM_019814842.2;refGene_gn=HSCB;refGene_tx=XM_019814843.2;refGene_gn=HSCB;refGene_tx=XM_019814844.1;refGene_gn=HSCB;refGene_tx=XM_023241622.1
D3 23993950 23994047
refGene_gn=HSCB;refGene_tx=XM_003994765.4;refGene_gn=HSCB;refGene_tx=XM_019814841.1;refGene_gn=HSCB;refGene_tx=XM_019814842.2;refGene_gn=HSCB;refGene_tx=XM_019814843.2;refGene_gn=HSCB;refGene_tx=XM_019814844.1;

refGene_gn=HSCB;refGene_tx=XM_023241622.1
D3 23995637 23995669 refGene_gn=HSCB;refGene_tx=XM_023241622.1
D3 23995746 23995982
refGene_gn=HSCB;refGene_tx=XM_003994765.4;refGene_gn=HSCB;refGene_tx=XM_019814841.1;refGene_gn=HSCB;refGene_tx=XM_019814842.2;refGene_gn=HSCB;refGene_tx=XM_019814843.2;refGene_gn=HSCB;refGene_tx=XM_019814844.1
D3 23996243 23996314 refGene_gn=CHEK2;refGene_tx=XM_019814836.2
D3 23997225 23997233 refGene_gn=CHEK2;refGene_tx=XM_023241620.1
D3 23998482 23998807
refGene_gn=CHEK2;refGene_tx=XM_003994763.4;refGene_gn=CHEK2;refGene_tx=XM_011287724.3;refGene_gn=CHEK2;refGene_tx=XM_019814836.2;refGene_gn=CHEK2;refGene_tx=XM_023241616.1;refGene_gn=CHEK2;refGene_tx=XM_023241617.1;refGene_gn=CHEK2;refGene_tx=XM_023241618.1;refGene_gn=CHEK2;refGene_tx=XM_023241619.1;refGene_gn=CHEK2;refGene_tx=XM_023241620.1;refGene_gn=CHEK2;refGene_tx=XM_023241621.1
D3 24016180 24016305
refGene_gn=CHEK2;refGene_tx=XM_003994763.4;refGene_gn=CHEK2;refGene_tx=XM_011287724.3;refGene_gn=CHEK2;refGene_tx=XM_019814836.2;refGene_gn=CHEK2;refGene_tx=XM_023241616.1;refGene_gn=CHEK2;refGene_tx=XM_023241617.1;refGene_gn=CHEK2;refGene_tx=XM_023241618.1;refGene_gn=CHEK2;refGene_tx=XM_023241619.1;refGene_gn=CHEK2;refGene_tx=XM_023241620.1
D3 24016422 24016570
refGene_gn=CHEK2;refGene_tx=XM_003994763.4;refGene_gn=CHEK2;refGene_tx=XM_011287724.3;refGene_gn=CHEK2;refGene_tx=XM_019814836.2;refGene_gn=CHEK2;refGene_tx=XM_019814838.2;refGene_gn=CHEK2;refGene_tx=XM_023241616.1;refGene_gn=CHEK2;refGene_tx=XM_023241617.1;refGene_gn=CHEK2;refGene_tx=XM_023241618.1;refGene_gn=CHEK2;refGene_tx=XM_023241619.1;refGene_gn=CHEK2;refGene_tx=XM_023241620.1
D3 24019692 24019783
refGene_gn=CHEK2;refGene_tx=XM_003994763.4;refGene_gn=CHEK2;refGene_tx=XM_011287724.3;refGene_gn=CHEK2;refGene_tx=XM_019814836.2;refGene_gn=CHEK2;refGene_tx=XM_019814838.2;refGene_gn=CHEK2;refGene_tx=XM_019814839.2;refGene_gn=CHEK2;refGene_tx=XM_023241616.1;refGene_gn=CHEK2;refGene_tx=XM_023241617.1;refGene_gn=CHEK2;refGene_tx=XM_023241618.1;refGene_gn=CHEK2;refGene_tx=XM_023241619.1;refGene_gn=CHEK2;refGene_tx=XM_023241620.1;refGene_gn=CHEK2;refGene_tx=XM_023241621.1
D3 24024377 24024486
refGene_gn=CHEK2;refGene_tx=XM_003994763.4;refGene_gn=CHEK2;refGene_tx=XM_011287724.3;refGene_gn=CHEK2;refGene_tx=XM_019814836.2;refGene_gn=CHEK2;refGene_tx=XM_019814838.2;refGene_gn=CHEK2;refGene_tx=XM_019814839.2;refGene_gn=CHEK2;refGene_tx=XM_023241616.1;refGene_gn=CHEK2;refGene_tx=XM_023241617.1;refGene_gn=CHEK2;refGene_tx=XM_023241618.1;refGene_gn=CHEK2;refGene_tx=XM_023241619.1;refGene_gn=CHEK2;refGene_tx=XM_023241620.1;refGene_gn=CHEK2;refGene_tx=XM_023241621.1
D3 24027388 24027442
refGene_gn=CHEK2;refGene_tx=XM_003994763.4;refGene_gn=CHEK2;refGene_tx=XM_011287724.3;refGene_gn=CHEK2;refGene_tx=XM_019814836.2;refGene_gn=CHEK2;refGene_tx=XM_019814838.2;refGene_gn=CHEK2;refGene_tx=XM_019814839.2;refGene_gn=CHEK2;refGene_tx=XM_023241616.1;refGene_gn=CHEK2;refGene_tx=XM_023241617.1;refGene_gn=CHEK2;refGene_tx=XM_023241618.1;refGen

e_gn=CHEK2;refGene_tx=XM_023241620.1;refGene_gn=CHEK2;refGene_tx=XM_023241621.1
D3 24032127 24032189
refGene_gn=CHEK2;refGene_tx=XM_003994763.4;refGene_gn=CHEK2;refGene_tx=XM_011287724.3;refGene_gn=CHEK2;refGene_tx=XM_019814836.2;refGene_gn=CHEK2;refGene_tx=XM_019814838.2;refGene_gn=CHEK2;refGene_tx=XM_019814839.2;refGene_gn=CHEK2;refGene_tx=XM_023241616.1;refGene_gn=CHEK2;refGene_tx=XM_023241617.1;refGene_gn=CHEK2;refGene_tx=XM_023241618.1;refGene_gn=CHEK2;refGene_tx=XM_023241619.1;refGene_gn=CHEK2;refGene_tx=XM_023241620.1;refGene_gn=CHEK2;refGene_tx=XM_023241621.1
D3 24034425 24034525
refGene_gn=CHEK2;refGene_tx=XM_003994763.4;refGene_gn=CHEK2;refGene_tx=XM_011287724.3;refGene_gn=CHEK2;refGene_tx=XM_019814836.2;refGene_gn=CHEK2;refGene_tx=XM_019814838.2;refGene_gn=CHEK2;refGene_tx=XM_019814839.2;refGene_gn=CHEK2;refGene_tx=XM_023241616.1;refGene_gn=CHEK2;refGene_tx=XM_023241617.1;refGene_gn=CHEK2;refGene_tx=XM_023241618.1;refGene_gn=CHEK2;refGene_tx=XM_023241619.1;refGene_gn=CHEK2;refGene_tx=XM_023241620.1;refGene_gn=CHEK2;refGene_tx=XM_023241621.1
D3 24035858 24035945
refGene_gn=CHEK2;refGene_tx=XM_003994763.4;refGene_gn=CHEK2;refGene_tx=XM_011287724.3;refGene_gn=CHEK2;refGene_tx=XM_019814836.2;refGene_gn=CHEK2;refGene_tx=XM_019814838.2;refGene_gn=CHEK2;refGene_tx=XM_019814839.2;refGene_gn=CHEK2;refGene_tx=XM_023241616.1;refGene_gn=CHEK2;refGene_tx=XM_023241617.1;refGene_gn=CHEK2;refGene_tx=XM_023241618.1;refGene_gn=CHEK2;refGene_tx=XM_023241619.1;refGene_gn=CHEK2;refGene_tx=XM_023241620.1;refGene_gn=CHEK2;refGene_tx=XM_023241621.1
D3 24036674 24036838
refGene_gn=CHEK2;refGene_tx=XM_003994763.4;refGene_gn=CHEK2;refGene_tx=XM_011287724.3;refGene_gn=CHEK2;refGene_tx=XM_019814836.2;refGene_gn=CHEK2;refGene_tx=XM_019814838.2;refGene_gn=CHEK2;refGene_tx=XM_019814839.2;refGene_gn=CHEK2;refGene_tx=XM_023241616.1;refGene_gn=CHEK2;refGene_tx=XM_023241617.1;refGene_gn=CHEK2;refGene_tx=XM_023241618.1;refGene_gn=CHEK2;refGene_tx=XM_023241619.1;refGene_gn=CHEK2;refGene_tx=XM_023241620.1;refGene_gn=CHEK2;refGene_tx=XM_023241621.1
D3 24037036 24037152
refGene_gn=CHEK2;refGene_tx=XM_003994763.4;refGene_gn=CHEK2;refGene_tx=XM_011287724.3;refGene_gn=CHEK2;refGene_tx=XM_019814836.2;refGene_gn=CHEK2;refGene_tx=XM_019814838.2;refGene_gn=CHEK2;refGene_tx=XM_019814839.2;refGene_gn=CHEK2;refGene_tx=XM_023241616.1;refGene_gn=CHEK2;refGene_tx=XM_023241617.1;refGene_gn=CHEK2;refGene_tx=XM_023241618.1;refGene_gn=CHEK2;refGene_tx=XM_023241619.1;refGene_gn=CHEK2;refGene_tx=XM_023241620.1;refGene_gn=CHEK2;refGene_tx=XM_023241621.1
D3 24038164 24038250
refGene_gn=CHEK2;refGene_tx=XM_003994763.4;refGene_gn=CHEK2;refGene_tx=XM_011287724.3;refGene_gn=CHEK2;refGene_tx=XM_019814836.2;refGene_gn=CHEK2;refGene_tx=XM_019814838.2;refGene_gn=CHEK2;refGene_tx=XM_019814839.2;refGene_gn=CHEK2;refGene_tx=XM_023241616.1;refGene_gn=CHEK2;refGene_tx=XM_023241617.1;refGene_gn=CHEK2;refGene_tx=XM_023241618.1;refGene_gn=CHEK2;refGene_tx=XM_023241619.1;refGene_gn=CHEK2;refGene_tx=XM_023241620.1;refGene_gn=CHEK2;refGene_tx=XM_023241621.1

D3 24042464 24042545
refGene_gn=CHEK2;refGene_tx=XM_003994763.4;refGene_gn=CHEK2;refGene_tx=XM_011287724.3;refGene_gn=CHEK2;refGene_tx=XM_019814836.2;refGene_gn=CHEK2;refGene_tx=XM_019814838.2;refGene_gn=CHEK2;refGene_tx=XM_019814839.2;refGene_gn=CHEK2;refGene_tx=XM_023241616.1;refGene_gn=CHEK2;refGene_tx=XM_023241617.1;refGene_gn=CHEK2;refGene_tx=XM_023241618.1;refGene_gn=CHEK2;refGene_tx=XM_023241619.1;refGene_gn=CHEK2;refGene_tx=XM_023241620.1;refGene_gn=CHEK2;refGene_tx=XM_023241621.1
D3 24043372 24043462
refGene_gn=CHEK2;refGene_tx=XM_003994763.4;refGene_gn=CHEK2;refGene_tx=XM_011287724.3;refGene_gn=CHEK2;refGene_tx=XM_019814836.2;refGene_gn=CHEK2;refGene_tx=XM_019814838.2;refGene_gn=CHEK2;refGene_tx=XM_019814839.2;refGene_gn=CHEK2;refGene_tx=XM_023241616.1;refGene_gn=CHEK2;refGene_tx=XM_023241617.1;refGene_gn=CHEK2;refGene_tx=XM_023241618.1;refGene_gn=CHEK2;refGene_tx=XM_023241619.1;refGene_gn=CHEK2;refGene_tx=XM_023241620.1;refGene_gn=CHEK2;refGene_tx=XM_023241621.1
D3 24049337 24049439 refGene_gn=TTC28;refGene_tx=XM_023241623.1
D3 24088191 24088470 refGene_gn=TTC28;refGene_tx=XM_023241623.1
D3 24365427 24365575 refGene_gn=TTC28;refGene_tx=XM_023241623.1
D3 24375774 24376047 refGene_gn=TTC28;refGene_tx=XM_023241623.1
D3 24377009 24377140 refGene_gn=TTC28;refGene_tx=XM_023241623.1
D3 24495419 24495927 refGene_gn=TTC28;refGene_tx=XM_023241623.1
D3 24559144 24560486 refGene_gn=TTC28;refGene_tx=XM_023241623.1
D3 24561764 24562288 refGene_gn=TTC28;refGene_tx=XM_023241623.1
D3 24566068 24566178 refGene_gn=TTC28;refGene_tx=XM_023241623.1
D3 24568411 24568541 refGene_gn=TTC28;refGene_tx=XM_023241623.1
D3 24570959 24571178 refGene_gn=TTC28;refGene_tx=XM_023241623.1
D3 24573143 24573309 refGene_gn=TTC28;refGene_tx=XM_023241623.1
D3 24630551 24630692 refGene_gn=TTC28;refGene_tx=XM_023241623.1
D3 24646553 24646698 refGene_gn=TTC28;refGene_tx=XM_023241623.1
D3 24656504 24656684 refGene_gn=TTC28;refGene_tx=XM_023241623.1
D3 24658861 24659582 refGene_gn=TTC28;refGene_tx=XM_023241623.1
D3 24661710 24661835 refGene_gn=TTC28;refGene_tx=XM_023241623.1
D3 24664218 24664450 refGene_gn=TTC28;refGene_tx=XM_023241623.1
D3 24665018 24665095 refGene_gn=TTC28;refGene_tx=XM_023241623.1
D3 24666676 24666700 refGene_gn=TTC28;refGene_tx=XM_023241623.1
D3 24667416 24667546 refGene_gn=TTC28;refGene_tx=XM_023241623.1
D3 24670692 24670800 refGene_gn=TTC28;refGene_tx=XM_023241623.1
D3 24671956 24673527 refGene_gn=TTC28;refGene_tx=XM_023241623.1
D3 24673528 24673591 refGene_gn=TTC28;refGene_tx=XM_023241623.1
D3 24742064 24742084
refGene_gn=PITPNB;refGene_tx=XM_003994761.5;refGene_gn=PITPNB;refGene_tx=XM_003994762.5
D3 24746796 24746827
refGene_gn=PITPNB;refGene_tx=XM_003994761.5;refGene_gn=PITPNB;refGene_tx=XM_003994762.5
D3 24749833 24749979
refGene_gn=PITPNB;refGene_tx=XM_003994761.5;refGene_gn=PITPNB;refGene_tx=XM_003994762.5
D3 24760889 24760981

refGene_gn=PITPNB;refGene_tx=XM_003994761.5;refGene_gn=PITPNB;refGene_tx=XM_003994762.5
D3 24761620 24761628
refGene_gn=PITPNB;refGene_tx=XM_003994761.5;refGene_gn=PITPNB;refGene_tx=XM_003994762.5
D3 24762135 24762210
refGene_gn=PITPNB;refGene_tx=XM_003994761.5;refGene_gn=PITPNB;refGene_tx=XM_003994762.5
D3 24763794 24763878
refGene_gn=PITPNB;refGene_tx=XM_003994761.5;refGene_gn=PITPNB;refGene_tx=XM_003994762.5
D3 24783922 24784000
refGene_gn=PITPNB;refGene_tx=XM_003994761.5;refGene_gn=PITPNB;refGene_tx=XM_003994762.5
D3 24797161 24797272
refGene_gn=PITPNB;refGene_tx=XM_003994761.5;refGene_gn=PITPNB;refGene_tx=XM_003994762.5
D3 24798893 24799016
refGene_gn=PITPNB;refGene_tx=XM_003994761.5;refGene_gn=PITPNB;refGene_tx=XM_003994762.5
D3 24802332 24802380 refGene_gn=PITPNB;refGene_tx=XM_003994761.5
D3 24803613 24803664 refGene_gn=PITPNB;refGene_tx=XM_003994762.5
D3 24849501 24849592 refGene_gn=MN1;refGene_tx=XM_023241624.1
D3 24849594 24850414 refGene_gn=MN1;refGene_tx=XM_023241624.1
D3 24850470 24851320 refGene_gn=MN1;refGene_tx=XM_023241624.1
D3 24851321 24853311 refGene_gn=MN1;refGene_tx=XM_023241624.1
D3 24894448 24894630 refGene_gn=MN1;refGene_tx=XM_023241624.1
D3 25499570 25499812
refGene_gn=LOC109493003;refGene_tx=XM_019814335.1
D3 25508938 25509059
refGene_gn=LOC109493003;refGene_tx=XM_019814335.1
D3 25857626 25857774
refGene_gn=CRYBA4;refGene_tx=XM_006938543.2;refGene_gn=CRYBA4;refGene_tx=XM_019814849.1
D3 25859320 25859463
refGene_gn=CRYBA4;refGene_tx=XM_006938543.2;refGene_gn=CRYBA4;refGene_tx=XM_019814849.1
D3 25861900 25862042
refGene_gn=CRYBA4;refGene_tx=XM_006938543.2;refGene_gn=CRYBA4;refGene_tx=XM_019814849.1
D3 25863668 25863787
refGene_gn=CRYBA4;refGene_tx=XM_006938543.2;refGene_gn=CRYBA4;refGene_tx=XM_019814849.1
D3 25864412 25864459
refGene_gn=CRYBA4;refGene_tx=XM_006938543.2;refGene_gn=CRYBA4;refGene_tx=XM_019814849.1
D3 25865038 25865069 refGene_gn=CRYBA4;refGene_tx=XM_006938543.2
D3 25870884 25871052
refGene_gn=CRYBB1;refGene_tx=XM_011287721.3;refGene_gn=CRYBB1;refGene_tx=XM_019814850.2;refGene_gn=CRYBB1;refGene_tx=XM_019814851.2

D3 25872955 25873074
refGene_gn=CRYBB1;refGene_tx=XM_011287721.3;refGene_gn=CRYBB1;refGene_tx=XM_019814850.2;refGene_gn=CRYBB1;refGene_tx=XM_019814851.2
D3 25876719 25876852
refGene_gn=CRYBB1;refGene_tx=XM_011287721.3;refGene_gn=CRYBB1;refGene_tx=XM_019814850.2;refGene_gn=CRYBB1;refGene_tx=XM_019814851.2
D3 25879066 25879209
refGene_gn=CRYBB1;refGene_tx=XM_011287721.3;refGene_gn=CRYBB1;refGene_tx=XM_019814850.2;refGene_gn=CRYBB1;refGene_tx=XM_019814851.2
D3 25879976 25880160
refGene_gn=CRYBB1;refGene_tx=XM_011287721.3;refGene_gn=CRYBB1;refGene_tx=XM_019814850.2;refGene_gn=CRYBB1;refGene_tx=XM_019814851.2
D3 25924418 25925260
refGene_gn=TPST2;refGene_tx=XM_023241625.1;refGene_gn=TPST2;refGene_tx=XM_023241626.1
D3 25928119 25928318
refGene_gn=TPST2;refGene_tx=XM_023241625.1;refGene_gn=TPST2;refGene_tx=XM_023241626.1
D3 25933488 25933539
refGene_gn=TPST2;refGene_tx=XM_023241625.1;refGene_gn=TPST2;refGene_tx=XM_023241626.1
D3 25936275 25936317
refGene_gn=TPST2;refGene_tx=XM_023241625.1;refGene_gn=TPST2;refGene_tx=XM_023241626.1
D3 25951309 25951518
refGene_gn=TFIP11;refGene_tx=XM_003994758.5;refGene_gn=TFIP11;refGene_tx=XM_019814852.2;refGene_gn=TFIP11;refGene_tx=XM_019814853.2
D3 25953556 25953710
refGene_gn=TFIP11;refGene_tx=XM_003994758.5;refGene_gn=TFIP11;refGene_tx=XM_019814852.2;refGene_gn=TFIP11;refGene_tx=XM_019814853.2
D3 25954019 25954176
refGene_gn=TFIP11;refGene_tx=XM_003994758.5;refGene_gn=TFIP11;refGene_tx=XM_019814852.2;refGene_gn=TFIP11;refGene_tx=XM_019814853.2
D3 25955872 25956000
refGene_gn=TFIP11;refGene_tx=XM_003994758.5;refGene_gn=TFIP11;refGene_tx=XM_019814852.2;refGene_gn=TFIP11;refGene_tx=XM_019814853.2
D3 25957450 25957603
refGene_gn=TFIP11;refGene_tx=XM_003994758.5;refGene_gn=TFIP11;refGene_tx=XM_019814852.2;refGene_gn=TFIP11;refGene_tx=XM_019814853.2
D3 25959642 25960170
refGene_gn=TFIP11;refGene_tx=XM_003994758.5;refGene_gn=TFIP11;refGene_tx=XM_019814852.2;refGene_gn=TFIP11;refGene_tx=XM_019814853.2
D3 25960251 25960358
refGene_gn=TFIP11;refGene_tx=XM_003994758.5;refGene_gn=TFIP11;refGene_tx=XM_019814852.2;refGene_gn=TFIP11;refGene_tx=XM_019814853.2
D3 25960957 25961126
refGene_gn=TFIP11;refGene_tx=XM_003994758.5;refGene_gn=TFIP11;refGene_tx=XM_019814852.2;refGene_gn=TFIP11;refGene_tx=XM_019814853.2
D3 25961465 25961709
refGene_gn=TFIP11;refGene_tx=XM_003994758.5;refGene_gn=TFIP11;refGene_

tx=XM_019814852.2;refGene_gn=TFIP11;refGene_tx=XM_019814853.2
D3 25962178 25962321
refGene_gn=TFIP11;refGene_tx=XM_003994758.5;refGene_gn=TFIP11;refGene_tx=XM_019814852.2;refGene_gn=TFIP11;refGene_tx=XM_019814853.2
D3 25962788 25962954
refGene_gn=TFIP11;refGene_tx=XM_003994758.5;refGene_gn=TFIP11;refGene_tx=XM_019814852.2;refGene_gn=TFIP11;refGene_tx=XM_019814853.2
D3 25964252 25964608
refGene_gn=TFIP11;refGene_tx=XM_003994758.5;refGene_gn=TFIP11;refGene_tx=XM_019814852.2;refGene_gn=TFIP11;refGene_tx=XM_019814853.2
D3 25964943 25965163 refGene_gn=SRRD;refGene_tx=XM_023241627.1
D3 25965547 25965595 refGene_gn=SRRD;refGene_tx=XM_023241627.1
D3 25966034 25966189 refGene_gn=SRRD;refGene_tx=XM_023241627.1
D3 25968145 25968244 refGene_gn=SRRD;refGene_tx=XM_023241627.1
D3 25968332 25968592 refGene_gn=SRRD;refGene_tx=XM_023241627.1
D3 25969908 25969949 refGene_gn=SRRD;refGene_tx=XM_023241627.1
D3 25980711 25980875 refGene_gn=SRRD;refGene_tx=XM_023241627.1
D3 25982921 25982962
refGene_gn=HPS4;refGene_tx=XM_003994757.5;refGene_gn=HPS4;refGene_tx=XM_019814854.2;refGene_gn=HPS4;refGene_tx=XM_019814855.2;refGene_gn=HPS4;refGene_tx=XM_019814856.2;refGene_gn=HPS4;refGene_tx=XM_019814858.2;refGene_gn=HPS4;refGene_tx=XM_019814859.2;refGene_gn=HPS4;refGene_tx=XM_019814861.2
D3 25984648 25984739
refGene_gn=HPS4;refGene_tx=XM_003994757.5;refGene_gn=HPS4;refGene_tx=XM_019814854.2;refGene_gn=HPS4;refGene_tx=XM_019814855.2;refGene_gn=HPS4;refGene_tx=XM_019814856.2;refGene_gn=HPS4;refGene_tx=XM_019814858.2;refGene_gn=HPS4;refGene_tx=XM_019814859.2;refGene_gn=HPS4;refGene_tx=XM_019814861.2
D3 25986715 25986859
refGene_gn=HPS4;refGene_tx=XM_003994757.5;refGene_gn=HPS4;refGene_tx=XM_019814854.2;refGene_gn=HPS4;refGene_tx=XM_019814855.2;refGene_gn=HPS4;refGene_tx=XM_019814856.2;refGene_gn=HPS4;refGene_tx=XM_019814858.2;refGene_gn=HPS4;refGene_tx=XM_019814859.2;refGene_gn=HPS4;refGene_tx=XM_019814861.2
D3 25990401 25990509
refGene_gn=HPS4;refGene_tx=XM_003994757.5;refGene_gn=HPS4;refGene_tx=XM_019814854.2;refGene_gn=HPS4;refGene_tx=XM_019814855.2;refGene_gn=HPS4;refGene_tx=XM_019814856.2;refGene_gn=HPS4;refGene_tx=XM_019814858.2;refGene_gn=HPS4;refGene_tx=XM_019814859.2
D3 25991144 25991261
refGene_gn=HPS4;refGene_tx=XM_003994757.5;refGene_gn=HPS4;refGene_tx=XM_019814854.2;refGene_gn=HPS4;refGene_tx=XM_019814855.2;refGene_gn=HPS4;refGene_tx=XM_019814856.2;refGene_gn=HPS4;refGene_tx=XM_019814858.2;refGene_gn=HPS4;refGene_tx=XM_019814859.2
D3 25993107 25993202
refGene_gn=HPS4;refGene_tx=XM_003994757.5;refGene_gn=HPS4;refGene_tx=XM_019814854.2;refGene_gn=HPS4;refGene_tx=XM_019814855.2;refGene_gn=HPS4;refGene_tx=XM_019814856.2;refGene_gn=HPS4;refGene_tx=XM_019814858.2;refGene_gn=HPS4;refGene_tx=XM_019814859.2;refGene_gn=HPS4;refGene_tx=X

M_019814861.2
D3 25995367 25995440
refGene_gn=HPS4;refGene_tx=XM_003994757.5;refGene_gn=HPS4;refGene_tx=XM_019814854.2;refGene_gn=HPS4;refGene_tx=XM_019814855.2;refGene_gn=HPS4;refGene_tx=XM_019814856.2;refGene_gn=HPS4;refGene_tx=XM_019814858.2;refGene_gn=HPS4;refGene_tx=XM_019814859.2;refGene_gn=HPS4;refGene_tx=XM_019814861.2
D3 25996350 25996387
refGene_gn=HPS4;refGene_tx=XM_003994757.5;refGene_gn=HPS4;refGene_tx=XM_019814854.2;refGene_gn=HPS4;refGene_tx=XM_019814855.2;refGene_gn=HPS4;refGene_tx=XM_019814856.2;refGene_gn=HPS4;refGene_tx=XM_019814858.2;refGene_gn=HPS4;refGene_tx=XM_019814859.2;refGene_gn=HPS4;refGene_tx=XM_019814861.2
D3 25997326 25997423
refGene_gn=HPS4;refGene_tx=XM_003994757.5;refGene_gn=HPS4;refGene_tx=XM_019814854.2;refGene_gn=HPS4;refGene_tx=XM_019814855.2;refGene_gn=HPS4;refGene_tx=XM_019814856.2;refGene_gn=HPS4;refGene_tx=XM_019814858.2;refGene_gn=HPS4;refGene_tx=XM_019814859.2;refGene_gn=HPS4;refGene_tx=XM_019814861.2
D3 25997941 25998833
refGene_gn=HPS4;refGene_tx=XM_003994757.5;refGene_gn=HPS4;refGene_tx=XM_019814854.2;refGene_gn=HPS4;refGene_tx=XM_019814855.2;refGene_gn=HPS4;refGene_tx=XM_019814856.2;refGene_gn=HPS4;refGene_tx=XM_019814858.2;refGene_gn=HPS4;refGene_tx=XM_019814859.2;refGene_gn=HPS4;refGene_tx=XM_019814861.2
D3 26004055 26004188
refGene_gn=HPS4;refGene_tx=XM_003994757.5;refGene_gn=HPS4;refGene_tx=XM_019814854.2;refGene_gn=HPS4;refGene_tx=XM_019814855.2;refGene_gn=HPS4;refGene_tx=XM_019814856.2;refGene_gn=HPS4;refGene_tx=XM_019814858.2;refGene_gn=HPS4;refGene_tx=XM_019814859.2;refGene_gn=HPS4;refGene_tx=XM_019814861.2
D3 26004681 26004790
refGene_gn=HPS4;refGene_tx=XM_003994757.5;refGene_gn=HPS4;refGene_tx=XM_019814854.2;refGene_gn=HPS4;refGene_tx=XM_019814855.2;refGene_gn=HPS4;refGene_tx=XM_019814856.2;refGene_gn=HPS4;refGene_tx=XM_019814858.2;refGene_gn=HPS4;refGene_tx=XM_019814859.2;refGene_gn=HPS4;refGene_tx=XM_019814861.2
D3 26005484 26005494
refGene_gn=HPS4;refGene_tx=XM_019814858.2;refGene_gn=HPS4;refGene_tx=XM_019814859.2
D3 26006993 26007144 refGene_gn=HPS4;refGene_tx=XM_019814856.2
D3 26007221 26007393
refGene_gn=HPS4;refGene_tx=XM_003994757.5;refGene_gn=HPS4;refGene_tx=XM_019814854.2;refGene_gn=HPS4;refGene_tx=XM_019814855.2;refGene_gn=HPS4;refGene_tx=XM_019814861.2
D3 26017078 26017188 refGene_gn=ASPHD2;refGene_tx=XM_003994756.5
D3 26017980 26018094 refGene_gn=ASPHD2;refGene_tx=XM_003994756.5
D3 26023087 26023904 refGene_gn=ASPHD2;refGene_tx=XM_003994756.5
D3 26023906 26023978 refGene_gn=ASPHD2;refGene_tx=XM_003994756.5
D3 26065750 26065780 refGene_gn=SEZ6L;refGene_tx=XM_023242157.1

D3 26067423 26067526 refGene_gn=SEZ6L;refGene_tx=XM_023242157.1
D3 26069273 26069388 refGene_gn=SEZ6L;refGene_tx=XM_023242157.1
D3 26071232 26071265 refGene_gn=SEZ6L;refGene_tx=XM_023242157.1
D3 26077068 26077263 refGene_gn=SEZ6L;refGene_tx=XM_023242157.1
D3 26084018 26084210 refGene_gn=SEZ6L;refGene_tx=XM_023242157.1
D3 26088724 26088919 refGene_gn=SEZ6L;refGene_tx=XM_023242157.1
D3 26094668 26094865 refGene_gn=SEZ6L;refGene_tx=XM_023242157.1
D3 26117530 26117669 refGene_gn=SEZ6L;refGene_tx=XM_023242157.1
D3 26119163 26119358 refGene_gn=SEZ6L;refGene_tx=XM_023242157.1
D3 26119764 26119931 refGene_gn=SEZ6L;refGene_tx=XM_023242157.1
D3 26122153 26122319 refGene_gn=SEZ6L;refGene_tx=XM_023242157.1
D3 26126487 26126673 refGene_gn=SEZ6L;refGene_tx=XM_023242157.1
D3 26128848 26129041 refGene_gn=SEZ6L;refGene_tx=XM_023242157.1
D3 26131336 26131470 refGene_gn=SEZ6L;refGene_tx=XM_023242157.1
D3 26133157 26133871 refGene_gn=SEZ6L;refGene_tx=XM_023242157.1
D3 26246634 26247272 refGene_gn=SEZ6L;refGene_tx=XM_023242157.1
D3 26248859 26248885 refGene_gn=SEZ6L;refGene_tx=XM_023242157.1
D3 26364112 26365349
refGene_gn=MYO18B;refGene_tx=XM_006938537.4;refGene_gn=MYO18B;refGene_tx=XM_023241628.1;refGene_gn=MYO18B;refGene_tx=XM_023241629.1;refGene_gn=MYO18B;refGene_tx=XM_023241630.1;refGene_gn=MYO18B;refGene_tx=XM_023241631.1
D3 26388767 26388905
refGene_gn=MYO18B;refGene_tx=XM_006938537.4;refGene_gn=MYO18B;refGene_tx=XM_023241628.1;refGene_gn=MYO18B;refGene_tx=XM_023241629.1;refGene_gn=MYO18B;refGene_tx=XM_023241630.1;refGene_gn=MYO18B;refGene_tx=XM_023241631.1
D3 26390278 26390323
refGene_gn=MYO18B;refGene_tx=XM_006938537.4;refGene_gn=MYO18B;refGene_tx=XM_023241628.1;refGene_gn=MYO18B;refGene_tx=XM_023241629.1;refGene_gn=MYO18B;refGene_tx=XM_023241630.1;refGene_gn=MYO18B;refGene_tx=XM_023241631.1
D3 26401747 26401878
refGene_gn=MYO18B;refGene_tx=XM_006938537.4;refGene_gn=MYO18B;refGene_tx=XM_023241628.1;refGene_gn=MYO18B;refGene_tx=XM_023241629.1;refGene_gn=MYO18B;refGene_tx=XM_023241630.1;refGene_gn=MYO18B;refGene_tx=XM_023241631.1
D3 26435599 26435785
refGene_gn=MYO18B;refGene_tx=XM_006938537.4;refGene_gn=MYO18B;refGene_tx=XM_023241628.1;refGene_gn=MYO18B;refGene_tx=XM_023241629.1;refGene_gn=MYO18B;refGene_tx=XM_023241630.1;refGene_gn=MYO18B;refGene_tx=XM_023241631.1
D3 26438545 26438683
refGene_gn=MYO18B;refGene_tx=XM_006938537.4;refGene_gn=MYO18B;refGene_tx=XM_023241628.1;refGene_gn=MYO18B;refGene_tx=XM_023241629.1;refGene_gn=MYO18B;refGene_tx=XM_023241630.1;refGene_gn=MYO18B;refGene_tx=XM_023241631.1
D3 26439771 26439855
refGene_gn=MYO18B;refGene_tx=XM_006938537.4;refGene_gn=MYO18B;refGene_tx=XM_023241628.1;refGene_gn=MYO18B;refGene_tx=XM_023241629.1;refGene_

gn=MYO18B;refGene_tx=XM_023241630.1;refGene_gn=MYO18B;refGene_tx=XM_023241631.1
D3	26442668	26442785
refGene_gn=MYO18B;refGene_tx=XM_006938537.4;refGene_gn=MYO18B;refGene_tx=XM_023241628.1;refGene_gn=MYO18B;refGene_tx=XM_023241629.1;refGene_gn=MYO18B;refGene_tx=XM_023241630.1;refGene_gn=MYO18B;refGene_tx=XM_023241631.1
D3	26444082	26444196
refGene_gn=MYO18B;refGene_tx=XM_006938537.4;refGene_gn=MYO18B;refGene_tx=XM_023241628.1;refGene_gn=MYO18B;refGene_tx=XM_023241629.1;refGene_gn=MYO18B;refGene_tx=XM_023241630.1;refGene_gn=MYO18B;refGene_tx=XM_023241631.1
D3	26464071	26464224
refGene_gn=MYO18B;refGene_tx=XM_006938537.4;refGene_gn=MYO18B;refGene_tx=XM_023241628.1;refGene_gn=MYO18B;refGene_tx=XM_023241629.1;refGene_gn=MYO18B;refGene_tx=XM_023241630.1;refGene_gn=MYO18B;refGene_tx=XM_023241631.1
D3	26471196	26471301
refGene_gn=MYO18B;refGene_tx=XM_006938537.4;refGene_gn=MYO18B;refGene_tx=XM_023241628.1;refGene_gn=MYO18B;refGene_tx=XM_023241629.1;refGene_gn=MYO18B;refGene_tx=XM_023241630.1;refGene_gn=MYO18B;refGene_tx=XM_023241631.1
D3	26475986	26476097
refGene_gn=MYO18B;refGene_tx=XM_006938537.4;refGene_gn=MYO18B;refGene_tx=XM_023241628.1;refGene_gn=MYO18B;refGene_tx=XM_023241629.1;refGene_gn=MYO18B;refGene_tx=XM_023241630.1;refGene_gn=MYO18B;refGene_tx=XM_023241631.1
D3	26480546	26480747
refGene_gn=MYO18B;refGene_tx=XM_006938537.4;refGene_gn=MYO18B;refGene_tx=XM_023241628.1;refGene_gn=MYO18B;refGene_tx=XM_023241629.1;refGene_gn=MYO18B;refGene_tx=XM_023241630.1;refGene_gn=MYO18B;refGene_tx=XM_023241631.1
D3	26481554	26481678
refGene_gn=MYO18B;refGene_tx=XM_006938537.4;refGene_gn=MYO18B;refGene_tx=XM_023241628.1;refGene_gn=MYO18B;refGene_tx=XM_023241629.1;refGene_gn=MYO18B;refGene_tx=XM_023241630.1;refGene_gn=MYO18B;refGene_tx=XM_023241631.1
D3	26485787	26485942
refGene_gn=MYO18B;refGene_tx=XM_006938537.4;refGene_gn=MYO18B;refGene_tx=XM_023241628.1;refGene_gn=MYO18B;refGene_tx=XM_023241629.1;refGene_gn=MYO18B;refGene_tx=XM_023241630.1;refGene_gn=MYO18B;refGene_tx=XM_023241631.1
D3	26490648	26490773
refGene_gn=MYO18B;refGene_tx=XM_006938537.4;refGene_gn=MYO18B;refGene_tx=XM_023241628.1;refGene_gn=MYO18B;refGene_tx=XM_023241629.1;refGene_gn=MYO18B;refGene_tx=XM_023241630.1;refGene_gn=MYO18B;refGene_tx=XM_023241631.1
D3	26495263	26495372
refGene_gn=MYO18B;refGene_tx=XM_006938537.4;refGene_gn=MYO18B;refGene_tx=XM_023241628.1;refGene_gn=MYO18B;refGene_tx=XM_023241629.1;refGene_

gn=MYO18B;refGene_tx=XM_023241630.1;refGene_gn=MYO18B;refGene_tx=XM_023241631.1
D3 26495688 26495808
refGene_gn=MYO18B;refGene_tx=XM_006938537.4;refGene_gn=MYO18B;refGene_tx=XM_023241628.1;refGene_gn=MYO18B;refGene_tx=XM_023241629.1;refGene_gn=MYO18B;refGene_tx=XM_023241630.1;refGene_gn=MYO18B;refGene_tx=XM_023241631.1
D3 26510252 26510342
refGene_gn=MYO18B;refGene_tx=XM_006938537.4;refGene_gn=MYO18B;refGene_tx=XM_023241628.1;refGene_gn=MYO18B;refGene_tx=XM_023241629.1;refGene_gn=MYO18B;refGene_tx=XM_023241630.1;refGene_gn=MYO18B;refGene_tx=XM_023241631.1
D3 26511304 26511448
refGene_gn=MYO18B;refGene_tx=XM_006938537.4;refGene_gn=MYO18B;refGene_tx=XM_023241628.1;refGene_gn=MYO18B;refGene_tx=XM_023241629.1;refGene_gn=MYO18B;refGene_tx=XM_023241630.1;refGene_gn=MYO18B;refGene_tx=XM_023241631.1
D3 26513434 26513563
refGene_gn=MYO18B;refGene_tx=XM_006938537.4;refGene_gn=MYO18B;refGene_tx=XM_023241628.1;refGene_gn=MYO18B;refGene_tx=XM_023241629.1;refGene_gn=MYO18B;refGene_tx=XM_023241630.1;refGene_gn=MYO18B;refGene_tx=XM_023241631.1
D3 26518454 26518520
refGene_gn=MYO18B;refGene_tx=XM_006938537.4;refGene_gn=MYO18B;refGene_tx=XM_023241628.1;refGene_gn=MYO18B;refGene_tx=XM_023241629.1;refGene_gn=MYO18B;refGene_tx=XM_023241630.1;refGene_gn=MYO18B;refGene_tx=XM_023241631.1
D3 26528946 26529056
refGene_gn=MYO18B;refGene_tx=XM_006938537.4;refGene_gn=MYO18B;refGene_tx=XM_023241628.1;refGene_gn=MYO18B;refGene_tx=XM_023241629.1;refGene_gn=MYO18B;refGene_tx=XM_023241630.1;refGene_gn=MYO18B;refGene_tx=XM_023241631.1
D3 26533631 26533854
refGene_gn=MYO18B;refGene_tx=XM_006938537.4;refGene_gn=MYO18B;refGene_tx=XM_023241628.1;refGene_gn=MYO18B;refGene_tx=XM_023241629.1;refGene_gn=MYO18B;refGene_tx=XM_023241630.1;refGene_gn=MYO18B;refGene_tx=XM_023241631.1
D3 26534696 26534880
refGene_gn=MYO18B;refGene_tx=XM_006938537.4;refGene_gn=MYO18B;refGene_tx=XM_023241628.1;refGene_gn=MYO18B;refGene_tx=XM_023241629.1;refGene_gn=MYO18B;refGene_tx=XM_023241630.1;refGene_gn=MYO18B;refGene_tx=XM_023241631.1
D3 26537834 26537994
refGene_gn=MYO18B;refGene_tx=XM_006938537.4;refGene_gn=MYO18B;refGene_tx=XM_023241628.1;refGene_gn=MYO18B;refGene_tx=XM_023241629.1;refGene_gn=MYO18B;refGene_tx=XM_023241630.1;refGene_gn=MYO18B;refGene_tx=XM_023241631.1
D3 26545090 26545238
refGene_gn=MYO18B;refGene_tx=XM_006938537.4;refGene_gn=MYO18B;refGene_tx=XM_023241628.1;refGene_gn=MYO18B;refGene_tx=XM_023241629.1;refGene_

gn=MYO18B;refGene_tx=XM_023241630.1;refGene_gn=MYO18B;refGene_tx=XM_023241631.1
D3 26547237 26547318
refGene_gn=MYO18B;refGene_tx=XM_006938537.4;refGene_gn=MYO18B;refGene_tx=XM_023241628.1;refGene_gn=MYO18B;refGene_tx=XM_023241629.1;refGene_gn=MYO18B;refGene_tx=XM_023241630.1;refGene_gn=MYO18B;refGene_tx=XM_023241631.1
D3 26550725 26550918
refGene_gn=MYO18B;refGene_tx=XM_006938537.4;refGene_gn=MYO18B;refGene_tx=XM_023241628.1;refGene_gn=MYO18B;refGene_tx=XM_023241629.1;refGene_gn=MYO18B;refGene_tx=XM_023241630.1;refGene_gn=MYO18B;refGene_tx=XM_023241631.1
D3 26552047 26552138
refGene_gn=MYO18B;refGene_tx=XM_006938537.4;refGene_gn=MYO18B;refGene_tx=XM_023241628.1;refGene_gn=MYO18B;refGene_tx=XM_023241629.1;refGene_gn=MYO18B;refGene_tx=XM_023241630.1;refGene_gn=MYO18B;refGene_tx=XM_023241631.1
D3 26555133 26555307
refGene_gn=MYO18B;refGene_tx=XM_006938537.4;refGene_gn=MYO18B;refGene_tx=XM_023241628.1;refGene_gn=MYO18B;refGene_tx=XM_023241629.1;refGene_gn=MYO18B;refGene_tx=XM_023241630.1;refGene_gn=MYO18B;refGene_tx=XM_023241631.1
D3 26580043 26580188
refGene_gn=MYO18B;refGene_tx=XM_006938537.4;refGene_gn=MYO18B;refGene_tx=XM_023241628.1;refGene_gn=MYO18B;refGene_tx=XM_023241629.1;refGene_gn=MYO18B;refGene_tx=XM_023241630.1;refGene_gn=MYO18B;refGene_tx=XM_023241631.1
D3 26590326 26590390
refGene_gn=MYO18B;refGene_tx=XM_006938537.4;refGene_gn=MYO18B;refGene_tx=XM_023241628.1;refGene_gn=MYO18B;refGene_tx=XM_023241629.1;refGene_gn=MYO18B;refGene_tx=XM_023241630.1;refGene_gn=MYO18B;refGene_tx=XM_023241631.1
D3 26590721 26590822
refGene_gn=MYO18B;refGene_tx=XM_006938537.4;refGene_gn=MYO18B;refGene_tx=XM_023241628.1;refGene_gn=MYO18B;refGene_tx=XM_023241629.1;refGene_gn=MYO18B;refGene_tx=XM_023241630.1;refGene_gn=MYO18B;refGene_tx=XM_023241631.1
D3 26591618 26591761
refGene_gn=MYO18B;refGene_tx=XM_006938537.4;refGene_gn=MYO18B;refGene_tx=XM_023241628.1;refGene_gn=MYO18B;refGene_tx=XM_023241629.1;refGene_gn=MYO18B;refGene_tx=XM_023241630.1;refGene_gn=MYO18B;refGene_tx=XM_023241631.1
D3 26593770 26593969
refGene_gn=MYO18B;refGene_tx=XM_006938537.4;refGene_gn=MYO18B;refGene_tx=XM_023241628.1;refGene_gn=MYO18B;refGene_tx=XM_023241629.1;refGene_gn=MYO18B;refGene_tx=XM_023241630.1;refGene_gn=MYO18B;refGene_tx=XM_023241631.1
D3 26598211 26598388
refGene_gn=MYO18B;refGene_tx=XM_006938537.4;refGene_gn=MYO18B;refGene_tx=XM_023241628.1;refGene_gn=MYO18B;refGene_tx=XM_023241629.1;refGene_

gn=MYO18B;refGene_tx=XM_023241630.1;refGene_gn=MYO18B;refGene_tx=XM_023241631.1
D3 26599668 26599781
refGene_gn=MYO18B;refGene_tx=XM_006938537.4;refGene_gn=MYO18B;refGene_tx=XM_023241628.1;refGene_gn=MYO18B;refGene_tx=XM_023241629.1;refGene_gn=MYO18B;refGene_tx=XM_023241630.1;refGene_gn=MYO18B;refGene_tx=XM_023241631.1
D3 26600065 26600132
refGene_gn=MYO18B;refGene_tx=XM_006938537.4;refGene_gn=MYO18B;refGene_tx=XM_023241628.1;refGene_gn=MYO18B;refGene_tx=XM_023241629.1;refGene_gn=MYO18B;refGene_tx=XM_023241630.1;refGene_gn=MYO18B;refGene_tx=XM_023241631.1
D3 26600636 26601101
refGene_gn=MYO18B;refGene_tx=XM_006938537.4;refGene_gn=MYO18B;refGene_tx=XM_023241628.1;refGene_gn=MYO18B;refGene_tx=XM_023241629.1;refGene_gn=MYO18B;refGene_tx=XM_023241630.1;refGene_gn=MYO18B;refGene_tx=XM_023241631.1
D3 26601110 26601998
refGene_gn=MYO18B;refGene_tx=XM_006938537.4;refGene_gn=MYO18B;refGene_tx=XM_023241628.1;refGene_gn=MYO18B;refGene_tx=XM_023241629.1;refGene_gn=MYO18B;refGene_tx=XM_023241630.1;refGene_gn=MYO18B;refGene_tx=XM_023241631.1
D3 26607496 26607655
refGene_gn=MYO18B;refGene_tx=XM_006938537.4;refGene_gn=MYO18B;refGene_tx=XM_023241628.1;refGene_gn=MYO18B;refGene_tx=XM_023241629.1;refGene_gn=MYO18B;refGene_tx=XM_023241630.1;refGene_gn=MYO18B;refGene_tx=XM_023241631.1
D3 26609065 26609104
refGene_gn=MYO18B;refGene_tx=XM_006938537.4;refGene_gn=MYO18B;refGene_tx=XM_023241628.1;refGene_gn=MYO18B;refGene_tx=XM_023241629.1;refGene_gn=MYO18B;refGene_tx=XM_023241630.1;refGene_gn=MYO18B;refGene_tx=XM_023241631.1
D3 26638908 26639070
refGene_gn=GRK3;refGene_tx=XM_019814870.2;refGene_gn=GRK3;refGene_tx=XM_023241632.1;refGene_gn=GRK3;refGene_tx=XM_023241633.1
D3 26640295 26640409
refGene_gn=GRK3;refGene_tx=XM_019814870.2;refGene_gn=GRK3;refGene_tx=XM_023241632.1;refGene_gn=GRK3;refGene_tx=XM_023241633.1
D3 26642816 26642953
refGene_gn=GRK3;refGene_tx=XM_019814870.2;refGene_gn=GRK3;refGene_tx=XM_023241632.1;refGene_gn=GRK3;refGene_tx=XM_023241633.1
D3 26643234 26643335 refGene_gn=GRK3;refGene_tx=XM_023241634.1
D3 26645661 26645824
refGene_gn=GRK3;refGene_tx=XM_019814870.2;refGene_gn=GRK3;refGene_tx=XM_023241632.1;refGene_gn=GRK3;refGene_tx=XM_023241633.1;refGene_gn=GRK3;refGene_tx=XM_023241634.1
D3 26647497 26647593
refGene_gn=GRK3;refGene_tx=XM_019814870.2;refGene_gn=GRK3;refGene_tx=XM_023241632.1;refGene_gn=GRK3;refGene_tx=XM_023241633.1;refGene_gn=GRK3;refGene_tx=XM_023241634.1

D3 26648624 26648691
refGene_gn=GRK3;refGene_tx=XM_019814870.2;refGene_gn=GRK3;refGene_tx=XM_023241632.1;refGene_gn=GRK3;refGene_tx=XM_023241633.1;refGene_gn=GRK3;refGene_tx=XM_023241634.1
D3 26653258 26653359
refGene_gn=GRK3;refGene_tx=XM_019814870.2;refGene_gn=GRK3;refGene_tx=XM_023241632.1;refGene_gn=GRK3;refGene_tx=XM_023241633.1;refGene_gn=GRK3;refGene_tx=XM_023241634.1
D3 26654167 26654234
refGene_gn=GRK3;refGene_tx=XM_019814870.2;refGene_gn=GRK3;refGene_tx=XM_023241632.1;refGene_gn=GRK3;refGene_tx=XM_023241633.1;refGene_gn=GRK3;refGene_tx=XM_023241634.1
D3 26662081 26662189
refGene_gn=GRK3;refGene_tx=XM_019814870.2;refGene_gn=GRK3;refGene_tx=XM_023241632.1;refGene_gn=GRK3;refGene_tx=XM_023241633.1;refGene_gn=GRK3;refGene_tx=XM_023241634.1
D3 26666688 26666783
refGene_gn=GRK3;refGene_tx=XM_019814870.2;refGene_gn=GRK3;refGene_tx=XM_023241632.1;refGene_gn=GRK3;refGene_tx=XM_023241633.1;refGene_gn=GRK3;refGene_tx=XM_023241634.1
D3 26668969 26669100
refGene_gn=GRK3;refGene_tx=XM_019814870.2;refGene_gn=GRK3;refGene_tx=XM_023241632.1;refGene_gn=GRK3;refGene_tx=XM_023241633.1;refGene_gn=GRK3;refGene_tx=XM_023241634.1
D3 26670690 26670769
refGene_gn=GRK3;refGene_tx=XM_019814870.2;refGene_gn=GRK3;refGene_tx=XM_023241632.1;refGene_gn=GRK3;refGene_tx=XM_023241633.1;refGene_gn=GRK3;refGene_tx=XM_023241634.1
D3 26678193 26678293
refGene_gn=GRK3;refGene_tx=XM_019814870.2;refGene_gn=GRK3;refGene_tx=XM_023241632.1;refGene_gn=GRK3;refGene_tx=XM_023241634.1
D3 26682605 26682697
refGene_gn=GRK3;refGene_tx=XM_019814870.2;refGene_gn=GRK3;refGene_tx=XM_023241632.1;refGene_gn=GRK3;refGene_tx=XM_023241634.1
D3 26684719 26684771
refGene_gn=GRK3;refGene_tx=XM_019814870.2;refGene_gn=GRK3;refGene_tx=XM_023241632.1;refGene_gn=GRK3;refGene_tx=XM_023241633.1;refGene_gn=GRK3;refGene_tx=XM_023241634.1
D3 26688750 26688812
refGene_gn=GRK3;refGene_tx=XM_019814870.2;refGene_gn=GRK3;refGene_tx=XM_023241632.1;refGene_gn=GRK3;refGene_tx=XM_023241633.1;refGene_gn=GRK3;refGene_tx=XM_023241634.1
D3 26689918 26689993
refGene_gn=GRK3;refGene_tx=XM_019814870.2;refGene_gn=GRK3;refGene_tx=XM_023241632.1;refGene_gn=GRK3;refGene_tx=XM_023241633.1;refGene_gn=GRK3;refGene_tx=XM_023241634.1
D3 26692186 26692288
refGene_gn=GRK3;refGene_tx=XM_019814870.2;refGene_gn=GRK3;refGene_tx=XM_023241632.1;refGene_gn=GRK3;refGene_tx=XM_023241633.1;refGene_gn=GRK3;refGene_tx=XM_023241634.1

D3 26702770 26702844
refGene_gn=GRK3;refGene_tx=XM_023241632.1;refGene_gn=GRK3;refGene_tx=XM_023241633.1;refGene_gn=GRK3;refGene_tx=XM_023241634.1
D3 26738845 26738922
refGene_gn=GRK3;refGene_tx=XM_023241632.1;refGene_gn=GRK3;refGene_tx=XM_023241633.1;refGene_gn=GRK3;refGene_tx=XM_023241634.1
D3 26764359 26764472
refGene_gn=GRK3;refGene_tx=XM_023241632.1;refGene_gn=GRK3;refGene_tx=XM_023241633.1;refGene_gn=GRK3;refGene_tx=XM_023241634.1
D3 26847317 26847486 refGene_gn=CRYBB2;refGene_tx=XM_006938536.2
D3 26849041 26849184 refGene_gn=CRYBB2;refGene_tx=XM_006938536.2
D3 26850347 26850480 refGene_gn=CRYBB2;refGene_tx=XM_006938536.2
D3 26851663 26851782 refGene_gn=CRYBB2;refGene_tx=XM_006938536.2
D3 26854622 26854676 refGene_gn=CRYBB2;refGene_tx=XM_006938536.2
D3 26865065 26865231 refGene_gn=CRYBB3;refGene_tx=XM_006938535.2
D3 26866343 26866486 refGene_gn=CRYBB3;refGene_tx=XM_006938535.2
D3 26867380 26867513 refGene_gn=CRYBB3;refGene_tx=XM_006938535.2
D3 26868048 26868167 refGene_gn=CRYBB3;refGene_tx=XM_006938535.2
D3 26869753 26869828 refGene_gn=CRYBB3;refGene_tx=XM_006938535.2
D3 26879853 26879932
refGene_gn=KIAA1671;refGene_tx=XM_003994871.5;refGene_gn=KIAA1671;refGene_tx=XM_019814871.1
D3 26883526 26883660
refGene_gn=KIAA1671;refGene_tx=XM_003994871.5;refGene_gn=KIAA1671;refGene_tx=XM_019814871.1
D3 26885088 26885213
refGene_gn=KIAA1671;refGene_tx=XM_003994871.5;refGene_gn=KIAA1671;refGene_tx=XM_019814871.1
D3 26887318 26887493
refGene_gn=KIAA1671;refGene_tx=XM_003994871.5;refGene_gn=KIAA1671;refGene_tx=XM_019814871.1
D3 26890272 26890528
refGene_gn=KIAA1671;refGene_tx=XM_003994871.5;refGene_gn=KIAA1671;refGene_tx=XM_019814871.1
D3 26894290 26894409
refGene_gn=KIAA1671;refGene_tx=XM_003994871.5;refGene_gn=KIAA1671;refGene_tx=XM_019814871.1
D3 27002973 27003108
refGene_gn=KIAA1671;refGene_tx=XM_003994871.5;refGene_gn=KIAA1671;refGene_tx=XM_019814871.1
D3 27010039 27012889
refGene_gn=KIAA1671;refGene_tx=XM_003994871.5;refGene_gn=KIAA1671;refGene_tx=XM_019814871.1
D3 27015520 27015608
refGene_gn=KIAA1671;refGene_tx=XM_003994871.5;refGene_gn=KIAA1671;refGene_tx=XM_019814871.1
D3 27017577 27019133
refGene_gn=KIAA1671;refGene_tx=XM_003994871.5;refGene_gn=KIAA1671;refGene_tx=XM_019814871.1
D3 27086237 27086302 refGene_gn=TMEM211;refGene_tx=XM_023242158.1

D3 27086408 27086550 refGene_gn=TMEM211;refGene_tx=XM_023242158.1
D3 27099592 27099709 refGene_gn=TMEM211;refGene_tx=XM_023242158.1
D3 27100461 27100600 refGene_gn=TMEM211;refGene_tx=XM_023242158.1
D3 27102675 27102938 refGene_gn=TMEM211;refGene_tx=XM_023242158.1
D3 27111002 27111091
refGene_gn=SGSM1;refGene_tx=XM_023241635.1;refGene_gn=SGSM1;refGene_tx=XM_023241636.1;refGene_gn=SGSM1;refGene_tx=XM_023241637.1;refGene_gn=SGSM1;refGene_tx=XM_023241638.1;refGene_gn=SGSM1;refGene_tx=XM_023241639.1;refGene_gn=SGSM1;refGene_tx=XM_023241640.1;refGene_gn=SGSM1;refGene_tx=XM_023241641.1;refGene_gn=SGSM1;refGene_tx=XM_023241644.1;refGene_gn=SGSM1;refGene_tx=XM_023241645.1
D3 27116504 27116672
refGene_gn=SGSM1;refGene_tx=XM_023241635.1;refGene_gn=SGSM1;refGene_tx=XM_023241636.1;refGene_gn=SGSM1;refGene_tx=XM_023241637.1;refGene_gn=SGSM1;refGene_tx=XM_023241638.1;refGene_gn=SGSM1;refGene_tx=XM_023241639.1;refGene_gn=SGSM1;refGene_tx=XM_023241640.1;refGene_gn=SGSM1;refGene_tx=XM_023241641.1;refGene_gn=SGSM1;refGene_tx=XM_023241644.1;refGene_gn=SGSM1;refGene_tx=XM_023241645.1
D3 27118071 27118168
refGene_gn=SGSM1;refGene_tx=XM_023241635.1;refGene_gn=SGSM1;refGene_tx=XM_023241636.1;refGene_gn=SGSM1;refGene_tx=XM_023241637.1;refGene_gn=SGSM1;refGene_tx=XM_023241638.1;refGene_gn=SGSM1;refGene_tx=XM_023241639.1;refGene_gn=SGSM1;refGene_tx=XM_023241640.1;refGene_gn=SGSM1;refGene_tx=XM_023241641.1;refGene_gn=SGSM1;refGene_tx=XM_023241644.1;refGene_gn=SGSM1;refGene_tx=XM_023241645.1
D3 27118820 27118930
refGene_gn=SGSM1;refGene_tx=XM_023241635.1;refGene_gn=SGSM1;refGene_tx=XM_023241636.1;refGene_gn=SGSM1;refGene_tx=XM_023241637.1;refGene_gn=SGSM1;refGene_tx=XM_023241638.1;refGene_gn=SGSM1;refGene_tx=XM_023241639.1;refGene_gn=SGSM1;refGene_tx=XM_023241640.1;refGene_gn=SGSM1;refGene_tx=XM_023241641.1;refGene_gn=SGSM1;refGene_tx=XM_023241644.1;refGene_gn=SGSM1;refGene_tx=XM_023241645.1
D3 27124886 27124969
refGene_gn=SGSM1;refGene_tx=XM_023241635.1;refGene_gn=SGSM1;refGene_tx=XM_023241636.1;refGene_gn=SGSM1;refGene_tx=XM_023241637.1;refGene_gn=SGSM1;refGene_tx=XM_023241638.1;refGene_gn=SGSM1;refGene_tx=XM_023241639.1;refGene_gn=SGSM1;refGene_tx=XM_023241640.1;refGene_gn=SGSM1;refGene_tx=XM_023241641.1;refGene_gn=SGSM1;refGene_tx=XM_023241644.1;refGene_gn=SGSM1;refGene_tx=XM_023241645.1
D3 27126012 27126022 refGene_gn=SGSM1;refGene_tx=XM_023241643.1
D3 27126646 27126683 refGene_gn=SGSM1;refGene_tx=XM_023241642.1
D3 27127095 27127220
refGene_gn=SGSM1;refGene_tx=XM_023241635.1;refGene_gn=SGSM1;refGene_tx=XM_023241636.1;refGene_gn=SGSM1;refGene_tx=XM_023241637.1;refGene_gn=SGSM1;refGene_tx=XM_023241638.1;refGene_gn=SGSM1;refGene_tx=XM_023241639.1;refGene_gn=SGSM1;refGene_tx=XM_023241640.1;refGene_gn=SGSM1;refGene_tx=XM_023241641.1;refGene_gn=SGSM1;refGene_tx=XM_023241642.1;refGene_gn=SGSM1;refGene_tx=XM_023241643.1;refGene_gn=SGSM1;refGene_tx=XM_023241644.1;refGene_gn=SGSM1;refGene_tx=XM_023241645.1
D3 27129187 27129778

refGene_gn=SGSM1;refGene_tx=XM_023241635.1;refGene_gn=SGSM1;refGene_tx
=XM_023241636.1;refGene_gn=SGSM1;refGene_tx=XM_023241637.1;refGene_gn=
SGSM1;refGene_tx=XM_023241638.1;refGene_gn=SGSM1;refGene_tx=XM_0232416
39.1;refGene_gn=SGSM1;refGene_tx=XM_023241640.1;refGene_gn=SGSM1;refGe
ne_tx=XM_023241641.1;refGene_gn=SGSM1;refGene_tx=XM_023241642.1;refGen
e_gn=SGSM1;refGene_tx=XM_023241643.1;refGene_gn=SGSM1;refGene_tx=XM_02
3241644.1;refGene_gn=SGSM1;refGene_tx=XM_023241645.1
D3 27131288 27131357
refGene_gn=SGSM1;refGene_tx=XM_023241635.1;refGene_gn=SGSM1;refGene_tx
=XM_023241636.1;refGene_gn=SGSM1;refGene_tx=XM_023241637.1;refGene_gn=
SGSM1;refGene_tx=XM_023241638.1;refGene_gn=SGSM1;refGene_tx=XM_0232416
39.1;refGene_gn=SGSM1;refGene_tx=XM_023241640.1;refGene_gn=SGSM1;refGe
ne_tx=XM_023241642.1;refGene_gn=SGSM1;refGene_tx=XM_023241643.1;refGen
e_gn=SGSM1;refGene_tx=XM_023241644.1;refGene_gn=SGSM1;refGene_tx=XM_02
3241645.1
D3 27132713 27132896
refGene_gn=SGSM1;refGene_tx=XM_023241635.1;refGene_gn=SGSM1;refGene_tx
=XM_023241636.1;refGene_gn=SGSM1;refGene_tx=XM_023241637.1;refGene_gn=
SGSM1;refGene_tx=XM_023241638.1;refGene_gn=SGSM1;refGene_tx=XM_0232416
39.1;refGene_gn=SGSM1;refGene_tx=XM_023241640.1;refGene_gn=SGSM1;refGe
ne_tx=XM_023241642.1;refGene_gn=SGSM1;refGene_tx=XM_023241643.1;refGen
e_gn=SGSM1;refGene_tx=XM_023241644.1;refGene_gn=SGSM1;refGene_tx=XM_02
3241645.1
D3 27140093 27140222
refGene_gn=SGSM1;refGene_tx=XM_023241635.1;refGene_gn=SGSM1;refGene_tx
=XM_023241636.1;refGene_gn=SGSM1;refGene_tx=XM_023241637.1;refGene_gn=
SGSM1;refGene_tx=XM_023241638.1;refGene_gn=SGSM1;refGene_tx=XM_0232416
39.1;refGene_gn=SGSM1;refGene_tx=XM_023241640.1;refGene_gn=SGSM1;refGe
ne_tx=XM_023241641.1;refGene_gn=SGSM1;refGene_tx=XM_023241642.1;refGen
e_gn=SGSM1;refGene_tx=XM_023241643.1;refGene_gn=SGSM1;refGene_tx=XM_02
3241644.1;refGene_gn=SGSM1;refGene_tx=XM_023241645.1
D3 27143349 27143495
refGene_gn=SGSM1;refGene_tx=XM_023241635.1;refGene_gn=SGSM1;refGene_tx
=XM_023241636.1;refGene_gn=SGSM1;refGene_tx=XM_023241637.1;refGene_gn=
SGSM1;refGene_tx=XM_023241638.1;refGene_gn=SGSM1;refGene_tx=XM_0232416
39.1;refGene_gn=SGSM1;refGene_tx=XM_023241640.1;refGene_gn=SGSM1;refGe
ne_tx=XM_023241641.1;refGene_gn=SGSM1;refGene_tx=XM_023241642.1;refGen
e_gn=SGSM1;refGene_tx=XM_023241643.1;refGene_gn=SGSM1;refGene_tx=XM_02
3241644.1;refGene_gn=SGSM1;refGene_tx=XM_023241645.1
D3 27145703 27145768
refGene_gn=SGSM1;refGene_tx=XM_023241635.1;refGene_gn=SGSM1;refGene_tx
=XM_023241636.1;refGene_gn=SGSM1;refGene_tx=XM_023241637.1;refGene_gn=
SGSM1;refGene_tx=XM_023241638.1;refGene_gn=SGSM1;refGene_tx=XM_0232416
39.1;refGene_gn=SGSM1;refGene_tx=XM_023241640.1;refGene_gn=SGSM1;refGe
ne_tx=XM_023241641.1;refGene_gn=SGSM1;refGene_tx=XM_023241642.1;refGen
e_gn=SGSM1;refGene_tx=XM_023241643.1;refGene_gn=SGSM1;refGene_tx=XM_02
3241644.1;refGene_gn=SGSM1;refGene_tx=XM_023241645.1
D3 27147183 27147322
refGene_gn=SGSM1;refGene_tx=XM_023241635.1;refGene_gn=SGSM1;refGene_tx
=XM_023241636.1;refGene_gn=SGSM1;refGene_tx=XM_023241637.1;refGene_gn=

SGSM1;refGene_tx=XM_023241638.1;refGene_gn=SGSM1;refGene_tx=XM_0232416
39.1;refGene_gn=SGSM1;refGene_tx=XM_023241640.1;refGene_gn=SGSM1;refGe
ne_tx=XM_023241641.1;refGene_gn=SGSM1;refGene_tx=XM_023241642.1;refGen
e_gn=SGSM1;refGene_tx=XM_023241643.1;refGene_gn=SGSM1;refGene_tx=XM_02
3241644.1;refGene_gn=SGSM1;refGene_tx=XM_023241645.1
D3 27149214 27149379
refGene_gn=SGSM1;refGene_tx=XM_023241635.1;refGene_gn=SGSM1;refGene_tx
=XM_023241636.1;refGene_gn=SGSM1;refGene_tx=XM_023241637.1;refGene_gn=
SGSM1;refGene_tx=XM_023241638.1;refGene_gn=SGSM1;refGene_tx=XM_0232416
40.1;refGene_gn=SGSM1;refGene_tx=XM_023241641.1;refGene_gn=SGSM1;refGe
ne_tx=XM_023241642.1;refGene_gn=SGSM1;refGene_tx=XM_023241643.1;refGen
e_gn=SGSM1;refGene_tx=XM_023241644.1;refGene_gn=SGSM1;refGene_tx=XM_02
3241645.1
D3 27153758 27153891
refGene_gn=SGSM1;refGene_tx=XM_023241635.1;refGene_gn=SGSM1;refGene_tx
=XM_023241636.1;refGene_gn=SGSM1;refGene_tx=XM_023241637.1;refGene_gn=
SGSM1;refGene_tx=XM_023241638.1;refGene_gn=SGSM1;refGene_tx=XM_0232416
39.1;refGene_gn=SGSM1;refGene_tx=XM_023241640.1;refGene_gn=SGSM1;refGe
ne_tx=XM_023241641.1;refGene_gn=SGSM1;refGene_tx=XM_023241642.1;refGen
e_gn=SGSM1;refGene_tx=XM_023241643.1;refGene_gn=SGSM1;refGene_tx=XM_02
3241644.1;refGene_gn=SGSM1;refGene_tx=XM_023241645.1
D3 27154302 27154466
refGene_gn=SGSM1;refGene_tx=XM_023241635.1;refGene_gn=SGSM1;refGene_tx
=XM_023241636.1;refGene_gn=SGSM1;refGene_tx=XM_023241637.1;refGene_gn=
SGSM1;refGene_tx=XM_023241638.1;refGene_gn=SGSM1;refGene_tx=XM_0232416
39.1;refGene_gn=SGSM1;refGene_tx=XM_023241640.1;refGene_gn=SGSM1;refGe
ne_tx=XM_023241641.1;refGene_gn=SGSM1;refGene_tx=XM_023241642.1;refGen
e_gn=SGSM1;refGene_tx=XM_023241643.1;refGene_gn=SGSM1;refGene_tx=XM_02
3241644.1;refGene_gn=SGSM1;refGene_tx=XM_023241645.1
D3 27155668 27155736
refGene_gn=SGSM1;refGene_tx=XM_023241635.1;refGene_gn=SGSM1;refGene_tx
=XM_023241636.1;refGene_gn=SGSM1;refGene_tx=XM_023241637.1;refGene_gn=
SGSM1;refGene_tx=XM_023241638.1;refGene_gn=SGSM1;refGene_tx=XM_0232416
39.1;refGene_gn=SGSM1;refGene_tx=XM_023241640.1;refGene_gn=SGSM1;refGe
ne_tx=XM_023241641.1;refGene_gn=SGSM1;refGene_tx=XM_023241642.1;refGen
e_gn=SGSM1;refGene_tx=XM_023241643.1;refGene_gn=SGSM1;refGene_tx=XM_02
3241644.1;refGene_gn=SGSM1;refGene_tx=XM_023241645.1
D3 27160868 27160993
refGene_gn=SGSM1;refGene_tx=XM_023241635.1;refGene_gn=SGSM1;refGene_tx
=XM_023241636.1;refGene_gn=SGSM1;refGene_tx=XM_023241637.1;refGene_gn=
SGSM1;refGene_tx=XM_023241638.1;refGene_gn=SGSM1;refGene_tx=XM_0232416
39.1;refGene_gn=SGSM1;refGene_tx=XM_023241640.1;refGene_gn=SGSM1;refGe
ne_tx=XM_023241641.1;refGene_gn=SGSM1;refGene_tx=XM_023241642.1;refGen
e_gn=SGSM1;refGene_tx=XM_023241643.1;refGene_gn=SGSM1;refGene_tx=XM_02
3241644.1;refGene_gn=SGSM1;refGene_tx=XM_023241645.1
D3 27165238 27165370
refGene_gn=SGSM1;refGene_tx=XM_023241635.1;refGene_gn=SGSM1;refGene_tx
=XM_023241636.1;refGene_gn=SGSM1;refGene_tx=XM_023241637.1;refGene_gn=
SGSM1;refGene_tx=XM_023241638.1;refGene_gn=SGSM1;refGene_tx=XM_0232416
39.1;refGene_gn=SGSM1;refGene_tx=XM_023241640.1;refGene_gn=SGSM1;refGe

ne_tx=XM_023241641.1;refGene_gn=SGSM1;refGene_tx=XM_023241642.1;refGene_gn=SGSM1;refGene_tx=XM_023241643.1;refGene_gn=SGSM1;refGene_tx=XM_023241644.1;refGene_gn=SGSM1;refGene_tx=XM_023241645.1
D3 27165503 27165649
refGene_gn=SGSM1;refGene_tx=XM_023241635.1;refGene_gn=SGSM1;refGene_tx=XM_023241636.1;refGene_gn=SGSM1;refGene_tx=XM_023241637.1;refGene_gn=SGSM1;refGene_tx=XM_023241638.1;refGene_gn=SGSM1;refGene_tx=XM_023241639.1;refGene_gn=SGSM1;refGene_tx=XM_023241640.1;refGene_gn=SGSM1;refGene_tx=XM_023241641.1;refGene_gn=SGSM1;refGene_tx=XM_023241642.1;refGene_gn=SGSM1;refGene_tx=XM_023241643.1;refGene_gn=SGSM1;refGene_tx=XM_023241644.1;refGene_gn=SGSM1;refGene_tx=XM_023241645.1
D3 27165929 27165997
refGene_gn=SGSM1;refGene_tx=XM_023241635.1;refGene_gn=SGSM1;refGene_tx=XM_023241636.1;refGene_gn=SGSM1;refGene_tx=XM_023241637.1;refGene_gn=SGSM1;refGene_tx=XM_023241638.1;refGene_gn=SGSM1;refGene_tx=XM_023241639.1;refGene_gn=SGSM1;refGene_tx=XM_023241640.1;refGene_gn=SGSM1;refGene_tx=XM_023241641.1;refGene_gn=SGSM1;refGene_tx=XM_023241642.1;refGene_gn=SGSM1;refGene_tx=XM_023241643.1;refGene_gn=SGSM1;refGene_tx=XM_023241644.1;refGene_gn=SGSM1;refGene_tx=XM_023241645.1
D3 27168256 27168261 refGene_gn=SGSM1;refGene_tx=XM_023241644.1
D3 27169239 27169392
refGene_gn=SGSM1;refGene_tx=XM_023241635.1;refGene_gn=SGSM1;refGene_tx=XM_023241636.1;refGene_gn=SGSM1;refGene_tx=XM_023241637.1;refGene_gn=SGSM1;refGene_tx=XM_023241638.1;refGene_gn=SGSM1;refGene_tx=XM_023241639.1;refGene_gn=SGSM1;refGene_tx=XM_023241640.1;refGene_gn=SGSM1;refGene_tx=XM_023241641.1;refGene_gn=SGSM1;refGene_tx=XM_023241642.1;refGene_gn=SGSM1;refGene_tx=XM_023241643.1
D3 27170950 27171113
refGene_gn=SGSM1;refGene_tx=XM_023241635.1;refGene_gn=SGSM1;refGene_tx=XM_023241636.1;refGene_gn=SGSM1;refGene_tx=XM_023241637.1;refGene_gn=SGSM1;refGene_tx=XM_023241638.1;refGene_gn=SGSM1;refGene_tx=XM_023241639.1;refGene_gn=SGSM1;refGene_tx=XM_023241640.1;refGene_gn=SGSM1;refGene_tx=XM_023241641.1;refGene_gn=SGSM1;refGene_tx=XM_023241642.1;refGene_gn=SGSM1;refGene_tx=XM_023241643.1
D3 27175394 27175470
refGene_gn=SGSM1;refGene_tx=XM_023241635.1;refGene_gn=SGSM1;refGene_tx=XM_023241636.1;refGene_gn=SGSM1;refGene_tx=XM_023241637.1;refGene_gn=SGSM1;refGene_tx=XM_023241638.1;refGene_gn=SGSM1;refGene_tx=XM_023241639.1;refGene_gn=SGSM1;refGene_tx=XM_023241640.1;refGene_gn=SGSM1;refGene_tx=XM_023241641.1;refGene_gn=SGSM1;refGene_tx=XM_023241642.1;refGene_gn=SGSM1;refGene_tx=XM_023241643.1
D3 27211773 27211817 refGene_gn=SGSM1;refGene_tx=XM_023241638.1
D3 27211914 27211933 refGene_gn=SGSM1;refGene_tx=XM_023241638.1
D3 27288448 27288651
refGene_gn=SGSM1;refGene_tx=XM_023241635.1;refGene_gn=SGSM1;refGene_tx=XM_023241636.1;refGene_gn=SGSM1;refGene_tx=XM_023241637.1;refGene_gn=SGSM1;refGene_tx=XM_023241639.1;refGene_gn=SGSM1;refGene_tx=XM_023241641.1;refGene_gn=SGSM1;refGene_tx=XM_023241642.1;refGene_gn=SGSM1;refGene_tx=XM_023241643.1
D3 27288763 27288764

refGene_gn=SGSM1;refGene_tx=XM_023241635.1;refGene_gn=SGSM1;refGene_tx=XM_023241636.1;refGene_gn=SGSM1;refGene_tx=XM_023241637.1;refGene_gn=SGSM1;refGene_tx=XM_023241639.1;refGene_gn=SGSM1;refGene_tx=XM_023241641.1;refGene_gn=SGSM1;refGene_tx=XM_023241642.1;refGene_gn=SGSM1;refGene_tx=XM_023241643.1
D3 27331279 27331513
refGene_gn=LOC101100160;refGene_tx=XM_023242160.1
D3 27332620 27332723
refGene_gn=LOC101100160;refGene_tx=XM_023242160.1
D3 27335591 27335720
refGene_gn=LOC101100160;refGene_tx=XM_023242160.1
D3 27336290 27336505
refGene_gn=LOC101100160;refGene_tx=XM_023242160.1
D3 27337791 27337910
refGene_gn=LOC101100160;refGene_tx=XM_023242160.1
D3 27338780 27338861
refGene_gn=LOC101100160;refGene_tx=XM_023242160.1
D3 27339508 27339703
refGene_gn=LOC101100160;refGene_tx=XM_023242160.1
D3 27341815 27341927
refGene_gn=LOC101100160;refGene_tx=XM_023242160.1
D3 27345264 27345351
refGene_gn=LOC101100160;refGene_tx=XM_023242160.1
D3 27345353 27345423
refGene_gn=LOC101100160;refGene_tx=XM_023242160.1
D3 27345645 27345733
refGene_gn=LOC101100160;refGene_tx=XM_023242160.1
D3 27346122 27346237
refGene_gn=LOC101100160;refGene_tx=XM_023242160.1
D3 27346738 27346926
refGene_gn=LOC101100160;refGene_tx=XM_023242160.1
D3 27347491 27347564
refGene_gn=LOC101100160;refGene_tx=XM_023242160.1
D3 27354286 27354487
refGene_gn=LOC101100160;refGene_tx=XM_023242160.1
D3 27354632 27354736
refGene_gn=LOC101100160;refGene_tx=XM_023242160.1
D3 27357044 27357115
refGene_gn=LOC101100160;refGene_tx=XM_023242160.1
D3 27357521 27357672
refGene_gn=LOC101100160;refGene_tx=XM_023242160.1
D3 27358239 27358365
refGene_gn=LOC101100160;refGene_tx=XM_023242160.1
D3 27359261 27359409
refGene_gn=LOC101100160;refGene_tx=XM_023242160.1
D3 27359571 27359688
refGene_gn=LOC101100160;refGene_tx=XM_023242160.1
D3 27986513 27986589
refGene_gn=LOC111557367;refGene_tx=XM_023242161.1
D3 27987155 27987228

refGene_gn=LOC111557367;refGene_tx=XM_023242161.1
D3 27987229 27987345
refGene_gn=LOC111557367;refGene_tx=XM_023242161.1
D3 27988554 27988575
refGene_gn=LOC111557367;refGene_tx=XM_023242161.1
D3 27988658 27988799
refGene_gn=LOC111557367;refGene_tx=XM_023242161.1
D3 27997166 27997297
refGene_gn=LOC111557367;refGene_tx=XM_023242161.1
D3 27997797 27997985
refGene_gn=LOC111557367;refGene_tx=XM_023242161.1
D3 27998554 27998627
refGene_gn=LOC111557367;refGene_tx=XM_023242161.1
D3 28005396 28005597
refGene_gn=LOC111557367;refGene_tx=XM_023242161.1
D3 28005742 28005846
refGene_gn=LOC111557367;refGene_tx=XM_023242161.1
D3 28008166 28008237
refGene_gn=LOC111557367;refGene_tx=XM_023242161.1
D3 28008643 28008794
refGene_gn=LOC111557367;refGene_tx=XM_023242161.1
D3 28009361 28009487
refGene_gn=LOC111557367;refGene_tx=XM_023242161.1
D3 28010384 28010532
refGene_gn=LOC111557367;refGene_tx=XM_023242161.1
D3 28010695 28010812
refGene_gn=LOC111557367;refGene_tx=XM_023242161.1
D3 28067527 28067730
refGene_gn=DGCR6L;refGene_tx=XM_003994924.5;refGene_gn=DGCR6L;refGene_tx=XM_019814874.2;refGene_gn=DGCR6L;refGene_tx=XM_019814875.2
D3 28067824 28067985
refGene_gn=DGCR6L;refGene_tx=XM_003994924.5;refGene_gn=DGCR6L;refGene_tx=XM_019814874.2;refGene_gn=DGCR6L;refGene_tx=XM_019814875.2
D3 28068367 28068540
refGene_gn=DGCR6L;refGene_tx=XM_019814874.2;refGene_gn=DGCR6L;refGene_tx=XM_019814875.2
D3 28070339 28070540
refGene_gn=DGCR6L;refGene_tx=XM_019814874.2;refGene_gn=DGCR6L;refGene_tx=XM_019814875.2
D3 28071682 28071783 refGene_gn=DGCR6L;refGene_tx=XM_003994924.5
D3 28072596 28072737 refGene_gn=DGCR6L;refGene_tx=XM_003994924.5
D3 28073201 28073285 refGene_gn=DGCR6L;refGene_tx=XM_003994924.5
D3 28074754 28074942
refGene_gn=LOC101100421;refGene_tx=XM_003994942.5;refGene_gn=LOC101100421;refGene_tx=XM_019814873.2
D3 28075019 28075108
refGene_gn=LOC101100421;refGene_tx=XM_003994942.5;refGene_gn=LOC101100421;refGene_tx=XM_019814873.2
D3 28077680 28077744
refGene_gn=LOC101100421;refGene_tx=XM_011287910.3

D3 28080354 28080453
refGene_gn=LOC101100421;refGene_tx=XM_003994942.5;refGene_gn=LOC101100
421;refGene_tx=XM_011287910.3;refGene_gn=LOC101100421;refGene_tx=XM_01
9814873.2
D3 28081637 28081813
refGene_gn=LOC101100421;refGene_tx=XM_003994942.5;refGene_gn=LOC101100
421;refGene_tx=XM_011287910.3;refGene_gn=LOC101100421;refGene_tx=XM_01
9814873.2
D3 28082621 28082768
refGene_gn=LOC101100421;refGene_tx=XM_003994942.5;refGene_gn=LOC101100
421;refGene_tx=XM_011287910.3;refGene_gn=LOC101100421;refGene_tx=XM_01
9814873.2
D3 28082860 28082953
refGene_gn=LOC101100421;refGene_tx=XM_003994942.5;refGene_gn=LOC101100
421;refGene_tx=XM_011287910.3;refGene_gn=LOC101100421;refGene_tx=XM_01
9814873.2
D3 28084354 28084436
refGene_gn=LOC101100421;refGene_tx=XM_003994942.5;refGene_gn=LOC101100
421;refGene_tx=XM_011287910.3;refGene_gn=LOC101100421;refGene_tx=XM_01
9814873.2
D3 28085175 28085255
refGene_gn=LOC101100421;refGene_tx=XM_003994942.5;refGene_gn=LOC101100
421;refGene_tx=XM_011287910.3;refGene_gn=LOC101100421;refGene_tx=XM_01
9814873.2
D3 28085632 28085749
refGene_gn=LOC101100421;refGene_tx=XM_003994942.5;refGene_gn=LOC101100
421;refGene_tx=XM_011287910.3;refGene_gn=LOC101100421;refGene_tx=XM_01
9814873.2
D3 28085953 28086018
refGene_gn=LOC101100421;refGene_tx=XM_003994942.5;refGene_gn=LOC101100
421;refGene_tx=XM_011287910.3;refGene_gn=LOC101100421;refGene_tx=XM_01
9814873.2
D3 28087483 28087633
refGene_gn=LOC101100421;refGene_tx=XM_003994942.5;refGene_gn=LOC101100
421;refGene_tx=XM_011287910.3
D3 28088129 28088164
refGene_gn=LOC101100421;refGene_tx=XM_003994942.5;refGene_gn=LOC101100
421;refGene_tx=XM_011287910.3;refGene_gn=LOC101100421;refGene_tx=XM_01
9814873.2
D3 28093580 28093789
refGene_gn=LOC101100421;refGene_tx=XM_003994942.5;refGene_gn=LOC101100
421;refGene_tx=XM_011287910.3;refGene_gn=LOC101100421;refGene_tx=XM_01
9814873.2
D3 28098149 28098440
refGene_gn=LOC101100421;refGene_tx=XM_003994942.5;refGene_gn=LOC101100
421;refGene_tx=XM_011287910.3;refGene_gn=LOC101100421;refGene_tx=XM_01
9814873.2
D3 28128087 28128109 refGene_gn=RTN4R;refGene_tx=XM_023241646.1
D3 28152192 28153595 refGene_gn=RTN4R;refGene_tx=XM_023241646.1
D3 28235560 28236094

refGene_gn=CCDC188;refGene_tx=XM_011287907.3;refGene_gn=CCDC188;refGene_tx=XM_019814880.2;refGene_gn=CCDC188;refGene_tx=XM_019814881.2;refGene_gn=CCDC188;refGene_tx=XM_019814882.2;refGene_gn=CCDC188;refGene_tx=XM_019814883.2;refGene_gn=CCDC188;refGene_tx=XM_019814884.2;refGene_gn=CCDC188;refGene_tx=XM_019814885.2;refGene_gn=CCDC188;refGene_tx=XM_023241647.1;refGene_gn=CCDC188;refGene_tx=XM_023241648.1
D3 28236269 28236377
refGene_gn=CCDC188;refGene_tx=XM_011287907.3;refGene_gn=CCDC188;refGene_tx=XM_019814880.2;refGene_gn=CCDC188;refGene_tx=XM_019814881.2;refGene_gn=CCDC188;refGene_tx=XM_019814882.2;refGene_gn=CCDC188;refGene_tx=XM_019814883.2;refGene_gn=CCDC188;refGene_tx=XM_019814884.2;refGene_gn=CCDC188;refGene_tx=XM_019814885.2;refGene_gn=CCDC188;refGene_tx=XM_023241647.1;refGene_gn=CCDC188;refGene_tx=XM_023241648.1
D3 28236476 28236641
refGene_gn=CCDC188;refGene_tx=XM_011287907.3;refGene_gn=CCDC188;refGene_tx=XM_019814880.2;refGene_gn=CCDC188;refGene_tx=XM_019814881.2;refGene_gn=CCDC188;refGene_tx=XM_019814882.2;refGene_gn=CCDC188;refGene_tx=XM_019814883.2;refGene_gn=CCDC188;refGene_tx=XM_019814884.2;refGene_gn=CCDC188;refGene_tx=XM_019814885.2;refGene_gn=CCDC188;refGene_tx=XM_023241647.1;refGene_gn=CCDC188;refGene_tx=XM_023241648.1
D3 28236825 28236882
refGene_gn=CCDC188;refGene_tx=XM_011287907.3;refGene_gn=CCDC188;refGene_tx=XM_019814880.2;refGene_gn=CCDC188;refGene_tx=XM_019814881.2;refGene_gn=CCDC188;refGene_tx=XM_019814882.2;refGene_gn=CCDC188;refGene_tx=XM_019814883.2;refGene_gn=CCDC188;refGene_tx=XM_019814884.2;refGene_gn=CCDC188;refGene_tx=XM_019814885.2;refGene_gn=CCDC188;refGene_tx=XM_023241647.1;refGene_gn=CCDC188;refGene_tx=XM_023241648.1
D3 28236989 28237049
refGene_gn=CCDC188;refGene_tx=XM_011287907.3;refGene_gn=CCDC188;refGene_tx=XM_019814880.2;refGene_gn=CCDC188;refGene_tx=XM_019814881.2;refGene_gn=CCDC188;refGene_tx=XM_019814882.2;refGene_gn=CCDC188;refGene_tx=XM_019814883.2;refGene_gn=CCDC188;refGene_tx=XM_019814884.2;refGene_gn=CCDC188;refGene_tx=XM_019814885.2;refGene_gn=CCDC188;refGene_tx=XM_023241647.1;refGene_gn=CCDC188;refGene_tx=XM_023241648.1
D3 28237206 28237271
refGene_gn=CCDC188;refGene_tx=XM_011287907.3;refGene_gn=CCDC188;refGene_tx=XM_019814880.2;refGene_gn=CCDC188;refGene_tx=XM_019814881.2;refGene_gn=CCDC188;refGene_tx=XM_019814882.2;refGene_gn=CCDC188;refGene_tx=XM_019814883.2;refGene_gn=CCDC188;refGene_tx=XM_019814884.2;refGene_gn=CCDC188;refGene_tx=XM_019814885.2;refGene_gn=CCDC188;refGene_tx=XM_023241647.1;refGene_gn=CCDC188;refGene_tx=XM_023241648.1
D3 28237369 28237508
refGene_gn=CCDC188;refGene_tx=XM_011287907.3;refGene_gn=CCDC188;refGene_tx=XM_019814880.2;refGene_gn=CCDC188;refGene_tx=XM_019814881.2;refGene_gn=CCDC188;refGene_tx=XM_019814882.2;refGene_gn=CCDC188;refGene_tx=XM_019814883.2;refGene_gn=CCDC188;refGene_tx=XM_019814884.2;refGene_gn=CCDC188;refGene_tx=XM_019814885.2;refGene_gn=CCDC188;refGene_tx=XM_023241647.1;refGene_gn=CCDC188;refGene_tx=XM_023241648.1
D3 28237754 28237804
refGene_gn=CCDC188;refGene_tx=XM_011287907.3;refGene_gn=CCDC188;refGen

e_tx=XM_019814880.2;refGene_gn=CCDC188;refGene_tx=XM_019814881.2;refGene_gn=CCDC188;refGene_tx=XM_019814882.2;refGene_gn=CCDC188;refGene_tx=XM_019814883.2;refGene_gn=CCDC188;refGene_tx=XM_019814884.2;refGene_gn=CCDC188;refGene_tx=XM_019814885.2;refGene_gn=CCDC188;refGene_tx=XM_023241647.1;refGene_gn=CCDC188;refGene_tx=XM_023241648.1
D3 28237894 28238232 refGene_gn=CCDC188;refGene_tx=XM_011287907.3;refGene_gn=CCDC188;refGene_tx=XM_019814880.2;refGene_gn=CCDC188;refGene_tx=XM_019814883.2;refGene_gn=CCDC188;refGene_tx=XM_023241647.1;refGene_gn=CCDC188;refGene_tx=XM_023241648.1
D3 28238711 28238817 refGene_gn=CCDC188;refGene_tx=XM_019814881.2;refGene_gn=CCDC188;refGene_tx=XM_019814882.2;refGene_gn=CCDC188;refGene_tx=XM_019814885.2
D3 28239564 28239790 refGene_gn=ZDHHC8;refGene_tx=XM_006938688.4;refGene_gn=ZDHHC8;refGene_tx=XM_011287906.3;refGene_gn=ZDHHC8;refGene_tx=XM_019814879.2
D3 28240216 28240304 refGene_gn=CCDC188;refGene_tx=XM_019814881.2;refGene_gn=CCDC188;refGene_tx=XM_019814884.2;refGene_gn=CCDC188;refGene_tx=XM_019814885.2
D3 28241505 28241677 refGene_gn=ZDHHC8;refGene_tx=XM_003994925.4
D3 28242811 28243812 refGene_gn=ZDHHC8;refGene_tx=XM_003994925.4;refGene_gn=ZDHHC8;refGene_tx=XM_006938688.4;refGene_gn=ZDHHC8;refGene_tx=XM_011287906.3;refGene_gn=ZDHHC8;refGene_tx=XM_019814879.2
D3 28245163 28245273 refGene_gn=ZDHHC8;refGene_tx=XM_003994925.4;refGene_gn=ZDHHC8;refGene_tx=XM_006938688.4
D3 28245340 28245466 refGene_gn=ZDHHC8;refGene_tx=XM_003994925.4;refGene_gn=ZDHHC8;refGene_tx=XM_006938688.4;refGene_gn=ZDHHC8;refGene_tx=XM_019814879.2
D3 28245675 28245817 refGene_gn=ZDHHC8;refGene_tx=XM_003994925.4;refGene_gn=ZDHHC8;refGene_tx=XM_006938688.4;refGene_gn=ZDHHC8;refGene_tx=XM_011287906.3;refGene_gn=ZDHHC8;refGene_tx=XM_019814879.2
D3 28245983 28246075 refGene_gn=ZDHHC8;refGene_tx=XM_003994925.4;refGene_gn=ZDHHC8;refGene_tx=XM_006938688.4;refGene_gn=ZDHHC8;refGene_tx=XM_011287906.3;refGene_gn=ZDHHC8;refGene_tx=XM_019814879.2
D3 28246525 28246628 refGene_gn=ZDHHC8;refGene_tx=XM_003994925.4;refGene_gn=ZDHHC8;refGene_tx=XM_006938688.4;refGene_gn=ZDHHC8;refGene_tx=XM_011287906.3;refGene_gn=ZDHHC8;refGene_tx=XM_019814879.2
D3 28246850 28247023 refGene_gn=ZDHHC8;refGene_tx=XM_003994925.4;refGene_gn=ZDHHC8;refGene_tx=XM_006938688.4;refGene_gn=ZDHHC8;refGene_tx=XM_011287906.3;refGene_gn=ZDHHC8;refGene_tx=XM_019814879.2
D3 28247110 28247268 refGene_gn=ZDHHC8;refGene_tx=XM_003994925.4;refGene_gn=ZDHHC8;refGene_tx=XM_006938688.4;refGene_gn=ZDHHC8;refGene_tx=XM_011287906.3;refGene_gn=ZDHHC8;refGene_tx=XM_019814879.2

D3 28247442 28247564
refGene_gn=ZDHHC8;refGene_tx=XM_003994925.4;refGene_gn=ZDHHC8;refGene_tx=XM_006938688.4;refGene_gn=ZDHHC8;refGene_tx=XM_011287906.3;refGene_gn=ZDHHC8;refGene_tx=XM_019814879.2
D3 28254577 28254681
refGene_gn=ZDHHC8;refGene_tx=XM_003994925.4;refGene_gn=ZDHHC8;refGene_tx=XM_006938688.4;refGene_gn=ZDHHC8;refGene_tx=XM_011287906.3;refGene_gn=ZDHHC8;refGene_tx=XM_019814879.2
D3 28258937 28259062
refGene_gn=RANBP1;refGene_tx=XM_023241651.1;refGene_gn=RANBP1;refGene_tx=XM_023241652.1
D3 28259569 28259635
refGene_gn=RANBP1;refGene_tx=XM_023241651.1;refGene_gn=RANBP1;refGene_tx=XM_023241652.1
D3 28260301 28260430
refGene_gn=RANBP1;refGene_tx=XM_023241651.1;refGene_gn=RANBP1;refGene_tx=XM_023241652.1
D3 28262548 28262706
refGene_gn=RANBP1;refGene_tx=XM_023241651.1;refGene_gn=RANBP1;refGene_tx=XM_023241652.1
D3 28264875 28265012
refGene_gn=RANBP1;refGene_tx=XM_023241651.1;refGene_gn=RANBP1;refGene_tx=XM_023241652.1
D3 28266338 28266353
refGene_gn=RANBP1;refGene_tx=XM_023241651.1;refGene_gn=RANBP1;refGene_tx=XM_023241652.1
D3 28267059 28267083
refGene_gn=TRMT2A;refGene_tx=XM_011287904.2;refGene_gn=TRMT2A;refGene_tx=XM_011287905.3;refGene_gn=TRMT2A;refGene_tx=XM_019814887.2;refGene_gn=TRMT2A;refGene_tx=XM_023241650.1
D3 28267321 28267884
refGene_gn=TRMT2A;refGene_tx=XM_011287904.2;refGene_gn=TRMT2A;refGene_tx=XM_011287905.3;refGene_gn=TRMT2A;refGene_tx=XM_019814887.2;refGene_gn=TRMT2A;refGene_tx=XM_023241650.1
D3 28268142 28268251
refGene_gn=TRMT2A;refGene_tx=XM_011287904.2;refGene_gn=TRMT2A;refGene_tx=XM_011287905.3;refGene_gn=TRMT2A;refGene_tx=XM_019814887.2;refGene_gn=TRMT2A;refGene_tx=XM_023241650.1
D3 28268538 28268720
refGene_gn=TRMT2A;refGene_tx=XM_011287904.2;refGene_gn=TRMT2A;refGene_tx=XM_011287905.3;refGene_gn=TRMT2A;refGene_tx=XM_019814887.2;refGene_gn=TRMT2A;refGene_tx=XM_023241650.1
D3 28268908 28269023
refGene_gn=TRMT2A;refGene_tx=XM_011287904.2;refGene_gn=TRMT2A;refGene_tx=XM_011287905.3;refGene_gn=TRMT2A;refGene_tx=XM_019814887.2;refGene_gn=TRMT2A;refGene_tx=XM_023241650.1
D3 28269115 28269231
refGene_gn=TRMT2A;refGene_tx=XM_011287904.2;refGene_gn=TRMT2A;refGene_tx=XM_011287905.3;refGene_gn=TRMT2A;refGene_tx=XM_019814887.2;refGene_gn=TRMT2A;refGene_tx=XM_023241650.1

D3 28269309 28269421
refGene_gn=TRMT2A;refGene_tx=XM_011287904.2;refGene_gn=TRMT2A;refGene_tx=XM_011287905.3;refGene_gn=TRMT2A;refGene_tx=XM_019814887.2;refGene_gn=TRMT2A;refGene_tx=XM_023241650.1
D3 28270031 28270154
refGene_gn=TRMT2A;refGene_tx=XM_011287904.2;refGene_gn=TRMT2A;refGene_tx=XM_011287905.3;refGene_gn=TRMT2A;refGene_tx=XM_019814887.2;refGene_gn=TRMT2A;refGene_tx=XM_023241650.1
D3 28270367 28270443
refGene_gn=TRMT2A;refGene_tx=XM_011287904.2;refGene_gn=TRMT2A;refGene_tx=XM_011287905.3;refGene_gn=TRMT2A;refGene_tx=XM_019814887.2;refGene_gn=TRMT2A;refGene_tx=XM_023241650.1
D3 28270682 28270877
refGene_gn=TRMT2A;refGene_tx=XM_011287904.2;refGene_gn=TRMT2A;refGene_tx=XM_011287905.3;refGene_gn=TRMT2A;refGene_tx=XM_019814887.2;refGene_gn=TRMT2A;refGene_tx=XM_023241650.1
D3 28270993 28271090
refGene_gn=TRMT2A;refGene_tx=XM_011287904.2;refGene_gn=TRMT2A;refGene_tx=XM_011287905.3;refGene_gn=TRMT2A;refGene_tx=XM_019814887.2;refGene_gn=TRMT2A;refGene_tx=XM_023241650.1
D3 28271209 28271441
refGene_gn=TRMT2A;refGene_tx=XM_011287904.2;refGene_gn=TRMT2A;refGene_tx=XM_011287905.3;refGene_gn=TRMT2A;refGene_tx=XM_019814887.2;refGene_gn=TRMT2A;refGene_tx=XM_023241650.1
D3 28274136 28274220
refGene_gn=DGCR8;refGene_tx=XM_019814888.2;refGene_gn=DGCR8;refGene_tx=XM_023241649.1
D3 28275202 28275316
refGene_gn=DGCR8;refGene_tx=XM_019814888.2;refGene_gn=DGCR8;refGene_tx=XM_023241649.1
D3 28276732 28276860
refGene_gn=DGCR8;refGene_tx=XM_019814888.2;refGene_gn=DGCR8;refGene_tx=XM_023241649.1
D3 28278605 28278712
refGene_gn=DGCR8;refGene_tx=XM_019814888.2;refGene_gn=DGCR8;refGene_tx=XM_023241649.1
D3 28279042 28279143
refGene_gn=DGCR8;refGene_tx=XM_019814888.2;refGene_gn=DGCR8;refGene_tx=XM_023241649.1
D3 28294525 28294608
refGene_gn=DGCR8;refGene_tx=XM_019814888.2;refGene_gn=DGCR8;refGene_tx=XM_023241649.1
D3 28296130 28296229
refGene_gn=DGCR8;refGene_tx=XM_019814888.2;refGene_gn=DGCR8;refGene_tx=XM_023241649.1
D3 28296991 28297093
refGene_gn=DGCR8;refGene_tx=XM_019814888.2;refGene_gn=DGCR8;refGene_tx=XM_023241649.1
D3 28297354 28297552
refGene_gn=DGCR8;refGene_tx=XM_019814888.2;refGene_gn=DGCR8;refGene_tx

=XM_023241649.1
D3 28299666 28299949
refGene_gn=DGCR8;refGene_tx=XM_019814888.2;refGene_gn=DGCR8;refGene_tx
=XM_023241649.1
D3 28300111 28300254
refGene_gn=DGCR8;refGene_tx=XM_019814888.2;refGene_gn=DGCR8;refGene_tx
=XM_023241649.1
D3 28302360 28302520
refGene_gn=DGCR8;refGene_tx=XM_019814888.2;refGene_gn=DGCR8;refGene_tx
=XM_023241649.1
D3 28302997 28303717
refGene_gn=DGCR8;refGene_tx=XM_019814888.2;refGene_gn=DGCR8;refGene_tx
=XM_023241649.1
D3 28315180 28315301
refGene_gn=TANGO2;refGene_tx=XM_011287897.3;refGene_gn=TANGO2;refGene_
tx=XM_011287900.2;refGene_gn=TANGO2;refGene_tx=XM_023241653.1
D3 28316894 28316999
refGene_gn=TANGO2;refGene_tx=XM_011287897.3;refGene_gn=TANGO2;refGene_
tx=XM_011287900.2;refGene_gn=TANGO2;refGene_tx=XM_023241653.1
D3 28318467 28318621
refGene_gn=TANGO2;refGene_tx=XM_011287897.3;refGene_gn=TANGO2;refGene_
tx=XM_011287900.2;refGene_gn=TANGO2;refGene_tx=XM_023241653.1
D3 28318963 28319005
refGene_gn=TANGO2;refGene_tx=XM_011287897.3;refGene_gn=TANGO2;refGene_
tx=XM_023241653.1
D3 28321701 28321772
refGene_gn=TANGO2;refGene_tx=XM_011287897.3;refGene_gn=TANGO2;refGene_
tx=XM_011287900.2;refGene_gn=TANGO2;refGene_tx=XM_023241653.1
D3 28325281 28325396
refGene_gn=TANGO2;refGene_tx=XM_011287897.3;refGene_gn=TANGO2;refGene_
tx=XM_011287900.2;refGene_gn=TANGO2;refGene_tx=XM_023241653.1
D3 28325938 28326058
refGene_gn=TANGO2;refGene_tx=XM_011287897.3;refGene_gn=TANGO2;refGene_
tx=XM_011287900.2;refGene_gn=TANGO2;refGene_tx=XM_023241653.1
D3 28329809 28329898
refGene_gn=TANGO2;refGene_tx=XM_011287897.3;refGene_gn=TANGO2;refGene_
tx=XM_011287900.2;refGene_gn=TANGO2;refGene_tx=XM_023241653.1
D3 28338407 28338504
refGene_gn=TANGO2;refGene_tx=XM_011287897.3;refGene_gn=TANGO2;refGene_
tx=XM_011287900.2;refGene_gn=TANGO2;refGene_tx=XM_023241653.1
D3 28350259 28350389 refGene_gn=TANGO2;refGene_tx=XM_011287897.3
D3 28383025 28383235
refGene_gn=ARVCF;refGene_tx=XM_006938701.2;refGene_gn=ARVCF;refGene_tx
=XM_011287895.3;refGene_gn=ARVCF;refGene_tx=XM_011287896.2;refGene_gn=
ARVCF;refGene_tx=XM_019814890.1;refGene_gn=ARVCF;refGene_tx=XM_0198148
91.1;refGene_gn=ARVCF;refGene_tx=XM_023241654.1
D3 28392403 28392562
refGene_gn=ARVCF;refGene_tx=XM_006938701.2;refGene_gn=ARVCF;refGene_tx
=XM_011287895.3;refGene_gn=ARVCF;refGene_tx=XM_011287896.2;refGene_gn=
ARVCF;refGene_tx=XM_019814890.1;refGene_gn=ARVCF;refGene_tx=XM_0198148

91.1;refGene_gn=ARVCF;refGene_tx=XM_023241654.1
D3 28392740 28393264
refGene_gn=ARVCF;refGene_tx=XM_006938701.2;refGene_gn=ARVCF;refGene_tx=XM_011287895.3;refGene_gn=ARVCF;refGene_tx=XM_011287896.2;refGene_gn=ARVCF;refGene_tx=XM_019814890.1;refGene_gn=ARVCF;refGene_tx=XM_019814891.1;refGene_gn=ARVCF;refGene_tx=XM_023241654.1
D3 28394286 28394786
refGene_gn=ARVCF;refGene_tx=XM_006938701.2;refGene_gn=ARVCF;refGene_tx=XM_011287895.3;refGene_gn=ARVCF;refGene_tx=XM_011287896.2;refGene_gn=ARVCF;refGene_tx=XM_019814890.1;refGene_gn=ARVCF;refGene_tx=XM_019814891.1;refGene_gn=ARVCF;refGene_tx=XM_023241654.1
D3 28395447 28395631
refGene_gn=ARVCF;refGene_tx=XM_006938701.2;refGene_gn=ARVCF;refGene_tx=XM_011287895.3;refGene_gn=ARVCF;refGene_tx=XM_011287896.2;refGene_gn=ARVCF;refGene_tx=XM_019814890.1;refGene_gn=ARVCF;refGene_tx=XM_019814891.1;refGene_gn=ARVCF;refGene_tx=XM_023241654.1
D3 28396462 28396580
refGene_gn=ARVCF;refGene_tx=XM_006938701.2;refGene_gn=ARVCF;refGene_tx=XM_011287895.3;refGene_gn=ARVCF;refGene_tx=XM_011287896.2;refGene_gn=ARVCF;refGene_tx=XM_019814890.1;refGene_gn=ARVCF;refGene_tx=XM_019814891.1;refGene_gn=ARVCF;refGene_tx=XM_023241654.1
D3 28397012 28397184
refGene_gn=ARVCF;refGene_tx=XM_006938701.2;refGene_gn=ARVCF;refGene_tx=XM_011287895.3;refGene_gn=ARVCF;refGene_tx=XM_011287896.2;refGene_gn=ARVCF;refGene_tx=XM_019814890.1;refGene_gn=ARVCF;refGene_tx=XM_019814891.1;refGene_gn=ARVCF;refGene_tx=XM_023241654.1
D3 28397890 28397908
refGene_gn=ARVCF;refGene_tx=XM_011287895.3;refGene_gn=ARVCF;refGene_tx=XM_011287896.2;refGene_gn=ARVCF;refGene_tx=XM_023241654.1
D3 28398946 28399018
refGene_gn=ARVCF;refGene_tx=XM_006938701.2;refGene_gn=ARVCF;refGene_tx=XM_011287895.3;refGene_gn=ARVCF;refGene_tx=XM_011287896.2;refGene_gn=ARVCF;refGene_tx=XM_019814890.1;refGene_gn=ARVCF;refGene_tx=XM_019814891.1;refGene_gn=ARVCF;refGene_tx=XM_023241654.1
D3 28400393 28400521
refGene_gn=ARVCF;refGene_tx=XM_006938701.2;refGene_gn=ARVCF;refGene_tx=XM_011287895.3;refGene_gn=ARVCF;refGene_tx=XM_011287896.2;refGene_gn=ARVCF;refGene_tx=XM_019814890.1;refGene_gn=ARVCF;refGene_tx=XM_019814891.1;refGene_gn=ARVCF;refGene_tx=XM_023241654.1
D3 28400839 28400990
refGene_gn=ARVCF;refGene_tx=XM_006938701.2;refGene_gn=ARVCF;refGene_tx=XM_011287895.3;refGene_gn=ARVCF;refGene_tx=XM_011287896.2;refGene_gn=ARVCF;refGene_tx=XM_019814890.1;refGene_gn=ARVCF;refGene_tx=XM_019814891.1;refGene_gn=ARVCF;refGene_tx=XM_023241654.1
D3 28401289 28401488
refGene_gn=ARVCF;refGene_tx=XM_006938701.2;refGene_gn=ARVCF;refGene_tx=XM_011287895.3;refGene_gn=ARVCF;refGene_tx=XM_011287896.2;refGene_gn=ARVCF;refGene_tx=XM_019814890.1;refGene_gn=ARVCF;refGene_tx=XM_019814891.1;refGene_gn=ARVCF;refGene_tx=XM_023241654.1
D3 28401576 28401688

refGene_gn=ARVCF;refGene_tx=XM_006938701.2;refGene_gn=ARVCF;refGene_tx=XM_011287895.3;refGene_gn=ARVCF;refGene_tx=XM_011287896.2;refGene_gn=ARVCF;refGene_tx=XM_019814890.1;refGene_gn=ARVCF;refGene_tx=XM_019814891.1;refGene_gn=ARVCF;refGene_tx=XM_023241654.1
D3 28401806 28401897
refGene_gn=ARVCF;refGene_tx=XM_006938701.2;refGene_gn=ARVCF;refGene_tx=XM_011287895.3;refGene_gn=ARVCF;refGene_tx=XM_011287896.2;refGene_gn=ARVCF;refGene_tx=XM_019814890.1;refGene_gn=ARVCF;refGene_tx=XM_019814891.1;refGene_gn=ARVCF;refGene_tx=XM_023241654.1
D3 28402214 28402268
refGene_gn=ARVCF;refGene_tx=XM_006938701.2;refGene_gn=ARVCF;refGene_tx=XM_011287895.3;refGene_gn=ARVCF;refGene_tx=XM_011287896.2;refGene_gn=ARVCF;refGene_tx=XM_019814890.1;refGene_gn=ARVCF;refGene_tx=XM_019814891.1;refGene_gn=ARVCF;refGene_tx=XM_023241654.1
D3 28402666 28402752
refGene_gn=ARVCF;refGene_tx=XM_006938701.2;refGene_gn=ARVCF;refGene_tx=XM_011287895.3;refGene_gn=ARVCF;refGene_tx=XM_011287896.2;refGene_gn=ARVCF;refGene_tx=XM_019814890.1;refGene_gn=ARVCF;refGene_tx=XM_019814891.1;refGene_gn=ARVCF;refGene_tx=XM_023241654.1
D3 28403352 28403457
refGene_gn=ARVCF;refGene_tx=XM_011287895.3;refGene_gn=ARVCF;refGene_tx=XM_019814891.1
D3 28403920 28403947
refGene_gn=ARVCF;refGene_tx=XM_006938701.2;refGene_gn=ARVCF;refGene_tx=XM_011287896.2
D3 28406148 28406392
refGene_gn=COMT;refGene_tx=XM_023241661.1;refGene_gn=COMT;refGene_tx=XM_023241662.1;refGene_gn=COMT;refGene_tx=XM_023241663.1;refGene_gn=COMT;refGene_tx=XM_023241664.1
D3 28409402 28409527
refGene_gn=COMT;refGene_tx=XM_023241661.1;refGene_gn=COMT;refGene_tx=XM_023241662.1;refGene_gn=COMT;refGene_tx=XM_023241663.1
D3 28409800 28409932
refGene_gn=COMT;refGene_tx=XM_023241661.1;refGene_gn=COMT;refGene_tx=XM_023241662.1;refGene_gn=COMT;refGene_tx=XM_023241663.1;refGene_gn=COMT;refGene_tx=XM_023241664.1
D3 28410290 28410484
refGene_gn=COMT;refGene_tx=XM_023241661.1;refGene_gn=COMT;refGene_tx=XM_023241662.1;refGene_gn=COMT;refGene_tx=XM_023241663.1;refGene_gn=COMT;refGene_tx=XM_023241664.1
D3 28411245 28411525
refGene_gn=COMT;refGene_tx=XM_023241661.1;refGene_gn=COMT;refGene_tx=XM_023241662.1;refGene_gn=COMT;refGene_tx=XM_023241663.1;refGene_gn=COMT;refGene_tx=XM_023241664.1
D3 28417138 28417354
refGene_gn=ARVCF;refGene_tx=XM_019814890.1;refGene_gn=ARVCF;refGene_tx=XM_023241654.1
D3 28425694 28425803
refGene_gn=TXNRD2;refGene_tx=XM_023241655.1;refGene_gn=TXNRD2;refGene_tx=XM_023241656.1;refGene_gn=TXNRD2;refGene_tx=XM_023241657.1;refGene_

gn=TXNRD2;refGene_tx=XM_023241658.1;refGene_gn=TXNRD2;refGene_tx=XM_023241659.1;refGene_gn=TXNRD2;refGene_tx=XM_023241660.1
D3 28429861 28429930
refGene_gn=TXNRD2;refGene_tx=XM_023241655.1;refGene_gn=TXNRD2;refGene_tx=XM_023241656.1;refGene_gn=TXNRD2;refGene_tx=XM_023241657.1;refGene_gn=TXNRD2;refGene_tx=XM_023241658.1;refGene_gn=TXNRD2;refGene_tx=XM_023241659.1;refGene_gn=TXNRD2;refGene_tx=XM_023241660.1
D3 28437594 28437651
refGene_gn=TXNRD2;refGene_tx=XM_023241655.1;refGene_gn=TXNRD2;refGene_tx=XM_023241656.1;refGene_gn=TXNRD2;refGene_tx=XM_023241657.1;refGene_gn=TXNRD2;refGene_tx=XM_023241658.1;refGene_gn=TXNRD2;refGene_tx=XM_023241659.1;refGene_gn=TXNRD2;refGene_tx=XM_023241660.1
D3 28437996 28438141
refGene_gn=TXNRD2;refGene_tx=XM_023241655.1;refGene_gn=TXNRD2;refGene_tx=XM_023241656.1;refGene_gn=TXNRD2;refGene_tx=XM_023241657.1;refGene_gn=TXNRD2;refGene_tx=XM_023241658.1;refGene_gn=TXNRD2;refGene_tx=XM_023241659.1;refGene_gn=TXNRD2;refGene_tx=XM_023241660.1
D3 28438759 28438834
refGene_gn=TXNRD2;refGene_tx=XM_023241655.1;refGene_gn=TXNRD2;refGene_tx=XM_023241656.1;refGene_gn=TXNRD2;refGene_tx=XM_023241657.1;refGene_gn=TXNRD2;refGene_tx=XM_023241658.1;refGene_gn=TXNRD2;refGene_tx=XM_023241659.1;refGene_gn=TXNRD2;refGene_tx=XM_023241660.1
D3 28440671 28440750
refGene_gn=TXNRD2;refGene_tx=XM_023241655.1;refGene_gn=TXNRD2;refGene_tx=XM_023241656.1;refGene_gn=TXNRD2;refGene_tx=XM_023241657.1;refGene_gn=TXNRD2;refGene_tx=XM_023241658.1;refGene_gn=TXNRD2;refGene_tx=XM_023241659.1;refGene_gn=TXNRD2;refGene_tx=XM_023241660.1
D3 28441308 28441371
refGene_gn=TXNRD2;refGene_tx=XM_023241655.1;refGene_gn=TXNRD2;refGene_tx=XM_023241656.1;refGene_gn=TXNRD2;refGene_tx=XM_023241657.1;refGene_gn=TXNRD2;refGene_tx=XM_023241658.1;refGene_gn=TXNRD2;refGene_tx=XM_023241659.1;refGene_gn=TXNRD2;refGene_tx=XM_023241660.1
D3 28445675 28445746
refGene_gn=TXNRD2;refGene_tx=XM_023241655.1;refGene_gn=TXNRD2;refGene_tx=XM_023241656.1;refGene_gn=TXNRD2;refGene_tx=XM_023241657.1;refGene_gn=TXNRD2;refGene_tx=XM_023241658.1;refGene_gn=TXNRD2;refGene_tx=XM_023241659.1;refGene_gn=TXNRD2;refGene_tx=XM_023241660.1
D3 28455025 28455045
refGene_gn=TXNRD2;refGene_tx=XM_023241655.1;refGene_gn=TXNRD2;refGene_tx=XM_023241656.1;refGene_gn=TXNRD2;refGene_tx=XM_023241657.1;refGene_gn=TXNRD2;refGene_tx=XM_023241658.1;refGene_gn=TXNRD2;refGene_tx=XM_023241659.1;refGene_gn=TXNRD2;refGene_tx=XM_023241660.1
D3 28455835 28455927
refGene_gn=TXNRD2;refGene_tx=XM_023241655.1;refGene_gn=TXNRD2;refGene_tx=XM_023241656.1;refGene_gn=TXNRD2;refGene_tx=XM_023241657.1;refGene_gn=TXNRD2;refGene_tx=XM_023241658.1;refGene_gn=TXNRD2;refGene_tx=XM_023241659.1;refGene_gn=TXNRD2;refGene_tx=XM_023241660.1
D3 28458191 28458366
refGene_gn=TXNRD2;refGene_tx=XM_023241655.1;refGene_gn=TXNRD2;refGene_tx=XM_023241656.1;refGene_gn=TXNRD2;refGene_tx=XM_023241657.1;refGene_

gn=TXNRD2;refGene_tx=XM_023241658.1;refGene_gn=TXNRD2;refGene_tx=XM_02
3241659.1;refGene_gn=TXNRD2;refGene_tx=XM_023241660.1
D3 28468316 28468453
refGene_gn=TXNRD2;refGene_tx=XM_023241655.1;refGene_gn=TXNRD2;refGene_
tx=XM_023241656.1;refGene_gn=TXNRD2;refGene_tx=XM_023241657.1;refGene_
gn=TXNRD2;refGene_tx=XM_023241658.1;refGene_gn=TXNRD2;refGene_tx=XM_02
3241659.1;refGene_gn=TXNRD2;refGene_tx=XM_023241660.1
D3 28471643 28471739
refGene_gn=TXNRD2;refGene_tx=XM_023241655.1;refGene_gn=TXNRD2;refGene_
tx=XM_023241656.1;refGene_gn=TXNRD2;refGene_tx=XM_023241657.1;refGene_
gn=TXNRD2;refGene_tx=XM_023241658.1;refGene_gn=TXNRD2;refGene_tx=XM_02
3241659.1;refGene_gn=TXNRD2;refGene_tx=XM_023241660.1
D3 28472051 28472144
refGene_gn=TXNRD2;refGene_tx=XM_023241655.1;refGene_gn=TXNRD2;refGene_
tx=XM_023241656.1;refGene_gn=TXNRD2;refGene_tx=XM_023241657.1;refGene_
gn=TXNRD2;refGene_tx=XM_023241658.1;refGene_gn=TXNRD2;refGene_tx=XM_02
3241659.1;refGene_gn=TXNRD2;refGene_tx=XM_023241660.1
D3 28473732 28473804
refGene_gn=TXNRD2;refGene_tx=XM_023241655.1;refGene_gn=TXNRD2;refGene_
tx=XM_023241656.1;refGene_gn=TXNRD2;refGene_tx=XM_023241657.1;refGene_
gn=TXNRD2;refGene_tx=XM_023241658.1;refGene_gn=TXNRD2;refGene_tx=XM_02
3241659.1;refGene_gn=TXNRD2;refGene_tx=XM_023241660.1
D3 28473912 28474079
refGene_gn=TXNRD2;refGene_tx=XM_023241655.1;refGene_gn=TXNRD2;refGene_
tx=XM_023241656.1;refGene_gn=TXNRD2;refGene_tx=XM_023241657.1;refGene_
gn=TXNRD2;refGene_tx=XM_023241658.1;refGene_gn=TXNRD2;refGene_tx=XM_02
3241660.1
D3 28474479 28474535 refGene_gn=TXNRD2;refGene_tx=XM_023241656.1
D3 28475836 28475960
refGene_gn=TXNRD2;refGene_tx=XM_023241655.1;refGene_gn=TXNRD2;refGene_
tx=XM_023241656.1;refGene_gn=TXNRD2;refGene_tx=XM_023241659.1
D3 28526594 28526719
refGene_gn=GNB1L;refGene_tx=XM_003994931.5;refGene_gn=GNB1L;refGene_tx
=XM_006938705.4
D3 28527213 28527339
refGene_gn=GNB1L;refGene_tx=XM_003994931.5;refGene_gn=GNB1L;refGene_tx
=XM_006938705.4
D3 28532945 28533108
refGene_gn=GNB1L;refGene_tx=XM_003994931.5;refGene_gn=GNB1L;refGene_tx
=XM_006938705.4
D3 28537223 28537322
refGene_gn=GNB1L;refGene_tx=XM_003994931.5;refGene_gn=GNB1L;refGene_tx
=XM_006938705.4
D3 28541924 28542140
refGene_gn=GNB1L;refGene_tx=XM_003994931.5;refGene_gn=GNB1L;refGene_tx
=XM_006938705.4
D3 28558665 28558983
refGene_gn=GNB1L;refGene_tx=XM_003994931.5;refGene_gn=GNB1L;refGene_tx
=XM_006938705.4
D3 28578889 28579377 refGene_gn=TBX1;refGene_tx=XM_023241665.1

D3 28579748 28579849 refGene_gn=TBX1;refGene_tx=XM_023241665.1
D3 28579933 28580001 refGene_gn=TBX1;refGene_tx=XM_023241665.1
D3 28580618 28580774 refGene_gn=TBX1;refGene_tx=XM_023241665.1
D3 28581353 28581525 refGene_gn=TBX1;refGene_tx=XM_023241665.1
D3 28582376 28582478 refGene_gn=TBX1;refGene_tx=XM_023241665.1
D3 28584474 28584844 refGene_gn=TBX1;refGene_tx=XM_023241665.1
D3 28586128 28586312 refGene_gn=TBX1;refGene_tx=XM_023241665.1
D3 28618651 28619517 refGene_gn=GP1BB;refGene_tx=XM_023242162.1
D3 28620372 28620400 refGene_gn=GP1BB;refGene_tx=XM_023242162.1
D3 28620770 28620827
refGene_gn=SEPT5;refGene_tx=XM_011279864.3;refGene_gn=SEPT5;refGene_tx=XM_023241666.1;refGene_gn=SEPT5;refGene_tx=XM_023241667.1
D3 28620925 28621028
refGene_gn=SEPT5;refGene_tx=XM_011279864.3;refGene_gn=SEPT5;refGene_tx=XM_023241666.1;refGene_gn=SEPT5;refGene_tx=XM_023241667.1
D3 28621264 28621400
refGene_gn=SEPT5;refGene_tx=XM_011279864.3;refGene_gn=SEPT5;refGene_tx=XM_023241666.1;refGene_gn=SEPT5;refGene_tx=XM_023241667.1
D3 28621485 28621582
refGene_gn=SEPT5;refGene_tx=XM_011279864.3;refGene_gn=SEPT5;refGene_tx=XM_023241666.1;refGene_gn=SEPT5;refGene_tx=XM_023241667.1
D3 28622388 28622490
refGene_gn=SEPT5;refGene_tx=XM_011279864.3;refGene_gn=SEPT5;refGene_tx=XM_023241666.1;refGene_gn=SEPT5;refGene_tx=XM_023241667.1
D3 28622622 28622740
refGene_gn=SEPT5;refGene_tx=XM_011279864.3;refGene_gn=SEPT5;refGene_tx=XM_023241666.1;refGene_gn=SEPT5;refGene_tx=XM_023241667.1
D3 28622936 28623071
refGene_gn=SEPT5;refGene_tx=XM_011279864.3;refGene_gn=SEPT5;refGene_tx=XM_023241666.1;refGene_gn=SEPT5;refGene_tx=XM_023241667.1
D3 28623158 28623282
refGene_gn=SEPT5;refGene_tx=XM_011279864.3;refGene_gn=SEPT5;refGene_tx=XM_023241666.1;refGene_gn=SEPT5;refGene_tx=XM_023241667.1
D3 28623553 28623640
refGene_gn=SEPT5;refGene_tx=XM_011279864.3;refGene_gn=SEPT5;refGene_tx=XM_023241666.1;refGene_gn=SEPT5;refGene_tx=XM_023241667.1
D3 28623746 28623843
refGene_gn=SEPT5;refGene_tx=XM_011279864.3;refGene_gn=SEPT5;refGene_tx=XM_023241666.1;refGene_gn=SEPT5;refGene_tx=XM_023241667.1
D3 28624495 28624576 refGene_gn=SEPT5;refGene_tx=XM_011279864.3
D3 28626971 28627166 refGene_gn=SEPT5;refGene_tx=XM_023241666.1
D3 28628241 28628252 refGene_gn=SEPT5;refGene_tx=XM_023241667.1
D3 28628385 28628428 refGene_gn=SEPT5;refGene_tx=XM_023241667.1
D3 28763406 28764063 refGene_gn=CLDN5;refGene_tx=XM_011287892.3
D3 28769074 28769139
refGene_gn=CDC45;refGene_tx=XM_003994932.4;refGene_gn=CDC45;refGene_tx=XM_023241671.1
D3 28771109 28771186
refGene_gn=CDC45;refGene_tx=XM_003994932.4;refGene_gn=CDC45;refGene_tx=XM_023241671.1

D3 28773718 28773837
refGene_gn=CDC45;refGene_tx=XM_003994932.4;refGene_gn=CDC45;refGene_tx
=XM_023241671.1
D3 28775431 28775515
refGene_gn=CDC45;refGene_tx=XM_003994932.4;refGene_gn=CDC45;refGene_tx
=XM_023241671.1
D3 28775598 28775737
refGene_gn=CDC45;refGene_tx=XM_003994932.4;refGene_gn=CDC45;refGene_tx
=XM_023241671.1
D3 28781214 28781376
refGene_gn=CDC45;refGene_tx=XM_003994932.4;refGene_gn=CDC45;refGene_tx
=XM_023241671.1
D3 28781968 28782067
refGene_gn=CDC45;refGene_tx=XM_003994932.4;refGene_gn=CDC45;refGene_tx
=XM_023241671.1
D3 28782337 28782469 refGene_gn=CDC45;refGene_tx=XM_003994932.4
D3 28784063 28784183
refGene_gn=CDC45;refGene_tx=XM_003994932.4;refGene_gn=CDC45;refGene_tx
=XM_023241671.1
D3 28785521 28785572
refGene_gn=CDC45;refGene_tx=XM_003994932.4;refGene_gn=CDC45;refGene_tx
=XM_023241671.1
D3 28787303 28787365
refGene_gn=CDC45;refGene_tx=XM_003994932.4;refGene_gn=CDC45;refGene_tx
=XM_023241671.1
D3 28789131 28789180
refGene_gn=CDC45;refGene_tx=XM_003994932.4;refGene_gn=CDC45;refGene_tx
=XM_023241671.1
D3 28790791 28790847
refGene_gn=CDC45;refGene_tx=XM_003994932.4;refGene_gn=CDC45;refGene_tx
=XM_023241671.1
D3 28796799 28796946
refGene_gn=CDC45;refGene_tx=XM_003994932.4;refGene_gn=CDC45;refGene_tx
=XM_023241671.1
D3 28797908 28798046
refGene_gn=CDC45;refGene_tx=XM_003994932.4;refGene_gn=CDC45;refGene_tx
=XM_023241671.1
D3 28799488 28799581
refGene_gn=CDC45;refGene_tx=XM_003994932.4;refGene_gn=CDC45;refGene_tx
=XM_023241671.1
D3 28800348 28800408
refGene_gn=CDC45;refGene_tx=XM_003994932.4;refGene_gn=CDC45;refGene_tx
=XM_023241671.1
D3 28800543 28800594
refGene_gn=CDC45;refGene_tx=XM_003994932.4;refGene_gn=CDC45;refGene_tx
=XM_023241671.1
D3 28801478 28801481 refGene_gn=UFD1;refGene_tx=XM_019814905.2
D3 28804788 28804921 refGene_gn=UFD1;refGene_tx=XM_019814905.2
D3 28805299 28805332 refGene_gn=UFD1;refGene_tx=XM_019814905.2
D3 28808846 28808968 refGene_gn=UFD1;refGene_tx=XM_019814905.2

D3 28812091 28812222 refGene_gn=UFD1;refGene_tx=XM_019814905.2
D3 28814565 28814638 refGene_gn=UFD1;refGene_tx=XM_019814905.2
D3 28818970 28819039 refGene_gn=UFD1;refGene_tx=XM_019814905.2
D3 28820341 28820407 refGene_gn=UFD1;refGene_tx=XM_019814905.2
D3 28820677 28820725 refGene_gn=UFD1;refGene_tx=XM_019814905.2
D3 28821502 28821591 refGene_gn=UFD1;refGene_tx=XM_019814905.2
D3 28823047 28823129 refGene_gn=UFD1;refGene_tx=XM_019814905.2
D3 28826589 28826664 refGene_gn=UFD1;refGene_tx=XM_019814905.2
D3 28829539 28829675
refGene_gn=LOC101098170;refGene_tx=XM_019814902.2;refGene_gn=LOC101098170;refGene_tx=XM_023241668.1
D3 28836992 28837317
refGene_gn=MRPL40;refGene_tx=XM_003994936.5;refGene_gn=MRPL40;refGene_tx=XM_019814906.2
D3 28838622 28838781
refGene_gn=MRPL40;refGene_tx=XM_003994936.5;refGene_gn=MRPL40;refGene_tx=XM_019814906.2
D3 28839723 28839807
refGene_gn=MRPL40;refGene_tx=XM_003994936.5;refGene_gn=MRPL40;refGene_tx=XM_019814906.2
D3 28840118 28840171 refGene_gn=MRPL40;refGene_tx=XM_003994936.5
D3 28841295 28841332
refGene_gn=LOC101098170;refGene_tx=XM_023241669.1
D3 28862285 28862348
refGene_gn=LOC101098170;refGene_tx=XM_019814902.2;refGene_gn=LOC101098170;refGene_tx=XM_023241668.1;refGene_gn=LOC101098170;refGene_tx=XM_023241669.1
D3 28864098 28864209
refGene_gn=LOC101098170;refGene_tx=XM_019814902.2;refGene_gn=LOC101098170;refGene_tx=XM_023241668.1;refGene_gn=LOC101098170;refGene_tx=XM_023241669.1
D3 28865448 28865539
refGene_gn=LOC101098170;refGene_tx=XM_019814902.2;refGene_gn=LOC101098170;refGene_tx=XM_019814903.2;refGene_gn=LOC101098170;refGene_tx=XM_023241668.1;refGene_gn=LOC101098170;refGene_tx=XM_023241669.1
D3 28866868 28866963
refGene_gn=LOC101098170;refGene_tx=XM_019814902.2;refGene_gn=LOC101098170;refGene_tx=XM_019814903.2;refGene_gn=LOC101098170;refGene_tx=XM_023241668.1;refGene_gn=LOC101098170;refGene_tx=XM_023241669.1
D3 28874787 28874883
refGene_gn=LOC101098170;refGene_tx=XM_019814902.2;refGene_gn=LOC101098170;refGene_tx=XM_019814903.2;refGene_gn=LOC101098170;refGene_tx=XM_023241668.1;refGene_gn=LOC101098170;refGene_tx=XM_023241669.1;refGene_gn=LOC101098170;refGene_tx=XM_023241670.1
D3 28876041 28876202
refGene_gn=LOC101098170;refGene_tx=XM_019814902.2;refGene_gn=LOC101098170;refGene_tx=XM_019814903.2;refGene_gn=LOC101098170;refGene_tx=XM_023241668.1;refGene_gn=LOC101098170;refGene_tx=XM_023241669.1;refGene_gn=LOC101098170;refGene_tx=XM_023241670.1
D3 28878020 28878188

refGene_gn=LOC101098170;refGene_tx=XM_019814902.2;refGene_gn=LOC101098170;refGene_tx=XM_019814903.2;refGene_gn=LOC101098170;refGene_tx=XM_023241668.1;refGene_gn=LOC101098170;refGene_tx=XM_023241669.1;refGene_gn=LOC101098170;refGene_tx=XM_023241670.1
D3 28879885 28879999
refGene_gn=LOC101098170;refGene_tx=XM_019814902.2;refGene_gn=LOC101098170;refGene_tx=XM_019814903.2;refGene_gn=LOC101098170;refGene_tx=XM_023241668.1;refGene_gn=LOC101098170;refGene_tx=XM_023241669.1;refGene_gn=LOC101098170;refGene_tx=XM_023241670.1
D3 28882222 28882293
refGene_gn=LOC101098170;refGene_tx=XM_019814902.2;refGene_gn=LOC101098170;refGene_tx=XM_019814903.2;refGene_gn=LOC101098170;refGene_tx=XM_023241668.1;refGene_gn=LOC101098170;refGene_tx=XM_023241669.1;refGene_gn=LOC101098170;refGene_tx=XM_023241670.1
D3 28882943 28883049
refGene_gn=LOC101098170;refGene_tx=XM_019814902.2;refGene_gn=LOC101098170;refGene_tx=XM_019814903.2;refGene_gn=LOC101098170;refGene_tx=XM_023241668.1;refGene_gn=LOC101098170;refGene_tx=XM_023241669.1;refGene_gn=LOC101098170;refGene_tx=XM_023241670.1
D3 28884908 28885124
refGene_gn=LOC101098170;refGene_tx=XM_019814902.2;refGene_gn=LOC101098170;refGene_tx=XM_019814903.2;refGene_gn=LOC101098170;refGene_tx=XM_023241668.1;refGene_gn=LOC101098170;refGene_tx=XM_023241669.1;refGene_gn=LOC101098170;refGene_tx=XM_023241670.1
D3 28886060 28886146
refGene_gn=LOC101098170;refGene_tx=XM_019814902.2;refGene_gn=LOC101098170;refGene_tx=XM_019814903.2;refGene_gn=LOC101098170;refGene_tx=XM_023241668.1;refGene_gn=LOC101098170;refGene_tx=XM_023241669.1;refGene_gn=LOC101098170;refGene_tx=XM_023241670.1
D3 28890549 28890747
refGene_gn=LOC101098170;refGene_tx=XM_019814902.2;refGene_gn=LOC101098170;refGene_tx=XM_019814903.2;refGene_gn=LOC101098170;refGene_tx=XM_023241668.1;refGene_gn=LOC101098170;refGene_tx=XM_023241669.1;refGene_gn=LOC101098170;refGene_tx=XM_023241670.1
D3 28891901 28892063
refGene_gn=LOC101098170;refGene_tx=XM_019814902.2;refGene_gn=LOC101098170;refGene_tx=XM_019814903.2;refGene_gn=LOC101098170;refGene_tx=XM_023241668.1;refGene_gn=LOC101098170;refGene_tx=XM_023241669.1;refGene_gn=LOC101098170;refGene_tx=XM_023241670.1
D3 28909430 28909635
refGene_gn=LOC101098170;refGene_tx=XM_019814902.2;refGene_gn=LOC101098170;refGene_tx=XM_019814903.2;refGene_gn=LOC101098170;refGene_tx=XM_023241668.1;refGene_gn=LOC101098170;refGene_tx=XM_023241669.1;refGene_gn=LOC101098170;refGene_tx=XM_023241670.1
D3 28910035 28910140
refGene_gn=LOC101098170;refGene_tx=XM_019814902.2;refGene_gn=LOC101098170;refGene_tx=XM_019814903.2;refGene_gn=LOC101098170;refGene_tx=XM_023241668.1;refGene_gn=LOC101098170;refGene_tx=XM_023241669.1;refGene_gn=LOC101098170;refGene_tx=XM_023241670.1
D3 28911394 28911543

refGene_gn=LOC101098170;refGene_tx=XM_019814902.2;refGene_gn=LOC101098170;refGene_tx=XM_019814903.2;refGene_gn=LOC101098170;refGene_tx=XM_023241668.1;refGene_gn=LOC101098170;refGene_tx=XM_023241669.1;refGene_gn=LOC101098170;refGene_tx=XM_023241670.1
D3 28912732 28912894
refGene_gn=LOC101098170;refGene_tx=XM_019814902.2;refGene_gn=LOC101098170;refGene_tx=XM_019814903.2;refGene_gn=LOC101098170;refGene_tx=XM_023241668.1;refGene_gn=LOC101098170;refGene_tx=XM_023241669.1;refGene_gn=LOC101098170;refGene_tx=XM_023241670.1
D3 28913486 28913545
refGene_gn=LOC101098170;refGene_tx=XM_019814902.2;refGene_gn=LOC101098170;refGene_tx=XM_019814903.2;refGene_gn=LOC101098170;refGene_tx=XM_023241668.1;refGene_gn=LOC101098170;refGene_tx=XM_023241669.1;refGene_gn=LOC101098170;refGene_tx=XM_023241670.1
D3 28913916 28914022
refGene_gn=LOC101098170;refGene_tx=XM_019814902.2;refGene_gn=LOC101098170;refGene_tx=XM_019814903.2;refGene_gn=LOC101098170;refGene_tx=XM_023241668.1;refGene_gn=LOC101098170;refGene_tx=XM_023241669.1;refGene_gn=LOC101098170;refGene_tx=XM_023241670.1
D3 28915890 28916013
refGene_gn=LOC101098170;refGene_tx=XM_019814902.2;refGene_gn=LOC101098170;refGene_tx=XM_019814903.2;refGene_gn=LOC101098170;refGene_tx=XM_023241668.1;refGene_gn=LOC101098170;refGene_tx=XM_023241669.1;refGene_gn=LOC101098170;refGene_tx=XM_023241670.1
D3 28916515 28916679
refGene_gn=LOC101098170;refGene_tx=XM_019814902.2;refGene_gn=LOC101098170;refGene_tx=XM_019814903.2;refGene_gn=LOC101098170;refGene_tx=XM_023241668.1;refGene_gn=LOC101098170;refGene_tx=XM_023241669.1;refGene_gn=LOC101098170;refGene_tx=XM_023241670.1
D3 28918800 28918889
refGene_gn=LOC101098170;refGene_tx=XM_019814902.2
D3 28934036 28934491
refGene_gn=LOC101098170;refGene_tx=XM_019814902.2;refGene_gn=LOC101098170;refGene_tx=XM_019814903.2;refGene_gn=LOC101098170;refGene_tx=XM_023241668.1;refGene_gn=LOC101098170;refGene_tx=XM_023241669.1;refGene_gn=LOC101098170;refGene_tx=XM_023241670.1
D3 28977340 28977382
refGene_gn=CLTCL1;refGene_tx=XM_011287889.3;refGene_gn=CLTCL1;refGene_tx=XM_023241672.1;refGene_gn=CLTCL1;refGene_tx=XM_023241673.1
D3 28988243 28988451
refGene_gn=CLTCL1;refGene_tx=XM_011287889.3;refGene_gn=CLTCL1;refGene_tx=XM_023241672.1;refGene_gn=CLTCL1;refGene_tx=XM_023241673.1
D3 29000544 29000813
refGene_gn=CLTCL1;refGene_tx=XM_011287889.3;refGene_gn=CLTCL1;refGene_tx=XM_023241672.1;refGene_gn=CLTCL1;refGene_tx=XM_023241673.1
D3 29018717 29018879
refGene_gn=CLTCL1;refGene_tx=XM_011287889.3;refGene_gn=CLTCL1;refGene_tx=XM_023241672.1;refGene_gn=CLTCL1;refGene_tx=XM_023241673.1
D3 29021236 29021350
refGene_gn=CLTCL1;refGene_tx=XM_011287889.3;refGene_gn=CLTCL1;refGene_

tx=XM_023241672.1;refGene_gn=CLTCL1;refGene_tx=XM_023241673.1
D3 29025519 29025693
refGene_gn=CLTCL1;refGene_tx=XM_011287889.3;refGene_gn=CLTCL1;refGene_tx=XM_023241672.1;refGene_gn=CLTCL1;refGene_tx=XM_023241673.1
D3 29026931 29027129
refGene_gn=CLTCL1;refGene_tx=XM_011287889.3;refGene_gn=CLTCL1;refGene_tx=XM_023241672.1;refGene_gn=CLTCL1;refGene_tx=XM_023241673.1
D3 29028146 29028347
refGene_gn=CLTCL1;refGene_tx=XM_011287889.3;refGene_gn=CLTCL1;refGene_tx=XM_023241672.1;refGene_gn=CLTCL1;refGene_tx=XM_023241673.1
D3 29028447 29028600
refGene_gn=CLTCL1;refGene_tx=XM_011287889.3;refGene_gn=CLTCL1;refGene_tx=XM_023241672.1;refGene_gn=CLTCL1;refGene_tx=XM_023241673.1
D3 29029209 29029332
refGene_gn=CLTCL1;refGene_tx=XM_011287889.3;refGene_gn=CLTCL1;refGene_tx=XM_023241672.1;refGene_gn=CLTCL1;refGene_tx=XM_023241673.1
D3 29030656 29030794
refGene_gn=CLTCL1;refGene_tx=XM_011287889.3;refGene_gn=CLTCL1;refGene_tx=XM_023241672.1;refGene_gn=CLTCL1;refGene_tx=XM_023241673.1
D3 29034309 29034474
refGene_gn=CLTCL1;refGene_tx=XM_011287889.3;refGene_gn=CLTCL1;refGene_tx=XM_023241672.1;refGene_gn=CLTCL1;refGene_tx=XM_023241673.1
D3 29035039 29035220
refGene_gn=CLTCL1;refGene_tx=XM_011287889.3;refGene_gn=CLTCL1;refGene_tx=XM_023241672.1;refGene_gn=CLTCL1;refGene_tx=XM_023241673.1
D3 29036562 29036726
refGene_gn=CLTCL1;refGene_tx=XM_011287889.3;refGene_gn=CLTCL1;refGene_tx=XM_023241672.1;refGene_gn=CLTCL1;refGene_tx=XM_023241673.1
D3 29038078 29038204
refGene_gn=CLTCL1;refGene_tx=XM_011287889.3;refGene_gn=CLTCL1;refGene_tx=XM_023241672.1;refGene_gn=CLTCL1;refGene_tx=XM_023241673.1
D3 29038742 29038885
refGene_gn=CLTCL1;refGene_tx=XM_011287889.3;refGene_gn=CLTCL1;refGene_tx=XM_023241672.1;refGene_gn=CLTCL1;refGene_tx=XM_023241673.1
D3 29039263 29039498
refGene_gn=CLTCL1;refGene_tx=XM_011287889.3;refGene_gn=CLTCL1;refGene_tx=XM_023241672.1;refGene_gn=CLTCL1;refGene_tx=XM_023241673.1
D3 29041489 29041612
refGene_gn=CLTCL1;refGene_tx=XM_011287889.3;refGene_gn=CLTCL1;refGene_tx=XM_023241672.1;refGene_gn=CLTCL1;refGene_tx=XM_023241673.1
D3 29044459 29044605
refGene_gn=CLTCL1;refGene_tx=XM_011287889.3;refGene_gn=CLTCL1;refGene_tx=XM_023241672.1;refGene_gn=CLTCL1;refGene_tx=XM_023241673.1
D3 29056493 29056677
refGene_gn=CLTCL1;refGene_tx=XM_011287889.3;refGene_gn=CLTCL1;refGene_tx=XM_023241672.1;refGene_gn=CLTCL1;refGene_tx=XM_023241673.1
D3 29058819 29059012
refGene_gn=CLTCL1;refGene_tx=XM_011287889.3;refGene_gn=CLTCL1;refGene_tx=XM_023241672.1;refGene_gn=CLTCL1;refGene_tx=XM_023241673.1
D3 29059584 29059742

refGene_gn=CLTCL1;refGene_tx=XM_011287889.3;refGene_gn=CLTCL1;refGene_tx=XM_023241672.1;refGene_gn=CLTCL1;refGene_tx=XM_023241673.1
D3 29065710 29065875
refGene_gn=CLTCL1;refGene_tx=XM_011287889.3;refGene_gn=CLTCL1;refGene_tx=XM_023241672.1;refGene_gn=CLTCL1;refGene_tx=XM_023241673.1
D3 29067223 29067331
refGene_gn=CLTCL1;refGene_tx=XM_011287889.3;refGene_gn=CLTCL1;refGene_tx=XM_023241672.1;refGene_gn=CLTCL1;refGene_tx=XM_023241673.1
D3 29070034 29070202
refGene_gn=CLTCL1;refGene_tx=XM_011287889.3;refGene_gn=CLTCL1;refGene_tx=XM_023241672.1;refGene_gn=CLTCL1;refGene_tx=XM_023241673.1
D3 29070278 29070428
refGene_gn=CLTCL1;refGene_tx=XM_011287889.3;refGene_gn=CLTCL1;refGene_tx=XM_023241672.1;refGene_gn=CLTCL1;refGene_tx=XM_023241673.1
D3 29073866 29073998
refGene_gn=CLTCL1;refGene_tx=XM_011287889.3;refGene_gn=CLTCL1;refGene_tx=XM_023241672.1;refGene_gn=CLTCL1;refGene_tx=XM_023241673.1
D3 29074366 29074477
refGene_gn=CLTCL1;refGene_tx=XM_011287889.3;refGene_gn=CLTCL1;refGene_tx=XM_023241672.1;refGene_gn=CLTCL1;refGene_tx=XM_023241673.1
D3 29074745 29074916
refGene_gn=CLTCL1;refGene_tx=XM_011287889.3;refGene_gn=CLTCL1;refGene_tx=XM_023241672.1;refGene_gn=CLTCL1;refGene_tx=XM_023241673.1
D3 29077595 29077817
refGene_gn=CLTCL1;refGene_tx=XM_011287889.3;refGene_gn=CLTCL1;refGene_tx=XM_023241672.1;refGene_gn=CLTCL1;refGene_tx=XM_023241673.1
D3 29080805 29080881
refGene_gn=CLTCL1;refGene_tx=XM_011287889.3;refGene_gn=CLTCL1;refGene_tx=XM_023241672.1;refGene_gn=CLTCL1;refGene_tx=XM_023241673.1
D3 29081430 29081450 refGene_gn=CLTCL1;refGene_tx=XM_023241673.1
D3 29081686 29081805 refGene_gn=CLTCL1;refGene_tx=XM_011287889.3
D3 29082591 29082994 refGene_gn=CLTCL1;refGene_tx=XM_023241672.1
D3 29083334 29083442 refGene_gn=CLTCL1;refGene_tx=XM_023241672.1
D3 29083601 29083701 refGene_gn=CLTCL1;refGene_tx=XM_023241672.1
D3 29083779 29083794 refGene_gn=CLTCL1;refGene_tx=XM_023241672.1
D3 29083995 29084010 refGene_gn=SLC25A1;refGene_tx=XM_023241674.1
D3 29084395 29084480 refGene_gn=SLC25A1;refGene_tx=XM_023241674.1
D3 29084622 29084727 refGene_gn=SLC25A1;refGene_tx=XM_023241674.1
D3 29084931 29085047 refGene_gn=SLC25A1;refGene_tx=XM_023241674.1
D3 29085134 29085208 refGene_gn=SLC25A1;refGene_tx=XM_023241674.1
D3 29085427 29085542 refGene_gn=SLC25A1;refGene_tx=XM_023241674.1
D3 29116242 29116498 refGene_gn=GSC2;refGene_tx=XM_023242163.1
D3 29116606 29116860 refGene_gn=GSC2;refGene_tx=XM_023242163.1
D3 29117428 29117872 refGene_gn=GSC2;refGene_tx=XM_023242163.1
D3 29121663 29121795
refGene_gn=ESS2;refGene_tx=XM_006938707.4;refGene_gn=ESS2;refGene_tx=XM_023241675.1
D3 29122570 29122739
refGene_gn=ESS2;refGene_tx=XM_006938707.4;refGene_gn=ESS2;refGene_tx=XM_023241675.1;refGene_gn=ESS2;refGene_tx=XM_023241676.1

D3 29122826 29122928
refGene_gn=ESS2;refGene_tx=XM_006938707.4;refGene_gn=ESS2;refGene_tx=XM_023241675.1;refGene_gn=ESS2;refGene_tx=XM_023241676.1
D3 29125076 29125246
refGene_gn=ESS2;refGene_tx=XM_006938707.4;refGene_gn=ESS2;refGene_tx=XM_023241675.1;refGene_gn=ESS2;refGene_tx=XM_023241676.1
D3 29125619 29125737
refGene_gn=ESS2;refGene_tx=XM_006938707.4;refGene_gn=ESS2;refGene_tx=XM_023241675.1;refGene_gn=ESS2;refGene_tx=XM_023241676.1
D3 29126040 29126174
refGene_gn=ESS2;refGene_tx=XM_006938707.4;refGene_gn=ESS2;refGene_tx=XM_023241675.1;refGene_gn=ESS2;refGene_tx=XM_023241676.1
D3 29126891 29126994
refGene_gn=ESS2;refGene_tx=XM_006938707.4;refGene_gn=ESS2;refGene_tx=XM_023241675.1;refGene_gn=ESS2;refGene_tx=XM_023241676.1
D3 29127962 29128072 refGene_gn=ESS2;refGene_tx=XM_006938707.4
D3 29130048 29130164
refGene_gn=ESS2;refGene_tx=XM_006938707.4;refGene_gn=ESS2;refGene_tx=XM_023241675.1;refGene_gn=ESS2;refGene_tx=XM_023241676.1
D3 29130667 29131495
refGene_gn=ESS2;refGene_tx=XM_006938707.4;refGene_gn=ESS2;refGene_tx=XM_023241675.1;refGene_gn=ESS2;refGene_tx=XM_023241676.1
D3 29133105 29134194 refGene_gn=TSSK2;refGene_tx=XM_003994938.4
D3 29138906 29140016 refGene_gn=TSSK1B;refGene_tx=XM_003994939.4
D3 29142759 29142838
refGene_gn=DGCR2;refGene_tx=XM_023241677.1;refGene_gn=DGCR2;refGene_tx=XM_023241678.1;refGene_gn=DGCR2;refGene_tx=XM_023241679.1;refGene_gn=DGCR2;refGene_tx=XM_023241680.1
D3 29177438 29177561
refGene_gn=DGCR2;refGene_tx=XM_023241677.1;refGene_gn=DGCR2;refGene_tx=XM_023241678.1
D3 29203609 29203735
refGene_gn=DGCR2;refGene_tx=XM_023241677.1;refGene_gn=DGCR2;refGene_tx=XM_023241678.1;refGene_gn=DGCR2;refGene_tx=XM_023241679.1;refGene_gn=DGCR2;refGene_tx=XM_023241680.1
D3 29206191 29206402
refGene_gn=DGCR2;refGene_tx=XM_023241677.1;refGene_gn=DGCR2;refGene_tx=XM_023241678.1;refGene_gn=DGCR2;refGene_tx=XM_023241679.1;refGene_gn=DGCR2;refGene_tx=XM_023241680.1
D3 29208341 29208418
refGene_gn=DGCR2;refGene_tx=XM_023241677.1;refGene_gn=DGCR2;refGene_tx=XM_023241678.1;refGene_gn=DGCR2;refGene_tx=XM_023241679.1;refGene_gn=DGCR2;refGene_tx=XM_023241680.1
D3 29216233 29216410
refGene_gn=DGCR2;refGene_tx=XM_023241677.1;refGene_gn=DGCR2;refGene_tx=XM_023241678.1;refGene_gn=DGCR2;refGene_tx=XM_023241679.1;refGene_gn=DGCR2;refGene_tx=XM_023241680.1
D3 29229372 29229576
refGene_gn=DGCR2;refGene_tx=XM_023241677.1;refGene_gn=DGCR2;refGene_tx=XM_023241678.1;refGene_gn=DGCR2;refGene_tx=XM_023241679.1;refGene_gn=

DGCR2;refGene_tx=XM_023241680.1
D3 29234468 29234564
refGene_gn=DGCR2;refGene_tx=XM_023241677.1;refGene_gn=DGCR2;refGene_tx=XM_023241679.1
D3 29234965 29235118
refGene_gn=DGCR2;refGene_tx=XM_023241677.1;refGene_gn=DGCR2;refGene_tx=XM_023241678.1;refGene_gn=DGCR2;refGene_tx=XM_023241679.1;refGene_gn=DGCR2;refGene_tx=XM_023241680.1
D3 29236083 29236320
refGene_gn=DGCR2;refGene_tx=XM_023241677.1;refGene_gn=DGCR2;refGene_tx=XM_023241678.1;refGene_gn=DGCR2;refGene_tx=XM_023241679.1;refGene_gn=DGCR2;refGene_tx=XM_023241680.1
D3 29237887 29238144
refGene_gn=DGCR2;refGene_tx=XM_023241677.1;refGene_gn=DGCR2;refGene_tx=XM_023241678.1;refGene_gn=DGCR2;refGene_tx=XM_023241679.1;refGene_gn=DGCR2;refGene_tx=XM_023241680.1
D3 29242211 29242278
refGene_gn=LOC101082268;refGene_tx=XM_003994953.5;refGene_gn=LOC101082268;refGene_tx=XM_019814908.2;refGene_gn=LOC101082268;refGene_tx=XM_019814909.2;refGene_gn=LOC101082268;refGene_tx=XM_019814910.2;refGene_gn=LOC101082268;refGene_tx=XM_019814911.2;refGene_gn=LOC101082268;refGene_tx=XM_019814913.2;refGene_gn=LOC101082268;refGene_tx=XM_019814914.2;refGene_gn=LOC101082268;refGene_tx=XM_019814916.2
D3 29242495 29242549
refGene_gn=LOC101082268;refGene_tx=XM_003994953.5;refGene_gn=LOC101082268;refGene_tx=XM_019814908.2;refGene_gn=LOC101082268;refGene_tx=XM_019814909.2;refGene_gn=LOC101082268;refGene_tx=XM_019814910.2;refGene_gn=LOC101082268;refGene_tx=XM_019814911.2;refGene_gn=LOC101082268;refGene_tx=XM_019814913.2;refGene_gn=LOC101082268;refGene_tx=XM_019814916.2
D3 29243188 29243359
refGene_gn=LOC101082268;refGene_tx=XM_003994953.5;refGene_gn=LOC101082268;refGene_tx=XM_019814908.2;refGene_gn=LOC101082268;refGene_tx=XM_019814909.2;refGene_gn=LOC101082268;refGene_tx=XM_019814910.2;refGene_gn=LOC101082268;refGene_tx=XM_019814911.2;refGene_gn=LOC101082268;refGene_tx=XM_019814913.2;refGene_gn=LOC101082268;refGene_tx=XM_019814914.2;refGene_gn=LOC101082268;refGene_tx=XM_019814916.2
D3 29243882 29244043
refGene_gn=LOC101082268;refGene_tx=XM_003994953.5;refGene_gn=LOC101082268;refGene_tx=XM_019814908.2;refGene_gn=LOC101082268;refGene_tx=XM_019814909.2;refGene_gn=LOC101082268;refGene_tx=XM_019814910.2;refGene_gn=LOC101082268;refGene_tx=XM_019814911.2;refGene_gn=LOC101082268;refGene_tx=XM_019814913.2;refGene_gn=LOC101082268;refGene_tx=XM_019814914.2;refGene_gn=LOC101082268;refGene_tx=XM_019814916.2
D3 29244125 29244218
refGene_gn=LOC101082268;refGene_tx=XM_003994953.5;refGene_gn=LOC101082268;refGene_tx=XM_019814908.2;refGene_gn=LOC101082268;refGene_tx=XM_019814909.2;refGene_gn=LOC101082268;refGene_tx=XM_019814910.2;refGene_gn=LOC101082268;refGene_tx=XM_019814911.2;refGene_gn=LOC101082268;refGene_tx=XM_019814913.2;refGene_gn=LOC101082268;refGene_tx=XM_019814914.2
D3 29244582 29244661

refGene_gn=LOC101082268;refGene_tx=XM_003994953.5;refGene_gn=LOC101082
268;refGene_tx=XM_019814908.2;refGene_gn=LOC101082268;refGene_tx=XM_01
9814909.2;refGene_gn=LOC101082268;refGene_tx=XM_019814910.2;refGene_gn
=LOC101082268;refGene_tx=XM_019814911.2;refGene_gn=LOC101082268;refGen
e_tx=XM_019814913.2;refGene_gn=LOC101082268;refGene_tx=XM_019814914.2;
refGene_gn=LOC101082268;refGene_tx=XM_019814916.2
D3 29244769 29244936
refGene_gn=LOC101082268;refGene_tx=XM_003994953.5;refGene_gn=LOC101082
268;refGene_tx=XM_019814908.2;refGene_gn=LOC101082268;refGene_tx=XM_01
9814909.2;refGene_gn=LOC101082268;refGene_tx=XM_019814910.2;refGene_gn
=LOC101082268;refGene_tx=XM_019814911.2;refGene_gn=LOC101082268;refGen
e_tx=XM_019814913.2;refGene_gn=LOC101082268;refGene_tx=XM_019814914.2;
refGene_gn=LOC101082268;refGene_tx=XM_019814916.2
D3 29245027 29245276
refGene_gn=LOC101082268;refGene_tx=XM_003994953.5;refGene_gn=LOC101082
268;refGene_tx=XM_019814908.2;refGene_gn=LOC101082268;refGene_tx=XM_01
9814909.2;refGene_gn=LOC101082268;refGene_tx=XM_019814910.2;refGene_gn
=LOC101082268;refGene_tx=XM_019814911.2;refGene_gn=LOC101082268;refGen
e_tx=XM_019814913.2;refGene_gn=LOC101082268;refGene_tx=XM_019814914.2;
refGene_gn=LOC101082268;refGene_tx=XM_019814916.2
D3 29312369 29312403
refGene_gn=ZNF74;refGene_tx=XM_006938708.4;refGene_gn=ZNF74;refGene_tx
=XM_019814917.2
D3 29312794 29312880
refGene_gn=ZNF74;refGene_tx=XM_006938708.4;refGene_gn=ZNF74;refGene_tx
=XM_019814917.2
D3 29323795 29323922
refGene_gn=ZNF74;refGene_tx=XM_006938708.4;refGene_gn=ZNF74;refGene_tx
=XM_019814917.2
D3 29324352 29324448 refGene_gn=ZNF74;refGene_tx=XM_006938708.4
D3 29325414 29327078
refGene_gn=ZNF74;refGene_tx=XM_006938708.4;refGene_gn=ZNF74;refGene_tx
=XM_019814917.2
D3 29351668 29352570
refGene_gn=SCARF2;refGene_tx=XM_023241681.1;refGene_gn=SCARF2;refGene_
tx=XM_023241682.1
D3 29353387 29353540
refGene_gn=SCARF2;refGene_tx=XM_023241681.1;refGene_gn=SCARF2;refGene_
tx=XM_023241682.1
D3 29354753 29354884
refGene_gn=SCARF2;refGene_tx=XM_023241681.1;refGene_gn=SCARF2;refGene_
tx=XM_023241682.1
D3 29355089 29355207
refGene_gn=SCARF2;refGene_tx=XM_023241681.1;refGene_gn=SCARF2;refGene_
tx=XM_023241682.1
D3 29355290 29355394
refGene_gn=SCARF2;refGene_tx=XM_023241681.1;refGene_gn=SCARF2;refGene_
tx=XM_023241682.1
D3 29356013 29356142
refGene_gn=SCARF2;refGene_tx=XM_023241681.1;refGene_gn=SCARF2;refGene_

tx=XM_023241682.1
D3 29356285 29356504
refGene_gn=SCARF2;refGene_tx=XM_023241681.1;refGene_gn=SCARF2;refGene_tx=XM_023241682.1
D3 29356610 29357130
refGene_gn=SCARF2;refGene_tx=XM_023241681.1;refGene_gn=SCARF2;refGene_tx=XM_023241682.1
D3 29357322 29357424
refGene_gn=SCARF2;refGene_tx=XM_023241681.1;refGene_gn=SCARF2;refGene_tx=XM_023241682.1
D3 29357531 29357590
refGene_gn=SCARF2;refGene_tx=XM_023241681.1;refGene_gn=SCARF2;refGene_tx=XM_023241682.1
D3 29362498 29362668
refGene_gn=SCARF2;refGene_tx=XM_023241681.1;refGene_gn=SCARF2;refGene_tx=XM_023241682.1
D3 29367122 29367488
refGene_gn=KLHL22;refGene_tx=XM_023241683.1;refGene_gn=KLHL22;refGene_tx=XM_023241684.1
D3 29370662 29370896
refGene_gn=KLHL22;refGene_tx=XM_023241683.1;refGene_gn=KLHL22;refGene_tx=XM_023241684.1
D3 29377810 29378003
refGene_gn=KLHL22;refGene_tx=XM_023241683.1;refGene_gn=KLHL22;refGene_tx=XM_023241684.1
D3 29385752 29386471
refGene_gn=KLHL22;refGene_tx=XM_023241683.1;refGene_gn=KLHL22;refGene_tx=XM_023241684.1
D3 29391498 29391664
refGene_gn=KLHL22;refGene_tx=XM_023241683.1;refGene_gn=KLHL22;refGene_tx=XM_023241684.1
D3 29400812 29401039
refGene_gn=KLHL22;refGene_tx=XM_023241683.1;refGene_gn=KLHL22;refGene_tx=XM_023241684.1
D3 29419198 29419266
refGene_gn=LOC101099909;refGene_tx=XM_023242164.1
D3 29442063 29442101
refGene_gn=LOC101099909;refGene_tx=XM_023242164.1
D3 29444350 29444433
refGene_gn=LOC101099909;refGene_tx=XM_023242164.1
D3 29457794 29457846
refGene_gn=LOC101099909;refGene_tx=XM_023242164.1
D3 29459588 29459618
refGene_gn=LOC101099909;refGene_tx=XM_023242164.1
D3 29461116 29461329
refGene_gn=LOC101099909;refGene_tx=XM_023242164.1
D3 29470100 29470348
refGene_gn=LOC101099909;refGene_tx=XM_023242164.1
D3 29471440 29471767
refGene_gn=LOC101099909;refGene_tx=XM_023242164.1

D3 29472843 29472969
refGene_gn=LOC101099909;refGene_tx=XM_023242164.1
D3 29477021 29477146
refGene_gn=LOC101099909;refGene_tx=XM_023242164.1
D3 29485053 29485185
refGene_gn=LOC101099909;refGene_tx=XM_023242164.1
D3 29485282 29485410
refGene_gn=LOC101099909;refGene_tx=XM_023242164.1
D3 29485626 29485761
refGene_gn=LOC101099909;refGene_tx=XM_023242164.1
D3 29485853 29485917
refGene_gn=LOC101099909;refGene_tx=XM_023242164.1
D3 29486791 29486858
refGene_gn=LOC101099909;refGene_tx=XM_023242164.1
D3 29487380 29487541
refGene_gn=LOC101099909;refGene_tx=XM_023242164.1
D3 29487639 29487806
refGene_gn=LOC101099909;refGene_tx=XM_023242164.1
D3 29488269 29488368
refGene_gn=LOC101099909;refGene_tx=XM_023242164.1
D3 29489053 29489190
refGene_gn=LOC101099909;refGene_tx=XM_023242164.1
D3 29491099 29491238
refGene_gn=LOC102900693;refGene_tx=XM_006938710.4
D3 29491391 29491463
refGene_gn=LOC102900693;refGene_tx=XM_006938710.4
D3 29491544 29491627
refGene_gn=LOC102900693;refGene_tx=XM_006938710.4
D3 29491701 29491879
refGene_gn=LOC102900693;refGene_tx=XM_006938710.4
D3 29492707 29492870
refGene_gn=LOC102900693;refGene_tx=XM_006938710.4
D3 29495020 29495062
refGene_gn=LOC102900693;refGene_tx=XM_006938710.4
D3 29496267 29496517
refGene_gn=LOC102900693;refGene_tx=XM_006938710.4
D3 29514486 29514816
refGene_gn=SMPD4;refGene_tx=XM_023241685.1;refGene_gn=SMPD4;refGene_tx=XM_023241686.1;refGene_gn=SMPD4;refGene_tx=XM_023241687.1;refGene_gn=SMPD4;refGene_tx=XM_023241688.1;refGene_gn=SMPD4;refGene_tx=XM_023241689.1;refGene_gn=SMPD4;refGene_tx=XM_023241690.1;refGene_gn=SMPD4;refGene_tx=XM_023241691.1
D3 29514930 29515059
refGene_gn=SMPD4;refGene_tx=XM_023241685.1;refGene_gn=SMPD4;refGene_tx=XM_023241686.1;refGene_gn=SMPD4;refGene_tx=XM_023241687.1;refGene_gn=SMPD4;refGene_tx=XM_023241688.1;refGene_gn=SMPD4;refGene_tx=XM_023241689.1;refGene_gn=SMPD4;refGene_tx=XM_023241690.1;refGene_gn=SMPD4;refGene_tx=XM_023241691.1
D3 29515255 29515387
refGene_gn=SMPD4;refGene_tx=XM_023241685.1;refGene_gn=SMPD4;refGene_tx

=XM_023241686.1;refGene_gn=SMPD4;refGene_tx=XM_023241687.1;refGene_gn=SMPD4;refGene_tx=XM_023241688.1;refGene_gn=SMPD4;refGene_tx=XM_023241689.1;refGene_gn=SMPD4;refGene_tx=XM_023241690.1;refGene_gn=SMPD4;refGene_tx=XM_023241691.1
D3 29515609 29515843
refGene_gn=SMPD4;refGene_tx=XM_023241685.1;refGene_gn=SMPD4;refGene_tx=XM_023241686.1;refGene_gn=SMPD4;refGene_tx=XM_023241687.1;refGene_gn=SMPD4;refGene_tx=XM_023241688.1;refGene_gn=SMPD4;refGene_tx=XM_023241689.1;refGene_gn=SMPD4;refGene_tx=XM_023241690.1;refGene_gn=SMPD4;refGene_tx=XM_023241691.1
D3 29516437 29516643
refGene_gn=SMPD4;refGene_tx=XM_023241685.1;refGene_gn=SMPD4;refGene_tx=XM_023241686.1;refGene_gn=SMPD4;refGene_tx=XM_023241687.1;refGene_gn=SMPD4;refGene_tx=XM_023241688.1;refGene_gn=SMPD4;refGene_tx=XM_023241689.1;refGene_gn=SMPD4;refGene_tx=XM_023241690.1;refGene_gn=SMPD4;refGene_tx=XM_023241691.1
D3 29517139 29517303
refGene_gn=SMPD4;refGene_tx=XM_023241685.1;refGene_gn=SMPD4;refGene_tx=XM_023241686.1;refGene_gn=SMPD4;refGene_tx=XM_023241687.1;refGene_gn=SMPD4;refGene_tx=XM_023241688.1;refGene_gn=SMPD4;refGene_tx=XM_023241689.1;refGene_gn=SMPD4;refGene_tx=XM_023241690.1;refGene_gn=SMPD4;refGene_tx=XM_023241691.1
D3 29517891 29517992
refGene_gn=SMPD4;refGene_tx=XM_023241685.1;refGene_gn=SMPD4;refGene_tx=XM_023241686.1;refGene_gn=SMPD4;refGene_tx=XM_023241687.1;refGene_gn=SMPD4;refGene_tx=XM_023241688.1;refGene_gn=SMPD4;refGene_tx=XM_023241689.1;refGene_gn=SMPD4;refGene_tx=XM_023241690.1;refGene_gn=SMPD4;refGene_tx=XM_023241691.1
D3 29518638 29518729
refGene_gn=SMPD4;refGene_tx=XM_023241685.1;refGene_gn=SMPD4;refGene_tx=XM_023241686.1;refGene_gn=SMPD4;refGene_tx=XM_023241687.1;refGene_gn=SMPD4;refGene_tx=XM_023241688.1;refGene_gn=SMPD4;refGene_tx=XM_023241689.1;refGene_gn=SMPD4;refGene_tx=XM_023241690.1;refGene_gn=SMPD4;refGene_tx=XM_023241691.1
D3 29519516 29519662
refGene_gn=SMPD4;refGene_tx=XM_023241685.1;refGene_gn=SMPD4;refGene_tx=XM_023241686.1;refGene_gn=SMPD4;refGene_tx=XM_023241687.1;refGene_gn=SMPD4;refGene_tx=XM_023241688.1;refGene_gn=SMPD4;refGene_tx=XM_023241689.1;refGene_gn=SMPD4;refGene_tx=XM_023241690.1;refGene_gn=SMPD4;refGene_tx=XM_023241691.1
D3 29522216 29522303
refGene_gn=SMPD4;refGene_tx=XM_023241685.1;refGene_gn=SMPD4;refGene_tx=XM_023241686.1;refGene_gn=SMPD4;refGene_tx=XM_023241687.1;refGene_gn=SMPD4;refGene_tx=XM_023241690.1;refGene_gn=SMPD4;refGene_tx=XM_023241691.1
D3 29525562 29525634
refGene_gn=SMPD4;refGene_tx=XM_023241685.1;refGene_gn=SMPD4;refGene_tx=XM_023241686.1;refGene_gn=SMPD4;refGene_tx=XM_023241687.1;refGene_gn=SMPD4;refGene_tx=XM_023241688.1;refGene_gn=SMPD4;refGene_tx=XM_023241689.1;refGene_gn=SMPD4;refGene_tx=XM_023241690.1;refGene_gn=SMPD4;refGe

ne_tx=XM_023241691.1
D3 29527186 29527319
refGene_gn=SMPD4;refGene_tx=XM_023241685.1;refGene_gn=SMPD4;refGene_tx=XM_023241686.1;refGene_gn=SMPD4;refGene_tx=XM_023241687.1;refGene_gn=SMPD4;refGene_tx=XM_023241688.1;refGene_gn=SMPD4;refGene_tx=XM_023241689.1;refGene_gn=SMPD4;refGene_tx=XM_023241690.1;refGene_gn=SMPD4;refGene_tx=XM_023241691.1
D3 29529250 29529402
refGene_gn=SMPD4;refGene_tx=XM_023241685.1;refGene_gn=SMPD4;refGene_tx=XM_023241686.1;refGene_gn=SMPD4;refGene_tx=XM_023241687.1;refGene_gn=SMPD4;refGene_tx=XM_023241688.1;refGene_gn=SMPD4;refGene_tx=XM_023241689.1;refGene_gn=SMPD4;refGene_tx=XM_023241690.1;refGene_gn=SMPD4;refGene_tx=XM_023241691.1
D3 29529916 29529969
refGene_gn=SMPD4;refGene_tx=XM_023241685.1;refGene_gn=SMPD4;refGene_tx=XM_023241686.1;refGene_gn=SMPD4;refGene_tx=XM_023241687.1;refGene_gn=SMPD4;refGene_tx=XM_023241688.1;refGene_gn=SMPD4;refGene_tx=XM_023241689.1;refGene_gn=SMPD4;refGene_tx=XM_023241690.1;refGene_gn=SMPD4;refGene_tx=XM_023241691.1
D3 29530077 29530186
refGene_gn=SMPD4;refGene_tx=XM_023241685.1;refGene_gn=SMPD4;refGene_tx=XM_023241686.1;refGene_gn=SMPD4;refGene_tx=XM_023241687.1;refGene_gn=SMPD4;refGene_tx=XM_023241688.1;refGene_gn=SMPD4;refGene_tx=XM_023241689.1;refGene_gn=SMPD4;refGene_tx=XM_023241690.1;refGene_gn=SMPD4;refGene_tx=XM_023241691.1
D3 29530778 29530854
refGene_gn=SMPD4;refGene_tx=XM_023241685.1;refGene_gn=SMPD4;refGene_tx=XM_023241686.1;refGene_gn=SMPD4;refGene_tx=XM_023241687.1;refGene_gn=SMPD4;refGene_tx=XM_023241688.1;refGene_gn=SMPD4;refGene_tx=XM_023241689.1;refGene_gn=SMPD4;refGene_tx=XM_023241690.1;refGene_gn=SMPD4;refGene_tx=XM_023241691.1
D3 29531004 29531147
refGene_gn=SMPD4;refGene_tx=XM_023241685.1;refGene_gn=SMPD4;refGene_tx=XM_023241686.1;refGene_gn=SMPD4;refGene_tx=XM_023241687.1;refGene_gn=SMPD4;refGene_tx=XM_023241688.1;refGene_gn=SMPD4;refGene_tx=XM_023241689.1;refGene_gn=SMPD4;refGene_tx=XM_023241690.1
D3 29532004 29532091
refGene_gn=SMPD4;refGene_tx=XM_023241685.1;refGene_gn=SMPD4;refGene_tx=XM_023241686.1;refGene_gn=SMPD4;refGene_tx=XM_023241687.1;refGene_gn=SMPD4;refGene_tx=XM_023241688.1;refGene_gn=SMPD4;refGene_tx=XM_023241689.1;refGene_gn=SMPD4;refGene_tx=XM_023241690.1
D3 29533546 29533630
refGene_gn=SMPD4;refGene_tx=XM_023241685.1;refGene_gn=SMPD4;refGene_tx=XM_023241686.1;refGene_gn=SMPD4;refGene_tx=XM_023241687.1;refGene_gn=SMPD4;refGene_tx=XM_023241688.1;refGene_gn=SMPD4;refGene_tx=XM_023241689.1;refGene_gn=SMPD4;refGene_tx=XM_023241690.1
D3 29535129 29535267
refGene_gn=SMPD4;refGene_tx=XM_023241685.1;refGene_gn=SMPD4;refGene_tx=XM_023241688.1
D3 29535671 29535734 refGene_gn=SMPD4;refGene_tx=XM_023241687.1

D3 29535910 29535985
refGene_gn=SMPD4;refGene_tx=XM_023241686.1;refGene_gn=SMPD4;refGene_tx
=XM_023241689.1
D3 29537070 29537234
refGene_gn=MZT2B;refGene_tx=XM_023241692.1;refGene_gn=MZT2B;refGene_tx
=XM_023241693.1
D3 29537453 29537602
refGene_gn=MZT2B;refGene_tx=XM_023241692.1;refGene_gn=MZT2B;refGene_tx
=XM_023241693.1
D3 29538520 29538700 refGene_gn=MZT2B;refGene_tx=XM_023241692.1
D3 29544228 29544386
refGene_gn=MZT2B;refGene_tx=XM_023241692.1;refGene_gn=MZT2B;refGene_tx
=XM_023241693.1
D3 29545735 29546032
refGene_gn=LOC101090556;refGene_tx=XM_011287925.3
D3 29547836 29548517
refGene_gn=LOC101090556;refGene_tx=XM_011287925.3
D3 29548829 29548978
refGene_gn=LOC101090556;refGene_tx=XM_011287925.3
D3 29549860 29550083
refGene_gn=LOC101090556;refGene_tx=XM_011287925.3
D3 29553808 29553811
refGene_gn=LOC101090556;refGene_tx=XM_011287925.3
D3 29558661 29558716
refGene_gn=LOC111557368;refGene_tx=XM_023242165.1
D3 29558717 29558815
refGene_gn=LOC111557368;refGene_tx=XM_023242165.1
D3 29561028 29561124
refGene_gn=LOC111557368;refGene_tx=XM_023242165.1
D3 29562910 29562989
refGene_gn=LOC111557368;refGene_tx=XM_023242165.1
D3 29563750 29563866
refGene_gn=LOC111557368;refGene_tx=XM_023242165.1
D3 29564853 29564963
refGene_gn=LOC111557368;refGene_tx=XM_023242165.1
D3 29569240 29569447
refGene_gn=LOC111557368;refGene_tx=XM_023242165.1
D3 29570373 29570506
refGene_gn=LOC111557368;refGene_tx=XM_023242165.1
D3 29570942 29571085
refGene_gn=LOC111557368;refGene_tx=XM_023242165.1
D3 29572539 29572645
refGene_gn=LOC111557368;refGene_tx=XM_023242165.1
D3 29572720 29572726
refGene_gn=LOC111557368;refGene_tx=XM_023242165.1
D3 29573911 29573967
refGene_gn=LOC105259651;refGene_tx=XM_023241694.1;refGene_gn=LOC105259
651;refGene_tx=XM_023241695.1;refGene_gn=LOC105259651;refGene_tx=XM_02
3241696.1
D3 29574733 29574884

refGene_gn=LOC105259651;refGene_tx=XM_023241694.1;refGene_gn=LOC105259651;refGene_tx=XM_023241695.1;refGene_gn=LOC105259651;refGene_tx=XM_023241696.1
D3 29576214 29576286
refGene_gn=LOC105259651;refGene_tx=XM_023241694.1;refGene_gn=LOC105259651;refGene_tx=XM_023241695.1;refGene_gn=LOC105259651;refGene_tx=XM_023241696.1
D3 29579903 29579979
refGene_gn=LOC105259651;refGene_tx=XM_023241694.1;refGene_gn=LOC105259651;refGene_tx=XM_023241695.1;refGene_gn=LOC105259651;refGene_tx=XM_023241696.1
D3 29580178 29580272
refGene_gn=LOC105259651;refGene_tx=XM_023241694.1;refGene_gn=LOC105259651;refGene_tx=XM_023241695.1;refGene_gn=LOC105259651;refGene_tx=XM_023241696.1
D3 29580352 29580436
refGene_gn=LOC105259651;refGene_tx=XM_023241694.1;refGene_gn=LOC105259651;refGene_tx=XM_023241695.1;refGene_gn=LOC105259651;refGene_tx=XM_023241696.1
D3 29580525 29580667
refGene_gn=LOC105259651;refGene_tx=XM_023241694.1;refGene_gn=LOC105259651;refGene_tx=XM_023241695.1;refGene_gn=LOC105259651;refGene_tx=XM_023241696.1
D3 29580761 29580871
refGene_gn=LOC105259651;refGene_tx=XM_023241694.1;refGene_gn=LOC105259651;refGene_tx=XM_023241695.1;refGene_gn=LOC105259651;refGene_tx=XM_023241696.1
D3 29582234 29582328
refGene_gn=LOC105259651;refGene_tx=XM_023241694.1;refGene_gn=LOC105259651;refGene_tx=XM_023241695.1;refGene_gn=LOC105259651;refGene_tx=XM_023241696.1
D3 29582417 29582483
refGene_gn=LOC105259651;refGene_tx=XM_023241694.1;refGene_gn=LOC105259651;refGene_tx=XM_023241695.1
D3 29582672 29582750
refGene_gn=LOC105259651;refGene_tx=XM_023241694.1;refGene_gn=LOC105259651;refGene_tx=XM_023241695.1;refGene_gn=LOC105259651;refGene_tx=XM_023241696.1
D3 29582838 29582922
refGene_gn=LOC105259651;refGene_tx=XM_023241694.1;refGene_gn=LOC105259651;refGene_tx=XM_023241695.1;refGene_gn=LOC105259651;refGene_tx=XM_023241696.1
D3 29605604 29606586
refGene_gn=SLC7A4;refGene_tx=XM_003994956.4;refGene_gn=SLC7A4;refGene_tx=XM_019814922.2
D3 29607041 29607682
refGene_gn=SLC7A4;refGene_tx=XM_003994956.4;refGene_gn=SLC7A4;refGene_tx=XM_019814922.2
D3 29607945 29608054
refGene_gn=SLC7A4;refGene_tx=XM_003994956.4;refGene_gn=SLC7A4;refGene_

tx=XM_019814922.2
D3 29608157 29608434
refGene_gn=SLC7A4;refGene_tx=XM_003994956.4;refGene_gn=SLC7A4;refGene_tx=XM_019814922.2
D3 29610982 29611066
refGene_gn=P2RX6;refGene_tx=XM_023241697.1;refGene_gn=P2RX6;refGene_tx=XM_023241698.1;refGene_gn=P2RX6;refGene_tx=XM_023241699.1;refGene_gn=P2RX6;refGene_tx=XM_023241700.1;refGene_gn=P2RX6;refGene_tx=XM_023241701.1
D3 29611154 29611232
refGene_gn=P2RX6;refGene_tx=XM_023241697.1;refGene_gn=P2RX6;refGene_tx=XM_023241698.1;refGene_gn=P2RX6;refGene_tx=XM_023241699.1;refGene_gn=P2RX6;refGene_tx=XM_023241700.1;refGene_gn=P2RX6;refGene_tx=XM_023241701.1
D3 29611421 29611487
refGene_gn=P2RX6;refGene_tx=XM_023241697.1;refGene_gn=P2RX6;refGene_tx=XM_023241698.1;refGene_gn=P2RX6;refGene_tx=XM_023241700.1;refGene_gn=P2RX6;refGene_tx=XM_023241701.1
D3 29611576 29611670
refGene_gn=P2RX6;refGene_tx=XM_023241697.1;refGene_gn=P2RX6;refGene_tx=XM_023241698.1;refGene_gn=P2RX6;refGene_tx=XM_023241699.1;refGene_gn=P2RX6;refGene_tx=XM_023241700.1;refGene_gn=P2RX6;refGene_tx=XM_023241701.1
D3 29613036 29613146
refGene_gn=P2RX6;refGene_tx=XM_023241697.1;refGene_gn=P2RX6;refGene_tx=XM_023241698.1;refGene_gn=P2RX6;refGene_tx=XM_023241699.1;refGene_gn=P2RX6;refGene_tx=XM_023241700.1;refGene_gn=P2RX6;refGene_tx=XM_023241701.1
D3 29613240 29613382
refGene_gn=P2RX6;refGene_tx=XM_023241697.1;refGene_gn=P2RX6;refGene_tx=XM_023241698.1;refGene_gn=P2RX6;refGene_tx=XM_023241699.1;refGene_gn=P2RX6;refGene_tx=XM_023241700.1;refGene_gn=P2RX6;refGene_tx=XM_023241701.1
D3 29613471 29613555
refGene_gn=P2RX6;refGene_tx=XM_023241697.1;refGene_gn=P2RX6;refGene_tx=XM_023241698.1;refGene_gn=P2RX6;refGene_tx=XM_023241699.1;refGene_gn=P2RX6;refGene_tx=XM_023241700.1;refGene_gn=P2RX6;refGene_tx=XM_023241701.1
D3 29613635 29613729
refGene_gn=P2RX6;refGene_tx=XM_023241697.1;refGene_gn=P2RX6;refGene_tx=XM_023241698.1;refGene_gn=P2RX6;refGene_tx=XM_023241699.1;refGene_gn=P2RX6;refGene_tx=XM_023241700.1;refGene_gn=P2RX6;refGene_tx=XM_023241701.1
D3 29613928 29614004
refGene_gn=P2RX6;refGene_tx=XM_023241697.1;refGene_gn=P2RX6;refGene_tx=XM_023241698.1;refGene_gn=P2RX6;refGene_tx=XM_023241699.1;refGene_gn=P2RX6;refGene_tx=XM_023241700.1;refGene_gn=P2RX6;refGene_tx=XM_023241701.1
D3 29617637 29617709
refGene_gn=P2RX6;refGene_tx=XM_023241697.1;refGene_gn=P2RX6;refGene_tx

=XM_023241698.1;refGene_gn=P2RX6;refGene_tx=XM_023241699.1;refGene_gn=P2RX6;refGene_tx=XM_023241700.1;refGene_gn=P2RX6;refGene_tx=XM_023241701.1
D3 29619036 29619187
refGene_gn=P2RX6;refGene_tx=XM_023241697.1;refGene_gn=P2RX6;refGene_tx=XM_023241698.1;refGene_gn=P2RX6;refGene_tx=XM_023241699.1;refGene_gn=P2RX6;refGene_tx=XM_023241700.1;refGene_gn=P2RX6;refGene_tx=XM_023241701.1
D3 29619869 29620003
refGene_gn=P2RX6;refGene_tx=XM_023241697.1;refGene_gn=P2RX6;refGene_tx=XM_023241698.1;refGene_gn=P2RX6;refGene_tx=XM_023241699.1;refGene_gn=P2RX6;refGene_tx=XM_023241700.1;refGene_gn=P2RX6;refGene_tx=XM_023241701.1
D3 29621278 29621384
refGene_gn=LOC105260043;refGene_tx=XM_023241702.1
D3 29622845 29622988
refGene_gn=LOC105260043;refGene_tx=XM_023241702.1
D3 29623422 29623555
refGene_gn=LOC105260043;refGene_tx=XM_023241702.1
D3 29624481 29624688
refGene_gn=LOC105260043;refGene_tx=XM_023241702.1
D3 29629934 29630044
refGene_gn=LOC105260043;refGene_tx=XM_023241702.1
D3 29631031 29631147
refGene_gn=LOC105260043;refGene_tx=XM_023241702.1
D3 29631881 29631960
refGene_gn=LOC105260043;refGene_tx=XM_023241702.1
D3 29633766 29633862
refGene_gn=LOC105260043;refGene_tx=XM_023241702.1
D3 29698280 29698376 refGene_gn=CCDC74B;refGene_tx=XM_023241703.1
D3 29698378 29698529 refGene_gn=CCDC74B;refGene_tx=XM_023241703.1
D3 29699733 29699775 refGene_gn=CCDC74B;refGene_tx=XM_023241703.1
D3 29700878 29700937 refGene_gn=CCDC74B;refGene_tx=XM_023241703.1
D3 29700968 29700969 refGene_gn=CCDC74B;refGene_tx=XM_023241703.1
D3 29701929 29702092 refGene_gn=CCDC74B;refGene_tx=XM_023241703.1
D3 29702920 29703098 refGene_gn=CCDC74B;refGene_tx=XM_023241703.1
D3 29703172 29703255 refGene_gn=CCDC74B;refGene_tx=XM_023241703.1
D3 29703336 29703408 refGene_gn=CCDC74B;refGene_tx=XM_023241703.1
D3 29703561 29703700 refGene_gn=CCDC74B;refGene_tx=XM_023241703.1
D3 29705586 29705723
refGene_gn=LOC105260067;refGene_tx=XM_011279895.3
D3 29706407 29706506
refGene_gn=LOC105260067;refGene_tx=XM_011279895.3
D3 29706969 29707136
refGene_gn=LOC105260067;refGene_tx=XM_011279895.3
D3 29707234 29707395
refGene_gn=LOC105260067;refGene_tx=XM_011279895.3
D3 29707917 29707926
refGene_gn=LOC105260067;refGene_tx=XM_011279895.3
D3 29710936 29711034

refGene_gn=LOC105260043;refGene_tx=XM_023241702.1
D3 29711035 29711090
refGene_gn=LOC105260043;refGene_tx=XM_023241702.1
D3 29726075 29726155
refGene_gn=THAP7;refGene_tx=XM_006938722.4;refGene_gn=THAP7;refGene_tx=XM_019814932.2
D3 29726575 29726731 refGene_gn=THAP7;refGene_tx=XM_006938722.4
D3 29727527 29727668
refGene_gn=THAP7;refGene_tx=XM_006938722.4;refGene_gn=THAP7;refGene_tx=XM_019814932.2
D3 29727859 29728424
refGene_gn=THAP7;refGene_tx=XM_006938722.4;refGene_gn=THAP7;refGene_tx=XM_019814932.2
D3 29730711 29730828 refGene_gn=LZTR1;refGene_tx=XM_011287923.3
D3 29731105 29731186 refGene_gn=LZTR1;refGene_tx=XM_011287923.3
D3 29731274 29731380 refGene_gn=LZTR1;refGene_tx=XM_011287923.3
D3 29732017 29732123 refGene_gn=LZTR1;refGene_tx=XM_011287923.3
D3 29732125 29732166 refGene_gn=LZTR1;refGene_tx=XM_011287923.3
D3 29732259 29732386 refGene_gn=LZTR1;refGene_tx=XM_011287923.3
D3 29733273 29733430 refGene_gn=LZTR1;refGene_tx=XM_011287923.3
D3 29733724 29733894 refGene_gn=LZTR1;refGene_tx=XM_011287923.3
D3 29734104 29734270 refGene_gn=LZTR1;refGene_tx=XM_011287923.3
D3 29734349 29734445 refGene_gn=LZTR1;refGene_tx=XM_011287923.3
D3 29734626 29734719 refGene_gn=LZTR1;refGene_tx=XM_011287923.3
D3 29735097 29735208 refGene_gn=LZTR1;refGene_tx=XM_011287923.3
D3 29735349 29735505 refGene_gn=LZTR1;refGene_tx=XM_011287923.3
D3 29735981 29736183 refGene_gn=LZTR1;refGene_tx=XM_011287923.3
D3 29738130 29738270 refGene_gn=LZTR1;refGene_tx=XM_011287923.3
D3 29738863 29738921 refGene_gn=LZTR1;refGene_tx=XM_011287923.3
D3 29739953 29740037 refGene_gn=LZTR1;refGene_tx=XM_011287923.3
D3 29740533 29740642 refGene_gn=LZTR1;refGene_tx=XM_011287923.3
D3 29741133 29741213 refGene_gn=LZTR1;refGene_tx=XM_011287923.3
D3 29742659 29742716 refGene_gn=LZTR1;refGene_tx=XM_011287923.3
D3 29746079 29746142 refGene_gn=LZTR1;refGene_tx=XM_011287923.3
D3 29746610 29746810 refGene_gn=LZTR1;refGene_tx=XM_011287923.3
D3 29746958 29747008 refGene_gn=AIFM3;refGene_tx=XM_019814931.2
D3 29748084 29748124
refGene_gn=AIFM3;refGene_tx=XM_019814924.2;refGene_gn=AIFM3;refGene_tx=XM_019814925.2;refGene_gn=AIFM3;refGene_tx=XM_019814926.2;refGene_gn=AIFM3;refGene_tx=XM_019814927.2;refGene_gn=AIFM3;refGene_tx=XM_019814928.2;refGene_gn=AIFM3;refGene_tx=XM_019814930.2;refGene_gn=AIFM3;refGene_tx=XM_023241704.1
D3 29748351 29748372
refGene_gn=AIFM3;refGene_tx=XM_019814924.2;refGene_gn=AIFM3;refGene_tx=XM_019814925.2;refGene_gn=AIFM3;refGene_tx=XM_019814927.2;refGene_gn=AIFM3;refGene_tx=XM_019814930.2;refGene_gn=AIFM3;refGene_tx=XM_023241704.1
D3 29749034 29749139
refGene_gn=AIFM3;refGene_tx=XM_019814924.2;refGene_gn=AIFM3;refGene_tx=XM_019814925.2;refGene_gn=AIFM3;refGene_tx=XM_019814926.2;refGene_gn=

AIFM3;refGene_tx=XM_019814927.2;refGene_gn=AIFM3;refGene_tx=XM_0198149
30.2;refGene_gn=AIFM3;refGene_tx=XM_023241704.1
D3 29749449 29749462 refGene_gn=AIFM3;refGene_tx=XM_019814929.2
D3 29749689 29749765
refGene_gn=AIFM3;refGene_tx=XM_019814924.2;refGene_gn=AIFM3;refGene_tx
=XM_019814925.2;refGene_gn=AIFM3;refGene_tx=XM_019814926.2;refGene_gn=
AIFM3;refGene_tx=XM_019814927.2;refGene_gn=AIFM3;refGene_tx=XM_0198149
28.2;refGene_gn=AIFM3;refGene_tx=XM_019814929.2;refGene_gn=AIFM3;refGe
ne_tx=XM_019814930.2;refGene_gn=AIFM3;refGene_tx=XM_023241704.1
D3 29750048 29750147
refGene_gn=AIFM3;refGene_tx=XM_019814924.2;refGene_gn=AIFM3;refGene_tx
=XM_019814925.2;refGene_gn=AIFM3;refGene_tx=XM_019814926.2;refGene_gn=
AIFM3;refGene_tx=XM_019814927.2;refGene_gn=AIFM3;refGene_tx=XM_0198149
28.2;refGene_gn=AIFM3;refGene_tx=XM_019814929.2;refGene_gn=AIFM3;refGe
ne_tx=XM_019814930.2;refGene_gn=AIFM3;refGene_tx=XM_019814931.2;refGen
e_gn=AIFM3;refGene_tx=XM_023241704.1
D3 29751554 29751672
refGene_gn=AIFM3;refGene_tx=XM_019814924.2;refGene_gn=AIFM3;refGene_tx
=XM_019814925.2;refGene_gn=AIFM3;refGene_tx=XM_019814926.2;refGene_gn=
AIFM3;refGene_tx=XM_019814927.2;refGene_gn=AIFM3;refGene_tx=XM_0198149
28.2;refGene_gn=AIFM3;refGene_tx=XM_019814929.2;refGene_gn=AIFM3;refGe
ne_tx=XM_019814930.2;refGene_gn=AIFM3;refGene_tx=XM_019814931.2;refGen
e_gn=AIFM3;refGene_tx=XM_023241704.1
D3 29751787 29751864
refGene_gn=AIFM3;refGene_tx=XM_019814924.2;refGene_gn=AIFM3;refGene_tx
=XM_019814925.2;refGene_gn=AIFM3;refGene_tx=XM_019814926.2;refGene_gn=
AIFM3;refGene_tx=XM_019814927.2;refGene_gn=AIFM3;refGene_tx=XM_0198149
28.2;refGene_gn=AIFM3;refGene_tx=XM_019814929.2;refGene_gn=AIFM3;refGe
ne_tx=XM_019814930.2;refGene_gn=AIFM3;refGene_tx=XM_019814931.2;refGen
e_gn=AIFM3;refGene_tx=XM_023241704.1
D3 29752454 29752518
refGene_gn=AIFM3;refGene_tx=XM_019814924.2;refGene_gn=AIFM3;refGene_tx
=XM_019814925.2;refGene_gn=AIFM3;refGene_tx=XM_019814926.2;refGene_gn=
AIFM3;refGene_tx=XM_019814927.2;refGene_gn=AIFM3;refGene_tx=XM_0198149
28.2;refGene_gn=AIFM3;refGene_tx=XM_019814929.2;refGene_gn=AIFM3;refGe
ne_tx=XM_019814930.2;refGene_gn=AIFM3;refGene_tx=XM_019814931.2;refGen
e_gn=AIFM3;refGene_tx=XM_023241704.1
D3 29752594 29752666
refGene_gn=AIFM3;refGene_tx=XM_019814924.2;refGene_gn=AIFM3;refGene_tx
=XM_019814925.2;refGene_gn=AIFM3;refGene_tx=XM_019814926.2;refGene_gn=
AIFM3;refGene_tx=XM_019814927.2;refGene_gn=AIFM3;refGene_tx=XM_0198149
28.2;refGene_gn=AIFM3;refGene_tx=XM_019814929.2;refGene_gn=AIFM3;refGe
ne_tx=XM_019814930.2;refGene_gn=AIFM3;refGene_tx=XM_019814931.2;refGen
e_gn=AIFM3;refGene_tx=XM_023241704.1
D3 29752774 29752890
refGene_gn=AIFM3;refGene_tx=XM_019814924.2;refGene_gn=AIFM3;refGene_tx
=XM_019814925.2;refGene_gn=AIFM3;refGene_tx=XM_019814926.2;refGene_gn=
AIFM3;refGene_tx=XM_019814927.2;refGene_gn=AIFM3;refGene_tx=XM_0198149
28.2;refGene_gn=AIFM3;refGene_tx=XM_019814929.2;refGene_gn=AIFM3;refGe
ne_tx=XM_019814931.2;refGene_gn=AIFM3;refGene_tx=XM_023241704.1

D3 29752981 29753112
refGene_gn=AIFM3;refGene_tx=XM_019814924.2;refGene_gn=AIFM3;refGene_tx=XM_019814925.2;refGene_gn=AIFM3;refGene_tx=XM_019814926.2;refGene_gn=AIFM3;refGene_tx=XM_019814927.2;refGene_gn=AIFM3;refGene_tx=XM_019814928.2;refGene_gn=AIFM3;refGene_tx=XM_019814929.2;refGene_gn=AIFM3;refGene_tx=XM_019814930.2;refGene_gn=AIFM3;refGene_tx=XM_019814931.2;refGene_gn=AIFM3;refGene_tx=XM_023241704.1
D3 29753220 29753312
refGene_gn=AIFM3;refGene_tx=XM_019814924.2;refGene_gn=AIFM3;refGene_tx=XM_019814925.2;refGene_gn=AIFM3;refGene_tx=XM_019814926.2;refGene_gn=AIFM3;refGene_tx=XM_019814927.2;refGene_gn=AIFM3;refGene_tx=XM_019814928.2;refGene_gn=AIFM3;refGene_tx=XM_019814929.2;refGene_gn=AIFM3;refGene_tx=XM_019814930.2;refGene_gn=AIFM3;refGene_tx=XM_019814931.2;refGene_gn=AIFM3;refGene_tx=XM_023241704.1
D3 29753743 29753830
refGene_gn=AIFM3;refGene_tx=XM_019814924.2;refGene_gn=AIFM3;refGene_tx=XM_019814925.2;refGene_gn=AIFM3;refGene_tx=XM_019814926.2;refGene_gn=AIFM3;refGene_tx=XM_019814927.2;refGene_gn=AIFM3;refGene_tx=XM_019814928.2;refGene_gn=AIFM3;refGene_tx=XM_019814929.2;refGene_gn=AIFM3;refGene_tx=XM_019814930.2;refGene_gn=AIFM3;refGene_tx=XM_019814931.2;refGene_gn=AIFM3;refGene_tx=XM_023241704.1
D3 29754121 29754234
refGene_gn=AIFM3;refGene_tx=XM_019814924.2;refGene_gn=AIFM3;refGene_tx=XM_019814925.2;refGene_gn=AIFM3;refGene_tx=XM_019814926.2;refGene_gn=AIFM3;refGene_tx=XM_019814927.2;refGene_gn=AIFM3;refGene_tx=XM_019814928.2;refGene_gn=AIFM3;refGene_tx=XM_019814929.2;refGene_gn=AIFM3;refGene_tx=XM_019814930.2;refGene_gn=AIFM3;refGene_tx=XM_019814931.2;refGene_gn=AIFM3;refGene_tx=XM_023241704.1
D3 29754310 29754407
refGene_gn=AIFM3;refGene_tx=XM_019814924.2;refGene_gn=AIFM3;refGene_tx=XM_019814925.2;refGene_gn=AIFM3;refGene_tx=XM_019814926.2;refGene_gn=AIFM3;refGene_tx=XM_019814927.2;refGene_gn=AIFM3;refGene_tx=XM_019814928.2;refGene_gn=AIFM3;refGene_tx=XM_019814929.2;refGene_gn=AIFM3;refGene_tx=XM_019814930.2;refGene_gn=AIFM3;refGene_tx=XM_019814931.2;refGene_gn=AIFM3;refGene_tx=XM_023241704.1
D3 29754635 29754680
refGene_gn=AIFM3;refGene_tx=XM_019814924.2;refGene_gn=AIFM3;refGene_tx=XM_019814925.2;refGene_gn=AIFM3;refGene_tx=XM_019814926.2;refGene_gn=AIFM3;refGene_tx=XM_019814928.2;refGene_gn=AIFM3;refGene_tx=XM_019814929.2;refGene_gn=AIFM3;refGene_tx=XM_019814930.2;refGene_gn=AIFM3;refGene_tx=XM_019814931.2;refGene_gn=AIFM3;refGene_tx=XM_023241704.1
D3 29754775 29754885
refGene_gn=AIFM3;refGene_tx=XM_019814924.2;refGene_gn=AIFM3;refGene_tx=XM_019814925.2;refGene_gn=AIFM3;refGene_tx=XM_019814926.2;refGene_gn=AIFM3;refGene_tx=XM_019814927.2;refGene_gn=AIFM3;refGene_tx=XM_019814928.2;refGene_gn=AIFM3;refGene_tx=XM_019814929.2;refGene_gn=AIFM3;refGene_tx=XM_019814930.2;refGene_gn=AIFM3;refGene_tx=XM_019814931.2;refGene_gn=AIFM3;refGene_tx=XM_023241704.1
D3 29755101 29755211
refGene_gn=AIFM3;refGene_tx=XM_019814924.2;refGene_gn=AIFM3;refGene_tx

=XM_019814925.2;refGene_gn=AIFM3;refGene_tx=XM_019814926.2;refGene_gn=AIFM3;refGene_tx=XM_019814927.2;refGene_gn=AIFM3;refGene_tx=XM_019814928.2;refGene_gn=AIFM3;refGene_tx=XM_019814929.2;refGene_gn=AIFM3;refGene_tx=XM_019814930.2;refGene_gn=AIFM3;refGene_tx=XM_019814931.2;refGene_gn=AIFM3;refGene_tx=XM_023241704.1
D3 29755459 29755673 refGene_gn=AIFM3;refGene_tx=XM_019814924.2;refGene_gn=AIFM3;refGene_tx=XM_019814925.2;refGene_gn=AIFM3;refGene_tx=XM_019814926.2;refGene_gn=AIFM3;refGene_tx=XM_019814927.2;refGene_gn=AIFM3;refGene_tx=XM_019814928.2;refGene_gn=AIFM3;refGene_tx=XM_019814929.2;refGene_gn=AIFM3;refGene_tx=XM_019814930.2;refGene_gn=AIFM3;refGene_tx=XM_019814931.2;refGene_gn=AIFM3;refGene_tx=XM_023241704.1
D3 29759803 29759834 refGene_gn=AIFM3;refGene_tx=XM_019814924.2;refGene_gn=AIFM3;refGene_tx=XM_019814925.2;refGene_gn=AIFM3;refGene_tx=XM_019814926.2;refGene_gn=AIFM3;refGene_tx=XM_019814927.2;refGene_gn=AIFM3;refGene_tx=XM_019814928.2;refGene_gn=AIFM3;refGene_tx=XM_019814929.2;refGene_gn=AIFM3;refGene_tx=XM_019814930.2;refGene_gn=AIFM3;refGene_tx=XM_019814931.2;refGene_gn=AIFM3;refGene_tx=XM_023241704.1
D3 29771784 29771919 refGene_gn=CRKL;refGene_tx=XM_003994962.5
D3 29796816 29797282 refGene_gn=CRKL;refGene_tx=XM_003994962.5
D3 29808147 29808458 refGene_gn=CRKL;refGene_tx=XM_003994962.5
D3 29832168 29832326 refGene_gn=SNAP29;refGene_tx=XM_003994964.5
D3 29837171 29837270 refGene_gn=SNAP29;refGene_tx=XM_003994964.5
D3 29839880 29839966 refGene_gn=SNAP29;refGene_tx=XM_003994964.5
D3 29845801 29845998 refGene_gn=SNAP29;refGene_tx=XM_003994964.5
D3 29854791 29855028 refGene_gn=SNAP29;refGene_tx=XM_003994964.5
D3 29855502 29855679 refGene_gn=PI4KA;refGene_tx=XM_006938723.4
D3 29870643 29870760 refGene_gn=PI4KA;refGene_tx=XM_006938723.4
D3 29875199 29875293 refGene_gn=PI4KA;refGene_tx=XM_006938723.4
D3 29877706 29877795 refGene_gn=PI4KA;refGene_tx=XM_006938723.4
D3 29879746 29879819 refGene_gn=PI4KA;refGene_tx=XM_006938723.4
D3 29880421 29880681 refGene_gn=PI4KA;refGene_tx=XM_006938723.4
D3 29881206 29881273 refGene_gn=PI4KA;refGene_tx=XM_006938723.4
D3 29884600 29884749 refGene_gn=PI4KA;refGene_tx=XM_006938723.4
D3 29886399 29886465 refGene_gn=PI4KA;refGene_tx=XM_006938723.4
D3 29887031 29887128 refGene_gn=PI4KA;refGene_tx=XM_006938723.4
D3 29888758 29888950 refGene_gn=PI4KA;refGene_tx=XM_006938723.4
D3 29889464 29889565 refGene_gn=PI4KA;refGene_tx=XM_006938723.4
D3 29890420 29890550 refGene_gn=PI4KA;refGene_tx=XM_006938723.4
D3 29891538 29891671 refGene_gn=PI4KA;refGene_tx=XM_006938723.4
D3 29892686 29892782 refGene_gn=PI4KA;refGene_tx=XM_006938723.4
D3 29893148 29893332 refGene_gn=PI4KA;refGene_tx=XM_006938723.4
D3 29893783 29893887 refGene_gn=PI4KA;refGene_tx=XM_006938723.4
D3 29895927 29896096 refGene_gn=PI4KA;refGene_tx=XM_006938723.4
D3 29898303 29898354 refGene_gn=PI4KA;refGene_tx=XM_006938723.4
D3 29903888 29904080 refGene_gn=SERPIND1;refGene_tx=XM_006938745.4
D3 29904992 29905137 refGene_gn=SERPIND1;refGene_tx=XM_006938745.4
D3 29907338 29907612 refGene_gn=SERPIND1;refGene_tx=XM_006938745.4
D3 29911632 29912512 refGene_gn=SERPIND1;refGene_tx=XM_006938745.4

D3 29920603 29920712 refGene_gn=PI4KA;refGene_tx=XM_006938723.4
D3 29921070 29921207 refGene_gn=PI4KA;refGene_tx=XM_006938723.4
D3 29921389 29921523 refGene_gn=PI4KA;refGene_tx=XM_006938723.4
D3 29924525 29924608 refGene_gn=PI4KA;refGene_tx=XM_006938723.4
D3 29931551 29931622 refGene_gn=PI4KA;refGene_tx=XM_006938723.4
D3 29931705 29931830 refGene_gn=PI4KA;refGene_tx=XM_006938723.4
D3 29933023 29933105 refGene_gn=PI4KA;refGene_tx=XM_006938723.4
D3 29933329 29933413 refGene_gn=PI4KA;refGene_tx=XM_006938723.4
D3 29934631 29934721 refGene_gn=PI4KA;refGene_tx=XM_006938723.4
D3 29936873 29936993 refGene_gn=PI4KA;refGene_tx=XM_006938723.4
D3 29940056 29940149 refGene_gn=PI4KA;refGene_tx=XM_006938723.4
D3 29941568 29941725 refGene_gn=PI4KA;refGene_tx=XM_006938723.4
D3 29941956 29942084 refGene_gn=PI4KA;refGene_tx=XM_006938723.4
D3 29947165 29947324 refGene_gn=PI4KA;refGene_tx=XM_006938723.4
D3 29947605 29947757 refGene_gn=PI4KA;refGene_tx=XM_006938723.4
D3 29949191 29949299 refGene_gn=PI4KA;refGene_tx=XM_006938723.4
D3 29949929 29950057 refGene_gn=PI4KA;refGene_tx=XM_006938723.4
D3 29951418 29951538 refGene_gn=PI4KA;refGene_tx=XM_006938723.4
D3 29951702 29951782 refGene_gn=PI4KA;refGene_tx=XM_006938723.4
D3 29951879 29952073 refGene_gn=PI4KA;refGene_tx=XM_006938723.4
D3 29953078 29953169 refGene_gn=PI4KA;refGene_tx=XM_006938723.4
D3 29953540 29953708 refGene_gn=PI4KA;refGene_tx=XM_006938723.4
D3 29954422 29954476 refGene_gn=PI4KA;refGene_tx=XM_006938723.4
D3 29957078 29957199 refGene_gn=PI4KA;refGene_tx=XM_006938723.4
D3 29958855 29958985 refGene_gn=PI4KA;refGene_tx=XM_006938723.4
D3 29959931 29960002 refGene_gn=PI4KA;refGene_tx=XM_006938723.4
D3 29961233 29961306 refGene_gn=PI4KA;refGene_tx=XM_006938723.4
D3 29961391 29961462 refGene_gn=PI4KA;refGene_tx=XM_006938723.4
D3 29962518 29962628 refGene_gn=PI4KA;refGene_tx=XM_006938723.4
D3 29963081 29963186 refGene_gn=PI4KA;refGene_tx=XM_006938723.4
D3 29963278 29963404 refGene_gn=PI4KA;refGene_tx=XM_006938723.4
D3 29963843 29963964 refGene_gn=PI4KA;refGene_tx=XM_006938723.4
D3 29964445 29964605 refGene_gn=PI4KA;refGene_tx=XM_006938723.4
D3 29965371 29965461 refGene_gn=PI4KA;refGene_tx=XM_006938723.4
D3 29965912 29965996 refGene_gn=PI4KA;refGene_tx=XM_006938723.4
D3 29967089 29967141 refGene_gn=PI4KA;refGene_tx=XM_006938723.4
D3 29975212 29975327 refGene_gn=TMEM191C;refGene_tx=XM_011287922.3
D3 29975828 29976033
refGene_gn=TMEM191C;refGene_tx=XM_006938730.3;refGene_gn=TMEM191C;refGene_tx=XM_011287921.3
D3 29976113 29976154
refGene_gn=TMEM191C;refGene_tx=XM_006938730.3;refGene_gn=TMEM191C;refGene_tx=XM_011287921.3;refGene_gn=TMEM191C;refGene_tx=XM_011287922.3
D3 29976390 29976463
refGene_gn=TMEM191C;refGene_tx=XM_006938730.3;refGene_gn=TMEM191C;refGene_tx=XM_011287921.3;refGene_gn=TMEM191C;refGene_tx=XM_011287922.3
D3 29976546 29976611
refGene_gn=TMEM191C;refGene_tx=XM_006938730.3;refGene_gn=TMEM191C;refGene_tx=XM_011287921.3;refGene_gn=TMEM191C;refGene_tx=XM_011287922.3
D3 29976763 29976823

refGene_gn=TMEM191C;refGene_tx=XM_006938730.3;refGene_gn=TMEM191C;refGene_tx=XM_011287921.3;refGene_gn=TMEM191C;refGene_tx=XM_011287922.3
D3 29976943 29977000
refGene_gn=TMEM191C;refGene_tx=XM_006938730.3;refGene_gn=TMEM191C;refGene_tx=XM_011287922.3
D3 29977133 29977211
refGene_gn=TMEM191C;refGene_tx=XM_006938730.3;refGene_gn=TMEM191C;refGene_tx=XM_011287921.3;refGene_gn=TMEM191C;refGene_tx=XM_011287922.3
D3 29977360 29977411
refGene_gn=TMEM191C;refGene_tx=XM_006938730.3;refGene_gn=TMEM191C;refGene_tx=XM_011287921.3;refGene_gn=TMEM191C;refGene_tx=XM_011287922.3
D3 29977599 29977723
refGene_gn=TMEM191C;refGene_tx=XM_006938730.3;refGene_gn=TMEM191C;refGene_tx=XM_011287921.3;refGene_gn=TMEM191C;refGene_tx=XM_011287922.3
D3 29978030 29978185
refGene_gn=TMEM191C;refGene_tx=XM_006938730.3;refGene_gn=TMEM191C;refGene_tx=XM_011287921.3;refGene_gn=TMEM191C;refGene_tx=XM_011287922.3
D3 29992719 29994544 refGene_gn=HIC2;refGene_tx=XM_023241706.1
D3 29996298 29996324 refGene_gn=HIC2;refGene_tx=XM_023241706.1
D3 30031153 30036238
refGene_gn=LOC101091053;refGene_tx=XM_019814344.2
D3 30045434 30045461
refGene_gn=UBE2L3;refGene_tx=XM_019814934.2;refGene_gn=UBE2L3;refGene_tx=XM_023241707.1
D3 30060806 30060902 refGene_gn=UBE2L3;refGene_tx=XM_019814934.2
D3 30094463 30094650
refGene_gn=UBE2L3;refGene_tx=XM_019814934.2;refGene_gn=UBE2L3;refGene_tx=XM_023241707.1
D3 30101165 30101320
refGene_gn=UBE2L3;refGene_tx=XM_019814934.2;refGene_gn=UBE2L3;refGene_tx=XM_023241707.1
D3 30107473 30107828 refGene_gn=YDJC;refGene_tx=XM_003994987.4
D3 30108024 30108202 refGene_gn=YDJC;refGene_tx=XM_003994987.4
D3 30108294 30108394 refGene_gn=YDJC;refGene_tx=XM_003994987.4
D3 30108595 30108755 refGene_gn=YDJC;refGene_tx=XM_003994987.4
D3 30108844 30109008 refGene_gn=YDJC;refGene_tx=XM_003994987.4
D3 30109147 30109218 refGene_gn=CCDC116;refGene_tx=XM_006938731.4
D3 30111347 30111486
refGene_gn=CCDC116;refGene_tx=XM_006938731.4;refGene_gn=CCDC116;refGene_tx=XM_006938733.4;refGene_gn=CCDC116;refGene_tx=XM_023241708.1
D3 30112328 30112886
refGene_gn=CCDC116;refGene_tx=XM_006938731.4;refGene_gn=CCDC116;refGene_tx=XM_006938733.4;refGene_gn=CCDC116;refGene_tx=XM_023241708.1
D3 30112996 30113608
refGene_gn=CCDC116;refGene_tx=XM_006938731.4;refGene_gn=CCDC116;refGene_tx=XM_006938733.4;refGene_gn=CCDC116;refGene_tx=XM_023241708.1
D3 30114743 30115520
refGene_gn=CCDC116;refGene_tx=XM_006938731.4;refGene_gn=CCDC116;refGene_tx=XM_006938733.4;refGene_gn=CCDC116;refGene_tx=XM_023241708.1
D3 30120247 30120467 refGene_gn=SDF2L1;refGene_tx=XM_003994969.4

D3	30120865 30121062 refGene_gn=SDF2L1;refGene_tx=XM_003994969.4
D3	30121849 30122131 refGene_gn=SDF2L1;refGene_tx=XM_003994969.4
D3	30128414 30128636
refGene_gn=LOC101086852;refGene_tx=XM_011287919.2;refGene_gn=LOC101086
852;refGene_tx=XM_023241709.1
D3	30128834 30128958
refGene_gn=LOC101086852;refGene_tx=XM_011287919.2;refGene_gn=LOC101086
852;refGene_tx=XM_023241709.1
D3	30131229 30131444
refGene_gn=LOC101086852;refGene_tx=XM_011287919.2;refGene_gn=LOC101086
852;refGene_tx=XM_023241709.1
D3	30132096 30132167
refGene_gn=LOC101086852;refGene_tx=XM_011287919.2;refGene_gn=LOC101086
852;refGene_tx=XM_023241709.1
D3	30132414 30132519
refGene_gn=LOC101086852;refGene_tx=XM_011287919.2;refGene_gn=LOC101086
852;refGene_tx=XM_023241709.1
D3	30137956 30138131
refGene_gn=LOC101086852;refGene_tx=XM_011287919.2;refGene_gn=LOC101086
852;refGene_tx=XM_023241709.1
D3	30139877 30139909
refGene_gn=PPIL2;refGene_tx=XM_003994971.5;refGene_gn=PPIL2;refGene_tx
=XM_006938737.4;refGene_gn=PPIL2;refGene_tx=XM_019814939.2;refGene_gn=
PPIL2;refGene_tx=XM_019814940.2
D3	30142642 30142692
refGene_gn=PPIL2;refGene_tx=XM_003994971.5;refGene_gn=PPIL2;refGene_tx
=XM_006938737.4;refGene_gn=PPIL2;refGene_tx=XM_019814939.2;refGene_gn=
PPIL2;refGene_tx=XM_019814940.2
D3	30143719 30143765
refGene_gn=PPIL2;refGene_tx=XM_003994971.5;refGene_gn=PPIL2;refGene_tx
=XM_006938737.4;refGene_gn=PPIL2;refGene_tx=XM_019814939.2;refGene_gn=
PPIL2;refGene_tx=XM_019814940.2
D3	30145910 30145973
refGene_gn=PPIL2;refGene_tx=XM_003994971.5;refGene_gn=PPIL2;refGene_tx
=XM_006938737.4;refGene_gn=PPIL2;refGene_tx=XM_019814939.2;refGene_gn=
PPIL2;refGene_tx=XM_019814940.2
D3	30147534 30147586
refGene_gn=PPIL2;refGene_tx=XM_003994971.5;refGene_gn=PPIL2;refGene_tx
=XM_006938737.4;refGene_gn=PPIL2;refGene_tx=XM_019814939.2;refGene_gn=
PPIL2;refGene_tx=XM_019814940.2
D3	30151569 30151621
refGene_gn=PPIL2;refGene_tx=XM_003994971.5;refGene_gn=PPIL2;refGene_tx
=XM_006938737.4;refGene_gn=PPIL2;refGene_tx=XM_019814939.2;refGene_gn=
PPIL2;refGene_tx=XM_019814940.2
D3	30156132 30156224
refGene_gn=PPIL2;refGene_tx=XM_003994971.5;refGene_gn=PPIL2;refGene_tx
=XM_006938737.4;refGene_gn=PPIL2;refGene_tx=XM_019814939.2;refGene_gn=
PPIL2;refGene_tx=XM_019814940.2
D3	30157085 30157175
refGene_gn=PPIL2;refGene_tx=XM_003994971.5;refGene_gn=PPIL2;refGene_tx

=XM_006938737.4;refGene_gn=PPIL2;refGene_tx=XM_019814939.2;refGene_gn=PPIL2;refGene_tx=XM_019814940.2
D3 30158027 30158103
refGene_gn=PPIL2;refGene_tx=XM_003994971.5;refGene_gn=PPIL2;refGene_tx=XM_006938737.4;refGene_gn=PPIL2;refGene_tx=XM_019814939.2;refGene_gn=PPIL2;refGene_tx=XM_019814940.2
D3 30158845 30159006
refGene_gn=PPIL2;refGene_tx=XM_003994971.5;refGene_gn=PPIL2;refGene_tx=XM_006938737.4;refGene_gn=PPIL2;refGene_tx=XM_019814939.2;refGene_gn=PPIL2;refGene_tx=XM_019814940.2
D3 30159835 30159911
refGene_gn=PPIL2;refGene_tx=XM_003994971.5;refGene_gn=PPIL2;refGene_tx=XM_006938737.4;refGene_gn=PPIL2;refGene_tx=XM_019814939.2;refGene_gn=PPIL2;refGene_tx=XM_019814940.2
D3 30160479 30160586
refGene_gn=PPIL2;refGene_tx=XM_003994971.5;refGene_gn=PPIL2;refGene_tx=XM_006938737.4;refGene_gn=PPIL2;refGene_tx=XM_019814939.2;refGene_gn=PPIL2;refGene_tx=XM_019814940.2
D3 30161039 30161129
refGene_gn=PPIL2;refGene_tx=XM_003994971.5;refGene_gn=PPIL2;refGene_tx=XM_006938737.4;refGene_gn=PPIL2;refGene_tx=XM_019814939.2;refGene_gn=PPIL2;refGene_tx=XM_019814940.2
D3 30161478 30161512
refGene_gn=PPIL2;refGene_tx=XM_003994971.5;refGene_gn=PPIL2;refGene_tx=XM_006938737.4;refGene_gn=PPIL2;refGene_tx=XM_019814939.2;refGene_gn=PPIL2;refGene_tx=XM_019814940.2
D3 30162065 30162183
refGene_gn=PPIL2;refGene_tx=XM_003994971.5;refGene_gn=PPIL2;refGene_tx=XM_006938737.4;refGene_gn=PPIL2;refGene_tx=XM_019814939.2;refGene_gn=PPIL2;refGene_tx=XM_019814940.2
D3 30167170 30167227
refGene_gn=PPIL2;refGene_tx=XM_003994971.5;refGene_gn=PPIL2;refGene_tx=XM_006938737.4;refGene_gn=PPIL2;refGene_tx=XM_019814939.2;refGene_gn=PPIL2;refGene_tx=XM_019814940.2
D3 30167813 30167886
refGene_gn=PPIL2;refGene_tx=XM_003994971.5;refGene_gn=PPIL2;refGene_tx=XM_006938737.4;refGene_gn=PPIL2;refGene_tx=XM_019814939.2;refGene_gn=PPIL2;refGene_tx=XM_019814940.2
D3 30168000 30168063
refGene_gn=PPIL2;refGene_tx=XM_003994971.5;refGene_gn=PPIL2;refGene_tx=XM_006938737.4;refGene_gn=PPIL2;refGene_tx=XM_019814939.2;refGene_gn=PPIL2;refGene_tx=XM_019814940.2
D3 30168157 30168297
refGene_gn=PPIL2;refGene_tx=XM_003994971.5;refGene_gn=PPIL2;refGene_tx=XM_006938737.4;refGene_gn=PPIL2;refGene_tx=XM_019814939.2;refGene_gn=PPIL2;refGene_tx=XM_019814940.2
D3 30168548 30168645
refGene_gn=PPIL2;refGene_tx=XM_003994971.5;refGene_gn=PPIL2;refGene_tx=XM_006938737.4;refGene_gn=PPIL2;refGene_tx=XM_019814939.2;refGene_gn=PPIL2;refGene_tx=XM_019814940.2

D3	30170670	30170868
refGene_gn=YPEL1;refGene_tx=XM_019814943.2;refGene_gn=YPEL1;refGene_tx=XM_023241710.1;refGene_gn=YPEL1;refGene_tx=XM_023241711.1;refGene_gn=YPEL1;refGene_tx=XM_023241712.1
D3	30173728	30173824	refGene_gn=YPEL1;refGene_tx=XM_023241713.1
D3	30175003	30175112
refGene_gn=YPEL1;refGene_tx=XM_019814943.2;refGene_gn=YPEL1;refGene_tx=XM_023241710.1;refGene_gn=YPEL1;refGene_tx=XM_023241711.1;refGene_gn=YPEL1;refGene_tx=XM_023241712.1;refGene_gn=YPEL1;refGene_tx=XM_023241713.1
D3	30176027	30176071
refGene_gn=YPEL1;refGene_tx=XM_019814943.2;refGene_gn=YPEL1;refGene_tx=XM_023241710.1;refGene_gn=YPEL1;refGene_tx=XM_023241711.1;refGene_gn=YPEL1;refGene_tx=XM_023241712.1;refGene_gn=YPEL1;refGene_tx=XM_023241713.1
D3	30178845	30178962
refGene_gn=YPEL1;refGene_tx=XM_019814943.2;refGene_gn=YPEL1;refGene_tx=XM_023241710.1;refGene_gn=YPEL1;refGene_tx=XM_023241711.1;refGene_gn=YPEL1;refGene_tx=XM_023241712.1;refGene_gn=YPEL1;refGene_tx=XM_023241713.1
D3	30226319	30226436	refGene_gn=MAPK1;refGene_tx=XM_003994973.4
D3	30230816	30230926	refGene_gn=MAPK1;refGene_tx=XM_003994973.4
D3	30243586	30243718	refGene_gn=MAPK1;refGene_tx=XM_003994973.4
D3	30244023	30244138	refGene_gn=MAPK1;refGene_tx=XM_003994973.4
D3	30250326	30250443	refGene_gn=MAPK1;refGene_tx=XM_003994973.4
D3	30261027	30261217	refGene_gn=MAPK1;refGene_tx=XM_003994973.4
D3	30262912	30263095	refGene_gn=MAPK1;refGene_tx=XM_003994973.4
D3	30327490	30327609	refGene_gn=MAPK1;refGene_tx=XM_003994973.4
D3	30360048	30360401
refGene_gn=PPM1F;refGene_tx=XM_003994974.5;refGene_gn=PPM1F;refGene_tx=XM_006938738.4
D3	30361568	30361662
refGene_gn=PPM1F;refGene_tx=XM_003994974.5;refGene_gn=PPM1F;refGene_tx=XM_006938738.4
D3	30365784	30365928
refGene_gn=PPM1F;refGene_tx=XM_003994974.5;refGene_gn=PPM1F;refGene_tx=XM_006938738.4
D3	30368909	30369098
refGene_gn=PPM1F;refGene_tx=XM_003994974.5;refGene_gn=PPM1F;refGene_tx=XM_006938738.4
D3	30369741	30369944
refGene_gn=PPM1F;refGene_tx=XM_003994974.5;refGene_gn=PPM1F;refGene_tx=XM_006938738.4
D3	30375150	30375308
refGene_gn=PPM1F;refGene_tx=XM_003994974.5;refGene_gn=PPM1F;refGene_tx=XM_006938738.4
D3	30379535	30379732
refGene_gn=PPM1F;refGene_tx=XM_003994974.5;refGene_gn=PPM1F;refGene_tx=XM_006938738.4
D3	30383186	30383227

refGene_gn=LOC109493009;refGene_tx=XM_019814345.2
D3 30383547 30383572
refGene_gn=LOC109493009;refGene_tx=XM_019814345.2
D3 30385257 30385321
refGene_gn=LOC109493009;refGene_tx=XM_019814345.2
D3 30386406 30386604
refGene_gn=LOC109493009;refGene_tx=XM_019814345.2
D3 30391490 30391972
refGene_gn=TOP3B;refGene_tx=XM_006938741.2;refGene_gn=TOP3B;refGene_tx=XM_011287918.2;refGene_gn=TOP3B;refGene_tx=XM_023241714.1
D3 30393012 30393214
refGene_gn=TOP3B;refGene_tx=XM_006938741.2;refGene_gn=TOP3B;refGene_tx=XM_011287918.2;refGene_gn=TOP3B;refGene_tx=XM_023241714.1
D3 30394051 30394152
refGene_gn=TOP3B;refGene_tx=XM_006938741.2;refGene_gn=TOP3B;refGene_tx=XM_011287918.2;refGene_gn=TOP3B;refGene_tx=XM_023241714.1
D3 30394770 30394920
refGene_gn=TOP3B;refGene_tx=XM_011287918.2;refGene_gn=TOP3B;refGene_tx=XM_023241714.1
D3 30395707 30395836
refGene_gn=TOP3B;refGene_tx=XM_006938741.2;refGene_gn=TOP3B;refGene_tx=XM_011287918.2;refGene_gn=TOP3B;refGene_tx=XM_023241714.1
D3 30397047 30397221
refGene_gn=TOP3B;refGene_tx=XM_006938741.2;refGene_gn=TOP3B;refGene_tx=XM_011287918.2;refGene_gn=TOP3B;refGene_tx=XM_023241714.1
D3 30397366 30397513
refGene_gn=TOP3B;refGene_tx=XM_006938741.2;refGene_gn=TOP3B;refGene_tx=XM_011287918.2;refGene_gn=TOP3B;refGene_tx=XM_023241714.1
D3 30398242 30398348
refGene_gn=TOP3B;refGene_tx=XM_006938741.2;refGene_gn=TOP3B;refGene_tx=XM_011287918.2;refGene_gn=TOP3B;refGene_tx=XM_023241714.1
D3 30398471 30398626
refGene_gn=TOP3B;refGene_tx=XM_006938741.2;refGene_gn=TOP3B;refGene_tx=XM_011287918.2;refGene_gn=TOP3B;refGene_tx=XM_023241714.1
D3 30400100 30400191
refGene_gn=TOP3B;refGene_tx=XM_006938741.2;refGene_gn=TOP3B;refGene_tx=XM_011287918.2;refGene_gn=TOP3B;refGene_tx=XM_023241714.1
D3 30401667 30401781
refGene_gn=TOP3B;refGene_tx=XM_006938741.2;refGene_gn=TOP3B;refGene_tx=XM_011287918.2;refGene_gn=TOP3B;refGene_tx=XM_023241714.1
D3 30402498 30402655
refGene_gn=TOP3B;refGene_tx=XM_006938741.2;refGene_gn=TOP3B;refGene_tx=XM_011287918.2;refGene_gn=TOP3B;refGene_tx=XM_023241714.1
D3 30403156 30403353
refGene_gn=TOP3B;refGene_tx=XM_006938741.2;refGene_gn=TOP3B;refGene_tx=XM_011287918.2;refGene_gn=TOP3B;refGene_tx=XM_023241714.1
D3 30404746 30404821
refGene_gn=TOP3B;refGene_tx=XM_006938741.2;refGene_gn=TOP3B;refGene_tx=XM_011287918.2;refGene_gn=TOP3B;refGene_tx=XM_023241714.1
D3 30405407 30405514

refGene_gn=TOP3B;refGene_tx=XM_006938741.2;refGene_gn=TOP3B;refGene_tx=XM_011287918.2;refGene_gn=TOP3B;refGene_tx=XM_023241714.1
D3 30407046 30407178
refGene_gn=TOP3B;refGene_tx=XM_006938741.2;refGene_gn=TOP3B;refGene_tx=XM_011287918.2;refGene_gn=TOP3B;refGene_tx=XM_023241714.1
D3 30408358 30408428
refGene_gn=TOP3B;refGene_tx=XM_006938741.2;refGene_gn=TOP3B;refGene_tx=XM_011287918.2
D3 30413112 30413708
refGene_gn=LOC109493009;refGene_tx=XM_019814345.2
D3 30510340 30510386
refGene_gn=LOC101091817;refGene_tx=XM_003994989.3
D3 30510470 30510768
refGene_gn=LOC101091817;refGene_tx=XM_003994989.3
D3 30510769 30510809
refGene_gn=LOC101091817;refGene_tx=XM_003994989.3
D3 30541063 30544939 refGene_gn=LOC101088367
D3 30574189 30575536
refGene_gn=ZNF70;refGene_tx=XM_011287927.3;refGene_gn=ZNF70;refGene_tx=XM_023241715.1
D3 30580717 30581022
refGene_gn=VPREB3;refGene_tx=XM_011287928.3;refGene_gn=VPREB3;refGene_tx=XM_019814370.2
D3 30581752 30581801
refGene_gn=VPREB3;refGene_tx=XM_011287928.3;refGene_gn=VPREB3;refGene_tx=XM_019814370.2
D3 30585449 30585474
refGene_gn=CD3H22orf15;refGene_tx=XM_023241716.1
D3 30586116 30586203
refGene_gn=CD3H22orf15;refGene_tx=XM_023241716.1
D3 30586297 30586423
refGene_gn=CD3H22orf15;refGene_tx=XM_023241716.1
D3 30586653 30586728
refGene_gn=CD3H22orf15;refGene_tx=XM_023241716.1
D3 30586808 30586948
refGene_gn=CD3H22orf15;refGene_tx=XM_023241716.1
D3 30588017 30588037 refGene_gn=CHCHD10;refGene_tx=XM_003994978.4
D3 30588133 30588281 refGene_gn=CHCHD10;refGene_tx=XM_003994978.4
D3 30589147 30589367 refGene_gn=CHCHD10;refGene_tx=XM_003994978.4
D3 30589630 30589671 refGene_gn=CHCHD10;refGene_tx=XM_003994978.4
D3 30592130 30592241 refGene_gn=MMP11;refGene_tx=XM_003994979.5
D3 30597412 30597639 refGene_gn=MMP11;refGene_tx=XM_003994979.5
D3 30598560 30598704 refGene_gn=MMP11;refGene_tx=XM_003994979.5
D3 30598782 30598916 refGene_gn=MMP11;refGene_tx=XM_003994979.5
D3 30599055 30599297 refGene_gn=MMP11;refGene_tx=XM_003994979.5
D3 30599383 30599600 refGene_gn=MMP11;refGene_tx=XM_003994979.5
D3 30600383 30600641 refGene_gn=MMP11;refGene_tx=XM_003994979.5
D3 30601603 30601737 refGene_gn=MMP11;refGene_tx=XM_003994979.5
D3 30603824 30603917
refGene_gn=SMARCB1;refGene_tx=XM_003994980.5;refGene_gn=SMARCB1;refGen

e_tx=XM_003994981.5
D3	30607199	30607338
refGene_gn=SMARCB1;refGene_tx=XM_003994980.5;refGene_gn=SMARCB1;refGene_tx=XM_003994981.5;refGene_gn=SMARCB1;refGene_tx=XM_023241717.1
D3	30609020	30609150
refGene_gn=SMARCB1;refGene_tx=XM_003994980.5;refGene_gn=SMARCB1;refGene_tx=XM_003994981.5;refGene_gn=SMARCB1;refGene_tx=XM_023241717.1
D3	30613278	30613416
refGene_gn=SMARCB1;refGene_tx=XM_003994980.5;refGene_gn=SMARCB1;refGene_tx=XM_003994981.5;refGene_gn=SMARCB1;refGene_tx=XM_023241717.1
D3	30615505	30615633
refGene_gn=SMARCB1;refGene_tx=XM_003994980.5;refGene_gn=SMARCB1;refGene_tx=XM_003994981.5;refGene_gn=SMARCB1;refGene_tx=XM_023241717.1
D3	30623545	30623712
refGene_gn=SMARCB1;refGene_tx=XM_003994980.5;refGene_gn=SMARCB1;refGene_tx=XM_003994981.5;refGene_gn=SMARCB1;refGene_tx=XM_023241717.1
D3	30631648	30631839
refGene_gn=SMARCB1;refGene_tx=XM_003994980.5;refGene_gn=SMARCB1;refGene_tx=XM_003994981.5;refGene_gn=SMARCB1;refGene_tx=XM_023241717.1
D3	30636362	30636408	refGene_gn=DERL3;refGene_tx=XM_019814949.2
D3	30637969	30638093	refGene_gn=DERL3;refGene_tx=XM_006938743.3
D3	30638978	30639110
refGene_gn=SMARCB1;refGene_tx=XM_003994980.5;refGene_gn=SMARCB1;refGene_tx=XM_003994981.5;refGene_gn=SMARCB1;refGene_tx=XM_023241717.1
D3	30639546	30639586
refGene_gn=SMARCB1;refGene_tx=XM_003994980.5;refGene_gn=SMARCB1;refGene_tx=XM_003994981.5;refGene_gn=SMARCB1;refGene_tx=XM_023241717.1
D3	30642206	30642288	refGene_gn=DERL3;refGene_tx=XM_003994982.3
D3	30642394	30642485
refGene_gn=DERL3;refGene_tx=XM_003994982.3;refGene_gn=DERL3;refGene_tx=XM_006938743.3;refGene_gn=DERL3;refGene_tx=XM_019814949.2
D3	30643034	30643230
refGene_gn=DERL3;refGene_tx=XM_003994982.3;refGene_gn=DERL3;refGene_tx=XM_006938743.3;refGene_gn=DERL3;refGene_tx=XM_019814949.2
D3	30643720	30643814
refGene_gn=DERL3;refGene_tx=XM_003994982.3;refGene_gn=DERL3;refGene_tx=XM_006938743.3;refGene_gn=DERL3;refGene_tx=XM_019814949.2
D3	30643963	30644037
refGene_gn=DERL3;refGene_tx=XM_003994982.3;refGene_gn=DERL3;refGene_tx=XM_006938743.3;refGene_gn=DERL3;refGene_tx=XM_019814949.2
D3	30644120	30644186
refGene_gn=DERL3;refGene_tx=XM_003994982.3;refGene_gn=DERL3;refGene_tx=XM_006938743.3;refGene_gn=DERL3;refGene_tx=XM_019814949.2
D3	30644299	30644392
refGene_gn=DERL3;refGene_tx=XM_003994982.3;refGene_gn=DERL3;refGene_tx=XM_006938743.3;refGene_gn=DERL3;refGene_tx=XM_019814949.2
D3	30660479	30660572	refGene_gn=SLC2A11;refGene_tx=XM_023241196.1
D3	30661166	30661259	refGene_gn=SLC2A11;refGene_tx=XM_023241196.1
D3	30664157	30664318	refGene_gn=SLC2A11;refGene_tx=XM_023241196.1
D3	30671645	30671770	refGene_gn=SLC2A11;refGene_tx=XM_023241196.1

D3 30672569 30672699 refGene_gn=SLC2A11;refGene_tx=XM_023241196.1
D3 30673251 30673400 refGene_gn=SLC2A11;refGene_tx=XM_023241196.1
D3 30676133 30676525 refGene_gn=SLC2A11;refGene_tx=XM_023241196.1
D3 30677066 30677168 refGene_gn=SLC2A11;refGene_tx=XM_023241196.1
D3 30677250 30677326 refGene_gn=SLC2A11;refGene_tx=XM_023241196.1
D3 30677561 30677689 refGene_gn=SLC2A11;refGene_tx=XM_023241196.1
D3 30677900 30678101 refGene_gn=SLC2A11;refGene_tx=XM_023241196.1
D3 30694817 30694925 refGene_gn=MIF;refGene_tx=XM_003994984.5
D3 30695103 30695276 refGene_gn=MIF;refGene_tx=XM_003994984.5
D3 30695400 30695467 refGene_gn=MIF;refGene_tx=XM_003994984.5
D3 33119731 33120625 refGene_gn=MC2R;refGene_tx=XM_003994992.3
D3 33203225 33204203 refGene_gn=MC5R;refGene_tx=XM_019814371.2
D3 33263155 33263193 refGene_gn=RNMT;refGene_tx=XM_003994994.5
D3 33266788 33266822 refGene_gn=RNMT;refGene_tx=XM_003994994.5
D3 33268192 33268294 refGene_gn=RNMT;refGene_tx=XM_003994994.5
D3 33277304 33277422 refGene_gn=RNMT;refGene_tx=XM_003994994.5
D3 33280642 33280807 refGene_gn=RNMT;refGene_tx=XM_003994994.5
D3 33281359 33281541 refGene_gn=RNMT;refGene_tx=XM_003994994.5
D3 33283462 33283578 refGene_gn=RNMT;refGene_tx=XM_003994994.5
D3 33286331 33286457 refGene_gn=RNMT;refGene_tx=XM_003994994.5
D3 33289457 33289593 refGene_gn=RNMT;refGene_tx=XM_003994994.5
D3 33291225 33291642 refGene_gn=RNMT;refGene_tx=XM_003994994.5
D3 33311209 33311664
refGene_gn=FAM210A;refGene_tx=XM_003994995.5;refGene_gn=FAM210A;refGene_tx=XM_006938748.4;refGene_gn=FAM210A;refGene_tx=XM_006938749.4;refGene_gn=FAM210A;refGene_tx=XM_006938750.4;refGene_gn=FAM210A;refGene_tx=XM_006938752.4;refGene_gn=FAM210A;refGene_tx=XM_006938753.4;refGene_gn=FAM210A;refGene_tx=XM_019814951.2;refGene_gn=FAM210A;refGene_tx=XM_019814952.2
D3 33317030 33317142
refGene_gn=FAM210A;refGene_tx=XM_003994995.5;refGene_gn=FAM210A;refGene_tx=XM_006938748.4;refGene_gn=FAM210A;refGene_tx=XM_006938749.4;refGene_gn=FAM210A;refGene_tx=XM_006938750.4;refGene_gn=FAM210A;refGene_tx=XM_006938752.4;refGene_gn=FAM210A;refGene_tx=XM_006938753.4;refGene_gn=FAM210A;refGene_tx=XM_019814951.2;refGene_gn=FAM210A;refGene_tx=XM_019814952.2
D3 33321173 33321407
refGene_gn=FAM210A;refGene_tx=XM_003994995.5;refGene_gn=FAM210A;refGene_tx=XM_006938748.4;refGene_gn=FAM210A;refGene_tx=XM_006938749.4;refGene_gn=FAM210A;refGene_tx=XM_006938750.4;refGene_gn=FAM210A;refGene_tx=XM_006938752.4;refGene_gn=FAM210A;refGene_tx=XM_006938753.4;refGene_gn=FAM210A;refGene_tx=XM_019814951.2;refGene_gn=FAM210A;refGene_tx=XM_019814952.2
D3 33348446 33348977
refGene_gn=LDLRAD4;refGene_tx=XM_003994996.5;refGene_gn=LDLRAD4;refGene_tx=XM_003994997.5;refGene_gn=LDLRAD4;refGene_tx=XM_006938755.4;refGene_gn=LDLRAD4;refGene_tx=XM_011287951.3;refGene_gn=LDLRAD4;refGene_tx=XM_019814958.2;refGene_gn=LDLRAD4;refGene_tx=XM_023241718.1;refGene_gn=LDLRAD4;refGene_tx=XM_023241719.1;refGene_gn=LDLRAD4;refGene_tx=XM_023241720.1;refGene_gn=LDLRAD4;refGene_tx=XM_023241721.1;refGene_gn=LDLR

AD4;refGene_tx=XM_023241722.1;refGene_gn=LDLRAD4;refGene_tx=XM_0232417
23.1;refGene_gn=LDLRAD4;refGene_tx=XM_023241724.1;refGene_gn=LDLRAD4;r
efGene_tx=XM_023241725.1
D3 33350490 33350544
refGene_gn=LDLRAD4;refGene_tx=XM_003994996.5;refGene_gn=LDLRAD4;refGen
e_tx=XM_006938755.4;refGene_gn=LDLRAD4;refGene_tx=XM_011287951.3;refGe
ne_gn=LDLRAD4;refGene_tx=XM_019814958.2;refGene_gn=LDLRAD4;refGene_tx=
XM_023241718.1;refGene_gn=LDLRAD4;refGene_tx=XM_023241719.1;refGene_gn
=LDLRAD4;refGene_tx=XM_023241720.1;refGene_gn=LDLRAD4;refGene_tx=XM_02
3241721.1;refGene_gn=LDLRAD4;refGene_tx=XM_023241722.1;refGene_gn=LDLR
AD4;refGene_tx=XM_023241724.1;refGene_gn=LDLRAD4;refGene_tx=XM_0232417
25.1
D3 33370145 33370300
refGene_gn=LDLRAD4;refGene_tx=XM_003994996.5;refGene_gn=LDLRAD4;refGen
e_tx=XM_003994997.5;refGene_gn=LDLRAD4;refGene_tx=XM_006938755.4;refGe
ne_gn=LDLRAD4;refGene_tx=XM_011287951.3;refGene_gn=LDLRAD4;refGene_tx=
XM_019814958.2;refGene_gn=LDLRAD4;refGene_tx=XM_023241718.1;refGene_gn
=LDLRAD4;refGene_tx=XM_023241719.1;refGene_gn=LDLRAD4;refGene_tx=XM_02
3241720.1;refGene_gn=LDLRAD4;refGene_tx=XM_023241721.1;refGene_gn=LDLR
AD4;refGene_tx=XM_023241722.1;refGene_gn=LDLRAD4;refGene_tx=XM_0232417
23.1;refGene_gn=LDLRAD4;refGene_tx=XM_023241724.1;refGene_gn=LDLRAD4;r
efGene_tx=XM_023241725.1
D3 33379871 33379941
refGene_gn=LDLRAD4;refGene_tx=XM_003994996.5;refGene_gn=LDLRAD4;refGen
e_tx=XM_003994997.5
D3 33539427 33539568
refGene_gn=LDLRAD4;refGene_tx=XM_006938755.4;refGene_gn=LDLRAD4;refGen
e_tx=XM_011287951.3;refGene_gn=LDLRAD4;refGene_tx=XM_023241718.1;refGe
ne_gn=LDLRAD4;refGene_tx=XM_023241719.1;refGene_gn=LDLRAD4;refGene_tx=
XM_023241720.1;refGene_gn=LDLRAD4;refGene_tx=XM_023241721.1;refGene_gn
=LDLRAD4;refGene_tx=XM_023241722.1;refGene_gn=LDLRAD4;refGene_tx=XM_02
3241723.1
D3 33597344 33597384
refGene_gn=LDLRAD4;refGene_tx=XM_006938755.4;refGene_gn=LDLRAD4;refGen
e_tx=XM_011287951.3;refGene_gn=LDLRAD4;refGene_tx=XM_023241718.1;refGe
ne_gn=LDLRAD4;refGene_tx=XM_023241719.1;refGene_gn=LDLRAD4;refGene_tx=
XM_023241720.1;refGene_gn=LDLRAD4;refGene_tx=XM_023241721.1;refGene_gn
=LDLRAD4;refGene_tx=XM_023241722.1;refGene_gn=LDLRAD4;refGene_tx=XM_02
3241723.1
D3 33901181 33901323
refGene_gn=CEP192;refGene_tx=XM_019814959.2;refGene_gn=CEP192;refGene_
tx=XM_023241726.1
D3 33906391 33906450
refGene_gn=CEP192;refGene_tx=XM_019814959.2;refGene_gn=CEP192;refGene_
tx=XM_023241726.1
D3 33906874 33907004
refGene_gn=CEP192;refGene_tx=XM_019814959.2;refGene_gn=CEP192;refGene_
tx=XM_023241726.1
D3 33909798 33909920
refGene_gn=CEP192;refGene_tx=XM_019814959.2;refGene_gn=CEP192;refGene_

tx=XM_023241726.1
D3	33910363	33910483
refGene_gn=CEP192;refGene_tx=XM_019814959.2;refGene_gn=CEP192;refGene_tx=XM_023241726.1
D3	33915076	33915172
refGene_gn=CEP192;refGene_tx=XM_019814959.2;refGene_gn=CEP192;refGene_tx=XM_023241726.1
D3	33916119	33916199
refGene_gn=CEP192;refGene_tx=XM_019814959.2;refGene_gn=CEP192;refGene_tx=XM_023241726.1
D3	33918864	33919072
refGene_gn=CEP192;refGene_tx=XM_019814959.2;refGene_gn=CEP192;refGene_tx=XM_023241726.1
D3	33919506	33919612
refGene_gn=CEP192;refGene_tx=XM_019814959.2;refGene_gn=CEP192;refGene_tx=XM_023241726.1
D3	33921599	33921723
refGene_gn=CEP192;refGene_tx=XM_019814959.2;refGene_gn=CEP192;refGene_tx=XM_023241726.1
D3	33922330	33922509
refGene_gn=CEP192;refGene_tx=XM_019814959.2;refGene_gn=CEP192;refGene_tx=XM_023241726.1
D3	33927664	33927815
refGene_gn=CEP192;refGene_tx=XM_019814959.2;refGene_gn=CEP192;refGene_tx=XM_023241726.1
D3	33930690	33930800
refGene_gn=CEP192;refGene_tx=XM_019814959.2;refGene_gn=CEP192;refGene_tx=XM_023241726.1
D3	33932584	33932700
refGene_gn=CEP192;refGene_tx=XM_019814959.2;refGene_gn=CEP192;refGene_tx=XM_023241726.1
D3	33932945	33933206
refGene_gn=CEP192;refGene_tx=XM_019814959.2;refGene_gn=CEP192;refGene_tx=XM_023241726.1
D3	33939416	33939593
refGene_gn=CEP192;refGene_tx=XM_019814959.2;refGene_gn=CEP192;refGene_tx=XM_023241726.1
D3	33939740	33939831
refGene_gn=CEP192;refGene_tx=XM_019814959.2;refGene_gn=CEP192;refGene_tx=XM_023241726.1
D3	33944714	33944891
refGene_gn=CEP192;refGene_tx=XM_019814959.2;refGene_gn=CEP192;refGene_tx=XM_023241726.1
D3	33945828	33945947
refGene_gn=CEP192;refGene_tx=XM_019814959.2;refGene_gn=CEP192;refGene_tx=XM_023241726.1
D3	33946454	33946547
refGene_gn=CEP192;refGene_tx=XM_019814959.2;refGene_gn=CEP192;refGene_tx=XM_023241726.1
D3	33946645	33946785

refGene_gn=CEP192;refGene_tx=XM_019814959.2;refGene_gn=CEP192;refGene_tx=XM_023241726.1
D3 33947435 33947499
refGene_gn=CEP192;refGene_tx=XM_019814959.2;refGene_gn=CEP192;refGene_tx=XM_023241726.1
D3 33947625 33947769
refGene_gn=CEP192;refGene_tx=XM_019814959.2;refGene_gn=CEP192;refGene_tx=XM_023241726.1
D3 33947960 33948086
refGene_gn=CEP192;refGene_tx=XM_019814959.2;refGene_gn=CEP192;refGene_tx=XM_023241726.1
D3 33952204 33952435
refGene_gn=CEP192;refGene_tx=XM_019814959.2;refGene_gn=CEP192;refGene_tx=XM_023241726.1
D3 33954287 33954436
refGene_gn=CEP192;refGene_tx=XM_019814959.2;refGene_gn=CEP192;refGene_tx=XM_023241726.1
D3 33955675 33956597
refGene_gn=CEP192;refGene_tx=XM_019814959.2;refGene_gn=CEP192;refGene_tx=XM_023241726.1
D3 33958716 33958885
refGene_gn=CEP192;refGene_tx=XM_019814959.2;refGene_gn=CEP192;refGene_tx=XM_023241726.1
D3 33967454 33967581
refGene_gn=CEP192;refGene_tx=XM_019814959.2;refGene_gn=CEP192;refGene_tx=XM_023241726.1
D3 33967675 33968498
refGene_gn=CEP192;refGene_tx=XM_019814959.2;refGene_gn=CEP192;refGene_tx=XM_023241726.1
D3 33974396 33974515
refGene_gn=CEP192;refGene_tx=XM_019814959.2;refGene_gn=CEP192;refGene_tx=XM_023241726.1
D3 33975909 33976036
refGene_gn=CEP192;refGene_tx=XM_019814959.2;refGene_gn=CEP192;refGene_tx=XM_023241726.1
D3 33978043 33978253
refGene_gn=CEP192;refGene_tx=XM_019814959.2;refGene_gn=CEP192;refGene_tx=XM_023241726.1
D3 33978904 33978969
refGene_gn=CEP192;refGene_tx=XM_019814959.2;refGene_gn=CEP192;refGene_tx=XM_023241726.1
D3 33986173 33986317
refGene_gn=CEP192;refGene_tx=XM_019814959.2;refGene_gn=CEP192;refGene_tx=XM_023241726.1
D3 33986771 33987132
refGene_gn=CEP192;refGene_tx=XM_019814959.2;refGene_gn=CEP192;refGene_tx=XM_023241726.1
D3 34000739 34000873
refGene_gn=CEP192;refGene_tx=XM_019814959.2;refGene_gn=CEP192;refGene_tx=XM_023241726.1

D3 34002196 34002341
refGene_gn=CEP192;refGene_tx=XM_019814959.2;refGene_gn=CEP192;refGene_
tx=XM_023241726.1
D3 34003489 34003638
refGene_gn=CEP192;refGene_tx=XM_019814959.2;refGene_gn=CEP192;refGene_
tx=XM_023241726.1
D3 34012005 34012126
refGene_gn=CEP192;refGene_tx=XM_019814959.2;refGene_gn=CEP192;refGene_
tx=XM_023241726.1
D3 34015362 34015415
refGene_gn=CEP192;refGene_tx=XM_019814959.2;refGene_gn=CEP192;refGene_
tx=XM_023241726.1
D3 34018583 34018759
refGene_gn=CEP192;refGene_tx=XM_019814959.2;refGene_gn=CEP192;refGene_
tx=XM_023241726.1
D3 34026413 34026539
refGene_gn=CEP192;refGene_tx=XM_019814959.2;refGene_gn=CEP192;refGene_
tx=XM_023241726.1
D3 34027257 34027421
refGene_gn=CEP192;refGene_tx=XM_019814959.2;refGene_gn=CEP192;refGene_
tx=XM_023241726.1
D3 34039212 34039408 refGene_gn=SEH1L;refGene_tx=XM_003994999.4
D3 34041039 34041052 refGene_gn=SEH1L;refGene_tx=XM_003994998.4
D3 34042120 34042271
refGene_gn=SEH1L;refGene_tx=XM_003994998.4;refGene_gn=SEH1L;refGene_tx
=XM_003994999.4
D3 34043567 34043725
refGene_gn=SEH1L;refGene_tx=XM_003994998.4;refGene_gn=SEH1L;refGene_tx
=XM_003994999.4
D3 34044454 34044595
refGene_gn=SEH1L;refGene_tx=XM_003994998.4;refGene_gn=SEH1L;refGene_tx
=XM_003994999.4
D3 34048988 34049087
refGene_gn=SEH1L;refGene_tx=XM_003994998.4;refGene_gn=SEH1L;refGene_tx
=XM_003994999.4
D3 34056900 34057112
refGene_gn=SEH1L;refGene_tx=XM_003994998.4;refGene_gn=SEH1L;refGene_tx
=XM_003994999.4
D3 34064472 34064619
refGene_gn=SEH1L;refGene_tx=XM_003994998.4;refGene_gn=SEH1L;refGene_tx
=XM_003994999.4
D3 34067215 34067266
refGene_gn=SEH1L;refGene_tx=XM_003994998.4;refGene_gn=SEH1L;refGene_tx
=XM_003994999.4
D3 34072404 34072515
refGene_gn=SEH1L;refGene_tx=XM_003994998.4;refGene_gn=SEH1L;refGene_tx
=XM_003994999.4
D3 34171935 34172004
refGene_gn=PTPN2;refGene_tx=XM_011287948.2;refGene_gn=PTPN2;refGene_tx
=XM_019814960.2;refGene_gn=PTPN2;refGene_tx=XM_019814962.2

D3 34195224 34195315
refGene_gn=PTPN2;refGene_tx=XM_011287948.2;refGene_gn=PTPN2;refGene_tx=XM_019814960.2;refGene_gn=PTPN2;refGene_tx=XM_019814962.2
D3 34204945 34204955 refGene_gn=PTPN2;refGene_tx=XM_019814961.2
D3 34218551 34218652
refGene_gn=PTPN2;refGene_tx=XM_011287948.2;refGene_gn=PTPN2;refGene_tx=XM_019814960.2;refGene_gn=PTPN2;refGene_tx=XM_019814961.2;refGene_gn=PTPN2;refGene_tx=XM_019814962.2
D3 34224367 34224466
refGene_gn=PTPN2;refGene_tx=XM_011287948.2;refGene_gn=PTPN2;refGene_tx=XM_019814960.2;refGene_gn=PTPN2;refGene_tx=XM_019814961.2;refGene_gn=PTPN2;refGene_tx=XM_019814962.2
D3 34228402 34228540
refGene_gn=PTPN2;refGene_tx=XM_011287948.2;refGene_gn=PTPN2;refGene_tx=XM_019814960.2;refGene_gn=PTPN2;refGene_tx=XM_019814961.2;refGene_gn=PTPN2;refGene_tx=XM_019814962.2
D3 34236556 34236766
refGene_gn=PTPN2;refGene_tx=XM_011287948.2;refGene_gn=PTPN2;refGene_tx=XM_019814960.2;refGene_gn=PTPN2;refGene_tx=XM_019814961.2;refGene_gn=PTPN2;refGene_tx=XM_019814962.2
D3 34238339 34238492
refGene_gn=PTPN2;refGene_tx=XM_011287948.2;refGene_gn=PTPN2;refGene_tx=XM_019814960.2;refGene_gn=PTPN2;refGene_tx=XM_019814961.2;refGene_gn=PTPN2;refGene_tx=XM_019814962.2
D3 34245156 34245338
refGene_gn=PTPN2;refGene_tx=XM_011287948.2;refGene_gn=PTPN2;refGene_tx=XM_019814960.2;refGene_gn=PTPN2;refGene_tx=XM_019814961.2;refGene_gn=PTPN2;refGene_tx=XM_019814962.2
D3 34249744 34249952
refGene_gn=PTPN2;refGene_tx=XM_011287948.2;refGene_gn=PTPN2;refGene_tx=XM_019814960.2;refGene_gn=PTPN2;refGene_tx=XM_019814961.2;refGene_gn=PTPN2;refGene_tx=XM_019814962.2
D3 34250642 34250694 refGene_gn=PTPN2;refGene_tx=XM_019814960.2
D3 34257370 34257392 refGene_gn=PTPN2;refGene_tx=XM_011287948.2
D3 34314483 34314582 refGene_gn=PSMG2;refGene_tx=XM_006938764.3
D3 34315409 34315530 refGene_gn=PSMG2;refGene_tx=XM_006938764.3
D3 34323342 34323516 refGene_gn=PSMG2;refGene_tx=XM_006938764.3
D3 34324371 34324490 refGene_gn=PSMG2;refGene_tx=XM_006938764.3
D3 34327301 34327360 refGene_gn=PSMG2;refGene_tx=XM_006938764.3
D3 34330934 34331106 refGene_gn=PSMG2;refGene_tx=XM_006938764.3
D3 34334424 34334481 refGene_gn=PSMG2;refGene_tx=XM_006938764.3
D3 34335062 34335125 refGene_gn=CEP76;refGene_tx=XM_023241197.1
D3 34336193 34336349 refGene_gn=CEP76;refGene_tx=XM_023241197.1
D3 34337510 34337592 refGene_gn=CEP76;refGene_tx=XM_023241197.1
D3 34338206 34338431 refGene_gn=CEP76;refGene_tx=XM_023241197.1
D3 34339692 34339878 refGene_gn=CEP76;refGene_tx=XM_023241197.1
D3 34341271 34341369 refGene_gn=CEP76;refGene_tx=XM_023241197.1
D3 34341945 34342074 refGene_gn=CEP76;refGene_tx=XM_023241197.1
D3 34346742 34347117 refGene_gn=CEP76;refGene_tx=XM_023241197.1
D3 34350544 34350892 refGene_gn=CEP76;refGene_tx=XM_023241197.1

D3 34354142 34354360 refGene_gn=CEP76;refGene_tx=XM_023241197.1
D3 34354766 34354905 refGene_gn=CEP76;refGene_tx=XM_023241197.1
D3 34371044 34371366
refGene_gn=SPIRE1;refGene_tx=XM_023241727.1;refGene_gn=SPIRE1;refGene_tx=XM_023241728.1;refGene_gn=SPIRE1;refGene_tx=XM_023241729.1;refGene_gn=SPIRE1;refGene_tx=XM_023241730.1;refGene_gn=SPIRE1;refGene_tx=XM_023241731.1
D3 34393899 34393934
refGene_gn=SPIRE1;refGene_tx=XM_023241727.1;refGene_gn=SPIRE1;refGene_tx=XM_023241728.1;refGene_gn=SPIRE1;refGene_tx=XM_023241729.1;refGene_gn=SPIRE1;refGene_tx=XM_023241730.1;refGene_gn=SPIRE1;refGene_tx=XM_023241731.1
D3 34405073 34405115
refGene_gn=SPIRE1;refGene_tx=XM_023241728.1;refGene_gn=SPIRE1;refGene_tx=XM_023241730.1
D3 34454488 34454722
refGene_gn=SPIRE1;refGene_tx=XM_023241727.1;refGene_gn=SPIRE1;refGene_tx=XM_023241728.1;refGene_gn=SPIRE1;refGene_tx=XM_023241729.1;refGene_gn=SPIRE1;refGene_tx=XM_023241730.1;refGene_gn=SPIRE1;refGene_tx=XM_023241731.1
D3 34474099 34474225
refGene_gn=SPIRE1;refGene_tx=XM_023241727.1;refGene_gn=SPIRE1;refGene_tx=XM_023241728.1;refGene_gn=SPIRE1;refGene_tx=XM_023241729.1;refGene_gn=SPIRE1;refGene_tx=XM_023241730.1;refGene_gn=SPIRE1;refGene_tx=XM_023241731.1
D3 34499138 34499216
refGene_gn=SPIRE1;refGene_tx=XM_023241727.1;refGene_gn=SPIRE1;refGene_tx=XM_023241728.1;refGene_gn=SPIRE1;refGene_tx=XM_023241729.1;refGene_gn=SPIRE1;refGene_tx=XM_023241730.1;refGene_gn=SPIRE1;refGene_tx=XM_023241731.1
D3 34503062 34503227
refGene_gn=SPIRE1;refGene_tx=XM_023241727.1;refGene_gn=SPIRE1;refGene_tx=XM_023241728.1;refGene_gn=SPIRE1;refGene_tx=XM_023241729.1;refGene_gn=SPIRE1;refGene_tx=XM_023241730.1;refGene_gn=SPIRE1;refGene_tx=XM_023241731.1
D3 34523420 34523507
refGene_gn=SPIRE1;refGene_tx=XM_023241727.1;refGene_gn=SPIRE1;refGene_tx=XM_023241728.1;refGene_gn=SPIRE1;refGene_tx=XM_023241729.1;refGene_gn=SPIRE1;refGene_tx=XM_023241730.1;refGene_gn=SPIRE1;refGene_tx=XM_023241731.1
D3 34524379 34524509
refGene_gn=SPIRE1;refGene_tx=XM_023241727.1;refGene_gn=SPIRE1;refGene_tx=XM_023241728.1;refGene_gn=SPIRE1;refGene_tx=XM_023241729.1;refGene_gn=SPIRE1;refGene_tx=XM_023241730.1;refGene_gn=SPIRE1;refGene_tx=XM_023241731.1
D3 34530818 34530860
refGene_gn=SPIRE1;refGene_tx=XM_023241727.1;refGene_gn=SPIRE1;refGene_tx=XM_023241729.1
D3 34538296 34538469
refGene_gn=SPIRE1;refGene_tx=XM_023241727.1;refGene_gn=SPIRE1;refGene_

tx=XM_023241728.1;refGene_gn=SPIRE1;refGene_tx=XM_023241729.1;refGene_gn=SPIRE1;refGene_tx=XM_023241730.1;refGene_gn=SPIRE1;refGene_tx=XM_023241731.1
D3 34541483 34541574 refGene_gn=SPIRE1;refGene_tx=XM_023241727.1;refGene_gn=SPIRE1;refGene_tx=XM_023241728.1;refGene_gn=SPIRE1;refGene_tx=XM_023241729.1;refGene_gn=SPIRE1;refGene_tx=XM_023241730.1;refGene_gn=SPIRE1;refGene_tx=XM_023241731.1
D3 34560196 34560339 refGene_gn=SPIRE1;refGene_tx=XM_023241727.1;refGene_gn=SPIRE1;refGene_tx=XM_023241728.1;refGene_gn=SPIRE1;refGene_tx=XM_023241729.1;refGene_gn=SPIRE1;refGene_tx=XM_023241730.1;refGene_gn=SPIRE1;refGene_tx=XM_023241731.1
D3 34562596 34562770 refGene_gn=SPIRE1;refGene_tx=XM_023241727.1;refGene_gn=SPIRE1;refGene_tx=XM_023241728.1
D3 34567033 34567171 refGene_gn=SPIRE1;refGene_tx=XM_023241727.1;refGene_gn=SPIRE1;refGene_tx=XM_023241728.1;refGene_gn=SPIRE1;refGene_tx=XM_023241729.1;refGene_gn=SPIRE1;refGene_tx=XM_023241730.1;refGene_gn=SPIRE1;refGene_tx=XM_023241731.1
D3 34567765 34567836 refGene_gn=SPIRE1;refGene_tx=XM_023241727.1;refGene_gn=SPIRE1;refGene_tx=XM_023241728.1;refGene_gn=SPIRE1;refGene_tx=XM_023241729.1;refGene_gn=SPIRE1;refGene_tx=XM_023241730.1;refGene_gn=SPIRE1;refGene_tx=XM_023241731.1
D3 34568373 34568401 refGene_gn=SPIRE1;refGene_tx=XM_023241727.1;refGene_gn=SPIRE1;refGene_tx=XM_023241728.1;refGene_gn=SPIRE1;refGene_tx=XM_023241729.1;refGene_gn=SPIRE1;refGene_tx=XM_023241730.1;refGene_gn=SPIRE1;refGene_tx=XM_023241731.1
D3 34568496 34568633 refGene_gn=SPIRE1;refGene_tx=XM_023241727.1;refGene_gn=SPIRE1;refGene_tx=XM_023241728.1;refGene_gn=SPIRE1;refGene_tx=XM_023241729.1;refGene_gn=SPIRE1;refGene_tx=XM_023241730.1;refGene_gn=SPIRE1;refGene_tx=XM_023241731.1
D3 34570177 34570436 refGene_gn=SPIRE1;refGene_tx=XM_023241727.1;refGene_gn=SPIRE1;refGene_tx=XM_023241728.1;refGene_gn=SPIRE1;refGene_tx=XM_023241729.1;refGene_gn=SPIRE1;refGene_tx=XM_023241730.1;refGene_gn=SPIRE1;refGene_tx=XM_023241731.1
D3 34582887 34582971 refGene_gn=PRELID3A;refGene_tx=XM_011287938.3;refGene_gn=PRELID3A;refGene_tx=XM_011287939.3;refGene_gn=PRELID3A;refGene_tx=XM_019814970.2;refGene_gn=PRELID3A;refGene_tx=XM_019814971.2
D3 34595974 34596058 refGene_gn=PRELID3A;refGene_tx=XM_019814972.2
D3 34597175 34597229 refGene_gn=PRELID3A;refGene_tx=XM_011287942.3;refGene_gn=PRELID3A;refGene_tx=XM_019814966.2;refGene_gn=PRELID3A;refGene_tx=XM_019814967.2;refGene_gn=PRELID3A;refGene_tx=XM_019814973.1;refGene_gn=PRELID3A;refGen

e_tx=XM_023241732.1
D3 34598266 34598456
refGene_gn=PRELID3A;refGene_tx=XM_011287940.2;refGene_gn=PRELID3A;refGene_tx=XM_019814966.2;refGene_gn=PRELID3A;refGene_tx=XM_019814967.2;refGene_gn=PRELID3A;refGene_tx=XM_023241732.1
D3 34598888 34598991
refGene_gn=PRELID3A;refGene_tx=XM_011287938.3;refGene_gn=PRELID3A;refGene_tx=XM_011287939.3;refGene_gn=PRELID3A;refGene_tx=XM_011287940.2;refGene_gn=PRELID3A;refGene_tx=XM_011287942.3;refGene_gn=PRELID3A;refGene_tx=XM_019814966.2;refGene_gn=PRELID3A;refGene_tx=XM_019814967.2;refGene_gn=PRELID3A;refGene_tx=XM_019814970.2;refGene_gn=PRELID3A;refGene_tx=XM_019814971.2;refGene_gn=PRELID3A;refGene_tx=XM_019814972.2;refGene_gn=PRELID3A;refGene_tx=XM_019814973.1;refGene_gn=PRELID3A;refGene_tx=XM_023241732.1
D3 34599099 34599170
refGene_gn=PRELID3A;refGene_tx=XM_011287938.3;refGene_gn=PRELID3A;refGene_tx=XM_011287939.3;refGene_gn=PRELID3A;refGene_tx=XM_011287940.2;refGene_gn=PRELID3A;refGene_tx=XM_011287942.3;refGene_gn=PRELID3A;refGene_tx=XM_019814966.2;refGene_gn=PRELID3A;refGene_tx=XM_019814967.2;refGene_gn=PRELID3A;refGene_tx=XM_019814970.2;refGene_gn=PRELID3A;refGene_tx=XM_019814971.2;refGene_gn=PRELID3A;refGene_tx=XM_019814972.2;refGene_gn=PRELID3A;refGene_tx=XM_019814973.1;refGene_gn=PRELID3A;refGene_tx=XM_023241732.1
D3 34600182 34600272
refGene_gn=PRELID3A;refGene_tx=XM_011287938.3;refGene_gn=PRELID3A;refGene_tx=XM_011287939.3;refGene_gn=PRELID3A;refGene_tx=XM_011287940.2;refGene_gn=PRELID3A;refGene_tx=XM_011287942.3;refGene_gn=PRELID3A;refGene_tx=XM_019814966.2;refGene_gn=PRELID3A;refGene_tx=XM_019814967.2;refGene_gn=PRELID3A;refGene_tx=XM_019814970.2;refGene_gn=PRELID3A;refGene_tx=XM_019814971.2;refGene_gn=PRELID3A;refGene_tx=XM_019814972.2;refGene_gn=PRELID3A;refGene_tx=XM_019814973.1;refGene_gn=PRELID3A;refGene_tx=XM_023241732.1
D3 34601771 34601940
refGene_gn=PRELID3A;refGene_tx=XM_011287938.3;refGene_gn=PRELID3A;refGene_tx=XM_011287939.3;refGene_gn=PRELID3A;refGene_tx=XM_011287940.2;refGene_gn=PRELID3A;refGene_tx=XM_011287942.3;refGene_gn=PRELID3A;refGene_tx=XM_019814966.2;refGene_gn=PRELID3A;refGene_tx=XM_019814967.2;refGene_gn=PRELID3A;refGene_tx=XM_019814970.2;refGene_gn=PRELID3A;refGene_tx=XM_019814971.2;refGene_gn=PRELID3A;refGene_tx=XM_019814972.2;refGene_gn=PRELID3A;refGene_tx=XM_019814973.1;refGene_gn=PRELID3A;refGene_tx=XM_023241732.1
D3 34612392 34612424
refGene_gn=PRELID3A;refGene_tx=XM_011287938.3;refGene_gn=PRELID3A;refGene_tx=XM_011287939.3;refGene_gn=PRELID3A;refGene_tx=XM_011287940.2;refGene_gn=PRELID3A;refGene_tx=XM_011287942.3;refGene_gn=PRELID3A;refGene_tx=XM_019814966.2;refGene_gn=PRELID3A;refGene_tx=XM_019814967.2;refGene_gn=PRELID3A;refGene_tx=XM_019814970.2;refGene_gn=PRELID3A;refGene_tx=XM_019814971.2;refGene_gn=PRELID3A;refGene_tx=XM_019814972.2;refGene_gn=PRELID3A;refGene_tx=XM_019814973.1
D3 34639649 34639766 refGene_gn=AFG3L2;refGene_tx=XM_023241733.1

D3 34644374 34644474 refGene_gn=AFG3L2;refGene_tx=XM_023241733.1
D3 34644888 34644966 refGene_gn=AFG3L2;refGene_tx=XM_023241733.1
D3 34647694 34647801 refGene_gn=AFG3L2;refGene_tx=XM_023241733.1
D3 34647939 34648092 refGene_gn=AFG3L2;refGene_tx=XM_023241733.1
D3 34650252 34650327 refGene_gn=AFG3L2;refGene_tx=XM_023241733.1
D3 34654143 34654268 refGene_gn=AFG3L2;refGene_tx=XM_023241733.1
D3 34656243 34656517 refGene_gn=AFG3L2;refGene_tx=XM_023241733.1
D3 34657910 34658048 refGene_gn=AFG3L2;refGene_tx=XM_023241733.1
D3 34660604 34660758 refGene_gn=AFG3L2;refGene_tx=XM_023241733.1
D3 34664457 34664565 refGene_gn=AFG3L2;refGene_tx=XM_023241733.1
D3 34664661 34664787 refGene_gn=AFG3L2;refGene_tx=XM_023241733.1
D3 34667260 34667371 refGene_gn=AFG3L2;refGene_tx=XM_023241733.1
D3 34670260 34670376 refGene_gn=AFG3L2;refGene_tx=XM_023241733.1
D3 34673710 34673911 refGene_gn=AFG3L2;refGene_tx=XM_023241733.1
D3 34675878 34676073 refGene_gn=AFG3L2;refGene_tx=XM_023241733.1
D3 34679339 34679582 refGene_gn=AFG3L2;refGene_tx=XM_023241733.1
D3 34682762 34683826
refGene_gn=TUBB6;refGene_tx=XM_003995004.4;refGene_gn=TUBB6;refGene_tx=XM_023241734.1
D3 34696032 34696143
refGene_gn=TUBB6;refGene_tx=XM_003995004.4;refGene_gn=TUBB6;refGene_tx=XM_023241734.1
D3 34698011 34698120 refGene_gn=TUBB6;refGene_tx=XM_003995004.4
D3 34698503 34698560 refGene_gn=TUBB6;refGene_tx=XM_003995004.4
D3 34722054 34722202
refGene_gn=CIDEA;refGene_tx=XM_003995005.5;refGene_gn=CIDEA;refGene_tx=XM_006938770.4;refGene_gn=CIDEA;refGene_tx=XM_011287932.2;refGene_gn=CIDEA;refGene_tx=XM_011287933.3;refGene_gn=CIDEA;refGene_tx=XM_011287934.3;refGene_gn=CIDEA;refGene_tx=XM_011287935.3;refGene_gn=CIDEA;refGene_tx=XM_023241735.1
D3 34724168 34724350
refGene_gn=CIDEA;refGene_tx=XM_003995005.5;refGene_gn=CIDEA;refGene_tx=XM_006938770.4;refGene_gn=CIDEA;refGene_tx=XM_011287932.2;refGene_gn=CIDEA;refGene_tx=XM_011287933.3;refGene_gn=CIDEA;refGene_tx=XM_011287934.3;refGene_gn=CIDEA;refGene_tx=XM_011287935.3;refGene_gn=CIDEA;refGene_tx=XM_023241735.1
D3 34730760 34730907
refGene_gn=CIDEA;refGene_tx=XM_003995005.5;refGene_gn=CIDEA;refGene_tx=XM_006938770.4;refGene_gn=CIDEA;refGene_tx=XM_011287932.2;refGene_gn=CIDEA;refGene_tx=XM_011287933.3;refGene_gn=CIDEA;refGene_tx=XM_011287934.3;refGene_gn=CIDEA;refGene_tx=XM_011287935.3;refGene_gn=CIDEA;refGene_tx=XM_023241735.1
D3 34731719 34731864
refGene_gn=CIDEA;refGene_tx=XM_003995005.5;refGene_gn=CIDEA;refGene_tx=XM_006938770.4;refGene_gn=CIDEA;refGene_tx=XM_011287932.2;refGene_gn=CIDEA;refGene_tx=XM_011287933.3;refGene_gn=CIDEA;refGene_tx=XM_011287934.3;refGene_gn=CIDEA;refGene_tx=XM_011287935.3;refGene_gn=CIDEA;refGene_tx=XM_023241735.1
D3 34740564 34740602 refGene_gn=CIDEA;refGene_tx=XM_003995005.5
D3 34740726 34740945

refGene_gn=CIDEA;refGene_tx=XM_011287932.2;refGene_gn=CIDEA;refGene_tx=XM_023241735.1
D3 34741811 34741816 refGene_gn=CIDEA;refGene_tx=XM_011287932.2
D3 34894606 34894875
refGene_gn=MPPE1;refGene_tx=XM_003995012.5;refGene_gn=MPPE1;refGene_tx=XM_006938773.4;refGene_gn=MPPE1;refGene_tx=XM_006938774.4;refGene_gn=MPPE1;refGene_tx=XM_006938775.4;refGene_gn=MPPE1;refGene_tx=XM_006938777.4;refGene_gn=MPPE1;refGene_tx=XM_006938778.4;refGene_gn=MPPE1;refGene_tx=XM_006938779.4;refGene_gn=MPPE1;refGene_tx=XM_019814975.2;refGene_gn=MPPE1;refGene_tx=XM_019814976.2
D3 34898060 34898169
refGene_gn=MPPE1;refGene_tx=XM_003995012.5;refGene_gn=MPPE1;refGene_tx=XM_006938773.4;refGene_gn=MPPE1;refGene_tx=XM_006938774.4;refGene_gn=MPPE1;refGene_tx=XM_006938775.4;refGene_gn=MPPE1;refGene_tx=XM_006938777.4;refGene_gn=MPPE1;refGene_tx=XM_006938778.4;refGene_gn=MPPE1;refGene_tx=XM_006938779.4;refGene_gn=MPPE1;refGene_tx=XM_019814975.2;refGene_gn=MPPE1;refGene_tx=XM_019814976.2
D3 34900721 34900825
refGene_gn=MPPE1;refGene_tx=XM_003995012.5;refGene_gn=MPPE1;refGene_tx=XM_006938773.4;refGene_gn=MPPE1;refGene_tx=XM_006938774.4;refGene_gn=MPPE1;refGene_tx=XM_006938775.4;refGene_gn=MPPE1;refGene_tx=XM_006938777.4;refGene_gn=MPPE1;refGene_tx=XM_006938778.4;refGene_gn=MPPE1;refGene_tx=XM_006938779.4;refGene_gn=MPPE1;refGene_tx=XM_019814975.2;refGene_gn=MPPE1;refGene_tx=XM_019814976.2
D3 34901467 34901542
refGene_gn=MPPE1;refGene_tx=XM_003995012.5;refGene_gn=MPPE1;refGene_tx=XM_006938773.4;refGene_gn=MPPE1;refGene_tx=XM_006938774.4;refGene_gn=MPPE1;refGene_tx=XM_006938775.4;refGene_gn=MPPE1;refGene_tx=XM_006938777.4;refGene_gn=MPPE1;refGene_tx=XM_006938778.4;refGene_gn=MPPE1;refGene_tx=XM_019814975.2;refGene_gn=MPPE1;refGene_tx=XM_019814976.2
D3 34903861 34903970
refGene_gn=MPPE1;refGene_tx=XM_003995012.5;refGene_gn=MPPE1;refGene_tx=XM_006938773.4;refGene_gn=MPPE1;refGene_tx=XM_006938774.4;refGene_gn=MPPE1;refGene_tx=XM_006938775.4;refGene_gn=MPPE1;refGene_tx=XM_006938777.4;refGene_gn=MPPE1;refGene_tx=XM_006938778.4;refGene_gn=MPPE1;refGene_tx=XM_006938779.4;refGene_gn=MPPE1;refGene_tx=XM_019814975.2;refGene_gn=MPPE1;refGene_tx=XM_019814976.2
D3 34904109 34904178
refGene_gn=MPPE1;refGene_tx=XM_003995012.5;refGene_gn=MPPE1;refGene_tx=XM_006938773.4;refGene_gn=MPPE1;refGene_tx=XM_006938774.4;refGene_gn=MPPE1;refGene_tx=XM_006938775.4;refGene_gn=MPPE1;refGene_tx=XM_006938777.4;refGene_gn=MPPE1;refGene_tx=XM_006938778.4;refGene_gn=MPPE1;refGene_tx=XM_006938779.4;refGene_gn=MPPE1;refGene_tx=XM_019814975.2;refGene_gn=MPPE1;refGene_tx=XM_019814976.2
D3 34904263 34904386
refGene_gn=MPPE1;refGene_tx=XM_003995012.5;refGene_gn=MPPE1;refGene_tx=XM_006938773.4;refGene_gn=MPPE1;refGene_tx=XM_006938774.4;refGene_gn=MPPE1;refGene_tx=XM_006938775.4;refGene_gn=MPPE1;refGene_tx=XM_006938777.4;refGene_gn=MPPE1;refGene_tx=XM_006938778.4;refGene_gn=MPPE1;refGene_tx=XM_006938779.4;refGene_gn=MPPE1;refGene_tx=XM_019814975.2;refGen

e_gn=MPPE1;refGene_tx=XM_019814976.2
D3 34905084 34905225
refGene_gn=MPPE1;refGene_tx=XM_003995012.5;refGene_gn=MPPE1;refGene_tx=XM_006938773.4;refGene_gn=MPPE1;refGene_tx=XM_006938774.4;refGene_gn=MPPE1;refGene_tx=XM_006938775.4;refGene_gn=MPPE1;refGene_tx=XM_006938777.4;refGene_gn=MPPE1;refGene_tx=XM_006938778.4;refGene_gn=MPPE1;refGene_tx=XM_006938779.4;refGene_gn=MPPE1;refGene_tx=XM_019814975.2;refGene_gn=MPPE1;refGene_tx=XM_019814976.2
D3 34906234 34906417
refGene_gn=MPPE1;refGene_tx=XM_003995012.5;refGene_gn=MPPE1;refGene_tx=XM_006938773.4;refGene_gn=MPPE1;refGene_tx=XM_006938774.4;refGene_gn=MPPE1;refGene_tx=XM_006938775.4;refGene_gn=MPPE1;refGene_tx=XM_006938777.4;refGene_gn=MPPE1;refGene_tx=XM_006938778.4;refGene_gn=MPPE1;refGene_tx=XM_006938779.4;refGene_gn=MPPE1;refGene_tx=XM_019814975.2;refGene_gn=MPPE1;refGene_tx=XM_019814976.2
D3 34909068 34909215
refGene_gn=GNAL;refGene_tx=XM_003995009.5;refGene_gn=GNAL;refGene_tx=XM_006938772.4;refGene_gn=GNAL;refGene_tx=XM_019814977.2
D3 34913168 34913236
refGene_gn=GNAL;refGene_tx=XM_003995009.5;refGene_gn=GNAL;refGene_tx=XM_006938772.4;refGene_gn=GNAL;refGene_tx=XM_019814977.2
D3 34915122 34915253
refGene_gn=GNAL;refGene_tx=XM_003995009.5;refGene_gn=GNAL;refGene_tx=XM_006938772.4;refGene_gn=GNAL;refGene_tx=XM_019814977.2
D3 34917161 34917282
refGene_gn=GNAL;refGene_tx=XM_003995009.5;refGene_gn=GNAL;refGene_tx=XM_006938772.4;refGene_gn=GNAL;refGene_tx=XM_019814977.2
D3 34917811 34917870
refGene_gn=GNAL;refGene_tx=XM_003995009.5;refGene_gn=GNAL;refGene_tx=XM_006938772.4;refGene_gn=GNAL;refGene_tx=XM_019814977.2
D3 34918418 34918492
refGene_gn=GNAL;refGene_tx=XM_003995009.5;refGene_gn=GNAL;refGene_tx=XM_006938772.4;refGene_gn=GNAL;refGene_tx=XM_019814977.2
D3 34919701 34919756
refGene_gn=GNAL;refGene_tx=XM_003995009.5;refGene_gn=GNAL;refGene_tx=XM_006938772.4;refGene_gn=GNAL;refGene_tx=XM_019814977.2
D3 34927388 34927988 refGene_gn=CHMP1B;refGene_tx=XM_006938780.3
D3 34950666 34950764
refGene_gn=GNAL;refGene_tx=XM_003995009.5;refGene_gn=GNAL;refGene_tx=XM_006938772.4;refGene_gn=GNAL;refGene_tx=XM_019814977.2
D3 35011614 35011734
refGene_gn=GNAL;refGene_tx=XM_003995009.5;refGene_gn=GNAL;refGene_tx=XM_006938772.4;refGene_gn=GNAL;refGene_tx=XM_019814977.2
D3 35011837 35011892
refGene_gn=GNAL;refGene_tx=XM_003995009.5;refGene_gn=GNAL;refGene_tx=XM_006938772.4;refGene_gn=GNAL;refGene_tx=XM_019814977.2
D3 35012590 35012663
refGene_gn=GNAL;refGene_tx=XM_003995009.5;refGene_gn=GNAL;refGene_tx=XM_006938772.4;refGene_gn=GNAL;refGene_tx=XM_019814977.2
D3 35012925 35013067

refGene_gn=GNAL;refGene_tx=XM_003995009.5;refGene_gn=GNAL;refGene_tx=XM_006938772.4
D3 35056842 35057209 refGene_gn=GNAL;refGene_tx=XM_019814977.2
D3 35094150 35094357
refGene_gn=LOC111557233;refGene_tx=XM_023241198.1
D3 35094455 35094603
refGene_gn=LOC111557233;refGene_tx=XM_023241198.1
D3 35096840 35096973
refGene_gn=LOC111557233;refGene_tx=XM_023241198.1
D3 35098198 35098451
refGene_gn=LOC111557233;refGene_tx=XM_023241198.1
D3 35101125 35101308
refGene_gn=LOC111557233;refGene_tx=XM_023241198.1
D3 35104389 35104648
refGene_gn=LOC111557233;refGene_tx=XM_023241198.1
D3 35104966 35105130
refGene_gn=LOC111557233;refGene_tx=XM_023241198.1
D3 35105207 35105370
refGene_gn=LOC111557233;refGene_tx=XM_023241198.1
D3 35107850 35108018
refGene_gn=LOC111557233;refGene_tx=XM_023241198.1
D3 35108860 35108924
refGene_gn=LOC111557233;refGene_tx=XM_023241198.1
D3 35109076 35109131
refGene_gn=LOC111557233;refGene_tx=XM_023241198.1
D3 35587101 35587165
refGene_gn=PIEZO2;refGene_tx=XM_023241737.1;refGene_gn=PIEZO2;refGene_tx=XM_023241738.1;refGene_gn=PIEZO2;refGene_tx=XM_023241739.1;refGene_gn=PIEZO2;refGene_tx=XM_023241740.1;refGene_gn=PIEZO2;refGene_tx=XM_023241741.1
D3 35670190 35670286
refGene_gn=PIEZO2;refGene_tx=XM_023241737.1;refGene_gn=PIEZO2;refGene_tx=XM_023241738.1;refGene_gn=PIEZO2;refGene_tx=XM_023241739.1;refGene_gn=PIEZO2;refGene_tx=XM_023241740.1;refGene_gn=PIEZO2;refGene_tx=XM_023241741.1
D3 35749117 35749243
refGene_gn=PIEZO2;refGene_tx=XM_023241737.1;refGene_gn=PIEZO2;refGene_tx=XM_023241738.1;refGene_gn=PIEZO2;refGene_tx=XM_023241739.1;refGene_gn=PIEZO2;refGene_tx=XM_023241740.1;refGene_gn=PIEZO2;refGene_tx=XM_023241741.1
D3 35809165 35809208
refGene_gn=PIEZO2;refGene_tx=XM_023241737.1;refGene_gn=PIEZO2;refGene_tx=XM_023241738.1;refGene_gn=PIEZO2;refGene_tx=XM_023241739.1;refGene_gn=PIEZO2;refGene_tx=XM_023241740.1;refGene_gn=PIEZO2;refGene_tx=XM_023241741.1
D3 35846301 35846464
refGene_gn=PIEZO2;refGene_tx=XM_023241737.1;refGene_gn=PIEZO2;refGene_tx=XM_023241738.1;refGene_gn=PIEZO2;refGene_tx=XM_023241739.1;refGene_gn=PIEZO2;refGene_tx=XM_023241740.1;refGene_gn=PIEZO2;refGene_tx=XM_023241741.1

D3 35860779 35860990
refGene_gn=PIEZO2;refGene_tx=XM_023241737.1;refGene_gn=PIEZO2;refGene_tx=XM_023241738.1;refGene_gn=PIEZO2;refGene_tx=XM_023241739.1;refGene_gn=PIEZO2;refGene_tx=XM_023241740.1;refGene_gn=PIEZO2;refGene_tx=XM_023241741.1
D3 35862721 35862935
refGene_gn=PIEZO2;refGene_tx=XM_023241737.1;refGene_gn=PIEZO2;refGene_tx=XM_023241738.1;refGene_gn=PIEZO2;refGene_tx=XM_023241739.1;refGene_gn=PIEZO2;refGene_tx=XM_023241740.1;refGene_gn=PIEZO2;refGene_tx=XM_023241741.1
D3 35921152 35921315
refGene_gn=PIEZO2;refGene_tx=XM_023241737.1;refGene_gn=PIEZO2;refGene_tx=XM_023241738.1;refGene_gn=PIEZO2;refGene_tx=XM_023241739.1;refGene_gn=PIEZO2;refGene_tx=XM_023241740.1;refGene_gn=PIEZO2;refGene_tx=XM_023241741.1
D3 35924138 35924258
refGene_gn=PIEZO2;refGene_tx=XM_023241737.1;refGene_gn=PIEZO2;refGene_tx=XM_023241738.1;refGene_gn=PIEZO2;refGene_tx=XM_023241739.1;refGene_gn=PIEZO2;refGene_tx=XM_023241740.1;refGene_gn=PIEZO2;refGene_tx=XM_023241741.1
D3 35927386 35927425
refGene_gn=PIEZO2;refGene_tx=XM_023241737.1;refGene_gn=PIEZO2;refGene_tx=XM_023241738.1;refGene_gn=PIEZO2;refGene_tx=XM_023241739.1;refGene_gn=PIEZO2;refGene_tx=XM_023241740.1;refGene_gn=PIEZO2;refGene_tx=XM_023241741.1
D3 35928363 35928502
refGene_gn=PIEZO2;refGene_tx=XM_023241737.1;refGene_gn=PIEZO2;refGene_tx=XM_023241738.1;refGene_gn=PIEZO2;refGene_tx=XM_023241739.1;refGene_gn=PIEZO2;refGene_tx=XM_023241740.1;refGene_gn=PIEZO2;refGene_tx=XM_023241741.1
D3 35931409 35931558
refGene_gn=PIEZO2;refGene_tx=XM_023241737.1;refGene_gn=PIEZO2;refGene_tx=XM_023241738.1;refGene_gn=PIEZO2;refGene_tx=XM_023241739.1;refGene_gn=PIEZO2;refGene_tx=XM_023241740.1;refGene_gn=PIEZO2;refGene_tx=XM_023241741.1
D3 35935021 35935252
refGene_gn=PIEZO2;refGene_tx=XM_023241737.1;refGene_gn=PIEZO2;refGene_tx=XM_023241738.1;refGene_gn=PIEZO2;refGene_tx=XM_023241739.1;refGene_gn=PIEZO2;refGene_tx=XM_023241740.1;refGene_gn=PIEZO2;refGene_tx=XM_023241741.1
D3 35939048 35939172
refGene_gn=PIEZO2;refGene_tx=XM_023241737.1;refGene_gn=PIEZO2;refGene_tx=XM_023241738.1;refGene_gn=PIEZO2;refGene_tx=XM_023241739.1;refGene_gn=PIEZO2;refGene_tx=XM_023241740.1;refGene_gn=PIEZO2;refGene_tx=XM_023241741.1
D3 35940467 35940751
refGene_gn=PIEZO2;refGene_tx=XM_023241737.1;refGene_gn=PIEZO2;refGene_tx=XM_023241738.1;refGene_gn=PIEZO2;refGene_tx=XM_023241739.1;refGene_gn=PIEZO2;refGene_tx=XM_023241740.1;refGene_gn=PIEZO2;refGene_tx=XM_023241741.1

D3 35942293 35942442
refGene_gn=PIEZO2;refGene_tx=XM_023241737.1;refGene_gn=PIEZO2;refGene_tx=XM_023241738.1;refGene_gn=PIEZO2;refGene_tx=XM_023241739.1;refGene_gn=PIEZO2;refGene_tx=XM_023241740.1;refGene_gn=PIEZO2;refGene_tx=XM_023241741.1
D3 35946280 35946454
refGene_gn=PIEZO2;refGene_tx=XM_023241737.1;refGene_gn=PIEZO2;refGene_tx=XM_023241738.1;refGene_gn=PIEZO2;refGene_tx=XM_023241739.1;refGene_gn=PIEZO2;refGene_tx=XM_023241740.1;refGene_gn=PIEZO2;refGene_tx=XM_023241741.1
D3 35950358 35950400
refGene_gn=PIEZO2;refGene_tx=XM_023241737.1;refGene_gn=PIEZO2;refGene_tx=XM_023241739.1;refGene_gn=PIEZO2;refGene_tx=XM_023241740.1
D3 35956748 35956781 refGene_gn=PIEZO2;refGene_tx=XM_023241739.1
D3 35957147 35957341
refGene_gn=PIEZO2;refGene_tx=XM_023241737.1;refGene_gn=PIEZO2;refGene_tx=XM_023241738.1;refGene_gn=PIEZO2;refGene_tx=XM_023241739.1;refGene_gn=PIEZO2;refGene_tx=XM_023241740.1;refGene_gn=PIEZO2;refGene_tx=XM_023241741.1
D3 35959368 35959529
refGene_gn=PIEZO2;refGene_tx=XM_023241737.1;refGene_gn=PIEZO2;refGene_tx=XM_023241738.1;refGene_gn=PIEZO2;refGene_tx=XM_023241739.1;refGene_gn=PIEZO2;refGene_tx=XM_023241740.1;refGene_gn=PIEZO2;refGene_tx=XM_023241741.1
D3 35968908 35969085
refGene_gn=PIEZO2;refGene_tx=XM_023241737.1;refGene_gn=PIEZO2;refGene_tx=XM_023241738.1;refGene_gn=PIEZO2;refGene_tx=XM_023241739.1;refGene_gn=PIEZO2;refGene_tx=XM_023241740.1;refGene_gn=PIEZO2;refGene_tx=XM_023241741.1
D3 35969344 35969470
refGene_gn=PIEZO2;refGene_tx=XM_023241737.1;refGene_gn=PIEZO2;refGene_tx=XM_023241738.1;refGene_gn=PIEZO2;refGene_tx=XM_023241739.1;refGene_gn=PIEZO2;refGene_tx=XM_023241740.1;refGene_gn=PIEZO2;refGene_tx=XM_023241741.1
D3 35970840 35971041
refGene_gn=PIEZO2;refGene_tx=XM_023241737.1;refGene_gn=PIEZO2;refGene_tx=XM_023241738.1;refGene_gn=PIEZO2;refGene_tx=XM_023241739.1;refGene_gn=PIEZO2;refGene_tx=XM_023241740.1;refGene_gn=PIEZO2;refGene_tx=XM_023241741.1
D3 35971753 35971958
refGene_gn=PIEZO2;refGene_tx=XM_023241737.1;refGene_gn=PIEZO2;refGene_tx=XM_023241738.1;refGene_gn=PIEZO2;refGene_tx=XM_023241739.1;refGene_gn=PIEZO2;refGene_tx=XM_023241740.1;refGene_gn=PIEZO2;refGene_tx=XM_023241741.1
D3 35972079 35972181
refGene_gn=PIEZO2;refGene_tx=XM_023241737.1;refGene_gn=PIEZO2;refGene_tx=XM_023241738.1;refGene_gn=PIEZO2;refGene_tx=XM_023241739.1;refGene_gn=PIEZO2;refGene_tx=XM_023241740.1;refGene_gn=PIEZO2;refGene_tx=XM_023241741.1
D3 35973290 35973456

refGene_gn=PIEZO2;refGene_tx=XM_023241737.1;refGene_gn=PIEZO2;refGene_tx=XM_023241738.1;refGene_gn=PIEZO2;refGene_tx=XM_023241739.1;refGene_gn=PIEZO2;refGene_tx=XM_023241740.1;refGene_gn=PIEZO2;refGene_tx=XM_023241741.1
D3 35977032 35977276
refGene_gn=PIEZO2;refGene_tx=XM_023241737.1;refGene_gn=PIEZO2;refGene_tx=XM_023241738.1;refGene_gn=PIEZO2;refGene_tx=XM_023241739.1;refGene_gn=PIEZO2;refGene_tx=XM_023241740.1;refGene_gn=PIEZO2;refGene_tx=XM_023241741.1
D3 35979631 35979728
refGene_gn=PIEZO2;refGene_tx=XM_023241737.1;refGene_gn=PIEZO2;refGene_tx=XM_023241738.1;refGene_gn=PIEZO2;refGene_tx=XM_023241739.1;refGene_gn=PIEZO2;refGene_tx=XM_023241740.1;refGene_gn=PIEZO2;refGene_tx=XM_023241741.1
D3 35981546 35981706
refGene_gn=PIEZO2;refGene_tx=XM_023241737.1;refGene_gn=PIEZO2;refGene_tx=XM_023241738.1;refGene_gn=PIEZO2;refGene_tx=XM_023241739.1;refGene_gn=PIEZO2;refGene_tx=XM_023241740.1;refGene_gn=PIEZO2;refGene_tx=XM_023241741.1
D3 35985756 35985846
refGene_gn=PIEZO2;refGene_tx=XM_023241737.1;refGene_gn=PIEZO2;refGene_tx=XM_023241738.1;refGene_gn=PIEZO2;refGene_tx=XM_023241739.1;refGene_gn=PIEZO2;refGene_tx=XM_023241740.1;refGene_gn=PIEZO2;refGene_tx=XM_023241741.1
D3 35987908 35988030
refGene_gn=PIEZO2;refGene_tx=XM_023241737.1;refGene_gn=PIEZO2;refGene_tx=XM_023241738.1;refGene_gn=PIEZO2;refGene_tx=XM_023241739.1;refGene_gn=PIEZO2;refGene_tx=XM_023241740.1;refGene_gn=PIEZO2;refGene_tx=XM_023241741.1
D3 35989119 35989191
refGene_gn=PIEZO2;refGene_tx=XM_023241737.1;refGene_gn=PIEZO2;refGene_tx=XM_023241738.1;refGene_gn=PIEZO2;refGene_tx=XM_023241739.1;refGene_gn=PIEZO2;refGene_tx=XM_023241740.1;refGene_gn=PIEZO2;refGene_tx=XM_023241741.1
D3 35993819 35993926
refGene_gn=PIEZO2;refGene_tx=XM_023241737.1;refGene_gn=PIEZO2;refGene_tx=XM_023241738.1;refGene_gn=PIEZO2;refGene_tx=XM_023241739.1;refGene_gn=PIEZO2;refGene_tx=XM_023241740.1;refGene_gn=PIEZO2;refGene_tx=XM_023241741.1
D3 35995204 35995303
refGene_gn=PIEZO2;refGene_tx=XM_023241737.1;refGene_gn=PIEZO2;refGene_tx=XM_023241738.1
D3 35999326 35999441
refGene_gn=PIEZO2;refGene_tx=XM_023241737.1;refGene_gn=PIEZO2;refGene_tx=XM_023241738.1;refGene_gn=PIEZO2;refGene_tx=XM_023241739.1;refGene_gn=PIEZO2;refGene_tx=XM_023241740.1;refGene_gn=PIEZO2;refGene_tx=XM_023241741.1
D3 36004693 36004753
refGene_gn=PIEZO2;refGene_tx=XM_023241737.1;refGene_gn=PIEZO2;refGene_tx=XM_023241738.1;refGene_gn=PIEZO2;refGene_tx=XM_023241739.1;refGene_

gn=PIEZO2;refGene_tx=XM_023241740.1;refGene_gn=PIEZO2;refGene_tx=XM_02
3241741.1
D3 36009444 36009611
refGene_gn=PIEZO2;refGene_tx=XM_023241737.1;refGene_gn=PIEZO2;refGene_
tx=XM_023241738.1;refGene_gn=PIEZO2;refGene_tx=XM_023241739.1;refGene_
gn=PIEZO2;refGene_tx=XM_023241740.1;refGene_gn=PIEZO2;refGene_tx=XM_02
3241741.1
D3 36010627 36010794
refGene_gn=PIEZO2;refGene_tx=XM_023241737.1;refGene_gn=PIEZO2;refGene_
tx=XM_023241738.1;refGene_gn=PIEZO2;refGene_tx=XM_023241739.1;refGene_
gn=PIEZO2;refGene_tx=XM_023241740.1;refGene_gn=PIEZO2;refGene_tx=XM_02
3241741.1
D3 36018208 36018376
refGene_gn=PIEZO2;refGene_tx=XM_023241737.1;refGene_gn=PIEZO2;refGene_
tx=XM_023241738.1;refGene_gn=PIEZO2;refGene_tx=XM_023241739.1;refGene_
gn=PIEZO2;refGene_tx=XM_023241740.1
D3 36020856 36021261
refGene_gn=PIEZO2;refGene_tx=XM_023241737.1;refGene_gn=PIEZO2;refGene_
tx=XM_023241738.1;refGene_gn=PIEZO2;refGene_tx=XM_023241739.1;refGene_
gn=PIEZO2;refGene_tx=XM_023241740.1;refGene_gn=PIEZO2;refGene_tx=XM_02
3241741.1
D3 36021578 36021837
refGene_gn=PIEZO2;refGene_tx=XM_023241737.1;refGene_gn=PIEZO2;refGene_
tx=XM_023241738.1;refGene_gn=PIEZO2;refGene_tx=XM_023241739.1;refGene_
gn=PIEZO2;refGene_tx=XM_023241740.1;refGene_gn=PIEZO2;refGene_tx=XM_02
3241741.1
D3 36024853 36025036
refGene_gn=PIEZO2;refGene_tx=XM_023241737.1;refGene_gn=PIEZO2;refGene_
tx=XM_023241738.1;refGene_gn=PIEZO2;refGene_tx=XM_023241739.1;refGene_
gn=PIEZO2;refGene_tx=XM_023241740.1;refGene_gn=PIEZO2;refGene_tx=XM_02
3241741.1
D3 36027762 36028015
refGene_gn=PIEZO2;refGene_tx=XM_023241737.1;refGene_gn=PIEZO2;refGene_
tx=XM_023241738.1;refGene_gn=PIEZO2;refGene_tx=XM_023241739.1;refGene_
gn=PIEZO2;refGene_tx=XM_023241740.1;refGene_gn=PIEZO2;refGene_tx=XM_02
3241741.1
D3 36029258 36029391
refGene_gn=PIEZO2;refGene_tx=XM_023241737.1;refGene_gn=PIEZO2;refGene_
tx=XM_023241738.1;refGene_gn=PIEZO2;refGene_tx=XM_023241739.1;refGene_
gn=PIEZO2;refGene_tx=XM_023241740.1;refGene_gn=PIEZO2;refGene_tx=XM_02
3241741.1
D3 36031572 36031720
refGene_gn=PIEZO2;refGene_tx=XM_023241737.1;refGene_gn=PIEZO2;refGene_
tx=XM_023241738.1;refGene_gn=PIEZO2;refGene_tx=XM_023241739.1;refGene_
gn=PIEZO2;refGene_tx=XM_023241740.1;refGene_gn=PIEZO2;refGene_tx=XM_02
3241741.1
D3 36031818 36032033
refGene_gn=PIEZO2;refGene_tx=XM_023241737.1;refGene_gn=PIEZO2;refGene_
tx=XM_023241738.1;refGene_gn=PIEZO2;refGene_tx=XM_023241739.1;refGene_
gn=PIEZO2;refGene_tx=XM_023241740.1;refGene_gn=PIEZO2;refGene_tx=XM_02

3241741.1
D3 36037674 36037833
refGene_gn=PIEZO2;refGene_tx=XM_023241737.1;refGene_gn=PIEZO2;refGene_tx=XM_023241738.1;refGene_gn=PIEZO2;refGene_tx=XM_023241739.1;refGene_gn=PIEZO2;refGene_tx=XM_023241740.1;refGene_gn=PIEZO2;refGene_tx=XM_023241741.1
D3 36039557 36039705
refGene_gn=PIEZO2;refGene_tx=XM_023241737.1;refGene_gn=PIEZO2;refGene_tx=XM_023241738.1;refGene_gn=PIEZO2;refGene_tx=XM_023241739.1;refGene_gn=PIEZO2;refGene_tx=XM_023241740.1;refGene_gn=PIEZO2;refGene_tx=XM_023241741.1
D3 36044724 36044913
refGene_gn=PIEZO2;refGene_tx=XM_023241737.1;refGene_gn=PIEZO2;refGene_tx=XM_023241738.1;refGene_gn=PIEZO2;refGene_tx=XM_023241739.1;refGene_gn=PIEZO2;refGene_tx=XM_023241740.1;refGene_gn=PIEZO2;refGene_tx=XM_023241741.1
D3 36045247 36045340
refGene_gn=PIEZO2;refGene_tx=XM_023241737.1;refGene_gn=PIEZO2;refGene_tx=XM_023241738.1;refGene_gn=PIEZO2;refGene_tx=XM_023241739.1;refGene_gn=PIEZO2;refGene_tx=XM_023241740.1;refGene_gn=PIEZO2;refGene_tx=XM_023241741.1
D3 36046909 36047082
refGene_gn=PIEZO2;refGene_tx=XM_023241737.1;refGene_gn=PIEZO2;refGene_tx=XM_023241738.1;refGene_gn=PIEZO2;refGene_tx=XM_023241739.1;refGene_gn=PIEZO2;refGene_tx=XM_023241740.1;refGene_gn=PIEZO2;refGene_tx=XM_023241741.1
D3 36048933 36049062
refGene_gn=PIEZO2;refGene_tx=XM_023241737.1;refGene_gn=PIEZO2;refGene_tx=XM_023241738.1;refGene_gn=PIEZO2;refGene_tx=XM_023241739.1;refGene_gn=PIEZO2;refGene_tx=XM_023241740.1;refGene_gn=PIEZO2;refGene_tx=XM_023241741.1
D3 36050249 36050329
refGene_gn=PIEZO2;refGene_tx=XM_023241737.1;refGene_gn=PIEZO2;refGene_tx=XM_023241738.1;refGene_gn=PIEZO2;refGene_tx=XM_023241739.1;refGene_gn=PIEZO2;refGene_tx=XM_023241740.1;refGene_gn=PIEZO2;refGene_tx=XM_023241741.1
D3 36052500 36052684
refGene_gn=PIEZO2;refGene_tx=XM_023241737.1;refGene_gn=PIEZO2;refGene_tx=XM_023241738.1;refGene_gn=PIEZO2;refGene_tx=XM_023241739.1;refGene_gn=PIEZO2;refGene_tx=XM_023241740.1;refGene_gn=PIEZO2;refGene_tx=XM_023241741.1
D3 36053770 36054023
refGene_gn=PIEZO2;refGene_tx=XM_023241737.1;refGene_gn=PIEZO2;refGene_tx=XM_023241738.1;refGene_gn=PIEZO2;refGene_tx=XM_023241739.1;refGene_gn=PIEZO2;refGene_tx=XM_023241740.1;refGene_gn=PIEZO2;refGene_tx=XM_023241741.1
D3 36108804 36108945
refGene_gn=NAPG;refGene_tx=XM_003995014.5;refGene_gn=NAPG;refGene_tx=XM_019814979.2
D3 36109990 36110120

refGene_gn=NAPG;refGene_tx=XM_003995014.5;refGene_gn=NAPG;refGene_tx=XM_019814979.2
D3 36110667 36110747
refGene_gn=NAPG;refGene_tx=XM_003995014.5;refGene_gn=NAPG;refGene_tx=XM_019814979.2
D3 36112937 36113016
refGene_gn=NAPG;refGene_tx=XM_003995014.5;refGene_gn=NAPG;refGene_tx=XM_019814979.2
D3 36118554 36118625
refGene_gn=NAPG;refGene_tx=XM_003995014.5;refGene_gn=NAPG;refGene_tx=XM_019814979.2
D3 36118908 36118975
refGene_gn=NAPG;refGene_tx=XM_003995014.5;refGene_gn=NAPG;refGene_tx=XM_019814979.2
D3 36119089 36119199
refGene_gn=NAPG;refGene_tx=XM_003995014.5;refGene_gn=NAPG;refGene_tx=XM_019814979.2
D3 36122347 36122378
refGene_gn=NAPG;refGene_tx=XM_003995014.5;refGene_gn=NAPG;refGene_tx=XM_019814979.2
D3 36123016 36123034
refGene_gn=NAPG;refGene_tx=XM_003995014.5;refGene_gn=NAPG;refGene_tx=XM_019814979.2
D3 36123730 36123815
refGene_gn=NAPG;refGene_tx=XM_003995014.5;refGene_gn=NAPG;refGene_tx=XM_019814979.2
D3 36125773 36125841
refGene_gn=NAPG;refGene_tx=XM_003995014.5;refGene_gn=NAPG;refGene_tx=XM_019814979.2
D3 36135354 36135410
refGene_gn=NAPG;refGene_tx=XM_003995014.5;refGene_gn=NAPG;refGene_tx=XM_019814979.2
D3 36174445 36174897 refGene_gn=APCDD1;refGene_tx=XM_023241742.1
D3 36176284 36176606 refGene_gn=APCDD1;refGene_tx=XM_023241742.1
D3 36189804 36190336 refGene_gn=APCDD1;refGene_tx=XM_023241742.1
D3 36193419 36193603 refGene_gn=APCDD1;refGene_tx=XM_023241742.1
D3 36206494 36206552 refGene_gn=APCDD1;refGene_tx=XM_023241742.1
D3 36663129 36663288 refGene_gn=VAPA;refGene_tx=XM_011287980.3
D3 36666409 36666439 refGene_gn=VAPA;refGene_tx=XM_011287980.3
D3 36666440 36666584 refGene_gn=VAPA;refGene_tx=XM_011287980.3
D3 36671345 36671480 refGene_gn=VAPA;refGene_tx=XM_011287980.3
D3 36680039 36680120 refGene_gn=VAPA;refGene_tx=XM_011287980.3
D3 36680527 36680631 refGene_gn=VAPA;refGene_tx=XM_011287980.3
D3 36684086 36684239 refGene_gn=VAPA;refGene_tx=XM_011287980.3
D3 36707905 36707984 refGene_gn=VAPA;refGene_tx=XM_011287980.3
D3 36729002 36730756
refGene_gn=TXNDC2;refGene_tx=XM_003995017.4;refGene_gn=TXNDC2;refGene_tx=XM_006938783.3
D3 36731072 36731172
refGene_gn=TXNDC2;refGene_tx=XM_003995017.4;refGene_gn=TXNDC2;refGene_

tx=XM_006938783.3
D3 36750327 36750425
refGene_gn=RAB31;refGene_tx=XM_019814984.2;refGene_gn=RAB31;refGene_tx
=XM_023241743.1
D3 36764411 36764521
refGene_gn=RAB31;refGene_tx=XM_019814984.2;refGene_gn=RAB31;refGene_tx
=XM_023241743.1
D3 36792209 36792316
refGene_gn=RAB31;refGene_tx=XM_019814984.2;refGene_gn=RAB31;refGene_tx
=XM_023241743.1
D3 36793369 36793441
refGene_gn=RAB31;refGene_tx=XM_019814984.2;refGene_gn=RAB31;refGene_tx
=XM_023241743.1
D3 36803562 36803755 refGene_gn=RAB31;refGene_tx=XM_019814984.2
D3 36814015 36814097
refGene_gn=RAB31;refGene_tx=XM_019814984.2;refGene_gn=RAB31;refGene_tx
=XM_023241743.1
D3 36823229 36823309 refGene_gn=RAB31;refGene_tx=XM_023241743.1
D3 36881991 36882030 refGene_gn=RAB31;refGene_tx=XM_023241743.1
D3 36972187 36972194
refGene_gn=PPP4R1;refGene_tx=XM_023241744.1;refGene_gn=PPP4R1;refGene_
tx=XM_023241746.1
D3 36972396 36972441
refGene_gn=PPP4R1;refGene_tx=XM_023241744.1;refGene_gn=PPP4R1;refGene_
tx=XM_023241746.1
D3 36984912 36984976 refGene_gn=PPP4R1;refGene_tx=XM_019814989.2
D3 36991906 36992042
refGene_gn=PPP4R1;refGene_tx=XM_019814989.2;refGene_gn=PPP4R1;refGene_
tx=XM_023241744.1;refGene_gn=PPP4R1;refGene_tx=XM_023241746.1;refGene_
gn=PPP4R1;refGene_tx=XM_023241747.1;refGene_gn=PPP4R1;refGene_tx=XM_02
3241748.1
D3 36992843 36992950
refGene_gn=PPP4R1;refGene_tx=XM_019814989.2;refGene_gn=PPP4R1;refGene_
tx=XM_023241744.1;refGene_gn=PPP4R1;refGene_tx=XM_023241746.1;refGene_
gn=PPP4R1;refGene_tx=XM_023241747.1;refGene_gn=PPP4R1;refGene_tx=XM_02
3241748.1
D3 36997525 36997668
refGene_gn=PPP4R1;refGene_tx=XM_019814989.2;refGene_gn=PPP4R1;refGene_
tx=XM_023241744.1;refGene_gn=PPP4R1;refGene_tx=XM_023241746.1;refGene_
gn=PPP4R1;refGene_tx=XM_023241747.1;refGene_gn=PPP4R1;refGene_tx=XM_02
3241748.1
D3 36998129 36998276
refGene_gn=PPP4R1;refGene_tx=XM_019814989.2;refGene_gn=PPP4R1;refGene_
tx=XM_023241744.1;refGene_gn=PPP4R1;refGene_tx=XM_023241746.1;refGene_
gn=PPP4R1;refGene_tx=XM_023241747.1;refGene_gn=PPP4R1;refGene_tx=XM_02
3241748.1
D3 37001147 37001255
refGene_gn=PPP4R1;refGene_tx=XM_019814989.2;refGene_gn=PPP4R1;refGene_
tx=XM_023241744.1;refGene_gn=PPP4R1;refGene_tx=XM_023241746.1;refGene_
gn=PPP4R1;refGene_tx=XM_023241747.1;refGene_gn=PPP4R1;refGene_tx=XM_02

3241748.1
D3	37001392	37001458
refGene_gn=PPP4R1;refGene_tx=XM_019814989.2;refGene_gn=PPP4R1;refGene_tx=XM_023241744.1;refGene_gn=PPP4R1;refGene_tx=XM_023241746.1;refGene_gn=PPP4R1;refGene_tx=XM_023241747.1;refGene_gn=PPP4R1;refGene_tx=XM_023241748.1
D3	37002133	37002292
refGene_gn=PPP4R1;refGene_tx=XM_019814989.2;refGene_gn=PPP4R1;refGene_tx=XM_023241744.1;refGene_gn=PPP4R1;refGene_tx=XM_023241746.1;refGene_gn=PPP4R1;refGene_tx=XM_023241747.1;refGene_gn=PPP4R1;refGene_tx=XM_023241748.1
D3	37006326	37006454
refGene_gn=PPP4R1;refGene_tx=XM_019814989.2;refGene_gn=PPP4R1;refGene_tx=XM_023241744.1;refGene_gn=PPP4R1;refGene_tx=XM_023241746.1;refGene_gn=PPP4R1;refGene_tx=XM_023241747.1;refGene_gn=PPP4R1;refGene_tx=XM_023241748.1
D3	37013638	37014162
refGene_gn=PPP4R1;refGene_tx=XM_019814989.2;refGene_gn=PPP4R1;refGene_tx=XM_023241744.1;refGene_gn=PPP4R1;refGene_tx=XM_023241746.1;refGene_gn=PPP4R1;refGene_tx=XM_023241747.1;refGene_gn=PPP4R1;refGene_tx=XM_023241748.1
D3	37020729	37020902
refGene_gn=PPP4R1;refGene_tx=XM_019814989.2;refGene_gn=PPP4R1;refGene_tx=XM_023241744.1;refGene_gn=PPP4R1;refGene_tx=XM_023241746.1;refGene_gn=PPP4R1;refGene_tx=XM_023241747.1;refGene_gn=PPP4R1;refGene_tx=XM_023241748.1
D3	37022228	37022324
refGene_gn=PPP4R1;refGene_tx=XM_019814989.2;refGene_gn=PPP4R1;refGene_tx=XM_023241744.1;refGene_gn=PPP4R1;refGene_tx=XM_023241746.1;refGene_gn=PPP4R1;refGene_tx=XM_023241747.1;refGene_gn=PPP4R1;refGene_tx=XM_023241748.1
D3	37024574	37024760
refGene_gn=PPP4R1;refGene_tx=XM_019814989.2;refGene_gn=PPP4R1;refGene_tx=XM_023241744.1;refGene_gn=PPP4R1;refGene_tx=XM_023241746.1;refGene_gn=PPP4R1;refGene_tx=XM_023241747.1;refGene_gn=PPP4R1;refGene_tx=XM_023241748.1
D3	37026442	37026604
refGene_gn=PPP4R1;refGene_tx=XM_019814989.2;refGene_gn=PPP4R1;refGene_tx=XM_023241744.1;refGene_gn=PPP4R1;refGene_tx=XM_023241746.1;refGene_gn=PPP4R1;refGene_tx=XM_023241747.1;refGene_gn=PPP4R1;refGene_tx=XM_023241748.1
D3	37029588	37029689
refGene_gn=PPP4R1;refGene_tx=XM_019814989.2;refGene_gn=PPP4R1;refGene_tx=XM_023241744.1;refGene_gn=PPP4R1;refGene_tx=XM_023241746.1;refGene_gn=PPP4R1;refGene_tx=XM_023241747.1;refGene_gn=PPP4R1;refGene_tx=XM_023241748.1
D3	37032466	37032587
refGene_gn=PPP4R1;refGene_tx=XM_019814989.2;refGene_gn=PPP4R1;refGene_tx=XM_023241744.1;refGene_gn=PPP4R1;refGene_tx=XM_023241746.1;refGene_gn=PPP4R1;refGene_tx=XM_023241747.1;refGene_gn=PPP4R1;refGene_tx=XM_02

3241748.1
D3 37032684 37032819
refGene_gn=PPP4R1;refGene_tx=XM_019814989.2;refGene_gn=PPP4R1;refGene_tx=XM_023241744.1;refGene_gn=PPP4R1;refGene_tx=XM_023241746.1;refGene_gn=PPP4R1;refGene_tx=XM_023241747.1;refGene_gn=PPP4R1;refGene_tx=XM_023241748.1
D3 37033281 37033423
refGene_gn=PPP4R1;refGene_tx=XM_019814989.2;refGene_gn=PPP4R1;refGene_tx=XM_023241744.1;refGene_gn=PPP4R1;refGene_tx=XM_023241746.1;refGene_gn=PPP4R1;refGene_tx=XM_023241747.1;refGene_gn=PPP4R1;refGene_tx=XM_023241748.1
D3 37034381 37034545
refGene_gn=PPP4R1;refGene_tx=XM_019814989.2;refGene_gn=PPP4R1;refGene_tx=XM_023241744.1;refGene_gn=PPP4R1;refGene_tx=XM_023241746.1;refGene_gn=PPP4R1;refGene_tx=XM_023241747.1;refGene_gn=PPP4R1;refGene_tx=XM_023241748.1
D3 37049428 37049671 refGene_gn=RALBP1;refGene_tx=XM_019814990.1
D3 37051430 37051554 refGene_gn=RALBP1;refGene_tx=XM_019814990.1
D3 37051800 37051922 refGene_gn=RALBP1;refGene_tx=XM_019814990.1
D3 37055975 37056083 refGene_gn=RALBP1;refGene_tx=XM_019814990.1
D3 37060331 37060463 refGene_gn=RALBP1;refGene_tx=XM_019814990.1
D3 37061467 37061629 refGene_gn=RALBP1;refGene_tx=XM_019814990.1
D3 37063162 37063512 refGene_gn=RALBP1;refGene_tx=XM_019814990.1
D3 37067701 37068160 refGene_gn=RALBP1;refGene_tx=XM_019814990.1
D3 37070128 37070372 refGene_gn=RALBP1;refGene_tx=XM_019814990.1
D3 37161481 37161663
refGene_gn=TWSG1;refGene_tx=XM_003995019.4;refGene_gn=TWSG1;refGene_tx=XM_023241749.1
D3 37164235 37164502
refGene_gn=TWSG1;refGene_tx=XM_003995019.4;refGene_gn=TWSG1;refGene_tx=XM_023241749.1
D3 37195138 37195238
refGene_gn=TWSG1;refGene_tx=XM_003995019.4;refGene_gn=TWSG1;refGene_tx=XM_023241749.1
D3 37228991 37229114
refGene_gn=TWSG1;refGene_tx=XM_003995019.4;refGene_gn=TWSG1;refGene_tx=XM_023241749.1
D3 37256940 37257126
refGene_gn=ANKRD12;refGene_tx=XM_011287973.3;refGene_gn=ANKRD12;refGene_tx=XM_023241750.1;refGene_gn=ANKRD12;refGene_tx=XM_023241751.1;refGene_gn=ANKRD12;refGene_tx=XM_023241752.1;refGene_gn=ANKRD12;refGene_tx=XM_023241753.1;refGene_gn=ANKRD12;refGene_tx=XM_023241754.1
D3 37257735 37257831
refGene_gn=ANKRD12;refGene_tx=XM_011287973.3;refGene_gn=ANKRD12;refGene_tx=XM_023241750.1;refGene_gn=ANKRD12;refGene_tx=XM_023241751.1;refGene_gn=ANKRD12;refGene_tx=XM_023241752.1;refGene_gn=ANKRD12;refGene_tx=XM_023241753.1;refGene_gn=ANKRD12;refGene_tx=XM_023241754.1
D3 37260583 37260727
refGene_gn=ANKRD12;refGene_tx=XM_011287973.3;refGene_gn=ANKRD12;refGene_tx=XM_023241750.1;refGene_gn=ANKRD12;refGene_tx=XM_023241751.1;refGe

ne_gn=ANKRD12;refGene_tx=XM_023241752.1;refGene_gn=ANKRD12;refGene_tx=XM_023241753.1;refGene_gn=ANKRD12;refGene_tx=XM_023241754.1
D3 37264186 37264285 refGene_gn=ANKRD12;refGene_tx=XM_011287973.3;refGene_gn=ANKRD12;refGene_tx=XM_023241750.1;refGene_gn=ANKRD12;refGene_tx=XM_023241751.1;refGene_gn=ANKRD12;refGene_tx=XM_023241752.1;refGene_gn=ANKRD12;refGene_tx=XM_023241753.1;refGene_gn=ANKRD12;refGene_tx=XM_023241754.1
D3 37272008 37276723 refGene_gn=ANKRD12;refGene_tx=XM_011287973.3;refGene_gn=ANKRD12;refGene_tx=XM_023241750.1;refGene_gn=ANKRD12;refGene_tx=XM_023241751.1;refGene_gn=ANKRD12;refGene_tx=XM_023241752.1;refGene_gn=ANKRD12;refGene_tx=XM_023241753.1;refGene_gn=ANKRD12;refGene_tx=XM_023241754.1
D3 37293402 37293466 refGene_gn=ANKRD12;refGene_tx=XM_023241753.1
D3 37302768 37302916 refGene_gn=ANKRD12;refGene_tx=XM_011287973.3;refGene_gn=ANKRD12;refGene_tx=XM_023241750.1;refGene_gn=ANKRD12;refGene_tx=XM_023241751.1;refGene_gn=ANKRD12;refGene_tx=XM_023241752.1
D3 37314921 37315064 refGene_gn=ANKRD12;refGene_tx=XM_011287973.3;refGene_gn=ANKRD12;refGene_tx=XM_023241750.1;refGene_gn=ANKRD12;refGene_tx=XM_023241751.1;refGene_gn=ANKRD12;refGene_tx=XM_023241752.1
D3 37317986 37318187 refGene_gn=ANKRD12;refGene_tx=XM_011287973.3;refGene_gn=ANKRD12;refGene_tx=XM_023241750.1;refGene_gn=ANKRD12;refGene_tx=XM_023241751.1;refGene_gn=ANKRD12;refGene_tx=XM_023241752.1
D3 37322688 37322835 refGene_gn=ANKRD12;refGene_tx=XM_011287973.3;refGene_gn=ANKRD12;refGene_tx=XM_023241750.1;refGene_gn=ANKRD12;refGene_tx=XM_023241751.1;refGene_gn=ANKRD12;refGene_tx=XM_023241752.1
D3 37326818 37326887 refGene_gn=ANKRD12;refGene_tx=XM_011287973.3;refGene_gn=ANKRD12;refGene_tx=XM_023241750.1
D3 37333688 37333836 refGene_gn=ANKRD12;refGene_tx=XM_011287973.3;refGene_gn=ANKRD12;refGene_tx=XM_023241750.1;refGene_gn=ANKRD12;refGene_tx=XM_023241751.1
D3 37345772 37345859 refGene_gn=ANKRD12;refGene_tx=XM_011287973.3;refGene_gn=ANKRD12;refGene_tx=XM_023241750.1;refGene_gn=ANKRD12;refGene_tx=XM_023241751.1
D3 37387325 37387419 refGene_gn=NDUFV2;refGene_tx=XM_003995021.5
D3 37391804 37391881 refGene_gn=NDUFV2;refGene_tx=XM_003995021.5
D3 37395634 37395744 refGene_gn=NDUFV2;refGene_tx=XM_003995021.5
D3 37397749 37397918 refGene_gn=NDUFV2;refGene_tx=XM_003995021.5
D3 37400920 37401037 refGene_gn=NDUFV2;refGene_tx=XM_003995021.5
D3 37401124 37401187 refGene_gn=NDUFV2;refGene_tx=XM_003995021.5
D3 37403018 37403084 refGene_gn=NDUFV2;refGene_tx=XM_003995021.5
D3 37418129 37418183 refGene_gn=NDUFV2;refGene_tx=XM_003995021.5
D3 37613490 37613685 refGene_gn=MTCL1;refGene_tx=XM_023241755.1;refGene_gn=MTCL1;refGene_tx=XM_023241756.1;refGene_gn=MTCL1;refGene_tx=XM_023241757.1;refGene_gn=MTCL1;refGene_tx=XM_023241758.1;refGene_gn=MTCL1;refGene_tx=XM_0232417

59.1;refGene_gn=MTCL1;refGene_tx=XM_023241761.1;refGene_gn=MTCL1;refGene_tx=XM_023241762.1;refGene_gn=MTCL1;refGene_tx=XM_023241763.1
D3 37616426 37616465
refGene_gn=MTCL1;refGene_tx=XM_019814991.2;refGene_gn=MTCL1;refGene_tx=XM_023241760.1;refGene_gn=MTCL1;refGene_tx=XM_023241764.1
D3 37620024 37621564
refGene_gn=MTCL1;refGene_tx=XM_019814991.2;refGene_gn=MTCL1;refGene_tx=XM_023241755.1;refGene_gn=MTCL1;refGene_tx=XM_023241756.1;refGene_gn=MTCL1;refGene_tx=XM_023241757.1;refGene_gn=MTCL1;refGene_tx=XM_023241758.1;refGene_gn=MTCL1;refGene_tx=XM_023241759.1;refGene_gn=MTCL1;refGene_tx=XM_023241760.1;refGene_gn=MTCL1;refGene_tx=XM_023241761.1;refGene_gn=MTCL1;refGene_tx=XM_023241762.1;refGene_gn=MTCL1;refGene_tx=XM_023241763.1;refGene_gn=MTCL1;refGene_tx=XM_023241764.1
D3 37626042 37626074
refGene_gn=MTCL1;refGene_tx=XM_019814991.2;refGene_gn=MTCL1;refGene_tx=XM_023241755.1;refGene_gn=MTCL1;refGene_tx=XM_023241756.1;refGene_gn=MTCL1;refGene_tx=XM_023241757.1;refGene_gn=MTCL1;refGene_tx=XM_023241758.1;refGene_gn=MTCL1;refGene_tx=XM_023241759.1;refGene_gn=MTCL1;refGene_tx=XM_023241760.1;refGene_gn=MTCL1;refGene_tx=XM_023241761.1;refGene_gn=MTCL1;refGene_tx=XM_023241762.1;refGene_gn=MTCL1;refGene_tx=XM_023241763.1;refGene_gn=MTCL1;refGene_tx=XM_023241764.1
D3 37627541 37627838
refGene_gn=MTCL1;refGene_tx=XM_019814991.2;refGene_gn=MTCL1;refGene_tx=XM_023241755.1;refGene_gn=MTCL1;refGene_tx=XM_023241756.1;refGene_gn=MTCL1;refGene_tx=XM_023241757.1;refGene_gn=MTCL1;refGene_tx=XM_023241758.1;refGene_gn=MTCL1;refGene_tx=XM_023241759.1;refGene_gn=MTCL1;refGene_tx=XM_023241760.1;refGene_gn=MTCL1;refGene_tx=XM_023241761.1;refGene_gn=MTCL1;refGene_tx=XM_023241762.1;refGene_gn=MTCL1;refGene_tx=XM_023241763.1;refGene_gn=MTCL1;refGene_tx=XM_023241764.1
D3 37633341 37633596
refGene_gn=MTCL1;refGene_tx=XM_019814991.2;refGene_gn=MTCL1;refGene_tx=XM_023241755.1;refGene_gn=MTCL1;refGene_tx=XM_023241756.1;refGene_gn=MTCL1;refGene_tx=XM_023241757.1;refGene_gn=MTCL1;refGene_tx=XM_023241758.1;refGene_gn=MTCL1;refGene_tx=XM_023241759.1;refGene_gn=MTCL1;refGene_tx=XM_023241760.1;refGene_gn=MTCL1;refGene_tx=XM_023241761.1;refGene_gn=MTCL1;refGene_tx=XM_023241762.1;refGene_gn=MTCL1;refGene_tx=XM_023241763.1;refGene_gn=MTCL1;refGene_tx=XM_023241764.1
D3 37637847 37638015
refGene_gn=MTCL1;refGene_tx=XM_019814991.2;refGene_gn=MTCL1;refGene_tx=XM_023241755.1;refGene_gn=MTCL1;refGene_tx=XM_023241756.1;refGene_gn=MTCL1;refGene_tx=XM_023241757.1;refGene_gn=MTCL1;refGene_tx=XM_023241758.1;refGene_gn=MTCL1;refGene_tx=XM_023241759.1;refGene_gn=MTCL1;refGene_tx=XM_023241760.1;refGene_gn=MTCL1;refGene_tx=XM_023241762.1;refGene_gn=MTCL1;refGene_tx=XM_023241763.1;refGene_gn=MTCL1;refGene_tx=XM_023241764.1
D3 37647644 37647842
refGene_gn=MTCL1;refGene_tx=XM_019814991.2;refGene_gn=MTCL1;refGene_tx=XM_023241755.1;refGene_gn=MTCL1;refGene_tx=XM_023241756.1;refGene_gn=MTCL1;refGene_tx=XM_023241757.1;refGene_gn=MTCL1;refGene_tx=XM_023241758.1;refGene_gn=MTCL1;refGene_tx=XM_023241759.1;refGene_gn=MTCL1;refGe

ne_tx=XM_023241760.1;refGene_gn=MTCL1;refGene_tx=XM_023241761.1;refGene_gn=MTCL1;refGene_tx=XM_023241763.1;refGene_gn=MTCL1;refGene_tx=XM_023241764.1
D3 37649487 37649718
refGene_gn=MTCL1;refGene_tx=XM_019814991.2;refGene_gn=MTCL1;refGene_tx=XM_023241755.1;refGene_gn=MTCL1;refGene_tx=XM_023241756.1;refGene_gn=MTCL1;refGene_tx=XM_023241757.1;refGene_gn=MTCL1;refGene_tx=XM_023241758.1;refGene_gn=MTCL1;refGene_tx=XM_023241759.1;refGene_gn=MTCL1;refGene_tx=XM_023241760.1;refGene_gn=MTCL1;refGene_tx=XM_023241761.1;refGene_gn=MTCL1;refGene_tx=XM_023241762.1;refGene_gn=MTCL1;refGene_tx=XM_023241763.1;refGene_gn=MTCL1;refGene_tx=XM_023241764.1
D3 37652063 37652186
refGene_gn=MTCL1;refGene_tx=XM_019814991.2;refGene_gn=MTCL1;refGene_tx=XM_023241755.1;refGene_gn=MTCL1;refGene_tx=XM_023241756.1;refGene_gn=MTCL1;refGene_tx=XM_023241757.1;refGene_gn=MTCL1;refGene_tx=XM_023241758.1;refGene_gn=MTCL1;refGene_tx=XM_023241760.1;refGene_gn=MTCL1;refGene_tx=XM_023241761.1;refGene_gn=MTCL1;refGene_tx=XM_023241762.1
D3 37660336 37660468
refGene_gn=MTCL1;refGene_tx=XM_019814991.2;refGene_gn=MTCL1;refGene_tx=XM_023241755.1;refGene_gn=MTCL1;refGene_tx=XM_023241756.1;refGene_gn=MTCL1;refGene_tx=XM_023241757.1;refGene_gn=MTCL1;refGene_tx=XM_023241758.1;refGene_gn=MTCL1;refGene_tx=XM_023241759.1;refGene_gn=MTCL1;refGene_tx=XM_023241760.1;refGene_gn=MTCL1;refGene_tx=XM_023241761.1;refGene_gn=MTCL1;refGene_tx=XM_023241762.1;refGene_gn=MTCL1;refGene_tx=XM_023241763.1;refGene_gn=MTCL1;refGene_tx=XM_023241764.1
D3 37661604 37662942
refGene_gn=MTCL1;refGene_tx=XM_019814991.2;refGene_gn=MTCL1;refGene_tx=XM_023241755.1;refGene_gn=MTCL1;refGene_tx=XM_023241756.1;refGene_gn=MTCL1;refGene_tx=XM_023241757.1;refGene_gn=MTCL1;refGene_tx=XM_023241758.1;refGene_gn=MTCL1;refGene_tx=XM_023241759.1;refGene_gn=MTCL1;refGene_tx=XM_023241760.1;refGene_gn=MTCL1;refGene_tx=XM_023241761.1;refGene_gn=MTCL1;refGene_tx=XM_023241762.1;refGene_gn=MTCL1;refGene_tx=XM_023241763.1;refGene_gn=MTCL1;refGene_tx=XM_023241764.1
D3 37667129 37667207
refGene_gn=MTCL1;refGene_tx=XM_023241755.1;refGene_gn=MTCL1;refGene_tx=XM_023241756.1;refGene_gn=MTCL1;refGene_tx=XM_023241757.1;refGene_gn=MTCL1;refGene_tx=XM_023241759.1;refGene_gn=MTCL1;refGene_tx=XM_023241760.1;refGene_gn=MTCL1;refGene_tx=XM_023241761.1;refGene_gn=MTCL1;refGene_tx=XM_023241762.1
D3 37668381 37668441
refGene_gn=MTCL1;refGene_tx=XM_019814991.2;refGene_gn=MTCL1;refGene_tx=XM_023241755.1;refGene_gn=MTCL1;refGene_tx=XM_023241756.1;refGene_gn=MTCL1;refGene_tx=XM_023241757.1;refGene_gn=MTCL1;refGene_tx=XM_023241758.1;refGene_gn=MTCL1;refGene_tx=XM_023241759.1;refGene_gn=MTCL1;refGene_tx=XM_023241760.1;refGene_gn=MTCL1;refGene_tx=XM_023241761.1;refGene_gn=MTCL1;refGene_tx=XM_023241762.1;refGene_gn=MTCL1;refGene_tx=XM_023241763.1;refGene_gn=MTCL1;refGene_tx=XM_023241764.1
D3 37731890 37732049
refGene_gn=MTCL1;refGene_tx=XM_019814991.2;refGene_gn=MTCL1;refGene_tx=XM_023241755.1;refGene_gn=MTCL1;refGene_tx=XM_023241756.1;refGene_gn=

MTCL1;refGene_tx=XM_023241757.1;refGene_gn=MTCL1;refGene_tx=XM_023241758.1;refGene_gn=MTCL1;refGene_tx=XM_023241759.1;refGene_gn=MTCL1;refGene_tx=XM_023241760.1;refGene_gn=MTCL1;refGene_tx=XM_023241761.1;refGene_gn=MTCL1;refGene_tx=XM_023241762.1;refGene_gn=MTCL1;refGene_tx=XM_023241763.1;refGene_gn=MTCL1;refGene_tx=XM_023241764.1
D3 37733455 37733680
refGene_gn=MTCL1;refGene_tx=XM_019814991.2;refGene_gn=MTCL1;refGene_tx=XM_023241755.1;refGene_gn=MTCL1;refGene_tx=XM_023241756.1;refGene_gn=MTCL1;refGene_tx=XM_023241757.1;refGene_gn=MTCL1;refGene_tx=XM_023241758.1;refGene_gn=MTCL1;refGene_tx=XM_023241759.1;refGene_gn=MTCL1;refGene_tx=XM_023241760.1;refGene_gn=MTCL1;refGene_tx=XM_023241761.1;refGene_gn=MTCL1;refGene_tx=XM_023241762.1;refGene_gn=MTCL1;refGene_tx=XM_023241763.1;refGene_gn=MTCL1;refGene_tx=XM_023241764.1
D3 37743369 37744440
refGene_gn=MTCL1;refGene_tx=XM_023241760.1;refGene_gn=MTCL1;refGene_tx=XM_023241761.1
D3 37794882 37794996 refGene_gn=RAB12;refGene_tx=XM_011287984.3
D3 37796433 37796538 refGene_gn=RAB12;refGene_tx=XM_011287984.3
D3 37797145 37797235 refGene_gn=RAB12;refGene_tx=XM_011287984.3
D3 37798493 37798632 refGene_gn=RAB12;refGene_tx=XM_011287984.3
D3 37805272 37805333 refGene_gn=RAB12;refGene_tx=XM_011287984.3
D3 37819901 37820127 refGene_gn=RAB12;refGene_tx=XM_011287984.3
D3 37997754 37997808
refGene_gn=PTPRM;refGene_tx=XM_003995022.5;refGene_gn=PTPRM;refGene_tx=XM_003995023.5;refGene_gn=PTPRM;refGene_tx=XM_011287970.3;refGene_gn=PTPRM;refGene_tx=XM_019814992.2;refGene_gn=PTPRM;refGene_tx=XM_019814993.2;refGene_gn=PTPRM;refGene_tx=XM_019814994.2;refGene_gn=PTPRM;refGene_tx=XM_019814995.2;refGene_gn=PTPRM;refGene_tx=XM_019814996.1
D3 38006476 38006612
refGene_gn=PTPRM;refGene_tx=XM_003995022.5;refGene_gn=PTPRM;refGene_tx=XM_003995023.5;refGene_gn=PTPRM;refGene_tx=XM_011287970.3;refGene_gn=PTPRM;refGene_tx=XM_019814992.2;refGene_gn=PTPRM;refGene_tx=XM_019814993.2;refGene_gn=PTPRM;refGene_tx=XM_019814994.2;refGene_gn=PTPRM;refGene_tx=XM_019814995.2;refGene_gn=PTPRM;refGene_tx=XM_019814996.1
D3 38013331 38013495
refGene_gn=PTPRM;refGene_tx=XM_003995022.5;refGene_gn=PTPRM;refGene_tx=XM_003995023.5;refGene_gn=PTPRM;refGene_tx=XM_011287970.3;refGene_gn=PTPRM;refGene_tx=XM_019814992.2;refGene_gn=PTPRM;refGene_tx=XM_019814993.2;refGene_gn=PTPRM;refGene_tx=XM_019814994.2;refGene_gn=PTPRM;refGene_tx=XM_019814995.2;refGene_gn=PTPRM;refGene_tx=XM_019814996.1
D3 38015429 38015555
refGene_gn=PTPRM;refGene_tx=XM_003995022.5;refGene_gn=PTPRM;refGene_tx=XM_003995023.5;refGene_gn=PTPRM;refGene_tx=XM_011287970.3;refGene_gn=PTPRM;refGene_tx=XM_019814992.2;refGene_gn=PTPRM;refGene_tx=XM_019814993.2;refGene_gn=PTPRM;refGene_tx=XM_019814994.2;refGene_gn=PTPRM;refGene_tx=XM_019814995.2;refGene_gn=PTPRM;refGene_tx=XM_019814996.1
D3 38018963 38019095
refGene_gn=PTPRM;refGene_tx=XM_003995022.5;refGene_gn=PTPRM;refGene_tx=XM_003995023.5;refGene_gn=PTPRM;refGene_tx=XM_011287970.3;refGene_gn=PTPRM;refGene_tx=XM_019814992.2;refGene_gn=PTPRM;refGene_tx=XM_0198149

93.2;refGene_gn=PTPRM;refGene_tx=XM_019814994.2;refGene_gn=PTPRM;refGene_tx=XM_019814995.2;refGene_gn=PTPRM;refGene_tx=XM_019814996.1
D3 38020110 38020284
refGene_gn=PTPRM;refGene_tx=XM_003995022.5;refGene_gn=PTPRM;refGene_tx=XM_003995023.5;refGene_gn=PTPRM;refGene_tx=XM_011287970.3;refGene_gn=PTPRM;refGene_tx=XM_019814992.2;refGene_gn=PTPRM;refGene_tx=XM_019814993.2;refGene_gn=PTPRM;refGene_tx=XM_019814994.2;refGene_gn=PTPRM;refGene_tx=XM_019814995.2;refGene_gn=PTPRM;refGene_tx=XM_019814996.1
D3 38021010 38021160
refGene_gn=PTPRM;refGene_tx=XM_003995022.5;refGene_gn=PTPRM;refGene_tx=XM_003995023.5;refGene_gn=PTPRM;refGene_tx=XM_011287970.3;refGene_gn=PTPRM;refGene_tx=XM_019814992.2;refGene_gn=PTPRM;refGene_tx=XM_019814993.2;refGene_gn=PTPRM;refGene_tx=XM_019814994.2;refGene_gn=PTPRM;refGene_tx=XM_019814995.2;refGene_gn=PTPRM;refGene_tx=XM_019814996.1
D3 38023104 38023240
refGene_gn=PTPRM;refGene_tx=XM_003995022.5;refGene_gn=PTPRM;refGene_tx=XM_003995023.5;refGene_gn=PTPRM;refGene_tx=XM_011287970.3;refGene_gn=PTPRM;refGene_tx=XM_019814992.2;refGene_gn=PTPRM;refGene_tx=XM_019814993.2;refGene_gn=PTPRM;refGene_tx=XM_019814994.2;refGene_gn=PTPRM;refGene_tx=XM_019814995.2;refGene_gn=PTPRM;refGene_tx=XM_019814996.1
D3 38023556 38023711
refGene_gn=PTPRM;refGene_tx=XM_003995022.5;refGene_gn=PTPRM;refGene_tx=XM_003995023.5;refGene_gn=PTPRM;refGene_tx=XM_011287970.3;refGene_gn=PTPRM;refGene_tx=XM_019814992.2;refGene_gn=PTPRM;refGene_tx=XM_019814993.2;refGene_gn=PTPRM;refGene_tx=XM_019814994.2;refGene_gn=PTPRM;refGene_tx=XM_019814995.2;refGene_gn=PTPRM;refGene_tx=XM_019814996.1
D3 38028002 38028053
refGene_gn=PTPRM;refGene_tx=XM_019814992.2;refGene_gn=PTPRM;refGene_tx=XM_019814993.2;refGene_gn=PTPRM;refGene_tx=XM_019814994.2;refGene_gn=PTPRM;refGene_tx=XM_019814995.2;refGene_gn=PTPRM;refGene_tx=XM_019814996.1
D3 38028773 38028890
refGene_gn=PTPRM;refGene_tx=XM_003995022.5;refGene_gn=PTPRM;refGene_tx=XM_003995023.5;refGene_gn=PTPRM;refGene_tx=XM_011287970.3;refGene_gn=PTPRM;refGene_tx=XM_019814992.2;refGene_gn=PTPRM;refGene_tx=XM_019814993.2;refGene_gn=PTPRM;refGene_tx=XM_019814994.2;refGene_gn=PTPRM;refGene_tx=XM_019814995.2;refGene_gn=PTPRM;refGene_tx=XM_019814996.1
D3 38055224 38055322
refGene_gn=PTPRM;refGene_tx=XM_003995022.5;refGene_gn=PTPRM;refGene_tx=XM_003995023.5;refGene_gn=PTPRM;refGene_tx=XM_011287970.3;refGene_gn=PTPRM;refGene_tx=XM_019814992.2;refGene_gn=PTPRM;refGene_tx=XM_019814993.2;refGene_gn=PTPRM;refGene_tx=XM_019814994.2;refGene_gn=PTPRM;refGene_tx=XM_019814995.2;refGene_gn=PTPRM;refGene_tx=XM_019814996.1
D3 38078493 38078530
refGene_gn=PTPRM;refGene_tx=XM_003995022.5;refGene_gn=PTPRM;refGene_tx=XM_003995023.5;refGene_gn=PTPRM;refGene_tx=XM_011287970.3;refGene_gn=PTPRM;refGene_tx=XM_019814992.2;refGene_gn=PTPRM;refGene_tx=XM_019814993.2;refGene_gn=PTPRM;refGene_tx=XM_019814994.2;refGene_gn=PTPRM;refGene_tx=XM_019814995.2;refGene_gn=PTPRM;refGene_tx=XM_019814996.1
D3 38081277 38081354

refGene_gn=PTPRM;refGene_tx=XM_003995022.5;refGene_gn=PTPRM;refGene_tx=XM_003995023.5;refGene_gn=PTPRM;refGene_tx=XM_011287970.3;refGene_gn=PTPRM;refGene_tx=XM_019814992.2;refGene_gn=PTPRM;refGene_tx=XM_019814993.2;refGene_gn=PTPRM;refGene_tx=XM_019814994.2;refGene_gn=PTPRM;refGene_tx=XM_019814995.2;refGene_gn=PTPRM;refGene_tx=XM_019814996.1
D3 38098103 38098191
refGene_gn=PTPRM;refGene_tx=XM_003995022.5;refGene_gn=PTPRM;refGene_tx=XM_003995023.5;refGene_gn=PTPRM;refGene_tx=XM_011287970.3;refGene_gn=PTPRM;refGene_tx=XM_019814992.2;refGene_gn=PTPRM;refGene_tx=XM_019814993.2;refGene_gn=PTPRM;refGene_tx=XM_019814994.2;refGene_gn=PTPRM;refGene_tx=XM_019814995.2;refGene_gn=PTPRM;refGene_tx=XM_019814996.1
D3 38135986 38136174
refGene_gn=PTPRM;refGene_tx=XM_003995022.5;refGene_gn=PTPRM;refGene_tx=XM_003995023.5;refGene_gn=PTPRM;refGene_tx=XM_011287970.3;refGene_gn=PTPRM;refGene_tx=XM_019814992.2;refGene_gn=PTPRM;refGene_tx=XM_019814993.2;refGene_gn=PTPRM;refGene_tx=XM_019814994.2;refGene_gn=PTPRM;refGene_tx=XM_019814995.2;refGene_gn=PTPRM;refGene_tx=XM_019814996.1
D3 38136884 38136896
refGene_gn=PTPRM;refGene_tx=XM_003995023.5;refGene_gn=PTPRM;refGene_tx=XM_011287970.3;refGene_gn=PTPRM;refGene_tx=XM_019814992.2;refGene_gn=PTPRM;refGene_tx=XM_019814993.2;refGene_gn=PTPRM;refGene_tx=XM_019814995.2;refGene_gn=PTPRM;refGene_tx=XM_019814996.1
D3 38141020 38141047
refGene_gn=PTPRM;refGene_tx=XM_003995023.5;refGene_gn=PTPRM;refGene_tx=XM_011287970.3;refGene_gn=PTPRM;refGene_tx=XM_019814992.2;refGene_gn=PTPRM;refGene_tx=XM_019814995.2;refGene_gn=PTPRM;refGene_tx=XM_019814996.1
D3 38141277 38141352
refGene_gn=PTPRM;refGene_tx=XM_003995022.5;refGene_gn=PTPRM;refGene_tx=XM_003995023.5;refGene_gn=PTPRM;refGene_tx=XM_011287970.3;refGene_gn=PTPRM;refGene_tx=XM_019814992.2;refGene_gn=PTPRM;refGene_tx=XM_019814993.2;refGene_gn=PTPRM;refGene_tx=XM_019814994.2;refGene_gn=PTPRM;refGene_tx=XM_019814995.2;refGene_gn=PTPRM;refGene_tx=XM_019814996.1
D3 38145745 38145897
refGene_gn=PTPRM;refGene_tx=XM_003995022.5;refGene_gn=PTPRM;refGene_tx=XM_003995023.5;refGene_gn=PTPRM;refGene_tx=XM_011287970.3;refGene_gn=PTPRM;refGene_tx=XM_019814992.2;refGene_gn=PTPRM;refGene_tx=XM_019814993.2;refGene_gn=PTPRM;refGene_tx=XM_019814994.2;refGene_gn=PTPRM;refGene_tx=XM_019814995.2;refGene_gn=PTPRM;refGene_tx=XM_019814996.1
D3 38227262 38227395
refGene_gn=PTPRM;refGene_tx=XM_003995022.5;refGene_gn=PTPRM;refGene_tx=XM_003995023.5;refGene_gn=PTPRM;refGene_tx=XM_011287970.3;refGene_gn=PTPRM;refGene_tx=XM_019814992.2;refGene_gn=PTPRM;refGene_tx=XM_019814993.2;refGene_gn=PTPRM;refGene_tx=XM_019814994.2;refGene_gn=PTPRM;refGene_tx=XM_019814995.2;refGene_gn=PTPRM;refGene_tx=XM_019814996.1
D3 38257834 38257909
refGene_gn=PTPRM;refGene_tx=XM_011287970.3;refGene_gn=PTPRM;refGene_tx=XM_019814992.2;refGene_gn=PTPRM;refGene_tx=XM_019814993.2;refGene_gn=PTPRM;refGene_tx=XM_019814994.2;refGene_gn=PTPRM;refGene_tx=XM_019814996.1

D3 38264405 38264442
refGene_gn=PTPRM;refGene_tx=XM_003995022.5;refGene_gn=PTPRM;refGene_tx=XM_003995023.5;refGene_gn=PTPRM;refGene_tx=XM_011287970.3;refGene_gn=PTPRM;refGene_tx=XM_019814992.2;refGene_gn=PTPRM;refGene_tx=XM_019814993.2;refGene_gn=PTPRM;refGene_tx=XM_019814994.2;refGene_gn=PTPRM;refGene_tx=XM_019814995.2;refGene_gn=PTPRM;refGene_tx=XM_019814996.1
D3 38265529 38265803
refGene_gn=PTPRM;refGene_tx=XM_003995022.5;refGene_gn=PTPRM;refGene_tx=XM_003995023.5;refGene_gn=PTPRM;refGene_tx=XM_011287970.3;refGene_gn=PTPRM;refGene_tx=XM_019814992.2;refGene_gn=PTPRM;refGene_tx=XM_019814993.2;refGene_gn=PTPRM;refGene_tx=XM_019814994.2;refGene_gn=PTPRM;refGene_tx=XM_019814995.2;refGene_gn=PTPRM;refGene_tx=XM_019814996.1
D3 38289828 38289931
refGene_gn=PTPRM;refGene_tx=XM_003995022.5;refGene_gn=PTPRM;refGene_tx=XM_003995023.5;refGene_gn=PTPRM;refGene_tx=XM_011287970.3;refGene_gn=PTPRM;refGene_tx=XM_019814992.2;refGene_gn=PTPRM;refGene_tx=XM_019814993.2;refGene_gn=PTPRM;refGene_tx=XM_019814994.2;refGene_gn=PTPRM;refGene_tx=XM_019814995.2;refGene_gn=PTPRM;refGene_tx=XM_019814996.1
D3 38293404 38293606
refGene_gn=PTPRM;refGene_tx=XM_003995022.5;refGene_gn=PTPRM;refGene_tx=XM_003995023.5;refGene_gn=PTPRM;refGene_tx=XM_011287970.3;refGene_gn=PTPRM;refGene_tx=XM_019814992.2;refGene_gn=PTPRM;refGene_tx=XM_019814993.2;refGene_gn=PTPRM;refGene_tx=XM_019814994.2;refGene_gn=PTPRM;refGene_tx=XM_019814995.2;refGene_gn=PTPRM;refGene_tx=XM_019814996.1
D3 38303208 38303318
refGene_gn=PTPRM;refGene_tx=XM_003995022.5;refGene_gn=PTPRM;refGene_tx=XM_003995023.5;refGene_gn=PTPRM;refGene_tx=XM_011287970.3;refGene_gn=PTPRM;refGene_tx=XM_019814992.2;refGene_gn=PTPRM;refGene_tx=XM_019814993.2;refGene_gn=PTPRM;refGene_tx=XM_019814994.2;refGene_gn=PTPRM;refGene_tx=XM_019814995.2;refGene_gn=PTPRM;refGene_tx=XM_019814996.1
D3 38312498 38312807
refGene_gn=PTPRM;refGene_tx=XM_003995022.5;refGene_gn=PTPRM;refGene_tx=XM_003995023.5;refGene_gn=PTPRM;refGene_tx=XM_011287970.3;refGene_gn=PTPRM;refGene_tx=XM_019814992.2;refGene_gn=PTPRM;refGene_tx=XM_019814993.2;refGene_gn=PTPRM;refGene_tx=XM_019814994.2;refGene_gn=PTPRM;refGene_tx=XM_019814995.2;refGene_gn=PTPRM;refGene_tx=XM_019814996.1
D3 38415185 38415479
refGene_gn=PTPRM;refGene_tx=XM_003995022.5;refGene_gn=PTPRM;refGene_tx=XM_003995023.5;refGene_gn=PTPRM;refGene_tx=XM_011287970.3;refGene_gn=PTPRM;refGene_tx=XM_019814992.2;refGene_gn=PTPRM;refGene_tx=XM_019814993.2;refGene_gn=PTPRM;refGene_tx=XM_019814994.2;refGene_gn=PTPRM;refGene_tx=XM_019814995.2;refGene_gn=PTPRM;refGene_tx=XM_019814996.1
D3 38421369 38421544
refGene_gn=PTPRM;refGene_tx=XM_003995022.5;refGene_gn=PTPRM;refGene_tx=XM_003995023.5;refGene_gn=PTPRM;refGene_tx=XM_011287970.3;refGene_gn=PTPRM;refGene_tx=XM_019814992.2;refGene_gn=PTPRM;refGene_tx=XM_019814993.2;refGene_gn=PTPRM;refGene_tx=XM_019814994.2;refGene_gn=PTPRM;refGene_tx=XM_019814995.2;refGene_gn=PTPRM;refGene_tx=XM_019814996.1
D3 38446028 38446144
refGene_gn=PTPRM;refGene_tx=XM_003995022.5;refGene_gn=PTPRM;refGene_tx

=XM_003995023.5;refGene_gn=PTPRM;refGene_tx=XM_011287970.3;refGene_gn=PTPRM;refGene_tx=XM_019814992.2;refGene_gn=PTPRM;refGene_tx=XM_019814993.2;refGene_gn=PTPRM;refGene_tx=XM_019814994.2;refGene_gn=PTPRM;refGene_tx=XM_019814995.2
D3 38466763 38466842
refGene_gn=PTPRM;refGene_tx=XM_003995022.5;refGene_gn=PTPRM;refGene_tx=XM_003995023.5;refGene_gn=PTPRM;refGene_tx=XM_011287970.3;refGene_gn=PTPRM;refGene_tx=XM_019814992.2;refGene_gn=PTPRM;refGene_tx=XM_019814993.2;refGene_gn=PTPRM;refGene_tx=XM_019814994.2;refGene_gn=PTPRM;refGene_tx=XM_019814995.2
D3 38495219 38495491
refGene_gn=PTPRM;refGene_tx=XM_003995022.5;refGene_gn=PTPRM;refGene_tx=XM_003995023.5;refGene_gn=PTPRM;refGene_tx=XM_011287970.3;refGene_gn=PTPRM;refGene_tx=XM_019814992.2;refGene_gn=PTPRM;refGene_tx=XM_019814993.2;refGene_gn=PTPRM;refGene_tx=XM_019814994.2;refGene_gn=PTPRM;refGene_tx=XM_019814995.2
D3 38599444 38599567
refGene_gn=PTPRM;refGene_tx=XM_003995022.5;refGene_gn=PTPRM;refGene_tx=XM_003995023.5;refGene_gn=PTPRM;refGene_tx=XM_011287970.3;refGene_gn=PTPRM;refGene_tx=XM_019814992.2;refGene_gn=PTPRM;refGene_tx=XM_019814993.2;refGene_gn=PTPRM;refGene_tx=XM_019814994.2;refGene_gn=PTPRM;refGene_tx=XM_019814995.2
D3 38798947 38799020
refGene_gn=PTPRM;refGene_tx=XM_003995022.5;refGene_gn=PTPRM;refGene_tx=XM_003995023.5;refGene_gn=PTPRM;refGene_tx=XM_011287970.3;refGene_gn=PTPRM;refGene_tx=XM_019814992.2;refGene_gn=PTPRM;refGene_tx=XM_019814993.2;refGene_gn=PTPRM;refGene_tx=XM_019814994.2;refGene_gn=PTPRM;refGene_tx=XM_019814995.2
D3 39087234 39088470 refGene_gn=LRRC30;refGene_tx=XM_006938789.4
D3 39166507 39166580 refGene_gn=LAMA1;refGene_tx=XM_019814998.2
D3 39196657 39196766 refGene_gn=LAMA1;refGene_tx=XM_023241765.1
D3 39197482 39197653
refGene_gn=LAMA1;refGene_tx=XM_019814998.2;refGene_gn=LAMA1;refGene_tx=XM_023241765.1
D3 39197842 39197955
refGene_gn=LAMA1;refGene_tx=XM_019814998.2;refGene_gn=LAMA1;refGene_tx=XM_023241765.1
D3 39224784 39225027
refGene_gn=LAMA1;refGene_tx=XM_019814998.2;refGene_gn=LAMA1;refGene_tx=XM_023241765.1
D3 39225929 39226109
refGene_gn=LAMA1;refGene_tx=XM_019814998.2;refGene_gn=LAMA1;refGene_tx=XM_023241765.1
D3 39228602 39228692
refGene_gn=LAMA1;refGene_tx=XM_019814998.2;refGene_gn=LAMA1;refGene_tx=XM_023241765.1
D3 39232268 39232386
refGene_gn=LAMA1;refGene_tx=XM_019814998.2;refGene_gn=LAMA1;refGene_tx=XM_023241765.1
D3 39234698 39234877

refGene_gn=LAMA1;refGene_tx=XM_019814998.2;refGene_gn=LAMA1;refGene_tx=XM_023241765.1
D3 39235559 39235665
refGene_gn=LAMA1;refGene_tx=XM_019814998.2;refGene_gn=LAMA1;refGene_tx=XM_023241765.1
D3 39237862 39238023
refGene_gn=LAMA1;refGene_tx=XM_019814998.2;refGene_gn=LAMA1;refGene_tx=XM_023241765.1
D3 39239689 39239830
refGene_gn=LAMA1;refGene_tx=XM_019814998.2;refGene_gn=LAMA1;refGene_tx=XM_023241765.1
D3 39241032 39241206
refGene_gn=LAMA1;refGene_tx=XM_019814998.2;refGene_gn=LAMA1;refGene_tx=XM_023241765.1
D3 39243369 39243471
refGene_gn=LAMA1;refGene_tx=XM_019814998.2;refGene_gn=LAMA1;refGene_tx=XM_023241765.1
D3 39244672 39244884
refGene_gn=LAMA1;refGene_tx=XM_019814998.2;refGene_gn=LAMA1;refGene_tx=XM_023241765.1
D3 39245665 39245777
refGene_gn=LAMA1;refGene_tx=XM_019814998.2;refGene_gn=LAMA1;refGene_tx=XM_023241765.1
D3 39249265 39249376
refGene_gn=LAMA1;refGene_tx=XM_019814998.2;refGene_gn=LAMA1;refGene_tx=XM_023241765.1
D3 39253002 39253130
refGene_gn=LAMA1;refGene_tx=XM_019814998.2;refGene_gn=LAMA1;refGene_tx=XM_023241765.1
D3 39255085 39255172
refGene_gn=LAMA1;refGene_tx=XM_019814998.2;refGene_gn=LAMA1;refGene_tx=XM_023241765.1
D3 39255870 39256082
refGene_gn=LAMA1;refGene_tx=XM_019814998.2;refGene_gn=LAMA1;refGene_tx=XM_023241765.1
D3 39262733 39262840
refGene_gn=LAMA1;refGene_tx=XM_019814998.2;refGene_gn=LAMA1;refGene_tx=XM_023241765.1
D3 39263157 39263338
refGene_gn=LAMA1;refGene_tx=XM_019814998.2;refGene_gn=LAMA1;refGene_tx=XM_023241765.1
D3 39263919 39264056
refGene_gn=LAMA1;refGene_tx=XM_019814998.2;refGene_gn=LAMA1;refGene_tx=XM_023241765.1
D3 39264953 39265190
refGene_gn=LAMA1;refGene_tx=XM_019814998.2;refGene_gn=LAMA1;refGene_tx=XM_023241765.1
D3 39266841 39266985
refGene_gn=LAMA1;refGene_tx=XM_019814998.2;refGene_gn=LAMA1;refGene_tx=XM_023241765.1

D3 39267501 39267681
refGene_gn=LAMA1;refGene_tx=XM_019814998.2;refGene_gn=LAMA1;refGene_tx=XM_023241765.1
D3 39268951 39269137
refGene_gn=LAMA1;refGene_tx=XM_019814998.2;refGene_gn=LAMA1;refGene_tx=XM_023241765.1
D3 39269679 39269807
refGene_gn=LAMA1;refGene_tx=XM_019814998.2;refGene_gn=LAMA1;refGene_tx=XM_023241765.1
D3 39270447 39270568
refGene_gn=LAMA1;refGene_tx=XM_019814998.2;refGene_gn=LAMA1;refGene_tx=XM_023241765.1
D3 39272516 39272654
refGene_gn=LAMA1;refGene_tx=XM_019814998.2;refGene_gn=LAMA1;refGene_tx=XM_023241765.1
D3 39280445 39280567
refGene_gn=LAMA1;refGene_tx=XM_019814998.2;refGene_gn=LAMA1;refGene_tx=XM_023241765.1
D3 39284436 39284523
refGene_gn=LAMA1;refGene_tx=XM_019814998.2;refGene_gn=LAMA1;refGene_tx=XM_023241765.1
D3 39284816 39285010
refGene_gn=LAMA1;refGene_tx=XM_019814998.2;refGene_gn=LAMA1;refGene_tx=XM_023241765.1
D3 39285484 39285627
refGene_gn=LAMA1;refGene_tx=XM_019814998.2;refGene_gn=LAMA1;refGene_tx=XM_023241765.1
D3 39287236 39287326
refGene_gn=LAMA1;refGene_tx=XM_019814998.2;refGene_gn=LAMA1;refGene_tx=XM_023241765.1
D3 39288818 39288930
refGene_gn=LAMA1;refGene_tx=XM_019814998.2;refGene_gn=LAMA1;refGene_tx=XM_023241765.1
D3 39290403 39290566
refGene_gn=LAMA1;refGene_tx=XM_019814998.2;refGene_gn=LAMA1;refGene_tx=XM_023241765.1
D3 39297072 39297283
refGene_gn=LAMA1;refGene_tx=XM_019814998.2;refGene_gn=LAMA1;refGene_tx=XM_023241765.1
D3 39297691 39297808
refGene_gn=LAMA1;refGene_tx=XM_019814998.2;refGene_gn=LAMA1;refGene_tx=XM_023241765.1
D3 39297933 39298097
refGene_gn=LAMA1;refGene_tx=XM_019814998.2;refGene_gn=LAMA1;refGene_tx=XM_023241765.1
D3 39299856 39299992
refGene_gn=LAMA1;refGene_tx=XM_019814998.2;refGene_gn=LAMA1;refGene_tx=XM_023241765.1
D3 39300530 39300624
refGene_gn=LAMA1;refGene_tx=XM_019814998.2;refGene_gn=LAMA1;refGene_tx

=XM_023241765.1
D3 39302228 39302345
refGene_gn=LAMA1;refGene_tx=XM_019814998.2;refGene_gn=LAMA1;refGene_tx=XM_023241765.1
D3 39303897 39304080
refGene_gn=LAMA1;refGene_tx=XM_019814998.2;refGene_gn=LAMA1;refGene_tx=XM_023241765.1
D3 39304722 39304877
refGene_gn=LAMA1;refGene_tx=XM_019814998.2;refGene_gn=LAMA1;refGene_tx=XM_023241765.1
D3 39306378 39306522
refGene_gn=LAMA1;refGene_tx=XM_019814998.2;refGene_gn=LAMA1;refGene_tx=XM_023241765.1
D3 39307505 39307639
refGene_gn=LAMA1;refGene_tx=XM_019814998.2;refGene_gn=LAMA1;refGene_tx=XM_023241765.1
D3 39308752 39308903
refGene_gn=LAMA1;refGene_tx=XM_019814998.2;refGene_gn=LAMA1;refGene_tx=XM_023241765.1
D3 39309663 39309788
refGene_gn=LAMA1;refGene_tx=XM_019814998.2;refGene_gn=LAMA1;refGene_tx=XM_023241765.1
D3 39315310 39315461
refGene_gn=LAMA1;refGene_tx=XM_019814998.2;refGene_gn=LAMA1;refGene_tx=XM_023241765.1
D3 39316119 39316264
refGene_gn=LAMA1;refGene_tx=XM_019814998.2;refGene_gn=LAMA1;refGene_tx=XM_023241765.1
D3 39316734 39316876
refGene_gn=LAMA1;refGene_tx=XM_019814998.2;refGene_gn=LAMA1;refGene_tx=XM_023241765.1
D3 39319489 39319604
refGene_gn=LAMA1;refGene_tx=XM_019814998.2;refGene_gn=LAMA1;refGene_tx=XM_023241765.1
D3 39319799 39319970
refGene_gn=LAMA1;refGene_tx=XM_019814998.2;refGene_gn=LAMA1;refGene_tx=XM_023241765.1
D3 39322176 39322340
refGene_gn=LAMA1;refGene_tx=XM_019814998.2;refGene_gn=LAMA1;refGene_tx=XM_023241765.1
D3 39323054 39323240
refGene_gn=LAMA1;refGene_tx=XM_019814998.2;refGene_gn=LAMA1;refGene_tx=XM_023241765.1
D3 39324628 39324764
refGene_gn=LAMA1;refGene_tx=XM_019814998.2;refGene_gn=LAMA1;refGene_tx=XM_023241765.1
D3 39326776 39326889
refGene_gn=LAMA1;refGene_tx=XM_019814998.2;refGene_gn=LAMA1;refGene_tx=XM_023241765.1
D3 39328957 39329147

refGene_gn=LAMA1;refGene_tx=XM_019814998.2;refGene_gn=LAMA1;refGene_tx=XM_023241765.1
D3 39329683 39329842
refGene_gn=LAMA1;refGene_tx=XM_019814998.2;refGene_gn=LAMA1;refGene_tx=XM_023241765.1
D3 39330363 39330517
refGene_gn=LAMA1;refGene_tx=XM_019814998.2;refGene_gn=LAMA1;refGene_tx=XM_023241765.1
D3 39331764 39331898
refGene_gn=LAMA1;refGene_tx=XM_019814998.2;refGene_gn=LAMA1;refGene_tx=XM_023241765.1
D3 39334898 39335121
refGene_gn=LAMA1;refGene_tx=XM_019814998.2;refGene_gn=LAMA1;refGene_tx=XM_023241765.1
D3 39335726 39335887
refGene_gn=LAMA1;refGene_tx=XM_019814998.2;refGene_gn=LAMA1;refGene_tx=XM_023241765.1
D3 39396376 39396471
refGene_gn=ARHGAP28;refGene_tx=XM_003995025.4;refGene_gn=ARHGAP28;refGene_tx=XM_019815000.2;refGene_gn=ARHGAP28;refGene_tx=XM_023241766.1;refGene_gn=ARHGAP28;refGene_tx=XM_023241767.1;refGene_gn=ARHGAP28;refGene_tx=XM_023241768.1;refGene_gn=ARHGAP28;refGene_tx=XM_023241769.1;refGene_gn=ARHGAP28;refGene_tx=XM_023241770.1;refGene_gn=ARHGAP28;refGene_tx=XM_023241771.1;refGene_gn=ARHGAP28;refGene_tx=XM_023241772.1
D3 39398638 39398703
refGene_gn=ARHGAP28;refGene_tx=XM_003995025.4;refGene_gn=ARHGAP28;refGene_tx=XM_019815000.2;refGene_gn=ARHGAP28;refGene_tx=XM_023241766.1;refGene_gn=ARHGAP28;refGene_tx=XM_023241767.1;refGene_gn=ARHGAP28;refGene_tx=XM_023241768.1;refGene_gn=ARHGAP28;refGene_tx=XM_023241769.1;refGene_gn=ARHGAP28;refGene_tx=XM_023241770.1;refGene_gn=ARHGAP28;refGene_tx=XM_023241771.1;refGene_gn=ARHGAP28;refGene_tx=XM_023241772.1
D3 39408523 39408648
refGene_gn=ARHGAP28;refGene_tx=XM_003995025.4;refGene_gn=ARHGAP28;refGene_tx=XM_019815000.2;refGene_gn=ARHGAP28;refGene_tx=XM_023241766.1;refGene_gn=ARHGAP28;refGene_tx=XM_023241767.1;refGene_gn=ARHGAP28;refGene_tx=XM_023241768.1;refGene_gn=ARHGAP28;refGene_tx=XM_023241769.1;refGene_gn=ARHGAP28;refGene_tx=XM_023241770.1;refGene_gn=ARHGAP28;refGene_tx=XM_023241771.1;refGene_gn=ARHGAP28;refGene_tx=XM_023241772.1
D3 39410682 39410739
refGene_gn=ARHGAP28;refGene_tx=XM_003995025.4;refGene_gn=ARHGAP28;refGene_tx=XM_019815000.2;refGene_gn=ARHGAP28;refGene_tx=XM_023241766.1;refGene_gn=ARHGAP28;refGene_tx=XM_023241768.1;refGene_gn=ARHGAP28;refGene_tx=XM_023241769.1;refGene_gn=ARHGAP28;refGene_tx=XM_023241770.1;refGene_gn=ARHGAP28;refGene_tx=XM_023241771.1;refGene_gn=ARHGAP28;refGene_tx=XM_023241772.1
D3 39419694 39419808
refGene_gn=ARHGAP28;refGene_tx=XM_003995025.4;refGene_gn=ARHGAP28;refGene_tx=XM_019815000.2;refGene_gn=ARHGAP28;refGene_tx=XM_023241766.1;refGene_gn=ARHGAP28;refGene_tx=XM_023241767.1;refGene_gn=ARHGAP28;refGene_tx=XM_023241768.1;refGene_gn=ARHGAP28;refGene_tx=XM_023241769.1;refG

ene_gn=ARHGAP28;refGene_tx=XM_023241770.1;refGene_gn=ARHGAP28;refGene_tx=XM_023241771.1;refGene_gn=ARHGAP28;refGene_tx=XM_023241772.1
D3 39420157 39420355
refGene_gn=ARHGAP28;refGene_tx=XM_003995025.4;refGene_gn=ARHGAP28;refGene_tx=XM_019815000.2;refGene_gn=ARHGAP28;refGene_tx=XM_023241766.1;refGene_gn=ARHGAP28;refGene_tx=XM_023241767.1;refGene_gn=ARHGAP28;refGene_tx=XM_023241768.1;refGene_gn=ARHGAP28;refGene_tx=XM_023241769.1;refGene_gn=ARHGAP28;refGene_tx=XM_023241770.1;refGene_gn=ARHGAP28;refGene_tx=XM_023241771.1;refGene_gn=ARHGAP28;refGene_tx=XM_023241772.1
D3 39422985 39423068
refGene_gn=ARHGAP28;refGene_tx=XM_003995025.4;refGene_gn=ARHGAP28;refGene_tx=XM_019815000.2;refGene_gn=ARHGAP28;refGene_tx=XM_023241766.1;refGene_gn=ARHGAP28;refGene_tx=XM_023241767.1;refGene_gn=ARHGAP28;refGene_tx=XM_023241768.1;refGene_gn=ARHGAP28;refGene_tx=XM_023241769.1;refGene_gn=ARHGAP28;refGene_tx=XM_023241770.1;refGene_gn=ARHGAP28;refGene_tx=XM_023241771.1;refGene_gn=ARHGAP28;refGene_tx=XM_023241772.1
D3 39429160 39429323
refGene_gn=ARHGAP28;refGene_tx=XM_003995025.4;refGene_gn=ARHGAP28;refGene_tx=XM_019815000.2;refGene_gn=ARHGAP28;refGene_tx=XM_023241766.1;refGene_gn=ARHGAP28;refGene_tx=XM_023241767.1;refGene_gn=ARHGAP28;refGene_tx=XM_023241768.1;refGene_gn=ARHGAP28;refGene_tx=XM_023241769.1;refGene_gn=ARHGAP28;refGene_tx=XM_023241770.1;refGene_gn=ARHGAP28;refGene_tx=XM_023241771.1;refGene_gn=ARHGAP28;refGene_tx=XM_023241772.1
D3 39435235 39435313
refGene_gn=ARHGAP28;refGene_tx=XM_003995025.4;refGene_gn=ARHGAP28;refGene_tx=XM_019815000.2;refGene_gn=ARHGAP28;refGene_tx=XM_023241766.1;refGene_gn=ARHGAP28;refGene_tx=XM_023241767.1;refGene_gn=ARHGAP28;refGene_tx=XM_023241768.1;refGene_gn=ARHGAP28;refGene_tx=XM_023241769.1;refGene_gn=ARHGAP28;refGene_tx=XM_023241770.1;refGene_gn=ARHGAP28;refGene_tx=XM_023241771.1;refGene_gn=ARHGAP28;refGene_tx=XM_023241772.1
D3 39437631 39437723
refGene_gn=ARHGAP28;refGene_tx=XM_003995025.4;refGene_gn=ARHGAP28;refGene_tx=XM_019815000.2;refGene_gn=ARHGAP28;refGene_tx=XM_023241766.1;refGene_gn=ARHGAP28;refGene_tx=XM_023241767.1;refGene_gn=ARHGAP28;refGene_tx=XM_023241768.1;refGene_gn=ARHGAP28;refGene_tx=XM_023241769.1;refGene_gn=ARHGAP28;refGene_tx=XM_023241770.1;refGene_gn=ARHGAP28;refGene_tx=XM_023241771.1;refGene_gn=ARHGAP28;refGene_tx=XM_023241772.1
D3 39437824 39437990
refGene_gn=ARHGAP28;refGene_tx=XM_003995025.4;refGene_gn=ARHGAP28;refGene_tx=XM_019815000.2;refGene_gn=ARHGAP28;refGene_tx=XM_023241766.1;refGene_gn=ARHGAP28;refGene_tx=XM_023241767.1;refGene_gn=ARHGAP28;refGene_tx=XM_023241768.1;refGene_gn=ARHGAP28;refGene_tx=XM_023241769.1;refGene_gn=ARHGAP28;refGene_tx=XM_023241770.1;refGene_gn=ARHGAP28;refGene_tx=XM_023241771.1;refGene_gn=ARHGAP28;refGene_tx=XM_023241772.1
D3 39440546 39440689
refGene_gn=ARHGAP28;refGene_tx=XM_003995025.4;refGene_gn=ARHGAP28;refGene_tx=XM_019815000.2;refGene_gn=ARHGAP28;refGene_tx=XM_023241766.1;refGene_gn=ARHGAP28;refGene_tx=XM_023241767.1;refGene_gn=ARHGAP28;refGene_tx=XM_023241768.1;refGene_gn=ARHGAP28;refGene_tx=XM_023241769.1;refGene_gn=ARHGAP28;refGene_tx=XM_023241770.1;refGene_gn=ARHGAP28;refGene_

tx=XM_023241771.1;refGene_gn=ARHGAP28;refGene_tx=XM_023241772.1
D3 39442580 39442665
refGene_gn=ARHGAP28;refGene_tx=XM_003995025.4;refGene_gn=ARHGAP28;refGene_tx=XM_019815000.2;refGene_gn=ARHGAP28;refGene_tx=XM_023241766.1;refGene_gn=ARHGAP28;refGene_tx=XM_023241767.1;refGene_gn=ARHGAP28;refGene_tx=XM_023241768.1;refGene_gn=ARHGAP28;refGene_tx=XM_023241769.1;refGene_gn=ARHGAP28;refGene_tx=XM_023241770.1;refGene_gn=ARHGAP28;refGene_tx=XM_023241771.1;refGene_gn=ARHGAP28;refGene_tx=XM_023241772.1
D3 39452140 39452230
refGene_gn=ARHGAP28;refGene_tx=XM_003995025.4;refGene_gn=ARHGAP28;refGene_tx=XM_023241766.1;refGene_gn=ARHGAP28;refGene_tx=XM_023241767.1;refGene_gn=ARHGAP28;refGene_tx=XM_023241769.1;refGene_gn=ARHGAP28;refGene_tx=XM_023241770.1;refGene_gn=ARHGAP28;refGene_tx=XM_023241771.1;refGene_gn=ARHGAP28;refGene_tx=XM_023241772.1
D3 39459909 39460002
refGene_gn=ARHGAP28;refGene_tx=XM_003995025.4;refGene_gn=ARHGAP28;refGene_tx=XM_019815000.2;refGene_gn=ARHGAP28;refGene_tx=XM_023241766.1;refGene_gn=ARHGAP28;refGene_tx=XM_023241767.1;refGene_gn=ARHGAP28;refGene_tx=XM_023241768.1;refGene_gn=ARHGAP28;refGene_tx=XM_023241769.1;refGene_gn=ARHGAP28;refGene_tx=XM_023241770.1;refGene_gn=ARHGAP28;refGene_tx=XM_023241771.1;refGene_gn=ARHGAP28;refGene_tx=XM_023241772.1
D3 39465522 39465615
refGene_gn=ARHGAP28;refGene_tx=XM_023241766.1;refGene_gn=ARHGAP28;refGene_tx=XM_023241767.1;refGene_gn=ARHGAP28;refGene_tx=XM_023241768.1;refGene_gn=ARHGAP28;refGene_tx=XM_023241769.1;refGene_gn=ARHGAP28;refGene_tx=XM_023241770.1;refGene_gn=ARHGAP28;refGene_tx=XM_023241771.1;refGene_gn=ARHGAP28;refGene_tx=XM_023241772.1
D3 39471083 39471301
refGene_gn=ARHGAP28;refGene_tx=XM_003995025.4;refGene_gn=ARHGAP28;refGene_tx=XM_019815000.2;refGene_gn=ARHGAP28;refGene_tx=XM_023241766.1;refGene_gn=ARHGAP28;refGene_tx=XM_023241767.1;refGene_gn=ARHGAP28;refGene_tx=XM_023241768.1;refGene_gn=ARHGAP28;refGene_tx=XM_023241769.1;refGene_gn=ARHGAP28;refGene_tx=XM_023241770.1;refGene_gn=ARHGAP28;refGene_tx=XM_023241771.1;refGene_gn=ARHGAP28;refGene_tx=XM_023241772.1
D3 39481417 39481620
refGene_gn=ARHGAP28;refGene_tx=XM_003995025.4;refGene_gn=ARHGAP28;refGene_tx=XM_019815000.2;refGene_gn=ARHGAP28;refGene_tx=XM_023241766.1;refGene_gn=ARHGAP28;refGene_tx=XM_023241767.1;refGene_gn=ARHGAP28;refGene_tx=XM_023241768.1;refGene_gn=ARHGAP28;refGene_tx=XM_023241769.1;refGene_gn=ARHGAP28;refGene_tx=XM_023241770.1;refGene_gn=ARHGAP28;refGene_tx=XM_023241771.1;refGene_gn=ARHGAP28;refGene_tx=XM_023241772.1
D3 39587025 39587201
refGene_gn=ARHGAP28;refGene_tx=XM_003995025.4;refGene_gn=ARHGAP28;refGene_tx=XM_019815000.2;refGene_gn=ARHGAP28;refGene_tx=XM_023241766.1;refGene_gn=ARHGAP28;refGene_tx=XM_023241767.1;refGene_gn=ARHGAP28;refGene_tx=XM_023241768.1
D3 39976186 39976255
refGene_gn=L3MBTL4;refGene_tx=XM_023241773.1;refGene_gn=L3MBTL4;refGene_tx=XM_023241775.1;refGene_gn=L3MBTL4;refGene_tx=XM_023241776.1
D3 39976629 39976734

refGene_gn=L3MBTL4;refGene_tx=XM_023241773.1;refGene_gn=L3MBTL4;refGen
e_tx=XM_023241774.1;refGene_gn=L3MBTL4;refGene_tx=XM_023241775.1;refGe
ne_gn=L3MBTL4;refGene_tx=XM_023241776.1
D3 39999080 39999135
refGene_gn=L3MBTL4;refGene_tx=XM_023241773.1;refGene_gn=L3MBTL4;refGen
e_tx=XM_023241774.1;refGene_gn=L3MBTL4;refGene_tx=XM_023241775.1;refGe
ne_gn=L3MBTL4;refGene_tx=XM_023241776.1
D3 40037413 40037505
refGene_gn=L3MBTL4;refGene_tx=XM_023241773.1;refGene_gn=L3MBTL4;refGen
e_tx=XM_023241774.1;refGene_gn=L3MBTL4;refGene_tx=XM_023241775.1;refGe
ne_gn=L3MBTL4;refGene_tx=XM_023241776.1
D3 40058212 40058317
refGene_gn=L3MBTL4;refGene_tx=XM_023241773.1;refGene_gn=L3MBTL4;refGen
e_tx=XM_023241774.1;refGene_gn=L3MBTL4;refGene_tx=XM_023241775.1;refGe
ne_gn=L3MBTL4;refGene_tx=XM_023241776.1
D3 40059611 40059747
refGene_gn=L3MBTL4;refGene_tx=XM_023241773.1;refGene_gn=L3MBTL4;refGen
e_tx=XM_023241774.1;refGene_gn=L3MBTL4;refGene_tx=XM_023241775.1;refGe
ne_gn=L3MBTL4;refGene_tx=XM_023241776.1
D3 40061515 40061607
refGene_gn=L3MBTL4;refGene_tx=XM_023241773.1;refGene_gn=L3MBTL4;refGen
e_tx=XM_023241774.1;refGene_gn=L3MBTL4;refGene_tx=XM_023241775.1;refGe
ne_gn=L3MBTL4;refGene_tx=XM_023241776.1
D3 40062813 40062968
refGene_gn=L3MBTL4;refGene_tx=XM_023241773.1;refGene_gn=L3MBTL4;refGen
e_tx=XM_023241774.1;refGene_gn=L3MBTL4;refGene_tx=XM_023241775.1;refGe
ne_gn=L3MBTL4;refGene_tx=XM_023241776.1
D3 40064040 40064117
refGene_gn=L3MBTL4;refGene_tx=XM_023241773.1;refGene_gn=L3MBTL4;refGen
e_tx=XM_023241774.1;refGene_gn=L3MBTL4;refGene_tx=XM_023241775.1;refGe
ne_gn=L3MBTL4;refGene_tx=XM_023241776.1
D3 40099100 40099186
refGene_gn=L3MBTL4;refGene_tx=XM_023241773.1;refGene_gn=L3MBTL4;refGen
e_tx=XM_023241774.1;refGene_gn=L3MBTL4;refGene_tx=XM_023241775.1;refGe
ne_gn=L3MBTL4;refGene_tx=XM_023241776.1
D3 40101265 40101376
refGene_gn=L3MBTL4;refGene_tx=XM_023241773.1;refGene_gn=L3MBTL4;refGen
e_tx=XM_023241774.1;refGene_gn=L3MBTL4;refGene_tx=XM_023241775.1;refGe
ne_gn=L3MBTL4;refGene_tx=XM_023241776.1
D3 40144534 40144649
refGene_gn=L3MBTL4;refGene_tx=XM_023241773.1;refGene_gn=L3MBTL4;refGen
e_tx=XM_023241774.1;refGene_gn=L3MBTL4;refGene_tx=XM_023241775.1;refGe
ne_gn=L3MBTL4;refGene_tx=XM_023241776.1
D3 40173800 40173903
refGene_gn=L3MBTL4;refGene_tx=XM_023241773.1;refGene_gn=L3MBTL4;refGen
e_tx=XM_023241774.1;refGene_gn=L3MBTL4;refGene_tx=XM_023241775.1;refGe
ne_gn=L3MBTL4;refGene_tx=XM_023241776.1
D3 40219331 40219505
refGene_gn=L3MBTL4;refGene_tx=XM_023241773.1;refGene_gn=L3MBTL4;refGen
e_tx=XM_023241774.1;refGene_gn=L3MBTL4;refGene_tx=XM_023241775.1;refGe

ne_gn=L3MBTL4;refGene_tx=XM_023241776.1
D3 40231370 40231441
refGene_gn=L3MBTL4;refGene_tx=XM_023241773.1;refGene_gn=L3MBTL4;refGene_tx=XM_023241774.1;refGene_gn=L3MBTL4;refGene_tx=XM_023241775.1;refGene_gn=L3MBTL4;refGene_tx=XM_023241776.1
D3 40297621 40297689 refGene_gn=L3MBTL4;refGene_tx=XM_023241776.1
D3 40317128 40317208 refGene_gn=L3MBTL4;refGene_tx=XM_023241775.1
D3 40324590 40324760
refGene_gn=L3MBTL4;refGene_tx=XM_023241773.1;refGene_gn=L3MBTL4;refGene_tx=XM_023241774.1
D3 40334588 40334651
refGene_gn=L3MBTL4;refGene_tx=XM_023241773.1;refGene_gn=L3MBTL4;refGene_tx=XM_023241774.1
D3 40336156 40336300 refGene_gn=L3MBTL4;refGene_tx=XM_023241773.1
D3 40337094 40337262 refGene_gn=L3MBTL4;refGene_tx=XM_023241774.1
D3 40405214 40407119 refGene_gn=TMEM200C;refGene_tx=XM_023241777.1
D3 40805172 40805358 refGene_gn=EPB41L3;refGene_tx=XM_019815027.2
D3 40809907 40810105 refGene_gn=EPB41L3;refGene_tx=XM_019815027.2
D3 40846915 40847020 refGene_gn=EPB41L3;refGene_tx=XM_019815027.2
D3 40848433 40848476 refGene_gn=EPB41L3;refGene_tx=XM_019815027.2
D3 40856741 40856817 refGene_gn=EPB41L3;refGene_tx=XM_019815027.2
D3 40858902 40859121 refGene_gn=EPB41L3;refGene_tx=XM_019815027.2
D3 40859455 40859543 refGene_gn=EPB41L3;refGene_tx=XM_019815027.2
D3 40864361 40864514 refGene_gn=EPB41L3;refGene_tx=XM_019815027.2
D3 40868123 40868221 refGene_gn=EPB41L3;refGene_tx=XM_019815027.2
D3 40869098 40869274 refGene_gn=EPB41L3;refGene_tx=XM_019815027.2
D3 40872914 40873081 refGene_gn=EPB41L3;refGene_tx=XM_019815027.2
D3 40876462 40876487 refGene_gn=EPB41L3;refGene_tx=XM_019815027.2
D3 40876489 40877025 refGene_gn=EPB41L3;refGene_tx=XM_019815027.2
D3 40881924 40881978 refGene_gn=EPB41L3;refGene_tx=XM_019815027.2
D3 40884691 40884727 refGene_gn=EPB41L3;refGene_tx=XM_019815027.2
D3 40885463 40885655 refGene_gn=EPB41L3;refGene_tx=XM_019815027.2
D3 40893182 40893305 refGene_gn=EPB41L3;refGene_tx=XM_019815027.2
D3 40893921 40894305 refGene_gn=EPB41L3;refGene_tx=XM_019815027.2
D3 40895126 40895258 refGene_gn=EPB41L3;refGene_tx=XM_019815027.2
D3 40895758 40895857 refGene_gn=EPB41L3;refGene_tx=XM_019815027.2
D3 40896514 40896595 refGene_gn=EPB41L3;refGene_tx=XM_019815027.2
D3 40896868 40896979 refGene_gn=EPB41L3;refGene_tx=XM_019815027.2
D3 40985589 40985592
refGene_gn=ZBTB14;refGene_tx=XM_003995033.5;refGene_gn=ZBTB14;refGene_tx=XM_019815041.2;refGene_gn=ZBTB14;refGene_tx=XM_023241778.1;refGene_gn=ZBTB14;refGene_tx=XM_023241779.1
D3 40986760 40988107
refGene_gn=ZBTB14;refGene_tx=XM_003995033.5;refGene_gn=ZBTB14;refGene_tx=XM_019815041.2;refGene_gn=ZBTB14;refGene_tx=XM_023241778.1;refGene_gn=ZBTB14;refGene_tx=XM_023241779.1
D3 41071260 41071276 refGene_gn=AKAIN1;refGene_tx=XM_019815043.2
D3 41125037 41125231 refGene_gn=AKAIN1;refGene_tx=XM_019815043.2
D3 42324025 42324982
refGene_gn=DLGAP1;refGene_tx=XM_023241780.1;refGene_gn=DLGAP1;refGene_

tx=XM_023241781.1;refGene_gn=DLGAP1;refGene_tx=XM_023241782.1;refGene_gn=DLGAP1;refGene_tx=XM_023241783.1
D3 42329078 42329171
refGene_gn=DLGAP1;refGene_tx=XM_023241785.1;refGene_gn=DLGAP1;refGene_tx=XM_023241790.1
D3 42329551 42329632
refGene_gn=DLGAP1;refGene_tx=XM_023241786.1;refGene_gn=DLGAP1;refGene_tx=XM_023241787.1;refGene_gn=DLGAP1;refGene_tx=XM_023241789.1
D3 42357337 42357388
refGene_gn=DLGAP1;refGene_tx=XM_023241784.1;refGene_gn=DLGAP1;refGene_tx=XM_023241788.1
D3 42380609 42380723 refGene_gn=DLGAP1;refGene_tx=XM_023241784.1
D3 42389835 42390050
refGene_gn=DLGAP1;refGene_tx=XM_023241780.1;refGene_gn=DLGAP1;refGene_tx=XM_023241781.1;refGene_gn=DLGAP1;refGene_tx=XM_023241782.1;refGene_gn=DLGAP1;refGene_tx=XM_023241783.1;refGene_gn=DLGAP1;refGene_tx=XM_023241784.1;refGene_gn=DLGAP1;refGene_tx=XM_023241785.1;refGene_gn=DLGAP1;refGene_tx=XM_023241786.1;refGene_gn=DLGAP1;refGene_tx=XM_023241787.1;refGene_gn=DLGAP1;refGene_tx=XM_023241788.1;refGene_gn=DLGAP1;refGene_tx=XM_023241789.1;refGene_gn=DLGAP1;refGene_tx=XM_023241790.1
D3 42455209 42455387
refGene_gn=DLGAP1;refGene_tx=XM_023241780.1;refGene_gn=DLGAP1;refGene_tx=XM_023241781.1;refGene_gn=DLGAP1;refGene_tx=XM_023241782.1;refGene_gn=DLGAP1;refGene_tx=XM_023241783.1;refGene_gn=DLGAP1;refGene_tx=XM_023241784.1;refGene_gn=DLGAP1;refGene_tx=XM_023241785.1;refGene_gn=DLGAP1;refGene_tx=XM_023241786.1;refGene_gn=DLGAP1;refGene_tx=XM_023241787.1;refGene_gn=DLGAP1;refGene_tx=XM_023241788.1;refGene_gn=DLGAP1;refGene_tx=XM_023241789.1;refGene_gn=DLGAP1;refGene_tx=XM_023241790.1
D3 42467130 42467371
refGene_gn=DLGAP1;refGene_tx=XM_023241780.1;refGene_gn=DLGAP1;refGene_tx=XM_023241781.1;refGene_gn=DLGAP1;refGene_tx=XM_023241782.1;refGene_gn=DLGAP1;refGene_tx=XM_023241783.1;refGene_gn=DLGAP1;refGene_tx=XM_023241784.1;refGene_gn=DLGAP1;refGene_tx=XM_023241785.1;refGene_gn=DLGAP1;refGene_tx=XM_023241786.1;refGene_gn=DLGAP1;refGene_tx=XM_023241787.1;refGene_gn=DLGAP1;refGene_tx=XM_023241788.1;refGene_gn=DLGAP1;refGene_tx=XM_023241789.1;refGene_gn=DLGAP1;refGene_tx=XM_023241790.1
D3 42535809 42535839
refGene_gn=DLGAP1;refGene_tx=XM_023241780.1;refGene_gn=DLGAP1;refGene_tx=XM_023241782.1;refGene_gn=DLGAP1;refGene_tx=XM_023241784.1;refGene_gn=DLGAP1;refGene_tx=XM_023241785.1;refGene_gn=DLGAP1;refGene_tx=XM_023241786.1;refGene_gn=DLGAP1;refGene_tx=XM_023241787.1;refGene_gn=DLGAP1;refGene_tx=XM_023241788.1
D3 42574000 42574205
refGene_gn=LOC102899787;refGene_tx=XM_019815046.2
D3 42580365 42580817
refGene_gn=LOC102899787;refGene_tx=XM_019815046.2
D3 42589714 42590088
refGene_gn=DLGAP1;refGene_tx=XM_023241780.1;refGene_gn=DLGAP1;refGene_tx=XM_023241781.1;refGene_gn=DLGAP1;refGene_tx=XM_023241782.1;refGene_gn=DLGAP1;refGene_tx=XM_023241783.1;refGene_gn=DLGAP1;refGene_tx=XM_02

3241784.1;refGene_gn=DLGAP1;refGene_tx=XM_023241785.1;refGene_gn=DLGAP1;refGene_tx=XM_023241786.1;refGene_gn=DLGAP1;refGene_tx=XM_023241787.1;refGene_gn=DLGAP1;refGene_tx=XM_023241788.1;refGene_gn=DLGAP1;refGene_tx=XM_023241789.1;refGene_gn=DLGAP1;refGene_tx=XM_023241790.1
D3 42602050 42602142
refGene_gn=DLGAP1;refGene_tx=XM_023241780.1;refGene_gn=DLGAP1;refGene_tx=XM_023241781.1;refGene_gn=DLGAP1;refGene_tx=XM_023241782.1;refGene_gn=DLGAP1;refGene_tx=XM_023241783.1;refGene_gn=DLGAP1;refGene_tx=XM_023241784.1;refGene_gn=DLGAP1;refGene_tx=XM_023241785.1;refGene_gn=DLGAP1;refGene_tx=XM_023241786.1;refGene_gn=DLGAP1;refGene_tx=XM_023241787.1;refGene_gn=DLGAP1;refGene_tx=XM_023241788.1;refGene_gn=DLGAP1;refGene_tx=XM_023241789.1;refGene_gn=DLGAP1;refGene_tx=XM_023241790.1
D3 42628376 42628798
refGene_gn=DLGAP1;refGene_tx=XM_023241780.1;refGene_gn=DLGAP1;refGene_tx=XM_023241781.1;refGene_gn=DLGAP1;refGene_tx=XM_023241782.1;refGene_gn=DLGAP1;refGene_tx=XM_023241783.1;refGene_gn=DLGAP1;refGene_tx=XM_023241784.1;refGene_gn=DLGAP1;refGene_tx=XM_023241785.1;refGene_gn=DLGAP1;refGene_tx=XM_023241786.1;refGene_gn=DLGAP1;refGene_tx=XM_023241787.1;refGene_gn=DLGAP1;refGene_tx=XM_023241788.1;refGene_gn=DLGAP1;refGene_tx=XM_023241789.1;refGene_gn=DLGAP1;refGene_tx=XM_023241790.1
D3 42646629 42646721
refGene_gn=DLGAP1;refGene_tx=XM_023241780.1;refGene_gn=DLGAP1;refGene_tx=XM_023241781.1;refGene_gn=DLGAP1;refGene_tx=XM_023241782.1;refGene_gn=DLGAP1;refGene_tx=XM_023241783.1;refGene_gn=DLGAP1;refGene_tx=XM_023241784.1;refGene_gn=DLGAP1;refGene_tx=XM_023241785.1;refGene_gn=DLGAP1;refGene_tx=XM_023241786.1;refGene_gn=DLGAP1;refGene_tx=XM_023241787.1;refGene_gn=DLGAP1;refGene_tx=XM_023241788.1;refGene_gn=DLGAP1;refGene_tx=XM_023241789.1;refGene_gn=DLGAP1;refGene_tx=XM_023241790.1
D3 42652198 42652351
refGene_gn=DLGAP1;refGene_tx=XM_023241780.1;refGene_gn=DLGAP1;refGene_tx=XM_023241781.1;refGene_gn=DLGAP1;refGene_tx=XM_023241782.1;refGene_gn=DLGAP1;refGene_tx=XM_023241783.1;refGene_gn=DLGAP1;refGene_tx=XM_023241784.1;refGene_gn=DLGAP1;refGene_tx=XM_023241785.1;refGene_gn=DLGAP1;refGene_tx=XM_023241786.1;refGene_gn=DLGAP1;refGene_tx=XM_023241787.1;refGene_gn=DLGAP1;refGene_tx=XM_023241788.1;refGene_gn=DLGAP1;refGene_tx=XM_023241789.1;refGene_gn=DLGAP1;refGene_tx=XM_023241790.1
D3 42655271 42655481
refGene_gn=DLGAP1;refGene_tx=XM_023241780.1;refGene_gn=DLGAP1;refGene_tx=XM_023241781.1;refGene_gn=DLGAP1;refGene_tx=XM_023241782.1;refGene_gn=DLGAP1;refGene_tx=XM_023241783.1;refGene_gn=DLGAP1;refGene_tx=XM_023241784.1;refGene_gn=DLGAP1;refGene_tx=XM_023241785.1;refGene_gn=DLGAP1;refGene_tx=XM_023241786.1;refGene_gn=DLGAP1;refGene_tx=XM_023241787.1;refGene_gn=DLGAP1;refGene_tx=XM_023241788.1;refGene_gn=DLGAP1;refGene_tx=XM_023241789.1;refGene_gn=DLGAP1;refGene_tx=XM_023241790.1
D3 42689902 42690478 refGene_gn=TGIF1;refGene_tx=XM_003995036.4
D3 42691277 42691504 refGene_gn=TGIF1;refGene_tx=XM_003995036.4
D3 42696924 42696940 refGene_gn=TGIF1;refGene_tx=XM_003995036.4
D3 42846393 42846566
refGene_gn=LOC101085247;refGene_tx=XM_011288088.2
D3 42847640 42847802

refGene_gn=LOC101085247;refGene_tx=XM_011288088.2
D3 42850796 42850980
refGene_gn=LOC101085247;refGene_tx=XM_011288088.2
D3 42869723 42869896
refGene_gn=LOC101085837;refGene_tx=XM_003995040.4;refGene_gn=LOC101085837;refGene_tx=XM_003995041.4;refGene_gn=LOC101085837;refGene_tx=XM_019815049.2
D3 42872726 42872888
refGene_gn=LOC101085837;refGene_tx=XM_003995040.4;refGene_gn=LOC101085837;refGene_tx=XM_003995041.4;refGene_gn=LOC101085837;refGene_tx=XM_019815049.2
D3 42873355 42873554
refGene_gn=LOC101085837;refGene_tx=XM_003995040.4;refGene_gn=LOC101085837;refGene_tx=XM_003995041.4;refGene_gn=LOC101085837;refGene_tx=XM_019815049.2
D3 42874569 42874572
refGene_gn=LOC101085837;refGene_tx=XM_003995041.4
D3 42925526 42925825 refGene_gn=MYOM1;refGene_tx=XM_023241791.1
D3 42948784 42948925 refGene_gn=MYOM1;refGene_tx=XM_023241791.1
D3 42954223 42954581 refGene_gn=MYOM1;refGene_tx=XM_023241791.1
D3 42955607 42955765 refGene_gn=MYOM1;refGene_tx=XM_023241791.1
D3 42964057 42964150 refGene_gn=MYOM1;refGene_tx=XM_023241791.1
D3 42966299 42966388 refGene_gn=MYOM1;refGene_tx=XM_023241791.1
D3 42966497 42966560 refGene_gn=MYOM1;refGene_tx=XM_023241791.1
D3 42970247 42970412 refGene_gn=MYOM1;refGene_tx=XM_023241791.1
D3 42973258 42973420 refGene_gn=MYOM1;refGene_tx=XM_023241791.1
D3 42978147 42978289 refGene_gn=MYOM1;refGene_tx=XM_023241791.1
D3 42981375 42981427 refGene_gn=MYOM1;refGene_tx=XM_023241791.1
D3 42981428 42981576 refGene_gn=MYOM1;refGene_tx=XM_023241791.1
D3 42983947 42984004 refGene_gn=MYOM1;refGene_tx=XM_023241791.1
D3 42989742 42989867 refGene_gn=MYOM1;refGene_tx=XM_023241791.1
D3 42994593 42994777 refGene_gn=MYOM1;refGene_tx=XM_023241791.1
D3 42995265 42995440 refGene_gn=MYOM1;refGene_tx=XM_023241791.1
D3 42997473 42997595 refGene_gn=MYOM1;refGene_tx=XM_023241791.1
D3 43001031 43001228 refGene_gn=MYOM1;refGene_tx=XM_023241791.1
D3 43007495 43007622 refGene_gn=MYOM1;refGene_tx=XM_023241791.1
D3 43010761 43010946 refGene_gn=MYOM1;refGene_tx=XM_023241791.1
D3 43013610 43013725 refGene_gn=MYOM1;refGene_tx=XM_023241791.1
D3 43031753 43031910 refGene_gn=MYOM1;refGene_tx=XM_023241791.1
D3 43033500 43033607 refGene_gn=MYOM1;refGene_tx=XM_023241791.1
D3 43033724 43033769 refGene_gn=MYOM1;refGene_tx=XM_023241791.1
D3 43039226 43039363 refGene_gn=MYOM1;refGene_tx=XM_023241791.1
D3 43041498 43041643 refGene_gn=MYOM1;refGene_tx=XM_023241791.1
D3 43042365 43042425 refGene_gn=MYOM1;refGene_tx=XM_023241791.1
D3 43042720 43042788 refGene_gn=MYOM1;refGene_tx=XM_023241791.1
D3 43046314 43046428 refGene_gn=MYOM1;refGene_tx=XM_023241791.1
D3 43047120 43047208 refGene_gn=MYOM1;refGene_tx=XM_023241791.1
D3 43048355 43048394 refGene_gn=MYOM1;refGene_tx=XM_023241791.1
D3 43048484 43048590 refGene_gn=MYOM1;refGene_tx=XM_023241791.1
D3 43051838 43052002 refGene_gn=MYOM1;refGene_tx=XM_023241791.1

D3 43054204 43054241 refGene_gn=MYOM1;refGene_tx=XM_023241791.1
D3 43054488 43054511 refGene_gn=MYOM1;refGene_tx=XM_023241791.1
D3 43057029 43057085 refGene_gn=MYOM1;refGene_tx=XM_023241791.1
D3 43060234 43060528 refGene_gn=MYOM1;refGene_tx=XM_023241791.1
D3 43123417 43123525 refGene_gn=LPIN2;refGene_tx=XM_019815051.2
D3 43148877 43149078
refGene_gn=LPIN2;refGene_tx=XM_019815051.2;refGene_gn=LPIN2;refGene_tx=XM_023241793.1
D3 43156758 43156854
refGene_gn=LPIN2;refGene_tx=XM_019815051.2;refGene_gn=LPIN2;refGene_tx=XM_023241793.1
D3 43159910 43160212
refGene_gn=LPIN2;refGene_tx=XM_019815051.2;refGene_gn=LPIN2;refGene_tx=XM_023241793.1
D3 43166389 43166497
refGene_gn=LPIN2;refGene_tx=XM_019815051.2;refGene_gn=LPIN2;refGene_tx=XM_023241793.1
D3 43167177 43167304
refGene_gn=LPIN2;refGene_tx=XM_019815051.2;refGene_gn=LPIN2;refGene_tx=XM_023241793.1
D3 43168139 43168485
refGene_gn=LPIN2;refGene_tx=XM_019815051.2;refGene_gn=LPIN2;refGene_tx=XM_023241793.1
D3 43172290 43172390
refGene_gn=LPIN2;refGene_tx=XM_019815051.2;refGene_gn=LPIN2;refGene_tx=XM_023241793.1
D3 43175325 43175513
refGene_gn=LPIN2;refGene_tx=XM_019815051.2;refGene_gn=LPIN2;refGene_tx=XM_023241793.1
D3 43177245 43177339
refGene_gn=LPIN2;refGene_tx=XM_019815051.2;refGene_gn=LPIN2;refGene_tx=XM_023241793.1
D3 43177778 43177848
refGene_gn=LPIN2;refGene_tx=XM_019815051.2;refGene_gn=LPIN2;refGene_tx=XM_023241793.1
D3 43178828 43178918
refGene_gn=LPIN2;refGene_tx=XM_019815051.2;refGene_gn=LPIN2;refGene_tx=XM_023241793.1
D3 43179907 43179990
refGene_gn=LPIN2;refGene_tx=XM_019815051.2;refGene_gn=LPIN2;refGene_tx=XM_023241793.1
D3 43181464 43181609
refGene_gn=LPIN2;refGene_tx=XM_019815051.2;refGene_gn=LPIN2;refGene_tx=XM_023241793.1
D3 43182496 43182645
refGene_gn=LPIN2;refGene_tx=XM_019815051.2;refGene_gn=LPIN2;refGene_tx=XM_023241793.1
D3 43183205 43183292
refGene_gn=LPIN2;refGene_tx=XM_019815051.2;refGene_gn=LPIN2;refGene_tx=XM_023241793.1

D3	43184682	43184835
refGene_gn=LPIN2;refGene_tx=XM_019815051.2;refGene_gn=LPIN2;refGene_tx
=XM_023241793.1
D3	43185268	43185383
refGene_gn=LPIN2;refGene_tx=XM_019815051.2;refGene_gn=LPIN2;refGene_tx
=XM_023241793.1
D3	43186040	43186144
refGene_gn=LPIN2;refGene_tx=XM_019815051.2;refGene_gn=LPIN2;refGene_tx
=XM_023241793.1
D3	43186489	43186634
refGene_gn=LPIN2;refGene_tx=XM_019815051.2;refGene_gn=LPIN2;refGene_tx
=XM_023241793.1
D3	43193129	43193467	refGene_gn=EMILIN2;refGene_tx=XM_023241792.1
D3	43196055	43196184
refGene_gn=EMILIN2;refGene_tx=XM_006938804.4;refGene_gn=EMILIN2;refGen
e_tx=XM_023241792.1
D3	43196742	43196775
refGene_gn=EMILIN2;refGene_tx=XM_006938804.4;refGene_gn=EMILIN2;refGen
e_tx=XM_023241792.1
D3	43197371	43197787
refGene_gn=EMILIN2;refGene_tx=XM_006938804.4;refGene_gn=EMILIN2;refGen
e_tx=XM_023241792.1
D3	43210239	43212168
refGene_gn=EMILIN2;refGene_tx=XM_006938804.4;refGene_gn=EMILIN2;refGen
e_tx=XM_023241792.1
D3	43216720	43216896
refGene_gn=EMILIN2;refGene_tx=XM_006938804.4;refGene_gn=EMILIN2;refGen
e_tx=XM_023241792.1
D3	43242181	43242304
refGene_gn=EMILIN2;refGene_tx=XM_006938804.4;refGene_gn=EMILIN2;refGen
e_tx=XM_023241792.1
D3	43242847	43242981
refGene_gn=EMILIN2;refGene_tx=XM_006938804.4;refGene_gn=EMILIN2;refGen
e_tx=XM_023241792.1
D3	43263447	43263472	refGene_gn=SMCHD1;refGene_tx=XM_003995046.5
D3	43270838	43270953	refGene_gn=SMCHD1;refGene_tx=XM_003995046.5
D3	43271211	43271370	refGene_gn=SMCHD1;refGene_tx=XM_003995046.5
D3	43281854	43282026	refGene_gn=SMCHD1;refGene_tx=XM_003995046.5
D3	43289142	43289213	refGene_gn=SMCHD1;refGene_tx=XM_003995046.5
D3	43289674	43289784	refGene_gn=SMCHD1;refGene_tx=XM_003995046.5
D3	43291459	43291650	refGene_gn=SMCHD1;refGene_tx=XM_003995046.5
D3	43295382	43295505	refGene_gn=SMCHD1;refGene_tx=XM_003995046.5
D3	43297569	43297655	refGene_gn=SMCHD1;refGene_tx=XM_003995046.5
D3	43298827	43298947	refGene_gn=SMCHD1;refGene_tx=XM_003995046.5
D3	43299115	43299242	refGene_gn=SMCHD1;refGene_tx=XM_003995046.5
D3	43307307	43307460	refGene_gn=SMCHD1;refGene_tx=XM_003995046.5
D3	43308322	43308454	refGene_gn=SMCHD1;refGene_tx=XM_003995046.5
D3	43309298	43309386	refGene_gn=SMCHD1;refGene_tx=XM_003995046.5
D3	43326029	43326094	refGene_gn=SMCHD1;refGene_tx=XM_003995046.5
D3	43326978	43327094	refGene_gn=SMCHD1;refGene_tx=XM_003995046.5

| | | | |
|---|---|---|---|
| D3 | 43328631 | 43328789 | refGene_gn=SMCHD1;refGene_tx=XM_003995046.5 |
| D3 | 43329023 | 43329103 | refGene_gn=SMCHD1;refGene_tx=XM_003995046.5 |
| D3 | 43331087 | 43331213 | refGene_gn=SMCHD1;refGene_tx=XM_003995046.5 |
| D3 | 43338234 | 43338402 | refGene_gn=SMCHD1;refGene_tx=XM_003995046.5 |
| D3 | 43341854 | 43341973 | refGene_gn=SMCHD1;refGene_tx=XM_003995046.5 |
| D3 | 43342541 | 43342630 | refGene_gn=SMCHD1;refGene_tx=XM_003995046.5 |
| D3 | 43343544 | 43343693 | refGene_gn=SMCHD1;refGene_tx=XM_003995046.5 |
| D3 | 43349761 | 43349989 | refGene_gn=SMCHD1;refGene_tx=XM_003995046.5 |
| D3 | 43351798 | 43351933 | refGene_gn=SMCHD1;refGene_tx=XM_003995046.5 |
| D3 | 43352594 | 43352734 | refGene_gn=SMCHD1;refGene_tx=XM_003995046.5 |
| D3 | 43355283 | 43355356 | refGene_gn=SMCHD1;refGene_tx=XM_003995046.5 |
| D3 | 43356689 | 43356786 | refGene_gn=SMCHD1;refGene_tx=XM_003995046.5 |
| D3 | 43362406 | 43362551 | refGene_gn=SMCHD1;refGene_tx=XM_003995046.5 |
| D3 | 43366616 | 43366736 | refGene_gn=SMCHD1;refGene_tx=XM_003995046.5 |
| D3 | 43366815 | 43366893 | refGene_gn=SMCHD1;refGene_tx=XM_003995046.5 |
| D3 | 43373585 | 43373699 | refGene_gn=SMCHD1;refGene_tx=XM_003995046.5 |
| D3 | 43374299 | 43374382 | refGene_gn=SMCHD1;refGene_tx=XM_003995046.5 |
| D3 | 43375101 | 43375208 | refGene_gn=SMCHD1;refGene_tx=XM_003995046.5 |
| D3 | 43375759 | 43375873 | refGene_gn=SMCHD1;refGene_tx=XM_003995046.5 |
| D3 | 43377411 | 43377606 | refGene_gn=SMCHD1;refGene_tx=XM_003995046.5 |
| D3 | 43378930 | 43379114 | refGene_gn=SMCHD1;refGene_tx=XM_003995046.5 |
| D3 | 43379192 | 43379313 | refGene_gn=SMCHD1;refGene_tx=XM_003995046.5 |
| D3 | 43381371 | 43381582 | refGene_gn=SMCHD1;refGene_tx=XM_003995046.5 |
| D3 | 43382270 | 43382361 | refGene_gn=SMCHD1;refGene_tx=XM_003995046.5 |
| D3 | 43384505 | 43384672 | refGene_gn=SMCHD1;refGene_tx=XM_003995046.5 |
| D3 | 43389479 | 43389599 | refGene_gn=SMCHD1;refGene_tx=XM_003995046.5 |
| D3 | 43389719 | 43389834 | refGene_gn=SMCHD1;refGene_tx=XM_003995046.5 |
| D3 | 43403384 | 43403515 | refGene_gn=SMCHD1;refGene_tx=XM_003995046.5 |
| D3 | 43404158 | 43404241 | refGene_gn=SMCHD1;refGene_tx=XM_003995046.5 |
| D3 | 43408813 | 43408975 | refGene_gn=SMCHD1;refGene_tx=XM_003995046.5 |
| D3 | 43409831 | 43409907 | refGene_gn=SMCHD1;refGene_tx=XM_003995046.5 |
| D3 | 43416820 | 43416997 | refGene_gn=SMCHD1;refGene_tx=XM_003995046.5 |
| D3 | 43461369 | 43461507 | refGene_gn=NDC80;refGene_tx=XM_003995047.4 |
| D3 | 43474992 | 43475095 | refGene_gn=NDC80;refGene_tx=XM_003995047.4 |
| D3 | 43477592 | 43477723 | refGene_gn=NDC80;refGene_tx=XM_003995047.4 |
| D3 | 43478962 | 43479055 | refGene_gn=NDC80;refGene_tx=XM_003995047.4 |
| D3 | 43493279 | 43493369 | refGene_gn=NDC80;refGene_tx=XM_003995047.4 |
| D3 | 43495678 | 43495831 | refGene_gn=NDC80;refGene_tx=XM_003995047.4 |
| D3 | 43502268 | 43502474 | refGene_gn=NDC80;refGene_tx=XM_003995047.4 |
| D3 | 43505311 | 43505456 | refGene_gn=NDC80;refGene_tx=XM_003995047.4 |
| D3 | 43506200 | 43506307 | refGene_gn=NDC80;refGene_tx=XM_003995047.4 |
| D3 | 43507525 | 43507619 | refGene_gn=NDC80;refGene_tx=XM_003995047.4 |
| D3 | 43510450 | 43510540 | refGene_gn=NDC80;refGene_tx=XM_003995047.4 |
| D3 | 43520193 | 43520296 | refGene_gn=NDC80;refGene_tx=XM_003995047.4 |
| D3 | 43520983 | 43521156 | refGene_gn=NDC80;refGene_tx=XM_003995047.4 |
| D3 | 43521234 | 43521358 | refGene_gn=NDC80;refGene_tx=XM_003995047.4 |
| D3 | 43523306 | 43523384 | refGene_gn=NDC80;refGene_tx=XM_003995047.4 |
| D3 | 43524554 | 43524655 | refGene_gn=NDC80;refGene_tx=XM_003995047.4 |
| D3 | 43527971 | 43528508 | refGene_gn=METTL4;refGene_tx=XM_019815053.2 |
| D3 | 43530317 | 43530380 | refGene_gn=METTL4;refGene_tx=XM_019815053.2 |

D3 43547356 43547650 refGene_gn=METTL4;refGene_tx=XM_019815053.2
D3 43547651 43547727 refGene_gn=METTL4;refGene_tx=XM_019815053.2
D3 43550157 43550227 refGene_gn=METTL4;refGene_tx=XM_019815053.2
D3 43565482 43565657 refGene_gn=METTL4;refGene_tx=XM_019815053.2
D3 43568424 43568531 refGene_gn=METTL4;refGene_tx=XM_019815053.2
D3 43568924 43569016 refGene_gn=METTL4;refGene_tx=XM_019815053.2
D3 43574593 43574739 refGene_gn=METTL4;refGene_tx=XM_019815053.2
D3 45225844 45226034
refGene_gn=ADCYAP1;refGene_tx=XM_003995051.4;refGene_gn=ADCYAP1;refGen
e_tx=XM_023241794.1
D3 45227143 45227242 refGene_gn=ADCYAP1;refGene_tx=XM_003995051.4
D3 45227715 45227847
refGene_gn=ADCYAP1;refGene_tx=XM_003995051.4;refGene_gn=ADCYAP1;refGen
e_tx=XM_023241794.1
D3 45230045 45230155
refGene_gn=ADCYAP1;refGene_tx=XM_003995051.4;refGene_gn=ADCYAP1;refGen
e_tx=XM_023241794.1
D3 45365555 45365820
refGene_gn=YES1;refGene_tx=XM_003995052.5;refGene_gn=YES1;refGene_tx=X
M_023241795.1;refGene_gn=YES1;refGene_tx=XM_023241796.1
D3 45368318 45368418
refGene_gn=YES1;refGene_tx=XM_003995052.5;refGene_gn=YES1;refGene_tx=X
M_023241795.1;refGene_gn=YES1;refGene_tx=XM_023241796.1
D3 45371459 45371558
refGene_gn=YES1;refGene_tx=XM_003995052.5;refGene_gn=YES1;refGene_tx=X
M_023241795.1;refGene_gn=YES1;refGene_tx=XM_023241796.1
D3 45372819 45372923
refGene_gn=YES1;refGene_tx=XM_003995052.5;refGene_gn=YES1;refGene_tx=X
M_023241795.1;refGene_gn=YES1;refGene_tx=XM_023241796.1
D3 45373009 45373159
refGene_gn=YES1;refGene_tx=XM_003995052.5;refGene_gn=YES1;refGene_tx=X
M_023241795.1;refGene_gn=YES1;refGene_tx=XM_023241796.1
D3 45374483 45374639
refGene_gn=YES1;refGene_tx=XM_003995052.5;refGene_gn=YES1;refGene_tx=X
M_023241795.1;refGene_gn=YES1;refGene_tx=XM_023241796.1
D3 45374815 45374995
refGene_gn=YES1;refGene_tx=XM_003995052.5;refGene_gn=YES1;refGene_tx=X
M_023241795.1;refGene_gn=YES1;refGene_tx=XM_023241796.1
D3 45379816 45379893
refGene_gn=YES1;refGene_tx=XM_003995052.5;refGene_gn=YES1;refGene_tx=X
M_023241795.1;refGene_gn=YES1;refGene_tx=XM_023241796.1
D3 45381422 45381576
refGene_gn=YES1;refGene_tx=XM_003995052.5;refGene_gn=YES1;refGene_tx=X
M_023241795.1;refGene_gn=YES1;refGene_tx=XM_023241796.1
D3 45383220 45383352
refGene_gn=YES1;refGene_tx=XM_003995052.5;refGene_gn=YES1;refGene_tx=X
M_023241795.1;refGene_gn=YES1;refGene_tx=XM_023241796.1
D3 45384640 45384849
refGene_gn=YES1;refGene_tx=XM_003995052.5;refGene_gn=YES1;refGene_tx=X
M_023241795.1;refGene_gn=YES1;refGene_tx=XM_023241796.1

D3 45398829 45398913
refGene_gn=ENOSF1;refGene_tx=XM_023241797.1;refGene_gn=ENOSF1;refGene_tx=XM_023241798.1
D3 45404327 45404436
refGene_gn=ENOSF1;refGene_tx=XM_003995053.4;refGene_gn=ENOSF1;refGene_tx=XM_023241797.1;refGene_gn=ENOSF1;refGene_tx=XM_023241798.1
D3 45409781 45409897
refGene_gn=ENOSF1;refGene_tx=XM_023241797.1;refGene_gn=ENOSF1;refGene_tx=XM_023241798.1
D3 45411813 45411900
refGene_gn=ENOSF1;refGene_tx=XM_003995053.4;refGene_gn=ENOSF1;refGene_tx=XM_023241797.1;refGene_gn=ENOSF1;refGene_tx=XM_023241798.1
D3 45412263 45412290
refGene_gn=ENOSF1;refGene_tx=XM_003995053.4;refGene_gn=ENOSF1;refGene_tx=XM_023241797.1;refGene_gn=ENOSF1;refGene_tx=XM_023241798.1
D3 45414403 45414476
refGene_gn=ENOSF1;refGene_tx=XM_003995053.4;refGene_gn=ENOSF1;refGene_tx=XM_023241797.1;refGene_gn=ENOSF1;refGene_tx=XM_023241798.1
D3 45414598 45414637
refGene_gn=ENOSF1;refGene_tx=XM_003995053.4;refGene_gn=ENOSF1;refGene_tx=XM_023241797.1;refGene_gn=ENOSF1;refGene_tx=XM_023241798.1
D3 45414999 45415082
refGene_gn=ENOSF1;refGene_tx=XM_003995053.4;refGene_gn=ENOSF1;refGene_tx=XM_023241797.1;refGene_gn=ENOSF1;refGene_tx=XM_023241798.1
D3 45415862 45415897
refGene_gn=ENOSF1;refGene_tx=XM_003995053.4;refGene_gn=ENOSF1;refGene_tx=XM_023241797.1;refGene_gn=ENOSF1;refGene_tx=XM_023241798.1
D3 45418001 45418089
refGene_gn=ENOSF1;refGene_tx=XM_003995053.4;refGene_gn=ENOSF1;refGene_tx=XM_023241797.1;refGene_gn=ENOSF1;refGene_tx=XM_023241798.1
D3 45419765 45419900
refGene_gn=ENOSF1;refGene_tx=XM_003995053.4;refGene_gn=ENOSF1;refGene_tx=XM_023241797.1;refGene_gn=ENOSF1;refGene_tx=XM_023241798.1
D3 45422641 45422683
refGene_gn=ENOSF1;refGene_tx=XM_003995053.4;refGene_gn=ENOSF1;refGene_tx=XM_023241797.1;refGene_gn=ENOSF1;refGene_tx=XM_023241798.1
D3 45423959 45424089
refGene_gn=ENOSF1;refGene_tx=XM_003995053.4;refGene_gn=ENOSF1;refGene_tx=XM_023241797.1;refGene_gn=ENOSF1;refGene_tx=XM_023241798.1
D3 45424322 45424422
refGene_gn=ENOSF1;refGene_tx=XM_003995053.4;refGene_gn=ENOSF1;refGene_tx=XM_023241797.1;refGene_gn=ENOSF1;refGene_tx=XM_023241798.1
D3 45426660 45426742
refGene_gn=ENOSF1;refGene_tx=XM_003995053.4;refGene_gn=ENOSF1;refGene_tx=XM_023241797.1
D3 45427323 45427425
refGene_gn=ENOSF1;refGene_tx=XM_003995053.4;refGene_gn=ENOSF1;refGene_tx=XM_023241797.1;refGene_gn=ENOSF1;refGene_tx=XM_023241798.1
D3 45428718 45428856 refGene_gn=TYMS;refGene_tx=XM_003995054.5
D3 45430153 45430225 refGene_gn=TYMS;refGene_tx=XM_003995054.5

D3 45430616 45430792 refGene_gn=TYMS;refGene_tx=XM_003995054.5
D3 45431597 45431699 refGene_gn=TYMS;refGene_tx=XM_003995054.5
D3 45433337 45433512 refGene_gn=TYMS;refGene_tx=XM_003995054.5
D3 45436047 45436121 refGene_gn=TYMS;refGene_tx=XM_003995054.5
D3 45437875 45438089 refGene_gn=TYMS;refGene_tx=XM_003995054.5
D3 45447590 45447770
refGene_gn=CLUL1;refGene_tx=XM_006938806.2;refGene_gn=CLUL1;refGene_tx
=XM_019815058.1
D3 45451597 45451812
refGene_gn=CLUL1;refGene_tx=XM_006938806.2;refGene_gn=CLUL1;refGene_tx
=XM_019815058.1
D3 45459887 45460025
refGene_gn=CLUL1;refGene_tx=XM_006938806.2;refGene_gn=CLUL1;refGene_tx
=XM_019815058.1
D3 45465673 45466106
refGene_gn=CLUL1;refGene_tx=XM_006938806.2;refGene_gn=CLUL1;refGene_tx
=XM_019815058.1
D3 45468320 45468488
refGene_gn=CLUL1;refGene_tx=XM_006938806.2;refGene_gn=CLUL1;refGene_tx
=XM_019815058.1
D3 45474979 45475128
refGene_gn=CLUL1;refGene_tx=XM_006938806.2;refGene_gn=CLUL1;refGene_tx
=XM_019815058.1
D3 45476781 45476887
refGene_gn=CLUL1;refGene_tx=XM_006938806.2;refGene_gn=CLUL1;refGene_tx
=XM_019815058.1
D3 45504662 45505181 refGene_gn=CETN1;refGene_tx=XM_003995056.4
D3 45556300 45556307 refGene_gn=COLEC12;refGene_tx=XM_023241800.1
D3 45575874 45575925 refGene_gn=COLEC12;refGene_tx=XM_023241800.1
D3 45703394 45703517 refGene_gn=COLEC12;refGene_tx=XM_023241800.1
D3 45710128 45710227 refGene_gn=COLEC12;refGene_tx=XM_023241800.1
D3 45711442 45712489 refGene_gn=COLEC12;refGene_tx=XM_023241800.1
D3 45725096 45725585 refGene_gn=COLEC12;refGene_tx=XM_023241800.1
D3 45726804 45726941 refGene_gn=COLEC12;refGene_tx=XM_023241800.1
D3 45728191 45728301 refGene_gn=COLEC12;refGene_tx=XM_023241800.1
D3 45738041 45738187 refGene_gn=COLEC12;refGene_tx=XM_023241800.1
D3 45750781 45750801 refGene_gn=COLEC12;refGene_tx=XM_023241800.1
D3 45816949 45817003 refGene_gn=THOC1;refGene_tx=XM_011288080.3
D3 45819427 45819501 refGene_gn=THOC1;refGene_tx=XM_011288080.3
D3 45819596 45819657 refGene_gn=THOC1;refGene_tx=XM_011288080.3
D3 45820868 45820935 refGene_gn=THOC1;refGene_tx=XM_011288080.3
D3 45824811 45824930 refGene_gn=THOC1;refGene_tx=XM_011288080.3
D3 45826063 45826112 refGene_gn=THOC1;refGene_tx=XM_011288080.3
D3 45826519 45826615 refGene_gn=THOC1;refGene_tx=XM_011288080.3
D3 45834331 45834414 refGene_gn=THOC1;refGene_tx=XM_011288080.3
D3 45837210 45837284 refGene_gn=THOC1;refGene_tx=XM_011288080.3
D3 45845828 45845937 refGene_gn=THOC1;refGene_tx=XM_011288080.3
D3 45848070 45848202 refGene_gn=THOC1;refGene_tx=XM_011288080.3
D3 45857074 45857175 refGene_gn=THOC1;refGene_tx=XM_011288080.3
D3 45858621 45858688 refGene_gn=THOC1;refGene_tx=XM_011288080.3

D3 45858886 45858937 refGene_gn=THOC1;refGene_tx=XM_011288080.3
D3 45859028 45859099 refGene_gn=THOC1;refGene_tx=XM_011288080.3
D3 45859953 45860049 refGene_gn=THOC1;refGene_tx=XM_011288080.3
D3 45860619 45860685 refGene_gn=THOC1;refGene_tx=XM_011288080.3
D3 45864632 45864716 refGene_gn=THOC1;refGene_tx=XM_011288080.3
D3 45867623 45867771 refGene_gn=THOC1;refGene_tx=XM_011288080.3
D3 45868503 45868579 refGene_gn=THOC1;refGene_tx=XM_011288080.3
D3 45869313 45869609 refGene_gn=THOC1;refGene_tx=XM_011288080.3
D3 45872564 45872716 refGene_gn=USP14;refGene_tx=XM_003995057.5
D3 45873358 45873466 refGene_gn=USP14;refGene_tx=XM_003995057.5
D3 45873857 45873918 refGene_gn=USP14;refGene_tx=XM_003995057.5
D3 45877825 45877954 refGene_gn=USP14;refGene_tx=XM_003995057.5
D3 45879540 45879633 refGene_gn=USP14;refGene_tx=XM_003995057.5
D3 45879781 45879847 refGene_gn=USP14;refGene_tx=XM_003995057.5
D3 45883266 45883381 refGene_gn=USP14;refGene_tx=XM_003995057.5
D3 45884971 45885057 refGene_gn=USP14;refGene_tx=XM_003995057.5
D3 45885429 45885510 refGene_gn=USP14;refGene_tx=XM_003995057.5
D3 45886438 45886569 refGene_gn=USP14;refGene_tx=XM_003995057.5
D3 45890350 45890409 refGene_gn=USP14;refGene_tx=XM_003995057.5
D3 45899170 45899274 refGene_gn=USP14;refGene_tx=XM_003995057.5
D3 45899990 45900095 refGene_gn=USP14;refGene_tx=XM_003995057.5
D3 45912776 45912809 refGene_gn=USP14;refGene_tx=XM_003995057.5
D3 45915935 45916081 refGene_gn=USP14;refGene_tx=XM_003995057.5
D3 45921316 45921332 refGene_gn=USP14;refGene_tx=XM_003995057.5
D3 45956990 45956997
refGene_gn=ROCK1;refGene_tx=XM_011288079.3;refGene_gn=ROCK1;refGene_tx
=XM_023241801.1;refGene_gn=ROCK1;refGene_tx=XM_023241802.1
D3 45958192 45958400
refGene_gn=ROCK1;refGene_tx=XM_011288079.3;refGene_gn=ROCK1;refGene_tx
=XM_023241801.1;refGene_gn=ROCK1;refGene_tx=XM_023241802.1
D3 45959322 45959584
refGene_gn=ROCK1;refGene_tx=XM_011288079.3;refGene_gn=ROCK1;refGene_tx
=XM_023241801.1;refGene_gn=ROCK1;refGene_tx=XM_023241802.1
D3 45959738 45959817
refGene_gn=ROCK1;refGene_tx=XM_011288079.3;refGene_gn=ROCK1;refGene_tx
=XM_023241801.1;refGene_gn=ROCK1;refGene_tx=XM_023241802.1
D3 45965028 45965117
refGene_gn=ROCK1;refGene_tx=XM_011288079.3;refGene_gn=ROCK1;refGene_tx
=XM_023241801.1;refGene_gn=ROCK1;refGene_tx=XM_023241802.1
D3 45965317 45965388
refGene_gn=ROCK1;refGene_tx=XM_011288079.3;refGene_gn=ROCK1;refGene_tx
=XM_023241801.1;refGene_gn=ROCK1;refGene_tx=XM_023241802.1
D3 45971944 45972104
refGene_gn=ROCK1;refGene_tx=XM_011288079.3;refGene_gn=ROCK1;refGene_tx
=XM_023241801.1;refGene_gn=ROCK1;refGene_tx=XM_023241802.1
D3 45972954 45973143
refGene_gn=ROCK1;refGene_tx=XM_011288079.3;refGene_gn=ROCK1;refGene_tx
=XM_023241801.1;refGene_gn=ROCK1;refGene_tx=XM_023241802.1
D3 45976073 45976162
refGene_gn=ROCK1;refGene_tx=XM_011288079.3;refGene_gn=ROCK1;refGene_tx

=XM_023241801.1;refGene_gn=ROCK1;refGene_tx=XM_023241802.1
D3 45976608 45976702
refGene_gn=ROCK1;refGene_tx=XM_011288079.3;refGene_gn=ROCK1;refGene_tx=XM_023241801.1;refGene_gn=ROCK1;refGene_tx=XM_023241802.1
D3 45977924 45978090
refGene_gn=ROCK1;refGene_tx=XM_011288079.3;refGene_gn=ROCK1;refGene_tx=XM_023241801.1;refGene_gn=ROCK1;refGene_tx=XM_023241802.1
D3 45986171 45986266
refGene_gn=ROCK1;refGene_tx=XM_011288079.3;refGene_gn=ROCK1;refGene_tx=XM_023241801.1;refGene_gn=ROCK1;refGene_tx=XM_023241802.1
D3 45989806 45989876
refGene_gn=ROCK1;refGene_tx=XM_011288079.3;refGene_gn=ROCK1;refGene_tx=XM_023241801.1;refGene_gn=ROCK1;refGene_tx=XM_023241802.1
D3 45991432 45991617
refGene_gn=ROCK1;refGene_tx=XM_011288079.3;refGene_gn=ROCK1;refGene_tx=XM_023241801.1;refGene_gn=ROCK1;refGene_tx=XM_023241802.1
D3 45993204 45993365
refGene_gn=ROCK1;refGene_tx=XM_011288079.3;refGene_gn=ROCK1;refGene_tx=XM_023241801.1;refGene_gn=ROCK1;refGene_tx=XM_023241802.1
D3 45997764 45997915
refGene_gn=ROCK1;refGene_tx=XM_011288079.3;refGene_gn=ROCK1;refGene_tx=XM_023241801.1;refGene_gn=ROCK1;refGene_tx=XM_023241802.1
D3 45999407 45999514
refGene_gn=ROCK1;refGene_tx=XM_011288079.3;refGene_gn=ROCK1;refGene_tx=XM_023241801.1;refGene_gn=ROCK1;refGene_tx=XM_023241802.1
D3 46018032 46018279
refGene_gn=ROCK1;refGene_tx=XM_011288079.3;refGene_gn=ROCK1;refGene_tx=XM_023241801.1;refGene_gn=ROCK1;refGene_tx=XM_023241802.1
D3 46018392 46018484
refGene_gn=ROCK1;refGene_tx=XM_011288079.3;refGene_gn=ROCK1;refGene_tx=XM_023241801.1;refGene_gn=ROCK1;refGene_tx=XM_023241802.1
D3 46019849 46019985
refGene_gn=ROCK1;refGene_tx=XM_011288079.3;refGene_gn=ROCK1;refGene_tx=XM_023241801.1;refGene_gn=ROCK1;refGene_tx=XM_023241802.1
D3 46025076 46025125
refGene_gn=ROCK1;refGene_tx=XM_011288079.3;refGene_gn=ROCK1;refGene_tx=XM_023241801.1;refGene_gn=ROCK1;refGene_tx=XM_023241802.1
D3 46027129 46027218
refGene_gn=ROCK1;refGene_tx=XM_011288079.3;refGene_gn=ROCK1;refGene_tx=XM_023241801.1;refGene_gn=ROCK1;refGene_tx=XM_023241802.1
D3 46032716 46032777
refGene_gn=ROCK1;refGene_tx=XM_011288079.3;refGene_gn=ROCK1;refGene_tx=XM_023241801.1;refGene_gn=ROCK1;refGene_tx=XM_023241802.1
D3 46036318 46036478
refGene_gn=ROCK1;refGene_tx=XM_011288079.3;refGene_gn=ROCK1;refGene_tx=XM_023241801.1;refGene_gn=ROCK1;refGene_tx=XM_023241802.1
D3 46047508 46047600
refGene_gn=ROCK1;refGene_tx=XM_011288079.3;refGene_gn=ROCK1;refGene_tx=XM_023241801.1;refGene_gn=ROCK1;refGene_tx=XM_023241802.1
D3 46051005 46051144

refGene_gn=ROCK1;refGene_tx=XM_011288079.3;refGene_gn=ROCK1;refGene_tx=XM_023241801.1;refGene_gn=ROCK1;refGene_tx=XM_023241802.1
D3 46051478 46051623
refGene_gn=ROCK1;refGene_tx=XM_011288079.3;refGene_gn=ROCK1;refGene_tx=XM_023241801.1;refGene_gn=ROCK1;refGene_tx=XM_023241802.1
D3 46052867 46052952
refGene_gn=ROCK1;refGene_tx=XM_011288079.3;refGene_gn=ROCK1;refGene_tx=XM_023241801.1;refGene_gn=ROCK1;refGene_tx=XM_023241802.1
D3 46053803 46053979
refGene_gn=ROCK1;refGene_tx=XM_011288079.3;refGene_gn=ROCK1;refGene_tx=XM_023241801.1;refGene_gn=ROCK1;refGene_tx=XM_023241802.1
D3 46056628 46056766
refGene_gn=ROCK1;refGene_tx=XM_011288079.3;refGene_gn=ROCK1;refGene_tx=XM_023241801.1;refGene_gn=ROCK1;refGene_tx=XM_023241802.1
D3 46057472 46057573
refGene_gn=ROCK1;refGene_tx=XM_011288079.3;refGene_gn=ROCK1;refGene_tx=XM_023241801.1;refGene_gn=ROCK1;refGene_tx=XM_023241802.1
D3 46076559 46076641
refGene_gn=ROCK1;refGene_tx=XM_011288079.3;refGene_gn=ROCK1;refGene_tx=XM_023241801.1;refGene_gn=ROCK1;refGene_tx=XM_023241802.1
D3 46114022 46114115
refGene_gn=ROCK1;refGene_tx=XM_011288079.3;refGene_gn=ROCK1;refGene_tx=XM_023241801.1;refGene_gn=ROCK1;refGene_tx=XM_023241802.1
D3 46361627 46361784
refGene_gn=GREB1L;refGene_tx=XM_023241804.1;refGene_gn=GREB1L;refGene_tx=XM_023241805.1;refGene_gn=GREB1L;refGene_tx=XM_023241806.1;refGene_gn=GREB1L;refGene_tx=XM_023241807.1;refGene_gn=GREB1L;refGene_tx=XM_023241808.1
D3 46361994 46362192
refGene_gn=GREB1L;refGene_tx=XM_023241804.1;refGene_gn=GREB1L;refGene_tx=XM_023241805.1;refGene_gn=GREB1L;refGene_tx=XM_023241806.1;refGene_gn=GREB1L;refGene_tx=XM_023241807.1;refGene_gn=GREB1L;refGene_tx=XM_023241808.1
D3 46373018 46373195
refGene_gn=GREB1L;refGene_tx=XM_023241804.1;refGene_gn=GREB1L;refGene_tx=XM_023241805.1;refGene_gn=GREB1L;refGene_tx=XM_023241806.1;refGene_gn=GREB1L;refGene_tx=XM_023241807.1;refGene_gn=GREB1L;refGene_tx=XM_023241808.1
D3 46378655 46378832
refGene_gn=GREB1L;refGene_tx=XM_023241804.1;refGene_gn=GREB1L;refGene_tx=XM_023241805.1;refGene_gn=GREB1L;refGene_tx=XM_023241806.1;refGene_gn=GREB1L;refGene_tx=XM_023241807.1;refGene_gn=GREB1L;refGene_tx=XM_023241808.1
D3 46381610 46381733
refGene_gn=GREB1L;refGene_tx=XM_023241804.1;refGene_gn=GREB1L;refGene_tx=XM_023241805.1;refGene_gn=GREB1L;refGene_tx=XM_023241806.1;refGene_gn=GREB1L;refGene_tx=XM_023241807.1;refGene_gn=GREB1L;refGene_tx=XM_023241808.1
D3 46420572 46420701
refGene_gn=GREB1L;refGene_tx=XM_023241804.1;refGene_gn=GREB1L;refGene_

tx=XM_023241805.1;refGene_gn=GREB1L;refGene_tx=XM_023241806.1;refGene_gn=GREB1L;refGene_tx=XM_023241807.1;refGene_gn=GREB1L;refGene_tx=XM_023241808.1
D3 46424060 46424177
refGene_gn=GREB1L;refGene_tx=XM_023241804.1;refGene_gn=GREB1L;refGene_tx=XM_023241805.1;refGene_gn=GREB1L;refGene_tx=XM_023241806.1;refGene_gn=GREB1L;refGene_tx=XM_023241807.1;refGene_gn=GREB1L;refGene_tx=XM_023241808.1
D3 46424776 46424896
refGene_gn=GREB1L;refGene_tx=XM_023241804.1;refGene_gn=GREB1L;refGene_tx=XM_023241805.1;refGene_gn=GREB1L;refGene_tx=XM_023241806.1;refGene_gn=GREB1L;refGene_tx=XM_023241807.1;refGene_gn=GREB1L;refGene_tx=XM_023241808.1
D3 46425923 46426061
refGene_gn=GREB1L;refGene_tx=XM_023241804.1;refGene_gn=GREB1L;refGene_tx=XM_023241805.1;refGene_gn=GREB1L;refGene_tx=XM_023241806.1;refGene_gn=GREB1L;refGene_tx=XM_023241807.1;refGene_gn=GREB1L;refGene_tx=XM_023241808.1
D3 46427695 46427881
refGene_gn=GREB1L;refGene_tx=XM_023241804.1;refGene_gn=GREB1L;refGene_tx=XM_023241805.1;refGene_gn=GREB1L;refGene_tx=XM_023241806.1;refGene_gn=GREB1L;refGene_tx=XM_023241807.1;refGene_gn=GREB1L;refGene_tx=XM_023241808.1
D3 46431766 46432093
refGene_gn=GREB1L;refGene_tx=XM_023241804.1;refGene_gn=GREB1L;refGene_tx=XM_023241805.1;refGene_gn=GREB1L;refGene_tx=XM_023241806.1;refGene_gn=GREB1L;refGene_tx=XM_023241807.1;refGene_gn=GREB1L;refGene_tx=XM_023241808.1
D3 46433134 46433263
refGene_gn=GREB1L;refGene_tx=XM_023241804.1;refGene_gn=GREB1L;refGene_tx=XM_023241805.1;refGene_gn=GREB1L;refGene_tx=XM_023241806.1;refGene_gn=GREB1L;refGene_tx=XM_023241807.1;refGene_gn=GREB1L;refGene_tx=XM_023241808.1
D3 46433846 46433981
refGene_gn=GREB1L;refGene_tx=XM_023241804.1;refGene_gn=GREB1L;refGene_tx=XM_023241805.1;refGene_gn=GREB1L;refGene_tx=XM_023241806.1;refGene_gn=GREB1L;refGene_tx=XM_023241807.1;refGene_gn=GREB1L;refGene_tx=XM_023241808.1
D3 46436178 46436376
refGene_gn=GREB1L;refGene_tx=XM_023241804.1;refGene_gn=GREB1L;refGene_tx=XM_023241805.1;refGene_gn=GREB1L;refGene_tx=XM_023241806.1;refGene_gn=GREB1L;refGene_tx=XM_023241807.1;refGene_gn=GREB1L;refGene_tx=XM_023241808.1
D3 46459467 46459648
refGene_gn=GREB1L;refGene_tx=XM_023241804.1;refGene_gn=GREB1L;refGene_tx=XM_023241805.1;refGene_gn=GREB1L;refGene_tx=XM_023241806.1;refGene_gn=GREB1L;refGene_tx=XM_023241807.1;refGene_gn=GREB1L;refGene_tx=XM_023241808.1
D3 46462116 46462309
refGene_gn=GREB1L;refGene_tx=XM_023241804.1;refGene_gn=GREB1L;refGene_

tx=XM_023241805.1;refGene_gn=GREB1L;refGene_tx=XM_023241806.1;refGene_gn=GREB1L;refGene_tx=XM_023241807.1;refGene_gn=GREB1L;refGene_tx=XM_023241808.1
D3	46477410	46477544
refGene_gn=GREB1L;refGene_tx=XM_023241804.1;refGene_gn=GREB1L;refGene_tx=XM_023241805.1;refGene_gn=GREB1L;refGene_tx=XM_023241806.1;refGene_gn=GREB1L;refGene_tx=XM_023241807.1;refGene_gn=GREB1L;refGene_tx=XM_023241808.1
D3	46480444	46480784
refGene_gn=GREB1L;refGene_tx=XM_023241804.1;refGene_gn=GREB1L;refGene_tx=XM_023241805.1;refGene_gn=GREB1L;refGene_tx=XM_023241806.1;refGene_gn=GREB1L;refGene_tx=XM_023241807.1;refGene_gn=GREB1L;refGene_tx=XM_023241808.1
D3	46486235	46486351
refGene_gn=GREB1L;refGene_tx=XM_023241804.1;refGene_gn=GREB1L;refGene_tx=XM_023241805.1;refGene_gn=GREB1L;refGene_tx=XM_023241806.1;refGene_gn=GREB1L;refGene_tx=XM_023241807.1;refGene_gn=GREB1L;refGene_tx=XM_023241808.1
D3	46486942	46487187
refGene_gn=GREB1L;refGene_tx=XM_023241804.1;refGene_gn=GREB1L;refGene_tx=XM_023241805.1;refGene_gn=GREB1L;refGene_tx=XM_023241806.1;refGene_gn=GREB1L;refGene_tx=XM_023241807.1;refGene_gn=GREB1L;refGene_tx=XM_023241808.1
D3	46490173	46490757
refGene_gn=GREB1L;refGene_tx=XM_023241804.1;refGene_gn=GREB1L;refGene_tx=XM_023241805.1;refGene_gn=GREB1L;refGene_tx=XM_023241806.1;refGene_gn=GREB1L;refGene_tx=XM_023241807.1;refGene_gn=GREB1L;refGene_tx=XM_023241808.1
D3	46490950	46491053
refGene_gn=GREB1L;refGene_tx=XM_023241804.1;refGene_gn=GREB1L;refGene_tx=XM_023241805.1;refGene_gn=GREB1L;refGene_tx=XM_023241806.1;refGene_gn=GREB1L;refGene_tx=XM_023241807.1;refGene_gn=GREB1L;refGene_tx=XM_023241808.1
D3	46493934	46494090
refGene_gn=GREB1L;refGene_tx=XM_023241804.1;refGene_gn=GREB1L;refGene_tx=XM_023241805.1;refGene_gn=GREB1L;refGene_tx=XM_023241806.1;refGene_gn=GREB1L;refGene_tx=XM_023241807.1;refGene_gn=GREB1L;refGene_tx=XM_023241808.1
D3	46494362	46494502
refGene_gn=GREB1L;refGene_tx=XM_023241804.1;refGene_gn=GREB1L;refGene_tx=XM_023241805.1;refGene_gn=GREB1L;refGene_tx=XM_023241806.1;refGene_gn=GREB1L;refGene_tx=XM_023241807.1;refGene_gn=GREB1L;refGene_tx=XM_023241808.1
D3	46496644	46496806
refGene_gn=GREB1L;refGene_tx=XM_023241804.1;refGene_gn=GREB1L;refGene_tx=XM_023241805.1;refGene_gn=GREB1L;refGene_tx=XM_023241806.1;refGene_gn=GREB1L;refGene_tx=XM_023241807.1;refGene_gn=GREB1L;refGene_tx=XM_023241808.1
D3	46496956	46497164
refGene_gn=GREB1L;refGene_tx=XM_023241804.1;refGene_gn=GREB1L;refGene_

tx=XM_023241805.1;refGene_gn=GREB1L;refGene_tx=XM_023241806.1;refGene_gn=GREB1L;refGene_tx=XM_023241807.1;refGene_gn=GREB1L;refGene_tx=XM_023241808.1
D3 46503060 46503226
refGene_gn=GREB1L;refGene_tx=XM_023241804.1;refGene_gn=GREB1L;refGene_tx=XM_023241805.1;refGene_gn=GREB1L;refGene_tx=XM_023241806.1;refGene_gn=GREB1L;refGene_tx=XM_023241807.1;refGene_gn=GREB1L;refGene_tx=XM_023241808.1
D3 46504137 46504365
refGene_gn=GREB1L;refGene_tx=XM_023241804.1;refGene_gn=GREB1L;refGene_tx=XM_023241805.1;refGene_gn=GREB1L;refGene_tx=XM_023241806.1;refGene_gn=GREB1L;refGene_tx=XM_023241807.1;refGene_gn=GREB1L;refGene_tx=XM_023241808.1
D3 46505498 46505640
refGene_gn=GREB1L;refGene_tx=XM_023241804.1;refGene_gn=GREB1L;refGene_tx=XM_023241805.1;refGene_gn=GREB1L;refGene_tx=XM_023241806.1;refGene_gn=GREB1L;refGene_tx=XM_023241807.1;refGene_gn=GREB1L;refGene_tx=XM_023241808.1
D3 46506478 46506679
refGene_gn=GREB1L;refGene_tx=XM_023241804.1;refGene_gn=GREB1L;refGene_tx=XM_023241805.1;refGene_gn=GREB1L;refGene_tx=XM_023241806.1;refGene_gn=GREB1L;refGene_tx=XM_023241807.1;refGene_gn=GREB1L;refGene_tx=XM_023241808.1
D3 46508444 46508583
refGene_gn=GREB1L;refGene_tx=XM_023241804.1;refGene_gn=GREB1L;refGene_tx=XM_023241805.1;refGene_gn=GREB1L;refGene_tx=XM_023241806.1;refGene_gn=GREB1L;refGene_tx=XM_023241807.1;refGene_gn=GREB1L;refGene_tx=XM_023241808.1
D3 46510142 46510306
refGene_gn=GREB1L;refGene_tx=XM_023241804.1;refGene_gn=GREB1L;refGene_tx=XM_023241805.1;refGene_gn=GREB1L;refGene_tx=XM_023241806.1;refGene_gn=GREB1L;refGene_tx=XM_023241807.1;refGene_gn=GREB1L;refGene_tx=XM_023241808.1
D3 46517565 46517713
refGene_gn=ESCO1;refGene_tx=XM_003995061.4;refGene_gn=ESCO1;refGene_tx=XM_011288077.2;refGene_gn=ESCO1;refGene_tx=XM_019815059.2
D3 46522843 46523031
refGene_gn=ESCO1;refGene_tx=XM_003995061.4;refGene_gn=ESCO1;refGene_tx=XM_011288077.2;refGene_gn=ESCO1;refGene_tx=XM_019815059.2
D3 46524813 46524957
refGene_gn=ESCO1;refGene_tx=XM_003995061.4;refGene_gn=ESCO1;refGene_tx=XM_011288077.2;refGene_gn=ESCO1;refGene_tx=XM_019815059.2
D3 46528036 46528126
refGene_gn=ESCO1;refGene_tx=XM_003995061.4;refGene_gn=ESCO1;refGene_tx=XM_011288077.2;refGene_gn=ESCO1;refGene_tx=XM_019815059.2
D3 46541591 46541723
refGene_gn=ESCO1;refGene_tx=XM_003995061.4;refGene_gn=ESCO1;refGene_tx=XM_011288077.2;refGene_gn=ESCO1;refGene_tx=XM_019815059.2
D3 46544817 46544932
refGene_gn=ESCO1;refGene_tx=XM_003995061.4;refGene_gn=ESCO1;refGene_tx

=XM_011288077.2;refGene_gn=ESCO1;refGene_tx=XM_019815059.2
D3 46546222 46546283
refGene_gn=ESCO1;refGene_tx=XM_003995061.4;refGene_gn=ESCO1;refGene_tx
=XM_011288077.2;refGene_gn=ESCO1;refGene_tx=XM_019815059.2
D3 46548807 46548922
refGene_gn=ESCO1;refGene_tx=XM_003995061.4;refGene_gn=ESCO1;refGene_tx
=XM_011288077.2;refGene_gn=ESCO1;refGene_tx=XM_019815059.2
D3 46553422 46554973
refGene_gn=ESCO1;refGene_tx=XM_003995061.4;refGene_gn=ESCO1;refGene_tx
=XM_011288077.2;refGene_gn=ESCO1;refGene_tx=XM_019815059.2
D3 46598364 46598378
refGene_gn=SNRPD1;refGene_tx=XM_003995062.4;refGene_gn=SNRPD1;refGene_
tx=XM_019815062.2
D3 46606230 46606307
refGene_gn=SNRPD1;refGene_tx=XM_003995062.4;refGene_gn=SNRPD1;refGene_
tx=XM_006938808.4;refGene_gn=SNRPD1;refGene_tx=XM_019815062.2
D3 46608525 46608717
refGene_gn=SNRPD1;refGene_tx=XM_003995062.4;refGene_gn=SNRPD1;refGene_
tx=XM_006938808.4;refGene_gn=SNRPD1;refGene_tx=XM_019815062.2
D3 46611621 46611698
refGene_gn=SNRPD1;refGene_tx=XM_003995062.4;refGene_gn=SNRPD1;refGene_
tx=XM_006938808.4;refGene_gn=SNRPD1;refGene_tx=XM_019815062.2
D3 46642568 46642747 refGene_gn=ABHD3;refGene_tx=XM_011288076.2
D3 46646265 46646431 refGene_gn=ABHD3;refGene_tx=XM_011288076.2
D3 46646510 46646559 refGene_gn=ABHD3;refGene_tx=XM_011288076.2
D3 46648099 46648273 refGene_gn=ABHD3;refGene_tx=XM_011288076.2
D3 46652381 46652494 refGene_gn=ABHD3;refGene_tx=XM_011288076.2
D3 46665705 46665751 refGene_gn=ABHD3;refGene_tx=XM_011288076.2
D3 46678049 46678232 refGene_gn=ABHD3;refGene_tx=XM_011288076.2
D3 46679513 46679677 refGene_gn=ABHD3;refGene_tx=XM_011288076.2
D3 46680220 46680382 refGene_gn=ABHD3;refGene_tx=XM_011288076.2
D3 46710541 46710770 refGene_gn=MIB1;refGene_tx=XM_011288075.3
D3 46732101 46732273 refGene_gn=MIB1;refGene_tx=XM_011288075.3
D3 46734078 46734208 refGene_gn=MIB1;refGene_tx=XM_011288075.3
D3 46735912 46736017 refGene_gn=MIB1;refGene_tx=XM_011288075.3
D3 46738835 46738902 refGene_gn=MIB1;refGene_tx=XM_011288075.3
D3 46740666 46740871 refGene_gn=MIB1;refGene_tx=XM_011288075.3
D3 46750058 46750242 refGene_gn=MIB1;refGene_tx=XM_011288075.3
D3 46759015 46759160 refGene_gn=MIB1;refGene_tx=XM_011288075.3
D3 46760693 46760827 refGene_gn=MIB1;refGene_tx=XM_011288075.3
D3 46764506 46764614 refGene_gn=MIB1;refGene_tx=XM_011288075.3
D3 46780975 46781173 refGene_gn=MIB1;refGene_tx=XM_011288075.3
D3 46783230 46783382 refGene_gn=MIB1;refGene_tx=XM_011288075.3
D3 46805785 46805918 refGene_gn=MIB1;refGene_tx=XM_011288075.3
D3 46809267 46809354 refGene_gn=MIB1;refGene_tx=XM_011288075.3
D3 46810965 46811127 refGene_gn=MIB1;refGene_tx=XM_011288075.3
D3 46813183 46813365 refGene_gn=MIB1;refGene_tx=XM_011288075.3
D3 46815528 46815721 refGene_gn=MIB1;refGene_tx=XM_011288075.3
D3 46819448 46819527 refGene_gn=MIB1;refGene_tx=XM_011288075.3
D3 46825376 46825490 refGene_gn=MIB1;refGene_tx=XM_011288075.3

D3 46827902 46828003 refGene_gn=MIB1;refGene_tx=XM_011288075.3
D3 46832162 46832303 refGene_gn=MIB1;refGene_tx=XM_011288075.3
D3 47049035 47050164
refGene_gn=GATA6;refGene_tx=XM_023241809.1;refGene_gn=GATA6;refGene_tx=XM_023241810.1;refGene_gn=GATA6;refGene_tx=XM_023241811.1
D3 47054698 47054865
refGene_gn=GATA6;refGene_tx=XM_023241809.1;refGene_gn=GATA6;refGene_tx=XM_023241810.1;refGene_gn=GATA6;refGene_tx=XM_023241811.1
D3 47058445 47058571
refGene_gn=GATA6;refGene_tx=XM_023241809.1;refGene_gn=GATA6;refGene_tx=XM_023241810.1;refGene_gn=GATA6;refGene_tx=XM_023241811.1
D3 47059431 47059519
refGene_gn=GATA6;refGene_tx=XM_023241809.1;refGene_gn=GATA6;refGene_tx=XM_023241810.1;refGene_gn=GATA6;refGene_tx=XM_023241811.1
D3 47059612 47059716
refGene_gn=GATA6;refGene_tx=XM_023241809.1;refGene_gn=GATA6;refGene_tx=XM_023241810.1;refGene_gn=GATA6;refGene_tx=XM_023241811.1
D3 47077912 47078080
refGene_gn=GATA6;refGene_tx=XM_023241809.1;refGene_gn=GATA6;refGene_tx=XM_023241810.1
D3 47171064 47171190 refGene_gn=GATA6;refGene_tx=XM_023241811.1
D3 47707412 47707521
refGene_gn=RBBP8;refGene_tx=XM_011288074.3;refGene_gn=RBBP8;refGene_tx=XM_019815064.2;refGene_gn=RBBP8;refGene_tx=XM_019815065.2;refGene_gn=RBBP8;refGene_tx=XM_019815066.2
D3 47722498 47722541
refGene_gn=RBBP8;refGene_tx=XM_011288074.3;refGene_gn=RBBP8;refGene_tx=XM_019815064.2;refGene_gn=RBBP8;refGene_tx=XM_019815065.2;refGene_gn=RBBP8;refGene_tx=XM_019815066.2
D3 47731203 47731299
refGene_gn=RBBP8;refGene_tx=XM_011288074.3;refGene_gn=RBBP8;refGene_tx=XM_019815064.2;refGene_gn=RBBP8;refGene_tx=XM_019815065.2;refGene_gn=RBBP8;refGene_tx=XM_019815066.2
D3 47742125 47742238
refGene_gn=RBBP8;refGene_tx=XM_011288074.3;refGene_gn=RBBP8;refGene_tx=XM_019815064.2;refGene_gn=RBBP8;refGene_tx=XM_019815065.2;refGene_gn=RBBP8;refGene_tx=XM_019815066.2
D3 47749605 47749672
refGene_gn=RBBP8;refGene_tx=XM_011288074.3;refGene_gn=RBBP8;refGene_tx=XM_019815064.2;refGene_gn=RBBP8;refGene_tx=XM_019815065.2;refGene_gn=RBBP8;refGene_tx=XM_019815066.2
D3 47761174 47761350
refGene_gn=RBBP8;refGene_tx=XM_011288074.3;refGene_gn=RBBP8;refGene_tx=XM_019815064.2;refGene_gn=RBBP8;refGene_tx=XM_019815065.2;refGene_gn=RBBP8;refGene_tx=XM_019815066.2
D3 47764710 47764815
refGene_gn=RBBP8;refGene_tx=XM_011288074.3;refGene_gn=RBBP8;refGene_tx=XM_019815064.2;refGene_gn=RBBP8;refGene_tx=XM_019815065.2;refGene_gn=RBBP8;refGene_tx=XM_019815066.2
D3 47769013 47769111

refGene_gn=RBBP8;refGene_tx=XM_011288074.3;refGene_gn=RBBP8;refGene_tx=XM_019815064.2;refGene_gn=RBBP8;refGene_tx=XM_019815065.2;refGene_gn=RBBP8;refGene_tx=XM_019815066.2
D3 47770940 47771047
refGene_gn=RBBP8;refGene_tx=XM_011288074.3;refGene_gn=RBBP8;refGene_tx=XM_019815064.2;refGene_gn=RBBP8;refGene_tx=XM_019815065.2;refGene_gn=RBBP8;refGene_tx=XM_019815066.2
D3 47772595 47773484
refGene_gn=RBBP8;refGene_tx=XM_011288074.3;refGene_gn=RBBP8;refGene_tx=XM_019815064.2;refGene_gn=RBBP8;refGene_tx=XM_019815065.2;refGene_gn=RBBP8;refGene_tx=XM_019815066.2
D3 47773566 47773690
refGene_gn=RBBP8;refGene_tx=XM_011288074.3;refGene_gn=RBBP8;refGene_tx=XM_019815064.2;refGene_gn=RBBP8;refGene_tx=XM_019815065.2;refGene_gn=RBBP8;refGene_tx=XM_019815066.2
D3 47777686 47777775
refGene_gn=RBBP8;refGene_tx=XM_011288074.3;refGene_gn=RBBP8;refGene_tx=XM_019815064.2;refGene_gn=RBBP8;refGene_tx=XM_019815065.2;refGene_gn=RBBP8;refGene_tx=XM_019815066.2
D3 47778414 47778532
refGene_gn=RBBP8;refGene_tx=XM_011288074.3;refGene_gn=RBBP8;refGene_tx=XM_019815064.2;refGene_gn=RBBP8;refGene_tx=XM_019815065.2;refGene_gn=RBBP8;refGene_tx=XM_019815066.2
D3 47782062 47782206
refGene_gn=RBBP8;refGene_tx=XM_011288074.3;refGene_gn=RBBP8;refGene_tx=XM_019815064.2;refGene_gn=RBBP8;refGene_tx=XM_019815065.2;refGene_gn=RBBP8;refGene_tx=XM_019815066.2
D3 47786066 47786136
refGene_gn=RBBP8;refGene_tx=XM_011288074.3;refGene_gn=RBBP8;refGene_tx=XM_019815064.2;refGene_gn=RBBP8;refGene_tx=XM_019815065.2;refGene_gn=RBBP8;refGene_tx=XM_019815066.2
D3 47790222 47790319
refGene_gn=RBBP8;refGene_tx=XM_011288074.3;refGene_gn=RBBP8;refGene_tx=XM_019815064.2;refGene_gn=RBBP8;refGene_tx=XM_019815065.2;refGene_gn=RBBP8;refGene_tx=XM_019815066.2
D3 47793758 47793900
refGene_gn=RBBP8;refGene_tx=XM_011288074.3;refGene_gn=RBBP8;refGene_tx=XM_019815064.2;refGene_gn=RBBP8;refGene_tx=XM_019815065.2;refGene_gn=RBBP8;refGene_tx=XM_019815066.2
D3 47795753 47795851
refGene_gn=RBBP8;refGene_tx=XM_011288074.3;refGene_gn=RBBP8;refGene_tx=XM_019815064.2;refGene_gn=RBBP8;refGene_tx=XM_019815065.2;refGene_gn=RBBP8;refGene_tx=XM_019815066.2
D3 47897026 47897883 refGene_gn=CABLES1;refGene_tx=XM_023241812.1
D3 47944651 47944723 refGene_gn=CABLES1;refGene_tx=XM_023241812.1
D3 47950104 47950197
refGene_gn=CABLES1;refGene_tx=XM_006938812.4;refGene_gn=CABLES1;refGene_tx=XM_023241812.1
D3 47969007 47969085
refGene_gn=CABLES1;refGene_tx=XM_006938812.4;refGene_gn=CABLES1;refGen

e_tx=XM_023241812.1
D3 47987931 47988028
refGene_gn=CABLES1;refGene_tx=XM_006938812.4;refGene_gn=CABLES1;refGene_tx=XM_023241812.1
D3 47989333 47989490
refGene_gn=CABLES1;refGene_tx=XM_006938812.4;refGene_gn=CABLES1;refGene_tx=XM_023241812.1
D3 47990639 47990743
refGene_gn=CABLES1;refGene_tx=XM_006938812.4;refGene_gn=CABLES1;refGene_tx=XM_023241812.1
D3 48005288 48005395
refGene_gn=CABLES1;refGene_tx=XM_006938812.4;refGene_gn=CABLES1;refGene_tx=XM_023241812.1
D3 48006605 48006813
refGene_gn=CABLES1;refGene_tx=XM_006938812.4;refGene_gn=CABLES1;refGene_tx=XM_023241812.1
D3 48010018 48010159
refGene_gn=CABLES1;refGene_tx=XM_006938812.4;refGene_gn=CABLES1;refGene_tx=XM_023241812.1
D3 48042825 48042886 refGene_gn=TMEM241;refGene_tx=XM_019815072.2
D3 48050131 48050196 refGene_gn=TMEM241;refGene_tx=XM_019815072.2
D3 48081272 48081366 refGene_gn=TMEM241;refGene_tx=XM_019815072.2
D3 48085863 48085933 refGene_gn=TMEM241;refGene_tx=XM_019815072.2
D3 48097580 48097615 refGene_gn=TMEM241;refGene_tx=XM_019815072.2
D3 48099950 48099986 refGene_gn=TMEM241;refGene_tx=XM_019815072.2
D3 48101848 48101894 refGene_gn=TMEM241;refGene_tx=XM_019815072.2
D3 48102780 48102829 refGene_gn=TMEM241;refGene_tx=XM_019815072.2
D3 48104666 48104718 refGene_gn=TMEM241;refGene_tx=XM_019815072.2
D3 48106499 48106525 refGene_gn=TMEM241;refGene_tx=XM_019815072.2
D3 48116114 48116192 refGene_gn=TMEM241;refGene_tx=XM_019815072.2
D3 48131715 48131797 refGene_gn=TMEM241;refGene_tx=XM_019815072.2
D3 48147557 48147635 refGene_gn=TMEM241;refGene_tx=XM_019815072.2
D3 48150646 48150693 refGene_gn=TMEM241;refGene_tx=XM_019815072.2
D3 48156315 48156387 refGene_gn=TMEM241;refGene_tx=XM_019815072.2
D3 48166153 48166216 refGene_gn=RIOK3;refGene_tx=XM_003995071.5
D3 48174112 48174228 refGene_gn=RIOK3;refGene_tx=XM_003995071.5
D3 48174813 48174959 refGene_gn=RIOK3;refGene_tx=XM_003995071.5
D3 48175053 48175161 refGene_gn=RIOK3;refGene_tx=XM_003995071.5
D3 48175378 48175488 refGene_gn=RIOK3;refGene_tx=XM_003995071.5
D3 48177062 48177206 refGene_gn=RIOK3;refGene_tx=XM_003995071.5
D3 48177963 48178091 refGene_gn=RIOK3;refGene_tx=XM_003995071.5
D3 48185200 48185398 refGene_gn=RIOK3;refGene_tx=XM_003995071.5
D3 48187060 48187220 refGene_gn=RIOK3;refGene_tx=XM_003995071.5
D3 48188616 48188697 refGene_gn=RIOK3;refGene_tx=XM_003995071.5
D3 48188783 48188873 refGene_gn=RIOK3;refGene_tx=XM_003995071.5
D3 48190444 48190552 refGene_gn=RIOK3;refGene_tx=XM_003995071.5
D3 48191489 48191597 refGene_gn=RIOK3;refGene_tx=XM_003995071.5
D3 48209199 48209301 refGene_gn=RMC1;refGene_tx=XM_023241813.1
D3 48210040 48210117 refGene_gn=RMC1;refGene_tx=XM_023241813.1
D3 48212219 48212304

refGene_gn=RMC1;refGene_tx=XM_023241813.1;refGene_gn=RMC1;refGene_tx=XM_023241814.1
D3 48213707 48213764
refGene_gn=RMC1;refGene_tx=XM_023241813.1;refGene_gn=RMC1;refGene_tx=XM_023241814.1
D3 48215092 48215179
refGene_gn=RMC1;refGene_tx=XM_023241813.1;refGene_gn=RMC1;refGene_tx=XM_023241814.1
D3 48217094 48217235
refGene_gn=RMC1;refGene_tx=XM_023241813.1;refGene_gn=RMC1;refGene_tx=XM_023241814.1
D3 48217564 48217668
refGene_gn=RMC1;refGene_tx=XM_023241813.1;refGene_gn=RMC1;refGene_tx=XM_023241814.1
D3 48218951 48219041
refGene_gn=RMC1;refGene_tx=XM_023241813.1;refGene_gn=RMC1;refGene_tx=XM_023241814.1
D3 48219129 48219235
refGene_gn=RMC1;refGene_tx=XM_023241813.1;refGene_gn=RMC1;refGene_tx=XM_023241814.1
D3 48220213 48220325
refGene_gn=RMC1;refGene_tx=XM_023241813.1;refGene_gn=RMC1;refGene_tx=XM_023241814.1
D3 48221724 48221769
refGene_gn=RMC1;refGene_tx=XM_023241813.1;refGene_gn=RMC1;refGene_tx=XM_023241814.1
D3 48222019 48222073
refGene_gn=RMC1;refGene_tx=XM_023241813.1;refGene_gn=RMC1;refGene_tx=XM_023241814.1
D3 48223458 48223587
refGene_gn=RMC1;refGene_tx=XM_023241813.1;refGene_gn=RMC1;refGene_tx=XM_023241814.1
D3 48224118 48224225
refGene_gn=RMC1;refGene_tx=XM_023241813.1;refGene_gn=RMC1;refGene_tx=XM_023241814.1
D3 48225614 48225734
refGene_gn=RMC1;refGene_tx=XM_023241813.1;refGene_gn=RMC1;refGene_tx=XM_023241814.1
D3 48226037 48226115
refGene_gn=RMC1;refGene_tx=XM_023241813.1;refGene_gn=RMC1;refGene_tx=XM_023241814.1
D3 48226370 48226475
refGene_gn=RMC1;refGene_tx=XM_023241813.1;refGene_gn=RMC1;refGene_tx=XM_023241814.1
D3 48226571 48226640
refGene_gn=RMC1;refGene_tx=XM_023241813.1;refGene_gn=RMC1;refGene_tx=XM_023241814.1
D3 48226725 48226951
refGene_gn=RMC1;refGene_tx=XM_023241813.1;refGene_gn=RMC1;refGene_tx=XM_023241814.1

D3 48227633 48227713
refGene_gn=RMC1;refGene_tx=XM_023241813.1;refGene_gn=RMC1;refGene_tx=XM_023241814.1
D3 48228235 48228318
refGene_gn=NPC1;refGene_tx=NM_001009829.2;refGene_gn=NPC1;refGene_tx=XM_019814307.2;refGene_gn=NPC1;refGene_tx=XM_019814308.1
D3 48228973 48229136
refGene_gn=NPC1;refGene_tx=NM_001009829.2;refGene_gn=NPC1;refGene_tx=XM_019814307.2;refGene_gn=NPC1;refGene_tx=XM_019814308.1
D3 48229867 48229981
refGene_gn=NPC1;refGene_tx=NM_001009829.2;refGene_gn=NPC1;refGene_tx=XM_019814307.2;refGene_gn=NPC1;refGene_tx=XM_019814308.1
D3 48230729 48230961
refGene_gn=NPC1;refGene_tx=NM_001009829.2;refGene_gn=NPC1;refGene_tx=XM_019814307.2;refGene_gn=NPC1;refGene_tx=XM_019814308.1
D3 48231926 48232127
refGene_gn=NPC1;refGene_tx=NM_001009829.2;refGene_gn=NPC1;refGene_tx=XM_019814307.2;refGene_gn=NPC1;refGene_tx=XM_019814308.1
D3 48233426 48233556
refGene_gn=NPC1;refGene_tx=NM_001009829.2;refGene_gn=NPC1;refGene_tx=XM_019814307.2;refGene_gn=NPC1;refGene_tx=XM_019814308.1
D3 48234172 48234288
refGene_gn=NPC1;refGene_tx=NM_001009829.2;refGene_gn=NPC1;refGene_tx=XM_019814307.2;refGene_gn=NPC1;refGene_tx=XM_019814308.1
D3 48234615 48234803
refGene_gn=NPC1;refGene_tx=NM_001009829.2;refGene_gn=NPC1;refGene_tx=XM_019814307.2;refGene_gn=NPC1;refGene_tx=XM_019814308.1
D3 48235239 48235329
refGene_gn=NPC1;refGene_tx=NM_001009829.2;refGene_gn=NPC1;refGene_tx=XM_019814307.2;refGene_gn=NPC1;refGene_tx=XM_019814308.1
D3 48236472 48236613
refGene_gn=NPC1;refGene_tx=NM_001009829.2;refGene_gn=NPC1;refGene_tx=XM_019814307.2;refGene_gn=NPC1;refGene_tx=XM_019814308.1
D3 48236706 48236834
refGene_gn=NPC1;refGene_tx=NM_001009829.2;refGene_gn=NPC1;refGene_tx=XM_019814307.2;refGene_gn=NPC1;refGene_tx=XM_019814308.1
D3 48238591 48238706
refGene_gn=NPC1;refGene_tx=NM_001009829.2;refGene_gn=NPC1;refGene_tx=XM_019814307.2;refGene_gn=NPC1;refGene_tx=XM_019814308.1
D3 48239416 48239599
refGene_gn=NPC1;refGene_tx=NM_001009829.2;refGene_gn=NPC1;refGene_tx=XM_019814307.2;refGene_gn=NPC1;refGene_tx=XM_019814308.1
D3 48240027 48240217
refGene_gn=NPC1;refGene_tx=NM_001009829.2;refGene_gn=NPC1;refGene_tx=XM_019814307.2;refGene_gn=NPC1;refGene_tx=XM_019814308.1
D3 48243203 48243306
refGene_gn=NPC1;refGene_tx=NM_001009829.2;refGene_gn=NPC1;refGene_tx=XM_019814307.2;refGene_gn=NPC1;refGene_tx=XM_019814308.1
D3 48246161 48246262
refGene_gn=NPC1;refGene_tx=NM_001009829.2;refGene_gn=NPC1;refGene_tx=X

M_019814307.2;refGene_gn=NPC1;refGene_tx=XM_019814308.1
D3 48248585 48248812
refGene_gn=NPC1;refGene_tx=NM_001009829.2;refGene_gn=NPC1;refGene_tx=XM_019814307.2;refGene_gn=NPC1;refGene_tx=XM_019814308.1
D3 48250285 48250707
refGene_gn=NPC1;refGene_tx=NM_001009829.2;refGene_gn=NPC1;refGene_tx=XM_019814307.2;refGene_gn=NPC1;refGene_tx=XM_019814308.1
D3 48251185 48251259
refGene_gn=NPC1;refGene_tx=NM_001009829.2;refGene_gn=NPC1;refGene_tx=XM_019814307.2;refGene_gn=NPC1;refGene_tx=XM_019814308.1
D3 48253103 48253353
refGene_gn=NPC1;refGene_tx=NM_001009829.2;refGene_gn=NPC1;refGene_tx=XM_019814307.2;refGene_gn=NPC1;refGene_tx=XM_019814308.1
D3 48253937 48254105
refGene_gn=NPC1;refGene_tx=NM_001009829.2;refGene_gn=NPC1;refGene_tx=XM_019814307.2;refGene_gn=NPC1;refGene_tx=XM_019814308.1
D3 48261357 48261533
refGene_gn=NPC1;refGene_tx=NM_001009829.2;refGene_gn=NPC1;refGene_tx=XM_019814307.2;refGene_gn=NPC1;refGene_tx=XM_019814308.1
D3 48264011 48264118
refGene_gn=NPC1;refGene_tx=NM_001009829.2;refGene_gn=NPC1;refGene_tx=XM_019814307.2;refGene_gn=NPC1;refGene_tx=XM_019814308.1
D3 48265075 48265198
refGene_gn=NPC1;refGene_tx=NM_001009829.2;refGene_gn=NPC1;refGene_tx=XM_019814307.2;refGene_gn=NPC1;refGene_tx=XM_019814308.1
D3 48276276 48276306 refGene_gn=NPC1;refGene_tx=XM_019814308.1
D3 48283886 48283943
refGene_gn=NPC1;refGene_tx=NM_001009829.2;refGene_gn=NPC1;refGene_tx=XM_019814307.2
D3 48299177 48299261
refGene_gn=ANKRD29;refGene_tx=XM_023241815.1;refGene_gn=ANKRD29;refGene_tx=XM_023241816.1;refGene_gn=ANKRD29;refGene_tx=XM_023241817.1;refGene_gn=ANKRD29;refGene_tx=XM_023241818.1;refGene_gn=ANKRD29;refGene_tx=XM_023241819.1;refGene_gn=ANKRD29;refGene_tx=XM_023241820.1
D3 48305081 48305180
refGene_gn=ANKRD29;refGene_tx=XM_023241815.1;refGene_gn=ANKRD29;refGene_tx=XM_023241816.1;refGene_gn=ANKRD29;refGene_tx=XM_023241817.1;refGene_gn=ANKRD29;refGene_tx=XM_023241818.1;refGene_gn=ANKRD29;refGene_tx=XM_023241819.1;refGene_gn=ANKRD29;refGene_tx=XM_023241820.1
D3 48307413 48307509
refGene_gn=ANKRD29;refGene_tx=XM_023241815.1;refGene_gn=ANKRD29;refGene_tx=XM_023241816.1;refGene_gn=ANKRD29;refGene_tx=XM_023241817.1;refGene_gn=ANKRD29;refGene_tx=XM_023241818.1;refGene_gn=ANKRD29;refGene_tx=XM_023241819.1;refGene_gn=ANKRD29;refGene_tx=XM_023241820.1
D3 48309162 48309261
refGene_gn=ANKRD29;refGene_tx=XM_023241815.1;refGene_gn=ANKRD29;refGene_tx=XM_023241816.1;refGene_gn=ANKRD29;refGene_tx=XM_023241817.1;refGene_gn=ANKRD29;refGene_tx=XM_023241818.1;refGene_gn=ANKRD29;refGene_tx=XM_023241819.1;refGene_gn=ANKRD29;refGene_tx=XM_023241820.1
D3 48319132 48319231

refGene_gn=ANKRD29;refGene_tx=XM_023241815.1;refGene_gn=ANKRD29;refGene_tx=XM_023241816.1;refGene_gn=ANKRD29;refGene_tx=XM_023241817.1;refGene_gn=ANKRD29;refGene_tx=XM_023241818.1;refGene_gn=ANKRD29;refGene_tx=XM_023241819.1;refGene_gn=ANKRD29;refGene_tx=XM_023241820.1
D3 48322758 48322857
refGene_gn=ANKRD29;refGene_tx=XM_023241815.1;refGene_gn=ANKRD29;refGene_tx=XM_023241816.1;refGene_gn=ANKRD29;refGene_tx=XM_023241817.1;refGene_gn=ANKRD29;refGene_tx=XM_023241818.1;refGene_gn=ANKRD29;refGene_tx=XM_023241819.1;refGene_gn=ANKRD29;refGene_tx=XM_023241820.1
D3 48325849 48325948
refGene_gn=ANKRD29;refGene_tx=XM_023241815.1;refGene_gn=ANKRD29;refGene_tx=XM_023241816.1;refGene_gn=ANKRD29;refGene_tx=XM_023241817.1;refGene_gn=ANKRD29;refGene_tx=XM_023241818.1;refGene_gn=ANKRD29;refGene_tx=XM_023241819.1;refGene_gn=ANKRD29;refGene_tx=XM_023241820.1
D3 48330581 48330680
refGene_gn=ANKRD29;refGene_tx=XM_023241815.1;refGene_gn=ANKRD29;refGene_tx=XM_023241816.1;refGene_gn=ANKRD29;refGene_tx=XM_023241817.1;refGene_gn=ANKRD29;refGene_tx=XM_023241818.1;refGene_gn=ANKRD29;refGene_tx=XM_023241819.1;refGene_gn=ANKRD29;refGene_tx=XM_023241820.1
D3 48339033 48339144 refGene_gn=ANKRD29;refGene_tx=XM_023241815.1
D3 48350971 48350992 refGene_gn=ANKRD29;refGene_tx=XM_023241815.1
D3 48384512 48384806
refGene_gn=LAMA3;refGene_tx=XM_023241821.1;refGene_gn=LAMA3;refGene_tx=XM_023241822.1;refGene_gn=LAMA3;refGene_tx=XM_023241823.1;refGene_gn=LAMA3;refGene_tx=XM_023241824.1
D3 48420349 48420502
refGene_gn=LAMA3;refGene_tx=XM_023241821.1;refGene_gn=LAMA3;refGene_tx=XM_023241822.1;refGene_gn=LAMA3;refGene_tx=XM_023241824.1
D3 48458599 48458717
refGene_gn=LAMA3;refGene_tx=XM_023241821.1;refGene_gn=LAMA3;refGene_tx=XM_023241822.1;refGene_gn=LAMA3;refGene_tx=XM_023241823.1;refGene_gn=LAMA3;refGene_tx=XM_023241824.1
D3 48459646 48459765
refGene_gn=LAMA3;refGene_tx=XM_023241821.1;refGene_gn=LAMA3;refGene_tx=XM_023241822.1;refGene_gn=LAMA3;refGene_tx=XM_023241823.1;refGene_gn=LAMA3;refGene_tx=XM_023241824.1
D3 48461272 48461443
refGene_gn=LAMA3;refGene_tx=XM_023241821.1;refGene_gn=LAMA3;refGene_tx=XM_023241822.1;refGene_gn=LAMA3;refGene_tx=XM_023241823.1;refGene_gn=LAMA3;refGene_tx=XM_023241824.1
D3 48463977 48464069
refGene_gn=LAMA3;refGene_tx=XM_023241821.1;refGene_gn=LAMA3;refGene_tx=XM_023241822.1;refGene_gn=LAMA3;refGene_tx=XM_023241823.1;refGene_gn=LAMA3;refGene_tx=XM_023241824.1
D3 48467257 48467373
refGene_gn=LAMA3;refGene_tx=XM_023241821.1;refGene_gn=LAMA3;refGene_tx=XM_023241822.1;refGene_gn=LAMA3;refGene_tx=XM_023241823.1;refGene_gn=LAMA3;refGene_tx=XM_023241824.1
D3 48470439 48470558
refGene_gn=LAMA3;refGene_tx=XM_023241821.1;refGene_gn=LAMA3;refGene_tx

=XM_023241822.1;refGene_gn=LAMA3;refGene_tx=XM_023241823.1;refGene_gn=LAMA3;refGene_tx=XM_023241824.1
D3 48478226 48478317 refGene_gn=LAMA3;refGene_tx=XM_023241821.1;refGene_gn=LAMA3;refGene_tx=XM_023241822.1;refGene_gn=LAMA3;refGene_tx=XM_023241823.1;refGene_gn=LAMA3;refGene_tx=XM_023241824.1
D3 48480678 48480810 refGene_gn=LAMA3;refGene_tx=XM_023241821.1;refGene_gn=LAMA3;refGene_tx=XM_023241822.1;refGene_gn=LAMA3;refGene_tx=XM_023241823.1;refGene_gn=LAMA3;refGene_tx=XM_023241824.1
D3 48482393 48482456 refGene_gn=LAMA3;refGene_tx=XM_023241821.1;refGene_gn=LAMA3;refGene_tx=XM_023241822.1;refGene_gn=LAMA3;refGene_tx=XM_023241823.1;refGene_gn=LAMA3;refGene_tx=XM_023241824.1
D3 48489875 48490010 refGene_gn=LAMA3;refGene_tx=XM_023241821.1;refGene_gn=LAMA3;refGene_tx=XM_023241822.1;refGene_gn=LAMA3;refGene_tx=XM_023241823.1;refGene_gn=LAMA3;refGene_tx=XM_023241824.1
D3 48504469 48504607 refGene_gn=LAMA3;refGene_tx=XM_023241821.1;refGene_gn=LAMA3;refGene_tx=XM_023241822.1;refGene_gn=LAMA3;refGene_tx=XM_023241823.1;refGene_gn=LAMA3;refGene_tx=XM_023241824.1
D3 48508654 48508701 refGene_gn=LAMA3;refGene_tx=XM_023241821.1;refGene_gn=LAMA3;refGene_tx=XM_023241822.1;refGene_gn=LAMA3;refGene_tx=XM_023241823.1;refGene_gn=LAMA3;refGene_tx=XM_023241824.1
D3 48510577 48510677 refGene_gn=LAMA3;refGene_tx=XM_023241821.1;refGene_gn=LAMA3;refGene_tx=XM_023241822.1;refGene_gn=LAMA3;refGene_tx=XM_023241823.1;refGene_gn=LAMA3;refGene_tx=XM_023241824.1
D3 48511348 48511401 refGene_gn=LAMA3;refGene_tx=XM_023241821.1;refGene_gn=LAMA3;refGene_tx=XM_023241822.1;refGene_gn=LAMA3;refGene_tx=XM_023241823.1;refGene_gn=LAMA3;refGene_tx=XM_023241824.1
D3 48511631 48511737 refGene_gn=LAMA3;refGene_tx=XM_023241821.1;refGene_gn=LAMA3;refGene_tx=XM_023241822.1;refGene_gn=LAMA3;refGene_tx=XM_023241823.1;refGene_gn=LAMA3;refGene_tx=XM_023241824.1
D3 48512460 48512560 refGene_gn=LAMA3;refGene_tx=XM_023241821.1;refGene_gn=LAMA3;refGene_tx=XM_023241822.1;refGene_gn=LAMA3;refGene_tx=XM_023241823.1;refGene_gn=LAMA3;refGene_tx=XM_023241824.1
D3 48519210 48519367 refGene_gn=LAMA3;refGene_tx=XM_023241821.1;refGene_gn=LAMA3;refGene_tx=XM_023241822.1;refGene_gn=LAMA3;refGene_tx=XM_023241823.1;refGene_gn=LAMA3;refGene_tx=XM_023241824.1
D3 48523814 48523947 refGene_gn=LAMA3;refGene_tx=XM_023241821.1;refGene_gn=LAMA3;refGene_tx=XM_023241822.1;refGene_gn=LAMA3;refGene_tx=XM_023241823.1;refGene_gn=LAMA3;refGene_tx=XM_023241824.1

D3 48525951 48526094
refGene_gn=LAMA3;refGene_tx=XM_023241821.1;refGene_gn=LAMA3;refGene_tx=XM_023241822.1;refGene_gn=LAMA3;refGene_tx=XM_023241823.1;refGene_gn=LAMA3;refGene_tx=XM_023241824.1
D3 48528219 48528317
refGene_gn=LAMA3;refGene_tx=XM_023241821.1;refGene_gn=LAMA3;refGene_tx=XM_023241822.1;refGene_gn=LAMA3;refGene_tx=XM_023241823.1;refGene_gn=LAMA3;refGene_tx=XM_023241824.1
D3 48528658 48528812
refGene_gn=LAMA3;refGene_tx=XM_023241821.1;refGene_gn=LAMA3;refGene_tx=XM_023241822.1;refGene_gn=LAMA3;refGene_tx=XM_023241823.1;refGene_gn=LAMA3;refGene_tx=XM_023241824.1
D3 48539017 48539178
refGene_gn=LAMA3;refGene_tx=XM_023241821.1;refGene_gn=LAMA3;refGene_tx=XM_023241822.1;refGene_gn=LAMA3;refGene_tx=XM_023241823.1;refGene_gn=LAMA3;refGene_tx=XM_023241824.1
D3 48542615 48542724
refGene_gn=LAMA3;refGene_tx=XM_023241821.1;refGene_gn=LAMA3;refGene_tx=XM_023241822.1;refGene_gn=LAMA3;refGene_tx=XM_023241823.1;refGene_gn=LAMA3;refGene_tx=XM_023241824.1
D3 48544918 48545016
refGene_gn=LAMA3;refGene_tx=XM_023241821.1;refGene_gn=LAMA3;refGene_tx=XM_023241822.1;refGene_gn=LAMA3;refGene_tx=XM_023241823.1;refGene_gn=LAMA3;refGene_tx=XM_023241824.1
D3 48546151 48546293
refGene_gn=LAMA3;refGene_tx=XM_023241821.1;refGene_gn=LAMA3;refGene_tx=XM_023241822.1;refGene_gn=LAMA3;refGene_tx=XM_023241823.1;refGene_gn=LAMA3;refGene_tx=XM_023241824.1
D3 48547974 48548101
refGene_gn=LAMA3;refGene_tx=XM_023241821.1;refGene_gn=LAMA3;refGene_tx=XM_023241822.1;refGene_gn=LAMA3;refGene_tx=XM_023241823.1;refGene_gn=LAMA3;refGene_tx=XM_023241824.1
D3 48548190 48548330
refGene_gn=LAMA3;refGene_tx=XM_023241821.1;refGene_gn=LAMA3;refGene_tx=XM_023241822.1;refGene_gn=LAMA3;refGene_tx=XM_023241823.1;refGene_gn=LAMA3;refGene_tx=XM_023241824.1
D3 48549863 48549979
refGene_gn=LAMA3;refGene_tx=XM_023241821.1;refGene_gn=LAMA3;refGene_tx=XM_023241822.1;refGene_gn=LAMA3;refGene_tx=XM_023241823.1;refGene_gn=LAMA3;refGene_tx=XM_023241824.1
D3 48551156 48551368
refGene_gn=LAMA3;refGene_tx=XM_023241821.1;refGene_gn=LAMA3;refGene_tx=XM_023241822.1;refGene_gn=LAMA3;refGene_tx=XM_023241823.1;refGene_gn=LAMA3;refGene_tx=XM_023241824.1
D3 48552808 48553013
refGene_gn=LAMA3;refGene_tx=XM_023241821.1;refGene_gn=LAMA3;refGene_tx=XM_023241822.1;refGene_gn=LAMA3;refGene_tx=XM_023241823.1;refGene_gn=LAMA3;refGene_tx=XM_023241824.1
D3 48565757 48565902
refGene_gn=LAMA3;refGene_tx=XM_023241821.1;refGene_gn=LAMA3;refGene_tx

=XM_023241822.1;refGene_gn=LAMA3;refGene_tx=XM_023241823.1;refGene_gn=LAMA3;refGene_tx=XM_023241824.1
D3 48566605 48566746
refGene_gn=LAMA3;refGene_tx=XM_023241821.1;refGene_gn=LAMA3;refGene_tx=XM_023241822.1;refGene_gn=LAMA3;refGene_tx=XM_023241823.1;refGene_gn=LAMA3;refGene_tx=XM_023241824.1
D3 48568013 48568175
refGene_gn=LAMA3;refGene_tx=XM_023241821.1;refGene_gn=LAMA3;refGene_tx=XM_023241822.1;refGene_gn=LAMA3;refGene_tx=XM_023241823.1;refGene_gn=LAMA3;refGene_tx=XM_023241824.1
D3 48570670 48570769
refGene_gn=LAMA3;refGene_tx=XM_023241821.1;refGene_gn=LAMA3;refGene_tx=XM_023241822.1;refGene_gn=LAMA3;refGene_tx=XM_023241823.1;refGene_gn=LAMA3;refGene_tx=XM_023241824.1
D3 48573118 48573202
refGene_gn=LAMA3;refGene_tx=XM_023241821.1;refGene_gn=LAMA3;refGene_tx=XM_023241822.1;refGene_gn=LAMA3;refGene_tx=XM_023241823.1;refGene_gn=LAMA3;refGene_tx=XM_023241824.1
D3 48581710 48581941
refGene_gn=LAMA3;refGene_tx=XM_023241821.1;refGene_gn=LAMA3;refGene_tx=XM_023241822.1;refGene_gn=LAMA3;refGene_tx=XM_023241823.1
D3 48583190 48583361 refGene_gn=LAMA3;refGene_tx=XM_019815081.2
D3 48586396 48586510
refGene_gn=LAMA3;refGene_tx=XM_019815081.2;refGene_gn=LAMA3;refGene_tx=XM_023241821.1;refGene_gn=LAMA3;refGene_tx=XM_023241822.1;refGene_gn=LAMA3;refGene_tx=XM_023241823.1;refGene_gn=LAMA3;refGene_tx=XM_023241824.1
D3 48591286 48591396
refGene_gn=LAMA3;refGene_tx=XM_019815081.2;refGene_gn=LAMA3;refGene_tx=XM_023241821.1;refGene_gn=LAMA3;refGene_tx=XM_023241822.1;refGene_gn=LAMA3;refGene_tx=XM_023241823.1;refGene_gn=LAMA3;refGene_tx=XM_023241824.1
D3 48593993 48594074
refGene_gn=LAMA3;refGene_tx=XM_019815081.2;refGene_gn=LAMA3;refGene_tx=XM_023241821.1;refGene_gn=LAMA3;refGene_tx=XM_023241822.1;refGene_gn=LAMA3;refGene_tx=XM_023241823.1;refGene_gn=LAMA3;refGene_tx=XM_023241824.1
D3 48600217 48600324
refGene_gn=LAMA3;refGene_tx=XM_019815081.2;refGene_gn=LAMA3;refGene_tx=XM_023241821.1;refGene_gn=LAMA3;refGene_tx=XM_023241822.1;refGene_gn=LAMA3;refGene_tx=XM_023241823.1;refGene_gn=LAMA3;refGene_tx=XM_023241824.1
D3 48603754 48603805
refGene_gn=LAMA3;refGene_tx=XM_019815081.2;refGene_gn=LAMA3;refGene_tx=XM_023241821.1;refGene_gn=LAMA3;refGene_tx=XM_023241822.1;refGene_gn=LAMA3;refGene_tx=XM_023241823.1;refGene_gn=LAMA3;refGene_tx=XM_023241824.1
D3 48604694 48604846
refGene_gn=LAMA3;refGene_tx=XM_019815081.2;refGene_gn=LAMA3;refGene_tx=XM_023241821.1;refGene_gn=LAMA3;refGene_tx=XM_023241822.1;refGene_gn=

LAMA3;refGene_tx=XM_023241823.1;refGene_gn=LAMA3;refGene_tx=XM_0232418
24.1
D3 48608762 48608873
refGene_gn=LAMA3;refGene_tx=XM_019815081.2;refGene_gn=LAMA3;refGene_tx
=XM_023241821.1;refGene_gn=LAMA3;refGene_tx=XM_023241822.1;refGene_gn=
LAMA3;refGene_tx=XM_023241823.1;refGene_gn=LAMA3;refGene_tx=XM_0232418
24.1
D3 48608998 48609110
refGene_gn=LAMA3;refGene_tx=XM_019815081.2;refGene_gn=LAMA3;refGene_tx
=XM_023241821.1;refGene_gn=LAMA3;refGene_tx=XM_023241822.1;refGene_gn=
LAMA3;refGene_tx=XM_023241823.1;refGene_gn=LAMA3;refGene_tx=XM_0232418
24.1
D3 48609331 48609499
refGene_gn=LAMA3;refGene_tx=XM_019815081.2;refGene_gn=LAMA3;refGene_tx
=XM_023241821.1;refGene_gn=LAMA3;refGene_tx=XM_023241822.1;refGene_gn=
LAMA3;refGene_tx=XM_023241823.1;refGene_gn=LAMA3;refGene_tx=XM_0232418
24.1
D3 48610753 48610950
refGene_gn=LAMA3;refGene_tx=XM_019815081.2;refGene_gn=LAMA3;refGene_tx
=XM_023241821.1;refGene_gn=LAMA3;refGene_tx=XM_023241822.1;refGene_gn=
LAMA3;refGene_tx=XM_023241823.1;refGene_gn=LAMA3;refGene_tx=XM_0232418
24.1
D3 48612148 48612265
refGene_gn=LAMA3;refGene_tx=XM_019815081.2;refGene_gn=LAMA3;refGene_tx
=XM_023241821.1;refGene_gn=LAMA3;refGene_tx=XM_023241822.1;refGene_gn=
LAMA3;refGene_tx=XM_023241823.1;refGene_gn=LAMA3;refGene_tx=XM_0232418
24.1
D3 48612898 48613053
refGene_gn=LAMA3;refGene_tx=XM_019815081.2;refGene_gn=LAMA3;refGene_tx
=XM_023241821.1;refGene_gn=LAMA3;refGene_tx=XM_023241822.1;refGene_gn=
LAMA3;refGene_tx=XM_023241823.1;refGene_gn=LAMA3;refGene_tx=XM_0232418
24.1
D3 48613335 48613477
refGene_gn=LAMA3;refGene_tx=XM_019815081.2;refGene_gn=LAMA3;refGene_tx
=XM_023241821.1;refGene_gn=LAMA3;refGene_tx=XM_023241822.1;refGene_gn=
LAMA3;refGene_tx=XM_023241823.1;refGene_gn=LAMA3;refGene_tx=XM_0232418
24.1
D3 48614551 48614654
refGene_gn=LAMA3;refGene_tx=XM_019815081.2;refGene_gn=LAMA3;refGene_tx
=XM_023241821.1;refGene_gn=LAMA3;refGene_tx=XM_023241822.1;refGene_gn=
LAMA3;refGene_tx=XM_023241823.1;refGene_gn=LAMA3;refGene_tx=XM_0232418
24.1
D3 48616212 48616329
refGene_gn=LAMA3;refGene_tx=XM_019815081.2;refGene_gn=LAMA3;refGene_tx
=XM_023241821.1;refGene_gn=LAMA3;refGene_tx=XM_023241822.1;refGene_gn=
LAMA3;refGene_tx=XM_023241823.1;refGene_gn=LAMA3;refGene_tx=XM_0232418
24.1
D3 48616413 48616593
refGene_gn=LAMA3;refGene_tx=XM_019815081.2;refGene_gn=LAMA3;refGene_tx
=XM_023241821.1;refGene_gn=LAMA3;refGene_tx=XM_023241822.1;refGene_gn=

LAMA3;refGene_tx=XM_023241823.1;refGene_gn=LAMA3;refGene_tx=XM_0232418
24.1
D3 48617109 48617252
refGene_gn=LAMA3;refGene_tx=XM_019815081.2;refGene_gn=LAMA3;refGene_tx
=XM_023241821.1;refGene_gn=LAMA3;refGene_tx=XM_023241822.1;refGene_gn=
LAMA3;refGene_tx=XM_023241823.1;refGene_gn=LAMA3;refGene_tx=XM_0232418
24.1
D3 48618428 48618599
refGene_gn=LAMA3;refGene_tx=XM_019815081.2;refGene_gn=LAMA3;refGene_tx
=XM_023241821.1;refGene_gn=LAMA3;refGene_tx=XM_023241822.1;refGene_gn=
LAMA3;refGene_tx=XM_023241823.1;refGene_gn=LAMA3;refGene_tx=XM_0232418
24.1
D3 48620470 48620622
refGene_gn=LAMA3;refGene_tx=XM_019815081.2;refGene_gn=LAMA3;refGene_tx
=XM_023241821.1;refGene_gn=LAMA3;refGene_tx=XM_023241822.1;refGene_gn=
LAMA3;refGene_tx=XM_023241823.1;refGene_gn=LAMA3;refGene_tx=XM_0232418
24.1
D3 48620768 48620931
refGene_gn=LAMA3;refGene_tx=XM_019815081.2;refGene_gn=LAMA3;refGene_tx
=XM_023241821.1;refGene_gn=LAMA3;refGene_tx=XM_023241822.1;refGene_gn=
LAMA3;refGene_tx=XM_023241823.1;refGene_gn=LAMA3;refGene_tx=XM_0232418
24.1
D3 48621596 48621730
refGene_gn=LAMA3;refGene_tx=XM_019815081.2;refGene_gn=LAMA3;refGene_tx
=XM_023241821.1;refGene_gn=LAMA3;refGene_tx=XM_023241822.1;refGene_gn=
LAMA3;refGene_tx=XM_023241823.1;refGene_gn=LAMA3;refGene_tx=XM_0232418
24.1
D3 48622797 48622942
refGene_gn=LAMA3;refGene_tx=XM_019815081.2;refGene_gn=LAMA3;refGene_tx
=XM_023241821.1;refGene_gn=LAMA3;refGene_tx=XM_023241822.1;refGene_gn=
LAMA3;refGene_tx=XM_023241823.1;refGene_gn=LAMA3;refGene_tx=XM_0232418
24.1
D3 48625975 48626095
refGene_gn=LAMA3;refGene_tx=XM_019815081.2;refGene_gn=LAMA3;refGene_tx
=XM_023241821.1;refGene_gn=LAMA3;refGene_tx=XM_023241822.1;refGene_gn=
LAMA3;refGene_tx=XM_023241823.1;refGene_gn=LAMA3;refGene_tx=XM_0232418
24.1
D3 48626478 48626612
refGene_gn=LAMA3;refGene_tx=XM_019815081.2;refGene_gn=LAMA3;refGene_tx
=XM_023241821.1;refGene_gn=LAMA3;refGene_tx=XM_023241822.1;refGene_gn=
LAMA3;refGene_tx=XM_023241823.1;refGene_gn=LAMA3;refGene_tx=XM_0232418
24.1
D3 48633078 48633196
refGene_gn=LAMA3;refGene_tx=XM_019815081.2;refGene_gn=LAMA3;refGene_tx
=XM_023241821.1;refGene_gn=LAMA3;refGene_tx=XM_023241822.1;refGene_gn=
LAMA3;refGene_tx=XM_023241823.1;refGene_gn=LAMA3;refGene_tx=XM_0232418
24.1
D3 48633576 48633717
refGene_gn=LAMA3;refGene_tx=XM_019815081.2;refGene_gn=LAMA3;refGene_tx
=XM_023241821.1;refGene_gn=LAMA3;refGene_tx=XM_023241822.1;refGene_gn=

LAMA3;refGene_tx=XM_023241823.1;refGene_gn=LAMA3;refGene_tx=XM_0232418
24.1
D3 48635605 48635745
refGene_gn=LAMA3;refGene_tx=XM_019815081.2;refGene_gn=LAMA3;refGene_tx
=XM_023241821.1;refGene_gn=LAMA3;refGene_tx=XM_023241822.1;refGene_gn=
LAMA3;refGene_tx=XM_023241823.1;refGene_gn=LAMA3;refGene_tx=XM_0232418
24.1
D3 48636495 48636627
refGene_gn=LAMA3;refGene_tx=XM_019815081.2;refGene_gn=LAMA3;refGene_tx
=XM_023241821.1;refGene_gn=LAMA3;refGene_tx=XM_023241822.1;refGene_gn=
LAMA3;refGene_tx=XM_023241823.1;refGene_gn=LAMA3;refGene_tx=XM_0232418
24.1
D3 48638397 48638551
refGene_gn=LAMA3;refGene_tx=XM_019815081.2;refGene_gn=LAMA3;refGene_tx
=XM_023241821.1;refGene_gn=LAMA3;refGene_tx=XM_023241822.1;refGene_gn=
LAMA3;refGene_tx=XM_023241823.1;refGene_gn=LAMA3;refGene_tx=XM_0232418
24.1
D3 48642817 48642987
refGene_gn=LAMA3;refGene_tx=XM_019815081.2;refGene_gn=LAMA3;refGene_tx
=XM_023241821.1;refGene_gn=LAMA3;refGene_tx=XM_023241822.1;refGene_gn=
LAMA3;refGene_tx=XM_023241823.1;refGene_gn=LAMA3;refGene_tx=XM_0232418
24.1
D3 48646772 48646956
refGene_gn=LAMA3;refGene_tx=XM_019815081.2;refGene_gn=LAMA3;refGene_tx
=XM_023241821.1;refGene_gn=LAMA3;refGene_tx=XM_023241822.1;refGene_gn=
LAMA3;refGene_tx=XM_023241823.1;refGene_gn=LAMA3;refGene_tx=XM_0232418
24.1
D3 48648835 48648976
refGene_gn=LAMA3;refGene_tx=XM_019815081.2;refGene_gn=LAMA3;refGene_tx
=XM_023241821.1;refGene_gn=LAMA3;refGene_tx=XM_023241822.1;refGene_gn=
LAMA3;refGene_tx=XM_023241823.1;refGene_gn=LAMA3;refGene_tx=XM_0232418
24.1
D3 48652459 48652619
refGene_gn=LAMA3;refGene_tx=XM_019815081.2;refGene_gn=LAMA3;refGene_tx
=XM_023241821.1;refGene_gn=LAMA3;refGene_tx=XM_023241822.1;refGene_gn=
LAMA3;refGene_tx=XM_023241823.1;refGene_gn=LAMA3;refGene_tx=XM_0232418
24.1
D3 48652718 48652849
refGene_gn=LAMA3;refGene_tx=XM_019815081.2;refGene_gn=LAMA3;refGene_tx
=XM_023241821.1;refGene_gn=LAMA3;refGene_tx=XM_023241822.1;refGene_gn=
LAMA3;refGene_tx=XM_023241823.1;refGene_gn=LAMA3;refGene_tx=XM_0232418
24.1
D3 48655258 48655352
refGene_gn=LAMA3;refGene_tx=XM_019815081.2;refGene_gn=LAMA3;refGene_tx
=XM_023241821.1;refGene_gn=LAMA3;refGene_tx=XM_023241822.1;refGene_gn=
LAMA3;refGene_tx=XM_023241823.1;refGene_gn=LAMA3;refGene_tx=XM_0232418
24.1
D3 48656540 48656660
refGene_gn=LAMA3;refGene_tx=XM_019815081.2;refGene_gn=LAMA3;refGene_tx
=XM_023241821.1;refGene_gn=LAMA3;refGene_tx=XM_023241822.1;refGene_gn=

LAMA3;refGene_tx=XM_023241823.1;refGene_gn=LAMA3;refGene_tx=XM_0232418
24.1
D3 48657390 48657536
refGene_gn=LAMA3;refGene_tx=XM_019815081.2;refGene_gn=LAMA3;refGene_tx
=XM_023241821.1;refGene_gn=LAMA3;refGene_tx=XM_023241822.1;refGene_gn=
LAMA3;refGene_tx=XM_023241823.1;refGene_gn=LAMA3;refGene_tx=XM_0232418
24.1
D3 48709801 48709959 refGene_gn=TTC39C;refGene_tx=XM_003995073.4
D3 48756191 48756240 refGene_gn=TTC39C;refGene_tx=XM_003995073.4
D3 48758069 48758198 refGene_gn=TTC39C;refGene_tx=XM_003995073.4
D3 48760659 48760774 refGene_gn=TTC39C;refGene_tx=XM_003995073.4
D3 48770612 48770967 refGene_gn=TTC39C;refGene_tx=XM_003995073.4
D3 48772680 48772849 refGene_gn=TTC39C;refGene_tx=XM_003995073.4
D3 48802691 48802785 refGene_gn=TTC39C;refGene_tx=XM_003995073.4
D3 48805655 48805763 refGene_gn=TTC39C;refGene_tx=XM_003995073.4
D3 48809132 48809242 refGene_gn=TTC39C;refGene_tx=XM_003995073.4
D3 48810267 48810391 refGene_gn=TTC39C;refGene_tx=XM_003995073.4
D3 48813936 48814034 refGene_gn=TTC39C;refGene_tx=XM_003995073.4
D3 48815115 48815220 refGene_gn=TTC39C;refGene_tx=XM_003995073.4
D3 48816481 48816520 refGene_gn=TTC39C;refGene_tx=XM_003995073.4
D3 48817186 48817276 refGene_gn=TTC39C;refGene_tx=XM_003995073.4
D3 48825072 48825220
refGene_gn=CABYR;refGene_tx=XM_011288071.3;refGene_gn=CABYR;refGene_tx
=XM_019815084.2;refGene_gn=CABYR;refGene_tx=XM_023241825.1;refGene_gn=
CABYR;refGene_tx=XM_023241826.1;refGene_gn=CABYR;refGene_tx=XM_0232418
27.1;refGene_gn=CABYR;refGene_tx=XM_023241828.1;refGene_gn=CABYR;refGe
ne_tx=XM_023241829.1;refGene_gn=CABYR;refGene_tx=XM_023241830.1
D3 48825319 48825373
refGene_gn=CABYR;refGene_tx=XM_011288071.3;refGene_gn=CABYR;refGene_tx
=XM_019815084.2;refGene_gn=CABYR;refGene_tx=XM_023241825.1;refGene_gn=
CABYR;refGene_tx=XM_023241826.1;refGene_gn=CABYR;refGene_tx=XM_0232418
27.1;refGene_gn=CABYR;refGene_tx=XM_023241828.1;refGene_gn=CABYR;refGe
ne_tx=XM_023241829.1;refGene_gn=CABYR;refGene_tx=XM_023241830.1
D3 48836092 48836443
refGene_gn=CABYR;refGene_tx=XM_011288071.3;refGene_gn=CABYR;refGene_tx
=XM_019815084.2;refGene_gn=CABYR;refGene_tx=XM_023241825.1;refGene_gn=
CABYR;refGene_tx=XM_023241826.1;refGene_gn=CABYR;refGene_tx=XM_0232418
27.1;refGene_gn=CABYR;refGene_tx=XM_023241828.1;refGene_gn=CABYR;refGe
ne_tx=XM_023241829.1;refGene_gn=CABYR;refGene_tx=XM_023241830.1
D3 48837295 48837364
refGene_gn=CABYR;refGene_tx=XM_019815084.2;refGene_gn=CABYR;refGene_tx
=XM_023241825.1;refGene_gn=CABYR;refGene_tx=XM_023241826.1;refGene_gn=
CABYR;refGene_tx=XM_023241827.1;refGene_gn=CABYR;refGene_tx=XM_0232418
28.1;refGene_gn=CABYR;refGene_tx=XM_023241829.1;refGene_gn=CABYR;refGe
ne_tx=XM_023241830.1
D3 48839559 48840167
refGene_gn=CABYR;refGene_tx=XM_011288071.3;refGene_gn=CABYR;refGene_tx
=XM_019815084.2;refGene_gn=CABYR;refGene_tx=XM_023241825.1;refGene_gn=
CABYR;refGene_tx=XM_023241826.1;refGene_gn=CABYR;refGene_tx=XM_0232418
27.1;refGene_gn=CABYR;refGene_tx=XM_023241828.1;refGene_gn=CABYR;refGe

ne_tx=XM_023241829.1
D3	48841549	48841572
refGene_gn=CABYR;refGene_tx=XM_019815084.2;refGene_gn=CABYR;refGene_tx=XM_023241830.1
D3	48843122	48843225
refGene_gn=OSBPL1A;refGene_tx=XM_003995075.4;refGene_gn=OSBPL1A;refGene_tx=XM_011288065.2;refGene_gn=OSBPL1A;refGene_tx=XM_011288066.3;refGene_gn=OSBPL1A;refGene_tx=XM_011288067.2
D3	48846726	48846817
refGene_gn=OSBPL1A;refGene_tx=XM_003995075.4;refGene_gn=OSBPL1A;refGene_tx=XM_011288065.2;refGene_gn=OSBPL1A;refGene_tx=XM_011288066.3;refGene_gn=OSBPL1A;refGene_tx=XM_011288067.2
D3	48848360	48848484
refGene_gn=OSBPL1A;refGene_tx=XM_003995075.4;refGene_gn=OSBPL1A;refGene_tx=XM_011288065.2;refGene_gn=OSBPL1A;refGene_tx=XM_011288066.3;refGene_gn=OSBPL1A;refGene_tx=XM_011288067.2
D3	48849093	48849210
refGene_gn=OSBPL1A;refGene_tx=XM_003995075.4;refGene_gn=OSBPL1A;refGene_tx=XM_011288065.2;refGene_gn=OSBPL1A;refGene_tx=XM_011288066.3;refGene_gn=OSBPL1A;refGene_tx=XM_011288067.2
D3	48851799	48851926
refGene_gn=OSBPL1A;refGene_tx=XM_003995075.4;refGene_gn=OSBPL1A;refGene_tx=XM_011288065.2;refGene_gn=OSBPL1A;refGene_tx=XM_011288066.3;refGene_gn=OSBPL1A;refGene_tx=XM_011288067.2
D3	48852162	48852252
refGene_gn=OSBPL1A;refGene_tx=XM_003995075.4;refGene_gn=OSBPL1A;refGene_tx=XM_011288065.2;refGene_gn=OSBPL1A;refGene_tx=XM_011288066.3;refGene_gn=OSBPL1A;refGene_tx=XM_011288067.2
D3	48853146	48853254
refGene_gn=OSBPL1A;refGene_tx=XM_003995075.4;refGene_gn=OSBPL1A;refGene_tx=XM_011288065.2;refGene_gn=OSBPL1A;refGene_tx=XM_011288066.3;refGene_gn=OSBPL1A;refGene_tx=XM_011288067.2
D3	48861360	48861543
refGene_gn=OSBPL1A;refGene_tx=XM_003995075.4;refGene_gn=OSBPL1A;refGene_tx=XM_011288065.2;refGene_gn=OSBPL1A;refGene_tx=XM_011288066.3;refGene_gn=OSBPL1A;refGene_tx=XM_011288067.2
D3	48862499	48862597
refGene_gn=OSBPL1A;refGene_tx=XM_003995075.4;refGene_gn=OSBPL1A;refGene_tx=XM_011288065.2;refGene_gn=OSBPL1A;refGene_tx=XM_011288066.3;refGene_gn=OSBPL1A;refGene_tx=XM_011288067.2
D3	48863569	48863704
refGene_gn=OSBPL1A;refGene_tx=XM_003995075.4;refGene_gn=OSBPL1A;refGene_tx=XM_011288065.2;refGene_gn=OSBPL1A;refGene_tx=XM_011288066.3;refGene_gn=OSBPL1A;refGene_tx=XM_011288067.2
D3	48881405	48881481
refGene_gn=OSBPL1A;refGene_tx=XM_003995075.4;refGene_gn=OSBPL1A;refGene_tx=XM_011288065.2;refGene_gn=OSBPL1A;refGene_tx=XM_011288066.3;refGene_gn=OSBPL1A;refGene_tx=XM_011288067.2
D3	48925460	48925617
refGene_gn=OSBPL1A;refGene_tx=XM_003995075.4;refGene_gn=OSBPL1A;refGen

e_tx=XM_011288065.2;refGene_gn=OSBPL1A;refGene_tx=XM_011288066.3;refGene_gn=OSBPL1A;refGene_tx=XM_011288067.2
D3 48949098 48949261 refGene_gn=OSBPL1A;refGene_tx=XM_003995075.4;refGene_gn=OSBPL1A;refGene_tx=XM_011288065.2
D3 48988648 48988755 refGene_gn=OSBPL1A;refGene_tx=XM_003995075.4
D3 48997395 48997477 refGene_gn=OSBPL1A;refGene_tx=XM_003995075.4
D3 49003342 49003465 refGene_gn=OSBPL1A;refGene_tx=XM_003995075.4;refGene_gn=OSBPL1A;refGene_tx=XM_011288065.2
D3 49004692 49004791 refGene_gn=OSBPL1A;refGene_tx=XM_003995075.4;refGene_gn=OSBPL1A;refGene_tx=XM_011288065.2
D3 49014766 49014830 refGene_gn=OSBPL1A;refGene_tx=XM_003995075.4;refGene_gn=OSBPL1A;refGene_tx=XM_011288065.2
D3 49014947 49015021 refGene_gn=OSBPL1A;refGene_tx=XM_003995075.4;refGene_gn=OSBPL1A;refGene_tx=XM_011288065.2
D3 49016200 49016245 refGene_gn=OSBPL1A;refGene_tx=XM_003995075.4;refGene_gn=OSBPL1A;refGene_tx=XM_011288065.2
D3 49016347 49016409 refGene_gn=OSBPL1A;refGene_tx=XM_003995075.4;refGene_gn=OSBPL1A;refGene_tx=XM_011288065.2
D3 49032988 49033133 refGene_gn=OSBPL1A;refGene_tx=XM_003995075.4;refGene_gn=OSBPL1A;refGene_tx=XM_011288065.2
D3 49034278 49034364 refGene_gn=OSBPL1A;refGene_tx=XM_003995075.4;refGene_gn=OSBPL1A;refGene_tx=XM_011288065.2
D3 49039542 49039654 refGene_gn=OSBPL1A;refGene_tx=XM_003995075.4;refGene_gn=OSBPL1A;refGene_tx=XM_011288065.2
D3 49065922 49065997 refGene_gn=OSBPL1A;refGene_tx=XM_003995075.4;refGene_gn=OSBPL1A;refGene_tx=XM_011288065.2
D3 49068414 49068500 refGene_gn=OSBPL1A;refGene_tx=XM_003995075.4;refGene_gn=OSBPL1A;refGene_tx=XM_011288065.2
D3 49075772 49075893 refGene_gn=OSBPL1A;refGene_tx=XM_003995075.4;refGene_gn=OSBPL1A;refGene_tx=XM_011288065.2
D3 49121325 49121361 refGene_gn=IMPACT;refGene_tx=XM_023241831.1;refGene_gn=IMPACT;refGene_tx=XM_023241832.1;refGene_gn=IMPACT;refGene_tx=XM_023241833.1
D3 49122433 49122562 refGene_gn=IMPACT;refGene_tx=XM_023241831.1;refGene_gn=IMPACT;refGene_tx=XM_023241832.1;refGene_gn=IMPACT;refGene_tx=XM_023241833.1;refGene_gn=IMPACT;refGene_tx=XM_023241834.1;refGene_gn=IMPACT;refGene_tx=XM_02

3241836.1;refGene_gn=IMPACT;refGene_tx=XM_023241837.1
D3 49123534 49123587
refGene_gn=IMPACT;refGene_tx=XM_023241831.1;refGene_gn=IMPACT;refGene_tx=XM_023241832.1;refGene_gn=IMPACT;refGene_tx=XM_023241833.1;refGene_gn=IMPACT;refGene_tx=XM_023241834.1;refGene_gn=IMPACT;refGene_tx=XM_023241836.1;refGene_gn=IMPACT;refGene_tx=XM_023241837.1
D3 49125429 49125492
refGene_gn=IMPACT;refGene_tx=XM_023241831.1;refGene_gn=IMPACT;refGene_tx=XM_023241832.1;refGene_gn=IMPACT;refGene_tx=XM_023241833.1;refGene_gn=IMPACT;refGene_tx=XM_023241834.1;refGene_gn=IMPACT;refGene_tx=XM_023241836.1;refGene_gn=IMPACT;refGene_tx=XM_023241837.1
D3 49134685 49134771
refGene_gn=IMPACT;refGene_tx=XM_023241831.1;refGene_gn=IMPACT;refGene_tx=XM_023241832.1;refGene_gn=IMPACT;refGene_tx=XM_023241833.1;refGene_gn=IMPACT;refGene_tx=XM_023241834.1;refGene_gn=IMPACT;refGene_tx=XM_023241836.1;refGene_gn=IMPACT;refGene_tx=XM_023241837.1
D3 49138148 49138271
refGene_gn=IMPACT;refGene_tx=XM_023241831.1;refGene_gn=IMPACT;refGene_tx=XM_023241832.1;refGene_gn=IMPACT;refGene_tx=XM_023241833.1;refGene_gn=IMPACT;refGene_tx=XM_023241834.1;refGene_gn=IMPACT;refGene_tx=XM_023241836.1;refGene_gn=IMPACT;refGene_tx=XM_023241837.1
D3 49140120 49140218
refGene_gn=IMPACT;refGene_tx=XM_023241831.1;refGene_gn=IMPACT;refGene_tx=XM_023241832.1;refGene_gn=IMPACT;refGene_tx=XM_023241833.1;refGene_gn=IMPACT;refGene_tx=XM_023241834.1;refGene_gn=IMPACT;refGene_tx=XM_023241836.1;refGene_gn=IMPACT;refGene_tx=XM_023241837.1
D3 49142329 49142403
refGene_gn=IMPACT;refGene_tx=XM_023241831.1;refGene_gn=IMPACT;refGene_tx=XM_023241832.1;refGene_gn=IMPACT;refGene_tx=XM_023241833.1;refGene_gn=IMPACT;refGene_tx=XM_023241834.1;refGene_gn=IMPACT;refGene_tx=XM_023241836.1;refGene_gn=IMPACT;refGene_tx=XM_023241837.1
D3 49145721 49145812
refGene_gn=IMPACT;refGene_tx=XM_023241831.1;refGene_gn=IMPACT;refGene_tx=XM_023241832.1;refGene_gn=IMPACT;refGene_tx=XM_023241833.1;refGene_gn=IMPACT;refGene_tx=XM_023241834.1;refGene_gn=IMPACT;refGene_tx=XM_023241836.1;refGene_gn=IMPACT;refGene_tx=XM_023241837.1
D3 49146592 49146610
refGene_gn=IMPACT;refGene_tx=XM_023241831.1;refGene_gn=IMPACT;refGene_tx=XM_023241833.1;refGene_gn=IMPACT;refGene_tx=XM_023241834.1;refGene_gn=IMPACT;refGene_tx=XM_023241836.1;refGene_gn=IMPACT;refGene_tx=XM_023241837.1
D3 49147139 49147274
refGene_gn=IMPACT;refGene_tx=XM_023241831.1;refGene_gn=IMPACT;refGene_tx=XM_023241832.1;refGene_gn=IMPACT;refGene_tx=XM_023241833.1;refGene_gn=IMPACT;refGene_tx=XM_023241834.1;refGene_gn=IMPACT;refGene_tx=XM_023241836.1;refGene_gn=IMPACT;refGene_tx=XM_023241837.1
D3 49148226 49148295
refGene_gn=IMPACT;refGene_tx=XM_023241831.1;refGene_gn=IMPACT;refGene_tx=XM_023241832.1;refGene_gn=IMPACT;refGene_tx=XM_023241833.1;refGene_gn=IMPACT;refGene_tx=XM_023241834.1;refGene_gn=IMPACT;refGene_tx=XM_02

3241836.1;refGene_gn=IMPACT;refGene_tx=XM_023241837.1
D3 49165501 49165727 refGene_gn=HRH4;refGene_tx=XM_003995081.3
D3 49171966 49172130 refGene_gn=HRH4;refGene_tx=XM_003995081.3
D3 49177585 49178419 refGene_gn=HRH4;refGene_tx=XM_003995081.3
D3 49702001 49702031
refGene_gn=ZNF521;refGene_tx=XM_023241838.1;refGene_gn=ZNF521;refGene_tx=XM_023241839.1;refGene_gn=ZNF521;refGene_tx=XM_023241840.1
D3 49710706 49710870
refGene_gn=ZNF521;refGene_tx=XM_023241838.1;refGene_gn=ZNF521;refGene_tx=XM_023241840.1
D3 49727691 49727807
refGene_gn=ZNF521;refGene_tx=XM_023241838.1;refGene_gn=ZNF521;refGene_tx=XM_023241839.1;refGene_gn=ZNF521;refGene_tx=XM_023241840.1
D3 49730441 49730573
refGene_gn=ZNF521;refGene_tx=XM_023241838.1;refGene_gn=ZNF521;refGene_tx=XM_023241839.1;refGene_gn=ZNF521;refGene_tx=XM_023241840.1
D3 49831417 49831502
refGene_gn=ZNF521;refGene_tx=XM_023241838.1;refGene_gn=ZNF521;refGene_tx=XM_023241839.1;refGene_gn=ZNF521;refGene_tx=XM_023241840.1
D3 49860653 49864006
refGene_gn=ZNF521;refGene_tx=XM_023241838.1;refGene_gn=ZNF521;refGene_tx=XM_023241839.1;refGene_gn=ZNF521;refGene_tx=XM_023241840.1
D3 49951363 49951543
refGene_gn=ZNF521;refGene_tx=XM_023241838.1;refGene_gn=ZNF521;refGene_tx=XM_023241839.1
D3 49979066 49979106
refGene_gn=ZNF521;refGene_tx=XM_023241838.1;refGene_gn=ZNF521;refGene_tx=XM_023241839.1
D3 50605391 50605418
refGene_gn=SS18;refGene_tx=XM_011288062.3;refGene_gn=SS18;refGene_tx=XM_019815093.2;refGene_gn=SS18;refGene_tx=XM_019815095.2;refGene_gn=SS18;refGene_tx=XM_023241841.1;refGene_gn=SS18;refGene_tx=XM_023241843.1;refGene_gn=SS18;refGene_tx=XM_023241844.1
D3 50624386 50624520
refGene_gn=SS18;refGene_tx=XM_011288062.3;refGene_gn=SS18;refGene_tx=XM_019815093.2;refGene_gn=SS18;refGene_tx=XM_019815095.2;refGene_gn=SS18;refGene_tx=XM_023241841.1;refGene_gn=SS18;refGene_tx=XM_023241843.1;refGene_gn=SS18;refGene_tx=XM_023241844.1
D3 50627320 50627443
refGene_gn=SS18;refGene_tx=XM_011288062.3;refGene_gn=SS18;refGene_tx=XM_019815093.2;refGene_gn=SS18;refGene_tx=XM_019815095.2;refGene_gn=SS18;refGene_tx=XM_023241841.1;refGene_gn=SS18;refGene_tx=XM_023241843.1;refGene_gn=SS18;refGene_tx=XM_023241844.1
D3 50628155 50628248
refGene_gn=SS18;refGene_tx=XM_011288062.3;refGene_gn=SS18;refGene_tx=XM_019815093.2;refGene_gn=SS18;refGene_tx=XM_023241841.1;refGene_gn=SS18;refGene_tx=XM_023241843.1;refGene_gn=SS18;refGene_tx=XM_023241844.1
D3 50631044 50631149
refGene_gn=SS18;refGene_tx=XM_011288062.3;refGene_gn=SS18;refGene_tx=XM_019815093.2;refGene_gn=SS18;refGene_tx=XM_019815095.2;refGene_gn=SS1

8;refGene_tx=XM_023241841.1;refGene_gn=SS18;refGene_tx=XM_023241843.1;
refGene_gn=SS18;refGene_tx=XM_023241844.1
D3 50631808 50631976
refGene_gn=SS18;refGene_tx=XM_011288062.3;refGene_gn=SS18;refGene_tx=X
M_019815093.2;refGene_gn=SS18;refGene_tx=XM_019815095.2;refGene_gn=SS1
8;refGene_tx=XM_023241841.1;refGene_gn=SS18;refGene_tx=XM_023241843.1;
refGene_gn=SS18;refGene_tx=XM_023241844.1
D3 50647209 50647431
refGene_gn=SS18;refGene_tx=XM_011288062.3;refGene_gn=SS18;refGene_tx=X
M_019815093.2;refGene_gn=SS18;refGene_tx=XM_019815095.2;refGene_gn=SS1
8;refGene_tx=XM_023241841.1;refGene_gn=SS18;refGene_tx=XM_023241843.1;
refGene_gn=SS18;refGene_tx=XM_023241844.1
D3 50651256 50651410
refGene_gn=SS18;refGene_tx=XM_011288062.3;refGene_gn=SS18;refGene_tx=X
M_019815093.2;refGene_gn=SS18;refGene_tx=XM_019815095.2;refGene_gn=SS1
8;refGene_tx=XM_023241841.1;refGene_gn=SS18;refGene_tx=XM_023241843.1;
refGene_gn=SS18;refGene_tx=XM_023241844.1
D3 50681691 50681776
refGene_gn=SS18;refGene_tx=XM_011288062.3;refGene_gn=SS18;refGene_tx=X
M_019815093.2;refGene_gn=SS18;refGene_tx=XM_019815095.2;refGene_gn=SS1
8;refGene_tx=XM_023241841.1;refGene_gn=SS18;refGene_tx=XM_023241843.1
D3 50681884 50681892 refGene_gn=SS18;refGene_tx=XM_023241843.1
D3 50690761 50690838
refGene_gn=SS18;refGene_tx=XM_011288062.3;refGene_gn=SS18;refGene_tx=X
M_019815093.2;refGene_gn=SS18;refGene_tx=XM_019815095.2;refGene_gn=SS1
8;refGene_tx=XM_023241841.1
D3 50693433 50693502
refGene_gn=SS18;refGene_tx=XM_011288062.3;refGene_gn=SS18;refGene_tx=X
M_019815095.2
D3 50713189 50713291 refGene_gn=PSMA8;refGene_tx=XM_003995084.4
D3 50727317 50727444 refGene_gn=PSMA8;refGene_tx=XM_003995084.4
D3 50746590 50746715 refGene_gn=PSMA8;refGene_tx=XM_003995084.4
D3 50749262 50749385 refGene_gn=PSMA8;refGene_tx=XM_003995084.4
D3 50757965 50758085 refGene_gn=PSMA8;refGene_tx=XM_003995084.4
D3 50758198 50758261 refGene_gn=PSMA8;refGene_tx=XM_003995084.4
D3 50761989 50762082 refGene_gn=PSMA8;refGene_tx=XM_003995084.4
D3 50789895 50790238 refGene_gn=TAF4B;refGene_tx=XM_006938826.4
D3 50803731 50803877 refGene_gn=TAF4B;refGene_tx=XM_006938826.4
D3 50806724 50806832 refGene_gn=TAF4B;refGene_tx=XM_006938826.4
D3 50810669 50810834 refGene_gn=TAF4B;refGene_tx=XM_006938826.4
D3 50810936 50811059 refGene_gn=TAF4B;refGene_tx=XM_006938826.4
D3 50814496 50814586 refGene_gn=TAF4B;refGene_tx=XM_006938826.4
D3 50818822 50819437 refGene_gn=TAF4B;refGene_tx=XM_006938826.4
D3 50831124 50831260 refGene_gn=TAF4B;refGene_tx=XM_006938826.4
D3 50831924 50832030 refGene_gn=TAF4B;refGene_tx=XM_006938826.4
D3 50852051 50852221 refGene_gn=TAF4B;refGene_tx=XM_006938826.4
D3 50856995 50857126 refGene_gn=TAF4B;refGene_tx=XM_006938826.4
D3 50861974 50862100 refGene_gn=TAF4B;refGene_tx=XM_006938826.4
D3 50868823 50868880 refGene_gn=TAF4B;refGene_tx=XM_006938826.4
D3 50883828 50883939 refGene_gn=TAF4B;refGene_tx=XM_006938826.4

D3 50914671 50914839 refGene_gn=TAF4B;refGene_tx=XM_006938826.4
D3 50984565 50984724
refGene_gn=KCTD1;refGene_tx=XM_019815099.2;refGene_gn=KCTD1;refGene_tx=XM_019815100.2;refGene_gn=KCTD1;refGene_tx=XM_019815101.2;refGene_gn=KCTD1;refGene_tx=XM_023241846.1
D3 50989074 50989380
refGene_gn=KCTD1;refGene_tx=XM_019815099.2;refGene_gn=KCTD1;refGene_tx=XM_019815100.2;refGene_gn=KCTD1;refGene_tx=XM_019815101.2;refGene_gn=KCTD1;refGene_tx=XM_023241846.1
D3 51005726 51005871
refGene_gn=KCTD1;refGene_tx=XM_019815099.2;refGene_gn=KCTD1;refGene_tx=XM_019815100.2;refGene_gn=KCTD1;refGene_tx=XM_019815101.2;refGene_gn=KCTD1;refGene_tx=XM_023241846.1
D3 51029268 51029447
refGene_gn=KCTD1;refGene_tx=XM_019815099.2;refGene_gn=KCTD1;refGene_tx=XM_019815100.2;refGene_gn=KCTD1;refGene_tx=XM_019815101.2;refGene_gn=KCTD1;refGene_tx=XM_023241846.1
D3 51038366 51038372 refGene_gn=KCTD1;refGene_tx=XM_019815101.2
D3 51073643 51075452 refGene_gn=KCTD1;refGene_tx=XM_019815099.2
D3 51172592 51172601 refGene_gn=KCTD1;refGene_tx=XM_019815100.2
D3 51346307 51346586
refGene_gn=AQP4;refGene_tx=XM_003995088.4;refGene_gn=AQP4;refGene_tx=XM_011288061.2;refGene_gn=AQP4;refGene_tx=XM_019815105.1
D3 51351270 51351351
refGene_gn=AQP4;refGene_tx=XM_003995088.4;refGene_gn=AQP4;refGene_tx=XM_011288061.2;refGene_gn=AQP4;refGene_tx=XM_019815105.1
D3 51351649 51351814
refGene_gn=AQP4;refGene_tx=XM_003995088.4;refGene_gn=AQP4;refGene_tx=XM_011288061.2;refGene_gn=AQP4;refGene_tx=XM_019815105.1
D3 51352740 51353155
refGene_gn=AQP4;refGene_tx=XM_003995088.4;refGene_gn=AQP4;refGene_tx=XM_011288061.2;refGene_gn=AQP4;refGene_tx=XM_019815105.1
D3 51356244 51356276 refGene_gn=AQP4;refGene_tx=XM_003995088.4
D3 51408301 51409393
refGene_gn=CHST9;refGene_tx=XM_006938827.4;refGene_gn=CHST9;refGene_tx=XM_019815102.2;refGene_gn=CHST9;refGene_tx=XM_019815103.2;refGene_gn=CHST9;refGene_tx=XM_019815104.2
D3 51435896 51435934
refGene_gn=CHST9;refGene_tx=XM_006938827.4;refGene_gn=CHST9;refGene_tx=XM_019815102.2;refGene_gn=CHST9;refGene_tx=XM_019815103.2;refGene_gn=CHST9;refGene_tx=XM_019815104.2
D3 51515315 51515357
refGene_gn=CHST9;refGene_tx=XM_006938827.4;refGene_gn=CHST9;refGene_tx=XM_019815102.2
D3 51541853 51541892
refGene_gn=CHST9;refGene_tx=XM_019815102.2;refGene_gn=CHST9;refGene_tx=XM_019815103.2
D3 51612167 51612288
refGene_gn=CHST9;refGene_tx=XM_006938827.4;refGene_gn=CHST9;refGene_tx=XM_019815102.2;refGene_gn=CHST9;refGene_tx=XM_019815103.2;refGene_gn=

CHST9;refGene_tx=XM_019815104.2
D3 52356167 52356374
refGene_gn=CDH2;refGene_tx=XM_006938831.4;refGene_gn=CDH2;refGene_tx=XM_023241848.1
D3 52367214 52367379
refGene_gn=CDH2;refGene_tx=XM_006938831.4;refGene_gn=CDH2;refGene_tx=XM_023241848.1
D3 52387141 52387281
refGene_gn=CDH2;refGene_tx=XM_006938831.4;refGene_gn=CDH2;refGene_tx=XM_023241848.1
D3 52389726 52389960
refGene_gn=CDH2;refGene_tx=XM_006938831.4;refGene_gn=CDH2;refGene_tx=XM_023241848.1
D3 52390254 52390488
refGene_gn=CDH2;refGene_tx=XM_006938831.4;refGene_gn=CDH2;refGene_tx=XM_023241848.1
D3 52393653 52393796
refGene_gn=CDH2;refGene_tx=XM_006938831.4;refGene_gn=CDH2;refGene_tx=XM_023241848.1
D3 52395117 52395371
refGene_gn=CDH2;refGene_tx=XM_006938831.4;refGene_gn=CDH2;refGene_tx=XM_023241848.1
D3 52397799 52397985
refGene_gn=CDH2;refGene_tx=XM_006938831.4;refGene_gn=CDH2;refGene_tx=XM_023241848.1
D3 52398660 52398798
refGene_gn=CDH2;refGene_tx=XM_006938831.4;refGene_gn=CDH2;refGene_tx=XM_023241848.1
D3 52406562 52406735
refGene_gn=CDH2;refGene_tx=XM_006938831.4;refGene_gn=CDH2;refGene_tx=XM_023241848.1
D3 52409335 52409480
refGene_gn=CDH2;refGene_tx=XM_006938831.4;refGene_gn=CDH2;refGene_tx=XM_023241848.1
D3 52413543 52413699
refGene_gn=CDH2;refGene_tx=XM_006938831.4;refGene_gn=CDH2;refGene_tx=XM_023241848.1
D3 52415424 52415571
refGene_gn=CDH2;refGene_tx=XM_006938831.4;refGene_gn=CDH2;refGene_tx=XM_023241848.1
D3 52417300 52417527
refGene_gn=CDH2;refGene_tx=XM_006938831.4;refGene_gn=CDH2;refGene_tx=XM_023241848.1
D3 52541262 52541374
refGene_gn=CDH2;refGene_tx=XM_006938831.4;refGene_gn=CDH2;refGene_tx=XM_023241848.1
D3 52547171 52547216 refGene_gn=CDH2;refGene_tx=XM_006938831.4
D3 52569582 52569642 refGene_gn=CDH2;refGene_tx=XM_023241848.1
D3 54938090 54938288 refGene_gn=DSC3;refGene_tx=XM_006938833.3
D3 54938398 54938425 refGene_gn=DSC3;refGene_tx=XM_019815113.2

D3 54940954 54941212
refGene_gn=DSC3;refGene_tx=XM_006938833.3;refGene_gn=DSC3;refGene_tx=XM_019815113.2
D3 54945853 54945975
refGene_gn=DSC3;refGene_tx=XM_006938833.3;refGene_gn=DSC3;refGene_tx=XM_019815113.2
D3 54948692 54948917
refGene_gn=DSC3;refGene_tx=XM_006938833.3;refGene_gn=DSC3;refGene_tx=XM_019815113.2
D3 54951103 54951328
refGene_gn=DSC3;refGene_tx=XM_006938833.3;refGene_gn=DSC3;refGene_tx=XM_019815113.2
D3 54952427 54952570
refGene_gn=DSC3;refGene_tx=XM_006938833.3;refGene_gn=DSC3;refGene_tx=XM_019815113.2
D3 54952680 54952937
refGene_gn=DSC3;refGene_tx=XM_006938833.3;refGene_gn=DSC3;refGene_tx=XM_019815113.2
D3 54965319 54965505
refGene_gn=DSC3;refGene_tx=XM_006938833.3;refGene_gn=DSC3;refGene_tx=XM_019815113.2
D3 54965872 54966007
refGene_gn=DSC3;refGene_tx=XM_006938833.3;refGene_gn=DSC3;refGene_tx=XM_019815113.2
D3 54969042 54969209
refGene_gn=DSC3;refGene_tx=XM_006938833.3;refGene_gn=DSC3;refGene_tx=XM_019815113.2
D3 54972191 54972336
refGene_gn=DSC3;refGene_tx=XM_006938833.3;refGene_gn=DSC3;refGene_tx=XM_019815113.2
D3 54973644 54973800
refGene_gn=DSC3;refGene_tx=XM_006938833.3;refGene_gn=DSC3;refGene_tx=XM_019815113.2
D3 54976862 54976982
refGene_gn=DSC3;refGene_tx=XM_006938833.3;refGene_gn=DSC3;refGene_tx=XM_019815113.2
D3 54978650 54978850
refGene_gn=DSC3;refGene_tx=XM_006938833.3;refGene_gn=DSC3;refGene_tx=XM_019815113.2
D3 54980894 54980979
refGene_gn=DSC3;refGene_tx=XM_006938833.3;refGene_gn=DSC3;refGene_tx=XM_019815113.2
D3 54991786 54991852
refGene_gn=DSC3;refGene_tx=XM_006938833.3;refGene_gn=DSC3;refGene_tx=XM_019815113.2
D3 55010022 55010220 refGene_gn=DSC2;refGene_tx=XM_011288060.3
D3 55010316 55010352 refGene_gn=DSC2;refGene_tx=XM_019815115.2
D3 55011012 55011270
refGene_gn=DSC2;refGene_tx=XM_011288060.3;refGene_gn=DSC2;refGene_tx=XM_019815115.2

D3 55013550 55013675
refGene_gn=DSC2;refGene_tx=XM_011288060.3;refGene_gn=DSC2;refGene_tx=XM_019815115.2
D3 55014042 55014279
refGene_gn=DSC2;refGene_tx=XM_011288060.3;refGene_gn=DSC2;refGene_tx=XM_019815115.2
D3 55017378 55017606
refGene_gn=DSC2;refGene_tx=XM_011288060.3;refGene_gn=DSC2;refGene_tx=XM_019815115.2
D3 55022217 55022360
refGene_gn=DSC2;refGene_tx=XM_011288060.3;refGene_gn=DSC2;refGene_tx=XM_019815115.2
D3 55022481 55022738
refGene_gn=DSC2;refGene_tx=XM_011288060.3;refGene_gn=DSC2;refGene_tx=XM_019815115.2
D3 55024306 55024492
refGene_gn=DSC2;refGene_tx=XM_011288060.3;refGene_gn=DSC2;refGene_tx=XM_019815115.2
D3 55024733 55024868
refGene_gn=DSC2;refGene_tx=XM_011288060.3;refGene_gn=DSC2;refGene_tx=XM_019815115.2
D3 55029654 55029821
refGene_gn=DSC2;refGene_tx=XM_011288060.3;refGene_gn=DSC2;refGene_tx=XM_019815115.2
D3 55030765 55030910
refGene_gn=DSC2;refGene_tx=XM_011288060.3;refGene_gn=DSC2;refGene_tx=XM_019815115.2
D3 55033931 55034087
refGene_gn=DSC2;refGene_tx=XM_011288060.3;refGene_gn=DSC2;refGene_tx=XM_019815115.2
D3 55035260 55035380
refGene_gn=DSC2;refGene_tx=XM_011288060.3;refGene_gn=DSC2;refGene_tx=XM_019815115.2
D3 55036328 55036528
refGene_gn=DSC2;refGene_tx=XM_011288060.3;refGene_gn=DSC2;refGene_tx=XM_019815115.2
D3 55037412 55037497
refGene_gn=DSC2;refGene_tx=XM_011288060.3;refGene_gn=DSC2;refGene_tx=XM_019815115.2
D3 55045070 55045139
refGene_gn=DSC2;refGene_tx=XM_011288060.3;refGene_gn=DSC2;refGene_tx=XM_019815115.2
D3 55068514 55068712
refGene_gn=DSC1;refGene_tx=XM_003995095.4;refGene_gn=DSC1;refGene_tx=XM_019815118.1
D3 55068804 55068840
refGene_gn=DSC1;refGene_tx=XM_006938832.2;refGene_gn=DSC1;refGene_tx=XM_019815119.2
D3 55069584 55069830
refGene_gn=DSC1;refGene_tx=XM_003995095.4;refGene_gn=DSC1;refGene_tx=X

M_006938832.2;refGene_gn=DSC1;refGene_tx=XM_019815118.1;refGene_gn=DSC1;refGene_tx=XM_019815119.2
D3 55070327 55070446
refGene_gn=DSC1;refGene_tx=XM_003995095.4;refGene_gn=DSC1;refGene_tx=XM_006938832.2;refGene_gn=DSC1;refGene_tx=XM_019815118.1;refGene_gn=DSC1;refGene_tx=XM_019815119.2
D3 55072075 55072318
refGene_gn=DSC1;refGene_tx=XM_003995095.4;refGene_gn=DSC1;refGene_tx=XM_006938832.2;refGene_gn=DSC1;refGene_tx=XM_019815118.1;refGene_gn=DSC1;refGene_tx=XM_019815119.2
D3 55073757 55073970
refGene_gn=DSC1;refGene_tx=XM_003995095.4;refGene_gn=DSC1;refGene_tx=XM_006938832.2;refGene_gn=DSC1;refGene_tx=XM_019815118.1;refGene_gn=DSC1;refGene_tx=XM_019815119.2
D3 55077841 55077984
refGene_gn=DSC1;refGene_tx=XM_003995095.4;refGene_gn=DSC1;refGene_tx=XM_006938832.2;refGene_gn=DSC1;refGene_tx=XM_019815118.1;refGene_gn=DSC1;refGene_tx=XM_019815119.2
D3 55078067 55078327
refGene_gn=DSC1;refGene_tx=XM_003995095.4;refGene_gn=DSC1;refGene_tx=XM_006938832.2;refGene_gn=DSC1;refGene_tx=XM_019815118.1;refGene_gn=DSC1;refGene_tx=XM_019815119.2
D3 55080630 55080816
refGene_gn=DSC1;refGene_tx=XM_003995095.4;refGene_gn=DSC1;refGene_tx=XM_006938832.2;refGene_gn=DSC1;refGene_tx=XM_019815118.1;refGene_gn=DSC1;refGene_tx=XM_019815119.2
D3 55082227 55082362
refGene_gn=DSC1;refGene_tx=XM_003995095.4;refGene_gn=DSC1;refGene_tx=XM_006938832.2;refGene_gn=DSC1;refGene_tx=XM_019815118.1;refGene_gn=DSC1;refGene_tx=XM_019815119.2
D3 55082858 55083025
refGene_gn=DSC1;refGene_tx=XM_003995095.4;refGene_gn=DSC1;refGene_tx=XM_006938832.2;refGene_gn=DSC1;refGene_tx=XM_019815118.1;refGene_gn=DSC1;refGene_tx=XM_019815119.2
D3 55084563 55084708
refGene_gn=DSC1;refGene_tx=XM_003995095.4;refGene_gn=DSC1;refGene_tx=XM_006938832.2;refGene_gn=DSC1;refGene_tx=XM_019815118.1;refGene_gn=DSC1;refGene_tx=XM_019815119.2
D3 55089925 55090081
refGene_gn=DSC1;refGene_tx=XM_003995095.4;refGene_gn=DSC1;refGene_tx=XM_006938832.2;refGene_gn=DSC1;refGene_tx=XM_019815118.1;refGene_gn=DSC1;refGene_tx=XM_019815119.2
D3 55091090 55091210
refGene_gn=DSC1;refGene_tx=XM_003995095.4;refGene_gn=DSC1;refGene_tx=XM_006938832.2;refGene_gn=DSC1;refGene_tx=XM_019815118.1;refGene_gn=DSC1;refGene_tx=XM_019815119.2
D3 55092416 55092619
refGene_gn=DSC1;refGene_tx=XM_003995095.4;refGene_gn=DSC1;refGene_tx=XM_006938832.2;refGene_gn=DSC1;refGene_tx=XM_019815118.1;refGene_gn=DSC1;refGene_tx=XM_019815119.2

D3 55093929 55094014
refGene_gn=DSC1;refGene_tx=XM_003995095.4;refGene_gn=DSC1;refGene_tx=XM_006938832.2;refGene_gn=DSC1;refGene_tx=XM_019815118.1;refGene_gn=DSC1;refGene_tx=XM_019815119.2
D3 55096725 55096788
refGene_gn=DSC1;refGene_tx=XM_003995095.4;refGene_gn=DSC1;refGene_tx=XM_006938832.2
D3 55243084 55243132 refGene_gn=DSG1;refGene_tx=XM_011288059.3
D3 55253521 55253557 refGene_gn=DSG1;refGene_tx=XM_011288059.3
D3 55253812 55253944 refGene_gn=DSG1;refGene_tx=XM_011288059.3
D3 55255617 55255773 refGene_gn=DSG1;refGene_tx=XM_011288059.3
D3 55258131 55258276 refGene_gn=DSG1;refGene_tx=XM_011288059.3
D3 55260142 55260309 refGene_gn=DSG1;refGene_tx=XM_011288059.3
D3 55262533 55262668 refGene_gn=DSG1;refGene_tx=XM_011288059.3
D3 55262971 55263157 refGene_gn=DSG1;refGene_tx=XM_011288059.3
D3 55265690 55265950 refGene_gn=DSG1;refGene_tx=XM_011288059.3
D3 55267263 55267403 refGene_gn=DSG1;refGene_tx=XM_011288059.3
D3 55270793 55271150 refGene_gn=DSG1;refGene_tx=XM_011288059.3
D3 55273023 55273157 refGene_gn=DSG1;refGene_tx=XM_011288059.3
D3 55273490 55273563 refGene_gn=DSG1;refGene_tx=XM_011288059.3
D3 55276414 55276623 refGene_gn=DSG1;refGene_tx=XM_011288059.3
D3 55278618 55279665 refGene_gn=DSG1;refGene_tx=XM_011288059.3
D3 55300789 55300837 refGene_gn=DSG4;refGene_tx=XM_019815116.1
D3 55306761 55306812
refGene_gn=DSC1;refGene_tx=XM_019815118.1;refGene_gn=DSC1;refGene_tx=XM_019815119.2
D3 55314982 55315018 refGene_gn=DSG4;refGene_tx=XM_019815116.1
D3 55316550 55316682 refGene_gn=DSG4;refGene_tx=XM_019815116.1
D3 55318348 55318504 refGene_gn=DSG4;refGene_tx=XM_019815116.1
D3 55318806 55318951 refGene_gn=DSG4;refGene_tx=XM_019815116.1
D3 55321135 55321302 refGene_gn=DSG4;refGene_tx=XM_019815116.1
D3 55321571 55321706 refGene_gn=DSG4;refGene_tx=XM_019815116.1
D3 55322738 55322924 refGene_gn=DSG4;refGene_tx=XM_019815116.1
D3 55328948 55329220 refGene_gn=DSG4;refGene_tx=XM_019815116.1
D3 55330434 55330574 refGene_gn=DSG4;refGene_tx=XM_019815116.1
D3 55332446 55332665 refGene_gn=DSG4;refGene_tx=XM_019815116.1
D3 55335986 55336292 refGene_gn=DSG4;refGene_tx=XM_019815116.1
D3 55339019 55339159 refGene_gn=DSG4;refGene_tx=XM_019815116.1
D3 55339311 55339375 refGene_gn=DSG4;refGene_tx=XM_019815116.1
D3 55340722 55340931 refGene_gn=DSG4;refGene_tx=XM_019815116.1
D3 55343299 55344067 refGene_gn=DSG4;refGene_tx=XM_019815116.1
D3 55372539 55372587 refGene_gn=DSG3;refGene_tx=XM_019815117.2
D3 55383506 55383542 refGene_gn=DSG3;refGene_tx=XM_019815117.2
D3 55384064 55384193 refGene_gn=DSG3;refGene_tx=XM_019815117.2
D3 55385058 55385214 refGene_gn=DSG3;refGene_tx=XM_019815117.2
D3 55385674 55385819 refGene_gn=DSG3;refGene_tx=XM_019815117.2
D3 55386474 55386641 refGene_gn=DSG3;refGene_tx=XM_019815117.2
D3 55387524 55387653 refGene_gn=DSG3;refGene_tx=XM_019815117.2
D3 55387956 55388142 refGene_gn=DSG3;refGene_tx=XM_019815117.2
D3 55391151 55391423 refGene_gn=DSG3;refGene_tx=XM_019815117.2

D3 55392036 55392176 refGene_gn=DSG3;refGene_tx=XM_019815117.2
D3 55393600 55393825 refGene_gn=DSG3;refGene_tx=XM_019815117.2
D3 55396737 55396995 refGene_gn=DSG3;refGene_tx=XM_019815117.2
D3 55400848 55400988 refGene_gn=DSG3;refGene_tx=XM_019815117.2
D3 55401510 55401574 refGene_gn=DSG3;refGene_tx=XM_019815117.2
D3 55402953 55403201 refGene_gn=DSG3;refGene_tx=XM_019815117.2
D3 55404337 55404904 refGene_gn=DSG3;refGene_tx=XM_019815117.2
D3 55414866 55414926
refGene_gn=DSC1;refGene_tx=XM_019815118.1;refGene_gn=DSC1;refGene_tx=XM_019815119.2
D3 55423101 55423731 refGene_gn=DSG2;refGene_tx=XM_023241850.1
D3 55444542 55444578 refGene_gn=DSG2;refGene_tx=XM_023241850.1
D3 55446276 55446411 refGene_gn=DSG2;refGene_tx=XM_023241850.1
D3 55449995 55450157 refGene_gn=DSG2;refGene_tx=XM_023241850.1
D3 55450300 55450445 refGene_gn=DSG2;refGene_tx=XM_023241850.1
D3 55451301 55451468 refGene_gn=DSG2;refGene_tx=XM_023241850.1
D3 55453395 55453533 refGene_gn=DSG2;refGene_tx=XM_023241850.1
D3 55453650 55453839 refGene_gn=DSG2;refGene_tx=XM_023241850.1
D3 55459787 55460053 refGene_gn=DSG2;refGene_tx=XM_023241850.1
D3 55461058 55461201 refGene_gn=DSG2;refGene_tx=XM_023241850.1
D3 55461514 55461742 refGene_gn=DSG2;refGene_tx=XM_023241850.1
D3 55467369 55467597 refGene_gn=DSG2;refGene_tx=XM_023241850.1
D3 55469589 55469711 refGene_gn=DSG2;refGene_tx=XM_023241850.1
D3 55470861 55471194 refGene_gn=DSG2;refGene_tx=XM_023241850.1
D3 55473046 55474117 refGene_gn=DSG2;refGene_tx=XM_023241850.1
D3 55510473 55510539 refGene_gn=TTR;refGene_tx=NM_001287556.1
D3 55511444 55511575 refGene_gn=TTR;refGene_tx=NM_001287556.1
D3 55514133 55514269 refGene_gn=TTR;refGene_tx=NM_001287556.1
D3 55517290 55517398 refGene_gn=TTR;refGene_tx=NM_001287556.1
D3 55553173 55553321 refGene_gn=B4GALT6;refGene_tx=XM_011288058.2
D3 55554347 55554449 refGene_gn=B4GALT6;refGene_tx=XM_011288058.2
D3 55555142 55555265 refGene_gn=B4GALT6;refGene_tx=XM_011288058.2
D3 55558862 55559050 refGene_gn=B4GALT6;refGene_tx=XM_011288058.2
D3 55564296 55564413 refGene_gn=B4GALT6;refGene_tx=XM_011288058.2
D3 55570595 55570720 refGene_gn=B4GALT6;refGene_tx=XM_011288058.2
D3 55587635 55587749 refGene_gn=B4GALT6;refGene_tx=XM_011288058.2
D3 55594940 55595057 refGene_gn=B4GALT6;refGene_tx=XM_011288058.2
D3 55617707 55617822 refGene_gn=B4GALT6;refGene_tx=XM_011288058.2
D3 55618458 55619319
refGene_gn=LOC109493397;refGene_tx=XM_023241851.1;refGene_gn=LOC109493397;refGene_tx=XM_023241852.1
D3 55621070 55621092
refGene_gn=LOC109493397;refGene_tx=XM_023241852.1
D3 55703433 55703668 refGene_gn=TRAPPC8;refGene_tx=XM_011288057.3
D3 55704619 55704709 refGene_gn=TRAPPC8;refGene_tx=XM_011288057.3
D3 55710480 55710626 refGene_gn=TRAPPC8;refGene_tx=XM_011288057.3
D3 55714915 55715017 refGene_gn=TRAPPC8;refGene_tx=XM_011288057.3
D3 55717848 55718022 refGene_gn=TRAPPC8;refGene_tx=XM_011288057.3
D3 55720315 55720374 refGene_gn=TRAPPC8;refGene_tx=XM_011288057.3
D3 55720462 55720531 refGene_gn=TRAPPC8;refGene_tx=XM_011288057.3

D3 55721819 55721916 refGene_gn=TRAPPC8;refGene_tx=XM_011288057.3
D3 55723246 55723394 refGene_gn=TRAPPC8;refGene_tx=XM_011288057.3
D3 55727142 55727585 refGene_gn=TRAPPC8;refGene_tx=XM_011288057.3
D3 55732640 55732795 refGene_gn=TRAPPC8;refGene_tx=XM_011288057.3
D3 55733219 55733346 refGene_gn=TRAPPC8;refGene_tx=XM_011288057.3
D3 55733872 55733947 refGene_gn=TRAPPC8;refGene_tx=XM_011288057.3
D3 55736552 55736683 refGene_gn=TRAPPC8;refGene_tx=XM_011288057.3
D3 55737101 55737296 refGene_gn=TRAPPC8;refGene_tx=XM_011288057.3
D3 55739365 55739474 refGene_gn=TRAPPC8;refGene_tx=XM_011288057.3
D3 55740287 55740512 refGene_gn=TRAPPC8;refGene_tx=XM_011288057.3
D3 55744189 55744321 refGene_gn=TRAPPC8;refGene_tx=XM_011288057.3
D3 55751818 55751924 refGene_gn=TRAPPC8;refGene_tx=XM_011288057.3
D3 55757868 55757969 refGene_gn=TRAPPC8;refGene_tx=XM_011288057.3
D3 55762745 55762896 refGene_gn=TRAPPC8;refGene_tx=XM_011288057.3
D3 55763590 55763706 refGene_gn=TRAPPC8;refGene_tx=XM_011288057.3
D3 55764055 55764312 refGene_gn=TRAPPC8;refGene_tx=XM_011288057.3
D3 55765616 55765671 refGene_gn=TRAPPC8;refGene_tx=XM_011288057.3
D3 55765672 55765711 refGene_gn=TRAPPC8;refGene_tx=XM_011288057.3
D3 55768242 55768396 refGene_gn=TRAPPC8;refGene_tx=XM_011288057.3
D3 55770954 55771129 refGene_gn=TRAPPC8;refGene_tx=XM_011288057.3
D3 55772642 55772732 refGene_gn=TRAPPC8;refGene_tx=XM_011288057.3
D3 55776721 55776916 refGene_gn=TRAPPC8;refGene_tx=XM_011288057.3
D3 55785291 55785448 refGene_gn=TRAPPC8;refGene_tx=XM_011288057.3
D3 55842552 55842716 refGene_gn=RNF125;refGene_tx=XM_003995101.5
D3 55858559 55858713 refGene_gn=RNF125;refGene_tx=XM_003995101.5
D3 55862606 55862701 refGene_gn=RNF125;refGene_tx=XM_003995101.5
D3 55864785 55864876 refGene_gn=RNF125;refGene_tx=XM_003995101.5
D3 55875280 55875388 refGene_gn=RNF125;refGene_tx=XM_003995101.5
D3 55878405 55878492 refGene_gn=RNF125;refGene_tx=XM_003995101.5
D3 55897524 55897527
refGene_gn=RNF138;refGene_tx=XM_023241853.1;refGene_gn=RNF138;refGene_tx=XM_023241858.1
D3 55898990 55899184
refGene_gn=RNF138;refGene_tx=XM_023241853.1;refGene_gn=RNF138;refGene_tx=XM_023241854.1;refGene_gn=RNF138;refGene_tx=XM_023241855.1;refGene_gn=RNF138;refGene_tx=XM_023241858.1
D3 55910763 55910929
refGene_gn=RNF138;refGene_tx=XM_023241853.1;refGene_gn=RNF138;refGene_tx=XM_023241854.1;refGene_gn=RNF138;refGene_tx=XM_023241855.1;refGene_gn=RNF138;refGene_tx=XM_023241856.1;refGene_gn=RNF138;refGene_tx=XM_023241857.1
D3 55912106 55912222
refGene_gn=RNF138;refGene_tx=XM_023241853.1;refGene_gn=RNF138;refGene_tx=XM_023241854.1;refGene_gn=RNF138;refGene_tx=XM_023241855.1;refGene_gn=RNF138;refGene_tx=XM_023241856.1;refGene_gn=RNF138;refGene_tx=XM_023241857.1
D3 55923724 55923781
refGene_gn=RNF138;refGene_tx=XM_023241853.1;refGene_gn=RNF138;refGene_tx=XM_023241854.1;refGene_gn=RNF138;refGene_tx=XM_023241855.1;refGene_gn=RNF138;refGene_tx=XM_023241856.1;refGene_gn=RNF138;refGene_tx=XM_02

3241857.1;refGene_gn=RNF138;refGene_tx=XM_023241858.1
D3 55924613 55924725
refGene_gn=RNF138;refGene_tx=XM_023241853.1;refGene_gn=RNF138;refGene_tx=XM_023241854.1;refGene_gn=RNF138;refGene_tx=XM_023241855.1;refGene_gn=RNF138;refGene_tx=XM_023241856.1;refGene_gn=RNF138;refGene_tx=XM_023241857.1;refGene_gn=RNF138;refGene_tx=XM_023241858.1
D3 55926267 55926375
refGene_gn=RNF138;refGene_tx=XM_023241853.1;refGene_gn=RNF138;refGene_tx=XM_023241854.1;refGene_gn=RNF138;refGene_tx=XM_023241855.1;refGene_gn=RNF138;refGene_tx=XM_023241856.1;refGene_gn=RNF138;refGene_tx=XM_023241857.1;refGene_gn=RNF138;refGene_tx=XM_023241858.1
D3 55928450 55928519
refGene_gn=RNF138;refGene_tx=XM_023241853.1;refGene_gn=RNF138;refGene_tx=XM_023241854.1;refGene_gn=RNF138;refGene_tx=XM_023241855.1;refGene_gn=RNF138;refGene_tx=XM_023241856.1;refGene_gn=RNF138;refGene_tx=XM_023241857.1;refGene_gn=RNF138;refGene_tx=XM_023241858.1
D3 55970952 55971015 refGene_gn=MEP1B;refGene_tx=XM_023241859.1
D3 55973474 55973493 refGene_gn=MEP1B;refGene_tx=XM_023241859.1
D3 55974342 55974393 refGene_gn=MEP1B;refGene_tx=XM_023241859.1
D3 55974479 55974523 refGene_gn=MEP1B;refGene_tx=XM_023241859.1
D3 55976094 55976173 refGene_gn=MEP1B;refGene_tx=XM_023241859.1
D3 55983541 55983659 refGene_gn=MEP1B;refGene_tx=XM_023241859.1
D3 55984711 55984890 refGene_gn=MEP1B;refGene_tx=XM_023241859.1
D3 55986235 55986454 refGene_gn=MEP1B;refGene_tx=XM_023241859.1
D3 55987003 55987156 refGene_gn=MEP1B;refGene_tx=XM_023241859.1
D3 55988980 55989196 refGene_gn=MEP1B;refGene_tx=XM_023241859.1
D3 55991063 55991507 refGene_gn=MEP1B;refGene_tx=XM_023241859.1
D3 55993000 55993180 refGene_gn=MEP1B;refGene_tx=XM_023241859.1
D3 55995381 55995508 refGene_gn=MEP1B;refGene_tx=XM_023241859.1
D3 55996142 55996347 refGene_gn=MEP1B;refGene_tx=XM_023241859.1
D3 55998364 55998379 refGene_gn=MEP1B;refGene_tx=XM_023241859.1
D3 56042190 56043088
refGene_gn=GAREM1;refGene_tx=XM_023241860.1;refGene_gn=GAREM1;refGene_tx=XM_023241861.1;refGene_gn=GAREM1;refGene_tx=XM_023241862.1
D3 56044539 56044706
refGene_gn=GAREM1;refGene_tx=XM_023241860.1;refGene_gn=GAREM1;refGene_tx=XM_023241861.1;refGene_gn=GAREM1;refGene_tx=XM_023241862.1
D3 56059841 56061014
refGene_gn=GAREM1;refGene_tx=XM_023241860.1;refGene_gn=GAREM1;refGene_tx=XM_023241861.1;refGene_gn=GAREM1;refGene_tx=XM_023241862.1
D3 56081616 56081747
refGene_gn=GAREM1;refGene_tx=XM_023241860.1;refGene_gn=GAREM1;refGene_tx=XM_023241861.1;refGene_gn=GAREM1;refGene_tx=XM_023241862.1
D3 56164912 56165053
refGene_gn=GAREM1;refGene_tx=XM_023241860.1;refGene_gn=GAREM1;refGene_tx=XM_023241861.1;refGene_gn=GAREM1;refGene_tx=XM_023241862.1
D3 56239829 56239863 refGene_gn=GAREM1;refGene_tx=XM_023241862.1
D3 56241926 56242047
refGene_gn=GAREM1;refGene_tx=XM_023241860.1;refGene_gn=GAREM1;refGene_tx=XM_023241861.1

D3 56408425 56408566 refGene_gn=KLHL14;refGene_tx=XM_006938836.3
D3 56412285 56412443 refGene_gn=KLHL14;refGene_tx=XM_006938836.3
D3 56416382 56416541 refGene_gn=KLHL14;refGene_tx=XM_006938836.3
D3 56416622 56416813 refGene_gn=KLHL14;refGene_tx=XM_006938836.3
D3 56423568 56423647 refGene_gn=KLHL14;refGene_tx=XM_006938836.3
D3 56432003 56432093 refGene_gn=KLHL14;refGene_tx=XM_006938836.3
D3 56480704 56480826 refGene_gn=KLHL14;refGene_tx=XM_006938836.3
D3 56508961 56509911 refGene_gn=KLHL14;refGene_tx=XM_006938836.3
D3 56655817 56655898 refGene_gn=CCDC178;refGene_tx=XM_023241866.1
D3 56698874 56698985
refGene_gn=CCDC178;refGene_tx=XM_019815136.2;refGene_gn=CCDC178;refGene_tx=XM_023241863.1;refGene_gn=CCDC178;refGene_tx=XM_023241864.1;refGene_gn=CCDC178;refGene_tx=XM_023241865.1
D3 56699263 56699398
refGene_gn=CCDC178;refGene_tx=XM_019815136.2;refGene_gn=CCDC178;refGene_tx=XM_023241863.1;refGene_gn=CCDC178;refGene_tx=XM_023241864.1;refGene_gn=CCDC178;refGene_tx=XM_023241865.1;refGene_gn=CCDC178;refGene_tx=XM_023241866.1
D3 56791834 56791984
refGene_gn=CCDC178;refGene_tx=XM_019815136.2;refGene_gn=CCDC178;refGene_tx=XM_023241863.1;refGene_gn=CCDC178;refGene_tx=XM_023241864.1;refGene_gn=CCDC178;refGene_tx=XM_023241865.1;refGene_gn=CCDC178;refGene_tx=XM_023241866.1
D3 56903282 56903442
refGene_gn=CCDC178;refGene_tx=XM_019815136.2;refGene_gn=CCDC178;refGene_tx=XM_023241863.1;refGene_gn=CCDC178;refGene_tx=XM_023241864.1;refGene_gn=CCDC178;refGene_tx=XM_023241865.1;refGene_gn=CCDC178;refGene_tx=XM_023241866.1
D3 56908238 56908384
refGene_gn=CCDC178;refGene_tx=XM_019815136.2;refGene_gn=CCDC178;refGene_tx=XM_023241863.1;refGene_gn=CCDC178;refGene_tx=XM_023241864.1;refGene_gn=CCDC178;refGene_tx=XM_023241865.1;refGene_gn=CCDC178;refGene_tx=XM_023241866.1
D3 56914746 56914860
refGene_gn=CCDC178;refGene_tx=XM_019815136.2;refGene_gn=CCDC178;refGene_tx=XM_023241863.1;refGene_gn=CCDC178;refGene_tx=XM_023241864.1;refGene_gn=CCDC178;refGene_tx=XM_023241865.1;refGene_gn=CCDC178;refGene_tx=XM_023241866.1
D3 56916441 56916603
refGene_gn=CCDC178;refGene_tx=XM_019815136.2;refGene_gn=CCDC178;refGene_tx=XM_023241863.1;refGene_gn=CCDC178;refGene_tx=XM_023241864.1;refGene_gn=CCDC178;refGene_tx=XM_023241865.1;refGene_gn=CCDC178;refGene_tx=XM_023241866.1
D3 56917873 56917936
refGene_gn=CCDC178;refGene_tx=XM_019815136.2;refGene_gn=CCDC178;refGene_tx=XM_023241863.1;refGene_gn=CCDC178;refGene_tx=XM_023241864.1;refGene_gn=CCDC178;refGene_tx=XM_023241865.1;refGene_gn=CCDC178;refGene_tx=XM_023241866.1
D3 56943046 56943230
refGene_gn=CCDC178;refGene_tx=XM_019815136.2;refGene_gn=CCDC178;refGen

e_tx=XM_023241863.1;refGene_gn=CCDC178;refGene_tx=XM_023241864.1;refGene_gn=CCDC178;refGene_tx=XM_023241865.1;refGene_gn=CCDC178;refGene_tx=XM_023241866.1
D3 56963059 56963196
refGene_gn=CCDC178;refGene_tx=XM_019815136.2;refGene_gn=CCDC178;refGene_tx=XM_023241863.1;refGene_gn=CCDC178;refGene_tx=XM_023241864.1;refGene_gn=CCDC178;refGene_tx=XM_023241865.1;refGene_gn=CCDC178;refGene_tx=XM_023241866.1
D3 56963350 56963446
refGene_gn=CCDC178;refGene_tx=XM_019815136.2;refGene_gn=CCDC178;refGene_tx=XM_023241863.1;refGene_gn=CCDC178;refGene_tx=XM_023241864.1;refGene_gn=CCDC178;refGene_tx=XM_023241865.1;refGene_gn=CCDC178;refGene_tx=XM_023241866.1
D3 56979132 56979286
refGene_gn=CCDC178;refGene_tx=XM_019815136.2;refGene_gn=CCDC178;refGene_tx=XM_023241863.1;refGene_gn=CCDC178;refGene_tx=XM_023241864.1;refGene_gn=CCDC178;refGene_tx=XM_023241865.1;refGene_gn=CCDC178;refGene_tx=XM_023241866.1
D3 56991561 56991704
refGene_gn=CCDC178;refGene_tx=XM_019815136.2;refGene_gn=CCDC178;refGene_tx=XM_023241863.1;refGene_gn=CCDC178;refGene_tx=XM_023241864.1;refGene_gn=CCDC178;refGene_tx=XM_023241865.1;refGene_gn=CCDC178;refGene_tx=XM_023241866.1
D3 56996949 56997170
refGene_gn=CCDC178;refGene_tx=XM_019815136.2;refGene_gn=CCDC178;refGene_tx=XM_023241863.1;refGene_gn=CCDC178;refGene_tx=XM_023241864.1;refGene_gn=CCDC178;refGene_tx=XM_023241865.1;refGene_gn=CCDC178;refGene_tx=XM_023241866.1
D3 57001633 57001834
refGene_gn=CCDC178;refGene_tx=XM_019815136.2;refGene_gn=CCDC178;refGene_tx=XM_023241863.1;refGene_gn=CCDC178;refGene_tx=XM_023241864.1;refGene_gn=CCDC178;refGene_tx=XM_023241865.1;refGene_gn=CCDC178;refGene_tx=XM_023241866.1
D3 57005110 57005196
refGene_gn=CCDC178;refGene_tx=XM_019815136.2;refGene_gn=CCDC178;refGene_tx=XM_023241863.1;refGene_gn=CCDC178;refGene_tx=XM_023241864.1;refGene_gn=CCDC178;refGene_tx=XM_023241865.1;refGene_gn=CCDC178;refGene_tx=XM_023241866.1
D3 57017064 57017087
refGene_gn=CCDC178;refGene_tx=XM_023241863.1;refGene_gn=CCDC178;refGene_tx=XM_023241864.1;refGene_gn=CCDC178;refGene_tx=XM_023241865.1;refGene_gn=CCDC178;refGene_tx=XM_023241866.1
D3 57032736 57032876
refGene_gn=CCDC178;refGene_tx=XM_023241863.1;refGene_gn=CCDC178;refGene_tx=XM_023241864.1;refGene_gn=CCDC178;refGene_tx=XM_023241865.1;refGene_gn=CCDC178;refGene_tx=XM_023241866.1
D3 57039931 57040021
refGene_gn=CCDC178;refGene_tx=XM_023241863.1;refGene_gn=CCDC178;refGene_tx=XM_023241864.1;refGene_gn=CCDC178;refGene_tx=XM_023241865.1;refGene_gn=CCDC178;refGene_tx=XM_023241866.1

D3	57059380 57059438
refGene_gn=CCDC178;refGene_tx=XM_023241863.1;refGene_gn=CCDC178;refGene_tx=XM_023241864.1;refGene_gn=CCDC178;refGene_tx=XM_023241865.1;refGene_gn=CCDC178;refGene_tx=XM_023241866.1
D3	57198649 57198703 refGene_gn=ASXL3;refGene_tx=XM_023241867.1
D3	57231611 57231694
refGene_gn=ASXL3;refGene_tx=XM_023241867.1;refGene_gn=ASXL3;refGene_tx=XM_023241868.1
D3	57242683 57242686
refGene_gn=ASXL3;refGene_tx=XM_023241867.1;refGene_gn=ASXL3;refGene_tx=XM_023241868.1
D3	57266919 57267028
refGene_gn=ASXL3;refGene_tx=XM_023241867.1;refGene_gn=ASXL3;refGene_tx=XM_023241868.1;refGene_gn=ASXL3;refGene_tx=XM_023241869.1;refGene_gn=ASXL3;refGene_tx=XM_023241870.1
D3	57268266 57268375
refGene_gn=ASXL3;refGene_tx=XM_023241867.1;refGene_gn=ASXL3;refGene_tx=XM_023241868.1;refGene_gn=ASXL3;refGene_tx=XM_023241869.1;refGene_gn=ASXL3;refGene_tx=XM_023241870.1
D3	57284073 57284195
refGene_gn=ASXL3;refGene_tx=XM_023241867.1;refGene_gn=ASXL3;refGene_tx=XM_023241868.1;refGene_gn=ASXL3;refGene_tx=XM_023241869.1;refGene_gn=ASXL3;refGene_tx=XM_023241870.1
D3	57295835 57295953
refGene_gn=ASXL3;refGene_tx=XM_023241867.1;refGene_gn=ASXL3;refGene_tx=XM_023241868.1;refGene_gn=ASXL3;refGene_tx=XM_023241869.1;refGene_gn=ASXL3;refGene_tx=XM_023241870.1
D3	57296911 57297031
refGene_gn=ASXL3;refGene_tx=XM_023241867.1;refGene_gn=ASXL3;refGene_tx=XM_023241868.1;refGene_gn=ASXL3;refGene_tx=XM_023241869.1;refGene_gn=ASXL3;refGene_tx=XM_023241870.1
D3	57309370 57309534
refGene_gn=ASXL3;refGene_tx=XM_023241867.1;refGene_gn=ASXL3;refGene_tx=XM_023241868.1;refGene_gn=ASXL3;refGene_tx=XM_023241869.1;refGene_gn=ASXL3;refGene_tx=XM_023241870.1
D3	57360785 57360882
refGene_gn=ASXL3;refGene_tx=XM_023241867.1;refGene_gn=ASXL3;refGene_tx=XM_023241868.1;refGene_gn=ASXL3;refGene_tx=XM_023241869.1;refGene_gn=ASXL3;refGene_tx=XM_023241870.1
D3	57362979 57363085
refGene_gn=ASXL3;refGene_tx=XM_023241867.1;refGene_gn=ASXL3;refGene_tx=XM_023241868.1;refGene_gn=ASXL3;refGene_tx=XM_023241869.1;refGene_gn=ASXL3;refGene_tx=XM_023241870.1
D3	57368130 57370072
refGene_gn=ASXL3;refGene_tx=XM_023241867.1;refGene_gn=ASXL3;refGene_tx=XM_023241868.1;refGene_gn=ASXL3;refGene_tx=XM_023241869.1;refGene_gn=ASXL3;refGene_tx=XM_023241870.1
D3	57372911 57376625
refGene_gn=ASXL3;refGene_tx=XM_023241867.1;refGene_gn=ASXL3;refGene_tx=XM_023241868.1;refGene_gn=ASXL3;refGene_tx=XM_023241869.1;refGene_gn=

ASXL3;refGene_tx=XM_023241870.1
D3 57475097 57475291
refGene_gn=NOL4;refGene_tx=XM_003995112.5;refGene_gn=NOL4;refGene_tx=XM_003995114.5;refGene_gn=NOL4;refGene_tx=XM_019815138.2;refGene_gn=NOL4;refGene_tx=XM_019815140.1;refGene_gn=NOL4;refGene_tx=XM_019815141.2;refGene_gn=NOL4;refGene_tx=XM_019815142.2
D3 57544740 57544921
refGene_gn=NOL4;refGene_tx=XM_003995112.5;refGene_gn=NOL4;refGene_tx=XM_003995114.5;refGene_gn=NOL4;refGene_tx=XM_019815138.2;refGene_gn=NOL4;refGene_tx=XM_019815140.1;refGene_gn=NOL4;refGene_tx=XM_019815141.2;refGene_gn=NOL4;refGene_tx=XM_019815142.2
D3 57582729 57582843
refGene_gn=NOL4;refGene_tx=XM_003995112.5;refGene_gn=NOL4;refGene_tx=XM_003995114.5;refGene_gn=NOL4;refGene_tx=XM_019815138.2;refGene_gn=NOL4;refGene_tx=XM_019815140.1;refGene_gn=NOL4;refGene_tx=XM_019815141.2;refGene_gn=NOL4;refGene_tx=XM_019815142.2
D3 57601969 57602161
refGene_gn=NOL4;refGene_tx=XM_003995112.5;refGene_gn=NOL4;refGene_tx=XM_019815138.2;refGene_gn=NOL4;refGene_tx=XM_019815140.1;refGene_gn=NOL4;refGene_tx=XM_019815141.2;refGene_gn=NOL4;refGene_tx=XM_019815142.2
D3 57602900 57603080
refGene_gn=NOL4;refGene_tx=XM_003995112.5;refGene_gn=NOL4;refGene_tx=XM_003995114.5;refGene_gn=NOL4;refGene_tx=XM_019815138.2;refGene_gn=NOL4;refGene_tx=XM_019815140.1;refGene_gn=NOL4;refGene_tx=XM_019815141.2;refGene_gn=NOL4;refGene_tx=XM_019815142.2
D3 57690784 57691068
refGene_gn=NOL4;refGene_tx=XM_003995112.5;refGene_gn=NOL4;refGene_tx=XM_003995114.5;refGene_gn=NOL4;refGene_tx=XM_019815138.2;refGene_gn=NOL4;refGene_tx=XM_019815140.1;refGene_gn=NOL4;refGene_tx=XM_019815141.2;refGene_gn=NOL4;refGene_tx=XM_019815142.2
D3 57696011 57696030 refGene_gn=NOL4;refGene_tx=XM_019815141.2
D3 57774433 57774566
refGene_gn=NOL4;refGene_tx=XM_003995112.5;refGene_gn=NOL4;refGene_tx=XM_003995114.5;refGene_gn=NOL4;refGene_tx=XM_019815138.2;refGene_gn=NOL4;refGene_tx=XM_019815140.1
D3 57780540 57780653
refGene_gn=NOL4;refGene_tx=XM_003995112.5;refGene_gn=NOL4;refGene_tx=XM_003995114.5;refGene_gn=NOL4;refGene_tx=XM_019815138.2;refGene_gn=NOL4;refGene_tx=XM_019815140.1
D3 57781563 57781675
refGene_gn=NOL4;refGene_tx=XM_003995112.5;refGene_gn=NOL4;refGene_tx=XM_003995114.5;refGene_gn=NOL4;refGene_tx=XM_019815138.2;refGene_gn=NOL4;refGene_tx=XM_019815140.1
D3 57802931 57803081
refGene_gn=NOL4;refGene_tx=XM_003995112.5;refGene_gn=NOL4;refGene_tx=XM_003995114.5;refGene_gn=NOL4;refGene_tx=XM_019815138.2
D3 57901296 57901560
refGene_gn=NOL4;refGene_tx=XM_003995112.5;refGene_gn=NOL4;refGene_tx=XM_003995114.5
D3 58340440 58340507

refGene_gn=DTNA;refGene_tx=XM_011288048.3;refGene_gn=DTNA;refGene_tx=XM_011288054.2;refGene_gn=DTNA;refGene_tx=XM_019815143.2;refGene_gn=DTNA;refGene_tx=XM_019815144.2;refGene_gn=DTNA;refGene_tx=XM_019815145.2;refGene_gn=DTNA;refGene_tx=XM_019815146.2;refGene_gn=DTNA;refGene_tx=XM_019815147.2;refGene_gn=DTNA;refGene_tx=XM_019815149.2;refGene_gn=DTNA;refGene_tx=XM_019815150.2;refGene_gn=DTNA;refGene_tx=XM_019815151.2;refGene_gn=DTNA;refGene_tx=XM_019815152.2;refGene_gn=DTNA;refGene_tx=XM_019815153.2;refGene_gn=DTNA;refGene_tx=XM_019815154.2;refGene_gn=DTNA;refGene_tx=XM_019815155.2;refGene_gn=DTNA;refGene_tx=XM_019815156.2;refGene_gn=DTNA;refGene_tx=XM_019815157.2;refGene_gn=DTNA;refGene_tx=XM_019815158.2;refGene_gn=DTNA;refGene_tx=XM_019815159.2;refGene_gn=DTNA;refGene_tx=XM_019815160.2;refGene_gn=DTNA;refGene_tx=XM_019815161.2;refGene_gn=DTNA;refGene_tx=XM_019815162.1;refGene_gn=DTNA;refGene_tx=XM_019815163.2;refGene_gn=DTNA;refGene_tx=XM_019815164.2;refGene_gn=DTNA;refGene_tx=XM_019815165.2;refGene_gn=DTNA;refGene_tx=XM_019815166.2;refGene_gn=DTNA;refGene_tx=XM_019815167.2;refGene_gn=DTNA;refGene_tx=XM_019815168.1;refGene_gn=DTNA;refGene_tx=XM_019815169.2;refGene_gn=DTNA;refGene_tx=XM_019815170.1;refGene_gn=DTNA;refGene_tx=XM_019815171.1;refGene_gn=DTNA;refGene_tx=XM_019815172.1;refGene_gn=DTNA;refGene_tx=XM_019815173.1;refGene_gn=DTNA;refGene_tx=XM_023241871.1

D3 58350253 58350334

refGene_gn=DTNA;refGene_tx=XM_011288048.3;refGene_gn=DTNA;refGene_tx=XM_011288054.2;refGene_gn=DTNA;refGene_tx=XM_019815143.2;refGene_gn=DTNA;refGene_tx=XM_019815144.2;refGene_gn=DTNA;refGene_tx=XM_019815145.2;refGene_gn=DTNA;refGene_tx=XM_019815146.2;refGene_gn=DTNA;refGene_tx=XM_019815147.2;refGene_gn=DTNA;refGene_tx=XM_019815149.2;refGene_gn=DTNA;refGene_tx=XM_019815150.2;refGene_gn=DTNA;refGene_tx=XM_019815151.2;refGene_gn=DTNA;refGene_tx=XM_019815152.2;refGene_gn=DTNA;refGene_tx=XM_019815153.2;refGene_gn=DTNA;refGene_tx=XM_019815154.2;refGene_gn=DTNA;refGene_tx=XM_019815155.2;refGene_gn=DTNA;refGene_tx=XM_019815156.2;refGene_gn=DTNA;refGene_tx=XM_019815157.2;refGene_gn=DTNA;refGene_tx=XM_019815158.2;refGene_gn=DTNA;refGene_tx=XM_019815159.2;refGene_gn=DTNA;refGene_tx=XM_019815160.2;refGene_gn=DTNA;refGene_tx=XM_019815161.2;refGene_gn=DTNA;refGene_tx=XM_019815162.1;refGene_gn=DTNA;refGene_tx=XM_019815163.2;refGene_gn=DTNA;refGene_tx=XM_019815164.2;refGene_gn=DTNA;refGene_tx=XM_019815165.2;refGene_gn=DTNA;refGene_tx=XM_019815166.2;refGene_gn=DTNA;refGene_tx=XM_019815167.2;refGene_gn=DTNA;refGene_tx=XM_019815168.1;refGene_gn=DTNA;refGene_tx=XM_019815169.2;refGene_gn=DTNA;refGene_tx=XM_019815170.1;refGene_gn=DTNA;refGene_tx=XM_019815171.1;refGene_gn=DTNA;refGene_tx=XM_019815172.1;refGene_gn=DTNA;refGene_tx=XM_019815173.1;refGene_gn=DTNA;refGene_tx=XM_023241871.1

D3 58370649 58370863

refGene_gn=DTNA;refGene_tx=XM_011288048.3;refGene_gn=DTNA;refGene_tx=XM_011288054.2;refGene_gn=DTNA;refGene_tx=XM_019815143.2;refGene_gn=DTNA;refGene_tx=XM_019815144.2;refGene_gn=DTNA;refGene_tx=XM_019815145.2;refGene_gn=DTNA;refGene_tx=XM_019815146.2;refGene_gn=DTNA;refGene_tx=XM_019815147.2;refGene_gn=DTNA;refGene_tx=XM_019815149.2;refGene_gn=DTNA;refGene_tx=XM_019815150.2;refGene_gn=DTNA;refGene_tx=XM_019815151.2;refGene_gn=DTNA;refGene_tx=XM_019815152.2;refGene_gn=DTNA;refGene_tx=XM_019815153.2;refGene_gn=DTNA;refGene_tx=XM_019815154.2;refGene_gn=DTN

A;refGene_tx=XM_019815155.2;refGene_gn=DTNA;refGene_tx=XM_019815156.2;
refGene_gn=DTNA;refGene_tx=XM_019815157.2;refGene_gn=DTNA;refGene_tx=X
M_019815158.2;refGene_gn=DTNA;refGene_tx=XM_019815159.2;refGene_gn=DTN
A;refGene_tx=XM_019815160.2;refGene_gn=DTNA;refGene_tx=XM_019815161.2;
refGene_gn=DTNA;refGene_tx=XM_019815162.1;refGene_gn=DTNA;refGene_tx=X
M_019815163.2;refGene_gn=DTNA;refGene_tx=XM_019815164.2;refGene_gn=DTN
A;refGene_tx=XM_019815165.2;refGene_gn=DTNA;refGene_tx=XM_019815166.2;
refGene_gn=DTNA;refGene_tx=XM_019815167.2;refGene_gn=DTNA;refGene_tx=X
M_019815168.1;refGene_gn=DTNA;refGene_tx=XM_019815169.2;refGene_gn=DTN
A;refGene_tx=XM_019815170.1;refGene_gn=DTNA;refGene_tx=XM_019815171.1;
refGene_gn=DTNA;refGene_tx=XM_019815172.1;refGene_gn=DTNA;refGene_tx=X
M_019815173.1;refGene_gn=DTNA;refGene_tx=XM_023241871.1
D3 58382847 58382933
refGene_gn=DTNA;refGene_tx=XM_011288048.3;refGene_gn=DTNA;refGene_tx=X
M_011288054.2;refGene_gn=DTNA;refGene_tx=XM_019815143.2;refGene_gn=DTN
A;refGene_tx=XM_019815144.2;refGene_gn=DTNA;refGene_tx=XM_019815145.2;
refGene_gn=DTNA;refGene_tx=XM_019815146.2;refGene_gn=DTNA;refGene_tx=X
M_019815147.2;refGene_gn=DTNA;refGene_tx=XM_019815149.2;refGene_gn=DTN
A;refGene_tx=XM_019815150.2;refGene_gn=DTNA;refGene_tx=XM_019815151.2;
refGene_gn=DTNA;refGene_tx=XM_019815152.2;refGene_gn=DTNA;refGene_tx=X
M_019815153.2;refGene_gn=DTNA;refGene_tx=XM_019815154.2;refGene_gn=DTN
A;refGene_tx=XM_019815155.2;refGene_gn=DTNA;refGene_tx=XM_019815156.2;
refGene_gn=DTNA;refGene_tx=XM_019815157.2;refGene_gn=DTNA;refGene_tx=X
M_019815158.2;refGene_gn=DTNA;refGene_tx=XM_019815159.2;refGene_gn=DTN
A;refGene_tx=XM_019815160.2;refGene_gn=DTNA;refGene_tx=XM_019815161.2;
refGene_gn=DTNA;refGene_tx=XM_019815162.1;refGene_gn=DTNA;refGene_tx=X
M_019815163.2;refGene_gn=DTNA;refGene_tx=XM_019815164.2;refGene_gn=DTN
A;refGene_tx=XM_019815165.2;refGene_gn=DTNA;refGene_tx=XM_019815166.2;
refGene_gn=DTNA;refGene_tx=XM_019815167.2;refGene_gn=DTNA;refGene_tx=X
M_019815168.1;refGene_gn=DTNA;refGene_tx=XM_019815169.2;refGene_gn=DTN
A;refGene_tx=XM_019815170.1;refGene_gn=DTNA;refGene_tx=XM_019815171.1;
refGene_gn=DTNA;refGene_tx=XM_019815172.1;refGene_gn=DTNA;refGene_tx=X
M_019815173.1;refGene_gn=DTNA;refGene_tx=XM_023241871.1
D3 58389851 58390006
refGene_gn=DTNA;refGene_tx=XM_011288048.3;refGene_gn=DTNA;refGene_tx=X
M_011288054.2;refGene_gn=DTNA;refGene_tx=XM_019815143.2;refGene_gn=DTN
A;refGene_tx=XM_019815144.2;refGene_gn=DTNA;refGene_tx=XM_019815145.2;
refGene_gn=DTNA;refGene_tx=XM_019815146.2;refGene_gn=DTNA;refGene_tx=X
M_019815147.2;refGene_gn=DTNA;refGene_tx=XM_019815149.2;refGene_gn=DTN
A;refGene_tx=XM_019815150.2;refGene_gn=DTNA;refGene_tx=XM_019815151.2;
refGene_gn=DTNA;refGene_tx=XM_019815152.2;refGene_gn=DTNA;refGene_tx=X
M_019815153.2;refGene_gn=DTNA;refGene_tx=XM_019815154.2;refGene_gn=DTN
A;refGene_tx=XM_019815155.2;refGene_gn=DTNA;refGene_tx=XM_019815156.2;
refGene_gn=DTNA;refGene_tx=XM_019815157.2;refGene_gn=DTNA;refGene_tx=X
M_019815158.2;refGene_gn=DTNA;refGene_tx=XM_019815159.2;refGene_gn=DTN
A;refGene_tx=XM_019815160.2;refGene_gn=DTNA;refGene_tx=XM_019815161.2;
refGene_gn=DTNA;refGene_tx=XM_019815162.1;refGene_gn=DTNA;refGene_tx=X
M_019815163.2;refGene_gn=DTNA;refGene_tx=XM_019815164.2;refGene_gn=DTN
A;refGene_tx=XM_019815165.2;refGene_gn=DTNA;refGene_tx=XM_019815166.2;
refGene_gn=DTNA;refGene_tx=XM_019815167.2;refGene_gn=DTNA;refGene_tx=X

M_019815168.1;refGene_gn=DTNA;refGene_tx=XM_019815169.2;refGene_gn=DTNA;refGene_tx=XM_019815170.1;refGene_gn=DTNA;refGene_tx=XM_019815171.1;refGene_gn=DTNA;refGene_tx=XM_019815172.1;refGene_gn=DTNA;refGene_tx=XM_019815173.1;refGene_gn=DTNA;refGene_tx=XM_023241871.1
D3 58395126 58395232
refGene_gn=DTNA;refGene_tx=XM_011288048.3;refGene_gn=DTNA;refGene_tx=XM_011288054.2;refGene_gn=DTNA;refGene_tx=XM_019815143.2;refGene_gn=DTNA;refGene_tx=XM_019815144.2;refGene_gn=DTNA;refGene_tx=XM_019815145.2;refGene_gn=DTNA;refGene_tx=XM_019815146.2;refGene_gn=DTNA;refGene_tx=XM_019815147.2;refGene_gn=DTNA;refGene_tx=XM_019815149.2;refGene_gn=DTNA;refGene_tx=XM_019815150.2;refGene_gn=DTNA;refGene_tx=XM_019815151.2;refGene_gn=DTNA;refGene_tx=XM_019815152.2;refGene_gn=DTNA;refGene_tx=XM_019815153.2;refGene_gn=DTNA;refGene_tx=XM_019815154.2;refGene_gn=DTNA;refGene_tx=XM_019815155.2;refGene_gn=DTNA;refGene_tx=XM_019815156.2;refGene_gn=DTNA;refGene_tx=XM_019815157.2;refGene_gn=DTNA;refGene_tx=XM_019815158.2;refGene_gn=DTNA;refGene_tx=XM_019815159.2;refGene_gn=DTNA;refGene_tx=XM_019815160.2;refGene_gn=DTNA;refGene_tx=XM_019815161.2;refGene_gn=DTNA;refGene_tx=XM_019815162.1;refGene_gn=DTNA;refGene_tx=XM_019815163.2;refGene_gn=DTNA;refGene_tx=XM_019815164.2;refGene_gn=DTNA;refGene_tx=XM_019815165.2;refGene_gn=DTNA;refGene_tx=XM_019815166.2;refGene_gn=DTNA;refGene_tx=XM_019815167.2;refGene_gn=DTNA;refGene_tx=XM_019815168.1;refGene_gn=DTNA;refGene_tx=XM_019815169.2;refGene_gn=DTNA;refGene_tx=XM_019815170.1;refGene_gn=DTNA;refGene_tx=XM_019815171.1;refGene_gn=DTNA;refGene_tx=XM_019815172.1;refGene_gn=DTNA;refGene_tx=XM_019815173.1;refGene_gn=DTNA;refGene_tx=XM_023241871.1
D3 58396488 58396655
refGene_gn=DTNA;refGene_tx=XM_011288048.3;refGene_gn=DTNA;refGene_tx=XM_011288054.2;refGene_gn=DTNA;refGene_tx=XM_019815143.2;refGene_gn=DTNA;refGene_tx=XM_019815144.2;refGene_gn=DTNA;refGene_tx=XM_019815145.2;refGene_gn=DTNA;refGene_tx=XM_019815146.2;refGene_gn=DTNA;refGene_tx=XM_019815147.2;refGene_gn=DTNA;refGene_tx=XM_019815149.2;refGene_gn=DTNA;refGene_tx=XM_019815150.2;refGene_gn=DTNA;refGene_tx=XM_019815151.2;refGene_gn=DTNA;refGene_tx=XM_019815152.2;refGene_gn=DTNA;refGene_tx=XM_019815153.2;refGene_gn=DTNA;refGene_tx=XM_019815154.2;refGene_gn=DTNA;refGene_tx=XM_019815155.2;refGene_gn=DTNA;refGene_tx=XM_019815156.2;refGene_gn=DTNA;refGene_tx=XM_019815157.2;refGene_gn=DTNA;refGene_tx=XM_019815158.2;refGene_gn=DTNA;refGene_tx=XM_019815159.2;refGene_gn=DTNA;refGene_tx=XM_019815160.2;refGene_gn=DTNA;refGene_tx=XM_019815161.2;refGene_gn=DTNA;refGene_tx=XM_019815162.1;refGene_gn=DTNA;refGene_tx=XM_019815163.2;refGene_gn=DTNA;refGene_tx=XM_019815164.2;refGene_gn=DTNA;refGene_tx=XM_019815165.2;refGene_gn=DTNA;refGene_tx=XM_019815166.2;refGene_gn=DTNA;refGene_tx=XM_019815167.2;refGene_gn=DTNA;refGene_tx=XM_019815168.1;refGene_gn=DTNA;refGene_tx=XM_019815169.2;refGene_gn=DTNA;refGene_tx=XM_019815170.1;refGene_gn=DTNA;refGene_tx=XM_019815171.1;refGene_gn=DTNA;refGene_tx=XM_019815172.1;refGene_gn=DTNA;refGene_tx=XM_019815173.1;refGene_gn=DTNA;refGene_tx=XM_023241871.1
D3 58399285 58399410
refGene_gn=DTNA;refGene_tx=XM_011288048.3;refGene_gn=DTNA;refGene_tx=XM_011288054.2;refGene_gn=DTNA;refGene_tx=XM_019815143.2;refGene_gn=DTNA;refGene_tx=XM_019815144.2;refGene_gn=DTNA;refGene_tx=XM_019815145.2;

refGene_gn=DTNA;refGene_tx=XM_019815146.2;refGene_gn=DTNA;refGene_tx=XM_019815147.2;refGene_gn=DTNA;refGene_tx=XM_019815149.2;refGene_gn=DTNA;refGene_tx=XM_019815150.2;refGene_gn=DTNA;refGene_tx=XM_019815151.2;refGene_gn=DTNA;refGene_tx=XM_019815152.2;refGene_gn=DTNA;refGene_tx=XM_019815153.2;refGene_gn=DTNA;refGene_tx=XM_019815154.2;refGene_gn=DTNA;refGene_tx=XM_019815155.2;refGene_gn=DTNA;refGene_tx=XM_019815156.2;refGene_gn=DTNA;refGene_tx=XM_019815157.2;refGene_gn=DTNA;refGene_tx=XM_019815158.2;refGene_gn=DTNA;refGene_tx=XM_019815159.2;refGene_gn=DTNA;refGene_tx=XM_019815160.2;refGene_gn=DTNA;refGene_tx=XM_019815161.2;refGene_gn=DTNA;refGene_tx=XM_019815162.1;refGene_gn=DTNA;refGene_tx=XM_019815163.2;refGene_gn=DTNA;refGene_tx=XM_019815164.2;refGene_gn=DTNA;refGene_tx=XM_019815165.2;refGene_gn=DTNA;refGene_tx=XM_019815166.2;refGene_gn=DTNA;refGene_tx=XM_019815167.2;refGene_gn=DTNA;refGene_tx=XM_019815168.1;refGene_gn=DTNA;refGene_tx=XM_019815169.2;refGene_gn=DTNA;refGene_tx=XM_019815170.1;refGene_gn=DTNA;refGene_tx=XM_019815171.1;refGene_gn=DTNA;refGene_tx=XM_019815172.1;refGene_gn=DTNA;refGene_tx=XM_019815173.1;refGene_gn=DTNA;refGene_tx=XM_023241871.1

D3 58401831 58401840

refGene_gn=DTNA;refGene_tx=XM_019815143.2;refGene_gn=DTNA;refGene_tx=XM_019815144.2;refGene_gn=DTNA;refGene_tx=XM_019815145.2;refGene_gn=DTNA;refGene_tx=XM_019815146.2;refGene_gn=DTNA;refGene_tx=XM_019815149.2;refGene_gn=DTNA;refGene_tx=XM_019815150.2;refGene_gn=DTNA;refGene_tx=XM_019815151.2;refGene_gn=DTNA;refGene_tx=XM_019815152.2;refGene_gn=DTNA;refGene_tx=XM_019815154.2;refGene_gn=DTNA;refGene_tx=XM_019815155.2;refGene_gn=DTNA;refGene_tx=XM_019815156.2;refGene_gn=DTNA;refGene_tx=XM_019815157.2;refGene_gn=DTNA;refGene_tx=XM_019815158.2;refGene_gn=DTNA;refGene_tx=XM_019815159.2;refGene_gn=DTNA;refGene_tx=XM_019815161.2;refGene_gn=DTNA;refGene_tx=XM_019815164.2;refGene_gn=DTNA;refGene_tx=XM_019815168.1;refGene_gn=DTNA;refGene_tx=XM_019815170.1;refGene_gn=DTNA;refGene_tx=XM_019815171.1;refGene_gn=DTNA;refGene_tx=XM_019815172.1;refGene_gn=DTNA;refGene_tx=XM_019815173.1;refGene_gn=DTNA;refGene_tx=XM_023241871.1

D3 58403959 58404043

refGene_gn=DTNA;refGene_tx=XM_011288048.3;refGene_gn=DTNA;refGene_tx=XM_011288054.2;refGene_gn=DTNA;refGene_tx=XM_019815143.2;refGene_gn=DTNA;refGene_tx=XM_019815144.2;refGene_gn=DTNA;refGene_tx=XM_019815145.2;refGene_gn=DTNA;refGene_tx=XM_019815146.2;refGene_gn=DTNA;refGene_tx=XM_019815147.2;refGene_gn=DTNA;refGene_tx=XM_019815149.2;refGene_gn=DTNA;refGene_tx=XM_019815150.2;refGene_gn=DTNA;refGene_tx=XM_019815151.2;refGene_gn=DTNA;refGene_tx=XM_019815152.2;refGene_gn=DTNA;refGene_tx=XM_019815153.2;refGene_gn=DTNA;refGene_tx=XM_019815154.2;refGene_gn=DTNA;refGene_tx=XM_019815155.2;refGene_gn=DTNA;refGene_tx=XM_019815156.2;refGene_gn=DTNA;refGene_tx=XM_019815157.2;refGene_gn=DTNA;refGene_tx=XM_019815158.2;refGene_gn=DTNA;refGene_tx=XM_019815159.2;refGene_gn=DTNA;refGene_tx=XM_019815160.2;refGene_gn=DTNA;refGene_tx=XM_019815161.2;refGene_gn=DTNA;refGene_tx=XM_019815162.1;refGene_gn=DTNA;refGene_tx=XM_019815163.2;refGene_gn=DTNA;refGene_tx=XM_019815164.2;refGene_gn=DTNA;refGene_tx=XM_019815165.2;refGene_gn=DTNA;refGene_tx=XM_019815166.2;refGene_gn=DTNA;refGene_tx=XM_019815167.2;refGene_gn=DTNA;refGene_tx=XM_019815168.1;refGene_gn=DTNA;refGene_tx=XM_019815169.2;refGene_gn=DTN

A;refGene_tx=XM_019815170.1;refGene_gn=DTNA;refGene_tx=XM_019815171.1;
refGene_gn=DTNA;refGene_tx=XM_019815172.1;refGene_gn=DTNA;refGene_tx=X
M_019815173.1;refGene_gn=DTNA;refGene_tx=XM_023241871.1
D3 58405398 58405429
refGene_gn=DTNA;refGene_tx=XM_011288054.2;refGene_gn=DTNA;refGene_tx=X
M_019815173.1
D3 58405760 58405850
refGene_gn=DTNA;refGene_tx=XM_019815143.2;refGene_gn=DTNA;refGene_tx=X
M_019815144.2;refGene_gn=DTNA;refGene_tx=XM_019815145.2;refGene_gn=DTN
A;refGene_tx=XM_019815146.2;refGene_gn=DTNA;refGene_tx=XM_019815147.2;
refGene_gn=DTNA;refGene_tx=XM_019815149.2;refGene_gn=DTNA;refGene_tx=X
M_019815150.2;refGene_gn=DTNA;refGene_tx=XM_019815152.2;refGene_gn=DTN
A;refGene_tx=XM_019815154.2;refGene_gn=DTNA;refGene_tx=XM_019815155.2;
refGene_gn=DTNA;refGene_tx=XM_019815156.2;refGene_gn=DTNA;refGene_tx=X
M_019815157.2;refGene_gn=DTNA;refGene_tx=XM_019815158.2;refGene_gn=DTN
A;refGene_tx=XM_019815159.2;refGene_gn=DTNA;refGene_tx=XM_019815160.2;
refGene_gn=DTNA;refGene_tx=XM_019815165.2;refGene_gn=DTNA;refGene_tx=X
M_019815170.1;refGene_gn=DTNA;refGene_tx=XM_019815171.1;refGene_gn=DTN
A;refGene_tx=XM_019815172.1;refGene_gn=DTNA;refGene_tx=XM_023241871.1
D3 58406066 58406133 refGene_gn=DTNA;refGene_tx=XM_019815172.1
D3 58413475 58413553
refGene_gn=DTNA;refGene_tx=XM_019815143.2;refGene_gn=DTNA;refGene_tx=X
M_019815144.2;refGene_gn=DTNA;refGene_tx=XM_019815145.2;refGene_gn=DTN
A;refGene_tx=XM_019815146.2;refGene_gn=DTNA;refGene_tx=XM_019815147.2;
refGene_gn=DTNA;refGene_tx=XM_019815149.2;refGene_gn=DTNA;refGene_tx=X
M_019815150.2;refGene_gn=DTNA;refGene_tx=XM_019815151.2;refGene_gn=DTN
A;refGene_tx=XM_019815152.2;refGene_gn=DTNA;refGene_tx=XM_019815153.2;
refGene_gn=DTNA;refGene_tx=XM_019815154.2;refGene_gn=DTNA;refGene_tx=X
M_019815155.2;refGene_gn=DTNA;refGene_tx=XM_019815156.2;refGene_gn=DTN
A;refGene_tx=XM_019815157.2;refGene_gn=DTNA;refGene_tx=XM_019815159.2;
refGene_gn=DTNA;refGene_tx=XM_019815160.2;refGene_gn=DTNA;refGene_tx=X
M_019815162.1;refGene_gn=DTNA;refGene_tx=XM_019815164.2;refGene_gn=DTN
A;refGene_tx=XM_019815166.2;refGene_gn=DTNA;refGene_tx=XM_019815170.1;
refGene_gn=DTNA;refGene_tx=XM_019815171.1;refGene_gn=DTNA;refGene_tx=X
M_023241871.1
D3 58414113 58414206
refGene_gn=DTNA;refGene_tx=XM_019815143.2;refGene_gn=DTNA;refGene_tx=X
M_019815144.2;refGene_gn=DTNA;refGene_tx=XM_019815145.2;refGene_gn=DTN
A;refGene_tx=XM_019815146.2;refGene_gn=DTNA;refGene_tx=XM_019815147.2;
refGene_gn=DTNA;refGene_tx=XM_019815149.2;refGene_gn=DTNA;refGene_tx=X
M_019815150.2;refGene_gn=DTNA;refGene_tx=XM_019815151.2;refGene_gn=DTN
A;refGene_tx=XM_019815153.2;refGene_gn=DTNA;refGene_tx=XM_019815154.2;
refGene_gn=DTNA;refGene_tx=XM_019815155.2;refGene_gn=DTNA;refGene_tx=X
M_019815156.2;refGene_gn=DTNA;refGene_tx=XM_019815157.2;refGene_gn=DTN
A;refGene_tx=XM_019815159.2;refGene_gn=DTNA;refGene_tx=XM_019815160.2;
refGene_gn=DTNA;refGene_tx=XM_019815162.1;refGene_gn=DTNA;refGene_tx=X
M_019815164.2;refGene_gn=DTNA;refGene_tx=XM_019815170.1;refGene_gn=DTN
A;refGene_tx=XM_019815171.1;refGene_gn=DTNA;refGene_tx=XM_023241871.1
D3 58426085 58426173
refGene_gn=DTNA;refGene_tx=XM_011288048.3;refGene_gn=DTNA;refGene_tx=X

M_019815143.2;refGene_gn=DTNA;refGene_tx=XM_019815144.2;refGene_gn=DTN
A;refGene_tx=XM_019815145.2;refGene_gn=DTNA;refGene_tx=XM_019815146.2;
refGene_gn=DTNA;refGene_tx=XM_019815147.2;refGene_gn=DTNA;refGene_tx=X
M_019815149.2;refGene_gn=DTNA;refGene_tx=XM_019815150.2;refGene_gn=DTN
A;refGene_tx=XM_019815151.2;refGene_gn=DTNA;refGene_tx=XM_019815152.2;
refGene_gn=DTNA;refGene_tx=XM_019815153.2;refGene_gn=DTNA;refGene_tx=X
M_019815154.2;refGene_gn=DTNA;refGene_tx=XM_019815155.2;refGene_gn=DTN
A;refGene_tx=XM_019815156.2;refGene_gn=DTNA;refGene_tx=XM_019815157.2;
refGene_gn=DTNA;refGene_tx=XM_019815158.2;refGene_gn=DTNA;refGene_tx=X
M_019815159.2;refGene_gn=DTNA;refGene_tx=XM_019815160.2;refGene_gn=DTN
A;refGene_tx=XM_019815161.2;refGene_gn=DTNA;refGene_tx=XM_019815162.1;
refGene_gn=DTNA;refGene_tx=XM_019815163.2;refGene_gn=DTNA;refGene_tx=X
M_019815164.2;refGene_gn=DTNA;refGene_tx=XM_019815165.2;refGene_gn=DTN
A;refGene_tx=XM_019815166.2;refGene_gn=DTNA;refGene_tx=XM_019815167.2;
refGene_gn=DTNA;refGene_tx=XM_019815168.1;refGene_gn=DTNA;refGene_tx=X
M_019815169.2;refGene_gn=DTNA;refGene_tx=XM_019815170.1;refGene_gn=DTN
A;refGene_tx=XM_019815171.1;refGene_gn=DTNA;refGene_tx=XM_023241871.1
D3 58428147 58428245
refGene_gn=DTNA;refGene_tx=XM_011288048.3;refGene_gn=DTNA;refGene_tx=X
M_019815143.2;refGene_gn=DTNA;refGene_tx=XM_019815144.2;refGene_gn=DTN
A;refGene_tx=XM_019815145.2;refGene_gn=DTNA;refGene_tx=XM_019815146.2;
refGene_gn=DTNA;refGene_tx=XM_019815147.2;refGene_gn=DTNA;refGene_tx=X
M_019815149.2;refGene_gn=DTNA;refGene_tx=XM_019815150.2;refGene_gn=DTN
A;refGene_tx=XM_019815151.2;refGene_gn=DTNA;refGene_tx=XM_019815152.2;
refGene_gn=DTNA;refGene_tx=XM_019815153.2;refGene_gn=DTNA;refGene_tx=X
M_019815154.2;refGene_gn=DTNA;refGene_tx=XM_019815155.2;refGene_gn=DTN
A;refGene_tx=XM_019815156.2;refGene_gn=DTNA;refGene_tx=XM_019815157.2;
refGene_gn=DTNA;refGene_tx=XM_019815158.2;refGene_gn=DTNA;refGene_tx=X
M_019815159.2;refGene_gn=DTNA;refGene_tx=XM_019815160.2;refGene_gn=DTN
A;refGene_tx=XM_019815161.2;refGene_gn=DTNA;refGene_tx=XM_019815162.1;
refGene_gn=DTNA;refGene_tx=XM_019815163.2;refGene_gn=DTNA;refGene_tx=X
M_019815164.2;refGene_gn=DTNA;refGene_tx=XM_019815165.2;refGene_gn=DTN
A;refGene_tx=XM_019815166.2;refGene_gn=DTNA;refGene_tx=XM_019815167.2;
refGene_gn=DTNA;refGene_tx=XM_019815168.1;refGene_gn=DTNA;refGene_tx=X
M_019815169.2;refGene_gn=DTNA;refGene_tx=XM_019815170.1;refGene_gn=DTN
A;refGene_tx=XM_019815171.1;refGene_gn=DTNA;refGene_tx=XM_023241871.1
D3 58432686 58432800
refGene_gn=DTNA;refGene_tx=XM_011288048.3;refGene_gn=DTNA;refGene_tx=X
M_019815143.2;refGene_gn=DTNA;refGene_tx=XM_019815144.2;refGene_gn=DTN
A;refGene_tx=XM_019815145.2;refGene_gn=DTNA;refGene_tx=XM_019815146.2;
refGene_gn=DTNA;refGene_tx=XM_019815147.2;refGene_gn=DTNA;refGene_tx=X
M_019815149.2;refGene_gn=DTNA;refGene_tx=XM_019815150.2;refGene_gn=DTN
A;refGene_tx=XM_019815151.2;refGene_gn=DTNA;refGene_tx=XM_019815152.2;
refGene_gn=DTNA;refGene_tx=XM_019815153.2;refGene_gn=DTNA;refGene_tx=X
M_019815154.2;refGene_gn=DTNA;refGene_tx=XM_019815155.2;refGene_gn=DTN
A;refGene_tx=XM_019815156.2;refGene_gn=DTNA;refGene_tx=XM_019815157.2;
refGene_gn=DTNA;refGene_tx=XM_019815158.2;refGene_gn=DTNA;refGene_tx=X
M_019815159.2;refGene_gn=DTNA;refGene_tx=XM_019815160.2;refGene_gn=DTN
A;refGene_tx=XM_019815161.2;refGene_gn=DTNA;refGene_tx=XM_019815162.1;
refGene_gn=DTNA;refGene_tx=XM_019815163.2;refGene_gn=DTNA;refGene_tx=X

M_019815164.2;refGene_gn=DTNA;refGene_tx=XM_019815165.2;refGene_gn=DTN
A;refGene_tx=XM_019815166.2;refGene_gn=DTNA;refGene_tx=XM_019815167.2;
refGene_gn=DTNA;refGene_tx=XM_019815168.1;refGene_gn=DTNA;refGene_tx=X
M_019815169.2;refGene_gn=DTNA;refGene_tx=XM_019815170.1;refGene_gn=DTN
A;refGene_tx=XM_019815171.1;refGene_gn=DTNA;refGene_tx=XM_023241871.1
D3 58440773 58440870
refGene_gn=DTNA;refGene_tx=XM_011288048.3;refGene_gn=DTNA;refGene_tx=X
M_019815143.2;refGene_gn=DTNA;refGene_tx=XM_019815144.2;refGene_gn=DTN
A;refGene_tx=XM_019815145.2;refGene_gn=DTNA;refGene_tx=XM_019815146.2;
refGene_gn=DTNA;refGene_tx=XM_019815147.2;refGene_gn=DTNA;refGene_tx=X
M_019815149.2;refGene_gn=DTNA;refGene_tx=XM_019815150.2;refGene_gn=DTN
A;refGene_tx=XM_019815151.2;refGene_gn=DTNA;refGene_tx=XM_019815152.2;
refGene_gn=DTNA;refGene_tx=XM_019815153.2;refGene_gn=DTNA;refGene_tx=X
M_019815154.2;refGene_gn=DTNA;refGene_tx=XM_019815155.2;refGene_gn=DTN
A;refGene_tx=XM_019815156.2;refGene_gn=DTNA;refGene_tx=XM_019815157.2;
refGene_gn=DTNA;refGene_tx=XM_019815158.2;refGene_gn=DTNA;refGene_tx=X
M_019815159.2;refGene_gn=DTNA;refGene_tx=XM_019815160.2;refGene_gn=DTN
A;refGene_tx=XM_019815161.2;refGene_gn=DTNA;refGene_tx=XM_019815162.1;
refGene_gn=DTNA;refGene_tx=XM_019815163.2;refGene_gn=DTNA;refGene_tx=X
M_019815164.2;refGene_gn=DTNA;refGene_tx=XM_019815165.2;refGene_gn=DTN
A;refGene_tx=XM_019815166.2;refGene_gn=DTNA;refGene_tx=XM_019815167.2;
refGene_gn=DTNA;refGene_tx=XM_019815168.1;refGene_gn=DTNA;refGene_tx=X
M_019815169.2;refGene_gn=DTNA;refGene_tx=XM_019815170.1;refGene_gn=DTN
A;refGene_tx=XM_019815171.1;refGene_gn=DTNA;refGene_tx=XM_023241871.1
D3 58442655 58442676
refGene_gn=DTNA;refGene_tx=XM_019815143.2;refGene_gn=DTNA;refGene_tx=X
M_019815144.2;refGene_gn=DTNA;refGene_tx=XM_019815145.2;refGene_gn=DTN
A;refGene_tx=XM_019815146.2;refGene_gn=DTNA;refGene_tx=XM_019815147.2;
refGene_gn=DTNA;refGene_tx=XM_019815150.2;refGene_gn=DTNA;refGene_tx=X
M_019815151.2;refGene_gn=DTNA;refGene_tx=XM_019815152.2;refGene_gn=DTN
A;refGene_tx=XM_019815153.2;refGene_gn=DTNA;refGene_tx=XM_019815154.2;
refGene_gn=DTNA;refGene_tx=XM_019815155.2;refGene_gn=DTNA;refGene_tx=X
M_019815156.2;refGene_gn=DTNA;refGene_tx=XM_019815157.2;refGene_gn=DTN
A;refGene_tx=XM_019815158.2;refGene_gn=DTNA;refGene_tx=XM_019815161.2;
refGene_gn=DTNA;refGene_tx=XM_019815162.1;refGene_gn=DTNA;refGene_tx=X
M_019815163.2;refGene_gn=DTNA;refGene_tx=XM_019815167.2;refGene_gn=DTN
A;refGene_tx=XM_019815170.1;refGene_gn=DTNA;refGene_tx=XM_023241871.1
D3 58443554 58443578
refGene_gn=DTNA;refGene_tx=XM_019815170.1;refGene_gn=DTNA;refGene_tx=X
M_019815171.1
D3 58450594 58450754
refGene_gn=DTNA;refGene_tx=XM_011288048.3;refGene_gn=DTNA;refGene_tx=X
M_019815143.2;refGene_gn=DTNA;refGene_tx=XM_019815144.2;refGene_gn=DTN
A;refGene_tx=XM_019815145.2;refGene_gn=DTNA;refGene_tx=XM_019815146.2;
refGene_gn=DTNA;refGene_tx=XM_019815147.2;refGene_gn=DTNA;refGene_tx=X
M_019815149.2;refGene_gn=DTNA;refGene_tx=XM_019815150.2;refGene_gn=DTN
A;refGene_tx=XM_019815151.2;refGene_gn=DTNA;refGene_tx=XM_019815152.2;
refGene_gn=DTNA;refGene_tx=XM_019815153.2;refGene_gn=DTNA;refGene_tx=X
M_019815154.2;refGene_gn=DTNA;refGene_tx=XM_019815155.2;refGene_gn=DTN
A;refGene_tx=XM_019815156.2;refGene_gn=DTNA;refGene_tx=XM_019815157.2;

refGene_gn=DTNA;refGene_tx=XM_019815158.2;refGene_gn=DTNA;refGene_tx=XM_019815159.2;refGene_gn=DTNA;refGene_tx=XM_019815160.2;refGene_gn=DTNA;refGene_tx=XM_019815161.2;refGene_gn=DTNA;refGene_tx=XM_019815162.1;refGene_gn=DTNA;refGene_tx=XM_019815163.2;refGene_gn=DTNA;refGene_tx=XM_019815164.2;refGene_gn=DTNA;refGene_tx=XM_019815165.2;refGene_gn=DTNA;refGene_tx=XM_019815166.2;refGene_gn=DTNA;refGene_tx=XM_019815167.2;refGene_gn=DTNA;refGene_tx=XM_019815168.1;refGene_gn=DTNA;refGene_tx=XM_019815169.2;refGene_gn=DTNA;refGene_tx=XM_023241871.1

D3 58453434 58453524

refGene_gn=DTNA;refGene_tx=XM_011288048.3;refGene_gn=DTNA;refGene_tx=XM_019815143.2;refGene_gn=DTNA;refGene_tx=XM_019815144.2;refGene_gn=DTNA;refGene_tx=XM_019815145.2;refGene_gn=DTNA;refGene_tx=XM_019815146.2;refGene_gn=DTNA;refGene_tx=XM_019815147.2;refGene_gn=DTNA;refGene_tx=XM_019815149.2;refGene_gn=DTNA;refGene_tx=XM_019815150.2;refGene_gn=DTNA;refGene_tx=XM_019815151.2;refGene_gn=DTNA;refGene_tx=XM_019815152.2;refGene_gn=DTNA;refGene_tx=XM_019815153.2;refGene_gn=DTNA;refGene_tx=XM_019815154.2;refGene_gn=DTNA;refGene_tx=XM_019815155.2;refGene_gn=DTNA;refGene_tx=XM_019815156.2;refGene_gn=DTNA;refGene_tx=XM_019815157.2;refGene_gn=DTNA;refGene_tx=XM_019815158.2;refGene_gn=DTNA;refGene_tx=XM_019815159.2;refGene_gn=DTNA;refGene_tx=XM_019815160.2;refGene_gn=DTNA;refGene_tx=XM_019815161.2;refGene_gn=DTNA;refGene_tx=XM_019815162.1;refGene_gn=DTNA;refGene_tx=XM_019815163.2;refGene_gn=DTNA;refGene_tx=XM_019815164.2;refGene_gn=DTNA;refGene_tx=XM_019815165.2;refGene_gn=DTNA;refGene_tx=XM_019815166.2;refGene_gn=DTNA;refGene_tx=XM_019815167.2;refGene_gn=DTNA;refGene_tx=XM_019815168.1;refGene_gn=DTNA;refGene_tx=XM_019815169.2;refGene_gn=DTNA;refGene_tx=XM_023241871.1

D3 58454812 58454981

refGene_gn=DTNA;refGene_tx=XM_011288048.3;refGene_gn=DTNA;refGene_tx=XM_019815143.2;refGene_gn=DTNA;refGene_tx=XM_019815144.2;refGene_gn=DTNA;refGene_tx=XM_019815145.2;refGene_gn=DTNA;refGene_tx=XM_019815146.2;refGene_gn=DTNA;refGene_tx=XM_019815147.2;refGene_gn=DTNA;refGene_tx=XM_019815149.2;refGene_gn=DTNA;refGene_tx=XM_019815150.2;refGene_gn=DTNA;refGene_tx=XM_019815151.2;refGene_gn=DTNA;refGene_tx=XM_019815152.2;refGene_gn=DTNA;refGene_tx=XM_019815153.2;refGene_gn=DTNA;refGene_tx=XM_019815154.2;refGene_gn=DTNA;refGene_tx=XM_019815155.2;refGene_gn=DTNA;refGene_tx=XM_019815156.2;refGene_gn=DTNA;refGene_tx=XM_019815157.2;refGene_gn=DTNA;refGene_tx=XM_019815158.2;refGene_gn=DTNA;refGene_tx=XM_019815159.2;refGene_gn=DTNA;refGene_tx=XM_019815160.2;refGene_gn=DTNA;refGene_tx=XM_019815161.2;refGene_gn=DTNA;refGene_tx=XM_019815162.1;refGene_gn=DTNA;refGene_tx=XM_019815163.2;refGene_gn=DTNA;refGene_tx=XM_019815164.2;refGene_gn=DTNA;refGene_tx=XM_019815165.2;refGene_gn=DTNA;refGene_tx=XM_019815166.2;refGene_gn=DTNA;refGene_tx=XM_019815167.2;refGene_gn=DTNA;refGene_tx=XM_019815168.1;refGene_gn=DTNA;refGene_tx=XM_019815169.2;refGene_gn=DTNA;refGene_tx=XM_023241871.1

D3 58457685 58457854

refGene_gn=DTNA;refGene_tx=XM_011288048.3;refGene_gn=DTNA;refGene_tx=XM_019815143.2;refGene_gn=DTNA;refGene_tx=XM_019815144.2;refGene_gn=DTNA;refGene_tx=XM_019815145.2;refGene_gn=DTNA;refGene_tx=XM_019815146.2;refGene_gn=DTNA;refGene_tx=XM_019815147.2;refGene_gn=DTNA;refGene_tx=XM_019815149.2;refGene_gn=DTNA;refGene_tx=XM_019815150.2;refGene_gn=DTN

A;refGene_tx=XM_019815151.2;refGene_gn=DTNA;refGene_tx=XM_019815152.2;refGene_gn=DTNA;refGene_tx=XM_019815153.2;refGene_gn=DTNA;refGene_tx=XM_019815154.2;refGene_gn=DTNA;refGene_tx=XM_019815155.2;refGene_gn=DTNA;refGene_tx=XM_019815156.2;refGene_gn=DTNA;refGene_tx=XM_019815157.2;refGene_gn=DTNA;refGene_tx=XM_019815158.2;refGene_gn=DTNA;refGene_tx=XM_019815159.2;refGene_gn=DTNA;refGene_tx=XM_019815160.2;refGene_gn=DTNA;refGene_tx=XM_019815161.2;refGene_gn=DTNA;refGene_tx=XM_019815162.1;refGene_gn=DTNA;refGene_tx=XM_019815163.2;refGene_gn=DTNA;refGene_tx=XM_019815164.2;refGene_gn=DTNA;refGene_tx=XM_019815165.2;refGene_gn=DTNA;refGene_tx=XM_019815166.2;refGene_gn=DTNA;refGene_tx=XM_019815167.2;refGene_gn=DTNA;refGene_tx=XM_019815168.1;refGene_gn=DTNA;refGene_tx=XM_019815169.2;refGene_gn=DTNA;refGene_tx=XM_023241871.1
D3 58460001 58460019
refGene_gn=DTNA;refGene_tx=XM_019815154.2;refGene_gn=DTNA;refGene_tx=XM_019815155.2;refGene_gn=DTNA;refGene_tx=XM_019815156.2;refGene_gn=DTNA;refGene_tx=XM_019815157.2;refGene_gn=DTNA;refGene_tx=XM_019815159.2;refGene_gn=DTNA;refGene_tx=XM_019815160.2;refGene_gn=DTNA;refGene_tx=XM_019815162.1;refGene_gn=DTNA;refGene_tx=XM_019815164.2;refGene_gn=DTNA;refGene_tx=XM_019815165.2;refGene_gn=DTNA;refGene_tx=XM_019815166.2;refGene_gn=DTNA;refGene_tx=XM_019815167.2;refGene_gn=DTNA;refGene_tx=XM_019815168.1;refGene_gn=DTNA;refGene_tx=XM_019815169.2
D3 58463027 58463171
refGene_gn=DTNA;refGene_tx=XM_019815143.2;refGene_gn=DTNA;refGene_tx=XM_019815144.2;refGene_gn=DTNA;refGene_tx=XM_019815145.2;refGene_gn=DTNA;refGene_tx=XM_019815146.2;refGene_gn=DTNA;refGene_tx=XM_019815147.2;refGene_gn=DTNA;refGene_tx=XM_019815149.2;refGene_gn=DTNA;refGene_tx=XM_019815150.2;refGene_gn=DTNA;refGene_tx=XM_019815151.2;refGene_gn=DTNA;refGene_tx=XM_019815152.2;refGene_gn=DTNA;refGene_tx=XM_019815153.2;refGene_gn=DTNA;refGene_tx=XM_019815158.2;refGene_gn=DTNA;refGene_tx=XM_019815161.2;refGene_gn=DTNA;refGene_tx=XM_019815163.2;refGene_gn=DTNA;refGene_tx=XM_023241871.1
D3 58465679 58465820 refGene_gn=DTNA;refGene_tx=XM_011288048.3
D3 58531985 58532009 refGene_gn=MAPRE2;refGene_tx=XM_006938845.3
D3 58558337 58558399 refGene_gn=MAPRE2;refGene_tx=XM_006938845.3
D3 58591352 58591471 refGene_gn=MAPRE2;refGene_tx=XM_003995130.4
D3 58622288 58622416
refGene_gn=MAPRE2;refGene_tx=XM_003995130.4;refGene_gn=MAPRE2;refGene_tx=XM_006938845.3;refGene_gn=MAPRE2;refGene_tx=XM_006938846.3;refGene_gn=MAPRE2;refGene_tx=XM_019815174.2
D3 58650200 58650346
refGene_gn=MAPRE2;refGene_tx=XM_003995130.4;refGene_gn=MAPRE2;refGene_tx=XM_006938845.3;refGene_gn=MAPRE2;refGene_tx=XM_006938846.3;refGene_gn=MAPRE2;refGene_tx=XM_019815174.2
D3 58653992 58654206
refGene_gn=MAPRE2;refGene_tx=XM_003995130.4;refGene_gn=MAPRE2;refGene_tx=XM_006938845.3;refGene_gn=MAPRE2;refGene_tx=XM_006938846.3;refGene_gn=MAPRE2;refGene_tx=XM_019815174.2
D3 58678990 58679130
refGene_gn=MAPRE2;refGene_tx=XM_003995130.4;refGene_gn=MAPRE2;refGene_tx=XM_006938845.3;refGene_gn=MAPRE2;refGene_tx=XM_006938846.3;refGene_

gn=MAPRE2;refGene_tx=XM_019815174.2
D3 58683224 58683383
refGene_gn=MAPRE2;refGene_tx=XM_003995130.4;refGene_gn=MAPRE2;refGene_tx=XM_006938845.3;refGene_gn=MAPRE2;refGene_tx=XM_006938846.3;refGene_gn=MAPRE2;refGene_tx=XM_019815174.2
D3 58688936 58689011
refGene_gn=MAPRE2;refGene_tx=XM_003995130.4;refGene_gn=MAPRE2;refGene_tx=XM_006938845.3;refGene_gn=MAPRE2;refGene_tx=XM_006938846.3;refGene_gn=MAPRE2;refGene_tx=XM_019815174.2
D3 58788923 58789337
refGene_gn=ZNF397;refGene_tx=XM_011288034.3;refGene_gn=ZNF397;refGene_tx=XM_019815177.2
D3 58789600 58789724
refGene_gn=ZNF397;refGene_tx=XM_011288034.3;refGene_gn=ZNF397;refGene_tx=XM_019815177.2
D3 58792942 58793991
refGene_gn=ZNF397;refGene_tx=XM_011288034.3;refGene_gn=ZNF397;refGene_tx=XM_019815177.2
D3 58800049 58800981
refGene_gn=LOC101093719;refGene_tx=XM_023241199.1
D3 58803955 58804094
refGene_gn=LOC101093719;refGene_tx=XM_023241199.1
D3 58804393 58804864
refGene_gn=LOC101093719;refGene_tx=XM_023241199.1
D3 58811242 58811391
refGene_gn=LOC101093719;refGene_tx=XM_023241199.1
D3 58820815 58820915
refGene_gn=LOC109493406;refGene_tx=XM_019815175.2;refGene_gn=LOC109493406;refGene_tx=XM_019815176.2
D3 58830802 58832814
refGene_gn=LOC109493406;refGene_tx=XM_019815175.2;refGene_gn=LOC109493406;refGene_tx=XM_019815176.2
D3 58851514 58852053
refGene_gn=LOC101093719;refGene_tx=XM_023241199.1;refGene_gn=ZNF24;refGene_tx=XM_006938853.4;refGene_gn=ZNF24;refGene_tx=XM_011288033.3
D3 58854195 58854343
refGene_gn=LOC101093719;refGene_tx=XM_023241199.1;refGene_gn=ZNF24;refGene_tx=XM_006938853.4;refGene_gn=ZNF24;refGene_tx=XM_011288033.3
D3 58854590 58855011
refGene_gn=LOC101093719;refGene_tx=XM_023241199.1;refGene_gn=ZNF24;refGene_tx=XM_006938853.4;refGene_gn=ZNF24;refGene_tx=XM_011288033.3
D3 58856474 58856971
refGene_gn=LOC101093719;refGene_tx=XM_023241199.1
D3 58857040 58857283
refGene_gn=LOC101093719;refGene_tx=XM_023241199.1
D3 58859496 58859518
refGene_gn=LOC101093719;refGene_tx=XM_023241199.1
D3 58875979 58876430
refGene_gn=LOC101093719;refGene_tx=XM_023241199.1
D3 58882574 58882716

refGene_gn=LOC101093719;refGene_tx=XM_023241199.1
D3 58882995 58883412
refGene_gn=LOC101093719;refGene_tx=XM_023241199.1
D3 58962274 58962406
refGene_gn=INO80C;refGene_tx=XM_003995136.4;refGene_gn=INO80C;refGene_tx=XM_019815178.2
D3 58968781 58968849
refGene_gn=INO80C;refGene_tx=XM_003995136.4;refGene_gn=INO80C;refGene_tx=XM_019815178.2
D3 58969781 58969893
refGene_gn=INO80C;refGene_tx=XM_003995136.4;refGene_gn=INO80C;refGene_tx=XM_019815178.2
D3 58970821 58970932
refGene_gn=INO80C;refGene_tx=XM_003995136.4;refGene_gn=INO80C;refGene_tx=XM_019815178.2
D3 58983376 58983532 refGene_gn=INO80C;refGene_tx=XM_003995136.4
D3 59117834 59117973 refGene_gn=GALNT1;refGene_tx=XM_003995137.5
D3 59127030 59127205 refGene_gn=GALNT1;refGene_tx=XM_003995137.5
D3 59156335 59156502 refGene_gn=GALNT1;refGene_tx=XM_003995137.5
D3 59161899 59162107 refGene_gn=GALNT1;refGene_tx=XM_003995137.5
D3 59166141 59166312 refGene_gn=GALNT1;refGene_tx=XM_003995137.5
D3 59168532 59168650 refGene_gn=GALNT1;refGene_tx=XM_003995137.5
D3 59170380 59170561 refGene_gn=GALNT1;refGene_tx=XM_003995137.5
D3 59171432 59171572 refGene_gn=GALNT1;refGene_tx=XM_003995137.5
D3 59176201 59176300 refGene_gn=GALNT1;refGene_tx=XM_003995137.5
D3 59176986 59177121 refGene_gn=GALNT1;refGene_tx=XM_003995137.5
D3 59180506 59180653 refGene_gn=GALNT1;refGene_tx=XM_003995137.5
D3 59382772 59382922
refGene_gn=RPRD1A;refGene_tx=XM_006938854.3;refGene_gn=RPRD1A;refGene_tx=XM_011288029.3
D3 59404920 59404959 refGene_gn=RPRD1A;refGene_tx=XM_006938855.4
D3 59416395 59416515 refGene_gn=RPRD1A;refGene_tx=XM_019815179.2
D3 59417786 59417962
refGene_gn=RPRD1A;refGene_tx=XM_006938854.3;refGene_gn=RPRD1A;refGene_tx=XM_006938855.4;refGene_gn=RPRD1A;refGene_tx=XM_011288029.3;refGene_gn=RPRD1A;refGene_tx=XM_019815179.2
D3 59418070 59418197
refGene_gn=RPRD1A;refGene_tx=XM_006938854.3;refGene_gn=RPRD1A;refGene_tx=XM_006938855.4;refGene_gn=RPRD1A;refGene_tx=XM_011288029.3;refGene_gn=RPRD1A;refGene_tx=XM_019815179.2
D3 59421123 59421221
refGene_gn=RPRD1A;refGene_tx=XM_006938854.3;refGene_gn=RPRD1A;refGene_tx=XM_006938855.4;refGene_gn=RPRD1A;refGene_tx=XM_011288029.3;refGene_gn=RPRD1A;refGene_tx=XM_019815179.2
D3 59421301 59421408
refGene_gn=RPRD1A;refGene_tx=XM_006938854.3;refGene_gn=RPRD1A;refGene_tx=XM_006938855.4;refGene_gn=RPRD1A;refGene_tx=XM_011288029.3;refGene_gn=RPRD1A;refGene_tx=XM_019815179.2
D3 59424598 59424728
refGene_gn=RPRD1A;refGene_tx=XM_006938854.3;refGene_gn=RPRD1A;refGene_

tx=XM_006938855.4;refGene_gn=RPRD1A;refGene_tx=XM_011288029.3;refGene_gn=RPRD1A;refGene_tx=XM_019815179.2
D3 59448672 59448823
refGene_gn=RPRD1A;refGene_tx=XM_006938854.3;refGene_gn=RPRD1A;refGene_tx=XM_006938855.4;refGene_gn=RPRD1A;refGene_tx=XM_011288029.3;refGene_gn=RPRD1A;refGene_tx=XM_019815179.2
D3 59482627 59482780
refGene_gn=SLC39A6;refGene_tx=XM_023241874.1;refGene_gn=SLC39A6;refGene_tx=XM_023241875.1
D3 59483904 59484095
refGene_gn=SLC39A6;refGene_tx=XM_023241874.1;refGene_gn=SLC39A6;refGene_tx=XM_023241875.1
D3 59485420 59485502
refGene_gn=SLC39A6;refGene_tx=XM_023241874.1;refGene_gn=SLC39A6;refGene_tx=XM_023241875.1
D3 59486694 59487071
refGene_gn=SLC39A6;refGene_tx=XM_023241874.1;refGene_gn=SLC39A6;refGene_tx=XM_023241875.1
D3 59490487 59490593
refGene_gn=SLC39A6;refGene_tx=XM_023241874.1;refGene_gn=SLC39A6;refGene_tx=XM_023241875.1
D3 59495819 59496038
refGene_gn=SLC39A6;refGene_tx=XM_023241874.1;refGene_gn=SLC39A6;refGene_tx=XM_023241875.1
D3 59499526 59499696
refGene_gn=SLC39A6;refGene_tx=XM_023241874.1;refGene_gn=SLC39A6;refGene_tx=XM_023241875.1
D3 59500982 59501166
refGene_gn=SLC39A6;refGene_tx=XM_023241874.1;refGene_gn=SLC39A6;refGene_tx=XM_023241875.1
D3 59502638 59503364
refGene_gn=SLC39A6;refGene_tx=XM_023241874.1;refGene_gn=SLC39A6;refGene_tx=XM_023241875.1
D3 59506107 59506245 refGene_gn=ELP2;refGene_tx=XM_003995141.5
D3 59508821 59508900 refGene_gn=ELP2;refGene_tx=XM_003995141.5
D3 59511097 59511168 refGene_gn=ELP2;refGene_tx=XM_003995141.5
D3 59515029 59515186 refGene_gn=ELP2;refGene_tx=XM_003995141.5
D3 59515446 59515524 refGene_gn=ELP2;refGene_tx=XM_003995141.5
D3 59517977 59518042 refGene_gn=ELP2;refGene_tx=XM_003995141.5
D3 59518801 59518868 refGene_gn=ELP2;refGene_tx=XM_003995141.5
D3 59519584 59519725 refGene_gn=ELP2;refGene_tx=XM_003995141.5
D3 59520809 59520905 refGene_gn=ELP2;refGene_tx=XM_003995141.5
D3 59522326 59522427 refGene_gn=ELP2;refGene_tx=XM_003995141.5
D3 59522620 59522752 refGene_gn=ELP2;refGene_tx=XM_003995141.5
D3 59526306 59526456 refGene_gn=ELP2;refGene_tx=XM_003995141.5
D3 59528546 59528735 refGene_gn=ELP2;refGene_tx=XM_003995141.5
D3 59531043 59531113 refGene_gn=ELP2;refGene_tx=XM_003995141.5
D3 59531653 59531749 refGene_gn=ELP2;refGene_tx=XM_003995141.5
D3 59531893 59531951 refGene_gn=ELP2;refGene_tx=XM_003995141.5
D3 59532686 59532759 refGene_gn=ELP2;refGene_tx=XM_003995141.5

D3 59540202 59540395 refGene_gn=ELP2;refGene_tx=XM_003995141.5
D3 59542042 59542164 refGene_gn=ELP2;refGene_tx=XM_003995141.5
D3 59544367 59544501 refGene_gn=ELP2;refGene_tx=XM_003995141.5
D3 59545466 59545476 refGene_gn=ELP2;refGene_tx=XM_003995141.5
D3 59545477 59545581 refGene_gn=ELP2;refGene_tx=XM_003995141.5
D3 59549445 59549602 refGene_gn=ELP2;refGene_tx=XM_003995141.5
D3 59560063 59560205
refGene_gn=MOCOS;refGene_tx=XM_023241876.1;refGene_gn=MOCOS;refGene_tx=XM_023241877.1
D3 59566104 59566194
refGene_gn=MOCOS;refGene_tx=XM_023241876.1;refGene_gn=MOCOS;refGene_tx=XM_023241877.1;refGene_gn=MOCOS;refGene_tx=XM_023241878.1
D3 59569320 59569387
refGene_gn=MOCOS;refGene_tx=XM_023241876.1;refGene_gn=MOCOS;refGene_tx=XM_023241877.1;refGene_gn=MOCOS;refGene_tx=XM_023241878.1
D3 59570278 59570920
refGene_gn=MOCOS;refGene_tx=XM_023241876.1;refGene_gn=MOCOS;refGene_tx=XM_023241877.1;refGene_gn=MOCOS;refGene_tx=XM_023241878.1;refGene_gn=MOCOS;refGene_tx=XM_023241879.1
D3 59573081 59573158
refGene_gn=MOCOS;refGene_tx=XM_023241876.1;refGene_gn=MOCOS;refGene_tx=XM_023241877.1;refGene_gn=MOCOS;refGene_tx=XM_023241878.1;refGene_gn=MOCOS;refGene_tx=XM_023241879.1
D3 59575284 59575484
refGene_gn=MOCOS;refGene_tx=XM_023241876.1;refGene_gn=MOCOS;refGene_tx=XM_023241877.1;refGene_gn=MOCOS;refGene_tx=XM_023241878.1;refGene_gn=MOCOS;refGene_tx=XM_023241879.1
D3 59581627 59581744
refGene_gn=MOCOS;refGene_tx=XM_023241876.1;refGene_gn=MOCOS;refGene_tx=XM_023241877.1;refGene_gn=MOCOS;refGene_tx=XM_023241878.1;refGene_gn=MOCOS;refGene_tx=XM_023241879.1
D3 59583056 59583515
refGene_gn=MOCOS;refGene_tx=XM_023241876.1;refGene_gn=MOCOS;refGene_tx=XM_023241877.1;refGene_gn=MOCOS;refGene_tx=XM_023241878.1;refGene_gn=MOCOS;refGene_tx=XM_023241879.1
D3 59590693 59590856
refGene_gn=MOCOS;refGene_tx=XM_023241876.1;refGene_gn=MOCOS;refGene_tx=XM_023241877.1;refGene_gn=MOCOS;refGene_tx=XM_023241878.1;refGene_gn=MOCOS;refGene_tx=XM_023241879.1
D3 59597445 59597524
refGene_gn=MOCOS;refGene_tx=XM_023241876.1;refGene_gn=MOCOS;refGene_tx=XM_023241877.1;refGene_gn=MOCOS;refGene_tx=XM_023241878.1;refGene_gn=MOCOS;refGene_tx=XM_023241879.1
D3 59599006 59599128
refGene_gn=MOCOS;refGene_tx=XM_023241876.1;refGene_gn=MOCOS;refGene_tx=XM_023241877.1;refGene_gn=MOCOS;refGene_tx=XM_023241878.1;refGene_gn=MOCOS;refGene_tx=XM_023241879.1
D3 59604765 59604871
refGene_gn=MOCOS;refGene_tx=XM_023241876.1;refGene_gn=MOCOS;refGene_tx=XM_023241877.1;refGene_gn=MOCOS;refGene_tx=XM_023241878.1;refGene_gn=

MOCOS;refGene_tx=XM_023241879.1
D3 59608800 59608939
refGene_gn=MOCOS;refGene_tx=XM_023241876.1;refGene_gn=MOCOS;refGene_tx=XM_023241877.1;refGene_gn=MOCOS;refGene_tx=XM_023241878.1;refGene_gn=MOCOS;refGene_tx=XM_023241879.1
D3 59613874 59613979
refGene_gn=MOCOS;refGene_tx=XM_023241876.1;refGene_gn=MOCOS;refGene_tx=XM_023241877.1;refGene_gn=MOCOS;refGene_tx=XM_023241878.1;refGene_gn=MOCOS;refGene_tx=XM_023241879.1
D3 59615845 59615998
refGene_gn=MOCOS;refGene_tx=XM_023241876.1;refGene_gn=MOCOS;refGene_tx=XM_023241877.1;refGene_gn=MOCOS;refGene_tx=XM_023241878.1;refGene_gn=MOCOS;refGene_tx=XM_023241879.1
D3 59631546 59631711
refGene_gn=FHOD3;refGene_tx=XM_023241880.1;refGene_gn=FHOD3;refGene_tx=XM_023241881.1;refGene_gn=FHOD3;refGene_tx=XM_023241882.1;refGene_gn=FHOD3;refGene_tx=XM_023241883.1;refGene_gn=FHOD3;refGene_tx=XM_023241884.1;refGene_gn=FHOD3;refGene_tx=XM_023241885.1;refGene_gn=FHOD3;refGene_tx=XM_023241886.1;refGene_gn=FHOD3;refGene_tx=XM_023241887.1;refGene_gn=FHOD3;refGene_tx=XM_023241888.1;refGene_gn=FHOD3;refGene_tx=XM_023241889.1;refGene_gn=FHOD3;refGene_tx=XM_023241890.1;refGene_gn=FHOD3;refGene_tx=XM_023241892.1;refGene_gn=FHOD3;refGene_tx=XM_023241893.1
D3 59691454 59691561
refGene_gn=FHOD3;refGene_tx=XM_023241880.1;refGene_gn=FHOD3;refGene_tx=XM_023241881.1;refGene_gn=FHOD3;refGene_tx=XM_023241882.1;refGene_gn=FHOD3;refGene_tx=XM_023241883.1;refGene_gn=FHOD3;refGene_tx=XM_023241884.1;refGene_gn=FHOD3;refGene_tx=XM_023241885.1;refGene_gn=FHOD3;refGene_tx=XM_023241886.1;refGene_gn=FHOD3;refGene_tx=XM_023241887.1;refGene_gn=FHOD3;refGene_tx=XM_023241888.1;refGene_gn=FHOD3;refGene_tx=XM_023241889.1;refGene_gn=FHOD3;refGene_tx=XM_023241890.1;refGene_gn=FHOD3;refGene_tx=XM_023241892.1;refGene_gn=FHOD3;refGene_tx=XM_023241893.1
D3 59703723 59703788
refGene_gn=FHOD3;refGene_tx=XM_023241880.1;refGene_gn=FHOD3;refGene_tx=XM_023241881.1;refGene_gn=FHOD3;refGene_tx=XM_023241882.1;refGene_gn=FHOD3;refGene_tx=XM_023241883.1;refGene_gn=FHOD3;refGene_tx=XM_023241884.1;refGene_gn=FHOD3;refGene_tx=XM_023241885.1;refGene_gn=FHOD3;refGene_tx=XM_023241886.1;refGene_gn=FHOD3;refGene_tx=XM_023241887.1;refGene_gn=FHOD3;refGene_tx=XM_023241888.1;refGene_gn=FHOD3;refGene_tx=XM_023241889.1;refGene_gn=FHOD3;refGene_tx=XM_023241890.1;refGene_gn=FHOD3;refGene_tx=XM_023241892.1;refGene_gn=FHOD3;refGene_tx=XM_023241893.1
D3 59831318 59831386
refGene_gn=FHOD3;refGene_tx=XM_023241880.1;refGene_gn=FHOD3;refGene_tx=XM_023241881.1;refGene_gn=FHOD3;refGene_tx=XM_023241882.1;refGene_gn=FHOD3;refGene_tx=XM_023241883.1;refGene_gn=FHOD3;refGene_tx=XM_023241884.1;refGene_gn=FHOD3;refGene_tx=XM_023241885.1;refGene_gn=FHOD3;refGene_tx=XM_023241886.1;refGene_gn=FHOD3;refGene_tx=XM_023241887.1;refGene_gn=FHOD3;refGene_tx=XM_023241888.1;refGene_gn=FHOD3;refGene_tx=XM_023241889.1;refGene_gn=FHOD3;refGene_tx=XM_023241890.1;refGene_gn=FHOD3;refGene_tx=XM_023241892.1;refGene_gn=FHOD3;refGene_tx=XM_023241893.1
D3 59839240 59839346

refGene_gn=FHOD3;refGene_tx=XM_023241880.1;refGene_gn=FHOD3;refGene_tx=XM_023241881.1;refGene_gn=FHOD3;refGene_tx=XM_023241882.1;refGene_gn=FHOD3;refGene_tx=XM_023241883.1;refGene_gn=FHOD3;refGene_tx=XM_023241884.1;refGene_gn=FHOD3;refGene_tx=XM_023241885.1;refGene_gn=FHOD3;refGene_tx=XM_023241886.1;refGene_gn=FHOD3;refGene_tx=XM_023241887.1;refGene_gn=FHOD3;refGene_tx=XM_023241888.1;refGene_gn=FHOD3;refGene_tx=XM_023241889.1;refGene_gn=FHOD3;refGene_tx=XM_023241890.1;refGene_gn=FHOD3;refGene_tx=XM_023241892.1;refGene_gn=FHOD3;refGene_tx=XM_023241893.1
D3 59907726 59907821
refGene_gn=FHOD3;refGene_tx=XM_023241880.1;refGene_gn=FHOD3;refGene_tx=XM_023241881.1;refGene_gn=FHOD3;refGene_tx=XM_023241882.1;refGene_gn=FHOD3;refGene_tx=XM_023241883.1;refGene_gn=FHOD3;refGene_tx=XM_023241884.1;refGene_gn=FHOD3;refGene_tx=XM_023241885.1;refGene_gn=FHOD3;refGene_tx=XM_023241886.1;refGene_gn=FHOD3;refGene_tx=XM_023241887.1;refGene_gn=FHOD3;refGene_tx=XM_023241888.1;refGene_gn=FHOD3;refGene_tx=XM_023241889.1;refGene_gn=FHOD3;refGene_tx=XM_023241890.1;refGene_gn=FHOD3;refGene_tx=XM_023241891.1;refGene_gn=FHOD3;refGene_tx=XM_023241892.1;refGene_gn=FHOD3;refGene_tx=XM_023241893.1
D3 59923245 59923357
refGene_gn=FHOD3;refGene_tx=XM_023241880.1;refGene_gn=FHOD3;refGene_tx=XM_023241881.1;refGene_gn=FHOD3;refGene_tx=XM_023241882.1;refGene_gn=FHOD3;refGene_tx=XM_023241883.1;refGene_gn=FHOD3;refGene_tx=XM_023241884.1;refGene_gn=FHOD3;refGene_tx=XM_023241885.1;refGene_gn=FHOD3;refGene_tx=XM_023241886.1;refGene_gn=FHOD3;refGene_tx=XM_023241887.1;refGene_gn=FHOD3;refGene_tx=XM_023241888.1;refGene_gn=FHOD3;refGene_tx=XM_023241889.1;refGene_gn=FHOD3;refGene_tx=XM_023241890.1;refGene_gn=FHOD3;refGene_tx=XM_023241891.1;refGene_gn=FHOD3;refGene_tx=XM_023241892.1;refGene_gn=FHOD3;refGene_tx=XM_023241893.1
D3 59930509 59930604
refGene_gn=FHOD3;refGene_tx=XM_023241880.1;refGene_gn=FHOD3;refGene_tx=XM_023241881.1;refGene_gn=FHOD3;refGene_tx=XM_023241882.1;refGene_gn=FHOD3;refGene_tx=XM_023241883.1;refGene_gn=FHOD3;refGene_tx=XM_023241884.1;refGene_gn=FHOD3;refGene_tx=XM_023241885.1;refGene_gn=FHOD3;refGene_tx=XM_023241886.1;refGene_gn=FHOD3;refGene_tx=XM_023241887.1;refGene_gn=FHOD3;refGene_tx=XM_023241888.1;refGene_gn=FHOD3;refGene_tx=XM_023241889.1;refGene_gn=FHOD3;refGene_tx=XM_023241890.1;refGene_gn=FHOD3;refGene_tx=XM_023241891.1;refGene_gn=FHOD3;refGene_tx=XM_023241892.1;refGene_gn=FHOD3;refGene_tx=XM_023241893.1
D3 59936895 59937039
refGene_gn=FHOD3;refGene_tx=XM_023241880.1;refGene_gn=FHOD3;refGene_tx=XM_023241881.1;refGene_gn=FHOD3;refGene_tx=XM_023241882.1;refGene_gn=FHOD3;refGene_tx=XM_023241883.1;refGene_gn=FHOD3;refGene_tx=XM_023241884.1;refGene_gn=FHOD3;refGene_tx=XM_023241885.1;refGene_gn=FHOD3;refGene_tx=XM_023241886.1;refGene_gn=FHOD3;refGene_tx=XM_023241887.1;refGene_gn=FHOD3;refGene_tx=XM_023241888.1;refGene_gn=FHOD3;refGene_tx=XM_023241889.1;refGene_gn=FHOD3;refGene_tx=XM_023241890.1;refGene_gn=FHOD3;refGene_tx=XM_023241891.1;refGene_gn=FHOD3;refGene_tx=XM_023241892.1;refGene_gn=FHOD3;refGene_tx=XM_023241893.1
D3 59952031 59952157 refGene_gn=FHOD3;refGene_tx=XM_023241894.1
D3 59952442 59952690

refGene_gn=FHOD3;refGene_tx=XM_023241880.1;refGene_gn=FHOD3;refGene_tx=XM_023241881.1;refGene_gn=FHOD3;refGene_tx=XM_023241882.1;refGene_gn=FHOD3;refGene_tx=XM_023241883.1;refGene_gn=FHOD3;refGene_tx=XM_023241884.1;refGene_gn=FHOD3;refGene_tx=XM_023241885.1;refGene_gn=FHOD3;refGene_tx=XM_023241886.1;refGene_gn=FHOD3;refGene_tx=XM_023241887.1;refGene_gn=FHOD3;refGene_tx=XM_023241888.1;refGene_gn=FHOD3;refGene_tx=XM_023241889.1;refGene_gn=FHOD3;refGene_tx=XM_023241890.1;refGene_gn=FHOD3;refGene_tx=XM_023241891.1;refGene_gn=FHOD3;refGene_tx=XM_023241892.1;refGene_gn=FHOD3;refGene_tx=XM_023241893.1;refGene_gn=FHOD3;refGene_tx=XM_023241894.1

D3 59976538 59976634

refGene_gn=FHOD3;refGene_tx=XM_023241880.1;refGene_gn=FHOD3;refGene_tx=XM_023241881.1;refGene_gn=FHOD3;refGene_tx=XM_023241882.1;refGene_gn=FHOD3;refGene_tx=XM_023241883.1;refGene_gn=FHOD3;refGene_tx=XM_023241884.1;refGene_gn=FHOD3;refGene_tx=XM_023241885.1;refGene_gn=FHOD3;refGene_tx=XM_023241887.1;refGene_gn=FHOD3;refGene_tx=XM_023241888.1;refGene_gn=FHOD3;refGene_tx=XM_023241891.1;refGene_gn=FHOD3;refGene_tx=XM_023241894.1

D3 59979616 59979970

refGene_gn=FHOD3;refGene_tx=XM_023241880.1;refGene_gn=FHOD3;refGene_tx=XM_023241881.1;refGene_gn=FHOD3;refGene_tx=XM_023241882.1;refGene_gn=FHOD3;refGene_tx=XM_023241883.1;refGene_gn=FHOD3;refGene_tx=XM_023241884.1;refGene_gn=FHOD3;refGene_tx=XM_023241885.1;refGene_gn=FHOD3;refGene_tx=XM_023241886.1;refGene_gn=FHOD3;refGene_tx=XM_023241891.1;refGene_gn=FHOD3;refGene_tx=XM_023241894.1

D3 59980399 59980474

refGene_gn=FHOD3;refGene_tx=XM_023241880.1;refGene_gn=FHOD3;refGene_tx=XM_023241881.1;refGene_gn=FHOD3;refGene_tx=XM_023241882.1;refGene_gn=FHOD3;refGene_tx=XM_023241883.1;refGene_gn=FHOD3;refGene_tx=XM_023241885.1;refGene_gn=FHOD3;refGene_tx=XM_023241886.1;refGene_gn=FHOD3;refGene_tx=XM_023241887.1;refGene_gn=FHOD3;refGene_tx=XM_023241889.1;refGene_gn=FHOD3;refGene_tx=XM_023241891.1;refGene_gn=FHOD3;refGene_tx=XM_023241894.1

D3 59986377 59986485

refGene_gn=FHOD3;refGene_tx=XM_023241880.1;refGene_gn=FHOD3;refGene_tx=XM_023241881.1;refGene_gn=FHOD3;refGene_tx=XM_023241882.1;refGene_gn=FHOD3;refGene_tx=XM_023241883.1;refGene_gn=FHOD3;refGene_tx=XM_023241884.1;refGene_gn=FHOD3;refGene_tx=XM_023241885.1;refGene_gn=FHOD3;refGene_tx=XM_023241886.1;refGene_gn=FHOD3;refGene_tx=XM_023241887.1;refGene_gn=FHOD3;refGene_tx=XM_023241888.1;refGene_gn=FHOD3;refGene_tx=XM_023241889.1;refGene_gn=FHOD3;refGene_tx=XM_023241890.1;refGene_gn=FHOD3;refGene_tx=XM_023241891.1;refGene_gn=FHOD3;refGene_tx=XM_023241892.1;refGene_gn=FHOD3;refGene_tx=XM_023241893.1;refGene_gn=FHOD3;refGene_tx=XM_023241894.1

D3 60005587 60005722

refGene_gn=FHOD3;refGene_tx=XM_023241880.1;refGene_gn=FHOD3;refGene_tx=XM_023241881.1;refGene_gn=FHOD3;refGene_tx=XM_023241882.1;refGene_gn=FHOD3;refGene_tx=XM_023241883.1;refGene_gn=FHOD3;refGene_tx=XM_023241884.1;refGene_gn=FHOD3;refGene_tx=XM_023241885.1;refGene_gn=FHOD3;refGene_tx=XM_023241886.1;refGene_gn=FHOD3;refGene_tx=XM_023241887.1;refGen

e_gn=FHOD3;refGene_tx=XM_023241888.1;refGene_gn=FHOD3;refGene_tx=XM_023241889.1;refGene_gn=FHOD3;refGene_tx=XM_023241890.1;refGene_gn=FHOD3;refGene_tx=XM_023241891.1;refGene_gn=FHOD3;refGene_tx=XM_023241892.1;refGene_gn=FHOD3;refGene_tx=XM_023241893.1;refGene_gn=FHOD3;refGene_tx=XM_023241894.1
D3 60009886 60009937
refGene_gn=FHOD3;refGene_tx=XM_023241880.1;refGene_gn=FHOD3;refGene_tx=XM_023241881.1;refGene_gn=FHOD3;refGene_tx=XM_023241882.1;refGene_gn=FHOD3;refGene_tx=XM_023241884.1;refGene_gn=FHOD3;refGene_tx=XM_023241886.1;refGene_gn=FHOD3;refGene_tx=XM_023241887.1;refGene_gn=FHOD3;refGene_tx=XM_023241888.1;refGene_gn=FHOD3;refGene_tx=XM_023241889.1;refGene_gn=FHOD3;refGene_tx=XM_023241890.1;refGene_gn=FHOD3;refGene_tx=XM_023241891.1;refGene_gn=FHOD3;refGene_tx=XM_023241892.1;refGene_gn=FHOD3;refGene_tx=XM_023241894.1
D3 60015860 60016075
refGene_gn=FHOD3;refGene_tx=XM_023241880.1;refGene_gn=FHOD3;refGene_tx=XM_023241881.1;refGene_gn=FHOD3;refGene_tx=XM_023241882.1;refGene_gn=FHOD3;refGene_tx=XM_023241883.1;refGene_gn=FHOD3;refGene_tx=XM_023241884.1;refGene_gn=FHOD3;refGene_tx=XM_023241885.1;refGene_gn=FHOD3;refGene_tx=XM_023241886.1;refGene_gn=FHOD3;refGene_tx=XM_023241887.1;refGene_gn=FHOD3;refGene_tx=XM_023241888.1;refGene_gn=FHOD3;refGene_tx=XM_023241889.1;refGene_gn=FHOD3;refGene_tx=XM_023241890.1;refGene_gn=FHOD3;refGene_tx=XM_023241891.1;refGene_gn=FHOD3;refGene_tx=XM_023241892.1;refGene_gn=FHOD3;refGene_tx=XM_023241893.1;refGene_gn=FHOD3;refGene_tx=XM_023241894.1
D3 60033899 60034178
refGene_gn=FHOD3;refGene_tx=XM_023241880.1;refGene_gn=FHOD3;refGene_tx=XM_023241881.1;refGene_gn=FHOD3;refGene_tx=XM_023241882.1;refGene_gn=FHOD3;refGene_tx=XM_023241883.1;refGene_gn=FHOD3;refGene_tx=XM_023241884.1;refGene_gn=FHOD3;refGene_tx=XM_023241885.1;refGene_gn=FHOD3;refGene_tx=XM_023241886.1;refGene_gn=FHOD3;refGene_tx=XM_023241887.1;refGene_gn=FHOD3;refGene_tx=XM_023241888.1;refGene_gn=FHOD3;refGene_tx=XM_023241889.1;refGene_gn=FHOD3;refGene_tx=XM_023241890.1;refGene_gn=FHOD3;refGene_tx=XM_023241891.1;refGene_gn=FHOD3;refGene_tx=XM_023241892.1;refGene_gn=FHOD3;refGene_tx=XM_023241893.1;refGene_gn=FHOD3;refGene_tx=XM_023241894.1
D3 60042489 60043373
refGene_gn=FHOD3;refGene_tx=XM_023241880.1;refGene_gn=FHOD3;refGene_tx=XM_023241881.1;refGene_gn=FHOD3;refGene_tx=XM_023241882.1;refGene_gn=FHOD3;refGene_tx=XM_023241883.1;refGene_gn=FHOD3;refGene_tx=XM_023241884.1;refGene_gn=FHOD3;refGene_tx=XM_023241885.1;refGene_gn=FHOD3;refGene_tx=XM_023241886.1;refGene_gn=FHOD3;refGene_tx=XM_023241887.1;refGene_gn=FHOD3;refGene_tx=XM_023241888.1;refGene_gn=FHOD3;refGene_tx=XM_023241889.1;refGene_gn=FHOD3;refGene_tx=XM_023241890.1;refGene_gn=FHOD3;refGene_tx=XM_023241891.1;refGene_gn=FHOD3;refGene_tx=XM_023241892.1;refGene_gn=FHOD3;refGene_tx=XM_023241893.1;refGene_gn=FHOD3;refGene_tx=XM_023241894.1
D3 60053048 60053207
refGene_gn=FHOD3;refGene_tx=XM_023241880.1;refGene_gn=FHOD3;refGene_tx=XM_023241881.1;refGene_gn=FHOD3;refGene_tx=XM_023241882.1;refGene_gn=

FHOD3;refGene_tx=XM_023241883.1;refGene_gn=FHOD3;refGene_tx=XM_0232418
84.1;refGene_gn=FHOD3;refGene_tx=XM_023241885.1;refGene_gn=FHOD3;refGe
ne_tx=XM_023241886.1;refGene_gn=FHOD3;refGene_tx=XM_023241887.1;refGen
e_gn=FHOD3;refGene_tx=XM_023241888.1;refGene_gn=FHOD3;refGene_tx=XM_02
3241889.1;refGene_gn=FHOD3;refGene_tx=XM_023241890.1;refGene_gn=FHOD3;
refGene_tx=XM_023241891.1;refGene_gn=FHOD3;refGene_tx=XM_023241892.1;r
efGene_gn=FHOD3;refGene_tx=XM_023241893.1;refGene_gn=FHOD3;refGene_tx=
XM_023241894.1
D3 60063476 60063659
refGene_gn=FHOD3;refGene_tx=XM_023241880.1;refGene_gn=FHOD3;refGene_tx
=XM_023241881.1;refGene_gn=FHOD3;refGene_tx=XM_023241882.1;refGene_gn=
FHOD3;refGene_tx=XM_023241883.1;refGene_gn=FHOD3;refGene_tx=XM_0232418
84.1;refGene_gn=FHOD3;refGene_tx=XM_023241885.1;refGene_gn=FHOD3;refGe
ne_tx=XM_023241886.1;refGene_gn=FHOD3;refGene_tx=XM_023241887.1;refGen
e_gn=FHOD3;refGene_tx=XM_023241888.1;refGene_gn=FHOD3;refGene_tx=XM_02
3241889.1;refGene_gn=FHOD3;refGene_tx=XM_023241890.1;refGene_gn=FHOD3;
refGene_tx=XM_023241891.1;refGene_gn=FHOD3;refGene_tx=XM_023241892.1;r
efGene_gn=FHOD3;refGene_tx=XM_023241893.1;refGene_gn=FHOD3;refGene_tx=
XM_023241894.1
D3 60065068 60065188
refGene_gn=FHOD3;refGene_tx=XM_023241880.1;refGene_gn=FHOD3;refGene_tx
=XM_023241881.1;refGene_gn=FHOD3;refGene_tx=XM_023241882.1;refGene_gn=
FHOD3;refGene_tx=XM_023241883.1;refGene_gn=FHOD3;refGene_tx=XM_0232418
84.1;refGene_gn=FHOD3;refGene_tx=XM_023241885.1;refGene_gn=FHOD3;refGe
ne_tx=XM_023241886.1;refGene_gn=FHOD3;refGene_tx=XM_023241887.1;refGen
e_gn=FHOD3;refGene_tx=XM_023241888.1;refGene_gn=FHOD3;refGene_tx=XM_02
3241889.1;refGene_gn=FHOD3;refGene_tx=XM_023241890.1;refGene_gn=FHOD3;
refGene_tx=XM_023241891.1;refGene_gn=FHOD3;refGene_tx=XM_023241892.1;r
efGene_gn=FHOD3;refGene_tx=XM_023241893.1;refGene_gn=FHOD3;refGene_tx=
XM_023241894.1
D3 60065875 60066037
refGene_gn=FHOD3;refGene_tx=XM_023241880.1;refGene_gn=FHOD3;refGene_tx
=XM_023241881.1;refGene_gn=FHOD3;refGene_tx=XM_023241882.1;refGene_gn=
FHOD3;refGene_tx=XM_023241883.1;refGene_gn=FHOD3;refGene_tx=XM_0232418
84.1;refGene_gn=FHOD3;refGene_tx=XM_023241885.1;refGene_gn=FHOD3;refGe
ne_tx=XM_023241886.1;refGene_gn=FHOD3;refGene_tx=XM_023241887.1;refGen
e_gn=FHOD3;refGene_tx=XM_023241888.1;refGene_gn=FHOD3;refGene_tx=XM_02
3241889.1;refGene_gn=FHOD3;refGene_tx=XM_023241890.1;refGene_gn=FHOD3;
refGene_tx=XM_023241891.1;refGene_gn=FHOD3;refGene_tx=XM_023241892.1;r
efGene_gn=FHOD3;refGene_tx=XM_023241893.1;refGene_gn=FHOD3;refGene_tx=
XM_023241894.1
D3 60070590 60070781
refGene_gn=FHOD3;refGene_tx=XM_023241880.1;refGene_gn=FHOD3;refGene_tx
=XM_023241881.1;refGene_gn=FHOD3;refGene_tx=XM_023241882.1;refGene_gn=
FHOD3;refGene_tx=XM_023241883.1;refGene_gn=FHOD3;refGene_tx=XM_0232418
84.1;refGene_gn=FHOD3;refGene_tx=XM_023241885.1;refGene_gn=FHOD3;refGe
ne_tx=XM_023241886.1;refGene_gn=FHOD3;refGene_tx=XM_023241887.1;refGen
e_gn=FHOD3;refGene_tx=XM_023241888.1;refGene_gn=FHOD3;refGene_tx=XM_02
3241889.1;refGene_gn=FHOD3;refGene_tx=XM_023241890.1;refGene_gn=FHOD3;
refGene_tx=XM_023241891.1;refGene_gn=FHOD3;refGene_tx=XM_023241892.1;r

efGene_gn=FHOD3;refGene_tx=XM_023241893.1;refGene_gn=FHOD3;refGene_tx=XM_023241894.1
D3 60077236 60077429
refGene_gn=FHOD3;refGene_tx=XM_023241880.1;refGene_gn=FHOD3;refGene_tx=XM_023241881.1;refGene_gn=FHOD3;refGene_tx=XM_023241882.1;refGene_gn=FHOD3;refGene_tx=XM_023241883.1;refGene_gn=FHOD3;refGene_tx=XM_023241884.1;refGene_gn=FHOD3;refGene_tx=XM_023241885.1;refGene_gn=FHOD3;refGene_tx=XM_023241886.1;refGene_gn=FHOD3;refGene_tx=XM_023241887.1;refGene_gn=FHOD3;refGene_tx=XM_023241888.1;refGene_gn=FHOD3;refGene_tx=XM_023241889.1;refGene_gn=FHOD3;refGene_tx=XM_023241890.1;refGene_gn=FHOD3;refGene_tx=XM_023241891.1;refGene_gn=FHOD3;refGene_tx=XM_023241892.1;refGene_gn=FHOD3;refGene_tx=XM_023241893.1;refGene_gn=FHOD3;refGene_tx=XM_023241894.1
D3 60082568 60082592
refGene_gn=FHOD3;refGene_tx=XM_023241880.1;refGene_gn=FHOD3;refGene_tx=XM_023241882.1;refGene_gn=FHOD3;refGene_tx=XM_023241883.1;refGene_gn=FHOD3;refGene_tx=XM_023241884.1;refGene_gn=FHOD3;refGene_tx=XM_023241885.1;refGene_gn=FHOD3;refGene_tx=XM_023241886.1;refGene_gn=FHOD3;refGene_tx=XM_023241887.1;refGene_gn=FHOD3;refGene_tx=XM_023241888.1;refGene_gn=FHOD3;refGene_tx=XM_023241889.1;refGene_gn=FHOD3;refGene_tx=XM_023241890.1;refGene_gn=FHOD3;refGene_tx=XM_023241891.1;refGene_gn=FHOD3;refGene_tx=XM_023241894.1
D3 60084140 60084315
refGene_gn=FHOD3;refGene_tx=XM_023241880.1;refGene_gn=FHOD3;refGene_tx=XM_023241881.1;refGene_gn=FHOD3;refGene_tx=XM_023241882.1;refGene_gn=FHOD3;refGene_tx=XM_023241883.1;refGene_gn=FHOD3;refGene_tx=XM_023241884.1;refGene_gn=FHOD3;refGene_tx=XM_023241885.1;refGene_gn=FHOD3;refGene_tx=XM_023241886.1;refGene_gn=FHOD3;refGene_tx=XM_023241887.1;refGene_gn=FHOD3;refGene_tx=XM_023241888.1;refGene_gn=FHOD3;refGene_tx=XM_023241889.1;refGene_gn=FHOD3;refGene_tx=XM_023241890.1;refGene_gn=FHOD3;refGene_tx=XM_023241891.1;refGene_gn=FHOD3;refGene_tx=XM_023241892.1;refGene_gn=FHOD3;refGene_tx=XM_023241893.1;refGene_gn=FHOD3;refGene_tx=XM_023241894.1
D3 60091312 60091474
refGene_gn=FHOD3;refGene_tx=XM_023241880.1;refGene_gn=FHOD3;refGene_tx=XM_023241881.1;refGene_gn=FHOD3;refGene_tx=XM_023241882.1;refGene_gn=FHOD3;refGene_tx=XM_023241883.1;refGene_gn=FHOD3;refGene_tx=XM_023241884.1;refGene_gn=FHOD3;refGene_tx=XM_023241885.1;refGene_gn=FHOD3;refGene_tx=XM_023241886.1;refGene_gn=FHOD3;refGene_tx=XM_023241887.1;refGene_gn=FHOD3;refGene_tx=XM_023241888.1;refGene_gn=FHOD3;refGene_tx=XM_023241889.1;refGene_gn=FHOD3;refGene_tx=XM_023241890.1;refGene_gn=FHOD3;refGene_tx=XM_023241891.1;refGene_gn=FHOD3;refGene_tx=XM_023241892.1;refGene_gn=FHOD3;refGene_tx=XM_023241893.1;refGene_gn=FHOD3;refGene_tx=XM_023241894.1
D3 60101599 60101682
refGene_gn=FHOD3;refGene_tx=XM_023241880.1;refGene_gn=FHOD3;refGene_tx=XM_023241881.1;refGene_gn=FHOD3;refGene_tx=XM_023241882.1;refGene_gn=FHOD3;refGene_tx=XM_023241883.1;refGene_gn=FHOD3;refGene_tx=XM_023241884.1;refGene_gn=FHOD3;refGene_tx=XM_023241885.1;refGene_gn=FHOD3;refGene_tx=XM_023241886.1;refGene_gn=FHOD3;refGene_tx=XM_023241887.1;refGen

e_gn=FHOD3;refGene_tx=XM_023241888.1;refGene_gn=FHOD3;refGene_tx=XM_023241889.1;refGene_gn=FHOD3;refGene_tx=XM_023241890.1;refGene_gn=FHOD3;refGene_tx=XM_023241891.1;refGene_gn=FHOD3;refGene_tx=XM_023241892.1;refGene_gn=FHOD3;refGene_tx=XM_023241893.1;refGene_gn=FHOD3;refGene_tx=XM_023241894.1
D3 60113975 60114206
refGene_gn=TPGS2;refGene_tx=XM_003995142.4;refGene_gn=TPGS2;refGene_tx=XM_006938858.3;refGene_gn=TPGS2;refGene_tx=XM_006938859.3
D3 60115469 60115630
refGene_gn=TPGS2;refGene_tx=XM_003995142.4;refGene_gn=TPGS2;refGene_tx=XM_006938858.3;refGene_gn=TPGS2;refGene_tx=XM_006938859.3
D3 60117382 60117496
refGene_gn=TPGS2;refGene_tx=XM_003995142.4;refGene_gn=TPGS2;refGene_tx=XM_006938858.3;refGene_gn=TPGS2;refGene_tx=XM_006938859.3
D3 60121451 60121522 refGene_gn=TPGS2;refGene_tx=XM_006938860.3
D3 60129891 60130020
refGene_gn=TPGS2;refGene_tx=XM_003995142.4;refGene_gn=TPGS2;refGene_tx=XM_006938859.3;refGene_gn=TPGS2;refGene_tx=XM_006938860.3
D3 60132094 60132182
refGene_gn=TPGS2;refGene_tx=XM_003995142.4;refGene_gn=TPGS2;refGene_tx=XM_006938858.3;refGene_gn=TPGS2;refGene_tx=XM_006938859.3;refGene_gn=TPGS2;refGene_tx=XM_006938860.3
D3 60166547 60166627
refGene_gn=TPGS2;refGene_tx=XM_003995142.4;refGene_gn=TPGS2;refGene_tx=XM_006938858.3;refGene_gn=TPGS2;refGene_tx=XM_006938860.3
D3 60181384 60181460
refGene_gn=TPGS2;refGene_tx=XM_003995142.4;refGene_gn=TPGS2;refGene_tx=XM_006938858.3;refGene_gn=TPGS2;refGene_tx=XM_006938860.3
D3 60181622 60181923
refGene_gn=KIAA1328;refGene_tx=XM_003995143.5;refGene_gn=KIAA1328;refGene_tx=XM_006938856.4;refGene_gn=KIAA1328;refGene_tx=XM_019815186.2;refGene_gn=KIAA1328;refGene_tx=XM_019815187.2
D3 60188302 60188338
refGene_gn=KIAA1328;refGene_tx=XM_003995143.5;refGene_gn=KIAA1328;refGene_tx=XM_006938856.4;refGene_gn=KIAA1328;refGene_tx=XM_019815186.2;refGene_gn=KIAA1328;refGene_tx=XM_019815187.2
D3 60189547 60189687
refGene_gn=KIAA1328;refGene_tx=XM_003995143.5;refGene_gn=KIAA1328;refGene_tx=XM_006938856.4;refGene_gn=KIAA1328;refGene_tx=XM_006938857.3;refGene_gn=KIAA1328;refGene_tx=XM_019815186.2;refGene_gn=KIAA1328;refGene_tx=XM_019815187.2;refGene_gn=KIAA1328;refGene_tx=XM_019815192.2;refGene_gn=KIAA1328;refGene_tx=XM_019815193.2;refGene_gn=KIAA1328;refGene_tx=XM_023241895.1;refGene_gn=KIAA1328;refGene_tx=XM_023241896.1
D3 60199437 60199532
refGene_gn=KIAA1328;refGene_tx=XM_003995143.5;refGene_gn=KIAA1328;refGene_tx=XM_006938856.4;refGene_gn=KIAA1328;refGene_tx=XM_006938857.3;refGene_gn=KIAA1328;refGene_tx=XM_019815186.2;refGene_gn=KIAA1328;refGene_tx=XM_019815187.2;refGene_gn=KIAA1328;refGene_tx=XM_019815192.2;refGene_gn=KIAA1328;refGene_tx=XM_019815193.2;refGene_gn=KIAA1328;refGene_tx=XM_023241895.1

D3 60228460 60228576
refGene_gn=KIAA1328;refGene_tx=XM_003995143.5;refGene_gn=KIAA1328;refGene_tx=XM_006938856.4;refGene_gn=KIAA1328;refGene_tx=XM_006938857.3;refGene_gn=KIAA1328;refGene_tx=XM_019815186.2;refGene_gn=KIAA1328;refGene_tx=XM_019815187.2;refGene_gn=KIAA1328;refGene_tx=XM_019815192.2;refGene_gn=KIAA1328;refGene_tx=XM_019815193.2;refGene_gn=KIAA1328;refGene_tx=XM_023241895.1;refGene_gn=KIAA1328;refGene_tx=XM_023241896.1
D3 60306304 60306432
refGene_gn=KIAA1328;refGene_tx=XM_003995143.5;refGene_gn=KIAA1328;refGene_tx=XM_006938856.4;refGene_gn=KIAA1328;refGene_tx=XM_006938857.3;refGene_gn=KIAA1328;refGene_tx=XM_019815186.2;refGene_gn=KIAA1328;refGene_tx=XM_019815187.2;refGene_gn=KIAA1328;refGene_tx=XM_019815192.2;refGene_gn=KIAA1328;refGene_tx=XM_019815193.2;refGene_gn=KIAA1328;refGene_tx=XM_023241895.1;refGene_gn=KIAA1328;refGene_tx=XM_023241896.1
D3 60392511 60393185
refGene_gn=KIAA1328;refGene_tx=XM_003995143.5;refGene_gn=KIAA1328;refGene_tx=XM_006938856.4;refGene_gn=KIAA1328;refGene_tx=XM_006938857.3;refGene_gn=KIAA1328;refGene_tx=XM_019815186.2;refGene_gn=KIAA1328;refGene_tx=XM_019815187.2;refGene_gn=KIAA1328;refGene_tx=XM_019815192.2;refGene_gn=KIAA1328;refGene_tx=XM_019815193.2;refGene_gn=KIAA1328;refGene_tx=XM_023241895.1;refGene_gn=KIAA1328;refGene_tx=XM_023241896.1
D3 60401242 60401350
refGene_gn=KIAA1328;refGene_tx=XM_006938856.4;refGene_gn=KIAA1328;refGene_tx=XM_006938857.3;refGene_gn=KIAA1328;refGene_tx=XM_019815186.2;refGene_gn=KIAA1328;refGene_tx=XM_019815187.2;refGene_gn=KIAA1328;refGene_tx=XM_019815192.2;refGene_gn=KIAA1328;refGene_tx=XM_019815193.2;refGene_gn=KIAA1328;refGene_tx=XM_023241895.1;refGene_gn=KIAA1328;refGene_tx=XM_023241896.1
D3 60454146 60454328
refGene_gn=KIAA1328;refGene_tx=XM_003995143.5;refGene_gn=KIAA1328;refGene_tx=XM_006938856.4;refGene_gn=KIAA1328;refGene_tx=XM_006938857.3;refGene_gn=KIAA1328;refGene_tx=XM_019815186.2;refGene_gn=KIAA1328;refGene_tx=XM_019815187.2;refGene_gn=KIAA1328;refGene_tx=XM_019815192.2;refGene_gn=KIAA1328;refGene_tx=XM_019815193.2;refGene_gn=KIAA1328;refGene_tx=XM_023241895.1;refGene_gn=KIAA1328;refGene_tx=XM_023241896.1
D3 60470398 60470507
refGene_gn=KIAA1328;refGene_tx=XM_003995143.5;refGene_gn=KIAA1328;refGene_tx=XM_006938856.4;refGene_gn=KIAA1328;refGene_tx=XM_006938857.3;refGene_gn=KIAA1328;refGene_tx=XM_019815186.2;refGene_gn=KIAA1328;refGene_tx=XM_019815187.2;refGene_gn=KIAA1328;refGene_tx=XM_019815192.2;refGene_gn=KIAA1328;refGene_tx=XM_019815193.2;refGene_gn=KIAA1328;refGene_tx=XM_023241895.1;refGene_gn=KIAA1328;refGene_tx=XM_023241896.1
D3 60519415 60519626
refGene_gn=KIAA1328;refGene_tx=XM_003995143.5;refGene_gn=KIAA1328;refGene_tx=XM_006938856.4;refGene_gn=KIAA1328;refGene_tx=XM_006938857.3;refGene_gn=KIAA1328;refGene_tx=XM_019815192.2;refGene_gn=KIAA1328;refGene_tx=XM_019815193.2;refGene_gn=KIAA1328;refGene_tx=XM_023241895.1;refGene_gn=KIAA1328;refGene_tx=XM_023241896.1
D3 60530186 60530364
refGene_gn=KIAA1328;refGene_tx=XM_019815186.2;refGene_gn=KIAA1328;refG

ene_tx=XM_019815187.2
D3 60537644 60537671
refGene_gn=CELF4;refGene_tx=XM_023241901.1;refGene_gn=CELF4;refGene_tx=XM_023241902.1;refGene_gn=CELF4;refGene_tx=XM_023241904.1;refGene_gn=CELF4;refGene_tx=XM_023241905.1;refGene_gn=CELF4;refGene_tx=XM_023241906.1
D3 60539707 60539761
refGene_gn=CELF4;refGene_tx=XM_023241897.1;refGene_gn=CELF4;refGene_tx=XM_023241898.1;refGene_gn=CELF4;refGene_tx=XM_023241899.1;refGene_gn=CELF4;refGene_tx=XM_023241900.1;refGene_gn=CELF4;refGene_tx=XM_023241903.1;refGene_gn=CELF4;refGene_tx=XM_023241943.1
D3 60546222 60546394
refGene_gn=CELF4;refGene_tx=XM_023241897.1;refGene_gn=CELF4;refGene_tx=XM_023241898.1;refGene_gn=CELF4;refGene_tx=XM_023241899.1;refGene_gn=CELF4;refGene_tx=XM_023241900.1;refGene_gn=CELF4;refGene_tx=XM_023241901.1;refGene_gn=CELF4;refGene_tx=XM_023241902.1;refGene_gn=CELF4;refGene_tx=XM_023241903.1;refGene_gn=CELF4;refGene_tx=XM_023241904.1;refGene_gn=CELF4;refGene_tx=XM_023241905.1;refGene_gn=CELF4;refGene_tx=XM_023241906.1;refGene_gn=CELF4;refGene_tx=XM_023241907.1;refGene_gn=CELF4;refGene_tx=XM_023241908.1;refGene_gn=CELF4;refGene_tx=XM_023241909.1;refGene_gn=CELF4;refGene_tx=XM_023241910.1;refGene_gn=CELF4;refGene_tx=XM_023241911.1;refGene_gn=CELF4;refGene_tx=XM_023241912.1;refGene_gn=CELF4;refGene_tx=XM_023241913.1;refGene_gn=CELF4;refGene_tx=XM_023241914.1;refGene_gn=CELF4;refGene_tx=XM_023241915.1;refGene_gn=CELF4;refGene_tx=XM_023241916.1;refGene_gn=CELF4;refGene_tx=XM_023241917.1;refGene_gn=CELF4;refGene_tx=XM_023241918.1;refGene_gn=CELF4;refGene_tx=XM_023241919.1;refGene_gn=CELF4;refGene_tx=XM_023241920.1;refGene_gn=CELF4;refGene_tx=XM_023241921.1;refGene_gn=CELF4;refGene_tx=XM_023241922.1;refGene_gn=CELF4;refGene_tx=XM_023241923.1;refGene_gn=CELF4;refGene_tx=XM_023241924.1;refGene_gn=CELF4;refGene_tx=XM_023241925.1;refGene_gn=CELF4;refGene_tx=XM_023241926.1;refGene_gn=CELF4;refGene_tx=XM_023241927.1;refGene_gn=CELF4;refGene_tx=XM_023241928.1;refGene_gn=CELF4;refGene_tx=XM_023241929.1;refGene_gn=CELF4;refGene_tx=XM_023241930.1;refGene_gn=CELF4;refGene_tx=XM_023241931.1;refGene_gn=CELF4;refGene_tx=XM_023241932.1;refGene_gn=CELF4;refGene_tx=XM_023241933.1;refGene_gn=CELF4;refGene_tx=XM_023241934.1;refGene_gn=CELF4;refGene_tx=XM_023241935.1;refGene_gn=CELF4;refGene_tx=XM_023241936.1;refGene_gn=CELF4;refGene_tx=XM_023241937.1;refGene_gn=CELF4;refGene_tx=XM_023241938.1;refGene_gn=CELF4;refGene_tx=XM_023241939.1;refGene_gn=CELF4;refGene_tx=XM_023241940.1;refGene_gn=CELF4;refGene_tx=XM_023241941.1;refGene_gn=CELF4;refGene_tx=XM_023241942.1;refGene_gn=CELF4;refGene_tx=XM_023241943.1;refGene_gn=CELF4;refGene_tx=XM_023241944.1;refGene_gn=CELF4;refGene_tx=XM_023241945.1;refGene_gn=CELF4;refGene_tx=XM_023241946.1
D3 60546791 60546870
refGene_gn=CELF4;refGene_tx=XM_023241898.1;refGene_gn=CELF4;refGene_tx=XM_023241899.1;refGene_gn=CELF4;refGene_tx=XM_023241900.1;refGene_gn=CELF4;refGene_tx=XM_023241901.1;refGene_gn=CELF4;refGene_tx=XM_023241902.1;refGene_gn=CELF4;refGene_tx=XM_023241903.1;refGene_gn=CELF4;refGene_tx=XM_023241904.1;refGene_gn=CELF4;refGene_tx=XM_023241905.1;refGen

e_gn=CELF4;refGene_tx=XM_023241906.1;refGene_gn=CELF4;refGene_tx=XM_02
3241943.1
D3 60550921 60551066 refGene_gn=CELF4;refGene_tx=XM_023241897.1
D3 60551615 60551759
refGene_gn=CELF4;refGene_tx=XM_023241897.1;refGene_gn=CELF4;refGene_tx
=XM_023241898.1;refGene_gn=CELF4;refGene_tx=XM_023241899.1;refGene_gn=
CELF4;refGene_tx=XM_023241900.1;refGene_gn=CELF4;refGene_tx=XM_0232419
01.1;refGene_gn=CELF4;refGene_tx=XM_023241902.1;refGene_gn=CELF4;refGe
ne_tx=XM_023241903.1;refGene_gn=CELF4;refGene_tx=XM_023241904.1;refGen
e_gn=CELF4;refGene_tx=XM_023241905.1;refGene_gn=CELF4;refGene_tx=XM_02
3241906.1;refGene_gn=CELF4;refGene_tx=XM_023241907.1;refGene_gn=CELF4;
refGene_tx=XM_023241908.1;refGene_gn=CELF4;refGene_tx=XM_023241909.1;r
efGene_gn=CELF4;refGene_tx=XM_023241910.1;refGene_gn=CELF4;refGene_tx=
XM_023241911.1;refGene_gn=CELF4;refGene_tx=XM_023241912.1;refGene_gn=C
ELF4;refGene_tx=XM_023241913.1;refGene_gn=CELF4;refGene_tx=XM_02324191
4.1;refGene_gn=CELF4;refGene_tx=XM_023241915.1;refGene_gn=CELF4;refGen
e_tx=XM_023241916.1;refGene_gn=CELF4;refGene_tx=XM_023241917.1;refGene
_gn=CELF4;refGene_tx=XM_023241918.1;refGene_gn=CELF4;refGene_tx=XM_023
241919.1;refGene_gn=CELF4;refGene_tx=XM_023241920.1;refGene_gn=CELF4;r
efGene_tx=XM_023241921.1;refGene_gn=CELF4;refGene_tx=XM_023241922.1;re
fGene_gn=CELF4;refGene_tx=XM_023241923.1;refGene_gn=CELF4;refGene_tx=X
M_023241924.1;refGene_gn=CELF4;refGene_tx=XM_023241925.1;refGene_gn=CE
LF4;refGene_tx=XM_023241926.1;refGene_gn=CELF4;refGene_tx=XM_023241927
.
1;refGene_gn=CELF4;refGene_tx=XM_023241943.1;refGene_gn=CELF4;refGene_
tx=XM_023241944.1;refGene_gn=CELF4;refGene_tx=XM_023241945.1;refGene_g
n=CELF4;refGene_tx=XM_023241946.1
D3 60556780 60556864
refGene_gn=CELF4;refGene_tx=XM_023241897.1;refGene_gn=CELF4;refGene_tx
=XM_023241898.1;refGene_gn=CELF4;refGene_tx=XM_023241899.1;refGene_gn=
CELF4;refGene_tx=XM_023241900.1;refGene_gn=CELF4;refGene_tx=XM_0232419
01.1;refGene_gn=CELF4;refGene_tx=XM_023241902.1;refGene_gn=CELF4;refGe
ne_tx=XM_023241903.1;refGene_gn=CELF4;refGene_tx=XM_023241904.1;refGen
e_gn=CELF4;refGene_tx=XM_023241905.1;refGene_gn=CELF4;refGene_tx=XM_02
3241906.1;refGene_gn=CELF4;refGene_tx=XM_023241907.1;refGene_gn=CELF4;
refGene_tx=XM_023241908.1;refGene_gn=CELF4;refGene_tx=XM_023241909.1;r
efGene_gn=CELF4;refGene_tx=XM_023241910.1;refGene_gn=CELF4;refGene_tx=
XM_023241911.1;refGene_gn=CELF4;refGene_tx=XM_023241912.1;refGene_gn=C
ELF4;refGene_tx=XM_023241913.1;refGene_gn=CELF4;refGene_tx=XM_02324191
4.1;refGene_gn=CELF4;refGene_tx=XM_023241915.1;refGene_gn=CELF4;refGen
e_tx=XM_023241916.1;refGene_gn=CELF4;refGene_tx=XM_023241917.1;refGene
_gn=CELF4;refGene_tx=XM_023241918.1;refGene_gn=CELF4;refGene_tx=XM_023
241919.1;refGene_gn=CELF4;refGene_tx=XM_023241920.1;refGene_gn=CELF4;r
efGene_tx=XM_023241921.1;refGene_gn=CELF4;refGene_tx=XM_023241922.1;re
fGene_gn=CELF4;refGene_tx=XM_023241923.1;refGene_gn=CELF4;refGene_tx=X
M_023241924.1;refGene_gn=CELF4;refGene_tx=XM_023241925.1;refGene_gn=CE
LF4;refGene_tx=XM_023241926.1;refGene_gn=CELF4;refGene_tx=XM_023241927
.
1;refGene_gn=CELF4;refGene_tx=XM_023241928.1;refGene_gn=CELF4;refGene_
tx=XM_023241929.1;refGene_gn=CELF4;refGene_tx=XM_023241930.1;refGene_g

n=CELF4;refGene_tx=XM_023241931.1;refGene_gn=CELF4;refGene_tx=XM_02324
1932.1;refGene_gn=CELF4;refGene_tx=XM_023241933.1;refGene_gn=CELF4;ref
Gene_tx=XM_023241934.1;refGene_gn=CELF4;refGene_tx=XM_023241935.1;refG
ene_gn=CELF4;refGene_tx=XM_023241936.1;refGene_gn=CELF4;refGene_tx=XM_
023241937.1;refGene_gn=CELF4;refGene_tx=XM_023241938.1;refGene_gn=CELF
4;refGene_tx=XM_023241939.1;refGene_gn=CELF4;refGene_tx=XM_023241940.1
;refGene_gn=CELF4;refGene_tx=XM_023241941.1;refGene_gn=CELF4;refGene_t
x=XM_023241942.1;refGene_gn=CELF4;refGene_tx=XM_023241943.1;refGene_gn
=CELF4;refGene_tx=XM_023241944.1;refGene_gn=CELF4;refGene_tx=XM_023241
945.1;refGene_gn=CELF4;refGene_tx=XM_023241946.1
D3 60558266 60558332
refGene_gn=CELF4;refGene_tx=XM_023241897.1;refGene_gn=CELF4;refGene_tx
=XM_023241898.1;refGene_gn=CELF4;refGene_tx=XM_023241899.1;refGene_gn=
CELF4;refGene_tx=XM_023241900.1;refGene_gn=CELF4;refGene_tx=XM_0232419
01.1;refGene_gn=CELF4;refGene_tx=XM_023241902.1;refGene_gn=CELF4;refGe
ne_tx=XM_023241903.1;refGene_gn=CELF4;refGene_tx=XM_023241904.1;refGen
e_gn=CELF4;refGene_tx=XM_023241905.1;refGene_gn=CELF4;refGene_tx=XM_02
3241906.1;refGene_gn=CELF4;refGene_tx=XM_023241907.1;refGene_gn=CELF4;
refGene_tx=XM_023241908.1;refGene_gn=CELF4;refGene_tx=XM_023241909.1;r
efGene_gn=CELF4;refGene_tx=XM_023241910.1;refGene_gn=CELF4;refGene_tx=
XM_023241911.1;refGene_gn=CELF4;refGene_tx=XM_023241912.1;refGene_gn=C
ELF4;refGene_tx=XM_023241913.1;refGene_gn=CELF4;refGene_tx=XM_02324191
4.1;refGene_gn=CELF4;refGene_tx=XM_023241915.1;refGene_gn=CELF4;refGen
e_tx=XM_023241916.1;refGene_gn=CELF4;refGene_tx=XM_023241917.1;refGene
_gn=CELF4;refGene_tx=XM_023241918.1;refGene_gn=CELF4;refGene_tx=XM_023
241919.1;refGene_gn=CELF4;refGene_tx=XM_023241920.1;refGene_gn=CELF4;r
efGene_tx=XM_023241921.1;refGene_gn=CELF4;refGene_tx=XM_023241922.1;re
fGene_gn=CELF4;refGene_tx=XM_023241923.1;refGene_gn=CELF4;refGene_tx=X
M_023241924.1;refGene_gn=CELF4;refGene_tx=XM_023241925.1;refGene_gn=CE
LF4;refGene_tx=XM_023241926.1;refGene_gn=CELF4;refGene_tx=XM_023241927
.
1;refGene_gn=CELF4;refGene_tx=XM_023241928.1;refGene_gn=CELF4;refGene_
tx=XM_023241929.1;refGene_gn=CELF4;refGene_tx=XM_023241930.1;refGene_g
n=CELF4;refGene_tx=XM_023241931.1;refGene_gn=CELF4;refGene_tx=XM_02324
1932.1;refGene_gn=CELF4;refGene_tx=XM_023241933.1;refGene_gn=CELF4;ref
Gene_tx=XM_023241934.1;refGene_gn=CELF4;refGene_tx=XM_023241935.1;refG
ene_gn=CELF4;refGene_tx=XM_023241936.1;refGene_gn=CELF4;refGene_tx=XM_
023241937.1;refGene_gn=CELF4;refGene_tx=XM_023241938.1;refGene_gn=CELF
4;refGene_tx=XM_023241939.1;refGene_gn=CELF4;refGene_tx=XM_023241940.1
;refGene_gn=CELF4;refGene_tx=XM_023241941.1;refGene_gn=CELF4;refGene_t
x=XM_023241942.1;refGene_gn=CELF4;refGene_tx=XM_023241943.1;refGene_gn
=CELF4;refGene_tx=XM_023241944.1;refGene_gn=CELF4;refGene_tx=XM_023241
945.1;refGene_gn=CELF4;refGene_tx=XM_023241946.1
D3 60562072 60562222
refGene_gn=CELF4;refGene_tx=XM_023241897.1;refGene_gn=CELF4;refGene_tx
=XM_023241898.1;refGene_gn=CELF4;refGene_tx=XM_023241899.1;refGene_gn=
CELF4;refGene_tx=XM_023241900.1;refGene_gn=CELF4;refGene_tx=XM_0232419
01.1;refGene_gn=CELF4;refGene_tx=XM_023241902.1;refGene_gn=CELF4;refGe
ne_tx=XM_023241903.1;refGene_gn=CELF4;refGene_tx=XM_023241904.1;refGen
e_gn=CELF4;refGene_tx=XM_023241905.1;refGene_gn=CELF4;refGene_tx=XM_02

3241906.1;refGene_gn=CELF4;refGene_tx=XM_023241907.1;refGene_gn=CELF4;
refGene_tx=XM_023241908.1;refGene_gn=CELF4;refGene_tx=XM_023241909.1;r
efGene_gn=CELF4;refGene_tx=XM_023241910.1;refGene_gn=CELF4;refGene_tx=
XM_023241911.1;refGene_gn=CELF4;refGene_tx=XM_023241912.1;refGene_gn=C
ELF4;refGene_tx=XM_023241913.1;refGene_gn=CELF4;refGene_tx=XM_02324191
4.1;refGene_gn=CELF4;refGene_tx=XM_023241915.1;refGene_gn=CELF4;refGen
e_tx=XM_023241916.1;refGene_gn=CELF4;refGene_tx=XM_023241917.1;refGene
_gn=CELF4;refGene_tx=XM_023241918.1;refGene_gn=CELF4;refGene_tx=XM_023
241919.1;refGene_gn=CELF4;refGene_tx=XM_023241920.1;refGene_gn=CELF4;r
efGene_tx=XM_023241921.1;refGene_gn=CELF4;refGene_tx=XM_023241922.1;re
fGene_gn=CELF4;refGene_tx=XM_023241923.1;refGene_gn=CELF4;refGene_tx=X
M_023241924.1;refGene_gn=CELF4;refGene_tx=XM_023241925.1;refGene_gn=CE
LF4;refGene_tx=XM_023241926.1;refGene_gn=CELF4;refGene_tx=XM_023241927
.
1;refGene_gn=CELF4;refGene_tx=XM_023241928.1;refGene_gn=CELF4;refGene_
tx=XM_023241929.1;refGene_gn=CELF4;refGene_tx=XM_023241930.1;refGene_g
n=CELF4;refGene_tx=XM_023241931.1;refGene_gn=CELF4;refGene_tx=XM_02324
1932.1;refGene_gn=CELF4;refGene_tx=XM_023241933.1;refGene_gn=CELF4;ref
Gene_tx=XM_023241934.1;refGene_gn=CELF4;refGene_tx=XM_023241935.1;refG
ene_gn=CELF4;refGene_tx=XM_023241936.1;refGene_gn=CELF4;refGene_tx=XM_
023241937.1;refGene_gn=CELF4;refGene_tx=XM_023241938.1;refGene_gn=CELF
4;refGene_tx=XM_023241939.1;refGene_gn=CELF4;refGene_tx=XM_023241940.1
;refGene_gn=CELF4;refGene_tx=XM_023241941.1;refGene_gn=CELF4;refGene_t
x=XM_023241942.1;refGene_gn=CELF4;refGene_tx=XM_023241943.1;refGene_gn
=CELF4;refGene_tx=XM_023241944.1;refGene_gn=CELF4;refGene_tx=XM_023241
945.1;refGene_gn=CELF4;refGene_tx=XM_023241946.1
D3 60564355 60564503
refGene_gn=CELF4;refGene_tx=XM_023241897.1;refGene_gn=CELF4;refGene_tx
=XM_023241898.1;refGene_gn=CELF4;refGene_tx=XM_023241899.1;refGene_gn=
CELF4;refGene_tx=XM_023241900.1;refGene_gn=CELF4;refGene_tx=XM_0232419
01.1;refGene_gn=CELF4;refGene_tx=XM_023241902.1;refGene_gn=CELF4;refGe
ne_tx=XM_023241903.1;refGene_gn=CELF4;refGene_tx=XM_023241904.1;refGen
e_gn=CELF4;refGene_tx=XM_023241905.1;refGene_gn=CELF4;refGene_tx=XM_02
3241906.1;refGene_gn=CELF4;refGene_tx=XM_023241907.1;refGene_gn=CELF4;
refGene_tx=XM_023241908.1;refGene_gn=CELF4;refGene_tx=XM_023241909.1;r
efGene_gn=CELF4;refGene_tx=XM_023241910.1;refGene_gn=CELF4;refGene_tx=
XM_023241911.1;refGene_gn=CELF4;refGene_tx=XM_023241912.1;refGene_gn=C
ELF4;refGene_tx=XM_023241913.1;refGene_gn=CELF4;refGene_tx=XM_02324191
4.1;refGene_gn=CELF4;refGene_tx=XM_023241915.1;refGene_gn=CELF4;refGen
e_tx=XM_023241916.1;refGene_gn=CELF4;refGene_tx=XM_023241917.1;refGene
_gn=CELF4;refGene_tx=XM_023241918.1;refGene_gn=CELF4;refGene_tx=XM_023
241919.1;refGene_gn=CELF4;refGene_tx=XM_023241920.1;refGene_gn=CELF4;r
efGene_tx=XM_023241921.1;refGene_gn=CELF4;refGene_tx=XM_023241922.1;re
fGene_gn=CELF4;refGene_tx=XM_023241923.1;refGene_gn=CELF4;refGene_tx=X
M_023241924.1;refGene_gn=CELF4;refGene_tx=XM_023241925.1;refGene_gn=CE
LF4;refGene_tx=XM_023241926.1;refGene_gn=CELF4;refGene_tx=XM_023241927
.
1;refGene_gn=CELF4;refGene_tx=XM_023241928.1;refGene_gn=CELF4;refGene_
tx=XM_023241929.1;refGene_gn=CELF4;refGene_tx=XM_023241930.1;refGene_g
n=CELF4;refGene_tx=XM_023241931.1;refGene_gn=CELF4;refGene_tx=XM_02324

1932.1;refGene_gn=CELF4;refGene_tx=XM_023241933.1;refGene_gn=CELF4;ref
Gene_tx=XM_023241934.1;refGene_gn=CELF4;refGene_tx=XM_023241935.1;refG
ene_gn=CELF4;refGene_tx=XM_023241936.1;refGene_gn=CELF4;refGene_tx=XM_
023241937.1;refGene_gn=CELF4;refGene_tx=XM_023241938.1;refGene_gn=CELF
4;refGene_tx=XM_023241939.1;refGene_gn=CELF4;refGene_tx=XM_023241940.1
;refGene_gn=CELF4;refGene_tx=XM_023241941.1;refGene_gn=CELF4;refGene_t
x=XM_023241942.1;refGene_gn=CELF4;refGene_tx=XM_023241943.1;refGene_gn
=CELF4;refGene_tx=XM_023241944.1;refGene_gn=CELF4;refGene_tx=XM_023241
945.1;refGene_gn=CELF4;refGene_tx=XM_023241946.1
D3 60565630 60565774
refGene_gn=CELF4;refGene_tx=XM_023241897.1;refGene_gn=CELF4;refGene_tx
=XM_023241898.1;refGene_gn=CELF4;refGene_tx=XM_023241899.1;refGene_gn=
CELF4;refGene_tx=XM_023241900.1;refGene_gn=CELF4;refGene_tx=XM_0232419
01.1;refGene_gn=CELF4;refGene_tx=XM_023241902.1;refGene_gn=CELF4;refGe
ne_tx=XM_023241903.1;refGene_gn=CELF4;refGene_tx=XM_023241904.1;refGen
e_gn=CELF4;refGene_tx=XM_023241905.1;refGene_gn=CELF4;refGene_tx=XM_02
3241906.1;refGene_gn=CELF4;refGene_tx=XM_023241907.1;refGene_gn=CELF4;
refGene_tx=XM_023241908.1;refGene_gn=CELF4;refGene_tx=XM_023241909.1;r
efGene_gn=CELF4;refGene_tx=XM_023241910.1;refGene_gn=CELF4;refGene_tx=
XM_023241911.1;refGene_gn=CELF4;refGene_tx=XM_023241912.1;refGene_gn=C
ELF4;refGene_tx=XM_023241913.1;refGene_gn=CELF4;refGene_tx=XM_02324191
4.1;refGene_gn=CELF4;refGene_tx=XM_023241915.1;refGene_gn=CELF4;refGen
e_tx=XM_023241916.1;refGene_gn=CELF4;refGene_tx=XM_023241917.1;refGene
_gn=CELF4;refGene_tx=XM_023241918.1;refGene_gn=CELF4;refGene_tx=XM_023
241919.1;refGene_gn=CELF4;refGene_tx=XM_023241920.1;refGene_gn=CELF4;r
efGene_tx=XM_023241921.1;refGene_gn=CELF4;refGene_tx=XM_023241922.1;re
fGene_gn=CELF4;refGene_tx=XM_023241923.1;refGene_gn=CELF4;refGene_tx=X
M_023241924.1;refGene_gn=CELF4;refGene_tx=XM_023241925.1;refGene_gn=CE
LF4;refGene_tx=XM_023241926.1;refGene_gn=CELF4;refGene_tx=XM_023241927
.
1;refGene_gn=CELF4;refGene_tx=XM_023241928.1;refGene_gn=CELF4;refGene_
tx=XM_023241929.1;refGene_gn=CELF4;refGene_tx=XM_023241930.1;refGene_g
n=CELF4;refGene_tx=XM_023241931.1;refGene_gn=CELF4;refGene_tx=XM_02324
1932.1;refGene_gn=CELF4;refGene_tx=XM_023241933.1;refGene_gn=CELF4;ref
Gene_tx=XM_023241934.1;refGene_gn=CELF4;refGene_tx=XM_023241935.1;refG
ene_gn=CELF4;refGene_tx=XM_023241936.1;refGene_gn=CELF4;refGene_tx=XM_
023241937.1;refGene_gn=CELF4;refGene_tx=XM_023241938.1;refGene_gn=CELF
4;refGene_tx=XM_023241939.1;refGene_gn=CELF4;refGene_tx=XM_023241940.1
;refGene_gn=CELF4;refGene_tx=XM_023241941.1;refGene_gn=CELF4;refGene_t
x=XM_023241942.1;refGene_gn=CELF4;refGene_tx=XM_023241943.1;refGene_gn
=CELF4;refGene_tx=XM_023241944.1;refGene_gn=CELF4;refGene_tx=XM_023241
945.1;refGene_gn=CELF4;refGene_tx=XM_023241946.1
D3 60566188 60566268
refGene_gn=CELF4;refGene_tx=XM_023241897.1;refGene_gn=CELF4;refGene_tx
=XM_023241898.1;refGene_gn=CELF4;refGene_tx=XM_023241899.1;refGene_gn=
CELF4;refGene_tx=XM_023241900.1;refGene_gn=CELF4;refGene_tx=XM_0232419
01.1;refGene_gn=CELF4;refGene_tx=XM_023241902.1;refGene_gn=CELF4;refGe
ne_tx=XM_023241903.1;refGene_gn=CELF4;refGene_tx=XM_023241904.1;refGen
e_gn=CELF4;refGene_tx=XM_023241905.1;refGene_gn=CELF4;refGene_tx=XM_02
3241906.1;refGene_gn=CELF4;refGene_tx=XM_023241907.1;refGene_gn=CELF4;

refGene_tx=XM_023241908.1;refGene_gn=CELF4;refGene_tx=XM_023241909.1;r
efGene_gn=CELF4;refGene_tx=XM_023241910.1;refGene_gn=CELF4;refGene_tx=
XM_023241911.1;refGene_gn=CELF4;refGene_tx=XM_023241912.1;refGene_gn=C
ELF4;refGene_tx=XM_023241913.1;refGene_gn=CELF4;refGene_tx=XM_02324191
4.1;refGene_gn=CELF4;refGene_tx=XM_023241915.1;refGene_gn=CELF4;refGen
e_tx=XM_023241916.1;refGene_gn=CELF4;refGene_tx=XM_023241917.1;refGene
_gn=CELF4;refGene_tx=XM_023241918.1;refGene_gn=CELF4;refGene_tx=XM_023
241919.1;refGene_gn=CELF4;refGene_tx=XM_023241920.1;refGene_gn=CELF4;r
efGene_tx=XM_023241921.1;refGene_gn=CELF4;refGene_tx=XM_023241922.1;re
fGene_gn=CELF4;refGene_tx=XM_023241923.1;refGene_gn=CELF4;refGene_tx=X
M_023241924.1;refGene_gn=CELF4;refGene_tx=XM_023241925.1;refGene_gn=CE
LF4;refGene_tx=XM_023241926.1;refGene_gn=CELF4;refGene_tx=XM_023241927
.
1;refGene_gn=CELF4;refGene_tx=XM_023241928.1;refGene_gn=CELF4;refGene_
tx=XM_023241929.1;refGene_gn=CELF4;refGene_tx=XM_023241930.1;refGene_g
n=CELF4;refGene_tx=XM_023241931.1;refGene_gn=CELF4;refGene_tx=XM_02324
1932.1;refGene_gn=CELF4;refGene_tx=XM_023241933.1;refGene_gn=CELF4;ref
Gene_tx=XM_023241934.1;refGene_gn=CELF4;refGene_tx=XM_023241935.1;refG
ene_gn=CELF4;refGene_tx=XM_023241936.1;refGene_gn=CELF4;refGene_tx=XM_
023241937.1;refGene_gn=CELF4;refGene_tx=XM_023241938.1;refGene_gn=CELF
4;refGene_tx=XM_023241939.1;refGene_gn=CELF4;refGene_tx=XM_023241940.1
;refGene_gn=CELF4;refGene_tx=XM_023241941.1;refGene_gn=CELF4;refGene_t
x=XM_023241942.1;refGene_gn=CELF4;refGene_tx=XM_023241943.1;refGene_gn
=CELF4;refGene_tx=XM_023241944.1;refGene_gn=CELF4;refGene_tx=XM_023241
945.1;refGene_gn=CELF4;refGene_tx=XM_023241946.1
D3 60566716 60566845
refGene_gn=CELF4;refGene_tx=XM_023241897.1;refGene_gn=CELF4;refGene_tx
=XM_023241898.1;refGene_gn=CELF4;refGene_tx=XM_023241899.1;refGene_gn=
CELF4;refGene_tx=XM_023241900.1;refGene_gn=CELF4;refGene_tx=XM_0232419
01.1;refGene_gn=CELF4;refGene_tx=XM_023241902.1;refGene_gn=CELF4;refGe
ne_tx=XM_023241903.1;refGene_gn=CELF4;refGene_tx=XM_023241904.1;refGen
e_gn=CELF4;refGene_tx=XM_023241905.1;refGene_gn=CELF4;refGene_tx=XM_02
3241906.1;refGene_gn=CELF4;refGene_tx=XM_023241907.1;refGene_gn=CELF4;
refGene_tx=XM_023241908.1;refGene_gn=CELF4;refGene_tx=XM_023241909.1;r
efGene_gn=CELF4;refGene_tx=XM_023241910.1;refGene_gn=CELF4;refGene_tx=
XM_023241911.1;refGene_gn=CELF4;refGene_tx=XM_023241912.1;refGene_gn=C
ELF4;refGene_tx=XM_023241913.1;refGene_gn=CELF4;refGene_tx=XM_02324191
4.1;refGene_gn=CELF4;refGene_tx=XM_023241915.1;refGene_gn=CELF4;refGen
e_tx=XM_023241916.1;refGene_gn=CELF4;refGene_tx=XM_023241917.1;refGene
_gn=CELF4;refGene_tx=XM_023241918.1;refGene_gn=CELF4;refGene_tx=XM_023
241919.1;refGene_gn=CELF4;refGene_tx=XM_023241920.1;refGene_gn=CELF4;r
efGene_tx=XM_023241921.1;refGene_gn=CELF4;refGene_tx=XM_023241922.1;re
fGene_gn=CELF4;refGene_tx=XM_023241923.1;refGene_gn=CELF4;refGene_tx=X
M_023241924.1;refGene_gn=CELF4;refGene_tx=XM_023241925.1;refGene_gn=CE
LF4;refGene_tx=XM_023241926.1;refGene_gn=CELF4;refGene_tx=XM_023241927
.
1;refGene_gn=CELF4;refGene_tx=XM_023241928.1;refGene_gn=CELF4;refGene_
tx=XM_023241929.1;refGene_gn=CELF4;refGene_tx=XM_023241930.1;refGene_g
n=CELF4;refGene_tx=XM_023241931.1;refGene_gn=CELF4;refGene_tx=XM_02324
1932.1;refGene_gn=CELF4;refGene_tx=XM_023241933.1;refGene_gn=CELF4;ref

Gene_tx=XM_023241934.1;refGene_gn=CELF4;refGene_tx=XM_023241935.1;refGene_gn=CELF4;refGene_tx=XM_023241936.1;refGene_gn=CELF4;refGene_tx=XM_023241937.1;refGene_gn=CELF4;refGene_tx=XM_023241938.1;refGene_gn=CELF4;refGene_tx=XM_023241939.1;refGene_gn=CELF4;refGene_tx=XM_023241940.1;refGene_gn=CELF4;refGene_tx=XM_023241941.1;refGene_gn=CELF4;refGene_tx=XM_023241942.1;refGene_gn=CELF4;refGene_tx=XM_023241943.1;refGene_gn=CELF4;refGene_tx=XM_023241944.1;refGene_gn=CELF4;refGene_tx=XM_023241945.1;refGene_gn=CELF4;refGene_tx=XM_023241946.1
D3 60609306 60609385
refGene_gn=CELF4;refGene_tx=XM_023241897.1;refGene_gn=CELF4;refGene_tx=XM_023241898.1;refGene_gn=CELF4;refGene_tx=XM_023241899.1;refGene_gn=CELF4;refGene_tx=XM_023241900.1;refGene_gn=CELF4;refGene_tx=XM_023241901.1;refGene_gn=CELF4;refGene_tx=XM_023241902.1;refGene_gn=CELF4;refGene_tx=XM_023241903.1;refGene_gn=CELF4;refGene_tx=XM_023241904.1;refGene_gn=CELF4;refGene_tx=XM_023241905.1;refGene_gn=CELF4;refGene_tx=XM_023241906.1;refGene_gn=CELF4;refGene_tx=XM_023241907.1;refGene_gn=CELF4;refGene_tx=XM_023241908.1;refGene_gn=CELF4;refGene_tx=XM_023241909.1;refGene_gn=CELF4;refGene_tx=XM_023241910.1;refGene_gn=CELF4;refGene_tx=XM_023241911.1;refGene_gn=CELF4;refGene_tx=XM_023241912.1;refGene_gn=CELF4;refGene_tx=XM_023241913.1;refGene_gn=CELF4;refGene_tx=XM_023241914.1;refGene_gn=CELF4;refGene_tx=XM_023241915.1;refGene_gn=CELF4;refGene_tx=XM_023241916.1;refGene_gn=CELF4;refGene_tx=XM_023241917.1;refGene_gn=CELF4;refGene_tx=XM_023241918.1;refGene_gn=CELF4;refGene_tx=XM_023241919.1;refGene_gn=CELF4;refGene_tx=XM_023241920.1;refGene_gn=CELF4;refGene_tx=XM_023241921.1;refGene_gn=CELF4;refGene_tx=XM_023241922.1;refGene_gn=CELF4;refGene_tx=XM_023241923.1;refGene_gn=CELF4;refGene_tx=XM_023241924.1;refGene_gn=CELF4;refGene_tx=XM_023241925.1;refGene_gn=CELF4;refGene_tx=XM_023241926.1;refGene_gn=CELF4;refGene_tx=XM_023241927.1;refGene_gn=CELF4;refGene_tx=XM_023241928.1;refGene_gn=CELF4;refGene_tx=XM_023241929.1;refGene_gn=CELF4;refGene_tx=XM_023241930.1;refGene_gn=CELF4;refGene_tx=XM_023241931.1;refGene_gn=CELF4;refGene_tx=XM_023241932.1;refGene_gn=CELF4;refGene_tx=XM_023241933.1;refGene_gn=CELF4;refGene_tx=XM_023241934.1;refGene_gn=CELF4;refGene_tx=XM_023241935.1;refGene_gn=CELF4;refGene_tx=XM_023241936.1;refGene_gn=CELF4;refGene_tx=XM_023241937.1;refGene_gn=CELF4;refGene_tx=XM_023241938.1;refGene_gn=CELF4;refGene_tx=XM_023241939.1;refGene_gn=CELF4;refGene_tx=XM_023241940.1;refGene_gn=CELF4;refGene_tx=XM_023241941.1;refGene_gn=CELF4;refGene_tx=XM_023241942.1;refGene_gn=CELF4;refGene_tx=XM_023241943.1;refGene_gn=CELF4;refGene_tx=XM_023241944.1;refGene_gn=CELF4;refGene_tx=XM_023241945.1;refGene_gn=CELF4;refGene_tx=XM_023241946.1
D3 60757321 60757404
refGene_gn=CELF4;refGene_tx=XM_023241897.1;refGene_gn=CELF4;refGene_tx=XM_023241898.1;refGene_gn=CELF4;refGene_tx=XM_023241899.1;refGene_gn=CELF4;refGene_tx=XM_023241900.1;refGene_gn=CELF4;refGene_tx=XM_023241901.1;refGene_gn=CELF4;refGene_tx=XM_023241902.1;refGene_gn=CELF4;refGene_tx=XM_023241903.1;refGene_gn=CELF4;refGene_tx=XM_023241904.1;refGene_gn=CELF4;refGene_tx=XM_023241905.1;refGene_gn=CELF4;refGene_tx=XM_023241906.1;refGene_gn=CELF4;refGene_tx=XM_023241907.1;refGene_gn=CELF4;refGene_tx=XM_023241908.1;refGene_gn=CELF4;refGene_tx=XM_023241909.1;r

efGene_gn=CELF4;refGene_tx=XM_023241910.1;refGene_gn=CELF4;refGene_tx=
XM_023241911.1;refGene_gn=CELF4;refGene_tx=XM_023241912.1;refGene_gn=C
ELF4;refGene_tx=XM_023241913.1;refGene_gn=CELF4;refGene_tx=XM_02324191
4.1;refGene_gn=CELF4;refGene_tx=XM_023241915.1;refGene_gn=CELF4;refGen
e_tx=XM_023241916.1;refGene_gn=CELF4;refGene_tx=XM_023241917.1;refGene
_gn=CELF4;refGene_tx=XM_023241918.1;refGene_gn=CELF4;refGene_tx=XM_023
241919.1;refGene_gn=CELF4;refGene_tx=XM_023241920.1;refGene_gn=CELF4;r
efGene_tx=XM_023241921.1;refGene_gn=CELF4;refGene_tx=XM_023241922.1;re
fGene_gn=CELF4;refGene_tx=XM_023241923.1;refGene_gn=CELF4;refGene_tx=X
M_023241924.1;refGene_gn=CELF4;refGene_tx=XM_023241925.1;refGene_gn=CE
LF4;refGene_tx=XM_023241926.1;refGene_gn=CELF4;refGene_tx=XM_023241927
.
1;refGene_gn=CELF4;refGene_tx=XM_023241928.1;refGene_gn=CELF4;refGene_
tx=XM_023241929.1;refGene_gn=CELF4;refGene_tx=XM_023241930.1;refGene_g
n=CELF4;refGene_tx=XM_023241931.1;refGene_gn=CELF4;refGene_tx=XM_02324
1932.1;refGene_gn=CELF4;refGene_tx=XM_023241933.1;refGene_gn=CELF4;ref
Gene_tx=XM_023241934.1;refGene_gn=CELF4;refGene_tx=XM_023241935.1;refG
ene_gn=CELF4;refGene_tx=XM_023241936.1;refGene_gn=CELF4;refGene_tx=XM_
023241937.1;refGene_gn=CELF4;refGene_tx=XM_023241938.1;refGene_gn=CELF
4;refGene_tx=XM_023241939.1;refGene_gn=CELF4;refGene_tx=XM_023241940.1
;refGene_gn=CELF4;refGene_tx=XM_023241941.1;refGene_gn=CELF4;refGene_t
x=XM_023241942.1;refGene_gn=CELF4;refGene_tx=XM_023241943.1;refGene_gn
=CELF4;refGene_tx=XM_023241944.1;refGene_gn=CELF4;refGene_tx=XM_023241
945.1;refGene_gn=CELF4;refGene_tx=XM_023241946.1
D3 60833042 60833328
refGene_gn=CELF4;refGene_tx=XM_023241897.1;refGene_gn=CELF4;refGene_tx
=XM_023241898.1;refGene_gn=CELF4;refGene_tx=XM_023241899.1;refGene_gn=
CELF4;refGene_tx=XM_023241900.1;refGene_gn=CELF4;refGene_tx=XM_0232419
01.1;refGene_gn=CELF4;refGene_tx=XM_023241902.1;refGene_gn=CELF4;refGe
ne_tx=XM_023241903.1;refGene_gn=CELF4;refGene_tx=XM_023241904.1;refGen
e_gn=CELF4;refGene_tx=XM_023241905.1;refGene_gn=CELF4;refGene_tx=XM_02
3241906.1;refGene_gn=CELF4;refGene_tx=XM_023241907.1;refGene_gn=CELF4;
refGene_tx=XM_023241908.1;refGene_gn=CELF4;refGene_tx=XM_023241909.1;r
efGene_gn=CELF4;refGene_tx=XM_023241910.1;refGene_gn=CELF4;refGene_tx=
XM_023241911.1;refGene_gn=CELF4;refGene_tx=XM_023241912.1;refGene_gn=C
ELF4;refGene_tx=XM_023241913.1;refGene_gn=CELF4;refGene_tx=XM_02324191
4.1;refGene_gn=CELF4;refGene_tx=XM_023241915.1;refGene_gn=CELF4;refGen
e_tx=XM_023241916.1;refGene_gn=CELF4;refGene_tx=XM_023241917.1;refGene
_gn=CELF4;refGene_tx=XM_023241918.1;refGene_gn=CELF4;refGene_tx=XM_023
241919.1;refGene_gn=CELF4;refGene_tx=XM_023241920.1;refGene_gn=CELF4;r
efGene_tx=XM_023241921.1;refGene_gn=CELF4;refGene_tx=XM_023241922.1;re
fGene_gn=CELF4;refGene_tx=XM_023241923.1;refGene_gn=CELF4;refGene_tx=X
M_023241924.1;refGene_gn=CELF4;refGene_tx=XM_023241925.1;refGene_gn=CE
LF4;refGene_tx=XM_023241926.1;refGene_gn=CELF4;refGene_tx=XM_023241927
.
1;refGene_gn=CELF4;refGene_tx=XM_023241928.1;refGene_gn=CELF4;refGene_
tx=XM_023241929.1;refGene_gn=CELF4;refGene_tx=XM_023241930.1;refGene_g
n=CELF4;refGene_tx=XM_023241931.1;refGene_gn=CELF4;refGene_tx=XM_02324
1932.1;refGene_gn=CELF4;refGene_tx=XM_023241933.1;refGene_gn=CELF4;ref
Gene_tx=XM_023241934.1;refGene_gn=CELF4;refGene_tx=XM_023241935.1;refG

ene_gn=CELF4;refGene_tx=XM_023241936.1;refGene_gn=CELF4;refGene_tx=XM_023241937.1;refGene_gn=CELF4;refGene_tx=XM_023241938.1;refGene_gn=CELF4;refGene_tx=XM_023241939.1;refGene_gn=CELF4;refGene_tx=XM_023241940.1;refGene_gn=CELF4;refGene_tx=XM_023241941.1;refGene_gn=CELF4;refGene_tx=XM_023241942.1;refGene_gn=CELF4;refGene_tx=XM_023241943.1;refGene_gn=CELF4;refGene_tx=XM_023241944.1;refGene_gn=CELF4;refGene_tx=XM_023241945.1;refGene_gn=CELF4;refGene_tx=XM_023241946.1
D3 64813573 64813641 refGene_gn=PIK3C3;refGene_tx=XM_003995149.5
D3 64817124 64817313 refGene_gn=PIK3C3;refGene_tx=XM_003995149.5
D3 64821792 64821820 refGene_gn=PIK3C3;refGene_tx=XM_003995149.5
D3 64821821 64821937 refGene_gn=PIK3C3;refGene_tx=XM_003995149.5
D3 64829213 64829343 refGene_gn=PIK3C3;refGene_tx=XM_003995149.5
D3 64858047 64858134 refGene_gn=PIK3C3;refGene_tx=XM_003995149.5
D3 64860237 64860333 refGene_gn=PIK3C3;refGene_tx=XM_003995149.5
D3 64862940 64863012 refGene_gn=PIK3C3;refGene_tx=XM_003995149.5
D3 64866372 64866477 refGene_gn=PIK3C3;refGene_tx=XM_003995149.5
D3 64867197 64867290 refGene_gn=PIK3C3;refGene_tx=XM_003995149.5
D3 64876207 64876393 refGene_gn=PIK3C3;refGene_tx=XM_003995149.5
D3 64880963 64881118 refGene_gn=PIK3C3;refGene_tx=XM_003995149.5
D3 64882705 64882796 refGene_gn=PIK3C3;refGene_tx=XM_003995149.5
D3 64888935 64889003 refGene_gn=PIK3C3;refGene_tx=XM_003995149.5
D3 64895383 64895489 refGene_gn=PIK3C3;refGene_tx=XM_003995149.5
D3 64896995 64897112 refGene_gn=PIK3C3;refGene_tx=XM_003995149.5
D3 64900163 64900295 refGene_gn=PIK3C3;refGene_tx=XM_003995149.5
D3 64903762 64903891 refGene_gn=PIK3C3;refGene_tx=XM_003995149.5
D3 64904749 64904819 refGene_gn=PIK3C3;refGene_tx=XM_003995149.5
D3 64906571 64906636 refGene_gn=PIK3C3;refGene_tx=XM_003995149.5
D3 64915841 64915926 refGene_gn=PIK3C3;refGene_tx=XM_003995149.5
D3 64922385 64922460 refGene_gn=PIK3C3;refGene_tx=XM_003995149.5
D3 64936947 64937116 refGene_gn=PIK3C3;refGene_tx=XM_003995149.5
D3 64942452 64942543 refGene_gn=PIK3C3;refGene_tx=XM_003995149.5
D3 64952099 64952225 refGene_gn=PIK3C3;refGene_tx=XM_003995149.5
D3 64961776 64961791 refGene_gn=PIK3C3;refGene_tx=XM_003995149.5
D3 65564399 65564627
refGene_gn=RIT2;refGene_tx=XM_003995151.4;refGene_gn=RIT2;refGene_tx=XM_019815210.2
D3 65755718 65755910
refGene_gn=RIT2;refGene_tx=XM_003995151.4;refGene_gn=RIT2;refGene_tx=XM_019815210.2
D3 65809877 65809951
refGene_gn=RIT2;refGene_tx=XM_003995151.4;refGene_gn=RIT2;refGene_tx=XM_019815210.2
D3 65866945 65867002
refGene_gn=RIT2;refGene_tx=XM_003995151.4;refGene_gn=RIT2;refGene_tx=XM_019815210.2
D3 65939374 65939477 refGene_gn=RIT2;refGene_tx=XM_003995151.4
D3 66049251 66049559
refGene_gn=SYT4;refGene_tx=XM_003995152.5;refGene_gn=SYT4;refGene_tx=XM_011288025.3;refGene_gn=SYT4;refGene_tx=XM_011288026.3
D3 66050894 66051015

refGene_gn=SYT4;refGene_tx=XM_003995152.5;refGene_gn=SYT4;refGene_tx=XM_011288025.3;refGene_gn=SYT4;refGene_tx=XM_011288026.3
D3 66052652 66053467
refGene_gn=SYT4;refGene_tx=XM_003995152.5;refGene_gn=SYT4;refGene_tx=XM_011288025.3;refGene_gn=SYT4;refGene_tx=XM_011288026.3
D3 66056447 66056481
refGene_gn=SYT4;refGene_tx=XM_003995152.5;refGene_gn=SYT4;refGene_tx=XM_011288025.3;refGene_gn=SYT4;refGene_tx=XM_011288026.3
D3 67282387 67282873 refGene_gn=SETBP1;refGene_tx=XM_011288024.3
D3 67450777 67450831 refGene_gn=SETBP1;refGene_tx=XM_011288024.3
D3 67524848 67528308 refGene_gn=SETBP1;refGene_tx=XM_011288024.3
D3 67607654 67607825 refGene_gn=SETBP1;refGene_tx=XM_011288024.3
D3 67633803 67634239 refGene_gn=SETBP1;refGene_tx=XM_011288024.3
D3 67634240 67634421 refGene_gn=SETBP1;refGene_tx=XM_011288024.3
D3 68165709 68165862
refGene_gn=LOC101084395;refGene_tx=XM_023241947.1;refGene_gn=LOC101084395;refGene_tx=XM_023241948.1
D3 68171534 68171715
refGene_gn=LOC101084395;refGene_tx=XM_023241947.1;refGene_gn=LOC101084395;refGene_tx=XM_023241948.1
D3 68172695 68172885
refGene_gn=LOC101084395;refGene_tx=XM_023241947.1;refGene_gn=LOC101084395;refGene_tx=XM_023241948.1
D3 68177290 68177419
refGene_gn=LOC101084395;refGene_tx=XM_023241947.1;refGene_gn=LOC101084395;refGene_tx=XM_023241948.1
D3 68181048 68181241
refGene_gn=LOC101084395;refGene_tx=XM_023241947.1;refGene_gn=LOC101084395;refGene_tx=XM_023241948.1
D3 68185179 68185327
refGene_gn=LOC101084395;refGene_tx=XM_023241947.1;refGene_gn=LOC101084395;refGene_tx=XM_023241948.1
D3 68186386 68186521
refGene_gn=LOC101084395;refGene_tx=XM_023241947.1;refGene_gn=LOC101084395;refGene_tx=XM_023241948.1
D3 68188247 68188297
refGene_gn=LOC101084395;refGene_tx=XM_023241947.1
D3 68188800 68188983
refGene_gn=LOC101084395;refGene_tx=XM_023241947.1;refGene_gn=LOC101084395;refGene_tx=XM_023241948.1
D3 68206970 68207118
refGene_gn=LOC111556206;refGene_tx=XM_019815213.1;refGene_gn=LOC111556206;refGene_tx=XM_019815214.2
D3 68207846 68208036
refGene_gn=LOC111556206;refGene_tx=XM_019815213.1;refGene_gn=LOC111556206;refGene_tx=XM_019815214.2
D3 68208389 68208518
refGene_gn=LOC111556206;refGene_tx=XM_019815213.1;refGene_gn=LOC111556206;refGene_tx=XM_019815214.2
D3 68209603 68209796

refGene_gn=LOC111556206;refGene_tx=XM_019815213.1;refGene_gn=LOC111556206;refGene_tx=XM_019815214.2
D3 68212778 68212926
refGene_gn=LOC111556206;refGene_tx=XM_019815213.1;refGene_gn=LOC111556206;refGene_tx=XM_019815214.2
D3 68213773 68213908
refGene_gn=LOC111556206;refGene_tx=XM_019815213.1;refGene_gn=LOC111556206;refGene_tx=XM_019815214.2
D3 68215015 68215065
refGene_gn=LOC111556206;refGene_tx=XM_019815213.1;refGene_gn=LOC111556206;refGene_tx=XM_019815214.2
D3 68218547 68218748
refGene_gn=LOC111556206;refGene_tx=XM_019815213.1;refGene_gn=LOC111556206;refGene_tx=XM_019815214.2
D3 68252553 68252700
refGene_gn=LOC101101172;refGene_tx=XM_019815215.1;refGene_gn=LOC101101172;refGene_tx=XM_019815216.2;refGene_gn=LOC101101172;refGene_tx=XM_019815217.2
D3 68262931 68263088
refGene_gn=LOC101101172;refGene_tx=XM_006938866.3;refGene_gn=LOC101101172;refGene_tx=XM_019815215.1;refGene_gn=LOC101101172;refGene_tx=XM_019815216.2;refGene_gn=LOC101101172;refGene_tx=XM_019815217.2;refGene_gn=LOC101101172;refGene_tx=XM_019815218.1;refGene_gn=LOC101101172;refGene_tx=XM_019815219.2
D3 68263777 68263967
refGene_gn=LOC101101172;refGene_tx=XM_006938866.3;refGene_gn=LOC101101172;refGene_tx=XM_019815215.1;refGene_gn=LOC101101172;refGene_tx=XM_019815216.2;refGene_gn=LOC101101172;refGene_tx=XM_019815217.2;refGene_gn=LOC101101172;refGene_tx=XM_019815218.1;refGene_gn=LOC101101172;refGene_tx=XM_019815219.2
D3 68266369 68266498
refGene_gn=LOC101101172;refGene_tx=XM_006938866.3;refGene_gn=LOC101101172;refGene_tx=XM_019815215.1;refGene_gn=LOC101101172;refGene_tx=XM_019815216.2;refGene_gn=LOC101101172;refGene_tx=XM_019815217.2;refGene_gn=LOC101101172;refGene_tx=XM_019815218.1;refGene_gn=LOC101101172;refGene_tx=XM_019815219.2
D3 68267967 68268160
refGene_gn=LOC101101172;refGene_tx=XM_006938866.3;refGene_gn=LOC101101172;refGene_tx=XM_019815215.1;refGene_gn=LOC101101172;refGene_tx=XM_019815216.2;refGene_gn=LOC101101172;refGene_tx=XM_019815217.2;refGene_gn=LOC101101172;refGene_tx=XM_019815218.1;refGene_gn=LOC101101172;refGene_tx=XM_019815219.2
D3 68272110 68272258
refGene_gn=LOC101101172;refGene_tx=XM_006938866.3;refGene_gn=LOC101101172;refGene_tx=XM_019815215.1;refGene_gn=LOC101101172;refGene_tx=XM_019815216.2;refGene_gn=LOC101101172;refGene_tx=XM_019815217.2;refGene_gn=LOC101101172;refGene_tx=XM_019815218.1;refGene_gn=LOC101101172;refGene_tx=XM_019815219.2
D3 68272454 68272589
refGene_gn=LOC101101172;refGene_tx=XM_006938866.3;refGene_gn=LOC101101

172;refGene_tx=XM_019815215.1;refGene_gn=LOC101101172;refGene_tx=XM_019815216.2;refGene_gn=LOC101101172;refGene_tx=XM_019815217.2;refGene_gn=LOC101101172;refGene_tx=XM_019815218.1;refGene_gn=LOC101101172;refGene_tx=XM_019815219.2
D3 68278488 68278538
refGene_gn=LOC101101172;refGene_tx=XM_006938866.3;refGene_gn=LOC101101172;refGene_tx=XM_019815215.1;refGene_gn=LOC101101172;refGene_tx=XM_019815216.2;refGene_gn=LOC101101172;refGene_tx=XM_019815217.2;refGene_gn=LOC101101172;refGene_tx=XM_019815218.1;refGene_gn=LOC101101172;refGene_tx=XM_019815219.2
D3 68280040 68280217
refGene_gn=LOC101101172;refGene_tx=XM_006938866.3;refGene_gn=LOC101101172;refGene_tx=XM_019815215.1;refGene_gn=LOC101101172;refGene_tx=XM_019815216.2;refGene_gn=LOC101101172;refGene_tx=XM_019815217.2;refGene_gn=LOC101101172;refGene_tx=XM_019815218.1;refGene_gn=LOC101101172;refGene_tx=XM_019815219.2
D3 68328554 68328606 refGene_gn=SIGLEC15;refGene_tx=XM_019815220.2
D3 68335828 68335888 refGene_gn=SIGLEC15;refGene_tx=XM_019815220.2
D3 68336262 68336646
refGene_gn=SIGLEC15;refGene_tx=XM_019815220.2;refGene_gn=SIGLEC15;refGene_tx=XM_023241949.1;refGene_gn=SIGLEC15;refGene_tx=XM_023241950.1;refGene_gn=SIGLEC15;refGene_tx=XM_023241951.1
D3 68337735 68338122
refGene_gn=SIGLEC15;refGene_tx=XM_019815220.2;refGene_gn=SIGLEC15;refGene_tx=XM_023241949.1;refGene_gn=SIGLEC15;refGene_tx=XM_023241950.1;refGene_gn=SIGLEC15;refGene_tx=XM_023241951.1
D3 68339204 68339235
refGene_gn=SIGLEC15;refGene_tx=XM_019815220.2;refGene_gn=SIGLEC15;refGene_tx=XM_023241949.1;refGene_gn=SIGLEC15;refGene_tx=XM_023241950.1;refGene_gn=SIGLEC15;refGene_tx=XM_023241951.1
D3 68340976 68341058
refGene_gn=SIGLEC15;refGene_tx=XM_019815220.2;refGene_gn=SIGLEC15;refGene_tx=XM_023241949.1;refGene_gn=SIGLEC15;refGene_tx=XM_023241950.1;refGene_gn=SIGLEC15;refGene_tx=XM_023241951.1
D3 68351253 68351436 refGene_gn=EPG5;refGene_tx=XM_011288022.2
D3 68353644 68353759 refGene_gn=EPG5;refGene_tx=XM_011288022.2
D3 68355576 68355792 refGene_gn=EPG5;refGene_tx=XM_011288022.2
D3 68355979 68356196 refGene_gn=EPG5;refGene_tx=XM_011288022.2
D3 68357463 68357706 refGene_gn=EPG5;refGene_tx=XM_011288022.2
D3 68364252 68364397 refGene_gn=EPG5;refGene_tx=XM_011288022.2
D3 68365337 68365547 refGene_gn=EPG5;refGene_tx=XM_011288022.2
D3 68366042 68366228 refGene_gn=EPG5;refGene_tx=XM_011288022.2
D3 68369163 68369339 refGene_gn=EPG5;refGene_tx=XM_011288022.2
D3 68373865 68373969 refGene_gn=EPG5;refGene_tx=XM_011288022.2
D3 68376007 68376080 refGene_gn=EPG5;refGene_tx=XM_011288022.2
D3 68376646 68376848 refGene_gn=EPG5;refGene_tx=XM_011288022.2
D3 68377366 68377515 refGene_gn=EPG5;refGene_tx=XM_011288022.2
D3 68380567 68380781 refGene_gn=EPG5;refGene_tx=XM_011288022.2
D3 68381781 68381976 refGene_gn=EPG5;refGene_tx=XM_011288022.2
D3 68384649 68384806 refGene_gn=EPG5;refGene_tx=XM_011288022.2

D3 68386519 68386662 refGene_gn=EPG5;refGene_tx=XM_011288022.2
D3 68395579 68395742 refGene_gn=EPG5;refGene_tx=XM_011288022.2
D3 68397288 68397460 refGene_gn=EPG5;refGene_tx=XM_011288022.2
D3 68399086 68399231 refGene_gn=EPG5;refGene_tx=XM_011288022.2
D3 68402701 68402825 refGene_gn=EPG5;refGene_tx=XM_011288022.2
D3 68404634 68404856 refGene_gn=EPG5;refGene_tx=XM_011288022.2
D3 68405707 68405874 refGene_gn=EPG5;refGene_tx=XM_011288022.2
D3 68406789 68406912 refGene_gn=EPG5;refGene_tx=XM_011288022.2
D3 68408409 68408520 refGene_gn=EPG5;refGene_tx=XM_011288022.2
D3 68408908 68409106 refGene_gn=EPG5;refGene_tx=XM_011288022.2
D3 68409331 68409476 refGene_gn=EPG5;refGene_tx=XM_011288022.2
D3 68412300 68412441 refGene_gn=EPG5;refGene_tx=XM_011288022.2
D3 68415560 68415820 refGene_gn=EPG5;refGene_tx=XM_011288022.2
D3 68416470 68416590 refGene_gn=EPG5;refGene_tx=XM_011288022.2
D3 68417963 68418128 refGene_gn=EPG5;refGene_tx=XM_011288022.2
D3 68422847 68422988 refGene_gn=EPG5;refGene_tx=XM_011288022.2
D3 68424883 68425038 refGene_gn=EPG5;refGene_tx=XM_011288022.2
D3 68426186 68426344 refGene_gn=EPG5;refGene_tx=XM_011288022.2
D3 68430045 68430201 refGene_gn=EPG5;refGene_tx=XM_011288022.2
D3 68433440 68433591 refGene_gn=EPG5;refGene_tx=XM_011288022.2
D3 68433942 68434057 refGene_gn=EPG5;refGene_tx=XM_011288022.2
D3 68436085 68436191 refGene_gn=EPG5;refGene_tx=XM_011288022.2
D3 68438109 68438183 refGene_gn=EPG5;refGene_tx=XM_011288022.2
D3 68439092 68439200 refGene_gn=EPG5;refGene_tx=XM_011288022.2
D3 68440091 68440228 refGene_gn=EPG5;refGene_tx=XM_011288022.2
D3 68441721 68441959 refGene_gn=EPG5;refGene_tx=XM_011288022.2
D3 68443212 68444157 refGene_gn=EPG5;refGene_tx=XM_011288022.2
D3 68454761 68454824 refGene_gn=EPG5;refGene_tx=XM_011288022.2
D3 68468974 68469024
refGene_gn=PSTPIP2;refGene_tx=XM_023241952.1;refGene_gn=PSTPIP2;refGene_tx=XM_023241953.1
D3 68470349 68470384
refGene_gn=PSTPIP2;refGene_tx=XM_023241952.1;refGene_gn=PSTPIP2;refGene_tx=XM_023241953.1
D3 68471848 68471930
refGene_gn=PSTPIP2;refGene_tx=XM_023241952.1;refGene_gn=PSTPIP2;refGene_tx=XM_023241953.1
D3 68472047 68472144
refGene_gn=PSTPIP2;refGene_tx=XM_023241952.1;refGene_gn=PSTPIP2;refGene_tx=XM_023241953.1
D3 68473029 68473128
refGene_gn=PSTPIP2;refGene_tx=XM_023241952.1;refGene_gn=PSTPIP2;refGene_tx=XM_023241953.1
D3 68475724 68475804
refGene_gn=PSTPIP2;refGene_tx=XM_023241952.1;refGene_gn=PSTPIP2;refGene_tx=XM_023241953.1
D3 68476733 68476779
refGene_gn=PSTPIP2;refGene_tx=XM_023241952.1;refGene_gn=PSTPIP2;refGene_tx=XM_023241953.1
D3 68477357 68477456 refGene_gn=PSTPIP2;refGene_tx=XM_023241952.1

D3 68483437 68483500
refGene_gn=PSTPIP2;refGene_tx=XM_023241952.1;refGene_gn=PSTPIP2;refGene_tx=XM_023241953.1
D3 68488002 68488109
refGene_gn=PSTPIP2;refGene_tx=XM_023241952.1;refGene_gn=PSTPIP2;refGene_tx=XM_023241953.1
D3 68491388 68491423
refGene_gn=PSTPIP2;refGene_tx=XM_023241952.1;refGene_gn=PSTPIP2;refGene_tx=XM_023241953.1
D3 68496613 68496691
refGene_gn=PSTPIP2;refGene_tx=XM_023241952.1;refGene_gn=PSTPIP2;refGene_tx=XM_023241953.1
D3 68500078 68500179
refGene_gn=PSTPIP2;refGene_tx=XM_023241952.1;refGene_gn=PSTPIP2;refGene_tx=XM_023241953.1
D3 68539698 68539731
refGene_gn=PSTPIP2;refGene_tx=XM_023241952.1;refGene_gn=PSTPIP2;refGene_tx=XM_023241953.1
D3 68555674 68555756 refGene_gn=ATP5F1A;refGene_tx=XM_003995155.5
D3 68555893 68556044 refGene_gn=ATP5F1A;refGene_tx=XM_003995155.5
D3 68556892 68557037 refGene_gn=ATP5F1A;refGene_tx=XM_003995155.5
D3 68557160 68557268 refGene_gn=ATP5F1A;refGene_tx=XM_003995155.5
D3 68558013 68558238 refGene_gn=ATP5F1A;refGene_tx=XM_003995155.5
D3 68558342 68558494 refGene_gn=ATP5F1A;refGene_tx=XM_003995155.5
D3 68559181 68559330 refGene_gn=ATP5F1A;refGene_tx=XM_003995155.5
D3 68559516 68559683 refGene_gn=ATP5F1A;refGene_tx=XM_003995155.5
D3 68559770 68559944 refGene_gn=ATP5F1A;refGene_tx=XM_003995155.5
D3 68561840 68562010 refGene_gn=ATP5F1A;refGene_tx=XM_003995155.5
D3 68563350 68563429 refGene_gn=ATP5F1A;refGene_tx=XM_003995155.5
D3 68565073 68565133 refGene_gn=ATP5F1A;refGene_tx=XM_003995155.5
D3 68565465 68565543
refGene_gn=HAUS1;refGene_tx=XM_019815222.2;refGene_gn=HAUS1;refGene_tx=XM_019815224.2
D3 68568877 68568907 refGene_gn=HAUS1;refGene_tx=XM_006938867.3
D3 68569680 68569855
refGene_gn=HAUS1;refGene_tx=XM_006938867.3;refGene_gn=HAUS1;refGene_tx=XM_019815222.2;refGene_gn=HAUS1;refGene_tx=XM_019815224.2
D3 68575050 68575186
refGene_gn=HAUS1;refGene_tx=XM_006938867.3;refGene_gn=HAUS1;refGene_tx=XM_019815222.2;refGene_gn=HAUS1;refGene_tx=XM_019815224.2
D3 68576801 68576936
refGene_gn=HAUS1;refGene_tx=XM_006938867.3;refGene_gn=HAUS1;refGene_tx=XM_019815222.2;refGene_gn=HAUS1;refGene_tx=XM_019815224.2
D3 68578169 68578293
refGene_gn=HAUS1;refGene_tx=XM_006938867.3;refGene_gn=HAUS1;refGene_tx=XM_019815222.2;refGene_gn=HAUS1;refGene_tx=XM_019815224.2
D3 68578694 68578760
refGene_gn=HAUS1;refGene_tx=XM_006938867.3;refGene_gn=HAUS1;refGene_tx=XM_019815222.2;refGene_gn=HAUS1;refGene_tx=XM_019815224.2
D3 68579956 68580028

refGene_gn=HAUS1;refGene_tx=XM_006938867.3;refGene_gn=HAUS1;refGene_tx=XM_019815222.2;refGene_gn=HAUS1;refGene_tx=XM_019815224.2
D3 68580267 68580315
refGene_gn=HAUS1;refGene_tx=XM_006938867.3;refGene_gn=HAUS1;refGene_tx=XM_019815222.2;refGene_gn=HAUS1;refGene_tx=XM_019815224.2
D3 68581552 68581603
refGene_gn=HAUS1;refGene_tx=XM_006938867.3;refGene_gn=HAUS1;refGene_tx=XM_019815222.2;refGene_gn=HAUS1;refGene_tx=XM_019815224.2
D3 68651939 68652654
refGene_gn=CD3H18orf25;refGene_tx=XM_023241954.1;refGene_gn=CD3H18orf25;refGene_tx=XM_023241955.1;refGene_gn=CD3H18orf25;refGene_tx=XM_023241956.1;refGene_gn=CD3H18orf25;refGene_tx=XM_023241957.1;refGene_gn=CD3H18orf25;refGene_tx=XM_023241958.1;refGene_gn=CD3H18orf25;refGene_tx=XM_023241959.1
D3 68674187 68674370
refGene_gn=CD3H18orf25;refGene_tx=XM_023241954.1;refGene_gn=CD3H18orf25;refGene_tx=XM_023241955.1;refGene_gn=CD3H18orf25;refGene_tx=XM_023241956.1;refGene_gn=CD3H18orf25;refGene_tx=XM_023241957.1
D3 68687204 68687331
refGene_gn=CD3H18orf25;refGene_tx=XM_023241954.1;refGene_gn=CD3H18orf25;refGene_tx=XM_023241955.1;refGene_gn=CD3H18orf25;refGene_tx=XM_023241956.1;refGene_gn=CD3H18orf25;refGene_tx=XM_023241957.1;refGene_gn=CD3H18orf25;refGene_tx=XM_023241958.1;refGene_gn=CD3H18orf25;refGene_tx=XM_023241959.1
D3 68688413 68688467
refGene_gn=CD3H18orf25;refGene_tx=XM_023241954.1;refGene_gn=CD3H18orf25;refGene_tx=XM_023241956.1;refGene_gn=CD3H18orf25;refGene_tx=XM_023241957.1;refGene_gn=CD3H18orf25;refGene_tx=XM_023241958.1
D3 68690447 68690475
refGene_gn=CD3H18orf25;refGene_tx=XM_023241956.1;refGene_gn=CD3H18orf25;refGene_tx=XM_023241957.1
D3 68695758 68695948
refGene_gn=CD3H18orf25;refGene_tx=XM_023241954.1;refGene_gn=CD3H18orf25;refGene_tx=XM_023241955.1;refGene_gn=CD3H18orf25;refGene_tx=XM_023241958.1;refGene_gn=CD3H18orf25;refGene_tx=XM_023241959.1
D3 68754894 68754955 refGene_gn=RNF165;refGene_tx=XM_023241960.1
D3 68845681 68845997 refGene_gn=RNF165;refGene_tx=XM_023241960.1
D3 68847871 68847982 refGene_gn=RNF165;refGene_tx=XM_023241960.1
D3 68853133 68853263 refGene_gn=RNF165;refGene_tx=XM_023241960.1
D3 68855442 68855547 refGene_gn=RNF165;refGene_tx=XM_023241960.1
D3 68855882 68855978 refGene_gn=RNF165;refGene_tx=XM_023241960.1
D3 68860821 68860928 refGene_gn=RNF165;refGene_tx=XM_023241960.1
D3 68861416 68861534 refGene_gn=RNF165;refGene_tx=XM_023241960.1
D3 68875638 68876119 refGene_gn=LOXHD1;refGene_tx=XM_023241200.1
D3 68881084 68881243 refGene_gn=LOXHD1;refGene_tx=XM_023241200.1
D3 68882564 68882697 refGene_gn=LOXHD1;refGene_tx=XM_023241200.1
D3 68888576 68888747 refGene_gn=LOXHD1;refGene_tx=XM_023241200.1
D3 68894076 68894262 refGene_gn=LOXHD1;refGene_tx=XM_023241200.1
D3 68895877 68896052 refGene_gn=LOXHD1;refGene_tx=XM_023241200.1
D3 68896773 68896891 refGene_gn=LOXHD1;refGene_tx=XM_023241200.1

D3 68903372 68903500 refGene_gn=LOXHD1;refGene_tx=XM_023241200.1
D3 68907341 68907550 refGene_gn=LOXHD1;refGene_tx=XM_023241200.1
D3 68909448 68909584 refGene_gn=LOXHD1;refGene_tx=XM_023241200.1
D3 68909689 68909899 refGene_gn=LOXHD1;refGene_tx=XM_023241200.1
D3 68913755 68913910 refGene_gn=LOXHD1;refGene_tx=XM_023241200.1
D3 68917385 68917548 refGene_gn=LOXHD1;refGene_tx=XM_023241200.1
D3 68918474 68918591 refGene_gn=LOXHD1;refGene_tx=XM_023241200.1
D3 68922767 68922949 refGene_gn=LOXHD1;refGene_tx=XM_023241200.1
D3 68927140 68927305 refGene_gn=LOXHD1;refGene_tx=XM_023241200.1
D3 68928465 68928594 refGene_gn=LOXHD1;refGene_tx=XM_023241200.1
D3 68931189 68931294 refGene_gn=LOXHD1;refGene_tx=XM_023241200.1
D3 68932183 68932353 refGene_gn=LOXHD1;refGene_tx=XM_023241200.1
D3 68941722 68941856 refGene_gn=LOXHD1;refGene_tx=XM_023241200.1
D3 68943781 68943936 refGene_gn=LOXHD1;refGene_tx=XM_023241200.1
D3 68944388 68944860 refGene_gn=LOXHD1;refGene_tx=XM_023241200.1
D3 68947221 68947382 refGene_gn=LOXHD1;refGene_tx=XM_023241200.1
D3 68950909 68951102 refGene_gn=LOXHD1;refGene_tx=XM_023241200.1
D3 68953487 68953684 refGene_gn=LOXHD1;refGene_tx=XM_023241200.1
D3 68954569 68954646 refGene_gn=LOXHD1;refGene_tx=XM_023241200.1
D3 68959195 68959356 refGene_gn=LOXHD1;refGene_tx=XM_023241200.1
D3 68961461 68961616 refGene_gn=LOXHD1;refGene_tx=XM_023241200.1
D3 68973495 68973631 refGene_gn=LOXHD1;refGene_tx=XM_023241200.1
D3 68974025 68974112 refGene_gn=LOXHD1;refGene_tx=XM_023241200.1
D3 68975096 68975257 refGene_gn=LOXHD1;refGene_tx=XM_023241200.1
D3 68975816 68975952 refGene_gn=LOXHD1;refGene_tx=XM_023241200.1
D3 68984827 68985078 refGene_gn=LOXHD1;refGene_tx=XM_023241200.1
D3 68987579 68987703 refGene_gn=LOXHD1;refGene_tx=XM_023241200.1
D3 68997405 68997554 refGene_gn=LOXHD1;refGene_tx=XM_023241200.1
D3 69004314 69004413 refGene_gn=LOXHD1;refGene_tx=XM_023241200.1
D3 69027179 69027364 refGene_gn=LOXHD1;refGene_tx=XM_023241200.1
D3 69029228 69029309 refGene_gn=LOXHD1;refGene_tx=XM_023241200.1
D3 69036049 69036164 refGene_gn=LOXHD1;refGene_tx=XM_023241200.1
D3 69036434 69036630 refGene_gn=LOXHD1;refGene_tx=XM_023241200.1
D3 69059563 69060032
refGene_gn=ST8SIA5;refGene_tx=XM_011288019.3;refGene_gn=ST8SIA5;refGene_tx=XM_011288020.3;refGene_gn=ST8SIA5;refGene_tx=XM_019815227.2;refGene_gn=ST8SIA5;refGene_tx=XM_019815228.2
D3 69061388 69061481
refGene_gn=ST8SIA5;refGene_tx=XM_011288019.3;refGene_gn=ST8SIA5;refGene_tx=XM_011288020.3;refGene_gn=ST8SIA5;refGene_tx=XM_019815227.2;refGene_gn=ST8SIA5;refGene_tx=XM_019815228.2
D3 69062276 69062389
refGene_gn=ST8SIA5;refGene_tx=XM_011288019.3;refGene_gn=ST8SIA5;refGene_tx=XM_011288020.3
D3 69066279 69066480
refGene_gn=ST8SIA5;refGene_tx=XM_011288019.3;refGene_gn=ST8SIA5;refGene_tx=XM_011288020.3;refGene_gn=ST8SIA5;refGene_tx=XM_019815227.2;refGene_gn=ST8SIA5;refGene_tx=XM_019815228.2
D3 69069904 69069991
refGene_gn=ST8SIA5;refGene_tx=XM_011288019.3;refGene_gn=ST8SIA5;refGen

e_tx=XM_011288020.3;refGene_gn=ST8SIA5;refGene_tx=XM_019815227.2;refGene_gn=ST8SIA5;refGene_tx=XM_019815228.2
D3 69081296 69081389
refGene_gn=ST8SIA5;refGene_tx=XM_011288019.3;refGene_gn=ST8SIA5;refGene_tx=XM_011288020.3;refGene_gn=ST8SIA5;refGene_tx=XM_019815227.2;refGene_gn=ST8SIA5;refGene_tx=XM_019815228.2
D3 69095378 69095486
refGene_gn=ST8SIA5;refGene_tx=XM_011288019.3;refGene_gn=ST8SIA5;refGene_tx=XM_019815227.2;refGene_gn=ST8SIA5;refGene_tx=XM_019815228.2
D3 69112678 69112809
refGene_gn=ST8SIA5;refGene_tx=XM_011288019.3;refGene_gn=ST8SIA5;refGene_tx=XM_011288020.3;refGene_gn=ST8SIA5;refGene_tx=XM_019815227.2;refGene_gn=ST8SIA5;refGene_tx=XM_019815228.2
D3 69112932 69113697
refGene_gn=LOC111557234;refGene_tx=XM_023241201.1
D3 69114025 69114157
refGene_gn=LOC111557234;refGene_tx=XM_023241201.1
D3 69115290 69115377
refGene_gn=LOC111557234;refGene_tx=XM_023241201.1
D3 69170889 69171072 refGene_gn=PIAS2;refGene_tx=XM_023241963.1
D3 69173739 69173777 refGene_gn=PIAS2;refGene_tx=XM_023241963.1
D3 69178491 69178631 refGene_gn=PIAS2;refGene_tx=XM_023241963.1
D3 69185224 69185396 refGene_gn=PIAS2;refGene_tx=XM_023241963.1
D3 69187011 69187145 refGene_gn=PIAS2;refGene_tx=XM_023241963.1
D3 69188344 69188505 refGene_gn=PIAS2;refGene_tx=XM_023241963.1
D3 69198171 69198245 refGene_gn=PIAS2;refGene_tx=XM_023241963.1
D3 69199612 69199718 refGene_gn=PIAS2;refGene_tx=XM_023241963.1
D3 69202110 69202176 refGene_gn=PIAS2;refGene_tx=XM_023241963.1
D3 69202177 69202228 refGene_gn=PIAS2;refGene_tx=XM_023241963.1
D3 69202229 69202247 refGene_gn=PIAS2;refGene_tx=XM_023241963.1
D3 69227779 69227870 refGene_gn=PIAS2;refGene_tx=XM_023241963.1
D3 69227992 69228043 refGene_gn=PIAS2;refGene_tx=XM_023241963.1
D3 69237083 69237168 refGene_gn=PIAS2;refGene_tx=XM_023241963.1
D3 69248392 69248867 refGene_gn=PIAS2;refGene_tx=XM_023241963.1
D3 69252478 69252526 refGene_gn=PIAS2;refGene_tx=XM_023241963.1
D3 69277474 69277498 refGene_gn=PIAS2;refGene_tx=XM_023241963.1
D3 69298600 69298651
refGene_gn=KATNAL2;refGene_tx=XM_006938879.3;refGene_gn=KATNAL2;refGene_tx=XM_006938880.3;refGene_gn=KATNAL2;refGene_tx=XM_023241962.1
D3 69323111 69323162 refGene_gn=KATNAL2;refGene_tx=XM_023241961.1
D3 69324915 69324986
refGene_gn=KATNAL2;refGene_tx=XM_006938879.3;refGene_gn=KATNAL2;refGene_tx=XM_006938880.3;refGene_gn=KATNAL2;refGene_tx=XM_023241961.1;refGene_gn=KATNAL2;refGene_tx=XM_023241962.1
D3 69328596 69328763
refGene_gn=KATNAL2;refGene_tx=XM_006938879.3;refGene_gn=KATNAL2;refGene_tx=XM_006938880.3;refGene_gn=KATNAL2;refGene_tx=XM_023241961.1;refGene_gn=KATNAL2;refGene_tx=XM_023241962.1
D3 69330095 69330138
refGene_gn=KATNAL2;refGene_tx=XM_006938879.3;refGene_gn=KATNAL2;refGen

e_tx=XM_006938880.3;refGene_gn=KATNAL2;refGene_tx=XM_023241961.1;refGene_gn=KATNAL2;refGene_tx=XM_023241962.1
D3 69333200 69333318
refGene_gn=KATNAL2;refGene_tx=XM_006938879.3;refGene_gn=KATNAL2;refGene_tx=XM_006938880.3;refGene_gn=KATNAL2;refGene_tx=XM_023241961.1;refGene_gn=KATNAL2;refGene_tx=XM_023241962.1
D3 69334690 69334789
refGene_gn=KATNAL2;refGene_tx=XM_006938879.3;refGene_gn=KATNAL2;refGene_tx=XM_006938880.3;refGene_gn=KATNAL2;refGene_tx=XM_023241961.1;refGene_gn=KATNAL2;refGene_tx=XM_023241962.1
D3 69336784 69336883
refGene_gn=KATNAL2;refGene_tx=XM_006938879.3;refGene_gn=KATNAL2;refGene_tx=XM_023241961.1;refGene_gn=KATNAL2;refGene_tx=XM_023241962.1
D3 69337091 69337169
refGene_gn=KATNAL2;refGene_tx=XM_006938879.3;refGene_gn=KATNAL2;refGene_tx=XM_006938880.3;refGene_gn=KATNAL2;refGene_tx=XM_023241961.1;refGene_gn=KATNAL2;refGene_tx=XM_023241962.1
D3 69339969 69340068
refGene_gn=KATNAL2;refGene_tx=XM_006938879.3;refGene_gn=KATNAL2;refGene_tx=XM_006938880.3;refGene_gn=KATNAL2;refGene_tx=XM_023241961.1
D3 69342077 69342141
refGene_gn=KATNAL2;refGene_tx=XM_006938879.3;refGene_gn=KATNAL2;refGene_tx=XM_006938880.3;refGene_gn=KATNAL2;refGene_tx=XM_023241961.1;refGene_gn=KATNAL2;refGene_tx=XM_023241962.1
D3 69342342 69342461
refGene_gn=KATNAL2;refGene_tx=XM_006938879.3;refGene_gn=KATNAL2;refGene_tx=XM_006938880.3;refGene_gn=KATNAL2;refGene_tx=XM_023241961.1;refGene_gn=KATNAL2;refGene_tx=XM_023241962.1
D3 69347363 69347455
refGene_gn=KATNAL2;refGene_tx=XM_006938879.3;refGene_gn=KATNAL2;refGene_tx=XM_006938880.3;refGene_gn=KATNAL2;refGene_tx=XM_023241961.1;refGene_gn=KATNAL2;refGene_tx=XM_023241962.1
D3 69349633 69349744
refGene_gn=KATNAL2;refGene_tx=XM_006938879.3;refGene_gn=KATNAL2;refGene_tx=XM_006938880.3;refGene_gn=KATNAL2;refGene_tx=XM_023241961.1;refGene_gn=KATNAL2;refGene_tx=XM_023241962.1
D3 69364736 69364899
refGene_gn=KATNAL2;refGene_tx=XM_006938879.3;refGene_gn=KATNAL2;refGene_tx=XM_006938880.3;refGene_gn=KATNAL2;refGene_tx=XM_023241961.1;refGene_gn=KATNAL2;refGene_tx=XM_023241962.1
D3 69365730 69365833
refGene_gn=KATNAL2;refGene_tx=XM_006938879.3;refGene_gn=KATNAL2;refGene_tx=XM_006938880.3;refGene_gn=KATNAL2;refGene_tx=XM_023241961.1;refGene_gn=KATNAL2;refGene_tx=XM_023241962.1
D3 69366343 69366483
refGene_gn=KATNAL2;refGene_tx=XM_006938879.3;refGene_gn=KATNAL2;refGene_tx=XM_006938880.3;refGene_gn=KATNAL2;refGene_tx=XM_023241961.1;refGene_gn=KATNAL2;refGene_tx=XM_023241962.1
D3 69373898 69374002
refGene_gn=HDHD2;refGene_tx=XM_011288013.3;refGene_gn=HDHD2;refGene_tx

=XM_011288014.2;refGene_gn=HDHD2;refGene_tx=XM_011288015.3;refGene_gn=
HDHD2;refGene_tx=XM_011288016.3
D3 69377860 69377924
refGene_gn=HDHD2;refGene_tx=XM_011288013.3;refGene_gn=HDHD2;refGene_tx
=XM_011288014.2;refGene_gn=HDHD2;refGene_tx=XM_011288015.3;refGene_gn=
HDHD2;refGene_tx=XM_011288016.3
D3 69380358 69380575
refGene_gn=HDHD2;refGene_tx=XM_011288013.3;refGene_gn=HDHD2;refGene_tx
=XM_011288014.2;refGene_gn=HDHD2;refGene_tx=XM_011288015.3;refGene_gn=
HDHD2;refGene_tx=XM_011288016.3
D3 69382190 69382345 refGene_gn=HDHD2;refGene_tx=XM_019815237.2
D3 69389159 69389239 refGene_gn=HDHD2;refGene_tx=XM_019815237.2
D3 69393883 69393968
refGene_gn=HDHD2;refGene_tx=XM_011288013.3;refGene_gn=HDHD2;refGene_tx
=XM_011288014.2;refGene_gn=HDHD2;refGene_tx=XM_011288015.3;refGene_gn=
HDHD2;refGene_tx=XM_011288016.3;refGene_gn=HDHD2;refGene_tx=XM_0198152
37.2
D3 69397747 69397956
refGene_gn=HDHD2;refGene_tx=XM_011288013.3;refGene_gn=HDHD2;refGene_tx
=XM_011288014.2;refGene_gn=HDHD2;refGene_tx=XM_011288015.3;refGene_gn=
HDHD2;refGene_tx=XM_011288016.3;refGene_gn=HDHD2;refGene_tx=XM_0198152
37.2
D3 69401663 69401764
refGene_gn=HDHD2;refGene_tx=XM_011288013.3;refGene_gn=HDHD2;refGene_tx
=XM_011288014.2;refGene_gn=HDHD2;refGene_tx=XM_011288015.3;refGene_gn=
HDHD2;refGene_tx=XM_011288016.3;refGene_gn=HDHD2;refGene_tx=XM_0198152
37.2
D3 69418034 69418090 refGene_gn=IER3IP1;refGene_tx=XM_003995172.5
D3 69419273 69419375 refGene_gn=IER3IP1;refGene_tx=XM_003995172.5
D3 69433201 69433292 refGene_gn=IER3IP1;refGene_tx=XM_003995172.5
D3 69457592 69457655
refGene_gn=SKOR2;refGene_tx=XM_023241964.1;refGene_gn=SKOR2;refGene_tx
=XM_023241965.1
D3 69466870 69466938
refGene_gn=SKOR2;refGene_tx=XM_023241964.1;refGene_gn=SKOR2;refGene_tx
=XM_023241965.1
D3 69478108 69478207
refGene_gn=SKOR2;refGene_tx=XM_023241964.1;refGene_gn=SKOR2;refGene_tx
=XM_023241965.1
D3 69478586 69478652
refGene_gn=SKOR2;refGene_tx=XM_023241964.1;refGene_gn=SKOR2;refGene_tx
=XM_023241965.1
D3 69490100 69490175
refGene_gn=SKOR2;refGene_tx=XM_023241964.1;refGene_gn=SKOR2;refGene_tx
=XM_023241965.1
D3 69490573 69490637
refGene_gn=SKOR2;refGene_tx=XM_023241964.1;refGene_gn=SKOR2;refGene_tx
=XM_023241965.1
D3 69491665 69494263
refGene_gn=SKOR2;refGene_tx=XM_023241964.1;refGene_gn=SKOR2;refGene_tx

=XM_023241965.1
D3 69980007 69980131
refGene_gn=SMAD2;refGene_tx=XM_023241966.1;refGene_gn=SMAD2;refGene_tx=XM_023241967.1;refGene_gn=SMAD2;refGene_tx=XM_023241968.1;refGene_gn=SMAD2;refGene_tx=XM_023241969.1;refGene_gn=SMAD2;refGene_tx=XM_023241970.1;refGene_gn=SMAD2;refGene_tx=XM_023241971.1
D3 69985012 69985157
refGene_gn=SMAD2;refGene_tx=XM_023241966.1;refGene_gn=SMAD2;refGene_tx=XM_023241967.1;refGene_gn=SMAD2;refGene_tx=XM_023241968.1;refGene_gn=SMAD2;refGene_tx=XM_023241969.1;refGene_gn=SMAD2;refGene_tx=XM_023241970.1;refGene_gn=SMAD2;refGene_tx=XM_023241971.1
D3 69985373 69985511
refGene_gn=SMAD2;refGene_tx=XM_023241966.1;refGene_gn=SMAD2;refGene_tx=XM_023241967.1;refGene_gn=SMAD2;refGene_tx=XM_023241968.1;refGene_gn=SMAD2;refGene_tx=XM_023241969.1;refGene_gn=SMAD2;refGene_tx=XM_023241970.1;refGene_gn=SMAD2;refGene_tx=XM_023241971.1
D3 69988464 69988677
refGene_gn=SMAD2;refGene_tx=XM_023241966.1;refGene_gn=SMAD2;refGene_tx=XM_023241967.1;refGene_gn=SMAD2;refGene_tx=XM_023241968.1;refGene_gn=SMAD2;refGene_tx=XM_023241969.1;refGene_gn=SMAD2;refGene_tx=XM_023241970.1;refGene_gn=SMAD2;refGene_tx=XM_023241971.1
D3 69989869 69989923
refGene_gn=SMAD2;refGene_tx=XM_023241966.1;refGene_gn=SMAD2;refGene_tx=XM_023241967.1;refGene_gn=SMAD2;refGene_tx=XM_023241968.1;refGene_gn=SMAD2;refGene_tx=XM_023241969.1;refGene_gn=SMAD2;refGene_tx=XM_023241970.1;refGene_gn=SMAD2;refGene_tx=XM_023241971.1
D3 69999877 69999952
refGene_gn=SMAD2;refGene_tx=XM_023241966.1;refGene_gn=SMAD2;refGene_tx=XM_023241967.1;refGene_gn=SMAD2;refGene_tx=XM_023241968.1;refGene_gn=SMAD2;refGene_tx=XM_023241969.1;refGene_gn=SMAD2;refGene_tx=XM_023241970.1;refGene_gn=SMAD2;refGene_tx=XM_023241971.1
D3 70002753 70002888
refGene_gn=SMAD2;refGene_tx=XM_023241966.1;refGene_gn=SMAD2;refGene_tx=XM_023241967.1;refGene_gn=SMAD2;refGene_tx=XM_023241968.1;refGene_gn=SMAD2;refGene_tx=XM_023241969.1;refGene_gn=SMAD2;refGene_tx=XM_023241970.1;refGene_gn=SMAD2;refGene_tx=XM_023241971.1
D3 70003979 70004173
refGene_gn=SMAD2;refGene_tx=XM_023241966.1;refGene_gn=SMAD2;refGene_tx=XM_023241967.1;refGene_gn=SMAD2;refGene_tx=XM_023241968.1;refGene_gn=SMAD2;refGene_tx=XM_023241969.1;refGene_gn=SMAD2;refGene_tx=XM_023241970.1;refGene_gn=SMAD2;refGene_tx=XM_023241971.1
D3 70005231 70005321
refGene_gn=SMAD2;refGene_tx=XM_023241966.1;refGene_gn=SMAD2;refGene_tx=XM_023241967.1;refGene_gn=SMAD2;refGene_tx=XM_023241968.1;refGene_gn=SMAD2;refGene_tx=XM_023241969.1;refGene_gn=SMAD2;refGene_tx=XM_023241970.1
D3 70032829 70033065
refGene_gn=SMAD2;refGene_tx=XM_023241966.1;refGene_gn=SMAD2;refGene_tx=XM_023241967.1;refGene_gn=SMAD2;refGene_tx=XM_023241968.1;refGene_gn=SMAD2;refGene_tx=XM_023241969.1;refGene_gn=SMAD2;refGene_tx=XM_0232419

70.1;refGene_gn=SMAD2;refGene_tx=XM_023241971.1
D3 70145470 70146122
refGene_gn=ZBTB7C;refGene_tx=XM_003995176.5;refGene_gn=ZBTB7C;refGene_tx=XM_019815241.2;refGene_gn=ZBTB7C;refGene_tx=XM_023241972.1;refGene_gn=ZBTB7C;refGene_tx=XM_023241973.1;refGene_gn=ZBTB7C;refGene_tx=XM_023241974.1;refGene_gn=ZBTB7C;refGene_tx=XM_023241975.1;refGene_gn=ZBTB7C;refGene_tx=XM_023241976.1;refGene_gn=ZBTB7C;refGene_tx=XM_023241977.1
D3 70155859 70157070
refGene_gn=ZBTB7C;refGene_tx=XM_003995176.5;refGene_gn=ZBTB7C;refGene_tx=XM_019815241.2;refGene_gn=ZBTB7C;refGene_tx=XM_023241972.1;refGene_gn=ZBTB7C;refGene_tx=XM_023241973.1;refGene_gn=ZBTB7C;refGene_tx=XM_023241974.1;refGene_gn=ZBTB7C;refGene_tx=XM_023241975.1;refGene_gn=ZBTB7C;refGene_tx=XM_023241976.1;refGene_gn=ZBTB7C;refGene_tx=XM_023241977.1
D3 70689141 70689321
refGene_gn=CTIF;refGene_tx=XM_003995177.5;refGene_gn=CTIF;refGene_tx=XM_006938888.4;refGene_gn=CTIF;refGene_tx=XM_019815242.2;refGene_gn=CTIF;refGene_tx=XM_019815243.1;refGene_gn=CTIF;refGene_tx=XM_019815244.2;refGene_gn=CTIF;refGene_tx=XM_019815245.2;refGene_gn=CTIF;refGene_tx=XM_019815246.2
D3 70694168 70694216
refGene_gn=CTIF;refGene_tx=XM_019815242.2;refGene_gn=CTIF;refGene_tx=XM_019815243.1;refGene_gn=CTIF;refGene_tx=XM_019815244.2;refGene_gn=CTIF;refGene_tx=XM_019815246.2
D3 70706121 70706193
refGene_gn=CTIF;refGene_tx=XM_003995177.5;refGene_gn=CTIF;refGene_tx=XM_006938888.4;refGene_gn=CTIF;refGene_tx=XM_019815242.2;refGene_gn=CTIF;refGene_tx=XM_019815243.1;refGene_gn=CTIF;refGene_tx=XM_019815244.2;refGene_gn=CTIF;refGene_tx=XM_019815246.2
D3 70725908 70725982
refGene_gn=CTIF;refGene_tx=XM_003995177.5;refGene_gn=CTIF;refGene_tx=XM_006938888.4;refGene_gn=CTIF;refGene_tx=XM_019815242.2;refGene_gn=CTIF;refGene_tx=XM_019815243.1;refGene_gn=CTIF;refGene_tx=XM_019815244.2;refGene_gn=CTIF;refGene_tx=XM_019815245.2;refGene_gn=CTIF;refGene_tx=XM_019815246.2
D3 70727209 70727314
refGene_gn=CTIF;refGene_tx=XM_003995177.5;refGene_gn=CTIF;refGene_tx=XM_006938888.4;refGene_gn=CTIF;refGene_tx=XM_019815242.2;refGene_gn=CTIF;refGene_tx=XM_019815243.1;refGene_gn=CTIF;refGene_tx=XM_019815244.2;refGene_gn=CTIF;refGene_tx=XM_019815245.2;refGene_gn=CTIF;refGene_tx=XM_019815246.2
D3 70732537 70732613
refGene_gn=CTIF;refGene_tx=XM_003995177.5;refGene_gn=CTIF;refGene_tx=XM_006938888.4;refGene_gn=CTIF;refGene_tx=XM_019815242.2;refGene_gn=CTIF;refGene_tx=XM_019815243.1;refGene_gn=CTIF;refGene_tx=XM_019815244.2;refGene_gn=CTIF;refGene_tx=XM_019815245.2;refGene_gn=CTIF;refGene_tx=XM_019815246.2
D3 70775567 70775644
refGene_gn=CTIF;refGene_tx=XM_003995177.5;refGene_gn=CTIF;refGene_tx=X

M_006938888.4;refGene_gn=CTIF;refGene_tx=XM_019815242.2;refGene_gn=CTIF;refGene_tx=XM_019815243.1;refGene_gn=CTIF;refGene_tx=XM_019815244.2;refGene_gn=CTIF;refGene_tx=XM_019815245.2;refGene_gn=CTIF;refGene_tx=XM_019815246.2
D3 70817943 70818436
refGene_gn=CTIF;refGene_tx=XM_003995177.5;refGene_gn=CTIF;refGene_tx=XM_006938888.4;refGene_gn=CTIF;refGene_tx=XM_019815242.2;refGene_gn=CTIF;refGene_tx=XM_019815243.1;refGene_gn=CTIF;refGene_tx=XM_019815244.2;refGene_gn=CTIF;refGene_tx=XM_019815245.2;refGene_gn=CTIF;refGene_tx=XM_019815246.2
D3 70821049 70821349
refGene_gn=CTIF;refGene_tx=XM_003995177.5;refGene_gn=CTIF;refGene_tx=XM_006938888.4;refGene_gn=CTIF;refGene_tx=XM_019815242.2;refGene_gn=CTIF;refGene_tx=XM_019815243.1;refGene_gn=CTIF;refGene_tx=XM_019815244.2;refGene_gn=CTIF;refGene_tx=XM_019815245.2;refGene_gn=CTIF;refGene_tx=XM_019815246.2
D3 70870331 70870487
refGene_gn=CTIF;refGene_tx=XM_003995177.5;refGene_gn=CTIF;refGene_tx=XM_006938888.4;refGene_gn=CTIF;refGene_tx=XM_019815242.2;refGene_gn=CTIF;refGene_tx=XM_019815243.1;refGene_gn=CTIF;refGene_tx=XM_019815244.2;refGene_gn=CTIF;refGene_tx=XM_019815245.2
D3 70905999 70906053
refGene_gn=CTIF;refGene_tx=XM_003995177.5;refGene_gn=CTIF;refGene_tx=XM_006938888.4;refGene_gn=CTIF;refGene_tx=XM_019815242.2;refGene_gn=CTIF;refGene_tx=XM_019815243.1;refGene_gn=CTIF;refGene_tx=XM_019815244.2;refGene_gn=CTIF;refGene_tx=XM_019815245.2;refGene_gn=CTIF;refGene_tx=XM_019815246.2
D3 70907548 70907764
refGene_gn=CTIF;refGene_tx=XM_003995177.5;refGene_gn=CTIF;refGene_tx=XM_006938888.4;refGene_gn=CTIF;refGene_tx=XM_019815242.2;refGene_gn=CTIF;refGene_tx=XM_019815243.1;refGene_gn=CTIF;refGene_tx=XM_019815244.2;refGene_gn=CTIF;refGene_tx=XM_019815245.2;refGene_gn=CTIF;refGene_tx=XM_019815246.2
D3 70958007 70958546
refGene_gn=SMAD7;refGene_tx=XM_023241979.1;refGene_gn=SMAD7;refGene_tx=XM_023241980.1
D3 70978070 70978145
refGene_gn=SMAD7;refGene_tx=XM_023241979.1;refGene_gn=SMAD7;refGene_tx=XM_023241980.1
D3 70984010 70984064
refGene_gn=SMAD7;refGene_tx=XM_023241979.1;refGene_gn=SMAD7;refGene_tx=XM_023241980.1
D3 70985379 70985992
refGene_gn=SMAD7;refGene_tx=XM_023241979.1;refGene_gn=SMAD7;refGene_tx=XM_023241980.1
D3 71067716 71067866 refGene_gn=DYM;refGene_tx=XM_019815247.1
D3 71118967 71119081 refGene_gn=DYM;refGene_tx=XM_019815247.1
D3 71132675 71132858 refGene_gn=DYM;refGene_tx=XM_019815247.1
D3 71176134 71176237 refGene_gn=DYM;refGene_tx=XM_019815247.1
D3 71259936 71260031 refGene_gn=DYM;refGene_tx=XM_019815247.1

D3 71261429 71261543 refGene_gn=DYM;refGene_tx=XM_019815247.1
D3 71272460 71272586 refGene_gn=DYM;refGene_tx=XM_019815247.1
D3 71274394 71274573 refGene_gn=DYM;refGene_tx=XM_019815247.1
D3 71279192 71279375 refGene_gn=DYM;refGene_tx=XM_019815247.1
D3 71310013 71310156 refGene_gn=DYM;refGene_tx=XM_019815247.1
D3 71311573 71311699 refGene_gn=DYM;refGene_tx=XM_019815247.1
D3 71336744 71336817 refGene_gn=DYM;refGene_tx=XM_019815247.1
D3 71352568 71352702 refGene_gn=DYM;refGene_tx=XM_019815247.1
D3 71353662 71353756 refGene_gn=DYM;refGene_tx=XM_019815247.1
D3 71365036 71365089 refGene_gn=DYM;refGene_tx=XM_019815247.1
D3 71397469 71397495 refGene_gn=DYM;refGene_tx=XM_019815247.1
D3 71397496 71397610 refGene_gn=DYM;refGene_tx=XM_019815247.1
D3 71437665 71437731
refGene_gn=CD3H18orf32;refGene_tx=XM_003995180.4;refGene_gn=CD3H18orf32;refGene_tx=XM_019815252.2
D3 71438741 71438906
refGene_gn=CD3H18orf32;refGene_tx=XM_003995180.4;refGene_gn=CD3H18orf32;refGene_tx=XM_019815252.2
D3 71443171 71443219 refGene_gn=RPL17;refGene_tx=NM_001128842.1
D3 71443717 71443909 refGene_gn=RPL17;refGene_tx=NM_001128842.1
D3 71444755 71444854 refGene_gn=RPL17;refGene_tx=NM_001128842.1
D3 71445092 71445195 refGene_gn=RPL17;refGene_tx=NM_001128842.1
D3 71445993 71446034 refGene_gn=RPL17;refGene_tx=NM_001128842.1
D3 71446125 71446165 refGene_gn=RPL17;refGene_tx=NM_001128842.1
D3 71484147 71484235 refGene_gn=LIPG;refGene_tx=XM_019815255.2
D3 71485301 71485398 refGene_gn=LIPG;refGene_tx=XM_003995203.5
D3 71487491 71487660
refGene_gn=LIPG;refGene_tx=XM_019815253.2;refGene_gn=LIPG;refGene_tx=XM_019815254.2
D3 71488161 71488343
refGene_gn=LIPG;refGene_tx=XM_003995203.5;refGene_gn=LIPG;refGene_tx=XM_019815253.2;refGene_gn=LIPG;refGene_tx=XM_019815254.2;refGene_gn=LIPG;refGene_tx=XM_019815255.2
D3 71489983 71490163
refGene_gn=LIPG;refGene_tx=XM_003995203.5;refGene_gn=LIPG;refGene_tx=XM_019815253.2;refGene_gn=LIPG;refGene_tx=XM_019815254.2;refGene_gn=LIPG;refGene_tx=XM_019815255.2
D3 71491288 71491400
refGene_gn=LIPG;refGene_tx=XM_003995203.5;refGene_gn=LIPG;refGene_tx=XM_019815253.2;refGene_gn=LIPG;refGene_tx=XM_019815254.2;refGene_gn=LIPG;refGene_tx=XM_019815255.2
D3 71496420 71496642
refGene_gn=LIPG;refGene_tx=XM_003995203.5;refGene_gn=LIPG;refGene_tx=XM_019815253.2;refGene_gn=LIPG;refGene_tx=XM_019815254.2;refGene_gn=LIPG;refGene_tx=XM_019815255.2
D3 71498220 71498463
refGene_gn=LIPG;refGene_tx=XM_003995203.5;refGene_gn=LIPG;refGene_tx=XM_019815253.2;refGene_gn=LIPG;refGene_tx=XM_019815254.2;refGene_gn=LIPG;refGene_tx=XM_019815255.2
D3 71499179 71499300

refGene_gn=LIPG;refGene_tx=XM_003995203.5;refGene_gn=LIPG;refGene_tx=XM_019815253.2;refGene_gn=LIPG;refGene_tx=XM_019815254.2;refGene_gn=LIPG;refGene_tx=XM_019815255.2
D3 71499995 71500214
refGene_gn=LIPG;refGene_tx=XM_003995203.5;refGene_gn=LIPG;refGene_tx=XM_019815253.2;refGene_gn=LIPG;refGene_tx=XM_019815254.2;refGene_gn=LIPG;refGene_tx=XM_019815255.2
D3 71502218 71502323
refGene_gn=LIPG;refGene_tx=XM_003995203.5;refGene_gn=LIPG;refGene_tx=XM_019815253.2;refGene_gn=LIPG;refGene_tx=XM_019815254.2;refGene_gn=LIPG;refGene_tx=XM_019815255.2
D3 71504464 71504486
refGene_gn=LIPG;refGene_tx=XM_003995203.5;refGene_gn=LIPG;refGene_tx=XM_019815253.2;refGene_gn=LIPG;refGene_tx=XM_019815254.2;refGene_gn=LIPG;refGene_tx=XM_019815255.2
D3 71665533 71665618 refGene_gn=ACAA2;refGene_tx=XM_003995181.5
D3 71666861 71667016 refGene_gn=ACAA2;refGene_tx=XM_003995181.5
D3 71669198 71669269 refGene_gn=ACAA2;refGene_tx=XM_003995181.5
D3 71673770 71673809 refGene_gn=ACAA2;refGene_tx=XM_003995181.5
D3 71673811 71673899 refGene_gn=ACAA2;refGene_tx=XM_003995181.5
D3 71674729 71674783 refGene_gn=ACAA2;refGene_tx=XM_003995181.5
D3 71674784 71674906 refGene_gn=ACAA2;refGene_tx=XM_003995181.5
D3 71677120 71677268 refGene_gn=ACAA2;refGene_tx=XM_003995181.5
D3 71678601 71678718 refGene_gn=ACAA2;refGene_tx=XM_003995181.5
D3 71688505 71688588 refGene_gn=ACAA2;refGene_tx=XM_003995181.5
D3 71688590 71688633 refGene_gn=ACAA2;refGene_tx=XM_003995181.5
D3 71692502 71692669 refGene_gn=ACAA2;refGene_tx=XM_003995181.5
D3 71698856 71698872 refGene_gn=ACAA2;refGene_tx=XM_003995181.5
D3 71710640 71710793
refGene_gn=MYO5B;refGene_tx=XM_023241982.1;refGene_gn=MYO5B;refGene_tx=XM_023241983.1
D3 71715602 71715683
refGene_gn=MYO5B;refGene_tx=XM_023241982.1;refGene_gn=MYO5B;refGene_tx=XM_023241983.1
D3 71717736 71717911
refGene_gn=MYO5B;refGene_tx=XM_023241982.1;refGene_gn=MYO5B;refGene_tx=XM_023241983.1
D3 71718500 71718786
refGene_gn=MYO5B;refGene_tx=XM_023241982.1;refGene_gn=MYO5B;refGene_tx=XM_023241983.1
D3 71720079 71720230
refGene_gn=MYO5B;refGene_tx=XM_023241982.1;refGene_gn=MYO5B;refGene_tx=XM_023241983.1
D3 71722991 71723081
refGene_gn=MYO5B;refGene_tx=XM_023241982.1;refGene_gn=MYO5B;refGene_tx=XM_023241983.1
D3 71724812 71724964
refGene_gn=MYO5B;refGene_tx=XM_023241982.1;refGene_gn=MYO5B;refGene_tx=XM_023241983.1
D3 71727006 71727150

refGene_gn=MYO5B;refGene_tx=XM_023241982.1;refGene_gn=MYO5B;refGene_tx=XM_023241983.1
D3 71729294 71729388
refGene_gn=MYO5B;refGene_tx=XM_023241982.1;refGene_gn=MYO5B;refGene_tx=XM_023241983.1
D3 71729547 71729746
refGene_gn=MYO5B;refGene_tx=XM_023241982.1;refGene_gn=MYO5B;refGene_tx=XM_023241983.1
D3 71732495 71732573 refGene_gn=MYO5B;refGene_tx=XM_023241982.1
D3 71738913 71739014
refGene_gn=MYO5B;refGene_tx=XM_023241982.1;refGene_gn=MYO5B;refGene_tx=XM_023241983.1
D3 71740081 71740321
refGene_gn=MYO5B;refGene_tx=XM_023241982.1;refGene_gn=MYO5B;refGene_tx=XM_023241983.1
D3 71744100 71744166
refGene_gn=MYO5B;refGene_tx=XM_023241982.1;refGene_gn=MYO5B;refGene_tx=XM_023241983.1
D3 71747793 71747934
refGene_gn=MYO5B;refGene_tx=XM_023241982.1;refGene_gn=MYO5B;refGene_tx=XM_023241983.1
D3 71750295 71750415
refGene_gn=MYO5B;refGene_tx=XM_023241982.1;refGene_gn=MYO5B;refGene_tx=XM_023241983.1
D3 71751765 71751911
refGene_gn=MYO5B;refGene_tx=XM_023241982.1;refGene_gn=MYO5B;refGene_tx=XM_023241983.1
D3 71753168 71753253
refGene_gn=MYO5B;refGene_tx=XM_023241982.1;refGene_gn=MYO5B;refGene_tx=XM_023241983.1
D3 71765749 71765983
refGene_gn=MYO5B;refGene_tx=XM_023241982.1;refGene_gn=MYO5B;refGene_tx=XM_023241983.1
D3 71773363 71773603
refGene_gn=MYO5B;refGene_tx=XM_023241982.1;refGene_gn=MYO5B;refGene_tx=XM_023241983.1
D3 71775461 71775618
refGene_gn=MYO5B;refGene_tx=XM_023241982.1;refGene_gn=MYO5B;refGene_tx=XM_023241983.1
D3 71777446 71777658
refGene_gn=MYO5B;refGene_tx=XM_023241982.1;refGene_gn=MYO5B;refGene_tx=XM_023241983.1
D3 71781739 71781851
refGene_gn=MYO5B;refGene_tx=XM_023241982.1;refGene_gn=MYO5B;refGene_tx=XM_023241983.1
D3 71795748 71795835
refGene_gn=MYO5B;refGene_tx=XM_023241982.1;refGene_gn=MYO5B;refGene_tx=XM_023241983.1
D3 71801384 71801482
refGene_gn=MYO5B;refGene_tx=XM_023241982.1;refGene_gn=MYO5B;refGene_tx

=XM_023241983.1
D3	71802371	71802527
refGene_gn=MYO5B;refGene_tx=XM_023241982.1;refGene_gn=MYO5B;refGene_tx
=XM_023241983.1
D3	71815082	71815166
refGene_gn=MYO5B;refGene_tx=XM_023241982.1;refGene_gn=MYO5B;refGene_tx
=XM_023241983.1
D3	71815923	71816046
refGene_gn=MYO5B;refGene_tx=XM_023241982.1;refGene_gn=MYO5B;refGene_tx
=XM_023241983.1
D3	71824078	71824219
refGene_gn=MYO5B;refGene_tx=XM_023241982.1;refGene_gn=MYO5B;refGene_tx
=XM_023241983.1
D3	71824677	71824759
refGene_gn=MYO5B;refGene_tx=XM_023241982.1;refGene_gn=MYO5B;refGene_tx
=XM_023241983.1
D3	71834987	71835253
refGene_gn=MYO5B;refGene_tx=XM_023241982.1;refGene_gn=MYO5B;refGene_tx
=XM_023241983.1
D3	71840657	71840767
refGene_gn=MYO5B;refGene_tx=XM_023241982.1;refGene_gn=MYO5B;refGene_tx
=XM_023241983.1
D3	71844301	71844409
refGene_gn=MYO5B;refGene_tx=XM_023241982.1;refGene_gn=MYO5B;refGene_tx
=XM_023241983.1
D3	71849580	71849662
refGene_gn=MYO5B;refGene_tx=XM_023241982.1;refGene_gn=MYO5B;refGene_tx
=XM_023241983.1
D3	71850949	71851093
refGene_gn=MYO5B;refGene_tx=XM_023241982.1;refGene_gn=MYO5B;refGene_tx
=XM_023241983.1
D3	71858402	71858559
refGene_gn=MYO5B;refGene_tx=XM_023241982.1;refGene_gn=MYO5B;refGene_tx
=XM_023241983.1
D3	71892427	71892572
refGene_gn=MYO5B;refGene_tx=XM_023241982.1;refGene_gn=MYO5B;refGene_tx
=XM_023241983.1
D3	71896908	71897080
refGene_gn=MYO5B;refGene_tx=XM_023241982.1;refGene_gn=MYO5B;refGene_tx
=XM_023241983.1
D3	71906829	71906940
refGene_gn=MYO5B;refGene_tx=XM_023241982.1;refGene_gn=MYO5B;refGene_tx
=XM_023241983.1
D3	72050983	72051010
refGene_gn=MYO5B;refGene_tx=XM_023241982.1;refGene_gn=MYO5B;refGene_tx
=XM_023241983.1
D3	72077590	72077819	refGene_gn=CFAP53;refGene_tx=XM_003995182.4
D3	72080791	72080894	refGene_gn=CFAP53;refGene_tx=XM_003995182.4
D3	72082739	72082956	refGene_gn=CFAP53;refGene_tx=XM_003995182.4
D3	72089972	72090191	refGene_gn=CFAP53;refGene_tx=XM_003995182.4

D3 72093809 72094113 refGene_gn=CFAP53;refGene_tx=XM_003995182.4
D3 72108568 72108742 refGene_gn=CFAP53;refGene_tx=XM_003995182.4
D3 72109214 72109444 refGene_gn=CFAP53;refGene_tx=XM_003995182.4
D3 72113264 72113498 refGene_gn=CFAP53;refGene_tx=XM_003995182.4
D3 72114643 72114716 refGene_gn=MBD1;refGene_tx=XM_023241986.1
D3 72116313 72116521
refGene_gn=MBD1;refGene_tx=XM_011287995.1;refGene_gn=MBD1;refGene_tx=XM_011288004.1;refGene_gn=MBD1;refGene_tx=XM_011288008.1;refGene_gn=MBD1;refGene_tx=XM_019815259.1;refGene_gn=MBD1;refGene_tx=XM_019815260.1;refGene_gn=MBD1;refGene_tx=XM_023241984.1;refGene_gn=MBD1;refGene_tx=XM_023241985.1;refGene_gn=MBD1;refGene_tx=XM_023241991.1
D3 72118214 72118254
refGene_gn=MBD1;refGene_tx=XM_011288001.2;refGene_gn=MBD1;refGene_tx=XM_011288002.2;refGene_gn=MBD1;refGene_tx=XM_011288006.2;refGene_gn=MBD1;refGene_tx=XM_011288009.2;refGene_gn=MBD1;refGene_tx=XM_019815267.1;refGene_gn=MBD1;refGene_tx=XM_019815269.1;refGene_gn=MBD1;refGene_tx=XM_023241987.1;refGene_gn=MBD1;refGene_tx=XM_023241988.1;refGene_gn=MBD1;refGene_tx=XM_023241989.1;refGene_gn=MBD1;refGene_tx=XM_023241990.1;refGene_gn=MBD1;refGene_tx=XM_023241992.1
D3 72119400 72119462
refGene_gn=MBD1;refGene_tx=XM_011287995.1;refGene_gn=MBD1;refGene_tx=XM_011288001.2;refGene_gn=MBD1;refGene_tx=XM_011288002.2;refGene_gn=MBD1;refGene_tx=XM_011288004.1;refGene_gn=MBD1;refGene_tx=XM_011288006.2;refGene_gn=MBD1;refGene_tx=XM_011288008.1;refGene_gn=MBD1;refGene_tx=XM_011288009.2;refGene_gn=MBD1;refGene_tx=XM_019815259.1;refGene_gn=MBD1;refGene_tx=XM_019815260.1;refGene_gn=MBD1;refGene_tx=XM_019815267.1;refGene_gn=MBD1;refGene_tx=XM_019815269.1;refGene_gn=MBD1;refGene_tx=XM_023241984.1;refGene_gn=MBD1;refGene_tx=XM_023241985.1;refGene_gn=MBD1;refGene_tx=XM_023241986.1;refGene_gn=MBD1;refGene_tx=XM_023241987.1;refGene_gn=MBD1;refGene_tx=XM_023241988.1;refGene_gn=MBD1;refGene_tx=XM_023241989.1;refGene_gn=MBD1;refGene_tx=XM_023241990.1;refGene_gn=MBD1;refGene_tx=XM_023241991.1;refGene_gn=MBD1;refGene_tx=XM_023241992.1
D3 72119548 72119683
refGene_gn=MBD1;refGene_tx=XM_011287995.1;refGene_gn=MBD1;refGene_tx=XM_011288001.2;refGene_gn=MBD1;refGene_tx=XM_011288002.2;refGene_gn=MBD1;refGene_tx=XM_011288004.1;refGene_gn=MBD1;refGene_tx=XM_011288006.2;refGene_gn=MBD1;refGene_tx=XM_011288008.1;refGene_gn=MBD1;refGene_tx=XM_011288009.2;refGene_gn=MBD1;refGene_tx=XM_019815259.1;refGene_gn=MBD1;refGene_tx=XM_019815260.1;refGene_gn=MBD1;refGene_tx=XM_019815267.1;refGene_gn=MBD1;refGene_tx=XM_019815269.1;refGene_gn=MBD1;refGene_tx=XM_023241984.1;refGene_gn=MBD1;refGene_tx=XM_023241985.1;refGene_gn=MBD1;refGene_tx=XM_023241986.1;refGene_gn=MBD1;refGene_tx=XM_023241987.1;refGene_gn=MBD1;refGene_tx=XM_023241988.1;refGene_gn=MBD1;refGene_tx=XM_023241989.1;refGene_gn=MBD1;refGene_tx=XM_023241990.1;refGene_gn=MBD1;refGene_tx=XM_023241991.1;refGene_gn=MBD1;refGene_tx=XM_023241992.1
D3 72120034 72120172
refGene_gn=MBD1;refGene_tx=XM_011287995.1;refGene_gn=MBD1;refGene_tx=XM_011288001.2;refGene_gn=MBD1;refGene_tx=XM_011288002.2;refGene_gn=MBD1;refGene_tx=XM_019815259.1;refGene_gn=MBD1;refGene_tx=XM_019815260.1;refGene_gn=MBD1;refGene_tx=XM_019815267.1;refGene_gn=MBD1;refGene_tx=X

M_019815269.1;refGene_gn=MBD1;refGene_tx=XM_023241984.1;refGene_gn=MBD1;refGene_tx=XM_023241985.1;refGene_gn=MBD1;refGene_tx=XM_023241986.1;refGene_gn=MBD1;refGene_tx=XM_023241987.1;refGene_gn=MBD1;refGene_tx=XM_023241988.1;refGene_gn=MBD1;refGene_tx=XM_023241989.1;refGene_gn=MBD1;refGene_tx=XM_023241990.1
D3 72120265 72120565
refGene_gn=MBD1;refGene_tx=XM_011287995.1;refGene_gn=MBD1;refGene_tx=XM_011288001.2;refGene_gn=MBD1;refGene_tx=XM_011288002.2;refGene_gn=MBD1;refGene_tx=XM_011288004.1;refGene_gn=MBD1;refGene_tx=XM_011288006.2;refGene_gn=MBD1;refGene_tx=XM_011288008.1;refGene_gn=MBD1;refGene_tx=XM_011288009.2;refGene_gn=MBD1;refGene_tx=XM_019815259.1;refGene_gn=MBD1;refGene_tx=XM_019815260.1;refGene_gn=MBD1;refGene_tx=XM_019815267.1;refGene_gn=MBD1;refGene_tx=XM_019815269.1;refGene_gn=MBD1;refGene_tx=XM_023241984.1;refGene_gn=MBD1;refGene_tx=XM_023241985.1;refGene_gn=MBD1;refGene_tx=XM_023241986.1;refGene_gn=MBD1;refGene_tx=XM_023241987.1;refGene_gn=MBD1;refGene_tx=XM_023241988.1;refGene_gn=MBD1;refGene_tx=XM_023241989.1;refGene_gn=MBD1;refGene_tx=XM_023241990.1;refGene_gn=MBD1;refGene_tx=XM_023241991.1;refGene_gn=MBD1;refGene_tx=XM_023241992.1
D3 72120916 72121084
refGene_gn=MBD1;refGene_tx=XM_011287995.1;refGene_gn=MBD1;refGene_tx=XM_011288001.2;refGene_gn=MBD1;refGene_tx=XM_011288002.2;refGene_gn=MBD1;refGene_tx=XM_011288004.1;refGene_gn=MBD1;refGene_tx=XM_011288006.2;refGene_gn=MBD1;refGene_tx=XM_011288008.1;refGene_gn=MBD1;refGene_tx=XM_011288009.2;refGene_gn=MBD1;refGene_tx=XM_019815259.1;refGene_gn=MBD1;refGene_tx=XM_019815260.1;refGene_gn=MBD1;refGene_tx=XM_019815267.1;refGene_gn=MBD1;refGene_tx=XM_019815269.1;refGene_gn=MBD1;refGene_tx=XM_023241984.1;refGene_gn=MBD1;refGene_tx=XM_023241985.1;refGene_gn=MBD1;refGene_tx=XM_023241986.1;refGene_gn=MBD1;refGene_tx=XM_023241987.1;refGene_gn=MBD1;refGene_tx=XM_023241988.1;refGene_gn=MBD1;refGene_tx=XM_023241989.1;refGene_gn=MBD1;refGene_tx=XM_023241990.1;refGene_gn=MBD1;refGene_tx=XM_023241991.1;refGene_gn=MBD1;refGene_tx=XM_023241992.1
D3 72121660 72121729
refGene_gn=MBD1;refGene_tx=XM_011287995.1;refGene_gn=MBD1;refGene_tx=XM_011288001.2;refGene_gn=MBD1;refGene_tx=XM_011288002.2;refGene_gn=MBD1;refGene_tx=XM_011288008.1;refGene_gn=MBD1;refGene_tx=XM_011288009.2;refGene_gn=MBD1;refGene_tx=XM_019815259.1;refGene_gn=MBD1;refGene_tx=XM_019815260.1;refGene_gn=MBD1;refGene_tx=XM_019815267.1;refGene_gn=MBD1;refGene_tx=XM_019815269.1;refGene_gn=MBD1;refGene_tx=XM_023241984.1;refGene_gn=MBD1;refGene_tx=XM_023241985.1;refGene_gn=MBD1;refGene_tx=XM_023241986.1;refGene_gn=MBD1;refGene_tx=XM_023241987.1;refGene_gn=MBD1;refGene_tx=XM_023241988.1;refGene_gn=MBD1;refGene_tx=XM_023241989.1;refGene_gn=MBD1;refGene_tx=XM_023241990.1;refGene_gn=MBD1;refGene_tx=XM_023241991.1;refGene_gn=MBD1;refGene_tx=XM_023241992.1
D3 72121811 72121931
refGene_gn=MBD1;refGene_tx=XM_011287995.1;refGene_gn=MBD1;refGene_tx=XM_011288001.2;refGene_gn=MBD1;refGene_tx=XM_011288002.2;refGene_gn=MBD1;refGene_tx=XM_011288004.1;refGene_gn=MBD1;refGene_tx=XM_011288006.2;refGene_gn=MBD1;refGene_tx=XM_011288008.1;refGene_gn=MBD1;refGene_tx=XM_011288009.2;refGene_gn=MBD1;refGene_tx=XM_019815259.1;refGene_gn=MBD1;refGene_tx=XM_019815260.1;refGene_gn=MBD1;refGene_tx=XM_019815267.1;

refGene_gn=MBD1;refGene_tx=XM_019815269.1;refGene_gn=MBD1;refGene_tx=X
M_023241984.1;refGene_gn=MBD1;refGene_tx=XM_023241985.1;refGene_gn=MBD
1;refGene_tx=XM_023241986.1;refGene_gn=MBD1;refGene_tx=XM_023241987.1;
refGene_gn=MBD1;refGene_tx=XM_023241988.1;refGene_gn=MBD1;refGene_tx=X
M_023241989.1;refGene_gn=MBD1;refGene_tx=XM_023241990.1;refGene_gn=MBD
1;refGene_tx=XM_023241991.1;refGene_gn=MBD1;refGene_tx=XM_023241992.1
D3 72122081 72122156
refGene_gn=MBD1;refGene_tx=XM_011288001.2;refGene_gn=MBD1;refGene_tx=X
M_019815259.1;refGene_gn=MBD1;refGene_tx=XM_019815260.1;refGene_gn=MBD
1;refGene_tx=XM_023241984.1;refGene_gn=MBD1;refGene_tx=XM_023241985.1;
refGene_gn=MBD1;refGene_tx=XM_023241986.1;refGene_gn=MBD1;refGene_tx=X
M_023241987.1;refGene_gn=MBD1;refGene_tx=XM_023241988.1;refGene_gn=MBD
1;refGene_tx=XM_023241989.1;refGene_gn=MBD1;refGene_tx=XM_023241990.1;
refGene_gn=MBD1;refGene_tx=XM_023241991.1;refGene_gn=MBD1;refGene_tx=X
M_023241992.1
D3 72122328 72122457
refGene_gn=MBD1;refGene_tx=XM_011287995.1;refGene_gn=MBD1;refGene_tx=X
M_011288001.2;refGene_gn=MBD1;refGene_tx=XM_011288002.2;refGene_gn=MBD
1;refGene_tx=XM_011288004.1;refGene_gn=MBD1;refGene_tx=XM_011288006.2;
refGene_gn=MBD1;refGene_tx=XM_011288008.1;refGene_gn=MBD1;refGene_tx=X
M_011288009.2;refGene_gn=MBD1;refGene_tx=XM_019815259.1;refGene_gn=MBD
1;refGene_tx=XM_019815260.1;refGene_gn=MBD1;refGene_tx=XM_019815267.1;
refGene_gn=MBD1;refGene_tx=XM_019815269.1;refGene_gn=MBD1;refGene_tx=X
M_023241984.1;refGene_gn=MBD1;refGene_tx=XM_023241985.1;refGene_gn=MBD
1;refGene_tx=XM_023241986.1;refGene_gn=MBD1;refGene_tx=XM_023241987.1;
refGene_gn=MBD1;refGene_tx=XM_023241988.1;refGene_gn=MBD1;refGene_tx=X
M_023241989.1;refGene_gn=MBD1;refGene_tx=XM_023241990.1;refGene_gn=MBD
1;refGene_tx=XM_023241991.1;refGene_gn=MBD1;refGene_tx=XM_023241992.1
D3 72122552 72122699
refGene_gn=MBD1;refGene_tx=XM_011287995.1;refGene_gn=MBD1;refGene_tx=X
M_011288001.2;refGene_gn=MBD1;refGene_tx=XM_011288002.2;refGene_gn=MBD
1;refGene_tx=XM_011288004.1;refGene_gn=MBD1;refGene_tx=XM_011288006.2;
refGene_gn=MBD1;refGene_tx=XM_011288008.1;refGene_gn=MBD1;refGene_tx=X
M_011288009.2;refGene_gn=MBD1;refGene_tx=XM_019815259.1;refGene_gn=MBD
1;refGene_tx=XM_019815260.1;refGene_gn=MBD1;refGene_tx=XM_019815267.1;
refGene_gn=MBD1;refGene_tx=XM_023241984.1;refGene_gn=MBD1;refGene_tx=X
M_023241985.1;refGene_gn=MBD1;refGene_tx=XM_023241986.1;refGene_gn=MBD
1;refGene_tx=XM_023241987.1;refGene_gn=MBD1;refGene_tx=XM_023241988.1;
refGene_gn=MBD1;refGene_tx=XM_023241989.1;refGene_gn=MBD1;refGene_tx=X
M_023241990.1;refGene_gn=MBD1;refGene_tx=XM_023241991.1;refGene_gn=MBD
1;refGene_tx=XM_023241992.1
D3 72123071 72123112
refGene_gn=MBD1;refGene_tx=XM_011287995.1;refGene_gn=MBD1;refGene_tx=X
M_011288001.2;refGene_gn=MBD1;refGene_tx=XM_011288002.2;refGene_gn=MBD
1;refGene_tx=XM_011288004.1;refGene_gn=MBD1;refGene_tx=XM_011288006.2;
refGene_gn=MBD1;refGene_tx=XM_011288008.1;refGene_gn=MBD1;refGene_tx=X
M_011288009.2;refGene_gn=MBD1;refGene_tx=XM_019815259.1;refGene_gn=MBD
1;refGene_tx=XM_019815260.1;refGene_gn=MBD1;refGene_tx=XM_019815267.1;
refGene_gn=MBD1;refGene_tx=XM_019815269.1;refGene_gn=MBD1;refGene_tx=X
M_023241984.1;refGene_gn=MBD1;refGene_tx=XM_023241985.1;refGene_gn=MBD

1;refGene_tx=XM_023241986.1;refGene_gn=MBD1;refGene_tx=XM_023241987.1;
refGene_gn=MBD1;refGene_tx=XM_023241988.1;refGene_gn=MBD1;refGene_tx=X
M_023241989.1;refGene_gn=MBD1;refGene_tx=XM_023241990.1;refGene_gn=MBD
1;refGene_tx=XM_023241991.1;refGene_gn=MBD1;refGene_tx=XM_023241992.1
D3 72123262 72123345
refGene_gn=MBD1;refGene_tx=XM_011287995.1;refGene_gn=MBD1;refGene_tx=X
M_011288001.2;refGene_gn=MBD1;refGene_tx=XM_011288002.2;refGene_gn=MBD
1;refGene_tx=XM_011288004.1;refGene_gn=MBD1;refGene_tx=XM_011288006.2;
refGene_gn=MBD1;refGene_tx=XM_011288008.1;refGene_gn=MBD1;refGene_tx=X
M_011288009.2;refGene_gn=MBD1;refGene_tx=XM_019815259.1;refGene_gn=MBD
1;refGene_tx=XM_019815260.1;refGene_gn=MBD1;refGene_tx=XM_019815267.1;
refGene_gn=MBD1;refGene_tx=XM_019815269.1;refGene_gn=MBD1;refGene_tx=X
M_023241984.1;refGene_gn=MBD1;refGene_tx=XM_023241985.1;refGene_gn=MBD
1;refGene_tx=XM_023241986.1;refGene_gn=MBD1;refGene_tx=XM_023241987.1;
refGene_gn=MBD1;refGene_tx=XM_023241988.1;refGene_gn=MBD1;refGene_tx=X
M_023241989.1;refGene_gn=MBD1;refGene_tx=XM_023241990.1;refGene_gn=MBD
1;refGene_tx=XM_023241991.1;refGene_gn=MBD1;refGene_tx=XM_023241992.1
D3 72123432 72123629
refGene_gn=MBD1;refGene_tx=XM_011287995.1;refGene_gn=MBD1;refGene_tx=X
M_011288001.2;refGene_gn=MBD1;refGene_tx=XM_011288002.2;refGene_gn=MBD
1;refGene_tx=XM_011288004.1;refGene_gn=MBD1;refGene_tx=XM_011288006.2;
refGene_gn=MBD1;refGene_tx=XM_011288008.1;refGene_gn=MBD1;refGene_tx=X
M_011288009.2;refGene_gn=MBD1;refGene_tx=XM_019815259.1;refGene_gn=MBD
1;refGene_tx=XM_019815260.1;refGene_gn=MBD1;refGene_tx=XM_019815267.1;
refGene_gn=MBD1;refGene_tx=XM_019815269.1;refGene_gn=MBD1;refGene_tx=X
M_023241984.1;refGene_gn=MBD1;refGene_tx=XM_023241985.1;refGene_gn=MBD
1;refGene_tx=XM_023241986.1;refGene_gn=MBD1;refGene_tx=XM_023241987.1;
refGene_gn=MBD1;refGene_tx=XM_023241988.1;refGene_gn=MBD1;refGene_tx=X
M_023241989.1;refGene_gn=MBD1;refGene_tx=XM_023241990.1;refGene_gn=MBD
1;refGene_tx=XM_023241991.1;refGene_gn=MBD1;refGene_tx=XM_023241992.1
D3 72123690 72123805
refGene_gn=MBD1;refGene_tx=XM_011287995.1;refGene_gn=MBD1;refGene_tx=X
M_011288001.2;refGene_gn=MBD1;refGene_tx=XM_011288002.2;refGene_gn=MBD
1;refGene_tx=XM_011288004.1;refGene_gn=MBD1;refGene_tx=XM_011288006.2;
refGene_gn=MBD1;refGene_tx=XM_011288008.1;refGene_gn=MBD1;refGene_tx=X
M_011288009.2;refGene_gn=MBD1;refGene_tx=XM_019815259.1;refGene_gn=MBD
1;refGene_tx=XM_019815260.1;refGene_gn=MBD1;refGene_tx=XM_019815267.1;
refGene_gn=MBD1;refGene_tx=XM_019815269.1;refGene_gn=MBD1;refGene_tx=X
M_023241984.1;refGene_gn=MBD1;refGene_tx=XM_023241985.1;refGene_gn=MBD
1;refGene_tx=XM_023241986.1;refGene_gn=MBD1;refGene_tx=XM_023241987.1;
refGene_gn=MBD1;refGene_tx=XM_023241988.1;refGene_gn=MBD1;refGene_tx=X
M_023241989.1;refGene_gn=MBD1;refGene_tx=XM_023241990.1;refGene_gn=MBD
1;refGene_tx=XM_023241991.1;refGene_gn=MBD1;refGene_tx=XM_023241992.1
D3 72126243 72126353
refGene_gn=MBD1;refGene_tx=XM_011287995.1;refGene_gn=MBD1;refGene_tx=X
M_011288001.2;refGene_gn=MBD1;refGene_tx=XM_011288002.2;refGene_gn=MBD
1;refGene_tx=XM_011288004.1;refGene_gn=MBD1;refGene_tx=XM_011288006.2;
refGene_gn=MBD1;refGene_tx=XM_011288008.1;refGene_gn=MBD1;refGene_tx=X
M_011288009.2;refGene_gn=MBD1;refGene_tx=XM_019815259.1;refGene_gn=MBD
1;refGene_tx=XM_019815260.1;refGene_gn=MBD1;refGene_tx=XM_019815267.1;

refGene_gn=MBD1;refGene_tx=XM_019815269.1;refGene_gn=MBD1;refGene_tx=XM_023241984.1;refGene_gn=MBD1;refGene_tx=XM_023241985.1;refGene_gn=MBD1;refGene_tx=XM_023241986.1;refGene_gn=MBD1;refGene_tx=XM_023241987.1;refGene_gn=MBD1;refGene_tx=XM_023241988.1;refGene_gn=MBD1;refGene_tx=XM_023241989.1;refGene_gn=MBD1;refGene_tx=XM_023241990.1;refGene_gn=MBD1;refGene_tx=XM_023241991.1;refGene_gn=MBD1;refGene_tx=XM_023241992.1
D3 72129308 72129455
refGene_gn=CXXC1;refGene_tx=XM_006938895.4;refGene_gn=CXXC1;refGene_tx=XM_019815270.2
D3 72129538 72129691
refGene_gn=CXXC1;refGene_tx=XM_006938895.4;refGene_gn=CXXC1;refGene_tx=XM_019815270.2
D3 72130009 72130106
refGene_gn=CXXC1;refGene_tx=XM_006938895.4;refGene_gn=CXXC1;refGene_tx=XM_019815270.2
D3 72130244 72130294
refGene_gn=CXXC1;refGene_tx=XM_006938895.4;refGene_gn=CXXC1;refGene_tx=XM_019815270.2
D3 72130418 72130529
refGene_gn=CXXC1;refGene_tx=XM_006938895.4;refGene_gn=CXXC1;refGene_tx=XM_019815270.2
D3 72130608 72130816
refGene_gn=CXXC1;refGene_tx=XM_006938895.4;refGene_gn=CXXC1;refGene_tx=XM_019815270.2
D3 72131102 72131287
refGene_gn=CXXC1;refGene_tx=XM_006938895.4;refGene_gn=CXXC1;refGene_tx=XM_019815270.2
D3 72131459 72131567
refGene_gn=CXXC1;refGene_tx=XM_006938895.4;refGene_gn=CXXC1;refGene_tx=XM_019815270.2
D3 72131731 72131977
refGene_gn=CXXC1;refGene_tx=XM_006938895.4;refGene_gn=CXXC1;refGene_tx=XM_019815270.2
D3 72132054 72132081 refGene_gn=CXXC1;refGene_tx=XM_006938895.4
D3 72132496 72132700
refGene_gn=CXXC1;refGene_tx=XM_006938895.4;refGene_gn=CXXC1;refGene_tx=XM_019815270.2
D3 72132791 72133027
refGene_gn=CXXC1;refGene_tx=XM_006938895.4;refGene_gn=CXXC1;refGene_tx=XM_019815270.2
D3 72133318 72133419
refGene_gn=CXXC1;refGene_tx=XM_006938895.4;refGene_gn=CXXC1;refGene_tx=XM_019815270.2
D3 72133519 72133638
refGene_gn=CXXC1;refGene_tx=XM_006938895.4;refGene_gn=CXXC1;refGene_tx=XM_019815270.2
D3 72134420 72134423
refGene_gn=CXXC1;refGene_tx=XM_006938895.4;refGene_gn=CXXC1;refGene_tx=XM_019815270.2
D3 72166185 72166273

refGene_gn=SKA1;refGene_tx=XM_006938908.4;refGene_gn=SKA1;refGene_tx=XM_023241993.1
D3 72169058 72169183
refGene_gn=SKA1;refGene_tx=XM_006938908.4;refGene_gn=SKA1;refGene_tx=XM_023241993.1
D3 72170722 72170820
refGene_gn=SKA1;refGene_tx=XM_006938908.4;refGene_gn=SKA1;refGene_tx=XM_023241993.1
D3 72173693 72173831
refGene_gn=SKA1;refGene_tx=XM_006938908.4;refGene_gn=SKA1;refGene_tx=XM_023241993.1
D3 72177020 72177190
refGene_gn=SKA1;refGene_tx=XM_006938908.4;refGene_gn=SKA1;refGene_tx=XM_023241993.1
D3 72177978 72178127
refGene_gn=SKA1;refGene_tx=XM_006938908.4;refGene_gn=SKA1;refGene_tx=XM_023241993.1
D3 72371491 72372037
refGene_gn=MAPK4;refGene_tx=XM_011287991.3;refGene_gn=MAPK4;refGene_tx=XM_019815272.2
D3 72411082 72411227
refGene_gn=MAPK4;refGene_tx=XM_011287991.3;refGene_gn=MAPK4;refGene_tx=XM_019815272.2
D3 72412948 72413110
refGene_gn=MAPK4;refGene_tx=XM_011287991.3;refGene_gn=MAPK4;refGene_tx=XM_019815272.2
D3 72418596 72418798
refGene_gn=MAPK4;refGene_tx=XM_011287991.3;refGene_gn=MAPK4;refGene_tx=XM_019815272.2
D3 72421553 72422250
refGene_gn=MAPK4;refGene_tx=XM_011287991.3;refGene_gn=MAPK4;refGene_tx=XM_019815272.2
D3 72469987 72470002 refGene_gn=MRO;refGene_tx=XM_023241996.1
D3 72471181 72471214 refGene_gn=MRO;refGene_tx=XM_023241995.1
D3 72471917 72471971
refGene_gn=MRO;refGene_tx=XM_011287988.3;refGene_gn=MRO;refGene_tx=XM_023241994.1;refGene_gn=MRO;refGene_tx=XM_023241997.1;refGene_gn=MRO;refGene_tx=XM_023241998.1;refGene_gn=MRO;refGene_tx=XM_023241999.1;refGene_gn=MRO;refGene_tx=XM_023242001.1
D3 72472319 72472427
refGene_gn=MRO;refGene_tx=XM_011287988.3;refGene_gn=MRO;refGene_tx=XM_023241994.1;refGene_gn=MRO;refGene_tx=XM_023241995.1;refGene_gn=MRO;refGene_tx=XM_023241996.1;refGene_gn=MRO;refGene_tx=XM_023241997.1;refGene_gn=MRO;refGene_tx=XM_023241998.1;refGene_gn=MRO;refGene_tx=XM_023241999.1;refGene_gn=MRO;refGene_tx=XM_023242001.1
D3 72473755 72473911
refGene_gn=MRO;refGene_tx=XM_011287988.3;refGene_gn=MRO;refGene_tx=XM_023241994.1;refGene_gn=MRO;refGene_tx=XM_023241995.1;refGene_gn=MRO;refGene_tx=XM_023241996.1;refGene_gn=MRO;refGene_tx=XM_023241997.1;refGene_gn=MRO;refGene_tx=XM_023241998.1;refGene_gn=MRO;refGene_tx=XM_02324

1999.1;refGene_gn=MRO;refGene_tx=XM_023242001.1
D3 72476257 72476440
refGene_gn=MRO;refGene_tx=XM_011287988.3;refGene_gn=MRO;refGene_tx=XM_023241994.1;refGene_gn=MRO;refGene_tx=XM_023241995.1;refGene_gn=MRO;refGene_tx=XM_023241996.1;refGene_gn=MRO;refGene_tx=XM_023241997.1;refGene_gn=MRO;refGene_tx=XM_023241998.1;refGene_gn=MRO;refGene_tx=XM_023241999.1;refGene_gn=MRO;refGene_tx=XM_023242001.1
D3 72477401 72477548
refGene_gn=MRO;refGene_tx=XM_011287988.3;refGene_gn=MRO;refGene_tx=XM_023241994.1;refGene_gn=MRO;refGene_tx=XM_023241995.1;refGene_gn=MRO;refGene_tx=XM_023241996.1;refGene_gn=MRO;refGene_tx=XM_023241997.1;refGene_gn=MRO;refGene_tx=XM_023241998.1;refGene_gn=MRO;refGene_tx=XM_023241999.1;refGene_gn=MRO;refGene_tx=XM_023242001.1
D3 72478781 72478884
refGene_gn=MRO;refGene_tx=XM_011287988.3;refGene_gn=MRO;refGene_tx=XM_023241994.1;refGene_gn=MRO;refGene_tx=XM_023241995.1;refGene_gn=MRO;refGene_tx=XM_023241996.1;refGene_gn=MRO;refGene_tx=XM_023241997.1;refGene_gn=MRO;refGene_tx=XM_023241998.1;refGene_gn=MRO;refGene_tx=XM_023241999.1;refGene_gn=MRO;refGene_tx=XM_023242001.1
D3 72486306 72486410
refGene_gn=MRO;refGene_tx=XM_023241994.1;refGene_gn=MRO;refGene_tx=XM_023241995.1;refGene_gn=MRO;refGene_tx=XM_023241996.1
D3 72542022 72542130 refGene_gn=ME2;refGene_tx=XM_003995198.4
D3 72551714 72551848 refGene_gn=ME2;refGene_tx=XM_003995198.4
D3 72555845 72555995 refGene_gn=ME2;refGene_tx=XM_003995198.4
D3 72558426 72558502 refGene_gn=ME2;refGene_tx=XM_003995198.4
D3 72559595 72559757 refGene_gn=ME2;refGene_tx=XM_003995198.4
D3 72560071 72560175 refGene_gn=ME2;refGene_tx=XM_003995198.4
D3 72563056 72563166 refGene_gn=ME2;refGene_tx=XM_003995198.4
D3 72563268 72563366 refGene_gn=ME2;refGene_tx=XM_003995198.4
D3 72563663 72563777 refGene_gn=ME2;refGene_tx=XM_003995198.4
D3 72566491 72566606 refGene_gn=ME2;refGene_tx=XM_003995198.4
D3 72568826 72568969 refGene_gn=ME2;refGene_tx=XM_003995198.4
D3 72572243 72572346 refGene_gn=ME2;refGene_tx=XM_003995198.4
D3 72580776 72580847 refGene_gn=ME2;refGene_tx=XM_003995198.4
D3 72581606 72581705 refGene_gn=ME2;refGene_tx=XM_003995198.4
D3 72586219 72586387 refGene_gn=ME2;refGene_tx=XM_003995198.4
D3 72606178 72606335 refGene_gn=ELAC1;refGene_tx=XM_003995200.3
D3 72612560 72613028 refGene_gn=ELAC1;refGene_tx=XM_003995200.3
D3 72614776 72615243 refGene_gn=ELAC1;refGene_tx=XM_003995200.3
D3 72647028 72647053
refGene_gn=LOC111557333;refGene_tx=XM_023242004.1
D3 72653400 72653816
refGene_gn=LOC111557333;refGene_tx=XM_023242004.1
D3 72673837 72674086
refGene_gn=SMAD4;refGene_tx=XM_011287987.3;refGene_gn=SMAD4;refGene_tx=XM_019815278.2;refGene_gn=SMAD4;refGene_tx=XM_023242002.1;refGene_gn=SMAD4;refGene_tx=XM_023242003.1
D3 72674927 72675102
refGene_gn=SMAD4;refGene_tx=XM_011287987.3;refGene_gn=SMAD4;refGene_tx

=XM_019815278.2;refGene_gn=SMAD4;refGene_tx=XM_023242002.1;refGene_gn=SMAD4;refGene_tx=XM_023242003.1
D3 72675537 72675567 refGene_gn=SMAD4;refGene_tx=XM_011287987.3;refGene_gn=SMAD4;refGene_tx=XM_019815278.2;refGene_gn=SMAD4;refGene_tx=XM_023242002.1;refGene_gn=SMAD4;refGene_tx=XM_023242003.1
D3 72680475 72680688 refGene_gn=SMAD4;refGene_tx=XM_011287987.3;refGene_gn=SMAD4;refGene_tx=XM_019815278.2;refGene_gn=SMAD4;refGene_tx=XM_023242002.1;refGene_gn=SMAD4;refGene_tx=XM_023242003.1
D3 72684657 72684777 refGene_gn=SMAD4;refGene_tx=XM_011287987.3;refGene_gn=SMAD4;refGene_tx=XM_019815278.2;refGene_gn=SMAD4;refGene_tx=XM_023242002.1;refGene_gn=SMAD4;refGene_tx=XM_023242003.1
D3 72684871 72684988 refGene_gn=SMAD4;refGene_tx=XM_011287987.3;refGene_gn=SMAD4;refGene_tx=XM_019815278.2;refGene_gn=SMAD4;refGene_tx=XM_023242002.1;refGene_gn=SMAD4;refGene_tx=XM_023242003.1
D3 72686378 72686429 refGene_gn=SMAD4;refGene_tx=XM_011287987.3;refGene_gn=SMAD4;refGene_tx=XM_019815278.2;refGene_gn=SMAD4;refGene_tx=XM_023242002.1;refGene_gn=SMAD4;refGene_tx=XM_023242003.1
D3 72696244 72696428 refGene_gn=SMAD4;refGene_tx=XM_011287987.3;refGene_gn=SMAD4;refGene_tx=XM_019815278.2;refGene_gn=SMAD4;refGene_tx=XM_023242002.1;refGene_gn=SMAD4;refGene_tx=XM_023242003.1
D3 72697727 72697896 refGene_gn=SMAD4;refGene_tx=XM_011287987.3;refGene_gn=SMAD4;refGene_tx=XM_019815278.2;refGene_gn=SMAD4;refGene_tx=XM_023242002.1;refGene_gn=SMAD4;refGene_tx=XM_023242003.1
D3 72699468 72699537 refGene_gn=SMAD4;refGene_tx=XM_023242003.1
D3 72709306 72709445 refGene_gn=SMAD4;refGene_tx=XM_011287987.3;refGene_gn=SMAD4;refGene_tx=XM_019815278.2;refGene_gn=SMAD4;refGene_tx=XM_023242002.1
D3 72710931 72711143 refGene_gn=SMAD4;refGene_tx=XM_011287987.3;refGene_gn=SMAD4;refGene_tx=XM_019815278.2;refGene_gn=SMAD4;refGene_tx=XM_023242002.1
D3 72769110 72770336 refGene_gn=MEX3C;refGene_tx=XM_023242005.1
D3 72787505 72788253 refGene_gn=MEX3C;refGene_tx=XM_023242005.1
D3 73132352 73132368 refGene_gn=LOC101095864;refGene_tx=XM_011287986.2
D3 73133837 73134253 refGene_gn=LOC101095864;refGene_tx=XM_011287986.2
D3 73790194 73790285 refGene_gn=DCC;refGene_tx=XM_019815281.2
D3 74192447 74192768 refGene_gn=DCC;refGene_tx=XM_019815281.2
D3 74314687 74314972 refGene_gn=DCC;refGene_tx=XM_019815281.2
D3 74332553 74332704 refGene_gn=DCC;refGene_tx=XM_019815281.2
D3 74333896 74334033 refGene_gn=DCC;refGene_tx=XM_019815281.2
D3 74489466 74489621 refGene_gn=DCC;refGene_tx=XM_019815281.2
D3 74492261 74492382 refGene_gn=DCC;refGene_tx=XM_019815281.2

D3 74595354 74595511 refGene_gn=DCC;refGene_tx=XM_019815281.2
D3 74614006 74614161 refGene_gn=DCC;refGene_tx=XM_019815281.2
D3 74640944 74641093 refGene_gn=DCC;refGene_tx=XM_019815281.2
D3 74642764 74642903 refGene_gn=DCC;refGene_tx=XM_019815281.2
D3 74649783 74649833 refGene_gn=DCC;refGene_tx=XM_019815281.2
D3 74732677 74732819 refGene_gn=DCC;refGene_tx=XM_019815281.2
D3 74744640 74744751 refGene_gn=DCC;refGene_tx=XM_019815281.2
D3 74767016 74767211 refGene_gn=DCC;refGene_tx=XM_019815281.2
D3 74809512 74809608 refGene_gn=DCC;refGene_tx=XM_019815281.2
D3 74815098 74815331 refGene_gn=DCC;refGene_tx=XM_019815281.2
D3 74820072 74820211 refGene_gn=DCC;refGene_tx=XM_019815281.2
D3 74825318 74825426 refGene_gn=DCC;refGene_tx=XM_019815281.2
D3 74831669 74831864 refGene_gn=DCC;refGene_tx=XM_019815281.2
D3 74836759 74836792 refGene_gn=DCC;refGene_tx=XM_019815281.2
D3 74859873 74859939 refGene_gn=DCC;refGene_tx=XM_019815281.2
D3 74871994 74872114 refGene_gn=DCC;refGene_tx=XM_019815281.2
D3 74872116 74872156 refGene_gn=DCC;refGene_tx=XM_019815281.2
D3 74880478 74880705 refGene_gn=DCC;refGene_tx=XM_019815281.2
D3 74885187 74885304 refGene_gn=DCC;refGene_tx=XM_019815281.2
D3 74898642 74898798 refGene_gn=DCC;refGene_tx=XM_019815281.2
D3 74908209 74908422 refGene_gn=DCC;refGene_tx=XM_019815281.2
D3 74934762 74934905 refGene_gn=DCC;refGene_tx=XM_019815281.2
D3 74937522 74937612 refGene_gn=DCC;refGene_tx=XM_019815281.2
D3 75468452 75468579 refGene_gn=MBD2;refGene_tx=XM_019815286.2
D3 75473252 75473430 refGene_gn=MBD2;refGene_tx=XM_019815286.2
D3 75474743 75474834 refGene_gn=MBD2;refGene_tx=XM_019815286.2
D3 75496870 75497008 refGene_gn=MBD2;refGene_tx=XM_019815286.2
D3 75513014 75513174 refGene_gn=MBD2;refGene_tx=XM_019815286.2
D3 75529450 75529536 refGene_gn=MBD2;refGene_tx=XM_019815286.2
D3 75529537 75529990 refGene_gn=MBD2;refGene_tx=XM_019815286.2
D3 75566892 75567040
refGene_gn=POLI;refGene_tx=XM_011288112.3;refGene_gn=POLI;refGene_tx=XM_019815287.2;refGene_gn=POLI;refGene_tx=XM_023242008.1
D3 75568399 75568510
refGene_gn=POLI;refGene_tx=XM_011288112.3;refGene_gn=POLI;refGene_tx=XM_019815287.2;refGene_gn=POLI;refGene_tx=XM_023242008.1
D3 75570941 75571106
refGene_gn=POLI;refGene_tx=XM_011288112.3;refGene_gn=POLI;refGene_tx=XM_019815287.2;refGene_gn=POLI;refGene_tx=XM_023242008.1
D3 75575901 75576054
refGene_gn=POLI;refGene_tx=XM_011288112.3;refGene_gn=POLI;refGene_tx=XM_023242006.1;refGene_gn=POLI;refGene_tx=XM_023242007.1;refGene_gn=POLI;refGene_tx=XM_023242008.1
D3 75578202 75578439
refGene_gn=POLI;refGene_tx=XM_011288112.3;refGene_gn=POLI;refGene_tx=XM_019815287.2;refGene_gn=POLI;refGene_tx=XM_023242006.1;refGene_gn=POLI;refGene_tx=XM_023242007.1;refGene_gn=POLI;refGene_tx=XM_023242008.1
D3 75580669 75580848
refGene_gn=POLI;refGene_tx=XM_011288112.3;refGene_gn=POLI;refGene_tx=XM_019815287.2;refGene_gn=POLI;refGene_tx=XM_023242006.1;refGene_gn=POL

I;refGene_tx=XM_023242007.1;refGene_gn=POLI;refGene_tx=XM_023242008.1
D3 75582395 75582487
refGene_gn=POLI;refGene_tx=XM_011288112.3;refGene_gn=POLI;refGene_tx=XM_019815287.2;refGene_gn=POLI;refGene_tx=XM_023242006.1;refGene_gn=POLI;refGene_tx=XM_023242007.1;refGene_gn=POLI;refGene_tx=XM_023242008.1
D3 75585501 75585632
refGene_gn=POLI;refGene_tx=XM_011288112.3;refGene_gn=POLI;refGene_tx=XM_019815287.2;refGene_gn=POLI;refGene_tx=XM_023242006.1;refGene_gn=POLI;refGene_tx=XM_023242007.1;refGene_gn=POLI;refGene_tx=XM_023242008.1
D3 75593408 75593614
refGene_gn=POLI;refGene_tx=XM_011288112.3;refGene_gn=POLI;refGene_tx=XM_019815287.2;refGene_gn=POLI;refGene_tx=XM_023242006.1;refGene_gn=POLI;refGene_tx=XM_023242007.1
D3 75595805 75596621
refGene_gn=POLI;refGene_tx=XM_011288112.3;refGene_gn=POLI;refGene_tx=XM_019815287.2;refGene_gn=POLI;refGene_tx=XM_023242006.1;refGene_gn=POLI;refGene_tx=XM_023242007.1;refGene_gn=POLI;refGene_tx=XM_023242008.1
D3 75615441 75615643
refGene_gn=STARD6;refGene_tx=XM_006938918.4;refGene_gn=STARD6;refGene_tx=XM_011288113.2;refGene_gn=STARD6;refGene_tx=XM_019815293.1;refGene_gn=STARD6;refGene_tx=XM_019815294.1;refGene_gn=STARD6;refGene_tx=XM_019815296.2;refGene_gn=STARD6;refGene_tx=XM_019815297.1;refGene_gn=STARD6;refGene_tx=XM_019815299.2
D3 75619791 75619885
refGene_gn=STARD6;refGene_tx=XM_006938918.4;refGene_gn=STARD6;refGene_tx=XM_011288113.2;refGene_gn=STARD6;refGene_tx=XM_019815293.1;refGene_gn=STARD6;refGene_tx=XM_019815294.1;refGene_gn=STARD6;refGene_tx=XM_019815296.2;refGene_gn=STARD6;refGene_tx=XM_019815297.1;refGene_gn=STARD6;refGene_tx=XM_019815299.2
D3 75622839 75622957
refGene_gn=STARD6;refGene_tx=XM_006938918.4;refGene_gn=STARD6;refGene_tx=XM_011288113.2;refGene_gn=STARD6;refGene_tx=XM_019815293.1;refGene_gn=STARD6;refGene_tx=XM_019815294.1;refGene_gn=STARD6;refGene_tx=XM_019815296.2;refGene_gn=STARD6;refGene_tx=XM_019815297.1;refGene_gn=STARD6;refGene_tx=XM_019815299.2
D3 75625933 75626060
refGene_gn=STARD6;refGene_tx=XM_006938918.4;refGene_gn=STARD6;refGene_tx=XM_011288113.2;refGene_gn=STARD6;refGene_tx=XM_019815293.1;refGene_gn=STARD6;refGene_tx=XM_019815294.1;refGene_gn=STARD6;refGene_tx=XM_019815296.2;refGene_gn=STARD6;refGene_tx=XM_019815297.1;refGene_gn=STARD6;refGene_tx=XM_019815299.2
D3 75635941 75635991
refGene_gn=STARD6;refGene_tx=XM_006938918.4;refGene_gn=STARD6;refGene_tx=XM_011288113.2;refGene_gn=STARD6;refGene_tx=XM_019815293.1;refGene_gn=STARD6;refGene_tx=XM_019815294.1;refGene_gn=STARD6;refGene_tx=XM_019815296.2;refGene_gn=STARD6;refGene_tx=XM_019815297.1;refGene_gn=STARD6;refGene_tx=XM_019815299.2
D3 75636188 75636278
refGene_gn=STARD6;refGene_tx=XM_006938918.4;refGene_gn=STARD6;refGene_tx=XM_011288113.2;refGene_gn=STARD6;refGene_tx=XM_019815293.1;refGene_

gn=STARD6;refGene_tx=XM_019815294.1;refGene_gn=STARD6;refGene_tx=XM_01
9815296.2;refGene_gn=STARD6;refGene_tx=XM_019815297.1;refGene_gn=STARD
6;refGene_tx=XM_019815299.2
D3 75643175 75643458
refGene_gn=CD3H18orf54;refGene_tx=XM_006938916.4;refGene_gn=CD3H18orf5
4;refGene_tx=XM_019815288.1;refGene_gn=CD3H18orf54;refGene_tx=XM_01981
5289.1;refGene_gn=CD3H18orf54;refGene_tx=XM_019815290.1;refGene_gn=CD3
H18orf54;refGene_tx=XM_019815291.1
D3 75644531 75645320
refGene_gn=CD3H18orf54;refGene_tx=XM_006938916.4;refGene_gn=CD3H18orf5
4;refGene_tx=XM_019815288.1;refGene_gn=CD3H18orf54;refGene_tx=XM_01981
5289.1;refGene_gn=CD3H18orf54;refGene_tx=XM_019815290.1;refGene_gn=CD3
H18orf54;refGene_tx=XM_019815291.1
D3 75645638 75645793
refGene_gn=CD3H18orf54;refGene_tx=XM_006938916.4;refGene_gn=CD3H18orf5
4;refGene_tx=XM_019815288.1;refGene_gn=CD3H18orf54;refGene_tx=XM_01981
5289.1;refGene_gn=CD3H18orf54;refGene_tx=XM_019815290.1;refGene_gn=CD3
H18orf54;refGene_tx=XM_019815291.1
D3 75648934 75649037
refGene_gn=CD3H18orf54;refGene_tx=XM_006938916.4;refGene_gn=CD3H18orf5
4;refGene_tx=XM_019815288.1;refGene_gn=CD3H18orf54;refGene_tx=XM_01981
5289.1;refGene_gn=CD3H18orf54;refGene_tx=XM_019815290.1
D3 75655079 75655211
refGene_gn=CD3H18orf54;refGene_tx=XM_006938916.4;refGene_gn=CD3H18orf5
4;refGene_tx=XM_019815288.1;refGene_gn=CD3H18orf54;refGene_tx=XM_01981
5289.1;refGene_gn=CD3H18orf54;refGene_tx=XM_019815290.1;refGene_gn=CD3
H18orf54;refGene_tx=XM_019815291.1
D3 75656472 75656543
refGene_gn=CD3H18orf54;refGene_tx=XM_006938916.4;refGene_gn=CD3H18orf5
4;refGene_tx=XM_019815288.1;refGene_gn=CD3H18orf54;refGene_tx=XM_01981
5289.1;refGene_gn=CD3H18orf54;refGene_tx=XM_019815290.1;refGene_gn=CD3
H18orf54;refGene_tx=XM_019815291.1
D3 75665176 75665249
refGene_gn=CD3H18orf54;refGene_tx=XM_006938916.4
D3 75688545 75688653
refGene_gn=CD3H18orf54;refGene_tx=XM_019815288.1;refGene_gn=CD3H18orf5
4;refGene_tx=XM_019815289.1;refGene_gn=CD3H18orf54;refGene_tx=XM_01981
5290.1;refGene_gn=CD3H18orf54;refGene_tx=XM_019815291.1;refGene_gn=CD3
H18orf54;refGene_tx=XM_023242009.1
D3 75689806 75690037
refGene_gn=CD3H18orf54;refGene_tx=XM_023242009.1
D3 75690533 75690740
refGene_gn=CD3H18orf54;refGene_tx=XM_023242009.1
D3 75693809 75694040
refGene_gn=CD3H18orf54;refGene_tx=XM_023242009.1
D3 75694968 75695122
refGene_gn=CD3H18orf54;refGene_tx=XM_023242009.1
D3 75696721 75696831
refGene_gn=CD3H18orf54;refGene_tx=XM_023242009.1
D3 75697614 75697728

refGene_gn=CD3H18orf54;refGene_tx=XM_023242009.1
D3 75699428 75699575
refGene_gn=CD3H18orf54;refGene_tx=XM_023242009.1
D3 75700447 75700582
refGene_gn=CD3H18orf54;refGene_tx=XM_023242009.1
D3 75782988 75783081
refGene_gn=LOC102900861;refGene_tx=XM_023242010.1
D3 75786532 75786706
refGene_gn=LOC102900861;refGene_tx=XM_023242010.1
D3 75789699 75790578
refGene_gn=LOC102900861;refGene_tx=XM_023242010.1
D3 75819696 75819752 refGene_gn=DYNAP;refGene_tx=XM_011288197.3
D3 75826004 75826068 refGene_gn=DYNAP;refGene_tx=XM_011288197.3
D3 75829738 75829903 refGene_gn=DYNAP;refGene_tx=XM_011288197.3
D3 75834848 75835508 refGene_gn=DYNAP;refGene_tx=XM_011288197.3
D3 75860891 75860948
refGene_gn=LOC102900145;refGene_tx=XM_006938920.4
D3 75863444 75863525
refGene_gn=LOC102900145;refGene_tx=XM_006938920.4
D3 75864290 75864608
refGene_gn=LOC102900145;refGene_tx=XM_006938920.4
D3 76054854 76054913 refGene_gn=RAB27B;refGene_tx=XM_019815307.1
D3 76074205 76074377
refGene_gn=RAB27B;refGene_tx=XM_011288174.2;refGene_gn=RAB27B;refGene_tx=XM_019815307.1;refGene_gn=RAB27B;refGene_tx=XM_019815308.1;refGene_gn=RAB27B;refGene_tx=XM_019815309.1;refGene_gn=RAB27B;refGene_tx=XM_019815310.2;refGene_gn=RAB27B;refGene_tx=XM_019815311.2
D3 76075992 76076078
refGene_gn=RAB27B;refGene_tx=XM_011288174.2;refGene_gn=RAB27B;refGene_tx=XM_019815307.1;refGene_gn=RAB27B;refGene_tx=XM_019815308.1;refGene_gn=RAB27B;refGene_tx=XM_019815309.1;refGene_gn=RAB27B;refGene_tx=XM_019815310.2;refGene_gn=RAB27B;refGene_tx=XM_019815311.2
D3 76080546 76080650
refGene_gn=RAB27B;refGene_tx=XM_011288174.2;refGene_gn=RAB27B;refGene_tx=XM_019815307.1;refGene_gn=RAB27B;refGene_tx=XM_019815308.1;refGene_gn=RAB27B;refGene_tx=XM_019815309.1;refGene_gn=RAB27B;refGene_tx=XM_019815310.2;refGene_gn=RAB27B;refGene_tx=XM_019815311.2
D3 76083473 76083597
refGene_gn=RAB27B;refGene_tx=XM_011288174.2;refGene_gn=RAB27B;refGene_tx=XM_019815307.1;refGene_gn=RAB27B;refGene_tx=XM_019815308.1;refGene_gn=RAB27B;refGene_tx=XM_019815309.1;refGene_gn=RAB27B;refGene_tx=XM_019815310.2;refGene_gn=RAB27B;refGene_tx=XM_019815311.2
D3 76084689 76084879
refGene_gn=RAB27B;refGene_tx=XM_011288174.2;refGene_gn=RAB27B;refGene_tx=XM_019815307.1;refGene_gn=RAB27B;refGene_tx=XM_019815308.1;refGene_gn=RAB27B;refGene_tx=XM_019815309.1;refGene_gn=RAB27B;refGene_tx=XM_019815310.2;refGene_gn=RAB27B;refGene_tx=XM_019815311.2
D3 76099734 76099792
refGene_gn=CCDC68;refGene_tx=XM_006938921.2;refGene_gn=CCDC68;refGene_tx=XM_011288173.1

D3 76102088 76102165
refGene_gn=CCDC68;refGene_tx=XM_006938921.2;refGene_gn=CCDC68;refGene_tx=XM_011288173.1
D3 76111349 76111433
refGene_gn=CCDC68;refGene_tx=XM_006938921.2;refGene_gn=CCDC68;refGene_tx=XM_011288173.1
D3 76112756 76112862
refGene_gn=CCDC68;refGene_tx=XM_006938921.2;refGene_gn=CCDC68;refGene_tx=XM_011288173.1
D3 76119923 76120006
refGene_gn=CCDC68;refGene_tx=XM_006938921.2;refGene_gn=CCDC68;refGene_tx=XM_011288173.1
D3 76125305 76125434
refGene_gn=CCDC68;refGene_tx=XM_006938921.2;refGene_gn=CCDC68;refGene_tx=XM_011288173.1
D3 76127156 76127282
refGene_gn=CCDC68;refGene_tx=XM_006938921.2;refGene_gn=CCDC68;refGene_tx=XM_011288173.1
D3 76128501 76128642
refGene_gn=CCDC68;refGene_tx=XM_006938921.2;refGene_gn=CCDC68;refGene_tx=XM_011288173.1
D3 76132131 76132218
refGene_gn=CCDC68;refGene_tx=XM_006938921.2;refGene_gn=CCDC68;refGene_tx=XM_011288173.1
D3 76133322 76133439
refGene_gn=CCDC68;refGene_tx=XM_006938921.2;refGene_gn=CCDC68;refGene_tx=XM_011288173.1
D3 76373754 76373891
refGene_gn=TCF4;refGene_tx=XM_006938933.4;refGene_gn=TCF4;refGene_tx=XM_006938934.4;refGene_gn=TCF4;refGene_tx=XM_006938935.4;refGene_gn=TCF4;refGene_tx=XM_011288171.3;refGene_gn=TCF4;refGene_tx=XM_019815315.2;refGene_gn=TCF4;refGene_tx=XM_019815321.2;refGene_gn=TCF4;refGene_tx=XM_019815322.2;refGene_gn=TCF4;refGene_tx=XM_019815323.2;refGene_gn=TCF4;refGene_tx=XM_019815324.2;refGene_gn=TCF4;refGene_tx=XM_019815325.2;refGene_gn=TCF4;refGene_tx=XM_019815326.2;refGene_gn=TCF4;refGene_tx=XM_019815327.2;refGene_gn=TCF4;refGene_tx=XM_019815328.2;refGene_gn=TCF4;refGene_tx=XM_023242011.1;refGene_gn=TCF4;refGene_tx=XM_023242012.1;refGene_gn=TCF4;refGene_tx=XM_023242013.1;refGene_gn=TCF4;refGene_tx=XM_023242014.1;refGene_gn=TCF4;refGene_tx=XM_023242016.1;refGene_gn=TCF4;refGene_tx=XM_023242017.1;refGene_gn=TCF4;refGene_tx=XM_023242018.1;refGene_gn=TCF4;refGene_tx=XM_023242019.1;refGene_gn=TCF4;refGene_tx=XM_023242020.1;refGene_gn=TCF4;refGene_tx=XM_023242021.1;refGene_gn=TCF4;refGene_tx=XM_023242022.1;refGene_gn=TCF4;refGene_tx=XM_023242023.1;refGene_gn=TCF4;refGene_tx=XM_023242024.1;refGene_gn=TCF4;refGene_tx=XM_023242025.1;refGene_gn=TCF4;refGene_tx=XM_023242026.1;refGene_gn=TCF4;refGene_tx=XM_023242027.1;refGene_gn=TCF4;refGene_tx=XM_023242028.1;refGene_gn=TCF4;refGene_tx=XM_023242029.1
D3 76374357 76374587
refGene_gn=TCF4;refGene_tx=XM_006938933.4;refGene_gn=TCF4;refGene_tx=XM_006938934.4;refGene_gn=TCF4;refGene_tx=XM_006938935.4;refGene_gn=TCF

4;refGene_tx=XM_011288171.3;refGene_gn=TCF4;refGene_tx=XM_019815315.2;
refGene_gn=TCF4;refGene_tx=XM_019815321.2;refGene_gn=TCF4;refGene_tx=X
M_019815322.2;refGene_gn=TCF4;refGene_tx=XM_019815323.2;refGene_gn=TCF
4;refGene_tx=XM_019815324.2;refGene_gn=TCF4;refGene_tx=XM_019815325.2;
refGene_gn=TCF4;refGene_tx=XM_019815326.2;refGene_gn=TCF4;refGene_tx=X
M_019815327.2;refGene_gn=TCF4;refGene_tx=XM_019815328.2;refGene_gn=TCF
4;refGene_tx=XM_023242011.1;refGene_gn=TCF4;refGene_tx=XM_023242012.1;
refGene_gn=TCF4;refGene_tx=XM_023242013.1;refGene_gn=TCF4;refGene_tx=X
M_023242014.1;refGene_gn=TCF4;refGene_tx=XM_023242016.1;refGene_gn=TCF
4;refGene_tx=XM_023242017.1;refGene_gn=TCF4;refGene_tx=XM_023242018.1;
refGene_gn=TCF4;refGene_tx=XM_023242019.1;refGene_gn=TCF4;refGene_tx=X
M_023242020.1;refGene_gn=TCF4;refGene_tx=XM_023242021.1;refGene_gn=TCF
4;refGene_tx=XM_023242022.1;refGene_gn=TCF4;refGene_tx=XM_023242023.1;
refGene_gn=TCF4;refGene_tx=XM_023242024.1;refGene_gn=TCF4;refGene_tx=X
M_023242025.1;refGene_gn=TCF4;refGene_tx=XM_023242026.1;refGene_gn=TCF
4;refGene_tx=XM_023242027.1;refGene_gn=TCF4;refGene_tx=XM_023242028.1;
refGene_gn=TCF4;refGene_tx=XM_023242029.1
D3 76377131 76377176
refGene_gn=TCF4;refGene_tx=XM_019815315.2;refGene_gn=TCF4;refGene_tx=X
M_019815321.2;refGene_gn=TCF4;refGene_tx=XM_019815322.2;refGene_gn=TCF
4;refGene_tx=XM_019815323.2;refGene_gn=TCF4;refGene_tx=XM_019815324.2;
refGene_gn=TCF4;refGene_tx=XM_019815325.2;refGene_gn=TCF4;refGene_tx=X
M_019815326.2;refGene_gn=TCF4;refGene_tx=XM_019815327.2;refGene_gn=TCF
4;refGene_tx=XM_019815328.2;refGene_gn=TCF4;refGene_tx=XM_023242011.1;
refGene_gn=TCF4;refGene_tx=XM_023242012.1;refGene_gn=TCF4;refGene_tx=X
M_023242013.1;refGene_gn=TCF4;refGene_tx=XM_023242020.1;refGene_gn=TCF
4;refGene_tx=XM_023242021.1;refGene_gn=TCF4;refGene_tx=XM_023242022.1;
refGene_gn=TCF4;refGene_tx=XM_023242027.1;refGene_gn=TCF4;refGene_tx=X
M_023242028.1
D3 76377667 76377830
refGene_gn=TCF4;refGene_tx=XM_006938933.4;refGene_gn=TCF4;refGene_tx=X
M_006938934.4;refGene_gn=TCF4;refGene_tx=XM_006938935.4;refGene_gn=TCF
4;refGene_tx=XM_011288171.3;refGene_gn=TCF4;refGene_tx=XM_019815315.2;
refGene_gn=TCF4;refGene_tx=XM_019815321.2;refGene_gn=TCF4;refGene_tx=X
M_019815322.2;refGene_gn=TCF4;refGene_tx=XM_019815323.2;refGene_gn=TCF
4;refGene_tx=XM_019815324.2;refGene_gn=TCF4;refGene_tx=XM_019815325.2;
refGene_gn=TCF4;refGene_tx=XM_019815326.2;refGene_gn=TCF4;refGene_tx=X
M_019815327.2;refGene_gn=TCF4;refGene_tx=XM_019815328.2;refGene_gn=TCF
4;refGene_tx=XM_023242011.1;refGene_gn=TCF4;refGene_tx=XM_023242012.1;
refGene_gn=TCF4;refGene_tx=XM_023242013.1;refGene_gn=TCF4;refGene_tx=X
M_023242014.1;refGene_gn=TCF4;refGene_tx=XM_023242016.1;refGene_gn=TCF
4;refGene_tx=XM_023242017.1;refGene_gn=TCF4;refGene_tx=XM_023242018.1;
refGene_gn=TCF4;refGene_tx=XM_023242019.1;refGene_gn=TCF4;refGene_tx=X
M_023242020.1;refGene_gn=TCF4;refGene_tx=XM_023242021.1;refGene_gn=TCF
4;refGene_tx=XM_023242022.1;refGene_gn=TCF4;refGene_tx=XM_023242023.1;
refGene_gn=TCF4;refGene_tx=XM_023242024.1;refGene_gn=TCF4;refGene_tx=X
M_023242025.1;refGene_gn=TCF4;refGene_tx=XM_023242026.1;refGene_gn=TCF
4;refGene_tx=XM_023242027.1;refGene_gn=TCF4;refGene_tx=XM_023242028.1;
refGene_gn=TCF4;refGene_tx=XM_023242029.1
D3 76379432 76379568

refGene_gn=TCF4;refGene_tx=XM_006938933.4;refGene_gn=TCF4;refGene_tx=X
M_006938934.4;refGene_gn=TCF4;refGene_tx=XM_006938935.4;refGene_gn=TCF
4;refGene_tx=XM_011288171.3;refGene_gn=TCF4;refGene_tx=XM_019815315.2;
refGene_gn=TCF4;refGene_tx=XM_019815321.2;refGene_gn=TCF4;refGene_tx=X
M_019815322.2;refGene_gn=TCF4;refGene_tx=XM_019815323.2;refGene_gn=TCF
4;refGene_tx=XM_019815324.2;refGene_gn=TCF4;refGene_tx=XM_019815325.2;
refGene_gn=TCF4;refGene_tx=XM_019815326.2;refGene_gn=TCF4;refGene_tx=X
M_019815327.2;refGene_gn=TCF4;refGene_tx=XM_019815328.2;refGene_gn=TCF
4;refGene_tx=XM_023242011.1;refGene_gn=TCF4;refGene_tx=XM_023242012.1;
refGene_gn=TCF4;refGene_tx=XM_023242013.1;refGene_gn=TCF4;refGene_tx=X
M_023242014.1;refGene_gn=TCF4;refGene_tx=XM_023242016.1;refGene_gn=TCF
4;refGene_tx=XM_023242017.1;refGene_gn=TCF4;refGene_tx=XM_023242018.1;
refGene_gn=TCF4;refGene_tx=XM_023242019.1;refGene_gn=TCF4;refGene_tx=X
M_023242020.1;refGene_gn=TCF4;refGene_tx=XM_023242021.1;refGene_gn=TCF
4;refGene_tx=XM_023242022.1;refGene_gn=TCF4;refGene_tx=XM_023242023.1;
refGene_gn=TCF4;refGene_tx=XM_023242024.1;refGene_gn=TCF4;refGene_tx=X
M_023242025.1;refGene_gn=TCF4;refGene_tx=XM_023242026.1;refGene_gn=TCF
4;refGene_tx=XM_023242027.1;refGene_gn=TCF4;refGene_tx=XM_023242028.1;
refGene_gn=TCF4;refGene_tx=XM_023242029.1
D3 76399370 76399574
refGene_gn=TCF4;refGene_tx=XM_006938933.4;refGene_gn=TCF4;refGene_tx=X
M_006938934.4;refGene_gn=TCF4;refGene_tx=XM_006938935.4;refGene_gn=TCF
4;refGene_tx=XM_011288171.3;refGene_gn=TCF4;refGene_tx=XM_019815315.2;
refGene_gn=TCF4;refGene_tx=XM_019815321.2;refGene_gn=TCF4;refGene_tx=X
M_019815322.2;refGene_gn=TCF4;refGene_tx=XM_019815323.2;refGene_gn=TCF
4;refGene_tx=XM_019815324.2;refGene_gn=TCF4;refGene_tx=XM_019815325.2;
refGene_gn=TCF4;refGene_tx=XM_019815326.2;refGene_gn=TCF4;refGene_tx=X
M_019815327.2;refGene_gn=TCF4;refGene_tx=XM_019815328.2;refGene_gn=TCF
4;refGene_tx=XM_023242011.1;refGene_gn=TCF4;refGene_tx=XM_023242012.1;
refGene_gn=TCF4;refGene_tx=XM_023242013.1;refGene_gn=TCF4;refGene_tx=X
M_023242014.1;refGene_gn=TCF4;refGene_tx=XM_023242016.1;refGene_gn=TCF
4;refGene_tx=XM_023242017.1;refGene_gn=TCF4;refGene_tx=XM_023242018.1;
refGene_gn=TCF4;refGene_tx=XM_023242019.1;refGene_gn=TCF4;refGene_tx=X
M_023242020.1;refGene_gn=TCF4;refGene_tx=XM_023242021.1;refGene_gn=TCF
4;refGene_tx=XM_023242022.1;refGene_gn=TCF4;refGene_tx=XM_023242023.1;
refGene_gn=TCF4;refGene_tx=XM_023242024.1;refGene_gn=TCF4;refGene_tx=X
M_023242025.1;refGene_gn=TCF4;refGene_tx=XM_023242026.1;refGene_gn=TCF
4;refGene_tx=XM_023242027.1;refGene_gn=TCF4;refGene_tx=XM_023242028.1;
refGene_gn=TCF4;refGene_tx=XM_023242029.1
D3 76402290 76402367
refGene_gn=TCF4;refGene_tx=XM_006938933.4;refGene_gn=TCF4;refGene_tx=X
M_006938934.4;refGene_gn=TCF4;refGene_tx=XM_006938935.4;refGene_gn=TCF
4;refGene_tx=XM_011288171.3;refGene_gn=TCF4;refGene_tx=XM_019815315.2;
refGene_gn=TCF4;refGene_tx=XM_019815321.2;refGene_gn=TCF4;refGene_tx=X
M_019815322.2;refGene_gn=TCF4;refGene_tx=XM_019815323.2;refGene_gn=TCF
4;refGene_tx=XM_019815324.2;refGene_gn=TCF4;refGene_tx=XM_019815325.2;
refGene_gn=TCF4;refGene_tx=XM_019815326.2;refGene_gn=TCF4;refGene_tx=X
M_019815327.2;refGene_gn=TCF4;refGene_tx=XM_019815328.2;refGene_gn=TCF
4;refGene_tx=XM_023242011.1;refGene_gn=TCF4;refGene_tx=XM_023242012.1;
refGene_gn=TCF4;refGene_tx=XM_023242013.1;refGene_gn=TCF4;refGene_tx=X

M_023242014.1;refGene_gn=TCF4;refGene_tx=XM_023242016.1;refGene_gn=TCF4;refGene_tx=XM_023242017.1;refGene_gn=TCF4;refGene_tx=XM_023242018.1;refGene_gn=TCF4;refGene_tx=XM_023242019.1;refGene_gn=TCF4;refGene_tx=XM_023242020.1;refGene_gn=TCF4;refGene_tx=XM_023242021.1;refGene_gn=TCF4;refGene_tx=XM_023242022.1;refGene_gn=TCF4;refGene_tx=XM_023242023.1;refGene_gn=TCF4;refGene_tx=XM_023242024.1;refGene_gn=TCF4;refGene_tx=XM_023242025.1;refGene_gn=TCF4;refGene_tx=XM_023242026.1;refGene_gn=TCF4;refGene_tx=XM_023242027.1;refGene_gn=TCF4;refGene_tx=XM_023242028.1;refGene_gn=TCF4;refGene_tx=XM_023242029.1

D3 76404951 76405030

refGene_gn=TCF4;refGene_tx=XM_006938933.4;refGene_gn=TCF4;refGene_tx=XM_006938934.4;refGene_gn=TCF4;refGene_tx=XM_006938935.4;refGene_gn=TCF4;refGene_tx=XM_011288171.3;refGene_gn=TCF4;refGene_tx=XM_019815315.2;refGene_gn=TCF4;refGene_tx=XM_019815321.2;refGene_gn=TCF4;refGene_tx=XM_019815322.2;refGene_gn=TCF4;refGene_tx=XM_019815323.2;refGene_gn=TCF4;refGene_tx=XM_019815324.2;refGene_gn=TCF4;refGene_tx=XM_019815325.2;refGene_gn=TCF4;refGene_tx=XM_019815326.2;refGene_gn=TCF4;refGene_tx=XM_019815327.2;refGene_gn=TCF4;refGene_tx=XM_019815328.2;refGene_gn=TCF4;refGene_tx=XM_023242011.1;refGene_gn=TCF4;refGene_tx=XM_023242012.1;refGene_gn=TCF4;refGene_tx=XM_023242013.1;refGene_gn=TCF4;refGene_tx=XM_023242014.1;refGene_gn=TCF4;refGene_tx=XM_023242016.1;refGene_gn=TCF4;refGene_tx=XM_023242017.1;refGene_gn=TCF4;refGene_tx=XM_023242018.1;refGene_gn=TCF4;refGene_tx=XM_023242019.1;refGene_gn=TCF4;refGene_tx=XM_023242020.1;refGene_gn=TCF4;refGene_tx=XM_023242021.1;refGene_gn=TCF4;refGene_tx=XM_023242022.1;refGene_gn=TCF4;refGene_tx=XM_023242023.1;refGene_gn=TCF4;refGene_tx=XM_023242024.1;refGene_gn=TCF4;refGene_tx=XM_023242025.1;refGene_gn=TCF4;refGene_tx=XM_023242026.1;refGene_gn=TCF4;refGene_tx=XM_023242027.1;refGene_gn=TCF4;refGene_tx=XM_023242028.1;refGene_gn=TCF4;refGene_tx=XM_023242029.1

D3 76406179 76406247

refGene_gn=TCF4;refGene_tx=XM_006938933.4;refGene_gn=TCF4;refGene_tx=XM_006938934.4;refGene_gn=TCF4;refGene_tx=XM_006938935.4;refGene_gn=TCF4;refGene_tx=XM_011288171.3;refGene_gn=TCF4;refGene_tx=XM_019815315.2;refGene_gn=TCF4;refGene_tx=XM_019815321.2;refGene_gn=TCF4;refGene_tx=XM_019815322.2;refGene_gn=TCF4;refGene_tx=XM_019815323.2;refGene_gn=TCF4;refGene_tx=XM_019815324.2;refGene_gn=TCF4;refGene_tx=XM_019815325.2;refGene_gn=TCF4;refGene_tx=XM_019815326.2;refGene_gn=TCF4;refGene_tx=XM_019815327.2;refGene_gn=TCF4;refGene_tx=XM_019815328.2;refGene_gn=TCF4;refGene_tx=XM_023242011.1;refGene_gn=TCF4;refGene_tx=XM_023242012.1;refGene_gn=TCF4;refGene_tx=XM_023242013.1;refGene_gn=TCF4;refGene_tx=XM_023242014.1;refGene_gn=TCF4;refGene_tx=XM_023242016.1;refGene_gn=TCF4;refGene_tx=XM_023242017.1;refGene_gn=TCF4;refGene_tx=XM_023242018.1;refGene_gn=TCF4;refGene_tx=XM_023242019.1;refGene_gn=TCF4;refGene_tx=XM_023242020.1;refGene_gn=TCF4;refGene_tx=XM_023242021.1;refGene_gn=TCF4;refGene_tx=XM_023242022.1;refGene_gn=TCF4;refGene_tx=XM_023242023.1;refGene_gn=TCF4;refGene_tx=XM_023242024.1;refGene_gn=TCF4;refGene_tx=XM_023242025.1;refGene_gn=TCF4;refGene_tx=XM_023242026.1;refGene_gn=TCF4;refGene_tx=XM_023242027.1;refGene_gn=TCF4;refGene_tx=XM_023242028.1;refGene_gn=TCF4;refGene_tx=XM_023242029.1

D3 76414013 76414146

refGene_gn=TCF4;refGene_tx=XM_006938933.4;refGene_gn=TCF4;refGene_tx=XM_006938934.4;refGene_gn=TCF4;refGene_tx=XM_006938935.4;refGene_gn=TCF4;refGene_tx=XM_011288171.3;refGene_gn=TCF4;refGene_tx=XM_019815315.2;refGene_gn=TCF4;refGene_tx=XM_019815321.2;refGene_gn=TCF4;refGene_tx=XM_019815322.2;refGene_gn=TCF4;refGene_tx=XM_019815323.2;refGene_gn=TCF4;refGene_tx=XM_019815324.2;refGene_gn=TCF4;refGene_tx=XM_019815325.2;refGene_gn=TCF4;refGene_tx=XM_019815326.2;refGene_gn=TCF4;refGene_tx=XM_019815327.2;refGene_gn=TCF4;refGene_tx=XM_019815328.2;refGene_gn=TCF4;refGene_tx=XM_023242011.1;refGene_gn=TCF4;refGene_tx=XM_023242012.1;refGene_gn=TCF4;refGene_tx=XM_023242013.1;refGene_gn=TCF4;refGene_tx=XM_023242014.1;refGene_gn=TCF4;refGene_tx=XM_023242016.1;refGene_gn=TCF4;refGene_tx=XM_023242017.1;refGene_gn=TCF4;refGene_tx=XM_023242018.1;refGene_gn=TCF4;refGene_tx=XM_023242019.1;refGene_gn=TCF4;refGene_tx=XM_023242020.1;refGene_gn=TCF4;refGene_tx=XM_023242021.1;refGene_gn=TCF4;refGene_tx=XM_023242022.1;refGene_gn=TCF4;refGene_tx=XM_023242023.1;refGene_gn=TCF4;refGene_tx=XM_023242024.1;refGene_gn=TCF4;refGene_tx=XM_023242025.1;refGene_gn=TCF4;refGene_tx=XM_023242026.1;refGene_gn=TCF4;refGene_tx=XM_023242027.1;refGene_gn=TCF4;refGene_tx=XM_023242028.1;refGene_gn=TCF4;refGene_tx=XM_023242029.1
D3 76419750 76419884
refGene_gn=TCF4;refGene_tx=XM_006938933.4;refGene_gn=TCF4;refGene_tx=XM_006938934.4;refGene_gn=TCF4;refGene_tx=XM_006938935.4;refGene_gn=TCF4;refGene_tx=XM_011288171.3;refGene_gn=TCF4;refGene_tx=XM_019815315.2;refGene_gn=TCF4;refGene_tx=XM_019815321.2;refGene_gn=TCF4;refGene_tx=XM_019815322.2;refGene_gn=TCF4;refGene_tx=XM_019815323.2;refGene_gn=TCF4;refGene_tx=XM_019815324.2;refGene_gn=TCF4;refGene_tx=XM_019815325.2;refGene_gn=TCF4;refGene_tx=XM_019815326.2;refGene_gn=TCF4;refGene_tx=XM_019815327.2;refGene_gn=TCF4;refGene_tx=XM_019815328.2;refGene_gn=TCF4;refGene_tx=XM_023242011.1;refGene_gn=TCF4;refGene_tx=XM_023242012.1;refGene_gn=TCF4;refGene_tx=XM_023242013.1;refGene_gn=TCF4;refGene_tx=XM_023242014.1;refGene_gn=TCF4;refGene_tx=XM_023242016.1;refGene_gn=TCF4;refGene_tx=XM_023242017.1;refGene_gn=TCF4;refGene_tx=XM_023242018.1;refGene_gn=TCF4;refGene_tx=XM_023242019.1;refGene_gn=TCF4;refGene_tx=XM_023242020.1;refGene_gn=TCF4;refGene_tx=XM_023242021.1;refGene_gn=TCF4;refGene_tx=XM_023242022.1;refGene_gn=TCF4;refGene_tx=XM_023242023.1;refGene_gn=TCF4;refGene_tx=XM_023242024.1;refGene_gn=TCF4;refGene_tx=XM_023242025.1;refGene_gn=TCF4;refGene_tx=XM_023242026.1;refGene_gn=TCF4;refGene_tx=XM_023242027.1;refGene_gn=TCF4;refGene_tx=XM_023242028.1;refGene_gn=TCF4;refGene_tx=XM_023242029.1
D3 76423377 76423483
refGene_gn=TCF4;refGene_tx=XM_006938933.4;refGene_gn=TCF4;refGene_tx=XM_006938934.4;refGene_gn=TCF4;refGene_tx=XM_006938935.4;refGene_gn=TCF4;refGene_tx=XM_011288171.3;refGene_gn=TCF4;refGene_tx=XM_019815315.2;refGene_gn=TCF4;refGene_tx=XM_019815321.2;refGene_gn=TCF4;refGene_tx=XM_019815322.2;refGene_gn=TCF4;refGene_tx=XM_019815323.2;refGene_gn=TCF4;refGene_tx=XM_019815324.2;refGene_gn=TCF4;refGene_tx=XM_019815325.2;refGene_gn=TCF4;refGene_tx=XM_019815326.2;refGene_gn=TCF4;refGene_tx=XM_019815327.2;refGene_gn=TCF4;refGene_tx=XM_019815328.2;refGene_gn=TCF4;refGene_tx=XM_023242011.1;refGene_gn=TCF4;refGene_tx=XM_023242012.1;refGene_gn=TCF4;refGene_tx=XM_023242013.1;refGene_gn=TCF4;refGene_tx=X

M_023242014.1;refGene_gn=TCF4;refGene_tx=XM_023242016.1;refGene_gn=TCF4;refGene_tx=XM_023242017.1;refGene_gn=TCF4;refGene_tx=XM_023242018.1;refGene_gn=TCF4;refGene_tx=XM_023242019.1;refGene_gn=TCF4;refGene_tx=XM_023242020.1;refGene_gn=TCF4;refGene_tx=XM_023242021.1;refGene_gn=TCF4;refGene_tx=XM_023242022.1;refGene_gn=TCF4;refGene_tx=XM_023242023.1;refGene_gn=TCF4;refGene_tx=XM_023242024.1;refGene_gn=TCF4;refGene_tx=XM_023242025.1;refGene_gn=TCF4;refGene_tx=XM_023242026.1;refGene_gn=TCF4;refGene_tx=XM_023242027.1;refGene_gn=TCF4;refGene_tx=XM_023242028.1;refGene_gn=TCF4;refGene_tx=XM_023242029.1

D3 76465720 76465789

refGene_gn=TCF4;refGene_tx=XM_006938933.4;refGene_gn=TCF4;refGene_tx=XM_006938935.4;refGene_gn=TCF4;refGene_tx=XM_011288171.3;refGene_gn=TCF4;refGene_tx=XM_023242029.1

D3 76494883 76494933

refGene_gn=TCF4;refGene_tx=XM_006938934.4;refGene_gn=TCF4;refGene_tx=XM_019815315.2;refGene_gn=TCF4;refGene_tx=XM_019815321.2;refGene_gn=TCF4;refGene_tx=XM_019815322.2;refGene_gn=TCF4;refGene_tx=XM_019815323.2;refGene_gn=TCF4;refGene_tx=XM_019815324.2;refGene_gn=TCF4;refGene_tx=XM_019815325.2;refGene_gn=TCF4;refGene_tx=XM_019815326.2;refGene_gn=TCF4;refGene_tx=XM_019815327.2;refGene_gn=TCF4;refGene_tx=XM_019815328.2;refGene_gn=TCF4;refGene_tx=XM_023242011.1;refGene_gn=TCF4;refGene_tx=XM_023242012.1;refGene_gn=TCF4;refGene_tx=XM_023242013.1;refGene_gn=TCF4;refGene_tx=XM_023242014.1;refGene_gn=TCF4;refGene_tx=XM_023242016.1;refGene_gn=TCF4;refGene_tx=XM_023242017.1;refGene_gn=TCF4;refGene_tx=XM_023242018.1;refGene_gn=TCF4;refGene_tx=XM_023242019.1;refGene_gn=TCF4;refGene_tx=XM_023242020.1;refGene_gn=TCF4;refGene_tx=XM_023242021.1;refGene_gn=TCF4;refGene_tx=XM_023242022.1;refGene_gn=TCF4;refGene_tx=XM_023242023.1;refGene_gn=TCF4;refGene_tx=XM_023242024.1;refGene_gn=TCF4;refGene_tx=XM_023242025.1;refGene_gn=TCF4;refGene_tx=XM_023242026.1;refGene_gn=TCF4;refGene_tx=XM_023242027.1;refGene_gn=TCF4;refGene_tx=XM_023242028.1

D3 76495413 76495543

refGene_gn=TCF4;refGene_tx=XM_006938934.4;refGene_gn=TCF4;refGene_tx=XM_019815315.2;refGene_gn=TCF4;refGene_tx=XM_019815321.2;refGene_gn=TCF4;refGene_tx=XM_019815322.2;refGene_gn=TCF4;refGene_tx=XM_019815323.2;refGene_gn=TCF4;refGene_tx=XM_019815324.2;refGene_gn=TCF4;refGene_tx=XM_019815325.2;refGene_gn=TCF4;refGene_tx=XM_019815326.2;refGene_gn=TCF4;refGene_tx=XM_019815327.2;refGene_gn=TCF4;refGene_tx=XM_019815328.2;refGene_gn=TCF4;refGene_tx=XM_023242011.1;refGene_gn=TCF4;refGene_tx=XM_023242012.1;refGene_gn=TCF4;refGene_tx=XM_023242013.1;refGene_gn=TCF4;refGene_tx=XM_023242014.1;refGene_gn=TCF4;refGene_tx=XM_023242016.1;refGene_gn=TCF4;refGene_tx=XM_023242017.1;refGene_gn=TCF4;refGene_tx=XM_023242018.1;refGene_gn=TCF4;refGene_tx=XM_023242019.1;refGene_gn=TCF4;refGene_tx=XM_023242020.1;refGene_gn=TCF4;refGene_tx=XM_023242021.1;refGene_gn=TCF4;refGene_tx=XM_023242022.1;refGene_gn=TCF4;refGene_tx=XM_023242023.1;refGene_gn=TCF4;refGene_tx=XM_023242024.1;refGene_gn=TCF4;refGene_tx=XM_023242025.1;refGene_gn=TCF4;refGene_tx=XM_023242026.1;refGene_gn=TCF4;refGene_tx=XM_023242027.1;refGene_gn=TCF4;refGene_tx=XM_023242028.1

D3 76545466 76545531

refGene_gn=TCF4;refGene_tx=XM_006938934.4;refGene_gn=TCF4;refGene_tx=XM_019815315.2;refGene_gn=TCF4;refGene_tx=XM_019815321.2;refGene_gn=TCF4;refGene_tx=XM_019815322.2;refGene_gn=TCF4;refGene_tx=XM_019815323.2;refGene_gn=TCF4;refGene_tx=XM_023242011.1;refGene_gn=TCF4;refGene_tx=XM_023242012.1;refGene_gn=TCF4;refGene_tx=XM_023242013.1;refGene_gn=TCF4;refGene_tx=XM_023242014.1;refGene_gn=TCF4;refGene_tx=XM_023242016.1;refGene_gn=TCF4;refGene_tx=XM_023242017.1;refGene_gn=TCF4;refGene_tx=XM_023242018.1;refGene_gn=TCF4;refGene_tx=XM_023242019.1;refGene_gn=TCF4;refGene_tx=XM_023242020.1;refGene_gn=TCF4;refGene_tx=XM_023242021.1;refGene_gn=TCF4;refGene_tx=XM_023242022.1;refGene_gn=TCF4;refGene_tx=XM_023242023.1;refGene_gn=TCF4;refGene_tx=XM_023242024.1;refGene_gn=TCF4;refGene_tx=XM_023242025.1;refGene_gn=TCF4;refGene_tx=XM_023242026.1;refGene_gn=TCF4;refGene_tx=XM_023242027.1;refGene_gn=TCF4;refGene_tx=XM_023242028.1

D3 76600662 76600759

refGene_gn=TCF4;refGene_tx=XM_006938934.4;refGene_gn=TCF4;refGene_tx=XM_019815315.2;refGene_gn=TCF4;refGene_tx=XM_019815321.2;refGene_gn=TCF4;refGene_tx=XM_019815322.2;refGene_gn=TCF4;refGene_tx=XM_019815323.2;refGene_gn=TCF4;refGene_tx=XM_023242011.1;refGene_gn=TCF4;refGene_tx=XM_023242012.1;refGene_gn=TCF4;refGene_tx=XM_023242013.1;refGene_gn=TCF4;refGene_tx=XM_023242014.1;refGene_gn=TCF4;refGene_tx=XM_023242016.1;refGene_gn=TCF4;refGene_tx=XM_023242017.1;refGene_gn=TCF4;refGene_tx=XM_023242018.1;refGene_gn=TCF4;refGene_tx=XM_023242019.1;refGene_gn=TCF4;refGene_tx=XM_023242020.1;refGene_gn=TCF4;refGene_tx=XM_023242021.1;refGene_gn=TCF4;refGene_tx=XM_023242022.1;refGene_gn=TCF4;refGene_tx=XM_023242023.1;refGene_gn=TCF4;refGene_tx=XM_023242024.1;refGene_gn=TCF4;refGene_tx=XM_023242025.1;refGene_gn=TCF4;refGene_tx=XM_023242026.1;refGene_gn=TCF4;refGene_tx=XM_023242027.1;refGene_gn=TCF4;refGene_tx=XM_023242028.1

D3 76603585 76603647

refGene_gn=TCF4;refGene_tx=XM_006938934.4;refGene_gn=TCF4;refGene_tx=XM_019815315.2;refGene_gn=TCF4;refGene_tx=XM_019815321.2;refGene_gn=TCF4;refGene_tx=XM_019815322.2;refGene_gn=TCF4;refGene_tx=XM_023242011.1;refGene_gn=TCF4;refGene_tx=XM_023242012.1;refGene_gn=TCF4;refGene_tx=XM_023242013.1;refGene_gn=TCF4;refGene_tx=XM_023242014.1;refGene_gn=TCF4;refGene_tx=XM_023242016.1;refGene_gn=TCF4;refGene_tx=XM_023242017.1;refGene_gn=TCF4;refGene_tx=XM_023242018.1;refGene_gn=TCF4;refGene_tx=XM_023242019.1;refGene_gn=TCF4;refGene_tx=XM_023242020.1;refGene_gn=TCF4;refGene_tx=XM_023242021.1;refGene_gn=TCF4;refGene_tx=XM_023242022.1;refGene_gn=TCF4;refGene_tx=XM_023242023.1;refGene_gn=TCF4;refGene_tx=XM_023242024.1;refGene_gn=TCF4;refGene_tx=XM_023242025.1;refGene_gn=TCF4;refGene_tx=XM_023242026.1;refGene_gn=TCF4;refGene_tx=XM_023242027.1;refGene_gn=TCF4;refGene_tx=XM_023242028.1

D3 76719825 76719898

refGene_gn=TCF4;refGene_tx=XM_006938934.4;refGene_gn=TCF4;refGene_tx=XM_019815315.2;refGene_gn=TCF4;refGene_tx=XM_019815321.2;refGene_gn=TCF4;refGene_tx=XM_019815322.2;refGene_gn=TCF4;refGene_tx=XM_023242011.1;refGene_gn=TCF4;refGene_tx=XM_023242012.1;refGene_gn=TCF4;refGene_tx=XM_023242013.1;refGene_gn=TCF4;refGene_tx=XM_023242014.1;refGene_gn=TCF4;refGene_tx=XM_023242016.1;refGene_gn=TCF4;refGene_tx=XM_023242017.1;

refGene_gn=TCF4;refGene_tx=XM_023242018.1;refGene_gn=TCF4;refGene_tx=XM_023242019.1;refGene_gn=TCF4;refGene_tx=XM_023242020.1;refGene_gn=TCF4;refGene_tx=XM_023242021.1;refGene_gn=TCF4;refGene_tx=XM_023242022.1;refGene_gn=TCF4;refGene_tx=XM_023242023.1;refGene_gn=TCF4;refGene_tx=XM_023242024.1;refGene_gn=TCF4;refGene_tx=XM_023242025.1;refGene_gn=TCF4;refGene_tx=XM_023242026.1;refGene_gn=TCF4;refGene_tx=XM_023242027.1;refGene_gn=TCF4;refGene_tx=XM_023242028.1
D3 76721601 76721673
refGene_gn=TCF4;refGene_tx=XM_006938934.4;refGene_gn=TCF4;refGene_tx=XM_019815315.2;refGene_gn=TCF4;refGene_tx=XM_023242020.1;refGene_gn=TCF4;refGene_tx=XM_023242021.1;refGene_gn=TCF4;refGene_tx=XM_023242023.1;refGene_gn=TCF4;refGene_tx=XM_023242024.1;refGene_gn=TCF4;refGene_tx=XM_023242025.1
D3 76723047 76723113
refGene_gn=TCF4;refGene_tx=XM_023242022.1;refGene_gn=TCF4;refGene_tx=XM_023242026.1
D3 77624550 77624580 refGene_gn=TXNL1;refGene_tx=XM_011288158.3
D3 77625978 77626017 refGene_gn=TXNL1;refGene_tx=XM_011288157.3
D3 77626906 77626915 refGene_gn=TXNL1;refGene_tx=XM_011288159.3
D3 77630765 77630870
refGene_gn=TXNL1;refGene_tx=XM_011288157.3;refGene_gn=TXNL1;refGene_tx=XM_011288158.3;refGene_gn=TXNL1;refGene_tx=XM_011288159.3
D3 77634041 77634214
refGene_gn=TXNL1;refGene_tx=XM_011288157.3;refGene_gn=TXNL1;refGene_tx=XM_011288158.3;refGene_gn=TXNL1;refGene_tx=XM_011288159.3
D3 77635599 77635669
refGene_gn=TXNL1;refGene_tx=XM_011288157.3;refGene_gn=TXNL1;refGene_tx=XM_011288158.3;refGene_gn=TXNL1;refGene_tx=XM_011288159.3
D3 77638313 77638436
refGene_gn=TXNL1;refGene_tx=XM_011288157.3;refGene_gn=TXNL1;refGene_tx=XM_011288158.3;refGene_gn=TXNL1;refGene_tx=XM_011288159.3
D3 77644297 77644471
refGene_gn=TXNL1;refGene_tx=XM_011288157.3;refGene_gn=TXNL1;refGene_tx=XM_011288158.3;refGene_gn=TXNL1;refGene_tx=XM_011288159.3
D3 77646590 77646687
refGene_gn=TXNL1;refGene_tx=XM_011288157.3;refGene_gn=TXNL1;refGene_tx=XM_011288158.3;refGene_gn=TXNL1;refGene_tx=XM_011288159.3
D3 77657623 77657721
refGene_gn=TXNL1;refGene_tx=XM_011288157.3;refGene_gn=TXNL1;refGene_tx=XM_011288158.3;refGene_gn=TXNL1;refGene_tx=XM_011288159.3
D3 77695540 77695699
refGene_gn=WDR7;refGene_tx=XM_006938939.3;refGene_gn=WDR7;refGene_tx=XM_019815330.2;refGene_gn=WDR7;refGene_tx=XM_019815331.2;refGene_gn=WDR7;refGene_tx=XM_019815332.2;refGene_gn=WDR7;refGene_tx=XM_019815334.2;refGene_gn=WDR7;refGene_tx=XM_019815335.2
D3 77702539 77702646
refGene_gn=WDR7;refGene_tx=XM_006938939.3;refGene_gn=WDR7;refGene_tx=XM_019815330.2;refGene_gn=WDR7;refGene_tx=XM_019815331.2;refGene_gn=WDR7;refGene_tx=XM_019815332.2;refGene_gn=WDR7;refGene_tx=XM_019815334.2;refGene_gn=WDR7;refGene_tx=XM_019815335.2

D3 77704186 77704265
refGene_gn=WDR7;refGene_tx=XM_006938939.3;refGene_gn=WDR7;refGene_tx=XM_019815330.2;refGene_gn=WDR7;refGene_tx=XM_019815331.2;refGene_gn=WDR7;refGene_tx=XM_019815332.2;refGene_gn=WDR7;refGene_tx=XM_019815334.2;refGene_gn=WDR7;refGene_tx=XM_019815335.2
D3 77707356 77707531
refGene_gn=WDR7;refGene_tx=XM_006938939.3;refGene_gn=WDR7;refGene_tx=XM_019815330.2;refGene_gn=WDR7;refGene_tx=XM_019815331.2;refGene_gn=WDR7;refGene_tx=XM_019815332.2;refGene_gn=WDR7;refGene_tx=XM_019815334.2;refGene_gn=WDR7;refGene_tx=XM_019815335.2
D3 77711438 77711515
refGene_gn=WDR7;refGene_tx=XM_006938939.3;refGene_gn=WDR7;refGene_tx=XM_019815330.2;refGene_gn=WDR7;refGene_tx=XM_019815331.2;refGene_gn=WDR7;refGene_tx=XM_019815332.2;refGene_gn=WDR7;refGene_tx=XM_019815334.2;refGene_gn=WDR7;refGene_tx=XM_019815335.2
D3 77712800 77712920
refGene_gn=WDR7;refGene_tx=XM_006938939.3;refGene_gn=WDR7;refGene_tx=XM_019815330.2;refGene_gn=WDR7;refGene_tx=XM_019815331.2;refGene_gn=WDR7;refGene_tx=XM_019815332.2;refGene_gn=WDR7;refGene_tx=XM_019815334.2;refGene_gn=WDR7;refGene_tx=XM_019815335.2
D3 77716029 77716175
refGene_gn=WDR7;refGene_tx=XM_006938939.3;refGene_gn=WDR7;refGene_tx=XM_019815330.2;refGene_gn=WDR7;refGene_tx=XM_019815331.2;refGene_gn=WDR7;refGene_tx=XM_019815332.2;refGene_gn=WDR7;refGene_tx=XM_019815334.2;refGene_gn=WDR7;refGene_tx=XM_019815335.2
D3 77716500 77716603
refGene_gn=WDR7;refGene_tx=XM_006938939.3;refGene_gn=WDR7;refGene_tx=XM_019815330.2;refGene_gn=WDR7;refGene_tx=XM_019815331.2;refGene_gn=WDR7;refGene_tx=XM_019815332.2;refGene_gn=WDR7;refGene_tx=XM_019815334.2;refGene_gn=WDR7;refGene_tx=XM_019815335.2
D3 77719450 77719583
refGene_gn=WDR7;refGene_tx=XM_006938939.3;refGene_gn=WDR7;refGene_tx=XM_019815330.2;refGene_gn=WDR7;refGene_tx=XM_019815331.2;refGene_gn=WDR7;refGene_tx=XM_019815332.2;refGene_gn=WDR7;refGene_tx=XM_019815334.2;refGene_gn=WDR7;refGene_tx=XM_019815335.2
D3 77719827 77720076
refGene_gn=WDR7;refGene_tx=XM_006938939.3;refGene_gn=WDR7;refGene_tx=XM_019815330.2;refGene_gn=WDR7;refGene_tx=XM_019815331.2;refGene_gn=WDR7;refGene_tx=XM_019815332.2;refGene_gn=WDR7;refGene_tx=XM_019815334.2;refGene_gn=WDR7;refGene_tx=XM_019815335.2
D3 77720869 77721090
refGene_gn=WDR7;refGene_tx=XM_006938939.3;refGene_gn=WDR7;refGene_tx=XM_019815330.2;refGene_gn=WDR7;refGene_tx=XM_019815331.2;refGene_gn=WDR7;refGene_tx=XM_019815332.2;refGene_gn=WDR7;refGene_tx=XM_019815334.2;refGene_gn=WDR7;refGene_tx=XM_019815335.2
D3 77739955 77740151
refGene_gn=WDR7;refGene_tx=XM_006938939.3;refGene_gn=WDR7;refGene_tx=XM_019815330.2;refGene_gn=WDR7;refGene_tx=XM_019815331.2;refGene_gn=WDR7;refGene_tx=XM_019815332.2;refGene_gn=WDR7;refGene_tx=XM_019815334.2;refGene_gn=WDR7;refGene_tx=XM_019815335.2

D3 77749432 77749647
refGene_gn=WDR7;refGene_tx=XM_006938939.3;refGene_gn=WDR7;refGene_tx=XM_019815330.2;refGene_gn=WDR7;refGene_tx=XM_019815331.2;refGene_gn=WDR7;refGene_tx=XM_019815332.2;refGene_gn=WDR7;refGene_tx=XM_019815334.2;refGene_gn=WDR7;refGene_tx=XM_019815335.2
D3 77774221 77774991
refGene_gn=WDR7;refGene_tx=XM_006938939.3;refGene_gn=WDR7;refGene_tx=XM_019815330.2;refGene_gn=WDR7;refGene_tx=XM_019815331.2;refGene_gn=WDR7;refGene_tx=XM_019815332.2;refGene_gn=WDR7;refGene_tx=XM_019815334.2;refGene_gn=WDR7;refGene_tx=XM_019815335.2
D3 77776259 77776348
refGene_gn=WDR7;refGene_tx=XM_006938939.3;refGene_gn=WDR7;refGene_tx=XM_019815330.2;refGene_gn=WDR7;refGene_tx=XM_019815331.2;refGene_gn=WDR7;refGene_tx=XM_019815332.2;refGene_gn=WDR7;refGene_tx=XM_019815334.2;refGene_gn=WDR7;refGene_tx=XM_019815335.2
D3 77790651 77790750
refGene_gn=WDR7;refGene_tx=XM_006938939.3;refGene_gn=WDR7;refGene_tx=XM_019815330.2;refGene_gn=WDR7;refGene_tx=XM_019815331.2;refGene_gn=WDR7;refGene_tx=XM_019815332.2;refGene_gn=WDR7;refGene_tx=XM_019815335.2
D3 77793507 77793626
refGene_gn=WDR7;refGene_tx=XM_006938939.3;refGene_gn=WDR7;refGene_tx=XM_019815330.2;refGene_gn=WDR7;refGene_tx=XM_019815331.2;refGene_gn=WDR7;refGene_tx=XM_019815332.2;refGene_gn=WDR7;refGene_tx=XM_019815334.2;refGene_gn=WDR7;refGene_tx=XM_019815335.2
D3 77795548 77795672
refGene_gn=WDR7;refGene_tx=XM_006938939.3;refGene_gn=WDR7;refGene_tx=XM_019815330.2;refGene_gn=WDR7;refGene_tx=XM_019815331.2;refGene_gn=WDR7;refGene_tx=XM_019815332.2;refGene_gn=WDR7;refGene_tx=XM_019815334.2;refGene_gn=WDR7;refGene_tx=XM_019815335.2
D3 77830142 77830256
refGene_gn=WDR7;refGene_tx=XM_006938939.3;refGene_gn=WDR7;refGene_tx=XM_019815330.2;refGene_gn=WDR7;refGene_tx=XM_019815331.2;refGene_gn=WDR7;refGene_tx=XM_019815332.2;refGene_gn=WDR7;refGene_tx=XM_019815334.2;refGene_gn=WDR7;refGene_tx=XM_019815335.2
D3 77877472 77877694
refGene_gn=WDR7;refGene_tx=XM_006938939.3;refGene_gn=WDR7;refGene_tx=XM_019815330.2;refGene_gn=WDR7;refGene_tx=XM_019815331.2;refGene_gn=WDR7;refGene_tx=XM_019815332.2;refGene_gn=WDR7;refGene_tx=XM_019815334.2
D3 77918371 77918558
refGene_gn=WDR7;refGene_tx=XM_006938939.3;refGene_gn=WDR7;refGene_tx=XM_019815330.2;refGene_gn=WDR7;refGene_tx=XM_019815331.2;refGene_gn=WDR7;refGene_tx=XM_019815332.2;refGene_gn=WDR7;refGene_tx=XM_019815334.2;refGene_gn=WDR7;refGene_tx=XM_019815335.2
D3 77929614 77929732
refGene_gn=WDR7;refGene_tx=XM_006938939.3;refGene_gn=WDR7;refGene_tx=XM_019815330.2;refGene_gn=WDR7;refGene_tx=XM_019815331.2;refGene_gn=WDR7;refGene_tx=XM_019815332.2;refGene_gn=WDR7;refGene_tx=XM_019815334.2;refGene_gn=WDR7;refGene_tx=XM_019815335.2
D3 77932049 77932199
refGene_gn=WDR7;refGene_tx=XM_006938939.3;refGene_gn=WDR7;refGene_tx=X

M_019815330.2;refGene_gn=WDR7;refGene_tx=XM_019815331.2;refGene_gn=WDR7;refGene_tx=XM_019815332.2;refGene_gn=WDR7;refGene_tx=XM_019815334.2;refGene_gn=WDR7;refGene_tx=XM_019815335.2
D3	77932780	77932863
refGene_gn=WDR7;refGene_tx=XM_006938939.3;refGene_gn=WDR7;refGene_tx=XM_019815330.2;refGene_gn=WDR7;refGene_tx=XM_019815331.2;refGene_gn=WDR7;refGene_tx=XM_019815332.2;refGene_gn=WDR7;refGene_tx=XM_019815334.2;refGene_gn=WDR7;refGene_tx=XM_019815335.2
D3	77958326	77958426
refGene_gn=WDR7;refGene_tx=XM_006938939.3;refGene_gn=WDR7;refGene_tx=XM_019815330.2;refGene_gn=WDR7;refGene_tx=XM_019815331.2;refGene_gn=WDR7;refGene_tx=XM_019815332.2;refGene_gn=WDR7;refGene_tx=XM_019815334.2;refGene_gn=WDR7;refGene_tx=XM_019815335.2
D3	78020300	78020405
refGene_gn=WDR7;refGene_tx=XM_006938939.3;refGene_gn=WDR7;refGene_tx=XM_019815330.2;refGene_gn=WDR7;refGene_tx=XM_019815331.2;refGene_gn=WDR7;refGene_tx=XM_019815332.2;refGene_gn=WDR7;refGene_tx=XM_019815334.2;refGene_gn=WDR7;refGene_tx=XM_019815335.2
D3	78026572	78026776
refGene_gn=WDR7;refGene_tx=XM_006938939.3;refGene_gn=WDR7;refGene_tx=XM_019815330.2;refGene_gn=WDR7;refGene_tx=XM_019815331.2;refGene_gn=WDR7;refGene_tx=XM_019815332.2;refGene_gn=WDR7;refGene_tx=XM_019815334.2;refGene_gn=WDR7;refGene_tx=XM_019815335.2
D3	78293644	78293823	refGene_gn=ST8SIA3;refGene_tx=XM_019815338.2
D3	78295214	78295351
refGene_gn=ST8SIA3;refGene_tx=XM_019815338.2;refGene_gn=ST8SIA3;refGene_tx=XM_019815339.2
D3	78297793	78298351
refGene_gn=ST8SIA3;refGene_tx=XM_019815338.2;refGene_gn=ST8SIA3;refGene_tx=XM_019815339.2
D3	78300934	78301217
refGene_gn=ST8SIA3;refGene_tx=XM_019815338.2;refGene_gn=ST8SIA3;refGene_tx=XM_019815339.2
D3	78369545	78370773	refGene_gn=ONECUT2;refGene_tx=XM_003995254.4
D3	78413569	78413856	refGene_gn=ONECUT2;refGene_tx=XM_003995254.4
D3	78475713	78475848	refGene_gn=FECH;refGene_tx=XM_006938940.3
D3	78476327	78476387	refGene_gn=FECH;refGene_tx=XM_006938940.3
D3	78479005	78479044	refGene_gn=FECH;refGene_tx=XM_006938941.3
D3	78479468	78479633
refGene_gn=FECH;refGene_tx=XM_006938940.3;refGene_gn=FECH;refGene_tx=XM_006938941.3
D3	78480064	78480172
refGene_gn=FECH;refGene_tx=XM_006938940.3;refGene_gn=FECH;refGene_tx=XM_006938941.3
D3	78483870	78483969
refGene_gn=FECH;refGene_tx=XM_006938940.3;refGene_gn=FECH;refGene_tx=XM_006938941.3
D3	78487235	78487342
refGene_gn=FECH;refGene_tx=XM_006938940.3;refGene_gn=FECH;refGene_tx=XM_006938941.3

D3 78491108 78491243
refGene_gn=FECH;refGene_tx=XM_006938940.3;refGene_gn=FECH;refGene_tx=XM_006938941.3
D3 78498592 78498741
refGene_gn=FECH;refGene_tx=XM_006938940.3;refGene_gn=FECH;refGene_tx=XM_006938941.3
D3 78500315 78500435
refGene_gn=FECH;refGene_tx=XM_006938940.3;refGene_gn=FECH;refGene_tx=XM_006938941.3
D3 78505675 78505802
refGene_gn=FECH;refGene_tx=XM_006938940.3;refGene_gn=FECH;refGene_tx=XM_006938941.3
D3 78511528 78511574
refGene_gn=FECH;refGene_tx=XM_006938940.3;refGene_gn=FECH;refGene_tx=XM_006938941.3
D3 78522893 78523025 refGene_gn=NARS;refGene_tx=XM_003995225.5
D3 78523757 78523889 refGene_gn=NARS;refGene_tx=XM_003995225.5
D3 78524183 78524315 refGene_gn=NARS;refGene_tx=XM_003995225.5
D3 78527535 78527649 refGene_gn=NARS;refGene_tx=XM_003995225.5
D3 78528286 78528422 refGene_gn=NARS;refGene_tx=XM_003995225.5
D3 78528837 78529037 refGene_gn=NARS;refGene_tx=XM_003995225.5
D3 78529142 78529364 refGene_gn=NARS;refGene_tx=XM_003995225.5
D3 78530560 78530647 refGene_gn=NARS;refGene_tx=XM_003995225.5
D3 78533003 78533074 refGene_gn=NARS;refGene_tx=XM_003995225.5
D3 78534347 78534426 refGene_gn=NARS;refGene_tx=XM_003995225.5
D3 78535558 78535648 refGene_gn=NARS;refGene_tx=XM_003995225.5
D3 78535734 78535827 refGene_gn=NARS;refGene_tx=XM_003995225.5
D3 78535920 78535986 refGene_gn=NARS;refGene_tx=XM_003995225.5
D3 78539349 78539429 refGene_gn=NARS;refGene_tx=XM_003995225.5
D3 78540517 78540644 refGene_gn=NARS;refGene_tx=XM_003995225.5
D3 78581890 78582115 refGene_gn=ATP8B1;refGene_tx=XM_011288155.3
D3 78583288 78583419 refGene_gn=ATP8B1;refGene_tx=XM_011288155.3
D3 78585368 78585507 refGene_gn=ATP8B1;refGene_tx=XM_011288155.3
D3 78585822 78586068 refGene_gn=ATP8B1;refGene_tx=XM_011288155.3
D3 78587796 78587880 refGene_gn=ATP8B1;refGene_tx=XM_011288155.3
D3 78589664 78589888 refGene_gn=ATP8B1;refGene_tx=XM_011288155.3
D3 78593846 78594135 refGene_gn=ATP8B1;refGene_tx=XM_011288155.3
D3 78594539 78594672 refGene_gn=ATP8B1;refGene_tx=XM_011288155.3
D3 78597880 78597956 refGene_gn=ATP8B1;refGene_tx=XM_011288155.3
D3 78599159 78599271 refGene_gn=ATP8B1;refGene_tx=XM_011288155.3
D3 78600016 78600181 refGene_gn=ATP8B1;refGene_tx=XM_011288155.3
D3 78602263 78602376 refGene_gn=ATP8B1;refGene_tx=XM_011288155.3
D3 78604088 78604277 refGene_gn=ATP8B1;refGene_tx=XM_011288155.3
D3 78610340 78610497 refGene_gn=ATP8B1;refGene_tx=XM_011288155.3
D3 78611204 78611248 refGene_gn=ATP8B1;refGene_tx=XM_011288155.3
D3 78613818 78614027 refGene_gn=ATP8B1;refGene_tx=XM_011288155.3
D3 78616312 78616503 refGene_gn=ATP8B1;refGene_tx=XM_011288155.3
D3 78618693 78618782 refGene_gn=ATP8B1;refGene_tx=XM_011288155.3
D3 78619143 78619302 refGene_gn=ATP8B1;refGene_tx=XM_011288155.3
D3 78619420 78619503 refGene_gn=ATP8B1;refGene_tx=XM_011288155.3

D3 78621257 78621328 refGene_gn=ATP8B1;refGene_tx=XM_011288155.3
D3 78621436 78621509 refGene_gn=ATP8B1;refGene_tx=XM_011288155.3
D3 78624261 78624323 refGene_gn=ATP8B1;refGene_tx=XM_011288155.3
D3 78624437 78624536 refGene_gn=ATP8B1;refGene_tx=XM_011288155.3
D3 78626581 78626695 refGene_gn=ATP8B1;refGene_tx=XM_011288155.3
D3 78628984 78629082 refGene_gn=ATP8B1;refGene_tx=XM_011288155.3
D3 78647771 78647952 refGene_gn=ATP8B1;refGene_tx=XM_011288155.3
D3 78900854 78900902
refGene_gn=NEDD4L;refGene_tx=XM_023242040.1;refGene_gn=NEDD4L;refGene_tx=XM_023242041.1;refGene_gn=NEDD4L;refGene_tx=XM_023242042.1
D3 79009497 79009571
refGene_gn=NEDD4L;refGene_tx=XM_023242040.1;refGene_gn=NEDD4L;refGene_tx=XM_023242041.1;refGene_gn=NEDD4L;refGene_tx=XM_023242042.1
D3 79061380 79061403 refGene_gn=NEDD4L;refGene_tx=XM_019815349.2
D3 79089552 79089634
refGene_gn=NEDD4L;refGene_tx=XM_019815349.2;refGene_gn=NEDD4L;refGene_tx=XM_023242040.1;refGene_gn=NEDD4L;refGene_tx=XM_023242041.1;refGene_gn=NEDD4L;refGene_tx=XM_023242042.1
D3 79092312 79092351
refGene_gn=NEDD4L;refGene_tx=XM_019815349.2;refGene_gn=NEDD4L;refGene_tx=XM_023242040.1;refGene_gn=NEDD4L;refGene_tx=XM_023242041.1;refGene_gn=NEDD4L;refGene_tx=XM_023242042.1
D3 79095037 79095091
refGene_gn=NEDD4L;refGene_tx=XM_019815349.2;refGene_gn=NEDD4L;refGene_tx=XM_023242040.1;refGene_gn=NEDD4L;refGene_tx=XM_023242041.1;refGene_gn=NEDD4L;refGene_tx=XM_023242042.1
D3 79162541 79162592
refGene_gn=NEDD4L;refGene_tx=XM_019815349.2;refGene_gn=NEDD4L;refGene_tx=XM_023242040.1;refGene_gn=NEDD4L;refGene_tx=XM_023242041.1;refGene_gn=NEDD4L;refGene_tx=XM_023242042.1
D3 79171113 79171175
refGene_gn=NEDD4L;refGene_tx=XM_019815349.2;refGene_gn=NEDD4L;refGene_tx=XM_019815353.2;refGene_gn=NEDD4L;refGene_tx=XM_019815354.2;refGene_gn=NEDD4L;refGene_tx=XM_023242031.1;refGene_gn=NEDD4L;refGene_tx=XM_023242032.1;refGene_gn=NEDD4L;refGene_tx=XM_023242033.1;refGene_gn=NEDD4L;refGene_tx=XM_023242034.1;refGene_gn=NEDD4L;refGene_tx=XM_023242035.1;refGene_gn=NEDD4L;refGene_tx=XM_023242036.1;refGene_gn=NEDD4L;refGene_tx=XM_023242037.1;refGene_gn=NEDD4L;refGene_tx=XM_023242038.1;refGene_gn=NEDD4L;refGene_tx=XM_023242039.1;refGene_gn=NEDD4L;refGene_tx=XM_023242040.1;refGene_gn=NEDD4L;refGene_tx=XM_023242041.1;refGene_gn=NEDD4L;refGene_tx=XM_023242042.1
D3 79171359 79171462
refGene_gn=NEDD4L;refGene_tx=XM_019815349.2;refGene_gn=NEDD4L;refGene_tx=XM_019815353.2;refGene_gn=NEDD4L;refGene_tx=XM_019815354.2;refGene_gn=NEDD4L;refGene_tx=XM_023242031.1;refGene_gn=NEDD4L;refGene_tx=XM_023242032.1;refGene_gn=NEDD4L;refGene_tx=XM_023242033.1;refGene_gn=NEDD4L;refGene_tx=XM_023242034.1;refGene_gn=NEDD4L;refGene_tx=XM_023242035.1;refGene_gn=NEDD4L;refGene_tx=XM_023242036.1;refGene_gn=NEDD4L;refGene_tx=XM_023242037.1;refGene_gn=NEDD4L;refGene_tx=XM_023242038.1;refGene_gn=NEDD4L;refGene_tx=XM_023242039.1;refGene_gn=NEDD4L;refGene_tx=XM_

023242040.1;refGene_gn=NEDD4L;refGene_tx=XM_023242041.1;refGene_gn=NED
D4L;refGene_tx=XM_023242042.1
D3 79173156 79173323
refGene_gn=NEDD4L;refGene_tx=XM_019815349.2;refGene_gn=NEDD4L;refGene_
tx=XM_019815353.2;refGene_gn=NEDD4L;refGene_tx=XM_019815354.2;refGene_
gn=NEDD4L;refGene_tx=XM_023242031.1;refGene_gn=NEDD4L;refGene_tx=XM_02
3242032.1;refGene_gn=NEDD4L;refGene_tx=XM_023242033.1;refGene_gn=NEDD4
L;refGene_tx=XM_023242034.1;refGene_gn=NEDD4L;refGene_tx=XM_023242035.
1;refGene_gn=NEDD4L;refGene_tx=XM_023242036.1;refGene_gn=NEDD4L;refGen
e_tx=XM_023242037.1;refGene_gn=NEDD4L;refGene_tx=XM_023242038.1;refGen
e_gn=NEDD4L;refGene_tx=XM_023242039.1;refGene_gn=NEDD4L;refGene_tx=XM_
023242040.1;refGene_gn=NEDD4L;refGene_tx=XM_023242041.1;refGene_gn=NED
D4L;refGene_tx=XM_023242042.1
D3 79180844 79180977
refGene_gn=NEDD4L;refGene_tx=XM_019815349.2;refGene_gn=NEDD4L;refGene_
tx=XM_019815353.2;refGene_gn=NEDD4L;refGene_tx=XM_019815354.2;refGene_
gn=NEDD4L;refGene_tx=XM_023242031.1;refGene_gn=NEDD4L;refGene_tx=XM_02
3242032.1;refGene_gn=NEDD4L;refGene_tx=XM_023242033.1;refGene_gn=NEDD4
L;refGene_tx=XM_023242034.1;refGene_gn=NEDD4L;refGene_tx=XM_023242035.
1;refGene_gn=NEDD4L;refGene_tx=XM_023242036.1;refGene_gn=NEDD4L;refGen
e_tx=XM_023242037.1;refGene_gn=NEDD4L;refGene_tx=XM_023242038.1;refGen
e_gn=NEDD4L;refGene_tx=XM_023242039.1;refGene_gn=NEDD4L;refGene_tx=XM_
023242040.1;refGene_gn=NEDD4L;refGene_tx=XM_023242041.1;refGene_gn=NED
D4L;refGene_tx=XM_023242042.1
D3 79182368 79182545
refGene_gn=NEDD4L;refGene_tx=XM_019815349.2;refGene_gn=NEDD4L;refGene_
tx=XM_019815353.2;refGene_gn=NEDD4L;refGene_tx=XM_019815354.2;refGene_
gn=NEDD4L;refGene_tx=XM_023242031.1;refGene_gn=NEDD4L;refGene_tx=XM_02
3242032.1;refGene_gn=NEDD4L;refGene_tx=XM_023242033.1;refGene_gn=NEDD4
L;refGene_tx=XM_023242034.1;refGene_gn=NEDD4L;refGene_tx=XM_023242035.
1;refGene_gn=NEDD4L;refGene_tx=XM_023242037.1;refGene_gn=NEDD4L;refGen
e_tx=XM_023242039.1;refGene_gn=NEDD4L;refGene_tx=XM_023242040.1;refGen
e_gn=NEDD4L;refGene_tx=XM_023242041.1;refGene_gn=NEDD4L;refGene_tx=XM_
023242042.1
D3 79185765 79185840
refGene_gn=NEDD4L;refGene_tx=XM_019815349.2;refGene_gn=NEDD4L;refGene_
tx=XM_019815353.2;refGene_gn=NEDD4L;refGene_tx=XM_019815354.2;refGene_
gn=NEDD4L;refGene_tx=XM_023242031.1;refGene_gn=NEDD4L;refGene_tx=XM_02
3242032.1;refGene_gn=NEDD4L;refGene_tx=XM_023242033.1;refGene_gn=NEDD4
L;refGene_tx=XM_023242034.1;refGene_gn=NEDD4L;refGene_tx=XM_023242035.
1;refGene_gn=NEDD4L;refGene_tx=XM_023242036.1;refGene_gn=NEDD4L;refGen
e_tx=XM_023242037.1;refGene_gn=NEDD4L;refGene_tx=XM_023242038.1;refGen
e_gn=NEDD4L;refGene_tx=XM_023242039.1;refGene_gn=NEDD4L;refGene_tx=XM_
023242040.1;refGene_gn=NEDD4L;refGene_tx=XM_023242041.1;refGene_gn=NED
D4L;refGene_tx=XM_023242042.1
D3 79187480 79187540
refGene_gn=NEDD4L;refGene_tx=XM_019815349.2;refGene_gn=NEDD4L;refGene_
tx=XM_019815353.2;refGene_gn=NEDD4L;refGene_tx=XM_019815354.2;refGene_
gn=NEDD4L;refGene_tx=XM_023242031.1;refGene_gn=NEDD4L;refGene_tx=XM_02
3242032.1;refGene_gn=NEDD4L;refGene_tx=XM_023242033.1;refGene_gn=NEDD4

L;refGene_tx=XM_023242040.1;refGene_gn=NEDD4L;refGene_tx=XM_023242041.
1
D3 79192921 79193053
refGene_gn=NEDD4L;refGene_tx=XM_019815349.2;refGene_gn=NEDD4L;refGene_
tx=XM_019815353.2;refGene_gn=NEDD4L;refGene_tx=XM_019815354.2;refGene_
gn=NEDD4L;refGene_tx=XM_023242031.1;refGene_gn=NEDD4L;refGene_tx=XM_02
3242032.1;refGene_gn=NEDD4L;refGene_tx=XM_023242033.1;refGene_gn=NEDD4
L;refGene_tx=XM_023242034.1;refGene_gn=NEDD4L;refGene_tx=XM_023242035.
1;refGene_gn=NEDD4L;refGene_tx=XM_023242036.1;refGene_gn=NEDD4L;refGen
e_tx=XM_023242040.1;refGene_gn=NEDD4L;refGene_tx=XM_023242041.1;refGen
e_gn=NEDD4L;refGene_tx=XM_023242042.1
D3 79193554 79193674
refGene_gn=NEDD4L;refGene_tx=XM_019815349.2;refGene_gn=NEDD4L;refGene_
tx=XM_019815353.2;refGene_gn=NEDD4L;refGene_tx=XM_019815354.2;refGene_
gn=NEDD4L;refGene_tx=XM_023242031.1;refGene_gn=NEDD4L;refGene_tx=XM_02
3242032.1;refGene_gn=NEDD4L;refGene_tx=XM_023242033.1;refGene_gn=NEDD4
L;refGene_tx=XM_023242034.1;refGene_gn=NEDD4L;refGene_tx=XM_023242035.
1;refGene_gn=NEDD4L;refGene_tx=XM_023242036.1;refGene_gn=NEDD4L;refGen
e_tx=XM_023242040.1;refGene_gn=NEDD4L;refGene_tx=XM_023242041.1;refGen
e_gn=NEDD4L;refGene_tx=XM_023242042.1
D3 79194769 79194967
refGene_gn=NEDD4L;refGene_tx=XM_019815349.2;refGene_gn=NEDD4L;refGene_
tx=XM_019815353.2;refGene_gn=NEDD4L;refGene_tx=XM_019815354.2;refGene_
gn=NEDD4L;refGene_tx=XM_023242031.1;refGene_gn=NEDD4L;refGene_tx=XM_02
3242032.1;refGene_gn=NEDD4L;refGene_tx=XM_023242033.1;refGene_gn=NEDD4
L;refGene_tx=XM_023242034.1;refGene_gn=NEDD4L;refGene_tx=XM_023242035.
1;refGene_gn=NEDD4L;refGene_tx=XM_023242036.1;refGene_gn=NEDD4L;refGen
e_tx=XM_023242037.1;refGene_gn=NEDD4L;refGene_tx=XM_023242038.1;refGen
e_gn=NEDD4L;refGene_tx=XM_023242040.1;refGene_gn=NEDD4L;refGene_tx=XM_
023242041.1;refGene_gn=NEDD4L;refGene_tx=XM_023242042.1
D3 79200170 79200248
refGene_gn=NEDD4L;refGene_tx=XM_019815349.2;refGene_gn=NEDD4L;refGene_
tx=XM_019815353.2;refGene_gn=NEDD4L;refGene_tx=XM_019815354.2;refGene_
gn=NEDD4L;refGene_tx=XM_023242031.1;refGene_gn=NEDD4L;refGene_tx=XM_02
3242032.1;refGene_gn=NEDD4L;refGene_tx=XM_023242033.1;refGene_gn=NEDD4
L;refGene_tx=XM_023242034.1;refGene_gn=NEDD4L;refGene_tx=XM_023242035.
1;refGene_gn=NEDD4L;refGene_tx=XM_023242036.1;refGene_gn=NEDD4L;refGen
e_tx=XM_023242037.1;refGene_gn=NEDD4L;refGene_tx=XM_023242038.1;refGen
e_gn=NEDD4L;refGene_tx=XM_023242039.1;refGene_gn=NEDD4L;refGene_tx=XM_
023242040.1;refGene_gn=NEDD4L;refGene_tx=XM_023242041.1;refGene_gn=NED
D4L;refGene_tx=XM_023242042.1
D3 79201814 79201869
refGene_gn=NEDD4L;refGene_tx=XM_019815349.2;refGene_gn=NEDD4L;refGene_
tx=XM_019815353.2;refGene_gn=NEDD4L;refGene_tx=XM_019815354.2;refGene_
gn=NEDD4L;refGene_tx=XM_023242031.1;refGene_gn=NEDD4L;refGene_tx=XM_02
3242032.1;refGene_gn=NEDD4L;refGene_tx=XM_023242033.1;refGene_gn=NEDD4
L;refGene_tx=XM_023242034.1;refGene_gn=NEDD4L;refGene_tx=XM_023242035.
1;refGene_gn=NEDD4L;refGene_tx=XM_023242036.1;refGene_gn=NEDD4L;refGen
e_tx=XM_023242037.1;refGene_gn=NEDD4L;refGene_tx=XM_023242038.1;refGen
e_gn=NEDD4L;refGene_tx=XM_023242039.1;refGene_gn=NEDD4L;refGene_tx=XM_

023242040.1;refGene_gn=NEDD4L;refGene_tx=XM_023242041.1;refGene_gn=NEDD4L;refGene_tx=XM_023242042.1
D3 79201978 79202032
refGene_gn=NEDD4L;refGene_tx=XM_019815349.2;refGene_gn=NEDD4L;refGene_tx=XM_019815353.2;refGene_gn=NEDD4L;refGene_tx=XM_019815354.2;refGene_gn=NEDD4L;refGene_tx=XM_023242031.1;refGene_gn=NEDD4L;refGene_tx=XM_023242034.1;refGene_gn=NEDD4L;refGene_tx=XM_023242036.1;refGene_gn=NEDD4L;refGene_tx=XM_023242037.1;refGene_gn=NEDD4L;refGene_tx=XM_023242038.1;refGene_gn=NEDD4L;refGene_tx=XM_023242039.1;refGene_gn=NEDD4L;refGene_tx=XM_023242040.1
D3 79207191 79207250
refGene_gn=NEDD4L;refGene_tx=XM_019815349.2;refGene_gn=NEDD4L;refGene_tx=XM_019815353.2;refGene_gn=NEDD4L;refGene_tx=XM_019815354.2;refGene_gn=NEDD4L;refGene_tx=XM_023242031.1;refGene_gn=NEDD4L;refGene_tx=XM_023242032.1;refGene_gn=NEDD4L;refGene_tx=XM_023242033.1;refGene_gn=NEDD4L;refGene_tx=XM_023242034.1;refGene_gn=NEDD4L;refGene_tx=XM_023242035.1;refGene_gn=NEDD4L;refGene_tx=XM_023242036.1;refGene_gn=NEDD4L;refGene_tx=XM_023242037.1;refGene_gn=NEDD4L;refGene_tx=XM_023242038.1;refGene_gn=NEDD4L;refGene_tx=XM_023242039.1;refGene_gn=NEDD4L;refGene_tx=XM_023242040.1;refGene_gn=NEDD4L;refGene_tx=XM_023242041.1;refGene_gn=NEDD4L;refGene_tx=XM_023242042.1
D3 79208418 79208421
refGene_gn=NEDD4L;refGene_tx=XM_019815349.2;refGene_gn=NEDD4L;refGene_tx=XM_019815353.2;refGene_gn=NEDD4L;refGene_tx=XM_019815354.2;refGene_gn=NEDD4L;refGene_tx=XM_023242032.1;refGene_gn=NEDD4L;refGene_tx=XM_023242034.1;refGene_gn=NEDD4L;refGene_tx=XM_023242036.1;refGene_gn=NEDD4L;refGene_tx=XM_023242037.1;refGene_gn=NEDD4L;refGene_tx=XM_023242038.1;refGene_gn=NEDD4L;refGene_tx=XM_023242039.1;refGene_gn=NEDD4L;refGene_tx=XM_023242040.1
D3 79214494 79214560
refGene_gn=NEDD4L;refGene_tx=XM_019815349.2;refGene_gn=NEDD4L;refGene_tx=XM_019815353.2;refGene_gn=NEDD4L;refGene_tx=XM_019815354.2;refGene_gn=NEDD4L;refGene_tx=XM_023242031.1;refGene_gn=NEDD4L;refGene_tx=XM_023242032.1;refGene_gn=NEDD4L;refGene_tx=XM_023242033.1;refGene_gn=NEDD4L;refGene_tx=XM_023242034.1;refGene_gn=NEDD4L;refGene_tx=XM_023242035.1;refGene_gn=NEDD4L;refGene_tx=XM_023242036.1;refGene_gn=NEDD4L;refGene_tx=XM_023242037.1;refGene_gn=NEDD4L;refGene_tx=XM_023242038.1;refGene_gn=NEDD4L;refGene_tx=XM_023242039.1;refGene_gn=NEDD4L;refGene_tx=XM_023242040.1;refGene_gn=NEDD4L;refGene_tx=XM_023242041.1;refGene_gn=NEDD4L;refGene_tx=XM_023242042.1
D3 79216248 79216478
refGene_gn=NEDD4L;refGene_tx=XM_019815349.2;refGene_gn=NEDD4L;refGene_tx=XM_019815353.2;refGene_gn=NEDD4L;refGene_tx=XM_019815354.2;refGene_gn=NEDD4L;refGene_tx=XM_023242031.1;refGene_gn=NEDD4L;refGene_tx=XM_023242032.1;refGene_gn=NEDD4L;refGene_tx=XM_023242033.1;refGene_gn=NEDD4L;refGene_tx=XM_023242034.1;refGene_gn=NEDD4L;refGene_tx=XM_023242035.1;refGene_gn=NEDD4L;refGene_tx=XM_023242036.1;refGene_gn=NEDD4L;refGene_tx=XM_023242037.1;refGene_gn=NEDD4L;refGene_tx=XM_023242038.1;refGene_gn=NEDD4L;refGene_tx=XM_023242039.1;refGene_gn=NEDD4L;refGene_tx=XM_023242040.1;refGene_gn=NEDD4L;refGene_tx=XM_023242041.1;refGene_gn=NED

D4L;refGene_tx=XM_023242042.1
D3 79218003 79218125
refGene_gn=NEDD4L;refGene_tx=XM_019815349.2;refGene_gn=NEDD4L;refGene_tx=XM_019815353.2;refGene_gn=NEDD4L;refGene_tx=XM_019815354.2;refGene_gn=NEDD4L;refGene_tx=XM_023242031.1;refGene_gn=NEDD4L;refGene_tx=XM_023242032.1;refGene_gn=NEDD4L;refGene_tx=XM_023242033.1;refGene_gn=NEDD4L;refGene_tx=XM_023242034.1;refGene_gn=NEDD4L;refGene_tx=XM_023242035.1;refGene_gn=NEDD4L;refGene_tx=XM_023242036.1;refGene_gn=NEDD4L;refGene_tx=XM_023242037.1;refGene_gn=NEDD4L;refGene_tx=XM_023242038.1;refGene_gn=NEDD4L;refGene_tx=XM_023242039.1;refGene_gn=NEDD4L;refGene_tx=XM_023242040.1;refGene_gn=NEDD4L;refGene_tx=XM_023242041.1;refGene_gn=NEDD4L;refGene_tx=XM_023242042.1
D3 79220469 79220540
refGene_gn=NEDD4L;refGene_tx=XM_019815349.2;refGene_gn=NEDD4L;refGene_tx=XM_019815353.2;refGene_gn=NEDD4L;refGene_tx=XM_019815354.2;refGene_gn=NEDD4L;refGene_tx=XM_023242031.1;refGene_gn=NEDD4L;refGene_tx=XM_023242032.1;refGene_gn=NEDD4L;refGene_tx=XM_023242033.1;refGene_gn=NEDD4L;refGene_tx=XM_023242034.1;refGene_gn=NEDD4L;refGene_tx=XM_023242035.1;refGene_gn=NEDD4L;refGene_tx=XM_023242036.1;refGene_gn=NEDD4L;refGene_tx=XM_023242037.1;refGene_gn=NEDD4L;refGene_tx=XM_023242038.1;refGene_gn=NEDD4L;refGene_tx=XM_023242039.1;refGene_gn=NEDD4L;refGene_tx=XM_023242040.1;refGene_gn=NEDD4L;refGene_tx=XM_023242041.1;refGene_gn=NEDD4L;refGene_tx=XM_023242042.1
D3 79222483 79222579
refGene_gn=NEDD4L;refGene_tx=XM_019815349.2;refGene_gn=NEDD4L;refGene_tx=XM_019815353.2;refGene_gn=NEDD4L;refGene_tx=XM_019815354.2;refGene_gn=NEDD4L;refGene_tx=XM_023242031.1;refGene_gn=NEDD4L;refGene_tx=XM_023242032.1;refGene_gn=NEDD4L;refGene_tx=XM_023242033.1;refGene_gn=NEDD4L;refGene_tx=XM_023242034.1;refGene_gn=NEDD4L;refGene_tx=XM_023242035.1;refGene_gn=NEDD4L;refGene_tx=XM_023242036.1;refGene_gn=NEDD4L;refGene_tx=XM_023242037.1;refGene_gn=NEDD4L;refGene_tx=XM_023242038.1;refGene_gn=NEDD4L;refGene_tx=XM_023242039.1;refGene_gn=NEDD4L;refGene_tx=XM_023242040.1;refGene_gn=NEDD4L;refGene_tx=XM_023242041.1;refGene_gn=NEDD4L;refGene_tx=XM_023242042.1
D3 79229754 79229828
refGene_gn=NEDD4L;refGene_tx=XM_019815349.2;refGene_gn=NEDD4L;refGene_tx=XM_019815353.2;refGene_gn=NEDD4L;refGene_tx=XM_019815354.2;refGene_gn=NEDD4L;refGene_tx=XM_023242031.1;refGene_gn=NEDD4L;refGene_tx=XM_023242032.1;refGene_gn=NEDD4L;refGene_tx=XM_023242033.1;refGene_gn=NEDD4L;refGene_tx=XM_023242034.1;refGene_gn=NEDD4L;refGene_tx=XM_023242035.1;refGene_gn=NEDD4L;refGene_tx=XM_023242036.1;refGene_gn=NEDD4L;refGene_tx=XM_023242037.1;refGene_gn=NEDD4L;refGene_tx=XM_023242038.1;refGene_gn=NEDD4L;refGene_tx=XM_023242039.1;refGene_gn=NEDD4L;refGene_tx=XM_023242040.1;refGene_gn=NEDD4L;refGene_tx=XM_023242041.1;refGene_gn=NEDD4L;refGene_tx=XM_023242042.1
D3 79232214 79232275
refGene_gn=NEDD4L;refGene_tx=XM_019815349.2;refGene_gn=NEDD4L;refGene_tx=XM_019815353.2;refGene_gn=NEDD4L;refGene_tx=XM_019815354.2;refGene_gn=NEDD4L;refGene_tx=XM_023242031.1;refGene_gn=NEDD4L;refGene_tx=XM_023242032.1;refGene_gn=NEDD4L;refGene_tx=XM_023242033.1;refGene_gn=NEDD4

L;refGene_tx=XM_023242034.1;refGene_gn=NEDD4L;refGene_tx=XM_023242035.1;refGene_gn=NEDD4L;refGene_tx=XM_023242036.1;refGene_gn=NEDD4L;refGene_tx=XM_023242037.1;refGene_gn=NEDD4L;refGene_tx=XM_023242038.1;refGene_gn=NEDD4L;refGene_tx=XM_023242039.1;refGene_gn=NEDD4L;refGene_tx=XM_023242040.1;refGene_gn=NEDD4L;refGene_tx=XM_023242041.1;refGene_gn=NEDD4L;refGene_tx=XM_023242042.1
D3 79233321 79233381
refGene_gn=NEDD4L;refGene_tx=XM_019815349.2;refGene_gn=NEDD4L;refGene_tx=XM_019815353.2;refGene_gn=NEDD4L;refGene_tx=XM_019815354.2;refGene_gn=NEDD4L;refGene_tx=XM_023242031.1;refGene_gn=NEDD4L;refGene_tx=XM_023242032.1;refGene_gn=NEDD4L;refGene_tx=XM_023242033.1;refGene_gn=NEDD4L;refGene_tx=XM_023242034.1;refGene_gn=NEDD4L;refGene_tx=XM_023242035.1;refGene_gn=NEDD4L;refGene_tx=XM_023242036.1;refGene_gn=NEDD4L;refGene_tx=XM_023242037.1;refGene_gn=NEDD4L;refGene_tx=XM_023242038.1;refGene_gn=NEDD4L;refGene_tx=XM_023242039.1;refGene_gn=NEDD4L;refGene_tx=XM_023242040.1;refGene_gn=NEDD4L;refGene_tx=XM_023242041.1;refGene_gn=NEDD4L;refGene_tx=XM_023242042.1
D3 79234818 79234926
refGene_gn=NEDD4L;refGene_tx=XM_019815349.2;refGene_gn=NEDD4L;refGene_tx=XM_019815353.2;refGene_gn=NEDD4L;refGene_tx=XM_019815354.2;refGene_gn=NEDD4L;refGene_tx=XM_023242031.1;refGene_gn=NEDD4L;refGene_tx=XM_023242032.1;refGene_gn=NEDD4L;refGene_tx=XM_023242033.1;refGene_gn=NEDD4L;refGene_tx=XM_023242034.1;refGene_gn=NEDD4L;refGene_tx=XM_023242035.1;refGene_gn=NEDD4L;refGene_tx=XM_023242036.1;refGene_gn=NEDD4L;refGene_tx=XM_023242037.1;refGene_gn=NEDD4L;refGene_tx=XM_023242038.1;refGene_gn=NEDD4L;refGene_tx=XM_023242039.1;refGene_gn=NEDD4L;refGene_tx=XM_023242040.1;refGene_gn=NEDD4L;refGene_tx=XM_023242041.1;refGene_gn=NEDD4L;refGene_tx=XM_023242042.1
D3 79236174 79236271
refGene_gn=NEDD4L;refGene_tx=XM_019815349.2;refGene_gn=NEDD4L;refGene_tx=XM_019815353.2;refGene_gn=NEDD4L;refGene_tx=XM_019815354.2;refGene_gn=NEDD4L;refGene_tx=XM_023242031.1;refGene_gn=NEDD4L;refGene_tx=XM_023242032.1;refGene_gn=NEDD4L;refGene_tx=XM_023242033.1;refGene_gn=NEDD4L;refGene_tx=XM_023242034.1;refGene_gn=NEDD4L;refGene_tx=XM_023242035.1;refGene_gn=NEDD4L;refGene_tx=XM_023242036.1;refGene_gn=NEDD4L;refGene_tx=XM_023242037.1;refGene_gn=NEDD4L;refGene_tx=XM_023242038.1;refGene_gn=NEDD4L;refGene_tx=XM_023242039.1;refGene_gn=NEDD4L;refGene_tx=XM_023242040.1;refGene_gn=NEDD4L;refGene_tx=XM_023242041.1;refGene_gn=NEDD4L;refGene_tx=XM_023242042.1
D3 79237119 79237192
refGene_gn=NEDD4L;refGene_tx=XM_019815349.2;refGene_gn=NEDD4L;refGene_tx=XM_019815353.2;refGene_gn=NEDD4L;refGene_tx=XM_019815354.2;refGene_gn=NEDD4L;refGene_tx=XM_023242031.1;refGene_gn=NEDD4L;refGene_tx=XM_023242032.1;refGene_gn=NEDD4L;refGene_tx=XM_023242033.1;refGene_gn=NEDD4L;refGene_tx=XM_023242034.1;refGene_gn=NEDD4L;refGene_tx=XM_023242035.1;refGene_gn=NEDD4L;refGene_tx=XM_023242036.1;refGene_gn=NEDD4L;refGene_tx=XM_023242037.1;refGene_gn=NEDD4L;refGene_tx=XM_023242038.1;refGene_gn=NEDD4L;refGene_tx=XM_023242039.1;refGene_gn=NEDD4L;refGene_tx=XM_023242040.1;refGene_gn=NEDD4L;refGene_tx=XM_023242041.1;refGene_gn=NEDD4L;refGene_tx=XM_023242042.1

D3 79241985 79242088
refGene_gn=NEDD4L;refGene_tx=XM_019815349.2;refGene_gn=NEDD4L;refGene_tx=XM_019815353.2;refGene_gn=NEDD4L;refGene_tx=XM_019815354.2;refGene_gn=NEDD4L;refGene_tx=XM_023242031.1;refGene_gn=NEDD4L;refGene_tx=XM_023242032.1;refGene_gn=NEDD4L;refGene_tx=XM_023242033.1;refGene_gn=NEDD4L;refGene_tx=XM_023242034.1;refGene_gn=NEDD4L;refGene_tx=XM_023242035.1;refGene_gn=NEDD4L;refGene_tx=XM_023242036.1;refGene_gn=NEDD4L;refGene_tx=XM_023242037.1;refGene_gn=NEDD4L;refGene_tx=XM_023242038.1;refGene_gn=NEDD4L;refGene_tx=XM_023242039.1;refGene_gn=NEDD4L;refGene_tx=XM_023242040.1;refGene_gn=NEDD4L;refGene_tx=XM_023242041.1;refGene_gn=NEDD4L;refGene_tx=XM_023242042.1
D3 79310605 79310834 refGene_gn=ALPK2;refGene_tx=XM_023242043.1
D3 79323918 79323967 refGene_gn=ALPK2;refGene_tx=XM_023242043.1
D3 79329397 79329615 refGene_gn=ALPK2;refGene_tx=XM_023242043.1
D3 79335878 79335967 refGene_gn=ALPK2;refGene_tx=XM_023242043.1
D3 79337426 79337701 refGene_gn=ALPK2;refGene_tx=XM_023242043.1
D3 79341023 79341059 refGene_gn=ALPK2;refGene_tx=XM_023242043.1
D3 79341172 79341300 refGene_gn=ALPK2;refGene_tx=XM_023242043.1
D3 79343543 79343691 refGene_gn=ALPK2;refGene_tx=XM_023242043.1
D3 79347472 79348279 refGene_gn=ALPK2;refGene_tx=XM_023242043.1
D3 79348281 79348392 refGene_gn=ALPK2;refGene_tx=XM_023242043.1
D3 79348394 79350696 refGene_gn=ALPK2;refGene_tx=XM_023242043.1
D3 79386144 79387879 refGene_gn=ALPK2;refGene_tx=XM_023242043.1
D3 79410742 79410860 refGene_gn=ALPK2;refGene_tx=XM_023242043.1
D3 79414541 79414650 refGene_gn=ALPK2;refGene_tx=XM_023242043.1
D3 79464402 79464674
refGene_gn=MALT1;refGene_tx=XM_003995258.5;refGene_gn=MALT1;refGene_tx=XM_011288150.3
D3 79474243 79474410
refGene_gn=MALT1;refGene_tx=XM_003995258.5;refGene_gn=MALT1;refGene_tx=XM_011288150.3
D3 79485418 79485540
refGene_gn=MALT1;refGene_tx=XM_003995258.5;refGene_gn=MALT1;refGene_tx=XM_011288150.3
D3 79492682 79492842
refGene_gn=MALT1;refGene_tx=XM_003995258.5;refGene_gn=MALT1;refGene_tx=XM_011288150.3
D3 79495142 79495321
refGene_gn=MALT1;refGene_tx=XM_003995258.5;refGene_gn=MALT1;refGene_tx=XM_011288150.3
D3 79495750 79495847
refGene_gn=MALT1;refGene_tx=XM_003995258.5;refGene_gn=MALT1;refGene_tx=XM_011288150.3
D3 79496367 79496397 refGene_gn=MALT1;refGene_tx=XM_003995258.5
D3 79499398 79499425
refGene_gn=MALT1;refGene_tx=XM_003995258.5;refGene_gn=MALT1;refGene_tx=XM_011288150.3
D3 79501270 79501303
refGene_gn=MALT1;refGene_tx=XM_003995258.5;refGene_gn=MALT1;refGene_tx=XM_011288150.3

D3 79506522 79506726
refGene_gn=MALT1;refGene_tx=XM_003995258.5;refGene_gn=MALT1;refGene_tx=XM_011288150.3
D3 79515463 79515641
refGene_gn=MALT1;refGene_tx=XM_003995258.5;refGene_gn=MALT1;refGene_tx=XM_011288150.3
D3 79516913 79516988
refGene_gn=MALT1;refGene_tx=XM_003995258.5;refGene_gn=MALT1;refGene_tx=XM_011288150.3
D3 79517332 79517460
refGene_gn=MALT1;refGene_tx=XM_003995258.5;refGene_gn=MALT1;refGene_tx=XM_011288150.3
D3 79519801 79519951
refGene_gn=MALT1;refGene_tx=XM_003995258.5;refGene_gn=MALT1;refGene_tx=XM_011288150.3
D3 79522518 79522676
refGene_gn=MALT1;refGene_tx=XM_003995258.5;refGene_gn=MALT1;refGene_tx=XM_011288150.3
D3 79523769 79523895
refGene_gn=MALT1;refGene_tx=XM_003995258.5;refGene_gn=MALT1;refGene_tx=XM_011288150.3
D3 79524967 79525408
refGene_gn=MALT1;refGene_tx=XM_003995258.5;refGene_gn=MALT1;refGene_tx=XM_011288150.3
D3 79673005 79675351
refGene_gn=ZNF532;refGene_tx=XM_011288142.3;refGene_gn=ZNF532;refGene_tx=XM_011288144.3;refGene_gn=ZNF532;refGene_tx=XM_011288145.3;refGene_gn=ZNF532;refGene_tx=XM_011288146.3;refGene_gn=ZNF532;refGene_tx=XM_011288147.3;refGene_gn=ZNF532;refGene_tx=XM_011288148.3;refGene_gn=ZNF532;refGene_tx=XM_019815359.2;refGene_gn=ZNF532;refGene_tx=XM_019815360.2;refGene_gn=ZNF532;refGene_tx=XM_019815361.2;refGene_gn=ZNF532;refGene_tx=XM_019815362.2;refGene_gn=ZNF532;refGene_tx=XM_023242046.1
D3 79688227 79688409
refGene_gn=ZNF532;refGene_tx=XM_011288142.3;refGene_gn=ZNF532;refGene_tx=XM_011288144.3;refGene_gn=ZNF532;refGene_tx=XM_011288145.3;refGene_gn=ZNF532;refGene_tx=XM_011288146.3;refGene_gn=ZNF532;refGene_tx=XM_011288147.3;refGene_gn=ZNF532;refGene_tx=XM_011288148.3;refGene_gn=ZNF532;refGene_tx=XM_019815359.2;refGene_gn=ZNF532;refGene_tx=XM_019815360.2;refGene_gn=ZNF532;refGene_tx=XM_019815361.2
D3 79692908 79693085
refGene_gn=ZNF532;refGene_tx=XM_011288142.3;refGene_gn=ZNF532;refGene_tx=XM_011288144.3;refGene_gn=ZNF532;refGene_tx=XM_011288145.3;refGene_gn=ZNF532;refGene_tx=XM_011288146.3;refGene_gn=ZNF532;refGene_tx=XM_011288147.3;refGene_gn=ZNF532;refGene_tx=XM_011288148.3;refGene_gn=ZNF532;refGene_tx=XM_019815360.2;refGene_gn=ZNF532;refGene_tx=XM_019815361.2
D3 79698915 79699078
refGene_gn=ZNF532;refGene_tx=XM_011288142.3;refGene_gn=ZNF532;refGene_tx=XM_011288144.3;refGene_gn=ZNF532;refGene_tx=XM_011288145.3;refGene_gn=ZNF532;refGene_tx=XM_011288146.3;refGene_gn=ZNF532;refGene_tx=XM_01

1288147.3;refGene_gn=ZNF532;refGene_tx=XM_011288148.3;refGene_gn=ZNF532;refGene_tx=XM_019815359.2;refGene_gn=ZNF532;refGene_tx=XM_019815360.2;refGene_gn=ZNF532;refGene_tx=XM_019815361.2
D3 79702998 79703280
refGene_gn=ZNF532;refGene_tx=XM_011288142.3;refGene_gn=ZNF532;refGene_tx=XM_011288144.3;refGene_gn=ZNF532;refGene_tx=XM_011288145.3;refGene_gn=ZNF532;refGene_tx=XM_011288146.3;refGene_gn=ZNF532;refGene_tx=XM_011288147.3;refGene_gn=ZNF532;refGene_tx=XM_011288148.3;refGene_gn=ZNF532;refGene_tx=XM_019815359.2;refGene_gn=ZNF532;refGene_tx=XM_019815360.2;refGene_gn=ZNF532;refGene_tx=XM_019815361.2;refGene_gn=ZNF532;refGene_tx=XM_019815362.2
D3 79728326 79728439
refGene_gn=ZNF532;refGene_tx=XM_011288142.3;refGene_gn=ZNF532;refGene_tx=XM_011288144.3;refGene_gn=ZNF532;refGene_tx=XM_011288145.3;refGene_gn=ZNF532;refGene_tx=XM_011288146.3;refGene_gn=ZNF532;refGene_tx=XM_011288147.3;refGene_gn=ZNF532;refGene_tx=XM_011288148.3;refGene_gn=ZNF532;refGene_tx=XM_019815359.2;refGene_gn=ZNF532;refGene_tx=XM_019815360.2;refGene_gn=ZNF532;refGene_tx=XM_019815361.2;refGene_gn=ZNF532;refGene_tx=XM_019815362.2;refGene_gn=ZNF532;refGene_tx=XM_023242046.1
D3 79730773 79730921
refGene_gn=ZNF532;refGene_tx=XM_011288142.3;refGene_gn=ZNF532;refGene_tx=XM_011288144.3;refGene_gn=ZNF532;refGene_tx=XM_011288145.3;refGene_gn=ZNF532;refGene_tx=XM_011288146.3;refGene_gn=ZNF532;refGene_tx=XM_011288147.3;refGene_gn=ZNF532;refGene_tx=XM_011288148.3;refGene_gn=ZNF532;refGene_tx=XM_019815359.2;refGene_gn=ZNF532;refGene_tx=XM_019815360.2;refGene_gn=ZNF532;refGene_tx=XM_019815361.2;refGene_gn=ZNF532;refGene_tx=XM_019815362.2;refGene_gn=ZNF532;refGene_tx=XM_023242046.1
D3 79732331 79732826
refGene_gn=ZNF532;refGene_tx=XM_011288142.3;refGene_gn=ZNF532;refGene_tx=XM_011288144.3;refGene_gn=ZNF532;refGene_tx=XM_011288145.3;refGene_gn=ZNF532;refGene_tx=XM_011288146.3;refGene_gn=ZNF532;refGene_tx=XM_011288147.3;refGene_gn=ZNF532;refGene_tx=XM_011288148.3;refGene_gn=ZNF532;refGene_tx=XM_019815359.2;refGene_gn=ZNF532;refGene_tx=XM_019815360.2;refGene_gn=ZNF532;refGene_tx=XM_019815361.2;refGene_gn=ZNF532;refGene_tx=XM_019815362.2;refGene_gn=ZNF532;refGene_tx=XM_023242046.1
D3 79744386 79744540
refGene_gn=LOC101089150;refGene_tx=XM_023241203.1
D3 79762981 79763121
refGene_gn=LOC101089150;refGene_tx=XM_023241203.1
D3 79764454 79764533
refGene_gn=LOC101089150;refGene_tx=XM_023241203.1
D3 79772480 79772622
refGene_gn=LOC101089150;refGene_tx=XM_023241203.1
D3 79773620 79773838
refGene_gn=LOC101089150;refGene_tx=XM_023241203.1
D3 79777660 79777840
refGene_gn=LOC101089150;refGene_tx=XM_023241203.1
D3 79778822 79778925
refGene_gn=LOC101089150;refGene_tx=XM_023241203.1
D3 79783533 79783635

refGene_gn=LOC101089150;refGene_tx=XM_023241203.1
D3 79790100 79790266
refGene_gn=LOC101089150;refGene_tx=XM_023241203.1
D3 79791682 79791762
refGene_gn=LOC101089150;refGene_tx=XM_023241203.1
D3 79796711 79796825
refGene_gn=LOC101089150;refGene_tx=XM_023241203.1
D3 79798389 79798522
refGene_gn=LOC101089150;refGene_tx=XM_023241203.1
D3 79801667 79801854
refGene_gn=LOC101089150;refGene_tx=XM_023241203.1
D3 79807389 79807532
refGene_gn=LOC101089150;refGene_tx=XM_023241203.1
D3 79808297 79808423
refGene_gn=LOC101089150;refGene_tx=XM_023241203.1
D3 79845949 79846036 refGene_gn=SEC11C;refGene_tx=XM_003995228.5
D3 79854218 79854328 refGene_gn=SEC11C;refGene_tx=XM_003995228.5
D3 79857274 79857424 refGene_gn=SEC11C;refGene_tx=XM_003995228.5
D3 79859407 79859527 refGene_gn=SEC11C;refGene_tx=XM_003995228.5
D3 79861456 79861514 refGene_gn=SEC11C;refGene_tx=XM_003995228.5
D3 79862153 79862207 refGene_gn=SEC11C;refGene_tx=XM_003995228.5
D3 79897215 79897354
refGene_gn=GRP;refGene_tx=XM_023242047.1;refGene_gn=GRP;refGene_tx=XM_023242048.1
D3 79902586 79902826
refGene_gn=GRP;refGene_tx=XM_023242047.1;refGene_gn=GRP;refGene_tx=XM_023242048.1
D3 79907179 79907236
refGene_gn=GRP;refGene_tx=XM_023242047.1;refGene_gn=GRP;refGene_tx=XM_023242048.1
D3 79928762 79929254 refGene_gn=RAX;refGene_tx=XM_023242049.1
D3 79932311 79932565 refGene_gn=RAX;refGene_tx=XM_023242049.1
D3 79932874 79933163 refGene_gn=RAX;refGene_tx=XM_023242049.1
D3 79948163 79948391 refGene_gn=CPLX4;refGene_tx=XM_003995229.4
D3 79963923 79964011 refGene_gn=CPLX4;refGene_tx=XM_003995229.4
D3 79970011 79970178 refGene_gn=CPLX4;refGene_tx=XM_003995229.4
D3 79976378 79976415
refGene_gn=LMAN1;refGene_tx=XM_003995230.5;refGene_gn=LMAN1;refGene_tx=XM_023242050.1;refGene_gn=LMAN1;refGene_tx=XM_023242051.1
D3 79976713 79976835
refGene_gn=LMAN1;refGene_tx=XM_003995230.5;refGene_gn=LMAN1;refGene_tx=XM_023242050.1;refGene_gn=LMAN1;refGene_tx=XM_023242051.1
D3 79981666 79981820
refGene_gn=LMAN1;refGene_tx=XM_003995230.5;refGene_gn=LMAN1;refGene_tx=XM_023242050.1;refGene_gn=LMAN1;refGene_tx=XM_023242051.1
D3 79992968 79993039
refGene_gn=LMAN1;refGene_tx=XM_003995230.5;refGene_gn=LMAN1;refGene_tx=XM_023242050.1;refGene_gn=LMAN1;refGene_tx=XM_023242051.1
D3 79993158 79993352
refGene_gn=LMAN1;refGene_tx=XM_003995230.5;refGene_gn=LMAN1;refGene_tx

=XM_023242050.1;refGene_gn=LMAN1;refGene_tx=XM_023242051.1
D3 79994900 79995033
refGene_gn=LMAN1;refGene_tx=XM_003995230.5;refGene_gn=LMAN1;refGene_tx
=XM_023242050.1;refGene_gn=LMAN1;refGene_tx=XM_023242051.1
D3 79995729 79995788
refGene_gn=LMAN1;refGene_tx=XM_003995230.5;refGene_gn=LMAN1;refGene_tx
=XM_023242050.1;refGene_gn=LMAN1;refGene_tx=XM_023242051.1
D3 79997632 79997756
refGene_gn=LMAN1;refGene_tx=XM_003995230.5;refGene_gn=LMAN1;refGene_tx
=XM_023242050.1;refGene_gn=LMAN1;refGene_tx=XM_023242051.1
D3 80002111 80002211
refGene_gn=LMAN1;refGene_tx=XM_003995230.5;refGene_gn=LMAN1;refGene_tx
=XM_023242050.1;refGene_gn=LMAN1;refGene_tx=XM_023242051.1
D3 80003321 80003383
refGene_gn=LMAN1;refGene_tx=XM_003995230.5;refGene_gn=LMAN1;refGene_tx
=XM_023242050.1;refGene_gn=LMAN1;refGene_tx=XM_023242051.1
D3 80004360 80004468
refGene_gn=LMAN1;refGene_tx=XM_003995230.5;refGene_gn=LMAN1;refGene_tx
=XM_023242050.1;refGene_gn=LMAN1;refGene_tx=XM_023242051.1
D3 80004549 80004704
refGene_gn=LMAN1;refGene_tx=XM_003995230.5;refGene_gn=LMAN1;refGene_tx
=XM_023242050.1;refGene_gn=LMAN1;refGene_tx=XM_023242051.1
D3 80008102 80008340
refGene_gn=LMAN1;refGene_tx=XM_003995230.5;refGene_gn=LMAN1;refGene_tx
=XM_023242050.1;refGene_gn=LMAN1;refGene_tx=XM_023242051.1
D3 80058997 80059231 refGene_gn=CCBE1;refGene_tx=XM_023242052.1
D3 80061105 80061141 refGene_gn=CCBE1;refGene_tx=XM_023242052.1
D3 80062170 80062206 refGene_gn=CCBE1;refGene_tx=XM_023242052.1
D3 80062307 80062447 refGene_gn=CCBE1;refGene_tx=XM_023242052.1
D3 80068266 80068387 refGene_gn=CCBE1;refGene_tx=XM_023242052.1
D3 80079505 80079609 refGene_gn=CCBE1;refGene_tx=XM_023242052.1
D3 80089693 80089846 refGene_gn=CCBE1;refGene_tx=XM_023242052.1
D3 80091966 80092101 refGene_gn=CCBE1;refGene_tx=XM_023242052.1
D3 80096359 80096412 refGene_gn=CCBE1;refGene_tx=XM_023242052.1
D3 80292747 80292828 refGene_gn=CCBE1;refGene_tx=XM_023242052.1
D3 80293188 80293319 refGene_gn=CCBE1;refGene_tx=XM_023242052.1
D3 80419371 80419429 refGene_gn=PMAIP1;refGene_tx=XM_023242053.1
D3 80421753 80421860 refGene_gn=PMAIP1;refGene_tx=XM_023242053.1
D3 80793681 80794680 refGene_gn=MC4R;refGene_tx=XM_019815373.2
D3 80929375 80929406
refGene_gn=LOC101089901;refGene_tx=XM_023242054.1
D3 80934183 80934274
refGene_gn=LOC101089901;refGene_tx=XM_023242054.1
D3 80937849 80937962
refGene_gn=LOC101089901;refGene_tx=XM_023242054.1
D3 80938282 80938398
refGene_gn=LOC101089901;refGene_tx=XM_023242054.1
D3 80939941 80940039
refGene_gn=LOC101089901;refGene_tx=XM_023242054.1
D3 80941814 80941869

refGene_gn=LOC101089901;refGene_tx=XM_023242054.1
D3 80942794 80942911
refGene_gn=LOC101089901;refGene_tx=XM_023242054.1
D3 80944519 80944584
refGene_gn=LOC101089901;refGene_tx=XM_023242054.1
D3 80944687 80944766
refGene_gn=LOC101089901;refGene_tx=XM_023242054.1
D3 80949799 80949949
refGene_gn=LOC101089901;refGene_tx=XM_023242054.1
D3 80955974 80956082
refGene_gn=LOC101089901;refGene_tx=XM_023242054.1
D3 80957450 80957548
refGene_gn=LOC101089901;refGene_tx=XM_023242054.1
D3 80960145 80960311
refGene_gn=LOC101089901;refGene_tx=XM_023242054.1
D3 81737686 81737932
refGene_gn=CDH20;refGene_tx=XM_003995232.5;refGene_gn=CDH20;refGene_tx=XM_023242055.1;refGene_gn=CDH20;refGene_tx=XM_023242056.1;refGene_gn=CDH20;refGene_tx=XM_023242057.1;refGene_gn=CDH20;refGene_tx=XM_023242058.1
D3 81745849 81746144
refGene_gn=CDH20;refGene_tx=XM_003995232.5;refGene_gn=CDH20;refGene_tx=XM_023242055.1;refGene_gn=CDH20;refGene_tx=XM_023242056.1;refGene_gn=CDH20;refGene_tx=XM_023242057.1;refGene_gn=CDH20;refGene_tx=XM_023242058.1
D3 81746659 81746779
refGene_gn=CDH20;refGene_tx=XM_003995232.5;refGene_gn=CDH20;refGene_tx=XM_023242055.1;refGene_gn=CDH20;refGene_tx=XM_023242056.1;refGene_gn=CDH20;refGene_tx=XM_023242057.1;refGene_gn=CDH20;refGene_tx=XM_023242058.1
D3 81749437 81749605
refGene_gn=CDH20;refGene_tx=XM_003995232.5;refGene_gn=CDH20;refGene_tx=XM_023242055.1;refGene_gn=CDH20;refGene_tx=XM_023242057.1;refGene_gn=CDH20;refGene_tx=XM_023242058.1
D3 81754207 81754395
refGene_gn=CDH20;refGene_tx=XM_003995232.5;refGene_gn=CDH20;refGene_tx=XM_023242055.1;refGene_gn=CDH20;refGene_tx=XM_023242056.1;refGene_gn=CDH20;refGene_tx=XM_023242057.1;refGene_gn=CDH20;refGene_tx=XM_023242058.1
D3 81763653 81763907
refGene_gn=CDH20;refGene_tx=XM_003995232.5;refGene_gn=CDH20;refGene_tx=XM_023242055.1;refGene_gn=CDH20;refGene_tx=XM_023242056.1;refGene_gn=CDH20;refGene_tx=XM_023242057.1;refGene_gn=CDH20;refGene_tx=XM_023242058.1
D3 81770241 81770378
refGene_gn=CDH20;refGene_tx=XM_003995232.5;refGene_gn=CDH20;refGene_tx=XM_023242055.1;refGene_gn=CDH20;refGene_tx=XM_023242056.1;refGene_gn=CDH20;refGene_tx=XM_023242057.1;refGene_gn=CDH20;refGene_tx=XM_023242058.1
D3 81773071 81773193

refGene_gn=CDH20;refGene_tx=XM_003995232.5;refGene_gn=CDH20;refGene_tx=XM_023242055.1;refGene_gn=CDH20;refGene_tx=XM_023242056.1;refGene_gn=CDH20;refGene_tx=XM_023242057.1;refGene_gn=CDH20;refGene_tx=XM_023242058.1
D3 81777721 81777839 refGene_gn=CDH20;refGene_tx=XM_003995232.5;refGene_gn=CDH20;refGene_tx=XM_023242055.1;refGene_gn=CDH20;refGene_tx=XM_023242056.1;refGene_gn=CDH20;refGene_tx=XM_023242057.1;refGene_gn=CDH20;refGene_tx=XM_023242058.1
D3 81779184 81779270 refGene_gn=CDH20;refGene_tx=XM_023242058.1
D3 81782736 81782988 refGene_gn=CDH20;refGene_tx=XM_003995232.5;refGene_gn=CDH20;refGene_tx=XM_023242055.1;refGene_gn=CDH20;refGene_tx=XM_023242056.1;refGene_gn=CDH20;refGene_tx=XM_023242057.1
D3 81786640 81786678 refGene_gn=CDH20;refGene_tx=XM_023242057.1
D3 81787443 81787949 refGene_gn=CDH20;refGene_tx=XM_003995232.5;refGene_gn=CDH20;refGene_tx=XM_023242055.1;refGene_gn=CDH20;refGene_tx=XM_023242056.1
D3 81989808 81990420 refGene_gn=RNF152;refGene_tx=XM_023242059.1;refGene_gn=RNF152;refGene_tx=XM_023242060.1;refGene_gn=RNF152;refGene_tx=XM_023242061.1;refGene_gn=RNF152;refGene_tx=XM_023242062.1;refGene_gn=RNF152;refGene_tx=XM_023242063.1;refGene_gn=RNF152;refGene_tx=XM_023242064.1;refGene_gn=RNF152;refGene_tx=XM_023242065.1
D3 82177292 82177416 refGene_gn=PIGN;refGene_tx=XM_006938956.3;refGene_gn=PIGN;refGene_tx=XM_006938957.3;refGene_gn=PIGN;refGene_tx=XM_006938958.3;refGene_gn=PIGN;refGene_tx=XM_006938959.3;refGene_gn=PIGN;refGene_tx=XM_006938960.3;refGene_gn=PIGN;refGene_tx=XM_011288141.3;refGene_gn=PIGN;refGene_tx=XM_019815378.2;refGene_gn=PIGN;refGene_tx=XM_019815379.2;refGene_gn=PIGN;refGene_tx=XM_019815380.2;refGene_gn=PIGN;refGene_tx=XM_019815381.2;refGene_gn=PIGN;refGene_tx=XM_019815382.2;refGene_gn=PIGN;refGene_tx=XM_023242066.1
D3 82185771 82185824 refGene_gn=PIGN;refGene_tx=XM_006938956.3;refGene_gn=PIGN;refGene_tx=XM_006938957.3;refGene_gn=PIGN;refGene_tx=XM_006938958.3;refGene_gn=PIGN;refGene_tx=XM_006938959.3;refGene_gn=PIGN;refGene_tx=XM_006938960.3;refGene_gn=PIGN;refGene_tx=XM_011288141.3;refGene_gn=PIGN;refGene_tx=XM_019815378.2;refGene_gn=PIGN;refGene_tx=XM_019815379.2;refGene_gn=PIGN;refGene_tx=XM_019815380.2;refGene_gn=PIGN;refGene_tx=XM_019815381.2;refGene_gn=PIGN;refGene_tx=XM_023242066.1
D3 82188025 82188068 refGene_gn=PIGN;refGene_tx=XM_006938956.3;refGene_gn=PIGN;refGene_tx=XM_006938957.3;refGene_gn=PIGN;refGene_tx=XM_006938958.3;refGene_gn=PIGN;refGene_tx=XM_006938959.3;refGene_gn=PIGN;refGene_tx=XM_006938960.3;refGene_gn=PIGN;refGene_tx=XM_011288141.3;refGene_gn=PIGN;refGene_tx=XM_019815378.2;refGene_gn=PIGN;refGene_tx=XM_019815379.2;refGene_gn=PIGN;refGene_tx=XM_019815380.2;refGene_gn=PIGN;refGene_tx=XM_019815381.2;refGene_gn=PIGN;refGene_tx=XM_023242066.1
D3 82196185 82196259

refGene_gn=PIGN;refGene_tx=XM_006938956.3;refGene_gn=PIGN;refGene_tx=XM_006938957.3;refGene_gn=PIGN;refGene_tx=XM_006938958.3;refGene_gn=PIGN;refGene_tx=XM_006938959.3;refGene_gn=PIGN;refGene_tx=XM_006938960.3;refGene_gn=PIGN;refGene_tx=XM_011288141.3;refGene_gn=PIGN;refGene_tx=XM_019815378.2;refGene_gn=PIGN;refGene_tx=XM_019815379.2;refGene_gn=PIGN;refGene_tx=XM_019815380.2;refGene_gn=PIGN;refGene_tx=XM_019815381.2;refGene_gn=PIGN;refGene_tx=XM_019815382.2;refGene_gn=PIGN;refGene_tx=XM_023242066.1

D3 82198784 82198860 refGene_gn=PIGN;refGene_tx=XM_006938956.3;refGene_gn=PIGN;refGene_tx=XM_006938957.3;refGene_gn=PIGN;refGene_tx=XM_006938958.3;refGene_gn=PIGN;refGene_tx=XM_006938959.3;refGene_gn=PIGN;refGene_tx=XM_006938960.3;refGene_gn=PIGN;refGene_tx=XM_011288141.3;refGene_gn=PIGN;refGene_tx=XM_019815378.2;refGene_gn=PIGN;refGene_tx=XM_019815379.2;refGene_gn=PIGN;refGene_tx=XM_019815380.2;refGene_gn=PIGN;refGene_tx=XM_019815381.2;refGene_gn=PIGN;refGene_tx=XM_019815382.2;refGene_gn=PIGN;refGene_tx=XM_023242066.1

D3 82199380 82199436 refGene_gn=PIGN;refGene_tx=XM_006938956.3;refGene_gn=PIGN;refGene_tx=XM_006938957.3;refGene_gn=PIGN;refGene_tx=XM_006938958.3;refGene_gn=PIGN;refGene_tx=XM_006938959.3;refGene_gn=PIGN;refGene_tx=XM_006938960.3;refGene_gn=PIGN;refGene_tx=XM_011288141.3;refGene_gn=PIGN;refGene_tx=XM_019815378.2;refGene_gn=PIGN;refGene_tx=XM_019815379.2;refGene_gn=PIGN;refGene_tx=XM_019815380.2;refGene_gn=PIGN;refGene_tx=XM_019815381.2;refGene_gn=PIGN;refGene_tx=XM_019815382.2;refGene_gn=PIGN;refGene_tx=XM_023242066.1

D3 82202190 82202277 refGene_gn=PIGN;refGene_tx=XM_006938956.3;refGene_gn=PIGN;refGene_tx=XM_006938957.3;refGene_gn=PIGN;refGene_tx=XM_006938958.3;refGene_gn=PIGN;refGene_tx=XM_006938959.3;refGene_gn=PIGN;refGene_tx=XM_006938960.3;refGene_gn=PIGN;refGene_tx=XM_011288141.3;refGene_gn=PIGN;refGene_tx=XM_019815378.2;refGene_gn=PIGN;refGene_tx=XM_019815379.2;refGene_gn=PIGN;refGene_tx=XM_019815380.2;refGene_gn=PIGN;refGene_tx=XM_019815381.2;refGene_gn=PIGN;refGene_tx=XM_019815382.2;refGene_gn=PIGN;refGene_tx=XM_023242066.1

D3 82203949 82204052 refGene_gn=PIGN;refGene_tx=XM_006938956.3;refGene_gn=PIGN;refGene_tx=XM_006938957.3;refGene_gn=PIGN;refGene_tx=XM_006938958.3;refGene_gn=PIGN;refGene_tx=XM_006938959.3;refGene_gn=PIGN;refGene_tx=XM_006938960.3;refGene_gn=PIGN;refGene_tx=XM_011288141.3;refGene_gn=PIGN;refGene_tx=XM_019815378.2;refGene_gn=PIGN;refGene_tx=XM_019815379.2;refGene_gn=PIGN;refGene_tx=XM_019815380.2;refGene_gn=PIGN;refGene_tx=XM_019815381.2;refGene_gn=PIGN;refGene_tx=XM_019815382.2;refGene_gn=PIGN;refGene_tx=XM_023242066.1

D3 82210241 82210344 refGene_gn=PIGN;refGene_tx=XM_006938956.3;refGene_gn=PIGN;refGene_tx=XM_006938957.3;refGene_gn=PIGN;refGene_tx=XM_006938958.3;refGene_gn=PIGN;refGene_tx=XM_006938959.3;refGene_gn=PIGN;refGene_tx=XM_006938960.3;refGene_gn=PIGN;refGene_tx=XM_011288141.3;refGene_gn=PIGN;refGene_tx=XM_019815378.2;refGene_gn=PIGN;refGene_tx=XM_019815379.2;refGene_gn=PIG

N;refGene_tx=XM_019815380.2;refGene_gn=PIGN;refGene_tx=XM_019815381.2;
refGene_gn=PIGN;refGene_tx=XM_019815382.2;refGene_gn=PIGN;refGene_tx=X
M_023242066.1
D3 82212904 82213013
refGene_gn=PIGN;refGene_tx=XM_006938956.3;refGene_gn=PIGN;refGene_tx=X
M_006938957.3;refGene_gn=PIGN;refGene_tx=XM_006938958.3;refGene_gn=PIG
N;refGene_tx=XM_006938959.3;refGene_gn=PIGN;refGene_tx=XM_006938960.3;
refGene_gn=PIGN;refGene_tx=XM_011288141.3;refGene_gn=PIGN;refGene_tx=X
M_019815378.2;refGene_gn=PIGN;refGene_tx=XM_019815379.2;refGene_gn=PIG
N;refGene_tx=XM_019815380.2;refGene_gn=PIGN;refGene_tx=XM_019815381.2;
refGene_gn=PIGN;refGene_tx=XM_019815382.2;refGene_gn=PIGN;refGene_tx=X
M_023242066.1
D3 82215570 82215679
refGene_gn=PIGN;refGene_tx=XM_006938956.3;refGene_gn=PIGN;refGene_tx=X
M_006938957.3;refGene_gn=PIGN;refGene_tx=XM_006938958.3;refGene_gn=PIG
N;refGene_tx=XM_006938959.3;refGene_gn=PIGN;refGene_tx=XM_006938960.3;
refGene_gn=PIGN;refGene_tx=XM_011288141.3;refGene_gn=PIGN;refGene_tx=X
M_019815378.2;refGene_gn=PIGN;refGene_tx=XM_019815379.2;refGene_gn=PIG
N;refGene_tx=XM_019815380.2;refGene_gn=PIGN;refGene_tx=XM_019815381.2;
refGene_gn=PIGN;refGene_tx=XM_019815382.2;refGene_gn=PIGN;refGene_tx=X
M_023242066.1
D3 82218642 82218734
refGene_gn=PIGN;refGene_tx=XM_006938956.3;refGene_gn=PIGN;refGene_tx=X
M_006938957.3;refGene_gn=PIGN;refGene_tx=XM_006938958.3;refGene_gn=PIG
N;refGene_tx=XM_006938959.3;refGene_gn=PIGN;refGene_tx=XM_006938960.3;
refGene_gn=PIGN;refGene_tx=XM_011288141.3;refGene_gn=PIGN;refGene_tx=X
M_019815378.2;refGene_gn=PIGN;refGene_tx=XM_019815379.2;refGene_gn=PIG
N;refGene_tx=XM_019815380.2;refGene_gn=PIGN;refGene_tx=XM_019815381.2;
refGene_gn=PIGN;refGene_tx=XM_019815382.2;refGene_gn=PIGN;refGene_tx=X
M_023242066.1
D3 82219645 82219738
refGene_gn=PIGN;refGene_tx=XM_006938956.3;refGene_gn=PIGN;refGene_tx=X
M_006938957.3;refGene_gn=PIGN;refGene_tx=XM_006938958.3;refGene_gn=PIG
N;refGene_tx=XM_006938959.3;refGene_gn=PIGN;refGene_tx=XM_006938960.3;
refGene_gn=PIGN;refGene_tx=XM_011288141.3;refGene_gn=PIGN;refGene_tx=X
M_019815378.2;refGene_gn=PIGN;refGene_tx=XM_019815379.2;refGene_gn=PIG
N;refGene_tx=XM_019815380.2;refGene_gn=PIGN;refGene_tx=XM_019815381.2;
refGene_gn=PIGN;refGene_tx=XM_019815382.2;refGene_gn=PIGN;refGene_tx=X
M_023242066.1
D3 82219842 82219942
refGene_gn=PIGN;refGene_tx=XM_006938956.3;refGene_gn=PIGN;refGene_tx=X
M_006938957.3;refGene_gn=PIGN;refGene_tx=XM_006938958.3;refGene_gn=PIG
N;refGene_tx=XM_006938959.3;refGene_gn=PIGN;refGene_tx=XM_006938960.3;
refGene_gn=PIGN;refGene_tx=XM_011288141.3;refGene_gn=PIGN;refGene_tx=X
M_019815378.2;refGene_gn=PIGN;refGene_tx=XM_019815379.2;refGene_gn=PIG
N;refGene_tx=XM_019815380.2;refGene_gn=PIGN;refGene_tx=XM_019815381.2;
refGene_gn=PIGN;refGene_tx=XM_019815382.2;refGene_gn=PIGN;refGene_tx=X
M_023242066.1
D3 82222550 82222690
refGene_gn=PIGN;refGene_tx=XM_006938956.3;refGene_gn=PIGN;refGene_tx=X

M_006938957.3;refGene_gn=PIGN;refGene_tx=XM_006938958.3;refGene_gn=PIG
N;refGene_tx=XM_006938959.3;refGene_gn=PIGN;refGene_tx=XM_006938960.3;
refGene_gn=PIGN;refGene_tx=XM_011288141.3;refGene_gn=PIGN;refGene_tx=X
M_019815378.2;refGene_gn=PIGN;refGene_tx=XM_019815379.2;refGene_gn=PIG
N;refGene_tx=XM_019815380.2;refGene_gn=PIGN;refGene_tx=XM_019815381.2;
refGene_gn=PIGN;refGene_tx=XM_019815382.2;refGene_gn=PIGN;refGene_tx=X
M_023242066.1
D3 82228324 82228507
refGene_gn=PIGN;refGene_tx=XM_006938956.3;refGene_gn=PIGN;refGene_tx=X
M_006938957.3;refGene_gn=PIGN;refGene_tx=XM_006938958.3;refGene_gn=PIG
N;refGene_tx=XM_006938959.3;refGene_gn=PIGN;refGene_tx=XM_011288141.3;
refGene_gn=PIGN;refGene_tx=XM_019815378.2;refGene_gn=PIGN;refGene_tx=X
M_019815379.2;refGene_gn=PIGN;refGene_tx=XM_019815380.2;refGene_gn=PIG
N;refGene_tx=XM_019815381.2;refGene_gn=PIGN;refGene_tx=XM_019815382.2;
refGene_gn=PIGN;refGene_tx=XM_023242066.1
D3 82229434 82229513
refGene_gn=PIGN;refGene_tx=XM_006938956.3;refGene_gn=PIGN;refGene_tx=X
M_006938957.3;refGene_gn=PIGN;refGene_tx=XM_006938958.3;refGene_gn=PIG
N;refGene_tx=XM_006938959.3;refGene_gn=PIGN;refGene_tx=XM_006938960.3;
refGene_gn=PIGN;refGene_tx=XM_011288141.3;refGene_gn=PIGN;refGene_tx=X
M_019815378.2;refGene_gn=PIGN;refGene_tx=XM_019815379.2;refGene_gn=PIG
N;refGene_tx=XM_019815380.2;refGene_gn=PIGN;refGene_tx=XM_019815381.2;
refGene_gn=PIGN;refGene_tx=XM_019815382.2;refGene_gn=PIGN;refGene_tx=X
M_023242066.1
D3 82240816 82240872
refGene_gn=PIGN;refGene_tx=XM_006938956.3;refGene_gn=PIGN;refGene_tx=X
M_006938957.3;refGene_gn=PIGN;refGene_tx=XM_006938958.3;refGene_gn=PIG
N;refGene_tx=XM_006938959.3;refGene_gn=PIGN;refGene_tx=XM_006938960.3;
refGene_gn=PIGN;refGene_tx=XM_011288141.3;refGene_gn=PIGN;refGene_tx=X
M_019815378.2;refGene_gn=PIGN;refGene_tx=XM_019815379.2;refGene_gn=PIG
N;refGene_tx=XM_019815380.2;refGene_gn=PIGN;refGene_tx=XM_019815381.2;
refGene_gn=PIGN;refGene_tx=XM_019815382.2;refGene_gn=PIGN;refGene_tx=X
M_023242066.1
D3 82241515 82241608
refGene_gn=PIGN;refGene_tx=XM_006938956.3;refGene_gn=PIGN;refGene_tx=X
M_006938957.3;refGene_gn=PIGN;refGene_tx=XM_006938958.3;refGene_gn=PIG
N;refGene_tx=XM_006938959.3;refGene_gn=PIGN;refGene_tx=XM_006938960.3;
refGene_gn=PIGN;refGene_tx=XM_011288141.3;refGene_gn=PIGN;refGene_tx=X
M_019815378.2;refGene_gn=PIGN;refGene_tx=XM_019815379.2;refGene_gn=PIG
N;refGene_tx=XM_019815380.2;refGene_gn=PIGN;refGene_tx=XM_019815381.2;
refGene_gn=PIGN;refGene_tx=XM_019815382.2;refGene_gn=PIGN;refGene_tx=X
M_023242066.1
D3 82243443 82243503
refGene_gn=PIGN;refGene_tx=XM_006938956.3;refGene_gn=PIGN;refGene_tx=X
M_006938957.3;refGene_gn=PIGN;refGene_tx=XM_006938958.3;refGene_gn=PIG
N;refGene_tx=XM_006938959.3;refGene_gn=PIGN;refGene_tx=XM_006938960.3;
refGene_gn=PIGN;refGene_tx=XM_011288141.3;refGene_gn=PIGN;refGene_tx=X
M_019815378.2;refGene_gn=PIGN;refGene_tx=XM_019815379.2;refGene_gn=PIG
N;refGene_tx=XM_019815380.2;refGene_gn=PIGN;refGene_tx=XM_019815381.2;
refGene_gn=PIGN;refGene_tx=XM_019815382.2;refGene_gn=PIGN;refGene_tx=X

M_023242066.1
D3 82246208 82246249
refGene_gn=PIGN;refGene_tx=XM_006938956.3;refGene_gn=PIGN;refGene_tx=XM_006938957.3;refGene_gn=PIGN;refGene_tx=XM_006938958.3;refGene_gn=PIGN;refGene_tx=XM_006938959.3;refGene_gn=PIGN;refGene_tx=XM_006938960.3;refGene_gn=PIGN;refGene_tx=XM_011288141.3;refGene_gn=PIGN;refGene_tx=XM_019815378.2;refGene_gn=PIGN;refGene_tx=XM_019815379.2;refGene_gn=PIGN;refGene_tx=XM_019815380.2;refGene_gn=PIGN;refGene_tx=XM_019815381.2;refGene_gn=PIGN;refGene_tx=XM_019815382.2;refGene_gn=PIGN;refGene_tx=XM_023242066.1
D3 82249192 82249309
refGene_gn=PIGN;refGene_tx=XM_006938956.3;refGene_gn=PIGN;refGene_tx=XM_006938957.3;refGene_gn=PIGN;refGene_tx=XM_006938958.3;refGene_gn=PIGN;refGene_tx=XM_006938959.3;refGene_gn=PIGN;refGene_tx=XM_006938960.3;refGene_gn=PIGN;refGene_tx=XM_011288141.3;refGene_gn=PIGN;refGene_tx=XM_019815378.2;refGene_gn=PIGN;refGene_tx=XM_019815379.2;refGene_gn=PIGN;refGene_tx=XM_019815380.2;refGene_gn=PIGN;refGene_tx=XM_019815381.2;refGene_gn=PIGN;refGene_tx=XM_019815382.2;refGene_gn=PIGN;refGene_tx=XM_023242066.1
D3 82250040 82250171
refGene_gn=PIGN;refGene_tx=XM_006938956.3;refGene_gn=PIGN;refGene_tx=XM_006938957.3;refGene_gn=PIGN;refGene_tx=XM_006938958.3;refGene_gn=PIGN;refGene_tx=XM_006938959.3;refGene_gn=PIGN;refGene_tx=XM_006938960.3;refGene_gn=PIGN;refGene_tx=XM_011288141.3;refGene_gn=PIGN;refGene_tx=XM_019815378.2;refGene_gn=PIGN;refGene_tx=XM_019815379.2;refGene_gn=PIGN;refGene_tx=XM_019815380.2;refGene_gn=PIGN;refGene_tx=XM_019815381.2;refGene_gn=PIGN;refGene_tx=XM_019815382.2;refGene_gn=PIGN;refGene_tx=XM_023242066.1
D3 82251455 82251580
refGene_gn=PIGN;refGene_tx=XM_006938956.3;refGene_gn=PIGN;refGene_tx=XM_006938957.3;refGene_gn=PIGN;refGene_tx=XM_006938958.3;refGene_gn=PIGN;refGene_tx=XM_006938959.3;refGene_gn=PIGN;refGene_tx=XM_006938960.3;refGene_gn=PIGN;refGene_tx=XM_011288141.3;refGene_gn=PIGN;refGene_tx=XM_019815378.2;refGene_gn=PIGN;refGene_tx=XM_019815379.2;refGene_gn=PIGN;refGene_tx=XM_019815380.2;refGene_gn=PIGN;refGene_tx=XM_019815381.2;refGene_gn=PIGN;refGene_tx=XM_019815382.2;refGene_gn=PIGN;refGene_tx=XM_023242066.1
D3 82256450 82256557
refGene_gn=PIGN;refGene_tx=XM_006938956.3;refGene_gn=PIGN;refGene_tx=XM_006938957.3;refGene_gn=PIGN;refGene_tx=XM_006938958.3;refGene_gn=PIGN;refGene_tx=XM_006938959.3;refGene_gn=PIGN;refGene_tx=XM_006938960.3;refGene_gn=PIGN;refGene_tx=XM_011288141.3;refGene_gn=PIGN;refGene_tx=XM_019815378.2;refGene_gn=PIGN;refGene_tx=XM_019815379.2;refGene_gn=PIGN;refGene_tx=XM_019815380.2;refGene_gn=PIGN;refGene_tx=XM_019815381.2;refGene_gn=PIGN;refGene_tx=XM_019815382.2;refGene_gn=PIGN;refGene_tx=XM_023242066.1
D3 82258592 82258691
refGene_gn=PIGN;refGene_tx=XM_006938956.3;refGene_gn=PIGN;refGene_tx=XM_006938957.3;refGene_gn=PIGN;refGene_tx=XM_006938958.3;refGene_gn=PIGN;refGene_tx=XM_006938959.3;refGene_gn=PIGN;refGene_tx=XM_006938960.3;

refGene_gn=PIGN;refGene_tx=XM_011288141.3;refGene_gn=PIGN;refGene_tx=XM_019815378.2;refGene_gn=PIGN;refGene_tx=XM_019815379.2;refGene_gn=PIGN;refGene_tx=XM_019815380.2;refGene_gn=PIGN;refGene_tx=XM_019815381.2;refGene_gn=PIGN;refGene_tx=XM_019815382.2;refGene_gn=PIGN;refGene_tx=XM_023242066.1
D3 82259207 82259329
refGene_gn=PIGN;refGene_tx=XM_006938956.3;refGene_gn=PIGN;refGene_tx=XM_006938957.3;refGene_gn=PIGN;refGene_tx=XM_006938958.3;refGene_gn=PIGN;refGene_tx=XM_006938959.3;refGene_gn=PIGN;refGene_tx=XM_006938960.3;refGene_gn=PIGN;refGene_tx=XM_011288141.3;refGene_gn=PIGN;refGene_tx=XM_019815378.2;refGene_gn=PIGN;refGene_tx=XM_019815379.2;refGene_gn=PIGN;refGene_tx=XM_019815380.2;refGene_gn=PIGN;refGene_tx=XM_019815381.2;refGene_gn=PIGN;refGene_tx=XM_019815382.2;refGene_gn=PIGN;refGene_tx=XM_023242066.1
D3 82262377 82262598
refGene_gn=PIGN;refGene_tx=XM_006938956.3;refGene_gn=PIGN;refGene_tx=XM_006938957.3;refGene_gn=PIGN;refGene_tx=XM_006938958.3;refGene_gn=PIGN;refGene_tx=XM_006938959.3;refGene_gn=PIGN;refGene_tx=XM_006938960.3;refGene_gn=PIGN;refGene_tx=XM_011288141.3;refGene_gn=PIGN;refGene_tx=XM_019815378.2;refGene_gn=PIGN;refGene_tx=XM_019815379.2;refGene_gn=PIGN;refGene_tx=XM_019815380.2;refGene_gn=PIGN;refGene_tx=XM_019815381.2;refGene_gn=PIGN;refGene_tx=XM_019815382.2;refGene_gn=PIGN;refGene_tx=XM_023242066.1
D3 82282167 82282696
refGene_gn=KIAA1468;refGene_tx=XM_003995235.5;refGene_gn=KIAA1468;refGene_tx=XM_006938961.4;refGene_gn=KIAA1468;refGene_tx=XM_006938962.4;refGene_gn=KIAA1468;refGene_tx=XM_011288140.3;refGene_gn=KIAA1468;refGene_tx=XM_019815385.2;refGene_gn=KIAA1468;refGene_tx=XM_019815386.2;refGene_gn=KIAA1468;refGene_tx=XM_019815387.2;refGene_gn=KIAA1468;refGene_tx=XM_019815388.2;refGene_gn=KIAA1468;refGene_tx=XM_019815389.2
D3 82306657 82306747
refGene_gn=KIAA1468;refGene_tx=XM_003995235.5;refGene_gn=KIAA1468;refGene_tx=XM_006938961.4;refGene_gn=KIAA1468;refGene_tx=XM_006938962.4;refGene_gn=KIAA1468;refGene_tx=XM_011288140.3;refGene_gn=KIAA1468;refGene_tx=XM_019815385.2;refGene_gn=KIAA1468;refGene_tx=XM_019815386.2;refGene_gn=KIAA1468;refGene_tx=XM_019815387.2;refGene_gn=KIAA1468;refGene_tx=XM_019815388.2;refGene_gn=KIAA1468;refGene_tx=XM_019815389.2
D3 82314832 82314904
refGene_gn=KIAA1468;refGene_tx=XM_003995235.5;refGene_gn=KIAA1468;refGene_tx=XM_006938961.4;refGene_gn=KIAA1468;refGene_tx=XM_019815385.2;refGene_gn=KIAA1468;refGene_tx=XM_019815386.2;refGene_gn=KIAA1468;refGene_tx=XM_019815387.2;refGene_gn=KIAA1468;refGene_tx=XM_019815388.2;refGene_gn=KIAA1468;refGene_tx=XM_019815389.2
D3 82315015 82315071
refGene_gn=KIAA1468;refGene_tx=XM_003995235.5;refGene_gn=KIAA1468;refGene_tx=XM_006938961.4;refGene_gn=KIAA1468;refGene_tx=XM_006938962.4;refGene_gn=KIAA1468;refGene_tx=XM_011288140.3;refGene_gn=KIAA1468;refGene_tx=XM_019815385.2;refGene_gn=KIAA1468;refGene_tx=XM_019815386.2;refGene_gn=KIAA1468;refGene_tx=XM_019815387.2;refGene_gn=KIAA1468;refGene_tx=XM_019815388.2;refGene_gn=KIAA1468;refGene_tx=XM_019815389.2

D3 82315172 82315286
refGene_gn=KIAA1468;refGene_tx=XM_003995235.5;refGene_gn=KIAA1468;refGene_tx=XM_006938961.4;refGene_gn=KIAA1468;refGene_tx=XM_006938962.4;refGene_gn=KIAA1468;refGene_tx=XM_011288140.3;refGene_gn=KIAA1468;refGene_tx=XM_019815385.2;refGene_gn=KIAA1468;refGene_tx=XM_019815386.2;refGene_gn=KIAA1468;refGene_tx=XM_019815387.2;refGene_gn=KIAA1468;refGene_tx=XM_019815388.2;refGene_gn=KIAA1468;refGene_tx=XM_019815389.2
D3 82317567 82317771
refGene_gn=KIAA1468;refGene_tx=XM_003995235.5;refGene_gn=KIAA1468;refGene_tx=XM_006938961.4;refGene_gn=KIAA1468;refGene_tx=XM_006938962.4;refGene_gn=KIAA1468;refGene_tx=XM_011288140.3;refGene_gn=KIAA1468;refGene_tx=XM_019815385.2;refGene_gn=KIAA1468;refGene_tx=XM_019815386.2;refGene_gn=KIAA1468;refGene_tx=XM_019815387.2;refGene_gn=KIAA1468;refGene_tx=XM_019815388.2;refGene_gn=KIAA1468;refGene_tx=XM_019815389.2
D3 82317876 82317968
refGene_gn=KIAA1468;refGene_tx=XM_003995235.5;refGene_gn=KIAA1468;refGene_tx=XM_006938961.4;refGene_gn=KIAA1468;refGene_tx=XM_006938962.4;refGene_gn=KIAA1468;refGene_tx=XM_011288140.3;refGene_gn=KIAA1468;refGene_tx=XM_019815385.2;refGene_gn=KIAA1468;refGene_tx=XM_019815386.2;refGene_gn=KIAA1468;refGene_tx=XM_019815387.2;refGene_gn=KIAA1468;refGene_tx=XM_019815388.2;refGene_gn=KIAA1468;refGene_tx=XM_019815389.2
D3 82318587 82318872
refGene_gn=KIAA1468;refGene_tx=XM_003995235.5;refGene_gn=KIAA1468;refGene_tx=XM_006938961.4;refGene_gn=KIAA1468;refGene_tx=XM_006938962.4;refGene_gn=KIAA1468;refGene_tx=XM_011288140.3;refGene_gn=KIAA1468;refGene_tx=XM_019815385.2;refGene_gn=KIAA1468;refGene_tx=XM_019815386.2;refGene_gn=KIAA1468;refGene_tx=XM_019815387.2;refGene_gn=KIAA1468;refGene_tx=XM_019815388.2;refGene_gn=KIAA1468;refGene_tx=XM_019815389.2
D3 82321743 82321819
refGene_gn=KIAA1468;refGene_tx=XM_003995235.5;refGene_gn=KIAA1468;refGene_tx=XM_006938961.4;refGene_gn=KIAA1468;refGene_tx=XM_006938962.4;refGene_gn=KIAA1468;refGene_tx=XM_011288140.3;refGene_gn=KIAA1468;refGene_tx=XM_019815385.2;refGene_gn=KIAA1468;refGene_tx=XM_019815386.2;refGene_gn=KIAA1468;refGene_tx=XM_019815387.2;refGene_gn=KIAA1468;refGene_tx=XM_019815388.2;refGene_gn=KIAA1468;refGene_tx=XM_019815389.2
D3 82322891 82322987
refGene_gn=KIAA1468;refGene_tx=XM_003995235.5;refGene_gn=KIAA1468;refGene_tx=XM_006938961.4;refGene_gn=KIAA1468;refGene_tx=XM_006938962.4;refGene_gn=KIAA1468;refGene_tx=XM_011288140.3;refGene_gn=KIAA1468;refGene_tx=XM_019815385.2;refGene_gn=KIAA1468;refGene_tx=XM_019815386.2;refGene_gn=KIAA1468;refGene_tx=XM_019815387.2;refGene_gn=KIAA1468;refGene_tx=XM_019815388.2;refGene_gn=KIAA1468;refGene_tx=XM_019815389.2
D3 82323136 82323160
refGene_gn=KIAA1468;refGene_tx=XM_011288140.3;refGene_gn=KIAA1468;refGene_tx=XM_019815385.2;refGene_gn=KIAA1468;refGene_tx=XM_019815386.2;refGene_gn=KIAA1468;refGene_tx=XM_019815387.2;refGene_gn=KIAA1468;refGene_tx=XM_019815388.2;refGene_gn=KIAA1468;refGene_tx=XM_019815389.2
D3 82334754 82334867
refGene_gn=KIAA1468;refGene_tx=XM_003995235.5;refGene_gn=KIAA1468;refGene_tx=XM_006938961.4;refGene_gn=KIAA1468;refGene_tx=XM_006938962.4;re

fGene_gn=KIAA1468;refGene_tx=XM_011288140.3;refGene_gn=KIAA1468;refGene_tx=XM_019815385.2;refGene_gn=KIAA1468;refGene_tx=XM_019815386.2;refGene_gn=KIAA1468;refGene_tx=XM_019815387.2;refGene_gn=KIAA1468;refGene_tx=XM_019815388.2;refGene_gn=KIAA1468;refGene_tx=XM_019815389.2
D3 82344145 82344236
refGene_gn=KIAA1468;refGene_tx=XM_003995235.5;refGene_gn=KIAA1468;refGene_tx=XM_006938961.4;refGene_gn=KIAA1468;refGene_tx=XM_006938962.4;refGene_gn=KIAA1468;refGene_tx=XM_011288140.3;refGene_gn=KIAA1468;refGene_tx=XM_019815385.2;refGene_gn=KIAA1468;refGene_tx=XM_019815386.2;refGene_gn=KIAA1468;refGene_tx=XM_019815387.2;refGene_gn=KIAA1468;refGene_tx=XM_019815388.2;refGene_gn=KIAA1468;refGene_tx=XM_019815389.2
D3 82346918 82346990
refGene_gn=KIAA1468;refGene_tx=XM_003995235.5;refGene_gn=KIAA1468;refGene_tx=XM_006938961.4;refGene_gn=KIAA1468;refGene_tx=XM_006938962.4;refGene_gn=KIAA1468;refGene_tx=XM_011288140.3;refGene_gn=KIAA1468;refGene_tx=XM_019815385.2;refGene_gn=KIAA1468;refGene_tx=XM_019815386.2;refGene_gn=KIAA1468;refGene_tx=XM_019815387.2;refGene_gn=KIAA1468;refGene_tx=XM_019815388.2;refGene_gn=KIAA1468;refGene_tx=XM_019815389.2
D3 82349470 82349611
refGene_gn=KIAA1468;refGene_tx=XM_003995235.5;refGene_gn=KIAA1468;refGene_tx=XM_006938961.4;refGene_gn=KIAA1468;refGene_tx=XM_006938962.4;refGene_gn=KIAA1468;refGene_tx=XM_011288140.3;refGene_gn=KIAA1468;refGene_tx=XM_019815385.2;refGene_gn=KIAA1468;refGene_tx=XM_019815386.2;refGene_gn=KIAA1468;refGene_tx=XM_019815387.2;refGene_gn=KIAA1468;refGene_tx=XM_019815388.2;refGene_gn=KIAA1468;refGene_tx=XM_019815389.2
D3 82350063 82350228
refGene_gn=KIAA1468;refGene_tx=XM_003995235.5;refGene_gn=KIAA1468;refGene_tx=XM_006938961.4;refGene_gn=KIAA1468;refGene_tx=XM_006938962.4;refGene_gn=KIAA1468;refGene_tx=XM_011288140.3;refGene_gn=KIAA1468;refGene_tx=XM_019815385.2;refGene_gn=KIAA1468;refGene_tx=XM_019815386.2;refGene_gn=KIAA1468;refGene_tx=XM_019815387.2;refGene_gn=KIAA1468;refGene_tx=XM_019815388.2;refGene_gn=KIAA1468;refGene_tx=XM_019815389.2
D3 82352274 82352422
refGene_gn=KIAA1468;refGene_tx=XM_003995235.5;refGene_gn=KIAA1468;refGene_tx=XM_006938961.4;refGene_gn=KIAA1468;refGene_tx=XM_006938962.4;refGene_gn=KIAA1468;refGene_tx=XM_011288140.3;refGene_gn=KIAA1468;refGene_tx=XM_019815385.2;refGene_gn=KIAA1468;refGene_tx=XM_019815386.2;refGene_gn=KIAA1468;refGene_tx=XM_019815387.2;refGene_gn=KIAA1468;refGene_tx=XM_019815388.2;refGene_gn=KIAA1468;refGene_tx=XM_019815389.2
D3 82355818 82355975
refGene_gn=KIAA1468;refGene_tx=XM_003995235.5;refGene_gn=KIAA1468;refGene_tx=XM_006938961.4;refGene_gn=KIAA1468;refGene_tx=XM_006938962.4;refGene_gn=KIAA1468;refGene_tx=XM_011288140.3;refGene_gn=KIAA1468;refGene_tx=XM_019815385.2;refGene_gn=KIAA1468;refGene_tx=XM_019815386.2;refGene_gn=KIAA1468;refGene_tx=XM_019815387.2;refGene_gn=KIAA1468;refGene_tx=XM_019815388.2;refGene_gn=KIAA1468;refGene_tx=XM_019815389.2
D3 82356599 82356723
refGene_gn=KIAA1468;refGene_tx=XM_003995235.5;refGene_gn=KIAA1468;refGene_tx=XM_006938961.4;refGene_gn=KIAA1468;refGene_tx=XM_006938962.4;refGene_gn=KIAA1468;refGene_tx=XM_011288140.3;refGene_gn=KIAA1468;refGen

e_tx=XM_019815385.2;refGene_gn=KIAA1468;refGene_tx=XM_019815386.2;refGene_gn=KIAA1468;refGene_tx=XM_019815387.2;refGene_gn=KIAA1468;refGene_tx=XM_019815388.2;refGene_gn=KIAA1468;refGene_tx=XM_019815389.2
D3 82358721 82358770
refGene_gn=KIAA1468;refGene_tx=XM_003995235.5;refGene_gn=KIAA1468;refGene_tx=XM_006938961.4;refGene_gn=KIAA1468;refGene_tx=XM_006938962.4;refGene_gn=KIAA1468;refGene_tx=XM_011288140.3;refGene_gn=KIAA1468;refGene_tx=XM_019815385.2;refGene_gn=KIAA1468;refGene_tx=XM_019815386.2;refGene_gn=KIAA1468;refGene_tx=XM_019815387.2;refGene_gn=KIAA1468;refGene_tx=XM_019815388.2;refGene_gn=KIAA1468;refGene_tx=XM_019815389.2
D3 82361227 82361307
refGene_gn=KIAA1468;refGene_tx=XM_003995235.5;refGene_gn=KIAA1468;refGene_tx=XM_006938961.4;refGene_gn=KIAA1468;refGene_tx=XM_006938962.4;refGene_gn=KIAA1468;refGene_tx=XM_011288140.3;refGene_gn=KIAA1468;refGene_tx=XM_019815385.2;refGene_gn=KIAA1468;refGene_tx=XM_019815386.2;refGene_gn=KIAA1468;refGene_tx=XM_019815387.2;refGene_gn=KIAA1468;refGene_tx=XM_019815388.2;refGene_gn=KIAA1468;refGene_tx=XM_019815389.2
D3 82364864 82364971
refGene_gn=KIAA1468;refGene_tx=XM_003995235.5;refGene_gn=KIAA1468;refGene_tx=XM_006938961.4;refGene_gn=KIAA1468;refGene_tx=XM_006938962.4;refGene_gn=KIAA1468;refGene_tx=XM_011288140.3;refGene_gn=KIAA1468;refGene_tx=XM_019815385.2;refGene_gn=KIAA1468;refGene_tx=XM_019815386.2;refGene_gn=KIAA1468;refGene_tx=XM_019815387.2;refGene_gn=KIAA1468;refGene_tx=XM_019815388.2;refGene_gn=KIAA1468;refGene_tx=XM_019815389.2
D3 82365912 82366012
refGene_gn=KIAA1468;refGene_tx=XM_003995235.5;refGene_gn=KIAA1468;refGene_tx=XM_006938961.4;refGene_gn=KIAA1468;refGene_tx=XM_006938962.4;refGene_gn=KIAA1468;refGene_tx=XM_011288140.3;refGene_gn=KIAA1468;refGene_tx=XM_019815385.2;refGene_gn=KIAA1468;refGene_tx=XM_019815386.2;refGene_gn=KIAA1468;refGene_tx=XM_019815387.2;refGene_gn=KIAA1468;refGene_tx=XM_019815388.2;refGene_gn=KIAA1468;refGene_tx=XM_019815389.2
D3 82370832 82370915
refGene_gn=KIAA1468;refGene_tx=XM_006938961.4;refGene_gn=KIAA1468;refGene_tx=XM_006938962.4;refGene_gn=KIAA1468;refGene_tx=XM_011288140.3;refGene_gn=KIAA1468;refGene_tx=XM_019815385.2;refGene_gn=KIAA1468;refGene_tx=XM_019815387.2;refGene_gn=KIAA1468;refGene_tx=XM_019815388.2;refGene_gn=KIAA1468;refGene_tx=XM_019815389.2
D3 82371405 82371488
refGene_gn=KIAA1468;refGene_tx=XM_003995235.5;refGene_gn=KIAA1468;refGene_tx=XM_019815386.2;refGene_gn=KIAA1468;refGene_tx=XM_019815389.2
D3 82371691 82371755
refGene_gn=KIAA1468;refGene_tx=XM_003995235.5;refGene_gn=KIAA1468;refGene_tx=XM_006938961.4;refGene_gn=KIAA1468;refGene_tx=XM_006938962.4;refGene_gn=KIAA1468;refGene_tx=XM_011288140.3;refGene_gn=KIAA1468;refGene_tx=XM_019815385.2;refGene_gn=KIAA1468;refGene_tx=XM_019815386.2;refGene_gn=KIAA1468;refGene_tx=XM_019815387.2;refGene_gn=KIAA1468;refGene_tx=XM_019815388.2
D3 82373530 82373669
refGene_gn=KIAA1468;refGene_tx=XM_003995235.5;refGene_gn=KIAA1468;refGene_tx=XM_006938961.4;refGene_gn=KIAA1468;refGene_tx=XM_006938962.4;re

fGene_gn=KIAA1468;refGene_tx=XM_011288140.3;refGene_gn=KIAA1468;refGene_tx=XM_019815385.2;refGene_gn=KIAA1468;refGene_tx=XM_019815386.2;refGene_gn=KIAA1468;refGene_tx=XM_019815387.2;refGene_gn=KIAA1468;refGene_tx=XM_019815388.2
D3 82377593 82377710
refGene_gn=KIAA1468;refGene_tx=XM_003995235.5;refGene_gn=KIAA1468;refGene_tx=XM_006938961.4;refGene_gn=KIAA1468;refGene_tx=XM_006938962.4;refGene_gn=KIAA1468;refGene_tx=XM_011288140.3;refGene_gn=KIAA1468;refGene_tx=XM_019815385.2;refGene_gn=KIAA1468;refGene_tx=XM_019815386.2;refGene_gn=KIAA1468;refGene_tx=XM_019815387.2;refGene_gn=KIAA1468;refGene_tx=XM_019815388.2
D3 82382085 82382174
refGene_gn=KIAA1468;refGene_tx=XM_003995235.5;refGene_gn=KIAA1468;refGene_tx=XM_006938961.4;refGene_gn=KIAA1468;refGene_tx=XM_006938962.4;refGene_gn=KIAA1468;refGene_tx=XM_011288140.3;refGene_gn=KIAA1468;refGene_tx=XM_019815385.2;refGene_gn=KIAA1468;refGene_tx=XM_019815386.2;refGene_gn=KIAA1468;refGene_tx=XM_019815387.2;refGene_gn=KIAA1468;refGene_tx=XM_019815388.2
D3 82385027 82385098
refGene_gn=KIAA1468;refGene_tx=XM_003995235.5;refGene_gn=KIAA1468;refGene_tx=XM_006938961.4;refGene_gn=KIAA1468;refGene_tx=XM_006938962.4;refGene_gn=KIAA1468;refGene_tx=XM_011288140.3;refGene_gn=KIAA1468;refGene_tx=XM_019815385.2;refGene_gn=KIAA1468;refGene_tx=XM_019815386.2;refGene_gn=KIAA1468;refGene_tx=XM_019815387.2;refGene_gn=KIAA1468;refGene_tx=XM_019815388.2
D3 82390377 82390498
refGene_gn=KIAA1468;refGene_tx=XM_003995235.5;refGene_gn=KIAA1468;refGene_tx=XM_006938961.4;refGene_gn=KIAA1468;refGene_tx=XM_006938962.4;refGene_gn=KIAA1468;refGene_tx=XM_011288140.3;refGene_gn=KIAA1468;refGene_tx=XM_019815385.2;refGene_gn=KIAA1468;refGene_tx=XM_019815386.2;refGene_gn=KIAA1468;refGene_tx=XM_019815387.2
D3 82395805 82395902 refGene_gn=KIAA1468;refGene_tx=XM_019815388.2
D3 82412400 82412475
refGene_gn=TNFRSF11A;refGene_tx=XM_023241204.1
D3 82435967 82436049
refGene_gn=TNFRSF11A;refGene_tx=XM_023241204.1
D3 82437282 82437408
refGene_gn=TNFRSF11A;refGene_tx=XM_023241204.1
D3 82441543 82441687
refGene_gn=TNFRSF11A;refGene_tx=XM_023241204.1
D3 82446414 82446508
refGene_gn=TNFRSF11A;refGene_tx=XM_023241204.1
D3 82447533 82447628
refGene_gn=TNFRSF11A;refGene_tx=XM_023241204.1
D3 82449239 82449353
refGene_gn=TNFRSF11A;refGene_tx=XM_023241204.1
D3 82452413 82452466
refGene_gn=TNFRSF11A;refGene_tx=XM_023241204.1
D3 82453807 82454570
refGene_gn=TNFRSF11A;refGene_tx=XM_023241204.1

D3 82467469 82467834
refGene_gn=TNFRSF11A;refGene_tx=XM_023241204.1
D3 82583581 82584538
refGene_gn=ZCCHC2;refGene_tx=XM_011288139.3;refGene_gn=ZCCHC2;refGene_tx=XM_023242067.1;refGene_gn=ZCCHC2;refGene_tx=XM_023242068.1;refGene_gn=ZCCHC2;refGene_tx=XM_023242069.1;refGene_gn=ZCCHC2;refGene_tx=XM_023242070.1
D3 82599326 82599438
refGene_gn=ZCCHC2;refGene_tx=XM_011288139.3;refGene_gn=ZCCHC2;refGene_tx=XM_023242067.1;refGene_gn=ZCCHC2;refGene_tx=XM_023242068.1;refGene_gn=ZCCHC2;refGene_tx=XM_023242069.1;refGene_gn=ZCCHC2;refGene_tx=XM_023242070.1
D3 82602251 82602328
refGene_gn=ZCCHC2;refGene_tx=XM_011288139.3;refGene_gn=ZCCHC2;refGene_tx=XM_023242067.1;refGene_gn=ZCCHC2;refGene_tx=XM_023242068.1;refGene_gn=ZCCHC2;refGene_tx=XM_023242069.1;refGene_gn=ZCCHC2;refGene_tx=XM_023242070.1
D3 82604606 82604678
refGene_gn=ZCCHC2;refGene_tx=XM_011288139.3;refGene_gn=ZCCHC2;refGene_tx=XM_023242067.1;refGene_gn=ZCCHC2;refGene_tx=XM_023242068.1;refGene_gn=ZCCHC2;refGene_tx=XM_023242069.1
D3 82608756 82608869
refGene_gn=ZCCHC2;refGene_tx=XM_011288139.3;refGene_gn=ZCCHC2;refGene_tx=XM_023242067.1;refGene_gn=ZCCHC2;refGene_tx=XM_023242068.1;refGene_gn=ZCCHC2;refGene_tx=XM_023242069.1;refGene_gn=ZCCHC2;refGene_tx=XM_023242070.1
D3 82621167 82621262
refGene_gn=ZCCHC2;refGene_tx=XM_011288139.3;refGene_gn=ZCCHC2;refGene_tx=XM_023242067.1;refGene_gn=ZCCHC2;refGene_tx=XM_023242068.1;refGene_gn=ZCCHC2;refGene_tx=XM_023242069.1;refGene_gn=ZCCHC2;refGene_tx=XM_023242070.1
D3 82623667 82623751
refGene_gn=ZCCHC2;refGene_tx=XM_011288139.3;refGene_gn=ZCCHC2;refGene_tx=XM_023242067.1;refGene_gn=ZCCHC2;refGene_tx=XM_023242068.1;refGene_gn=ZCCHC2;refGene_tx=XM_023242069.1;refGene_gn=ZCCHC2;refGene_tx=XM_023242070.1
D3 82625208 82625266
refGene_gn=ZCCHC2;refGene_tx=XM_011288139.3;refGene_gn=ZCCHC2;refGene_tx=XM_023242067.1;refGene_gn=ZCCHC2;refGene_tx=XM_023242068.1;refGene_gn=ZCCHC2;refGene_tx=XM_023242069.1;refGene_gn=ZCCHC2;refGene_tx=XM_023242070.1
D3 82627565 82627701
refGene_gn=ZCCHC2;refGene_tx=XM_011288139.3;refGene_gn=ZCCHC2;refGene_tx=XM_023242067.1;refGene_gn=ZCCHC2;refGene_tx=XM_023242068.1;refGene_gn=ZCCHC2;refGene_tx=XM_023242069.1;refGene_gn=ZCCHC2;refGene_tx=XM_023242070.1
D3 82628749 82628814
refGene_gn=ZCCHC2;refGene_tx=XM_011288139.3;refGene_gn=ZCCHC2;refGene_tx=XM_023242067.1;refGene_gn=ZCCHC2;refGene_tx=XM_023242068.1;refGene_gn=ZCCHC2;refGene_tx=XM_023242069.1;refGene_gn=ZCCHC2;refGene_tx=XM_02

3242070.1
D3 82629184 82629279
refGene_gn=ZCCHC2;refGene_tx=XM_011288139.3;refGene_gn=ZCCHC2;refGene_tx=XM_023242067.1;refGene_gn=ZCCHC2;refGene_tx=XM_023242068.1;refGene_gn=ZCCHC2;refGene_tx=XM_023242069.1;refGene_gn=ZCCHC2;refGene_tx=XM_023242070.1
D3 82634112 82634241
refGene_gn=ZCCHC2;refGene_tx=XM_023242067.1;refGene_gn=ZCCHC2;refGene_tx=XM_023242068.1;refGene_gn=ZCCHC2;refGene_tx=XM_023242069.1;refGene_gn=ZCCHC2;refGene_tx=XM_023242070.1
D3 82640693 82642184
refGene_gn=ZCCHC2;refGene_tx=XM_011288139.3;refGene_gn=ZCCHC2;refGene_tx=XM_023242067.1;refGene_gn=ZCCHC2;refGene_tx=XM_023242068.1;refGene_gn=ZCCHC2;refGene_tx=XM_023242069.1;refGene_gn=ZCCHC2;refGene_tx=XM_023242070.1
D3 82643145 82643213
refGene_gn=ZCCHC2;refGene_tx=XM_011288139.3;refGene_gn=ZCCHC2;refGene_tx=XM_023242067.1;refGene_gn=ZCCHC2;refGene_tx=XM_023242070.1
D3 82647773 82647784 refGene_gn=ZCCHC2;refGene_tx=XM_023242068.1
D3 82648682 82648693 refGene_gn=ZCCHC2;refGene_tx=XM_023242069.1
D3 82659230 82659753
refGene_gn=LOC102900632;refGene_tx=XM_023242071.1
D3 82660622 82660686
refGene_gn=LOC102900632;refGene_tx=XM_023242071.1
D3 82660861 82661150
refGene_gn=LOC102900632;refGene_tx=XM_023242071.1
D3 82751502 82753036 refGene_gn=PHLPP1;refGene_tx=XM_023242072.1
D3 82856899 82857096 refGene_gn=PHLPP1;refGene_tx=XM_023242072.1
D3 82863690 82863816 refGene_gn=PHLPP1;refGene_tx=XM_023242072.1
D3 82885273 82885440 refGene_gn=PHLPP1;refGene_tx=XM_023242072.1
D3 82911462 82911609 refGene_gn=PHLPP1;refGene_tx=XM_023242072.1
D3 82912434 82912665 refGene_gn=PHLPP1;refGene_tx=XM_023242072.1
D3 82916913 82917116 refGene_gn=PHLPP1;refGene_tx=XM_023242072.1
D3 82918921 82918982 refGene_gn=PHLPP1;refGene_tx=XM_023242072.1
D3 82921775 82921871 refGene_gn=PHLPP1;refGene_tx=XM_023242072.1
D3 82925217 82925373 refGene_gn=PHLPP1;refGene_tx=XM_023242072.1
D3 82935083 82935284 refGene_gn=PHLPP1;refGene_tx=XM_023242072.1
D3 82938769 82938932 refGene_gn=PHLPP1;refGene_tx=XM_023242072.1
D3 82949542 82949673 refGene_gn=PHLPP1;refGene_tx=XM_023242072.1
D3 82955000 82955105 refGene_gn=PHLPP1;refGene_tx=XM_023242072.1
D3 82962397 82962592 refGene_gn=PHLPP1;refGene_tx=XM_023242072.1
D3 82964884 82965113 refGene_gn=PHLPP1;refGene_tx=XM_023242072.1
D3 82966026 82967157 refGene_gn=PHLPP1;refGene_tx=XM_023242072.1
D3 83102813 83102948
refGene_gn=BCL2;refGene_tx=NM_001009340.1;refGene_gn=BCL2;refGene_tx=XM_023241191.1
D3 83275585 83276164
refGene_gn=BCL2;refGene_tx=NM_001009340.1;refGene_gn=BCL2;refGene_tx=XM_023241191.1
D3 83287757 83287877 refGene_gn=KDSR;refGene_tx=XM_023242075.1

D3 83290095 83290539
refGene_gn=KDSR;refGene_tx=XM_023242074.1;refGene_gn=KDSR;refGene_tx=XM_023242076.1
D3 83291955 83292057
refGene_gn=KDSR;refGene_tx=XM_023242074.1;refGene_gn=KDSR;refGene_tx=XM_023242075.1;refGene_gn=KDSR;refGene_tx=XM_023242076.1
D3 83295974 83296058
refGene_gn=KDSR;refGene_tx=XM_023242074.1;refGene_gn=KDSR;refGene_tx=XM_023242075.1;refGene_gn=KDSR;refGene_tx=XM_023242076.1
D3 83299767 83299851
refGene_gn=KDSR;refGene_tx=XM_023242074.1;refGene_gn=KDSR;refGene_tx=XM_023242075.1;refGene_gn=KDSR;refGene_tx=XM_023242076.1
D3 83305431 83305623
refGene_gn=KDSR;refGene_tx=XM_023242074.1;refGene_gn=KDSR;refGene_tx=XM_023242075.1;refGene_gn=KDSR;refGene_tx=XM_023242076.1
D3 83311477 83311573
refGene_gn=KDSR;refGene_tx=XM_023242074.1;refGene_gn=KDSR;refGene_tx=XM_023242075.1
D3 83311762 83311828
refGene_gn=KDSR;refGene_tx=XM_023242074.1;refGene_gn=KDSR;refGene_tx=XM_023242075.1
D3 83316163 83316220
refGene_gn=KDSR;refGene_tx=XM_023242074.1;refGene_gn=KDSR;refGene_tx=XM_023242075.1
D3 83318512 83318602
refGene_gn=KDSR;refGene_tx=XM_023242074.1;refGene_gn=KDSR;refGene_tx=XM_023242075.1
D3 83323676 83323784
refGene_gn=KDSR;refGene_tx=XM_023242074.1;refGene_gn=KDSR;refGene_tx=XM_023242075.1
D3 83343711 83343813 refGene_gn=VPS4B;refGene_tx=XM_003995236.5
D3 83345271 83345412 refGene_gn=VPS4B;refGene_tx=XM_003995236.5
D3 83348308 83348528 refGene_gn=VPS4B;refGene_tx=XM_003995236.5
D3 83350566 83350648 refGene_gn=VPS4B;refGene_tx=XM_003995236.5
D3 83351139 83351288 refGene_gn=VPS4B;refGene_tx=XM_003995236.5
D3 83351724 83351881 refGene_gn=VPS4B;refGene_tx=XM_003995236.5
D3 83355007 83355127 refGene_gn=VPS4B;refGene_tx=XM_003995236.5
D3 83358899 83358967 refGene_gn=VPS4B;refGene_tx=XM_003995236.5
D3 83361103 83361260 refGene_gn=VPS4B;refGene_tx=XM_003995236.5
D3 83362428 83362540 refGene_gn=VPS4B;refGene_tx=XM_003995236.5
D3 83374167 83374194 refGene_gn=VPS4B;refGene_tx=XM_003995236.5
D3 83423907 83424075 refGene_gn=SERPINB5;refGene_tx=XM_003995237.4
D3 83427922 83428060 refGene_gn=SERPINB5;refGene_tx=XM_003995237.4
D3 83430850 83430968 refGene_gn=SERPINB5;refGene_tx=XM_003995237.4
D3 83433494 83433637 refGene_gn=SERPINB5;refGene_tx=XM_003995237.4
D3 83436271 83436439 refGene_gn=SERPINB5;refGene_tx=XM_003995237.4
D3 83439544 83439937 refGene_gn=SERPINB5;refGene_tx=XM_003995237.4
D3 83485617 83485785
refGene_gn=SERPINB12;refGene_tx=XM_003995238.3;refGene_gn=SERPINB12;refGene_tx=XM_019815392.2

D3 83488542 83488677
refGene_gn=SERPINB12;refGene_tx=XM_003995238.3;refGene_gn=SERPINB12;re
fGene_tx=XM_019815392.2
D3 83489441 83489582
refGene_gn=SERPINB12;refGene_tx=XM_003995238.3;refGene_gn=SERPINB12;re
fGene_tx=XM_019815392.2
D3 83491328 83491446
refGene_gn=SERPINB12;refGene_tx=XM_003995238.3;refGene_gn=SERPINB12;re
fGene_tx=XM_019815392.2
D3 83493637 83493780
refGene_gn=SERPINB12;refGene_tx=XM_003995238.3;refGene_gn=SERPINB12;re
fGene_tx=XM_019815392.2
D3 83494738 83494906
refGene_gn=SERPINB12;refGene_tx=XM_003995238.3;refGene_gn=SERPINB12;re
fGene_tx=XM_019815392.2
D3 83495887 83496292
refGene_gn=SERPINB12;refGene_tx=XM_003995238.3;refGene_gn=SERPINB12;re
fGene_tx=XM_019815392.2
D3 83510838 83510920
refGene_gn=SERPINB13;refGene_tx=XM_019815393.1;refGene_gn=SERPINB13;re
fGene_tx=XM_019815395.2;refGene_gn=SERPINB13;refGene_tx=XM_019815396.1
D3 83511080 83511262
refGene_gn=SERPINB13;refGene_tx=XM_019815393.1;refGene_gn=SERPINB13;re
fGene_tx=XM_019815394.1;refGene_gn=SERPINB13;refGene_tx=XM_019815395.2
;refGene_gn=SERPINB13;refGene_tx=XM_019815396.1
D3 83511753 83511837
refGene_gn=SERPINB13;refGene_tx=XM_019815393.1;refGene_gn=SERPINB13;re
fGene_tx=XM_019815394.1;refGene_gn=SERPINB13;refGene_tx=XM_019815395.2
;refGene_gn=SERPINB13;refGene_tx=XM_019815396.1
D3 83514260 83514263
refGene_gn=SERPINB13;refGene_tx=XM_019815396.1
D3 83515745 83515874
refGene_gn=SERPINB13;refGene_tx=XM_019815393.1;refGene_gn=SERPINB13;re
fGene_tx=XM_019815394.1;refGene_gn=SERPINB13;refGene_tx=XM_019815396.1
D3 83516290 83516408
refGene_gn=SERPINB13;refGene_tx=XM_019815393.1;refGene_gn=SERPINB13;re
fGene_tx=XM_019815394.1;refGene_gn=SERPINB13;refGene_tx=XM_019815395.2
;refGene_gn=SERPINB13;refGene_tx=XM_019815396.1
D3 83517808 83517951
refGene_gn=SERPINB13;refGene_tx=XM_019815393.1;refGene_gn=SERPINB13;re
fGene_tx=XM_019815394.1;refGene_gn=SERPINB13;refGene_tx=XM_019815395.2
;refGene_gn=SERPINB13;refGene_tx=XM_019815396.1
D3 83518160 83518316
refGene_gn=SERPINB13;refGene_tx=XM_019815393.1;refGene_gn=SERPINB13;re
fGene_tx=XM_019815394.1;refGene_gn=SERPINB13;refGene_tx=XM_019815395.2
;refGene_gn=SERPINB13;refGene_tx=XM_019815396.1
D3 83520136 83520550
refGene_gn=SERPINB13;refGene_tx=XM_019815393.1;refGene_gn=SERPINB13;re
fGene_tx=XM_019815394.1;refGene_gn=SERPINB13;refGene_tx=XM_019815395.2
;refGene_gn=SERPINB13;refGene_tx=XM_019815396.1

D3 83538861 83539263 refGene_gn=SERPINB3;refGene_tx=XM_003995242.2
D3 83541125 83541281 refGene_gn=SERPINB3;refGene_tx=XM_003995242.2
D3 83541773 83541913 refGene_gn=SERPINB3;refGene_tx=XM_003995242.2
D3 83542733 83542851 refGene_gn=SERPINB3;refGene_tx=XM_003995242.2
D3 83543600 83543729 refGene_gn=SERPINB3;refGene_tx=XM_003995242.2
D3 83544961 83545018 refGene_gn=SERPINB3;refGene_tx=XM_003995242.2
D3 83545564 83545732 refGene_gn=SERPINB3;refGene_tx=XM_003995242.2
D3 83584867 83585035
refGene_gn=SERPINB11;refGene_tx=XM_003995243.5;refGene_gn=SERPINB11;refGene_tx=XM_006938963.4;refGene_gn=SERPINB11;refGene_tx=XM_023242077.1
D3 83586065 83586125
refGene_gn=SERPINB11;refGene_tx=XM_003995243.5;refGene_gn=SERPINB11;refGene_tx=XM_006938963.4;refGene_gn=SERPINB11;refGene_tx=XM_023242077.1
D3 83587296 83587425
refGene_gn=SERPINB11;refGene_tx=XM_003995243.5;refGene_gn=SERPINB11;refGene_tx=XM_006938963.4;refGene_gn=SERPINB11;refGene_tx=XM_023242077.1
D3 83590237 83590355
refGene_gn=SERPINB11;refGene_tx=XM_003995243.5;refGene_gn=SERPINB11;refGene_tx=XM_006938963.4;refGene_gn=SERPINB11;refGene_tx=XM_023242077.1
D3 83599000 83599143
refGene_gn=SERPINB11;refGene_tx=XM_003995243.5;refGene_gn=SERPINB11;refGene_tx=XM_006938963.4;refGene_gn=SERPINB11;refGene_tx=XM_023242077.1
D3 83600281 83600437
refGene_gn=SERPINB11;refGene_tx=XM_003995243.5;refGene_gn=SERPINB11;refGene_tx=XM_006938963.4;refGene_gn=SERPINB11;refGene_tx=XM_023242077.1
D3 83602545 83602950
refGene_gn=SERPINB11;refGene_tx=XM_003995243.5;refGene_gn=SERPINB11;refGene_tx=XM_006938963.4;refGene_gn=SERPINB11;refGene_tx=XM_023242077.1
D3 83611609 83611618 refGene_gn=SERPINB7;refGene_tx=XM_023242078.1
D3 83648251 83648437
refGene_gn=SERPINB7;refGene_tx=XM_003995244.4;refGene_gn=SERPINB7;refGene_tx=XM_019815398.2;refGene_gn=SERPINB7;refGene_tx=XM_019815399.2;refGene_gn=SERPINB7;refGene_tx=XM_023242078.1
D3 83654428 83654479
refGene_gn=SERPINB7;refGene_tx=XM_003995244.4;refGene_gn=SERPINB7;refGene_tx=XM_019815398.2;refGene_gn=SERPINB7;refGene_tx=XM_019815399.2;refGene_gn=SERPINB7;refGene_tx=XM_023242078.1
D3 83655050 83655167
refGene_gn=SERPINB7;refGene_tx=XM_003995244.4;refGene_gn=SERPINB7;refGene_tx=XM_019815398.2;refGene_gn=SERPINB7;refGene_tx=XM_019815399.2;refGene_gn=SERPINB7;refGene_tx=XM_023242078.1
D3 83660315 83660433
refGene_gn=SERPINB7;refGene_tx=XM_003995244.4;refGene_gn=SERPINB7;refGene_tx=XM_019815398.2;refGene_gn=SERPINB7;refGene_tx=XM_019815399.2;refGene_gn=SERPINB7;refGene_tx=XM_023242078.1
D3 83663998 83664141
refGene_gn=SERPINB7;refGene_tx=XM_003995244.4;refGene_gn=SERPINB7;refGene_tx=XM_019815398.2;refGene_gn=SERPINB7;refGene_tx=XM_019815399.2;refGene_gn=SERPINB7;refGene_tx=XM_023242078.1
D3 83665461 83665608

refGene_gn=SERPINB7;refGene_tx=XM_003995244.4;refGene_gn=SERPINB7;refGene_tx=XM_019815398.2;refGene_gn=SERPINB7;refGene_tx=XM_019815399.2;refGene_gn=SERPINB7;refGene_tx=XM_023242078.1
D3 83668325 83668724
refGene_gn=SERPINB7;refGene_tx=XM_003995244.4;refGene_gn=SERPINB7;refGene_tx=XM_019815398.2;refGene_gn=SERPINB7;refGene_tx=XM_019815399.2;refGene_gn=SERPINB7;refGene_tx=XM_023242078.1
D3 83726900 83727068 refGene_gn=SERPINB2;refGene_tx=XM_003995246.3
D3 83731110 83731230 refGene_gn=SERPINB2;refGene_tx=XM_003995246.3
D3 83732531 83732660 refGene_gn=SERPINB2;refGene_tx=XM_003995246.3
D3 83733168 83733286 refGene_gn=SERPINB2;refGene_tx=XM_003995246.3
D3 83735693 83735836 refGene_gn=SERPINB2;refGene_tx=XM_003995246.3
D3 83736707 83736872 refGene_gn=SERPINB2;refGene_tx=XM_003995246.3
D3 83737452 83737857 refGene_gn=SERPINB2;refGene_tx=XM_003995246.3
D3 83746674 83746842
refGene_gn=SERPINB10;refGene_tx=XM_003995247.3
D3 83748918 83748981
refGene_gn=SERPINB10;refGene_tx=XM_003995247.3
D3 83749669 83749807
refGene_gn=SERPINB10;refGene_tx=XM_003995247.3
D3 83751540 83751658
refGene_gn=SERPINB10;refGene_tx=XM_003995247.3
D3 83760516 83760659
refGene_gn=SERPINB10;refGene_tx=XM_003995247.3
D3 83763511 83763667
refGene_gn=SERPINB10;refGene_tx=XM_003995247.3
D3 83765679 83766084
refGene_gn=SERPINB10;refGene_tx=XM_003995247.3
D3 83780113 83780281
refGene_gn=LOC101085929;refGene_tx=XM_023242080.1;refGene_gn=LOC101085929;refGene_tx=XM_023242081.1
D3 83781782 83781920
refGene_gn=LOC101085929;refGene_tx=XM_023242080.1;refGene_gn=LOC101085929;refGene_tx=XM_023242081.1
D3 83782935 83782995
refGene_gn=LOC101085929;refGene_tx=XM_019815401.1
D3 83783666 83783784
refGene_gn=LOC101085929;refGene_tx=XM_019815401.1;refGene_gn=LOC101085929;refGene_tx=XM_023242080.1;refGene_gn=LOC101085929;refGene_tx=XM_023242081.1
D3 83785361 83785504
refGene_gn=LOC101085929;refGene_tx=XM_019815401.1;refGene_gn=LOC101085929;refGene_tx=XM_019815402.1;refGene_gn=LOC101085929;refGene_tx=XM_023242080.1;refGene_gn=LOC101085929;refGene_tx=XM_023242081.1
D3 83787105 83787258
refGene_gn=LOC101085929;refGene_tx=XM_019815401.1;refGene_gn=LOC101085929;refGene_tx=XM_019815402.1;refGene_gn=LOC101085929;refGene_tx=XM_023242080.1;refGene_gn=LOC101085929;refGene_tx=XM_023242081.1
D3 83789030 83789435
refGene_gn=LOC101085929;refGene_tx=XM_019815401.1;refGene_gn=LOC101085

929;refGene_tx=XM_019815402.1;refGene_gn=LOC101085929;refGene_tx=XM_023242080.1;refGene_gn=LOC101085929;refGene_tx=XM_023242081.1
D3 85195946 85196003 refGene_gn=CDH7;refGene_tx=XM_023242084.1
D3 85207901 85208111 refGene_gn=CDH7;refGene_tx=XM_003995250.4;refGene_gn=CDH7;refGene_tx=XM_019815403.2;refGene_gn=CDH7;refGene_tx=XM_023242082.1;refGene_gn=CDH7;refGene_tx=XM_023242083.1
D3 85240911 85240995 refGene_gn=CDH7;refGene_tx=XM_019815404.2
D3 85251894 85252189 refGene_gn=CDH7;refGene_tx=XM_003995250.4;refGene_gn=CDH7;refGene_tx=XM_019815403.2;refGene_gn=CDH7;refGene_tx=XM_019815404.2;refGene_gn=CDH7;refGene_tx=XM_023242082.1;refGene_gn=CDH7;refGene_tx=XM_023242083.1;refGene_gn=CDH7;refGene_tx=XM_023242084.1
D3 85255985 85256105 refGene_gn=CDH7;refGene_tx=XM_003995250.4;refGene_gn=CDH7;refGene_tx=XM_019815403.2;refGene_gn=CDH7;refGene_tx=XM_019815404.2;refGene_gn=CDH7;refGene_tx=XM_023242082.1;refGene_gn=CDH7;refGene_tx=XM_023242083.1;refGene_gn=CDH7;refGene_tx=XM_023242084.1
D3 85263306 85263474 refGene_gn=CDH7;refGene_tx=XM_003995250.4;refGene_gn=CDH7;refGene_tx=XM_019815403.2;refGene_gn=CDH7;refGene_tx=XM_019815404.2;refGene_gn=CDH7;refGene_tx=XM_023242082.1;refGene_gn=CDH7;refGene_tx=XM_023242083.1;refGene_gn=CDH7;refGene_tx=XM_023242084.1
D3 85266103 85266291 refGene_gn=CDH7;refGene_tx=XM_003995250.4;refGene_gn=CDH7;refGene_tx=XM_019815403.2;refGene_gn=CDH7;refGene_tx=XM_019815404.2;refGene_gn=CDH7;refGene_tx=XM_023242082.1;refGene_gn=CDH7;refGene_tx=XM_023242083.1;refGene_gn=CDH7;refGene_tx=XM_023242084.1
D3 85281749 85282003 refGene_gn=CDH7;refGene_tx=XM_003995250.4;refGene_gn=CDH7;refGene_tx=XM_019815403.2;refGene_gn=CDH7;refGene_tx=XM_019815404.2;refGene_gn=CDH7;refGene_tx=XM_023242082.1;refGene_gn=CDH7;refGene_tx=XM_023242083.1;refGene_gn=CDH7;refGene_tx=XM_023242084.1
D3 85294370 85294507 refGene_gn=CDH7;refGene_tx=XM_003995250.4;refGene_gn=CDH7;refGene_tx=XM_019815403.2;refGene_gn=CDH7;refGene_tx=XM_019815404.2;refGene_gn=CDH7;refGene_tx=XM_023242082.1;refGene_gn=CDH7;refGene_tx=XM_023242083.1;refGene_gn=CDH7;refGene_tx=XM_023242084.1
D3 85295315 85295437 refGene_gn=CDH7;refGene_tx=XM_003995250.4;refGene_gn=CDH7;refGene_tx=XM_019815403.2;refGene_gn=CDH7;refGene_tx=XM_019815404.2;refGene_gn=CDH7;refGene_tx=XM_023242082.1;refGene_gn=CDH7;refGene_tx=XM_023242083.1;refGene_gn=CDH7;refGene_tx=XM_023242084.1
D3 85296091 85296209 refGene_gn=CDH7;refGene_tx=XM_003995250.4;refGene_gn=CDH7;refGene_tx=XM_019815403.2;refGene_gn=CDH7;refGene_tx=XM_019815404.2;refGene_gn=CDH7;refGene_tx=XM_023242082.1;refGene_gn=CDH7;refGene_tx=XM_023242083.1;refGene_gn=CDH7;refGene_tx=XM_023242084.1
D3 85299284 85299536 refGene_gn=CDH7;refGene_tx=XM_003995250.4;refGene_gn=CDH7;refGene_tx=X

M_019815403.2;refGene_gn=CDH7;refGene_tx=XM_019815404.2;refGene_gn=CDH7;refGene_tx=XM_023242082.1;refGene_gn=CDH7;refGene_tx=XM_023242083.1;refGene_gn=CDH7;refGene_tx=XM_023242084.1
D3 85316221 85316715 refGene_gn=CDH7;refGene_tx=XM_003995250.4;refGene_gn=CDH7;refGene_tx=XM_019815403.2;refGene_gn=CDH7;refGene_tx=XM_019815404.2;refGene_gn=CDH7;refGene_tx=XM_023242082.1;refGene_gn=CDH7;refGene_tx=XM_023242083.1;refGene_gn=CDH7;refGene_tx=XM_023242084.1
D3 85840968 85841459 refGene_gn=CDH19;refGene_tx=XM_023242085.1;refGene_gn=CDH19;refGene_tx=XM_023242086.1
D3 85844086 85844338 refGene_gn=CDH19;refGene_tx=XM_023242085.1;refGene_gn=CDH19;refGene_tx=XM_023242086.1;refGene_gn=CDH19;refGene_tx=XM_023242087.1
D3 85846316 85846434 refGene_gn=CDH19;refGene_tx=XM_023242085.1;refGene_gn=CDH19;refGene_tx=XM_023242086.1;refGene_gn=CDH19;refGene_tx=XM_023242087.1
D3 85868089 85868134 refGene_gn=CDH19;refGene_tx=XM_023242088.1
D3 85868339 85868461 refGene_gn=CDH19;refGene_tx=XM_023242085.1;refGene_gn=CDH19;refGene_tx=XM_023242086.1;refGene_gn=CDH19;refGene_tx=XM_023242087.1;refGene_gn=CDH19;refGene_tx=XM_023242088.1
D3 85877370 85877492 refGene_gn=CDH19;refGene_tx=XM_023242085.1;refGene_gn=CDH19;refGene_tx=XM_023242086.1;refGene_gn=CDH19;refGene_tx=XM_023242087.1;refGene_gn=CDH19;refGene_tx=XM_023242088.1
D3 85882829 85883083 refGene_gn=CDH19;refGene_tx=XM_023242085.1;refGene_gn=CDH19;refGene_tx=XM_023242086.1;refGene_gn=CDH19;refGene_tx=XM_023242087.1;refGene_gn=CDH19;refGene_tx=XM_023242088.1
D3 85883622 85883804 refGene_gn=CDH19;refGene_tx=XM_023242085.1;refGene_gn=CDH19;refGene_tx=XM_023242086.1;refGene_gn=CDH19;refGene_tx=XM_023242087.1;refGene_gn=CDH19;refGene_tx=XM_023242088.1
D3 85891612 85891777 refGene_gn=CDH19;refGene_tx=XM_023242085.1;refGene_gn=CDH19;refGene_tx=XM_023242086.1;refGene_gn=CDH19;refGene_tx=XM_023242087.1;refGene_gn=CDH19;refGene_tx=XM_023242088.1
D3 85893154 85893274 refGene_gn=CDH19;refGene_tx=XM_023242085.1;refGene_gn=CDH19;refGene_tx=XM_023242086.1;refGene_gn=CDH19;refGene_tx=XM_023242087.1;refGene_gn=CDH19;refGene_tx=XM_023242088.1
D3 85908669 85908964 refGene_gn=CDH19;refGene_tx=XM_023242085.1;refGene_gn=CDH19;refGene_tx=XM_023242086.1;refGene_gn=CDH19;refGene_tx=XM_023242087.1;refGene_gn=CDH19;refGene_tx=XM_023242088.1
D3 85912518 85912713 refGene_gn=CDH19;refGene_tx=XM_023242085.1;refGene_gn=CDH19;refGene_tx=XM_023242086.1;refGene_gn=CDH19;refGene_tx=XM_023242087.1;refGene_gn=CDH19;refGene_tx=XM_023242088.1

D3 86691588 86695227
refGene_gn=DSEL;refGene_tx=XM_003995252.5;refGene_gn=DSEL;refGene_tx=XM_019815405.2;refGene_gn=DSEL;refGene_tx=XM_019815406.2
D3 87598439 87600144
refGene_gn=LOC111557345;refGene_tx=XM_023242089.1
D3 87600146 87600374
refGene_gn=LOC111557345;refGene_tx=XM_023242089.1
D3 87600376 87601767
refGene_gn=LOC111557345;refGene_tx=XM_023242089.1
D3 87601768 87602337
refGene_gn=LOC111557345;refGene_tx=XM_023242089.1
D3 87602384 87603299
refGene_gn=LOC111557345;refGene_tx=XM_023242089.1
D3 87603300 87603655
refGene_gn=LOC111557345;refGene_tx=XM_023242089.1
D3 87616824 87617082
refGene_gn=TMX3;refGene_tx=XM_003995253.4;refGene_gn=TMX3;refGene_tx=XM_023242091.1;refGene_gn=TMX3;refGene_tx=XM_023242092.1
D3 87618676 87618745
refGene_gn=TMX3;refGene_tx=XM_003995253.4;refGene_gn=TMX3;refGene_tx=XM_023242092.1
D3 87620112 87620242
refGene_gn=TMX3;refGene_tx=XM_003995253.4;refGene_gn=TMX3;refGene_tx=XM_023242091.1;refGene_gn=TMX3;refGene_tx=XM_023242092.1
D3 87624840 87624897
refGene_gn=TMX3;refGene_tx=XM_003995253.4;refGene_gn=TMX3;refGene_tx=XM_023242091.1;refGene_gn=TMX3;refGene_tx=XM_023242092.1
D3 87625737 87625791
refGene_gn=TMX3;refGene_tx=XM_003995253.4;refGene_gn=TMX3;refGene_tx=XM_023242091.1;refGene_gn=TMX3;refGene_tx=XM_023242092.1
D3 87625955 87626013
refGene_gn=TMX3;refGene_tx=XM_003995253.4;refGene_gn=TMX3;refGene_tx=XM_023242091.1;refGene_gn=TMX3;refGene_tx=XM_023242092.1
D3 87628545 87628644
refGene_gn=TMX3;refGene_tx=XM_003995253.4;refGene_gn=TMX3;refGene_tx=XM_023242091.1;refGene_gn=TMX3;refGene_tx=XM_023242092.1
D3 87632914 87632981
refGene_gn=TMX3;refGene_tx=XM_003995253.4;refGene_gn=TMX3;refGene_tx=XM_023242091.1;refGene_gn=TMX3;refGene_tx=XM_023242092.1
D3 87638415 87638493
refGene_gn=TMX3;refGene_tx=XM_003995253.4;refGene_gn=TMX3;refGene_tx=XM_023242091.1;refGene_gn=TMX3;refGene_tx=XM_023242092.1
D3 87639129 87639229
refGene_gn=TMX3;refGene_tx=XM_003995253.4;refGene_gn=TMX3;refGene_tx=XM_023242091.1;refGene_gn=TMX3;refGene_tx=XM_023242092.1
D3 87641407 87641488
refGene_gn=TMX3;refGene_tx=XM_003995253.4;refGene_gn=TMX3;refGene_tx=XM_023242091.1;refGene_gn=TMX3;refGene_tx=XM_023242092.1
D3 87642849 87642895
refGene_gn=TMX3;refGene_tx=XM_003995253.4;refGene_gn=TMX3;refGene_tx=X

M_023242091.1;refGene_gn=TMX3;refGene_tx=XM_023242092.1
D3 87649929 87650053
refGene_gn=TMX3;refGene_tx=XM_003995253.4;refGene_gn=TMX3;refGene_tx=XM_023242091.1;refGene_gn=TMX3;refGene_tx=XM_023242092.1
D3 87651406 87651446
refGene_gn=TMX3;refGene_tx=XM_003995253.4;refGene_gn=TMX3;refGene_tx=XM_023242091.1
D3 87658235 87658290
refGene_gn=TMX3;refGene_tx=XM_003995253.4;refGene_gn=TMX3;refGene_tx=XM_023242091.1
D3 87659332 87659582
refGene_gn=TMX3;refGene_tx=XM_003995253.4;refGene_gn=TMX3;refGene_tx=XM_023242091.1
D3 87769980 87770580
refGene_gn=CCDC102B;refGene_tx=XM_019815409.2;refGene_gn=CCDC102B;refGene_tx=XM_019815410.2;refGene_gn=CCDC102B;refGene_tx=XM_019815412.2;refGene_gn=CCDC102B;refGene_tx=XM_019815415.2;refGene_gn=CCDC102B;refGene_tx=XM_019815417.2;refGene_gn=CCDC102B;refGene_tx=XM_023242090.1
D3 87771114 87771335
refGene_gn=CCDC102B;refGene_tx=XM_019815409.2;refGene_gn=CCDC102B;refGene_tx=XM_019815410.2;refGene_gn=CCDC102B;refGene_tx=XM_019815412.2;refGene_gn=CCDC102B;refGene_tx=XM_019815415.2;refGene_gn=CCDC102B;refGene_tx=XM_019815417.2;refGene_gn=CCDC102B;refGene_tx=XM_023242090.1
D3 87782802 87782911
refGene_gn=CCDC102B;refGene_tx=XM_019815409.2;refGene_gn=CCDC102B;refGene_tx=XM_019815410.2;refGene_gn=CCDC102B;refGene_tx=XM_019815412.2;refGene_gn=CCDC102B;refGene_tx=XM_019815415.2;refGene_gn=CCDC102B;refGene_tx=XM_019815417.2;refGene_gn=CCDC102B;refGene_tx=XM_023242090.1
D3 87814182 87814299
refGene_gn=CCDC102B;refGene_tx=XM_019815409.2;refGene_gn=CCDC102B;refGene_tx=XM_019815410.2;refGene_gn=CCDC102B;refGene_tx=XM_019815412.2;refGene_gn=CCDC102B;refGene_tx=XM_019815415.2;refGene_gn=CCDC102B;refGene_tx=XM_019815417.2;refGene_gn=CCDC102B;refGene_tx=XM_023242090.1
D3 87853205 87853415
refGene_gn=CCDC102B;refGene_tx=XM_019815409.2;refGene_gn=CCDC102B;refGene_tx=XM_019815410.2;refGene_gn=CCDC102B;refGene_tx=XM_019815412.2;refGene_gn=CCDC102B;refGene_tx=XM_019815415.2;refGene_gn=CCDC102B;refGene_tx=XM_019815417.2;refGene_gn=CCDC102B;refGene_tx=XM_023242090.1
D3 87857282 87857585
refGene_gn=CCDC102B;refGene_tx=XM_019815409.2;refGene_gn=CCDC102B;refGene_tx=XM_019815410.2;refGene_gn=CCDC102B;refGene_tx=XM_019815412.2;refGene_gn=CCDC102B;refGene_tx=XM_019815415.2;refGene_gn=CCDC102B;refGene_tx=XM_019815417.2
D3 87865388 87865427 refGene_gn=CCDC102B;refGene_tx=XM_023242090.1
D3 88280620 88280686 refGene_gn=DOK6;refGene_tx=XM_023242093.1
D3 88397681 88397789 refGene_gn=DOK6;refGene_tx=XM_023242093.1
D3 88422152 88422267 refGene_gn=DOK6;refGene_tx=XM_023242093.1
D3 88489667 88489787 refGene_gn=DOK6;refGene_tx=XM_023242093.1
D3 88510685 88510875 refGene_gn=DOK6;refGene_tx=XM_023242093.1
D3 88568902 88569041 refGene_gn=DOK6;refGene_tx=XM_023242093.1

D3 88582728 88582846 refGene_gn=DOK6;refGene_tx=XM_023242093.1
D3 88644211 88644351 refGene_gn=DOK6;refGene_tx=XM_023242093.1
D3 88661296 88661410 refGene_gn=CD226;refGene_tx=XM_023242094.1
D3 88663680 88663741 refGene_gn=CD226;refGene_tx=XM_023242094.1
D3 88668295 88668398 refGene_gn=CD226;refGene_tx=XM_023242094.1
D3 88689339 88689687 refGene_gn=CD226;refGene_tx=XM_023242094.1
D3 88734218 88734557 refGene_gn=CD226;refGene_tx=XM_023242094.1
D3 88734867 88734913 refGene_gn=CD226;refGene_tx=XM_023242094.1
D3 88778805 88778891
refGene_gn=RTTN;refGene_tx=XM_011288133.3;refGene_gn=RTTN;refGene_tx=XM_019815422.2;refGene_gn=RTTN;refGene_tx=XM_019815424.2
D3 88779810 88779880
refGene_gn=RTTN;refGene_tx=XM_011288133.3;refGene_gn=RTTN;refGene_tx=XM_019815424.2
D3 88781000 88781104
refGene_gn=RTTN;refGene_tx=XM_011288133.3;refGene_gn=RTTN;refGene_tx=XM_019815422.2;refGene_gn=RTTN;refGene_tx=XM_019815424.2
D3 88788104 88788372
refGene_gn=RTTN;refGene_tx=XM_011288133.3;refGene_gn=RTTN;refGene_tx=XM_019815422.2;refGene_gn=RTTN;refGene_tx=XM_019815424.2;refGene_gn=RTTN;refGene_tx=XM_023242095.1
D3 88791548 88791751
refGene_gn=RTTN;refGene_tx=XM_011288133.3;refGene_gn=RTTN;refGene_tx=XM_019815422.2;refGene_gn=RTTN;refGene_tx=XM_019815424.2
D3 88794545 88794672
refGene_gn=RTTN;refGene_tx=XM_011288133.3;refGene_gn=RTTN;refGene_tx=XM_019815422.2;refGene_gn=RTTN;refGene_tx=XM_019815424.2;refGene_gn=RTTN;refGene_tx=XM_023242095.1
D3 88798347 88798425
refGene_gn=RTTN;refGene_tx=XM_011288133.3;refGene_gn=RTTN;refGene_tx=XM_019815422.2;refGene_gn=RTTN;refGene_tx=XM_019815424.2;refGene_gn=RTTN;refGene_tx=XM_023242095.1
D3 88799692 88799790
refGene_gn=RTTN;refGene_tx=XM_011288133.3;refGene_gn=RTTN;refGene_tx=XM_019815422.2;refGene_gn=RTTN;refGene_tx=XM_019815424.2;refGene_gn=RTTN;refGene_tx=XM_023242095.1
D3 88800795 88800901
refGene_gn=RTTN;refGene_tx=XM_011288133.3;refGene_gn=RTTN;refGene_tx=XM_019815422.2;refGene_gn=RTTN;refGene_tx=XM_019815424.2;refGene_gn=RTTN;refGene_tx=XM_023242095.1
D3 88810418 88810636
refGene_gn=RTTN;refGene_tx=XM_011288133.3;refGene_gn=RTTN;refGene_tx=XM_019815422.2;refGene_gn=RTTN;refGene_tx=XM_019815424.2;refGene_gn=RTTN;refGene_tx=XM_023242095.1
D3 88813787 88813925
refGene_gn=RTTN;refGene_tx=XM_011288133.3;refGene_gn=RTTN;refGene_tx=XM_019815422.2;refGene_gn=RTTN;refGene_tx=XM_019815424.2;refGene_gn=RTTN;refGene_tx=XM_023242095.1
D3 88816546 88816700
refGene_gn=RTTN;refGene_tx=XM_011288133.3;refGene_gn=RTTN;refGene_tx=X

M_019815422.2;refGene_gn=RTTN;refGene_tx=XM_019815424.2;refGene_gn=RTTN;refGene_tx=XM_023242095.1
D3 88819202 88819293 refGene_gn=RTTN;refGene_tx=XM_011288133.3;refGene_gn=RTTN;refGene_tx=XM_019815422.2;refGene_gn=RTTN;refGene_tx=XM_019815424.2;refGene_gn=RTTN;refGene_tx=XM_023242095.1
D3 88821696 88821892 refGene_gn=RTTN;refGene_tx=XM_011288133.3;refGene_gn=RTTN;refGene_tx=XM_019815422.2;refGene_gn=RTTN;refGene_tx=XM_019815424.2;refGene_gn=RTTN;refGene_tx=XM_023242095.1
D3 88826399 88826493 refGene_gn=RTTN;refGene_tx=XM_011288133.3;refGene_gn=RTTN;refGene_tx=XM_019815422.2;refGene_gn=RTTN;refGene_tx=XM_019815424.2;refGene_gn=RTTN;refGene_tx=XM_023242095.1
D3 88835325 88835417 refGene_gn=RTTN;refGene_tx=XM_011288133.3;refGene_gn=RTTN;refGene_tx=XM_019815422.2;refGene_gn=RTTN;refGene_tx=XM_019815424.2;refGene_gn=RTTN;refGene_tx=XM_023242095.1
D3 88836870 88837060 refGene_gn=RTTN;refGene_tx=XM_011288133.3;refGene_gn=RTTN;refGene_tx=XM_019815422.2;refGene_gn=RTTN;refGene_tx=XM_019815424.2;refGene_gn=RTTN;refGene_tx=XM_023242095.1
D3 88849291 88849363 refGene_gn=RTTN;refGene_tx=XM_011288133.3;refGene_gn=RTTN;refGene_tx=XM_019815422.2;refGene_gn=RTTN;refGene_tx=XM_019815424.2;refGene_gn=RTTN;refGene_tx=XM_023242095.1
D3 88850893 88851052 refGene_gn=RTTN;refGene_tx=XM_011288133.3;refGene_gn=RTTN;refGene_tx=XM_019815422.2;refGene_gn=RTTN;refGene_tx=XM_019815424.2;refGene_gn=RTTN;refGene_tx=XM_023242095.1
D3 88852834 88852945 refGene_gn=RTTN;refGene_tx=XM_011288133.3;refGene_gn=RTTN;refGene_tx=XM_019815422.2;refGene_gn=RTTN;refGene_tx=XM_019815424.2;refGene_gn=RTTN;refGene_tx=XM_023242095.1
D3 88853411 88853540 refGene_gn=RTTN;refGene_tx=XM_011288133.3;refGene_gn=RTTN;refGene_tx=XM_019815422.2;refGene_gn=RTTN;refGene_tx=XM_019815424.2;refGene_gn=RTTN;refGene_tx=XM_023242095.1
D3 88861106 88861326 refGene_gn=RTTN;refGene_tx=XM_011288133.3;refGene_gn=RTTN;refGene_tx=XM_019815422.2;refGene_gn=RTTN;refGene_tx=XM_019815424.2;refGene_gn=RTTN;refGene_tx=XM_023242095.1
D3 88865719 88865874 refGene_gn=RTTN;refGene_tx=XM_011288133.3;refGene_gn=RTTN;refGene_tx=XM_019815422.2;refGene_gn=RTTN;refGene_tx=XM_019815424.2;refGene_gn=RTTN;refGene_tx=XM_023242095.1
D3 88875250 88875395 refGene_gn=RTTN;refGene_tx=XM_011288133.3;refGene_gn=RTTN;refGene_tx=XM_019815422.2;refGene_gn=RTTN;refGene_tx=XM_019815424.2;refGene_gn=RTTN;refGene_tx=XM_023242095.1

D3 88876589 88876829
refGene_gn=RTTN;refGene_tx=XM_011288133.3;refGene_gn=RTTN;refGene_tx=XM_019815422.2;refGene_gn=RTTN;refGene_tx=XM_019815424.2;refGene_gn=RTTN;refGene_tx=XM_023242095.1
D3 88877959 88878148
refGene_gn=RTTN;refGene_tx=XM_011288133.3;refGene_gn=RTTN;refGene_tx=XM_019815422.2;refGene_gn=RTTN;refGene_tx=XM_019815424.2;refGene_gn=RTTN;refGene_tx=XM_023242095.1
D3 88880775 88880844
refGene_gn=RTTN;refGene_tx=XM_011288133.3;refGene_gn=RTTN;refGene_tx=XM_019815422.2;refGene_gn=RTTN;refGene_tx=XM_019815424.2;refGene_gn=RTTN;refGene_tx=XM_023242095.1
D3 88881872 88881969
refGene_gn=RTTN;refGene_tx=XM_011288133.3;refGene_gn=RTTN;refGene_tx=XM_019815422.2;refGene_gn=RTTN;refGene_tx=XM_019815424.2;refGene_gn=RTTN;refGene_tx=XM_023242095.1
D3 88886876 88886994
refGene_gn=RTTN;refGene_tx=XM_011288133.3;refGene_gn=RTTN;refGene_tx=XM_019815422.2;refGene_gn=RTTN;refGene_tx=XM_019815424.2;refGene_gn=RTTN;refGene_tx=XM_023242095.1
D3 88887588 88887677
refGene_gn=RTTN;refGene_tx=XM_011288133.3;refGene_gn=RTTN;refGene_tx=XM_019815422.2;refGene_gn=RTTN;refGene_tx=XM_019815424.2;refGene_gn=RTTN;refGene_tx=XM_023242095.1
D3 88890206 88890306
refGene_gn=RTTN;refGene_tx=XM_011288133.3;refGene_gn=RTTN;refGene_tx=XM_019815422.2;refGene_gn=RTTN;refGene_tx=XM_019815424.2;refGene_gn=RTTN;refGene_tx=XM_023242095.1
D3 88891927 88892099
refGene_gn=RTTN;refGene_tx=XM_011288133.3;refGene_gn=RTTN;refGene_tx=XM_019815422.2;refGene_gn=RTTN;refGene_tx=XM_023242095.1
D3 88895013 88895153
refGene_gn=RTTN;refGene_tx=XM_011288133.3;refGene_gn=RTTN;refGene_tx=XM_019815422.2;refGene_gn=RTTN;refGene_tx=XM_023242095.1
D3 88895964 88896081
refGene_gn=RTTN;refGene_tx=XM_011288133.3;refGene_gn=RTTN;refGene_tx=XM_019815422.2;refGene_gn=RTTN;refGene_tx=XM_023242095.1
D3 88896639 88896765
refGene_gn=RTTN;refGene_tx=XM_011288133.3;refGene_gn=RTTN;refGene_tx=XM_019815422.2;refGene_gn=RTTN;refGene_tx=XM_023242095.1
D3 88905180 88905307
refGene_gn=RTTN;refGene_tx=XM_011288133.3;refGene_gn=RTTN;refGene_tx=XM_019815422.2;refGene_gn=RTTN;refGene_tx=XM_023242095.1
D3 88907499 88907612
refGene_gn=RTTN;refGene_tx=XM_011288133.3;refGene_gn=RTTN;refGene_tx=XM_019815422.2;refGene_gn=RTTN;refGene_tx=XM_023242095.1
D3 88908818 88909031
refGene_gn=RTTN;refGene_tx=XM_011288133.3;refGene_gn=RTTN;refGene_tx=XM_019815422.2;refGene_gn=RTTN;refGene_tx=XM_023242095.1
D3 88914149 88914320

refGene_gn=RTTN;refGene_tx=XM_011288133.3;refGene_gn=RTTN;refGene_tx=XM_019815422.2;refGene_gn=RTTN;refGene_tx=XM_023242095.1
D3 88925549 88925665
refGene_gn=RTTN;refGene_tx=XM_011288133.3;refGene_gn=RTTN;refGene_tx=XM_019815422.2;refGene_gn=RTTN;refGene_tx=XM_023242095.1
D3 88927886 88928068
refGene_gn=RTTN;refGene_tx=XM_011288133.3;refGene_gn=RTTN;refGene_tx=XM_019815422.2;refGene_gn=RTTN;refGene_tx=XM_023242095.1
D3 88930710 88930876
refGene_gn=RTTN;refGene_tx=XM_011288133.3;refGene_gn=RTTN;refGene_tx=XM_019815422.2;refGene_gn=RTTN;refGene_tx=XM_023242095.1
D3 88933879 88934030
refGene_gn=RTTN;refGene_tx=XM_011288133.3;refGene_gn=RTTN;refGene_tx=XM_019815422.2;refGene_gn=RTTN;refGene_tx=XM_023242095.1
D3 88934951 88935066
refGene_gn=RTTN;refGene_tx=XM_011288133.3;refGene_gn=RTTN;refGene_tx=XM_019815422.2;refGene_gn=RTTN;refGene_tx=XM_023242095.1
D3 88936221 88936312
refGene_gn=RTTN;refGene_tx=XM_011288133.3;refGene_gn=RTTN;refGene_tx=XM_019815422.2;refGene_gn=RTTN;refGene_tx=XM_023242095.1
D3 88938388 88938478
refGene_gn=RTTN;refGene_tx=XM_011288133.3;refGene_gn=RTTN;refGene_tx=XM_019815422.2;refGene_gn=RTTN;refGene_tx=XM_023242095.1
D3 88940116 88940294
refGene_gn=RTTN;refGene_tx=XM_011288133.3;refGene_gn=RTTN;refGene_tx=XM_019815422.2;refGene_gn=RTTN;refGene_tx=XM_023242095.1
D3 88941353 88941541
refGene_gn=RTTN;refGene_tx=XM_011288133.3;refGene_gn=RTTN;refGene_tx=XM_019815422.2;refGene_gn=RTTN;refGene_tx=XM_023242095.1
D3 88941897 88941928
refGene_gn=RTTN;refGene_tx=XM_011288133.3;refGene_gn=RTTN;refGene_tx=XM_019815422.2;refGene_gn=RTTN;refGene_tx=XM_023242095.1
D3 89032540 89034148
refGene_gn=SOCS6;refGene_tx=XM_023242096.1;refGene_gn=SOCS6;refGene_tx=XM_023242097.1;refGene_gn=SOCS6;refGene_tx=XM_023242098.1;refGene_gn=SOCS6;refGene_tx=XM_023242099.1
D3 90762265 90762463
refGene_gn=CBLN2;refGene_tx=XM_019815426.2;refGene_gn=CBLN2;refGene_tx=XM_023242100.1;refGene_gn=CBLN2;refGene_tx=XM_023242101.1;refGene_gn=CBLN2;refGene_tx=XM_023242102.1
D3 90762727 90762847
refGene_gn=CBLN2;refGene_tx=XM_019815426.2;refGene_gn=CBLN2;refGene_tx=XM_023242100.1;refGene_gn=CBLN2;refGene_tx=XM_023242101.1;refGene_gn=CBLN2;refGene_tx=XM_023242102.1
D3 90765765 90766290
refGene_gn=CBLN2;refGene_tx=XM_019815426.2;refGene_gn=CBLN2;refGene_tx=XM_023242100.1;refGene_gn=CBLN2;refGene_tx=XM_023242101.1;refGene_gn=CBLN2;refGene_tx=XM_023242102.1
D3 90767488 90767497
refGene_gn=CBLN2;refGene_tx=XM_023242100.1;refGene_gn=CBLN2;refGene_tx

=XM_023242101.1;refGene_gn=CBLN2;refGene_tx=XM_023242102.1
D3 90929736 90929797
refGene_gn=NETO1;refGene_tx=XM_011288131.3;refGene_gn=NETO1;refGene_tx=XM_019815427.2;refGene_gn=NETO1;refGene_tx=XM_019815428.2;refGene_gn=NETO1;refGene_tx=XM_019815429.2;refGene_gn=NETO1;refGene_tx=XM_023242104.1;refGene_gn=NETO1;refGene_tx=XM_023242105.1
D3 90930804 90931363
refGene_gn=NETO1;refGene_tx=XM_011288131.3;refGene_gn=NETO1;refGene_tx=XM_019815427.2;refGene_gn=NETO1;refGene_tx=XM_019815428.2;refGene_gn=NETO1;refGene_tx=XM_019815429.2;refGene_gn=NETO1;refGene_tx=XM_019815431.2;refGene_gn=NETO1;refGene_tx=XM_023242104.1;refGene_gn=NETO1;refGene_tx=XM_023242105.1
D3 90938023 90938137
refGene_gn=NETO1;refGene_tx=XM_011288131.3;refGene_gn=NETO1;refGene_tx=XM_019815427.2;refGene_gn=NETO1;refGene_tx=XM_019815428.2;refGene_gn=NETO1;refGene_tx=XM_019815429.2;refGene_gn=NETO1;refGene_tx=XM_019815431.2;refGene_gn=NETO1;refGene_tx=XM_023242104.1;refGene_gn=NETO1;refGene_tx=XM_023242105.1
D3 90945546 90945569 refGene_gn=NETO1;refGene_tx=XM_019815433.2
D3 90947668 90947760 refGene_gn=NETO1;refGene_tx=XM_019815432.2
D3 90962631 90962860
refGene_gn=NETO1;refGene_tx=XM_011288131.3;refGene_gn=NETO1;refGene_tx=XM_019815427.2;refGene_gn=NETO1;refGene_tx=XM_019815428.2;refGene_gn=NETO1;refGene_tx=XM_019815429.2;refGene_gn=NETO1;refGene_tx=XM_019815431.2;refGene_gn=NETO1;refGene_tx=XM_019815432.2;refGene_gn=NETO1;refGene_tx=XM_019815433.2;refGene_gn=NETO1;refGene_tx=XM_019815434.2;refGene_gn=NETO1;refGene_tx=XM_023242104.1;refGene_gn=NETO1;refGene_tx=XM_023242105.1
D3 90973245 90973326
refGene_gn=NETO1;refGene_tx=XM_019815427.2;refGene_gn=NETO1;refGene_tx=XM_019815428.2;refGene_gn=NETO1;refGene_tx=XM_019815429.2;refGene_gn=NETO1;refGene_tx=XM_019815431.2;refGene_gn=NETO1;refGene_tx=XM_019815432.2;refGene_gn=NETO1;refGene_tx=XM_019815433.2;refGene_gn=NETO1;refGene_tx=XM_023242104.1;refGene_gn=NETO1;refGene_tx=XM_023242105.1
D3 90973487 90973615
refGene_gn=NETO1;refGene_tx=XM_011288131.3;refGene_gn=NETO1;refGene_tx=XM_019815427.2;refGene_gn=NETO1;refGene_tx=XM_019815428.2;refGene_gn=NETO1;refGene_tx=XM_019815429.2;refGene_gn=NETO1;refGene_tx=XM_019815431.2;refGene_gn=NETO1;refGene_tx=XM_019815432.2;refGene_gn=NETO1;refGene_tx=XM_019815433.2;refGene_gn=NETO1;refGene_tx=XM_023242104.1;refGene_gn=NETO1;refGene_tx=XM_023242105.1
D3 90973733 90973775
refGene_gn=NETO1;refGene_tx=XM_011288131.3;refGene_gn=NETO1;refGene_tx=XM_019815427.2;refGene_gn=NETO1;refGene_tx=XM_019815428.2;refGene_gn=NETO1;refGene_tx=XM_019815429.2;refGene_gn=NETO1;refGene_tx=XM_019815431.2;refGene_gn=NETO1;refGene_tx=XM_019815432.2;refGene_gn=NETO1;refGene_tx=XM_019815433.2;refGene_gn=NETO1;refGene_tx=XM_023242104.1;refGene_gn=NETO1;refGene_tx=XM_023242105.1
D3 91031823 91032072
refGene_gn=NETO1;refGene_tx=XM_011288131.3;refGene_gn=NETO1;refGene_tx

=XM_019815427.2;refGene_gn=NETO1;refGene_tx=XM_019815428.2;refGene_gn=NETO1;refGene_tx=XM_019815429.2;refGene_gn=NETO1;refGene_tx=XM_019815431.2;refGene_gn=NETO1;refGene_tx=XM_019815432.2;refGene_gn=NETO1;refGene_tx=XM_019815433.2;refGene_gn=NETO1;refGene_tx=XM_019815434.2;refGene_gn=NETO1;refGene_tx=XM_023242104.1;refGene_gn=NETO1;refGene_tx=XM_023242105.1
D3 91038851 91038989
refGene_gn=NETO1;refGene_tx=XM_011288131.3;refGene_gn=NETO1;refGene_tx=XM_019815427.2;refGene_gn=NETO1;refGene_tx=XM_019815428.2;refGene_gn=NETO1;refGene_tx=XM_019815429.2;refGene_gn=NETO1;refGene_tx=XM_019815431.2;refGene_gn=NETO1;refGene_tx=XM_019815432.2;refGene_gn=NETO1;refGene_tx=XM_019815433.2;refGene_gn=NETO1;refGene_tx=XM_019815434.2;refGene_gn=NETO1;refGene_tx=XM_023242104.1;refGene_gn=NETO1;refGene_tx=XM_023242105.1
D3 91039222 91039276
refGene_gn=NETO1;refGene_tx=XM_011288131.3;refGene_gn=NETO1;refGene_tx=XM_019815427.2;refGene_gn=NETO1;refGene_tx=XM_019815428.2;refGene_gn=NETO1;refGene_tx=XM_019815429.2;refGene_gn=NETO1;refGene_tx=XM_019815431.2;refGene_gn=NETO1;refGene_tx=XM_019815432.2;refGene_gn=NETO1;refGene_tx=XM_019815433.2;refGene_gn=NETO1;refGene_tx=XM_019815434.2;refGene_gn=NETO1;refGene_tx=XM_023242104.1;refGene_gn=NETO1;refGene_tx=XM_023242105.1
D3 91041286 91041314
refGene_gn=NETO1;refGene_tx=XM_011288131.3;refGene_gn=NETO1;refGene_tx=XM_019815427.2;refGene_gn=NETO1;refGene_tx=XM_019815428.2;refGene_gn=NETO1;refGene_tx=XM_019815429.2;refGene_gn=NETO1;refGene_tx=XM_019815431.2;refGene_gn=NETO1;refGene_tx=XM_019815432.2;refGene_gn=NETO1;refGene_tx=XM_019815433.2;refGene_gn=NETO1;refGene_tx=XM_019815434.2;refGene_gn=NETO1;refGene_tx=XM_023242104.1;refGene_gn=NETO1;refGene_tx=XM_023242105.1
D3 91949154 91949421
refGene_gn=FBXO15;refGene_tx=XM_011288129.3;refGene_gn=FBXO15;refGene_tx=XM_011288130.3;refGene_gn=FBXO15;refGene_tx=XM_023242106.1
D3 91951934 91952059
refGene_gn=FBXO15;refGene_tx=XM_011288129.3;refGene_gn=FBXO15;refGene_tx=XM_011288130.3;refGene_gn=FBXO15;refGene_tx=XM_023242106.1
D3 91975488 91975631
refGene_gn=FBXO15;refGene_tx=XM_011288129.3;refGene_gn=FBXO15;refGene_tx=XM_011288130.3;refGene_gn=FBXO15;refGene_tx=XM_023242106.1
D3 91976574 91976657
refGene_gn=FBXO15;refGene_tx=XM_011288129.3;refGene_gn=FBXO15;refGene_tx=XM_011288130.3;refGene_gn=FBXO15;refGene_tx=XM_023242106.1
D3 91978056 91978183
refGene_gn=FBXO15;refGene_tx=XM_011288129.3;refGene_gn=FBXO15;refGene_tx=XM_011288130.3;refGene_gn=FBXO15;refGene_tx=XM_023242106.1
D3 91982517 91982727
refGene_gn=FBXO15;refGene_tx=XM_011288129.3;refGene_gn=FBXO15;refGene_tx=XM_011288130.3;refGene_gn=FBXO15;refGene_tx=XM_023242106.1
D3 91983281 91983524
refGene_gn=FBXO15;refGene_tx=XM_011288129.3;refGene_gn=FBXO15;refGene_

tx=XM_011288130.3;refGene_gn=FBXO15;refGene_tx=XM_023242106.1
D3	91986873 91986978
refGene_gn=FBXO15;refGene_tx=XM_011288129.3;refGene_gn=FBXO15;refGene_tx=XM_011288130.3;refGene_gn=FBXO15;refGene_tx=XM_023242106.1
D3	91989725 91989797
refGene_gn=FBXO15;refGene_tx=XM_011288129.3;refGene_gn=FBXO15;refGene_tx=XM_011288130.3
D3	91997025 91997132
refGene_gn=FBXO15;refGene_tx=XM_011288129.3;refGene_gn=FBXO15;refGene_tx=XM_011288130.3
D3	91998061 91998359
refGene_gn=TIMM21;refGene_tx=XM_003995272.5;refGene_gn=TIMM21;refGene_tx=XM_006938975.3;refGene_gn=TIMM21;refGene_tx=XM_011288128.3
D3	92004202 92004265
refGene_gn=TIMM21;refGene_tx=XM_003995272.5;refGene_gn=TIMM21;refGene_tx=XM_006938975.3;refGene_gn=TIMM21;refGene_tx=XM_011288128.3
D3	92004360 92004458
refGene_gn=TIMM21;refGene_tx=XM_003995272.5;refGene_gn=TIMM21;refGene_tx=XM_006938975.3;refGene_gn=TIMM21;refGene_tx=XM_011288128.3
D3	92006225 92006299
refGene_gn=TIMM21;refGene_tx=XM_003995272.5;refGene_gn=TIMM21;refGene_tx=XM_006938975.3;refGene_gn=TIMM21;refGene_tx=XM_011288128.3
D3	92006384 92006490
refGene_gn=TIMM21;refGene_tx=XM_003995272.5;refGene_gn=TIMM21;refGene_tx=XM_006938975.3;refGene_gn=TIMM21;refGene_tx=XM_011288128.3
D3	92006595 92006691
refGene_gn=TIMM21;refGene_tx=XM_003995272.5;refGene_gn=TIMM21;refGene_tx=XM_006938975.3;refGene_gn=TIMM21;refGene_tx=XM_011288128.3
D3	92086128 92086210
refGene_gn=LOC101098747;refGene_tx=XM_011288127.3
D3	92087937 92087972
refGene_gn=LOC101098747;refGene_tx=XM_011288127.3
D3	92092542 92092572
refGene_gn=LOC101098747;refGene_tx=XM_011288127.3
D3	92094671 92094800
refGene_gn=LOC101098747;refGene_tx=XM_011288127.3
D3	92121822 92121951
refGene_gn=LOC101098747;refGene_tx=XM_011288127.3
D3	92137639 92137773
refGene_gn=CD3H18orf63;refGene_tx=XM_023242107.1
D3	92139774 92139853
refGene_gn=CD3H18orf63;refGene_tx=XM_023242107.1
D3	92141631 92141688
refGene_gn=CD3H18orf63;refGene_tx=XM_023242107.1
D3	92143790 92143902
refGene_gn=CD3H18orf63;refGene_tx=XM_023242107.1
D3	92144481 92144523
refGene_gn=CD3H18orf63;refGene_tx=XM_023242107.1
D3	92146321 92146398
refGene_gn=CD3H18orf63;refGene_tx=XM_023242107.1

D3 92151604 92151714
refGene_gn=CD3H18orf63;refGene_tx=XM_023242107.1
D3 92152531 92152628
refGene_gn=CD3H18orf63;refGene_tx=XM_023242107.1
D3 92154149 92154203
refGene_gn=CD3H18orf63;refGene_tx=XM_023242107.1
D3 92163360 92164386
refGene_gn=CD3H18orf63;refGene_tx=XM_023242107.1
D3 92164574 92164631
refGene_gn=CD3H18orf63;refGene_tx=XM_023242107.1
D3 92234279 92234498 refGene_gn=FAM69C;refGene_tx=XM_023241205.1
D3 92237303 92237468 refGene_gn=FAM69C;refGene_tx=XM_023241205.1
D3 92241928 92242612 refGene_gn=FAM69C;refGene_tx=XM_023241205.1
D3 92251723 92251924 refGene_gn=FAM69C;refGene_tx=XM_023241205.1
D3 92286836 92286896
refGene_gn=CNDP2;refGene_tx=XM_003995273.5;refGene_gn=CNDP2;refGene_tx=XM_023242108.1
D3 92288515 92288659
refGene_gn=CNDP2;refGene_tx=XM_003995273.5;refGene_gn=CNDP2;refGene_tx=XM_023242108.1
D3 92292203 92292366
refGene_gn=CNDP2;refGene_tx=XM_003995273.5;refGene_gn=CNDP2;refGene_tx=XM_023242108.1
D3 92294773 92294862
refGene_gn=CNDP2;refGene_tx=XM_003995273.5;refGene_gn=CNDP2;refGene_tx=XM_023242108.1
D3 92296449 92296650
refGene_gn=CNDP2;refGene_tx=XM_003995273.5;refGene_gn=CNDP2;refGene_tx=XM_023242108.1
D3 92297714 92297799
refGene_gn=CNDP2;refGene_tx=XM_003995273.5;refGene_gn=CNDP2;refGene_tx=XM_023242108.1
D3 92298984 92299145
refGene_gn=CNDP2;refGene_tx=XM_003995273.5;refGene_gn=CNDP2;refGene_tx=XM_023242108.1
D3 92300937 92301102
refGene_gn=CNDP2;refGene_tx=XM_003995273.5;refGene_gn=CNDP2;refGene_tx=XM_023242108.1
D3 92301999 92302141
refGene_gn=CNDP2;refGene_tx=XM_003995273.5;refGene_gn=CNDP2;refGene_tx=XM_023242108.1
D3 92302320 92302468
refGene_gn=CNDP2;refGene_tx=XM_003995273.5;refGene_gn=CNDP2;refGene_tx=XM_023242108.1
D3 92303275 92303345
refGene_gn=CNDP2;refGene_tx=XM_003995273.5;refGene_gn=CNDP2;refGene_tx=XM_023242108.1
D3 92320213 92320297
refGene_gn=CNDP1;refGene_tx=XM_006938977.4;refGene_gn=CNDP1;refGene_tx=XM_019815453.1

D3 92323352 92323502
refGene_gn=CNDP1;refGene_tx=XM_006938977.4;refGene_gn=CNDP1;refGene_tx=XM_019815453.1
D3 92324228 92324391
refGene_gn=CNDP1;refGene_tx=XM_006938977.4;refGene_gn=CNDP1;refGene_tx=XM_019815453.1
D3 92325106 92325195
refGene_gn=CNDP1;refGene_tx=XM_006938977.4;refGene_gn=CNDP1;refGene_tx=XM_019815453.1
D3 92330456 92330657
refGene_gn=CNDP1;refGene_tx=XM_006938977.4;refGene_gn=CNDP1;refGene_tx=XM_019815453.1
D3 92333322 92333407
refGene_gn=CNDP1;refGene_tx=XM_006938977.4;refGene_gn=CNDP1;refGene_tx=XM_019815453.1
D3 92338339 92338500
refGene_gn=CNDP1;refGene_tx=XM_006938977.4;refGene_gn=CNDP1;refGene_tx=XM_019815453.1
D3 92339546 92339711
refGene_gn=CNDP1;refGene_tx=XM_006938977.4;refGene_gn=CNDP1;refGene_tx=XM_019815453.1
D3 92340952 92341094
refGene_gn=CNDP1;refGene_tx=XM_006938977.4;refGene_gn=CNDP1;refGene_tx=XM_019815453.1
D3 92344233 92344381
refGene_gn=CNDP1;refGene_tx=XM_006938977.4;refGene_gn=CNDP1;refGene_tx=XM_019815453.1
D3 92344572 92344642
refGene_gn=CNDP1;refGene_tx=XM_006938977.4;refGene_gn=CNDP1;refGene_tx=XM_019815453.1
D3 92379980 92384658 refGene_gn=ZNF407;refGene_tx=XM_011288125.3
D3 92390529 92390644 refGene_gn=ZNF407;refGene_tx=XM_011288125.3
D3 92528254 92528329 refGene_gn=ZNF407;refGene_tx=XM_011288125.3
D3 92624136 92624303 refGene_gn=ZNF407;refGene_tx=XM_011288125.3
D3 92627987 92628071 refGene_gn=ZNF407;refGene_tx=XM_011288125.3
D3 92638627 92638710 refGene_gn=ZNF407;refGene_tx=XM_011288125.3
D3 92638711 92638749 refGene_gn=ZNF407;refGene_tx=XM_011288125.3
D3 92665640 92665819 refGene_gn=ZNF407;refGene_tx=XM_011288125.3
D3 92806998 92808317 refGene_gn=ZNF407;refGene_tx=XM_011288125.3
D3 92921247 92922172
refGene_gn=ZADH2;refGene_tx=XM_019815454.2;refGene_gn=ZADH2;refGene_tx=XM_019815455.2;refGene_gn=ZADH2;refGene_tx=XM_023242109.1
D3 92928713 92928922 refGene_gn=ZADH2;refGene_tx=XM_023242109.1
D3 92931760 92931800 refGene_gn=TSHZ1;refGene_tx=XM_023242110.1
D3 93002861 93006043 refGene_gn=TSHZ1;refGene_tx=XM_023242110.1
D3 93865493 93865553 refGene_gn=ZNF516;refGene_tx=XM_023242111.1
D3 93872708 93872776 refGene_gn=ZNF516;refGene_tx=XM_023242111.1
D3 93882022 93883102 refGene_gn=ZNF516;refGene_tx=XM_023242111.1
D3 93883104 93883482 refGene_gn=ZNF516;refGene_tx=XM_023242111.1
D3 93932612 93934374 refGene_gn=ZNF516;refGene_tx=XM_023242111.1

D3 94184936 94184991
refGene_gn=ZNF236;refGene_tx=XM_023242112.1;refGene_gn=ZNF236;refGene_tx=XM_023242113.1;refGene_gn=ZNF236;refGene_tx=XM_023242119.1
D3 94200465 94200608
refGene_gn=ZNF236;refGene_tx=XM_023242112.1;refGene_gn=ZNF236;refGene_tx=XM_023242113.1;refGene_gn=ZNF236;refGene_tx=XM_023242114.1;refGene_gn=ZNF236;refGene_tx=XM_023242115.1;refGene_gn=ZNF236;refGene_tx=XM_023242116.1;refGene_gn=ZNF236;refGene_tx=XM_023242119.1
D3 94202349 94202514
refGene_gn=ZNF236;refGene_tx=XM_023242112.1;refGene_gn=ZNF236;refGene_tx=XM_023242113.1;refGene_gn=ZNF236;refGene_tx=XM_023242114.1;refGene_gn=ZNF236;refGene_tx=XM_023242115.1;refGene_gn=ZNF236;refGene_tx=XM_023242116.1;refGene_gn=ZNF236;refGene_tx=XM_023242119.1
D3 94202790 94202928 refGene_gn=ZNF236;refGene_tx=XM_023242117.1
D3 94211667 94211846
refGene_gn=ZNF236;refGene_tx=XM_023242112.1;refGene_gn=ZNF236;refGene_tx=XM_023242113.1;refGene_gn=ZNF236;refGene_tx=XM_023242114.1;refGene_gn=ZNF236;refGene_tx=XM_023242115.1;refGene_gn=ZNF236;refGene_tx=XM_023242116.1;refGene_gn=ZNF236;refGene_tx=XM_023242117.1;refGene_gn=ZNF236;refGene_tx=XM_023242118.1;refGene_gn=ZNF236;refGene_tx=XM_023242119.1
D3 94213846 94213971
refGene_gn=ZNF236;refGene_tx=XM_023242112.1;refGene_gn=ZNF236;refGene_tx=XM_023242113.1;refGene_gn=ZNF236;refGene_tx=XM_023242114.1;refGene_gn=ZNF236;refGene_tx=XM_023242115.1;refGene_gn=ZNF236;refGene_tx=XM_023242116.1;refGene_gn=ZNF236;refGene_tx=XM_023242117.1;refGene_gn=ZNF236;refGene_tx=XM_023242118.1;refGene_gn=ZNF236;refGene_tx=XM_023242119.1
D3 94214267 94214440
refGene_gn=ZNF236;refGene_tx=XM_023242112.1;refGene_gn=ZNF236;refGene_tx=XM_023242113.1;refGene_gn=ZNF236;refGene_tx=XM_023242114.1;refGene_gn=ZNF236;refGene_tx=XM_023242115.1;refGene_gn=ZNF236;refGene_tx=XM_023242116.1;refGene_gn=ZNF236;refGene_tx=XM_023242117.1;refGene_gn=ZNF236;refGene_tx=XM_023242118.1;refGene_gn=ZNF236;refGene_tx=XM_023242119.1
D3 94216501 94216645
refGene_gn=ZNF236;refGene_tx=XM_023242112.1;refGene_gn=ZNF236;refGene_tx=XM_023242113.1;refGene_gn=ZNF236;refGene_tx=XM_023242114.1;refGene_gn=ZNF236;refGene_tx=XM_023242115.1;refGene_gn=ZNF236;refGene_tx=XM_023242116.1;refGene_gn=ZNF236;refGene_tx=XM_023242117.1;refGene_gn=ZNF236;refGene_tx=XM_023242118.1;refGene_gn=ZNF236;refGene_tx=XM_023242119.1
D3 94219156 94219360
refGene_gn=ZNF236;refGene_tx=XM_023242112.1;refGene_gn=ZNF236;refGene_tx=XM_023242113.1;refGene_gn=ZNF236;refGene_tx=XM_023242114.1;refGene_gn=ZNF236;refGene_tx=XM_023242115.1;refGene_gn=ZNF236;refGene_tx=XM_023242116.1;refGene_gn=ZNF236;refGene_tx=XM_023242117.1;refGene_gn=ZNF236;refGene_tx=XM_023242118.1;refGene_gn=ZNF236;refGene_tx=XM_023242119.1
D3 94220328 94220557

refGene_gn=ZNF236;refGene_tx=XM_023242112.1;refGene_gn=ZNF236;refGene_tx=XM_023242113.1;refGene_gn=ZNF236;refGene_tx=XM_023242114.1;refGene_gn=ZNF236;refGene_tx=XM_023242115.1;refGene_gn=ZNF236;refGene_tx=XM_023242116.1;refGene_gn=ZNF236;refGene_tx=XM_023242117.1;refGene_gn=ZNF236;refGene_tx=XM_023242118.1;refGene_gn=ZNF236;refGene_tx=XM_023242119.1
D3 94233382 94233655
refGene_gn=ZNF236;refGene_tx=XM_023242112.1;refGene_gn=ZNF236;refGene_tx=XM_023242113.1;refGene_gn=ZNF236;refGene_tx=XM_023242114.1;refGene_gn=ZNF236;refGene_tx=XM_023242115.1;refGene_gn=ZNF236;refGene_tx=XM_023242116.1;refGene_gn=ZNF236;refGene_tx=XM_023242117.1;refGene_gn=ZNF236;refGene_tx=XM_023242118.1;refGene_gn=ZNF236;refGene_tx=XM_023242119.1
D3 94234910 94235114
refGene_gn=ZNF236;refGene_tx=XM_023242112.1;refGene_gn=ZNF236;refGene_tx=XM_023242113.1;refGene_gn=ZNF236;refGene_tx=XM_023242114.1;refGene_gn=ZNF236;refGene_tx=XM_023242115.1;refGene_gn=ZNF236;refGene_tx=XM_023242116.1;refGene_gn=ZNF236;refGene_tx=XM_023242117.1;refGene_gn=ZNF236;refGene_tx=XM_023242118.1;refGene_gn=ZNF236;refGene_tx=XM_023242119.1
D3 94246420 94246562
refGene_gn=ZNF236;refGene_tx=XM_023242112.1;refGene_gn=ZNF236;refGene_tx=XM_023242113.1;refGene_gn=ZNF236;refGene_tx=XM_023242114.1;refGene_gn=ZNF236;refGene_tx=XM_023242115.1;refGene_gn=ZNF236;refGene_tx=XM_023242116.1;refGene_gn=ZNF236;refGene_tx=XM_023242117.1;refGene_gn=ZNF236;refGene_tx=XM_023242118.1;refGene_gn=ZNF236;refGene_tx=XM_023242119.1
D3 94247445 94247706
refGene_gn=ZNF236;refGene_tx=XM_023242112.1;refGene_gn=ZNF236;refGene_tx=XM_023242113.1;refGene_gn=ZNF236;refGene_tx=XM_023242114.1;refGene_gn=ZNF236;refGene_tx=XM_023242115.1;refGene_gn=ZNF236;refGene_tx=XM_023242116.1;refGene_gn=ZNF236;refGene_tx=XM_023242117.1;refGene_gn=ZNF236;refGene_tx=XM_023242118.1;refGene_gn=ZNF236;refGene_tx=XM_023242119.1
D3 94250330 94250590
refGene_gn=ZNF236;refGene_tx=XM_023242112.1;refGene_gn=ZNF236;refGene_tx=XM_023242113.1;refGene_gn=ZNF236;refGene_tx=XM_023242114.1;refGene_gn=ZNF236;refGene_tx=XM_023242115.1;refGene_gn=ZNF236;refGene_tx=XM_023242116.1;refGene_gn=ZNF236;refGene_tx=XM_023242117.1;refGene_gn=ZNF236;refGene_tx=XM_023242118.1;refGene_gn=ZNF236;refGene_tx=XM_023242119.1
D3 94251736 94251835
refGene_gn=ZNF236;refGene_tx=XM_023242112.1;refGene_gn=ZNF236;refGene_tx=XM_023242113.1;refGene_gn=ZNF236;refGene_tx=XM_023242114.1;refGene_gn=ZNF236;refGene_tx=XM_023242115.1;refGene_gn=ZNF236;refGene_tx=XM_023242116.1;refGene_gn=ZNF236;refGene_tx=XM_023242117.1;refGene_gn=ZNF236;refGene_tx=XM_023242118.1;refGene_gn=ZNF236;refGene_tx=XM_023242119.1
D3 94252296 94252448
refGene_gn=ZNF236;refGene_tx=XM_023242112.1;refGene_gn=ZNF236;refGene_

tx=XM_023242113.1;refGene_gn=ZNF236;refGene_tx=XM_023242114.1;refGene_gn=ZNF236;refGene_tx=XM_023242115.1;refGene_gn=ZNF236;refGene_tx=XM_023242116.1;refGene_gn=ZNF236;refGene_tx=XM_023242117.1;refGene_gn=ZNF236;refGene_tx=XM_023242118.1;refGene_gn=ZNF236;refGene_tx=XM_023242119.1
D3 94253911 94254015
refGene_gn=ZNF236;refGene_tx=XM_023242112.1;refGene_gn=ZNF236;refGene_tx=XM_023242113.1;refGene_gn=ZNF236;refGene_tx=XM_023242114.1;refGene_gn=ZNF236;refGene_tx=XM_023242115.1;refGene_gn=ZNF236;refGene_tx=XM_023242116.1;refGene_gn=ZNF236;refGene_tx=XM_023242117.1;refGene_gn=ZNF236;refGene_tx=XM_023242118.1;refGene_gn=ZNF236;refGene_tx=XM_023242119.1
D3 94256875 94257027
refGene_gn=ZNF236;refGene_tx=XM_023242112.1;refGene_gn=ZNF236;refGene_tx=XM_023242113.1;refGene_gn=ZNF236;refGene_tx=XM_023242114.1;refGene_gn=ZNF236;refGene_tx=XM_023242115.1;refGene_gn=ZNF236;refGene_tx=XM_023242116.1;refGene_gn=ZNF236;refGene_tx=XM_023242117.1;refGene_gn=ZNF236;refGene_tx=XM_023242118.1;refGene_gn=ZNF236;refGene_tx=XM_023242119.1
D3 94258535 94258748
refGene_gn=ZNF236;refGene_tx=XM_023242112.1;refGene_gn=ZNF236;refGene_tx=XM_023242113.1;refGene_gn=ZNF236;refGene_tx=XM_023242114.1;refGene_gn=ZNF236;refGene_tx=XM_023242115.1;refGene_gn=ZNF236;refGene_tx=XM_023242116.1;refGene_gn=ZNF236;refGene_tx=XM_023242117.1;refGene_gn=ZNF236;refGene_tx=XM_023242118.1;refGene_gn=ZNF236;refGene_tx=XM_023242119.1
D3 94260826 94261109
refGene_gn=ZNF236;refGene_tx=XM_023242112.1;refGene_gn=ZNF236;refGene_tx=XM_023242113.1;refGene_gn=ZNF236;refGene_tx=XM_023242114.1;refGene_gn=ZNF236;refGene_tx=XM_023242115.1;refGene_gn=ZNF236;refGene_tx=XM_023242116.1;refGene_gn=ZNF236;refGene_tx=XM_023242117.1;refGene_gn=ZNF236;refGene_tx=XM_023242118.1;refGene_gn=ZNF236;refGene_tx=XM_023242119.1
D3 94263462 94263566
refGene_gn=ZNF236;refGene_tx=XM_023242112.1;refGene_gn=ZNF236;refGene_tx=XM_023242113.1;refGene_gn=ZNF236;refGene_tx=XM_023242114.1;refGene_gn=ZNF236;refGene_tx=XM_023242115.1;refGene_gn=ZNF236;refGene_tx=XM_023242116.1;refGene_gn=ZNF236;refGene_tx=XM_023242117.1;refGene_gn=ZNF236;refGene_tx=XM_023242118.1;refGene_gn=ZNF236;refGene_tx=XM_023242119.1
D3 94265419 94265785
refGene_gn=ZNF236;refGene_tx=XM_023242112.1;refGene_gn=ZNF236;refGene_tx=XM_023242113.1;refGene_gn=ZNF236;refGene_tx=XM_023242114.1;refGene_gn=ZNF236;refGene_tx=XM_023242115.1;refGene_gn=ZNF236;refGene_tx=XM_023242116.1;refGene_gn=ZNF236;refGene_tx=XM_023242117.1;refGene_gn=ZNF236;refGene_tx=XM_023242118.1;refGene_gn=ZNF236;refGene_tx=XM_023242119.1
D3 94266547 94266693
refGene_gn=ZNF236;refGene_tx=XM_023242112.1;refGene_gn=ZNF236;refGene_tx=XM_023242113.1;refGene_gn=ZNF236;refGene_tx=XM_023242114.1;refGene_

gn=ZNF236;refGene_tx=XM_023242115.1;refGene_gn=ZNF236;refGene_tx=XM_02
3242116.1;refGene_gn=ZNF236;refGene_tx=XM_023242117.1;refGene_gn=ZNF23
6;refGene_tx=XM_023242118.1;refGene_gn=ZNF236;refGene_tx=XM_023242119.
1
D3 94266829 94267067
refGene_gn=ZNF236;refGene_tx=XM_023242112.1;refGene_gn=ZNF236;refGene_
tx=XM_023242113.1;refGene_gn=ZNF236;refGene_tx=XM_023242114.1;refGene_
gn=ZNF236;refGene_tx=XM_023242115.1;refGene_gn=ZNF236;refGene_tx=XM_02
3242116.1;refGene_gn=ZNF236;refGene_tx=XM_023242117.1;refGene_gn=ZNF23
6;refGene_tx=XM_023242118.1;refGene_gn=ZNF236;refGene_tx=XM_023242119.
1
D3 94267473 94267653
refGene_gn=ZNF236;refGene_tx=XM_023242112.1;refGene_gn=ZNF236;refGene_
tx=XM_023242113.1;refGene_gn=ZNF236;refGene_tx=XM_023242114.1;refGene_
gn=ZNF236;refGene_tx=XM_023242115.1;refGene_gn=ZNF236;refGene_tx=XM_02
3242116.1;refGene_gn=ZNF236;refGene_tx=XM_023242117.1;refGene_gn=ZNF23
6;refGene_tx=XM_023242118.1;refGene_gn=ZNF236;refGene_tx=XM_023242119.
1
D3 94274864 94274946 refGene_gn=ZNF236;refGene_tx=XM_023242119.1
D3 94275992 94276180
refGene_gn=ZNF236;refGene_tx=XM_023242112.1;refGene_gn=ZNF236;refGene_
tx=XM_023242113.1;refGene_gn=ZNF236;refGene_tx=XM_023242114.1;refGene_
gn=ZNF236;refGene_tx=XM_023242115.1;refGene_gn=ZNF236;refGene_tx=XM_02
3242116.1;refGene_gn=ZNF236;refGene_tx=XM_023242117.1;refGene_gn=ZNF23
6;refGene_tx=XM_023242118.1;refGene_gn=ZNF236;refGene_tx=XM_023242119.
1
D3 94283439 94283574
refGene_gn=ZNF236;refGene_tx=XM_023242112.1;refGene_gn=ZNF236;refGene_
tx=XM_023242113.1;refGene_gn=ZNF236;refGene_tx=XM_023242114.1;refGene_
gn=ZNF236;refGene_tx=XM_023242115.1;refGene_gn=ZNF236;refGene_tx=XM_02
3242116.1;refGene_gn=ZNF236;refGene_tx=XM_023242117.1;refGene_gn=ZNF23
6;refGene_tx=XM_023242118.1
D3 94286071 94286269
refGene_gn=ZNF236;refGene_tx=XM_023242112.1;refGene_gn=ZNF236;refGene_
tx=XM_023242113.1;refGene_gn=ZNF236;refGene_tx=XM_023242114.1;refGene_
gn=ZNF236;refGene_tx=XM_023242115.1;refGene_gn=ZNF236;refGene_tx=XM_02
3242116.1;refGene_gn=ZNF236;refGene_tx=XM_023242117.1;refGene_gn=ZNF23
6;refGene_tx=XM_023242118.1
D3 94287084 94287214
refGene_gn=ZNF236;refGene_tx=XM_023242112.1;refGene_gn=ZNF236;refGene_
tx=XM_023242113.1;refGene_gn=ZNF236;refGene_tx=XM_023242114.1;refGene_
gn=ZNF236;refGene_tx=XM_023242115.1;refGene_gn=ZNF236;refGene_tx=XM_02
3242116.1;refGene_gn=ZNF236;refGene_tx=XM_023242117.1;refGene_gn=ZNF23
6;refGene_tx=XM_023242118.1
D3 94289108 94289285
refGene_gn=ZNF236;refGene_tx=XM_023242112.1;refGene_gn=ZNF236;refGene_
tx=XM_023242113.1;refGene_gn=ZNF236;refGene_tx=XM_023242114.1;refGene_
gn=ZNF236;refGene_tx=XM_023242115.1;refGene_gn=ZNF236;refGene_tx=XM_02
3242116.1;refGene_gn=ZNF236;refGene_tx=XM_023242117.1;refGene_gn=ZNF23
6;refGene_tx=XM_023242118.1

D3 94290427 94290552
refGene_gn=ZNF236;refGene_tx=XM_023242112.1;refGene_gn=ZNF236;refGene_tx=XM_023242113.1;refGene_gn=ZNF236;refGene_tx=XM_023242114.1;refGene_gn=ZNF236;refGene_tx=XM_023242115.1;refGene_gn=ZNF236;refGene_tx=XM_023242116.1;refGene_gn=ZNF236;refGene_tx=XM_023242117.1;refGene_gn=ZNF236;refGene_tx=XM_023242118.1
D3 94300282 94300327
refGene_gn=MBP;refGene_tx=XM_019815459.2;refGene_gn=MBP;refGene_tx=XM_019815460.2;refGene_gn=MBP;refGene_tx=XM_019815461.2;refGene_gn=MBP;refGene_tx=XM_019815464.1;refGene_gn=MBP;refGene_tx=XM_023242120.1
D3 94305746 94305854
refGene_gn=MBP;refGene_tx=XM_019815459.2;refGene_gn=MBP;refGene_tx=XM_019815460.2;refGene_gn=MBP;refGene_tx=XM_019815461.2;refGene_gn=MBP;refGene_tx=XM_019815464.1;refGene_gn=MBP;refGene_tx=XM_023242120.1
D3 94308965 94308998
refGene_gn=MBP;refGene_tx=XM_019815460.2;refGene_gn=MBP;refGene_tx=XM_019815464.1;refGene_gn=MBP;refGene_tx=XM_023242120.1
D3 94309360 94309396
refGene_gn=MBP;refGene_tx=XM_019815459.2;refGene_gn=MBP;refGene_tx=XM_019815460.2;refGene_gn=MBP;refGene_tx=XM_019815461.2;refGene_gn=MBP;refGene_tx=XM_019815464.1;refGene_gn=MBP;refGene_tx=XM_023242120.1
D3 94310280 94310388
refGene_gn=MBP;refGene_tx=XM_019815459.2;refGene_gn=MBP;refGene_tx=XM_019815460.2;refGene_gn=MBP;refGene_tx=XM_019815461.2;refGene_gn=MBP;refGene_tx=XM_019815464.1;refGene_gn=MBP;refGene_tx=XM_023242120.1
D3 94325088 94325166
refGene_gn=MBP;refGene_tx=XM_019815459.2;refGene_gn=MBP;refGene_tx=XM_019815464.1;refGene_gn=MBP;refGene_tx=XM_023242120.1
D3 94331594 94332049
refGene_gn=MBP;refGene_tx=XM_019815459.2;refGene_gn=MBP;refGene_tx=XM_019815460.2;refGene_gn=MBP;refGene_tx=XM_019815461.2;refGene_gn=MBP;refGene_tx=XM_019815462.2;refGene_gn=MBP;refGene_tx=XM_019815464.1;refGene_gn=MBP;refGene_tx=XM_023242120.1
D3 94365468 94365556 refGene_gn=MBP;refGene_tx=XM_023242120.1
D3 94391560 94391636
refGene_gn=MBP;refGene_tx=XM_019815462.2;refGene_gn=MBP;refGene_tx=XM_019815464.1;refGene_gn=MBP;refGene_tx=XM_023242120.1
D3 94393276 94393327 refGene_gn=MBP;refGene_tx=XM_019815464.1
D3 94399936 94400464 refGene_gn=MBP;refGene_tx=XM_019815462.2
D3 94506983 94507649 refGene_gn=GALR1;refGene_tx=XM_011288124.3
D3 94514034 94514100 refGene_gn=GALR1;refGene_tx=XM_011288124.3
D3 94521492 94521810 refGene_gn=GALR1;refGene_tx=XM_011288124.3
D3 95802277 95802359 refGene_gn=SALL3;refGene_tx=XM_023242121.1
D3 95813343 95816825 refGene_gn=SALL3;refGene_tx=XM_023242121.1
D3 95819102 95819534 refGene_gn=SALL3;refGene_tx=XM_023242121.1
D3 95861246 95861362
refGene_gn=ATP9B;refGene_tx=XM_023242122.1;refGene_gn=ATP9B;refGene_tx=XM_023242123.1;refGene_gn=ATP9B;refGene_tx=XM_023242124.1;refGene_gn=ATP9B;refGene_tx=XM_023242125.1;refGene_gn=ATP9B;refGene_tx=XM_023242126.1;refGene_gn=ATP9B;refGene_tx=XM_023242127.1;refGene_gn=ATP9B;refGe

ne_tx=XM_023242128.1
D3	95881853	95882030
refGene_gn=ATP9B;refGene_tx=XM_023242122.1;refGene_gn=ATP9B;refGene_tx=XM_023242123.1;refGene_gn=ATP9B;refGene_tx=XM_023242124.1;refGene_gn=ATP9B;refGene_tx=XM_023242125.1;refGene_gn=ATP9B;refGene_tx=XM_023242126.1;refGene_gn=ATP9B;refGene_tx=XM_023242127.1;refGene_gn=ATP9B;refGene_tx=XM_023242128.1
D3	95899714	95899865
refGene_gn=ATP9B;refGene_tx=XM_023242122.1;refGene_gn=ATP9B;refGene_tx=XM_023242123.1;refGene_gn=ATP9B;refGene_tx=XM_023242124.1;refGene_gn=ATP9B;refGene_tx=XM_023242125.1;refGene_gn=ATP9B;refGene_tx=XM_023242126.1;refGene_gn=ATP9B;refGene_tx=XM_023242127.1;refGene_gn=ATP9B;refGene_tx=XM_023242128.1
D3	95902781	95902895
refGene_gn=ATP9B;refGene_tx=XM_023242122.1;refGene_gn=ATP9B;refGene_tx=XM_023242123.1;refGene_gn=ATP9B;refGene_tx=XM_023242124.1;refGene_gn=ATP9B;refGene_tx=XM_023242125.1;refGene_gn=ATP9B;refGene_tx=XM_023242127.1;refGene_gn=ATP9B;refGene_tx=XM_023242128.1
D3	95911856	95911965
refGene_gn=ATP9B;refGene_tx=XM_023242122.1;refGene_gn=ATP9B;refGene_tx=XM_023242123.1;refGene_gn=ATP9B;refGene_tx=XM_023242124.1;refGene_gn=ATP9B;refGene_tx=XM_023242125.1;refGene_gn=ATP9B;refGene_tx=XM_023242126.1;refGene_gn=ATP9B;refGene_tx=XM_023242127.1;refGene_gn=ATP9B;refGene_tx=XM_023242128.1
D3	95926261	95926320
refGene_gn=ATP9B;refGene_tx=XM_023242122.1;refGene_gn=ATP9B;refGene_tx=XM_023242123.1;refGene_gn=ATP9B;refGene_tx=XM_023242124.1;refGene_gn=ATP9B;refGene_tx=XM_023242125.1;refGene_gn=ATP9B;refGene_tx=XM_023242126.1;refGene_gn=ATP9B;refGene_tx=XM_023242127.1;refGene_gn=ATP9B;refGene_tx=XM_023242128.1
D3	95936779	95936831
refGene_gn=ATP9B;refGene_tx=XM_023242122.1;refGene_gn=ATP9B;refGene_tx=XM_023242123.1;refGene_gn=ATP9B;refGene_tx=XM_023242124.1;refGene_gn=ATP9B;refGene_tx=XM_023242125.1;refGene_gn=ATP9B;refGene_tx=XM_023242126.1;refGene_gn=ATP9B;refGene_tx=XM_023242127.1;refGene_gn=ATP9B;refGene_tx=XM_023242128.1
D3	95955117	95955212
refGene_gn=ATP9B;refGene_tx=XM_023242122.1;refGene_gn=ATP9B;refGene_tx=XM_023242123.1;refGene_gn=ATP9B;refGene_tx=XM_023242124.1;refGene_gn=ATP9B;refGene_tx=XM_023242125.1;refGene_gn=ATP9B;refGene_tx=XM_023242126.1;refGene_gn=ATP9B;refGene_tx=XM_023242127.1;refGene_gn=ATP9B;refGene_tx=XM_023242128.1
D3	95976232	95976313
refGene_gn=ATP9B;refGene_tx=XM_023242122.1;refGene_gn=ATP9B;refGene_tx=XM_023242123.1;refGene_gn=ATP9B;refGene_tx=XM_023242124.1;refGene_gn=ATP9B;refGene_tx=XM_023242126.1;refGene_gn=ATP9B;refGene_tx=XM_023242127.1;refGene_gn=ATP9B;refGene_tx=XM_023242128.1
D3	95993710	95993786
refGene_gn=ATP9B;refGene_tx=XM_023242122.1;refGene_gn=ATP9B;refGene_tx=XM_023242123.1;refGene_gn=ATP9B;refGene_tx=XM_023242124.1;refGene_gn=

ATP9B;refGene_tx=XM_023242125.1;refGene_gn=ATP9B;refGene_tx=XM_0232421
26.1;refGene_gn=ATP9B;refGene_tx=XM_023242127.1;refGene_gn=ATP9B;refGe
ne_tx=XM_023242128.1
D3 96000224 96000301
refGene_gn=ATP9B;refGene_tx=XM_023242122.1;refGene_gn=ATP9B;refGene_tx
=XM_023242123.1;refGene_gn=ATP9B;refGene_tx=XM_023242124.1;refGene_gn=
ATP9B;refGene_tx=XM_023242125.1;refGene_gn=ATP9B;refGene_tx=XM_0232421
26.1;refGene_gn=ATP9B;refGene_tx=XM_023242127.1
D3 96033905 96034066
refGene_gn=ATP9B;refGene_tx=XM_023242122.1;refGene_gn=ATP9B;refGene_tx
=XM_023242123.1;refGene_gn=ATP9B;refGene_tx=XM_023242124.1;refGene_gn=
ATP9B;refGene_tx=XM_023242125.1;refGene_gn=ATP9B;refGene_tx=XM_0232421
26.1;refGene_gn=ATP9B;refGene_tx=XM_023242127.1
D3 96050138 96050281
refGene_gn=ATP9B;refGene_tx=XM_023242122.1;refGene_gn=ATP9B;refGene_tx
=XM_023242123.1;refGene_gn=ATP9B;refGene_tx=XM_023242124.1;refGene_gn=
ATP9B;refGene_tx=XM_023242125.1;refGene_gn=ATP9B;refGene_tx=XM_0232421
26.1;refGene_gn=ATP9B;refGene_tx=XM_023242127.1
D3 96067709 96067822
refGene_gn=ATP9B;refGene_tx=XM_023242122.1;refGene_gn=ATP9B;refGene_tx
=XM_023242123.1;refGene_gn=ATP9B;refGene_tx=XM_023242124.1;refGene_gn=
ATP9B;refGene_tx=XM_023242125.1;refGene_gn=ATP9B;refGene_tx=XM_0232421
26.1;refGene_gn=ATP9B;refGene_tx=XM_023242127.1;refGene_gn=ATP9B;refGe
ne_tx=XM_023242128.1
D3 96071584 96071833
refGene_gn=ATP9B;refGene_tx=XM_023242122.1;refGene_gn=ATP9B;refGene_tx
=XM_023242123.1;refGene_gn=ATP9B;refGene_tx=XM_023242124.1;refGene_gn=
ATP9B;refGene_tx=XM_023242125.1;refGene_gn=ATP9B;refGene_tx=XM_0232421
26.1;refGene_gn=ATP9B;refGene_tx=XM_023242127.1;refGene_gn=ATP9B;refGe
ne_tx=XM_023242128.1
D3 96087027 96087189
refGene_gn=ATP9B;refGene_tx=XM_023242122.1;refGene_gn=ATP9B;refGene_tx
=XM_023242123.1;refGene_gn=ATP9B;refGene_tx=XM_023242124.1;refGene_gn=
ATP9B;refGene_tx=XM_023242125.1;refGene_gn=ATP9B;refGene_tx=XM_0232421
26.1;refGene_gn=ATP9B;refGene_tx=XM_023242127.1;refGene_gn=ATP9B;refGe
ne_tx=XM_023242128.1
D3 96087881 96087974
refGene_gn=ATP9B;refGene_tx=XM_023242122.1;refGene_gn=ATP9B;refGene_tx
=XM_023242123.1;refGene_gn=ATP9B;refGene_tx=XM_023242124.1;refGene_gn=
ATP9B;refGene_tx=XM_023242125.1;refGene_gn=ATP9B;refGene_tx=XM_0232421
26.1;refGene_gn=ATP9B;refGene_tx=XM_023242127.1;refGene_gn=ATP9B;refGe
ne_tx=XM_023242128.1
D3 96092450 96092534
refGene_gn=ATP9B;refGene_tx=XM_023242122.1;refGene_gn=ATP9B;refGene_tx
=XM_023242123.1;refGene_gn=ATP9B;refGene_tx=XM_023242124.1;refGene_gn=
ATP9B;refGene_tx=XM_023242125.1;refGene_gn=ATP9B;refGene_tx=XM_0232421
26.1;refGene_gn=ATP9B;refGene_tx=XM_023242128.1
D3 96092623 96092794
refGene_gn=ATP9B;refGene_tx=XM_023242122.1;refGene_gn=ATP9B;refGene_tx
=XM_023242123.1;refGene_gn=ATP9B;refGene_tx=XM_023242124.1;refGene_gn=

ATP9B;refGene_tx=XM_023242125.1;refGene_gn=ATP9B;refGene_tx=XM_0232421
26.1;refGene_gn=ATP9B;refGene_tx=XM_023242128.1
D3 96095998 96096097
refGene_gn=ATP9B;refGene_tx=XM_023242122.1;refGene_gn=ATP9B;refGene_tx
=XM_023242123.1;refGene_gn=ATP9B;refGene_tx=XM_023242124.1;refGene_gn=
ATP9B;refGene_tx=XM_023242125.1;refGene_gn=ATP9B;refGene_tx=XM_0232421
26.1;refGene_gn=ATP9B;refGene_tx=XM_023242127.1;refGene_gn=ATP9B;refGe
ne_tx=XM_023242128.1
D3 96096731 96096821
refGene_gn=ATP9B;refGene_tx=XM_023242122.1;refGene_gn=ATP9B;refGene_tx
=XM_023242123.1;refGene_gn=ATP9B;refGene_tx=XM_023242124.1;refGene_gn=
ATP9B;refGene_tx=XM_023242125.1;refGene_gn=ATP9B;refGene_tx=XM_0232421
26.1;refGene_gn=ATP9B;refGene_tx=XM_023242127.1;refGene_gn=ATP9B;refGe
ne_tx=XM_023242128.1
D3 96097571 96097716
refGene_gn=ATP9B;refGene_tx=XM_023242122.1;refGene_gn=ATP9B;refGene_tx
=XM_023242123.1;refGene_gn=ATP9B;refGene_tx=XM_023242124.1;refGene_gn=
ATP9B;refGene_tx=XM_023242125.1;refGene_gn=ATP9B;refGene_tx=XM_0232421
26.1;refGene_gn=ATP9B;refGene_tx=XM_023242127.1;refGene_gn=ATP9B;refGe
ne_tx=XM_023242128.1
D3 96097815 96097880
refGene_gn=ATP9B;refGene_tx=XM_023242122.1;refGene_gn=ATP9B;refGene_tx
=XM_023242123.1;refGene_gn=ATP9B;refGene_tx=XM_023242124.1;refGene_gn=
ATP9B;refGene_tx=XM_023242125.1;refGene_gn=ATP9B;refGene_tx=XM_0232421
26.1;refGene_gn=ATP9B;refGene_tx=XM_023242127.1;refGene_gn=ATP9B;refGe
ne_tx=XM_023242128.1
D3 96098401 96098557
refGene_gn=ATP9B;refGene_tx=XM_023242122.1;refGene_gn=ATP9B;refGene_tx
=XM_023242123.1;refGene_gn=ATP9B;refGene_tx=XM_023242124.1;refGene_gn=
ATP9B;refGene_tx=XM_023242125.1;refGene_gn=ATP9B;refGene_tx=XM_0232421
26.1;refGene_gn=ATP9B;refGene_tx=XM_023242127.1;refGene_gn=ATP9B;refGe
ne_tx=XM_023242128.1
D3 96098668 96098733
refGene_gn=ATP9B;refGene_tx=XM_023242122.1;refGene_gn=ATP9B;refGene_tx
=XM_023242123.1;refGene_gn=ATP9B;refGene_tx=XM_023242124.1;refGene_gn=
ATP9B;refGene_tx=XM_023242125.1;refGene_gn=ATP9B;refGene_tx=XM_0232421
26.1;refGene_gn=ATP9B;refGene_tx=XM_023242127.1;refGene_gn=ATP9B;refGe
ne_tx=XM_023242128.1
D3 96103038 96103147
refGene_gn=ATP9B;refGene_tx=XM_023242122.1;refGene_gn=ATP9B;refGene_tx
=XM_023242123.1;refGene_gn=ATP9B;refGene_tx=XM_023242124.1;refGene_gn=
ATP9B;refGene_tx=XM_023242125.1;refGene_gn=ATP9B;refGene_tx=XM_0232421
26.1;refGene_gn=ATP9B;refGene_tx=XM_023242127.1;refGene_gn=ATP9B;refGe
ne_tx=XM_023242128.1
D3 96111695 96111753
refGene_gn=ATP9B;refGene_tx=XM_023242122.1;refGene_gn=ATP9B;refGene_tx
=XM_023242123.1;refGene_gn=ATP9B;refGene_tx=XM_023242124.1;refGene_gn=
ATP9B;refGene_tx=XM_023242125.1;refGene_gn=ATP9B;refGene_tx=XM_0232421
26.1;refGene_gn=ATP9B;refGene_tx=XM_023242127.1;refGene_gn=ATP9B;refGe
ne_tx=XM_023242128.1

D3 96112362 96112566
refGene_gn=ATP9B;refGene_tx=XM_023242122.1;refGene_gn=ATP9B;refGene_tx=XM_023242123.1;refGene_gn=ATP9B;refGene_tx=XM_023242124.1;refGene_gn=ATP9B;refGene_tx=XM_023242125.1;refGene_gn=ATP9B;refGene_tx=XM_023242126.1;refGene_gn=ATP9B;refGene_tx=XM_023242127.1;refGene_gn=ATP9B;refGene_tx=XM_023242128.1
D3 96112652 96112717 refGene_gn=ATP9B;refGene_tx=XM_023242124.1
D3 96117097 96117234
refGene_gn=ATP9B;refGene_tx=XM_023242122.1;refGene_gn=ATP9B;refGene_tx=XM_023242123.1;refGene_gn=ATP9B;refGene_tx=XM_023242125.1;refGene_gn=ATP9B;refGene_tx=XM_023242126.1;refGene_gn=ATP9B;refGene_tx=XM_023242127.1;refGene_gn=ATP9B;refGene_tx=XM_023242128.1
D3 96135462 96135589
refGene_gn=NFATC1;refGene_tx=XM_023242129.1;refGene_gn=NFATC1;refGene_tx=XM_023242130.1;refGene_gn=NFATC1;refGene_tx=XM_023242131.1;refGene_gn=NFATC1;refGene_tx=XM_023242132.1;refGene_gn=NFATC1;refGene_tx=XM_023242133.1
D3 96139847 96139935 refGene_gn=NFATC1;refGene_tx=XM_006938981.4
D3 96148874 96149970
refGene_gn=NFATC1;refGene_tx=XM_006938981.4;refGene_gn=NFATC1;refGene_tx=XM_023242129.1;refGene_gn=NFATC1;refGene_tx=XM_023242130.1;refGene_gn=NFATC1;refGene_tx=XM_023242131.1;refGene_gn=NFATC1;refGene_tx=XM_023242132.1;refGene_gn=NFATC1;refGene_tx=XM_023242133.1
D3 96163668 96163828
refGene_gn=NFATC1;refGene_tx=XM_006938981.4;refGene_gn=NFATC1;refGene_tx=XM_023242129.1;refGene_gn=NFATC1;refGene_tx=XM_023242130.1;refGene_gn=NFATC1;refGene_tx=XM_023242131.1;refGene_gn=NFATC1;refGene_tx=XM_023242132.1;refGene_gn=NFATC1;refGene_tx=XM_023242133.1
D3 96179823 96180026
refGene_gn=NFATC1;refGene_tx=XM_006938981.4;refGene_gn=NFATC1;refGene_tx=XM_023242129.1;refGene_gn=NFATC1;refGene_tx=XM_023242130.1;refGene_gn=NFATC1;refGene_tx=XM_023242131.1;refGene_gn=NFATC1;refGene_tx=XM_023242132.1;refGene_gn=NFATC1;refGene_tx=XM_023242133.1
D3 96180522 96180695
refGene_gn=NFATC1;refGene_tx=XM_006938981.4;refGene_gn=NFATC1;refGene_tx=XM_023242129.1;refGene_gn=NFATC1;refGene_tx=XM_023242130.1;refGene_gn=NFATC1;refGene_tx=XM_023242131.1;refGene_gn=NFATC1;refGene_tx=XM_023242132.1;refGene_gn=NFATC1;refGene_tx=XM_023242133.1
D3 96182285 96182426
refGene_gn=NFATC1;refGene_tx=XM_006938981.4;refGene_gn=NFATC1;refGene_tx=XM_023242129.1;refGene_gn=NFATC1;refGene_tx=XM_023242130.1;refGene_gn=NFATC1;refGene_tx=XM_023242131.1;refGene_gn=NFATC1;refGene_tx=XM_023242132.1;refGene_gn=NFATC1;refGene_tx=XM_023242133.1
D3 96192585 96192641
refGene_gn=NFATC1;refGene_tx=XM_006938981.4;refGene_gn=NFATC1;refGene_tx=XM_023242129.1;refGene_gn=NFATC1;refGene_tx=XM_023242130.1;refGene_gn=NFATC1;refGene_tx=XM_023242131.1;refGene_gn=NFATC1;refGene_tx=XM_023242132.1;refGene_gn=NFATC1;refGene_tx=XM_023242133.1
D3 96196644 96196836
refGene_gn=NFATC1;refGene_tx=XM_006938981.4;refGene_gn=NFATC1;refGene_

tx=XM_023242129.1;refGene_gn=NFATC1;refGene_tx=XM_023242130.1;refGene_gn=NFATC1;refGene_tx=XM_023242131.1;refGene_gn=NFATC1;refGene_tx=XM_023242132.1;refGene_gn=NFATC1;refGene_tx=XM_023242133.1
D3 96198191 96198253 refGene_gn=NFATC1;refGene_tx=XM_023242131.1
D3 96209343 96209726
refGene_gn=NFATC1;refGene_tx=XM_006938981.4;refGene_gn=NFATC1;refGene_tx=XM_023242129.1;refGene_gn=NFATC1;refGene_tx=XM_023242130.1
D3 96209837 96210033
refGene_gn=NFATC1;refGene_tx=XM_006938981.4;refGene_gn=NFATC1;refGene_tx=XM_023242129.1
D3 96239866 96239910
refGene_gn=NFATC1;refGene_tx=XM_006938981.4;refGene_gn=NFATC1;refGene_tx=XM_023242129.1;refGene_gn=NFATC1;refGene_tx=XM_023242130.1;refGene_gn=NFATC1;refGene_tx=XM_023242133.1
D3 96347177 96347482
refGene_gn=CTDP1;refGene_tx=XM_006938983.4;refGene_gn=CTDP1;refGene_tx=XM_011288118.3;refGene_gn=CTDP1;refGene_tx=XM_019815469.2;refGene_gn=CTDP1;refGene_tx=XM_019815470.2
D3 96357407 96357491
refGene_gn=CTDP1;refGene_tx=XM_006938983.4;refGene_gn=CTDP1;refGene_tx=XM_011288118.3;refGene_gn=CTDP1;refGene_tx=XM_019815469.2;refGene_gn=CTDP1;refGene_tx=XM_019815470.2
D3 96358120 96358214
refGene_gn=CTDP1;refGene_tx=XM_006938983.4;refGene_gn=CTDP1;refGene_tx=XM_011288118.3;refGene_gn=CTDP1;refGene_tx=XM_019815469.2;refGene_gn=CTDP1;refGene_tx=XM_019815470.2
D3 96359462 96359591
refGene_gn=CTDP1;refGene_tx=XM_006938983.4;refGene_gn=CTDP1;refGene_tx=XM_011288118.3;refGene_gn=CTDP1;refGene_tx=XM_019815469.2;refGene_gn=CTDP1;refGene_tx=XM_019815470.2
D3 96362324 96362475
refGene_gn=CTDP1;refGene_tx=XM_006938983.4;refGene_gn=CTDP1;refGene_tx=XM_011288118.3;refGene_gn=CTDP1;refGene_tx=XM_019815469.2;refGene_gn=CTDP1;refGene_tx=XM_019815470.2
D3 96364895 96364986
refGene_gn=CTDP1;refGene_tx=XM_006938983.4;refGene_gn=CTDP1;refGene_tx=XM_011288118.3;refGene_gn=CTDP1;refGene_tx=XM_019815469.2;refGene_gn=CTDP1;refGene_tx=XM_019815470.2
D3 96366754 96366921
refGene_gn=CTDP1;refGene_tx=XM_006938983.4;refGene_gn=CTDP1;refGene_tx=XM_011288118.3;refGene_gn=CTDP1;refGene_tx=XM_019815469.2;refGene_gn=CTDP1;refGene_tx=XM_019815470.2
D3 96368013 96369003
refGene_gn=CTDP1;refGene_tx=XM_006938983.4;refGene_gn=CTDP1;refGene_tx=XM_011288118.3;refGene_gn=CTDP1;refGene_tx=XM_019815469.2;refGene_gn=CTDP1;refGene_tx=XM_019815470.2
D3 96370377 96370519
refGene_gn=CTDP1;refGene_tx=XM_006938983.4;refGene_gn=CTDP1;refGene_tx=XM_011288118.3;refGene_gn=CTDP1;refGene_tx=XM_019815469.2;refGene_gn=CTDP1;refGene_tx=XM_019815470.2

D3 96371063 96371298
refGene_gn=CTDP1;refGene_tx=XM_006938983.4;refGene_gn=CTDP1;refGene_tx=XM_011288118.3;refGene_gn=CTDP1;refGene_tx=XM_019815469.2;refGene_gn=CTDP1;refGene_tx=XM_019815470.2
D3 96379813 96379973
refGene_gn=CTDP1;refGene_tx=XM_006938983.4;refGene_gn=CTDP1;refGene_tx=XM_011288118.3;refGene_gn=CTDP1;refGene_tx=XM_019815469.2;refGene_gn=CTDP1;refGene_tx=XM_019815470.2
D3 96391685 96391846 refGene_gn=CTDP1;refGene_tx=XM_006938983.4
D3 96403334 96403571
refGene_gn=CTDP1;refGene_tx=XM_006938983.4;refGene_gn=CTDP1;refGene_tx=XM_011288118.3
D3 96478750 96479305 refGene_gn=KCNG2;refGene_tx=XM_023242134.1
D3 96505283 96506093 refGene_gn=KCNG2;refGene_tx=XM_023242134.1
D3 96511734 96511782 refGene_gn=PQLC1;refGene_tx=XM_023242137.1
D3 96512073 96512281
refGene_gn=PQLC1;refGene_tx=XM_019815472.2;refGene_gn=PQLC1;refGene_tx=XM_023242135.1;refGene_gn=PQLC1;refGene_tx=XM_023242136.1;refGene_gn=PQLC1;refGene_tx=XM_023242137.1
D3 96519406 96519488 refGene_gn=PQLC1;refGene_tx=XM_023242138.1
D3 96520302 96520519
refGene_gn=PQLC1;refGene_tx=XM_019815472.2;refGene_gn=PQLC1;refGene_tx=XM_023242135.1;refGene_gn=PQLC1;refGene_tx=XM_023242136.1;refGene_gn=PQLC1;refGene_tx=XM_023242137.1;refGene_gn=PQLC1;refGene_tx=XM_023242138.1
D3 96547146 96547200
refGene_gn=PQLC1;refGene_tx=XM_023242135.1;refGene_gn=PQLC1;refGene_tx=XM_023242136.1;refGene_gn=PQLC1;refGene_tx=XM_023242137.1;refGene_gn=PQLC1;refGene_tx=XM_023242138.1
D3 96555269 96555403
refGene_gn=PQLC1;refGene_tx=XM_019815472.2;refGene_gn=PQLC1;refGene_tx=XM_023242135.1;refGene_gn=PQLC1;refGene_tx=XM_023242136.1;refGene_gn=PQLC1;refGene_tx=XM_023242137.1;refGene_gn=PQLC1;refGene_tx=XM_023242138.1
D3 96563570 96563773
refGene_gn=PQLC1;refGene_tx=XM_019815472.2;refGene_gn=PQLC1;refGene_tx=XM_023242135.1;refGene_gn=PQLC1;refGene_tx=XM_023242136.1;refGene_gn=PQLC1;refGene_tx=XM_023242137.1;refGene_gn=PQLC1;refGene_tx=XM_023242138.1
D3 96564129 96564387
refGene_gn=LOC111557235;refGene_tx=XM_023241206.1
D3 96564826 96564967
refGene_gn=LOC111557235;refGene_tx=XM_023241206.1
D3 96565247 96565411
refGene_gn=LOC111557235;refGene_tx=XM_023241206.1
D3 96567473 96567918
refGene_gn=LOC111557235;refGene_tx=XM_023241206.1
D3 96581082 96581133
refGene_gn=HSBP1L1;refGene_tx=XM_019815475.2;refGene_gn=HSBP1L1;refGene_tx=XM_019815476.2;refGene_gn=HSBP1L1;refGene_tx=XM_019815477.2

D3	96582104 96582171
refGene_gn=HSBP1L1;refGene_tx=XM_019815475.2;refGene_gn=HSBP1L1;refGene_tx=XM_019815476.2;refGene_gn=HSBP1L1;refGene_tx=XM_019815477.2
D3	96583160 96583255
refGene_gn=HSBP1L1;refGene_tx=XM_019815475.2;refGene_gn=HSBP1L1;refGene_tx=XM_019815476.2;refGene_gn=HSBP1L1;refGene_tx=XM_019815477.2
D3	96585744 96585750 refGene_gn=HSBP1L1;refGene_tx=XM_019815477.2
D3	96590906 96590927 refGene_gn=HSBP1L1;refGene_tx=XM_019815475.2
D3	96592398 96592570
refGene_gn=TXNL4A;refGene_tx=XM_003995280.4;refGene_gn=TXNL4A;refGene_tx=XM_019815478.1;refGene_gn=TXNL4A;refGene_tx=XM_023242144.1
D3	96595534 96595638
refGene_gn=TXNL4A;refGene_tx=XM_003995280.4;refGene_gn=TXNL4A;refGene_tx=XM_019815478.1;refGene_gn=TXNL4A;refGene_tx=XM_023242144.1
D3	96603834 96603987
refGene_gn=TXNL4A;refGene_tx=XM_003995280.4;refGene_gn=TXNL4A;refGene_tx=XM_023242144.1
D3	96619369 96619381
refGene_gn=LOC102902085;refGene_tx=XM_011288115.3;refGene_gn=LOC102902085;refGene_tx=XM_019815479.2;refGene_gn=LOC102902085;refGene_tx=XM_019815480.2;refGene_gn=LOC102902085;refGene_tx=XM_019815481.2
D3	96619712 96619802
refGene_gn=LOC102902085;refGene_tx=XM_006938986.4;refGene_gn=LOC102902085;refGene_tx=XM_023242140.1
D3	96620230 96620615
refGene_gn=LOC102902085;refGene_tx=XM_006938986.4;refGene_gn=LOC102902085;refGene_tx=XM_011288115.3;refGene_gn=LOC102902085;refGene_tx=XM_019815479.2;refGene_gn=LOC102902085;refGene_tx=XM_019815480.2;refGene_gn=LOC102902085;refGene_tx=XM_019815481.2;refGene_gn=LOC102902085;refGene_tx=XM_019815483.2;refGene_gn=LOC102902085;refGene_tx=XM_019815484.2;refGene_gn=LOC102902085;refGene_tx=XM_019815485.2;refGene_gn=LOC102902085;refGene_tx=XM_019815486.2;refGene_gn=LOC102902085;refGene_tx=XM_023242139.1;refGene_gn=LOC102902085;refGene_tx=XM_023242140.1
D3	96621470 96621642
refGene_gn=LOC102902085;refGene_tx=XM_006938986.4;refGene_gn=LOC102902085;refGene_tx=XM_011288115.3;refGene_gn=LOC102902085;refGene_tx=XM_019815479.2;refGene_gn=LOC102902085;refGene_tx=XM_019815480.2;refGene_gn=LOC102902085;refGene_tx=XM_019815481.2;refGene_gn=LOC102902085;refGene_tx=XM_019815483.2;refGene_gn=LOC102902085;refGene_tx=XM_019815484.2;refGene_gn=LOC102902085;refGene_tx=XM_019815485.2;refGene_gn=LOC102902085;refGene_tx=XM_019815486.2;refGene_gn=LOC102902085;refGene_tx=XM_023242139.1;refGene_gn=LOC102902085;refGene_tx=XM_023242140.1
D3	96623100 96623279
refGene_gn=LOC102902085;refGene_tx=XM_006938986.4;refGene_gn=LOC102902085;refGene_tx=XM_011288115.3;refGene_gn=LOC102902085;refGene_tx=XM_019815479.2;refGene_gn=LOC102902085;refGene_tx=XM_019815480.2;refGene_gn=LOC102902085;refGene_tx=XM_019815481.2;refGene_gn=LOC102902085;refGene_tx=XM_019815483.2;refGene_gn=LOC102902085;refGene_tx=XM_019815484.2;refGene_gn=LOC102902085;refGene_tx=XM_019815485.2;refGene_gn=LOC102902085;refGene_tx=XM_019815486.2;refGene_gn=LOC102902085;refGene_tx=XM_02

3242139.1;refGene_gn=LOC102902085;refGene_tx=XM_023242140.1
D3 96624276 96624302
refGene_gn=LOC102902085;refGene_tx=XM_023242140.1
D3 96626041 96626138
refGene_gn=LOC102902085;refGene_tx=XM_006938986.4;refGene_gn=LOC102902085;refGene_tx=XM_011288115.3;refGene_gn=LOC102902085;refGene_tx=XM_019815479.2;refGene_gn=LOC102902085;refGene_tx=XM_019815480.2;refGene_gn=LOC102902085;refGene_tx=XM_019815481.2;refGene_gn=LOC102902085;refGene_tx=XM_019815483.2;refGene_gn=LOC102902085;refGene_tx=XM_019815484.2;refGene_gn=LOC102902085;refGene_tx=XM_019815485.2;refGene_gn=LOC102902085;refGene_tx=XM_019815486.2;refGene_gn=LOC102902085;refGene_tx=XM_023242139.1
D3 96629101 96629276
refGene_gn=LOC102902085;refGene_tx=XM_006938986.4;refGene_gn=LOC102902085;refGene_tx=XM_011288115.3;refGene_gn=LOC102902085;refGene_tx=XM_019815479.2;refGene_gn=LOC102902085;refGene_tx=XM_019815480.2;refGene_gn=LOC102902085;refGene_tx=XM_019815481.2;refGene_gn=LOC102902085;refGene_tx=XM_019815483.2;refGene_gn=LOC102902085;refGene_tx=XM_019815484.2;refGene_gn=LOC102902085;refGene_tx=XM_019815485.2;refGene_gn=LOC102902085;refGene_tx=XM_019815486.2;refGene_gn=LOC102902085;refGene_tx=XM_023242139.1
D3 96630335 96630496
refGene_gn=RBFA;refGene_tx=XM_023242141.1;refGene_gn=RBFA;refGene_tx=XM_023242142.1;refGene_gn=RBFA;refGene_tx=XM_023242143.1
D3 96633132 96633175
refGene_gn=RBFA;refGene_tx=XM_023242141.1;refGene_gn=RBFA;refGene_tx=XM_023242142.1;refGene_gn=RBFA;refGene_tx=XM_023242143.1
D3 96633799 96633976
refGene_gn=RBFA;refGene_tx=XM_023242141.1;refGene_gn=RBFA;refGene_tx=XM_023242142.1;refGene_gn=RBFA;refGene_tx=XM_023242143.1
D3 96634555 96634668
refGene_gn=RBFA;refGene_tx=XM_023242141.1;refGene_gn=RBFA;refGene_tx=XM_023242142.1;refGene_gn=RBFA;refGene_tx=XM_023242143.1
D3 96637445 96637530
refGene_gn=RBFA;refGene_tx=XM_023242141.1;refGene_gn=RBFA;refGene_tx=XM_023242142.1;refGene_gn=RBFA;refGene_tx=XM_023242143.1
D3 96641319 96641393 refGene_gn=RBFA;refGene_tx=XM_023242141.1
D3 96642457 96642749 refGene_gn=RBFA;refGene_tx=XM_023242141.1
D3 96679801 96680005
refGene_gn=RBFA;refGene_tx=XM_023242142.1;refGene_gn=RBFA;refGene_tx=XM_023242143.1
D3 96680019 96680100 refGene_gn=RBFA;refGene_tx=XM_023242143.1
D3 96713962 96714070
refGene_gn=ADNP2;refGene_tx=XM_003995299.5;refGene_gn=ADNP2;refGene_tx=XM_019815488.2;refGene_gn=ADNP2;refGene_tx=XM_023242145.1
D3 96728206 96728296
refGene_gn=ADNP2;refGene_tx=XM_003995299.5;refGene_gn=ADNP2;refGene_tx=XM_019815488.2;refGene_gn=ADNP2;refGene_tx=XM_023242145.1
D3 96733134 96736410
refGene_gn=ADNP2;refGene_tx=XM_003995299.5;refGene_gn=ADNP2;refGene_tx

=XM_019815488.2;refGene_gn=ADNP2;refGene_tx=XM_019815489.2;refGene_gn=ADNP2;refGene_tx=XM_023242145.1
D3 96771785 96772582 refGene_gn=PARD6G;refGene_tx=XM_023242146.1
D3 96830733 96830956 refGene_gn=PARD6G;refGene_tx=XM_023242146.1
D3 96858352 96858424 refGene_gn=PARD6G;refGene_tx=XM_023242146.1
D4 4958 5147
refGene_gn=LOC111557576;refGene_tx=XM_023243177.1
D4 7812 7936
refGene_gn=LOC111557576;refGene_tx=XM_023243178.1
D4 8208 8919
refGene_gn=LOC111557576;refGene_tx=XM_023243176.1;refGene_gn=LOC111557576;refGene_tx=XM_023243178.1
D4 73636 74390
refGene_gn=LOC105260391;refGene_tx=XM_011280555.3
D4 74391 75509
refGene_gn=LOC105260391;refGene_tx=XM_011280555.3
D4 75511 75931
refGene_gn=LOC105260391;refGene_tx=XM_011280555.3
D4 92187 92657
refGene_gn=LOC111557577;refGene_tx=XM_023243182.1
D4 96033 96217
refGene_gn=LOC111557577;refGene_tx=XM_023243182.1
D4 127280 127388
refGene_gn=MFSD14B;refGene_tx=XM_003995301.5;refGene_gn=MFSD14B;refGene_tx=XM_023243180.1
D4 157036 157132
refGene_gn=MFSD14B;refGene_tx=XM_003995301.5;refGene_gn=MFSD14B;refGene_tx=XM_023243180.1;refGene_gn=MFSD14B;refGene_tx=XM_023243181.1
D4 170976 171039
refGene_gn=MFSD14B;refGene_tx=XM_003995301.5;refGene_gn=MFSD14B;refGene_tx=XM_023243181.1
D4 183407 183535
refGene_gn=MFSD14B;refGene_tx=XM_003995301.5;refGene_gn=MFSD14B;refGene_tx=XM_023243180.1;refGene_gn=MFSD14B;refGene_tx=XM_023243181.1
D4 185093 185208
refGene_gn=MFSD14B;refGene_tx=XM_003995301.5;refGene_gn=MFSD14B;refGene_tx=XM_023243180.1;refGene_gn=MFSD14B;refGene_tx=XM_023243181.1
D4 190239 190458
refGene_gn=MFSD14B;refGene_tx=XM_003995301.5;refGene_gn=MFSD14B;refGene_tx=XM_023243180.1;refGene_gn=MFSD14B;refGene_tx=XM_023243181.1
D4 192004 192118
refGene_gn=MFSD14B;refGene_tx=XM_003995301.5;refGene_gn=MFSD14B;refGene_tx=XM_023243180.1;refGene_gn=MFSD14B;refGene_tx=XM_023243181.1
D4 194704 194776
refGene_gn=MFSD14B;refGene_tx=XM_003995301.5;refGene_gn=MFSD14B;refGene_tx=XM_023243180.1;refGene_gn=MFSD14B;refGene_tx=XM_023243181.1
D4 196366 196479
refGene_gn=MFSD14B;refGene_tx=XM_003995301.5;refGene_gn=MFSD14B;refGene_tx=XM_023243180.1;refGene_gn=MFSD14B;refGene_tx=XM_023243181.1
D4 197767 197865

refGene_gn=MFSD14B;refGene_tx=XM_003995301.5;refGene_gn=MFSD14B;refGene_tx=XM_023243180.1;refGene_gn=MFSD14B;refGene_tx=XM_023243181.1
D4	199845	199988
refGene_gn=MFSD14B;refGene_tx=XM_003995301.5;refGene_gn=MFSD14B;refGene_tx=XM_023243180.1;refGene_gn=MFSD14B;refGene_tx=XM_023243181.1
D4	200673	200925
refGene_gn=MFSD14B;refGene_tx=XM_003995301.5;refGene_gn=MFSD14B;refGene_tx=XM_023243180.1;refGene_gn=MFSD14B;refGene_tx=XM_023243181.1
D4	261327	261512
refGene_gn=LOC102899568;refGene_tx=XM_023243185.1
D4	297640	297739
refGene_gn=LOC102899568;refGene_tx=XM_023243185.1
D4	298013	298047
refGene_gn=LOC102899568;refGene_tx=XM_023243185.1
D4	335389	337704
refGene_gn=ZNF484;refGene_tx=XM_011288210.3;refGene_gn=ZNF484;refGene_tx=XM_019815630.2;refGene_gn=ZNF484;refGene_tx=XM_019815631.2;refGene_gn=ZNF484;refGene_tx=XM_023243183.1
D4	344520	344613
refGene_gn=ZNF484;refGene_tx=XM_011288210.3;refGene_gn=ZNF484;refGene_tx=XM_019815630.2;refGene_gn=ZNF484;refGene_tx=XM_019815631.2;refGene_gn=ZNF484;refGene_tx=XM_023243183.1
D4	351893	352020
refGene_gn=ZNF484;refGene_tx=XM_011288210.3;refGene_gn=ZNF484;refGene_tx=XM_019815630.2;refGene_gn=ZNF484;refGene_tx=XM_019815631.2;refGene_gn=ZNF484;refGene_tx=XM_023243183.1
D4	352537	352582	refGene_gn=ZNF484;refGene_tx=XM_023243183.1
D4	360840	360855
refGene_gn=ZNF484;refGene_tx=XM_011288210.3;refGene_gn=ZNF484;refGene_tx=XM_019815630.2
D4	397616	397699
refGene_gn=IARS;refGene_tx=XM_006939004.4;refGene_gn=IARS;refGene_tx=XM_011288208.3;refGene_gn=IARS;refGene_tx=XM_011288209.3;refGene_gn=IARS;refGene_tx=XM_023243186.1
D4	401287	401440
refGene_gn=IARS;refGene_tx=XM_006939004.4;refGene_gn=IARS;refGene_tx=XM_011288208.3;refGene_gn=IARS;refGene_tx=XM_011288209.3;refGene_gn=IARS;refGene_tx=XM_023243186.1
D4	402022	402166
refGene_gn=IARS;refGene_tx=XM_006939004.4;refGene_gn=IARS;refGene_tx=XM_011288208.3;refGene_gn=IARS;refGene_tx=XM_011288209.3;refGene_gn=IARS;refGene_tx=XM_023243186.1
D4	403573	403699
refGene_gn=IARS;refGene_tx=XM_006939004.4;refGene_gn=IARS;refGene_tx=XM_011288208.3;refGene_gn=IARS;refGene_tx=XM_011288209.3;refGene_gn=IARS;refGene_tx=XM_023243186.1
D4	410508	410614
refGene_gn=IARS;refGene_tx=XM_006939004.4;refGene_gn=IARS;refGene_tx=XM_011288208.3;refGene_gn=IARS;refGene_tx=XM_011288209.3;refGene_gn=IARS;refGene_tx=XM_023243186.1

D4 411417 411594
refGene_gn=IARS;refGene_tx=XM_006939004.4;refGene_gn=IARS;refGene_tx=XM_011288208.3;refGene_gn=IARS;refGene_tx=XM_011288209.3;refGene_gn=IARS;refGene_tx=XM_023243186.1
D4 412440 412536
refGene_gn=IARS;refGene_tx=XM_006939004.4;refGene_gn=IARS;refGene_tx=XM_011288208.3;refGene_gn=IARS;refGene_tx=XM_011288209.3;refGene_gn=IARS;refGene_tx=XM_023243186.1
D4 414237 414350
refGene_gn=IARS;refGene_tx=XM_006939004.4;refGene_gn=IARS;refGene_tx=XM_011288208.3;refGene_gn=IARS;refGene_tx=XM_011288209.3;refGene_gn=IARS;refGene_tx=XM_023243186.1
D4 416674 416849
refGene_gn=IARS;refGene_tx=XM_006939004.4;refGene_gn=IARS;refGene_tx=XM_011288208.3;refGene_gn=IARS;refGene_tx=XM_011288209.3;refGene_gn=IARS;refGene_tx=XM_023243186.1
D4 418586 418670
refGene_gn=IARS;refGene_tx=XM_006939004.4;refGene_gn=IARS;refGene_tx=XM_011288208.3;refGene_gn=IARS;refGene_tx=XM_011288209.3;refGene_gn=IARS;refGene_tx=XM_023243186.1
D4 418911 419014
refGene_gn=IARS;refGene_tx=XM_006939004.4;refGene_gn=IARS;refGene_tx=XM_011288208.3;refGene_gn=IARS;refGene_tx=XM_011288209.3;refGene_gn=IARS;refGene_tx=XM_023243186.1
D4 419251 419373
refGene_gn=IARS;refGene_tx=XM_006939004.4;refGene_gn=IARS;refGene_tx=XM_011288208.3;refGene_gn=IARS;refGene_tx=XM_011288209.3;refGene_gn=IARS;refGene_tx=XM_023243186.1
D4 420181 420259
refGene_gn=IARS;refGene_tx=XM_006939004.4;refGene_gn=IARS;refGene_tx=XM_011288208.3;refGene_gn=IARS;refGene_tx=XM_011288209.3;refGene_gn=IARS;refGene_tx=XM_023243186.1
D4 421238 421330
refGene_gn=IARS;refGene_tx=XM_006939004.4;refGene_gn=IARS;refGene_tx=XM_011288208.3;refGene_gn=IARS;refGene_tx=XM_011288209.3;refGene_gn=IARS;refGene_tx=XM_023243186.1
D4 423819 423940
refGene_gn=IARS;refGene_tx=XM_006939004.4;refGene_gn=IARS;refGene_tx=XM_011288208.3;refGene_gn=IARS;refGene_tx=XM_011288209.3;refGene_gn=IARS;refGene_tx=XM_023243186.1
D4 425608 425753
refGene_gn=IARS;refGene_tx=XM_006939004.4;refGene_gn=IARS;refGene_tx=XM_011288208.3;refGene_gn=IARS;refGene_tx=XM_011288209.3;refGene_gn=IARS;refGene_tx=XM_023243186.1
D4 426712 426796
refGene_gn=IARS;refGene_tx=XM_006939004.4;refGene_gn=IARS;refGene_tx=XM_011288208.3;refGene_gn=IARS;refGene_tx=XM_011288209.3;refGene_gn=IARS;refGene_tx=XM_023243186.1
D4 428201 428288
refGene_gn=IARS;refGene_tx=XM_006939004.4;refGene_gn=IARS;refGene_tx=X

M_011288208.3;refGene_gn=IARS;refGene_tx=XM_011288209.3;refGene_gn=IARS;refGene_tx=XM_023243186.1
D4 430051 430246
refGene_gn=IARS;refGene_tx=XM_006939004.4;refGene_gn=IARS;refGene_tx=XM_011288208.3;refGene_gn=IARS;refGene_tx=XM_011288209.3;refGene_gn=IARS;refGene_tx=XM_023243186.1
D4 430616 430690
refGene_gn=IARS;refGene_tx=XM_006939004.4;refGene_gn=IARS;refGene_tx=XM_011288208.3;refGene_gn=IARS;refGene_tx=XM_011288209.3;refGene_gn=IARS;refGene_tx=XM_023243186.1
D4 431893 432020
refGene_gn=IARS;refGene_tx=XM_006939004.4;refGene_gn=IARS;refGene_tx=XM_011288208.3;refGene_gn=IARS;refGene_tx=XM_011288209.3;refGene_gn=IARS;refGene_tx=XM_023243186.1
D4 432383 432482
refGene_gn=IARS;refGene_tx=XM_006939004.4;refGene_gn=IARS;refGene_tx=XM_011288208.3;refGene_gn=IARS;refGene_tx=XM_011288209.3;refGene_gn=IARS;refGene_tx=XM_023243186.1
D4 433815 433907
refGene_gn=IARS;refGene_tx=XM_006939004.4;refGene_gn=IARS;refGene_tx=XM_011288208.3;refGene_gn=IARS;refGene_tx=XM_011288209.3;refGene_gn=IARS;refGene_tx=XM_023243186.1
D4 434417 434540
refGene_gn=IARS;refGene_tx=XM_006939004.4;refGene_gn=IARS;refGene_tx=XM_011288208.3;refGene_gn=IARS;refGene_tx=XM_011288209.3;refGene_gn=IARS;refGene_tx=XM_023243186.1
D4 437449 437545
refGene_gn=IARS;refGene_tx=XM_006939004.4;refGene_gn=IARS;refGene_tx=XM_011288208.3;refGene_gn=IARS;refGene_tx=XM_011288209.3;refGene_gn=IARS;refGene_tx=XM_023243186.1
D4 442466 442527
refGene_gn=IARS;refGene_tx=XM_006939004.4;refGene_gn=IARS;refGene_tx=XM_011288208.3;refGene_gn=IARS;refGene_tx=XM_011288209.3;refGene_gn=IARS;refGene_tx=XM_023243186.1
D4 442786 442874
refGene_gn=IARS;refGene_tx=XM_006939004.4;refGene_gn=IARS;refGene_tx=XM_011288208.3;refGene_gn=IARS;refGene_tx=XM_011288209.3;refGene_gn=IARS;refGene_tx=XM_023243186.1
D4 445878 446026
refGene_gn=IARS;refGene_tx=XM_006939004.4;refGene_gn=IARS;refGene_tx=XM_011288208.3;refGene_gn=IARS;refGene_tx=XM_011288209.3;refGene_gn=IARS;refGene_tx=XM_023243186.1
D4 450546 450664
refGene_gn=IARS;refGene_tx=XM_006939004.4;refGene_gn=IARS;refGene_tx=XM_011288208.3;refGene_gn=IARS;refGene_tx=XM_011288209.3;refGene_gn=IARS;refGene_tx=XM_023243186.1
D4 454998 455081
refGene_gn=IARS;refGene_tx=XM_006939004.4;refGene_gn=IARS;refGene_tx=XM_011288208.3;refGene_gn=IARS;refGene_tx=XM_011288209.3;refGene_gn=IARS;refGene_tx=XM_023243186.1

D4 457394 457514
refGene_gn=IARS;refGene_tx=XM_006939004.4;refGene_gn=IARS;refGene_tx=XM_011288208.3;refGene_gn=IARS;refGene_tx=XM_011288209.3;refGene_gn=IARS;refGene_tx=XM_023243186.1
D4 457762 457919
refGene_gn=IARS;refGene_tx=XM_006939004.4;refGene_gn=IARS;refGene_tx=XM_011288208.3;refGene_gn=IARS;refGene_tx=XM_023243186.1
D4 458596 458715
refGene_gn=IARS;refGene_tx=XM_006939004.4;refGene_gn=IARS;refGene_tx=XM_011288208.3;refGene_gn=IARS;refGene_tx=XM_023243186.1
D4 468094 468145
refGene_gn=NOL8;refGene_tx=XM_003995302.5;refGene_gn=NOL8;refGene_tx=XM_006939005.4;refGene_gn=NOL8;refGene_tx=XM_019815641.2;refGene_gn=NOL8;refGene_tx=XM_023243188.1;refGene_gn=NOL8;refGene_tx=XM_023243189.1
D4 468609 468689
refGene_gn=NOL8;refGene_tx=XM_003995302.5;refGene_gn=NOL8;refGene_tx=XM_006939005.4;refGene_gn=NOL8;refGene_tx=XM_019815641.2;refGene_gn=NOL8;refGene_tx=XM_023243188.1;refGene_gn=NOL8;refGene_tx=XM_023243189.1
D4 469860 469934
refGene_gn=NOL8;refGene_tx=XM_003995302.5;refGene_gn=NOL8;refGene_tx=XM_006939005.4;refGene_gn=NOL8;refGene_tx=XM_019815641.2;refGene_gn=NOL8;refGene_tx=XM_023243188.1;refGene_gn=NOL8;refGene_tx=XM_023243189.1
D4 470427 470554
refGene_gn=NOL8;refGene_tx=XM_003995302.5;refGene_gn=NOL8;refGene_tx=XM_006939005.4;refGene_gn=NOL8;refGene_tx=XM_019815641.2;refGene_gn=NOL8;refGene_tx=XM_023243188.1;refGene_gn=NOL8;refGene_tx=XM_023243189.1
D4 471974 472243
refGene_gn=NOL8;refGene_tx=XM_003995302.5;refGene_gn=NOL8;refGene_tx=XM_006939005.4;refGene_gn=NOL8;refGene_tx=XM_019815641.2;refGene_gn=NOL8;refGene_tx=XM_023243188.1;refGene_gn=NOL8;refGene_tx=XM_023243189.1
D4 477183 477265
refGene_gn=NOL8;refGene_tx=XM_003995302.5;refGene_gn=NOL8;refGene_tx=XM_006939005.4;refGene_gn=NOL8;refGene_tx=XM_019815641.2;refGene_gn=NOL8;refGene_tx=XM_019815642.2;refGene_gn=NOL8;refGene_tx=XM_023243188.1;refGene_gn=NOL8;refGene_tx=XM_023243189.1
D4 478859 478998
refGene_gn=NOL8;refGene_tx=XM_003995302.5;refGene_gn=NOL8;refGene_tx=XM_006939005.4;refGene_gn=NOL8;refGene_tx=XM_019815641.2;refGene_gn=NOL8;refGene_tx=XM_019815642.2;refGene_gn=NOL8;refGene_tx=XM_023243188.1;refGene_gn=NOL8;refGene_tx=XM_023243189.1
D4 481564 481655
refGene_gn=NOL8;refGene_tx=XM_003995302.5;refGene_gn=NOL8;refGene_tx=XM_006939005.4;refGene_gn=NOL8;refGene_tx=XM_019815641.2;refGene_gn=NOL8;refGene_tx=XM_019815642.2;refGene_gn=NOL8;refGene_tx=XM_023243188.1;refGene_gn=NOL8;refGene_tx=XM_023243189.1
D4 481883 482003
refGene_gn=NOL8;refGene_tx=XM_003995302.5;refGene_gn=NOL8;refGene_tx=XM_006939005.4;refGene_gn=NOL8;refGene_tx=XM_019815641.2;refGene_gn=NOL8;refGene_tx=XM_019815642.2;refGene_gn=NOL8;refGene_tx=XM_023243188.1;refGene_gn=NOL8;refGene_tx=XM_023243189.1

D4 482730 482838
refGene_gn=NOL8;refGene_tx=XM_003995302.5;refGene_gn=NOL8;refGene_tx=XM_006939005.4;refGene_gn=NOL8;refGene_tx=XM_019815641.2;refGene_gn=NOL8;refGene_tx=XM_019815642.2;refGene_gn=NOL8;refGene_tx=XM_023243188.1;refGene_gn=NOL8;refGene_tx=XM_023243189.1
D4 484337 486170
refGene_gn=NOL8;refGene_tx=XM_003995302.5;refGene_gn=NOL8;refGene_tx=XM_006939005.4;refGene_gn=NOL8;refGene_tx=XM_019815641.2;refGene_gn=NOL8;refGene_tx=XM_019815642.2;refGene_gn=NOL8;refGene_tx=XM_023243188.1;refGene_gn=NOL8;refGene_tx=XM_023243189.1
D4 488145 488214
refGene_gn=NOL8;refGene_tx=XM_003995302.5;refGene_gn=NOL8;refGene_tx=XM_006939005.4;refGene_gn=NOL8;refGene_tx=XM_019815641.2;refGene_gn=NOL8;refGene_tx=XM_019815642.2;refGene_gn=NOL8;refGene_tx=XM_023243188.1;refGene_gn=NOL8;refGene_tx=XM_023243189.1
D4 488727 488863
refGene_gn=NOL8;refGene_tx=XM_003995302.5;refGene_gn=NOL8;refGene_tx=XM_006939005.4;refGene_gn=NOL8;refGene_tx=XM_019815641.2;refGene_gn=NOL8;refGene_tx=XM_019815642.2;refGene_gn=NOL8;refGene_tx=XM_023243188.1;refGene_gn=NOL8;refGene_tx=XM_023243189.1
D4 490508 490587
refGene_gn=NOL8;refGene_tx=XM_003995302.5;refGene_gn=NOL8;refGene_tx=XM_006939005.4;refGene_gn=NOL8;refGene_tx=XM_019815641.2;refGene_gn=NOL8;refGene_tx=XM_019815642.2;refGene_gn=NOL8;refGene_tx=XM_023243188.1;refGene_gn=NOL8;refGene_tx=XM_023243189.1
D4 491934 491997
refGene_gn=NOL8;refGene_tx=XM_003995302.5;refGene_gn=NOL8;refGene_tx=XM_006939005.4;refGene_gn=NOL8;refGene_tx=XM_019815642.2;refGene_gn=NOL8;refGene_tx=XM_023243188.1;refGene_gn=NOL8;refGene_tx=XM_023243189.1
D4 492576 492715
refGene_gn=NOL8;refGene_tx=XM_003995302.5;refGene_gn=NOL8;refGene_tx=XM_006939005.4;refGene_gn=NOL8;refGene_tx=XM_019815641.2;refGene_gn=NOL8;refGene_tx=XM_019815642.2;refGene_gn=NOL8;refGene_tx=XM_023243188.1;refGene_gn=NOL8;refGene_tx=XM_023243189.1
D4 494449 494556
refGene_gn=CENPP;refGene_tx=XM_023243190.1;refGene_gn=CENPP;refGene_tx=XM_023243191.1;refGene_gn=CENPP;refGene_tx=XM_023243192.1;refGene_gn=CENPP;refGene_tx=XM_023243193.1;refGene_gn=CENPP;refGene_tx=XM_023243194.1
D4 513495 513677
refGene_gn=CENPP;refGene_tx=XM_023243190.1;refGene_gn=CENPP;refGene_tx=XM_023243191.1;refGene_gn=CENPP;refGene_tx=XM_023243192.1;refGene_gn=CENPP;refGene_tx=XM_023243193.1;refGene_gn=CENPP;refGene_tx=XM_023243194.1;refGene_gn=CENPP;refGene_tx=XM_023243195.1;refGene_gn=CENPP;refGene_tx=XM_023243196.1;refGene_gn=CENPP;refGene_tx=XM_023243197.1
D4 515229 515318
refGene_gn=CENPP;refGene_tx=XM_023243190.1;refGene_gn=CENPP;refGene_tx=XM_023243191.1;refGene_gn=CENPP;refGene_tx=XM_023243192.1;refGene_gn=CENPP;refGene_tx=XM_023243193.1;refGene_gn=CENPP;refGene_tx=XM_023243194.1;refGene_gn=CENPP;refGene_tx=XM_023243195.1;refGene_gn=CENPP;refGe

ne_tx=XM_023243196.1;refGene_gn=CENPP;refGene_tx=XM_023243197.1
D4	526860	526949
refGene_gn=CENPP;refGene_tx=XM_023243190.1;refGene_gn=CENPP;refGene_tx=XM_023243191.1;refGene_gn=CENPP;refGene_tx=XM_023243192.1;refGene_gn=CENPP;refGene_tx=XM_023243193.1;refGene_gn=CENPP;refGene_tx=XM_023243194.1;refGene_gn=CENPP;refGene_tx=XM_023243195.1;refGene_gn=CENPP;refGene_tx=XM_023243196.1;refGene_gn=CENPP;refGene_tx=XM_023243197.1
D4	551969	552066
refGene_gn=CENPP;refGene_tx=XM_023243190.1;refGene_gn=CENPP;refGene_tx=XM_023243191.1;refGene_gn=CENPP;refGene_tx=XM_023243192.1;refGene_gn=CENPP;refGene_tx=XM_023243193.1;refGene_gn=CENPP;refGene_tx=XM_023243194.1;refGene_gn=CENPP;refGene_tx=XM_023243195.1;refGene_gn=CENPP;refGene_tx=XM_023243196.1;refGene_gn=CENPP;refGene_tx=XM_023243197.1
D4	553141	553276	refGene_gn=CENPP;refGene_tx=XM_023243194.1
D4	557756	557927	refGene_gn=OGN;refGene_tx=XM_023243198.1
D4	558720	558816	refGene_gn=OGN;refGene_tx=XM_023243198.1
D4	560772	560975	refGene_gn=OGN;refGene_tx=XM_023243198.1
D4	563459	563618	refGene_gn=OGN;refGene_tx=XM_023243198.1
D4	566951	566958	refGene_gn=OGN;refGene_tx=XM_023243198.1
D4	573964	574051	refGene_gn=OGN;refGene_tx=XM_023243198.1
D4	576324	576498	refGene_gn=OGN;refGene_tx=XM_023243198.1
D4	587808	588116	refGene_gn=OMD;refGene_tx=XM_006939009.4
D4	589021	589961	refGene_gn=OMD;refGene_tx=XM_006939009.4
D4	611878	612076
refGene_gn=ASPN;refGene_tx=XM_003995305.5;refGene_gn=ASPN;refGene_tx=XM_019815645.2
D4	614189	614328
refGene_gn=ASPN;refGene_tx=XM_003995305.5;refGene_gn=ASPN;refGene_tx=XM_019815645.2
D4	614954	615048
refGene_gn=ASPN;refGene_tx=XM_003995305.5;refGene_gn=ASPN;refGene_tx=XM_019815645.2
D4	618976	619087
refGene_gn=ASPN;refGene_tx=XM_003995305.5;refGene_gn=ASPN;refGene_tx=XM_019815645.2
D4	620196	620410
refGene_gn=ASPN;refGene_tx=XM_003995305.5;refGene_gn=ASPN;refGene_tx=XM_019815645.2
D4	624262	624375
refGene_gn=ASPN;refGene_tx=XM_003995305.5;refGene_gn=ASPN;refGene_tx=XM_019815645.2
D4	627348	627598	refGene_gn=ASPN;refGene_tx=XM_003995305.5
D4	633948	634026	refGene_gn=ASPN;refGene_tx=XM_019815645.2
D4	634489	634541	refGene_gn=ASPN;refGene_tx=XM_019815645.2
D4	650415	650584
refGene_gn=ECM2;refGene_tx=XM_003995303.5;refGene_gn=ECM2;refGene_tx=XM_006939006.4;refGene_gn=ECM2;refGene_tx=XM_006939007.4;refGene_gn=ECM2;refGene_tx=XM_011288207.3
D4	655686	656013
refGene_gn=ECM2;refGene_tx=XM_003995303.5;refGene_gn=ECM2;refGene_tx=X

M_006939006.4;refGene_gn=ECM2;refGene_tx=XM_006939007.4;refGene_gn=ECM2;refGene_tx=XM_011288207.3
D4 657000 657140
refGene_gn=ECM2;refGene_tx=XM_003995303.5;refGene_gn=ECM2;refGene_tx=XM_006939006.4;refGene_gn=ECM2;refGene_tx=XM_006939007.4;refGene_gn=ECM2;refGene_tx=XM_011288207.3
D4 659411 659569
refGene_gn=ECM2;refGene_tx=XM_003995303.5;refGene_gn=ECM2;refGene_tx=XM_006939006.4;refGene_gn=ECM2;refGene_tx=XM_006939007.4;refGene_gn=ECM2;refGene_tx=XM_011288207.3
D4 662272 662408
refGene_gn=ECM2;refGene_tx=XM_003995303.5;refGene_gn=ECM2;refGene_tx=XM_006939006.4;refGene_gn=ECM2;refGene_tx=XM_006939007.4;refGene_gn=ECM2;refGene_tx=XM_011288207.3
D4 664081 664197
refGene_gn=ECM2;refGene_tx=XM_003995303.5;refGene_gn=ECM2;refGene_tx=XM_006939006.4;refGene_gn=ECM2;refGene_tx=XM_006939007.4;refGene_gn=ECM2;refGene_tx=XM_011288207.3
D4 665751 666363
refGene_gn=ECM2;refGene_tx=XM_003995303.5;refGene_gn=ECM2;refGene_tx=XM_006939006.4;refGene_gn=ECM2;refGene_tx=XM_006939007.4;refGene_gn=ECM2;refGene_tx=XM_011288207.3
D4 668253 668442
refGene_gn=ECM2;refGene_tx=XM_003995303.5;refGene_gn=ECM2;refGene_tx=XM_006939006.4;refGene_gn=ECM2;refGene_tx=XM_006939007.4;refGene_gn=ECM2;refGene_tx=XM_011288207.3
D4 669753 670039
refGene_gn=ECM2;refGene_tx=XM_003995303.5;refGene_gn=ECM2;refGene_tx=XM_006939006.4;refGene_gn=ECM2;refGene_tx=XM_006939007.4;refGene_gn=ECM2;refGene_tx=XM_011288207.3
D4 720990 721070
refGene_gn=CENPP;refGene_tx=XM_023243190.1;refGene_gn=CENPP;refGene_tx=XM_023243191.1;refGene_gn=CENPP;refGene_tx=XM_023243192.1;refGene_gn=CENPP;refGene_tx=XM_023243193.1;refGene_gn=CENPP;refGene_tx=XM_023243195.1;refGene_gn=CENPP;refGene_tx=XM_023243196.1;refGene_gn=CENPP;refGene_tx=XM_023243197.1
D4 721396 721503 refGene_gn=CENPP;refGene_tx=XM_023243193.1
D4 721594 721674 refGene_gn=CENPP;refGene_tx=XM_023243193.1
D4 721953 722045
refGene_gn=CENPP;refGene_tx=XM_023243190.1;refGene_gn=CENPP;refGene_tx=XM_023243191.1;refGene_gn=CENPP;refGene_tx=XM_023243192.1;refGene_gn=CENPP;refGene_tx=XM_023243195.1;refGene_gn=CENPP;refGene_tx=XM_023243196.1;refGene_gn=CENPP;refGene_tx=XM_023243197.1
D4 722352 722483
refGene_gn=CENPP;refGene_tx=XM_023243190.1;refGene_gn=CENPP;refGene_tx=XM_023243191.1;refGene_gn=CENPP;refGene_tx=XM_023243192.1;refGene_gn=CENPP;refGene_tx=XM_023243195.1;refGene_gn=CENPP;refGene_tx=XM_023243196.1;refGene_gn=CENPP;refGene_tx=XM_023243197.1
D4 724422 724648
refGene_gn=LOC101083117;refGene_tx=XM_003995306.4;refGene_gn=LOC101083

117;refGene_tx=XM_019815636.2
D4 726625 726705
refGene_gn=LOC101083117;refGene_tx=XM_003995306.4;refGene_gn=LOC101083
117;refGene_tx=XM_019815636.2
D4 739809 739912
refGene_gn=LOC101083117;refGene_tx=XM_003995306.4;refGene_gn=LOC101083
117;refGene_tx=XM_019815636.2
D4 740486 740637
refGene_gn=LOC101083117;refGene_tx=XM_003995306.4;refGene_gn=LOC101083
117;refGene_tx=XM_019815636.2;refGene_gn=LOC101083117;refGene_tx=XM_01
9815637.2
D4 741749 742029
refGene_gn=LOC101083117;refGene_tx=XM_003995306.4;refGene_gn=LOC101083
117;refGene_tx=XM_019815636.2;refGene_gn=LOC101083117;refGene_tx=XM_01
9815637.2
D4 748931 749004
refGene_gn=LOC101083117;refGene_tx=XM_003995306.4;refGene_gn=LOC101083
117;refGene_tx=XM_019815636.2;refGene_gn=LOC101083117;refGene_tx=XM_01
9815637.2
D4 749416 749475
refGene_gn=LOC101083117;refGene_tx=XM_003995306.4;refGene_gn=LOC101083
117;refGene_tx=XM_019815636.2;refGene_gn=LOC101083117;refGene_tx=XM_01
9815637.2
D4 751249 751264
refGene_gn=LOC101083117;refGene_tx=XM_019815639.2
D4 752347 752398
refGene_gn=LOC101083117;refGene_tx=XM_019815638.2
D4 757622 757712
refGene_gn=LOC101083117;refGene_tx=XM_003995306.4;refGene_gn=LOC101083
117;refGene_tx=XM_019815636.2;refGene_gn=LOC101083117;refGene_tx=XM_01
9815637.2;refGene_gn=LOC101083117;refGene_tx=XM_019815638.2;refGene_gn
=LOC101083117;refGene_tx=XM_019815639.2
D4 758799 758921
refGene_gn=LOC101083117;refGene_tx=XM_003995306.4;refGene_gn=LOC101083
117;refGene_tx=XM_019815636.2;refGene_gn=LOC101083117;refGene_tx=XM_01
9815637.2;refGene_gn=LOC101083117;refGene_tx=XM_019815638.2;refGene_gn
=LOC101083117;refGene_tx=XM_019815639.2
D4 760578 760645
refGene_gn=LOC101083117;refGene_tx=XM_003995306.4;refGene_gn=LOC101083
117;refGene_tx=XM_019815636.2;refGene_gn=LOC101083117;refGene_tx=XM_01
9815637.2;refGene_gn=LOC101083117;refGene_tx=XM_019815638.2;refGene_gn
=LOC101083117;refGene_tx=XM_019815639.2
D4 763456 763552
refGene_gn=LOC101083117;refGene_tx=XM_003995306.4;refGene_gn=LOC101083
117;refGene_tx=XM_019815636.2;refGene_gn=LOC101083117;refGene_tx=XM_01
9815637.2;refGene_gn=LOC101083117;refGene_tx=XM_019815638.2;refGene_gn
=LOC101083117;refGene_tx=XM_019815639.2
D4 765130 765178
refGene_gn=LOC101083117;refGene_tx=XM_003995306.4;refGene_gn=LOC101083
117;refGene_tx=XM_019815636.2;refGene_gn=LOC101083117;refGene_tx=XM_01

9815637.2;refGene_gn=LOC101083117;refGene_tx=XM_019815638.2;refGene_gn=LOC101083117;refGene_tx=XM_019815639.2
D4 771605 771686
refGene_gn=LOC101083117;refGene_tx=XM_003995306.4
D4 808239 808245
refGene_gn=LOC101083117;refGene_tx=XM_003995307.5
D4 809916 810127
refGene_gn=LOC101083117;refGene_tx=XM_003995307.5;refGene_gn=LOC101083117;refGene_tx=XM_019815636.2;refGene_gn=LOC101083117;refGene_tx=XM_019815637.2;refGene_gn=LOC101083117;refGene_tx=XM_019815638.2;refGene_gn=LOC101083117;refGene_tx=XM_019815639.2
D4 811224 811376
refGene_gn=LOC101083117;refGene_tx=XM_003995307.5;refGene_gn=LOC101083117;refGene_tx=XM_019815636.2;refGene_gn=LOC101083117;refGene_tx=XM_019815637.2;refGene_gn=LOC101083117;refGene_tx=XM_019815638.2;refGene_gn=LOC101083117;refGene_tx=XM_019815639.2
D4 811500 812508
refGene_gn=LOC101083117;refGene_tx=XM_003995307.5;refGene_gn=LOC101083117;refGene_tx=XM_019815636.2;refGene_gn=LOC101083117;refGene_tx=XM_019815637.2;refGene_gn=LOC101083117;refGene_tx=XM_019815638.2;refGene_gn=LOC101083117;refGene_tx=XM_019815639.2
D4 813007 813442
refGene_gn=LOC101083117;refGene_tx=XM_003995307.5;refGene_gn=LOC101083117;refGene_tx=XM_019815636.2;refGene_gn=LOC101083117;refGene_tx=XM_019815637.2;refGene_gn=LOC101083117;refGene_tx=XM_019815638.2;refGene_gn=LOC101083117;refGene_tx=XM_019815639.2
D4 814362 814515
refGene_gn=LOC101083117;refGene_tx=XM_003995307.5;refGene_gn=LOC101083117;refGene_tx=XM_019815636.2;refGene_gn=LOC101083117;refGene_tx=XM_019815637.2;refGene_gn=LOC101083117;refGene_tx=XM_019815638.2;refGene_gn=LOC101083117;refGene_tx=XM_019815639.2
D4 820525 820738
refGene_gn=LOC101083117;refGene_tx=XM_003995307.5;refGene_gn=LOC101083117;refGene_tx=XM_019815636.2;refGene_gn=LOC101083117;refGene_tx=XM_019815637.2;refGene_gn=LOC101083117;refGene_tx=XM_019815638.2;refGene_gn=LOC101083117;refGene_tx=XM_019815639.2
D4 851650 851887
refGene_gn=LOC101083117;refGene_tx=XM_003995307.5;refGene_gn=LOC101083117;refGene_tx=XM_019815636.2;refGene_gn=LOC101083117;refGene_tx=XM_019815637.2;refGene_gn=LOC101083117;refGene_tx=XM_019815638.2;refGene_gn=LOC101083117;refGene_tx=XM_019815639.2
D4 919539 920022
refGene_gn=FGD3;refGene_tx=XM_003995335.5;refGene_gn=FGD3;refGene_tx=XM_006939011.4;refGene_gn=FGD3;refGene_tx=XM_006939012.4;refGene_gn=FGD3;refGene_tx=XM_019815650.2;refGene_gn=FGD3;refGene_tx=XM_019815651.2;refGene_gn=FGD3;refGene_tx=XM_019815652.2
D4 924489 924579
refGene_gn=FGD3;refGene_tx=XM_003995335.5;refGene_gn=FGD3;refGene_tx=XM_006939011.4;refGene_gn=FGD3;refGene_tx=XM_006939012.4;refGene_gn=FGD3;refGene_tx=XM_019815650.2;refGene_gn=FGD3;refGene_tx=XM_019815651.2;

refGene_gn=FGD3;refGene_tx=XM_019815652.2
D4 925218 925355
refGene_gn=FGD3;refGene_tx=XM_003995335.5;refGene_gn=FGD3;refGene_tx=XM_006939011.4;refGene_gn=FGD3;refGene_tx=XM_006939012.4;refGene_gn=FGD3;refGene_tx=XM_019815650.2;refGene_gn=FGD3;refGene_tx=XM_019815651.2;refGene_gn=FGD3;refGene_tx=XM_019815652.2
D4 926766 926923
refGene_gn=FGD3;refGene_tx=XM_003995335.5;refGene_gn=FGD3;refGene_tx=XM_006939011.4;refGene_gn=FGD3;refGene_tx=XM_006939012.4;refGene_gn=FGD3;refGene_tx=XM_019815650.2;refGene_gn=FGD3;refGene_tx=XM_019815651.2;refGene_gn=FGD3;refGene_tx=XM_019815652.2
D4 929388 929527
refGene_gn=FGD3;refGene_tx=XM_003995335.5;refGene_gn=FGD3;refGene_tx=XM_006939011.4;refGene_gn=FGD3;refGene_tx=XM_006939012.4;refGene_gn=FGD3;refGene_tx=XM_019815650.2;refGene_gn=FGD3;refGene_tx=XM_019815651.2;refGene_gn=FGD3;refGene_tx=XM_019815652.2
D4 929976 930035
refGene_gn=FGD3;refGene_tx=XM_003995335.5;refGene_gn=FGD3;refGene_tx=XM_006939011.4;refGene_gn=FGD3;refGene_tx=XM_006939012.4;refGene_gn=FGD3;refGene_tx=XM_019815650.2;refGene_gn=FGD3;refGene_tx=XM_019815651.2;refGene_gn=FGD3;refGene_tx=XM_019815652.2
D4 932572 932719
refGene_gn=FGD3;refGene_tx=XM_003995335.5;refGene_gn=FGD3;refGene_tx=XM_006939011.4;refGene_gn=FGD3;refGene_tx=XM_006939012.4;refGene_gn=FGD3;refGene_tx=XM_019815650.2;refGene_gn=FGD3;refGene_tx=XM_019815651.2;refGene_gn=FGD3;refGene_tx=XM_019815652.2
D4 933715 933808
refGene_gn=FGD3;refGene_tx=XM_003995335.5;refGene_gn=FGD3;refGene_tx=XM_006939011.4;refGene_gn=FGD3;refGene_tx=XM_006939012.4;refGene_gn=FGD3;refGene_tx=XM_019815650.2;refGene_gn=FGD3;refGene_tx=XM_019815651.2;refGene_gn=FGD3;refGene_tx=XM_019815652.2
D4 935546 935626
refGene_gn=FGD3;refGene_tx=XM_003995335.5;refGene_gn=FGD3;refGene_tx=XM_006939011.4;refGene_gn=FGD3;refGene_tx=XM_006939012.4;refGene_gn=FGD3;refGene_tx=XM_019815650.2;refGene_gn=FGD3;refGene_tx=XM_019815651.2;refGene_gn=FGD3;refGene_tx=XM_019815652.2
D4 936849 936880
refGene_gn=FGD3;refGene_tx=XM_003995335.5;refGene_gn=FGD3;refGene_tx=XM_006939011.4;refGene_gn=FGD3;refGene_tx=XM_006939012.4;refGene_gn=FGD3;refGene_tx=XM_019815650.2;refGene_gn=FGD3;refGene_tx=XM_019815651.2;refGene_gn=FGD3;refGene_tx=XM_019815652.2
D4 937723 937831
refGene_gn=FGD3;refGene_tx=XM_003995335.5;refGene_gn=FGD3;refGene_tx=XM_006939011.4;refGene_gn=FGD3;refGene_tx=XM_006939012.4;refGene_gn=FGD3;refGene_tx=XM_019815650.2;refGene_gn=FGD3;refGene_tx=XM_019815651.2;refGene_gn=FGD3;refGene_tx=XM_019815652.2
D4 938703 938766
refGene_gn=FGD3;refGene_tx=XM_003995335.5;refGene_gn=FGD3;refGene_tx=XM_006939011.4;refGene_gn=FGD3;refGene_tx=XM_006939012.4;refGene_gn=FGD3;refGene_tx=XM_019815650.2;refGene_gn=FGD3;refGene_tx=XM_019815651.2;

refGene_gn=FGD3;refGene_tx=XM_019815652.2
D4 943244 943367
refGene_gn=FGD3;refGene_tx=XM_003995335.5;refGene_gn=FGD3;refGene_tx=XM_006939011.4;refGene_gn=FGD3;refGene_tx=XM_006939012.4;refGene_gn=FGD3;refGene_tx=XM_019815650.2;refGene_gn=FGD3;refGene_tx=XM_019815651.2;refGene_gn=FGD3;refGene_tx=XM_019815652.2
D4 944785 944929
refGene_gn=FGD3;refGene_tx=XM_003995335.5;refGene_gn=FGD3;refGene_tx=XM_006939011.4;refGene_gn=FGD3;refGene_tx=XM_006939012.4;refGene_gn=FGD3;refGene_tx=XM_019815650.2;refGene_gn=FGD3;refGene_tx=XM_019815651.2
D4 945395 945542
refGene_gn=FGD3;refGene_tx=XM_003995335.5;refGene_gn=FGD3;refGene_tx=XM_006939011.4;refGene_gn=FGD3;refGene_tx=XM_006939012.4;refGene_gn=FGD3;refGene_tx=XM_019815650.2;refGene_gn=FGD3;refGene_tx=XM_019815651.2;refGene_gn=FGD3;refGene_tx=XM_019815652.2
D4 945786 946005
refGene_gn=FGD3;refGene_tx=XM_003995335.5;refGene_gn=FGD3;refGene_tx=XM_006939011.4;refGene_gn=FGD3;refGene_tx=XM_006939012.4;refGene_gn=FGD3;refGene_tx=XM_019815650.2;refGene_gn=FGD3;refGene_tx=XM_019815651.2;refGene_gn=FGD3;refGene_tx=XM_019815652.2
D4 958701 958789 refGene_gn=SUSD3;refGene_tx=XM_023243199.1
D4 959199 959233 refGene_gn=SUSD3;refGene_tx=XM_019815655.2
D4 963204 963250 refGene_gn=SUSD3;refGene_tx=XM_023243200.1
D4 963308 963360 refGene_gn=SUSD3;refGene_tx=XM_019815653.2
D4 969565 969754
refGene_gn=SUSD3;refGene_tx=XM_019815653.2;refGene_gn=SUSD3;refGene_tx=XM_019815655.2;refGene_gn=SUSD3;refGene_tx=XM_023243199.1;refGene_gn=SUSD3;refGene_tx=XM_023243200.1
D4 971236 971381
refGene_gn=SUSD3;refGene_tx=XM_019815653.2;refGene_gn=SUSD3;refGene_tx=XM_019815655.2;refGene_gn=SUSD3;refGene_tx=XM_023243199.1;refGene_gn=SUSD3;refGene_tx=XM_023243200.1
D4 971761 971914
refGene_gn=SUSD3;refGene_tx=XM_019815653.2;refGene_gn=SUSD3;refGene_tx=XM_019815655.2;refGene_gn=SUSD3;refGene_tx=XM_023243199.1;refGene_gn=SUSD3;refGene_tx=XM_023243200.1
D4 974764 974957
refGene_gn=SUSD3;refGene_tx=XM_019815653.2;refGene_gn=SUSD3;refGene_tx=XM_019815655.2;refGene_gn=SUSD3;refGene_tx=XM_023243199.1;refGene_gn=SUSD3;refGene_tx=XM_023243200.1
D4 981833 981840
refGene_gn=CARD19;refGene_tx=XM_023243201.1;refGene_gn=CARD19;refGene_tx=XM_023243202.1;refGene_gn=CARD19;refGene_tx=XM_023243203.1
D4 990932 991075
refGene_gn=CARD19;refGene_tx=XM_023243201.1;refGene_gn=CARD19;refGene_tx=XM_023243202.1;refGene_gn=CARD19;refGene_tx=XM_023243203.1
D4 992516 992670
refGene_gn=CARD19;refGene_tx=XM_023243201.1;refGene_gn=CARD19;refGene_tx=XM_023243202.1;refGene_gn=CARD19;refGene_tx=XM_023243203.1
D4 993728 993788

refGene_gn=CARD19;refGene_tx=XM_023243201.1;refGene_gn=CARD19;refGene_tx=XM_023243203.1
D4	993972	994044
refGene_gn=CARD19;refGene_tx=XM_023243201.1;refGene_gn=CARD19;refGene_tx=XM_023243202.1
D4	994494	994610
refGene_gn=CARD19;refGene_tx=XM_023243201.1;refGene_gn=CARD19;refGene_tx=XM_023243202.1;refGene_gn=CARD19;refGene_tx=XM_023243203.1
D4	1003371	1003526
refGene_gn=NINJ1;refGene_tx=XM_023243204.1;refGene_gn=NINJ1;refGene_tx=XM_023243205.1
D4	1004051	1004279
refGene_gn=NINJ1;refGene_tx=XM_023243204.1;refGene_gn=NINJ1;refGene_tx=XM_023243205.1
D4	1011422	1011497
refGene_gn=NINJ1;refGene_tx=XM_023243204.1;refGene_gn=NINJ1;refGene_tx=XM_023243205.1
D4	1055076	1055754	refGene_gn=WNK2;refGene_tx=XM_023243243.1
D4	1093337	1093510	refGene_gn=WNK2;refGene_tx=XM_023243243.1
D4	1093680	1093901	refGene_gn=WNK2;refGene_tx=XM_023243243.1
D4	1097595	1097932	refGene_gn=WNK2;refGene_tx=XM_023243243.1
D4	1098302	1098412	refGene_gn=WNK2;refGene_tx=XM_023243243.1
D4	1121980	1122289	refGene_gn=WNK2;refGene_tx=XM_023243243.1
D4	1126356	1126559	refGene_gn=WNK2;refGene_tx=XM_023243243.1
D4	1128253	1128385	refGene_gn=WNK2;refGene_tx=XM_023243243.1
D4	1128669	1128861	refGene_gn=WNK2;refGene_tx=XM_023243243.1
D4	1130125	1130695	refGene_gn=WNK2;refGene_tx=XM_023243243.1
D4	1132575	1132869	refGene_gn=WNK2;refGene_tx=XM_023243243.1
D4	1133272	1133322	refGene_gn=WNK2;refGene_tx=XM_023243243.1
D4	1134049	1134218	refGene_gn=WNK2;refGene_tx=XM_023243243.1
D4	1134304	1134421	refGene_gn=WNK2;refGene_tx=XM_023243243.1
D4	1134616	1134787	refGene_gn=WNK2;refGene_tx=XM_023243243.1
D4	1134981	1135027	refGene_gn=WNK2;refGene_tx=XM_023243243.1
D4	1135501	1135621	refGene_gn=WNK2;refGene_tx=XM_023243243.1
D4	1166169	1166936	refGene_gn=WNK2;refGene_tx=XM_023243243.1
D4	1167203	1167273	refGene_gn=WNK2;refGene_tx=XM_023243243.1
D4	1169119	1169181	refGene_gn=WNK2;refGene_tx=XM_023243243.1
D4	1169270	1169950	refGene_gn=WNK2;refGene_tx=XM_023243243.1
D4	1171816	1172028	refGene_gn=WNK2;refGene_tx=XM_023243243.1
D4	1172876	1173065	refGene_gn=WNK2;refGene_tx=XM_023243243.1
D4	1173789	1173888	refGene_gn=WNK2;refGene_tx=XM_023243243.1
D4	1179429	1179474	refGene_gn=WNK2;refGene_tx=XM_023243243.1
D4	1180902	1181162	refGene_gn=WNK2;refGene_tx=XM_023243243.1
D4	1201755	1201861	refGene_gn=WNK2;refGene_tx=XM_023243243.1
D4	1203240	1203419	refGene_gn=WNK2;refGene_tx=XM_023243243.1
D4	1339684	1340158
refGene_gn=LOC101083883;refGene_tx=XM_023243206.1;refGene_gn=LOC101083883;refGene_tx=XM_023243207.1
D4	1358074	1358321
refGene_gn=LOC101083883;refGene_tx=XM_023243206.1;refGene_gn=LOC101083

883;refGene_tx=XM_023243207.1
D4 1362071 1362154
refGene_gn=LOC101083883;refGene_tx=XM_023243206.1;refGene_gn=LOC101083
883;refGene_tx=XM_023243207.1
D4 1376853 1376982
refGene_gn=LOC101083883;refGene_tx=XM_023243206.1;refGene_gn=LOC101083
883;refGene_tx=XM_023243207.1
D4 1377513 1377610
refGene_gn=LOC101083883;refGene_tx=XM_023243206.1;refGene_gn=LOC101083
883;refGene_tx=XM_023243207.1
D4 1389091 1389189
refGene_gn=LOC101083883;refGene_tx=XM_023243206.1;refGene_gn=LOC101083
883;refGene_tx=XM_023243207.1
D4 1389414 1389695
refGene_gn=LOC101083883;refGene_tx=XM_023243206.1;refGene_gn=LOC101083
883;refGene_tx=XM_023243207.1
D4 1398332 1398420
refGene_gn=LOC101083883;refGene_tx=XM_023243206.1;refGene_gn=LOC101083
883;refGene_tx=XM_023243207.1
D4 1400302 1400530
refGene_gn=LOC101083883;refGene_tx=XM_023243206.1;refGene_gn=LOC101083
883;refGene_tx=XM_023243207.1
D4 1402792 1402967
refGene_gn=LOC101083883;refGene_tx=XM_023243206.1;refGene_gn=LOC101083
883;refGene_tx=XM_023243207.1
D4 1412300 1412550
refGene_gn=LOC101083883;refGene_tx=XM_023243206.1;refGene_gn=LOC101083
883;refGene_tx=XM_023243207.1
D4 1418978 1419093
refGene_gn=LOC101083883;refGene_tx=XM_023243206.1;refGene_gn=LOC101083
883;refGene_tx=XM_023243207.1
D4 1424131 1424341
refGene_gn=LOC101083883;refGene_tx=XM_023243206.1;refGene_gn=LOC101083
883;refGene_tx=XM_023243207.1
D4 1425767 1425951
refGene_gn=LOC101083883;refGene_tx=XM_023243206.1;refGene_gn=LOC101083
883;refGene_tx=XM_023243207.1
D4 1426448 1426586
refGene_gn=LOC101083883;refGene_tx=XM_023243206.1
D4 1428558 1428700
refGene_gn=LOC101083883;refGene_tx=XM_023243206.1;refGene_gn=LOC101083
883;refGene_tx=XM_023243207.1
D4 1429138 1429235
refGene_gn=LOC101083883;refGene_tx=XM_023243206.1;refGene_gn=LOC101083
883;refGene_tx=XM_023243207.1
D4 1435028 1435340
refGene_gn=LOC101083883;refGene_tx=XM_023243206.1;refGene_gn=LOC101083
883;refGene_tx=XM_023243207.1
D4 1446103 1446201 refGene_gn=PHF2;refGene_tx=XM_023243208.1
D4 1483375 1483461 refGene_gn=PHF2;refGene_tx=XM_023243208.1

D4	1488932	1489047	refGene_gn=PHF2;refGene_tx=XM_023243208.1
D4	1498210	1498371	refGene_gn=PHF2;refGene_tx=XM_023243208.1
D4	1500334	1500476	refGene_gn=PHF2;refGene_tx=XM_023243208.1
D4	1503314	1503501	refGene_gn=PHF2;refGene_tx=XM_023243208.1
D4	1504339	1504502	refGene_gn=PHF2;refGene_tx=XM_023243208.1
D4	1505123	1505211	refGene_gn=PHF2;refGene_tx=XM_023243208.1
D4	1505572	1505679	refGene_gn=PHF2;refGene_tx=XM_023243208.1
D4	1506848	1506940	refGene_gn=PHF2;refGene_tx=XM_023243208.1
D4	1507745	1507835	refGene_gn=PHF2;refGene_tx=XM_023243208.1
D4	1508708	1508768	refGene_gn=PHF2;refGene_tx=XM_023243208.1
D4	1508769	1509078	refGene_gn=PHF2;refGene_tx=XM_023243208.1
D4	1510035	1510155	refGene_gn=PHF2;refGene_tx=XM_023243208.1
D4	1510326	1510442	refGene_gn=PHF2;refGene_tx=XM_023243208.1
D4	1511860	1512039	refGene_gn=PHF2;refGene_tx=XM_023243208.1
D4	1512116	1512187	refGene_gn=PHF2;refGene_tx=XM_023243208.1
D4	1512902	1513063	refGene_gn=PHF2;refGene_tx=XM_023243208.1
D4	1517213	1517488	refGene_gn=PHF2;refGene_tx=XM_023243208.1
D4	1518670	1518766	refGene_gn=PHF2;refGene_tx=XM_023243208.1
D4	1519032	1519142	refGene_gn=PHF2;refGene_tx=XM_023243208.1
D4	1520721	1521073	refGene_gn=PHF2;refGene_tx=XM_023243208.1
D4	1521625	1521714	refGene_gn=PHF2;refGene_tx=XM_023243208.1
D4	1674083	1674248	refGene_gn=BARX1;refGene_tx=XM_023242740.1
D4	1674667	1674752	refGene_gn=BARX1;refGene_tx=XM_023242740.1
D4	1674833	1675125	refGene_gn=BARX1;refGene_tx=XM_023242740.1
D4	1676731	1676969	refGene_gn=BARX1;refGene_tx=XM_023242740.1
D4	1730741	1730744
refGene_gn=PTPDC1;refGene_tx=XM_019815663.2;refGene_gn=PTPDC1;refGene_tx=XM_019815664.2
D4	1751626	1751714
refGene_gn=PTPDC1;refGene_tx=XM_011288200.3;refGene_gn=PTPDC1;refGene_tx=XM_019815663.2;refGene_gn=PTPDC1;refGene_tx=XM_019815664.2;refGene_gn=PTPDC1;refGene_tx=XM_019815665.2;refGene_gn=PTPDC1;refGene_tx=XM_019815666.2;refGene_gn=PTPDC1;refGene_tx=XM_019815667.2;refGene_gn=PTPDC1;refGene_tx=XM_023243209.1;refGene_gn=PTPDC1;refGene_tx=XM_023243210.1
D4	1764705	1764923	refGene_gn=PTPDC1;refGene_tx=XM_023243211.1
D4	1765384	1765558
refGene_gn=PTPDC1;refGene_tx=XM_011288200.3;refGene_gn=PTPDC1;refGene_tx=XM_019815663.2;refGene_gn=PTPDC1;refGene_tx=XM_019815664.2;refGene_gn=PTPDC1;refGene_tx=XM_019815665.2;refGene_gn=PTPDC1;refGene_tx=XM_019815666.2;refGene_gn=PTPDC1;refGene_tx=XM_019815667.2;refGene_gn=PTPDC1;refGene_tx=XM_023243209.1;refGene_gn=PTPDC1;refGene_tx=XM_023243210.1;refGene_gn=PTPDC1;refGene_tx=XM_023243211.1
D4	1767351	1767432
refGene_gn=PTPDC1;refGene_tx=XM_011288200.3;refGene_gn=PTPDC1;refGene_tx=XM_019815663.2;refGene_gn=PTPDC1;refGene_tx=XM_019815664.2;refGene_gn=PTPDC1;refGene_tx=XM_019815665.2;refGene_gn=PTPDC1;refGene_tx=XM_019815666.2;refGene_gn=PTPDC1;refGene_tx=XM_019815667.2;refGene_gn=PTPDC1;refGene_tx=XM_019815669.2;refGene_gn=PTPDC1;refGene_tx=XM_023243209.1;refGene_gn=PTPDC1;refGene_tx=XM_023243210.1;refGene_gn=PTPDC1;refGen

e_tx=XM_023243211.1
D4 1767653 1767772
refGene_gn=PTPDC1;refGene_tx=XM_011288200.3;refGene_gn=PTPDC1;refGene_tx=XM_019815663.2;refGene_gn=PTPDC1;refGene_tx=XM_019815664.2;refGene_gn=PTPDC1;refGene_tx=XM_019815665.2;refGene_gn=PTPDC1;refGene_tx=XM_019815666.2;refGene_gn=PTPDC1;refGene_tx=XM_019815667.2;refGene_gn=PTPDC1;refGene_tx=XM_019815669.2;refGene_gn=PTPDC1;refGene_tx=XM_023243209.1;refGene_gn=PTPDC1;refGene_tx=XM_023243210.1;refGene_gn=PTPDC1;refGene_tx=XM_023243211.1
D4 1768738 1768876
refGene_gn=PTPDC1;refGene_tx=XM_011288200.3;refGene_gn=PTPDC1;refGene_tx=XM_019815663.2;refGene_gn=PTPDC1;refGene_tx=XM_019815664.2;refGene_gn=PTPDC1;refGene_tx=XM_019815665.2;refGene_gn=PTPDC1;refGene_tx=XM_019815666.2;refGene_gn=PTPDC1;refGene_tx=XM_019815667.2;refGene_gn=PTPDC1;refGene_tx=XM_019815669.2;refGene_gn=PTPDC1;refGene_tx=XM_023243209.1;refGene_gn=PTPDC1;refGene_tx=XM_023243211.1
D4 1770667 1771902
refGene_gn=PTPDC1;refGene_tx=XM_011288200.3;refGene_gn=PTPDC1;refGene_tx=XM_019815663.2;refGene_gn=PTPDC1;refGene_tx=XM_019815664.2;refGene_gn=PTPDC1;refGene_tx=XM_019815665.2;refGene_gn=PTPDC1;refGene_tx=XM_019815666.2;refGene_gn=PTPDC1;refGene_tx=XM_019815667.2;refGene_gn=PTPDC1;refGene_tx=XM_019815669.2;refGene_gn=PTPDC1;refGene_tx=XM_023243209.1;refGene_gn=PTPDC1;refGene_tx=XM_023243210.1;refGene_gn=PTPDC1;refGene_tx=XM_023243211.1
D4 1776273 1776459
refGene_gn=PTPDC1;refGene_tx=XM_011288200.3;refGene_gn=PTPDC1;refGene_tx=XM_019815663.2;refGene_gn=PTPDC1;refGene_tx=XM_019815664.2;refGene_gn=PTPDC1;refGene_tx=XM_019815665.2;refGene_gn=PTPDC1;refGene_tx=XM_019815666.2;refGene_gn=PTPDC1;refGene_tx=XM_019815667.2;refGene_gn=PTPDC1;refGene_tx=XM_019815669.2;refGene_gn=PTPDC1;refGene_tx=XM_023243209.1;refGene_gn=PTPDC1;refGene_tx=XM_023243210.1;refGene_gn=PTPDC1;refGene_tx=XM_023243211.1
D4 1778014 1778125
refGene_gn=PTPDC1;refGene_tx=XM_011288200.3;refGene_gn=PTPDC1;refGene_tx=XM_019815663.2;refGene_gn=PTPDC1;refGene_tx=XM_019815664.2;refGene_gn=PTPDC1;refGene_tx=XM_019815665.2;refGene_gn=PTPDC1;refGene_tx=XM_019815666.2;refGene_gn=PTPDC1;refGene_tx=XM_019815667.2;refGene_gn=PTPDC1;refGene_tx=XM_019815669.2;refGene_gn=PTPDC1;refGene_tx=XM_023243209.1;refGene_gn=PTPDC1;refGene_tx=XM_023243210.1;refGene_gn=PTPDC1;refGene_tx=XM_023243211.1
D4 1780713 1780830
refGene_gn=PTPDC1;refGene_tx=XM_011288200.3;refGene_gn=PTPDC1;refGene_tx=XM_019815663.2;refGene_gn=PTPDC1;refGene_tx=XM_019815664.2;refGene_gn=PTPDC1;refGene_tx=XM_019815665.2;refGene_gn=PTPDC1;refGene_tx=XM_019815666.2;refGene_gn=PTPDC1;refGene_tx=XM_019815667.2;refGene_gn=PTPDC1;refGene_tx=XM_019815669.2;refGene_gn=PTPDC1;refGene_tx=XM_023243209.1;refGene_gn=PTPDC1;refGene_tx=XM_023243210.1;refGene_gn=PTPDC1;refGene_tx=XM_023243211.1
D4 1810096 1810213
refGene_gn=LOC109493659;refGene_tx=XM_023243244.1

D4	1810850	1810948
refGene_gn=LOC109493659;refGene_tx=XM_023243244.1
D4	1811198	1811463
refGene_gn=LOC109493659;refGene_tx=XM_023243244.1
D4	1811994	1812072
refGene_gn=LOC109493659;refGene_tx=XM_023243244.1
D4	1816740	1816876
refGene_gn=LOC109493659;refGene_tx=XM_023243244.1
D4	1817080	1818036
refGene_gn=LOC109493659;refGene_tx=XM_023243244.1
D4	1891344	1891377
refGene_gn=ZNF169;refGene_tx=XM_006939021.2;refGene_gn=ZNF169;refGene_tx=XM_006939022.3;refGene_gn=ZNF169;refGene_tx=XM_006939023.3;refGene_gn=ZNF169;refGene_tx=XM_006939024.4;refGene_gn=ZNF169;refGene_tx=XM_019815672.2;refGene_gn=ZNF169;refGene_tx=XM_019815675.2;refGene_gn=ZNF169;refGene_tx=XM_019815682.2;refGene_gn=ZNF169;refGene_tx=XM_019815683.2
D4	1914654	1914781
refGene_gn=ZNF169;refGene_tx=XM_006939021.2;refGene_gn=ZNF169;refGene_tx=XM_006939022.3;refGene_gn=ZNF169;refGene_tx=XM_006939023.3;refGene_gn=ZNF169;refGene_tx=XM_006939024.4;refGene_gn=ZNF169;refGene_tx=XM_006939025.4;refGene_gn=ZNF169;refGene_tx=XM_006939026.3;refGene_gn=ZNF169;refGene_tx=XM_019815672.2;refGene_gn=ZNF169;refGene_tx=XM_019815675.2;refGene_gn=ZNF169;refGene_tx=XM_019815678.1;refGene_gn=ZNF169;refGene_tx=XM_019815679.2;refGene_gn=ZNF169;refGene_tx=XM_019815682.2;refGene_gn=ZNF169;refGene_tx=XM_019815683.2
D4	1915264	1915360
refGene_gn=ZNF169;refGene_tx=XM_006939021.2;refGene_gn=ZNF169;refGene_tx=XM_006939022.3;refGene_gn=ZNF169;refGene_tx=XM_006939023.3;refGene_gn=ZNF169;refGene_tx=XM_006939024.4;refGene_gn=ZNF169;refGene_tx=XM_006939025.4;refGene_gn=ZNF169;refGene_tx=XM_006939026.3;refGene_gn=ZNF169;refGene_tx=XM_019815672.2;refGene_gn=ZNF169;refGene_tx=XM_019815675.2;refGene_gn=ZNF169;refGene_tx=XM_019815678.1;refGene_gn=ZNF169;refGene_tx=XM_019815679.2
D4	1928887	1930404
refGene_gn=ZNF169;refGene_tx=XM_006939021.2;refGene_gn=ZNF169;refGene_tx=XM_006939022.3;refGene_gn=ZNF169;refGene_tx=XM_006939023.3;refGene_gn=ZNF169;refGene_tx=XM_006939024.4;refGene_gn=ZNF169;refGene_tx=XM_006939025.4;refGene_gn=ZNF169;refGene_tx=XM_006939026.3;refGene_gn=ZNF169;refGene_tx=XM_019815672.2;refGene_gn=ZNF169;refGene_tx=XM_019815675.2;refGene_gn=ZNF169;refGene_tx=XM_019815678.1;refGene_gn=ZNF169;refGene_tx=XM_019815679.2;refGene_gn=ZNF169;refGene_tx=XM_019815682.2;refGene_gn=ZNF169;refGene_tx=XM_019815683.2
D4	1978322	1978967
refGene_gn=LOC111557589;refGene_tx=XM_023243245.1
D4	2105585	2105712
refGene_gn=LOC101094887;refGene_tx=XM_011288220.3;refGene_gn=LOC101094887;refGene_tx=XM_011288221.2;refGene_gn=LOC101094887;refGene_tx=XM_019815687.2
D4	2109741	2109869

refGene_gn=LOC101094887;refGene_tx=XM_011288220.3;refGene_gn=LOC101094887;refGene_tx=XM_011288221.2;refGene_gn=LOC101094887;refGene_tx=XM_019815687.2
D4 2109946 2110125
refGene_gn=LOC101094887;refGene_tx=XM_011288220.3;refGene_gn=LOC101094887;refGene_tx=XM_011288221.2;refGene_gn=LOC101094887;refGene_tx=XM_019815687.2
D4 2111721 2111932
refGene_gn=LOC101094887;refGene_tx=XM_011288220.3;refGene_gn=LOC101094887;refGene_tx=XM_011288221.2;refGene_gn=LOC101094887;refGene_tx=XM_019815687.2
D4 2112387 2112546
refGene_gn=LOC101094887;refGene_tx=XM_011288220.3;refGene_gn=LOC101094887;refGene_tx=XM_011288221.2;refGene_gn=LOC101094887;refGene_tx=XM_019815687.2
D4 2136042 2136252
refGene_gn=LOC101094887;refGene_tx=XM_011288220.3;refGene_gn=LOC101094887;refGene_tx=XM_011288221.2;refGene_gn=LOC101094887;refGene_tx=XM_019815687.2
D4 2139951 2140070
refGene_gn=LOC101094887;refGene_tx=XM_011288220.3;refGene_gn=LOC101094887;refGene_tx=XM_011288221.2;refGene_gn=LOC101094887;refGene_tx=XM_019815687.2
D4 2140505 2140669
refGene_gn=LOC101094887;refGene_tx=XM_011288220.3;refGene_gn=LOC101094887;refGene_tx=XM_011288221.2;refGene_gn=LOC101094887;refGene_tx=XM_019815687.2
D4 2148063 2148101
refGene_gn=LOC101094887;refGene_tx=XM_011288220.3;refGene_gn=LOC101094887;refGene_tx=XM_011288221.2;refGene_gn=LOC101094887;refGene_tx=XM_019815687.2
D4 2150800 2151007
refGene_gn=LOC101094887;refGene_tx=XM_011288221.2
D4 2152264 2152702
refGene_gn=LOC101094887;refGene_tx=XM_011288220.3;refGene_gn=LOC101094887;refGene_tx=XM_019815687.2
D4 2152993 2153191 refGene_gn=CDK20;refGene_tx=XM_003995311.5
D4 2153852 2153858 refGene_gn=CDK20;refGene_tx=XM_006939028.3
D4 2154287 2154443
refGene_gn=CDK20;refGene_tx=XM_003995311.5;refGene_gn=CDK20;refGene_tx=XM_006939028.3
D4 2154764 2154888
refGene_gn=CDK20;refGene_tx=XM_003995311.5;refGene_gn=CDK20;refGene_tx=XM_006939028.3
D4 2155555 2155618
refGene_gn=CDK20;refGene_tx=XM_003995311.5;refGene_gn=CDK20;refGene_tx=XM_006939028.3
D4 2155802 2155924
refGene_gn=CDK20;refGene_tx=XM_003995311.5;refGene_gn=CDK20;refGene_tx=XM_006939028.3

D4	2156148	2156337
refGene_gn=CDK20;refGene_tx=XM_003995311.5;refGene_gn=CDK20;refGene_tx
=XM_006939028.3
D4	2158460	2158574
refGene_gn=CDK20;refGene_tx=XM_003995311.5;refGene_gn=CDK20;refGene_tx
=XM_006939028.3
D4	2159009	2159084
refGene_gn=CDK20;refGene_tx=XM_003995311.5;refGene_gn=CDK20;refGene_tx
=XM_006939028.3
D4	2413169	2413225
refGene_gn=SPIN1;refGene_tx=XM_023243215.1;refGene_gn=SPIN1;refGene_tx
=XM_023243216.1;refGene_gn=SPIN1;refGene_tx=XM_023243217.1
D4	2429504	2429553
refGene_gn=SPIN1;refGene_tx=XM_023243215.1;refGene_gn=SPIN1;refGene_tx
=XM_023243216.1;refGene_gn=SPIN1;refGene_tx=XM_023243217.1
D4	2439297	2439551
refGene_gn=SPIN1;refGene_tx=XM_023243215.1;refGene_gn=SPIN1;refGene_tx
=XM_023243216.1;refGene_gn=SPIN1;refGene_tx=XM_023243217.1
D4	2444271	2444505
refGene_gn=SPIN1;refGene_tx=XM_023243215.1;refGene_gn=SPIN1;refGene_tx
=XM_023243216.1;refGene_gn=SPIN1;refGene_tx=XM_023243217.1
D4	2449577	2449777
refGene_gn=SPIN1;refGene_tx=XM_023243215.1;refGene_gn=SPIN1;refGene_tx
=XM_023243216.1;refGene_gn=SPIN1;refGene_tx=XM_023243217.1
D4	2483339	2483641
refGene_gn=NXNL2;refGene_tx=XM_006939029.3;refGene_gn=NXNL2;refGene_tx
=XM_023243218.1
D4	2490261	2490430	refGene_gn=NXNL2;refGene_tx=XM_006939029.3
D4	2672312	2672510	refGene_gn=NXNL2;refGene_tx=XM_023243218.1
D4	2703254	2703330	refGene_gn=NXNL2;refGene_tx=XM_023243218.1
D4	2719686	2719808
refGene_gn=LOC111557590;refGene_tx=XM_023243247.1
D4	2719962	2720217
refGene_gn=LOC111557590;refGene_tx=XM_023243247.1
D4	2727182	2727312
refGene_gn=LOC111557590;refGene_tx=XM_023243247.1
D4	2727324	2728464	refGene_gn=S1PR3;refGene_tx=XM_006939030.3
D4	2738584	2738713	refGene_gn=SHC3;refGene_tx=XM_023243220.1
D4	2753930	2754226	refGene_gn=SHC3;refGene_tx=XM_023243220.1
D4	2758483	2758639	refGene_gn=SHC3;refGene_tx=XM_023243220.1
D4	2761425	2761513	refGene_gn=SHC3;refGene_tx=XM_023243220.1
D4	2762476	2762627	refGene_gn=SHC3;refGene_tx=XM_023243220.1
D4	2766038	2766165	refGene_gn=SHC3;refGene_tx=XM_023243220.1
D4	2771097	2771149	refGene_gn=SHC3;refGene_tx=XM_023243220.1
D4	2776375	2776429	refGene_gn=SHC3;refGene_tx=XM_023243220.1
D4	2779210	2779330	refGene_gn=SHC3;refGene_tx=XM_023243220.1
D4	2781604	2781668	refGene_gn=SHC3;refGene_tx=XM_023243220.1
D4	2812900	2812971	refGene_gn=SHC3;refGene_tx=XM_023243220.1
D4	2838487	2838943	refGene_gn=SHC3;refGene_tx=XM_023243220.1
D4	2902931	2902990	refGene_gn=CKS2;refGene_tx=XM_003995316.5

D4 2905874 2906002 refGene_gn=CKS2;refGene_tx=XM_003995316.5
D4 2907261 2907314 refGene_gn=CKS2;refGene_tx=XM_003995316.5
D4 2909151 2909187
refGene_gn=SECISBP2;refGene_tx=XM_023243228.1;refGene_gn=SECISBP2;refGene_tx=XM_023243230.1;refGene_gn=SECISBP2;refGene_tx=XM_023243231.1;refGene_gn=SECISBP2;refGene_tx=XM_023243232.1;refGene_gn=SECISBP2;refGene_tx=XM_023243233.1;refGene_gn=SECISBP2;refGene_tx=XM_023243234.1;refGene_gn=SECISBP2;refGene_tx=XM_023243237.1
D4 2910024 2910170
refGene_gn=SECISBP2;refGene_tx=XM_023243228.1;refGene_gn=SECISBP2;refGene_tx=XM_023243230.1;refGene_gn=SECISBP2;refGene_tx=XM_023243231.1;refGene_gn=SECISBP2;refGene_tx=XM_023243232.1;refGene_gn=SECISBP2;refGene_tx=XM_023243233.1;refGene_gn=SECISBP2;refGene_tx=XM_023243234.1;refGene_gn=SECISBP2;refGene_tx=XM_023243235.1;refGene_gn=SECISBP2;refGene_tx=XM_023243236.1;refGene_gn=SECISBP2;refGene_tx=XM_023243237.1
D4 2914885 2915135
refGene_gn=SECISBP2;refGene_tx=XM_023243228.1;refGene_gn=SECISBP2;refGene_tx=XM_023243230.1;refGene_gn=SECISBP2;refGene_tx=XM_023243231.1;refGene_gn=SECISBP2;refGene_tx=XM_023243232.1;refGene_gn=SECISBP2;refGene_tx=XM_023243233.1;refGene_gn=SECISBP2;refGene_tx=XM_023243234.1;refGene_gn=SECISBP2;refGene_tx=XM_023243237.1
D4 2915366 2915490
refGene_gn=SECISBP2;refGene_tx=XM_023243228.1;refGene_gn=SECISBP2;refGene_tx=XM_023243230.1;refGene_gn=SECISBP2;refGene_tx=XM_023243231.1;refGene_gn=SECISBP2;refGene_tx=XM_023243232.1;refGene_gn=SECISBP2;refGene_tx=XM_023243233.1;refGene_gn=SECISBP2;refGene_tx=XM_023243234.1;refGene_gn=SECISBP2;refGene_tx=XM_023243237.1
D4 2916921 2917151
refGene_gn=SECISBP2;refGene_tx=XM_023243228.1;refGene_gn=SECISBP2;refGene_tx=XM_023243230.1;refGene_gn=SECISBP2;refGene_tx=XM_023243231.1;refGene_gn=SECISBP2;refGene_tx=XM_023243232.1;refGene_gn=SECISBP2;refGene_tx=XM_023243233.1;refGene_gn=SECISBP2;refGene_tx=XM_023243234.1;refGene_gn=SECISBP2;refGene_tx=XM_023243235.1;refGene_gn=SECISBP2;refGene_tx=XM_023243236.1;refGene_gn=SECISBP2;refGene_tx=XM_023243237.1
D4 2919818 2919989
refGene_gn=SECISBP2;refGene_tx=XM_023243228.1;refGene_gn=SECISBP2;refGene_tx=XM_023243230.1;refGene_gn=SECISBP2;refGene_tx=XM_023243231.1;refGene_gn=SECISBP2;refGene_tx=XM_023243232.1;refGene_gn=SECISBP2;refGene_tx=XM_023243233.1;refGene_gn=SECISBP2;refGene_tx=XM_023243235.1;refGene_gn=SECISBP2;refGene_tx=XM_023243236.1
D4 2921363 2921566
refGene_gn=SECISBP2;refGene_tx=XM_023243228.1;refGene_gn=SECISBP2;refGene_tx=XM_023243230.1;refGene_gn=SECISBP2;refGene_tx=XM_023243231.1;refGene_gn=SECISBP2;refGene_tx=XM_023243232.1;refGene_gn=SECISBP2;refGene_tx=XM_023243233.1;refGene_gn=SECISBP2;refGene_tx=XM_023243235.1;refGene_gn=SECISBP2;refGene_tx=XM_023243236.1
D4 2923785 2923908
refGene_gn=SECISBP2;refGene_tx=XM_023243228.1;refGene_gn=SECISBP2;refGene_tx=XM_023243230.1;refGene_gn=SECISBP2;refGene_tx=XM_023243231.1;refGene_gn=SECISBP2;refGene_tx=XM_023243232.1;refGene_gn=SECISBP2;refGen

e_tx=XM_023243233.1;refGene_gn=SECISBP2;refGene_tx=XM_023243234.1;refGene_gn=SECISBP2;refGene_tx=XM_023243235.1;refGene_gn=SECISBP2;refGene_tx=XM_023243236.1
D4	2925075	2925165	refGene_gn=SECISBP2;refGene_tx=XM_023243228.1;refGene_gn=SECISBP2;refGene_tx=XM_023243230.1;refGene_gn=SECISBP2;refGene_tx=XM_023243231.1;refGene_gn=SECISBP2;refGene_tx=XM_023243232.1;refGene_gn=SECISBP2;refGene_tx=XM_023243233.1;refGene_gn=SECISBP2;refGene_tx=XM_023243234.1;refGene_gn=SECISBP2;refGene_tx=XM_023243235.1;refGene_gn=SECISBP2;refGene_tx=XM_023243236.1
D4	2926561	2926694	refGene_gn=SECISBP2;refGene_tx=XM_023243228.1;refGene_gn=SECISBP2;refGene_tx=XM_023243230.1;refGene_gn=SECISBP2;refGene_tx=XM_023243231.1;refGene_gn=SECISBP2;refGene_tx=XM_023243232.1;refGene_gn=SECISBP2;refGene_tx=XM_023243233.1;refGene_gn=SECISBP2;refGene_tx=XM_023243234.1;refGene_gn=SECISBP2;refGene_tx=XM_023243235.1;refGene_gn=SECISBP2;refGene_tx=XM_023243236.1;refGene_gn=SECISBP2;refGene_tx=XM_023243237.1
D4	2941073	2941237	refGene_gn=SECISBP2;refGene_tx=XM_023243228.1;refGene_gn=SECISBP2;refGene_tx=XM_023243230.1;refGene_gn=SECISBP2;refGene_tx=XM_023243231.1;refGene_gn=SECISBP2;refGene_tx=XM_023243232.1;refGene_gn=SECISBP2;refGene_tx=XM_023243233.1;refGene_gn=SECISBP2;refGene_tx=XM_023243234.1;refGene_gn=SECISBP2;refGene_tx=XM_023243235.1;refGene_gn=SECISBP2;refGene_tx=XM_023243236.1;refGene_gn=SECISBP2;refGene_tx=XM_023243237.1
D4	2941703	2941839	refGene_gn=SECISBP2;refGene_tx=XM_023243228.1;refGene_gn=SECISBP2;refGene_tx=XM_023243230.1;refGene_gn=SECISBP2;refGene_tx=XM_023243231.1;refGene_gn=SECISBP2;refGene_tx=XM_023243232.1;refGene_gn=SECISBP2;refGene_tx=XM_023243233.1;refGene_gn=SECISBP2;refGene_tx=XM_023243234.1;refGene_gn=SECISBP2;refGene_tx=XM_023243235.1;refGene_gn=SECISBP2;refGene_tx=XM_023243236.1;refGene_gn=SECISBP2;refGene_tx=XM_023243237.1
D4	2943122	2943273	refGene_gn=SECISBP2;refGene_tx=XM_023243228.1;refGene_gn=SECISBP2;refGene_tx=XM_023243230.1;refGene_gn=SECISBP2;refGene_tx=XM_023243231.1;refGene_gn=SECISBP2;refGene_tx=XM_023243232.1;refGene_gn=SECISBP2;refGene_tx=XM_023243233.1;refGene_gn=SECISBP2;refGene_tx=XM_023243234.1;refGene_gn=SECISBP2;refGene_tx=XM_023243235.1;refGene_gn=SECISBP2;refGene_tx=XM_023243236.1;refGene_gn=SECISBP2;refGene_tx=XM_023243237.1
D4	2943746	2943967	refGene_gn=SECISBP2;refGene_tx=XM_023243228.1;refGene_gn=SECISBP2;refGene_tx=XM_023243230.1;refGene_gn=SECISBP2;refGene_tx=XM_023243231.1;refGene_gn=SECISBP2;refGene_tx=XM_023243232.1;refGene_gn=SECISBP2;refGene_tx=XM_023243233.1;refGene_gn=SECISBP2;refGene_tx=XM_023243234.1;refGene_gn=SECISBP2;refGene_tx=XM_023243235.1;refGene_gn=SECISBP2;refGene_tx=XM_023243236.1;refGene_gn=SECISBP2;refGene_tx=XM_023243237.1
D4	2948238	2948393	refGene_gn=SECISBP2;refGene_tx=XM_023243228.1;refGene_gn=SECISBP2;refGene_tx=XM_023243230.1;refGene_gn=SECISBP2;refGene_tx=XM_023243231.1;refGene_gn=SECISBP2;refGene_tx=XM_023243232.1;refGene_gn=SECISBP2;refGene_tx=XM_023243233.1;refGene_gn=SECISBP2;refGene_tx=XM_023243234.1;refG

ene_gn=SECISBP2;refGene_tx=XM_023243235.1;refGene_gn=SECISBP2;refGene_tx=XM_023243236.1;refGene_gn=SECISBP2;refGene_tx=XM_023243237.1
D4 2948630 2948802 refGene_gn=SECISBP2;refGene_tx=XM_023243228.1;refGene_gn=SECISBP2;refGene_tx=XM_023243230.1;refGene_gn=SECISBP2;refGene_tx=XM_023243231.1;refGene_gn=SECISBP2;refGene_tx=XM_023243232.1;refGene_gn=SECISBP2;refGene_tx=XM_023243233.1;refGene_gn=SECISBP2;refGene_tx=XM_023243234.1;refGene_gn=SECISBP2;refGene_tx=XM_023243235.1;refGene_gn=SECISBP2;refGene_tx=XM_023243236.1;refGene_gn=SECISBP2;refGene_tx=XM_023243237.1
D4 2949163 2949267 refGene_gn=SECISBP2;refGene_tx=XM_023243228.1;refGene_gn=SECISBP2;refGene_tx=XM_023243230.1;refGene_gn=SECISBP2;refGene_tx=XM_023243231.1;refGene_gn=SECISBP2;refGene_tx=XM_023243232.1;refGene_gn=SECISBP2;refGene_tx=XM_023243233.1;refGene_gn=SECISBP2;refGene_tx=XM_023243234.1;refGene_gn=SECISBP2;refGene_tx=XM_023243235.1;refGene_gn=SECISBP2;refGene_tx=XM_023243236.1;refGene_gn=SECISBP2;refGene_tx=XM_023243237.1
D4 2952199 2952226 refGene_gn=SEMA4D;refGene_tx=XM_023243227.1
D4 2952789 2952887 refGene_gn=SEMA4D;refGene_tx=XM_023243227.1
D4 2954455 2954665 refGene_gn=SEMA4D;refGene_tx=XM_023243227.1
D4 2960337 2960544 refGene_gn=SEMA4D;refGene_tx=XM_023243227.1
D4 2961977 2962903 refGene_gn=SEMA4D;refGene_tx=XM_023243222.1;refGene_gn=SEMA4D;refGene_tx=XM_023243223.1;refGene_gn=SEMA4D;refGene_tx=XM_023243224.1;refGene_gn=SEMA4D;refGene_tx=XM_023243225.1;refGene_gn=SEMA4D;refGene_tx=XM_023243226.1
D4 2963702 2963746 refGene_gn=SEMA4D;refGene_tx=XM_023243222.1;refGene_gn=SEMA4D;refGene_tx=XM_023243223.1;refGene_gn=SEMA4D;refGene_tx=XM_023243224.1;refGene_gn=SEMA4D;refGene_tx=XM_023243225.1;refGene_gn=SEMA4D;refGene_tx=XM_023243226.1;refGene_gn=SEMA4D;refGene_tx=XM_023243227.1
D4 2963829 2964008 refGene_gn=SEMA4D;refGene_tx=XM_023243222.1;refGene_gn=SEMA4D;refGene_tx=XM_023243223.1;refGene_gn=SEMA4D;refGene_tx=XM_023243224.1;refGene_gn=SEMA4D;refGene_tx=XM_023243225.1;refGene_gn=SEMA4D;refGene_tx=XM_023243226.1;refGene_gn=SEMA4D;refGene_tx=XM_023243227.1
D4 2967453 2967569 refGene_gn=SEMA4D;refGene_tx=XM_023243222.1;refGene_gn=SEMA4D;refGene_tx=XM_023243223.1;refGene_gn=SEMA4D;refGene_tx=XM_023243224.1;refGene_gn=SEMA4D;refGene_tx=XM_023243225.1;refGene_gn=SEMA4D;refGene_tx=XM_023243226.1;refGene_gn=SEMA4D;refGene_tx=XM_023243227.1
D4 2968333 2968556 refGene_gn=SEMA4D;refGene_tx=XM_023243222.1;refGene_gn=SEMA4D;refGene_tx=XM_023243223.1;refGene_gn=SEMA4D;refGene_tx=XM_023243224.1;refGene_gn=SEMA4D;refGene_tx=XM_023243225.1;refGene_gn=SEMA4D;refGene_tx=XM_023243226.1;refGene_gn=SEMA4D;refGene_tx=XM_023243227.1
D4 2969393 2969550 refGene_gn=SEMA4D;refGene_tx=XM_023243222.1;refGene_gn=SEMA4D;refGene_tx=XM_023243223.1;refGene_gn=SEMA4D;refGene_tx=XM_023243224.1;refGene_gn=SEMA4D;refGene_tx=XM_023243225.1;refGene_gn=SEMA4D;refGene_tx=XM_023243226.1;refGene_gn=SEMA4D;refGene_tx=XM_023243227.1

D4 2969638 2969814
refGene_gn=SEMA4D;refGene_tx=XM_023243222.1;refGene_gn=SEMA4D;refGene_tx=XM_023243223.1;refGene_gn=SEMA4D;refGene_tx=XM_023243224.1;refGene_gn=SEMA4D;refGene_tx=XM_023243225.1;refGene_gn=SEMA4D;refGene_tx=XM_023243226.1;refGene_gn=SEMA4D;refGene_tx=XM_023243227.1
D4 2970777 2970929
refGene_gn=SEMA4D;refGene_tx=XM_023243222.1;refGene_gn=SEMA4D;refGene_tx=XM_023243223.1;refGene_gn=SEMA4D;refGene_tx=XM_023243224.1;refGene_gn=SEMA4D;refGene_tx=XM_023243225.1;refGene_gn=SEMA4D;refGene_tx=XM_023243226.1;refGene_gn=SEMA4D;refGene_tx=XM_023243227.1
D4 2971633 2971747
refGene_gn=SEMA4D;refGene_tx=XM_023243222.1;refGene_gn=SEMA4D;refGene_tx=XM_023243223.1;refGene_gn=SEMA4D;refGene_tx=XM_023243224.1;refGene_gn=SEMA4D;refGene_tx=XM_023243225.1;refGene_gn=SEMA4D;refGene_tx=XM_023243226.1;refGene_gn=SEMA4D;refGene_tx=XM_023243227.1
D4 2972180 2972274
refGene_gn=SEMA4D;refGene_tx=XM_023243222.1;refGene_gn=SEMA4D;refGene_tx=XM_023243223.1;refGene_gn=SEMA4D;refGene_tx=XM_023243224.1;refGene_gn=SEMA4D;refGene_tx=XM_023243225.1;refGene_gn=SEMA4D;refGene_tx=XM_023243226.1;refGene_gn=SEMA4D;refGene_tx=XM_023243227.1
D4 2974876 2974975
refGene_gn=SEMA4D;refGene_tx=XM_023243222.1;refGene_gn=SEMA4D;refGene_tx=XM_023243223.1;refGene_gn=SEMA4D;refGene_tx=XM_023243224.1;refGene_gn=SEMA4D;refGene_tx=XM_023243225.1;refGene_gn=SEMA4D;refGene_tx=XM_023243226.1;refGene_gn=SEMA4D;refGene_tx=XM_023243227.1
D4 2976416 2976479
refGene_gn=SEMA4D;refGene_tx=XM_023243222.1;refGene_gn=SEMA4D;refGene_tx=XM_023243223.1;refGene_gn=SEMA4D;refGene_tx=XM_023243224.1;refGene_gn=SEMA4D;refGene_tx=XM_023243225.1;refGene_gn=SEMA4D;refGene_tx=XM_023243226.1;refGene_gn=SEMA4D;refGene_tx=XM_023243227.1
D4 2978684 2978830
refGene_gn=SEMA4D;refGene_tx=XM_023243222.1;refGene_gn=SEMA4D;refGene_tx=XM_023243223.1;refGene_gn=SEMA4D;refGene_tx=XM_023243224.1;refGene_gn=SEMA4D;refGene_tx=XM_023243225.1;refGene_gn=SEMA4D;refGene_tx=XM_023243226.1;refGene_gn=SEMA4D;refGene_tx=XM_023243227.1
D4 2980252 2980352
refGene_gn=SEMA4D;refGene_tx=XM_023243222.1;refGene_gn=SEMA4D;refGene_tx=XM_023243223.1;refGene_gn=SEMA4D;refGene_tx=XM_023243224.1;refGene_gn=SEMA4D;refGene_tx=XM_023243225.1;refGene_gn=SEMA4D;refGene_tx=XM_023243226.1;refGene_gn=SEMA4D;refGene_tx=XM_023243227.1
D4 3105355 3105408 refGene_gn=GADD45G;refGene_tx=XM_023243239.1
D4 3105643 3105745 refGene_gn=GADD45G;refGene_tx=XM_023243239.1
D4 3105854 3106068 refGene_gn=GADD45G;refGene_tx=XM_023243239.1
D4 3106160 3106271 refGene_gn=GADD45G;refGene_tx=XM_023243239.1
D4 3437898 3438358
refGene_gn=LOC109493572;refGene_tx=XM_019815516.1
D4 3439415 3439890
refGene_gn=LOC109493572;refGene_tx=XM_019815516.1
D4 3442086 3442138
refGene_gn=LOC109493572;refGene_tx=XM_019815516.1

D4	3484462	3484573
refGene_gn=LOC109493572;refGene_tx=XM_019815516.1
D4	3751553	3752153
refGene_gn=DIRAS2;refGene_tx=XM_003995319.4;refGene_gn=DIRAS2;refGene_tx=XM_023243248.1
D4	3867656	3868073
refGene_gn=SYK;refGene_tx=XM_023243249.1;refGene_gn=SYK;refGene_tx=XM_023243250.1
D4	3870136	3870297
refGene_gn=SYK;refGene_tx=XM_023243249.1;refGene_gn=SYK;refGene_tx=XM_023243250.1
D4	3882194	3882333
refGene_gn=SYK;refGene_tx=XM_023243249.1;refGene_gn=SYK;refGene_tx=XM_023243250.1
D4	3886089	3886171
refGene_gn=SYK;refGene_tx=XM_023243249.1;refGene_gn=SYK;refGene_tx=XM_023243250.1
D4	3886524	3886574
refGene_gn=SYK;refGene_tx=XM_023243249.1;refGene_gn=SYK;refGene_tx=XM_023243250.1
D4	3888199	3888268
refGene_gn=SYK;refGene_tx=XM_019815699.1;refGene_gn=SYK;refGene_tx=XM_023243249.1
D4	3896904	3896992
refGene_gn=SYK;refGene_tx=XM_019815699.1;refGene_gn=SYK;refGene_tx=XM_023243249.1;refGene_gn=SYK;refGene_tx=XM_023243250.1
D4	3898251	3898455
refGene_gn=SYK;refGene_tx=XM_019815699.1;refGene_gn=SYK;refGene_tx=XM_023243249.1;refGene_gn=SYK;refGene_tx=XM_023243250.1
D4	3901640	3901850
refGene_gn=SYK;refGene_tx=XM_019815699.1;refGene_gn=SYK;refGene_tx=XM_023243249.1;refGene_gn=SYK;refGene_tx=XM_023243250.1
D4	3903000	3903190
refGene_gn=SYK;refGene_tx=XM_019815699.1;refGene_gn=SYK;refGene_tx=XM_023243249.1;refGene_gn=SYK;refGene_tx=XM_023243250.1
D4	3909884	3910025
refGene_gn=SYK;refGene_tx=XM_019815699.1;refGene_gn=SYK;refGene_tx=XM_023243249.1;refGene_gn=SYK;refGene_tx=XM_023243250.1
D4	3910720	3910833
refGene_gn=SYK;refGene_tx=XM_019815699.1;refGene_gn=SYK;refGene_tx=XM_023243249.1;refGene_gn=SYK;refGene_tx=XM_023243250.1
D4	3916551	3916624
refGene_gn=SYK;refGene_tx=XM_019815699.1;refGene_gn=SYK;refGene_tx=XM_023243249.1;refGene_gn=SYK;refGene_tx=XM_023243250.1
D4	3925180	3925350
refGene_gn=AUH;refGene_tx=XM_023243254.1;refGene_gn=AUH;refGene_tx=XM_023243255.1;refGene_gn=AUH;refGene_tx=XM_023243257.1;refGene_gn=AUH;refGene_tx=XM_023243258.1
D4	3926268	3926350
refGene_gn=AUH;refGene_tx=XM_023243254.1;refGene_gn=AUH;refGene_tx=XM_

023243255.1;refGene_gn=AUH;refGene_tx=XM_023243257.1;refGene_gn=AUH;refGene_tx=XM_023243258.1
D4 4086246 4086360 refGene_gn=AUH;refGene_tx=XM_023243256.1
D4 4114334 4114475
refGene_gn=AUH;refGene_tx=XM_023243254.1;refGene_gn=AUH;refGene_tx=XM_023243255.1;refGene_gn=AUH;refGene_tx=XM_023243256.1;refGene_gn=AUH;refGene_tx=XM_023243257.1;refGene_gn=AUH;refGene_tx=XM_023243258.1
D4 4117666 4117744
refGene_gn=AUH;refGene_tx=XM_019815701.2;refGene_gn=AUH;refGene_tx=XM_019815702.2
D4 4122426 4122474
refGene_gn=AUH;refGene_tx=XM_019815701.2;refGene_gn=AUH;refGene_tx=XM_019815702.2;refGene_gn=AUH;refGene_tx=XM_023243254.1;refGene_gn=AUH;refGene_tx=XM_023243255.1;refGene_gn=AUH;refGene_tx=XM_023243256.1;refGene_gn=AUH;refGene_tx=XM_023243257.1;refGene_gn=AUH;refGene_tx=XM_023243258.1
D4 4123603 4123654
refGene_gn=AUH;refGene_tx=XM_019815701.2;refGene_gn=AUH;refGene_tx=XM_019815702.2;refGene_gn=AUH;refGene_tx=XM_023243254.1;refGene_gn=AUH;refGene_tx=XM_023243255.1;refGene_gn=AUH;refGene_tx=XM_023243256.1;refGene_gn=AUH;refGene_tx=XM_023243257.1;refGene_gn=AUH;refGene_tx=XM_023243258.1
D4 4126538 4126726
refGene_gn=AUH;refGene_tx=XM_019815701.2;refGene_gn=AUH;refGene_tx=XM_019815702.2;refGene_gn=AUH;refGene_tx=XM_023243254.1;refGene_gn=AUH;refGene_tx=XM_023243255.1;refGene_gn=AUH;refGene_tx=XM_023243256.1;refGene_gn=AUH;refGene_tx=XM_023243257.1;refGene_gn=AUH;refGene_tx=XM_023243258.1
D4 4208974 4209031
refGene_gn=AUH;refGene_tx=XM_019815701.2;refGene_gn=AUH;refGene_tx=XM_019815702.2;refGene_gn=AUH;refGene_tx=XM_023243254.1;refGene_gn=AUH;refGene_tx=XM_023243255.1;refGene_gn=AUH;refGene_tx=XM_023243256.1;refGene_gn=AUH;refGene_tx=XM_023243257.1;refGene_gn=AUH;refGene_tx=XM_023243258.1
D4 4210603 4210696
refGene_gn=AUH;refGene_tx=XM_019815701.2;refGene_gn=AUH;refGene_tx=XM_019815702.2;refGene_gn=AUH;refGene_tx=XM_023243254.1;refGene_gn=AUH;refGene_tx=XM_023243255.1;refGene_gn=AUH;refGene_tx=XM_023243256.1;refGene_gn=AUH;refGene_tx=XM_023243257.1;refGene_gn=AUH;refGene_tx=XM_023243258.1
D4 4240734 4240821
refGene_gn=AUH;refGene_tx=XM_019815701.2;refGene_gn=AUH;refGene_tx=XM_023243254.1;refGene_gn=AUH;refGene_tx=XM_023243256.1;refGene_gn=AUH;refGene_tx=XM_023243257.1;refGene_gn=AUH;refGene_tx=XM_023243258.1
D4 4267741 4267829
refGene_gn=AUH;refGene_tx=XM_019815701.2;refGene_gn=AUH;refGene_tx=XM_019815702.2;refGene_gn=AUH;refGene_tx=XM_023243254.1;refGene_gn=AUH;refGene_tx=XM_023243255.1;refGene_gn=AUH;refGene_tx=XM_023243256.1;refGene_gn=AUH;refGene_tx=XM_023243258.1
D4 4267940 4268008

refGene_gn=AUH;refGene_tx=XM_019815701.2;refGene_gn=AUH;refGene_tx=XM_019815702.2;refGene_gn=AUH;refGene_tx=XM_023243254.1;refGene_gn=AUH;refGene_tx=XM_023243255.1;refGene_gn=AUH;refGene_tx=XM_023243256.1;refGene_gn=AUH;refGene_tx=XM_023243258.1
D4 4277242 4277507
refGene_gn=AUH;refGene_tx=XM_019815701.2;refGene_gn=AUH;refGene_tx=XM_019815702.2;refGene_gn=AUH;refGene_tx=XM_023243254.1;refGene_gn=AUH;refGene_tx=XM_023243255.1;refGene_gn=AUH;refGene_tx=XM_023243256.1;refGene_gn=AUH;refGene_tx=XM_023243257.1
D4 4317299 4318691
refGene_gn=NFIL3;refGene_tx=XM_006939033.4;refGene_gn=NFIL3;refGene_tx=XM_023243252.1;refGene_gn=NFIL3;refGene_tx=XM_023243253.1
D4 4523182 4524625
refGene_gn=ROR2;refGene_tx=XM_003995323.4;refGene_gn=ROR2;refGene_tx=XM_023243262.1
D4 4526142 4526345
refGene_gn=ROR2;refGene_tx=XM_003995323.4;refGene_gn=ROR2;refGene_tx=XM_023243262.1
D4 4529510 4529756
refGene_gn=ROR2;refGene_tx=XM_003995323.4;refGene_gn=ROR2;refGene_tx=XM_023243262.1
D4 4531288 4531603
refGene_gn=ROR2;refGene_tx=XM_003995323.4;refGene_gn=ROR2;refGene_tx=XM_023243262.1
D4 4534946 4535074
refGene_gn=ROR2;refGene_tx=XM_003995323.4;refGene_gn=ROR2;refGene_tx=XM_023243262.1
D4 4550175 4550206
refGene_gn=ROR2;refGene_tx=XM_003995323.4;refGene_gn=ROR2;refGene_tx=XM_023243262.1
D4 4551048 4551336
refGene_gn=ROR2;refGene_tx=XM_003995323.4;refGene_gn=ROR2;refGene_tx=XM_023243262.1
D4 4567934 4568012
refGene_gn=ROR2;refGene_tx=XM_003995323.4;refGene_gn=ROR2;refGene_tx=XM_023243262.1
D4 4719786 4719874 refGene_gn=ROR2;refGene_tx=XM_023243262.1
D4 4769157 4769251 refGene_gn=SPTLC1;refGene_tx=XM_003995324.5
D4 4770692 4770766 refGene_gn=SPTLC1;refGene_tx=XM_003995324.5
D4 4774013 4774131 refGene_gn=SPTLC1;refGene_tx=XM_003995324.5
D4 4777410 4777465 refGene_gn=SPTLC1;refGene_tx=XM_003995324.5
D4 4778501 4778598 refGene_gn=SPTLC1;refGene_tx=XM_003995324.5
D4 4779678 4779774 refGene_gn=SPTLC1;refGene_tx=XM_003995324.5
D4 4781725 4781833 refGene_gn=SPTLC1;refGene_tx=XM_003995324.5
D4 4786235 4786325 refGene_gn=SPTLC1;refGene_tx=XM_003995324.5
D4 4787796 4787926 refGene_gn=SPTLC1;refGene_tx=XM_003995324.5
D4 4795772 4795905 refGene_gn=SPTLC1;refGene_tx=XM_003995324.5
D4 4804954 4805027 refGene_gn=SPTLC1;refGene_tx=XM_003995324.5
D4 4805592 4805686 refGene_gn=SPTLC1;refGene_tx=XM_003995324.5
D4 4816316 4816411 refGene_gn=SPTLC1;refGene_tx=XM_003995324.5

D4 4818379 4818487 refGene_gn=SPTLC1;refGene_tx=XM_003995324.5
D4 4821061 4821118 refGene_gn=SPTLC1;refGene_tx=XM_003995324.5
D4 4981874 4985162
refGene_gn=LOC101096613;refGene_tx=XM_023243265.1
D4 4985164 4985873
refGene_gn=LOC101096613;refGene_tx=XM_023243265.1
D4 4986102 4986163
refGene_gn=LOC101096613;refGene_tx=XM_023243265.1
D4 4986672 4986727
refGene_gn=LOC101096613;refGene_tx=XM_023243265.1
D4 4987482 4987671
refGene_gn=LOC101096613;refGene_tx=XM_023243265.1
D4 5086433 5086518
refGene_gn=LOC101096853;refGene_tx=XM_023243266.1
D4 5118606 5118717
refGene_gn=LOC101096853;refGene_tx=XM_023243266.1
D4 5132141 5132305
refGene_gn=LOC101096853;refGene_tx=XM_023243266.1
D4 5132306 5132336
refGene_gn=LOC101096853;refGene_tx=XM_023243266.1
D4 5175861 5176009
refGene_gn=LOC101096853;refGene_tx=XM_023243266.1
D4 5186459 5186663
refGene_gn=LOC101096853;refGene_tx=XM_023243266.1
D4 5187770 5187955
refGene_gn=LOC101096853;refGene_tx=XM_023243266.1
D4 5189022 5189166
refGene_gn=LOC101096853;refGene_tx=XM_023243266.1
D4 5192172 5192434
refGene_gn=LOC101096853;refGene_tx=XM_023243266.1
D4 5198665 5198812
refGene_gn=LOC101096853;refGene_tx=XM_023243266.1
D4 5208276 5208448
refGene_gn=LOC101096853;refGene_tx=XM_023243266.1
D4 5216860 5216967
refGene_gn=LOC101096853;refGene_tx=XM_023243266.1
D4 5220812 5220932
refGene_gn=LOC101096853;refGene_tx=XM_023243266.1
D4 5235038 5235236
refGene_gn=LOC101096853;refGene_tx=XM_023243266.1
D4 5238973 5239130
refGene_gn=LOC101096853;refGene_tx=XM_023243266.1
D4 5244956 5245084
refGene_gn=LOC101096853;refGene_tx=XM_023243266.1
D4 5252754 5252925
refGene_gn=LOC101096853;refGene_tx=XM_023243266.1
D4 5253899 5254118
refGene_gn=LOC101096853;refGene_tx=XM_023243266.1
D4 5256309 5256549
refGene_gn=LOC101096853;refGene_tx=XM_023243266.1

D4 5257316 5257547
refGene_gn=LOC101096853;refGene_tx=XM_023243266.1
D4 5259061 5259195
refGene_gn=LOC101096853;refGene_tx=XM_023243266.1
D4 5259999 5260087
refGene_gn=LOC101096853;refGene_tx=XM_023243266.1
D4 5273515 5273746
refGene_gn=LOC101096853;refGene_tx=XM_023243266.1
D4 5273964 5274036
refGene_gn=LOC101096853;refGene_tx=XM_023243266.1
D4 5278883 5279077
refGene_gn=LOC101096853;refGene_tx=XM_023243266.1
D4 5721379 5721568
refGene_gn=LOC101097105;refGene_tx=XM_023243270.1
D4 5722329 5722384
refGene_gn=LOC101097105;refGene_tx=XM_023243270.1
D4 5722893 5722954
refGene_gn=LOC101097105;refGene_tx=XM_023243270.1
D4 5723178 5726723
refGene_gn=LOC101097105;refGene_tx=XM_023243270.1
D4 5726725 5726791
refGene_gn=LOC101097105;refGene_tx=XM_023243270.1
D4 5726793 5727149
refGene_gn=LOC101097105;refGene_tx=XM_023243270.1
D4 5840940 5841957
refGene_gn=LOC111557592;refGene_tx=XM_023243260.1
D4 5939901 5939921 refGene_gn=ERMP1;refGene_tx=XM_023243275.1
D4 5941041 5941379
refGene_gn=ERMP1;refGene_tx=XM_023243273.1;refGene_gn=ERMP1;refGene_tx
=XM_023243274.1
D4 5942748 5943050
refGene_gn=ERMP1;refGene_tx=XM_023243273.1;refGene_gn=ERMP1;refGene_tx
=XM_023243274.1;refGene_gn=ERMP1;refGene_tx=XM_023243275.1
D4 5947470 5947598
refGene_gn=ERMP1;refGene_tx=XM_023243273.1;refGene_gn=ERMP1;refGene_tx
=XM_023243275.1
D4 5948199 5948305
refGene_gn=ERMP1;refGene_tx=XM_023243273.1;refGene_gn=ERMP1;refGene_tx
=XM_023243275.1
D4 5959911 5960058
refGene_gn=ERMP1;refGene_tx=XM_023243273.1;refGene_gn=ERMP1;refGene_tx
=XM_023243274.1;refGene_gn=ERMP1;refGene_tx=XM_023243275.1
D4 5960756 5960849
refGene_gn=ERMP1;refGene_tx=XM_023243273.1;refGene_gn=ERMP1;refGene_tx
=XM_023243274.1;refGene_gn=ERMP1;refGene_tx=XM_023243275.1
D4 5962614 5962827
refGene_gn=ERMP1;refGene_tx=XM_023243273.1;refGene_gn=ERMP1;refGene_tx
=XM_023243274.1;refGene_gn=ERMP1;refGene_tx=XM_023243275.1
D4 5963655 5963876
refGene_gn=ERMP1;refGene_tx=XM_023243273.1;refGene_gn=ERMP1;refGene_tx

=XM_023243274.1;refGene_gn=ERMP1;refGene_tx=XM_023243275.1
D4	5966778	5966953
refGene_gn=ERMP1;refGene_tx=XM_023243273.1;refGene_gn=ERMP1;refGene_tx
=XM_023243274.1;refGene_gn=ERMP1;refGene_tx=XM_023243275.1
D4	5967318	5967509
refGene_gn=ERMP1;refGene_tx=XM_023243273.1;refGene_gn=ERMP1;refGene_tx
=XM_023243274.1;refGene_gn=ERMP1;refGene_tx=XM_023243275.1
D4	5971448	5971601
refGene_gn=ERMP1;refGene_tx=XM_023243273.1;refGene_gn=ERMP1;refGene_tx
=XM_023243274.1;refGene_gn=ERMP1;refGene_tx=XM_023243275.1
D4	5973755	5973958
refGene_gn=ERMP1;refGene_tx=XM_023243273.1;refGene_gn=ERMP1;refGene_tx
=XM_023243274.1;refGene_gn=ERMP1;refGene_tx=XM_023243275.1
D4	5974685	5974801
refGene_gn=ERMP1;refGene_tx=XM_023243273.1;refGene_gn=ERMP1;refGene_tx
=XM_023243274.1;refGene_gn=ERMP1;refGene_tx=XM_023243275.1
D4	5982523	5982687
refGene_gn=ERMP1;refGene_tx=XM_023243273.1;refGene_gn=ERMP1;refGene_tx
=XM_023243274.1;refGene_gn=ERMP1;refGene_tx=XM_023243275.1
D4	5982802	5982967
refGene_gn=ERMP1;refGene_tx=XM_023243273.1;refGene_gn=ERMP1;refGene_tx
=XM_023243274.1;refGene_gn=ERMP1;refGene_tx=XM_023243275.1
D4	5990213	5990502
refGene_gn=RIC1;refGene_tx=XM_019815710.2;refGene_gn=RIC1;refGene_tx=X
M_023243271.1;refGene_gn=RIC1;refGene_tx=XM_023243272.1
D4	5991934	5992123
refGene_gn=RIC1;refGene_tx=XM_019815710.2;refGene_gn=RIC1;refGene_tx=X
M_023243271.1;refGene_gn=RIC1;refGene_tx=XM_023243272.1
D4	5992268	5992446
refGene_gn=RIC1;refGene_tx=XM_019815710.2;refGene_gn=RIC1;refGene_tx=X
M_023243271.1;refGene_gn=RIC1;refGene_tx=XM_023243272.1
D4	5994531	5994720
refGene_gn=RIC1;refGene_tx=XM_019815710.2;refGene_gn=RIC1;refGene_tx=X
M_023243271.1;refGene_gn=RIC1;refGene_tx=XM_023243272.1
D4	5995906	5996193
refGene_gn=RIC1;refGene_tx=XM_019815710.2;refGene_gn=RIC1;refGene_tx=X
M_023243271.1;refGene_gn=RIC1;refGene_tx=XM_023243272.1
D4	5997808	5997945
refGene_gn=RIC1;refGene_tx=XM_019815710.2;refGene_gn=RIC1;refGene_tx=X
M_023243271.1;refGene_gn=RIC1;refGene_tx=XM_023243272.1
D4	5998033	5998192
refGene_gn=RIC1;refGene_tx=XM_019815710.2;refGene_gn=RIC1;refGene_tx=X
M_023243271.1;refGene_gn=RIC1;refGene_tx=XM_023243272.1
D4	6000433	6001162
refGene_gn=RIC1;refGene_tx=XM_019815710.2;refGene_gn=RIC1;refGene_tx=X
M_023243271.1;refGene_gn=RIC1;refGene_tx=XM_023243272.1
D4	6001627	6001744
refGene_gn=RIC1;refGene_tx=XM_019815710.2;refGene_gn=RIC1;refGene_tx=X
M_023243271.1;refGene_gn=RIC1;refGene_tx=XM_023243272.1
D4	6007910	6008049

refGene_gn=RIC1;refGene_tx=XM_019815710.2;refGene_gn=RIC1;refGene_tx=XM_023243271.1;refGene_gn=RIC1;refGene_tx=XM_023243272.1
D4 6009010 6009171
refGene_gn=RIC1;refGene_tx=XM_019815710.2;refGene_gn=RIC1;refGene_tx=XM_023243271.1;refGene_gn=RIC1;refGene_tx=XM_023243272.1
D4 6010528 6010618
refGene_gn=RIC1;refGene_tx=XM_019815710.2;refGene_gn=RIC1;refGene_tx=XM_023243271.1;refGene_gn=RIC1;refGene_tx=XM_023243272.1
D4 6011424 6011535
refGene_gn=RIC1;refGene_tx=XM_019815710.2;refGene_gn=RIC1;refGene_tx=XM_023243271.1;refGene_gn=RIC1;refGene_tx=XM_023243272.1
D4 6011875 6011914
refGene_gn=RIC1;refGene_tx=XM_019815710.2;refGene_gn=RIC1;refGene_tx=XM_023243271.1;refGene_gn=RIC1;refGene_tx=XM_023243272.1
D4 6017106 6017310
refGene_gn=RIC1;refGene_tx=XM_019815710.2;refGene_gn=RIC1;refGene_tx=XM_023243271.1;refGene_gn=RIC1;refGene_tx=XM_023243272.1
D4 6018973 6019126
refGene_gn=RIC1;refGene_tx=XM_019815710.2;refGene_gn=RIC1;refGene_tx=XM_023243271.1;refGene_gn=RIC1;refGene_tx=XM_023243272.1
D4 6020694 6020743
refGene_gn=RIC1;refGene_tx=XM_019815710.2;refGene_gn=RIC1;refGene_tx=XM_023243271.1;refGene_gn=RIC1;refGene_tx=XM_023243272.1
D4 6021402 6021547
refGene_gn=RIC1;refGene_tx=XM_019815710.2;refGene_gn=RIC1;refGene_tx=XM_023243271.1;refGene_gn=RIC1;refGene_tx=XM_023243272.1
D4 6028037 6028126
refGene_gn=RIC1;refGene_tx=XM_019815710.2;refGene_gn=RIC1;refGene_tx=XM_023243271.1;refGene_gn=RIC1;refGene_tx=XM_023243272.1
D4 6033603 6033695
refGene_gn=RIC1;refGene_tx=XM_019815710.2;refGene_gn=RIC1;refGene_tx=XM_023243271.1;refGene_gn=RIC1;refGene_tx=XM_023243272.1
D4 6038031 6038168
refGene_gn=RIC1;refGene_tx=XM_019815710.2;refGene_gn=RIC1;refGene_tx=XM_023243271.1;refGene_gn=RIC1;refGene_tx=XM_023243272.1
D4 6038758 6038901
refGene_gn=RIC1;refGene_tx=XM_019815710.2;refGene_gn=RIC1;refGene_tx=XM_023243271.1;refGene_gn=RIC1;refGene_tx=XM_023243272.1
D4 6045070 6045178
refGene_gn=RIC1;refGene_tx=XM_019815710.2;refGene_gn=RIC1;refGene_tx=XM_023243271.1;refGene_gn=RIC1;refGene_tx=XM_023243272.1
D4 6063957 6064037
refGene_gn=RIC1;refGene_tx=XM_023243271.1;refGene_gn=RIC1;refGene_tx=XM_023243272.1
D4 6093189 6093297
refGene_gn=RIC1;refGene_tx=XM_023243271.1;refGene_gn=RIC1;refGene_tx=XM_023243272.1
D4 6135336 6135480
refGene_gn=RIC1;refGene_tx=XM_023243271.1;refGene_gn=RIC1;refGene_tx=XM_023243272.1

D4 6179168 6179195
refGene_gn=PDCD1LG2;refGene_tx=XM_006939037.3;refGene_gn=PDCD1LG2;refGene_tx=XM_019815494.2;refGene_gn=PDCD1LG2;refGene_tx=XM_023242193.1
D4 6190109 6190115
refGene_gn=PDCD1LG2;refGene_tx=NM_001290244.1;refGene_gn=PDCD1LG2;refGene_tx=XM_006939038.3;refGene_gn=PDCD1LG2;refGene_tx=XM_023242194.1
D4 6202094 6202144
refGene_gn=PDCD1LG2;refGene_tx=NM_001290244.1;refGene_gn=PDCD1LG2;refGene_tx=XM_006939037.3;refGene_gn=PDCD1LG2;refGene_tx=XM_006939038.3;refGene_gn=PDCD1LG2;refGene_tx=XM_019815494.2;refGene_gn=PDCD1LG2;refGene_tx=XM_023242193.1;refGene_gn=PDCD1LG2;refGene_tx=XM_023242194.1
D4 6207456 6207591
refGene_gn=PDCD1LG2;refGene_tx=NM_001290244.1;refGene_gn=PDCD1LG2;refGene_tx=XM_006939037.3;refGene_gn=PDCD1LG2;refGene_tx=XM_006939038.3;refGene_gn=PDCD1LG2;refGene_tx=XM_019815494.2;refGene_gn=PDCD1LG2;refGene_tx=XM_023242193.1;refGene_gn=PDCD1LG2;refGene_tx=XM_023242194.1
D4 6213562 6213657 refGene_gn=PDCD1LG2;refGene_tx=XM_023242195.1
D4 6214567 6214837
refGene_gn=PDCD1LG2;refGene_tx=NM_001290244.1;refGene_gn=PDCD1LG2;refGene_tx=XM_006939037.3;refGene_gn=PDCD1LG2;refGene_tx=XM_006939038.3;refGene_gn=PDCD1LG2;refGene_tx=XM_019815494.2;refGene_gn=PDCD1LG2;refGene_tx=XM_023242193.1;refGene_gn=PDCD1LG2;refGene_tx=XM_023242194.1;refGene_gn=PDCD1LG2;refGene_tx=XM_023242195.1
D4 6223469 6223775
refGene_gn=PDCD1LG2;refGene_tx=NM_001290244.1;refGene_gn=PDCD1LG2;refGene_tx=XM_006939037.3;refGene_gn=PDCD1LG2;refGene_tx=XM_006939038.3;refGene_gn=PDCD1LG2;refGene_tx=XM_019815494.2;refGene_gn=PDCD1LG2;refGene_tx=XM_023242193.1;refGene_gn=PDCD1LG2;refGene_tx=XM_023242194.1;refGene_gn=PDCD1LG2;refGene_tx=XM_023242195.1
D4 6232953 6233019
refGene_gn=PDCD1LG2;refGene_tx=NM_001290244.1;refGene_gn=PDCD1LG2;refGene_tx=XM_006939037.3;refGene_gn=PDCD1LG2;refGene_tx=XM_006939038.3;refGene_gn=PDCD1LG2;refGene_tx=XM_019815494.2;refGene_gn=PDCD1LG2;refGene_tx=XM_023242193.1;refGene_gn=PDCD1LG2;refGene_tx=XM_023242194.1;refGene_gn=PDCD1LG2;refGene_tx=XM_023242195.1
D4 6245060 6245082
refGene_gn=PDCD1LG2;refGene_tx=XM_006939037.3;refGene_gn=PDCD1LG2;refGene_tx=XM_006939038.3
D4 6255332 6255378
refGene_gn=PDCD1LG2;refGene_tx=XM_023242193.1;refGene_gn=PDCD1LG2;refGene_tx=XM_023242194.1;refGene_gn=PDCD1LG2;refGene_tx=XM_023242195.1
D4 6280847 6280870 refGene_gn=CD274;refGene_tx=XM_006939039.2
D4 6282295 6282355 refGene_gn=CD274;refGene_tx=XM_006939039.2
D4 6284060 6284171 refGene_gn=CD274;refGene_tx=XM_006939039.2
D4 6286324 6286612 refGene_gn=CD274;refGene_tx=XM_006939039.2
D4 6294906 6295248 refGene_gn=CD274;refGene_tx=XM_006939039.2
D4 6296129 6296181 refGene_gn=CD274;refGene_tx=XM_006939039.2
D4 6315770 6315851
refGene_gn=PLGRKT;refGene_tx=XM_011288234.3;refGene_gn=PLGRKT;refGene_tx=XM_011288235.3;refGene_gn=PLGRKT;refGene_tx=XM_011288236.3

D4 6351689 6351820
refGene_gn=PLGRKT;refGene_tx=XM_011288234.3;refGene_gn=PLGRKT;refGene_
tx=XM_011288235.3;refGene_gn=PLGRKT;refGene_tx=XM_011288236.3
D4 6352393 6352503
refGene_gn=PLGRKT;refGene_tx=XM_011288234.3;refGene_gn=PLGRKT;refGene_
tx=XM_011288235.3;refGene_gn=PLGRKT;refGene_tx=XM_011288236.3
D4 6356242 6356364
refGene_gn=PLGRKT;refGene_tx=XM_011288234.3;refGene_gn=PLGRKT;refGene_
tx=XM_011288235.3;refGene_gn=PLGRKT;refGene_tx=XM_011288236.3
D4 6375806 6376011 refGene_gn=RLN;refGene_tx=NM_001009317.1
D4 6381026 6381364 refGene_gn=RLN;refGene_tx=NM_001009317.1
D4 6421342 6421625 refGene_gn=INSL6;refGene_tx=XM_003995326.3
D4 6434584 6434943 refGene_gn=INSL6;refGene_tx=XM_003995326.3
D4 6463126 6463234
refGene_gn=JAK2;refGene_tx=XM_019815715.2;refGene_gn=JAK2;refGene_tx=X
M_023243278.1;refGene_gn=JAK2;refGene_tx=XM_023243279.1
D4 6463680 6463794
refGene_gn=JAK2;refGene_tx=XM_019815715.2;refGene_gn=JAK2;refGene_tx=X
M_023243278.1;refGene_gn=JAK2;refGene_tx=XM_023243279.1
D4 6465124 6465242
refGene_gn=JAK2;refGene_tx=XM_019815715.2;refGene_gn=JAK2;refGene_tx=X
M_023243278.1;refGene_gn=JAK2;refGene_tx=XM_023243279.1
D4 6471069 6471145 refGene_gn=JAK2;refGene_tx=XM_019815713.2
D4 6471649 6471662 refGene_gn=JAK2;refGene_tx=XM_019815714.2
D4 6474065 6474238
refGene_gn=JAK2;refGene_tx=XM_019815713.2;refGene_gn=JAK2;refGene_tx=X
M_019815714.2;refGene_gn=JAK2;refGene_tx=XM_019815715.2;refGene_gn=JAK
2;refGene_tx=XM_023243278.1;refGene_gn=JAK2;refGene_tx=XM_023243279.1
D4 6474410 6474535
refGene_gn=JAK2;refGene_tx=XM_019815713.2;refGene_gn=JAK2;refGene_tx=X
M_019815714.2;refGene_gn=JAK2;refGene_tx=XM_019815715.2;refGene_gn=JAK
2;refGene_tx=XM_023243278.1;refGene_gn=JAK2;refGene_tx=XM_023243279.1
D4 6475384 6475574
refGene_gn=JAK2;refGene_tx=XM_019815713.2;refGene_gn=JAK2;refGene_tx=X
M_019815714.2;refGene_gn=JAK2;refGene_tx=XM_019815715.2;refGene_gn=JAK
2;refGene_tx=XM_023243278.1;refGene_gn=JAK2;refGene_tx=XM_023243279.1
D4 6483152 6483286
refGene_gn=JAK2;refGene_tx=XM_019815713.2;refGene_gn=JAK2;refGene_tx=X
M_019815714.2;refGene_gn=JAK2;refGene_tx=XM_019815715.2;refGene_gn=JAK
2;refGene_tx=XM_023243278.1;refGene_gn=JAK2;refGene_tx=XM_023243279.1
D4 6484363 6484514
refGene_gn=JAK2;refGene_tx=XM_019815713.2;refGene_gn=JAK2;refGene_tx=X
M_019815714.2;refGene_gn=JAK2;refGene_tx=XM_019815715.2;refGene_gn=JAK
2;refGene_tx=XM_023243278.1;refGene_gn=JAK2;refGene_tx=XM_023243279.1
D4 6484668 6484820
refGene_gn=JAK2;refGene_tx=XM_019815713.2;refGene_gn=JAK2;refGene_tx=X
M_019815714.2;refGene_gn=JAK2;refGene_tx=XM_019815715.2;refGene_gn=JAK
2;refGene_tx=XM_023243278.1;refGene_gn=JAK2;refGene_tx=XM_023243279.1
D4 6486528 6486667
refGene_gn=JAK2;refGene_tx=XM_019815713.2;refGene_gn=JAK2;refGene_tx=X

M_019815714.2;refGene_gn=JAK2;refGene_tx=XM_019815715.2;refGene_gn=JAK2;refGene_tx=XM_023243278.1;refGene_gn=JAK2;refGene_tx=XM_023243279.1
D4 6487402 6487530
refGene_gn=JAK2;refGene_tx=XM_019815713.2;refGene_gn=JAK2;refGene_tx=XM_019815714.2;refGene_gn=JAK2;refGene_tx=XM_019815715.2;refGene_gn=JAK2;refGene_tx=XM_023243278.1;refGene_gn=JAK2;refGene_tx=XM_023243279.1
D4 6490627 6490715
refGene_gn=JAK2;refGene_tx=XM_019815713.2;refGene_gn=JAK2;refGene_tx=XM_019815714.2;refGene_gn=JAK2;refGene_tx=XM_019815715.2;refGene_gn=JAK2;refGene_tx=XM_023243278.1;refGene_gn=JAK2;refGene_tx=XM_023243279.1
D4 6491806 6491941
refGene_gn=JAK2;refGene_tx=XM_019815713.2;refGene_gn=JAK2;refGene_tx=XM_019815714.2;refGene_gn=JAK2;refGene_tx=XM_019815715.2;refGene_gn=JAK2;refGene_tx=XM_023243278.1;refGene_gn=JAK2;refGene_tx=XM_023243279.1
D4 6494383 6494511
refGene_gn=JAK2;refGene_tx=XM_019815713.2;refGene_gn=JAK2;refGene_tx=XM_019815714.2;refGene_gn=JAK2;refGene_tx=XM_019815715.2;refGene_gn=JAK2;refGene_tx=XM_023243278.1;refGene_gn=JAK2;refGene_tx=XM_023243279.1
D4 6495153 6495340
refGene_gn=JAK2;refGene_tx=XM_019815713.2;refGene_gn=JAK2;refGene_tx=XM_019815714.2;refGene_gn=JAK2;refGene_tx=XM_019815715.2;refGene_gn=JAK2;refGene_tx=XM_023243278.1;refGene_gn=JAK2;refGene_tx=XM_023243279.1
D4 6498771 6498883
refGene_gn=JAK2;refGene_tx=XM_019815713.2;refGene_gn=JAK2;refGene_tx=XM_019815714.2;refGene_gn=JAK2;refGene_tx=XM_019815715.2;refGene_gn=JAK2;refGene_tx=XM_023243278.1;refGene_gn=JAK2;refGene_tx=XM_023243279.1
D4 6500615 6500773
refGene_gn=JAK2;refGene_tx=XM_019815713.2;refGene_gn=JAK2;refGene_tx=XM_019815714.2;refGene_gn=JAK2;refGene_tx=XM_019815715.2;refGene_gn=JAK2;refGene_tx=XM_023243278.1;refGene_gn=JAK2;refGene_tx=XM_023243279.1
D4 6503531 6503651
refGene_gn=JAK2;refGene_tx=XM_019815713.2;refGene_gn=JAK2;refGene_tx=XM_019815714.2;refGene_gn=JAK2;refGene_tx=XM_019815715.2;refGene_gn=JAK2;refGene_tx=XM_023243278.1;refGene_gn=JAK2;refGene_tx=XM_023243279.1
D4 6504380 6504702
refGene_gn=JAK2;refGene_tx=XM_019815713.2;refGene_gn=JAK2;refGene_tx=XM_019815714.2;refGene_gn=JAK2;refGene_tx=XM_019815715.2;refGene_gn=JAK2;refGene_tx=XM_023243278.1;refGene_gn=JAK2;refGene_tx=XM_023243279.1
D4 6511541 6511687
refGene_gn=JAK2;refGene_tx=XM_019815713.2;refGene_gn=JAK2;refGene_tx=XM_019815714.2;refGene_gn=JAK2;refGene_tx=XM_019815715.2;refGene_gn=JAK2;refGene_tx=XM_023243278.1;refGene_gn=JAK2;refGene_tx=XM_023243279.1
D4 6516745 6516863
refGene_gn=JAK2;refGene_tx=XM_019815713.2;refGene_gn=JAK2;refGene_tx=XM_019815714.2;refGene_gn=JAK2;refGene_tx=XM_023243278.1;refGene_gn=JAK2;refGene_tx=XM_023243279.1
D4 6523202 6523326
refGene_gn=JAK2;refGene_tx=XM_019815713.2;refGene_gn=JAK2;refGene_tx=XM_019815714.2;refGene_gn=JAK2;refGene_tx=XM_023243278.1;refGene_gn=JAK2;refGene_tx=XM_023243279.1

D4 6537418 6537644
refGene_gn=JAK2;refGene_tx=XM_019815713.2;refGene_gn=JAK2;refGene_tx=XM_019815714.2;refGene_gn=JAK2;refGene_tx=XM_023243278.1;refGene_gn=JAK2;refGene_tx=XM_023243279.1
D4 6714695 6714846
refGene_gn=RCL1;refGene_tx=XM_003995328.5;refGene_gn=RCL1;refGene_tx=XM_019815716.2;refGene_gn=RCL1;refGene_tx=XM_019815717.2
D4 6724412 6724516
refGene_gn=RCL1;refGene_tx=XM_003995328.5;refGene_gn=RCL1;refGene_tx=XM_019815717.2
D4 6728373 6728514 refGene_gn=RCL1;refGene_tx=XM_023243281.1
D4 6729532 6729689
refGene_gn=RCL1;refGene_tx=XM_003995328.5;refGene_gn=RCL1;refGene_tx=XM_019815716.2;refGene_gn=RCL1;refGene_tx=XM_019815717.2;refGene_gn=RCL1;refGene_tx=XM_023243281.1
D4 6733621 6733831
refGene_gn=RCL1;refGene_tx=XM_003995328.5;refGene_gn=RCL1;refGene_tx=XM_019815716.2;refGene_gn=RCL1;refGene_tx=XM_019815717.2;refGene_gn=RCL1;refGene_tx=XM_023243281.1
D4 6735165 6735182 refGene_gn=RCL1;refGene_tx=XM_019815717.2
D4 6746738 6746863
refGene_gn=RCL1;refGene_tx=XM_003995328.5;refGene_gn=RCL1;refGene_tx=XM_019815716.2;refGene_gn=RCL1;refGene_tx=XM_023243281.1
D4 6747758 6747833
refGene_gn=RCL1;refGene_tx=XM_003995328.5;refGene_gn=RCL1;refGene_tx=XM_019815716.2;refGene_gn=RCL1;refGene_tx=XM_023243281.1
D4 6752981 6753157
refGene_gn=RCL1;refGene_tx=XM_003995328.5;refGene_gn=RCL1;refGene_tx=XM_019815716.2;refGene_gn=RCL1;refGene_tx=XM_023243281.1
D4 6755729 6755801
refGene_gn=RCL1;refGene_tx=XM_003995328.5;refGene_gn=RCL1;refGene_tx=XM_019815716.2;refGene_gn=RCL1;refGene_tx=XM_023243281.1
D4 6771668 6771804
refGene_gn=RCL1;refGene_tx=XM_003995328.5;refGene_gn=RCL1;refGene_tx=XM_019815716.2;refGene_gn=RCL1;refGene_tx=XM_023243281.1
D4 6813366 6813512 refGene_gn=AK3;refGene_tx=XM_023243284.1
D4 6819958 6820083
refGene_gn=AK3;refGene_tx=XM_019815521.2;refGene_gn=AK3;refGene_tx=XM_023243282.1;refGene_gn=AK3;refGene_tx=XM_023243284.1
D4 6821092 6821265
refGene_gn=AK3;refGene_tx=XM_019815521.2;refGene_gn=AK3;refGene_tx=XM_023243282.1;refGene_gn=AK3;refGene_tx=XM_023243284.1
D4 6821810 6821929
refGene_gn=AK3;refGene_tx=XM_019815521.2;refGene_gn=AK3;refGene_tx=XM_023243282.1;refGene_gn=AK3;refGene_tx=XM_023243284.1
D4 6826388 6826509
refGene_gn=AK3;refGene_tx=XM_019815521.2;refGene_gn=AK3;refGene_tx=XM_023243282.1;refGene_gn=AK3;refGene_tx=XM_023243284.1
D4 6832948 6833050 refGene_gn=CDC37L1;refGene_tx=XM_023243285.1
D4 6837052 6837217 refGene_gn=CDC37L1;refGene_tx=XM_023243285.1

D4 6840682 6840805 refGene_gn=CDC37L1;refGene_tx=XM_023243285.1
D4 6841157 6841273 refGene_gn=CDC37L1;refGene_tx=XM_023243285.1
D4 6847701 6847795 refGene_gn=CDC37L1;refGene_tx=XM_023243285.1
D4 6850669 6850951 refGene_gn=CDC37L1;refGene_tx=XM_023243285.1
D4 6855078 6855210 refGene_gn=CDC37L1;refGene_tx=XM_023243285.1
D4 6867803 6867842
refGene_gn=SPATA6L;refGene_tx=XM_006939045.4;refGene_gn=SPATA6L;refGene_tx=XM_006939047.3;refGene_gn=SPATA6L;refGene_tx=XM_023243286.1;refGene_gn=SPATA6L;refGene_tx=XM_023243287.1
D4 6871006 6871909
refGene_gn=PLPP6;refGene_tx=XM_023243289.1;refGene_gn=PLPP6;refGene_tx=XM_023243290.1;refGene_gn=PLPP6;refGene_tx=XM_023243291.1;refGene_gn=PLPP6;refGene_tx=XM_023243292.1
D4 6872224 6872362
refGene_gn=SPATA6L;refGene_tx=XM_006939045.4;refGene_gn=SPATA6L;refGene_tx=XM_006939047.3;refGene_gn=SPATA6L;refGene_tx=XM_023243286.1;refGene_gn=SPATA6L;refGene_tx=XM_023243287.1
D4 6875020 6875069
refGene_gn=SPATA6L;refGene_tx=XM_006939045.4;refGene_gn=SPATA6L;refGene_tx=XM_006939047.3;refGene_gn=SPATA6L;refGene_tx=XM_023243286.1;refGene_gn=SPATA6L;refGene_tx=XM_023243287.1
D4 6878444 6878486
refGene_gn=SPATA6L;refGene_tx=XM_006939045.4;refGene_gn=SPATA6L;refGene_tx=XM_006939047.3;refGene_gn=SPATA6L;refGene_tx=XM_023243286.1;refGene_gn=SPATA6L;refGene_tx=XM_023243287.1
D4 6890173 6890298
refGene_gn=SPATA6L;refGene_tx=XM_006939045.4;refGene_gn=SPATA6L;refGene_tx=XM_006939047.3;refGene_gn=SPATA6L;refGene_tx=XM_023243286.1;refGene_gn=SPATA6L;refGene_tx=XM_023243287.1;refGene_gn=SPATA6L;refGene_tx=XM_023243288.1
D4 6895733 6895811
refGene_gn=SPATA6L;refGene_tx=XM_006939045.4;refGene_gn=SPATA6L;refGene_tx=XM_023243286.1;refGene_gn=SPATA6L;refGene_tx=XM_023243287.1;refGene_gn=SPATA6L;refGene_tx=XM_023243288.1
D4 6899107 6899326
refGene_gn=SPATA6L;refGene_tx=XM_006939045.4;refGene_gn=SPATA6L;refGene_tx=XM_006939047.3;refGene_gn=SPATA6L;refGene_tx=XM_023243286.1;refGene_gn=SPATA6L;refGene_tx=XM_023243288.1
D4 6902097 6902200
refGene_gn=SPATA6L;refGene_tx=XM_006939045.4;refGene_gn=SPATA6L;refGene_tx=XM_006939047.3;refGene_gn=SPATA6L;refGene_tx=XM_023243286.1;refGene_gn=SPATA6L;refGene_tx=XM_023243287.1;refGene_gn=SPATA6L;refGene_tx=XM_023243288.1
D4 6905424 6905459
refGene_gn=SPATA6L;refGene_tx=XM_006939045.4;refGene_gn=SPATA6L;refGene_tx=XM_006939047.3;refGene_gn=SPATA6L;refGene_tx=XM_023243286.1;refGene_gn=SPATA6L;refGene_tx=XM_023243287.1;refGene_gn=SPATA6L;refGene_tx=XM_023243288.1
D4 6906211 6906408
refGene_gn=SPATA6L;refGene_tx=XM_006939045.4;refGene_gn=SPATA6L;refGen

e_tx=XM_006939047.3;refGene_gn=SPATA6L;refGene_tx=XM_023243286.1;refGene_gn=SPATA6L;refGene_tx=XM_023243287.1;refGene_gn=SPATA6L;refGene_tx=XM_023243288.1
D4 6909963 6910057
refGene_gn=SPATA6L;refGene_tx=XM_006939045.4;refGene_gn=SPATA6L;refGene_tx=XM_006939047.3;refGene_gn=SPATA6L;refGene_tx=XM_023243286.1;refGene_gn=SPATA6L;refGene_tx=XM_023243287.1;refGene_gn=SPATA6L;refGene_tx=XM_023243288.1
D4 6911631 6911721
refGene_gn=SPATA6L;refGene_tx=XM_006939047.3;refGene_gn=SPATA6L;refGene_tx=XM_023243286.1;refGene_gn=SPATA6L;refGene_tx=XM_023243287.1;refGene_gn=SPATA6L;refGene_tx=XM_023243288.1
D4 6915829 6915913 refGene_gn=SPATA6L;refGene_tx=XM_006939045.4
D4 6936789 6937036 refGene_gn=SLC1A1;refGene_tx=XM_023243293.1
D4 6939104 6939239 refGene_gn=SLC1A1;refGene_tx=XM_023243293.1
D4 6946471 6946666 refGene_gn=SLC1A1;refGene_tx=XM_023243293.1
D4 6947328 6947451 refGene_gn=SLC1A1;refGene_tx=XM_023243293.1
D4 6949348 6949456 refGene_gn=SLC1A1;refGene_tx=XM_023243293.1
D4 6951002 6951187 refGene_gn=SLC1A1;refGene_tx=XM_023243293.1
D4 6954144 6954243 refGene_gn=SLC1A1;refGene_tx=XM_023243293.1
D4 6955803 6955846 refGene_gn=SLC1A1;refGene_tx=XM_023243293.1
D4 6957303 6957418 refGene_gn=SLC1A1;refGene_tx=XM_023243293.1
D4 6959052 6959145 refGene_gn=SLC1A1;refGene_tx=XM_023243293.1
D4 6978517 6978548 refGene_gn=SLC1A1;refGene_tx=XM_023243293.1
D4 6978549 6978659 refGene_gn=SLC1A1;refGene_tx=XM_023243293.1
D4 7018441 7018532 refGene_gn=SLC1A1;refGene_tx=XM_023243293.1
D4 7205087 7205400 refGene_gn=GLIS3;refGene_tx=XM_023243295.1
D4 7218610 7218998
refGene_gn=GLIS3;refGene_tx=XM_023243294.1;refGene_gn=GLIS3;refGene_tx=XM_023243296.1
D4 7351376 7351584
refGene_gn=GLIS3;refGene_tx=XM_023243294.1;refGene_gn=GLIS3;refGene_tx=XM_023243295.1;refGene_gn=GLIS3;refGene_tx=XM_023243296.1
D4 7358618 7359744
refGene_gn=GLIS3;refGene_tx=XM_023243294.1;refGene_gn=GLIS3;refGene_tx=XM_023243295.1;refGene_gn=GLIS3;refGene_tx=XM_023243296.1
D4 7550230 7550392
refGene_gn=GLIS3;refGene_tx=XM_023243294.1;refGene_gn=GLIS3;refGene_tx=XM_023243295.1;refGene_gn=GLIS3;refGene_tx=XM_023243296.1
D4 7554016 7554142
refGene_gn=GLIS3;refGene_tx=XM_023243294.1;refGene_gn=GLIS3;refGene_tx=XM_023243295.1
D4 7554448 7554559
refGene_gn=GLIS3;refGene_tx=XM_023243294.1;refGene_gn=GLIS3;refGene_tx=XM_023243295.1;refGene_gn=GLIS3;refGene_tx=XM_023243296.1
D4 7584651 7584796
refGene_gn=GLIS3;refGene_tx=XM_023243294.1;refGene_gn=GLIS3;refGene_tx=XM_023243295.1;refGene_gn=GLIS3;refGene_tx=XM_023243296.1
D4 7603854 7604002
refGene_gn=GLIS3;refGene_tx=XM_023243294.1;refGene_gn=GLIS3;refGene_tx

=XM_023243295.1;refGene_gn=GLIS3;refGene_tx=XM_023243296.1
D4 7621715 7621891
refGene_gn=GLIS3;refGene_tx=XM_023243294.1;refGene_gn=GLIS3;refGene_tx=XM_023243295.1;refGene_gn=GLIS3;refGene_tx=XM_023243296.1
D4 7647070 7647253
refGene_gn=GLIS3;refGene_tx=XM_023243294.1;refGene_gn=GLIS3;refGene_tx=XM_023243295.1;refGene_gn=GLIS3;refGene_tx=XM_023243296.1
D4 7648107 7648244
refGene_gn=GLIS3;refGene_tx=XM_023243294.1;refGene_gn=GLIS3;refGene_tx=XM_023243295.1;refGene_gn=GLIS3;refGene_tx=XM_023243296.1
D4 7928913 7928947
refGene_gn=RFX3;refGene_tx=XM_019815720.2;refGene_gn=RFX3;refGene_tx=XM_019815721.2;refGene_gn=RFX3;refGene_tx=XM_019815727.2;refGene_gn=RFX3;refGene_tx=XM_023243297.1;refGene_gn=RFX3;refGene_tx=XM_023243299.1;refGene_gn=RFX3;refGene_tx=XM_023243300.1;refGene_gn=RFX3;refGene_tx=XM_023243301.1
D4 8006558 8006683
refGene_gn=RFX3;refGene_tx=XM_019815720.2;refGene_gn=RFX3;refGene_tx=XM_019815721.2;refGene_gn=RFX3;refGene_tx=XM_019815723.2;refGene_gn=RFX3;refGene_tx=XM_019815724.2;refGene_gn=RFX3;refGene_tx=XM_019815726.2;refGene_gn=RFX3;refGene_tx=XM_019815727.2;refGene_gn=RFX3;refGene_tx=XM_023243297.1;refGene_gn=RFX3;refGene_tx=XM_023243298.1;refGene_gn=RFX3;refGene_tx=XM_023243299.1;refGene_gn=RFX3;refGene_tx=XM_023243300.1;refGene_gn=RFX3;refGene_tx=XM_023243301.1
D4 8045233 8045331
refGene_gn=RFX3;refGene_tx=XM_019815720.2;refGene_gn=RFX3;refGene_tx=XM_019815721.2;refGene_gn=RFX3;refGene_tx=XM_019815723.2;refGene_gn=RFX3;refGene_tx=XM_019815724.2;refGene_gn=RFX3;refGene_tx=XM_019815726.2;refGene_gn=RFX3;refGene_tx=XM_019815727.2;refGene_gn=RFX3;refGene_tx=XM_023243297.1;refGene_gn=RFX3;refGene_tx=XM_023243298.1;refGene_gn=RFX3;refGene_tx=XM_023243299.1;refGene_gn=RFX3;refGene_tx=XM_023243300.1;refGene_gn=RFX3;refGene_tx=XM_023243301.1
D4 8055642 8055681
refGene_gn=RFX3;refGene_tx=XM_019815720.2;refGene_gn=RFX3;refGene_tx=XM_019815721.2;refGene_gn=RFX3;refGene_tx=XM_019815723.2;refGene_gn=RFX3;refGene_tx=XM_019815724.2;refGene_gn=RFX3;refGene_tx=XM_019815726.2;refGene_gn=RFX3;refGene_tx=XM_019815727.2;refGene_gn=RFX3;refGene_tx=XM_023243298.1;refGene_gn=RFX3;refGene_tx=XM_023243299.1;refGene_gn=RFX3;refGene_tx=XM_023243301.1
D4 8064788 8065047
refGene_gn=RFX3;refGene_tx=XM_019815720.2;refGene_gn=RFX3;refGene_tx=XM_019815721.2;refGene_gn=RFX3;refGene_tx=XM_019815723.2;refGene_gn=RFX3;refGene_tx=XM_019815724.2;refGene_gn=RFX3;refGene_tx=XM_019815726.2;refGene_gn=RFX3;refGene_tx=XM_019815727.2;refGene_gn=RFX3;refGene_tx=XM_023243297.1;refGene_gn=RFX3;refGene_tx=XM_023243298.1;refGene_gn=RFX3;refGene_tx=XM_023243299.1;refGene_gn=RFX3;refGene_tx=XM_023243300.1;refGene_gn=RFX3;refGene_tx=XM_023243301.1
D4 8083548 8083623
refGene_gn=RFX3;refGene_tx=XM_019815720.2;refGene_gn=RFX3;refGene_tx=XM_019815721.2;refGene_gn=RFX3;refGene_tx=XM_019815723.2;refGene_gn=RFX

3;refGene_tx=XM_019815724.2;refGene_gn=RFX3;refGene_tx=XM_019815726.2;
refGene_gn=RFX3;refGene_tx=XM_019815727.2;refGene_gn=RFX3;refGene_tx=X
M_023243297.1;refGene_gn=RFX3;refGene_tx=XM_023243298.1;refGene_gn=RFX
3;refGene_tx=XM_023243301.1
D4 8093048 8093230
refGene_gn=RFX3;refGene_tx=XM_019815720.2;refGene_gn=RFX3;refGene_tx=X
M_019815721.2;refGene_gn=RFX3;refGene_tx=XM_019815723.2;refGene_gn=RFX
3;refGene_tx=XM_019815724.2;refGene_gn=RFX3;refGene_tx=XM_019815726.2;
refGene_gn=RFX3;refGene_tx=XM_019815727.2;refGene_gn=RFX3;refGene_tx=X
M_023243297.1;refGene_gn=RFX3;refGene_tx=XM_023243298.1;refGene_gn=RFX
3;refGene_tx=XM_023243299.1;refGene_gn=RFX3;refGene_tx=XM_023243300.1;
refGene_gn=RFX3;refGene_tx=XM_023243301.1
D4 8096674 8096794
refGene_gn=RFX3;refGene_tx=XM_019815720.2;refGene_gn=RFX3;refGene_tx=X
M_019815721.2;refGene_gn=RFX3;refGene_tx=XM_019815723.2;refGene_gn=RFX
3;refGene_tx=XM_019815724.2;refGene_gn=RFX3;refGene_tx=XM_019815726.2;
refGene_gn=RFX3;refGene_tx=XM_019815727.2;refGene_gn=RFX3;refGene_tx=X
M_023243297.1;refGene_gn=RFX3;refGene_tx=XM_023243298.1;refGene_gn=RFX
3;refGene_tx=XM_023243299.1;refGene_gn=RFX3;refGene_tx=XM_023243300.1
D4 8109021 8109143
refGene_gn=RFX3;refGene_tx=XM_019815720.2;refGene_gn=RFX3;refGene_tx=X
M_019815721.2;refGene_gn=RFX3;refGene_tx=XM_019815723.2;refGene_gn=RFX
3;refGene_tx=XM_019815724.2;refGene_gn=RFX3;refGene_tx=XM_019815726.2;
refGene_gn=RFX3;refGene_tx=XM_019815727.2;refGene_gn=RFX3;refGene_tx=X
M_023243297.1;refGene_gn=RFX3;refGene_tx=XM_023243298.1;refGene_gn=RFX
3;refGene_tx=XM_023243299.1;refGene_gn=RFX3;refGene_tx=XM_023243300.1;
refGene_gn=RFX3;refGene_tx=XM_023243301.1
D4 8110909 8111022
refGene_gn=RFX3;refGene_tx=XM_019815720.2;refGene_gn=RFX3;refGene_tx=X
M_019815721.2;refGene_gn=RFX3;refGene_tx=XM_019815723.2;refGene_gn=RFX
3;refGene_tx=XM_019815724.2;refGene_gn=RFX3;refGene_tx=XM_019815726.2;
refGene_gn=RFX3;refGene_tx=XM_019815727.2;refGene_gn=RFX3;refGene_tx=X
M_023243297.1;refGene_gn=RFX3;refGene_tx=XM_023243298.1;refGene_gn=RFX
3;refGene_tx=XM_023243299.1;refGene_gn=RFX3;refGene_tx=XM_023243300.1;
refGene_gn=RFX3;refGene_tx=XM_023243301.1
D4 8115446 8115562
refGene_gn=RFX3;refGene_tx=XM_019815720.2;refGene_gn=RFX3;refGene_tx=X
M_019815721.2;refGene_gn=RFX3;refGene_tx=XM_019815723.2;refGene_gn=RFX
3;refGene_tx=XM_019815724.2;refGene_gn=RFX3;refGene_tx=XM_019815726.2;
refGene_gn=RFX3;refGene_tx=XM_019815727.2;refGene_gn=RFX3;refGene_tx=X
M_023243297.1;refGene_gn=RFX3;refGene_tx=XM_023243298.1;refGene_gn=RFX
3;refGene_tx=XM_023243299.1;refGene_gn=RFX3;refGene_tx=XM_023243300.1;
refGene_gn=RFX3;refGene_tx=XM_023243301.1
D4 8115999 8116154
refGene_gn=RFX3;refGene_tx=XM_019815720.2;refGene_gn=RFX3;refGene_tx=X
M_019815721.2;refGene_gn=RFX3;refGene_tx=XM_019815723.2;refGene_gn=RFX
3;refGene_tx=XM_019815724.2;refGene_gn=RFX3;refGene_tx=XM_019815726.2;
refGene_gn=RFX3;refGene_tx=XM_019815727.2;refGene_gn=RFX3;refGene_tx=X
M_023243297.1;refGene_gn=RFX3;refGene_tx=XM_023243298.1;refGene_gn=RFX
3;refGene_tx=XM_023243299.1;refGene_gn=RFX3;refGene_tx=XM_023243300.1;

refGene_gn=RFX3;refGene_tx=XM_023243301.1
D4 8120071 8120169
refGene_gn=RFX3;refGene_tx=XM_019815720.2;refGene_gn=RFX3;refGene_tx=XM_019815721.2;refGene_gn=RFX3;refGene_tx=XM_019815723.2;refGene_gn=RFX3;refGene_tx=XM_019815724.2;refGene_gn=RFX3;refGene_tx=XM_019815726.2;refGene_gn=RFX3;refGene_tx=XM_019815727.2;refGene_gn=RFX3;refGene_tx=XM_023243297.1;refGene_gn=RFX3;refGene_tx=XM_023243298.1;refGene_gn=RFX3;refGene_tx=XM_023243299.1;refGene_gn=RFX3;refGene_tx=XM_023243300.1;refGene_gn=RFX3;refGene_tx=XM_023243301.1
D4 8123239 8123389
refGene_gn=RFX3;refGene_tx=XM_019815720.2;refGene_gn=RFX3;refGene_tx=XM_019815721.2;refGene_gn=RFX3;refGene_tx=XM_019815723.2;refGene_gn=RFX3;refGene_tx=XM_019815724.2;refGene_gn=RFX3;refGene_tx=XM_019815726.2;refGene_gn=RFX3;refGene_tx=XM_019815727.2;refGene_gn=RFX3;refGene_tx=XM_023243297.1;refGene_gn=RFX3;refGene_tx=XM_023243298.1;refGene_gn=RFX3;refGene_tx=XM_023243299.1;refGene_gn=RFX3;refGene_tx=XM_023243300.1;refGene_gn=RFX3;refGene_tx=XM_023243301.1
D4 8130446 8130655
refGene_gn=RFX3;refGene_tx=XM_019815720.2;refGene_gn=RFX3;refGene_tx=XM_019815721.2;refGene_gn=RFX3;refGene_tx=XM_019815723.2;refGene_gn=RFX3;refGene_tx=XM_019815724.2;refGene_gn=RFX3;refGene_tx=XM_019815726.2;refGene_gn=RFX3;refGene_tx=XM_019815727.2;refGene_gn=RFX3;refGene_tx=XM_023243297.1;refGene_gn=RFX3;refGene_tx=XM_023243298.1;refGene_gn=RFX3;refGene_tx=XM_023243299.1;refGene_gn=RFX3;refGene_tx=XM_023243300.1;refGene_gn=RFX3;refGene_tx=XM_023243301.1
D4 8142050 8142204
refGene_gn=RFX3;refGene_tx=XM_019815720.2;refGene_gn=RFX3;refGene_tx=XM_019815721.2;refGene_gn=RFX3;refGene_tx=XM_019815723.2;refGene_gn=RFX3;refGene_tx=XM_019815724.2;refGene_gn=RFX3;refGene_tx=XM_019815726.2;refGene_gn=RFX3;refGene_tx=XM_019815727.2;refGene_gn=RFX3;refGene_tx=XM_023243297.1;refGene_gn=RFX3;refGene_tx=XM_023243298.1;refGene_gn=RFX3;refGene_tx=XM_023243299.1;refGene_gn=RFX3;refGene_tx=XM_023243300.1;refGene_gn=RFX3;refGene_tx=XM_023243301.1
D4 8160626 8160629 refGene_gn=RFX3;refGene_tx=XM_019815727.2
D4 8161599 8161642
refGene_gn=RFX3;refGene_tx=XM_019815720.2;refGene_gn=RFX3;refGene_tx=XM_019815721.2;refGene_gn=RFX3;refGene_tx=XM_019815723.2;refGene_gn=RFX3;refGene_tx=XM_019815724.2;refGene_gn=RFX3;refGene_tx=XM_019815726.2;refGene_gn=RFX3;refGene_tx=XM_023243297.1;refGene_gn=RFX3;refGene_tx=XM_023243298.1;refGene_gn=RFX3;refGene_tx=XM_023243299.1;refGene_gn=RFX3;refGene_tx=XM_023243300.1;refGene_gn=RFX3;refGene_tx=XM_023243301.1
D4 8164720 8164959
refGene_gn=RFX3;refGene_tx=XM_019815720.2;refGene_gn=RFX3;refGene_tx=XM_019815721.2;refGene_gn=RFX3;refGene_tx=XM_019815723.2;refGene_gn=RFX3;refGene_tx=XM_019815724.2;refGene_gn=RFX3;refGene_tx=XM_019815726.2;refGene_gn=RFX3;refGene_tx=XM_023243297.1;refGene_gn=RFX3;refGene_tx=XM_023243298.1;refGene_gn=RFX3;refGene_tx=XM_023243299.1;refGene_gn=RFX3;refGene_tx=XM_023243300.1;refGene_gn=RFX3;refGene_tx=XM_023243301.1
D4 8199624 8199901
refGene_gn=LOC101100603;refGene_tx=XM_023243261.1

D4 8202948 8203710
refGene_gn=LOC101100603;refGene_tx=XM_023243261.1
D4 8299400 8299526
refGene_gn=LOC101100603;refGene_tx=XM_023243261.1
D4 8348419 8348526
refGene_gn=LOC101100603;refGene_tx=XM_023243261.1
D4 8349043 8349148
refGene_gn=LOC101100603;refGene_tx=XM_023243261.1
D4 8369925 8370036
refGene_gn=LOC101100603;refGene_tx=XM_023243261.1
D4 8408486 8408664
refGene_gn=LOC101100603;refGene_tx=XM_023243261.1
D4 8413832 8413923
refGene_gn=LOC101100603;refGene_tx=XM_023243261.1
D4 8428895 8428977
refGene_gn=LOC101100603;refGene_tx=XM_023243261.1
D4 8430154 8430240
refGene_gn=LOC101100603;refGene_tx=XM_023243261.1
D4 8433762 8433852
refGene_gn=LOC101100603;refGene_tx=XM_023243261.1
D4 8436053 8436191
refGene_gn=LOC101100603;refGene_tx=XM_023243261.1
D4 8445167 8445257
refGene_gn=LOC101100603;refGene_tx=XM_023243261.1
D4 8451009 8451019
refGene_gn=LOC101100603;refGene_tx=XM_023243261.1
D4 8506901 8506983 refGene_gn=PUM3;refGene_tx=XM_003995332.5
D4 8508288 8508507 refGene_gn=PUM3;refGene_tx=XM_003995332.5
D4 8511503 8511639 refGene_gn=PUM3;refGene_tx=XM_003995332.5
D4 8512243 8512319 refGene_gn=PUM3;refGene_tx=XM_003995332.5
D4 8513557 8513651 refGene_gn=PUM3;refGene_tx=XM_003995332.5
D4 8513916 8513983 refGene_gn=PUM3;refGene_tx=XM_003995332.5
D4 8515211 8515371 refGene_gn=PUM3;refGene_tx=XM_003995332.5
D4 8515372 8515387 refGene_gn=PUM3;refGene_tx=XM_003995332.5
D4 8516335 8516439 refGene_gn=PUM3;refGene_tx=XM_003995332.5
D4 8517445 8517524 refGene_gn=PUM3;refGene_tx=XM_003995332.5
D4 8519441 8519540 refGene_gn=PUM3;refGene_tx=XM_003995332.5
D4 8520421 8520475 refGene_gn=PUM3;refGene_tx=XM_003995332.5
D4 8524167 8524248 refGene_gn=PUM3;refGene_tx=XM_003995332.5
D4 8529424 8529567 refGene_gn=PUM3;refGene_tx=XM_003995332.5
D4 8537281 8537504 refGene_gn=PUM3;refGene_tx=XM_003995332.5
D4 8539028 8539116 refGene_gn=PUM3;refGene_tx=XM_003995332.5
D4 8541942 8542033 refGene_gn=PUM3;refGene_tx=XM_003995332.5
D4 8545374 8545507 refGene_gn=PUM3;refGene_tx=XM_003995332.5
D4 8592223 8592505 refGene_gn=KCNV2;refGene_tx=XM_023243303.1
D4 8603003 8604404 refGene_gn=KCNV2;refGene_tx=XM_023243303.1
D4 8679509 8679545
refGene_gn=VLDLR;refGene_tx=XM_023243304.1;refGene_gn=VLDLR;refGene_tx=XM_023243305.1;refGene_gn=VLDLR;refGene_tx=XM_023243306.1;refGene_gn=VLDLR;refGene_tx=XM_023243307.1

D4 8680425 8680595
refGene_gn=VLDLR;refGene_tx=XM_023243304.1;refGene_gn=VLDLR;refGene_tx=XM_023243305.1;refGene_gn=VLDLR;refGene_tx=XM_023243306.1;refGene_gn=VLDLR;refGene_tx=XM_023243307.1
D4 8681448 8681529
refGene_gn=VLDLR;refGene_tx=XM_023243304.1;refGene_gn=VLDLR;refGene_tx=XM_023243305.1;refGene_gn=VLDLR;refGene_tx=XM_023243306.1;refGene_gn=VLDLR;refGene_tx=XM_023243307.1
D4 8681905 8681989
refGene_gn=VLDLR;refGene_tx=XM_023243304.1;refGene_gn=VLDLR;refGene_tx=XM_023243306.1
D4 8682845 8682992
refGene_gn=VLDLR;refGene_tx=XM_023243304.1;refGene_gn=VLDLR;refGene_tx=XM_023243305.1;refGene_gn=VLDLR;refGene_tx=XM_023243306.1;refGene_gn=VLDLR;refGene_tx=XM_023243307.1
D4 8683821 8683963
refGene_gn=VLDLR;refGene_tx=XM_023243304.1;refGene_gn=VLDLR;refGene_tx=XM_023243305.1;refGene_gn=VLDLR;refGene_tx=XM_023243306.1;refGene_gn=VLDLR;refGene_tx=XM_023243307.1
D4 8684264 8684404
refGene_gn=VLDLR;refGene_tx=XM_023243304.1;refGene_gn=VLDLR;refGene_tx=XM_023243305.1;refGene_gn=VLDLR;refGene_tx=XM_023243306.1;refGene_gn=VLDLR;refGene_tx=XM_023243307.1
D4 8684937 8685056
refGene_gn=VLDLR;refGene_tx=XM_023243304.1;refGene_gn=VLDLR;refGene_tx=XM_023243305.1;refGene_gn=VLDLR;refGene_tx=XM_023243306.1;refGene_gn=VLDLR;refGene_tx=XM_023243307.1
D4 8685987 8686206
refGene_gn=VLDLR;refGene_tx=XM_023243304.1;refGene_gn=VLDLR;refGene_tx=XM_023243305.1;refGene_gn=VLDLR;refGene_tx=XM_023243306.1;refGene_gn=VLDLR;refGene_tx=XM_023243307.1
D4 8686809 8686981
refGene_gn=VLDLR;refGene_tx=XM_023243304.1;refGene_gn=VLDLR;refGene_tx=XM_023243305.1;refGene_gn=VLDLR;refGene_tx=XM_023243306.1;refGene_gn=VLDLR;refGene_tx=XM_023243307.1
D4 8687972 8688098
refGene_gn=VLDLR;refGene_tx=XM_023243304.1;refGene_gn=VLDLR;refGene_tx=XM_023243305.1;refGene_gn=VLDLR;refGene_tx=XM_023243306.1;refGene_gn=VLDLR;refGene_tx=XM_023243307.1
D4 8688225 8688345
refGene_gn=VLDLR;refGene_tx=XM_023243304.1;refGene_gn=VLDLR;refGene_tx=XM_023243305.1;refGene_gn=VLDLR;refGene_tx=XM_023243306.1;refGene_gn=VLDLR;refGene_tx=XM_023243307.1
D4 8688804 8688927
refGene_gn=VLDLR;refGene_tx=XM_023243304.1;refGene_gn=VLDLR;refGene_tx=XM_023243305.1;refGene_gn=VLDLR;refGene_tx=XM_023243306.1;refGene_gn=VLDLR;refGene_tx=XM_023243307.1
D4 8689014 8689137
refGene_gn=VLDLR;refGene_tx=XM_023243304.1;refGene_gn=VLDLR;refGene_tx=XM_023243305.1;refGene_gn=VLDLR;refGene_tx=XM_023243306.1;refGene_gn=

VLDLR;refGene_tx=XM_023243307.1
D4	8689250	8689622
refGene_gn=VLDLR;refGene_tx=XM_023243304.1;refGene_gn=VLDLR;refGene_tx=XM_023243305.1;refGene_gn=VLDLR;refGene_tx=XM_023243306.1;refGene_gn=VLDLR;refGene_tx=XM_023243307.1
D4	8691298	8691421
refGene_gn=VLDLR;refGene_tx=XM_023243304.1;refGene_gn=VLDLR;refGene_tx=XM_023243305.1
D4	8692791	8692914
refGene_gn=VLDLR;refGene_tx=XM_023243304.1;refGene_gn=VLDLR;refGene_tx=XM_023243305.1;refGene_gn=VLDLR;refGene_tx=XM_023243306.1;refGene_gn=VLDLR;refGene_tx=XM_023243307.1
D4	8697149	8697269
refGene_gn=VLDLR;refGene_tx=XM_023243304.1;refGene_gn=VLDLR;refGene_tx=XM_023243305.1;refGene_gn=VLDLR;refGene_tx=XM_023243306.1;refGene_gn=VLDLR;refGene_tx=XM_023243307.1
D4	8709012	8709146
refGene_gn=VLDLR;refGene_tx=XM_023243304.1;refGene_gn=VLDLR;refGene_tx=XM_023243305.1;refGene_gn=VLDLR;refGene_tx=XM_023243306.1;refGene_gn=VLDLR;refGene_tx=XM_023243307.1
D4	8709311	8710027
refGene_gn=VLDLR;refGene_tx=XM_023243304.1;refGene_gn=VLDLR;refGene_tx=XM_023243305.1;refGene_gn=VLDLR;refGene_tx=XM_023243306.1;refGene_gn=VLDLR;refGene_tx=XM_023243307.1
D4	9064459	9064495	refGene_gn=SMARCA2;refGene_tx=XM_023243309.1
D4	9065593	9065736	refGene_gn=SMARCA2;refGene_tx=XM_023243309.1
D4	9081851	9081984	refGene_gn=SMARCA2;refGene_tx=XM_023243309.1
D4	9084916	9085018	refGene_gn=SMARCA2;refGene_tx=XM_023243309.1
D4	9085496	9085602	refGene_gn=SMARCA2;refGene_tx=XM_023243309.1
D4	9095014	9095068	refGene_gn=SMARCA2;refGene_tx=XM_023243309.1
D4	9103772	9103990	refGene_gn=SMARCA2;refGene_tx=XM_023243309.1
D4	9135158	9135377	refGene_gn=SMARCA2;refGene_tx=XM_023243309.1
D4	9140838	9140916	refGene_gn=SMARCA2;refGene_tx=XM_023243309.1
D4	9144443	9144671	refGene_gn=SMARCA2;refGene_tx=XM_023243309.1
D4	9152572	9152736	refGene_gn=SMARCA2;refGene_tx=XM_023243309.1
D4	9158515	9158682	refGene_gn=SMARCA2;refGene_tx=XM_023243309.1
D4	9161004	9161051	refGene_gn=SMARCA2;refGene_tx=XM_023243309.1
D4	9164773	9164860	refGene_gn=SMARCA2;refGene_tx=XM_023243309.1
D4	9165452	9165560	refGene_gn=SMARCA2;refGene_tx=XM_023243309.1
D4	9175451	9175565	refGene_gn=SMARCA2;refGene_tx=XM_023243309.1
D4	9176964	9177207	refGene_gn=SMARCA2;refGene_tx=XM_023243309.1
D4	9179865	9179976	refGene_gn=SMARCA2;refGene_tx=XM_023243309.1
D4	9180646	9180713	refGene_gn=SMARCA2;refGene_tx=XM_023243309.1
D4	9182040	9182204	refGene_gn=SMARCA2;refGene_tx=XM_023243309.1
D4	9185977	9186125	refGene_gn=SMARCA2;refGene_tx=XM_023243309.1
D4	9188011	9188112	refGene_gn=SMARCA2;refGene_tx=XM_023243309.1
D4	9190248	9190306	refGene_gn=SMARCA2;refGene_tx=XM_023243309.1
D4	9190775	9190906	refGene_gn=SMARCA2;refGene_tx=XM_023243309.1
D4	9194086	9194140	refGene_gn=SMARCA2;refGene_tx=XM_023243309.1
D4	9202862	9203033	refGene_gn=SMARCA2;refGene_tx=XM_023243309.1

D4 9205376 9205550 refGene_gn=SMARCA2;refGene_tx=XM_023243309.1
D4 9207051 9207078 refGene_gn=SMARCA2;refGene_tx=XM_023243309.1
D4 9207080 9207227 refGene_gn=SMARCA2;refGene_tx=XM_023243309.1
D4 9209210 9209337 refGene_gn=SMARCA2;refGene_tx=XM_023243309.1
D4 9216162 9216418 refGene_gn=SMARCA2;refGene_tx=XM_023243309.1
D4 9223620 9224007 refGene_gn=SMARCA2;refGene_tx=XM_023243309.1
D4 9230410 9230540 refGene_gn=SMARCA2;refGene_tx=XM_023243309.1
D4 9234060 9234285 refGene_gn=SMARCA2;refGene_tx=XM_023243309.1
D4 10083944 10085002
refGene_gn=DMRT2;refGene_tx=XM_023243310.1;refGene_gn=DMRT2;refGene_tx
=XM_023243311.1
D4 10085402 10085481 refGene_gn=DMRT2;refGene_tx=XM_023243311.1
D4 10087068 10087171
refGene_gn=DMRT2;refGene_tx=XM_023243310.1;refGene_gn=DMRT2;refGene_tx
=XM_023243311.1
D4 10088696 10089218
refGene_gn=DMRT2;refGene_tx=XM_023243310.1;refGene_gn=DMRT2;refGene_tx
=XM_023243311.1
D4 10151633 10152601
refGene_gn=DMRT3;refGene_tx=XM_023243312.1;refGene_gn=DMRT3;refGene_tx
=XM_023243313.1
D4 10165978 10166444
refGene_gn=DMRT3;refGene_tx=XM_023243312.1;refGene_gn=DMRT3;refGene_tx
=XM_023243313.1
D4 10175100 10175255
refGene_gn=DMRT1;refGene_tx=XM_023243314.1;refGene_gn=DMRT1;refGene_tx
=XM_023243315.1
D4 10219124 10219272
refGene_gn=DMRT1;refGene_tx=XM_023243314.1;refGene_gn=DMRT1;refGene_tx
=XM_023243315.1
D4 10240309 10240593
refGene_gn=DMRT1;refGene_tx=XM_023243314.1;refGene_gn=DMRT1;refGene_tx
=XM_023243315.1
D4 10281986 10282170
refGene_gn=DMRT1;refGene_tx=XM_023243314.1;refGene_gn=DMRT1;refGene_tx
=XM_023243315.1
D4 10285839 10286208
refGene_gn=DMRT1;refGene_tx=XM_023243314.1;refGene_gn=DMRT1;refGene_tx
=XM_023243315.1
D4 10348244 10348307
refGene_gn=KANK1;refGene_tx=XM_006939071.3;refGene_gn=KANK1;refGene_tx
=XM_023243316.1;refGene_gn=KANK1;refGene_tx=XM_023243318.1;refGene_gn=
KANK1;refGene_tx=XM_023243319.1
D4 10348894 10348993
refGene_gn=KANK1;refGene_tx=XM_006939071.3;refGene_gn=KANK1;refGene_tx
=XM_023243316.1;refGene_gn=KANK1;refGene_tx=XM_023243318.1;refGene_gn=
KANK1;refGene_tx=XM_023243319.1
D4 10351253 10351454
refGene_gn=KANK1;refGene_tx=XM_006939071.3;refGene_gn=KANK1;refGene_tx
=XM_023243316.1;refGene_gn=KANK1;refGene_tx=XM_023243318.1;refGene_gn=

KANK1;refGene_tx=XM_023243319.1
D4 10352235 10352378
refGene_gn=KANK1;refGene_tx=XM_006939071.3;refGene_gn=KANK1;refGene_tx=XM_023243316.1;refGene_gn=KANK1;refGene_tx=XM_023243318.1;refGene_gn=KANK1;refGene_tx=XM_023243319.1
D4 10355235 10355455
refGene_gn=KANK1;refGene_tx=XM_006939071.3;refGene_gn=KANK1;refGene_tx=XM_023243316.1;refGene_gn=KANK1;refGene_tx=XM_023243318.1;refGene_gn=KANK1;refGene_tx=XM_023243319.1
D4 10358657 10358745
refGene_gn=KANK1;refGene_tx=XM_006939071.3;refGene_gn=KANK1;refGene_tx=XM_023243316.1;refGene_gn=KANK1;refGene_tx=XM_023243318.1;refGene_gn=KANK1;refGene_tx=XM_023243319.1
D4 10360130 10360373
refGene_gn=KANK1;refGene_tx=XM_006939071.3;refGene_gn=KANK1;refGene_tx=XM_023243316.1;refGene_gn=KANK1;refGene_tx=XM_023243318.1
D4 10361553 10361662
refGene_gn=KANK1;refGene_tx=XM_006939071.3;refGene_gn=KANK1;refGene_tx=XM_023243316.1;refGene_gn=KANK1;refGene_tx=XM_023243318.1;refGene_gn=KANK1;refGene_tx=XM_023243319.1
D4 10362314 10362512
refGene_gn=KANK1;refGene_tx=XM_006939071.3;refGene_gn=KANK1;refGene_tx=XM_023243316.1;refGene_gn=KANK1;refGene_tx=XM_023243318.1;refGene_gn=KANK1;refGene_tx=XM_023243319.1
D4 10376492 10379159
refGene_gn=KANK1;refGene_tx=XM_006939071.3;refGene_gn=KANK1;refGene_tx=XM_023243316.1;refGene_gn=KANK1;refGene_tx=XM_023243318.1;refGene_gn=KANK1;refGene_tx=XM_023243319.1
D4 10413983 10414020
refGene_gn=KANK1;refGene_tx=XM_023243316.1;refGene_gn=KANK1;refGene_tx=XM_023243318.1;refGene_gn=KANK1;refGene_tx=XM_023243319.1
D4 10584748 10584809
refGene_gn=DOCK8;refGene_tx=XM_006939072.4;refGene_gn=DOCK8;refGene_tx=XM_006939075.4;refGene_gn=DOCK8;refGene_tx=XM_023243320.1;refGene_gn=DOCK8;refGene_tx=XM_023243321.1
D4 10585122 10585293
refGene_gn=DOCK8;refGene_tx=XM_006939072.4;refGene_gn=DOCK8;refGene_tx=XM_006939075.4;refGene_gn=DOCK8;refGene_tx=XM_023243320.1;refGene_gn=DOCK8;refGene_tx=XM_023243321.1
D4 10597603 10597710
refGene_gn=DOCK8;refGene_tx=XM_006939072.4;refGene_gn=DOCK8;refGene_tx=XM_006939075.4;refGene_gn=DOCK8;refGene_tx=XM_023243320.1;refGene_gn=DOCK8;refGene_tx=XM_023243321.1
D4 10598982 10599126
refGene_gn=DOCK8;refGene_tx=XM_006939072.4;refGene_gn=DOCK8;refGene_tx=XM_006939075.4;refGene_gn=DOCK8;refGene_tx=XM_023243320.1;refGene_gn=DOCK8;refGene_tx=XM_023243321.1
D4 10601515 10601752
refGene_gn=DOCK8;refGene_tx=XM_006939072.4;refGene_gn=DOCK8;refGene_tx=XM_006939075.4;refGene_gn=DOCK8;refGene_tx=XM_023243320.1;refGene_gn=

DOCK8;refGene_tx=XM_023243321.1
D4 10606193 10606283
refGene_gn=DOCK8;refGene_tx=XM_006939072.4;refGene_gn=DOCK8;refGene_tx=XM_006939075.4;refGene_gn=DOCK8;refGene_tx=XM_023243320.1;refGene_gn=DOCK8;refGene_tx=XM_023243321.1
D4 10607933 10608068
refGene_gn=DOCK8;refGene_tx=XM_006939072.4;refGene_gn=DOCK8;refGene_tx=XM_006939075.4;refGene_gn=DOCK8;refGene_tx=XM_023243320.1;refGene_gn=DOCK8;refGene_tx=XM_023243321.1
D4 10608528 10608660
refGene_gn=DOCK8;refGene_tx=XM_006939072.4;refGene_gn=DOCK8;refGene_tx=XM_006939075.4;refGene_gn=DOCK8;refGene_tx=XM_023243320.1;refGene_gn=DOCK8;refGene_tx=XM_023243321.1
D4 10609902 10610046
refGene_gn=DOCK8;refGene_tx=XM_006939072.4;refGene_gn=DOCK8;refGene_tx=XM_006939075.4;refGene_gn=DOCK8;refGene_tx=XM_023243320.1;refGene_gn=DOCK8;refGene_tx=XM_023243321.1
D4 10614079 10614265
refGene_gn=DOCK8;refGene_tx=XM_023243323.1;refGene_gn=DOCK8;refGene_tx=XM_023243324.1
D4 10615254 10615447
refGene_gn=DOCK8;refGene_tx=XM_006939072.4;refGene_gn=DOCK8;refGene_tx=XM_006939075.4;refGene_gn=DOCK8;refGene_tx=XM_023243320.1;refGene_gn=DOCK8;refGene_tx=XM_023243321.1;refGene_gn=DOCK8;refGene_tx=XM_023243323.1;refGene_gn=DOCK8;refGene_tx=XM_023243324.1
D4 10616221 10616322
refGene_gn=DOCK8;refGene_tx=XM_006939072.4;refGene_gn=DOCK8;refGene_tx=XM_006939075.4;refGene_gn=DOCK8;refGene_tx=XM_023243320.1;refGene_gn=DOCK8;refGene_tx=XM_023243321.1;refGene_gn=DOCK8;refGene_tx=XM_023243323.1;refGene_gn=DOCK8;refGene_tx=XM_023243324.1
D4 10616959 10617118
refGene_gn=DOCK8;refGene_tx=XM_006939072.4;refGene_gn=DOCK8;refGene_tx=XM_006939075.4;refGene_gn=DOCK8;refGene_tx=XM_023243320.1;refGene_gn=DOCK8;refGene_tx=XM_023243321.1;refGene_gn=DOCK8;refGene_tx=XM_023243323.1;refGene_gn=DOCK8;refGene_tx=XM_023243324.1
D4 10618193 10618346
refGene_gn=DOCK8;refGene_tx=XM_006939072.4;refGene_gn=DOCK8;refGene_tx=XM_006939075.4;refGene_gn=DOCK8;refGene_tx=XM_023243320.1;refGene_gn=DOCK8;refGene_tx=XM_023243321.1;refGene_gn=DOCK8;refGene_tx=XM_023243323.1;refGene_gn=DOCK8;refGene_tx=XM_023243324.1
D4 10619336 10619471
refGene_gn=DOCK8;refGene_tx=XM_006939072.4;refGene_gn=DOCK8;refGene_tx=XM_006939075.4;refGene_gn=DOCK8;refGene_tx=XM_023243320.1;refGene_gn=DOCK8;refGene_tx=XM_023243321.1;refGene_gn=DOCK8;refGene_tx=XM_023243323.1;refGene_gn=DOCK8;refGene_tx=XM_023243324.1
D4 10621099 10621196
refGene_gn=DOCK8;refGene_tx=XM_006939072.4;refGene_gn=DOCK8;refGene_tx=XM_006939075.4;refGene_gn=DOCK8;refGene_tx=XM_023243320.1;refGene_gn=DOCK8;refGene_tx=XM_023243321.1;refGene_gn=DOCK8;refGene_tx=XM_023243323.1;refGene_gn=DOCK8;refGene_tx=XM_023243324.1

D4 10625437 10625525
refGene_gn=DOCK8;refGene_tx=XM_006939072.4;refGene_gn=DOCK8;refGene_tx=XM_006939075.4;refGene_gn=DOCK8;refGene_tx=XM_023243320.1;refGene_gn=DOCK8;refGene_tx=XM_023243321.1;refGene_gn=DOCK8;refGene_tx=XM_023243323.1;refGene_gn=DOCK8;refGene_tx=XM_023243324.1
D4 10626489 10626622
refGene_gn=DOCK8;refGene_tx=XM_006939072.4;refGene_gn=DOCK8;refGene_tx=XM_006939075.4;refGene_gn=DOCK8;refGene_tx=XM_023243320.1;refGene_gn=DOCK8;refGene_tx=XM_023243321.1;refGene_gn=DOCK8;refGene_tx=XM_023243323.1;refGene_gn=DOCK8;refGene_tx=XM_023243324.1
D4 10626990 10627173
refGene_gn=DOCK8;refGene_tx=XM_006939072.4;refGene_gn=DOCK8;refGene_tx=XM_006939075.4;refGene_gn=DOCK8;refGene_tx=XM_023243320.1;refGene_gn=DOCK8;refGene_tx=XM_023243321.1;refGene_gn=DOCK8;refGene_tx=XM_023243323.1;refGene_gn=DOCK8;refGene_tx=XM_023243324.1
D4 10629066 10629212
refGene_gn=DOCK8;refGene_tx=XM_006939072.4;refGene_gn=DOCK8;refGene_tx=XM_006939075.4;refGene_gn=DOCK8;refGene_tx=XM_023243320.1;refGene_gn=DOCK8;refGene_tx=XM_023243321.1;refGene_gn=DOCK8;refGene_tx=XM_023243323.1;refGene_gn=DOCK8;refGene_tx=XM_023243324.1
D4 10633380 10633550
refGene_gn=DOCK8;refGene_tx=XM_006939072.4;refGene_gn=DOCK8;refGene_tx=XM_006939075.4;refGene_gn=DOCK8;refGene_tx=XM_023243320.1;refGene_gn=DOCK8;refGene_tx=XM_023243321.1;refGene_gn=DOCK8;refGene_tx=XM_023243323.1;refGene_gn=DOCK8;refGene_tx=XM_023243324.1
D4 10638934 10639074
refGene_gn=DOCK8;refGene_tx=XM_006939072.4;refGene_gn=DOCK8;refGene_tx=XM_006939075.4;refGene_gn=DOCK8;refGene_tx=XM_023243320.1;refGene_gn=DOCK8;refGene_tx=XM_023243321.1;refGene_gn=DOCK8;refGene_tx=XM_023243323.1;refGene_gn=DOCK8;refGene_tx=XM_023243324.1
D4 10641191 10641347
refGene_gn=DOCK8;refGene_tx=XM_006939072.4;refGene_gn=DOCK8;refGene_tx=XM_006939075.4;refGene_gn=DOCK8;refGene_tx=XM_023243320.1;refGene_gn=DOCK8;refGene_tx=XM_023243321.1;refGene_gn=DOCK8;refGene_tx=XM_023243323.1;refGene_gn=DOCK8;refGene_tx=XM_023243324.1
D4 10645766 10645880
refGene_gn=DOCK8;refGene_tx=XM_006939072.4;refGene_gn=DOCK8;refGene_tx=XM_006939075.4;refGene_gn=DOCK8;refGene_tx=XM_023243320.1;refGene_gn=DOCK8;refGene_tx=XM_023243321.1;refGene_gn=DOCK8;refGene_tx=XM_023243323.1;refGene_gn=DOCK8;refGene_tx=XM_023243324.1
D4 10647891 10648041
refGene_gn=DOCK8;refGene_tx=XM_006939072.4;refGene_gn=DOCK8;refGene_tx=XM_006939075.4;refGene_gn=DOCK8;refGene_tx=XM_023243320.1;refGene_gn=DOCK8;refGene_tx=XM_023243321.1;refGene_gn=DOCK8;refGene_tx=XM_023243323.1;refGene_gn=DOCK8;refGene_tx=XM_023243324.1
D4 10654595 10654691
refGene_gn=DOCK8;refGene_tx=XM_006939072.4;refGene_gn=DOCK8;refGene_tx=XM_006939075.4;refGene_gn=DOCK8;refGene_tx=XM_023243320.1;refGene_gn=DOCK8;refGene_tx=XM_023243321.1;refGene_gn=DOCK8;refGene_tx=XM_023243323.1;refGene_gn=DOCK8;refGene_tx=XM_023243324.1

D4 10659147 10659243
refGene_gn=DOCK8;refGene_tx=XM_006939072.4;refGene_gn=DOCK8;refGene_tx=XM_006939075.4;refGene_gn=DOCK8;refGene_tx=XM_023243320.1;refGene_gn=DOCK8;refGene_tx=XM_023243321.1;refGene_gn=DOCK8;refGene_tx=XM_023243323.1;refGene_gn=DOCK8;refGene_tx=XM_023243324.1
D4 10662089 10662259
refGene_gn=DOCK8;refGene_tx=XM_006939072.4;refGene_gn=DOCK8;refGene_tx=XM_006939075.4;refGene_gn=DOCK8;refGene_tx=XM_023243320.1;refGene_gn=DOCK8;refGene_tx=XM_023243321.1;refGene_gn=DOCK8;refGene_tx=XM_023243323.1;refGene_gn=DOCK8;refGene_tx=XM_023243324.1
D4 10663645 10663810
refGene_gn=DOCK8;refGene_tx=XM_006939072.4;refGene_gn=DOCK8;refGene_tx=XM_006939075.4;refGene_gn=DOCK8;refGene_tx=XM_023243320.1;refGene_gn=DOCK8;refGene_tx=XM_023243321.1;refGene_gn=DOCK8;refGene_tx=XM_023243323.1;refGene_gn=DOCK8;refGene_tx=XM_023243324.1
D4 10668196 10668431
refGene_gn=DOCK8;refGene_tx=XM_006939072.4;refGene_gn=DOCK8;refGene_tx=XM_006939075.4;refGene_gn=DOCK8;refGene_tx=XM_023243320.1;refGene_gn=DOCK8;refGene_tx=XM_023243321.1;refGene_gn=DOCK8;refGene_tx=XM_023243323.1;refGene_gn=DOCK8;refGene_tx=XM_023243324.1
D4 10669349 10669445
refGene_gn=DOCK8;refGene_tx=XM_006939072.4;refGene_gn=DOCK8;refGene_tx=XM_006939075.4;refGene_gn=DOCK8;refGene_tx=XM_023243320.1;refGene_gn=DOCK8;refGene_tx=XM_023243321.1;refGene_gn=DOCK8;refGene_tx=XM_023243323.1;refGene_gn=DOCK8;refGene_tx=XM_023243324.1
D4 10671408 10671510
refGene_gn=DOCK8;refGene_tx=XM_006939072.4;refGene_gn=DOCK8;refGene_tx=XM_006939075.4;refGene_gn=DOCK8;refGene_tx=XM_023243320.1;refGene_gn=DOCK8;refGene_tx=XM_023243321.1;refGene_gn=DOCK8;refGene_tx=XM_023243323.1;refGene_gn=DOCK8;refGene_tx=XM_023243324.1
D4 10672141 10672280
refGene_gn=DOCK8;refGene_tx=XM_006939072.4;refGene_gn=DOCK8;refGene_tx=XM_006939075.4;refGene_gn=DOCK8;refGene_tx=XM_023243320.1;refGene_gn=DOCK8;refGene_tx=XM_023243321.1;refGene_gn=DOCK8;refGene_tx=XM_023243323.1;refGene_gn=DOCK8;refGene_tx=XM_023243324.1
D4 10674029 10674100
refGene_gn=DOCK8;refGene_tx=XM_006939072.4;refGene_gn=DOCK8;refGene_tx=XM_006939075.4;refGene_gn=DOCK8;refGene_tx=XM_023243320.1;refGene_gn=DOCK8;refGene_tx=XM_023243321.1;refGene_gn=DOCK8;refGene_tx=XM_023243323.1;refGene_gn=DOCK8;refGene_tx=XM_023243324.1
D4 10675501 10675619
refGene_gn=DOCK8;refGene_tx=XM_006939072.4;refGene_gn=DOCK8;refGene_tx=XM_006939075.4;refGene_gn=DOCK8;refGene_tx=XM_023243320.1;refGene_gn=DOCK8;refGene_tx=XM_023243321.1;refGene_gn=DOCK8;refGene_tx=XM_023243323.1;refGene_gn=DOCK8;refGene_tx=XM_023243324.1
D4 10705330 10705493
refGene_gn=DOCK8;refGene_tx=XM_006939072.4;refGene_gn=DOCK8;refGene_tx=XM_006939075.4;refGene_gn=DOCK8;refGene_tx=XM_023243320.1;refGene_gn=DOCK8;refGene_tx=XM_023243321.1;refGene_gn=DOCK8;refGene_tx=XM_023243323.1;refGene_gn=DOCK8;refGene_tx=XM_023243324.1

D4 10706239 10706333
refGene_gn=DOCK8;refGene_tx=XM_006939072.4;refGene_gn=DOCK8;refGene_tx=XM_006939075.4;refGene_gn=DOCK8;refGene_tx=XM_023243320.1;refGene_gn=DOCK8;refGene_tx=XM_023243321.1;refGene_gn=DOCK8;refGene_tx=XM_023243323.1;refGene_gn=DOCK8;refGene_tx=XM_023243324.1
D4 10709471 10709608
refGene_gn=DOCK8;refGene_tx=XM_006939072.4;refGene_gn=DOCK8;refGene_tx=XM_006939075.4;refGene_gn=DOCK8;refGene_tx=XM_023243320.1;refGene_gn=DOCK8;refGene_tx=XM_023243321.1;refGene_gn=DOCK8;refGene_tx=XM_023243323.1;refGene_gn=DOCK8;refGene_tx=XM_023243324.1
D4 10711845 10712005
refGene_gn=DOCK8;refGene_tx=XM_006939072.4;refGene_gn=DOCK8;refGene_tx=XM_006939075.4;refGene_gn=DOCK8;refGene_tx=XM_023243320.1;refGene_gn=DOCK8;refGene_tx=XM_023243321.1;refGene_gn=DOCK8;refGene_tx=XM_023243323.1;refGene_gn=DOCK8;refGene_tx=XM_023243324.1
D4 10713610 10713691
refGene_gn=DOCK8;refGene_tx=XM_006939072.4;refGene_gn=DOCK8;refGene_tx=XM_006939075.4;refGene_gn=DOCK8;refGene_tx=XM_023243320.1;refGene_gn=DOCK8;refGene_tx=XM_023243321.1;refGene_gn=DOCK8;refGene_tx=XM_023243323.1;refGene_gn=DOCK8;refGene_tx=XM_023243324.1
D4 10717752 10717902
refGene_gn=DOCK8;refGene_tx=XM_006939072.4;refGene_gn=DOCK8;refGene_tx=XM_006939075.4;refGene_gn=DOCK8;refGene_tx=XM_023243320.1;refGene_gn=DOCK8;refGene_tx=XM_023243321.1;refGene_gn=DOCK8;refGene_tx=XM_023243323.1;refGene_gn=DOCK8;refGene_tx=XM_023243324.1
D4 10718868 10718935
refGene_gn=DOCK8;refGene_tx=XM_006939072.4;refGene_gn=DOCK8;refGene_tx=XM_006939075.4;refGene_gn=DOCK8;refGene_tx=XM_023243320.1;refGene_gn=DOCK8;refGene_tx=XM_023243321.1;refGene_gn=DOCK8;refGene_tx=XM_023243323.1;refGene_gn=DOCK8;refGene_tx=XM_023243324.1
D4 10723639 10723725
refGene_gn=DOCK8;refGene_tx=XM_006939072.4;refGene_gn=DOCK8;refGene_tx=XM_006939075.4;refGene_gn=DOCK8;refGene_tx=XM_023243320.1;refGene_gn=DOCK8;refGene_tx=XM_023243321.1;refGene_gn=DOCK8;refGene_tx=XM_023243323.1;refGene_gn=DOCK8;refGene_tx=XM_023243324.1
D4 10730621 10730834
refGene_gn=DOCK8;refGene_tx=XM_006939072.4;refGene_gn=DOCK8;refGene_tx=XM_006939075.4;refGene_gn=DOCK8;refGene_tx=XM_023243320.1;refGene_gn=DOCK8;refGene_tx=XM_023243321.1;refGene_gn=DOCK8;refGene_tx=XM_023243323.1;refGene_gn=DOCK8;refGene_tx=XM_023243324.1
D4 10735818 10735942
refGene_gn=DOCK8;refGene_tx=XM_006939072.4;refGene_gn=DOCK8;refGene_tx=XM_006939075.4;refGene_gn=DOCK8;refGene_tx=XM_023243320.1;refGene_gn=DOCK8;refGene_tx=XM_023243321.1;refGene_gn=DOCK8;refGene_tx=XM_023243323.1;refGene_gn=DOCK8;refGene_tx=XM_023243324.1
D4 10750948 10751020
refGene_gn=DOCK8;refGene_tx=XM_006939072.4;refGene_gn=DOCK8;refGene_tx=XM_006939075.4;refGene_gn=DOCK8;refGene_tx=XM_023243320.1;refGene_gn=DOCK8;refGene_tx=XM_023243321.1;refGene_gn=DOCK8;refGene_tx=XM_023243323.1;refGene_gn=DOCK8;refGene_tx=XM_023243324.1

D4 10753563 10753739
refGene_gn=DOCK8;refGene_tx=XM_006939072.4;refGene_gn=DOCK8;refGene_tx
=XM_006939075.4;refGene_gn=DOCK8;refGene_tx=XM_023243320.1;refGene_gn=
DOCK8;refGene_tx=XM_023243321.1;refGene_gn=DOCK8;refGene_tx=XM_0232433
23.1;refGene_gn=DOCK8;refGene_tx=XM_023243324.1
D4 10763972 10764075
refGene_gn=DOCK8;refGene_tx=XM_023243320.1;refGene_gn=DOCK8;refGene_tx
=XM_023243321.1;refGene_gn=DOCK8;refGene_tx=XM_023243323.1;refGene_gn=
DOCK8;refGene_tx=XM_023243324.1
D4 10816393 10816446
refGene_gn=DOCK8;refGene_tx=XM_023243320.1;refGene_gn=DOCK8;refGene_tx
=XM_023243321.1;refGene_gn=DOCK8;refGene_tx=XM_023243323.1;refGene_gn=
DOCK8;refGene_tx=XM_023243324.1
D4 10846944 10847098
refGene_gn=LOC101080413;refGene_tx=XM_003995366.4
D4 10851340 10851427
refGene_gn=LOC101080413;refGene_tx=XM_003995366.4
D4 10854421 10854518
refGene_gn=LOC101080413;refGene_tx=XM_003995366.4
D4 10855863 10855955
refGene_gn=LOC101080413;refGene_tx=XM_003995366.4
D4 10870557 10870617
refGene_gn=LOC101080413;refGene_tx=XM_003995366.4
D4 10872160 10872198
refGene_gn=LOC101080413;refGene_tx=XM_003995366.4
D4 10877004 10877051
refGene_gn=LOC101080413;refGene_tx=XM_003995366.4
D4 10878793 10878880
refGene_gn=LOC101080413;refGene_tx=XM_003995366.4
D4 10881909 10881954
refGene_gn=LOC101080413;refGene_tx=XM_003995366.4
D4 10887068 10887125
refGene_gn=LOC101080413;refGene_tx=XM_003995366.4
D4 10890255 10890307
refGene_gn=LOC101080413;refGene_tx=XM_003995366.4
D4 10895264 10895333
refGene_gn=LOC101080413;refGene_tx=XM_003995366.4
D4 10895436 10895502
refGene_gn=LOC101080413;refGene_tx=XM_003995366.4
D4 10897787 10897917
refGene_gn=LOC101080413;refGene_tx=XM_003995366.4
D4 10898294 10898401
refGene_gn=LOC101080413;refGene_tx=XM_003995366.4
D4 10909249 10910605
refGene_gn=LOC101084143;refGene_tx=XM_019815527.2
D4 10952710 10952971 refGene_gn=PGM5;refGene_tx=XM_011288248.3
D4 10973482 10973645
refGene_gn=PGM5;refGene_tx=XM_011288248.3;refGene_gn=PGM5;refGene_tx=X
M_019815739.1
D4 10979933 10980080

refGene_gn=PGM5;refGene_tx=XM_011288248.3;refGene_gn=PGM5;refGene_tx=XM_019815739.1
D4 10984008 10984134
refGene_gn=PGM5;refGene_tx=XM_011288248.3;refGene_gn=PGM5;refGene_tx=XM_019815739.1
D4 10987948 10988139
refGene_gn=PGM5;refGene_tx=XM_011288248.3;refGene_gn=PGM5;refGene_tx=XM_019815739.1
D4 10988744 10988899
refGene_gn=PGM5;refGene_tx=XM_011288248.3;refGene_gn=PGM5;refGene_tx=XM_019815739.1
D4 11065558 11065674
refGene_gn=PGM5;refGene_tx=XM_011288248.3;refGene_gn=PGM5;refGene_tx=XM_019815739.1
D4 11082681 11082817
refGene_gn=PGM5;refGene_tx=XM_011288248.3;refGene_gn=PGM5;refGene_tx=XM_019815739.1
D4 11086834 11087018
refGene_gn=PGM5;refGene_tx=XM_011288248.3;refGene_gn=PGM5;refGene_tx=XM_019815739.1
D4 11101505 11101640
refGene_gn=PGM5;refGene_tx=XM_011288248.3;refGene_gn=PGM5;refGene_tx=XM_019815739.1
D4 11135770 11135860
refGene_gn=PGM5;refGene_tx=XM_011288248.3;refGene_gn=PGM5;refGene_tx=XM_019815739.1
D4 11142590 11142816 refGene_gn=TMEM252;refGene_tx=XM_003995382.3
D4 11146608 11146886 refGene_gn=TMEM252;refGene_tx=XM_003995382.3
D4 11373893 11373993 refGene_gn=FAM122A;refGene_tx=XM_003995370.5
D4 11373995 11374882 refGene_gn=FAM122A;refGene_tx=XM_003995370.5
D4 11419460 11419529 refGene_gn=PIP5K1B;refGene_tx=XM_019815741.2
D4 11433271 11433364
refGene_gn=PIP5K1B;refGene_tx=XM_023243327.1;refGene_gn=PIP5K1B;refGene_tx=XM_023243328.1
D4 11467593 11467724
refGene_gn=PIP5K1B;refGene_tx=XM_019815741.2;refGene_gn=PIP5K1B;refGene_tx=XM_023243327.1;refGene_gn=PIP5K1B;refGene_tx=XM_023243328.1
D4 11477898 11478016
refGene_gn=PIP5K1B;refGene_tx=XM_019815741.2;refGene_gn=PIP5K1B;refGene_tx=XM_023243327.1;refGene_gn=PIP5K1B;refGene_tx=XM_023243328.1
D4 11486761 11486914
refGene_gn=PIP5K1B;refGene_tx=XM_019815741.2;refGene_gn=PIP5K1B;refGene_tx=XM_023243327.1;refGene_gn=PIP5K1B;refGene_tx=XM_023243328.1
D4 11492801 11493101
refGene_gn=PIP5K1B;refGene_tx=XM_019815741.2;refGene_gn=PIP5K1B;refGene_tx=XM_023243327.1;refGene_gn=PIP5K1B;refGene_tx=XM_023243328.1
D4 11516356 11516568
refGene_gn=PIP5K1B;refGene_tx=XM_019815741.2;refGene_gn=PIP5K1B;refGene_tx=XM_023243327.1;refGene_gn=PIP5K1B;refGene_tx=XM_023243328.1
D4 11518267 11518351

refGene_gn=PIP5K1B;refGene_tx=XM_019815741.2;refGene_gn=PIP5K1B;refGene_tx=XM_023243327.1;refGene_gn=PIP5K1B;refGene_tx=XM_023243328.1
D4 11518472 11518521
refGene_gn=PIP5K1B;refGene_tx=XM_019815741.2;refGene_gn=PIP5K1B;refGene_tx=XM_023243327.1;refGene_gn=PIP5K1B;refGene_tx=XM_023243328.1
D4 11521830 11521915
refGene_gn=PIP5K1B;refGene_tx=XM_019815741.2;refGene_gn=PIP5K1B;refGene_tx=XM_023243327.1;refGene_gn=PIP5K1B;refGene_tx=XM_023243328.1
D4 11525065 11525221
refGene_gn=PIP5K1B;refGene_tx=XM_019815741.2;refGene_gn=PIP5K1B;refGene_tx=XM_023243327.1;refGene_gn=PIP5K1B;refGene_tx=XM_023243328.1
D4 11531454 11531599
refGene_gn=PIP5K1B;refGene_tx=XM_019815741.2;refGene_gn=PIP5K1B;refGene_tx=XM_023243327.1;refGene_gn=PIP5K1B;refGene_tx=XM_023243328.1
D4 11583395 11583513
refGene_gn=PIP5K1B;refGene_tx=XM_019815741.2;refGene_gn=PIP5K1B;refGene_tx=XM_023243327.1;refGene_gn=PIP5K1B;refGene_tx=XM_023243328.1
D4 11604592 11604595 refGene_gn=PIP5K1B;refGene_tx=XM_023243328.1
D4 11611629 11611665
refGene_gn=PIP5K1B;refGene_tx=XM_019815741.2;refGene_gn=PIP5K1B;refGene_tx=XM_023243327.1
D4 11663346 11663514 refGene_gn=FXN;refGene_tx=XM_003995383.5
D4 11672705 11672803 refGene_gn=FXN;refGene_tx=XM_003995383.5
D4 11674982 11675103 refGene_gn=FXN;refGene_tx=XM_003995383.5
D4 11683605 11683703 refGene_gn=FXN;refGene_tx=XM_003995383.5
D4 11687322 11687473 refGene_gn=FXN;refGene_tx=XM_003995383.5
D4 11746598 11746714
refGene_gn=TJP2;refGene_tx=XM_006939079.4;refGene_gn=TJP2;refGene_tx=XM_019815742.2
D4 11757813 11757934
refGene_gn=TJP2;refGene_tx=XM_006939079.4;refGene_gn=TJP2;refGene_tx=XM_019815742.2
D4 11777966 11778026 refGene_gn=TJP2;refGene_tx=XM_006939078.4
D4 11805695 11805767 refGene_gn=TJP2;refGene_tx=XM_006939077.4
D4 11815958 11816012
refGene_gn=TJP2;refGene_tx=XM_006939077.4;refGene_gn=TJP2;refGene_tx=XM_006939078.4;refGene_gn=TJP2;refGene_tx=XM_006939079.4;refGene_gn=TJP2;refGene_tx=XM_019815742.2;refGene_gn=TJP2;refGene_tx=XM_023243331.1
D4 11818353 11818478
refGene_gn=TJP2;refGene_tx=XM_006939077.4;refGene_gn=TJP2;refGene_tx=XM_006939078.4;refGene_gn=TJP2;refGene_tx=XM_006939079.4;refGene_gn=TJP2;refGene_tx=XM_019815742.2;refGene_gn=TJP2;refGene_tx=XM_023243331.1
D4 11819905 11820008
refGene_gn=TJP2;refGene_tx=XM_006939077.4;refGene_gn=TJP2;refGene_tx=XM_006939078.4;refGene_gn=TJP2;refGene_tx=XM_006939079.4;refGene_gn=TJP2;refGene_tx=XM_019815742.2;refGene_gn=TJP2;refGene_tx=XM_023243331.1
D4 11822217 11822797
refGene_gn=TJP2;refGene_tx=XM_006939077.4;refGene_gn=TJP2;refGene_tx=XM_006939078.4;refGene_gn=TJP2;refGene_tx=XM_006939079.4;refGene_gn=TJP2;refGene_tx=XM_019815742.2;refGene_gn=TJP2;refGene_tx=XM_023243331.1

D4 11826476 11826580
refGene_gn=TJP2;refGene_tx=XM_006939077.4;refGene_gn=TJP2;refGene_tx=XM_006939078.4;refGene_gn=TJP2;refGene_tx=XM_006939079.4;refGene_gn=TJP2;refGene_tx=XM_019815742.2;refGene_gn=TJP2;refGene_tx=XM_023243331.1
D4 11827211 11827365
refGene_gn=TJP2;refGene_tx=XM_006939077.4;refGene_gn=TJP2;refGene_tx=XM_006939078.4;refGene_gn=TJP2;refGene_tx=XM_006939079.4;refGene_gn=TJP2;refGene_tx=XM_019815742.2;refGene_gn=TJP2;refGene_tx=XM_023243331.1
D4 11828848 11828957
refGene_gn=TJP2;refGene_tx=XM_006939077.4;refGene_gn=TJP2;refGene_tx=XM_006939078.4;refGene_gn=TJP2;refGene_tx=XM_006939079.4;refGene_gn=TJP2;refGene_tx=XM_019815742.2;refGene_gn=TJP2;refGene_tx=XM_023243331.1
D4 11829061 11829195
refGene_gn=TJP2;refGene_tx=XM_006939077.4;refGene_gn=TJP2;refGene_tx=XM_006939078.4;refGene_gn=TJP2;refGene_tx=XM_006939079.4;refGene_gn=TJP2;refGene_tx=XM_019815742.2;refGene_gn=TJP2;refGene_tx=XM_023243331.1
D4 11831232 11831299
refGene_gn=TJP2;refGene_tx=XM_006939077.4;refGene_gn=TJP2;refGene_tx=XM_006939078.4;refGene_gn=TJP2;refGene_tx=XM_006939079.4;refGene_gn=TJP2;refGene_tx=XM_019815742.2;refGene_gn=TJP2;refGene_tx=XM_023243331.1
D4 11832106 11832257
refGene_gn=TJP2;refGene_tx=XM_006939077.4;refGene_gn=TJP2;refGene_tx=XM_006939078.4;refGene_gn=TJP2;refGene_tx=XM_006939079.4;refGene_gn=TJP2;refGene_tx=XM_019815742.2;refGene_gn=TJP2;refGene_tx=XM_023243331.1
D4 11838032 11838141
refGene_gn=TJP2;refGene_tx=XM_006939077.4;refGene_gn=TJP2;refGene_tx=XM_006939078.4;refGene_gn=TJP2;refGene_tx=XM_006939079.4;refGene_gn=TJP2;refGene_tx=XM_019815742.2;refGene_gn=TJP2;refGene_tx=XM_023243331.1
D4 11839391 11839602
refGene_gn=TJP2;refGene_tx=XM_006939077.4;refGene_gn=TJP2;refGene_tx=XM_006939078.4;refGene_gn=TJP2;refGene_tx=XM_006939079.4;refGene_gn=TJP2;refGene_tx=XM_019815742.2;refGene_gn=TJP2;refGene_tx=XM_023243331.1
D4 11840329 11840517
refGene_gn=TJP2;refGene_tx=XM_006939077.4;refGene_gn=TJP2;refGene_tx=XM_006939078.4;refGene_gn=TJP2;refGene_tx=XM_006939079.4;refGene_gn=TJP2;refGene_tx=XM_019815742.2;refGene_gn=TJP2;refGene_tx=XM_023243331.1
D4 11841378 11841474
refGene_gn=TJP2;refGene_tx=XM_006939077.4;refGene_gn=TJP2;refGene_tx=XM_006939078.4;refGene_gn=TJP2;refGene_tx=XM_006939079.4;refGene_gn=TJP2;refGene_tx=XM_019815742.2;refGene_gn=TJP2;refGene_tx=XM_023243331.1
D4 11842050 11842130
refGene_gn=TJP2;refGene_tx=XM_006939077.4;refGene_gn=TJP2;refGene_tx=XM_006939078.4;refGene_gn=TJP2;refGene_tx=XM_006939079.4;refGene_gn=TJP2;refGene_tx=XM_019815742.2;refGene_gn=TJP2;refGene_tx=XM_023243331.1
D4 11842537 11842748
refGene_gn=TJP2;refGene_tx=XM_006939077.4;refGene_gn=TJP2;refGene_tx=XM_006939078.4;refGene_gn=TJP2;refGene_tx=XM_006939079.4;refGene_gn=TJP2;refGene_tx=XM_019815742.2;refGene_gn=TJP2;refGene_tx=XM_023243331.1
D4 11849171 11849272
refGene_gn=TJP2;refGene_tx=XM_006939077.4;refGene_gn=TJP2;refGene_tx=X

M_006939078.4;refGene_gn=TJP2;refGene_tx=XM_006939079.4;refGene_gn=TJP2;refGene_tx=XM_019815742.2;refGene_gn=TJP2;refGene_tx=XM_023243331.1
D4 11850491 11850704
refGene_gn=TJP2;refGene_tx=XM_006939077.4;refGene_gn=TJP2;refGene_tx=XM_006939078.4;refGene_gn=TJP2;refGene_tx=XM_006939079.4;refGene_gn=TJP2;refGene_tx=XM_019815742.2;refGene_gn=TJP2;refGene_tx=XM_023243331.1
D4 11851894 11852005
refGene_gn=TJP2;refGene_tx=XM_006939077.4;refGene_gn=TJP2;refGene_tx=XM_006939078.4;refGene_gn=TJP2;refGene_tx=XM_006939079.4;refGene_gn=TJP2;refGene_tx=XM_023243331.1
D4 11853369 11853699
refGene_gn=TJP2;refGene_tx=XM_006939077.4;refGene_gn=TJP2;refGene_tx=XM_006939078.4;refGene_gn=TJP2;refGene_tx=XM_006939079.4;refGene_gn=TJP2;refGene_tx=XM_019815742.2;refGene_gn=TJP2;refGene_tx=XM_023243331.1
D4 11855654 11855740
refGene_gn=TJP2;refGene_tx=XM_006939077.4;refGene_gn=TJP2;refGene_tx=XM_006939078.4;refGene_gn=TJP2;refGene_tx=XM_006939079.4;refGene_gn=TJP2;refGene_tx=XM_019815742.2;refGene_gn=TJP2;refGene_tx=XM_023243331.1
D4 11856646 11856812
refGene_gn=TJP2;refGene_tx=XM_006939077.4;refGene_gn=TJP2;refGene_tx=XM_006939078.4;refGene_gn=TJP2;refGene_tx=XM_006939079.4;refGene_gn=TJP2;refGene_tx=XM_019815742.2;refGene_gn=TJP2;refGene_tx=XM_023243331.1
D4 11911510 11911528 refGene_gn=FAM189A2;refGene_tx=XM_006939080.4
D4 11912283 11912712 refGene_gn=FAM189A2;refGene_tx=XM_023243332.1
D4 11922606 11922663
refGene_gn=FAM189A2;refGene_tx=XM_006939080.4;refGene_gn=FAM189A2;refGene_tx=XM_023243332.1
D4 11952583 11952694
refGene_gn=FAM189A2;refGene_tx=XM_006939080.4;refGene_gn=FAM189A2;refGene_tx=XM_023243332.1
D4 11961304 11961433
refGene_gn=FAM189A2;refGene_tx=XM_006939080.4;refGene_gn=FAM189A2;refGene_tx=XM_023243332.1
D4 11963274 11963382
refGene_gn=FAM189A2;refGene_tx=XM_006939080.4;refGene_gn=FAM189A2;refGene_tx=XM_023243332.1
D4 11963470 11963604
refGene_gn=FAM189A2;refGene_tx=XM_006939080.4;refGene_gn=FAM189A2;refGene_tx=XM_023243332.1
D4 11969505 11969635
refGene_gn=FAM189A2;refGene_tx=XM_006939080.4;refGene_gn=FAM189A2;refGene_tx=XM_023243332.1
D4 11969804 11969883
refGene_gn=FAM189A2;refGene_tx=XM_006939080.4;refGene_gn=FAM189A2;refGene_tx=XM_023243332.1
D4 11971503 11971662
refGene_gn=FAM189A2;refGene_tx=XM_006939080.4;refGene_gn=FAM189A2;refGene_tx=XM_023243332.1
D4 11974118 11974411
refGene_gn=FAM189A2;refGene_tx=XM_006939080.4;refGene_gn=FAM189A2;refG

ene_tx=XM_023243332.1
D4 11977451 11977655
refGene_gn=FAM189A2;refGene_tx=XM_006939080.4;refGene_gn=FAM189A2;refGene_tx=XM_023243332.1
D4 12000067 12000139
refGene_gn=APBA1;refGene_tx=XM_023243333.1;refGene_gn=APBA1;refGene_tx=XM_023243334.1;refGene_gn=APBA1;refGene_tx=XM_023243335.1;refGene_gn=APBA1;refGene_tx=XM_023243336.1;refGene_gn=APBA1;refGene_tx=XM_023243337.1;refGene_gn=APBA1;refGene_tx=XM_023243338.1
D4 12000917 12001058
refGene_gn=APBA1;refGene_tx=XM_023243333.1;refGene_gn=APBA1;refGene_tx=XM_023243334.1;refGene_gn=APBA1;refGene_tx=XM_023243335.1;refGene_gn=APBA1;refGene_tx=XM_023243336.1;refGene_gn=APBA1;refGene_tx=XM_023243337.1;refGene_gn=APBA1;refGene_tx=XM_023243338.1
D4 12004921 12005041
refGene_gn=APBA1;refGene_tx=XM_023243333.1;refGene_gn=APBA1;refGene_tx=XM_023243334.1;refGene_gn=APBA1;refGene_tx=XM_023243335.1;refGene_gn=APBA1;refGene_tx=XM_023243336.1;refGene_gn=APBA1;refGene_tx=XM_023243337.1;refGene_gn=APBA1;refGene_tx=XM_023243338.1
D4 12012442 12012655
refGene_gn=APBA1;refGene_tx=XM_023243333.1;refGene_gn=APBA1;refGene_tx=XM_023243334.1;refGene_gn=APBA1;refGene_tx=XM_023243335.1;refGene_gn=APBA1;refGene_tx=XM_023243336.1;refGene_gn=APBA1;refGene_tx=XM_023243337.1;refGene_gn=APBA1;refGene_tx=XM_023243338.1
D4 12015379 12015559
refGene_gn=APBA1;refGene_tx=XM_023243333.1;refGene_gn=APBA1;refGene_tx=XM_023243334.1;refGene_gn=APBA1;refGene_tx=XM_023243335.1;refGene_gn=APBA1;refGene_tx=XM_023243336.1;refGene_gn=APBA1;refGene_tx=XM_023243337.1;refGene_gn=APBA1;refGene_tx=XM_023243338.1
D4 12019153 12019339
refGene_gn=APBA1;refGene_tx=XM_023243333.1;refGene_gn=APBA1;refGene_tx=XM_023243334.1;refGene_gn=APBA1;refGene_tx=XM_023243335.1;refGene_gn=APBA1;refGene_tx=XM_023243336.1;refGene_gn=APBA1;refGene_tx=XM_023243337.1;refGene_gn=APBA1;refGene_tx=XM_023243338.1
D4 12019971 12020058
refGene_gn=APBA1;refGene_tx=XM_023243333.1;refGene_gn=APBA1;refGene_tx=XM_023243334.1;refGene_gn=APBA1;refGene_tx=XM_023243335.1;refGene_gn=APBA1;refGene_tx=XM_023243336.1;refGene_gn=APBA1;refGene_tx=XM_023243337.1;refGene_gn=APBA1;refGene_tx=XM_023243338.1
D4 12021057 12021093
refGene_gn=APBA1;refGene_tx=XM_023243333.1;refGene_gn=APBA1;refGene_tx=XM_023243334.1;refGene_gn=APBA1;refGene_tx=XM_023243335.1;refGene_gn=APBA1;refGene_tx=XM_023243336.1;refGene_gn=APBA1;refGene_tx=XM_023243338.1
D4 12028457 12028603
refGene_gn=APBA1;refGene_tx=XM_023243333.1;refGene_gn=APBA1;refGene_tx=XM_023243334.1;refGene_gn=APBA1;refGene_tx=XM_023243335.1;refGene_gn=APBA1;refGene_tx=XM_023243336.1;refGene_gn=APBA1;refGene_tx=XM_023243337.1;refGene_gn=APBA1;refGene_tx=XM_023243338.1
D4 12031989 12032029

refGene_gn=APBA1;refGene_tx=XM_023243333.1;refGene_gn=APBA1;refGene_tx=XM_023243334.1;refGene_gn=APBA1;refGene_tx=XM_023243335.1;refGene_gn=APBA1;refGene_tx=XM_023243336.1;refGene_gn=APBA1;refGene_tx=XM_023243337.1;refGene_gn=APBA1;refGene_tx=XM_023243338.1
D4 12036617 12036713
refGene_gn=APBA1;refGene_tx=XM_023243333.1;refGene_gn=APBA1;refGene_tx=XM_023243334.1;refGene_gn=APBA1;refGene_tx=XM_023243335.1;refGene_gn=APBA1;refGene_tx=XM_023243336.1;refGene_gn=APBA1;refGene_tx=XM_023243337.1
D4 12066116 12067298
refGene_gn=APBA1;refGene_tx=XM_023243333.1;refGene_gn=APBA1;refGene_tx=XM_023243334.1;refGene_gn=APBA1;refGene_tx=XM_023243335.1;refGene_gn=APBA1;refGene_tx=XM_023243336.1;refGene_gn=APBA1;refGene_tx=XM_023243337.1;refGene_gn=APBA1;refGene_tx=XM_023243338.1
D4 12241780 12242123 refGene_gn=PTAR1;refGene_tx=XM_023243339.1
D4 12244056 12244361 refGene_gn=PTAR1;refGene_tx=XM_023243339.1
D4 12252621 12252835 refGene_gn=PTAR1;refGene_tx=XM_023243339.1
D4 12254870 12254975 refGene_gn=PTAR1;refGene_tx=XM_023243339.1
D4 12264399 12264466 refGene_gn=PTAR1;refGene_tx=XM_023243339.1
D4 12275052 12275222 refGene_gn=PTAR1;refGene_tx=XM_023243339.1
D4 12283950 12284036 refGene_gn=PTAR1;refGene_tx=XM_023243339.1
D4 12327753 12327794 refGene_gn=MAMDC2;refGene_tx=XM_011288225.3
D4 12335341 12335508
refGene_gn=MAMDC2;refGene_tx=XM_003995375.4;refGene_gn=MAMDC2;refGene_tx=XM_011288224.3;refGene_gn=MAMDC2;refGene_tx=XM_019815750.1
D4 12359205 12359314
refGene_gn=MAMDC2;refGene_tx=XM_003995375.4;refGene_gn=MAMDC2;refGene_tx=XM_011288224.3;refGene_gn=MAMDC2;refGene_tx=XM_011288225.3;refGene_gn=MAMDC2;refGene_tx=XM_019815750.1
D4 12372097 12372231
refGene_gn=MAMDC2;refGene_tx=XM_003995375.4;refGene_gn=MAMDC2;refGene_tx=XM_011288224.3;refGene_gn=MAMDC2;refGene_tx=XM_011288225.3;refGene_gn=MAMDC2;refGene_tx=XM_019815747.2;refGene_gn=MAMDC2;refGene_tx=XM_019815749.2;refGene_gn=MAMDC2;refGene_tx=XM_019815750.1
D4 12374736 12374805
refGene_gn=MAMDC2;refGene_tx=XM_003995375.4;refGene_gn=MAMDC2;refGene_tx=XM_011288224.3;refGene_gn=MAMDC2;refGene_tx=XM_011288225.3;refGene_gn=MAMDC2;refGene_tx=XM_019815747.2;refGene_gn=MAMDC2;refGene_tx=XM_019815749.2;refGene_gn=MAMDC2;refGene_tx=XM_019815750.1
D4 12395806 12395864
refGene_gn=MAMDC2;refGene_tx=XM_003995375.4;refGene_gn=MAMDC2;refGene_tx=XM_011288224.3;refGene_gn=MAMDC2;refGene_tx=XM_011288225.3;refGene_gn=MAMDC2;refGene_tx=XM_019815747.2;refGene_gn=MAMDC2;refGene_tx=XM_019815749.2;refGene_gn=MAMDC2;refGene_tx=XM_019815750.1
D4 12412595 12412784
refGene_gn=MAMDC2;refGene_tx=XM_003995375.4;refGene_gn=MAMDC2;refGene_tx=XM_019815750.1
D4 12536922 12537217
refGene_gn=MAMDC2;refGene_tx=XM_011288224.3;refGene_gn=MAMDC2;refGene_tx=XM_011288225.3;refGene_gn=MAMDC2;refGene_tx=XM_019815747.2;refGene_

gn=MAMDC2;refGene_tx=XM_019815749.2
D4	12537554	12537668
refGene_gn=MAMDC2;refGene_tx=XM_011288224.3;refGene_gn=MAMDC2;refGene_tx=XM_011288225.3;refGene_gn=MAMDC2;refGene_tx=XM_019815747.2;refGene_gn=MAMDC2;refGene_tx=XM_019815749.2
D4	12600259	12600531
refGene_gn=MAMDC2;refGene_tx=XM_011288224.3;refGene_gn=MAMDC2;refGene_tx=XM_011288225.3;refGene_gn=MAMDC2;refGene_tx=XM_019815747.2;refGene_gn=MAMDC2;refGene_tx=XM_019815749.2
D4	12602717	12602802
refGene_gn=MAMDC2;refGene_tx=XM_011288224.3;refGene_gn=MAMDC2;refGene_tx=XM_011288225.3;refGene_gn=MAMDC2;refGene_tx=XM_019815747.2;refGene_gn=MAMDC2;refGene_tx=XM_019815749.2
D4	12605775	12605913
refGene_gn=MAMDC2;refGene_tx=XM_011288224.3;refGene_gn=MAMDC2;refGene_tx=XM_011288225.3;refGene_gn=MAMDC2;refGene_tx=XM_019815747.2;refGene_gn=MAMDC2;refGene_tx=XM_019815749.2
D4	12619069	12619326
refGene_gn=MAMDC2;refGene_tx=XM_011288224.3;refGene_gn=MAMDC2;refGene_tx=XM_011288225.3;refGene_gn=MAMDC2;refGene_tx=XM_019815747.2;refGene_gn=MAMDC2;refGene_tx=XM_019815749.2
D4	12623092	12623186
refGene_gn=MAMDC2;refGene_tx=XM_011288224.3;refGene_gn=MAMDC2;refGene_tx=XM_011288225.3;refGene_gn=MAMDC2;refGene_tx=XM_019815747.2;refGene_gn=MAMDC2;refGene_tx=XM_019815749.2
D4	12629948	12630092
refGene_gn=MAMDC2;refGene_tx=XM_011288224.3;refGene_gn=MAMDC2;refGene_tx=XM_011288225.3;refGene_gn=MAMDC2;refGene_tx=XM_019815747.2;refGene_gn=MAMDC2;refGene_tx=XM_019815749.2
D4	12634341	12634607
refGene_gn=MAMDC2;refGene_tx=XM_011288224.3;refGene_gn=MAMDC2;refGene_tx=XM_011288225.3;refGene_gn=MAMDC2;refGene_tx=XM_019815747.2;refGene_gn=MAMDC2;refGene_tx=XM_019815749.2
D4	12656309	12656403
refGene_gn=MAMDC2;refGene_tx=XM_011288224.3;refGene_gn=MAMDC2;refGene_tx=XM_011288225.3;refGene_gn=MAMDC2;refGene_tx=XM_019815747.2;refGene_gn=MAMDC2;refGene_tx=XM_019815749.2
D4	12659283	12659436
refGene_gn=MAMDC2;refGene_tx=XM_011288224.3;refGene_gn=MAMDC2;refGene_tx=XM_011288225.3;refGene_gn=MAMDC2;refGene_tx=XM_019815747.2;refGene_gn=MAMDC2;refGene_tx=XM_019815749.2
D4	12679226	12679486
refGene_gn=MAMDC2;refGene_tx=XM_011288224.3;refGene_gn=MAMDC2;refGene_tx=XM_011288225.3;refGene_gn=MAMDC2;refGene_tx=XM_019815747.2;refGene_gn=MAMDC2;refGene_tx=XM_019815749.2
D4	12683878	12683963
refGene_gn=MAMDC2;refGene_tx=XM_011288224.3;refGene_gn=MAMDC2;refGene_tx=XM_011288225.3;refGene_gn=MAMDC2;refGene_tx=XM_019815747.2;refGene_gn=MAMDC2;refGene_tx=XM_019815749.2
D4	12684110	12684175

refGene_gn=MAMDC2;refGene_tx=XM_011288224.3;refGene_gn=MAMDC2;refGene_tx=XM_011288225.3;refGene_gn=MAMDC2;refGene_tx=XM_019815747.2;refGene_gn=MAMDC2;refGene_tx=XM_019815749.2
D4 12701483 12701677 refGene_gn=SMC5;refGene_tx=XM_003995387.5
D4 12704041 12704183 refGene_gn=SMC5;refGene_tx=XM_003995387.5
D4 12718219 12718272 refGene_gn=SMC5;refGene_tx=XM_003995387.5
D4 12725754 12725917 refGene_gn=SMC5;refGene_tx=XM_003995387.5
D4 12727382 12727517 refGene_gn=SMC5;refGene_tx=XM_003995387.5
D4 12729250 12729391 refGene_gn=SMC5;refGene_tx=XM_003995387.5
D4 12730753 12730915 refGene_gn=SMC5;refGene_tx=XM_003995387.5
D4 12731939 12732011 refGene_gn=SMC5;refGene_tx=XM_003995387.5
D4 12749530 12749786 refGene_gn=SMC5;refGene_tx=XM_003995387.5
D4 12752143 12752298 refGene_gn=SMC5;refGene_tx=XM_003995387.5
D4 12755908 12756022 refGene_gn=SMC5;refGene_tx=XM_003995387.5
D4 12758490 12758585 refGene_gn=SMC5;refGene_tx=XM_003995387.5
D4 12759213 12759346 refGene_gn=SMC5;refGene_tx=XM_003995387.5
D4 12761940 12762114 refGene_gn=SMC5;refGene_tx=XM_003995387.5
D4 12762208 12762242 refGene_gn=SMC5;refGene_tx=XM_003995387.5
D4 12762244 12762377 refGene_gn=SMC5;refGene_tx=XM_003995387.5
D4 12773670 12773794 refGene_gn=SMC5;refGene_tx=XM_003995387.5
D4 12774157 12774280 refGene_gn=SMC5;refGene_tx=XM_003995387.5
D4 12786653 12786779 refGene_gn=SMC5;refGene_tx=XM_003995387.5
D4 12789495 12789540 refGene_gn=SMC5;refGene_tx=XM_003995387.5
D4 12789955 12790051 refGene_gn=SMC5;refGene_tx=XM_003995387.5
D4 12790492 12790597 refGene_gn=SMC5;refGene_tx=XM_003995387.5
D4 12790799 12790919 refGene_gn=SMC5;refGene_tx=XM_003995387.5
D4 12791746 12791926 refGene_gn=SMC5;refGene_tx=XM_003995387.5
D4 12792005 12792101 refGene_gn=SMC5;refGene_tx=XM_003995387.5
D4 12793249 12793390 refGene_gn=SMC5;refGene_tx=XM_003995387.5
D4 12872137 12872367 refGene_gn=KLF9;refGene_tx=XM_003995409.5
D4 12893627 12894132 refGene_gn=KLF9;refGene_tx=XM_003995409.5
D4 12987728 12989181
refGene_gn=TRPM3;refGene_tx=XM_003995388.5;refGene_gn=TRPM3;refGene_tx=XM_003995389.5;refGene_gn=TRPM3;refGene_tx=XM_003995390.5;refGene_gn=TRPM3;refGene_tx=XM_003995398.5;refGene_gn=TRPM3;refGene_tx=XM_003995399.5;refGene_gn=TRPM3;refGene_tx=XM_019815753.1;refGene_gn=TRPM3;refGene_tx=XM_019815754.1
D4 12997262 12997395
refGene_gn=TRPM3;refGene_tx=XM_003995388.5;refGene_gn=TRPM3;refGene_tx=XM_003995389.5;refGene_gn=TRPM3;refGene_tx=XM_003995390.5;refGene_gn=TRPM3;refGene_tx=XM_003995398.5;refGene_gn=TRPM3;refGene_tx=XM_003995399.5;refGene_gn=TRPM3;refGene_tx=XM_019815753.1;refGene_gn=TRPM3;refGene_tx=XM_019815754.1
D4 12999282 12999482
refGene_gn=TRPM3;refGene_tx=XM_003995388.5;refGene_gn=TRPM3;refGene_tx=XM_003995389.5;refGene_gn=TRPM3;refGene_tx=XM_003995390.5;refGene_gn=TRPM3;refGene_tx=XM_003995398.5;refGene_gn=TRPM3;refGene_tx=XM_003995399.5;refGene_gn=TRPM3;refGene_tx=XM_019815753.1;refGene_gn=TRPM3;refGene_tx=XM_019815754.1
D4 12999608 12999759

refGene_gn=TRPM3;refGene_tx=XM_003995388.5;refGene_gn=TRPM3;refGene_tx=XM_003995389.5;refGene_gn=TRPM3;refGene_tx=XM_003995390.5;refGene_gn=TRPM3;refGene_tx=XM_003995398.5;refGene_gn=TRPM3;refGene_tx=XM_003995399.5;refGene_gn=TRPM3;refGene_tx=XM_019815753.1;refGene_gn=TRPM3;refGene_tx=XM_019815754.1
D4 13036561 13036736
refGene_gn=TRPM3;refGene_tx=XM_003995388.5;refGene_gn=TRPM3;refGene_tx=XM_003995389.5;refGene_gn=TRPM3;refGene_tx=XM_003995390.5;refGene_gn=TRPM3;refGene_tx=XM_003995398.5;refGene_gn=TRPM3;refGene_tx=XM_003995399.5;refGene_gn=TRPM3;refGene_tx=XM_019815753.1;refGene_gn=TRPM3;refGene_tx=XM_019815754.1
D4 13043086 13043338
refGene_gn=TRPM3;refGene_tx=XM_003995388.5;refGene_gn=TRPM3;refGene_tx=XM_003995389.5;refGene_gn=TRPM3;refGene_tx=XM_003995390.5;refGene_gn=TRPM3;refGene_tx=XM_003995398.5;refGene_gn=TRPM3;refGene_tx=XM_003995399.5;refGene_gn=TRPM3;refGene_tx=XM_019815753.1;refGene_gn=TRPM3;refGene_tx=XM_019815754.1
D4 13047813 13047942
refGene_gn=TRPM3;refGene_tx=XM_003995388.5;refGene_gn=TRPM3;refGene_tx=XM_003995389.5;refGene_gn=TRPM3;refGene_tx=XM_003995390.5;refGene_gn=TRPM3;refGene_tx=XM_003995398.5;refGene_gn=TRPM3;refGene_tx=XM_003995399.5;refGene_gn=TRPM3;refGene_tx=XM_019815753.1;refGene_gn=TRPM3;refGene_tx=XM_019815754.1
D4 13055165 13055306
refGene_gn=TRPM3;refGene_tx=XM_003995388.5;refGene_gn=TRPM3;refGene_tx=XM_003995389.5;refGene_gn=TRPM3;refGene_tx=XM_003995390.5;refGene_gn=TRPM3;refGene_tx=XM_003995398.5;refGene_gn=TRPM3;refGene_tx=XM_003995399.5;refGene_gn=TRPM3;refGene_tx=XM_019815753.1;refGene_gn=TRPM3;refGene_tx=XM_019815754.1
D4 13060164 13060332
refGene_gn=TRPM3;refGene_tx=XM_003995388.5;refGene_gn=TRPM3;refGene_tx=XM_003995389.5;refGene_gn=TRPM3;refGene_tx=XM_003995390.5;refGene_gn=TRPM3;refGene_tx=XM_003995398.5;refGene_gn=TRPM3;refGene_tx=XM_003995399.5;refGene_gn=TRPM3;refGene_tx=XM_019815753.1;refGene_gn=TRPM3;refGene_tx=XM_019815754.1
D4 13062721 13062950
refGene_gn=TRPM3;refGene_tx=XM_003995388.5;refGene_gn=TRPM3;refGene_tx=XM_003995389.5;refGene_gn=TRPM3;refGene_tx=XM_003995390.5;refGene_gn=TRPM3;refGene_tx=XM_003995398.5;refGene_gn=TRPM3;refGene_tx=XM_003995399.5;refGene_gn=TRPM3;refGene_tx=XM_019815753.1;refGene_gn=TRPM3;refGene_tx=XM_019815754.1
D4 13063853 13064143
refGene_gn=TRPM3;refGene_tx=XM_003995388.5;refGene_gn=TRPM3;refGene_tx=XM_003995389.5;refGene_gn=TRPM3;refGene_tx=XM_003995390.5;refGene_gn=TRPM3;refGene_tx=XM_003995398.5;refGene_gn=TRPM3;refGene_tx=XM_003995399.5;refGene_gn=TRPM3;refGene_tx=XM_019815753.1;refGene_gn=TRPM3;refGene_tx=XM_019815754.1
D4 13065015 13065045
refGene_gn=TRPM3;refGene_tx=XM_003995388.5;refGene_gn=TRPM3;refGene_tx=XM_003995389.5;refGene_gn=TRPM3;refGene_tx=XM_003995390.5;refGene_gn=

TRPM3;refGene_tx=XM_003995398.5;refGene_gn=TRPM3;refGene_tx=XM_0039953
99.5;refGene_gn=TRPM3;refGene_tx=XM_019815753.1;refGene_gn=TRPM3;refGe
ne_tx=XM_019815754.1
D4 13068047 13068188
refGene_gn=TRPM3;refGene_tx=XM_003995388.5;refGene_gn=TRPM3;refGene_tx
=XM_003995389.5;refGene_gn=TRPM3;refGene_tx=XM_003995390.5;refGene_gn=
TRPM3;refGene_tx=XM_003995398.5;refGene_gn=TRPM3;refGene_tx=XM_0039953
99.5;refGene_gn=TRPM3;refGene_tx=XM_019815753.1;refGene_gn=TRPM3;refGe
ne_tx=XM_019815754.1
D4 13068347 13068383
refGene_gn=TRPM3;refGene_tx=XM_003995398.5;refGene_gn=TRPM3;refGene_tx
=XM_003995399.5;refGene_gn=TRPM3;refGene_tx=XM_019815753.1
D4 13076364 13076415
refGene_gn=TRPM3;refGene_tx=XM_003995388.5;refGene_gn=TRPM3;refGene_tx
=XM_003995389.5;refGene_gn=TRPM3;refGene_tx=XM_003995390.5;refGene_gn=
TRPM3;refGene_tx=XM_003995398.5;refGene_gn=TRPM3;refGene_tx=XM_0039953
99.5;refGene_gn=TRPM3;refGene_tx=XM_019815753.1;refGene_gn=TRPM3;refGe
ne_tx=XM_019815754.1
D4 13080387 13080522
refGene_gn=TRPM3;refGene_tx=XM_003995388.5;refGene_gn=TRPM3;refGene_tx
=XM_003995389.5;refGene_gn=TRPM3;refGene_tx=XM_003995390.5;refGene_gn=
TRPM3;refGene_tx=XM_003995398.5;refGene_gn=TRPM3;refGene_tx=XM_0039953
99.5;refGene_gn=TRPM3;refGene_tx=XM_019815753.1;refGene_gn=TRPM3;refGe
ne_tx=XM_019815754.1
D4 13082144 13082245
refGene_gn=TRPM3;refGene_tx=XM_003995388.5;refGene_gn=TRPM3;refGene_tx
=XM_003995389.5;refGene_gn=TRPM3;refGene_tx=XM_003995390.5;refGene_gn=
TRPM3;refGene_tx=XM_003995398.5;refGene_gn=TRPM3;refGene_tx=XM_0039953
99.5;refGene_gn=TRPM3;refGene_tx=XM_019815753.1;refGene_gn=TRPM3;refGe
ne_tx=XM_019815754.1
D4 13105967 13106040
refGene_gn=TRPM3;refGene_tx=XM_003995388.5;refGene_gn=TRPM3;refGene_tx
=XM_003995389.5;refGene_gn=TRPM3;refGene_tx=XM_003995390.5;refGene_gn=
TRPM3;refGene_tx=XM_003995398.5;refGene_gn=TRPM3;refGene_tx=XM_0039953
99.5;refGene_gn=TRPM3;refGene_tx=XM_019815753.1;refGene_gn=TRPM3;refGe
ne_tx=XM_019815754.1
D4 13149257 13149381
refGene_gn=TRPM3;refGene_tx=XM_003995388.5;refGene_gn=TRPM3;refGene_tx
=XM_003995389.5;refGene_gn=TRPM3;refGene_tx=XM_003995390.5;refGene_gn=
TRPM3;refGene_tx=XM_003995398.5;refGene_gn=TRPM3;refGene_tx=XM_0039953
99.5;refGene_gn=TRPM3;refGene_tx=XM_019815753.1;refGene_gn=TRPM3;refGe
ne_tx=XM_019815754.1
D4 13168811 13168986
refGene_gn=TRPM3;refGene_tx=XM_003995388.5;refGene_gn=TRPM3;refGene_tx
=XM_003995389.5;refGene_gn=TRPM3;refGene_tx=XM_003995390.5;refGene_gn=
TRPM3;refGene_tx=XM_003995398.5;refGene_gn=TRPM3;refGene_tx=XM_0039953
99.5;refGene_gn=TRPM3;refGene_tx=XM_019815753.1;refGene_gn=TRPM3;refGe
ne_tx=XM_019815754.1
D4 13189195 13189270
refGene_gn=TRPM3;refGene_tx=XM_003995390.5;refGene_gn=TRPM3;refGene_tx

=XM_019815753.1;refGene_gn=TRPM3;refGene_tx=XM_019815754.1
D4 13206838 13207010
refGene_gn=TRPM3;refGene_tx=XM_003995388.5;refGene_gn=TRPM3;refGene_tx=XM_003995389.5;refGene_gn=TRPM3;refGene_tx=XM_003995390.5;refGene_gn=TRPM3;refGene_tx=XM_003995398.5;refGene_gn=TRPM3;refGene_tx=XM_003995399.5;refGene_gn=TRPM3;refGene_tx=XM_019815753.1;refGene_gn=TRPM3;refGene_tx=XM_019815754.1
D4 13221142 13221267
refGene_gn=TRPM3;refGene_tx=XM_003995388.5;refGene_gn=TRPM3;refGene_tx=XM_003995389.5;refGene_gn=TRPM3;refGene_tx=XM_003995390.5;refGene_gn=TRPM3;refGene_tx=XM_003995398.5;refGene_gn=TRPM3;refGene_tx=XM_003995399.5;refGene_gn=TRPM3;refGene_tx=XM_019815753.1;refGene_gn=TRPM3;refGene_tx=XM_019815754.1
D4 13224742 13224956
refGene_gn=TRPM3;refGene_tx=XM_003995388.5;refGene_gn=TRPM3;refGene_tx=XM_003995389.5;refGene_gn=TRPM3;refGene_tx=XM_003995390.5;refGene_gn=TRPM3;refGene_tx=XM_003995398.5;refGene_gn=TRPM3;refGene_tx=XM_003995399.5;refGene_gn=TRPM3;refGene_tx=XM_019815753.1;refGene_gn=TRPM3;refGene_tx=XM_019815754.1
D4 13238523 13238728
refGene_gn=TRPM3;refGene_tx=XM_003995388.5;refGene_gn=TRPM3;refGene_tx=XM_003995389.5;refGene_gn=TRPM3;refGene_tx=XM_003995390.5;refGene_gn=TRPM3;refGene_tx=XM_003995398.5;refGene_gn=TRPM3;refGene_tx=XM_003995399.5;refGene_gn=TRPM3;refGene_tx=XM_019815753.1;refGene_gn=TRPM3;refGene_tx=XM_019815754.1
D4 13240079 13240159
refGene_gn=TRPM3;refGene_tx=XM_003995388.5;refGene_gn=TRPM3;refGene_tx=XM_003995389.5;refGene_gn=TRPM3;refGene_tx=XM_003995390.5;refGene_gn=TRPM3;refGene_tx=XM_003995398.5;refGene_gn=TRPM3;refGene_tx=XM_003995399.5;refGene_gn=TRPM3;refGene_tx=XM_019815753.1;refGene_gn=TRPM3;refGene_tx=XM_019815754.1
D4 13479235 13479412
refGene_gn=TRPM3;refGene_tx=XM_003995388.5;refGene_gn=TRPM3;refGene_tx=XM_003995390.5;refGene_gn=TRPM3;refGene_tx=XM_003995399.5;refGene_gn=TRPM3;refGene_tx=XM_019815753.1
D4 13786469 13786652
refGene_gn=TRPM3;refGene_tx=XM_003995389.5;refGene_gn=TRPM3;refGene_tx=XM_003995398.5;refGene_gn=TRPM3;refGene_tx=XM_019815754.1
D4 14020297 14020494
refGene_gn=TMEM2;refGene_tx=XM_003995401.5;refGene_gn=TMEM2;refGene_tx=XM_023243341.1
D4 14020847 14020951
refGene_gn=TMEM2;refGene_tx=XM_003995401.5;refGene_gn=TMEM2;refGene_tx=XM_023243341.1
D4 14023264 14023419
refGene_gn=TMEM2;refGene_tx=XM_003995401.5;refGene_gn=TMEM2;refGene_tx=XM_023243341.1
D4 14027138 14027237
refGene_gn=TMEM2;refGene_tx=XM_003995401.5;refGene_gn=TMEM2;refGene_tx=XM_023243341.1

D4 14029664 14029884
refGene_gn=TMEM2;refGene_tx=XM_003995401.5;refGene_gn=TMEM2;refGene_tx
=XM_023243341.1
D4 14033966 14034149
refGene_gn=TMEM2;refGene_tx=XM_003995401.5;refGene_gn=TMEM2;refGene_tx
=XM_023243341.1
D4 14038429 14038638
refGene_gn=TMEM2;refGene_tx=XM_003995401.5;refGene_gn=TMEM2;refGene_tx
=XM_023243341.1
D4 14042581 14042797
refGene_gn=TMEM2;refGene_tx=XM_003995401.5;refGene_gn=TMEM2;refGene_tx
=XM_023243341.1
D4 14045366 14045544
refGene_gn=TMEM2;refGene_tx=XM_003995401.5;refGene_gn=TMEM2;refGene_tx
=XM_023243341.1
D4 14048160 14048316
refGene_gn=TMEM2;refGene_tx=XM_003995401.5;refGene_gn=TMEM2;refGene_tx
=XM_023243341.1
D4 14049303 14049339
refGene_gn=TMEM2;refGene_tx=XM_003995401.5;refGene_gn=TMEM2;refGene_tx
=XM_023243341.1
D4 14050568 14050700
refGene_gn=TMEM2;refGene_tx=XM_003995401.5;refGene_gn=TMEM2;refGene_tx
=XM_023243341.1
D4 14055078 14055167
refGene_gn=TMEM2;refGene_tx=XM_003995401.5;refGene_gn=TMEM2;refGene_tx
=XM_023243341.1
D4 14057723 14057852
refGene_gn=TMEM2;refGene_tx=XM_003995401.5;refGene_gn=TMEM2;refGene_tx
=XM_023243341.1
D4 14061513 14061583
refGene_gn=TMEM2;refGene_tx=XM_003995401.5;refGene_gn=TMEM2;refGene_tx
=XM_023243341.1
D4 14061726 14061932
refGene_gn=TMEM2;refGene_tx=XM_003995401.5;refGene_gn=TMEM2;refGene_tx
=XM_023243341.1
D4 14062241 14062400 refGene_gn=TMEM2;refGene_tx=XM_023243341.1
D4 14063603 14063813 refGene_gn=TMEM2;refGene_tx=XM_003995401.5
D4 14065972 14066142 refGene_gn=TMEM2;refGene_tx=XM_003995401.5
D4 14067971 14068160 refGene_gn=TMEM2;refGene_tx=XM_003995401.5
D4 14073650 14073820 refGene_gn=TMEM2;refGene_tx=XM_003995401.5
D4 14079673 14080235 refGene_gn=TMEM2;refGene_tx=XM_003995401.5
D4 14080680 14080821 refGene_gn=TMEM2;refGene_tx=XM_003995401.5
D4 14086009 14086340 refGene_gn=TMEM2;refGene_tx=XM_003995401.5
D4 14185756 14185976
refGene_gn=ABHD17B;refGene_tx=XM_023243342.1;refGene_gn=ABHD17B;refGen
e_tx=XM_023243343.1;refGene_gn=ABHD17B;refGene_tx=XM_023243344.1
D4 14188845 14189025
refGene_gn=ABHD17B;refGene_tx=XM_023243342.1;refGene_gn=ABHD17B;refGen
e_tx=XM_023243343.1;refGene_gn=ABHD17B;refGene_tx=XM_023243344.1

D4 14192829 14193299
refGene_gn=ABHD17B;refGene_tx=XM_023243342.1;refGene_gn=ABHD17B;refGene_tx=XM_023243343.1;refGene_gn=ABHD17B;refGene_tx=XM_023243344.1
D4 14225229 14225283 refGene_gn=ABHD17B;refGene_tx=XM_023243342.1
D4 14237991 14238093
refGene_gn=CD4H9orf85;refGene_tx=XM_003995404.5
D4 14275265 14275372
refGene_gn=CD4H9orf85;refGene_tx=XM_003995404.5
D4 14302118 14302232
refGene_gn=CD4H9orf85;refGene_tx=XM_003995404.5
D4 14302748 14302896
refGene_gn=CD4H9orf85;refGene_tx=XM_003995404.5
D4 14455037 14455103 refGene_gn=GDA;refGene_tx=XM_003995405.5
D4 14455104 14455161 refGene_gn=GDA;refGene_tx=XM_003995405.5
D4 14489768 14489857 refGene_gn=GDA;refGene_tx=XM_003995405.5
D4 14496692 14496864 refGene_gn=GDA;refGene_tx=XM_003995405.5
D4 14504408 14504496 refGene_gn=GDA;refGene_tx=XM_003995405.5
D4 14505978 14506084 refGene_gn=GDA;refGene_tx=XM_003995405.5
D4 14507515 14507543 refGene_gn=GDA;refGene_tx=XM_003995405.5
D4 14508696 14508804 refGene_gn=GDA;refGene_tx=XM_003995405.5
D4 14511324 14511432 refGene_gn=GDA;refGene_tx=XM_003995405.5
D4 14514215 14514313 refGene_gn=GDA;refGene_tx=XM_003995405.5
D4 14515159 14515227 refGene_gn=GDA;refGene_tx=XM_003995405.5
D4 14524871 14525018 refGene_gn=GDA;refGene_tx=XM_003995405.5
D4 14527835 14527966 refGene_gn=GDA;refGene_tx=XM_003995405.5
D4 14529522 14529550 refGene_gn=GDA;refGene_tx=XM_003995405.5
D4 14530442 14530513 refGene_gn=GDA;refGene_tx=XM_003995405.5
D4 14611289 14611438
refGene_gn=ZFAND5;refGene_tx=XM_003995406.5;refGene_gn=ZFAND5;refGene_tx=XM_023243345.1
D4 14612158 14612284
refGene_gn=ZFAND5;refGene_tx=XM_003995406.5;refGene_gn=ZFAND5;refGene_tx=XM_023243345.1
D4 14614698 14614802
refGene_gn=ZFAND5;refGene_tx=XM_003995406.5;refGene_gn=ZFAND5;refGene_tx=XM_023243345.1
D4 14615432 14615544
refGene_gn=ZFAND5;refGene_tx=XM_003995406.5;refGene_gn=ZFAND5;refGene_tx=XM_023243345.1
D4 14615975 14616126
refGene_gn=ZFAND5;refGene_tx=XM_003995406.5;refGene_gn=ZFAND5;refGene_tx=XM_023243345.1
D4 14864545 14864909 refGene_gn=TMC1;refGene_tx=XM_023243263.1
D4 14909934 14909982 refGene_gn=TMC1;refGene_tx=XM_023243263.1
D4 14913736 14913908 refGene_gn=TMC1;refGene_tx=XM_023243263.1
D4 14922588 14922717 refGene_gn=TMC1;refGene_tx=XM_023243263.1
D4 14942841 14942932 refGene_gn=TMC1;refGene_tx=XM_023243263.1
D4 14945789 14945871 refGene_gn=TMC1;refGene_tx=XM_023243263.1
D4 14953230 14953337 refGene_gn=TMC1;refGene_tx=XM_023243263.1
D4 14956239 14956338 refGene_gn=TMC1;refGene_tx=XM_023243263.1

D4 14972498 14972641 refGene_gn=TMC1;refGene_tx=XM_023243263.1
D4 14976871 14977016 refGene_gn=TMC1;refGene_tx=XM_023243263.1
D4 14977628 14977823 refGene_gn=TMC1;refGene_tx=XM_023243263.1
D4 14980537 14980717 refGene_gn=TMC1;refGene_tx=XM_023243263.1
D4 14980843 14981005 refGene_gn=TMC1;refGene_tx=XM_023243263.1
D4 14988056 14988185 refGene_gn=TMC1;refGene_tx=XM_023243263.1
D4 14997998 14998066 refGene_gn=TMC1;refGene_tx=XM_023243263.1
D4 15003186 15003426 refGene_gn=TMC1;refGene_tx=XM_023243263.1
D4 15008436 15008562 refGene_gn=TMC1;refGene_tx=XM_023243263.1
D4 15012205 15012284 refGene_gn=TMC1;refGene_tx=XM_023243263.1
D4 15012386 15012438 refGene_gn=TMC1;refGene_tx=XM_023243263.1
D4 15018485 15018540 refGene_gn=TMC1;refGene_tx=XM_023243263.1
D4 15027594 15027688 refGene_gn=TMC1;refGene_tx=XM_023243263.1
D4 15057584 15057657 refGene_gn=ALDH1A1;refGene_tx=XM_003995407.3
D4 15062410 15062485 refGene_gn=ALDH1A1;refGene_tx=XM_003995407.3
D4 15069587 15069745 refGene_gn=ALDH1A1;refGene_tx=XM_003995407.3
D4 15074753 15074918 refGene_gn=ALDH1A1;refGene_tx=XM_003995407.3
D4 15079499 15079684 refGene_gn=ALDH1A1;refGene_tx=XM_003995407.3
D4 15081245 15081348 refGene_gn=ALDH1A1;refGene_tx=XM_003995407.3
D4 15086025 15086139 refGene_gn=ALDH1A1;refGene_tx=XM_003995407.3
D4 15087581 15087710 refGene_gn=ALDH1A1;refGene_tx=XM_003995407.3
D4 15089483 15089545 refGene_gn=ALDH1A1;refGene_tx=XM_003995407.3
D4 15091673 15091803 refGene_gn=ALDH1A1;refGene_tx=XM_003995407.3
D4 15093974 15094115 refGene_gn=ALDH1A1;refGene_tx=XM_003995407.3
D4 15102248 15102353 refGene_gn=ALDH1A1;refGene_tx=XM_003995407.3
D4 15115895 15115961 refGene_gn=ALDH1A1;refGene_tx=XM_003995407.3
D4 15308784 15308850
refGene_gn=ANXA1;refGene_tx=NM_001009857.1;refGene_gn=ANXA1;refGene_tx
=XM_006939090.4
D4 15308943 15309052
refGene_gn=ANXA1;refGene_tx=NM_001009857.1;refGene_gn=ANXA1;refGene_tx
=XM_006939090.4
D4 15309575 15309670
refGene_gn=ANXA1;refGene_tx=NM_001009857.1;refGene_gn=ANXA1;refGene_tx
=XM_006939090.4
D4 15310707 15310821
refGene_gn=ANXA1;refGene_tx=NM_001009857.1;refGene_gn=ANXA1;refGene_tx
=XM_006939090.4
D4 15311247 15311338
refGene_gn=ANXA1;refGene_tx=NM_001009857.1;refGene_gn=ANXA1;refGene_tx
=XM_006939090.4
D4 15313214 15313294
refGene_gn=ANXA1;refGene_tx=NM_001009857.1;refGene_gn=ANXA1;refGene_tx
=XM_006939090.4
D4 15313753 15313810
refGene_gn=ANXA1;refGene_tx=NM_001009857.1;refGene_gn=ANXA1;refGene_tx
=XM_006939090.4
D4 15315668 15315762
refGene_gn=ANXA1;refGene_tx=NM_001009857.1;refGene_gn=ANXA1;refGene_tx
=XM_006939090.4

D4 15316626 15316722
refGene_gn=ANXA1;refGene_tx=NM_001009857.1;refGene_gn=ANXA1;refGene_tx=XM_006939090.4
D4 15317802 15317861 refGene_gn=ANXA1;refGene_tx=XM_006939090.4
D4 15319276 15319399 refGene_gn=ANXA1;refGene_tx=XM_006939090.4
D4 15320301 15320355 refGene_gn=ANXA1;refGene_tx=XM_006939090.4
D4 16427350 16427357 refGene_gn=RORB;refGene_tx=XM_019815772.2
D4 16547397 16547437 refGene_gn=RORB;refGene_tx=XM_019815771.2
D4 16563071 16563157
refGene_gn=RORB;refGene_tx=XM_019815771.2;refGene_gn=RORB;refGene_tx=XM_019815772.2
D4 16568418 16568560
refGene_gn=RORB;refGene_tx=XM_019815771.2;refGene_gn=RORB;refGene_tx=XM_019815772.2
D4 16576991 16577393
refGene_gn=RORB;refGene_tx=XM_019815771.2;refGene_gn=RORB;refGene_tx=XM_019815772.2
D4 16596546 16596668
refGene_gn=RORB;refGene_tx=XM_019815771.2;refGene_gn=RORB;refGene_tx=XM_019815772.2
D4 16598059 16598192
refGene_gn=RORB;refGene_tx=XM_019815771.2;refGene_gn=RORB;refGene_tx=XM_019815772.2
D4 16601734 16601842
refGene_gn=RORB;refGene_tx=XM_019815771.2;refGene_gn=RORB;refGene_tx=XM_019815772.2
D4 16603351 16603462
refGene_gn=RORB;refGene_tx=XM_019815771.2;refGene_gn=RORB;refGene_tx=XM_019815772.2
D4 16606677 16606790
refGene_gn=RORB;refGene_tx=XM_019815771.2;refGene_gn=RORB;refGene_tx=XM_019815772.2
D4 16618826 16618982
refGene_gn=RORB;refGene_tx=XM_019815771.2;refGene_gn=RORB;refGene_tx=XM_019815772.2
D4 16662257 16662391
refGene_gn=TRPM6;refGene_tx=XM_019815765.2;refGene_gn=TRPM6;refGene_tx=XM_023243347.1;refGene_gn=TRPM6;refGene_tx=XM_023243348.1
D4 16664992 16665099
refGene_gn=TRPM6;refGene_tx=XM_019815765.2;refGene_gn=TRPM6;refGene_tx=XM_023243347.1;refGene_gn=TRPM6;refGene_tx=XM_023243348.1
D4 16669538 16669590
refGene_gn=TRPM6;refGene_tx=XM_019815765.2;refGene_gn=TRPM6;refGene_tx=XM_023243347.1;refGene_gn=TRPM6;refGene_tx=XM_023243348.1
D4 16676668 16676874
refGene_gn=TRPM6;refGene_tx=XM_019815765.2;refGene_gn=TRPM6;refGene_tx=XM_023243347.1;refGene_gn=TRPM6;refGene_tx=XM_023243348.1
D4 16677642 16677725
refGene_gn=TRPM6;refGene_tx=XM_019815765.2;refGene_gn=TRPM6;refGene_tx=XM_023243347.1;refGene_gn=TRPM6;refGene_tx=XM_023243348.1

D4 16678016 16678303
refGene_gn=TRPM6;refGene_tx=XM_019815765.2;refGene_gn=TRPM6;refGene_tx=XM_023243347.1;refGene_gn=TRPM6;refGene_tx=XM_023243348.1
D4 16680290 16680356
refGene_gn=TRPM6;refGene_tx=XM_019815765.2;refGene_gn=TRPM6;refGene_tx=XM_023243347.1;refGene_gn=TRPM6;refGene_tx=XM_023243348.1
D4 16681817 16681868
refGene_gn=TRPM6;refGene_tx=XM_019815765.2;refGene_gn=TRPM6;refGene_tx=XM_023243347.1;refGene_gn=TRPM6;refGene_tx=XM_023243348.1
D4 16685665 16685691
refGene_gn=TRPM6;refGene_tx=XM_019815765.2;refGene_gn=TRPM6;refGene_tx=XM_023243347.1;refGene_gn=TRPM6;refGene_tx=XM_023243348.1
D4 16687231 16687290
refGene_gn=TRPM6;refGene_tx=XM_019815765.2;refGene_gn=TRPM6;refGene_tx=XM_023243347.1;refGene_gn=TRPM6;refGene_tx=XM_023243348.1
D4 16688808 16688900
refGene_gn=TRPM6;refGene_tx=XM_019815765.2;refGene_gn=TRPM6;refGene_tx=XM_023243347.1;refGene_gn=TRPM6;refGene_tx=XM_023243348.1
D4 16692211 16692335
refGene_gn=TRPM6;refGene_tx=XM_019815765.2;refGene_gn=TRPM6;refGene_tx=XM_023243347.1;refGene_gn=TRPM6;refGene_tx=XM_023243348.1
D4 16696787 16696900
refGene_gn=TRPM6;refGene_tx=XM_019815765.2;refGene_gn=TRPM6;refGene_tx=XM_023243347.1;refGene_gn=TRPM6;refGene_tx=XM_023243348.1
D4 16697083 16698219
refGene_gn=TRPM6;refGene_tx=XM_019815765.2;refGene_gn=TRPM6;refGene_tx=XM_023243347.1;refGene_gn=TRPM6;refGene_tx=XM_023243348.1
D4 16715133 16715266
refGene_gn=TRPM6;refGene_tx=XM_019815765.2;refGene_gn=TRPM6;refGene_tx=XM_023243347.1;refGene_gn=TRPM6;refGene_tx=XM_023243348.1
D4 16718410 16718604
refGene_gn=TRPM6;refGene_tx=XM_019815765.2;refGene_gn=TRPM6;refGene_tx=XM_023243347.1;refGene_gn=TRPM6;refGene_tx=XM_023243348.1
D4 16722437 16722552
refGene_gn=TRPM6;refGene_tx=XM_019815765.2;refGene_gn=TRPM6;refGene_tx=XM_023243347.1;refGene_gn=TRPM6;refGene_tx=XM_023243348.1
D4 16722756 16722931
refGene_gn=TRPM6;refGene_tx=XM_019815765.2;refGene_gn=TRPM6;refGene_tx=XM_023243347.1;refGene_gn=TRPM6;refGene_tx=XM_023243348.1
D4 16724542 16724794
refGene_gn=TRPM6;refGene_tx=XM_019815765.2;refGene_gn=TRPM6;refGene_tx=XM_023243347.1;refGene_gn=TRPM6;refGene_tx=XM_023243348.1
D4 16726133 16726262
refGene_gn=TRPM6;refGene_tx=XM_019815765.2;refGene_gn=TRPM6;refGene_tx=XM_023243347.1
D4 16729097 16729244
refGene_gn=TRPM6;refGene_tx=XM_019815765.2;refGene_gn=TRPM6;refGene_tx=XM_023243347.1;refGene_gn=TRPM6;refGene_tx=XM_023243348.1
D4 16734170 16734323
refGene_gn=TRPM6;refGene_tx=XM_019815765.2;refGene_gn=TRPM6;refGene_tx

=XM_023243347.1;refGene_gn=TRPM6;refGene_tx=XM_023243348.1
D4 16738787 16739016
refGene_gn=TRPM6;refGene_tx=XM_019815765.2;refGene_gn=TRPM6;refGene_tx=XM_023243347.1;refGene_gn=TRPM6;refGene_tx=XM_023243348.1
D4 16740488 16740769
refGene_gn=TRPM6;refGene_tx=XM_019815765.2;refGene_gn=TRPM6;refGene_tx=XM_023243347.1;refGene_gn=TRPM6;refGene_tx=XM_023243348.1
D4 16742220 16742355
refGene_gn=TRPM6;refGene_tx=XM_019815765.2;refGene_gn=TRPM6;refGene_tx=XM_023243347.1;refGene_gn=TRPM6;refGene_tx=XM_023243348.1
D4 16744513 16744666
refGene_gn=TRPM6;refGene_tx=XM_019815765.2;refGene_gn=TRPM6;refGene_tx=XM_023243347.1;refGene_gn=TRPM6;refGene_tx=XM_023243348.1
D4 16747729 16747783
refGene_gn=TRPM6;refGene_tx=XM_019815765.2;refGene_gn=TRPM6;refGene_tx=XM_023243347.1;refGene_gn=TRPM6;refGene_tx=XM_023243348.1
D4 16749092 16749227
refGene_gn=TRPM6;refGene_tx=XM_019815765.2;refGene_gn=TRPM6;refGene_tx=XM_023243347.1;refGene_gn=TRPM6;refGene_tx=XM_023243348.1
D4 16752514 16752615
refGene_gn=TRPM6;refGene_tx=XM_019815765.2;refGene_gn=TRPM6;refGene_tx=XM_023243347.1;refGene_gn=TRPM6;refGene_tx=XM_023243348.1
D4 16752736 16752809
refGene_gn=TRPM6;refGene_tx=XM_019815765.2;refGene_gn=TRPM6;refGene_tx=XM_023243347.1;refGene_gn=TRPM6;refGene_tx=XM_023243348.1
D4 16755417 16755541
refGene_gn=TRPM6;refGene_tx=XM_019815765.2;refGene_gn=TRPM6;refGene_tx=XM_023243347.1;refGene_gn=TRPM6;refGene_tx=XM_023243348.1
D4 16756646 16756815
refGene_gn=TRPM6;refGene_tx=XM_019815765.2;refGene_gn=TRPM6;refGene_tx=XM_023243347.1;refGene_gn=TRPM6;refGene_tx=XM_023243348.1
D4 16762217 16762389
refGene_gn=TRPM6;refGene_tx=XM_019815765.2;refGene_gn=TRPM6;refGene_tx=XM_023243347.1;refGene_gn=TRPM6;refGene_tx=XM_023243348.1
D4 16767721 16767846
refGene_gn=TRPM6;refGene_tx=XM_019815765.2;refGene_gn=TRPM6;refGene_tx=XM_023243347.1;refGene_gn=TRPM6;refGene_tx=XM_023243348.1
D4 16771078 16771292
refGene_gn=TRPM6;refGene_tx=XM_019815765.2;refGene_gn=TRPM6;refGene_tx=XM_023243347.1;refGene_gn=TRPM6;refGene_tx=XM_023243348.1
D4 16774413 16774591
refGene_gn=TRPM6;refGene_tx=XM_019815765.2;refGene_gn=TRPM6;refGene_tx=XM_023243347.1;refGene_gn=TRPM6;refGene_tx=XM_023243348.1
D4 16780946 16780985
refGene_gn=TRPM6;refGene_tx=XM_019815765.2;refGene_gn=TRPM6;refGene_tx=XM_023243347.1;refGene_gn=TRPM6;refGene_tx=XM_023243348.1
D4 16784599 16784679
refGene_gn=TRPM6;refGene_tx=XM_019815765.2;refGene_gn=TRPM6;refGene_tx=XM_023243347.1;refGene_gn=TRPM6;refGene_tx=XM_023243348.1
D4 16810281 16810299

refGene_gn=TRPM6;refGene_tx=XM_019815765.2;refGene_gn=TRPM6;refGene_tx=XM_023243348.1
D4 16834421 16834430 refGene_gn=TRPM6;refGene_tx=XM_023243347.1
D4 16892129 16892231 refGene_gn=CARNMT1;refGene_tx=XM_011288282.3;refGene_gn=CARNMT1;refGene_tx=XM_011288284.2
D4 16893165 16893269 refGene_gn=CARNMT1;refGene_tx=XM_011288282.3;refGene_gn=CARNMT1;refGene_tx=XM_011288284.2
D4 16903431 16903545 refGene_gn=CARNMT1;refGene_tx=XM_011288282.3;refGene_gn=CARNMT1;refGene_tx=XM_011288284.2
D4 16905784 16905963 refGene_gn=CARNMT1;refGene_tx=XM_011288282.3;refGene_gn=CARNMT1;refGene_tx=XM_011288284.2
D4 16906927 16907068 refGene_gn=CARNMT1;refGene_tx=XM_011288282.3;refGene_gn=CARNMT1;refGene_tx=XM_011288284.2
D4 16924745 16924909 refGene_gn=CARNMT1;refGene_tx=XM_011288282.3;refGene_gn=CARNMT1;refGene_tx=XM_011288284.2
D4 16925709 16925905 refGene_gn=CARNMT1;refGene_tx=XM_011288282.3;refGene_gn=CARNMT1;refGene_tx=XM_011288284.2
D4 16926403 16926417 refGene_gn=CARNMT1;refGene_tx=XM_011288284.2
D4 16933808 16934044 refGene_gn=CARNMT1;refGene_tx=XM_011288282.3
D4 16959125 16959145 refGene_gn=NMRK1;refGene_tx=XM_011288280.3;refGene_gn=NMRK1;refGene_tx=XM_011288281.3
D4 16964277 16964361 refGene_gn=NMRK1;refGene_tx=XM_011288280.3;refGene_gn=NMRK1;refGene_tx=XM_011288281.3
D4 16966648 16966782 refGene_gn=NMRK1;refGene_tx=XM_011288280.3;refGene_gn=NMRK1;refGene_tx=XM_011288281.3
D4 16967840 16967912 refGene_gn=NMRK1;refGene_tx=XM_011288280.3;refGene_gn=NMRK1;refGene_tx=XM_011288281.3
D4 16968003 16968151 refGene_gn=NMRK1;refGene_tx=XM_011288280.3;refGene_gn=NMRK1;refGene_tx=XM_011288281.3
D4 16974589 16974638 refGene_gn=NMRK1;refGene_tx=XM_011288280.3;refGene_gn=NMRK1;refGene_tx=XM_011288281.3
D4 16974926 16975017 refGene_gn=NMRK1;refGene_tx=XM_011288280.3;refGene_gn=NMRK1;refGene_tx=XM_011288281.3
D4 16980630 16980659 refGene_gn=NMRK1;refGene_tx=XM_011288280.3;refGene_gn=NMRK1;refGene_tx=XM_011288281.3

D4 16984665 16984699 refGene_gn=OSTF1;refGene_tx=XM_003995417.5
D4 17010577 17010624 refGene_gn=OSTF1;refGene_tx=XM_003995417.5
D4 17019647 17019698 refGene_gn=OSTF1;refGene_tx=XM_003995417.5
D4 17023099 17023163 refGene_gn=OSTF1;refGene_tx=XM_003995417.5
D4 17025982 17026036 refGene_gn=OSTF1;refGene_tx=XM_003995417.5
D4 17028616 17028724 refGene_gn=OSTF1;refGene_tx=XM_003995417.5
D4 17029665 17029715 refGene_gn=OSTF1;refGene_tx=XM_003995417.5
D4 17033580 17033659 refGene_gn=OSTF1;refGene_tx=XM_003995417.5
D4 17036280 17036379 refGene_gn=OSTF1;refGene_tx=XM_003995417.5
D4 17042286 17042345 refGene_gn=OSTF1;refGene_tx=XM_003995417.5
D4 17623103 17623820
refGene_gn=PCSK5;refGene_tx=XM_023243354.1;refGene_gn=PCSK5;refGene_tx
=XM_023243355.1;refGene_gn=PCSK5;refGene_tx=XM_023243356.1
D4 17665055 17665160
refGene_gn=PCSK5;refGene_tx=XM_023243354.1;refGene_gn=PCSK5;refGene_tx
=XM_023243355.1;refGene_gn=PCSK5;refGene_tx=XM_023243356.1
D4 17713971 17714085
refGene_gn=PCSK5;refGene_tx=XM_023243354.1;refGene_gn=PCSK5;refGene_tx
=XM_023243355.1;refGene_gn=PCSK5;refGene_tx=XM_023243356.1
D4 17754637 17754781
refGene_gn=PCSK5;refGene_tx=XM_023243354.1;refGene_gn=PCSK5;refGene_tx
=XM_023243355.1;refGene_gn=PCSK5;refGene_tx=XM_023243356.1
D4 17759077 17759154
refGene_gn=PCSK5;refGene_tx=XM_023243354.1;refGene_gn=PCSK5;refGene_tx
=XM_023243355.1;refGene_gn=PCSK5;refGene_tx=XM_023243356.1
D4 17794852 17794941
refGene_gn=PCSK5;refGene_tx=XM_023243354.1;refGene_gn=PCSK5;refGene_tx
=XM_023243355.1;refGene_gn=PCSK5;refGene_tx=XM_023243356.1
D4 17798622 17798795
refGene_gn=PCSK5;refGene_tx=XM_023243354.1;refGene_gn=PCSK5;refGene_tx
=XM_023243355.1;refGene_gn=PCSK5;refGene_tx=XM_023243356.1
D4 17822960 17823173
refGene_gn=PCSK5;refGene_tx=XM_023243354.1;refGene_gn=PCSK5;refGene_tx
=XM_023243355.1;refGene_gn=PCSK5;refGene_tx=XM_023243356.1
D4 17833809 17833910
refGene_gn=PCSK5;refGene_tx=XM_023243354.1;refGene_gn=PCSK5;refGene_tx
=XM_023243355.1;refGene_gn=PCSK5;refGene_tx=XM_023243356.1
D4 17863135 17863239
refGene_gn=PCSK5;refGene_tx=XM_023243354.1;refGene_gn=PCSK5;refGene_tx
=XM_023243355.1;refGene_gn=PCSK5;refGene_tx=XM_023243356.1
D4 17886652 17886770
refGene_gn=PCSK5;refGene_tx=XM_023243354.1;refGene_gn=PCSK5;refGene_tx
=XM_023243355.1;refGene_gn=PCSK5;refGene_tx=XM_023243356.1
D4 17887961 17888150
refGene_gn=PCSK5;refGene_tx=XM_023243354.1;refGene_gn=PCSK5;refGene_tx
=XM_023243355.1;refGene_gn=PCSK5;refGene_tx=XM_023243356.1
D4 17897608 17897745
refGene_gn=PCSK5;refGene_tx=XM_023243354.1;refGene_gn=PCSK5;refGene_tx
=XM_023243355.1;refGene_gn=PCSK5;refGene_tx=XM_023243356.1
D4 17911606 17911750

refGene_gn=PCSK5;refGene_tx=XM_023243354.1;refGene_gn=PCSK5;refGene_tx=XM_023243355.1;refGene_gn=PCSK5;refGene_tx=XM_023243356.1
D4 17916319 17916422 refGene_gn=PCSK5;refGene_tx=XM_023243354.1;refGene_gn=PCSK5;refGene_tx=XM_023243355.1;refGene_gn=PCSK5;refGene_tx=XM_023243356.1
D4 17917686 17917880 refGene_gn=PCSK5;refGene_tx=XM_023243354.1;refGene_gn=PCSK5;refGene_tx=XM_023243355.1;refGene_gn=PCSK5;refGene_tx=XM_023243356.1
D4 17923651 17923736 refGene_gn=PCSK5;refGene_tx=XM_023243354.1;refGene_gn=PCSK5;refGene_tx=XM_023243355.1;refGene_gn=PCSK5;refGene_tx=XM_023243356.1
D4 17928263 17928361 refGene_gn=PCSK5;refGene_tx=XM_023243354.1;refGene_gn=PCSK5;refGene_tx=XM_023243355.1;refGene_gn=PCSK5;refGene_tx=XM_023243356.1
D4 17928789 17928919 refGene_gn=PCSK5;refGene_tx=XM_023243354.1;refGene_gn=PCSK5;refGene_tx=XM_023243355.1;refGene_gn=PCSK5;refGene_tx=XM_023243356.1
D4 17929814 17929930 refGene_gn=PCSK5;refGene_tx=XM_023243354.1;refGene_gn=PCSK5;refGene_tx=XM_023243355.1;refGene_gn=PCSK5;refGene_tx=XM_023243356.1
D4 17933365 17933481 refGene_gn=PCSK5;refGene_tx=XM_023243356.1
D4 17977680 17977783 refGene_gn=PCSK5;refGene_tx=XM_023243354.1;refGene_gn=PCSK5;refGene_tx=XM_023243355.1
D4 17983688 17983825 refGene_gn=PCSK5;refGene_tx=XM_023243354.1;refGene_gn=PCSK5;refGene_tx=XM_023243355.1
D4 17990213 17990420 refGene_gn=PCSK5;refGene_tx=XM_023243354.1;refGene_gn=PCSK5;refGene_tx=XM_023243355.1
D4 17993906 17993975 refGene_gn=PCSK5;refGene_tx=XM_023243354.1;refGene_gn=PCSK5;refGene_tx=XM_023243355.1
D4 18032963 18033006 refGene_gn=PCSK5;refGene_tx=XM_023243354.1;refGene_gn=PCSK5;refGene_tx=XM_023243355.1
D4 18035564 18035701 refGene_gn=PCSK5;refGene_tx=XM_023243354.1;refGene_gn=PCSK5;refGene_tx=XM_023243355.1
D4 18036812 18037013 refGene_gn=PCSK5;refGene_tx=XM_023243354.1;refGene_gn=PCSK5;refGene_tx=XM_023243355.1
D4 18040971 18041049 refGene_gn=PCSK5;refGene_tx=XM_023243354.1
D4 18045003 18045087 refGene_gn=PCSK5;refGene_tx=XM_023243354.1;refGene_gn=PCSK5;refGene_tx=XM_023243355.1
D4 18046153 18046349 refGene_gn=PCSK5;refGene_tx=XM_023243354.1;refGene_gn=PCSK5;refGene_tx=XM_023243355.1
D4 18052878 18053096

refGene_gn=PCSK5;refGene_tx=XM_023243354.1;refGene_gn=PCSK5;refGene_tx=XM_023243355.1
D4 18054143 18054380
refGene_gn=PCSK5;refGene_tx=XM_023243354.1;refGene_gn=PCSK5;refGene_tx=XM_023243355.1
D4 18057288 18057516
refGene_gn=PCSK5;refGene_tx=XM_023243354.1;refGene_gn=PCSK5;refGene_tx=XM_023243355.1
D4 18058709 18058887
refGene_gn=PCSK5;refGene_tx=XM_023243354.1;refGene_gn=PCSK5;refGene_tx=XM_023243355.1
D4 18063454 18063672
refGene_gn=PCSK5;refGene_tx=XM_023243354.1;refGene_gn=PCSK5;refGene_tx=XM_023243355.1
D4 18075521 18075622
refGene_gn=PCSK5;refGene_tx=XM_023243354.1;refGene_gn=PCSK5;refGene_tx=XM_023243355.1
D4 18078116 18078303
refGene_gn=PCSK5;refGene_tx=XM_023243354.1;refGene_gn=PCSK5;refGene_tx=XM_023243355.1
D4 18081445 18081849
refGene_gn=PCSK5;refGene_tx=XM_023243354.1;refGene_gn=PCSK5;refGene_tx=XM_023243355.1
D4 18103650 18103781 refGene_gn=RFK;refGene_tx=XM_011288290.3
D4 18104060 18104215 refGene_gn=RFK;refGene_tx=XM_023243357.1
D4 18104537 18104640
refGene_gn=RFK;refGene_tx=XM_011288290.3;refGene_gn=RFK;refGene_tx=XM_023243357.1
D4 18106177 18106329
refGene_gn=RFK;refGene_tx=XM_011288290.3;refGene_gn=RFK;refGene_tx=XM_023243357.1
D4 18107211 18107293
refGene_gn=RFK;refGene_tx=XM_011288290.3;refGene_gn=RFK;refGene_tx=XM_023243357.1
D4 18184114 18185401
refGene_gn=GCNT1;refGene_tx=XM_006939108.4;refGene_gn=GCNT1;refGene_tx=XM_011288277.3;refGene_gn=GCNT1;refGene_tx=XM_011288278.3;refGene_gn=GCNT1;refGene_tx=XM_011288279.3;refGene_gn=GCNT1;refGene_tx=XM_019815774.2;refGene_gn=GCNT1;refGene_tx=XM_019815775.2;refGene_gn=GCNT1;refGene_tx=XM_019815776.2;refGene_gn=GCNT1;refGene_tx=XM_019815777.2;refGene_gn=GCNT1;refGene_tx=XM_023243358.1;refGene_gn=GCNT1;refGene_tx=XM_023243359.1;refGene_gn=GCNT1;refGene_tx=XM_023243360.1
D4 18277509 18277540
refGene_gn=PRUNE2;refGene_tx=XM_019815778.1;refGene_gn=PRUNE2;refGene_tx=XM_019815779.2;refGene_gn=PRUNE2;refGene_tx=XM_019815780.2;refGene_gn=PRUNE2;refGene_tx=XM_019815781.1;refGene_gn=PRUNE2;refGene_tx=XM_019815782.1;refGene_gn=PRUNE2;refGene_tx=XM_019815784.1;refGene_gn=PRUNE2;refGene_tx=XM_019815785.1;refGene_gn=PRUNE2;refGene_tx=XM_019815787.2
D4 18281652 18281700

refGene_gn=PRUNE2;refGene_tx=XM_019815778.1;refGene_gn=PRUNE2;refGene_tx=XM_019815779.2;refGene_gn=PRUNE2;refGene_tx=XM_019815780.2;refGene_gn=PRUNE2;refGene_tx=XM_019815781.1;refGene_gn=PRUNE2;refGene_tx=XM_019815785.1;refGene_gn=PRUNE2;refGene_tx=XM_019815787.2
D4 18286296 18286335
refGene_gn=PRUNE2;refGene_tx=XM_019815779.2;refGene_gn=PRUNE2;refGene_tx=XM_019815785.1;refGene_gn=PRUNE2;refGene_tx=XM_019815787.2
D4 18286872 18286908
refGene_gn=PRUNE2;refGene_tx=XM_019815778.1;refGene_gn=PRUNE2;refGene_tx=XM_019815780.2;refGene_gn=PRUNE2;refGene_tx=XM_019815782.1
D4 18290143 18290242
refGene_gn=PRUNE2;refGene_tx=XM_019815778.1;refGene_gn=PRUNE2;refGene_tx=XM_019815779.2;refGene_gn=PRUNE2;refGene_tx=XM_019815780.2;refGene_gn=PRUNE2;refGene_tx=XM_019815781.1;refGene_gn=PRUNE2;refGene_tx=XM_019815782.1;refGene_gn=PRUNE2;refGene_tx=XM_019815784.1;refGene_gn=PRUNE2;refGene_tx=XM_019815785.1;refGene_gn=PRUNE2;refGene_tx=XM_019815787.2
D4 18295704 18295791
refGene_gn=PRUNE2;refGene_tx=XM_019815778.1;refGene_gn=PRUNE2;refGene_tx=XM_019815779.2;refGene_gn=PRUNE2;refGene_tx=XM_019815780.2;refGene_gn=PRUNE2;refGene_tx=XM_019815781.1;refGene_gn=PRUNE2;refGene_tx=XM_019815782.1;refGene_gn=PRUNE2;refGene_tx=XM_019815784.1;refGene_gn=PRUNE2;refGene_tx=XM_019815785.1;refGene_gn=PRUNE2;refGene_tx=XM_019815787.2
D4 18297405 18297537
refGene_gn=PRUNE2;refGene_tx=XM_019815778.1;refGene_gn=PRUNE2;refGene_tx=XM_019815779.2;refGene_gn=PRUNE2;refGene_tx=XM_019815780.2;refGene_gn=PRUNE2;refGene_tx=XM_019815781.1;refGene_gn=PRUNE2;refGene_tx=XM_019815782.1;refGene_gn=PRUNE2;refGene_tx=XM_019815784.1;refGene_gn=PRUNE2;refGene_tx=XM_019815785.1;refGene_gn=PRUNE2;refGene_tx=XM_019815787.2
D4 18298166 18298269
refGene_gn=PRUNE2;refGene_tx=XM_019815778.1;refGene_gn=PRUNE2;refGene_tx=XM_019815779.2;refGene_gn=PRUNE2;refGene_tx=XM_019815780.2;refGene_gn=PRUNE2;refGene_tx=XM_019815781.1;refGene_gn=PRUNE2;refGene_tx=XM_019815782.1;refGene_gn=PRUNE2;refGene_tx=XM_019815784.1;refGene_gn=PRUNE2;refGene_tx=XM_019815785.1;refGene_gn=PRUNE2;refGene_tx=XM_019815787.2
D4 18301095 18301104
refGene_gn=PRUNE2;refGene_tx=XM_019815778.1;refGene_gn=PRUNE2;refGene_tx=XM_019815781.1;refGene_gn=PRUNE2;refGene_tx=XM_019815782.1;refGene_gn=PRUNE2;refGene_tx=XM_019815784.1;refGene_gn=PRUNE2;refGene_tx=XM_019815785.1;refGene_gn=PRUNE2;refGene_tx=XM_019815787.2
D4 18304129 18304303
refGene_gn=PRUNE2;refGene_tx=XM_019815778.1;refGene_gn=PRUNE2;refGene_tx=XM_019815779.2;refGene_gn=PRUNE2;refGene_tx=XM_019815780.2;refGene_gn=PRUNE2;refGene_tx=XM_019815781.1;refGene_gn=PRUNE2;refGene_tx=XM_019815782.1;refGene_gn=PRUNE2;refGene_tx=XM_019815784.1;refGene_gn=PRUNE2;refGene_tx=XM_019815785.1;refGene_gn=PRUNE2;refGene_tx=XM_019815787.2

D4 18313530 18313734
refGene_gn=PRUNE2;refGene_tx=XM_019815778.1;refGene_gn=PRUNE2;refGene_tx=XM_019815779.2;refGene_gn=PRUNE2;refGene_tx=XM_019815780.2;refGene_gn=PRUNE2;refGene_tx=XM_019815781.1;refGene_gn=PRUNE2;refGene_tx=XM_019815782.1;refGene_gn=PRUNE2;refGene_tx=XM_019815784.1;refGene_gn=PRUNE2;refGene_tx=XM_019815785.1;refGene_gn=PRUNE2;refGene_tx=XM_019815787.2
D4 18315339 18315419
refGene_gn=PRUNE2;refGene_tx=XM_019815778.1;refGene_gn=PRUNE2;refGene_tx=XM_019815779.2;refGene_gn=PRUNE2;refGene_tx=XM_019815780.2;refGene_gn=PRUNE2;refGene_tx=XM_019815781.1;refGene_gn=PRUNE2;refGene_tx=XM_019815782.1;refGene_gn=PRUNE2;refGene_tx=XM_019815784.1;refGene_gn=PRUNE2;refGene_tx=XM_019815785.1;refGene_gn=PRUNE2;refGene_tx=XM_019815787.2
D4 18353258 18354024
refGene_gn=PRUNE2;refGene_tx=XM_019815778.1;refGene_gn=PRUNE2;refGene_tx=XM_019815779.2;refGene_gn=PRUNE2;refGene_tx=XM_019815780.2;refGene_gn=PRUNE2;refGene_tx=XM_019815781.1;refGene_gn=PRUNE2;refGene_tx=XM_019815782.1;refGene_gn=PRUNE2;refGene_tx=XM_019815784.1;refGene_gn=PRUNE2;refGene_tx=XM_019815785.1;refGene_gn=PRUNE2;refGene_tx=XM_019815787.2
D4 18354366 18360934
refGene_gn=PRUNE2;refGene_tx=XM_019815778.1;refGene_gn=PRUNE2;refGene_tx=XM_019815779.2;refGene_gn=PRUNE2;refGene_tx=XM_019815780.2;refGene_gn=PRUNE2;refGene_tx=XM_019815781.1;refGene_gn=PRUNE2;refGene_tx=XM_019815782.1;refGene_gn=PRUNE2;refGene_tx=XM_019815784.1;refGene_gn=PRUNE2;refGene_tx=XM_019815785.1;refGene_gn=PRUNE2;refGene_tx=XM_019815787.2
D4 18364080 18364239
refGene_gn=PRUNE2;refGene_tx=XM_019815778.1;refGene_gn=PRUNE2;refGene_tx=XM_019815779.2;refGene_gn=PRUNE2;refGene_tx=XM_019815780.2;refGene_gn=PRUNE2;refGene_tx=XM_019815781.1;refGene_gn=PRUNE2;refGene_tx=XM_019815782.1;refGene_gn=PRUNE2;refGene_tx=XM_019815784.1;refGene_gn=PRUNE2;refGene_tx=XM_019815785.1;refGene_gn=PRUNE2;refGene_tx=XM_019815787.2
D4 18461474 18461569
refGene_gn=PRUNE2;refGene_tx=XM_019815778.1;refGene_gn=PRUNE2;refGene_tx=XM_019815779.2;refGene_gn=PRUNE2;refGene_tx=XM_019815780.2;refGene_gn=PRUNE2;refGene_tx=XM_019815781.1;refGene_gn=PRUNE2;refGene_tx=XM_019815782.1;refGene_gn=PRUNE2;refGene_tx=XM_019815784.1;refGene_gn=PRUNE2;refGene_tx=XM_019815785.1;refGene_gn=PRUNE2;refGene_tx=XM_019815787.2
D4 18464124 18464277
refGene_gn=PRUNE2;refGene_tx=XM_019815778.1;refGene_gn=PRUNE2;refGene_tx=XM_019815779.2;refGene_gn=PRUNE2;refGene_tx=XM_019815780.2;refGene_gn=PRUNE2;refGene_tx=XM_019815781.1;refGene_gn=PRUNE2;refGene_tx=XM_019815782.1;refGene_gn=PRUNE2;refGene_tx=XM_019815784.1;refGene_gn=PRUNE2;refGene_tx=XM_019815785.1;refGene_gn=PRUNE2;refGene_tx=XM_019815787.2
D4 18480454 18480618

refGene_gn=PRUNE2;refGene_tx=XM_019815778.1;refGene_gn=PRUNE2;refGene_tx=XM_019815779.2;refGene_gn=PRUNE2;refGene_tx=XM_019815780.2;refGene_gn=PRUNE2;refGene_tx=XM_019815781.1;refGene_gn=PRUNE2;refGene_tx=XM_019815782.1;refGene_gn=PRUNE2;refGene_tx=XM_019815784.1;refGene_gn=PRUNE2;refGene_tx=XM_019815785.1;refGene_gn=PRUNE2;refGene_tx=XM_019815787.2
D4 18483802 18484005
refGene_gn=PRUNE2;refGene_tx=XM_019815778.1;refGene_gn=PRUNE2;refGene_tx=XM_019815779.2;refGene_gn=PRUNE2;refGene_tx=XM_019815780.2;refGene_gn=PRUNE2;refGene_tx=XM_019815781.1;refGene_gn=PRUNE2;refGene_tx=XM_019815782.1;refGene_gn=PRUNE2;refGene_tx=XM_019815784.1;refGene_gn=PRUNE2;refGene_tx=XM_019815785.1;refGene_gn=PRUNE2;refGene_tx=XM_019815787.2
D4 18488462 18488567
refGene_gn=PRUNE2;refGene_tx=XM_019815778.1;refGene_gn=PRUNE2;refGene_tx=XM_019815779.2;refGene_gn=PRUNE2;refGene_tx=XM_019815780.2;refGene_gn=PRUNE2;refGene_tx=XM_019815781.1;refGene_gn=PRUNE2;refGene_tx=XM_019815782.1;refGene_gn=PRUNE2;refGene_tx=XM_019815784.1;refGene_gn=PRUNE2;refGene_tx=XM_019815785.1;refGene_gn=PRUNE2;refGene_tx=XM_019815787.2
D4 18550226 18550262
refGene_gn=PRUNE2;refGene_tx=XM_019815778.1;refGene_gn=PRUNE2;refGene_tx=XM_019815779.2;refGene_gn=PRUNE2;refGene_tx=XM_019815780.2;refGene_gn=PRUNE2;refGene_tx=XM_019815781.1;refGene_gn=PRUNE2;refGene_tx=XM_019815782.1;refGene_gn=PRUNE2;refGene_tx=XM_019815784.1;refGene_gn=PRUNE2;refGene_tx=XM_019815787.2
D4 18650116 18651412 refGene_gn=FOXB2;refGene_tx=XM_023242453.1
D4 18772743 18772843
refGene_gn=VPS13A;refGene_tx=XM_023243366.1;refGene_gn=VPS13A;refGene_tx=XM_023243367.1;refGene_gn=VPS13A;refGene_tx=XM_023243368.1
D4 18785771 18785815
refGene_gn=VPS13A;refGene_tx=XM_023243366.1;refGene_gn=VPS13A;refGene_tx=XM_023243367.1;refGene_gn=VPS13A;refGene_tx=XM_023243368.1
D4 18787840 18787883
refGene_gn=VPS13A;refGene_tx=XM_023243366.1;refGene_gn=VPS13A;refGene_tx=XM_023243367.1;refGene_gn=VPS13A;refGene_tx=XM_023243368.1
D4 18790355 18790451
refGene_gn=VPS13A;refGene_tx=XM_023243366.1;refGene_gn=VPS13A;refGene_tx=XM_023243367.1;refGene_gn=VPS13A;refGene_tx=XM_023243368.1
D4 18790719 18790821
refGene_gn=VPS13A;refGene_tx=XM_023243366.1;refGene_gn=VPS13A;refGene_tx=XM_023243367.1;refGene_gn=VPS13A;refGene_tx=XM_023243368.1
D4 18804246 18804356
refGene_gn=VPS13A;refGene_tx=XM_023243366.1;refGene_gn=VPS13A;refGene_tx=XM_023243367.1;refGene_gn=VPS13A;refGene_tx=XM_023243368.1
D4 18805031 18805091
refGene_gn=VPS13A;refGene_tx=XM_023243366.1;refGene_gn=VPS13A;refGene_tx=XM_023243367.1;refGene_gn=VPS13A;refGene_tx=XM_023243368.1
D4 18807016 18807076
refGene_gn=VPS13A;refGene_tx=XM_023243366.1;refGene_gn=VPS13A;refGene_

tx=XM_023243367.1;refGene_gn=VPS13A;refGene_tx=XM_023243368.1
D4 18807255 18807336
refGene_gn=VPS13A;refGene_tx=XM_023243366.1;refGene_gn=VPS13A;refGene_tx=XM_023243367.1;refGene_gn=VPS13A;refGene_tx=XM_023243368.1
D4 18811025 18811083
refGene_gn=VPS13A;refGene_tx=XM_023243366.1;refGene_gn=VPS13A;refGene_tx=XM_023243367.1;refGene_gn=VPS13A;refGene_tx=XM_023243368.1
D4 18815557 18815685
refGene_gn=VPS13A;refGene_tx=XM_023243366.1;refGene_gn=VPS13A;refGene_tx=XM_023243367.1;refGene_gn=VPS13A;refGene_tx=XM_023243368.1
D4 18815888 18815995
refGene_gn=VPS13A;refGene_tx=XM_023243366.1;refGene_gn=VPS13A;refGene_tx=XM_023243367.1;refGene_gn=VPS13A;refGene_tx=XM_023243368.1
D4 18818110 18818282
refGene_gn=VPS13A;refGene_tx=XM_023243366.1;refGene_gn=VPS13A;refGene_tx=XM_023243367.1;refGene_gn=VPS13A;refGene_tx=XM_023243368.1
D4 18820203 18820266
refGene_gn=VPS13A;refGene_tx=XM_023243366.1;refGene_gn=VPS13A;refGene_tx=XM_023243367.1;refGene_gn=VPS13A;refGene_tx=XM_023243368.1
D4 18820748 18820878
refGene_gn=VPS13A;refGene_tx=XM_023243366.1;refGene_gn=VPS13A;refGene_tx=XM_023243367.1;refGene_gn=VPS13A;refGene_tx=XM_023243368.1
D4 18821943 18822045
refGene_gn=VPS13A;refGene_tx=XM_023243363.1;refGene_gn=VPS13A;refGene_tx=XM_023243364.1;refGene_gn=VPS13A;refGene_tx=XM_023243365.1;refGene_gn=VPS13A;refGene_tx=XM_023243366.1;refGene_gn=VPS13A;refGene_tx=XM_023243367.1;refGene_gn=VPS13A;refGene_tx=XM_023243368.1
D4 18822341 18822485
refGene_gn=VPS13A;refGene_tx=XM_023243363.1;refGene_gn=VPS13A;refGene_tx=XM_023243364.1;refGene_gn=VPS13A;refGene_tx=XM_023243365.1;refGene_gn=VPS13A;refGene_tx=XM_023243366.1;refGene_gn=VPS13A;refGene_tx=XM_023243367.1;refGene_gn=VPS13A;refGene_tx=XM_023243368.1
D4 18833511 18833701
refGene_gn=VPS13A;refGene_tx=XM_023243363.1;refGene_gn=VPS13A;refGene_tx=XM_023243364.1;refGene_gn=VPS13A;refGene_tx=XM_023243365.1;refGene_gn=VPS13A;refGene_tx=XM_023243366.1;refGene_gn=VPS13A;refGene_tx=XM_023243367.1;refGene_gn=VPS13A;refGene_tx=XM_023243368.1
D4 18833786 18833901
refGene_gn=VPS13A;refGene_tx=XM_023243363.1;refGene_gn=VPS13A;refGene_tx=XM_023243364.1;refGene_gn=VPS13A;refGene_tx=XM_023243365.1;refGene_gn=VPS13A;refGene_tx=XM_023243366.1;refGene_gn=VPS13A;refGene_tx=XM_023243367.1;refGene_gn=VPS13A;refGene_tx=XM_023243368.1
D4 18837466 18837603
refGene_gn=VPS13A;refGene_tx=XM_023243363.1;refGene_gn=VPS13A;refGene_tx=XM_023243364.1;refGene_gn=VPS13A;refGene_tx=XM_023243365.1;refGene_gn=VPS13A;refGene_tx=XM_023243366.1;refGene_gn=VPS13A;refGene_tx=XM_023243367.1;refGene_gn=VPS13A;refGene_tx=XM_023243368.1
D4 18839530 18839663
refGene_gn=VPS13A;refGene_tx=XM_023243363.1;refGene_gn=VPS13A;refGene_tx=XM_023243364.1;refGene_gn=VPS13A;refGene_tx=XM_023243365.1;refGene_

gn=VPS13A;refGene_tx=XM_023243366.1;refGene_gn=VPS13A;refGene_tx=XM_023243367.1;refGene_gn=VPS13A;refGene_tx=XM_023243368.1
D4 18842465 18842583
refGene_gn=VPS13A;refGene_tx=XM_023243363.1;refGene_gn=VPS13A;refGene_tx=XM_023243364.1;refGene_gn=VPS13A;refGene_tx=XM_023243365.1;refGene_gn=VPS13A;refGene_tx=XM_023243366.1;refGene_gn=VPS13A;refGene_tx=XM_023243367.1;refGene_gn=VPS13A;refGene_tx=XM_023243368.1
D4 18855158 18855297
refGene_gn=VPS13A;refGene_tx=XM_023243363.1;refGene_gn=VPS13A;refGene_tx=XM_023243364.1;refGene_gn=VPS13A;refGene_tx=XM_023243365.1;refGene_gn=VPS13A;refGene_tx=XM_023243366.1;refGene_gn=VPS13A;refGene_tx=XM_023243367.1;refGene_gn=VPS13A;refGene_tx=XM_023243368.1
D4 18873545 18873627
refGene_gn=VPS13A;refGene_tx=XM_023243363.1;refGene_gn=VPS13A;refGene_tx=XM_023243364.1;refGene_gn=VPS13A;refGene_tx=XM_023243365.1;refGene_gn=VPS13A;refGene_tx=XM_023243366.1;refGene_gn=VPS13A;refGene_tx=XM_023243367.1;refGene_gn=VPS13A;refGene_tx=XM_023243368.1
D4 18874872 18875027
refGene_gn=VPS13A;refGene_tx=XM_023243363.1;refGene_gn=VPS13A;refGene_tx=XM_023243364.1;refGene_gn=VPS13A;refGene_tx=XM_023243365.1;refGene_gn=VPS13A;refGene_tx=XM_023243366.1;refGene_gn=VPS13A;refGene_tx=XM_023243367.1;refGene_gn=VPS13A;refGene_tx=XM_023243368.1
D4 18875407 18875564
refGene_gn=VPS13A;refGene_tx=XM_023243363.1;refGene_gn=VPS13A;refGene_tx=XM_023243364.1;refGene_gn=VPS13A;refGene_tx=XM_023243365.1;refGene_gn=VPS13A;refGene_tx=XM_023243366.1;refGene_gn=VPS13A;refGene_tx=XM_023243367.1;refGene_gn=VPS13A;refGene_tx=XM_023243368.1
D4 18887128 18887208
refGene_gn=VPS13A;refGene_tx=XM_023243363.1;refGene_gn=VPS13A;refGene_tx=XM_023243364.1;refGene_gn=VPS13A;refGene_tx=XM_023243365.1;refGene_gn=VPS13A;refGene_tx=XM_023243366.1;refGene_gn=VPS13A;refGene_tx=XM_023243367.1;refGene_gn=VPS13A;refGene_tx=XM_023243368.1
D4 18887931 18887991
refGene_gn=VPS13A;refGene_tx=XM_023243363.1;refGene_gn=VPS13A;refGene_tx=XM_023243364.1;refGene_gn=VPS13A;refGene_tx=XM_023243365.1;refGene_gn=VPS13A;refGene_tx=XM_023243366.1;refGene_gn=VPS13A;refGene_tx=XM_023243367.1;refGene_gn=VPS13A;refGene_tx=XM_023243368.1
D4 18888441 18888595
refGene_gn=VPS13A;refGene_tx=XM_023243363.1;refGene_gn=VPS13A;refGene_tx=XM_023243364.1;refGene_gn=VPS13A;refGene_tx=XM_023243365.1;refGene_gn=VPS13A;refGene_tx=XM_023243366.1;refGene_gn=VPS13A;refGene_tx=XM_023243367.1;refGene_gn=VPS13A;refGene_tx=XM_023243368.1
D4 18889307 18889424
refGene_gn=VPS13A;refGene_tx=XM_023243363.1;refGene_gn=VPS13A;refGene_tx=XM_023243364.1;refGene_gn=VPS13A;refGene_tx=XM_023243365.1;refGene_gn=VPS13A;refGene_tx=XM_023243366.1;refGene_gn=VPS13A;refGene_tx=XM_023243367.1;refGene_gn=VPS13A;refGene_tx=XM_023243368.1
D4 18889499 18889603
refGene_gn=VPS13A;refGene_tx=XM_023243363.1;refGene_gn=VPS13A;refGene_tx=XM_023243364.1;refGene_gn=VPS13A;refGene_tx=XM_023243365.1;refGene_

gn=VPS13A;refGene_tx=XM_023243366.1;refGene_gn=VPS13A;refGene_tx=XM_02
3243367.1;refGene_gn=VPS13A;refGene_tx=XM_023243368.1
D4 18898751 18898919
refGene_gn=VPS13A;refGene_tx=XM_023243363.1;refGene_gn=VPS13A;refGene_
tx=XM_023243364.1;refGene_gn=VPS13A;refGene_tx=XM_023243365.1;refGene_
gn=VPS13A;refGene_tx=XM_023243366.1;refGene_gn=VPS13A;refGene_tx=XM_02
3243367.1;refGene_gn=VPS13A;refGene_tx=XM_023243368.1
D4 18901289 18901594
refGene_gn=VPS13A;refGene_tx=XM_023243363.1;refGene_gn=VPS13A;refGene_
tx=XM_023243364.1;refGene_gn=VPS13A;refGene_tx=XM_023243365.1;refGene_
gn=VPS13A;refGene_tx=XM_023243366.1;refGene_gn=VPS13A;refGene_tx=XM_02
3243367.1;refGene_gn=VPS13A;refGene_tx=XM_023243368.1
D4 18909353 18909501
refGene_gn=VPS13A;refGene_tx=XM_023243363.1;refGene_gn=VPS13A;refGene_
tx=XM_023243364.1;refGene_gn=VPS13A;refGene_tx=XM_023243365.1;refGene_
gn=VPS13A;refGene_tx=XM_023243366.1;refGene_gn=VPS13A;refGene_tx=XM_02
3243367.1;refGene_gn=VPS13A;refGene_tx=XM_023243368.1
D4 18912310 18912461
refGene_gn=VPS13A;refGene_tx=XM_023243363.1;refGene_gn=VPS13A;refGene_
tx=XM_023243364.1;refGene_gn=VPS13A;refGene_tx=XM_023243365.1;refGene_
gn=VPS13A;refGene_tx=XM_023243366.1;refGene_gn=VPS13A;refGene_tx=XM_02
3243367.1;refGene_gn=VPS13A;refGene_tx=XM_023243368.1
D4 18920144 18920272
refGene_gn=VPS13A;refGene_tx=XM_023243363.1;refGene_gn=VPS13A;refGene_
tx=XM_023243364.1;refGene_gn=VPS13A;refGene_tx=XM_023243365.1;refGene_
gn=VPS13A;refGene_tx=XM_023243366.1;refGene_gn=VPS13A;refGene_tx=XM_02
3243367.1;refGene_gn=VPS13A;refGene_tx=XM_023243368.1
D4 18920848 18921018
refGene_gn=VPS13A;refGene_tx=XM_023243363.1;refGene_gn=VPS13A;refGene_
tx=XM_023243364.1;refGene_gn=VPS13A;refGene_tx=XM_023243365.1;refGene_
gn=VPS13A;refGene_tx=XM_023243366.1;refGene_gn=VPS13A;refGene_tx=XM_02
3243367.1;refGene_gn=VPS13A;refGene_tx=XM_023243368.1
D4 18921547 18921765
refGene_gn=VPS13A;refGene_tx=XM_023243363.1;refGene_gn=VPS13A;refGene_
tx=XM_023243364.1;refGene_gn=VPS13A;refGene_tx=XM_023243365.1;refGene_
gn=VPS13A;refGene_tx=XM_023243366.1;refGene_gn=VPS13A;refGene_tx=XM_02
3243367.1;refGene_gn=VPS13A;refGene_tx=XM_023243368.1
D4 18922460 18922693
refGene_gn=VPS13A;refGene_tx=XM_023243363.1;refGene_gn=VPS13A;refGene_
tx=XM_023243364.1;refGene_gn=VPS13A;refGene_tx=XM_023243365.1;refGene_
gn=VPS13A;refGene_tx=XM_023243366.1;refGene_gn=VPS13A;refGene_tx=XM_02
3243367.1;refGene_gn=VPS13A;refGene_tx=XM_023243368.1
D4 18932840 18932933
refGene_gn=VPS13A;refGene_tx=XM_023243363.1;refGene_gn=VPS13A;refGene_
tx=XM_023243364.1;refGene_gn=VPS13A;refGene_tx=XM_023243365.1;refGene_
gn=VPS13A;refGene_tx=XM_023243366.1;refGene_gn=VPS13A;refGene_tx=XM_02
3243367.1;refGene_gn=VPS13A;refGene_tx=XM_023243368.1
D4 18933427 18933784
refGene_gn=VPS13A;refGene_tx=XM_023243363.1;refGene_gn=VPS13A;refGene_
tx=XM_023243364.1;refGene_gn=VPS13A;refGene_tx=XM_023243365.1;refGene_

gn=VPS13A;refGene_tx=XM_023243366.1;refGene_gn=VPS13A;refGene_tx=XM_023243367.1;refGene_gn=VPS13A;refGene_tx=XM_023243368.1
D4 18935109 18935211
refGene_gn=VPS13A;refGene_tx=XM_023243363.1;refGene_gn=VPS13A;refGene_tx=XM_023243364.1;refGene_gn=VPS13A;refGene_tx=XM_023243365.1;refGene_gn=VPS13A;refGene_tx=XM_023243366.1;refGene_gn=VPS13A;refGene_tx=XM_023243367.1;refGene_gn=VPS13A;refGene_tx=XM_023243368.1
D4 18936500 18936659
refGene_gn=VPS13A;refGene_tx=XM_023243363.1;refGene_gn=VPS13A;refGene_tx=XM_023243364.1;refGene_gn=VPS13A;refGene_tx=XM_023243365.1;refGene_gn=VPS13A;refGene_tx=XM_023243366.1;refGene_gn=VPS13A;refGene_tx=XM_023243367.1;refGene_gn=VPS13A;refGene_tx=XM_023243368.1
D4 18936821 18937077
refGene_gn=VPS13A;refGene_tx=XM_023243363.1;refGene_gn=VPS13A;refGene_tx=XM_023243364.1;refGene_gn=VPS13A;refGene_tx=XM_023243365.1;refGene_gn=VPS13A;refGene_tx=XM_023243366.1;refGene_gn=VPS13A;refGene_tx=XM_023243367.1;refGene_gn=VPS13A;refGene_tx=XM_023243368.1
D4 18938408 18938569
refGene_gn=VPS13A;refGene_tx=XM_023243363.1;refGene_gn=VPS13A;refGene_tx=XM_023243364.1;refGene_gn=VPS13A;refGene_tx=XM_023243365.1;refGene_gn=VPS13A;refGene_tx=XM_023243366.1;refGene_gn=VPS13A;refGene_tx=XM_023243367.1;refGene_gn=VPS13A;refGene_tx=XM_023243368.1
D4 18950178 18950282
refGene_gn=VPS13A;refGene_tx=XM_023243363.1;refGene_gn=VPS13A;refGene_tx=XM_023243364.1;refGene_gn=VPS13A;refGene_tx=XM_023243365.1;refGene_gn=VPS13A;refGene_tx=XM_023243366.1;refGene_gn=VPS13A;refGene_tx=XM_023243367.1;refGene_gn=VPS13A;refGene_tx=XM_023243368.1
D4 18953849 18954132
refGene_gn=VPS13A;refGene_tx=XM_023243363.1;refGene_gn=VPS13A;refGene_tx=XM_023243364.1;refGene_gn=VPS13A;refGene_tx=XM_023243365.1;refGene_gn=VPS13A;refGene_tx=XM_023243366.1;refGene_gn=VPS13A;refGene_tx=XM_023243367.1;refGene_gn=VPS13A;refGene_tx=XM_023243368.1
D4 18957479 18957875
refGene_gn=VPS13A;refGene_tx=XM_023243363.1;refGene_gn=VPS13A;refGene_tx=XM_023243364.1;refGene_gn=VPS13A;refGene_tx=XM_023243365.1;refGene_gn=VPS13A;refGene_tx=XM_023243366.1;refGene_gn=VPS13A;refGene_tx=XM_023243367.1;refGene_gn=VPS13A;refGene_tx=XM_023243368.1
D4 18958124 18958229
refGene_gn=VPS13A;refGene_tx=XM_023243363.1;refGene_gn=VPS13A;refGene_tx=XM_023243364.1;refGene_gn=VPS13A;refGene_tx=XM_023243365.1;refGene_gn=VPS13A;refGene_tx=XM_023243366.1;refGene_gn=VPS13A;refGene_tx=XM_023243367.1;refGene_gn=VPS13A;refGene_tx=XM_023243368.1
D4 18958349 18958496
refGene_gn=VPS13A;refGene_tx=XM_023243363.1;refGene_gn=VPS13A;refGene_tx=XM_023243364.1;refGene_gn=VPS13A;refGene_tx=XM_023243365.1;refGene_gn=VPS13A;refGene_tx=XM_023243366.1;refGene_gn=VPS13A;refGene_tx=XM_023243367.1;refGene_gn=VPS13A;refGene_tx=XM_023243368.1
D4 18963037 18963166
refGene_gn=VPS13A;refGene_tx=XM_023243363.1;refGene_gn=VPS13A;refGene_tx=XM_023243364.1;refGene_gn=VPS13A;refGene_tx=XM_023243365.1;refGene_

gn=VPS13A;refGene_tx=XM_023243366.1;refGene_gn=VPS13A;refGene_tx=XM_023243367.1;refGene_gn=VPS13A;refGene_tx=XM_023243368.1
D4 18963575 18963709
refGene_gn=VPS13A;refGene_tx=XM_023243363.1;refGene_gn=VPS13A;refGene_tx=XM_023243364.1;refGene_gn=VPS13A;refGene_tx=XM_023243365.1;refGene_gn=VPS13A;refGene_tx=XM_023243366.1;refGene_gn=VPS13A;refGene_tx=XM_023243367.1;refGene_gn=VPS13A;refGene_tx=XM_023243368.1
D4 18969060 18969190
refGene_gn=VPS13A;refGene_tx=XM_023243363.1;refGene_gn=VPS13A;refGene_tx=XM_023243364.1;refGene_gn=VPS13A;refGene_tx=XM_023243365.1;refGene_gn=VPS13A;refGene_tx=XM_023243366.1;refGene_gn=VPS13A;refGene_tx=XM_023243367.1;refGene_gn=VPS13A;refGene_tx=XM_023243368.1
D4 18977108 18977341
refGene_gn=VPS13A;refGene_tx=XM_023243363.1;refGene_gn=VPS13A;refGene_tx=XM_023243364.1;refGene_gn=VPS13A;refGene_tx=XM_023243365.1;refGene_gn=VPS13A;refGene_tx=XM_023243366.1;refGene_gn=VPS13A;refGene_tx=XM_023243367.1;refGene_gn=VPS13A;refGene_tx=XM_023243368.1
D4 18980612 18980766
refGene_gn=VPS13A;refGene_tx=XM_023243363.1;refGene_gn=VPS13A;refGene_tx=XM_023243364.1;refGene_gn=VPS13A;refGene_tx=XM_023243365.1;refGene_gn=VPS13A;refGene_tx=XM_023243366.1;refGene_gn=VPS13A;refGene_tx=XM_023243367.1;refGene_gn=VPS13A;refGene_tx=XM_023243368.1
D4 18981375 18981522
refGene_gn=VPS13A;refGene_tx=XM_023243363.1;refGene_gn=VPS13A;refGene_tx=XM_023243364.1;refGene_gn=VPS13A;refGene_tx=XM_023243365.1;refGene_gn=VPS13A;refGene_tx=XM_023243366.1;refGene_gn=VPS13A;refGene_tx=XM_023243367.1;refGene_gn=VPS13A;refGene_tx=XM_023243368.1
D4 18981724 18981806
refGene_gn=VPS13A;refGene_tx=XM_023243363.1;refGene_gn=VPS13A;refGene_tx=XM_023243364.1;refGene_gn=VPS13A;refGene_tx=XM_023243365.1;refGene_gn=VPS13A;refGene_tx=XM_023243366.1;refGene_gn=VPS13A;refGene_tx=XM_023243367.1;refGene_gn=VPS13A;refGene_tx=XM_023243368.1
D4 18982674 18982744
refGene_gn=VPS13A;refGene_tx=XM_023243363.1;refGene_gn=VPS13A;refGene_tx=XM_023243364.1;refGene_gn=VPS13A;refGene_tx=XM_023243365.1;refGene_gn=VPS13A;refGene_tx=XM_023243366.1;refGene_gn=VPS13A;refGene_tx=XM_023243367.1;refGene_gn=VPS13A;refGene_tx=XM_023243368.1
D4 18983552 18983658
refGene_gn=VPS13A;refGene_tx=XM_023243363.1;refGene_gn=VPS13A;refGene_tx=XM_023243364.1;refGene_gn=VPS13A;refGene_tx=XM_023243365.1;refGene_gn=VPS13A;refGene_tx=XM_023243366.1;refGene_gn=VPS13A;refGene_tx=XM_023243367.1;refGene_gn=VPS13A;refGene_tx=XM_023243368.1
D4 18989959 18990073
refGene_gn=VPS13A;refGene_tx=XM_023243363.1;refGene_gn=VPS13A;refGene_tx=XM_023243364.1;refGene_gn=VPS13A;refGene_tx=XM_023243365.1;refGene_gn=VPS13A;refGene_tx=XM_023243366.1;refGene_gn=VPS13A;refGene_tx=XM_023243367.1;refGene_gn=VPS13A;refGene_tx=XM_023243368.1
D4 18991157 18991303
refGene_gn=VPS13A;refGene_tx=XM_023243363.1;refGene_gn=VPS13A;refGene_tx=XM_023243364.1;refGene_gn=VPS13A;refGene_tx=XM_023243365.1;refGene_

gn=VPS13A;refGene_tx=XM_023243366.1;refGene_gn=VPS13A;refGene_tx=XM_023243367.1;refGene_gn=VPS13A;refGene_tx=XM_023243368.1
D4 18992497 18992579
refGene_gn=VPS13A;refGene_tx=XM_023243363.1;refGene_gn=VPS13A;refGene_tx=XM_023243364.1;refGene_gn=VPS13A;refGene_tx=XM_023243365.1;refGene_gn=VPS13A;refGene_tx=XM_023243366.1;refGene_gn=VPS13A;refGene_tx=XM_023243367.1;refGene_gn=VPS13A;refGene_tx=XM_023243368.1
D4 18993411 18993525
refGene_gn=VPS13A;refGene_tx=XM_023243363.1;refGene_gn=VPS13A;refGene_tx=XM_023243364.1;refGene_gn=VPS13A;refGene_tx=XM_023243365.1;refGene_gn=VPS13A;refGene_tx=XM_023243366.1;refGene_gn=VPS13A;refGene_tx=XM_023243367.1;refGene_gn=VPS13A;refGene_tx=XM_023243368.1
D4 18994345 18994421
refGene_gn=VPS13A;refGene_tx=XM_023243363.1;refGene_gn=VPS13A;refGene_tx=XM_023243364.1;refGene_gn=VPS13A;refGene_tx=XM_023243365.1;refGene_gn=VPS13A;refGene_tx=XM_023243366.1;refGene_gn=VPS13A;refGene_tx=XM_023243367.1;refGene_gn=VPS13A;refGene_tx=XM_023243368.1
D4 18994506 18994670
refGene_gn=VPS13A;refGene_tx=XM_023243363.1;refGene_gn=VPS13A;refGene_tx=XM_023243364.1;refGene_gn=VPS13A;refGene_tx=XM_023243365.1;refGene_gn=VPS13A;refGene_tx=XM_023243366.1;refGene_gn=VPS13A;refGene_tx=XM_023243367.1;refGene_gn=VPS13A;refGene_tx=XM_023243368.1
D4 18995207 18995253
refGene_gn=VPS13A;refGene_tx=XM_023243363.1;refGene_gn=VPS13A;refGene_tx=XM_023243364.1;refGene_gn=VPS13A;refGene_tx=XM_023243365.1;refGene_gn=VPS13A;refGene_tx=XM_023243366.1;refGene_gn=VPS13A;refGene_tx=XM_023243367.1;refGene_gn=VPS13A;refGene_tx=XM_023243368.1
D4 18995358 18995482
refGene_gn=VPS13A;refGene_tx=XM_023243363.1;refGene_gn=VPS13A;refGene_tx=XM_023243364.1;refGene_gn=VPS13A;refGene_tx=XM_023243365.1;refGene_gn=VPS13A;refGene_tx=XM_023243366.1;refGene_gn=VPS13A;refGene_tx=XM_023243367.1;refGene_gn=VPS13A;refGene_tx=XM_023243368.1
D4 19004867 19004982
refGene_gn=VPS13A;refGene_tx=XM_023243363.1;refGene_gn=VPS13A;refGene_tx=XM_023243364.1;refGene_gn=VPS13A;refGene_tx=XM_023243365.1;refGene_gn=VPS13A;refGene_tx=XM_023243366.1;refGene_gn=VPS13A;refGene_tx=XM_023243367.1;refGene_gn=VPS13A;refGene_tx=XM_023243368.1
D4 19005181 19005202
refGene_gn=VPS13A;refGene_tx=XM_023243365.1;refGene_gn=VPS13A;refGene_tx=XM_023243368.1
D4 19007552 19007645
refGene_gn=VPS13A;refGene_tx=XM_023243364.1;refGene_gn=VPS13A;refGene_tx=XM_023243367.1
D4 19021065 19021151
refGene_gn=VPS13A;refGene_tx=XM_023243363.1;refGene_gn=VPS13A;refGene_tx=XM_023243366.1
D4 19023799 19023923
refGene_gn=VPS13A;refGene_tx=XM_023243363.1;refGene_gn=VPS13A;refGene_tx=XM_023243366.1
D4 19026388 19026463

refGene_gn=VPS13A;refGene_tx=XM_023243363.1;refGene_gn=VPS13A;refGene_tx=XM_023243366.1
D4 19026557 19026608
refGene_gn=VPS13A;refGene_tx=XM_023243363.1;refGene_gn=VPS13A;refGene_tx=XM_023243366.1
D4 19050799 19050990 refGene_gn=GNA14;refGene_tx=XM_003995421.5
D4 19051701 19051844 refGene_gn=GNA14;refGene_tx=XM_019815788.2
D4 19051951 19052105
refGene_gn=GNA14;refGene_tx=XM_003995421.5;refGene_gn=GNA14;refGene_tx=XM_019815788.2
D4 19055704 19055834
refGene_gn=GNA14;refGene_tx=XM_003995421.5;refGene_gn=GNA14;refGene_tx=XM_019815788.2
D4 19057957 19058086
refGene_gn=GNA14;refGene_tx=XM_003995421.5;refGene_gn=GNA14;refGene_tx=XM_019815788.2
D4 19060854 19061009
refGene_gn=GNA14;refGene_tx=XM_003995421.5;refGene_gn=GNA14;refGene_tx=XM_019815788.2
D4 19140295 19140480
refGene_gn=GNA14;refGene_tx=XM_003995421.5;refGene_gn=GNA14;refGene_tx=XM_019815788.2
D4 19257150 19257274
refGene_gn=GNA14;refGene_tx=XM_003995421.5;refGene_gn=GNA14;refGene_tx=XM_019815788.2
D4 19327664 19327855
refGene_gn=GNAQ;refGene_tx=XM_019815794.2;refGene_gn=GNAQ;refGene_tx=XM_023243369.1;refGene_gn=GNAQ;refGene_tx=XM_023243370.1
D4 19341484 19341638
refGene_gn=GNAQ;refGene_tx=XM_019815794.2;refGene_gn=GNAQ;refGene_tx=XM_023243369.1;refGene_gn=GNAQ;refGene_tx=XM_023243370.1
D4 19405992 19406122
refGene_gn=GNAQ;refGene_tx=XM_019815794.2;refGene_gn=GNAQ;refGene_tx=XM_023243369.1;refGene_gn=GNAQ;refGene_tx=XM_023243370.1
D4 19408665 19408794
refGene_gn=GNAQ;refGene_tx=XM_019815794.2;refGene_gn=GNAQ;refGene_tx=XM_023243369.1;refGene_gn=GNAQ;refGene_tx=XM_023243370.1
D4 19426550 19426705
refGene_gn=GNAQ;refGene_tx=XM_019815794.2;refGene_gn=GNAQ;refGene_tx=XM_023243369.1;refGene_gn=GNAQ;refGene_tx=XM_023243370.1
D4 19533418 19533603
refGene_gn=GNAQ;refGene_tx=XM_019815794.2;refGene_gn=GNAQ;refGene_tx=XM_023243369.1;refGene_gn=GNAQ;refGene_tx=XM_023243370.1
D4 19639925 19639935 refGene_gn=GNAQ;refGene_tx=XM_023243370.1
D4 19640336 19640361 refGene_gn=GNAQ;refGene_tx=XM_019815794.2
D4 19640756 19640892 refGene_gn=GNAQ;refGene_tx=XM_023243369.1
D4 19806287 19806540 refGene_gn=CEP78;refGene_tx=XM_023243372.1
D4 19809778 19809951 refGene_gn=CEP78;refGene_tx=XM_023243372.1
D4 19810046 19810119 refGene_gn=CEP78;refGene_tx=XM_023243372.1
D4 19811408 19811512 refGene_gn=CEP78;refGene_tx=XM_023243372.1

D4 19812942 19813117 refGene_gn=CEP78;refGene_tx=XM_023243372.1
D4 19815066 19815180 refGene_gn=CEP78;refGene_tx=XM_023243372.1
D4 19816722 19816787 refGene_gn=CEP78;refGene_tx=XM_023243372.1
D4 19817070 19817182 refGene_gn=CEP78;refGene_tx=XM_023243372.1
D4 19820006 19820142 refGene_gn=CEP78;refGene_tx=XM_023243372.1
D4 19821610 19821656 refGene_gn=CEP78;refGene_tx=XM_023243372.1
D4 19823277 19823406 refGene_gn=CEP78;refGene_tx=XM_023243372.1
D4 19830646 19830724 refGene_gn=CEP78;refGene_tx=XM_023243372.1
D4 19831917 19831935 refGene_gn=CEP78;refGene_tx=XM_023243372.1
D4 19831936 19832085 refGene_gn=CEP78;refGene_tx=XM_023243372.1
D4 19833350 19833522 refGene_gn=CEP78;refGene_tx=XM_023243372.1
D4 19833842 19833890 refGene_gn=CEP78;refGene_tx=XM_023243372.1
D4 19834656 19835220 refGene_gn=CEP78;refGene_tx=XM_023243372.1
D4 19868490 19868550 refGene_gn=PSAT1;refGene_tx=XM_003995426.4
D4 19873810 19873871 refGene_gn=PSAT1;refGene_tx=XM_003995426.4
D4 19874832 19874902 refGene_gn=PSAT1;refGene_tx=XM_003995426.4
D4 19877433 19877639 refGene_gn=PSAT1;refGene_tx=XM_003995426.4
D4 19878451 19878624 refGene_gn=PSAT1;refGene_tx=XM_003995426.4
D4 19880830 19881000 refGene_gn=PSAT1;refGene_tx=XM_003995426.4
D4 19888400 19888529 refGene_gn=PSAT1;refGene_tx=XM_003995426.4
D4 19898859 19898997 refGene_gn=PSAT1;refGene_tx=XM_003995426.4
D4 19899943 19900049 refGene_gn=PSAT1;refGene_tx=XM_003995426.4
D4 20183953 20184274
refGene_gn=LOC101095924;refGene_tx=XM_011288275.2
D4 21039422 21039467
refGene_gn=TLE4;refGene_tx=XM_023243373.1;refGene_gn=TLE4;refGene_tx=XM_023243374.1;refGene_gn=TLE4;refGene_tx=XM_023243378.1;refGene_gn=TLE4;refGene_tx=XM_023243379.1;refGene_gn=TLE4;refGene_tx=XM_023243381.1;refGene_gn=TLE4;refGene_tx=XM_023243382.1;refGene_gn=TLE4;refGene_tx=XM_023243385.1
D4 21039969 21040008
refGene_gn=TLE4;refGene_tx=XM_023243376.1;refGene_gn=TLE4;refGene_tx=XM_023243377.1;refGene_gn=TLE4;refGene_tx=XM_023243380.1;refGene_gn=TLE4;refGene_tx=XM_023243383.1;refGene_gn=TLE4;refGene_tx=XM_023243384.1
D4 21040327 21040425
refGene_gn=TLE4;refGene_tx=XM_023243373.1;refGene_gn=TLE4;refGene_tx=XM_023243374.1;refGene_gn=TLE4;refGene_tx=XM_023243376.1;refGene_gn=TLE4;refGene_tx=XM_023243377.1;refGene_gn=TLE4;refGene_tx=XM_023243378.1;refGene_gn=TLE4;refGene_tx=XM_023243379.1;refGene_gn=TLE4;refGene_tx=XM_023243380.1;refGene_gn=TLE4;refGene_tx=XM_023243381.1;refGene_gn=TLE4;refGene_tx=XM_023243382.1;refGene_gn=TLE4;refGene_tx=XM_023243383.1;refGene_gn=TLE4;refGene_tx=XM_023243384.1;refGene_gn=TLE4;refGene_tx=XM_023243385.1
D4 21041522 21041586
refGene_gn=TLE4;refGene_tx=XM_023243373.1;refGene_gn=TLE4;refGene_tx=XM_023243374.1;refGene_gn=TLE4;refGene_tx=XM_023243376.1;refGene_gn=TLE4;refGene_tx=XM_023243377.1;refGene_gn=TLE4;refGene_tx=XM_023243378.1;refGene_gn=TLE4;refGene_tx=XM_023243379.1;refGene_gn=TLE4;refGene_tx=XM_023243380.1;refGene_gn=TLE4;refGene_tx=XM_023243381.1;refGene_gn=TLE4;refGene_tx=XM_023243382.1;refGene_gn=TLE4;refGene_tx=XM_023243383.1;

refGene_gn=TLE4;refGene_tx=XM_023243384.1;refGene_gn=TLE4;refGene_tx=XM_023243385.1
D4 21042789 21042834
refGene_gn=TLE4;refGene_tx=XM_023243373.1;refGene_gn=TLE4;refGene_tx=XM_023243374.1;refGene_gn=TLE4;refGene_tx=XM_023243376.1;refGene_gn=TLE4;refGene_tx=XM_023243377.1;refGene_gn=TLE4;refGene_tx=XM_023243378.1;refGene_gn=TLE4;refGene_tx=XM_023243379.1;refGene_gn=TLE4;refGene_tx=XM_023243380.1;refGene_gn=TLE4;refGene_tx=XM_023243381.1;refGene_gn=TLE4;refGene_tx=XM_023243382.1;refGene_gn=TLE4;refGene_tx=XM_023243383.1;refGene_gn=TLE4;refGene_tx=XM_023243384.1;refGene_gn=TLE4;refGene_tx=XM_023243385.1
D4 21073426 21073489
refGene_gn=TLE4;refGene_tx=XM_023243373.1;refGene_gn=TLE4;refGene_tx=XM_023243374.1;refGene_gn=TLE4;refGene_tx=XM_023243376.1;refGene_gn=TLE4;refGene_tx=XM_023243377.1;refGene_gn=TLE4;refGene_tx=XM_023243381.1;refGene_gn=TLE4;refGene_tx=XM_023243382.1;refGene_gn=TLE4;refGene_tx=XM_023243383.1;refGene_gn=TLE4;refGene_tx=XM_023243384.1
D4 21085197 21085272
refGene_gn=TLE4;refGene_tx=XM_023243373.1;refGene_gn=TLE4;refGene_tx=XM_023243374.1;refGene_gn=TLE4;refGene_tx=XM_023243376.1;refGene_gn=TLE4;refGene_tx=XM_023243377.1;refGene_gn=TLE4;refGene_tx=XM_023243378.1;refGene_gn=TLE4;refGene_tx=XM_023243379.1;refGene_gn=TLE4;refGene_tx=XM_023243380.1
D4 21108932 21109137
refGene_gn=TLE4;refGene_tx=XM_023243373.1;refGene_gn=TLE4;refGene_tx=XM_023243374.1;refGene_gn=TLE4;refGene_tx=XM_023243376.1;refGene_gn=TLE4;refGene_tx=XM_023243377.1;refGene_gn=TLE4;refGene_tx=XM_023243378.1;refGene_gn=TLE4;refGene_tx=XM_023243379.1;refGene_gn=TLE4;refGene_tx=XM_023243380.1;refGene_gn=TLE4;refGene_tx=XM_023243381.1;refGene_gn=TLE4;refGene_tx=XM_023243382.1;refGene_gn=TLE4;refGene_tx=XM_023243383.1;refGene_gn=TLE4;refGene_tx=XM_023243384.1;refGene_gn=TLE4;refGene_tx=XM_023243385.1
D4 21110399 21110416
refGene_gn=TLE4;refGene_tx=XM_023243373.1;refGene_gn=TLE4;refGene_tx=XM_023243374.1;refGene_gn=TLE4;refGene_tx=XM_023243376.1;refGene_gn=TLE4;refGene_tx=XM_023243377.1;refGene_gn=TLE4;refGene_tx=XM_023243378.1;refGene_gn=TLE4;refGene_tx=XM_023243379.1;refGene_gn=TLE4;refGene_tx=XM_023243380.1;refGene_gn=TLE4;refGene_tx=XM_023243381.1;refGene_gn=TLE4;refGene_tx=XM_023243382.1;refGene_gn=TLE4;refGene_tx=XM_023243383.1;refGene_gn=TLE4;refGene_tx=XM_023243384.1;refGene_gn=TLE4;refGene_tx=XM_023243385.1
D4 21156898 21157018
refGene_gn=TLE4;refGene_tx=XM_023243373.1;refGene_gn=TLE4;refGene_tx=XM_023243374.1;refGene_gn=TLE4;refGene_tx=XM_023243376.1;refGene_gn=TLE4;refGene_tx=XM_023243377.1;refGene_gn=TLE4;refGene_tx=XM_023243378.1;refGene_gn=TLE4;refGene_tx=XM_023243379.1;refGene_gn=TLE4;refGene_tx=XM_023243380.1;refGene_gn=TLE4;refGene_tx=XM_023243381.1;refGene_gn=TLE4;refGene_tx=XM_023243382.1;refGene_gn=TLE4;refGene_tx=XM_023243383.1;refGene_gn=TLE4;refGene_tx=XM_023243384.1;refGene_gn=TLE4;refGene_tx=XM_023243385.1

D4 21158663 21158717
refGene_gn=TLE4;refGene_tx=XM_023243373.1;refGene_gn=TLE4;refGene_tx=XM_023243374.1;refGene_gn=TLE4;refGene_tx=XM_023243376.1;refGene_gn=TLE4;refGene_tx=XM_023243377.1;refGene_gn=TLE4;refGene_tx=XM_023243378.1;refGene_gn=TLE4;refGene_tx=XM_023243379.1;refGene_gn=TLE4;refGene_tx=XM_023243380.1;refGene_gn=TLE4;refGene_tx=XM_023243381.1;refGene_gn=TLE4;refGene_tx=XM_023243382.1;refGene_gn=TLE4;refGene_tx=XM_023243383.1;refGene_gn=TLE4;refGene_tx=XM_023243384.1;refGene_gn=TLE4;refGene_tx=XM_023243385.1
D4 21159644 21159797
refGene_gn=TLE4;refGene_tx=XM_023243373.1;refGene_gn=TLE4;refGene_tx=XM_023243374.1;refGene_gn=TLE4;refGene_tx=XM_023243376.1;refGene_gn=TLE4;refGene_tx=XM_023243377.1;refGene_gn=TLE4;refGene_tx=XM_023243378.1;refGene_gn=TLE4;refGene_tx=XM_023243379.1;refGene_gn=TLE4;refGene_tx=XM_023243380.1;refGene_gn=TLE4;refGene_tx=XM_023243381.1;refGene_gn=TLE4;refGene_tx=XM_023243382.1;refGene_gn=TLE4;refGene_tx=XM_023243383.1;refGene_gn=TLE4;refGene_tx=XM_023243384.1;refGene_gn=TLE4;refGene_tx=XM_023243385.1
D4 21161093 21161226
refGene_gn=TLE4;refGene_tx=XM_023243373.1;refGene_gn=TLE4;refGene_tx=XM_023243374.1;refGene_gn=TLE4;refGene_tx=XM_023243376.1;refGene_gn=TLE4;refGene_tx=XM_023243377.1;refGene_gn=TLE4;refGene_tx=XM_023243378.1;refGene_gn=TLE4;refGene_tx=XM_023243379.1;refGene_gn=TLE4;refGene_tx=XM_023243380.1;refGene_gn=TLE4;refGene_tx=XM_023243381.1;refGene_gn=TLE4;refGene_tx=XM_023243382.1;refGene_gn=TLE4;refGene_tx=XM_023243383.1;refGene_gn=TLE4;refGene_tx=XM_023243384.1;refGene_gn=TLE4;refGene_tx=XM_023243385.1
D4 21161502 21161696
refGene_gn=TLE4;refGene_tx=XM_023243373.1;refGene_gn=TLE4;refGene_tx=XM_023243374.1;refGene_gn=TLE4;refGene_tx=XM_023243376.1;refGene_gn=TLE4;refGene_tx=XM_023243377.1;refGene_gn=TLE4;refGene_tx=XM_023243378.1;refGene_gn=TLE4;refGene_tx=XM_023243379.1;refGene_gn=TLE4;refGene_tx=XM_023243380.1;refGene_gn=TLE4;refGene_tx=XM_023243381.1;refGene_gn=TLE4;refGene_tx=XM_023243382.1;refGene_gn=TLE4;refGene_tx=XM_023243383.1;refGene_gn=TLE4;refGene_tx=XM_023243384.1;refGene_gn=TLE4;refGene_tx=XM_023243385.1
D4 21162206 21162283
refGene_gn=TLE4;refGene_tx=XM_023243373.1;refGene_gn=TLE4;refGene_tx=XM_023243374.1;refGene_gn=TLE4;refGene_tx=XM_023243376.1;refGene_gn=TLE4;refGene_tx=XM_023243377.1;refGene_gn=TLE4;refGene_tx=XM_023243378.1;refGene_gn=TLE4;refGene_tx=XM_023243379.1;refGene_gn=TLE4;refGene_tx=XM_023243380.1;refGene_gn=TLE4;refGene_tx=XM_023243381.1;refGene_gn=TLE4;refGene_tx=XM_023243382.1;refGene_gn=TLE4;refGene_tx=XM_023243383.1;refGene_gn=TLE4;refGene_tx=XM_023243384.1;refGene_gn=TLE4;refGene_tx=XM_023243385.1
D4 21171467 21171717
refGene_gn=TLE4;refGene_tx=XM_023243373.1;refGene_gn=TLE4;refGene_tx=XM_023243374.1;refGene_gn=TLE4;refGene_tx=XM_023243376.1;refGene_gn=TLE4;refGene_tx=XM_023243377.1;refGene_gn=TLE4;refGene_tx=XM_023243378.1;refGene_gn=TLE4;refGene_tx=XM_023243379.1;refGene_gn=TLE4;refGene_tx=X

M_023243380.1;refGene_gn=TLE4;refGene_tx=XM_023243381.1;refGene_gn=TLE4;refGene_tx=XM_023243382.1;refGene_gn=TLE4;refGene_tx=XM_023243383.1;refGene_gn=TLE4;refGene_tx=XM_023243384.1;refGene_gn=TLE4;refGene_tx=XM_023243385.1
D4 21172784 21173032
refGene_gn=TLE4;refGene_tx=XM_023243373.1;refGene_gn=TLE4;refGene_tx=XM_023243374.1;refGene_gn=TLE4;refGene_tx=XM_023243376.1;refGene_gn=TLE4;refGene_tx=XM_023243377.1;refGene_gn=TLE4;refGene_tx=XM_023243378.1;refGene_gn=TLE4;refGene_tx=XM_023243379.1;refGene_gn=TLE4;refGene_tx=XM_023243380.1;refGene_gn=TLE4;refGene_tx=XM_023243381.1;refGene_gn=TLE4;refGene_tx=XM_023243382.1;refGene_gn=TLE4;refGene_tx=XM_023243383.1;refGene_gn=TLE4;refGene_tx=XM_023243384.1;refGene_gn=TLE4;refGene_tx=XM_023243385.1
D4 21175341 21175489
refGene_gn=TLE4;refGene_tx=XM_023243373.1;refGene_gn=TLE4;refGene_tx=XM_023243374.1;refGene_gn=TLE4;refGene_tx=XM_023243376.1;refGene_gn=TLE4;refGene_tx=XM_023243377.1;refGene_gn=TLE4;refGene_tx=XM_023243378.1;refGene_gn=TLE4;refGene_tx=XM_023243379.1;refGene_gn=TLE4;refGene_tx=XM_023243380.1;refGene_gn=TLE4;refGene_tx=XM_023243381.1;refGene_gn=TLE4;refGene_tx=XM_023243382.1;refGene_gn=TLE4;refGene_tx=XM_023243383.1;refGene_gn=TLE4;refGene_tx=XM_023243384.1;refGene_gn=TLE4;refGene_tx=XM_023243385.1
D4 21175877 21176028
refGene_gn=TLE4;refGene_tx=XM_023243373.1;refGene_gn=TLE4;refGene_tx=XM_023243374.1;refGene_gn=TLE4;refGene_tx=XM_023243376.1;refGene_gn=TLE4;refGene_tx=XM_023243377.1;refGene_gn=TLE4;refGene_tx=XM_023243378.1;refGene_gn=TLE4;refGene_tx=XM_023243379.1;refGene_gn=TLE4;refGene_tx=XM_023243380.1;refGene_gn=TLE4;refGene_tx=XM_023243381.1;refGene_gn=TLE4;refGene_tx=XM_023243382.1;refGene_gn=TLE4;refGene_tx=XM_023243383.1;refGene_gn=TLE4;refGene_tx=XM_023243384.1;refGene_gn=TLE4;refGene_tx=XM_023243385.1
D4 21176364 21176441
refGene_gn=TLE4;refGene_tx=XM_023243373.1;refGene_gn=TLE4;refGene_tx=XM_023243374.1;refGene_gn=TLE4;refGene_tx=XM_023243376.1;refGene_gn=TLE4;refGene_tx=XM_023243377.1;refGene_gn=TLE4;refGene_tx=XM_023243378.1;refGene_gn=TLE4;refGene_tx=XM_023243379.1;refGene_gn=TLE4;refGene_tx=XM_023243380.1;refGene_gn=TLE4;refGene_tx=XM_023243381.1;refGene_gn=TLE4;refGene_tx=XM_023243382.1;refGene_gn=TLE4;refGene_tx=XM_023243383.1;refGene_gn=TLE4;refGene_tx=XM_023243384.1;refGene_gn=TLE4;refGene_tx=XM_023243385.1
D4 21178703 21178811
refGene_gn=TLE4;refGene_tx=XM_023243373.1;refGene_gn=TLE4;refGene_tx=XM_023243374.1;refGene_gn=TLE4;refGene_tx=XM_023243376.1;refGene_gn=TLE4;refGene_tx=XM_023243377.1;refGene_gn=TLE4;refGene_tx=XM_023243378.1;refGene_gn=TLE4;refGene_tx=XM_023243379.1;refGene_gn=TLE4;refGene_tx=XM_023243380.1;refGene_gn=TLE4;refGene_tx=XM_023243381.1;refGene_gn=TLE4;refGene_tx=XM_023243382.1;refGene_gn=TLE4;refGene_tx=XM_023243383.1;refGene_gn=TLE4;refGene_tx=XM_023243384.1;refGene_gn=TLE4;refGene_tx=XM_023243385.1
D4 22513967 22514465

refGene_gn=LOC105261088;refGene_tx=XM_023243391.1
D4 22774991 22775099
refGene_gn=TLE1;refGene_tx=XM_011288270.3;refGene_gn=TLE1;refGene_tx=XM_011288272.3;refGene_gn=TLE1;refGene_tx=XM_011288273.3;refGene_gn=TLE1;refGene_tx=XM_011288274.3;refGene_gn=TLE1;refGene_tx=XM_019815802.2;refGene_gn=TLE1;refGene_tx=XM_019815803.2
D4 22775251 22775328
refGene_gn=TLE1;refGene_tx=XM_011288270.3;refGene_gn=TLE1;refGene_tx=XM_011288272.3;refGene_gn=TLE1;refGene_tx=XM_011288273.3;refGene_gn=TLE1;refGene_tx=XM_011288274.3;refGene_gn=TLE1;refGene_tx=XM_019815802.2;refGene_gn=TLE1;refGene_tx=XM_019815803.2
D4 22776249 22776400
refGene_gn=TLE1;refGene_tx=XM_011288270.3;refGene_gn=TLE1;refGene_tx=XM_011288272.3;refGene_gn=TLE1;refGene_tx=XM_011288273.3;refGene_gn=TLE1;refGene_tx=XM_011288274.3;refGene_gn=TLE1;refGene_tx=XM_019815802.2;refGene_gn=TLE1;refGene_tx=XM_019815803.2
D4 22777835 22777983
refGene_gn=TLE1;refGene_tx=XM_011288270.3;refGene_gn=TLE1;refGene_tx=XM_011288272.3;refGene_gn=TLE1;refGene_tx=XM_011288273.3;refGene_gn=TLE1;refGene_tx=XM_011288274.3;refGene_gn=TLE1;refGene_tx=XM_019815802.2;refGene_gn=TLE1;refGene_tx=XM_019815803.2
D4 22781717 22781965
refGene_gn=TLE1;refGene_tx=XM_011288270.3;refGene_gn=TLE1;refGene_tx=XM_011288272.3;refGene_gn=TLE1;refGene_tx=XM_011288273.3;refGene_gn=TLE1;refGene_tx=XM_011288274.3;refGene_gn=TLE1;refGene_tx=XM_019815802.2;refGene_gn=TLE1;refGene_tx=XM_019815803.2
D4 22784062 22784312
refGene_gn=TLE1;refGene_tx=XM_011288270.3;refGene_gn=TLE1;refGene_tx=XM_011288272.3;refGene_gn=TLE1;refGene_tx=XM_011288273.3;refGene_gn=TLE1;refGene_tx=XM_011288274.3;refGene_gn=TLE1;refGene_tx=XM_019815802.2;refGene_gn=TLE1;refGene_tx=XM_019815803.2
D4 22798824 22798901
refGene_gn=TLE1;refGene_tx=XM_011288270.3;refGene_gn=TLE1;refGene_tx=XM_011288272.3;refGene_gn=TLE1;refGene_tx=XM_011288273.3;refGene_gn=TLE1;refGene_tx=XM_011288274.3;refGene_gn=TLE1;refGene_tx=XM_019815802.2;refGene_gn=TLE1;refGene_tx=XM_019815803.2
D4 22800459 22800650
refGene_gn=TLE1;refGene_tx=XM_011288270.3;refGene_gn=TLE1;refGene_tx=XM_011288272.3;refGene_gn=TLE1;refGene_tx=XM_011288273.3;refGene_gn=TLE1;refGene_tx=XM_011288274.3;refGene_gn=TLE1;refGene_tx=XM_019815802.2;refGene_gn=TLE1;refGene_tx=XM_019815803.2
D4 22801652 22801797
refGene_gn=TLE1;refGene_tx=XM_011288270.3;refGene_gn=TLE1;refGene_tx=XM_011288272.3;refGene_gn=TLE1;refGene_tx=XM_011288273.3;refGene_gn=TLE1;refGene_tx=XM_011288274.3;refGene_gn=TLE1;refGene_tx=XM_019815802.2;refGene_gn=TLE1;refGene_tx=XM_019815803.2
D4 22803139 22803292
refGene_gn=TLE1;refGene_tx=XM_011288270.3;refGene_gn=TLE1;refGene_tx=XM_011288272.3;refGene_gn=TLE1;refGene_tx=XM_011288273.3;refGene_gn=TLE1;refGene_tx=XM_011288274.3;refGene_gn=TLE1;refGene_tx=XM_019815802.2;

refGene_gn=TLE1;refGene_tx=XM_019815803.2
D4 22803788 22803842
refGene_gn=TLE1;refGene_tx=XM_011288270.3;refGene_gn=TLE1;refGene_tx=XM_011288272.3;refGene_gn=TLE1;refGene_tx=XM_011288273.3;refGene_gn=TLE1;refGene_tx=XM_011288274.3;refGene_gn=TLE1;refGene_tx=XM_019815802.2;refGene_gn=TLE1;refGene_tx=XM_019815803.2
D4 22807148 22807265
refGene_gn=TLE1;refGene_tx=XM_011288270.3;refGene_gn=TLE1;refGene_tx=XM_011288272.3;refGene_gn=TLE1;refGene_tx=XM_011288273.3;refGene_gn=TLE1;refGene_tx=XM_011288274.3;refGene_gn=TLE1;refGene_tx=XM_019815802.2;refGene_gn=TLE1;refGene_tx=XM_019815803.2
D4 22820458 22820475
refGene_gn=TLE1;refGene_tx=XM_011288270.3;refGene_gn=TLE1;refGene_tx=XM_011288272.3;refGene_gn=TLE1;refGene_tx=XM_011288273.3;refGene_gn=TLE1;refGene_tx=XM_011288274.3;refGene_gn=TLE1;refGene_tx=XM_019815802.2;refGene_gn=TLE1;refGene_tx=XM_019815803.2
D4 22821208 22821413
refGene_gn=TLE1;refGene_tx=XM_011288270.3;refGene_gn=TLE1;refGene_tx=XM_011288272.3;refGene_gn=TLE1;refGene_tx=XM_011288273.3;refGene_gn=TLE1;refGene_tx=XM_011288274.3;refGene_gn=TLE1;refGene_tx=XM_019815802.2;refGene_gn=TLE1;refGene_tx=XM_019815803.2
D4 22835194 22835299
refGene_gn=TLE1;refGene_tx=XM_011288270.3;refGene_gn=TLE1;refGene_tx=XM_011288272.3;refGene_gn=TLE1;refGene_tx=XM_011288273.3;refGene_gn=TLE1;refGene_tx=XM_011288274.3;refGene_gn=TLE1;refGene_tx=XM_019815802.2
D4 22836681 22836744
refGene_gn=TLE1;refGene_tx=XM_011288270.3;refGene_gn=TLE1;refGene_tx=XM_011288272.3;refGene_gn=TLE1;refGene_tx=XM_011288273.3;refGene_gn=TLE1;refGene_tx=XM_011288274.3;refGene_gn=TLE1;refGene_tx=XM_019815802.2
D4 22862446 22862491
refGene_gn=TLE1;refGene_tx=XM_011288270.3;refGene_gn=TLE1;refGene_tx=XM_011288272.3;refGene_gn=TLE1;refGene_tx=XM_011288273.3;refGene_gn=TLE1;refGene_tx=XM_011288274.3;refGene_gn=TLE1;refGene_tx=XM_019815802.2;refGene_gn=TLE1;refGene_tx=XM_019815803.2
D4 22862602 22862666
refGene_gn=TLE1;refGene_tx=XM_011288270.3;refGene_gn=TLE1;refGene_tx=XM_011288272.3;refGene_gn=TLE1;refGene_tx=XM_011288273.3;refGene_gn=TLE1;refGene_tx=XM_011288274.3;refGene_gn=TLE1;refGene_tx=XM_019815802.2;refGene_gn=TLE1;refGene_tx=XM_019815803.2
D4 22864112 22864213
refGene_gn=TLE1;refGene_tx=XM_011288270.3;refGene_gn=TLE1;refGene_tx=XM_011288272.3;refGene_gn=TLE1;refGene_tx=XM_011288273.3;refGene_gn=TLE1;refGene_tx=XM_011288274.3;refGene_gn=TLE1;refGene_tx=XM_019815803.2
D4 22865013 22865037
refGene_gn=TLE1;refGene_tx=XM_011288270.3;refGene_gn=TLE1;refGene_tx=XM_011288272.3;refGene_gn=TLE1;refGene_tx=XM_011288273.3;refGene_gn=TLE1;refGene_tx=XM_011288274.3;refGene_gn=TLE1;refGene_tx=XM_019815803.2
D4 23133463 23133601
refGene_gn=LOC111556292;refGene_tx=XM_023243394.1
D4 23134437 23134483

refGene_gn=LOC111556292;refGene_tx=XM_023243394.1
D4 23135034 23135095
refGene_gn=LOC111556292;refGene_tx=XM_023243394.1
D4 23135378 23138622
refGene_gn=LOC111556292;refGene_tx=XM_023243394.1
D4 24068490 24068596
refGene_gn=RASEF;refGene_tx=XM_003995449.5;refGene_gn=RASEF;refGene_tx=XM_019815810.2;refGene_gn=RASEF;refGene_tx=XM_019815811.2
D4 24075070 24075147
refGene_gn=RASEF;refGene_tx=XM_003995449.5;refGene_gn=RASEF;refGene_tx=XM_019815810.2;refGene_gn=RASEF;refGene_tx=XM_019815811.2
D4 24077733 24077853
refGene_gn=RASEF;refGene_tx=XM_003995449.5;refGene_gn=RASEF;refGene_tx=XM_019815810.2
D4 24082586 24082701
refGene_gn=RASEF;refGene_tx=XM_003995449.5;refGene_gn=RASEF;refGene_tx=XM_019815810.2;refGene_gn=RASEF;refGene_tx=XM_019815811.2
D4 24085120 24085202
refGene_gn=RASEF;refGene_tx=XM_003995449.5;refGene_gn=RASEF;refGene_tx=XM_019815810.2;refGene_gn=RASEF;refGene_tx=XM_019815811.2
D4 24086497 24086645
refGene_gn=RASEF;refGene_tx=XM_003995449.5;refGene_gn=RASEF;refGene_tx=XM_019815810.2;refGene_gn=RASEF;refGene_tx=XM_019815811.2
D4 24086761 24086899
refGene_gn=RASEF;refGene_tx=XM_003995449.5;refGene_gn=RASEF;refGene_tx=XM_019815810.2;refGene_gn=RASEF;refGene_tx=XM_019815811.2
D4 24087202 24087437
refGene_gn=RASEF;refGene_tx=XM_003995449.5;refGene_gn=RASEF;refGene_tx=XM_019815810.2;refGene_gn=RASEF;refGene_tx=XM_019815811.2
D4 24090689 24090778
refGene_gn=RASEF;refGene_tx=XM_003995449.5;refGene_gn=RASEF;refGene_tx=XM_019815810.2;refGene_gn=RASEF;refGene_tx=XM_019815811.2
D4 24092175 24092260
refGene_gn=RASEF;refGene_tx=XM_003995449.5;refGene_gn=RASEF;refGene_tx=XM_019815810.2;refGene_gn=RASEF;refGene_tx=XM_019815811.2
D4 24094096 24094165
refGene_gn=RASEF;refGene_tx=XM_003995449.5;refGene_gn=RASEF;refGene_tx=XM_019815811.2
D4 24096231 24096347
refGene_gn=RASEF;refGene_tx=XM_003995449.5;refGene_gn=RASEF;refGene_tx=XM_019815810.2;refGene_gn=RASEF;refGene_tx=XM_019815811.2
D4 24098911 24098989
refGene_gn=RASEF;refGene_tx=XM_003995449.5;refGene_gn=RASEF;refGene_tx=XM_019815810.2;refGene_gn=RASEF;refGene_tx=XM_019815811.2
D4 24102449 24102545
refGene_gn=RASEF;refGene_tx=XM_003995449.5;refGene_gn=RASEF;refGene_tx=XM_019815810.2;refGene_gn=RASEF;refGene_tx=XM_019815811.2
D4 24107787 24107878
refGene_gn=RASEF;refGene_tx=XM_003995449.5;refGene_gn=RASEF;refGene_tx=XM_019815810.2;refGene_gn=RASEF;refGene_tx=XM_019815811.2

D4 24111042 24111189
refGene_gn=RASEF;refGene_tx=XM_003995449.5;refGene_gn=RASEF;refGene_tx=XM_019815810.2;refGene_gn=RASEF;refGene_tx=XM_019815811.2
D4 24144292 24144714
refGene_gn=RASEF;refGene_tx=XM_003995449.5;refGene_gn=RASEF;refGene_tx=XM_019815810.2;refGene_gn=RASEF;refGene_tx=XM_019815811.2
D4 24349678 24349692 refGene_gn=FRMD3;refGene_tx=XM_019815813.2
D4 24354318 24354914
refGene_gn=FRMD3;refGene_tx=XM_006939122.4;refGene_gn=FRMD3;refGene_tx=XM_019815813.2
D4 24421287 24421412
refGene_gn=FRMD3;refGene_tx=XM_006939122.4;refGene_gn=FRMD3;refGene_tx=XM_019815813.2
D4 24429570 24429639
refGene_gn=FRMD3;refGene_tx=XM_006939122.4;refGene_gn=FRMD3;refGene_tx=XM_019815813.2
D4 24429949 24430024
refGene_gn=FRMD3;refGene_tx=XM_006939122.4;refGene_gn=FRMD3;refGene_tx=XM_019815813.2
D4 24441663 24441752
refGene_gn=FRMD3;refGene_tx=XM_006939122.4;refGene_gn=FRMD3;refGene_tx=XM_019815813.2
D4 24442886 24442950
refGene_gn=FRMD3;refGene_tx=XM_006939122.4;refGene_gn=FRMD3;refGene_tx=XM_019815813.2
D4 24446072 24446161
refGene_gn=FRMD3;refGene_tx=XM_006939122.4;refGene_gn=FRMD3;refGene_tx=XM_019815813.2
D4 24447714 24447802
refGene_gn=FRMD3;refGene_tx=XM_006939122.4;refGene_gn=FRMD3;refGene_tx=XM_019815813.2
D4 24464158 24464282
refGene_gn=FRMD3;refGene_tx=XM_006939122.4;refGene_gn=FRMD3;refGene_tx=XM_019815813.2
D4 24472565 24472663
refGene_gn=FRMD3;refGene_tx=XM_006939122.4;refGene_gn=FRMD3;refGene_tx=XM_019815813.2
D4 24478887 24478966
refGene_gn=FRMD3;refGene_tx=XM_006939122.4;refGene_gn=FRMD3;refGene_tx=XM_019815813.2
D4 24499098 24499141
refGene_gn=FRMD3;refGene_tx=XM_006939122.4;refGene_gn=FRMD3;refGene_tx=XM_019815813.2
D4 24515746 24515851
refGene_gn=FRMD3;refGene_tx=XM_006939122.4;refGene_gn=FRMD3;refGene_tx=XM_019815813.2
D4 24665870 24666017
refGene_gn=FRMD3;refGene_tx=XM_006939122.4;refGene_gn=FRMD3;refGene_tx=XM_019815813.2
D4 24731992 24732042

refGene_gn=IDNK;refGene_tx=XM_023242202.1;refGene_gn=IDNK;refGene_tx=XM_023242203.1
D4 24737649 24737680 refGene_gn=IDNK;refGene_tx=XM_023242202.1
D4 24738125 24738212
refGene_gn=IDNK;refGene_tx=XM_019815815.2;refGene_gn=IDNK;refGene_tx=XM_023242202.1;refGene_gn=IDNK;refGene_tx=XM_023242203.1;refGene_gn=IDNK;refGene_tx=XM_023242204.1;refGene_gn=IDNK;refGene_tx=XM_023242205.1;refGene_gn=IDNK;refGene_tx=XM_023242206.1;refGene_gn=IDNK;refGene_tx=XM_023242207.1;refGene_gn=IDNK;refGene_tx=XM_023242208.1;refGene_gn=IDNK;refGene_tx=XM_023242209.1;refGene_gn=IDNK;refGene_tx=XM_023242210.1
D4 24742870 24742914
refGene_gn=IDNK;refGene_tx=XM_019815815.2;refGene_gn=IDNK;refGene_tx=XM_023242202.1;refGene_gn=IDNK;refGene_tx=XM_023242203.1;refGene_gn=IDNK;refGene_tx=XM_023242204.1;refGene_gn=IDNK;refGene_tx=XM_023242205.1;refGene_gn=IDNK;refGene_tx=XM_023242206.1;refGene_gn=IDNK;refGene_tx=XM_023242207.1;refGene_gn=IDNK;refGene_tx=XM_023242208.1;refGene_gn=IDNK;refGene_tx=XM_023242209.1;refGene_gn=IDNK;refGene_tx=XM_023242210.1
D4 24744778 24745124
refGene_gn=IDNK;refGene_tx=XM_019815815.2;refGene_gn=IDNK;refGene_tx=XM_023242202.1;refGene_gn=IDNK;refGene_tx=XM_023242203.1;refGene_gn=IDNK;refGene_tx=XM_023242204.1;refGene_gn=IDNK;refGene_tx=XM_023242205.1;refGene_gn=IDNK;refGene_tx=XM_023242206.1;refGene_gn=IDNK;refGene_tx=XM_023242207.1;refGene_gn=IDNK;refGene_tx=XM_023242208.1;refGene_gn=IDNK;refGene_tx=XM_023242209.1;refGene_gn=IDNK;refGene_tx=XM_023242210.1
D4 24758439 24758592
refGene_gn=UBQLN1;refGene_tx=XM_011288266.2;refGene_gn=UBQLN1;refGene_tx=XM_011288267.2
D4 24759974 24760143
refGene_gn=UBQLN1;refGene_tx=XM_011288266.2;refGene_gn=UBQLN1;refGene_tx=XM_011288267.2
D4 24760371 24760487
refGene_gn=UBQLN1;refGene_tx=XM_011288266.2;refGene_gn=UBQLN1;refGene_tx=XM_011288267.2
D4 24761718 24761802 refGene_gn=UBQLN1;refGene_tx=XM_011288266.2
D4 24765122 24765265
refGene_gn=UBQLN1;refGene_tx=XM_011288266.2;refGene_gn=UBQLN1;refGene_tx=XM_011288267.2
D4 24770519 24770754
refGene_gn=UBQLN1;refGene_tx=XM_011288266.2;refGene_gn=UBQLN1;refGene_tx=XM_011288267.2
D4 24770933 24771092
refGene_gn=UBQLN1;refGene_tx=XM_011288266.2;refGene_gn=UBQLN1;refGene_tx=XM_011288267.2
D4 24771565 24771828
refGene_gn=UBQLN1;refGene_tx=XM_011288266.2;refGene_gn=UBQLN1;refGene_tx=XM_011288267.2
D4 24775738 24775854
refGene_gn=UBQLN1;refGene_tx=XM_011288266.2;refGene_gn=UBQLN1;refGene_tx=XM_011288267.2
D4 24778243 24778395

refGene_gn=UBQLN1;refGene_tx=XM_011288266.2;refGene_gn=UBQLN1;refGene_tx=XM_011288267.2
D4 24801295 24801475
refGene_gn=UBQLN1;refGene_tx=XM_011288266.2;refGene_gn=UBQLN1;refGene_tx=XM_011288267.2
D4 24834318 24834366
refGene_gn=GKAP1;refGene_tx=XM_006939126.4;refGene_gn=GKAP1;refGene_tx=XM_006939127.4
D4 24836559 24836637
refGene_gn=GKAP1;refGene_tx=XM_006939126.4;refGene_gn=GKAP1;refGene_tx=XM_006939127.4
D4 24837428 24837499
refGene_gn=GKAP1;refGene_tx=XM_006939126.4;refGene_gn=GKAP1;refGene_tx=XM_006939127.4
D4 24841672 24841736
refGene_gn=GKAP1;refGene_tx=XM_006939126.4;refGene_gn=GKAP1;refGene_tx=XM_006939127.4
D4 24846533 24846635
refGene_gn=GKAP1;refGene_tx=XM_006939126.4;refGene_gn=GKAP1;refGene_tx=XM_006939127.4
D4 24858176 24858326 refGene_gn=GKAP1;refGene_tx=XM_006939126.4
D4 24886956 24886979
refGene_gn=GKAP1;refGene_tx=XM_006939126.4;refGene_gn=GKAP1;refGene_tx=XM_006939127.4
D4 24889356 24889480
refGene_gn=GKAP1;refGene_tx=XM_006939126.4;refGene_gn=GKAP1;refGene_tx=XM_006939127.4
D4 24893493 24893571
refGene_gn=GKAP1;refGene_tx=XM_006939126.4;refGene_gn=GKAP1;refGene_tx=XM_006939127.4
D4 24903381 24903525
refGene_gn=GKAP1;refGene_tx=XM_006939126.4;refGene_gn=GKAP1;refGene_tx=XM_006939127.4
D4 24904955 24905165
refGene_gn=GKAP1;refGene_tx=XM_006939126.4;refGene_gn=GKAP1;refGene_tx=XM_006939127.4
D4 24937722 24937779
refGene_gn=KIF27;refGene_tx=XM_023242214.1;refGene_gn=KIF27;refGene_tx=XM_023242215.1
D4 24940534 24941019
refGene_gn=KIF27;refGene_tx=XM_023242211.1;refGene_gn=KIF27;refGene_tx=XM_023242212.1;refGene_gn=KIF27;refGene_tx=XM_023242213.1;refGene_gn=KIF27;refGene_tx=XM_023242214.1;refGene_gn=KIF27;refGene_tx=XM_023242215.1
D4 24945675 24945840
refGene_gn=KIF27;refGene_tx=XM_023242211.1;refGene_gn=KIF27;refGene_tx=XM_023242212.1;refGene_gn=KIF27;refGene_tx=XM_023242214.1;refGene_gn=KIF27;refGene_tx=XM_023242215.1
D4 24949057 24949256
refGene_gn=KIF27;refGene_tx=XM_023242211.1;refGene_gn=KIF27;refGene_tx

=XM_023242212.1;refGene_gn=KIF27;refGene_tx=XM_023242213.1;refGene_gn=KIF27;refGene_tx=XM_023242214.1;refGene_gn=KIF27;refGene_tx=XM_023242215.1
D4 24950615 24950822
refGene_gn=KIF27;refGene_tx=XM_023242211.1;refGene_gn=KIF27;refGene_tx=XM_023242212.1;refGene_gn=KIF27;refGene_tx=XM_023242213.1;refGene_gn=KIF27;refGene_tx=XM_023242214.1;refGene_gn=KIF27;refGene_tx=XM_023242215.1
D4 24955420 24955636
refGene_gn=KIF27;refGene_tx=XM_023242211.1;refGene_gn=KIF27;refGene_tx=XM_023242212.1;refGene_gn=KIF27;refGene_tx=XM_023242213.1;refGene_gn=KIF27;refGene_tx=XM_023242214.1;refGene_gn=KIF27;refGene_tx=XM_023242215.1
D4 24963228 24963405
refGene_gn=KIF27;refGene_tx=XM_023242211.1;refGene_gn=KIF27;refGene_tx=XM_023242212.1;refGene_gn=KIF27;refGene_tx=XM_023242213.1;refGene_gn=KIF27;refGene_tx=XM_023242214.1;refGene_gn=KIF27;refGene_tx=XM_023242215.1
D4 24966815 24966929
refGene_gn=KIF27;refGene_tx=XM_023242211.1;refGene_gn=KIF27;refGene_tx=XM_023242212.1;refGene_gn=KIF27;refGene_tx=XM_023242213.1;refGene_gn=KIF27;refGene_tx=XM_023242214.1;refGene_gn=KIF27;refGene_tx=XM_023242215.1
D4 24977656 24977854
refGene_gn=KIF27;refGene_tx=XM_023242211.1;refGene_gn=KIF27;refGene_tx=XM_023242212.1;refGene_gn=KIF27;refGene_tx=XM_023242213.1;refGene_gn=KIF27;refGene_tx=XM_023242214.1;refGene_gn=KIF27;refGene_tx=XM_023242215.1
D4 24982755 24982961
refGene_gn=KIF27;refGene_tx=XM_023242211.1;refGene_gn=KIF27;refGene_tx=XM_023242212.1;refGene_gn=KIF27;refGene_tx=XM_023242213.1;refGene_gn=KIF27;refGene_tx=XM_023242214.1;refGene_gn=KIF27;refGene_tx=XM_023242215.1
D4 24987209 24987365
refGene_gn=KIF27;refGene_tx=XM_023242211.1;refGene_gn=KIF27;refGene_tx=XM_023242212.1;refGene_gn=KIF27;refGene_tx=XM_023242213.1;refGene_gn=KIF27;refGene_tx=XM_023242214.1;refGene_gn=KIF27;refGene_tx=XM_023242215.1
D4 24989076 24989180
refGene_gn=KIF27;refGene_tx=XM_023242211.1;refGene_gn=KIF27;refGene_tx=XM_023242212.1;refGene_gn=KIF27;refGene_tx=XM_023242213.1;refGene_gn=KIF27;refGene_tx=XM_023242214.1;refGene_gn=KIF27;refGene_tx=XM_023242215.1
D4 24991065 24991232
refGene_gn=KIF27;refGene_tx=XM_023242211.1;refGene_gn=KIF27;refGene_tx=XM_023242212.1;refGene_gn=KIF27;refGene_tx=XM_023242213.1;refGene_gn=KIF27;refGene_tx=XM_023242214.1;refGene_gn=KIF27;refGene_tx=XM_023242215.1
D4 24996724 24996931
refGene_gn=KIF27;refGene_tx=XM_023242211.1;refGene_gn=KIF27;refGene_tx

=XM_023242212.1;refGene_gn=KIF27;refGene_tx=XM_023242213.1;refGene_gn=KIF27;refGene_tx=XM_023242214.1;refGene_gn=KIF27;refGene_tx=XM_023242215.1
D4 25008727 25008871
refGene_gn=KIF27;refGene_tx=XM_023242211.1;refGene_gn=KIF27;refGene_tx=XM_023242212.1;refGene_gn=KIF27;refGene_tx=XM_023242213.1;refGene_gn=KIF27;refGene_tx=XM_023242214.1;refGene_gn=KIF27;refGene_tx=XM_023242215.1
D4 25016495 25017454
refGene_gn=KIF27;refGene_tx=XM_023242211.1;refGene_gn=KIF27;refGene_tx=XM_023242212.1;refGene_gn=KIF27;refGene_tx=XM_023242213.1;refGene_gn=KIF27;refGene_tx=XM_023242214.1;refGene_gn=KIF27;refGene_tx=XM_023242215.1
D4 25020656 25020857
refGene_gn=KIF27;refGene_tx=XM_023242211.1;refGene_gn=KIF27;refGene_tx=XM_023242212.1;refGene_gn=KIF27;refGene_tx=XM_023242213.1;refGene_gn=KIF27;refGene_tx=XM_023242214.1;refGene_gn=KIF27;refGene_tx=XM_023242215.1
D4 25029624 25029922
refGene_gn=KIF27;refGene_tx=XM_023242211.1;refGene_gn=KIF27;refGene_tx=XM_023242212.1;refGene_gn=KIF27;refGene_tx=XM_023242213.1;refGene_gn=KIF27;refGene_tx=XM_023242214.1;refGene_gn=KIF27;refGene_tx=XM_023242215.1
D4 25053623 25053856
refGene_gn=CD4H9orf64;refGene_tx=XM_003995439.4;refGene_gn=CD4H9orf64;refGene_tx=XM_019815823.2
D4 25057778 25057974
refGene_gn=CD4H9orf64;refGene_tx=XM_003995439.4;refGene_gn=CD4H9orf64;refGene_tx=XM_019815823.2
D4 25065203 25065463
refGene_gn=CD4H9orf64;refGene_tx=XM_003995439.4;refGene_gn=CD4H9orf64;refGene_tx=XM_019815823.2
D4 25065975 25066312
refGene_gn=CD4H9orf64;refGene_tx=XM_003995439.4
D4 25081959 25081990
refGene_gn=HNRNPK;refGene_tx=XM_011288263.3;refGene_gn=HNRNPK;refGene_tx=XM_019815822.2
D4 25082017 25082051
refGene_gn=HNRNPK;refGene_tx=XM_006939129.4;refGene_gn=HNRNPK;refGene_tx=XM_006939130.3;refGene_gn=HNRNPK;refGene_tx=XM_011288260.3;refGene_gn=HNRNPK;refGene_tx=XM_011288261.3;refGene_gn=HNRNPK;refGene_tx=XM_011288262.3;refGene_gn=HNRNPK;refGene_tx=XM_011288264.3;refGene_gn=HNRNPK;refGene_tx=XM_019815821.2
D4 25082766 25082936
refGene_gn=HNRNPK;refGene_tx=XM_006939129.4;refGene_gn=HNRNPK;refGene_tx=XM_006939130.3;refGene_gn=HNRNPK;refGene_tx=XM_011288260.3;refGene_gn=HNRNPK;refGene_tx=XM_011288261.3;refGene_gn=HNRNPK;refGene_tx=XM_011288262.3;refGene_gn=HNRNPK;refGene_tx=XM_011288263.3;refGene_gn=HNRNPK;refGene_tx=XM_011288264.3;refGene_gn=HNRNPK;refGene_tx=XM_019815821.2;refGene_gn=HNRNPK;refGene_tx=XM_019815822.2

D4 25083340 25083423
refGene_gn=HNRNPK;refGene_tx=XM_006939129.4;refGene_gn=HNRNPK;refGene_tx=XM_006939130.3;refGene_gn=HNRNPK;refGene_tx=XM_011288260.3;refGene_gn=HNRNPK;refGene_tx=XM_011288261.3;refGene_gn=HNRNPK;refGene_tx=XM_011288262.3;refGene_gn=HNRNPK;refGene_tx=XM_011288263.3;refGene_gn=HNRNPK;refGene_tx=XM_011288264.3;refGene_gn=HNRNPK;refGene_tx=XM_019815821.2;refGene_gn=HNRNPK;refGene_tx=XM_019815822.2
D4 25083500 25083516
refGene_gn=HNRNPK;refGene_tx=XM_006939129.4;refGene_gn=HNRNPK;refGene_tx=XM_006939130.3;refGene_gn=HNRNPK;refGene_tx=XM_011288260.3;refGene_gn=HNRNPK;refGene_tx=XM_011288261.3;refGene_gn=HNRNPK;refGene_tx=XM_011288262.3;refGene_gn=HNRNPK;refGene_tx=XM_011288263.3;refGene_gn=HNRNPK;refGene_tx=XM_011288264.3;refGene_gn=HNRNPK;refGene_tx=XM_019815821.2;refGene_gn=HNRNPK;refGene_tx=XM_019815822.2
D4 25083780 25083864
refGene_gn=HNRNPK;refGene_tx=XM_006939129.4;refGene_gn=HNRNPK;refGene_tx=XM_006939130.3;refGene_gn=HNRNPK;refGene_tx=XM_011288260.3;refGene_gn=HNRNPK;refGene_tx=XM_011288261.3;refGene_gn=HNRNPK;refGene_tx=XM_011288262.3;refGene_gn=HNRNPK;refGene_tx=XM_011288263.3;refGene_gn=HNRNPK;refGene_tx=XM_011288264.3;refGene_gn=HNRNPK;refGene_tx=XM_019815821.2;refGene_gn=HNRNPK;refGene_tx=XM_019815822.2
D4 25084176 25084231
refGene_gn=HNRNPK;refGene_tx=XM_006939129.4;refGene_gn=HNRNPK;refGene_tx=XM_006939130.3;refGene_gn=HNRNPK;refGene_tx=XM_011288260.3;refGene_gn=HNRNPK;refGene_tx=XM_011288261.3;refGene_gn=HNRNPK;refGene_tx=XM_011288262.3;refGene_gn=HNRNPK;refGene_tx=XM_011288263.3;refGene_gn=HNRNPK;refGene_tx=XM_011288264.3;refGene_gn=HNRNPK;refGene_tx=XM_019815821.2;refGene_gn=HNRNPK;refGene_tx=XM_019815822.2
D4 25084343 25084651
refGene_gn=HNRNPK;refGene_tx=XM_006939129.4;refGene_gn=HNRNPK;refGene_tx=XM_006939130.3;refGene_gn=HNRNPK;refGene_tx=XM_011288260.3;refGene_gn=HNRNPK;refGene_tx=XM_011288261.3;refGene_gn=HNRNPK;refGene_tx=XM_011288262.3;refGene_gn=HNRNPK;refGene_tx=XM_011288263.3;refGene_gn=HNRNPK;refGene_tx=XM_011288264.3;refGene_gn=HNRNPK;refGene_tx=XM_019815821.2;refGene_gn=HNRNPK;refGene_tx=XM_019815822.2
D4 25085268 25085397
refGene_gn=HNRNPK;refGene_tx=XM_006939129.4;refGene_gn=HNRNPK;refGene_tx=XM_006939130.3;refGene_gn=HNRNPK;refGene_tx=XM_011288260.3;refGene_gn=HNRNPK;refGene_tx=XM_011288261.3;refGene_gn=HNRNPK;refGene_tx=XM_011288262.3;refGene_gn=HNRNPK;refGene_tx=XM_011288263.3;refGene_gn=HNRNPK;refGene_tx=XM_011288264.3;refGene_gn=HNRNPK;refGene_tx=XM_019815821.2;refGene_gn=HNRNPK;refGene_tx=XM_019815822.2
D4 25085769 25085883
refGene_gn=HNRNPK;refGene_tx=XM_006939129.4;refGene_gn=HNRNPK;refGene_tx=XM_006939130.3;refGene_gn=HNRNPK;refGene_tx=XM_011288260.3;refGene_gn=HNRNPK;refGene_tx=XM_011288261.3;refGene_gn=HNRNPK;refGene_tx=XM_011288262.3;refGene_gn=HNRNPK;refGene_tx=XM_011288263.3;refGene_gn=HNRNPK;refGene_tx=XM_011288264.3;refGene_gn=HNRNPK;refGene_tx=XM_019815821.2;refGene_gn=HNRNPK;refGene_tx=XM_019815822.2
D4 25086382 25086454

refGene_gn=HNRNPK;refGene_tx=XM_006939129.4;refGene_gn=HNRNPK;refGene_tx=XM_006939130.3;refGene_gn=HNRNPK;refGene_tx=XM_011288260.3;refGene_gn=HNRNPK;refGene_tx=XM_011288261.3;refGene_gn=HNRNPK;refGene_tx=XM_011288262.3;refGene_gn=HNRNPK;refGene_tx=XM_011288263.3;refGene_gn=HNRNPK;refGene_tx=XM_019815821.2
D4 25086972 25087045
refGene_gn=HNRNPK;refGene_tx=XM_006939129.4;refGene_gn=HNRNPK;refGene_tx=XM_006939130.3;refGene_gn=HNRNPK;refGene_tx=XM_011288260.3;refGene_gn=HNRNPK;refGene_tx=XM_011288261.3;refGene_gn=HNRNPK;refGene_tx=XM_011288262.3;refGene_gn=HNRNPK;refGene_tx=XM_011288263.3;refGene_gn=HNRNPK;refGene_tx=XM_011288264.3;refGene_gn=HNRNPK;refGene_tx=XM_019815821.2;refGene_gn=HNRNPK;refGene_tx=XM_019815822.2
D4 25087924 25087968
refGene_gn=HNRNPK;refGene_tx=XM_006939129.4;refGene_gn=HNRNPK;refGene_tx=XM_006939130.3;refGene_gn=HNRNPK;refGene_tx=XM_011288260.3;refGene_gn=HNRNPK;refGene_tx=XM_011288261.3;refGene_gn=HNRNPK;refGene_tx=XM_011288262.3;refGene_gn=HNRNPK;refGene_tx=XM_011288263.3;refGene_gn=HNRNPK;refGene_tx=XM_011288264.3;refGene_gn=HNRNPK;refGene_tx=XM_019815821.2;refGene_gn=HNRNPK;refGene_tx=XM_019815822.2
D4 25089422 25089479
refGene_gn=HNRNPK;refGene_tx=XM_006939129.4;refGene_gn=HNRNPK;refGene_tx=XM_006939130.3;refGene_gn=HNRNPK;refGene_tx=XM_011288260.3;refGene_gn=HNRNPK;refGene_tx=XM_011288261.3;refGene_gn=HNRNPK;refGene_tx=XM_011288262.3;refGene_gn=HNRNPK;refGene_tx=XM_011288263.3;refGene_gn=HNRNPK;refGene_tx=XM_011288264.3;refGene_gn=HNRNPK;refGene_tx=XM_019815821.2;refGene_gn=HNRNPK;refGene_tx=XM_019815822.2
D4 25090141 25090239
refGene_gn=HNRNPK;refGene_tx=XM_006939129.4;refGene_gn=HNRNPK;refGene_tx=XM_006939130.3;refGene_gn=HNRNPK;refGene_tx=XM_011288260.3;refGene_gn=HNRNPK;refGene_tx=XM_011288261.3;refGene_gn=HNRNPK;refGene_tx=XM_011288262.3;refGene_gn=HNRNPK;refGene_tx=XM_011288263.3;refGene_gn=HNRNPK;refGene_tx=XM_011288264.3;refGene_gn=HNRNPK;refGene_tx=XM_019815821.2;refGene_gn=HNRNPK;refGene_tx=XM_019815822.2
D4 25090679 25090737
refGene_gn=HNRNPK;refGene_tx=XM_006939129.4;refGene_gn=HNRNPK;refGene_tx=XM_006939130.3;refGene_gn=HNRNPK;refGene_tx=XM_011288260.3;refGene_gn=HNRNPK;refGene_tx=XM_011288261.3;refGene_gn=HNRNPK;refGene_tx=XM_011288262.3;refGene_gn=HNRNPK;refGene_tx=XM_011288263.3;refGene_gn=HNRNPK;refGene_tx=XM_011288264.3;refGene_gn=HNRNPK;refGene_tx=XM_019815821.2;refGene_gn=HNRNPK;refGene_tx=XM_019815822.2
D4 25111490 25113398
refGene_gn=RMI1;refGene_tx=XM_023242216.1;refGene_gn=RMI1;refGene_tx=XM_023242217.1
D4 25364286 25364413 refGene_gn=SLC28A3;refGene_tx=XM_003995451.3
D4 25365495 25365616 refGene_gn=SLC28A3;refGene_tx=XM_003995451.3
D4 25365864 25365963 refGene_gn=SLC28A3;refGene_tx=XM_003995451.3
D4 25366918 25367000 refGene_gn=SLC28A3;refGene_tx=XM_003995451.3
D4 25373210 25373408 refGene_gn=SLC28A3;refGene_tx=XM_003995451.3
D4 25373786 25373955 refGene_gn=SLC28A3;refGene_tx=XM_003995451.3
D4 25375113 25375244 refGene_gn=SLC28A3;refGene_tx=XM_003995451.3

D4 25376876 25377002 refGene_gn=SLC28A3;refGene_tx=XM_003995451.3
D4 25378953 25379034 refGene_gn=SLC28A3;refGene_tx=XM_003995451.3
D4 25380479 25380560 refGene_gn=SLC28A3;refGene_tx=XM_003995451.3
D4 25382730 25382808 refGene_gn=SLC28A3;refGene_tx=XM_003995451.3
D4 25383897 25384011 refGene_gn=SLC28A3;refGene_tx=XM_003995451.3
D4 25385245 25385390 refGene_gn=SLC28A3;refGene_tx=XM_003995451.3
D4 25386478 25386668 refGene_gn=SLC28A3;refGene_tx=XM_003995451.3
D4 25388086 25388178 refGene_gn=SLC28A3;refGene_tx=XM_003995451.3
D4 25391299 25391382 refGene_gn=SLC28A3;refGene_tx=XM_003995451.3
D4 25396174 25396258 refGene_gn=SLC28A3;refGene_tx=XM_003995451.3
D4 25418119 25418197 refGene_gn=SLC28A3;refGene_tx=XM_003995451.3
D4 25709021 25709233
refGene_gn=NTRK2;refGene_tx=XM_023242218.1;refGene_gn=NTRK2;refGene_tx=XM_023242219.1;refGene_gn=NTRK2;refGene_tx=XM_023242220.1;refGene_gn=NTRK2;refGene_tx=XM_023242221.1;refGene_gn=NTRK2;refGene_tx=XM_023242222.1
D4 25736801 25736876
refGene_gn=NTRK2;refGene_tx=XM_023242218.1;refGene_gn=NTRK2;refGene_tx=XM_023242219.1;refGene_gn=NTRK2;refGene_tx=XM_023242220.1;refGene_gn=NTRK2;refGene_tx=XM_023242221.1;refGene_gn=NTRK2;refGene_tx=XM_023242222.1
D4 25736998 25737070
refGene_gn=NTRK2;refGene_tx=XM_023242218.1;refGene_gn=NTRK2;refGene_tx=XM_023242219.1;refGene_gn=NTRK2;refGene_tx=XM_023242220.1;refGene_gn=NTRK2;refGene_tx=XM_023242221.1;refGene_gn=NTRK2;refGene_tx=XM_023242222.1
D4 25741794 25741863
refGene_gn=NTRK2;refGene_tx=XM_023242218.1;refGene_gn=NTRK2;refGene_tx=XM_023242219.1;refGene_gn=NTRK2;refGene_tx=XM_023242220.1;refGene_gn=NTRK2;refGene_tx=XM_023242221.1;refGene_gn=NTRK2;refGene_tx=XM_023242222.1
D4 25744628 25744783
refGene_gn=NTRK2;refGene_tx=XM_023242218.1;refGene_gn=NTRK2;refGene_tx=XM_023242219.1;refGene_gn=NTRK2;refGene_tx=XM_023242220.1;refGene_gn=NTRK2;refGene_tx=XM_023242221.1;refGene_gn=NTRK2;refGene_tx=XM_023242222.1
D4 25757557 25757694
refGene_gn=NTRK2;refGene_tx=XM_023242218.1;refGene_gn=NTRK2;refGene_tx=XM_023242219.1;refGene_gn=NTRK2;refGene_tx=XM_023242220.1;refGene_gn=NTRK2;refGene_tx=XM_023242221.1;refGene_gn=NTRK2;refGene_tx=XM_023242222.1
D4 25758201 25758334
refGene_gn=NTRK2;refGene_tx=XM_023242218.1;refGene_gn=NTRK2;refGene_tx=XM_023242219.1;refGene_gn=NTRK2;refGene_tx=XM_023242220.1;refGene_gn=NTRK2;refGene_tx=XM_023242221.1;refGene_gn=NTRK2;refGene_tx=XM_023242222.1
D4 25760799 25761105
refGene_gn=NTRK2;refGene_tx=XM_023242218.1;refGene_gn=NTRK2;refGene_tx=XM_023242219.1;refGene_gn=NTRK2;refGene_tx=XM_023242220.1;refGene_gn=NTRK2;refGene_tx=XM_023242221.1;refGene_gn=NTRK2;refGene_tx=XM_0232422

22.1
D4	25772644	25772680
refGene_gn=NTRK2;refGene_tx=XM_023242218.1;refGene_gn=NTRK2;refGene_tx
=XM_023242219.1;refGene_gn=NTRK2;refGene_tx=XM_023242220.1;refGene_gn=
NTRK2;refGene_tx=XM_023242221.1;refGene_gn=NTRK2;refGene_tx=XM_0232422
22.1
D4	25775719	25775820
refGene_gn=NTRK2;refGene_tx=XM_023242218.1;refGene_gn=NTRK2;refGene_tx
=XM_023242219.1;refGene_gn=NTRK2;refGene_tx=XM_023242220.1;refGene_gn=
NTRK2;refGene_tx=XM_023242221.1;refGene_gn=NTRK2;refGene_tx=XM_0232422
22.1
D4	25782281	25782381
refGene_gn=NTRK2;refGene_tx=XM_023242218.1;refGene_gn=NTRK2;refGene_tx
=XM_023242219.1;refGene_gn=NTRK2;refGene_tx=XM_023242220.1;refGene_gn=
NTRK2;refGene_tx=XM_023242221.1;refGene_gn=NTRK2;refGene_tx=XM_0232422
22.1
D4	25841102	25841140	refGene_gn=NTRK2;refGene_tx=XM_023242222.1
D4	25890785	25890833
refGene_gn=NTRK2;refGene_tx=XM_023242218.1;refGene_gn=NTRK2;refGene_tx
=XM_023242220.1
D4	25896367	25896556
refGene_gn=NTRK2;refGene_tx=XM_023242218.1;refGene_gn=NTRK2;refGene_tx
=XM_023242219.1;refGene_gn=NTRK2;refGene_tx=XM_023242220.1;refGene_gn=
NTRK2;refGene_tx=XM_023242221.1
D4	25900698	25900727
refGene_gn=NTRK2;refGene_tx=XM_023242220.1;refGene_gn=NTRK2;refGene_tx
=XM_023242221.1
D4	25958767	25958898
refGene_gn=NTRK2;refGene_tx=XM_023242218.1;refGene_gn=NTRK2;refGene_tx
=XM_023242219.1
D4	25973475	25973648
refGene_gn=NTRK2;refGene_tx=XM_023242218.1;refGene_gn=NTRK2;refGene_tx
=XM_023242219.1
D4	25978575	25978810
refGene_gn=NTRK2;refGene_tx=XM_023242218.1;refGene_gn=NTRK2;refGene_tx
=XM_023242219.1
D4	26040495	26040654
refGene_gn=NTRK2;refGene_tx=XM_023242218.1;refGene_gn=NTRK2;refGene_tx
=XM_023242219.1
D4	26041435	26041621
refGene_gn=NTRK2;refGene_tx=XM_023242218.1;refGene_gn=NTRK2;refGene_tx
=XM_023242219.1
D4	26450919	26451097
refGene_gn=AGTPBP1;refGene_tx=XM_023242223.1;refGene_gn=AGTPBP1;refGen
e_tx=XM_023242224.1
D4	26456849	26456895	refGene_gn=AGTPBP1;refGene_tx=XM_023242226.1
D4	26457163	26457329	refGene_gn=AGTPBP1;refGene_tx=XM_023242225.1
D4	26465034	26465195
refGene_gn=AGTPBP1;refGene_tx=XM_023242223.1;refGene_gn=AGTPBP1;refGen
e_tx=XM_023242224.1;refGene_gn=AGTPBP1;refGene_tx=XM_023242225.1;refGe

ne_gn=AGTPBP1;refGene_tx=XM_023242226.1
D4 26468840 26469017
refGene_gn=AGTPBP1;refGene_tx=XM_023242223.1;refGene_gn=AGTPBP1;refGen
e_tx=XM_023242224.1;refGene_gn=AGTPBP1;refGene_tx=XM_023242225.1;refGe
ne_gn=AGTPBP1;refGene_tx=XM_023242226.1
D4 26471034 26471166
refGene_gn=AGTPBP1;refGene_tx=XM_023242223.1;refGene_gn=AGTPBP1;refGen
e_tx=XM_023242224.1;refGene_gn=AGTPBP1;refGene_tx=XM_023242225.1;refGe
ne_gn=AGTPBP1;refGene_tx=XM_023242226.1
D4 26472379 26472509
refGene_gn=AGTPBP1;refGene_tx=XM_023242223.1;refGene_gn=AGTPBP1;refGen
e_tx=XM_023242224.1;refGene_gn=AGTPBP1;refGene_tx=XM_023242225.1;refGe
ne_gn=AGTPBP1;refGene_tx=XM_023242226.1
D4 26473996 26474177
refGene_gn=AGTPBP1;refGene_tx=XM_023242223.1;refGene_gn=AGTPBP1;refGen
e_tx=XM_023242224.1;refGene_gn=AGTPBP1;refGene_tx=XM_023242225.1;refGe
ne_gn=AGTPBP1;refGene_tx=XM_023242226.1
D4 26475209 26475363
refGene_gn=AGTPBP1;refGene_tx=XM_023242223.1;refGene_gn=AGTPBP1;refGen
e_tx=XM_023242224.1;refGene_gn=AGTPBP1;refGene_tx=XM_023242225.1;refGe
ne_gn=AGTPBP1;refGene_tx=XM_023242226.1
D4 26477888 26478033
refGene_gn=AGTPBP1;refGene_tx=XM_023242223.1;refGene_gn=AGTPBP1;refGen
e_tx=XM_023242224.1;refGene_gn=AGTPBP1;refGene_tx=XM_023242225.1;refGe
ne_gn=AGTPBP1;refGene_tx=XM_023242226.1
D4 26481424 26481512
refGene_gn=AGTPBP1;refGene_tx=XM_023242223.1;refGene_gn=AGTPBP1;refGen
e_tx=XM_023242224.1;refGene_gn=AGTPBP1;refGene_tx=XM_023242225.1;refGe
ne_gn=AGTPBP1;refGene_tx=XM_023242226.1
D4 26499538 26499687
refGene_gn=AGTPBP1;refGene_tx=XM_023242223.1;refGene_gn=AGTPBP1;refGen
e_tx=XM_023242224.1;refGene_gn=AGTPBP1;refGene_tx=XM_023242225.1;refGe
ne_gn=AGTPBP1;refGene_tx=XM_023242226.1
D4 26499771 26499858
refGene_gn=AGTPBP1;refGene_tx=XM_023242223.1;refGene_gn=AGTPBP1;refGen
e_tx=XM_023242224.1;refGene_gn=AGTPBP1;refGene_tx=XM_023242225.1;refGe
ne_gn=AGTPBP1;refGene_tx=XM_023242226.1
D4 26501437 26501521
refGene_gn=AGTPBP1;refGene_tx=XM_023242223.1;refGene_gn=AGTPBP1;refGen
e_tx=XM_023242224.1;refGene_gn=AGTPBP1;refGene_tx=XM_023242225.1;refGe
ne_gn=AGTPBP1;refGene_tx=XM_023242226.1
D4 26517164 26517880
refGene_gn=AGTPBP1;refGene_tx=XM_023242223.1;refGene_gn=AGTPBP1;refGen
e_tx=XM_023242224.1;refGene_gn=AGTPBP1;refGene_tx=XM_023242225.1;refGe
ne_gn=AGTPBP1;refGene_tx=XM_023242226.1
D4 26527749 26527866
refGene_gn=AGTPBP1;refGene_tx=XM_023242223.1;refGene_gn=AGTPBP1;refGen
e_tx=XM_023242224.1;refGene_gn=AGTPBP1;refGene_tx=XM_023242225.1;refGe
ne_gn=AGTPBP1;refGene_tx=XM_023242226.1
D4 26532631 26532729

refGene_gn=AGTPBP1;refGene_tx=XM_023242223.1;refGene_gn=AGTPBP1;refGene_tx=XM_023242224.1;refGene_gn=AGTPBP1;refGene_tx=XM_023242225.1;refGene_gn=AGTPBP1;refGene_tx=XM_023242226.1
D4 26549575 26549753
refGene_gn=AGTPBP1;refGene_tx=XM_023242223.1;refGene_gn=AGTPBP1;refGene_tx=XM_023242224.1;refGene_gn=AGTPBP1;refGene_tx=XM_023242225.1;refGene_gn=AGTPBP1;refGene_tx=XM_023242226.1
D4 26551804 26552013
refGene_gn=AGTPBP1;refGene_tx=XM_023242223.1;refGene_gn=AGTPBP1;refGene_tx=XM_023242224.1;refGene_gn=AGTPBP1;refGene_tx=XM_023242225.1;refGene_gn=AGTPBP1;refGene_tx=XM_023242226.1
D4 26555603 26555641
refGene_gn=AGTPBP1;refGene_tx=XM_023242223.1;refGene_gn=AGTPBP1;refGene_tx=XM_023242224.1;refGene_gn=AGTPBP1;refGene_tx=XM_023242225.1;refGene_gn=AGTPBP1;refGene_tx=XM_023242226.1
D4 26562412 26562506
refGene_gn=AGTPBP1;refGene_tx=XM_023242223.1;refGene_gn=AGTPBP1;refGene_tx=XM_023242224.1;refGene_gn=AGTPBP1;refGene_tx=XM_023242225.1;refGene_gn=AGTPBP1;refGene_tx=XM_023242226.1
D4 26565332 26565464
refGene_gn=AGTPBP1;refGene_tx=XM_023242223.1;refGene_gn=AGTPBP1;refGene_tx=XM_023242224.1;refGene_gn=AGTPBP1;refGene_tx=XM_023242225.1;refGene_gn=AGTPBP1;refGene_tx=XM_023242226.1
D4 26567630 26567777
refGene_gn=AGTPBP1;refGene_tx=XM_023242223.1;refGene_gn=AGTPBP1;refGene_tx=XM_023242224.1;refGene_gn=AGTPBP1;refGene_tx=XM_023242225.1;refGene_gn=AGTPBP1;refGene_tx=XM_023242226.1
D4 26568220 26568284
refGene_gn=AGTPBP1;refGene_tx=XM_023242223.1;refGene_gn=AGTPBP1;refGene_tx=XM_023242224.1;refGene_gn=AGTPBP1;refGene_tx=XM_023242225.1;refGene_gn=AGTPBP1;refGene_tx=XM_023242226.1
D4 26571417 26571485
refGene_gn=AGTPBP1;refGene_tx=XM_023242223.1;refGene_gn=AGTPBP1;refGene_tx=XM_023242224.1;refGene_gn=AGTPBP1;refGene_tx=XM_023242225.1;refGene_gn=AGTPBP1;refGene_tx=XM_023242226.1
D4 26582268 26582393
refGene_gn=AGTPBP1;refGene_tx=XM_023242223.1;refGene_gn=AGTPBP1;refGene_tx=XM_023242224.1;refGene_gn=AGTPBP1;refGene_tx=XM_023242225.1;refGene_gn=AGTPBP1;refGene_tx=XM_023242226.1
D4 26592232 26592264
refGene_gn=AGTPBP1;refGene_tx=XM_023242223.1;refGene_gn=AGTPBP1;refGene_tx=XM_023242224.1;refGene_gn=AGTPBP1;refGene_tx=XM_023242225.1;refGene_gn=AGTPBP1;refGene_tx=XM_023242226.1
D4 26699083 26699207
refGene_gn=NAA35;refGene_tx=XM_011288304.3;refGene_gn=NAA35;refGene_tx=XM_023242227.1;refGene_gn=NAA35;refGene_tx=XM_023242228.1
D4 26710442 26710476
refGene_gn=NAA35;refGene_tx=XM_011288304.3;refGene_gn=NAA35;refGene_tx=XM_023242227.1;refGene_gn=NAA35;refGene_tx=XM_023242228.1
D4 26714338 26714453

refGene_gn=NAA35;refGene_tx=XM_011288304.3;refGene_gn=NAA35;refGene_tx=XM_023242227.1;refGene_gn=NAA35;refGene_tx=XM_023242228.1
D4 26715723 26715798
refGene_gn=NAA35;refGene_tx=XM_011288304.3;refGene_gn=NAA35;refGene_tx=XM_023242227.1;refGene_gn=NAA35;refGene_tx=XM_023242228.1
D4 26716381 26716549
refGene_gn=NAA35;refGene_tx=XM_011288304.3;refGene_gn=NAA35;refGene_tx=XM_023242227.1;refGene_gn=NAA35;refGene_tx=XM_023242228.1
D4 26733640 26733707
refGene_gn=NAA35;refGene_tx=XM_011288304.3;refGene_gn=NAA35;refGene_tx=XM_023242227.1;refGene_gn=NAA35;refGene_tx=XM_023242228.1
D4 26733786 26733830
refGene_gn=NAA35;refGene_tx=XM_011288304.3;refGene_gn=NAA35;refGene_tx=XM_023242227.1;refGene_gn=NAA35;refGene_tx=XM_023242228.1
D4 26735066 26735117
refGene_gn=NAA35;refGene_tx=XM_011288304.3;refGene_gn=NAA35;refGene_tx=XM_023242227.1;refGene_gn=NAA35;refGene_tx=XM_023242228.1
D4 26736002 26736086
refGene_gn=NAA35;refGene_tx=XM_011288304.3;refGene_gn=NAA35;refGene_tx=XM_023242227.1;refGene_gn=NAA35;refGene_tx=XM_023242228.1
D4 26737049 26737164
refGene_gn=NAA35;refGene_tx=XM_011288304.3;refGene_gn=NAA35;refGene_tx=XM_023242227.1;refGene_gn=NAA35;refGene_tx=XM_023242228.1
D4 26762470 26762649
refGene_gn=NAA35;refGene_tx=XM_011288304.3;refGene_gn=NAA35;refGene_tx=XM_023242227.1;refGene_gn=NAA35;refGene_tx=XM_023242228.1
D4 26766750 26766810
refGene_gn=NAA35;refGene_tx=XM_011288304.3;refGene_gn=NAA35;refGene_tx=XM_023242227.1;refGene_gn=NAA35;refGene_tx=XM_023242228.1
D4 26771627 26771734
refGene_gn=NAA35;refGene_tx=XM_011288304.3;refGene_gn=NAA35;refGene_tx=XM_023242227.1;refGene_gn=NAA35;refGene_tx=XM_023242228.1
D4 26779966 26780033
refGene_gn=NAA35;refGene_tx=XM_011288304.3;refGene_gn=NAA35;refGene_tx=XM_023242227.1;refGene_gn=NAA35;refGene_tx=XM_023242228.1
D4 26781914 26782013
refGene_gn=NAA35;refGene_tx=XM_011288304.3;refGene_gn=NAA35;refGene_tx=XM_023242227.1;refGene_gn=NAA35;refGene_tx=XM_023242228.1
D4 26782826 26783005
refGene_gn=NAA35;refGene_tx=XM_011288304.3;refGene_gn=NAA35;refGene_tx=XM_023242227.1;refGene_gn=NAA35;refGene_tx=XM_023242228.1
D4 26785982 26786119
refGene_gn=NAA35;refGene_tx=XM_011288304.3;refGene_gn=NAA35;refGene_tx=XM_023242227.1;refGene_gn=NAA35;refGene_tx=XM_023242228.1
D4 26786983 26787051
refGene_gn=NAA35;refGene_tx=XM_011288304.3;refGene_gn=NAA35;refGene_tx=XM_023242227.1;refGene_gn=NAA35;refGene_tx=XM_023242228.1
D4 26787993 26788134
refGene_gn=NAA35;refGene_tx=XM_011288304.3;refGene_gn=NAA35;refGene_tx=XM_023242227.1;refGene_gn=NAA35;refGene_tx=XM_023242228.1

D4 26788528 26788651
refGene_gn=NAA35;refGene_tx=XM_011288304.3;refGene_gn=NAA35;refGene_tx=XM_023242227.1;refGene_gn=NAA35;refGene_tx=XM_023242228.1
D4 26791714 26791795
refGene_gn=NAA35;refGene_tx=XM_011288304.3;refGene_gn=NAA35;refGene_tx=XM_023242227.1;refGene_gn=NAA35;refGene_tx=XM_023242228.1
D4 26792518 26792578
refGene_gn=NAA35;refGene_tx=XM_011288304.3;refGene_gn=NAA35;refGene_tx=XM_023242227.1;refGene_gn=NAA35;refGene_tx=XM_023242228.1
D4 26796272 26796349 refGene_gn=GOLM1;refGene_tx=XM_023242229.1
D4 26802933 26803047 refGene_gn=GOLM1;refGene_tx=XM_023242229.1
D4 26804564 26804837 refGene_gn=GOLM1;refGene_tx=XM_023242229.1
D4 26805526 26805686 refGene_gn=GOLM1;refGene_tx=XM_023242229.1
D4 26810051 26810181 refGene_gn=GOLM1;refGene_tx=XM_023242229.1
D4 26817855 26817958 refGene_gn=GOLM1;refGene_tx=XM_023242229.1
D4 26821241 26821296 refGene_gn=GOLM1;refGene_tx=XM_023242229.1
D4 26837563 26837743 refGene_gn=GOLM1;refGene_tx=XM_023242229.1
D4 26839553 26839682 refGene_gn=GOLM1;refGene_tx=XM_023242229.1
D4 26926699 26926923
refGene_gn=CD4H9orf153;refGene_tx=XM_019815828.2;refGene_gn=CD4H9orf153;refGene_tx=XM_019815829.2;refGene_gn=CD4H9orf153;refGene_tx=XM_019815830.2;refGene_gn=CD4H9orf153;refGene_tx=XM_019815831.2
D4 26930140 26930636
refGene_gn=CD4H9orf153;refGene_tx=XM_019815828.2;refGene_gn=CD4H9orf153;refGene_tx=XM_019815829.2;refGene_gn=CD4H9orf153;refGene_tx=XM_019815830.2;refGene_gn=CD4H9orf153;refGene_tx=XM_019815831.2
D4 26932105 26932171
refGene_gn=CD4H9orf153;refGene_tx=XM_019815829.2;refGene_gn=CD4H9orf153;refGene_tx=XM_019815830.2;refGene_gn=CD4H9orf153;refGene_tx=XM_019815831.2
D4 26954951 26955086
refGene_gn=CD4H9orf153;refGene_tx=XM_019815828.2
D4 26962380 26962529 refGene_gn=ISCA1;refGene_tx=XM_003995454.4
D4 26964136 26964242 refGene_gn=ISCA1;refGene_tx=XM_003995454.4
D4 26967386 26967440 refGene_gn=ISCA1;refGene_tx=XM_003995454.4
D4 26974943 26975024 refGene_gn=ISCA1;refGene_tx=XM_003995454.4
D4 26980401 26980469
refGene_gn=ZCCHC6;refGene_tx=XM_003995455.5;refGene_gn=ZCCHC6;refGene_tx=XM_019815832.2;refGene_gn=ZCCHC6;refGene_tx=XM_019815833.2;refGene_gn=ZCCHC6;refGene_tx=XM_019815834.2;refGene_gn=ZCCHC6;refGene_tx=XM_019815835.2;refGene_gn=ZCCHC6;refGene_tx=XM_019815837.2
D4 26991556 26991620
refGene_gn=ZCCHC6;refGene_tx=XM_019815832.2;refGene_gn=ZCCHC6;refGene_tx=XM_019815833.2;refGene_gn=ZCCHC6;refGene_tx=XM_019815834.2;refGene_gn=ZCCHC6;refGene_tx=XM_019815835.2;refGene_gn=ZCCHC6;refGene_tx=XM_019815837.2;refGene_gn=ZCCHC6;refGene_tx=XM_019815838.2
D4 26991978 26992055
refGene_gn=ZCCHC6;refGene_tx=XM_019815832.2;refGene_gn=ZCCHC6;refGene_tx=XM_019815833.2;refGene_gn=ZCCHC6;refGene_tx=XM_019815834.2;refGene_gn=ZCCHC6;refGene_tx=XM_019815835.2;refGene_gn=ZCCHC6;refGene_tx=XM_01

9815837.2
D4	26994011	26994337
refGene_gn=ZCCHC6;refGene_tx=XM_003995455.5;refGene_gn=ZCCHC6;refGene_tx=XM_019815832.2;refGene_gn=ZCCHC6;refGene_tx=XM_019815833.2;refGene_gn=ZCCHC6;refGene_tx=XM_019815834.2;refGene_gn=ZCCHC6;refGene_tx=XM_019815835.2;refGene_gn=ZCCHC6;refGene_tx=XM_019815837.2;refGene_gn=ZCCHC6;refGene_tx=XM_019815838.2
D4	26995448	26995564
refGene_gn=ZCCHC6;refGene_tx=XM_003995455.5;refGene_gn=ZCCHC6;refGene_tx=XM_019815832.2;refGene_gn=ZCCHC6;refGene_tx=XM_019815833.2;refGene_gn=ZCCHC6;refGene_tx=XM_019815834.2;refGene_gn=ZCCHC6;refGene_tx=XM_019815835.2;refGene_gn=ZCCHC6;refGene_tx=XM_019815837.2;refGene_gn=ZCCHC6;refGene_tx=XM_019815838.2
D4	26997425	26997517
refGene_gn=ZCCHC6;refGene_tx=XM_003995455.5;refGene_gn=ZCCHC6;refGene_tx=XM_019815832.2;refGene_gn=ZCCHC6;refGene_tx=XM_019815833.2;refGene_gn=ZCCHC6;refGene_tx=XM_019815834.2;refGene_gn=ZCCHC6;refGene_tx=XM_019815835.2;refGene_gn=ZCCHC6;refGene_tx=XM_019815837.2;refGene_gn=ZCCHC6;refGene_tx=XM_019815838.2
D4	26997603	26997651
refGene_gn=ZCCHC6;refGene_tx=XM_003995455.5;refGene_gn=ZCCHC6;refGene_tx=XM_019815832.2;refGene_gn=ZCCHC6;refGene_tx=XM_019815833.2;refGene_gn=ZCCHC6;refGene_tx=XM_019815834.2;refGene_gn=ZCCHC6;refGene_tx=XM_019815835.2;refGene_gn=ZCCHC6;refGene_tx=XM_019815837.2;refGene_gn=ZCCHC6;refGene_tx=XM_019815838.2
D4	27001973	27002151
refGene_gn=ZCCHC6;refGene_tx=XM_003995455.5;refGene_gn=ZCCHC6;refGene_tx=XM_019815832.2;refGene_gn=ZCCHC6;refGene_tx=XM_019815833.2;refGene_gn=ZCCHC6;refGene_tx=XM_019815834.2;refGene_gn=ZCCHC6;refGene_tx=XM_019815835.2;refGene_gn=ZCCHC6;refGene_tx=XM_019815837.2;refGene_gn=ZCCHC6;refGene_tx=XM_019815838.2
D4	27002796	27002874
refGene_gn=ZCCHC6;refGene_tx=XM_003995455.5;refGene_gn=ZCCHC6;refGene_tx=XM_019815832.2;refGene_gn=ZCCHC6;refGene_tx=XM_019815833.2;refGene_gn=ZCCHC6;refGene_tx=XM_019815834.2;refGene_gn=ZCCHC6;refGene_tx=XM_019815835.2;refGene_gn=ZCCHC6;refGene_tx=XM_019815837.2;refGene_gn=ZCCHC6;refGene_tx=XM_019815838.2
D4	27003048	27003162
refGene_gn=ZCCHC6;refGene_tx=XM_003995455.5;refGene_gn=ZCCHC6;refGene_tx=XM_019815832.2;refGene_gn=ZCCHC6;refGene_tx=XM_019815833.2;refGene_gn=ZCCHC6;refGene_tx=XM_019815834.2;refGene_gn=ZCCHC6;refGene_tx=XM_019815835.2;refGene_gn=ZCCHC6;refGene_tx=XM_019815837.2;refGene_gn=ZCCHC6;refGene_tx=XM_019815838.2
D4	27003528	27003618
refGene_gn=ZCCHC6;refGene_tx=XM_003995455.5;refGene_gn=ZCCHC6;refGene_tx=XM_019815832.2;refGene_gn=ZCCHC6;refGene_tx=XM_019815833.2;refGene_gn=ZCCHC6;refGene_tx=XM_019815834.2;refGene_gn=ZCCHC6;refGene_tx=XM_019815835.2;refGene_gn=ZCCHC6;refGene_tx=XM_019815837.2;refGene_gn=ZCCHC6;refGene_tx=XM_019815838.2
D4	27004232	27004336

refGene_gn=ZCCHC6;refGene_tx=XM_003995455.5;refGene_gn=ZCCHC6;refGene_tx=XM_019815832.2;refGene_gn=ZCCHC6;refGene_tx=XM_019815833.2;refGene_gn=ZCCHC6;refGene_tx=XM_019815834.2;refGene_gn=ZCCHC6;refGene_tx=XM_019815835.2;refGene_gn=ZCCHC6;refGene_tx=XM_019815837.2;refGene_gn=ZCCHC6;refGene_tx=XM_019815838.2
D4 27009226 27009284
refGene_gn=ZCCHC6;refGene_tx=XM_003995455.5;refGene_gn=ZCCHC6;refGene_tx=XM_019815832.2;refGene_gn=ZCCHC6;refGene_tx=XM_019815833.2;refGene_gn=ZCCHC6;refGene_tx=XM_019815834.2;refGene_gn=ZCCHC6;refGene_tx=XM_019815835.2;refGene_gn=ZCCHC6;refGene_tx=XM_019815837.2;refGene_gn=ZCCHC6;refGene_tx=XM_019815838.2
D4 27010924 27011025
refGene_gn=ZCCHC6;refGene_tx=XM_003995455.5;refGene_gn=ZCCHC6;refGene_tx=XM_019815832.2;refGene_gn=ZCCHC6;refGene_tx=XM_019815833.2;refGene_gn=ZCCHC6;refGene_tx=XM_019815834.2;refGene_gn=ZCCHC6;refGene_tx=XM_019815835.2;refGene_gn=ZCCHC6;refGene_tx=XM_019815837.2;refGene_gn=ZCCHC6;refGene_tx=XM_019815838.2
D4 27011890 27011977
refGene_gn=ZCCHC6;refGene_tx=XM_003995455.5;refGene_gn=ZCCHC6;refGene_tx=XM_019815832.2;refGene_gn=ZCCHC6;refGene_tx=XM_019815833.2;refGene_gn=ZCCHC6;refGene_tx=XM_019815834.2;refGene_gn=ZCCHC6;refGene_tx=XM_019815835.2;refGene_gn=ZCCHC6;refGene_tx=XM_019815837.2;refGene_gn=ZCCHC6;refGene_tx=XM_019815838.2
D4 27013169 27013320
refGene_gn=ZCCHC6;refGene_tx=XM_003995455.5;refGene_gn=ZCCHC6;refGene_tx=XM_019815832.2;refGene_gn=ZCCHC6;refGene_tx=XM_019815833.2;refGene_gn=ZCCHC6;refGene_tx=XM_019815834.2;refGene_gn=ZCCHC6;refGene_tx=XM_019815835.2;refGene_gn=ZCCHC6;refGene_tx=XM_019815837.2;refGene_gn=ZCCHC6;refGene_tx=XM_019815838.2
D4 27013690 27014769
refGene_gn=ZCCHC6;refGene_tx=XM_003995455.5;refGene_gn=ZCCHC6;refGene_tx=XM_019815832.2;refGene_gn=ZCCHC6;refGene_tx=XM_019815833.2;refGene_gn=ZCCHC6;refGene_tx=XM_019815834.2;refGene_gn=ZCCHC6;refGene_tx=XM_019815835.2;refGene_gn=ZCCHC6;refGene_tx=XM_019815837.2;refGene_gn=ZCCHC6;refGene_tx=XM_019815838.2
D4 27016167 27016348
refGene_gn=ZCCHC6;refGene_tx=XM_003995455.5;refGene_gn=ZCCHC6;refGene_tx=XM_019815832.2;refGene_gn=ZCCHC6;refGene_tx=XM_019815833.2;refGene_gn=ZCCHC6;refGene_tx=XM_019815834.2;refGene_gn=ZCCHC6;refGene_tx=XM_019815835.2;refGene_gn=ZCCHC6;refGene_tx=XM_019815837.2;refGene_gn=ZCCHC6;refGene_tx=XM_019815838.2
D4 27019748 27019901
refGene_gn=ZCCHC6;refGene_tx=XM_003995455.5;refGene_gn=ZCCHC6;refGene_tx=XM_019815832.2;refGene_gn=ZCCHC6;refGene_tx=XM_019815833.2;refGene_gn=ZCCHC6;refGene_tx=XM_019815834.2;refGene_gn=ZCCHC6;refGene_tx=XM_019815835.2;refGene_gn=ZCCHC6;refGene_tx=XM_019815837.2;refGene_gn=ZCCHC6;refGene_tx=XM_019815838.2
D4 27025393 27025513
refGene_gn=ZCCHC6;refGene_tx=XM_003995455.5;refGene_gn=ZCCHC6;refGene_tx=XM_019815832.2;refGene_gn=ZCCHC6;refGene_tx=XM_019815833.2;refGene_

gn=ZCCHC6;refGene_tx=XM_019815834.2;refGene_gn=ZCCHC6;refGene_tx=XM_019815835.2;refGene_gn=ZCCHC6;refGene_tx=XM_019815837.2;refGene_gn=ZCCHC6;refGene_tx=XM_019815838.2
D4 27028659 27028786
refGene_gn=ZCCHC6;refGene_tx=XM_003995455.5;refGene_gn=ZCCHC6;refGene_tx=XM_019815832.2;refGene_gn=ZCCHC6;refGene_tx=XM_019815833.2;refGene_gn=ZCCHC6;refGene_tx=XM_019815834.2;refGene_gn=ZCCHC6;refGene_tx=XM_019815835.2;refGene_gn=ZCCHC6;refGene_tx=XM_019815837.2;refGene_gn=ZCCHC6;refGene_tx=XM_019815838.2
D4 27029583 27029653
refGene_gn=ZCCHC6;refGene_tx=XM_003995455.5;refGene_gn=ZCCHC6;refGene_tx=XM_019815832.2;refGene_gn=ZCCHC6;refGene_tx=XM_019815833.2;refGene_gn=ZCCHC6;refGene_tx=XM_019815834.2;refGene_gn=ZCCHC6;refGene_tx=XM_019815835.2;refGene_gn=ZCCHC6;refGene_tx=XM_019815837.2;refGene_gn=ZCCHC6;refGene_tx=XM_019815838.2
D4 27030581 27030633
refGene_gn=ZCCHC6;refGene_tx=XM_003995455.5;refGene_gn=ZCCHC6;refGene_tx=XM_019815832.2;refGene_gn=ZCCHC6;refGene_tx=XM_019815833.2;refGene_gn=ZCCHC6;refGene_tx=XM_019815834.2;refGene_gn=ZCCHC6;refGene_tx=XM_019815835.2;refGene_gn=ZCCHC6;refGene_tx=XM_019815837.2;refGene_gn=ZCCHC6;refGene_tx=XM_019815838.2
D4 27032797 27032886
refGene_gn=ZCCHC6;refGene_tx=XM_003995455.5;refGene_gn=ZCCHC6;refGene_tx=XM_019815832.2;refGene_gn=ZCCHC6;refGene_tx=XM_019815833.2;refGene_gn=ZCCHC6;refGene_tx=XM_019815834.2;refGene_gn=ZCCHC6;refGene_tx=XM_019815835.2;refGene_gn=ZCCHC6;refGene_tx=XM_019815837.2;refGene_gn=ZCCHC6;refGene_tx=XM_019815838.2
D4 27034485 27034663
refGene_gn=ZCCHC6;refGene_tx=XM_003995455.5;refGene_gn=ZCCHC6;refGene_tx=XM_019815832.2;refGene_gn=ZCCHC6;refGene_tx=XM_019815833.2;refGene_gn=ZCCHC6;refGene_tx=XM_019815834.2;refGene_gn=ZCCHC6;refGene_tx=XM_019815835.2;refGene_gn=ZCCHC6;refGene_tx=XM_019815837.2;refGene_gn=ZCCHC6;refGene_tx=XM_019815838.2
D4 27035164 27035281
refGene_gn=ZCCHC6;refGene_tx=XM_003995455.5;refGene_gn=ZCCHC6;refGene_tx=XM_019815832.2;refGene_gn=ZCCHC6;refGene_tx=XM_019815833.2;refGene_gn=ZCCHC6;refGene_tx=XM_019815834.2;refGene_gn=ZCCHC6;refGene_tx=XM_019815835.2;refGene_gn=ZCCHC6;refGene_tx=XM_019815837.2;refGene_gn=ZCCHC6;refGene_tx=XM_019815838.2
D4 27035594 27035776
refGene_gn=ZCCHC6;refGene_tx=XM_003995455.5;refGene_gn=ZCCHC6;refGene_tx=XM_019815832.2;refGene_gn=ZCCHC6;refGene_tx=XM_019815833.2;refGene_gn=ZCCHC6;refGene_tx=XM_019815834.2;refGene_gn=ZCCHC6;refGene_tx=XM_019815835.2;refGene_gn=ZCCHC6;refGene_tx=XM_019815837.2;refGene_gn=ZCCHC6;refGene_tx=XM_019815838.2
D4 27040834 27041354
refGene_gn=ZCCHC6;refGene_tx=XM_003995455.5;refGene_gn=ZCCHC6;refGene_tx=XM_019815832.2;refGene_gn=ZCCHC6;refGene_tx=XM_019815833.2;refGene_gn=ZCCHC6;refGene_tx=XM_019815834.2;refGene_gn=ZCCHC6;refGene_tx=XM_019815835.2;refGene_gn=ZCCHC6;refGene_tx=XM_019815837.2;refGene_gn=ZCCHC

6;refGene_tx=XM_019815838.2
D4 27349834 27349856
refGene_gn=LOC109493773;refGene_tx=XM_019815840.2;refGene_gn=LOC109493773;refGene_tx=XM_019815841.2;refGene_gn=LOC109493773;refGene_tx=XM_019815842.2;refGene_gn=LOC109493773;refGene_tx=XM_019815843.2;refGene_gn=LOC109493773;refGene_tx=XM_019815844.2;refGene_gn=LOC109493773;refGene_tx=XM_023242230.1;refGene_gn=LOC109493773;refGene_tx=XM_023242231.1;refGene_gn=LOC109493773;refGene_tx=XM_023242232.1;refGene_gn=LOC109493773;refGene_tx=XM_023242233.1;refGene_gn=LOC109493773;refGene_tx=XM_023242234.1
D4 27350209 27350367
refGene_gn=LOC109493773;refGene_tx=XM_019815840.2;refGene_gn=LOC109493773;refGene_tx=XM_019815841.2;refGene_gn=LOC109493773;refGene_tx=XM_019815842.2;refGene_gn=LOC109493773;refGene_tx=XM_019815843.2;refGene_gn=LOC109493773;refGene_tx=XM_019815844.2;refGene_gn=LOC109493773;refGene_tx=XM_023242230.1;refGene_gn=LOC109493773;refGene_tx=XM_023242231.1;refGene_gn=LOC109493773;refGene_tx=XM_023242232.1;refGene_gn=LOC109493773;refGene_tx=XM_023242233.1;refGene_gn=LOC109493773;refGene_tx=XM_023242234.1
D4 27353430 27353567
refGene_gn=LOC109493773;refGene_tx=XM_019815840.2;refGene_gn=LOC109493773;refGene_tx=XM_019815841.2;refGene_gn=LOC109493773;refGene_tx=XM_019815842.2;refGene_gn=LOC109493773;refGene_tx=XM_019815843.2;refGene_gn=LOC109493773;refGene_tx=XM_019815844.2;refGene_gn=LOC109493773;refGene_tx=XM_023242230.1;refGene_gn=LOC109493773;refGene_tx=XM_023242231.1;refGene_gn=LOC109493773;refGene_tx=XM_023242232.1;refGene_gn=LOC109493773;refGene_tx=XM_023242233.1;refGene_gn=LOC109493773;refGene_tx=XM_023242234.1
D4 27354054 27354201
refGene_gn=LOC109493773;refGene_tx=XM_019815840.2;refGene_gn=LOC109493773;refGene_tx=XM_019815841.2;refGene_gn=LOC109493773;refGene_tx=XM_019815842.2;refGene_gn=LOC109493773;refGene_tx=XM_019815843.2;refGene_gn=LOC109493773;refGene_tx=XM_019815844.2;refGene_gn=LOC109493773;refGene_tx=XM_023242230.1;refGene_gn=LOC109493773;refGene_tx=XM_023242231.1;refGene_gn=LOC109493773;refGene_tx=XM_023242232.1;refGene_gn=LOC109493773;refGene_tx=XM_023242233.1;refGene_gn=LOC109493773;refGene_tx=XM_023242234.1
D4 27358322 27358381
refGene_gn=LOC109493773;refGene_tx=XM_019815840.2;refGene_gn=LOC109493773;refGene_tx=XM_019815841.2;refGene_gn=LOC109493773;refGene_tx=XM_019815842.2;refGene_gn=LOC109493773;refGene_tx=XM_019815843.2;refGene_gn=LOC109493773;refGene_tx=XM_019815844.2;refGene_gn=LOC109493773;refGene_tx=XM_023242230.1;refGene_gn=LOC109493773;refGene_tx=XM_023242231.1;refGene_gn=LOC109493773;refGene_tx=XM_023242232.1;refGene_gn=LOC109493773;refGene_tx=XM_023242233.1;refGene_gn=LOC109493773;refGene_tx=XM_023242234.1
D4 27360454 27360674
refGene_gn=LOC109493773;refGene_tx=XM_019815840.2;refGene_gn=LOC109493773;refGene_tx=XM_019815841.2;refGene_gn=LOC109493773;refGene_tx=XM_019815842.2;refGene_gn=LOC109493773;refGene_tx=XM_019815843.2;refGene_gn

=LOC109493773;refGene_tx=XM_019815844.2;refGene_gn=LOC109493773;refGene_tx=XM_023242230.1;refGene_gn=LOC109493773;refGene_tx=XM_023242231.1;refGene_gn=LOC109493773;refGene_tx=XM_023242232.1;refGene_gn=LOC109493773;refGene_tx=XM_023242233.1;refGene_gn=LOC109493773;refGene_tx=XM_023242234.1
D4 27361086 27361179 refGene_gn=LOC109493773;refGene_tx=XM_019815840.2;refGene_gn=LOC109493773;refGene_tx=XM_019815841.2;refGene_gn=LOC109493773;refGene_tx=XM_019815842.2;refGene_gn=LOC109493773;refGene_tx=XM_019815843.2;refGene_gn=LOC109493773;refGene_tx=XM_019815844.2;refGene_gn=LOC109493773;refGene_tx=XM_023242230.1;refGene_gn=LOC109493773;refGene_tx=XM_023242231.1;refGene_gn=LOC109493773;refGene_tx=XM_023242232.1;refGene_gn=LOC109493773;refGene_tx=XM_023242233.1;refGene_gn=LOC109493773;refGene_tx=XM_023242234.1
D4 27362888 27362896 refGene_gn=LOC109493773;refGene_tx=XM_019815840.2;refGene_gn=LOC109493773;refGene_tx=XM_019815841.2;refGene_gn=LOC109493773;refGene_tx=XM_019815842.2;refGene_gn=LOC109493773;refGene_tx=XM_019815843.2;refGene_gn=LOC109493773;refGene_tx=XM_019815844.2;refGene_gn=LOC109493773;refGene_tx=XM_023242230.1;refGene_gn=LOC109493773;refGene_tx=XM_023242231.1;refGene_gn=LOC109493773;refGene_tx=XM_023242232.1;refGene_gn=LOC109493773;refGene_tx=XM_023242233.1;refGene_gn=LOC109493773;refGene_tx=XM_023242234.1
D4 27549281 27550316 refGene_gn=GAS1;refGene_tx=XM_011288313.3
D4 28034085 28034147 refGene_gn=DAPK1;refGene_tx=XM_023242235.1
D4 28129806 28130028 refGene_gn=DAPK1;refGene_tx=XM_023242235.1
D4 28158308 28158447 refGene_gn=DAPK1;refGene_tx=XM_023242235.1
D4 28160383 28160513 refGene_gn=DAPK1;refGene_tx=XM_023242235.1
D4 28160694 28160743 refGene_gn=DAPK1;refGene_tx=XM_023242235.1
D4 28160830 28160857 refGene_gn=DAPK1;refGene_tx=XM_023242235.1
D4 28161555 28161708 refGene_gn=DAPK1;refGene_tx=XM_023242235.1
D4 28162074 28162120 refGene_gn=DAPK1;refGene_tx=XM_023242235.1
D4 28163290 28163380 refGene_gn=DAPK1;refGene_tx=XM_023242235.1
D4 28164751 28164844 refGene_gn=DAPK1;refGene_tx=XM_023242235.1
D4 28167079 28167199 refGene_gn=DAPK1;refGene_tx=XM_023242235.1
D4 28167644 28167743 refGene_gn=DAPK1;refGene_tx=XM_023242235.1
D4 28168495 28168594 refGene_gn=DAPK1;refGene_tx=XM_023242235.1
D4 28169950 28170049 refGene_gn=DAPK1;refGene_tx=XM_023242235.1
D4 28171044 28171242 refGene_gn=DAPK1;refGene_tx=XM_023242235.1
D4 28173096 28173294 refGene_gn=DAPK1;refGene_tx=XM_023242235.1
D4 28177533 28177632 refGene_gn=DAPK1;refGene_tx=XM_023242235.1
D4 28184824 28184902 refGene_gn=DAPK1;refGene_tx=XM_023242235.1
D4 28197436 28197659 refGene_gn=DAPK1;refGene_tx=XM_023242235.1
D4 28204968 28205157 refGene_gn=DAPK1;refGene_tx=XM_023242235.1
D4 28207019 28207217 refGene_gn=DAPK1;refGene_tx=XM_023242235.1
D4 28208394 28208533 refGene_gn=DAPK1;refGene_tx=XM_023242235.1
D4 28209940 28210061 refGene_gn=DAPK1;refGene_tx=XM_023242235.1
D4 28212845 28213034 refGene_gn=DAPK1;refGene_tx=XM_023242235.1
D4 28216217 28216566 refGene_gn=DAPK1;refGene_tx=XM_023242235.1
D4 28216568 28217449 refGene_gn=DAPK1;refGene_tx=XM_023242235.1

D4 28237197 28237281
refGene_gn=LOC101101202;refGene_tx=XM_023242236.1
D4 28237377 28237471
refGene_gn=LOC101101202;refGene_tx=XM_023242236.1
D4 28237473 28237523
refGene_gn=LOC101101202;refGene_tx=XM_023242236.1
D4 28237700 28237925
refGene_gn=LOC101101202;refGene_tx=XM_023242236.1
D4 28238098 28238261
refGene_gn=LOC101101202;refGene_tx=XM_023242236.1
D4 28239142 28239260
refGene_gn=LOC101101202;refGene_tx=XM_023242236.1
D4 28239677 28239777
refGene_gn=LOC101101202;refGene_tx=XM_023242236.1
D4 28267916 28268079
refGene_gn=LOC101080443;refGene_tx=XM_023242238.1
D4 28268700 28268818
refGene_gn=LOC101080443;refGene_tx=XM_023242238.1
D4 28269532 28269548
refGene_gn=LOC101080443;refGene_tx=XM_023242238.1
D4 28269549 28269633
refGene_gn=LOC101080443;refGene_tx=XM_023242238.1
D4 28477953 28478053
refGene_gn=LOC101082891;refGene_tx=XM_019815855.1
D4 28478536 28478654
refGene_gn=LOC101082891;refGene_tx=XM_019815855.1
D4 28479548 28479711
refGene_gn=LOC101082891;refGene_tx=XM_019815855.1
D4 28480415 28480640
refGene_gn=LOC101082891;refGene_tx=XM_019815855.1
D4 28480828 28480963
refGene_gn=LOC101082891;refGene_tx=XM_019815855.1
D4 28481066 28481186
refGene_gn=LOC101082891;refGene_tx=XM_019815855.1
D4 28481478 28481604
refGene_gn=LOC101082891;refGene_tx=XM_019815855.1
D4 28566818 28566918 refGene_gn=PDP;refGene_tx=XM_023242241.1
D4 28567250 28567525
refGene_gn=PDP;refGene_tx=XM_023242239.1;refGene_gn=PDP;refGene_tx=XM_
023242240.1;refGene_gn=PDP;refGene_tx=XM_023242241.1
D4 28568199 28568362
refGene_gn=PDP;refGene_tx=XM_023242239.1;refGene_gn=PDP;refGene_tx=XM_
023242240.1;refGene_gn=PDP;refGene_tx=XM_023242241.1
D4 28568822 28569047
refGene_gn=PDP;refGene_tx=XM_023242239.1;refGene_gn=PDP;refGene_tx=XM_
023242240.1;refGene_gn=PDP;refGene_tx=XM_023242241.1
D4 28569453 28569600
refGene_gn=PDP;refGene_tx=XM_023242239.1;refGene_gn=PDP;refGene_tx=XM_
023242240.1;refGene_gn=PDP;refGene_tx=XM_023242241.1
D4 28569766 28569886

refGene_gn=PDP;refGene_tx=XM_023242239.1;refGene_gn=PDP;refGene_tx=XM_023242240.1;refGene_gn=PDP;refGene_tx=XM_023242241.1
D4 28570187 28570313
refGene_gn=PDP;refGene_tx=XM_023242239.1;refGene_gn=PDP;refGene_tx=XM_023242240.1;refGene_gn=PDP;refGene_tx=XM_023242241.1
D4 28627527 28627653
refGene_gn=LOC101087880;refGene_tx=XM_003995465.5
D4 28627949 28628072
refGene_gn=LOC101087880;refGene_tx=XM_003995465.5
D4 28628173 28628320
refGene_gn=LOC101087880;refGene_tx=XM_003995465.5
D4 28628500 28628725
refGene_gn=LOC101087880;refGene_tx=XM_003995465.5
D4 28629411 28629574
refGene_gn=LOC101087880;refGene_tx=XM_003995465.5
D4 28630211 28630329
refGene_gn=LOC101087880;refGene_tx=XM_003995465.5
D4 28630804 28630904
refGene_gn=LOC101087880;refGene_tx=XM_003995465.5
D4 28635565 28635760 refGene_gn=FBP2;refGene_tx=XM_003995457.5
D4 28637224 28637344 refGene_gn=FBP2;refGene_tx=XM_003995457.5
D4 28643635 28643773 refGene_gn=FBP2;refGene_tx=XM_003995457.5
D4 28645912 28646053 refGene_gn=FBP2;refGene_tx=XM_003995457.5
D4 28657220 28657313 refGene_gn=FBP2;refGene_tx=XM_003995457.5
D4 28659391 28659554 refGene_gn=FBP2;refGene_tx=XM_003995457.5
D4 28664172 28664342 refGene_gn=FBP2;refGene_tx=XM_003995457.5
D4 28671908 28672100 refGene_gn=FBP1;refGene_tx=XM_003995458.5
D4 28673726 28673846 refGene_gn=FBP1;refGene_tx=XM_003995458.5
D4 28674920 28675058 refGene_gn=FBP1;refGene_tx=XM_003995458.5
D4 28678311 28678452 refGene_gn=FBP1;refGene_tx=XM_003995458.5
D4 28684009 28684102 refGene_gn=FBP1;refGene_tx=XM_003995458.5
D4 28686371 28686534 refGene_gn=FBP1;refGene_tx=XM_003995458.5
D4 28700047 28700217 refGene_gn=FBP1;refGene_tx=XM_003995458.5
D4 28832119 28832907
refGene_gn=CD4H9orf3;refGene_tx=XM_023242243.1
D4 28847623 28847790
refGene_gn=CD4H9orf3;refGene_tx=XM_023242243.1
D4 28867722 28867876
refGene_gn=CD4H9orf3;refGene_tx=XM_023242243.1
D4 28875276 28875522
refGene_gn=CD4H9orf3;refGene_tx=XM_023242243.1
D4 28986131 28986321
refGene_gn=CD4H9orf3;refGene_tx=XM_023242243.1
D4 28990333 28990440
refGene_gn=CD4H9orf3;refGene_tx=XM_023242243.1
D4 29014523 29014626
refGene_gn=CD4H9orf3;refGene_tx=XM_023242243.1
D4 29015259 29015367
refGene_gn=CD4H9orf3;refGene_tx=XM_023242243.1
D4 29026794 29026806

refGene_gn=CD4H9orf3;refGene_tx=XM_023242243.1
D4	29026807	29026839
refGene_gn=CD4H9orf3;refGene_tx=XM_023242243.1
D4	29037436	29037497
refGene_gn=CD4H9orf3;refGene_tx=XM_023242243.1
D4	29063433	29063496
refGene_gn=CD4H9orf3;refGene_tx=XM_023242243.1
D4	29063914	29063989
refGene_gn=CD4H9orf3;refGene_tx=XM_023242243.1
D4	29115496	29115613
refGene_gn=CD4H9orf3;refGene_tx=XM_023242243.1
D4	29133747	29133834
refGene_gn=CD4H9orf3;refGene_tx=XM_023242243.1
D4	29135729	29135870
refGene_gn=CD4H9orf3;refGene_tx=XM_023242243.1
D4	29155629	29155884
refGene_gn=FANCC;refGene_tx=XM_003995467.3;refGene_gn=FANCC;refGene_tx
=XM_019815867.1;refGene_gn=FANCC;refGene_tx=XM_019815868.1;refGene_gn=
FANCC;refGene_tx=XM_019815869.1;refGene_gn=FANCC;refGene_tx=XM_0232422
44.1
D4	29170992	29171199
refGene_gn=FANCC;refGene_tx=XM_003995467.3;refGene_gn=FANCC;refGene_tx
=XM_019815867.1;refGene_gn=FANCC;refGene_tx=XM_019815868.1;refGene_gn=
FANCC;refGene_tx=XM_019815869.1;refGene_gn=FANCC;refGene_tx=XM_0232422
44.1
D4	29175520	29175695
refGene_gn=FANCC;refGene_tx=XM_003995467.3;refGene_gn=FANCC;refGene_tx
=XM_019815867.1;refGene_gn=FANCC;refGene_tx=XM_019815868.1;refGene_gn=
FANCC;refGene_tx=XM_023242244.1
D4	29178401	29178483
refGene_gn=FANCC;refGene_tx=XM_003995467.3;refGene_gn=FANCC;refGene_tx
=XM_019815867.1;refGene_gn=FANCC;refGene_tx=XM_019815868.1;refGene_gn=
FANCC;refGene_tx=XM_023242244.1
D4	29180990	29181066
refGene_gn=FANCC;refGene_tx=XM_003995467.3;refGene_gn=FANCC;refGene_tx
=XM_019815867.1;refGene_gn=FANCC;refGene_tx=XM_019815868.1;refGene_gn=
FANCC;refGene_tx=XM_023242244.1
D4	29188722	29188822
refGene_gn=FANCC;refGene_tx=XM_003995467.3;refGene_gn=FANCC;refGene_tx
=XM_019815867.1;refGene_gn=FANCC;refGene_tx=XM_019815868.1;refGene_gn=
FANCC;refGene_tx=XM_019815869.1;refGene_gn=FANCC;refGene_tx=XM_0232422
44.1
D4	29190093	29190146
refGene_gn=FANCC;refGene_tx=XM_003995467.3;refGene_gn=FANCC;refGene_tx
=XM_019815867.1;refGene_gn=FANCC;refGene_tx=XM_019815868.1;refGene_gn=
FANCC;refGene_tx=XM_019815869.1;refGene_gn=FANCC;refGene_tx=XM_0232422
44.1
D4	29201453	29201610
refGene_gn=FANCC;refGene_tx=XM_003995467.3;refGene_gn=FANCC;refGene_tx
=XM_019815867.1;refGene_gn=FANCC;refGene_tx=XM_019815868.1;refGene_gn=

FANCC;refGene_tx=XM_019815869.1;refGene_gn=FANCC;refGene_tx=XM_0232422
44.1
D4 29221479 29221644
refGene_gn=FANCC;refGene_tx=XM_003995467.3;refGene_gn=FANCC;refGene_tx
=XM_019815867.1;refGene_gn=FANCC;refGene_tx=XM_019815868.1;refGene_gn=
FANCC;refGene_tx=XM_019815869.1;refGene_gn=FANCC;refGene_tx=XM_0232422
44.1
D4 29240784 29240849
refGene_gn=FANCC;refGene_tx=XM_003995467.3;refGene_gn=FANCC;refGene_tx
=XM_019815867.1;refGene_gn=FANCC;refGene_tx=XM_019815868.1;refGene_gn=
FANCC;refGene_tx=XM_019815869.1;refGene_gn=FANCC;refGene_tx=XM_0232422
44.1
D4 29241735 29241849
refGene_gn=FANCC;refGene_tx=XM_003995467.3;refGene_gn=FANCC;refGene_tx
=XM_019815867.1;refGene_gn=FANCC;refGene_tx=XM_019815868.1;refGene_gn=
FANCC;refGene_tx=XM_019815869.1;refGene_gn=FANCC;refGene_tx=XM_0232422
44.1
D4 29312163 29312258
refGene_gn=FANCC;refGene_tx=XM_003995467.3;refGene_gn=FANCC;refGene_tx
=XM_019815867.1;refGene_gn=FANCC;refGene_tx=XM_019815868.1;refGene_gn=
FANCC;refGene_tx=XM_019815869.1;refGene_gn=FANCC;refGene_tx=XM_0232422
44.1
D4 29317995 29318080
refGene_gn=FANCC;refGene_tx=XM_003995467.3;refGene_gn=FANCC;refGene_tx
=XM_019815867.1;refGene_gn=FANCC;refGene_tx=XM_019815868.1;refGene_gn=
FANCC;refGene_tx=XM_019815869.1;refGene_gn=FANCC;refGene_tx=XM_0232422
44.1
D4 29319665 29319830
refGene_gn=FANCC;refGene_tx=XM_003995467.3;refGene_gn=FANCC;refGene_tx
=XM_019815867.1;refGene_gn=FANCC;refGene_tx=XM_019815868.1;refGene_gn=
FANCC;refGene_tx=XM_019815869.1;refGene_gn=FANCC;refGene_tx=XM_0232422
44.1
D4 29506815 29507370
refGene_gn=PTCH1;refGene_tx=XM_011288296.3;refGene_gn=PTCH1;refGene_tx
=XM_019815872.2;refGene_gn=PTCH1;refGene_tx=XM_023242245.1;refGene_gn=
PTCH1;refGene_tx=XM_023242246.1;refGene_gn=PTCH1;refGene_tx=XM_0232422
47.1
D4 29508954 29509209
refGene_gn=PTCH1;refGene_tx=XM_011288296.3;refGene_gn=PTCH1;refGene_tx
=XM_019815872.2;refGene_gn=PTCH1;refGene_tx=XM_023242245.1;refGene_gn=
PTCH1;refGene_tx=XM_023242246.1;refGene_gn=PTCH1;refGene_tx=XM_0232422
47.1
D4 29509760 29509860
refGene_gn=PTCH1;refGene_tx=XM_011288296.3;refGene_gn=PTCH1;refGene_tx
=XM_019815872.2;refGene_gn=PTCH1;refGene_tx=XM_023242245.1;refGene_gn=
PTCH1;refGene_tx=XM_023242246.1;refGene_gn=PTCH1;refGene_tx=XM_0232422
47.1
D4 29513206 29513349
refGene_gn=PTCH1;refGene_tx=XM_011288296.3;refGene_gn=PTCH1;refGene_tx
=XM_019815872.2;refGene_gn=PTCH1;refGene_tx=XM_023242245.1;refGene_gn=

PTCH1;refGene_tx=XM_023242246.1;refGene_gn=PTCH1;refGene_tx=XM_0232422
47.1
D4 29517106 29517244
refGene_gn=PTCH1;refGene_tx=XM_011288296.3;refGene_gn=PTCH1;refGene_tx
=XM_019815872.2;refGene_gn=PTCH1;refGene_tx=XM_023242245.1;refGene_gn=
PTCH1;refGene_tx=XM_023242246.1;refGene_gn=PTCH1;refGene_tx=XM_0232422
47.1
D4 29518993 29519274
refGene_gn=PTCH1;refGene_tx=XM_011288296.3;refGene_gn=PTCH1;refGene_tx
=XM_019815872.2;refGene_gn=PTCH1;refGene_tx=XM_023242245.1;refGene_gn=
PTCH1;refGene_tx=XM_023242246.1;refGene_gn=PTCH1;refGene_tx=XM_0232422
47.1
D4 29520693 29520877
refGene_gn=PTCH1;refGene_tx=XM_011288296.3;refGene_gn=PTCH1;refGene_tx
=XM_019815872.2;refGene_gn=PTCH1;refGene_tx=XM_023242245.1;refGene_gn=
PTCH1;refGene_tx=XM_023242246.1;refGene_gn=PTCH1;refGene_tx=XM_0232422
47.1
D4 29522932 29523075
refGene_gn=PTCH1;refGene_tx=XM_011288296.3;refGene_gn=PTCH1;refGene_tx
=XM_019815872.2;refGene_gn=PTCH1;refGene_tx=XM_023242245.1;refGene_gn=
PTCH1;refGene_tx=XM_023242246.1;refGene_gn=PTCH1;refGene_tx=XM_0232422
47.1
D4 29527682 29527992
refGene_gn=PTCH1;refGene_tx=XM_011288296.3;refGene_gn=PTCH1;refGene_tx
=XM_019815872.2;refGene_gn=PTCH1;refGene_tx=XM_023242245.1;refGene_gn=
PTCH1;refGene_tx=XM_023242246.1;refGene_gn=PTCH1;refGene_tx=XM_0232422
47.1
D4 29528412 29528815
refGene_gn=PTCH1;refGene_tx=XM_011288296.3;refGene_gn=PTCH1;refGene_tx
=XM_019815872.2;refGene_gn=PTCH1;refGene_tx=XM_023242245.1;refGene_gn=
PTCH1;refGene_tx=XM_023242246.1;refGene_gn=PTCH1;refGene_tx=XM_0232422
47.1
D4 29529490 29529609
refGene_gn=PTCH1;refGene_tx=XM_011288296.3;refGene_gn=PTCH1;refGene_tx
=XM_019815872.2;refGene_gn=PTCH1;refGene_tx=XM_023242245.1;refGene_gn=
PTCH1;refGene_tx=XM_023242246.1;refGene_gn=PTCH1;refGene_tx=XM_0232422
47.1
D4 29535812 29535938
refGene_gn=PTCH1;refGene_tx=XM_011288296.3;refGene_gn=PTCH1;refGene_tx
=XM_019815872.2;refGene_gn=PTCH1;refGene_tx=XM_023242245.1;refGene_gn=
PTCH1;refGene_tx=XM_023242246.1;refGene_gn=PTCH1;refGene_tx=XM_0232422
47.1
D4 29536715 29536814
refGene_gn=PTCH1;refGene_tx=XM_011288296.3;refGene_gn=PTCH1;refGene_tx
=XM_019815872.2;refGene_gn=PTCH1;refGene_tx=XM_023242245.1;refGene_gn=
PTCH1;refGene_tx=XM_023242246.1;refGene_gn=PTCH1;refGene_tx=XM_0232422
47.1
D4 29537473 29537629
refGene_gn=PTCH1;refGene_tx=XM_011288296.3;refGene_gn=PTCH1;refGene_tx
=XM_019815872.2;refGene_gn=PTCH1;refGene_tx=XM_023242245.1;refGene_gn=

PTCH1;refGene_tx=XM_023242247.1
D4 29538037 29538169
refGene_gn=PTCH1;refGene_tx=XM_011288296.3;refGene_gn=PTCH1;refGene_tx=XM_019815872.2;refGene_gn=PTCH1;refGene_tx=XM_023242245.1;refGene_gn=PTCH1;refGene_tx=XM_023242246.1;refGene_gn=PTCH1;refGene_tx=XM_023242247.1
D4 29538996 29539144
refGene_gn=PTCH1;refGene_tx=XM_011288296.3;refGene_gn=PTCH1;refGene_tx=XM_019815872.2;refGene_gn=PTCH1;refGene_tx=XM_023242245.1;refGene_gn=PTCH1;refGene_tx=XM_023242246.1;refGene_gn=PTCH1;refGene_tx=XM_023242247.1
D4 29539918 29540040
refGene_gn=PTCH1;refGene_tx=XM_011288296.3;refGene_gn=PTCH1;refGene_tx=XM_019815872.2;refGene_gn=PTCH1;refGene_tx=XM_023242245.1;refGene_gn=PTCH1;refGene_tx=XM_023242246.1;refGene_gn=PTCH1;refGene_tx=XM_023242247.1
D4 29540331 29540530
refGene_gn=PTCH1;refGene_tx=XM_011288296.3;refGene_gn=PTCH1;refGene_tx=XM_019815872.2;refGene_gn=PTCH1;refGene_tx=XM_023242245.1;refGene_gn=PTCH1;refGene_tx=XM_023242246.1;refGene_gn=PTCH1;refGene_tx=XM_023242247.1
D4 29541938 29542030
refGene_gn=PTCH1;refGene_tx=XM_011288296.3;refGene_gn=PTCH1;refGene_tx=XM_019815872.2;refGene_gn=PTCH1;refGene_tx=XM_023242245.1;refGene_gn=PTCH1;refGene_tx=XM_023242246.1;refGene_gn=PTCH1;refGene_tx=XM_023242247.1
D4 29542127 29542197
refGene_gn=PTCH1;refGene_tx=XM_011288296.3;refGene_gn=PTCH1;refGene_tx=XM_019815872.2;refGene_gn=PTCH1;refGene_tx=XM_023242245.1;refGene_gn=PTCH1;refGene_tx=XM_023242246.1;refGene_gn=PTCH1;refGene_tx=XM_023242247.1
D4 29545294 29545484
refGene_gn=PTCH1;refGene_tx=XM_011288296.3;refGene_gn=PTCH1;refGene_tx=XM_019815872.2;refGene_gn=PTCH1;refGene_tx=XM_023242245.1;refGene_gn=PTCH1;refGene_tx=XM_023242246.1;refGene_gn=PTCH1;refGene_tx=XM_023242247.1
D4 29564412 29564605
refGene_gn=PTCH1;refGene_tx=XM_019815872.2;refGene_gn=PTCH1;refGene_tx=XM_023242245.1;refGene_gn=PTCH1;refGene_tx=XM_023242246.1
D4 29566262 29566460
refGene_gn=PTCH1;refGene_tx=XM_023242245.1;refGene_gn=PTCH1;refGene_tx=XM_023242246.1
D4 29574596 29574599 refGene_gn=PTCH1;refGene_tx=XM_019815872.2
D4 29922767 29922813
refGene_gn=ERCC6L2;refGene_tx=XM_011288295.3;refGene_gn=ERCC6L2;refGene_tx=XM_019815875.2
D4 29926552 29926977
refGene_gn=ERCC6L2;refGene_tx=XM_011288295.3;refGene_gn=ERCC6L2;refGene_tx=XM_019815875.2
D4 29960958 29961078

refGene_gn=ERCC6L2;refGene_tx=XM_011288295.3;refGene_gn=ERCC6L2;refGene_tx=XM_019815875.2
D4 29975570 29975764
refGene_gn=ERCC6L2;refGene_tx=XM_011288295.3;refGene_gn=ERCC6L2;refGene_tx=XM_019815875.2
D4 29979561 29979723
refGene_gn=ERCC6L2;refGene_tx=XM_011288295.3;refGene_gn=ERCC6L2;refGene_tx=XM_019815875.2
D4 29980134 29980342
refGene_gn=ERCC6L2;refGene_tx=XM_011288295.3;refGene_gn=ERCC6L2;refGene_tx=XM_019815875.2
D4 29983207 29983348
refGene_gn=ERCC6L2;refGene_tx=XM_011288295.3;refGene_gn=ERCC6L2;refGene_tx=XM_019815875.2
D4 29984380 29984494
refGene_gn=ERCC6L2;refGene_tx=XM_011288295.3;refGene_gn=ERCC6L2;refGene_tx=XM_019815875.2
D4 29985397 29985517
refGene_gn=ERCC6L2;refGene_tx=XM_011288295.3;refGene_gn=ERCC6L2;refGene_tx=XM_019815875.2
D4 29991699 29991771
refGene_gn=ERCC6L2;refGene_tx=XM_011288295.3;refGene_gn=ERCC6L2;refGene_tx=XM_019815875.2
D4 29992377 29992523
refGene_gn=ERCC6L2;refGene_tx=XM_011288295.3;refGene_gn=ERCC6L2;refGene_tx=XM_019815875.2
D4 29995215 29995311
refGene_gn=ERCC6L2;refGene_tx=XM_011288295.3;refGene_gn=ERCC6L2;refGene_tx=XM_019815875.2
D4 30005529 30005629
refGene_gn=ERCC6L2;refGene_tx=XM_011288295.3;refGene_gn=ERCC6L2;refGene_tx=XM_019815875.2
D4 30012618 30012771 refGene_gn=ERCC6L2;refGene_tx=XM_011288295.3
D4 30017556 30017637
refGene_gn=ERCC6L2;refGene_tx=XM_011288295.3;refGene_gn=ERCC6L2;refGene_tx=XM_019815875.2
D4 30019197 30020362
refGene_gn=ERCC6L2;refGene_tx=XM_011288295.3;refGene_gn=ERCC6L2;refGene_tx=XM_019815875.2
D4 30023812 30023967
refGene_gn=ERCC6L2;refGene_tx=XM_011288295.3;refGene_gn=ERCC6L2;refGene_tx=XM_019815875.2
D4 30048853 30049035
refGene_gn=ERCC6L2;refGene_tx=XM_011288295.3;refGene_gn=ERCC6L2;refGene_tx=XM_019815875.2
D4 30055363 30056336
refGene_gn=ERCC6L2;refGene_tx=XM_011288295.3;refGene_gn=ERCC6L2;refGene_tx=XM_019815875.2
D4 30286894 30286993 refGene_gn=HSD17B3;refGene_tx=XM_003995460.5
D4 30290909 30291059 refGene_gn=HSD17B3;refGene_tx=XM_003995460.5

D4 30291658 30291805 refGene_gn=HSD17B3;refGene_tx=XM_023242249.1
D4 30294338 30294404
refGene_gn=HSD17B3;refGene_tx=XM_003995460.5;refGene_gn=HSD17B3;refGene_tx=XM_023242249.1
D4 30295055 30295137
refGene_gn=HSD17B3;refGene_tx=XM_003995460.5;refGene_gn=HSD17B3;refGene_tx=XM_023242249.1
D4 30296438 30296473
refGene_gn=HSD17B3;refGene_tx=XM_003995460.5;refGene_gn=HSD17B3;refGene_tx=XM_023242249.1
D4 30299580 30299616
refGene_gn=HSD17B3;refGene_tx=XM_003995460.5;refGene_gn=HSD17B3;refGene_tx=XM_023242249.1
D4 30300913 30300981
refGene_gn=HSD17B3;refGene_tx=XM_003995460.5;refGene_gn=HSD17B3;refGene_tx=XM_023242249.1
D4 30302282 30302390
refGene_gn=HSD17B3;refGene_tx=XM_003995460.5;refGene_gn=HSD17B3;refGene_tx=XM_023242249.1
D4 30304061 30304137
refGene_gn=HSD17B3;refGene_tx=XM_003995460.5;refGene_gn=HSD17B3;refGene_tx=XM_023242249.1
D4 30331346 30331393
refGene_gn=HSD17B3;refGene_tx=XM_003995460.5;refGene_gn=HSD17B3;refGene_tx=XM_023242249.1
D4 30334387 30334541
refGene_gn=HSD17B3;refGene_tx=XM_003995460.5;refGene_gn=HSD17B3;refGene_tx=XM_023242249.1
D4 30359280 30359383
refGene_gn=SLC35D2;refGene_tx=XM_023242250.1;refGene_gn=SLC35D2;refGene_tx=XM_023242252.1;refGene_gn=SLC35D2;refGene_tx=XM_023242253.1
D4 30360015 30360098
refGene_gn=SLC35D2;refGene_tx=XM_023242250.1;refGene_gn=SLC35D2;refGene_tx=XM_023242252.1;refGene_gn=SLC35D2;refGene_tx=XM_023242253.1
D4 30361603 30361682
refGene_gn=SLC35D2;refGene_tx=XM_023242250.1;refGene_gn=SLC35D2;refGene_tx=XM_023242252.1;refGene_gn=SLC35D2;refGene_tx=XM_023242253.1
D4 30386525 30386667 refGene_gn=SLC35D2;refGene_tx=XM_023242251.1
D4 30386778 30386846
refGene_gn=SLC35D2;refGene_tx=XM_023242250.1;refGene_gn=SLC35D2;refGene_tx=XM_023242251.1;refGene_gn=SLC35D2;refGene_tx=XM_023242252.1;refGene_gn=SLC35D2;refGene_tx=XM_023242253.1
D4 30391486 30391579
refGene_gn=SLC35D2;refGene_tx=XM_023242250.1;refGene_gn=SLC35D2;refGene_tx=XM_023242251.1;refGene_gn=SLC35D2;refGene_tx=XM_023242252.1;refGene_gn=SLC35D2;refGene_tx=XM_023242253.1
D4 30392864 30392967
refGene_gn=SLC35D2;refGene_tx=XM_023242250.1;refGene_gn=SLC35D2;refGene_tx=XM_023242251.1;refGene_gn=SLC35D2;refGene_tx=XM_023242252.1;refGene_gn=SLC35D2;refGene_tx=XM_023242253.1

D4 30397614 30397683
refGene_gn=SLC35D2;refGene_tx=XM_023242250.1;refGene_gn=SLC35D2;refGene_tx=XM_023242251.1;refGene_gn=SLC35D2;refGene_tx=XM_023242253.1
D4 30398578 30398650
refGene_gn=SLC35D2;refGene_tx=XM_023242250.1;refGene_gn=SLC35D2;refGene_tx=XM_023242251.1;refGene_gn=SLC35D2;refGene_tx=XM_023242253.1
D4 30407654 30407722
refGene_gn=SLC35D2;refGene_tx=XM_023242250.1;refGene_gn=SLC35D2;refGene_tx=XM_023242251.1;refGene_gn=SLC35D2;refGene_tx=XM_023242252.1;refGene_gn=SLC35D2;refGene_tx=XM_023242253.1
D4 30412926 30413013
refGene_gn=SLC35D2;refGene_tx=XM_023242250.1;refGene_gn=SLC35D2;refGene_tx=XM_023242251.1;refGene_gn=SLC35D2;refGene_tx=XM_023242252.1;refGene_gn=SLC35D2;refGene_tx=XM_023242253.1
D4 30415015 30415049
refGene_gn=SLC35D2;refGene_tx=XM_023242250.1;refGene_gn=SLC35D2;refGene_tx=XM_023242251.1;refGene_gn=SLC35D2;refGene_tx=XM_023242252.1
D4 30422405 30422557
refGene_gn=SLC35D2;refGene_tx=XM_023242250.1;refGene_gn=SLC35D2;refGene_tx=XM_023242251.1;refGene_gn=SLC35D2;refGene_tx=XM_023242252.1
D4 30427421 30427644 refGene_gn=ZNF367;refGene_tx=XM_003995461.5
D4 30430225 30430364 refGene_gn=ZNF367;refGene_tx=XM_003995461.5
D4 30431465 30431585 refGene_gn=ZNF367;refGene_tx=XM_003995461.5
D4 30434876 30435027 refGene_gn=ZNF367;refGene_tx=XM_003995461.5
D4 30450728 30451142 refGene_gn=ZNF367;refGene_tx=XM_003995461.5
D4 30524184 30524578
refGene_gn=HABP4;refGene_tx=XM_023242254.1;refGene_gn=HABP4;refGene_tx=XM_023242255.1;refGene_gn=HABP4;refGene_tx=XM_023242256.1;refGene_gn=HABP4;refGene_tx=XM_023242257.1;refGene_gn=HABP4;refGene_tx=XM_023242258.1
D4 30529934 30530097
refGene_gn=HABP4;refGene_tx=XM_023242254.1;refGene_gn=HABP4;refGene_tx=XM_023242255.1;refGene_gn=HABP4;refGene_tx=XM_023242256.1;refGene_gn=HABP4;refGene_tx=XM_023242257.1;refGene_gn=HABP4;refGene_tx=XM_023242258.1
D4 30537714 30537855
refGene_gn=HABP4;refGene_tx=XM_023242254.1;refGene_gn=HABP4;refGene_tx=XM_023242255.1;refGene_gn=HABP4;refGene_tx=XM_023242256.1;refGene_gn=HABP4;refGene_tx=XM_023242257.1;refGene_gn=HABP4;refGene_tx=XM_023242258.1
D4 30538090 30538159
refGene_gn=HABP4;refGene_tx=XM_023242254.1;refGene_gn=HABP4;refGene_tx=XM_023242255.1;refGene_gn=HABP4;refGene_tx=XM_023242256.1;refGene_gn=HABP4;refGene_tx=XM_023242257.1;refGene_gn=HABP4;refGene_tx=XM_023242258.1
D4 30547566 30547650
refGene_gn=HABP4;refGene_tx=XM_023242254.1;refGene_gn=HABP4;refGene_tx=XM_023242255.1;refGene_gn=HABP4;refGene_tx=XM_023242256.1;refGene_gn=HABP4;refGene_tx=XM_023242257.1;refGene_gn=HABP4;refGene_tx=XM_023242258.1

D4 30559232 30559404
refGene_gn=HABP4;refGene_tx=XM_023242254.1;refGene_gn=HABP4;refGene_tx=XM_023242255.1;refGene_gn=HABP4;refGene_tx=XM_023242256.1;refGene_gn=HABP4;refGene_tx=XM_023242257.1;refGene_gn=HABP4;refGene_tx=XM_023242258.1
D4 30562338 30562524
refGene_gn=HABP4;refGene_tx=XM_023242254.1;refGene_gn=HABP4;refGene_tx=XM_023242255.1;refGene_gn=HABP4;refGene_tx=XM_023242256.1;refGene_gn=HABP4;refGene_tx=XM_023242257.1;refGene_gn=HABP4;refGene_tx=XM_023242258.1
D4 30562998 30563195
refGene_gn=HABP4;refGene_tx=XM_023242254.1;refGene_gn=HABP4;refGene_tx=XM_023242255.1
D4 30564001 30564058
refGene_gn=HABP4;refGene_tx=XM_023242256.1;refGene_gn=HABP4;refGene_tx=XM_023242257.1;refGene_gn=HABP4;refGene_tx=XM_023242258.1
D4 30576971 30577008
refGene_gn=CDC14B;refGene_tx=XM_003995462.5;refGene_gn=CDC14B;refGene_tx=XM_023242262.1;refGene_gn=CDC14B;refGene_tx=XM_023242263.1;refGene_gn=CDC14B;refGene_tx=XM_023242264.1;refGene_gn=CDC14B;refGene_tx=XM_023242265.1;refGene_gn=CDC14B;refGene_tx=XM_023242268.1
D4 30577067 30577371
refGene_gn=CDC14B;refGene_tx=XM_019815883.2;refGene_gn=CDC14B;refGene_tx=XM_023242260.1;refGene_gn=CDC14B;refGene_tx=XM_023242261.1;refGene_gn=CDC14B;refGene_tx=XM_023242262.1;refGene_gn=CDC14B;refGene_tx=XM_023242263.1
D4 30581980 30582097
refGene_gn=CDC14B;refGene_tx=XM_003995462.5;refGene_gn=CDC14B;refGene_tx=XM_019815883.2;refGene_gn=CDC14B;refGene_tx=XM_023242260.1;refGene_gn=CDC14B;refGene_tx=XM_023242261.1;refGene_gn=CDC14B;refGene_tx=XM_023242262.1;refGene_gn=CDC14B;refGene_tx=XM_023242263.1;refGene_gn=CDC14B;refGene_tx=XM_023242264.1;refGene_gn=CDC14B;refGene_tx=XM_023242265.1
D4 30586873 30586988
refGene_gn=CDC14B;refGene_tx=XM_019815885.2;refGene_gn=CDC14B;refGene_tx=XM_023242266.1;refGene_gn=CDC14B;refGene_tx=XM_023242267.1
D4 30593041 30593139
refGene_gn=CDC14B;refGene_tx=XM_003995462.5;refGene_gn=CDC14B;refGene_tx=XM_019815883.2;refGene_gn=CDC14B;refGene_tx=XM_019815885.2;refGene_gn=CDC14B;refGene_tx=XM_023242260.1;refGene_gn=CDC14B;refGene_tx=XM_023242261.1;refGene_gn=CDC14B;refGene_tx=XM_023242262.1;refGene_gn=CDC14B;refGene_tx=XM_023242263.1;refGene_gn=CDC14B;refGene_tx=XM_023242264.1;refGene_gn=CDC14B;refGene_tx=XM_023242265.1;refGene_gn=CDC14B;refGene_tx=XM_023242266.1;refGene_gn=CDC14B;refGene_tx=XM_023242267.1;refGene_gn=CDC14B;refGene_tx=XM_023242268.1
D4 30593873 30594033
refGene_gn=CDC14B;refGene_tx=XM_003995462.5;refGene_gn=CDC14B;refGene_tx=XM_019815883.2;refGene_gn=CDC14B;refGene_tx=XM_019815885.2;refGene_gn=CDC14B;refGene_tx=XM_023242260.1;refGene_gn=CDC14B;refGene_tx=XM_023242261.1;refGene_gn=CDC14B;refGene_tx=XM_023242262.1;refGene_gn=CDC14

B;refGene_tx=XM_023242263.1;refGene_gn=CDC14B;refGene_tx=XM_023242264.1;refGene_gn=CDC14B;refGene_tx=XM_023242265.1;refGene_gn=CDC14B;refGene_tx=XM_023242266.1;refGene_gn=CDC14B;refGene_tx=XM_023242267.1;refGene_gn=CDC14B;refGene_tx=XM_023242268.1
D4 30594196 30594335
refGene_gn=CDC14B;refGene_tx=XM_003995462.5;refGene_gn=CDC14B;refGene_tx=XM_019815883.2;refGene_gn=CDC14B;refGene_tx=XM_019815885.2;refGene_gn=CDC14B;refGene_tx=XM_023242260.1;refGene_gn=CDC14B;refGene_tx=XM_023242261.1;refGene_gn=CDC14B;refGene_tx=XM_023242262.1;refGene_gn=CDC14B;refGene_tx=XM_023242263.1;refGene_gn=CDC14B;refGene_tx=XM_023242264.1;refGene_gn=CDC14B;refGene_tx=XM_023242265.1;refGene_gn=CDC14B;refGene_tx=XM_023242266.1;refGene_gn=CDC14B;refGene_tx=XM_023242267.1;refGene_gn=CDC14B;refGene_tx=XM_023242268.1
D4 30599323 30599554
refGene_gn=CDC14B;refGene_tx=XM_003995462.5;refGene_gn=CDC14B;refGene_tx=XM_019815883.2;refGene_gn=CDC14B;refGene_tx=XM_019815885.2;refGene_gn=CDC14B;refGene_tx=XM_023242260.1;refGene_gn=CDC14B;refGene_tx=XM_023242261.1;refGene_gn=CDC14B;refGene_tx=XM_023242262.1;refGene_gn=CDC14B;refGene_tx=XM_023242263.1;refGene_gn=CDC14B;refGene_tx=XM_023242264.1;refGene_gn=CDC14B;refGene_tx=XM_023242265.1;refGene_gn=CDC14B;refGene_tx=XM_023242266.1;refGene_gn=CDC14B;refGene_tx=XM_023242267.1;refGene_gn=CDC14B;refGene_tx=XM_023242268.1
D4 30599834 30599922
refGene_gn=CDC14B;refGene_tx=XM_003995462.5;refGene_gn=CDC14B;refGene_tx=XM_019815883.2;refGene_gn=CDC14B;refGene_tx=XM_019815885.2;refGene_gn=CDC14B;refGene_tx=XM_023242260.1;refGene_gn=CDC14B;refGene_tx=XM_023242261.1;refGene_gn=CDC14B;refGene_tx=XM_023242262.1;refGene_gn=CDC14B;refGene_tx=XM_023242263.1;refGene_gn=CDC14B;refGene_tx=XM_023242264.1;refGene_gn=CDC14B;refGene_tx=XM_023242265.1;refGene_gn=CDC14B;refGene_tx=XM_023242266.1;refGene_gn=CDC14B;refGene_tx=XM_023242267.1;refGene_gn=CDC14B;refGene_tx=XM_023242268.1
D4 30602949 30603012
refGene_gn=CDC14B;refGene_tx=XM_003995462.5;refGene_gn=CDC14B;refGene_tx=XM_019815883.2;refGene_gn=CDC14B;refGene_tx=XM_019815885.2;refGene_gn=CDC14B;refGene_tx=XM_023242260.1;refGene_gn=CDC14B;refGene_tx=XM_023242261.1;refGene_gn=CDC14B;refGene_tx=XM_023242262.1;refGene_gn=CDC14B;refGene_tx=XM_023242263.1;refGene_gn=CDC14B;refGene_tx=XM_023242264.1;refGene_gn=CDC14B;refGene_tx=XM_023242265.1;refGene_gn=CDC14B;refGene_tx=XM_023242266.1;refGene_gn=CDC14B;refGene_tx=XM_023242267.1;refGene_gn=CDC14B;refGene_tx=XM_023242268.1
D4 30605577 30605644
refGene_gn=CDC14B;refGene_tx=XM_003995462.5;refGene_gn=CDC14B;refGene_tx=XM_019815883.2;refGene_gn=CDC14B;refGene_tx=XM_019815885.2;refGene_gn=CDC14B;refGene_tx=XM_023242260.1;refGene_gn=CDC14B;refGene_tx=XM_023242261.1;refGene_gn=CDC14B;refGene_tx=XM_023242262.1;refGene_gn=CDC14B;refGene_tx=XM_023242263.1;refGene_gn=CDC14B;refGene_tx=XM_023242264.1;refGene_gn=CDC14B;refGene_tx=XM_023242265.1;refGene_gn=CDC14B;refGene_tx=XM_023242266.1;refGene_gn=CDC14B;refGene_tx=XM_023242267.1;refGene_gn=CDC14B;refGene_tx=XM_023242268.1
D4 30610818 30610895

refGene_gn=CDC14B;refGene_tx=XM_003995462.5;refGene_gn=CDC14B;refGene_tx=XM_019815883.2;refGene_gn=CDC14B;refGene_tx=XM_019815885.2;refGene_gn=CDC14B;refGene_tx=XM_023242260.1;refGene_gn=CDC14B;refGene_tx=XM_023242261.1;refGene_gn=CDC14B;refGene_tx=XM_023242262.1;refGene_gn=CDC14B;refGene_tx=XM_023242263.1;refGene_gn=CDC14B;refGene_tx=XM_023242264.1;refGene_gn=CDC14B;refGene_tx=XM_023242265.1;refGene_gn=CDC14B;refGene_tx=XM_023242266.1;refGene_gn=CDC14B;refGene_tx=XM_023242267.1;refGene_gn=CDC14B;refGene_tx=XM_023242268.1
D4 30623595 30623688
refGene_gn=CDC14B;refGene_tx=XM_003995462.5;refGene_gn=CDC14B;refGene_tx=XM_019815883.2;refGene_gn=CDC14B;refGene_tx=XM_019815885.2;refGene_gn=CDC14B;refGene_tx=XM_023242260.1;refGene_gn=CDC14B;refGene_tx=XM_023242261.1;refGene_gn=CDC14B;refGene_tx=XM_023242262.1;refGene_gn=CDC14B;refGene_tx=XM_023242263.1;refGene_gn=CDC14B;refGene_tx=XM_023242264.1;refGene_gn=CDC14B;refGene_tx=XM_023242265.1;refGene_gn=CDC14B;refGene_tx=XM_023242266.1;refGene_gn=CDC14B;refGene_tx=XM_023242267.1;refGene_gn=CDC14B;refGene_tx=XM_023242268.1
D4 30626261 30626337
refGene_gn=CDC14B;refGene_tx=XM_003995462.5;refGene_gn=CDC14B;refGene_tx=XM_019815883.2;refGene_gn=CDC14B;refGene_tx=XM_019815885.2;refGene_gn=CDC14B;refGene_tx=XM_023242260.1;refGene_gn=CDC14B;refGene_tx=XM_023242261.1;refGene_gn=CDC14B;refGene_tx=XM_023242262.1;refGene_gn=CDC14B;refGene_tx=XM_023242263.1;refGene_gn=CDC14B;refGene_tx=XM_023242264.1;refGene_gn=CDC14B;refGene_tx=XM_023242265.1;refGene_gn=CDC14B;refGene_tx=XM_023242266.1;refGene_gn=CDC14B;refGene_tx=XM_023242267.1;refGene_gn=CDC14B;refGene_tx=XM_023242268.1
D4 30626875 30626966
refGene_gn=CDC14B;refGene_tx=XM_003995462.5;refGene_gn=CDC14B;refGene_tx=XM_019815883.2;refGene_gn=CDC14B;refGene_tx=XM_019815885.2;refGene_gn=CDC14B;refGene_tx=XM_023242260.1;refGene_gn=CDC14B;refGene_tx=XM_023242261.1;refGene_gn=CDC14B;refGene_tx=XM_023242262.1;refGene_gn=CDC14B;refGene_tx=XM_023242263.1;refGene_gn=CDC14B;refGene_tx=XM_023242264.1;refGene_gn=CDC14B;refGene_tx=XM_023242265.1;refGene_gn=CDC14B;refGene_tx=XM_023242266.1;refGene_gn=CDC14B;refGene_tx=XM_023242267.1;refGene_gn=CDC14B;refGene_tx=XM_023242268.1
D4 30628344 30628405
refGene_gn=CDC14B;refGene_tx=XM_003995462.5;refGene_gn=CDC14B;refGene_tx=XM_019815883.2;refGene_gn=CDC14B;refGene_tx=XM_019815885.2
D4 30671273 30671300
refGene_gn=CDC14B;refGene_tx=XM_023242260.1;refGene_gn=CDC14B;refGene_tx=XM_023242262.1;refGene_gn=CDC14B;refGene_tx=XM_023242264.1;refGene_gn=CDC14B;refGene_tx=XM_023242266.1;refGene_gn=CDC14B;refGene_tx=XM_023242268.1
D4 30678725 30678885
refGene_gn=CDC14B;refGene_tx=XM_023242260.1;refGene_gn=CDC14B;refGene_tx=XM_023242261.1;refGene_gn=CDC14B;refGene_tx=XM_023242262.1;refGene_gn=CDC14B;refGene_tx=XM_023242263.1;refGene_gn=CDC14B;refGene_tx=XM_023242264.1;refGene_gn=CDC14B;refGene_tx=XM_023242265.1;refGene_gn=CDC14B;refGene_tx=XM_023242266.1;refGene_gn=CDC14B;refGene_tx=XM_023242267.1;refGene_gn=CDC14B;refGene_tx=XM_023242268.1

D4 30706842 30706970 refGene_gn=AAED1;refGene_tx=XM_023242269.1
D4 30708071 30708238 refGene_gn=AAED1;refGene_tx=XM_023242270.1
D4 30709334 30709466
refGene_gn=AAED1;refGene_tx=XM_023242269.1;refGene_gn=AAED1;refGene_tx=XM_023242270.1
D4 30714753 30714859
refGene_gn=AAED1;refGene_tx=XM_023242269.1;refGene_gn=AAED1;refGene_tx=XM_023242270.1
D4 30715032 30715086
refGene_gn=AAED1;refGene_tx=XM_023242269.1;refGene_gn=AAED1;refGene_tx=XM_023242270.1
D4 30715973 30716042
refGene_gn=AAED1;refGene_tx=XM_023242269.1;refGene_gn=AAED1;refGene_tx=XM_023242270.1
D4 30716441 30716651
refGene_gn=AAED1;refGene_tx=XM_023242269.1;refGene_gn=AAED1;refGene_tx=XM_023242270.1
D4 30785819 30785884
refGene_gn=LOC101090147;refGene_tx=XM_023242273.1
D4 30808361 30808468
refGene_gn=LOC101090147;refGene_tx=XM_023242273.1
D4 30808881 30808947
refGene_gn=LOC101090147;refGene_tx=XM_023242273.1
D4 30829514 30829610
refGene_gn=LOC101090147;refGene_tx=XM_023242273.1
D4 30847602 30848036
refGene_gn=LOC101090147;refGene_tx=XM_023242273.1
D4 30848037 30849087
refGene_gn=LOC101090147;refGene_tx=XM_023242273.1
D4 30849401 30850287
refGene_gn=LOC102900842;refGene_tx=XM_019815890.1
D4 30850289 30850776
refGene_gn=LOC102900842;refGene_tx=XM_019815890.1
D4 30850777 30851129
refGene_gn=LOC102900842;refGene_tx=XM_019815890.1
D4 30880381 30881994 refGene_gn=ZNF510;refGene_tx=XM_023242272.1
D4 30884124 30884220 refGene_gn=ZNF510;refGene_tx=XM_023242272.1
D4 30884517 30884644 refGene_gn=ZNF510;refGene_tx=XM_023242272.1
D4 30911698 30911757 refGene_gn=ZNF510;refGene_tx=XM_023242272.1
D4 30912169 30912236 refGene_gn=ZNF510;refGene_tx=XM_023242272.1
D4 30914040 30914498
refGene_gn=LOC109493787;refGene_tx=XM_023242274.1;refGene_gn=LOC109493787;refGene_tx=XM_023242275.1
D4 30916991 30917298
refGene_gn=LOC109493787;refGene_tx=XM_023242274.1;refGene_gn=LOC109493787;refGene_tx=XM_023242275.1
D4 30937878 30939749
refGene_gn=LOC101090653;refGene_tx=XM_019815888.1
D4 30945512 30945608
refGene_gn=LOC101090653;refGene_tx=XM_019815888.1

D4 31006059 31006186
refGene_gn=LOC101090653;refGene_tx=XM_019815888.1
D4 31011050 31011106
refGene_gn=LOC101090653;refGene_tx=XM_019815888.1
D4 31013232 31013245
refGene_gn=LOC101090653;refGene_tx=XM_019815888.1
D4 31091689 31091789
refGene_gn=LOC101086605;refGene_tx=XM_011288293.2
D4 31092544 31092659
refGene_gn=LOC101086605;refGene_tx=XM_011288293.2
D4 31093310 31093476
refGene_gn=LOC101086605;refGene_tx=XM_011288293.2
D4 31093923 31094148
refGene_gn=LOC101086605;refGene_tx=XM_011288293.2
D4 31094330 31094477
refGene_gn=LOC101086605;refGene_tx=XM_011288293.2
D4 31094581 31094704
refGene_gn=LOC101086605;refGene_tx=XM_011288293.2
D4 31094998 31095124
refGene_gn=LOC101086605;refGene_tx=XM_011288293.2
D4 31300642 31300825
refGene_gn=LOC101091410;refGene_tx=XM_003995475.5
D4 31301556 31301614
refGene_gn=LOC101091410;refGene_tx=XM_003995475.5
D4 31302197 31302258
refGene_gn=LOC101091410;refGene_tx=XM_003995475.5
D4 31302531 31306444
refGene_gn=LOC101091410;refGene_tx=XM_003995475.5
D4 31369404 31370541
refGene_gn=LOC101091908;refGene_tx=XM_006939139.4
D4 31370542 31372299
refGene_gn=LOC101091908;refGene_tx=XM_006939139.4
D4 31378975 31379071
refGene_gn=LOC101091908;refGene_tx=XM_006939139.4
D4 31379365 31379492
refGene_gn=LOC101091908;refGene_tx=XM_006939139.4
D4 31384537 31384567
refGene_gn=LOC101091908;refGene_tx=XM_006939139.4
D4 33593847 33593924
refGene_gn=MLANA;refGene_tx=XM_003995477.4;refGene_gn=MLANA;refGene_tx
=XM_019815897.1
D4 33598348 33598445
refGene_gn=MLANA;refGene_tx=XM_003995477.4;refGene_gn=MLANA;refGene_tx
=XM_019815897.1
D4 33603672 33603786
refGene_gn=MLANA;refGene_tx=XM_003995477.4;refGene_gn=MLANA;refGene_tx
=XM_019815897.1
D4 33605068 33605137
refGene_gn=MLANA;refGene_tx=XM_003995477.4;refGene_gn=MLANA;refGene_tx
=XM_019815897.1

D4 33759087 33762405
refGene_gn=RANBP6;refGene_tx=XM_003995478.4;refGene_gn=RANBP6;refGene_
tx=XM_006939153.4
D4 33831567 33831664
refGene_gn=IL33;refGene_tx=XM_003995479.5;refGene_gn=IL33;refGene_tx=X
M_006939154.3;refGene_gn=IL33;refGene_tx=XM_006939155.3;refGene_gn=IL3
3;refGene_tx=XM_006939156.4;refGene_gn=IL33;refGene_tx=XM_006939157.3;
refGene_gn=IL33;refGene_tx=XM_006939158.3;refGene_gn=IL33;refGene_tx=X
M_006939159.3;refGene_gn=IL33;refGene_tx=XM_019815899.2;refGene_gn=IL3
3;refGene_tx=XM_019815901.2;refGene_gn=IL33;refGene_tx=XM_019815902.2
D4 33838525 33838648
refGene_gn=IL33;refGene_tx=XM_003995479.5;refGene_gn=IL33;refGene_tx=X
M_006939154.3;refGene_gn=IL33;refGene_tx=XM_006939155.3;refGene_gn=IL3
3;refGene_tx=XM_006939156.4;refGene_gn=IL33;refGene_tx=XM_006939157.3;
refGene_gn=IL33;refGene_tx=XM_006939158.3;refGene_gn=IL33;refGene_tx=X
M_006939159.3;refGene_gn=IL33;refGene_tx=XM_019815899.2;refGene_gn=IL3
3;refGene_tx=XM_019815901.2;refGene_gn=IL33;refGene_tx=XM_019815902.2
D4 33839189 33839321
refGene_gn=IL33;refGene_tx=XM_003995479.5;refGene_gn=IL33;refGene_tx=X
M_006939154.3;refGene_gn=IL33;refGene_tx=XM_006939155.3;refGene_gn=IL3
3;refGene_tx=XM_006939156.4;refGene_gn=IL33;refGene_tx=XM_006939157.3;
refGene_gn=IL33;refGene_tx=XM_006939158.3;refGene_gn=IL33;refGene_tx=X
M_006939159.3;refGene_gn=IL33;refGene_tx=XM_019815899.2;refGene_gn=IL3
3;refGene_tx=XM_019815901.2;refGene_gn=IL33;refGene_tx=XM_019815902.2
D4 33840914 33841022
refGene_gn=IL33;refGene_tx=XM_003995479.5;refGene_gn=IL33;refGene_tx=X
M_006939154.3;refGene_gn=IL33;refGene_tx=XM_006939155.3;refGene_gn=IL3
3;refGene_tx=XM_006939156.4;refGene_gn=IL33;refGene_tx=XM_006939157.3;
refGene_gn=IL33;refGene_tx=XM_006939158.3;refGene_gn=IL33;refGene_tx=X
M_006939159.3;refGene_gn=IL33;refGene_tx=XM_019815899.2;refGene_gn=IL3
3;refGene_tx=XM_019815901.2;refGene_gn=IL33;refGene_tx=XM_019815902.2
D4 33841799 33841850
refGene_gn=IL33;refGene_tx=XM_003995479.5;refGene_gn=IL33;refGene_tx=X
M_006939154.3;refGene_gn=IL33;refGene_tx=XM_006939155.3;refGene_gn=IL3
3;refGene_tx=XM_006939156.4;refGene_gn=IL33;refGene_tx=XM_006939157.3;
refGene_gn=IL33;refGene_tx=XM_006939158.3;refGene_gn=IL33;refGene_tx=X
M_006939159.3;refGene_gn=IL33;refGene_tx=XM_019815899.2;refGene_gn=IL3
3;refGene_tx=XM_019815901.2;refGene_gn=IL33;refGene_tx=XM_019815902.2
D4 33842791 33842889
refGene_gn=IL33;refGene_tx=XM_003995479.5;refGene_gn=IL33;refGene_tx=X
M_006939154.3;refGene_gn=IL33;refGene_tx=XM_006939155.3;refGene_gn=IL3
3;refGene_tx=XM_006939156.4;refGene_gn=IL33;refGene_tx=XM_006939157.3;
refGene_gn=IL33;refGene_tx=XM_006939158.3;refGene_gn=IL33;refGene_tx=X
M_006939159.3;refGene_gn=IL33;refGene_tx=XM_019815899.2;refGene_gn=IL3
3;refGene_tx=XM_019815901.2;refGene_gn=IL33;refGene_tx=XM_019815902.2
D4 33844239 33844437
refGene_gn=IL33;refGene_tx=XM_003995479.5;refGene_gn=IL33;refGene_tx=X
M_006939154.3;refGene_gn=IL33;refGene_tx=XM_006939155.3;refGene_gn=IL3
3;refGene_tx=XM_006939156.4;refGene_gn=IL33;refGene_tx=XM_006939157.3;
refGene_gn=IL33;refGene_tx=XM_006939158.3;refGene_gn=IL33;refGene_tx=X

M_006939159.3;refGene_gn=IL33;refGene_tx=XM_019815899.2;refGene_gn=IL33;refGene_tx=XM_019815901.2;refGene_gn=IL33;refGene_tx=XM_019815902.2
D4 33967100 33967952 refGene_gn=TPD52L3;refGene_tx=XM_006944340.3
D4 34014840 34014993 refGene_gn=UHRF2;refGene_tx=XM_003995480.5;refGene_gn=UHRF2;refGene_tx=XM_019815903.2
D4 34021493 34021727 refGene_gn=UHRF2;refGene_tx=XM_003995480.5;refGene_gn=UHRF2;refGene_tx=XM_019815903.2
D4 34033509 34033769 refGene_gn=UHRF2;refGene_tx=XM_003995480.5;refGene_gn=UHRF2;refGene_tx=XM_019815903.2
D4 34051123 34051342 refGene_gn=UHRF2;refGene_tx=XM_003995480.5;refGene_gn=UHRF2;refGene_tx=XM_019815903.2
D4 34063074 34063184 refGene_gn=UHRF2;refGene_tx=XM_003995480.5;refGene_gn=UHRF2;refGene_tx=XM_019815903.2
D4 34066629 34066816 refGene_gn=UHRF2;refGene_tx=XM_003995480.5;refGene_gn=UHRF2;refGene_tx=XM_019815903.2
D4 34071058 34071182 refGene_gn=UHRF2;refGene_tx=XM_003995480.5;refGene_gn=UHRF2;refGene_tx=XM_019815903.2
D4 34071476 34071584 refGene_gn=UHRF2;refGene_tx=XM_003995480.5;refGene_gn=UHRF2;refGene_tx=XM_019815903.2
D4 34075031 34075136 refGene_gn=UHRF2;refGene_tx=XM_003995480.5;refGene_gn=UHRF2;refGene_tx=XM_019815903.2
D4 34080782 34080889 refGene_gn=UHRF2;refGene_tx=XM_003995480.5;refGene_gn=UHRF2;refGene_tx=XM_019815903.2
D4 34084151 34084314 refGene_gn=UHRF2;refGene_tx=XM_003995480.5;refGene_gn=UHRF2;refGene_tx=XM_019815903.2
D4 34085001 34085142 refGene_gn=UHRF2;refGene_tx=XM_003995480.5;refGene_gn=UHRF2;refGene_tx=XM_019815903.2
D4 34085597 34085669 refGene_gn=UHRF2;refGene_tx=XM_019815903.2
D4 34086840 34086937 refGene_gn=UHRF2;refGene_tx=XM_003995480.5
D4 34087269 34087427 refGene_gn=UHRF2;refGene_tx=XM_003995480.5
D4 34091245 34091344 refGene_gn=UHRF2;refGene_tx=XM_003995480.5
D4 34092327 34092474 refGene_gn=UHRF2;refGene_tx=XM_003995480.5
D4 34111533 34111677 refGene_gn=GLDC;refGene_tx=XM_011288328.3
D4 34118772 34118853 refGene_gn=GLDC;refGene_tx=XM_011288328.3
D4 34120347 34120520 refGene_gn=GLDC;refGene_tx=XM_011288328.3
D4 34125042 34125138 refGene_gn=GLDC;refGene_tx=XM_011288328.3
D4 34132138 34132250 refGene_gn=GLDC;refGene_tx=XM_011288328.3
D4 34134673 34134815 refGene_gn=GLDC;refGene_tx=XM_011288328.3

D4 34136073 34136186 refGene_gn=GLDC;refGene_tx=XM_011288328.3
D4 34137408 34137558 refGene_gn=GLDC;refGene_tx=XM_011288328.3
D4 34138986 34139112 refGene_gn=GLDC;refGene_tx=XM_011288328.3
D4 34144914 34144990 refGene_gn=GLDC;refGene_tx=XM_011288328.3
D4 34160002 34160145 refGene_gn=GLDC;refGene_tx=XM_011288328.3
D4 34161272 34161314 refGene_gn=GLDC;refGene_tx=XM_011288328.3
D4 34161476 34161561 refGene_gn=GLDC;refGene_tx=XM_011288328.3
D4 34162062 34162160 refGene_gn=GLDC;refGene_tx=XM_011288328.3
D4 34163788 34163869 refGene_gn=GLDC;refGene_tx=XM_011288328.3
D4 34164474 34164614 refGene_gn=GLDC;refGene_tx=XM_011288328.3
D4 34166052 34166158 refGene_gn=GLDC;refGene_tx=XM_011288328.3
D4 34174987 34175084 refGene_gn=GLDC;refGene_tx=XM_011288328.3
D4 34176924 34177121 refGene_gn=GLDC;refGene_tx=XM_011288328.3
D4 34177444 34177592 refGene_gn=GLDC;refGene_tx=XM_011288328.3
D4 34179272 34179331 refGene_gn=GLDC;refGene_tx=XM_011288328.3
D4 34179332 34179351 refGene_gn=GLDC;refGene_tx=XM_011288328.3
D4 34184642 34184807 refGene_gn=GLDC;refGene_tx=XM_011288328.3
D4 34195086 34195222 refGene_gn=GLDC;refGene_tx=XM_011288328.3
D4 34215417 34215496 refGene_gn=GLDC;refGene_tx=XM_011288328.3
D4 34216038 34216290 refGene_gn=GLDC;refGene_tx=XM_011288328.3
D4 34377647 34377791
refGene_gn=KDM4C;refGene_tx=XM_023242276.1;refGene_gn=KDM4C;refGene_tx=XM_023242277.1;refGene_gn=KDM4C;refGene_tx=XM_023242278.1;refGene_gn=KDM4C;refGene_tx=XM_023242279.1;refGene_gn=KDM4C;refGene_tx=XM_023242280.1;refGene_gn=KDM4C;refGene_tx=XM_023242283.1;refGene_gn=KDM4C;refGene_tx=XM_023242285.1;refGene_gn=KDM4C;refGene_tx=XM_023242286.1;refGene_gn=KDM4C;refGene_tx=XM_023242287.1;refGene_gn=KDM4C;refGene_tx=XM_023242288.1;refGene_gn=KDM4C;refGene_tx=XM_023242289.1
D4 34385146 34385322
refGene_gn=KDM4C;refGene_tx=XM_023242276.1;refGene_gn=KDM4C;refGene_tx=XM_023242277.1;refGene_gn=KDM4C;refGene_tx=XM_023242278.1;refGene_gn=KDM4C;refGene_tx=XM_023242279.1;refGene_gn=KDM4C;refGene_tx=XM_023242280.1;refGene_gn=KDM4C;refGene_tx=XM_023242281.1;refGene_gn=KDM4C;refGene_tx=XM_023242282.1;refGene_gn=KDM4C;refGene_tx=XM_023242283.1;refGene_gn=KDM4C;refGene_tx=XM_023242285.1;refGene_gn=KDM4C;refGene_tx=XM_023242286.1;refGene_gn=KDM4C;refGene_tx=XM_023242287.1;refGene_gn=KDM4C;refGene_tx=XM_023242288.1;refGene_gn=KDM4C;refGene_tx=XM_023242289.1
D4 34393040 34393155
refGene_gn=KDM4C;refGene_tx=XM_023242276.1;refGene_gn=KDM4C;refGene_tx=XM_023242277.1;refGene_gn=KDM4C;refGene_tx=XM_023242278.1;refGene_gn=KDM4C;refGene_tx=XM_023242279.1;refGene_gn=KDM4C;refGene_tx=XM_023242280.1;refGene_gn=KDM4C;refGene_tx=XM_023242281.1;refGene_gn=KDM4C;refGene_tx=XM_023242282.1;refGene_gn=KDM4C;refGene_tx=XM_023242283.1;refGene_gn=KDM4C;refGene_tx=XM_023242284.1;refGene_gn=KDM4C;refGene_tx=XM_023242285.1;refGene_gn=KDM4C;refGene_tx=XM_023242286.1;refGene_gn=KDM4C;refGene_tx=XM_023242287.1;refGene_gn=KDM4C;refGene_tx=XM_023242288.1;refGene_gn=KDM4C;refGene_tx=XM_023242289.1
D4 34414353 34414399 refGene_gn=KDM4C;refGene_tx=XM_023242287.1
D4 34424725 34424919
refGene_gn=KDM4C;refGene_tx=XM_023242276.1;refGene_gn=KDM4C;refGene_tx

=XM_023242277.1;refGene_gn=KDM4C;refGene_tx=XM_023242278.1;refGene_gn=KDM4C;refGene_tx=XM_023242279.1;refGene_gn=KDM4C;refGene_tx=XM_023242280.1;refGene_gn=KDM4C;refGene_tx=XM_023242281.1;refGene_gn=KDM4C;refGene_tx=XM_023242282.1;refGene_gn=KDM4C;refGene_tx=XM_023242283.1;refGene_gn=KDM4C;refGene_tx=XM_023242285.1;refGene_gn=KDM4C;refGene_tx=XM_023242286.1;refGene_gn=KDM4C;refGene_tx=XM_023242287.1;refGene_gn=KDM4C;refGene_tx=XM_023242289.1
D4 34451166 34451216
refGene_gn=KDM4C;refGene_tx=XM_023242276.1;refGene_gn=KDM4C;refGene_tx=XM_023242277.1;refGene_gn=KDM4C;refGene_tx=XM_023242278.1;refGene_gn=KDM4C;refGene_tx=XM_023242279.1;refGene_gn=KDM4C;refGene_tx=XM_023242280.1;refGene_gn=KDM4C;refGene_tx=XM_023242281.1;refGene_gn=KDM4C;refGene_tx=XM_023242282.1;refGene_gn=KDM4C;refGene_tx=XM_023242283.1;refGene_gn=KDM4C;refGene_tx=XM_023242284.1;refGene_gn=KDM4C;refGene_tx=XM_023242285.1;refGene_gn=KDM4C;refGene_tx=XM_023242286.1;refGene_gn=KDM4C;refGene_tx=XM_023242288.1;refGene_gn=KDM4C;refGene_tx=XM_023242289.1
D4 34466602 34466706
refGene_gn=KDM4C;refGene_tx=XM_023242276.1;refGene_gn=KDM4C;refGene_tx=XM_023242277.1;refGene_gn=KDM4C;refGene_tx=XM_023242278.1;refGene_gn=KDM4C;refGene_tx=XM_023242279.1;refGene_gn=KDM4C;refGene_tx=XM_023242280.1;refGene_gn=KDM4C;refGene_tx=XM_023242281.1;refGene_gn=KDM4C;refGene_tx=XM_023242282.1;refGene_gn=KDM4C;refGene_tx=XM_023242283.1;refGene_gn=KDM4C;refGene_tx=XM_023242284.1;refGene_gn=KDM4C;refGene_tx=XM_023242285.1;refGene_gn=KDM4C;refGene_tx=XM_023242286.1;refGene_gn=KDM4C;refGene_tx=XM_023242289.1
D4 34470721 34470859
refGene_gn=KDM4C;refGene_tx=XM_023242276.1;refGene_gn=KDM4C;refGene_tx=XM_023242277.1;refGene_gn=KDM4C;refGene_tx=XM_023242278.1;refGene_gn=KDM4C;refGene_tx=XM_023242279.1;refGene_gn=KDM4C;refGene_tx=XM_023242280.1;refGene_gn=KDM4C;refGene_tx=XM_023242281.1;refGene_gn=KDM4C;refGene_tx=XM_023242282.1;refGene_gn=KDM4C;refGene_tx=XM_023242283.1;refGene_gn=KDM4C;refGene_tx=XM_023242284.1;refGene_gn=KDM4C;refGene_tx=XM_023242285.1;refGene_gn=KDM4C;refGene_tx=XM_023242286.1;refGene_gn=KDM4C;refGene_tx=XM_023242289.1
D4 34582670 34582864
refGene_gn=KDM4C;refGene_tx=XM_023242276.1;refGene_gn=KDM4C;refGene_tx=XM_023242277.1;refGene_gn=KDM4C;refGene_tx=XM_023242278.1;refGene_gn=KDM4C;refGene_tx=XM_023242279.1;refGene_gn=KDM4C;refGene_tx=XM_023242280.1;refGene_gn=KDM4C;refGene_tx=XM_023242281.1;refGene_gn=KDM4C;refGene_tx=XM_023242282.1;refGene_gn=KDM4C;refGene_tx=XM_023242283.1;refGene_gn=KDM4C;refGene_tx=XM_023242284.1;refGene_gn=KDM4C;refGene_tx=XM_023242285.1;refGene_gn=KDM4C;refGene_tx=XM_023242286.1;refGene_gn=KDM4C;refGene_tx=XM_023242289.1
D4 34585585 34585863
refGene_gn=KDM4C;refGene_tx=XM_023242276.1;refGene_gn=KDM4C;refGene_tx=XM_023242277.1;refGene_gn=KDM4C;refGene_tx=XM_023242278.1;refGene_gn=KDM4C;refGene_tx=XM_023242279.1;refGene_gn=KDM4C;refGene_tx=XM_023242280.1;refGene_gn=KDM4C;refGene_tx=XM_023242281.1;refGene_gn=KDM4C;refGene_tx=XM_023242282.1;refGene_gn=KDM4C;refGene_tx=XM_023242283.1;refGene_gn=KDM4C;refGene_tx=XM_023242284.1;refGene_gn=KDM4C;refGene_tx=XM_02

3242285.1;refGene_gn=KDM4C;refGene_tx=XM_023242286.1;refGene_gn=KDM4C;refGene_tx=XM_023242289.1
D4 34588259 34588573
refGene_gn=KDM4C;refGene_tx=XM_023242276.1;refGene_gn=KDM4C;refGene_tx=XM_023242277.1;refGene_gn=KDM4C;refGene_tx=XM_023242278.1;refGene_gn=KDM4C;refGene_tx=XM_023242279.1;refGene_gn=KDM4C;refGene_tx=XM_023242280.1;refGene_gn=KDM4C;refGene_tx=XM_023242281.1;refGene_gn=KDM4C;refGene_tx=XM_023242282.1;refGene_gn=KDM4C;refGene_tx=XM_023242283.1;refGene_gn=KDM4C;refGene_tx=XM_023242284.1;refGene_gn=KDM4C;refGene_tx=XM_023242285.1;refGene_gn=KDM4C;refGene_tx=XM_023242286.1;refGene_gn=KDM4C;refGene_tx=XM_023242289.1
D4 34591934 34592043
refGene_gn=KDM4C;refGene_tx=XM_023242276.1;refGene_gn=KDM4C;refGene_tx=XM_023242277.1;refGene_gn=KDM4C;refGene_tx=XM_023242278.1;refGene_gn=KDM4C;refGene_tx=XM_023242279.1;refGene_gn=KDM4C;refGene_tx=XM_023242280.1;refGene_gn=KDM4C;refGene_tx=XM_023242281.1;refGene_gn=KDM4C;refGene_tx=XM_023242282.1;refGene_gn=KDM4C;refGene_tx=XM_023242283.1;refGene_gn=KDM4C;refGene_tx=XM_023242284.1;refGene_gn=KDM4C;refGene_tx=XM_023242285.1;refGene_gn=KDM4C;refGene_tx=XM_023242286.1;refGene_gn=KDM4C;refGene_tx=XM_023242289.1
D4 34617893 34618075
refGene_gn=KDM4C;refGene_tx=XM_023242276.1;refGene_gn=KDM4C;refGene_tx=XM_023242277.1;refGene_gn=KDM4C;refGene_tx=XM_023242278.1;refGene_gn=KDM4C;refGene_tx=XM_023242279.1;refGene_gn=KDM4C;refGene_tx=XM_023242280.1;refGene_gn=KDM4C;refGene_tx=XM_023242281.1;refGene_gn=KDM4C;refGene_tx=XM_023242282.1;refGene_gn=KDM4C;refGene_tx=XM_023242283.1;refGene_gn=KDM4C;refGene_tx=XM_023242284.1;refGene_gn=KDM4C;refGene_tx=XM_023242285.1;refGene_gn=KDM4C;refGene_tx=XM_023242286.1;refGene_gn=KDM4C;refGene_tx=XM_023242289.1
D4 34619959 34620173
refGene_gn=KDM4C;refGene_tx=XM_023242276.1;refGene_gn=KDM4C;refGene_tx=XM_023242277.1;refGene_gn=KDM4C;refGene_tx=XM_023242278.1;refGene_gn=KDM4C;refGene_tx=XM_023242279.1;refGene_gn=KDM4C;refGene_tx=XM_023242280.1;refGene_gn=KDM4C;refGene_tx=XM_023242281.1;refGene_gn=KDM4C;refGene_tx=XM_023242282.1;refGene_gn=KDM4C;refGene_tx=XM_023242283.1;refGene_gn=KDM4C;refGene_tx=XM_023242284.1;refGene_gn=KDM4C;refGene_tx=XM_023242285.1;refGene_gn=KDM4C;refGene_tx=XM_023242286.1;refGene_gn=KDM4C;refGene_tx=XM_023242289.1
D4 34621993 34622070
refGene_gn=KDM4C;refGene_tx=XM_023242276.1;refGene_gn=KDM4C;refGene_tx=XM_023242277.1;refGene_gn=KDM4C;refGene_tx=XM_023242278.1;refGene_gn=KDM4C;refGene_tx=XM_023242279.1;refGene_gn=KDM4C;refGene_tx=XM_023242280.1;refGene_gn=KDM4C;refGene_tx=XM_023242281.1;refGene_gn=KDM4C;refGene_tx=XM_023242282.1;refGene_gn=KDM4C;refGene_tx=XM_023242283.1;refGene_gn=KDM4C;refGene_tx=XM_023242284.1;refGene_gn=KDM4C;refGene_tx=XM_023242285.1;refGene_gn=KDM4C;refGene_tx=XM_023242286.1;refGene_gn=KDM4C;refGene_tx=XM_023242289.1
D4 34644693 34644749
refGene_gn=KDM4C;refGene_tx=XM_023242276.1;refGene_gn=KDM4C;refGene_tx=XM_023242277.1;refGene_gn=KDM4C;refGene_tx=XM_023242278.1;refGene_gn=

KDM4C;refGene_tx=XM_023242279.1;refGene_gn=KDM4C;refGene_tx=XM_0232422
80.1;refGene_gn=KDM4C;refGene_tx=XM_023242281.1;refGene_gn=KDM4C;refGe
ne_tx=XM_023242282.1;refGene_gn=KDM4C;refGene_tx=XM_023242283.1;refGen
e_gn=KDM4C;refGene_tx=XM_023242284.1;refGene_gn=KDM4C;refGene_tx=XM_02
3242285.1;refGene_gn=KDM4C;refGene_tx=XM_023242286.1;refGene_gn=KDM4C;
refGene_tx=XM_023242289.1
D4 34647187 34647296
refGene_gn=KDM4C;refGene_tx=XM_023242276.1;refGene_gn=KDM4C;refGene_tx
=XM_023242277.1;refGene_gn=KDM4C;refGene_tx=XM_023242278.1;refGene_gn=
KDM4C;refGene_tx=XM_023242279.1;refGene_gn=KDM4C;refGene_tx=XM_0232422
80.1;refGene_gn=KDM4C;refGene_tx=XM_023242281.1;refGene_gn=KDM4C;refGe
ne_tx=XM_023242282.1;refGene_gn=KDM4C;refGene_tx=XM_023242283.1;refGen
e_gn=KDM4C;refGene_tx=XM_023242284.1;refGene_gn=KDM4C;refGene_tx=XM_02
3242285.1;refGene_gn=KDM4C;refGene_tx=XM_023242286.1;refGene_gn=KDM4C;
refGene_tx=XM_023242289.1
D4 34713627 34713813
refGene_gn=KDM4C;refGene_tx=XM_023242276.1;refGene_gn=KDM4C;refGene_tx
=XM_023242277.1;refGene_gn=KDM4C;refGene_tx=XM_023242278.1;refGene_gn=
KDM4C;refGene_tx=XM_023242279.1;refGene_gn=KDM4C;refGene_tx=XM_0232422
80.1;refGene_gn=KDM4C;refGene_tx=XM_023242281.1;refGene_gn=KDM4C;refGe
ne_tx=XM_023242282.1;refGene_gn=KDM4C;refGene_tx=XM_023242283.1;refGen
e_gn=KDM4C;refGene_tx=XM_023242284.1;refGene_gn=KDM4C;refGene_tx=XM_02
3242285.1;refGene_gn=KDM4C;refGene_tx=XM_023242286.1;refGene_gn=KDM4C;
refGene_tx=XM_023242289.1
D4 34719659 34719797 refGene_gn=KDM4C;refGene_tx=XM_023242283.1
D4 34727986 34727995 refGene_gn=KDM4C;refGene_tx=XM_023242285.1
D4 34734125 34734176
refGene_gn=KDM4C;refGene_tx=XM_023242276.1;refGene_gn=KDM4C;refGene_tx
=XM_023242277.1;refGene_gn=KDM4C;refGene_tx=XM_023242278.1;refGene_gn=
KDM4C;refGene_tx=XM_023242279.1;refGene_gn=KDM4C;refGene_tx=XM_0232422
81.1;refGene_gn=KDM4C;refGene_tx=XM_023242282.1;refGene_gn=KDM4C;refGe
ne_tx=XM_023242284.1;refGene_gn=KDM4C;refGene_tx=XM_023242286.1
D4 34736435 34736606
refGene_gn=KDM4C;refGene_tx=XM_023242276.1;refGene_gn=KDM4C;refGene_tx
=XM_023242277.1;refGene_gn=KDM4C;refGene_tx=XM_023242278.1;refGene_gn=
KDM4C;refGene_tx=XM_023242279.1;refGene_gn=KDM4C;refGene_tx=XM_0232422
80.1;refGene_gn=KDM4C;refGene_tx=XM_023242281.1;refGene_gn=KDM4C;refGe
ne_tx=XM_023242282.1;refGene_gn=KDM4C;refGene_tx=XM_023242284.1;refGen
e_gn=KDM4C;refGene_tx=XM_023242286.1;refGene_gn=KDM4C;refGene_tx=XM_02
3242289.1
D4 34770061 34770181
refGene_gn=KDM4C;refGene_tx=XM_023242276.1;refGene_gn=KDM4C;refGene_tx
=XM_023242277.1;refGene_gn=KDM4C;refGene_tx=XM_023242278.1;refGene_gn=
KDM4C;refGene_tx=XM_023242279.1;refGene_gn=KDM4C;refGene_tx=XM_0232422
80.1;refGene_gn=KDM4C;refGene_tx=XM_023242281.1;refGene_gn=KDM4C;refGe
ne_tx=XM_023242282.1;refGene_gn=KDM4C;refGene_tx=XM_023242284.1;refGen
e_gn=KDM4C;refGene_tx=XM_023242286.1;refGene_gn=KDM4C;refGene_tx=XM_02
3242289.1
D4 34774309 34774402
refGene_gn=KDM4C;refGene_tx=XM_023242276.1;refGene_gn=KDM4C;refGene_tx

=XM_023242277.1;refGene_gn=KDM4C;refGene_tx=XM_023242278.1;refGene_gn=KDM4C;refGene_tx=XM_023242279.1;refGene_gn=KDM4C;refGene_tx=XM_023242280.1;refGene_gn=KDM4C;refGene_tx=XM_023242281.1;refGene_gn=KDM4C;refGene_tx=XM_023242282.1;refGene_gn=KDM4C;refGene_tx=XM_023242284.1;refGene_gn=KDM4C;refGene_tx=XM_023242286.1;refGene_gn=KDM4C;refGene_tx=XM_023242289.1
D4 34778775 34778952
refGene_gn=KDM4C;refGene_tx=XM_023242276.1;refGene_gn=KDM4C;refGene_tx=XM_023242277.1;refGene_gn=KDM4C;refGene_tx=XM_023242278.1;refGene_gn=KDM4C;refGene_tx=XM_023242279.1;refGene_gn=KDM4C;refGene_tx=XM_023242280.1;refGene_gn=KDM4C;refGene_tx=XM_023242281.1;refGene_gn=KDM4C;refGene_tx=XM_023242282.1;refGene_gn=KDM4C;refGene_tx=XM_023242284.1;refGene_gn=KDM4C;refGene_tx=XM_023242286.1;refGene_gn=KDM4C;refGene_tx=XM_023242289.1
D4 35348067 35348132 refGene_gn=DMAC1;refGene_tx=XM_003995481.3
D4 35349036 35349310 refGene_gn=DMAC1;refGene_tx=XM_003995481.3
D4 35805492 35805561
refGene_gn=PTPRD;refGene_tx=XM_019815913.2;refGene_gn=PTPRD;refGene_tx=XM_019815914.2;refGene_gn=PTPRD;refGene_tx=XM_019815915.2;refGene_gn=PTPRD;refGene_tx=XM_019815916.2;refGene_gn=PTPRD;refGene_tx=XM_019815917.2;refGene_gn=PTPRD;refGene_tx=XM_019815918.2;refGene_gn=PTPRD;refGene_tx=XM_019815919.2;refGene_gn=PTPRD;refGene_tx=XM_019815920.2;refGene_gn=PTPRD;refGene_tx=XM_019815922.2;refGene_gn=PTPRD;refGene_tx=XM_019815923.2;refGene_gn=PTPRD;refGene_tx=XM_019815924.2;refGene_gn=PTPRD;refGene_tx=XM_019815925.2;refGene_gn=PTPRD;refGene_tx=XM_019815926.2;refGene_gn=PTPRD;refGene_tx=XM_019815927.2;refGene_gn=PTPRD;refGene_tx=XM_019815928.2;refGene_gn=PTPRD;refGene_tx=XM_019815929.2;refGene_gn=PTPRD;refGene_tx=XM_019815930.2;refGene_gn=PTPRD;refGene_tx=XM_019815931.2;refGene_gn=PTPRD;refGene_tx=XM_019815932.2;refGene_gn=PTPRD;refGene_tx=XM_019815933.2;refGene_gn=PTPRD;refGene_tx=XM_019815934.2;refGene_gn=PTPRD;refGene_tx=XM_019815935.2;refGene_gn=PTPRD;refGene_tx=XM_019815937.2;refGene_gn=PTPRD;refGene_tx=XM_019815939.2;refGene_gn=PTPRD;refGene_tx=XM_019815942.2;refGene_gn=PTPRD;refGene_tx=XM_019815943.2;refGene_gn=PTPRD;refGene_tx=XM_019815944.2;refGene_gn=PTPRD;refGene_tx=XM_019815945.2;refGene_gn=PTPRD;refGene_tx=XM_019815946.2;refGene_gn=PTPRD;refGene_tx=XM_019815947.2;refGene_gn=PTPRD;refGene_tx=XM_019815948.2;refGene_gn=PTPRD;refGene_tx=XM_019815949.2;refGene_gn=PTPRD;refGene_tx=XM_019815951.2;refGene_gn=PTPRD;refGene_tx=XM_019815952.2;refGene_gn=PTPRD;refGene_tx=XM_023242291.1;refGene_gn=PTPRD;refGene_tx=XM_023242292.1;refGene_gn=PTPRD;refGene_tx=XM_023242293.1;refGene_gn=PTPRD;refGene_tx=XM_023242294.1;refGene_gn=PTPRD;refGene_tx=XM_023242296.1
D4 35807078 35807214
refGene_gn=PTPRD;refGene_tx=XM_019815913.2;refGene_gn=PTPRD;refGene_tx=XM_019815914.2;refGene_gn=PTPRD;refGene_tx=XM_019815915.2;refGene_gn=PTPRD;refGene_tx=XM_019815916.2;refGene_gn=PTPRD;refGene_tx=XM_019815917.2;refGene_gn=PTPRD;refGene_tx=XM_019815918.2;refGene_gn=PTPRD;refGene_tx=XM_019815919.2;refGene_gn=PTPRD;refGene_tx=XM_019815920.2;refGene_gn=PTPRD;refGene_tx=XM_019815922.2;refGene_gn=PTPRD;refGene_tx=XM_019815923.2;refGene_gn=PTPRD;refGene_tx=XM_019815924.2;refGene_gn=PTPRD;

refGene_tx=XM_019815925.2;refGene_gn=PTPRD;refGene_tx=XM_019815926.2;r
efGene_gn=PTPRD;refGene_tx=XM_019815927.2;refGene_gn=PTPRD;refGene_tx=
XM_019815928.2;refGene_gn=PTPRD;refGene_tx=XM_019815929.2;refGene_gn=P
TPRD;refGene_tx=XM_019815930.2;refGene_gn=PTPRD;refGene_tx=XM_01981593
1.2;refGene_gn=PTPRD;refGene_tx=XM_019815932.2;refGene_gn=PTPRD;refGen
e_tx=XM_019815933.2;refGene_gn=PTPRD;refGene_tx=XM_019815934.2;refGene
_gn=PTPRD;refGene_tx=XM_019815935.2;refGene_gn=PTPRD;refGene_tx=XM_019
815937.2;refGene_gn=PTPRD;refGene_tx=XM_019815939.2;refGene_gn=PTPRD;r
efGene_tx=XM_019815942.2;refGene_gn=PTPRD;refGene_tx=XM_019815943.2;re
fGene_gn=PTPRD;refGene_tx=XM_019815944.2;refGene_gn=PTPRD;refGene_tx=X
M_019815945.2;refGene_gn=PTPRD;refGene_tx=XM_019815946.2;refGene_gn=PT
PRD;refGene_tx=XM_019815947.2;refGene_gn=PTPRD;refGene_tx=XM_019815948
.
2;refGene_gn=PTPRD;refGene_tx=XM_019815949.2;refGene_gn=PTPRD;refGene_
tx=XM_019815951.2;refGene_gn=PTPRD;refGene_tx=XM_019815952.2;refGene_g
n=PTPRD;refGene_tx=XM_023242291.1;refGene_gn=PTPRD;refGene_tx=XM_02324
2292.1;refGene_gn=PTPRD;refGene_tx=XM_023242293.1;refGene_gn=PTPRD;ref
Gene_tx=XM_023242294.1;refGene_gn=PTPRD;refGene_tx=XM_023242296.1
D4 35810251 35810406
refGene_gn=PTPRD;refGene_tx=XM_019815913.2;refGene_gn=PTPRD;refGene_tx
=XM_019815914.2;refGene_gn=PTPRD;refGene_tx=XM_019815915.2;refGene_gn=
PTPRD;refGene_tx=XM_019815916.2;refGene_gn=PTPRD;refGene_tx=XM_0198159
17.2;refGene_gn=PTPRD;refGene_tx=XM_019815918.2;refGene_gn=PTPRD;refGe
ne_tx=XM_019815919.2;refGene_gn=PTPRD;refGene_tx=XM_019815920.2;refGen
e_gn=PTPRD;refGene_tx=XM_019815922.2;refGene_gn=PTPRD;refGene_tx=XM_01
9815923.2;refGene_gn=PTPRD;refGene_tx=XM_019815924.2;refGene_gn=PTPRD;
refGene_tx=XM_019815925.2;refGene_gn=PTPRD;refGene_tx=XM_019815926.2;r
efGene_gn=PTPRD;refGene_tx=XM_019815927.2;refGene_gn=PTPRD;refGene_tx=
XM_019815928.2;refGene_gn=PTPRD;refGene_tx=XM_019815929.2;refGene_gn=P
TPRD;refGene_tx=XM_019815930.2;refGene_gn=PTPRD;refGene_tx=XM_01981593
1.2;refGene_gn=PTPRD;refGene_tx=XM_019815932.2;refGene_gn=PTPRD;refGen
e_tx=XM_019815933.2;refGene_gn=PTPRD;refGene_tx=XM_019815934.2;refGene
_gn=PTPRD;refGene_tx=XM_019815935.2;refGene_gn=PTPRD;refGene_tx=XM_019
815937.2;refGene_gn=PTPRD;refGene_tx=XM_019815939.2;refGene_gn=PTPRD;r
efGene_tx=XM_019815942.2;refGene_gn=PTPRD;refGene_tx=XM_019815943.2;re
fGene_gn=PTPRD;refGene_tx=XM_019815944.2;refGene_gn=PTPRD;refGene_tx=X
M_019815945.2;refGene_gn=PTPRD;refGene_tx=XM_019815946.2;refGene_gn=PT
PRD;refGene_tx=XM_019815947.2;refGene_gn=PTPRD;refGene_tx=XM_019815948
.
2;refGene_gn=PTPRD;refGene_tx=XM_019815949.2;refGene_gn=PTPRD;refGene_
tx=XM_019815951.2;refGene_gn=PTPRD;refGene_tx=XM_019815952.2;refGene_g
n=PTPRD;refGene_tx=XM_023242291.1;refGene_gn=PTPRD;refGene_tx=XM_02324
2292.1;refGene_gn=PTPRD;refGene_tx=XM_023242293.1;refGene_gn=PTPRD;ref
Gene_tx=XM_023242294.1;refGene_gn=PTPRD;refGene_tx=XM_023242296.1
D4 35813641 35813767
refGene_gn=PTPRD;refGene_tx=XM_019815913.2;refGene_gn=PTPRD;refGene_tx
=XM_019815914.2;refGene_gn=PTPRD;refGene_tx=XM_019815915.2;refGene_gn=
PTPRD;refGene_tx=XM_019815916.2;refGene_gn=PTPRD;refGene_tx=XM_0198159
17.2;refGene_gn=PTPRD;refGene_tx=XM_019815918.2;refGene_gn=PTPRD;refGe
ne_tx=XM_019815919.2;refGene_gn=PTPRD;refGene_tx=XM_019815920.2;refGen

e_gn=PTPRD;refGene_tx=XM_019815922.2;refGene_gn=PTPRD;refGene_tx=XM_01
9815923.2;refGene_gn=PTPRD;refGene_tx=XM_019815924.2;refGene_gn=PTPRD;
refGene_tx=XM_019815925.2;refGene_gn=PTPRD;refGene_tx=XM_019815926.2;r
efGene_gn=PTPRD;refGene_tx=XM_019815927.2;refGene_gn=PTPRD;refGene_tx=
XM_019815928.2;refGene_gn=PTPRD;refGene_tx=XM_019815929.2;refGene_gn=P
TPRD;refGene_tx=XM_019815930.2;refGene_gn=PTPRD;refGene_tx=XM_01981593
1.2;refGene_gn=PTPRD;refGene_tx=XM_019815932.2;refGene_gn=PTPRD;refGen
e_tx=XM_019815933.2;refGene_gn=PTPRD;refGene_tx=XM_019815934.2;refGene
_gn=PTPRD;refGene_tx=XM_019815935.2;refGene_gn=PTPRD;refGene_tx=XM_019
815937.2;refGene_gn=PTPRD;refGene_tx=XM_019815939.2;refGene_gn=PTPRD;r
efGene_tx=XM_019815942.2;refGene_gn=PTPRD;refGene_tx=XM_019815943.2;re
fGene_gn=PTPRD;refGene_tx=XM_019815944.2;refGene_gn=PTPRD;refGene_tx=X
M_019815945.2;refGene_gn=PTPRD;refGene_tx=XM_019815946.2;refGene_gn=PT
PRD;refGene_tx=XM_019815947.2;refGene_gn=PTPRD;refGene_tx=XM_019815948
.
2;refGene_gn=PTPRD;refGene_tx=XM_019815949.2;refGene_gn=PTPRD;refGene_
tx=XM_019815951.2;refGene_gn=PTPRD;refGene_tx=XM_019815952.2;refGene_g
n=PTPRD;refGene_tx=XM_023242291.1;refGene_gn=PTPRD;refGene_tx=XM_02324
2292.1;refGene_gn=PTPRD;refGene_tx=XM_023242293.1;refGene_gn=PTPRD;ref
Gene_tx=XM_023242294.1;refGene_gn=PTPRD;refGene_tx=XM_023242296.1
D4 35815117 35815244
refGene_gn=PTPRD;refGene_tx=XM_019815913.2;refGene_gn=PTPRD;refGene_tx
=XM_019815914.2;refGene_gn=PTPRD;refGene_tx=XM_019815915.2;refGene_gn=
PTPRD;refGene_tx=XM_019815916.2;refGene_gn=PTPRD;refGene_tx=XM_0198159
17.2;refGene_gn=PTPRD;refGene_tx=XM_019815918.2;refGene_gn=PTPRD;refGe
ne_tx=XM_019815919.2;refGene_gn=PTPRD;refGene_tx=XM_019815920.2;refGen
e_gn=PTPRD;refGene_tx=XM_019815922.2;refGene_gn=PTPRD;refGene_tx=XM_01
9815923.2;refGene_gn=PTPRD;refGene_tx=XM_019815924.2;refGene_gn=PTPRD;
refGene_tx=XM_019815925.2;refGene_gn=PTPRD;refGene_tx=XM_019815926.2;r
efGene_gn=PTPRD;refGene_tx=XM_019815927.2;refGene_gn=PTPRD;refGene_tx=
XM_019815928.2;refGene_gn=PTPRD;refGene_tx=XM_019815929.2;refGene_gn=P
TPRD;refGene_tx=XM_019815930.2;refGene_gn=PTPRD;refGene_tx=XM_01981593
1.2;refGene_gn=PTPRD;refGene_tx=XM_019815932.2;refGene_gn=PTPRD;refGen
e_tx=XM_019815933.2;refGene_gn=PTPRD;refGene_tx=XM_019815934.2;refGene
_gn=PTPRD;refGene_tx=XM_019815935.2;refGene_gn=PTPRD;refGene_tx=XM_019
815937.2;refGene_gn=PTPRD;refGene_tx=XM_019815939.2;refGene_gn=PTPRD;r
efGene_tx=XM_019815942.2;refGene_gn=PTPRD;refGene_tx=XM_019815943.2;re
fGene_gn=PTPRD;refGene_tx=XM_019815944.2;refGene_gn=PTPRD;refGene_tx=X
M_019815945.2;refGene_gn=PTPRD;refGene_tx=XM_019815946.2;refGene_gn=PT
PRD;refGene_tx=XM_019815947.2;refGene_gn=PTPRD;refGene_tx=XM_019815948
.
2;refGene_gn=PTPRD;refGene_tx=XM_019815949.2;refGene_gn=PTPRD;refGene_
tx=XM_019815951.2;refGene_gn=PTPRD;refGene_tx=XM_019815952.2;refGene_g
n=PTPRD;refGene_tx=XM_023242291.1;refGene_gn=PTPRD;refGene_tx=XM_02324
2292.1;refGene_gn=PTPRD;refGene_tx=XM_023242293.1;refGene_gn=PTPRD;ref
Gene_tx=XM_023242294.1;refGene_gn=PTPRD;refGene_tx=XM_023242296.1
D4 35815861 35816040
refGene_gn=PTPRD;refGene_tx=XM_019815913.2;refGene_gn=PTPRD;refGene_tx
=XM_019815914.2;refGene_gn=PTPRD;refGene_tx=XM_019815915.2;refGene_gn=
PTPRD;refGene_tx=XM_019815916.2;refGene_gn=PTPRD;refGene_tx=XM_0198159

17.2;refGene_gn=PTPRD;refGene_tx=XM_019815918.2;refGene_gn=PTPRD;refGene_tx=XM_019815919.2;refGene_gn=PTPRD;refGene_tx=XM_019815920.2;refGene_gn=PTPRD;refGene_tx=XM_019815922.2;refGene_gn=PTPRD;refGene_tx=XM_019815923.2;refGene_gn=PTPRD;refGene_tx=XM_019815924.2;refGene_gn=PTPRD;refGene_tx=XM_019815925.2;refGene_gn=PTPRD;refGene_tx=XM_019815926.2;refGene_gn=PTPRD;refGene_tx=XM_019815927.2;refGene_gn=PTPRD;refGene_tx=XM_019815928.2;refGene_gn=PTPRD;refGene_tx=XM_019815929.2;refGene_gn=PTPRD;refGene_tx=XM_019815930.2;refGene_gn=PTPRD;refGene_tx=XM_019815931.2;refGene_gn=PTPRD;refGene_tx=XM_019815932.2;refGene_gn=PTPRD;refGene_tx=XM_019815933.2;refGene_gn=PTPRD;refGene_tx=XM_019815934.2;refGene_gn=PTPRD;refGene_tx=XM_019815935.2;refGene_gn=PTPRD;refGene_tx=XM_019815937.2;refGene_gn=PTPRD;refGene_tx=XM_019815939.2;refGene_gn=PTPRD;refGene_tx=XM_019815942.2;refGene_gn=PTPRD;refGene_tx=XM_019815943.2;refGene_gn=PTPRD;refGene_tx=XM_019815944.2;refGene_gn=PTPRD;refGene_tx=XM_019815945.2;refGene_gn=PTPRD;refGene_tx=XM_019815946.2;refGene_gn=PTPRD;refGene_tx=XM_019815947.2;refGene_gn=PTPRD;refGene_tx=XM_019815948.
2;refGene_gn=PTPRD;refGene_tx=XM_019815949.2;refGene_gn=PTPRD;refGene_tx=XM_019815951.2;refGene_gn=PTPRD;refGene_tx=XM_019815952.2;refGene_gn=PTPRD;refGene_tx=XM_023242291.1;refGene_gn=PTPRD;refGene_tx=XM_023242292.1;refGene_gn=PTPRD;refGene_tx=XM_023242293.1;refGene_gn=PTPRD;refGene_tx=XM_023242294.1;refGene_gn=PTPRD;refGene_tx=XM_023242296.1

D4 35816480 35816766

refGene_gn=PTPRD;refGene_tx=XM_019815913.2;refGene_gn=PTPRD;refGene_tx=XM_019815914.2;refGene_gn=PTPRD;refGene_tx=XM_019815915.2;refGene_gn=PTPRD;refGene_tx=XM_019815916.2;refGene_gn=PTPRD;refGene_tx=XM_019815917.2;refGene_gn=PTPRD;refGene_tx=XM_019815918.2;refGene_gn=PTPRD;refGene_tx=XM_019815919.2;refGene_gn=PTPRD;refGene_tx=XM_019815920.2;refGene_gn=PTPRD;refGene_tx=XM_019815922.2;refGene_gn=PTPRD;refGene_tx=XM_019815923.2;refGene_gn=PTPRD;refGene_tx=XM_019815924.2;refGene_gn=PTPRD;refGene_tx=XM_019815925.2;refGene_gn=PTPRD;refGene_tx=XM_019815926.2;refGene_gn=PTPRD;refGene_tx=XM_019815927.2;refGene_gn=PTPRD;refGene_tx=XM_019815928.2;refGene_gn=PTPRD;refGene_tx=XM_019815929.2;refGene_gn=PTPRD;refGene_tx=XM_019815930.2;refGene_gn=PTPRD;refGene_tx=XM_019815931.2;refGene_gn=PTPRD;refGene_tx=XM_019815932.2;refGene_gn=PTPRD;refGene_tx=XM_019815933.2;refGene_gn=PTPRD;refGene_tx=XM_019815934.2;refGene_gn=PTPRD;refGene_tx=XM_019815935.2;refGene_gn=PTPRD;refGene_tx=XM_019815937.2;refGene_gn=PTPRD;refGene_tx=XM_019815939.2;refGene_gn=PTPRD;refGene_tx=XM_019815942.2;refGene_gn=PTPRD;refGene_tx=XM_019815943.2;refGene_gn=PTPRD;refGene_tx=XM_019815944.2;refGene_gn=PTPRD;refGene_tx=XM_019815945.2;refGene_gn=PTPRD;refGene_tx=XM_019815946.2;refGene_gn=PTPRD;refGene_tx=XM_019815947.2;refGene_gn=PTPRD;refGene_tx=XM_019815948.
2;refGene_gn=PTPRD;refGene_tx=XM_019815949.2;refGene_gn=PTPRD;refGene_tx=XM_019815951.2;refGene_gn=PTPRD;refGene_tx=XM_019815952.2;refGene_gn=PTPRD;refGene_tx=XM_023242291.1;refGene_gn=PTPRD;refGene_tx=XM_023242292.1;refGene_gn=PTPRD;refGene_tx=XM_023242293.1;refGene_gn=PTPRD;refGene_tx=XM_023242294.1;refGene_gn=PTPRD;refGene_tx=XM_023242296.1

D4 35852597 35852752

refGene_gn=PTPRD;refGene_tx=XM_019815913.2;refGene_gn=PTPRD;refGene_tx

=XM_019815914.2;refGene_gn=PTPRD;refGene_tx=XM_019815915.2;refGene_gn=
PTPRD;refGene_tx=XM_019815916.2;refGene_gn=PTPRD;refGene_tx=XM_0198159
17.2;refGene_gn=PTPRD;refGene_tx=XM_019815918.2;refGene_gn=PTPRD;refGe
ne_tx=XM_019815919.2;refGene_gn=PTPRD;refGene_tx=XM_019815920.2;refGen
e_gn=PTPRD;refGene_tx=XM_019815922.2;refGene_gn=PTPRD;refGene_tx=XM_01
9815923.2;refGene_gn=PTPRD;refGene_tx=XM_019815924.2;refGene_gn=PTPRD;
refGene_tx=XM_019815925.2;refGene_gn=PTPRD;refGene_tx=XM_019815926.2;r
efGene_gn=PTPRD;refGene_tx=XM_019815927.2;refGene_gn=PTPRD;refGene_tx=
XM_019815928.2;refGene_gn=PTPRD;refGene_tx=XM_019815929.2;refGene_gn=P
TPRD;refGene_tx=XM_019815930.2;refGene_gn=PTPRD;refGene_tx=XM_01981593
1.2;refGene_gn=PTPRD;refGene_tx=XM_019815932.2;refGene_gn=PTPRD;refGen
e_tx=XM_019815933.2;refGene_gn=PTPRD;refGene_tx=XM_019815934.2;refGene
_gn=PTPRD;refGene_tx=XM_019815935.2;refGene_gn=PTPRD;refGene_tx=XM_019
815937.2;refGene_gn=PTPRD;refGene_tx=XM_019815939.2;refGene_gn=PTPRD;r
efGene_tx=XM_019815942.2;refGene_gn=PTPRD;refGene_tx=XM_019815943.2;re
fGene_gn=PTPRD;refGene_tx=XM_019815944.2;refGene_gn=PTPRD;refGene_tx=X
M_019815945.2;refGene_gn=PTPRD;refGene_tx=XM_019815946.2;refGene_gn=PT
PRD;refGene_tx=XM_019815947.2;refGene_gn=PTPRD;refGene_tx=XM_019815948
.
2;refGene_gn=PTPRD;refGene_tx=XM_019815949.2;refGene_gn=PTPRD;refGene_
tx=XM_019815951.2;refGene_gn=PTPRD;refGene_tx=XM_019815952.2;refGene_g
n=PTPRD;refGene_tx=XM_023242291.1;refGene_gn=PTPRD;refGene_tx=XM_02324
2292.1;refGene_gn=PTPRD;refGene_tx=XM_023242293.1;refGene_gn=PTPRD;ref
Gene_tx=XM_023242294.1;refGene_gn=PTPRD;refGene_tx=XM_023242296.1
D4 35853256 35853376
refGene_gn=PTPRD;refGene_tx=XM_019815913.2;refGene_gn=PTPRD;refGene_tx
=XM_019815914.2;refGene_gn=PTPRD;refGene_tx=XM_019815915.2;refGene_gn=
PTPRD;refGene_tx=XM_019815916.2;refGene_gn=PTPRD;refGene_tx=XM_0198159
17.2;refGene_gn=PTPRD;refGene_tx=XM_019815918.2;refGene_gn=PTPRD;refGe
ne_tx=XM_019815919.2;refGene_gn=PTPRD;refGene_tx=XM_019815920.2;refGen
e_gn=PTPRD;refGene_tx=XM_019815922.2;refGene_gn=PTPRD;refGene_tx=XM_01
9815923.2;refGene_gn=PTPRD;refGene_tx=XM_019815924.2;refGene_gn=PTPRD;
refGene_tx=XM_019815925.2;refGene_gn=PTPRD;refGene_tx=XM_019815926.2;r
efGene_gn=PTPRD;refGene_tx=XM_019815927.2;refGene_gn=PTPRD;refGene_tx=
XM_019815928.2;refGene_gn=PTPRD;refGene_tx=XM_019815929.2;refGene_gn=P
TPRD;refGene_tx=XM_019815930.2;refGene_gn=PTPRD;refGene_tx=XM_01981593
1.2;refGene_gn=PTPRD;refGene_tx=XM_019815932.2;refGene_gn=PTPRD;refGen
e_tx=XM_019815933.2;refGene_gn=PTPRD;refGene_tx=XM_019815934.2;refGene
_gn=PTPRD;refGene_tx=XM_019815935.2;refGene_gn=PTPRD;refGene_tx=XM_019
815937.2;refGene_gn=PTPRD;refGene_tx=XM_019815939.2;refGene_gn=PTPRD;r
efGene_tx=XM_019815942.2;refGene_gn=PTPRD;refGene_tx=XM_019815943.2;re
fGene_gn=PTPRD;refGene_tx=XM_019815944.2;refGene_gn=PTPRD;refGene_tx=X
M_019815945.2;refGene_gn=PTPRD;refGene_tx=XM_019815946.2;refGene_gn=PT
PRD;refGene_tx=XM_019815947.2;refGene_gn=PTPRD;refGene_tx=XM_019815948
.
2;refGene_gn=PTPRD;refGene_tx=XM_019815949.2;refGene_gn=PTPRD;refGene_
tx=XM_019815951.2;refGene_gn=PTPRD;refGene_tx=XM_019815952.2;refGene_g
n=PTPRD;refGene_tx=XM_023242291.1;refGene_gn=PTPRD;refGene_tx=XM_02324
2292.1;refGene_gn=PTPRD;refGene_tx=XM_023242293.1;refGene_gn=PTPRD;ref
Gene_tx=XM_023242294.1;refGene_gn=PTPRD;refGene_tx=XM_023242296.1

D4 35867402 35867578
refGene_gn=PTPRD;refGene_tx=XM_019815913.2;refGene_gn=PTPRD;refGene_tx=XM_019815914.2;refGene_gn=PTPRD;refGene_tx=XM_019815915.2;refGene_gn=PTPRD;refGene_tx=XM_019815916.2;refGene_gn=PTPRD;refGene_tx=XM_019815917.2;refGene_gn=PTPRD;refGene_tx=XM_019815918.2;refGene_gn=PTPRD;refGene_tx=XM_019815919.2;refGene_gn=PTPRD;refGene_tx=XM_019815920.2;refGene_gn=PTPRD;refGene_tx=XM_019815922.2;refGene_gn=PTPRD;refGene_tx=XM_019815923.2;refGene_gn=PTPRD;refGene_tx=XM_019815924.2;refGene_gn=PTPRD;refGene_tx=XM_019815925.2;refGene_gn=PTPRD;refGene_tx=XM_019815926.2;refGene_gn=PTPRD;refGene_tx=XM_019815927.2;refGene_gn=PTPRD;refGene_tx=XM_019815928.2;refGene_gn=PTPRD;refGene_tx=XM_019815929.2;refGene_gn=PTPRD;refGene_tx=XM_019815930.2;refGene_gn=PTPRD;refGene_tx=XM_019815931.2;refGene_gn=PTPRD;refGene_tx=XM_019815932.2;refGene_gn=PTPRD;refGene_tx=XM_019815933.2;refGene_gn=PTPRD;refGene_tx=XM_019815934.2;refGene_gn=PTPRD;refGene_tx=XM_019815935.2;refGene_gn=PTPRD;refGene_tx=XM_019815937.2;refGene_gn=PTPRD;refGene_tx=XM_019815939.2;refGene_gn=PTPRD;refGene_tx=XM_019815942.2;refGene_gn=PTPRD;refGene_tx=XM_019815943.2;refGene_gn=PTPRD;refGene_tx=XM_019815944.2;refGene_gn=PTPRD;refGene_tx=XM_019815945.2;refGene_gn=PTPRD;refGene_tx=XM_019815946.2;refGene_gn=PTPRD;refGene_tx=XM_019815947.2;refGene_gn=PTPRD;refGene_tx=XM_019815948.
2;refGene_gn=PTPRD;refGene_tx=XM_019815949.2;refGene_gn=PTPRD;refGene_tx=XM_019815951.2;refGene_gn=PTPRD;refGene_tx=XM_019815952.2;refGene_gn=PTPRD;refGene_tx=XM_023242291.1;refGene_gn=PTPRD;refGene_tx=XM_023242292.1;refGene_gn=PTPRD;refGene_tx=XM_023242293.1;refGene_gn=PTPRD;refGene_tx=XM_023242294.1;refGene_gn=PTPRD;refGene_tx=XM_023242296.1
D4 35883323 35883447
refGene_gn=PTPRD;refGene_tx=XM_019815913.2;refGene_gn=PTPRD;refGene_tx=XM_019815914.2;refGene_gn=PTPRD;refGene_tx=XM_019815915.2;refGene_gn=PTPRD;refGene_tx=XM_019815916.2;refGene_gn=PTPRD;refGene_tx=XM_019815917.2;refGene_gn=PTPRD;refGene_tx=XM_019815918.2;refGene_gn=PTPRD;refGene_tx=XM_019815919.2;refGene_gn=PTPRD;refGene_tx=XM_019815920.2;refGene_gn=PTPRD;refGene_tx=XM_019815922.2;refGene_gn=PTPRD;refGene_tx=XM_019815923.2;refGene_gn=PTPRD;refGene_tx=XM_019815924.2;refGene_gn=PTPRD;refGene_tx=XM_019815925.2;refGene_gn=PTPRD;refGene_tx=XM_019815926.2;refGene_gn=PTPRD;refGene_tx=XM_019815927.2;refGene_gn=PTPRD;refGene_tx=XM_019815928.2;refGene_gn=PTPRD;refGene_tx=XM_019815929.2;refGene_gn=PTPRD;refGene_tx=XM_019815930.2;refGene_gn=PTPRD;refGene_tx=XM_019815931.2;refGene_gn=PTPRD;refGene_tx=XM_019815932.2;refGene_gn=PTPRD;refGene_tx=XM_019815933.2;refGene_gn=PTPRD;refGene_tx=XM_019815934.2;refGene_gn=PTPRD;refGene_tx=XM_019815935.2;refGene_gn=PTPRD;refGene_tx=XM_019815937.2;refGene_gn=PTPRD;refGene_tx=XM_019815939.2;refGene_gn=PTPRD;refGene_tx=XM_019815942.2;refGene_gn=PTPRD;refGene_tx=XM_019815943.2;refGene_gn=PTPRD;refGene_tx=XM_019815944.2;refGene_gn=PTPRD;refGene_tx=XM_019815945.2;refGene_gn=PTPRD;refGene_tx=XM_019815946.2;refGene_gn=PTPRD;refGene_tx=XM_019815947.2;refGene_gn=PTPRD;refGene_tx=XM_019815948.
2;refGene_gn=PTPRD;refGene_tx=XM_019815949.2;refGene_gn=PTPRD;refGene_tx=XM_019815951.2;refGene_gn=PTPRD;refGene_tx=XM_019815952.2;refGene_gn=PTPRD;refGene_tx=XM_023242291.1;refGene_gn=PTPRD;refGene_tx=XM_02324

2292.1;refGene_gn=PTPRD;refGene_tx=XM_023242293.1;refGene_gn=PTPRD;refGene_tx=XM_023242294.1;refGene_gn=PTPRD;refGene_tx=XM_023242296.1
D4 35915623 35915721
refGene_gn=PTPRD;refGene_tx=XM_019815913.2;refGene_gn=PTPRD;refGene_tx=XM_019815914.2;refGene_gn=PTPRD;refGene_tx=XM_019815915.2;refGene_gn=PTPRD;refGene_tx=XM_019815916.2;refGene_gn=PTPRD;refGene_tx=XM_019815917.2;refGene_gn=PTPRD;refGene_tx=XM_019815918.2;refGene_gn=PTPRD;refGene_tx=XM_019815919.2;refGene_gn=PTPRD;refGene_tx=XM_019815920.2;refGene_gn=PTPRD;refGene_tx=XM_019815922.2;refGene_gn=PTPRD;refGene_tx=XM_019815923.2;refGene_gn=PTPRD;refGene_tx=XM_019815924.2;refGene_gn=PTPRD;refGene_tx=XM_019815925.2;refGene_gn=PTPRD;refGene_tx=XM_019815926.2;refGene_gn=PTPRD;refGene_tx=XM_019815927.2;refGene_gn=PTPRD;refGene_tx=XM_019815928.2;refGene_gn=PTPRD;refGene_tx=XM_019815929.2;refGene_gn=PTPRD;refGene_tx=XM_019815930.2;refGene_gn=PTPRD;refGene_tx=XM_019815931.2;refGene_gn=PTPRD;refGene_tx=XM_019815932.2;refGene_gn=PTPRD;refGene_tx=XM_019815933.2;refGene_gn=PTPRD;refGene_tx=XM_019815934.2;refGene_gn=PTPRD;refGene_tx=XM_019815935.2;refGene_gn=PTPRD;refGene_tx=XM_019815937.2;refGene_gn=PTPRD;refGene_tx=XM_019815939.2;refGene_gn=PTPRD;refGene_tx=XM_019815942.2;refGene_gn=PTPRD;refGene_tx=XM_019815943.2;refGene_gn=PTPRD;refGene_tx=XM_019815944.2;refGene_gn=PTPRD;refGene_tx=XM_019815945.2;refGene_gn=PTPRD;refGene_tx=XM_019815946.2;refGene_gn=PTPRD;refGene_tx=XM_019815947.2;refGene_gn=PTPRD;refGene_tx=XM_019815948.
2;refGene_gn=PTPRD;refGene_tx=XM_019815949.2;refGene_gn=PTPRD;refGene_tx=XM_019815951.2;refGene_gn=PTPRD;refGene_tx=XM_019815952.2;refGene_gn=PTPRD;refGene_tx=XM_023242291.1;refGene_gn=PTPRD;refGene_tx=XM_023242292.1;refGene_gn=PTPRD;refGene_tx=XM_023242293.1;refGene_gn=PTPRD;refGene_tx=XM_023242294.1;refGene_gn=PTPRD;refGene_tx=XM_023242296.1
D4 35916226 35916268
refGene_gn=PTPRD;refGene_tx=XM_019815913.2;refGene_gn=PTPRD;refGene_tx=XM_019815914.2;refGene_gn=PTPRD;refGene_tx=XM_019815915.2;refGene_gn=PTPRD;refGene_tx=XM_019815916.2;refGene_gn=PTPRD;refGene_tx=XM_019815917.2;refGene_gn=PTPRD;refGene_tx=XM_019815918.2;refGene_gn=PTPRD;refGene_tx=XM_019815919.2;refGene_gn=PTPRD;refGene_tx=XM_019815920.2;refGene_gn=PTPRD;refGene_tx=XM_019815922.2;refGene_gn=PTPRD;refGene_tx=XM_019815923.2;refGene_gn=PTPRD;refGene_tx=XM_019815924.2;refGene_gn=PTPRD;refGene_tx=XM_019815925.2;refGene_gn=PTPRD;refGene_tx=XM_019815926.2;refGene_gn=PTPRD;refGene_tx=XM_019815928.2;refGene_gn=PTPRD;refGene_tx=XM_019815933.2;refGene_gn=PTPRD;refGene_tx=XM_019815934.2;refGene_gn=PTPRD;refGene_tx=XM_019815935.2;refGene_gn=PTPRD;refGene_tx=XM_019815937.2;refGene_gn=PTPRD;refGene_tx=XM_019815939.2;refGene_gn=PTPRD;refGene_tx=XM_019815944.2;refGene_gn=PTPRD;refGene_tx=XM_023242291.1;refGene_gn=PTPRD;refGene_tx=XM_023242292.1;refGene_gn=PTPRD;refGene_tx=XM_023242293.1
D4 35927961 35928074
refGene_gn=PTPRD;refGene_tx=XM_019815913.2;refGene_gn=PTPRD;refGene_tx=XM_019815914.2;refGene_gn=PTPRD;refGene_tx=XM_019815915.2;refGene_gn=PTPRD;refGene_tx=XM_019815916.2;refGene_gn=PTPRD;refGene_tx=XM_019815917.2;refGene_gn=PTPRD;refGene_tx=XM_019815918.2;refGene_gn=PTPRD;refGene_tx=XM_019815919.2;refGene_gn=PTPRD;refGene_tx=XM_019815920.2;refGen

e_gn=PTPRD;refGene_tx=XM_019815922.2;refGene_gn=PTPRD;refGene_tx=XM_01
9815923.2;refGene_gn=PTPRD;refGene_tx=XM_019815924.2;refGene_gn=PTPRD;
refGene_tx=XM_019815925.2;refGene_gn=PTPRD;refGene_tx=XM_019815926.2;r
efGene_gn=PTPRD;refGene_tx=XM_019815927.2;refGene_gn=PTPRD;refGene_tx=
XM_019815928.2;refGene_gn=PTPRD;refGene_tx=XM_019815929.2;refGene_gn=P
TPRD;refGene_tx=XM_019815930.2;refGene_gn=PTPRD;refGene_tx=XM_01981593
1.2;refGene_gn=PTPRD;refGene_tx=XM_019815932.2;refGene_gn=PTPRD;refGen
e_tx=XM_019815933.2;refGene_gn=PTPRD;refGene_tx=XM_019815934.2;refGene
_gn=PTPRD;refGene_tx=XM_019815935.2;refGene_gn=PTPRD;refGene_tx=XM_019
815937.2;refGene_gn=PTPRD;refGene_tx=XM_019815939.2;refGene_gn=PTPRD;r
efGene_tx=XM_019815942.2;refGene_gn=PTPRD;refGene_tx=XM_019815943.2;re
fGene_gn=PTPRD;refGene_tx=XM_019815944.2;refGene_gn=PTPRD;refGene_tx=X
M_019815945.2;refGene_gn=PTPRD;refGene_tx=XM_019815946.2;refGene_gn=PT
PRD;refGene_tx=XM_019815947.2;refGene_gn=PTPRD;refGene_tx=XM_019815948
.
2;refGene_gn=PTPRD;refGene_tx=XM_019815949.2;refGene_gn=PTPRD;refGene_
tx=XM_019815951.2;refGene_gn=PTPRD;refGene_tx=XM_019815952.2;refGene_g
n=PTPRD;refGene_tx=XM_023242291.1;refGene_gn=PTPRD;refGene_tx=XM_02324
2292.1;refGene_gn=PTPRD;refGene_tx=XM_023242293.1;refGene_gn=PTPRD;ref
Gene_tx=XM_023242294.1;refGene_gn=PTPRD;refGene_tx=XM_023242296.1
D4 35932098 35932113
refGene_gn=PTPRD;refGene_tx=XM_019815913.2;refGene_gn=PTPRD;refGene_tx
=XM_019815914.2;refGene_gn=PTPRD;refGene_tx=XM_019815915.2;refGene_gn=
PTPRD;refGene_tx=XM_019815916.2;refGene_gn=PTPRD;refGene_tx=XM_0198159
17.2;refGene_gn=PTPRD;refGene_tx=XM_019815918.2;refGene_gn=PTPRD;refGe
ne_tx=XM_019815919.2;refGene_gn=PTPRD;refGene_tx=XM_019815920.2;refGen
e_gn=PTPRD;refGene_tx=XM_019815922.2;refGene_gn=PTPRD;refGene_tx=XM_01
9815923.2;refGene_gn=PTPRD;refGene_tx=XM_019815924.2;refGene_gn=PTPRD;
refGene_tx=XM_019815925.2;refGene_gn=PTPRD;refGene_tx=XM_019815926.2;r
efGene_gn=PTPRD;refGene_tx=XM_019815927.2;refGene_gn=PTPRD;refGene_tx=
XM_019815928.2;refGene_gn=PTPRD;refGene_tx=XM_019815929.2;refGene_gn=P
TPRD;refGene_tx=XM_019815933.2;refGene_gn=PTPRD;refGene_tx=XM_01981593
4.2;refGene_gn=PTPRD;refGene_tx=XM_019815935.2;refGene_gn=PTPRD;refGen
e_tx=XM_019815937.2;refGene_gn=PTPRD;refGene_tx=XM_019815939.2;refGene
_gn=PTPRD;refGene_tx=XM_019815942.2;refGene_gn=PTPRD;refGene_tx=XM_019
815943.2;refGene_gn=PTPRD;refGene_tx=XM_019815945.2;refGene_gn=PTPRD;r
efGene_tx=XM_019815949.2;refGene_gn=PTPRD;refGene_tx=XM_019815951.2;re
fGene_gn=PTPRD;refGene_tx=XM_019815952.2;refGene_gn=PTPRD;refGene_tx=X
M_023242292.1;refGene_gn=PTPRD;refGene_tx=XM_023242293.1
D4 35938379 35938540
refGene_gn=PTPRD;refGene_tx=XM_019815913.2;refGene_gn=PTPRD;refGene_tx
=XM_019815914.2;refGene_gn=PTPRD;refGene_tx=XM_019815915.2;refGene_gn=
PTPRD;refGene_tx=XM_019815916.2;refGene_gn=PTPRD;refGene_tx=XM_0198159
17.2;refGene_gn=PTPRD;refGene_tx=XM_019815918.2;refGene_gn=PTPRD;refGe
ne_tx=XM_019815919.2;refGene_gn=PTPRD;refGene_tx=XM_019815920.2;refGen
e_gn=PTPRD;refGene_tx=XM_019815922.2;refGene_gn=PTPRD;refGene_tx=XM_01
9815923.2;refGene_gn=PTPRD;refGene_tx=XM_019815924.2;refGene_gn=PTPRD;
refGene_tx=XM_019815925.2;refGene_gn=PTPRD;refGene_tx=XM_019815926.2;r
efGene_gn=PTPRD;refGene_tx=XM_019815927.2;refGene_gn=PTPRD;refGene_tx=
XM_019815928.2;refGene_gn=PTPRD;refGene_tx=XM_019815929.2;refGene_gn=P

TPRD;refGene_tx=XM_019815930.2;refGene_gn=PTPRD;refGene_tx=XM_01981593
1.2;refGene_gn=PTPRD;refGene_tx=XM_019815932.2;refGene_gn=PTPRD;refGen
e_tx=XM_019815933.2;refGene_gn=PTPRD;refGene_tx=XM_019815934.2;refGene
_gn=PTPRD;refGene_tx=XM_019815935.2;refGene_gn=PTPRD;refGene_tx=XM_019
815937.2;refGene_gn=PTPRD;refGene_tx=XM_019815939.2;refGene_gn=PTPRD;r
efGene_tx=XM_019815942.2;refGene_gn=PTPRD;refGene_tx=XM_019815943.2;re
fGene_gn=PTPRD;refGene_tx=XM_019815944.2;refGene_gn=PTPRD;refGene_tx=X
M_019815945.2;refGene_gn=PTPRD;refGene_tx=XM_019815946.2;refGene_gn=PT
PRD;refGene_tx=XM_019815947.2;refGene_gn=PTPRD;refGene_tx=XM_019815948
.
2;refGene_gn=PTPRD;refGene_tx=XM_019815949.2;refGene_gn=PTPRD;refGene_
tx=XM_019815951.2;refGene_gn=PTPRD;refGene_tx=XM_019815952.2;refGene_g
n=PTPRD;refGene_tx=XM_023242291.1;refGene_gn=PTPRD;refGene_tx=XM_02324
2292.1;refGene_gn=PTPRD;refGene_tx=XM_023242293.1;refGene_gn=PTPRD;ref
Gene_tx=XM_023242294.1;refGene_gn=PTPRD;refGene_tx=XM_023242296.1
D4 35943190 35943400
refGene_gn=PTPRD;refGene_tx=XM_019815913.2;refGene_gn=PTPRD;refGene_tx
=XM_019815914.2;refGene_gn=PTPRD;refGene_tx=XM_019815915.2;refGene_gn=
PTPRD;refGene_tx=XM_019815916.2;refGene_gn=PTPRD;refGene_tx=XM_0198159
17.2;refGene_gn=PTPRD;refGene_tx=XM_019815918.2;refGene_gn=PTPRD;refGe
ne_tx=XM_019815919.2;refGene_gn=PTPRD;refGene_tx=XM_019815920.2;refGen
e_gn=PTPRD;refGene_tx=XM_019815922.2;refGene_gn=PTPRD;refGene_tx=XM_01
9815923.2;refGene_gn=PTPRD;refGene_tx=XM_019815924.2;refGene_gn=PTPRD;
refGene_tx=XM_019815925.2;refGene_gn=PTPRD;refGene_tx=XM_019815926.2;r
efGene_gn=PTPRD;refGene_tx=XM_019815927.2;refGene_gn=PTPRD;refGene_tx=
XM_019815928.2;refGene_gn=PTPRD;refGene_tx=XM_019815929.2;refGene_gn=P
TPRD;refGene_tx=XM_019815930.2;refGene_gn=PTPRD;refGene_tx=XM_01981593
1.2;refGene_gn=PTPRD;refGene_tx=XM_019815932.2;refGene_gn=PTPRD;refGen
e_tx=XM_019815933.2;refGene_gn=PTPRD;refGene_tx=XM_019815934.2;refGene
_gn=PTPRD;refGene_tx=XM_019815935.2;refGene_gn=PTPRD;refGene_tx=XM_019
815937.2;refGene_gn=PTPRD;refGene_tx=XM_019815939.2;refGene_gn=PTPRD;r
efGene_tx=XM_019815942.2;refGene_gn=PTPRD;refGene_tx=XM_019815943.2;re
fGene_gn=PTPRD;refGene_tx=XM_019815944.2;refGene_gn=PTPRD;refGene_tx=X
M_019815945.2;refGene_gn=PTPRD;refGene_tx=XM_019815946.2;refGene_gn=PT
PRD;refGene_tx=XM_019815947.2;refGene_gn=PTPRD;refGene_tx=XM_019815948
.
2;refGene_gn=PTPRD;refGene_tx=XM_019815949.2;refGene_gn=PTPRD;refGene_
tx=XM_019815951.2;refGene_gn=PTPRD;refGene_tx=XM_019815952.2;refGene_g
n=PTPRD;refGene_tx=XM_023242291.1;refGene_gn=PTPRD;refGene_tx=XM_02324
2292.1;refGene_gn=PTPRD;refGene_tx=XM_023242293.1;refGene_gn=PTPRD;ref
Gene_tx=XM_023242294.1;refGene_gn=PTPRD;refGene_tx=XM_023242296.1
D4 35948815 35948906
refGene_gn=PTPRD;refGene_tx=XM_019815913.2;refGene_gn=PTPRD;refGene_tx
=XM_019815914.2;refGene_gn=PTPRD;refGene_tx=XM_019815915.2;refGene_gn=
PTPRD;refGene_tx=XM_019815916.2;refGene_gn=PTPRD;refGene_tx=XM_0198159
17.2;refGene_gn=PTPRD;refGene_tx=XM_019815918.2;refGene_gn=PTPRD;refGe
ne_tx=XM_019815919.2;refGene_gn=PTPRD;refGene_tx=XM_019815920.2;refGen
e_gn=PTPRD;refGene_tx=XM_019815922.2;refGene_gn=PTPRD;refGene_tx=XM_01
9815923.2;refGene_gn=PTPRD;refGene_tx=XM_019815924.2;refGene_gn=PTPRD;
refGene_tx=XM_019815925.2;refGene_gn=PTPRD;refGene_tx=XM_019815926.2;r

efGene_gn=PTPRD;refGene_tx=XM_019815927.2;refGene_gn=PTPRD;refGene_tx=XM_019815928.2;refGene_gn=PTPRD;refGene_tx=XM_019815929.2;refGene_gn=PTPRD;refGene_tx=XM_019815930.2;refGene_gn=PTPRD;refGene_tx=XM_019815931.2;refGene_gn=PTPRD;refGene_tx=XM_019815932.2;refGene_gn=PTPRD;refGene_tx=XM_019815933.2;refGene_gn=PTPRD;refGene_tx=XM_019815934.2;refGene_gn=PTPRD;refGene_tx=XM_019815935.2;refGene_gn=PTPRD;refGene_tx=XM_019815937.2;refGene_gn=PTPRD;refGene_tx=XM_019815939.2;refGene_gn=PTPRD;refGene_tx=XM_019815942.2;refGene_gn=PTPRD;refGene_tx=XM_019815943.2;refGene_gn=PTPRD;refGene_tx=XM_019815944.2;refGene_gn=PTPRD;refGene_tx=XM_019815945.2;refGene_gn=PTPRD;refGene_tx=XM_019815946.2;refGene_gn=PTPRD;refGene_tx=XM_019815947.2;refGene_gn=PTPRD;refGene_tx=XM_019815948.
2;refGene_gn=PTPRD;refGene_tx=XM_019815949.2;refGene_gn=PTPRD;refGene_tx=XM_019815951.2;refGene_gn=PTPRD;refGene_tx=XM_019815952.2;refGene_gn=PTPRD;refGene_tx=XM_023242291.1;refGene_gn=PTPRD;refGene_tx=XM_023242292.1;refGene_gn=PTPRD;refGene_tx=XM_023242293.1;refGene_gn=PTPRD;refGene_tx=XM_023242294.1;refGene_gn=PTPRD;refGene_tx=XM_023242296.1

D4 35967583 35967843

refGene_gn=PTPRD;refGene_tx=XM_019815913.2;refGene_gn=PTPRD;refGene_tx=XM_019815914.2;refGene_gn=PTPRD;refGene_tx=XM_019815915.2;refGene_gn=PTPRD;refGene_tx=XM_019815916.2;refGene_gn=PTPRD;refGene_tx=XM_019815917.2;refGene_gn=PTPRD;refGene_tx=XM_019815918.2;refGene_gn=PTPRD;refGene_tx=XM_019815919.2;refGene_gn=PTPRD;refGene_tx=XM_019815920.2;refGene_gn=PTPRD;refGene_tx=XM_019815922.2;refGene_gn=PTPRD;refGene_tx=XM_019815923.2;refGene_gn=PTPRD;refGene_tx=XM_019815924.2;refGene_gn=PTPRD;refGene_tx=XM_019815925.2;refGene_gn=PTPRD;refGene_tx=XM_019815926.2;refGene_gn=PTPRD;refGene_tx=XM_019815927.2;refGene_gn=PTPRD;refGene_tx=XM_019815928.2;refGene_gn=PTPRD;refGene_tx=XM_019815929.2;refGene_gn=PTPRD;refGene_tx=XM_019815930.2;refGene_gn=PTPRD;refGene_tx=XM_019815931.2;refGene_gn=PTPRD;refGene_tx=XM_019815932.2;refGene_gn=PTPRD;refGene_tx=XM_019815933.2;refGene_gn=PTPRD;refGene_tx=XM_019815934.2;refGene_gn=PTPRD;refGene_tx=XM_019815935.2;refGene_gn=PTPRD;refGene_tx=XM_019815937.2;refGene_gn=PTPRD;refGene_tx=XM_019815939.2;refGene_gn=PTPRD;refGene_tx=XM_019815942.2;refGene_gn=PTPRD;refGene_tx=XM_019815943.2;refGene_gn=PTPRD;refGene_tx=XM_019815944.2;refGene_gn=PTPRD;refGene_tx=XM_019815945.2;refGene_gn=PTPRD;refGene_tx=XM_019815946.2;refGene_gn=PTPRD;refGene_tx=XM_019815947.2;refGene_gn=PTPRD;refGene_tx=XM_019815948.
2;refGene_gn=PTPRD;refGene_tx=XM_019815949.2;refGene_gn=PTPRD;refGene_tx=XM_019815951.2;refGene_gn=PTPRD;refGene_tx=XM_019815952.2;refGene_gn=PTPRD;refGene_tx=XM_023242291.1;refGene_gn=PTPRD;refGene_tx=XM_023242292.1;refGene_gn=PTPRD;refGene_tx=XM_023242293.1;refGene_gn=PTPRD;refGene_tx=XM_023242294.1;refGene_gn=PTPRD;refGene_tx=XM_023242296.1

D4 35968640 35968741

refGene_gn=PTPRD;refGene_tx=XM_019815913.2;refGene_gn=PTPRD;refGene_tx=XM_019815914.2;refGene_gn=PTPRD;refGene_tx=XM_019815915.2;refGene_gn=PTPRD;refGene_tx=XM_019815916.2;refGene_gn=PTPRD;refGene_tx=XM_019815917.2;refGene_gn=PTPRD;refGene_tx=XM_019815918.2;refGene_gn=PTPRD;refGene_tx=XM_019815919.2;refGene_gn=PTPRD;refGene_tx=XM_019815920.2;refGene_gn=PTPRD;refGene_tx=XM_019815922.2;refGene_gn=PTPRD;refGene_tx=XM_01

9815923.2;refGene_gn=PTPRD;refGene_tx=XM_019815924.2;refGene_gn=PTPRD;
refGene_tx=XM_019815925.2;refGene_gn=PTPRD;refGene_tx=XM_019815926.2;r
efGene_gn=PTPRD;refGene_tx=XM_019815927.2;refGene_gn=PTPRD;refGene_tx=
XM_019815928.2;refGene_gn=PTPRD;refGene_tx=XM_019815929.2;refGene_gn=P
TPRD;refGene_tx=XM_019815930.2;refGene_gn=PTPRD;refGene_tx=XM_01981593
1.2;refGene_gn=PTPRD;refGene_tx=XM_019815932.2;refGene_gn=PTPRD;refGen
e_tx=XM_019815933.2;refGene_gn=PTPRD;refGene_tx=XM_019815934.2;refGene
_gn=PTPRD;refGene_tx=XM_019815935.2;refGene_gn=PTPRD;refGene_tx=XM_019
815937.2;refGene_gn=PTPRD;refGene_tx=XM_019815939.2;refGene_gn=PTPRD;r
efGene_tx=XM_019815942.2;refGene_gn=PTPRD;refGene_tx=XM_019815943.2;re
fGene_gn=PTPRD;refGene_tx=XM_019815944.2;refGene_gn=PTPRD;refGene_tx=X
M_019815945.2;refGene_gn=PTPRD;refGene_tx=XM_019815946.2;refGene_gn=PT
PRD;refGene_tx=XM_019815947.2;refGene_gn=PTPRD;refGene_tx=XM_019815948
.
2;refGene_gn=PTPRD;refGene_tx=XM_019815949.2;refGene_gn=PTPRD;refGene_
tx=XM_019815951.2;refGene_gn=PTPRD;refGene_tx=XM_019815952.2;refGene_g
n=PTPRD;refGene_tx=XM_023242291.1;refGene_gn=PTPRD;refGene_tx=XM_02324
2292.1;refGene_gn=PTPRD;refGene_tx=XM_023242293.1;refGene_gn=PTPRD;ref
Gene_tx=XM_023242294.1;refGene_gn=PTPRD;refGene_tx=XM_023242296.1
D4 35969177 35969765
refGene_gn=PTPRD;refGene_tx=XM_019815913.2;refGene_gn=PTPRD;refGene_tx
=XM_019815914.2;refGene_gn=PTPRD;refGene_tx=XM_019815915.2;refGene_gn=
PTPRD;refGene_tx=XM_019815916.2;refGene_gn=PTPRD;refGene_tx=XM_0198159
17.2;refGene_gn=PTPRD;refGene_tx=XM_019815918.2;refGene_gn=PTPRD;refGe
ne_tx=XM_019815919.2;refGene_gn=PTPRD;refGene_tx=XM_019815920.2;refGen
e_gn=PTPRD;refGene_tx=XM_019815922.2;refGene_gn=PTPRD;refGene_tx=XM_01
9815923.2;refGene_gn=PTPRD;refGene_tx=XM_019815924.2;refGene_gn=PTPRD;
refGene_tx=XM_019815925.2;refGene_gn=PTPRD;refGene_tx=XM_019815926.2;r
efGene_gn=PTPRD;refGene_tx=XM_019815927.2;refGene_gn=PTPRD;refGene_tx=
XM_019815928.2;refGene_gn=PTPRD;refGene_tx=XM_019815929.2;refGene_gn=P
TPRD;refGene_tx=XM_019815930.2;refGene_gn=PTPRD;refGene_tx=XM_01981593
1.2;refGene_gn=PTPRD;refGene_tx=XM_019815932.2;refGene_gn=PTPRD;refGen
e_tx=XM_019815933.2;refGene_gn=PTPRD;refGene_tx=XM_019815934.2;refGene
_gn=PTPRD;refGene_tx=XM_019815935.2;refGene_gn=PTPRD;refGene_tx=XM_019
815937.2;refGene_gn=PTPRD;refGene_tx=XM_019815939.2;refGene_gn=PTPRD;r
efGene_tx=XM_019815948.2;refGene_gn=PTPRD;refGene_tx=XM_019815949.2;re
fGene_gn=PTPRD;refGene_tx=XM_023242291.1;refGene_gn=PTPRD;refGene_tx=X
M_023242292.1;refGene_gn=PTPRD;refGene_tx=XM_023242293.1;refGene_gn=PT
PRD;refGene_tx=XM_023242296.1
D4 35972391 35972509
refGene_gn=PTPRD;refGene_tx=XM_019815913.2;refGene_gn=PTPRD;refGene_tx
=XM_019815914.2;refGene_gn=PTPRD;refGene_tx=XM_019815915.2;refGene_gn=
PTPRD;refGene_tx=XM_019815916.2;refGene_gn=PTPRD;refGene_tx=XM_0198159
17.2;refGene_gn=PTPRD;refGene_tx=XM_019815918.2;refGene_gn=PTPRD;refGe
ne_tx=XM_019815919.2;refGene_gn=PTPRD;refGene_tx=XM_019815920.2;refGen
e_gn=PTPRD;refGene_tx=XM_019815922.2;refGene_gn=PTPRD;refGene_tx=XM_01
9815923.2;refGene_gn=PTPRD;refGene_tx=XM_019815924.2;refGene_gn=PTPRD;
refGene_tx=XM_019815925.2;refGene_gn=PTPRD;refGene_tx=XM_019815926.2;r
efGene_gn=PTPRD;refGene_tx=XM_019815927.2;refGene_gn=PTPRD;refGene_tx=
XM_019815928.2;refGene_gn=PTPRD;refGene_tx=XM_019815929.2;refGene_gn=P

TPRD;refGene_tx=XM_019815930.2;refGene_gn=PTPRD;refGene_tx=XM_01981593
1.2;refGene_gn=PTPRD;refGene_tx=XM_019815932.2;refGene_gn=PTPRD;refGen
e_tx=XM_019815933.2;refGene_gn=PTPRD;refGene_tx=XM_019815934.2;refGene
_gn=PTPRD;refGene_tx=XM_019815935.2;refGene_gn=PTPRD;refGene_tx=XM_019
815937.2;refGene_gn=PTPRD;refGene_tx=XM_019815939.2;refGene_gn=PTPRD;r
efGene_tx=XM_019815948.2;refGene_gn=PTPRD;refGene_tx=XM_019815949.2;re
fGene_gn=PTPRD;refGene_tx=XM_023242291.1;refGene_gn=PTPRD;refGene_tx=X
M_023242292.1;refGene_gn=PTPRD;refGene_tx=XM_023242293.1;refGene_gn=PT
PRD;refGene_tx=XM_023242296.1
D4 35976442 35976469
refGene_gn=PTPRD;refGene_tx=XM_019815913.2;refGene_gn=PTPRD;refGene_tx
=XM_019815914.2;refGene_gn=PTPRD;refGene_tx=XM_019815915.2;refGene_gn=
PTPRD;refGene_tx=XM_019815916.2;refGene_gn=PTPRD;refGene_tx=XM_0198159
17.2;refGene_gn=PTPRD;refGene_tx=XM_019815918.2;refGene_gn=PTPRD;refGe
ne_tx=XM_019815919.2;refGene_gn=PTPRD;refGene_tx=XM_019815920.2;refGen
e_gn=PTPRD;refGene_tx=XM_019815922.2;refGene_gn=PTPRD;refGene_tx=XM_01
9815923.2;refGene_gn=PTPRD;refGene_tx=XM_019815925.2;refGene_gn=PTPRD;
refGene_tx=XM_019815927.2;refGene_gn=PTPRD;refGene_tx=XM_019815928.2;r
efGene_gn=PTPRD;refGene_tx=XM_019815929.2;refGene_gn=PTPRD;refGene_tx=
XM_019815930.2;refGene_gn=PTPRD;refGene_tx=XM_019815931.2;refGene_gn=P
TPRD;refGene_tx=XM_019815932.2;refGene_gn=PTPRD;refGene_tx=XM_01981593
4.2;refGene_gn=PTPRD;refGene_tx=XM_019815935.2;refGene_gn=PTPRD;refGen
e_tx=XM_019815937.2;refGene_gn=PTPRD;refGene_tx=XM_023242291.1;refGene
_gn=PTPRD;refGene_tx=XM_023242292.1;refGene_gn=PTPRD;refGene_tx=XM_023
242293.1
D4 35978584 35978778
refGene_gn=PTPRD;refGene_tx=XM_019815913.2;refGene_gn=PTPRD;refGene_tx
=XM_019815914.2;refGene_gn=PTPRD;refGene_tx=XM_019815915.2;refGene_gn=
PTPRD;refGene_tx=XM_019815916.2;refGene_gn=PTPRD;refGene_tx=XM_0198159
17.2;refGene_gn=PTPRD;refGene_tx=XM_019815918.2;refGene_gn=PTPRD;refGe
ne_tx=XM_019815919.2;refGene_gn=PTPRD;refGene_tx=XM_019815920.2;refGen
e_gn=PTPRD;refGene_tx=XM_019815922.2;refGene_gn=PTPRD;refGene_tx=XM_01
9815923.2;refGene_gn=PTPRD;refGene_tx=XM_019815924.2;refGene_gn=PTPRD;
refGene_tx=XM_019815925.2;refGene_gn=PTPRD;refGene_tx=XM_019815926.2;r
efGene_gn=PTPRD;refGene_tx=XM_019815927.2;refGene_gn=PTPRD;refGene_tx=
XM_019815928.2;refGene_gn=PTPRD;refGene_tx=XM_019815929.2;refGene_gn=P
TPRD;refGene_tx=XM_019815930.2;refGene_gn=PTPRD;refGene_tx=XM_01981593
1.2;refGene_gn=PTPRD;refGene_tx=XM_019815932.2;refGene_gn=PTPRD;refGen
e_tx=XM_019815933.2;refGene_gn=PTPRD;refGene_tx=XM_019815934.2;refGene
_gn=PTPRD;refGene_tx=XM_019815935.2;refGene_gn=PTPRD;refGene_tx=XM_019
815937.2;refGene_gn=PTPRD;refGene_tx=XM_019815939.2;refGene_gn=PTPRD;r
efGene_tx=XM_019815948.2;refGene_gn=PTPRD;refGene_tx=XM_019815949.2;re
fGene_gn=PTPRD;refGene_tx=XM_023242291.1;refGene_gn=PTPRD;refGene_tx=X
M_023242292.1;refGene_gn=PTPRD;refGene_tx=XM_023242293.1;refGene_gn=PT
PRD;refGene_tx=XM_023242296.1
D4 35979671 35979977
refGene_gn=PTPRD;refGene_tx=XM_019815913.2;refGene_gn=PTPRD;refGene_tx
=XM_019815914.2;refGene_gn=PTPRD;refGene_tx=XM_019815915.2;refGene_gn=
PTPRD;refGene_tx=XM_019815916.2;refGene_gn=PTPRD;refGene_tx=XM_0198159
17.2;refGene_gn=PTPRD;refGene_tx=XM_019815918.2;refGene_gn=PTPRD;refGe

ne_tx=XM_019815919.2;refGene_gn=PTPRD;refGene_tx=XM_019815920.2;refGene_gn=PTPRD;refGene_tx=XM_019815922.2;refGene_gn=PTPRD;refGene_tx=XM_019815923.2;refGene_gn=PTPRD;refGene_tx=XM_019815924.2;refGene_gn=PTPRD;refGene_tx=XM_019815925.2;refGene_gn=PTPRD;refGene_tx=XM_019815926.2;refGene_gn=PTPRD;refGene_tx=XM_019815927.2;refGene_gn=PTPRD;refGene_tx=XM_019815928.2;refGene_gn=PTPRD;refGene_tx=XM_019815929.2;refGene_gn=PTPRD;refGene_tx=XM_019815930.2;refGene_gn=PTPRD;refGene_tx=XM_019815931.2;refGene_gn=PTPRD;refGene_tx=XM_019815932.2;refGene_gn=PTPRD;refGene_tx=XM_019815933.2;refGene_gn=PTPRD;refGene_tx=XM_019815948.2;refGene_gn=PTPRD;refGene_tx=XM_019815949.2;refGene_gn=PTPRD;refGene_tx=XM_023242291.1;refGene_gn=PTPRD;refGene_tx=XM_023242292.1

D4 35983205 35983350

refGene_gn=PTPRD;refGene_tx=XM_019815913.2;refGene_gn=PTPRD;refGene_tx=XM_019815914.2;refGene_gn=PTPRD;refGene_tx=XM_019815915.2;refGene_gn=PTPRD;refGene_tx=XM_019815916.2;refGene_gn=PTPRD;refGene_tx=XM_019815917.2;refGene_gn=PTPRD;refGene_tx=XM_019815918.2;refGene_gn=PTPRD;refGene_tx=XM_019815919.2;refGene_gn=PTPRD;refGene_tx=XM_019815920.2;refGene_gn=PTPRD;refGene_tx=XM_019815922.2;refGene_gn=PTPRD;refGene_tx=XM_019815923.2;refGene_gn=PTPRD;refGene_tx=XM_019815924.2;refGene_gn=PTPRD;refGene_tx=XM_019815925.2;refGene_gn=PTPRD;refGene_tx=XM_019815926.2;refGene_gn=PTPRD;refGene_tx=XM_019815927.2;refGene_gn=PTPRD;refGene_tx=XM_019815928.2;refGene_gn=PTPRD;refGene_tx=XM_019815929.2;refGene_gn=PTPRD;refGene_tx=XM_019815930.2;refGene_gn=PTPRD;refGene_tx=XM_019815931.2;refGene_gn=PTPRD;refGene_tx=XM_019815932.2;refGene_gn=PTPRD;refGene_tx=XM_019815933.2;refGene_gn=PTPRD;refGene_tx=XM_019815934.2;refGene_gn=PTPRD;refGene_tx=XM_019815935.2;refGene_gn=PTPRD;refGene_tx=XM_019815937.2;refGene_gn=PTPRD;refGene_tx=XM_019815939.2;refGene_gn=PTPRD;refGene_tx=XM_019815942.2;refGene_gn=PTPRD;refGene_tx=XM_019815943.2;refGene_gn=PTPRD;refGene_tx=XM_019815944.2;refGene_gn=PTPRD;refGene_tx=XM_019815945.2;refGene_gn=PTPRD;refGene_tx=XM_019815946.2;refGene_gn=PTPRD;refGene_tx=XM_019815947.2;refGene_gn=PTPRD;refGene_tx=XM_019815948.2;refGene_gn=PTPRD;refGene_tx=XM_019815949.2;refGene_gn=PTPRD;refGene_tx=XM_019815951.2;refGene_gn=PTPRD;refGene_tx=XM_019815952.2;refGene_gn=PTPRD;refGene_tx=XM_023242291.1;refGene_gn=PTPRD;refGene_tx=XM_023242292.1;refGene_gn=PTPRD;refGene_tx=XM_023242293.1;refGene_gn=PTPRD;refGene_tx=XM_023242294.1;refGene_gn=PTPRD;refGene_tx=XM_023242296.1

D4 35985863 35985997

refGene_gn=PTPRD;refGene_tx=XM_019815913.2;refGene_gn=PTPRD;refGene_tx=XM_019815914.2;refGene_gn=PTPRD;refGene_tx=XM_019815915.2;refGene_gn=PTPRD;refGene_tx=XM_019815916.2;refGene_gn=PTPRD;refGene_tx=XM_019815917.2;refGene_gn=PTPRD;refGene_tx=XM_019815918.2;refGene_gn=PTPRD;refGene_tx=XM_019815919.2;refGene_gn=PTPRD;refGene_tx=XM_019815920.2;refGene_gn=PTPRD;refGene_tx=XM_019815922.2;refGene_gn=PTPRD;refGene_tx=XM_019815923.2;refGene_gn=PTPRD;refGene_tx=XM_019815924.2;refGene_gn=PTPRD;refGene_tx=XM_019815925.2;refGene_gn=PTPRD;refGene_tx=XM_019815926.2;refGene_gn=PTPRD;refGene_tx=XM_019815927.2;refGene_gn=PTPRD;refGene_tx=XM_019815928.2;refGene_gn=PTPRD;refGene_tx=XM_019815929.2;refGene_gn=PTPRD;refGene_tx=XM_019815930.2;refGene_gn=PTPRD;refGene_tx=XM_019815931.2;refGene_gn=PTPRD;refGene_tx=XM_019815932.2;refGene_gn=PTPRD;refGen

e_tx=XM_019815933.2;refGene_gn=PTPRD;refGene_tx=XM_019815934.2;refGene_gn=PTPRD;refGene_tx=XM_019815935.2;refGene_gn=PTPRD;refGene_tx=XM_019815937.2;refGene_gn=PTPRD;refGene_tx=XM_019815939.2;refGene_gn=PTPRD;refGene_tx=XM_019815942.2;refGene_gn=PTPRD;refGene_tx=XM_019815943.2;refGene_gn=PTPRD;refGene_tx=XM_019815944.2;refGene_gn=PTPRD;refGene_tx=XM_019815945.2;refGene_gn=PTPRD;refGene_tx=XM_019815946.2;refGene_gn=PTPRD;refGene_tx=XM_019815947.2;refGene_gn=PTPRD;refGene_tx=XM_019815948.
2;refGene_gn=PTPRD;refGene_tx=XM_019815949.2;refGene_gn=PTPRD;refGene_tx=XM_019815951.2;refGene_gn=PTPRD;refGene_tx=XM_019815952.2;refGene_gn=PTPRD;refGene_tx=XM_023242291.1;refGene_gn=PTPRD;refGene_tx=XM_023242292.1;refGene_gn=PTPRD;refGene_tx=XM_023242293.1;refGene_gn=PTPRD;refGene_tx=XM_023242294.1;refGene_gn=PTPRD;refGene_tx=XM_023242296.1
D4 35995711 35996293
refGene_gn=PTPRD;refGene_tx=XM_019815913.2;refGene_gn=PTPRD;refGene_tx=XM_019815914.2;refGene_gn=PTPRD;refGene_tx=XM_019815915.2;refGene_gn=PTPRD;refGene_tx=XM_019815916.2;refGene_gn=PTPRD;refGene_tx=XM_019815917.2;refGene_gn=PTPRD;refGene_tx=XM_019815918.2;refGene_gn=PTPRD;refGene_tx=XM_019815919.2;refGene_gn=PTPRD;refGene_tx=XM_019815920.2;refGene_gn=PTPRD;refGene_tx=XM_019815922.2;refGene_gn=PTPRD;refGene_tx=XM_019815923.2;refGene_gn=PTPRD;refGene_tx=XM_019815924.2;refGene_gn=PTPRD;refGene_tx=XM_019815925.2;refGene_gn=PTPRD;refGene_tx=XM_019815926.2;refGene_gn=PTPRD;refGene_tx=XM_019815927.2;refGene_gn=PTPRD;refGene_tx=XM_019815928.2;refGene_gn=PTPRD;refGene_tx=XM_019815929.2;refGene_gn=PTPRD;refGene_tx=XM_019815930.2;refGene_gn=PTPRD;refGene_tx=XM_019815931.2;refGene_gn=PTPRD;refGene_tx=XM_019815932.2;refGene_gn=PTPRD;refGene_tx=XM_019815933.2;refGene_gn=PTPRD;refGene_tx=XM_019815934.2;refGene_gn=PTPRD;refGene_tx=XM_019815935.2;refGene_gn=PTPRD;refGene_tx=XM_019815937.2;refGene_gn=PTPRD;refGene_tx=XM_019815939.2;refGene_gn=PTPRD;refGene_tx=XM_019815942.2;refGene_gn=PTPRD;refGene_tx=XM_019815943.2;refGene_gn=PTPRD;refGene_tx=XM_019815944.2;refGene_gn=PTPRD;refGene_tx=XM_019815945.2;refGene_gn=PTPRD;refGene_tx=XM_019815946.2;refGene_gn=PTPRD;refGene_tx=XM_019815947.2;refGene_gn=PTPRD;refGene_tx=XM_019815948.
2;refGene_gn=PTPRD;refGene_tx=XM_019815949.2;refGene_gn=PTPRD;refGene_tx=XM_019815951.2;refGene_gn=PTPRD;refGene_tx=XM_019815952.2;refGene_gn=PTPRD;refGene_tx=XM_023242291.1;refGene_gn=PTPRD;refGene_tx=XM_023242292.1;refGene_gn=PTPRD;refGene_tx=XM_023242293.1;refGene_gn=PTPRD;refGene_tx=XM_023242294.1;refGene_gn=PTPRD;refGene_tx=XM_023242296.1
D4 35999228 35999498
refGene_gn=PTPRD;refGene_tx=XM_019815913.2;refGene_gn=PTPRD;refGene_tx=XM_019815914.2;refGene_gn=PTPRD;refGene_tx=XM_019815915.2;refGene_gn=PTPRD;refGene_tx=XM_019815916.2;refGene_gn=PTPRD;refGene_tx=XM_019815917.2;refGene_gn=PTPRD;refGene_tx=XM_019815918.2;refGene_gn=PTPRD;refGene_tx=XM_019815919.2;refGene_gn=PTPRD;refGene_tx=XM_019815920.2;refGene_gn=PTPRD;refGene_tx=XM_019815922.2;refGene_gn=PTPRD;refGene_tx=XM_019815923.2;refGene_gn=PTPRD;refGene_tx=XM_019815924.2;refGene_gn=PTPRD;refGene_tx=XM_019815925.2;refGene_gn=PTPRD;refGene_tx=XM_019815926.2;refGene_gn=PTPRD;refGene_tx=XM_019815927.2;refGene_gn=PTPRD;refGene_tx=XM_019815928.2;refGene_gn=PTPRD;refGene_tx=XM_019815929.2;refGene_gn=P

TPRD;refGene_tx=XM_019815930.2;refGene_gn=PTPRD;refGene_tx=XM_01981593
1.2;refGene_gn=PTPRD;refGene_tx=XM_019815932.2;refGene_gn=PTPRD;refGen
e_tx=XM_019815933.2;refGene_gn=PTPRD;refGene_tx=XM_019815934.2;refGene
_gn=PTPRD;refGene_tx=XM_019815935.2;refGene_gn=PTPRD;refGene_tx=XM_019
815937.2;refGene_gn=PTPRD;refGene_tx=XM_019815939.2;refGene_gn=PTPRD;r
efGene_tx=XM_019815942.2;refGene_gn=PTPRD;refGene_tx=XM_019815943.2;re
fGene_gn=PTPRD;refGene_tx=XM_019815944.2;refGene_gn=PTPRD;refGene_tx=X
M_019815945.2;refGene_gn=PTPRD;refGene_tx=XM_019815946.2;refGene_gn=PT
PRD;refGene_tx=XM_019815947.2;refGene_gn=PTPRD;refGene_tx=XM_019815948
.
2;refGene_gn=PTPRD;refGene_tx=XM_019815949.2;refGene_gn=PTPRD;refGene_
tx=XM_019815951.2;refGene_gn=PTPRD;refGene_tx=XM_019815952.2;refGene_g
n=PTPRD;refGene_tx=XM_023242291.1;refGene_gn=PTPRD;refGene_tx=XM_02324
2292.1;refGene_gn=PTPRD;refGene_tx=XM_023242293.1;refGene_gn=PTPRD;ref
Gene_tx=XM_023242294.1;refGene_gn=PTPRD;refGene_tx=XM_023242296.1
D4 36001480 36001492
refGene_gn=PTPRD;refGene_tx=XM_019815913.2;refGene_gn=PTPRD;refGene_tx
=XM_019815914.2;refGene_gn=PTPRD;refGene_tx=XM_019815915.2;refGene_gn=
PTPRD;refGene_tx=XM_019815916.2;refGene_gn=PTPRD;refGene_tx=XM_0198159
17.2;refGene_gn=PTPRD;refGene_tx=XM_019815918.2;refGene_gn=PTPRD;refGe
ne_tx=XM_019815919.2;refGene_gn=PTPRD;refGene_tx=XM_019815922.2;refGen
e_gn=PTPRD;refGene_tx=XM_019815923.2;refGene_gn=PTPRD;refGene_tx=XM_01
9815924.2;refGene_gn=PTPRD;refGene_tx=XM_019815925.2;refGene_gn=PTPRD;
refGene_tx=XM_019815926.2;refGene_gn=PTPRD;refGene_tx=XM_019815927.2;r
efGene_gn=PTPRD;refGene_tx=XM_019815929.2;refGene_gn=PTPRD;refGene_tx=
XM_019815930.2;refGene_gn=PTPRD;refGene_tx=XM_019815931.2;refGene_gn=P
TPRD;refGene_tx=XM_019815932.2;refGene_gn=PTPRD;refGene_tx=XM_01981593
4.2;refGene_gn=PTPRD;refGene_tx=XM_019815935.2;refGene_gn=PTPRD;refGen
e_tx=XM_019815942.2;refGene_gn=PTPRD;refGene_tx=XM_019815943.2;refGene
_gn=PTPRD;refGene_tx=XM_019815945.2;refGene_gn=PTPRD;refGene_tx=XM_019
815946.2;refGene_gn=PTPRD;refGene_tx=XM_019815947.2;refGene_gn=PTPRD;r
efGene_tx=XM_019815951.2;refGene_gn=PTPRD;refGene_tx=XM_023242291.1;re
fGene_gn=PTPRD;refGene_tx=XM_023242292.1;refGene_gn=PTPRD;refGene_tx=X
M_023242294.1
D4 36002894 36003005
refGene_gn=PTPRD;refGene_tx=XM_019815913.2;refGene_gn=PTPRD;refGene_tx
=XM_019815914.2;refGene_gn=PTPRD;refGene_tx=XM_019815915.2;refGene_gn=
PTPRD;refGene_tx=XM_019815916.2;refGene_gn=PTPRD;refGene_tx=XM_0198159
17.2;refGene_gn=PTPRD;refGene_tx=XM_019815918.2;refGene_gn=PTPRD;refGe
ne_tx=XM_019815919.2;refGene_gn=PTPRD;refGene_tx=XM_019815920.2;refGen
e_gn=PTPRD;refGene_tx=XM_019815922.2;refGene_gn=PTPRD;refGene_tx=XM_01
9815923.2;refGene_gn=PTPRD;refGene_tx=XM_019815924.2;refGene_gn=PTPRD;
refGene_tx=XM_019815925.2;refGene_gn=PTPRD;refGene_tx=XM_019815926.2;r
efGene_gn=PTPRD;refGene_tx=XM_019815927.2;refGene_gn=PTPRD;refGene_tx=
XM_019815928.2;refGene_gn=PTPRD;refGene_tx=XM_019815929.2;refGene_gn=P
TPRD;refGene_tx=XM_019815930.2;refGene_gn=PTPRD;refGene_tx=XM_01981593
1.2;refGene_gn=PTPRD;refGene_tx=XM_019815932.2;refGene_gn=PTPRD;refGen
e_tx=XM_019815933.2;refGene_gn=PTPRD;refGene_tx=XM_019815934.2;refGene
_gn=PTPRD;refGene_tx=XM_019815935.2;refGene_gn=PTPRD;refGene_tx=XM_019
815937.2;refGene_gn=PTPRD;refGene_tx=XM_019815939.2;refGene_gn=PTPRD;r

efGene_tx=XM_019815942.2;refGene_gn=PTPRD;refGene_tx=XM_019815943.2;refGene_gn=PTPRD;refGene_tx=XM_019815944.2;refGene_gn=PTPRD;refGene_tx=XM_019815945.2;refGene_gn=PTPRD;refGene_tx=XM_019815946.2;refGene_gn=PTPRD;refGene_tx=XM_019815947.2;refGene_gn=PTPRD;refGene_tx=XM_019815948.
2;refGene_gn=PTPRD;refGene_tx=XM_019815949.2;refGene_gn=PTPRD;refGene_tx=XM_019815951.2;refGene_gn=PTPRD;refGene_tx=XM_019815952.2;refGene_gn=PTPRD;refGene_tx=XM_023242291.1;refGene_gn=PTPRD;refGene_tx=XM_023242292.1;refGene_gn=PTPRD;refGene_tx=XM_023242293.1;refGene_gn=PTPRD;refGene_tx=XM_023242294.1;refGene_gn=PTPRD;refGene_tx=XM_023242296.1
D4 36004606 36004624
refGene_gn=PTPRD;refGene_tx=XM_019815913.2;refGene_gn=PTPRD;refGene_tx=XM_019815914.2;refGene_gn=PTPRD;refGene_tx=XM_019815915.2;refGene_gn=PTPRD;refGene_tx=XM_019815916.2;refGene_gn=PTPRD;refGene_tx=XM_019815917.2;refGene_gn=PTPRD;refGene_tx=XM_019815918.2;refGene_gn=PTPRD;refGene_tx=XM_019815919.2;refGene_gn=PTPRD;refGene_tx=XM_019815920.2;refGene_gn=PTPRD;refGene_tx=XM_019815924.2;refGene_gn=PTPRD;refGene_tx=XM_019815926.2;refGene_gn=PTPRD;refGene_tx=XM_019815927.2;refGene_gn=PTPRD;refGene_tx=XM_019815929.2;refGene_gn=PTPRD;refGene_tx=XM_019815930.2;refGene_gn=PTPRD;refGene_tx=XM_019815931.2;refGene_gn=PTPRD;refGene_tx=XM_019815932.2;refGene_gn=PTPRD;refGene_tx=XM_019815934.2;refGene_gn=PTPRD;refGene_tx=XM_019815935.2;refGene_gn=PTPRD;refGene_tx=XM_019815942.2;refGene_gn=PTPRD;refGene_tx=XM_019815943.2;refGene_gn=PTPRD;refGene_tx=XM_019815945.2;refGene_gn=PTPRD;refGene_tx=XM_019815946.2;refGene_gn=PTPRD;refGene_tx=XM_023242291.1;refGene_gn=PTPRD;refGene_tx=XM_023242293.1;refGene_gn=PTPRD;refGene_tx=XM_023242294.1
D4 36005355 36005364
refGene_gn=PTPRD;refGene_tx=XM_019815913.2;refGene_gn=PTPRD;refGene_tx=XM_019815914.2;refGene_gn=PTPRD;refGene_tx=XM_019815915.2;refGene_gn=PTPRD;refGene_tx=XM_019815916.2;refGene_gn=PTPRD;refGene_tx=XM_019815917.2;refGene_gn=PTPRD;refGene_tx=XM_019815918.2;refGene_gn=PTPRD;refGene_tx=XM_019815920.2;refGene_gn=PTPRD;refGene_tx=XM_019815922.2;refGene_gn=PTPRD;refGene_tx=XM_019815924.2;refGene_gn=PTPRD;refGene_tx=XM_019815926.2;refGene_gn=PTPRD;refGene_tx=XM_019815927.2;refGene_gn=PTPRD;refGene_tx=XM_019815929.2;refGene_gn=PTPRD;refGene_tx=XM_019815930.2;refGene_gn=PTPRD;refGene_tx=XM_019815931.2;refGene_gn=PTPRD;refGene_tx=XM_019815932.2;refGene_gn=PTPRD;refGene_tx=XM_019815934.2;refGene_gn=PTPRD;refGene_tx=XM_019815942.2;refGene_gn=PTPRD;refGene_tx=XM_019815943.2;refGene_gn=PTPRD;refGene_tx=XM_019815944.2;refGene_gn=PTPRD;refGene_tx=XM_019815946.2;refGene_gn=PTPRD;refGene_tx=XM_019815947.2;refGene_gn=PTPRD;refGene_tx=XM_023242291.1;refGene_gn=PTPRD;refGene_tx=XM_023242292.1;refGene_gn=PTPRD;refGene_tx=XM_023242294.1
D4 36006616 36006805
refGene_gn=PTPRD;refGene_tx=XM_019815913.2;refGene_gn=PTPRD;refGene_tx=XM_019815914.2;refGene_gn=PTPRD;refGene_tx=XM_019815915.2;refGene_gn=PTPRD;refGene_tx=XM_019815916.2;refGene_gn=PTPRD;refGene_tx=XM_019815917.2;refGene_gn=PTPRD;refGene_tx=XM_019815918.2;refGene_gn=PTPRD;refGene_tx=XM_019815919.2;refGene_gn=PTPRD;refGene_tx=XM_019815920.2;refGene_gn=PTPRD;refGene_tx=XM_019815922.2;refGene_gn=PTPRD;refGene_tx=XM_019815923.2;refGene_gn=PTPRD;refGene_tx=XM_019815924.2;refGene_gn=PTPRD;

refGene_tx=XM_019815925.2;refGene_gn=PTPRD;refGene_tx=XM_019815926.2;refGene_gn=PTPRD;refGene_tx=XM_019815927.2;refGene_gn=PTPRD;refGene_tx=XM_019815928.2;refGene_gn=PTPRD;refGene_tx=XM_019815929.2;refGene_gn=PTPRD;refGene_tx=XM_019815930.2;refGene_gn=PTPRD;refGene_tx=XM_019815931.2;refGene_gn=PTPRD;refGene_tx=XM_019815932.2;refGene_gn=PTPRD;refGene_tx=XM_019815933.2;refGene_gn=PTPRD;refGene_tx=XM_019815934.2;refGene_gn=PTPRD;refGene_tx=XM_019815935.2;refGene_gn=PTPRD;refGene_tx=XM_019815937.2;refGene_gn=PTPRD;refGene_tx=XM_019815939.2;refGene_gn=PTPRD;refGene_tx=XM_019815942.2;refGene_gn=PTPRD;refGene_tx=XM_019815943.2;refGene_gn=PTPRD;refGene_tx=XM_019815944.2;refGene_gn=PTPRD;refGene_tx=XM_019815945.2;refGene_gn=PTPRD;refGene_tx=XM_019815946.2;refGene_gn=PTPRD;refGene_tx=XM_019815947.2;refGene_gn=PTPRD;refGene_tx=XM_019815948.
2;refGene_gn=PTPRD;refGene_tx=XM_019815949.2;refGene_gn=PTPRD;refGene_tx=XM_019815951.2;refGene_gn=PTPRD;refGene_tx=XM_019815952.2;refGene_gn=PTPRD;refGene_tx=XM_023242291.1;refGene_gn=PTPRD;refGene_tx=XM_023242292.1;refGene_gn=PTPRD;refGene_tx=XM_023242293.1;refGene_gn=PTPRD;refGene_tx=XM_023242294.1;refGene_gn=PTPRD;refGene_tx=XM_023242296.1

D4 36124795 36124937
refGene_gn=PTPRD;refGene_tx=XM_019815913.2;refGene_gn=PTPRD;refGene_tx=XM_019815914.2;refGene_gn=PTPRD;refGene_tx=XM_019815915.2;refGene_gn=PTPRD;refGene_tx=XM_019815916.2;refGene_gn=PTPRD;refGene_tx=XM_019815917.2;refGene_gn=PTPRD;refGene_tx=XM_019815918.2;refGene_gn=PTPRD;refGene_tx=XM_019815919.2;refGene_gn=PTPRD;refGene_tx=XM_019815920.2;refGene_gn=PTPRD;refGene_tx=XM_019815922.2;refGene_gn=PTPRD;refGene_tx=XM_019815923.2;refGene_gn=PTPRD;refGene_tx=XM_019815924.2;refGene_gn=PTPRD;refGene_tx=XM_019815925.2;refGene_gn=PTPRD;refGene_tx=XM_019815926.2;refGene_gn=PTPRD;refGene_tx=XM_019815927.2;refGene_gn=PTPRD;refGene_tx=XM_019815928.2;refGene_gn=PTPRD;refGene_tx=XM_019815929.2;refGene_gn=PTPRD;refGene_tx=XM_019815930.2;refGene_gn=PTPRD;refGene_tx=XM_019815931.2;refGene_gn=PTPRD;refGene_tx=XM_019815932.2;refGene_gn=PTPRD;refGene_tx=XM_019815933.2;refGene_gn=PTPRD;refGene_tx=XM_019815934.2;refGene_gn=PTPRD;refGene_tx=XM_019815935.2;refGene_gn=PTPRD;refGene_tx=XM_019815937.2;refGene_gn=PTPRD;refGene_tx=XM_019815939.2;refGene_gn=PTPRD;refGene_tx=XM_019815942.2;refGene_gn=PTPRD;refGene_tx=XM_019815943.2;refGene_gn=PTPRD;refGene_tx=XM_019815944.2;refGene_gn=PTPRD;refGene_tx=XM_019815945.2;refGene_gn=PTPRD;refGene_tx=XM_019815946.2;refGene_gn=PTPRD;refGene_tx=XM_019815947.2;refGene_gn=PTPRD;refGene_tx=XM_019815948.
2;refGene_gn=PTPRD;refGene_tx=XM_019815949.2;refGene_gn=PTPRD;refGene_tx=XM_019815951.2;refGene_gn=PTPRD;refGene_tx=XM_019815952.2;refGene_gn=PTPRD;refGene_tx=XM_023242291.1;refGene_gn=PTPRD;refGene_tx=XM_023242292.1;refGene_gn=PTPRD;refGene_tx=XM_023242293.1;refGene_gn=PTPRD;refGene_tx=XM_023242294.1;refGene_gn=PTPRD;refGene_tx=XM_023242296.1

D4 36128292 36128441
refGene_gn=PTPRD;refGene_tx=XM_019815913.2;refGene_gn=PTPRD;refGene_tx=XM_019815914.2;refGene_gn=PTPRD;refGene_tx=XM_019815915.2;refGene_gn=PTPRD;refGene_tx=XM_019815916.2;refGene_gn=PTPRD;refGene_tx=XM_019815917.2;refGene_gn=PTPRD;refGene_tx=XM_019815918.2;refGene_gn=PTPRD;refGene_tx=XM_019815919.2;refGene_gn=PTPRD;refGene_tx=XM_019815920.2;refGen

e_gn=PTPRD;refGene_tx=XM_019815922.2;refGene_gn=PTPRD;refGene_tx=XM_01
9815923.2;refGene_gn=PTPRD;refGene_tx=XM_019815924.2;refGene_gn=PTPRD;
refGene_tx=XM_019815925.2;refGene_gn=PTPRD;refGene_tx=XM_019815926.2;r
efGene_gn=PTPRD;refGene_tx=XM_019815927.2;refGene_gn=PTPRD;refGene_tx=
XM_019815928.2;refGene_gn=PTPRD;refGene_tx=XM_019815929.2;refGene_gn=P
TPRD;refGene_tx=XM_019815930.2;refGene_gn=PTPRD;refGene_tx=XM_01981593
1.2;refGene_gn=PTPRD;refGene_tx=XM_019815932.2;refGene_gn=PTPRD;refGen
e_tx=XM_019815933.2;refGene_gn=PTPRD;refGene_tx=XM_019815934.2;refGene
_gn=PTPRD;refGene_tx=XM_019815935.2;refGene_gn=PTPRD;refGene_tx=XM_019
815937.2;refGene_gn=PTPRD;refGene_tx=XM_019815939.2;refGene_gn=PTPRD;r
efGene_tx=XM_019815942.2;refGene_gn=PTPRD;refGene_tx=XM_019815943.2;re
fGene_gn=PTPRD;refGene_tx=XM_019815944.2;refGene_gn=PTPRD;refGene_tx=X
M_019815945.2;refGene_gn=PTPRD;refGene_tx=XM_019815946.2;refGene_gn=PT
PRD;refGene_tx=XM_019815947.2;refGene_gn=PTPRD;refGene_tx=XM_019815948
.
2;refGene_gn=PTPRD;refGene_tx=XM_019815949.2;refGene_gn=PTPRD;refGene_
tx=XM_019815951.2;refGene_gn=PTPRD;refGene_tx=XM_019815952.2;refGene_g
n=PTPRD;refGene_tx=XM_023242291.1;refGene_gn=PTPRD;refGene_tx=XM_02324
2292.1;refGene_gn=PTPRD;refGene_tx=XM_023242293.1;refGene_gn=PTPRD;ref
Gene_tx=XM_023242294.1;refGene_gn=PTPRD;refGene_tx=XM_023242296.1
D4 36223558 36223622
refGene_gn=PTPRD;refGene_tx=XM_019815913.2;refGene_gn=PTPRD;refGene_tx
=XM_019815914.2;refGene_gn=PTPRD;refGene_tx=XM_019815915.2;refGene_gn=
PTPRD;refGene_tx=XM_019815916.2;refGene_gn=PTPRD;refGene_tx=XM_0198159
17.2;refGene_gn=PTPRD;refGene_tx=XM_019815918.2;refGene_gn=PTPRD;refGe
ne_tx=XM_019815919.2;refGene_gn=PTPRD;refGene_tx=XM_019815920.2;refGen
e_gn=PTPRD;refGene_tx=XM_019815922.2;refGene_gn=PTPRD;refGene_tx=XM_01
9815923.2;refGene_gn=PTPRD;refGene_tx=XM_019815924.2;refGene_gn=PTPRD;
refGene_tx=XM_019815925.2;refGene_gn=PTPRD;refGene_tx=XM_019815926.2;r
efGene_gn=PTPRD;refGene_tx=XM_019815927.2;refGene_gn=PTPRD;refGene_tx=
XM_019815928.2;refGene_gn=PTPRD;refGene_tx=XM_019815929.2;refGene_gn=P
TPRD;refGene_tx=XM_019815930.2;refGene_gn=PTPRD;refGene_tx=XM_01981593
1.2;refGene_gn=PTPRD;refGene_tx=XM_019815932.2;refGene_gn=PTPRD;refGen
e_tx=XM_019815933.2;refGene_gn=PTPRD;refGene_tx=XM_019815934.2;refGene
_gn=PTPRD;refGene_tx=XM_019815935.2;refGene_gn=PTPRD;refGene_tx=XM_019
815937.2;refGene_gn=PTPRD;refGene_tx=XM_019815939.2;refGene_gn=PTPRD;r
efGene_tx=XM_019815942.2;refGene_gn=PTPRD;refGene_tx=XM_019815943.2;re
fGene_gn=PTPRD;refGene_tx=XM_019815944.2;refGene_gn=PTPRD;refGene_tx=X
M_019815945.2;refGene_gn=PTPRD;refGene_tx=XM_019815946.2;refGene_gn=PT
PRD;refGene_tx=XM_019815947.2;refGene_gn=PTPRD;refGene_tx=XM_019815948
.
2;refGene_gn=PTPRD;refGene_tx=XM_019815949.2;refGene_gn=PTPRD;refGene_
tx=XM_019815951.2;refGene_gn=PTPRD;refGene_tx=XM_019815952.2;refGene_g
n=PTPRD;refGene_tx=XM_023242291.1;refGene_gn=PTPRD;refGene_tx=XM_02324
2292.1;refGene_gn=PTPRD;refGene_tx=XM_023242293.1;refGene_gn=PTPRD;ref
Gene_tx=XM_023242294.1;refGene_gn=PTPRD;refGene_tx=XM_023242296.1
D4 40068863 40069248 refGene_gn=TYRP1;refGene_tx=NM_001042560.2
D4 40070349 40070672 refGene_gn=TYRP1;refGene_tx=NM_001042560.2
D4 40073738 40073943 refGene_gn=TYRP1;refGene_tx=NM_001042560.2
D4 40078022 40078190 refGene_gn=TYRP1;refGene_tx=NM_001042560.2

| | | | |
|---|---|---|---|
| D4 | 40081044 | 40081224 | refGene_gn=TYRP1;refGene_tx=NM_001042560.2 |
| D4 | 40085942 | 40086089 | refGene_gn=TYRP1;refGene_tx=NM_001042560.2 |
| D4 | 40087195 | 40087401 | refGene_gn=TYRP1;refGene_tx=NM_001042560.2 |
| D4 | 40147586 | 40147892 | refGene_gn=LURAP1L;refGene_tx=XM_019815953.2 |
| D4 | 40198771 | 40198912 | refGene_gn=LURAP1L;refGene_tx=XM_019815953.2 |
| D4 | 40198913 | 40199147 | refGene_gn=LURAP1L;refGene_tx=XM_019815953.2 |
| D4 | 40449560 | 40449707 | refGene_gn=MPDZ;refGene_tx=XM_023242297.1 |
| D4 | 40451499 | 40451623 | refGene_gn=MPDZ;refGene_tx=XM_023242297.1 |
| D4 | 40452858 | 40452971 | refGene_gn=MPDZ;refGene_tx=XM_023242297.1 |
| D4 | 40453537 | 40453642 | refGene_gn=MPDZ;refGene_tx=XM_023242297.1 |
| D4 | 40454926 | 40455049 | refGene_gn=MPDZ;refGene_tx=XM_023242297.1 |
| D4 | 40456153 | 40456197 | refGene_gn=MPDZ;refGene_tx=XM_023242297.1 |
| D4 | 40457336 | 40457427 | refGene_gn=MPDZ;refGene_tx=XM_023242297.1 |
| D4 | 40458260 | 40458347 | refGene_gn=MPDZ;refGene_tx=XM_023242297.1 |
| D4 | 40459206 | 40459354 | refGene_gn=MPDZ;refGene_tx=XM_023242297.1 |
| D4 | 40461870 | 40462064 | refGene_gn=MPDZ;refGene_tx=XM_023242297.1 |
| D4 | 40462235 | 40462319 | refGene_gn=MPDZ;refGene_tx=XM_023242297.1 |
| D4 | 40463263 | 40463409 | refGene_gn=MPDZ;refGene_tx=XM_023242297.1 |
| D4 | 40465848 | 40466020 | refGene_gn=MPDZ;refGene_tx=XM_023242297.1 |
| D4 | 40466749 | 40466824 | refGene_gn=MPDZ;refGene_tx=XM_023242297.1 |
| D4 | 40466908 | 40467001 | refGene_gn=MPDZ;refGene_tx=XM_023242297.1 |
| D4 | 40475371 | 40475452 | refGene_gn=MPDZ;refGene_tx=XM_023242297.1 |
| D4 | 40477460 | 40477551 | refGene_gn=MPDZ;refGene_tx=XM_023242297.1 |
| D4 | 40477771 | 40477863 | refGene_gn=MPDZ;refGene_tx=XM_023242297.1 |
| D4 | 40478989 | 40479186 | refGene_gn=MPDZ;refGene_tx=XM_023242297.1 |
| D4 | 40480994 | 40481157 | refGene_gn=MPDZ;refGene_tx=XM_023242297.1 |
| D4 | 40484204 | 40484303 | refGene_gn=MPDZ;refGene_tx=XM_023242297.1 |
| D4 | 40488828 | 40488939 | refGene_gn=MPDZ;refGene_tx=XM_023242297.1 |
| D4 | 40492009 | 40492187 | refGene_gn=MPDZ;refGene_tx=XM_023242297.1 |
| D4 | 40499338 | 40499431 | refGene_gn=MPDZ;refGene_tx=XM_023242297.1 |
| D4 | 40503146 | 40503251 | refGene_gn=MPDZ;refGene_tx=XM_023242297.1 |
| D4 | 40509054 | 40509253 | refGene_gn=MPDZ;refGene_tx=XM_023242297.1 |
| D4 | 40519860 | 40519984 | refGene_gn=MPDZ;refGene_tx=XM_023242297.1 |
| D4 | 40520255 | 40520537 | refGene_gn=MPDZ;refGene_tx=XM_023242297.1 |
| D4 | 40528140 | 40528308 | refGene_gn=MPDZ;refGene_tx=XM_023242297.1 |
| D4 | 40530848 | 40530965 | refGene_gn=MPDZ;refGene_tx=XM_023242297.1 |
| D4 | 40532625 | 40532835 | refGene_gn=MPDZ;refGene_tx=XM_023242297.1 |
| D4 | 40533989 | 40534117 | refGene_gn=MPDZ;refGene_tx=XM_023242297.1 |
| D4 | 40534119 | 40534174 | refGene_gn=MPDZ;refGene_tx=XM_023242297.1 |
| D4 | 40535384 | 40535549 | refGene_gn=MPDZ;refGene_tx=XM_023242297.1 |
| D4 | 40536398 | 40536545 | refGene_gn=MPDZ;refGene_tx=XM_023242297.1 |
| D4 | 40538980 | 40539092 | refGene_gn=MPDZ;refGene_tx=XM_023242297.1 |
| D4 | 40541043 | 40541113 | refGene_gn=MPDZ;refGene_tx=XM_023242297.1 |
| D4 | 40541867 | 40542051 | refGene_gn=MPDZ;refGene_tx=XM_023242297.1 |
| D4 | 40551940 | 40552029 | refGene_gn=MPDZ;refGene_tx=XM_023242297.1 |
| D4 | 40552378 | 40552493 | refGene_gn=MPDZ;refGene_tx=XM_023242297.1 |
| D4 | 40554882 | 40555089 | refGene_gn=MPDZ;refGene_tx=XM_023242297.1 |
| D4 | 40557344 | 40557473 | refGene_gn=MPDZ;refGene_tx=XM_023242297.1 |
| D4 | 40558194 | 40558408 | refGene_gn=MPDZ;refGene_tx=XM_023242297.1 |
| D4 | 40559036 | 40559176 | refGene_gn=MPDZ;refGene_tx=XM_023242297.1 |

D4 40559927 40560137 refGene_gn=MPDZ;refGene_tx=XM_023242297.1
D4 40580507 40580674 refGene_gn=MPDZ;refGene_tx=XM_023242297.1
D4 40583488 40583504 refGene_gn=MPDZ;refGene_tx=XM_023242297.1
D4 41369567 41369697
refGene_gn=NFIB;refGene_tx=XM_011288336.3;refGene_gn=NFIB;refGene_tx=XM_011288337.3;refGene_gn=NFIB;refGene_tx=XM_011288339.3;refGene_gn=NFIB;refGene_tx=XM_019815963.2;refGene_gn=NFIB;refGene_tx=XM_019815964.2;refGene_gn=NFIB;refGene_tx=XM_019815965.2;refGene_gn=NFIB;refGene_tx=XM_019815966.2;refGene_gn=NFIB;refGene_tx=XM_019815967.2;refGene_gn=NFIB;refGene_tx=XM_019815968.2;refGene_gn=NFIB;refGene_tx=XM_019815971.2
D4 41384274 41384363
refGene_gn=NFIB;refGene_tx=XM_011288336.3;refGene_gn=NFIB;refGene_tx=XM_011288337.3;refGene_gn=NFIB;refGene_tx=XM_011288339.3;refGene_gn=NFIB;refGene_tx=XM_019815963.2;refGene_gn=NFIB;refGene_tx=XM_019815964.2;refGene_gn=NFIB;refGene_tx=XM_019815965.2;refGene_gn=NFIB;refGene_tx=XM_019815968.2
D4 41394281 41394364
refGene_gn=NFIB;refGene_tx=XM_011288336.3;refGene_gn=NFIB;refGene_tx=XM_011288337.3;refGene_gn=NFIB;refGene_tx=XM_019815963.2;refGene_gn=NFIB;refGene_tx=XM_019815964.2;refGene_gn=NFIB;refGene_tx=XM_019815965.2;refGene_gn=NFIB;refGene_tx=XM_019815966.2;refGene_gn=NFIB;refGene_tx=XM_019815967.2;refGene_gn=NFIB;refGene_tx=XM_019815969.1;refGene_gn=NFIB;refGene_tx=XM_019815970.1
D4 41397786 41397952
refGene_gn=NFIB;refGene_tx=XM_011288336.3;refGene_gn=NFIB;refGene_tx=XM_011288337.3;refGene_gn=NFIB;refGene_tx=XM_019815963.2;refGene_gn=NFIB;refGene_tx=XM_019815964.2;refGene_gn=NFIB;refGene_tx=XM_019815965.2;refGene_gn=NFIB;refGene_tx=XM_019815966.2;refGene_gn=NFIB;refGene_tx=XM_019815967.2
D4 41402405 41402590
refGene_gn=NFIB;refGene_tx=XM_011288336.3;refGene_gn=NFIB;refGene_tx=XM_011288337.3;refGene_gn=NFIB;refGene_tx=XM_011288339.3;refGene_gn=NFIB;refGene_tx=XM_019815963.2;refGene_gn=NFIB;refGene_tx=XM_019815964.2;refGene_gn=NFIB;refGene_tx=XM_019815965.2;refGene_gn=NFIB;refGene_tx=XM_019815966.2;refGene_gn=NFIB;refGene_tx=XM_019815967.2;refGene_gn=NFIB;refGene_tx=XM_019815968.2;refGene_gn=NFIB;refGene_tx=XM_019815969.1;refGene_gn=NFIB;refGene_tx=XM_019815970.1;refGene_gn=NFIB;refGene_tx=XM_019815971.2
D4 41406446 41406581
refGene_gn=NFIB;refGene_tx=XM_011288336.3;refGene_gn=NFIB;refGene_tx=XM_011288337.3;refGene_gn=NFIB;refGene_tx=XM_011288339.3;refGene_gn=NFIB;refGene_tx=XM_019815963.2;refGene_gn=NFIB;refGene_tx=XM_019815964.2;refGene_gn=NFIB;refGene_tx=XM_019815965.2;refGene_gn=NFIB;refGene_tx=XM_019815966.2;refGene_gn=NFIB;refGene_tx=XM_019815967.2;refGene_gn=NFIB;refGene_tx=XM_019815968.2;refGene_gn=NFIB;refGene_tx=XM_019815969.1;refGene_gn=NFIB;refGene_tx=XM_019815970.1;refGene_gn=NFIB;refGene_tx=XM_019815971.2
D4 41428812 41428931
refGene_gn=NFIB;refGene_tx=XM_011288336.3;refGene_gn=NFIB;refGene_tx=XM_011288337.3;refGene_gn=NFIB;refGene_tx=XM_011288339.3;refGene_gn=NFI

B;refGene_tx=XM_019815963.2;refGene_gn=NFIB;refGene_tx=XM_019815964.2;refGene_gn=NFIB;refGene_tx=XM_019815965.2;refGene_gn=NFIB;refGene_tx=XM_019815966.2;refGene_gn=NFIB;refGene_tx=XM_019815967.2;refGene_gn=NFIB;refGene_tx=XM_019815968.2;refGene_gn=NFIB;refGene_tx=XM_019815969.1;refGene_gn=NFIB;refGene_tx=XM_019815970.1;refGene_gn=NFIB;refGene_tx=XM_019815971.2
D4 41431699 41431820
refGene_gn=NFIB;refGene_tx=XM_011288336.3;refGene_gn=NFIB;refGene_tx=XM_011288337.3;refGene_gn=NFIB;refGene_tx=XM_011288339.3;refGene_gn=NFIB;refGene_tx=XM_019815963.2;refGene_gn=NFIB;refGene_tx=XM_019815964.2;refGene_gn=NFIB;refGene_tx=XM_019815965.2;refGene_gn=NFIB;refGene_tx=XM_019815966.2;refGene_gn=NFIB;refGene_tx=XM_019815967.2;refGene_gn=NFIB;refGene_tx=XM_019815968.2;refGene_gn=NFIB;refGene_tx=XM_019815969.1;refGene_gn=NFIB;refGene_tx=XM_019815970.1;refGene_gn=NFIB;refGene_tx=XM_019815971.2
D4 41437454 41437523
refGene_gn=NFIB;refGene_tx=XM_011288336.3;refGene_gn=NFIB;refGene_tx=XM_011288337.3;refGene_gn=NFIB;refGene_tx=XM_011288339.3;refGene_gn=NFIB;refGene_tx=XM_019815963.2;refGene_gn=NFIB;refGene_tx=XM_019815964.2;refGene_gn=NFIB;refGene_tx=XM_019815965.2;refGene_gn=NFIB;refGene_tx=XM_019815966.2;refGene_gn=NFIB;refGene_tx=XM_019815967.2;refGene_gn=NFIB;refGene_tx=XM_019815968.2;refGene_gn=NFIB;refGene_tx=XM_019815969.1;refGene_gn=NFIB;refGene_tx=XM_019815970.1;refGene_gn=NFIB;refGene_tx=XM_019815971.2
D4 41470635 41470689
refGene_gn=NFIB;refGene_tx=XM_011288336.3;refGene_gn=NFIB;refGene_tx=XM_011288337.3;refGene_gn=NFIB;refGene_tx=XM_011288339.3;refGene_gn=NFIB;refGene_tx=XM_019815963.2;refGene_gn=NFIB;refGene_tx=XM_019815964.2;refGene_gn=NFIB;refGene_tx=XM_019815965.2;refGene_gn=NFIB;refGene_tx=XM_019815966.2;refGene_gn=NFIB;refGene_tx=XM_019815967.2;refGene_gn=NFIB;refGene_tx=XM_019815968.2;refGene_gn=NFIB;refGene_tx=XM_019815969.1;refGene_gn=NFIB;refGene_tx=XM_019815970.1;refGene_gn=NFIB;refGene_tx=XM_019815971.2
D4 41598916 41599448
refGene_gn=NFIB;refGene_tx=XM_011288336.3;refGene_gn=NFIB;refGene_tx=XM_011288337.3;refGene_gn=NFIB;refGene_tx=XM_011288339.3;refGene_gn=NFIB;refGene_tx=XM_019815963.2;refGene_gn=NFIB;refGene_tx=XM_019815964.2;refGene_gn=NFIB;refGene_tx=XM_019815965.2;refGene_gn=NFIB;refGene_tx=XM_019815966.2;refGene_gn=NFIB;refGene_tx=XM_019815967.2;refGene_gn=NFIB;refGene_tx=XM_019815968.2;refGene_gn=NFIB;refGene_tx=XM_019815969.1;refGene_gn=NFIB;refGene_tx=XM_019815970.1;refGene_gn=NFIB;refGene_tx=XM_019815971.2
D4 41599845 41599848 refGene_gn=NFIB;refGene_tx=XM_019815965.2
D4 41605587 41605617
refGene_gn=NFIB;refGene_tx=XM_011288336.3;refGene_gn=NFIB;refGene_tx=XM_011288337.3;refGene_gn=NFIB;refGene_tx=XM_011288339.3
D4 41824323 41824419
refGene_gn=NFIB;refGene_tx=XM_019815963.2;refGene_gn=NFIB;refGene_tx=XM_019815964.2;refGene_gn=NFIB;refGene_tx=XM_019815966.2;refGene_gn=NFIB;refGene_tx=XM_019815967.2;refGene_gn=NFIB;refGene_tx=XM_019815968.2;

refGene_gn=NFIB;refGene_tx=XM_019815969.1;refGene_gn=NFIB;refGene_tx=XM_019815970.1;refGene_gn=NFIB;refGene_tx=XM_019815971.2
D4 41902640 41902773
refGene_gn=ZDHHC21;refGene_tx=XM_019815972.2;refGene_gn=ZDHHC21;refGene_tx=XM_019815974.2;refGene_gn=ZDHHC21;refGene_tx=XM_023242330.1;refGene_gn=ZDHHC21;refGene_tx=XM_023242331.1;refGene_gn=ZDHHC21;refGene_tx=XM_023242332.1;refGene_gn=ZDHHC21;refGene_tx=XM_023242333.1;refGene_gn=ZDHHC21;refGene_tx=XM_023242334.1
D4 41903320 41903364
refGene_gn=ZDHHC21;refGene_tx=XM_019815972.2;refGene_gn=ZDHHC21;refGene_tx=XM_019815974.2;refGene_gn=ZDHHC21;refGene_tx=XM_023242330.1;refGene_gn=ZDHHC21;refGene_tx=XM_023242331.1;refGene_gn=ZDHHC21;refGene_tx=XM_023242332.1;refGene_gn=ZDHHC21;refGene_tx=XM_023242333.1;refGene_gn=ZDHHC21;refGene_tx=XM_023242334.1
D4 41915878 41915995
refGene_gn=ZDHHC21;refGene_tx=XM_019815972.2;refGene_gn=ZDHHC21;refGene_tx=XM_019815974.2;refGene_gn=ZDHHC21;refGene_tx=XM_023242330.1;refGene_gn=ZDHHC21;refGene_tx=XM_023242331.1;refGene_gn=ZDHHC21;refGene_tx=XM_023242332.1;refGene_gn=ZDHHC21;refGene_tx=XM_023242333.1;refGene_gn=ZDHHC21;refGene_tx=XM_023242334.1
D4 41936841 41936980
refGene_gn=ZDHHC21;refGene_tx=XM_019815972.2;refGene_gn=ZDHHC21;refGene_tx=XM_019815974.2;refGene_gn=ZDHHC21;refGene_tx=XM_023242330.1;refGene_gn=ZDHHC21;refGene_tx=XM_023242331.1;refGene_gn=ZDHHC21;refGene_tx=XM_023242332.1;refGene_gn=ZDHHC21;refGene_tx=XM_023242333.1;refGene_gn=ZDHHC21;refGene_tx=XM_023242334.1
D4 41939803 41939915
refGene_gn=ZDHHC21;refGene_tx=XM_019815972.2;refGene_gn=ZDHHC21;refGene_tx=XM_019815974.2;refGene_gn=ZDHHC21;refGene_tx=XM_023242330.1;refGene_gn=ZDHHC21;refGene_tx=XM_023242331.1;refGene_gn=ZDHHC21;refGene_tx=XM_023242332.1;refGene_gn=ZDHHC21;refGene_tx=XM_023242333.1;refGene_gn=ZDHHC21;refGene_tx=XM_023242334.1
D4 41946946 41947045
refGene_gn=ZDHHC21;refGene_tx=XM_019815972.2;refGene_gn=ZDHHC21;refGene_tx=XM_019815974.2;refGene_gn=ZDHHC21;refGene_tx=XM_023242330.1;refGene_gn=ZDHHC21;refGene_tx=XM_023242331.1;refGene_gn=ZDHHC21;refGene_tx=XM_023242332.1;refGene_gn=ZDHHC21;refGene_tx=XM_023242333.1;refGene_gn=ZDHHC21;refGene_tx=XM_023242334.1
D4 41948321 41948475
refGene_gn=ZDHHC21;refGene_tx=XM_019815972.2;refGene_gn=ZDHHC21;refGene_tx=XM_019815974.2;refGene_gn=ZDHHC21;refGene_tx=XM_023242330.1;refGene_gn=ZDHHC21;refGene_tx=XM_023242331.1;refGene_gn=ZDHHC21;refGene_tx=XM_023242332.1;refGene_gn=ZDHHC21;refGene_tx=XM_023242333.1;refGene_gn=ZDHHC21;refGene_tx=XM_023242334.1
D4 41981328 41981643 refGene_gn=CER1;refGene_tx=XM_023242335.1
D4 41983606 41984104 refGene_gn=CER1;refGene_tx=XM_023242335.1
D4 42002241 42002447
refGene_gn=FREM1;refGene_tx=XM_019815975.2;refGene_gn=FREM1;refGene_tx=XM_019815979.2;refGene_gn=FREM1;refGene_tx=XM_019815981.2;refGene_gn=FREM1;refGene_tx=XM_019815982.1;refGene_gn=FREM1;refGene_tx=XM_0198159

84.2;refGene_gn=FREM1;refGene_tx=XM_019815985.2;refGene_gn=FREM1;refGene_tx=XM_019815986.2;refGene_gn=FREM1;refGene_tx=XM_019815987.2;refGene_gn=FREM1;refGene_tx=XM_019815988.1;refGene_gn=FREM1;refGene_tx=XM_019815989.1;refGene_gn=FREM1;refGene_tx=XM_023242336.1
D4 42004263 42004349
refGene_gn=FREM1;refGene_tx=XM_019815975.2;refGene_gn=FREM1;refGene_tx=XM_019815979.2;refGene_gn=FREM1;refGene_tx=XM_019815981.2;refGene_gn=FREM1;refGene_tx=XM_019815982.1;refGene_gn=FREM1;refGene_tx=XM_019815984.2;refGene_gn=FREM1;refGene_tx=XM_019815985.2;refGene_gn=FREM1;refGene_tx=XM_019815986.2;refGene_gn=FREM1;refGene_tx=XM_019815987.2;refGene_gn=FREM1;refGene_tx=XM_019815988.1;refGene_gn=FREM1;refGene_tx=XM_019815989.1;refGene_gn=FREM1;refGene_tx=XM_023242336.1
D4 42016157 42016273
refGene_gn=FREM1;refGene_tx=XM_019815975.2;refGene_gn=FREM1;refGene_tx=XM_019815979.2;refGene_gn=FREM1;refGene_tx=XM_019815981.2;refGene_gn=FREM1;refGene_tx=XM_019815982.1;refGene_gn=FREM1;refGene_tx=XM_019815984.2;refGene_gn=FREM1;refGene_tx=XM_019815985.2;refGene_gn=FREM1;refGene_tx=XM_019815986.2;refGene_gn=FREM1;refGene_tx=XM_019815987.2;refGene_gn=FREM1;refGene_tx=XM_019815988.1;refGene_gn=FREM1;refGene_tx=XM_019815989.1;refGene_gn=FREM1;refGene_tx=XM_023242336.1
D4 42016680 42016833
refGene_gn=FREM1;refGene_tx=XM_019815975.2;refGene_gn=FREM1;refGene_tx=XM_019815979.2;refGene_gn=FREM1;refGene_tx=XM_019815981.2;refGene_gn=FREM1;refGene_tx=XM_019815982.1;refGene_gn=FREM1;refGene_tx=XM_019815984.2;refGene_gn=FREM1;refGene_tx=XM_019815985.2;refGene_gn=FREM1;refGene_tx=XM_019815986.2;refGene_gn=FREM1;refGene_tx=XM_019815987.2;refGene_gn=FREM1;refGene_tx=XM_019815988.1;refGene_gn=FREM1;refGene_tx=XM_019815989.1;refGene_gn=FREM1;refGene_tx=XM_023242336.1
D4 42017017 42017182
refGene_gn=FREM1;refGene_tx=XM_019815975.2;refGene_gn=FREM1;refGene_tx=XM_019815979.2;refGene_gn=FREM1;refGene_tx=XM_019815981.2;refGene_gn=FREM1;refGene_tx=XM_019815982.1;refGene_gn=FREM1;refGene_tx=XM_019815984.2;refGene_gn=FREM1;refGene_tx=XM_019815985.2;refGene_gn=FREM1;refGene_tx=XM_019815986.2;refGene_gn=FREM1;refGene_tx=XM_019815987.2;refGene_gn=FREM1;refGene_tx=XM_019815988.1;refGene_gn=FREM1;refGene_tx=XM_019815989.1;refGene_gn=FREM1;refGene_tx=XM_023242336.1
D4 42017464 42017512
refGene_gn=FREM1;refGene_tx=XM_019815975.2;refGene_gn=FREM1;refGene_tx=XM_019815979.2;refGene_gn=FREM1;refGene_tx=XM_019815981.2;refGene_gn=FREM1;refGene_tx=XM_019815982.1;refGene_gn=FREM1;refGene_tx=XM_019815984.2;refGene_gn=FREM1;refGene_tx=XM_019815985.2;refGene_gn=FREM1;refGene_tx=XM_019815986.2;refGene_gn=FREM1;refGene_tx=XM_019815987.2;refGene_gn=FREM1;refGene_tx=XM_019815988.1;refGene_gn=FREM1;refGene_tx=XM_019815989.1;refGene_gn=FREM1;refGene_tx=XM_023242336.1
D4 42018638 42018877
refGene_gn=FREM1;refGene_tx=XM_019815975.2;refGene_gn=FREM1;refGene_tx=XM_019815979.2;refGene_gn=FREM1;refGene_tx=XM_019815981.2;refGene_gn=FREM1;refGene_tx=XM_019815982.1;refGene_gn=FREM1;refGene_tx=XM_019815984.2;refGene_gn=FREM1;refGene_tx=XM_019815985.2;refGene_gn=FREM1;refGene_tx=XM_019815986.2;refGene_gn=FREM1;refGene_tx=XM_019815987.2;refGen

e_gn=FREM1;refGene_tx=XM_019815988.1;refGene_gn=FREM1;refGene_tx=XM_01
9815989.1;refGene_gn=FREM1;refGene_tx=XM_023242336.1
D4	42020625	42020775
refGene_gn=FREM1;refGene_tx=XM_019815975.2;refGene_gn=FREM1;refGene_tx
=XM_019815979.2;refGene_gn=FREM1;refGene_tx=XM_019815981.2;refGene_gn=
FREM1;refGene_tx=XM_019815982.1;refGene_gn=FREM1;refGene_tx=XM_0198159
84.2;refGene_gn=FREM1;refGene_tx=XM_019815985.2;refGene_gn=FREM1;refGe
ne_tx=XM_019815986.2;refGene_gn=FREM1;refGene_tx=XM_019815987.2;refGen
e_gn=FREM1;refGene_tx=XM_019815988.1;refGene_gn=FREM1;refGene_tx=XM_01
9815989.1;refGene_gn=FREM1;refGene_tx=XM_023242336.1
D4	42027295	42027368
refGene_gn=FREM1;refGene_tx=XM_019815975.2;refGene_gn=FREM1;refGene_tx
=XM_019815979.2;refGene_gn=FREM1;refGene_tx=XM_019815981.2;refGene_gn=
FREM1;refGene_tx=XM_019815982.1;refGene_gn=FREM1;refGene_tx=XM_0198159
84.2;refGene_gn=FREM1;refGene_tx=XM_019815985.2;refGene_gn=FREM1;refGe
ne_tx=XM_019815986.2;refGene_gn=FREM1;refGene_tx=XM_019815987.2;refGen
e_gn=FREM1;refGene_tx=XM_019815988.1;refGene_gn=FREM1;refGene_tx=XM_01
9815989.1;refGene_gn=FREM1;refGene_tx=XM_023242336.1
D4	42030465	42030595
refGene_gn=FREM1;refGene_tx=XM_019815975.2;refGene_gn=FREM1;refGene_tx
=XM_019815979.2;refGene_gn=FREM1;refGene_tx=XM_019815981.2;refGene_gn=
FREM1;refGene_tx=XM_019815982.1;refGene_gn=FREM1;refGene_tx=XM_0198159
84.2;refGene_gn=FREM1;refGene_tx=XM_019815985.2;refGene_gn=FREM1;refGe
ne_tx=XM_019815986.2;refGene_gn=FREM1;refGene_tx=XM_019815987.2;refGen
e_gn=FREM1;refGene_tx=XM_019815988.1;refGene_gn=FREM1;refGene_tx=XM_01
9815989.1;refGene_gn=FREM1;refGene_tx=XM_023242336.1
D4	42041201	42041346
refGene_gn=FREM1;refGene_tx=XM_019815975.2;refGene_gn=FREM1;refGene_tx
=XM_019815979.2;refGene_gn=FREM1;refGene_tx=XM_019815981.2;refGene_gn=
FREM1;refGene_tx=XM_019815982.1;refGene_gn=FREM1;refGene_tx=XM_0198159
84.2;refGene_gn=FREM1;refGene_tx=XM_019815985.2;refGene_gn=FREM1;refGe
ne_tx=XM_019815986.2;refGene_gn=FREM1;refGene_tx=XM_019815987.2;refGen
e_gn=FREM1;refGene_tx=XM_019815988.1;refGene_gn=FREM1;refGene_tx=XM_01
9815989.1;refGene_gn=FREM1;refGene_tx=XM_023242336.1
D4	42042071	42042273
refGene_gn=FREM1;refGene_tx=XM_019815975.2;refGene_gn=FREM1;refGene_tx
=XM_019815979.2;refGene_gn=FREM1;refGene_tx=XM_019815981.2;refGene_gn=
FREM1;refGene_tx=XM_019815982.1;refGene_gn=FREM1;refGene_tx=XM_0198159
84.2;refGene_gn=FREM1;refGene_tx=XM_019815985.2;refGene_gn=FREM1;refGe
ne_tx=XM_019815986.2;refGene_gn=FREM1;refGene_tx=XM_019815987.2;refGen
e_gn=FREM1;refGene_tx=XM_019815988.1;refGene_gn=FREM1;refGene_tx=XM_01
9815989.1;refGene_gn=FREM1;refGene_tx=XM_023242336.1
D4	42048454	42048919
refGene_gn=FREM1;refGene_tx=XM_019815975.2;refGene_gn=FREM1;refGene_tx
=XM_019815979.2;refGene_gn=FREM1;refGene_tx=XM_019815981.2;refGene_gn=
FREM1;refGene_tx=XM_019815982.1;refGene_gn=FREM1;refGene_tx=XM_0198159
84.2;refGene_gn=FREM1;refGene_tx=XM_019815985.2;refGene_gn=FREM1;refGe
ne_tx=XM_019815986.2;refGene_gn=FREM1;refGene_tx=XM_019815987.2;refGen
e_gn=FREM1;refGene_tx=XM_019815988.1;refGene_gn=FREM1;refGene_tx=XM_01
9815989.1;refGene_gn=FREM1;refGene_tx=XM_023242336.1

D4 42056836 42057101
refGene_gn=FREM1;refGene_tx=XM_019815975.2;refGene_gn=FREM1;refGene_tx=XM_019815979.2;refGene_gn=FREM1;refGene_tx=XM_019815981.2;refGene_gn=FREM1;refGene_tx=XM_019815982.1;refGene_gn=FREM1;refGene_tx=XM_019815984.2;refGene_gn=FREM1;refGene_tx=XM_019815985.2;refGene_gn=FREM1;refGene_tx=XM_019815986.2;refGene_gn=FREM1;refGene_tx=XM_019815987.2;refGene_gn=FREM1;refGene_tx=XM_019815988.1;refGene_gn=FREM1;refGene_tx=XM_023242336.1
D4 42062708 42062904
refGene_gn=FREM1;refGene_tx=XM_019815975.2;refGene_gn=FREM1;refGene_tx=XM_019815979.2;refGene_gn=FREM1;refGene_tx=XM_019815981.2;refGene_gn=FREM1;refGene_tx=XM_019815982.1;refGene_gn=FREM1;refGene_tx=XM_019815984.2;refGene_gn=FREM1;refGene_tx=XM_019815985.2;refGene_gn=FREM1;refGene_tx=XM_019815986.2;refGene_gn=FREM1;refGene_tx=XM_019815987.2;refGene_gn=FREM1;refGene_tx=XM_019815988.1;refGene_gn=FREM1;refGene_tx=XM_023242336.1
D4 42065263 42065405
refGene_gn=FREM1;refGene_tx=XM_019815975.2;refGene_gn=FREM1;refGene_tx=XM_019815979.2;refGene_gn=FREM1;refGene_tx=XM_019815981.2;refGene_gn=FREM1;refGene_tx=XM_019815982.1;refGene_gn=FREM1;refGene_tx=XM_019815984.2;refGene_gn=FREM1;refGene_tx=XM_019815985.2;refGene_gn=FREM1;refGene_tx=XM_019815986.2;refGene_gn=FREM1;refGene_tx=XM_019815987.2;refGene_gn=FREM1;refGene_tx=XM_019815988.1;refGene_gn=FREM1;refGene_tx=XM_023242336.1
D4 42069497 42069642
refGene_gn=FREM1;refGene_tx=XM_019815975.2;refGene_gn=FREM1;refGene_tx=XM_019815979.2;refGene_gn=FREM1;refGene_tx=XM_019815981.2;refGene_gn=FREM1;refGene_tx=XM_019815982.1;refGene_gn=FREM1;refGene_tx=XM_019815984.2;refGene_gn=FREM1;refGene_tx=XM_019815985.2;refGene_gn=FREM1;refGene_tx=XM_019815986.2;refGene_gn=FREM1;refGene_tx=XM_019815987.2;refGene_gn=FREM1;refGene_tx=XM_019815988.1;refGene_gn=FREM1;refGene_tx=XM_023242336.1
D4 42073838 42074061
refGene_gn=FREM1;refGene_tx=XM_019815975.2;refGene_gn=FREM1;refGene_tx=XM_019815979.2;refGene_gn=FREM1;refGene_tx=XM_019815981.2;refGene_gn=FREM1;refGene_tx=XM_019815982.1;refGene_gn=FREM1;refGene_tx=XM_019815984.2;refGene_gn=FREM1;refGene_tx=XM_019815985.2;refGene_gn=FREM1;refGene_tx=XM_019815986.2;refGene_gn=FREM1;refGene_tx=XM_019815987.2;refGene_gn=FREM1;refGene_tx=XM_019815988.1;refGene_gn=FREM1;refGene_tx=XM_023242336.1
D4 42078811 42079008
refGene_gn=FREM1;refGene_tx=XM_019815975.2;refGene_gn=FREM1;refGene_tx=XM_019815979.2;refGene_gn=FREM1;refGene_tx=XM_019815981.2;refGene_gn=FREM1;refGene_tx=XM_019815982.1;refGene_gn=FREM1;refGene_tx=XM_019815984.2;refGene_gn=FREM1;refGene_tx=XM_019815985.2;refGene_gn=FREM1;refGene_tx=XM_019815986.2;refGene_gn=FREM1;refGene_tx=XM_019815987.2;refGene_gn=FREM1;refGene_tx=XM_019815988.1;refGene_gn=FREM1;refGene_tx=XM_023242336.1
D4 42085907 42086093
refGene_gn=FREM1;refGene_tx=XM_019815975.2;refGene_gn=FREM1;refGene_tx

=XM_019815979.2;refGene_gn=FREM1;refGene_tx=XM_019815981.2;refGene_gn=FREM1;refGene_tx=XM_019815982.1;refGene_gn=FREM1;refGene_tx=XM_019815984.2;refGene_gn=FREM1;refGene_tx=XM_019815985.2;refGene_gn=FREM1;refGene_tx=XM_019815986.2;refGene_gn=FREM1;refGene_tx=XM_019815987.2;refGene_gn=FREM1;refGene_tx=XM_019815988.1;refGene_gn=FREM1;refGene_tx=XM_023242336.1
D4 42087193 42087388
refGene_gn=FREM1;refGene_tx=XM_019815975.2;refGene_gn=FREM1;refGene_tx=XM_019815979.2;refGene_gn=FREM1;refGene_tx=XM_019815981.2;refGene_gn=FREM1;refGene_tx=XM_019815982.1;refGene_gn=FREM1;refGene_tx=XM_019815984.2;refGene_gn=FREM1;refGene_tx=XM_019815985.2;refGene_gn=FREM1;refGene_tx=XM_019815986.2;refGene_gn=FREM1;refGene_tx=XM_019815987.2;refGene_gn=FREM1;refGene_tx=XM_019815988.1;refGene_gn=FREM1;refGene_tx=XM_023242336.1
D4 42093129 42093382
refGene_gn=FREM1;refGene_tx=XM_019815975.2;refGene_gn=FREM1;refGene_tx=XM_019815979.2;refGene_gn=FREM1;refGene_tx=XM_019815981.2;refGene_gn=FREM1;refGene_tx=XM_019815982.1;refGene_gn=FREM1;refGene_tx=XM_019815984.2;refGene_gn=FREM1;refGene_tx=XM_019815985.2;refGene_gn=FREM1;refGene_tx=XM_019815986.2;refGene_gn=FREM1;refGene_tx=XM_019815987.2;refGene_gn=FREM1;refGene_tx=XM_019815988.1;refGene_gn=FREM1;refGene_tx=XM_023242336.1
D4 42096562 42096656
refGene_gn=FREM1;refGene_tx=XM_019815975.2;refGene_gn=FREM1;refGene_tx=XM_019815979.2;refGene_gn=FREM1;refGene_tx=XM_019815981.2;refGene_gn=FREM1;refGene_tx=XM_019815982.1;refGene_gn=FREM1;refGene_tx=XM_019815984.2;refGene_gn=FREM1;refGene_tx=XM_019815985.2;refGene_gn=FREM1;refGene_tx=XM_019815986.2;refGene_gn=FREM1;refGene_tx=XM_019815987.2;refGene_gn=FREM1;refGene_tx=XM_019815988.1;refGene_gn=FREM1;refGene_tx=XM_023242336.1
D4 42099953 42100162
refGene_gn=FREM1;refGene_tx=XM_019815975.2;refGene_gn=FREM1;refGene_tx=XM_019815979.2;refGene_gn=FREM1;refGene_tx=XM_019815981.2;refGene_gn=FREM1;refGene_tx=XM_019815982.1;refGene_gn=FREM1;refGene_tx=XM_019815984.2;refGene_gn=FREM1;refGene_tx=XM_019815985.2;refGene_gn=FREM1;refGene_tx=XM_019815986.2;refGene_gn=FREM1;refGene_tx=XM_019815987.2;refGene_gn=FREM1;refGene_tx=XM_019815988.1;refGene_gn=FREM1;refGene_tx=XM_023242336.1
D4 42105221 42105389
refGene_gn=FREM1;refGene_tx=XM_019815975.2;refGene_gn=FREM1;refGene_tx=XM_019815979.2;refGene_gn=FREM1;refGene_tx=XM_019815981.2;refGene_gn=FREM1;refGene_tx=XM_019815982.1;refGene_gn=FREM1;refGene_tx=XM_019815984.2;refGene_gn=FREM1;refGene_tx=XM_019815985.2;refGene_gn=FREM1;refGene_tx=XM_019815986.2;refGene_gn=FREM1;refGene_tx=XM_019815987.2;refGene_gn=FREM1;refGene_tx=XM_019815988.1;refGene_gn=FREM1;refGene_tx=XM_023242336.1
D4 42108449 42108540
refGene_gn=FREM1;refGene_tx=XM_019815975.2;refGene_gn=FREM1;refGene_tx=XM_019815979.2;refGene_gn=FREM1;refGene_tx=XM_019815981.2;refGene_gn=FREM1;refGene_tx=XM_019815982.1;refGene_gn=FREM1;refGene_tx=XM_0198159

84.2;refGene_gn=FREM1;refGene_tx=XM_019815985.2;refGene_gn=FREM1;refGene_tx=XM_019815986.2;refGene_gn=FREM1;refGene_tx=XM_019815987.2;refGene_gn=FREM1;refGene_tx=XM_019815988.1;refGene_gn=FREM1;refGene_tx=XM_023242336.1
D4 42109235 42109429
refGene_gn=FREM1;refGene_tx=XM_019815975.2;refGene_gn=FREM1;refGene_tx=XM_019815979.2;refGene_gn=FREM1;refGene_tx=XM_019815981.2;refGene_gn=FREM1;refGene_tx=XM_019815982.1;refGene_gn=FREM1;refGene_tx=XM_019815984.2;refGene_gn=FREM1;refGene_tx=XM_019815985.2;refGene_gn=FREM1;refGene_tx=XM_019815986.2;refGene_gn=FREM1;refGene_tx=XM_019815987.2;refGene_gn=FREM1;refGene_tx=XM_019815988.1;refGene_gn=FREM1;refGene_tx=XM_023242336.1
D4 42117276 42117419
refGene_gn=FREM1;refGene_tx=XM_019815975.2;refGene_gn=FREM1;refGene_tx=XM_019815979.2;refGene_gn=FREM1;refGene_tx=XM_019815981.2;refGene_gn=FREM1;refGene_tx=XM_019815982.1;refGene_gn=FREM1;refGene_tx=XM_019815984.2;refGene_gn=FREM1;refGene_tx=XM_019815985.2;refGene_gn=FREM1;refGene_tx=XM_019815986.2;refGene_gn=FREM1;refGene_tx=XM_019815987.2;refGene_gn=FREM1;refGene_tx=XM_019815988.1;refGene_gn=FREM1;refGene_tx=XM_023242336.1
D4 42118038 42118383
refGene_gn=FREM1;refGene_tx=XM_019815975.2;refGene_gn=FREM1;refGene_tx=XM_019815979.2;refGene_gn=FREM1;refGene_tx=XM_019815981.2;refGene_gn=FREM1;refGene_tx=XM_019815982.1;refGene_gn=FREM1;refGene_tx=XM_019815984.2;refGene_gn=FREM1;refGene_tx=XM_019815985.2;refGene_gn=FREM1;refGene_tx=XM_019815986.2;refGene_gn=FREM1;refGene_tx=XM_019815987.2;refGene_gn=FREM1;refGene_tx=XM_019815988.1;refGene_gn=FREM1;refGene_tx=XM_023242336.1
D4 42120625 42120757
refGene_gn=FREM1;refGene_tx=XM_019815975.2;refGene_gn=FREM1;refGene_tx=XM_019815979.2;refGene_gn=FREM1;refGene_tx=XM_019815981.2;refGene_gn=FREM1;refGene_tx=XM_019815982.1;refGene_gn=FREM1;refGene_tx=XM_019815984.2;refGene_gn=FREM1;refGene_tx=XM_019815985.2;refGene_gn=FREM1;refGene_tx=XM_019815987.2;refGene_gn=FREM1;refGene_tx=XM_019815988.1;refGene_gn=FREM1;refGene_tx=XM_023242336.1
D4 42123211 42123221
refGene_gn=FREM1;refGene_tx=XM_019815987.2;refGene_gn=FREM1;refGene_tx=XM_019815988.1
D4 42123712 42123821
refGene_gn=FREM1;refGene_tx=XM_019815975.2;refGene_gn=FREM1;refGene_tx=XM_019815979.2;refGene_gn=FREM1;refGene_tx=XM_019815981.2;refGene_gn=FREM1;refGene_tx=XM_019815982.1;refGene_gn=FREM1;refGene_tx=XM_019815984.2;refGene_gn=FREM1;refGene_tx=XM_019815985.2;refGene_gn=FREM1;refGene_tx=XM_019815986.2;refGene_gn=FREM1;refGene_tx=XM_023242336.1
D4 42126534 42126858
refGene_gn=FREM1;refGene_tx=XM_019815975.2;refGene_gn=FREM1;refGene_tx=XM_019815979.2;refGene_gn=FREM1;refGene_tx=XM_019815981.2;refGene_gn=FREM1;refGene_tx=XM_019815982.1;refGene_gn=FREM1;refGene_tx=XM_019815984.2;refGene_gn=FREM1;refGene_tx=XM_019815985.2;refGene_gn=FREM1;refGene_tx=XM_019815986.2;refGene_gn=FREM1;refGene_tx=XM_023242336.1

D4 42133956 42134153
refGene_gn=FREM1;refGene_tx=XM_019815975.2;refGene_gn=FREM1;refGene_tx=XM_019815979.2;refGene_gn=FREM1;refGene_tx=XM_019815981.2;refGene_gn=FREM1;refGene_tx=XM_019815982.1;refGene_gn=FREM1;refGene_tx=XM_019815984.2;refGene_gn=FREM1;refGene_tx=XM_019815985.2;refGene_gn=FREM1;refGene_tx=XM_019815986.2;refGene_gn=FREM1;refGene_tx=XM_023242336.1
D4 42135519 42135821
refGene_gn=FREM1;refGene_tx=XM_019815975.2;refGene_gn=FREM1;refGene_tx=XM_019815979.2;refGene_gn=FREM1;refGene_tx=XM_019815981.2;refGene_gn=FREM1;refGene_tx=XM_019815982.1;refGene_gn=FREM1;refGene_tx=XM_019815984.2;refGene_gn=FREM1;refGene_tx=XM_019815985.2;refGene_gn=FREM1;refGene_tx=XM_019815986.2;refGene_gn=FREM1;refGene_tx=XM_023242336.1
D4 42137944 42138039
refGene_gn=FREM1;refGene_tx=XM_019815975.2;refGene_gn=FREM1;refGene_tx=XM_019815979.2;refGene_gn=FREM1;refGene_tx=XM_019815981.2;refGene_gn=FREM1;refGene_tx=XM_019815982.1;refGene_gn=FREM1;refGene_tx=XM_019815984.2;refGene_gn=FREM1;refGene_tx=XM_019815985.2;refGene_gn=FREM1;refGene_tx=XM_019815986.2;refGene_gn=FREM1;refGene_tx=XM_023242336.1
D4 42142643 42142934
refGene_gn=FREM1;refGene_tx=XM_019815975.2;refGene_gn=FREM1;refGene_tx=XM_019815979.2;refGene_gn=FREM1;refGene_tx=XM_019815981.2;refGene_gn=FREM1;refGene_tx=XM_019815982.1;refGene_gn=FREM1;refGene_tx=XM_019815984.2;refGene_gn=FREM1;refGene_tx=XM_019815985.2;refGene_gn=FREM1;refGene_tx=XM_019815986.2;refGene_gn=FREM1;refGene_tx=XM_023242336.1
D4 42332042 42332133 refGene_gn=TTC39B;refGene_tx=XM_023242337.1
D4 42335340 42335457 refGene_gn=TTC39B;refGene_tx=XM_023242337.1
D4 42337256 42337374 refGene_gn=TTC39B;refGene_tx=XM_023242337.1
D4 42348047 42348156 refGene_gn=TTC39B;refGene_tx=XM_023242337.1
D4 42352128 42352255 refGene_gn=TTC39B;refGene_tx=XM_023242337.1
D4 42356753 42356845 refGene_gn=TTC39B;refGene_tx=XM_023242337.1
D4 42357712 42357874 refGene_gn=TTC39B;refGene_tx=XM_023242337.1
D4 42359305 42359365 refGene_gn=TTC39B;refGene_tx=XM_023242337.1
D4 42359455 42359523 refGene_gn=TTC39B;refGene_tx=XM_023242337.1
D4 42360414 42360523 refGene_gn=TTC39B;refGene_tx=XM_023242337.1
D4 42362043 42362109 refGene_gn=TTC39B;refGene_tx=XM_023242337.1
D4 42363489 42363595 refGene_gn=TTC39B;refGene_tx=XM_023242337.1
D4 42373439 42373504 refGene_gn=TTC39B;refGene_tx=XM_023242337.1
D4 42377402 42377470 refGene_gn=TTC39B;refGene_tx=XM_023242337.1
D4 42380887 42380964 refGene_gn=TTC39B;refGene_tx=XM_023242337.1
D4 42381936 42382068 refGene_gn=TTC39B;refGene_tx=XM_023242337.1
D4 42384567 42384678 refGene_gn=TTC39B;refGene_tx=XM_023242337.1
D4 42388728 42388824 refGene_gn=TTC39B;refGene_tx=XM_023242337.1
D4 42423091 42423126 refGene_gn=TTC39B;refGene_tx=XM_023242337.1
D4 42456123 42456324 refGene_gn=TTC39B;refGene_tx=XM_023242337.1
D4 42456325 42456367 refGene_gn=TTC39B;refGene_tx=XM_023242337.1
D4 42456538 42456658
refGene_gn=LOC111557468;refGene_tx=XM_023242298.1
D4 42461449 42461530
refGene_gn=LOC111557468;refGene_tx=XM_023242298.1
D4 42462116 42462194

refGene_gn=LOC111557468;refGene_tx=XM_023242298.1
D4	42464358	42464498
refGene_gn=LOC111557468;refGene_tx=XM_023242298.1
D4	42493459	42493700
refGene_gn=LOC111557468;refGene_tx=XM_023242298.1
D4	42572172	42572489	refGene_gn=SNAPC3;refGene_tx=XM_003995489.4
D4	42573218	42573296	refGene_gn=SNAPC3;refGene_tx=XM_003995489.4
D4	42590365	42590450	refGene_gn=SNAPC3;refGene_tx=XM_003995489.4
D4	42594453	42594558	refGene_gn=SNAPC3;refGene_tx=XM_003995489.4
D4	42603334	42603484	refGene_gn=SNAPC3;refGene_tx=XM_003995489.4
D4	42609829	42609912	refGene_gn=SNAPC3;refGene_tx=XM_003995489.4
D4	42612197	42612362	refGene_gn=SNAPC3;refGene_tx=XM_003995489.4
D4	42615268	42615376	refGene_gn=SNAPC3;refGene_tx=XM_003995489.4
D4	42616591	42616739	refGene_gn=SNAPC3;refGene_tx=XM_003995489.4
D4	42621842	42621903
refGene_gn=PSIP1;refGene_tx=NM_001009372.1;refGene_gn=PSIP1;refGene_tx
=XM_023242197.1
D4	42622717	42622829
refGene_gn=PSIP1;refGene_tx=NM_001009372.1;refGene_gn=PSIP1;refGene_tx
=XM_023242197.1
D4	42624477	42624691
refGene_gn=PSIP1;refGene_tx=NM_001009372.1;refGene_gn=PSIP1;refGene_tx
=XM_023242197.1
D4	42624808	42624910
refGene_gn=PSIP1;refGene_tx=NM_001009372.1;refGene_gn=PSIP1;refGene_tx
=XM_023242197.1
D4	42625119	42625190
refGene_gn=PSIP1;refGene_tx=NM_001009372.1;refGene_gn=PSIP1;refGene_tx
=XM_023242197.1
D4	42625801	42625857
refGene_gn=PSIP1;refGene_tx=NM_001009372.1;refGene_gn=PSIP1;refGene_tx
=XM_023242197.1
D4	42627037	42627116	refGene_gn=PSIP1;refGene_tx=XM_023242198.1
D4	42627146	42627171	refGene_gn=PSIP1;refGene_tx=XM_023242199.1
D4	42627238	42627251	refGene_gn=PSIP1;refGene_tx=XM_023242200.1
D4	42628453	42628460	refGene_gn=PSIP1;refGene_tx=XM_023242201.1
D4	42628537	42628656
refGene_gn=PSIP1;refGene_tx=NM_001009372.1;refGene_gn=PSIP1;refGene_tx
=XM_023242197.1;refGene_gn=PSIP1;refGene_tx=XM_023242198.1;refGene_gn=
PSIP1;refGene_tx=XM_023242199.1;refGene_gn=PSIP1;refGene_tx=XM_0232422
00.1;refGene_gn=PSIP1;refGene_tx=XM_023242201.1
D4	42629491	42629720
refGene_gn=PSIP1;refGene_tx=NM_001009372.1;refGene_gn=PSIP1;refGene_tx
=XM_023242197.1;refGene_gn=PSIP1;refGene_tx=XM_023242198.1;refGene_gn=
PSIP1;refGene_tx=XM_023242199.1;refGene_gn=PSIP1;refGene_tx=XM_0232422
00.1;refGene_gn=PSIP1;refGene_tx=XM_023242201.1
D4	42634534	42634610
refGene_gn=PSIP1;refGene_tx=NM_001009372.1;refGene_gn=PSIP1;refGene_tx
=XM_023242197.1;refGene_gn=PSIP1;refGene_tx=XM_023242198.1;refGene_gn=
PSIP1;refGene_tx=XM_023242199.1;refGene_gn=PSIP1;refGene_tx=XM_0232422

00.1;refGene_gn=PSIP1;refGene_tx=XM_023242201.1
D4 42635523 42635620
refGene_gn=PSIP1;refGene_tx=NM_001009372.1;refGene_gn=PSIP1;refGene_tx=XM_023242197.1;refGene_gn=PSIP1;refGene_tx=XM_023242198.1;refGene_gn=PSIP1;refGene_tx=XM_023242199.1;refGene_gn=PSIP1;refGene_tx=XM_023242200.1;refGene_gn=PSIP1;refGene_tx=XM_023242201.1
D4 42641474 42641537
refGene_gn=PSIP1;refGene_tx=NM_001009372.1;refGene_gn=PSIP1;refGene_tx=XM_023242197.1;refGene_gn=PSIP1;refGene_tx=XM_023242198.1;refGene_gn=PSIP1;refGene_tx=XM_023242199.1;refGene_gn=PSIP1;refGene_tx=XM_023242200.1;refGene_gn=PSIP1;refGene_tx=XM_023242201.1
D4 42642417 42642522
refGene_gn=PSIP1;refGene_tx=NM_001009372.1;refGene_gn=PSIP1;refGene_tx=XM_023242197.1;refGene_gn=PSIP1;refGene_tx=XM_023242198.1;refGene_gn=PSIP1;refGene_tx=XM_023242199.1;refGene_gn=PSIP1;refGene_tx=XM_023242200.1;refGene_gn=PSIP1;refGene_tx=XM_023242201.1
D4 42645010 42645149
refGene_gn=PSIP1;refGene_tx=NM_001009372.1;refGene_gn=PSIP1;refGene_tx=XM_023242197.1;refGene_gn=PSIP1;refGene_tx=XM_023242198.1;refGene_gn=PSIP1;refGene_tx=XM_023242199.1;refGene_gn=PSIP1;refGene_tx=XM_023242200.1;refGene_gn=PSIP1;refGene_tx=XM_023242201.1
D4 42655858 42655935
refGene_gn=PSIP1;refGene_tx=NM_001009372.1;refGene_gn=PSIP1;refGene_tx=XM_023242197.1;refGene_gn=PSIP1;refGene_tx=XM_023242198.1;refGene_gn=PSIP1;refGene_tx=XM_023242199.1;refGene_gn=PSIP1;refGene_tx=XM_023242200.1;refGene_gn=PSIP1;refGene_tx=XM_023242201.1
D4 42659004 42659076
refGene_gn=PSIP1;refGene_tx=NM_001009372.1;refGene_gn=PSIP1;refGene_tx=XM_023242197.1;refGene_gn=PSIP1;refGene_tx=XM_023242198.1;refGene_gn=PSIP1;refGene_tx=XM_023242199.1;refGene_gn=PSIP1;refGene_tx=XM_023242200.1;refGene_gn=PSIP1;refGene_tx=XM_023242201.1
D4 42705632 42705676
refGene_gn=CCDC171;refGene_tx=XM_011288345.3;refGene_gn=CCDC171;refGene_tx=XM_011288346.2;refGene_gn=CCDC171;refGene_tx=XM_011288347.3;refGene_gn=CCDC171;refGene_tx=XM_011288348.3;refGene_gn=CCDC171;refGene_tx=XM_011288349.3;refGene_gn=CCDC171;refGene_tx=XM_019815993.2;refGene_gn=CCDC171;refGene_tx=XM_019815994.2;refGene_gn=CCDC171;refGene_tx=XM_019815997.2
D4 42716891 42717027
refGene_gn=CCDC171;refGene_tx=XM_011288345.3;refGene_gn=CCDC171;refGene_tx=XM_011288346.2;refGene_gn=CCDC171;refGene_tx=XM_011288347.3;refGene_gn=CCDC171;refGene_tx=XM_011288348.3;refGene_gn=CCDC171;refGene_tx=XM_011288349.3;refGene_gn=CCDC171;refGene_tx=XM_019815993.2;refGene_gn=CCDC171;refGene_tx=XM_019815994.2;refGene_gn=CCDC171;refGene_tx=XM_019815997.2
D4 42726926 42727101
refGene_gn=CCDC171;refGene_tx=XM_011288345.3;refGene_gn=CCDC171;refGene_tx=XM_011288346.2;refGene_gn=CCDC171;refGene_tx=XM_011288347.3;refGene_gn=CCDC171;refGene_tx=XM_011288348.3;refGene_gn=CCDC171;refGene_tx=XM_011288349.3;refGene_gn=CCDC171;refGene_tx=XM_019815993.2;refGene_gn

=CCDC171;refGene_tx=XM_019815994.2;refGene_gn=CCDC171;refGene_tx=XM_019815997.2
D4	42730189	42730380
refGene_gn=CCDC171;refGene_tx=XM_011288345.3;refGene_gn=CCDC171;refGene_tx=XM_011288346.2;refGene_gn=CCDC171;refGene_tx=XM_011288347.3;refGene_gn=CCDC171;refGene_tx=XM_011288348.3;refGene_gn=CCDC171;refGene_tx=XM_011288349.3;refGene_gn=CCDC171;refGene_tx=XM_019815993.2;refGene_gn=CCDC171;refGene_tx=XM_019815994.2;refGene_gn=CCDC171;refGene_tx=XM_019815997.2
D4	42732415	42732547
refGene_gn=CCDC171;refGene_tx=XM_011288345.3;refGene_gn=CCDC171;refGene_tx=XM_011288346.2;refGene_gn=CCDC171;refGene_tx=XM_011288347.3;refGene_gn=CCDC171;refGene_tx=XM_011288348.3;refGene_gn=CCDC171;refGene_tx=XM_019815993.2;refGene_gn=CCDC171;refGene_tx=XM_019815994.2;refGene_gn=CCDC171;refGene_tx=XM_019815997.2
D4	42741934	42742081
refGene_gn=CCDC171;refGene_tx=XM_011288345.3;refGene_gn=CCDC171;refGene_tx=XM_011288346.2;refGene_gn=CCDC171;refGene_tx=XM_011288347.3;refGene_gn=CCDC171;refGene_tx=XM_011288348.3;refGene_gn=CCDC171;refGene_tx=XM_011288349.3;refGene_gn=CCDC171;refGene_tx=XM_019815993.2;refGene_gn=CCDC171;refGene_tx=XM_019815994.2;refGene_gn=CCDC171;refGene_tx=XM_019815997.2
D4	42763183	42763276
refGene_gn=CCDC171;refGene_tx=XM_011288345.3;refGene_gn=CCDC171;refGene_tx=XM_011288346.2;refGene_gn=CCDC171;refGene_tx=XM_011288347.3;refGene_gn=CCDC171;refGene_tx=XM_011288348.3;refGene_gn=CCDC171;refGene_tx=XM_011288349.3;refGene_gn=CCDC171;refGene_tx=XM_019815993.2;refGene_gn=CCDC171;refGene_tx=XM_019815994.2
D4	42768396	42768557
refGene_gn=CCDC171;refGene_tx=XM_011288345.3;refGene_gn=CCDC171;refGene_tx=XM_011288346.2;refGene_gn=CCDC171;refGene_tx=XM_011288347.3;refGene_gn=CCDC171;refGene_tx=XM_011288348.3;refGene_gn=CCDC171;refGene_tx=XM_011288349.3;refGene_gn=CCDC171;refGene_tx=XM_019815993.2;refGene_gn=CCDC171;refGene_tx=XM_019815994.2
D4	42782856	42782995
refGene_gn=CCDC171;refGene_tx=XM_011288345.3;refGene_gn=CCDC171;refGene_tx=XM_011288346.2;refGene_gn=CCDC171;refGene_tx=XM_011288347.3;refGene_gn=CCDC171;refGene_tx=XM_011288348.3;refGene_gn=CCDC171;refGene_tx=XM_011288349.3;refGene_gn=CCDC171;refGene_tx=XM_019815993.2;refGene_gn=CCDC171;refGene_tx=XM_019815994.2
D4	42804604	42804707
refGene_gn=CCDC171;refGene_tx=XM_011288345.3;refGene_gn=CCDC171;refGene_tx=XM_011288346.2;refGene_gn=CCDC171;refGene_tx=XM_011288347.3;refGene_gn=CCDC171;refGene_tx=XM_011288348.3;refGene_gn=CCDC171;refGene_tx=XM_011288349.3;refGene_gn=CCDC171;refGene_tx=XM_019815993.2;refGene_gn=CCDC171;refGene_tx=XM_019815994.2;refGene_gn=CCDC171;refGene_tx=XM_019815997.2
D4	42820646	42820753
refGene_gn=CCDC171;refGene_tx=XM_011288345.3;refGene_gn=CCDC171;refGene_tx=XM_011288346.2;refGene_gn=CCDC171;refGene_tx=XM_011288347.3;refGe

ne_gn=CCDC171;refGene_tx=XM_011288348.3;refGene_gn=CCDC171;refGene_tx=XM_011288349.3;refGene_gn=CCDC171;refGene_tx=XM_019815993.2;refGene_gn=CCDC171;refGene_tx=XM_019815994.2;refGene_gn=CCDC171;refGene_tx=XM_019815997.2
D4 42822525 42822615
refGene_gn=CCDC171;refGene_tx=XM_011288345.3;refGene_gn=CCDC171;refGene_tx=XM_011288346.2;refGene_gn=CCDC171;refGene_tx=XM_011288347.3;refGene_gn=CCDC171;refGene_tx=XM_011288348.3;refGene_gn=CCDC171;refGene_tx=XM_011288349.3;refGene_gn=CCDC171;refGene_tx=XM_019815993.2;refGene_gn=CCDC171;refGene_tx=XM_019815994.2;refGene_gn=CCDC171;refGene_tx=XM_019815997.2
D4 42823671 42823872
refGene_gn=CCDC171;refGene_tx=XM_011288345.3;refGene_gn=CCDC171;refGene_tx=XM_011288346.2;refGene_gn=CCDC171;refGene_tx=XM_011288347.3;refGene_gn=CCDC171;refGene_tx=XM_011288348.3;refGene_gn=CCDC171;refGene_tx=XM_011288349.3;refGene_gn=CCDC171;refGene_tx=XM_019815993.2;refGene_gn=CCDC171;refGene_tx=XM_019815994.2;refGene_gn=CCDC171;refGene_tx=XM_019815997.2
D4 42826436 42826604
refGene_gn=CCDC171;refGene_tx=XM_011288345.3;refGene_gn=CCDC171;refGene_tx=XM_011288346.2;refGene_gn=CCDC171;refGene_tx=XM_011288347.3;refGene_gn=CCDC171;refGene_tx=XM_011288348.3;refGene_gn=CCDC171;refGene_tx=XM_011288349.3;refGene_gn=CCDC171;refGene_tx=XM_019815993.2;refGene_gn=CCDC171;refGene_tx=XM_019815994.2;refGene_gn=CCDC171;refGene_tx=XM_019815997.2
D4 42828186 42828375
refGene_gn=CCDC171;refGene_tx=XM_011288345.3;refGene_gn=CCDC171;refGene_tx=XM_011288346.2;refGene_gn=CCDC171;refGene_tx=XM_011288347.3;refGene_gn=CCDC171;refGene_tx=XM_011288348.3;refGene_gn=CCDC171;refGene_tx=XM_011288349.3;refGene_gn=CCDC171;refGene_tx=XM_019815993.2;refGene_gn=CCDC171;refGene_tx=XM_019815994.2;refGene_gn=CCDC171;refGene_tx=XM_019815997.2
D4 42843971 42844479
refGene_gn=CCDC171;refGene_tx=XM_011288345.3;refGene_gn=CCDC171;refGene_tx=XM_011288346.2;refGene_gn=CCDC171;refGene_tx=XM_011288347.3;refGene_gn=CCDC171;refGene_tx=XM_011288348.3;refGene_gn=CCDC171;refGene_tx=XM_011288349.3;refGene_gn=CCDC171;refGene_tx=XM_019815993.2;refGene_gn=CCDC171;refGene_tx=XM_019815994.2;refGene_gn=CCDC171;refGene_tx=XM_019815997.2
D4 42845560 42845677
refGene_gn=CCDC171;refGene_tx=XM_011288345.3;refGene_gn=CCDC171;refGene_tx=XM_011288346.2;refGene_gn=CCDC171;refGene_tx=XM_011288347.3;refGene_gn=CCDC171;refGene_tx=XM_011288348.3;refGene_gn=CCDC171;refGene_tx=XM_011288349.3;refGene_gn=CCDC171;refGene_tx=XM_019815993.2;refGene_gn=CCDC171;refGene_tx=XM_019815994.2;refGene_gn=CCDC171;refGene_tx=XM_019815997.2
D4 42864226 42864453
refGene_gn=CCDC171;refGene_tx=XM_011288345.3;refGene_gn=CCDC171;refGene_tx=XM_011288346.2;refGene_gn=CCDC171;refGene_tx=XM_011288347.3;refGene_gn=CCDC171;refGene_tx=XM_011288348.3;refGene_gn=CCDC171;refGene_tx=

XM_011288349.3;refGene_gn=CCDC171;refGene_tx=XM_019815993.2;refGene_gn=CCDC171;refGene_tx=XM_019815994.2;refGene_gn=CCDC171;refGene_tx=XM_019815997.2
D4 42865172 42865355
refGene_gn=CCDC171;refGene_tx=XM_011288345.3;refGene_gn=CCDC171;refGene_tx=XM_011288346.2;refGene_gn=CCDC171;refGene_tx=XM_011288347.3;refGene_gn=CCDC171;refGene_tx=XM_011288348.3;refGene_gn=CCDC171;refGene_tx=XM_011288349.3;refGene_gn=CCDC171;refGene_tx=XM_019815993.2;refGene_gn=CCDC171;refGene_tx=XM_019815994.2;refGene_gn=CCDC171;refGene_tx=XM_019815997.2
D4 42872201 42872387
refGene_gn=CCDC171;refGene_tx=XM_011288345.3;refGene_gn=CCDC171;refGene_tx=XM_011288346.2;refGene_gn=CCDC171;refGene_tx=XM_011288347.3;refGene_gn=CCDC171;refGene_tx=XM_011288348.3;refGene_gn=CCDC171;refGene_tx=XM_011288349.3;refGene_gn=CCDC171;refGene_tx=XM_019815993.2;refGene_gn=CCDC171;refGene_tx=XM_019815994.2;refGene_gn=CCDC171;refGene_tx=XM_019815997.2
D4 42911695 42911841
refGene_gn=CCDC171;refGene_tx=XM_011288345.3;refGene_gn=CCDC171;refGene_tx=XM_011288346.2;refGene_gn=CCDC171;refGene_tx=XM_011288347.3;refGene_gn=CCDC171;refGene_tx=XM_011288348.3;refGene_gn=CCDC171;refGene_tx=XM_011288349.3;refGene_gn=CCDC171;refGene_tx=XM_019815993.2;refGene_gn=CCDC171;refGene_tx=XM_019815994.2;refGene_gn=CCDC171;refGene_tx=XM_019815997.2
D4 42913939 42913994
refGene_gn=CCDC171;refGene_tx=XM_011288345.3;refGene_gn=CCDC171;refGene_tx=XM_011288346.2;refGene_gn=CCDC171;refGene_tx=XM_011288347.3;refGene_gn=CCDC171;refGene_tx=XM_011288348.3;refGene_gn=CCDC171;refGene_tx=XM_011288349.3;refGene_gn=CCDC171;refGene_tx=XM_019815993.2;refGene_gn=CCDC171;refGene_tx=XM_019815994.2;refGene_gn=CCDC171;refGene_tx=XM_019815997.2
D4 42931585 42931717
refGene_gn=CCDC171;refGene_tx=XM_011288345.3;refGene_gn=CCDC171;refGene_tx=XM_011288346.2;refGene_gn=CCDC171;refGene_tx=XM_011288347.3;refGene_gn=CCDC171;refGene_tx=XM_011288348.3;refGene_gn=CCDC171;refGene_tx=XM_011288349.3;refGene_gn=CCDC171;refGene_tx=XM_019815993.2;refGene_gn=CCDC171;refGene_tx=XM_019815997.2
D4 42965422 42965575
refGene_gn=CCDC171;refGene_tx=XM_011288345.3;refGene_gn=CCDC171;refGene_tx=XM_011288346.2;refGene_gn=CCDC171;refGene_tx=XM_011288347.3;refGene_gn=CCDC171;refGene_tx=XM_011288348.3;refGene_gn=CCDC171;refGene_tx=XM_011288349.3;refGene_gn=CCDC171;refGene_tx=XM_019815993.2;refGene_gn=CCDC171;refGene_tx=XM_019815994.2;refGene_gn=CCDC171;refGene_tx=XM_019815997.2
D4 43007883 43008096
refGene_gn=CCDC171;refGene_tx=XM_011288345.3;refGene_gn=CCDC171;refGene_tx=XM_011288346.2;refGene_gn=CCDC171;refGene_tx=XM_011288347.3;refGene_gn=CCDC171;refGene_tx=XM_011288348.3;refGene_gn=CCDC171;refGene_tx=XM_011288349.3;refGene_gn=CCDC171;refGene_tx=XM_019815993.2;refGene_gn=CCDC171;refGene_tx=XM_019815994.2;refGene_gn=CCDC171;refGene_tx=XM_01

9815997.2
D4 43396777 43397438
refGene_gn=BNC2;refGene_tx=XM_003995490.4;refGene_gn=BNC2;refGene_tx=XM_006939167.3;refGene_gn=BNC2;refGene_tx=XM_011288344.3;refGene_gn=BNC2;refGene_tx=XM_019815998.2;refGene_gn=BNC2;refGene_tx=XM_019815999.2;refGene_gn=BNC2;refGene_tx=XM_019816000.2;refGene_gn=BNC2;refGene_tx=XM_019816005.2;refGene_gn=BNC2;refGene_tx=XM_019816006.2;refGene_gn=BNC2;refGene_tx=XM_019816007.2;refGene_gn=BNC2;refGene_tx=XM_019816008.2;refGene_gn=BNC2;refGene_tx=XM_019816009.2;refGene_gn=BNC2;refGene_tx=XM_019816011.2;refGene_gn=BNC2;refGene_tx=XM_019816012.2;refGene_gn=BNC2;refGene_tx=XM_019816013.2;refGene_gn=BNC2;refGene_tx=XM_019816014.2;refGene_gn=BNC2;refGene_tx=XM_019816018.1;refGene_gn=BNC2;refGene_tx=XM_019816019.2;refGene_gn=BNC2;refGene_tx=XM_019816022.2
D4 43399030 43399049
refGene_gn=BNC2;refGene_tx=XM_019816017.2;refGene_gn=BNC2;refGene_tx=XM_019816020.2;refGene_gn=BNC2;refGene_tx=XM_019816021.2
D4 43412788 43412854
refGene_gn=BNC2;refGene_tx=XM_011288344.3;refGene_gn=BNC2;refGene_tx=XM_019815998.2;refGene_gn=BNC2;refGene_tx=XM_019815999.2;refGene_gn=BNC2;refGene_tx=XM_019816000.2;refGene_gn=BNC2;refGene_tx=XM_019816005.2;refGene_gn=BNC2;refGene_tx=XM_019816006.2;refGene_gn=BNC2;refGene_tx=XM_019816007.2;refGene_gn=BNC2;refGene_tx=XM_019816008.2;refGene_gn=BNC2;refGene_tx=XM_019816009.2;refGene_gn=BNC2;refGene_tx=XM_019816011.2;refGene_gn=BNC2;refGene_tx=XM_019816012.2;refGene_gn=BNC2;refGene_tx=XM_019816013.2;refGene_gn=BNC2;refGene_tx=XM_019816014.2;refGene_gn=BNC2;refGene_tx=XM_019816017.2;refGene_gn=BNC2;refGene_tx=XM_019816018.1;refGene_gn=BNC2;refGene_tx=XM_019816019.2;refGene_gn=BNC2;refGene_tx=XM_019816020.2;refGene_gn=BNC2;refGene_tx=XM_019816022.2
D4 43413285 43415255
refGene_gn=BNC2;refGene_tx=XM_003995490.4;refGene_gn=BNC2;refGene_tx=XM_006939167.3;refGene_gn=BNC2;refGene_tx=XM_011288344.3;refGene_gn=BNC2;refGene_tx=XM_019815998.2;refGene_gn=BNC2;refGene_tx=XM_019815999.2;refGene_gn=BNC2;refGene_tx=XM_019816000.2;refGene_gn=BNC2;refGene_tx=XM_019816005.2;refGene_gn=BNC2;refGene_tx=XM_019816006.2;refGene_gn=BNC2;refGene_tx=XM_019816007.2;refGene_gn=BNC2;refGene_tx=XM_019816008.2;refGene_gn=BNC2;refGene_tx=XM_019816009.2;refGene_gn=BNC2;refGene_tx=XM_019816011.2;refGene_gn=BNC2;refGene_tx=XM_019816012.2;refGene_gn=BNC2;refGene_tx=XM_019816013.2;refGene_gn=BNC2;refGene_tx=XM_019816014.2;refGene_gn=BNC2;refGene_tx=XM_019816017.2;refGene_gn=BNC2;refGene_tx=XM_019816018.1;refGene_gn=BNC2;refGene_tx=XM_019816019.2;refGene_gn=BNC2;refGene_tx=XM_019816020.2;refGene_gn=BNC2;refGene_tx=XM_019816021.2;refGene_gn=BNC2;refGene_tx=XM_019816022.2
D4 43521380 43521616
refGene_gn=BNC2;refGene_tx=XM_003995490.4;refGene_gn=BNC2;refGene_tx=XM_006939167.3;refGene_gn=BNC2;refGene_tx=XM_011288344.3;refGene_gn=BNC2;refGene_tx=XM_019815998.2;refGene_gn=BNC2;refGene_tx=XM_019815999.2;refGene_gn=BNC2;refGene_tx=XM_019816000.2;refGene_gn=BNC2;refGene_tx=XM_019816005.2;refGene_gn=BNC2;refGene_tx=XM_019816006.2;refGene_gn=BNC2;refGene_tx=XM_019816007.2;refGene_gn=BNC2;refGene_tx=XM_019816008.2;refGene_gn=BNC2;refGene_tx=XM_019816009.2;refGene_gn=BNC2;refGene_tx=X

M_019816011.2;refGene_gn=BNC2;refGene_tx=XM_019816012.2;refGene_gn=BNC2;refGene_tx=XM_019816013.2;refGene_gn=BNC2;refGene_tx=XM_019816014.2;refGene_gn=BNC2;refGene_tx=XM_019816017.2;refGene_gn=BNC2;refGene_tx=XM_019816018.1;refGene_gn=BNC2;refGene_tx=XM_019816019.2;refGene_gn=BNC2;refGene_tx=XM_019816020.2;refGene_gn=BNC2;refGene_tx=XM_019816021.2;refGene_gn=BNC2;refGene_tx=XM_019816022.2
D4 43546948 43546958 refGene_gn=BNC2;refGene_tx=XM_019816009.2
D4 43547386 43547489
refGene_gn=BNC2;refGene_tx=XM_003995490.4;refGene_gn=BNC2;refGene_tx=XM_006939167.3;refGene_gn=BNC2;refGene_tx=XM_011288344.3;refGene_gn=BNC2;refGene_tx=XM_019815998.2;refGene_gn=BNC2;refGene_tx=XM_019815999.2;refGene_gn=BNC2;refGene_tx=XM_019816000.2;refGene_gn=BNC2;refGene_tx=XM_019816005.2;refGene_gn=BNC2;refGene_tx=XM_019816006.2;refGene_gn=BNC2;refGene_tx=XM_019816007.2;refGene_gn=BNC2;refGene_tx=XM_019816008.2;refGene_gn=BNC2;refGene_tx=XM_019816017.2;refGene_gn=BNC2;refGene_tx=XM_019816018.1;refGene_gn=BNC2;refGene_tx=XM_019816019.2;refGene_gn=BNC2;refGene_tx=XM_019816020.2;refGene_gn=BNC2;refGene_tx=XM_019816021.2;refGene_gn=BNC2;refGene_tx=XM_019816022.2
D4 43550675 43550708 refGene_gn=BNC2;refGene_tx=XM_019816006.2
D4 43645042 43645126
refGene_gn=BNC2;refGene_tx=XM_006939167.3;refGene_gn=BNC2;refGene_tx=XM_019815998.2;refGene_gn=BNC2;refGene_tx=XM_019815999.2;refGene_gn=BNC2;refGene_tx=XM_019816000.2;refGene_gn=BNC2;refGene_tx=XM_019816005.2;refGene_gn=BNC2;refGene_tx=XM_019816017.2;refGene_gn=BNC2;refGene_tx=XM_019816018.1;refGene_gn=BNC2;refGene_tx=XM_019816019.2;refGene_gn=BNC2;refGene_tx=XM_019816020.2;refGene_gn=BNC2;refGene_tx=XM_019816021.2;refGene_gn=BNC2;refGene_tx=XM_019816022.2
D4 43662058 43662061
refGene_gn=BNC2;refGene_tx=XM_019816008.2;refGene_gn=BNC2;refGene_tx=XM_019816022.2
D4 43684382 43684583
refGene_gn=BNC2;refGene_tx=XM_003995490.4;refGene_gn=BNC2;refGene_tx=XM_006939167.3;refGene_gn=BNC2;refGene_tx=XM_011288344.3;refGene_gn=BNC2;refGene_tx=XM_019815998.2;refGene_gn=BNC2;refGene_tx=XM_019815999.2;refGene_gn=BNC2;refGene_tx=XM_019816000.2;refGene_gn=BNC2;refGene_tx=XM_019816017.2;refGene_gn=BNC2;refGene_tx=XM_019816018.1;refGene_gn=BNC2;refGene_tx=XM_019816019.2;refGene_gn=BNC2;refGene_tx=XM_019816020.2;refGene_gn=BNC2;refGene_tx=XM_019816021.2
D4 43748557 43748656
refGene_gn=BNC2;refGene_tx=XM_003995490.4;refGene_gn=BNC2;refGene_tx=XM_006939167.3;refGene_gn=BNC2;refGene_tx=XM_011288344.3;refGene_gn=BNC2;refGene_tx=XM_019815998.2;refGene_gn=BNC2;refGene_tx=XM_019815999.2;refGene_gn=BNC2;refGene_tx=XM_019816017.2;refGene_gn=BNC2;refGene_tx=XM_019816019.2;refGene_gn=BNC2;refGene_tx=XM_019816020.2;refGene_gn=BNC2;refGene_tx=XM_019816021.2
D4 43825395 43825398
refGene_gn=BNC2;refGene_tx=XM_019815999.2;refGene_gn=BNC2;refGene_tx=XM_019816000.2;refGene_gn=BNC2;refGene_tx=XM_019816005.2;refGene_gn=BNC2;refGene_tx=XM_019816007.2
D4 44050835 44051168

refGene_gn=CNTLN;refGene_tx=XM_011288695.3;refGene_gn=CNTLN;refGene_tx=XM_019816024.2
D4 44059996 44060085
refGene_gn=CNTLN;refGene_tx=XM_011288695.3;refGene_gn=CNTLN;refGene_tx=XM_019816024.2
D4 44137931 44138016
refGene_gn=CNTLN;refGene_tx=XM_011288695.3;refGene_gn=CNTLN;refGene_tx=XM_019816024.2
D4 44157833 44157968
refGene_gn=CNTLN;refGene_tx=XM_011288695.3;refGene_gn=CNTLN;refGene_tx=XM_019816024.2
D4 44158578 44158758
refGene_gn=CNTLN;refGene_tx=XM_011288695.3;refGene_gn=CNTLN;refGene_tx=XM_019816024.2
D4 44169453 44169587
refGene_gn=CNTLN;refGene_tx=XM_011288695.3;refGene_gn=CNTLN;refGene_tx=XM_019816024.2
D4 44179573 44179736
refGene_gn=CNTLN;refGene_tx=XM_011288695.3;refGene_gn=CNTLN;refGene_tx=XM_019816024.2
D4 44190637 44190832
refGene_gn=CNTLN;refGene_tx=XM_011288695.3;refGene_gn=CNTLN;refGene_tx=XM_019816024.2
D4 44214607 44214784
refGene_gn=CNTLN;refGene_tx=XM_011288695.3;refGene_gn=CNTLN;refGene_tx=XM_019816024.2
D4 44216614 44216740
refGene_gn=CNTLN;refGene_tx=XM_011288695.3;refGene_gn=CNTLN;refGene_tx=XM_019816024.2
D4 44226737 44226859
refGene_gn=CNTLN;refGene_tx=XM_011288695.3;refGene_gn=CNTLN;refGene_tx=XM_019816024.2
D4 44228206 44228326
refGene_gn=CNTLN;refGene_tx=XM_011288695.3;refGene_gn=CNTLN;refGene_tx=XM_019816024.2
D4 44244573 44244674
refGene_gn=CNTLN;refGene_tx=XM_011288695.3;refGene_gn=CNTLN;refGene_tx=XM_019816024.2
D4 44253453 44253545
refGene_gn=CNTLN;refGene_tx=XM_011288695.3;refGene_gn=CNTLN;refGene_tx=XM_019816024.2
D4 44258594 44259130
refGene_gn=CNTLN;refGene_tx=XM_011288695.3;refGene_gn=CNTLN;refGene_tx=XM_019816024.2
D4 44272632 44272813
refGene_gn=CNTLN;refGene_tx=XM_011288695.3;refGene_gn=CNTLN;refGene_tx=XM_019816024.2
D4 44276798 44276892
refGene_gn=CNTLN;refGene_tx=XM_011288695.3;refGene_gn=CNTLN;refGene_tx=XM_019816024.2

D4 44276972 44277196
refGene_gn=CNTLN;refGene_tx=XM_011288695.3;refGene_gn=CNTLN;refGene_tx=XM_019816024.2
D4 44312486 44312678
refGene_gn=CNTLN;refGene_tx=XM_011288695.3;refGene_gn=CNTLN;refGene_tx=XM_019816024.2
D4 44318743 44318841
refGene_gn=CNTLN;refGene_tx=XM_011288695.3;refGene_gn=CNTLN;refGene_tx=XM_019816024.2
D4 44320521 44320648
refGene_gn=CNTLN;refGene_tx=XM_011288695.3;refGene_gn=CNTLN;refGene_tx=XM_019816024.2
D4 44322047 44322185
refGene_gn=CNTLN;refGene_tx=XM_011288695.3;refGene_gn=CNTLN;refGene_tx=XM_019816024.2
D4 44322962 44323148
refGene_gn=CNTLN;refGene_tx=XM_011288695.3;refGene_gn=CNTLN;refGene_tx=XM_019816024.2
D4 44352515 44352701
refGene_gn=CNTLN;refGene_tx=XM_011288695.3;refGene_gn=CNTLN;refGene_tx=XM_019816024.2
D4 44355409 44355487
refGene_gn=CNTLN;refGene_tx=XM_011288695.3;refGene_gn=CNTLN;refGene_tx=XM_019816024.2
D4 44366410 44366512
refGene_gn=CNTLN;refGene_tx=XM_011288695.3;refGene_gn=CNTLN;refGene_tx=XM_019816024.2
D4 44439590 44439635 refGene_gn=SH3GL2;refGene_tx=XM_023242339.1
D4 44596594 44596663 refGene_gn=SH3GL2;refGene_tx=XM_023242339.1
D4 44606028 44606101 refGene_gn=SH3GL2;refGene_tx=XM_023242339.1
D4 44640201 44640345 refGene_gn=SH3GL2;refGene_tx=XM_023242339.1
D4 44641168 44641302 refGene_gn=SH3GL2;refGene_tx=XM_023242339.1
D4 44643136 44643295 refGene_gn=SH3GL2;refGene_tx=XM_023242339.1
D4 44644749 44644853 refGene_gn=SH3GL2;refGene_tx=XM_023242339.1
D4 44646400 44646531 refGene_gn=SH3GL2;refGene_tx=XM_023242339.1
D4 44648597 44648797 refGene_gn=SH3GL2;refGene_tx=XM_023242339.1
D4 44667151 44667696
refGene_gn=LOC101081765;refGene_tx=XM_019815533.1
D4 44674332 44674406
refGene_gn=LOC101081765;refGene_tx=XM_019815533.1
D4 44704566 44704982
refGene_gn=LOC101081765;refGene_tx=XM_019815533.1
D4 44766049 44766270
refGene_gn=LOC111557469;refGene_tx=XM_023242299.1
D4 44767022 44767568
refGene_gn=LOC111557469;refGene_tx=XM_023242299.1
D4 44768834 44768949
refGene_gn=LOC111557469;refGene_tx=XM_023242299.1
D4 45276144 45276207
refGene_gn=ADAMTSL1;refGene_tx=XM_011288618.2;refGene_gn=ADAMTSL1;refG

ene_tx=XM_023242340.1;refGene_gn=ADAMTSL1;refGene_tx=XM_023242341.1;re
fGene_gn=ADAMTSL1;refGene_tx=XM_023242342.1;refGene_gn=ADAMTSL1;refGen
e_tx=XM_023242343.1;refGene_gn=ADAMTSL1;refGene_tx=XM_023242344.1
D4 45308272 45308400
refGene_gn=ADAMTSL1;refGene_tx=XM_011288618.2;refGene_gn=ADAMTSL1;refG
ene_tx=XM_023242340.1;refGene_gn=ADAMTSL1;refGene_tx=XM_023242341.1;re
fGene_gn=ADAMTSL1;refGene_tx=XM_023242342.1;refGene_gn=ADAMTSL1;refGen
e_tx=XM_023242343.1;refGene_gn=ADAMTSL1;refGene_tx=XM_023242344.1
D4 45331315 45331361
refGene_gn=ADAMTSL1;refGene_tx=XM_011288618.2;refGene_gn=ADAMTSL1;refG
ene_tx=XM_023242340.1;refGene_gn=ADAMTSL1;refGene_tx=XM_023242341.1;re
fGene_gn=ADAMTSL1;refGene_tx=XM_023242342.1;refGene_gn=ADAMTSL1;refGen
e_tx=XM_023242343.1;refGene_gn=ADAMTSL1;refGene_tx=XM_023242344.1
D4 45363984 45364221
refGene_gn=ADAMTSL1;refGene_tx=XM_011288618.2;refGene_gn=ADAMTSL1;refG
ene_tx=XM_023242340.1;refGene_gn=ADAMTSL1;refGene_tx=XM_023242341.1;re
fGene_gn=ADAMTSL1;refGene_tx=XM_023242342.1;refGene_gn=ADAMTSL1;refGen
e_tx=XM_023242343.1;refGene_gn=ADAMTSL1;refGene_tx=XM_023242344.1
D4 45404904 45405031
refGene_gn=ADAMTSL1;refGene_tx=XM_011288618.2;refGene_gn=ADAMTSL1;refG
ene_tx=XM_023242340.1;refGene_gn=ADAMTSL1;refGene_tx=XM_023242341.1;re
fGene_gn=ADAMTSL1;refGene_tx=XM_023242342.1;refGene_gn=ADAMTSL1;refGen
e_tx=XM_023242343.1;refGene_gn=ADAMTSL1;refGene_tx=XM_023242344.1
D4 45420190 45420265
refGene_gn=ADAMTSL1;refGene_tx=XM_011288618.2;refGene_gn=ADAMTSL1;refG
ene_tx=XM_023242340.1;refGene_gn=ADAMTSL1;refGene_tx=XM_023242342.1;re
fGene_gn=ADAMTSL1;refGene_tx=XM_023242343.1;refGene_gn=ADAMTSL1;refGen
e_tx=XM_023242344.1
D4 45423691 45423849
refGene_gn=ADAMTSL1;refGene_tx=XM_011288618.2;refGene_gn=ADAMTSL1;refG
ene_tx=XM_023242340.1;refGene_gn=ADAMTSL1;refGene_tx=XM_023242341.1;re
fGene_gn=ADAMTSL1;refGene_tx=XM_023242342.1;refGene_gn=ADAMTSL1;refGen
e_tx=XM_023242343.1;refGene_gn=ADAMTSL1;refGene_tx=XM_023242344.1
D4 45429688 45429800
refGene_gn=ADAMTSL1;refGene_tx=XM_011288618.2;refGene_gn=ADAMTSL1;refG
ene_tx=XM_023242340.1;refGene_gn=ADAMTSL1;refGene_tx=XM_023242341.1;re
fGene_gn=ADAMTSL1;refGene_tx=XM_023242342.1;refGene_gn=ADAMTSL1;refGen
e_tx=XM_023242343.1;refGene_gn=ADAMTSL1;refGene_tx=XM_023242344.1
D4 45433316 45433455
refGene_gn=ADAMTSL1;refGene_tx=XM_011288618.2;refGene_gn=ADAMTSL1;refG
ene_tx=XM_023242340.1;refGene_gn=ADAMTSL1;refGene_tx=XM_023242341.1;re
fGene_gn=ADAMTSL1;refGene_tx=XM_023242342.1;refGene_gn=ADAMTSL1;refGen
e_tx=XM_023242343.1;refGene_gn=ADAMTSL1;refGene_tx=XM_023242344.1
D4 45446092 45446143
refGene_gn=ADAMTSL1;refGene_tx=XM_011288618.2;refGene_gn=ADAMTSL1;refG
ene_tx=XM_023242341.1;refGene_gn=ADAMTSL1;refGene_tx=XM_023242342.1;re
fGene_gn=ADAMTSL1;refGene_tx=XM_023242343.1;refGene_gn=ADAMTSL1;refGen
e_tx=XM_023242344.1
D4 45450681 45450886
refGene_gn=ADAMTSL1;refGene_tx=XM_011288618.2;refGene_gn=ADAMTSL1;refG

ene_tx=XM_023242340.1;refGene_gn=ADAMTSL1;refGene_tx=XM_023242341.1;refGene_gn=ADAMTSL1;refGene_tx=XM_023242342.1;refGene_gn=ADAMTSL1;refGene_tx=XM_023242343.1;refGene_gn=ADAMTSL1;refGene_tx=XM_023242344.1
D4 45452111 45452259
refGene_gn=ADAMTSL1;refGene_tx=XM_011288618.2;refGene_gn=ADAMTSL1;refGene_tx=XM_023242340.1;refGene_gn=ADAMTSL1;refGene_tx=XM_023242341.1;refGene_gn=ADAMTSL1;refGene_tx=XM_023242342.1;refGene_gn=ADAMTSL1;refGene_tx=XM_023242343.1;refGene_gn=ADAMTSL1;refGene_tx=XM_023242344.1
D4 45455222 45455307
refGene_gn=ADAMTSL1;refGene_tx=XM_011288618.2;refGene_gn=ADAMTSL1;refGene_tx=XM_023242340.1;refGene_gn=ADAMTSL1;refGene_tx=XM_023242341.1;refGene_gn=ADAMTSL1;refGene_tx=XM_023242342.1;refGene_gn=ADAMTSL1;refGene_tx=XM_023242343.1;refGene_gn=ADAMTSL1;refGene_tx=XM_023242344.1
D4 45473045 45473347
refGene_gn=ADAMTSL1;refGene_tx=XM_011288618.2;refGene_gn=ADAMTSL1;refGene_tx=XM_023242340.1;refGene_gn=ADAMTSL1;refGene_tx=XM_023242341.1;refGene_gn=ADAMTSL1;refGene_tx=XM_023242342.1;refGene_gn=ADAMTSL1;refGene_tx=XM_023242343.1;refGene_gn=ADAMTSL1;refGene_tx=XM_023242344.1
D4 45480223 45480353
refGene_gn=ADAMTSL1;refGene_tx=XM_011288618.2;refGene_gn=ADAMTSL1;refGene_tx=XM_023242340.1;refGene_gn=ADAMTSL1;refGene_tx=XM_023242341.1;refGene_gn=ADAMTSL1;refGene_tx=XM_023242342.1;refGene_gn=ADAMTSL1;refGene_tx=XM_023242343.1;refGene_gn=ADAMTSL1;refGene_tx=XM_023242344.1
D4 45512285 45512496
refGene_gn=ADAMTSL1;refGene_tx=XM_011288618.2;refGene_gn=ADAMTSL1;refGene_tx=XM_023242340.1;refGene_gn=ADAMTSL1;refGene_tx=XM_023242341.1;refGene_gn=ADAMTSL1;refGene_tx=XM_023242342.1;refGene_gn=ADAMTSL1;refGene_tx=XM_023242343.1;refGene_gn=ADAMTSL1;refGene_tx=XM_023242344.1
D4 45525586 45525766
refGene_gn=ADAMTSL1;refGene_tx=XM_011288618.2;refGene_gn=ADAMTSL1;refGene_tx=XM_023242340.1;refGene_gn=ADAMTSL1;refGene_tx=XM_023242341.1;refGene_gn=ADAMTSL1;refGene_tx=XM_023242342.1;refGene_gn=ADAMTSL1;refGene_tx=XM_023242343.1;refGene_gn=ADAMTSL1;refGene_tx=XM_023242344.1
D4 45531229 45531383
refGene_gn=ADAMTSL1;refGene_tx=XM_011288618.2;refGene_gn=ADAMTSL1;refGene_tx=XM_023242340.1;refGene_gn=ADAMTSL1;refGene_tx=XM_023242341.1;refGene_gn=ADAMTSL1;refGene_tx=XM_023242342.1;refGene_gn=ADAMTSL1;refGene_tx=XM_023242343.1;refGene_gn=ADAMTSL1;refGene_tx=XM_023242344.1
D4 45532344 45533470
refGene_gn=ADAMTSL1;refGene_tx=XM_011288618.2;refGene_gn=ADAMTSL1;refGene_tx=XM_023242340.1;refGene_gn=ADAMTSL1;refGene_tx=XM_023242341.1;refGene_gn=ADAMTSL1;refGene_tx=XM_023242342.1;refGene_gn=ADAMTSL1;refGene_tx=XM_023242343.1;refGene_gn=ADAMTSL1;refGene_tx=XM_023242344.1
D4 45553433 45553561
refGene_gn=ADAMTSL1;refGene_tx=XM_011288618.2;refGene_gn=ADAMTSL1;refGene_tx=XM_023242340.1;refGene_gn=ADAMTSL1;refGene_tx=XM_023242341.1;refGene_gn=ADAMTSL1;refGene_tx=XM_023242342.1;refGene_gn=ADAMTSL1;refGene_tx=XM_023242343.1;refGene_gn=ADAMTSL1;refGene_tx=XM_023242344.1
D4 45571038 45571167
refGene_gn=ADAMTSL1;refGene_tx=XM_011288618.2;refGene_gn=ADAMTSL1;refG

ene_tx=XM_023242340.1;refGene_gn=ADAMTSL1;refGene_tx=XM_023242341.1;refGene_gn=ADAMTSL1;refGene_tx=XM_023242342.1;refGene_gn=ADAMTSL1;refGene_tx=XM_023242343.1;refGene_gn=ADAMTSL1;refGene_tx=XM_023242344.1
D4	45581214	45581394	refGene_gn=ADAMTSL1;refGene_tx=XM_011288618.2;refGene_gn=ADAMTSL1;refGene_tx=XM_023242340.1;refGene_gn=ADAMTSL1;refGene_tx=XM_023242341.1;refGene_gn=ADAMTSL1;refGene_tx=XM_023242342.1;refGene_gn=ADAMTSL1;refGene_tx=XM_023242343.1;refGene_gn=ADAMTSL1;refGene_tx=XM_023242344.1
D4	45585351	45585486	refGene_gn=ADAMTSL1;refGene_tx=XM_011288618.2;refGene_gn=ADAMTSL1;refGene_tx=XM_023242340.1;refGene_gn=ADAMTSL1;refGene_tx=XM_023242341.1;refGene_gn=ADAMTSL1;refGene_tx=XM_023242342.1;refGene_gn=ADAMTSL1;refGene_tx=XM_023242343.1;refGene_gn=ADAMTSL1;refGene_tx=XM_023242344.1
D4	45588075	45588188	refGene_gn=ADAMTSL1;refGene_tx=XM_023242343.1
D4	45601655	45601792	refGene_gn=ADAMTSL1;refGene_tx=XM_023242344.1
D4	45621471	45621684	refGene_gn=ADAMTSL1;refGene_tx=XM_011288618.2;refGene_gn=ADAMTSL1;refGene_tx=XM_023242340.1;refGene_gn=ADAMTSL1;refGene_tx=XM_023242341.1;refGene_gn=ADAMTSL1;refGene_tx=XM_023242342.1;refGene_gn=ADAMTSL1;refGene_tx=XM_023242343.1
D4	45621788	45621805	refGene_gn=ADAMTSL1;refGene_tx=XM_023242342.1
D4	45623294	45623475	refGene_gn=ADAMTSL1;refGene_tx=XM_011288618.2;refGene_gn=ADAMTSL1;refGene_tx=XM_023242340.1;refGene_gn=ADAMTSL1;refGene_tx=XM_023242341.1
D4	45626425	45626633	refGene_gn=ADAMTSL1;refGene_tx=XM_011288618.2;refGene_gn=ADAMTSL1;refGene_tx=XM_023242340.1;refGene_gn=ADAMTSL1;refGene_tx=XM_023242341.1
D4	45650815	45650925	refGene_gn=ADAMTSL1;refGene_tx=XM_011288618.2;refGene_gn=ADAMTSL1;refGene_tx=XM_023242340.1;refGene_gn=ADAMTSL1;refGene_tx=XM_023242341.1
D4	45651689	45651910	refGene_gn=ADAMTSL1;refGene_tx=XM_011288618.2;refGene_gn=ADAMTSL1;refGene_tx=XM_023242340.1;refGene_gn=ADAMTSL1;refGene_tx=XM_023242341.1
D4	45653411	45653518	refGene_gn=ADAMTSL1;refGene_tx=XM_011288618.2;refGene_gn=ADAMTSL1;refGene_tx=XM_023242340.1;refGene_gn=ADAMTSL1;refGene_tx=XM_023242341.1
D4	45676024	45677028	refGene_gn=SAXO1;refGene_tx=XM_006939172.4
D4	45690504	45690707	refGene_gn=SAXO1;refGene_tx=XM_006939172.4
D4	45700259	45700439	refGene_gn=SAXO1;refGene_tx=XM_006939172.4
D4	45774257	45774304	refGene_gn=SAXO1;refGene_tx=XM_006939172.4
D4	45792968	45793910	refGene_gn=RRAGA;refGene_tx=XM_003995492.5
D4	45799696	45799749	refGene_gn=HAUS6;refGene_tx=XM_011288616.3
D4	45801310	45802360	refGene_gn=HAUS6;refGene_tx=XM_011288616.3
D4	45803754	45803890	refGene_gn=HAUS6;refGene_tx=XM_011288616.3
D4	45805263	45805449	refGene_gn=HAUS6;refGene_tx=XM_011288616.3
D4	45805662	45805729	refGene_gn=HAUS6;refGene_tx=XM_011288616.3
D4	45812534	45812616	refGene_gn=HAUS6;refGene_tx=XM_011288616.3
D4	45814364	45814467	refGene_gn=HAUS6;refGene_tx=XM_011288616.3
D4	45814940	45815067	refGene_gn=HAUS6;refGene_tx=XM_011288616.3
D4	45815686	45815880	refGene_gn=HAUS6;refGene_tx=XM_011288616.3

D4 45817966 45818137 refGene_gn=HAUS6;refGene_tx=XM_011288616.3
D4 45821446 45821495 refGene_gn=HAUS6;refGene_tx=XM_011288616.3
D4 45822038 45822104 refGene_gn=HAUS6;refGene_tx=XM_011288616.3
D4 45824642 45824790 refGene_gn=HAUS6;refGene_tx=XM_011288616.3
D4 45828302 45828435 refGene_gn=HAUS6;refGene_tx=XM_011288616.3
D4 45828959 45829038 refGene_gn=HAUS6;refGene_tx=XM_011288616.3
D4 45830726 45830822 refGene_gn=HAUS6;refGene_tx=XM_011288616.3
D4 45835425 45835553 refGene_gn=HAUS6;refGene_tx=XM_011288616.3
D4 45840303 45840444 refGene_gn=PLIN2;refGene_tx=XM_023242345.1
D4 45848189 45848588
refGene_gn=PLIN2;refGene_tx=XM_023242345.1;refGene_gn=PLIN2;refGene_tx=XM_023242346.1
D4 45849276 45849411
refGene_gn=PLIN2;refGene_tx=XM_023242345.1;refGene_gn=PLIN2;refGene_tx=XM_023242346.1
D4 45850537 45850719
refGene_gn=PLIN2;refGene_tx=XM_023242345.1;refGene_gn=PLIN2;refGene_tx=XM_023242346.1
D4 45851261 45851547
refGene_gn=PLIN2;refGene_tx=XM_023242345.1;refGene_gn=PLIN2;refGene_tx=XM_023242346.1
D4 45852611 45852694
refGene_gn=PLIN2;refGene_tx=XM_023242345.1;refGene_gn=PLIN2;refGene_tx=XM_023242346.1
D4 45854746 45854942
refGene_gn=PLIN2;refGene_tx=XM_023242345.1;refGene_gn=PLIN2;refGene_tx=XM_023242346.1
D4 45855026 45855056
refGene_gn=PLIN2;refGene_tx=XM_023242345.1;refGene_gn=PLIN2;refGene_tx=XM_023242346.1
D4 45967439 45967744
refGene_gn=DENND4C;refGene_tx=XM_003995496.5;refGene_gn=DENND4C;refGene_tx=XM_019816032.2;refGene_gn=DENND4C;refGene_tx=XM_019816033.2;refGene_gn=DENND4C;refGene_tx=XM_019816034.2
D4 45977895 45978148
refGene_gn=DENND4C;refGene_tx=XM_003995496.5;refGene_gn=DENND4C;refGene_tx=XM_019816032.2;refGene_gn=DENND4C;refGene_tx=XM_019816033.2;refGene_gn=DENND4C;refGene_tx=XM_019816034.2
D4 45979265 45979335
refGene_gn=DENND4C;refGene_tx=XM_003995496.5;refGene_gn=DENND4C;refGene_tx=XM_019816032.2;refGene_gn=DENND4C;refGene_tx=XM_019816033.2;refGene_gn=DENND4C;refGene_tx=XM_019816034.2
D4 45981396 45981566
refGene_gn=DENND4C;refGene_tx=XM_003995496.5;refGene_gn=DENND4C;refGene_tx=XM_019816032.2;refGene_gn=DENND4C;refGene_tx=XM_019816033.2;refGene_gn=DENND4C;refGene_tx=XM_019816034.2;refGene_gn=DENND4C;refGene_tx=XM_019816035.2
D4 45991854 45992093
refGene_gn=DENND4C;refGene_tx=XM_003995496.5;refGene_gn=DENND4C;refGene_tx=XM_019816032.2;refGene_gn=DENND4C;refGene_tx=XM_019816033.2;refGe

ne_gn=DENND4C;refGene_tx=XM_019816034.2;refGene_gn=DENND4C;refGene_tx=XM_019816035.2
D4 45993795 45993862 refGene_gn=DENND4C;refGene_tx=XM_003995496.5;refGene_gn=DENND4C;refGene_tx=XM_019816032.2;refGene_gn=DENND4C;refGene_tx=XM_019816033.2;refGene_gn=DENND4C;refGene_tx=XM_019816034.2;refGene_gn=DENND4C;refGene_tx=XM_019816035.2
D4 45994935 45994994 refGene_gn=DENND4C;refGene_tx=XM_003995496.5;refGene_gn=DENND4C;refGene_tx=XM_019816032.2;refGene_gn=DENND4C;refGene_tx=XM_019816033.2;refGene_gn=DENND4C;refGene_tx=XM_019816034.2;refGene_gn=DENND4C;refGene_tx=XM_019816035.2
D4 45995820 45995965 refGene_gn=DENND4C;refGene_tx=XM_003995496.5;refGene_gn=DENND4C;refGene_tx=XM_019816032.2;refGene_gn=DENND4C;refGene_tx=XM_019816033.2;refGene_gn=DENND4C;refGene_tx=XM_019816034.2;refGene_gn=DENND4C;refGene_tx=XM_019816035.2
D4 45999936 46000112 refGene_gn=DENND4C;refGene_tx=XM_003995496.5;refGene_gn=DENND4C;refGene_tx=XM_019816032.2;refGene_gn=DENND4C;refGene_tx=XM_019816033.2;refGene_gn=DENND4C;refGene_tx=XM_019816034.2;refGene_gn=DENND4C;refGene_tx=XM_019816035.2
D4 46003616 46003723 refGene_gn=DENND4C;refGene_tx=XM_003995496.5;refGene_gn=DENND4C;refGene_tx=XM_019816032.2;refGene_gn=DENND4C;refGene_tx=XM_019816033.2;refGene_gn=DENND4C;refGene_tx=XM_019816034.2;refGene_gn=DENND4C;refGene_tx=XM_019816035.2
D4 46004040 46004259 refGene_gn=DENND4C;refGene_tx=XM_003995496.5;refGene_gn=DENND4C;refGene_tx=XM_019816032.2;refGene_gn=DENND4C;refGene_tx=XM_019816033.2;refGene_gn=DENND4C;refGene_tx=XM_019816034.2;refGene_gn=DENND4C;refGene_tx=XM_019816035.2
D4 46010555 46010701 refGene_gn=DENND4C;refGene_tx=XM_003995496.5;refGene_gn=DENND4C;refGene_tx=XM_019816032.2;refGene_gn=DENND4C;refGene_tx=XM_019816033.2;refGene_gn=DENND4C;refGene_tx=XM_019816034.2;refGene_gn=DENND4C;refGene_tx=XM_019816035.2
D4 46012072 46012108 refGene_gn=DENND4C;refGene_tx=XM_003995496.5;refGene_gn=DENND4C;refGene_tx=XM_019816032.2;refGene_gn=DENND4C;refGene_tx=XM_019816033.2;refGene_gn=DENND4C;refGene_tx=XM_019816034.2;refGene_gn=DENND4C;refGene_tx=XM_019816035.2
D4 46012188 46012319 refGene_gn=DENND4C;refGene_tx=XM_003995496.5;refGene_gn=DENND4C;refGene_tx=XM_019816032.2;refGene_gn=DENND4C;refGene_tx=XM_019816033.2;refGene_gn=DENND4C;refGene_tx=XM_019816034.2;refGene_gn=DENND4C;refGene_tx=XM_019816035.2
D4 46014346 46014479 refGene_gn=DENND4C;refGene_tx=XM_003995496.5;refGene_gn=DENND4C;refGene_tx=XM_019816032.2;refGene_gn=DENND4C;refGene_tx=XM_019816033.2;refGe

ne_gn=DENND4C;refGene_tx=XM_019816034.2;refGene_gn=DENND4C;refGene_tx=XM_019816035.2
D4 46016885 46017092
refGene_gn=DENND4C;refGene_tx=XM_003995496.5;refGene_gn=DENND4C;refGene_tx=XM_019816032.2;refGene_gn=DENND4C;refGene_tx=XM_019816033.2;refGene_gn=DENND4C;refGene_tx=XM_019816034.2;refGene_gn=DENND4C;refGene_tx=XM_019816035.2
D4 46021780 46021909
refGene_gn=DENND4C;refGene_tx=XM_003995496.5;refGene_gn=DENND4C;refGene_tx=XM_019816032.2;refGene_gn=DENND4C;refGene_tx=XM_019816033.2;refGene_gn=DENND4C;refGene_tx=XM_019816034.2;refGene_gn=DENND4C;refGene_tx=XM_019816035.2
D4 46022103 46022248
refGene_gn=DENND4C;refGene_tx=XM_003995496.5;refGene_gn=DENND4C;refGene_tx=XM_019816032.2;refGene_gn=DENND4C;refGene_tx=XM_019816033.2;refGene_gn=DENND4C;refGene_tx=XM_019816034.2;refGene_gn=DENND4C;refGene_tx=XM_019816035.2
D4 46022513 46022660
refGene_gn=DENND4C;refGene_tx=XM_019816032.2;refGene_gn=DENND4C;refGene_tx=XM_019816033.2;refGene_gn=DENND4C;refGene_tx=XM_019816035.2
D4 46027895 46028018
refGene_gn=DENND4C;refGene_tx=XM_003995496.5;refGene_gn=DENND4C;refGene_tx=XM_019816032.2;refGene_gn=DENND4C;refGene_tx=XM_019816033.2;refGene_gn=DENND4C;refGene_tx=XM_019816034.2;refGene_gn=DENND4C;refGene_tx=XM_019816035.2
D4 46029032 46029179
refGene_gn=DENND4C;refGene_tx=XM_003995496.5;refGene_gn=DENND4C;refGene_tx=XM_019816032.2;refGene_gn=DENND4C;refGene_tx=XM_019816033.2;refGene_gn=DENND4C;refGene_tx=XM_019816034.2;refGene_gn=DENND4C;refGene_tx=XM_019816035.2
D4 46030696 46031862
refGene_gn=DENND4C;refGene_tx=XM_003995496.5;refGene_gn=DENND4C;refGene_tx=XM_019816032.2;refGene_gn=DENND4C;refGene_tx=XM_019816033.2;refGene_gn=DENND4C;refGene_tx=XM_019816034.2;refGene_gn=DENND4C;refGene_tx=XM_019816035.2
D4 46035307 46035485
refGene_gn=DENND4C;refGene_tx=XM_003995496.5;refGene_gn=DENND4C;refGene_tx=XM_019816032.2;refGene_gn=DENND4C;refGene_tx=XM_019816033.2;refGene_gn=DENND4C;refGene_tx=XM_019816034.2;refGene_gn=DENND4C;refGene_tx=XM_019816035.2
D4 46036223 46036333
refGene_gn=DENND4C;refGene_tx=XM_003995496.5;refGene_gn=DENND4C;refGene_tx=XM_019816032.2;refGene_gn=DENND4C;refGene_tx=XM_019816033.2;refGene_gn=DENND4C;refGene_tx=XM_019816034.2;refGene_gn=DENND4C;refGene_tx=XM_019816035.2
D4 46036630 46036806
refGene_gn=DENND4C;refGene_tx=XM_003995496.5;refGene_gn=DENND4C;refGene_tx=XM_019816032.2;refGene_gn=DENND4C;refGene_tx=XM_019816033.2;refGene_gn=DENND4C;refGene_tx=XM_019816034.2;refGene_gn=DENND4C;refGene_tx=XM_019816035.2

D4 46040577 46040760
refGene_gn=DENND4C;refGene_tx=XM_003995496.5;refGene_gn=DENND4C;refGene_tx=XM_019816032.2;refGene_gn=DENND4C;refGene_tx=XM_019816033.2;refGene_gn=DENND4C;refGene_tx=XM_019816034.2;refGene_gn=DENND4C;refGene_tx=XM_019816035.2
D4 46041691 46041887
refGene_gn=DENND4C;refGene_tx=XM_003995496.5;refGene_gn=DENND4C;refGene_tx=XM_019816032.2;refGene_gn=DENND4C;refGene_tx=XM_019816033.2;refGene_gn=DENND4C;refGene_tx=XM_019816034.2;refGene_gn=DENND4C;refGene_tx=XM_019816035.2
D4 46044620 46044866
refGene_gn=DENND4C;refGene_tx=XM_003995496.5;refGene_gn=DENND4C;refGene_tx=XM_019816032.2;refGene_gn=DENND4C;refGene_tx=XM_019816033.2;refGene_gn=DENND4C;refGene_tx=XM_019816034.2;refGene_gn=DENND4C;refGene_tx=XM_019816035.2
D4 46045911 46046029
refGene_gn=DENND4C;refGene_tx=XM_003995496.5;refGene_gn=DENND4C;refGene_tx=XM_019816032.2;refGene_gn=DENND4C;refGene_tx=XM_019816033.2;refGene_gn=DENND4C;refGene_tx=XM_019816034.2;refGene_gn=DENND4C;refGene_tx=XM_019816035.2
D4 46052346 46052497
refGene_gn=DENND4C;refGene_tx=XM_003995496.5;refGene_gn=DENND4C;refGene_tx=XM_019816032.2;refGene_gn=DENND4C;refGene_tx=XM_019816033.2;refGene_gn=DENND4C;refGene_tx=XM_019816034.2;refGene_gn=DENND4C;refGene_tx=XM_019816035.2
D4 46054289 46054354
refGene_gn=DENND4C;refGene_tx=XM_003995496.5;refGene_gn=DENND4C;refGene_tx=XM_019816032.2;refGene_gn=DENND4C;refGene_tx=XM_019816033.2;refGene_gn=DENND4C;refGene_tx=XM_019816034.2;refGene_gn=DENND4C;refGene_tx=XM_019816035.2
D4 46054571 46054708
refGene_gn=DENND4C;refGene_tx=XM_003995496.5;refGene_gn=DENND4C;refGene_tx=XM_019816032.2;refGene_gn=DENND4C;refGene_tx=XM_019816033.2;refGene_gn=DENND4C;refGene_tx=XM_019816034.2;refGene_gn=DENND4C;refGene_tx=XM_019816035.2
D4 46058188 46058284 refGene_gn=RPS6;refGene_tx=XM_003995497.3
D4 46058378 46058536 refGene_gn=RPS6;refGene_tx=XM_003995497.3
D4 46060815 46060962 refGene_gn=RPS6;refGene_tx=XM_003995497.3
D4 46061157 46061368 refGene_gn=RPS6;refGene_tx=XM_003995497.3
D4 46061883 46062015 refGene_gn=RPS6;refGene_tx=XM_003995497.3
D4 46062546 46062750 refGene_gn=RPS6;refGene_tx=XM_003995497.3
D4 46088871 46088979 refGene_gn=ACER2;refGene_tx=XM_003995498.4
D4 46098940 46099055 refGene_gn=ACER2;refGene_tx=XM_003995498.4
D4 46099796 46099938 refGene_gn=ACER2;refGene_tx=XM_003995498.4
D4 46107719 46107857 refGene_gn=ACER2;refGene_tx=XM_003995498.4
D4 46114973 46115111 refGene_gn=ACER2;refGene_tx=XM_003995498.4
D4 46118862 46119049 refGene_gn=ACER2;refGene_tx=XM_003995498.4
D4 46178366 46178636
refGene_gn=SLC24A2;refGene_tx=XM_019816036.1;refGene_gn=SLC24A2;refGene_tx=XM_023242348.1;refGene_gn=SLC24A2;refGene_tx=XM_023242349.1;refGe

ne_gn=SLC24A2;refGene_tx=XM_023242350.1;refGene_gn=SLC24A2;refGene_tx=XM_023242351.1
D4 46182815 46182982
refGene_gn=SLC24A2;refGene_tx=XM_019816036.1;refGene_gn=SLC24A2;refGene_tx=XM_023242348.1;refGene_gn=SLC24A2;refGene_tx=XM_023242349.1;refGene_gn=SLC24A2;refGene_tx=XM_023242350.1
D4 46189386 46189476
refGene_gn=SLC24A2;refGene_tx=XM_019816036.1;refGene_gn=SLC24A2;refGene_tx=XM_023242348.1;refGene_gn=SLC24A2;refGene_tx=XM_023242349.1;refGene_gn=SLC24A2;refGene_tx=XM_023242350.1;refGene_gn=SLC24A2;refGene_tx=XM_023242351.1
D4 46198683 46198821
refGene_gn=SLC24A2;refGene_tx=XM_019816036.1;refGene_gn=SLC24A2;refGene_tx=XM_023242348.1;refGene_gn=SLC24A2;refGene_tx=XM_023242349.1;refGene_gn=SLC24A2;refGene_tx=XM_023242350.1;refGene_gn=SLC24A2;refGene_tx=XM_023242351.1
D4 46217900 46218019
refGene_gn=SLC24A2;refGene_tx=XM_019816036.1;refGene_gn=SLC24A2;refGene_tx=XM_023242348.1;refGene_gn=SLC24A2;refGene_tx=XM_023242349.1;refGene_gn=SLC24A2;refGene_tx=XM_023242350.1;refGene_gn=SLC24A2;refGene_tx=XM_023242351.1
D4 46221596 46221695
refGene_gn=SLC24A2;refGene_tx=XM_019816036.1;refGene_gn=SLC24A2;refGene_tx=XM_023242348.1;refGene_gn=SLC24A2;refGene_tx=XM_023242349.1;refGene_gn=SLC24A2;refGene_tx=XM_023242350.1;refGene_gn=SLC24A2;refGene_tx=XM_023242351.1
D4 46241208 46241259
refGene_gn=SLC24A2;refGene_tx=XM_019816036.1;refGene_gn=SLC24A2;refGene_tx=XM_023242348.1;refGene_gn=SLC24A2;refGene_tx=XM_023242349.1;refGene_gn=SLC24A2;refGene_tx=XM_023242351.1
D4 46266651 46266760
refGene_gn=SLC24A2;refGene_tx=XM_019816036.1;refGene_gn=SLC24A2;refGene_tx=XM_023242348.1;refGene_gn=SLC24A2;refGene_tx=XM_023242349.1;refGene_gn=SLC24A2;refGene_tx=XM_023242350.1;refGene_gn=SLC24A2;refGene_tx=XM_023242351.1
D4 46269353 46269392
refGene_gn=SLC24A2;refGene_tx=XM_019816036.1;refGene_gn=SLC24A2;refGene_tx=XM_023242348.1;refGene_gn=SLC24A2;refGene_tx=XM_023242349.1;refGene_gn=SLC24A2;refGene_tx=XM_023242350.1;refGene_gn=SLC24A2;refGene_tx=XM_023242351.1
D4 46418247 46419177
refGene_gn=SLC24A2;refGene_tx=XM_019816036.1;refGene_gn=SLC24A2;refGene_tx=XM_023242348.1;refGene_gn=SLC24A2;refGene_tx=XM_023242349.1;refGene_gn=SLC24A2;refGene_tx=XM_023242350.1;refGene_gn=SLC24A2;refGene_tx=XM_023242351.1
D4 46846656 46846788 refGene_gn=MLLT3;refGene_tx=XM_023242352.1
D4 46853567 46853639 refGene_gn=MLLT3;refGene_tx=XM_023242352.1
D4 46854808 46854880 refGene_gn=MLLT3;refGene_tx=XM_023242352.1
D4 46861017 46861117 refGene_gn=MLLT3;refGene_tx=XM_023242352.1
D4 46863742 46863872 refGene_gn=MLLT3;refGene_tx=XM_023242352.1

D4 46865739 46865815 refGene_gn=MLLT3;refGene_tx=XM_023242352.1
D4 46921213 46921918 refGene_gn=MLLT3;refGene_tx=XM_023242352.1
D4 46965162 46965306 refGene_gn=MLLT3;refGene_tx=XM_023242352.1
D4 46975819 46975864 refGene_gn=MLLT3;refGene_tx=XM_023242352.1
D4 47233316 47233373
refGene_gn=FOCAD;refGene_tx=XM_023242353.1;refGene_gn=FOCAD;refGene_tx=XM_023242354.1
D4 47236582 47236657 refGene_gn=FOCAD;refGene_tx=XM_023242353.1
D4 47238959 47239114 refGene_gn=FOCAD;refGene_tx=XM_023242353.1
D4 47260898 47261003
refGene_gn=FOCAD;refGene_tx=XM_023242353.1;refGene_gn=FOCAD;refGene_tx=XM_023242354.1;refGene_gn=FOCAD;refGene_tx=XM_023242355.1
D4 47280233 47280335
refGene_gn=FOCAD;refGene_tx=XM_023242353.1;refGene_gn=FOCAD;refGene_tx=XM_023242354.1;refGene_gn=FOCAD;refGene_tx=XM_023242355.1
D4 47284985 47285190
refGene_gn=FOCAD;refGene_tx=XM_023242353.1;refGene_gn=FOCAD;refGene_tx=XM_023242354.1;refGene_gn=FOCAD;refGene_tx=XM_023242355.1
D4 47289947 47290154
refGene_gn=FOCAD;refGene_tx=XM_023242353.1;refGene_gn=FOCAD;refGene_tx=XM_023242354.1;refGene_gn=FOCAD;refGene_tx=XM_023242355.1
D4 47295751 47295839
refGene_gn=FOCAD;refGene_tx=XM_023242353.1;refGene_gn=FOCAD;refGene_tx=XM_023242354.1;refGene_gn=FOCAD;refGene_tx=XM_023242355.1
D4 47301195 47301398
refGene_gn=FOCAD;refGene_tx=XM_023242353.1;refGene_gn=FOCAD;refGene_tx=XM_023242354.1;refGene_gn=FOCAD;refGene_tx=XM_023242355.1
D4 47310314 47310572
refGene_gn=FOCAD;refGene_tx=XM_023242353.1;refGene_gn=FOCAD;refGene_tx=XM_023242354.1;refGene_gn=FOCAD;refGene_tx=XM_023242355.1
D4 47353150 47353255
refGene_gn=FOCAD;refGene_tx=XM_023242353.1;refGene_gn=FOCAD;refGene_tx=XM_023242354.1;refGene_gn=FOCAD;refGene_tx=XM_023242355.1
D4 47354116 47354218
refGene_gn=FOCAD;refGene_tx=XM_023242353.1;refGene_gn=FOCAD;refGene_tx=XM_023242354.1;refGene_gn=FOCAD;refGene_tx=XM_023242355.1
D4 47354740 47354871
refGene_gn=FOCAD;refGene_tx=XM_023242353.1;refGene_gn=FOCAD;refGene_tx=XM_023242354.1;refGene_gn=FOCAD;refGene_tx=XM_023242355.1
D4 47357013 47357140
refGene_gn=FOCAD;refGene_tx=XM_023242353.1;refGene_gn=FOCAD;refGene_tx=XM_023242354.1;refGene_gn=FOCAD;refGene_tx=XM_023242355.1
D4 47393800 47393935
refGene_gn=FOCAD;refGene_tx=XM_023242353.1;refGene_gn=FOCAD;refGene_tx=XM_023242354.1;refGene_gn=FOCAD;refGene_tx=XM_023242355.1
D4 47400510 47400561
refGene_gn=FOCAD;refGene_tx=XM_023242353.1;refGene_gn=FOCAD;refGene_tx=XM_023242354.1;refGene_gn=FOCAD;refGene_tx=XM_023242355.1
D4 47401583 47401667
refGene_gn=FOCAD;refGene_tx=XM_023242353.1;refGene_gn=FOCAD;refGene_tx

=XM_023242354.1;refGene_gn=FOCAD;refGene_tx=XM_023242355.1
D4 47408875 47409014
refGene_gn=FOCAD;refGene_tx=XM_023242353.1;refGene_gn=FOCAD;refGene_tx=XM_023242354.1;refGene_gn=FOCAD;refGene_tx=XM_023242355.1
D4 47413203 47413389
refGene_gn=FOCAD;refGene_tx=XM_023242353.1;refGene_gn=FOCAD;refGene_tx=XM_023242354.1;refGene_gn=FOCAD;refGene_tx=XM_023242355.1
D4 47422921 47423043
refGene_gn=FOCAD;refGene_tx=XM_023242353.1;refGene_gn=FOCAD;refGene_tx=XM_023242354.1;refGene_gn=FOCAD;refGene_tx=XM_023242355.1
D4 47440374 47440467
refGene_gn=FOCAD;refGene_tx=XM_023242353.1;refGene_gn=FOCAD;refGene_tx=XM_023242354.1;refGene_gn=FOCAD;refGene_tx=XM_023242355.1
D4 47446797 47446886
refGene_gn=FOCAD;refGene_tx=XM_023242353.1;refGene_gn=FOCAD;refGene_tx=XM_023242354.1;refGene_gn=FOCAD;refGene_tx=XM_023242355.1
D4 47449555 47449600
refGene_gn=FOCAD;refGene_tx=XM_023242353.1;refGene_gn=FOCAD;refGene_tx=XM_023242354.1;refGene_gn=FOCAD;refGene_tx=XM_023242355.1
D4 47454944 47455053
refGene_gn=FOCAD;refGene_tx=XM_023242353.1;refGene_gn=FOCAD;refGene_tx=XM_023242354.1;refGene_gn=FOCAD;refGene_tx=XM_023242355.1
D4 47457663 47457780
refGene_gn=FOCAD;refGene_tx=XM_023242353.1;refGene_gn=FOCAD;refGene_tx=XM_023242354.1;refGene_gn=FOCAD;refGene_tx=XM_023242355.1
D4 47460725 47460964
refGene_gn=FOCAD;refGene_tx=XM_023242353.1;refGene_gn=FOCAD;refGene_tx=XM_023242354.1;refGene_gn=FOCAD;refGene_tx=XM_023242355.1
D4 47464089 47464179
refGene_gn=FOCAD;refGene_tx=XM_023242353.1;refGene_gn=FOCAD;refGene_tx=XM_023242354.1;refGene_gn=FOCAD;refGene_tx=XM_023242355.1
D4 47470186 47470334
refGene_gn=FOCAD;refGene_tx=XM_023242353.1;refGene_gn=FOCAD;refGene_tx=XM_023242354.1;refGene_gn=FOCAD;refGene_tx=XM_023242355.1
D4 47472521 47472641
refGene_gn=FOCAD;refGene_tx=XM_023242353.1;refGene_gn=FOCAD;refGene_tx=XM_023242354.1;refGene_gn=FOCAD;refGene_tx=XM_023242355.1
D4 47474271 47474394
refGene_gn=FOCAD;refGene_tx=XM_023242353.1;refGene_gn=FOCAD;refGene_tx=XM_023242354.1;refGene_gn=FOCAD;refGene_tx=XM_023242355.1
D4 47474864 47474942
refGene_gn=FOCAD;refGene_tx=XM_023242353.1;refGene_gn=FOCAD;refGene_tx=XM_023242354.1;refGene_gn=FOCAD;refGene_tx=XM_023242355.1
D4 47475569 47475641
refGene_gn=FOCAD;refGene_tx=XM_023242353.1;refGene_gn=FOCAD;refGene_tx=XM_023242354.1;refGene_gn=FOCAD;refGene_tx=XM_023242355.1
D4 47477681 47477784
refGene_gn=FOCAD;refGene_tx=XM_023242353.1;refGene_gn=FOCAD;refGene_tx=XM_023242354.1;refGene_gn=FOCAD;refGene_tx=XM_023242355.1
D4 47479920 47480001

refGene_gn=FOCAD;refGene_tx=XM_023242353.1;refGene_gn=FOCAD;refGene_tx=XM_023242354.1;refGene_gn=FOCAD;refGene_tx=XM_023242355.1
D4 47501761 47501890
refGene_gn=FOCAD;refGene_tx=XM_023242353.1;refGene_gn=FOCAD;refGene_tx=XM_023242354.1;refGene_gn=FOCAD;refGene_tx=XM_023242355.1
D4 47504499 47504615
refGene_gn=FOCAD;refGene_tx=XM_023242353.1;refGene_gn=FOCAD;refGene_tx=XM_023242354.1;refGene_gn=FOCAD;refGene_tx=XM_023242355.1
D4 47508453 47508714
refGene_gn=FOCAD;refGene_tx=XM_023242353.1;refGene_gn=FOCAD;refGene_tx=XM_023242354.1;refGene_gn=FOCAD;refGene_tx=XM_023242355.1
D4 47509555 47509645
refGene_gn=FOCAD;refGene_tx=XM_023242353.1;refGene_gn=FOCAD;refGene_tx=XM_023242354.1;refGene_gn=FOCAD;refGene_tx=XM_023242355.1
D4 47512874 47513052
refGene_gn=FOCAD;refGene_tx=XM_023242353.1;refGene_gn=FOCAD;refGene_tx=XM_023242354.1;refGene_gn=FOCAD;refGene_tx=XM_023242355.1
D4 47514683 47514781
refGene_gn=FOCAD;refGene_tx=XM_023242353.1;refGene_gn=FOCAD;refGene_tx=XM_023242354.1;refGene_gn=FOCAD;refGene_tx=XM_023242355.1
D4 47522107 47522359
refGene_gn=FOCAD;refGene_tx=XM_023242353.1;refGene_gn=FOCAD;refGene_tx=XM_023242354.1;refGene_gn=FOCAD;refGene_tx=XM_023242355.1
D4 47522587 47522688
refGene_gn=HACD4;refGene_tx=XM_019816041.2;refGene_gn=HACD4;refGene_tx=XM_019816042.1;refGene_gn=HACD4;refGene_tx=XM_019816043.2;refGene_gn=HACD4;refGene_tx=XM_023242356.1;refGene_gn=HACD4;refGene_tx=XM_023242359.1;refGene_gn=HACD4;refGene_tx=XM_023242360.1;refGene_gn=HACD4;refGene_tx=XM_023242361.1
D4 47524387 47524463
refGene_gn=FOCAD;refGene_tx=XM_023242353.1;refGene_gn=FOCAD;refGene_tx=XM_023242354.1;refGene_gn=FOCAD;refGene_tx=XM_023242355.1
D4 47526712 47526786
refGene_gn=FOCAD;refGene_tx=XM_023242353.1;refGene_gn=FOCAD;refGene_tx=XM_023242354.1;refGene_gn=FOCAD;refGene_tx=XM_023242355.1
D4 47536387 47536470 refGene_gn=HACD4;refGene_tx=XM_023242358.1
D4 47537540 47537666
refGene_gn=HACD4;refGene_tx=XM_019816041.2;refGene_gn=HACD4;refGene_tx=XM_019816042.1;refGene_gn=HACD4;refGene_tx=XM_019816043.2;refGene_gn=HACD4;refGene_tx=XM_023242356.1;refGene_gn=HACD4;refGene_tx=XM_023242358.1;refGene_gn=HACD4;refGene_tx=XM_023242359.1;refGene_gn=HACD4;refGene_tx=XM_023242360.1;refGene_gn=HACD4;refGene_tx=XM_023242361.1
D4 47549218 47549325
refGene_gn=HACD4;refGene_tx=XM_019816041.2;refGene_gn=HACD4;refGene_tx=XM_019816042.1;refGene_gn=HACD4;refGene_tx=XM_019816043.2;refGene_gn=HACD4;refGene_tx=XM_023242356.1;refGene_gn=HACD4;refGene_tx=XM_023242358.1;refGene_gn=HACD4;refGene_tx=XM_023242359.1;refGene_gn=HACD4;refGene_tx=XM_023242360.1;refGene_gn=HACD4;refGene_tx=XM_023242361.1
D4 47554083 47554196
refGene_gn=HACD4;refGene_tx=XM_019816041.2;refGene_gn=HACD4;refGene_tx

=XM_019816042.1;refGene_gn=HACD4;refGene_tx=XM_019816043.2;refGene_gn=HACD4;refGene_tx=XM_023242356.1;refGene_gn=HACD4;refGene_tx=XM_023242358.1;refGene_gn=HACD4;refGene_tx=XM_023242359.1;refGene_gn=HACD4;refGene_tx=XM_023242360.1;refGene_gn=HACD4;refGene_tx=XM_023242361.1
D4 47569458 47569586
refGene_gn=HACD4;refGene_tx=XM_019816041.2;refGene_gn=HACD4;refGene_tx=XM_019816042.1;refGene_gn=HACD4;refGene_tx=XM_019816043.2;refGene_gn=HACD4;refGene_tx=XM_023242356.1;refGene_gn=HACD4;refGene_tx=XM_023242358.1;refGene_gn=HACD4;refGene_tx=XM_023242359.1;refGene_gn=HACD4;refGene_tx=XM_023242360.1;refGene_gn=HACD4;refGene_tx=XM_023242361.1
D4 47572160 47572189 refGene_gn=HACD4;refGene_tx=XM_023242360.1
D4 47572349 47572453
refGene_gn=HACD4;refGene_tx=XM_019816041.2;refGene_gn=HACD4;refGene_tx=XM_023242356.1;refGene_gn=HACD4;refGene_tx=XM_023242358.1;refGene_gn=HACD4;refGene_tx=XM_023242359.1;refGene_gn=HACD4;refGene_tx=XM_023242360.1
D4 47573729 47573758
refGene_gn=HACD4;refGene_tx=XM_023242356.1;refGene_gn=HACD4;refGene_tx=XM_023242358.1;refGene_gn=HACD4;refGene_tx=XM_023242359.1
D4 47624643 47625204 refGene_gn=IFNB1;refGene_tx=NM_001009297.1
D4 47654165 47654717
refGene_gn=LOC101088440;refGene_tx=XM_003995521.1
D4 47659495 47659681
refGene_gn=LOC101088699;refGene_tx=XM_011288694.1
D4 47659682 47660039
refGene_gn=LOC101088699;refGene_tx=XM_011288694.1
D4 47665812 47666016
refGene_gn=LOC101088699;refGene_tx=XM_011288694.1
D4 47677855 47678446 refGene_gn=IFNW4;refGene_tx=NM_001048159.1
D4 47683814 47684381 refGene_gn=IFNA8;refGene_tx=NM_001009193.1
D4 47700704 47700806 refGene_gn=IFNW11;refGene_tx=XM_023243267.1
D4 47700807 47701062 refGene_gn=IFNW11;refGene_tx=XM_023243267.1
D4 47701063 47701133 refGene_gn=IFNW11;refGene_tx=XM_023243267.1
D4 47701134 47701171 refGene_gn=IFNW11;refGene_tx=XM_023243267.1
D4 47701173 47701294 refGene_gn=IFNW11;refGene_tx=XM_023243267.1
D4 47706574 47707093 refGene_gn=IFNA5;refGene_tx=XM_023243268.1
D4 47727699 47728281
refGene_gn=LOC101081371;refGene_tx=XM_004001268.2
D4 47735292 47736033
refGene_gn=LOC111557471;refGene_tx=XM_023242300.1
D4 47742430 47742637
refGene_gn=LOC105259601;refGene_tx=XM_023242362.1
D4 47742638 47743022
refGene_gn=LOC105259601;refGene_tx=XM_023242362.1
D4 47764224 47766078 refGene_gn=KLHL9;refGene_tx=XM_003995514.5
D4 47789806 47790388
refGene_gn=LOC111557596;refGene_tx=XM_023243269.1
D4 47798899 47799484 refGene_gn=IFNA7;refGene_tx=NM_001009197.2
D4 47806520 47807111 refGene_gn=IFNW12;refGene_tx=NM_001089309.1
D4 47829020 47829647 refGene_gn=IFNE;refGene_tx=NM_001278829.1

D4	48103268	48103301
refGene_gn=MTAP;refGene_tx=XM_011288609.3;refGene_gn=MTAP;refGene_tx=XM_011288610.3;refGene_gn=MTAP;refGene_tx=XM_019816044.2
D4	48112586	48112673
refGene_gn=MTAP;refGene_tx=XM_011288609.3;refGene_gn=MTAP;refGene_tx=XM_011288610.3;refGene_gn=MTAP;refGene_tx=XM_019816044.2
D4	48113894	48113953
refGene_gn=MTAP;refGene_tx=XM_011288609.3;refGene_gn=MTAP;refGene_tx=XM_011288610.3;refGene_gn=MTAP;refGene_tx=XM_019816044.2
D4	48117640	48117808
refGene_gn=MTAP;refGene_tx=XM_011288609.3;refGene_gn=MTAP;refGene_tx=XM_011288610.3;refGene_gn=MTAP;refGene_tx=XM_019816044.2
D4	48137805	48137908
refGene_gn=MTAP;refGene_tx=XM_011288609.3;refGene_gn=MTAP;refGene_tx=XM_011288610.3;refGene_gn=MTAP;refGene_tx=XM_019816044.2
D4	48142497	48142737
refGene_gn=MTAP;refGene_tx=XM_011288609.3;refGene_gn=MTAP;refGene_tx=XM_011288610.3;refGene_gn=MTAP;refGene_tx=XM_019816044.2
D4	48147902	48148025
refGene_gn=MTAP;refGene_tx=XM_011288609.3;refGene_gn=MTAP;refGene_tx=XM_011288610.3;refGene_gn=MTAP;refGene_tx=XM_019816044.2
D4	48148738	48148786	refGene_gn=MTAP;refGene_tx=XM_019816044.2
D4	48149442	48149466	refGene_gn=MTAP;refGene_tx=XM_011288610.3
D4	48150580	48150619	refGene_gn=MTAP;refGene_tx=XM_011288609.3
D4	48177668	48177682	refGene_gn=CDKN2A;refGene_tx=NM_001290248.1
D4	48179848	48180155	refGene_gn=CDKN2A;refGene_tx=NM_001290248.1
D4	48183665	48183791	refGene_gn=CDKN2A;refGene_tx=NM_001290248.1
D4	48221547	48221808	refGene_gn=CDKN2B;refGene_tx=XM_003995516.5
D4	48224047	48224203	refGene_gn=CDKN2B;refGene_tx=XM_003995516.5
D4	48534731	48535371	refGene_gn=DMRTA1;refGene_tx=XM_003995524.3
D4	48538474	48538597	refGene_gn=DMRTA1;refGene_tx=XM_003995524.3
D4	48538599	48539321	refGene_gn=DMRTA1;refGene_tx=XM_003995524.3
D4	49515758	49516089
refGene_gn=ELAVL2;refGene_tx=XM_011288604.3;refGene_gn=ELAVL2;refGene_tx=XM_011288606.3;refGene_gn=ELAVL2;refGene_tx=XM_011288607.3;refGene_gn=ELAVL2;refGene_tx=XM_019816046.2;refGene_gn=ELAVL2;refGene_tx=XM_019816047.2;refGene_gn=ELAVL2;refGene_tx=XM_019816049.2;refGene_gn=ELAVL2;refGene_tx=XM_019816051.2;refGene_gn=ELAVL2;refGene_tx=XM_019816052.2;refGene_gn=ELAVL2;refGene_tx=XM_019816053.2;refGene_gn=ELAVL2;refGene_tx=XM_019816054.2;refGene_gn=ELAVL2;refGene_tx=XM_023242363.1;refGene_gn=ELAVL2;refGene_tx=XM_023242364.1;refGene_gn=ELAVL2;refGene_tx=XM_023242365.1
D4	49516710	49516749
refGene_gn=ELAVL2;refGene_tx=XM_011288604.3;refGene_gn=ELAVL2;refGene_tx=XM_011288606.3;refGene_gn=ELAVL2;refGene_tx=XM_011288607.3;refGene_gn=ELAVL2;refGene_tx=XM_019816046.2;refGene_gn=ELAVL2;refGene_tx=XM_019816047.2;refGene_gn=ELAVL2;refGene_tx=XM_019816049.2;refGene_gn=ELAVL2;refGene_tx=XM_019816051.2;refGene_gn=ELAVL2;refGene_tx=XM_019816052.2;refGene_gn=ELAVL2;refGene_tx=XM_019816053.2;refGene_gn=ELAVL2;refGene_tx=XM_019816054.2;refGene_gn=ELAVL2;refGene_tx=XM_023242363.1;refGen

e_gn=ELAVL2;refGene_tx=XM_023242364.1;refGene_gn=ELAVL2;refGene_tx=XM_023242365.1
D4 49524666 49524892
refGene_gn=ELAVL2;refGene_tx=XM_011288604.3;refGene_gn=ELAVL2;refGene_tx=XM_011288606.3;refGene_gn=ELAVL2;refGene_tx=XM_011288607.3;refGene_gn=ELAVL2;refGene_tx=XM_019816046.2;refGene_gn=ELAVL2;refGene_tx=XM_019816047.2;refGene_gn=ELAVL2;refGene_tx=XM_019816049.2;refGene_gn=ELAVL2;refGene_tx=XM_019816051.2;refGene_gn=ELAVL2;refGene_tx=XM_019816052.2;refGene_gn=ELAVL2;refGene_tx=XM_019816053.2;refGene_gn=ELAVL2;refGene_tx=XM_019816054.2;refGene_gn=ELAVL2;refGene_tx=XM_023242363.1;refGene_gn=ELAVL2;refGene_tx=XM_023242364.1;refGene_gn=ELAVL2;refGene_tx=XM_023242365.1
D4 49528468 49528622
refGene_gn=ELAVL2;refGene_tx=XM_011288604.3;refGene_gn=ELAVL2;refGene_tx=XM_011288606.3;refGene_gn=ELAVL2;refGene_tx=XM_011288607.3;refGene_gn=ELAVL2;refGene_tx=XM_019816046.2;refGene_gn=ELAVL2;refGene_tx=XM_019816047.2;refGene_gn=ELAVL2;refGene_tx=XM_019816049.2;refGene_gn=ELAVL2;refGene_tx=XM_019816051.2;refGene_gn=ELAVL2;refGene_tx=XM_019816052.2;refGene_gn=ELAVL2;refGene_tx=XM_019816053.2;refGene_gn=ELAVL2;refGene_tx=XM_019816054.2;refGene_gn=ELAVL2;refGene_tx=XM_023242363.1;refGene_gn=ELAVL2;refGene_tx=XM_023242364.1;refGene_gn=ELAVL2;refGene_tx=XM_023242365.1
D4 49553383 49553487
refGene_gn=ELAVL2;refGene_tx=XM_011288604.3;refGene_gn=ELAVL2;refGene_tx=XM_011288606.3;refGene_gn=ELAVL2;refGene_tx=XM_011288607.3;refGene_gn=ELAVL2;refGene_tx=XM_019816046.2;refGene_gn=ELAVL2;refGene_tx=XM_019816047.2;refGene_gn=ELAVL2;refGene_tx=XM_019816049.2;refGene_gn=ELAVL2;refGene_tx=XM_019816051.2;refGene_gn=ELAVL2;refGene_tx=XM_019816052.2;refGene_gn=ELAVL2;refGene_tx=XM_019816053.2;refGene_gn=ELAVL2;refGene_tx=XM_019816054.2;refGene_gn=ELAVL2;refGene_tx=XM_023242363.1;refGene_gn=ELAVL2;refGene_tx=XM_023242364.1;refGene_gn=ELAVL2;refGene_tx=XM_023242365.1
D4 49579615 49579859
refGene_gn=ELAVL2;refGene_tx=XM_011288604.3;refGene_gn=ELAVL2;refGene_tx=XM_011288606.3;refGene_gn=ELAVL2;refGene_tx=XM_011288607.3;refGene_gn=ELAVL2;refGene_tx=XM_019816046.2;refGene_gn=ELAVL2;refGene_tx=XM_019816047.2;refGene_gn=ELAVL2;refGene_tx=XM_019816049.2;refGene_gn=ELAVL2;refGene_tx=XM_019816051.2;refGene_gn=ELAVL2;refGene_tx=XM_019816052.2;refGene_gn=ELAVL2;refGene_tx=XM_019816053.2;refGene_gn=ELAVL2;refGene_tx=XM_019816054.2;refGene_gn=ELAVL2;refGene_tx=XM_023242363.1;refGene_gn=ELAVL2;refGene_tx=XM_023242364.1;refGene_gn=ELAVL2;refGene_tx=XM_023242365.1
D4 49582684 49582756
refGene_gn=ELAVL2;refGene_tx=XM_011288604.3;refGene_gn=ELAVL2;refGene_tx=XM_011288606.3;refGene_gn=ELAVL2;refGene_tx=XM_019816046.2;refGene_gn=ELAVL2;refGene_tx=XM_019816047.2;refGene_gn=ELAVL2;refGene_tx=XM_019816049.2;refGene_gn=ELAVL2;refGene_tx=XM_023242363.1;refGene_gn=ELAVL2;refGene_tx=XM_023242364.1
D4 49678305 49678332 refGene_gn=ELAVL2;refGene_tx=XM_019816051.2
D4 50188315 50188421

refGene_gn=IZUMO3;refGene_tx=XM_006939186.3;refGene_gn=IZUMO3;refGene_tx=XM_019816055.2
D4 50188762 50188853 refGene_gn=IZUMO3;refGene_tx=XM_006939186.3;refGene_gn=IZUMO3;refGene_tx=XM_019816055.2
D4 50189217 50189298 refGene_gn=IZUMO3;refGene_tx=XM_006939186.3;refGene_gn=IZUMO3;refGene_tx=XM_019816055.2
D4 50189764 50189782 refGene_gn=IZUMO3;refGene_tx=XM_006939186.3;refGene_gn=IZUMO3;refGene_tx=XM_019816055.2
D4 50189997 50190087 refGene_gn=IZUMO3;refGene_tx=XM_006939186.3;refGene_gn=IZUMO3;refGene_tx=XM_019816055.2
D4 50190236 50190311 refGene_gn=IZUMO3;refGene_tx=XM_006939186.3;refGene_gn=IZUMO3;refGene_tx=XM_019816055.2
D4 50190447 50190679 refGene_gn=IZUMO3;refGene_tx=XM_006939186.3
D4 50190901 50191034 refGene_gn=IZUMO3;refGene_tx=XM_019816055.2
D4 51136951 51137974 refGene_gn=TUSC1;refGene_tx=XM_023242366.1
D4 52113031 52113378 refGene_gn=CAAP1;refGene_tx=XM_003995527.4;refGene_gn=CAAP1;refGene_tx=XM_006939188.3
D4 52130387 52130461 refGene_gn=CAAP1;refGene_tx=XM_003995527.4;refGene_gn=CAAP1;refGene_tx=XM_006939188.3
D4 52156777 52156853 refGene_gn=CAAP1;refGene_tx=XM_003995527.4;refGene_gn=CAAP1;refGene_tx=XM_006939188.3
D4 52157896 52157981 refGene_gn=CAAP1;refGene_tx=XM_003995527.4;refGene_gn=CAAP1;refGene_tx=XM_006939188.3
D4 52158950 52159151 refGene_gn=CAAP1;refGene_tx=XM_003995527.4;refGene_gn=CAAP1;refGene_tx=XM_006939188.3
D4 52164894 52164900 refGene_gn=CAAP1;refGene_tx=XM_006939188.3
D4 52164981 52165287 refGene_gn=CAAP1;refGene_tx=XM_003995527.4
D4 52179985 52180551 refGene_gn=PLAA;refGene_tx=XM_003995528.5;refGene_gn=PLAA;refGene_tx=XM_006939189.4;refGene_gn=PLAA;refGene_tx=XM_019816056.2
D4 52181862 52182027 refGene_gn=PLAA;refGene_tx=XM_003995528.5;refGene_gn=PLAA;refGene_tx=XM_006939189.4;refGene_gn=PLAA;refGene_tx=XM_019816056.2
D4 52183623 52183725 refGene_gn=PLAA;refGene_tx=XM_003995528.5;refGene_gn=PLAA;refGene_tx=XM_006939189.4;refGene_gn=PLAA;refGene_tx=XM_019816056.2
D4 52185266 52185335 refGene_gn=PLAA;refGene_tx=XM_003995528.5;refGene_gn=PLAA;refGene_tx=XM_006939189.4
D4 52190589 52190661 refGene_gn=PLAA;refGene_tx=XM_003995528.5

D4 52193199 52193419
refGene_gn=PLAA;refGene_tx=XM_003995528.5;refGene_gn=PLAA;refGene_tx=XM_006939189.4;refGene_gn=PLAA;refGene_tx=XM_019816056.2
D4 52194160 52194318
refGene_gn=PLAA;refGene_tx=XM_003995528.5;refGene_gn=PLAA;refGene_tx=XM_006939189.4;refGene_gn=PLAA;refGene_tx=XM_019816056.2
D4 52198671 52198841
refGene_gn=PLAA;refGene_tx=XM_003995528.5;refGene_gn=PLAA;refGene_tx=XM_006939189.4;refGene_gn=PLAA;refGene_tx=XM_019816056.2
D4 52200503 52200639
refGene_gn=PLAA;refGene_tx=XM_003995528.5;refGene_gn=PLAA;refGene_tx=XM_006939189.4;refGene_gn=PLAA;refGene_tx=XM_019816056.2
D4 52201102 52201270
refGene_gn=PLAA;refGene_tx=XM_003995528.5;refGene_gn=PLAA;refGene_tx=XM_006939189.4;refGene_gn=PLAA;refGene_tx=XM_019816056.2
D4 52204857 52204978
refGene_gn=PLAA;refGene_tx=XM_003995528.5;refGene_gn=PLAA;refGene_tx=XM_006939189.4;refGene_gn=PLAA;refGene_tx=XM_019816056.2
D4 52205068 52205169
refGene_gn=PLAA;refGene_tx=XM_003995528.5;refGene_gn=PLAA;refGene_tx=XM_006939189.4;refGene_gn=PLAA;refGene_tx=XM_019816056.2
D4 52216616 52216810
refGene_gn=PLAA;refGene_tx=XM_003995528.5;refGene_gn=PLAA;refGene_tx=XM_006939189.4;refGene_gn=PLAA;refGene_tx=XM_019816056.2
D4 52222630 52222779
refGene_gn=PLAA;refGene_tx=XM_003995528.5;refGene_gn=PLAA;refGene_tx=XM_006939189.4;refGene_gn=PLAA;refGene_tx=XM_019816056.2
D4 52228559 52228679
refGene_gn=IFT74;refGene_tx=XM_023242367.1;refGene_gn=IFT74;refGene_tx=XM_023242368.1
D4 52240243 52240379
refGene_gn=IFT74;refGene_tx=XM_023242367.1;refGene_gn=IFT74;refGene_tx=XM_023242368.1
D4 52242655 52242704
refGene_gn=IFT74;refGene_tx=XM_023242367.1;refGene_gn=IFT74;refGene_tx=XM_023242368.1
D4 52244161 52244260
refGene_gn=IFT74;refGene_tx=XM_023242367.1;refGene_gn=IFT74;refGene_tx=XM_023242368.1
D4 52244419 52244480
refGene_gn=IFT74;refGene_tx=XM_023242367.1;refGene_gn=IFT74;refGene_tx=XM_023242368.1
D4 52251055 52251115
refGene_gn=IFT74;refGene_tx=XM_023242367.1;refGene_gn=IFT74;refGene_tx=XM_023242368.1
D4 52252666 52252728
refGene_gn=IFT74;refGene_tx=XM_023242367.1;refGene_gn=IFT74;refGene_tx=XM_023242368.1
D4 52260874 52261203 refGene_gn=LRRC19;refGene_tx=XM_023242369.1
D4 52261698 52261884 refGene_gn=LRRC19;refGene_tx=XM_023242369.1

D4 52262547 52263061 refGene_gn=LRRC19;refGene_tx=XM_023242369.1
D4 52264730 52264820 refGene_gn=LRRC19;refGene_tx=XM_023242369.1
D4 52268786 52268879 refGene_gn=LRRC19;refGene_tx=XM_023242369.1
D4 52271421 52271560
refGene_gn=IFT74;refGene_tx=XM_023242367.1;refGene_gn=IFT74;refGene_tx=XM_023242368.1
D4 52272915 52272978
refGene_gn=IFT74;refGene_tx=XM_023242367.1;refGene_gn=IFT74;refGene_tx=XM_023242368.1
D4 52281799 52281943
refGene_gn=IFT74;refGene_tx=XM_023242367.1;refGene_gn=IFT74;refGene_tx=XM_023242368.1
D4 52283843 52283884
refGene_gn=IFT74;refGene_tx=XM_023242367.1;refGene_gn=IFT74;refGene_tx=XM_023242368.1
D4 52286798 52286878
refGene_gn=IFT74;refGene_tx=XM_023242367.1;refGene_gn=IFT74;refGene_tx=XM_023242368.1
D4 52300272 52300326
refGene_gn=IFT74;refGene_tx=XM_023242367.1;refGene_gn=IFT74;refGene_tx=XM_023242368.1
D4 52302946 52303044 refGene_gn=IFT74;refGene_tx=XM_023242367.1
D4 52304108 52304235 refGene_gn=IFT74;refGene_tx=XM_023242367.1
D4 52318063 52318227
refGene_gn=IFT74;refGene_tx=XM_023242367.1;refGene_gn=IFT74;refGene_tx=XM_023242368.1
D4 52318775 52318901
refGene_gn=IFT74;refGene_tx=XM_023242367.1;refGene_gn=IFT74;refGene_tx=XM_023242368.1
D4 52322526 52322587
refGene_gn=IFT74;refGene_tx=XM_023242367.1;refGene_gn=IFT74;refGene_tx=XM_023242368.1
D4 52324113 52324232
refGene_gn=IFT74;refGene_tx=XM_023242367.1;refGene_gn=IFT74;refGene_tx=XM_023242368.1
D4 52357835 52357887
refGene_gn=TEK;refGene_tx=XM_003995529.5;refGene_gn=TEK;refGene_tx=XM_006939192.3;refGene_gn=TEK;refGene_tx=XM_019816057.2
D4 52402321 52402633
refGene_gn=TEK;refGene_tx=XM_003995529.5;refGene_gn=TEK;refGene_tx=XM_006939192.3;refGene_gn=TEK;refGene_tx=XM_019816057.2
D4 52412255 52412366
refGene_gn=TEK;refGene_tx=XM_003995529.5;refGene_gn=TEK;refGene_tx=XM_006939192.3;refGene_gn=TEK;refGene_tx=XM_019816057.2
D4 52413242 52413395
refGene_gn=TEK;refGene_tx=XM_003995529.5;refGene_gn=TEK;refGene_tx=XM_006939192.3;refGene_gn=TEK;refGene_tx=XM_019816057.2
D4 52416964 52417096
refGene_gn=TEK;refGene_tx=XM_003995529.5;refGene_gn=TEK;refGene_tx=XM_006939192.3;refGene_gn=TEK;refGene_tx=XM_019816057.2

D4 52417416 52417557
refGene_gn=TEK;refGene_tx=XM_003995529.5;refGene_gn=TEK;refGene_tx=XM_006939192.3;refGene_gn=TEK;refGene_tx=XM_019816057.2
D4 52418531 52418660
refGene_gn=TEK;refGene_tx=XM_003995529.5;refGene_gn=TEK;refGene_tx=XM_006939192.3;refGene_gn=TEK;refGene_tx=XM_019816057.2
D4 52422326 52422478
refGene_gn=TEK;refGene_tx=XM_003995529.5;refGene_gn=TEK;refGene_tx=XM_006939192.3;refGene_gn=TEK;refGene_tx=XM_019816057.2
D4 52424089 52424234
refGene_gn=TEK;refGene_tx=XM_003995529.5;refGene_gn=TEK;refGene_tx=XM_006939192.3;refGene_gn=TEK;refGene_tx=XM_019816057.2
D4 52430132 52430294
refGene_gn=TEK;refGene_tx=XM_003995529.5;refGene_gn=TEK;refGene_tx=XM_006939192.3;refGene_gn=TEK;refGene_tx=XM_019816057.2
D4 52432054 52432189
refGene_gn=TEK;refGene_tx=XM_003995529.5;refGene_gn=TEK;refGene_tx=XM_019816057.2
D4 52436631 52436919
refGene_gn=TEK;refGene_tx=XM_003995529.5;refGene_gn=TEK;refGene_tx=XM_006939192.3;refGene_gn=TEK;refGene_tx=XM_019816057.2
D4 52442987 52443287
refGene_gn=TEK;refGene_tx=XM_003995529.5;refGene_gn=TEK;refGene_tx=XM_006939192.3;refGene_gn=TEK;refGene_tx=XM_019816057.2
D4 52445176 52445331
refGene_gn=TEK;refGene_tx=XM_003995529.5;refGene_gn=TEK;refGene_tx=XM_006939192.3;refGene_gn=TEK;refGene_tx=XM_019816057.2
D4 52446839 52447050
refGene_gn=TEK;refGene_tx=XM_003995529.5;refGene_gn=TEK;refGene_tx=XM_006939192.3;refGene_gn=TEK;refGene_tx=XM_019816057.2
D4 52449451 52449562
refGene_gn=TEK;refGene_tx=XM_003995529.5;refGene_gn=TEK;refGene_tx=XM_006939192.3;refGene_gn=TEK;refGene_tx=XM_019816057.2
D4 52451819 52452010
refGene_gn=TEK;refGene_tx=XM_003995529.5;refGene_gn=TEK;refGene_tx=XM_006939192.3;refGene_gn=TEK;refGene_tx=XM_019816057.2
D4 52452557 52452671
refGene_gn=TEK;refGene_tx=XM_003995529.5;refGene_gn=TEK;refGene_tx=XM_006939192.3;refGene_gn=TEK;refGene_tx=XM_019816057.2
D4 52457298 52457369
refGene_gn=TEK;refGene_tx=XM_003995529.5;refGene_gn=TEK;refGene_tx=XM_006939192.3;refGene_gn=TEK;refGene_tx=XM_019816057.2
D4 52458415 52458456
refGene_gn=TEK;refGene_tx=XM_003995529.5;refGene_gn=TEK;refGene_tx=XM_006939192.3;refGene_gn=TEK;refGene_tx=XM_019816057.2
D4 52459388 52459485
refGene_gn=TEK;refGene_tx=XM_003995529.5;refGene_gn=TEK;refGene_tx=XM_006939192.3;refGene_gn=TEK;refGene_tx=XM_019816057.2
D4 52461300 52461400
refGene_gn=TEK;refGene_tx=XM_003995529.5;refGene_gn=TEK;refGene_tx=XM_

006939192.3;refGene_gn=TEK;refGene_tx=XM_019816057.2
D4 52462194 52462269
refGene_gn=TEK;refGene_tx=XM_003995529.5;refGene_gn=TEK;refGene_tx=XM_006939192.3;refGene_gn=TEK;refGene_tx=XM_019816057.2
D4 52464666 52464749
refGene_gn=LOC101093554;refGene_tx=XM_023242371.1
D4 52466128 52466422
refGene_gn=LOC101093554;refGene_tx=XM_023242370.1;refGene_gn=LOC101093554;refGene_tx=XM_023242371.1
D4 52468573 52468712
refGene_gn=LOC101093554;refGene_tx=XM_023242370.1;refGene_gn=LOC101093554;refGene_tx=XM_023242371.1
D4 52474317 52474433
refGene_gn=LOC101093554;refGene_tx=XM_023242370.1;refGene_gn=LOC101093554;refGene_tx=XM_023242371.1
D4 52475708 52475816
refGene_gn=LOC101093554;refGene_tx=XM_023242370.1;refGene_gn=LOC101093554;refGene_tx=XM_023242371.1
D4 52475988 52476062
refGene_gn=LOC101093554;refGene_tx=XM_023242370.1;refGene_gn=LOC101093554;refGene_tx=XM_023242371.1
D4 52478708 52478822
refGene_gn=LOC101093554;refGene_tx=XM_023242370.1;refGene_gn=LOC101093554;refGene_tx=XM_023242371.1
D4 52480319 52480469
refGene_gn=LOC101093554;refGene_tx=XM_023242370.1;refGene_gn=LOC101093554;refGene_tx=XM_023242371.1
D4 52481875 52481938
refGene_gn=LOC101093554;refGene_tx=XM_023242370.1;refGene_gn=LOC101093554;refGene_tx=XM_023242371.1
D4 52485755 52485852
refGene_gn=LOC101093554;refGene_tx=XM_023242370.1;refGene_gn=LOC101093554;refGene_tx=XM_023242371.1
D4 52495976 52496129
refGene_gn=LOC101093554;refGene_tx=XM_023242370.1;refGene_gn=LOC101093554;refGene_tx=XM_023242371.1
D4 52497533 52497652
refGene_gn=LOC101093554;refGene_tx=XM_023242370.1;refGene_gn=LOC101093554;refGene_tx=XM_023242371.1
D4 52498094 52498302
refGene_gn=LOC101093554;refGene_tx=XM_023242370.1;refGene_gn=LOC101093554;refGene_tx=XM_023242371.1
D4 52500641 52500731
refGene_gn=LOC101093554;refGene_tx=XM_023242370.1;refGene_gn=LOC101093554;refGene_tx=XM_023242371.1
D4 52501166 52502327
refGene_gn=LOC101093554;refGene_tx=XM_023242370.1;refGene_gn=LOC101093554;refGene_tx=XM_023242371.1
D4 52506480 52506760
refGene_gn=EQTN;refGene_tx=XM_006939193.3;refGene_gn=EQTN;refGene_tx=X

M_019816060.2;refGene_gn=EQTN;refGene_tx=XM_019816061.2;refGene_gn=EQTN;refGene_tx=XM_023242372.1
D4 52507234 52507388 refGene_gn=EQTN;refGene_tx=XM_006939193.3;refGene_gn=EQTN;refGene_tx=XM_019816060.2;refGene_gn=EQTN;refGene_tx=XM_019816061.2;refGene_gn=EQTN;refGene_tx=XM_023242372.1
D4 52510152 52510218 refGene_gn=EQTN;refGene_tx=XM_006939193.3;refGene_gn=EQTN;refGene_tx=XM_019816060.2;refGene_gn=EQTN;refGene_tx=XM_019816061.2;refGene_gn=EQTN;refGene_tx=XM_023242372.1
D4 52511599 52511650 refGene_gn=EQTN;refGene_tx=XM_006939193.3;refGene_gn=EQTN;refGene_tx=XM_019816060.2;refGene_gn=EQTN;refGene_tx=XM_019816061.2;refGene_gn=EQTN;refGene_tx=XM_023242372.1
D4 52512962 52513052 refGene_gn=EQTN;refGene_tx=XM_006939193.3;refGene_gn=EQTN;refGene_tx=XM_019816060.2;refGene_gn=EQTN;refGene_tx=XM_019816061.2;refGene_gn=EQTN;refGene_tx=XM_023242372.1
D4 52515016 52515103 refGene_gn=EQTN;refGene_tx=XM_006939193.3;refGene_gn=EQTN;refGene_tx=XM_019816060.2;refGene_gn=EQTN;refGene_tx=XM_019816061.2;refGene_gn=EQTN;refGene_tx=XM_023242372.1
D4 52518796 52519096 refGene_gn=EQTN;refGene_tx=XM_006939193.3;refGene_gn=EQTN;refGene_tx=XM_019816060.2;refGene_gn=EQTN;refGene_tx=XM_019816061.2;refGene_gn=EQTN;refGene_tx=XM_023242372.1
D4 52519317 52519399 refGene_gn=EQTN;refGene_tx=XM_019816060.2;refGene_gn=EQTN;refGene_tx=XM_019816061.2
D4 52526415 52526565 refGene_gn=EQTN;refGene_tx=XM_006939193.3;refGene_gn=EQTN;refGene_tx=XM_023242372.1
D4 52529827 52529885 refGene_gn=EQTN;refGene_tx=XM_006939193.3
D4 52532119 52532135 refGene_gn=EQTN;refGene_tx=XM_023242372.1
D4 52548733 52548894 refGene_gn=MOB3B;refGene_tx=XM_006939194.4;refGene_gn=MOB3B;refGene_tx=XM_006939195.4;refGene_gn=MOB3B;refGene_tx=XM_006939196.4;refGene_gn=MOB3B;refGene_tx=XM_011288600.3;refGene_gn=MOB3B;refGene_tx=XM_019816063.2;refGene_gn=MOB3B;refGene_tx=XM_023242374.1
D4 52572764 52572773 refGene_gn=MOB3B;refGene_tx=XM_019816065.2
D4 52579835 52580038 refGene_gn=MOB3B;refGene_tx=XM_006939194.4;refGene_gn=MOB3B;refGene_tx=XM_006939195.4;refGene_gn=MOB3B;refGene_tx=XM_006939196.4;refGene_gn=MOB3B;refGene_tx=XM_011288600.3;refGene_gn=MOB3B;refGene_tx=XM_019816063.2;refGene_gn=MOB3B;refGene_tx=XM_019816065.2
D4 52667224 52667642 refGene_gn=MOB3B;refGene_tx=XM_006939194.4;refGene_gn=MOB3B;refGene_tx=XM_006939195.4;refGene_gn=MOB3B;refGene_tx=XM_006939196.4;refGene_gn=MOB3B;refGene_tx=XM_011288600.3;refGene_gn=MOB3B;refGene_tx=XM_019816063.2;refGene_gn=MOB3B;refGene_tx=XM_019816065.2;refGene_gn=MOB3B;refGe

ne_tx=XM_023242374.1
D4	52741254 52741911 refGene_gn=IFNK;refGene_tx=XM_019816062.2
D4	52773409 52773596
refGene_gn=CD4H9orf72;refGene_tx=XM_023242375.1;refGene_gn=CD4H9orf72;
refGene_tx=XM_023242376.1;refGene_gn=CD4H9orf72;refGene_tx=XM_02324237
7.1;refGene_gn=CD4H9orf72;refGene_tx=XM_023242379.1;refGene_gn=CD4H9or
f72;refGene_tx=XM_023242380.1;refGene_gn=CD4H9orf72;refGene_tx=XM_0232
42381.1;refGene_gn=CD4H9orf72;refGene_tx=XM_023242382.1
D4	52773726 52773836
refGene_gn=CD4H9orf72;refGene_tx=XM_023242375.1;refGene_gn=CD4H9orf72;
refGene_tx=XM_023242376.1;refGene_gn=CD4H9orf72;refGene_tx=XM_02324237
7.1;refGene_gn=CD4H9orf72;refGene_tx=XM_023242379.1;refGene_gn=CD4H9or
f72;refGene_tx=XM_023242380.1;refGene_gn=CD4H9orf72;refGene_tx=XM_0232
42381.1;refGene_gn=CD4H9orf72;refGene_tx=XM_023242382.1
D4	52774937 52774995
refGene_gn=CD4H9orf72;refGene_tx=XM_023242375.1;refGene_gn=CD4H9orf72;
refGene_tx=XM_023242376.1;refGene_gn=CD4H9orf72;refGene_tx=XM_02324237
7.1;refGene_gn=CD4H9orf72;refGene_tx=XM_023242379.1;refGene_gn=CD4H9or
f72;refGene_tx=XM_023242380.1;refGene_gn=CD4H9orf72;refGene_tx=XM_0232
42381.1;refGene_gn=CD4H9orf72;refGene_tx=XM_023242382.1
D4	52778611 52778847
refGene_gn=CD4H9orf72;refGene_tx=XM_023242375.1;refGene_gn=CD4H9orf72;
refGene_tx=XM_023242376.1;refGene_gn=CD4H9orf72;refGene_tx=XM_02324237
7.1;refGene_gn=CD4H9orf72;refGene_tx=XM_023242379.1;refGene_gn=CD4H9or
f72;refGene_tx=XM_023242380.1;refGene_gn=CD4H9orf72;refGene_tx=XM_0232
42381.1;refGene_gn=CD4H9orf72;refGene_tx=XM_023242382.1
D4	52782572 52782689
refGene_gn=CD4H9orf72;refGene_tx=XM_023242375.1;refGene_gn=CD4H9orf72;
refGene_tx=XM_023242376.1;refGene_gn=CD4H9orf72;refGene_tx=XM_02324237
7.1;refGene_gn=CD4H9orf72;refGene_tx=XM_023242379.1;refGene_gn=CD4H9or
f72;refGene_tx=XM_023242380.1;refGene_gn=CD4H9orf72;refGene_tx=XM_0232
42381.1;refGene_gn=CD4H9orf72;refGene_tx=XM_023242382.1
D4	52784423 52784496
refGene_gn=CD4H9orf72;refGene_tx=XM_023242375.1;refGene_gn=CD4H9orf72;
refGene_tx=XM_023242376.1;refGene_gn=CD4H9orf72;refGene_tx=XM_02324237
7.1;refGene_gn=CD4H9orf72;refGene_tx=XM_023242379.1;refGene_gn=CD4H9or
f72;refGene_tx=XM_023242380.1;refGene_gn=CD4H9orf72;refGene_tx=XM_0232
42381.1;refGene_gn=CD4H9orf72;refGene_tx=XM_023242382.1
D4	52786646 52786711
refGene_gn=CD4H9orf72;refGene_tx=XM_023242375.1;refGene_gn=CD4H9orf72;
refGene_tx=XM_023242376.1;refGene_gn=CD4H9orf72;refGene_tx=XM_02324237
7.1;refGene_gn=CD4H9orf72;refGene_tx=XM_023242379.1;refGene_gn=CD4H9or
f72;refGene_tx=XM_023242380.1;refGene_gn=CD4H9orf72;refGene_tx=XM_0232
42381.1;refGene_gn=CD4H9orf72;refGene_tx=XM_023242382.1
D4	52787421 52787517
refGene_gn=CD4H9orf72;refGene_tx=XM_023242375.1;refGene_gn=CD4H9orf72;
refGene_tx=XM_023242376.1;refGene_gn=CD4H9orf72;refGene_tx=XM_02324237
7.1;refGene_gn=CD4H9orf72;refGene_tx=XM_023242379.1;refGene_gn=CD4H9or
f72;refGene_tx=XM_023242380.1;refGene_gn=CD4H9orf72;refGene_tx=XM_0232
42381.1;refGene_gn=CD4H9orf72;refGene_tx=XM_023242382.1

D4 52792710 52792770
refGene_gn=CD4H9orf72;refGene_tx=XM_023242375.1;refGene_gn=CD4H9orf72;
refGene_tx=XM_023242376.1;refGene_gn=CD4H9orf72;refGene_tx=XM_02324237
7.1;refGene_gn=CD4H9orf72;refGene_tx=XM_023242379.1;refGene_gn=CD4H9or
f72;refGene_tx=XM_023242380.1;refGene_gn=CD4H9orf72;refGene_tx=XM_0232
42381.1;refGene_gn=CD4H9orf72;refGene_tx=XM_023242382.1
D4 52794122 52794566
refGene_gn=CD4H9orf72;refGene_tx=XM_023242375.1;refGene_gn=CD4H9orf72;
refGene_tx=XM_023242376.1;refGene_gn=CD4H9orf72;refGene_tx=XM_02324237
7.1;refGene_gn=CD4H9orf72;refGene_tx=XM_023242379.1;refGene_gn=CD4H9or
f72;refGene_tx=XM_023242380.1;refGene_gn=CD4H9orf72;refGene_tx=XM_0232
42381.1;refGene_gn=CD4H9orf72;refGene_tx=XM_023242382.1
D4 53121427 53123248
refGene_gn=LINGO2;refGene_tx=XM_006939197.4;refGene_gn=LINGO2;refGene_
tx=XM_019816066.2;refGene_gn=LINGO2;refGene_tx=XM_019816067.2;refGene_
gn=LINGO2;refGene_tx=XM_023242383.1
D4 56832714 56832811 refGene_gn=ACO1;refGene_tx=XM_006939201.3
D4 56833863 56834032 refGene_gn=ACO1;refGene_tx=XM_006939201.3
D4 56835057 56835195 refGene_gn=ACO1;refGene_tx=XM_006939201.3
D4 56841943 56842013 refGene_gn=ACO1;refGene_tx=XM_006939201.3
D4 56842139 56842323 refGene_gn=ACO1;refGene_tx=XM_006939201.3
D4 56842794 56842934 refGene_gn=ACO1;refGene_tx=XM_006939201.3
D4 56846399 56846571 refGene_gn=ACO1;refGene_tx=XM_006939201.3
D4 56849810 56849911 refGene_gn=ACO1;refGene_tx=XM_006939201.3
D4 56850860 56850977 refGene_gn=ACO1;refGene_tx=XM_006939201.3
D4 56852096 56852256 refGene_gn=ACO1;refGene_tx=XM_006939201.3
D4 56854201 56854337 refGene_gn=ACO1;refGene_tx=XM_006939201.3
D4 56855652 56855737 refGene_gn=ACO1;refGene_tx=XM_006939201.3
D4 56856514 56856671 refGene_gn=ACO1;refGene_tx=XM_006939201.3
D4 56858364 56858489 refGene_gn=ACO1;refGene_tx=XM_006939201.3
D4 56860537 56860642 refGene_gn=ACO1;refGene_tx=XM_006939201.3
D4 56861374 56861517 refGene_gn=ACO1;refGene_tx=XM_006939201.3
D4 56864146 56864294 refGene_gn=ACO1;refGene_tx=XM_006939201.3
D4 56868204 56868327 refGene_gn=ACO1;refGene_tx=XM_006939201.3
D4 56870868 56871054 refGene_gn=ACO1;refGene_tx=XM_006939201.3
D4 56872001 56872115 refGene_gn=ACO1;refGene_tx=XM_006939201.3
D4 56874472 56874769
refGene_gn=DDX58;refGene_tx=XM_006939199.4;refGene_gn=DDX58;refGene_tx
=XM_006939200.4;refGene_gn=DDX58;refGene_tx=XM_019816078.2
D4 56876383 56876527
refGene_gn=DDX58;refGene_tx=XM_006939199.4;refGene_gn=DDX58;refGene_tx
=XM_006939200.4;refGene_gn=DDX58;refGene_tx=XM_019816078.2
D4 56879947 56880099
refGene_gn=DDX58;refGene_tx=XM_006939199.4;refGene_gn=DDX58;refGene_tx
=XM_006939200.4;refGene_gn=DDX58;refGene_tx=XM_019816078.2
D4 56881319 56881490
refGene_gn=DDX58;refGene_tx=XM_006939199.4;refGene_gn=DDX58;refGene_tx
=XM_006939200.4;refGene_gn=DDX58;refGene_tx=XM_019816078.2
D4 56884263 56884354
refGene_gn=DDX58;refGene_tx=XM_006939199.4;refGene_gn=DDX58;refGene_tx

=XM_006939200.4;refGene_gn=DDX58;refGene_tx=XM_019816078.2
D4	56886510	56886659
refGene_gn=DDX58;refGene_tx=XM_006939199.4;refGene_gn=DDX58;refGene_tx
=XM_006939200.4;refGene_gn=DDX58;refGene_tx=XM_019816078.2
D4	56887378	56887514
refGene_gn=DDX58;refGene_tx=XM_006939199.4;refGene_gn=DDX58;refGene_tx
=XM_006939200.4;refGene_gn=DDX58;refGene_tx=XM_019816078.2
D4	56888504	56888662
refGene_gn=DDX58;refGene_tx=XM_006939199.4;refGene_gn=DDX58;refGene_tx
=XM_006939200.4;refGene_gn=DDX58;refGene_tx=XM_019816078.2
D4	56892230	56892350
refGene_gn=DDX58;refGene_tx=XM_006939199.4;refGene_gn=DDX58;refGene_tx
=XM_006939200.4;refGene_gn=DDX58;refGene_tx=XM_019816078.2
D4	56894065	56894228
refGene_gn=DDX58;refGene_tx=XM_006939199.4;refGene_gn=DDX58;refGene_tx
=XM_006939200.4;refGene_gn=DDX58;refGene_tx=XM_019816078.2
D4	56896207	56896466
refGene_gn=DDX58;refGene_tx=XM_006939199.4;refGene_gn=DDX58;refGene_tx
=XM_006939200.4;refGene_gn=DDX58;refGene_tx=XM_019816078.2
D4	56897001	56897155
refGene_gn=DDX58;refGene_tx=XM_006939199.4;refGene_gn=DDX58;refGene_tx
=XM_006939200.4;refGene_gn=DDX58;refGene_tx=XM_019816078.2
D4	56897577	56897679
refGene_gn=DDX58;refGene_tx=XM_006939199.4;refGene_gn=DDX58;refGene_tx
=XM_006939200.4;refGene_gn=DDX58;refGene_tx=XM_019816078.2
D4	56898768	56898891
refGene_gn=DDX58;refGene_tx=XM_006939199.4;refGene_gn=DDX58;refGene_tx
=XM_019816078.2
D4	56899955	56900103
refGene_gn=DDX58;refGene_tx=XM_006939199.4;refGene_gn=DDX58;refGene_tx
=XM_006939200.4;refGene_gn=DDX58;refGene_tx=XM_019816078.2
D4	56901192	56901374
refGene_gn=DDX58;refGene_tx=XM_006939199.4;refGene_gn=DDX58;refGene_tx
=XM_006939200.4;refGene_gn=DDX58;refGene_tx=XM_019816078.2
D4	56905283	56905418
refGene_gn=DDX58;refGene_tx=XM_006939199.4;refGene_gn=DDX58;refGene_tx
=XM_006939200.4
D4	56911321	56911427
refGene_gn=DDX58;refGene_tx=XM_006939199.4;refGene_gn=DDX58;refGene_tx
=XM_006939200.4
D4	56922667	56925598	refGene_gn=TOPORS;refGene_tx=XM_011288597.3
D4	56931771	56931966	refGene_gn=TOPORS;refGene_tx=XM_011288597.3
D4	56933494	56933497	refGene_gn=TOPORS;refGene_tx=XM_011288597.3
D4	56934660	56934729	refGene_gn=NDUFB6;refGene_tx=XM_003995543.5
D4	56938122	56938167	refGene_gn=NDUFB6;refGene_tx=XM_003995543.5
D4	56944313	56944406	refGene_gn=NDUFB6;refGene_tx=XM_003995543.5
D4	56946523	56946703	refGene_gn=NDUFB6;refGene_tx=XM_003995543.5
D4	57098489	57099197
refGene_gn=TMEM215;refGene_tx=XM_019816080.2;refGene_gn=TMEM215;refGen
e_tx=XM_019816081.2

D4 57245072 57245227 refGene_gn=APTX;refGene_tx=XM_006939202.3
D4 57245990 57246094 refGene_gn=APTX;refGene_tx=XM_006939202.3
D4 57251901 57252128 refGene_gn=APTX;refGene_tx=XM_006939202.3
D4 57252942 57253002 refGene_gn=APTX;refGene_tx=XM_006939202.3
D4 57254353 57254656 refGene_gn=APTX;refGene_tx=XM_006939202.3
D4 57254870 57254917 refGene_gn=APTX;refGene_tx=XM_006939202.3
D4 57256483 57256616 refGene_gn=APTX;refGene_tx=XM_006939202.3
D4 57290855 57290987
refGene_gn=DNAJA1;refGene_tx=XM_003995550.5;refGene_gn=DNAJA1;refGene_tx=XM_019816079.2
D4 57291157 57291335
refGene_gn=DNAJA1;refGene_tx=XM_003995550.5;refGene_gn=DNAJA1;refGene_tx=XM_019816079.2
D4 57293130 57293235
refGene_gn=DNAJA1;refGene_tx=XM_003995550.5;refGene_gn=DNAJA1;refGene_tx=XM_019816079.2
D4 57294175 57294403
refGene_gn=DNAJA1;refGene_tx=XM_003995550.5;refGene_gn=DNAJA1;refGene_tx=XM_019816079.2
D4 57296386 57296501
refGene_gn=DNAJA1;refGene_tx=XM_003995550.5;refGene_gn=DNAJA1;refGene_tx=XM_019816079.2
D4 57298173 57298289
refGene_gn=DNAJA1;refGene_tx=XM_003995550.5;refGene_gn=DNAJA1;refGene_tx=XM_019816079.2
D4 57298576 57298677
refGene_gn=DNAJA1;refGene_tx=XM_003995550.5;refGene_gn=DNAJA1;refGene_tx=XM_019816079.2
D4 57299548 57299767
refGene_gn=DNAJA1;refGene_tx=XM_003995550.5;refGene_gn=DNAJA1;refGene_tx=XM_019816079.2
D4 57306958 57307057
refGene_gn=SMU1;refGene_tx=XM_003995551.5;refGene_gn=SMU1;refGene_tx=XM_006939205.4
D4 57308307 57308460
refGene_gn=SMU1;refGene_tx=XM_003995551.5;refGene_gn=SMU1;refGene_tx=XM_006939205.4
D4 57310973 57311141
refGene_gn=SMU1;refGene_tx=XM_003995551.5;refGene_gn=SMU1;refGene_tx=XM_006939205.4
D4 57313130 57313257
refGene_gn=SMU1;refGene_tx=XM_003995551.5;refGene_gn=SMU1;refGene_tx=XM_006939205.4
D4 57313763 57313891
refGene_gn=SMU1;refGene_tx=XM_003995551.5;refGene_gn=SMU1;refGene_tx=XM_006939205.4
D4 57314511 57314628
refGene_gn=SMU1;refGene_tx=XM_003995551.5;refGene_gn=SMU1;refGene_tx=XM_006939205.4
D4 57316153 57316273

refGene_gn=SMU1;refGene_tx=XM_003995551.5;refGene_gn=SMU1;refGene_tx=X
M_006939205.4
D4	57317705	57317834
refGene_gn=SMU1;refGene_tx=XM_003995551.5;refGene_gn=SMU1;refGene_tx=X
M_006939205.4
D4	57320947	57321058
refGene_gn=SMU1;refGene_tx=XM_003995551.5;refGene_gn=SMU1;refGene_tx=X
M_006939205.4
D4	57322071	57322224
refGene_gn=SMU1;refGene_tx=XM_003995551.5;refGene_gn=SMU1;refGene_tx=X
M_006939205.4
D4	57324000	57324211
refGene_gn=SMU1;refGene_tx=XM_003995551.5;refGene_gn=SMU1;refGene_tx=X
M_006939205.4
D4	57327589	57327615	refGene_gn=SMU1;refGene_tx=XM_003995551.5
D4	57356744	57356877
refGene_gn=B4GALT1;refGene_tx=XM_006939208.3;refGene_gn=B4GALT1;refGen
e_tx=XM_023242387.1
D4	57357118	57357223
refGene_gn=B4GALT1;refGene_tx=XM_006939208.3;refGene_gn=B4GALT1;refGen
e_tx=XM_023242387.1
D4	57359277	57359400	refGene_gn=B4GALT1;refGene_tx=XM_006939208.3
D4	57361981	57362169
refGene_gn=B4GALT1;refGene_tx=XM_006939208.3;refGene_gn=B4GALT1;refGen
e_tx=XM_023242387.1
D4	57378693	57378929
refGene_gn=B4GALT1;refGene_tx=XM_006939208.3;refGene_gn=B4GALT1;refGen
e_tx=XM_023242387.1
D4	57409122	57409540
refGene_gn=B4GALT1;refGene_tx=XM_006939208.3;refGene_gn=B4GALT1;refGen
e_tx=XM_023242387.1
D4	57468687	57468868	refGene_gn=SPINK4;refGene_tx=XM_003995552.4
D4	57473303	57473347	refGene_gn=SPINK4;refGene_tx=XM_003995552.4
D4	57474698	57474811	refGene_gn=SPINK4;refGene_tx=XM_003995552.4
D4	57476274	57476320	refGene_gn=SPINK4;refGene_tx=XM_003995552.4
D4	57481546	57481636	refGene_gn=BAG1;refGene_tx=XM_023242386.1
D4	57482139	57482202	refGene_gn=BAG1;refGene_tx=XM_023242386.1
D4	57483259	57483367	refGene_gn=BAG1;refGene_tx=XM_023242386.1
D4	57485466	57485580	refGene_gn=BAG1;refGene_tx=XM_023242386.1
D4	57487180	57487263	refGene_gn=BAG1;refGene_tx=XM_023242386.1
D4	57489921	57490050	refGene_gn=BAG1;refGene_tx=XM_023242386.1
D4	57491450	57491478	refGene_gn=BAG1;refGene_tx=XM_023242386.1
D4	57491479	57492151	refGene_gn=BAG1;refGene_tx=XM_023242386.1
D4	57492257	57492326	refGene_gn=CHMP5;refGene_tx=XM_003995553.5
D4	57493106	57493211	refGene_gn=CHMP5;refGene_tx=XM_003995553.5
D4	57494631	57494678	refGene_gn=CHMP5;refGene_tx=XM_003995553.5
D4	57496481	57496575	refGene_gn=CHMP5;refGene_tx=XM_003995553.5
D4	57496704	57496776	refGene_gn=CHMP5;refGene_tx=XM_003995553.5
D4	57501503	57501612	refGene_gn=CHMP5;refGene_tx=XM_003995553.5
D4	57504268	57504381	refGene_gn=CHMP5;refGene_tx=XM_003995553.5

D4 57506858 57506909 refGene_gn=CHMP5;refGene_tx=XM_003995553.5
D4 57516567 57516592
refGene_gn=NFX1;refGene_tx=XM_011288594.3;refGene_gn=NFX1;refGene_tx=XM_019816083.2
D4 57521163 57522165
refGene_gn=NFX1;refGene_tx=XM_011288594.3;refGene_gn=NFX1;refGene_tx=XM_019816083.2
D4 57533391 57533550
refGene_gn=NFX1;refGene_tx=XM_011288594.3;refGene_gn=NFX1;refGene_tx=XM_019816083.2
D4 57535791 57535869
refGene_gn=NFX1;refGene_tx=XM_011288594.3;refGene_gn=NFX1;refGene_tx=XM_019816083.2
D4 57540632 57540738
refGene_gn=NFX1;refGene_tx=XM_011288594.3;refGene_gn=NFX1;refGene_tx=XM_019816083.2
D4 57544305 57544377
refGene_gn=NFX1;refGene_tx=XM_011288594.3;refGene_gn=NFX1;refGene_tx=XM_019816083.2
D4 57546070 57546210
refGene_gn=NFX1;refGene_tx=XM_011288594.3;refGene_gn=NFX1;refGene_tx=XM_019816083.2
D4 57555397 57555497
refGene_gn=NFX1;refGene_tx=XM_011288594.3;refGene_gn=NFX1;refGene_tx=XM_019816083.2
D4 57555580 57555798
refGene_gn=NFX1;refGene_tx=XM_011288594.3;refGene_gn=NFX1;refGene_tx=XM_019816083.2
D4 57562975 57563073
refGene_gn=NFX1;refGene_tx=XM_011288594.3;refGene_gn=NFX1;refGene_tx=XM_019816083.2
D4 57566187 57566218
refGene_gn=NFX1;refGene_tx=XM_011288594.3;refGene_gn=NFX1;refGene_tx=XM_019816083.2
D4 57570392 57570472
refGene_gn=NFX1;refGene_tx=XM_011288594.3;refGene_gn=NFX1;refGene_tx=XM_019816083.2
D4 57571624 57571733
refGene_gn=NFX1;refGene_tx=XM_011288594.3;refGene_gn=NFX1;refGene_tx=XM_019816083.2
D4 57575799 57575919
refGene_gn=NFX1;refGene_tx=XM_011288594.3;refGene_gn=NFX1;refGene_tx=XM_019816083.2
D4 57577995 57578075
refGene_gn=NFX1;refGene_tx=XM_011288594.3;refGene_gn=NFX1;refGene_tx=XM_019816083.2
D4 57581758 57581989
refGene_gn=NFX1;refGene_tx=XM_011288594.3;refGene_gn=NFX1;refGene_tx=XM_019816083.2
D4 57582847 57582921

refGene_gn=NFX1;refGene_tx=XM_011288594.3;refGene_gn=NFX1;refGene_tx=XM_019816083.2
D4 57583734 57583836
refGene_gn=NFX1;refGene_tx=XM_011288594.3;refGene_gn=NFX1;refGene_tx=XM_019816083.2
D4 57584355 57584397
refGene_gn=NFX1;refGene_tx=XM_011288594.3;refGene_gn=NFX1;refGene_tx=XM_019816083.2
D4 57588961 57589060
refGene_gn=NFX1;refGene_tx=XM_011288594.3;refGene_gn=NFX1;refGene_tx=XM_019816083.2
D4 57589685 57589788
refGene_gn=NFX1;refGene_tx=XM_011288594.3;refGene_gn=NFX1;refGene_tx=XM_019816083.2
D4 57590631 57590777 refGene_gn=NFX1;refGene_tx=XM_011288594.3
D4 57591516 57591621 refGene_gn=NFX1;refGene_tx=XM_011288594.3
D4 57592847 57592920 refGene_gn=NFX1;refGene_tx=XM_011288594.3
D4 57605531 57605808
refGene_gn=AQP7;refGene_tx=XM_006939211.4;refGene_gn=AQP7;refGene_tx=XM_006939215.3;refGene_gn=AQP7;refGene_tx=XM_011288589.3;refGene_gn=AQP7;refGene_tx=XM_011288590.3;refGene_gn=AQP7;refGene_tx=XM_011288593.3;refGene_gn=AQP7;refGene_tx=XM_019816084.2;refGene_gn=AQP7;refGene_tx=XM_019816087.2;refGene_gn=AQP7;refGene_tx=XM_019816088.1
D4 57606188 57606406
refGene_gn=AQP7;refGene_tx=XM_006939211.4;refGene_gn=AQP7;refGene_tx=XM_006939215.3;refGene_gn=AQP7;refGene_tx=XM_011288589.3;refGene_gn=AQP7;refGene_tx=XM_011288590.3;refGene_gn=AQP7;refGene_tx=XM_011288593.3;refGene_gn=AQP7;refGene_tx=XM_019816084.2;refGene_gn=AQP7;refGene_tx=XM_019816087.2;refGene_gn=AQP7;refGene_tx=XM_019816088.1
D4 57606602 57606721
refGene_gn=AQP7;refGene_tx=XM_006939211.4;refGene_gn=AQP7;refGene_tx=XM_006939215.3;refGene_gn=AQP7;refGene_tx=XM_011288589.3;refGene_gn=AQP7;refGene_tx=XM_011288590.3;refGene_gn=AQP7;refGene_tx=XM_011288593.3;refGene_gn=AQP7;refGene_tx=XM_019816084.2;refGene_gn=AQP7;refGene_tx=XM_019816087.2;refGene_gn=AQP7;refGene_tx=XM_019816088.1
D4 57606930 57607068
refGene_gn=AQP7;refGene_tx=XM_006939211.4;refGene_gn=AQP7;refGene_tx=XM_006939215.3;refGene_gn=AQP7;refGene_tx=XM_011288589.3;refGene_gn=AQP7;refGene_tx=XM_011288590.3;refGene_gn=AQP7;refGene_tx=XM_011288593.3;refGene_gn=AQP7;refGene_tx=XM_019816084.2;refGene_gn=AQP7;refGene_tx=XM_019816087.2;refGene_gn=AQP7;refGene_tx=XM_019816088.1
D4 57607486 57607610
refGene_gn=AQP7;refGene_tx=XM_006939211.4;refGene_gn=AQP7;refGene_tx=XM_006939215.3;refGene_gn=AQP7;refGene_tx=XM_011288589.3;refGene_gn=AQP7;refGene_tx=XM_011288590.3;refGene_gn=AQP7;refGene_tx=XM_011288593.3;refGene_gn=AQP7;refGene_tx=XM_019816084.2;refGene_gn=AQP7;refGene_tx=XM_019816087.2;refGene_gn=AQP7;refGene_tx=XM_019816088.1
D4 57615404 57615522
refGene_gn=AQP7;refGene_tx=XM_006939211.4;refGene_gn=AQP7;refGene_tx=XM_011288589.3;refGene_gn=AQP7;refGene_tx=XM_011288590.3;refGene_gn=AQP

7;refGene_tx=XM_011288593.3;refGene_gn=AQP7;refGene_tx=XM_019816084.2
D4 57619328 57619354
refGene_gn=AQP7;refGene_tx=XM_006939211.4;refGene_gn=AQP7;refGene_tx=XM_011288589.3;refGene_gn=AQP7;refGene_tx=XM_011288590.3;refGene_gn=AQP7;refGene_tx=XM_011288593.3;refGene_gn=AQP7;refGene_tx=XM_019816084.2
D4 57654040 57654209 refGene_gn=AQP3;refGene_tx=XM_011288588.3
D4 57654297 57654515 refGene_gn=AQP3;refGene_tx=XM_011288588.3
D4 57654843 57654962 refGene_gn=AQP3;refGene_tx=XM_011288588.3
D4 57655339 57655477 refGene_gn=AQP3;refGene_tx=XM_011288588.3
D4 57655838 57655965 refGene_gn=AQP3;refGene_tx=XM_011288588.3
D4 57659027 57659135 refGene_gn=AQP3;refGene_tx=XM_011288588.3
D4 57669746 57669896 refGene_gn=NOL6;refGene_tx=XM_023242389.1
D4 57670119 57670221 refGene_gn=NOL6;refGene_tx=XM_023242389.1
D4 57670337 57670532 refGene_gn=NOL6;refGene_tx=XM_023242389.1
D4 57670931 57671021 refGene_gn=NOL6;refGene_tx=XM_023242389.1
D4 57671143 57671268 refGene_gn=NOL6;refGene_tx=XM_023242389.1
D4 57672041 57672139 refGene_gn=NOL6;refGene_tx=XM_023242389.1
D4 57672374 57672527 refGene_gn=NOL6;refGene_tx=XM_023242389.1
D4 57673238 57673402 refGene_gn=NOL6;refGene_tx=XM_023242389.1
D4 57673505 57673660 refGene_gn=NOL6;refGene_tx=XM_023242389.1
D4 57673738 57673856 refGene_gn=NOL6;refGene_tx=XM_023242389.1
D4 57673999 57674140 refGene_gn=NOL6;refGene_tx=XM_023242389.1
D4 57674368 57674444 refGene_gn=NOL6;refGene_tx=XM_023242389.1
D4 57674594 57674743 refGene_gn=NOL6;refGene_tx=XM_023242389.1
D4 57674916 57675039 refGene_gn=NOL6;refGene_tx=XM_023242389.1
D4 57675215 57675393 refGene_gn=NOL6;refGene_tx=XM_023242389.1
D4 57675541 57675657 refGene_gn=NOL6;refGene_tx=XM_023242389.1
D4 57675860 57675970 refGene_gn=NOL6;refGene_tx=XM_023242389.1
D4 57676055 57676106 refGene_gn=NOL6;refGene_tx=XM_023242389.1
D4 57676311 57676432 refGene_gn=NOL6;refGene_tx=XM_023242389.1
D4 57676520 57676684 refGene_gn=NOL6;refGene_tx=XM_023242389.1
D4 57676768 57676903 refGene_gn=NOL6;refGene_tx=XM_023242389.1
D4 57677135 57677304 refGene_gn=NOL6;refGene_tx=XM_023242389.1
D4 57677672 57677852 refGene_gn=NOL6;refGene_tx=XM_023242389.1
D4 57680739 57680856 refGene_gn=NOL6;refGene_tx=XM_023242389.1
D4 57680941 57681148 refGene_gn=NOL6;refGene_tx=XM_023242389.1
D4 57682194 57682248 refGene_gn=NOL6;refGene_tx=XM_023242389.1
D4 57692775 57692952 refGene_gn=UBE2R2;refGene_tx=XM_011288587.3
D4 57777018 57777105 refGene_gn=UBE2R2;refGene_tx=XM_011288587.3
D4 57789417 57789515 refGene_gn=UBE2R2;refGene_tx=XM_011288587.3
D4 57803548 57803683 refGene_gn=UBE2R2;refGene_tx=XM_011288587.3
D4 57806140 57806360 refGene_gn=UBE2R2;refGene_tx=XM_011288587.3
D4 57811797 57811893
refGene_gn=UBAP2;refGene_tx=XM_006939216.3;refGene_gn=UBAP2;refGene_tx=XM_006939217.3;refGene_gn=UBAP2;refGene_tx=XM_006939218.3;refGene_gn=UBAP2;refGene_tx=XM_006939219.3;refGene_gn=UBAP2;refGene_tx=XM_019816090.1
D4 57811983 57812175
refGene_gn=UBAP2;refGene_tx=XM_006939216.3;refGene_gn=UBAP2;refGene_tx=XM_006939217.3;refGene_gn=UBAP2;refGene_tx=XM_006939218.3;refGene_gn=

UBAP2;refGene_tx=XM_006939219.3;refGene_gn=UBAP2;refGene_tx=XM_0198160
90.1
D4 57812260 57812328
refGene_gn=UBAP2;refGene_tx=XM_006939216.3;refGene_gn=UBAP2;refGene_tx
=XM_006939217.3;refGene_gn=UBAP2;refGene_tx=XM_006939218.3;refGene_gn=
UBAP2;refGene_tx=XM_006939219.3;refGene_gn=UBAP2;refGene_tx=XM_0198160
90.1
D4 57812483 57812591
refGene_gn=UBAP2;refGene_tx=XM_006939216.3;refGene_gn=UBAP2;refGene_tx
=XM_006939217.3;refGene_gn=UBAP2;refGene_tx=XM_006939218.3;refGene_gn=
UBAP2;refGene_tx=XM_006939219.3
D4 57812675 57812775
refGene_gn=UBAP2;refGene_tx=XM_006939216.3;refGene_gn=UBAP2;refGene_tx
=XM_006939217.3;refGene_gn=UBAP2;refGene_tx=XM_006939218.3;refGene_gn=
UBAP2;refGene_tx=XM_006939219.3;refGene_gn=UBAP2;refGene_tx=XM_0198160
90.1
D4 57813097 57813303
refGene_gn=UBAP2;refGene_tx=XM_006939216.3;refGene_gn=UBAP2;refGene_tx
=XM_006939217.3;refGene_gn=UBAP2;refGene_tx=XM_006939218.3;refGene_gn=
UBAP2;refGene_tx=XM_006939219.3;refGene_gn=UBAP2;refGene_tx=XM_0198160
90.1
D4 57813523 57813602
refGene_gn=UBAP2;refGene_tx=XM_006939216.3;refGene_gn=UBAP2;refGene_tx
=XM_006939217.3;refGene_gn=UBAP2;refGene_tx=XM_006939218.3;refGene_gn=
UBAP2;refGene_tx=XM_006939219.3;refGene_gn=UBAP2;refGene_tx=XM_0198160
90.1
D4 57815944 57815992
refGene_gn=UBAP2;refGene_tx=XM_006939216.3;refGene_gn=UBAP2;refGene_tx
=XM_006939217.3;refGene_gn=UBAP2;refGene_tx=XM_006939218.3;refGene_gn=
UBAP2;refGene_tx=XM_006939219.3;refGene_gn=UBAP2;refGene_tx=XM_0198160
90.1
D4 57816272 57816364
refGene_gn=UBAP2;refGene_tx=XM_006939216.3;refGene_gn=UBAP2;refGene_tx
=XM_006939217.3;refGene_gn=UBAP2;refGene_tx=XM_006939218.3;refGene_gn=
UBAP2;refGene_tx=XM_006939219.3;refGene_gn=UBAP2;refGene_tx=XM_0198160
90.1
D4 57817207 57817403
refGene_gn=UBAP2;refGene_tx=XM_006939216.3;refGene_gn=UBAP2;refGene_tx
=XM_006939217.3;refGene_gn=UBAP2;refGene_tx=XM_006939218.3;refGene_gn=
UBAP2;refGene_tx=XM_006939219.3;refGene_gn=UBAP2;refGene_tx=XM_0198160
90.1
D4 57821851 57821921
refGene_gn=UBAP2;refGene_tx=XM_006939216.3;refGene_gn=UBAP2;refGene_tx
=XM_006939217.3;refGene_gn=UBAP2;refGene_tx=XM_006939218.3;refGene_gn=
UBAP2;refGene_tx=XM_006939219.3;refGene_gn=UBAP2;refGene_tx=XM_0198160
90.1
D4 57823099 57823283
refGene_gn=UBAP2;refGene_tx=XM_006939216.3;refGene_gn=UBAP2;refGene_tx
=XM_006939217.3;refGene_gn=UBAP2;refGene_tx=XM_006939218.3;refGene_gn=
UBAP2;refGene_tx=XM_006939219.3;refGene_gn=UBAP2;refGene_tx=XM_0198160

90.1
D4	57825417 57825457
refGene_gn=UBAP2;refGene_tx=XM_006939216.3;refGene_gn=UBAP2;refGene_tx=XM_006939217.3;refGene_gn=UBAP2;refGene_tx=XM_006939218.3;refGene_gn=UBAP2;refGene_tx=XM_006939219.3;refGene_gn=UBAP2;refGene_tx=XM_019816090.1
D4	57829596 57829810
refGene_gn=UBAP2;refGene_tx=XM_006939216.3;refGene_gn=UBAP2;refGene_tx=XM_006939217.3;refGene_gn=UBAP2;refGene_tx=XM_006939218.3;refGene_gn=UBAP2;refGene_tx=XM_006939219.3;refGene_gn=UBAP2;refGene_tx=XM_019816090.1
D4	57831195 57831365
refGene_gn=UBAP2;refGene_tx=XM_006939216.3;refGene_gn=UBAP2;refGene_tx=XM_006939217.3;refGene_gn=UBAP2;refGene_tx=XM_006939218.3;refGene_gn=UBAP2;refGene_tx=XM_006939219.3;refGene_gn=UBAP2;refGene_tx=XM_019816090.1
D4	57831976 57832251
refGene_gn=UBAP2;refGene_tx=XM_006939216.3;refGene_gn=UBAP2;refGene_tx=XM_006939217.3;refGene_gn=UBAP2;refGene_tx=XM_006939218.3;refGene_gn=UBAP2;refGene_tx=XM_006939219.3;refGene_gn=UBAP2;refGene_tx=XM_019816090.1
D4	57835345 57835559
refGene_gn=UBAP2;refGene_tx=XM_006939216.3;refGene_gn=UBAP2;refGene_tx=XM_006939217.3;refGene_gn=UBAP2;refGene_tx=XM_006939218.3;refGene_gn=UBAP2;refGene_tx=XM_006939219.3;refGene_gn=UBAP2;refGene_tx=XM_019816090.1
D4	57838899 57839089
refGene_gn=UBAP2;refGene_tx=XM_006939216.3;refGene_gn=UBAP2;refGene_tx=XM_006939217.3;refGene_gn=UBAP2;refGene_tx=XM_006939218.3;refGene_gn=UBAP2;refGene_tx=XM_006939219.3;refGene_gn=UBAP2;refGene_tx=XM_019816090.1
D4	57841448 57841516
refGene_gn=UBAP2;refGene_tx=XM_006939216.3;refGene_gn=UBAP2;refGene_tx=XM_006939217.3;refGene_gn=UBAP2;refGene_tx=XM_006939218.3;refGene_gn=UBAP2;refGene_tx=XM_006939219.3;refGene_gn=UBAP2;refGene_tx=XM_019816090.1
D4	57846939 57846995
refGene_gn=UBAP2;refGene_tx=XM_006939216.3;refGene_gn=UBAP2;refGene_tx=XM_006939217.3;refGene_gn=UBAP2;refGene_tx=XM_006939218.3;refGene_gn=UBAP2;refGene_tx=XM_006939219.3;refGene_gn=UBAP2;refGene_tx=XM_019816090.1
D4	57852291 57852357
refGene_gn=UBAP2;refGene_tx=XM_006939216.3;refGene_gn=UBAP2;refGene_tx=XM_006939217.3;refGene_gn=UBAP2;refGene_tx=XM_019816090.1
D4	57858647 57858751
refGene_gn=UBAP2;refGene_tx=XM_006939216.3;refGene_gn=UBAP2;refGene_tx=XM_006939217.3;refGene_gn=UBAP2;refGene_tx=XM_006939218.3;refGene_gn=UBAP2;refGene_tx=XM_006939219.3;refGene_gn=UBAP2;refGene_tx=XM_019816090.1
D4	57861564 57861619

refGene_gn=UBAP2;refGene_tx=XM_006939216.3;refGene_gn=UBAP2;refGene_tx=XM_006939217.3;refGene_gn=UBAP2;refGene_tx=XM_006939218.3;refGene_gn=UBAP2;refGene_tx=XM_006939219.3;refGene_gn=UBAP2;refGene_tx=XM_019816090.1
D4 57875645 57875723
refGene_gn=UBAP2;refGene_tx=XM_006939216.3;refGene_gn=UBAP2;refGene_tx=XM_006939217.3;refGene_gn=UBAP2;refGene_tx=XM_006939218.3;refGene_gn=UBAP2;refGene_tx=XM_006939219.3;refGene_gn=UBAP2;refGene_tx=XM_019816090.1
D4 57877580 57877734
refGene_gn=UBAP2;refGene_tx=XM_006939216.3;refGene_gn=UBAP2;refGene_tx=XM_006939217.3;refGene_gn=UBAP2;refGene_tx=XM_006939218.3;refGene_gn=UBAP2;refGene_tx=XM_006939219.3;refGene_gn=UBAP2;refGene_tx=XM_019816090.1
D4 57883007 57883118
refGene_gn=UBAP2;refGene_tx=XM_006939216.3;refGene_gn=UBAP2;refGene_tx=XM_006939217.3;refGene_gn=UBAP2;refGene_tx=XM_006939218.3;refGene_gn=UBAP2;refGene_tx=XM_006939219.3;refGene_gn=UBAP2;refGene_tx=XM_019816090.1
D4 57886368 57886446
refGene_gn=UBAP2;refGene_tx=XM_006939216.3;refGene_gn=UBAP2;refGene_tx=XM_006939217.3;refGene_gn=UBAP2;refGene_tx=XM_006939218.3;refGene_gn=UBAP2;refGene_tx=XM_006939219.3;refGene_gn=UBAP2;refGene_tx=XM_019816090.1
D4 57892744 57892843
refGene_gn=UBAP2;refGene_tx=XM_006939216.3;refGene_gn=UBAP2;refGene_tx=XM_006939217.3;refGene_gn=UBAP2;refGene_tx=XM_006939218.3;refGene_gn=UBAP2;refGene_tx=XM_006939219.3;refGene_gn=UBAP2;refGene_tx=XM_019816090.1
D4 57961397 57961556
refGene_gn=DCAF12;refGene_tx=XM_003995562.4;refGene_gn=DCAF12;refGene_tx=XM_019816092.1
D4 57962099 57962278
refGene_gn=DCAF12;refGene_tx=XM_003995562.4;refGene_gn=DCAF12;refGene_tx=XM_019816092.1
D4 57964736 57964899
refGene_gn=DCAF12;refGene_tx=XM_003995562.4;refGene_gn=DCAF12;refGene_tx=XM_019816092.1
D4 57966034 57966100
refGene_gn=DCAF12;refGene_tx=XM_003995562.4;refGene_gn=DCAF12;refGene_tx=XM_019816092.1
D4 57967071 57967265
refGene_gn=DCAF12;refGene_tx=XM_003995562.4;refGene_gn=DCAF12;refGene_tx=XM_019816092.1
D4 57972222 57972283
refGene_gn=DCAF12;refGene_tx=XM_003995562.4;refGene_gn=DCAF12;refGene_tx=XM_019816092.1
D4 57973372 57973579
refGene_gn=DCAF12;refGene_tx=XM_003995562.4;refGene_gn=DCAF12;refGene_tx=XM_019816092.1

D4 57996785 57997040
refGene_gn=DCAF12;refGene_tx=XM_003995562.4;refGene_gn=DCAF12;refGene_tx=XM_019816092.1
D4 57997376 57997436 refGene_gn=DCAF12;refGene_tx=XM_019816092.1
D4 57998136 57998214 refGene_gn=DCAF12;refGene_tx=XM_003995562.4
D4 58083577 58083611
refGene_gn=UBAP1;refGene_tx=XM_006939220.4;refGene_gn=UBAP1;refGene_tx=XM_006939221.4;refGene_gn=UBAP1;refGene_tx=XM_019816104.2
D4 58094318 58094443
refGene_gn=UBAP1;refGene_tx=XM_006939220.4;refGene_gn=UBAP1;refGene_tx=XM_006939221.4;refGene_gn=UBAP1;refGene_tx=XM_019816104.2
D4 58099599 58100535
refGene_gn=UBAP1;refGene_tx=XM_006939220.4;refGene_gn=UBAP1;refGene_tx=XM_006939221.4;refGene_gn=UBAP1;refGene_tx=XM_019816104.2
D4 58109284 58109467
refGene_gn=UBAP1;refGene_tx=XM_006939220.4;refGene_gn=UBAP1;refGene_tx=XM_006939221.4;refGene_gn=UBAP1;refGene_tx=XM_019816104.2
D4 58110190 58110292
refGene_gn=UBAP1;refGene_tx=XM_006939220.4;refGene_gn=UBAP1;refGene_tx=XM_006939221.4
D4 58110878 58111019
refGene_gn=UBAP1;refGene_tx=XM_006939220.4;refGene_gn=UBAP1;refGene_tx=XM_006939221.4;refGene_gn=UBAP1;refGene_tx=XM_019816104.2
D4 58113734 58113875
refGene_gn=KIF24;refGene_tx=XM_011288585.2;refGene_gn=KIF24;refGene_tx=XM_019816094.2;refGene_gn=KIF24;refGene_tx=XM_019816096.2;refGene_gn=KIF24;refGene_tx=XM_019816100.2;refGene_gn=KIF24;refGene_tx=XM_023242392.1
D4 58114946 58115040
refGene_gn=KIF24;refGene_tx=XM_011288585.2;refGene_gn=KIF24;refGene_tx=XM_019816094.2;refGene_gn=KIF24;refGene_tx=XM_019816096.2;refGene_gn=KIF24;refGene_tx=XM_019816100.2;refGene_gn=KIF24;refGene_tx=XM_023242392.1
D4 58116191 58118420
refGene_gn=KIF24;refGene_tx=XM_011288585.2;refGene_gn=KIF24;refGene_tx=XM_019816094.2;refGene_gn=KIF24;refGene_tx=XM_019816096.2;refGene_gn=KIF24;refGene_tx=XM_019816100.2;refGene_gn=KIF24;refGene_tx=XM_023242392.1
D4 58122377 58122487
refGene_gn=KIF24;refGene_tx=XM_011288585.2;refGene_gn=KIF24;refGene_tx=XM_019816094.2;refGene_gn=KIF24;refGene_tx=XM_019816096.2;refGene_gn=KIF24;refGene_tx=XM_019816100.2;refGene_gn=KIF24;refGene_tx=XM_023242392.1
D4 58125228 58125300
refGene_gn=KIF24;refGene_tx=XM_011288585.2;refGene_gn=KIF24;refGene_tx=XM_019816094.2;refGene_gn=KIF24;refGene_tx=XM_019816096.2;refGene_gn=KIF24;refGene_tx=XM_019816100.2;refGene_gn=KIF24;refGene_tx=XM_023242392.1
D4 58132949 58133055
refGene_gn=KIF24;refGene_tx=XM_011288585.2;refGene_gn=KIF24;refGene_tx

=XM_019816094.2;refGene_gn=KIF24;refGene_tx=XM_019816096.2;refGene_gn=KIF24;refGene_tx=XM_019816100.2;refGene_gn=KIF24;refGene_tx=XM_023242392.1
D4 58133940 58134062
refGene_gn=KIF24;refGene_tx=XM_011288585.2;refGene_gn=KIF24;refGene_tx=XM_019816094.2;refGene_gn=KIF24;refGene_tx=XM_019816096.2;refGene_gn=KIF24;refGene_tx=XM_019816100.2;refGene_gn=KIF24;refGene_tx=XM_023242392.1
D4 58142969 58143057
refGene_gn=KIF24;refGene_tx=XM_011288585.2;refGene_gn=KIF24;refGene_tx=XM_019816094.2;refGene_gn=KIF24;refGene_tx=XM_019816096.2;refGene_gn=KIF24;refGene_tx=XM_019816100.2;refGene_gn=KIF24;refGene_tx=XM_023242392.1
D4 58146899 58147115
refGene_gn=KIF24;refGene_tx=XM_011288585.2;refGene_gn=KIF24;refGene_tx=XM_019816094.2;refGene_gn=KIF24;refGene_tx=XM_019816100.2;refGene_gn=KIF24;refGene_tx=XM_023242392.1
D4 58150901 58150999
refGene_gn=KIF24;refGene_tx=XM_011288585.2;refGene_gn=KIF24;refGene_tx=XM_019816094.2;refGene_gn=KIF24;refGene_tx=XM_019816096.2;refGene_gn=KIF24;refGene_tx=XM_019816100.2;refGene_gn=KIF24;refGene_tx=XM_023242392.1
D4 58155998 58156188
refGene_gn=KIF24;refGene_tx=XM_011288585.2;refGene_gn=KIF24;refGene_tx=XM_019816094.2;refGene_gn=KIF24;refGene_tx=XM_019816096.2;refGene_gn=KIF24;refGene_tx=XM_019816100.2;refGene_gn=KIF24;refGene_tx=XM_023242392.1
D4 58163257 58163906
refGene_gn=KIF24;refGene_tx=XM_011288585.2;refGene_gn=KIF24;refGene_tx=XM_019816094.2;refGene_gn=KIF24;refGene_tx=XM_019816096.2;refGene_gn=KIF24;refGene_tx=XM_023242392.1
D4 58183571 58183638
refGene_gn=KIF24;refGene_tx=XM_019816096.2;refGene_gn=KIF24;refGene_tx=XM_023242392.1
D4 58189113 58189215
refGene_gn=KIF24;refGene_tx=XM_019816096.2;refGene_gn=KIF24;refGene_tx=XM_023242392.1
D4 58199062 58199189
refGene_gn=NUDT2;refGene_tx=XM_006939223.4;refGene_gn=NUDT2;refGene_tx=XM_019816105.1
D4 58201329 58201646
refGene_gn=NUDT2;refGene_tx=XM_006939223.4;refGene_gn=NUDT2;refGene_tx=XM_019816105.1
D4 58209763 58211908
refGene_gn=MYORG;refGene_tx=XM_003995566.5;refGene_gn=MYORG;refGene_tx=XM_006939224.4
D4 58216822 58216928
refGene_gn=CD4H9orf24;refGene_tx=XM_006939225.3;refGene_gn=CD4H9orf24;refGene_tx=XM_019816106.2;refGene_gn=CD4H9orf24;refGene_tx=XM_019816107.2;refGene_gn=CD4H9orf24;refGene_tx=XM_019816108.2;refGene_gn=CD4H9or

f24;refGene_tx=XM_019816109.2;refGene_gn=CD4H9orf24;refGene_tx=XM_019816110.2;refGene_gn=CD4H9orf24;refGene_tx=XM_023242394.1;refGene_gn=CD4H9orf24;refGene_tx=XM_023242395.1;refGene_gn=CD4H9orf24;refGene_tx=XM_023242396.1
D4 58217352 58217439
refGene_gn=CD4H9orf24;refGene_tx=XM_003995567.4;refGene_gn=CD4H9orf24;refGene_tx=XM_006939225.3;refGene_gn=CD4H9orf24;refGene_tx=XM_019816106.2;refGene_gn=CD4H9orf24;refGene_tx=XM_019816107.2;refGene_gn=CD4H9orf24;refGene_tx=XM_019816108.2;refGene_gn=CD4H9orf24;refGene_tx=XM_019816109.2;refGene_gn=CD4H9orf24;refGene_tx=XM_019816110.2;refGene_gn=CD4H9orf24;refGene_tx=XM_023242396.1
D4 58218732 58218967
refGene_gn=CD4H9orf24;refGene_tx=XM_003995567.4;refGene_gn=CD4H9orf24;refGene_tx=XM_006939225.3;refGene_gn=CD4H9orf24;refGene_tx=XM_019816106.2;refGene_gn=CD4H9orf24;refGene_tx=XM_019816107.2;refGene_gn=CD4H9orf24;refGene_tx=XM_019816108.2;refGene_gn=CD4H9orf24;refGene_tx=XM_019816109.2;refGene_gn=CD4H9orf24;refGene_tx=XM_019816110.2;refGene_gn=CD4H9orf24;refGene_tx=XM_023242394.1;refGene_gn=CD4H9orf24;refGene_tx=XM_023242395.1;refGene_gn=CD4H9orf24;refGene_tx=XM_023242396.1
D4 58219178 58219254
refGene_gn=CD4H9orf24;refGene_tx=XM_003995567.4;refGene_gn=CD4H9orf24;refGene_tx=XM_006939225.3;refGene_gn=CD4H9orf24;refGene_tx=XM_019816106.2;refGene_gn=CD4H9orf24;refGene_tx=XM_019816107.2;refGene_gn=CD4H9orf24;refGene_tx=XM_019816108.2;refGene_gn=CD4H9orf24;refGene_tx=XM_019816109.2;refGene_gn=CD4H9orf24;refGene_tx=XM_019816110.2;refGene_gn=CD4H9orf24;refGene_tx=XM_023242394.1;refGene_gn=CD4H9orf24;refGene_tx=XM_023242395.1;refGene_gn=CD4H9orf24;refGene_tx=XM_023242396.1
D4 58220490 58220606
refGene_gn=CD4H9orf24;refGene_tx=XM_003995567.4;refGene_gn=CD4H9orf24;refGene_tx=XM_006939225.3;refGene_gn=CD4H9orf24;refGene_tx=XM_019816106.2;refGene_gn=CD4H9orf24;refGene_tx=XM_019816107.2;refGene_gn=CD4H9orf24;refGene_tx=XM_019816108.2;refGene_gn=CD4H9orf24;refGene_tx=XM_023242394.1;refGene_gn=CD4H9orf24;refGene_tx=XM_023242395.1;refGene_gn=CD4H9orf24;refGene_tx=XM_023242396.1
D4 58222696 58222841
refGene_gn=CD4H9orf24;refGene_tx=XM_003995567.4;refGene_gn=CD4H9orf24;refGene_tx=XM_006939225.3;refGene_gn=CD4H9orf24;refGene_tx=XM_019816106.2;refGene_gn=CD4H9orf24;refGene_tx=XM_019816107.2;refGene_gn=CD4H9orf24;refGene_tx=XM_019816108.2;refGene_gn=CD4H9orf24;refGene_tx=XM_023242394.1;refGene_gn=CD4H9orf24;refGene_tx=XM_023242395.1;refGene_gn=CD4H9orf24;refGene_tx=XM_023242396.1
D4 58235180 58235315
refGene_gn=CD4H9orf24;refGene_tx=XM_003995567.4;refGene_gn=CD4H9orf24;refGene_tx=XM_006939225.3;refGene_gn=CD4H9orf24;refGene_tx=XM_019816106.2;refGene_gn=CD4H9orf24;refGene_tx=XM_019816107.2;refGene_gn=CD4H9orf24;refGene_tx=XM_023242394.1;refGene_gn=CD4H9orf24;refGene_tx=XM_023242395.1
D4 58235353 58235373
refGene_gn=CD4H9orf24;refGene_tx=XM_019816107.2
D4 58238900 58239059

refGene_gn=FAM219A;refGene_tx=XM_011288579.2;refGene_gn=FAM219A;refGene_tx=XM_011288580.2;refGene_gn=FAM219A;refGene_tx=XM_011288582.2;refGene_gn=FAM219A;refGene_tx=XM_011288584.2
D4 58239612 58239667
refGene_gn=FAM219A;refGene_tx=XM_011288579.2;refGene_gn=FAM219A;refGene_tx=XM_011288580.2;refGene_gn=FAM219A;refGene_tx=XM_011288582.2;refGene_gn=FAM219A;refGene_tx=XM_011288584.2
D4 58240275 58240356
refGene_gn=FAM219A;refGene_tx=XM_011288579.2;refGene_gn=FAM219A;refGene_tx=XM_011288580.2;refGene_gn=FAM219A;refGene_tx=XM_011288582.2;refGene_gn=FAM219A;refGene_tx=XM_011288584.2
D4 58240580 58240683
refGene_gn=FAM219A;refGene_tx=XM_011288579.2;refGene_gn=FAM219A;refGene_tx=XM_011288580.2;refGene_gn=FAM219A;refGene_tx=XM_011288582.2;refGene_gn=FAM219A;refGene_tx=XM_011288584.2
D4 58243688 58243788
refGene_gn=FAM219A;refGene_tx=XM_011288579.2;refGene_gn=FAM219A;refGene_tx=XM_011288580.2;refGene_gn=FAM219A;refGene_tx=XM_011288582.2;refGene_gn=FAM219A;refGene_tx=XM_011288584.2
D4 58289967 58290027
refGene_gn=FAM219A;refGene_tx=XM_011288579.2;refGene_gn=FAM219A;refGene_tx=XM_011288580.2;refGene_gn=FAM219A;refGene_tx=XM_011288582.2;refGene_gn=FAM219A;refGene_tx=XM_011288584.2
D4 58290798 58290828 refGene_gn=DNAI1;refGene_tx=XM_003995569.5
D4 58311163 58311196 refGene_gn=DNAI1;refGene_tx=XM_003995569.5
D4 58312867 58312966 refGene_gn=DNAI1;refGene_tx=XM_003995569.5
D4 58313154 58313235 refGene_gn=DNAI1;refGene_tx=XM_003995569.5
D4 58315084 58315211 refGene_gn=DNAI1;refGene_tx=XM_003995569.5
D4 58315479 58315607 refGene_gn=DNAI1;refGene_tx=XM_003995569.5
D4 58315851 58315971 refGene_gn=DNAI1;refGene_tx=XM_003995569.5
D4 58316951 58317011 refGene_gn=DNAI1;refGene_tx=XM_003995569.5
D4 58318221 58318356 refGene_gn=DNAI1;refGene_tx=XM_003995569.5
D4 58321261 58321346 refGene_gn=DNAI1;refGene_tx=XM_003995569.5
D4 58324667 58324785 refGene_gn=DNAI1;refGene_tx=XM_003995569.5
D4 58325070 58325114 refGene_gn=DNAI1;refGene_tx=XM_003995569.5
D4 58334424 58334672 refGene_gn=DNAI1;refGene_tx=XM_003995569.5
D4 58339498 58339588 refGene_gn=DNAI1;refGene_tx=XM_003995569.5
D4 58339718 58339806 refGene_gn=DNAI1;refGene_tx=XM_003995569.5
D4 58340439 58340519 refGene_gn=DNAI1;refGene_tx=XM_003995569.5
D4 58341983 58342132 refGene_gn=DNAI1;refGene_tx=XM_003995569.5
D4 58342238 58342338 refGene_gn=DNAI1;refGene_tx=XM_003995569.5
D4 58344723 58344909 refGene_gn=DNAI1;refGene_tx=XM_003995569.5
D4 58347936 58348035 refGene_gn=DNAI1;refGene_tx=XM_003995569.5
D4 58348739 58348970 refGene_gn=ENHO;refGene_tx=XM_023242405.1
D4 58368197 58368270 refGene_gn=CNTFR;refGene_tx=XM_023242403.1
D4 58369365 58369376 refGene_gn=CNTFR;refGene_tx=XM_023242403.1
D4 58370093 58370175
refGene_gn=CNTFR;refGene_tx=XM_023242398.1;refGene_gn=CNTFR;refGene_tx=XM_023242399.1;refGene_gn=CNTFR;refGene_tx=XM_023242400.1;refGene_gn=CNTFR;refGene_tx=XM_023242401.1;refGene_gn=CNTFR;refGene_tx=XM_0232424

04.1
D4 58370245 58370414
refGene_gn=CNTFR;refGene_tx=XM_023242398.1;refGene_gn=CNTFR;refGene_tx=XM_023242399.1;refGene_gn=CNTFR;refGene_tx=XM_023242400.1;refGene_gn=CNTFR;refGene_tx=XM_023242401.1;refGene_gn=CNTFR;refGene_tx=XM_023242402.1;refGene_gn=CNTFR;refGene_tx=XM_023242404.1
D4 58370815 58370996
refGene_gn=CNTFR;refGene_tx=XM_023242398.1;refGene_gn=CNTFR;refGene_tx=XM_023242399.1;refGene_gn=CNTFR;refGene_tx=XM_023242400.1;refGene_gn=CNTFR;refGene_tx=XM_023242401.1;refGene_gn=CNTFR;refGene_tx=XM_023242402.1;refGene_gn=CNTFR;refGene_tx=XM_023242403.1;refGene_gn=CNTFR;refGene_tx=XM_023242404.1
D4 58374458 58374622
refGene_gn=CNTFR;refGene_tx=XM_023242398.1;refGene_gn=CNTFR;refGene_tx=XM_023242399.1;refGene_gn=CNTFR;refGene_tx=XM_023242400.1;refGene_gn=CNTFR;refGene_tx=XM_023242401.1;refGene_gn=CNTFR;refGene_tx=XM_023242402.1;refGene_gn=CNTFR;refGene_tx=XM_023242403.1;refGene_gn=CNTFR;refGene_tx=XM_023242404.1
D4 58375861 58376028
refGene_gn=CNTFR;refGene_tx=XM_023242398.1;refGene_gn=CNTFR;refGene_tx=XM_023242399.1;refGene_gn=CNTFR;refGene_tx=XM_023242400.1;refGene_gn=CNTFR;refGene_tx=XM_023242401.1;refGene_gn=CNTFR;refGene_tx=XM_023242402.1;refGene_gn=CNTFR;refGene_tx=XM_023242403.1;refGene_gn=CNTFR;refGene_tx=XM_023242404.1
D4 58376202 58376320
refGene_gn=CNTFR;refGene_tx=XM_023242398.1;refGene_gn=CNTFR;refGene_tx=XM_023242399.1;refGene_gn=CNTFR;refGene_tx=XM_023242400.1;refGene_gn=CNTFR;refGene_tx=XM_023242401.1;refGene_gn=CNTFR;refGene_tx=XM_023242402.1;refGene_gn=CNTFR;refGene_tx=XM_023242403.1;refGene_gn=CNTFR;refGene_tx=XM_023242404.1
D4 58383172 58383406
refGene_gn=CNTFR;refGene_tx=XM_023242398.1;refGene_gn=CNTFR;refGene_tx=XM_023242399.1;refGene_gn=CNTFR;refGene_tx=XM_023242400.1;refGene_gn=CNTFR;refGene_tx=XM_023242401.1;refGene_gn=CNTFR;refGene_tx=XM_023242402.1;refGene_gn=CNTFR;refGene_tx=XM_023242403.1
D4 58387627 58387712
refGene_gn=CNTFR;refGene_tx=XM_023242398.1;refGene_gn=CNTFR;refGene_tx=XM_023242399.1;refGene_gn=CNTFR;refGene_tx=XM_023242400.1;refGene_gn=CNTFR;refGene_tx=XM_023242401.1;refGene_gn=CNTFR;refGene_tx=XM_023242402.1;refGene_gn=CNTFR;refGene_tx=XM_023242403.1;refGene_gn=CNTFR;refGene_tx=XM_023242404.1
D4 58426669 58427161
refGene_gn=RPP25L;refGene_tx=XM_003995575.4;refGene_gn=RPP25L;refGene_tx=XM_006939231.4;refGene_gn=RPP25L;refGene_tx=XM_006939232.4;refGene_gn=RPP25L;refGene_tx=XM_011288578.2
D4 58430617 58430707
refGene_gn=DCTN3;refGene_tx=XM_003995573.5;refGene_gn=DCTN3;refGene_tx=XM_011288575.3;refGene_gn=DCTN3;refGene_tx=XM_011288576.3;refGene_gn=DCTN3;refGene_tx=XM_019816118.2
D4 58430896 58430956

refGene_gn=DCTN3;refGene_tx=XM_003995573.5;refGene_gn=DCTN3;refGene_tx
=XM_011288575.3;refGene_gn=DCTN3;refGene_tx=XM_011288576.3;refGene_gn=
DCTN3;refGene_tx=XM_019816118.2
D4 58431576 58431635
refGene_gn=DCTN3;refGene_tx=XM_003995573.5;refGene_gn=DCTN3;refGene_tx
=XM_011288575.3;refGene_gn=DCTN3;refGene_tx=XM_011288576.3;refGene_gn=
DCTN3;refGene_tx=XM_019816118.2
D4 58431773 58432013
refGene_gn=DCTN3;refGene_tx=XM_019816115.2;refGene_gn=DCTN3;refGene_tx
=XM_019816116.2
D4 58432405 58432458
refGene_gn=DCTN3;refGene_tx=XM_011288575.3;refGene_gn=DCTN3;refGene_tx
=XM_019816115.2;refGene_gn=DCTN3;refGene_tx=XM_019816116.2;refGene_gn=
DCTN3;refGene_tx=XM_019816118.2
D4 58432609 58432705
refGene_gn=DCTN3;refGene_tx=XM_003995573.5;refGene_gn=DCTN3;refGene_tx
=XM_011288575.3;refGene_gn=DCTN3;refGene_tx=XM_011288576.3;refGene_gn=
DCTN3;refGene_tx=XM_019816115.2;refGene_gn=DCTN3;refGene_tx=XM_0198161
16.2;refGene_gn=DCTN3;refGene_tx=XM_019816118.2
D4 58433708 58433795
refGene_gn=DCTN3;refGene_tx=XM_003995573.5;refGene_gn=DCTN3;refGene_tx
=XM_011288575.3;refGene_gn=DCTN3;refGene_tx=XM_011288576.3;refGene_gn=
DCTN3;refGene_tx=XM_019816115.2;refGene_gn=DCTN3;refGene_tx=XM_0198161
16.2;refGene_gn=DCTN3;refGene_tx=XM_019816118.2
D4 58434492 58434577
refGene_gn=DCTN3;refGene_tx=XM_003995573.5;refGene_gn=DCTN3;refGene_tx
=XM_011288575.3;refGene_gn=DCTN3;refGene_tx=XM_011288576.3;refGene_gn=
DCTN3;refGene_tx=XM_019816115.2;refGene_gn=DCTN3;refGene_tx=XM_0198161
16.2;refGene_gn=DCTN3;refGene_tx=XM_019816118.2
D4 58436061 58436157
refGene_gn=DCTN3;refGene_tx=XM_003995573.5;refGene_gn=DCTN3;refGene_tx
=XM_011288575.3;refGene_gn=DCTN3;refGene_tx=XM_011288576.3;refGene_gn=
DCTN3;refGene_tx=XM_019816115.2;refGene_gn=DCTN3;refGene_tx=XM_0198161
16.2;refGene_gn=DCTN3;refGene_tx=XM_019816118.2
D4 58437203 58437304
refGene_gn=ARID3C;refGene_tx=XM_003995576.4;refGene_gn=ARID3C;refGene_
tx=XM_023242406.1
D4 58437764 58437854 refGene_gn=ARID3C;refGene_tx=XM_003995576.4
D4 58438085 58438268
refGene_gn=ARID3C;refGene_tx=XM_003995576.4;refGene_gn=ARID3C;refGene_
tx=XM_023242406.1
D4 58438797 58439081
refGene_gn=ARID3C;refGene_tx=XM_003995576.4;refGene_gn=ARID3C;refGene_
tx=XM_023242406.1
D4 58439226 58439410
refGene_gn=ARID3C;refGene_tx=XM_003995576.4;refGene_gn=ARID3C;refGene_
tx=XM_023242406.1
D4 58441246 58441319
refGene_gn=ARID3C;refGene_tx=XM_003995576.4;refGene_gn=ARID3C;refGene_
tx=XM_023242406.1

D4 58443177 58443564
refGene_gn=ARID3C;refGene_tx=XM_003995576.4;refGene_gn=ARID3C;refGene_tx=XM_023242406.1
D4 58444421 58444778
refGene_gn=ARID3C;refGene_tx=XM_003995576.4;refGene_gn=ARID3C;refGene_tx=XM_023242406.1
D4 58452434 58452661 refGene_gn=SIGMAR1;refGene_tx=XM_003995577.4
D4 58452784 58452804 refGene_gn=SIGMAR1;refGene_tx=XM_019816131.1
D4 58452980 58452997 refGene_gn=SIGMAR1;refGene_tx=XM_019816132.1
D4 58453478 58453571
refGene_gn=SIGMAR1;refGene_tx=XM_003995577.4;refGene_gn=SIGMAR1;refGene_tx=XM_019816131.1;refGene_gn=SIGMAR1;refGene_tx=XM_019816132.1
D4 58453704 58453905
refGene_gn=SIGMAR1;refGene_tx=XM_003995577.4;refGene_gn=SIGMAR1;refGene_tx=XM_019816131.1;refGene_gn=SIGMAR1;refGene_tx=XM_019816132.1
D4 58454049 58454200
refGene_gn=SIGMAR1;refGene_tx=XM_003995577.4;refGene_gn=SIGMAR1;refGene_tx=XM_019816131.1;refGene_gn=SIGMAR1;refGene_tx=XM_019816132.1
D4 58459886 58459968
refGene_gn=GALT;refGene_tx=XM_003995580.4;refGene_gn=GALT;refGene_tx=XM_006939234.3;refGene_gn=GALT;refGene_tx=XM_019816130.1
D4 58460267 58460437
refGene_gn=GALT;refGene_tx=XM_003995580.4;refGene_gn=GALT;refGene_tx=XM_006939234.3;refGene_gn=GALT;refGene_tx=XM_006939235.4;refGene_gn=GALT;refGene_tx=XM_019816130.1
D4 58460667 58460743
refGene_gn=GALT;refGene_tx=XM_003995580.4;refGene_gn=GALT;refGene_tx=XM_006939234.3;refGene_gn=GALT;refGene_tx=XM_006939235.4;refGene_gn=GALT;refGene_tx=XM_019816130.1
D4 58460829 58460878
refGene_gn=GALT;refGene_tx=XM_003995580.4;refGene_gn=GALT;refGene_tx=XM_006939234.3;refGene_gn=GALT;refGene_tx=XM_006939235.4;refGene_gn=GALT;refGene_tx=XM_019816130.1
D4 58460998 58461128
refGene_gn=GALT;refGene_tx=XM_003995580.4;refGene_gn=GALT;refGene_tx=XM_006939234.3;refGene_gn=GALT;refGene_tx=XM_006939235.4;refGene_gn=GALT;refGene_tx=XM_019816130.1
D4 58461281 58461338
refGene_gn=GALT;refGene_tx=XM_003995580.4;refGene_gn=GALT;refGene_tx=XM_006939235.4;refGene_gn=GALT;refGene_tx=XM_019816130.1
D4 58461496 58461619
refGene_gn=GALT;refGene_tx=XM_003995580.4;refGene_gn=GALT;refGene_tx=XM_006939234.3;refGene_gn=GALT;refGene_tx=XM_006939235.4;refGene_gn=GALT;refGene_tx=XM_019816130.1
D4 58461897 58462030
refGene_gn=GALT;refGene_tx=XM_003995580.4;refGene_gn=GALT;refGene_tx=XM_006939234.3;refGene_gn=GALT;refGene_tx=XM_006939235.4;refGene_gn=GALT;refGene_tx=XM_019816130.1
D4 58462133 58462217
refGene_gn=GALT;refGene_tx=XM_003995580.4;refGene_gn=GALT;refGene_tx=X

M_006939234.3;refGene_gn=GALT;refGene_tx=XM_006939235.4
D4 58462535 58462690
refGene_gn=GALT;refGene_tx=XM_003995580.4;refGene_gn=GALT;refGene_tx=XM_006939234.3;refGene_gn=GALT;refGene_tx=XM_006939235.4;refGene_gn=GALT;refGene_tx=XM_019816130.1
D4 58462985 58463066
refGene_gn=GALT;refGene_tx=XM_003995580.4;refGene_gn=GALT;refGene_tx=XM_006939234.3;refGene_gn=GALT;refGene_tx=XM_006939235.4;refGene_gn=GALT;refGene_tx=XM_019816130.1
D4 58467788 58467888
refGene_gn=IL11RA;refGene_tx=XM_003995583.3;refGene_gn=IL11RA;refGene_tx=XM_006939236.2;refGene_gn=IL11RA;refGene_tx=XM_006939237.3;refGene_gn=IL11RA;refGene_tx=XM_006939238.2;refGene_gn=IL11RA;refGene_tx=XM_006939239.3;refGene_gn=IL11RA;refGene_tx=XM_019816124.1
D4 58468200 58468261
refGene_gn=IL11RA;refGene_tx=XM_003995583.3;refGene_gn=IL11RA;refGene_tx=XM_006939236.2;refGene_gn=IL11RA;refGene_tx=XM_006939237.3;refGene_gn=IL11RA;refGene_tx=XM_006939238.2;refGene_gn=IL11RA;refGene_tx=XM_006939239.3;refGene_gn=IL11RA;refGene_tx=XM_019816124.1
D4 58468985 58469155
refGene_gn=IL11RA;refGene_tx=XM_003995583.3;refGene_gn=IL11RA;refGene_tx=XM_006939236.2;refGene_gn=IL11RA;refGene_tx=XM_006939237.3;refGene_gn=IL11RA;refGene_tx=XM_006939238.2;refGene_gn=IL11RA;refGene_tx=XM_006939239.3;refGene_gn=IL11RA;refGene_tx=XM_019816124.1
D4 58469291 58469406
refGene_gn=IL11RA;refGene_tx=XM_003995583.3;refGene_gn=IL11RA;refGene_tx=XM_006939236.2;refGene_gn=IL11RA;refGene_tx=XM_006939237.3;refGene_gn=IL11RA;refGene_tx=XM_006939238.2;refGene_gn=IL11RA;refGene_tx=XM_006939239.3;refGene_gn=IL11RA;refGene_tx=XM_019816124.1
D4 58469792 58469825
refGene_gn=IL11RA;refGene_tx=XM_003995583.3;refGene_gn=IL11RA;refGene_tx=XM_006939236.2;refGene_gn=IL11RA;refGene_tx=XM_006939237.3;refGene_gn=IL11RA;refGene_tx=XM_006939238.2;refGene_gn=IL11RA;refGene_tx=XM_006939239.3;refGene_gn=IL11RA;refGene_tx=XM_019816124.1
D4 58469909 58470076
refGene_gn=IL11RA;refGene_tx=XM_003995583.3;refGene_gn=IL11RA;refGene_tx=XM_006939236.2;refGene_gn=IL11RA;refGene_tx=XM_006939237.3;refGene_gn=IL11RA;refGene_tx=XM_006939238.2;refGene_gn=IL11RA;refGene_tx=XM_006939239.3;refGene_gn=IL11RA;refGene_tx=XM_019816124.1
D4 58470749 58470913
refGene_gn=IL11RA;refGene_tx=XM_003995583.3;refGene_gn=IL11RA;refGene_tx=XM_006939236.2;refGene_gn=IL11RA;refGene_tx=XM_006939237.3;refGene_gn=IL11RA;refGene_tx=XM_006939238.2;refGene_gn=IL11RA;refGene_tx=XM_006939239.3;refGene_gn=IL11RA;refGene_tx=XM_019816124.1
D4 58471708 58471850
refGene_gn=IL11RA;refGene_tx=XM_003995583.3;refGene_gn=IL11RA;refGene_tx=XM_006939236.2;refGene_gn=IL11RA;refGene_tx=XM_006939237.3;refGene_gn=IL11RA;refGene_tx=XM_006939238.2;refGene_gn=IL11RA;refGene_tx=XM_006939239.3;refGene_gn=IL11RA;refGene_tx=XM_019816124.1
D4 58472116 58472236

refGene_gn=IL11RA;refGene_tx=XM_003995583.3;refGene_gn=IL11RA;refGene_tx=XM_006939236.2;refGene_gn=IL11RA;refGene_tx=XM_006939237.3;refGene_gn=IL11RA;refGene_tx=XM_006939239.3;refGene_gn=IL11RA;refGene_tx=XM_019816124.1
D4 58472348 58472445
refGene_gn=IL11RA;refGene_tx=XM_003995583.3;refGene_gn=IL11RA;refGene_tx=XM_006939236.2;refGene_gn=IL11RA;refGene_tx=XM_006939237.3;refGene_gn=IL11RA;refGene_tx=XM_006939238.2;refGene_gn=IL11RA;refGene_tx=XM_006939239.3;refGene_gn=IL11RA;refGene_tx=XM_019816124.1
D4 58472671 58472754
refGene_gn=IL11RA;refGene_tx=XM_003995583.3;refGene_gn=IL11RA;refGene_tx=XM_006939236.2;refGene_gn=IL11RA;refGene_tx=XM_006939237.3;refGene_gn=IL11RA;refGene_tx=XM_006939238.2;refGene_gn=IL11RA;refGene_tx=XM_006939239.3;refGene_gn=IL11RA;refGene_tx=XM_019816124.1
D4 58473259 58473276 refGene_gn=IL11RA;refGene_tx=XM_003995583.3
D4 58473799 58473944 refGene_gn=CCL27;refGene_tx=XM_003995585.4
D4 58474148 58474281 refGene_gn=CCL27;refGene_tx=XM_003995585.4
D4 58474424 58474494 refGene_gn=CCL27;refGene_tx=XM_003995585.4
D4 58474653 58474744 refGene_gn=IL11RA;refGene_tx=XM_019816124.1
D4 58476367 58476461
refGene_gn=IL11RA;refGene_tx=XM_006939236.2;refGene_gn=IL11RA;refGene_tx=XM_006939237.3;refGene_gn=IL11RA;refGene_tx=XM_006939238.2
D4 58477338 58477495
refGene_gn=LOC109493846;refGene_tx=XM_019816134.2;refGene_gn=LOC109493846;refGene_tx=XM_019816135.2;refGene_gn=LOC109493846;refGene_tx=XM_019816136.2;refGene_gn=LOC109493846;refGene_tx=XM_019816137.1;refGene_gn=LOC109493846;refGene_tx=XM_019816138.2
D4 58477615 58477675
refGene_gn=LOC109493846;refGene_tx=XM_019816134.2;refGene_gn=LOC109493846;refGene_tx=XM_019816135.2;refGene_gn=LOC109493846;refGene_tx=XM_019816136.2;refGene_gn=LOC109493846;refGene_tx=XM_019816137.1;refGene_gn=LOC109493846;refGene_tx=XM_019816138.2
D4 58477904 58477951
refGene_gn=LOC109493846;refGene_tx=XM_019816134.2;refGene_gn=LOC109493846;refGene_tx=XM_019816135.2;refGene_gn=LOC109493846;refGene_tx=XM_019816136.2;refGene_gn=LOC109493846;refGene_tx=XM_019816137.1;refGene_gn=LOC109493846;refGene_tx=XM_019816138.2
D4 58480497 58480657
refGene_gn=IL11RA;refGene_tx=XM_006939236.2;refGene_gn=IL11RA;refGene_tx=XM_006939237.3;refGene_gn=IL11RA;refGene_tx=XM_006939238.2;refGene_gn=IL11RA;refGene_tx=XM_006939239.3;refGene_gn=IL11RA;refGene_tx=XM_019816124.1
D4 58501421 58501442 refGene_gn=CCL19;refGene_tx=XM_003995586.5
D4 58501532 58501626 refGene_gn=CCL19;refGene_tx=XM_003995586.5
D4 58501859 58501986 refGene_gn=CCL19;refGene_tx=XM_003995586.5
D4 58502698 58502759 refGene_gn=CCL19;refGene_tx=XM_003995586.5
D4 58521044 58521078 refGene_gn=CCL21;refGene_tx=XM_003995587.5
D4 58521159 58521342 refGene_gn=CCL21;refGene_tx=XM_003995587.5
D4 58521430 58521551 refGene_gn=CCL21;refGene_tx=XM_003995587.5
D4 58521645 58521712 refGene_gn=CCL21;refGene_tx=XM_003995587.5

D4 58594118 58597944 refGene_gn=FAM205A;refGene_tx=XM_006939241.3
D4 58598966 58599027 refGene_gn=FAM205A;refGene_tx=XM_006939241.3
D4 58599574 58599614 refGene_gn=FAM205A;refGene_tx=XM_006939241.3
D4 58600477 58600621 refGene_gn=FAM205A;refGene_tx=XM_006939241.3
D4 58665575 58666039 refGene_gn=FAM205C;refGene_tx=XM_023242410.1
D4 58669430 58669917
refGene_gn=FAM205C;refGene_tx=XM_023242410.1;refGene_gn=FAM205C;refGene_tx=XM_023242411.1;refGene_gn=FAM205C;refGene_tx=XM_023242413.1;refGene_gn=FAM205C;refGene_tx=XM_023242414.1
D4 58670944 58671005
refGene_gn=FAM205C;refGene_tx=XM_023242410.1;refGene_gn=FAM205C;refGene_tx=XM_023242411.1;refGene_gn=FAM205C;refGene_tx=XM_023242413.1;refGene_gn=FAM205C;refGene_tx=XM_023242414.1
D4 58671503 58671549
refGene_gn=FAM205C;refGene_tx=XM_023242410.1;refGene_gn=FAM205C;refGene_tx=XM_023242411.1;refGene_gn=FAM205C;refGene_tx=XM_023242413.1;refGene_gn=FAM205C;refGene_tx=XM_023242414.1
D4 58672048 58672192
refGene_gn=FAM205C;refGene_tx=XM_023242410.1;refGene_gn=FAM205C;refGene_tx=XM_023242411.1;refGene_gn=FAM205C;refGene_tx=XM_023242413.1;refGene_gn=FAM205C;refGene_tx=XM_023242414.1
D4 58726802 58726922
refGene_gn=LOC111557473;refGene_tx=XM_023242301.1
D4 58767030 58767424
refGene_gn=LOC111557473;refGene_tx=XM_023242301.1
D4 58767473 58767853
refGene_gn=LOC111557473;refGene_tx=XM_023242301.1
D4 58781022 58781400
refGene_gn=PHF24;refGene_tx=XM_006939244.4;refGene_gn=PHF24;refGene_tx=XM_019816129.2;refGene_gn=PHF24;refGene_tx=XM_023242409.1
D4 58782042 58782228
refGene_gn=PHF24;refGene_tx=XM_006939244.4;refGene_gn=PHF24;refGene_tx=XM_019816129.2;refGene_gn=PHF24;refGene_tx=XM_023242409.1
D4 58785590 58785669
refGene_gn=PHF24;refGene_tx=XM_006939244.4;refGene_gn=PHF24;refGene_tx=XM_019816129.2;refGene_gn=PHF24;refGene_tx=XM_023242409.1
D4 58785983 58786189
refGene_gn=PHF24;refGene_tx=XM_006939244.4;refGene_gn=PHF24;refGene_tx=XM_019816129.2;refGene_gn=PHF24;refGene_tx=XM_023242409.1
D4 58786536 58786697
refGene_gn=PHF24;refGene_tx=XM_006939244.4;refGene_gn=PHF24;refGene_tx=XM_019816129.2;refGene_gn=PHF24;refGene_tx=XM_023242409.1
D4 58787211 58787307
refGene_gn=PHF24;refGene_tx=XM_006939244.4;refGene_gn=PHF24;refGene_tx=XM_019816129.2;refGene_gn=PHF24;refGene_tx=XM_023242409.1
D4 58787627 58787724
refGene_gn=PHF24;refGene_tx=XM_006939244.4;refGene_gn=PHF24;refGene_tx=XM_019816129.2;refGene_gn=PHF24;refGene_tx=XM_023242409.1
D4 58798105 58798122 refGene_gn=DNAJB5;refGene_tx=XM_023242407.1
D4 58799426 58799589

refGene_gn=DNAJB5;refGene_tx=XM_006939245.4;refGene_gn=DNAJB5;refGene_tx=XM_019816125.2
D4 58800452 58800597
refGene_gn=DNAJB5;refGene_tx=XM_006939245.4;refGene_gn=DNAJB5;refGene_tx=XM_019816125.2
D4 58802949 58803194
refGene_gn=DNAJB5;refGene_tx=XM_003995590.5;refGene_gn=DNAJB5;refGene_tx=XM_006939245.4;refGene_gn=DNAJB5;refGene_tx=XM_006939246.4;refGene_gn=DNAJB5;refGene_tx=XM_019816125.2;refGene_gn=DNAJB5;refGene_tx=XM_019816126.2;refGene_gn=DNAJB5;refGene_tx=XM_019816127.2;refGene_gn=DNAJB5;refGene_tx=XM_023242407.1;refGene_gn=DNAJB5;refGene_tx=XM_023242408.1
D4 58806049 58806651
refGene_gn=DNAJB5;refGene_tx=XM_003995590.5;refGene_gn=DNAJB5;refGene_tx=XM_006939245.4;refGene_gn=DNAJB5;refGene_tx=XM_006939246.4;refGene_gn=DNAJB5;refGene_tx=XM_019816125.2;refGene_gn=DNAJB5;refGene_tx=XM_019816126.2;refGene_gn=DNAJB5;refGene_tx=XM_019816127.2;refGene_gn=DNAJB5;refGene_tx=XM_023242407.1;refGene_gn=DNAJB5;refGene_tx=XM_023242408.1
D4 58806789 58807023
refGene_gn=DNAJB5;refGene_tx=XM_003995590.5;refGene_gn=DNAJB5;refGene_tx=XM_006939245.4;refGene_gn=DNAJB5;refGene_tx=XM_006939246.4;refGene_gn=DNAJB5;refGene_tx=XM_019816125.2;refGene_gn=DNAJB5;refGene_tx=XM_019816126.2;refGene_gn=DNAJB5;refGene_tx=XM_019816127.2;refGene_gn=DNAJB5;refGene_tx=XM_023242407.1;refGene_gn=DNAJB5;refGene_tx=XM_023242408.1
D4 58841221 58841452
refGene_gn=CD4H9orf131;refGene_tx=XM_006939259.3
D4 58841855 58845335
refGene_gn=CD4H9orf131;refGene_tx=XM_006939259.3
D4 58852847 58852953
refGene_gn=VCP;refGene_tx=XM_019816140.2;refGene_gn=VCP;refGene_tx=XM_023242416.1;refGene_gn=VCP;refGene_tx=XM_023242417.1
D4 58853106 58853261
refGene_gn=VCP;refGene_tx=XM_019816140.2;refGene_gn=VCP;refGene_tx=XM_023242416.1;refGene_gn=VCP;refGene_tx=XM_023242417.1
D4 58855239 58855395
refGene_gn=VCP;refGene_tx=XM_019816140.2;refGene_gn=VCP;refGene_tx=XM_023242416.1;refGene_gn=VCP;refGene_tx=XM_023242417.1
D4 58855616 58855925
refGene_gn=VCP;refGene_tx=XM_019816140.2;refGene_gn=VCP;refGene_tx=XM_023242416.1;refGene_gn=VCP;refGene_tx=XM_023242417.1
D4 58856114 58856327
refGene_gn=VCP;refGene_tx=XM_019816140.2;refGene_gn=VCP;refGene_tx=XM_023242416.1;refGene_gn=VCP;refGene_tx=XM_023242417.1
D4 58856587 58856710
refGene_gn=VCP;refGene_tx=XM_019816140.2;refGene_gn=VCP;refGene_tx=XM_023242416.1;refGene_gn=VCP;refGene_tx=XM_023242417.1
D4 58856828 58856993
refGene_gn=VCP;refGene_tx=XM_019816140.2;refGene_gn=VCP;refGene_tx=XM_

023242416.1;refGene_gn=VCP;refGene_tx=XM_023242417.1
D4 58857400 58857513
refGene_gn=VCP;refGene_tx=XM_019816140.2;refGene_gn=VCP;refGene_tx=XM_023242416.1;refGene_gn=VCP;refGene_tx=XM_023242417.1
D4 58858074 58858210
refGene_gn=VCP;refGene_tx=XM_019816140.2;refGene_gn=VCP;refGene_tx=XM_023242416.1;refGene_gn=VCP;refGene_tx=XM_023242417.1
D4 58858287 58858421
refGene_gn=VCP;refGene_tx=XM_019816140.2;refGene_gn=VCP;refGene_tx=XM_023242416.1;refGene_gn=VCP;refGene_tx=XM_023242417.1
D4 58859082 58859185
refGene_gn=VCP;refGene_tx=XM_019816140.2;refGene_gn=VCP;refGene_tx=XM_023242416.1;refGene_gn=VCP;refGene_tx=XM_023242417.1
D4 58860248 58860380
refGene_gn=VCP;refGene_tx=XM_019816140.2;refGene_gn=VCP;refGene_tx=XM_023242416.1;refGene_gn=VCP;refGene_tx=XM_023242417.1
D4 58860903 58861034
refGene_gn=VCP;refGene_tx=XM_019816140.2;refGene_gn=VCP;refGene_tx=XM_023242416.1;refGene_gn=VCP;refGene_tx=XM_023242417.1
D4 58861850 58861993
refGene_gn=VCP;refGene_tx=XM_019816140.2;refGene_gn=VCP;refGene_tx=XM_023242416.1;refGene_gn=VCP;refGene_tx=XM_023242417.1
D4 58863525 58863698
refGene_gn=VCP;refGene_tx=XM_019816140.2;refGene_gn=VCP;refGene_tx=XM_023242416.1;refGene_gn=VCP;refGene_tx=XM_023242417.1
D4 58863885 58863997 refGene_gn=VCP;refGene_tx=XM_023242416.1
D4 58867540 58867557 refGene_gn=VCP;refGene_tx=XM_023242416.1
D4 58869376 58869485 refGene_gn=FANCG;refGene_tx=XM_003995603.3
D4 58869638 58869762 refGene_gn=FANCG;refGene_tx=XM_003995603.3
D4 58870216 58870372 refGene_gn=FANCG;refGene_tx=XM_003995603.3
D4 58870562 58870609 refGene_gn=FANCG;refGene_tx=XM_003995603.3
D4 58870760 58871050 refGene_gn=FANCG;refGene_tx=XM_003995603.3
D4 58871311 58871378 refGene_gn=FANCG;refGene_tx=XM_003995603.3
D4 58871761 58871913 refGene_gn=FANCG;refGene_tx=XM_003995603.3
D4 58872037 58872184 refGene_gn=FANCG;refGene_tx=XM_003995603.3
D4 58872282 58872413 refGene_gn=FANCG;refGene_tx=XM_003995603.3
D4 58872576 58872712 refGene_gn=FANCG;refGene_tx=XM_003995603.3
D4 58873169 58873372 refGene_gn=FANCG;refGene_tx=XM_003995603.3
D4 58873604 58873736 refGene_gn=FANCG;refGene_tx=XM_003995603.3
D4 58874125 58874216 refGene_gn=FANCG;refGene_tx=XM_003995603.3
D4 58874421 58874622 refGene_gn=FANCG;refGene_tx=XM_003995603.3
D4 58880609 58880739
refGene_gn=PIGO;refGene_tx=XM_003995592.4;refGene_gn=PIGO;refGene_tx=XM_006939248.3;refGene_gn=PIGO;refGene_tx=XM_019816142.2
D4 58880891 58880962
refGene_gn=PIGO;refGene_tx=XM_003995592.4;refGene_gn=PIGO;refGene_tx=XM_006939248.3;refGene_gn=PIGO;refGene_tx=XM_019816142.2
D4 58881574 58881789
refGene_gn=PIGO;refGene_tx=XM_003995592.4;refGene_gn=PIGO;refGene_tx=XM_006939248.3;refGene_gn=PIGO;refGene_tx=XM_019816142.2

D4 58882006 58882213
refGene_gn=PIGO;refGene_tx=XM_003995592.4;refGene_gn=PIGO;refGene_tx=XM_006939248.3;refGene_gn=PIGO;refGene_tx=XM_019816142.2
D4 58882777 58884346
refGene_gn=PIGO;refGene_tx=XM_003995592.4;refGene_gn=PIGO;refGene_tx=XM_006939248.3;refGene_gn=PIGO;refGene_tx=XM_006939249.3;refGene_gn=PIGO;refGene_tx=XM_019816142.2
D4 58884607 58884787
refGene_gn=PIGO;refGene_tx=XM_003995592.4;refGene_gn=PIGO;refGene_tx=XM_006939248.3;refGene_gn=PIGO;refGene_tx=XM_006939249.3;refGene_gn=PIGO;refGene_tx=XM_019816142.2
D4 58884993 58885153
refGene_gn=PIGO;refGene_tx=XM_003995592.4;refGene_gn=PIGO;refGene_tx=XM_006939248.3;refGene_gn=PIGO;refGene_tx=XM_006939249.3;refGene_gn=PIGO;refGene_tx=XM_019816142.2
D4 58885472 58885596
refGene_gn=PIGO;refGene_tx=XM_003995592.4;refGene_gn=PIGO;refGene_tx=XM_006939248.3;refGene_gn=PIGO;refGene_tx=XM_006939249.3;refGene_gn=PIGO;refGene_tx=XM_019816142.2
D4 58885783 58885927
refGene_gn=PIGO;refGene_tx=XM_003995592.4;refGene_gn=PIGO;refGene_tx=XM_006939248.3;refGene_gn=PIGO;refGene_tx=XM_006939249.3;refGene_gn=PIGO;refGene_tx=XM_019816142.2
D4 58886554 58887365
refGene_gn=PIGO;refGene_tx=XM_003995592.4;refGene_gn=PIGO;refGene_tx=XM_006939248.3;refGene_gn=PIGO;refGene_tx=XM_006939249.3;refGene_gn=PIGO;refGene_tx=XM_019816142.2
D4 58887842 58887942 refGene_gn=PIGO;refGene_tx=XM_003995592.4
D4 58891860 58891998 refGene_gn=STOML2;refGene_tx=XM_006939250.3
D4 58892419 58892548 refGene_gn=STOML2;refGene_tx=XM_006939250.3
D4 58892797 58892877 refGene_gn=STOML2;refGene_tx=XM_006939250.3
D4 58892995 58893140 refGene_gn=STOML2;refGene_tx=XM_006939250.3
D4 58893293 58893428 refGene_gn=STOML2;refGene_tx=XM_006939250.3
D4 58893578 58893680 refGene_gn=STOML2;refGene_tx=XM_006939250.3
D4 58893771 58893830 refGene_gn=STOML2;refGene_tx=XM_006939250.3
D4 58893968 58894068 refGene_gn=STOML2;refGene_tx=XM_006939250.3
D4 58894521 58894659 refGene_gn=STOML2;refGene_tx=XM_006939250.3
D4 58894899 58894944 refGene_gn=STOML2;refGene_tx=XM_006939250.3
D4 58897198 58897352
refGene_gn=FAM214B;refGene_tx=XM_003995595.5;refGene_gn=FAM214B;refGene_tx=XM_006939252.3;refGene_gn=FAM214B;refGene_tx=XM_006939253.3;refGene_gn=FAM214B;refGene_tx=XM_006939254.3;refGene_gn=FAM214B;refGene_tx=XM_019816164.2
D4 58897469 58897645
refGene_gn=FAM214B;refGene_tx=XM_003995595.5;refGene_gn=FAM214B;refGene_tx=XM_006939252.3;refGene_gn=FAM214B;refGene_tx=XM_006939253.3;refGene_gn=FAM214B;refGene_tx=XM_006939254.3;refGene_gn=FAM214B;refGene_tx=XM_019816164.2
D4 58897728 58897797
refGene_gn=FAM214B;refGene_tx=XM_003995595.5;refGene_gn=FAM214B;refGen

e_tx=XM_006939252.3;refGene_gn=FAM214B;refGene_tx=XM_006939253.3;refGene_gn=FAM214B;refGene_tx=XM_006939254.3;refGene_gn=FAM214B;refGene_tx=XM_019816164.2
D4 58898020 58898179
refGene_gn=FAM214B;refGene_tx=XM_003995595.5;refGene_gn=FAM214B;refGene_tx=XM_006939252.3;refGene_gn=FAM214B;refGene_tx=XM_006939253.3;refGene_gn=FAM214B;refGene_tx=XM_006939254.3;refGene_gn=FAM214B;refGene_tx=XM_019816164.2
D4 58898304 58898410
refGene_gn=FAM214B;refGene_tx=XM_003995595.5;refGene_gn=FAM214B;refGene_tx=XM_006939252.3;refGene_gn=FAM214B;refGene_tx=XM_006939253.3;refGene_gn=FAM214B;refGene_tx=XM_006939254.3;refGene_gn=FAM214B;refGene_tx=XM_019816164.2
D4 58898599 58898658
refGene_gn=FAM214B;refGene_tx=XM_003995595.5;refGene_gn=FAM214B;refGene_tx=XM_006939252.3;refGene_gn=FAM214B;refGene_tx=XM_006939253.3;refGene_gn=FAM214B;refGene_tx=XM_006939254.3;refGene_gn=FAM214B;refGene_tx=XM_019816164.2
D4 58898841 58899735
refGene_gn=FAM214B;refGene_tx=XM_003995595.5;refGene_gn=FAM214B;refGene_tx=XM_006939252.3;refGene_gn=FAM214B;refGene_tx=XM_006939253.3;refGene_gn=FAM214B;refGene_tx=XM_006939254.3;refGene_gn=FAM214B;refGene_tx=XM_019816164.2
D4 58907748 58907755 refGene_gn=UNC13B;refGene_tx=XM_019816146.1
D4 58924588 58924610
refGene_gn=UNC13B;refGene_tx=XM_003995596.3;refGene_gn=UNC13B;refGene_tx=XM_019816143.1;refGene_gn=UNC13B;refGene_tx=XM_019816144.1;refGene_gn=UNC13B;refGene_tx=XM_019816145.1;refGene_gn=UNC13B;refGene_tx=XM_019816147.1;refGene_gn=UNC13B;refGene_tx=XM_019816148.2;refGene_gn=UNC13B;refGene_tx=XM_019816149.2;refGene_gn=UNC13B;refGene_tx=XM_019816150.1;refGene_gn=UNC13B;refGene_tx=XM_019816151.1;refGene_gn=UNC13B;refGene_tx=XM_019816152.2;refGene_gn=UNC13B;refGene_tx=XM_019816153.1
D4 58980972 58981002
refGene_gn=UNC13B;refGene_tx=XM_003995596.3;refGene_gn=UNC13B;refGene_tx=XM_019816143.1;refGene_gn=UNC13B;refGene_tx=XM_019816144.1;refGene_gn=UNC13B;refGene_tx=XM_019816145.1;refGene_gn=UNC13B;refGene_tx=XM_019816146.1;refGene_gn=UNC13B;refGene_tx=XM_019816147.1;refGene_gn=UNC13B;refGene_tx=XM_019816148.2;refGene_gn=UNC13B;refGene_tx=XM_019816149.2;refGene_gn=UNC13B;refGene_tx=XM_019816150.1;refGene_gn=UNC13B;refGene_tx=XM_019816151.1;refGene_gn=UNC13B;refGene_tx=XM_019816152.2;refGene_gn=UNC13B;refGene_tx=XM_019816153.1
D4 58985278 58985378
refGene_gn=UNC13B;refGene_tx=XM_003995596.3;refGene_gn=UNC13B;refGene_tx=XM_019816143.1;refGene_gn=UNC13B;refGene_tx=XM_019816144.1;refGene_gn=UNC13B;refGene_tx=XM_019816145.1;refGene_gn=UNC13B;refGene_tx=XM_019816146.1;refGene_gn=UNC13B;refGene_tx=XM_019816147.1;refGene_gn=UNC13B;refGene_tx=XM_019816148.2;refGene_gn=UNC13B;refGene_tx=XM_019816149.2;refGene_gn=UNC13B;refGene_tx=XM_019816150.1;refGene_gn=UNC13B;refGene_tx=XM_019816151.1;refGene_gn=UNC13B;refGene_tx=XM_019816152.2;refGene_gn=UNC13B;refGene_tx=XM_019816153.1

D4 59003870 59003988
refGene_gn=UNC13B;refGene_tx=XM_003995596.3;refGene_gn=UNC13B;refGene_tx=XM_019816143.1;refGene_gn=UNC13B;refGene_tx=XM_019816144.1;refGene_gn=UNC13B;refGene_tx=XM_019816145.1;refGene_gn=UNC13B;refGene_tx=XM_019816146.1;refGene_gn=UNC13B;refGene_tx=XM_019816147.1;refGene_gn=UNC13B;refGene_tx=XM_019816148.2;refGene_gn=UNC13B;refGene_tx=XM_019816149.2;refGene_gn=UNC13B;refGene_tx=XM_019816150.1;refGene_gn=UNC13B;refGene_tx=XM_019816151.1;refGene_gn=UNC13B;refGene_tx=XM_019816152.2;refGene_gn=UNC13B;refGene_tx=XM_019816153.1
D4 59004922 59005046
refGene_gn=UNC13B;refGene_tx=XM_003995596.3;refGene_gn=UNC13B;refGene_tx=XM_019816143.1;refGene_gn=UNC13B;refGene_tx=XM_019816144.1;refGene_gn=UNC13B;refGene_tx=XM_019816145.1;refGene_gn=UNC13B;refGene_tx=XM_019816146.1;refGene_gn=UNC13B;refGene_tx=XM_019816147.1;refGene_gn=UNC13B;refGene_tx=XM_019816148.2;refGene_gn=UNC13B;refGene_tx=XM_019816149.2;refGene_gn=UNC13B;refGene_tx=XM_019816150.1;refGene_gn=UNC13B;refGene_tx=XM_019816151.1;refGene_gn=UNC13B;refGene_tx=XM_019816152.2;refGene_gn=UNC13B;refGene_tx=XM_019816153.1
D4 59010356 59010430
refGene_gn=UNC13B;refGene_tx=XM_003995596.3;refGene_gn=UNC13B;refGene_tx=XM_019816143.1;refGene_gn=UNC13B;refGene_tx=XM_019816144.1;refGene_gn=UNC13B;refGene_tx=XM_019816145.1;refGene_gn=UNC13B;refGene_tx=XM_019816146.1;refGene_gn=UNC13B;refGene_tx=XM_019816147.1;refGene_gn=UNC13B;refGene_tx=XM_019816148.2;refGene_gn=UNC13B;refGene_tx=XM_019816149.2;refGene_gn=UNC13B;refGene_tx=XM_019816150.1;refGene_gn=UNC13B;refGene_tx=XM_019816151.1;refGene_gn=UNC13B;refGene_tx=XM_019816152.2;refGene_gn=UNC13B;refGene_tx=XM_019816153.1
D4 59024768 59024826
refGene_gn=UNC13B;refGene_tx=XM_003995596.3;refGene_gn=UNC13B;refGene_tx=XM_019816143.1;refGene_gn=UNC13B;refGene_tx=XM_019816144.1;refGene_gn=UNC13B;refGene_tx=XM_019816145.1;refGene_gn=UNC13B;refGene_tx=XM_019816146.1;refGene_gn=UNC13B;refGene_tx=XM_019816147.1;refGene_gn=UNC13B;refGene_tx=XM_019816148.2;refGene_gn=UNC13B;refGene_tx=XM_019816149.2;refGene_gn=UNC13B;refGene_tx=XM_019816150.1;refGene_gn=UNC13B;refGene_tx=XM_019816151.1;refGene_gn=UNC13B;refGene_tx=XM_019816152.2;refGene_gn=UNC13B;refGene_tx=XM_019816153.1
D4 59053665 59053701
refGene_gn=UNC13B;refGene_tx=XM_019816143.1;refGene_gn=UNC13B;refGene_tx=XM_019816144.1;refGene_gn=UNC13B;refGene_tx=XM_019816145.1;refGene_gn=UNC13B;refGene_tx=XM_019816146.1;refGene_gn=UNC13B;refGene_tx=XM_019816148.2;refGene_gn=UNC13B;refGene_tx=XM_019816149.2;refGene_gn=UNC13B;refGene_tx=XM_019816150.1;refGene_gn=UNC13B;refGene_tx=XM_019816151.1;refGene_gn=UNC13B;refGene_tx=XM_019816153.1
D4 59057742 59057977
refGene_gn=UNC13B;refGene_tx=XM_003995596.3;refGene_gn=UNC13B;refGene_tx=XM_019816143.1;refGene_gn=UNC13B;refGene_tx=XM_019816144.1;refGene_gn=UNC13B;refGene_tx=XM_019816145.1;refGene_gn=UNC13B;refGene_tx=XM_019816146.1;refGene_gn=UNC13B;refGene_tx=XM_019816147.1;refGene_gn=UNC13B;refGene_tx=XM_019816148.2;refGene_gn=UNC13B;refGene_tx=XM_019816149.2;refGene_gn=UNC13B;refGene_tx=XM_019816150.1;refGene_gn=UNC13B;refGen

e_tx=XM_019816151.1;refGene_gn=UNC13B;refGene_tx=XM_019816152.2;refGene_gn=UNC13B;refGene_tx=XM_019816153.1
D4 59062155 59070609
refGene_gn=UNC13B;refGene_tx=XM_019816143.1;refGene_gn=UNC13B;refGene_tx=XM_019816144.1;refGene_gn=UNC13B;refGene_tx=XM_019816145.1;refGene_gn=UNC13B;refGene_tx=XM_019816146.1;refGene_gn=UNC13B;refGene_tx=XM_019816147.1;refGene_gn=UNC13B;refGene_tx=XM_019816148.2;refGene_gn=UNC13B;refGene_tx=XM_019816149.2
D4 59073859 59074174
refGene_gn=UNC13B;refGene_tx=XM_003995596.3;refGene_gn=UNC13B;refGene_tx=XM_019816143.1;refGene_gn=UNC13B;refGene_tx=XM_019816144.1;refGene_gn=UNC13B;refGene_tx=XM_019816145.1;refGene_gn=UNC13B;refGene_tx=XM_019816146.1;refGene_gn=UNC13B;refGene_tx=XM_019816147.1;refGene_gn=UNC13B;refGene_tx=XM_019816148.2;refGene_gn=UNC13B;refGene_tx=XM_019816149.2;refGene_gn=UNC13B;refGene_tx=XM_019816150.1;refGene_gn=UNC13B;refGene_tx=XM_019816151.1;refGene_gn=UNC13B;refGene_tx=XM_019816152.2;refGene_gn=UNC13B;refGene_tx=XM_019816153.1
D4 59076211 59076302
refGene_gn=UNC13B;refGene_tx=XM_003995596.3;refGene_gn=UNC13B;refGene_tx=XM_019816143.1;refGene_gn=UNC13B;refGene_tx=XM_019816144.1;refGene_gn=UNC13B;refGene_tx=XM_019816145.1;refGene_gn=UNC13B;refGene_tx=XM_019816146.1;refGene_gn=UNC13B;refGene_tx=XM_019816147.1;refGene_gn=UNC13B;refGene_tx=XM_019816148.2;refGene_gn=UNC13B;refGene_tx=XM_019816149.2;refGene_gn=UNC13B;refGene_tx=XM_019816150.1;refGene_gn=UNC13B;refGene_tx=XM_019816151.1;refGene_gn=UNC13B;refGene_tx=XM_019816152.2;refGene_gn=UNC13B;refGene_tx=XM_019816153.1
D4 59120216 59120263
refGene_gn=UNC13B;refGene_tx=XM_003995596.3;refGene_gn=UNC13B;refGene_tx=XM_019816143.1;refGene_gn=UNC13B;refGene_tx=XM_019816144.1;refGene_gn=UNC13B;refGene_tx=XM_019816145.1;refGene_gn=UNC13B;refGene_tx=XM_019816146.1;refGene_gn=UNC13B;refGene_tx=XM_019816147.1;refGene_gn=UNC13B;refGene_tx=XM_019816148.2;refGene_gn=UNC13B;refGene_tx=XM_019816149.2;refGene_gn=UNC13B;refGene_tx=XM_019816150.1;refGene_gn=UNC13B;refGene_tx=XM_019816151.1;refGene_gn=UNC13B;refGene_tx=XM_019816152.2;refGene_gn=UNC13B;refGene_tx=XM_019816153.1
D4 59122653 59122732
refGene_gn=UNC13B;refGene_tx=XM_003995596.3;refGene_gn=UNC13B;refGene_tx=XM_019816143.1;refGene_gn=UNC13B;refGene_tx=XM_019816144.1;refGene_gn=UNC13B;refGene_tx=XM_019816145.1;refGene_gn=UNC13B;refGene_tx=XM_019816146.1;refGene_gn=UNC13B;refGene_tx=XM_019816147.1;refGene_gn=UNC13B;refGene_tx=XM_019816148.2;refGene_gn=UNC13B;refGene_tx=XM_019816149.2;refGene_gn=UNC13B;refGene_tx=XM_019816150.1;refGene_gn=UNC13B;refGene_tx=XM_019816151.1;refGene_gn=UNC13B;refGene_tx=XM_019816152.2;refGene_gn=UNC13B;refGene_tx=XM_019816153.1
D4 59124146 59124152
refGene_gn=UNC13B;refGene_tx=XM_019816143.1;refGene_gn=UNC13B;refGene_tx=XM_019816145.1;refGene_gn=UNC13B;refGene_tx=XM_019816146.1;refGene_gn=UNC13B;refGene_tx=XM_019816147.1;refGene_gn=UNC13B;refGene_tx=XM_019816148.2;refGene_gn=UNC13B;refGene_tx=XM_019816149.2;refGene_gn=UNC13B;refGene_tx=XM_019816150.1;refGene_gn=UNC13B;refGene_tx=XM_019816151.

1
D4	59125756 59125831
refGene_gn=UNC13B;refGene_tx=XM_003995596.3;refGene_gn=UNC13B;refGene_tx=XM_019816143.1;refGene_gn=UNC13B;refGene_tx=XM_019816144.1;refGene_gn=UNC13B;refGene_tx=XM_019816145.1;refGene_gn=UNC13B;refGene_tx=XM_019816146.1;refGene_gn=UNC13B;refGene_tx=XM_019816147.1;refGene_gn=UNC13B;refGene_tx=XM_019816148.2;refGene_gn=UNC13B;refGene_tx=XM_019816149.2;refGene_gn=UNC13B;refGene_tx=XM_019816150.1;refGene_gn=UNC13B;refGene_tx=XM_019816151.1;refGene_gn=UNC13B;refGene_tx=XM_019816152.2;refGene_gn=UNC13B;refGene_tx=XM_019816153.1
D4	59126321 59126541
refGene_gn=UNC13B;refGene_tx=XM_003995596.3;refGene_gn=UNC13B;refGene_tx=XM_019816143.1;refGene_gn=UNC13B;refGene_tx=XM_019816144.1;refGene_gn=UNC13B;refGene_tx=XM_019816145.1;refGene_gn=UNC13B;refGene_tx=XM_019816146.1;refGene_gn=UNC13B;refGene_tx=XM_019816147.1;refGene_gn=UNC13B;refGene_tx=XM_019816148.2;refGene_gn=UNC13B;refGene_tx=XM_019816149.2;refGene_gn=UNC13B;refGene_tx=XM_019816150.1;refGene_gn=UNC13B;refGene_tx=XM_019816151.1;refGene_gn=UNC13B;refGene_tx=XM_019816152.2;refGene_gn=UNC13B;refGene_tx=XM_019816153.1
D4	59128650 59128878
refGene_gn=UNC13B;refGene_tx=XM_003995596.3;refGene_gn=UNC13B;refGene_tx=XM_019816143.1;refGene_gn=UNC13B;refGene_tx=XM_019816144.1;refGene_gn=UNC13B;refGene_tx=XM_019816145.1;refGene_gn=UNC13B;refGene_tx=XM_019816146.1;refGene_gn=UNC13B;refGene_tx=XM_019816147.1;refGene_gn=UNC13B;refGene_tx=XM_019816148.2;refGene_gn=UNC13B;refGene_tx=XM_019816149.2;refGene_gn=UNC13B;refGene_tx=XM_019816150.1;refGene_gn=UNC13B;refGene_tx=XM_019816151.1;refGene_gn=UNC13B;refGene_tx=XM_019816152.2;refGene_gn=UNC13B;refGene_tx=XM_019816153.1
D4	59129489 59129631
refGene_gn=UNC13B;refGene_tx=XM_003995596.3;refGene_gn=UNC13B;refGene_tx=XM_019816143.1;refGene_gn=UNC13B;refGene_tx=XM_019816144.1;refGene_gn=UNC13B;refGene_tx=XM_019816145.1;refGene_gn=UNC13B;refGene_tx=XM_019816146.1;refGene_gn=UNC13B;refGene_tx=XM_019816147.1;refGene_gn=UNC13B;refGene_tx=XM_019816148.2;refGene_gn=UNC13B;refGene_tx=XM_019816149.2;refGene_gn=UNC13B;refGene_tx=XM_019816150.1;refGene_gn=UNC13B;refGene_tx=XM_019816151.1;refGene_gn=UNC13B;refGene_tx=XM_019816152.2;refGene_gn=UNC13B;refGene_tx=XM_019816153.1
D4	59131391 59131561
refGene_gn=UNC13B;refGene_tx=XM_003995596.3;refGene_gn=UNC13B;refGene_tx=XM_019816143.1;refGene_gn=UNC13B;refGene_tx=XM_019816144.1;refGene_gn=UNC13B;refGene_tx=XM_019816145.1;refGene_gn=UNC13B;refGene_tx=XM_019816146.1;refGene_gn=UNC13B;refGene_tx=XM_019816147.1;refGene_gn=UNC13B;refGene_tx=XM_019816148.2;refGene_gn=UNC13B;refGene_tx=XM_019816149.2;refGene_gn=UNC13B;refGene_tx=XM_019816150.1;refGene_gn=UNC13B;refGene_tx=XM_019816151.1;refGene_gn=UNC13B;refGene_tx=XM_019816152.2;refGene_gn=UNC13B;refGene_tx=XM_019816153.1
D4	59132061 59132177
refGene_gn=UNC13B;refGene_tx=XM_003995596.3;refGene_gn=UNC13B;refGene_tx=XM_019816143.1;refGene_gn=UNC13B;refGene_tx=XM_019816144.1;refGene_gn=UNC13B;refGene_tx=XM_019816145.1;refGene_gn=UNC13B;refGene_tx=XM_01

9816146.1;refGene_gn=UNC13B;refGene_tx=XM_019816147.1;refGene_gn=UNC13B;refGene_tx=XM_019816148.2;refGene_gn=UNC13B;refGene_tx=XM_019816149.2;refGene_gn=UNC13B;refGene_tx=XM_019816150.1;refGene_gn=UNC13B;refGene_tx=XM_019816151.1;refGene_gn=UNC13B;refGene_tx=XM_019816152.2;refGene_gn=UNC13B;refGene_tx=XM_019816153.1
D4 59132490 59132654
refGene_gn=UNC13B;refGene_tx=XM_003995596.3;refGene_gn=UNC13B;refGene_tx=XM_019816143.1;refGene_gn=UNC13B;refGene_tx=XM_019816144.1;refGene_gn=UNC13B;refGene_tx=XM_019816145.1;refGene_gn=UNC13B;refGene_tx=XM_019816146.1;refGene_gn=UNC13B;refGene_tx=XM_019816147.1;refGene_gn=UNC13B;refGene_tx=XM_019816148.2;refGene_gn=UNC13B;refGene_tx=XM_019816149.2;refGene_gn=UNC13B;refGene_tx=XM_019816150.1;refGene_gn=UNC13B;refGene_tx=XM_019816151.1;refGene_gn=UNC13B;refGene_tx=XM_019816152.2;refGene_gn=UNC13B;refGene_tx=XM_019816153.1
D4 59134966 59135117
refGene_gn=UNC13B;refGene_tx=XM_003995596.3;refGene_gn=UNC13B;refGene_tx=XM_019816143.1;refGene_gn=UNC13B;refGene_tx=XM_019816144.1;refGene_gn=UNC13B;refGene_tx=XM_019816145.1;refGene_gn=UNC13B;refGene_tx=XM_019816146.1;refGene_gn=UNC13B;refGene_tx=XM_019816147.1;refGene_gn=UNC13B;refGene_tx=XM_019816148.2;refGene_gn=UNC13B;refGene_tx=XM_019816149.2;refGene_gn=UNC13B;refGene_tx=XM_019816150.1;refGene_gn=UNC13B;refGene_tx=XM_019816151.1;refGene_gn=UNC13B;refGene_tx=XM_019816152.2;refGene_gn=UNC13B;refGene_tx=XM_019816153.1
D4 59136546 59136615
refGene_gn=UNC13B;refGene_tx=XM_003995596.3;refGene_gn=UNC13B;refGene_tx=XM_019816143.1;refGene_gn=UNC13B;refGene_tx=XM_019816144.1;refGene_gn=UNC13B;refGene_tx=XM_019816145.1;refGene_gn=UNC13B;refGene_tx=XM_019816146.1;refGene_gn=UNC13B;refGene_tx=XM_019816147.1;refGene_gn=UNC13B;refGene_tx=XM_019816148.2;refGene_gn=UNC13B;refGene_tx=XM_019816149.2;refGene_gn=UNC13B;refGene_tx=XM_019816150.1;refGene_gn=UNC13B;refGene_tx=XM_019816151.1;refGene_gn=UNC13B;refGene_tx=XM_019816152.2;refGene_gn=UNC13B;refGene_tx=XM_019816153.1
D4 59138010 59138100
refGene_gn=UNC13B;refGene_tx=XM_003995596.3;refGene_gn=UNC13B;refGene_tx=XM_019816143.1;refGene_gn=UNC13B;refGene_tx=XM_019816144.1;refGene_gn=UNC13B;refGene_tx=XM_019816145.1;refGene_gn=UNC13B;refGene_tx=XM_019816146.1;refGene_gn=UNC13B;refGene_tx=XM_019816147.1;refGene_gn=UNC13B;refGene_tx=XM_019816148.2;refGene_gn=UNC13B;refGene_tx=XM_019816149.2;refGene_gn=UNC13B;refGene_tx=XM_019816150.1;refGene_gn=UNC13B;refGene_tx=XM_019816151.1;refGene_gn=UNC13B;refGene_tx=XM_019816152.2;refGene_gn=UNC13B;refGene_tx=XM_019816153.1
D4 59138441 59138570
refGene_gn=UNC13B;refGene_tx=XM_003995596.3;refGene_gn=UNC13B;refGene_tx=XM_019816143.1;refGene_gn=UNC13B;refGene_tx=XM_019816144.1;refGene_gn=UNC13B;refGene_tx=XM_019816145.1;refGene_gn=UNC13B;refGene_tx=XM_019816146.1;refGene_gn=UNC13B;refGene_tx=XM_019816147.1;refGene_gn=UNC13B;refGene_tx=XM_019816148.2;refGene_gn=UNC13B;refGene_tx=XM_019816149.2;refGene_gn=UNC13B;refGene_tx=XM_019816150.1;refGene_gn=UNC13B;refGene_tx=XM_019816151.1;refGene_gn=UNC13B;refGene_tx=XM_019816152.2;refGene_gn=UNC13B;refGene_tx=XM_019816153.1

D4 59144179 59144307
refGene_gn=UNC13B;refGene_tx=XM_003995596.3;refGene_gn=UNC13B;refGene_tx=XM_019816143.1;refGene_gn=UNC13B;refGene_tx=XM_019816144.1;refGene_gn=UNC13B;refGene_tx=XM_019816145.1;refGene_gn=UNC13B;refGene_tx=XM_019816146.1;refGene_gn=UNC13B;refGene_tx=XM_019816147.1;refGene_gn=UNC13B;refGene_tx=XM_019816148.2;refGene_gn=UNC13B;refGene_tx=XM_019816149.2;refGene_gn=UNC13B;refGene_tx=XM_019816150.1;refGene_gn=UNC13B;refGene_tx=XM_019816151.1;refGene_gn=UNC13B;refGene_tx=XM_019816152.2;refGene_gn=UNC13B;refGene_tx=XM_019816153.1
D4 59144942 59145028
refGene_gn=UNC13B;refGene_tx=XM_003995596.3;refGene_gn=UNC13B;refGene_tx=XM_019816143.1;refGene_gn=UNC13B;refGene_tx=XM_019816144.1;refGene_gn=UNC13B;refGene_tx=XM_019816145.1;refGene_gn=UNC13B;refGene_tx=XM_019816146.1;refGene_gn=UNC13B;refGene_tx=XM_019816147.1;refGene_gn=UNC13B;refGene_tx=XM_019816148.2;refGene_gn=UNC13B;refGene_tx=XM_019816149.2;refGene_gn=UNC13B;refGene_tx=XM_019816150.1;refGene_gn=UNC13B;refGene_tx=XM_019816151.1;refGene_gn=UNC13B;refGene_tx=XM_019816152.2;refGene_gn=UNC13B;refGene_tx=XM_019816153.1
D4 59150254 59150381
refGene_gn=UNC13B;refGene_tx=XM_003995596.3;refGene_gn=UNC13B;refGene_tx=XM_019816143.1;refGene_gn=UNC13B;refGene_tx=XM_019816144.1;refGene_gn=UNC13B;refGene_tx=XM_019816145.1;refGene_gn=UNC13B;refGene_tx=XM_019816146.1;refGene_gn=UNC13B;refGene_tx=XM_019816147.1;refGene_gn=UNC13B;refGene_tx=XM_019816148.2;refGene_gn=UNC13B;refGene_tx=XM_019816149.2;refGene_gn=UNC13B;refGene_tx=XM_019816150.1;refGene_gn=UNC13B;refGene_tx=XM_019816151.1;refGene_gn=UNC13B;refGene_tx=XM_019816152.2;refGene_gn=UNC13B;refGene_tx=XM_019816153.1
D4 59150632 59150729
refGene_gn=UNC13B;refGene_tx=XM_003995596.3;refGene_gn=UNC13B;refGene_tx=XM_019816143.1;refGene_gn=UNC13B;refGene_tx=XM_019816144.1;refGene_gn=UNC13B;refGene_tx=XM_019816145.1;refGene_gn=UNC13B;refGene_tx=XM_019816146.1;refGene_gn=UNC13B;refGene_tx=XM_019816147.1;refGene_gn=UNC13B;refGene_tx=XM_019816148.2;refGene_gn=UNC13B;refGene_tx=XM_019816149.2;refGene_gn=UNC13B;refGene_tx=XM_019816150.1;refGene_gn=UNC13B;refGene_tx=XM_019816151.1;refGene_gn=UNC13B;refGene_tx=XM_019816152.2;refGene_gn=UNC13B;refGene_tx=XM_019816153.1
D4 59150949 59151093
refGene_gn=UNC13B;refGene_tx=XM_003995596.3;refGene_gn=UNC13B;refGene_tx=XM_019816143.1;refGene_gn=UNC13B;refGene_tx=XM_019816144.1;refGene_gn=UNC13B;refGene_tx=XM_019816145.1;refGene_gn=UNC13B;refGene_tx=XM_019816146.1;refGene_gn=UNC13B;refGene_tx=XM_019816147.1;refGene_gn=UNC13B;refGene_tx=XM_019816148.2;refGene_gn=UNC13B;refGene_tx=XM_019816149.2;refGene_gn=UNC13B;refGene_tx=XM_019816150.1;refGene_gn=UNC13B;refGene_tx=XM_019816151.1;refGene_gn=UNC13B;refGene_tx=XM_019816152.2;refGene_gn=UNC13B;refGene_tx=XM_019816153.1
D4 59151415 59151493
refGene_gn=UNC13B;refGene_tx=XM_003995596.3;refGene_gn=UNC13B;refGene_tx=XM_019816143.1;refGene_gn=UNC13B;refGene_tx=XM_019816144.1;refGene_gn=UNC13B;refGene_tx=XM_019816145.1;refGene_gn=UNC13B;refGene_tx=XM_019816146.1;refGene_gn=UNC13B;refGene_tx=XM_019816147.1;refGene_gn=UNC13

B;refGene_tx=XM_019816148.2;refGene_gn=UNC13B;refGene_tx=XM_019816149.2;refGene_gn=UNC13B;refGene_tx=XM_019816150.1;refGene_gn=UNC13B;refGene_tx=XM_019816151.1;refGene_gn=UNC13B;refGene_tx=XM_019816152.2;refGene_gn=UNC13B;refGene_tx=XM_019816153.1
D4 59152026 59152104
refGene_gn=UNC13B;refGene_tx=XM_003995596.3;refGene_gn=UNC13B;refGene_tx=XM_019816143.1;refGene_gn=UNC13B;refGene_tx=XM_019816144.1;refGene_gn=UNC13B;refGene_tx=XM_019816145.1;refGene_gn=UNC13B;refGene_tx=XM_019816146.1;refGene_gn=UNC13B;refGene_tx=XM_019816147.1;refGene_gn=UNC13B;refGene_tx=XM_019816148.2;refGene_gn=UNC13B;refGene_tx=XM_019816149.2;refGene_gn=UNC13B;refGene_tx=XM_019816150.1;refGene_gn=UNC13B;refGene_tx=XM_019816151.1;refGene_gn=UNC13B;refGene_tx=XM_019816152.2;refGene_gn=UNC13B;refGene_tx=XM_019816153.1
D4 59152346 59152435
refGene_gn=UNC13B;refGene_tx=XM_003995596.3;refGene_gn=UNC13B;refGene_tx=XM_019816143.1;refGene_gn=UNC13B;refGene_tx=XM_019816144.1;refGene_gn=UNC13B;refGene_tx=XM_019816145.1;refGene_gn=UNC13B;refGene_tx=XM_019816146.1;refGene_gn=UNC13B;refGene_tx=XM_019816147.1;refGene_gn=UNC13B;refGene_tx=XM_019816148.2;refGene_gn=UNC13B;refGene_tx=XM_019816149.2;refGene_gn=UNC13B;refGene_tx=XM_019816150.1;refGene_gn=UNC13B;refGene_tx=XM_019816151.1;refGene_gn=UNC13B;refGene_tx=XM_019816152.2;refGene_gn=UNC13B;refGene_tx=XM_019816153.1
D4 59152673 59152826
refGene_gn=UNC13B;refGene_tx=XM_003995596.3;refGene_gn=UNC13B;refGene_tx=XM_019816143.1;refGene_gn=UNC13B;refGene_tx=XM_019816144.1;refGene_gn=UNC13B;refGene_tx=XM_019816145.1;refGene_gn=UNC13B;refGene_tx=XM_019816146.1;refGene_gn=UNC13B;refGene_tx=XM_019816147.1;refGene_gn=UNC13B;refGene_tx=XM_019816148.2;refGene_gn=UNC13B;refGene_tx=XM_019816149.2;refGene_gn=UNC13B;refGene_tx=XM_019816150.1;refGene_gn=UNC13B;refGene_tx=XM_019816151.1;refGene_gn=UNC13B;refGene_tx=XM_019816152.2;refGene_gn=UNC13B;refGene_tx=XM_019816153.1
D4 59152943 59153232
refGene_gn=UNC13B;refGene_tx=XM_003995596.3;refGene_gn=UNC13B;refGene_tx=XM_019816143.1;refGene_gn=UNC13B;refGene_tx=XM_019816144.1;refGene_gn=UNC13B;refGene_tx=XM_019816145.1;refGene_gn=UNC13B;refGene_tx=XM_019816146.1;refGene_gn=UNC13B;refGene_tx=XM_019816147.1;refGene_gn=UNC13B;refGene_tx=XM_019816148.2;refGene_gn=UNC13B;refGene_tx=XM_019816149.2;refGene_gn=UNC13B;refGene_tx=XM_019816150.1;refGene_gn=UNC13B;refGene_tx=XM_019816151.1;refGene_gn=UNC13B;refGene_tx=XM_019816152.2;refGene_gn=UNC13B;refGene_tx=XM_019816153.1
D4 59153420 59153501
refGene_gn=UNC13B;refGene_tx=XM_003995596.3;refGene_gn=UNC13B;refGene_tx=XM_019816143.1;refGene_gn=UNC13B;refGene_tx=XM_019816144.1;refGene_gn=UNC13B;refGene_tx=XM_019816145.1;refGene_gn=UNC13B;refGene_tx=XM_019816146.1;refGene_gn=UNC13B;refGene_tx=XM_019816147.1;refGene_gn=UNC13B;refGene_tx=XM_019816148.2;refGene_gn=UNC13B;refGene_tx=XM_019816149.2;refGene_gn=UNC13B;refGene_tx=XM_019816150.1;refGene_gn=UNC13B;refGene_tx=XM_019816151.1;refGene_gn=UNC13B;refGene_tx=XM_019816152.2;refGene_gn=UNC13B;refGene_tx=XM_019816153.1
D4 59154445 59154593

refGene_gn=UNC13B;refGene_tx=XM_003995596.3;refGene_gn=UNC13B;refGene_tx=XM_019816143.1;refGene_gn=UNC13B;refGene_tx=XM_019816144.1;refGene_gn=UNC13B;refGene_tx=XM_019816145.1;refGene_gn=UNC13B;refGene_tx=XM_019816146.1;refGene_gn=UNC13B;refGene_tx=XM_019816147.1;refGene_gn=UNC13B;refGene_tx=XM_019816148.2;refGene_gn=UNC13B;refGene_tx=XM_019816149.2;refGene_gn=UNC13B;refGene_tx=XM_019816150.1;refGene_gn=UNC13B;refGene_tx=XM_019816151.1;refGene_gn=UNC13B;refGene_tx=XM_019816152.2;refGene_gn=UNC13B;refGene_tx=XM_019816153.1
D4 59156127 59156184
refGene_gn=UNC13B;refGene_tx=XM_019816148.2;refGene_gn=UNC13B;refGene_tx=XM_019816151.1;refGene_gn=UNC13B;refGene_tx=XM_019816152.2
D4 59157009 59157102
refGene_gn=UNC13B;refGene_tx=XM_003995596.3;refGene_gn=UNC13B;refGene_tx=XM_019816143.1;refGene_gn=UNC13B;refGene_tx=XM_019816144.1;refGene_gn=UNC13B;refGene_tx=XM_019816145.1;refGene_gn=UNC13B;refGene_tx=XM_019816146.1;refGene_gn=UNC13B;refGene_tx=XM_019816147.1;refGene_gn=UNC13B;refGene_tx=XM_019816148.2;refGene_gn=UNC13B;refGene_tx=XM_019816149.2;refGene_gn=UNC13B;refGene_tx=XM_019816150.1;refGene_gn=UNC13B;refGene_tx=XM_019816151.1;refGene_gn=UNC13B;refGene_tx=XM_019816152.2;refGene_gn=UNC13B;refGene_tx=XM_019816153.1
D4 59157282 59157442
refGene_gn=UNC13B;refGene_tx=XM_003995596.3;refGene_gn=UNC13B;refGene_tx=XM_019816143.1;refGene_gn=UNC13B;refGene_tx=XM_019816144.1;refGene_gn=UNC13B;refGene_tx=XM_019816145.1;refGene_gn=UNC13B;refGene_tx=XM_019816146.1;refGene_gn=UNC13B;refGene_tx=XM_019816147.1;refGene_gn=UNC13B;refGene_tx=XM_019816148.2;refGene_gn=UNC13B;refGene_tx=XM_019816149.2;refGene_gn=UNC13B;refGene_tx=XM_019816150.1;refGene_gn=UNC13B;refGene_tx=XM_019816151.1;refGene_gn=UNC13B;refGene_tx=XM_019816152.2;refGene_gn=UNC13B;refGene_tx=XM_019816153.1
D4 59157606 59157892
refGene_gn=UNC13B;refGene_tx=XM_003995596.3;refGene_gn=UNC13B;refGene_tx=XM_019816143.1;refGene_gn=UNC13B;refGene_tx=XM_019816144.1;refGene_gn=UNC13B;refGene_tx=XM_019816145.1;refGene_gn=UNC13B;refGene_tx=XM_019816146.1;refGene_gn=UNC13B;refGene_tx=XM_019816147.1;refGene_gn=UNC13B;refGene_tx=XM_019816148.2;refGene_gn=UNC13B;refGene_tx=XM_019816149.2;refGene_gn=UNC13B;refGene_tx=XM_019816150.1;refGene_gn=UNC13B;refGene_tx=XM_019816151.1;refGene_gn=UNC13B;refGene_tx=XM_019816152.2;refGene_gn=UNC13B;refGene_tx=XM_019816153.1
D4 59167819 59167847
refGene_gn=LOC101086263;refGene_tx=XM_006939264.3;refGene_gn=LOC101086263;refGene_tx=XM_019816159.2;refGene_gn=LOC101086263;refGene_tx=XM_019816163.2
D4 59188343 59188406
refGene_gn=LOC101086263;refGene_tx=XM_006939264.3;refGene_gn=LOC101086263;refGene_tx=XM_019816158.2;refGene_gn=LOC101086263;refGene_tx=XM_019816159.2;refGene_gn=LOC101086263;refGene_tx=XM_019816163.2
D4 59203286 59203345
refGene_gn=LOC101086263;refGene_tx=XM_006939264.3;refGene_gn=LOC101086263;refGene_tx=XM_019816158.2;refGene_gn=LOC101086263;refGene_tx=XM_019816159.2;refGene_gn=LOC101086263;refGene_tx=XM_019816163.2

D4 59207544 59207658
refGene_gn=LOC101086263;refGene_tx=XM_006939264.3;refGene_gn=LOC101086263;refGene_tx=XM_019816158.2;refGene_gn=LOC101086263;refGene_tx=XM_019816159.2;refGene_gn=LOC101086263;refGene_tx=XM_019816163.2
D4 59208447 59208546
refGene_gn=LOC101086263;refGene_tx=XM_006939264.3;refGene_gn=LOC101086263;refGene_tx=XM_019816158.2;refGene_gn=LOC101086263;refGene_tx=XM_019816159.2;refGene_gn=LOC101086263;refGene_tx=XM_019816163.2
D4 59210375 59210437
refGene_gn=LOC101086263;refGene_tx=XM_006939264.3;refGene_gn=LOC101086263;refGene_tx=XM_019816158.2;refGene_gn=LOC101086263;refGene_tx=XM_019816159.2;refGene_gn=LOC101086263;refGene_tx=XM_019816163.2
D4 59215677 59215750
refGene_gn=LOC101086263;refGene_tx=XM_006939264.3;refGene_gn=LOC101086263;refGene_tx=XM_019816158.2;refGene_gn=LOC101086263;refGene_tx=XM_019816159.2;refGene_gn=LOC101086263;refGene_tx=XM_019816163.2
D4 59216120 59216191
refGene_gn=LOC101086263;refGene_tx=XM_006939264.3;refGene_gn=LOC101086263;refGene_tx=XM_019816158.2;refGene_gn=LOC101086263;refGene_tx=XM_019816159.2;refGene_gn=LOC101086263;refGene_tx=XM_019816163.2
D4 59217248 59217331
refGene_gn=LOC101086263;refGene_tx=XM_006939264.3;refGene_gn=LOC101086263;refGene_tx=XM_019816158.2;refGene_gn=LOC101086263;refGene_tx=XM_019816159.2;refGene_gn=LOC101086263;refGene_tx=XM_019816163.2
D4 59218106 59218265
refGene_gn=LOC101086263;refGene_tx=XM_006939264.3;refGene_gn=LOC101086263;refGene_tx=XM_019816158.2;refGene_gn=LOC101086263;refGene_tx=XM_019816159.2;refGene_gn=LOC101086263;refGene_tx=XM_019816163.2
D4 59220693 59220782
refGene_gn=LOC101086263;refGene_tx=XM_006939264.3;refGene_gn=LOC101086263;refGene_tx=XM_019816158.2;refGene_gn=LOC101086263;refGene_tx=XM_019816159.2;refGene_gn=LOC101086263;refGene_tx=XM_019816163.2
D4 59227554 59227751
refGene_gn=LOC101086263;refGene_tx=XM_006939264.3;refGene_gn=LOC101086263;refGene_tx=XM_019816158.2;refGene_gn=LOC101086263;refGene_tx=XM_019816159.2;refGene_gn=LOC101086263;refGene_tx=XM_019816163.2
D4 59231174 59231383
refGene_gn=LOC101086263;refGene_tx=XM_006939264.3;refGene_gn=LOC101086263;refGene_tx=XM_019816158.2;refGene_gn=LOC101086263;refGene_tx=XM_019816159.2;refGene_gn=LOC101086263;refGene_tx=XM_019816163.2
D4 59233696 59233740
refGene_gn=LOC101086263;refGene_tx=XM_006939264.3;refGene_gn=LOC101086263;refGene_tx=XM_019816158.2;refGene_gn=LOC101086263;refGene_tx=XM_019816159.2;refGene_gn=LOC101086263;refGene_tx=XM_019816163.2
D4 59240620 59240786
refGene_gn=LOC101086263;refGene_tx=XM_006939264.3;refGene_gn=LOC101086263;refGene_tx=XM_019816158.2;refGene_gn=LOC101086263;refGene_tx=XM_019816159.2;refGene_gn=LOC101086263;refGene_tx=XM_019816163.2
D4 59243298 59243487
refGene_gn=LOC101086263;refGene_tx=XM_006939264.3;refGene_gn=LOC101086

263;refGene_tx=XM_019816158.2;refGene_gn=LOC101086263;refGene_tx=XM_019816159.2;refGene_gn=LOC101086263;refGene_tx=XM_019816163.2
D4 59244194 59244310
refGene_gn=LOC101086263;refGene_tx=XM_006939264.3;refGene_gn=LOC101086263;refGene_tx=XM_019816158.2;refGene_gn=LOC101086263;refGene_tx=XM_019816159.2;refGene_gn=LOC101086263;refGene_tx=XM_019816163.2
D4 59248144 59248309
refGene_gn=LOC101086263;refGene_tx=XM_006939264.3;refGene_gn=LOC101086263;refGene_tx=XM_019816158.2;refGene_gn=LOC101086263;refGene_tx=XM_019816159.2;refGene_gn=LOC101086263;refGene_tx=XM_019816163.2
D4 59249963 59250075
refGene_gn=LOC101086263;refGene_tx=XM_006939264.3;refGene_gn=LOC101086263;refGene_tx=XM_019816158.2;refGene_gn=LOC101086263;refGene_tx=XM_019816159.2;refGene_gn=LOC101086263;refGene_tx=XM_019816163.2
D4 59251489 59251595
refGene_gn=LOC101086263;refGene_tx=XM_006939264.3;refGene_gn=LOC101086263;refGene_tx=XM_019816158.2;refGene_gn=LOC101086263;refGene_tx=XM_019816159.2;refGene_gn=LOC101086263;refGene_tx=XM_019816163.2
D4 59251953 59252098
refGene_gn=LOC101086263;refGene_tx=XM_006939264.3;refGene_gn=LOC101086263;refGene_tx=XM_019816158.2;refGene_gn=LOC101086263;refGene_tx=XM_019816159.2;refGene_gn=LOC101086263;refGene_tx=XM_019816163.2
D4 59252634 59252818
refGene_gn=LOC101086263;refGene_tx=XM_006939264.3;refGene_gn=LOC101086263;refGene_tx=XM_019816158.2;refGene_gn=LOC101086263;refGene_tx=XM_019816159.2;refGene_gn=LOC101086263;refGene_tx=XM_019816163.2
D4 59255035 59255259
refGene_gn=LOC101086263;refGene_tx=XM_006939264.3;refGene_gn=LOC101086263;refGene_tx=XM_019816158.2;refGene_gn=LOC101086263;refGene_tx=XM_019816159.2;refGene_gn=LOC101086263;refGene_tx=XM_019816163.2
D4 59256582 59256666
refGene_gn=LOC101086263;refGene_tx=XM_006939264.3;refGene_gn=LOC101086263;refGene_tx=XM_019816158.2;refGene_gn=LOC101086263;refGene_tx=XM_019816159.2;refGene_gn=LOC101086263;refGene_tx=XM_019816163.2
D4 59257886 59257889
refGene_gn=LOC101086263;refGene_tx=XM_019816163.2
D4 59258016 59258262
refGene_gn=LOC101086263;refGene_tx=XM_006939264.3;refGene_gn=LOC101086263;refGene_tx=XM_019816158.2;refGene_gn=LOC101086263;refGene_tx=XM_019816159.2
D4 59258448 59258587
refGene_gn=LOC101086263;refGene_tx=XM_006939264.3;refGene_gn=LOC101086263;refGene_tx=XM_019816158.2;refGene_gn=LOC101086263;refGene_tx=XM_019816159.2
D4 59259387 59259518
refGene_gn=LOC101086263;refGene_tx=XM_006939264.3;refGene_gn=LOC101086263;refGene_tx=XM_019816158.2;refGene_gn=LOC101086263;refGene_tx=XM_019816159.2
D4 59259614 59259806
refGene_gn=LOC101086263;refGene_tx=XM_006939264.3;refGene_gn=LOC101086

263;refGene_tx=XM_019816158.2;refGene_gn=LOC101086263;refGene_tx=XM_019816159.2
D4 59323646 59325675
refGene_gn=RUSC2;refGene_tx=XM_003995604.5;refGene_gn=RUSC2;refGene_tx=XM_006939262.3;refGene_gn=RUSC2;refGene_tx=XM_019816155.2
D4 59330824 59331454
refGene_gn=RUSC2;refGene_tx=XM_003995604.5;refGene_gn=RUSC2;refGene_tx=XM_006939262.3;refGene_gn=RUSC2;refGene_tx=XM_019816155.2;refGene_gn=RUSC2;refGene_tx=XM_019816156.1
D4 59331705 59331891
refGene_gn=RUSC2;refGene_tx=XM_003995604.5;refGene_gn=RUSC2;refGene_tx=XM_006939262.3;refGene_gn=RUSC2;refGene_tx=XM_019816155.2;refGene_gn=RUSC2;refGene_tx=XM_019816156.1
D4 59332038 59332179
refGene_gn=RUSC2;refGene_tx=XM_003995604.5;refGene_gn=RUSC2;refGene_tx=XM_006939262.3;refGene_gn=RUSC2;refGene_tx=XM_019816155.2;refGene_gn=RUSC2;refGene_tx=XM_019816156.1
D4 59333270 59333347
refGene_gn=RUSC2;refGene_tx=XM_003995604.5;refGene_gn=RUSC2;refGene_tx=XM_006939262.3;refGene_gn=RUSC2;refGene_tx=XM_019816155.2;refGene_gn=RUSC2;refGene_tx=XM_019816156.1
D4 59333556 59333731
refGene_gn=RUSC2;refGene_tx=XM_003995604.5;refGene_gn=RUSC2;refGene_tx=XM_006939262.3;refGene_gn=RUSC2;refGene_tx=XM_019816155.2;refGene_gn=RUSC2;refGene_tx=XM_019816156.1
D4 59333821 59333927
refGene_gn=RUSC2;refGene_tx=XM_003995604.5;refGene_gn=RUSC2;refGene_tx=XM_006939262.3;refGene_gn=RUSC2;refGene_tx=XM_019816155.2;refGene_gn=RUSC2;refGene_tx=XM_019816156.1
D4 59334648 59334695
refGene_gn=RUSC2;refGene_tx=XM_003995604.5;refGene_gn=RUSC2;refGene_tx=XM_006939262.3;refGene_gn=RUSC2;refGene_tx=XM_019816155.2;refGene_gn=RUSC2;refGene_tx=XM_019816156.1
D4 59335147 59335955
refGene_gn=RUSC2;refGene_tx=XM_003995604.5;refGene_gn=RUSC2;refGene_tx=XM_006939262.3;refGene_gn=RUSC2;refGene_tx=XM_019816155.2;refGene_gn=RUSC2;refGene_tx=XM_019816156.1
D4 59336049 59336187
refGene_gn=RUSC2;refGene_tx=XM_003995604.5;refGene_gn=RUSC2;refGene_tx=XM_006939262.3;refGene_gn=RUSC2;refGene_tx=XM_019816155.2;refGene_gn=RUSC2;refGene_tx=XM_019816156.1
D4 59336268 59336470
refGene_gn=RUSC2;refGene_tx=XM_003995604.5;refGene_gn=RUSC2;refGene_tx=XM_006939262.3;refGene_gn=RUSC2;refGene_tx=XM_019816155.2;refGene_gn=RUSC2;refGene_tx=XM_019816156.1
D4 59336939 59337161
refGene_gn=FAM166B;refGene_tx=XM_003995605.5;refGene_gn=FAM166B;refGene_tx=XM_019816165.2
D4 59337430 59337628
refGene_gn=FAM166B;refGene_tx=XM_003995605.5;refGene_gn=FAM166B;refGen

e_tx=XM_019816165.2
D4	59337705 59337757
refGene_gn=FAM166B;refGene_tx=XM_003995605.5;refGene_gn=FAM166B;refGene_tx=XM_019816165.2
D4	59337954 59338172
refGene_gn=FAM166B;refGene_tx=XM_003995605.5;refGene_gn=FAM166B;refGene_tx=XM_019816165.2
D4	59338261 59338485
refGene_gn=FAM166B;refGene_tx=XM_003995605.5;refGene_gn=FAM166B;refGene_tx=XM_019816165.2
D4	59338865 59338927
refGene_gn=FAM166B;refGene_tx=XM_003995605.5;refGene_gn=FAM166B;refGene_tx=XM_019816165.2
D4	59359087 59359306 refGene_gn=TESK1;refGene_tx=XM_019815537.2
D4	59359439 59359561 refGene_gn=TESK1;refGene_tx=XM_019815537.2
D4	59359690 59359739 refGene_gn=TESK1;refGene_tx=XM_019815537.2
D4	59360243 59360390 refGene_gn=TESK1;refGene_tx=XM_019815537.2
D4	59360725 59360808 refGene_gn=TESK1;refGene_tx=XM_019815537.2
D4	59360976 59361067 refGene_gn=TESK1;refGene_tx=XM_019815537.2
D4	59361319 59361403 refGene_gn=TESK1;refGene_tx=XM_019815537.2
D4	59361534 59361624 refGene_gn=TESK1;refGene_tx=XM_019815537.2
D4	59361764 59361879 refGene_gn=TESK1;refGene_tx=XM_019815537.2
D4	59362227 59363114 refGene_gn=TESK1;refGene_tx=XM_019815537.2
D4	59363982 59364124
refGene_gn=CD72;refGene_tx=XM_023242421.1;refGene_gn=CD72;refGene_tx=XM_023242423.1
D4	59364231 59364322 refGene_gn=CD72;refGene_tx=XM_023242422.1
D4	59365064 59365183
refGene_gn=CD72;refGene_tx=XM_023242421.1;refGene_gn=CD72;refGene_tx=XM_023242422.1;refGene_gn=CD72;refGene_tx=XM_023242423.1
D4	59365820 59365963
refGene_gn=CD72;refGene_tx=XM_023242421.1;refGene_gn=CD72;refGene_tx=XM_023242422.1;refGene_gn=CD72;refGene_tx=XM_023242423.1
D4	59368848 59369184
refGene_gn=CD72;refGene_tx=XM_023242421.1;refGene_gn=CD72;refGene_tx=XM_023242422.1;refGene_gn=CD72;refGene_tx=XM_023242423.1
D4	59369509 59369599
refGene_gn=CD72;refGene_tx=XM_023242421.1;refGene_gn=CD72;refGene_tx=XM_023242422.1
D4	59370224 59370320
refGene_gn=CD72;refGene_tx=XM_023242421.1;refGene_gn=CD72;refGene_tx=XM_023242422.1;refGene_gn=CD72;refGene_tx=XM_023242423.1
D4	59370930 59371038
refGene_gn=CD72;refGene_tx=XM_023242421.1;refGene_gn=CD72;refGene_tx=XM_023242422.1;refGene_gn=CD72;refGene_tx=XM_023242423.1
D4	59371132 59371214
refGene_gn=CD72;refGene_tx=XM_023242421.1;refGene_gn=CD72;refGene_tx=XM_023242422.1;refGene_gn=CD72;refGene_tx=XM_023242423.1
D4	59382597 59382831
refGene_gn=SIT1;refGene_tx=XM_011288566.3;refGene_gn=SIT1;refGene_tx=X

M_023242424.1
D4 59382921 59382983
refGene_gn=SIT1;refGene_tx=XM_011288566.3;refGene_gn=SIT1;refGene_tx=XM_023242424.1
D4 59383100 59383159
refGene_gn=SIT1;refGene_tx=XM_011288566.3;refGene_gn=SIT1;refGene_tx=XM_023242424.1
D4 59383244 59383380
refGene_gn=SIT1;refGene_tx=XM_011288566.3;refGene_gn=SIT1;refGene_tx=XM_023242424.1
D4 59383494 59383540
refGene_gn=SIT1;refGene_tx=XM_011288566.3;refGene_gn=SIT1;refGene_tx=XM_023242424.1
D4 59387636 59387742
refGene_gn=CCDC107;refGene_tx=XM_003995672.4;refGene_gn=CCDC107;refGene_tx=XM_019816167.2
D4 59387824 59387976
refGene_gn=CCDC107;refGene_tx=XM_003995672.4;refGene_gn=CCDC107;refGene_tx=XM_019816167.2
D4 59389325 59389389
refGene_gn=CCDC107;refGene_tx=XM_003995672.4;refGene_gn=CCDC107;refGene_tx=XM_019816167.2
D4 59389480 59389571
refGene_gn=CCDC107;refGene_tx=XM_003995672.4;refGene_gn=CCDC107;refGene_tx=XM_019816167.2
D4 59389667 59389780
refGene_gn=CCDC107;refGene_tx=XM_003995672.4;refGene_gn=CCDC107;refGene_tx=XM_019816167.2
D4 59389861 59390085
refGene_gn=CCDC107;refGene_tx=XM_003995672.4;refGene_gn=CCDC107;refGene_tx=XM_019816167.2
D4 59390735 59390751 refGene_gn=ARHGEF39;refGene_tx=XM_003995673.4
D4 59390918 59391007 refGene_gn=ARHGEF39;refGene_tx=XM_003995673.4
D4 59391256 59391486 refGene_gn=ARHGEF39;refGene_tx=XM_003995673.4
D4 59391665 59391794 refGene_gn=ARHGEF39;refGene_tx=XM_003995673.4
D4 59392041 59392112 refGene_gn=ARHGEF39;refGene_tx=XM_003995673.4
D4 59392717 59392836 refGene_gn=ARHGEF39;refGene_tx=XM_003995673.4
D4 59393089 59393210 refGene_gn=ARHGEF39;refGene_tx=XM_003995673.4
D4 59393471 59393566 refGene_gn=ARHGEF39;refGene_tx=XM_003995673.4
D4 59393769 59393907 refGene_gn=ARHGEF39;refGene_tx=XM_003995673.4
D4 59399082 59399464
refGene_gn=CA9;refGene_tx=XM_003995606.5;refGene_gn=CA9;refGene_tx=XM_019816173.2;refGene_gn=CA9;refGene_tx=XM_019816174.2
D4 59400316 59400346
refGene_gn=CA9;refGene_tx=XM_003995606.5;refGene_gn=CA9;refGene_tx=XM_019816173.2;refGene_gn=CA9;refGene_tx=XM_019816174.2
D4 59400624 59400795
refGene_gn=CA9;refGene_tx=XM_003995606.5;refGene_gn=CA9;refGene_tx=XM_019816173.2;refGene_gn=CA9;refGene_tx=XM_019816174.2
D4 59400939 59401082

refGene_gn=CA9;refGene_tx=XM_003995606.5;refGene_gn=CA9;refGene_tx=XM_019816173.2;refGene_gn=CA9;refGene_tx=XM_019816174.2
D4 59401159 59401252 refGene_gn=CA9;refGene_tx=XM_003995606.5;refGene_gn=CA9;refGene_tx=XM_019816173.2;refGene_gn=CA9;refGene_tx=XM_019816174.2
D4 59402326 59402393 refGene_gn=CA9;refGene_tx=XM_003995606.5;refGene_gn=CA9;refGene_tx=XM_019816173.2
D4 59403248 59403406 refGene_gn=CA9;refGene_tx=XM_003995606.5;refGene_gn=CA9;refGene_tx=XM_019816173.2
D4 59403610 59403755 refGene_gn=CA9;refGene_tx=XM_003995606.5;refGene_gn=CA9;refGene_tx=XM_019816173.2;refGene_gn=CA9;refGene_tx=XM_019816174.2
D4 59403893 59403920 refGene_gn=CA9;refGene_tx=XM_003995606.5;refGene_gn=CA9;refGene_tx=XM_019816173.2;refGene_gn=CA9;refGene_tx=XM_019816174.2
D4 59404331 59404413 refGene_gn=CA9;refGene_tx=XM_003995606.5
D4 59404556 59404747 refGene_gn=CA9;refGene_tx=XM_003995606.5;refGene_gn=CA9;refGene_tx=XM_019816173.2;refGene_gn=CA9;refGene_tx=XM_019816174.2
D4 59405831 59405914 refGene_gn=TPM2;refGene_tx=XM_023242426.1;refGene_gn=TPM2;refGene_tx=XM_023242428.1
D4 59406951 59407034 refGene_gn=TPM2;refGene_tx=XM_023242425.1;refGene_gn=TPM2;refGene_tx=XM_023242427.1
D4 59408063 59408133 refGene_gn=TPM2;refGene_tx=XM_023242425.1;refGene_gn=TPM2;refGene_tx=XM_023242426.1;refGene_gn=TPM2;refGene_tx=XM_023242427.1;refGene_gn=TPM2;refGene_tx=XM_023242428.1
D4 59408540 59408603 refGene_gn=TPM2;refGene_tx=XM_023242425.1;refGene_gn=TPM2;refGene_tx=XM_023242426.1;refGene_gn=TPM2;refGene_tx=XM_023242427.1;refGene_gn=TPM2;refGene_tx=XM_023242428.1
D4 59408779 59408855 refGene_gn=TPM2;refGene_tx=XM_023242425.1;refGene_gn=TPM2;refGene_tx=XM_023242426.1
D4 59409099 59409175 refGene_gn=TPM2;refGene_tx=XM_023242427.1;refGene_gn=TPM2;refGene_tx=XM_023242428.1
D4 59409313 59409384 refGene_gn=TPM2;refGene_tx=XM_023242425.1;refGene_gn=TPM2;refGene_tx=XM_023242426.1;refGene_gn=TPM2;refGene_tx=XM_023242427.1;refGene_gn=TPM2;refGene_tx=XM_023242428.1
D4 59409478 59409596 refGene_gn=TPM2;refGene_tx=XM_023242425.1;refGene_gn=TPM2;refGene_tx=XM_023242426.1;refGene_gn=TPM2;refGene_tx=XM_023242427.1;refGene_gn=TPM2;refGene_tx=XM_023242428.1
D4 59409693 59409827

refGene_gn=TPM2;refGene_tx=XM_023242425.1;refGene_gn=TPM2;refGene_tx=XM_023242426.1;refGene_gn=TPM2;refGene_tx=XM_023242427.1;refGene_gn=TPM2;refGene_tx=XM_023242428.1
D4 59413155 59413281
refGene_gn=TPM2;refGene_tx=XM_023242425.1;refGene_gn=TPM2;refGene_tx=XM_023242426.1;refGene_gn=TPM2;refGene_tx=XM_023242427.1;refGene_gn=TPM2;refGene_tx=XM_023242428.1
D4 59413727 59413841
refGene_gn=TPM2;refGene_tx=XM_023242425.1;refGene_gn=TPM2;refGene_tx=XM_023242426.1;refGene_gn=TPM2;refGene_tx=XM_023242427.1;refGene_gn=TPM2;refGene_tx=XM_023242428.1
D4 59422181 59422307
refGene_gn=TLN1;refGene_tx=XM_011288561.2;refGene_gn=TLN1;refGene_tx=XM_019816168.1;refGene_gn=TLN1;refGene_tx=XM_019816169.1;refGene_gn=TLN1;refGene_tx=XM_019816170.1;refGene_gn=TLN1;refGene_tx=XM_019816171.1
D4 59422436 59422565
refGene_gn=TLN1;refGene_tx=XM_011288561.2;refGene_gn=TLN1;refGene_tx=XM_019816168.1;refGene_gn=TLN1;refGene_tx=XM_019816169.1;refGene_gn=TLN1;refGene_tx=XM_019816170.1;refGene_gn=TLN1;refGene_tx=XM_019816171.1
D4 59422703 59422886
refGene_gn=TLN1;refGene_tx=XM_011288561.2;refGene_gn=TLN1;refGene_tx=XM_019816168.1;refGene_gn=TLN1;refGene_tx=XM_019816169.1;refGene_gn=TLN1;refGene_tx=XM_019816170.1;refGene_gn=TLN1;refGene_tx=XM_019816171.1
D4 59422982 59423045
refGene_gn=TLN1;refGene_tx=XM_011288561.2;refGene_gn=TLN1;refGene_tx=XM_019816168.1;refGene_gn=TLN1;refGene_tx=XM_019816169.1;refGene_gn=TLN1;refGene_tx=XM_019816170.1;refGene_gn=TLN1;refGene_tx=XM_019816171.1
D4 59423167 59423293
refGene_gn=TLN1;refGene_tx=XM_011288561.2;refGene_gn=TLN1;refGene_tx=XM_019816168.1;refGene_gn=TLN1;refGene_tx=XM_019816169.1;refGene_gn=TLN1;refGene_tx=XM_019816170.1;refGene_gn=TLN1;refGene_tx=XM_019816171.1
D4 59423389 59423514
refGene_gn=TLN1;refGene_tx=XM_011288561.2;refGene_gn=TLN1;refGene_tx=XM_019816168.1;refGene_gn=TLN1;refGene_tx=XM_019816169.1;refGene_gn=TLN1;refGene_tx=XM_019816170.1;refGene_gn=TLN1;refGene_tx=XM_019816171.1
D4 59423714 59423820
refGene_gn=TLN1;refGene_tx=XM_011288561.2;refGene_gn=TLN1;refGene_tx=XM_019816168.1;refGene_gn=TLN1;refGene_tx=XM_019816169.1;refGene_gn=TLN1;refGene_tx=XM_019816170.1;refGene_gn=TLN1;refGene_tx=XM_019816171.1
D4 59424048 59424051
refGene_gn=TLN1;refGene_tx=XM_011288561.2;refGene_gn=TLN1;refGene_tx=XM_019816168.1;refGene_gn=TLN1;refGene_tx=XM_019816169.1
D4 59424366 59424474
refGene_gn=TLN1;refGene_tx=XM_011288561.2;refGene_gn=TLN1;refGene_tx=XM_019816168.1;refGene_gn=TLN1;refGene_tx=XM_019816169.1;refGene_gn=TLN1;refGene_tx=XM_019816170.1;refGene_gn=TLN1;refGene_tx=XM_019816171.1
D4 59424601 59424787
refGene_gn=TLN1;refGene_tx=XM_011288561.2;refGene_gn=TLN1;refGene_tx=XM_019816168.1;refGene_gn=TLN1;refGene_tx=XM_019816169.1;refGene_gn=TLN1;refGene_tx=XM_019816170.1;refGene_gn=TLN1;refGene_tx=XM_019816171.1

D4 59427115 59427232
refGene_gn=TLN1;refGene_tx=XM_011288561.2;refGene_gn=TLN1;refGene_tx=XM_019816168.1;refGene_gn=TLN1;refGene_tx=XM_019816169.1;refGene_gn=TLN1;refGene_tx=XM_019816170.1;refGene_gn=TLN1;refGene_tx=XM_019816171.1
D4 59427327 59427453
refGene_gn=TLN1;refGene_tx=XM_011288561.2;refGene_gn=TLN1;refGene_tx=XM_019816168.1;refGene_gn=TLN1;refGene_tx=XM_019816169.1;refGene_gn=TLN1;refGene_tx=XM_019816170.1;refGene_gn=TLN1;refGene_tx=XM_019816171.1
D4 59427543 59427727
refGene_gn=TLN1;refGene_tx=XM_011288561.2;refGene_gn=TLN1;refGene_tx=XM_019816168.1;refGene_gn=TLN1;refGene_tx=XM_019816169.1;refGene_gn=TLN1;refGene_tx=XM_019816170.1;refGene_gn=TLN1;refGene_tx=XM_019816171.1
D4 59427841 59428008
refGene_gn=TLN1;refGene_tx=XM_011288561.2;refGene_gn=TLN1;refGene_tx=XM_019816168.1;refGene_gn=TLN1;refGene_tx=XM_019816169.1;refGene_gn=TLN1;refGene_tx=XM_019816170.1;refGene_gn=TLN1;refGene_tx=XM_019816171.1
D4 59428181 59428328
refGene_gn=TLN1;refGene_tx=XM_011288561.2;refGene_gn=TLN1;refGene_tx=XM_019816168.1;refGene_gn=TLN1;refGene_tx=XM_019816169.1;refGene_gn=TLN1;refGene_tx=XM_019816170.1;refGene_gn=TLN1;refGene_tx=XM_019816171.1
D4 59429183 59429303
refGene_gn=TLN1;refGene_tx=XM_011288561.2;refGene_gn=TLN1;refGene_tx=XM_019816168.1;refGene_gn=TLN1;refGene_tx=XM_019816169.1;refGene_gn=TLN1;refGene_tx=XM_019816170.1;refGene_gn=TLN1;refGene_tx=XM_019816171.1
D4 59429379 59429481
refGene_gn=TLN1;refGene_tx=XM_011288561.2;refGene_gn=TLN1;refGene_tx=XM_019816168.1;refGene_gn=TLN1;refGene_tx=XM_019816169.1;refGene_gn=TLN1;refGene_tx=XM_019816170.1;refGene_gn=TLN1;refGene_tx=XM_019816171.1
D4 59429590 59429740
refGene_gn=TLN1;refGene_tx=XM_011288561.2;refGene_gn=TLN1;refGene_tx=XM_019816168.1;refGene_gn=TLN1;refGene_tx=XM_019816169.1;refGene_gn=TLN1;refGene_tx=XM_019816170.1;refGene_gn=TLN1;refGene_tx=XM_019816171.1
D4 59429831 59430002
refGene_gn=TLN1;refGene_tx=XM_011288561.2;refGene_gn=TLN1;refGene_tx=XM_019816168.1;refGene_gn=TLN1;refGene_tx=XM_019816169.1;refGene_gn=TLN1;refGene_tx=XM_019816170.1;refGene_gn=TLN1;refGene_tx=XM_019816171.1
D4 59430091 59430193
refGene_gn=TLN1;refGene_tx=XM_011288561.2;refGene_gn=TLN1;refGene_tx=XM_019816168.1;refGene_gn=TLN1;refGene_tx=XM_019816169.1;refGene_gn=TLN1;refGene_tx=XM_019816170.1;refGene_gn=TLN1;refGene_tx=XM_019816171.1
D4 59430399 59430532
refGene_gn=TLN1;refGene_tx=XM_011288561.2;refGene_gn=TLN1;refGene_tx=XM_019816168.1;refGene_gn=TLN1;refGene_tx=XM_019816169.1;refGene_gn=TLN1;refGene_tx=XM_019816170.1;refGene_gn=TLN1;refGene_tx=XM_019816171.1
D4 59430723 59430905
refGene_gn=TLN1;refGene_tx=XM_011288561.2;refGene_gn=TLN1;refGene_tx=XM_019816168.1;refGene_gn=TLN1;refGene_tx=XM_019816169.1;refGene_gn=TLN1;refGene_tx=XM_019816170.1;refGene_gn=TLN1;refGene_tx=XM_019816171.1
D4 59430998 59431139
refGene_gn=TLN1;refGene_tx=XM_011288561.2;refGene_gn=TLN1;refGene_tx=X

M_019816168.1;refGene_gn=TLN1;refGene_tx=XM_019816169.1;refGene_gn=TLN1;refGene_tx=XM_019816170.1;refGene_gn=TLN1;refGene_tx=XM_019816171.1
D4 59431373 59431535 refGene_gn=TLN1;refGene_tx=XM_011288561.2;refGene_gn=TLN1;refGene_tx=XM_019816168.1;refGene_gn=TLN1;refGene_tx=XM_019816169.1;refGene_gn=TLN1;refGene_tx=XM_019816170.1;refGene_gn=TLN1;refGene_tx=XM_019816171.1
D4 59432007 59432151 refGene_gn=TLN1;refGene_tx=XM_011288561.2;refGene_gn=TLN1;refGene_tx=XM_019816168.1;refGene_gn=TLN1;refGene_tx=XM_019816169.1;refGene_gn=TLN1;refGene_tx=XM_019816170.1;refGene_gn=TLN1;refGene_tx=XM_019816171.1
D4 59433302 59433425 refGene_gn=TLN1;refGene_tx=XM_011288561.2;refGene_gn=TLN1;refGene_tx=XM_019816168.1;refGene_gn=TLN1;refGene_tx=XM_019816169.1;refGene_gn=TLN1;refGene_tx=XM_019816170.1;refGene_gn=TLN1;refGene_tx=XM_019816171.1
D4 59433534 59433624 refGene_gn=TLN1;refGene_tx=XM_011288561.2;refGene_gn=TLN1;refGene_tx=XM_019816168.1;refGene_gn=TLN1;refGene_tx=XM_019816169.1;refGene_gn=TLN1;refGene_tx=XM_019816170.1;refGene_gn=TLN1;refGene_tx=XM_019816171.1
D4 59433726 59433820 refGene_gn=TLN1;refGene_tx=XM_011288561.2;refGene_gn=TLN1;refGene_tx=XM_019816168.1;refGene_gn=TLN1;refGene_tx=XM_019816169.1;refGene_gn=TLN1;refGene_tx=XM_019816170.1;refGene_gn=TLN1;refGene_tx=XM_019816171.1
D4 59434002 59434142 refGene_gn=TLN1;refGene_tx=XM_011288561.2;refGene_gn=TLN1;refGene_tx=XM_019816168.1;refGene_gn=TLN1;refGene_tx=XM_019816169.1;refGene_gn=TLN1;refGene_tx=XM_019816170.1;refGene_gn=TLN1;refGene_tx=XM_019816171.1
D4 59434338 59434536 refGene_gn=TLN1;refGene_tx=XM_011288561.2;refGene_gn=TLN1;refGene_tx=XM_019816168.1;refGene_gn=TLN1;refGene_tx=XM_019816169.1;refGene_gn=TLN1;refGene_tx=XM_019816170.1;refGene_gn=TLN1;refGene_tx=XM_019816171.1
D4 59434743 59434863 refGene_gn=TLN1;refGene_tx=XM_011288561.2;refGene_gn=TLN1;refGene_tx=XM_019816168.1;refGene_gn=TLN1;refGene_tx=XM_019816169.1;refGene_gn=TLN1;refGene_tx=XM_019816170.1;refGene_gn=TLN1;refGene_tx=XM_019816171.1
D4 59435221 59435430 refGene_gn=TLN1;refGene_tx=XM_011288561.2;refGene_gn=TLN1;refGene_tx=XM_019816168.1;refGene_gn=TLN1;refGene_tx=XM_019816169.1;refGene_gn=TLN1;refGene_tx=XM_019816170.1;refGene_gn=TLN1;refGene_tx=XM_019816171.1
D4 59435553 59435656 refGene_gn=TLN1;refGene_tx=XM_011288561.2;refGene_gn=TLN1;refGene_tx=XM_019816168.1;refGene_gn=TLN1;refGene_tx=XM_019816169.1;refGene_gn=TLN1;refGene_tx=XM_019816170.1;refGene_gn=TLN1;refGene_tx=XM_019816171.1
D4 59436254 59436383 refGene_gn=TLN1;refGene_tx=XM_011288561.2;refGene_gn=TLN1;refGene_tx=XM_019816168.1;refGene_gn=TLN1;refGene_tx=XM_019816169.1;refGene_gn=TLN1;refGene_tx=XM_019816170.1;refGene_gn=TLN1;refGene_tx=XM_019816171.1
D4 59436535 59436670 refGene_gn=TLN1;refGene_tx=XM_011288561.2;refGene_gn=TLN1;refGene_tx=XM_019816168.1;refGene_gn=TLN1;refGene_tx=XM_019816169.1;refGene_gn=TLN1;refGene_tx=XM_019816170.1;refGene_gn=TLN1;refGene_tx=XM_019816171.1

D4 59436861 59436975
refGene_gn=TLN1;refGene_tx=XM_011288561.2;refGene_gn=TLN1;refGene_tx=XM_019816168.1;refGene_gn=TLN1;refGene_tx=XM_019816169.1;refGene_gn=TLN1;refGene_tx=XM_019816170.1;refGene_gn=TLN1;refGene_tx=XM_019816171.1
D4 59437062 59437179
refGene_gn=TLN1;refGene_tx=XM_011288561.2;refGene_gn=TLN1;refGene_tx=XM_019816168.1;refGene_gn=TLN1;refGene_tx=XM_019816169.1;refGene_gn=TLN1;refGene_tx=XM_019816170.1;refGene_gn=TLN1;refGene_tx=XM_019816171.1
D4 59437357 59437486
refGene_gn=TLN1;refGene_tx=XM_011288561.2;refGene_gn=TLN1;refGene_tx=XM_019816168.1;refGene_gn=TLN1;refGene_tx=XM_019816169.1;refGene_gn=TLN1;refGene_tx=XM_019816170.1;refGene_gn=TLN1;refGene_tx=XM_019816171.1
D4 59438296 59438463
refGene_gn=TLN1;refGene_tx=XM_011288561.2;refGene_gn=TLN1;refGene_tx=XM_019816168.1;refGene_gn=TLN1;refGene_tx=XM_019816169.1;refGene_gn=TLN1;refGene_tx=XM_019816170.1;refGene_gn=TLN1;refGene_tx=XM_019816171.1
D4 59439033 59439328
refGene_gn=TLN1;refGene_tx=XM_011288561.2;refGene_gn=TLN1;refGene_tx=XM_019816168.1;refGene_gn=TLN1;refGene_tx=XM_019816169.1;refGene_gn=TLN1;refGene_tx=XM_019816170.1;refGene_gn=TLN1;refGene_tx=XM_019816171.1
D4 59439492 59439660
refGene_gn=TLN1;refGene_tx=XM_011288561.2;refGene_gn=TLN1;refGene_tx=XM_019816168.1;refGene_gn=TLN1;refGene_tx=XM_019816169.1;refGene_gn=TLN1;refGene_tx=XM_019816170.1;refGene_gn=TLN1;refGene_tx=XM_019816171.1
D4 59440213 59440264
refGene_gn=TLN1;refGene_tx=XM_019816168.1;refGene_gn=TLN1;refGene_tx=XM_019816169.1;refGene_gn=TLN1;refGene_tx=XM_019816170.1
D4 59440659 59440758
refGene_gn=TLN1;refGene_tx=XM_011288561.2;refGene_gn=TLN1;refGene_tx=XM_019816168.1;refGene_gn=TLN1;refGene_tx=XM_019816169.1;refGene_gn=TLN1;refGene_tx=XM_019816170.1;refGene_gn=TLN1;refGene_tx=XM_019816171.1
D4 59440876 59441085
refGene_gn=TLN1;refGene_tx=XM_011288561.2;refGene_gn=TLN1;refGene_tx=XM_019816168.1;refGene_gn=TLN1;refGene_tx=XM_019816169.1;refGene_gn=TLN1;refGene_tx=XM_019816170.1;refGene_gn=TLN1;refGene_tx=XM_019816171.1
D4 59441304 59441413
refGene_gn=TLN1;refGene_tx=XM_011288561.2;refGene_gn=TLN1;refGene_tx=XM_019816168.1;refGene_gn=TLN1;refGene_tx=XM_019816169.1;refGene_gn=TLN1;refGene_tx=XM_019816170.1;refGene_gn=TLN1;refGene_tx=XM_019816171.1
D4 59441547 59441661
refGene_gn=TLN1;refGene_tx=XM_011288561.2;refGene_gn=TLN1;refGene_tx=XM_019816168.1;refGene_gn=TLN1;refGene_tx=XM_019816169.1;refGene_gn=TLN1;refGene_tx=XM_019816170.1;refGene_gn=TLN1;refGene_tx=XM_019816171.1
D4 59441809 59441990
refGene_gn=TLN1;refGene_tx=XM_011288561.2;refGene_gn=TLN1;refGene_tx=XM_019816168.1;refGene_gn=TLN1;refGene_tx=XM_019816169.1;refGene_gn=TLN1;refGene_tx=XM_019816170.1;refGene_gn=TLN1;refGene_tx=XM_019816171.1
D4 59442213 59442290
refGene_gn=TLN1;refGene_tx=XM_011288561.2;refGene_gn=TLN1;refGene_tx=XM_019816168.1;refGene_gn=TLN1;refGene_tx=XM_019816169.1;refGene_gn=TLN

1;refGene_tx=XM_019816170.1;refGene_gn=TLN1;refGene_tx=XM_019816171.1
D4 59442453 59442555
refGene_gn=TLN1;refGene_tx=XM_011288561.2;refGene_gn=TLN1;refGene_tx=XM_019816168.1;refGene_gn=TLN1;refGene_tx=XM_019816169.1;refGene_gn=TLN1;refGene_tx=XM_019816170.1;refGene_gn=TLN1;refGene_tx=XM_019816171.1
D4 59443303 59443459
refGene_gn=TLN1;refGene_tx=XM_011288561.2;refGene_gn=TLN1;refGene_tx=XM_019816168.1;refGene_gn=TLN1;refGene_tx=XM_019816169.1;refGene_gn=TLN1;refGene_tx=XM_019816170.1;refGene_gn=TLN1;refGene_tx=XM_019816171.1
D4 59443789 59443894
refGene_gn=TLN1;refGene_tx=XM_011288561.2;refGene_gn=TLN1;refGene_tx=XM_019816168.1;refGene_gn=TLN1;refGene_tx=XM_019816169.1;refGene_gn=TLN1;refGene_tx=XM_019816170.1;refGene_gn=TLN1;refGene_tx=XM_019816171.1
D4 59444535 59444596
refGene_gn=TLN1;refGene_tx=XM_011288561.2;refGene_gn=TLN1;refGene_tx=XM_019816168.1;refGene_gn=TLN1;refGene_tx=XM_019816169.1;refGene_gn=TLN1;refGene_tx=XM_019816170.1;refGene_gn=TLN1;refGene_tx=XM_019816171.1
D4 59445314 59445442
refGene_gn=TLN1;refGene_tx=XM_011288561.2;refGene_gn=TLN1;refGene_tx=XM_019816168.1;refGene_gn=TLN1;refGene_tx=XM_019816169.1;refGene_gn=TLN1;refGene_tx=XM_019816170.1;refGene_gn=TLN1;refGene_tx=XM_019816171.1
D4 59445555 59445698
refGene_gn=TLN1;refGene_tx=XM_011288561.2;refGene_gn=TLN1;refGene_tx=XM_019816168.1;refGene_gn=TLN1;refGene_tx=XM_019816169.1;refGene_gn=TLN1;refGene_tx=XM_019816170.1;refGene_gn=TLN1;refGene_tx=XM_019816171.1
D4 59445958 59446111
refGene_gn=TLN1;refGene_tx=XM_011288561.2;refGene_gn=TLN1;refGene_tx=XM_019816168.1;refGene_gn=TLN1;refGene_tx=XM_019816169.1;refGene_gn=TLN1;refGene_tx=XM_019816170.1;refGene_gn=TLN1;refGene_tx=XM_019816171.1
D4 59446217 59446347
refGene_gn=TLN1;refGene_tx=XM_011288561.2;refGene_gn=TLN1;refGene_tx=XM_019816168.1;refGene_gn=TLN1;refGene_tx=XM_019816169.1;refGene_gn=TLN1;refGene_tx=XM_019816170.1;refGene_gn=TLN1;refGene_tx=XM_019816171.1
D4 59446601 59446699
refGene_gn=TLN1;refGene_tx=XM_011288561.2;refGene_gn=TLN1;refGene_tx=XM_019816168.1;refGene_gn=TLN1;refGene_tx=XM_019816169.1;refGene_gn=TLN1;refGene_tx=XM_019816170.1;refGene_gn=TLN1;refGene_tx=XM_019816171.1
D4 59446943 59447073
refGene_gn=TLN1;refGene_tx=XM_011288561.2;refGene_gn=TLN1;refGene_tx=XM_019816168.1;refGene_gn=TLN1;refGene_tx=XM_019816169.1;refGene_gn=TLN1;refGene_tx=XM_019816170.1;refGene_gn=TLN1;refGene_tx=XM_019816171.1
D4 59453769 59453862 refGene_gn=CREB3;refGene_tx=XM_003995609.4
D4 59453977 59454125 refGene_gn=CREB3;refGene_tx=XM_003995609.4
D4 59454201 59454269 refGene_gn=CREB3;refGene_tx=XM_003995609.4
D4 59454365 59454455 refGene_gn=CREB3;refGene_tx=XM_003995609.4
D4 59456637 59456744 refGene_gn=CREB3;refGene_tx=XM_003995609.4
D4 59456853 59456922 refGene_gn=CREB3;refGene_tx=XM_003995609.4
D4 59457770 59457855 refGene_gn=CREB3;refGene_tx=XM_003995609.4
D4 59457952 59458037 refGene_gn=CREB3;refGene_tx=XM_003995609.4
D4 59458115 59458432 refGene_gn=CREB3;refGene_tx=XM_003995609.4

D4 59458888 59459161
refGene_gn=GBA2;refGene_tx=XM_003995608.3;refGene_gn=GBA2;refGene_tx=XM_019816172.2
D4 59459439 59459631
refGene_gn=GBA2;refGene_tx=XM_003995608.3;refGene_gn=GBA2;refGene_tx=XM_019816172.2
D4 59459730 59459846
refGene_gn=GBA2;refGene_tx=XM_003995608.3;refGene_gn=GBA2;refGene_tx=XM_019816172.2
D4 59459924 59460067
refGene_gn=GBA2;refGene_tx=XM_003995608.3;refGene_gn=GBA2;refGene_tx=XM_019816172.2
D4 59460214 59460321
refGene_gn=GBA2;refGene_tx=XM_003995608.3;refGene_gn=GBA2;refGene_tx=XM_019816172.2
D4 59460446 59460598
refGene_gn=GBA2;refGene_tx=XM_003995608.3;refGene_gn=GBA2;refGene_tx=XM_019816172.2
D4 59460700 59460808
refGene_gn=GBA2;refGene_tx=XM_003995608.3;refGene_gn=GBA2;refGene_tx=XM_019816172.2
D4 59461018 59461123
refGene_gn=GBA2;refGene_tx=XM_003995608.3;refGene_gn=GBA2;refGene_tx=XM_019816172.2
D4 59461360 59461533
refGene_gn=GBA2;refGene_tx=XM_003995608.3;refGene_gn=GBA2;refGene_tx=XM_019816172.2
D4 59461731 59461857
refGene_gn=GBA2;refGene_tx=XM_003995608.3;refGene_gn=GBA2;refGene_tx=XM_019816172.2
D4 59461944 59462098
refGene_gn=GBA2;refGene_tx=XM_003995608.3;refGene_gn=GBA2;refGene_tx=XM_019816172.2
D4 59462259 59462362
refGene_gn=GBA2;refGene_tx=XM_003995608.3;refGene_gn=GBA2;refGene_tx=XM_019816172.2
D4 59462594 59462834
refGene_gn=GBA2;refGene_tx=XM_003995608.3;refGene_gn=GBA2;refGene_tx=XM_019816172.2
D4 59463108 59463327
refGene_gn=GBA2;refGene_tx=XM_003995608.3;refGene_gn=GBA2;refGene_tx=XM_019816172.2
D4 59465425 59465541
refGene_gn=GBA2;refGene_tx=XM_003995608.3;refGene_gn=GBA2;refGene_tx=XM_019816172.2
D4 59465841 59465933
refGene_gn=GBA2;refGene_tx=XM_003995608.3;refGene_gn=GBA2;refGene_tx=XM_019816172.2
D4 59469295 59469880
refGene_gn=GBA2;refGene_tx=XM_003995608.3;refGene_gn=GBA2;refGene_tx=X

M_019816172.2
D4	59470665	59470826
refGene_gn=GBA2;refGene_tx=XM_003995608.3;refGene_gn=RGP1;refGene_tx=XM_006939269.3;refGene_gn=RGP1;refGene_tx=XM_019816175.2
D4	59471210	59471347
refGene_gn=RGP1;refGene_tx=XM_006939269.3;refGene_gn=RGP1;refGene_tx=XM_019816175.2
D4	59471846	59471930
refGene_gn=RGP1;refGene_tx=XM_006939269.3;refGene_gn=RGP1;refGene_tx=XM_019816175.2
D4	59472034	59472184
refGene_gn=RGP1;refGene_tx=XM_006939269.3;refGene_gn=RGP1;refGene_tx=XM_019816175.2
D4	59472458	59472605
refGene_gn=RGP1;refGene_tx=XM_006939269.3;refGene_gn=RGP1;refGene_tx=XM_019816175.2
D4	59472809	59472937
refGene_gn=RGP1;refGene_tx=XM_006939269.3;refGene_gn=RGP1;refGene_tx=XM_019816175.2
D4	59473148	59473338
refGene_gn=RGP1;refGene_tx=XM_006939269.3;refGene_gn=RGP1;refGene_tx=XM_019816175.2
D4	59473841	59474065
refGene_gn=RGP1;refGene_tx=XM_006939269.3;refGene_gn=RGP1;refGene_tx=XM_019816175.2
D4	59474292	59474473	refGene_gn=MSMP;refGene_tx=XM_003995611.4
D4	59474809	59474918	refGene_gn=MSMP;refGene_tx=XM_003995611.4
D4	59475144	59475274	refGene_gn=MSMP;refGene_tx=XM_003995611.4
D4	59514918	59515585
refGene_gn=NPR2;refGene_tx=XM_003995612.3;refGene_gn=NPR2;refGene_tx=XM_006939272.2;refGene_gn=NPR2;refGene_tx=XM_006939273.2;refGene_gn=NPR2;refGene_tx=XM_006939274.2
D4	59516409	59516615
refGene_gn=NPR2;refGene_tx=XM_003995612.3;refGene_gn=NPR2;refGene_tx=XM_006939272.2;refGene_gn=NPR2;refGene_tx=XM_006939273.2;refGene_gn=NPR2;refGene_tx=XM_006939274.2
D4	59521615	59521729
refGene_gn=NPR2;refGene_tx=XM_003995612.3;refGene_gn=NPR2;refGene_tx=XM_006939272.2;refGene_gn=NPR2;refGene_tx=XM_006939273.2;refGene_gn=NPR2;refGene_tx=XM_006939274.2
D4	59522030	59522166
refGene_gn=NPR2;refGene_tx=XM_003995612.3;refGene_gn=NPR2;refGene_tx=XM_006939272.2;refGene_gn=NPR2;refGene_tx=XM_006939273.2;refGene_gn=NPR2;refGene_tx=XM_006939274.2
D4	59522408	59522503
refGene_gn=NPR2;refGene_tx=XM_003995612.3;refGene_gn=NPR2;refGene_tx=XM_006939272.2;refGene_gn=NPR2;refGene_tx=XM_006939273.2;refGene_gn=NPR2;refGene_tx=XM_006939274.2
D4	59522729	59522862
refGene_gn=NPR2;refGene_tx=XM_003995612.3;refGene_gn=NPR2;refGene_tx=X

M_006939272.2;refGene_gn=NPR2;refGene_tx=XM_006939273.2;refGene_gn=NPR2;refGene_tx=XM_006939274.2
D4 59523091 59523176
refGene_gn=NPR2;refGene_tx=XM_003995612.3;refGene_gn=NPR2;refGene_tx=XM_006939272.2;refGene_gn=NPR2;refGene_tx=XM_006939273.2;refGene_gn=NPR2;refGene_tx=XM_006939274.2
D4 59523643 59523773
refGene_gn=NPR2;refGene_tx=XM_003995612.3;refGene_gn=NPR2;refGene_tx=XM_006939272.2;refGene_gn=NPR2;refGene_tx=XM_006939273.2;refGene_gn=NPR2;refGene_tx=XM_006939274.2
D4 59523935 59524010
refGene_gn=NPR2;refGene_tx=XM_003995612.3;refGene_gn=NPR2;refGene_tx=XM_006939272.2
D4 59524207 59524285
refGene_gn=NPR2;refGene_tx=XM_003995612.3;refGene_gn=NPR2;refGene_tx=XM_006939272.2;refGene_gn=NPR2;refGene_tx=XM_006939273.2;refGene_gn=NPR2;refGene_tx=XM_006939274.2
D4 59524517 59524622
refGene_gn=NPR2;refGene_tx=XM_003995612.3;refGene_gn=NPR2;refGene_tx=XM_006939272.2;refGene_gn=NPR2;refGene_tx=XM_006939273.2;refGene_gn=NPR2;refGene_tx=XM_006939274.2
D4 59524719 59524791
refGene_gn=NPR2;refGene_tx=XM_003995612.3;refGene_gn=NPR2;refGene_tx=XM_006939272.2;refGene_gn=NPR2;refGene_tx=XM_006939273.2;refGene_gn=NPR2;refGene_tx=XM_006939274.2
D4 59526530 59526690
refGene_gn=NPR2;refGene_tx=XM_003995612.3;refGene_gn=NPR2;refGene_tx=XM_006939272.2;refGene_gn=NPR2;refGene_tx=XM_006939273.2;refGene_gn=NPR2;refGene_tx=XM_006939274.2
D4 59526844 59527000
refGene_gn=NPR2;refGene_tx=XM_003995612.3;refGene_gn=NPR2;refGene_tx=XM_006939272.2;refGene_gn=NPR2;refGene_tx=XM_006939273.2;refGene_gn=NPR2;refGene_tx=XM_006939274.2
D4 59527078 59527247
refGene_gn=NPR2;refGene_tx=XM_003995612.3;refGene_gn=NPR2;refGene_tx=XM_006939272.2;refGene_gn=NPR2;refGene_tx=XM_006939273.2;refGene_gn=NPR2;refGene_tx=XM_006939274.2
D4 59527411 59527558
refGene_gn=NPR2;refGene_tx=XM_003995612.3;refGene_gn=NPR2;refGene_tx=XM_006939272.2;refGene_gn=NPR2;refGene_tx=XM_006939273.2;refGene_gn=NPR2;refGene_tx=XM_006939274.2
D4 59528041 59528165
refGene_gn=NPR2;refGene_tx=XM_003995612.3;refGene_gn=NPR2;refGene_tx=XM_006939272.2;refGene_gn=NPR2;refGene_tx=XM_006939273.2;refGene_gn=NPR2;refGene_tx=XM_006939274.2
D4 59528354 59528423
refGene_gn=NPR2;refGene_tx=XM_003995612.3;refGene_gn=NPR2;refGene_tx=XM_006939272.2;refGene_gn=NPR2;refGene_tx=XM_006939273.2;refGene_gn=NPR2;refGene_tx=XM_006939274.2
D4 59529682 59529857

refGene_gn=NPR2;refGene_tx=XM_003995612.3;refGene_gn=NPR2;refGene_tx=XM_006939272.2;refGene_gn=NPR2;refGene_tx=XM_006939273.2;refGene_gn=NPR2;refGene_tx=XM_006939274.2
D4 59529928 59530027
refGene_gn=NPR2;refGene_tx=XM_003995612.3;refGene_gn=NPR2;refGene_tx=XM_006939272.2;refGene_gn=NPR2;refGene_tx=XM_006939273.2;refGene_gn=NPR2;refGene_tx=XM_006939274.2
D4 59530300 59530392
refGene_gn=NPR2;refGene_tx=XM_003995612.3;refGene_gn=NPR2;refGene_tx=XM_006939272.2;refGene_gn=NPR2;refGene_tx=XM_006939273.2;refGene_gn=NPR2;refGene_tx=XM_006939274.2
D4 59530523 59530589
refGene_gn=NPR2;refGene_tx=XM_003995612.3;refGene_gn=NPR2;refGene_tx=XM_006939272.2;refGene_gn=NPR2;refGene_tx=XM_006939273.2;refGene_gn=NPR2;refGene_tx=XM_006939274.2
D4 59531066 59531442
refGene_gn=SPAG8;refGene_tx=XM_006939270.4;refGene_gn=SPAG8;refGene_tx=XM_006939271.4;refGene_gn=SPAG8;refGene_tx=XM_019816178.2;refGene_gn=SPAG8;refGene_tx=XM_019816179.2;refGene_gn=SPAG8;refGene_tx=XM_023242429.1;refGene_gn=SPAG8;refGene_tx=XM_023242430.1
D4 59531542 59531667
refGene_gn=SPAG8;refGene_tx=XM_006939270.4;refGene_gn=SPAG8;refGene_tx=XM_006939271.4;refGene_gn=SPAG8;refGene_tx=XM_019816178.2;refGene_gn=SPAG8;refGene_tx=XM_019816179.2;refGene_gn=SPAG8;refGene_tx=XM_023242429.1;refGene_gn=SPAG8;refGene_tx=XM_023242430.1
D4 59531742 59531788
refGene_gn=SPAG8;refGene_tx=XM_006939270.4;refGene_gn=SPAG8;refGene_tx=XM_006939271.4;refGene_gn=SPAG8;refGene_tx=XM_019816178.2;refGene_gn=SPAG8;refGene_tx=XM_019816179.2;refGene_gn=SPAG8;refGene_tx=XM_023242429.1;refGene_gn=SPAG8;refGene_tx=XM_023242430.1
D4 59531984 59532159
refGene_gn=SPAG8;refGene_tx=XM_006939270.4;refGene_gn=SPAG8;refGene_tx=XM_006939271.4;refGene_gn=SPAG8;refGene_tx=XM_019816179.2;refGene_gn=SPAG8;refGene_tx=XM_023242429.1;refGene_gn=SPAG8;refGene_tx=XM_023242430.1
D4 59532284 59532978
refGene_gn=SPAG8;refGene_tx=XM_006939270.4;refGene_gn=SPAG8;refGene_tx=XM_006939271.4;refGene_gn=SPAG8;refGene_tx=XM_019816178.2;refGene_gn=SPAG8;refGene_tx=XM_019816179.2;refGene_gn=SPAG8;refGene_tx=XM_023242429.1;refGene_gn=SPAG8;refGene_tx=XM_023242430.1
D4 59533075 59533113
refGene_gn=SPAG8;refGene_tx=XM_006939270.4;refGene_gn=SPAG8;refGene_tx=XM_006939271.4;refGene_gn=SPAG8;refGene_tx=XM_019816178.2;refGene_gn=SPAG8;refGene_tx=XM_023242429.1;refGene_gn=SPAG8;refGene_tx=XM_023242430.1
D4 59534166 59534269
refGene_gn=HINT2;refGene_tx=XM_003995613.5;refGene_gn=HINT2;refGene_tx=XM_011288559.3
D4 59534379 59534452
refGene_gn=HINT2;refGene_tx=XM_003995613.5;refGene_gn=HINT2;refGene_tx

=XM_011288559.3
D4	59534558	59534663
refGene_gn=HINT2;refGene_tx=XM_003995613.5;refGene_gn=HINT2;refGene_tx
=XM_011288559.3
D4	59534759	59534900
refGene_gn=HINT2;refGene_tx=XM_003995613.5;refGene_gn=HINT2;refGene_tx
=XM_011288559.3
D4	59536032	59536242
refGene_gn=HINT2;refGene_tx=XM_003995613.5;refGene_gn=HINT2;refGene_tx
=XM_011288559.3
D4	59540091	59540129
refGene_gn=FAM221B;refGene_tx=XM_023242432.1;refGene_gn=FAM221B;refGen
e_tx=XM_023242433.1
D4	59540493	59540613
refGene_gn=FAM221B;refGene_tx=XM_023242432.1;refGene_gn=FAM221B;refGen
e_tx=XM_023242433.1
D4	59540799	59540997
refGene_gn=FAM221B;refGene_tx=XM_023242432.1;refGene_gn=FAM221B;refGen
e_tx=XM_023242433.1
D4	59541172	59541283
refGene_gn=FAM221B;refGene_tx=XM_023242432.1;refGene_gn=FAM221B;refGen
e_tx=XM_023242433.1
D4	59546143	59546287	refGene_gn=FAM221B;refGene_tx=XM_023242432.1
D4	59548414	59549201
refGene_gn=FAM221B;refGene_tx=XM_023242432.1;refGene_gn=FAM221B;refGen
e_tx=XM_023242433.1
D4	59551765	59552585
refGene_gn=TMEM8B;refGene_tx=XM_003995615.5;refGene_gn=TMEM8B;refGene_
tx=XM_006939275.4;refGene_gn=TMEM8B;refGene_tx=XM_019816180.2;refGene_
gn=TMEM8B;refGene_tx=XM_019816181.2;refGene_gn=TMEM8B;refGene_tx=XM_01
9816182.2;refGene_gn=TMEM8B;refGene_tx=XM_019816183.2;refGene_gn=TMEM8
B;refGene_tx=XM_019816184.2;refGene_gn=TMEM8B;refGene_tx=XM_019816185.
2
D4	59556859	59557049
refGene_gn=TMEM8B;refGene_tx=XM_003995615.5;refGene_gn=TMEM8B;refGene_
tx=XM_006939275.4;refGene_gn=TMEM8B;refGene_tx=XM_019816180.2;refGene_
gn=TMEM8B;refGene_tx=XM_019816181.2;refGene_gn=TMEM8B;refGene_tx=XM_01
9816182.2;refGene_gn=TMEM8B;refGene_tx=XM_019816183.2;refGene_gn=TMEM8
B;refGene_tx=XM_019816184.2;refGene_gn=TMEM8B;refGene_tx=XM_019816185.
2
D4	59557424	59557632
refGene_gn=TMEM8B;refGene_tx=XM_003995615.5;refGene_gn=TMEM8B;refGene_
tx=XM_006939275.4;refGene_gn=TMEM8B;refGene_tx=XM_019816180.2;refGene_
gn=TMEM8B;refGene_tx=XM_019816181.2;refGene_gn=TMEM8B;refGene_tx=XM_01
9816182.2;refGene_gn=TMEM8B;refGene_tx=XM_019816183.2;refGene_gn=TMEM8
B;refGene_tx=XM_019816184.2;refGene_gn=TMEM8B;refGene_tx=XM_019816185.
2
D4	59563681	59563815
refGene_gn=TMEM8B;refGene_tx=XM_003995615.5;refGene_gn=TMEM8B;refGene_
tx=XM_006939275.4;refGene_gn=TMEM8B;refGene_tx=XM_019816180.2;refGene_

gn=TMEM8B;refGene_tx=XM_019816181.2;refGene_gn=TMEM8B;refGene_tx=XM_01
9816182.2;refGene_gn=TMEM8B;refGene_tx=XM_019816183.2;refGene_gn=TMEM8
B;refGene_tx=XM_019816184.2;refGene_gn=TMEM8B;refGene_tx=XM_019816185.
2
D4 59564130 59564399
refGene_gn=TMEM8B;refGene_tx=XM_003995615.5;refGene_gn=TMEM8B;refGene_
tx=XM_006939275.4;refGene_gn=TMEM8B;refGene_tx=XM_019816180.2;refGene_
gn=TMEM8B;refGene_tx=XM_019816181.2;refGene_gn=TMEM8B;refGene_tx=XM_01
9816182.2;refGene_gn=TMEM8B;refGene_tx=XM_019816183.2;refGene_gn=TMEM8
B;refGene_tx=XM_019816184.2;refGene_gn=TMEM8B;refGene_tx=XM_019816185.
2
D4 59565005 59565331
refGene_gn=TMEM8B;refGene_tx=XM_003995615.5;refGene_gn=TMEM8B;refGene_
tx=XM_006939275.4;refGene_gn=TMEM8B;refGene_tx=XM_019816180.2;refGene_
gn=TMEM8B;refGene_tx=XM_019816181.2;refGene_gn=TMEM8B;refGene_tx=XM_01
9816182.2;refGene_gn=TMEM8B;refGene_tx=XM_019816183.2;refGene_gn=TMEM8
B;refGene_tx=XM_019816184.2;refGene_gn=TMEM8B;refGene_tx=XM_019816185.
2
D4 59568320 59568414
refGene_gn=TMEM8B;refGene_tx=XM_003995615.5;refGene_gn=TMEM8B;refGene_
tx=XM_006939275.4;refGene_gn=TMEM8B;refGene_tx=XM_019816180.2;refGene_
gn=TMEM8B;refGene_tx=XM_019816181.2;refGene_gn=TMEM8B;refGene_tx=XM_01
9816182.2;refGene_gn=TMEM8B;refGene_tx=XM_019816183.2;refGene_gn=TMEM8
B;refGene_tx=XM_019816184.2;refGene_gn=TMEM8B;refGene_tx=XM_019816185.
2
D4 59568598 59568722
refGene_gn=TMEM8B;refGene_tx=XM_003995615.5;refGene_gn=TMEM8B;refGene_
tx=XM_006939275.4;refGene_gn=TMEM8B;refGene_tx=XM_019816180.2;refGene_
gn=TMEM8B;refGene_tx=XM_019816181.2;refGene_gn=TMEM8B;refGene_tx=XM_01
9816182.2;refGene_gn=TMEM8B;refGene_tx=XM_019816183.2;refGene_gn=TMEM8
B;refGene_tx=XM_019816184.2;refGene_gn=TMEM8B;refGene_tx=XM_019816185.
2
D4 59568814 59568957
refGene_gn=TMEM8B;refGene_tx=XM_003995615.5;refGene_gn=TMEM8B;refGene_
tx=XM_006939275.4;refGene_gn=TMEM8B;refGene_tx=XM_019816180.2;refGene_
gn=TMEM8B;refGene_tx=XM_019816181.2;refGene_gn=TMEM8B;refGene_tx=XM_01
9816182.2;refGene_gn=TMEM8B;refGene_tx=XM_019816183.2;refGene_gn=TMEM8
B;refGene_tx=XM_019816184.2;refGene_gn=TMEM8B;refGene_tx=XM_019816185.
2
D4 59569226 59569405
refGene_gn=TMEM8B;refGene_tx=XM_003995615.5;refGene_gn=TMEM8B;refGene_
tx=XM_006939275.4;refGene_gn=TMEM8B;refGene_tx=XM_019816180.2;refGene_
gn=TMEM8B;refGene_tx=XM_019816181.2;refGene_gn=TMEM8B;refGene_tx=XM_01
9816182.2;refGene_gn=TMEM8B;refGene_tx=XM_019816183.2;refGene_gn=TMEM8
B;refGene_tx=XM_019816184.2;refGene_gn=TMEM8B;refGene_tx=XM_019816185.
2
D4 59576999 59577146
refGene_gn=TMEM8B;refGene_tx=XM_003995615.5;refGene_gn=TMEM8B;refGene_
tx=XM_006939275.4;refGene_gn=TMEM8B;refGene_tx=XM_019816180.2;refGene_
gn=TMEM8B;refGene_tx=XM_019816181.2;refGene_gn=TMEM8B;refGene_tx=XM_01

9816182.2;refGene_gn=TMEM8B;refGene_tx=XM_019816183.2;refGene_gn=TMEM8B;refGene_tx=XM_019816184.2;refGene_gn=TMEM8B;refGene_tx=XM_019816185.2
D4 59577256 59577396 refGene_gn=TMEM8B;refGene_tx=XM_003995615.5;refGene_gn=TMEM8B;refGene_tx=XM_006939275.4;refGene_gn=TMEM8B;refGene_tx=XM_019816180.2;refGene_gn=TMEM8B;refGene_tx=XM_019816181.2;refGene_gn=TMEM8B;refGene_tx=XM_019816182.2;refGene_gn=TMEM8B;refGene_tx=XM_019816183.2;refGene_gn=TMEM8B;refGene_tx=XM_019816184.2;refGene_gn=TMEM8B;refGene_tx=XM_019816185.2
D4 59577646 59578028 refGene_gn=TMEM8B;refGene_tx=XM_003995615.5;refGene_gn=TMEM8B;refGene_tx=XM_006939275.4;refGene_gn=TMEM8B;refGene_tx=XM_019816180.2;refGene_gn=TMEM8B;refGene_tx=XM_019816181.2;refGene_gn=TMEM8B;refGene_tx=XM_019816182.2;refGene_gn=TMEM8B;refGene_tx=XM_019816183.2
D4 59580880 59581897 refGene_gn=LOC101088278;refGene_tx=XM_003995676.2
D4 59584849 59584985 refGene_gn=TMEM8B;refGene_tx=XM_019816184.2;refGene_gn=TMEM8B;refGene_tx=XM_019816185.2
D4 59590834 59591773 refGene_gn=LOC101095289;refGene_tx=XM_003995616.3
D4 59616440 59616782 refGene_gn=HRCT1;refGene_tx=XM_003995677.5;refGene_gn=HRCT1;refGene_tx=XM_019816188.2
D4 59621615 59621843 refGene_gn=LOC111557510;refGene_tx=XM_023242435.1
D4 59649057 59650017 refGene_gn=LOC105261116;refGene_tx=XM_011288557.2
D4 59664700 59665660 refGene_gn=LOC101095775;refGene_tx=XM_011288555.1
D4 59674675 59675635 refGene_gn=LOC101096016;refGene_tx=XM_003995617.1
D4 59688007 59688961 refGene_gn=LOC101096269;refGene_tx=XM_003995618.2
D4 59694333 59695290 refGene_gn=LOC101096518;refGene_tx=XM_003995619.3
D4 59706294 59707239 refGene_gn=LOC101096768;refGene_tx=XM_011288554.2
D4 59720542 59720642 refGene_gn=RECK;refGene_tx=XM_011288552.3
D4 59732589 59732648 refGene_gn=RECK;refGene_tx=XM_011288552.3
D4 59735447 59735522 refGene_gn=RECK;refGene_tx=XM_011288552.3
D4 59736438 59736475 refGene_gn=RECK;refGene_tx=XM_011288552.3
D4 59738931 59739017 refGene_gn=RECK;refGene_tx=XM_011288552.3
D4 59740429 59740477 refGene_gn=RECK;refGene_tx=XM_011288552.3
D4 59741776 59741797 refGene_gn=RECK;refGene_tx=XM_019816189.2
D4 59753444 59753478 refGene_gn=RECK;refGene_tx=XM_011288552.3;refGene_gn=RECK;refGene_tx=XM_019816189.2
D4 59755879 59756077

refGene_gn=RECK;refGene_tx=XM_011288552.3;refGene_gn=RECK;refGene_tx=XM_019816189.2
D4 59757983 59758251
refGene_gn=RECK;refGene_tx=XM_011288552.3;refGene_gn=RECK;refGene_tx=XM_019816189.2
D4 59760455 59760635
refGene_gn=RECK;refGene_tx=XM_011288552.3;refGene_gn=RECK;refGene_tx=XM_019816189.2
D4 59766398 59766611
refGene_gn=RECK;refGene_tx=XM_011288552.3;refGene_gn=RECK;refGene_tx=XM_019816189.2
D4 59768333 59768470
refGene_gn=RECK;refGene_tx=XM_011288552.3;refGene_gn=RECK;refGene_tx=XM_019816189.2
D4 59770431 59770572
refGene_gn=RECK;refGene_tx=XM_011288552.3;refGene_gn=RECK;refGene_tx=XM_019816189.2
D4 59771990 59772179
refGene_gn=RECK;refGene_tx=XM_011288552.3;refGene_gn=RECK;refGene_tx=XM_019816189.2
D4 59774958 59775081
refGene_gn=RECK;refGene_tx=XM_011288552.3;refGene_gn=RECK;refGene_tx=XM_019816189.2
D4 59777369 59777541
refGene_gn=RECK;refGene_tx=XM_011288552.3;refGene_gn=RECK;refGene_tx=XM_019816189.2
D4 59798953 59799146
refGene_gn=RECK;refGene_tx=XM_011288552.3;refGene_gn=RECK;refGene_tx=XM_019816189.2
D4 59800326 59800537
refGene_gn=RECK;refGene_tx=XM_011288552.3;refGene_gn=RECK;refGene_tx=XM_019816189.2
D4 59802084 59802158
refGene_gn=RECK;refGene_tx=XM_011288552.3;refGene_gn=RECK;refGene_tx=XM_019816189.2
D4 59802974 59803130
refGene_gn=RECK;refGene_tx=XM_011288552.3;refGene_gn=RECK;refGene_tx=XM_019816189.2
D4 59804015 59804237
refGene_gn=RECK;refGene_tx=XM_011288552.3;refGene_gn=RECK;refGene_tx=XM_019816189.2
D4 59823954 59823967 refGene_gn=GLIPR2;refGene_tx=XM_003995621.4
D4 59833139 59833248 refGene_gn=GLIPR2;refGene_tx=XM_003995621.4
D4 59833596 59833700 refGene_gn=GLIPR2;refGene_tx=XM_003995621.4
D4 59835605 59835683 refGene_gn=GLIPR2;refGene_tx=XM_003995621.4
D4 59842354 59842515 refGene_gn=GLIPR2;refGene_tx=XM_003995621.4
D4 59847626 59849393 refGene_gn=CCIN;refGene_tx=XM_003995622.5
D4 59870829 59871046
refGene_gn=CLTA;refGene_tx=XM_003995623.4;refGene_gn=CLTA;refGene_tx=XM_003995626.4;refGene_gn=CLTA;refGene_tx=XM_011288550.3;refGene_gn=CLT

A;refGene_tx=XM_019816191.2;refGene_gn=CLTA;refGene_tx=XM_019816192.2;refGene_gn=CLTA;refGene_tx=XM_019816193.2;refGene_gn=CLTA;refGene_tx=XM_019816194.2
D4 59879225 59879263
refGene_gn=CLTA;refGene_tx=XM_003995623.4;refGene_gn=CLTA;refGene_tx=XM_003995626.4;refGene_gn=CLTA;refGene_tx=XM_011288550.3;refGene_gn=CLTA;refGene_tx=XM_019816191.2
D4 59881291 59881409
refGene_gn=CLTA;refGene_tx=XM_003995623.4;refGene_gn=CLTA;refGene_tx=XM_003995626.4;refGene_gn=CLTA;refGene_tx=XM_011288550.3;refGene_gn=CLTA;refGene_tx=XM_019816191.2
D4 59886014 59886126
refGene_gn=CLTA;refGene_tx=XM_003995623.4;refGene_gn=CLTA;refGene_tx=XM_003995626.4;refGene_gn=CLTA;refGene_tx=XM_011288550.3;refGene_gn=CLTA;refGene_tx=XM_019816191.2;refGene_gn=CLTA;refGene_tx=XM_019816192.2;refGene_gn=CLTA;refGene_tx=XM_019816193.2;refGene_gn=CLTA;refGene_tx=XM_019816194.2
D4 59890442 59890496
refGene_gn=CLTA;refGene_tx=XM_011288550.3;refGene_gn=CLTA;refGene_tx=XM_019816191.2;refGene_gn=CLTA;refGene_tx=XM_019816192.2
D4 59891664 59891700
refGene_gn=CLTA;refGene_tx=XM_003995626.4;refGene_gn=CLTA;refGene_tx=XM_011288550.3;refGene_gn=CLTA;refGene_tx=XM_019816192.2;refGene_gn=CLTA;refGene_tx=XM_019816193.2
D4 59892657 59892829
refGene_gn=CLTA;refGene_tx=XM_003995623.4;refGene_gn=CLTA;refGene_tx=XM_003995626.4;refGene_gn=CLTA;refGene_tx=XM_011288550.3;refGene_gn=CLTA;refGene_tx=XM_019816191.2;refGene_gn=CLTA;refGene_tx=XM_019816192.2;refGene_gn=CLTA;refGene_tx=XM_019816193.2;refGene_gn=CLTA;refGene_tx=XM_019816194.2
D4 59902277 59902513
refGene_gn=GNE;refGene_tx=XM_003995628.4;refGene_gn=GNE;refGene_tx=XM_006939279.4;refGene_gn=GNE;refGene_tx=XM_006939281.3;refGene_gn=GNE;refGene_tx=XM_006939282.4;refGene_gn=GNE;refGene_tx=XM_019816195.1
D4 59903732 59903849
refGene_gn=GNE;refGene_tx=XM_003995628.4;refGene_gn=GNE;refGene_tx=XM_006939279.4;refGene_gn=GNE;refGene_tx=XM_006939281.3;refGene_gn=GNE;refGene_tx=XM_006939282.4;refGene_gn=GNE;refGene_tx=XM_019816195.1
D4 59912846 59913029
refGene_gn=GNE;refGene_tx=XM_003995628.4;refGene_gn=GNE;refGene_tx=XM_006939279.4;refGene_gn=GNE;refGene_tx=XM_006939281.3;refGene_gn=GNE;refGene_tx=XM_006939282.4;refGene_gn=GNE;refGene_tx=XM_019816195.1
D4 59914921 59915143
refGene_gn=GNE;refGene_tx=XM_003995628.4;refGene_gn=GNE;refGene_tx=XM_006939279.4;refGene_gn=GNE;refGene_tx=XM_006939281.3;refGene_gn=GNE;refGene_tx=XM_006939282.4;refGene_gn=GNE;refGene_tx=XM_019816195.1
D4 59916188 59916318
refGene_gn=GNE;refGene_tx=XM_003995628.4;refGene_gn=GNE;refGene_tx=XM_006939279.4;refGene_gn=GNE;refGene_tx=XM_006939281.3;refGene_gn=GNE;refGene_tx=XM_006939282.4;refGene_gn=GNE;refGene_tx=XM_019816195.1

D4 59919405 59919616
refGene_gn=GNE;refGene_tx=XM_003995628.4;refGene_gn=GNE;refGene_tx=XM_006939279.4;refGene_gn=GNE;refGene_tx=XM_006939281.3;refGene_gn=GNE;refGene_tx=XM_006939282.4;refGene_gn=GNE;refGene_tx=XM_019816195.1
D4 59920357 59920445
refGene_gn=GNE;refGene_tx=XM_003995628.4;refGene_gn=GNE;refGene_tx=XM_006939279.4;refGene_gn=GNE;refGene_tx=XM_006939281.3;refGene_gn=GNE;refGene_tx=XM_006939282.4;refGene_gn=GNE;refGene_tx=XM_019816195.1
D4 59925346 59925559
refGene_gn=GNE;refGene_tx=XM_003995628.4;refGene_gn=GNE;refGene_tx=XM_006939279.4;refGene_gn=GNE;refGene_tx=XM_006939281.3;refGene_gn=GNE;refGene_tx=XM_006939282.4;refGene_gn=GNE;refGene_tx=XM_019816195.1
D4 59927661 59927814
refGene_gn=GNE;refGene_tx=XM_003995628.4;refGene_gn=GNE;refGene_tx=XM_006939279.4;refGene_gn=GNE;refGene_tx=XM_006939281.3;refGene_gn=GNE;refGene_tx=XM_006939282.4;refGene_gn=GNE;refGene_tx=XM_019816195.1
D4 59929410 59929862
refGene_gn=GNE;refGene_tx=XM_003995628.4;refGene_gn=GNE;refGene_tx=XM_006939279.4;refGene_gn=GNE;refGene_tx=XM_006939281.3;refGene_gn=GNE;refGene_tx=XM_006939282.4;refGene_gn=GNE;refGene_tx=XM_019816195.1
D4 59935944 59936150
refGene_gn=GNE;refGene_tx=XM_003995628.4;refGene_gn=GNE;refGene_tx=XM_006939279.4;refGene_gn=GNE;refGene_tx=XM_006939281.3;refGene_gn=GNE;refGene_tx=XM_019816195.1
D4 59940993 59941023 refGene_gn=GNE;refGene_tx=XM_006939279.4
D4 59996610 59996673
refGene_gn=RNF38;refGene_tx=XM_023242438.1;refGene_gn=RNF38;refGene_tx=XM_023242439.1;refGene_gn=RNF38;refGene_tx=XM_023242440.1;refGene_gn=RNF38;refGene_tx=XM_023242441.1;refGene_gn=RNF38;refGene_tx=XM_023242442.1;refGene_gn=RNF38;refGene_tx=XM_023242443.1;refGene_gn=RNF38;refGene_tx=XM_023242444.1;refGene_gn=RNF38;refGene_tx=XM_023242445.1;refGene_gn=RNF38;refGene_tx=XM_023242447.1;refGene_gn=RNF38;refGene_tx=XM_023242448.1;refGene_gn=RNF38;refGene_tx=XM_023242449.1;refGene_gn=RNF38;refGene_tx=XM_023242450.1
D4 59998060 59998160
refGene_gn=RNF38;refGene_tx=XM_023242438.1;refGene_gn=RNF38;refGene_tx=XM_023242439.1;refGene_gn=RNF38;refGene_tx=XM_023242440.1;refGene_gn=RNF38;refGene_tx=XM_023242441.1;refGene_gn=RNF38;refGene_tx=XM_023242442.1;refGene_gn=RNF38;refGene_tx=XM_023242443.1;refGene_gn=RNF38;refGene_tx=XM_023242444.1;refGene_gn=RNF38;refGene_tx=XM_023242445.1;refGene_gn=RNF38;refGene_tx=XM_023242447.1;refGene_gn=RNF38;refGene_tx=XM_023242448.1;refGene_gn=RNF38;refGene_tx=XM_023242449.1;refGene_gn=RNF38;refGene_tx=XM_023242450.1
D4 59999425 59999547
refGene_gn=RNF38;refGene_tx=XM_023242438.1;refGene_gn=RNF38;refGene_tx=XM_023242439.1;refGene_gn=RNF38;refGene_tx=XM_023242440.1;refGene_gn=RNF38;refGene_tx=XM_023242441.1;refGene_gn=RNF38;refGene_tx=XM_023242442.1;refGene_gn=RNF38;refGene_tx=XM_023242443.1;refGene_gn=RNF38;refGene_tx=XM_023242444.1;refGene_gn=RNF38;refGene_tx=XM_023242445.1;refGene_gn=RNF38;refGene_tx=XM_023242447.1;refGene_gn=RNF38;refGene_tx=XM_02

3242448.1;refGene_gn=RNF38;refGene_tx=XM_023242449.1;refGene_gn=RNF38;refGene_tx=XM_023242450.1
D4 60003059 60003144
refGene_gn=RNF38;refGene_tx=XM_023242438.1;refGene_gn=RNF38;refGene_tx=XM_023242439.1;refGene_gn=RNF38;refGene_tx=XM_023242440.1;refGene_gn=RNF38;refGene_tx=XM_023242441.1;refGene_gn=RNF38;refGene_tx=XM_023242442.1;refGene_gn=RNF38;refGene_tx=XM_023242443.1;refGene_gn=RNF38;refGene_tx=XM_023242444.1;refGene_gn=RNF38;refGene_tx=XM_023242445.1;refGene_gn=RNF38;refGene_tx=XM_023242447.1;refGene_gn=RNF38;refGene_tx=XM_023242448.1;refGene_gn=RNF38;refGene_tx=XM_023242449.1;refGene_gn=RNF38;refGene_tx=XM_023242450.1
D4 60004613 60004720
refGene_gn=RNF38;refGene_tx=XM_023242438.1;refGene_gn=RNF38;refGene_tx=XM_023242439.1;refGene_gn=RNF38;refGene_tx=XM_023242440.1;refGene_gn=RNF38;refGene_tx=XM_023242441.1;refGene_gn=RNF38;refGene_tx=XM_023242442.1;refGene_gn=RNF38;refGene_tx=XM_023242443.1;refGene_gn=RNF38;refGene_tx=XM_023242444.1;refGene_gn=RNF38;refGene_tx=XM_023242445.1;refGene_gn=RNF38;refGene_tx=XM_023242447.1;refGene_gn=RNF38;refGene_tx=XM_023242448.1;refGene_gn=RNF38;refGene_tx=XM_023242449.1;refGene_gn=RNF38;refGene_tx=XM_023242450.1
D4 60004999 60005182
refGene_gn=RNF38;refGene_tx=XM_023242438.1;refGene_gn=RNF38;refGene_tx=XM_023242439.1;refGene_gn=RNF38;refGene_tx=XM_023242440.1;refGene_gn=RNF38;refGene_tx=XM_023242441.1;refGene_gn=RNF38;refGene_tx=XM_023242442.1;refGene_gn=RNF38;refGene_tx=XM_023242443.1;refGene_gn=RNF38;refGene_tx=XM_023242444.1;refGene_gn=RNF38;refGene_tx=XM_023242445.1;refGene_gn=RNF38;refGene_tx=XM_023242447.1;refGene_gn=RNF38;refGene_tx=XM_023242448.1;refGene_gn=RNF38;refGene_tx=XM_023242449.1;refGene_gn=RNF38;refGene_tx=XM_023242450.1
D4 60007820 60007991
refGene_gn=RNF38;refGene_tx=XM_023242438.1;refGene_gn=RNF38;refGene_tx=XM_023242439.1;refGene_gn=RNF38;refGene_tx=XM_023242440.1;refGene_gn=RNF38;refGene_tx=XM_023242441.1;refGene_gn=RNF38;refGene_tx=XM_023242442.1;refGene_gn=RNF38;refGene_tx=XM_023242443.1;refGene_gn=RNF38;refGene_tx=XM_023242444.1;refGene_gn=RNF38;refGene_tx=XM_023242445.1;refGene_gn=RNF38;refGene_tx=XM_023242447.1;refGene_gn=RNF38;refGene_tx=XM_023242448.1;refGene_gn=RNF38;refGene_tx=XM_023242449.1;refGene_gn=RNF38;refGene_tx=XM_023242450.1
D4 60009251 60009419
refGene_gn=RNF38;refGene_tx=XM_023242438.1;refGene_gn=RNF38;refGene_tx=XM_023242439.1;refGene_gn=RNF38;refGene_tx=XM_023242440.1;refGene_gn=RNF38;refGene_tx=XM_023242441.1;refGene_gn=RNF38;refGene_tx=XM_023242442.1;refGene_gn=RNF38;refGene_tx=XM_023242443.1;refGene_gn=RNF38;refGene_tx=XM_023242444.1;refGene_gn=RNF38;refGene_tx=XM_023242445.1;refGene_gn=RNF38;refGene_tx=XM_023242447.1;refGene_gn=RNF38;refGene_tx=XM_023242448.1;refGene_gn=RNF38;refGene_tx=XM_023242449.1;refGene_gn=RNF38;refGene_tx=XM_023242450.1
D4 60016426 60016640
refGene_gn=RNF38;refGene_tx=XM_023242438.1;refGene_gn=RNF38;refGene_tx=XM_023242439.1;refGene_gn=RNF38;refGene_tx=XM_023242440.1;refGene_gn=

RNF38;refGene_tx=XM_023242441.1;refGene_gn=RNF38;refGene_tx=XM_0232424
42.1;refGene_gn=RNF38;refGene_tx=XM_023242443.1;refGene_gn=RNF38;refGe
ne_tx=XM_023242444.1;refGene_gn=RNF38;refGene_tx=XM_023242445.1;refGen
e_gn=RNF38;refGene_tx=XM_023242447.1;refGene_gn=RNF38;refGene_tx=XM_02
3242448.1;refGene_gn=RNF38;refGene_tx=XM_023242449.1;refGene_gn=RNF38;
refGene_tx=XM_023242450.1
D4 60023768 60023962
refGene_gn=RNF38;refGene_tx=XM_023242438.1;refGene_gn=RNF38;refGene_tx
=XM_023242439.1;refGene_gn=RNF38;refGene_tx=XM_023242440.1;refGene_gn=
RNF38;refGene_tx=XM_023242441.1;refGene_gn=RNF38;refGene_tx=XM_0232424
42.1;refGene_gn=RNF38;refGene_tx=XM_023242443.1;refGene_gn=RNF38;refGe
ne_tx=XM_023242444.1;refGene_gn=RNF38;refGene_tx=XM_023242445.1;refGen
e_gn=RNF38;refGene_tx=XM_023242447.1;refGene_gn=RNF38;refGene_tx=XM_02
3242448.1;refGene_gn=RNF38;refGene_tx=XM_023242449.1;refGene_gn=RNF38;
refGene_tx=XM_023242450.1
D4 60035244 60035394
refGene_gn=RNF38;refGene_tx=XM_023242440.1;refGene_gn=RNF38;refGene_tx
=XM_023242441.1;refGene_gn=RNF38;refGene_tx=XM_023242442.1;refGene_gn=
RNF38;refGene_tx=XM_023242443.1
D4 60042870 60042879
refGene_gn=RNF38;refGene_tx=XM_023242440.1;refGene_gn=RNF38;refGene_tx
=XM_023242442.1;refGene_gn=RNF38;refGene_tx=XM_023242444.1
D4 60056279 60056282
refGene_gn=RNF38;refGene_tx=XM_023242441.1;refGene_gn=RNF38;refGene_tx
=XM_023242443.1;refGene_gn=RNF38;refGene_tx=XM_023242445.1
D4 60201475 60201533
refGene_gn=MELK;refGene_tx=XM_023242451.1;refGene_gn=MELK;refGene_tx=X
M_023242452.1
D4 60204008 60204094
refGene_gn=MELK;refGene_tx=XM_023242451.1;refGene_gn=MELK;refGene_tx=X
M_023242452.1
D4 60206032 60206149
refGene_gn=MELK;refGene_tx=XM_023242451.1;refGene_gn=MELK;refGene_tx=X
M_023242452.1
D4 60209627 60209771
refGene_gn=MELK;refGene_tx=XM_023242451.1;refGene_gn=MELK;refGene_tx=X
M_023242452.1
D4 60211696 60211765
refGene_gn=MELK;refGene_tx=XM_023242451.1;refGene_gn=MELK;refGene_tx=X
M_023242452.1
D4 60213550 60213643
refGene_gn=MELK;refGene_tx=XM_023242451.1;refGene_gn=MELK;refGene_tx=X
M_023242452.1
D4 60218218 60218317
refGene_gn=MELK;refGene_tx=XM_023242451.1;refGene_gn=MELK;refGene_tx=X
M_023242452.1
D4 60240413 60240482
refGene_gn=MELK;refGene_tx=XM_023242451.1;refGene_gn=MELK;refGene_tx=X
M_023242452.1
D4 60242065 60242164

refGene_gn=MELK;refGene_tx=XM_023242451.1;refGene_gn=MELK;refGene_tx=XM_023242452.1
D4 60253045 60253132
refGene_gn=MELK;refGene_tx=XM_023242451.1;refGene_gn=MELK;refGene_tx=XM_023242452.1
D4 60260630 60260762
refGene_gn=MELK;refGene_tx=XM_023242451.1;refGene_gn=MELK;refGene_tx=XM_023242452.1
D4 60264293 60264416
refGene_gn=MELK;refGene_tx=XM_023242451.1;refGene_gn=MELK;refGene_tx=XM_023242452.1
D4 60269333 60269565
refGene_gn=MELK;refGene_tx=XM_023242451.1;refGene_gn=MELK;refGene_tx=XM_023242452.1
D4 60272941 60273038
refGene_gn=MELK;refGene_tx=XM_023242451.1;refGene_gn=MELK;refGene_tx=XM_023242452.1
D4 60274637 60274806
refGene_gn=MELK;refGene_tx=XM_023242451.1;refGene_gn=MELK;refGene_tx=XM_023242452.1
D4 60283885 60283989
refGene_gn=MELK;refGene_tx=XM_023242451.1;refGene_gn=MELK;refGene_tx=XM_023242452.1
D4 60285087 60285145 refGene_gn=MELK;refGene_tx=XM_023242452.1
D4 60286126 60286304 refGene_gn=MELK;refGene_tx=XM_023242451.1
D4 60419256 60419333
refGene_gn=PAX5;refGene_tx=XM_011288547.2;refGene_gn=PAX5;refGene_tx=XM_019816199.1;refGene_gn=PAX5;refGene_tx=XM_019816200.2
D4 60425596 60425683
refGene_gn=PAX5;refGene_tx=XM_011288547.2;refGene_gn=PAX5;refGene_tx=XM_019816199.1;refGene_gn=PAX5;refGene_tx=XM_019816200.2
D4 60458518 60458620
refGene_gn=PAX5;refGene_tx=XM_011288547.2;refGene_gn=PAX5;refGene_tx=XM_019816199.1;refGene_gn=PAX5;refGene_tx=XM_019816200.2
D4 60497817 60497947
refGene_gn=PAX5;refGene_tx=XM_011288547.2;refGene_gn=PAX5;refGene_tx=XM_019816199.1;refGene_gn=PAX5;refGene_tx=XM_019816200.2
D4 60503922 60503979
refGene_gn=PAX5;refGene_tx=XM_019816199.1;refGene_gn=PAX5;refGene_tx=XM_019816200.2
D4 60541790 60541966
refGene_gn=PAX5;refGene_tx=XM_011288547.2;refGene_gn=PAX5;refGene_tx=XM_019816199.1;refGene_gn=PAX5;refGene_tx=XM_019816200.2
D4 60574407 60574536
refGene_gn=PAX5;refGene_tx=XM_011288547.2;refGene_gn=PAX5;refGene_tx=XM_019816199.1;refGene_gn=PAX5;refGene_tx=XM_019816200.2
D4 60577997 60578062
refGene_gn=PAX5;refGene_tx=XM_011288547.2;refGene_gn=PAX5;refGene_tx=XM_019816199.1;refGene_gn=PAX5;refGene_tx=XM_019816200.2
D4 60585683 60585881

refGene_gn=PAX5;refGene_tx=XM_011288547.2;refGene_gn=PAX5;refGene_tx=XM_019816199.1;refGene_gn=PAX5;refGene_tx=XM_019816200.2
D4 60591338 60591504
refGene_gn=PAX5;refGene_tx=XM_011288547.2;refGene_gn=PAX5;refGene_tx=XM_019816199.1;refGene_gn=PAX5;refGene_tx=XM_019816200.2
D4 60597293 60597336 refGene_gn=PAX5;refGene_tx=XM_019816200.2
D4 60604897 60604943
refGene_gn=PAX5;refGene_tx=XM_011288547.2;refGene_gn=PAX5;refGene_tx=XM_019816199.1
D4 60651910 60652520
refGene_gn=ZCCHC7;refGene_tx=XM_006939289.3;refGene_gn=ZCCHC7;refGene_tx=XM_006939290.4;refGene_gn=ZCCHC7;refGene_tx=XM_006939291.4
D4 60846235 60846279
refGene_gn=ZCCHC7;refGene_tx=XM_006939289.3;refGene_gn=ZCCHC7;refGene_tx=XM_006939290.4;refGene_gn=ZCCHC7;refGene_tx=XM_006939291.4
D4 60847526 60847652
refGene_gn=ZCCHC7;refGene_tx=XM_006939289.3;refGene_gn=ZCCHC7;refGene_tx=XM_006939290.4;refGene_gn=ZCCHC7;refGene_tx=XM_006939291.4
D4 60848579 60848750
refGene_gn=ZCCHC7;refGene_tx=XM_006939289.3;refGene_gn=ZCCHC7;refGene_tx=XM_006939290.4;refGene_gn=ZCCHC7;refGene_tx=XM_006939291.4
D4 60871077 60871113
refGene_gn=ZCCHC7;refGene_tx=XM_006939289.3;refGene_gn=ZCCHC7;refGene_tx=XM_006939290.4;refGene_gn=ZCCHC7;refGene_tx=XM_006939291.4
D4 60891703 60891799
refGene_gn=ZCCHC7;refGene_tx=XM_006939289.3;refGene_gn=ZCCHC7;refGene_tx=XM_006939290.4;refGene_gn=ZCCHC7;refGene_tx=XM_006939291.4
D4 60897220 60897338
refGene_gn=ZCCHC7;refGene_tx=XM_006939289.3;refGene_gn=ZCCHC7;refGene_tx=XM_006939290.4;refGene_gn=ZCCHC7;refGene_tx=XM_006939291.4
D4 60899791 60900228
refGene_gn=ZCCHC7;refGene_tx=XM_006939289.3;refGene_gn=ZCCHC7;refGene_tx=XM_006939290.4;refGene_gn=ZCCHC7;refGene_tx=XM_006939291.4
D4 60963761 60963844
refGene_gn=GRHPR;refGene_tx=XM_003995641.3;refGene_gn=GRHPR;refGene_tx=XM_006939292.3
D4 60965000 60965102 refGene_gn=GRHPR;refGene_tx=XM_006939292.3
D4 60965414 60965545
refGene_gn=GRHPR;refGene_tx=XM_003995641.3;refGene_gn=GRHPR;refGene_tx=XM_006939292.3
D4 60966460 60966533
refGene_gn=GRHPR;refGene_tx=XM_003995641.3;refGene_gn=GRHPR;refGene_tx=XM_006939292.3
D4 60967076 60967193
refGene_gn=GRHPR;refGene_tx=XM_003995641.3;refGene_gn=GRHPR;refGene_tx=XM_006939292.3
D4 60968927 60969016
refGene_gn=GRHPR;refGene_tx=XM_003995641.3;refGene_gn=GRHPR;refGene_tx=XM_006939292.3
D4 60970097 60970202

refGene_gn=GRHPR;refGene_tx=XM_003995641.3;refGene_gn=GRHPR;refGene_tx=XM_006939292.3
D4 60970990 60971126
refGene_gn=GRHPR;refGene_tx=XM_003995641.3;refGene_gn=GRHPR;refGene_tx=XM_006939292.3
D4 60972744 60972875
refGene_gn=GRHPR;refGene_tx=XM_003995641.3;refGene_gn=GRHPR;refGene_tx=XM_006939292.3
D4 60975204 60975326
refGene_gn=GRHPR;refGene_tx=XM_003995641.3;refGene_gn=GRHPR;refGene_tx=XM_006939292.3
D4 60978820 60980854
refGene_gn=ZBTB5;refGene_tx=XM_003995642.5;refGene_gn=ZBTB5;refGene_tx=XM_006939293.4
D4 61005061 61005137 refGene_gn=POLR1E;refGene_tx=XM_011288546.3
D4 61005702 61005806 refGene_gn=POLR1E;refGene_tx=XM_011288546.3
D4 61007023 61007100 refGene_gn=POLR1E;refGene_tx=XM_011288546.3
D4 61008788 61008874 refGene_gn=POLR1E;refGene_tx=XM_011288546.3
D4 61011053 61011112 refGene_gn=POLR1E;refGene_tx=XM_011288546.3
D4 61011895 61012040 refGene_gn=POLR1E;refGene_tx=XM_011288546.3
D4 61013862 61013970 refGene_gn=POLR1E;refGene_tx=XM_011288546.3
D4 61014567 61014664 refGene_gn=POLR1E;refGene_tx=XM_011288546.3
D4 61016833 61016967 refGene_gn=POLR1E;refGene_tx=XM_011288546.3
D4 61017857 61017939 refGene_gn=POLR1E;refGene_tx=XM_011288546.3
D4 61018613 61018745 refGene_gn=POLR1E;refGene_tx=XM_011288546.3
D4 61020074 61020234 refGene_gn=POLR1E;refGene_tx=XM_011288546.3
D4 61038303 61038478
refGene_gn=FBXO10;refGene_tx=XM_003995643.5;refGene_gn=FBXO10;refGene_tx=XM_006939295.4
D4 61040453 61040635
refGene_gn=FBXO10;refGene_tx=XM_003995643.5;refGene_gn=FBXO10;refGene_tx=XM_006939295.4
D4 61041754 61042068
refGene_gn=FBXO10;refGene_tx=XM_003995643.5;refGene_gn=FBXO10;refGene_tx=XM_006939295.4
D4 61043711 61043981
refGene_gn=FBXO10;refGene_tx=XM_003995643.5;refGene_gn=FBXO10;refGene_tx=XM_006939295.4
D4 61044878 61045031
refGene_gn=FBXO10;refGene_tx=XM_003995643.5;refGene_gn=FBXO10;refGene_tx=XM_006939295.4
D4 61046778 61046849
refGene_gn=FBXO10;refGene_tx=XM_003995643.5;refGene_gn=FBXO10;refGene_tx=XM_006939295.4
D4 61051191 61051328
refGene_gn=FBXO10;refGene_tx=XM_003995643.5;refGene_gn=FBXO10;refGene_tx=XM_006939295.4
D4 61053609 61053759
refGene_gn=FBXO10;refGene_tx=XM_003995643.5;refGene_gn=FBXO10;refGene_tx=XM_006939295.4

D4 61059209 61060043
refGene_gn=FBXO10;refGene_tx=XM_003995643.5;refGene_gn=FBXO10;refGene_tx=XM_006939295.4
D4 61063066 61063657
refGene_gn=FBXO10;refGene_tx=XM_003995643.5;refGene_gn=FBXO10;refGene_tx=XM_006939295.4
D4 61089156 61089231 refGene_gn=FBXO10;refGene_tx=XM_006939295.4
D4 61104566 61104601 refGene_gn=TOMM5;refGene_tx=XM_003995644.4
D4 61108427 61108548 refGene_gn=TOMM5;refGene_tx=XM_003995644.4
D4 61213133 61213234
refGene_gn=FRMPD1;refGene_tx=XM_003995646.5;refGene_gn=FRMPD1;refGene_tx=XM_023242454.1;refGene_gn=FRMPD1;refGene_tx=XM_023242455.1;refGene_gn=FRMPD1;refGene_tx=XM_023242456.1;refGene_gn=FRMPD1;refGene_tx=XM_023242457.1;refGene_gn=FRMPD1;refGene_tx=XM_023242458.1;refGene_gn=FRMPD1;refGene_tx=XM_023242459.1
D4 61219896 61219898
refGene_gn=FRMPD1;refGene_tx=XM_023242460.1;refGene_gn=FRMPD1;refGene_tx=XM_023242461.1
D4 61223710 61223868
refGene_gn=FRMPD1;refGene_tx=XM_003995646.5;refGene_gn=FRMPD1;refGene_tx=XM_023242454.1;refGene_gn=FRMPD1;refGene_tx=XM_023242455.1;refGene_gn=FRMPD1;refGene_tx=XM_023242456.1;refGene_gn=FRMPD1;refGene_tx=XM_023242457.1;refGene_gn=FRMPD1;refGene_tx=XM_023242458.1;refGene_gn=FRMPD1;refGene_tx=XM_023242459.1;refGene_gn=FRMPD1;refGene_tx=XM_023242460.1;refGene_gn=FRMPD1;refGene_tx=XM_023242461.1
D4 61224746 61224849
refGene_gn=FRMPD1;refGene_tx=XM_003995646.5;refGene_gn=FRMPD1;refGene_tx=XM_023242454.1;refGene_gn=FRMPD1;refGene_tx=XM_023242455.1;refGene_gn=FRMPD1;refGene_tx=XM_023242456.1;refGene_gn=FRMPD1;refGene_tx=XM_023242457.1;refGene_gn=FRMPD1;refGene_tx=XM_023242458.1;refGene_gn=FRMPD1;refGene_tx=XM_023242459.1;refGene_gn=FRMPD1;refGene_tx=XM_023242460.1;refGene_gn=FRMPD1;refGene_tx=XM_023242461.1
D4 61227727 61227773
refGene_gn=FRMPD1;refGene_tx=XM_003995646.5;refGene_gn=FRMPD1;refGene_tx=XM_023242454.1;refGene_gn=FRMPD1;refGene_tx=XM_023242455.1;refGene_gn=FRMPD1;refGene_tx=XM_023242456.1;refGene_gn=FRMPD1;refGene_tx=XM_023242457.1;refGene_gn=FRMPD1;refGene_tx=XM_023242458.1;refGene_gn=FRMPD1;refGene_tx=XM_023242459.1;refGene_gn=FRMPD1;refGene_tx=XM_023242460.1;refGene_gn=FRMPD1;refGene_tx=XM_023242461.1
D4 61235980 61236115
refGene_gn=FRMPD1;refGene_tx=XM_023242454.1;refGene_gn=FRMPD1;refGene_tx=XM_023242455.1;refGene_gn=FRMPD1;refGene_tx=XM_023242456.1;refGene_gn=FRMPD1;refGene_tx=XM_023242457.1;refGene_gn=FRMPD1;refGene_tx=XM_023242458.1;refGene_gn=FRMPD1;refGene_tx=XM_023242459.1;refGene_gn=FRMPD1;refGene_tx=XM_023242460.1;refGene_gn=FRMPD1;refGene_tx=XM_023242461.1
D4 61237826 61237934
refGene_gn=FRMPD1;refGene_tx=XM_003995646.5;refGene_gn=FRMPD1;refGene_tx=XM_023242454.1;refGene_gn=FRMPD1;refGene_tx=XM_023242455.1;refGene_gn=FRMPD1;refGene_tx=XM_023242456.1;refGene_gn=FRMPD1;refGene_tx=XM_02

3242457.1;refGene_gn=FRMPD1;refGene_tx=XM_023242458.1;refGene_gn=FRMPD
1;refGene_tx=XM_023242459.1;refGene_gn=FRMPD1;refGene_tx=XM_023242460.
1;refGene_gn=FRMPD1;refGene_tx=XM_023242461.1
D4 61241976 61242072
refGene_gn=FRMPD1;refGene_tx=XM_003995646.5;refGene_gn=FRMPD1;refGene_
tx=XM_023242454.1;refGene_gn=FRMPD1;refGene_tx=XM_023242455.1;refGene_
gn=FRMPD1;refGene_tx=XM_023242456.1;refGene_gn=FRMPD1;refGene_tx=XM_02
3242457.1;refGene_gn=FRMPD1;refGene_tx=XM_023242458.1;refGene_gn=FRMPD
1;refGene_tx=XM_023242459.1;refGene_gn=FRMPD1;refGene_tx=XM_023242460.
1;refGene_gn=FRMPD1;refGene_tx=XM_023242461.1
D4 61246865 61246991
refGene_gn=FRMPD1;refGene_tx=XM_003995646.5;refGene_gn=FRMPD1;refGene_
tx=XM_023242454.1;refGene_gn=FRMPD1;refGene_tx=XM_023242455.1;refGene_
gn=FRMPD1;refGene_tx=XM_023242456.1;refGene_gn=FRMPD1;refGene_tx=XM_02
3242457.1;refGene_gn=FRMPD1;refGene_tx=XM_023242458.1;refGene_gn=FRMPD
1;refGene_tx=XM_023242459.1;refGene_gn=FRMPD1;refGene_tx=XM_023242460.
1;refGene_gn=FRMPD1;refGene_tx=XM_023242461.1
D4 61248103 61248223
refGene_gn=FRMPD1;refGene_tx=XM_003995646.5;refGene_gn=FRMPD1;refGene_
tx=XM_023242454.1;refGene_gn=FRMPD1;refGene_tx=XM_023242455.1;refGene_
gn=FRMPD1;refGene_tx=XM_023242456.1;refGene_gn=FRMPD1;refGene_tx=XM_02
3242457.1;refGene_gn=FRMPD1;refGene_tx=XM_023242458.1;refGene_gn=FRMPD
1;refGene_tx=XM_023242459.1;refGene_gn=FRMPD1;refGene_tx=XM_023242460.
1;refGene_gn=FRMPD1;refGene_tx=XM_023242461.1
D4 61249261 61249398
refGene_gn=FRMPD1;refGene_tx=XM_003995646.5;refGene_gn=FRMPD1;refGene_
tx=XM_023242454.1;refGene_gn=FRMPD1;refGene_tx=XM_023242455.1;refGene_
gn=FRMPD1;refGene_tx=XM_023242456.1;refGene_gn=FRMPD1;refGene_tx=XM_02
3242457.1;refGene_gn=FRMPD1;refGene_tx=XM_023242458.1;refGene_gn=FRMPD
1;refGene_tx=XM_023242459.1;refGene_gn=FRMPD1;refGene_tx=XM_023242460.
1;refGene_gn=FRMPD1;refGene_tx=XM_023242461.1
D4 61250738 61250865
refGene_gn=FRMPD1;refGene_tx=XM_003995646.5;refGene_gn=FRMPD1;refGene_
tx=XM_023242454.1;refGene_gn=FRMPD1;refGene_tx=XM_023242455.1;refGene_
gn=FRMPD1;refGene_tx=XM_023242456.1;refGene_gn=FRMPD1;refGene_tx=XM_02
3242457.1;refGene_gn=FRMPD1;refGene_tx=XM_023242458.1;refGene_gn=FRMPD
1;refGene_tx=XM_023242459.1;refGene_gn=FRMPD1;refGene_tx=XM_023242460.
1;refGene_gn=FRMPD1;refGene_tx=XM_023242461.1
D4 61251009 61251105
refGene_gn=FRMPD1;refGene_tx=XM_003995646.5;refGene_gn=FRMPD1;refGene_
tx=XM_023242454.1;refGene_gn=FRMPD1;refGene_tx=XM_023242455.1;refGene_
gn=FRMPD1;refGene_tx=XM_023242456.1;refGene_gn=FRMPD1;refGene_tx=XM_02
3242457.1;refGene_gn=FRMPD1;refGene_tx=XM_023242458.1;refGene_gn=FRMPD
1;refGene_tx=XM_023242459.1;refGene_gn=FRMPD1;refGene_tx=XM_023242460.
1;refGene_gn=FRMPD1;refGene_tx=XM_023242461.1
D4 61253203 61253386
refGene_gn=FRMPD1;refGene_tx=XM_003995646.5;refGene_gn=FRMPD1;refGene_
tx=XM_023242454.1;refGene_gn=FRMPD1;refGene_tx=XM_023242455.1;refGene_
gn=FRMPD1;refGene_tx=XM_023242456.1;refGene_gn=FRMPD1;refGene_tx=XM_02
3242457.1;refGene_gn=FRMPD1;refGene_tx=XM_023242458.1;refGene_gn=FRMPD

1;refGene_tx=XM_023242459.1;refGene_gn=FRMPD1;refGene_tx=XM_023242460.
1;refGene_gn=FRMPD1;refGene_tx=XM_023242461.1
D4 61254229 61254377
refGene_gn=FRMPD1;refGene_tx=XM_003995646.5;refGene_gn=FRMPD1;refGene_
tx=XM_023242454.1;refGene_gn=FRMPD1;refGene_tx=XM_023242455.1;refGene_
gn=FRMPD1;refGene_tx=XM_023242456.1;refGene_gn=FRMPD1;refGene_tx=XM_02
3242457.1;refGene_gn=FRMPD1;refGene_tx=XM_023242458.1;refGene_gn=FRMPD
1;refGene_tx=XM_023242459.1;refGene_gn=FRMPD1;refGene_tx=XM_023242460.
1;refGene_gn=FRMPD1;refGene_tx=XM_023242461.1
D4 61256177 61256984
refGene_gn=FRMPD1;refGene_tx=XM_003995646.5;refGene_gn=FRMPD1;refGene_
tx=XM_023242454.1;refGene_gn=FRMPD1;refGene_tx=XM_023242455.1;refGene_
gn=FRMPD1;refGene_tx=XM_023242456.1;refGene_gn=FRMPD1;refGene_tx=XM_02
3242457.1;refGene_gn=FRMPD1;refGene_tx=XM_023242458.1;refGene_gn=FRMPD
1;refGene_tx=XM_023242459.1;refGene_gn=FRMPD1;refGene_tx=XM_023242460.
1;refGene_gn=FRMPD1;refGene_tx=XM_023242461.1
D4 61259918 61262305
refGene_gn=FRMPD1;refGene_tx=XM_003995646.5;refGene_gn=FRMPD1;refGene_
tx=XM_023242454.1;refGene_gn=FRMPD1;refGene_tx=XM_023242455.1;refGene_
gn=FRMPD1;refGene_tx=XM_023242456.1;refGene_gn=FRMPD1;refGene_tx=XM_02
3242457.1;refGene_gn=FRMPD1;refGene_tx=XM_023242458.1;refGene_gn=FRMPD
1;refGene_tx=XM_023242459.1;refGene_gn=FRMPD1;refGene_tx=XM_023242460.
1;refGene_gn=FRMPD1;refGene_tx=XM_023242461.1
D4 61273177 61273363
refGene_gn=TRMT10B;refGene_tx=XM_003995647.4;refGene_gn=TRMT10B;refGen
e_tx=XM_006939296.3;refGene_gn=TRMT10B;refGene_tx=XM_006939298.3;refGe
ne_gn=TRMT10B;refGene_tx=XM_006939299.4;refGene_gn=TRMT10B;refGene_tx=
XM_011288543.3;refGene_gn=TRMT10B;refGene_tx=XM_019816208.2
D4 61274049 61274158
refGene_gn=TRMT10B;refGene_tx=XM_003995647.4;refGene_gn=TRMT10B;refGen
e_tx=XM_006939296.3;refGene_gn=TRMT10B;refGene_tx=XM_006939298.3;refGe
ne_gn=TRMT10B;refGene_tx=XM_006939299.4;refGene_gn=TRMT10B;refGene_tx=
XM_011288543.3;refGene_gn=TRMT10B;refGene_tx=XM_019816208.2
D4 61274765 61274890
refGene_gn=TRMT10B;refGene_tx=XM_003995647.4;refGene_gn=TRMT10B;refGen
e_tx=XM_006939296.3;refGene_gn=TRMT10B;refGene_tx=XM_006939298.3;refGe
ne_gn=TRMT10B;refGene_tx=XM_006939299.4;refGene_gn=TRMT10B;refGene_tx=
XM_011288543.3;refGene_gn=TRMT10B;refGene_tx=XM_019816208.2
D4 61277138 61277291
refGene_gn=TRMT10B;refGene_tx=XM_003995647.4;refGene_gn=TRMT10B;refGen
e_tx=XM_006939296.3;refGene_gn=TRMT10B;refGene_tx=XM_006939298.3;refGe
ne_gn=TRMT10B;refGene_tx=XM_006939299.4;refGene_gn=TRMT10B;refGene_tx=
XM_011288543.3;refGene_gn=TRMT10B;refGene_tx=XM_019816208.2
D4 61278428 61278507
refGene_gn=TRMT10B;refGene_tx=XM_003995647.4;refGene_gn=TRMT10B;refGen
e_tx=XM_006939296.3;refGene_gn=TRMT10B;refGene_tx=XM_006939299.4;refGe
ne_gn=TRMT10B;refGene_tx=XM_011288543.3;refGene_gn=TRMT10B;refGene_tx=
XM_019816208.2
D4 61279151 61279219
refGene_gn=TRMT10B;refGene_tx=XM_003995647.4;refGene_gn=TRMT10B;refGen

e_tx=XM_006939296.3;refGene_gn=TRMT10B;refGene_tx=XM_006939299.4;refGene_gn=TRMT10B;refGene_tx=XM_011288543.3;refGene_gn=TRMT10B;refGene_tx=XM_019816208.2
D4 61283348 61283472 refGene_gn=TRMT10B;refGene_tx=XM_003995647.4;refGene_gn=TRMT10B;refGene_tx=XM_006939296.3;refGene_gn=TRMT10B;refGene_tx=XM_006939298.3;refGene_gn=TRMT10B;refGene_tx=XM_006939299.4;refGene_gn=TRMT10B;refGene_tx=XM_011288543.3;refGene_gn=TRMT10B;refGene_tx=XM_019816208.2
D4 61284266 61284373 refGene_gn=TRMT10B;refGene_tx=XM_003995647.4;refGene_gn=TRMT10B;refGene_tx=XM_006939296.3;refGene_gn=TRMT10B;refGene_tx=XM_006939298.3;refGene_gn=TRMT10B;refGene_tx=XM_006939299.4;refGene_gn=TRMT10B;refGene_tx=XM_011288543.3;refGene_gn=TRMT10B;refGene_tx=XM_019816208.2
D4 61286327 61286529 refGene_gn=EXOSC3;refGene_tx=XM_003995648.4
D4 61287611 61287763 refGene_gn=EXOSC3;refGene_tx=XM_003995648.4
D4 61290214 61290364 refGene_gn=EXOSC3;refGene_tx=XM_003995648.4
D4 61291104 61291431 refGene_gn=EXOSC3;refGene_tx=XM_003995648.4
D4 61314603 61315133 refGene_gn=DCAF10;refGene_tx=XM_003995649.5;refGene_gn=DCAF10;refGene_tx=XM_019816207.2
D4 61325741 61325855 refGene_gn=DCAF10;refGene_tx=XM_003995649.5;refGene_gn=DCAF10;refGene_tx=XM_019816207.2
D4 61338046 61338244 refGene_gn=DCAF10;refGene_tx=XM_003995649.5;refGene_gn=DCAF10;refGene_tx=XM_019816207.2
D4 61346815 61347018 refGene_gn=DCAF10;refGene_tx=XM_003995649.5;refGene_gn=DCAF10;refGene_tx=XM_019816207.2
D4 61349256 61349367 refGene_gn=DCAF10;refGene_tx=XM_003995649.5
D4 61352243 61352389 refGene_gn=DCAF10;refGene_tx=XM_003995649.5;refGene_gn=DCAF10;refGene_tx=XM_019816207.2
D4 61353170 61353539 refGene_gn=DCAF10;refGene_tx=XM_003995649.5;refGene_gn=DCAF10;refGene_tx=XM_019816207.2
D4 61370999 61371893 refGene_gn=SLC25A51;refGene_tx=XM_006939300.4;refGene_gn=SLC25A51;refGene_tx=XM_023242463.1;refGene_gn=SLC25A51;refGene_tx=XM_023242464.1;refGene_gn=SLC25A51;refGene_tx=XM_023242465.1;refGene_gn=SLC25A51;refGene_tx=XM_023242466.1
D4 61399918 61400102 refGene_gn=SHB;refGene_tx=XM_003995682.5;refGene_gn=SHB;refGene_tx=XM_023242467.1
D4 61424974 61425094 refGene_gn=SHB;refGene_tx=XM_003995682.5
D4 61430325 61430497 refGene_gn=SHB;refGene_tx=XM_003995682.5;refGene_gn=SHB;refGene_tx=XM_023242467.1
D4 61447094 61447310 refGene_gn=SHB;refGene_tx=XM_003995682.5;refGene_gn=SHB;refGene_tx=XM_

023242467.1
D4 61485403 61485524
refGene_gn=SHB;refGene_tx=XM_003995682.5;refGene_gn=SHB;refGene_tx=XM_023242467.1
D4 61534463 61535177
refGene_gn=SHB;refGene_tx=XM_003995682.5;refGene_gn=SHB;refGene_tx=XM_023242467.1
D4 61773917 61775477
refGene_gn=ALDH1B1;refGene_tx=XM_003995651.3;refGene_gn=ALDH1B1;refGene_tx=XM_019816209.2;refGene_gn=ALDH1B1;refGene_tx=XM_019816210.2
D4 61787607 61787757
refGene_gn=IGFBPL1;refGene_tx=XM_003995683.5;refGene_gn=IGFBPL1;refGene_tx=XM_023242470.1
D4 61789595 61789712
refGene_gn=IGFBPL1;refGene_tx=XM_003995683.5;refGene_gn=IGFBPL1;refGene_tx=XM_023242470.1
D4 61790442 61790552
refGene_gn=IGFBPL1;refGene_tx=XM_003995683.5;refGene_gn=IGFBPL1;refGene_tx=XM_023242470.1
D4 61799940 61800388
refGene_gn=IGFBPL1;refGene_tx=XM_003995683.5;refGene_gn=IGFBPL1;refGene_tx=XM_023242470.1
D4 61810635 61810655
refGene_gn=LOC102901088;refGene_tx=XM_023242469.1
D4 61826566 61826642
refGene_gn=LOC102901088;refGene_tx=XM_023242469.1
D4 61828323 61828424
refGene_gn=LOC102901088;refGene_tx=XM_023242469.1
D4 61831123 61831254
refGene_gn=LOC102901088;refGene_tx=XM_023242469.1
D4 61835908 61836029
refGene_gn=LOC102901088;refGene_tx=XM_023242469.1
D4 61837589 61837635
refGene_gn=LOC102901088;refGene_tx=XM_023242469.1
D4 61841206 61841335
refGene_gn=LOC102901088;refGene_tx=XM_023242469.1
D4 61841672 61841743
refGene_gn=LOC102901088;refGene_tx=XM_023242469.1
D4 61843164 61843276
refGene_gn=LOC102901088;refGene_tx=XM_023242469.1
D4 61848102 61848256
refGene_gn=LOC102901088;refGene_tx=XM_023242469.1
D4 61854992 61855068
refGene_gn=LOC102901088;refGene_tx=XM_023242469.1
D4 61858345 61858427
refGene_gn=LOC102901088;refGene_tx=XM_023242469.1
D4 61858761 61858810
refGene_gn=LOC102901088;refGene_tx=XM_023242469.1
D4 61859263 61859381
refGene_gn=LOC102901088;refGene_tx=XM_023242469.1

D4 61860335 61860455
refGene_gn=LOC102901088;refGene_tx=XM_023242469.1
D4 61861712 61861876
refGene_gn=LOC102901088;refGene_tx=XM_023242469.1
D4 61862838 61862985
refGene_gn=LOC102901088;refGene_tx=XM_023242469.1
D4 61863696 61863845
refGene_gn=LOC102901088;refGene_tx=XM_023242469.1
D4 61971097 61971304
refGene_gn=TDRD7;refGene_tx=XM_023242472.1;refGene_gn=TDRD7;refGene_tx=XM_023242473.1;refGene_gn=TDRD7;refGene_tx=XM_023242474.1;refGene_gn=TDRD7;refGene_tx=XM_023242475.1
D4 61974320 61974462
refGene_gn=TDRD7;refGene_tx=XM_019816214.1;refGene_gn=TDRD7;refGene_tx=XM_023242472.1;refGene_gn=TDRD7;refGene_tx=XM_023242473.1;refGene_gn=TDRD7;refGene_tx=XM_023242474.1;refGene_gn=TDRD7;refGene_tx=XM_023242475.1;refGene_gn=TDRD7;refGene_tx=XM_023242476.1
D4 61975426 61975640
refGene_gn=TDRD7;refGene_tx=XM_019816214.1;refGene_gn=TDRD7;refGene_tx=XM_023242472.1;refGene_gn=TDRD7;refGene_tx=XM_023242473.1;refGene_gn=TDRD7;refGene_tx=XM_023242474.1;refGene_gn=TDRD7;refGene_tx=XM_023242475.1;refGene_gn=TDRD7;refGene_tx=XM_023242476.1
D4 61986555 61986629
refGene_gn=TDRD7;refGene_tx=XM_019816214.1;refGene_gn=TDRD7;refGene_tx=XM_023242472.1;refGene_gn=TDRD7;refGene_tx=XM_023242473.1;refGene_gn=TDRD7;refGene_tx=XM_023242474.1;refGene_gn=TDRD7;refGene_tx=XM_023242475.1;refGene_gn=TDRD7;refGene_tx=XM_023242476.1
D4 61988852 61989070
refGene_gn=TDRD7;refGene_tx=XM_019816214.1;refGene_gn=TDRD7;refGene_tx=XM_023242472.1;refGene_gn=TDRD7;refGene_tx=XM_023242473.1;refGene_gn=TDRD7;refGene_tx=XM_023242474.1;refGene_gn=TDRD7;refGene_tx=XM_023242475.1;refGene_gn=TDRD7;refGene_tx=XM_023242476.1
D4 62001236 62001823
refGene_gn=TDRD7;refGene_tx=XM_019816214.1;refGene_gn=TDRD7;refGene_tx=XM_023242472.1;refGene_gn=TDRD7;refGene_tx=XM_023242473.1;refGene_gn=TDRD7;refGene_tx=XM_023242474.1;refGene_gn=TDRD7;refGene_tx=XM_023242475.1;refGene_gn=TDRD7;refGene_tx=XM_023242476.1
D4 62005280 62005467
refGene_gn=TDRD7;refGene_tx=XM_019816214.1;refGene_gn=TDRD7;refGene_tx=XM_023242472.1;refGene_gn=TDRD7;refGene_tx=XM_023242473.1;refGene_gn=TDRD7;refGene_tx=XM_023242474.1;refGene_gn=TDRD7;refGene_tx=XM_023242475.1;refGene_gn=TDRD7;refGene_tx=XM_023242476.1
D4 62009798 62009910
refGene_gn=TDRD7;refGene_tx=XM_019816214.1;refGene_gn=TDRD7;refGene_tx=XM_023242472.1;refGene_gn=TDRD7;refGene_tx=XM_023242473.1;refGene_gn=TDRD7;refGene_tx=XM_023242474.1;refGene_gn=TDRD7;refGene_tx=XM_023242475.1;refGene_gn=TDRD7;refGene_tx=XM_023242476.1
D4 62011574 62011777
refGene_gn=TDRD7;refGene_tx=XM_019816214.1;refGene_gn=TDRD7;refGene_tx=XM_023242472.1;refGene_gn=TDRD7;refGene_tx=XM_023242473.1;refGene_gn=

TDRD7;refGene_tx=XM_023242474.1;refGene_gn=TDRD7;refGene_tx=XM_023242475.1;refGene_gn=TDRD7;refGene_tx=XM_023242476.1
D4 62013311 62013446
refGene_gn=TDRD7;refGene_tx=XM_019816214.1;refGene_gn=TDRD7;refGene_tx=XM_023242472.1;refGene_gn=TDRD7;refGene_tx=XM_023242473.1;refGene_gn=TDRD7;refGene_tx=XM_023242474.1;refGene_gn=TDRD7;refGene_tx=XM_023242476.1
D4 62015208 62015295
refGene_gn=TDRD7;refGene_tx=XM_019816214.1;refGene_gn=TDRD7;refGene_tx=XM_023242472.1;refGene_gn=TDRD7;refGene_tx=XM_023242473.1;refGene_gn=TDRD7;refGene_tx=XM_023242474.1;refGene_gn=TDRD7;refGene_tx=XM_023242475.1;refGene_gn=TDRD7;refGene_tx=XM_023242476.1
D4 62021649 62021784
refGene_gn=TDRD7;refGene_tx=XM_019816214.1;refGene_gn=TDRD7;refGene_tx=XM_023242472.1;refGene_gn=TDRD7;refGene_tx=XM_023242473.1;refGene_gn=TDRD7;refGene_tx=XM_023242474.1;refGene_gn=TDRD7;refGene_tx=XM_023242475.1;refGene_gn=TDRD7;refGene_tx=XM_023242476.1
D4 62022296 62022407
refGene_gn=TDRD7;refGene_tx=XM_019816214.1;refGene_gn=TDRD7;refGene_tx=XM_023242472.1;refGene_gn=TDRD7;refGene_tx=XM_023242473.1;refGene_gn=TDRD7;refGene_tx=XM_023242474.1;refGene_gn=TDRD7;refGene_tx=XM_023242475.1;refGene_gn=TDRD7;refGene_tx=XM_023242476.1
D4 62024218 62024721
refGene_gn=TDRD7;refGene_tx=XM_019816214.1;refGene_gn=TDRD7;refGene_tx=XM_023242472.1;refGene_gn=TDRD7;refGene_tx=XM_023242473.1;refGene_gn=TDRD7;refGene_tx=XM_023242474.1;refGene_gn=TDRD7;refGene_tx=XM_023242475.1;refGene_gn=TDRD7;refGene_tx=XM_023242476.1
D4 62028845 62029006
refGene_gn=TDRD7;refGene_tx=XM_019816214.1;refGene_gn=TDRD7;refGene_tx=XM_023242472.1;refGene_gn=TDRD7;refGene_tx=XM_023242473.1;refGene_gn=TDRD7;refGene_tx=XM_023242474.1;refGene_gn=TDRD7;refGene_tx=XM_023242475.1;refGene_gn=TDRD7;refGene_tx=XM_023242476.1
D4 62031834 62032055
refGene_gn=TDRD7;refGene_tx=XM_019816214.1;refGene_gn=TDRD7;refGene_tx=XM_023242472.1;refGene_gn=TDRD7;refGene_tx=XM_023242473.1;refGene_gn=TDRD7;refGene_tx=XM_023242474.1;refGene_gn=TDRD7;refGene_tx=XM_023242475.1;refGene_gn=TDRD7;refGene_tx=XM_023242476.1
D4 62058139 62058220
refGene_gn=TMOD1;refGene_tx=XM_003995653.5;refGene_gn=TMOD1;refGene_tx=XM_023242477.1
D4 62078318 62078475
refGene_gn=TMOD1;refGene_tx=XM_003995653.5;refGene_gn=TMOD1;refGene_tx=XM_023242477.1
D4 62085608 62085728
refGene_gn=TMOD1;refGene_tx=XM_003995653.5;refGene_gn=TMOD1;refGene_tx=XM_023242477.1
D4 62094709 62094799
refGene_gn=TMOD1;refGene_tx=XM_003995653.5;refGene_gn=TMOD1;refGene_tx=XM_023242477.1
D4 62095749 62095880

refGene_gn=TMOD1;refGene_tx=XM_003995653.5;refGene_gn=TMOD1;refGene_tx=XM_023242477.1
D4 62097565 62097673
refGene_gn=TMOD1;refGene_tx=XM_003995653.5;refGene_gn=TMOD1;refGene_tx=XM_023242477.1
D4 62103220 62103364
refGene_gn=TMOD1;refGene_tx=XM_003995653.5;refGene_gn=TMOD1;refGene_tx=XM_023242477.1
D4 62115760 62115905
refGene_gn=TMOD1;refGene_tx=XM_003995653.5;refGene_gn=TMOD1;refGene_tx=XM_023242477.1
D4 62122507 62122572
refGene_gn=TMOD1;refGene_tx=XM_003995653.5;refGene_gn=TMOD1;refGene_tx=XM_023242477.1
D4 62125320 62125559
refGene_gn=TSTD2;refGene_tx=XM_011288536.3;refGene_gn=TSTD2;refGene_tx=XM_011288537.3;refGene_gn=TSTD2;refGene_tx=XM_023242479.1;refGene_gn=TSTD2;refGene_tx=XM_023242480.1;refGene_gn=TSTD2;refGene_tx=XM_023242481.1
D4 62127419 62127558
refGene_gn=TSTD2;refGene_tx=XM_011288536.3;refGene_gn=TSTD2;refGene_tx=XM_011288537.3;refGene_gn=TSTD2;refGene_tx=XM_023242479.1;refGene_gn=TSTD2;refGene_tx=XM_023242480.1;refGene_gn=TSTD2;refGene_tx=XM_023242481.1
D4 62128028 62128187
refGene_gn=TSTD2;refGene_tx=XM_011288536.3;refGene_gn=TSTD2;refGene_tx=XM_011288537.3;refGene_gn=TSTD2;refGene_tx=XM_023242479.1;refGene_gn=TSTD2;refGene_tx=XM_023242480.1;refGene_gn=TSTD2;refGene_tx=XM_023242481.1
D4 62128699 62128818
refGene_gn=TSTD2;refGene_tx=XM_011288536.3;refGene_gn=TSTD2;refGene_tx=XM_011288537.3;refGene_gn=TSTD2;refGene_tx=XM_023242479.1;refGene_gn=TSTD2;refGene_tx=XM_023242480.1;refGene_gn=TSTD2;refGene_tx=XM_023242481.1
D4 62132086 62132192
refGene_gn=TSTD2;refGene_tx=XM_011288536.3;refGene_gn=TSTD2;refGene_tx=XM_011288537.3;refGene_gn=TSTD2;refGene_tx=XM_023242479.1;refGene_gn=TSTD2;refGene_tx=XM_023242480.1;refGene_gn=TSTD2;refGene_tx=XM_023242481.1
D4 62133030 62133156
refGene_gn=TSTD2;refGene_tx=XM_011288536.3;refGene_gn=TSTD2;refGene_tx=XM_023242479.1;refGene_gn=TSTD2;refGene_tx=XM_023242480.1;refGene_gn=TSTD2;refGene_tx=XM_023242481.1
D4 62138116 62138237
refGene_gn=TSTD2;refGene_tx=XM_011288536.3;refGene_gn=TSTD2;refGene_tx=XM_011288537.3;refGene_gn=TSTD2;refGene_tx=XM_023242479.1;refGene_gn=TSTD2;refGene_tx=XM_023242480.1;refGene_gn=TSTD2;refGene_tx=XM_023242481.1
D4 62140397 62140711
refGene_gn=TSTD2;refGene_tx=XM_011288536.3;refGene_gn=TSTD2;refGene_tx

=XM_011288537.3;refGene_gn=TSTD2;refGene_tx=XM_023242479.1;refGene_gn=TSTD2;refGene_tx=XM_023242480.1;refGene_gn=TSTD2;refGene_tx=XM_023242481.1
D4 62144291 62144456 refGene_gn=TSTD2;refGene_tx=XM_011288536.3;refGene_gn=TSTD2;refGene_tx=XM_011288537.3;refGene_gn=TSTD2;refGene_tx=XM_023242479.1;refGene_gn=TSTD2;refGene_tx=XM_023242480.1;refGene_gn=TSTD2;refGene_tx=XM_023242481.1
D4 62147358 62147392 refGene_gn=NCBP1;refGene_tx=XM_003995654.4
D4 62153995 62154084 refGene_gn=NCBP1;refGene_tx=XM_003995654.4
D4 62154625 62154726 refGene_gn=NCBP1;refGene_tx=XM_003995654.4
D4 62156791 62156948 refGene_gn=NCBP1;refGene_tx=XM_003995654.4
D4 62159317 62159425 refGene_gn=NCBP1;refGene_tx=XM_003995654.4
D4 62159755 62159877 refGene_gn=NCBP1;refGene_tx=XM_003995654.4
D4 62162205 62162275 refGene_gn=NCBP1;refGene_tx=XM_003995654.4
D4 62162732 62162922 refGene_gn=NCBP1;refGene_tx=XM_003995654.4
D4 62162924 62162947 refGene_gn=NCBP1;refGene_tx=XM_003995654.4
D4 62165785 62165883 refGene_gn=NCBP1;refGene_tx=XM_003995654.4
D4 62166584 62166648 refGene_gn=NCBP1;refGene_tx=XM_003995654.4
D4 62168325 62168436 refGene_gn=NCBP1;refGene_tx=XM_003995654.4
D4 62169192 62169257 refGene_gn=NCBP1;refGene_tx=XM_003995654.4
D4 62170878 62170941 refGene_gn=NCBP1;refGene_tx=XM_003995654.4
D4 62171175 62171250 refGene_gn=NCBP1;refGene_tx=XM_003995654.4
D4 62173424 62173528 refGene_gn=NCBP1;refGene_tx=XM_003995654.4
D4 62175506 62175629 refGene_gn=NCBP1;refGene_tx=XM_003995654.4
D4 62176801 62176904 refGene_gn=NCBP1;refGene_tx=XM_003995654.4
D4 62177700 62177794 refGene_gn=NCBP1;refGene_tx=XM_003995654.4
D4 62178063 62178167 refGene_gn=NCBP1;refGene_tx=XM_003995654.4
D4 62180214 62180329 refGene_gn=NCBP1;refGene_tx=XM_003995654.4
D4 62183286 62183415 refGene_gn=NCBP1;refGene_tx=XM_003995654.4
D4 62183952 62184066 refGene_gn=NCBP1;refGene_tx=XM_003995654.4
D4 62184946 62185060 refGene_gn=NCBP1;refGene_tx=XM_003995654.4
D4 62188317 62188466 refGene_gn=XPA;refGene_tx=XM_023242483.1
D4 62193475 62193645 refGene_gn=XPA;refGene_tx=XM_023242482.1
D4 62197035 62197153 refGene_gn=XPA;refGene_tx=XM_023242482.1;refGene_gn=XPA;refGene_tx=XM_023242483.1
D4 62200203 62200369 refGene_gn=XPA;refGene_tx=XM_023242482.1;refGene_gn=XPA;refGene_tx=XM_023242483.1
D4 62203018 62203124 refGene_gn=XPA;refGene_tx=XM_023242482.1;refGene_gn=XPA;refGene_tx=XM_023242483.1
D4 62207660 62207771 refGene_gn=XPA;refGene_tx=XM_023242482.1;refGene_gn=XPA;refGene_tx=XM_023242483.1
D4 62209727 62209899 refGene_gn=XPA;refGene_tx=XM_023242482.1;refGene_gn=XPA;refGene_tx=XM_023242483.1
D4 62350040 62351159 refGene_gn=FOXE1;refGene_tx=XM_023242484.1

D4 62391348 62391608 refGene_gn=TRMO;refGene_tx=XM_006939305.2
D4 62394147 62394197
refGene_gn=TRMO;refGene_tx=XM_006939306.3;refGene_gn=TRMO;refGene_tx=XM_019816221.2
D4 62394758 62395424
refGene_gn=TRMO;refGene_tx=XM_006939305.2;refGene_gn=TRMO;refGene_tx=XM_006939306.3;refGene_gn=TRMO;refGene_tx=XM_019816221.2
D4 62397684 62397842
refGene_gn=TRMO;refGene_tx=XM_006939305.2;refGene_gn=TRMO;refGene_tx=XM_006939306.3;refGene_gn=TRMO;refGene_tx=XM_019816221.2
D4 62400269 62400444
refGene_gn=TRMO;refGene_tx=XM_006939305.2;refGene_gn=TRMO;refGene_tx=XM_006939306.3;refGene_gn=TRMO;refGene_tx=XM_019816221.2
D4 62404803 62404879
refGene_gn=TRMO;refGene_tx=XM_006939305.2;refGene_gn=TRMO;refGene_tx=XM_006939306.3;refGene_gn=TRMO;refGene_tx=XM_019816221.2
D4 62410016 62410123
refGene_gn=HEMGN;refGene_tx=XM_019816217.2;refGene_gn=HEMGN;refGene_tx=XM_019816218.2;refGene_gn=HEMGN;refGene_tx=XM_023242486.1
D4 62412947 62414167
refGene_gn=HEMGN;refGene_tx=XM_019816217.2;refGene_gn=HEMGN;refGene_tx=XM_019816218.2;refGene_gn=HEMGN;refGene_tx=XM_023242486.1
D4 62418669 62418763
refGene_gn=HEMGN;refGene_tx=XM_019816217.2;refGene_gn=HEMGN;refGene_tx=XM_019816218.2;refGene_gn=HEMGN;refGene_tx=XM_023242486.1
D4 62420518 62420597
refGene_gn=HEMGN;refGene_tx=XM_019816217.2;refGene_gn=HEMGN;refGene_tx=XM_019816218.2;refGene_gn=HEMGN;refGene_tx=XM_023242486.1
D4 62454210 62454264
refGene_gn=ANP32B;refGene_tx=XM_023242491.1;refGene_gn=ANP32B;refGene_tx=XM_023242492.1
D4 62463528 62463678 refGene_gn=ANP32B;refGene_tx=XM_023242491.1
D4 62467215 62467338
refGene_gn=ANP32B;refGene_tx=XM_023242491.1;refGene_gn=ANP32B;refGene_tx=XM_023242492.1
D4 62472709 62472899
refGene_gn=ANP32B;refGene_tx=XM_023242491.1;refGene_gn=ANP32B;refGene_tx=XM_023242492.1
D4 62475507 62475632
refGene_gn=ANP32B;refGene_tx=XM_023242491.1;refGene_gn=ANP32B;refGene_tx=XM_023242492.1
D4 62476860 62476930
refGene_gn=ANP32B;refGene_tx=XM_023242491.1;refGene_gn=ANP32B;refGene_tx=XM_023242492.1
D4 62477916 62477984
refGene_gn=ANP32B;refGene_tx=XM_023242491.1;refGene_gn=ANP32B;refGene_tx=XM_023242492.1
D4 62492326 62492458
refGene_gn=NANS;refGene_tx=XM_023242488.1;refGene_gn=NANS;refGene_tx=XM_023242490.1

D4	62495657 62495663 refGene_gn=NANS;refGene_tx=XM_023242489.1
D4	62496638 62496854
refGene_gn=NANS;refGene_tx=XM_023242488.1;refGene_gn=NANS;refGene_tx=XM_023242489.1;refGene_gn=NANS;refGene_tx=XM_023242490.1
D4	62508133 62508233
refGene_gn=NANS;refGene_tx=XM_023242488.1;refGene_gn=NANS;refGene_tx=XM_023242489.1;refGene_gn=NANS;refGene_tx=XM_023242490.1
D4	62509044 62509199
refGene_gn=NANS;refGene_tx=XM_023242488.1;refGene_gn=NANS;refGene_tx=XM_023242489.1;refGene_gn=NANS;refGene_tx=XM_023242490.1
D4	62511675 62511942
refGene_gn=NANS;refGene_tx=XM_023242488.1;refGene_gn=NANS;refGene_tx=XM_023242489.1;refGene_gn=NANS;refGene_tx=XM_023242490.1
D4	62512670 62512697 refGene_gn=NANS;refGene_tx=XM_023242490.1
D4	62513684 62513894
refGene_gn=NANS;refGene_tx=XM_023242488.1;refGene_gn=NANS;refGene_tx=XM_023242489.1
D4	62519247 62519783
refGene_gn=TRIM14;refGene_tx=XM_019816223.2;refGene_gn=TRIM14;refGene_tx=XM_023242493.1;refGene_gn=TRIM14;refGene_tx=XM_023242494.1;refGene_gn=TRIM14;refGene_tx=XM_023242495.1
D4	62522951 62523044
refGene_gn=TRIM14;refGene_tx=XM_019816223.2;refGene_gn=TRIM14;refGene_tx=XM_023242493.1;refGene_gn=TRIM14;refGene_tx=XM_023242494.1;refGene_gn=TRIM14;refGene_tx=XM_023242495.1
D4	62527128 62527291
refGene_gn=TRIM14;refGene_tx=XM_019816223.2;refGene_gn=TRIM14;refGene_tx=XM_023242493.1;refGene_gn=TRIM14;refGene_tx=XM_023242494.1;refGene_gn=TRIM14;refGene_tx=XM_023242495.1
D4	62532439 62532673
refGene_gn=TRIM14;refGene_tx=XM_019816223.2;refGene_gn=TRIM14;refGene_tx=XM_023242493.1;refGene_gn=TRIM14;refGene_tx=XM_023242494.1;refGene_gn=TRIM14;refGene_tx=XM_023242495.1
D4	62541190 62541301
refGene_gn=TRIM14;refGene_tx=XM_019816223.2;refGene_gn=TRIM14;refGene_tx=XM_023242493.1;refGene_gn=TRIM14;refGene_tx=XM_023242494.1;refGene_gn=TRIM14;refGene_tx=XM_023242495.1
D4	62543554 62543578
refGene_gn=TRIM14;refGene_tx=XM_019816223.2;refGene_gn=TRIM14;refGene_tx=XM_023242495.1
D4	62543790 62544006
refGene_gn=TRIM14;refGene_tx=XM_023242493.1;refGene_gn=TRIM14;refGene_tx=XM_023242494.1
D4	62548700 62548832 refGene_gn=CORO2A;refGene_tx=XM_003995689.3
D4	62550078 62550323 refGene_gn=CORO2A;refGene_tx=XM_003995689.3
D4	62551246 62551337 refGene_gn=CORO2A;refGene_tx=XM_003995689.3
D4	62551685 62551798 refGene_gn=CORO2A;refGene_tx=XM_003995689.3
D4	62552564 62552661 refGene_gn=CORO2A;refGene_tx=XM_003995689.3
D4	62553391 62553496 refGene_gn=CORO2A;refGene_tx=XM_003995689.3
D4	62554290 62554407 refGene_gn=CORO2A;refGene_tx=XM_003995689.3

D4 62555163 62555343 refGene_gn=CORO2A;refGene_tx=XM_003995689.3
D4 62556895 62557045 refGene_gn=CORO2A;refGene_tx=XM_003995689.3
D4 62559489 62559606 refGene_gn=CORO2A;refGene_tx=XM_003995689.3
D4 62574565 62574766 refGene_gn=CORO2A;refGene_tx=XM_003995689.3
D4 62607874 62608082
refGene_gn=TBC1D2;refGene_tx=XM_006939308.3;refGene_gn=TBC1D2;refGene_tx=XM_006939310.2
D4 62608780 62608902
refGene_gn=TBC1D2;refGene_tx=XM_006939308.3;refGene_gn=TBC1D2;refGene_tx=XM_006939310.2
D4 62609656 62609842
refGene_gn=TBC1D2;refGene_tx=XM_006939308.3;refGene_gn=TBC1D2;refGene_tx=XM_006939310.2
D4 62611242 62611363
refGene_gn=TBC1D2;refGene_tx=XM_006939308.3;refGene_gn=TBC1D2;refGene_tx=XM_006939310.2
D4 62615246 62615723
refGene_gn=TBC1D2;refGene_tx=XM_006939308.3;refGene_gn=TBC1D2;refGene_tx=XM_006939310.2
D4 62616851 62617039
refGene_gn=TBC1D2;refGene_tx=XM_006939308.3;refGene_gn=TBC1D2;refGene_tx=XM_006939310.2
D4 62622485 62622596
refGene_gn=TBC1D2;refGene_tx=XM_006939308.3;refGene_gn=TBC1D2;refGene_tx=XM_006939310.2
D4 62629733 62630126
refGene_gn=TBC1D2;refGene_tx=XM_006939308.3;refGene_gn=TBC1D2;refGene_tx=XM_006939310.2
D4 62638095 62638286
refGene_gn=TBC1D2;refGene_tx=XM_006939308.3;refGene_gn=TBC1D2;refGene_tx=XM_006939310.2
D4 62642911 62643045
refGene_gn=TBC1D2;refGene_tx=XM_006939308.3;refGene_gn=TBC1D2;refGene_tx=XM_006939310.2
D4 62652602 62652750
refGene_gn=TBC1D2;refGene_tx=XM_006939308.3;refGene_gn=TBC1D2;refGene_tx=XM_006939310.2
D4 62658633 62658775 refGene_gn=TBC1D2;refGene_tx=XM_006939308.3
D4 62663132 62663498 refGene_gn=TBC1D2;refGene_tx=XM_006939308.3
D4 62681102 62681268 refGene_gn=GABBR2;refGene_tx=XM_003995690.5
D4 62683903 62684021 refGene_gn=GABBR2;refGene_tx=XM_003995690.5
D4 62687799 62687929 refGene_gn=GABBR2;refGene_tx=XM_003995690.5
D4 62690481 62690664 refGene_gn=GABBR2;refGene_tx=XM_003995690.5
D4 62692825 62693050 refGene_gn=GABBR2;refGene_tx=XM_003995690.5
D4 62697392 62697503 refGene_gn=GABBR2;refGene_tx=XM_003995690.5
D4 62737048 62737171 refGene_gn=GABBR2;refGene_tx=XM_003995690.5
D4 62746152 62746260 refGene_gn=GABBR2;refGene_tx=XM_003995690.5
D4 62759815 62759948 refGene_gn=GABBR2;refGene_tx=XM_003995690.5
D4 62763053 62763204 refGene_gn=GABBR2;refGene_tx=XM_003995690.5
D4 62766640 62766721 refGene_gn=GABBR2;refGene_tx=XM_003995690.5

| | | | |
|---|---|---|---|
| D4 | 62780725 | 62780786 | refGene_gn=GABBR2;refGene_tx=XM_003995690.5 |
| D4 | 62821031 | 62821268 | refGene_gn=GABBR2;refGene_tx=XM_003995690.5 |
| D4 | 62837047 | 62837248 | refGene_gn=GABBR2;refGene_tx=XM_003995690.5 |
| D4 | 62843915 | 62843981 | refGene_gn=GABBR2;refGene_tx=XM_003995690.5 |
| D4 | 62856997 | 62857099 | refGene_gn=GABBR2;refGene_tx=XM_003995690.5 |
| D4 | 62891139 | 62891310 | refGene_gn=GABBR2;refGene_tx=XM_003995690.5 |
| D4 | 62922458 | 62922596 | refGene_gn=GABBR2;refGene_tx=XM_003995690.5 |
| D4 | 63032721 | 63033042 | refGene_gn=GABBR2;refGene_tx=XM_003995690.5 |
| D4 | 63061159 | 63061264 | refGene_gn=ANKS6;refGene_tx=XM_023242497.1 |
| D4 | 63068416 | 63068533 | refGene_gn=ANKS6;refGene_tx=XM_023242497.1 |
| D4 | 63074337 | 63074405 | refGene_gn=ANKS6;refGene_tx=XM_023242497.1 |
| D4 | 63079360 | 63079541 | refGene_gn=ANKS6;refGene_tx=XM_023242497.1 |
| D4 | 63086198 | 63086368 | refGene_gn=ANKS6;refGene_tx=XM_023242497.1 |
| D4 | 63087186 | 63087340 | refGene_gn=ANKS6;refGene_tx=XM_023242497.1 |
| D4 | 63088416 | 63088626 | refGene_gn=ANKS6;refGene_tx=XM_023242497.1 |
| D4 | 63091568 | 63091618 | refGene_gn=ANKS6;refGene_tx=XM_023242497.1 |
| D4 | 63092380 | 63092579 | refGene_gn=ANKS6;refGene_tx=XM_023242497.1 |
| D4 | 63093845 | 63093994 | refGene_gn=ANKS6;refGene_tx=XM_023242497.1 |
| D4 | 63096038 | 63096145 | refGene_gn=ANKS6;refGene_tx=XM_023242497.1 |
| D4 | 63097125 | 63097330 | refGene_gn=ANKS6;refGene_tx=XM_023242497.1 |
| D4 | 63098002 | 63098047 | refGene_gn=ANKS6;refGene_tx=XM_023242497.1 |
| D4 | 63100702 | 63101199 | refGene_gn=ANKS6;refGene_tx=XM_023242497.1 |
| D4 | 63105910 | 63106269 | refGene_gn=ANKS6;refGene_tx=XM_023242497.1 |
| D4 | 63119646 | 63120011 | refGene_gn=GALNT12;refGene_tx=XM_011288676.2 |
| D4 | 63129894 | 63130064 | refGene_gn=GALNT12;refGene_tx=XM_011288676.2 |
| D4 | 63134709 | 63134899 | refGene_gn=GALNT12;refGene_tx=XM_011288676.2 |
| D4 | 63138737 | 63138923 | refGene_gn=GALNT12;refGene_tx=XM_011288676.2 |
| D4 | 63142497 | 63142615 | refGene_gn=GALNT12;refGene_tx=XM_011288676.2 |
| D4 | 63144045 | 63144222 | refGene_gn=GALNT12;refGene_tx=XM_011288676.2 |
| D4 | 63146616 | 63146748 | refGene_gn=GALNT12;refGene_tx=XM_011288676.2 |
| D4 | 63151075 | 63151189 | refGene_gn=GALNT12;refGene_tx=XM_011288676.2 |
| D4 | 63152421 | 63152568 | refGene_gn=GALNT12;refGene_tx=XM_011288676.2 |
| D4 | 63154524 | 63154662 | refGene_gn=GALNT12;refGene_tx=XM_011288676.2 |
| D4 | 63231508 | 63231519 | refGene_gn=COL15A1;refGene_tx=XM_006939312.2 |
| D4 | 63231594 | 63231680 | refGene_gn=COL15A1;refGene_tx=XM_006939312.2 |
| D4 | 63269398 | 63269946 | refGene_gn=COL15A1;refGene_tx=XM_006939312.2 |
| D4 | 63271584 | 63271659 | refGene_gn=COL15A1;refGene_tx=XM_006939312.2 |
| D4 | 63273184 | 63273265 | refGene_gn=COL15A1;refGene_tx=XM_006939312.2 |
| D4 | 63278733 | 63278881 | refGene_gn=COL15A1;refGene_tx=XM_006939312.2 |
| D4 | 63282828 | 63282938 | refGene_gn=COL15A1;refGene_tx=XM_006939312.2 |
| D4 | 63284881 | 63285037 | refGene_gn=COL15A1;refGene_tx=XM_006939312.2 |
| D4 | 63286104 | 63286257 | refGene_gn=COL15A1;refGene_tx=XM_006939312.2 |
| D4 | 63288273 | 63288423 | refGene_gn=COL15A1;refGene_tx=XM_006939312.2 |
| D4 | 63288902 | 63289046 | refGene_gn=COL15A1;refGene_tx=XM_006939312.2 |
| D4 | 63294709 | 63294769 | refGene_gn=COL15A1;refGene_tx=XM_006939312.2 |
| D4 | 63296160 | 63296253 | refGene_gn=COL15A1;refGene_tx=XM_006939312.2 |
| D4 | 63297737 | 63297863 | refGene_gn=COL15A1;refGene_tx=XM_006939312.2 |
| D4 | 63298437 | 63298500 | refGene_gn=COL15A1;refGene_tx=XM_006939312.2 |
| D4 | 63304519 | 63304555 | refGene_gn=COL15A1;refGene_tx=XM_006939312.2 |
| D4 | 63304981 | 63305122 | refGene_gn=COL15A1;refGene_tx=XM_006939312.2 |

D4 63305326 63305395 refGene_gn=COL15A1;refGene_tx=XM_006939312.2
D4 63306208 63306244 refGene_gn=COL15A1;refGene_tx=XM_006939312.2
D4 63306349 63306433 refGene_gn=COL15A1;refGene_tx=XM_006939312.2
D4 63307950 63308016 refGene_gn=COL15A1;refGene_tx=XM_006939312.2
D4 63309974 63310010 refGene_gn=COL15A1;refGene_tx=XM_006939312.2
D4 63312836 63312899 refGene_gn=COL15A1;refGene_tx=XM_006939312.2
D4 63315168 63315237 refGene_gn=COL15A1;refGene_tx=XM_006939312.2
D4 63315347 63315383 refGene_gn=COL15A1;refGene_tx=XM_006939312.2
D4 63318008 63318062 refGene_gn=COL15A1;refGene_tx=XM_006939312.2
D4 63318163 63318223 refGene_gn=COL15A1;refGene_tx=XM_006939312.2
D4 63319677 63319746 refGene_gn=COL15A1;refGene_tx=XM_006939312.2
D4 63319868 63319910 refGene_gn=COL15A1;refGene_tx=XM_006939312.2
D4 63322403 63322449 refGene_gn=COL15A1;refGene_tx=XM_006939312.2
D4 63324572 63324653 refGene_gn=COL15A1;refGene_tx=XM_006939312.2
D4 63325059 63325109 refGene_gn=COL15A1;refGene_tx=XM_006939312.2
D4 63325211 63325313 refGene_gn=COL15A1;refGene_tx=XM_006939312.2
D4 63326230 63326375 refGene_gn=COL15A1;refGene_tx=XM_006939312.2
D4 63329466 63329531 refGene_gn=COL15A1;refGene_tx=XM_006939312.2
D4 63330886 63331015 refGene_gn=COL15A1;refGene_tx=XM_006939312.2
D4 63331162 63331222 refGene_gn=COL15A1;refGene_tx=XM_006939312.2
D4 63331962 63332022 refGene_gn=COL15A1;refGene_tx=XM_006939312.2
D4 63335780 63335966 refGene_gn=COL15A1;refGene_tx=XM_006939312.2
D4 63337271 63337387 refGene_gn=COL15A1;refGene_tx=XM_006939312.2
D4 63338392 63338606 refGene_gn=COL15A1;refGene_tx=XM_006939312.2
D4 63370963 63371048
refGene_gn=TGFBR1;refGene_tx=XM_023242501.1;refGene_gn=TGFBR1;refGene_tx=XM_023242502.1
D4 63393995 63394098
refGene_gn=TGFBR1;refGene_tx=XM_023242499.1;refGene_gn=TGFBR1;refGene_tx=XM_023242500.1
D4 63400048 63400294
refGene_gn=TGFBR1;refGene_tx=XM_023242499.1;refGene_gn=TGFBR1;refGene_tx=XM_023242500.1;refGene_gn=TGFBR1;refGene_tx=XM_023242501.1;refGene_gn=TGFBR1;refGene_tx=XM_023242502.1
D4 63404379 63404622
refGene_gn=TGFBR1;refGene_tx=XM_023242499.1;refGene_gn=TGFBR1;refGene_tx=XM_023242500.1;refGene_gn=TGFBR1;refGene_tx=XM_023242501.1;refGene_gn=TGFBR1;refGene_tx=XM_023242502.1
D4 63409609 63409840
refGene_gn=TGFBR1;refGene_tx=XM_023242499.1;refGene_gn=TGFBR1;refGene_tx=XM_023242500.1;refGene_gn=TGFBR1;refGene_tx=XM_023242501.1;refGene_gn=TGFBR1;refGene_tx=XM_023242502.1
D4 63414016 63414184
refGene_gn=TGFBR1;refGene_tx=XM_023242499.1;refGene_gn=TGFBR1;refGene_tx=XM_023242500.1;refGene_gn=TGFBR1;refGene_tx=XM_023242501.1;refGene_gn=TGFBR1;refGene_tx=XM_023242502.1
D4 63415807 63415964
refGene_gn=TGFBR1;refGene_tx=XM_023242499.1;refGene_gn=TGFBR1;refGene_tx=XM_023242500.1;refGene_gn=TGFBR1;refGene_tx=XM_023242501.1;refGene_gn=TGFBR1;refGene_tx=XM_023242502.1

D4 63418051 63418176
refGene_gn=TGFBR1;refGene_tx=XM_023242499.1;refGene_gn=TGFBR1;refGene_tx=XM_023242500.1;refGene_gn=TGFBR1;refGene_tx=XM_023242501.1;refGene_gn=TGFBR1;refGene_tx=XM_023242502.1
D4 63419464 63419595
refGene_gn=TGFBR1;refGene_tx=XM_023242499.1;refGene_gn=TGFBR1;refGene_tx=XM_023242500.1;refGene_gn=TGFBR1;refGene_tx=XM_023242501.1;refGene_gn=TGFBR1;refGene_tx=XM_023242502.1
D4 63424387 63424513
refGene_gn=TGFBR1;refGene_tx=XM_023242499.1;refGene_gn=TGFBR1;refGene_tx=XM_023242500.1;refGene_gn=TGFBR1;refGene_tx=XM_023242501.1;refGene_gn=TGFBR1;refGene_tx=XM_023242502.1
D4 63476993 63477896 refGene_gn=ALG2;refGene_tx=XM_003995659.4
D4 63481134 63481740 refGene_gn=ALG2;refGene_tx=XM_003995659.4
D4 63481946 63481949 refGene_gn=SEC61B;refGene_tx=XM_003995660.4
D4 63482144 63482242 refGene_gn=SEC61B;refGene_tx=XM_003995660.4
D4 63484871 63484973 refGene_gn=SEC61B;refGene_tx=XM_003995660.4
D4 63488394 63488482 refGene_gn=SEC61B;refGene_tx=XM_003995660.4
D4 64019336 64019861
refGene_gn=LOC109493876;refGene_tx=XM_019816230.2
D4 64025540 64026509 refGene_gn=NR4A3;refGene_tx=XM_023242504.1
D4 64030030 64030160 refGene_gn=NR4A3;refGene_tx=XM_023242504.1
D4 64030624 64030797 refGene_gn=NR4A3;refGene_tx=XM_023242504.1
D4 64040800 64041000 refGene_gn=NR4A3;refGene_tx=XM_023242504.1
D4 64043254 64043433 refGene_gn=NR4A3;refGene_tx=XM_023242504.1
D4 64056567 64056743 refGene_gn=NR4A3;refGene_tx=XM_023242504.1
D4 64056744 64056816 refGene_gn=NR4A3;refGene_tx=XM_023242504.1
D4 64109331 64109454
refGene_gn=STX17;refGene_tx=XM_023242506.1;refGene_gn=STX17;refGene_tx=XM_023242507.1
D4 64121486 64121552
refGene_gn=STX17;refGene_tx=XM_023242506.1;refGene_gn=STX17;refGene_tx=XM_023242507.1
D4 64144745 64144971
refGene_gn=STX17;refGene_tx=XM_023242506.1;refGene_gn=STX17;refGene_tx=XM_023242507.1
D4 64153055 64153171
refGene_gn=STX17;refGene_tx=XM_023242506.1;refGene_gn=STX17;refGene_tx=XM_023242507.1
D4 64153250 64153301
refGene_gn=STX17;refGene_tx=XM_023242506.1;refGene_gn=STX17;refGene_tx=XM_023242507.1
D4 64161359 64161446
refGene_gn=STX17;refGene_tx=XM_023242506.1;refGene_gn=STX17;refGene_tx=XM_023242507.1
D4 64162183 64162423
refGene_gn=STX17;refGene_tx=XM_023242506.1;refGene_gn=STX17;refGene_tx=XM_023242507.1
D4 64175966 64176068 refGene_gn=ERP44;refGene_tx=XM_011288533.3
D4 64178198 64178301 refGene_gn=ERP44;refGene_tx=XM_011288533.3

D4	64195167	64195309	refGene_gn=ERP44;refGene_tx=XM_011288533.3
D4	64196259	64196371	refGene_gn=ERP44;refGene_tx=XM_011288533.3
D4	64200372	64200489	refGene_gn=ERP44;refGene_tx=XM_011288533.3
D4	64202060	64202118	refGene_gn=ERP44;refGene_tx=XM_011288533.3
D4	64204760	64204876	refGene_gn=ERP44;refGene_tx=XM_011288533.3
D4	64206088	64206273	refGene_gn=ERP44;refGene_tx=XM_011288533.3
D4	64226218	64226334	refGene_gn=ERP44;refGene_tx=XM_011288533.3
D4	64230430	64230470	refGene_gn=ERP44;refGene_tx=XM_011288533.3
D4	64232406	64232479	refGene_gn=ERP44;refGene_tx=XM_011288533.3
D4	64270281	64270575	refGene_gn=ERP44;refGene_tx=XM_011288533.3
D4	64273691	64273797
refGene_gn=INVS;refGene_tx=XM_003995663.5;refGene_gn=INVS;refGene_tx=XM_006939315.3;refGene_gn=INVS;refGene_tx=XM_006939320.3
D4	64291165	64291332
refGene_gn=INVS;refGene_tx=XM_003995663.5;refGene_gn=INVS;refGene_tx=XM_006939315.3;refGene_gn=INVS;refGene_tx=XM_006939316.3;refGene_gn=INVS;refGene_tx=XM_006939320.3
D4	64350942	64351116
refGene_gn=INVS;refGene_tx=XM_003995663.5;refGene_gn=INVS;refGene_tx=XM_006939315.3;refGene_gn=INVS;refGene_tx=XM_006939316.3;refGene_gn=INVS;refGene_tx=XM_006939317.4;refGene_gn=INVS;refGene_tx=XM_006939318.4;refGene_gn=INVS;refGene_tx=XM_006939320.3;refGene_gn=INVS;refGene_tx=XM_019816232.2;refGene_gn=INVS;refGene_tx=XM_023242508.1
D4	64355734	64355902
refGene_gn=INVS;refGene_tx=XM_003995663.5;refGene_gn=INVS;refGene_tx=XM_006939315.3;refGene_gn=INVS;refGene_tx=XM_006939316.3;refGene_gn=INVS;refGene_tx=XM_006939317.4;refGene_gn=INVS;refGene_tx=XM_006939318.4;refGene_gn=INVS;refGene_tx=XM_006939320.3;refGene_gn=INVS;refGene_tx=XM_019816232.2;refGene_gn=INVS;refGene_tx=XM_023242508.1
D4	64360497	64360678
refGene_gn=INVS;refGene_tx=XM_003995663.5;refGene_gn=INVS;refGene_tx=XM_006939315.3;refGene_gn=INVS;refGene_tx=XM_006939316.3;refGene_gn=INVS;refGene_tx=XM_006939317.4;refGene_gn=INVS;refGene_tx=XM_006939318.4;refGene_gn=INVS;refGene_tx=XM_006939320.3;refGene_gn=INVS;refGene_tx=XM_019816232.2;refGene_gn=INVS;refGene_tx=XM_023242508.1
D4	64363102	64363212
refGene_gn=INVS;refGene_tx=XM_003995663.5;refGene_gn=INVS;refGene_tx=XM_006939315.3;refGene_gn=INVS;refGene_tx=XM_006939316.3;refGene_gn=INVS;refGene_tx=XM_006939317.4;refGene_gn=INVS;refGene_tx=XM_006939318.4;refGene_gn=INVS;refGene_tx=XM_006939320.3;refGene_gn=INVS;refGene_tx=XM_019816232.2;refGene_gn=INVS;refGene_tx=XM_023242508.1
D4	64366546	64366718
refGene_gn=INVS;refGene_tx=XM_003995663.5;refGene_gn=INVS;refGene_tx=XM_006939315.3;refGene_gn=INVS;refGene_tx=XM_006939316.3;refGene_gn=INVS;refGene_tx=XM_006939317.4;refGene_gn=INVS;refGene_tx=XM_006939318.4;refGene_gn=INVS;refGene_tx=XM_006939320.3;refGene_gn=INVS;refGene_tx=XM_019816232.2;refGene_gn=INVS;refGene_tx=XM_019816233.2;refGene_gn=INVS;refGene_tx=XM_023242508.1
D4	64370709	64370865
refGene_gn=INVS;refGene_tx=XM_003995663.5;refGene_gn=INVS;refGene_tx=X

M_006939315.3;refGene_gn=INVS;refGene_tx=XM_006939316.3;refGene_gn=INVS;refGene_tx=XM_006939317.4;refGene_gn=INVS;refGene_tx=XM_006939318.4;refGene_gn=INVS;refGene_tx=XM_006939320.3;refGene_gn=INVS;refGene_tx=XM_019816232.2;refGene_gn=INVS;refGene_tx=XM_019816233.2;refGene_gn=INVS;refGene_tx=XM_023242508.1
D4 64371348 64371578
refGene_gn=INVS;refGene_tx=XM_003995663.5;refGene_gn=INVS;refGene_tx=XM_006939315.3;refGene_gn=INVS;refGene_tx=XM_006939316.3;refGene_gn=INVS;refGene_tx=XM_006939317.4;refGene_gn=INVS;refGene_tx=XM_006939318.4;refGene_gn=INVS;refGene_tx=XM_006939320.3;refGene_gn=INVS;refGene_tx=XM_019816232.2;refGene_gn=INVS;refGene_tx=XM_019816233.2;refGene_gn=INVS;refGene_tx=XM_023242508.1
D4 64375623 64375730
refGene_gn=INVS;refGene_tx=XM_003995663.5;refGene_gn=INVS;refGene_tx=XM_006939315.3;refGene_gn=INVS;refGene_tx=XM_006939316.3;refGene_gn=INVS;refGene_tx=XM_006939317.4;refGene_gn=INVS;refGene_tx=XM_006939318.4;refGene_gn=INVS;refGene_tx=XM_006939320.3;refGene_gn=INVS;refGene_tx=XM_019816232.2;refGene_gn=INVS;refGene_tx=XM_019816233.2;refGene_gn=INVS;refGene_tx=XM_023242508.1
D4 64382879 64383092
refGene_gn=INVS;refGene_tx=XM_003995663.5;refGene_gn=INVS;refGene_tx=XM_006939315.3;refGene_gn=INVS;refGene_tx=XM_006939316.3;refGene_gn=INVS;refGene_tx=XM_006939317.4;refGene_gn=INVS;refGene_tx=XM_006939318.4;refGene_gn=INVS;refGene_tx=XM_006939320.3;refGene_gn=INVS;refGene_tx=XM_019816232.2;refGene_gn=INVS;refGene_tx=XM_019816233.2;refGene_gn=INVS;refGene_tx=XM_023242508.1
D4 64395759 64396034
refGene_gn=INVS;refGene_tx=XM_003995663.5;refGene_gn=INVS;refGene_tx=XM_006939315.3;refGene_gn=INVS;refGene_tx=XM_006939316.3;refGene_gn=INVS;refGene_tx=XM_006939317.4;refGene_gn=INVS;refGene_tx=XM_006939318.4;refGene_gn=INVS;refGene_tx=XM_006939320.3;refGene_gn=INVS;refGene_tx=XM_019816232.2;refGene_gn=INVS;refGene_tx=XM_019816233.2;refGene_gn=INVS;refGene_tx=XM_023242508.1
D4 64411392 64412110
refGene_gn=INVS;refGene_tx=XM_003995663.5;refGene_gn=INVS;refGene_tx=XM_006939315.3;refGene_gn=INVS;refGene_tx=XM_006939316.3;refGene_gn=INVS;refGene_tx=XM_006939317.4;refGene_gn=INVS;refGene_tx=XM_006939318.4;refGene_gn=INVS;refGene_tx=XM_006939320.3;refGene_gn=INVS;refGene_tx=XM_019816232.2;refGene_gn=INVS;refGene_tx=XM_019816233.2;refGene_gn=INVS;refGene_tx=XM_023242508.1
D4 64422422 64422652
refGene_gn=INVS;refGene_tx=XM_003995663.5;refGene_gn=INVS;refGene_tx=XM_006939315.3;refGene_gn=INVS;refGene_tx=XM_006939316.3;refGene_gn=INVS;refGene_tx=XM_006939317.4;refGene_gn=INVS;refGene_tx=XM_006939318.4;refGene_gn=INVS;refGene_tx=XM_006939320.3;refGene_gn=INVS;refGene_tx=XM_019816232.2;refGene_gn=INVS;refGene_tx=XM_019816233.2;refGene_gn=INVS;refGene_tx=XM_023242508.1
D4 64423283 64423324 refGene_gn=INVS;refGene_tx=XM_006939315.3
D4 64423386 64423461
refGene_gn=INVS;refGene_tx=XM_003995663.5;refGene_gn=INVS;refGene_tx=X

M_006939316.3;refGene_gn=INVS;refGene_tx=XM_006939317.4;refGene_gn=INVS;refGene_tx=XM_006939318.4;refGene_gn=INVS;refGene_tx=XM_006939320.3;refGene_gn=INVS;refGene_tx=XM_019816232.2;refGene_gn=INVS;refGene_tx=XM_019816233.2;refGene_gn=INVS;refGene_tx=XM_023242508.1
D4 64426610 64426762 refGene_gn=INVS;refGene_tx=XM_003995663.5;refGene_gn=INVS;refGene_tx=XM_006939316.3;refGene_gn=INVS;refGene_tx=XM_006939317.4;refGene_gn=INVS;refGene_tx=XM_006939318.4;refGene_gn=INVS;refGene_tx=XM_006939320.3;refGene_gn=INVS;refGene_tx=XM_019816232.2;refGene_gn=INVS;refGene_tx=XM_019816233.2;refGene_gn=INVS;refGene_tx=XM_023242508.1
D4 64428959 64429073 refGene_gn=TEX10;refGene_tx=XM_003995664.4
D4 64430546 64430757 refGene_gn=TEX10;refGene_tx=XM_003995664.4
D4 64435108 64435290 refGene_gn=TEX10;refGene_tx=XM_003995664.4
D4 64437241 64437322 refGene_gn=TEX10;refGene_tx=XM_003995664.4
D4 64447279 64447413 refGene_gn=TEX10;refGene_tx=XM_003995664.4
D4 64449015 64449104 refGene_gn=TEX10;refGene_tx=XM_003995664.4
D4 64453831 64454009 refGene_gn=TEX10;refGene_tx=XM_003995664.4
D4 64455547 64455723 refGene_gn=TEX10;refGene_tx=XM_003995664.4
D4 64457524 64457660 refGene_gn=TEX10;refGene_tx=XM_003995664.4
D4 64458416 64458655 refGene_gn=TEX10;refGene_tx=XM_003995664.4
D4 64474454 64474477 refGene_gn=TEX10;refGene_tx=XM_003995664.4
D4 64474478 64474568 refGene_gn=TEX10;refGene_tx=XM_003995664.4
D4 64480335 64480579 refGene_gn=TEX10;refGene_tx=XM_003995664.4
D4 64480968 64481681 refGene_gn=TEX10;refGene_tx=XM_003995664.4
D4 64483189 64483369 refGene_gn=TEX10;refGene_tx=XM_003995664.4
D4 64571509 64571927 refGene_gn=MSANTD3;refGene_tx=XM_023242512.1;refGene_gn=MSANTD3;refGene_tx=XM_023242513.1;refGene_gn=MSANTD3;refGene_tx=XM_023242514.1;refGene_gn=MSANTD3;refGene_tx=XM_023242516.1
D4 64579700 64580110 refGene_gn=MSANTD3;refGene_tx=XM_023242512.1;refGene_gn=MSANTD3;refGene_tx=XM_023242513.1;refGene_gn=MSANTD3;refGene_tx=XM_023242514.1;refGene_gn=MSANTD3;refGene_tx=XM_023242516.1
D4 64601295 64601491 refGene_gn=TMEFF1;refGene_tx=XM_023242510.1;refGene_gn=TMEFF1;refGene_tx=XM_023242511.1
D4 64622815 64622925 refGene_gn=TMEFF1;refGene_tx=XM_023242510.1;refGene_gn=TMEFF1;refGene_tx=XM_023242511.1
D4 64629685 64629815 refGene_gn=TMEFF1;refGene_tx=XM_023242510.1;refGene_gn=TMEFF1;refGene_tx=XM_023242511.1
D4 64634346 64634373 refGene_gn=TMEFF1;refGene_tx=XM_023242510.1;refGene_gn=TMEFF1;refGene_tx=XM_023242511.1
D4 64636665 64636765 refGene_gn=TMEFF1;refGene_tx=XM_023242510.1;refGene_gn=TMEFF1;refGene_tx=XM_023242511.1
D4 64653249 64653398 refGene_gn=TMEFF1;refGene_tx=XM_023242510.1;refGene_gn=TMEFF1;refGene_

tx=XM_023242511.1
D4 64655733 64655799
refGene_gn=TMEFF1;refGene_tx=XM_023242510.1;refGene_gn=TMEFF1;refGene_
tx=XM_023242511.1
D4 64669619 64669743
refGene_gn=TMEFF1;refGene_tx=XM_023242510.1;refGene_gn=TMEFF1;refGene_
tx=XM_023242511.1
D4 64678820 64678979
refGene_gn=TMEFF1;refGene_tx=XM_023242510.1;refGene_gn=TMEFF1;refGene_
tx=XM_023242511.1
D4 64682506 64682591
refGene_gn=TMEFF1;refGene_tx=XM_023242510.1;refGene_gn=TMEFF1;refGene_
tx=XM_023242511.1
D4 64684139 64684547 refGene_gn=CAVIN4;refGene_tx=XM_019816237.1
D4 64691961 64692639 refGene_gn=CAVIN4;refGene_tx=XM_019816237.1
D4 65185990 65186053
refGene_gn=PLPPR1;refGene_tx=XM_006939332.4;refGene_gn=PLPPR1;refGene_
tx=XM_019816239.2;refGene_gn=PLPPR1;refGene_tx=XM_019816240.2;refGene_
gn=PLPPR1;refGene_tx=XM_023242517.1
D4 65268529 65268718
refGene_gn=PLPPR1;refGene_tx=XM_006939332.4;refGene_gn=PLPPR1;refGene_
tx=XM_019816239.2;refGene_gn=PLPPR1;refGene_tx=XM_019816240.2;refGene_
gn=PLPPR1;refGene_tx=XM_019816241.2;refGene_gn=PLPPR1;refGene_tx=XM_02
3242517.1
D4 65282332 65282465
refGene_gn=PLPPR1;refGene_tx=XM_006939332.4;refGene_gn=PLPPR1;refGene_
tx=XM_019816239.2;refGene_gn=PLPPR1;refGene_tx=XM_019816240.2;refGene_
gn=PLPPR1;refGene_tx=XM_019816241.2;refGene_gn=PLPPR1;refGene_tx=XM_02
3242517.1
D4 65300777 65301028
refGene_gn=PLPPR1;refGene_tx=XM_006939332.4;refGene_gn=PLPPR1;refGene_
tx=XM_019816239.2;refGene_gn=PLPPR1;refGene_tx=XM_019816240.2;refGene_
gn=PLPPR1;refGene_tx=XM_019816241.2;refGene_gn=PLPPR1;refGene_tx=XM_02
3242517.1
D4 65301292 65301481 refGene_gn=PLPPR1;refGene_tx=XM_023242517.1
D4 65304524 65304701
refGene_gn=PLPPR1;refGene_tx=XM_006939332.4;refGene_gn=PLPPR1;refGene_
tx=XM_019816239.2;refGene_gn=PLPPR1;refGene_tx=XM_019816240.2;refGene_
gn=PLPPR1;refGene_tx=XM_019816241.2
D4 65308192 65308324
refGene_gn=PLPPR1;refGene_tx=XM_006939332.4;refGene_gn=PLPPR1;refGene_
tx=XM_019816239.2;refGene_gn=PLPPR1;refGene_tx=XM_019816240.2;refGene_
gn=PLPPR1;refGene_tx=XM_019816241.2
D4 65313967 65314000
refGene_gn=PLPPR1;refGene_tx=XM_006939332.4;refGene_gn=PLPPR1;refGene_
tx=XM_019816239.2;refGene_gn=PLPPR1;refGene_tx=XM_019816240.2;refGene_
gn=PLPPR1;refGene_tx=XM_019816241.2
D4 65328672 65329266
refGene_gn=BAAT;refGene_tx=XM_003995698.3;refGene_gn=BAAT;refGene_tx=X
M_019816244.1;refGene_gn=BAAT;refGene_tx=XM_019816245.2

D4 65332512 65332715
refGene_gn=BAAT;refGene_tx=XM_003995698.3;refGene_gn=BAAT;refGene_tx=XM_019816244.1;refGene_gn=BAAT;refGene_tx=XM_019816245.2
D4 65332953 65333447
refGene_gn=BAAT;refGene_tx=XM_003995698.3;refGene_gn=BAAT;refGene_tx=XM_019816244.1;refGene_gn=BAAT;refGene_tx=XM_019816245.2
D4 65349586 65350183
refGene_gn=BAAT;refGene_tx=XM_019816244.1;refGene_gn=LOC101096116;refGene_tx=XM_003995700.4
D4 65353297 65353500
refGene_gn=LOC101096116;refGene_tx=XM_003995700.4
D4 65357047 65357513
refGene_gn=LOC101096116;refGene_tx=XM_003995700.4
D4 65372370 65372758 refGene_gn=MRPL50;refGene_tx=XM_003995701.4
D4 65374918 65375010 refGene_gn=MRPL50;refGene_tx=XM_003995701.4
D4 65375579 65375612
refGene_gn=ZNF189;refGene_tx=XM_003995702.4;refGene_gn=ZNF189;refGene_tx=XM_003995703.4;refGene_gn=ZNF189;refGene_tx=XM_006939333.3
D4 65376316 65376443
refGene_gn=ZNF189;refGene_tx=XM_003995702.4;refGene_gn=ZNF189;refGene_tx=XM_003995703.4;refGene_gn=ZNF189;refGene_tx=XM_006939333.3
D4 65385191 65386912
refGene_gn=ZNF189;refGene_tx=XM_003995702.4;refGene_gn=ZNF189;refGene_tx=XM_003995703.4;refGene_gn=ZNF189;refGene_tx=XM_006939333.3
D4 65396068 65396164 refGene_gn=ALDOB;refGene_tx=XM_006939334.3
D4 65398744 65398944 refGene_gn=ALDOB;refGene_tx=XM_006939334.3
D4 65399352 65399527 refGene_gn=ALDOB;refGene_tx=XM_006939334.3
D4 65400423 65400507 refGene_gn=ALDOB;refGene_tx=XM_006939334.3
D4 65401617 65401778 refGene_gn=ALDOB;refGene_tx=XM_006939334.3
D4 65402654 65402709 refGene_gn=ALDOB;refGene_tx=XM_006939334.3
D4 65403745 65403957 refGene_gn=ALDOB;refGene_tx=XM_006939334.3
D4 65405010 65405122 refGene_gn=ALDOB;refGene_tx=XM_006939334.3
D4 65445378 65446590
refGene_gn=TMEM246;refGene_tx=XM_006939336.4;refGene_gn=TMEM246;refGene_tx=XM_023242519.1
D4 65489639 65489642 refGene_gn=RNF20;refGene_tx=XM_019816251.2
D4 65490164 65490319
refGene_gn=RNF20;refGene_tx=XM_003995706.3;refGene_gn=RNF20;refGene_tx=XM_019816251.2;refGene_gn=RNF20;refGene_tx=XM_019816252.2
D4 65493272 65493345 refGene_gn=RNF20;refGene_tx=XM_019816251.2
D4 65495357 65495525
refGene_gn=RNF20;refGene_tx=XM_003995706.3;refGene_gn=RNF20;refGene_tx=XM_019816251.2;refGene_gn=RNF20;refGene_tx=XM_019816252.2
D4 65495644 65495792
refGene_gn=RNF20;refGene_tx=XM_003995706.3;refGene_gn=RNF20;refGene_tx=XM_019816251.2;refGene_gn=RNF20;refGene_tx=XM_019816252.2
D4 65495941 65496124
refGene_gn=RNF20;refGene_tx=XM_003995706.3;refGene_gn=RNF20;refGene_tx=XM_019816251.2;refGene_gn=RNF20;refGene_tx=XM_019816252.2
D4 65499776 65499895

refGene_gn=RNF20;refGene_tx=XM_003995706.3;refGene_gn=RNF20;refGene_tx=XM_019816251.2;refGene_gn=RNF20;refGene_tx=XM_019816252.2
D4 65502676 65502823
refGene_gn=RNF20;refGene_tx=XM_003995706.3;refGene_gn=RNF20;refGene_tx=XM_019816251.2;refGene_gn=RNF20;refGene_tx=XM_019816252.2
D4 65503220 65503298
refGene_gn=RNF20;refGene_tx=XM_003995706.3;refGene_gn=RNF20;refGene_tx=XM_019816251.2;refGene_gn=RNF20;refGene_tx=XM_019816252.2
D4 65503491 65503611
refGene_gn=RNF20;refGene_tx=XM_003995706.3;refGene_gn=RNF20;refGene_tx=XM_019816251.2;refGene_gn=RNF20;refGene_tx=XM_019816252.2
D4 65504699 65504879
refGene_gn=RNF20;refGene_tx=XM_003995706.3;refGene_gn=RNF20;refGene_tx=XM_019816251.2;refGene_gn=RNF20;refGene_tx=XM_019816252.2
D4 65505796 65505932
refGene_gn=RNF20;refGene_tx=XM_003995706.3;refGene_gn=RNF20;refGene_tx=XM_019816251.2;refGene_gn=RNF20;refGene_tx=XM_019816252.2
D4 65506251 65506373
refGene_gn=RNF20;refGene_tx=XM_003995706.3;refGene_gn=RNF20;refGene_tx=XM_019816251.2;refGene_gn=RNF20;refGene_tx=XM_019816252.2
D4 65506491 65506862
refGene_gn=RNF20;refGene_tx=XM_003995706.3;refGene_gn=RNF20;refGene_tx=XM_019816251.2;refGene_gn=RNF20;refGene_tx=XM_019816252.2
D4 65508073 65508191
refGene_gn=RNF20;refGene_tx=XM_003995706.3;refGene_gn=RNF20;refGene_tx=XM_019816251.2;refGene_gn=RNF20;refGene_tx=XM_019816252.2
D4 65508757 65508907
refGene_gn=RNF20;refGene_tx=XM_003995706.3;refGene_gn=RNF20;refGene_tx=XM_019816251.2;refGene_gn=RNF20;refGene_tx=XM_019816252.2
D4 65512299 65512512
refGene_gn=RNF20;refGene_tx=XM_003995706.3;refGene_gn=RNF20;refGene_tx=XM_019816251.2;refGene_gn=RNF20;refGene_tx=XM_019816252.2
D4 65513354 65513480
refGene_gn=RNF20;refGene_tx=XM_003995706.3;refGene_gn=RNF20;refGene_tx=XM_019816251.2;refGene_gn=RNF20;refGene_tx=XM_019816252.2
D4 65513636 65513777
refGene_gn=RNF20;refGene_tx=XM_003995706.3;refGene_gn=RNF20;refGene_tx=XM_019816251.2;refGene_gn=RNF20;refGene_tx=XM_019816252.2
D4 65514458 65514560
refGene_gn=RNF20;refGene_tx=XM_003995706.3;refGene_gn=RNF20;refGene_tx=XM_019816251.2;refGene_gn=RNF20;refGene_tx=XM_019816252.2
D4 65515003 65515180
refGene_gn=RNF20;refGene_tx=XM_003995706.3;refGene_gn=RNF20;refGene_tx=XM_019816251.2;refGene_gn=RNF20;refGene_tx=XM_019816252.2
D4 65523902 65524245 refGene_gn=GRIN3A;refGene_tx=XM_019816253.2
D4 65528910 65528987 refGene_gn=GRIN3A;refGene_tx=XM_019816253.2
D4 65530546 65530711 refGene_gn=GRIN3A;refGene_tx=XM_019816253.2
D4 65547852 65548365 refGene_gn=PPP3R2;refGene_tx=XM_003995707.4
D4 65564108 65564260 refGene_gn=GRIN3A;refGene_tx=XM_019816253.2
D4 65572264 65572380 refGene_gn=GRIN3A;refGene_tx=XM_019816253.2

D4 65577893 65578039 refGene_gn=GRIN3A;refGene_tx=XM_019816253.2
D4 65619354 65620402 refGene_gn=GRIN3A;refGene_tx=XM_019816253.2
D4 65636584 65637189 refGene_gn=GRIN3A;refGene_tx=XM_019816253.2
D4 65689678 65690377 refGene_gn=GRIN3A;refGene_tx=XM_019816253.2
D4 66504336 66504353 refGene_gn=CYLC2;refGene_tx=XM_023242522.1
D4 66514045 66514086 refGene_gn=CYLC2;refGene_tx=XM_023242522.1
D4 66515458 66515580 refGene_gn=CYLC2;refGene_tx=XM_023242522.1
D4 66517005 66517444 refGene_gn=CYLC2;refGene_tx=XM_023242522.1
D4 66517761 66518231 refGene_gn=CYLC2;refGene_tx=XM_023242522.1
D4 67423127 67423295 refGene_gn=SMC2;refGene_tx=XM_023242302.1
D4 67424247 67424397 refGene_gn=SMC2;refGene_tx=XM_023242302.1
D4 67426334 67426457 refGene_gn=SMC2;refGene_tx=XM_023242302.1
D4 67427706 67427745 refGene_gn=SMC2;refGene_tx=XM_023242302.1
D4 67430751 67430862 refGene_gn=SMC2;refGene_tx=XM_023242302.1
D4 67431036 67431081 refGene_gn=SMC2;refGene_tx=XM_023242302.1
D4 67432136 67432370 refGene_gn=SMC2;refGene_tx=XM_023242302.1
D4 67432814 67432964 refGene_gn=SMC2;refGene_tx=XM_023242302.1
D4 67435596 67435830 refGene_gn=SMC2;refGene_tx=XM_023242302.1
D4 67437461 67437629 refGene_gn=SMC2;refGene_tx=XM_023242302.1
D4 67447803 67447992 refGene_gn=SMC2;refGene_tx=XM_023242302.1
D4 67448348 67448468 refGene_gn=SMC2;refGene_tx=XM_023242302.1
D4 67450488 67450693 refGene_gn=SMC2;refGene_tx=XM_023242302.1
D4 67460181 67460317 refGene_gn=SMC2;refGene_tx=XM_023242302.1
D4 67462609 67462734 refGene_gn=SMC2;refGene_tx=XM_023242302.1
D4 67466544 67466738 refGene_gn=SMC2;refGene_tx=XM_023242302.1
D4 67467704 67467848 refGene_gn=SMC2;refGene_tx=XM_023242302.1
D4 67471077 67471272 refGene_gn=SMC2;refGene_tx=XM_023242302.1
D4 67473732 67473933 refGene_gn=SMC2;refGene_tx=XM_023242302.1
D4 67476096 67476213 refGene_gn=SMC2;refGene_tx=XM_023242302.1
D4 67482262 67482423 refGene_gn=SMC2;refGene_tx=XM_023242302.1
D4 67486125 67486273 refGene_gn=SMC2;refGene_tx=XM_023242302.1
D4 67487280 67487439 refGene_gn=SMC2;refGene_tx=XM_023242302.1
D4 67625646 67627821
refGene_gn=LOC102902407;refGene_tx=XM_006939339.3
D4 67827890 67828874
refGene_gn=LOC101080583;refGene_tx=XM_003995717.1
D4 67858144 67859101
refGene_gn=LOC101100006;refGene_tx=XM_011288527.3
D4 67877735 67878689
refGene_gn=LOC111556266;refGene_tx=XM_023242524.1
D4 67898693 67899653
refGene_gn=LOC101100254;refGene_tx=XM_003995711.3
D4 67954227 67955166
refGene_gn=LOC101081336;refGene_tx=XM_011288671.2
D4 67981193 67981270
refGene_gn=LOC101101005;refGene_tx=XM_003995714.4
D4 67981272 67981392
refGene_gn=LOC101101005;refGene_tx=XM_003995714.4
D4 67981394 67982145
refGene_gn=LOC101101005;refGene_tx=XM_003995714.4

D4 67993021 67994020
refGene_gn=LOC111556284;refGene_tx=XM_023242523.1
D4 68011782 68012684
refGene_gn=LOC101081581;refGene_tx=XM_023242303.1
D4 68013127 68013170
refGene_gn=LOC101081581;refGene_tx=XM_023242303.1
D4 68017938 68018901
refGene_gn=LOC111557474;refGene_tx=XM_023242304.1
D4 68031248 68032205
refGene_gn=LOC111556257;refGene_tx=XM_023243326.1
D4 68068128 68069067
refGene_gn=LOC101082351;refGene_tx=XM_003995721.3
D4 68074851 68075715
refGene_gn=LOC111557476;refGene_tx=XM_023242305.1
D4 68095870 68096842
refGene_gn=LOC111557478;refGene_tx=XM_023242307.1
D4 68149953 68150652
refGene_gn=LOC101083628;refGene_tx=XM_023243330.1
D4 68150653 68150908
refGene_gn=LOC101083628;refGene_tx=XM_023243330.1
D4 68163253 68164198
refGene_gn=LOC101101260;refGene_tx=XM_003995715.4
D4 68193168 68193228
refGene_gn=NIPSNAP3A;refGene_tx=XM_011288525.3;refGene_gn=NIPSNAP3A;refGene_tx=XM_019816267.2
D4 68196246 68196457
refGene_gn=NIPSNAP3A;refGene_tx=XM_011288525.3;refGene_gn=NIPSNAP3A;refGene_tx=XM_019816267.2
D4 68198182 68198341
refGene_gn=NIPSNAP3A;refGene_tx=XM_011288525.3;refGene_gn=NIPSNAP3A;refGene_tx=XM_019816267.2
D4 68199859 68200009
refGene_gn=NIPSNAP3A;refGene_tx=XM_011288525.3;refGene_gn=NIPSNAP3A;refGene_tx=XM_019816267.2
D4 68207501 68207569
refGene_gn=NIPSNAP3A;refGene_tx=XM_019816267.2
D4 68208707 68208794
refGene_gn=NIPSNAP3A;refGene_tx=XM_011288525.3
D4 68208898 68208975
refGene_gn=NIPSNAP3A;refGene_tx=XM_011288525.3
D4 68232712 68232772
refGene_gn=LOC101080332;refGene_tx=XM_006939340.3
D4 68235627 68235838
refGene_gn=LOC101080332;refGene_tx=XM_006939340.3;refGene_gn=LOC101080332;refGene_tx=XM_023242527.1
D4 68237373 68237529
refGene_gn=LOC101080332;refGene_tx=XM_006939340.3;refGene_gn=LOC101080332;refGene_tx=XM_023242527.1
D4 68238602 68238752
refGene_gn=LOC101080332;refGene_tx=XM_006939340.3;refGene_gn=LOC101080

332;refGene_tx=XM_023242527.1
D4	68243882	68243969
refGene_gn=LOC101080332;refGene_tx=XM_006939340.3;refGene_gn=LOC101080332;refGene_tx=XM_023242527.1
D4	68244061	68244138
refGene_gn=LOC101080332;refGene_tx=XM_006939340.3;refGene_gn=LOC101080332;refGene_tx=XM_023242527.1
D4	68250616	68250757
refGene_gn=ABCA1;refGene_tx=XM_011288524.3;refGene_gn=ABCA1;refGene_tx=XM_019816256.2;refGene_gn=ABCA1;refGene_tx=XM_019816257.2
D4	68251573	68251817
refGene_gn=ABCA1;refGene_tx=XM_011288524.3;refGene_gn=ABCA1;refGene_tx=XM_019816256.2;refGene_gn=ABCA1;refGene_tx=XM_019816257.2
D4	68252472	68252565
refGene_gn=ABCA1;refGene_tx=XM_011288524.3;refGene_gn=ABCA1;refGene_tx=XM_019816256.2;refGene_gn=ABCA1;refGene_tx=XM_019816257.2
D4	68253230	68253334
refGene_gn=ABCA1;refGene_tx=XM_011288524.3;refGene_gn=ABCA1;refGene_tx=XM_019816256.2;refGene_gn=ABCA1;refGene_tx=XM_019816257.2
D4	68254986	68255121
refGene_gn=ABCA1;refGene_tx=XM_011288524.3;refGene_gn=ABCA1;refGene_tx=XM_019816256.2;refGene_gn=ABCA1;refGene_tx=XM_019816257.2
D4	68255540	68255682
refGene_gn=ABCA1;refGene_tx=XM_011288524.3;refGene_gn=ABCA1;refGene_tx=XM_019816256.2;refGene_gn=ABCA1;refGene_tx=XM_019816257.2
D4	68258089	68258196
refGene_gn=ABCA1;refGene_tx=XM_011288524.3;refGene_gn=ABCA1;refGene_tx=XM_019816256.2;refGene_gn=ABCA1;refGene_tx=XM_019816257.2
D4	68259055	68259118
refGene_gn=ABCA1;refGene_tx=XM_011288524.3;refGene_gn=ABCA1;refGene_tx=XM_019816256.2;refGene_gn=ABCA1;refGene_tx=XM_019816257.2
D4	68260349	68260470
refGene_gn=ABCA1;refGene_tx=XM_011288524.3;refGene_gn=ABCA1;refGene_tx=XM_019816256.2;refGene_gn=ABCA1;refGene_tx=XM_019816257.2
D4	68260736	68260866
refGene_gn=ABCA1;refGene_tx=XM_011288524.3;refGene_gn=ABCA1;refGene_tx=XM_019816256.2;refGene_gn=ABCA1;refGene_tx=XM_019816257.2
D4	68261920	68262044
refGene_gn=ABCA1;refGene_tx=XM_011288524.3;refGene_gn=ABCA1;refGene_tx=XM_019816256.2;refGene_gn=ABCA1;refGene_tx=XM_019816257.2
D4	68263804	68263949
refGene_gn=ABCA1;refGene_tx=XM_011288524.3;refGene_gn=ABCA1;refGene_tx=XM_019816256.2;refGene_gn=ABCA1;refGene_tx=XM_019816257.2
D4	68264045	68264161
refGene_gn=ABCA1;refGene_tx=XM_011288524.3;refGene_gn=ABCA1;refGene_tx=XM_019816256.2;refGene_gn=ABCA1;refGene_tx=XM_019816257.2
D4	68266124	68266302
refGene_gn=ABCA1;refGene_tx=XM_011288524.3;refGene_gn=ABCA1;refGene_tx=XM_019816256.2;refGene_gn=ABCA1;refGene_tx=XM_019816257.2
D4	68267748	68267918

refGene_gn=ABCA1;refGene_tx=XM_011288524.3;refGene_gn=ABCA1;refGene_tx
=XM_019816256.2;refGene_gn=ABCA1;refGene_tx=XM_019816257.2
D4 68268647 68268722
refGene_gn=ABCA1;refGene_tx=XM_011288524.3;refGene_gn=ABCA1;refGene_tx
=XM_019816256.2;refGene_gn=ABCA1;refGene_tx=XM_019816257.2
D4 68270020 68270126
refGene_gn=ABCA1;refGene_tx=XM_011288524.3;refGene_gn=ABCA1;refGene_tx
=XM_019816256.2;refGene_gn=ABCA1;refGene_tx=XM_019816257.2
D4 68271138 68271171
refGene_gn=ABCA1;refGene_tx=XM_011288524.3;refGene_gn=ABCA1;refGene_tx
=XM_019816256.2;refGene_gn=ABCA1;refGene_tx=XM_019816257.2
D4 68272478 68272573
refGene_gn=ABCA1;refGene_tx=XM_011288524.3;refGene_gn=ABCA1;refGene_tx
=XM_019816256.2;refGene_gn=ABCA1;refGene_tx=XM_019816257.2
D4 68273585 68273775
refGene_gn=ABCA1;refGene_tx=XM_011288524.3;refGene_gn=ABCA1;refGene_tx
=XM_019816256.2;refGene_gn=ABCA1;refGene_tx=XM_019816257.2
D4 68276336 68276435
refGene_gn=ABCA1;refGene_tx=XM_011288524.3;refGene_gn=ABCA1;refGene_tx
=XM_019816256.2;refGene_gn=ABCA1;refGene_tx=XM_019816257.2
D4 68277956 68278081
refGene_gn=ABCA1;refGene_tx=XM_011288524.3;refGene_gn=ABCA1;refGene_tx
=XM_019816256.2;refGene_gn=ABCA1;refGene_tx=XM_019816257.2
D4 68279527 68279676
refGene_gn=ABCA1;refGene_tx=XM_011288524.3;refGene_gn=ABCA1;refGene_tx
=XM_019816256.2;refGene_gn=ABCA1;refGene_tx=XM_019816257.2
D4 68280740 68280854
refGene_gn=ABCA1;refGene_tx=XM_011288524.3;refGene_gn=ABCA1;refGene_tx
=XM_019816256.2;refGene_gn=ABCA1;refGene_tx=XM_019816257.2
D4 68281042 68281091
refGene_gn=ABCA1;refGene_tx=XM_011288524.3;refGene_gn=ABCA1;refGene_tx
=XM_019816256.2;refGene_gn=ABCA1;refGene_tx=XM_019816257.2
D4 68282150 68282353
refGene_gn=ABCA1;refGene_tx=XM_011288524.3;refGene_gn=ABCA1;refGene_tx
=XM_019816256.2;refGene_gn=ABCA1;refGene_tx=XM_019816257.2
D4 68283173 68283252
refGene_gn=ABCA1;refGene_tx=XM_019816256.2;refGene_gn=ABCA1;refGene_tx
=XM_019816257.2
D4 68283279 68283352 refGene_gn=ABCA1;refGene_tx=XM_011288524.3
D4 68284606 68284827
refGene_gn=ABCA1;refGene_tx=XM_011288524.3;refGene_gn=ABCA1;refGene_tx
=XM_019816256.2;refGene_gn=ABCA1;refGene_tx=XM_019816257.2
D4 68285847 68285985
refGene_gn=ABCA1;refGene_tx=XM_011288524.3;refGene_gn=ABCA1;refGene_tx
=XM_019816256.2;refGene_gn=ABCA1;refGene_tx=XM_019816257.2
D4 68286188 68286331
refGene_gn=ABCA1;refGene_tx=XM_011288524.3;refGene_gn=ABCA1;refGene_tx
=XM_019816256.2;refGene_gn=ABCA1;refGene_tx=XM_019816257.2
D4 68287261 68287393
refGene_gn=ABCA1;refGene_tx=XM_011288524.3;refGene_gn=ABCA1;refGene_tx

=XM_019816256.2;refGene_gn=ABCA1;refGene_tx=XM_019816257.2
D4 68288906 68289078
refGene_gn=ABCA1;refGene_tx=XM_011288524.3;refGene_gn=ABCA1;refGene_tx=XM_019816256.2;refGene_gn=ABCA1;refGene_tx=XM_019816257.2
D4 68291193 68291307
refGene_gn=ABCA1;refGene_tx=XM_011288524.3;refGene_gn=ABCA1;refGene_tx=XM_019816256.2;refGene_gn=ABCA1;refGene_tx=XM_019816257.2
D4 68292371 68292576
refGene_gn=ABCA1;refGene_tx=XM_011288524.3;refGene_gn=ABCA1;refGene_tx=XM_019816256.2;refGene_gn=ABCA1;refGene_tx=XM_019816257.2
D4 68292987 68293209
refGene_gn=ABCA1;refGene_tx=XM_011288524.3;refGene_gn=ABCA1;refGene_tx=XM_019816256.2;refGene_gn=ABCA1;refGene_tx=XM_019816257.2
D4 68294930 68295153
refGene_gn=ABCA1;refGene_tx=XM_011288524.3;refGene_gn=ABCA1;refGene_tx=XM_019816256.2;refGene_gn=ABCA1;refGene_tx=XM_019816257.2
D4 68297114 68297291
refGene_gn=ABCA1;refGene_tx=XM_011288524.3;refGene_gn=ABCA1;refGene_tx=XM_019816256.2;refGene_gn=ABCA1;refGene_tx=XM_019816257.2
D4 68298246 68298452
refGene_gn=ABCA1;refGene_tx=XM_011288524.3;refGene_gn=ABCA1;refGene_tx=XM_019816256.2;refGene_gn=ABCA1;refGene_tx=XM_019816257.2
D4 68299260 68299458
refGene_gn=ABCA1;refGene_tx=XM_011288524.3;refGene_gn=ABCA1;refGene_tx=XM_019816256.2;refGene_gn=ABCA1;refGene_tx=XM_019816257.2
D4 68303916 68304033
refGene_gn=ABCA1;refGene_tx=XM_011288524.3;refGene_gn=ABCA1;refGene_tx=XM_019816256.2;refGene_gn=ABCA1;refGene_tx=XM_019816257.2
D4 68304364 68304504
refGene_gn=ABCA1;refGene_tx=XM_011288524.3;refGene_gn=ABCA1;refGene_tx=XM_019816256.2;refGene_gn=ABCA1;refGene_tx=XM_019816257.2
D4 68306255 68306496
refGene_gn=ABCA1;refGene_tx=XM_011288524.3;refGene_gn=ABCA1;refGene_tx=XM_019816256.2;refGene_gn=ABCA1;refGene_tx=XM_019816257.2
D4 68310888 68310981
refGene_gn=ABCA1;refGene_tx=XM_011288524.3;refGene_gn=ABCA1;refGene_tx=XM_019816256.2;refGene_gn=ABCA1;refGene_tx=XM_019816257.2
D4 68325026 68325203
refGene_gn=ABCA1;refGene_tx=XM_011288524.3;refGene_gn=ABCA1;refGene_tx=XM_019816256.2;refGene_gn=ABCA1;refGene_tx=XM_019816257.2
D4 68327232 68327354
refGene_gn=ABCA1;refGene_tx=XM_011288524.3;refGene_gn=ABCA1;refGene_tx=XM_019816256.2;refGene_gn=ABCA1;refGene_tx=XM_019816257.2
D4 68344167 68344286
refGene_gn=ABCA1;refGene_tx=XM_011288524.3;refGene_gn=ABCA1;refGene_tx=XM_019816256.2;refGene_gn=ABCA1;refGene_tx=XM_019816257.2
D4 68345574 68345716
refGene_gn=ABCA1;refGene_tx=XM_011288524.3;refGene_gn=ABCA1;refGene_tx=XM_019816256.2;refGene_gn=ABCA1;refGene_tx=XM_019816257.2
D4 68349961 68350055

refGene_gn=ABCA1;refGene_tx=XM_011288524.3;refGene_gn=ABCA1;refGene_tx=XM_019816256.2;refGene_gn=ABCA1;refGene_tx=XM_019816257.2
D4 68359798 68359864
refGene_gn=ABCA1;refGene_tx=XM_011288524.3;refGene_gn=ABCA1;refGene_tx=XM_019816256.2;refGene_gn=ABCA1;refGene_tx=XM_019816257.2
D4 68645929 68645965
refGene_gn=SLC44A1;refGene_tx=XM_023242528.1;refGene_gn=SLC44A1;refGene_tx=XM_023242529.1;refGene_gn=SLC44A1;refGene_tx=XM_023242530.1
D4 68700690 68700780
refGene_gn=SLC44A1;refGene_tx=XM_023242528.1;refGene_gn=SLC44A1;refGene_tx=XM_023242529.1;refGene_gn=SLC44A1;refGene_tx=XM_023242530.1
D4 68711836 68711979
refGene_gn=SLC44A1;refGene_tx=XM_023242528.1;refGene_gn=SLC44A1;refGene_tx=XM_023242529.1;refGene_gn=SLC44A1;refGene_tx=XM_023242530.1
D4 68739819 68739956
refGene_gn=SLC44A1;refGene_tx=XM_023242528.1;refGene_gn=SLC44A1;refGene_tx=XM_023242529.1;refGene_gn=SLC44A1;refGene_tx=XM_023242530.1
D4 68754642 68754736
refGene_gn=SLC44A1;refGene_tx=XM_023242528.1;refGene_gn=SLC44A1;refGene_tx=XM_023242529.1;refGene_gn=SLC44A1;refGene_tx=XM_023242530.1
D4 68762591 68762761
refGene_gn=SLC44A1;refGene_tx=XM_023242528.1;refGene_gn=SLC44A1;refGene_tx=XM_023242529.1;refGene_gn=SLC44A1;refGene_tx=XM_023242530.1
D4 68764711 68764801
refGene_gn=SLC44A1;refGene_tx=XM_023242528.1;refGene_gn=SLC44A1;refGene_tx=XM_023242529.1;refGene_gn=SLC44A1;refGene_tx=XM_023242530.1
D4 68770370 68770510
refGene_gn=SLC44A1;refGene_tx=XM_023242528.1;refGene_gn=SLC44A1;refGene_tx=XM_023242529.1;refGene_gn=SLC44A1;refGene_tx=XM_023242530.1
D4 68772404 68772591
refGene_gn=SLC44A1;refGene_tx=XM_023242528.1;refGene_gn=SLC44A1;refGene_tx=XM_023242529.1;refGene_gn=SLC44A1;refGene_tx=XM_023242530.1
D4 68773767 68773933
refGene_gn=SLC44A1;refGene_tx=XM_023242528.1;refGene_gn=SLC44A1;refGene_tx=XM_023242529.1;refGene_gn=SLC44A1;refGene_tx=XM_023242530.1
D4 68775478 68775635
refGene_gn=SLC44A1;refGene_tx=XM_023242528.1;refGene_gn=SLC44A1;refGene_tx=XM_023242529.1;refGene_gn=SLC44A1;refGene_tx=XM_023242530.1
D4 68776337 68776421
refGene_gn=SLC44A1;refGene_tx=XM_023242528.1;refGene_gn=SLC44A1;refGene_tx=XM_023242529.1;refGene_gn=SLC44A1;refGene_tx=XM_023242530.1
D4 68787301 68787439
refGene_gn=SLC44A1;refGene_tx=XM_023242528.1;refGene_gn=SLC44A1;refGene_tx=XM_023242529.1;refGene_gn=SLC44A1;refGene_tx=XM_023242530.1
D4 68795393 68795630
refGene_gn=SLC44A1;refGene_tx=XM_023242528.1;refGene_gn=SLC44A1;refGene_tx=XM_023242529.1;refGene_gn=SLC44A1;refGene_tx=XM_023242530.1
D4 68797644 68797935
refGene_gn=SLC44A1;refGene_tx=XM_023242528.1;refGene_gn=SLC44A1;refGene_tx=XM_023242529.1;refGene_gn=SLC44A1;refGene_tx=XM_023242530.1

D4 68800902 68800926 refGene_gn=SLC44A1;refGene_tx=XM_023242528.1
D4 68845865 68845880 refGene_gn=SLC44A1;refGene_tx=XM_023242529.1
D4 68849881 68849935 refGene_gn=FSD1L;refGene_tx=XM_019816275.2
D4 68850385 68850505
refGene_gn=FSD1L;refGene_tx=XM_006939343.4;refGene_gn=FSD1L;refGene_tx=XM_006939344.4;refGene_gn=FSD1L;refGene_tx=XM_019816273.2;refGene_gn=FSD1L;refGene_tx=XM_019816274.2
D4 68854742 68854757 refGene_gn=FSD1L;refGene_tx=XM_023242532.1
D4 68869290 68869386
refGene_gn=FSD1L;refGene_tx=XM_006939343.4;refGene_gn=FSD1L;refGene_tx=XM_006939344.4;refGene_gn=FSD1L;refGene_tx=XM_019816273.2;refGene_gn=FSD1L;refGene_tx=XM_019816274.2;refGene_gn=FSD1L;refGene_tx=XM_019816275.2;refGene_gn=FSD1L;refGene_tx=XM_023242532.1
D4 68871027 68871159
refGene_gn=FSD1L;refGene_tx=XM_006939343.4;refGene_gn=FSD1L;refGene_tx=XM_006939344.4;refGene_gn=FSD1L;refGene_tx=XM_019816273.2;refGene_gn=FSD1L;refGene_tx=XM_019816274.2;refGene_gn=FSD1L;refGene_tx=XM_019816275.2;refGene_gn=FSD1L;refGene_tx=XM_023242532.1
D4 68877145 68877247
refGene_gn=FSD1L;refGene_tx=XM_006939343.4;refGene_gn=FSD1L;refGene_tx=XM_006939344.4;refGene_gn=FSD1L;refGene_tx=XM_019816273.2;refGene_gn=FSD1L;refGene_tx=XM_019816274.2;refGene_gn=FSD1L;refGene_tx=XM_019816275.2;refGene_gn=FSD1L;refGene_tx=XM_023242532.1
D4 68886581 68886604
refGene_gn=FSD1L;refGene_tx=XM_006939343.4;refGene_gn=FSD1L;refGene_tx=XM_006939344.4;refGene_gn=FSD1L;refGene_tx=XM_019816273.2;refGene_gn=FSD1L;refGene_tx=XM_019816274.2;refGene_gn=FSD1L;refGene_tx=XM_019816275.2;refGene_gn=FSD1L;refGene_tx=XM_023242532.1
D4 68891140 68891262
refGene_gn=FSD1L;refGene_tx=XM_006939343.4;refGene_gn=FSD1L;refGene_tx=XM_006939344.4;refGene_gn=FSD1L;refGene_tx=XM_019816273.2;refGene_gn=FSD1L;refGene_tx=XM_019816274.2;refGene_gn=FSD1L;refGene_tx=XM_019816275.2;refGene_gn=FSD1L;refGene_tx=XM_023242532.1
D4 68893280 68893490
refGene_gn=FSD1L;refGene_tx=XM_006939343.4;refGene_gn=FSD1L;refGene_tx=XM_006939344.4;refGene_gn=FSD1L;refGene_tx=XM_019816273.2;refGene_gn=FSD1L;refGene_tx=XM_019816274.2;refGene_gn=FSD1L;refGene_tx=XM_019816275.2;refGene_gn=FSD1L;refGene_tx=XM_023242532.1
D4 68895465 68895564
refGene_gn=FSD1L;refGene_tx=XM_006939343.4;refGene_gn=FSD1L;refGene_tx=XM_006939344.4;refGene_gn=FSD1L;refGene_tx=XM_019816273.2;refGene_gn=FSD1L;refGene_tx=XM_019816274.2;refGene_gn=FSD1L;refGene_tx=XM_019816275.2;refGene_gn=FSD1L;refGene_tx=XM_023242532.1
D4 68900920 68901050
refGene_gn=FSD1L;refGene_tx=XM_006939343.4;refGene_gn=FSD1L;refGene_tx=XM_006939344.4;refGene_gn=FSD1L;refGene_tx=XM_019816273.2;refGene_gn=FSD1L;refGene_tx=XM_019816274.2;refGene_gn=FSD1L;refGene_tx=XM_019816275.2;refGene_gn=FSD1L;refGene_tx=XM_023242532.1
D4 68901668 68901701
refGene_gn=FSD1L;refGene_tx=XM_006939343.4;refGene_gn=FSD1L;refGene_tx

=XM_019816273.2;refGene_gn=FSD1L;refGene_tx=XM_019816275.2;refGene_gn=FSD1L;refGene_tx=XM_023242532.1
D4 68904767 68904868
refGene_gn=FSD1L;refGene_tx=XM_006939343.4;refGene_gn=FSD1L;refGene_tx=XM_006939344.4;refGene_gn=FSD1L;refGene_tx=XM_019816273.2;refGene_gn=FSD1L;refGene_tx=XM_019816274.2;refGene_gn=FSD1L;refGene_tx=XM_019816275.2;refGene_gn=FSD1L;refGene_tx=XM_023242532.1
D4 68905315 68905567
refGene_gn=FSD1L;refGene_tx=XM_006939343.4;refGene_gn=FSD1L;refGene_tx=XM_006939344.4;refGene_gn=FSD1L;refGene_tx=XM_019816273.2;refGene_gn=FSD1L;refGene_tx=XM_019816274.2;refGene_gn=FSD1L;refGene_tx=XM_019816275.2;refGene_gn=FSD1L;refGene_tx=XM_023242532.1
D4 68908863 68908952
refGene_gn=FSD1L;refGene_tx=XM_006939343.4;refGene_gn=FSD1L;refGene_tx=XM_006939344.4;refGene_gn=FSD1L;refGene_tx=XM_019816273.2;refGene_gn=FSD1L;refGene_tx=XM_019816274.2;refGene_gn=FSD1L;refGene_tx=XM_019816275.2;refGene_gn=FSD1L;refGene_tx=XM_023242532.1
D4 68922927 68923050
refGene_gn=FSD1L;refGene_tx=XM_006939343.4;refGene_gn=FSD1L;refGene_tx=XM_006939344.4;refGene_gn=FSD1L;refGene_tx=XM_019816273.2;refGene_gn=FSD1L;refGene_tx=XM_019816274.2;refGene_gn=FSD1L;refGene_tx=XM_019816275.2;refGene_gn=FSD1L;refGene_tx=XM_023242532.1
D4 68947435 68947540
refGene_gn=FKTN;refGene_tx=XM_011288523.3;refGene_gn=FKTN;refGene_tx=XM_023242533.1;refGene_gn=FKTN;refGene_tx=XM_023242534.1
D4 68958124 68958184
refGene_gn=FKTN;refGene_tx=XM_011288523.3;refGene_gn=FKTN;refGene_tx=XM_023242533.1
D4 68963948 68964152
refGene_gn=FKTN;refGene_tx=XM_011288523.3;refGene_gn=FKTN;refGene_tx=XM_023242533.1;refGene_gn=FKTN;refGene_tx=XM_023242534.1
D4 68970762 68971040
refGene_gn=FKTN;refGene_tx=XM_011288523.3;refGene_gn=FKTN;refGene_tx=XM_023242533.1;refGene_gn=FKTN;refGene_tx=XM_023242534.1
D4 68971693 68971826
refGene_gn=FKTN;refGene_tx=XM_011288523.3;refGene_gn=FKTN;refGene_tx=XM_023242533.1;refGene_gn=FKTN;refGene_tx=XM_023242534.1
D4 68978988 68979118
refGene_gn=FKTN;refGene_tx=XM_011288523.3;refGene_gn=FKTN;refGene_tx=XM_023242533.1;refGene_gn=FKTN;refGene_tx=XM_023242534.1
D4 68984030 68984164
refGene_gn=FKTN;refGene_tx=XM_011288523.3;refGene_gn=FKTN;refGene_tx=XM_023242533.1;refGene_gn=FKTN;refGene_tx=XM_023242534.1
D4 68985698 68985826
refGene_gn=FKTN;refGene_tx=XM_011288523.3;refGene_gn=FKTN;refGene_tx=XM_023242533.1;refGene_gn=FKTN;refGene_tx=XM_023242534.1
D4 69004911 69005009
refGene_gn=FKTN;refGene_tx=XM_011288523.3;refGene_gn=FKTN;refGene_tx=XM_023242533.1;refGene_gn=FKTN;refGene_tx=XM_023242534.1
D4 69014623 69014838

refGene_gn=FKTN;refGene_tx=XM_011288523.3;refGene_gn=FKTN;refGene_tx=XM_023242533.1;refGene_gn=FKTN;refGene_tx=XM_023242534.1
D4 69031510 69031828 refGene_gn=TAL2;refGene_tx=XM_023242536.1
D4 69063791 69063903 refGene_gn=TMEM38B;refGene_tx=XM_023242537.1
D4 69064318 69064462 refGene_gn=TMEM38B;refGene_tx=XM_006939345.4
D4 69074202 69074350 refGene_gn=TMEM38B;refGene_tx=XM_006939345.4;refGene_gn=TMEM38B;refGene_tx=XM_023242538.1
D4 69076358 69076515 refGene_gn=TMEM38B;refGene_tx=XM_006939345.4;refGene_gn=TMEM38B;refGene_tx=XM_023242537.1;refGene_gn=TMEM38B;refGene_tx=XM_023242538.1
D4 69089239 69089424 refGene_gn=TMEM38B;refGene_tx=XM_006939345.4;refGene_gn=TMEM38B;refGene_tx=XM_023242537.1;refGene_gn=TMEM38B;refGene_tx=XM_023242538.1
D4 69090236 69090324 refGene_gn=TMEM38B;refGene_tx=XM_006939345.4;refGene_gn=TMEM38B;refGene_tx=XM_023242537.1;refGene_gn=TMEM38B;refGene_tx=XM_023242538.1
D4 69094001 69094119 refGene_gn=TMEM38B;refGene_tx=XM_006939345.4;refGene_gn=TMEM38B;refGene_tx=XM_023242537.1;refGene_gn=TMEM38B;refGene_tx=XM_023242538.1
D4 69113897 69114110 refGene_gn=TMEM38B;refGene_tx=XM_006939345.4;refGene_gn=TMEM38B;refGene_tx=XM_023242537.1;refGene_gn=TMEM38B;refGene_tx=XM_023242538.1
D4 70055060 70055280 refGene_gn=ZNF462;refGene_tx=XM_019816283.2;refGene_gn=ZNF462;refGene_tx=XM_023242539.1;refGene_gn=ZNF462;refGene_tx=XM_023242541.1
D4 70055823 70061441 refGene_gn=ZNF462;refGene_tx=XM_019816283.2;refGene_gn=ZNF462;refGene_tx=XM_023242539.1;refGene_gn=ZNF462;refGene_tx=XM_023242541.1
D4 70062212 70062377 refGene_gn=ZNF462;refGene_tx=XM_019816283.2;refGene_gn=ZNF462;refGene_tx=XM_023242539.1;refGene_gn=ZNF462;refGene_tx=XM_023242541.1
D4 70064143 70064247 refGene_gn=ZNF462;refGene_tx=XM_019816283.2;refGene_gn=ZNF462;refGene_tx=XM_023242539.1;refGene_gn=ZNF462;refGene_tx=XM_023242541.1
D4 70067238 70067357 refGene_gn=ZNF462;refGene_tx=XM_019816283.2;refGene_gn=ZNF462;refGene_tx=XM_023242539.1;refGene_gn=ZNF462;refGene_tx=XM_023242541.1
D4 70071460 70071652 refGene_gn=ZNF462;refGene_tx=XM_019816283.2;refGene_gn=ZNF462;refGene_tx=XM_023242539.1;refGene_gn=ZNF462;refGene_tx=XM_023242541.1
D4 70103327 70103595 refGene_gn=ZNF462;refGene_tx=XM_019816283.2;refGene_gn=ZNF462;refGene_tx=XM_023242539.1;refGene_gn=ZNF462;refGene_tx=XM_023242541.1
D4 70105515 70105652 refGene_gn=ZNF462;refGene_tx=XM_019816283.2;refGene_gn=ZNF462;refGene_tx=XM_023242539.1;refGene_gn=ZNF462;refGene_tx=XM_023242541.1
D4 70114788 70115012 refGene_gn=ZNF462;refGene_tx=XM_019816283.2;refGene_gn=ZNF462;refGene_tx=XM_023242539.1;refGene_gn=ZNF462;refGene_tx=XM_023242541.1

D4 70129088 70129221
refGene_gn=ZNF462;refGene_tx=XM_019816283.2;refGene_gn=ZNF462;refGene_tx=XM_023242539.1;refGene_gn=ZNF462;refGene_tx=XM_023242541.1
D4 70133577 70133701
refGene_gn=ZNF462;refGene_tx=XM_019816283.2;refGene_gn=ZNF462;refGene_tx=XM_023242539.1;refGene_gn=ZNF462;refGene_tx=XM_023242541.1
D4 70134865 70135073
refGene_gn=ZNF462;refGene_tx=XM_019816283.2;refGene_gn=ZNF462;refGene_tx=XM_023242539.1;refGene_gn=ZNF462;refGene_tx=XM_023242541.1
D4 70347140 70347206 refGene_gn=RAD23B;refGene_tx=XM_003995729.5
D4 70366296 70366378 refGene_gn=RAD23B;refGene_tx=XM_003995729.5
D4 70366672 70366752 refGene_gn=RAD23B;refGene_tx=XM_003995729.5
D4 70370153 70370422 refGene_gn=RAD23B;refGene_tx=XM_003995729.5
D4 70375449 70375505 refGene_gn=RAD23B;refGene_tx=XM_003995729.5
D4 70380268 70380396 refGene_gn=RAD23B;refGene_tx=XM_003995729.5
D4 70384211 70384344 refGene_gn=RAD23B;refGene_tx=XM_003995729.5
D4 70386084 70386212 refGene_gn=RAD23B;refGene_tx=XM_003995729.5
D4 70386887 70387058 refGene_gn=RAD23B;refGene_tx=XM_003995729.5
D4 70392182 70392296 refGene_gn=RAD23B;refGene_tx=XM_003995729.5
D4 70502020 70502196
refGene_gn=KLF4;refGene_tx=NM_001173444.2;refGene_gn=KLF4;refGene_tx=XM_023242190.1
D4 70503626 70504881
refGene_gn=KLF4;refGene_tx=NM_001173444.2;refGene_gn=KLF4;refGene_tx=XM_023242190.1
D4 70505553 70505674
refGene_gn=KLF4;refGene_tx=NM_001173444.2;refGene_gn=KLF4;refGene_tx=XM_023242190.1
D4 70505791 70505796
refGene_gn=KLF4;refGene_tx=NM_001173444.2;refGene_gn=KLF4;refGene_tx=XM_023242190.1
D4 71592740 71592837
refGene_gn=LOC101088887;refGene_tx=XM_019815542.1
D4 71604540 71605001
refGene_gn=LOC101088887;refGene_tx=XM_019815542.1
D4 71657867 71659121 refGene_gn=ACTL7B;refGene_tx=XM_019816285.2
D4 71664335 71665658 refGene_gn=ACTL7A;refGene_tx=XM_003995735.5
D4 71671224 71671292
refGene_gn=ELP1;refGene_tx=XM_006939347.4;refGene_gn=ELP1;refGene_tx=XM_011288516.3
D4 71674578 71674654
refGene_gn=ELP1;refGene_tx=XM_006939347.4;refGene_gn=ELP1;refGene_tx=XM_011288516.3
D4 71677203 71677358
refGene_gn=ELP1;refGene_tx=XM_006939347.4;refGene_gn=ELP1;refGene_tx=XM_011288516.3
D4 71677818 71677946
refGene_gn=ELP1;refGene_tx=XM_006939347.4;refGene_gn=ELP1;refGene_tx=XM_011288516.3
D4 71679245 71679357

refGene_gn=ELP1;refGene_tx=XM_006939347.4;refGene_gn=ELP1;refGene_tx=XM_011288516.3
D4 71679880 71679994
refGene_gn=ELP1;refGene_tx=XM_006939347.4;refGene_gn=ELP1;refGene_tx=XM_011288516.3
D4 71681515 71681576
refGene_gn=ELP1;refGene_tx=XM_006939347.4;refGene_gn=ELP1;refGene_tx=XM_011288516.3
D4 71681948 71682011
refGene_gn=ELP1;refGene_tx=XM_006939347.4;refGene_gn=ELP1;refGene_tx=XM_011288516.3
D4 71688895 71688957
refGene_gn=ELP1;refGene_tx=XM_006939347.4;refGene_gn=ELP1;refGene_tx=XM_011288516.3
D4 71690717 71690919
refGene_gn=ELP1;refGene_tx=XM_006939347.4;refGene_gn=ELP1;refGene_tx=XM_011288516.3
D4 71692668 71692766
refGene_gn=ELP1;refGene_tx=XM_006939347.4;refGene_gn=ELP1;refGene_tx=XM_011288516.3
D4 71693647 71693771
refGene_gn=ELP1;refGene_tx=XM_006939347.4;refGene_gn=ELP1;refGene_tx=XM_011288516.3
D4 71695998 71696147
refGene_gn=ELP1;refGene_tx=XM_006939347.4;refGene_gn=ELP1;refGene_tx=XM_011288516.3
D4 71696459 71696545
refGene_gn=ELP1;refGene_tx=XM_006939347.4;refGene_gn=ELP1;refGene_tx=XM_011288516.3
D4 71696668 71696806
refGene_gn=ELP1;refGene_tx=XM_006939347.4;refGene_gn=ELP1;refGene_tx=XM_011288516.3
D4 71698201 71698281
refGene_gn=ELP1;refGene_tx=XM_006939347.4;refGene_gn=ELP1;refGene_tx=XM_011288516.3
D4 71698378 71698457
refGene_gn=ELP1;refGene_tx=XM_006939347.4;refGene_gn=ELP1;refGene_tx=XM_011288516.3
D4 71699187 71699261
refGene_gn=ELP1;refGene_tx=XM_006939347.4;refGene_gn=ELP1;refGene_tx=XM_011288516.3
D4 71699593 71699709
refGene_gn=ELP1;refGene_tx=XM_006939347.4;refGene_gn=ELP1;refGene_tx=XM_011288516.3
D4 71700535 71700641
refGene_gn=ELP1;refGene_tx=XM_006939347.4;refGene_gn=ELP1;refGene_tx=XM_011288516.3
D4 71700725 71700779
refGene_gn=ELP1;refGene_tx=XM_006939347.4;refGene_gn=ELP1;refGene_tx=XM_011288516.3

D4 71701900 71702019
refGene_gn=ELP1;refGene_tx=XM_006939347.4;refGene_gn=ELP1;refGene_tx=XM_011288516.3
D4 71702324 71702431
refGene_gn=ELP1;refGene_tx=XM_006939347.4;refGene_gn=ELP1;refGene_tx=XM_011288516.3
D4 71704932 71705115
refGene_gn=ELP1;refGene_tx=XM_006939347.4;refGene_gn=ELP1;refGene_tx=XM_011288516.3
D4 71707275 71707375
refGene_gn=ELP1;refGene_tx=XM_006939347.4;refGene_gn=ELP1;refGene_tx=XM_011288516.3
D4 71708169 71708340
refGene_gn=ELP1;refGene_tx=XM_006939347.4;refGene_gn=ELP1;refGene_tx=XM_011288516.3
D4 71709389 71709620
refGene_gn=ELP1;refGene_tx=XM_006939347.4;refGene_gn=ELP1;refGene_tx=XM_011288516.3
D4 71713313 71713407
refGene_gn=ELP1;refGene_tx=XM_006939347.4;refGene_gn=ELP1;refGene_tx=XM_011288516.3
D4 71714696 71714820
refGene_gn=ELP1;refGene_tx=XM_006939347.4;refGene_gn=ELP1;refGene_tx=XM_011288516.3
D4 71716118 71716209
refGene_gn=ELP1;refGene_tx=XM_006939347.4;refGene_gn=ELP1;refGene_tx=XM_011288516.3
D4 71716529 71716626
refGene_gn=ELP1;refGene_tx=XM_006939347.4;refGene_gn=ELP1;refGene_tx=XM_011288516.3
D4 71718311 71718397
refGene_gn=ELP1;refGene_tx=XM_006939347.4;refGene_gn=ELP1;refGene_tx=XM_011288516.3
D4 71721630 71721711
refGene_gn=ELP1;refGene_tx=XM_006939347.4;refGene_gn=ELP1;refGene_tx=XM_011288516.3
D4 71722450 71722532
refGene_gn=ELP1;refGene_tx=XM_006939347.4;refGene_gn=ELP1;refGene_tx=XM_011288516.3
D4 71722753 71722906
refGene_gn=ELP1;refGene_tx=XM_006939347.4;refGene_gn=ELP1;refGene_tx=XM_011288516.3
D4 71723716 71723866
refGene_gn=ELP1;refGene_tx=XM_006939347.4;refGene_gn=ELP1;refGene_tx=XM_011288516.3
D4 71728239 71728318 refGene_gn=FAM206A;refGene_tx=XM_003995736.5
D4 71729650 71729742
refGene_gn=FAM206A;refGene_tx=XM_003995736.5;refGene_gn=FAM206A;refGene_tx=XM_006939348.3;refGene_gn=FAM206A;refGene_tx=XM_019816287.2
D4 71730450 71730580

refGene_gn=FAM206A;refGene_tx=XM_003995736.5;refGene_gn=FAM206A;refGene_tx=XM_006939348.3;refGene_gn=FAM206A;refGene_tx=XM_019816287.2
D4 71733243 71733320
refGene_gn=FAM206A;refGene_tx=XM_003995736.5;refGene_gn=FAM206A;refGene_tx=XM_006939348.3;refGene_gn=FAM206A;refGene_tx=XM_019816287.2
D4 71733400 71733470
refGene_gn=FAM206A;refGene_tx=XM_003995736.5;refGene_gn=FAM206A;refGene_tx=XM_006939348.3;refGene_gn=FAM206A;refGene_tx=XM_019816287.2
D4 71733563 71733701
refGene_gn=FAM206A;refGene_tx=XM_003995736.5;refGene_gn=FAM206A;refGene_tx=XM_006939348.3;refGene_gn=FAM206A;refGene_tx=XM_019816287.2
D4 71736530 71736596 refGene_gn=CTNNAL1;refGene_tx=XM_023242544.1
D4 71736732 71736816 refGene_gn=CTNNAL1;refGene_tx=XM_023242544.1
D4 71737482 71737596 refGene_gn=CTNNAL1;refGene_tx=XM_023242544.1
D4 71737741 71737798 refGene_gn=CTNNAL1;refGene_tx=XM_023242544.1
D4 71740568 71740617 refGene_gn=CTNNAL1;refGene_tx=XM_023242544.1
D4 71744112 71744267 refGene_gn=CTNNAL1;refGene_tx=XM_023242544.1
D4 71744359 71744410 refGene_gn=CTNNAL1;refGene_tx=XM_023242544.1
D4 71746126 71746164 refGene_gn=CTNNAL1;refGene_tx=XM_023242544.1
D4 71751897 71752048 refGene_gn=CTNNAL1;refGene_tx=XM_023242544.1
D4 71755823 71755916 refGene_gn=CTNNAL1;refGene_tx=XM_023242544.1
D4 71757509 71757668 refGene_gn=CTNNAL1;refGene_tx=XM_023242544.1
D4 71760716 71760803 refGene_gn=CTNNAL1;refGene_tx=XM_023242544.1
D4 71763302 71763503 refGene_gn=CTNNAL1;refGene_tx=XM_023242544.1
D4 71767068 71767239 refGene_gn=CTNNAL1;refGene_tx=XM_023242544.1
D4 71768240 71768330 refGene_gn=CTNNAL1;refGene_tx=XM_023242544.1
D4 71770400 71770520 refGene_gn=CTNNAL1;refGene_tx=XM_023242544.1
D4 71772169 71772357 refGene_gn=CTNNAL1;refGene_tx=XM_023242544.1
D4 71780733 71780923 refGene_gn=CTNNAL1;refGene_tx=XM_023242544.1
D4 71800184 71800307 refGene_gn=CTNNAL1;refGene_tx=XM_023242544.1
D4 71807444 71807490 refGene_gn=TMEM245;refGene_tx=XM_003995737.5
D4 71812344 71812813
refGene_gn=TMEM245;refGene_tx=XM_023242545.1;refGene_gn=TMEM245;refGene_tx=XM_023242546.1;refGene_gn=TMEM245;refGene_tx=XM_023242547.1
D4 71820478 71820673
refGene_gn=TMEM245;refGene_tx=XM_003995737.5;refGene_gn=TMEM245;refGene_tx=XM_023242545.1;refGene_gn=TMEM245;refGene_tx=XM_023242546.1;refGene_gn=TMEM245;refGene_tx=XM_023242547.1
D4 71822922 71823097
refGene_gn=TMEM245;refGene_tx=XM_003995737.5;refGene_gn=TMEM245;refGene_tx=XM_023242545.1;refGene_gn=TMEM245;refGene_tx=XM_023242546.1;refGene_gn=TMEM245;refGene_tx=XM_023242547.1
D4 71824949 71825050
refGene_gn=TMEM245;refGene_tx=XM_003995737.5;refGene_gn=TMEM245;refGene_tx=XM_023242545.1;refGene_gn=TMEM245;refGene_tx=XM_023242546.1;refGene_gn=TMEM245;refGene_tx=XM_023242547.1
D4 71838250 71838396
refGene_gn=TMEM245;refGene_tx=XM_003995737.5;refGene_gn=TMEM245;refGene_tx=XM_023242545.1;refGene_gn=TMEM245;refGene_tx=XM_023242546.1;refGene_gn=TMEM245;refGene_tx=XM_023242547.1

D4 71838525 71838648
refGene_gn=TMEM245;refGene_tx=XM_003995737.5;refGene_gn=TMEM245;refGene_tx=XM_023242545.1;refGene_gn=TMEM245;refGene_tx=XM_023242546.1;refGene_gn=TMEM245;refGene_tx=XM_023242547.1
D4 71842374 71842506
refGene_gn=TMEM245;refGene_tx=XM_003995737.5;refGene_gn=TMEM245;refGene_tx=XM_023242545.1;refGene_gn=TMEM245;refGene_tx=XM_023242546.1;refGene_gn=TMEM245;refGene_tx=XM_023242547.1
D4 71844692 71844791
refGene_gn=TMEM245;refGene_tx=XM_003995737.5;refGene_gn=TMEM245;refGene_tx=XM_023242545.1;refGene_gn=TMEM245;refGene_tx=XM_023242546.1;refGene_gn=TMEM245;refGene_tx=XM_023242547.1
D4 71851757 71851848
refGene_gn=TMEM245;refGene_tx=XM_003995737.5;refGene_gn=TMEM245;refGene_tx=XM_023242545.1;refGene_gn=TMEM245;refGene_tx=XM_023242546.1;refGene_gn=TMEM245;refGene_tx=XM_023242547.1
D4 71857298 71857381
refGene_gn=TMEM245;refGene_tx=XM_003995737.5;refGene_gn=TMEM245;refGene_tx=XM_023242545.1;refGene_gn=TMEM245;refGene_tx=XM_023242546.1;refGene_gn=TMEM245;refGene_tx=XM_023242547.1
D4 71863282 71863387
refGene_gn=TMEM245;refGene_tx=XM_003995737.5;refGene_gn=TMEM245;refGene_tx=XM_023242545.1;refGene_gn=TMEM245;refGene_tx=XM_023242546.1;refGene_gn=TMEM245;refGene_tx=XM_023242547.1
D4 71869179 71869203
refGene_gn=TMEM245;refGene_tx=XM_003995737.5;refGene_gn=TMEM245;refGene_tx=XM_023242545.1;refGene_gn=TMEM245;refGene_tx=XM_023242547.1
D4 71872147 71872317
refGene_gn=TMEM245;refGene_tx=XM_003995737.5;refGene_gn=TMEM245;refGene_tx=XM_023242545.1;refGene_gn=TMEM245;refGene_tx=XM_023242546.1;refGene_gn=TMEM245;refGene_tx=XM_023242547.1
D4 71876064 71876283
refGene_gn=TMEM245;refGene_tx=XM_003995737.5;refGene_gn=TMEM245;refGene_tx=XM_023242545.1;refGene_gn=TMEM245;refGene_tx=XM_023242546.1;refGene_gn=TMEM245;refGene_tx=XM_023242547.1
D4 71878795 71878912
refGene_gn=TMEM245;refGene_tx=XM_003995737.5;refGene_gn=TMEM245;refGene_tx=XM_023242545.1;refGene_gn=TMEM245;refGene_tx=XM_023242546.1
D4 71891732 71891834
refGene_gn=TMEM245;refGene_tx=XM_003995737.5;refGene_gn=TMEM245;refGene_tx=XM_023242545.1;refGene_gn=TMEM245;refGene_tx=XM_023242546.1;refGene_gn=TMEM245;refGene_tx=XM_023242547.1
D4 71893795 71893913
refGene_gn=TMEM245;refGene_tx=XM_003995737.5;refGene_gn=TMEM245;refGene_tx=XM_023242545.1;refGene_gn=TMEM245;refGene_tx=XM_023242546.1;refGene_gn=TMEM245;refGene_tx=XM_023242547.1
D4 71904255 71904840
refGene_gn=TMEM245;refGene_tx=XM_003995737.5;refGene_gn=TMEM245;refGene_tx=XM_023242545.1;refGene_gn=TMEM245;refGene_tx=XM_023242546.1;refGene_gn=TMEM245;refGene_tx=XM_023242547.1

D4 71918161 71918334 refGene_gn=FRRS1L;refGene_tx=XM_023242308.1
D4 71921839 71922086 refGene_gn=FRRS1L;refGene_tx=XM_023242308.1
D4 71924626 71924765 refGene_gn=FRRS1L;refGene_tx=XM_023242308.1
D4 71926834 71926919 refGene_gn=FRRS1L;refGene_tx=XM_023242308.1
D4 71937715 71938010 refGene_gn=FRRS1L;refGene_tx=XM_023242308.1
D4 71938011 71938134 refGene_gn=FRRS1L;refGene_tx=XM_023242308.1
D4 71945391 71945461 refGene_gn=EPB41L4B;refGene_tx=XM_023242549.1
D4 71946414 71946560 refGene_gn=EPB41L4B;refGene_tx=XM_023242549.1
D4 71949098 71949167 refGene_gn=EPB41L4B;refGene_tx=XM_023242549.1
D4 71951786 71951903 refGene_gn=EPB41L4B;refGene_tx=XM_023242549.1
D4 71958132 71958210 refGene_gn=EPB41L4B;refGene_tx=XM_023242549.1
D4 71960669 71960681 refGene_gn=EPB41L4B;refGene_tx=XM_023242549.1
D4 71960682 71960848 refGene_gn=EPB41L4B;refGene_tx=XM_023242549.1
D4 71966199 71966298 refGene_gn=EPB41L4B;refGene_tx=XM_023242549.1
D4 71968931 71968999 refGene_gn=EPB41L4B;refGene_tx=XM_023242549.1
D4 71969811 71969937 refGene_gn=EPB41L4B;refGene_tx=XM_023242549.1
D4 71976841 71976960 refGene_gn=EPB41L4B;refGene_tx=XM_023242549.1
D4 71979117 71979332 refGene_gn=EPB41L4B;refGene_tx=XM_023242549.1
D4 72001664 72001729 refGene_gn=EPB41L4B;refGene_tx=XM_023242549.1
D4 72005466 72005500 refGene_gn=EPB41L4B;refGene_tx=XM_023242549.1
D4 72008854 72008885 refGene_gn=EPB41L4B;refGene_tx=XM_023242549.1
D4 72011833 72011943 refGene_gn=EPB41L4B;refGene_tx=XM_023242549.1
D4 72013723 72013893 refGene_gn=EPB41L4B;refGene_tx=XM_023242549.1
D4 72013990 72014060 refGene_gn=EPB41L4B;refGene_tx=XM_023242549.1
D4 72014351 72014440 refGene_gn=EPB41L4B;refGene_tx=XM_023242549.1
D4 72014607 72014695 refGene_gn=EPB41L4B;refGene_tx=XM_023242549.1
D4 72015734 72015855 refGene_gn=EPB41L4B;refGene_tx=XM_023242549.1
D4 72020500 72020553 refGene_gn=EPB41L4B;refGene_tx=XM_023242549.1
D4 72022378 72022423 refGene_gn=EPB41L4B;refGene_tx=XM_023242549.1
D4 72024609 72024688 refGene_gn=EPB41L4B;refGene_tx=XM_023242549.1
D4 72025534 72025577 refGene_gn=EPB41L4B;refGene_tx=XM_023242549.1
D4 72036953 72037058 refGene_gn=EPB41L4B;refGene_tx=XM_023242549.1
D4 72075978 72076287 refGene_gn=EPB41L4B;refGene_tx=XM_023242549.1
D4 72118825 72118903
refGene_gn=PTPN3;refGene_tx=XM_011288512.3;refGene_gn=PTPN3;refGene_tx=XM_011288513.3;refGene_gn=PTPN3;refGene_tx=XM_011288514.3;refGene_gn=PTPN3;refGene_tx=XM_019816304.2;refGene_gn=PTPN3;refGene_tx=XM_019816305.2;refGene_gn=PTPN3;refGene_tx=XM_023242550.1
D4 72120590 72120726
refGene_gn=PTPN3;refGene_tx=XM_011288512.3;refGene_gn=PTPN3;refGene_tx=XM_011288513.3;refGene_gn=PTPN3;refGene_tx=XM_011288514.3;refGene_gn=PTPN3;refGene_tx=XM_019816304.2;refGene_gn=PTPN3;refGene_tx=XM_019816305.2;refGene_gn=PTPN3;refGene_tx=XM_023242550.1
D4 72121241 72121387
refGene_gn=PTPN3;refGene_tx=XM_011288512.3;refGene_gn=PTPN3;refGene_tx=XM_011288513.3;refGene_gn=PTPN3;refGene_tx=XM_011288514.3;refGene_gn=PTPN3;refGene_tx=XM_019816304.2;refGene_gn=PTPN3;refGene_tx=XM_019816305.2;refGene_gn=PTPN3;refGene_tx=XM_023242550.1
D4 72122236 72122365
refGene_gn=PTPN3;refGene_tx=XM_011288512.3;refGene_gn=PTPN3;refGene_tx

=XM_011288513.3;refGene_gn=PTPN3;refGene_tx=XM_011288514.3;refGene_gn=PTPN3;refGene_tx=XM_019816304.2;refGene_gn=PTPN3;refGene_tx=XM_019816305.2;refGene_gn=PTPN3;refGene_tx=XM_023242550.1
D4 72127570 72127717
refGene_gn=PTPN3;refGene_tx=XM_011288512.3;refGene_gn=PTPN3;refGene_tx=XM_011288513.3;refGene_gn=PTPN3;refGene_tx=XM_011288514.3;refGene_gn=PTPN3;refGene_tx=XM_019816304.2;refGene_gn=PTPN3;refGene_tx=XM_019816305.2;refGene_gn=PTPN3;refGene_tx=XM_023242550.1
D4 72129589 72129651
refGene_gn=PTPN3;refGene_tx=XM_011288512.3;refGene_gn=PTPN3;refGene_tx=XM_011288513.3;refGene_gn=PTPN3;refGene_tx=XM_011288514.3;refGene_gn=PTPN3;refGene_tx=XM_019816304.2;refGene_gn=PTPN3;refGene_tx=XM_019816305.2;refGene_gn=PTPN3;refGene_tx=XM_023242550.1
D4 72129940 72130031
refGene_gn=PTPN3;refGene_tx=XM_011288512.3;refGene_gn=PTPN3;refGene_tx=XM_011288513.3;refGene_gn=PTPN3;refGene_tx=XM_011288514.3;refGene_gn=PTPN3;refGene_tx=XM_019816304.2;refGene_gn=PTPN3;refGene_tx=XM_019816305.2;refGene_gn=PTPN3;refGene_tx=XM_023242550.1
D4 72142202 72142363
refGene_gn=PTPN3;refGene_tx=XM_011288512.3;refGene_gn=PTPN3;refGene_tx=XM_011288513.3;refGene_gn=PTPN3;refGene_tx=XM_011288514.3;refGene_gn=PTPN3;refGene_tx=XM_019816304.2;refGene_gn=PTPN3;refGene_tx=XM_019816305.2;refGene_gn=PTPN3;refGene_tx=XM_023242550.1
D4 72145255 72145412
refGene_gn=PTPN3;refGene_tx=XM_011288512.3;refGene_gn=PTPN3;refGene_tx=XM_011288513.3;refGene_gn=PTPN3;refGene_tx=XM_011288514.3;refGene_gn=PTPN3;refGene_tx=XM_019816304.2;refGene_gn=PTPN3;refGene_tx=XM_019816305.2;refGene_gn=PTPN3;refGene_tx=XM_023242550.1
D4 72147138 72147195
refGene_gn=PTPN3;refGene_tx=XM_011288512.3;refGene_gn=PTPN3;refGene_tx=XM_011288513.3;refGene_gn=PTPN3;refGene_tx=XM_011288514.3;refGene_gn=PTPN3;refGene_tx=XM_019816304.2;refGene_gn=PTPN3;refGene_tx=XM_019816305.2;refGene_gn=PTPN3;refGene_tx=XM_023242550.1
D4 72148274 72148352
refGene_gn=PTPN3;refGene_tx=XM_011288512.3;refGene_gn=PTPN3;refGene_tx=XM_011288513.3;refGene_gn=PTPN3;refGene_tx=XM_011288514.3;refGene_gn=PTPN3;refGene_tx=XM_019816304.2;refGene_gn=PTPN3;refGene_tx=XM_019816305.2;refGene_gn=PTPN3;refGene_tx=XM_023242550.1
D4 72148507 72148694
refGene_gn=PTPN3;refGene_tx=XM_011288512.3;refGene_gn=PTPN3;refGene_tx=XM_011288513.3;refGene_gn=PTPN3;refGene_tx=XM_011288514.3;refGene_gn=PTPN3;refGene_tx=XM_019816304.2;refGene_gn=PTPN3;refGene_tx=XM_019816305.2;refGene_gn=PTPN3;refGene_tx=XM_023242550.1
D4 72157803 72157980
refGene_gn=PTPN3;refGene_tx=XM_011288512.3;refGene_gn=PTPN3;refGene_tx=XM_011288513.3;refGene_gn=PTPN3;refGene_tx=XM_011288514.3;refGene_gn=PTPN3;refGene_tx=XM_019816304.2;refGene_gn=PTPN3;refGene_tx=XM_019816305.2;refGene_gn=PTPN3;refGene_tx=XM_023242550.1
D4 72160791 72160926
refGene_gn=PTPN3;refGene_tx=XM_011288512.3;refGene_gn=PTPN3;refGene_tx

=XM_011288513.3;refGene_gn=PTPN3;refGene_tx=XM_011288514.3;refGene_gn=PTPN3;refGene_tx=XM_019816304.2;refGene_gn=PTPN3;refGene_tx=XM_019816305.2;refGene_gn=PTPN3;refGene_tx=XM_023242550.1
D4 72164539 72164712
refGene_gn=PTPN3;refGene_tx=XM_011288512.3;refGene_gn=PTPN3;refGene_tx=XM_011288513.3;refGene_gn=PTPN3;refGene_tx=XM_011288514.3;refGene_gn=PTPN3;refGene_tx=XM_019816304.2;refGene_gn=PTPN3;refGene_tx=XM_019816305.2;refGene_gn=PTPN3;refGene_tx=XM_023242550.1
D4 72166232 72166296
refGene_gn=PTPN3;refGene_tx=XM_011288512.3;refGene_gn=PTPN3;refGene_tx=XM_011288513.3;refGene_gn=PTPN3;refGene_tx=XM_011288514.3;refGene_gn=PTPN3;refGene_tx=XM_019816304.2;refGene_gn=PTPN3;refGene_tx=XM_019816305.2;refGene_gn=PTPN3;refGene_tx=XM_023242550.1
D4 72167291 72167380
refGene_gn=PTPN3;refGene_tx=XM_011288512.3;refGene_gn=PTPN3;refGene_tx=XM_011288513.3;refGene_gn=PTPN3;refGene_tx=XM_011288514.3;refGene_gn=PTPN3;refGene_tx=XM_019816304.2;refGene_gn=PTPN3;refGene_tx=XM_019816305.2;refGene_gn=PTPN3;refGene_tx=XM_023242550.1
D4 72171836 72171924
refGene_gn=PTPN3;refGene_tx=XM_011288512.3;refGene_gn=PTPN3;refGene_tx=XM_019816304.2;refGene_gn=PTPN3;refGene_tx=XM_019816305.2;refGene_gn=PTPN3;refGene_tx=XM_023242550.1
D4 72173055 72173176
refGene_gn=PTPN3;refGene_tx=XM_011288512.3;refGene_gn=PTPN3;refGene_tx=XM_019816304.2;refGene_gn=PTPN3;refGene_tx=XM_019816305.2;refGene_gn=PTPN3;refGene_tx=XM_023242550.1
D4 72179640 72179693
refGene_gn=PTPN3;refGene_tx=XM_011288512.3;refGene_gn=PTPN3;refGene_tx=XM_019816304.2;refGene_gn=PTPN3;refGene_tx=XM_019816305.2;refGene_gn=PTPN3;refGene_tx=XM_023242550.1
D4 72183340 72183385
refGene_gn=PTPN3;refGene_tx=XM_011288512.3;refGene_gn=PTPN3;refGene_tx=XM_019816304.2;refGene_gn=PTPN3;refGene_tx=XM_019816305.2;refGene_gn=PTPN3;refGene_tx=XM_023242550.1
D4 72187968 72188047
refGene_gn=PTPN3;refGene_tx=XM_019816304.2;refGene_gn=PTPN3;refGene_tx=XM_023242550.1
D4 72190607 72190650
refGene_gn=PTPN3;refGene_tx=XM_019816304.2;refGene_gn=PTPN3;refGene_tx=XM_023242550.1
D4 72190727 72190835
refGene_gn=PTPN3;refGene_tx=XM_019816304.2;refGene_gn=PTPN3;refGene_tx=XM_023242550.1
D4 72196435 72196590
refGene_gn=PTPN3;refGene_tx=XM_019816304.2;refGene_gn=PTPN3;refGene_tx=XM_023242550.1
D4 72265117 72265166 refGene_gn=PTPN3;refGene_tx=XM_019816304.2
D4 72344769 72344774
refGene_gn=LOC105261111;refGene_tx=XM_019816297.2;refGene_gn=LOC105261111;refGene_tx=XM_023242552.1;refGene_gn=LOC105261111;refGene_tx=XM_02

3242553.1;refGene_gn=LOC105261111;refGene_tx=XM_023242554.1;refGene_gn=LOC105261111;refGene_tx=XM_023242555.1;refGene_gn=LOC105261111;refGene_tx=XM_023242556.1
D4 72463862 72463908
refGene_gn=LOC105261111;refGene_tx=XM_019816297.2;refGene_gn=LOC105261111;refGene_tx=XM_023242552.1;refGene_gn=LOC105261111;refGene_tx=XM_023242553.1;refGene_gn=LOC105261111;refGene_tx=XM_023242554.1;refGene_gn=LOC105261111;refGene_tx=XM_023242555.1;refGene_gn=LOC105261111;refGene_tx=XM_023242556.1
D4 72530981 72531062
refGene_gn=LOC105261111;refGene_tx=XM_019816297.2;refGene_gn=LOC105261111;refGene_tx=XM_023242552.1;refGene_gn=LOC105261111;refGene_tx=XM_023242553.1;refGene_gn=LOC105261111;refGene_tx=XM_023242554.1;refGene_gn=LOC105261111;refGene_tx=XM_023242555.1;refGene_gn=LOC105261111;refGene_tx=XM_023242556.1
D4 72544462 72544593
refGene_gn=LOC105261111;refGene_tx=XM_019816297.2;refGene_gn=LOC105261111;refGene_tx=XM_023242552.1;refGene_gn=LOC105261111;refGene_tx=XM_023242553.1;refGene_gn=LOC105261111;refGene_tx=XM_023242554.1;refGene_gn=LOC105261111;refGene_tx=XM_023242555.1;refGene_gn=LOC105261111;refGene_tx=XM_023242556.1
D4 72584183 72584298
refGene_gn=LOC105261111;refGene_tx=XM_019816297.2;refGene_gn=LOC105261111;refGene_tx=XM_023242552.1;refGene_gn=LOC105261111;refGene_tx=XM_023242553.1;refGene_gn=LOC105261111;refGene_tx=XM_023242554.1;refGene_gn=LOC105261111;refGene_tx=XM_023242555.1;refGene_gn=LOC105261111;refGene_tx=XM_023242556.1
D4 72585146 72585168
refGene_gn=LOC105261111;refGene_tx=XM_019816297.2;refGene_gn=LOC105261111;refGene_tx=XM_023242552.1;refGene_gn=LOC105261111;refGene_tx=XM_023242553.1;refGene_gn=LOC105261111;refGene_tx=XM_023242554.1;refGene_gn=LOC105261111;refGene_tx=XM_023242555.1;refGene_gn=LOC105261111;refGene_tx=XM_023242556.1
D4 72590755 72590857
refGene_gn=LOC105261111;refGene_tx=XM_019816297.2;refGene_gn=LOC105261111;refGene_tx=XM_023242552.1;refGene_gn=LOC105261111;refGene_tx=XM_023242554.1;refGene_gn=LOC105261111;refGene_tx=XM_023242555.1
D4 72601045 72601741
refGene_gn=LOC105261111;refGene_tx=XM_019816297.2;refGene_gn=LOC105261111;refGene_tx=XM_023242552.1;refGene_gn=LOC105261111;refGene_tx=XM_023242553.1;refGene_gn=LOC105261111;refGene_tx=XM_023242554.1
D4 72665105 72665191
refGene_gn=LOC105261111;refGene_tx=XM_019816297.2;refGene_gn=LOC105261111;refGene_tx=XM_023242552.1;refGene_gn=LOC105261111;refGene_tx=XM_023242553.1;refGene_gn=LOC105261111;refGene_tx=XM_023242554.1;refGene_gn=LOC105261111;refGene_tx=XM_023242555.1;refGene_gn=LOC105261111;refGene_tx=XM_023242556.1
D4 72761424 72763828
refGene_gn=LOC105261111;refGene_tx=XM_006939352.2;refGene_gn=LOC105261111;refGene_tx=XM_006939353.4;refGene_gn=LOC105261111;refGene_tx=XM_01

9816297.2;refGene_gn=LOC105261111;refGene_tx=XM_019816301.1;refGene_gn=LOC105261111;refGene_tx=XM_023242552.1;refGene_gn=LOC105261111;refGene_tx=XM_023242553.1;refGene_gn=LOC105261111;refGene_tx=XM_023242554.1;refGene_gn=LOC105261111;refGene_tx=XM_023242555.1;refGene_gn=LOC105261111;refGene_tx=XM_023242556.1
D4 72781783 72781962
refGene_gn=LOC105261111;refGene_tx=XM_006939352.2;refGene_gn=LOC105261111;refGene_tx=XM_006939353.4;refGene_gn=LOC105261111;refGene_tx=XM_019816297.2;refGene_gn=LOC105261111;refGene_tx=XM_019816301.1;refGene_gn=LOC105261111;refGene_tx=XM_023242552.1;refGene_gn=LOC105261111;refGene_tx=XM_023242553.1;refGene_gn=LOC105261111;refGene_tx=XM_023242555.1;refGene_gn=LOC105261111;refGene_tx=XM_023242556.1
D4 72787121 72787160
refGene_gn=LOC105261111;refGene_tx=XM_006939352.2;refGene_gn=LOC105261111;refGene_tx=XM_006939353.4;refGene_gn=LOC105261111;refGene_tx=XM_019816297.2;refGene_gn=LOC105261111;refGene_tx=XM_019816301.1;refGene_gn=LOC105261111;refGene_tx=XM_023242553.1;refGene_gn=LOC105261111;refGene_tx=XM_023242555.1;refGene_gn=LOC105261111;refGene_tx=XM_023242556.1
D4 72792818 72792917
refGene_gn=LOC105261111;refGene_tx=XM_006939352.2;refGene_gn=LOC105261111;refGene_tx=XM_006939353.4;refGene_gn=LOC105261111;refGene_tx=XM_019816297.2;refGene_gn=LOC105261111;refGene_tx=XM_019816301.1;refGene_gn=LOC105261111;refGene_tx=XM_023242552.1;refGene_gn=LOC105261111;refGene_tx=XM_023242553.1;refGene_gn=LOC105261111;refGene_tx=XM_023242554.1;refGene_gn=LOC105261111;refGene_tx=XM_023242555.1;refGene_gn=LOC105261111;refGene_tx=XM_023242556.1
D4 72813796 72814323
refGene_gn=CD4H9orf152;refGene_tx=XM_023242557.1
D4 72817020 72817213
refGene_gn=CD4H9orf152;refGene_tx=XM_023242557.1
D4 72864468 72864531 refGene_gn=TXN;refGene_tx=XM_023242558.1
D4 72865008 72865074 refGene_gn=TXN;refGene_tx=XM_023242558.1
D4 72872899 72872959 refGene_gn=TXN;refGene_tx=XM_023242558.1
D4 72873406 72873511 refGene_gn=TXN;refGene_tx=XM_023242558.1
D4 72877884 72877908 refGene_gn=TXN;refGene_tx=XM_023242558.1
D4 72904117 72904180
refGene_gn=TXNDC8;refGene_tx=XM_006939354.3;refGene_gn=TXNDC8;refGene_tx=XM_023242559.1
D4 72905110 72905176 refGene_gn=TXNDC8;refGene_tx=XM_006939354.3
D4 72914410 72914668 refGene_gn=TXNDC8;refGene_tx=XM_019816309.2
D4 72916188 72916254
refGene_gn=TXNDC8;refGene_tx=XM_006939354.3;refGene_gn=TXNDC8;refGene_tx=XM_019816309.2;refGene_gn=TXNDC8;refGene_tx=XM_023242559.1
D4 72919274 72919334
refGene_gn=TXNDC8;refGene_tx=XM_006939354.3;refGene_gn=TXNDC8;refGene_tx=XM_023242559.1
D4 72925033 72925138
refGene_gn=TXNDC8;refGene_tx=XM_006939354.3;refGene_gn=TXNDC8;refGene_tx=XM_019816309.2;refGene_gn=TXNDC8;refGene_tx=XM_023242559.1
D4 72927940 72927964

refGene_gn=TXNDC8;refGene_tx=XM_006939354.3;refGene_gn=TXNDC8;refGene_tx=XM_019816309.2;refGene_gn=TXNDC8;refGene_tx=XM_023242559.1
D4 72955971 72955993 refGene_gn=SVEP1;refGene_tx=XM_011288665.2;refGene_gn=SVEP1;refGene_tx=XM_023242560.1
D4 72960567 72960661 refGene_gn=SVEP1;refGene_tx=XM_011288665.2;refGene_gn=SVEP1;refGene_tx=XM_023242560.1
D4 72965467 72965563 refGene_gn=SVEP1;refGene_tx=XM_011288665.2;refGene_gn=SVEP1;refGene_tx=XM_023242560.1
D4 72967493 72967589 refGene_gn=SVEP1;refGene_tx=XM_011288665.2;refGene_gn=SVEP1;refGene_tx=XM_023242560.1
D4 72969406 72969577 refGene_gn=SVEP1;refGene_tx=XM_011288665.2;refGene_gn=SVEP1;refGene_tx=XM_023242560.1
D4 72973285 72973462 refGene_gn=SVEP1;refGene_tx=XM_011288665.2
D4 72975334 72975508 refGene_gn=SVEP1;refGene_tx=XM_011288665.2;refGene_gn=SVEP1;refGene_tx=XM_023242560.1
D4 72978723 72978787 refGene_gn=SVEP1;refGene_tx=XM_011288665.2;refGene_gn=SVEP1;refGene_tx=XM_023242560.1
D4 72991746 72991902 refGene_gn=SVEP1;refGene_tx=XM_011288665.2;refGene_gn=SVEP1;refGene_tx=XM_023242560.1
D4 72994419 72994645 refGene_gn=SVEP1;refGene_tx=XM_011288665.2;refGene_gn=SVEP1;refGene_tx=XM_023242560.1
D4 72996566 72999358 refGene_gn=SVEP1;refGene_tx=XM_011288665.2;refGene_gn=SVEP1;refGene_tx=XM_023242560.1
D4 73002062 73002735 refGene_gn=SVEP1;refGene_tx=XM_011288665.2;refGene_gn=SVEP1;refGene_tx=XM_023242560.1
D4 73020309 73020477 refGene_gn=SVEP1;refGene_tx=XM_011288665.2;refGene_gn=SVEP1;refGene_tx=XM_023242560.1
D4 73021422 73021614 refGene_gn=SVEP1;refGene_tx=XM_011288665.2;refGene_gn=SVEP1;refGene_tx=XM_023242560.1
D4 73022498 73022583 refGene_gn=SVEP1;refGene_tx=XM_011288665.2;refGene_gn=SVEP1;refGene_tx=XM_023242560.1
D4 73022809 73022986 refGene_gn=SVEP1;refGene_tx=XM_011288665.2;refGene_gn=SVEP1;refGene_tx=XM_023242560.1
D4 73024384 73024504 refGene_gn=SVEP1;refGene_tx=XM_011288665.2;refGene_gn=SVEP1;refGene_tx

=XM_023242560.1
D4 73025515 73025689
refGene_gn=SVEP1;refGene_tx=XM_011288665.2;refGene_gn=SVEP1;refGene_tx
=XM_023242560.1
D4 73026685 73026856
refGene_gn=SVEP1;refGene_tx=XM_011288665.2;refGene_gn=SVEP1;refGene_tx
=XM_023242560.1
D4 73028075 73028199
refGene_gn=SVEP1;refGene_tx=XM_011288665.2;refGene_gn=SVEP1;refGene_tx
=XM_023242560.1
D4 73029256 73029381
refGene_gn=SVEP1;refGene_tx=XM_011288665.2;refGene_gn=SVEP1;refGene_tx
=XM_023242560.1
D4 73032680 73032856
refGene_gn=SVEP1;refGene_tx=XM_011288665.2;refGene_gn=SVEP1;refGene_tx
=XM_023242560.1
D4 73034679 73034881
refGene_gn=SVEP1;refGene_tx=XM_011288665.2;refGene_gn=SVEP1;refGene_tx
=XM_023242560.1
D4 73035734 73035892
refGene_gn=SVEP1;refGene_tx=XM_011288665.2;refGene_gn=SVEP1;refGene_tx
=XM_023242560.1
D4 73039368 73039570
refGene_gn=SVEP1;refGene_tx=XM_011288665.2;refGene_gn=SVEP1;refGene_tx
=XM_023242560.1
D4 73040377 73040491
refGene_gn=SVEP1;refGene_tx=XM_011288665.2;refGene_gn=SVEP1;refGene_tx
=XM_023242560.1
D4 73041537 73041651
refGene_gn=SVEP1;refGene_tx=XM_011288665.2;refGene_gn=SVEP1;refGene_tx
=XM_023242560.1
D4 73043282 73043379
refGene_gn=SVEP1;refGene_tx=XM_011288665.2;refGene_gn=SVEP1;refGene_tx
=XM_023242560.1
D4 73044459 73044551
refGene_gn=SVEP1;refGene_tx=XM_011288665.2;refGene_gn=SVEP1;refGene_tx
=XM_023242560.1
D4 73044956 73045118
refGene_gn=SVEP1;refGene_tx=XM_011288665.2;refGene_gn=SVEP1;refGene_tx
=XM_023242560.1
D4 73051867 73052029
refGene_gn=SVEP1;refGene_tx=XM_011288665.2;refGene_gn=SVEP1;refGene_tx
=XM_023242560.1
D4 73053496 73053658
refGene_gn=SVEP1;refGene_tx=XM_011288665.2;refGene_gn=SVEP1;refGene_tx
=XM_023242560.1
D4 73055791 73056025
refGene_gn=SVEP1;refGene_tx=XM_011288665.2;refGene_gn=SVEP1;refGene_tx
=XM_023242560.1
D4 73056585 73056744

refGene_gn=SVEP1;refGene_tx=XM_011288665.2;refGene_gn=SVEP1;refGene_tx=XM_023242560.1
D4 73060723 73060832 refGene_gn=SVEP1;refGene_tx=XM_011288665.2;refGene_gn=SVEP1;refGene_tx=XM_023242560.1
D4 73062707 73062829 refGene_gn=SVEP1;refGene_tx=XM_011288665.2;refGene_gn=SVEP1;refGene_tx=XM_023242560.1
D4 73066165 73066360 refGene_gn=SVEP1;refGene_tx=XM_011288665.2;refGene_gn=SVEP1;refGene_tx=XM_023242560.1
D4 73067447 73067579 refGene_gn=SVEP1;refGene_tx=XM_011288665.2;refGene_gn=SVEP1;refGene_tx=XM_023242560.1
D4 73068557 73068665 refGene_gn=SVEP1;refGene_tx=XM_011288665.2;refGene_gn=SVEP1;refGene_tx=XM_023242560.1
D4 73071590 73071720 refGene_gn=SVEP1;refGene_tx=XM_011288665.2;refGene_gn=SVEP1;refGene_tx=XM_023242560.1
D4 73074667 73074786 refGene_gn=SVEP1;refGene_tx=XM_011288665.2;refGene_gn=SVEP1;refGene_tx=XM_023242560.1
D4 73080415 73080613 refGene_gn=SVEP1;refGene_tx=XM_011288665.2;refGene_gn=SVEP1;refGene_tx=XM_023242560.1
D4 73086542 73086722 refGene_gn=SVEP1;refGene_tx=XM_011288665.2;refGene_gn=SVEP1;refGene_tx=XM_023242560.1
D4 73095309 73095489 refGene_gn=SVEP1;refGene_tx=XM_011288665.2;refGene_gn=SVEP1;refGene_tx=XM_023242560.1
D4 73096837 73096996 refGene_gn=SVEP1;refGene_tx=XM_011288665.2;refGene_gn=SVEP1;refGene_tx=XM_023242560.1
D4 73124923 73125100 refGene_gn=SVEP1;refGene_tx=XM_011288665.2;refGene_gn=SVEP1;refGene_tx=XM_023242560.1
D4 73128096 73128352 refGene_gn=SVEP1;refGene_tx=XM_011288665.2;refGene_gn=SVEP1;refGene_tx=XM_023242560.1
D4 73153876 73154407 refGene_gn=SVEP1;refGene_tx=XM_011288665.2;refGene_gn=SVEP1;refGene_tx=XM_023242560.1
D4 73221194 73221273 refGene_gn=MUSK;refGene_tx=XM_011288501.2
D4 73231468 73231595 refGene_gn=MUSK;refGene_tx=XM_011288501.2
D4 73236753 73236905 refGene_gn=MUSK;refGene_tx=XM_011288501.2
D4 73241753 73241881 refGene_gn=MUSK;refGene_tx=XM_011288501.2
D4 73243141 73243283 refGene_gn=MUSK;refGene_tx=XM_011288501.2
D4 73269789 73269914 refGene_gn=MUSK;refGene_tx=XM_011288501.2

D4 73280184 73280344 refGene_gn=MUSK;refGene_tx=XM_011288501.2
D4 73298786 73299050 refGene_gn=MUSK;refGene_tx=XM_011288501.2
D4 73302244 73302420 refGene_gn=MUSK;refGene_tx=XM_011288501.2
D4 73309090 73309292 refGene_gn=MUSK;refGene_tx=XM_011288501.2
D4 73310001 73310193 refGene_gn=MUSK;refGene_tx=XM_011288501.2
D4 73311890 73312039 refGene_gn=MUSK;refGene_tx=XM_011288501.2
D4 73313182 73313865 refGene_gn=MUSK;refGene_tx=XM_011288501.2
D4 73368638 73368940
refGene_gn=LPAR1;refGene_tx=XM_003995751.4;refGene_gn=LPAR1;refGene_tx=XM_006939364.3;refGene_gn=LPAR1;refGene_tx=XM_011288500.3;refGene_gn=LPAR1;refGene_tx=XM_019816314.1;refGene_gn=LPAR1;refGene_tx=XM_019816316.1;refGene_gn=LPAR1;refGene_tx=XM_019816317.1;refGene_gn=LPAR1;refGene_tx=XM_023242561.1
D4 73423671 73424419
refGene_gn=LPAR1;refGene_tx=XM_003995751.4;refGene_gn=LPAR1;refGene_tx=XM_006939364.3;refGene_gn=LPAR1;refGene_tx=XM_011288500.3;refGene_gn=LPAR1;refGene_tx=XM_019816314.1;refGene_gn=LPAR1;refGene_tx=XM_019816316.1;refGene_gn=LPAR1;refGene_tx=XM_019816317.1;refGene_gn=LPAR1;refGene_tx=XM_023242561.1
D4 73447386 73447431
refGene_gn=LPAR1;refGene_tx=XM_003995751.4;refGene_gn=LPAR1;refGene_tx=XM_006939364.3;refGene_gn=LPAR1;refGene_tx=XM_011288500.3;refGene_gn=LPAR1;refGene_tx=XM_019816314.1;refGene_gn=LPAR1;refGene_tx=XM_019816316.1;refGene_gn=LPAR1;refGene_tx=XM_019816317.1;refGene_gn=LPAR1;refGene_tx=XM_023242561.1
D4 73724749 73725727
refGene_gn=LOC101092965;refGene_tx=XM_003995753.4
D4 73742807 73742947 refGene_gn=KIAA0368;refGene_tx=XM_023242563.1
D4 73744264 73744336 refGene_gn=KIAA0368;refGene_tx=XM_023242563.1
D4 73746810 73746899 refGene_gn=KIAA0368;refGene_tx=XM_023242563.1
D4 73747348 73747454 refGene_gn=KIAA0368;refGene_tx=XM_023242563.1
D4 73749409 73749548 refGene_gn=KIAA0368;refGene_tx=XM_023242563.1
D4 73752522 73752715 refGene_gn=KIAA0368;refGene_tx=XM_023242563.1
D4 73752811 73752855 refGene_gn=KIAA0368;refGene_tx=XM_023242563.1
D4 73753662 73753871 refGene_gn=KIAA0368;refGene_tx=XM_023242563.1
D4 73754470 73754662 refGene_gn=KIAA0368;refGene_tx=XM_023242563.1
D4 73754898 73754965 refGene_gn=KIAA0368;refGene_tx=XM_023242563.1
D4 73755045 73755137 refGene_gn=KIAA0368;refGene_tx=XM_023242563.1
D4 73755953 73756020 refGene_gn=KIAA0368;refGene_tx=XM_023242563.1
D4 73757099 73757189 refGene_gn=KIAA0368;refGene_tx=XM_023242563.1
D4 73758588 73758654 refGene_gn=KIAA0368;refGene_tx=XM_023242563.1
D4 73759587 73759738 refGene_gn=KIAA0368;refGene_tx=XM_023242563.1
D4 73761607 73761729 refGene_gn=KIAA0368;refGene_tx=XM_023242563.1
D4 73762789 73762837 refGene_gn=KIAA0368;refGene_tx=XM_023242563.1
D4 73763597 73763703 refGene_gn=KIAA0368;refGene_tx=XM_023242563.1
D4 73764306 73764386 refGene_gn=KIAA0368;refGene_tx=XM_023242563.1
D4 73767131 73767299 refGene_gn=KIAA0368;refGene_tx=XM_023242563.1
D4 73767583 73767628 refGene_gn=KIAA0368;refGene_tx=XM_023242563.1
D4 73767629 73767702 refGene_gn=KIAA0368;refGene_tx=XM_023242563.1
D4 73770708 73770777 refGene_gn=KIAA0368;refGene_tx=XM_023242563.1

D4 73771479 73771594 refGene_gn=KIAA0368;refGene_tx=XM_023242563.1
D4 73772415 73772470 refGene_gn=KIAA0368;refGene_tx=XM_023242563.1
D4 73772880 73773026 refGene_gn=KIAA0368;refGene_tx=XM_023242563.1
D4 73778102 73778226 refGene_gn=KIAA0368;refGene_tx=XM_023242563.1
D4 73781851 73781953 refGene_gn=KIAA0368;refGene_tx=XM_023242563.1
D4 73783834 73784007 refGene_gn=KIAA0368;refGene_tx=XM_023242563.1
D4 73784712 73784875 refGene_gn=KIAA0368;refGene_tx=XM_023242563.1
D4 73785640 73785775 refGene_gn=KIAA0368;refGene_tx=XM_023242563.1
D4 73788578 73788670 refGene_gn=KIAA0368;refGene_tx=XM_023242563.1
D4 73789064 73789287 refGene_gn=KIAA0368;refGene_tx=XM_023242563.1
D4 73790531 73790612 refGene_gn=KIAA0368;refGene_tx=XM_023242563.1
D4 73791594 73791718 refGene_gn=KIAA0368;refGene_tx=XM_023242563.1
D4 73792113 73792217 refGene_gn=KIAA0368;refGene_tx=XM_023242563.1
D4 73794300 73794423 refGene_gn=KIAA0368;refGene_tx=XM_023242563.1
D4 73794515 73794582 refGene_gn=KIAA0368;refGene_tx=XM_023242563.1
D4 73795577 73795627 refGene_gn=KIAA0368;refGene_tx=XM_023242563.1
D4 73797424 73797503 refGene_gn=KIAA0368;refGene_tx=XM_023242563.1
D4 73797757 73797843 refGene_gn=KIAA0368;refGene_tx=XM_023242563.1
D4 73800857 73800977 refGene_gn=KIAA0368;refGene_tx=XM_023242563.1
D4 73803133 73803215 refGene_gn=KIAA0368;refGene_tx=XM_023242563.1
D4 73806925 73807065 refGene_gn=KIAA0368;refGene_tx=XM_023242563.1
D4 73809949 73810118 refGene_gn=KIAA0368;refGene_tx=XM_023242563.1
D4 73813211 73813361 refGene_gn=KIAA0368;refGene_tx=XM_023242563.1
D4 73814493 73814612 refGene_gn=KIAA0368;refGene_tx=XM_023242563.1
D4 73816318 73816435 refGene_gn=KIAA0368;refGene_tx=XM_023242563.1
D4 73822437 73822568 refGene_gn=KIAA0368;refGene_tx=XM_023242563.1
D4 73846034 73846074 refGene_gn=KIAA0368;refGene_tx=XM_023242563.1
D4 73862753 73863141 refGene_gn=ZNF483;refGene_tx=XM_011288499.3
D4 73864807 73864896 refGene_gn=ZNF483;refGene_tx=XM_011288499.3
D4 73865836 73865963 refGene_gn=ZNF483;refGene_tx=XM_011288499.3
D4 73866342 73866438 refGene_gn=ZNF483;refGene_tx=XM_011288499.3
D4 73870192 73871742 refGene_gn=ZNF483;refGene_tx=XM_011288499.3
D4 73889059 73889170
refGene_gn=PTGR1;refGene_tx=XM_003995755.5;refGene_gn=PTGR1;refGene_tx
=XM_006939366.3;refGene_gn=PTGR1;refGene_tx=XM_006939367.4;refGene_gn=
PTGR1;refGene_tx=XM_019816320.2;refGene_gn=PTGR1;refGene_tx=XM_0232425
65.1;refGene_gn=PTGR1;refGene_tx=XM_023242566.1
D4 73892926 73893045
refGene_gn=PTGR1;refGene_tx=XM_003995755.5;refGene_gn=PTGR1;refGene_tx
=XM_006939366.3;refGene_gn=PTGR1;refGene_tx=XM_006939367.4;refGene_gn=
PTGR1;refGene_tx=XM_019816320.2;refGene_gn=PTGR1;refGene_tx=XM_0232425
65.1;refGene_gn=PTGR1;refGene_tx=XM_023242566.1
D4 73896630 73896739
refGene_gn=PTGR1;refGene_tx=XM_003995755.5;refGene_gn=PTGR1;refGene_tx
=XM_006939366.3;refGene_gn=PTGR1;refGene_tx=XM_006939367.4;refGene_gn=
PTGR1;refGene_tx=XM_019816320.2;refGene_gn=PTGR1;refGene_tx=XM_0232425
65.1;refGene_gn=PTGR1;refGene_tx=XM_023242566.1
D4 73899700 73899856
refGene_gn=PTGR1;refGene_tx=XM_003995755.5;refGene_gn=PTGR1;refGene_tx
=XM_006939366.3;refGene_gn=PTGR1;refGene_tx=XM_006939367.4;refGene_gn=

PTGR1;refGene_tx=XM_019816320.2;refGene_gn=PTGR1;refGene_tx=XM_0232425
65.1;refGene_gn=PTGR1;refGene_tx=XM_023242566.1
D4 73900581 73900699
refGene_gn=PTGR1;refGene_tx=XM_003995755.5;refGene_gn=PTGR1;refGene_tx
=XM_006939366.3;refGene_gn=PTGR1;refGene_tx=XM_006939367.4;refGene_gn=
PTGR1;refGene_tx=XM_019816320.2;refGene_gn=PTGR1;refGene_tx=XM_0232425
65.1;refGene_gn=PTGR1;refGene_tx=XM_023242566.1
D4 73904753 73904921
refGene_gn=PTGR1;refGene_tx=XM_003995755.5;refGene_gn=PTGR1;refGene_tx
=XM_006939366.3;refGene_gn=PTGR1;refGene_tx=XM_006939367.4;refGene_gn=
PTGR1;refGene_tx=XM_019816320.2;refGene_gn=PTGR1;refGene_tx=XM_0232425
65.1;refGene_gn=PTGR1;refGene_tx=XM_023242566.1
D4 73909214 73909271
refGene_gn=PTGR1;refGene_tx=XM_003995755.5;refGene_gn=PTGR1;refGene_tx
=XM_006939366.3;refGene_gn=PTGR1;refGene_tx=XM_006939367.4;refGene_gn=
PTGR1;refGene_tx=XM_019816320.2;refGene_gn=PTGR1;refGene_tx=XM_0232425
65.1;refGene_gn=PTGR1;refGene_tx=XM_023242566.1
D4 73911220 73911266
refGene_gn=PTGR1;refGene_tx=XM_003995755.5;refGene_gn=PTGR1;refGene_tx
=XM_006939366.3;refGene_gn=PTGR1;refGene_tx=XM_006939367.4;refGene_gn=
PTGR1;refGene_tx=XM_019816320.2;refGene_gn=PTGR1;refGene_tx=XM_0232425
65.1;refGene_gn=PTGR1;refGene_tx=XM_023242566.1
D4 73912688 73912794
refGene_gn=PTGR1;refGene_tx=XM_003995755.5;refGene_gn=PTGR1;refGene_tx
=XM_006939366.3;refGene_gn=PTGR1;refGene_tx=XM_006939367.4;refGene_gn=
PTGR1;refGene_tx=XM_019816320.2;refGene_gn=PTGR1;refGene_tx=XM_0232425
65.1
D4 73947279 73947612
refGene_gn=LOC101093721;refGene_tx=XM_023242564.1
D4 73960340 73960493
refGene_gn=LOC101093721;refGene_tx=XM_023242564.1
D4 73962780 73963251
refGene_gn=LOC101093721;refGene_tx=XM_023242564.1
D4 73965825 73965948
refGene_gn=LOC101093721;refGene_tx=XM_023242564.1
D4 73976359 73976440 refGene_gn=GNG10;refGene_tx=XM_023242567.1
D4 73980659 73980785 refGene_gn=GNG10;refGene_tx=XM_023242567.1
D4 73990569 73990670
refGene_gn=CD4H9orf84;refGene_tx=XM_011288498.2;refGene_gn=CD4H9orf84;
refGene_tx=XM_019816321.1
D4 73993868 73994829
refGene_gn=CD4H9orf84;refGene_tx=XM_011288498.2;refGene_gn=CD4H9orf84;
refGene_tx=XM_019816321.1
D4 73995696 73995846
refGene_gn=CD4H9orf84;refGene_tx=XM_011288498.2;refGene_gn=CD4H9orf84;
refGene_tx=XM_019816321.1
D4 73998315 73998447
refGene_gn=CD4H9orf84;refGene_tx=XM_011288498.2;refGene_gn=CD4H9orf84;
refGene_tx=XM_019816321.1
D4 74001534 74001628

refGene_gn=CD4H9orf84;refGene_tx=XM_011288498.2;refGene_gn=CD4H9orf84;
refGene_tx=XM_019816321.1
D4 74005708 74005830
refGene_gn=CD4H9orf84;refGene_tx=XM_011288498.2;refGene_gn=CD4H9orf84;
refGene_tx=XM_019816321.1
D4 74007098 74007210
refGene_gn=CD4H9orf84;refGene_tx=XM_011288498.2;refGene_gn=CD4H9orf84;
refGene_tx=XM_019816321.1
D4 74008528 74008646
refGene_gn=CD4H9orf84;refGene_tx=XM_011288498.2;refGene_gn=CD4H9orf84;
refGene_tx=XM_019816321.1
D4 74010169 74010348
refGene_gn=CD4H9orf84;refGene_tx=XM_011288498.2;refGene_gn=CD4H9orf84;
refGene_tx=XM_019816321.1
D4 74011571 74011638
refGene_gn=CD4H9orf84;refGene_tx=XM_011288498.2;refGene_gn=CD4H9orf84;
refGene_tx=XM_019816321.1
D4 74017390 74017463
refGene_gn=CD4H9orf84;refGene_tx=XM_011288498.2;refGene_gn=CD4H9orf84;
refGene_tx=XM_019816321.1
D4 74018915 74019094
refGene_gn=CD4H9orf84;refGene_tx=XM_011288498.2;refGene_gn=CD4H9orf84;
refGene_tx=XM_019816321.1
D4 74022444 74022549
refGene_gn=CD4H9orf84;refGene_tx=XM_011288498.2;refGene_gn=CD4H9orf84;
refGene_tx=XM_019816321.1
D4 74025524 74025701
refGene_gn=CD4H9orf84;refGene_tx=XM_011288498.2;refGene_gn=CD4H9orf84;
refGene_tx=XM_019816321.1
D4 74028525 74028654
refGene_gn=CD4H9orf84;refGene_tx=XM_011288498.2;refGene_gn=CD4H9orf84;
refGene_tx=XM_019816321.1
D4 74030345 74030762
refGene_gn=CD4H9orf84;refGene_tx=XM_011288498.2;refGene_gn=CD4H9orf84;
refGene_tx=XM_019816321.1
D4 74035329 74035572
refGene_gn=CD4H9orf84;refGene_tx=XM_011288498.2;refGene_gn=CD4H9orf84;
refGene_tx=XM_019816321.1
D4 74036301 74036387
refGene_gn=CD4H9orf84;refGene_tx=XM_011288498.2;refGene_gn=CD4H9orf84;
refGene_tx=XM_019816321.1
D4 74041077 74041183
refGene_gn=CD4H9orf84;refGene_tx=XM_011288498.2;refGene_gn=CD4H9orf84;
refGene_tx=XM_019816321.1
D4 74042245 74042347
refGene_gn=CD4H9orf84;refGene_tx=XM_011288498.2;refGene_gn=CD4H9orf84;
refGene_tx=XM_019816321.1
D4 74043291 74043442
refGene_gn=CD4H9orf84;refGene_tx=XM_011288498.2;refGene_gn=CD4H9orf84;
refGene_tx=XM_019816321.1

D4 74045253 74045350
refGene_gn=CD4H9orf84;refGene_tx=XM_011288498.2;refGene_gn=CD4H9orf84;
refGene_tx=XM_019816321.1
D4 74048791 74048954
refGene_gn=CD4H9orf84;refGene_tx=XM_011288498.2;refGene_gn=CD4H9orf84;
refGene_tx=XM_019816321.1
D4 74058312 74058497
refGene_gn=CD4H9orf84;refGene_tx=XM_011288498.2;refGene_gn=CD4H9orf84;
refGene_tx=XM_019816321.1
D4 74065516 74065604
refGene_gn=CD4H9orf84;refGene_tx=XM_011288498.2;refGene_gn=CD4H9orf84;
refGene_tx=XM_019816321.1
D4 74069953 74070077
refGene_gn=CD4H9orf84;refGene_tx=XM_011288498.2;refGene_gn=CD4H9orf84;
refGene_tx=XM_019816321.1
D4 74072412 74072493
refGene_gn=CD4H9orf84;refGene_tx=XM_011288498.2;refGene_gn=CD4H9orf84;
refGene_tx=XM_019816321.1
D4 74075395 74075464
refGene_gn=CD4H9orf84;refGene_tx=XM_011288498.2;refGene_gn=CD4H9orf84;
refGene_tx=XM_019816321.1
D4 74148229 74148327 refGene_gn=UGCG;refGene_tx=XM_011288497.3
D4 74168101 74168243 refGene_gn=UGCG;refGene_tx=XM_011288497.3
D4 74176648 74176751 refGene_gn=UGCG;refGene_tx=XM_011288497.3
D4 74177479 74177581 refGene_gn=UGCG;refGene_tx=XM_011288497.3
D4 74179064 74179177 refGene_gn=UGCG;refGene_tx=XM_011288497.3
D4 74182842 74183021 refGene_gn=UGCG;refGene_tx=XM_011288497.3
D4 74184542 74184629 refGene_gn=UGCG;refGene_tx=XM_011288497.3
D4 74185131 74185321 refGene_gn=UGCG;refGene_tx=XM_011288497.3
D4 74185682 74185853 refGene_gn=UGCG;refGene_tx=XM_011288497.3
D4 74332194 74332195 refGene_gn=SUSD1;refGene_tx=XM_023242309.1
D4 74332520 74332614 refGene_gn=SUSD1;refGene_tx=XM_023242309.1
D4 74342952 74342992 refGene_gn=SUSD1;refGene_tx=XM_023242309.1
D4 74347505 74347764 refGene_gn=SUSD1;refGene_tx=XM_023242309.1
D4 74351702 74351799 refGene_gn=SUSD1;refGene_tx=XM_023242309.1
D4 74362459 74362646 refGene_gn=SUSD1;refGene_tx=XM_023242309.1
D4 74364098 74364190 refGene_gn=SUSD1;refGene_tx=XM_023242309.1
D4 74386003 74386196 refGene_gn=SUSD1;refGene_tx=XM_023242309.1
D4 74390861 74390971 refGene_gn=SUSD1;refGene_tx=XM_023242309.1
D4 74405822 74406009 refGene_gn=SUSD1;refGene_tx=XM_023242309.1
D4 74406787 74406885 refGene_gn=SUSD1;refGene_tx=XM_023242309.1
D4 74415719 74415899 refGene_gn=SUSD1;refGene_tx=XM_023242309.1
D4 74430557 74430737 refGene_gn=SUSD1;refGene_tx=XM_023242309.1
D4 74431398 74431551 refGene_gn=SUSD1;refGene_tx=XM_023242309.1
D4 74446640 74446710 refGene_gn=SUSD1;refGene_tx=XM_023242309.1
D4 74446711 74446797 refGene_gn=SUSD1;refGene_tx=XM_023242309.1
D4 74449445 74449559 refGene_gn=SUSD1;refGene_tx=XM_023242309.1
D4 74463664 74463961 refGene_gn=SUSD1;refGene_tx=XM_023242309.1
D4 74471173 74471312 refGene_gn=SUSD1;refGene_tx=XM_023242309.1
D4 74502907 74503040 refGene_gn=PTBP3;refGene_tx=XM_023242569.1

D4 74503182 74503260 refGene_gn=PTBP3;refGene_tx=XM_023242569.1
D4 74507180 74507397 refGene_gn=PTBP3;refGene_tx=XM_023242569.1
D4 74508496 74508589 refGene_gn=PTBP3;refGene_tx=XM_023242569.1
D4 74511366 74511400 refGene_gn=PTBP3;refGene_tx=XM_023242569.1
D4 74511650 74511790 refGene_gn=PTBP3;refGene_tx=XM_023242569.1
D4 74514659 74514737 refGene_gn=PTBP3;refGene_tx=XM_023242569.1
D4 74540097 74540272 refGene_gn=PTBP3;refGene_tx=XM_023242569.1
D4 74541685 74541786 refGene_gn=PTBP3;refGene_tx=XM_023242569.1
D4 74541788 74541795 refGene_gn=PTBP3;refGene_tx=XM_023242569.1
D4 74552063 74552228 refGene_gn=PTBP3;refGene_tx=XM_023242569.1
D4 74561743 74561890 refGene_gn=PTBP3;refGene_tx=XM_023242569.1
D4 74565305 74565475 refGene_gn=PTBP3;refGene_tx=XM_023242569.1
D4 74587842 74587924 refGene_gn=PTBP3;refGene_tx=XM_023242569.1
D4 74624260 74624294 refGene_gn=PTBP3;refGene_tx=XM_023242569.1
D4 74627146 74627385 refGene_gn=PTBP3;refGene_tx=XM_023242569.1
D4 74659040 74659057 refGene_gn=HSDL2;refGene_tx=XM_003995758.5
D4 74672116 74672280 refGene_gn=HSDL2;refGene_tx=XM_003995758.5
D4 74673683 74673782 refGene_gn=HSDL2;refGene_tx=XM_003995758.5
D4 74679418 74679533 refGene_gn=HSDL2;refGene_tx=XM_003995758.5
D4 74685497 74685601 refGene_gn=HSDL2;refGene_tx=XM_003995758.5
D4 74686290 74686389 refGene_gn=HSDL2;refGene_tx=XM_003995758.5
D4 74697774 74697969 refGene_gn=HSDL2;refGene_tx=XM_003995758.5
D4 74702129 74702201 refGene_gn=HSDL2;refGene_tx=XM_003995758.5
D4 74706220 74706370 refGene_gn=HSDL2;refGene_tx=XM_003995758.5
D4 74710581 74710710 refGene_gn=HSDL2;refGene_tx=XM_003995758.5
D4 74725228 74725341 refGene_gn=HSDL2;refGene_tx=XM_003995758.5
D4 74810647 74811818
refGene_gn=KIAA1958;refGene_tx=XM_023242571.1;refGene_gn=KIAA1958;refGene_tx=XM_023242572.1
D4 74851718 74851802 refGene_gn=KIAA1958;refGene_tx=XM_023242572.1
D4 74877699 74877872
refGene_gn=KIAA1958;refGene_tx=XM_023242571.1;refGene_gn=KIAA1958;refGene_tx=XM_023242572.1
D4 74889652 74890459
refGene_gn=KIAA1958;refGene_tx=XM_023242571.1;refGene_gn=KIAA1958;refGene_tx=XM_023242572.1
D4 74914090 74914186
refGene_gn=INIP;refGene_tx=XM_006939370.4;refGene_gn=INIP;refGene_tx=XM_019816332.2;refGene_gn=INIP;refGene_tx=XM_019816333.1;refGene_gn=INIP;refGene_tx=XM_019816334.1;refGene_gn=INIP;refGene_tx=XM_019816335.2;refGene_gn=INIP;refGene_tx=XM_019816336.2
D4 74915296 74915387
refGene_gn=INIP;refGene_tx=XM_006939370.4;refGene_gn=INIP;refGene_tx=XM_019816332.2;refGene_gn=INIP;refGene_tx=XM_019816333.1;refGene_gn=INIP;refGene_tx=XM_019816334.1;refGene_gn=INIP;refGene_tx=XM_019816335.2;refGene_gn=INIP;refGene_tx=XM_019816336.2
D4 74917768 74917871
refGene_gn=INIP;refGene_tx=XM_006939370.4;refGene_gn=INIP;refGene_tx=XM_019816332.2;refGene_gn=INIP;refGene_tx=XM_019816333.1;refGene_gn=INIP;refGene_tx=XM_019816334.1;refGene_gn=INIP;refGene_tx=XM_019816335.2;

refGene_gn=INIP;refGene_tx=XM_019816336.2
D4	74923778	74923884	refGene_gn=INIP;refGene_tx=XM_019816332.2
D4	74930742	74930767	refGene_gn=INIP;refGene_tx=XM_006939370.4
D4	74962862	74963018
refGene_gn=SNX30;refGene_tx=XM_023242574.1;refGene_gn=SNX30;refGene_tx=XM_023242575.1
D4	75017432	75017624
refGene_gn=SNX30;refGene_tx=XM_023242574.1;refGene_gn=SNX30;refGene_tx=XM_023242575.1
D4	75029168	75029279
refGene_gn=SNX30;refGene_tx=XM_023242574.1;refGene_gn=SNX30;refGene_tx=XM_023242575.1
D4	75042874	75043033
refGene_gn=SNX30;refGene_tx=XM_023242574.1;refGene_gn=SNX30;refGene_tx=XM_023242575.1
D4	75047263	75047459
refGene_gn=SNX30;refGene_tx=XM_023242574.1;refGene_gn=SNX30;refGene_tx=XM_023242575.1
D4	75048903	75049103
refGene_gn=SNX30;refGene_tx=XM_023242574.1;refGene_gn=SNX30;refGene_tx=XM_023242575.1
D4	75062046	75062133
refGene_gn=SNX30;refGene_tx=XM_023242574.1;refGene_gn=SNX30;refGene_tx=XM_023242575.1
D4	75072657	75072810
refGene_gn=SNX30;refGene_tx=XM_023242574.1;refGene_gn=SNX30;refGene_tx=XM_023242575.1
D4	75074282	75074342
refGene_gn=SNX30;refGene_tx=XM_023242574.1;refGene_gn=SNX30;refGene_tx=XM_023242575.1
D4	75074507	75074547	refGene_gn=SNX30;refGene_tx=XM_023242574.1
D4	75084519	75084806
refGene_gn=SLC46A2;refGene_tx=XM_011288491.3;refGene_gn=SLC46A2;refGene_tx=XM_019816339.2
D4	75090777	75090934	refGene_gn=SLC46A2;refGene_tx=XM_011288491.3
D4	75091333	75091417	refGene_gn=SLC46A2;refGene_tx=XM_011288491.3
D4	75092859	75093991
refGene_gn=SLC46A2;refGene_tx=XM_011288491.3;refGene_gn=SLC46A2;refGene_tx=XM_019816339.2
D4	75147238	75147248
refGene_gn=LOC493739;refGene_tx=NM_001009233.1
D4	75148177	75148223
refGene_gn=LOC493739;refGene_tx=NM_001009233.1;refGene_gn=LOC493739;refGene_tx=XM_019815497.2
D4	75148423	75148519
refGene_gn=LOC493739;refGene_tx=NM_001009233.1;refGene_gn=LOC493739;refGene_tx=XM_019815497.2
D4	75148843	75148954
refGene_gn=LOC493739;refGene_tx=NM_001009233.1;refGene_gn=LOC493739;refGene_tx=XM_019815497.2

D4	75149673	75149744
refGene_gn=LOC493739;refGene_tx=NM_001009233.1;refGene_gn=LOC493739;re
fGene_tx=XM_019815497.2
D4	75150709	75150846
refGene_gn=LOC493739;refGene_tx=NM_001009233.1;refGene_gn=LOC493739;re
fGene_tx=XM_019815497.2
D4	75151203	75151293
refGene_gn=LOC493739;refGene_tx=NM_001009233.1;refGene_gn=LOC493739;re
fGene_tx=XM_019815497.2
D4	75170825	75171474	refGene_gn=ZNF883;refGene_tx=XM_019816358.2
D4	75171475	75171798	refGene_gn=ZNF883;refGene_tx=XM_019816358.2
D4	75191286	75192536
refGene_gn=ZFP37;refGene_tx=XM_019816341.2;refGene_gn=ZFP37;refGene_tx
=XM_019816342.2;refGene_gn=ZFP37;refGene_tx=XM_019816348.2;refGene_gn=
ZFP37;refGene_tx=XM_019816349.2;refGene_gn=ZFP37;refGene_tx=XM_0198163
50.2;refGene_gn=ZFP37;refGene_tx=XM_019816352.2;refGene_gn=ZFP37;refGe
ne_tx=XM_019816353.2;refGene_gn=ZFP37;refGene_tx=XM_019816354.1;refGen
e_gn=ZFP37;refGene_tx=XM_023242578.1;refGene_gn=ZFP37;refGene_tx=XM_02
3242579.1
D4	75194542	75194632
refGene_gn=ZFP37;refGene_tx=XM_019816341.2;refGene_gn=ZFP37;refGene_tx
=XM_019816342.2;refGene_gn=ZFP37;refGene_tx=XM_019816348.2;refGene_gn=
ZFP37;refGene_tx=XM_019816350.2;refGene_gn=ZFP37;refGene_tx=XM_0198163
52.2;refGene_gn=ZFP37;refGene_tx=XM_019816353.2;refGene_gn=ZFP37;refGe
ne_tx=XM_019816354.1;refGene_gn=ZFP37;refGene_tx=XM_023242578.1
D4	75204167	75204318
refGene_gn=ZFP37;refGene_tx=XM_019816341.2;refGene_gn=ZFP37;refGene_tx
=XM_019816342.2;refGene_gn=ZFP37;refGene_tx=XM_019816348.2;refGene_gn=
ZFP37;refGene_tx=XM_019816349.2;refGene_gn=ZFP37;refGene_tx=XM_0198163
50.2;refGene_gn=ZFP37;refGene_tx=XM_019816352.2;refGene_gn=ZFP37;refGe
ne_tx=XM_019816353.2;refGene_gn=ZFP37;refGene_tx=XM_019816354.1;refGen
e_gn=ZFP37;refGene_tx=XM_023242578.1
D4	75210061	75210106
refGene_gn=ZFP37;refGene_tx=XM_019816341.2;refGene_gn=ZFP37;refGene_tx
=XM_019816342.2;refGene_gn=ZFP37;refGene_tx=XM_019816348.2;refGene_gn=
ZFP37;refGene_tx=XM_019816349.2;refGene_gn=ZFP37;refGene_tx=XM_0198163
50.2;refGene_gn=ZFP37;refGene_tx=XM_023242578.1
D4	75210294	75210354
refGene_gn=ZFP37;refGene_tx=XM_019816341.2;refGene_gn=ZFP37;refGene_tx
=XM_019816342.2;refGene_gn=ZFP37;refGene_tx=XM_019816349.2;refGene_gn=
ZFP37;refGene_tx=XM_023242578.1
D4	75224393	75225934
refGene_gn=ZFP37;refGene_tx=XM_023242576.1;refGene_gn=ZFP37;refGene_tx
=XM_023242579.1
D4	75231092	75231227
refGene_gn=ZFP37;refGene_tx=XM_023242576.1;refGene_gn=ZFP37;refGene_tx
=XM_023242579.1
D4	75231535	75231662
refGene_gn=ZFP37;refGene_tx=XM_023242576.1;refGene_gn=ZFP37;refGene_tx
=XM_023242579.1

D4 75236770 75236965 refGene_gn=ZFP37;refGene_tx=XM_023242579.1
D4 75296450 75296764 refGene_gn=SLC31A2;refGene_tx=XM_023242584.1
D4 75300705 75300772
refGene_gn=SLC31A2;refGene_tx=XM_011288489.2;refGene_gn=SLC31A2;refGene_tx=XM_023242581.1;refGene_gn=SLC31A2;refGene_tx=XM_023242582.1;refGene_gn=SLC31A2;refGene_tx=XM_023242584.1
D4 75303292 75303482
refGene_gn=SLC31A2;refGene_tx=XM_011288489.2;refGene_gn=SLC31A2;refGene_tx=XM_023242581.1;refGene_gn=SLC31A2;refGene_tx=XM_023242582.1
D4 75304505 75304674 refGene_gn=SLC31A2;refGene_tx=XM_011288489.2
D4 75306253 75306428
refGene_gn=SLC31A2;refGene_tx=XM_023242581.1;refGene_gn=SLC31A2;refGene_tx=XM_023242582.1;refGene_gn=SLC31A2;refGene_tx=XM_023242584.1
D4 75308615 75308693
refGene_gn=FKBP15;refGene_tx=XM_003995763.5;refGene_gn=FKBP15;refGene_tx=XM_006939372.4;refGene_gn=FKBP15;refGene_tx=XM_011288490.3
D4 75310939 75311036
refGene_gn=FKBP15;refGene_tx=XM_003995763.5;refGene_gn=FKBP15;refGene_tx=XM_006939372.4;refGene_gn=FKBP15;refGene_tx=XM_011288490.3
D4 75311864 75312535
refGene_gn=FKBP15;refGene_tx=XM_003995763.5;refGene_gn=FKBP15;refGene_tx=XM_006939372.4;refGene_gn=FKBP15;refGene_tx=XM_011288490.3
D4 75312818 75312926
refGene_gn=FKBP15;refGene_tx=XM_003995763.5;refGene_gn=FKBP15;refGene_tx=XM_006939372.4;refGene_gn=FKBP15;refGene_tx=XM_011288490.3
D4 75313517 75313640
refGene_gn=FKBP15;refGene_tx=XM_003995763.5;refGene_gn=FKBP15;refGene_tx=XM_006939372.4;refGene_gn=FKBP15;refGene_tx=XM_011288490.3
D4 75314616 75314769
refGene_gn=FKBP15;refGene_tx=XM_003995763.5;refGene_gn=FKBP15;refGene_tx=XM_006939372.4;refGene_gn=FKBP15;refGene_tx=XM_011288490.3
D4 75315327 75315483
refGene_gn=FKBP15;refGene_tx=XM_003995763.5;refGene_gn=FKBP15;refGene_tx=XM_006939372.4;refGene_gn=FKBP15;refGene_tx=XM_011288490.3
D4 75317452 75317589
refGene_gn=FKBP15;refGene_tx=XM_003995763.5;refGene_gn=FKBP15;refGene_tx=XM_006939372.4;refGene_gn=FKBP15;refGene_tx=XM_011288490.3
D4 75319872 75320044
refGene_gn=FKBP15;refGene_tx=XM_003995763.5;refGene_gn=FKBP15;refGene_tx=XM_006939372.4;refGene_gn=FKBP15;refGene_tx=XM_011288490.3
D4 75325121 75325224
refGene_gn=FKBP15;refGene_tx=XM_003995763.5;refGene_gn=FKBP15;refGene_tx=XM_006939372.4;refGene_gn=FKBP15;refGene_tx=XM_011288490.3
D4 75326035 75326130
refGene_gn=FKBP15;refGene_tx=XM_003995763.5;refGene_gn=FKBP15;refGene_tx=XM_006939372.4;refGene_gn=FKBP15;refGene_tx=XM_011288490.3
D4 75326578 75326686
refGene_gn=FKBP15;refGene_tx=XM_003995763.5;refGene_gn=FKBP15;refGene_tx=XM_006939372.4;refGene_gn=FKBP15;refGene_tx=XM_011288490.3
D4 75326977 75327087

refGene_gn=FKBP15;refGene_tx=XM_003995763.5;refGene_gn=FKBP15;refGene_tx=XM_006939372.4;refGene_gn=FKBP15;refGene_tx=XM_011288490.3
D4 75328704 75328819
refGene_gn=FKBP15;refGene_tx=XM_003995763.5;refGene_gn=FKBP15;refGene_tx=XM_006939372.4;refGene_gn=FKBP15;refGene_tx=XM_011288490.3
D4 75330280 75330387
refGene_gn=FKBP15;refGene_tx=XM_003995763.5;refGene_gn=FKBP15;refGene_tx=XM_006939372.4;refGene_gn=FKBP15;refGene_tx=XM_011288490.3
D4 75330869 75330972
refGene_gn=FKBP15;refGene_tx=XM_003995763.5;refGene_gn=FKBP15;refGene_tx=XM_006939372.4;refGene_gn=FKBP15;refGene_tx=XM_011288490.3
D4 75333149 75333257
refGene_gn=FKBP15;refGene_tx=XM_003995763.5;refGene_gn=FKBP15;refGene_tx=XM_006939372.4;refGene_gn=FKBP15;refGene_tx=XM_011288490.3
D4 75335753 75335811
refGene_gn=FKBP15;refGene_tx=XM_003995763.5;refGene_gn=FKBP15;refGene_tx=XM_006939372.4;refGene_gn=FKBP15;refGene_tx=XM_011288490.3
D4 75336307 75336450
refGene_gn=FKBP15;refGene_tx=XM_003995763.5;refGene_gn=FKBP15;refGene_tx=XM_006939372.4;refGene_gn=FKBP15;refGene_tx=XM_011288490.3
D4 75339204 75339351
refGene_gn=FKBP15;refGene_tx=XM_003995763.5;refGene_gn=FKBP15;refGene_tx=XM_006939372.4;refGene_gn=FKBP15;refGene_tx=XM_011288490.3
D4 75341096 75341165
refGene_gn=FKBP15;refGene_tx=XM_003995763.5;refGene_gn=FKBP15;refGene_tx=XM_006939372.4;refGene_gn=FKBP15;refGene_tx=XM_011288490.3
D4 75341942 75342092
refGene_gn=FKBP15;refGene_tx=XM_003995763.5;refGene_gn=FKBP15;refGene_tx=XM_006939372.4;refGene_gn=FKBP15;refGene_tx=XM_011288490.3
D4 75344416 75344515
refGene_gn=FKBP15;refGene_tx=XM_003995763.5;refGene_gn=FKBP15;refGene_tx=XM_006939372.4;refGene_gn=FKBP15;refGene_tx=XM_011288490.3
D4 75344843 75344918
refGene_gn=FKBP15;refGene_tx=XM_003995763.5;refGene_gn=FKBP15;refGene_tx=XM_006939372.4;refGene_gn=FKBP15;refGene_tx=XM_011288490.3
D4 75348654 75348724
refGene_gn=FKBP15;refGene_tx=XM_003995763.5;refGene_gn=FKBP15;refGene_tx=XM_006939372.4;refGene_gn=FKBP15;refGene_tx=XM_011288490.3
D4 75349172 75349257
refGene_gn=FKBP15;refGene_tx=XM_003995763.5;refGene_gn=FKBP15;refGene_tx=XM_006939372.4;refGene_gn=FKBP15;refGene_tx=XM_011288490.3
D4 75356234 75356350
refGene_gn=FKBP15;refGene_tx=XM_003995763.5;refGene_gn=FKBP15;refGene_tx=XM_006939372.4;refGene_gn=FKBP15;refGene_tx=XM_011288490.3
D4 75365562 75365615
refGene_gn=FKBP15;refGene_tx=XM_003995763.5;refGene_gn=FKBP15;refGene_tx=XM_006939372.4;refGene_gn=FKBP15;refGene_tx=XM_011288490.3
D4 75365687 75366052 refGene_gn=SLC31A1;refGene_tx=XM_023242310.1
D4 75398716 75398882 refGene_gn=SLC31A1;refGene_tx=XM_023242310.1
D4 75399441 75399514 refGene_gn=SLC31A1;refGene_tx=XM_023242310.1

D4 75401898 75402067 refGene_gn=SLC31A1;refGene_tx=XM_023242310.1
D4 75403139 75403341 refGene_gn=SLC31A1;refGene_tx=XM_023242310.1
D4 75409651 75409828
refGene_gn=CDC26;refGene_tx=XM_003995764.4;refGene_gn=CDC26;refGene_tx=XM_006939376.4;refGene_gn=CDC26;refGene_tx=XM_006939377.4;refGene_gn=CDC26;refGene_tx=XM_006939378.3
D4 75413204 75413326
refGene_gn=CDC26;refGene_tx=XM_003995764.4;refGene_gn=CDC26;refGene_tx=XM_006939376.4;refGene_gn=CDC26;refGene_tx=XM_006939377.4;refGene_gn=CDC26;refGene_tx=XM_006939378.3
D4 75413842 75413894
refGene_gn=CDC26;refGene_tx=XM_006939376.4;refGene_gn=CDC26;refGene_tx=XM_006939377.4
D4 75416164 75416191
refGene_gn=PRPF4;refGene_tx=XM_003995765.4;refGene_gn=PRPF4;refGene_tx=XM_019816360.2
D4 75416949 75417130
refGene_gn=PRPF4;refGene_tx=XM_003995765.4;refGene_gn=PRPF4;refGene_tx=XM_019816360.2
D4 75418177 75418364
refGene_gn=PRPF4;refGene_tx=XM_003995765.4;refGene_gn=PRPF4;refGene_tx=XM_019816360.2
D4 75421916 75422004
refGene_gn=PRPF4;refGene_tx=XM_003995765.4;refGene_gn=PRPF4;refGene_tx=XM_019816360.2
D4 75422524 75422604
refGene_gn=PRPF4;refGene_tx=XM_003995765.4;refGene_gn=PRPF4;refGene_tx=XM_019816360.2
D4 75422776 75422870
refGene_gn=PRPF4;refGene_tx=XM_003995765.4;refGene_gn=PRPF4;refGene_tx=XM_019816360.2
D4 75423336 75423431
refGene_gn=PRPF4;refGene_tx=XM_003995765.4;refGene_gn=PRPF4;refGene_tx=XM_019816360.2
D4 75424572 75424654
refGene_gn=PRPF4;refGene_tx=XM_003995765.4;refGene_gn=PRPF4;refGene_tx=XM_019816360.2;refGene_gn=PRPF4;refGene_tx=XM_019816361.1
D4 75425570 75425694
refGene_gn=PRPF4;refGene_tx=XM_003995765.4;refGene_gn=PRPF4;refGene_tx=XM_019816360.2;refGene_gn=PRPF4;refGene_tx=XM_019816361.1
D4 75427295 75427385
refGene_gn=PRPF4;refGene_tx=XM_003995765.4;refGene_gn=PRPF4;refGene_tx=XM_019816360.2;refGene_gn=PRPF4;refGene_tx=XM_019816361.1
D4 75429795 75429918
refGene_gn=PRPF4;refGene_tx=XM_003995765.4;refGene_gn=PRPF4;refGene_tx=XM_019816360.2;refGene_gn=PRPF4;refGene_tx=XM_019816361.1
D4 75430028 75430136
refGene_gn=PRPF4;refGene_tx=XM_003995765.4;refGene_gn=PRPF4;refGene_tx=XM_019816360.2;refGene_gn=PRPF4;refGene_tx=XM_019816361.1
D4 75430226 75430345

refGene_gn=PRPF4;refGene_tx=XM_003995765.4;refGene_gn=PRPF4;refGene_tx=XM_019816360.2;refGene_gn=PRPF4;refGene_tx=XM_019816361.1
D4 75430862 75431056
refGene_gn=PRPF4;refGene_tx=XM_003995765.4;refGene_gn=PRPF4;refGene_tx=XM_019816360.2;refGene_gn=PRPF4;refGene_tx=XM_019816361.1
D4 75435448 75436021 refGene_gn=RNF183;refGene_tx=XM_003995766.4
D4 75457274 75457567
refGene_gn=WDR31;refGene_tx=XM_023242585.1;refGene_gn=WDR31;refGene_tx=XM_023242587.1;refGene_gn=WDR31;refGene_tx=XM_023242588.1;refGene_gn=WDR31;refGene_tx=XM_023242589.1;refGene_gn=WDR31;refGene_tx=XM_023242590.1;refGene_gn=WDR31;refGene_tx=XM_023242591.1;refGene_gn=WDR31;refGene_tx=XM_023242592.1;refGene_gn=WDR31;refGene_tx=XM_023242593.1
D4 75459204 75459367
refGene_gn=WDR31;refGene_tx=XM_023242585.1;refGene_gn=WDR31;refGene_tx=XM_023242587.1;refGene_gn=WDR31;refGene_tx=XM_023242588.1;refGene_gn=WDR31;refGene_tx=XM_023242589.1;refGene_gn=WDR31;refGene_tx=XM_023242590.1;refGene_gn=WDR31;refGene_tx=XM_023242591.1;refGene_gn=WDR31;refGene_tx=XM_023242592.1;refGene_gn=WDR31;refGene_tx=XM_023242593.1
D4 75461512 75461654
refGene_gn=WDR31;refGene_tx=XM_023242585.1;refGene_gn=WDR31;refGene_tx=XM_023242587.1;refGene_gn=WDR31;refGene_tx=XM_023242588.1;refGene_gn=WDR31;refGene_tx=XM_023242589.1;refGene_gn=WDR31;refGene_tx=XM_023242590.1;refGene_gn=WDR31;refGene_tx=XM_023242591.1;refGene_gn=WDR31;refGene_tx=XM_023242592.1;refGene_gn=WDR31;refGene_tx=XM_023242593.1
D4 75462640 75462708
refGene_gn=WDR31;refGene_tx=XM_023242585.1;refGene_gn=WDR31;refGene_tx=XM_023242587.1;refGene_gn=WDR31;refGene_tx=XM_023242588.1;refGene_gn=WDR31;refGene_tx=XM_023242589.1;refGene_gn=WDR31;refGene_tx=XM_023242590.1;refGene_gn=WDR31;refGene_tx=XM_023242591.1;refGene_gn=WDR31;refGene_tx=XM_023242592.1;refGene_gn=WDR31;refGene_tx=XM_023242593.1
D4 75464191 75464292
refGene_gn=WDR31;refGene_tx=XM_023242585.1;refGene_gn=WDR31;refGene_tx=XM_023242587.1;refGene_gn=WDR31;refGene_tx=XM_023242588.1;refGene_gn=WDR31;refGene_tx=XM_023242589.1;refGene_gn=WDR31;refGene_tx=XM_023242590.1;refGene_gn=WDR31;refGene_tx=XM_023242591.1;refGene_gn=WDR31;refGene_tx=XM_023242592.1;refGene_gn=WDR31;refGene_tx=XM_023242593.1
D4 75464389 75464534
refGene_gn=WDR31;refGene_tx=XM_023242585.1;refGene_gn=WDR31;refGene_tx=XM_023242587.1;refGene_gn=WDR31;refGene_tx=XM_023242588.1;refGene_gn=WDR31;refGene_tx=XM_023242589.1;refGene_gn=WDR31;refGene_tx=XM_023242590.1;refGene_gn=WDR31;refGene_tx=XM_023242591.1;refGene_gn=WDR31;refGene_tx=XM_023242592.1;refGene_gn=WDR31;refGene_tx=XM_023242593.1
D4 75465946 75466021
refGene_gn=WDR31;refGene_tx=XM_023242585.1;refGene_gn=WDR31;refGene_tx=XM_023242587.1;refGene_gn=WDR31;refGene_tx=XM_023242588.1;refGene_gn=WDR31;refGene_tx=XM_023242589.1;refGene_gn=WDR31;refGene_tx=XM_023242590.1;refGene_gn=WDR31;refGene_tx=XM_023242591.1;refGene_gn=WDR31;refGene_tx=XM_023242592.1;refGene_gn=WDR31;refGene_tx=XM_023242593.1
D4 75468124 75468257
refGene_gn=WDR31;refGene_tx=XM_023242585.1;refGene_gn=WDR31;refGene_tx

=XM_023242587.1;refGene_gn=WDR31;refGene_tx=XM_023242588.1;refGene_gn=WDR31;refGene_tx=XM_023242589.1;refGene_gn=WDR31;refGene_tx=XM_023242590.1;refGene_gn=WDR31;refGene_tx=XM_023242591.1;refGene_gn=WDR31;refGene_tx=XM_023242592.1;refGene_gn=WDR31;refGene_tx=XM_023242593.1
D4	75468816	75468948	refGene_gn=WDR31;refGene_tx=XM_023242585.1;refGene_gn=WDR31;refGene_tx=XM_023242587.1;refGene_gn=WDR31;refGene_tx=XM_023242588.1;refGene_gn=WDR31;refGene_tx=XM_023242589.1;refGene_gn=WDR31;refGene_tx=XM_023242590.1;refGene_gn=WDR31;refGene_tx=XM_023242591.1;refGene_gn=WDR31;refGene_tx=XM_023242592.1;refGene_gn=WDR31;refGene_tx=XM_023242593.1
D4	75479157	75479243	refGene_gn=WDR31;refGene_tx=XM_023242585.1;refGene_gn=WDR31;refGene_tx=XM_023242587.1;refGene_gn=WDR31;refGene_tx=XM_023242592.1;refGene_gn=WDR31;refGene_tx=XM_023242593.1
D4	75486715	75486892	refGene_gn=BSPRY;refGene_tx=XM_023242594.1
D4	75491196	75491295	refGene_gn=BSPRY;refGene_tx=XM_023242594.1
D4	75498163	75498394	refGene_gn=BSPRY;refGene_tx=XM_023242594.1
D4	75499497	75499523	refGene_gn=BSPRY;refGene_tx=XM_023242594.1
D4	75504977	75504999	refGene_gn=BSPRY;refGene_tx=XM_023242594.1
D4	75505000	75505103	refGene_gn=BSPRY;refGene_tx=XM_023242594.1
D4	75506340	75506867	refGene_gn=BSPRY;refGene_tx=XM_023242594.1
D4	75510035	75510785	refGene_gn=HDHD3;refGene_tx=XM_003995767.5
D4	75521740	75521802	refGene_gn=ALAD;refGene_tx=XM_006939387.3;refGene_gn=ALAD;refGene_tx=XM_019816383.1;refGene_gn=ALAD;refGene_tx=XM_019816384.1
D4	75522453	75522583	refGene_gn=ALAD;refGene_tx=XM_006939387.3;refGene_gn=ALAD;refGene_tx=XM_019816383.1;refGene_gn=ALAD;refGene_tx=XM_019816384.1
D4	75523181	75523268	refGene_gn=ALAD;refGene_tx=XM_006939387.3;refGene_gn=ALAD;refGene_tx=XM_019816383.1;refGene_gn=ALAD;refGene_tx=XM_019816384.1
D4	75523356	75523444	refGene_gn=ALAD;refGene_tx=XM_006939387.3;refGene_gn=ALAD;refGene_tx=XM_019816383.1;refGene_gn=ALAD;refGene_tx=XM_019816384.1
D4	75523531	75523587	refGene_gn=ALAD;refGene_tx=XM_006939387.3;refGene_gn=ALAD;refGene_tx=XM_019816383.1;refGene_gn=ALAD;refGene_tx=XM_019816384.1
D4	75524121	75524210	refGene_gn=ALAD;refGene_tx=XM_006939387.3;refGene_gn=ALAD;refGene_tx=XM_019816383.1;refGene_gn=ALAD;refGene_tx=XM_019816384.1
D4	75524310	75524394	refGene_gn=ALAD;refGene_tx=XM_006939387.3;refGene_gn=ALAD;refGene_tx=XM_019816383.1;refGene_gn=ALAD;refGene_tx=XM_019816384.1
D4	75524498	75524634	refGene_gn=ALAD;refGene_tx=XM_006939387.3;refGene_gn=ALAD;refGene_tx=XM_019816383.1;refGene_gn=ALAD;refGene_tx=XM_019816384.1
D4	75524974	75525071	refGene_gn=ALAD;refGene_tx=XM_006939387.3;refGene_gn=ALAD;refGene_tx=XM_019816383.1;refGene_gn=ALAD;refGene_tx=XM_019816384.1
D4	75525543	75525594

refGene_gn=ALAD;refGene_tx=XM_006939387.3;refGene_gn=ALAD;refGene_tx=XM_019816383.1;refGene_gn=ALAD;refGene_tx=XM_019816384.1
D4 75526574 75526687
refGene_gn=ALAD;refGene_tx=XM_006939387.3;refGene_gn=ALAD;refGene_tx=XM_019816383.1;refGene_gn=ALAD;refGene_tx=XM_019816384.1
D4 75537969 75538142
refGene_gn=POLE3;refGene_tx=XM_003995771.5;refGene_gn=POLE3;refGene_tx=XM_011288488.3
D4 75538407 75538526
refGene_gn=POLE3;refGene_tx=XM_003995771.5;refGene_gn=POLE3;refGene_tx=XM_011288488.3
D4 75538894 75538980
refGene_gn=POLE3;refGene_tx=XM_003995771.5;refGene_gn=POLE3;refGene_tx=XM_011288488.3
D4 75539082 75539148
refGene_gn=POLE3;refGene_tx=XM_003995771.5;refGene_gn=POLE3;refGene_tx=XM_011288488.3
D4 75542111 75542262
refGene_gn=CD4H9orf43;refGene_tx=XM_006939388.3;refGene_gn=CD4H9orf43;refGene_tx=XM_019816375.2;refGene_gn=CD4H9orf43;refGene_tx=XM_019816378.2;refGene_gn=CD4H9orf43;refGene_tx=XM_019816379.2;refGene_gn=CD4H9orf43;refGene_tx=XM_019816380.2
D4 75542362 75542513
refGene_gn=CD4H9orf43;refGene_tx=XM_006939388.3;refGene_gn=CD4H9orf43;refGene_tx=XM_019816375.2;refGene_gn=CD4H9orf43;refGene_tx=XM_019816378.2;refGene_gn=CD4H9orf43;refGene_tx=XM_019816379.2;refGene_gn=CD4H9orf43;refGene_tx=XM_019816380.2
D4 75546565 75546623
refGene_gn=CD4H9orf43;refGene_tx=XM_006939388.3;refGene_gn=CD4H9orf43;refGene_tx=XM_019816375.2;refGene_gn=CD4H9orf43;refGene_tx=XM_019816378.2;refGene_gn=CD4H9orf43;refGene_tx=XM_019816379.2;refGene_gn=CD4H9orf43;refGene_tx=XM_019816380.2
D4 75547507 75547608
refGene_gn=CD4H9orf43;refGene_tx=XM_006939388.3;refGene_gn=CD4H9orf43;refGene_tx=XM_019816375.2;refGene_gn=CD4H9orf43;refGene_tx=XM_019816378.2;refGene_gn=CD4H9orf43;refGene_tx=XM_019816379.2;refGene_gn=CD4H9orf43;refGene_tx=XM_019816380.2
D4 75549814 75549872
refGene_gn=CD4H9orf43;refGene_tx=XM_006939388.3;refGene_gn=CD4H9orf43;refGene_tx=XM_019816375.2;refGene_gn=CD4H9orf43;refGene_tx=XM_019816378.2;refGene_gn=CD4H9orf43;refGene_tx=XM_019816380.2
D4 75550607 75550777
refGene_gn=CD4H9orf43;refGene_tx=XM_006939388.3;refGene_gn=CD4H9orf43;refGene_tx=XM_019816375.2;refGene_gn=CD4H9orf43;refGene_tx=XM_019816378.2;refGene_gn=CD4H9orf43;refGene_tx=XM_019816380.2
D4 75551665 75551816
refGene_gn=CD4H9orf43;refGene_tx=XM_006939388.3;refGene_gn=CD4H9orf43;refGene_tx=XM_019816375.2;refGene_gn=CD4H9orf43;refGene_tx=XM_019816378.2;refGene_gn=CD4H9orf43;refGene_tx=XM_019816379.2;refGene_gn=CD4H9orf43;refGene_tx=XM_019816380.2

D4 75553320 75553411
refGene_gn=CD4H9orf43;refGene_tx=XM_006939388.3;refGene_gn=CD4H9orf43;
refGene_tx=XM_019816375.2;refGene_gn=CD4H9orf43;refGene_tx=XM_01981637
8.2;refGene_gn=CD4H9orf43;refGene_tx=XM_019816379.2;refGene_gn=CD4H9or
f43;refGene_tx=XM_019816380.2
D4 75553678 75553761
refGene_gn=CD4H9orf43;refGene_tx=XM_006939388.3;refGene_gn=CD4H9orf43;
refGene_tx=XM_019816375.2;refGene_gn=CD4H9orf43;refGene_tx=XM_01981637
8.2;refGene_gn=CD4H9orf43;refGene_tx=XM_019816379.2;refGene_gn=CD4H9or
f43;refGene_tx=XM_019816380.2
D4 75553963 75554051
refGene_gn=CD4H9orf43;refGene_tx=XM_006939388.3;refGene_gn=CD4H9orf43;
refGene_tx=XM_019816375.2;refGene_gn=CD4H9orf43;refGene_tx=XM_01981637
8.2;refGene_gn=CD4H9orf43;refGene_tx=XM_019816379.2;refGene_gn=CD4H9or
f43;refGene_tx=XM_019816380.2
D4 75556424 75556504
refGene_gn=CD4H9orf43;refGene_tx=XM_006939388.3;refGene_gn=CD4H9orf43;
refGene_tx=XM_019816375.2;refGene_gn=CD4H9orf43;refGene_tx=XM_01981637
8.2;refGene_gn=CD4H9orf43;refGene_tx=XM_019816379.2;refGene_gn=CD4H9or
f43;refGene_tx=XM_019816380.2
D4 75557634 75557692
refGene_gn=CD4H9orf43;refGene_tx=XM_006939388.3;refGene_gn=CD4H9orf43;
refGene_tx=XM_019816375.2;refGene_gn=CD4H9orf43;refGene_tx=XM_01981637
8.2;refGene_gn=CD4H9orf43;refGene_tx=XM_019816379.2;refGene_gn=CD4H9or
f43;refGene_tx=XM_019816380.2
D4 75557901 75558086
refGene_gn=CD4H9orf43;refGene_tx=XM_006939388.3;refGene_gn=CD4H9orf43;
refGene_tx=XM_019816375.2;refGene_gn=CD4H9orf43;refGene_tx=XM_01981637
8.2;refGene_gn=CD4H9orf43;refGene_tx=XM_019816379.2;refGene_gn=CD4H9or
f43;refGene_tx=XM_019816380.2
D4 75576495 75576613 refGene_gn=RGS3;refGene_tx=XM_019816366.2
D4 75578230 75578309
refGene_gn=RGS3;refGene_tx=XM_011288486.3;refGene_gn=RGS3;refGene_tx=X
M_019816367.2
D4 75592307 75592358
refGene_gn=RGS3;refGene_tx=XM_011288486.3;refGene_gn=RGS3;refGene_tx=X
M_019816366.2;refGene_gn=RGS3;refGene_tx=XM_019816367.2
D4 75594462 75594521
refGene_gn=RGS3;refGene_tx=XM_011288486.3;refGene_gn=RGS3;refGene_tx=X
M_019816366.2;refGene_gn=RGS3;refGene_tx=XM_019816367.2
D4 75595556 75595654
refGene_gn=RGS3;refGene_tx=XM_011288486.3;refGene_gn=RGS3;refGene_tx=X
M_019816366.2;refGene_gn=RGS3;refGene_tx=XM_019816367.2
D4 75596648 75596717
refGene_gn=RGS3;refGene_tx=XM_011288486.3;refGene_gn=RGS3;refGene_tx=X
M_019816366.2;refGene_gn=RGS3;refGene_tx=XM_019816367.2
D4 75605052 75605113
refGene_gn=RGS3;refGene_tx=XM_011288486.3;refGene_gn=RGS3;refGene_tx=X
M_019816366.2;refGene_gn=RGS3;refGene_tx=XM_019816367.2
D4 75606289 75606380

refGene_gn=RGS3;refGene_tx=XM_011288486.3;refGene_gn=RGS3;refGene_tx=XM_019816366.2;refGene_gn=RGS3;refGene_tx=XM_019816367.2
D4 75607018 75607074
refGene_gn=RGS3;refGene_tx=XM_011288486.3;refGene_gn=RGS3;refGene_tx=XM_019816366.2;refGene_gn=RGS3;refGene_tx=XM_019816367.2
D4 75609809 75609863
refGene_gn=RGS3;refGene_tx=XM_011288482.3;refGene_gn=RGS3;refGene_tx=XM_019816372.2
D4 75614436 75614518
refGene_gn=RGS3;refGene_tx=XM_011288482.3;refGene_gn=RGS3;refGene_tx=XM_011288486.3;refGene_gn=RGS3;refGene_tx=XM_019816366.2;refGene_gn=RGS3;refGene_tx=XM_019816367.2;refGene_gn=RGS3;refGene_tx=XM_019816372.2
D4 75615346 75615452
refGene_gn=RGS3;refGene_tx=XM_011288482.3;refGene_gn=RGS3;refGene_tx=XM_011288486.3;refGene_gn=RGS3;refGene_tx=XM_019816366.2;refGene_gn=RGS3;refGene_tx=XM_019816367.2;refGene_gn=RGS3;refGene_tx=XM_019816372.2
D4 75616181 75616533
refGene_gn=RGS3;refGene_tx=XM_011288482.3;refGene_gn=RGS3;refGene_tx=XM_011288486.3;refGene_gn=RGS3;refGene_tx=XM_019816366.2;refGene_gn=RGS3;refGene_tx=XM_019816367.2;refGene_gn=RGS3;refGene_tx=XM_019816372.2
D4 75617402 75617442
refGene_gn=RGS3;refGene_tx=XM_011288482.3;refGene_gn=RGS3;refGene_tx=XM_011288486.3;refGene_gn=RGS3;refGene_tx=XM_019816366.2;refGene_gn=RGS3;refGene_tx=XM_019816367.2;refGene_gn=RGS3;refGene_tx=XM_019816372.2
D4 75622991 75623188
refGene_gn=RGS3;refGene_tx=XM_011288482.3;refGene_gn=RGS3;refGene_tx=XM_011288486.3;refGene_gn=RGS3;refGene_tx=XM_019816366.2;refGene_gn=RGS3;refGene_tx=XM_019816367.2;refGene_gn=RGS3;refGene_tx=XM_019816372.2
D4 75625696 75625780
refGene_gn=RGS3;refGene_tx=XM_011288482.3;refGene_gn=RGS3;refGene_tx=XM_011288486.3;refGene_gn=RGS3;refGene_tx=XM_019816366.2;refGene_gn=RGS3;refGene_tx=XM_019816367.2;refGene_gn=RGS3;refGene_tx=XM_019816372.2
D4 75631132 75631244
refGene_gn=RGS3;refGene_tx=XM_011288482.3;refGene_gn=RGS3;refGene_tx=XM_011288486.3;refGene_gn=RGS3;refGene_tx=XM_019816366.2;refGene_gn=RGS3;refGene_tx=XM_019816367.2;refGene_gn=RGS3;refGene_tx=XM_019816372.2
D4 75637742 75637786
refGene_gn=RGS3;refGene_tx=XM_011288482.3;refGene_gn=RGS3;refGene_tx=XM_011288486.3;refGene_gn=RGS3;refGene_tx=XM_019816366.2;refGene_gn=RGS3;refGene_tx=XM_019816367.2;refGene_gn=RGS3;refGene_tx=XM_019816372.2
D4 75644173 75644296
refGene_gn=RGS3;refGene_tx=XM_011288482.3;refGene_gn=RGS3;refGene_tx=XM_011288486.3;refGene_gn=RGS3;refGene_tx=XM_019816366.2;refGene_gn=RGS3;refGene_tx=XM_019816367.2;refGene_gn=RGS3;refGene_tx=XM_019816372.2
D4 75649006 75649114
refGene_gn=RGS3;refGene_tx=XM_011288486.3;refGene_gn=RGS3;refGene_tx=XM_019816372.2
D4 75684833 75685772
refGene_gn=RGS3;refGene_tx=XM_011288482.3;refGene_gn=RGS3;refGene_tx=XM_011288483.3;refGene_gn=RGS3;refGene_tx=XM_011288484.3;refGene_gn=RGS

3;refGene_tx=XM_011288485.3;refGene_gn=RGS3;refGene_tx=XM_019816366.2;
refGene_gn=RGS3;refGene_tx=XM_019816367.2;refGene_gn=RGS3;refGene_tx=X
M_019816371.2
D4 75693374 75693439
refGene_gn=RGS3;refGene_tx=XM_011288482.3;refGene_gn=RGS3;refGene_tx=X
M_011288483.3;refGene_gn=RGS3;refGene_tx=XM_011288484.3;refGene_gn=RGS
3;refGene_tx=XM_011288485.3;refGene_gn=RGS3;refGene_tx=XM_019816366.2;
refGene_gn=RGS3;refGene_tx=XM_019816367.2;refGene_gn=RGS3;refGene_tx=X
M_019816371.2
D4 75697021 75697083 refGene_gn=RGS3;refGene_tx=XM_019816373.1
D4 75697182 75697284
refGene_gn=RGS3;refGene_tx=XM_011288482.3;refGene_gn=RGS3;refGene_tx=X
M_011288483.3;refGene_gn=RGS3;refGene_tx=XM_011288484.3;refGene_gn=RGS
3;refGene_tx=XM_011288485.3;refGene_gn=RGS3;refGene_tx=XM_019816366.2;
refGene_gn=RGS3;refGene_tx=XM_019816367.2;refGene_gn=RGS3;refGene_tx=X
M_019816371.2;refGene_gn=RGS3;refGene_tx=XM_019816373.1
D4 75697688 75697750
refGene_gn=RGS3;refGene_tx=XM_011288482.3;refGene_gn=RGS3;refGene_tx=X
M_011288483.3;refGene_gn=RGS3;refGene_tx=XM_011288484.3;refGene_gn=RGS
3;refGene_tx=XM_011288485.3;refGene_gn=RGS3;refGene_tx=XM_019816366.2;
refGene_gn=RGS3;refGene_tx=XM_019816367.2;refGene_gn=RGS3;refGene_tx=X
M_019816371.2;refGene_gn=RGS3;refGene_tx=XM_019816373.1
D4 75698361 75698528
refGene_gn=RGS3;refGene_tx=XM_011288482.3;refGene_gn=RGS3;refGene_tx=X
M_011288483.3;refGene_gn=RGS3;refGene_tx=XM_011288484.3;refGene_gn=RGS
3;refGene_tx=XM_011288485.3;refGene_gn=RGS3;refGene_tx=XM_019816366.2;
refGene_gn=RGS3;refGene_tx=XM_019816367.2;refGene_gn=RGS3;refGene_tx=X
M_019816371.2;refGene_gn=RGS3;refGene_tx=XM_019816373.1
D4 75699548 75699734
refGene_gn=RGS3;refGene_tx=XM_011288482.3;refGene_gn=RGS3;refGene_tx=X
M_011288483.3;refGene_gn=RGS3;refGene_tx=XM_011288484.3;refGene_gn=RGS
3;refGene_tx=XM_011288485.3;refGene_gn=RGS3;refGene_tx=XM_019816366.2;
refGene_gn=RGS3;refGene_tx=XM_019816367.2;refGene_gn=RGS3;refGene_tx=X
M_019816371.2;refGene_gn=RGS3;refGene_tx=XM_019816373.1
D4 75945025 75945430
refGene_gn=LOC111557480;refGene_tx=XM_023242311.1
D4 75945572 75945605
refGene_gn=ZNF618;refGene_tx=XM_023242597.1;refGene_gn=ZNF618;refGene_
tx=XM_023242599.1;refGene_gn=ZNF618;refGene_tx=XM_023242600.1;refGene_
gn=ZNF618;refGene_tx=XM_023242601.1;refGene_gn=ZNF618;refGene_tx=XM_02
3242602.1;refGene_gn=ZNF618;refGene_tx=XM_023242603.1;refGene_gn=ZNF61
8;refGene_tx=XM_023242604.1;refGene_gn=ZNF618;refGene_tx=XM_023242605.
1;refGene_gn=ZNF618;refGene_tx=XM_023242606.1;refGene_gn=ZNF618;refGen
e_tx=XM_023242607.1;refGene_gn=ZNF618;refGene_tx=XM_023242608.1;refGen
e_gn=ZNF618;refGene_tx=XM_023242610.1;refGene_gn=ZNF618;refGene_tx=XM_
023242611.1;refGene_gn=ZNF618;refGene_tx=XM_023242612.1;refGene_gn=ZNF
618;refGene_tx=XM_023242613.1;refGene_gn=ZNF618;refGene_tx=XM_02324261
4.1;refGene_gn=ZNF618;refGene_tx=XM_023242615.1;refGene_gn=ZNF618;refG
ene_tx=XM_023242616.1;refGene_gn=ZNF618;refGene_tx=XM_023242617.1;refG
ene_gn=ZNF618;refGene_tx=XM_023242618.1

D4 76045946 76045990
refGene_gn=ZNF618;refGene_tx=XM_023242597.1;refGene_gn=ZNF618;refGene_tx=XM_023242599.1;refGene_gn=ZNF618;refGene_tx=XM_023242600.1;refGene_gn=ZNF618;refGene_tx=XM_023242601.1;refGene_gn=ZNF618;refGene_tx=XM_023242602.1;refGene_gn=ZNF618;refGene_tx=XM_023242603.1;refGene_gn=ZNF618;refGene_tx=XM_023242604.1;refGene_gn=ZNF618;refGene_tx=XM_023242605.1;refGene_gn=ZNF618;refGene_tx=XM_023242606.1;refGene_gn=ZNF618;refGene_tx=XM_023242607.1;refGene_gn=ZNF618;refGene_tx=XM_023242608.1;refGene_gn=ZNF618;refGene_tx=XM_023242610.1;refGene_gn=ZNF618;refGene_tx=XM_023242611.1;refGene_gn=ZNF618;refGene_tx=XM_023242612.1;refGene_gn=ZNF618;refGene_tx=XM_023242613.1;refGene_gn=ZNF618;refGene_tx=XM_023242614.1;refGene_gn=ZNF618;refGene_tx=XM_023242615.1;refGene_gn=ZNF618;refGene_tx=XM_023242616.1;refGene_gn=ZNF618;refGene_tx=XM_023242617.1;refGene_gn=ZNF618;refGene_tx=XM_023242618.1
D4 76063998 76064258
refGene_gn=ZNF618;refGene_tx=XM_023242597.1;refGene_gn=ZNF618;refGene_tx=XM_023242599.1;refGene_gn=ZNF618;refGene_tx=XM_023242600.1;refGene_gn=ZNF618;refGene_tx=XM_023242601.1;refGene_gn=ZNF618;refGene_tx=XM_023242602.1;refGene_gn=ZNF618;refGene_tx=XM_023242603.1;refGene_gn=ZNF618;refGene_tx=XM_023242604.1;refGene_gn=ZNF618;refGene_tx=XM_023242605.1;refGene_gn=ZNF618;refGene_tx=XM_023242606.1;refGene_gn=ZNF618;refGene_tx=XM_023242607.1;refGene_gn=ZNF618;refGene_tx=XM_023242608.1;refGene_gn=ZNF618;refGene_tx=XM_023242610.1;refGene_gn=ZNF618;refGene_tx=XM_023242611.1;refGene_gn=ZNF618;refGene_tx=XM_023242612.1;refGene_gn=ZNF618;refGene_tx=XM_023242613.1;refGene_gn=ZNF618;refGene_tx=XM_023242614.1;refGene_gn=ZNF618;refGene_tx=XM_023242615.1;refGene_gn=ZNF618;refGene_tx=XM_023242616.1;refGene_gn=ZNF618;refGene_tx=XM_023242617.1;refGene_gn=ZNF618;refGene_tx=XM_023242618.1
D4 76073574 76073670
refGene_gn=ZNF618;refGene_tx=XM_023242597.1;refGene_gn=ZNF618;refGene_tx=XM_023242599.1;refGene_gn=ZNF618;refGene_tx=XM_023242600.1;refGene_gn=ZNF618;refGene_tx=XM_023242601.1;refGene_gn=ZNF618;refGene_tx=XM_023242602.1;refGene_gn=ZNF618;refGene_tx=XM_023242603.1;refGene_gn=ZNF618;refGene_tx=XM_023242605.1;refGene_gn=ZNF618;refGene_tx=XM_023242606.1;refGene_gn=ZNF618;refGene_tx=XM_023242610.1;refGene_gn=ZNF618;refGene_tx=XM_023242611.1;refGene_gn=ZNF618;refGene_tx=XM_023242612.1;refGene_gn=ZNF618;refGene_tx=XM_023242613.1;refGene_gn=ZNF618;refGene_tx=XM_023242614.1;refGene_gn=ZNF618;refGene_tx=XM_023242615.1;refGene_gn=ZNF618;refGene_tx=XM_023242617.1;refGene_gn=ZNF618;refGene_tx=XM_023242618.1
D4 76077088 76077166
refGene_gn=ZNF618;refGene_tx=XM_023242597.1;refGene_gn=ZNF618;refGene_tx=XM_023242599.1;refGene_gn=ZNF618;refGene_tx=XM_023242600.1;refGene_gn=ZNF618;refGene_tx=XM_023242601.1;refGene_gn=ZNF618;refGene_tx=XM_023242602.1;refGene_gn=ZNF618;refGene_tx=XM_023242603.1;refGene_gn=ZNF618;refGene_tx=XM_023242604.1;refGene_gn=ZNF618;refGene_tx=XM_023242605.1;refGene_gn=ZNF618;refGene_tx=XM_023242606.1;refGene_gn=ZNF618;refGene_tx=XM_023242607.1;refGene_gn=ZNF618;refGene_tx=XM_023242608.1;refGene_gn=ZNF618;refGene_tx=XM_023242610.1;refGene_gn=ZNF618;refGene_tx=XM_023242611.1;refGene_gn=ZNF618;refGene_tx=XM_023242612.1;refGene_gn=ZNF

618;refGene_tx=XM_023242613.1;refGene_gn=ZNF618;refGene_tx=XM_02324261
4.1;refGene_gn=ZNF618;refGene_tx=XM_023242615.1;refGene_gn=ZNF618;refG
ene_tx=XM_023242616.1;refGene_gn=ZNF618;refGene_tx=XM_023242617.1;refG
ene_gn=ZNF618;refGene_tx=XM_023242618.1
D4 76077784 76077823
refGene_gn=ZNF618;refGene_tx=XM_023242597.1;refGene_gn=ZNF618;refGene_
tx=XM_023242599.1;refGene_gn=ZNF618;refGene_tx=XM_023242600.1;refGene_
gn=ZNF618;refGene_tx=XM_023242601.1;refGene_gn=ZNF618;refGene_tx=XM_02
3242602.1;refGene_gn=ZNF618;refGene_tx=XM_023242603.1;refGene_gn=ZNF61
8;refGene_tx=XM_023242604.1;refGene_gn=ZNF618;refGene_tx=XM_023242605.
1;refGene_gn=ZNF618;refGene_tx=XM_023242606.1;refGene_gn=ZNF618;refGen
e_tx=XM_023242607.1;refGene_gn=ZNF618;refGene_tx=XM_023242608.1;refGen
e_gn=ZNF618;refGene_tx=XM_023242610.1;refGene_gn=ZNF618;refGene_tx=XM_
023242611.1;refGene_gn=ZNF618;refGene_tx=XM_023242612.1;refGene_gn=ZNF
618;refGene_tx=XM_023242613.1;refGene_gn=ZNF618;refGene_tx=XM_02324261
4.1;refGene_gn=ZNF618;refGene_tx=XM_023242615.1;refGene_gn=ZNF618;refG
ene_tx=XM_023242616.1;refGene_gn=ZNF618;refGene_tx=XM_023242617.1;refG
ene_gn=ZNF618;refGene_tx=XM_023242618.1
D4 76082948 76083038
refGene_gn=ZNF618;refGene_tx=XM_023242597.1;refGene_gn=ZNF618;refGene_
tx=XM_023242599.1;refGene_gn=ZNF618;refGene_tx=XM_023242600.1;refGene_
gn=ZNF618;refGene_tx=XM_023242601.1;refGene_gn=ZNF618;refGene_tx=XM_02
3242602.1;refGene_gn=ZNF618;refGene_tx=XM_023242603.1;refGene_gn=ZNF61
8;refGene_tx=XM_023242604.1;refGene_gn=ZNF618;refGene_tx=XM_023242605.
1;refGene_gn=ZNF618;refGene_tx=XM_023242606.1;refGene_gn=ZNF618;refGen
e_tx=XM_023242607.1;refGene_gn=ZNF618;refGene_tx=XM_023242608.1;refGen
e_gn=ZNF618;refGene_tx=XM_023242610.1;refGene_gn=ZNF618;refGene_tx=XM_
023242611.1;refGene_gn=ZNF618;refGene_tx=XM_023242612.1;refGene_gn=ZNF
618;refGene_tx=XM_023242613.1;refGene_gn=ZNF618;refGene_tx=XM_02324261
4.1;refGene_gn=ZNF618;refGene_tx=XM_023242615.1;refGene_gn=ZNF618;refG
ene_tx=XM_023242616.1;refGene_gn=ZNF618;refGene_tx=XM_023242617.1;refG
ene_gn=ZNF618;refGene_tx=XM_023242618.1
D4 76083951 76083987
refGene_gn=ZNF618;refGene_tx=XM_023242597.1;refGene_gn=ZNF618;refGene_
tx=XM_023242599.1;refGene_gn=ZNF618;refGene_tx=XM_023242601.1;refGene_
gn=ZNF618;refGene_tx=XM_023242602.1;refGene_gn=ZNF618;refGene_tx=XM_02
3242603.1;refGene_gn=ZNF618;refGene_tx=XM_023242604.1;refGene_gn=ZNF61
8;refGene_tx=XM_023242606.1;refGene_gn=ZNF618;refGene_tx=XM_023242607.
1;refGene_gn=ZNF618;refGene_tx=XM_023242608.1;refGene_gn=ZNF618;refGen
e_tx=XM_023242610.1;refGene_gn=ZNF618;refGene_tx=XM_023242611.1;refGen
e_gn=ZNF618;refGene_tx=XM_023242612.1;refGene_gn=ZNF618;refGene_tx=XM_
023242613.1;refGene_gn=ZNF618;refGene_tx=XM_023242614.1;refGene_gn=ZNF
618;refGene_tx=XM_023242615.1;refGene_gn=ZNF618;refGene_tx=XM_02324261
6.1;refGene_gn=ZNF618;refGene_tx=XM_023242617.1;refGene_gn=ZNF618;refG
ene_tx=XM_023242618.1
D4 76084079 76084157
refGene_gn=ZNF618;refGene_tx=XM_023242597.1;refGene_gn=ZNF618;refGene_
tx=XM_023242599.1;refGene_gn=ZNF618;refGene_tx=XM_023242600.1;refGene_
gn=ZNF618;refGene_tx=XM_023242601.1;refGene_gn=ZNF618;refGene_tx=XM_02
3242602.1;refGene_gn=ZNF618;refGene_tx=XM_023242603.1;refGene_gn=ZNF61

8;refGene_tx=XM_023242604.1;refGene_gn=ZNF618;refGene_tx=XM_023242605.
1;refGene_gn=ZNF618;refGene_tx=XM_023242606.1;refGene_gn=ZNF618;refGen
e_tx=XM_023242607.1;refGene_gn=ZNF618;refGene_tx=XM_023242608.1;refGen
e_gn=ZNF618;refGene_tx=XM_023242610.1;refGene_gn=ZNF618;refGene_tx=XM_
023242611.1;refGene_gn=ZNF618;refGene_tx=XM_023242612.1;refGene_gn=ZNF
618;refGene_tx=XM_023242613.1;refGene_gn=ZNF618;refGene_tx=XM_02324261
4.1;refGene_gn=ZNF618;refGene_tx=XM_023242615.1;refGene_gn=ZNF618;refG
ene_tx=XM_023242616.1;refGene_gn=ZNF618;refGene_tx=XM_023242617.1;refG
ene_gn=ZNF618;refGene_tx=XM_023242618.1
D4 76091198 76091258
refGene_gn=ZNF618;refGene_tx=XM_023242597.1;refGene_gn=ZNF618;refGene_
tx=XM_023242599.1;refGene_gn=ZNF618;refGene_tx=XM_023242600.1;refGene_
gn=ZNF618;refGene_tx=XM_023242601.1;refGene_gn=ZNF618;refGene_tx=XM_02
3242602.1;refGene_gn=ZNF618;refGene_tx=XM_023242604.1;refGene_gn=ZNF61
8;refGene_tx=XM_023242610.1;refGene_gn=ZNF618;refGene_tx=XM_023242611.
1;refGene_gn=ZNF618;refGene_tx=XM_023242612.1;refGene_gn=ZNF618;refGen
e_tx=XM_023242613.1;refGene_gn=ZNF618;refGene_tx=XM_023242616.1;refGen
e_gn=ZNF618;refGene_tx=XM_023242617.1
D4 76091772 76091862
refGene_gn=ZNF618;refGene_tx=XM_023242597.1;refGene_gn=ZNF618;refGene_
tx=XM_023242599.1;refGene_gn=ZNF618;refGene_tx=XM_023242600.1;refGene_
gn=ZNF618;refGene_tx=XM_023242601.1;refGene_gn=ZNF618;refGene_tx=XM_02
3242602.1;refGene_gn=ZNF618;refGene_tx=XM_023242603.1;refGene_gn=ZNF61
8;refGene_tx=XM_023242604.1;refGene_gn=ZNF618;refGene_tx=XM_023242605.
1;refGene_gn=ZNF618;refGene_tx=XM_023242606.1;refGene_gn=ZNF618;refGen
e_tx=XM_023242607.1;refGene_gn=ZNF618;refGene_tx=XM_023242608.1;refGen
e_gn=ZNF618;refGene_tx=XM_023242610.1;refGene_gn=ZNF618;refGene_tx=XM_
023242611.1;refGene_gn=ZNF618;refGene_tx=XM_023242612.1;refGene_gn=ZNF
618;refGene_tx=XM_023242613.1;refGene_gn=ZNF618;refGene_tx=XM_02324261
4.1;refGene_gn=ZNF618;refGene_tx=XM_023242615.1;refGene_gn=ZNF618;refG
ene_tx=XM_023242616.1;refGene_gn=ZNF618;refGene_tx=XM_023242617.1;refG
ene_gn=ZNF618;refGene_tx=XM_023242618.1
D4 76103010 76103250
refGene_gn=ZNF618;refGene_tx=XM_023242597.1;refGene_gn=ZNF618;refGene_
tx=XM_023242599.1;refGene_gn=ZNF618;refGene_tx=XM_023242600.1;refGene_
gn=ZNF618;refGene_tx=XM_023242601.1;refGene_gn=ZNF618;refGene_tx=XM_02
3242602.1;refGene_gn=ZNF618;refGene_tx=XM_023242603.1;refGene_gn=ZNF61
8;refGene_tx=XM_023242604.1;refGene_gn=ZNF618;refGene_tx=XM_023242605.
1;refGene_gn=ZNF618;refGene_tx=XM_023242606.1;refGene_gn=ZNF618;refGen
e_tx=XM_023242607.1;refGene_gn=ZNF618;refGene_tx=XM_023242608.1
D4 76106926 76107010
refGene_gn=ZNF618;refGene_tx=XM_023242597.1;refGene_gn=ZNF618;refGene_
tx=XM_023242599.1;refGene_gn=ZNF618;refGene_tx=XM_023242600.1;refGene_
gn=ZNF618;refGene_tx=XM_023242601.1;refGene_gn=ZNF618;refGene_tx=XM_02
3242602.1;refGene_gn=ZNF618;refGene_tx=XM_023242603.1;refGene_gn=ZNF61
8;refGene_tx=XM_023242604.1;refGene_gn=ZNF618;refGene_tx=XM_023242605.
1;refGene_gn=ZNF618;refGene_tx=XM_023242606.1;refGene_gn=ZNF618;refGen
e_tx=XM_023242607.1;refGene_gn=ZNF618;refGene_tx=XM_023242608.1;refGen
e_gn=ZNF618;refGene_tx=XM_023242610.1;refGene_gn=ZNF618;refGene_tx=XM_
023242611.1;refGene_gn=ZNF618;refGene_tx=XM_023242612.1;refGene_gn=ZNF

618;refGene_tx=XM_023242613.1;refGene_gn=ZNF618;refGene_tx=XM_02324261
4.1;refGene_gn=ZNF618;refGene_tx=XM_023242615.1;refGene_gn=ZNF618;refG
ene_tx=XM_023242616.1
D4 76109027 76109072
refGene_gn=ZNF618;refGene_tx=XM_023242597.1;refGene_gn=ZNF618;refGene_
tx=XM_023242599.1;refGene_gn=ZNF618;refGene_tx=XM_023242600.1;refGene_
gn=ZNF618;refGene_tx=XM_023242603.1;refGene_gn=ZNF618;refGene_tx=XM_02
3242604.1;refGene_gn=ZNF618;refGene_tx=XM_023242605.1;refGene_gn=ZNF61
8;refGene_tx=XM_023242610.1;refGene_gn=ZNF618;refGene_tx=XM_023242611.
1;refGene_gn=ZNF618;refGene_tx=XM_023242614.1;refGene_gn=ZNF618;refGen
e_tx=XM_023242615.1
D4 76110048 76110126
refGene_gn=ZNF618;refGene_tx=XM_023242597.1;refGene_gn=ZNF618;refGene_
tx=XM_023242599.1;refGene_gn=ZNF618;refGene_tx=XM_023242600.1;refGene_
gn=ZNF618;refGene_tx=XM_023242601.1;refGene_gn=ZNF618;refGene_tx=XM_02
3242602.1;refGene_gn=ZNF618;refGene_tx=XM_023242603.1;refGene_gn=ZNF61
8;refGene_tx=XM_023242604.1;refGene_gn=ZNF618;refGene_tx=XM_023242605.
1;refGene_gn=ZNF618;refGene_tx=XM_023242606.1;refGene_gn=ZNF618;refGen
e_tx=XM_023242607.1;refGene_gn=ZNF618;refGene_tx=XM_023242608.1;refGen
e_gn=ZNF618;refGene_tx=XM_023242610.1;refGene_gn=ZNF618;refGene_tx=XM_
023242611.1;refGene_gn=ZNF618;refGene_tx=XM_023242612.1;refGene_gn=ZNF
618;refGene_tx=XM_023242613.1;refGene_gn=ZNF618;refGene_tx=XM_02324261
4.1;refGene_gn=ZNF618;refGene_tx=XM_023242615.1;refGene_gn=ZNF618;refG
ene_tx=XM_023242616.1
D4 76120994 76121096
refGene_gn=ZNF618;refGene_tx=XM_023242597.1;refGene_gn=ZNF618;refGene_
tx=XM_023242599.1;refGene_gn=ZNF618;refGene_tx=XM_023242600.1;refGene_
gn=ZNF618;refGene_tx=XM_023242601.1;refGene_gn=ZNF618;refGene_tx=XM_02
3242602.1;refGene_gn=ZNF618;refGene_tx=XM_023242603.1;refGene_gn=ZNF61
8;refGene_tx=XM_023242604.1;refGene_gn=ZNF618;refGene_tx=XM_023242605.
1;refGene_gn=ZNF618;refGene_tx=XM_023242606.1;refGene_gn=ZNF618;refGen
e_tx=XM_023242607.1;refGene_gn=ZNF618;refGene_tx=XM_023242608.1;refGen
e_gn=ZNF618;refGene_tx=XM_023242610.1;refGene_gn=ZNF618;refGene_tx=XM_
023242611.1;refGene_gn=ZNF618;refGene_tx=XM_023242612.1;refGene_gn=ZNF
618;refGene_tx=XM_023242613.1;refGene_gn=ZNF618;refGene_tx=XM_02324261
4.1;refGene_gn=ZNF618;refGene_tx=XM_023242615.1;refGene_gn=ZNF618;refG
ene_tx=XM_023242616.1;refGene_gn=ZNF618;refGene_tx=XM_023242617.1;refG
ene_gn=ZNF618;refGene_tx=XM_023242618.1
D4 76121763 76123280
refGene_gn=ZNF618;refGene_tx=XM_023242597.1;refGene_gn=ZNF618;refGene_
tx=XM_023242599.1;refGene_gn=ZNF618;refGene_tx=XM_023242600.1;refGene_
gn=ZNF618;refGene_tx=XM_023242601.1;refGene_gn=ZNF618;refGene_tx=XM_02
3242602.1;refGene_gn=ZNF618;refGene_tx=XM_023242603.1;refGene_gn=ZNF61
8;refGene_tx=XM_023242604.1;refGene_gn=ZNF618;refGene_tx=XM_023242605.
1;refGene_gn=ZNF618;refGene_tx=XM_023242606.1;refGene_gn=ZNF618;refGen
e_tx=XM_023242607.1;refGene_gn=ZNF618;refGene_tx=XM_023242608.1;refGen
e_gn=ZNF618;refGene_tx=XM_023242610.1;refGene_gn=ZNF618;refGene_tx=XM_
023242611.1;refGene_gn=ZNF618;refGene_tx=XM_023242612.1;refGene_gn=ZNF
618;refGene_tx=XM_023242613.1;refGene_gn=ZNF618;refGene_tx=XM_02324261
4.1;refGene_gn=ZNF618;refGene_tx=XM_023242615.1;refGene_gn=ZNF618;refG

ene_tx=XM_023242616.1;refGene_gn=ZNF618;refGene_tx=XM_023242617.1;refGene_gn=ZNF618;refGene_tx=XM_023242618.1
D4 76132918 76132950 refGene_gn=AMBP;refGene_tx=NM_001198554.1
D4 76133620 76133794 refGene_gn=AMBP;refGene_tx=NM_001198554.1
D4 76134115 76134283 refGene_gn=AMBP;refGene_tx=NM_001198554.1
D4 76135352 76135434 refGene_gn=AMBP;refGene_tx=NM_001198554.1
D4 76140861 76140908 refGene_gn=AMBP;refGene_tx=NM_001198554.1
D4 76141667 76141769 refGene_gn=AMBP;refGene_tx=NM_001198554.1
D4 76142237 76142354 refGene_gn=AMBP;refGene_tx=NM_001198554.1
D4 76143168 76143245 refGene_gn=AMBP;refGene_tx=NM_001198554.1
D4 76144949 76145092 refGene_gn=AMBP;refGene_tx=NM_001198554.1
D4 76146341 76146458 refGene_gn=AMBP;refGene_tx=NM_001198554.1
D4 76155954 76156094
refGene_gn=KIF12;refGene_tx=XM_023242622.1;refGene_gn=KIF12;refGene_tx=XM_023242623.1;refGene_gn=KIF12;refGene_tx=XM_023242629.1
D4 76156154 76156530
refGene_gn=KIF12;refGene_tx=XM_019816393.2;refGene_gn=KIF12;refGene_tx=XM_023242619.1;refGene_gn=KIF12;refGene_tx=XM_023242620.1;refGene_gn=KIF12;refGene_tx=XM_023242621.1;refGene_gn=KIF12;refGene_tx=XM_023242622.1;refGene_gn=KIF12;refGene_tx=XM_023242623.1;refGene_gn=KIF12;refGene_tx=XM_023242625.1;refGene_gn=KIF12;refGene_tx=XM_023242626.1;refGene_gn=KIF12;refGene_tx=XM_023242629.1
D4 76156625 76156726
refGene_gn=KIF12;refGene_tx=XM_019816393.2;refGene_gn=KIF12;refGene_tx=XM_023242619.1;refGene_gn=KIF12;refGene_tx=XM_023242620.1;refGene_gn=KIF12;refGene_tx=XM_023242621.1;refGene_gn=KIF12;refGene_tx=XM_023242622.1;refGene_gn=KIF12;refGene_tx=XM_023242623.1;refGene_gn=KIF12;refGene_tx=XM_023242625.1;refGene_gn=KIF12;refGene_tx=XM_023242626.1;refGene_gn=KIF12;refGene_tx=XM_023242629.1
D4 76157321 76157426
refGene_gn=KIF12;refGene_tx=XM_019816393.2;refGene_gn=KIF12;refGene_tx=XM_023242619.1;refGene_gn=KIF12;refGene_tx=XM_023242620.1;refGene_gn=KIF12;refGene_tx=XM_023242621.1;refGene_gn=KIF12;refGene_tx=XM_023242622.1;refGene_gn=KIF12;refGene_tx=XM_023242623.1;refGene_gn=KIF12;refGene_tx=XM_023242625.1;refGene_gn=KIF12;refGene_tx=XM_023242626.1;refGene_gn=KIF12;refGene_tx=XM_023242629.1
D4 76157497 76157588
refGene_gn=KIF12;refGene_tx=XM_019816393.2;refGene_gn=KIF12;refGene_tx=XM_023242619.1;refGene_gn=KIF12;refGene_tx=XM_023242620.1;refGene_gn=KIF12;refGene_tx=XM_023242621.1;refGene_gn=KIF12;refGene_tx=XM_023242622.1;refGene_gn=KIF12;refGene_tx=XM_023242623.1;refGene_gn=KIF12;refGene_tx=XM_023242625.1;refGene_gn=KIF12;refGene_tx=XM_023242626.1;refGene_gn=KIF12;refGene_tx=XM_023242627.1;refGene_gn=KIF12;refGene_tx=XM_023242628.1;refGene_gn=KIF12;refGene_tx=XM_023242629.1
D4 76157967 76158054
refGene_gn=KIF12;refGene_tx=XM_019816393.2;refGene_gn=KIF12;refGene_tx=XM_023242619.1;refGene_gn=KIF12;refGene_tx=XM_023242620.1;refGene_gn=KIF12;refGene_tx=XM_023242622.1;refGene_gn=KIF12;refGene_tx=XM_023242623.1;refGene_gn=KIF12;refGene_tx=XM_023242625.1;refGene_gn=KIF12;refGene_tx=XM_023242626.1;refGene_gn=KIF12;refGene_tx=XM_023242629.1

D4 76158263 76158354
refGene_gn=KIF12;refGene_tx=XM_019816393.2;refGene_gn=KIF12;refGene_tx=XM_023242619.1;refGene_gn=KIF12;refGene_tx=XM_023242620.1;refGene_gn=KIF12;refGene_tx=XM_023242621.1;refGene_gn=KIF12;refGene_tx=XM_023242622.1;refGene_gn=KIF12;refGene_tx=XM_023242623.1;refGene_gn=KIF12;refGene_tx=XM_023242625.1;refGene_gn=KIF12;refGene_tx=XM_023242626.1;refGene_gn=KIF12;refGene_tx=XM_023242629.1
D4 76158434 76158537
refGene_gn=KIF12;refGene_tx=XM_019816393.2;refGene_gn=KIF12;refGene_tx=XM_023242619.1;refGene_gn=KIF12;refGene_tx=XM_023242620.1;refGene_gn=KIF12;refGene_tx=XM_023242621.1;refGene_gn=KIF12;refGene_tx=XM_023242622.1;refGene_gn=KIF12;refGene_tx=XM_023242623.1;refGene_gn=KIF12;refGene_tx=XM_023242625.1;refGene_gn=KIF12;refGene_tx=XM_023242626.1;refGene_gn=KIF12;refGene_tx=XM_023242627.1;refGene_gn=KIF12;refGene_tx=XM_023242628.1;refGene_gn=KIF12;refGene_tx=XM_023242629.1
D4 76159123 76159228
refGene_gn=KIF12;refGene_tx=XM_019816393.2;refGene_gn=KIF12;refGene_tx=XM_023242619.1;refGene_gn=KIF12;refGene_tx=XM_023242620.1;refGene_gn=KIF12;refGene_tx=XM_023242621.1;refGene_gn=KIF12;refGene_tx=XM_023242622.1;refGene_gn=KIF12;refGene_tx=XM_023242623.1;refGene_gn=KIF12;refGene_tx=XM_023242625.1;refGene_gn=KIF12;refGene_tx=XM_023242626.1;refGene_gn=KIF12;refGene_tx=XM_023242627.1;refGene_gn=KIF12;refGene_tx=XM_023242628.1;refGene_gn=KIF12;refGene_tx=XM_023242629.1
D4 76159315 76159434
refGene_gn=KIF12;refGene_tx=XM_019816393.2;refGene_gn=KIF12;refGene_tx=XM_023242619.1;refGene_gn=KIF12;refGene_tx=XM_023242620.1;refGene_gn=KIF12;refGene_tx=XM_023242621.1;refGene_gn=KIF12;refGene_tx=XM_023242622.1;refGene_gn=KIF12;refGene_tx=XM_023242623.1;refGene_gn=KIF12;refGene_tx=XM_023242625.1;refGene_gn=KIF12;refGene_tx=XM_023242626.1;refGene_gn=KIF12;refGene_tx=XM_023242627.1;refGene_gn=KIF12;refGene_tx=XM_023242628.1;refGene_gn=KIF12;refGene_tx=XM_023242629.1
D4 76160154 76160302
refGene_gn=KIF12;refGene_tx=XM_019816393.2;refGene_gn=KIF12;refGene_tx=XM_023242619.1;refGene_gn=KIF12;refGene_tx=XM_023242620.1;refGene_gn=KIF12;refGene_tx=XM_023242621.1;refGene_gn=KIF12;refGene_tx=XM_023242622.1;refGene_gn=KIF12;refGene_tx=XM_023242623.1;refGene_gn=KIF12;refGene_tx=XM_023242625.1;refGene_gn=KIF12;refGene_tx=XM_023242626.1;refGene_gn=KIF12;refGene_tx=XM_023242627.1;refGene_gn=KIF12;refGene_tx=XM_023242628.1;refGene_gn=KIF12;refGene_tx=XM_023242629.1
D4 76160481 76160573
refGene_gn=KIF12;refGene_tx=XM_019816393.2;refGene_gn=KIF12;refGene_tx=XM_023242619.1;refGene_gn=KIF12;refGene_tx=XM_023242620.1;refGene_gn=KIF12;refGene_tx=XM_023242621.1;refGene_gn=KIF12;refGene_tx=XM_023242622.1;refGene_gn=KIF12;refGene_tx=XM_023242623.1;refGene_gn=KIF12;refGene_tx=XM_023242625.1;refGene_gn=KIF12;refGene_tx=XM_023242626.1;refGene_gn=KIF12;refGene_tx=XM_023242627.1;refGene_gn=KIF12;refGene_tx=XM_023242628.1;refGene_gn=KIF12;refGene_tx=XM_023242629.1
D4 76160979 76161033
refGene_gn=KIF12;refGene_tx=XM_019816393.2;refGene_gn=KIF12;refGene_tx=XM_023242619.1;refGene_gn=KIF12;refGene_tx=XM_023242621.1;refGene_gn=

KIF12;refGene_tx=XM_023242622.1;refGene_gn=KIF12;refGene_tx=XM_0232426
23.1;refGene_gn=KIF12;refGene_tx=XM_023242625.1;refGene_gn=KIF12;refGe
ne_tx=XM_023242627.1;refGene_gn=KIF12;refGene_tx=XM_023242628.1
D4 76161410 76161546
refGene_gn=KIF12;refGene_tx=XM_019816393.2;refGene_gn=KIF12;refGene_tx
=XM_023242619.1;refGene_gn=KIF12;refGene_tx=XM_023242620.1;refGene_gn=
KIF12;refGene_tx=XM_023242621.1;refGene_gn=KIF12;refGene_tx=XM_0232426
22.1;refGene_gn=KIF12;refGene_tx=XM_023242623.1;refGene_gn=KIF12;refGe
ne_tx=XM_023242625.1;refGene_gn=KIF12;refGene_tx=XM_023242626.1;refGen
e_gn=KIF12;refGene_tx=XM_023242627.1;refGene_gn=KIF12;refGene_tx=XM_02
3242628.1;refGene_gn=KIF12;refGene_tx=XM_023242629.1
D4 76161713 76161848
refGene_gn=KIF12;refGene_tx=XM_019816393.2;refGene_gn=KIF12;refGene_tx
=XM_023242619.1;refGene_gn=KIF12;refGene_tx=XM_023242620.1;refGene_gn=
KIF12;refGene_tx=XM_023242621.1;refGene_gn=KIF12;refGene_tx=XM_0232426
22.1;refGene_gn=KIF12;refGene_tx=XM_023242623.1;refGene_gn=KIF12;refGe
ne_tx=XM_023242625.1;refGene_gn=KIF12;refGene_tx=XM_023242626.1;refGen
e_gn=KIF12;refGene_tx=XM_023242627.1;refGene_gn=KIF12;refGene_tx=XM_02
3242628.1;refGene_gn=KIF12;refGene_tx=XM_023242629.1
D4 76162223 76162299
refGene_gn=KIF12;refGene_tx=XM_023242619.1;refGene_gn=KIF12;refGene_tx
=XM_023242620.1;refGene_gn=KIF12;refGene_tx=XM_023242621.1;refGene_gn=
KIF12;refGene_tx=XM_023242622.1;refGene_gn=KIF12;refGene_tx=XM_0232426
23.1;refGene_gn=KIF12;refGene_tx=XM_023242625.1;refGene_gn=KIF12;refGe
ne_tx=XM_023242626.1;refGene_gn=KIF12;refGene_tx=XM_023242627.1;refGen
e_gn=KIF12;refGene_tx=XM_023242628.1;refGene_gn=KIF12;refGene_tx=XM_02
3242629.1
D4 76162388 76162516
refGene_gn=KIF12;refGene_tx=XM_023242619.1;refGene_gn=KIF12;refGene_tx
=XM_023242620.1;refGene_gn=KIF12;refGene_tx=XM_023242621.1;refGene_gn=
KIF12;refGene_tx=XM_023242622.1;refGene_gn=KIF12;refGene_tx=XM_0232426
23.1;refGene_gn=KIF12;refGene_tx=XM_023242625.1;refGene_gn=KIF12;refGe
ne_tx=XM_023242626.1;refGene_gn=KIF12;refGene_tx=XM_023242627.1;refGen
e_gn=KIF12;refGene_tx=XM_023242628.1;refGene_gn=KIF12;refGene_tx=XM_02
3242629.1
D4 76162728 76162974
refGene_gn=KIF12;refGene_tx=XM_023242619.1;refGene_gn=KIF12;refGene_tx
=XM_023242620.1;refGene_gn=KIF12;refGene_tx=XM_023242621.1;refGene_gn=
KIF12;refGene_tx=XM_023242622.1;refGene_gn=KIF12;refGene_tx=XM_0232426
23.1;refGene_gn=KIF12;refGene_tx=XM_023242625.1;refGene_gn=KIF12;refGe
ne_tx=XM_023242626.1;refGene_gn=KIF12;refGene_tx=XM_023242627.1;refGen
e_gn=KIF12;refGene_tx=XM_023242628.1;refGene_gn=KIF12;refGene_tx=XM_02
3242629.1
D4 76163044 76163070
refGene_gn=KIF12;refGene_tx=XM_023242625.1;refGene_gn=KIF12;refGene_tx
=XM_023242626.1;refGene_gn=KIF12;refGene_tx=XM_023242629.1
D4 76163267 76163441
refGene_gn=KIF12;refGene_tx=XM_023242619.1;refGene_gn=KIF12;refGene_tx
=XM_023242620.1;refGene_gn=KIF12;refGene_tx=XM_023242621.1;refGene_gn=
KIF12;refGene_tx=XM_023242622.1;refGene_gn=KIF12;refGene_tx=XM_0232426

23.1;refGene_gn=KIF12;refGene_tx=XM_023242627.1;refGene_gn=KIF12;refGene_tx=XM_023242628.1
D4 76209000 76209056
refGene_gn=COL27A1;refGene_tx=XM_019816400.2;refGene_gn=COL27A1;refGene_tx=XM_019816401.1;refGene_gn=COL27A1;refGene_tx=XM_019816402.1;refGene_gn=COL27A1;refGene_tx=XM_023242631.1
D4 76217260 76217331
refGene_gn=COL27A1;refGene_tx=XM_019816400.2;refGene_gn=COL27A1;refGene_tx=XM_019816401.1;refGene_gn=COL27A1;refGene_tx=XM_019816402.1;refGene_gn=COL27A1;refGene_tx=XM_023242631.1
D4 76220853 76222586
refGene_gn=COL27A1;refGene_tx=XM_019816400.2;refGene_gn=COL27A1;refGene_tx=XM_019816401.1;refGene_gn=COL27A1;refGene_tx=XM_019816402.1;refGene_gn=COL27A1;refGene_tx=XM_023242631.1
D4 76230102 76230156
refGene_gn=COL27A1;refGene_tx=XM_019816400.2;refGene_gn=COL27A1;refGene_tx=XM_019816401.1;refGene_gn=COL27A1;refGene_tx=XM_019816402.1;refGene_gn=COL27A1;refGene_tx=XM_023242631.1
D4 76234212 76234266
refGene_gn=COL27A1;refGene_tx=XM_019816400.2;refGene_gn=COL27A1;refGene_tx=XM_019816401.1;refGene_gn=COL27A1;refGene_tx=XM_019816402.1;refGene_gn=COL27A1;refGene_tx=XM_023242631.1
D4 76244217 76244271
refGene_gn=COL27A1;refGene_tx=XM_019816400.2;refGene_gn=COL27A1;refGene_tx=XM_019816401.1;refGene_gn=COL27A1;refGene_tx=XM_023242631.1
D4 76245711 76245765
refGene_gn=COL27A1;refGene_tx=XM_019816400.2;refGene_gn=COL27A1;refGene_tx=XM_019816401.1;refGene_gn=COL27A1;refGene_tx=XM_019816402.1;refGene_gn=COL27A1;refGene_tx=XM_023242631.1
D4 76254221 76254266
refGene_gn=COL27A1;refGene_tx=XM_019816400.2;refGene_gn=COL27A1;refGene_tx=XM_019816401.1;refGene_gn=COL27A1;refGene_tx=XM_019816402.1;refGene_gn=COL27A1;refGene_tx=XM_023242631.1
D4 76254897 76254951
refGene_gn=COL27A1;refGene_tx=XM_019816400.2;refGene_gn=COL27A1;refGene_tx=XM_019816401.1;refGene_gn=COL27A1;refGene_tx=XM_019816402.1;refGene_gn=COL27A1;refGene_tx=XM_023242631.1
D4 76255416 76255461
refGene_gn=COL27A1;refGene_tx=XM_019816400.2;refGene_gn=COL27A1;refGene_tx=XM_019816401.1;refGene_gn=COL27A1;refGene_tx=XM_019816402.1;refGene_gn=COL27A1;refGene_tx=XM_023242631.1
D4 76258319 76258373
refGene_gn=COL27A1;refGene_tx=XM_019816400.2;refGene_gn=COL27A1;refGene_tx=XM_019816401.1;refGene_gn=COL27A1;refGene_tx=XM_019816402.1;refGene_gn=COL27A1;refGene_tx=XM_023242631.1
D4 76259262 76259307
refGene_gn=COL27A1;refGene_tx=XM_019816400.2;refGene_gn=COL27A1;refGene_tx=XM_019816401.1;refGene_gn=COL27A1;refGene_tx=XM_019816402.1;refGene_gn=COL27A1;refGene_tx=XM_023242631.1
D4 76267608 76267662

refGene_gn=COL27A1;refGene_tx=XM_019816400.2;refGene_gn=COL27A1;refGene_tx=XM_019816401.1;refGene_gn=COL27A1;refGene_tx=XM_019816402.1;refGene_gn=COL27A1;refGene_tx=XM_023242631.1
D4 76270096 76270141
refGene_gn=COL27A1;refGene_tx=XM_019816400.2;refGene_gn=COL27A1;refGene_tx=XM_019816401.1;refGene_gn=COL27A1;refGene_tx=XM_019816402.1;refGene_gn=COL27A1;refGene_tx=XM_023242631.1
D4 76277584 76277638
refGene_gn=COL27A1;refGene_tx=XM_019816400.2;refGene_gn=COL27A1;refGene_tx=XM_019816401.1;refGene_gn=COL27A1;refGene_tx=XM_019816402.1;refGene_gn=COL27A1;refGene_tx=XM_023242631.1
D4 76278370 76278415
refGene_gn=COL27A1;refGene_tx=XM_019816400.2;refGene_gn=COL27A1;refGene_tx=XM_019816401.1;refGene_gn=COL27A1;refGene_tx=XM_019816402.1;refGene_gn=COL27A1;refGene_tx=XM_023242631.1
D4 76281733 76281787
refGene_gn=COL27A1;refGene_tx=XM_019816400.2;refGene_gn=COL27A1;refGene_tx=XM_019816401.1;refGene_gn=COL27A1;refGene_tx=XM_019816402.1;refGene_gn=COL27A1;refGene_tx=XM_023242631.1
D4 76283011 76283065
refGene_gn=COL27A1;refGene_tx=XM_019816400.2;refGene_gn=COL27A1;refGene_tx=XM_019816401.1;refGene_gn=COL27A1;refGene_tx=XM_019816402.1;refGene_gn=COL27A1;refGene_tx=XM_023242631.1
D4 76283703 76283757
refGene_gn=COL27A1;refGene_tx=XM_019816400.2;refGene_gn=COL27A1;refGene_tx=XM_019816401.1;refGene_gn=COL27A1;refGene_tx=XM_019816402.1;refGene_gn=COL27A1;refGene_tx=XM_023242631.1
D4 76286965 76287019
refGene_gn=COL27A1;refGene_tx=XM_019816400.2;refGene_gn=COL27A1;refGene_tx=XM_019816401.1;refGene_gn=COL27A1;refGene_tx=XM_019816402.1;refGene_gn=COL27A1;refGene_tx=XM_023242631.1
D4 76287172 76287226
refGene_gn=COL27A1;refGene_tx=XM_019816400.2;refGene_gn=COL27A1;refGene_tx=XM_019816401.1;refGene_gn=COL27A1;refGene_tx=XM_019816402.1;refGene_gn=COL27A1;refGene_tx=XM_023242631.1
D4 76288868 76288913
refGene_gn=COL27A1;refGene_tx=XM_019816400.2;refGene_gn=COL27A1;refGene_tx=XM_019816401.1;refGene_gn=COL27A1;refGene_tx=XM_019816402.1;refGene_gn=COL27A1;refGene_tx=XM_023242631.1
D4 76290116 76290170
refGene_gn=COL27A1;refGene_tx=XM_019816400.2;refGene_gn=COL27A1;refGene_tx=XM_019816401.1;refGene_gn=COL27A1;refGene_tx=XM_019816402.1;refGene_gn=COL27A1;refGene_tx=XM_023242631.1
D4 76291841 76291886
refGene_gn=COL27A1;refGene_tx=XM_019816400.2;refGene_gn=COL27A1;refGene_tx=XM_019816401.1;refGene_gn=COL27A1;refGene_tx=XM_019816402.1;refGene_gn=COL27A1;refGene_tx=XM_023242631.1
D4 76296451 76296505
refGene_gn=COL27A1;refGene_tx=XM_019816400.2;refGene_gn=COL27A1;refGene_tx=XM_019816401.1;refGene_gn=COL27A1;refGene_tx=XM_019816402.1;refGe

ne_gn=COL27A1;refGene_tx=XM_023242631.1
D4 76298342 76298396
refGene_gn=COL27A1;refGene_tx=XM_019816400.2;refGene_gn=COL27A1;refGene_tx=XM_019816401.1;refGene_gn=COL27A1;refGene_tx=XM_019816402.1;refGene_gn=COL27A1;refGene_tx=XM_023242631.1
D4 76298634 76298688
refGene_gn=COL27A1;refGene_tx=XM_019816400.2;refGene_gn=COL27A1;refGene_tx=XM_019816401.1;refGene_gn=COL27A1;refGene_tx=XM_019816402.1;refGene_gn=COL27A1;refGene_tx=XM_023242631.1
D4 76302758 76302812
refGene_gn=COL27A1;refGene_tx=XM_019816400.2;refGene_gn=COL27A1;refGene_tx=XM_019816401.1;refGene_gn=COL27A1;refGene_tx=XM_019816402.1;refGene_gn=COL27A1;refGene_tx=XM_023242631.1
D4 76308135 76308189
refGene_gn=COL27A1;refGene_tx=XM_019816400.2;refGene_gn=COL27A1;refGene_tx=XM_019816401.1;refGene_gn=COL27A1;refGene_tx=XM_019816402.1;refGene_gn=COL27A1;refGene_tx=XM_023242631.1
D4 76308703 76308748
refGene_gn=COL27A1;refGene_tx=XM_019816400.2;refGene_gn=COL27A1;refGene_tx=XM_019816401.1;refGene_gn=COL27A1;refGene_tx=XM_019816402.1;refGene_gn=COL27A1;refGene_tx=XM_023242631.1
D4 76308831 76308876
refGene_gn=COL27A1;refGene_tx=XM_019816400.2;refGene_gn=COL27A1;refGene_tx=XM_019816401.1;refGene_gn=COL27A1;refGene_tx=XM_019816402.1;refGene_gn=COL27A1;refGene_tx=XM_023242631.1
D4 76309137 76309191
refGene_gn=COL27A1;refGene_tx=XM_019816400.2;refGene_gn=COL27A1;refGene_tx=XM_019816401.1;refGene_gn=COL27A1;refGene_tx=XM_019816402.1;refGene_gn=COL27A1;refGene_tx=XM_023242631.1
D4 76310262 76310316
refGene_gn=COL27A1;refGene_tx=XM_019816400.2;refGene_gn=COL27A1;refGene_tx=XM_019816401.1;refGene_gn=COL27A1;refGene_tx=XM_019816402.1;refGene_gn=COL27A1;refGene_tx=XM_023242631.1
D4 76311211 76311265
refGene_gn=COL27A1;refGene_tx=XM_019816400.2;refGene_gn=COL27A1;refGene_tx=XM_019816401.1;refGene_gn=COL27A1;refGene_tx=XM_019816402.1;refGene_gn=COL27A1;refGene_tx=XM_023242631.1
D4 76312541 76312595
refGene_gn=COL27A1;refGene_tx=XM_019816400.2;refGene_gn=COL27A1;refGene_tx=XM_019816401.1;refGene_gn=COL27A1;refGene_tx=XM_019816402.1;refGene_gn=COL27A1;refGene_tx=XM_023242631.1
D4 76314504 76314558
refGene_gn=COL27A1;refGene_tx=XM_019816400.2;refGene_gn=COL27A1;refGene_tx=XM_019816401.1;refGene_gn=COL27A1;refGene_tx=XM_019816402.1;refGene_gn=COL27A1;refGene_tx=XM_023242631.1
D4 76318732 76318840
refGene_gn=COL27A1;refGene_tx=XM_019816400.2;refGene_gn=COL27A1;refGene_tx=XM_019816401.1;refGene_gn=COL27A1;refGene_tx=XM_019816402.1;refGene_gn=COL27A1;refGene_tx=XM_023242631.1
D4 76324049 76324103

refGene_gn=COL27A1;refGene_tx=XM_019816400.2;refGene_gn=COL27A1;refGene_tx=XM_019816401.1;refGene_gn=COL27A1;refGene_tx=XM_019816402.1
D4	76324233	76324341
refGene_gn=COL27A1;refGene_tx=XM_019816400.2;refGene_gn=COL27A1;refGene_tx=XM_019816401.1;refGene_gn=COL27A1;refGene_tx=XM_019816402.1;refGene_gn=COL27A1;refGene_tx=XM_023242631.1
D4	76325110	76325164
refGene_gn=COL27A1;refGene_tx=XM_019816400.2;refGene_gn=COL27A1;refGene_tx=XM_019816401.1;refGene_gn=COL27A1;refGene_tx=XM_019816402.1;refGene_gn=COL27A1;refGene_tx=XM_023242631.1
D4	76325711	76325765
refGene_gn=COL27A1;refGene_tx=XM_019816400.2;refGene_gn=COL27A1;refGene_tx=XM_019816401.1;refGene_gn=COL27A1;refGene_tx=XM_019816402.1;refGene_gn=COL27A1;refGene_tx=XM_023242631.1
D4	76329191	76329248
refGene_gn=COL27A1;refGene_tx=XM_019816400.2;refGene_gn=COL27A1;refGene_tx=XM_019816401.1;refGene_gn=COL27A1;refGene_tx=XM_019816402.1;refGene_gn=COL27A1;refGene_tx=XM_023242631.1
D4	76329438	76329492
refGene_gn=COL27A1;refGene_tx=XM_019816400.2;refGene_gn=COL27A1;refGene_tx=XM_019816401.1;refGene_gn=COL27A1;refGene_tx=XM_019816402.1;refGene_gn=COL27A1;refGene_tx=XM_023242631.1
D4	76330019	76330073
refGene_gn=COL27A1;refGene_tx=XM_019816400.2;refGene_gn=COL27A1;refGene_tx=XM_019816401.1;refGene_gn=COL27A1;refGene_tx=XM_019816402.1;refGene_gn=COL27A1;refGene_tx=XM_023242631.1
D4	76330350	76330404
refGene_gn=COL27A1;refGene_tx=XM_019816400.2;refGene_gn=COL27A1;refGene_tx=XM_019816401.1;refGene_gn=COL27A1;refGene_tx=XM_019816402.1;refGene_gn=COL27A1;refGene_tx=XM_023242631.1
D4	76331056	76331110
refGene_gn=COL27A1;refGene_tx=XM_019816400.2;refGene_gn=COL27A1;refGene_tx=XM_019816401.1;refGene_gn=COL27A1;refGene_tx=XM_019816402.1;refGene_gn=COL27A1;refGene_tx=XM_023242631.1
D4	76331426	76331534
refGene_gn=COL27A1;refGene_tx=XM_019816400.2;refGene_gn=COL27A1;refGene_tx=XM_019816401.1;refGene_gn=COL27A1;refGene_tx=XM_019816402.1;refGene_gn=COL27A1;refGene_tx=XM_023242631.1
D4	76331980	76332088
refGene_gn=COL27A1;refGene_tx=XM_019816400.2;refGene_gn=COL27A1;refGene_tx=XM_019816401.1;refGene_gn=COL27A1;refGene_tx=XM_019816402.1;refGene_gn=COL27A1;refGene_tx=XM_023242631.1
D4	76332953	76333061
refGene_gn=COL27A1;refGene_tx=XM_019816400.2;refGene_gn=COL27A1;refGene_tx=XM_019816401.1;refGene_gn=COL27A1;refGene_tx=XM_019816402.1;refGene_gn=COL27A1;refGene_tx=XM_023242631.1
D4	76336717	76336771
refGene_gn=COL27A1;refGene_tx=XM_019816400.2;refGene_gn=COL27A1;refGene_tx=XM_019816401.1;refGene_gn=COL27A1;refGene_tx=XM_019816402.1;refGene_gn=COL27A1;refGene_tx=XM_023242631.1

D4 76337158 76337221
refGene_gn=COL27A1;refGene_tx=XM_019816400.2;refGene_gn=COL27A1;refGen
e_tx=XM_019816401.1;refGene_gn=COL27A1;refGene_tx=XM_019816402.1;refGe
ne_gn=COL27A1;refGene_tx=XM_023242631.1
D4 76337608 76337662
refGene_gn=COL27A1;refGene_tx=XM_019816400.2;refGene_gn=COL27A1;refGen
e_tx=XM_019816401.1;refGene_gn=COL27A1;refGene_tx=XM_019816402.1;refGe
ne_gn=COL27A1;refGene_tx=XM_023242631.1
D4 76337804 76337840
refGene_gn=COL27A1;refGene_tx=XM_019816400.2;refGene_gn=COL27A1;refGen
e_tx=XM_019816401.1;refGene_gn=COL27A1;refGene_tx=XM_019816402.1;refGe
ne_gn=COL27A1;refGene_tx=XM_023242631.1
D4 76337960 76337978
refGene_gn=COL27A1;refGene_tx=XM_019816400.2;refGene_gn=COL27A1;refGen
e_tx=XM_019816401.1;refGene_gn=COL27A1;refGene_tx=XM_019816402.1;refGe
ne_gn=COL27A1;refGene_tx=XM_023242631.1
D4 76338209 76338245
refGene_gn=COL27A1;refGene_tx=XM_019816400.2;refGene_gn=COL27A1;refGen
e_tx=XM_019816401.1;refGene_gn=COL27A1;refGene_tx=XM_019816402.1;refGe
ne_gn=COL27A1;refGene_tx=XM_023242631.1
D4 76338532 76338559
refGene_gn=COL27A1;refGene_tx=XM_019816400.2;refGene_gn=COL27A1;refGen
e_tx=XM_019816401.1;refGene_gn=COL27A1;refGene_tx=XM_019816402.1;refGe
ne_gn=COL27A1;refGene_tx=XM_023242631.1
D4 76340913 76340979
refGene_gn=COL27A1;refGene_tx=XM_019816400.2;refGene_gn=COL27A1;refGen
e_tx=XM_019816401.1;refGene_gn=COL27A1;refGene_tx=XM_019816402.1;refGe
ne_gn=COL27A1;refGene_tx=XM_023242631.1
D4 76341329 76341498
refGene_gn=COL27A1;refGene_tx=XM_019816400.2;refGene_gn=COL27A1;refGen
e_tx=XM_019816401.1;refGene_gn=COL27A1;refGene_tx=XM_019816402.1;refGe
ne_gn=COL27A1;refGene_tx=XM_023242631.1
D4 76342397 76342507
refGene_gn=COL27A1;refGene_tx=XM_019816400.2;refGene_gn=COL27A1;refGen
e_tx=XM_019816401.1;refGene_gn=COL27A1;refGene_tx=XM_019816402.1;refGe
ne_gn=COL27A1;refGene_tx=XM_023242631.1
D4 76344204 76344423
refGene_gn=COL27A1;refGene_tx=XM_019816400.2;refGene_gn=COL27A1;refGen
e_tx=XM_019816401.1;refGene_gn=COL27A1;refGene_tx=XM_019816402.1;refGe
ne_gn=COL27A1;refGene_tx=XM_023242631.1
D4 76347903 76348050
refGene_gn=COL27A1;refGene_tx=XM_019816400.2;refGene_gn=COL27A1;refGen
e_tx=XM_019816401.1;refGene_gn=COL27A1;refGene_tx=XM_019816402.1;refGe
ne_gn=COL27A1;refGene_tx=XM_023242631.1
D4 76359973 76360087 refGene_gn=ORM2;refGene_tx=XM_003995775.3
D4 76360487 76360630 refGene_gn=ORM2;refGene_tx=XM_003995775.3
D4 76360859 76360930 refGene_gn=ORM2;refGene_tx=XM_003995775.3
D4 76361723 76361831 refGene_gn=ORM2;refGene_tx=XM_003995775.3
D4 76361969 76362073 refGene_gn=ORM2;refGene_tx=XM_003995775.3
D4 76363093 76363159 refGene_gn=ORM2;refGene_tx=XM_003995775.3

D4	76366204 76366469 refGene_gn=AKNA;refGene_tx=XM_006939392.4
D4	76370972 76371171 refGene_gn=AKNA;refGene_tx=XM_006939392.4
D4	76371451 76371568 refGene_gn=AKNA;refGene_tx=XM_006939392.4
D4	76372906 76373002 refGene_gn=AKNA;refGene_tx=XM_006939392.4
D4	76374219 76374366 refGene_gn=AKNA;refGene_tx=XM_006939392.4
D4	76375000 76375116 refGene_gn=AKNA;refGene_tx=XM_006939392.4
D4	76375677 76375854 refGene_gn=AKNA;refGene_tx=XM_006939392.4
D4	76378865 76379028 refGene_gn=AKNA;refGene_tx=XM_006939392.4
D4	76381026 76381201 refGene_gn=AKNA;refGene_tx=XM_006939392.4
D4	76381202 76381239 refGene_gn=AKNA;refGene_tx=XM_006939392.4
D4	76381918 76382025 refGene_gn=AKNA;refGene_tx=XM_006939392.4
D4	76383029 76383279 refGene_gn=AKNA;refGene_tx=XM_006939392.4
D4	76384110 76384305 refGene_gn=AKNA;refGene_tx=XM_006939392.4
D4	76384398 76384553 refGene_gn=AKNA;refGene_tx=XM_006939392.4
D4	76386334 76386542 refGene_gn=AKNA;refGene_tx=XM_006939392.4
D4	76387020 76387148 refGene_gn=AKNA;refGene_tx=XM_006939392.4
D4	76389101 76389161 refGene_gn=AKNA;refGene_tx=XM_006939392.4
D4	76392210 76392353 refGene_gn=AKNA;refGene_tx=XM_006939392.4
D4	76393102 76393259 refGene_gn=AKNA;refGene_tx=XM_006939392.4
D4	76398262 76398337 refGene_gn=AKNA;refGene_tx=XM_006939392.4
D4	76400339 76401400 refGene_gn=AKNA;refGene_tx=XM_006939392.4
D4	76404654 76404916 refGene_gn=AKNA;refGene_tx=XM_006939392.4
D4	76424877 76425060
refGene_gn=WHRN;refGene_tx=XM_019815545.2;refGene_gn=WHRN;refGene_tx=XM_023242632.1;refGene_gn=WHRN;refGene_tx=XM_023242633.1;refGene_gn=WHRN;refGene_tx=XM_023242635.1;refGene_gn=WHRN;refGene_tx=XM_023242636.1
D4	76425309 76425432
refGene_gn=WHRN;refGene_tx=XM_019815545.2;refGene_gn=WHRN;refGene_tx=XM_023242632.1;refGene_gn=WHRN;refGene_tx=XM_023242633.1;refGene_gn=WHRN;refGene_tx=XM_023242635.1;refGene_gn=WHRN;refGene_tx=XM_023242636.1
D4	76425961 76426143
refGene_gn=WHRN;refGene_tx=XM_019815545.2;refGene_gn=WHRN;refGene_tx=XM_023242632.1;refGene_gn=WHRN;refGene_tx=XM_023242633.1;refGene_gn=WHRN;refGene_tx=XM_023242635.1;refGene_gn=WHRN;refGene_tx=XM_023242636.1
D4	76428180 76428718
refGene_gn=WHRN;refGene_tx=XM_019815545.2;refGene_gn=WHRN;refGene_tx=XM_023242632.1;refGene_gn=WHRN;refGene_tx=XM_023242633.1;refGene_gn=WHRN;refGene_tx=XM_023242635.1;refGene_gn=WHRN;refGene_tx=XM_023242636.1
D4	76429960 76430032
refGene_gn=WHRN;refGene_tx=XM_019815545.2;refGene_gn=WHRN;refGene_tx=XM_023242632.1;refGene_gn=WHRN;refGene_tx=XM_023242633.1;refGene_gn=WHRN;refGene_tx=XM_023242635.1;refGene_gn=WHRN;refGene_tx=XM_023242636.1
D4	76438855 76438888
refGene_gn=WHRN;refGene_tx=XM_019815545.2;refGene_gn=WHRN;refGene_tx=XM_023242632.1
D4	76441444 76441654
refGene_gn=WHRN;refGene_tx=XM_019815545.2;refGene_gn=WHRN;refGene_tx=XM_023242632.1;refGene_gn=WHRN;refGene_tx=XM_023242633.1;refGene_gn=WHRN;refGene_tx=XM_023242635.1
D4	76442430 76442643

refGene_gn=WHRN;refGene_tx=XM_019815545.2;refGene_gn=WHRN;refGene_tx=XM_023242632.1;refGene_gn=WHRN;refGene_tx=XM_023242633.1;refGene_gn=WHRN;refGene_tx=XM_023242635.1
D4 76443861 76443898
refGene_gn=WHRN;refGene_tx=XM_019815545.2;refGene_gn=WHRN;refGene_tx=XM_023242632.1;refGene_gn=WHRN;refGene_tx=XM_023242633.1;refGene_gn=WHRN;refGene_tx=XM_023242635.1;refGene_gn=WHRN;refGene_tx=XM_023242636.1
D4 76444983 76445186
refGene_gn=WHRN;refGene_tx=XM_019815545.2;refGene_gn=WHRN;refGene_tx=XM_023242632.1;refGene_gn=WHRN;refGene_tx=XM_023242633.1;refGene_gn=WHRN;refGene_tx=XM_023242635.1;refGene_gn=WHRN;refGene_tx=XM_023242636.1
D4 76482508 76482634
refGene_gn=WHRN;refGene_tx=XM_019815545.2;refGene_gn=WHRN;refGene_tx=XM_023242632.1;refGene_gn=WHRN;refGene_tx=XM_023242633.1;refGene_gn=WHRN;refGene_tx=XM_023242635.1;refGene_gn=WHRN;refGene_tx=XM_023242636.1
D4 76494257 76494476
refGene_gn=WHRN;refGene_tx=XM_019815545.2;refGene_gn=WHRN;refGene_tx=XM_023242632.1;refGene_gn=WHRN;refGene_tx=XM_023242633.1;refGene_gn=WHRN;refGene_tx=XM_023242635.1;refGene_gn=WHRN;refGene_tx=XM_023242636.1
D4 76518878 76519496
refGene_gn=WHRN;refGene_tx=XM_019815545.2;refGene_gn=WHRN;refGene_tx=XM_023242632.1;refGene_gn=WHRN;refGene_tx=XM_023242633.1;refGene_gn=WHRN;refGene_tx=XM_023242635.1;refGene_gn=WHRN;refGene_tx=XM_023242636.1
D4 76573226 76573308 refGene_gn=ATP6V1G1;refGene_tx=XM_003995777.5
D4 76577171 76577272 refGene_gn=ATP6V1G1;refGene_tx=XM_003995777.5
D4 76579696 76579870 refGene_gn=ATP6V1G1;refGene_tx=XM_003995777.5
D4 76593230 76593339
refGene_gn=TMEM268;refGene_tx=XM_006939395.4;refGene_gn=TMEM268;refGene_tx=XM_006939396.4;refGene_gn=TMEM268;refGene_tx=XM_006939397.4;refGene_gn=TMEM268;refGene_tx=XM_019816415.2;refGene_gn=TMEM268;refGene_tx=XM_023242638.1
D4 76599341 76599451
refGene_gn=TMEM268;refGene_tx=XM_006939395.4;refGene_gn=TMEM268;refGene_tx=XM_006939396.4;refGene_gn=TMEM268;refGene_tx=XM_006939397.4;refGene_gn=TMEM268;refGene_tx=XM_019816415.2;refGene_gn=TMEM268;refGene_tx=XM_023242638.1
D4 76601340 76601448
refGene_gn=TMEM268;refGene_tx=XM_006939395.4;refGene_gn=TMEM268;refGene_tx=XM_006939396.4;refGene_gn=TMEM268;refGene_tx=XM_006939397.4;refGene_gn=TMEM268;refGene_tx=XM_019816415.2;refGene_gn=TMEM268;refGene_tx=XM_023242638.1
D4 76603751 76603901
refGene_gn=TMEM268;refGene_tx=XM_006939395.4;refGene_gn=TMEM268;refGene_tx=XM_006939396.4;refGene_gn=TMEM268;refGene_tx=XM_006939397.4;refGene_gn=TMEM268;refGene_tx=XM_019816415.2;refGene_gn=TMEM268;refGene_tx=XM_023242638.1
D4 76608132 76608243
refGene_gn=TMEM268;refGene_tx=XM_006939395.4;refGene_gn=TMEM268;refGene_tx=XM_006939396.4;refGene_gn=TMEM268;refGene_tx=XM_006939397.4;refGene_gn=TMEM268;refGene_tx=XM_019816415.2;refGene_gn=TMEM268;refGene_tx=

XM_023242638.1
D4 76610954 76611035
refGene_gn=TMEM268;refGene_tx=XM_006939395.4;refGene_gn=TMEM268;refGene_tx=XM_006939396.4;refGene_gn=TMEM268;refGene_tx=XM_006939397.4;refGene_gn=TMEM268;refGene_tx=XM_019816415.2;refGene_gn=TMEM268;refGene_tx=XM_023242638.1
D4 76611985 76612171
refGene_gn=TMEM268;refGene_tx=XM_006939395.4;refGene_gn=TMEM268;refGene_tx=XM_006939396.4;refGene_gn=TMEM268;refGene_tx=XM_006939397.4;refGene_gn=TMEM268;refGene_tx=XM_019816415.2;refGene_gn=TMEM268;refGene_tx=XM_023242638.1
D4 76615323 76615503
refGene_gn=TMEM268;refGene_tx=XM_006939395.4;refGene_gn=TMEM268;refGene_tx=XM_006939396.4;refGene_gn=TMEM268;refGene_tx=XM_006939397.4;refGene_gn=TMEM268;refGene_tx=XM_019816415.2;refGene_gn=TMEM268;refGene_tx=XM_023242638.1
D4 76631393 76631497
refGene_gn=TEX48;refGene_tx=XM_023242639.1;refGene_gn=TEX48;refGene_tx=XM_023242640.1
D4 76632835 76632964
refGene_gn=TEX48;refGene_tx=XM_023242639.1;refGene_gn=TEX48;refGene_tx=XM_023242640.1
D4 76635138 76635265
refGene_gn=TEX48;refGene_tx=XM_023242639.1;refGene_gn=TEX48;refGene_tx=XM_023242640.1
D4 76635495 76635603
refGene_gn=TEX48;refGene_tx=XM_023242639.1;refGene_gn=TEX48;refGene_tx=XM_023242640.1
D4 76641898 76642153 refGene_gn=TEX48;refGene_tx=XM_023242639.1
D4 76735570 76736025 refGene_gn=TNFSF15;refGene_tx=XM_003995779.4
D4 76739266 76739314 refGene_gn=TNFSF15;refGene_tx=XM_003995779.4
D4 76740037 76740080 refGene_gn=TNFSF15;refGene_tx=XM_003995779.4
D4 76751468 76751690 refGene_gn=TNFSF15;refGene_tx=XM_003995779.4
D4 76826307 76826702 refGene_gn=TNFSF8;refGene_tx=XM_003995780.4
D4 76828384 76828456 refGene_gn=TNFSF8;refGene_tx=XM_003995780.4
D4 76840624 76840667 refGene_gn=TNFSF8;refGene_tx=XM_003995780.4
D4 76851243 76851438 refGene_gn=TNFSF8;refGene_tx=XM_003995780.4
D4 76920246 76920357
refGene_gn=TNC;refGene_tx=XM_011288474.3;refGene_gn=TNC;refGene_tx=XM_023242642.1
D4 76922579 76922743
refGene_gn=TNC;refGene_tx=XM_011288474.3;refGene_gn=TNC;refGene_tx=XM_023242642.1
D4 76925255 76925417
refGene_gn=TNC;refGene_tx=XM_011288474.3;refGene_gn=TNC;refGene_tx=XM_023242642.1
D4 76926629 76926726
refGene_gn=TNC;refGene_tx=XM_011288474.3;refGene_gn=TNC;refGene_tx=XM_023242642.1
D4 76927487 76927639

refGene_gn=TNC;refGene_tx=XM_011288474.3;refGene_gn=TNC;refGene_tx=XM_023242642.1
D4	76929297	76929430
refGene_gn=TNC;refGene_tx=XM_011288474.3;refGene_gn=TNC;refGene_tx=XM_023242642.1
D4	76932712	76932843
refGene_gn=TNC;refGene_tx=XM_011288474.3;refGene_gn=TNC;refGene_tx=XM_023242642.1
D4	76933610	76933754
refGene_gn=TNC;refGene_tx=XM_011288474.3;refGene_gn=TNC;refGene_tx=XM_023242642.1
D4	76935827	76935947
refGene_gn=TNC;refGene_tx=XM_011288474.3;refGene_gn=TNC;refGene_tx=XM_023242642.1
D4	76938122	76938266
refGene_gn=TNC;refGene_tx=XM_011288474.3;refGene_gn=TNC;refGene_tx=XM_023242642.1
D4	76939632	76939755
refGene_gn=TNC;refGene_tx=XM_011288474.3;refGene_gn=TNC;refGene_tx=XM_023242642.1
D4	76945264	76945537
refGene_gn=TNC;refGene_tx=XM_011288474.3;refGene_gn=TNC;refGene_tx=XM_023242642.1
D4	76947102	76947375
refGene_gn=TNC;refGene_tx=XM_011288474.3;refGene_gn=TNC;refGene_tx=XM_023242642.1
D4	76954754	76955027
refGene_gn=TNC;refGene_tx=XM_011288474.3;refGene_gn=TNC;refGene_tx=XM_023242642.1
D4	76957315	76957588
refGene_gn=TNC;refGene_tx=XM_011288474.3;refGene_gn=TNC;refGene_tx=XM_023242642.1
D4	76960030	76960303
refGene_gn=TNC;refGene_tx=XM_011288474.3;refGene_gn=TNC;refGene_tx=XM_023242642.1
D4	76960907	76961180
refGene_gn=TNC;refGene_tx=XM_011288474.3;refGene_gn=TNC;refGene_tx=XM_023242642.1
D4	76961991	76962264
refGene_gn=TNC;refGene_tx=XM_011288474.3;refGene_gn=TNC;refGene_tx=XM_023242642.1
D4	76970468	76970732
refGene_gn=TNC;refGene_tx=XM_011288474.3;refGene_gn=TNC;refGene_tx=XM_023242642.1
D4	76972173	76972263
refGene_gn=TNC;refGene_tx=XM_011288474.3;refGene_gn=TNC;refGene_tx=XM_023242642.1
D4	76972519	76972705
refGene_gn=TNC;refGene_tx=XM_011288474.3;refGene_gn=TNC;refGene_tx=XM_023242642.1

D4 76973978 76974248
refGene_gn=TNC;refGene_tx=XM_011288474.3;refGene_gn=TNC;refGene_tx=XM_023242642.1
D4 76977635 76977792
refGene_gn=TNC;refGene_tx=XM_011288474.3;refGene_gn=TNC;refGene_tx=XM_023242642.1
D4 76978759 76978875
refGene_gn=TNC;refGene_tx=XM_011288474.3;refGene_gn=TNC;refGene_tx=XM_023242642.1
D4 76979880 76980144
refGene_gn=TNC;refGene_tx=XM_011288474.3;refGene_gn=TNC;refGene_tx=XM_023242642.1
D4 76981465 76982872
refGene_gn=TNC;refGene_tx=XM_011288474.3;refGene_gn=TNC;refGene_tx=XM_023242642.1
D4 76985546 76986003
refGene_gn=TNC;refGene_tx=XM_011288474.3;refGene_gn=TNC;refGene_tx=XM_023242642.1
D4 77867954 77868375 refGene_gn=PAPPA;refGene_tx=XM_023242643.1
D4 77900452 77901515 refGene_gn=PAPPA;refGene_tx=XM_023242643.1
D4 77922788 77922934 refGene_gn=PAPPA;refGene_tx=XM_023242643.1
D4 77927114 77927408 refGene_gn=PAPPA;refGene_tx=XM_023242643.1
D4 77934861 77935054 refGene_gn=PAPPA;refGene_tx=XM_023242643.1
D4 77941479 77941601 refGene_gn=PAPPA;refGene_tx=XM_023242643.1
D4 77949132 77949631 refGene_gn=PAPPA;refGene_tx=XM_023242643.1
D4 77977603 77977732 refGene_gn=PAPPA;refGene_tx=XM_023242643.1
D4 77982697 77982789 refGene_gn=PAPPA;refGene_tx=XM_023242643.1
D4 78012034 78012228 refGene_gn=PAPPA;refGene_tx=XM_023242643.1
D4 78038671 78038785 refGene_gn=PAPPA;refGene_tx=XM_023242643.1
D4 78039692 78039828 refGene_gn=PAPPA;refGene_tx=XM_023242643.1
D4 78042132 78042346 refGene_gn=PAPPA;refGene_tx=XM_023242643.1
D4 78051460 78051629 refGene_gn=PAPPA;refGene_tx=XM_023242643.1
D4 78053905 78054089 refGene_gn=PAPPA;refGene_tx=XM_023242643.1
D4 78063703 78063914 refGene_gn=PAPPA;refGene_tx=XM_023242643.1
D4 78064542 78064714 refGene_gn=PAPPA;refGene_tx=XM_023242643.1
D4 78073766 78073914 refGene_gn=PAPPA;refGene_tx=XM_023242643.1
D4 78079292 78079402 refGene_gn=PAPPA;refGene_tx=XM_023242643.1
D4 78088180 78088252 refGene_gn=PAPPA;refGene_tx=XM_023242643.1
D4 78093171 78093270 refGene_gn=PAPPA;refGene_tx=XM_023242643.1
D4 78104575 78104683 refGene_gn=PAPPA;refGene_tx=XM_023242643.1
D4 78137198 78137436
refGene_gn=ASTN2;refGene_tx=XM_011288658.3;refGene_gn=ASTN2;refGene_tx=XM_019816422.2;refGene_gn=ASTN2;refGene_tx=XM_019816424.1;refGene_gn=ASTN2;refGene_tx=XM_023242644.1;refGene_gn=ASTN2;refGene_tx=XM_023242645.1;refGene_gn=ASTN2;refGene_tx=XM_023242646.1
D4 78150623 78150821
refGene_gn=ASTN2;refGene_tx=XM_011288658.3;refGene_gn=ASTN2;refGene_tx=XM_019816422.2;refGene_gn=ASTN2;refGene_tx=XM_019816424.1;refGene_gn=ASTN2;refGene_tx=XM_023242644.1;refGene_gn=ASTN2;refGene_tx=XM_023242645.1;refGene_gn=ASTN2;refGene_tx=XM_023242646.1

D4 78152751 78152852
refGene_gn=ASTN2;refGene_tx=XM_011288658.3;refGene_gn=ASTN2;refGene_tx=XM_019816422.2;refGene_gn=ASTN2;refGene_tx=XM_019816424.1;refGene_gn=ASTN2;refGene_tx=XM_023242644.1;refGene_gn=ASTN2;refGene_tx=XM_023242645.1;refGene_gn=ASTN2;refGene_tx=XM_023242646.1
D4 78190501 78190643
refGene_gn=ASTN2;refGene_tx=XM_011288658.3;refGene_gn=ASTN2;refGene_tx=XM_019816422.2;refGene_gn=ASTN2;refGene_tx=XM_019816424.1;refGene_gn=ASTN2;refGene_tx=XM_023242644.1;refGene_gn=ASTN2;refGene_tx=XM_023242645.1;refGene_gn=ASTN2;refGene_tx=XM_023242646.1
D4 78288959 78289108
refGene_gn=ASTN2;refGene_tx=XM_011288658.3;refGene_gn=ASTN2;refGene_tx=XM_019816422.2;refGene_gn=ASTN2;refGene_tx=XM_019816424.1;refGene_gn=ASTN2;refGene_tx=XM_023242644.1;refGene_gn=ASTN2;refGene_tx=XM_023242645.1;refGene_gn=ASTN2;refGene_tx=XM_023242646.1
D4 78290605 78290739
refGene_gn=ASTN2;refGene_tx=XM_011288658.3;refGene_gn=ASTN2;refGene_tx=XM_019816422.2;refGene_gn=ASTN2;refGene_tx=XM_019816424.1;refGene_gn=ASTN2;refGene_tx=XM_023242644.1;refGene_gn=ASTN2;refGene_tx=XM_023242645.1;refGene_gn=ASTN2;refGene_tx=XM_023242646.1
D4 78321636 78321902
refGene_gn=ASTN2;refGene_tx=XM_011288658.3;refGene_gn=ASTN2;refGene_tx=XM_019816422.2;refGene_gn=ASTN2;refGene_tx=XM_019816424.1;refGene_gn=ASTN2;refGene_tx=XM_023242644.1;refGene_gn=ASTN2;refGene_tx=XM_023242645.1;refGene_gn=ASTN2;refGene_tx=XM_023242646.1
D4 78352083 78354102
refGene_gn=TRIM32;refGene_tx=NM_001309047.1;refGene_gn=TRIM32;refGene_tx=XM_019815492.2;refGene_gn=TRIM32;refGene_tx=XM_023242191.1;refGene_gn=TRIM32;refGene_tx=XM_023242192.1
D4 78373198 78373378
refGene_gn=ASTN2;refGene_tx=XM_011288658.3;refGene_gn=ASTN2;refGene_tx=XM_019816422.2;refGene_gn=ASTN2;refGene_tx=XM_019816424.1;refGene_gn=ASTN2;refGene_tx=XM_023242644.1;refGene_gn=ASTN2;refGene_tx=XM_023242645.1;refGene_gn=ASTN2;refGene_tx=XM_023242646.1
D4 78376476 78376581
refGene_gn=ASTN2;refGene_tx=XM_011288658.3;refGene_gn=ASTN2;refGene_tx=XM_019816422.2;refGene_gn=ASTN2;refGene_tx=XM_019816424.1;refGene_gn=ASTN2;refGene_tx=XM_023242644.1;refGene_gn=ASTN2;refGene_tx=XM_023242645.1;refGene_gn=ASTN2;refGene_tx=XM_023242646.1
D4 78381059 78381184
refGene_gn=ASTN2;refGene_tx=XM_011288658.3;refGene_gn=ASTN2;refGene_tx=XM_019816422.2;refGene_gn=ASTN2;refGene_tx=XM_019816424.1;refGene_gn=ASTN2;refGene_tx=XM_023242644.1;refGene_gn=ASTN2;refGene_tx=XM_023242645.1;refGene_gn=ASTN2;refGene_tx=XM_023242646.1
D4 78444323 78444512
refGene_gn=ASTN2;refGene_tx=XM_011288658.3;refGene_gn=ASTN2;refGene_tx=XM_019816422.2;refGene_gn=ASTN2;refGene_tx=XM_019816424.1;refGene_gn=ASTN2;refGene_tx=XM_023242644.1;refGene_gn=ASTN2;refGene_tx=XM_023242645.1;refGene_gn=ASTN2;refGene_tx=XM_023242646.1
D4 78460624 78460791

refGene_gn=ASTN2;refGene_tx=XM_011288658.3;refGene_gn=ASTN2;refGene_tx=XM_019816422.2;refGene_gn=ASTN2;refGene_tx=XM_019816424.1;refGene_gn=ASTN2;refGene_tx=XM_023242644.1;refGene_gn=ASTN2;refGene_tx=XM_023242645.1;refGene_gn=ASTN2;refGene_tx=XM_023242646.1
D4 78496567 78496718
refGene_gn=ASTN2;refGene_tx=XM_011288658.3;refGene_gn=ASTN2;refGene_tx=XM_019816422.2;refGene_gn=ASTN2;refGene_tx=XM_019816424.1;refGene_gn=ASTN2;refGene_tx=XM_023242644.1;refGene_gn=ASTN2;refGene_tx=XM_023242645.1;refGene_gn=ASTN2;refGene_tx=XM_023242646.1
D4 78594307 78594445
refGene_gn=ASTN2;refGene_tx=XM_011288658.3;refGene_gn=ASTN2;refGene_tx=XM_019816422.2;refGene_gn=ASTN2;refGene_tx=XM_023242644.1;refGene_gn=ASTN2;refGene_tx=XM_023242645.1;refGene_gn=ASTN2;refGene_tx=XM_023242646.1
D4 78595218 78595293
refGene_gn=ASTN2;refGene_tx=XM_011288658.3;refGene_gn=ASTN2;refGene_tx=XM_019816422.2;refGene_gn=ASTN2;refGene_tx=XM_023242644.1;refGene_gn=ASTN2;refGene_tx=XM_023242645.1;refGene_gn=ASTN2;refGene_tx=XM_023242646.1
D4 78596244 78596329
refGene_gn=ASTN2;refGene_tx=XM_011288658.3;refGene_gn=ASTN2;refGene_tx=XM_019816422.2;refGene_gn=ASTN2;refGene_tx=XM_023242644.1;refGene_gn=ASTN2;refGene_tx=XM_023242645.1;refGene_gn=ASTN2;refGene_tx=XM_023242646.1
D4 78625905 78626073
refGene_gn=ASTN2;refGene_tx=XM_011288658.3;refGene_gn=ASTN2;refGene_tx=XM_019816422.2;refGene_gn=ASTN2;refGene_tx=XM_023242644.1;refGene_gn=ASTN2;refGene_tx=XM_023242645.1;refGene_gn=ASTN2;refGene_tx=XM_023242646.1
D4 78659638 78659785
refGene_gn=ASTN2;refGene_tx=XM_011288658.3;refGene_gn=ASTN2;refGene_tx=XM_019816422.2;refGene_gn=ASTN2;refGene_tx=XM_023242644.1;refGene_gn=ASTN2;refGene_tx=XM_023242645.1;refGene_gn=ASTN2;refGene_tx=XM_023242646.1
D4 78706313 78706421
refGene_gn=ASTN2;refGene_tx=XM_011288658.3;refGene_gn=ASTN2;refGene_tx=XM_019816422.2;refGene_gn=ASTN2;refGene_tx=XM_023242644.1;refGene_gn=ASTN2;refGene_tx=XM_023242645.1;refGene_gn=ASTN2;refGene_tx=XM_023242646.1
D4 78752086 78752239
refGene_gn=ASTN2;refGene_tx=XM_011288658.3;refGene_gn=ASTN2;refGene_tx=XM_019816422.2;refGene_gn=ASTN2;refGene_tx=XM_023242645.1;refGene_gn=ASTN2;refGene_tx=XM_023242646.1
D4 78811016 78811401
refGene_gn=ASTN2;refGene_tx=XM_011288658.3;refGene_gn=ASTN2;refGene_tx=XM_019816422.2;refGene_gn=ASTN2;refGene_tx=XM_023242644.1;refGene_gn=ASTN2;refGene_tx=XM_023242645.1;refGene_gn=ASTN2;refGene_tx=XM_023242646.1
D4 78886607 78886795
refGene_gn=ASTN2;refGene_tx=XM_011288658.3;refGene_gn=ASTN2;refGene_tx

=XM_019816422.2;refGene_gn=ASTN2;refGene_tx=XM_023242644.1;refGene_gn=ASTN2;refGene_tx=XM_023242645.1;refGene_gn=ASTN2;refGene_tx=XM_023242646.1
D4 79013191 79013642
refGene_gn=ASTN2;refGene_tx=XM_011288658.3;refGene_gn=ASTN2;refGene_tx=XM_023242644.1
D4 79270747 79270840 refGene_gn=TLR4;refGene_tx=NM_001009223.1
D4 79275204 79275371 refGene_gn=TLR4;refGene_tx=NM_001009223.1
D4 79278651 79280893
refGene_gn=TLR4;refGene_tx=NM_001009223.1;refGene_gn=TLR4;refGene_tx=XM_019815496.1
D4 80613346 80614487
refGene_gn=BRINP1;refGene_tx=XM_006939410.4;refGene_gn=BRINP1;refGene_tx=XM_006939411.4;refGene_gn=BRINP1;refGene_tx=XM_006939412.4;refGene_gn=BRINP1;refGene_tx=XM_011288472.3;refGene_gn=BRINP1;refGene_tx=XM_019816425.2
D4 80640702 80640925
refGene_gn=BRINP1;refGene_tx=XM_006939410.4;refGene_gn=BRINP1;refGene_tx=XM_006939411.4;refGene_gn=BRINP1;refGene_tx=XM_006939412.4;refGene_gn=BRINP1;refGene_tx=XM_011288472.3;refGene_gn=BRINP1;refGene_tx=XM_019816425.2
D4 80645575 80645812
refGene_gn=BRINP1;refGene_tx=XM_006939410.4;refGene_gn=BRINP1;refGene_tx=XM_006939411.4;refGene_gn=BRINP1;refGene_tx=XM_006939412.4;refGene_gn=BRINP1;refGene_tx=XM_011288472.3
D4 80667514 80667620
refGene_gn=BRINP1;refGene_tx=XM_006939410.4;refGene_gn=BRINP1;refGene_tx=XM_006939411.4;refGene_gn=BRINP1;refGene_tx=XM_006939412.4;refGene_gn=BRINP1;refGene_tx=XM_011288472.3;refGene_gn=BRINP1;refGene_tx=XM_019816425.2
D4 80670950 80671120
refGene_gn=BRINP1;refGene_tx=XM_006939410.4;refGene_gn=BRINP1;refGene_tx=XM_006939411.4;refGene_gn=BRINP1;refGene_tx=XM_006939412.4;refGene_gn=BRINP1;refGene_tx=XM_011288472.3;refGene_gn=BRINP1;refGene_tx=XM_019816425.2
D4 80678379 80678570
refGene_gn=BRINP1;refGene_tx=XM_006939410.4;refGene_gn=BRINP1;refGene_tx=XM_006939411.4;refGene_gn=BRINP1;refGene_tx=XM_006939412.4;refGene_gn=BRINP1;refGene_tx=XM_011288472.3;refGene_gn=BRINP1;refGene_tx=XM_019816425.2
D4 80739443 80739661
refGene_gn=BRINP1;refGene_tx=XM_006939410.4;refGene_gn=BRINP1;refGene_tx=XM_006939411.4;refGene_gn=BRINP1;refGene_tx=XM_006939412.4;refGene_gn=BRINP1;refGene_tx=XM_011288472.3;refGene_gn=BRINP1;refGene_tx=XM_019816425.2
D4 81619477 81619534 refGene_gn=CDK5RAP2;refGene_tx=XM_023242647.1
D4 81620008 81620055 refGene_gn=CDK5RAP2;refGene_tx=XM_023242647.1
D4 81620556 81620699
refGene_gn=CDK5RAP2;refGene_tx=XM_019816426.2;refGene_gn=CDK5RAP2;refGene_tx=XM_019816427.2;refGene_gn=CDK5RAP2;refGene_tx=XM_019816428.2;re

fGene_gn=CDK5RAP2;refGene_tx=XM_023242648.1
D4 81624327 81624454
refGene_gn=CDK5RAP2;refGene_tx=XM_019816426.2;refGene_gn=CDK5RAP2;refGene_tx=XM_019816427.2;refGene_gn=CDK5RAP2;refGene_tx=XM_019816428.2;refGene_gn=CDK5RAP2;refGene_tx=XM_023242647.1;refGene_gn=CDK5RAP2;refGene_tx=XM_023242648.1
D4 81630588 81630732
refGene_gn=CDK5RAP2;refGene_tx=XM_019816426.2;refGene_gn=CDK5RAP2;refGene_tx=XM_019816427.2;refGene_gn=CDK5RAP2;refGene_tx=XM_019816428.2;refGene_gn=CDK5RAP2;refGene_tx=XM_023242647.1;refGene_gn=CDK5RAP2;refGene_tx=XM_023242648.1
D4 81632544 81632810
refGene_gn=CDK5RAP2;refGene_tx=XM_019816426.2;refGene_gn=CDK5RAP2;refGene_tx=XM_019816427.2;refGene_gn=CDK5RAP2;refGene_tx=XM_019816428.2;refGene_gn=CDK5RAP2;refGene_tx=XM_023242647.1;refGene_gn=CDK5RAP2;refGene_tx=XM_023242648.1
D4 81633831 81633909
refGene_gn=CDK5RAP2;refGene_tx=XM_019816426.2;refGene_gn=CDK5RAP2;refGene_tx=XM_019816427.2;refGene_gn=CDK5RAP2;refGene_tx=XM_019816428.2;refGene_gn=CDK5RAP2;refGene_tx=XM_023242647.1;refGene_gn=CDK5RAP2;refGene_tx=XM_023242648.1
D4 81636641 81636878
refGene_gn=CDK5RAP2;refGene_tx=XM_019816426.2;refGene_gn=CDK5RAP2;refGene_tx=XM_019816427.2;refGene_gn=CDK5RAP2;refGene_tx=XM_019816428.2;refGene_gn=CDK5RAP2;refGene_tx=XM_023242647.1;refGene_gn=CDK5RAP2;refGene_tx=XM_023242648.1
D4 81637957 81638079
refGene_gn=CDK5RAP2;refGene_tx=XM_019816426.2;refGene_gn=CDK5RAP2;refGene_tx=XM_019816427.2;refGene_gn=CDK5RAP2;refGene_tx=XM_019816428.2;refGene_gn=CDK5RAP2;refGene_tx=XM_023242647.1;refGene_gn=CDK5RAP2;refGene_tx=XM_023242648.1
D4 81638897 81639087
refGene_gn=CDK5RAP2;refGene_tx=XM_019816426.2;refGene_gn=CDK5RAP2;refGene_tx=XM_019816427.2;refGene_gn=CDK5RAP2;refGene_tx=XM_019816428.2;refGene_gn=CDK5RAP2;refGene_tx=XM_023242647.1;refGene_gn=CDK5RAP2;refGene_tx=XM_023242648.1
D4 81640970 81641087
refGene_gn=CDK5RAP2;refGene_tx=XM_019816426.2;refGene_gn=CDK5RAP2;refGene_tx=XM_019816427.2;refGene_gn=CDK5RAP2;refGene_tx=XM_019816428.2;refGene_gn=CDK5RAP2;refGene_tx=XM_023242647.1;refGene_gn=CDK5RAP2;refGene_tx=XM_023242648.1
D4 81645150 81645270
refGene_gn=CDK5RAP2;refGene_tx=XM_019816426.2;refGene_gn=CDK5RAP2;refGene_tx=XM_019816427.2;refGene_gn=CDK5RAP2;refGene_tx=XM_019816428.2;refGene_gn=CDK5RAP2;refGene_tx=XM_023242647.1;refGene_gn=CDK5RAP2;refGene_tx=XM_023242648.1
D4 81649490 81649666
refGene_gn=CDK5RAP2;refGene_tx=XM_019816426.2;refGene_gn=CDK5RAP2;refGene_tx=XM_019816427.2;refGene_gn=CDK5RAP2;refGene_tx=XM_019816428.2;refGene_gn=CDK5RAP2;refGene_tx=XM_023242647.1;refGene_gn=CDK5RAP2;refGen

e_tx=XM_023242648.1
D4 81652361 81652410
refGene_gn=CDK5RAP2;refGene_tx=XM_019816426.2;refGene_gn=CDK5RAP2;refGene_tx=XM_019816427.2;refGene_gn=CDK5RAP2;refGene_tx=XM_019816428.2;refGene_gn=CDK5RAP2;refGene_tx=XM_023242647.1;refGene_gn=CDK5RAP2;refGene_tx=XM_023242648.1
D4 81662898 81663131
refGene_gn=CDK5RAP2;refGene_tx=XM_019816426.2;refGene_gn=CDK5RAP2;refGene_tx=XM_019816427.2;refGene_gn=CDK5RAP2;refGene_tx=XM_019816428.2;refGene_gn=CDK5RAP2;refGene_tx=XM_023242647.1
D4 81664975 81665546
refGene_gn=CDK5RAP2;refGene_tx=XM_019816426.2;refGene_gn=CDK5RAP2;refGene_tx=XM_019816427.2;refGene_gn=CDK5RAP2;refGene_tx=XM_019816428.2;refGene_gn=CDK5RAP2;refGene_tx=XM_023242647.1
D4 81673850 81674082
refGene_gn=CDK5RAP2;refGene_tx=XM_019816426.2;refGene_gn=CDK5RAP2;refGene_tx=XM_019816427.2;refGene_gn=CDK5RAP2;refGene_tx=XM_019816428.2;refGene_gn=CDK5RAP2;refGene_tx=XM_023242647.1;refGene_gn=CDK5RAP2;refGene_tx=XM_023242648.1
D4 81679299 81679732
refGene_gn=CDK5RAP2;refGene_tx=XM_019816426.2;refGene_gn=CDK5RAP2;refGene_tx=XM_019816427.2;refGene_gn=CDK5RAP2;refGene_tx=XM_019816428.2;refGene_gn=CDK5RAP2;refGene_tx=XM_023242647.1;refGene_gn=CDK5RAP2;refGene_tx=XM_023242648.1
D4 81685972 81686160
refGene_gn=CDK5RAP2;refGene_tx=XM_019816426.2;refGene_gn=CDK5RAP2;refGene_tx=XM_019816427.2;refGene_gn=CDK5RAP2;refGene_tx=XM_019816428.2;refGene_gn=CDK5RAP2;refGene_tx=XM_023242647.1;refGene_gn=CDK5RAP2;refGene_tx=XM_023242648.1
D4 81687990 81688086
refGene_gn=CDK5RAP2;refGene_tx=XM_019816426.2;refGene_gn=CDK5RAP2;refGene_tx=XM_019816427.2;refGene_gn=CDK5RAP2;refGene_tx=XM_019816428.2;refGene_gn=CDK5RAP2;refGene_tx=XM_023242647.1;refGene_gn=CDK5RAP2;refGene_tx=XM_023242648.1
D4 81694815 81694944
refGene_gn=CDK5RAP2;refGene_tx=XM_019816426.2;refGene_gn=CDK5RAP2;refGene_tx=XM_019816427.2;refGene_gn=CDK5RAP2;refGene_tx=XM_019816428.2;refGene_gn=CDK5RAP2;refGene_tx=XM_023242647.1;refGene_gn=CDK5RAP2;refGene_tx=XM_023242648.1
D4 81697021 81697131
refGene_gn=CDK5RAP2;refGene_tx=XM_019816426.2;refGene_gn=CDK5RAP2;refGene_tx=XM_019816427.2;refGene_gn=CDK5RAP2;refGene_tx=XM_019816428.2;refGene_gn=CDK5RAP2;refGene_tx=XM_023242647.1;refGene_gn=CDK5RAP2;refGene_tx=XM_023242648.1
D4 81698223 81698354
refGene_gn=CDK5RAP2;refGene_tx=XM_019816426.2;refGene_gn=CDK5RAP2;refGene_tx=XM_019816427.2;refGene_gn=CDK5RAP2;refGene_tx=XM_019816428.2;refGene_gn=CDK5RAP2;refGene_tx=XM_023242647.1;refGene_gn=CDK5RAP2;refGene_tx=XM_023242648.1
D4 81704116 81704214

refGene_gn=CDK5RAP2;refGene_tx=XM_019816426.2;refGene_gn=CDK5RAP2;refGene_tx=XM_019816427.2;refGene_gn=CDK5RAP2;refGene_tx=XM_019816428.2;refGene_gn=CDK5RAP2;refGene_tx=XM_023242647.1;refGene_gn=CDK5RAP2;refGene_tx=XM_023242648.1
D4 81711980 81712118
refGene_gn=CDK5RAP2;refGene_tx=XM_019816426.2;refGene_gn=CDK5RAP2;refGene_tx=XM_019816427.2;refGene_gn=CDK5RAP2;refGene_tx=XM_019816428.2;refGene_gn=CDK5RAP2;refGene_tx=XM_023242647.1;refGene_gn=CDK5RAP2;refGene_tx=XM_023242648.1
D4 81715920 81716091
refGene_gn=CDK5RAP2;refGene_tx=XM_019816426.2;refGene_gn=CDK5RAP2;refGene_tx=XM_019816427.2;refGene_gn=CDK5RAP2;refGene_tx=XM_019816428.2;refGene_gn=CDK5RAP2;refGene_tx=XM_023242647.1;refGene_gn=CDK5RAP2;refGene_tx=XM_023242648.1
D4 81730882 81731101
refGene_gn=CDK5RAP2;refGene_tx=XM_019816426.2;refGene_gn=CDK5RAP2;refGene_tx=XM_019816427.2;refGene_gn=CDK5RAP2;refGene_tx=XM_019816428.2;refGene_gn=CDK5RAP2;refGene_tx=XM_023242647.1;refGene_gn=CDK5RAP2;refGene_tx=XM_023242648.1
D4 81739482 81739575
refGene_gn=CDK5RAP2;refGene_tx=XM_019816426.2;refGene_gn=CDK5RAP2;refGene_tx=XM_019816427.2;refGene_gn=CDK5RAP2;refGene_tx=XM_019816428.2;refGene_gn=CDK5RAP2;refGene_tx=XM_023242647.1;refGene_gn=CDK5RAP2;refGene_tx=XM_023242648.1
D4 81741942 81742062
refGene_gn=CDK5RAP2;refGene_tx=XM_019816426.2;refGene_gn=CDK5RAP2;refGene_tx=XM_019816427.2;refGene_gn=CDK5RAP2;refGene_tx=XM_019816428.2;refGene_gn=CDK5RAP2;refGene_tx=XM_023242647.1;refGene_gn=CDK5RAP2;refGene_tx=XM_023242648.1
D4 81742922 81742976
refGene_gn=CDK5RAP2;refGene_tx=XM_019816426.2;refGene_gn=CDK5RAP2;refGene_tx=XM_019816427.2;refGene_gn=CDK5RAP2;refGene_tx=XM_019816428.2;refGene_gn=CDK5RAP2;refGene_tx=XM_023242647.1;refGene_gn=CDK5RAP2;refGene_tx=XM_023242648.1
D4 81744268 81744431
refGene_gn=CDK5RAP2;refGene_tx=XM_019816426.2;refGene_gn=CDK5RAP2;refGene_tx=XM_019816427.2;refGene_gn=CDK5RAP2;refGene_tx=XM_019816428.2;refGene_gn=CDK5RAP2;refGene_tx=XM_023242647.1;refGene_gn=CDK5RAP2;refGene_tx=XM_023242648.1
D4 81750526 81750681
refGene_gn=CDK5RAP2;refGene_tx=XM_019816426.2;refGene_gn=CDK5RAP2;refGene_tx=XM_019816427.2;refGene_gn=CDK5RAP2;refGene_tx=XM_019816428.2;refGene_gn=CDK5RAP2;refGene_tx=XM_023242647.1;refGene_gn=CDK5RAP2;refGene_tx=XM_023242648.1
D4 81752370 81752494
refGene_gn=CDK5RAP2;refGene_tx=XM_019816426.2;refGene_gn=CDK5RAP2;refGene_tx=XM_019816427.2;refGene_gn=CDK5RAP2;refGene_tx=XM_019816428.2;refGene_gn=CDK5RAP2;refGene_tx=XM_023242647.1;refGene_gn=CDK5RAP2;refGene_tx=XM_023242648.1
D4 81759480 81759557

refGene_gn=CDK5RAP2;refGene_tx=XM_019816426.2;refGene_gn=CDK5RAP2;refGene_tx=XM_019816427.2;refGene_gn=CDK5RAP2;refGene_tx=XM_019816428.2;refGene_gn=CDK5RAP2;refGene_tx=XM_023242647.1;refGene_gn=CDK5RAP2;refGene_tx=XM_023242648.1
D4 81762815 81762827 refGene_gn=CDK5RAP2;refGene_tx=XM_019816428.2
D4 81762979 81763090
refGene_gn=CDK5RAP2;refGene_tx=XM_019816426.2;refGene_gn=CDK5RAP2;refGene_tx=XM_019816427.2;refGene_gn=CDK5RAP2;refGene_tx=XM_023242647.1;refGene_gn=CDK5RAP2;refGene_tx=XM_023242648.1
D4 81767728 81767776 refGene_gn=CDK5RAP2;refGene_tx=XM_019816427.2
D4 81778271 81778339
refGene_gn=CDK5RAP2;refGene_tx=XM_019816426.2;refGene_gn=CDK5RAP2;refGene_tx=XM_023242647.1;refGene_gn=CDK5RAP2;refGene_tx=XM_023242648.1
D4 81782046 81782114
refGene_gn=CDK5RAP2;refGene_tx=XM_019816426.2;refGene_gn=CDK5RAP2;refGene_tx=XM_023242647.1;refGene_gn=CDK5RAP2;refGene_tx=XM_023242648.1
D4 81791784 81791843
refGene_gn=CDK5RAP2;refGene_tx=XM_019816426.2;refGene_gn=CDK5RAP2;refGene_tx=XM_023242647.1;refGene_gn=CDK5RAP2;refGene_tx=XM_023242648.1
D4 81813879 81814331 refGene_gn=MEGF9;refGene_tx=XM_023242649.1
D4 81816136 81816406 refGene_gn=MEGF9;refGene_tx=XM_023242649.1
D4 81816896 81817040 refGene_gn=MEGF9;refGene_tx=XM_023242649.1
D4 81829095 81829235 refGene_gn=MEGF9;refGene_tx=XM_023242649.1
D4 81855841 81856043 refGene_gn=MEGF9;refGene_tx=XM_023242649.1
D4 81899681 81900276 refGene_gn=MEGF9;refGene_tx=XM_023242649.1
D4 81907447 81907628 refGene_gn=FBXW2;refGene_tx=XM_023242650.1
D4 81936496 81936785
refGene_gn=FBXW2;refGene_tx=XM_003995804.4;refGene_gn=FBXW2;refGene_tx=XM_019816430.2;refGene_gn=FBXW2;refGene_tx=XM_019816432.2
D4 81943108 81943278
refGene_gn=FBXW2;refGene_tx=XM_003995804.4;refGene_gn=FBXW2;refGene_tx=XM_019816430.2;refGene_gn=FBXW2;refGene_tx=XM_019816432.2;refGene_gn=FBXW2;refGene_tx=XM_023242650.1
D4 81944112 81944199
refGene_gn=FBXW2;refGene_tx=XM_003995804.4;refGene_gn=FBXW2;refGene_tx=XM_019816430.2;refGene_gn=FBXW2;refGene_tx=XM_019816432.2;refGene_gn=FBXW2;refGene_tx=XM_023242650.1
D4 81946521 81946655
refGene_gn=FBXW2;refGene_tx=XM_003995804.4;refGene_gn=FBXW2;refGene_tx=XM_019816430.2;refGene_gn=FBXW2;refGene_tx=XM_019816432.2;refGene_gn=FBXW2;refGene_tx=XM_023242650.1
D4 81948724 81948919
refGene_gn=FBXW2;refGene_tx=XM_003995804.4;refGene_gn=FBXW2;refGene_tx=XM_019816430.2;refGene_gn=FBXW2;refGene_tx=XM_023242650.1
D4 81957653 81958143
refGene_gn=FBXW2;refGene_tx=XM_003995804.4;refGene_gn=FBXW2;refGene_tx=XM_019816430.2;refGene_gn=FBXW2;refGene_tx=XM_019816432.2;refGene_gn=FBXW2;refGene_tx=XM_023242650.1
D4 81964373 81964388 refGene_gn=FBXW2;refGene_tx=XM_019816430.2
D4 81967719 81967725

refGene_gn=LOC101090150;refGene_tx=XM_003995818.4
D4	81968987	81970091
refGene_gn=LOC101090150;refGene_tx=XM_003995818.4
D4	81974076	81974334	refGene_gn=PSMD5;refGene_tx=XM_023242651.1
D4	81975886	81976027	refGene_gn=PSMD5;refGene_tx=XM_023242651.1
D4	81976908	81977018	refGene_gn=PSMD5;refGene_tx=XM_023242651.1
D4	81980234	81980426	refGene_gn=PSMD5;refGene_tx=XM_023242651.1
D4	81982121	81982264	refGene_gn=PSMD5;refGene_tx=XM_023242651.1
D4	81984151	81984261	refGene_gn=PSMD5;refGene_tx=XM_023242651.1
D4	81987176	81987305	refGene_gn=PSMD5;refGene_tx=XM_023242651.1
D4	81987744	81987858	refGene_gn=PSMD5;refGene_tx=XM_023242651.1
D4	81989044	81989189	refGene_gn=PSMD5;refGene_tx=XM_023242651.1
D4	81996629	81996802	refGene_gn=PSMD5;refGene_tx=XM_023242651.1
D4	82001246	82001315
refGene_gn=LOC102901472;refGene_tx=XM_006939417.3;refGene_gn=LOC102901
472;refGene_tx=XM_019816434.2
D4	82005201	82005350
refGene_gn=LOC102901472;refGene_tx=XM_006939417.3;refGene_gn=LOC102901
472;refGene_tx=XM_019816434.2
D4	82012262	82012322
refGene_gn=LOC102901472;refGene_tx=XM_006939417.3
D4	82014508	82014554
refGene_gn=LOC102901472;refGene_tx=XM_006939417.3;refGene_gn=LOC102901
472;refGene_tx=XM_019816434.2
D4	82014806	82014856
refGene_gn=LOC102901472;refGene_tx=XM_006939417.3;refGene_gn=LOC102901
472;refGene_tx=XM_019816434.2
D4	82017723	82017826
refGene_gn=LOC102901472;refGene_tx=XM_006939417.3;refGene_gn=LOC102901
472;refGene_tx=XM_019816434.2
D4	82024618	82024961
refGene_gn=PHF19;refGene_tx=XM_019816437.1;refGene_gn=PHF19;refGene_tx
=XM_023242653.1;refGene_gn=PHF19;refGene_tx=XM_023242654.1
D4	82026075	82026171
refGene_gn=PHF19;refGene_tx=XM_019816437.1;refGene_gn=PHF19;refGene_tx
=XM_023242653.1;refGene_gn=PHF19;refGene_tx=XM_023242654.1
D4	82027152	82027238
refGene_gn=PHF19;refGene_tx=XM_019816437.1;refGene_gn=PHF19;refGene_tx
=XM_023242653.1;refGene_gn=PHF19;refGene_tx=XM_023242654.1
D4	82028028	82028116
refGene_gn=PHF19;refGene_tx=XM_019816437.1;refGene_gn=PHF19;refGene_tx
=XM_023242653.1;refGene_gn=PHF19;refGene_tx=XM_023242654.1
D4	82028675	82028837
refGene_gn=PHF19;refGene_tx=XM_019816437.1;refGene_gn=PHF19;refGene_tx
=XM_023242653.1;refGene_gn=PHF19;refGene_tx=XM_023242654.1
D4	82030369	82030437
refGene_gn=PHF19;refGene_tx=XM_019816437.1;refGene_gn=PHF19;refGene_tx
=XM_023242653.1;refGene_gn=PHF19;refGene_tx=XM_023242654.1
D4	82031939	82032060
refGene_gn=PHF19;refGene_tx=XM_019816437.1;refGene_gn=PHF19;refGene_tx

=XM_023242653.1;refGene_gn=PHF19;refGene_tx=XM_023242654.1
D4 82032264 82032333
refGene_gn=PHF19;refGene_tx=XM_019816437.1;refGene_gn=PHF19;refGene_tx
=XM_023242653.1;refGene_gn=PHF19;refGene_tx=XM_023242654.1
D4 82033097 82033193
refGene_gn=PHF19;refGene_tx=XM_019816437.1;refGene_gn=PHF19;refGene_tx
=XM_023242653.1;refGene_gn=PHF19;refGene_tx=XM_023242654.1
D4 82034752 82034901
refGene_gn=PHF19;refGene_tx=XM_019816437.1;refGene_gn=PHF19;refGene_tx
=XM_023242653.1;refGene_gn=PHF19;refGene_tx=XM_023242654.1
D4 82035445 82035546
refGene_gn=PHF19;refGene_tx=XM_019816437.1;refGene_gn=PHF19;refGene_tx
=XM_023242653.1;refGene_gn=PHF19;refGene_tx=XM_023242654.1
D4 82036036 82036132
refGene_gn=PHF19;refGene_tx=XM_019816437.1;refGene_gn=PHF19;refGene_tx
=XM_023242653.1;refGene_gn=PHF19;refGene_tx=XM_023242654.1
D4 82038968 82039050
refGene_gn=PHF19;refGene_tx=XM_019816437.1;refGene_gn=PHF19;refGene_tx
=XM_023242653.1;refGene_gn=PHF19;refGene_tx=XM_023242654.1
D4 82039539 82039725
refGene_gn=PHF19;refGene_tx=XM_019816437.1;refGene_gn=PHF19;refGene_tx
=XM_023242653.1;refGene_gn=PHF19;refGene_tx=XM_023242654.1
D4 82072316 82072535
refGene_gn=TRAF1;refGene_tx=XM_006939418.3;refGene_gn=TRAF1;refGene_tx
=XM_006939419.3;refGene_gn=TRAF1;refGene_tx=XM_006939420.4;refGene_gn=
TRAF1;refGene_tx=XM_019816441.1;refGene_gn=TRAF1;refGene_tx=XM_0198164
42.2;refGene_gn=TRAF1;refGene_tx=XM_023242655.1;refGene_gn=TRAF1;refGe
ne_tx=XM_023242656.1
D4 82074753 82074902
refGene_gn=TRAF1;refGene_tx=XM_006939418.3;refGene_gn=TRAF1;refGene_tx
=XM_006939419.3;refGene_gn=TRAF1;refGene_tx=XM_006939420.4;refGene_gn=
TRAF1;refGene_tx=XM_019816441.1;refGene_gn=TRAF1;refGene_tx=XM_0198164
42.2;refGene_gn=TRAF1;refGene_tx=XM_023242655.1;refGene_gn=TRAF1;refGe
ne_tx=XM_023242656.1
D4 82076882 82077060
refGene_gn=TRAF1;refGene_tx=XM_006939418.3;refGene_gn=TRAF1;refGene_tx
=XM_006939419.3;refGene_gn=TRAF1;refGene_tx=XM_006939420.4;refGene_gn=
TRAF1;refGene_tx=XM_019816441.1;refGene_gn=TRAF1;refGene_tx=XM_0198164
42.2;refGene_gn=TRAF1;refGene_tx=XM_023242655.1;refGene_gn=TRAF1;refGe
ne_tx=XM_023242656.1
D4 82078588 82078999
refGene_gn=TRAF1;refGene_tx=XM_006939418.3;refGene_gn=TRAF1;refGene_tx
=XM_006939419.3;refGene_gn=TRAF1;refGene_tx=XM_006939420.4;refGene_gn=
TRAF1;refGene_tx=XM_019816441.1;refGene_gn=TRAF1;refGene_tx=XM_0198164
42.2;refGene_gn=TRAF1;refGene_tx=XM_023242655.1;refGene_gn=TRAF1;refGe
ne_tx=XM_023242656.1
D4 82079484 82079550
refGene_gn=TRAF1;refGene_tx=XM_006939418.3;refGene_gn=TRAF1;refGene_tx
=XM_006939419.3;refGene_gn=TRAF1;refGene_tx=XM_006939420.4;refGene_gn=
TRAF1;refGene_tx=XM_019816441.1;refGene_gn=TRAF1;refGene_tx=XM_0198164

42.2;refGene_gn=TRAF1;refGene_tx=XM_023242655.1;refGene_gn=TRAF1;refGene_tx=XM_023242656.1
D4 82088078 82088163
refGene_gn=TRAF1;refGene_tx=XM_006939418.3;refGene_gn=TRAF1;refGene_tx=XM_006939419.3;refGene_gn=TRAF1;refGene_tx=XM_006939420.4;refGene_gn=TRAF1;refGene_tx=XM_019816441.1;refGene_gn=TRAF1;refGene_tx=XM_019816442.2;refGene_gn=TRAF1;refGene_tx=XM_023242655.1;refGene_gn=TRAF1;refGene_tx=XM_023242656.1
D4 82090005 82090145
refGene_gn=TRAF1;refGene_tx=XM_006939418.3;refGene_gn=TRAF1;refGene_tx=XM_006939419.3;refGene_gn=TRAF1;refGene_tx=XM_006939420.4;refGene_gn=TRAF1;refGene_tx=XM_023242655.1;refGene_gn=TRAF1;refGene_tx=XM_023242656.1
D4 82115802 82115941 refGene_gn=C5;refGene_tx=XM_023242658.1
D4 82116522 82116661 refGene_gn=C5;refGene_tx=XM_023242658.1
D4 82122797 82122881 refGene_gn=C5;refGene_tx=XM_023242658.1
D4 82125073 82125163 refGene_gn=C5;refGene_tx=XM_023242658.1
D4 82126092 82126176 refGene_gn=C5;refGene_tx=XM_023242658.1
D4 82126950 82127056 refGene_gn=C5;refGene_tx=XM_023242658.1
D4 82127172 82127247 refGene_gn=C5;refGene_tx=XM_023242658.1
D4 82127645 82127748 refGene_gn=C5;refGene_tx=XM_023242658.1
D4 82132945 82133005 refGene_gn=C5;refGene_tx=XM_023242658.1
D4 82133529 82133609 refGene_gn=C5;refGene_tx=XM_023242658.1
D4 82135398 82135461 refGene_gn=C5;refGene_tx=XM_023242658.1
D4 82137858 82138011 refGene_gn=C5;refGene_tx=XM_023242658.1
D4 82139291 82139497 refGene_gn=C5;refGene_tx=XM_023242658.1
D4 82141957 82142129 refGene_gn=C5;refGene_tx=XM_023242658.1
D4 82143406 82143502 refGene_gn=C5;refGene_tx=XM_023242658.1
D4 82145704 82145864 refGene_gn=C5;refGene_tx=XM_023242658.1
D4 82149418 82149494 refGene_gn=C5;refGene_tx=XM_023242658.1
D4 82149960 82150173 refGene_gn=C5;refGene_tx=XM_023242658.1
D4 82152052 82152142 refGene_gn=C5;refGene_tx=XM_023242658.1
D4 82157009 82157070 refGene_gn=C5;refGene_tx=XM_023242658.1
D4 82158522 82158750 refGene_gn=C5;refGene_tx=XM_023242658.1
D4 82160736 82160876 refGene_gn=C5;refGene_tx=XM_023242658.1
D4 82161573 82161647 refGene_gn=C5;refGene_tx=XM_023242658.1
D4 82163656 82163747 refGene_gn=C5;refGene_tx=XM_023242658.1
D4 82166110 82166308 refGene_gn=C5;refGene_tx=XM_023242658.1
D4 82167714 82167777 refGene_gn=C5;refGene_tx=XM_023242658.1
D4 82168989 82169119 refGene_gn=C5;refGene_tx=XM_023242658.1
D4 82170181 82170334 refGene_gn=C5;refGene_tx=XM_023242658.1
D4 82170460 82170670 refGene_gn=C5;refGene_tx=XM_023242658.1
D4 82172429 82172633 refGene_gn=C5;refGene_tx=XM_023242658.1
D4 82173921 82174107 refGene_gn=C5;refGene_tx=XM_023242658.1
D4 82175512 82175628 refGene_gn=C5;refGene_tx=XM_023242658.1
D4 82177676 82177803 refGene_gn=C5;refGene_tx=XM_023242658.1
D4 82179597 82179712 refGene_gn=C5;refGene_tx=XM_023242658.1
D4 82181732 82181823 refGene_gn=C5;refGene_tx=XM_023242658.1
D4 82184494 82184577 refGene_gn=C5;refGene_tx=XM_023242658.1
D4 82187812 82187904 refGene_gn=C5;refGene_tx=XM_023242658.1

D4 82191322 82191393 refGene_gn=C5;refGene_tx=XM_023242658.1
D4 82195617 82195780 refGene_gn=C5;refGene_tx=XM_023242658.1
D4 82198636 82198829 refGene_gn=C5;refGene_tx=XM_023242658.1
D4 82203184 82203249 refGene_gn=C5;refGene_tx=XM_023242658.1
D4 82216115 82216201
refGene_gn=LOC102902597;refGene_tx=XM_019816453.1;refGene_gn=LOC102902
597;refGene_tx=XM_023242666.1
D4 82218295 82218369
refGene_gn=LOC102902597;refGene_tx=XM_019816453.1;refGene_gn=LOC102902
597;refGene_tx=XM_023242666.1
D4 82220290 82220375
refGene_gn=LOC102902597;refGene_tx=XM_019816453.1;refGene_gn=LOC102902
597;refGene_tx=XM_023242666.1
D4 82222961 82223156
refGene_gn=LOC102902597;refGene_tx=XM_019816453.1;refGene_gn=LOC102902
597;refGene_tx=XM_023242666.1
D4 82225614 82225677
refGene_gn=LOC102902597;refGene_tx=XM_019816453.1;refGene_gn=LOC102902
597;refGene_tx=XM_023242666.1
D4 82232955 82233085
refGene_gn=LOC102902597;refGene_tx=XM_019816453.1;refGene_gn=LOC102902
597;refGene_tx=XM_023242666.1
D4 82234504 82234654
refGene_gn=LOC102902597;refGene_tx=XM_019816453.1;refGene_gn=LOC102902
597;refGene_tx=XM_023242666.1
D4 82234748 82234958
refGene_gn=LOC102902597;refGene_tx=XM_019816453.1;refGene_gn=LOC102902
597;refGene_tx=XM_023242666.1
D4 82235963 82236167
refGene_gn=LOC102902597;refGene_tx=XM_019816453.1;refGene_gn=LOC102902
597;refGene_tx=XM_023242666.1
D4 82238531 82238717
refGene_gn=LOC102902597;refGene_tx=XM_019816453.1;refGene_gn=LOC102902
597;refGene_tx=XM_023242666.1
D4 82239159 82239275
refGene_gn=LOC102902597;refGene_tx=XM_019816453.1;refGene_gn=LOC102902
597;refGene_tx=XM_023242666.1
D4 82240052 82240179
refGene_gn=LOC102902597;refGene_tx=XM_019816453.1;refGene_gn=LOC102902
597;refGene_tx=XM_023242666.1
D4 82240708 82240823
refGene_gn=LOC102902597;refGene_tx=XM_019816453.1;refGene_gn=LOC102902
597;refGene_tx=XM_023242666.1
D4 82241750 82241853
refGene_gn=LOC102902597;refGene_tx=XM_019816453.1;refGene_gn=LOC102902
597;refGene_tx=XM_023242666.1
D4 82246653 82246736
refGene_gn=LOC102902597;refGene_tx=XM_019816453.1;refGene_gn=LOC102902
597;refGene_tx=XM_023242666.1
D4 82248394 82248486

refGene_gn=LOC102902597;refGene_tx=XM_019816453.1;refGene_gn=LOC102902597;refGene_tx=XM_023242666.1
D4 82249962 82250033
refGene_gn=LOC102902597;refGene_tx=XM_019816453.1;refGene_gn=LOC102902597;refGene_tx=XM_023242666.1
D4 82251628 82251791
refGene_gn=LOC102902597;refGene_tx=XM_019816453.1;refGene_gn=LOC102902597;refGene_tx=XM_023242666.1
D4 82252649 82252730
refGene_gn=LOC102902597;refGene_tx=XM_023242666.1
D4 82253186 82253379
refGene_gn=LOC102902597;refGene_tx=XM_019816453.1
D4 82253955 82254020
refGene_gn=LOC102902597;refGene_tx=XM_019816453.1
D4 82275795 82276012
refGene_gn=CNTRL;refGene_tx=XM_023242659.1;refGene_gn=CNTRL;refGene_tx=XM_023242660.1;refGene_gn=CNTRL;refGene_tx=XM_023242661.1;refGene_gn=CNTRL;refGene_tx=XM_023242662.1;refGene_gn=CNTRL;refGene_tx=XM_023242663.1;refGene_gn=CNTRL;refGene_tx=XM_023242664.1;refGene_gn=CNTRL;refGene_tx=XM_023242665.1
D4 82279507 82279638
refGene_gn=CNTRL;refGene_tx=XM_023242659.1;refGene_gn=CNTRL;refGene_tx=XM_023242660.1;refGene_gn=CNTRL;refGene_tx=XM_023242661.1;refGene_gn=CNTRL;refGene_tx=XM_023242662.1;refGene_gn=CNTRL;refGene_tx=XM_023242663.1;refGene_gn=CNTRL;refGene_tx=XM_023242664.1;refGene_gn=CNTRL;refGene_tx=XM_023242665.1
D4 82283574 82283705
refGene_gn=CNTRL;refGene_tx=XM_023242659.1;refGene_gn=CNTRL;refGene_tx=XM_023242660.1;refGene_gn=CNTRL;refGene_tx=XM_023242661.1;refGene_gn=CNTRL;refGene_tx=XM_023242662.1;refGene_gn=CNTRL;refGene_tx=XM_023242663.1;refGene_gn=CNTRL;refGene_tx=XM_023242664.1;refGene_gn=CNTRL;refGene_tx=XM_023242665.1
D4 82285005 82285147
refGene_gn=CNTRL;refGene_tx=XM_023242659.1;refGene_gn=CNTRL;refGene_tx=XM_023242660.1;refGene_gn=CNTRL;refGene_tx=XM_023242661.1;refGene_gn=CNTRL;refGene_tx=XM_023242662.1;refGene_gn=CNTRL;refGene_tx=XM_023242663.1;refGene_gn=CNTRL;refGene_tx=XM_023242664.1;refGene_gn=CNTRL;refGene_tx=XM_023242665.1
D4 82287529 82287716
refGene_gn=CNTRL;refGene_tx=XM_023242659.1;refGene_gn=CNTRL;refGene_tx=XM_023242660.1;refGene_gn=CNTRL;refGene_tx=XM_023242661.1;refGene_gn=CNTRL;refGene_tx=XM_023242662.1;refGene_gn=CNTRL;refGene_tx=XM_023242663.1;refGene_gn=CNTRL;refGene_tx=XM_023242664.1;refGene_gn=CNTRL;refGene_tx=XM_023242665.1
D4 82291096 82291290
refGene_gn=CNTRL;refGene_tx=XM_023242659.1;refGene_gn=CNTRL;refGene_tx=XM_023242660.1;refGene_gn=CNTRL;refGene_tx=XM_023242661.1;refGene_gn=CNTRL;refGene_tx=XM_023242662.1;refGene_gn=CNTRL;refGene_tx=XM_023242663.1;refGene_gn=CNTRL;refGene_tx=XM_023242664.1;refGene_gn=CNTRL;refGene_tx=XM_023242665.1

D4 82296736 82296793
refGene_gn=CNTRL;refGene_tx=XM_023242659.1;refGene_gn=CNTRL;refGene_tx=XM_023242660.1;refGene_gn=CNTRL;refGene_tx=XM_023242662.1;refGene_gn=CNTRL;refGene_tx=XM_023242663.1;refGene_gn=CNTRL;refGene_tx=XM_023242665.1
D4 82298713 82298833
refGene_gn=CNTRL;refGene_tx=XM_023242659.1;refGene_gn=CNTRL;refGene_tx=XM_023242660.1;refGene_gn=CNTRL;refGene_tx=XM_023242661.1;refGene_gn=CNTRL;refGene_tx=XM_023242662.1;refGene_gn=CNTRL;refGene_tx=XM_023242663.1;refGene_gn=CNTRL;refGene_tx=XM_023242664.1;refGene_gn=CNTRL;refGene_tx=XM_023242665.1
D4 82299370 82299599
refGene_gn=CNTRL;refGene_tx=XM_023242659.1;refGene_gn=CNTRL;refGene_tx=XM_023242660.1;refGene_gn=CNTRL;refGene_tx=XM_023242661.1;refGene_gn=CNTRL;refGene_tx=XM_023242662.1;refGene_gn=CNTRL;refGene_tx=XM_023242663.1;refGene_gn=CNTRL;refGene_tx=XM_023242664.1;refGene_gn=CNTRL;refGene_tx=XM_023242665.1
D4 82300279 82300402
refGene_gn=CNTRL;refGene_tx=XM_023242659.1;refGene_gn=CNTRL;refGene_tx=XM_023242660.1;refGene_gn=CNTRL;refGene_tx=XM_023242661.1;refGene_gn=CNTRL;refGene_tx=XM_023242663.1;refGene_gn=CNTRL;refGene_tx=XM_023242665.1
D4 82300944 82301054
refGene_gn=CNTRL;refGene_tx=XM_023242659.1;refGene_gn=CNTRL;refGene_tx=XM_023242660.1;refGene_gn=CNTRL;refGene_tx=XM_023242661.1;refGene_gn=CNTRL;refGene_tx=XM_023242662.1;refGene_gn=CNTRL;refGene_tx=XM_023242663.1;refGene_gn=CNTRL;refGene_tx=XM_023242664.1;refGene_gn=CNTRL;refGene_tx=XM_023242665.1
D4 82303038 82303233
refGene_gn=CNTRL;refGene_tx=XM_023242659.1;refGene_gn=CNTRL;refGene_tx=XM_023242660.1;refGene_gn=CNTRL;refGene_tx=XM_023242661.1;refGene_gn=CNTRL;refGene_tx=XM_023242662.1;refGene_gn=CNTRL;refGene_tx=XM_023242663.1;refGene_gn=CNTRL;refGene_tx=XM_023242664.1
D4 82307490 82307644
refGene_gn=CNTRL;refGene_tx=XM_019816451.2;refGene_gn=CNTRL;refGene_tx=XM_019816452.1;refGene_gn=CNTRL;refGene_tx=XM_023242659.1;refGene_gn=CNTRL;refGene_tx=XM_023242660.1;refGene_gn=CNTRL;refGene_tx=XM_023242661.1;refGene_gn=CNTRL;refGene_tx=XM_023242662.1;refGene_gn=CNTRL;refGene_tx=XM_023242663.1;refGene_gn=CNTRL;refGene_tx=XM_023242664.1;refGene_gn=CNTRL;refGene_tx=XM_023242665.1
D4 82308714 82308935
refGene_gn=CNTRL;refGene_tx=XM_019816451.2;refGene_gn=CNTRL;refGene_tx=XM_019816452.1;refGene_gn=CNTRL;refGene_tx=XM_023242659.1;refGene_gn=CNTRL;refGene_tx=XM_023242660.1;refGene_gn=CNTRL;refGene_tx=XM_023242661.1;refGene_gn=CNTRL;refGene_tx=XM_023242662.1;refGene_gn=CNTRL;refGene_tx=XM_023242663.1;refGene_gn=CNTRL;refGene_tx=XM_023242664.1;refGene_gn=CNTRL;refGene_tx=XM_023242665.1
D4 82311891 82312068
refGene_gn=CNTRL;refGene_tx=XM_019816451.2;refGene_gn=CNTRL;refGene_tx=XM_019816452.1;refGene_gn=CNTRL;refGene_tx=XM_023242659.1;refGene_gn=

CNTRL;refGene_tx=XM_023242660.1;refGene_gn=CNTRL;refGene_tx=XM_0232426
61.1;refGene_gn=CNTRL;refGene_tx=XM_023242662.1;refGene_gn=CNTRL;refGe
ne_tx=XM_023242664.1;refGene_gn=CNTRL;refGene_tx=XM_023242665.1
D4 82314200 82314335
refGene_gn=CNTRL;refGene_tx=XM_019816451.2;refGene_gn=CNTRL;refGene_tx
=XM_019816452.1;refGene_gn=CNTRL;refGene_tx=XM_023242659.1;refGene_gn=
CNTRL;refGene_tx=XM_023242660.1;refGene_gn=CNTRL;refGene_tx=XM_0232426
61.1;refGene_gn=CNTRL;refGene_tx=XM_023242662.1;refGene_gn=CNTRL;refGe
ne_tx=XM_023242663.1;refGene_gn=CNTRL;refGene_tx=XM_023242664.1;refGen
e_gn=CNTRL;refGene_tx=XM_023242665.1
D4 82317656 82317802
refGene_gn=CNTRL;refGene_tx=XM_019816451.2;refGene_gn=CNTRL;refGene_tx
=XM_019816452.1;refGene_gn=CNTRL;refGene_tx=XM_023242659.1;refGene_gn=
CNTRL;refGene_tx=XM_023242660.1;refGene_gn=CNTRL;refGene_tx=XM_0232426
61.1;refGene_gn=CNTRL;refGene_tx=XM_023242662.1;refGene_gn=CNTRL;refGe
ne_tx=XM_023242663.1;refGene_gn=CNTRL;refGene_tx=XM_023242664.1;refGen
e_gn=CNTRL;refGene_tx=XM_023242665.1
D4 82318450 82318658
refGene_gn=CNTRL;refGene_tx=XM_019816451.2;refGene_gn=CNTRL;refGene_tx
=XM_019816452.1;refGene_gn=CNTRL;refGene_tx=XM_023242659.1;refGene_gn=
CNTRL;refGene_tx=XM_023242660.1;refGene_gn=CNTRL;refGene_tx=XM_0232426
61.1;refGene_gn=CNTRL;refGene_tx=XM_023242662.1;refGene_gn=CNTRL;refGe
ne_tx=XM_023242663.1;refGene_gn=CNTRL;refGene_tx=XM_023242664.1;refGen
e_gn=CNTRL;refGene_tx=XM_023242665.1
D4 82319142 82319322
refGene_gn=CNTRL;refGene_tx=XM_019816451.2;refGene_gn=CNTRL;refGene_tx
=XM_019816452.1;refGene_gn=CNTRL;refGene_tx=XM_023242659.1;refGene_gn=
CNTRL;refGene_tx=XM_023242660.1;refGene_gn=CNTRL;refGene_tx=XM_0232426
61.1;refGene_gn=CNTRL;refGene_tx=XM_023242662.1;refGene_gn=CNTRL;refGe
ne_tx=XM_023242663.1;refGene_gn=CNTRL;refGene_tx=XM_023242664.1;refGen
e_gn=CNTRL;refGene_tx=XM_023242665.1
D4 82321962 82322142
refGene_gn=CNTRL;refGene_tx=XM_019816451.2;refGene_gn=CNTRL;refGene_tx
=XM_019816452.1;refGene_gn=CNTRL;refGene_tx=XM_023242659.1;refGene_gn=
CNTRL;refGene_tx=XM_023242660.1;refGene_gn=CNTRL;refGene_tx=XM_0232426
61.1;refGene_gn=CNTRL;refGene_tx=XM_023242662.1;refGene_gn=CNTRL;refGe
ne_tx=XM_023242663.1;refGene_gn=CNTRL;refGene_tx=XM_023242664.1;refGen
e_gn=CNTRL;refGene_tx=XM_023242665.1
D4 82322893 82323010
refGene_gn=CNTRL;refGene_tx=XM_019816451.2;refGene_gn=CNTRL;refGene_tx
=XM_019816452.1;refGene_gn=CNTRL;refGene_tx=XM_023242659.1;refGene_gn=
CNTRL;refGene_tx=XM_023242660.1;refGene_gn=CNTRL;refGene_tx=XM_0232426
61.1;refGene_gn=CNTRL;refGene_tx=XM_023242662.1;refGene_gn=CNTRL;refGe
ne_tx=XM_023242663.1;refGene_gn=CNTRL;refGene_tx=XM_023242664.1;refGen
e_gn=CNTRL;refGene_tx=XM_023242665.1
D4 82323275 82323420
refGene_gn=CNTRL;refGene_tx=XM_019816451.2;refGene_gn=CNTRL;refGene_tx
=XM_019816452.1;refGene_gn=CNTRL;refGene_tx=XM_023242659.1;refGene_gn=
CNTRL;refGene_tx=XM_023242660.1;refGene_gn=CNTRL;refGene_tx=XM_0232426
61.1;refGene_gn=CNTRL;refGene_tx=XM_023242662.1;refGene_gn=CNTRL;refGe

ne_tx=XM_023242663.1;refGene_gn=CNTRL;refGene_tx=XM_023242664.1;refGene_gn=CNTRL;refGene_tx=XM_023242665.1
D4 82324383 82324532
refGene_gn=CNTRL;refGene_tx=XM_019816451.2;refGene_gn=CNTRL;refGene_tx=XM_019816452.1;refGene_gn=CNTRL;refGene_tx=XM_023242659.1;refGene_gn=CNTRL;refGene_tx=XM_023242660.1;refGene_gn=CNTRL;refGene_tx=XM_023242661.1;refGene_gn=CNTRL;refGene_tx=XM_023242662.1;refGene_gn=CNTRL;refGene_tx=XM_023242663.1;refGene_gn=CNTRL;refGene_tx=XM_023242664.1;refGene_gn=CNTRL;refGene_tx=XM_023242665.1
D4 82326199 82326389
refGene_gn=CNTRL;refGene_tx=XM_019816451.2;refGene_gn=CNTRL;refGene_tx=XM_019816452.1;refGene_gn=CNTRL;refGene_tx=XM_023242659.1;refGene_gn=CNTRL;refGene_tx=XM_023242660.1;refGene_gn=CNTRL;refGene_tx=XM_023242661.1;refGene_gn=CNTRL;refGene_tx=XM_023242662.1;refGene_gn=CNTRL;refGene_tx=XM_023242663.1;refGene_gn=CNTRL;refGene_tx=XM_023242664.1;refGene_gn=CNTRL;refGene_tx=XM_023242665.1
D4 82327432 82327746
refGene_gn=CNTRL;refGene_tx=XM_019816451.2;refGene_gn=CNTRL;refGene_tx=XM_019816452.1;refGene_gn=CNTRL;refGene_tx=XM_023242659.1;refGene_gn=CNTRL;refGene_tx=XM_023242660.1;refGene_gn=CNTRL;refGene_tx=XM_023242661.1;refGene_gn=CNTRL;refGene_tx=XM_023242662.1;refGene_gn=CNTRL;refGene_tx=XM_023242663.1;refGene_gn=CNTRL;refGene_tx=XM_023242664.1;refGene_gn=CNTRL;refGene_tx=XM_023242665.1
D4 82329472 82329678
refGene_gn=CNTRL;refGene_tx=XM_019816451.2;refGene_gn=CNTRL;refGene_tx=XM_019816452.1;refGene_gn=CNTRL;refGene_tx=XM_023242659.1;refGene_gn=CNTRL;refGene_tx=XM_023242660.1;refGene_gn=CNTRL;refGene_tx=XM_023242661.1;refGene_gn=CNTRL;refGene_tx=XM_023242662.1;refGene_gn=CNTRL;refGene_tx=XM_023242663.1;refGene_gn=CNTRL;refGene_tx=XM_023242664.1;refGene_gn=CNTRL;refGene_tx=XM_023242665.1
D4 82331902 82332095
refGene_gn=CNTRL;refGene_tx=XM_019816451.2;refGene_gn=CNTRL;refGene_tx=XM_019816452.1;refGene_gn=CNTRL;refGene_tx=XM_023242659.1;refGene_gn=CNTRL;refGene_tx=XM_023242660.1;refGene_gn=CNTRL;refGene_tx=XM_023242661.1;refGene_gn=CNTRL;refGene_tx=XM_023242662.1;refGene_gn=CNTRL;refGene_tx=XM_023242663.1;refGene_gn=CNTRL;refGene_tx=XM_023242664.1;refGene_gn=CNTRL;refGene_tx=XM_023242665.1
D4 82334542 82334673
refGene_gn=CNTRL;refGene_tx=XM_019816451.2;refGene_gn=CNTRL;refGene_tx=XM_019816452.1;refGene_gn=CNTRL;refGene_tx=XM_023242659.1;refGene_gn=CNTRL;refGene_tx=XM_023242660.1;refGene_gn=CNTRL;refGene_tx=XM_023242661.1;refGene_gn=CNTRL;refGene_tx=XM_023242662.1;refGene_gn=CNTRL;refGene_tx=XM_023242663.1;refGene_gn=CNTRL;refGene_tx=XM_023242664.1;refGene_gn=CNTRL;refGene_tx=XM_023242665.1
D4 82334813 82334954
refGene_gn=CNTRL;refGene_tx=XM_019816451.2;refGene_gn=CNTRL;refGene_tx=XM_019816452.1;refGene_gn=CNTRL;refGene_tx=XM_023242659.1;refGene_gn=CNTRL;refGene_tx=XM_023242660.1;refGene_gn=CNTRL;refGene_tx=XM_023242661.1;refGene_gn=CNTRL;refGene_tx=XM_023242662.1;refGene_gn=CNTRL;refGene_tx=XM_023242663.1;refGene_gn=CNTRL;refGene_tx=XM_023242664.1;refGen

e_gn=CNTRL;refGene_tx=XM_023242665.1
D4 82335055 82335182
refGene_gn=CNTRL;refGene_tx=XM_019816451.2;refGene_gn=CNTRL;refGene_tx=XM_019816452.1;refGene_gn=CNTRL;refGene_tx=XM_023242659.1;refGene_gn=CNTRL;refGene_tx=XM_023242660.1;refGene_gn=CNTRL;refGene_tx=XM_023242661.1;refGene_gn=CNTRL;refGene_tx=XM_023242662.1;refGene_gn=CNTRL;refGene_tx=XM_023242663.1;refGene_gn=CNTRL;refGene_tx=XM_023242664.1;refGene_gn=CNTRL;refGene_tx=XM_023242665.1
D4 82336462 82336627
refGene_gn=CNTRL;refGene_tx=XM_019816451.2;refGene_gn=CNTRL;refGene_tx=XM_019816452.1;refGene_gn=CNTRL;refGene_tx=XM_023242659.1;refGene_gn=CNTRL;refGene_tx=XM_023242660.1;refGene_gn=CNTRL;refGene_tx=XM_023242661.1;refGene_gn=CNTRL;refGene_tx=XM_023242662.1;refGene_gn=CNTRL;refGene_tx=XM_023242663.1;refGene_gn=CNTRL;refGene_tx=XM_023242664.1;refGene_gn=CNTRL;refGene_tx=XM_023242665.1
D4 82337387 82337547
refGene_gn=CNTRL;refGene_tx=XM_019816451.2;refGene_gn=CNTRL;refGene_tx=XM_019816452.1;refGene_gn=CNTRL;refGene_tx=XM_023242659.1;refGene_gn=CNTRL;refGene_tx=XM_023242660.1;refGene_gn=CNTRL;refGene_tx=XM_023242661.1;refGene_gn=CNTRL;refGene_tx=XM_023242662.1;refGene_gn=CNTRL;refGene_tx=XM_023242663.1;refGene_gn=CNTRL;refGene_tx=XM_023242664.1;refGene_gn=CNTRL;refGene_tx=XM_023242665.1
D4 82338684 82338800
refGene_gn=CNTRL;refGene_tx=XM_019816451.2;refGene_gn=CNTRL;refGene_tx=XM_019816452.1;refGene_gn=CNTRL;refGene_tx=XM_023242659.1;refGene_gn=CNTRL;refGene_tx=XM_023242660.1;refGene_gn=CNTRL;refGene_tx=XM_023242661.1;refGene_gn=CNTRL;refGene_tx=XM_023242662.1;refGene_gn=CNTRL;refGene_tx=XM_023242663.1;refGene_gn=CNTRL;refGene_tx=XM_023242664.1;refGene_gn=CNTRL;refGene_tx=XM_023242665.1
D4 82338882 82339100
refGene_gn=CNTRL;refGene_tx=XM_019816451.2;refGene_gn=CNTRL;refGene_tx=XM_019816452.1;refGene_gn=CNTRL;refGene_tx=XM_023242659.1;refGene_gn=CNTRL;refGene_tx=XM_023242660.1;refGene_gn=CNTRL;refGene_tx=XM_023242661.1;refGene_gn=CNTRL;refGene_tx=XM_023242662.1;refGene_gn=CNTRL;refGene_tx=XM_023242663.1;refGene_gn=CNTRL;refGene_tx=XM_023242664.1;refGene_gn=CNTRL;refGene_tx=XM_023242665.1
D4 82342095 82342253
refGene_gn=CNTRL;refGene_tx=XM_019816451.2;refGene_gn=CNTRL;refGene_tx=XM_019816452.1;refGene_gn=CNTRL;refGene_tx=XM_023242659.1;refGene_gn=CNTRL;refGene_tx=XM_023242660.1;refGene_gn=CNTRL;refGene_tx=XM_023242661.1;refGene_gn=CNTRL;refGene_tx=XM_023242662.1;refGene_gn=CNTRL;refGene_tx=XM_023242663.1;refGene_gn=CNTRL;refGene_tx=XM_023242664.1;refGene_gn=CNTRL;refGene_tx=XM_023242665.1
D4 82343278 82343352
refGene_gn=CNTRL;refGene_tx=XM_019816451.2;refGene_gn=CNTRL;refGene_tx=XM_019816452.1;refGene_gn=CNTRL;refGene_tx=XM_023242659.1;refGene_gn=CNTRL;refGene_tx=XM_023242660.1;refGene_gn=CNTRL;refGene_tx=XM_023242661.1;refGene_gn=CNTRL;refGene_tx=XM_023242662.1;refGene_gn=CNTRL;refGene_tx=XM_023242663.1;refGene_gn=CNTRL;refGene_tx=XM_023242664.1;refGene_gn=CNTRL;refGene_tx=XM_023242665.1

D4 82344729 82344918
refGene_gn=CNTRL;refGene_tx=XM_019816451.2;refGene_gn=CNTRL;refGene_tx=XM_019816452.1;refGene_gn=CNTRL;refGene_tx=XM_023242659.1;refGene_gn=CNTRL;refGene_tx=XM_023242660.1;refGene_gn=CNTRL;refGene_tx=XM_023242661.1;refGene_gn=CNTRL;refGene_tx=XM_023242662.1;refGene_gn=CNTRL;refGene_tx=XM_023242663.1;refGene_gn=CNTRL;refGene_tx=XM_023242664.1;refGene_gn=CNTRL;refGene_tx=XM_023242665.1
D4 82345325 82345551
refGene_gn=CNTRL;refGene_tx=XM_019816451.2;refGene_gn=CNTRL;refGene_tx=XM_019816452.1;refGene_gn=CNTRL;refGene_tx=XM_023242659.1;refGene_gn=CNTRL;refGene_tx=XM_023242660.1;refGene_gn=CNTRL;refGene_tx=XM_023242661.1;refGene_gn=CNTRL;refGene_tx=XM_023242662.1;refGene_gn=CNTRL;refGene_tx=XM_023242663.1;refGene_gn=CNTRL;refGene_tx=XM_023242664.1;refGene_gn=CNTRL;refGene_tx=XM_023242665.1
D4 82347013 82347219
refGene_gn=CNTRL;refGene_tx=XM_019816451.2;refGene_gn=CNTRL;refGene_tx=XM_019816452.1;refGene_gn=CNTRL;refGene_tx=XM_023242659.1;refGene_gn=CNTRL;refGene_tx=XM_023242660.1;refGene_gn=CNTRL;refGene_tx=XM_023242661.1;refGene_gn=CNTRL;refGene_tx=XM_023242662.1;refGene_gn=CNTRL;refGene_tx=XM_023242663.1;refGene_gn=CNTRL;refGene_tx=XM_023242664.1;refGene_gn=CNTRL;refGene_tx=XM_023242665.1
D4 82348224 82348365
refGene_gn=CNTRL;refGene_tx=XM_019816451.2;refGene_gn=CNTRL;refGene_tx=XM_019816452.1;refGene_gn=CNTRL;refGene_tx=XM_023242659.1;refGene_gn=CNTRL;refGene_tx=XM_023242660.1;refGene_gn=CNTRL;refGene_tx=XM_023242661.1;refGene_gn=CNTRL;refGene_tx=XM_023242662.1;refGene_gn=CNTRL;refGene_tx=XM_023242663.1;refGene_gn=CNTRL;refGene_tx=XM_023242664.1;refGene_gn=CNTRL;refGene_tx=XM_023242665.1
D4 82348941 82349214
refGene_gn=CNTRL;refGene_tx=XM_019816451.2;refGene_gn=CNTRL;refGene_tx=XM_019816452.1;refGene_gn=CNTRL;refGene_tx=XM_023242659.1;refGene_gn=CNTRL;refGene_tx=XM_023242660.1;refGene_gn=CNTRL;refGene_tx=XM_023242661.1;refGene_gn=CNTRL;refGene_tx=XM_023242662.1;refGene_gn=CNTRL;refGene_tx=XM_023242663.1;refGene_gn=CNTRL;refGene_tx=XM_023242664.1;refGene_gn=CNTRL;refGene_tx=XM_023242665.1
D4 82349388 82349451
refGene_gn=CNTRL;refGene_tx=XM_019816451.2;refGene_gn=CNTRL;refGene_tx=XM_019816452.1;refGene_gn=CNTRL;refGene_tx=XM_023242659.1;refGene_gn=CNTRL;refGene_tx=XM_023242660.1;refGene_gn=CNTRL;refGene_tx=XM_023242661.1;refGene_gn=CNTRL;refGene_tx=XM_023242662.1;refGene_gn=CNTRL;refGene_tx=XM_023242663.1;refGene_gn=CNTRL;refGene_tx=XM_023242664.1;refGene_gn=CNTRL;refGene_tx=XM_023242665.1
D4 82351147 82351354
refGene_gn=CNTRL;refGene_tx=XM_019816451.2;refGene_gn=CNTRL;refGene_tx=XM_019816452.1;refGene_gn=CNTRL;refGene_tx=XM_023242659.1;refGene_gn=CNTRL;refGene_tx=XM_023242660.1;refGene_gn=CNTRL;refGene_tx=XM_023242661.1;refGene_gn=CNTRL;refGene_tx=XM_023242662.1;refGene_gn=CNTRL;refGene_tx=XM_023242663.1;refGene_gn=CNTRL;refGene_tx=XM_023242664.1;refGene_gn=CNTRL;refGene_tx=XM_023242665.1
D4 82352573 82352630

refGene_gn=CNTRL;refGene_tx=XM_019816451.2;refGene_gn=CNTRL;refGene_tx=XM_019816452.1;refGene_gn=CNTRL;refGene_tx=XM_023242659.1;refGene_gn=CNTRL;refGene_tx=XM_023242660.1;refGene_gn=CNTRL;refGene_tx=XM_023242661.1;refGene_gn=CNTRL;refGene_tx=XM_023242662.1;refGene_gn=CNTRL;refGene_tx=XM_023242663.1;refGene_gn=CNTRL;refGene_tx=XM_023242664.1;refGene_gn=CNTRL;refGene_tx=XM_023242665.1
D4 82357118 82357296 refGene_gn=RAB14;refGene_tx=XM_003995812.5
D4 82358279 82358310 refGene_gn=RAB14;refGene_tx=XM_003995812.5
D4 82358704 82358792 refGene_gn=RAB14;refGene_tx=XM_003995812.5
D4 82363562 82363629 refGene_gn=RAB14;refGene_tx=XM_003995812.5
D4 82368016 82368194 refGene_gn=RAB14;refGene_tx=XM_003995812.5
D4 82369700 82369754 refGene_gn=RAB14;refGene_tx=XM_003995812.5
D4 82370877 82370929 refGene_gn=RAB14;refGene_tx=XM_003995812.5
D4 82414477 82414759 refGene_gn=LOC101088122;refGene_tx=XM_011288466.2
D4 82463948 82464092 refGene_gn=GSN;refGene_tx=XM_003995813.4
D4 82464188 82464212 refGene_gn=GSN;refGene_tx=XM_023242670.1
D4 82465545 82465750 refGene_gn=GSN;refGene_tx=XM_003995813.4;refGene_gn=GSN;refGene_tx=XM_023242668.1;refGene_gn=GSN;refGene_tx=XM_023242669.1;refGene_gn=GSN;refGene_tx=XM_023242670.1
D4 82466463 82466618 refGene_gn=GSN;refGene_tx=XM_003995813.4;refGene_gn=GSN;refGene_tx=XM_023242668.1;refGene_gn=GSN;refGene_tx=XM_023242669.1;refGene_gn=GSN;refGene_tx=XM_023242670.1
D4 82472646 82472808 refGene_gn=GSN;refGene_tx=XM_003995813.4;refGene_gn=GSN;refGene_tx=XM_023242668.1;refGene_gn=GSN;refGene_tx=XM_023242669.1;refGene_gn=GSN;refGene_tx=XM_023242670.1
D4 82474336 82474486 refGene_gn=GSN;refGene_tx=XM_003995813.4;refGene_gn=GSN;refGene_tx=XM_023242668.1;refGene_gn=GSN;refGene_tx=XM_023242669.1;refGene_gn=GSN;refGene_tx=XM_023242670.1
D4 82475663 82475753 refGene_gn=GSN;refGene_tx=XM_003995813.4;refGene_gn=GSN;refGene_tx=XM_023242668.1;refGene_gn=GSN;refGene_tx=XM_023242669.1;refGene_gn=GSN;refGene_tx=XM_023242670.1
D4 82477990 82478123 refGene_gn=GSN;refGene_tx=XM_003995813.4;refGene_gn=GSN;refGene_tx=XM_023242668.1;refGene_gn=GSN;refGene_tx=XM_023242669.1;refGene_gn=GSN;refGene_tx=XM_023242670.1
D4 82479416 82479505 refGene_gn=GSN;refGene_tx=XM_003995813.4;refGene_gn=GSN;refGene_tx=XM_023242668.1;refGene_gn=GSN;refGene_tx=XM_023242669.1;refGene_gn=GSN;refGene_tx=XM_023242670.1
D4 82479692 82479908 refGene_gn=GSN;refGene_tx=XM_003995813.4;refGene_gn=GSN;refGene_tx=XM_023242668.1;refGene_gn=GSN;refGene_tx=XM_023242669.1;refGene_gn=GSN;refGene_tx=XM_023242670.1
D4 82481628 82481762

refGene_gn=GSN;refGene_tx=XM_003995813.4;refGene_gn=GSN;refGene_tx=XM_023242668.1;refGene_gn=GSN;refGene_tx=XM_023242669.1;refGene_gn=GSN;refGene_tx=XM_023242670.1
D4 82484600 82484691
refGene_gn=GSN;refGene_tx=XM_003995813.4;refGene_gn=GSN;refGene_tx=XM_023242668.1;refGene_gn=GSN;refGene_tx=XM_023242669.1;refGene_gn=GSN;refGene_tx=XM_023242670.1
D4 82485953 82486124
refGene_gn=GSN;refGene_tx=XM_003995813.4;refGene_gn=GSN;refGene_tx=XM_023242668.1;refGene_gn=GSN;refGene_tx=XM_023242669.1;refGene_gn=GSN;refGene_tx=XM_023242670.1
D4 82486745 82486920
refGene_gn=GSN;refGene_tx=XM_003995813.4;refGene_gn=GSN;refGene_tx=XM_023242668.1;refGene_gn=GSN;refGene_tx=XM_023242669.1;refGene_gn=GSN;refGene_tx=XM_023242670.1
D4 82488005 82488130
refGene_gn=GSN;refGene_tx=XM_003995813.4;refGene_gn=GSN;refGene_tx=XM_023242668.1;refGene_gn=GSN;refGene_tx=XM_023242669.1;refGene_gn=GSN;refGene_tx=XM_023242670.1
D4 82488340 82488418
refGene_gn=GSN;refGene_tx=XM_003995813.4;refGene_gn=GSN;refGene_tx=XM_023242668.1;refGene_gn=GSN;refGene_tx=XM_023242669.1;refGene_gn=GSN;refGene_tx=XM_023242670.1
D4 82490323 82490384
refGene_gn=GSN;refGene_tx=XM_003995813.4;refGene_gn=GSN;refGene_tx=XM_023242668.1;refGene_gn=GSN;refGene_tx=XM_023242669.1;refGene_gn=GSN;refGene_tx=XM_023242670.1
D4 82490970 82491140
refGene_gn=GSN;refGene_tx=XM_003995813.4;refGene_gn=GSN;refGene_tx=XM_023242668.1;refGene_gn=GSN;refGene_tx=XM_023242669.1;refGene_gn=GSN;refGene_tx=XM_023242670.1
D4 82497863 82498058 refGene_gn=STOM;refGene_tx=XM_003995814.5
D4 82505626 82505761 refGene_gn=STOM;refGene_tx=XM_003995814.5
D4 82506756 82506960 refGene_gn=STOM;refGene_tx=XM_003995814.5
D4 82511549 82511632 refGene_gn=STOM;refGene_tx=XM_003995814.5
D4 82512684 82512757 refGene_gn=STOM;refGene_tx=XM_003995814.5
D4 82514562 82514666 refGene_gn=STOM;refGene_tx=XM_003995814.5
D4 82526807 82526868 refGene_gn=STOM;refGene_tx=XM_003995814.5
D4 82565357 82566024
refGene_gn=LOC101088888;refGene_tx=XM_019816463.2;refGene_gn=LOC101088888;refGene_tx=XM_019816464.2;refGene_gn=LOC101088888;refGene_tx=XM_019816466.2;refGene_gn=LOC101088888;refGene_tx=XM_019816467.2
D4 82566456 82566594
refGene_gn=LOC101088888;refGene_tx=XM_019816463.2;refGene_gn=LOC101088888;refGene_tx=XM_019816464.2;refGene_gn=LOC101088888;refGene_tx=XM_019816466.2;refGene_gn=LOC101088888;refGene_tx=XM_019816467.2
D4 82576592 82576637
refGene_gn=LOC101088888;refGene_tx=XM_019816463.2;refGene_gn=LOC101088888;refGene_tx=XM_019816464.2
D4 82580355 82580391

refGene_gn=LOC101088888;refGene_tx=XM_019816463.2;refGene_gn=LOC101088
888;refGene_tx=XM_019816464.2
D4 82581552 82581632
refGene_gn=LOC101088888;refGene_tx=XM_019816463.2;refGene_gn=LOC101088
888;refGene_tx=XM_019816464.2;refGene_gn=LOC101088888;refGene_tx=XM_01
9816466.2;refGene_gn=LOC101088888;refGene_tx=XM_019816467.2
D4 82582885 82582888
refGene_gn=LOC101088888;refGene_tx=XM_019816464.2;refGene_gn=LOC101088
888;refGene_tx=XM_019816467.2
D4 82589403 82589478
refGene_gn=LOC101088888;refGene_tx=XM_019816463.2;refGene_gn=LOC101088
888;refGene_tx=XM_019816466.2
D4 82594023 82594717
refGene_gn=LOC105261108;refGene_tx=XM_019816456.2;refGene_gn=LOC105261
108;refGene_tx=XM_019816457.2;refGene_gn=LOC105261108;refGene_tx=XM_01
9816458.2
D4 82595891 82595910
refGene_gn=LOC105261108;refGene_tx=XM_019816460.2
D4 82597194 82597329
refGene_gn=LOC105261108;refGene_tx=XM_019816456.2;refGene_gn=LOC105261
108;refGene_tx=XM_019816457.2;refGene_gn=LOC105261108;refGene_tx=XM_01
9816458.2;refGene_gn=LOC105261108;refGene_tx=XM_019816460.2
D4 82604588 82604642
refGene_gn=LOC105261108;refGene_tx=XM_019816456.2;refGene_gn=LOC105261
108;refGene_tx=XM_019816457.2;refGene_gn=LOC105261108;refGene_tx=XM_01
9816458.2;refGene_gn=LOC105261108;refGene_tx=XM_019816460.2
D4 82606225 82606348
refGene_gn=LOC105261108;refGene_tx=XM_019816456.2;refGene_gn=LOC105261
108;refGene_tx=XM_019816460.2
D4 82607271 82607304
refGene_gn=LOC105261108;refGene_tx=XM_019816457.2
D4 82608627 82608713
refGene_gn=LOC105261108;refGene_tx=XM_019816456.2;refGene_gn=LOC105261
108;refGene_tx=XM_019816457.2;refGene_gn=LOC105261108;refGene_tx=XM_01
9816458.2;refGene_gn=LOC105261108;refGene_tx=XM_019816460.2
D4 82609422 82609443
refGene_gn=LOC105261108;refGene_tx=XM_019816456.2;refGene_gn=LOC105261
108;refGene_tx=XM_019816457.2;refGene_gn=LOC105261108;refGene_tx=XM_01
9816458.2;refGene_gn=LOC105261108;refGene_tx=XM_019816460.2
D4 82623516 82624210
refGene_gn=GGTA1;refGene_tx=NM_001009308.1;refGene_gn=GGTA1;refGene_tx
=XM_023242196.1
D4 82628121 82628259
refGene_gn=GGTA1;refGene_tx=NM_001009308.1;refGene_gn=GGTA1;refGene_tx
=XM_023242196.1
D4 82634801 82634900
refGene_gn=GGTA1;refGene_tx=NM_001009308.1;refGene_gn=GGTA1;refGene_tx
=XM_023242196.1
D4 82635444 82635509
refGene_gn=GGTA1;refGene_tx=NM_001009308.1;refGene_gn=GGTA1;refGene_tx

=XM_023242196.1
D4 82637949 82637989
refGene_gn=GGTA1;refGene_tx=NM_001009308.1;refGene_gn=GGTA1;refGene_tx
=XM_023242196.1
D4 82643673 82643753
refGene_gn=GGTA1;refGene_tx=NM_001009308.1;refGene_gn=GGTA1;refGene_tx
=XM_023242196.1
D4 82735637 82735804
refGene_gn=LOC102902330;refGene_tx=XM_019816469.2;refGene_gn=LOC102902
330;refGene_tx=XM_019816470.1;refGene_gn=LOC102902330;refGene_tx=XM_01
9816472.1
D4 82735946 82735951
refGene_gn=LOC102902330;refGene_tx=XM_019816474.1
D4 82736191 82736208
refGene_gn=LOC102902330;refGene_tx=XM_019816473.1
D4 82736302 82736486
refGene_gn=LOC102902330;refGene_tx=XM_019816469.2;refGene_gn=LOC102902
330;refGene_tx=XM_019816470.1;refGene_gn=LOC102902330;refGene_tx=XM_01
9816472.1;refGene_gn=LOC102902330;refGene_tx=XM_019816473.1;refGene_gn
=LOC102902330;refGene_tx=XM_019816474.1
D4 82739609 82739767
refGene_gn=LOC102902330;refGene_tx=XM_019816469.2;refGene_gn=LOC102902
330;refGene_tx=XM_019816470.1;refGene_gn=LOC102902330;refGene_tx=XM_01
9816472.1;refGene_gn=LOC102902330;refGene_tx=XM_019816473.1;refGene_gn
=LOC102902330;refGene_tx=XM_019816474.1
D4 82739981 82740210
refGene_gn=LOC102902330;refGene_tx=XM_019816469.2;refGene_gn=LOC102902
330;refGene_tx=XM_019816470.1;refGene_gn=LOC102902330;refGene_tx=XM_01
9816472.1;refGene_gn=LOC102902330;refGene_tx=XM_019816473.1;refGene_gn
=LOC102902330;refGene_tx=XM_019816474.1
D4 82740319 82740359 refGene_gn=DAB2IP;refGene_tx=XM_023242674.1
D4 82806506 82806630
refGene_gn=DAB2IP;refGene_tx=XM_023242672.1;refGene_gn=DAB2IP;refGene_
tx=XM_023242673.1;refGene_gn=DAB2IP;refGene_tx=XM_023242675.1;refGene_
gn=DAB2IP;refGene_tx=XM_023242676.1
D4 82828700 82828804
refGene_gn=DAB2IP;refGene_tx=XM_023242672.1;refGene_gn=DAB2IP;refGene_
tx=XM_023242673.1;refGene_gn=DAB2IP;refGene_tx=XM_023242674.1;refGene_
gn=DAB2IP;refGene_tx=XM_023242675.1;refGene_gn=DAB2IP;refGene_tx=XM_02
3242676.1;refGene_gn=DAB2IP;refGene_tx=XM_023242677.1;refGene_gn=DAB2I
P;refGene_tx=XM_023242678.1
D4 82846867 82847019
refGene_gn=DAB2IP;refGene_tx=XM_023242672.1;refGene_gn=DAB2IP;refGene_
tx=XM_023242673.1;refGene_gn=DAB2IP;refGene_tx=XM_023242674.1;refGene_
gn=DAB2IP;refGene_tx=XM_023242675.1;refGene_gn=DAB2IP;refGene_tx=XM_02
3242676.1;refGene_gn=DAB2IP;refGene_tx=XM_023242677.1;refGene_gn=DAB2I
P;refGene_tx=XM_023242678.1
D4 82884424 82884447 refGene_gn=DAB2IP;refGene_tx=XM_023242679.1
D4 82900731 82900885
refGene_gn=DAB2IP;refGene_tx=XM_023242672.1;refGene_gn=DAB2IP;refGene_

tx=XM_023242673.1;refGene_gn=DAB2IP;refGene_tx=XM_023242674.1;refGene_gn=DAB2IP;refGene_tx=XM_023242675.1;refGene_gn=DAB2IP;refGene_tx=XM_023242676.1;refGene_gn=DAB2IP;refGene_tx=XM_023242677.1;refGene_gn=DAB2IP;refGene_tx=XM_023242678.1;refGene_gn=DAB2IP;refGene_tx=XM_023242679.1;refGene_gn=DAB2IP;refGene_tx=XM_023242680.1;refGene_gn=DAB2IP;refGene_tx=XM_023242681.1;refGene_gn=DAB2IP;refGene_tx=XM_023242682.1;refGene_gn=DAB2IP;refGene_tx=XM_023242683.1
D4 82902462 82902561
refGene_gn=DAB2IP;refGene_tx=XM_023242672.1;refGene_gn=DAB2IP;refGene_tx=XM_023242673.1;refGene_gn=DAB2IP;refGene_tx=XM_023242674.1;refGene_gn=DAB2IP;refGene_tx=XM_023242675.1;refGene_gn=DAB2IP;refGene_tx=XM_023242676.1;refGene_gn=DAB2IP;refGene_tx=XM_023242677.1;refGene_gn=DAB2IP;refGene_tx=XM_023242678.1;refGene_gn=DAB2IP;refGene_tx=XM_023242679.1;refGene_gn=DAB2IP;refGene_tx=XM_023242680.1;refGene_gn=DAB2IP;refGene_tx=XM_023242681.1;refGene_gn=DAB2IP;refGene_tx=XM_023242682.1;refGene_gn=DAB2IP;refGene_tx=XM_023242683.1
D4 82903380 82903935
refGene_gn=DAB2IP;refGene_tx=XM_023242672.1;refGene_gn=DAB2IP;refGene_tx=XM_023242673.1;refGene_gn=DAB2IP;refGene_tx=XM_023242674.1;refGene_gn=DAB2IP;refGene_tx=XM_023242675.1;refGene_gn=DAB2IP;refGene_tx=XM_023242676.1;refGene_gn=DAB2IP;refGene_tx=XM_023242677.1;refGene_gn=DAB2IP;refGene_tx=XM_023242678.1;refGene_gn=DAB2IP;refGene_tx=XM_023242679.1;refGene_gn=DAB2IP;refGene_tx=XM_023242680.1;refGene_gn=DAB2IP;refGene_tx=XM_023242681.1;refGene_gn=DAB2IP;refGene_tx=XM_023242682.1;refGene_gn=DAB2IP;refGene_tx=XM_023242683.1
D4 82906850 82906995
refGene_gn=DAB2IP;refGene_tx=XM_023242672.1;refGene_gn=DAB2IP;refGene_tx=XM_023242673.1;refGene_gn=DAB2IP;refGene_tx=XM_023242674.1;refGene_gn=DAB2IP;refGene_tx=XM_023242675.1;refGene_gn=DAB2IP;refGene_tx=XM_023242676.1;refGene_gn=DAB2IP;refGene_tx=XM_023242677.1;refGene_gn=DAB2IP;refGene_tx=XM_023242678.1;refGene_gn=DAB2IP;refGene_tx=XM_023242679.1;refGene_gn=DAB2IP;refGene_tx=XM_023242680.1;refGene_gn=DAB2IP;refGene_tx=XM_023242681.1;refGene_gn=DAB2IP;refGene_tx=XM_023242682.1;refGene_gn=DAB2IP;refGene_tx=XM_023242683.1
D4 82907085 82907230
refGene_gn=DAB2IP;refGene_tx=XM_023242672.1;refGene_gn=DAB2IP;refGene_tx=XM_023242673.1;refGene_gn=DAB2IP;refGene_tx=XM_023242674.1;refGene_gn=DAB2IP;refGene_tx=XM_023242675.1;refGene_gn=DAB2IP;refGene_tx=XM_023242676.1;refGene_gn=DAB2IP;refGene_tx=XM_023242677.1;refGene_gn=DAB2IP;refGene_tx=XM_023242678.1;refGene_gn=DAB2IP;refGene_tx=XM_023242679.1;refGene_gn=DAB2IP;refGene_tx=XM_023242680.1;refGene_gn=DAB2IP;refGene_tx=XM_023242681.1;refGene_gn=DAB2IP;refGene_tx=XM_023242682.1;refGene_gn=DAB2IP;refGene_tx=XM_023242683.1
D4 82909457 82909694
refGene_gn=DAB2IP;refGene_tx=XM_023242672.1;refGene_gn=DAB2IP;refGene_tx=XM_023242673.1;refGene_gn=DAB2IP;refGene_tx=XM_023242674.1;refGene_gn=DAB2IP;refGene_tx=XM_023242675.1;refGene_gn=DAB2IP;refGene_tx=XM_023242676.1;refGene_gn=DAB2IP;refGene_tx=XM_023242677.1;refGene_gn=DAB2IP;refGene_tx=XM_023242678.1;refGene_gn=DAB2IP;refGene_tx=XM_023242679.1;refGene_gn=DAB2IP;refGene_tx=XM_023242680.1;refGene_gn=DAB2IP;refGen

e_tx=XM_023242681.1;refGene_gn=DAB2IP;refGene_tx=XM_023242682.1;refGene_gn=DAB2IP;refGene_tx=XM_023242683.1
D4 82911316 82911518
refGene_gn=DAB2IP;refGene_tx=XM_023242672.1;refGene_gn=DAB2IP;refGene_tx=XM_023242673.1;refGene_gn=DAB2IP;refGene_tx=XM_023242674.1;refGene_gn=DAB2IP;refGene_tx=XM_023242675.1;refGene_gn=DAB2IP;refGene_tx=XM_023242676.1;refGene_gn=DAB2IP;refGene_tx=XM_023242677.1;refGene_gn=DAB2IP;refGene_tx=XM_023242678.1;refGene_gn=DAB2IP;refGene_tx=XM_023242679.1;refGene_gn=DAB2IP;refGene_tx=XM_023242680.1;refGene_gn=DAB2IP;refGene_tx=XM_023242681.1;refGene_gn=DAB2IP;refGene_tx=XM_023242682.1;refGene_gn=DAB2IP;refGene_tx=XM_023242683.1
D4 82913202 82913381
refGene_gn=DAB2IP;refGene_tx=XM_023242672.1;refGene_gn=DAB2IP;refGene_tx=XM_023242673.1;refGene_gn=DAB2IP;refGene_tx=XM_023242674.1;refGene_gn=DAB2IP;refGene_tx=XM_023242675.1;refGene_gn=DAB2IP;refGene_tx=XM_023242676.1;refGene_gn=DAB2IP;refGene_tx=XM_023242677.1;refGene_gn=DAB2IP;refGene_tx=XM_023242678.1;refGene_gn=DAB2IP;refGene_tx=XM_023242679.1;refGene_gn=DAB2IP;refGene_tx=XM_023242680.1;refGene_gn=DAB2IP;refGene_tx=XM_023242681.1;refGene_gn=DAB2IP;refGene_tx=XM_023242682.1;refGene_gn=DAB2IP;refGene_tx=XM_023242683.1
D4 82915297 82916186
refGene_gn=DAB2IP;refGene_tx=XM_023242672.1;refGene_gn=DAB2IP;refGene_tx=XM_023242673.1;refGene_gn=DAB2IP;refGene_tx=XM_023242674.1;refGene_gn=DAB2IP;refGene_tx=XM_023242675.1;refGene_gn=DAB2IP;refGene_tx=XM_023242676.1;refGene_gn=DAB2IP;refGene_tx=XM_023242677.1;refGene_gn=DAB2IP;refGene_tx=XM_023242678.1;refGene_gn=DAB2IP;refGene_tx=XM_023242679.1;refGene_gn=DAB2IP;refGene_tx=XM_023242680.1;refGene_gn=DAB2IP;refGene_tx=XM_023242681.1;refGene_gn=DAB2IP;refGene_tx=XM_023242682.1;refGene_gn=DAB2IP;refGene_tx=XM_023242683.1
D4 82916905 82917058
refGene_gn=DAB2IP;refGene_tx=XM_023242672.1;refGene_gn=DAB2IP;refGene_tx=XM_023242673.1;refGene_gn=DAB2IP;refGene_tx=XM_023242674.1;refGene_gn=DAB2IP;refGene_tx=XM_023242675.1;refGene_gn=DAB2IP;refGene_tx=XM_023242677.1;refGene_gn=DAB2IP;refGene_tx=XM_023242678.1;refGene_gn=DAB2IP;refGene_tx=XM_023242679.1;refGene_gn=DAB2IP;refGene_tx=XM_023242680.1;refGene_gn=DAB2IP;refGene_tx=XM_023242681.1;refGene_gn=DAB2IP;refGene_tx=XM_023242682.1;refGene_gn=DAB2IP;refGene_tx=XM_023242683.1
D4 82919145 82919339
refGene_gn=DAB2IP;refGene_tx=XM_023242672.1;refGene_gn=DAB2IP;refGene_tx=XM_023242673.1;refGene_gn=DAB2IP;refGene_tx=XM_023242674.1;refGene_gn=DAB2IP;refGene_tx=XM_023242675.1;refGene_gn=DAB2IP;refGene_tx=XM_023242676.1;refGene_gn=DAB2IP;refGene_tx=XM_023242677.1;refGene_gn=DAB2IP;refGene_tx=XM_023242678.1;refGene_gn=DAB2IP;refGene_tx=XM_023242679.1;refGene_gn=DAB2IP;refGene_tx=XM_023242680.1;refGene_gn=DAB2IP;refGene_tx=XM_023242681.1;refGene_gn=DAB2IP;refGene_tx=XM_023242682.1;refGene_gn=DAB2IP;refGene_tx=XM_023242683.1
D4 82923896 82923984
refGene_gn=DAB2IP;refGene_tx=XM_023242672.1;refGene_gn=DAB2IP;refGene_tx=XM_023242673.1;refGene_gn=DAB2IP;refGene_tx=XM_023242674.1;refGene_gn=DAB2IP;refGene_tx=XM_023242675.1;refGene_gn=DAB2IP;refGene_tx=XM_02

3242676.1;refGene_gn=DAB2IP;refGene_tx=XM_023242677.1;refGene_gn=DAB2IP;refGene_tx=XM_023242678.1;refGene_gn=DAB2IP;refGene_tx=XM_023242679.1;refGene_gn=DAB2IP;refGene_tx=XM_023242680.1;refGene_gn=DAB2IP;refGene_tx=XM_023242681.1;refGene_gn=DAB2IP;refGene_tx=XM_023242682.1;refGene_gn=DAB2IP;refGene_tx=XM_023242683.1
D4 82925180 82925348
refGene_gn=DAB2IP;refGene_tx=XM_023242672.1;refGene_gn=DAB2IP;refGene_tx=XM_023242673.1;refGene_gn=DAB2IP;refGene_tx=XM_023242674.1;refGene_gn=DAB2IP;refGene_tx=XM_023242675.1;refGene_gn=DAB2IP;refGene_tx=XM_023242676.1;refGene_gn=DAB2IP;refGene_tx=XM_023242677.1;refGene_gn=DAB2IP;refGene_tx=XM_023242678.1;refGene_gn=DAB2IP;refGene_tx=XM_023242679.1;refGene_gn=DAB2IP;refGene_tx=XM_023242680.1;refGene_gn=DAB2IP;refGene_tx=XM_023242681.1;refGene_gn=DAB2IP;refGene_tx=XM_023242682.1;refGene_gn=DAB2IP;refGene_tx=XM_023242683.1
D4 82926340 82926350 refGene_gn=DAB2IP;refGene_tx=XM_023242675.1
D4 82965323 82965757 refGene_gn=TTLL11;refGene_tx=XM_023242684.1
D4 82986747 82986854 refGene_gn=TTLL11;refGene_tx=XM_023242684.1
D4 82995036 82995288 refGene_gn=TTLL11;refGene_tx=XM_023242684.1
D4 83018805 83018889 refGene_gn=TTLL11;refGene_tx=XM_023242684.1
D4 83082436 83082552 refGene_gn=TTLL11;refGene_tx=XM_023242684.1
D4 83083298 83083394 refGene_gn=TTLL11;refGene_tx=XM_023242684.1
D4 83095088 83095664 refGene_gn=TTLL11;refGene_tx=XM_023242684.1
D4 83135383 83135517 refGene_gn=TTLL11;refGene_tx=XM_023242684.1
D4 83194047 83194702 refGene_gn=TTLL11;refGene_tx=XM_023242684.1
D4 83240109 83240247 refGene_gn=NDUFA8;refGene_tx=XM_003995822.4
D4 83244951 83245117 refGene_gn=NDUFA8;refGene_tx=XM_003995822.4
D4 83249847 83250011 refGene_gn=NDUFA8;refGene_tx=XM_003995822.4
D4 83257013 83257166 refGene_gn=NDUFA8;refGene_tx=XM_003995822.4
D4 83257353 83257400
refGene_gn=MORN5;refGene_tx=XM_003995834.5;refGene_gn=MORN5;refGene_tx=XM_006939430.4
D4 83260905 83261053
refGene_gn=MORN5;refGene_tx=XM_003995834.5;refGene_gn=MORN5;refGene_tx=XM_006939430.4
D4 83262655 83262767 refGene_gn=MORN5;refGene_tx=XM_003995834.5
D4 83266415 83266547
refGene_gn=MORN5;refGene_tx=XM_003995834.5;refGene_gn=MORN5;refGene_tx=XM_006939430.4
D4 83287025 83287105 refGene_gn=MORN5;refGene_tx=XM_003995834.5
D4 83291530 83291551
refGene_gn=LHX6;refGene_tx=XM_019816479.2;refGene_gn=LHX6;refGene_tx=XM_023242685.1
D4 83296488 83296592
refGene_gn=LHX6;refGene_tx=XM_019816479.2;refGene_gn=LHX6;refGene_tx=XM_023242685.1
D4 83299679 83299854
refGene_gn=LHX6;refGene_tx=XM_019816479.2;refGene_gn=LHX6;refGene_tx=XM_023242685.1
D4 83300072 83300168
refGene_gn=LHX6;refGene_tx=XM_019816479.2;refGene_gn=LHX6;refGene_tx=X

M_023242685.1
D4 83300388 83300489
refGene_gn=LHX6;refGene_tx=XM_019816479.2;refGene_gn=LHX6;refGene_tx=X
M_023242685.1
D4 83303101 83303322
refGene_gn=LHX6;refGene_tx=XM_019816479.2;refGene_gn=LHX6;refGene_tx=X
M_023242685.1
D4 83312099 83312221 refGene_gn=LHX6;refGene_tx=XM_023242685.1
D4 83312580 83312763 refGene_gn=LHX6;refGene_tx=XM_023242685.1
D4 83313160 83313232 refGene_gn=LHX6;refGene_tx=XM_023242685.1
D4 83314394 83314478 refGene_gn=LHX6;refGene_tx=XM_023242685.1
D4 83326196 83326356 refGene_gn=RBM18;refGene_tx=XM_003995825.4
D4 83329896 83329982 refGene_gn=RBM18;refGene_tx=XM_003995825.4
D4 83331678 83331765 refGene_gn=RBM18;refGene_tx=XM_003995825.4
D4 83334717 83334844 refGene_gn=RBM18;refGene_tx=XM_003995825.4
D4 83342336 83342449 refGene_gn=RBM18;refGene_tx=XM_003995825.4
D4 83345320 83345514
refGene_gn=MRRF;refGene_tx=XM_006939432.4;refGene_gn=MRRF;refGene_tx=X
M_006939433.4;refGene_gn=MRRF;refGene_tx=XM_011288462.3;refGene_gn=MRR
F;refGene_tx=XM_019816480.2;refGene_gn=MRRF;refGene_tx=XM_019816482.2;
refGene_gn=MRRF;refGene_tx=XM_019816486.2
D4 83345817 83345828 refGene_gn=MRRF;refGene_tx=XM_019816481.2
D4 83350191 83350412
refGene_gn=MRRF;refGene_tx=XM_006939432.4;refGene_gn=MRRF;refGene_tx=X
M_006939433.4;refGene_gn=MRRF;refGene_tx=XM_011288462.3;refGene_gn=MRR
F;refGene_tx=XM_019816480.2;refGene_gn=MRRF;refGene_tx=XM_019816481.2;
refGene_gn=MRRF;refGene_tx=XM_019816482.2;refGene_gn=MRRF;refGene_tx=X
M_019816483.2;refGene_gn=MRRF;refGene_tx=XM_019816484.2;refGene_gn=MRR
F;refGene_tx=XM_019816485.2;refGene_gn=MRRF;refGene_tx=XM_019816486.2;
refGene_gn=MRRF;refGene_tx=XM_019816489.2;refGene_gn=MRRF;refGene_tx=X
M_019816490.2;refGene_gn=MRRF;refGene_tx=XM_023242686.1
D4 83358632 83358788
refGene_gn=MRRF;refGene_tx=XM_006939432.4;refGene_gn=MRRF;refGene_tx=X
M_006939433.4;refGene_gn=MRRF;refGene_tx=XM_011288462.3;refGene_gn=MRR
F;refGene_tx=XM_019816480.2;refGene_gn=MRRF;refGene_tx=XM_019816481.2;
refGene_gn=MRRF;refGene_tx=XM_019816482.2;refGene_gn=MRRF;refGene_tx=X
M_019816483.2;refGene_gn=MRRF;refGene_tx=XM_019816484.2;refGene_gn=MRR
F;refGene_tx=XM_019816485.2;refGene_gn=MRRF;refGene_tx=XM_019816486.2;
refGene_gn=MRRF;refGene_tx=XM_019816489.2;refGene_gn=MRRF;refGene_tx=X
M_019816490.2;refGene_gn=MRRF;refGene_tx=XM_023242686.1
D4 83362104 83362223
refGene_gn=MRRF;refGene_tx=XM_006939432.4;refGene_gn=MRRF;refGene_tx=X
M_006939433.4;refGene_gn=MRRF;refGene_tx=XM_011288462.3;refGene_gn=MRR
F;refGene_tx=XM_019816480.2;refGene_gn=MRRF;refGene_tx=XM_019816481.2;
refGene_gn=MRRF;refGene_tx=XM_019816482.2;refGene_gn=MRRF;refGene_tx=X
M_019816483.2;refGene_gn=MRRF;refGene_tx=XM_019816484.2;refGene_gn=MRR
F;refGene_tx=XM_019816485.2;refGene_gn=MRRF;refGene_tx=XM_019816486.2;
refGene_gn=MRRF;refGene_tx=XM_019816489.2;refGene_gn=MRRF;refGene_tx=X
M_019816490.2;refGene_gn=MRRF;refGene_tx=XM_023242686.1
D4 83368016 83368108

refGene_gn=MRRF;refGene_tx=XM_006939432.4;refGene_gn=MRRF;refGene_tx=XM_006939433.4;refGene_gn=MRRF;refGene_tx=XM_011288462.3;refGene_gn=MRRF;refGene_tx=XM_019816480.2;refGene_gn=MRRF;refGene_tx=XM_019816481.2;refGene_gn=MRRF;refGene_tx=XM_019816482.2;refGene_gn=MRRF;refGene_tx=XM_019816483.2;refGene_gn=MRRF;refGene_tx=XM_019816484.2;refGene_gn=MRRF;refGene_tx=XM_019816485.2;refGene_gn=MRRF;refGene_tx=XM_019816486.2;refGene_gn=MRRF;refGene_tx=XM_019816489.2;refGene_gn=MRRF;refGene_tx=XM_019816490.2;refGene_gn=MRRF;refGene_tx=XM_023242686.1
D4 83378524 83378588 refGene_gn=MRRF;refGene_tx=XM_019816486.2
D4 83388170 83388330
refGene_gn=MRRF;refGene_tx=XM_006939432.4;refGene_gn=MRRF;refGene_tx=XM_006939433.4;refGene_gn=MRRF;refGene_tx=XM_011288462.3;refGene_gn=MRRF;refGene_tx=XM_019816480.2;refGene_gn=MRRF;refGene_tx=XM_019816481.2;refGene_gn=MRRF;refGene_tx=XM_019816482.2;refGene_gn=MRRF;refGene_tx=XM_019816483.2;refGene_gn=MRRF;refGene_tx=XM_019816484.2;refGene_gn=MRRF;refGene_tx=XM_019816485.2;refGene_gn=MRRF;refGene_tx=XM_019816489.2;refGene_gn=MRRF;refGene_tx=XM_019816490.2;refGene_gn=MRRF;refGene_tx=XM_023242686.1
D4 83388566 83388611 refGene_gn=MRRF;refGene_tx=XM_011288462.3
D4 83396365 83396443 refGene_gn=MRRF;refGene_tx=XM_006939432.4
D4 83400568 83400625 refGene_gn=MRRF;refGene_tx=XM_019816482.2
D4 83403147 83403396
refGene_gn=MRRF;refGene_tx=XM_019816480.2;refGene_gn=MRRF;refGene_tx=XM_019816481.2;refGene_gn=MRRF;refGene_tx=XM_019816483.2;refGene_gn=MRRF;refGene_tx=XM_019816484.2;refGene_gn=MRRF;refGene_tx=XM_019816485.2;refGene_gn=MRRF;refGene_tx=XM_019816489.2;refGene_gn=MRRF;refGene_tx=XM_019816490.2;refGene_gn=MRRF;refGene_tx=XM_023242686.1
D4 83404854 83404911 refGene_gn=MRRF;refGene_tx=XM_006939433.4
D4 83438138 83438145 refGene_gn=PTGS1;refGene_tx=XM_006939439.2
D4 83438238 83438334 refGene_gn=PTGS1;refGene_tx=XM_006939439.2
D4 83444856 83444973 refGene_gn=PTGS1;refGene_tx=XM_006939439.2
D4 83445356 83445497 refGene_gn=PTGS1;refGene_tx=XM_006939439.2
D4 83445698 83445842 refGene_gn=PTGS1;refGene_tx=XM_006939439.2
D4 83447933 83448115 refGene_gn=PTGS1;refGene_tx=XM_006939439.2
D4 83448227 83448311 refGene_gn=PTGS1;refGene_tx=XM_006939439.2
D4 83450063 83450310 refGene_gn=PTGS1;refGene_tx=XM_006939439.2
D4 83452465 83452752 refGene_gn=PTGS1;refGene_tx=XM_006939439.2
D4 83456704 83456852 refGene_gn=PTGS1;refGene_tx=XM_006939439.2
D4 83458258 83458614 refGene_gn=PTGS1;refGene_tx=XM_006939439.2
D4 83503670 83504615
refGene_gn=LOC109496998;refGene_tx=XM_023243351.1
D4 83519812 83520772
refGene_gn=LOC109493961;refGene_tx=XM_019816492.2
D4 83534148 83535159
refGene_gn=LOC109493962;refGene_tx=XM_019816493.2
D4 83549947 83550187
refGene_gn=LOC111557539;refGene_tx=XM_023242687.1
D4 83550188 83550893
refGene_gn=LOC111557539;refGene_tx=XM_023242687.1
D4 83564389 83565319

refGene_gn=LOC109493581;refGene_tx=XM_023242312.1
D4 83569445 83569566
refGene_gn=LOC111556267;refGene_tx=XM_023243352.1
D4 83569568 83569597
refGene_gn=LOC111556267;refGene_tx=XM_023243352.1
D4 83569599 83569774
refGene_gn=LOC111556267;refGene_tx=XM_023243352.1
D4 83569776 83569942
refGene_gn=LOC111556267;refGene_tx=XM_023243352.1
D4 83583677 83584620
refGene_gn=LOC111556267;refGene_tx=XM_023243352.1
D4 83611583 83611988
refGene_gn=LOC101095928;refGene_tx=XM_011288653.1
D4 83611990 83612119
refGene_gn=LOC101095928;refGene_tx=XM_011288653.1
D4 83612120 83612651
refGene_gn=LOC101095928;refGene_tx=XM_011288653.1
D4 83631987 83632929
refGene_gn=LOC101096180;refGene_tx=XM_003995838.1
D4 83648995 83649937
refGene_gn=LOC101096425;refGene_tx=XM_011288652.1
D4 83667205 83667961
refGene_gn=LOC101092804;refGene_tx=XM_003995829.1
D4 83667962 83668148
refGene_gn=LOC101092804;refGene_tx=XM_003995829.1
D4 83673698 83674640
refGene_gn=LOC105261127;refGene_tx=XM_011288651.2
D4 83681336 83681450
refGene_gn=LOC105259623;refGene_tx=XM_011288460.2
D4 83681451 83681799
refGene_gn=LOC105259623;refGene_tx=XM_011288460.2
D4 83681800 83681967
refGene_gn=LOC105259623;refGene_tx=XM_011288460.2
D4 83681969 83682270
refGene_gn=LOC105259623;refGene_tx=XM_011288460.2
D4 83718230 83718727
refGene_gn=LOC101097432;refGene_tx=XM_003995842.2
D4 83724282 83725230
refGene_gn=LOC101097186;refGene_tx=XM_003995841.1
D4 83737873 83738816
refGene_gn=LOC101097432;refGene_tx=XM_003995842.2
D4 83747221 83748154
refGene_gn=LOC101097681;refGene_tx=XM_023242313.1
D4 83763682 83764594
refGene_gn=LOC101093469;refGene_tx=XM_003995830.3
D4 83768096 83768132
refGene_gn=LOC101093469;refGene_tx=XM_003995830.3
D4 83775533 83776490
refGene_gn=LOC101097928;refGene_tx=XM_023242314.1
D4 83787573 83788002

refGene_gn=LOC101098174;refGene_tx=XM_003995845.1
D4 83788003 83788549
refGene_gn=LOC101098174;refGene_tx=XM_003995845.1
D4 83815020 83815953
refGene_gn=LOC101093722;refGene_tx=XM_011288459.2
D4 83847035 83848013
refGene_gn=LOC101098412;refGene_tx=XM_003995846.3
D4 83865729 83866707
refGene_gn=LOC101093971;refGene_tx=XM_003995832.3
D4 83883224 83883268
refGene_gn=LOC105261126;refGene_tx=XM_019815552.1
D4 83883352 83883584
refGene_gn=LOC105261126;refGene_tx=XM_019815552.1
D4 83884002 83884041
refGene_gn=LOC105261126;refGene_tx=XM_019815552.1
D4 83897468 83898401
refGene_gn=LOC101098902;refGene_tx=XM_011288648.2
D4 83910818 83911751
refGene_gn=LOC101099158;refGene_tx=XM_011288457.2
D4 83924445 83925375
refGene_gn=LOC101099411;refGene_tx=XM_006939443.3
D4 83942376 83943306
refGene_gn=LOC101099668;refGene_tx=XM_003995848.3
D4 83949698 83950640
refGene_gn=LOC101084181;refGene_tx=XM_004001198.3
D4 84041276 84041357
refGene_gn=LOC101087883;refGene_tx=XM_019815553.2
D4 84043958 84044094
refGene_gn=LOC101087883;refGene_tx=XM_019815553.2
D4 84044096 84044896
refGene_gn=LOC101087883;refGene_tx=XM_019815553.2
D4 84060026 84060980
refGene_gn=LOC101099914;refGene_tx=XM_003995849.4
D4 84067265 84068216
refGene_gn=LOC101100165;refGene_tx=XM_003995850.2
D4 84082301 84082853
refGene_gn=PDCL;refGene_tx=XM_006939444.4;refGene_gn=PDCL;refGene_tx=XM_011288455.3
D4 84085246 84085428
refGene_gn=PDCL;refGene_tx=XM_006939444.4;refGene_gn=PDCL;refGene_tx=XM_011288455.3
D4 84088344 84088516
refGene_gn=PDCL;refGene_tx=XM_006939444.4;refGene_gn=PDCL;refGene_tx=XM_011288455.3
D4 84104698 84104894
refGene_gn=RC3H2;refGene_tx=XM_011288453.3;refGene_gn=RC3H2;refGene_tx=XM_011288454.2;refGene_gn=RC3H2;refGene_tx=XM_019816494.2;refGene_gn=RC3H2;refGene_tx=XM_019816495.2
D4 84106122 84106271
refGene_gn=RC3H2;refGene_tx=XM_011288453.3;refGene_gn=RC3H2;refGene_tx

=XM_011288454.2;refGene_gn=RC3H2;refGene_tx=XM_019816494.2;refGene_gn=RC3H2;refGene_tx=XM_019816495.2
D4 84106363 84106471
refGene_gn=RC3H2;refGene_tx=XM_011288453.3;refGene_gn=RC3H2;refGene_tx=XM_011288454.2;refGene_gn=RC3H2;refGene_tx=XM_019816494.2;refGene_gn=RC3H2;refGene_tx=XM_019816495.2
D4 84106720 84106855
refGene_gn=RC3H2;refGene_tx=XM_011288453.3;refGene_gn=RC3H2;refGene_tx=XM_011288454.2;refGene_gn=RC3H2;refGene_tx=XM_019816494.2;refGene_gn=RC3H2;refGene_tx=XM_019816495.2
D4 84106949 84107031
refGene_gn=RC3H2;refGene_tx=XM_011288453.3;refGene_gn=RC3H2;refGene_tx=XM_011288454.2;refGene_gn=RC3H2;refGene_tx=XM_019816494.2;refGene_gn=RC3H2;refGene_tx=XM_019816495.2
D4 84107292 84107377
refGene_gn=RC3H2;refGene_tx=XM_011288453.3;refGene_gn=RC3H2;refGene_tx=XM_011288454.2;refGene_gn=RC3H2;refGene_tx=XM_019816494.2;refGene_gn=RC3H2;refGene_tx=XM_019816495.2
D4 84107618 84107832
refGene_gn=RC3H2;refGene_tx=XM_011288453.3;refGene_gn=RC3H2;refGene_tx=XM_011288454.2;refGene_gn=RC3H2;refGene_tx=XM_019816494.2;refGene_gn=RC3H2;refGene_tx=XM_019816495.2
D4 84108166 84108313
refGene_gn=RC3H2;refGene_tx=XM_011288453.3;refGene_gn=RC3H2;refGene_tx=XM_011288454.2;refGene_gn=RC3H2;refGene_tx=XM_019816494.2;refGene_gn=RC3H2;refGene_tx=XM_019816495.2
D4 84110349 84110520
refGene_gn=RC3H2;refGene_tx=XM_011288453.3;refGene_gn=RC3H2;refGene_tx=XM_011288454.2;refGene_gn=RC3H2;refGene_tx=XM_019816494.2
D4 84111092 84111526
refGene_gn=RC3H2;refGene_tx=XM_011288453.3;refGene_gn=RC3H2;refGene_tx=XM_011288454.2;refGene_gn=RC3H2;refGene_tx=XM_019816494.2;refGene_gn=RC3H2;refGene_tx=XM_019816495.2
D4 84112167 84112382
refGene_gn=RC3H2;refGene_tx=XM_011288453.3;refGene_gn=RC3H2;refGene_tx=XM_011288454.2;refGene_gn=RC3H2;refGene_tx=XM_019816494.2;refGene_gn=RC3H2;refGene_tx=XM_019816495.2
D4 84116853 84117162
refGene_gn=RC3H2;refGene_tx=XM_011288453.3;refGene_gn=RC3H2;refGene_tx=XM_011288454.2;refGene_gn=RC3H2;refGene_tx=XM_019816494.2;refGene_gn=RC3H2;refGene_tx=XM_019816495.2
D4 84127137 84127250
refGene_gn=RC3H2;refGene_tx=XM_011288453.3;refGene_gn=RC3H2;refGene_tx=XM_011288454.2;refGene_gn=RC3H2;refGene_tx=XM_019816494.2;refGene_gn=RC3H2;refGene_tx=XM_019816495.2
D4 84129070 84129189
refGene_gn=RC3H2;refGene_tx=XM_011288453.3;refGene_gn=RC3H2;refGene_tx=XM_011288454.2;refGene_gn=RC3H2;refGene_tx=XM_019816494.2;refGene_gn=RC3H2;refGene_tx=XM_019816495.2
D4 84129299 84129432

refGene_gn=RC3H2;refGene_tx=XM_011288453.3;refGene_gn=RC3H2;refGene_tx=XM_011288454.2;refGene_gn=RC3H2;refGene_tx=XM_019816494.2;refGene_gn=RC3H2;refGene_tx=XM_019816495.2
D4 84130136 84130337 refGene_gn=RC3H2;refGene_tx=XM_011288453.3;refGene_gn=RC3H2;refGene_tx=XM_011288454.2;refGene_gn=RC3H2;refGene_tx=XM_019816494.2;refGene_gn=RC3H2;refGene_tx=XM_019816495.2
D4 84133516 84133692 refGene_gn=RC3H2;refGene_tx=XM_011288453.3;refGene_gn=RC3H2;refGene_tx=XM_011288454.2;refGene_gn=RC3H2;refGene_tx=XM_019816494.2;refGene_gn=RC3H2;refGene_tx=XM_019816495.2
D4 84138442 84138676 refGene_gn=RC3H2;refGene_tx=XM_011288453.3;refGene_gn=RC3H2;refGene_tx=XM_011288454.2;refGene_gn=RC3H2;refGene_tx=XM_019816494.2;refGene_gn=RC3H2;refGene_tx=XM_019816495.2
D4 84141719 84141837 refGene_gn=RC3H2;refGene_tx=XM_011288453.3;refGene_gn=RC3H2;refGene_tx=XM_011288454.2;refGene_gn=RC3H2;refGene_tx=XM_019816494.2;refGene_gn=RC3H2;refGene_tx=XM_019816495.2
D4 84145778 84146009 refGene_gn=RC3H2;refGene_tx=XM_011288453.3;refGene_gn=RC3H2;refGene_tx=XM_011288454.2;refGene_gn=RC3H2;refGene_tx=XM_019816494.2;refGene_gn=RC3H2;refGene_tx=XM_019816495.2
D4 84157547 84158819 refGene_gn=ZBTB6;refGene_tx=XM_003995853.5;refGene_gn=ZBTB6;refGene_tx=XM_019815615.2;refGene_gn=ZBTB6;refGene_tx=XM_019815616.2
D4 84164047 84165373 refGene_gn=ZBTB26;refGene_tx=XM_003995854.4;refGene_gn=ZBTB26;refGene_tx=XM_019815611.1;refGene_gn=ZBTB26;refGene_tx=XM_019815613.1
D4 84199670 84199820 refGene_gn=RABGAP1;refGene_tx=XM_023242688.1
D4 84222980 84223215 refGene_gn=RABGAP1;refGene_tx=XM_023242688.1
D4 84224101 84224306 refGene_gn=RABGAP1;refGene_tx=XM_023242688.1
D4 84227689 84227864 refGene_gn=RABGAP1;refGene_tx=XM_023242688.1
D4 84228438 84228596 refGene_gn=RABGAP1;refGene_tx=XM_023242688.1
D4 84234609 84234720 refGene_gn=RABGAP1;refGene_tx=XM_023242688.1
D4 84235365 84235432 refGene_gn=RABGAP1;refGene_tx=XM_023242688.1
D4 84242487 84242590 refGene_gn=RABGAP1;refGene_tx=XM_023242688.1
D4 84243484 84243654 refGene_gn=RABGAP1;refGene_tx=XM_023242688.1
D4 84254624 84254799 refGene_gn=RABGAP1;refGene_tx=XM_023242688.1
D4 84258913 84259007 refGene_gn=RABGAP1;refGene_tx=XM_023242688.1
D4 84262940 84263091 refGene_gn=RABGAP1;refGene_tx=XM_023242688.1
D4 84278370 84278607 refGene_gn=RABGAP1;refGene_tx=XM_023242689.1
D4 84312303 84312417 refGene_gn=RABGAP1;refGene_tx=XM_023242688.1;refGene_gn=RABGAP1;refGene_tx=XM_023242689.1
D4 84317763 84317838 refGene_gn=RABGAP1;refGene_tx=XM_023242688.1;refGene_gn=RABGAP1;refGene_tx=XM_023242689.1
D4 84321475 84321601 refGene_gn=RABGAP1;refGene_tx=XM_023242688.1;refGene_gn=RABGAP1;refGen

e_tx=XM_023242689.1
D4	84322202	84322346
refGene_gn=RABGAP1;refGene_tx=XM_023242688.1;refGene_gn=RABGAP1;refGene_tx=XM_023242689.1
D4	84324537	84324579
refGene_gn=RABGAP1;refGene_tx=XM_023242688.1;refGene_gn=RABGAP1;refGene_tx=XM_023242689.1
D4	84324829	84324958
refGene_gn=RABGAP1;refGene_tx=XM_023242688.1;refGene_gn=RABGAP1;refGene_tx=XM_023242689.1
D4	84328661	84328682	refGene_gn=RABGAP1;refGene_tx=XM_023242690.1
D4	84336523	84336616
refGene_gn=RABGAP1;refGene_tx=XM_023242688.1;refGene_gn=RABGAP1;refGene_tx=XM_023242689.1;refGene_gn=RABGAP1;refGene_tx=XM_023242690.1
D4	84337052	84337163
refGene_gn=RABGAP1;refGene_tx=XM_023242688.1;refGene_gn=RABGAP1;refGene_tx=XM_023242689.1;refGene_gn=RABGAP1;refGene_tx=XM_023242690.1
D4	84346816	84346921
refGene_gn=RABGAP1;refGene_tx=XM_023242688.1;refGene_gn=RABGAP1;refGene_tx=XM_023242689.1;refGene_gn=RABGAP1;refGene_tx=XM_023242690.1
D4	84347780	84347864
refGene_gn=RABGAP1;refGene_tx=XM_023242688.1;refGene_gn=RABGAP1;refGene_tx=XM_023242689.1;refGene_gn=RABGAP1;refGene_tx=XM_023242690.1
D4	84348800	84348872
refGene_gn=RABGAP1;refGene_tx=XM_023242688.1;refGene_gn=RABGAP1;refGene_tx=XM_023242689.1;refGene_gn=RABGAP1;refGene_tx=XM_023242690.1
D4	84350510	84350708
refGene_gn=RABGAP1;refGene_tx=XM_023242688.1;refGene_gn=RABGAP1;refGene_tx=XM_023242689.1;refGene_gn=RABGAP1;refGene_tx=XM_023242690.1
D4	84352042	84352165
refGene_gn=RABGAP1;refGene_tx=XM_023242688.1;refGene_gn=RABGAP1;refGene_tx=XM_023242689.1;refGene_gn=RABGAP1;refGene_tx=XM_023242690.1
D4	84369535	84369540	refGene_gn=STRBP;refGene_tx=XM_023242696.1
D4	84370317	84370412
refGene_gn=STRBP;refGene_tx=XM_019815606.2;refGene_gn=STRBP;refGene_tx=XM_019815607.2;refGene_gn=STRBP;refGene_tx=XM_023242692.1;refGene_gn=STRBP;refGene_tx=XM_023242693.1;refGene_gn=STRBP;refGene_tx=XM_023242697.1
D4	84372818	84372895	refGene_gn=STRBP;refGene_tx=XM_023242694.1
D4	84375039	84375050	refGene_gn=STRBP;refGene_tx=XM_023242695.1
D4	84375513	84375558
refGene_gn=STRBP;refGene_tx=XM_019815606.2;refGene_gn=STRBP;refGene_tx=XM_019815607.2;refGene_gn=STRBP;refGene_tx=XM_023242692.1;refGene_gn=STRBP;refGene_tx=XM_023242693.1;refGene_gn=STRBP;refGene_tx=XM_023242694.1;refGene_gn=STRBP;refGene_tx=XM_023242695.1;refGene_gn=STRBP;refGene_tx=XM_023242696.1;refGene_gn=STRBP;refGene_tx=XM_023242697.1
D4	84380960	84381084
refGene_gn=STRBP;refGene_tx=XM_019815606.2;refGene_gn=STRBP;refGene_tx=XM_019815607.2;refGene_gn=STRBP;refGene_tx=XM_023242692.1;refGene_gn=STRBP;refGene_tx=XM_023242693.1;refGene_gn=STRBP;refGene_tx=XM_0232426

94.1;refGene_gn=STRBP;refGene_tx=XM_023242695.1;refGene_gn=STRBP;refGene_tx=XM_023242696.1;refGene_gn=STRBP;refGene_tx=XM_023242697.1
D4 84383771 84383912
refGene_gn=STRBP;refGene_tx=XM_019815606.2;refGene_gn=STRBP;refGene_tx=XM_019815607.2;refGene_gn=STRBP;refGene_tx=XM_023242692.1;refGene_gn=STRBP;refGene_tx=XM_023242693.1;refGene_gn=STRBP;refGene_tx=XM_023242694.1;refGene_gn=STRBP;refGene_tx=XM_023242695.1;refGene_gn=STRBP;refGene_tx=XM_023242696.1;refGene_gn=STRBP;refGene_tx=XM_023242697.1
D4 84384111 84384246
refGene_gn=STRBP;refGene_tx=XM_019815606.2;refGene_gn=STRBP;refGene_tx=XM_019815607.2;refGene_gn=STRBP;refGene_tx=XM_023242692.1;refGene_gn=STRBP;refGene_tx=XM_023242693.1;refGene_gn=STRBP;refGene_tx=XM_023242694.1;refGene_gn=STRBP;refGene_tx=XM_023242695.1;refGene_gn=STRBP;refGene_tx=XM_023242696.1;refGene_gn=STRBP;refGene_tx=XM_023242697.1
D4 84387370 84387529
refGene_gn=STRBP;refGene_tx=XM_019815606.2;refGene_gn=STRBP;refGene_tx=XM_019815607.2;refGene_gn=STRBP;refGene_tx=XM_023242692.1;refGene_gn=STRBP;refGene_tx=XM_023242693.1;refGene_gn=STRBP;refGene_tx=XM_023242694.1;refGene_gn=STRBP;refGene_tx=XM_023242695.1;refGene_gn=STRBP;refGene_tx=XM_023242696.1;refGene_gn=STRBP;refGene_tx=XM_023242697.1
D4 84393432 84393632
refGene_gn=STRBP;refGene_tx=XM_019815606.2;refGene_gn=STRBP;refGene_tx=XM_019815607.2;refGene_gn=STRBP;refGene_tx=XM_023242692.1;refGene_gn=STRBP;refGene_tx=XM_023242693.1;refGene_gn=STRBP;refGene_tx=XM_023242694.1;refGene_gn=STRBP;refGene_tx=XM_023242695.1;refGene_gn=STRBP;refGene_tx=XM_023242696.1;refGene_gn=STRBP;refGene_tx=XM_023242697.1
D4 84394052 84394145
refGene_gn=STRBP;refGene_tx=XM_019815606.2;refGene_gn=STRBP;refGene_tx=XM_019815607.2;refGene_gn=STRBP;refGene_tx=XM_023242692.1;refGene_gn=STRBP;refGene_tx=XM_023242693.1;refGene_gn=STRBP;refGene_tx=XM_023242694.1;refGene_gn=STRBP;refGene_tx=XM_023242695.1;refGene_gn=STRBP;refGene_tx=XM_023242696.1;refGene_gn=STRBP;refGene_tx=XM_023242697.1
D4 84406445 84406557
refGene_gn=STRBP;refGene_tx=XM_019815606.2;refGene_gn=STRBP;refGene_tx=XM_019815607.2;refGene_gn=STRBP;refGene_tx=XM_023242692.1;refGene_gn=STRBP;refGene_tx=XM_023242693.1;refGene_gn=STRBP;refGene_tx=XM_023242694.1;refGene_gn=STRBP;refGene_tx=XM_023242695.1;refGene_gn=STRBP;refGene_tx=XM_023242696.1;refGene_gn=STRBP;refGene_tx=XM_023242697.1
D4 84406763 84406861
refGene_gn=STRBP;refGene_tx=XM_019815606.2;refGene_gn=STRBP;refGene_tx=XM_019815607.2;refGene_gn=STRBP;refGene_tx=XM_023242692.1;refGene_gn=STRBP;refGene_tx=XM_023242693.1;refGene_gn=STRBP;refGene_tx=XM_023242694.1;refGene_gn=STRBP;refGene_tx=XM_023242695.1;refGene_gn=STRBP;refGene_tx=XM_023242696.1;refGene_gn=STRBP;refGene_tx=XM_023242697.1
D4 84407521 84407633
refGene_gn=STRBP;refGene_tx=XM_019815606.2;refGene_gn=STRBP;refGene_tx=XM_019815607.2;refGene_gn=STRBP;refGene_tx=XM_023242692.1;refGene_gn=STRBP;refGene_tx=XM_023242693.1;refGene_gn=STRBP;refGene_tx=XM_023242694.1;refGene_gn=STRBP;refGene_tx=XM_023242695.1;refGene_gn=STRBP;refGene_tx=XM_023242696.1

D4 84408737 84408833
refGene_gn=STRBP;refGene_tx=XM_019815606.2;refGene_gn=STRBP;refGene_tx=XM_019815607.2;refGene_gn=STRBP;refGene_tx=XM_023242692.1;refGene_gn=STRBP;refGene_tx=XM_023242693.1;refGene_gn=STRBP;refGene_tx=XM_023242694.1;refGene_gn=STRBP;refGene_tx=XM_023242695.1;refGene_gn=STRBP;refGene_tx=XM_023242696.1
D4 84409373 84409465
refGene_gn=STRBP;refGene_tx=XM_019815606.2;refGene_gn=STRBP;refGene_tx=XM_019815607.2;refGene_gn=STRBP;refGene_tx=XM_023242692.1;refGene_gn=STRBP;refGene_tx=XM_023242693.1;refGene_gn=STRBP;refGene_tx=XM_023242694.1;refGene_gn=STRBP;refGene_tx=XM_023242695.1;refGene_gn=STRBP;refGene_tx=XM_023242696.1
D4 84417369 84417514
refGene_gn=STRBP;refGene_tx=XM_019815606.2;refGene_gn=STRBP;refGene_tx=XM_019815607.2;refGene_gn=STRBP;refGene_tx=XM_023242692.1;refGene_gn=STRBP;refGene_tx=XM_023242693.1;refGene_gn=STRBP;refGene_tx=XM_023242694.1;refGene_gn=STRBP;refGene_tx=XM_023242695.1;refGene_gn=STRBP;refGene_tx=XM_023242696.1;refGene_gn=STRBP;refGene_tx=XM_023242697.1
D4 84421018 84421184
refGene_gn=STRBP;refGene_tx=XM_019815606.2;refGene_gn=STRBP;refGene_tx=XM_019815607.2;refGene_gn=STRBP;refGene_tx=XM_023242692.1;refGene_gn=STRBP;refGene_tx=XM_023242693.1;refGene_gn=STRBP;refGene_tx=XM_023242694.1;refGene_gn=STRBP;refGene_tx=XM_023242695.1;refGene_gn=STRBP;refGene_tx=XM_023242696.1;refGene_gn=STRBP;refGene_tx=XM_023242697.1
D4 84426627 84426848
refGene_gn=STRBP;refGene_tx=XM_019815606.2;refGene_gn=STRBP;refGene_tx=XM_019815607.2;refGene_gn=STRBP;refGene_tx=XM_023242692.1;refGene_gn=STRBP;refGene_tx=XM_023242693.1;refGene_gn=STRBP;refGene_tx=XM_023242694.1;refGene_gn=STRBP;refGene_tx=XM_023242695.1;refGene_gn=STRBP;refGene_tx=XM_023242696.1;refGene_gn=STRBP;refGene_tx=XM_023242697.1
D4 84430742 84430745
refGene_gn=STRBP;refGene_tx=XM_019815606.2;refGene_gn=STRBP;refGene_tx=XM_019815607.2;refGene_gn=STRBP;refGene_tx=XM_023242692.1;refGene_gn=STRBP;refGene_tx=XM_023242693.1;refGene_gn=STRBP;refGene_tx=XM_023242694.1;refGene_gn=STRBP;refGene_tx=XM_023242695.1;refGene_gn=STRBP;refGene_tx=XM_023242696.1;refGene_gn=STRBP;refGene_tx=XM_023242697.1
D4 84598492 84598589
refGene_gn=CRB2;refGene_tx=XM_023242698.1;refGene_gn=CRB2;refGene_tx=XM_023242699.1;refGene_gn=CRB2;refGene_tx=XM_023242700.1
D4 84605870 84606194
refGene_gn=CRB2;refGene_tx=XM_023242698.1;refGene_gn=CRB2;refGene_tx=XM_023242699.1;refGene_gn=CRB2;refGene_tx=XM_023242700.1
D4 84608088 84608284
refGene_gn=CRB2;refGene_tx=XM_023242698.1;refGene_gn=CRB2;refGene_tx=XM_023242699.1;refGene_gn=CRB2;refGene_tx=XM_023242700.1
D4 84608401 84608541
refGene_gn=CRB2;refGene_tx=XM_023242698.1;refGene_gn=CRB2;refGene_tx=XM_023242699.1;refGene_gn=CRB2;refGene_tx=XM_023242700.1
D4 84608968 84609154
refGene_gn=CRB2;refGene_tx=XM_023242698.1;refGene_gn=CRB2;refGene_tx=X

M_023242699.1
D4	84609351 84609465 refGene_gn=CRB2;refGene_tx=XM_023242698.1
D4	84612173 84613049
refGene_gn=CRB2;refGene_tx=XM_023242698.1;refGene_gn=CRB2;refGene_tx=XM_023242699.1;refGene_gn=CRB2;refGene_tx=XM_023242700.1
D4	84613138 84613647
refGene_gn=CRB2;refGene_tx=XM_023242698.1;refGene_gn=CRB2;refGene_tx=XM_023242699.1;refGene_gn=CRB2;refGene_tx=XM_023242700.1
D4	84614179 84614345
refGene_gn=CRB2;refGene_tx=XM_023242698.1;refGene_gn=CRB2;refGene_tx=XM_023242699.1;refGene_gn=CRB2;refGene_tx=XM_023242700.1
D4	84615185 84615969
refGene_gn=CRB2;refGene_tx=XM_023242698.1;refGene_gn=CRB2;refGene_tx=XM_023242699.1;refGene_gn=CRB2;refGene_tx=XM_023242700.1
D4	84616784 84616901
refGene_gn=CRB2;refGene_tx=XM_023242698.1;refGene_gn=CRB2;refGene_tx=XM_023242699.1;refGene_gn=CRB2;refGene_tx=XM_023242700.1
D4	84617474 84617592
refGene_gn=CRB2;refGene_tx=XM_023242698.1;refGene_gn=CRB2;refGene_tx=XM_023242699.1;refGene_gn=CRB2;refGene_tx=XM_023242700.1
D4	84619273 84619498
refGene_gn=CRB2;refGene_tx=XM_023242698.1;refGene_gn=CRB2;refGene_tx=XM_023242699.1;refGene_gn=CRB2;refGene_tx=XM_023242700.1
D4	84623723 84624917
refGene_gn=DENND1A;refGene_tx=XM_019815603.2;refGene_gn=DENND1A;refGene_tx=XM_019815604.2
D4	84626071 84626330
refGene_gn=DENND1A;refGene_tx=XM_019815603.2;refGene_gn=DENND1A;refGene_tx=XM_019815604.2
D4	84629069 84629198
refGene_gn=DENND1A;refGene_tx=XM_019815603.2;refGene_gn=DENND1A;refGene_tx=XM_019815604.2
D4	84643219 84643308
refGene_gn=DENND1A;refGene_tx=XM_019815603.2;refGene_gn=DENND1A;refGene_tx=XM_019815604.2
D4	84650039 84650096
refGene_gn=DENND1A;refGene_tx=XM_019815603.2;refGene_gn=DENND1A;refGene_tx=XM_019815604.2
D4	84673765 84673897
refGene_gn=DENND1A;refGene_tx=XM_019815603.2;refGene_gn=DENND1A;refGene_tx=XM_019815604.2
D4	84682450 84682507
refGene_gn=DENND1A;refGene_tx=XM_019815603.2;refGene_gn=DENND1A;refGene_tx=XM_019815604.2
D4	84683753 84683825
refGene_gn=DENND1A;refGene_tx=XM_019815603.2;refGene_gn=DENND1A;refGene_tx=XM_019815604.2
D4	84686104 84686145
refGene_gn=DENND1A;refGene_tx=XM_019815603.2;refGene_gn=DENND1A;refGene_tx=XM_019815604.2

D4 84688030 84688118
refGene_gn=DENND1A;refGene_tx=XM_019815603.2;refGene_gn=DENND1A;refGene_tx=XM_019815604.2
D4 84688500 84688605
refGene_gn=DENND1A;refGene_tx=XM_019815603.2;refGene_gn=DENND1A;refGene_tx=XM_019815604.2
D4 84776920 84777046
refGene_gn=DENND1A;refGene_tx=XM_019815603.2;refGene_gn=DENND1A;refGene_tx=XM_019815604.2
D4 84801350 84801452
refGene_gn=DENND1A;refGene_tx=XM_019815603.2;refGene_gn=DENND1A;refGene_tx=XM_019815604.2
D4 84826240 84826286
refGene_gn=DENND1A;refGene_tx=XM_019815603.2;refGene_gn=DENND1A;refGene_tx=XM_019815604.2
D4 84842914 84843015
refGene_gn=DENND1A;refGene_tx=XM_019815603.2;refGene_gn=DENND1A;refGene_tx=XM_019815604.2
D4 84864180 84864291
refGene_gn=DENND1A;refGene_tx=XM_019815603.2;refGene_gn=DENND1A;refGene_tx=XM_019815604.2
D4 84890420 84890474 refGene_gn=DENND1A;refGene_tx=XM_019815604.2
D4 84895030 84895111 refGene_gn=DENND1A;refGene_tx=XM_019815604.2
D4 84900497 84900567 refGene_gn=DENND1A;refGene_tx=XM_019815604.2
D4 84977062 84977182 refGene_gn=DENND1A;refGene_tx=XM_019815604.2
D4 84988656 84988706 refGene_gn=DENND1A;refGene_tx=XM_019815604.2
D4 85013603 85013647 refGene_gn=DENND1A;refGene_tx=XM_019815604.2
D4 85100373 85100444 refGene_gn=DENND1A;refGene_tx=XM_019815604.2
D4 85144489 85144509 refGene_gn=DENND1A;refGene_tx=XM_019815604.2
D4 85215486 85215606 refGene_gn=LHX2;refGene_tx=XM_011288451.3
D4 85217115 85217318 refGene_gn=LHX2;refGene_tx=XM_011288451.3
D4 85218316 85218720 refGene_gn=LHX2;refGene_tx=XM_011288451.3
D4 85224365 85224571 refGene_gn=LHX2;refGene_tx=XM_011288451.3
D4 85234509 85234797 refGene_gn=LHX2;refGene_tx=XM_011288451.3
D4 85413327 85413425 refGene_gn=NEK6;refGene_tx=XM_003995858.4
D4 85420714 85420832
refGene_gn=NEK6;refGene_tx=XM_003995858.4;refGene_gn=NEK6;refGene_tx=XM_006939451.4;refGene_gn=NEK6;refGene_tx=XM_023242702.1
D4 85431698 85431839
refGene_gn=NEK6;refGene_tx=XM_003995858.4;refGene_gn=NEK6;refGene_tx=XM_006939451.4;refGene_gn=NEK6;refGene_tx=XM_023242702.1
D4 85433137 85433200
refGene_gn=NEK6;refGene_tx=XM_003995858.4;refGene_gn=NEK6;refGene_tx=XM_006939451.4;refGene_gn=NEK6;refGene_tx=XM_023242702.1
D4 85437851 85437962
refGene_gn=NEK6;refGene_tx=XM_003995858.4;refGene_gn=NEK6;refGene_tx=XM_006939451.4;refGene_gn=NEK6;refGene_tx=XM_023242702.1
D4 85442330 85442439
refGene_gn=NEK6;refGene_tx=XM_003995858.4;refGene_gn=NEK6;refGene_tx=XM_006939451.4;refGene_gn=NEK6;refGene_tx=XM_023242702.1

D4 85442788 85442896
refGene_gn=NEK6;refGene_tx=XM_003995858.4;refGene_gn=NEK6;refGene_tx=XM_006939451.4;refGene_gn=NEK6;refGene_tx=XM_023242702.1
D4 85452932 85453027
refGene_gn=NEK6;refGene_tx=XM_003995858.4;refGene_gn=NEK6;refGene_tx=XM_006939451.4;refGene_gn=NEK6;refGene_tx=XM_023242702.1
D4 85460888 85461002
refGene_gn=NEK6;refGene_tx=XM_003995858.4;refGene_gn=NEK6;refGene_tx=XM_006939451.4;refGene_gn=NEK6;refGene_tx=XM_023242702.1
D4 85464420 85464531
refGene_gn=NEK6;refGene_tx=XM_003995858.4;refGene_gn=NEK6;refGene_tx=XM_006939451.4;refGene_gn=NEK6;refGene_tx=XM_023242702.1
D4 85467159 85467271 refGene_gn=PSMB7;refGene_tx=XM_003995859.3
D4 85470238 85470390 refGene_gn=PSMB7;refGene_tx=XM_003995859.3
D4 85477527 85477866 refGene_gn=PSMB7;refGene_tx=XM_019816496.1
D4 85496359 85496418
refGene_gn=PSMB7;refGene_tx=XM_003995859.3;refGene_gn=PSMB7;refGene_tx=XM_019816496.1
D4 85516541 85516657
refGene_gn=PSMB7;refGene_tx=XM_003995859.3;refGene_gn=PSMB7;refGene_tx=XM_019816496.1
D4 85522856 85522997
refGene_gn=PSMB7;refGene_tx=XM_003995859.3;refGene_gn=PSMB7;refGene_tx=XM_019816496.1
D4 85524450 85524548
refGene_gn=PSMB7;refGene_tx=XM_003995859.3;refGene_gn=PSMB7;refGene_tx=XM_019816496.1
D4 85525273 85525367
refGene_gn=PSMB7;refGene_tx=XM_003995859.3;refGene_gn=PSMB7;refGene_tx=XM_019816496.1
D4 85525808 85525870
refGene_gn=PSMB7;refGene_tx=XM_003995859.3;refGene_gn=PSMB7;refGene_tx=XM_019816496.1
D4 85559740 85559837
refGene_gn=ADGRD2;refGene_tx=XM_019816497.2;refGene_gn=ADGRD2;refGene_tx=XM_019816498.2;refGene_gn=ADGRD2;refGene_tx=XM_019816499.2;refGene_gn=ADGRD2;refGene_tx=XM_019816500.2
D4 85560566 85560701
refGene_gn=ADGRD2;refGene_tx=XM_019816497.2;refGene_gn=ADGRD2;refGene_tx=XM_019816498.2;refGene_gn=ADGRD2;refGene_tx=XM_019816499.2;refGene_gn=ADGRD2;refGene_tx=XM_019816500.2
D4 85561433 85561557
refGene_gn=ADGRD2;refGene_tx=XM_019816497.2;refGene_gn=ADGRD2;refGene_tx=XM_019816498.2;refGene_gn=ADGRD2;refGene_tx=XM_019816499.2;refGene_gn=ADGRD2;refGene_tx=XM_019816500.2
D4 85562009 85562115
refGene_gn=ADGRD2;refGene_tx=XM_019816497.2;refGene_gn=ADGRD2;refGene_tx=XM_019816498.2;refGene_gn=ADGRD2;refGene_tx=XM_019816499.2;refGene_gn=ADGRD2;refGene_tx=XM_019816500.2
D4 85564875 85564931

refGene_gn=ADGRD2;refGene_tx=XM_019816497.2;refGene_gn=ADGRD2;refGene_tx=XM_019816498.2;refGene_gn=ADGRD2;refGene_tx=XM_019816499.2;refGene_gn=ADGRD2;refGene_tx=XM_019816500.2
D4 85565580 85565707
refGene_gn=ADGRD2;refGene_tx=XM_019816497.2;refGene_gn=ADGRD2;refGene_tx=XM_019816498.2;refGene_gn=ADGRD2;refGene_tx=XM_019816499.2;refGene_gn=ADGRD2;refGene_tx=XM_019816500.2
D4 85566942 85567046
refGene_gn=ADGRD2;refGene_tx=XM_019816497.2;refGene_gn=ADGRD2;refGene_tx=XM_019816498.2;refGene_gn=ADGRD2;refGene_tx=XM_019816499.2;refGene_gn=ADGRD2;refGene_tx=XM_019816500.2
D4 85567305 85567408
refGene_gn=ADGRD2;refGene_tx=XM_019816497.2;refGene_gn=ADGRD2;refGene_tx=XM_019816498.2;refGene_gn=ADGRD2;refGene_tx=XM_019816499.2;refGene_gn=ADGRD2;refGene_tx=XM_019816500.2
D4 85567670 85567824
refGene_gn=ADGRD2;refGene_tx=XM_019816497.2;refGene_gn=ADGRD2;refGene_tx=XM_019816498.2;refGene_gn=ADGRD2;refGene_tx=XM_019816499.2;refGene_gn=ADGRD2;refGene_tx=XM_019816500.2
D4 85568339 85568473
refGene_gn=ADGRD2;refGene_tx=XM_019816497.2;refGene_gn=ADGRD2;refGene_tx=XM_019816498.2;refGene_gn=ADGRD2;refGene_tx=XM_019816499.2;refGene_gn=ADGRD2;refGene_tx=XM_019816500.2
D4 85568559 85568675
refGene_gn=ADGRD2;refGene_tx=XM_019816497.2;refGene_gn=ADGRD2;refGene_tx=XM_019816498.2;refGene_gn=ADGRD2;refGene_tx=XM_019816499.2;refGene_gn=ADGRD2;refGene_tx=XM_019816500.2
D4 85569535 85569656
refGene_gn=ADGRD2;refGene_tx=XM_019816497.2;refGene_gn=ADGRD2;refGene_tx=XM_019816498.2;refGene_gn=ADGRD2;refGene_tx=XM_019816499.2;refGene_gn=ADGRD2;refGene_tx=XM_019816500.2
D4 85573665 85573707
refGene_gn=ADGRD2;refGene_tx=XM_019816497.2;refGene_gn=ADGRD2;refGene_tx=XM_019816498.2;refGene_gn=ADGRD2;refGene_tx=XM_019816499.2;refGene_gn=ADGRD2;refGene_tx=XM_019816500.2
D4 85573825 85573879
refGene_gn=ADGRD2;refGene_tx=XM_019816497.2;refGene_gn=ADGRD2;refGene_tx=XM_019816498.2;refGene_gn=ADGRD2;refGene_tx=XM_019816499.2;refGene_gn=ADGRD2;refGene_tx=XM_019816500.2
D4 85573963 85574234
refGene_gn=ADGRD2;refGene_tx=XM_019816497.2;refGene_gn=ADGRD2;refGene_tx=XM_019816498.2;refGene_gn=ADGRD2;refGene_tx=XM_019816499.2;refGene_gn=ADGRD2;refGene_tx=XM_019816500.2
D4 85574651 85574710
refGene_gn=ADGRD2;refGene_tx=XM_019816497.2;refGene_gn=ADGRD2;refGene_tx=XM_019816498.2;refGene_gn=ADGRD2;refGene_tx=XM_019816499.2;refGene_gn=ADGRD2;refGene_tx=XM_019816500.2
D4 85575006 85575144
refGene_gn=ADGRD2;refGene_tx=XM_019816497.2;refGene_gn=ADGRD2;refGene_tx=XM_019816498.2;refGene_gn=ADGRD2;refGene_tx=XM_019816499.2;refGene_

gn=ADGRD2;refGene_tx=XM_019816500.2
D4 85576574 85576582
refGene_gn=ADGRD2;refGene_tx=XM_019816498.2;refGene_gn=ADGRD2;refGene_
tx=XM_019816500.2
D4 85581085 85581333 refGene_gn=NR5A1;refGene_tx=XM_023242703.1
D4 85588937 85589085 refGene_gn=NR5A1;refGene_tx=XM_023242703.1
D4 85591011 85591131 refGene_gn=NR5A1;refGene_tx=XM_023242703.1
D4 85598325 85598951 refGene_gn=NR5A1;refGene_tx=XM_023242703.1
D4 85600991 85601133 refGene_gn=NR5A1;refGene_tx=XM_023242703.1
D4 85601306 85601408 refGene_gn=NR5A1;refGene_tx=XM_023242703.1
D4 85601799 85601841
refGene_gn=LOC111557482;refGene_tx=XM_023242316.1
D4 85602419 85602872
refGene_gn=LOC111557482;refGene_tx=XM_023242316.1
D4 85619542 85619631
refGene_gn=NR6A1;refGene_tx=XM_019816505.2;refGene_gn=NR6A1;refGene_tx
=XM_023242704.1;refGene_gn=NR6A1;refGene_tx=XM_023242705.1
D4 85621494 85621647
refGene_gn=NR6A1;refGene_tx=XM_019816505.2;refGene_gn=NR6A1;refGene_tx
=XM_023242704.1;refGene_gn=NR6A1;refGene_tx=XM_023242705.1
D4 85623317 85623439
refGene_gn=NR6A1;refGene_tx=XM_019816505.2;refGene_gn=NR6A1;refGene_tx
=XM_023242704.1;refGene_gn=NR6A1;refGene_tx=XM_023242705.1
D4 85630850 85631105
refGene_gn=NR6A1;refGene_tx=XM_019816505.2;refGene_gn=NR6A1;refGene_tx
=XM_023242704.1;refGene_gn=NR6A1;refGene_tx=XM_023242705.1
D4 85632431 85632659
refGene_gn=NR6A1;refGene_tx=XM_019816505.2;refGene_gn=NR6A1;refGene_tx
=XM_023242704.1;refGene_gn=NR6A1;refGene_tx=XM_023242705.1
D4 85634077 85634232
refGene_gn=NR6A1;refGene_tx=XM_019816505.2;refGene_gn=NR6A1;refGene_tx
=XM_023242704.1;refGene_gn=NR6A1;refGene_tx=XM_023242705.1
D4 85641118 85641174
refGene_gn=NR6A1;refGene_tx=XM_019816505.2;refGene_gn=NR6A1;refGene_tx
=XM_023242704.1;refGene_gn=NR6A1;refGene_tx=XM_023242705.1
D4 85652292 85652535
refGene_gn=NR6A1;refGene_tx=XM_019816505.2;refGene_gn=NR6A1;refGene_tx
=XM_023242704.1;refGene_gn=NR6A1;refGene_tx=XM_023242705.1
D4 85687165 85687181 refGene_gn=NR6A1;refGene_tx=XM_019816505.2
D4 85814415 85814457
refGene_gn=NR6A1;refGene_tx=XM_023242704.1;refGene_gn=NR6A1;refGene_tx
=XM_023242705.1
D4 85851329 85851429
refGene_gn=NR6A1;refGene_tx=XM_023242704.1;refGene_gn=NR6A1;refGene_tx
=XM_023242705.1
D4 85857259 85857376 refGene_gn=OLFML2A;refGene_tx=XM_003995860.5
D4 85865212 85865476 refGene_gn=OLFML2A;refGene_tx=XM_003995860.5
D4 85871291 85871399 refGene_gn=OLFML2A;refGene_tx=XM_003995860.5
D4 85872851 85873061 refGene_gn=OLFML2A;refGene_tx=XM_003995860.5
D4 85875332 85875582 refGene_gn=OLFML2A;refGene_tx=XM_003995860.5

D4 85877776 85878007 refGene_gn=OLFML2A;refGene_tx=XM_003995860.5
D4 85879041 85879227 refGene_gn=OLFML2A;refGene_tx=XM_003995860.5
D4 85880619 85881224 refGene_gn=OLFML2A;refGene_tx=XM_003995860.5
D4 85913982 85914132 refGene_gn=WDR38;refGene_tx=XM_003995861.5
D4 85914654 85914775 refGene_gn=WDR38;refGene_tx=XM_003995861.5
D4 85915831 85915948 refGene_gn=WDR38;refGene_tx=XM_003995861.5
D4 85916028 85916126 refGene_gn=WDR38;refGene_tx=XM_003995861.5
D4 85916530 85916611 refGene_gn=WDR38;refGene_tx=XM_003995861.5
D4 85916826 85916958 refGene_gn=WDR38;refGene_tx=XM_003995861.5
D4 85917037 85917187 refGene_gn=WDR38;refGene_tx=XM_003995861.5
D4 85917618 85917666 refGene_gn=WDR38;refGene_tx=XM_003995861.5
D4 85917778 85917907 refGene_gn=WDR38;refGene_tx=XM_003995861.5
D4 85918244 85918394 refGene_gn=RPL35;refGene_tx=XM_003995862.3
D4 85920185 85920267 refGene_gn=RPL35;refGene_tx=XM_003995862.3
D4 85921739 85921876 refGene_gn=RPL35;refGene_tx=XM_003995862.3
D4 85922314 85922317 refGene_gn=RPL35;refGene_tx=XM_003995862.3
D4 85927121 85927270 refGene_gn=ARPC5L;refGene_tx=XM_003995863.5
D4 85931332 85931405 refGene_gn=ARPC5L;refGene_tx=XM_003995863.5
D4 85932348 85932525 refGene_gn=ARPC5L;refGene_tx=XM_003995863.5
D4 85933820 85933883 refGene_gn=ARPC5L;refGene_tx=XM_003995863.5
D4 85937797 85937878
refGene_gn=GOLGA1;refGene_tx=XM_003995864.5;refGene_gn=GOLGA1;refGene_tx=XM_006939457.3;refGene_gn=GOLGA1;refGene_tx=XM_019816511.2
D4 85938898 85938985
refGene_gn=GOLGA1;refGene_tx=XM_003995864.5;refGene_gn=GOLGA1;refGene_tx=XM_006939457.3;refGene_gn=GOLGA1;refGene_tx=XM_019816511.2
D4 85939518 85939689
refGene_gn=GOLGA1;refGene_tx=XM_003995864.5;refGene_gn=GOLGA1;refGene_tx=XM_006939457.3;refGene_gn=GOLGA1;refGene_tx=XM_019816511.2
D4 85940281 85940341
refGene_gn=GOLGA1;refGene_tx=XM_003995864.5;refGene_gn=GOLGA1;refGene_tx=XM_006939457.3;refGene_gn=GOLGA1;refGene_tx=XM_019816511.2
D4 85943642 85943786
refGene_gn=GOLGA1;refGene_tx=XM_003995864.5;refGene_gn=GOLGA1;refGene_tx=XM_006939457.3;refGene_gn=GOLGA1;refGene_tx=XM_019816511.2
D4 85944221 85944379
refGene_gn=GOLGA1;refGene_tx=XM_003995864.5;refGene_gn=GOLGA1;refGene_tx=XM_006939457.3;refGene_gn=GOLGA1;refGene_tx=XM_019816511.2
D4 85944495 85944598
refGene_gn=GOLGA1;refGene_tx=XM_003995864.5;refGene_gn=GOLGA1;refGene_tx=XM_006939457.3;refGene_gn=GOLGA1;refGene_tx=XM_019816511.2
D4 85945440 85945530
refGene_gn=GOLGA1;refGene_tx=XM_003995864.5;refGene_gn=GOLGA1;refGene_tx=XM_006939457.3;refGene_gn=GOLGA1;refGene_tx=XM_019816511.2
D4 85951484 85951580
refGene_gn=GOLGA1;refGene_tx=XM_003995864.5;refGene_gn=GOLGA1;refGene_tx=XM_006939457.3;refGene_gn=GOLGA1;refGene_tx=XM_019816511.2
D4 85952085 85952235
refGene_gn=GOLGA1;refGene_tx=XM_003995864.5;refGene_gn=GOLGA1;refGene_tx=XM_006939457.3;refGene_gn=GOLGA1;refGene_tx=XM_019816511.2

D4 85954078 85954174
refGene_gn=GOLGA1;refGene_tx=XM_003995864.5;refGene_gn=GOLGA1;refGene_tx=XM_006939457.3;refGene_gn=GOLGA1;refGene_tx=XM_019816511.2
D4 85957304 85957400
refGene_gn=GOLGA1;refGene_tx=XM_003995864.5;refGene_gn=GOLGA1;refGene_tx=XM_006939457.3;refGene_gn=GOLGA1;refGene_tx=XM_019816511.2
D4 85959611 85959737
refGene_gn=GOLGA1;refGene_tx=XM_003995864.5;refGene_gn=GOLGA1;refGene_tx=XM_006939457.3;refGene_gn=GOLGA1;refGene_tx=XM_019816511.2
D4 85967247 85967359
refGene_gn=GOLGA1;refGene_tx=XM_003995864.5;refGene_gn=GOLGA1;refGene_tx=XM_006939457.3;refGene_gn=GOLGA1;refGene_tx=XM_019816511.2
D4 85967837 85968007
refGene_gn=GOLGA1;refGene_tx=XM_003995864.5;refGene_gn=GOLGA1;refGene_tx=XM_006939457.3;refGene_gn=GOLGA1;refGene_tx=XM_019816511.2
D4 85968335 85968464
refGene_gn=GOLGA1;refGene_tx=XM_003995864.5;refGene_gn=GOLGA1;refGene_tx=XM_006939457.3;refGene_gn=GOLGA1;refGene_tx=XM_019816511.2
D4 85971177 85971210
refGene_gn=GOLGA1;refGene_tx=XM_003995864.5;refGene_gn=GOLGA1;refGene_tx=XM_006939457.3;refGene_gn=GOLGA1;refGene_tx=XM_019816511.2
D4 85972570 85972668
refGene_gn=GOLGA1;refGene_tx=XM_003995864.5;refGene_gn=GOLGA1;refGene_tx=XM_006939457.3;refGene_gn=GOLGA1;refGene_tx=XM_019816511.2
D4 85973609 85973684
refGene_gn=GOLGA1;refGene_tx=XM_003995864.5;refGene_gn=GOLGA1;refGene_tx=XM_006939457.3
D4 85975700 85975791
refGene_gn=GOLGA1;refGene_tx=XM_003995864.5;refGene_gn=GOLGA1;refGene_tx=XM_006939457.3;refGene_gn=GOLGA1;refGene_tx=XM_019816511.2
D4 85981162 85981297
refGene_gn=GOLGA1;refGene_tx=XM_003995864.5;refGene_gn=GOLGA1;refGene_tx=XM_006939457.3;refGene_gn=GOLGA1;refGene_tx=XM_019816511.2
D4 85993366 85993513
refGene_gn=SCAI;refGene_tx=XM_023242707.1;refGene_gn=SCAI;refGene_tx=XM_023242708.1
D4 86008253 86008354
refGene_gn=SCAI;refGene_tx=XM_023242707.1;refGene_gn=SCAI;refGene_tx=XM_023242708.1
D4 86008540 86008714
refGene_gn=SCAI;refGene_tx=XM_023242707.1;refGene_gn=SCAI;refGene_tx=XM_023242708.1;refGene_gn=SCAI;refGene_tx=XM_023242710.1
D4 86012180 86012253
refGene_gn=SCAI;refGene_tx=XM_023242707.1;refGene_gn=SCAI;refGene_tx=XM_023242708.1;refGene_gn=SCAI;refGene_tx=XM_023242710.1
D4 86028839 86028921
refGene_gn=SCAI;refGene_tx=XM_023242707.1;refGene_gn=SCAI;refGene_tx=XM_023242708.1;refGene_gn=SCAI;refGene_tx=XM_023242710.1
D4 86030652 86030752
refGene_gn=SCAI;refGene_tx=XM_023242707.1;refGene_gn=SCAI;refGene_tx=X

M_023242708.1;refGene_gn=SCAI;refGene_tx=XM_023242710.1
D4 86032646 86032725
refGene_gn=SCAI;refGene_tx=XM_023242707.1;refGene_gn=SCAI;refGene_tx=XM_023242708.1;refGene_gn=SCAI;refGene_tx=XM_023242710.1
D4 86033428 86033530
refGene_gn=SCAI;refGene_tx=XM_023242707.1;refGene_gn=SCAI;refGene_tx=XM_023242708.1;refGene_gn=SCAI;refGene_tx=XM_023242710.1
D4 86033783 86033885
refGene_gn=SCAI;refGene_tx=XM_023242707.1;refGene_gn=SCAI;refGene_tx=XM_023242708.1;refGene_gn=SCAI;refGene_tx=XM_023242710.1
D4 86036407 86036560
refGene_gn=SCAI;refGene_tx=XM_023242707.1;refGene_gn=SCAI;refGene_tx=XM_023242708.1;refGene_gn=SCAI;refGene_tx=XM_023242710.1
D4 86036711 86036810
refGene_gn=SCAI;refGene_tx=XM_023242707.1;refGene_gn=SCAI;refGene_tx=XM_023242708.1;refGene_gn=SCAI;refGene_tx=XM_023242710.1
D4 86038082 86038179
refGene_gn=SCAI;refGene_tx=XM_023242707.1;refGene_gn=SCAI;refGene_tx=XM_023242708.1;refGene_gn=SCAI;refGene_tx=XM_023242710.1
D4 86045492 86045591
refGene_gn=SCAI;refGene_tx=XM_023242707.1;refGene_gn=SCAI;refGene_tx=XM_023242708.1;refGene_gn=SCAI;refGene_tx=XM_023242710.1
D4 86046603 86046690
refGene_gn=SCAI;refGene_tx=XM_023242707.1;refGene_gn=SCAI;refGene_tx=XM_023242708.1;refGene_gn=SCAI;refGene_tx=XM_023242710.1
D4 86047219 86047315
refGene_gn=SCAI;refGene_tx=XM_023242707.1;refGene_gn=SCAI;refGene_tx=XM_023242708.1;refGene_gn=SCAI;refGene_tx=XM_023242710.1
D4 86066333 86066465
refGene_gn=SCAI;refGene_tx=XM_023242707.1;refGene_gn=SCAI;refGene_tx=XM_023242708.1;refGene_gn=SCAI;refGene_tx=XM_023242710.1
D4 86142307 86142352
refGene_gn=SCAI;refGene_tx=XM_023242707.1;refGene_gn=SCAI;refGene_tx=XM_023242708.1;refGene_gn=SCAI;refGene_tx=XM_023242710.1
D4 86143076 86143129
refGene_gn=SCAI;refGene_tx=XM_023242707.1;refGene_gn=SCAI;refGene_tx=XM_023242708.1;refGene_gn=SCAI;refGene_tx=XM_023242710.1
D4 86149266 86149515 refGene_gn=PPP6C;refGene_tx=XM_003995865.5
D4 86150386 86150596 refGene_gn=PPP6C;refGene_tx=XM_003995865.5
D4 86150826 86150906 refGene_gn=PPP6C;refGene_tx=XM_003995865.5
D4 86156074 86156216 refGene_gn=PPP6C;refGene_tx=XM_003995865.5
D4 86157223 86157289 refGene_gn=PPP6C;refGene_tx=XM_003995865.5
D4 86161470 86161566 refGene_gn=PPP6C;refGene_tx=XM_003995865.5
D4 86181956 86182031 refGene_gn=PPP6C;refGene_tx=XM_003995865.5
D4 86189782 86189835
refGene_gn=RABEPK;refGene_tx=XM_006939460.3;refGene_gn=RABEPK;refGene_tx=XM_006939461.4;refGene_gn=RABEPK;refGene_tx=XM_019816513.2
D4 86192252 86192410
refGene_gn=RABEPK;refGene_tx=XM_006939460.3;refGene_gn=RABEPK;refGene_tx=XM_006939461.4;refGene_gn=RABEPK;refGene_tx=XM_019816513.2;refGene_

gn=RABEPK;refGene_tx=XM_019816515.2
D4 86196832 86196985
refGene_gn=RABEPK;refGene_tx=XM_006939460.3;refGene_gn=RABEPK;refGene_tx=XM_006939461.4;refGene_gn=RABEPK;refGene_tx=XM_019816513.2;refGene_gn=RABEPK;refGene_tx=XM_019816515.2
D4 86201542 86201704
refGene_gn=RABEPK;refGene_tx=XM_006939460.3;refGene_gn=RABEPK;refGene_tx=XM_006939461.4;refGene_gn=RABEPK;refGene_tx=XM_019816513.2;refGene_gn=RABEPK;refGene_tx=XM_019816515.2
D4 86205103 86205253
refGene_gn=RABEPK;refGene_tx=XM_006939460.3;refGene_gn=RABEPK;refGene_tx=XM_006939461.4;refGene_gn=RABEPK;refGene_tx=XM_019816513.2;refGene_gn=RABEPK;refGene_tx=XM_019816515.2
D4 86207372 86207522
refGene_gn=RABEPK;refGene_tx=XM_006939460.3;refGene_gn=RABEPK;refGene_tx=XM_006939461.4;refGene_gn=RABEPK;refGene_tx=XM_019816513.2;refGene_gn=RABEPK;refGene_tx=XM_019816515.2
D4 86208105 86208398
refGene_gn=RABEPK;refGene_tx=XM_006939460.3;refGene_gn=RABEPK;refGene_tx=XM_006939461.4;refGene_gn=RABEPK;refGene_tx=XM_019816513.2;refGene_gn=RABEPK;refGene_tx=XM_019816515.2
D4 86210011 86210574
refGene_gn=HSPA5;refGene_tx=XM_006939458.4;refGene_gn=HSPA5;refGene_tx=XM_006939459.4
D4 86211266 86211434
refGene_gn=HSPA5;refGene_tx=XM_006939458.4;refGene_gn=HSPA5;refGene_tx=XM_006939459.4
D4 86211755 86211993
refGene_gn=HSPA5;refGene_tx=XM_006939458.4;refGene_gn=HSPA5;refGene_tx=XM_006939459.4
D4 86212119 86212510
refGene_gn=HSPA5;refGene_tx=XM_006939458.4;refGene_gn=HSPA5;refGene_tx=XM_006939459.4
D4 86212610 86212723
refGene_gn=HSPA5;refGene_tx=XM_006939458.4;refGene_gn=HSPA5;refGene_tx=XM_006939459.4
D4 86213050 86213188
refGene_gn=HSPA5;refGene_tx=XM_006939458.4;refGene_gn=HSPA5;refGene_tx=XM_006939459.4
D4 86213607 86213839
refGene_gn=HSPA5;refGene_tx=XM_006939458.4;refGene_gn=HSPA5;refGene_tx=XM_006939459.4
D4 86213941 86214063
refGene_gn=HSPA5;refGene_tx=XM_006939458.4;refGene_gn=HSPA5;refGene_tx=XM_006939459.4
D4 86220015 86220672
refGene_gn=HPRT1;refGene_tx=XM_006939462.4;refGene_gn=HPRT1;refGene_tx=XM_019816516.2;refGene_gn=HPRT1;refGene_tx=XM_019816517.1;refGene_gn=HPRT1;refGene_tx=XM_023242712.1
D4 86253711 86253896

refGene_gn=GAPVD1;refGene_tx=XM_011288444.2;refGene_gn=GAPVD1;refGene_tx=XM_011288445.2;refGene_gn=GAPVD1;refGene_tx=XM_011288446.2;refGene_gn=GAPVD1;refGene_tx=XM_011288447.2;refGene_gn=GAPVD1;refGene_tx=XM_019816519.2;refGene_gn=GAPVD1;refGene_tx=XM_023242714.1;refGene_gn=GAPVD1;refGene_tx=XM_023242715.1
D4 86255703 86256547
refGene_gn=GAPVD1;refGene_tx=XM_011288444.2;refGene_gn=GAPVD1;refGene_tx=XM_011288445.2;refGene_gn=GAPVD1;refGene_tx=XM_011288446.2;refGene_gn=GAPVD1;refGene_tx=XM_011288447.2;refGene_gn=GAPVD1;refGene_tx=XM_019816519.2;refGene_gn=GAPVD1;refGene_tx=XM_023242714.1;refGene_gn=GAPVD1;refGene_tx=XM_023242715.1
D4 86258740 86258827
refGene_gn=GAPVD1;refGene_tx=XM_011288444.2;refGene_gn=GAPVD1;refGene_tx=XM_011288445.2;refGene_gn=GAPVD1;refGene_tx=XM_011288446.2;refGene_gn=GAPVD1;refGene_tx=XM_011288447.2;refGene_gn=GAPVD1;refGene_tx=XM_019816519.2;refGene_gn=GAPVD1;refGene_tx=XM_023242714.1;refGene_gn=GAPVD1;refGene_tx=XM_023242715.1
D4 86260848 86260983
refGene_gn=GAPVD1;refGene_tx=XM_011288444.2;refGene_gn=GAPVD1;refGene_tx=XM_011288445.2;refGene_gn=GAPVD1;refGene_tx=XM_011288446.2;refGene_gn=GAPVD1;refGene_tx=XM_011288447.2;refGene_gn=GAPVD1;refGene_tx=XM_019816519.2;refGene_gn=GAPVD1;refGene_tx=XM_023242714.1;refGene_gn=GAPVD1;refGene_tx=XM_023242715.1
D4 86261128 86261318
refGene_gn=GAPVD1;refGene_tx=XM_011288444.2;refGene_gn=GAPVD1;refGene_tx=XM_011288445.2;refGene_gn=GAPVD1;refGene_tx=XM_011288446.2;refGene_gn=GAPVD1;refGene_tx=XM_011288447.2;refGene_gn=GAPVD1;refGene_tx=XM_019816519.2;refGene_gn=GAPVD1;refGene_tx=XM_023242714.1;refGene_gn=GAPVD1;refGene_tx=XM_023242715.1
D4 86265544 86265705
refGene_gn=GAPVD1;refGene_tx=XM_011288444.2;refGene_gn=GAPVD1;refGene_tx=XM_011288445.2;refGene_gn=GAPVD1;refGene_tx=XM_011288446.2;refGene_gn=GAPVD1;refGene_tx=XM_011288447.2;refGene_gn=GAPVD1;refGene_tx=XM_019816519.2;refGene_gn=GAPVD1;refGene_tx=XM_023242714.1;refGene_gn=GAPVD1;refGene_tx=XM_023242715.1
D4 86268063 86268193
refGene_gn=GAPVD1;refGene_tx=XM_011288444.2;refGene_gn=GAPVD1;refGene_tx=XM_011288445.2;refGene_gn=GAPVD1;refGene_tx=XM_011288446.2;refGene_gn=GAPVD1;refGene_tx=XM_011288447.2;refGene_gn=GAPVD1;refGene_tx=XM_019816519.2;refGene_gn=GAPVD1;refGene_tx=XM_023242714.1;refGene_gn=GAPVD1;refGene_tx=XM_023242715.1
D4 86269331 86269457
refGene_gn=GAPVD1;refGene_tx=XM_011288444.2;refGene_gn=GAPVD1;refGene_tx=XM_011288445.2;refGene_gn=GAPVD1;refGene_tx=XM_011288446.2;refGene_gn=GAPVD1;refGene_tx=XM_011288447.2;refGene_gn=GAPVD1;refGene_tx=XM_019816519.2;refGene_gn=GAPVD1;refGene_tx=XM_023242714.1;refGene_gn=GAPVD1;refGene_tx=XM_023242715.1
D4 86272464 86272638
refGene_gn=GAPVD1;refGene_tx=XM_011288444.2;refGene_gn=GAPVD1;refGene_tx=XM_011288445.2;refGene_gn=GAPVD1;refGene_tx=XM_011288446.2;refGene_

gn=GAPVD1;refGene_tx=XM_011288447.2;refGene_gn=GAPVD1;refGene_tx=XM_01
9816519.2;refGene_gn=GAPVD1;refGene_tx=XM_023242714.1;refGene_gn=GAPVD
1;refGene_tx=XM_023242715.1
D4 86274264 86274405
refGene_gn=GAPVD1;refGene_tx=XM_011288444.2;refGene_gn=GAPVD1;refGene_
tx=XM_011288445.2;refGene_gn=GAPVD1;refGene_tx=XM_011288446.2;refGene_
gn=GAPVD1;refGene_tx=XM_011288447.2;refGene_gn=GAPVD1;refGene_tx=XM_02
3242714.1
D4 86275486 86275621
refGene_gn=GAPVD1;refGene_tx=XM_011288444.2;refGene_gn=GAPVD1;refGene_
tx=XM_011288445.2;refGene_gn=GAPVD1;refGene_tx=XM_011288446.2;refGene_
gn=GAPVD1;refGene_tx=XM_011288447.2;refGene_gn=GAPVD1;refGene_tx=XM_01
9816519.2;refGene_gn=GAPVD1;refGene_tx=XM_023242714.1;refGene_gn=GAPVD
1;refGene_tx=XM_023242715.1
D4 86276092 86276212
refGene_gn=GAPVD1;refGene_tx=XM_011288444.2;refGene_gn=GAPVD1;refGene_
tx=XM_011288445.2;refGene_gn=GAPVD1;refGene_tx=XM_011288446.2;refGene_
gn=GAPVD1;refGene_tx=XM_011288447.2;refGene_gn=GAPVD1;refGene_tx=XM_01
9816519.2;refGene_gn=GAPVD1;refGene_tx=XM_023242714.1;refGene_gn=GAPVD
1;refGene_tx=XM_023242715.1
D4 86280130 86280208
refGene_gn=GAPVD1;refGene_tx=XM_011288444.2;refGene_gn=GAPVD1;refGene_
tx=XM_011288445.2;refGene_gn=GAPVD1;refGene_tx=XM_011288446.2;refGene_
gn=GAPVD1;refGene_tx=XM_011288447.2;refGene_gn=GAPVD1;refGene_tx=XM_01
9816519.2
D4 86280343 86280714
refGene_gn=GAPVD1;refGene_tx=XM_011288444.2;refGene_gn=GAPVD1;refGene_
tx=XM_011288445.2;refGene_gn=GAPVD1;refGene_tx=XM_011288446.2;refGene_
gn=GAPVD1;refGene_tx=XM_011288447.2;refGene_gn=GAPVD1;refGene_tx=XM_01
9816519.2
D4 86281679 86281767
refGene_gn=GAPVD1;refGene_tx=XM_011288444.2;refGene_gn=GAPVD1;refGene_
tx=XM_011288445.2;refGene_gn=GAPVD1;refGene_tx=XM_011288446.2;refGene_
gn=GAPVD1;refGene_tx=XM_011288447.2;refGene_gn=GAPVD1;refGene_tx=XM_01
9816519.2
D4 86282772 86282853
refGene_gn=GAPVD1;refGene_tx=XM_011288444.2;refGene_gn=GAPVD1;refGene_
tx=XM_011288445.2;refGene_gn=GAPVD1;refGene_tx=XM_019816519.2
D4 86286398 86286521
refGene_gn=GAPVD1;refGene_tx=XM_011288444.2;refGene_gn=GAPVD1;refGene_
tx=XM_011288445.2;refGene_gn=GAPVD1;refGene_tx=XM_011288446.2;refGene_
gn=GAPVD1;refGene_tx=XM_011288447.2;refGene_gn=GAPVD1;refGene_tx=XM_01
9816519.2
D4 86288046 86288176
refGene_gn=GAPVD1;refGene_tx=XM_011288444.2;refGene_gn=GAPVD1;refGene_
tx=XM_011288445.2;refGene_gn=GAPVD1;refGene_tx=XM_011288446.2;refGene_
gn=GAPVD1;refGene_tx=XM_011288447.2;refGene_gn=GAPVD1;refGene_tx=XM_01
9816519.2
D4 86289193 86289303
refGene_gn=GAPVD1;refGene_tx=XM_011288444.2;refGene_gn=GAPVD1;refGene_

tx=XM_011288445.2;refGene_gn=GAPVD1;refGene_tx=XM_011288446.2;refGene_gn=GAPVD1;refGene_tx=XM_011288447.2;refGene_gn=GAPVD1;refGene_tx=XM_019816519.2
D4	86289498	86289658
refGene_gn=GAPVD1;refGene_tx=XM_011288444.2;refGene_gn=GAPVD1;refGene_tx=XM_011288445.2;refGene_gn=GAPVD1;refGene_tx=XM_011288446.2;refGene_gn=GAPVD1;refGene_tx=XM_011288447.2;refGene_gn=GAPVD1;refGene_tx=XM_019816519.2
D4	86290502	86290690
refGene_gn=GAPVD1;refGene_tx=XM_011288444.2;refGene_gn=GAPVD1;refGene_tx=XM_011288445.2;refGene_gn=GAPVD1;refGene_tx=XM_011288446.2;refGene_gn=GAPVD1;refGene_tx=XM_011288447.2;refGene_gn=GAPVD1;refGene_tx=XM_019816519.2
D4	86291386	86291600
refGene_gn=GAPVD1;refGene_tx=XM_011288444.2;refGene_gn=GAPVD1;refGene_tx=XM_011288445.2;refGene_gn=GAPVD1;refGene_tx=XM_011288446.2;refGene_gn=GAPVD1;refGene_tx=XM_011288447.2;refGene_gn=GAPVD1;refGene_tx=XM_019816519.2
D4	86293799	86293872
refGene_gn=GAPVD1;refGene_tx=XM_011288444.2;refGene_gn=GAPVD1;refGene_tx=XM_011288445.2;refGene_gn=GAPVD1;refGene_tx=XM_011288446.2;refGene_gn=GAPVD1;refGene_tx=XM_011288447.2;refGene_gn=GAPVD1;refGene_tx=XM_019816519.2
D4	86294799	86294997
refGene_gn=GAPVD1;refGene_tx=XM_011288444.2;refGene_gn=GAPVD1;refGene_tx=XM_011288445.2;refGene_gn=GAPVD1;refGene_tx=XM_011288446.2;refGene_gn=GAPVD1;refGene_tx=XM_011288447.2;refGene_gn=GAPVD1;refGene_tx=XM_019816519.2
D4	86296720	86296861
refGene_gn=GAPVD1;refGene_tx=XM_011288444.2;refGene_gn=GAPVD1;refGene_tx=XM_011288445.2;refGene_gn=GAPVD1;refGene_tx=XM_011288446.2;refGene_gn=GAPVD1;refGene_tx=XM_011288447.2;refGene_gn=GAPVD1;refGene_tx=XM_019816519.2
D4	86331459	86331581
refGene_gn=LOC109493970;refGene_tx=XM_023242317.1
D4	86331685	86332254
refGene_gn=LOC109493970;refGene_tx=XM_023242317.1
D4	86332594	86332676
refGene_gn=LOC109493970;refGene_tx=XM_023242317.1
D4	86333271	86333293
refGene_gn=LOC109493970;refGene_tx=XM_023242317.1
D4	86354507	86354633
refGene_gn=MAPKAP1;refGene_tx=XM_006939465.4;refGene_gn=MAPKAP1;refGene_tx=XM_011288442.3;refGene_gn=MAPKAP1;refGene_tx=XM_019816520.2;refGene_gn=MAPKAP1;refGene_tx=XM_019816521.2;refGene_gn=MAPKAP1;refGene_tx=XM_019816522.2;refGene_gn=MAPKAP1;refGene_tx=XM_019816525.2
D4	86359609	86359707
refGene_gn=MAPKAP1;refGene_tx=XM_006939465.4;refGene_gn=MAPKAP1;refGene_tx=XM_011288442.3;refGene_gn=MAPKAP1;refGene_tx=XM_019816520.2;refGene_gn=MAPKAP1;refGene_tx=XM_019816521.2;refGene_gn=MAPKAP1;refGene_tx=

XM_019816522.2;refGene_gn=MAPKAP1;refGene_tx=XM_019816525.2
D4 86381163 86381301
refGene_gn=MAPKAP1;refGene_tx=XM_006939465.4;refGene_gn=MAPKAP1;refGen
e_tx=XM_011288442.3;refGene_gn=MAPKAP1;refGene_tx=XM_019816520.2;refGe
ne_gn=MAPKAP1;refGene_tx=XM_019816521.2;refGene_gn=MAPKAP1;refGene_tx=
XM_019816525.2
D4 86398318 86398459
refGene_gn=MAPKAP1;refGene_tx=XM_006939465.4;refGene_gn=MAPKAP1;refGen
e_tx=XM_011288442.3;refGene_gn=MAPKAP1;refGene_tx=XM_019816520.2;refGe
ne_gn=MAPKAP1;refGene_tx=XM_019816521.2;refGene_gn=MAPKAP1;refGene_tx=
XM_019816522.2;refGene_gn=MAPKAP1;refGene_tx=XM_019816525.2
D4 86417841 86417949
refGene_gn=MAPKAP1;refGene_tx=XM_006939465.4;refGene_gn=MAPKAP1;refGen
e_tx=XM_019816520.2;refGene_gn=MAPKAP1;refGene_tx=XM_019816521.2;refGe
ne_gn=MAPKAP1;refGene_tx=XM_019816522.2;refGene_gn=MAPKAP1;refGene_tx=
XM_019816525.2
D4 86430170 86430329
refGene_gn=MAPKAP1;refGene_tx=XM_019816520.2;refGene_gn=MAPKAP1;refGen
e_tx=XM_019816521.2;refGene_gn=MAPKAP1;refGene_tx=XM_019816522.2;refGe
ne_gn=MAPKAP1;refGene_tx=XM_019816525.2
D4 86448359 86448409 refGene_gn=MAPKAP1;refGene_tx=XM_019816524.2
D4 86451103 86451213
refGene_gn=MAPKAP1;refGene_tx=XM_006939465.4;refGene_gn=MAPKAP1;refGen
e_tx=XM_011288442.3;refGene_gn=MAPKAP1;refGene_tx=XM_019816520.2;refGe
ne_gn=MAPKAP1;refGene_tx=XM_019816521.2;refGene_gn=MAPKAP1;refGene_tx=
XM_019816522.2;refGene_gn=MAPKAP1;refGene_tx=XM_019816524.2;refGene_gn
=MAPKAP1;refGene_tx=XM_019816525.2
D4 86466249 86466426
refGene_gn=MAPKAP1;refGene_tx=XM_006939465.4;refGene_gn=MAPKAP1;refGen
e_tx=XM_011288442.3;refGene_gn=MAPKAP1;refGene_tx=XM_019816520.2;refGe
ne_gn=MAPKAP1;refGene_tx=XM_019816521.2;refGene_gn=MAPKAP1;refGene_tx=
XM_019816522.2;refGene_gn=MAPKAP1;refGene_tx=XM_019816524.2;refGene_gn
=MAPKAP1;refGene_tx=XM_019816525.2
D4 86490105 86490278
refGene_gn=MAPKAP1;refGene_tx=XM_006939465.4;refGene_gn=MAPKAP1;refGen
e_tx=XM_011288442.3;refGene_gn=MAPKAP1;refGene_tx=XM_019816520.2;refGe
ne_gn=MAPKAP1;refGene_tx=XM_019816521.2;refGene_gn=MAPKAP1;refGene_tx=
XM_019816522.2;refGene_gn=MAPKAP1;refGene_tx=XM_019816524.2;refGene_gn
=MAPKAP1;refGene_tx=XM_019816525.2
D4 86554833 86554982
refGene_gn=MAPKAP1;refGene_tx=XM_006939465.4;refGene_gn=MAPKAP1;refGen
e_tx=XM_011288442.3;refGene_gn=MAPKAP1;refGene_tx=XM_019816520.2;refGe
ne_gn=MAPKAP1;refGene_tx=XM_019816521.2;refGene_gn=MAPKAP1;refGene_tx=
XM_019816522.2;refGene_gn=MAPKAP1;refGene_tx=XM_019816524.2
D4 86566774 86566864
refGene_gn=MAPKAP1;refGene_tx=XM_006939465.4;refGene_gn=MAPKAP1;refGen
e_tx=XM_011288442.3;refGene_gn=MAPKAP1;refGene_tx=XM_019816520.2;refGe
ne_gn=MAPKAP1;refGene_tx=XM_019816521.2;refGene_gn=MAPKAP1;refGene_tx=
XM_019816522.2;refGene_gn=MAPKAP1;refGene_tx=XM_019816524.2
D4 86569295 86569554

refGene_gn=MAPKAP1;refGene_tx=XM_006939465.4;refGene_gn=MAPKAP1;refGene_tx=XM_011288442.3;refGene_gn=MAPKAP1;refGene_tx=XM_019816520.2;refGene_gn=MAPKAP1;refGene_tx=XM_019816521.2;refGene_gn=MAPKAP1;refGene_tx=XM_019816522.2;refGene_gn=MAPKAP1;refGene_tx=XM_019816524.2
D4 86631119 86631319
refGene_gn=PBX3;refGene_tx=XM_023242717.1;refGene_gn=PBX3;refGene_tx=XM_023242719.1
D4 86632258 86632332
refGene_gn=PBX3;refGene_tx=XM_023242717.1;refGene_gn=PBX3;refGene_tx=XM_023242718.1;refGene_gn=PBX3;refGene_tx=XM_023242719.1
D4 86792567 86792809
refGene_gn=PBX3;refGene_tx=XM_023242717.1;refGene_gn=PBX3;refGene_tx=XM_023242718.1;refGene_gn=PBX3;refGene_tx=XM_023242719.1
D4 86803477 86803668
refGene_gn=PBX3;refGene_tx=XM_023242717.1;refGene_gn=PBX3;refGene_tx=XM_023242718.1;refGene_gn=PBX3;refGene_tx=XM_023242719.1
D4 86813802 86813938
refGene_gn=PBX3;refGene_tx=XM_023242717.1;refGene_gn=PBX3;refGene_tx=XM_023242718.1;refGene_gn=PBX3;refGene_tx=XM_023242719.1
D4 86843828 86843994
refGene_gn=PBX3;refGene_tx=XM_023242717.1;refGene_gn=PBX3;refGene_tx=XM_023242718.1;refGene_gn=PBX3;refGene_tx=XM_023242719.1
D4 86845249 86845362
refGene_gn=PBX3;refGene_tx=XM_023242717.1;refGene_gn=PBX3;refGene_tx=XM_023242718.1
D4 86846143 86846233
refGene_gn=PBX3;refGene_tx=XM_023242717.1;refGene_gn=PBX3;refGene_tx=XM_023242718.1;refGene_gn=PBX3;refGene_tx=XM_023242719.1
D4 86849280 86849373
refGene_gn=PBX3;refGene_tx=XM_023242717.1;refGene_gn=PBX3;refGene_tx=XM_023242718.1
D4 87154630 87154705
refGene_gn=MVB12B;refGene_tx=XM_023242720.1;refGene_gn=MVB12B;refGene_tx=XM_023242721.1;refGene_gn=MVB12B;refGene_tx=XM_023242723.1;refGene_gn=MVB12B;refGene_tx=XM_023242724.1;refGene_gn=MVB12B;refGene_tx=XM_023242725.1
D4 87167740 87167863
refGene_gn=MVB12B;refGene_tx=XM_023242720.1;refGene_gn=MVB12B;refGene_tx=XM_023242721.1;refGene_gn=MVB12B;refGene_tx=XM_023242723.1;refGene_gn=MVB12B;refGene_tx=XM_023242724.1;refGene_gn=MVB12B;refGene_tx=XM_023242725.1
D4 87218676 87218784
refGene_gn=MVB12B;refGene_tx=XM_023242720.1;refGene_gn=MVB12B;refGene_tx=XM_023242721.1;refGene_gn=MVB12B;refGene_tx=XM_023242723.1;refGene_gn=MVB12B;refGene_tx=XM_023242724.1;refGene_gn=MVB12B;refGene_tx=XM_023242725.1
D4 87223721 87223818
refGene_gn=MVB12B;refGene_tx=XM_023242720.1;refGene_gn=MVB12B;refGene_tx=XM_023242721.1;refGene_gn=MVB12B;refGene_tx=XM_023242723.1;refGene_gn=MVB12B;refGene_tx=XM_023242724.1;refGene_gn=MVB12B;refGene_tx=XM_02

3242725.1
D4 87231621 87231751
refGene_gn=MVB12B;refGene_tx=XM_023242720.1;refGene_gn=MVB12B;refGene_tx=XM_023242721.1;refGene_gn=MVB12B;refGene_tx=XM_023242723.1;refGene_gn=MVB12B;refGene_tx=XM_023242724.1;refGene_gn=MVB12B;refGene_tx=XM_023242725.1
D4 87235582 87235705
refGene_gn=MVB12B;refGene_tx=XM_023242720.1;refGene_gn=MVB12B;refGene_tx=XM_023242721.1;refGene_gn=MVB12B;refGene_tx=XM_023242723.1;refGene_gn=MVB12B;refGene_tx=XM_023242724.1;refGene_gn=MVB12B;refGene_tx=XM_023242725.1
D4 87265601 87265696
refGene_gn=MVB12B;refGene_tx=XM_023242720.1;refGene_gn=MVB12B;refGene_tx=XM_023242721.1;refGene_gn=MVB12B;refGene_tx=XM_023242723.1;refGene_gn=MVB12B;refGene_tx=XM_023242724.1;refGene_gn=MVB12B;refGene_tx=XM_023242725.1
D4 87303008 87303109 refGene_gn=MVB12B;refGene_tx=XM_023242725.1
D4 87324215 87324271
refGene_gn=MVB12B;refGene_tx=XM_023242720.1;refGene_gn=MVB12B;refGene_tx=XM_023242721.1;refGene_gn=MVB12B;refGene_tx=XM_023242723.1;refGene_gn=MVB12B;refGene_tx=XM_023242724.1
D4 87326846 87326906
refGene_gn=MVB12B;refGene_tx=XM_023242720.1;refGene_gn=MVB12B;refGene_tx=XM_023242721.1;refGene_gn=MVB12B;refGene_tx=XM_023242723.1;refGene_gn=MVB12B;refGene_tx=XM_023242724.1
D4 87329044 87329113 refGene_gn=MVB12B;refGene_tx=XM_023242723.1
D4 87344719 87344806 refGene_gn=MVB12B;refGene_tx=XM_023242721.1
D4 87362612 87362642 refGene_gn=MVB12B;refGene_tx=XM_023242724.1
D4 87375061 87375229 refGene_gn=MVB12B;refGene_tx=XM_023242720.1
D4 87436183 87436322
refGene_gn=LMX1B;refGene_tx=XM_003995881.4;refGene_gn=LMX1B;refGene_tx=XM_003995882.4
D4 87437147 87437334
refGene_gn=LMX1B;refGene_tx=XM_003995881.4;refGene_gn=LMX1B;refGene_tx=XM_003995882.4
D4 87509966 87510199
refGene_gn=LMX1B;refGene_tx=XM_003995881.4;refGene_gn=LMX1B;refGene_tx=XM_003995882.4
D4 87512127 87512309
refGene_gn=LMX1B;refGene_tx=XM_003995881.4;refGene_gn=LMX1B;refGene_tx=XM_003995882.4
D4 87512493 87512571
refGene_gn=LMX1B;refGene_tx=XM_003995881.4;refGene_gn=LMX1B;refGene_tx=XM_003995882.4
D4 87512712 87512779
refGene_gn=LMX1B;refGene_tx=XM_003995881.4;refGene_gn=LMX1B;refGene_tx=XM_003995882.4
D4 87514963 87515128
refGene_gn=LMX1B;refGene_tx=XM_003995881.4;refGene_gn=LMX1B;refGene_tx=XM_003995882.4

D4 87515438 87515596
refGene_gn=LMX1B;refGene_tx=XM_003995881.4;refGene_gn=LMX1B;refGene_tx=XM_003995882.4
D4 87618274 87619678
refGene_gn=ZBTB43;refGene_tx=XM_011288440.3;refGene_gn=ZBTB43;refGene_tx=XM_019816531.2;refGene_gn=ZBTB43;refGene_tx=XM_019816532.2;refGene_gn=ZBTB43;refGene_tx=XM_019816533.2;refGene_gn=ZBTB43;refGene_tx=XM_019816534.2
D4 87625296 87625331 refGene_gn=ZBTB34;refGene_tx=XM_019816535.2
D4 87646849 87646881 refGene_gn=ZBTB34;refGene_tx=XM_006939470.3
D4 87664429 87665954
refGene_gn=ZBTB34;refGene_tx=XM_006939470.3;refGene_gn=ZBTB34;refGene_tx=XM_019816535.2;refGene_gn=ZBTB34;refGene_tx=XM_019816536.2;refGene_gn=ZBTB34;refGene_tx=XM_023242727.1
D4 87732239 87732296
refGene_gn=RALGPS1;refGene_tx=XM_011288438.3;refGene_gn=RALGPS1;refGene_tx=XM_011288439.3;refGene_gn=RALGPS1;refGene_tx=XM_019816547.2;refGene_gn=RALGPS1;refGene_tx=XM_019816549.2;refGene_gn=RALGPS1;refGene_tx=XM_019816550.2;refGene_gn=RALGPS1;refGene_tx=XM_019816554.2;refGene_gn=RALGPS1;refGene_tx=XM_023242731.1;refGene_gn=RALGPS1;refGene_tx=XM_023242733.1;refGene_gn=RALGPS1;refGene_tx=XM_023242734.1;refGene_gn=RALGPS1;refGene_tx=XM_023242735.1
D4 87735636 87735744
refGene_gn=RALGPS1;refGene_tx=XM_011288438.3;refGene_gn=RALGPS1;refGene_tx=XM_011288439.3;refGene_gn=RALGPS1;refGene_tx=XM_019816547.2;refGene_gn=RALGPS1;refGene_tx=XM_019816549.2;refGene_gn=RALGPS1;refGene_tx=XM_019816550.2;refGene_gn=RALGPS1;refGene_tx=XM_019816554.2;refGene_gn=RALGPS1;refGene_tx=XM_023242731.1;refGene_gn=RALGPS1;refGene_tx=XM_023242733.1;refGene_gn=RALGPS1;refGene_tx=XM_023242734.1;refGene_gn=RALGPS1;refGene_tx=XM_023242735.1
D4 87749718 87749769
refGene_gn=RALGPS1;refGene_tx=XM_011288438.3;refGene_gn=RALGPS1;refGene_tx=XM_011288439.3;refGene_gn=RALGPS1;refGene_tx=XM_019816547.2;refGene_gn=RALGPS1;refGene_tx=XM_019816549.2;refGene_gn=RALGPS1;refGene_tx=XM_019816550.2;refGene_gn=RALGPS1;refGene_tx=XM_019816554.2;refGene_gn=RALGPS1;refGene_tx=XM_023242731.1;refGene_gn=RALGPS1;refGene_tx=XM_023242734.1;refGene_gn=RALGPS1;refGene_tx=XM_023242735.1
D4 87782566 87782650
refGene_gn=RALGPS1;refGene_tx=XM_011288438.3;refGene_gn=RALGPS1;refGene_tx=XM_011288439.3;refGene_gn=RALGPS1;refGene_tx=XM_019816547.2;refGene_gn=RALGPS1;refGene_tx=XM_019816549.2;refGene_gn=RALGPS1;refGene_tx=XM_019816550.2;refGene_gn=RALGPS1;refGene_tx=XM_019816554.2;refGene_gn=RALGPS1;refGene_tx=XM_023242731.1;refGene_gn=RALGPS1;refGene_tx=XM_023242733.1;refGene_gn=RALGPS1;refGene_tx=XM_023242734.1;refGene_gn=RALGPS1;refGene_tx=XM_023242735.1
D4 87792391 87792481
refGene_gn=RALGPS1;refGene_tx=XM_011288438.3;refGene_gn=RALGPS1;refGene_tx=XM_011288439.3;refGene_gn=RALGPS1;refGene_tx=XM_019816547.2;refGene_gn=RALGPS1;refGene_tx=XM_019816549.2;refGene_gn=RALGPS1;refGene_tx=XM_019816550.2;refGene_gn=RALGPS1;refGene_tx=XM_019816554.2;refGene_gn

=RALGPS1;refGene_tx=XM_023242731.1;refGene_gn=RALGPS1;refGene_tx=XM_023242733.1;refGene_gn=RALGPS1;refGene_tx=XM_023242734.1;refGene_gn=RALGPS1;refGene_tx=XM_023242735.1
D4 87795051 87795144
refGene_gn=RALGPS1;refGene_tx=XM_011288438.3;refGene_gn=RALGPS1;refGene_tx=XM_011288439.3;refGene_gn=RALGPS1;refGene_tx=XM_019816547.2;refGene_gn=RALGPS1;refGene_tx=XM_019816549.2;refGene_gn=RALGPS1;refGene_tx=XM_019816550.2;refGene_gn=RALGPS1;refGene_tx=XM_019816554.2;refGene_gn=RALGPS1;refGene_tx=XM_023242731.1;refGene_gn=RALGPS1;refGene_tx=XM_023242733.1;refGene_gn=RALGPS1;refGene_tx=XM_023242734.1;refGene_gn=RALGPS1;refGene_tx=XM_023242735.1
D4 87809948 87810075
refGene_gn=RALGPS1;refGene_tx=XM_011288438.3;refGene_gn=RALGPS1;refGene_tx=XM_011288439.3;refGene_gn=RALGPS1;refGene_tx=XM_019816547.2;refGene_gn=RALGPS1;refGene_tx=XM_019816549.2;refGene_gn=RALGPS1;refGene_tx=XM_019816550.2;refGene_gn=RALGPS1;refGene_tx=XM_019816554.2;refGene_gn=RALGPS1;refGene_tx=XM_023242731.1;refGene_gn=RALGPS1;refGene_tx=XM_023242733.1;refGene_gn=RALGPS1;refGene_tx=XM_023242734.1;refGene_gn=RALGPS1;refGene_tx=XM_023242735.1
D4 87828626 87828826
refGene_gn=ANGPTL2;refGene_tx=XM_003995887.5;refGene_gn=ANGPTL2;refGene_tx=XM_006939476.4;refGene_gn=ANGPTL2;refGene_tx=XM_006939478.4;refGene_gn=ANGPTL2;refGene_tx=XM_006939479.4;refGene_gn=ANGPTL2;refGene_tx=XM_006939480.4;refGene_gn=ANGPTL2;refGene_tx=XM_011288433.3;refGene_gn=ANGPTL2;refGene_tx=XM_011288435.3;refGene_gn=ANGPTL2;refGene_tx=XM_011288436.3;refGene_gn=ANGPTL2;refGene_tx=XM_019816556.2
D4 87831228 87831499
refGene_gn=ANGPTL2;refGene_tx=XM_003995887.5;refGene_gn=ANGPTL2;refGene_tx=XM_006939476.4;refGene_gn=ANGPTL2;refGene_tx=XM_006939478.4;refGene_gn=ANGPTL2;refGene_tx=XM_006939479.4;refGene_gn=ANGPTL2;refGene_tx=XM_006939480.4;refGene_gn=ANGPTL2;refGene_tx=XM_011288433.3;refGene_gn=ANGPTL2;refGene_tx=XM_011288435.3;refGene_gn=ANGPTL2;refGene_tx=XM_011288436.3;refGene_gn=ANGPTL2;refGene_tx=XM_019816556.2
D4 87833202 87833396
refGene_gn=ANGPTL2;refGene_tx=XM_003995887.5;refGene_gn=ANGPTL2;refGene_tx=XM_006939476.4;refGene_gn=ANGPTL2;refGene_tx=XM_006939478.4;refGene_gn=ANGPTL2;refGene_tx=XM_006939479.4;refGene_gn=ANGPTL2;refGene_tx=XM_006939480.4;refGene_gn=ANGPTL2;refGene_tx=XM_011288433.3;refGene_gn=ANGPTL2;refGene_tx=XM_011288435.3;refGene_gn=ANGPTL2;refGene_tx=XM_011288436.3;refGene_gn=ANGPTL2;refGene_tx=XM_019816556.2
D4 87847330 87848196
refGene_gn=ANGPTL2;refGene_tx=XM_003995887.5;refGene_gn=ANGPTL2;refGene_tx=XM_006939476.4;refGene_gn=ANGPTL2;refGene_tx=XM_006939478.4;refGene_gn=ANGPTL2;refGene_tx=XM_006939479.4;refGene_gn=ANGPTL2;refGene_tx=XM_006939480.4;refGene_gn=ANGPTL2;refGene_tx=XM_011288433.3;refGene_gn=ANGPTL2;refGene_tx=XM_011288435.3;refGene_gn=ANGPTL2;refGene_tx=XM_011288436.3;refGene_gn=ANGPTL2;refGene_tx=XM_019816556.2
D4 87850289 87850361 refGene_gn=ANGPTL2;refGene_tx=XM_006939476.4
D4 87861799 87861846
refGene_gn=ANGPTL2;refGene_tx=XM_006939476.4;refGene_gn=ANGPTL2;refGen

e_tx=XM_019816556.2
D4 87908440 87908578
refGene_gn=RALGPS1;refGene_tx=XM_011288438.3;refGene_gn=RALGPS1;refGene_tx=XM_011288439.3;refGene_gn=RALGPS1;refGene_tx=XM_019816547.2;refGene_gn=RALGPS1;refGene_tx=XM_019816549.2;refGene_gn=RALGPS1;refGene_tx=XM_019816550.2;refGene_gn=RALGPS1;refGene_tx=XM_019816554.2;refGene_gn=RALGPS1;refGene_tx=XM_023242731.1;refGene_gn=RALGPS1;refGene_tx=XM_023242733.1;refGene_gn=RALGPS1;refGene_tx=XM_023242734.1;refGene_gn=RALGPS1;refGene_tx=XM_023242735.1
D4 87911105 87911199
refGene_gn=RALGPS1;refGene_tx=XM_011288438.3;refGene_gn=RALGPS1;refGene_tx=XM_011288439.3;refGene_gn=RALGPS1;refGene_tx=XM_019816547.2;refGene_gn=RALGPS1;refGene_tx=XM_019816549.2;refGene_gn=RALGPS1;refGene_tx=XM_019816550.2;refGene_gn=RALGPS1;refGene_tx=XM_019816554.2;refGene_gn=RALGPS1;refGene_tx=XM_023242731.1;refGene_gn=RALGPS1;refGene_tx=XM_023242733.1;refGene_gn=RALGPS1;refGene_tx=XM_023242734.1;refGene_gn=RALGPS1;refGene_tx=XM_023242735.1
D4 87916484 87916552
refGene_gn=RALGPS1;refGene_tx=XM_011288438.3;refGene_gn=RALGPS1;refGene_tx=XM_011288439.3;refGene_gn=RALGPS1;refGene_tx=XM_019816547.2;refGene_gn=RALGPS1;refGene_tx=XM_019816549.2;refGene_gn=RALGPS1;refGene_tx=XM_019816550.2;refGene_gn=RALGPS1;refGene_tx=XM_019816554.2;refGene_gn=RALGPS1;refGene_tx=XM_023242731.1;refGene_gn=RALGPS1;refGene_tx=XM_023242733.1;refGene_gn=RALGPS1;refGene_tx=XM_023242734.1;refGene_gn=RALGPS1;refGene_tx=XM_023242735.1
D4 87922867 87922989
refGene_gn=RALGPS1;refGene_tx=XM_023242734.1;refGene_gn=RALGPS1;refGene_tx=XM_023242735.1
D4 87923668 87923697 refGene_gn=RALGPS1;refGene_tx=XM_023242735.1
D4 87934214 87934341
refGene_gn=RALGPS1;refGene_tx=XM_011288438.3;refGene_gn=RALGPS1;refGene_tx=XM_011288439.3;refGene_gn=RALGPS1;refGene_tx=XM_019816547.2;refGene_gn=RALGPS1;refGene_tx=XM_019816549.2;refGene_gn=RALGPS1;refGene_tx=XM_019816550.2;refGene_gn=RALGPS1;refGene_tx=XM_019816554.2;refGene_gn=RALGPS1;refGene_tx=XM_023242731.1;refGene_gn=RALGPS1;refGene_tx=XM_023242733.1
D4 87935590 87935748
refGene_gn=RALGPS1;refGene_tx=XM_011288438.3;refGene_gn=RALGPS1;refGene_tx=XM_019816547.2;refGene_gn=RALGPS1;refGene_tx=XM_019816549.2;refGene_gn=RALGPS1;refGene_tx=XM_019816550.2;refGene_gn=RALGPS1;refGene_tx=XM_023242731.1;refGene_gn=RALGPS1;refGene_tx=XM_023242733.1
D4 87937994 87938046
refGene_gn=RALGPS1;refGene_tx=XM_011288438.3;refGene_gn=RALGPS1;refGene_tx=XM_019816547.2;refGene_gn=RALGPS1;refGene_tx=XM_019816549.2;refGene_gn=RALGPS1;refGene_tx=XM_019816550.2;refGene_gn=RALGPS1;refGene_tx=XM_023242731.1;refGene_gn=RALGPS1;refGene_tx=XM_023242733.1
D4 87949211 87949295
refGene_gn=RALGPS1;refGene_tx=XM_011288438.3;refGene_gn=RALGPS1;refGene_tx=XM_011288439.3;refGene_gn=RALGPS1;refGene_tx=XM_019816547.2;refGene_gn=RALGPS1;refGene_tx=XM_019816549.2;refGene_gn=RALGPS1;refGene_tx=

XM_019816550.2;refGene_gn=RALGPS1;refGene_tx=XM_019816554.2;refGene_gn=RALGPS1;refGene_tx=XM_023242731.1;refGene_gn=RALGPS1;refGene_tx=XM_023242733.1
D4 87950624 87950730 refGene_gn=RALGPS1;refGene_tx=XM_011288438.3;refGene_gn=RALGPS1;refGene_tx=XM_011288439.3;refGene_gn=RALGPS1;refGene_tx=XM_019816547.2;refGene_gn=RALGPS1;refGene_tx=XM_019816549.2;refGene_gn=RALGPS1;refGene_tx=XM_019816550.2;refGene_gn=RALGPS1;refGene_tx=XM_019816554.2;refGene_gn=RALGPS1;refGene_tx=XM_023242731.1;refGene_gn=RALGPS1;refGene_tx=XM_023242733.1
D4 87951696 87951789 refGene_gn=RALGPS1;refGene_tx=XM_011288438.3;refGene_gn=RALGPS1;refGene_tx=XM_011288439.3;refGene_gn=RALGPS1;refGene_tx=XM_019816547.2;refGene_gn=RALGPS1;refGene_tx=XM_019816549.2;refGene_gn=RALGPS1;refGene_tx=XM_019816550.2;refGene_gn=RALGPS1;refGene_tx=XM_019816554.2;refGene_gn=RALGPS1;refGene_tx=XM_023242731.1;refGene_gn=RALGPS1;refGene_tx=XM_023242733.1
D4 87951979 87952085 refGene_gn=RALGPS1;refGene_tx=XM_011288438.3;refGene_gn=RALGPS1;refGene_tx=XM_011288439.3;refGene_gn=RALGPS1;refGene_tx=XM_019816547.2;refGene_gn=RALGPS1;refGene_tx=XM_019816549.2;refGene_gn=RALGPS1;refGene_tx=XM_019816550.2;refGene_gn=RALGPS1;refGene_tx=XM_019816554.2;refGene_gn=RALGPS1;refGene_tx=XM_023242731.1;refGene_gn=RALGPS1;refGene_tx=XM_023242733.1
D4 87954852 87954944 refGene_gn=RALGPS1;refGene_tx=XM_011288438.3;refGene_gn=RALGPS1;refGene_tx=XM_011288439.3;refGene_gn=RALGPS1;refGene_tx=XM_019816547.2;refGene_gn=RALGPS1;refGene_tx=XM_019816549.2;refGene_gn=RALGPS1;refGene_tx=XM_019816550.2;refGene_gn=RALGPS1;refGene_tx=XM_019816554.2;refGene_gn=RALGPS1;refGene_tx=XM_023242731.1;refGene_gn=RALGPS1;refGene_tx=XM_023242733.1
D4 87958930 87958960 refGene_gn=RALGPS1;refGene_tx=XM_011288438.3;refGene_gn=RALGPS1;refGene_tx=XM_011288439.3;refGene_gn=RALGPS1;refGene_tx=XM_019816547.2;refGene_gn=RALGPS1;refGene_tx=XM_019816549.2;refGene_gn=RALGPS1;refGene_tx=XM_019816550.2;refGene_gn=RALGPS1;refGene_tx=XM_019816554.2;refGene_gn=RALGPS1;refGene_tx=XM_023242731.1;refGene_gn=RALGPS1;refGene_tx=XM_023242733.1
D4 87965599 87965691 refGene_gn=GARNL3;refGene_tx=XM_023242728.1
D4 87982529 87982700 refGene_gn=GARNL3;refGene_tx=XM_011288430.2;refGene_gn=GARNL3;refGene_tx=XM_023242728.1;refGene_gn=GARNL3;refGene_tx=XM_023242730.1
D4 88003732 88003740 refGene_gn=GARNL3;refGene_tx=XM_019816540.2;refGene_gn=GARNL3;refGene_tx=XM_019816542.2
D4 88003995 88004000 refGene_gn=GARNL3;refGene_tx=XM_019816541.1
D4 88004728 88004798 refGene_gn=GARNL3;refGene_tx=XM_011288430.2;refGene_gn=GARNL3;refGene_tx=XM_019816540.2;refGene_gn=GARNL3;refGene_tx=XM_019816541.1;refGene_gn=GARNL3;refGene_tx=XM_019816542.2;refGene_gn=GARNL3;refGene_tx=XM_02

3242728.1;refGene_gn=GARNL3;refGene_tx=XM_023242730.1
D4 88032842 88032917
refGene_gn=GARNL3;refGene_tx=XM_011288430.2;refGene_gn=GARNL3;refGene_tx=XM_019816540.2;refGene_gn=GARNL3;refGene_tx=XM_019816541.1;refGene_gn=GARNL3;refGene_tx=XM_023242728.1;refGene_gn=GARNL3;refGene_tx=XM_023242730.1
D4 88046924 88047024
refGene_gn=GARNL3;refGene_tx=XM_011288430.2;refGene_gn=GARNL3;refGene_tx=XM_019816540.2;refGene_gn=GARNL3;refGene_tx=XM_019816541.1;refGene_gn=GARNL3;refGene_tx=XM_019816542.2;refGene_gn=GARNL3;refGene_tx=XM_023242728.1;refGene_gn=GARNL3;refGene_tx=XM_023242730.1
D4 88048596 88048715
refGene_gn=GARNL3;refGene_tx=XM_011288430.2;refGene_gn=GARNL3;refGene_tx=XM_019816540.2;refGene_gn=GARNL3;refGene_tx=XM_019816541.1;refGene_gn=GARNL3;refGene_tx=XM_019816542.2;refGene_gn=GARNL3;refGene_tx=XM_023242728.1;refGene_gn=GARNL3;refGene_tx=XM_023242730.1
D4 88055189 88055254
refGene_gn=GARNL3;refGene_tx=XM_011288430.2;refGene_gn=GARNL3;refGene_tx=XM_019816540.2;refGene_gn=GARNL3;refGene_tx=XM_019816541.1;refGene_gn=GARNL3;refGene_tx=XM_019816542.2;refGene_gn=GARNL3;refGene_tx=XM_023242728.1;refGene_gn=GARNL3;refGene_tx=XM_023242730.1
D4 88058068 88058132
refGene_gn=GARNL3;refGene_tx=XM_011288430.2;refGene_gn=GARNL3;refGene_tx=XM_019816540.2;refGene_gn=GARNL3;refGene_tx=XM_019816541.1;refGene_gn=GARNL3;refGene_tx=XM_019816542.2;refGene_gn=GARNL3;refGene_tx=XM_023242728.1;refGene_gn=GARNL3;refGene_tx=XM_023242730.1
D4 88061117 88061144
refGene_gn=GARNL3;refGene_tx=XM_011288430.2;refGene_gn=GARNL3;refGene_tx=XM_019816540.2;refGene_gn=GARNL3;refGene_tx=XM_019816541.1;refGene_gn=GARNL3;refGene_tx=XM_019816542.2;refGene_gn=GARNL3;refGene_tx=XM_023242728.1;refGene_gn=GARNL3;refGene_tx=XM_023242730.1
D4 88068281 88068357
refGene_gn=GARNL3;refGene_tx=XM_011288430.2;refGene_gn=GARNL3;refGene_tx=XM_019816540.2;refGene_gn=GARNL3;refGene_tx=XM_019816541.1;refGene_gn=GARNL3;refGene_tx=XM_019816542.2;refGene_gn=GARNL3;refGene_tx=XM_023242728.1;refGene_gn=GARNL3;refGene_tx=XM_023242730.1
D4 88069028 88069127
refGene_gn=GARNL3;refGene_tx=XM_011288430.2;refGene_gn=GARNL3;refGene_tx=XM_019816540.2;refGene_gn=GARNL3;refGene_tx=XM_019816541.1;refGene_gn=GARNL3;refGene_tx=XM_019816542.2;refGene_gn=GARNL3;refGene_tx=XM_023242728.1;refGene_gn=GARNL3;refGene_tx=XM_023242730.1
D4 88070561 88070665
refGene_gn=GARNL3;refGene_tx=XM_011288430.2;refGene_gn=GARNL3;refGene_tx=XM_019816540.2;refGene_gn=GARNL3;refGene_tx=XM_019816541.1;refGene_gn=GARNL3;refGene_tx=XM_019816542.2;refGene_gn=GARNL3;refGene_tx=XM_023242728.1;refGene_gn=GARNL3;refGene_tx=XM_023242730.1
D4 88074951 88075060
refGene_gn=GARNL3;refGene_tx=XM_011288430.2;refGene_gn=GARNL3;refGene_tx=XM_019816540.2;refGene_gn=GARNL3;refGene_tx=XM_019816541.1;refGene_gn=GARNL3;refGene_tx=XM_019816542.2;refGene_gn=GARNL3;refGene_tx=XM_02

3242728.1;refGene_gn=GARNL3;refGene_tx=XM_023242730.1
D4 88076966 88077012
refGene_gn=GARNL3;refGene_tx=XM_011288430.2;refGene_gn=GARNL3;refGene_tx=XM_019816540.2;refGene_gn=GARNL3;refGene_tx=XM_019816541.1;refGene_gn=GARNL3;refGene_tx=XM_019816542.2;refGene_gn=GARNL3;refGene_tx=XM_023242728.1;refGene_gn=GARNL3;refGene_tx=XM_023242730.1
D4 88077468 88077575
refGene_gn=GARNL3;refGene_tx=XM_011288430.2;refGene_gn=GARNL3;refGene_tx=XM_019816540.2;refGene_gn=GARNL3;refGene_tx=XM_019816541.1;refGene_gn=GARNL3;refGene_tx=XM_019816542.2;refGene_gn=GARNL3;refGene_tx=XM_023242728.1;refGene_gn=GARNL3;refGene_tx=XM_023242730.1
D4 88080093 88080209
refGene_gn=GARNL3;refGene_tx=XM_011288430.2;refGene_gn=GARNL3;refGene_tx=XM_019816540.2;refGene_gn=GARNL3;refGene_tx=XM_019816541.1;refGene_gn=GARNL3;refGene_tx=XM_019816542.2;refGene_gn=GARNL3;refGene_tx=XM_023242728.1;refGene_gn=GARNL3;refGene_tx=XM_023242730.1
D4 88082047 88082152
refGene_gn=GARNL3;refGene_tx=XM_011288430.2;refGene_gn=GARNL3;refGene_tx=XM_019816540.2;refGene_gn=GARNL3;refGene_tx=XM_019816541.1;refGene_gn=GARNL3;refGene_tx=XM_019816542.2;refGene_gn=GARNL3;refGene_tx=XM_023242728.1;refGene_gn=GARNL3;refGene_tx=XM_023242730.1
D4 88083178 88083253
refGene_gn=GARNL3;refGene_tx=XM_011288430.2;refGene_gn=GARNL3;refGene_tx=XM_019816540.2;refGene_gn=GARNL3;refGene_tx=XM_019816541.1;refGene_gn=GARNL3;refGene_tx=XM_019816542.2;refGene_gn=GARNL3;refGene_tx=XM_023242728.1;refGene_gn=GARNL3;refGene_tx=XM_023242730.1
D4 88086775 88086887
refGene_gn=GARNL3;refGene_tx=XM_011288430.2;refGene_gn=GARNL3;refGene_tx=XM_019816540.2;refGene_gn=GARNL3;refGene_tx=XM_019816541.1;refGene_gn=GARNL3;refGene_tx=XM_019816542.2;refGene_gn=GARNL3;refGene_tx=XM_023242728.1;refGene_gn=GARNL3;refGene_tx=XM_023242730.1
D4 88091429 88091528
refGene_gn=GARNL3;refGene_tx=XM_011288430.2;refGene_gn=GARNL3;refGene_tx=XM_019816540.2;refGene_gn=GARNL3;refGene_tx=XM_019816541.1;refGene_gn=GARNL3;refGene_tx=XM_019816542.2;refGene_gn=GARNL3;refGene_tx=XM_023242728.1;refGene_gn=GARNL3;refGene_tx=XM_023242730.1
D4 88091878 88091995
refGene_gn=GARNL3;refGene_tx=XM_011288430.2;refGene_gn=GARNL3;refGene_tx=XM_019816540.2;refGene_gn=GARNL3;refGene_tx=XM_019816541.1;refGene_gn=GARNL3;refGene_tx=XM_019816542.2;refGene_gn=GARNL3;refGene_tx=XM_023242728.1;refGene_gn=GARNL3;refGene_tx=XM_023242730.1
D4 88092742 88092918
refGene_gn=GARNL3;refGene_tx=XM_011288430.2;refGene_gn=GARNL3;refGene_tx=XM_019816540.2;refGene_gn=GARNL3;refGene_tx=XM_019816541.1;refGene_gn=GARNL3;refGene_tx=XM_019816542.2;refGene_gn=GARNL3;refGene_tx=XM_023242728.1;refGene_gn=GARNL3;refGene_tx=XM_023242730.1
D4 88094377 88094536
refGene_gn=GARNL3;refGene_tx=XM_011288430.2;refGene_gn=GARNL3;refGene_tx=XM_019816540.2;refGene_gn=GARNL3;refGene_tx=XM_019816541.1;refGene_gn=GARNL3;refGene_tx=XM_019816542.2;refGene_gn=GARNL3;refGene_tx=XM_02

3242728.1;refGene_gn=GARNL3;refGene_tx=XM_023242730.1
D4 88100792 88100859
refGene_gn=GARNL3;refGene_tx=XM_011288430.2;refGene_gn=GARNL3;refGene_tx=XM_019816540.2;refGene_gn=GARNL3;refGene_tx=XM_019816541.1;refGene_gn=GARNL3;refGene_tx=XM_019816542.2;refGene_gn=GARNL3;refGene_tx=XM_023242728.1;refGene_gn=GARNL3;refGene_tx=XM_023242730.1
D4 88106228 88106336
refGene_gn=GARNL3;refGene_tx=XM_011288430.2;refGene_gn=GARNL3;refGene_tx=XM_019816540.2;refGene_gn=GARNL3;refGene_tx=XM_019816541.1;refGene_gn=GARNL3;refGene_tx=XM_019816542.2;refGene_gn=GARNL3;refGene_tx=XM_023242728.1;refGene_gn=GARNL3;refGene_tx=XM_023242730.1
D4 88107834 88107953
refGene_gn=GARNL3;refGene_tx=XM_011288430.2;refGene_gn=GARNL3;refGene_tx=XM_019816540.2;refGene_gn=GARNL3;refGene_tx=XM_019816541.1;refGene_gn=GARNL3;refGene_tx=XM_019816542.2;refGene_gn=GARNL3;refGene_tx=XM_023242728.1;refGene_gn=GARNL3;refGene_tx=XM_023242730.1
D4 88110212 88110351
refGene_gn=GARNL3;refGene_tx=XM_011288430.2;refGene_gn=GARNL3;refGene_tx=XM_019816540.2;refGene_gn=GARNL3;refGene_tx=XM_019816541.1;refGene_gn=GARNL3;refGene_tx=XM_019816542.2;refGene_gn=GARNL3;refGene_tx=XM_023242728.1;refGene_gn=GARNL3;refGene_tx=XM_023242730.1
D4 88111146 88111362
refGene_gn=GARNL3;refGene_tx=XM_011288430.2;refGene_gn=GARNL3;refGene_tx=XM_019816540.2;refGene_gn=GARNL3;refGene_tx=XM_019816541.1;refGene_gn=GARNL3;refGene_tx=XM_019816542.2;refGene_gn=GARNL3;refGene_tx=XM_023242728.1;refGene_gn=GARNL3;refGene_tx=XM_023242730.1
D4 88112886 88113013
refGene_gn=GARNL3;refGene_tx=XM_011288430.2;refGene_gn=GARNL3;refGene_tx=XM_019816540.2;refGene_gn=GARNL3;refGene_tx=XM_019816541.1;refGene_gn=GARNL3;refGene_tx=XM_019816542.2;refGene_gn=GARNL3;refGene_tx=XM_023242728.1;refGene_gn=GARNL3;refGene_tx=XM_023242730.1
D4 88114514 88114692
refGene_gn=GARNL3;refGene_tx=XM_011288430.2;refGene_gn=GARNL3;refGene_tx=XM_019816540.2;refGene_gn=GARNL3;refGene_tx=XM_019816541.1;refGene_gn=GARNL3;refGene_tx=XM_019816542.2;refGene_gn=GARNL3;refGene_tx=XM_023242728.1;refGene_gn=GARNL3;refGene_tx=XM_023242730.1
D4 88118187 88118243
refGene_gn=SLC2A8;refGene_tx=XM_019816561.2;refGene_gn=SLC2A8;refGene_tx=XM_019816562.2
D4 88118339 88118502
refGene_gn=SLC2A8;refGene_tx=XM_019816561.2;refGene_gn=SLC2A8;refGene_tx=XM_019816562.2
D4 88118780 88118987
refGene_gn=SLC2A8;refGene_tx=XM_011288428.2;refGene_gn=SLC2A8;refGene_tx=XM_019816561.2;refGene_gn=SLC2A8;refGene_tx=XM_019816562.2
D4 88120585 88120685
refGene_gn=SLC2A8;refGene_tx=XM_011288428.2;refGene_gn=SLC2A8;refGene_tx=XM_019816561.2;refGene_gn=SLC2A8;refGene_tx=XM_019816562.2
D4 88122096 88122296
refGene_gn=SLC2A8;refGene_tx=XM_011288428.2;refGene_gn=SLC2A8;refGene_

tx=XM_019816561.2;refGene_gn=SLC2A8;refGene_tx=XM_019816562.2
D4 88123242 88123386
refGene_gn=SLC2A8;refGene_tx=XM_011288428.2;refGene_gn=SLC2A8;refGene_tx=XM_019816561.2;refGene_gn=SLC2A8;refGene_tx=XM_019816562.2
D4 88123485 88123594
refGene_gn=SLC2A8;refGene_tx=XM_011288428.2;refGene_gn=SLC2A8;refGene_tx=XM_019816561.2;refGene_gn=SLC2A8;refGene_tx=XM_019816562.2
D4 88124259 88124433
refGene_gn=SLC2A8;refGene_tx=XM_011288428.2;refGene_gn=SLC2A8;refGene_tx=XM_019816561.2;refGene_gn=SLC2A8;refGene_tx=XM_019816562.2
D4 88124832 88124978
refGene_gn=SLC2A8;refGene_tx=XM_011288428.2;refGene_gn=SLC2A8;refGene_tx=XM_019816561.2
D4 88126571 88126709
refGene_gn=SLC2A8;refGene_tx=XM_011288428.2;refGene_gn=SLC2A8;refGene_tx=XM_019816561.2;refGene_gn=SLC2A8;refGene_tx=XM_019816562.2
D4 88133022 88133038
refGene_gn=ZNF79;refGene_tx=XM_003995897.4;refGene_gn=ZNF79;refGene_tx=XM_023242737.1
D4 88138034 88138120
refGene_gn=ZNF79;refGene_tx=XM_003995897.4;refGene_gn=ZNF79;refGene_tx=XM_023242737.1
D4 88141113 88141116 refGene_gn=ZNF79;refGene_tx=XM_019816558.2
D4 88142175 88142302
refGene_gn=ZNF79;refGene_tx=XM_003995897.4;refGene_gn=ZNF79;refGene_tx=XM_019816558.2;refGene_gn=ZNF79;refGene_tx=XM_019816559.2;refGene_gn=ZNF79;refGene_tx=XM_023242737.1
D4 88142808 88142907
refGene_gn=ZNF79;refGene_tx=XM_003995897.4;refGene_gn=ZNF79;refGene_tx=XM_006939495.2;refGene_gn=ZNF79;refGene_tx=XM_011288427.2;refGene_gn=ZNF79;refGene_tx=XM_019816558.2;refGene_gn=ZNF79;refGene_tx=XM_019816559.2;refGene_gn=ZNF79;refGene_tx=XM_019816560.2;refGene_gn=ZNF79;refGene_tx=XM_023242737.1;refGene_gn=ZNF79;refGene_tx=XM_023242739.1
D4 88145291 88146463
refGene_gn=ZNF79;refGene_tx=XM_003995897.4;refGene_gn=ZNF79;refGene_tx=XM_006939495.2;refGene_gn=ZNF79;refGene_tx=XM_011288427.2;refGene_gn=ZNF79;refGene_tx=XM_019816558.2;refGene_gn=ZNF79;refGene_tx=XM_019816559.2;refGene_gn=ZNF79;refGene_tx=XM_019816560.2;refGene_gn=ZNF79;refGene_tx=XM_023242737.1;refGene_gn=ZNF79;refGene_tx=XM_023242739.1
D4 88148417 88148423 refGene_gn=RPL12;refGene_tx=XM_003995898.3
D4 88148590 88148703 refGene_gn=RPL12;refGene_tx=XM_003995898.3
D4 88149076 88149163 refGene_gn=RPL12;refGene_tx=XM_003995898.3
D4 88149753 88149835 refGene_gn=RPL12;refGene_tx=XM_003995898.3
D4 88150212 88150311 refGene_gn=RPL12;refGene_tx=XM_003995898.3
D4 88151398 88151472 refGene_gn=RPL12;refGene_tx=XM_003995898.3
D4 88152245 88152282 refGene_gn=RPL12;refGene_tx=XM_003995898.3
D4 88154361 88154433
refGene_gn=LRSAM1;refGene_tx=XM_011288422.2;refGene_gn=LRSAM1;refGene_tx=XM_011288424.2
D4 88154850 88154907

refGene_gn=LRSAM1;refGene_tx=XM_011288422.2;refGene_gn=LRSAM1;refGene_
tx=XM_011288424.2
D4 88155693 88155738
refGene_gn=LRSAM1;refGene_tx=XM_011288422.2;refGene_gn=LRSAM1;refGene_
tx=XM_011288424.2
D4 88156743 88156821
refGene_gn=LRSAM1;refGene_tx=XM_011288422.2;refGene_gn=LRSAM1;refGene_
tx=XM_011288424.2
D4 88158423 88158492
refGene_gn=LRSAM1;refGene_tx=XM_011288422.2;refGene_gn=LRSAM1;refGene_
tx=XM_011288424.2
D4 88160179 88160264
refGene_gn=LRSAM1;refGene_tx=XM_011288422.2;refGene_gn=LRSAM1;refGene_
tx=XM_011288424.2
D4 88161266 88161388
refGene_gn=LRSAM1;refGene_tx=XM_011288422.2;refGene_gn=LRSAM1;refGene_
tx=XM_011288424.2
D4 88164227 88164318
refGene_gn=LRSAM1;refGene_tx=XM_011288422.2;refGene_gn=LRSAM1;refGene_
tx=XM_011288424.2
D4 88167261 88167392
refGene_gn=LRSAM1;refGene_tx=XM_011288422.2;refGene_gn=LRSAM1;refGene_
tx=XM_011288424.2
D4 88171538 88171568
refGene_gn=LRSAM1;refGene_tx=XM_011288422.2;refGene_gn=LRSAM1;refGene_
tx=XM_011288424.2
D4 88172017 88172140
refGene_gn=LRSAM1;refGene_tx=XM_011288422.2;refGene_gn=LRSAM1;refGene_
tx=XM_011288424.2
D4 88172426 88172566
refGene_gn=LRSAM1;refGene_tx=XM_011288422.2;refGene_gn=LRSAM1;refGene_
tx=XM_011288424.2
D4 88173486 88173531
refGene_gn=LRSAM1;refGene_tx=XM_011288422.2;refGene_gn=LRSAM1;refGene_
tx=XM_011288424.2
D4 88175470 88175541
refGene_gn=LRSAM1;refGene_tx=XM_011288422.2;refGene_gn=LRSAM1;refGene_
tx=XM_011288424.2
D4 88178376 88178476
refGene_gn=LRSAM1;refGene_tx=XM_011288422.2;refGene_gn=LRSAM1;refGene_
tx=XM_011288424.2
D4 88179898 88179986
refGene_gn=LRSAM1;refGene_tx=XM_011288422.2;refGene_gn=LRSAM1;refGene_
tx=XM_011288424.2
D4 88180944 88181019
refGene_gn=LRSAM1;refGene_tx=XM_011288422.2;refGene_gn=LRSAM1;refGene_
tx=XM_011288424.2
D4 88182430 88182511 refGene_gn=LRSAM1;refGene_tx=XM_011288422.2
D4 88184045 88184141
refGene_gn=LRSAM1;refGene_tx=XM_011288422.2;refGene_gn=LRSAM1;refGene_

tx=XM_011288424.2
D4 88186082 88186181
refGene_gn=LRSAM1;refGene_tx=XM_011288422.2;refGene_gn=LRSAM1;refGene_tx=XM_011288424.2
D4 88186972 88187104
refGene_gn=LRSAM1;refGene_tx=XM_011288422.2;refGene_gn=LRSAM1;refGene_tx=XM_011288424.2
D4 88188223 88188305
refGene_gn=LRSAM1;refGene_tx=XM_011288422.2;refGene_gn=LRSAM1;refGene_tx=XM_011288424.2
D4 88190432 88190581
refGene_gn=LRSAM1;refGene_tx=XM_011288422.2;refGene_gn=LRSAM1;refGene_tx=XM_011288424.2
D4 88191520 88191646
refGene_gn=LRSAM1;refGene_tx=XM_011288422.2;refGene_gn=LRSAM1;refGene_tx=XM_011288424.2
D4 88195393 88195977
refGene_gn=FAM129B;refGene_tx=XM_006939496.4;refGene_gn=FAM129B;refGene_tx=XM_019816564.2
D4 88196388 88196500
refGene_gn=FAM129B;refGene_tx=XM_006939496.4;refGene_gn=FAM129B;refGene_tx=XM_019816564.2
D4 88196623 88196731
refGene_gn=FAM129B;refGene_tx=XM_006939496.4;refGene_gn=FAM129B;refGene_tx=XM_019816564.2
D4 88196877 88196994
refGene_gn=FAM129B;refGene_tx=XM_006939496.4;refGene_gn=FAM129B;refGene_tx=XM_019816564.2
D4 88197496 88197652
refGene_gn=FAM129B;refGene_tx=XM_006939496.4;refGene_gn=FAM129B;refGene_tx=XM_019816564.2
D4 88198448 88198636
refGene_gn=FAM129B;refGene_tx=XM_006939496.4;refGene_gn=FAM129B;refGene_tx=XM_019816564.2
D4 88199860 88200023
refGene_gn=FAM129B;refGene_tx=XM_006939496.4;refGene_gn=FAM129B;refGene_tx=XM_019816564.2
D4 88200101 88200206
refGene_gn=FAM129B;refGene_tx=XM_006939496.4;refGene_gn=FAM129B;refGene_tx=XM_019816564.2
D4 88200733 88200849
refGene_gn=FAM129B;refGene_tx=XM_006939496.4;refGene_gn=FAM129B;refGene_tx=XM_019816564.2
D4 88205557 88205725
refGene_gn=FAM129B;refGene_tx=XM_006939496.4;refGene_gn=FAM129B;refGene_tx=XM_019816564.2
D4 88206985 88207091
refGene_gn=FAM129B;refGene_tx=XM_006939496.4;refGene_gn=FAM129B;refGene_tx=XM_019816564.2
D4 88208983 88209112

refGene_gn=FAM129B;refGene_tx=XM_006939496.4;refGene_gn=FAM129B;refGene_tx=XM_019816564.2
D4 88214413 88214544
refGene_gn=FAM129B;refGene_tx=XM_006939496.4;refGene_gn=FAM129B;refGene_tx=XM_019816564.2
D4 88225917 88226077 refGene_gn=FAM129B;refGene_tx=XM_019816564.2
D4 88247210 88247265 refGene_gn=FAM129B;refGene_tx=XM_006939496.4
D4 88284964 88285001
refGene_gn=STXBP1;refGene_tx=XM_011288419.2;refGene_gn=STXBP1;refGene_tx=XM_011288421.2;refGene_gn=STXBP1;refGene_tx=XM_019816566.1
D4 88321648 88321698
refGene_gn=STXBP1;refGene_tx=XM_011288419.2;refGene_gn=STXBP1;refGene_tx=XM_011288421.2;refGene_gn=STXBP1;refGene_tx=XM_019816566.1
D4 88323315 88323397
refGene_gn=STXBP1;refGene_tx=XM_011288419.2;refGene_gn=STXBP1;refGene_tx=XM_011288421.2;refGene_gn=STXBP1;refGene_tx=XM_019816566.1
D4 88327913 88327990
refGene_gn=STXBP1;refGene_tx=XM_011288419.2;refGene_gn=STXBP1;refGene_tx=XM_011288421.2;refGene_gn=STXBP1;refGene_tx=XM_019816566.1
D4 88328847 88328926
refGene_gn=STXBP1;refGene_tx=XM_011288419.2;refGene_gn=STXBP1;refGene_tx=XM_011288421.2;refGene_gn=STXBP1;refGene_tx=XM_019816566.1
D4 88330112 88330216
refGene_gn=STXBP1;refGene_tx=XM_011288419.2;refGene_gn=STXBP1;refGene_tx=XM_011288421.2;refGene_gn=STXBP1;refGene_tx=XM_019816566.1
D4 88332161 88332310
refGene_gn=STXBP1;refGene_tx=XM_011288419.2;refGene_gn=STXBP1;refGene_tx=XM_011288421.2;refGene_gn=STXBP1;refGene_tx=XM_019816566.1
D4 88334698 88334783
refGene_gn=STXBP1;refGene_tx=XM_011288419.2;refGene_gn=STXBP1;refGene_tx=XM_011288421.2;refGene_gn=STXBP1;refGene_tx=XM_019816566.1
D4 88335571 88335702
refGene_gn=STXBP1;refGene_tx=XM_011288419.2;refGene_gn=STXBP1;refGene_tx=XM_011288421.2;refGene_gn=STXBP1;refGene_tx=XM_019816566.1
D4 88336792 88336900
refGene_gn=STXBP1;refGene_tx=XM_011288419.2;refGene_gn=STXBP1;refGene_tx=XM_011288421.2;refGene_gn=STXBP1;refGene_tx=XM_019816566.1
D4 88338362 88338423
refGene_gn=STXBP1;refGene_tx=XM_011288419.2;refGene_gn=STXBP1;refGene_tx=XM_011288421.2;refGene_gn=STXBP1;refGene_tx=XM_019816566.1
D4 88339788 88339854
refGene_gn=STXBP1;refGene_tx=XM_011288419.2;refGene_gn=STXBP1;refGene_tx=XM_011288421.2;refGene_gn=STXBP1;refGene_tx=XM_019816566.1
D4 88340594 88340675
refGene_gn=STXBP1;refGene_tx=XM_011288419.2;refGene_gn=STXBP1;refGene_tx=XM_011288421.2;refGene_gn=STXBP1;refGene_tx=XM_019816566.1
D4 88342818 88342957
refGene_gn=STXBP1;refGene_tx=XM_011288419.2;refGene_gn=STXBP1;refGene_tx=XM_011288421.2;refGene_gn=STXBP1;refGene_tx=XM_019816566.1
D4 88343637 88343747

refGene_gn=STXBP1;refGene_tx=XM_011288419.2;refGene_gn=STXBP1;refGene_tx=XM_011288421.2;refGene_gn=STXBP1;refGene_tx=XM_019816566.1
D4 88344777 88344879
refGene_gn=STXBP1;refGene_tx=XM_011288419.2;refGene_gn=STXBP1;refGene_tx=XM_011288421.2;refGene_gn=STXBP1;refGene_tx=XM_019816566.1
D4 88347261 88347347
refGene_gn=STXBP1;refGene_tx=XM_011288419.2;refGene_gn=STXBP1;refGene_tx=XM_011288421.2;refGene_gn=STXBP1;refGene_tx=XM_019816566.1
D4 88350128 88350283
refGene_gn=STXBP1;refGene_tx=XM_011288419.2;refGene_gn=STXBP1;refGene_tx=XM_011288421.2
D4 88351843 88351953 refGene_gn=STXBP1;refGene_tx=XM_011288419.2
D4 88357390 88357473
refGene_gn=STXBP1;refGene_tx=XM_011288421.2;refGene_gn=STXBP1;refGene_tx=XM_019816566.1
D4 88364728 88364901 refGene_gn=CFAP157;refGene_tx=XM_003995921.5
D4 88367283 88367555 refGene_gn=CFAP157;refGene_tx=XM_003995921.5
D4 88368293 88368447 refGene_gn=CFAP157;refGene_tx=XM_003995921.5
D4 88368866 88369134 refGene_gn=CFAP157;refGene_tx=XM_003995921.5
D4 88369540 88369671 refGene_gn=CFAP157;refGene_tx=XM_003995921.5
D4 88369880 88370031 refGene_gn=CFAP157;refGene_tx=XM_003995921.5
D4 88370403 88370570 refGene_gn=CFAP157;refGene_tx=XM_003995921.5
D4 88370736 88370923 refGene_gn=CFAP157;refGene_tx=XM_003995921.5
D4 88371212 88371284 refGene_gn=CFAP157;refGene_tx=XM_003995921.5
D4 88371455 88371662
refGene_gn=PTRH1;refGene_tx=XM_003995922.5;refGene_gn=PTRH1;refGene_tx=XM_019816584.1
D4 88371766 88371812
refGene_gn=PTRH1;refGene_tx=XM_003995922.5;refGene_gn=PTRH1;refGene_tx=XM_019816584.1
D4 88371992 88372208
refGene_gn=PTRH1;refGene_tx=XM_003995922.5;refGene_gn=PTRH1;refGene_tx=XM_019816584.1
D4 88372355 88372575
refGene_gn=PTRH1;refGene_tx=XM_003995922.5;refGene_gn=PTRH1;refGene_tx=XM_019816584.1
D4 88372937 88373103
refGene_gn=PTRH1;refGene_tx=XM_003995922.5;refGene_gn=PTRH1;refGene_tx=XM_019816584.1
D4 88374307 88374432
refGene_gn=TTC16;refGene_tx=XM_011288418.3;refGene_gn=TTC16;refGene_tx=XM_019816579.2
D4 88374572 88374852
refGene_gn=TTC16;refGene_tx=XM_011288418.3;refGene_gn=TTC16;refGene_tx=XM_019816576.1;refGene_gn=TTC16;refGene_tx=XM_019816579.2;refGene_gn=TTC16;refGene_tx=XM_019816580.1;refGene_gn=TTC16;refGene_tx=XM_019816583.1
D4 88375002 88375146
refGene_gn=TTC16;refGene_tx=XM_011288418.3;refGene_gn=TTC16;refGene_tx=XM_019816576.1;refGene_gn=TTC16;refGene_tx=XM_019816583.1

D4 88376115 88376216
refGene_gn=TTC16;refGene_tx=XM_011288418.3;refGene_gn=TTC16;refGene_tx=XM_019816576.1;refGene_gn=TTC16;refGene_tx=XM_019816579.2;refGene_gn=TTC16;refGene_tx=XM_019816580.1;refGene_gn=TTC16;refGene_tx=XM_019816583.1
D4 88376307 88376437
refGene_gn=TTC16;refGene_tx=XM_011288418.3;refGene_gn=TTC16;refGene_tx=XM_019816576.1;refGene_gn=TTC16;refGene_tx=XM_019816579.2;refGene_gn=TTC16;refGene_tx=XM_019816582.2;refGene_gn=TTC16;refGene_tx=XM_019816583.1
D4 88378551 88378766
refGene_gn=TTC16;refGene_tx=XM_011288418.3;refGene_gn=TTC16;refGene_tx=XM_019816576.1;refGene_gn=TTC16;refGene_tx=XM_019816579.2;refGene_gn=TTC16;refGene_tx=XM_019816580.1;refGene_gn=TTC16;refGene_tx=XM_019816582.2;refGene_gn=TTC16;refGene_tx=XM_019816583.1
D4 88379426 88379671
refGene_gn=TTC16;refGene_tx=XM_011288418.3;refGene_gn=TTC16;refGene_tx=XM_019816576.1;refGene_gn=TTC16;refGene_tx=XM_019816579.2;refGene_gn=TTC16;refGene_tx=XM_019816580.1;refGene_gn=TTC16;refGene_tx=XM_019816582.2;refGene_gn=TTC16;refGene_tx=XM_019816583.1
D4 88380025 88380167
refGene_gn=TTC16;refGene_tx=XM_011288418.3;refGene_gn=TTC16;refGene_tx=XM_019816576.1;refGene_gn=TTC16;refGene_tx=XM_019816579.2;refGene_gn=TTC16;refGene_tx=XM_019816580.1;refGene_gn=TTC16;refGene_tx=XM_019816582.2;refGene_gn=TTC16;refGene_tx=XM_019816583.1
D4 88382463 88382629
refGene_gn=TTC16;refGene_tx=XM_011288418.3;refGene_gn=TTC16;refGene_tx=XM_019816576.1;refGene_gn=TTC16;refGene_tx=XM_019816579.2;refGene_gn=TTC16;refGene_tx=XM_019816580.1;refGene_gn=TTC16;refGene_tx=XM_019816582.2;refGene_gn=TTC16;refGene_tx=XM_019816583.1
D4 88383160 88383303
refGene_gn=TTC16;refGene_tx=XM_011288418.3;refGene_gn=TTC16;refGene_tx=XM_019816576.1;refGene_gn=TTC16;refGene_tx=XM_019816579.2;refGene_gn=TTC16;refGene_tx=XM_019816580.1;refGene_gn=TTC16;refGene_tx=XM_019816582.2;refGene_gn=TTC16;refGene_tx=XM_019816583.1
D4 88383465 88383646
refGene_gn=TTC16;refGene_tx=XM_011288418.3;refGene_gn=TTC16;refGene_tx=XM_019816576.1;refGene_gn=TTC16;refGene_tx=XM_019816579.2;refGene_gn=TTC16;refGene_tx=XM_019816580.1;refGene_gn=TTC16;refGene_tx=XM_019816582.2;refGene_gn=TTC16;refGene_tx=XM_019816583.1
D4 88385070 88385158
refGene_gn=TTC16;refGene_tx=XM_011288418.3;refGene_gn=TTC16;refGene_tx=XM_019816576.1;refGene_gn=TTC16;refGene_tx=XM_019816579.2;refGene_gn=TTC16;refGene_tx=XM_019816580.1;refGene_gn=TTC16;refGene_tx=XM_019816582.2;refGene_gn=TTC16;refGene_tx=XM_019816583.1
D4 88385562 88386200
refGene_gn=TTC16;refGene_tx=XM_011288418.3;refGene_gn=TTC16;refGene_tx=XM_019816576.1;refGene_gn=TTC16;refGene_tx=XM_019816579.2;refGene_gn=TTC16;refGene_tx=XM_019816580.1;refGene_gn=TTC16;refGene_tx=XM_019816582.2;refGene_gn=TTC16;refGene_tx=XM_019816583.1

D4 88386951 88387196 refGene_gn=TOR2A;refGene_tx=XM_003995900.5
D4 88387372 88387500 refGene_gn=TOR2A;refGene_tx=XM_003995900.5
D4 88388189 88388365 refGene_gn=TOR2A;refGene_tx=XM_003995900.5
D4 88389133 88389399 refGene_gn=TOR2A;refGene_tx=XM_003995900.5
D4 88389954 88390105 refGene_gn=TOR2A;refGene_tx=XM_003995900.5
D4 88392554 88392730
refGene_gn=SH2D3C;refGene_tx=XM_003995901.5;refGene_gn=SH2D3C;refGene_tx=XM_019816568.2;refGene_gn=SH2D3C;refGene_tx=XM_019816569.1;refGene_gn=SH2D3C;refGene_tx=XM_019816570.2;refGene_gn=SH2D3C;refGene_tx=XM_019816572.2;refGene_gn=SH2D3C;refGene_tx=XM_019816573.2;refGene_gn=SH2D3C;refGene_tx=XM_019816574.2;refGene_gn=SH2D3C;refGene_tx=XM_019816575.1;refGene_gn=SH2D3C;refGene_tx=XM_023242741.1;refGene_gn=SH2D3C;refGene_tx=XM_023242742.1;refGene_gn=SH2D3C;refGene_tx=XM_023242743.1;refGene_gn=SH2D3C;refGene_tx=XM_023242744.1
D4 88393536 88393743
refGene_gn=SH2D3C;refGene_tx=XM_003995901.5;refGene_gn=SH2D3C;refGene_tx=XM_019816568.2;refGene_gn=SH2D3C;refGene_tx=XM_019816569.1;refGene_gn=SH2D3C;refGene_tx=XM_019816570.2;refGene_gn=SH2D3C;refGene_tx=XM_019816572.2;refGene_gn=SH2D3C;refGene_tx=XM_019816573.2;refGene_gn=SH2D3C;refGene_tx=XM_019816574.2;refGene_gn=SH2D3C;refGene_tx=XM_019816575.1;refGene_gn=SH2D3C;refGene_tx=XM_023242741.1;refGene_gn=SH2D3C;refGene_tx=XM_023242742.1;refGene_gn=SH2D3C;refGene_tx=XM_023242743.1;refGene_gn=SH2D3C;refGene_tx=XM_023242744.1
D4 88394112 88394224
refGene_gn=SH2D3C;refGene_tx=XM_003995901.5;refGene_gn=SH2D3C;refGene_tx=XM_019816568.2;refGene_gn=SH2D3C;refGene_tx=XM_019816569.1;refGene_gn=SH2D3C;refGene_tx=XM_019816570.2;refGene_gn=SH2D3C;refGene_tx=XM_019816572.2;refGene_gn=SH2D3C;refGene_tx=XM_019816573.2;refGene_gn=SH2D3C;refGene_tx=XM_019816574.2;refGene_gn=SH2D3C;refGene_tx=XM_019816575.1;refGene_gn=SH2D3C;refGene_tx=XM_023242741.1;refGene_gn=SH2D3C;refGene_tx=XM_023242742.1;refGene_gn=SH2D3C;refGene_tx=XM_023242743.1;refGene_gn=SH2D3C;refGene_tx=XM_023242744.1
D4 88395019 88395191
refGene_gn=SH2D3C;refGene_tx=XM_003995901.5;refGene_gn=SH2D3C;refGene_tx=XM_019816568.2;refGene_gn=SH2D3C;refGene_tx=XM_019816569.1;refGene_gn=SH2D3C;refGene_tx=XM_019816570.2;refGene_gn=SH2D3C;refGene_tx=XM_019816572.2;refGene_gn=SH2D3C;refGene_tx=XM_019816573.2;refGene_gn=SH2D3C;refGene_tx=XM_019816574.2;refGene_gn=SH2D3C;refGene_tx=XM_019816575.1;refGene_gn=SH2D3C;refGene_tx=XM_023242741.1;refGene_gn=SH2D3C;refGene_tx=XM_023242742.1;refGene_gn=SH2D3C;refGene_tx=XM_023242743.1;refGene_gn=SH2D3C;refGene_tx=XM_023242744.1
D4 88395745 88395861
refGene_gn=SH2D3C;refGene_tx=XM_003995901.5;refGene_gn=SH2D3C;refGene_tx=XM_019816568.2;refGene_gn=SH2D3C;refGene_tx=XM_019816569.1;refGene_gn=SH2D3C;refGene_tx=XM_019816570.2;refGene_gn=SH2D3C;refGene_tx=XM_019816572.2;refGene_gn=SH2D3C;refGene_tx=XM_019816573.2;refGene_gn=SH2D3C;refGene_tx=XM_019816574.2;refGene_gn=SH2D3C;refGene_tx=XM_019816575.1;refGene_gn=SH2D3C;refGene_tx=XM_023242741.1;refGene_gn=SH2D3C;refGene_tx=XM_023242742.1;refGene_gn=SH2D3C;refGene_tx=XM_023242743.1;refGene_gn=SH2D3C;refGene_tx=XM_023242744.1

D4 88396792 88397328
refGene_gn=SH2D3C;refGene_tx=XM_003995901.5;refGene_gn=SH2D3C;refGene_tx=XM_019816568.2;refGene_gn=SH2D3C;refGene_tx=XM_019816569.1;refGene_gn=SH2D3C;refGene_tx=XM_019816570.2;refGene_gn=SH2D3C;refGene_tx=XM_019816572.2;refGene_gn=SH2D3C;refGene_tx=XM_019816573.2;refGene_gn=SH2D3C;refGene_tx=XM_019816574.2;refGene_gn=SH2D3C;refGene_tx=XM_019816575.1;refGene_gn=SH2D3C;refGene_tx=XM_023242741.1;refGene_gn=SH2D3C;refGene_tx=XM_023242742.1;refGene_gn=SH2D3C;refGene_tx=XM_023242743.1;refGene_gn=SH2D3C;refGene_tx=XM_023242744.1
D4 88398895 88399035
refGene_gn=SH2D3C;refGene_tx=XM_003995901.5;refGene_gn=SH2D3C;refGene_tx=XM_019816568.2;refGene_gn=SH2D3C;refGene_tx=XM_019816569.1;refGene_gn=SH2D3C;refGene_tx=XM_019816570.2;refGene_gn=SH2D3C;refGene_tx=XM_019816572.2;refGene_gn=SH2D3C;refGene_tx=XM_019816573.2;refGene_gn=SH2D3C;refGene_tx=XM_019816574.2;refGene_gn=SH2D3C;refGene_tx=XM_019816575.1;refGene_gn=SH2D3C;refGene_tx=XM_023242741.1;refGene_gn=SH2D3C;refGene_tx=XM_023242742.1;refGene_gn=SH2D3C;refGene_tx=XM_023242743.1;refGene_gn=SH2D3C;refGene_tx=XM_023242744.1
D4 88400279 88400734
refGene_gn=SH2D3C;refGene_tx=XM_003995901.5;refGene_gn=SH2D3C;refGene_tx=XM_019816568.2;refGene_gn=SH2D3C;refGene_tx=XM_019816569.1;refGene_gn=SH2D3C;refGene_tx=XM_019816570.2;refGene_gn=SH2D3C;refGene_tx=XM_019816572.2;refGene_gn=SH2D3C;refGene_tx=XM_019816573.2;refGene_gn=SH2D3C;refGene_tx=XM_019816574.2;refGene_gn=SH2D3C;refGene_tx=XM_019816575.1;refGene_gn=SH2D3C;refGene_tx=XM_023242741.1;refGene_gn=SH2D3C;refGene_tx=XM_023242742.1;refGene_gn=SH2D3C;refGene_tx=XM_023242743.1;refGene_gn=SH2D3C;refGene_tx=XM_023242744.1
D4 88401851 88401980
refGene_gn=SH2D3C;refGene_tx=XM_003995901.5;refGene_gn=SH2D3C;refGene_tx=XM_019816568.2;refGene_gn=SH2D3C;refGene_tx=XM_019816569.1;refGene_gn=SH2D3C;refGene_tx=XM_019816570.2;refGene_gn=SH2D3C;refGene_tx=XM_019816572.2;refGene_gn=SH2D3C;refGene_tx=XM_019816573.2;refGene_gn=SH2D3C;refGene_tx=XM_019816574.2;refGene_gn=SH2D3C;refGene_tx=XM_019816575.1;refGene_gn=SH2D3C;refGene_tx=XM_023242741.1;refGene_gn=SH2D3C;refGene_tx=XM_023242742.1;refGene_gn=SH2D3C;refGene_tx=XM_023242743.1;refGene_gn=SH2D3C;refGene_tx=XM_023242744.1
D4 88405706 88405988
refGene_gn=SH2D3C;refGene_tx=XM_023242741.1;refGene_gn=SH2D3C;refGene_tx=XM_023242742.1;refGene_gn=SH2D3C;refGene_tx=XM_023242743.1;refGene_gn=SH2D3C;refGene_tx=XM_023242744.1
D4 88406506 88406596
refGene_gn=SH2D3C;refGene_tx=XM_019816569.1;refGene_gn=SH2D3C;refGene_tx=XM_019816570.2;refGene_gn=SH2D3C;refGene_tx=XM_019816572.2;refGene_gn=SH2D3C;refGene_tx=XM_019816573.2
D4 88408929 88408969
refGene_gn=SH2D3C;refGene_tx=XM_003995901.5;refGene_gn=SH2D3C;refGene_tx=XM_019816568.2;refGene_gn=SH2D3C;refGene_tx=XM_019816574.2
D4 88418169 88418213
refGene_gn=SH2D3C;refGene_tx=XM_019816574.2;refGene_gn=SH2D3C;refGene_tx=XM_019816575.1

D4	88420003	88420475
refGene_gn=SH2D3C;refGene_tx=XM_003995901.5;refGene_gn=SH2D3C;refGene_tx=XM_019816568.2
D4	88422372	88422409
refGene_gn=SH2D3C;refGene_tx=XM_003995901.5;refGene_gn=SH2D3C;refGene_tx=XM_019816568.2
D4	88428921	88429013	refGene_gn=CDK9;refGene_tx=XM_023242746.1
D4	88429486	88429568	refGene_gn=CDK9;refGene_tx=XM_023242746.1
D4	88430344	88430435	refGene_gn=CDK9;refGene_tx=XM_023242746.1
D4	88430793	88430960	refGene_gn=CDK9;refGene_tx=XM_023242746.1
D4	88431078	88431250	refGene_gn=CDK9;refGene_tx=XM_023242746.1
D4	88431407	88431556	refGene_gn=CDK9;refGene_tx=XM_023242746.1
D4	88432049	88432415	refGene_gn=CDK9;refGene_tx=XM_023242746.1
D4	88437667	88437814	refGene_gn=FPGS;refGene_tx=XM_023242747.1
D4	88446027	88446165
refGene_gn=FPGS;refGene_tx=XM_003995903.5;refGene_gn=FPGS;refGene_tx=XM_023242748.1
D4	88447513	88447642
refGene_gn=FPGS;refGene_tx=XM_003995903.5;refGene_gn=FPGS;refGene_tx=XM_023242747.1
D4	88447714	88447789
refGene_gn=FPGS;refGene_tx=XM_003995903.5;refGene_gn=FPGS;refGene_tx=XM_006939502.3;refGene_gn=FPGS;refGene_tx=XM_023242747.1;refGene_gn=FPGS;refGene_tx=XM_023242748.1
D4	88447873	88447938
refGene_gn=FPGS;refGene_tx=XM_003995903.5;refGene_gn=FPGS;refGene_tx=XM_006939502.3;refGene_gn=FPGS;refGene_tx=XM_023242747.1;refGene_gn=FPGS;refGene_tx=XM_023242748.1
D4	88449531	88449646
refGene_gn=FPGS;refGene_tx=XM_003995903.5;refGene_gn=FPGS;refGene_tx=XM_006939502.3;refGene_gn=FPGS;refGene_tx=XM_023242747.1;refGene_gn=FPGS;refGene_tx=XM_023242748.1
D4	88449781	88449862
refGene_gn=FPGS;refGene_tx=XM_003995903.5;refGene_gn=FPGS;refGene_tx=XM_006939502.3;refGene_gn=FPGS;refGene_tx=XM_023242747.1;refGene_gn=FPGS;refGene_tx=XM_023242748.1
D4	88449954	88450016
refGene_gn=FPGS;refGene_tx=XM_003995903.5;refGene_gn=FPGS;refGene_tx=XM_006939502.3;refGene_gn=FPGS;refGene_tx=XM_023242747.1;refGene_gn=FPGS;refGene_tx=XM_023242748.1
D4	88450102	88450205
refGene_gn=FPGS;refGene_tx=XM_003995903.5;refGene_gn=FPGS;refGene_tx=XM_006939502.3;refGene_gn=FPGS;refGene_tx=XM_023242747.1;refGene_gn=FPGS;refGene_tx=XM_023242748.1
D4	88450480	88450558
refGene_gn=FPGS;refGene_tx=XM_003995903.5;refGene_gn=FPGS;refGene_tx=XM_006939502.3;refGene_gn=FPGS;refGene_tx=XM_023242747.1;refGene_gn=FPGS;refGene_tx=XM_023242748.1
D4	88450819	88450967
refGene_gn=FPGS;refGene_tx=XM_003995903.5;refGene_gn=FPGS;refGene_tx=X

M_006939502.3;refGene_gn=FPGS;refGene_tx=XM_023242747.1;refGene_gn=FPGS;refGene_tx=XM_023242748.1
D4 88451092 88451182 refGene_gn=FPGS;refGene_tx=XM_003995903.5;refGene_gn=FPGS;refGene_tx=XM_006939502.3;refGene_gn=FPGS;refGene_tx=XM_023242747.1;refGene_gn=FPGS;refGene_tx=XM_023242748.1
D4 88451926 88452077 refGene_gn=FPGS;refGene_tx=XM_003995903.5;refGene_gn=FPGS;refGene_tx=XM_006939502.3;refGene_gn=FPGS;refGene_tx=XM_023242747.1;refGene_gn=FPGS;refGene_tx=XM_023242748.1
D4 88452330 88452406 refGene_gn=FPGS;refGene_tx=XM_003995903.5;refGene_gn=FPGS;refGene_tx=XM_006939502.3;refGene_gn=FPGS;refGene_tx=XM_023242747.1;refGene_gn=FPGS;refGene_tx=XM_023242748.1
D4 88453250 88453317 refGene_gn=FPGS;refGene_tx=XM_003995903.5;refGene_gn=FPGS;refGene_tx=XM_006939502.3;refGene_gn=FPGS;refGene_tx=XM_023242747.1;refGene_gn=FPGS;refGene_tx=XM_023242748.1
D4 88454056 88454442 refGene_gn=FPGS;refGene_tx=XM_003995903.5;refGene_gn=FPGS;refGene_tx=XM_006939502.3;refGene_gn=FPGS;refGene_tx=XM_023242747.1;refGene_gn=FPGS;refGene_tx=XM_023242748.1
D4 88455857 88455982 refGene_gn=ENG;refGene_tx=XM_006939503.3
D4 88456111 88456213 refGene_gn=ENG;refGene_tx=XM_006939503.3
D4 88457269 88457324 refGene_gn=ENG;refGene_tx=XM_006939503.3
D4 88458625 88458877 refGene_gn=ENG;refGene_tx=XM_006939503.3
D4 88459150 88459267 refGene_gn=ENG;refGene_tx=XM_006939503.3
D4 88459795 88459834 refGene_gn=ENG;refGene_tx=XM_006939503.3
D4 88460056 88460194 refGene_gn=ENG;refGene_tx=XM_006939503.3
D4 88463873 88464016 refGene_gn=ENG;refGene_tx=XM_006939503.3
D4 88464250 88464425 refGene_gn=ENG;refGene_tx=XM_006939503.3
D4 88464856 88464983 refGene_gn=ENG;refGene_tx=XM_006939503.3
D4 88465264 88465430 refGene_gn=ENG;refGene_tx=XM_006939503.3
D4 88466109 88466275 refGene_gn=ENG;refGene_tx=XM_006939503.3
D4 88468856 88468997 refGene_gn=ENG;refGene_tx=XM_006939503.3
D4 88477960 88478103 refGene_gn=ENG;refGene_tx=XM_006939503.3
D4 88486099 88486166 refGene_gn=ENG;refGene_tx=XM_006939503.3
D4 88496332 88496401 refGene_gn=AK1;refGene_tx=XM_003995906.4;refGene_gn=AK1;refGene_tx=XM_003995908.5;refGene_gn=AK1;refGene_tx=XM_006939504.4
D4 88496652 88496844 refGene_gn=AK1;refGene_tx=XM_003995906.4;refGene_gn=AK1;refGene_tx=XM_003995908.5;refGene_gn=AK1;refGene_tx=XM_006939504.4
D4 88499060 88499177 refGene_gn=AK1;refGene_tx=XM_003995906.4;refGene_gn=AK1;refGene_tx=XM_003995908.5;refGene_gn=AK1;refGene_tx=XM_006939504.4
D4 88499877 88500041 refGene_gn=AK1;refGene_tx=XM_003995906.4;refGene_gn=AK1;refGene_tx=XM_003995908.5;refGene_gn=AK1;refGene_tx=XM_006939504.4
D4 88500211 88500247

refGene_gn=AK1;refGene_tx=XM_003995906.4;refGene_gn=AK1;refGene_tx=XM_003995908.5;refGene_gn=AK1;refGene_tx=XM_006939504.4
D4 88500411 88500466 refGene_gn=AK1;refGene_tx=XM_003995908.5
D4 88501638 88501645
refGene_gn=AK1;refGene_tx=XM_003995906.4;refGene_gn=AK1;refGene_tx=XM_006939504.4
D4 88511508 88511906
refGene_gn=ST6GALNAC6;refGene_tx=XM_019816591.2;refGene_gn=ST6GALNAC6;refGene_tx=XM_019816593.1;refGene_gn=ST6GALNAC6;refGene_tx=XM_023242749.1;refGene_gn=ST6GALNAC6;refGene_tx=XM_023242750.1;refGene_gn=ST6GALNAC6;refGene_tx=XM_023242751.1;refGene_gn=ST6GALNAC6;refGene_tx=XM_023242752.1;refGene_gn=ST6GALNAC6;refGene_tx=XM_023242753.1;refGene_gn=ST6GALNAC6;refGene_tx=XM_023242754.1
D4 88512248 88512252
refGene_gn=ST6GALNAC6;refGene_tx=XM_023242755.1
D4 88512565 88512673
refGene_gn=ST6GALNAC6;refGene_tx=XM_019816591.2;refGene_gn=ST6GALNAC6;refGene_tx=XM_019816593.1;refGene_gn=ST6GALNAC6;refGene_tx=XM_023242749.1;refGene_gn=ST6GALNAC6;refGene_tx=XM_023242750.1;refGene_gn=ST6GALNAC6;refGene_tx=XM_023242751.1;refGene_gn=ST6GALNAC6;refGene_tx=XM_023242752.1;refGene_gn=ST6GALNAC6;refGene_tx=XM_023242753.1;refGene_gn=ST6GALNAC6;refGene_tx=XM_023242754.1;refGene_gn=ST6GALNAC6;refGene_tx=XM_023242755.1
D4 88515472 88515879
refGene_gn=ST6GALNAC6;refGene_tx=XM_019816591.2;refGene_gn=ST6GALNAC6;refGene_tx=XM_019816593.1;refGene_gn=ST6GALNAC6;refGene_tx=XM_023242749.1;refGene_gn=ST6GALNAC6;refGene_tx=XM_023242750.1;refGene_gn=ST6GALNAC6;refGene_tx=XM_023242751.1;refGene_gn=ST6GALNAC6;refGene_tx=XM_023242752.1;refGene_gn=ST6GALNAC6;refGene_tx=XM_023242753.1;refGene_gn=ST6GALNAC6;refGene_tx=XM_023242754.1;refGene_gn=ST6GALNAC6;refGene_tx=XM_023242755.1
D4 88518328 88518508
refGene_gn=ST6GALNAC6;refGene_tx=XM_019816591.2;refGene_gn=ST6GALNAC6;refGene_tx=XM_023242749.1;refGene_gn=ST6GALNAC6;refGene_tx=XM_023242750.1;refGene_gn=ST6GALNAC6;refGene_tx=XM_023242751.1;refGene_gn=ST6GALNAC6;refGene_tx=XM_023242752.1;refGene_gn=ST6GALNAC6;refGene_tx=XM_023242753.1;refGene_gn=ST6GALNAC6;refGene_tx=XM_023242754.1;refGene_gn=ST6GALNAC6;refGene_tx=XM_023242755.1
D4 88520337 88520428
refGene_gn=ST6GALNAC6;refGene_tx=XM_019816591.2;refGene_gn=ST6GALNAC6;refGene_tx=XM_023242749.1;refGene_gn=ST6GALNAC6;refGene_tx=XM_023242750.1;refGene_gn=ST6GALNAC6;refGene_tx=XM_023242751.1;refGene_gn=ST6GALNAC6;refGene_tx=XM_023242752.1;refGene_gn=ST6GALNAC6;refGene_tx=XM_023242753.1;refGene_gn=ST6GALNAC6;refGene_tx=XM_023242754.1;refGene_gn=ST6GALNAC6;refGene_tx=XM_023242755.1
D4 88522061 88522087
refGene_gn=ST6GALNAC6;refGene_tx=XM_023242749.1;refGene_gn=ST6GALNAC6;refGene_tx=XM_023242754.1;refGene_gn=ST6GALNAC6;refGene_tx=XM_023242755.1
D4 88529960 88530150

refGene_gn=ST6GALNAC4;refGene_tx=XM_003995910.5
D4 88531008 88531116
refGene_gn=ST6GALNAC4;refGene_tx=XM_003995910.5
D4 88532874 88533287
refGene_gn=ST6GALNAC4;refGene_tx=XM_003995910.5
D4 88536870 88537056
refGene_gn=ST6GALNAC4;refGene_tx=XM_003995910.5
D4 88538578 88538590
refGene_gn=ST6GALNAC4;refGene_tx=XM_003995910.5
D4 88543243 88543511 refGene_gn=PIP5KL1;refGene_tx=XM_023242757.1
D4 88543951 88544153 refGene_gn=PIP5KL1;refGene_tx=XM_023242758.1
D4 88545551 88545705
refGene_gn=PIP5KL1;refGene_tx=XM_023242757.1;refGene_gn=PIP5KL1;refGen
e_tx=XM_023242758.1
D4 88546055 88546168
refGene_gn=PIP5KL1;refGene_tx=XM_023242757.1;refGene_gn=PIP5KL1;refGen
e_tx=XM_023242758.1
D4 88547165 88547221
refGene_gn=PIP5KL1;refGene_tx=XM_023242757.1;refGene_gn=PIP5KL1;refGen
e_tx=XM_023242758.1
D4 88547311 88547346
refGene_gn=PIP5KL1;refGene_tx=XM_023242757.1;refGene_gn=PIP5KL1;refGen
e_tx=XM_023242758.1
D4 88547848 88547973
refGene_gn=PIP5KL1;refGene_tx=XM_023242757.1;refGene_gn=PIP5KL1;refGen
e_tx=XM_023242758.1
D4 88548245 88548400
refGene_gn=PIP5KL1;refGene_tx=XM_023242757.1;refGene_gn=PIP5KL1;refGen
e_tx=XM_023242758.1
D4 88548606 88548657
refGene_gn=PIP5KL1;refGene_tx=XM_023242757.1;refGene_gn=PIP5KL1;refGen
e_tx=XM_023242758.1
D4 88549755 88549959
refGene_gn=PIP5KL1;refGene_tx=XM_023242757.1;refGene_gn=PIP5KL1;refGen
e_tx=XM_023242758.1
D4 88550770 88550800
refGene_gn=PIP5KL1;refGene_tx=XM_023242757.1;refGene_gn=PIP5KL1;refGen
e_tx=XM_023242758.1
D4 88554192 88554251 refGene_gn=DPM2;refGene_tx=XM_006939507.3
D4 88555074 88555177 refGene_gn=DPM2;refGene_tx=XM_006939507.3
D4 88555963 88556053 refGene_gn=DPM2;refGene_tx=XM_006939507.3
D4 88556346 88556349 refGene_gn=DPM2;refGene_tx=XM_006939507.3
D4 88561022 88561068 refGene_gn=FAM102A;refGene_tx=XM_003995912.5
D4 88562475 88562545 refGene_gn=FAM102A;refGene_tx=XM_003995912.5
D4 88562651 88562773 refGene_gn=FAM102A;refGene_tx=XM_003995912.5
D4 88563225 88563421 refGene_gn=FAM102A;refGene_tx=XM_003995912.5
D4 88563504 88563609 refGene_gn=FAM102A;refGene_tx=XM_003995912.5
D4 88565646 88565801 refGene_gn=FAM102A;refGene_tx=XM_003995912.5
D4 88565928 88565975 refGene_gn=FAM102A;refGene_tx=XM_003995912.5
D4 88568170 88568287 refGene_gn=FAM102A;refGene_tx=XM_003995912.5

D4	88570258 88570288 refGene_gn=FAM102A;refGene_tx=XM_003995912.5
D4	88570550 88570671 refGene_gn=FAM102A;refGene_tx=XM_003995912.5
D4	88593363 88593509 refGene_gn=FAM102A;refGene_tx=XM_003995912.5
D4	88646242 88646715 refGene_gn=NAIF1;refGene_tx=XM_019816594.2
D4	88648402 88648913 refGene_gn=NAIF1;refGene_tx=XM_019816594.2
D4	88650114 88650369
refGene_gn=SLC25A25;refGene_tx=XM_023242759.1;refGene_gn=SLC25A25;refGene_tx=XM_023242760.1
D4	88667781 88668000
refGene_gn=SLC25A25;refGene_tx=XM_003995915.5;refGene_gn=SLC25A25;refGene_tx=XM_019816595.2
D4	88668597 88668615 refGene_gn=SLC25A25;refGene_tx=XM_019816597.2
D4	88674443 88674602
refGene_gn=SLC25A25;refGene_tx=XM_003995914.5;refGene_gn=SLC25A25;refGene_tx=XM_019816596.2
D4	88677014 88677141
refGene_gn=SLC25A25;refGene_tx=XM_003995914.5;refGene_gn=SLC25A25;refGene_tx=XM_003995915.5;refGene_gn=SLC25A25;refGene_tx=XM_019816595.2;refGene_gn=SLC25A25;refGene_tx=XM_019816596.2;refGene_gn=SLC25A25;refGene_tx=XM_019816597.2;refGene_gn=SLC25A25;refGene_tx=XM_023242759.1;refGene_gn=SLC25A25;refGene_tx=XM_023242760.1
D4	88677232 88677320
refGene_gn=SLC25A25;refGene_tx=XM_003995914.5;refGene_gn=SLC25A25;refGene_tx=XM_003995915.5;refGene_gn=SLC25A25;refGene_tx=XM_019816595.2;refGene_gn=SLC25A25;refGene_tx=XM_019816596.2;refGene_gn=SLC25A25;refGene_tx=XM_019816597.2;refGene_gn=SLC25A25;refGene_tx=XM_023242759.1;refGene_gn=SLC25A25;refGene_tx=XM_023242760.1
D4	88677968 88678004
refGene_gn=SLC25A25;refGene_tx=XM_019816595.2;refGene_gn=SLC25A25;refGene_tx=XM_019816596.2;refGene_gn=SLC25A25;refGene_tx=XM_019816597.2;refGene_gn=SLC25A25;refGene_tx=XM_023242759.1
D4	88678266 88678378
refGene_gn=SLC25A25;refGene_tx=XM_003995914.5;refGene_gn=SLC25A25;refGene_tx=XM_003995915.5;refGene_gn=SLC25A25;refGene_tx=XM_019816595.2;refGene_gn=SLC25A25;refGene_tx=XM_019816596.2;refGene_gn=SLC25A25;refGene_tx=XM_019816597.2;refGene_gn=SLC25A25;refGene_tx=XM_023242759.1;refGene_gn=SLC25A25;refGene_tx=XM_023242760.1
D4	88679548 88679707
refGene_gn=SLC25A25;refGene_tx=XM_003995914.5;refGene_gn=SLC25A25;refGene_tx=XM_003995915.5;refGene_gn=SLC25A25;refGene_tx=XM_019816595.2;refGene_gn=SLC25A25;refGene_tx=XM_019816596.2;refGene_gn=SLC25A25;refGene_tx=XM_019816597.2;refGene_gn=SLC25A25;refGene_tx=XM_023242759.1;refGene_gn=SLC25A25;refGene_tx=XM_023242760.1
D4	88681248 88681401
refGene_gn=SLC25A25;refGene_tx=XM_003995914.5;refGene_gn=SLC25A25;refGene_tx=XM_003995915.5;refGene_gn=SLC25A25;refGene_tx=XM_019816595.2;refGene_gn=SLC25A25;refGene_tx=XM_019816596.2;refGene_gn=SLC25A25;refGene_tx=XM_019816597.2;refGene_gn=SLC25A25;refGene_tx=XM_023242759.1;refGene_gn=SLC25A25;refGene_tx=XM_023242760.1
D4	88681680 88681788

refGene_gn=SLC25A25;refGene_tx=XM_003995914.5;refGene_gn=SLC25A25;refGene_tx=XM_003995915.5;refGene_gn=SLC25A25;refGene_tx=XM_019816595.2;refGene_gn=SLC25A25;refGene_tx=XM_019816596.2;refGene_gn=SLC25A25;refGene_tx=XM_019816597.2;refGene_gn=SLC25A25;refGene_tx=XM_023242759.1;refGene_gn=SLC25A25;refGene_tx=XM_023242760.1
D4 88681901 88682069
refGene_gn=SLC25A25;refGene_tx=XM_003995914.5;refGene_gn=SLC25A25;refGene_tx=XM_003995915.5;refGene_gn=SLC25A25;refGene_tx=XM_019816595.2;refGene_gn=SLC25A25;refGene_tx=XM_019816596.2;refGene_gn=SLC25A25;refGene_tx=XM_019816597.2;refGene_gn=SLC25A25;refGene_tx=XM_023242759.1;refGene_gn=SLC25A25;refGene_tx=XM_023242760.1
D4 88682585 88682736
refGene_gn=SLC25A25;refGene_tx=XM_003995914.5;refGene_gn=SLC25A25;refGene_tx=XM_003995915.5;refGene_gn=SLC25A25;refGene_tx=XM_019816595.2;refGene_gn=SLC25A25;refGene_tx=XM_019816596.2;refGene_gn=SLC25A25;refGene_tx=XM_019816597.2;refGene_gn=SLC25A25;refGene_tx=XM_023242759.1;refGene_gn=SLC25A25;refGene_tx=XM_023242760.1
D4 88682839 88683024
refGene_gn=SLC25A25;refGene_tx=XM_003995914.5;refGene_gn=SLC25A25;refGene_tx=XM_003995915.5;refGene_gn=SLC25A25;refGene_tx=XM_019816595.2;refGene_gn=SLC25A25;refGene_tx=XM_019816596.2;refGene_gn=SLC25A25;refGene_tx=XM_019816597.2;refGene_gn=SLC25A25;refGene_tx=XM_023242759.1;refGene_gn=SLC25A25;refGene_tx=XM_023242760.1
D4 88695391 88695514
refGene_gn=PTGES2;refGene_tx=XM_011288407.2;refGene_gn=PTGES2;refGene_tx=XM_011288408.3;refGene_gn=PTGES2;refGene_tx=XM_023242761.1
D4 88696183 88696301
refGene_gn=PTGES2;refGene_tx=XM_011288407.2;refGene_gn=PTGES2;refGene_tx=XM_011288408.3;refGene_gn=PTGES2;refGene_tx=XM_023242761.1
D4 88696689 88696890
refGene_gn=PTGES2;refGene_tx=XM_011288407.2;refGene_gn=PTGES2;refGene_tx=XM_011288408.3;refGene_gn=PTGES2;refGene_tx=XM_023242761.1
D4 88697170 88697320
refGene_gn=PTGES2;refGene_tx=XM_011288407.2;refGene_gn=PTGES2;refGene_tx=XM_011288408.3;refGene_gn=PTGES2;refGene_tx=XM_023242761.1
D4 88697711 88697770
refGene_gn=PTGES2;refGene_tx=XM_011288407.2;refGene_gn=PTGES2;refGene_tx=XM_011288408.3
D4 88698458 88698656
refGene_gn=PTGES2;refGene_tx=XM_011288407.2;refGene_gn=PTGES2;refGene_tx=XM_011288408.3
D4 88699892 88700162
refGene_gn=PTGES2;refGene_tx=XM_011288407.2;refGene_gn=PTGES2;refGene_tx=XM_011288408.3
D4 88709016 88709154
refGene_gn=LOC101098498;refGene_tx=XM_003995927.3;refGene_gn=LOC101098498;refGene_tx=XM_011288406.2
D4 88709666 88709803
refGene_gn=LOC101098498;refGene_tx=XM_003995927.3;refGene_gn=LOC101098498;refGene_tx=XM_011288406.2

D4 88711203 88711283
refGene_gn=LOC101098498;refGene_tx=XM_003995927.3;refGene_gn=LOC101098
498;refGene_tx=XM_011288406.2
D4 88711426 88711546
refGene_gn=LOC101098498;refGene_tx=XM_003995927.3;refGene_gn=LOC101098
498;refGene_tx=XM_011288406.2
D4 88711765 88711867
refGene_gn=LOC101098498;refGene_tx=XM_003995927.3;refGene_gn=LOC101098
498;refGene_tx=XM_011288406.2
D4 88712573 88712593
refGene_gn=LOC101098498;refGene_tx=XM_003995927.3
D4 88722816 88722954
refGene_gn=LOC101086640;refGene_tx=XM_019816608.1;refGene_gn=LOC101086
640;refGene_tx=XM_019816609.1
D4 88723472 88723609
refGene_gn=LOC101086640;refGene_tx=XM_019816608.1;refGene_gn=LOC101086
640;refGene_tx=XM_019816609.1
D4 88724570 88724650
refGene_gn=LOC101086640;refGene_tx=XM_019816608.1;refGene_gn=LOC101086
640;refGene_tx=XM_019816609.1
D4 88724793 88724913
refGene_gn=LOC101086640;refGene_tx=XM_019816608.1;refGene_gn=LOC101086
640;refGene_tx=XM_019816609.1
D4 88725222 88725324
refGene_gn=LOC101086640;refGene_tx=XM_019816608.1;refGene_gn=LOC101086
640;refGene_tx=XM_019816609.1
D4 88726018 88726038
refGene_gn=LOC101086640;refGene_tx=XM_019816608.1;refGene_gn=LOC101086
640;refGene_tx=XM_019816609.1
D4 88743222 88743360
refGene_gn=LOC101091057;refGene_tx=XM_023243242.1
D4 88743883 88744020
refGene_gn=LOC101091057;refGene_tx=XM_023243242.1
D4 88745394 88745478
refGene_gn=LOC101091057;refGene_tx=XM_023243242.1
D4 88745603 88745737
refGene_gn=LOC101091057;refGene_tx=XM_023243242.1
D4 88745949 88746104
refGene_gn=LOC101091057;refGene_tx=XM_023243242.1
D4 88746745 88746765
refGene_gn=LOC101098498;refGene_tx=XM_011288406.2
D4 88756636 88756774
refGene_gn=LOC101091317;refGene_tx=XM_006939511.3
D4 88757294 88757431
refGene_gn=LOC101091317;refGene_tx=XM_006939511.3
D4 88758843 88758923
refGene_gn=LOC101091317;refGene_tx=XM_006939511.3
D4 88759066 88759186
refGene_gn=LOC101091317;refGene_tx=XM_006939511.3
D4 88759406 88759505

refGene_gn=LOC101091317;refGene_tx=XM_006939511.3
D4 88760198 88760218
refGene_gn=LOC101091317;refGene_tx=XM_006939511.3
D4 88778900 88779038
refGene_gn=LOC101091569;refGene_tx=XM_003995991.3
D4 88779556 88779693
refGene_gn=LOC101091569;refGene_tx=XM_003995991.3
D4 88781087 88781167
refGene_gn=LOC101091569;refGene_tx=XM_003995991.3
D4 88781312 88781432
refGene_gn=LOC101091569;refGene_tx=XM_003995991.3
D4 88781652 88781754
refGene_gn=LOC101091569;refGene_tx=XM_003995991.3
D4 88782444 88782464
refGene_gn=LOC101091569;refGene_tx=XM_003995991.3
D4 88793749 88793887
refGene_gn=LOC101091822;refGene_tx=XM_019816614.2;refGene_gn=LOC101091822;refGene_tx=XM_019816615.2
D4 88794402 88794539
refGene_gn=LOC101091822;refGene_tx=XM_019816614.2;refGene_gn=LOC101091822;refGene_tx=XM_019816615.2
D4 88795937 88796017
refGene_gn=LOC101091822;refGene_tx=XM_019816614.2;refGene_gn=LOC101091822;refGene_tx=XM_019816615.2
D4 88796160 88796280
refGene_gn=LOC101091822;refGene_tx=XM_019816614.2;refGene_gn=LOC101091822;refGene_tx=XM_019816615.2
D4 88796501 88796603
refGene_gn=LOC101091822;refGene_tx=XM_019816614.2;refGene_gn=LOC101091822;refGene_tx=XM_019816615.2
D4 88797295 88797315
refGene_gn=LOC101091822;refGene_tx=XM_019816614.2;refGene_gn=LOC101091822;refGene_tx=XM_019816615.2
D4 88817323 88817461
refGene_gn=LOC101092062;refGene_tx=XM_003995993.3
D4 88817980 88818117
refGene_gn=LOC101092062;refGene_tx=XM_003995993.3
D4 88819511 88819591
refGene_gn=LOC101092062;refGene_tx=XM_003995993.3
D4 88819734 88819854
refGene_gn=LOC101092062;refGene_tx=XM_003995993.3
D4 88820071 88820173
refGene_gn=LOC101092062;refGene_tx=XM_003995993.3
D4 88820831 88820851
refGene_gn=LOC101092062;refGene_tx=XM_003995993.3
D4 88830791 88830929
refGene_gn=LOC101092306;refGene_tx=XM_011288638.3;refGene_gn=LOC101092306;refGene_tx=XM_019816610.2
D4 88831442 88831579
refGene_gn=LOC101092306;refGene_tx=XM_011288638.3;refGene_gn=LOC101092

306;refGene_tx=XM_019816610.2
D4 88832942 88833022
refGene_gn=LOC101092306;refGene_tx=XM_011288638.3;refGene_gn=LOC101092306;refGene_tx=XM_019816610.2
D4 88833164 88833284
refGene_gn=LOC101092306;refGene_tx=XM_011288638.3;refGene_gn=LOC101092306;refGene_tx=XM_019816610.2
D4 88833504 88833606
refGene_gn=LOC101092306;refGene_tx=XM_011288638.3;refGene_gn=LOC101092306;refGene_tx=XM_019816610.2
D4 88834299 88834319
refGene_gn=LOC101092306;refGene_tx=XM_011288638.3;refGene_gn=LOC101092306;refGene_tx=XM_019816610.2
D4 88851297 88851435
refGene_gn=LOC101092559;refGene_tx=XM_019816611.2;refGene_gn=LOC101092559;refGene_tx=XM_019816612.2;refGene_gn=LOC101092559;refGene_tx=XM_019816613.2
D4 88851955 88852092
refGene_gn=LOC101092559;refGene_tx=XM_019816611.2;refGene_gn=LOC101092559;refGene_tx=XM_019816612.2;refGene_gn=LOC101092559;refGene_tx=XM_019816613.2
D4 88853499 88853579
refGene_gn=LOC101092559;refGene_tx=XM_019816611.2;refGene_gn=LOC101092559;refGene_tx=XM_019816612.2;refGene_gn=LOC101092559;refGene_tx=XM_019816613.2
D4 88853716 88853836
refGene_gn=LOC101092559;refGene_tx=XM_019816611.2;refGene_gn=LOC101092559;refGene_tx=XM_019816612.2;refGene_gn=LOC101092559;refGene_tx=XM_019816613.2
D4 88854053 88854155
refGene_gn=LOC101092559;refGene_tx=XM_019816611.2;refGene_gn=LOC101092559;refGene_tx=XM_019816612.2;refGene_gn=LOC101092559;refGene_tx=XM_019816613.2
D4 88854813 88854833
refGene_gn=LOC101092559;refGene_tx=XM_019816611.2;refGene_gn=LOC101092559;refGene_tx=XM_019816612.2;refGene_gn=LOC101092559;refGene_tx=XM_019816613.2
D4 88858998 88859077
refGene_gn=CD4H9orf16;refGene_tx=XM_003995928.3
D4 88860870 88860981
refGene_gn=CIZ1;refGene_tx=XM_019816598.1;refGene_gn=CIZ1;refGene_tx=XM_019816606.2;refGene_gn=CIZ1;refGene_tx=XM_023242764.1;refGene_gn=CIZ1;refGene_tx=XM_023242765.1;refGene_gn=CIZ1;refGene_tx=XM_023242768.1;refGene_gn=CIZ1;refGene_tx=XM_023242769.1;refGene_gn=CIZ1;refGene_tx=XM_023242770.1
D4 88861889 88862062
refGene_gn=CD4H9orf16;refGene_tx=XM_003995928.3
D4 88863452 88863668
refGene_gn=CIZ1;refGene_tx=XM_011288402.1;refGene_gn=CIZ1;refGene_tx=XM_019816598.1;refGene_gn=CIZ1;refGene_tx=XM_019816606.2;refGene_gn=CIZ

1;refGene_tx=XM_023242764.1;refGene_gn=CIZ1;refGene_tx=XM_023242765.1;
refGene_gn=CIZ1;refGene_tx=XM_023242766.1;refGene_gn=CIZ1;refGene_tx=X
M_023242767.1;refGene_gn=CIZ1;refGene_tx=XM_023242768.1;refGene_gn=CIZ
1;refGene_tx=XM_023242769.1;refGene_gn=CIZ1;refGene_tx=XM_023242770.1;
refGene_gn=CIZ1;refGene_tx=XM_023242771.1;refGene_gn=CIZ1;refGene_tx=X
M_023242772.1;refGene_gn=CIZ1;refGene_tx=XM_023242774.1
D4 88864008 88864130
refGene_gn=CIZ1;refGene_tx=XM_011288402.1;refGene_gn=CIZ1;refGene_tx=X
M_019816598.1;refGene_gn=CIZ1;refGene_tx=XM_019816606.2;refGene_gn=CIZ
1;refGene_tx=XM_023242764.1;refGene_gn=CIZ1;refGene_tx=XM_023242765.1;
refGene_gn=CIZ1;refGene_tx=XM_023242766.1;refGene_gn=CIZ1;refGene_tx=X
M_023242767.1;refGene_gn=CIZ1;refGene_tx=XM_023242768.1;refGene_gn=CIZ
1;refGene_tx=XM_023242769.1;refGene_gn=CIZ1;refGene_tx=XM_023242770.1;
refGene_gn=CIZ1;refGene_tx=XM_023242771.1;refGene_gn=CIZ1;refGene_tx=X
M_023242772.1;refGene_gn=CIZ1;refGene_tx=XM_023242774.1
D4 88864259 88864329
refGene_gn=CIZ1;refGene_tx=XM_011288402.1;refGene_gn=CIZ1;refGene_tx=X
M_019816598.1;refGene_gn=CIZ1;refGene_tx=XM_019816606.2;refGene_gn=CIZ
1;refGene_tx=XM_023242764.1;refGene_gn=CIZ1;refGene_tx=XM_023242765.1;
refGene_gn=CIZ1;refGene_tx=XM_023242766.1;refGene_gn=CIZ1;refGene_tx=X
M_023242767.1;refGene_gn=CIZ1;refGene_tx=XM_023242768.1;refGene_gn=CIZ
1;refGene_tx=XM_023242769.1;refGene_gn=CIZ1;refGene_tx=XM_023242770.1;
refGene_gn=CIZ1;refGene_tx=XM_023242771.1;refGene_gn=CIZ1;refGene_tx=X
M_023242772.1;refGene_gn=CIZ1;refGene_tx=XM_023242774.1
D4 88865763 88865919
refGene_gn=CIZ1;refGene_tx=XM_011288402.1;refGene_gn=CIZ1;refGene_tx=X
M_019816598.1;refGene_gn=CIZ1;refGene_tx=XM_019816606.2;refGene_gn=CIZ
1;refGene_tx=XM_023242764.1;refGene_gn=CIZ1;refGene_tx=XM_023242765.1;
refGene_gn=CIZ1;refGene_tx=XM_023242766.1;refGene_gn=CIZ1;refGene_tx=X
M_023242767.1;refGene_gn=CIZ1;refGene_tx=XM_023242768.1;refGene_gn=CIZ
1;refGene_tx=XM_023242769.1;refGene_gn=CIZ1;refGene_tx=XM_023242770.1;
refGene_gn=CIZ1;refGene_tx=XM_023242771.1;refGene_gn=CIZ1;refGene_tx=X
M_023242772.1;refGene_gn=CIZ1;refGene_tx=XM_023242774.1
D4 88866120 88866234
refGene_gn=CIZ1;refGene_tx=XM_011288402.1;refGene_gn=CIZ1;refGene_tx=X
M_019816598.1;refGene_gn=CIZ1;refGene_tx=XM_019816606.2;refGene_gn=CIZ
1;refGene_tx=XM_023242764.1;refGene_gn=CIZ1;refGene_tx=XM_023242765.1;
refGene_gn=CIZ1;refGene_tx=XM_023242766.1;refGene_gn=CIZ1;refGene_tx=X
M_023242767.1;refGene_gn=CIZ1;refGene_tx=XM_023242768.1;refGene_gn=CIZ
1;refGene_tx=XM_023242769.1;refGene_gn=CIZ1;refGene_tx=XM_023242770.1;
refGene_gn=CIZ1;refGene_tx=XM_023242771.1;refGene_gn=CIZ1;refGene_tx=X
M_023242772.1;refGene_gn=CIZ1;refGene_tx=XM_023242774.1
D4 88866715 88866803
refGene_gn=CIZ1;refGene_tx=XM_011288402.1;refGene_gn=CIZ1;refGene_tx=X
M_019816598.1;refGene_gn=CIZ1;refGene_tx=XM_019816606.2;refGene_gn=CIZ
1;refGene_tx=XM_023242764.1;refGene_gn=CIZ1;refGene_tx=XM_023242765.1;
refGene_gn=CIZ1;refGene_tx=XM_023242766.1;refGene_gn=CIZ1;refGene_tx=X
M_023242767.1;refGene_gn=CIZ1;refGene_tx=XM_023242768.1;refGene_gn=CIZ
1;refGene_tx=XM_023242769.1;refGene_gn=CIZ1;refGene_tx=XM_023242770.1;
refGene_gn=CIZ1;refGene_tx=XM_023242771.1;refGene_gn=CIZ1;refGene_tx=X

M_023242772.1;refGene_gn=CIZ1;refGene_tx=XM_023242774.1
D4 88868408 88868560
refGene_gn=CIZ1;refGene_tx=XM_011288402.1;refGene_gn=CIZ1;refGene_tx=XM_019816598.1;refGene_gn=CIZ1;refGene_tx=XM_019816606.2;refGene_gn=CIZ1;refGene_tx=XM_023242764.1;refGene_gn=CIZ1;refGene_tx=XM_023242765.1;refGene_gn=CIZ1;refGene_tx=XM_023242766.1;refGene_gn=CIZ1;refGene_tx=XM_023242767.1;refGene_gn=CIZ1;refGene_tx=XM_023242768.1;refGene_gn=CIZ1;refGene_tx=XM_023242769.1;refGene_gn=CIZ1;refGene_tx=XM_023242770.1;refGene_gn=CIZ1;refGene_tx=XM_023242771.1;refGene_gn=CIZ1;refGene_tx=XM_023242772.1;refGene_gn=CIZ1;refGene_tx=XM_023242774.1
D4 88869264 88869462
refGene_gn=CIZ1;refGene_tx=XM_011288402.1;refGene_gn=CIZ1;refGene_tx=XM_019816598.1;refGene_gn=CIZ1;refGene_tx=XM_019816606.2;refGene_gn=CIZ1;refGene_tx=XM_023242764.1;refGene_gn=CIZ1;refGene_tx=XM_023242765.1;refGene_gn=CIZ1;refGene_tx=XM_023242766.1;refGene_gn=CIZ1;refGene_tx=XM_023242767.1;refGene_gn=CIZ1;refGene_tx=XM_023242768.1;refGene_gn=CIZ1;refGene_tx=XM_023242769.1;refGene_gn=CIZ1;refGene_tx=XM_023242770.1;refGene_gn=CIZ1;refGene_tx=XM_023242771.1;refGene_gn=CIZ1;refGene_tx=XM_023242772.1;refGene_gn=CIZ1;refGene_tx=XM_023242774.1
D4 88870071 88870190
refGene_gn=CIZ1;refGene_tx=XM_011288402.1;refGene_gn=CIZ1;refGene_tx=XM_019816598.1;refGene_gn=CIZ1;refGene_tx=XM_019816606.2;refGene_gn=CIZ1;refGene_tx=XM_023242764.1;refGene_gn=CIZ1;refGene_tx=XM_023242765.1;refGene_gn=CIZ1;refGene_tx=XM_023242766.1;refGene_gn=CIZ1;refGene_tx=XM_023242767.1;refGene_gn=CIZ1;refGene_tx=XM_023242768.1;refGene_gn=CIZ1;refGene_tx=XM_023242769.1;refGene_gn=CIZ1;refGene_tx=XM_023242770.1;refGene_gn=CIZ1;refGene_tx=XM_023242771.1;refGene_gn=CIZ1;refGene_tx=XM_023242772.1;refGene_gn=CIZ1;refGene_tx=XM_023242774.1
D4 88870370 88870972
refGene_gn=CIZ1;refGene_tx=XM_011288402.1;refGene_gn=CIZ1;refGene_tx=XM_019816598.1;refGene_gn=CIZ1;refGene_tx=XM_019816606.2;refGene_gn=CIZ1;refGene_tx=XM_023242764.1;refGene_gn=CIZ1;refGene_tx=XM_023242765.1;refGene_gn=CIZ1;refGene_tx=XM_023242766.1;refGene_gn=CIZ1;refGene_tx=XM_023242767.1;refGene_gn=CIZ1;refGene_tx=XM_023242768.1;refGene_gn=CIZ1;refGene_tx=XM_023242769.1;refGene_gn=CIZ1;refGene_tx=XM_023242770.1;refGene_gn=CIZ1;refGene_tx=XM_023242771.1;refGene_gn=CIZ1;refGene_tx=XM_023242772.1;refGene_gn=CIZ1;refGene_tx=XM_023242774.1
D4 88871483 88871592
refGene_gn=CIZ1;refGene_tx=XM_011288402.1;refGene_gn=CIZ1;refGene_tx=XM_019816598.1;refGene_gn=CIZ1;refGene_tx=XM_019816606.2;refGene_gn=CIZ1;refGene_tx=XM_023242764.1;refGene_gn=CIZ1;refGene_tx=XM_023242765.1;refGene_gn=CIZ1;refGene_tx=XM_023242766.1;refGene_gn=CIZ1;refGene_tx=XM_023242767.1;refGene_gn=CIZ1;refGene_tx=XM_023242768.1;refGene_gn=CIZ1;refGene_tx=XM_023242769.1;refGene_gn=CIZ1;refGene_tx=XM_023242770.1;refGene_gn=CIZ1;refGene_tx=XM_023242771.1;refGene_gn=CIZ1;refGene_tx=XM_023242772.1;refGene_gn=CIZ1;refGene_tx=XM_023242774.1
D4 88871785 88871879
refGene_gn=CIZ1;refGene_tx=XM_011288402.1;refGene_gn=CIZ1;refGene_tx=XM_019816598.1;refGene_gn=CIZ1;refGene_tx=XM_019816606.2;refGene_gn=CIZ1;refGene_tx=XM_023242764.1;refGene_gn=CIZ1;refGene_tx=XM_023242765.1;

refGene_gn=CIZ1;refGene_tx=XM_023242766.1;refGene_gn=CIZ1;refGene_tx=XM_023242767.1;refGene_gn=CIZ1;refGene_tx=XM_023242768.1;refGene_gn=CIZ1;refGene_tx=XM_023242769.1;refGene_gn=CIZ1;refGene_tx=XM_023242770.1;refGene_gn=CIZ1;refGene_tx=XM_023242771.1;refGene_gn=CIZ1;refGene_tx=XM_023242772.1;refGene_gn=CIZ1;refGene_tx=XM_023242774.1
D4 88874045 88874275
refGene_gn=CIZ1;refGene_tx=XM_011288402.1;refGene_gn=CIZ1;refGene_tx=XM_019816598.1;refGene_gn=CIZ1;refGene_tx=XM_019816606.2;refGene_gn=CIZ1;refGene_tx=XM_023242764.1;refGene_gn=CIZ1;refGene_tx=XM_023242765.1;refGene_gn=CIZ1;refGene_tx=XM_023242766.1;refGene_gn=CIZ1;refGene_tx=XM_023242767.1;refGene_gn=CIZ1;refGene_tx=XM_023242768.1;refGene_gn=CIZ1;refGene_tx=XM_023242769.1;refGene_gn=CIZ1;refGene_tx=XM_023242770.1;refGene_gn=CIZ1;refGene_tx=XM_023242771.1;refGene_gn=CIZ1;refGene_tx=XM_023242772.1;refGene_gn=CIZ1;refGene_tx=XM_023242774.1
D4 88875911 88875983
refGene_gn=CIZ1;refGene_tx=XM_011288402.1;refGene_gn=CIZ1;refGene_tx=XM_019816598.1;refGene_gn=CIZ1;refGene_tx=XM_019816606.2;refGene_gn=CIZ1;refGene_tx=XM_023242764.1;refGene_gn=CIZ1;refGene_tx=XM_023242765.1;refGene_gn=CIZ1;refGene_tx=XM_023242766.1;refGene_gn=CIZ1;refGene_tx=XM_023242767.1;refGene_gn=CIZ1;refGene_tx=XM_023242768.1;refGene_gn=CIZ1;refGene_tx=XM_023242771.1
D4 88877811 88877927
refGene_gn=CIZ1;refGene_tx=XM_011288402.1;refGene_gn=CIZ1;refGene_tx=XM_019816598.1;refGene_gn=CIZ1;refGene_tx=XM_023242769.1;refGene_gn=CIZ1;refGene_tx=XM_023242770.1;refGene_gn=CIZ1;refGene_tx=XM_023242771.1;refGene_gn=CIZ1;refGene_tx=XM_023242772.1;refGene_gn=CIZ1;refGene_tx=XM_023242774.1
D4 88878164 88878297
refGene_gn=CIZ1;refGene_tx=XM_011288402.1;refGene_gn=CIZ1;refGene_tx=XM_019816598.1;refGene_gn=CIZ1;refGene_tx=XM_023242769.1;refGene_gn=CIZ1;refGene_tx=XM_023242770.1;refGene_gn=CIZ1;refGene_tx=XM_023242771.1;refGene_gn=CIZ1;refGene_tx=XM_023242772.1;refGene_gn=CIZ1;refGene_tx=XM_023242774.1
D4 88879087 88879241
refGene_gn=CIZ1;refGene_tx=XM_023242769.1;refGene_gn=CIZ1;refGene_tx=XM_023242770.1;refGene_gn=CIZ1;refGene_tx=XM_023242771.1;refGene_gn=CIZ1;refGene_tx=XM_023242772.1;refGene_gn=CIZ1;refGene_tx=XM_023242774.1
D4 88885257 88885418
refGene_gn=DNM1;refGene_tx=XM_023242775.1;refGene_gn=DNM1;refGene_tx=XM_023242776.1;refGene_gn=DNM1;refGene_tx=XM_023242777.1;refGene_gn=DNM1;refGene_tx=XM_023242778.1;refGene_gn=DNM1;refGene_tx=XM_023242780.1;refGene_gn=DNM1;refGene_tx=XM_023242781.1;refGene_gn=DNM1;refGene_tx=XM_023242782.1;refGene_gn=DNM1;refGene_tx=XM_023242783.1;refGene_gn=DNM1;refGene_tx=XM_023242784.1;refGene_gn=DNM1;refGene_tx=XM_023242785.1;refGene_gn=DNM1;refGene_tx=XM_023242786.1;refGene_gn=DNM1;refGene_tx=XM_023242787.1;refGene_gn=DNM1;refGene_tx=XM_023242788.1;refGene_gn=DNM1;refGene_tx=XM_023242790.1;refGene_gn=DNM1;refGene_tx=XM_023242791.1
D4 88898110 88898184
refGene_gn=DNM1;refGene_tx=XM_023242775.1;refGene_gn=DNM1;refGene_tx=XM_023242776.1;refGene_gn=DNM1;refGene_tx=XM_023242777.1;refGene_gn=DNM

1;refGene_tx=XM_023242778.1;refGene_gn=DNM1;refGene_tx=XM_023242780.1;
refGene_gn=DNM1;refGene_tx=XM_023242781.1;refGene_gn=DNM1;refGene_tx=X
M_023242782.1;refGene_gn=DNM1;refGene_tx=XM_023242783.1;refGene_gn=DNM
1;refGene_tx=XM_023242784.1;refGene_gn=DNM1;refGene_tx=XM_023242785.1;
refGene_gn=DNM1;refGene_tx=XM_023242786.1;refGene_gn=DNM1;refGene_tx=X
M_023242787.1;refGene_gn=DNM1;refGene_tx=XM_023242788.1;refGene_gn=DNM
1;refGene_tx=XM_023242790.1;refGene_gn=DNM1;refGene_tx=XM_023242791.1
D4 88898500 88898650
refGene_gn=DNM1;refGene_tx=XM_023242775.1;refGene_gn=DNM1;refGene_tx=X
M_023242776.1;refGene_gn=DNM1;refGene_tx=XM_023242777.1;refGene_gn=DNM
1;refGene_tx=XM_023242778.1;refGene_gn=DNM1;refGene_tx=XM_023242780.1;
refGene_gn=DNM1;refGene_tx=XM_023242781.1;refGene_gn=DNM1;refGene_tx=X
M_023242782.1;refGene_gn=DNM1;refGene_tx=XM_023242783.1;refGene_gn=DNM
1;refGene_tx=XM_023242784.1;refGene_gn=DNM1;refGene_tx=XM_023242785.1;
refGene_gn=DNM1;refGene_tx=XM_023242786.1;refGene_gn=DNM1;refGene_tx=X
M_023242787.1;refGene_gn=DNM1;refGene_tx=XM_023242788.1;refGene_gn=DNM
1;refGene_tx=XM_023242790.1;refGene_gn=DNM1;refGene_tx=XM_023242791.1
D4 88899014 88899218
refGene_gn=DNM1;refGene_tx=XM_023242775.1;refGene_gn=DNM1;refGene_tx=X
M_023242776.1;refGene_gn=DNM1;refGene_tx=XM_023242777.1;refGene_gn=DNM
1;refGene_tx=XM_023242778.1;refGene_gn=DNM1;refGene_tx=XM_023242780.1;
refGene_gn=DNM1;refGene_tx=XM_023242781.1;refGene_gn=DNM1;refGene_tx=X
M_023242782.1;refGene_gn=DNM1;refGene_tx=XM_023242783.1;refGene_gn=DNM
1;refGene_tx=XM_023242784.1;refGene_gn=DNM1;refGene_tx=XM_023242785.1;
refGene_gn=DNM1;refGene_tx=XM_023242786.1;refGene_gn=DNM1;refGene_tx=X
M_023242787.1;refGene_gn=DNM1;refGene_tx=XM_023242788.1;refGene_gn=DNM
1;refGene_tx=XM_023242790.1;refGene_gn=DNM1;refGene_tx=XM_023242791.1
D4 88899452 88899551
refGene_gn=DNM1;refGene_tx=XM_023242775.1;refGene_gn=DNM1;refGene_tx=X
M_023242776.1;refGene_gn=DNM1;refGene_tx=XM_023242777.1;refGene_gn=DNM
1;refGene_tx=XM_023242778.1;refGene_gn=DNM1;refGene_tx=XM_023242780.1;
refGene_gn=DNM1;refGene_tx=XM_023242781.1;refGene_gn=DNM1;refGene_tx=X
M_023242782.1;refGene_gn=DNM1;refGene_tx=XM_023242783.1;refGene_gn=DNM
1;refGene_tx=XM_023242784.1;refGene_gn=DNM1;refGene_tx=XM_023242785.1;
refGene_gn=DNM1;refGene_tx=XM_023242786.1;refGene_gn=DNM1;refGene_tx=X
M_023242787.1;refGene_gn=DNM1;refGene_tx=XM_023242788.1;refGene_gn=DNM
1;refGene_tx=XM_023242790.1;refGene_gn=DNM1;refGene_tx=XM_023242791.1
D4 88899647 88899808
refGene_gn=DNM1;refGene_tx=XM_023242775.1;refGene_gn=DNM1;refGene_tx=X
M_023242776.1;refGene_gn=DNM1;refGene_tx=XM_023242777.1;refGene_gn=DNM
1;refGene_tx=XM_023242778.1;refGene_gn=DNM1;refGene_tx=XM_023242780.1;
refGene_gn=DNM1;refGene_tx=XM_023242781.1;refGene_gn=DNM1;refGene_tx=X
M_023242782.1;refGene_gn=DNM1;refGene_tx=XM_023242783.1;refGene_gn=DNM
1;refGene_tx=XM_023242784.1;refGene_gn=DNM1;refGene_tx=XM_023242785.1;
refGene_gn=DNM1;refGene_tx=XM_023242786.1;refGene_gn=DNM1;refGene_tx=X
M_023242787.1;refGene_gn=DNM1;refGene_tx=XM_023242788.1;refGene_gn=DNM
1;refGene_tx=XM_023242790.1;refGene_gn=DNM1;refGene_tx=XM_023242791.1
D4 88900880 88901023
refGene_gn=DNM1;refGene_tx=XM_023242775.1;refGene_gn=DNM1;refGene_tx=X
M_023242776.1;refGene_gn=DNM1;refGene_tx=XM_023242777.1;refGene_gn=DNM

1;refGene_tx=XM_023242778.1;refGene_gn=DNM1;refGene_tx=XM_023242780.1;
refGene_gn=DNM1;refGene_tx=XM_023242781.1;refGene_gn=DNM1;refGene_tx=X
M_023242782.1;refGene_gn=DNM1;refGene_tx=XM_023242783.1;refGene_gn=DNM
1;refGene_tx=XM_023242784.1;refGene_gn=DNM1;refGene_tx=XM_023242785.1;
refGene_gn=DNM1;refGene_tx=XM_023242786.1;refGene_gn=DNM1;refGene_tx=X
M_023242787.1;refGene_gn=DNM1;refGene_tx=XM_023242788.1;refGene_gn=DNM
1;refGene_tx=XM_023242790.1;refGene_gn=DNM1;refGene_tx=XM_023242791.1
D4 88901133 88901269
refGene_gn=DNM1;refGene_tx=XM_023242775.1;refGene_gn=DNM1;refGene_tx=X
M_023242776.1;refGene_gn=DNM1;refGene_tx=XM_023242777.1;refGene_gn=DNM
1;refGene_tx=XM_023242778.1;refGene_gn=DNM1;refGene_tx=XM_023242780.1;
refGene_gn=DNM1;refGene_tx=XM_023242781.1;refGene_gn=DNM1;refGene_tx=X
M_023242782.1;refGene_gn=DNM1;refGene_tx=XM_023242783.1;refGene_gn=DNM
1;refGene_tx=XM_023242784.1;refGene_gn=DNM1;refGene_tx=XM_023242785.1;
refGene_gn=DNM1;refGene_tx=XM_023242786.1;refGene_gn=DNM1;refGene_tx=X
M_023242787.1;refGene_gn=DNM1;refGene_tx=XM_023242788.1;refGene_gn=DNM
1;refGene_tx=XM_023242790.1;refGene_gn=DNM1;refGene_tx=XM_023242791.1
D4 88901451 88901519
refGene_gn=DNM1;refGene_tx=XM_023242775.1;refGene_gn=DNM1;refGene_tx=X
M_023242776.1;refGene_gn=DNM1;refGene_tx=XM_023242777.1;refGene_gn=DNM
1;refGene_tx=XM_023242778.1;refGene_gn=DNM1;refGene_tx=XM_023242780.1;
refGene_gn=DNM1;refGene_tx=XM_023242781.1;refGene_gn=DNM1;refGene_tx=X
M_023242782.1;refGene_gn=DNM1;refGene_tx=XM_023242783.1;refGene_gn=DNM
1;refGene_tx=XM_023242784.1;refGene_gn=DNM1;refGene_tx=XM_023242785.1;
refGene_gn=DNM1;refGene_tx=XM_023242786.1;refGene_gn=DNM1;refGene_tx=X
M_023242787.1;refGene_gn=DNM1;refGene_tx=XM_023242788.1;refGene_gn=DNM
1;refGene_tx=XM_023242790.1;refGene_gn=DNM1;refGene_tx=XM_023242791.1
D4 88902706 88902845
refGene_gn=DNM1;refGene_tx=XM_023242776.1;refGene_gn=DNM1;refGene_tx=X
M_023242780.1;refGene_gn=DNM1;refGene_tx=XM_023242782.1;refGene_gn=DNM
1;refGene_tx=XM_023242785.1;refGene_gn=DNM1;refGene_tx=XM_023242790.1;
refGene_gn=DNM1;refGene_tx=XM_023242791.1
D4 88904483 88904622
refGene_gn=DNM1;refGene_tx=XM_023242775.1;refGene_gn=DNM1;refGene_tx=X
M_023242777.1;refGene_gn=DNM1;refGene_tx=XM_023242778.1;refGene_gn=DNM
1;refGene_tx=XM_023242781.1;refGene_gn=DNM1;refGene_tx=XM_023242783.1;
refGene_gn=DNM1;refGene_tx=XM_023242784.1;refGene_gn=DNM1;refGene_tx=X
M_023242786.1;refGene_gn=DNM1;refGene_tx=XM_023242787.1;refGene_gn=DNM
1;refGene_tx=XM_023242788.1
D4 88910861 88910948
refGene_gn=DNM1;refGene_tx=XM_023242775.1;refGene_gn=DNM1;refGene_tx=X
M_023242776.1;refGene_gn=DNM1;refGene_tx=XM_023242777.1;refGene_gn=DNM
1;refGene_tx=XM_023242778.1;refGene_gn=DNM1;refGene_tx=XM_023242780.1;
refGene_gn=DNM1;refGene_tx=XM_023242781.1;refGene_gn=DNM1;refGene_tx=X
M_023242782.1;refGene_gn=DNM1;refGene_tx=XM_023242783.1;refGene_gn=DNM
1;refGene_tx=XM_023242784.1;refGene_gn=DNM1;refGene_tx=XM_023242785.1;
refGene_gn=DNM1;refGene_tx=XM_023242786.1;refGene_gn=DNM1;refGene_tx=X
M_023242787.1;refGene_gn=DNM1;refGene_tx=XM_023242788.1;refGene_gn=DNM
1;refGene_tx=XM_023242790.1;refGene_gn=DNM1;refGene_tx=XM_023242791.1
D4 88914366 88914437

refGene_gn=DNM1;refGene_tx=XM_023242775.1;refGene_gn=DNM1;refGene_tx=XM_023242776.1;refGene_gn=DNM1;refGene_tx=XM_023242777.1;refGene_gn=DNM1;refGene_tx=XM_023242778.1;refGene_gn=DNM1;refGene_tx=XM_023242780.1;refGene_gn=DNM1;refGene_tx=XM_023242781.1;refGene_gn=DNM1;refGene_tx=XM_023242782.1;refGene_gn=DNM1;refGene_tx=XM_023242783.1;refGene_gn=DNM1;refGene_tx=XM_023242784.1;refGene_gn=DNM1;refGene_tx=XM_023242785.1;refGene_gn=DNM1;refGene_tx=XM_023242786.1;refGene_gn=DNM1;refGene_tx=XM_023242787.1;refGene_gn=DNM1;refGene_tx=XM_023242788.1;refGene_gn=DNM1;refGene_tx=XM_023242790.1;refGene_gn=DNM1;refGene_tx=XM_023242791.1
D4 88914652 88914704
refGene_gn=DNM1;refGene_tx=XM_023242775.1;refGene_gn=DNM1;refGene_tx=XM_023242776.1;refGene_gn=DNM1;refGene_tx=XM_023242777.1;refGene_gn=DNM1;refGene_tx=XM_023242778.1;refGene_gn=DNM1;refGene_tx=XM_023242780.1;refGene_gn=DNM1;refGene_tx=XM_023242781.1;refGene_gn=DNM1;refGene_tx=XM_023242782.1;refGene_gn=DNM1;refGene_tx=XM_023242783.1;refGene_gn=DNM1;refGene_tx=XM_023242784.1;refGene_gn=DNM1;refGene_tx=XM_023242785.1;refGene_gn=DNM1;refGene_tx=XM_023242786.1;refGene_gn=DNM1;refGene_tx=XM_023242787.1;refGene_gn=DNM1;refGene_tx=XM_023242788.1;refGene_gn=DNM1;refGene_tx=XM_023242790.1;refGene_gn=DNM1;refGene_tx=XM_023242791.1
D4 88914910 88914922
refGene_gn=DNM1;refGene_tx=XM_023242775.1;refGene_gn=DNM1;refGene_tx=XM_023242776.1;refGene_gn=DNM1;refGene_tx=XM_023242777.1;refGene_gn=DNM1;refGene_tx=XM_023242778.1;refGene_gn=DNM1;refGene_tx=XM_023242780.1;refGene_gn=DNM1;refGene_tx=XM_023242783.1;refGene_gn=DNM1;refGene_tx=XM_023242784.1;refGene_gn=DNM1;refGene_tx=XM_023242785.1;refGene_gn=DNM1;refGene_tx=XM_023242786.1;refGene_gn=DNM1;refGene_tx=XM_023242787.1;refGene_gn=DNM1;refGene_tx=XM_023242788.1;refGene_gn=DNM1;refGene_tx=XM_023242791.1
D4 88917289 88917403
refGene_gn=DNM1;refGene_tx=XM_023242775.1;refGene_gn=DNM1;refGene_tx=XM_023242776.1;refGene_gn=DNM1;refGene_tx=XM_023242777.1;refGene_gn=DNM1;refGene_tx=XM_023242778.1;refGene_gn=DNM1;refGene_tx=XM_023242780.1;refGene_gn=DNM1;refGene_tx=XM_023242781.1;refGene_gn=DNM1;refGene_tx=XM_023242782.1;refGene_gn=DNM1;refGene_tx=XM_023242783.1;refGene_gn=DNM1;refGene_tx=XM_023242784.1;refGene_gn=DNM1;refGene_tx=XM_023242785.1;refGene_gn=DNM1;refGene_tx=XM_023242786.1;refGene_gn=DNM1;refGene_tx=XM_023242787.1;refGene_gn=DNM1;refGene_tx=XM_023242788.1;refGene_gn=DNM1;refGene_tx=XM_023242790.1;refGene_gn=DNM1;refGene_tx=XM_023242791.1
D4 88921072 88921182
refGene_gn=DNM1;refGene_tx=XM_023242775.1;refGene_gn=DNM1;refGene_tx=XM_023242776.1;refGene_gn=DNM1;refGene_tx=XM_023242777.1;refGene_gn=DNM1;refGene_tx=XM_023242778.1;refGene_gn=DNM1;refGene_tx=XM_023242780.1;refGene_gn=DNM1;refGene_tx=XM_023242781.1;refGene_gn=DNM1;refGene_tx=XM_023242782.1;refGene_gn=DNM1;refGene_tx=XM_023242783.1;refGene_gn=DNM1;refGene_tx=XM_023242784.1;refGene_gn=DNM1;refGene_tx=XM_023242785.1;refGene_gn=DNM1;refGene_tx=XM_023242786.1;refGene_gn=DNM1;refGene_tx=XM_023242787.1;refGene_gn=DNM1;refGene_tx=XM_023242788.1;refGene_gn=DNM1;refGene_tx=XM_023242790.1;refGene_gn=DNM1;refGene_tx=XM_023242791.1
D4 88921924 88922036
refGene_gn=DNM1;refGene_tx=XM_023242775.1;refGene_gn=DNM1;refGene_tx=X

M_023242776.1;refGene_gn=DNM1;refGene_tx=XM_023242777.1;refGene_gn=DNM1;refGene_tx=XM_023242778.1;refGene_gn=DNM1;refGene_tx=XM_023242780.1;refGene_gn=DNM1;refGene_tx=XM_023242781.1;refGene_gn=DNM1;refGene_tx=XM_023242782.1;refGene_gn=DNM1;refGene_tx=XM_023242783.1;refGene_gn=DNM1;refGene_tx=XM_023242784.1;refGene_gn=DNM1;refGene_tx=XM_023242785.1;refGene_gn=DNM1;refGene_tx=XM_023242786.1;refGene_gn=DNM1;refGene_tx=XM_023242787.1;refGene_gn=DNM1;refGene_tx=XM_023242788.1;refGene_gn=DNM1;refGene_tx=XM_023242790.1;refGene_gn=DNM1;refGene_tx=XM_023242791.1
D4 88922430 88922442 refGene_gn=DNM1;refGene_tx=XM_023242775.1;refGene_gn=DNM1;refGene_tx=XM_023242776.1;refGene_gn=DNM1;refGene_tx=XM_023242777.1;refGene_gn=DNM1;refGene_tx=XM_023242778.1;refGene_gn=DNM1;refGene_tx=XM_023242780.1;refGene_gn=DNM1;refGene_tx=XM_023242781.1;refGene_gn=DNM1;refGene_tx=XM_023242782.1;refGene_gn=DNM1;refGene_tx=XM_023242783.1;refGene_gn=DNM1;refGene_tx=XM_023242784.1;refGene_gn=DNM1;refGene_tx=XM_023242785.1;refGene_gn=DNM1;refGene_tx=XM_023242786.1;refGene_gn=DNM1;refGene_tx=XM_023242787.1;refGene_gn=DNM1;refGene_tx=XM_023242788.1;refGene_gn=DNM1;refGene_tx=XM_023242790.1;refGene_gn=DNM1;refGene_tx=XM_023242791.1
D4 88922811 88922982 refGene_gn=DNM1;refGene_tx=XM_023242775.1;refGene_gn=DNM1;refGene_tx=XM_023242776.1;refGene_gn=DNM1;refGene_tx=XM_023242777.1;refGene_gn=DNM1;refGene_tx=XM_023242778.1;refGene_gn=DNM1;refGene_tx=XM_023242780.1;refGene_gn=DNM1;refGene_tx=XM_023242781.1;refGene_gn=DNM1;refGene_tx=XM_023242782.1;refGene_gn=DNM1;refGene_tx=XM_023242783.1;refGene_gn=DNM1;refGene_tx=XM_023242784.1;refGene_gn=DNM1;refGene_tx=XM_023242785.1;refGene_gn=DNM1;refGene_tx=XM_023242786.1;refGene_gn=DNM1;refGene_tx=XM_023242787.1;refGene_gn=DNM1;refGene_tx=XM_023242788.1;refGene_gn=DNM1;refGene_tx=XM_023242790.1;refGene_gn=DNM1;refGene_tx=XM_023242791.1
D4 88924322 88924564 refGene_gn=DNM1;refGene_tx=XM_023242775.1;refGene_gn=DNM1;refGene_tx=XM_023242776.1;refGene_gn=DNM1;refGene_tx=XM_023242777.1;refGene_gn=DNM1;refGene_tx=XM_023242778.1;refGene_gn=DNM1;refGene_tx=XM_023242780.1;refGene_gn=DNM1;refGene_tx=XM_023242781.1;refGene_gn=DNM1;refGene_tx=XM_023242782.1;refGene_gn=DNM1;refGene_tx=XM_023242783.1;refGene_gn=DNM1;refGene_tx=XM_023242784.1;refGene_gn=DNM1;refGene_tx=XM_023242785.1;refGene_gn=DNM1;refGene_tx=XM_023242786.1;refGene_gn=DNM1;refGene_tx=XM_023242787.1;refGene_gn=DNM1;refGene_tx=XM_023242788.1;refGene_gn=DNM1;refGene_tx=XM_023242790.1;refGene_gn=DNM1;refGene_tx=XM_023242791.1
D4 88924931 88925147 refGene_gn=DNM1;refGene_tx=XM_023242775.1;refGene_gn=DNM1;refGene_tx=XM_023242776.1;refGene_gn=DNM1;refGene_tx=XM_023242777.1;refGene_gn=DNM1;refGene_tx=XM_023242778.1;refGene_gn=DNM1;refGene_tx=XM_023242780.1;refGene_gn=DNM1;refGene_tx=XM_023242781.1;refGene_gn=DNM1;refGene_tx=XM_023242782.1;refGene_gn=DNM1;refGene_tx=XM_023242783.1;refGene_gn=DNM1;refGene_tx=XM_023242784.1;refGene_gn=DNM1;refGene_tx=XM_023242785.1;refGene_gn=DNM1;refGene_tx=XM_023242786.1;refGene_gn=DNM1;refGene_tx=XM_023242787.1;refGene_gn=DNM1;refGene_tx=XM_023242788.1;refGene_gn=DNM1;refGene_tx=XM_023242790.1;refGene_gn=DNM1;refGene_tx=XM_023242791.1
D4 88925860 88925862 refGene_gn=DNM1;refGene_tx=XM_023242790.1
D4 88927274 88927296

refGene_gn=DNM1;refGene_tx=XM_023242784.1;refGene_gn=DNM1;refGene_tx=XM_023242785.1;refGene_gn=DNM1;refGene_tx=XM_023242786.1;refGene_gn=DNM1;refGene_tx=XM_023242787.1;refGene_gn=DNM1;refGene_tx=XM_023242788.1
D4 88928233 88928306 refGene_gn=DNM1;refGene_tx=XM_023242775.1;refGene_gn=DNM1;refGene_tx=XM_023242776.1;refGene_gn=DNM1;refGene_tx=XM_023242777.1;refGene_gn=DNM1;refGene_tx=XM_023242781.1;refGene_gn=DNM1;refGene_tx=XM_023242782.1;refGene_gn=DNM1;refGene_tx=XM_023242790.1;refGene_gn=DNM1;refGene_tx=XM_023242791.1
D4 88928902 88928967 refGene_gn=DNM1;refGene_tx=XM_023242778.1;refGene_gn=DNM1;refGene_tx=XM_023242780.1;refGene_gn=DNM1;refGene_tx=XM_023242783.1
D4 88930924 88931151 refGene_gn=GOLGA2;refGene_tx=XM_006939517.4
D4 88931240 88931394 refGene_gn=GOLGA2;refGene_tx=XM_006939517.4
D4 88931476 88931583 refGene_gn=GOLGA2;refGene_tx=XM_006939517.4
D4 88931666 88931769 refGene_gn=GOLGA2;refGene_tx=XM_006939517.4
D4 88931863 88932046 refGene_gn=GOLGA2;refGene_tx=XM_006939517.4
D4 88932302 88932391 refGene_gn=GOLGA2;refGene_tx=XM_006939517.4
D4 88932597 88932673 refGene_gn=GOLGA2;refGene_tx=XM_006939517.4
D4 88932750 88932996 refGene_gn=GOLGA2;refGene_tx=XM_006939517.4
D4 88933601 88933715 refGene_gn=GOLGA2;refGene_tx=XM_006939517.4
D4 88933993 88934331 refGene_gn=GOLGA2;refGene_tx=XM_006939517.4
D4 88935081 88935169 refGene_gn=GOLGA2;refGene_tx=XM_006939517.4
D4 88935412 88935520 refGene_gn=GOLGA2;refGene_tx=XM_006939517.4
D4 88935627 88935717 refGene_gn=GOLGA2;refGene_tx=XM_006939517.4
D4 88937095 88937237 refGene_gn=GOLGA2;refGene_tx=XM_006939517.4
D4 88937526 88937585 refGene_gn=GOLGA2;refGene_tx=XM_006939517.4
D4 88938741 88938848 refGene_gn=GOLGA2;refGene_tx=XM_006939517.4
D4 88938942 88939036 refGene_gn=GOLGA2;refGene_tx=XM_006939517.4
D4 88939122 88939185 refGene_gn=GOLGA2;refGene_tx=XM_006939517.4
D4 88939477 88939516 refGene_gn=GOLGA2;refGene_tx=XM_006939517.4
D4 88940325 88940406 refGene_gn=GOLGA2;refGene_tx=XM_006939517.4
D4 88940599 88940659 refGene_gn=GOLGA2;refGene_tx=XM_006939517.4
D4 88941059 88941123 refGene_gn=GOLGA2;refGene_tx=XM_006939517.4
D4 88941243 88941287 refGene_gn=GOLGA2;refGene_tx=XM_006939517.4
D4 88941576 88941693 refGene_gn=GOLGA2;refGene_tx=XM_006939517.4
D4 88945006 88945129 refGene_gn=GOLGA2;refGene_tx=XM_006939517.4
D4 88946847 88946917 refGene_gn=GOLGA2;refGene_tx=XM_006939517.4
D4 88946918 88946932 refGene_gn=GOLGA2;refGene_tx=XM_006939517.4
D4 88947075 88947362 refGene_gn=SWI5;refGene_tx=XM_019816617.2
D4 88947649 88947698 refGene_gn=SWI5;refGene_tx=XM_019816617.2
D4 88949800 88949922 refGene_gn=SWI5;refGene_tx=XM_019816617.2
D4 88950644 88950739 refGene_gn=SWI5;refGene_tx=XM_019816617.2
D4 88951582 88951647 refGene_gn=SWI5;refGene_tx=XM_019816617.2
D4 88977672 88977756 refGene_gn=TRUB2;refGene_tx=XM_003995997.5
D4 88977757 88977999 refGene_gn=TRUB2;refGene_tx=XM_003995997.5
D4 88979041 88979178 refGene_gn=TRUB2;refGene_tx=XM_003995997.5
D4 88979715 88979788 refGene_gn=TRUB2;refGene_tx=XM_003995997.5
D4 88983380 88983462 refGene_gn=TRUB2;refGene_tx=XM_003995997.5
D4 88985079 88985141 refGene_gn=TRUB2;refGene_tx=XM_003995997.5

D4 88986282 88986357 refGene_gn=TRUB2;refGene_tx=XM_003995997.5
D4 88988051 88988183 refGene_gn=TRUB2;refGene_tx=XM_003995997.5
D4 88988778 88988893 refGene_gn=TRUB2;refGene_tx=XM_003995997.5
D4 88989311 88989381
refGene_gn=COQ4;refGene_tx=XM_003995932.5;refGene_gn=COQ4;refGene_tx=XM_019816619.2
D4 88989472 88989604
refGene_gn=COQ4;refGene_tx=XM_003995932.5;refGene_gn=COQ4;refGene_tx=XM_019816619.2
D4 88991001 88991098
refGene_gn=COQ4;refGene_tx=XM_003995932.5;refGene_gn=COQ4;refGene_tx=XM_019816619.2
D4 88991616 88991719
refGene_gn=COQ4;refGene_tx=XM_003995932.5;refGene_gn=COQ4;refGene_tx=XM_019816619.2
D4 88992598 88992955 refGene_gn=COQ4;refGene_tx=XM_019816619.2
D4 88996783 88996913 refGene_gn=COQ4;refGene_tx=XM_003995932.5
D4 88997426 88997520 refGene_gn=COQ4;refGene_tx=XM_003995932.5
D4 88998036 88998208 refGene_gn=COQ4;refGene_tx=XM_003995932.5
D4 89008153 89008314 refGene_gn=SLC27A4;refGene_tx=XM_003995933.4
D4 89009919 89010314 refGene_gn=SLC27A4;refGene_tx=XM_003995933.4
D4 89011712 89011871 refGene_gn=SLC27A4;refGene_tx=XM_003995933.4
D4 89012970 89013040 refGene_gn=SLC27A4;refGene_tx=XM_003995933.4
D4 89013139 89013231 refGene_gn=SLC27A4;refGene_tx=XM_003995933.4
D4 89014039 89014149 refGene_gn=SLC27A4;refGene_tx=XM_003995933.4
D4 89014359 89014569 refGene_gn=SLC27A4;refGene_tx=XM_003995933.4
D4 89014805 89014932 refGene_gn=SLC27A4;refGene_tx=XM_003995933.4
D4 89015589 89015727 refGene_gn=SLC27A4;refGene_tx=XM_003995933.4
D4 89015905 89016070 refGene_gn=SLC27A4;refGene_tx=XM_003995933.4
D4 89016163 89016310 refGene_gn=SLC27A4;refGene_tx=XM_003995933.4
D4 89018862 89019020 refGene_gn=SLC27A4;refGene_tx=XM_003995933.4
D4 89026094 89026129 refGene_gn=URM1;refGene_tx=XM_003995934.4
D4 89030274 89030345 refGene_gn=URM1;refGene_tx=XM_003995934.4
D4 89038584 89038666 refGene_gn=URM1;refGene_tx=XM_003995934.4
D4 89039989 89040038 refGene_gn=URM1;refGene_tx=XM_003995934.4
D4 89040375 89040444 refGene_gn=URM1;refGene_tx=XM_003995934.4
D4 89042937 89043594
refGene_gn=LOC111557546;refGene_tx=XM_023242793.1
D4 89056844 89056944
refGene_gn=URM1;refGene_tx=XM_023242794.1;refGene_gn=URM1;refGene_tx=XM_023242796.1;refGene_gn=URM1;refGene_tx=XM_023242797.1
D4 89059039 89059134
refGene_gn=URM1;refGene_tx=XM_023242794.1;refGene_gn=URM1;refGene_tx=XM_023242797.1
D4 89060799 89060838 refGene_gn=URM1;refGene_tx=XM_023242794.1
D4 89061574 89061918
refGene_gn=URM1;refGene_tx=XM_023242794.1;refGene_gn=URM1;refGene_tx=XM_023242796.1;refGene_gn=URM1;refGene_tx=XM_023242797.1
D4 89062310 89062507 refGene_gn=CERCAM;refGene_tx=XM_011288393.2
D4 89063914 89064025

refGene_gn=CERCAM;refGene_tx=XM_011288391.3;refGene_gn=CERCAM;refGene_tx=XM_011288393.2;refGene_gn=CERCAM;refGene_tx=XM_019816620.2
D4 89064199 89064317
refGene_gn=CERCAM;refGene_tx=XM_011288391.3;refGene_gn=CERCAM;refGene_tx=XM_011288393.2;refGene_gn=CERCAM;refGene_tx=XM_019816620.2
D4 89065346 89065481
refGene_gn=CERCAM;refGene_tx=XM_011288391.3;refGene_gn=CERCAM;refGene_tx=XM_011288393.2;refGene_gn=CERCAM;refGene_tx=XM_019816620.2
D4 89065584 89065789
refGene_gn=CERCAM;refGene_tx=XM_011288391.3;refGene_gn=CERCAM;refGene_tx=XM_011288393.2;refGene_gn=CERCAM;refGene_tx=XM_019816620.2
D4 89067775 89067895
refGene_gn=CERCAM;refGene_tx=XM_011288391.3;refGene_gn=CERCAM;refGene_tx=XM_011288393.2;refGene_gn=CERCAM;refGene_tx=XM_019816620.2
D4 89068229 89068306
refGene_gn=CERCAM;refGene_tx=XM_011288391.3;refGene_gn=CERCAM;refGene_tx=XM_011288393.2;refGene_gn=CERCAM;refGene_tx=XM_019816620.2
D4 89068410 89068517
refGene_gn=CERCAM;refGene_tx=XM_011288391.3;refGene_gn=CERCAM;refGene_tx=XM_011288393.2;refGene_gn=CERCAM;refGene_tx=XM_019816620.2
D4 89069335 89069468
refGene_gn=CERCAM;refGene_tx=XM_011288391.3;refGene_gn=CERCAM;refGene_tx=XM_011288393.2;refGene_gn=CERCAM;refGene_tx=XM_019816620.2
D4 89071081 89071209
refGene_gn=CERCAM;refGene_tx=XM_011288391.3;refGene_gn=CERCAM;refGene_tx=XM_011288393.2;refGene_gn=CERCAM;refGene_tx=XM_019816620.2
D4 89071349 89071553
refGene_gn=CERCAM;refGene_tx=XM_011288391.3;refGene_gn=CERCAM;refGene_tx=XM_011288393.2;refGene_gn=CERCAM;refGene_tx=XM_019816620.2
D4 89071947 89072203
refGene_gn=CERCAM;refGene_tx=XM_011288391.3;refGene_gn=CERCAM;refGene_tx=XM_011288393.2;refGene_gn=CERCAM;refGene_tx=XM_019816620.2
D4 89081658 89081690
refGene_gn=ODF2;refGene_tx=XM_006939521.4;refGene_gn=ODF2;refGene_tx=XM_019816625.2;refGene_gn=ODF2;refGene_tx=XM_019816629.2
D4 89082210 89082374
refGene_gn=ODF2;refGene_tx=XM_023242798.1;refGene_gn=ODF2;refGene_tx=XM_023242799.1;refGene_gn=ODF2;refGene_tx=XM_023242800.1;refGene_gn=ODF2;refGene_tx=XM_023242801.1;refGene_gn=ODF2;refGene_tx=XM_023242802.1;refGene_gn=ODF2;refGene_tx=XM_023242808.1
D4 89082680 89082712
refGene_gn=ODF2;refGene_tx=XM_003995938.5;refGene_gn=ODF2;refGene_tx=XM_006939522.2;refGene_gn=ODF2;refGene_tx=XM_019816626.2;refGene_gn=ODF2;refGene_tx=XM_019816630.1
D4 89083675 89083707
refGene_gn=ODF2;refGene_tx=XM_006939523.3;refGene_gn=ODF2;refGene_tx=XM_019816621.2;refGene_gn=ODF2;refGene_tx=XM_019816622.2;refGene_gn=ODF2;refGene_tx=XM_019816623.2;refGene_gn=ODF2;refGene_tx=XM_019816624.2;refGene_gn=ODF2;refGene_tx=XM_019816627.2;refGene_gn=ODF2;refGene_tx=XM_019816628.2;refGene_gn=ODF2;refGene_tx=XM_023242803.1;refGene_gn=ODF

2;refGene_tx=XM_023242805.1;refGene_gn=ODF2;refGene_tx=XM_023242809.1
D4 89085797 89085888
refGene_gn=ODF2;refGene_tx=XM_003995938.5;refGene_gn=ODF2;refGene_tx=XM_006939521.4;refGene_gn=ODF2;refGene_tx=XM_006939522.2;refGene_gn=ODF2;refGene_tx=XM_006939523.3;refGene_gn=ODF2;refGene_tx=XM_019816621.2;refGene_gn=ODF2;refGene_tx=XM_019816622.2;refGene_gn=ODF2;refGene_tx=XM_019816623.2;refGene_gn=ODF2;refGene_tx=XM_019816624.2;refGene_gn=ODF2;refGene_tx=XM_019816625.2;refGene_gn=ODF2;refGene_tx=XM_019816626.2;refGene_gn=ODF2;refGene_tx=XM_019816627.2;refGene_gn=ODF2;refGene_tx=XM_019816628.2;refGene_gn=ODF2;refGene_tx=XM_019816629.2;refGene_gn=ODF2;refGene_tx=XM_019816630.1;refGene_gn=ODF2;refGene_tx=XM_023242798.1;refGene_gn=ODF2;refGene_tx=XM_023242799.1;refGene_gn=ODF2;refGene_tx=XM_023242800.1;refGene_gn=ODF2;refGene_tx=XM_023242801.1;refGene_gn=ODF2;refGene_tx=XM_023242802.1;refGene_gn=ODF2;refGene_tx=XM_023242803.1;refGene_gn=ODF2;refGene_tx=XM_023242805.1;refGene_gn=ODF2;refGene_tx=XM_023242808.1;refGene_gn=ODF2;refGene_tx=XM_023242809.1
D4 89086789 89086906
refGene_gn=ODF2;refGene_tx=XM_019816621.2;refGene_gn=ODF2;refGene_tx=XM_019816622.2;refGene_gn=ODF2;refGene_tx=XM_019816623.2;refGene_gn=ODF2;refGene_tx=XM_019816624.2;refGene_gn=ODF2;refGene_tx=XM_019816625.2;refGene_gn=ODF2;refGene_tx=XM_019816626.2;refGene_gn=ODF2;refGene_tx=XM_023242798.1;refGene_gn=ODF2;refGene_tx=XM_023242799.1;refGene_gn=ODF2;refGene_tx=XM_023242803.1;refGene_gn=ODF2;refGene_tx=XM_023242805.1;refGene_gn=ODF2;refGene_tx=XM_023242806.1
D4 89087185 89087311
refGene_gn=ODF2;refGene_tx=XM_003995938.5;refGene_gn=ODF2;refGene_tx=XM_006939521.4;refGene_gn=ODF2;refGene_tx=XM_006939522.2;refGene_gn=ODF2;refGene_tx=XM_006939523.3;refGene_gn=ODF2;refGene_tx=XM_019816621.2;refGene_gn=ODF2;refGene_tx=XM_019816622.2;refGene_gn=ODF2;refGene_tx=XM_019816623.2;refGene_gn=ODF2;refGene_tx=XM_019816624.2;refGene_gn=ODF2;refGene_tx=XM_019816625.2;refGene_gn=ODF2;refGene_tx=XM_019816626.2;refGene_gn=ODF2;refGene_tx=XM_019816627.2;refGene_gn=ODF2;refGene_tx=XM_023242798.1;refGene_gn=ODF2;refGene_tx=XM_023242799.1;refGene_gn=ODF2;refGene_tx=XM_023242800.1;refGene_gn=ODF2;refGene_tx=XM_023242801.1;refGene_gn=ODF2;refGene_tx=XM_023242803.1;refGene_gn=ODF2;refGene_tx=XM_023242805.1;refGene_gn=ODF2;refGene_tx=XM_023242806.1;refGene_gn=ODF2;refGene_tx=XM_023242808.1;refGene_gn=ODF2;refGene_tx=XM_023242809.1
D4 89094436 89094607
refGene_gn=ODF2;refGene_tx=XM_003995938.5;refGene_gn=ODF2;refGene_tx=XM_006939521.4;refGene_gn=ODF2;refGene_tx=XM_006939522.2;refGene_gn=ODF2;refGene_tx=XM_006939523.3;refGene_gn=ODF2;refGene_tx=XM_019816621.2;refGene_gn=ODF2;refGene_tx=XM_019816622.2;refGene_gn=ODF2;refGene_tx=XM_019816623.2;refGene_gn=ODF2;refGene_tx=XM_019816624.2;refGene_gn=ODF2;refGene_tx=XM_019816625.2;refGene_gn=ODF2;refGene_tx=XM_019816626.2;refGene_gn=ODF2;refGene_tx=XM_019816627.2;refGene_gn=ODF2;refGene_tx=XM_019816628.2;refGene_gn=ODF2;refGene_tx=XM_019816629.2;refGene_gn=ODF2;refGene_tx=XM_019816630.1;refGene_gn=ODF2;refGene_tx=XM_023242798.1;refGene_gn=ODF2;refGene_tx=XM_023242799.1;refGene_gn=ODF2;refGene_tx=XM_023242800.1;refGene_gn=ODF2;refGene_tx=XM_023242801.1;refGene_gn=ODF2;refGene_tx=XM_023242802.1;refGene_gn=ODF2;refGene_tx=XM_023242803.1;

refGene_gn=ODF2;refGene_tx=XM_023242805.1;refGene_gn=ODF2;refGene_tx=XM_023242806.1;refGene_gn=ODF2;refGene_tx=XM_023242808.1;refGene_gn=ODF2;refGene_tx=XM_023242809.1
D4 89095460 89095621
refGene_gn=ODF2;refGene_tx=XM_003995938.5;refGene_gn=ODF2;refGene_tx=XM_006939521.4;refGene_gn=ODF2;refGene_tx=XM_006939522.2;refGene_gn=ODF2;refGene_tx=XM_006939523.3;refGene_gn=ODF2;refGene_tx=XM_019816621.2;refGene_gn=ODF2;refGene_tx=XM_019816622.2;refGene_gn=ODF2;refGene_tx=XM_019816623.2;refGene_gn=ODF2;refGene_tx=XM_019816624.2;refGene_gn=ODF2;refGene_tx=XM_019816625.2;refGene_gn=ODF2;refGene_tx=XM_019816626.2;refGene_gn=ODF2;refGene_tx=XM_019816627.2;refGene_gn=ODF2;refGene_tx=XM_019816628.2;refGene_gn=ODF2;refGene_tx=XM_019816629.2;refGene_gn=ODF2;refGene_tx=XM_019816630.1;refGene_gn=ODF2;refGene_tx=XM_023242798.1;refGene_gn=ODF2;refGene_tx=XM_023242799.1;refGene_gn=ODF2;refGene_tx=XM_023242800.1;refGene_gn=ODF2;refGene_tx=XM_023242801.1;refGene_gn=ODF2;refGene_tx=XM_023242802.1;refGene_gn=ODF2;refGene_tx=XM_023242803.1;refGene_gn=ODF2;refGene_tx=XM_023242805.1;refGene_gn=ODF2;refGene_tx=XM_023242806.1;refGene_gn=ODF2;refGene_tx=XM_023242808.1;refGene_gn=ODF2;refGene_tx=XM_023242809.1
D4 89096396 89096526
refGene_gn=ODF2;refGene_tx=XM_003995938.5;refGene_gn=ODF2;refGene_tx=XM_006939521.4;refGene_gn=ODF2;refGene_tx=XM_006939522.2;refGene_gn=ODF2;refGene_tx=XM_006939523.3;refGene_gn=ODF2;refGene_tx=XM_019816621.2;refGene_gn=ODF2;refGene_tx=XM_019816622.2;refGene_gn=ODF2;refGene_tx=XM_019816623.2;refGene_gn=ODF2;refGene_tx=XM_019816624.2;refGene_gn=ODF2;refGene_tx=XM_019816625.2;refGene_gn=ODF2;refGene_tx=XM_019816626.2;refGene_gn=ODF2;refGene_tx=XM_019816627.2;refGene_gn=ODF2;refGene_tx=XM_019816628.2;refGene_gn=ODF2;refGene_tx=XM_019816629.2;refGene_gn=ODF2;refGene_tx=XM_019816630.1;refGene_gn=ODF2;refGene_tx=XM_023242798.1;refGene_gn=ODF2;refGene_tx=XM_023242799.1;refGene_gn=ODF2;refGene_tx=XM_023242800.1;refGene_gn=ODF2;refGene_tx=XM_023242801.1;refGene_gn=ODF2;refGene_tx=XM_023242802.1;refGene_gn=ODF2;refGene_tx=XM_023242803.1;refGene_gn=ODF2;refGene_tx=XM_023242805.1;refGene_gn=ODF2;refGene_tx=XM_023242806.1;refGene_gn=ODF2;refGene_tx=XM_023242808.1;refGene_gn=ODF2;refGene_tx=XM_023242809.1
D4 89097107 89097239
refGene_gn=ODF2;refGene_tx=XM_003995938.5;refGene_gn=ODF2;refGene_tx=XM_006939521.4;refGene_gn=ODF2;refGene_tx=XM_006939522.2;refGene_gn=ODF2;refGene_tx=XM_006939523.3;refGene_gn=ODF2;refGene_tx=XM_019816621.2;refGene_gn=ODF2;refGene_tx=XM_019816622.2;refGene_gn=ODF2;refGene_tx=XM_019816623.2;refGene_gn=ODF2;refGene_tx=XM_019816624.2;refGene_gn=ODF2;refGene_tx=XM_019816625.2;refGene_gn=ODF2;refGene_tx=XM_019816626.2;refGene_gn=ODF2;refGene_tx=XM_019816627.2;refGene_gn=ODF2;refGene_tx=XM_019816628.2;refGene_gn=ODF2;refGene_tx=XM_019816629.2;refGene_gn=ODF2;refGene_tx=XM_019816630.1;refGene_gn=ODF2;refGene_tx=XM_023242798.1;refGene_gn=ODF2;refGene_tx=XM_023242799.1;refGene_gn=ODF2;refGene_tx=XM_023242800.1;refGene_gn=ODF2;refGene_tx=XM_023242801.1;refGene_gn=ODF2;refGene_tx=XM_023242802.1;refGene_gn=ODF2;refGene_tx=XM_023242803.1;refGene_gn=ODF2;refGene_tx=XM_023242805.1;refGene_gn=ODF2;refGene_tx=XM_023242806.1;refGene_gn=ODF2;refGene_tx=XM_023242808.1;refGene_gn=ODF

2;refGene_tx=XM_023242809.1
D4 89101837 89101909
refGene_gn=ODF2;refGene_tx=XM_003995938.5;refGene_gn=ODF2;refGene_tx=XM_006939521.4;refGene_gn=ODF2;refGene_tx=XM_006939522.2;refGene_gn=ODF2;refGene_tx=XM_006939523.3;refGene_gn=ODF2;refGene_tx=XM_019816621.2;refGene_gn=ODF2;refGene_tx=XM_019816622.2;refGene_gn=ODF2;refGene_tx=XM_019816623.2;refGene_gn=ODF2;refGene_tx=XM_019816624.2;refGene_gn=ODF2;refGene_tx=XM_019816625.2;refGene_gn=ODF2;refGene_tx=XM_019816626.2;refGene_gn=ODF2;refGene_tx=XM_019816627.2;refGene_gn=ODF2;refGene_tx=XM_019816628.2;refGene_gn=ODF2;refGene_tx=XM_019816629.2;refGene_gn=ODF2;refGene_tx=XM_019816630.1;refGene_gn=ODF2;refGene_tx=XM_023242798.1;refGene_gn=ODF2;refGene_tx=XM_023242799.1;refGene_gn=ODF2;refGene_tx=XM_023242800.1;refGene_gn=ODF2;refGene_tx=XM_023242801.1;refGene_gn=ODF2;refGene_tx=XM_023242802.1;refGene_gn=ODF2;refGene_tx=XM_023242803.1;refGene_gn=ODF2;refGene_tx=XM_023242805.1;refGene_gn=ODF2;refGene_tx=XM_023242806.1;refGene_gn=ODF2;refGene_tx=XM_023242808.1;refGene_gn=ODF2;refGene_tx=XM_023242809.1
D4 89103457 89103529
refGene_gn=ODF2;refGene_tx=XM_003995938.5;refGene_gn=ODF2;refGene_tx=XM_006939521.4;refGene_gn=ODF2;refGene_tx=XM_006939522.2;refGene_gn=ODF2;refGene_tx=XM_006939523.3;refGene_gn=ODF2;refGene_tx=XM_019816621.2;refGene_gn=ODF2;refGene_tx=XM_019816622.2;refGene_gn=ODF2;refGene_tx=XM_019816623.2;refGene_gn=ODF2;refGene_tx=XM_019816624.2;refGene_gn=ODF2;refGene_tx=XM_019816625.2;refGene_gn=ODF2;refGene_tx=XM_019816626.2;refGene_gn=ODF2;refGene_tx=XM_019816627.2;refGene_gn=ODF2;refGene_tx=XM_019816628.2;refGene_gn=ODF2;refGene_tx=XM_019816629.2;refGene_gn=ODF2;refGene_tx=XM_019816630.1;refGene_gn=ODF2;refGene_tx=XM_023242798.1;refGene_gn=ODF2;refGene_tx=XM_023242799.1;refGene_gn=ODF2;refGene_tx=XM_023242800.1;refGene_gn=ODF2;refGene_tx=XM_023242801.1;refGene_gn=ODF2;refGene_tx=XM_023242802.1;refGene_gn=ODF2;refGene_tx=XM_023242803.1;refGene_gn=ODF2;refGene_tx=XM_023242805.1;refGene_gn=ODF2;refGene_tx=XM_023242806.1;refGene_gn=ODF2;refGene_tx=XM_023242808.1;refGene_gn=ODF2;refGene_tx=XM_023242809.1
D4 89103650 89103767
refGene_gn=ODF2;refGene_tx=XM_003995938.5;refGene_gn=ODF2;refGene_tx=XM_006939521.4;refGene_gn=ODF2;refGene_tx=XM_006939522.2;refGene_gn=ODF2;refGene_tx=XM_006939523.3;refGene_gn=ODF2;refGene_tx=XM_019816621.2;refGene_gn=ODF2;refGene_tx=XM_019816622.2;refGene_gn=ODF2;refGene_tx=XM_019816623.2;refGene_gn=ODF2;refGene_tx=XM_019816624.2;refGene_gn=ODF2;refGene_tx=XM_019816625.2;refGene_gn=ODF2;refGene_tx=XM_019816626.2;refGene_gn=ODF2;refGene_tx=XM_019816627.2;refGene_gn=ODF2;refGene_tx=XM_019816628.2;refGene_gn=ODF2;refGene_tx=XM_019816629.2;refGene_gn=ODF2;refGene_tx=XM_019816630.1;refGene_gn=ODF2;refGene_tx=XM_023242798.1;refGene_gn=ODF2;refGene_tx=XM_023242799.1;refGene_gn=ODF2;refGene_tx=XM_023242800.1;refGene_gn=ODF2;refGene_tx=XM_023242801.1;refGene_gn=ODF2;refGene_tx=XM_023242802.1;refGene_gn=ODF2;refGene_tx=XM_023242803.1;refGene_gn=ODF2;refGene_tx=XM_023242805.1;refGene_gn=ODF2;refGene_tx=XM_023242806.1;refGene_gn=ODF2;refGene_tx=XM_023242808.1;refGene_gn=ODF2;refGene_tx=XM_023242809.1
D4 89104411 89104597

refGene_gn=ODF2;refGene_tx=XM_003995938.5;refGene_gn=ODF2;refGene_tx=XM_006939521.4;refGene_gn=ODF2;refGene_tx=XM_006939522.2;refGene_gn=ODF2;refGene_tx=XM_006939523.3;refGene_gn=ODF2;refGene_tx=XM_019816621.2;refGene_gn=ODF2;refGene_tx=XM_019816622.2;refGene_gn=ODF2;refGene_tx=XM_019816623.2;refGene_gn=ODF2;refGene_tx=XM_019816624.2;refGene_gn=ODF2;refGene_tx=XM_019816625.2;refGene_gn=ODF2;refGene_tx=XM_019816626.2;refGene_gn=ODF2;refGene_tx=XM_019816627.2;refGene_gn=ODF2;refGene_tx=XM_019816628.2;refGene_gn=ODF2;refGene_tx=XM_019816629.2;refGene_gn=ODF2;refGene_tx=XM_019816630.1;refGene_gn=ODF2;refGene_tx=XM_023242798.1;refGene_gn=ODF2;refGene_tx=XM_023242799.1;refGene_gn=ODF2;refGene_tx=XM_023242800.1;refGene_gn=ODF2;refGene_tx=XM_023242801.1;refGene_gn=ODF2;refGene_tx=XM_023242802.1;refGene_gn=ODF2;refGene_tx=XM_023242803.1;refGene_gn=ODF2;refGene_tx=XM_023242805.1;refGene_gn=ODF2;refGene_tx=XM_023242806.1;refGene_gn=ODF2;refGene_tx=XM_023242808.1;refGene_gn=ODF2;refGene_tx=XM_023242809.1

D4 89105022 89105132 refGene_gn=ODF2;refGene_tx=XM_003995938.5;refGene_gn=ODF2;refGene_tx=XM_006939521.4;refGene_gn=ODF2;refGene_tx=XM_006939522.2;refGene_gn=ODF2;refGene_tx=XM_006939523.3;refGene_gn=ODF2;refGene_tx=XM_019816621.2;refGene_gn=ODF2;refGene_tx=XM_019816622.2;refGene_gn=ODF2;refGene_tx=XM_019816623.2;refGene_gn=ODF2;refGene_tx=XM_019816624.2;refGene_gn=ODF2;refGene_tx=XM_019816625.2;refGene_gn=ODF2;refGene_tx=XM_019816626.2;refGene_gn=ODF2;refGene_tx=XM_019816627.2;refGene_gn=ODF2;refGene_tx=XM_019816628.2;refGene_gn=ODF2;refGene_tx=XM_019816629.2;refGene_gn=ODF2;refGene_tx=XM_019816630.1;refGene_gn=ODF2;refGene_tx=XM_023242798.1;refGene_gn=ODF2;refGene_tx=XM_023242799.1;refGene_gn=ODF2;refGene_tx=XM_023242800.1;refGene_gn=ODF2;refGene_tx=XM_023242801.1;refGene_gn=ODF2;refGene_tx=XM_023242802.1;refGene_gn=ODF2;refGene_tx=XM_023242803.1;refGene_gn=ODF2;refGene_tx=XM_023242805.1;refGene_gn=ODF2;refGene_tx=XM_023242806.1;refGene_gn=ODF2;refGene_tx=XM_023242808.1;refGene_gn=ODF2;refGene_tx=XM_023242809.1

D4 89106686 89106822 refGene_gn=ODF2;refGene_tx=XM_003995938.5;refGene_gn=ODF2;refGene_tx=XM_006939521.4;refGene_gn=ODF2;refGene_tx=XM_006939522.2;refGene_gn=ODF2;refGene_tx=XM_006939523.3;refGene_gn=ODF2;refGene_tx=XM_019816621.2;refGene_gn=ODF2;refGene_tx=XM_019816622.2;refGene_gn=ODF2;refGene_tx=XM_019816623.2;refGene_gn=ODF2;refGene_tx=XM_019816624.2;refGene_gn=ODF2;refGene_tx=XM_019816625.2;refGene_gn=ODF2;refGene_tx=XM_019816626.2;refGene_gn=ODF2;refGene_tx=XM_019816627.2;refGene_gn=ODF2;refGene_tx=XM_019816628.2;refGene_gn=ODF2;refGene_tx=XM_019816629.2;refGene_gn=ODF2;refGene_tx=XM_019816630.1;refGene_gn=ODF2;refGene_tx=XM_023242798.1;refGene_gn=ODF2;refGene_tx=XM_023242799.1;refGene_gn=ODF2;refGene_tx=XM_023242800.1;refGene_gn=ODF2;refGene_tx=XM_023242801.1;refGene_gn=ODF2;refGene_tx=XM_023242802.1;refGene_gn=ODF2;refGene_tx=XM_023242803.1;refGene_gn=ODF2;refGene_tx=XM_023242805.1;refGene_gn=ODF2;refGene_tx=XM_023242806.1;refGene_gn=ODF2;refGene_tx=XM_023242808.1;refGene_gn=ODF2;refGene_tx=XM_023242809.1

D4 89109847 89109958 refGene_gn=ODF2;refGene_tx=XM_003995938.5;refGene_gn=ODF2;refGene_tx=XM_006939521.4;refGene_gn=ODF2;refGene_tx=XM_006939522.2;refGene_gn=ODF

2;refGene_tx=XM_006939523.3;refGene_gn=ODF2;refGene_tx=XM_019816621.2;
refGene_gn=ODF2;refGene_tx=XM_019816622.2;refGene_gn=ODF2;refGene_tx=X
M_019816623.2;refGene_gn=ODF2;refGene_tx=XM_019816624.2;refGene_gn=ODF
2;refGene_tx=XM_019816625.2;refGene_gn=ODF2;refGene_tx=XM_019816626.2;
refGene_gn=ODF2;refGene_tx=XM_019816627.2;refGene_gn=ODF2;refGene_tx=X
M_019816628.2;refGene_gn=ODF2;refGene_tx=XM_019816629.2;refGene_gn=ODF
2;refGene_tx=XM_019816630.1;refGene_gn=ODF2;refGene_tx=XM_023242798.1;
refGene_gn=ODF2;refGene_tx=XM_023242799.1;refGene_gn=ODF2;refGene_tx=X
M_023242800.1;refGene_gn=ODF2;refGene_tx=XM_023242801.1;refGene_gn=ODF
2;refGene_tx=XM_023242802.1;refGene_gn=ODF2;refGene_tx=XM_023242803.1;
refGene_gn=ODF2;refGene_tx=XM_023242805.1;refGene_gn=ODF2;refGene_tx=X
M_023242806.1;refGene_gn=ODF2;refGene_tx=XM_023242808.1;refGene_gn=ODF
2;refGene_tx=XM_023242809.1
D4 89110061 89110166
refGene_gn=ODF2;refGene_tx=XM_003995938.5;refGene_gn=ODF2;refGene_tx=X
M_006939521.4;refGene_gn=ODF2;refGene_tx=XM_006939522.2;refGene_gn=ODF
2;refGene_tx=XM_006939523.3;refGene_gn=ODF2;refGene_tx=XM_019816621.2;
refGene_gn=ODF2;refGene_tx=XM_019816622.2;refGene_gn=ODF2;refGene_tx=X
M_019816623.2;refGene_gn=ODF2;refGene_tx=XM_019816624.2;refGene_gn=ODF
2;refGene_tx=XM_019816625.2;refGene_gn=ODF2;refGene_tx=XM_019816626.2;
refGene_gn=ODF2;refGene_tx=XM_019816627.2;refGene_gn=ODF2;refGene_tx=X
M_019816628.2;refGene_gn=ODF2;refGene_tx=XM_019816629.2;refGene_gn=ODF
2;refGene_tx=XM_019816630.1;refGene_gn=ODF2;refGene_tx=XM_023242798.1;
refGene_gn=ODF2;refGene_tx=XM_023242799.1;refGene_gn=ODF2;refGene_tx=X
M_023242800.1;refGene_gn=ODF2;refGene_tx=XM_023242801.1;refGene_gn=ODF
2;refGene_tx=XM_023242802.1;refGene_gn=ODF2;refGene_tx=XM_023242803.1;
refGene_gn=ODF2;refGene_tx=XM_023242805.1;refGene_gn=ODF2;refGene_tx=X
M_023242806.1;refGene_gn=ODF2;refGene_tx=XM_023242808.1;refGene_gn=ODF
2;refGene_tx=XM_023242809.1
D4 89112311 89112533
refGene_gn=ODF2;refGene_tx=XM_003995938.5;refGene_gn=ODF2;refGene_tx=X
M_006939521.4;refGene_gn=ODF2;refGene_tx=XM_006939522.2;refGene_gn=ODF
2;refGene_tx=XM_006939523.3;refGene_gn=ODF2;refGene_tx=XM_019816621.2;
refGene_gn=ODF2;refGene_tx=XM_019816622.2;refGene_gn=ODF2;refGene_tx=X
M_019816623.2;refGene_gn=ODF2;refGene_tx=XM_019816624.2;refGene_gn=ODF
2;refGene_tx=XM_019816625.2;refGene_gn=ODF2;refGene_tx=XM_019816626.2;
refGene_gn=ODF2;refGene_tx=XM_019816627.2;refGene_gn=ODF2;refGene_tx=X
M_019816628.2;refGene_gn=ODF2;refGene_tx=XM_019816629.2;refGene_gn=ODF
2;refGene_tx=XM_019816630.1;refGene_gn=ODF2;refGene_tx=XM_023242798.1;
refGene_gn=ODF2;refGene_tx=XM_023242799.1;refGene_gn=ODF2;refGene_tx=X
M_023242800.1;refGene_gn=ODF2;refGene_tx=XM_023242801.1;refGene_gn=ODF
2;refGene_tx=XM_023242802.1;refGene_gn=ODF2;refGene_tx=XM_023242803.1;
refGene_gn=ODF2;refGene_tx=XM_023242805.1;refGene_gn=ODF2;refGene_tx=X
M_023242806.1;refGene_gn=ODF2;refGene_tx=XM_023242808.1;refGene_gn=ODF
2;refGene_tx=XM_023242809.1
D4 89113801 89113902
refGene_gn=ODF2;refGene_tx=XM_006939521.4;refGene_gn=ODF2;refGene_tx=X
M_006939522.2;refGene_gn=ODF2;refGene_tx=XM_006939523.3;refGene_gn=ODF
2;refGene_tx=XM_019816621.2;refGene_gn=ODF2;refGene_tx=XM_019816622.2;
refGene_gn=ODF2;refGene_tx=XM_019816623.2;refGene_gn=ODF2;refGene_tx=X

M_019816624.2;refGene_gn=ODF2;refGene_tx=XM_019816625.2;refGene_gn=ODF2;refGene_tx=XM_019816626.2;refGene_gn=ODF2;refGene_tx=XM_019816627.2;refGene_gn=ODF2;refGene_tx=XM_019816628.2;refGene_gn=ODF2;refGene_tx=XM_019816629.2;refGene_gn=ODF2;refGene_tx=XM_019816630.1;refGene_gn=ODF2;refGene_tx=XM_023242798.1;refGene_gn=ODF2;refGene_tx=XM_023242799.1;refGene_gn=ODF2;refGene_tx=XM_023242800.1;refGene_gn=ODF2;refGene_tx=XM_023242801.1;refGene_gn=ODF2;refGene_tx=XM_023242802.1;refGene_gn=ODF2;refGene_tx=XM_023242803.1;refGene_gn=ODF2;refGene_tx=XM_023242805.1;refGene_gn=ODF2;refGene_tx=XM_023242806.1
D4 89115670 89115833
refGene_gn=ODF2;refGene_tx=XM_006939521.4;refGene_gn=ODF2;refGene_tx=XM_006939522.2;refGene_gn=ODF2;refGene_tx=XM_006939523.3;refGene_gn=ODF2;refGene_tx=XM_019816621.2;refGene_gn=ODF2;refGene_tx=XM_019816622.2;refGene_gn=ODF2;refGene_tx=XM_019816623.2;refGene_gn=ODF2;refGene_tx=XM_019816624.2;refGene_gn=ODF2;refGene_tx=XM_019816625.2;refGene_gn=ODF2;refGene_tx=XM_019816626.2;refGene_gn=ODF2;refGene_tx=XM_019816627.2;refGene_gn=ODF2;refGene_tx=XM_019816628.2;refGene_gn=ODF2;refGene_tx=XM_019816629.2;refGene_gn=ODF2;refGene_tx=XM_019816630.1;refGene_gn=ODF2;refGene_tx=XM_023242798.1;refGene_gn=ODF2;refGene_tx=XM_023242799.1;refGene_gn=ODF2;refGene_tx=XM_023242800.1;refGene_gn=ODF2;refGene_tx=XM_023242801.1;refGene_gn=ODF2;refGene_tx=XM_023242802.1;refGene_gn=ODF2;refGene_tx=XM_023242803.1;refGene_gn=ODF2;refGene_tx=XM_023242805.1;refGene_gn=ODF2;refGene_tx=XM_023242806.1
D4 89116255 89116381
refGene_gn=ODF2;refGene_tx=XM_006939521.4;refGene_gn=ODF2;refGene_tx=XM_006939522.2;refGene_gn=ODF2;refGene_tx=XM_006939523.3;refGene_gn=ODF2;refGene_tx=XM_019816621.2;refGene_gn=ODF2;refGene_tx=XM_019816622.2;refGene_gn=ODF2;refGene_tx=XM_019816623.2;refGene_gn=ODF2;refGene_tx=XM_019816624.2;refGene_gn=ODF2;refGene_tx=XM_019816625.2;refGene_gn=ODF2;refGene_tx=XM_019816626.2;refGene_gn=ODF2;refGene_tx=XM_019816627.2;refGene_gn=ODF2;refGene_tx=XM_019816628.2;refGene_gn=ODF2;refGene_tx=XM_019816629.2;refGene_gn=ODF2;refGene_tx=XM_019816630.1;refGene_gn=ODF2;refGene_tx=XM_023242798.1;refGene_gn=ODF2;refGene_tx=XM_023242799.1;refGene_gn=ODF2;refGene_tx=XM_023242800.1;refGene_gn=ODF2;refGene_tx=XM_023242801.1;refGene_gn=ODF2;refGene_tx=XM_023242802.1;refGene_gn=ODF2;refGene_tx=XM_023242803.1;refGene_gn=ODF2;refGene_tx=XM_023242805.1;refGene_gn=ODF2;refGene_tx=XM_023242806.1
D4 89116829 89117018
refGene_gn=ODF2;refGene_tx=XM_006939521.4;refGene_gn=ODF2;refGene_tx=XM_006939522.2;refGene_gn=ODF2;refGene_tx=XM_006939523.3;refGene_gn=ODF2;refGene_tx=XM_019816621.2;refGene_gn=ODF2;refGene_tx=XM_019816622.2;refGene_gn=ODF2;refGene_tx=XM_019816623.2;refGene_gn=ODF2;refGene_tx=XM_019816624.2;refGene_gn=ODF2;refGene_tx=XM_019816625.2;refGene_gn=ODF2;refGene_tx=XM_019816626.2;refGene_gn=ODF2;refGene_tx=XM_019816627.2;refGene_gn=ODF2;refGene_tx=XM_019816628.2;refGene_gn=ODF2;refGene_tx=XM_019816629.2;refGene_gn=ODF2;refGene_tx=XM_019816630.1;refGene_gn=ODF2;refGene_tx=XM_023242798.1;refGene_gn=ODF2;refGene_tx=XM_023242799.1;refGene_gn=ODF2;refGene_tx=XM_023242800.1;refGene_gn=ODF2;refGene_tx=XM_023242801.1;refGene_gn=ODF2;refGene_tx=XM_023242802.1;refGene_gn=ODF2;refGene_tx=XM_023242803.1;refGene_gn=ODF2;refGene_tx=XM_023242805.1;

refGene_gn=ODF2;refGene_tx=XM_023242806.1
D4	89120363 89120462 refGene_gn=GLE1;refGene_tx=XM_003995943.5
D4	89126344 89126356 refGene_gn=GLE1;refGene_tx=XM_019816631.1
D4	89129835 89130057
refGene_gn=GLE1;refGene_tx=XM_003995943.5;refGene_gn=GLE1;refGene_tx=XM_019816631.1
D4	89133136 89133247
refGene_gn=GLE1;refGene_tx=XM_003995943.5;refGene_gn=GLE1;refGene_tx=XM_019816631.1
D4	89138324 89138473
refGene_gn=GLE1;refGene_tx=XM_003995943.5;refGene_gn=GLE1;refGene_tx=XM_019816631.1
D4	89138715 89138776
refGene_gn=GLE1;refGene_tx=XM_003995943.5;refGene_gn=GLE1;refGene_tx=XM_019816631.1
D4	89138970 89139225
refGene_gn=GLE1;refGene_tx=XM_003995943.5;refGene_gn=GLE1;refGene_tx=XM_019816631.1
D4	89142564 89142793
refGene_gn=GLE1;refGene_tx=XM_003995943.5;refGene_gn=GLE1;refGene_tx=XM_019816631.1
D4	89143341 89143454
refGene_gn=GLE1;refGene_tx=XM_003995943.5;refGene_gn=GLE1;refGene_tx=XM_019816631.1
D4	89143615 89143685
refGene_gn=GLE1;refGene_tx=XM_003995943.5;refGene_gn=GLE1;refGene_tx=XM_019816631.1
D4	89147046 89147189
refGene_gn=GLE1;refGene_tx=XM_003995943.5;refGene_gn=GLE1;refGene_tx=XM_019816631.1
D4	89147286 89147477
refGene_gn=GLE1;refGene_tx=XM_003995943.5;refGene_gn=GLE1;refGene_tx=XM_019816631.1
D4	89148656 89148786
refGene_gn=GLE1;refGene_tx=XM_003995943.5;refGene_gn=GLE1;refGene_tx=XM_019816631.1
D4	89150558 89150663
refGene_gn=GLE1;refGene_tx=XM_003995943.5;refGene_gn=GLE1;refGene_tx=XM_019816631.1
D4	89151837 89151920
refGene_gn=GLE1;refGene_tx=XM_003995943.5;refGene_gn=GLE1;refGene_tx=XM_019816631.1
D4	89152823 89152887
refGene_gn=GLE1;refGene_tx=XM_003995943.5;refGene_gn=GLE1;refGene_tx=XM_019816631.1
D4	89153846 89153915
refGene_gn=GLE1;refGene_tx=XM_003995943.5;refGene_gn=GLE1;refGene_tx=XM_019816631.1
D4	89176230 89176467
refGene_gn=SPTAN1;refGene_tx=XM_019816632.1;refGene_gn=SPTAN1;refGene_

tx=XM_019816633.1;refGene_gn=SPTAN1;refGene_tx=XM_019816634.2;refGene_gn=SPTAN1;refGene_tx=XM_019816635.1;refGene_gn=SPTAN1;refGene_tx=XM_019816636.1;refGene_gn=SPTAN1;refGene_tx=XM_019816637.2;refGene_gn=SPTAN1;refGene_tx=XM_019816638.2;refGene_gn=SPTAN1;refGene_tx=XM_019816639.2;refGene_gn=SPTAN1;refGene_tx=XM_019816640.2;refGene_gn=SPTAN1;refGene_tx=XM_019816641.2
D4 89177674 89177800
refGene_gn=SPTAN1;refGene_tx=XM_019816632.1;refGene_gn=SPTAN1;refGene_tx=XM_019816633.1;refGene_gn=SPTAN1;refGene_tx=XM_019816634.2;refGene_gn=SPTAN1;refGene_tx=XM_019816635.1;refGene_gn=SPTAN1;refGene_tx=XM_019816636.1;refGene_gn=SPTAN1;refGene_tx=XM_019816637.2;refGene_gn=SPTAN1;refGene_tx=XM_019816638.2;refGene_gn=SPTAN1;refGene_tx=XM_019816639.2;refGene_gn=SPTAN1;refGene_tx=XM_019816640.2;refGene_gn=SPTAN1;refGene_tx=XM_019816641.2
D4 89181532 89181673
refGene_gn=SPTAN1;refGene_tx=XM_019816632.1;refGene_gn=SPTAN1;refGene_tx=XM_019816633.1;refGene_gn=SPTAN1;refGene_tx=XM_019816634.2;refGene_gn=SPTAN1;refGene_tx=XM_019816635.1;refGene_gn=SPTAN1;refGene_tx=XM_019816636.1;refGene_gn=SPTAN1;refGene_tx=XM_019816637.2;refGene_gn=SPTAN1;refGene_tx=XM_019816638.2;refGene_gn=SPTAN1;refGene_tx=XM_019816639.2;refGene_gn=SPTAN1;refGene_tx=XM_019816640.2;refGene_gn=SPTAN1;refGene_tx=XM_019816641.2
D4 89182030 89182177
refGene_gn=SPTAN1;refGene_tx=XM_019816632.1;refGene_gn=SPTAN1;refGene_tx=XM_019816633.1;refGene_gn=SPTAN1;refGene_tx=XM_019816634.2;refGene_gn=SPTAN1;refGene_tx=XM_019816635.1;refGene_gn=SPTAN1;refGene_tx=XM_019816636.1;refGene_gn=SPTAN1;refGene_tx=XM_019816637.2;refGene_gn=SPTAN1;refGene_tx=XM_019816638.2;refGene_gn=SPTAN1;refGene_tx=XM_019816639.2;refGene_gn=SPTAN1;refGene_tx=XM_019816640.2;refGene_gn=SPTAN1;refGene_tx=XM_019816641.2
D4 89183608 89183742
refGene_gn=SPTAN1;refGene_tx=XM_019816632.1;refGene_gn=SPTAN1;refGene_tx=XM_019816633.1;refGene_gn=SPTAN1;refGene_tx=XM_019816634.2;refGene_gn=SPTAN1;refGene_tx=XM_019816635.1;refGene_gn=SPTAN1;refGene_tx=XM_019816636.1;refGene_gn=SPTAN1;refGene_tx=XM_019816637.2;refGene_gn=SPTAN1;refGene_tx=XM_019816638.2;refGene_gn=SPTAN1;refGene_tx=XM_019816639.2;refGene_gn=SPTAN1;refGene_tx=XM_019816640.2;refGene_gn=SPTAN1;refGene_tx=XM_019816641.2
D4 89183897 89184042
refGene_gn=SPTAN1;refGene_tx=XM_019816632.1;refGene_gn=SPTAN1;refGene_tx=XM_019816633.1;refGene_gn=SPTAN1;refGene_tx=XM_019816634.2;refGene_gn=SPTAN1;refGene_tx=XM_019816635.1;refGene_gn=SPTAN1;refGene_tx=XM_019816636.1;refGene_gn=SPTAN1;refGene_tx=XM_019816637.2;refGene_gn=SPTAN1;refGene_tx=XM_019816638.2;refGene_gn=SPTAN1;refGene_tx=XM_019816639.2;refGene_gn=SPTAN1;refGene_tx=XM_019816640.2;refGene_gn=SPTAN1;refGene_tx=XM_019816641.2
D4 89184119 89184274
refGene_gn=SPTAN1;refGene_tx=XM_019816632.1;refGene_gn=SPTAN1;refGene_tx=XM_019816633.1;refGene_gn=SPTAN1;refGene_tx=XM_019816634.2;refGene_gn=SPTAN1;refGene_tx=XM_019816635.1;refGene_gn=SPTAN1;refGene_tx=XM_01

9816636.1;refGene_gn=SPTAN1;refGene_tx=XM_019816637.2;refGene_gn=SPTAN1;refGene_tx=XM_019816638.2;refGene_gn=SPTAN1;refGene_tx=XM_019816639.2;refGene_gn=SPTAN1;refGene_tx=XM_019816640.2;refGene_gn=SPTAN1;refGene_tx=XM_019816641.2
D4 89184924 89185060
refGene_gn=SPTAN1;refGene_tx=XM_019816632.1;refGene_gn=SPTAN1;refGene_tx=XM_019816633.1;refGene_gn=SPTAN1;refGene_tx=XM_019816634.2;refGene_gn=SPTAN1;refGene_tx=XM_019816635.1;refGene_gn=SPTAN1;refGene_tx=XM_019816636.1;refGene_gn=SPTAN1;refGene_tx=XM_019816637.2;refGene_gn=SPTAN1;refGene_tx=XM_019816638.2;refGene_gn=SPTAN1;refGene_tx=XM_019816639.2;refGene_gn=SPTAN1;refGene_tx=XM_019816640.2;refGene_gn=SPTAN1;refGene_tx=XM_019816641.2
D4 89185880 89185982
refGene_gn=SPTAN1;refGene_tx=XM_019816632.1;refGene_gn=SPTAN1;refGene_tx=XM_019816633.1;refGene_gn=SPTAN1;refGene_tx=XM_019816634.2;refGene_gn=SPTAN1;refGene_tx=XM_019816635.1;refGene_gn=SPTAN1;refGene_tx=XM_019816636.1;refGene_gn=SPTAN1;refGene_tx=XM_019816637.2;refGene_gn=SPTAN1;refGene_tx=XM_019816638.2;refGene_gn=SPTAN1;refGene_tx=XM_019816639.2;refGene_gn=SPTAN1;refGene_tx=XM_019816640.2;refGene_gn=SPTAN1;refGene_tx=XM_019816641.2
D4 89186841 89186979
refGene_gn=SPTAN1;refGene_tx=XM_019816632.1;refGene_gn=SPTAN1;refGene_tx=XM_019816633.1;refGene_gn=SPTAN1;refGene_tx=XM_019816634.2;refGene_gn=SPTAN1;refGene_tx=XM_019816635.1;refGene_gn=SPTAN1;refGene_tx=XM_019816636.1;refGene_gn=SPTAN1;refGene_tx=XM_019816637.2;refGene_gn=SPTAN1;refGene_tx=XM_019816638.2;refGene_gn=SPTAN1;refGene_tx=XM_019816639.2;refGene_gn=SPTAN1;refGene_tx=XM_019816640.2;refGene_gn=SPTAN1;refGene_tx=XM_019816641.2
D4 89187424 89187535
refGene_gn=SPTAN1;refGene_tx=XM_019816632.1;refGene_gn=SPTAN1;refGene_tx=XM_019816633.1;refGene_gn=SPTAN1;refGene_tx=XM_019816634.2;refGene_gn=SPTAN1;refGene_tx=XM_019816635.1;refGene_gn=SPTAN1;refGene_tx=XM_019816636.1;refGene_gn=SPTAN1;refGene_tx=XM_019816637.2;refGene_gn=SPTAN1;refGene_tx=XM_019816638.2;refGene_gn=SPTAN1;refGene_tx=XM_019816639.2;refGene_gn=SPTAN1;refGene_tx=XM_019816640.2;refGene_gn=SPTAN1;refGene_tx=XM_019816641.2
D4 89188461 89188539
refGene_gn=SPTAN1;refGene_tx=XM_019816632.1;refGene_gn=SPTAN1;refGene_tx=XM_019816633.1;refGene_gn=SPTAN1;refGene_tx=XM_019816634.2;refGene_gn=SPTAN1;refGene_tx=XM_019816635.1;refGene_gn=SPTAN1;refGene_tx=XM_019816636.1;refGene_gn=SPTAN1;refGene_tx=XM_019816637.2;refGene_gn=SPTAN1;refGene_tx=XM_019816638.2;refGene_gn=SPTAN1;refGene_tx=XM_019816639.2;refGene_gn=SPTAN1;refGene_tx=XM_019816640.2;refGene_gn=SPTAN1;refGene_tx=XM_019816641.2
D4 89188653 89188809
refGene_gn=SPTAN1;refGene_tx=XM_019816632.1;refGene_gn=SPTAN1;refGene_tx=XM_019816633.1;refGene_gn=SPTAN1;refGene_tx=XM_019816634.2;refGene_gn=SPTAN1;refGene_tx=XM_019816635.1;refGene_gn=SPTAN1;refGene_tx=XM_019816636.1;refGene_gn=SPTAN1;refGene_tx=XM_019816637.2;refGene_gn=SPTAN1;refGene_tx=XM_019816638.2;refGene_gn=SPTAN1;refGene_tx=XM_019816639.

2;refGene_gn=SPTAN1;refGene_tx=XM_019816640.2;refGene_gn=SPTAN1;refGene_tx=XM_019816641.2
D4 89189043 89189248
refGene_gn=SPTAN1;refGene_tx=XM_019816632.1;refGene_gn=SPTAN1;refGene_tx=XM_019816633.1;refGene_gn=SPTAN1;refGene_tx=XM_019816634.2;refGene_gn=SPTAN1;refGene_tx=XM_019816635.1;refGene_gn=SPTAN1;refGene_tx=XM_019816636.1;refGene_gn=SPTAN1;refGene_tx=XM_019816637.2;refGene_gn=SPTAN1;refGene_tx=XM_019816638.2;refGene_gn=SPTAN1;refGene_tx=XM_019816639.2;refGene_gn=SPTAN1;refGene_tx=XM_019816640.2;refGene_gn=SPTAN1;refGene_tx=XM_019816641.2
D4 89189943 89190125
refGene_gn=SPTAN1;refGene_tx=XM_019816632.1;refGene_gn=SPTAN1;refGene_tx=XM_019816633.1;refGene_gn=SPTAN1;refGene_tx=XM_019816634.2;refGene_gn=SPTAN1;refGene_tx=XM_019816635.1;refGene_gn=SPTAN1;refGene_tx=XM_019816636.1;refGene_gn=SPTAN1;refGene_tx=XM_019816637.2;refGene_gn=SPTAN1;refGene_tx=XM_019816638.2;refGene_gn=SPTAN1;refGene_tx=XM_019816639.2;refGene_gn=SPTAN1;refGene_tx=XM_019816640.2;refGene_gn=SPTAN1;refGene_tx=XM_019816641.2
D4 89190464 89190708
refGene_gn=SPTAN1;refGene_tx=XM_019816632.1;refGene_gn=SPTAN1;refGene_tx=XM_019816633.1;refGene_gn=SPTAN1;refGene_tx=XM_019816634.2;refGene_gn=SPTAN1;refGene_tx=XM_019816635.1;refGene_gn=SPTAN1;refGene_tx=XM_019816636.1;refGene_gn=SPTAN1;refGene_tx=XM_019816637.2;refGene_gn=SPTAN1;refGene_tx=XM_019816638.2;refGene_gn=SPTAN1;refGene_tx=XM_019816639.2;refGene_gn=SPTAN1;refGene_tx=XM_019816640.2;refGene_gn=SPTAN1;refGene_tx=XM_019816641.2
D4 89190901 89191024
refGene_gn=SPTAN1;refGene_tx=XM_019816632.1;refGene_gn=SPTAN1;refGene_tx=XM_019816633.1;refGene_gn=SPTAN1;refGene_tx=XM_019816634.2;refGene_gn=SPTAN1;refGene_tx=XM_019816635.1;refGene_gn=SPTAN1;refGene_tx=XM_019816636.1;refGene_gn=SPTAN1;refGene_tx=XM_019816637.2;refGene_gn=SPTAN1;refGene_tx=XM_019816638.2;refGene_gn=SPTAN1;refGene_tx=XM_019816639.2;refGene_gn=SPTAN1;refGene_tx=XM_019816640.2;refGene_gn=SPTAN1;refGene_tx=XM_019816641.2
D4 89191575 89191793
refGene_gn=SPTAN1;refGene_tx=XM_019816632.1;refGene_gn=SPTAN1;refGene_tx=XM_019816633.1;refGene_gn=SPTAN1;refGene_tx=XM_019816634.2;refGene_gn=SPTAN1;refGene_tx=XM_019816635.1;refGene_gn=SPTAN1;refGene_tx=XM_019816636.1;refGene_gn=SPTAN1;refGene_tx=XM_019816637.2;refGene_gn=SPTAN1;refGene_tx=XM_019816638.2;refGene_gn=SPTAN1;refGene_tx=XM_019816639.2;refGene_gn=SPTAN1;refGene_tx=XM_019816640.2;refGene_gn=SPTAN1;refGene_tx=XM_019816641.2
D4 89192626 89192719
refGene_gn=SPTAN1;refGene_tx=XM_019816632.1;refGene_gn=SPTAN1;refGene_tx=XM_019816633.1;refGene_gn=SPTAN1;refGene_tx=XM_019816634.2;refGene_gn=SPTAN1;refGene_tx=XM_019816635.1;refGene_gn=SPTAN1;refGene_tx=XM_019816636.1;refGene_gn=SPTAN1;refGene_tx=XM_019816637.2;refGene_gn=SPTAN1;refGene_tx=XM_019816638.2;refGene_gn=SPTAN1;refGene_tx=XM_019816639.2;refGene_gn=SPTAN1;refGene_tx=XM_019816640.2;refGene_gn=SPTAN1;refGene_tx=XM_019816641.2

D4 89193079 89193214
refGene_gn=SPTAN1;refGene_tx=XM_019816632.1;refGene_gn=SPTAN1;refGene_tx=XM_019816633.1;refGene_gn=SPTAN1;refGene_tx=XM_019816634.2;refGene_gn=SPTAN1;refGene_tx=XM_019816635.1;refGene_gn=SPTAN1;refGene_tx=XM_019816636.1;refGene_gn=SPTAN1;refGene_tx=XM_019816637.2;refGene_gn=SPTAN1;refGene_tx=XM_019816638.2;refGene_gn=SPTAN1;refGene_tx=XM_019816639.2;refGene_gn=SPTAN1;refGene_tx=XM_019816640.2;refGene_gn=SPTAN1;refGene_tx=XM_019816641.2
D4 89194276 89194425
refGene_gn=SPTAN1;refGene_tx=XM_019816632.1;refGene_gn=SPTAN1;refGene_tx=XM_019816633.1;refGene_gn=SPTAN1;refGene_tx=XM_019816634.2;refGene_gn=SPTAN1;refGene_tx=XM_019816635.1;refGene_gn=SPTAN1;refGene_tx=XM_019816636.1;refGene_gn=SPTAN1;refGene_tx=XM_019816637.2;refGene_gn=SPTAN1;refGene_tx=XM_019816638.2;refGene_gn=SPTAN1;refGene_tx=XM_019816639.2;refGene_gn=SPTAN1;refGene_tx=XM_019816640.2;refGene_gn=SPTAN1;refGene_tx=XM_019816641.2
D4 89195485 89195545
refGene_gn=SPTAN1;refGene_tx=XM_019816632.1;refGene_gn=SPTAN1;refGene_tx=XM_019816633.1;refGene_gn=SPTAN1;refGene_tx=XM_019816634.2;refGene_gn=SPTAN1;refGene_tx=XM_019816635.1;refGene_gn=SPTAN1;refGene_tx=XM_019816637.2;refGene_gn=SPTAN1;refGene_tx=XM_019816638.2;refGene_gn=SPTAN1;refGene_tx=XM_019816639.2
D4 89196647 89196846
refGene_gn=SPTAN1;refGene_tx=XM_019816632.1;refGene_gn=SPTAN1;refGene_tx=XM_019816633.1;refGene_gn=SPTAN1;refGene_tx=XM_019816634.2;refGene_gn=SPTAN1;refGene_tx=XM_019816635.1;refGene_gn=SPTAN1;refGene_tx=XM_019816636.1;refGene_gn=SPTAN1;refGene_tx=XM_019816637.2;refGene_gn=SPTAN1;refGene_tx=XM_019816638.2;refGene_gn=SPTAN1;refGene_tx=XM_019816639.2;refGene_gn=SPTAN1;refGene_tx=XM_019816640.2;refGene_gn=SPTAN1;refGene_tx=XM_019816641.2
D4 89198775 89198880
refGene_gn=SPTAN1;refGene_tx=XM_019816632.1;refGene_gn=SPTAN1;refGene_tx=XM_019816633.1;refGene_gn=SPTAN1;refGene_tx=XM_019816634.2;refGene_gn=SPTAN1;refGene_tx=XM_019816635.1;refGene_gn=SPTAN1;refGene_tx=XM_019816636.1;refGene_gn=SPTAN1;refGene_tx=XM_019816637.2;refGene_gn=SPTAN1;refGene_tx=XM_019816638.2;refGene_gn=SPTAN1;refGene_tx=XM_019816639.2;refGene_gn=SPTAN1;refGene_tx=XM_019816640.2;refGene_gn=SPTAN1;refGene_tx=XM_019816641.2
D4 89199301 89199325
refGene_gn=SPTAN1;refGene_tx=XM_019816632.1;refGene_gn=SPTAN1;refGene_tx=XM_019816633.1;refGene_gn=SPTAN1;refGene_tx=XM_019816634.2;refGene_gn=SPTAN1;refGene_tx=XM_019816635.1;refGene_gn=SPTAN1;refGene_tx=XM_019816636.1;refGene_gn=SPTAN1;refGene_tx=XM_019816637.2;refGene_gn=SPTAN1;refGene_tx=XM_019816638.2;refGene_gn=SPTAN1;refGene_tx=XM_019816639.2;refGene_gn=SPTAN1;refGene_tx=XM_019816640.2;refGene_gn=SPTAN1;refGene_tx=XM_019816641.2
D4 89199865 89199901
refGene_gn=SPTAN1;refGene_tx=XM_019816632.1;refGene_gn=SPTAN1;refGene_tx=XM_019816633.1;refGene_gn=SPTAN1;refGene_tx=XM_019816634.2;refGene_gn=SPTAN1;refGene_tx=XM_019816635.1;refGene_gn=SPTAN1;refGene_tx=XM_01

9816636.1;refGene_gn=SPTAN1;refGene_tx=XM_019816637.2;refGene_gn=SPTAN1;refGene_tx=XM_019816638.2;refGene_gn=SPTAN1;refGene_tx=XM_019816639.2;refGene_gn=SPTAN1;refGene_tx=XM_019816640.2;refGene_gn=SPTAN1;refGene_tx=XM_019816641.2
D4 89201822 89201870
refGene_gn=SPTAN1;refGene_tx=XM_019816632.1;refGene_gn=SPTAN1;refGene_tx=XM_019816633.1;refGene_gn=SPTAN1;refGene_tx=XM_019816634.2;refGene_gn=SPTAN1;refGene_tx=XM_019816636.1;refGene_gn=SPTAN1;refGene_tx=XM_019816637.2;refGene_gn=SPTAN1;refGene_tx=XM_019816638.2;refGene_gn=SPTAN1;refGene_tx=XM_019816639.2;refGene_gn=SPTAN1;refGene_tx=XM_019816640.2;refGene_gn=SPTAN1;refGene_tx=XM_019816641.2
D4 89202644 89202736
refGene_gn=SPTAN1;refGene_tx=XM_019816632.1;refGene_gn=SPTAN1;refGene_tx=XM_019816633.1;refGene_gn=SPTAN1;refGene_tx=XM_019816634.2;refGene_gn=SPTAN1;refGene_tx=XM_019816635.1;refGene_gn=SPTAN1;refGene_tx=XM_019816636.1;refGene_gn=SPTAN1;refGene_tx=XM_019816637.2;refGene_gn=SPTAN1;refGene_tx=XM_019816638.2;refGene_gn=SPTAN1;refGene_tx=XM_019816639.2;refGene_gn=SPTAN1;refGene_tx=XM_019816640.2;refGene_gn=SPTAN1;refGene_tx=XM_019816641.2
D4 89203021 89203166
refGene_gn=SPTAN1;refGene_tx=XM_019816632.1;refGene_gn=SPTAN1;refGene_tx=XM_019816633.1;refGene_gn=SPTAN1;refGene_tx=XM_019816634.2;refGene_gn=SPTAN1;refGene_tx=XM_019816635.1;refGene_gn=SPTAN1;refGene_tx=XM_019816636.1;refGene_gn=SPTAN1;refGene_tx=XM_019816637.2;refGene_gn=SPTAN1;refGene_tx=XM_019816638.2;refGene_gn=SPTAN1;refGene_tx=XM_019816639.2;refGene_gn=SPTAN1;refGene_tx=XM_019816640.2;refGene_gn=SPTAN1;refGene_tx=XM_019816641.2
D4 89203273 89203455
refGene_gn=SPTAN1;refGene_tx=XM_019816632.1;refGene_gn=SPTAN1;refGene_tx=XM_019816633.1;refGene_gn=SPTAN1;refGene_tx=XM_019816634.2;refGene_gn=SPTAN1;refGene_tx=XM_019816635.1;refGene_gn=SPTAN1;refGene_tx=XM_019816636.1;refGene_gn=SPTAN1;refGene_tx=XM_019816637.2;refGene_gn=SPTAN1;refGene_tx=XM_019816638.2;refGene_gn=SPTAN1;refGene_tx=XM_019816639.2;refGene_gn=SPTAN1;refGene_tx=XM_019816640.2;refGene_gn=SPTAN1;refGene_tx=XM_019816641.2
D4 89205605 89205705
refGene_gn=SPTAN1;refGene_tx=XM_019816632.1;refGene_gn=SPTAN1;refGene_tx=XM_019816633.1;refGene_gn=SPTAN1;refGene_tx=XM_019816634.2;refGene_gn=SPTAN1;refGene_tx=XM_019816635.1;refGene_gn=SPTAN1;refGene_tx=XM_019816636.1;refGene_gn=SPTAN1;refGene_tx=XM_019816637.2;refGene_gn=SPTAN1;refGene_tx=XM_019816638.2;refGene_gn=SPTAN1;refGene_tx=XM_019816639.2;refGene_gn=SPTAN1;refGene_tx=XM_019816640.2;refGene_gn=SPTAN1;refGene_tx=XM_019816641.2
D4 89205865 89206063
refGene_gn=SPTAN1;refGene_tx=XM_019816632.1;refGene_gn=SPTAN1;refGene_tx=XM_019816633.1;refGene_gn=SPTAN1;refGene_tx=XM_019816634.2;refGene_gn=SPTAN1;refGene_tx=XM_019816635.1;refGene_gn=SPTAN1;refGene_tx=XM_019816636.1;refGene_gn=SPTAN1;refGene_tx=XM_019816637.2;refGene_gn=SPTAN1;refGene_tx=XM_019816638.2;refGene_gn=SPTAN1;refGene_tx=XM_019816639.2;refGene_gn=SPTAN1;refGene_tx=XM_019816640.2;refGene_gn=SPTAN1;refGen

e_tx=XM_019816641.2
D4 89206161 89206308
refGene_gn=SPTAN1;refGene_tx=XM_019816632.1;refGene_gn=SPTAN1;refGene_tx=XM_019816633.1;refGene_gn=SPTAN1;refGene_tx=XM_019816634.2;refGene_gn=SPTAN1;refGene_tx=XM_019816635.1;refGene_gn=SPTAN1;refGene_tx=XM_019816636.1;refGene_gn=SPTAN1;refGene_tx=XM_019816637.2;refGene_gn=SPTAN1;refGene_tx=XM_019816638.2;refGene_gn=SPTAN1;refGene_tx=XM_019816639.2;refGene_gn=SPTAN1;refGene_tx=XM_019816640.2;refGene_gn=SPTAN1;refGene_tx=XM_019816641.2
D4 89206997 89207101
refGene_gn=SPTAN1;refGene_tx=XM_019816632.1;refGene_gn=SPTAN1;refGene_tx=XM_019816633.1;refGene_gn=SPTAN1;refGene_tx=XM_019816634.2;refGene_gn=SPTAN1;refGene_tx=XM_019816635.1;refGene_gn=SPTAN1;refGene_tx=XM_019816636.1;refGene_gn=SPTAN1;refGene_tx=XM_019816637.2;refGene_gn=SPTAN1;refGene_tx=XM_019816638.2;refGene_gn=SPTAN1;refGene_tx=XM_019816639.2;refGene_gn=SPTAN1;refGene_tx=XM_019816640.2;refGene_gn=SPTAN1;refGene_tx=XM_019816641.2
D4 89207186 89207349
refGene_gn=SPTAN1;refGene_tx=XM_019816632.1;refGene_gn=SPTAN1;refGene_tx=XM_019816633.1;refGene_gn=SPTAN1;refGene_tx=XM_019816634.2;refGene_gn=SPTAN1;refGene_tx=XM_019816635.1;refGene_gn=SPTAN1;refGene_tx=XM_019816636.1;refGene_gn=SPTAN1;refGene_tx=XM_019816637.2;refGene_gn=SPTAN1;refGene_tx=XM_019816638.2;refGene_gn=SPTAN1;refGene_tx=XM_019816639.2;refGene_gn=SPTAN1;refGene_tx=XM_019816640.2;refGene_gn=SPTAN1;refGene_tx=XM_019816641.2
D4 89207759 89207774
refGene_gn=SPTAN1;refGene_tx=XM_019816632.1;refGene_gn=SPTAN1;refGene_tx=XM_019816634.2;refGene_gn=SPTAN1;refGene_tx=XM_019816635.1;refGene_gn=SPTAN1;refGene_tx=XM_019816636.1;refGene_gn=SPTAN1;refGene_tx=XM_019816637.2;refGene_gn=SPTAN1;refGene_tx=XM_019816639.2;refGene_gn=SPTAN1;refGene_tx=XM_019816640.2
D4 89209592 89209724
refGene_gn=SPTAN1;refGene_tx=XM_019816632.1;refGene_gn=SPTAN1;refGene_tx=XM_019816633.1;refGene_gn=SPTAN1;refGene_tx=XM_019816634.2;refGene_gn=SPTAN1;refGene_tx=XM_019816635.1;refGene_gn=SPTAN1;refGene_tx=XM_019816636.1;refGene_gn=SPTAN1;refGene_tx=XM_019816637.2;refGene_gn=SPTAN1;refGene_tx=XM_019816638.2;refGene_gn=SPTAN1;refGene_tx=XM_019816639.2;refGene_gn=SPTAN1;refGene_tx=XM_019816640.2;refGene_gn=SPTAN1;refGene_tx=XM_019816641.2
D4 89209901 89210039
refGene_gn=SPTAN1;refGene_tx=XM_019816632.1;refGene_gn=SPTAN1;refGene_tx=XM_019816633.1;refGene_gn=SPTAN1;refGene_tx=XM_019816634.2;refGene_gn=SPTAN1;refGene_tx=XM_019816635.1;refGene_gn=SPTAN1;refGene_tx=XM_019816636.1;refGene_gn=SPTAN1;refGene_tx=XM_019816637.2;refGene_gn=SPTAN1;refGene_tx=XM_019816638.2;refGene_gn=SPTAN1;refGene_tx=XM_019816639.2;refGene_gn=SPTAN1;refGene_tx=XM_019816640.2;refGene_gn=SPTAN1;refGene_tx=XM_019816641.2
D4 89211052 89211157
refGene_gn=SPTAN1;refGene_tx=XM_019816632.1;refGene_gn=SPTAN1;refGene_tx=XM_019816633.1;refGene_gn=SPTAN1;refGene_tx=XM_019816634.2;refGene_

gn=SPTAN1;refGene_tx=XM_019816635.1;refGene_gn=SPTAN1;refGene_tx=XM_01
9816636.1;refGene_gn=SPTAN1;refGene_tx=XM_019816637.2;refGene_gn=SPTAN
1;refGene_tx=XM_019816638.2;refGene_gn=SPTAN1;refGene_tx=XM_019816639.
2;refGene_gn=SPTAN1;refGene_tx=XM_019816640.2;refGene_gn=SPTAN1;refGen
e_tx=XM_019816641.2
D4 89213325 89213534
refGene_gn=SPTAN1;refGene_tx=XM_019816632.1;refGene_gn=SPTAN1;refGene_
tx=XM_019816633.1;refGene_gn=SPTAN1;refGene_tx=XM_019816634.2;refGene_
gn=SPTAN1;refGene_tx=XM_019816635.1;refGene_gn=SPTAN1;refGene_tx=XM_01
9816636.1;refGene_gn=SPTAN1;refGene_tx=XM_019816637.2;refGene_gn=SPTAN
1;refGene_tx=XM_019816638.2;refGene_gn=SPTAN1;refGene_tx=XM_019816639.
2;refGene_gn=SPTAN1;refGene_tx=XM_019816640.2;refGene_gn=SPTAN1;refGen
e_tx=XM_019816641.2
D4 89214226 89214347
refGene_gn=SPTAN1;refGene_tx=XM_019816632.1;refGene_gn=SPTAN1;refGene_
tx=XM_019816633.1;refGene_gn=SPTAN1;refGene_tx=XM_019816634.2;refGene_
gn=SPTAN1;refGene_tx=XM_019816635.1;refGene_gn=SPTAN1;refGene_tx=XM_01
9816636.1;refGene_gn=SPTAN1;refGene_tx=XM_019816637.2;refGene_gn=SPTAN
1;refGene_tx=XM_019816638.2;refGene_gn=SPTAN1;refGene_tx=XM_019816639.
2;refGene_gn=SPTAN1;refGene_tx=XM_019816640.2;refGene_gn=SPTAN1;refGen
e_tx=XM_019816641.2
D4 89214565 89214687
refGene_gn=SPTAN1;refGene_tx=XM_019816632.1;refGene_gn=SPTAN1;refGene_
tx=XM_019816633.1;refGene_gn=SPTAN1;refGene_tx=XM_019816634.2;refGene_
gn=SPTAN1;refGene_tx=XM_019816635.1;refGene_gn=SPTAN1;refGene_tx=XM_01
9816636.1;refGene_gn=SPTAN1;refGene_tx=XM_019816637.2;refGene_gn=SPTAN
1;refGene_tx=XM_019816638.2;refGene_gn=SPTAN1;refGene_tx=XM_019816639.
2;refGene_gn=SPTAN1;refGene_tx=XM_019816640.2;refGene_gn=SPTAN1;refGen
e_tx=XM_019816641.2
D4 89215050 89215183
refGene_gn=SPTAN1;refGene_tx=XM_019816632.1;refGene_gn=SPTAN1;refGene_
tx=XM_019816633.1;refGene_gn=SPTAN1;refGene_tx=XM_019816634.2;refGene_
gn=SPTAN1;refGene_tx=XM_019816635.1;refGene_gn=SPTAN1;refGene_tx=XM_01
9816636.1;refGene_gn=SPTAN1;refGene_tx=XM_019816637.2;refGene_gn=SPTAN
1;refGene_tx=XM_019816638.2;refGene_gn=SPTAN1;refGene_tx=XM_019816639.
2;refGene_gn=SPTAN1;refGene_tx=XM_019816640.2;refGene_gn=SPTAN1;refGen
e_tx=XM_019816641.2
D4 89216783 89216882
refGene_gn=SPTAN1;refGene_tx=XM_019816632.1;refGene_gn=SPTAN1;refGene_
tx=XM_019816633.1;refGene_gn=SPTAN1;refGene_tx=XM_019816634.2;refGene_
gn=SPTAN1;refGene_tx=XM_019816635.1;refGene_gn=SPTAN1;refGene_tx=XM_01
9816636.1;refGene_gn=SPTAN1;refGene_tx=XM_019816637.2;refGene_gn=SPTAN
1;refGene_tx=XM_019816638.2;refGene_gn=SPTAN1;refGene_tx=XM_019816639.
2;refGene_gn=SPTAN1;refGene_tx=XM_019816640.2;refGene_gn=SPTAN1;refGen
e_tx=XM_019816641.2
D4 89218818 89218978
refGene_gn=SPTAN1;refGene_tx=XM_019816632.1;refGene_gn=SPTAN1;refGene_
tx=XM_019816633.1;refGene_gn=SPTAN1;refGene_tx=XM_019816634.2;refGene_
gn=SPTAN1;refGene_tx=XM_019816635.1;refGene_gn=SPTAN1;refGene_tx=XM_01
9816636.1;refGene_gn=SPTAN1;refGene_tx=XM_019816637.2;refGene_gn=SPTAN

1;refGene_tx=XM_019816638.2;refGene_gn=SPTAN1;refGene_tx=XM_019816639.2;refGene_gn=SPTAN1;refGene_tx=XM_019816640.2;refGene_gn=SPTAN1;refGene_tx=XM_019816641.2
D4 89219591 89219668
refGene_gn=SPTAN1;refGene_tx=XM_019816632.1;refGene_gn=SPTAN1;refGene_tx=XM_019816633.1;refGene_gn=SPTAN1;refGene_tx=XM_019816634.2;refGene_gn=SPTAN1;refGene_tx=XM_019816635.1;refGene_gn=SPTAN1;refGene_tx=XM_019816636.1;refGene_gn=SPTAN1;refGene_tx=XM_019816637.2;refGene_gn=SPTAN1;refGene_tx=XM_019816638.2;refGene_gn=SPTAN1;refGene_tx=XM_019816639.2;refGene_gn=SPTAN1;refGene_tx=XM_019816640.2;refGene_gn=SPTAN1;refGene_tx=XM_019816641.2
D4 89220224 89220434
refGene_gn=SPTAN1;refGene_tx=XM_019816632.1;refGene_gn=SPTAN1;refGene_tx=XM_019816633.1;refGene_gn=SPTAN1;refGene_tx=XM_019816634.2;refGene_gn=SPTAN1;refGene_tx=XM_019816635.1;refGene_gn=SPTAN1;refGene_tx=XM_019816636.1;refGene_gn=SPTAN1;refGene_tx=XM_019816637.2;refGene_gn=SPTAN1;refGene_tx=XM_019816638.2;refGene_gn=SPTAN1;refGene_tx=XM_019816639.2;refGene_gn=SPTAN1;refGene_tx=XM_019816640.2;refGene_gn=SPTAN1;refGene_tx=XM_019816641.2
D4 89220733 89221030
refGene_gn=SPTAN1;refGene_tx=XM_019816632.1;refGene_gn=SPTAN1;refGene_tx=XM_019816633.1;refGene_gn=SPTAN1;refGene_tx=XM_019816634.2;refGene_gn=SPTAN1;refGene_tx=XM_019816635.1;refGene_gn=SPTAN1;refGene_tx=XM_019816636.1;refGene_gn=SPTAN1;refGene_tx=XM_019816637.2;refGene_gn=SPTAN1;refGene_tx=XM_019816638.2;refGene_gn=SPTAN1;refGene_tx=XM_019816639.2;refGene_gn=SPTAN1;refGene_tx=XM_019816640.2;refGene_gn=SPTAN1;refGene_tx=XM_019816641.2
D4 89221448 89221561
refGene_gn=SPTAN1;refGene_tx=XM_019816632.1;refGene_gn=SPTAN1;refGene_tx=XM_019816633.1;refGene_gn=SPTAN1;refGene_tx=XM_019816634.2;refGene_gn=SPTAN1;refGene_tx=XM_019816635.1;refGene_gn=SPTAN1;refGene_tx=XM_019816636.1;refGene_gn=SPTAN1;refGene_tx=XM_019816637.2;refGene_gn=SPTAN1;refGene_tx=XM_019816638.2;refGene_gn=SPTAN1;refGene_tx=XM_019816639.2;refGene_gn=SPTAN1;refGene_tx=XM_019816640.2;refGene_gn=SPTAN1;refGene_tx=XM_019816641.2
D4 89221976 89221994
refGene_gn=SPTAN1;refGene_tx=XM_019816632.1;refGene_gn=SPTAN1;refGene_tx=XM_019816633.1;refGene_gn=SPTAN1;refGene_tx=XM_019816634.2;refGene_gn=SPTAN1;refGene_tx=XM_019816635.1;refGene_gn=SPTAN1;refGene_tx=XM_019816636.1;refGene_gn=SPTAN1;refGene_tx=XM_019816637.2;refGene_gn=SPTAN1;refGene_tx=XM_019816638.2;refGene_gn=SPTAN1;refGene_tx=XM_019816640.2;refGene_gn=SPTAN1;refGene_tx=XM_019816641.2
D4 89223208 89223271
refGene_gn=SPTAN1;refGene_tx=XM_019816632.1;refGene_gn=SPTAN1;refGene_tx=XM_019816633.1;refGene_gn=SPTAN1;refGene_tx=XM_019816634.2;refGene_gn=SPTAN1;refGene_tx=XM_019816635.1;refGene_gn=SPTAN1;refGene_tx=XM_019816636.1
D4 89224511 89224566
refGene_gn=SPTAN1;refGene_tx=XM_019816632.1;refGene_gn=SPTAN1;refGene_tx=XM_019816633.1;refGene_gn=SPTAN1;refGene_tx=XM_019816634.2;refGene_

gn=SPTAN1;refGene_tx=XM_019816635.1;refGene_gn=SPTAN1;refGene_tx=XM_01
9816636.1;refGene_gn=SPTAN1;refGene_tx=XM_019816637.2;refGene_gn=SPTAN
1;refGene_tx=XM_019816638.2;refGene_gn=SPTAN1;refGene_tx=XM_019816639.
2;refGene_gn=SPTAN1;refGene_tx=XM_019816640.2;refGene_gn=SPTAN1;refGen
e_tx=XM_019816641.2
D4 89224968 89225165
refGene_gn=SPTAN1;refGene_tx=XM_019816632.1;refGene_gn=SPTAN1;refGene_
tx=XM_019816633.1;refGene_gn=SPTAN1;refGene_tx=XM_019816634.2;refGene_
gn=SPTAN1;refGene_tx=XM_019816635.1;refGene_gn=SPTAN1;refGene_tx=XM_01
9816636.1;refGene_gn=SPTAN1;refGene_tx=XM_019816637.2;refGene_gn=SPTAN
1;refGene_tx=XM_019816638.2;refGene_gn=SPTAN1;refGene_tx=XM_019816639.
2;refGene_gn=SPTAN1;refGene_tx=XM_019816640.2;refGene_gn=SPTAN1;refGen
e_tx=XM_019816641.2
D4 89225335 89225389
refGene_gn=SPTAN1;refGene_tx=XM_019816632.1;refGene_gn=SPTAN1;refGene_
tx=XM_019816633.1;refGene_gn=SPTAN1;refGene_tx=XM_019816634.2;refGene_
gn=SPTAN1;refGene_tx=XM_019816635.1;refGene_gn=SPTAN1;refGene_tx=XM_01
9816636.1;refGene_gn=SPTAN1;refGene_tx=XM_019816637.2;refGene_gn=SPTAN
1;refGene_tx=XM_019816638.2;refGene_gn=SPTAN1;refGene_tx=XM_019816639.
2;refGene_gn=SPTAN1;refGene_tx=XM_019816640.2;refGene_gn=SPTAN1;refGen
e_tx=XM_019816641.2
D4 89225469 89225616
refGene_gn=SPTAN1;refGene_tx=XM_019816632.1;refGene_gn=SPTAN1;refGene_
tx=XM_019816633.1;refGene_gn=SPTAN1;refGene_tx=XM_019816634.2;refGene_
gn=SPTAN1;refGene_tx=XM_019816635.1;refGene_gn=SPTAN1;refGene_tx=XM_01
9816636.1;refGene_gn=SPTAN1;refGene_tx=XM_019816637.2;refGene_gn=SPTAN
1;refGene_tx=XM_019816638.2;refGene_gn=SPTAN1;refGene_tx=XM_019816639.
2;refGene_gn=SPTAN1;refGene_tx=XM_019816640.2;refGene_gn=SPTAN1;refGen
e_tx=XM_019816641.2
D4 89225701 89225849
refGene_gn=SPTAN1;refGene_tx=XM_019816632.1;refGene_gn=SPTAN1;refGene_
tx=XM_019816633.1;refGene_gn=SPTAN1;refGene_tx=XM_019816634.2;refGene_
gn=SPTAN1;refGene_tx=XM_019816635.1;refGene_gn=SPTAN1;refGene_tx=XM_01
9816636.1;refGene_gn=SPTAN1;refGene_tx=XM_019816637.2;refGene_gn=SPTAN
1;refGene_tx=XM_019816638.2;refGene_gn=SPTAN1;refGene_tx=XM_019816639.
2;refGene_gn=SPTAN1;refGene_tx=XM_019816640.2;refGene_gn=SPTAN1;refGen
e_tx=XM_019816641.2
D4 89226151 89226277
refGene_gn=SPTAN1;refGene_tx=XM_019816632.1;refGene_gn=SPTAN1;refGene_
tx=XM_019816633.1;refGene_gn=SPTAN1;refGene_tx=XM_019816634.2;refGene_
gn=SPTAN1;refGene_tx=XM_019816635.1;refGene_gn=SPTAN1;refGene_tx=XM_01
9816636.1;refGene_gn=SPTAN1;refGene_tx=XM_019816637.2;refGene_gn=SPTAN
1;refGene_tx=XM_019816638.2;refGene_gn=SPTAN1;refGene_tx=XM_019816639.
2;refGene_gn=SPTAN1;refGene_tx=XM_019816640.2;refGene_gn=SPTAN1;refGen
e_tx=XM_019816641.2
D4 89226715 89226954
refGene_gn=WDR34;refGene_tx=XM_003995947.5;refGene_gn=WDR34;refGene_tx
=XM_006939526.3;refGene_gn=WDR34;refGene_tx=XM_019816642.2;refGene_gn=
WDR34;refGene_tx=XM_019816644.2;refGene_gn=WDR34;refGene_tx=XM_0198166
45.2

D4 89227154 89227312
refGene_gn=WDR34;refGene_tx=XM_003995947.5;refGene_gn=WDR34;refGene_tx
=XM_006939526.3;refGene_gn=WDR34;refGene_tx=XM_019816642.2;refGene_gn=
WDR34;refGene_tx=XM_019816644.2;refGene_gn=WDR34;refGene_tx=XM_0198166
45.2
D4 89227594 89227827
refGene_gn=WDR34;refGene_tx=XM_003995947.5;refGene_gn=WDR34;refGene_tx
=XM_006939526.3;refGene_gn=WDR34;refGene_tx=XM_019816642.2;refGene_gn=
WDR34;refGene_tx=XM_019816644.2;refGene_gn=WDR34;refGene_tx=XM_0198166
45.2
D4 89228029 89228197
refGene_gn=WDR34;refGene_tx=XM_003995947.5;refGene_gn=WDR34;refGene_tx
=XM_006939526.3;refGene_gn=WDR34;refGene_tx=XM_019816642.2;refGene_gn=
WDR34;refGene_tx=XM_019816644.2;refGene_gn=WDR34;refGene_tx=XM_0198166
45.2
D4 89228699 89228809
refGene_gn=WDR34;refGene_tx=XM_003995947.5;refGene_gn=WDR34;refGene_tx
=XM_006939526.3;refGene_gn=WDR34;refGene_tx=XM_019816642.2;refGene_gn=
WDR34;refGene_tx=XM_019816644.2;refGene_gn=WDR34;refGene_tx=XM_0198166
45.2
D4 89229403 89229561
refGene_gn=WDR34;refGene_tx=XM_003995947.5;refGene_gn=WDR34;refGene_tx
=XM_006939526.3;refGene_gn=WDR34;refGene_tx=XM_019816642.2;refGene_gn=
WDR34;refGene_tx=XM_019816644.2;refGene_gn=WDR34;refGene_tx=XM_0198166
45.2
D4 89230076 89230186
refGene_gn=WDR34;refGene_tx=XM_003995947.5;refGene_gn=WDR34;refGene_tx
=XM_006939526.3;refGene_gn=WDR34;refGene_tx=XM_019816642.2;refGene_gn=
WDR34;refGene_tx=XM_019816644.2;refGene_gn=WDR34;refGene_tx=XM_0198166
45.2
D4 89234911 89235160
refGene_gn=WDR34;refGene_tx=XM_003995947.5;refGene_gn=WDR34;refGene_tx
=XM_006939526.3;refGene_gn=WDR34;refGene_tx=XM_019816642.2;refGene_gn=
WDR34;refGene_tx=XM_019816644.2;refGene_gn=WDR34;refGene_tx=XM_0198166
45.2
D4 89236444 89236476 refGene_gn=WDR34;refGene_tx=XM_006939526.3
D4 89243580 89243655
refGene_gn=WDR34;refGene_tx=XM_019816642.2;refGene_gn=WDR34;refGene_tx
=XM_019816644.2;refGene_gn=WDR34;refGene_tx=XM_019816645.2
D4 89247519 89247705
refGene_gn=WDR34;refGene_tx=XM_003995947.5;refGene_gn=WDR34;refGene_tx
=XM_019816642.2;refGene_gn=WDR34;refGene_tx=XM_019816644.2;refGene_gn=
WDR34;refGene_tx=XM_019816645.2
D4 89270952 89271025 refGene_gn=SET;refGene_tx=XM_023242812.1
D4 89272589 89272647 refGene_gn=SET;refGene_tx=XM_023242812.1
D4 89273314 89273457 refGene_gn=SET;refGene_tx=XM_023242812.1
D4 89273814 89273918 refGene_gn=SET;refGene_tx=XM_023242812.1
D4 89274022 89274136 refGene_gn=SET;refGene_tx=XM_023242812.1
D4 89274759 89274930 refGene_gn=SET;refGene_tx=XM_023242812.1
D4 89275272 89275419 refGene_gn=SET;refGene_tx=XM_023242812.1

D4 89276038 89276062 refGene_gn=SET;refGene_tx=XM_023242812.1
D4 89281281 89281305
refGene_gn=PKN3;refGene_tx=XM_023242813.1;refGene_gn=PKN3;refGene_tx=XM_023242814.1;refGene_gn=PKN3;refGene_tx=XM_023242815.1
D4 89282539 89282554 refGene_gn=PKN3;refGene_tx=XM_006939532.4
D4 89283716 89283791
refGene_gn=PKN3;refGene_tx=XM_006939530.3;refGene_gn=PKN3;refGene_tx=XM_006939531.3;refGene_gn=PKN3;refGene_tx=XM_011288387.2;refGene_gn=PKN3;refGene_tx=XM_019816646.1;refGene_gn=PKN3;refGene_tx=XM_019816647.1;refGene_gn=PKN3;refGene_tx=XM_019816648.1
D4 89283905 89284140
refGene_gn=PKN3;refGene_tx=XM_006939530.3;refGene_gn=PKN3;refGene_tx=XM_006939531.3;refGene_gn=PKN3;refGene_tx=XM_006939532.4;refGene_gn=PKN3;refGene_tx=XM_011288387.2;refGene_gn=PKN3;refGene_tx=XM_019816646.1;refGene_gn=PKN3;refGene_tx=XM_019816647.1;refGene_gn=PKN3;refGene_tx=XM_019816648.1;refGene_gn=PKN3;refGene_tx=XM_023242813.1;refGene_gn=PKN3;refGene_tx=XM_023242814.1;refGene_gn=PKN3;refGene_tx=XM_023242815.1
D4 89284347 89284493
refGene_gn=PKN3;refGene_tx=XM_006939530.3;refGene_gn=PKN3;refGene_tx=XM_006939531.3;refGene_gn=PKN3;refGene_tx=XM_006939532.4;refGene_gn=PKN3;refGene_tx=XM_011288387.2;refGene_gn=PKN3;refGene_tx=XM_019816646.1;refGene_gn=PKN3;refGene_tx=XM_019816647.1;refGene_gn=PKN3;refGene_tx=XM_019816648.1;refGene_gn=PKN3;refGene_tx=XM_023242813.1;refGene_gn=PKN3;refGene_tx=XM_023242814.1;refGene_gn=PKN3;refGene_tx=XM_023242815.1
D4 89284984 89285096
refGene_gn=PKN3;refGene_tx=XM_006939530.3;refGene_gn=PKN3;refGene_tx=XM_006939531.3;refGene_gn=PKN3;refGene_tx=XM_006939532.4;refGene_gn=PKN3;refGene_tx=XM_011288387.2;refGene_gn=PKN3;refGene_tx=XM_019816646.1;refGene_gn=PKN3;refGene_tx=XM_019816647.1;refGene_gn=PKN3;refGene_tx=XM_019816648.1;refGene_gn=PKN3;refGene_tx=XM_023242813.1;refGene_gn=PKN3;refGene_tx=XM_023242814.1;refGene_gn=PKN3;refGene_tx=XM_023242815.1
D4 89285178 89285306
refGene_gn=PKN3;refGene_tx=XM_006939530.3;refGene_gn=PKN3;refGene_tx=XM_006939531.3;refGene_gn=PKN3;refGene_tx=XM_006939532.4;refGene_gn=PKN3;refGene_tx=XM_011288387.2;refGene_gn=PKN3;refGene_tx=XM_019816646.1;refGene_gn=PKN3;refGene_tx=XM_019816647.1;refGene_gn=PKN3;refGene_tx=XM_019816648.1;refGene_gn=PKN3;refGene_tx=XM_023242813.1;refGene_gn=PKN3;refGene_tx=XM_023242814.1;refGene_gn=PKN3;refGene_tx=XM_023242815.1
D4 89285516 89285700
refGene_gn=PKN3;refGene_tx=XM_006939530.3;refGene_gn=PKN3;refGene_tx=XM_006939531.3;refGene_gn=PKN3;refGene_tx=XM_006939532.4;refGene_gn=PKN3;refGene_tx=XM_011288387.2;refGene_gn=PKN3;refGene_tx=XM_019816646.1;refGene_gn=PKN3;refGene_tx=XM_019816647.1;refGene_gn=PKN3;refGene_tx=XM_019816648.1;refGene_gn=PKN3;refGene_tx=XM_023242813.1;refGene_gn=PKN3;refGene_tx=XM_023242814.1;refGene_gn=PKN3;refGene_tx=XM_023242815.1
D4 89286383 89286530
refGene_gn=PKN3;refGene_tx=XM_006939530.3;refGene_gn=PKN3;refGene_tx=XM_006939531.3;refGene_gn=PKN3;refGene_tx=XM_006939532.4;refGene_gn=PKN3;refGene_tx=XM_011288387.2;refGene_gn=PKN3;refGene_tx=XM_019816646.1;refGene_gn=PKN3;refGene_tx=XM_019816647.1;refGene_gn=PKN3;refGene_tx=X

M_019816648.1;refGene_gn=PKN3;refGene_tx=XM_023242813.1;refGene_gn=PKN3;refGene_tx=XM_023242814.1;refGene_gn=PKN3;refGene_tx=XM_023242815.1
D4 89286613 89286723
refGene_gn=PKN3;refGene_tx=XM_006939530.3;refGene_gn=PKN3;refGene_tx=XM_006939531.3;refGene_gn=PKN3;refGene_tx=XM_006939532.4;refGene_gn=PKN3;refGene_tx=XM_011288387.2;refGene_gn=PKN3;refGene_tx=XM_019816646.1;refGene_gn=PKN3;refGene_tx=XM_019816647.1;refGene_gn=PKN3;refGene_tx=XM_019816648.1;refGene_gn=PKN3;refGene_tx=XM_023242813.1;refGene_gn=PKN3;refGene_tx=XM_023242814.1;refGene_gn=PKN3;refGene_tx=XM_023242815.1
D4 89286829 89286973
refGene_gn=PKN3;refGene_tx=XM_006939530.3;refGene_gn=PKN3;refGene_tx=XM_006939531.3;refGene_gn=PKN3;refGene_tx=XM_006939532.4;refGene_gn=PKN3;refGene_tx=XM_011288387.2;refGene_gn=PKN3;refGene_tx=XM_019816646.1;refGene_gn=PKN3;refGene_tx=XM_019816647.1;refGene_gn=PKN3;refGene_tx=XM_019816648.1;refGene_gn=PKN3;refGene_tx=XM_023242813.1;refGene_gn=PKN3;refGene_tx=XM_023242814.1;refGene_gn=PKN3;refGene_tx=XM_023242815.1
D4 89287392 89287468
refGene_gn=PKN3;refGene_tx=XM_006939530.3;refGene_gn=PKN3;refGene_tx=XM_006939531.3;refGene_gn=PKN3;refGene_tx=XM_006939532.4;refGene_gn=PKN3;refGene_tx=XM_011288387.2;refGene_gn=PKN3;refGene_tx=XM_019816646.1;refGene_gn=PKN3;refGene_tx=XM_019816647.1;refGene_gn=PKN3;refGene_tx=XM_019816648.1;refGene_gn=PKN3;refGene_tx=XM_023242813.1;refGene_gn=PKN3;refGene_tx=XM_023242814.1;refGene_gn=PKN3;refGene_tx=XM_023242815.1
D4 89287555 89287724
refGene_gn=PKN3;refGene_tx=XM_006939530.3;refGene_gn=PKN3;refGene_tx=XM_006939531.3;refGene_gn=PKN3;refGene_tx=XM_006939532.4;refGene_gn=PKN3;refGene_tx=XM_011288387.2;refGene_gn=PKN3;refGene_tx=XM_019816646.1;refGene_gn=PKN3;refGene_tx=XM_019816647.1;refGene_gn=PKN3;refGene_tx=XM_019816648.1;refGene_gn=PKN3;refGene_tx=XM_023242813.1;refGene_gn=PKN3;refGene_tx=XM_023242814.1;refGene_gn=PKN3;refGene_tx=XM_023242815.1
D4 89287859 89288010
refGene_gn=PKN3;refGene_tx=XM_006939530.3;refGene_gn=PKN3;refGene_tx=XM_006939531.3;refGene_gn=PKN3;refGene_tx=XM_006939532.4;refGene_gn=PKN3;refGene_tx=XM_011288387.2;refGene_gn=PKN3;refGene_tx=XM_019816646.1;refGene_gn=PKN3;refGene_tx=XM_019816647.1;refGene_gn=PKN3;refGene_tx=XM_019816648.1;refGene_gn=PKN3;refGene_tx=XM_023242813.1;refGene_gn=PKN3;refGene_tx=XM_023242814.1;refGene_gn=PKN3;refGene_tx=XM_023242815.1
D4 89288117 89288185
refGene_gn=PKN3;refGene_tx=XM_006939530.3;refGene_gn=PKN3;refGene_tx=XM_006939531.3;refGene_gn=PKN3;refGene_tx=XM_006939532.4;refGene_gn=PKN3;refGene_tx=XM_011288387.2;refGene_gn=PKN3;refGene_tx=XM_019816646.1;refGene_gn=PKN3;refGene_tx=XM_019816647.1;refGene_gn=PKN3;refGene_tx=XM_019816648.1;refGene_gn=PKN3;refGene_tx=XM_023242813.1;refGene_gn=PKN3;refGene_tx=XM_023242814.1;refGene_gn=PKN3;refGene_tx=XM_023242815.1
D4 89288499 89288563
refGene_gn=PKN3;refGene_tx=XM_006939530.3;refGene_gn=PKN3;refGene_tx=XM_006939531.3;refGene_gn=PKN3;refGene_tx=XM_006939532.4;refGene_gn=PKN3;refGene_tx=XM_011288387.2;refGene_gn=PKN3;refGene_tx=XM_019816646.1;refGene_gn=PKN3;refGene_tx=XM_019816647.1;refGene_gn=PKN3;refGene_tx=XM_019816648.1;refGene_gn=PKN3;refGene_tx=XM_023242813.1;refGene_gn=PKN

3;refGene_tx=XM_023242814.1;refGene_gn=PKN3;refGene_tx=XM_023242815.1
D4 89288713 89288805
refGene_gn=PKN3;refGene_tx=XM_006939530.3;refGene_gn=PKN3;refGene_tx=XM_006939531.3;refGene_gn=PKN3;refGene_tx=XM_006939532.4;refGene_gn=PKN3;refGene_tx=XM_011288387.2;refGene_gn=PKN3;refGene_tx=XM_019816646.1;refGene_gn=PKN3;refGene_tx=XM_019816647.1;refGene_gn=PKN3;refGene_tx=XM_019816648.1;refGene_gn=PKN3;refGene_tx=XM_023242813.1;refGene_gn=PKN3;refGene_tx=XM_023242814.1;refGene_gn=PKN3;refGene_tx=XM_023242815.1
D4 89289139 89289316
refGene_gn=PKN3;refGene_tx=XM_006939530.3;refGene_gn=PKN3;refGene_tx=XM_006939531.3;refGene_gn=PKN3;refGene_tx=XM_006939532.4;refGene_gn=PKN3;refGene_tx=XM_011288387.2;refGene_gn=PKN3;refGene_tx=XM_019816646.1;refGene_gn=PKN3;refGene_tx=XM_019816647.1;refGene_gn=PKN3;refGene_tx=XM_019816648.1;refGene_gn=PKN3;refGene_tx=XM_023242813.1;refGene_gn=PKN3;refGene_tx=XM_023242814.1;refGene_gn=PKN3;refGene_tx=XM_023242815.1
D4 89290056 89290119
refGene_gn=PKN3;refGene_tx=XM_006939530.3;refGene_gn=PKN3;refGene_tx=XM_006939531.3;refGene_gn=PKN3;refGene_tx=XM_006939532.4;refGene_gn=PKN3;refGene_tx=XM_011288387.2;refGene_gn=PKN3;refGene_tx=XM_019816646.1;refGene_gn=PKN3;refGene_tx=XM_019816647.1;refGene_gn=PKN3;refGene_tx=XM_019816648.1;refGene_gn=PKN3;refGene_tx=XM_023242813.1;refGene_gn=PKN3;refGene_tx=XM_023242814.1;refGene_gn=PKN3;refGene_tx=XM_023242815.1
D4 89290212 89290289
refGene_gn=PKN3;refGene_tx=XM_006939530.3;refGene_gn=PKN3;refGene_tx=XM_006939531.3;refGene_gn=PKN3;refGene_tx=XM_006939532.4;refGene_gn=PKN3;refGene_tx=XM_011288387.2;refGene_gn=PKN3;refGene_tx=XM_019816646.1;refGene_gn=PKN3;refGene_tx=XM_019816647.1;refGene_gn=PKN3;refGene_tx=XM_019816648.1;refGene_gn=PKN3;refGene_tx=XM_023242813.1;refGene_gn=PKN3;refGene_tx=XM_023242814.1;refGene_gn=PKN3;refGene_tx=XM_023242815.1
D4 89290729 89290872
refGene_gn=PKN3;refGene_tx=XM_006939530.3;refGene_gn=PKN3;refGene_tx=XM_006939531.3;refGene_gn=PKN3;refGene_tx=XM_006939532.4;refGene_gn=PKN3;refGene_tx=XM_011288387.2;refGene_gn=PKN3;refGene_tx=XM_019816646.1;refGene_gn=PKN3;refGene_tx=XM_019816647.1;refGene_gn=PKN3;refGene_tx=XM_019816648.1;refGene_gn=PKN3;refGene_tx=XM_023242813.1;refGene_gn=PKN3;refGene_tx=XM_023242814.1;refGene_gn=PKN3;refGene_tx=XM_023242815.1
D4 89290960 89291068
refGene_gn=PKN3;refGene_tx=XM_006939530.3;refGene_gn=PKN3;refGene_tx=XM_006939531.3;refGene_gn=PKN3;refGene_tx=XM_006939532.4;refGene_gn=PKN3;refGene_tx=XM_011288387.2;refGene_gn=PKN3;refGene_tx=XM_019816646.1;refGene_gn=PKN3;refGene_tx=XM_019816647.1;refGene_gn=PKN3;refGene_tx=XM_019816648.1;refGene_gn=PKN3;refGene_tx=XM_023242813.1;refGene_gn=PKN3;refGene_tx=XM_023242814.1;refGene_gn=PKN3;refGene_tx=XM_023242815.1
D4 89291229 89291310
refGene_gn=PKN3;refGene_tx=XM_006939530.3;refGene_gn=PKN3;refGene_tx=XM_006939531.3;refGene_gn=PKN3;refGene_tx=XM_006939532.4;refGene_gn=PKN3;refGene_tx=XM_011288387.2;refGene_gn=PKN3;refGene_tx=XM_019816646.1;refGene_gn=PKN3;refGene_tx=XM_019816647.1;refGene_gn=PKN3;refGene_tx=XM_019816648.1;refGene_gn=PKN3;refGene_tx=XM_023242813.1;refGene_gn=PKN3;refGene_tx=XM_023242814.1;refGene_gn=PKN3;refGene_tx=XM_023242815.1

D4 89291405 89291618
refGene_gn=PKN3;refGene_tx=XM_006939530.3;refGene_gn=PKN3;refGene_tx=XM_006939531.3;refGene_gn=PKN3;refGene_tx=XM_006939532.4;refGene_gn=PKN3;refGene_tx=XM_011288387.2;refGene_gn=PKN3;refGene_tx=XM_019816646.1;refGene_gn=PKN3;refGene_tx=XM_019816647.1;refGene_gn=PKN3;refGene_tx=XM_019816648.1;refGene_gn=PKN3;refGene_tx=XM_023242813.1;refGene_gn=PKN3;refGene_tx=XM_023242814.1;refGene_gn=PKN3;refGene_tx=XM_023242815.1
D4 89292125 89292555
refGene_gn=ZDHHC12;refGene_tx=XM_011288388.3;refGene_gn=ZDHHC12;refGene_tx=XM_019816651.2;refGene_gn=ZDHHC12;refGene_tx=XM_023242816.1
D4 89292660 89292827
refGene_gn=ZDHHC12;refGene_tx=XM_011288388.3;refGene_gn=ZDHHC12;refGene_tx=XM_019816651.2;refGene_gn=ZDHHC12;refGene_tx=XM_023242816.1
D4 89292988 89293066
refGene_gn=ZDHHC12;refGene_tx=XM_011288388.3;refGene_gn=ZDHHC12;refGene_tx=XM_019816651.2;refGene_gn=ZDHHC12;refGene_tx=XM_023242816.1
D4 89293422 89293559
refGene_gn=ZDHHC12;refGene_tx=XM_011288388.3;refGene_gn=ZDHHC12;refGene_tx=XM_019816651.2;refGene_gn=ZDHHC12;refGene_tx=XM_023242816.1
D4 89295022 89295122
refGene_gn=ZDHHC12;refGene_tx=XM_011288388.3;refGene_gn=ZDHHC12;refGene_tx=XM_019816651.2;refGene_gn=ZDHHC12;refGene_tx=XM_023242816.1
D4 89299524 89299582
refGene_gn=ZER1;refGene_tx=XM_011288383.2;refGene_gn=ZER1;refGene_tx=XM_011288384.3;refGene_gn=ZER1;refGene_tx=XM_011288386.2
D4 89301581 89301684
refGene_gn=ZER1;refGene_tx=XM_011288383.2;refGene_gn=ZER1;refGene_tx=XM_011288384.3;refGene_gn=ZER1;refGene_tx=XM_011288386.2
D4 89303096 89303194
refGene_gn=ZER1;refGene_tx=XM_011288383.2;refGene_gn=ZER1;refGene_tx=XM_011288384.3;refGene_gn=ZER1;refGene_tx=XM_011288386.2
D4 89304376 89304565
refGene_gn=ZER1;refGene_tx=XM_011288383.2;refGene_gn=ZER1;refGene_tx=XM_011288384.3;refGene_gn=ZER1;refGene_tx=XM_011288386.2
D4 89304880 89304996
refGene_gn=ZER1;refGene_tx=XM_011288383.2;refGene_gn=ZER1;refGene_tx=XM_011288384.3;refGene_gn=ZER1;refGene_tx=XM_011288386.2
D4 89306833 89306953
refGene_gn=ZER1;refGene_tx=XM_011288383.2;refGene_gn=ZER1;refGene_tx=XM_011288384.3;refGene_gn=ZER1;refGene_tx=XM_011288386.2
D4 89307115 89307157
refGene_gn=ZER1;refGene_tx=XM_011288383.2;refGene_gn=ZER1;refGene_tx=XM_011288384.3;refGene_gn=ZER1;refGene_tx=XM_011288386.2
D4 89307893 89308109
refGene_gn=ZER1;refGene_tx=XM_011288383.2;refGene_gn=ZER1;refGene_tx=XM_011288384.3;refGene_gn=ZER1;refGene_tx=XM_011288386.2
D4 89312122 89312296
refGene_gn=ZER1;refGene_tx=XM_011288383.2;refGene_gn=ZER1;refGene_tx=XM_011288384.3;refGene_gn=ZER1;refGene_tx=XM_011288386.2
D4 89312664 89312811

refGene_gn=ZER1;refGene_tx=XM_011288383.2;refGene_gn=ZER1;refGene_tx=XM_011288384.3;refGene_gn=ZER1;refGene_tx=XM_011288386.2
D4 89312974 89313089
refGene_gn=ZER1;refGene_tx=XM_011288383.2;refGene_gn=ZER1;refGene_tx=XM_011288384.3;refGene_gn=ZER1;refGene_tx=XM_011288386.2
D4 89313957 89314134
refGene_gn=ZER1;refGene_tx=XM_011288383.2;refGene_gn=ZER1;refGene_tx=XM_011288384.3;refGene_gn=ZER1;refGene_tx=XM_011288386.2
D4 89314446 89314883
refGene_gn=ZER1;refGene_tx=XM_011288383.2;refGene_gn=ZER1;refGene_tx=XM_011288384.3;refGene_gn=ZER1;refGene_tx=XM_011288386.2
D4 89315085 89315236
refGene_gn=ZER1;refGene_tx=XM_011288383.2;refGene_gn=ZER1;refGene_tx=XM_011288384.3;refGene_gn=ZER1;refGene_tx=XM_011288386.2
D4 89317546 89317704
refGene_gn=ZER1;refGene_tx=XM_011288383.2;refGene_gn=ZER1;refGene_tx=XM_011288384.3;refGene_gn=ZER1;refGene_tx=XM_011288386.2
D4 89342234 89342257 refGene_gn=TBC1D13;refGene_tx=XM_003995951.5
D4 89343220 89343294 refGene_gn=TBC1D13;refGene_tx=XM_003995951.5
D4 89344843 89344884 refGene_gn=TBC1D13;refGene_tx=XM_003995951.5
D4 89345450 89345512 refGene_gn=TBC1D13;refGene_tx=XM_003995951.5
D4 89345663 89345763
refGene_gn=TBC1D13;refGene_tx=XM_003995951.5;refGene_gn=TBC1D13;refGene_tx=XM_019816650.2
D4 89346530 89346613
refGene_gn=TBC1D13;refGene_tx=XM_003995951.5;refGene_gn=TBC1D13;refGene_tx=XM_019816650.2
D4 89349132 89349292
refGene_gn=TBC1D13;refGene_tx=XM_003995951.5;refGene_gn=TBC1D13;refGene_tx=XM_019816650.2
D4 89352415 89352626
refGene_gn=TBC1D13;refGene_tx=XM_003995951.5;refGene_gn=TBC1D13;refGene_tx=XM_019816650.2
D4 89353191 89353355
refGene_gn=TBC1D13;refGene_tx=XM_003995951.5;refGene_gn=TBC1D13;refGene_tx=XM_019816650.2
D4 89355223 89355384
refGene_gn=TBC1D13;refGene_tx=XM_003995951.5;refGene_gn=TBC1D13;refGene_tx=XM_019816650.2
D4 89355800 89355858
refGene_gn=TBC1D13;refGene_tx=XM_003995951.5;refGene_gn=TBC1D13;refGene_tx=XM_019816650.2
D4 89356722 89356788
refGene_gn=TBC1D13;refGene_tx=XM_003995951.5;refGene_gn=TBC1D13;refGene_tx=XM_019816650.2
D4 89360654 89361173 refGene_gn=ENDOG;refGene_tx=XM_003996000.4
D4 89362212 89362322 refGene_gn=ENDOG;refGene_tx=XM_003996000.4
D4 89363494 89363777 refGene_gn=ENDOG;refGene_tx=XM_003996000.4
D4 89363842 89363977
refGene_gn=SPOUT1;refGene_tx=XM_003996001.4;refGene_gn=SPOUT1;refGene_

tx=XM_019816652.2
D4 89364888 89365036
refGene_gn=SPOUT1;refGene_tx=XM_003996001.4;refGene_gn=SPOUT1;refGene_tx=XM_019816652.2
D4 89365227 89365330
refGene_gn=SPOUT1;refGene_tx=XM_003996001.4;refGene_gn=SPOUT1;refGene_tx=XM_019816652.2
D4 89365778 89365877
refGene_gn=SPOUT1;refGene_tx=XM_003996001.4;refGene_gn=SPOUT1;refGene_tx=XM_019816652.2
D4 89365982 89366055
refGene_gn=SPOUT1;refGene_tx=XM_003996001.4;refGene_gn=SPOUT1;refGene_tx=XM_019816652.2
D4 89367001 89367132
refGene_gn=SPOUT1;refGene_tx=XM_003996001.4;refGene_gn=SPOUT1;refGene_tx=XM_019816652.2
D4 89367324 89367374
refGene_gn=SPOUT1;refGene_tx=XM_003996001.4;refGene_gn=SPOUT1;refGene_tx=XM_019816652.2
D4 89367494 89367584
refGene_gn=SPOUT1;refGene_tx=XM_003996001.4;refGene_gn=SPOUT1;refGene_tx=XM_019816652.2
D4 89367818 89367978
refGene_gn=SPOUT1;refGene_tx=XM_003996001.4;refGene_gn=SPOUT1;refGene_tx=XM_019816652.2
D4 89369859 89369985
refGene_gn=SPOUT1;refGene_tx=XM_003996001.4;refGene_gn=SPOUT1;refGene_tx=XM_019816652.2
D4 89370297 89370343 refGene_gn=SPOUT1;refGene_tx=XM_003996001.4
D4 89370909 89370945 refGene_gn=SPOUT1;refGene_tx=XM_003996001.4
D4 89376604 89376670
refGene_gn=KYAT1;refGene_tx=XM_003995952.4;refGene_gn=KYAT1;refGene_tx=XM_006939536.3;refGene_gn=KYAT1;refGene_tx=XM_006939537.3;refGene_gn=KYAT1;refGene_tx=XM_006939538.4;refGene_gn=KYAT1;refGene_tx=XM_011288382.3;refGene_gn=KYAT1;refGene_tx=XM_023242817.1;refGene_gn=KYAT1;refGene_tx=XM_023242818.1
D4 89376771 89376858
refGene_gn=KYAT1;refGene_tx=XM_003995952.4;refGene_gn=KYAT1;refGene_tx=XM_006939536.3;refGene_gn=KYAT1;refGene_tx=XM_006939537.3;refGene_gn=KYAT1;refGene_tx=XM_006939538.4;refGene_gn=KYAT1;refGene_tx=XM_011288382.3;refGene_gn=KYAT1;refGene_tx=XM_023242817.1;refGene_gn=KYAT1;refGene_tx=XM_023242818.1
D4 89377625 89377705
refGene_gn=KYAT1;refGene_tx=XM_003995952.4;refGene_gn=KYAT1;refGene_tx=XM_006939536.3;refGene_gn=KYAT1;refGene_tx=XM_006939537.3;refGene_gn=KYAT1;refGene_tx=XM_006939538.4;refGene_gn=KYAT1;refGene_tx=XM_011288382.3;refGene_gn=KYAT1;refGene_tx=XM_023242817.1;refGene_gn=KYAT1;refGene_tx=XM_023242818.1
D4 89377801 89377988
refGene_gn=KYAT1;refGene_tx=XM_003995952.4;refGene_gn=KYAT1;refGene_tx

=XM_006939536.3;refGene_gn=KYAT1;refGene_tx=XM_006939537.3;refGene_gn=KYAT1;refGene_tx=XM_006939538.4;refGene_gn=KYAT1;refGene_tx=XM_011288382.3;refGene_gn=KYAT1;refGene_tx=XM_023242817.1;refGene_gn=KYAT1;refGene_tx=XM_023242818.1
D4 89378098 89378188
refGene_gn=KYAT1;refGene_tx=XM_003995952.4;refGene_gn=KYAT1;refGene_tx=XM_006939536.3;refGene_gn=KYAT1;refGene_tx=XM_006939537.3;refGene_gn=KYAT1;refGene_tx=XM_006939538.4;refGene_gn=KYAT1;refGene_tx=XM_011288382.3;refGene_gn=KYAT1;refGene_tx=XM_023242817.1;refGene_gn=KYAT1;refGene_tx=XM_023242818.1
D4 89378326 89378403
refGene_gn=KYAT1;refGene_tx=XM_003995952.4;refGene_gn=KYAT1;refGene_tx=XM_006939536.3;refGene_gn=KYAT1;refGene_tx=XM_006939537.3;refGene_gn=KYAT1;refGene_tx=XM_006939538.4;refGene_gn=KYAT1;refGene_tx=XM_011288382.3;refGene_gn=KYAT1;refGene_tx=XM_023242817.1;refGene_gn=KYAT1;refGene_tx=XM_023242818.1
D4 89378747 89378868
refGene_gn=KYAT1;refGene_tx=XM_003995952.4;refGene_gn=KYAT1;refGene_tx=XM_006939536.3;refGene_gn=KYAT1;refGene_tx=XM_006939537.3;refGene_gn=KYAT1;refGene_tx=XM_006939538.4;refGene_gn=KYAT1;refGene_tx=XM_011288382.3;refGene_gn=KYAT1;refGene_tx=XM_023242817.1;refGene_gn=KYAT1;refGene_tx=XM_023242818.1
D4 89379732 89379861
refGene_gn=KYAT1;refGene_tx=XM_003995952.4;refGene_gn=KYAT1;refGene_tx=XM_006939536.3;refGene_gn=KYAT1;refGene_tx=XM_006939537.3;refGene_gn=KYAT1;refGene_tx=XM_006939538.4;refGene_gn=KYAT1;refGene_tx=XM_011288382.3;refGene_gn=KYAT1;refGene_tx=XM_023242817.1;refGene_gn=KYAT1;refGene_tx=XM_023242818.1
D4 89380087 89380174
refGene_gn=KYAT1;refGene_tx=XM_003995952.4;refGene_gn=KYAT1;refGene_tx=XM_006939536.3;refGene_gn=KYAT1;refGene_tx=XM_006939537.3;refGene_gn=KYAT1;refGene_tx=XM_006939538.4;refGene_gn=KYAT1;refGene_tx=XM_011288382.3;refGene_gn=KYAT1;refGene_tx=XM_023242817.1;refGene_gn=KYAT1;refGene_tx=XM_023242818.1
D4 89380254 89380404
refGene_gn=KYAT1;refGene_tx=XM_003995952.4;refGene_gn=KYAT1;refGene_tx=XM_006939536.3;refGene_gn=KYAT1;refGene_tx=XM_006939537.3;refGene_gn=KYAT1;refGene_tx=XM_006939538.4;refGene_gn=KYAT1;refGene_tx=XM_011288382.3;refGene_gn=KYAT1;refGene_tx=XM_023242817.1;refGene_gn=KYAT1;refGene_tx=XM_023242818.1
D4 89382526 89382674
refGene_gn=KYAT1;refGene_tx=XM_003995952.4;refGene_gn=KYAT1;refGene_tx=XM_006939536.3;refGene_gn=KYAT1;refGene_tx=XM_006939537.3;refGene_gn=KYAT1;refGene_tx=XM_006939538.4;refGene_gn=KYAT1;refGene_tx=XM_011288382.3;refGene_gn=KYAT1;refGene_tx=XM_023242817.1;refGene_gn=KYAT1;refGene_tx=XM_023242818.1
D4 89384772 89384831
refGene_gn=KYAT1;refGene_tx=XM_003995952.4;refGene_gn=KYAT1;refGene_tx=XM_006939536.3;refGene_gn=KYAT1;refGene_tx=XM_006939537.3;refGene_gn=KYAT1;refGene_tx=XM_006939538.4;refGene_gn=KYAT1;refGene_tx=XM_0112883

82.3;refGene_gn=KYAT1;refGene_tx=XM_023242817.1;refGene_gn=KYAT1;refGene_tx=XM_023242818.1
D4 89386870 89386966 refGene_gn=KYAT1;refGene_tx=XM_006939536.3;refGene_gn=KYAT1;refGene_tx=XM_006939537.3;refGene_gn=KYAT1;refGene_tx=XM_011288382.3
D4 89387746 89387842 refGene_gn=KYAT1;refGene_tx=XM_023242817.1;refGene_gn=KYAT1;refGene_tx=XM_023242818.1
D4 89432172 89434329 refGene_gn=LRRC8A;refGene_tx=XM_003995953.4;refGene_gn=LRRC8A;refGene_tx=XM_006939539.4;refGene_gn=LRRC8A;refGene_tx=XM_019816654.2;refGene_gn=LRRC8A;refGene_tx=XM_019816655.2
D4 89439638 89439914 refGene_gn=LRRC8A;refGene_tx=XM_003995953.4;refGene_gn=LRRC8A;refGene_tx=XM_006939539.4;refGene_gn=LRRC8A;refGene_tx=XM_019816654.2;refGene_gn=LRRC8A;refGene_tx=XM_019816655.2
D4 89444963 89444996 refGene_gn=PHYHD1;refGene_tx=XM_003995955.4;refGene_gn=PHYHD1;refGene_tx=XM_006939540.3;refGene_gn=PHYHD1;refGene_tx=XM_006939541.3
D4 89447885 89448041 refGene_gn=PHYHD1;refGene_tx=XM_003995955.4;refGene_gn=PHYHD1;refGene_tx=XM_006939540.3;refGene_gn=PHYHD1;refGene_tx=XM_006939541.3
D4 89452642 89452718 refGene_gn=PHYHD1;refGene_tx=XM_003995955.4;refGene_gn=PHYHD1;refGene_tx=XM_006939540.3;refGene_gn=PHYHD1;refGene_tx=XM_006939541.3
D4 89452857 89452905 refGene_gn=PHYHD1;refGene_tx=XM_003995955.4;refGene_gn=PHYHD1;refGene_tx=XM_006939540.3;refGene_gn=PHYHD1;refGene_tx=XM_006939541.3
D4 89453932 89453988 refGene_gn=PHYHD1;refGene_tx=XM_003995955.4;refGene_gn=PHYHD1;refGene_tx=XM_006939540.3;refGene_gn=PHYHD1;refGene_tx=XM_006939541.3
D4 89454086 89454149 refGene_gn=PHYHD1;refGene_tx=XM_003995955.4;refGene_gn=PHYHD1;refGene_tx=XM_006939540.3;refGene_gn=PHYHD1;refGene_tx=XM_006939541.3
D4 89454834 89454856 refGene_gn=PHYHD1;refGene_tx=XM_003995955.4;refGene_gn=PHYHD1;refGene_tx=XM_006939540.3;refGene_gn=PHYHD1;refGene_tx=XM_006939541.3
D4 89456363 89456492 refGene_gn=PHYHD1;refGene_tx=XM_003995955.4;refGene_gn=PHYHD1;refGene_tx=XM_006939541.3
D4 89456591 89456708 refGene_gn=PHYHD1;refGene_tx=XM_003995955.4;refGene_gn=PHYHD1;refGene_tx=XM_006939540.3;refGene_gn=PHYHD1;refGene_tx=XM_006939541.3
D4 89457444 89457449 refGene_gn=PHYHD1;refGene_tx=XM_006939541.3
D4 89457743 89457870 refGene_gn=PHYHD1;refGene_tx=XM_003995955.4;refGene_gn=PHYHD1;refGene_tx=XM_006939540.3
D4 89457974 89458020 refGene_gn=PHYHD1;refGene_tx=XM_003995955.4;refGene_gn=PHYHD1;refGene_tx=XM_006939540.3

D4 89461106 89462723 refGene_gn=DOLK;refGene_tx=XM_003995957.5
D4 89463138 89463170
refGene_gn=NUP188;refGene_tx=XM_023242820.1;refGene_gn=NUP188;refGene_tx=XM_023242821.1
D4 89475304 89475359
refGene_gn=NUP188;refGene_tx=XM_023242820.1;refGene_gn=NUP188;refGene_tx=XM_023242821.1
D4 89477092 89477166
refGene_gn=NUP188;refGene_tx=XM_023242820.1;refGene_gn=NUP188;refGene_tx=XM_023242821.1
D4 89479716 89479801
refGene_gn=NUP188;refGene_tx=XM_023242820.1;refGene_gn=NUP188;refGene_tx=XM_023242821.1
D4 89481867 89481948
refGene_gn=NUP188;refGene_tx=XM_023242820.1;refGene_gn=NUP188;refGene_tx=XM_023242821.1
D4 89482430 89482475
refGene_gn=NUP188;refGene_tx=XM_023242820.1;refGene_gn=NUP188;refGene_tx=XM_023242821.1
D4 89483474 89483567
refGene_gn=NUP188;refGene_tx=XM_023242820.1;refGene_gn=NUP188;refGene_tx=XM_023242821.1
D4 89483960 89484080
refGene_gn=NUP188;refGene_tx=XM_023242820.1;refGene_gn=NUP188;refGene_tx=XM_023242821.1
D4 89492663 89492875
refGene_gn=NUP188;refGene_tx=XM_023242820.1;refGene_gn=NUP188;refGene_tx=XM_023242821.1
D4 89494379 89494494
refGene_gn=NUP188;refGene_tx=XM_023242820.1;refGene_gn=NUP188;refGene_tx=XM_023242821.1
D4 89495134 89495335
refGene_gn=NUP188;refGene_tx=XM_023242820.1;refGene_gn=NUP188;refGene_tx=XM_023242821.1
D4 89496243 89496333
refGene_gn=NUP188;refGene_tx=XM_023242820.1;refGene_gn=NUP188;refGene_tx=XM_023242821.1
D4 89499110 89499176
refGene_gn=NUP188;refGene_tx=XM_023242820.1;refGene_gn=NUP188;refGene_tx=XM_023242821.1
D4 89500140 89500260
refGene_gn=NUP188;refGene_tx=XM_023242820.1;refGene_gn=NUP188;refGene_tx=XM_023242821.1
D4 89500755 89500882
refGene_gn=NUP188;refGene_tx=XM_023242820.1;refGene_gn=NUP188;refGene_tx=XM_023242821.1
D4 89501838 89501991
refGene_gn=NUP188;refGene_tx=XM_023242820.1;refGene_gn=NUP188;refGene_tx=XM_023242821.1
D4 89502166 89502293

refGene_gn=NUP188;refGene_tx=XM_023242820.1;refGene_gn=NUP188;refGene_tx=XM_023242821.1
D4	89502500	89502588	refGene_gn=NUP188;refGene_tx=XM_023242820.1;refGene_gn=NUP188;refGene_tx=XM_023242821.1
D4	89502681	89502758	refGene_gn=NUP188;refGene_tx=XM_023242820.1;refGene_gn=NUP188;refGene_tx=XM_023242821.1
D4	89503334	89503449	refGene_gn=NUP188;refGene_tx=XM_023242820.1;refGene_gn=NUP188;refGene_tx=XM_023242821.1
D4	89504816	89504937	refGene_gn=NUP188;refGene_tx=XM_023242820.1;refGene_gn=NUP188;refGene_tx=XM_023242821.1
D4	89505064	89505131	refGene_gn=NUP188;refGene_tx=XM_023242820.1;refGene_gn=NUP188;refGene_tx=XM_023242821.1
D4	89505822	89505951	refGene_gn=NUP188;refGene_tx=XM_023242820.1;refGene_gn=NUP188;refGene_tx=XM_023242821.1
D4	89506275	89506415	refGene_gn=NUP188;refGene_tx=XM_023242820.1;refGene_gn=NUP188;refGene_tx=XM_023242821.1
D4	89507087	89507093	refGene_gn=NUP188;refGene_tx=XM_023242820.1
D4	89507240	89507347	refGene_gn=NUP188;refGene_tx=XM_023242820.1;refGene_gn=NUP188;refGene_tx=XM_023242821.1
D4	89509001	89509208	refGene_gn=NUP188;refGene_tx=XM_023242820.1;refGene_gn=NUP188;refGene_tx=XM_023242821.1
D4	89509321	89509491	refGene_gn=NUP188;refGene_tx=XM_023242820.1;refGene_gn=NUP188;refGene_tx=XM_023242821.1
D4	89510295	89510365	refGene_gn=NUP188;refGene_tx=XM_023242820.1;refGene_gn=NUP188;refGene_tx=XM_023242821.1
D4	89510766	89510834	refGene_gn=NUP188;refGene_tx=XM_023242820.1;refGene_gn=NUP188;refGene_tx=XM_023242821.1
D4	89511261	89511457	refGene_gn=NUP188;refGene_tx=XM_023242820.1;refGene_gn=NUP188;refGene_tx=XM_023242821.1
D4	89513370	89513448	refGene_gn=NUP188;refGene_tx=XM_023242820.1;refGene_gn=NUP188;refGene_tx=XM_023242821.1
D4	89513706	89513792	refGene_gn=NUP188;refGene_tx=XM_023242820.1;refGene_gn=NUP188;refGene_tx=XM_023242821.1
D4	89514009	89514155	refGene_gn=NUP188;refGene_tx=XM_023242820.1;refGene_gn=NUP188;refGene_

tx=XM_023242821.1
D4 89514461 89514643
refGene_gn=NUP188;refGene_tx=XM_023242820.1;refGene_gn=NUP188;refGene_tx=XM_023242821.1
D4 89515800 89516001
refGene_gn=NUP188;refGene_tx=XM_023242820.1;refGene_gn=NUP188;refGene_tx=XM_023242821.1
D4 89516187 89516280
refGene_gn=NUP188;refGene_tx=XM_023242820.1;refGene_gn=NUP188;refGene_tx=XM_023242821.1
D4 89516714 89516873
refGene_gn=NUP188;refGene_tx=XM_023242820.1;refGene_gn=NUP188;refGene_tx=XM_023242821.1
D4 89517090 89517228
refGene_gn=NUP188;refGene_tx=XM_023242820.1;refGene_gn=NUP188;refGene_tx=XM_023242821.1
D4 89517910 89517985
refGene_gn=NUP188;refGene_tx=XM_023242820.1;refGene_gn=NUP188;refGene_tx=XM_023242821.1
D4 89518067 89518298
refGene_gn=NUP188;refGene_tx=XM_023242820.1;refGene_gn=NUP188;refGene_tx=XM_023242821.1
D4 89518433 89518565
refGene_gn=NUP188;refGene_tx=XM_023242820.1;refGene_gn=NUP188;refGene_tx=XM_023242821.1
D4 89518831 89518905
refGene_gn=NUP188;refGene_tx=XM_023242820.1;refGene_gn=NUP188;refGene_tx=XM_023242821.1
D4 89519038 89519168
refGene_gn=NUP188;refGene_tx=XM_023242820.1;refGene_gn=NUP188;refGene_tx=XM_023242821.1
D4 89519279 89519456
refGene_gn=NUP188;refGene_tx=XM_023242820.1;refGene_gn=NUP188;refGene_tx=XM_023242821.1
D4 89521429 89521537
refGene_gn=SH3GLB2;refGene_tx=XM_019815558.2;refGene_gn=SH3GLB2;refGene_tx=XM_023242822.1;refGene_gn=SH3GLB2;refGene_tx=XM_023242823.1;refGene_gn=SH3GLB2;refGene_tx=XM_023242824.1;refGene_gn=SH3GLB2;refGene_tx=XM_023242825.1;refGene_gn=SH3GLB2;refGene_tx=XM_023242826.1
D4 89521823 89522064
refGene_gn=SH3GLB2;refGene_tx=XM_019815558.2;refGene_gn=SH3GLB2;refGene_tx=XM_023242822.1;refGene_gn=SH3GLB2;refGene_tx=XM_023242823.1;refGene_gn=SH3GLB2;refGene_tx=XM_023242824.1;refGene_gn=SH3GLB2;refGene_tx=XM_023242825.1;refGene_gn=SH3GLB2;refGene_tx=XM_023242826.1
D4 89522158 89522173
refGene_gn=SH3GLB2;refGene_tx=XM_019815558.2;refGene_gn=SH3GLB2;refGene_tx=XM_023242822.1;refGene_gn=SH3GLB2;refGene_tx=XM_023242825.1;refGene_gn=SH3GLB2;refGene_tx=XM_023242826.1
D4 89522409 89522510
refGene_gn=SH3GLB2;refGene_tx=XM_019815558.2;refGene_gn=SH3GLB2;refGen

e_tx=XM_023242822.1;refGene_gn=SH3GLB2;refGene_tx=XM_023242823.1;refGene_gn=SH3GLB2;refGene_tx=XM_023242824.1;refGene_gn=SH3GLB2;refGene_tx=XM_023242825.1;refGene_gn=SH3GLB2;refGene_tx=XM_023242826.1
D4 89522827 89522917
refGene_gn=SH3GLB2;refGene_tx=XM_019815558.2;refGene_gn=SH3GLB2;refGene_tx=XM_023242822.1;refGene_gn=SH3GLB2;refGene_tx=XM_023242823.1;refGene_gn=SH3GLB2;refGene_tx=XM_023242824.1;refGene_gn=SH3GLB2;refGene_tx=XM_023242825.1;refGene_gn=SH3GLB2;refGene_tx=XM_023242826.1
D4 89523417 89523441
refGene_gn=SH3GLB2;refGene_tx=XM_019815558.2;refGene_gn=SH3GLB2;refGene_tx=XM_023242822.1;refGene_gn=SH3GLB2;refGene_tx=XM_023242823.1;refGene_gn=SH3GLB2;refGene_tx=XM_023242824.1;refGene_gn=SH3GLB2;refGene_tx=XM_023242825.1;refGene_gn=SH3GLB2;refGene_tx=XM_023242826.1
D4 89525008 89525071
refGene_gn=SH3GLB2;refGene_tx=XM_019815558.2;refGene_gn=SH3GLB2;refGene_tx=XM_023242822.1;refGene_gn=SH3GLB2;refGene_tx=XM_023242823.1;refGene_gn=SH3GLB2;refGene_tx=XM_023242824.1
D4 89525746 89525758
refGene_gn=SH3GLB2;refGene_tx=XM_019815558.2;refGene_gn=SH3GLB2;refGene_tx=XM_023242823.1;refGene_gn=SH3GLB2;refGene_tx=XM_023242825.1
D4 89527227 89527320
refGene_gn=SH3GLB2;refGene_tx=XM_019815558.2;refGene_gn=SH3GLB2;refGene_tx=XM_023242822.1;refGene_gn=SH3GLB2;refGene_tx=XM_023242823.1;refGene_gn=SH3GLB2;refGene_tx=XM_023242824.1;refGene_gn=SH3GLB2;refGene_tx=XM_023242825.1;refGene_gn=SH3GLB2;refGene_tx=XM_023242826.1
D4 89527575 89527709
refGene_gn=SH3GLB2;refGene_tx=XM_019815558.2;refGene_gn=SH3GLB2;refGene_tx=XM_023242822.1;refGene_gn=SH3GLB2;refGene_tx=XM_023242823.1;refGene_gn=SH3GLB2;refGene_tx=XM_023242824.1;refGene_gn=SH3GLB2;refGene_tx=XM_023242825.1;refGene_gn=SH3GLB2;refGene_tx=XM_023242826.1
D4 89534143 89534272
refGene_gn=SH3GLB2;refGene_tx=XM_019815558.2;refGene_gn=SH3GLB2;refGene_tx=XM_023242822.1;refGene_gn=SH3GLB2;refGene_tx=XM_023242823.1;refGene_gn=SH3GLB2;refGene_tx=XM_023242824.1;refGene_gn=SH3GLB2;refGene_tx=XM_023242825.1;refGene_gn=SH3GLB2;refGene_tx=XM_023242826.1
D4 89535103 89535245
refGene_gn=SH3GLB2;refGene_tx=XM_019815558.2;refGene_gn=SH3GLB2;refGene_tx=XM_023242822.1;refGene_gn=SH3GLB2;refGene_tx=XM_023242823.1;refGene_gn=SH3GLB2;refGene_tx=XM_023242824.1;refGene_gn=SH3GLB2;refGene_tx=XM_023242825.1;refGene_gn=SH3GLB2;refGene_tx=XM_023242826.1
D4 89539433 89539496
refGene_gn=SH3GLB2;refGene_tx=XM_019815558.2;refGene_gn=SH3GLB2;refGene_tx=XM_023242822.1;refGene_gn=SH3GLB2;refGene_tx=XM_023242823.1;refGene_gn=SH3GLB2;refGene_tx=XM_023242824.1;refGene_gn=SH3GLB2;refGene_tx=XM_023242825.1;refGene_gn=SH3GLB2;refGene_tx=XM_023242826.1
D4 89547938 89548034
refGene_gn=MIGA2;refGene_tx=XM_003995958.5;refGene_gn=MIGA2;refGene_tx=XM_006939543.4
D4 89548646 89548857
refGene_gn=MIGA2;refGene_tx=XM_003995958.5;refGene_gn=MIGA2;refGene_tx

=XM_006939543.4;refGene_gn=MIGA2;refGene_tx=XM_019816658.2
D4	89551256	89551369
refGene_gn=MIGA2;refGene_tx=XM_003995958.5;refGene_gn=MIGA2;refGene_tx=XM_006939543.4;refGene_gn=MIGA2;refGene_tx=XM_019816658.2
D4	89552515	89552645
refGene_gn=MIGA2;refGene_tx=XM_003995958.5;refGene_gn=MIGA2;refGene_tx=XM_006939543.4;refGene_gn=MIGA2;refGene_tx=XM_019816658.2
D4	89553043	89553180
refGene_gn=MIGA2;refGene_tx=XM_003995958.5;refGene_gn=MIGA2;refGene_tx=XM_006939543.4;refGene_gn=MIGA2;refGene_tx=XM_019816658.2
D4	89559674	89559792
refGene_gn=MIGA2;refGene_tx=XM_003995958.5;refGene_gn=MIGA2;refGene_tx=XM_006939543.4;refGene_gn=MIGA2;refGene_tx=XM_019816658.2
D4	89561749	89561850
refGene_gn=MIGA2;refGene_tx=XM_003995958.5;refGene_gn=MIGA2;refGene_tx=XM_006939543.4;refGene_gn=MIGA2;refGene_tx=XM_019816658.2
D4	89562450	89562566
refGene_gn=MIGA2;refGene_tx=XM_003995958.5;refGene_gn=MIGA2;refGene_tx=XM_006939543.4;refGene_gn=MIGA2;refGene_tx=XM_019816658.2
D4	89563759	89563832
refGene_gn=MIGA2;refGene_tx=XM_003995958.5;refGene_gn=MIGA2;refGene_tx=XM_006939543.4;refGene_gn=MIGA2;refGene_tx=XM_019816658.2
D4	89564025	89564112
refGene_gn=MIGA2;refGene_tx=XM_003995958.5;refGene_gn=MIGA2;refGene_tx=XM_006939543.4;refGene_gn=MIGA2;refGene_tx=XM_019816658.2
D4	89567830	89567929
refGene_gn=MIGA2;refGene_tx=XM_003995958.5;refGene_gn=MIGA2;refGene_tx=XM_006939543.4;refGene_gn=MIGA2;refGene_tx=XM_019816658.2
D4	89568235	89568370
refGene_gn=MIGA2;refGene_tx=XM_003995958.5;refGene_gn=MIGA2;refGene_tx=XM_006939543.4;refGene_gn=MIGA2;refGene_tx=XM_019816658.2
D4	89569105	89569159
refGene_gn=MIGA2;refGene_tx=XM_003995958.5;refGene_gn=MIGA2;refGene_tx=XM_006939543.4;refGene_gn=MIGA2;refGene_tx=XM_019816658.2
D4	89569933	89570050
refGene_gn=MIGA2;refGene_tx=XM_003995958.5;refGene_gn=MIGA2;refGene_tx=XM_006939543.4;refGene_gn=MIGA2;refGene_tx=XM_019816658.2
D4	89570275	89570482
refGene_gn=MIGA2;refGene_tx=XM_003995958.5;refGene_gn=MIGA2;refGene_tx=XM_006939543.4;refGene_gn=MIGA2;refGene_tx=XM_019816658.2
D4	89576957	89577033
refGene_gn=DOLPP1;refGene_tx=XM_003995959.4;refGene_gn=DOLPP1;refGene_tx=XM_019816660.1
D4	89579713	89579759 refGene_gn=DOLPP1;refGene_tx=XM_019816659.2
D4	89580539	89580640
refGene_gn=DOLPP1;refGene_tx=XM_003995959.4;refGene_gn=DOLPP1;refGene_tx=XM_019816659.2;refGene_gn=DOLPP1;refGene_tx=XM_019816660.1
D4	89580855	89580940
refGene_gn=DOLPP1;refGene_tx=XM_003995959.4;refGene_gn=DOLPP1;refGene_tx=XM_019816659.2;refGene_gn=DOLPP1;refGene_tx=XM_019816660.1

D4 89581028 89581128
refGene_gn=DOLPP1;refGene_tx=XM_003995959.4;refGene_gn=DOLPP1;refGene_
tx=XM_019816659.2;refGene_gn=DOLPP1;refGene_tx=XM_019816660.1
D4 89581358 89581457
refGene_gn=DOLPP1;refGene_tx=XM_003995959.4;refGene_gn=DOLPP1;refGene_
tx=XM_019816659.2;refGene_gn=DOLPP1;refGene_tx=XM_019816660.1
D4 89582008 89582137
refGene_gn=DOLPP1;refGene_tx=XM_003995959.4;refGene_gn=DOLPP1;refGene_
tx=XM_019816659.2
D4 89582572 89582662
refGene_gn=DOLPP1;refGene_tx=XM_003995959.4;refGene_gn=DOLPP1;refGene_
tx=XM_019816659.2;refGene_gn=DOLPP1;refGene_tx=XM_019816660.1
D4 89584657 89584694
refGene_gn=DOLPP1;refGene_tx=XM_003995959.4;refGene_gn=DOLPP1;refGene_
tx=XM_019816659.2;refGene_gn=DOLPP1;refGene_tx=XM_019816660.1
D4 89591011 89591227
refGene_gn=CRAT;refGene_tx=XM_003995961.5;refGene_gn=CRAT;refGene_tx=X
M_006939545.4
D4 89591892 89592030
refGene_gn=CRAT;refGene_tx=XM_003995961.5;refGene_gn=CRAT;refGene_tx=X
M_006939545.4
D4 89592790 89592853
refGene_gn=CRAT;refGene_tx=XM_003995961.5;refGene_gn=CRAT;refGene_tx=X
M_006939545.4
D4 89593220 89593356
refGene_gn=CRAT;refGene_tx=XM_003995961.5;refGene_gn=CRAT;refGene_tx=X
M_006939545.4
D4 89593460 89593583
refGene_gn=CRAT;refGene_tx=XM_003995961.5;refGene_gn=CRAT;refGene_tx=X
M_006939545.4
D4 89593782 89593902
refGene_gn=CRAT;refGene_tx=XM_003995961.5;refGene_gn=CRAT;refGene_tx=X
M_006939545.4
D4 89594301 89594402
refGene_gn=CRAT;refGene_tx=XM_003995961.5;refGene_gn=CRAT;refGene_tx=X
M_006939545.4
D4 89594918 89595097
refGene_gn=CRAT;refGene_tx=XM_003995961.5;refGene_gn=CRAT;refGene_tx=X
M_006939545.4
D4 89595821 89595996
refGene_gn=CRAT;refGene_tx=XM_003995961.5;refGene_gn=CRAT;refGene_tx=X
M_006939545.4
D4 89596364 89596530
refGene_gn=CRAT;refGene_tx=XM_003995961.5;refGene_gn=CRAT;refGene_tx=X
M_006939545.4
D4 89597038 89597092
refGene_gn=CRAT;refGene_tx=XM_003995961.5;refGene_gn=CRAT;refGene_tx=X
M_006939545.4
D4 89598296 89598415
refGene_gn=CRAT;refGene_tx=XM_003995961.5;refGene_gn=CRAT;refGene_tx=X

M_006939545.4
D4 89600872 89601136
refGene_gn=CRAT;refGene_tx=XM_003995961.5;refGene_gn=CRAT;refGene_tx=XM_006939545.4
D4 89601830 89601857 refGene_gn=CRAT;refGene_tx=XM_006939545.4
D4 89603547 89603574 refGene_gn=CRAT;refGene_tx=XM_003995961.5
D4 89604079 89604108 refGene_gn=PTPA;refGene_tx=XM_003995962.5
D4 89604354 89604512 refGene_gn=PTPA;refGene_tx=XM_003995962.5
D4 89609770 89609868 refGene_gn=PTPA;refGene_tx=XM_003995962.5
D4 89613405 89613492 refGene_gn=PTPA;refGene_tx=XM_003995962.5
D4 89618074 89618095 refGene_gn=PTPA;refGene_tx=XM_003995962.5
D4 89618096 89618201 refGene_gn=PTPA;refGene_tx=XM_003995962.5
D4 89619984 89620102 refGene_gn=PTPA;refGene_tx=XM_003995962.5
D4 89621964 89622064 refGene_gn=PTPA;refGene_tx=XM_003995962.5
D4 89622953 89623078 refGene_gn=PTPA;refGene_tx=XM_003995962.5
D4 89624100 89624201 refGene_gn=PTPA;refGene_tx=XM_003995962.5
D4 89629040 89629148 refGene_gn=PTPA;refGene_tx=XM_003995962.5
D4 89633216 89633294 refGene_gn=PTPA;refGene_tx=XM_003995962.5
D4 89654861 89656091 refGene_gn=IER5L;refGene_tx=XM_023242827.1
D4 89981939 89982101
refGene_gn=NTMT1;refGene_tx=XM_003995964.5;refGene_gn=NTMT1;refGene_tx=XM_006939548.4;refGene_gn=NTMT1;refGene_tx=XM_006939549.4;refGene_gn=NTMT1;refGene_tx=XM_023242830.1
D4 89982949 89983202
refGene_gn=NTMT1;refGene_tx=XM_003995964.5;refGene_gn=NTMT1;refGene_tx=XM_006939548.4;refGene_gn=NTMT1;refGene_tx=XM_006939549.4;refGene_gn=NTMT1;refGene_tx=XM_006939550.4;refGene_gn=NTMT1;refGene_tx=XM_023242830.1
D4 89984256 89984513
refGene_gn=NTMT1;refGene_tx=XM_003995964.5;refGene_gn=NTMT1;refGene_tx=XM_006939548.4;refGene_gn=NTMT1;refGene_tx=XM_006939549.4;refGene_gn=NTMT1;refGene_tx=XM_006939550.4;refGene_gn=NTMT1;refGene_tx=XM_023242830.1
D4 89986184 89986852
refGene_gn=ASB6;refGene_tx=XM_003995967.5;refGene_gn=ASB6;refGene_tx=XM_011288379.3
D4 89986961 89987048
refGene_gn=ASB6;refGene_tx=XM_003995967.5;refGene_gn=ASB6;refGene_tx=XM_011288379.3
D4 89988095 89988204 refGene_gn=ASB6;refGene_tx=XM_011288379.3
D4 89988314 89988421
refGene_gn=ASB6;refGene_tx=XM_003995967.5;refGene_gn=ASB6;refGene_tx=XM_011288379.3
D4 89989479 89989661
refGene_gn=ASB6;refGene_tx=XM_003995967.5;refGene_gn=ASB6;refGene_tx=XM_011288379.3
D4 89991141 89991254
refGene_gn=ASB6;refGene_tx=XM_003995967.5;refGene_gn=ASB6;refGene_tx=XM_011288379.3
D4 90008668 90008948 refGene_gn=PRRX2;refGene_tx=XM_023242831.1

D4 90046601 90046789 refGene_gn=PRRX2;refGene_tx=XM_023242831.1
D4 90047625 90047804 refGene_gn=PRRX2;refGene_tx=XM_023242831.1
D4 90048962 90049098 refGene_gn=PRRX2;refGene_tx=XM_023242831.1
D4 90060219 90060469 refGene_gn=PTGES;refGene_tx=XM_006939551.4
D4 90067066 90067149 refGene_gn=PTGES;refGene_tx=XM_006939551.4
D4 90070207 90070342 refGene_gn=PTGES;refGene_tx=XM_006939551.4
D4 90102148 90102626
refGene_gn=TOR1B;refGene_tx=XM_023242832.1;refGene_gn=TOR1B;refGene_tx
=XM_023242833.1
D4 90103239 90103505
refGene_gn=TOR1B;refGene_tx=XM_023242832.1;refGene_gn=TOR1B;refGene_tx
=XM_023242833.1
D4 90104923 90105099
refGene_gn=TOR1B;refGene_tx=XM_023242832.1;refGene_gn=TOR1B;refGene_tx
=XM_023242833.1
D4 90105668 90105796
refGene_gn=TOR1B;refGene_tx=XM_023242832.1;refGene_gn=TOR1B;refGene_tx
=XM_023242833.1
D4 90106109 90106351
refGene_gn=TOR1B;refGene_tx=XM_023242832.1;refGene_gn=TOR1B;refGene_tx
=XM_023242833.1
D4 90110766 90111017 refGene_gn=TOR1A;refGene_tx=XM_023242834.1
D4 90114948 90115076 refGene_gn=TOR1A;refGene_tx=XM_023242834.1
D4 90115173 90115349 refGene_gn=TOR1A;refGene_tx=XM_023242834.1
D4 90117804 90118070 refGene_gn=TOR1A;refGene_tx=XM_023242834.1
D4 90118560 90118738 refGene_gn=TOR1A;refGene_tx=XM_023242834.1
D4 90120887 90120979
refGene_gn=CD4H9orf78;refGene_tx=XM_011288375.3
D4 90121881 90121980
refGene_gn=CD4H9orf78;refGene_tx=XM_011288375.3
D4 90122088 90122225
refGene_gn=CD4H9orf78;refGene_tx=XM_011288375.3
D4 90123236 90123434
refGene_gn=CD4H9orf78;refGene_tx=XM_011288375.3
D4 90124019 90124097
refGene_gn=CD4H9orf78;refGene_tx=XM_011288375.3
D4 90124901 90124972
refGene_gn=CD4H9orf78;refGene_tx=XM_011288375.3
D4 90125382 90125434
refGene_gn=CD4H9orf78;refGene_tx=XM_011288375.3
D4 90126582 90126642
refGene_gn=CD4H9orf78;refGene_tx=XM_011288375.3
D4 90126983 90127066
refGene_gn=CD4H9orf78;refGene_tx=XM_011288375.3
D4 90141337 90141418
refGene_gn=USP20;refGene_tx=XM_003995969.4;refGene_gn=USP20;refGene_tx
=XM_023242835.1;refGene_gn=USP20;refGene_tx=XM_023242836.1;refGene_gn=
USP20;refGene_tx=XM_023242837.1
D4 90146884 90146938
refGene_gn=USP20;refGene_tx=XM_003995969.4;refGene_gn=USP20;refGene_tx

=XM_023242835.1;refGene_gn=USP20;refGene_tx=XM_023242836.1
D4	90148308 90148371
refGene_gn=USP20;refGene_tx=XM_003995969.4;refGene_gn=USP20;refGene_tx=XM_023242835.1;refGene_gn=USP20;refGene_tx=XM_023242836.1;refGene_gn=USP20;refGene_tx=XM_023242837.1
D4	90148713 90148845
refGene_gn=USP20;refGene_tx=XM_003995969.4;refGene_gn=USP20;refGene_tx=XM_023242835.1;refGene_gn=USP20;refGene_tx=XM_023242836.1;refGene_gn=USP20;refGene_tx=XM_023242837.1
D4	90149893 90149990
refGene_gn=USP20;refGene_tx=XM_003995969.4;refGene_gn=USP20;refGene_tx=XM_023242835.1;refGene_gn=USP20;refGene_tx=XM_023242836.1;refGene_gn=USP20;refGene_tx=XM_023242837.1
D4	90150448 90150518
refGene_gn=USP20;refGene_tx=XM_003995969.4;refGene_gn=USP20;refGene_tx=XM_023242835.1;refGene_gn=USP20;refGene_tx=XM_023242836.1;refGene_gn=USP20;refGene_tx=XM_023242837.1
D4	90152361 90152475
refGene_gn=USP20;refGene_tx=XM_003995969.4;refGene_gn=USP20;refGene_tx=XM_023242835.1;refGene_gn=USP20;refGene_tx=XM_023242836.1;refGene_gn=USP20;refGene_tx=XM_023242837.1
D4	90153486 90153565
refGene_gn=USP20;refGene_tx=XM_003995969.4;refGene_gn=USP20;refGene_tx=XM_023242835.1;refGene_gn=USP20;refGene_tx=XM_023242836.1;refGene_gn=USP20;refGene_tx=XM_023242837.1
D4	90155987 90156435
refGene_gn=USP20;refGene_tx=XM_003995969.4;refGene_gn=USP20;refGene_tx=XM_023242835.1;refGene_gn=USP20;refGene_tx=XM_023242836.1;refGene_gn=USP20;refGene_tx=XM_023242837.1
D4	90156797 90156938
refGene_gn=USP20;refGene_tx=XM_003995969.4;refGene_gn=USP20;refGene_tx=XM_023242835.1;refGene_gn=USP20;refGene_tx=XM_023242836.1;refGene_gn=USP20;refGene_tx=XM_023242837.1
D4	90157248 90157364
refGene_gn=USP20;refGene_tx=XM_003995969.4;refGene_gn=USP20;refGene_tx=XM_023242835.1;refGene_gn=USP20;refGene_tx=XM_023242836.1;refGene_gn=USP20;refGene_tx=XM_023242837.1
D4	90157617 90157784
refGene_gn=USP20;refGene_tx=XM_003995969.4;refGene_gn=USP20;refGene_tx=XM_023242835.1;refGene_gn=USP20;refGene_tx=XM_023242836.1;refGene_gn=USP20;refGene_tx=XM_023242837.1
D4	90158367 90158462
refGene_gn=USP20;refGene_tx=XM_003995969.4;refGene_gn=USP20;refGene_tx=XM_023242835.1;refGene_gn=USP20;refGene_tx=XM_023242836.1;refGene_gn=USP20;refGene_tx=XM_023242837.1
D4	90164912 90164946
refGene_gn=USP20;refGene_tx=XM_003995969.4;refGene_gn=USP20;refGene_tx=XM_023242835.1;refGene_gn=USP20;refGene_tx=XM_023242836.1;refGene_gn=USP20;refGene_tx=XM_023242837.1
D4	90165138 90165184

refGene_gn=USP20;refGene_tx=XM_003995969.4;refGene_gn=USP20;refGene_tx=XM_023242835.1;refGene_gn=USP20;refGene_tx=XM_023242836.1;refGene_gn=USP20;refGene_tx=XM_023242837.1
D4 90165955 90166136
refGene_gn=USP20;refGene_tx=XM_003995969.4;refGene_gn=USP20;refGene_tx=XM_023242835.1;refGene_gn=USP20;refGene_tx=XM_023242836.1;refGene_gn=USP20;refGene_tx=XM_023242837.1
D4 90166213 90166340
refGene_gn=USP20;refGene_tx=XM_003995969.4;refGene_gn=USP20;refGene_tx=XM_023242835.1;refGene_gn=USP20;refGene_tx=XM_023242836.1;refGene_gn=USP20;refGene_tx=XM_023242837.1
D4 90166766 90166936
refGene_gn=USP20;refGene_tx=XM_003995969.4;refGene_gn=USP20;refGene_tx=XM_023242835.1;refGene_gn=USP20;refGene_tx=XM_023242836.1;refGene_gn=USP20;refGene_tx=XM_023242837.1
D4 90167017 90167099
refGene_gn=USP20;refGene_tx=XM_003995969.4;refGene_gn=USP20;refGene_tx=XM_023242835.1;refGene_gn=USP20;refGene_tx=XM_023242836.1;refGene_gn=USP20;refGene_tx=XM_023242837.1
D4 90168044 90168153
refGene_gn=USP20;refGene_tx=XM_003995969.4;refGene_gn=USP20;refGene_tx=XM_023242835.1;refGene_gn=USP20;refGene_tx=XM_023242836.1;refGene_gn=USP20;refGene_tx=XM_023242837.1
D4 90169015 90169118
refGene_gn=USP20;refGene_tx=XM_003995969.4;refGene_gn=USP20;refGene_tx=XM_023242835.1;refGene_gn=USP20;refGene_tx=XM_023242836.1;refGene_gn=USP20;refGene_tx=XM_023242837.1
D4 90170425 90170497
refGene_gn=USP20;refGene_tx=XM_003995969.4;refGene_gn=USP20;refGene_tx=XM_023242835.1;refGene_gn=USP20;refGene_tx=XM_023242836.1;refGene_gn=USP20;refGene_tx=XM_023242837.1
D4 90170890 90171051
refGene_gn=USP20;refGene_tx=XM_003995969.4;refGene_gn=USP20;refGene_tx=XM_023242835.1;refGene_gn=USP20;refGene_tx=XM_023242836.1;refGene_gn=USP20;refGene_tx=XM_023242837.1
D4 90179275 90179283
refGene_gn=FNBP1;refGene_tx=XM_023242838.1;refGene_gn=FNBP1;refGene_tx=XM_023242839.1;refGene_gn=FNBP1;refGene_tx=XM_023242840.1;refGene_gn=FNBP1;refGene_tx=XM_023242842.1;refGene_gn=FNBP1;refGene_tx=XM_023242844.1;refGene_gn=FNBP1;refGene_tx=XM_023242849.1
D4 90182707 90182883
refGene_gn=FNBP1;refGene_tx=XM_023242838.1;refGene_gn=FNBP1;refGene_tx=XM_023242839.1;refGene_gn=FNBP1;refGene_tx=XM_023242840.1;refGene_gn=FNBP1;refGene_tx=XM_023242841.1;refGene_gn=FNBP1;refGene_tx=XM_023242842.1;refGene_gn=FNBP1;refGene_tx=XM_023242843.1;refGene_gn=FNBP1;refGene_tx=XM_023242844.1;refGene_gn=FNBP1;refGene_tx=XM_023242845.1;refGene_gn=FNBP1;refGene_tx=XM_023242846.1;refGene_gn=FNBP1;refGene_tx=XM_023242847.1;refGene_gn=FNBP1;refGene_tx=XM_023242848.1;refGene_gn=FNBP1;refGene_tx=XM_023242849.1
D4 90184555 90184695

refGene_gn=FNBP1;refGene_tx=XM_023242838.1;refGene_gn=FNBP1;refGene_tx
=XM_023242839.1;refGene_gn=FNBP1;refGene_tx=XM_023242840.1;refGene_gn=
FNBP1;refGene_tx=XM_023242841.1;refGene_gn=FNBP1;refGene_tx=XM_0232428
42.1;refGene_gn=FNBP1;refGene_tx=XM_023242843.1;refGene_gn=FNBP1;refGe
ne_tx=XM_023242844.1;refGene_gn=FNBP1;refGene_tx=XM_023242845.1;refGen
e_gn=FNBP1;refGene_tx=XM_023242846.1;refGene_gn=FNBP1;refGene_tx=XM_02
3242847.1;refGene_gn=FNBP1;refGene_tx=XM_023242848.1;refGene_gn=FNBP1;
refGene_tx=XM_023242849.1
D4 90184990 90185112
refGene_gn=FNBP1;refGene_tx=XM_023242838.1;refGene_gn=FNBP1;refGene_tx
=XM_023242839.1;refGene_gn=FNBP1;refGene_tx=XM_023242840.1;refGene_gn=
FNBP1;refGene_tx=XM_023242841.1;refGene_gn=FNBP1;refGene_tx=XM_0232428
42.1;refGene_gn=FNBP1;refGene_tx=XM_023242843.1;refGene_gn=FNBP1;refGe
ne_tx=XM_023242844.1;refGene_gn=FNBP1;refGene_tx=XM_023242845.1;refGen
e_gn=FNBP1;refGene_tx=XM_023242846.1;refGene_gn=FNBP1;refGene_tx=XM_02
3242847.1;refGene_gn=FNBP1;refGene_tx=XM_023242848.1;refGene_gn=FNBP1;
refGene_tx=XM_023242849.1
D4 90186359 90186492
refGene_gn=FNBP1;refGene_tx=XM_023242838.1;refGene_gn=FNBP1;refGene_tx
=XM_023242839.1;refGene_gn=FNBP1;refGene_tx=XM_023242840.1;refGene_gn=
FNBP1;refGene_tx=XM_023242841.1;refGene_gn=FNBP1;refGene_tx=XM_0232428
42.1;refGene_gn=FNBP1;refGene_tx=XM_023242843.1;refGene_gn=FNBP1;refGe
ne_tx=XM_023242844.1;refGene_gn=FNBP1;refGene_tx=XM_023242845.1;refGen
e_gn=FNBP1;refGene_tx=XM_023242846.1;refGene_gn=FNBP1;refGene_tx=XM_02
3242847.1;refGene_gn=FNBP1;refGene_tx=XM_023242848.1;refGene_gn=FNBP1;
refGene_tx=XM_023242849.1
D4 90191976 90192086
refGene_gn=FNBP1;refGene_tx=XM_023242838.1;refGene_gn=FNBP1;refGene_tx
=XM_023242839.1;refGene_gn=FNBP1;refGene_tx=XM_023242840.1;refGene_gn=
FNBP1;refGene_tx=XM_023242841.1;refGene_gn=FNBP1;refGene_tx=XM_0232428
42.1;refGene_gn=FNBP1;refGene_tx=XM_023242843.1;refGene_gn=FNBP1;refGe
ne_tx=XM_023242844.1;refGene_gn=FNBP1;refGene_tx=XM_023242845.1;refGen
e_gn=FNBP1;refGene_tx=XM_023242846.1;refGene_gn=FNBP1;refGene_tx=XM_02
3242847.1;refGene_gn=FNBP1;refGene_tx=XM_023242848.1;refGene_gn=FNBP1;
refGene_tx=XM_023242849.1
D4 90196772 90196787
refGene_gn=FNBP1;refGene_tx=XM_023242838.1;refGene_gn=FNBP1;refGene_tx
=XM_023242839.1;refGene_gn=FNBP1;refGene_tx=XM_023242840.1;refGene_gn=
FNBP1;refGene_tx=XM_023242841.1;refGene_gn=FNBP1;refGene_tx=XM_0232428
42.1;refGene_gn=FNBP1;refGene_tx=XM_023242844.1;refGene_gn=FNBP1;refGe
ne_tx=XM_023242845.1;refGene_gn=FNBP1;refGene_tx=XM_023242847.1
D4 90199333 90199417
refGene_gn=FNBP1;refGene_tx=XM_023242838.1;refGene_gn=FNBP1;refGene_tx
=XM_023242839.1;refGene_gn=FNBP1;refGene_tx=XM_023242840.1;refGene_gn=
FNBP1;refGene_tx=XM_023242842.1
D4 90202782 90202965
refGene_gn=FNBP1;refGene_tx=XM_023242838.1;refGene_gn=FNBP1;refGene_tx
=XM_023242839.1;refGene_gn=FNBP1;refGene_tx=XM_023242840.1;refGene_gn=
FNBP1;refGene_tx=XM_023242841.1;refGene_gn=FNBP1;refGene_tx=XM_0232428
42.1;refGene_gn=FNBP1;refGene_tx=XM_023242843.1;refGene_gn=FNBP1;refGe

ne_tx=XM_023242844.1;refGene_gn=FNBP1;refGene_tx=XM_023242845.1;refGene_gn=FNBP1;refGene_tx=XM_023242846.1
D4 90203893 90204091
refGene_gn=FNBP1;refGene_tx=XM_023242838.1;refGene_gn=FNBP1;refGene_tx=XM_023242839.1;refGene_gn=FNBP1;refGene_tx=XM_023242840.1;refGene_gn=FNBP1;refGene_tx=XM_023242841.1;refGene_gn=FNBP1;refGene_tx=XM_023242842.1;refGene_gn=FNBP1;refGene_tx=XM_023242843.1;refGene_gn=FNBP1;refGene_tx=XM_023242844.1;refGene_gn=FNBP1;refGene_tx=XM_023242845.1;refGene_gn=FNBP1;refGene_tx=XM_023242846.1;refGene_gn=FNBP1;refGene_tx=XM_023242847.1;refGene_gn=FNBP1;refGene_tx=XM_023242848.1;refGene_gn=FNBP1;refGene_tx=XM_023242849.1
D4 90211344 90211491
refGene_gn=FNBP1;refGene_tx=XM_023242838.1;refGene_gn=FNBP1;refGene_tx=XM_023242839.1;refGene_gn=FNBP1;refGene_tx=XM_023242840.1;refGene_gn=FNBP1;refGene_tx=XM_023242841.1;refGene_gn=FNBP1;refGene_tx=XM_023242842.1;refGene_gn=FNBP1;refGene_tx=XM_023242843.1;refGene_gn=FNBP1;refGene_tx=XM_023242844.1;refGene_gn=FNBP1;refGene_tx=XM_023242845.1;refGene_gn=FNBP1;refGene_tx=XM_023242846.1;refGene_gn=FNBP1;refGene_tx=XM_023242847.1;refGene_gn=FNBP1;refGene_tx=XM_023242848.1;refGene_gn=FNBP1;refGene_tx=XM_023242849.1
D4 90212680 90212809
refGene_gn=FNBP1;refGene_tx=XM_023242838.1;refGene_gn=FNBP1;refGene_tx=XM_023242839.1;refGene_gn=FNBP1;refGene_tx=XM_023242840.1;refGene_gn=FNBP1;refGene_tx=XM_023242841.1;refGene_gn=FNBP1;refGene_tx=XM_023242842.1;refGene_gn=FNBP1;refGene_tx=XM_023242843.1;refGene_gn=FNBP1;refGene_tx=XM_023242844.1;refGene_gn=FNBP1;refGene_tx=XM_023242845.1;refGene_gn=FNBP1;refGene_tx=XM_023242846.1;refGene_gn=FNBP1;refGene_tx=XM_023242847.1;refGene_gn=FNBP1;refGene_tx=XM_023242848.1;refGene_gn=FNBP1;refGene_tx=XM_023242849.1
D4 90239224 90239329
refGene_gn=FNBP1;refGene_tx=XM_023242838.1;refGene_gn=FNBP1;refGene_tx=XM_023242839.1;refGene_gn=FNBP1;refGene_tx=XM_023242840.1;refGene_gn=FNBP1;refGene_tx=XM_023242841.1;refGene_gn=FNBP1;refGene_tx=XM_023242842.1;refGene_gn=FNBP1;refGene_tx=XM_023242843.1;refGene_gn=FNBP1;refGene_tx=XM_023242844.1;refGene_gn=FNBP1;refGene_tx=XM_023242845.1;refGene_gn=FNBP1;refGene_tx=XM_023242846.1;refGene_gn=FNBP1;refGene_tx=XM_023242847.1;refGene_gn=FNBP1;refGene_tx=XM_023242848.1;refGene_gn=FNBP1;refGene_tx=XM_023242849.1
D4 90239748 90239811
refGene_gn=FNBP1;refGene_tx=XM_023242838.1;refGene_gn=FNBP1;refGene_tx=XM_023242839.1;refGene_gn=FNBP1;refGene_tx=XM_023242840.1;refGene_gn=FNBP1;refGene_tx=XM_023242841.1;refGene_gn=FNBP1;refGene_tx=XM_023242842.1;refGene_gn=FNBP1;refGene_tx=XM_023242843.1;refGene_gn=FNBP1;refGene_tx=XM_023242844.1;refGene_gn=FNBP1;refGene_tx=XM_023242845.1;refGene_gn=FNBP1;refGene_tx=XM_023242846.1;refGene_gn=FNBP1;refGene_tx=XM_023242847.1;refGene_gn=FNBP1;refGene_tx=XM_023242848.1;refGene_gn=FNBP1;refGene_tx=XM_023242849.1
D4 90253633 90253781
refGene_gn=FNBP1;refGene_tx=XM_023242838.1;refGene_gn=FNBP1;refGene_tx=XM_023242839.1;refGene_gn=FNBP1;refGene_tx=XM_023242840.1;refGene_gn=

FNBP1;refGene_tx=XM_023242841.1;refGene_gn=FNBP1;refGene_tx=XM_023242842.1;refGene_gn=FNBP1;refGene_tx=XM_023242843.1;refGene_gn=FNBP1;refGene_tx=XM_023242844.1;refGene_gn=FNBP1;refGene_tx=XM_023242845.1;refGene_gn=FNBP1;refGene_tx=XM_023242846.1;refGene_gn=FNBP1;refGene_tx=XM_023242847.1;refGene_gn=FNBP1;refGene_tx=XM_023242848.1;refGene_gn=FNBP1;refGene_tx=XM_023242849.1
D4	90254475	90254532
refGene_gn=FNBP1;refGene_tx=XM_023242838.1;refGene_gn=FNBP1;refGene_tx=XM_023242839.1;refGene_gn=FNBP1;refGene_tx=XM_023242840.1;refGene_gn=FNBP1;refGene_tx=XM_023242841.1;refGene_gn=FNBP1;refGene_tx=XM_023242842.1;refGene_gn=FNBP1;refGene_tx=XM_023242843.1;refGene_gn=FNBP1;refGene_tx=XM_023242844.1;refGene_gn=FNBP1;refGene_tx=XM_023242845.1;refGene_gn=FNBP1;refGene_tx=XM_023242846.1;refGene_gn=FNBP1;refGene_tx=XM_023242847.1;refGene_gn=FNBP1;refGene_tx=XM_023242848.1;refGene_gn=FNBP1;refGene_tx=XM_023242849.1
D4	90267620	90267736
refGene_gn=FNBP1;refGene_tx=XM_023242838.1;refGene_gn=FNBP1;refGene_tx=XM_023242839.1;refGene_gn=FNBP1;refGene_tx=XM_023242840.1;refGene_gn=FNBP1;refGene_tx=XM_023242841.1;refGene_gn=FNBP1;refGene_tx=XM_023242842.1;refGene_gn=FNBP1;refGene_tx=XM_023242843.1;refGene_gn=FNBP1;refGene_tx=XM_023242844.1;refGene_gn=FNBP1;refGene_tx=XM_023242845.1;refGene_gn=FNBP1;refGene_tx=XM_023242846.1;refGene_gn=FNBP1;refGene_tx=XM_023242847.1;refGene_gn=FNBP1;refGene_tx=XM_023242848.1;refGene_gn=FNBP1;refGene_tx=XM_023242849.1
D4	90321583	90321607
refGene_gn=FNBP1;refGene_tx=XM_023242838.1;refGene_gn=FNBP1;refGene_tx=XM_023242839.1;refGene_gn=FNBP1;refGene_tx=XM_023242840.1;refGene_gn=FNBP1;refGene_tx=XM_023242841.1;refGene_gn=FNBP1;refGene_tx=XM_023242842.1;refGene_gn=FNBP1;refGene_tx=XM_023242843.1;refGene_gn=FNBP1;refGene_tx=XM_023242844.1;refGene_gn=FNBP1;refGene_tx=XM_023242845.1;refGene_gn=FNBP1;refGene_tx=XM_023242846.1;refGene_gn=FNBP1;refGene_tx=XM_023242847.1;refGene_gn=FNBP1;refGene_tx=XM_023242848.1;refGene_gn=FNBP1;refGene_tx=XM_023242849.1
D4	90329601	90329742	refGene_gn=GPR107;refGene_tx=XM_023242851.1
D4	90351478	90351601	refGene_gn=GPR107;refGene_tx=XM_023242851.1
D4	90352176	90352227	refGene_gn=GPR107;refGene_tx=XM_023242851.1
D4	90352481	90352561	refGene_gn=GPR107;refGene_tx=XM_023242851.1
D4	90353317	90353457	refGene_gn=GPR107;refGene_tx=XM_023242851.1
D4	90356428	90356466	refGene_gn=GPR107;refGene_tx=XM_023242851.1
D4	90357912	90357969	refGene_gn=GPR107;refGene_tx=XM_023242851.1
D4	90361148	90361259	refGene_gn=GPR107;refGene_tx=XM_023242851.1
D4	90361584	90361718	refGene_gn=GPR107;refGene_tx=XM_023242851.1
D4	90365666	90365742	refGene_gn=GPR107;refGene_tx=XM_023242851.1
D4	90367796	90367870	refGene_gn=GPR107;refGene_tx=XM_023242851.1
D4	90368246	90368364	refGene_gn=GPR107;refGene_tx=XM_023242851.1
D4	90371179	90371310	refGene_gn=GPR107;refGene_tx=XM_023242851.1
D4	90373367	90373411	refGene_gn=GPR107;refGene_tx=XM_023242851.1
D4	90385900	90385950	refGene_gn=GPR107;refGene_tx=XM_023242851.1
D4	90387552	90387636	refGene_gn=GPR107;refGene_tx=XM_023242851.1
D4	90388441	90388560	refGene_gn=GPR107;refGene_tx=XM_023242851.1

| | | | |
|---|---|---|---|
| D4 | 90392657 | 90392754 | refGene_gn=GPR107;refGene_tx=XM_023242851.1 |
| D4 | 90418462 | 90418526 | refGene_gn=NCS1;refGene_tx=XM_023242852.1 |
| D4 | 90444062 | 90444087 | refGene_gn=NCS1;refGene_tx=XM_023242852.1 |
| D4 | 90455932 | 90456071 | refGene_gn=NCS1;refGene_tx=XM_023242852.1 |
| D4 | 90457438 | 90457517 | refGene_gn=NCS1;refGene_tx=XM_023242852.1 |
| D4 | 90459224 | 90459313 | refGene_gn=NCS1;refGene_tx=XM_023242852.1 |
| D4 | 90459602 | 90459680 | refGene_gn=NCS1;refGene_tx=XM_023242852.1 |
| D4 | 90461446 | 90461545 | refGene_gn=NCS1;refGene_tx=XM_023242852.1 |
| D4 | 90499488 | 90499750 | refGene_gn=HMCN2;refGene_tx=XM_023242853.1 |
| D4 | 90518786 | 90518857 | refGene_gn=HMCN2;refGene_tx=XM_023242853.1 |
| D4 | 90520010 | 90520169 | refGene_gn=HMCN2;refGene_tx=XM_023242853.1 |
| D4 | 90521452 | 90521575 | refGene_gn=HMCN2;refGene_tx=XM_023242853.1 |
| D4 | 90529420 | 90529592 | refGene_gn=HMCN2;refGene_tx=XM_023242853.1 |
| D4 | 90530475 | 90530582 | refGene_gn=HMCN2;refGene_tx=XM_023242853.1 |
| D4 | 90531466 | 90531587 | refGene_gn=HMCN2;refGene_tx=XM_023242853.1 |
| D4 | 90533665 | 90533929 | refGene_gn=HMCN2;refGene_tx=XM_023242853.1 |
| D4 | 90536558 | 90536703 | refGene_gn=HMCN2;refGene_tx=XM_023242853.1 |
| D4 | 90537188 | 90537310 | refGene_gn=HMCN2;refGene_tx=XM_023242853.1 |
| D4 | 90539233 | 90539506 | refGene_gn=HMCN2;refGene_tx=XM_023242853.1 |
| D4 | 90540763 | 90540905 | refGene_gn=HMCN2;refGene_tx=XM_023242853.1 |
| D4 | 90541393 | 90541521 | refGene_gn=HMCN2;refGene_tx=XM_023242853.1 |
| D4 | 90542213 | 90542327 | refGene_gn=HMCN2;refGene_tx=XM_023242853.1 |
| D4 | 90544478 | 90544628 | refGene_gn=HMCN2;refGene_tx=XM_023242853.1 |
| D4 | 90551654 | 90551831 | refGene_gn=HMCN2;refGene_tx=XM_023242853.1 |
| D4 | 90552265 | 90552361 | refGene_gn=HMCN2;refGene_tx=XM_023242853.1 |
| D4 | 90552661 | 90552789 | refGene_gn=HMCN2;refGene_tx=XM_023242853.1 |
| D4 | 90554532 | 90554677 | refGene_gn=HMCN2;refGene_tx=XM_023242853.1 |
| D4 | 90557764 | 90557877 | refGene_gn=HMCN2;refGene_tx=XM_023242853.1 |
| D4 | 90558002 | 90558162 | refGene_gn=HMCN2;refGene_tx=XM_023242853.1 |
| D4 | 90560017 | 90560183 | refGene_gn=HMCN2;refGene_tx=XM_023242853.1 |
| D4 | 90569487 | 90569615 | refGene_gn=HMCN2;refGene_tx=XM_023242853.1 |
| D4 | 90571276 | 90571531 | refGene_gn=HMCN2;refGene_tx=XM_023242853.1 |
| D4 | 90572647 | 90572734 | refGene_gn=HMCN2;refGene_tx=XM_023242853.1 |
| D4 | 90575402 | 90575597 | refGene_gn=HMCN2;refGene_tx=XM_023242853.1 |
| D4 | 90576558 | 90576689 | refGene_gn=HMCN2;refGene_tx=XM_023242853.1 |
| D4 | 90576926 | 90577074 | refGene_gn=HMCN2;refGene_tx=XM_023242853.1 |
| D4 | 90577423 | 90577553 | refGene_gn=HMCN2;refGene_tx=XM_023242853.1 |
| D4 | 90578905 | 90579060 | refGene_gn=HMCN2;refGene_tx=XM_023242853.1 |
| D4 | 90579862 | 90580141 | refGene_gn=HMCN2;refGene_tx=XM_023242853.1 |
| D4 | 90581097 | 90581379 | refGene_gn=HMCN2;refGene_tx=XM_023242853.1 |
| D4 | 90582061 | 90582170 | refGene_gn=HMCN2;refGene_tx=XM_023242853.1 |
| D4 | 90582351 | 90582521 | refGene_gn=HMCN2;refGene_tx=XM_023242853.1 |
| D4 | 90583418 | 90583573 | refGene_gn=HMCN2;refGene_tx=XM_023242853.1 |
| D4 | 90583937 | 90584034 | refGene_gn=HMCN2;refGene_tx=XM_023242853.1 |
| D4 | 90584516 | 90584612 | refGene_gn=HMCN2;refGene_tx=XM_023242853.1 |
| D4 | 90585335 | 90585512 | refGene_gn=HMCN2;refGene_tx=XM_023242853.1 |
| D4 | 90587051 | 90587209 | refGene_gn=HMCN2;refGene_tx=XM_023242853.1 |
| D4 | 90587833 | 90587957 | refGene_gn=HMCN2;refGene_tx=XM_023242853.1 |
| D4 | 90588927 | 90589103 | refGene_gn=HMCN2;refGene_tx=XM_023242853.1 |
| D4 | 90590150 | 90590247 | refGene_gn=HMCN2;refGene_tx=XM_023242853.1 |

| | | | |
|---|---|---|---|
| D4 | 90590421 | 90590541 | refGene_gn=HMCN2;refGene_tx=XM_023242853.1 |
| D4 | 90591500 | 90591662 | refGene_gn=HMCN2;refGene_tx=XM_023242853.1 |
| D4 | 90592451 | 90592733 | refGene_gn=HMCN2;refGene_tx=XM_023242853.1 |
| D4 | 90593763 | 90593931 | refGene_gn=HMCN2;refGene_tx=XM_023242853.1 |
| D4 | 90595319 | 90595433 | refGene_gn=HMCN2;refGene_tx=XM_023242853.1 |
| D4 | 90596062 | 90596149 | refGene_gn=HMCN2;refGene_tx=XM_023242853.1 |
| D4 | 90597056 | 90597248 | refGene_gn=HMCN2;refGene_tx=XM_023242853.1 |
| D4 | 90597973 | 90598147 | refGene_gn=HMCN2;refGene_tx=XM_023242853.1 |
| D4 | 90598237 | 90598351 | refGene_gn=HMCN2;refGene_tx=XM_023242853.1 |
| D4 | 90598900 | 90599043 | refGene_gn=HMCN2;refGene_tx=XM_023242853.1 |
| D4 | 90599347 | 90599498 | refGene_gn=HMCN2;refGene_tx=XM_023242853.1 |
| D4 | 90600118 | 90600337 | refGene_gn=HMCN2;refGene_tx=XM_023242853.1 |
| D4 | 90601875 | 90601989 | refGene_gn=HMCN2;refGene_tx=XM_023242853.1 |
| D4 | 90602351 | 90602539 | refGene_gn=HMCN2;refGene_tx=XM_023242853.1 |
| D4 | 90603153 | 90603250 | refGene_gn=HMCN2;refGene_tx=XM_023242853.1 |
| D4 | 90603511 | 90603673 | refGene_gn=HMCN2;refGene_tx=XM_023242853.1 |
| D4 | 90603837 | 90603951 | refGene_gn=HMCN2;refGene_tx=XM_023242853.1 |
| D4 | 90604862 | 90605065 | refGene_gn=HMCN2;refGene_tx=XM_023242853.1 |
| D4 | 90605765 | 90605847 | refGene_gn=HMCN2;refGene_tx=XM_023242853.1 |
| D4 | 90607389 | 90607521 | refGene_gn=HMCN2;refGene_tx=XM_023242853.1 |
| D4 | 90609903 | 90610047 | refGene_gn=HMCN2;refGene_tx=XM_023242853.1 |
| D4 | 90610139 | 90610299 | refGene_gn=HMCN2;refGene_tx=XM_023242853.1 |
| D4 | 90610396 | 90610521 | refGene_gn=HMCN2;refGene_tx=XM_023242853.1 |
| D4 | 90610850 | 90611034 | refGene_gn=HMCN2;refGene_tx=XM_023242853.1 |
| D4 | 90612035 | 90612133 | refGene_gn=HMCN2;refGene_tx=XM_023242853.1 |
| D4 | 90612638 | 90612905 | refGene_gn=HMCN2;refGene_tx=XM_023242853.1 |
| D4 | 90613364 | 90613546 | refGene_gn=HMCN2;refGene_tx=XM_023242853.1 |
| D4 | 90614010 | 90614092 | refGene_gn=HMCN2;refGene_tx=XM_023242853.1 |
| D4 | 90614185 | 90614233 | refGene_gn=HMCN2;refGene_tx=XM_023242853.1 |
| D4 | 90614235 | 90614321 | refGene_gn=HMCN2;refGene_tx=XM_023242853.1 |
| D4 | 90614815 | 90614957 | refGene_gn=HMCN2;refGene_tx=XM_023242853.1 |
| D4 | 90615049 | 90615194 | refGene_gn=HMCN2;refGene_tx=XM_023242853.1 |
| D4 | 90616136 | 90616264 | refGene_gn=HMCN2;refGene_tx=XM_023242853.1 |
| D4 | 90616841 | 90616998 | refGene_gn=HMCN2;refGene_tx=XM_023242853.1 |
| D4 | 90618337 | 90618459 | refGene_gn=HMCN2;refGene_tx=XM_023242853.1 |
| D4 | 90619226 | 90619391 | refGene_gn=HMCN2;refGene_tx=XM_023242853.1 |
| D4 | 90620970 | 90621078 | refGene_gn=HMCN2;refGene_tx=XM_023242853.1 |
| D4 | 90621326 | 90621461 | refGene_gn=HMCN2;refGene_tx=XM_023242853.1 |
| D4 | 90621804 | 90621939 | refGene_gn=HMCN2;refGene_tx=XM_023242853.1 |
| D4 | 90622424 | 90622615 | refGene_gn=HMCN2;refGene_tx=XM_023242853.1 |
| D4 | 90623472 | 90623686 | refGene_gn=HMCN2;refGene_tx=XM_023242853.1 |
| D4 | 90624720 | 90624855 | refGene_gn=HMCN2;refGene_tx=XM_023242853.1 |
| D4 | 90625836 | 90626027 | refGene_gn=HMCN2;refGene_tx=XM_023242853.1 |
| D4 | 90627290 | 90627372 | refGene_gn=HMCN2;refGene_tx=XM_023242853.1 |
| D4 | 90631604 | 90631874 | refGene_gn=HMCN2;refGene_tx=XM_023242853.1 |
| D4 | 90634317 | 90634467 | refGene_gn=HMCN2;refGene_tx=XM_023242853.1 |
| D4 | 90636057 | 90636195 | refGene_gn=HMCN2;refGene_tx=XM_023242853.1 |
| D4 | 90636294 | 90636416 | refGene_gn=HMCN2;refGene_tx=XM_023242853.1 |
| D4 | 90637483 | 90637721 | refGene_gn=HMCN2;refGene_tx=XM_023242853.1 |
| D4 | 90638912 | 90638975 | refGene_gn=HMCN2;refGene_tx=XM_023242853.1 |

D4 90639107 90639230 refGene_gn=HMCN2;refGene_tx=XM_023242853.1
D4 90639942 90640077 refGene_gn=HMCN2;refGene_tx=XM_023242853.1
D4 90640873 90641002 refGene_gn=HMCN2;refGene_tx=XM_023242853.1
D4 90641520 90641841 refGene_gn=HMCN2;refGene_tx=XM_023242853.1
D4 90642504 90642624 refGene_gn=HMCN2;refGene_tx=XM_023242853.1
D4 90643472 90643599 refGene_gn=HMCN2;refGene_tx=XM_023242853.1
D4 90644604 90644950 refGene_gn=HMCN2;refGene_tx=XM_023242853.1
D4 90660745 90660850
refGene_gn=ASS1;refGene_tx=XM_011288374.2;refGene_gn=ASS1;refGene_tx=XM_019816669.2
D4 90661667 90661736
refGene_gn=ASS1;refGene_tx=XM_011288374.2;refGene_gn=ASS1;refGene_tx=XM_019816669.2
D4 90666849 90667038
refGene_gn=ASS1;refGene_tx=XM_011288374.2;refGene_gn=ASS1;refGene_tx=XM_019816669.2
D4 90672308 90672365
refGene_gn=ASS1;refGene_tx=XM_011288374.2;refGene_gn=ASS1;refGene_tx=XM_019816669.2
D4 90675169 90675244
refGene_gn=ASS1;refGene_tx=XM_011288374.2;refGene_gn=ASS1;refGene_tx=XM_019816669.2
D4 90678427 90678498
refGene_gn=ASS1;refGene_tx=XM_011288374.2;refGene_gn=ASS1;refGene_tx=XM_019816669.2
D4 90679036 90679067
refGene_gn=ASS1;refGene_tx=XM_011288374.2;refGene_gn=ASS1;refGene_tx=XM_019816669.2
D4 90684229 90684320
refGene_gn=ASS1;refGene_tx=XM_011288374.2;refGene_gn=ASS1;refGene_tx=XM_019816669.2
D4 90687194 90687279
refGene_gn=ASS1;refGene_tx=XM_011288374.2;refGene_gn=ASS1;refGene_tx=XM_019816669.2
D4 90687814 90687879
refGene_gn=ASS1;refGene_tx=XM_011288374.2;refGene_gn=ASS1;refGene_tx=XM_019816669.2
D4 90696589 90696721
refGene_gn=ASS1;refGene_tx=XM_011288374.2;refGene_gn=ASS1;refGene_tx=XM_019816669.2
D4 90701418 90701575
refGene_gn=ASS1;refGene_tx=XM_011288374.2;refGene_gn=ASS1;refGene_tx=XM_019816669.2
D4 90705509 90705575
refGene_gn=ASS1;refGene_tx=XM_011288374.2;refGene_gn=ASS1;refGene_tx=XM_019816669.2
D4 90706337 90706374 refGene_gn=ASS1;refGene_tx=XM_019816669.2
D4 90707011 90707057 refGene_gn=ASS1;refGene_tx=XM_011288374.2
D4 90764587 90764671
refGene_gn=FUBP3;refGene_tx=XM_003995971.4;refGene_gn=FUBP3;refGene_tx

=XM_006939555.3
D4	90766408 90766439 refGene_gn=FUBP3;refGene_tx=XM_019816670.1
D4	90777003 90777109
refGene_gn=FUBP3;refGene_tx=XM_003995971.4;refGene_gn=FUBP3;refGene_tx
=XM_006939555.3
D4	90788754 90788788
refGene_gn=FUBP3;refGene_tx=XM_003995971.4;refGene_gn=FUBP3;refGene_tx
=XM_006939555.3;refGene_gn=FUBP3;refGene_tx=XM_019816670.1
D4	90791197 90791247
refGene_gn=FUBP3;refGene_tx=XM_003995971.4;refGene_gn=FUBP3;refGene_tx
=XM_006939555.3;refGene_gn=FUBP3;refGene_tx=XM_019816670.1
D4	90791628 90791700
refGene_gn=FUBP3;refGene_tx=XM_003995971.4;refGene_gn=FUBP3;refGene_tx
=XM_006939555.3;refGene_gn=FUBP3;refGene_tx=XM_019816670.1
D4	90792947 90793005
refGene_gn=FUBP3;refGene_tx=XM_003995971.4;refGene_gn=FUBP3;refGene_tx
=XM_006939555.3;refGene_gn=FUBP3;refGene_tx=XM_019816670.1
D4	90795196 90795359
refGene_gn=FUBP3;refGene_tx=XM_003995971.4;refGene_gn=FUBP3;refGene_tx
=XM_006939555.3;refGene_gn=FUBP3;refGene_tx=XM_019816670.1
D4	90796527 90796626
refGene_gn=FUBP3;refGene_tx=XM_003995971.4;refGene_gn=FUBP3;refGene_tx
=XM_006939555.3;refGene_gn=FUBP3;refGene_tx=XM_019816670.1
D4	90799809 90799914
refGene_gn=FUBP3;refGene_tx=XM_003995971.4;refGene_gn=FUBP3;refGene_tx
=XM_006939555.3;refGene_gn=FUBP3;refGene_tx=XM_019816670.1
D4	90800975 90801078
refGene_gn=FUBP3;refGene_tx=XM_003995971.4;refGene_gn=FUBP3;refGene_tx
=XM_006939555.3;refGene_gn=FUBP3;refGene_tx=XM_019816670.1
D4	90801919 90802020
refGene_gn=FUBP3;refGene_tx=XM_003995971.4;refGene_gn=FUBP3;refGene_tx
=XM_006939555.3;refGene_gn=FUBP3;refGene_tx=XM_019816670.1
D4	90804091 90804233
refGene_gn=FUBP3;refGene_tx=XM_003995971.4;refGene_gn=FUBP3;refGene_tx
=XM_006939555.3;refGene_gn=FUBP3;refGene_tx=XM_019816670.1
D4	90807832 90807993
refGene_gn=FUBP3;refGene_tx=XM_003995971.4;refGene_gn=FUBP3;refGene_tx
=XM_006939555.3;refGene_gn=FUBP3;refGene_tx=XM_019816670.1
D4	90808781 90808855
refGene_gn=FUBP3;refGene_tx=XM_003995971.4;refGene_gn=FUBP3;refGene_tx
=XM_006939555.3;refGene_gn=FUBP3;refGene_tx=XM_019816670.1
D4	90809117 90809198
refGene_gn=FUBP3;refGene_tx=XM_003995971.4;refGene_gn=FUBP3;refGene_tx
=XM_006939555.3;refGene_gn=FUBP3;refGene_tx=XM_019816670.1
D4	90809372 90809449
refGene_gn=FUBP3;refGene_tx=XM_003995971.4;refGene_gn=FUBP3;refGene_tx
=XM_006939555.3;refGene_gn=FUBP3;refGene_tx=XM_019816670.1
D4	90811802 90811874
refGene_gn=FUBP3;refGene_tx=XM_003995971.4;refGene_gn=FUBP3;refGene_tx
=XM_006939555.3;refGene_gn=FUBP3;refGene_tx=XM_019816670.1

D4 90813422 90813550
refGene_gn=FUBP3;refGene_tx=XM_003995971.4;refGene_gn=FUBP3;refGene_tx=XM_006939555.3;refGene_gn=FUBP3;refGene_tx=XM_019816670.1
D4 90814502 90814511
refGene_gn=FUBP3;refGene_tx=XM_003995971.4;refGene_gn=FUBP3;refGene_tx=XM_006939555.3;refGene_gn=FUBP3;refGene_tx=XM_019816670.1
D4 90830257 90830480 refGene_gn=PRDM12;refGene_tx=XM_023242854.1
D4 90832318 90832509 refGene_gn=PRDM12;refGene_tx=XM_023242854.1
D4 90833880 90834036 refGene_gn=PRDM12;refGene_tx=XM_023242854.1
D4 90842670 90842782 refGene_gn=PRDM12;refGene_tx=XM_023242854.1
D4 90844114 90844524 refGene_gn=PRDM12;refGene_tx=XM_023242854.1
D4 90852362 90852431 refGene_gn=EXOSC2;refGene_tx=XM_003995972.5
D4 90852432 90852485 refGene_gn=EXOSC2;refGene_tx=XM_003995972.5
D4 90853768 90853870 refGene_gn=EXOSC2;refGene_tx=XM_003995972.5
D4 90855573 90855619 refGene_gn=EXOSC2;refGene_tx=XM_003995972.5
D4 90855841 90855931 refGene_gn=EXOSC2;refGene_tx=XM_003995972.5
D4 90857220 90857286 refGene_gn=EXOSC2;refGene_tx=XM_003995972.5
D4 90857774 90857843 refGene_gn=EXOSC2;refGene_tx=XM_003995972.5
D4 90859230 90859407 refGene_gn=EXOSC2;refGene_tx=XM_003995972.5
D4 90860284 90860413 refGene_gn=EXOSC2;refGene_tx=XM_003995972.5
D4 90860966 90861047 refGene_gn=EXOSC2;refGene_tx=XM_003995972.5
D4 90866912 90867048 refGene_gn=ABL1;refGene_tx=XM_003995974.5
D4 90965571 90965650 refGene_gn=ABL1;refGene_tx=XM_023242856.1
D4 90980056 90980230
refGene_gn=ABL1;refGene_tx=XM_003995974.5;refGene_gn=ABL1;refGene_tx=XM_023242856.1
D4 90980776 90981072
refGene_gn=ABL1;refGene_tx=XM_003995974.5;refGene_gn=ABL1;refGene_tx=XM_023242856.1
D4 90990655 90990928
refGene_gn=ABL1;refGene_tx=XM_003995974.5;refGene_gn=ABL1;refGene_tx=XM_023242856.1
D4 90998300 90998385
refGene_gn=ABL1;refGene_tx=XM_003995974.5;refGene_gn=ABL1;refGene_tx=XM_023242856.1
D4 90999201 90999379
refGene_gn=ABL1;refGene_tx=XM_003995974.5;refGene_gn=ABL1;refGene_tx=XM_023242856.1
D4 91001096 91001281
refGene_gn=ABL1;refGene_tx=XM_003995974.5;refGene_gn=ABL1;refGene_tx=XM_023242856.1
D4 91003613 91003766
refGene_gn=ABL1;refGene_tx=XM_003995974.5;refGene_gn=ABL1;refGene_tx=XM_023242856.1
D4 91004978 91005068
refGene_gn=ABL1;refGene_tx=XM_003995974.5;refGene_gn=ABL1;refGene_tx=XM_023242856.1
D4 91005418 91005583
refGene_gn=ABL1;refGene_tx=XM_003995974.5;refGene_gn=ABL1;refGene_tx=XM_023242856.1

D4 91008000 91009718
refGene_gn=ABL1;refGene_tx=XM_003995974.5;refGene_gn=ABL1;refGene_tx=XM_023242856.1
D4 91017012 91017408 refGene_gn=QRFP;refGene_tx=XM_023242857.1
D4 91026650 91026910
refGene_gn=FIBCD1;refGene_tx=XM_023242858.1;refGene_gn=FIBCD1;refGene_tx=XM_023242859.1
D4 91027451 91027631
refGene_gn=FIBCD1;refGene_tx=XM_023242858.1;refGene_gn=FIBCD1;refGene_tx=XM_023242859.1
D4 91032660 91032757
refGene_gn=FIBCD1;refGene_tx=XM_023242858.1;refGene_gn=FIBCD1;refGene_tx=XM_023242859.1
D4 91044038 91044175
refGene_gn=FIBCD1;refGene_tx=XM_023242858.1;refGene_gn=FIBCD1;refGene_tx=XM_023242859.1
D4 91044548 91044708
refGene_gn=FIBCD1;refGene_tx=XM_023242858.1;refGene_gn=FIBCD1;refGene_tx=XM_023242859.1
D4 91048851 91049331
refGene_gn=FIBCD1;refGene_tx=XM_023242858.1;refGene_gn=FIBCD1;refGene_tx=XM_023242859.1
D4 91049907 91049982 refGene_gn=FIBCD1;refGene_tx=XM_023242858.1
D4 91058018 91058090
refGene_gn=FIBCD1;refGene_tx=XM_023242858.1;refGene_gn=FIBCD1;refGene_tx=XM_023242859.1
D4 91101652 91102025 refGene_gn=LAMC3;refGene_tx=XM_023242860.1
D4 91115321 91115626 refGene_gn=LAMC3;refGene_tx=XM_023242860.1
D4 91118907 91119038 refGene_gn=LAMC3;refGene_tx=XM_023242860.1
D4 91121107 91121274 refGene_gn=LAMC3;refGene_tx=XM_023242860.1
D4 91123165 91123354 refGene_gn=LAMC3;refGene_tx=XM_023242860.1
D4 91123439 91123557 refGene_gn=LAMC3;refGene_tx=XM_023242860.1
D4 91125238 91125337 refGene_gn=LAMC3;refGene_tx=XM_023242860.1
D4 91128231 91128368 refGene_gn=LAMC3;refGene_tx=XM_023242860.1
D4 91130929 91131040 refGene_gn=LAMC3;refGene_tx=XM_023242860.1
D4 91132945 91133138 refGene_gn=LAMC3;refGene_tx=XM_023242860.1
D4 91133318 91133437 refGene_gn=LAMC3;refGene_tx=XM_023242860.1
D4 91133528 91133579 refGene_gn=LAMC3;refGene_tx=XM_023242860.1
D4 91137766 91137985 refGene_gn=LAMC3;refGene_tx=XM_023242860.1
D4 91139889 91140078 refGene_gn=LAMC3;refGene_tx=XM_023242860.1
D4 91141823 91141850 refGene_gn=LAMC3;refGene_tx=XM_023242860.1
D4 91141852 91142068 refGene_gn=LAMC3;refGene_tx=XM_023242860.1
D4 91142924 91143078 refGene_gn=LAMC3;refGene_tx=XM_023242860.1
D4 91143720 91143863 refGene_gn=LAMC3;refGene_tx=XM_023242860.1
D4 91144356 91144535 refGene_gn=LAMC3;refGene_tx=XM_023242860.1
D4 91146001 91146140 refGene_gn=LAMC3;refGene_tx=XM_023242860.1
D4 91147167 91147373 refGene_gn=LAMC3;refGene_tx=XM_023242860.1
D4 91147741 91147818 refGene_gn=LAMC3;refGene_tx=XM_023242860.1
D4 91148287 91148422 refGene_gn=LAMC3;refGene_tx=XM_023242860.1
D4 91149565 91149713 refGene_gn=LAMC3;refGene_tx=XM_023242860.1

D4 91150730 91150874 refGene_gn=LAMC3;refGene_tx=XM_023242860.1
D4 91151033 91151136 refGene_gn=LAMC3;refGene_tx=XM_023242860.1
D4 91153637 91153837 refGene_gn=LAMC3;refGene_tx=XM_023242860.1
D4 91154410 91154557 refGene_gn=LAMC3;refGene_tx=XM_023242860.1
D4 91154659 91154759 refGene_gn=LAMC3;refGene_tx=XM_023242860.1
D4 91157155 91157406 refGene_gn=LAMC3;refGene_tx=XM_023242860.1
D4 91162197 91162228 refGene_gn=AIF1L;refGene_tx=XM_023242862.1
D4 91162380 91162442 refGene_gn=AIF1L;refGene_tx=XM_023242862.1
D4 91173137 91173204 refGene_gn=AIF1L;refGene_tx=XM_023242862.1
D4 91175650 91175692
refGene_gn=AIF1L;refGene_tx=XM_006939558.4;refGene_gn=AIF1L;refGene_tx
=XM_019816672.2;refGene_gn=AIF1L;refGene_tx=XM_023242862.1
D4 91178786 91178949
refGene_gn=AIF1L;refGene_tx=XM_006939558.4;refGene_gn=AIF1L;refGene_tx
=XM_019816672.2;refGene_gn=AIF1L;refGene_tx=XM_023242862.1
D4 91181844 91181929
refGene_gn=AIF1L;refGene_tx=XM_006939558.4;refGene_gn=AIF1L;refGene_tx
=XM_019816672.2;refGene_gn=AIF1L;refGene_tx=XM_023242862.1
D4 91185770 91185815 refGene_gn=NUP214;refGene_tx=XM_023242861.1
D4 91188006 91188144 refGene_gn=NUP214;refGene_tx=XM_023242861.1
D4 91188145 91188203 refGene_gn=NUP214;refGene_tx=XM_023242861.1
D4 91188501 91188653 refGene_gn=NUP214;refGene_tx=XM_023242861.1
D4 91189474 91189673 refGene_gn=NUP214;refGene_tx=XM_023242861.1
D4 91190908 91190979 refGene_gn=NUP214;refGene_tx=XM_023242861.1
D4 91192142 91192206 refGene_gn=NUP214;refGene_tx=XM_023242861.1
D4 91192914 91193018 refGene_gn=NUP214;refGene_tx=XM_023242861.1
D4 91194928 91195033 refGene_gn=NUP214;refGene_tx=XM_023242861.1
D4 91195034 91195036 refGene_gn=NUP214;refGene_tx=XM_023242861.1
D4 91195677 91195744 refGene_gn=NUP214;refGene_tx=XM_023242861.1
D4 91199030 91199157 refGene_gn=NUP214;refGene_tx=XM_023242861.1
D4 91200126 91200288 refGene_gn=NUP214;refGene_tx=XM_023242861.1
D4 91204486 91204931 refGene_gn=NUP214;refGene_tx=XM_023242861.1
D4 91206474 91206647 refGene_gn=NUP214;refGene_tx=XM_023242861.1
D4 91209234 91209329 refGene_gn=NUP214;refGene_tx=XM_023242861.1
D4 91212325 91212412 refGene_gn=NUP214;refGene_tx=XM_023242861.1
D4 91212621 91212771 refGene_gn=NUP214;refGene_tx=XM_023242861.1
D4 91213709 91213868 refGene_gn=NUP214;refGene_tx=XM_023242861.1
D4 91216963 91217067 refGene_gn=NUP214;refGene_tx=XM_023242861.1
D4 91219035 91219218 refGene_gn=NUP214;refGene_tx=XM_023242861.1
D4 91220083 91220169 refGene_gn=NUP214;refGene_tx=XM_023242861.1
D4 91220265 91220349 refGene_gn=NUP214;refGene_tx=XM_023242861.1
D4 91228629 91228890 refGene_gn=NUP214;refGene_tx=XM_023242861.1
D4 91230009 91230171 refGene_gn=NUP214;refGene_tx=XM_023242861.1
D4 91232309 91232409 refGene_gn=NUP214;refGene_tx=XM_023242861.1
D4 91240592 91240668 refGene_gn=NUP214;refGene_tx=XM_023242861.1
D4 91241994 91242073 refGene_gn=NUP214;refGene_tx=XM_023242861.1
D4 91244652 91244743 refGene_gn=NUP214;refGene_tx=XM_023242861.1
D4 91247016 91247078 refGene_gn=NUP214;refGene_tx=XM_023242861.1
D4 91249173 91251015 refGene_gn=NUP214;refGene_tx=XM_023242861.1
D4 91253061 91253132 refGene_gn=NUP214;refGene_tx=XM_023242861.1

D4 91264366 91264520 refGene_gn=NUP214;refGene_tx=XM_023242861.1
D4 91271287 91271440 refGene_gn=NUP214;refGene_tx=XM_023242861.1
D4 91276086 91276258 refGene_gn=NUP214;refGene_tx=XM_023242861.1
D4 91278215 91278355 refGene_gn=NUP214;refGene_tx=XM_023242861.1
D4 91279173 91279198 refGene_gn=NUP214;refGene_tx=XM_023242861.1
D4 91279724 91279758 refGene_gn=NUP214;refGene_tx=XM_023242861.1
D4 91304040 91304569
refGene_gn=FAM78A;refGene_tx=XM_023242864.1;refGene_gn=FAM78A;refGene_tx=XM_023242865.1
D4 91314789 91315112
refGene_gn=FAM78A;refGene_tx=XM_023242864.1;refGene_gn=FAM78A;refGene_tx=XM_023242865.1
D4 91315356 91315892
refGene_gn=LOC111557483;refGene_tx=XM_023242318.1
D4 91316508 91316671
refGene_gn=LOC111557483;refGene_tx=XM_023242318.1
D4 91326296 91326584
refGene_gn=LOC111557483;refGene_tx=XM_023242318.1
D4 91326807 91327258
refGene_gn=PLPP7;refGene_tx=XM_003995976.4;refGene_gn=PLPP7;refGene_tx=XM_006939559.4
D4 91337360 91337725
refGene_gn=PLPP7;refGene_tx=XM_003995976.4;refGene_gn=PLPP7;refGene_tx=XM_006939559.4
D4 91412782 91412897
refGene_gn=PRRC2B;refGene_tx=XM_019816673.2;refGene_gn=PRRC2B;refGene_tx=XM_023242866.1;refGene_gn=PRRC2B;refGene_tx=XM_023242867.1;refGene_gn=PRRC2B;refGene_tx=XM_023242868.1;refGene_gn=PRRC2B;refGene_tx=XM_023242869.1;refGene_gn=PRRC2B;refGene_tx=XM_023242870.1
D4 91413653 91413831
refGene_gn=PRRC2B;refGene_tx=XM_019816673.2;refGene_gn=PRRC2B;refGene_tx=XM_023242866.1;refGene_gn=PRRC2B;refGene_tx=XM_023242867.1;refGene_gn=PRRC2B;refGene_tx=XM_023242868.1;refGene_gn=PRRC2B;refGene_tx=XM_023242869.1;refGene_gn=PRRC2B;refGene_tx=XM_023242870.1
D4 91416590 91416693
refGene_gn=PRRC2B;refGene_tx=XM_019816673.2;refGene_gn=PRRC2B;refGene_tx=XM_023242866.1;refGene_gn=PRRC2B;refGene_tx=XM_023242867.1;refGene_gn=PRRC2B;refGene_tx=XM_023242868.1;refGene_gn=PRRC2B;refGene_tx=XM_023242869.1;refGene_gn=PRRC2B;refGene_tx=XM_023242870.1
D4 91419215 91419288
refGene_gn=PRRC2B;refGene_tx=XM_019816673.2;refGene_gn=PRRC2B;refGene_tx=XM_023242866.1;refGene_gn=PRRC2B;refGene_tx=XM_023242867.1;refGene_gn=PRRC2B;refGene_tx=XM_023242868.1;refGene_gn=PRRC2B;refGene_tx=XM_023242869.1;refGene_gn=PRRC2B;refGene_tx=XM_023242870.1
D4 91423357 91423501
refGene_gn=PRRC2B;refGene_tx=XM_019816673.2;refGene_gn=PRRC2B;refGene_tx=XM_023242866.1;refGene_gn=PRRC2B;refGene_tx=XM_023242867.1;refGene_gn=PRRC2B;refGene_tx=XM_023242868.1;refGene_gn=PRRC2B;refGene_tx=XM_023242869.1;refGene_gn=PRRC2B;refGene_tx=XM_023242870.1
D4 91425768 91426010

refGene_gn=PRRC2B;refGene_tx=XM_019816673.2;refGene_gn=PRRC2B;refGene_tx=XM_023242866.1;refGene_gn=PRRC2B;refGene_tx=XM_023242867.1;refGene_gn=PRRC2B;refGene_tx=XM_023242868.1;refGene_gn=PRRC2B;refGene_tx=XM_023242869.1;refGene_gn=PRRC2B;refGene_tx=XM_023242870.1
D4 91426474 91426596
refGene_gn=PRRC2B;refGene_tx=XM_019816673.2;refGene_gn=PRRC2B;refGene_tx=XM_023242866.1;refGene_gn=PRRC2B;refGene_tx=XM_023242867.1;refGene_gn=PRRC2B;refGene_tx=XM_023242868.1;refGene_gn=PRRC2B;refGene_tx=XM_023242869.1;refGene_gn=PRRC2B;refGene_tx=XM_023242870.1
D4 91427362 91427505
refGene_gn=PRRC2B;refGene_tx=XM_019816673.2;refGene_gn=PRRC2B;refGene_tx=XM_023242866.1;refGene_gn=PRRC2B;refGene_tx=XM_023242867.1;refGene_gn=PRRC2B;refGene_tx=XM_023242868.1;refGene_gn=PRRC2B;refGene_tx=XM_023242869.1;refGene_gn=PRRC2B;refGene_tx=XM_023242870.1
D4 91432774 91432865
refGene_gn=PRRC2B;refGene_tx=XM_019816673.2;refGene_gn=PRRC2B;refGene_tx=XM_023242866.1;refGene_gn=PRRC2B;refGene_tx=XM_023242867.1;refGene_gn=PRRC2B;refGene_tx=XM_023242868.1;refGene_gn=PRRC2B;refGene_tx=XM_023242869.1;refGene_gn=PRRC2B;refGene_tx=XM_023242870.1
D4 91436788 91436981
refGene_gn=PRRC2B;refGene_tx=XM_019816673.2;refGene_gn=PRRC2B;refGene_tx=XM_023242866.1;refGene_gn=PRRC2B;refGene_tx=XM_023242867.1;refGene_gn=PRRC2B;refGene_tx=XM_023242868.1;refGene_gn=PRRC2B;refGene_tx=XM_023242869.1;refGene_gn=PRRC2B;refGene_tx=XM_023242870.1
D4 91440642 91440958
refGene_gn=PRRC2B;refGene_tx=XM_019816673.2;refGene_gn=PRRC2B;refGene_tx=XM_023242866.1;refGene_gn=PRRC2B;refGene_tx=XM_023242867.1;refGene_gn=PRRC2B;refGene_tx=XM_023242868.1;refGene_gn=PRRC2B;refGene_tx=XM_023242869.1;refGene_gn=PRRC2B;refGene_tx=XM_023242870.1
D4 91442982 91443176
refGene_gn=PRRC2B;refGene_tx=XM_019816673.2;refGene_gn=PRRC2B;refGene_tx=XM_023242866.1;refGene_gn=PRRC2B;refGene_tx=XM_023242867.1;refGene_gn=PRRC2B;refGene_tx=XM_023242868.1;refGene_gn=PRRC2B;refGene_tx=XM_023242869.1;refGene_gn=PRRC2B;refGene_tx=XM_023242870.1
D4 91445337 91445533
refGene_gn=PRRC2B;refGene_tx=XM_019816673.2;refGene_gn=PRRC2B;refGene_tx=XM_023242866.1;refGene_gn=PRRC2B;refGene_tx=XM_023242867.1;refGene_gn=PRRC2B;refGene_tx=XM_023242868.1;refGene_gn=PRRC2B;refGene_tx=XM_023242869.1;refGene_gn=PRRC2B;refGene_tx=XM_023242870.1
D4 91450605 91450822
refGene_gn=PRRC2B;refGene_tx=XM_019816673.2;refGene_gn=PRRC2B;refGene_tx=XM_023242866.1;refGene_gn=PRRC2B;refGene_tx=XM_023242867.1;refGene_gn=PRRC2B;refGene_tx=XM_023242868.1;refGene_gn=PRRC2B;refGene_tx=XM_023242869.1;refGene_gn=PRRC2B;refGene_tx=XM_023242870.1
D4 91451472 91453536
refGene_gn=PRRC2B;refGene_tx=XM_019816673.2;refGene_gn=PRRC2B;refGene_tx=XM_023242866.1;refGene_gn=PRRC2B;refGene_tx=XM_023242867.1;refGene_gn=PRRC2B;refGene_tx=XM_023242868.1;refGene_gn=PRRC2B;refGene_tx=XM_023242869.1;refGene_gn=PRRC2B;refGene_tx=XM_023242870.1
D4 91454846 91455043

refGene_gn=PRRC2B;refGene_tx=XM_019816673.2;refGene_gn=PRRC2B;refGene_tx=XM_023242866.1;refGene_gn=PRRC2B;refGene_tx=XM_023242867.1;refGene_gn=PRRC2B;refGene_tx=XM_023242868.1;refGene_gn=PRRC2B;refGene_tx=XM_023242869.1;refGene_gn=PRRC2B;refGene_tx=XM_023242870.1
D4 91455633 91455779
refGene_gn=PRRC2B;refGene_tx=XM_019816673.2;refGene_gn=PRRC2B;refGene_tx=XM_023242866.1;refGene_gn=PRRC2B;refGene_tx=XM_023242867.1;refGene_gn=PRRC2B;refGene_tx=XM_023242868.1;refGene_gn=PRRC2B;refGene_tx=XM_023242869.1;refGene_gn=PRRC2B;refGene_tx=XM_023242870.1
D4 91456320 91456462
refGene_gn=PRRC2B;refGene_tx=XM_019816673.2;refGene_gn=PRRC2B;refGene_tx=XM_023242866.1;refGene_gn=PRRC2B;refGene_tx=XM_023242867.1;refGene_gn=PRRC2B;refGene_tx=XM_023242868.1;refGene_gn=PRRC2B;refGene_tx=XM_023242869.1;refGene_gn=PRRC2B;refGene_tx=XM_023242870.1
D4 91458203 91458286
refGene_gn=PRRC2B;refGene_tx=XM_019816673.2;refGene_gn=PRRC2B;refGene_tx=XM_023242866.1;refGene_gn=PRRC2B;refGene_tx=XM_023242867.1;refGene_gn=PRRC2B;refGene_tx=XM_023242868.1;refGene_gn=PRRC2B;refGene_tx=XM_023242869.1;refGene_gn=PRRC2B;refGene_tx=XM_023242870.1
D4 91458752 91458941
refGene_gn=PRRC2B;refGene_tx=XM_019816673.2;refGene_gn=PRRC2B;refGene_tx=XM_023242866.1;refGene_gn=PRRC2B;refGene_tx=XM_023242867.1;refGene_gn=PRRC2B;refGene_tx=XM_023242868.1;refGene_gn=PRRC2B;refGene_tx=XM_023242869.1;refGene_gn=PRRC2B;refGene_tx=XM_023242870.1
D4 91459079 91459277
refGene_gn=PRRC2B;refGene_tx=XM_019816673.2;refGene_gn=PRRC2B;refGene_tx=XM_023242866.1;refGene_gn=PRRC2B;refGene_tx=XM_023242867.1;refGene_gn=PRRC2B;refGene_tx=XM_023242868.1;refGene_gn=PRRC2B;refGene_tx=XM_023242869.1;refGene_gn=PRRC2B;refGene_tx=XM_023242870.1
D4 91459721 91459808
refGene_gn=PRRC2B;refGene_tx=XM_019816673.2;refGene_gn=PRRC2B;refGene_tx=XM_023242866.1;refGene_gn=PRRC2B;refGene_tx=XM_023242867.1;refGene_gn=PRRC2B;refGene_tx=XM_023242868.1;refGene_gn=PRRC2B;refGene_tx=XM_023242869.1;refGene_gn=PRRC2B;refGene_tx=XM_023242870.1
D4 91460505 91460610
refGene_gn=PRRC2B;refGene_tx=XM_019816673.2;refGene_gn=PRRC2B;refGene_tx=XM_023242866.1;refGene_gn=PRRC2B;refGene_tx=XM_023242867.1;refGene_gn=PRRC2B;refGene_tx=XM_023242868.1;refGene_gn=PRRC2B;refGene_tx=XM_023242869.1;refGene_gn=PRRC2B;refGene_tx=XM_023242870.1
D4 91460768 91460961
refGene_gn=PRRC2B;refGene_tx=XM_019816673.2;refGene_gn=PRRC2B;refGene_tx=XM_023242866.1;refGene_gn=PRRC2B;refGene_tx=XM_023242867.1;refGene_gn=PRRC2B;refGene_tx=XM_023242868.1;refGene_gn=PRRC2B;refGene_tx=XM_023242869.1;refGene_gn=PRRC2B;refGene_tx=XM_023242870.1
D4 91461807 91461905
refGene_gn=PRRC2B;refGene_tx=XM_019816673.2;refGene_gn=PRRC2B;refGene_tx=XM_023242866.1;refGene_gn=PRRC2B;refGene_tx=XM_023242867.1;refGene_gn=PRRC2B;refGene_tx=XM_023242868.1;refGene_gn=PRRC2B;refGene_tx=XM_023242869.1;refGene_gn=PRRC2B;refGene_tx=XM_023242870.1
D4 91462844 91462972

refGene_gn=PRRC2B;refGene_tx=XM_019816673.2;refGene_gn=PRRC2B;refGene_tx=XM_023242866.1;refGene_gn=PRRC2B;refGene_tx=XM_023242867.1;refGene_gn=PRRC2B;refGene_tx=XM_023242868.1;refGene_gn=PRRC2B;refGene_tx=XM_023242869.1;refGene_gn=PRRC2B;refGene_tx=XM_023242870.1
D4 91463549 91463790
refGene_gn=PRRC2B;refGene_tx=XM_019816673.2;refGene_gn=PRRC2B;refGene_tx=XM_023242866.1;refGene_gn=PRRC2B;refGene_tx=XM_023242867.1;refGene_gn=PRRC2B;refGene_tx=XM_023242868.1;refGene_gn=PRRC2B;refGene_tx=XM_023242869.1;refGene_gn=PRRC2B;refGene_tx=XM_023242870.1
D4 91466548 91466728
refGene_gn=PRRC2B;refGene_tx=XM_019816673.2;refGene_gn=PRRC2B;refGene_tx=XM_023242866.1;refGene_gn=PRRC2B;refGene_tx=XM_023242867.1;refGene_gn=PRRC2B;refGene_tx=XM_023242868.1;refGene_gn=PRRC2B;refGene_tx=XM_023242869.1;refGene_gn=PRRC2B;refGene_tx=XM_023242870.1
D4 91467309 91467401
refGene_gn=PRRC2B;refGene_tx=XM_019816673.2;refGene_gn=PRRC2B;refGene_tx=XM_023242866.1;refGene_gn=PRRC2B;refGene_tx=XM_023242867.1;refGene_gn=PRRC2B;refGene_tx=XM_023242868.1;refGene_gn=PRRC2B;refGene_tx=XM_023242869.1;refGene_gn=PRRC2B;refGene_tx=XM_023242870.1
D4 91469528 91469610
refGene_gn=PRRC2B;refGene_tx=XM_019816673.2;refGene_gn=PRRC2B;refGene_tx=XM_023242866.1;refGene_gn=PRRC2B;refGene_tx=XM_023242867.1;refGene_gn=PRRC2B;refGene_tx=XM_023242868.1;refGene_gn=PRRC2B;refGene_tx=XM_023242869.1
D4 91470928 91471063
refGene_gn=PRRC2B;refGene_tx=XM_019816673.2;refGene_gn=PRRC2B;refGene_tx=XM_023242866.1;refGene_gn=PRRC2B;refGene_tx=XM_023242867.1;refGene_gn=PRRC2B;refGene_tx=XM_023242868.1;refGene_gn=PRRC2B;refGene_tx=XM_023242869.1;refGene_gn=PRRC2B;refGene_tx=XM_023242870.1
D4 91477292 91477330
refGene_gn=POMT1;refGene_tx=XM_019816676.2;refGene_gn=POMT1;refGene_tx=XM_019816677.2
D4 91478208 91478331
refGene_gn=POMT1;refGene_tx=XM_011288369.3;refGene_gn=POMT1;refGene_tx=XM_011288370.2;refGene_gn=POMT1;refGene_tx=XM_019816676.2;refGene_gn=POMT1;refGene_tx=XM_019816677.2
D4 91479696 91479803
refGene_gn=POMT1;refGene_tx=XM_011288369.3;refGene_gn=POMT1;refGene_tx=XM_011288370.2;refGene_gn=POMT1;refGene_tx=XM_019816676.2;refGene_gn=POMT1;refGene_tx=XM_019816677.2
D4 91480220 91480271
refGene_gn=POMT1;refGene_tx=XM_011288369.3;refGene_gn=POMT1;refGene_tx=XM_011288370.2;refGene_gn=POMT1;refGene_tx=XM_019816676.2;refGene_gn=POMT1;refGene_tx=XM_019816677.2
D4 91480825 91480972
refGene_gn=POMT1;refGene_tx=XM_011288369.3;refGene_gn=POMT1;refGene_tx=XM_011288370.2;refGene_gn=POMT1;refGene_tx=XM_019816676.2;refGene_gn=POMT1;refGene_tx=XM_019816677.2
D4 91481678 91481790
refGene_gn=POMT1;refGene_tx=XM_011288369.3;refGene_gn=POMT1;refGene_tx

=XM_011288370.2;refGene_gn=POMT1;refGene_tx=XM_019816676.2;refGene_gn=POMT1;refGene_tx=XM_019816677.2
D4 91482170 91482236
refGene_gn=POMT1;refGene_tx=XM_011288369.3;refGene_gn=POMT1;refGene_tx=XM_011288370.2;refGene_gn=POMT1;refGene_tx=XM_019816676.2;refGene_gn=POMT1;refGene_tx=XM_019816677.2
D4 91482323 91482417
refGene_gn=POMT1;refGene_tx=XM_011288369.3;refGene_gn=POMT1;refGene_tx=XM_011288370.2;refGene_gn=POMT1;refGene_tx=XM_019816676.2;refGene_gn=POMT1;refGene_tx=XM_019816677.2
D4 91482618 91482774
refGene_gn=POMT1;refGene_tx=XM_011288369.3;refGene_gn=POMT1;refGene_tx=XM_011288370.2;refGene_gn=POMT1;refGene_tx=XM_019816676.2;refGene_gn=POMT1;refGene_tx=XM_019816677.2
D4 91483675 91483806
refGene_gn=POMT1;refGene_tx=XM_011288369.3;refGene_gn=POMT1;refGene_tx=XM_011288370.2;refGene_gn=POMT1;refGene_tx=XM_019816676.2;refGene_gn=POMT1;refGene_tx=XM_019816677.2
D4 91484297 91484393
refGene_gn=POMT1;refGene_tx=XM_011288369.3;refGene_gn=POMT1;refGene_tx=XM_011288370.2;refGene_gn=POMT1;refGene_tx=XM_019816676.2;refGene_gn=POMT1;refGene_tx=XM_019816677.2
D4 91485332 91485425
refGene_gn=POMT1;refGene_tx=XM_011288369.3;refGene_gn=POMT1;refGene_tx=XM_011288370.2;refGene_gn=POMT1;refGene_tx=XM_019816676.2;refGene_gn=POMT1;refGene_tx=XM_019816677.2
D4 91486905 91487002
refGene_gn=POMT1;refGene_tx=XM_011288369.3;refGene_gn=POMT1;refGene_tx=XM_011288370.2;refGene_gn=POMT1;refGene_tx=XM_019816676.2;refGene_gn=POMT1;refGene_tx=XM_019816677.2
D4 91488857 91488950
refGene_gn=POMT1;refGene_tx=XM_011288369.3;refGene_gn=POMT1;refGene_tx=XM_011288370.2;refGene_gn=POMT1;refGene_tx=XM_019816676.2;refGene_gn=POMT1;refGene_tx=XM_019816677.2
D4 91489258 91489379
refGene_gn=POMT1;refGene_tx=XM_011288369.3;refGene_gn=POMT1;refGene_tx=XM_011288370.2;refGene_gn=POMT1;refGene_tx=XM_019816676.2;refGene_gn=POMT1;refGene_tx=XM_019816677.2
D4 91489753 91489851
refGene_gn=POMT1;refGene_tx=XM_011288369.3;refGene_gn=POMT1;refGene_tx=XM_011288370.2;refGene_gn=POMT1;refGene_tx=XM_019816676.2;refGene_gn=POMT1;refGene_tx=XM_019816677.2
D4 91490342 91490456
refGene_gn=POMT1;refGene_tx=XM_011288369.3;refGene_gn=POMT1;refGene_tx=XM_011288370.2;refGene_gn=POMT1;refGene_tx=XM_019816676.2;refGene_gn=POMT1;refGene_tx=XM_019816677.2
D4 91491224 91491351
refGene_gn=POMT1;refGene_tx=XM_011288369.3;refGene_gn=POMT1;refGene_tx=XM_011288370.2;refGene_gn=POMT1;refGene_tx=XM_019816676.2;refGene_gn=POMT1;refGene_tx=XM_019816677.2

D4 91492352 91492530
refGene_gn=POMT1;refGene_tx=XM_011288369.3;refGene_gn=POMT1;refGene_tx
=XM_011288370.2;refGene_gn=POMT1;refGene_tx=XM_019816676.2;refGene_gn=
POMT1;refGene_tx=XM_019816677.2
D4 91492956 91493131
refGene_gn=POMT1;refGene_tx=XM_011288369.3;refGene_gn=POMT1;refGene_tx
=XM_011288370.2;refGene_gn=POMT1;refGene_tx=XM_019816676.2;refGene_gn=
POMT1;refGene_tx=XM_019816677.2
D4 91494930 91495109 refGene_gn=UCK1;refGene_tx=XM_023242873.1
D4 91495559 91495608 refGene_gn=UCK1;refGene_tx=XM_023242873.1
D4 91496350 91496479 refGene_gn=UCK1;refGene_tx=XM_023242874.1
D4 91496806 91497061 refGene_gn=UCK1;refGene_tx=XM_023242872.1
D4 91497370 91497465
refGene_gn=UCK1;refGene_tx=XM_023242872.1;refGene_gn=UCK1;refGene_tx=X
M_023242873.1;refGene_gn=UCK1;refGene_tx=XM_023242874.1
D4 91497542 91497685
refGene_gn=UCK1;refGene_tx=XM_023242872.1;refGene_gn=UCK1;refGene_tx=X
M_023242873.1;refGene_gn=UCK1;refGene_tx=XM_023242874.1
D4 91497903 91498000
refGene_gn=UCK1;refGene_tx=XM_023242872.1;refGene_gn=UCK1;refGene_tx=X
M_023242873.1;refGene_gn=UCK1;refGene_tx=XM_023242874.1
D4 91498986 91499146
refGene_gn=UCK1;refGene_tx=XM_023242872.1;refGene_gn=UCK1;refGene_tx=X
M_023242873.1;refGene_gn=UCK1;refGene_tx=XM_023242874.1
D4 91499587 91499695
refGene_gn=UCK1;refGene_tx=XM_023242872.1;refGene_gn=UCK1;refGene_tx=X
M_023242873.1;refGene_gn=UCK1;refGene_tx=XM_023242874.1
D4 91526264 91526337 refGene_gn=PRRT1B;refGene_tx=XM_019815566.2
D4 91526541 91527002 refGene_gn=PRRT1B;refGene_tx=XM_019815566.2
D4 91527751 91527895 refGene_gn=PRRT1B;refGene_tx=XM_019815566.2
D4 91528960 91529110 refGene_gn=PRRT1B;refGene_tx=XM_019815566.2
D4 91550300 91550451
refGene_gn=RAPGEF1;refGene_tx=XM_011288359.3;refGene_gn=RAPGEF1;refGen
e_tx=XM_011288361.3;refGene_gn=RAPGEF1;refGene_tx=XM_023242875.1;refGe
ne_gn=RAPGEF1;refGene_tx=XM_023242876.1;refGene_gn=RAPGEF1;refGene_tx=
XM_023242877.1;refGene_gn=RAPGEF1;refGene_tx=XM_023242878.1;refGene_gn
=RAPGEF1;refGene_tx=XM_023242879.1;refGene_gn=RAPGEF1;refGene_tx=XM_02
3242880.1;refGene_gn=RAPGEF1;refGene_tx=XM_023242881.1;refGene_gn=RAPG
EF1;refGene_tx=XM_023242882.1
D4 91550682 91550811
refGene_gn=RAPGEF1;refGene_tx=XM_011288359.3;refGene_gn=RAPGEF1;refGen
e_tx=XM_011288361.3;refGene_gn=RAPGEF1;refGene_tx=XM_023242875.1;refGe
ne_gn=RAPGEF1;refGene_tx=XM_023242876.1;refGene_gn=RAPGEF1;refGene_tx=
XM_023242877.1;refGene_gn=RAPGEF1;refGene_tx=XM_023242878.1;refGene_gn
=RAPGEF1;refGene_tx=XM_023242879.1;refGene_gn=RAPGEF1;refGene_tx=XM_02
3242880.1;refGene_gn=RAPGEF1;refGene_tx=XM_023242881.1;refGene_gn=RAPG
EF1;refGene_tx=XM_023242882.1
D4 91551842 91551940
refGene_gn=RAPGEF1;refGene_tx=XM_011288359.3;refGene_gn=RAPGEF1;refGen
e_tx=XM_011288361.3;refGene_gn=RAPGEF1;refGene_tx=XM_023242875.1;refGe

ne_gn=RAPGEF1;refGene_tx=XM_023242876.1;refGene_gn=RAPGEF1;refGene_tx=XM_023242877.1;refGene_gn=RAPGEF1;refGene_tx=XM_023242878.1;refGene_gn=RAPGEF1;refGene_tx=XM_023242879.1;refGene_gn=RAPGEF1;refGene_tx=XM_023242880.1;refGene_gn=RAPGEF1;refGene_tx=XM_023242881.1;refGene_gn=RAPGEF1;refGene_tx=XM_023242882.1
D4 91552926 91553028
refGene_gn=RAPGEF1;refGene_tx=XM_011288359.3;refGene_gn=RAPGEF1;refGene_tx=XM_011288361.3;refGene_gn=RAPGEF1;refGene_tx=XM_023242875.1;refGene_gn=RAPGEF1;refGene_tx=XM_023242876.1;refGene_gn=RAPGEF1;refGene_tx=XM_023242877.1;refGene_gn=RAPGEF1;refGene_tx=XM_023242878.1;refGene_gn=RAPGEF1;refGene_tx=XM_023242879.1;refGene_gn=RAPGEF1;refGene_tx=XM_023242880.1;refGene_gn=RAPGEF1;refGene_tx=XM_023242881.1;refGene_gn=RAPGEF1;refGene_tx=XM_023242882.1
D4 91553117 91553196
refGene_gn=RAPGEF1;refGene_tx=XM_011288359.3;refGene_gn=RAPGEF1;refGene_tx=XM_011288361.3;refGene_gn=RAPGEF1;refGene_tx=XM_023242875.1;refGene_gn=RAPGEF1;refGene_tx=XM_023242876.1;refGene_gn=RAPGEF1;refGene_tx=XM_023242877.1;refGene_gn=RAPGEF1;refGene_tx=XM_023242878.1;refGene_gn=RAPGEF1;refGene_tx=XM_023242879.1;refGene_gn=RAPGEF1;refGene_tx=XM_023242880.1;refGene_gn=RAPGEF1;refGene_tx=XM_023242881.1;refGene_gn=RAPGEF1;refGene_tx=XM_023242882.1
D4 91554384 91554479
refGene_gn=RAPGEF1;refGene_tx=XM_011288359.3;refGene_gn=RAPGEF1;refGene_tx=XM_011288361.3;refGene_gn=RAPGEF1;refGene_tx=XM_023242875.1;refGene_gn=RAPGEF1;refGene_tx=XM_023242876.1;refGene_gn=RAPGEF1;refGene_tx=XM_023242877.1;refGene_gn=RAPGEF1;refGene_tx=XM_023242878.1;refGene_gn=RAPGEF1;refGene_tx=XM_023242879.1;refGene_gn=RAPGEF1;refGene_tx=XM_023242880.1;refGene_gn=RAPGEF1;refGene_tx=XM_023242881.1;refGene_gn=RAPGEF1;refGene_tx=XM_023242882.1
D4 91554626 91554711
refGene_gn=RAPGEF1;refGene_tx=XM_011288359.3;refGene_gn=RAPGEF1;refGene_tx=XM_011288361.3;refGene_gn=RAPGEF1;refGene_tx=XM_023242875.1;refGene_gn=RAPGEF1;refGene_tx=XM_023242876.1;refGene_gn=RAPGEF1;refGene_tx=XM_023242877.1;refGene_gn=RAPGEF1;refGene_tx=XM_023242878.1;refGene_gn=RAPGEF1;refGene_tx=XM_023242879.1;refGene_gn=RAPGEF1;refGene_tx=XM_023242880.1;refGene_gn=RAPGEF1;refGene_tx=XM_023242881.1;refGene_gn=RAPGEF1;refGene_tx=XM_023242882.1
D4 91555452 91555638
refGene_gn=RAPGEF1;refGene_tx=XM_011288359.3;refGene_gn=RAPGEF1;refGene_tx=XM_011288361.3;refGene_gn=RAPGEF1;refGene_tx=XM_023242875.1;refGene_gn=RAPGEF1;refGene_tx=XM_023242876.1;refGene_gn=RAPGEF1;refGene_tx=XM_023242877.1;refGene_gn=RAPGEF1;refGene_tx=XM_023242878.1;refGene_gn=RAPGEF1;refGene_tx=XM_023242879.1;refGene_gn=RAPGEF1;refGene_tx=XM_023242880.1;refGene_gn=RAPGEF1;refGene_tx=XM_023242881.1;refGene_gn=RAPGEF1;refGene_tx=XM_023242882.1
D4 91556483 91556576
refGene_gn=RAPGEF1;refGene_tx=XM_011288359.3;refGene_gn=RAPGEF1;refGene_tx=XM_011288361.3;refGene_gn=RAPGEF1;refGene_tx=XM_023242875.1;refGene_gn=RAPGEF1;refGene_tx=XM_023242876.1;refGene_gn=RAPGEF1;refGene_tx=XM_023242877.1;refGene_gn=RAPGEF1;refGene_tx=XM_023242878.1;refGene_gn

=RAPGEF1;refGene_tx=XM_023242879.1;refGene_gn=RAPGEF1;refGene_tx=XM_023242880.1;refGene_gn=RAPGEF1;refGene_tx=XM_023242881.1;refGene_gn=RAPGEF1;refGene_tx=XM_023242882.1
D4 91558206 91558291
refGene_gn=RAPGEF1;refGene_tx=XM_011288359.3;refGene_gn=RAPGEF1;refGene_tx=XM_011288361.3;refGene_gn=RAPGEF1;refGene_tx=XM_023242875.1;refGene_gn=RAPGEF1;refGene_tx=XM_023242876.1;refGene_gn=RAPGEF1;refGene_tx=XM_023242877.1;refGene_gn=RAPGEF1;refGene_tx=XM_023242878.1;refGene_gn=RAPGEF1;refGene_tx=XM_023242879.1;refGene_gn=RAPGEF1;refGene_tx=XM_023242880.1;refGene_gn=RAPGEF1;refGene_tx=XM_023242881.1;refGene_gn=RAPGEF1;refGene_tx=XM_023242882.1
D4 91560828 91560904
refGene_gn=RAPGEF1;refGene_tx=XM_011288359.3;refGene_gn=RAPGEF1;refGene_tx=XM_011288361.3;refGene_gn=RAPGEF1;refGene_tx=XM_023242875.1;refGene_gn=RAPGEF1;refGene_tx=XM_023242876.1;refGene_gn=RAPGEF1;refGene_tx=XM_023242877.1;refGene_gn=RAPGEF1;refGene_tx=XM_023242878.1;refGene_gn=RAPGEF1;refGene_tx=XM_023242879.1;refGene_gn=RAPGEF1;refGene_tx=XM_023242880.1;refGene_gn=RAPGEF1;refGene_tx=XM_023242881.1;refGene_gn=RAPGEF1;refGene_tx=XM_023242882.1
D4 91562523 91562635
refGene_gn=RAPGEF1;refGene_tx=XM_011288359.3;refGene_gn=RAPGEF1;refGene_tx=XM_011288361.3;refGene_gn=RAPGEF1;refGene_tx=XM_023242875.1;refGene_gn=RAPGEF1;refGene_tx=XM_023242876.1;refGene_gn=RAPGEF1;refGene_tx=XM_023242877.1;refGene_gn=RAPGEF1;refGene_tx=XM_023242878.1;refGene_gn=RAPGEF1;refGene_tx=XM_023242879.1;refGene_gn=RAPGEF1;refGene_tx=XM_023242880.1;refGene_gn=RAPGEF1;refGene_tx=XM_023242881.1;refGene_gn=RAPGEF1;refGene_tx=XM_023242882.1
D4 91565140 91565226
refGene_gn=RAPGEF1;refGene_tx=XM_011288359.3;refGene_gn=RAPGEF1;refGene_tx=XM_011288361.3;refGene_gn=RAPGEF1;refGene_tx=XM_023242875.1;refGene_gn=RAPGEF1;refGene_tx=XM_023242876.1;refGene_gn=RAPGEF1;refGene_tx=XM_023242877.1;refGene_gn=RAPGEF1;refGene_tx=XM_023242878.1;refGene_gn=RAPGEF1;refGene_tx=XM_023242879.1;refGene_gn=RAPGEF1;refGene_tx=XM_023242880.1;refGene_gn=RAPGEF1;refGene_tx=XM_023242881.1;refGene_gn=RAPGEF1;refGene_tx=XM_023242882.1
D4 91566953 91567046
refGene_gn=RAPGEF1;refGene_tx=XM_011288359.3;refGene_gn=RAPGEF1;refGene_tx=XM_011288361.3;refGene_gn=RAPGEF1;refGene_tx=XM_023242875.1;refGene_gn=RAPGEF1;refGene_tx=XM_023242876.1;refGene_gn=RAPGEF1;refGene_tx=XM_023242878.1;refGene_gn=RAPGEF1;refGene_tx=XM_023242879.1
D4 91567873 91568146
refGene_gn=RAPGEF1;refGene_tx=XM_011288359.3;refGene_gn=RAPGEF1;refGene_tx=XM_011288361.3;refGene_gn=RAPGEF1;refGene_tx=XM_023242875.1;refGene_gn=RAPGEF1;refGene_tx=XM_023242876.1;refGene_gn=RAPGEF1;refGene_tx=XM_023242877.1;refGene_gn=RAPGEF1;refGene_tx=XM_023242878.1;refGene_gn=RAPGEF1;refGene_tx=XM_023242879.1
D4 91580345 91580504
refGene_gn=RAPGEF1;refGene_tx=XM_011288359.3;refGene_gn=RAPGEF1;refGene_tx=XM_011288361.3;refGene_gn=RAPGEF1;refGene_tx=XM_023242875.1;refGene_gn=RAPGEF1;refGene_tx=XM_023242876.1;refGene_gn=RAPGEF1;refGene_tx=

XM_023242877.1;refGene_gn=RAPGEF1;refGene_tx=XM_023242878.1;refGene_gn=RAPGEF1;refGene_tx=XM_023242880.1
D4	91582791 91582991
refGene_gn=RAPGEF1;refGene_tx=XM_011288359.3;refGene_gn=RAPGEF1;refGene_tx=XM_011288361.3;refGene_gn=RAPGEF1;refGene_tx=XM_023242875.1;refGene_gn=RAPGEF1;refGene_tx=XM_023242876.1;refGene_gn=RAPGEF1;refGene_tx=XM_023242877.1;refGene_gn=RAPGEF1;refGene_tx=XM_023242878.1;refGene_gn=RAPGEF1;refGene_tx=XM_023242879.1;refGene_gn=RAPGEF1;refGene_tx=XM_023242880.1;refGene_gn=RAPGEF1;refGene_tx=XM_023242881.1;refGene_gn=RAPGEF1;refGene_tx=XM_023242882.1
D4	91588373 91588880
refGene_gn=RAPGEF1;refGene_tx=XM_011288359.3;refGene_gn=RAPGEF1;refGene_tx=XM_011288361.3;refGene_gn=RAPGEF1;refGene_tx=XM_023242875.1;refGene_gn=RAPGEF1;refGene_tx=XM_023242876.1;refGene_gn=RAPGEF1;refGene_tx=XM_023242877.1;refGene_gn=RAPGEF1;refGene_tx=XM_023242878.1;refGene_gn=RAPGEF1;refGene_tx=XM_023242879.1;refGene_gn=RAPGEF1;refGene_tx=XM_023242880.1;refGene_gn=RAPGEF1;refGene_tx=XM_023242881.1;refGene_gn=RAPGEF1;refGene_tx=XM_023242882.1
D4	91590408 91590592
refGene_gn=RAPGEF1;refGene_tx=XM_011288359.3;refGene_gn=RAPGEF1;refGene_tx=XM_011288361.3;refGene_gn=RAPGEF1;refGene_tx=XM_023242875.1;refGene_gn=RAPGEF1;refGene_tx=XM_023242876.1;refGene_gn=RAPGEF1;refGene_tx=XM_023242877.1;refGene_gn=RAPGEF1;refGene_tx=XM_023242878.1;refGene_gn=RAPGEF1;refGene_tx=XM_023242879.1;refGene_gn=RAPGEF1;refGene_tx=XM_023242880.1;refGene_gn=RAPGEF1;refGene_tx=XM_023242881.1;refGene_gn=RAPGEF1;refGene_tx=XM_023242882.1
D4	91591033 91591157
refGene_gn=RAPGEF1;refGene_tx=XM_011288359.3;refGene_gn=RAPGEF1;refGene_tx=XM_011288361.3;refGene_gn=RAPGEF1;refGene_tx=XM_023242875.1;refGene_gn=RAPGEF1;refGene_tx=XM_023242876.1;refGene_gn=RAPGEF1;refGene_tx=XM_023242877.1;refGene_gn=RAPGEF1;refGene_tx=XM_023242878.1;refGene_gn=RAPGEF1;refGene_tx=XM_023242879.1;refGene_gn=RAPGEF1;refGene_tx=XM_023242880.1;refGene_gn=RAPGEF1;refGene_tx=XM_023242881.1;refGene_gn=RAPGEF1;refGene_tx=XM_023242882.1
D4	91591632 91591785
refGene_gn=RAPGEF1;refGene_tx=XM_011288359.3;refGene_gn=RAPGEF1;refGene_tx=XM_011288361.3;refGene_gn=RAPGEF1;refGene_tx=XM_023242875.1;refGene_gn=RAPGEF1;refGene_tx=XM_023242876.1;refGene_gn=RAPGEF1;refGene_tx=XM_023242877.1;refGene_gn=RAPGEF1;refGene_tx=XM_023242878.1;refGene_gn=RAPGEF1;refGene_tx=XM_023242879.1;refGene_gn=RAPGEF1;refGene_tx=XM_023242880.1;refGene_gn=RAPGEF1;refGene_tx=XM_023242881.1;refGene_gn=RAPGEF1;refGene_tx=XM_023242882.1
D4	91592461 91592550
refGene_gn=RAPGEF1;refGene_tx=XM_011288359.3;refGene_gn=RAPGEF1;refGene_tx=XM_011288361.3;refGene_gn=RAPGEF1;refGene_tx=XM_023242875.1;refGene_gn=RAPGEF1;refGene_tx=XM_023242876.1;refGene_gn=RAPGEF1;refGene_tx=XM_023242877.1;refGene_gn=RAPGEF1;refGene_tx=XM_023242878.1;refGene_gn=RAPGEF1;refGene_tx=XM_023242879.1;refGene_gn=RAPGEF1;refGene_tx=XM_023242880.1;refGene_gn=RAPGEF1;refGene_tx=XM_023242881.1;refGene_gn=RAPGEF1;refGene_tx=XM_023242882.1

D4 91604638 91604795
refGene_gn=RAPGEF1;refGene_tx=XM_011288359.3;refGene_gn=RAPGEF1;refGene_tx=XM_011288361.3;refGene_gn=RAPGEF1;refGene_tx=XM_023242875.1;refGene_gn=RAPGEF1;refGene_tx=XM_023242876.1;refGene_gn=RAPGEF1;refGene_tx=XM_023242877.1;refGene_gn=RAPGEF1;refGene_tx=XM_023242878.1;refGene_gn=RAPGEF1;refGene_tx=XM_023242879.1;refGene_gn=RAPGEF1;refGene_tx=XM_023242880.1;refGene_gn=RAPGEF1;refGene_tx=XM_023242881.1;refGene_gn=RAPGEF1;refGene_tx=XM_023242882.1
D4 91608682 91608861
refGene_gn=RAPGEF1;refGene_tx=XM_011288359.3;refGene_gn=RAPGEF1;refGene_tx=XM_011288361.3;refGene_gn=RAPGEF1;refGene_tx=XM_023242875.1;refGene_gn=RAPGEF1;refGene_tx=XM_023242876.1;refGene_gn=RAPGEF1;refGene_tx=XM_023242877.1;refGene_gn=RAPGEF1;refGene_tx=XM_023242878.1;refGene_gn=RAPGEF1;refGene_tx=XM_023242879.1;refGene_gn=RAPGEF1;refGene_tx=XM_023242880.1;refGene_gn=RAPGEF1;refGene_tx=XM_023242881.1;refGene_gn=RAPGEF1;refGene_tx=XM_023242882.1
D4 91613675 91613789
refGene_gn=RAPGEF1;refGene_tx=XM_011288359.3;refGene_gn=RAPGEF1;refGene_tx=XM_011288361.3;refGene_gn=RAPGEF1;refGene_tx=XM_023242875.1;refGene_gn=RAPGEF1;refGene_tx=XM_023242876.1;refGene_gn=RAPGEF1;refGene_tx=XM_023242877.1;refGene_gn=RAPGEF1;refGene_tx=XM_023242879.1;refGene_gn=RAPGEF1;refGene_tx=XM_023242880.1;refGene_gn=RAPGEF1;refGene_tx=XM_023242881.1;refGene_gn=RAPGEF1;refGene_tx=XM_023242882.1
D4 91614375 91614515
refGene_gn=RAPGEF1;refGene_tx=XM_011288359.3;refGene_gn=RAPGEF1;refGene_tx=XM_011288361.3;refGene_gn=RAPGEF1;refGene_tx=XM_023242875.1;refGene_gn=RAPGEF1;refGene_tx=XM_023242876.1;refGene_gn=RAPGEF1;refGene_tx=XM_023242877.1;refGene_gn=RAPGEF1;refGene_tx=XM_023242878.1;refGene_gn=RAPGEF1;refGene_tx=XM_023242879.1;refGene_gn=RAPGEF1;refGene_tx=XM_023242880.1;refGene_gn=RAPGEF1;refGene_tx=XM_023242881.1;refGene_gn=RAPGEF1;refGene_tx=XM_023242882.1
D4 91634705 91634745 refGene_gn=RAPGEF1;refGene_tx=XM_011288361.3
D4 91658681 91658745
refGene_gn=RAPGEF1;refGene_tx=XM_011288359.3;refGene_gn=RAPGEF1;refGene_tx=XM_023242876.1;refGene_gn=RAPGEF1;refGene_tx=XM_023242877.1;refGene_gn=RAPGEF1;refGene_tx=XM_023242878.1;refGene_gn=RAPGEF1;refGene_tx=XM_023242879.1;refGene_gn=RAPGEF1;refGene_tx=XM_023242880.1;refGene_gn=RAPGEF1;refGene_tx=XM_023242881.1
D4 91680289 91680350
refGene_gn=RAPGEF1;refGene_tx=XM_023242875.1;refGene_gn=RAPGEF1;refGene_tx=XM_023242882.1
D4 91739071 91739152
refGene_gn=MED27;refGene_tx=XM_019816686.2;refGene_gn=MED27;refGene_tx=XM_023242883.1
D4 91743405 91743462
refGene_gn=MED27;refGene_tx=XM_019816686.2;refGene_gn=MED27;refGene_tx=XM_023242883.1
D4 91779352 91779487
refGene_gn=MED27;refGene_tx=XM_003995979.4;refGene_gn=MED27;refGene_tx=XM_003995980.4

D4 91781273 91781359
refGene_gn=MED27;refGene_tx=XM_006939575.4;refGene_gn=MED27;refGene_tx=XM_019816690.2
D4 91781647 91781725
refGene_gn=MED27;refGene_tx=XM_003995979.4;refGene_gn=MED27;refGene_tx=XM_003995980.4;refGene_gn=MED27;refGene_tx=XM_006939575.4;refGene_gn=MED27;refGene_tx=XM_019816686.2;refGene_gn=MED27;refGene_tx=XM_019816690.2;refGene_gn=MED27;refGene_tx=XM_023242883.1
D4 91799420 91799462
refGene_gn=MED27;refGene_tx=XM_003995979.4;refGene_gn=MED27;refGene_tx=XM_003995980.4;refGene_gn=MED27;refGene_tx=XM_006939575.4;refGene_gn=MED27;refGene_tx=XM_019816686.2;refGene_gn=MED27;refGene_tx=XM_019816690.2;refGene_gn=MED27;refGene_tx=XM_023242883.1
D4 91809039 91809147
refGene_gn=MED27;refGene_tx=XM_003995979.4;refGene_gn=MED27;refGene_tx=XM_006939575.4;refGene_gn=MED27;refGene_tx=XM_019816686.2;refGene_gn=MED27;refGene_tx=XM_019816690.2
D4 91846088 91846182
refGene_gn=MED27;refGene_tx=XM_003995979.4;refGene_gn=MED27;refGene_tx=XM_003995980.4;refGene_gn=MED27;refGene_tx=XM_006939575.4;refGene_gn=MED27;refGene_tx=XM_019816686.2;refGene_gn=MED27;refGene_tx=XM_019816690.2;refGene_gn=MED27;refGene_tx=XM_023242883.1
D4 91916833 91916964
refGene_gn=MED27;refGene_tx=XM_003995979.4;refGene_gn=MED27;refGene_tx=XM_003995980.4;refGene_gn=MED27;refGene_tx=XM_006939575.4;refGene_gn=MED27;refGene_tx=XM_019816686.2;refGene_gn=MED27;refGene_tx=XM_019816690.2;refGene_gn=MED27;refGene_tx=XM_023242883.1
D4 91979517 91979662
refGene_gn=MED27;refGene_tx=XM_003995979.4;refGene_gn=MED27;refGene_tx=XM_003995980.4;refGene_gn=MED27;refGene_tx=XM_006939575.4;refGene_gn=MED27;refGene_tx=XM_019816686.2;refGene_gn=MED27;refGene_tx=XM_019816690.2;refGene_gn=MED27;refGene_tx=XM_023242883.1
D4 91981722 91981925
refGene_gn=MED27;refGene_tx=XM_003995979.4;refGene_gn=MED27;refGene_tx=XM_003995980.4;refGene_gn=MED27;refGene_tx=XM_006939575.4;refGene_gn=MED27;refGene_tx=XM_019816686.2;refGene_gn=MED27;refGene_tx=XM_019816690.2;refGene_gn=MED27;refGene_tx=XM_023242883.1
D4 92058826 92059039
refGene_gn=NTNG2;refGene_tx=XM_023242884.1;refGene_gn=NTNG2;refGene_tx=XM_023242885.1;refGene_gn=NTNG2;refGene_tx=XM_023242886.1;refGene_gn=NTNG2;refGene_tx=XM_023242887.1;refGene_gn=NTNG2;refGene_tx=XM_023242888.1;refGene_gn=NTNG2;refGene_tx=XM_023242889.1;refGene_gn=NTNG2;refGene_tx=XM_023242890.1;refGene_gn=NTNG2;refGene_tx=XM_023242891.1;refGene_gn=NTNG2;refGene_tx=XM_023242893.1;refGene_gn=NTNG2;refGene_tx=XM_023242894.1;refGene_gn=NTNG2;refGene_tx=XM_023242896.1;refGene_gn=NTNG2;refGene_tx=XM_023242897.1;refGene_gn=NTNG2;refGene_tx=XM_023242898.1;refGene_gn=NTNG2;refGene_tx=XM_023242899.1;refGene_gn=NTNG2;refGene_tx=XM_023242900.1;refGene_gn=NTNG2;refGene_tx=XM_023242901.1
D4 92071505 92071520 refGene_gn=NTNG2;refGene_tx=XM_023242892.1
D4 92082847 92083491

refGene_gn=NTNG2;refGene_tx=XM_023242884.1;refGene_gn=NTNG2;refGene_tx=XM_023242885.1;refGene_gn=NTNG2;refGene_tx=XM_023242886.1;refGene_gn=NTNG2;refGene_tx=XM_023242887.1;refGene_gn=NTNG2;refGene_tx=XM_023242888.1;refGene_gn=NTNG2;refGene_tx=XM_023242889.1;refGene_gn=NTNG2;refGene_tx=XM_023242890.1;refGene_gn=NTNG2;refGene_tx=XM_023242891.1;refGene_gn=NTNG2;refGene_tx=XM_023242892.1;refGene_gn=NTNG2;refGene_tx=XM_023242893.1;refGene_gn=NTNG2;refGene_tx=XM_023242894.1;refGene_gn=NTNG2;refGene_tx=XM_023242895.1;refGene_gn=NTNG2;refGene_tx=XM_023242896.1;refGene_gn=NTNG2;refGene_tx=XM_023242897.1;refGene_gn=NTNG2;refGene_tx=XM_023242898.1;refGene_gn=NTNG2;refGene_tx=XM_023242899.1;refGene_gn=NTNG2;refGene_tx=XM_023242900.1;refGene_gn=NTNG2;refGene_tx=XM_023242901.1
D4 92107398 92107571
refGene_gn=NTNG2;refGene_tx=XM_023242884.1;refGene_gn=NTNG2;refGene_tx=XM_023242885.1;refGene_gn=NTNG2;refGene_tx=XM_023242886.1;refGene_gn=NTNG2;refGene_tx=XM_023242887.1;refGene_gn=NTNG2;refGene_tx=XM_023242888.1;refGene_gn=NTNG2;refGene_tx=XM_023242889.1;refGene_gn=NTNG2;refGene_tx=XM_023242890.1;refGene_gn=NTNG2;refGene_tx=XM_023242891.1;refGene_gn=NTNG2;refGene_tx=XM_023242892.1;refGene_gn=NTNG2;refGene_tx=XM_023242893.1;refGene_gn=NTNG2;refGene_tx=XM_023242894.1;refGene_gn=NTNG2;refGene_tx=XM_023242895.1;refGene_gn=NTNG2;refGene_tx=XM_023242896.1;refGene_gn=NTNG2;refGene_tx=XM_023242897.1;refGene_gn=NTNG2;refGene_tx=XM_023242898.1;refGene_gn=NTNG2;refGene_tx=XM_023242899.1;refGene_gn=NTNG2;refGene_tx=XM_023242900.1;refGene_gn=NTNG2;refGene_tx=XM_023242901.1
D4 92110567 92110594
refGene_gn=NTNG2;refGene_tx=XM_023242884.1;refGene_gn=NTNG2;refGene_tx=XM_023242885.1;refGene_gn=NTNG2;refGene_tx=XM_023242886.1;refGene_gn=NTNG2;refGene_tx=XM_023242887.1;refGene_gn=NTNG2;refGene_tx=XM_023242888.1;refGene_gn=NTNG2;refGene_tx=XM_023242889.1;refGene_gn=NTNG2;refGene_tx=XM_023242890.1;refGene_gn=NTNG2;refGene_tx=XM_023242891.1;refGene_gn=NTNG2;refGene_tx=XM_023242892.1;refGene_gn=NTNG2;refGene_tx=XM_023242893.1;refGene_gn=NTNG2;refGene_tx=XM_023242894.1;refGene_gn=NTNG2;refGene_tx=XM_023242895.1;refGene_gn=NTNG2;refGene_tx=XM_023242896.1;refGene_gn=NTNG2;refGene_tx=XM_023242897.1;refGene_gn=NTNG2;refGene_tx=XM_023242898.1;refGene_gn=NTNG2;refGene_tx=XM_023242899.1;refGene_gn=NTNG2;refGene_tx=XM_023242900.1;refGene_gn=NTNG2;refGene_tx=XM_023242901.1
D4 92111448 92111586
refGene_gn=NTNG2;refGene_tx=XM_023242884.1;refGene_gn=NTNG2;refGene_tx=XM_023242885.1;refGene_gn=NTNG2;refGene_tx=XM_023242886.1;refGene_gn=NTNG2;refGene_tx=XM_023242887.1;refGene_gn=NTNG2;refGene_tx=XM_023242888.1;refGene_gn=NTNG2;refGene_tx=XM_023242889.1;refGene_gn=NTNG2;refGene_tx=XM_023242890.1;refGene_gn=NTNG2;refGene_tx=XM_023242892.1;refGene_gn=NTNG2;refGene_tx=XM_023242895.1;refGene_gn=NTNG2;refGene_tx=XM_023242898.1;refGene_gn=NTNG2;refGene_tx=XM_023242899.1;refGene_gn=NTNG2;refGene_tx=XM_023242900.1;refGene_gn=NTNG2;refGene_tx=XM_023242901.1
D4 92114442 92114512
refGene_gn=NTNG2;refGene_tx=XM_023242884.1;refGene_gn=NTNG2;refGene_tx=XM_023242885.1;refGene_gn=NTNG2;refGene_tx=XM_023242886.1;refGene_gn=

NTNG2;refGene_tx=XM_023242887.1;refGene_gn=NTNG2;refGene_tx=XM_0232428
92.1;refGene_gn=NTNG2;refGene_tx=XM_023242895.1;refGene_gn=NTNG2;refGe
ne_tx=XM_023242898.1;refGene_gn=NTNG2;refGene_tx=XM_023242899.1;refGen
e_gn=NTNG2;refGene_tx=XM_023242900.1
D4 92114822 92114897
refGene_gn=NTNG2;refGene_tx=XM_023242884.1;refGene_gn=NTNG2;refGene_tx
=XM_023242885.1;refGene_gn=NTNG2;refGene_tx=XM_023242886.1;refGene_gn=
NTNG2;refGene_tx=XM_023242887.1;refGene_gn=NTNG2;refGene_tx=XM_0232428
88.1;refGene_gn=NTNG2;refGene_tx=XM_023242889.1;refGene_gn=NTNG2;refGe
ne_tx=XM_023242891.1;refGene_gn=NTNG2;refGene_tx=XM_023242892.1;refGen
e_gn=NTNG2;refGene_tx=XM_023242893.1;refGene_gn=NTNG2;refGene_tx=XM_02
3242895.1;refGene_gn=NTNG2;refGene_tx=XM_023242898.1;refGene_gn=NTNG2;
refGene_tx=XM_023242899.1;refGene_gn=NTNG2;refGene_tx=XM_023242900.1;r
efGene_gn=NTNG2;refGene_tx=XM_023242901.1
D4 92116355 92116457
refGene_gn=NTNG2;refGene_tx=XM_023242884.1;refGene_gn=NTNG2;refGene_tx
=XM_023242885.1;refGene_gn=NTNG2;refGene_tx=XM_023242886.1;refGene_gn=
NTNG2;refGene_tx=XM_023242887.1;refGene_gn=NTNG2;refGene_tx=XM_0232428
88.1;refGene_gn=NTNG2;refGene_tx=XM_023242889.1;refGene_gn=NTNG2;refGe
ne_tx=XM_023242890.1;refGene_gn=NTNG2;refGene_tx=XM_023242891.1;refGen
e_gn=NTNG2;refGene_tx=XM_023242892.1;refGene_gn=NTNG2;refGene_tx=XM_02
3242893.1;refGene_gn=NTNG2;refGene_tx=XM_023242894.1;refGene_gn=NTNG2;
refGene_tx=XM_023242895.1;refGene_gn=NTNG2;refGene_tx=XM_023242896.1;r
efGene_gn=NTNG2;refGene_tx=XM_023242898.1;refGene_gn=NTNG2;refGene_tx=
XM_023242899.1;refGene_gn=NTNG2;refGene_tx=XM_023242900.1;refGene_gn=N
TNG2;refGene_tx=XM_023242901.1
D4 92117356 92117425
refGene_gn=NTNG2;refGene_tx=XM_023242888.1;refGene_gn=NTNG2;refGene_tx
=XM_023242890.1;refGene_gn=NTNG2;refGene_tx=XM_023242891.1;refGene_gn=
NTNG2;refGene_tx=XM_023242894.1
D4 92117864 92118102
refGene_gn=NTNG2;refGene_tx=XM_023242884.1;refGene_gn=NTNG2;refGene_tx
=XM_023242885.1;refGene_gn=NTNG2;refGene_tx=XM_023242886.1;refGene_gn=
NTNG2;refGene_tx=XM_023242887.1;refGene_gn=NTNG2;refGene_tx=XM_0232428
88.1;refGene_gn=NTNG2;refGene_tx=XM_023242889.1;refGene_gn=NTNG2;refGe
ne_tx=XM_023242890.1;refGene_gn=NTNG2;refGene_tx=XM_023242891.1;refGen
e_gn=NTNG2;refGene_tx=XM_023242892.1;refGene_gn=NTNG2;refGene_tx=XM_02
3242893.1;refGene_gn=NTNG2;refGene_tx=XM_023242894.1;refGene_gn=NTNG2;
refGene_tx=XM_023242895.1;refGene_gn=NTNG2;refGene_tx=XM_023242896.1;r
efGene_gn=NTNG2;refGene_tx=XM_023242897.1;refGene_gn=NTNG2;refGene_tx=
XM_023242898.1;refGene_gn=NTNG2;refGene_tx=XM_023242901.1
D4 92119889 92120024
refGene_gn=NTNG2;refGene_tx=XM_023242884.1;refGene_gn=NTNG2;refGene_tx
=XM_023242885.1;refGene_gn=NTNG2;refGene_tx=XM_023242886.1;refGene_gn=
NTNG2;refGene_tx=XM_023242887.1;refGene_gn=NTNG2;refGene_tx=XM_0232428
88.1;refGene_gn=NTNG2;refGene_tx=XM_023242889.1;refGene_gn=NTNG2;refGe
ne_tx=XM_023242890.1;refGene_gn=NTNG2;refGene_tx=XM_023242891.1;refGen
e_gn=NTNG2;refGene_tx=XM_023242892.1;refGene_gn=NTNG2;refGene_tx=XM_02
3242893.1;refGene_gn=NTNG2;refGene_tx=XM_023242894.1;refGene_gn=NTNG2;
refGene_tx=XM_023242895.1;refGene_gn=NTNG2;refGene_tx=XM_023242896.1;r

efGene_gn=NTNG2;refGene_tx=XM_023242897.1
D4 92120615 92120851
refGene_gn=NTNG2;refGene_tx=XM_023242884.1;refGene_gn=NTNG2;refGene_tx=XM_023242885.1;refGene_gn=NTNG2;refGene_tx=XM_023242886.1;refGene_gn=NTNG2;refGene_tx=XM_023242887.1;refGene_gn=NTNG2;refGene_tx=XM_023242888.1;refGene_gn=NTNG2;refGene_tx=XM_023242889.1;refGene_gn=NTNG2;refGene_tx=XM_023242890.1;refGene_gn=NTNG2;refGene_tx=XM_023242891.1;refGene_gn=NTNG2;refGene_tx=XM_023242892.1;refGene_gn=NTNG2;refGene_tx=XM_023242893.1;refGene_gn=NTNG2;refGene_tx=XM_023242894.1;refGene_gn=NTNG2;refGene_tx=XM_023242895.1;refGene_gn=NTNG2;refGene_tx=XM_023242896.1;refGene_gn=NTNG2;refGene_tx=XM_023242897.1
D4 92134343 92135090
refGene_gn=SETX;refGene_tx=XM_003996022.4;refGene_gn=SETX;refGene_tx=XM_006939577.3;refGene_gn=SETX;refGene_tx=XM_023242903.1;refGene_gn=SETX;refGene_tx=XM_023242904.1;refGene_gn=SETX;refGene_tx=XM_023242905.1;refGene_gn=SETX;refGene_tx=XM_023242906.1;refGene_gn=SETX;refGene_tx=XM_023242907.1;refGene_gn=SETX;refGene_tx=XM_023242908.1;refGene_gn=SETX;refGene_tx=XM_023242909.1;refGene_gn=SETX;refGene_tx=XM_023242910.1
D4 92138886 92138974
refGene_gn=SETX;refGene_tx=XM_003996022.4;refGene_gn=SETX;refGene_tx=XM_006939577.3;refGene_gn=SETX;refGene_tx=XM_023242903.1;refGene_gn=SETX;refGene_tx=XM_023242904.1;refGene_gn=SETX;refGene_tx=XM_023242905.1;refGene_gn=SETX;refGene_tx=XM_023242906.1;refGene_gn=SETX;refGene_tx=XM_023242907.1;refGene_gn=SETX;refGene_tx=XM_023242908.1;refGene_gn=SETX;refGene_tx=XM_023242909.1;refGene_gn=SETX;refGene_tx=XM_023242910.1
D4 92140260 92140359
refGene_gn=SETX;refGene_tx=XM_003996022.4;refGene_gn=SETX;refGene_tx=XM_006939577.3;refGene_gn=SETX;refGene_tx=XM_023242903.1;refGene_gn=SETX;refGene_tx=XM_023242904.1;refGene_gn=SETX;refGene_tx=XM_023242905.1;refGene_gn=SETX;refGene_tx=XM_023242906.1;refGene_gn=SETX;refGene_tx=XM_023242907.1;refGene_gn=SETX;refGene_tx=XM_023242908.1;refGene_gn=SETX;refGene_tx=XM_023242909.1;refGene_gn=SETX;refGene_tx=XM_023242910.1
D4 92143861 92144026
refGene_gn=SETX;refGene_tx=XM_003996022.4;refGene_gn=SETX;refGene_tx=XM_006939577.3;refGene_gn=SETX;refGene_tx=XM_023242903.1;refGene_gn=SETX;refGene_tx=XM_023242904.1;refGene_gn=SETX;refGene_tx=XM_023242905.1;refGene_gn=SETX;refGene_tx=XM_023242906.1;refGene_gn=SETX;refGene_tx=XM_023242907.1;refGene_gn=SETX;refGene_tx=XM_023242908.1;refGene_gn=SETX;refGene_tx=XM_023242909.1
D4 92145676 92145769
refGene_gn=SETX;refGene_tx=XM_003996022.4;refGene_gn=SETX;refGene_tx=XM_006939577.3;refGene_gn=SETX;refGene_tx=XM_023242903.1;refGene_gn=SETX;refGene_tx=XM_023242904.1;refGene_gn=SETX;refGene_tx=XM_023242905.1;refGene_gn=SETX;refGene_tx=XM_023242906.1;refGene_gn=SETX;refGene_tx=XM_023242907.1;refGene_gn=SETX;refGene_tx=XM_023242908.1;refGene_gn=SETX;refGene_tx=XM_023242909.1;refGene_gn=SETX;refGene_tx=XM_023242910.1
D4 92147611 92147799
refGene_gn=SETX;refGene_tx=XM_003996022.4;refGene_gn=SETX;refGene_tx=XM_006939577.3;refGene_gn=SETX;refGene_tx=XM_023242903.1;refGene_gn=SETX;refGene_tx=XM_023242904.1;refGene_gn=SETX;refGene_tx=XM_023242905.1;

refGene_gn=SETX;refGene_tx=XM_023242906.1;refGene_gn=SETX;refGene_tx=XM_023242907.1;refGene_gn=SETX;refGene_tx=XM_023242908.1;refGene_gn=SETX;refGene_tx=XM_023242909.1;refGene_gn=SETX;refGene_tx=XM_023242910.1
D4 92150600 92150708
refGene_gn=SETX;refGene_tx=XM_003996022.4;refGene_gn=SETX;refGene_tx=XM_006939577.3;refGene_gn=SETX;refGene_tx=XM_023242903.1;refGene_gn=SETX;refGene_tx=XM_023242904.1;refGene_gn=SETX;refGene_tx=XM_023242905.1;refGene_gn=SETX;refGene_tx=XM_023242906.1;refGene_gn=SETX;refGene_tx=XM_023242907.1;refGene_gn=SETX;refGene_tx=XM_023242908.1;refGene_gn=SETX;refGene_tx=XM_023242909.1;refGene_gn=SETX;refGene_tx=XM_023242910.1
D4 92151768 92151918
refGene_gn=SETX;refGene_tx=XM_003996022.4;refGene_gn=SETX;refGene_tx=XM_006939577.3;refGene_gn=SETX;refGene_tx=XM_023242903.1;refGene_gn=SETX;refGene_tx=XM_023242904.1;refGene_gn=SETX;refGene_tx=XM_023242905.1;refGene_gn=SETX;refGene_tx=XM_023242906.1;refGene_gn=SETX;refGene_tx=XM_023242907.1;refGene_gn=SETX;refGene_tx=XM_023242908.1;refGene_gn=SETX;refGene_tx=XM_023242909.1;refGene_gn=SETX;refGene_tx=XM_023242910.1
D4 92154088 92154160
refGene_gn=SETX;refGene_tx=XM_003996022.4;refGene_gn=SETX;refGene_tx=XM_006939577.3;refGene_gn=SETX;refGene_tx=XM_023242903.1;refGene_gn=SETX;refGene_tx=XM_023242904.1;refGene_gn=SETX;refGene_tx=XM_023242905.1;refGene_gn=SETX;refGene_tx=XM_023242906.1;refGene_gn=SETX;refGene_tx=XM_023242907.1;refGene_gn=SETX;refGene_tx=XM_023242908.1;refGene_gn=SETX;refGene_tx=XM_023242909.1;refGene_gn=SETX;refGene_tx=XM_023242910.1
D4 92155653 92155769
refGene_gn=SETX;refGene_tx=XM_003996022.4;refGene_gn=SETX;refGene_tx=XM_006939577.3;refGene_gn=SETX;refGene_tx=XM_023242903.1;refGene_gn=SETX;refGene_tx=XM_023242904.1;refGene_gn=SETX;refGene_tx=XM_023242905.1;refGene_gn=SETX;refGene_tx=XM_023242906.1;refGene_gn=SETX;refGene_tx=XM_023242907.1;refGene_gn=SETX;refGene_tx=XM_023242908.1;refGene_gn=SETX;refGene_tx=XM_023242909.1;refGene_gn=SETX;refGene_tx=XM_023242910.1
D4 92155966 92156068
refGene_gn=SETX;refGene_tx=XM_003996022.4;refGene_gn=SETX;refGene_tx=XM_006939577.3;refGene_gn=SETX;refGene_tx=XM_023242903.1;refGene_gn=SETX;refGene_tx=XM_023242904.1;refGene_gn=SETX;refGene_tx=XM_023242905.1;refGene_gn=SETX;refGene_tx=XM_023242906.1;refGene_gn=SETX;refGene_tx=XM_023242907.1;refGene_gn=SETX;refGene_tx=XM_023242908.1;refGene_gn=SETX;refGene_tx=XM_023242909.1;refGene_gn=SETX;refGene_tx=XM_023242910.1
D4 92160087 92160244
refGene_gn=SETX;refGene_tx=XM_003996022.4;refGene_gn=SETX;refGene_tx=XM_006939577.3;refGene_gn=SETX;refGene_tx=XM_023242903.1;refGene_gn=SETX;refGene_tx=XM_023242904.1;refGene_gn=SETX;refGene_tx=XM_023242905.1;refGene_gn=SETX;refGene_tx=XM_023242906.1;refGene_gn=SETX;refGene_tx=XM_023242907.1;refGene_gn=SETX;refGene_tx=XM_023242908.1;refGene_gn=SETX;refGene_tx=XM_023242909.1;refGene_gn=SETX;refGene_tx=XM_023242910.1
D4 92160726 92160894
refGene_gn=SETX;refGene_tx=XM_003996022.4;refGene_gn=SETX;refGene_tx=XM_006939577.3;refGene_gn=SETX;refGene_tx=XM_023242903.1;refGene_gn=SETX;refGene_tx=XM_023242904.1;refGene_gn=SETX;refGene_tx=XM_023242905.1;refGene_gn=SETX;refGene_tx=XM_023242906.1;refGene_gn=SETX;refGene_tx=X

M_023242907.1;refGene_gn=SETX;refGene_tx=XM_023242908.1;refGene_gn=SETX;refGene_tx=XM_023242909.1;refGene_gn=SETX;refGene_tx=XM_023242910.1
D4 92161924 92162157
refGene_gn=SETX;refGene_tx=XM_003996022.4;refGene_gn=SETX;refGene_tx=XM_006939577.3;refGene_gn=SETX;refGene_tx=XM_023242903.1;refGene_gn=SETX;refGene_tx=XM_023242904.1;refGene_gn=SETX;refGene_tx=XM_023242905.1;refGene_gn=SETX;refGene_tx=XM_023242906.1;refGene_gn=SETX;refGene_tx=XM_023242907.1;refGene_gn=SETX;refGene_tx=XM_023242908.1;refGene_gn=SETX;refGene_tx=XM_023242909.1;refGene_gn=SETX;refGene_tx=XM_023242910.1
D4 92164260 92164434
refGene_gn=SETX;refGene_tx=XM_003996022.4;refGene_gn=SETX;refGene_tx=XM_006939577.3;refGene_gn=SETX;refGene_tx=XM_023242903.1;refGene_gn=SETX;refGene_tx=XM_023242904.1;refGene_gn=SETX;refGene_tx=XM_023242905.1;refGene_gn=SETX;refGene_tx=XM_023242906.1;refGene_gn=SETX;refGene_tx=XM_023242907.1;refGene_gn=SETX;refGene_tx=XM_023242908.1;refGene_gn=SETX;refGene_tx=XM_023242909.1;refGene_gn=SETX;refGene_tx=XM_023242910.1
D4 92171311 92171411
refGene_gn=SETX;refGene_tx=XM_003996022.4;refGene_gn=SETX;refGene_tx=XM_006939577.3;refGene_gn=SETX;refGene_tx=XM_023242903.1;refGene_gn=SETX;refGene_tx=XM_023242904.1;refGene_gn=SETX;refGene_tx=XM_023242905.1;refGene_gn=SETX;refGene_tx=XM_023242906.1;refGene_gn=SETX;refGene_tx=XM_023242907.1;refGene_gn=SETX;refGene_tx=XM_023242908.1;refGene_gn=SETX;refGene_tx=XM_023242909.1;refGene_gn=SETX;refGene_tx=XM_023242910.1
D4 92173104 92177256
refGene_gn=SETX;refGene_tx=XM_003996022.4;refGene_gn=SETX;refGene_tx=XM_006939577.3;refGene_gn=SETX;refGene_tx=XM_023242903.1;refGene_gn=SETX;refGene_tx=XM_023242904.1;refGene_gn=SETX;refGene_tx=XM_023242905.1;refGene_gn=SETX;refGene_tx=XM_023242906.1;refGene_gn=SETX;refGene_tx=XM_023242907.1;refGene_gn=SETX;refGene_tx=XM_023242908.1;refGene_gn=SETX;refGene_tx=XM_023242909.1;refGene_gn=SETX;refGene_tx=XM_023242910.1
D4 92177739 92177827
refGene_gn=SETX;refGene_tx=XM_003996022.4;refGene_gn=SETX;refGene_tx=XM_006939577.3;refGene_gn=SETX;refGene_tx=XM_023242903.1;refGene_gn=SETX;refGene_tx=XM_023242904.1;refGene_gn=SETX;refGene_tx=XM_023242905.1;refGene_gn=SETX;refGene_tx=XM_023242906.1;refGene_gn=SETX;refGene_tx=XM_023242907.1;refGene_gn=SETX;refGene_tx=XM_023242908.1;refGene_gn=SETX;refGene_tx=XM_023242909.1;refGene_gn=SETX;refGene_tx=XM_023242910.1
D4 92178222 92178394
refGene_gn=SETX;refGene_tx=XM_003996022.4;refGene_gn=SETX;refGene_tx=XM_006939577.3;refGene_gn=SETX;refGene_tx=XM_023242903.1;refGene_gn=SETX;refGene_tx=XM_023242904.1;refGene_gn=SETX;refGene_tx=XM_023242905.1;refGene_gn=SETX;refGene_tx=XM_023242906.1;refGene_gn=SETX;refGene_tx=XM_023242907.1;refGene_gn=SETX;refGene_tx=XM_023242908.1;refGene_gn=SETX;refGene_tx=XM_023242909.1;refGene_gn=SETX;refGene_tx=XM_023242910.1
D4 92180495 92180615
refGene_gn=SETX;refGene_tx=XM_003996022.4;refGene_gn=SETX;refGene_tx=XM_006939577.3;refGene_gn=SETX;refGene_tx=XM_023242903.1;refGene_gn=SETX;refGene_tx=XM_023242904.1;refGene_gn=SETX;refGene_tx=XM_023242905.1;refGene_gn=SETX;refGene_tx=XM_023242906.1;refGene_gn=SETX;refGene_tx=XM_023242907.1;refGene_gn=SETX;refGene_tx=XM_023242908.1;refGene_gn=SET

X;refGene_tx=XM_023242909.1;refGene_gn=SETX;refGene_tx=XM_023242910.1
D4 92181354 92181574
refGene_gn=SETX;refGene_tx=XM_003996022.4;refGene_gn=SETX;refGene_tx=XM_006939577.3;refGene_gn=SETX;refGene_tx=XM_023242903.1;refGene_gn=SETX;refGene_tx=XM_023242904.1;refGene_gn=SETX;refGene_tx=XM_023242905.1;refGene_gn=SETX;refGene_tx=XM_023242906.1;refGene_gn=SETX;refGene_tx=XM_023242907.1;refGene_gn=SETX;refGene_tx=XM_023242908.1;refGene_gn=SETX;refGene_tx=XM_023242909.1;refGene_gn=SETX;refGene_tx=XM_023242910.1
D4 92186167 92186277
refGene_gn=SETX;refGene_tx=XM_003996022.4;refGene_gn=SETX;refGene_tx=XM_006939577.3;refGene_gn=SETX;refGene_tx=XM_023242903.1;refGene_gn=SETX;refGene_tx=XM_023242904.1;refGene_gn=SETX;refGene_tx=XM_023242905.1;refGene_gn=SETX;refGene_tx=XM_023242906.1;refGene_gn=SETX;refGene_tx=XM_023242907.1;refGene_gn=SETX;refGene_tx=XM_023242908.1;refGene_gn=SETX;refGene_tx=XM_023242909.1;refGene_gn=SETX;refGene_tx=XM_023242910.1
D4 92188479 92188690
refGene_gn=SETX;refGene_tx=XM_003996022.4;refGene_gn=SETX;refGene_tx=XM_006939577.3;refGene_gn=SETX;refGene_tx=XM_023242903.1;refGene_gn=SETX;refGene_tx=XM_023242904.1;refGene_gn=SETX;refGene_tx=XM_023242905.1;refGene_gn=SETX;refGene_tx=XM_023242906.1;refGene_gn=SETX;refGene_tx=XM_023242907.1;refGene_gn=SETX;refGene_tx=XM_023242908.1;refGene_gn=SETX;refGene_tx=XM_023242909.1;refGene_gn=SETX;refGene_tx=XM_023242910.1
D4 92189545 92189722
refGene_gn=SETX;refGene_tx=XM_003996022.4;refGene_gn=SETX;refGene_tx=XM_006939577.3;refGene_gn=SETX;refGene_tx=XM_023242903.1;refGene_gn=SETX;refGene_tx=XM_023242904.1;refGene_gn=SETX;refGene_tx=XM_023242905.1;refGene_gn=SETX;refGene_tx=XM_023242906.1;refGene_gn=SETX;refGene_tx=XM_023242907.1;refGene_gn=SETX;refGene_tx=XM_023242908.1;refGene_gn=SETX;refGene_tx=XM_023242909.1;refGene_gn=SETX;refGene_tx=XM_023242910.1
D4 92221341 92221523 refGene_gn=TTF1;refGene_tx=XM_023242911.1
D4 92222437 92222523 refGene_gn=TTF1;refGene_tx=XM_023242911.1
D4 92226266 92226332 refGene_gn=TTF1;refGene_tx=XM_023242911.1
D4 92226784 92226874 refGene_gn=TTF1;refGene_tx=XM_023242911.1
D4 92229035 92229270 refGene_gn=TTF1;refGene_tx=XM_023242911.1
D4 92231171 92231302 refGene_gn=TTF1;refGene_tx=XM_023242911.1
D4 92233344 92233423 refGene_gn=TTF1;refGene_tx=XM_023242911.1
D4 92234603 92234789 refGene_gn=TTF1;refGene_tx=XM_023242911.1
D4 92236615 92236842 refGene_gn=TTF1;refGene_tx=XM_023242911.1
D4 92237047 92238039 refGene_gn=TTF1;refGene_tx=XM_023242911.1
D4 92239922 92239934 refGene_gn=TTF1;refGene_tx=XM_023242911.1
D4 92243220 92243412 refGene_gn=CFAP77;refGene_tx=XM_023242912.1
D4 92315374 92315474 refGene_gn=CFAP77;refGene_tx=XM_023242912.1
D4 92315975 92316204 refGene_gn=CFAP77;refGene_tx=XM_023242912.1
D4 92344801 92344907 refGene_gn=CFAP77;refGene_tx=XM_023242912.1
D4 92348772 92348874 refGene_gn=CFAP77;refGene_tx=XM_023242912.1
D4 92373441 92373564 refGene_gn=CFAP77;refGene_tx=XM_023242912.1
D4 92384129 92384595 refGene_gn=BARHL1;refGene_tx=XM_023242914.1
D4 92388579 92388802 refGene_gn=BARHL1;refGene_tx=XM_023242914.1
D4 92390453 92390748 refGene_gn=BARHL1;refGene_tx=XM_023242914.1
D4 92396095 92396321

refGene_gn=DDX31;refGene_tx=XM_023242915.1;refGene_gn=DDX31;refGene_tx=XM_023242916.1;refGene_gn=DDX31;refGene_tx=XM_023242917.1
D4 92411974 92412143
refGene_gn=DDX31;refGene_tx=XM_023242915.1;refGene_gn=DDX31;refGene_tx=XM_023242916.1;refGene_gn=DDX31;refGene_tx=XM_023242917.1;refGene_gn=DDX31;refGene_tx=XM_023242918.1;refGene_gn=DDX31;refGene_tx=XM_023242919.1
D4 92419033 92419145
refGene_gn=DDX31;refGene_tx=XM_023242915.1;refGene_gn=DDX31;refGene_tx=XM_023242916.1;refGene_gn=DDX31;refGene_tx=XM_023242917.1;refGene_gn=DDX31;refGene_tx=XM_023242918.1;refGene_gn=DDX31;refGene_tx=XM_023242919.1
D4 92423779 92423911
refGene_gn=DDX31;refGene_tx=XM_023242915.1;refGene_gn=DDX31;refGene_tx=XM_023242916.1;refGene_gn=DDX31;refGene_tx=XM_023242920.1
D4 92424580 92424705
refGene_gn=DDX31;refGene_tx=XM_023242918.1;refGene_gn=DDX31;refGene_tx=XM_023242919.1;refGene_gn=DDX31;refGene_tx=XM_023242920.1
D4 92425251 92425302
refGene_gn=DDX31;refGene_tx=XM_023242918.1;refGene_gn=DDX31;refGene_tx=XM_023242920.1
D4 92425534 92425616
refGene_gn=DDX31;refGene_tx=XM_023242915.1;refGene_gn=DDX31;refGene_tx=XM_023242916.1;refGene_gn=DDX31;refGene_tx=XM_023242917.1;refGene_gn=DDX31;refGene_tx=XM_023242918.1;refGene_gn=DDX31;refGene_tx=XM_023242919.1;refGene_gn=DDX31;refGene_tx=XM_023242920.1
D4 92430936 92431076
refGene_gn=DDX31;refGene_tx=XM_023242915.1;refGene_gn=DDX31;refGene_tx=XM_023242916.1;refGene_gn=DDX31;refGene_tx=XM_023242917.1;refGene_gn=DDX31;refGene_tx=XM_023242918.1;refGene_gn=DDX31;refGene_tx=XM_023242919.1;refGene_gn=DDX31;refGene_tx=XM_023242920.1
D4 92432676 92432727
refGene_gn=DDX31;refGene_tx=XM_023242915.1;refGene_gn=DDX31;refGene_tx=XM_023242916.1;refGene_gn=DDX31;refGene_tx=XM_023242917.1;refGene_gn=DDX31;refGene_tx=XM_023242918.1;refGene_gn=DDX31;refGene_tx=XM_023242919.1;refGene_gn=DDX31;refGene_tx=XM_023242920.1
D4 92435980 92436223 refGene_gn=DDX31;refGene_tx=XM_023242921.1
D4 92441775 92441835
refGene_gn=DDX31;refGene_tx=XM_023242915.1;refGene_gn=DDX31;refGene_tx=XM_023242916.1;refGene_gn=DDX31;refGene_tx=XM_023242917.1;refGene_gn=DDX31;refGene_tx=XM_023242918.1;refGene_gn=DDX31;refGene_tx=XM_023242919.1;refGene_gn=DDX31;refGene_tx=XM_023242920.1;refGene_gn=DDX31;refGene_tx=XM_023242921.1
D4 92445708 92445885
refGene_gn=DDX31;refGene_tx=XM_023242915.1;refGene_gn=DDX31;refGene_tx=XM_023242916.1;refGene_gn=DDX31;refGene_tx=XM_023242917.1;refGene_gn=DDX31;refGene_tx=XM_023242918.1;refGene_gn=DDX31;refGene_tx=XM_023242919.1;refGene_gn=DDX31;refGene_tx=XM_023242920.1;refGene_gn=DDX31;refGene_tx=XM_023242921.1
D4 92446685 92446921

refGene_gn=DDX31;refGene_tx=XM_023242915.1;refGene_gn=DDX31;refGene_tx=XM_023242916.1;refGene_gn=DDX31;refGene_tx=XM_023242917.1;refGene_gn=DDX31;refGene_tx=XM_023242918.1;refGene_gn=DDX31;refGene_tx=XM_023242919.1;refGene_gn=DDX31;refGene_tx=XM_023242920.1;refGene_gn=DDX31;refGene_tx=XM_023242921.1
D4 92447950 92448057
refGene_gn=DDX31;refGene_tx=XM_023242915.1;refGene_gn=DDX31;refGene_tx=XM_023242916.1;refGene_gn=DDX31;refGene_tx=XM_023242917.1;refGene_gn=DDX31;refGene_tx=XM_023242918.1;refGene_gn=DDX31;refGene_tx=XM_023242919.1;refGene_gn=DDX31;refGene_tx=XM_023242920.1;refGene_gn=DDX31;refGene_tx=XM_023242921.1
D4 92448164 92448284
refGene_gn=DDX31;refGene_tx=XM_023242915.1;refGene_gn=DDX31;refGene_tx=XM_023242916.1;refGene_gn=DDX31;refGene_tx=XM_023242917.1;refGene_gn=DDX31;refGene_tx=XM_023242918.1;refGene_gn=DDX31;refGene_tx=XM_023242919.1;refGene_gn=DDX31;refGene_tx=XM_023242920.1;refGene_gn=DDX31;refGene_tx=XM_023242921.1
D4 92449857 92449922
refGene_gn=DDX31;refGene_tx=XM_023242915.1;refGene_gn=DDX31;refGene_tx=XM_023242916.1;refGene_gn=DDX31;refGene_tx=XM_023242917.1;refGene_gn=DDX31;refGene_tx=XM_023242918.1;refGene_gn=DDX31;refGene_tx=XM_023242919.1;refGene_gn=DDX31;refGene_tx=XM_023242920.1;refGene_gn=DDX31;refGene_tx=XM_023242921.1
D4 92450643 92450685
refGene_gn=DDX31;refGene_tx=XM_023242915.1;refGene_gn=DDX31;refGene_tx=XM_023242916.1;refGene_gn=DDX31;refGene_tx=XM_023242917.1;refGene_gn=DDX31;refGene_tx=XM_023242918.1;refGene_gn=DDX31;refGene_tx=XM_023242919.1;refGene_gn=DDX31;refGene_tx=XM_023242920.1;refGene_gn=DDX31;refGene_tx=XM_023242921.1
D4 92452038 92452083
refGene_gn=DDX31;refGene_tx=XM_023242915.1;refGene_gn=DDX31;refGene_tx=XM_023242916.1;refGene_gn=DDX31;refGene_tx=XM_023242917.1;refGene_gn=DDX31;refGene_tx=XM_023242918.1;refGene_gn=DDX31;refGene_tx=XM_023242919.1;refGene_gn=DDX31;refGene_tx=XM_023242920.1;refGene_gn=DDX31;refGene_tx=XM_023242921.1
D4 92458099 92458164
refGene_gn=DDX31;refGene_tx=XM_023242915.1;refGene_gn=DDX31;refGene_tx=XM_023242916.1;refGene_gn=DDX31;refGene_tx=XM_023242917.1;refGene_gn=DDX31;refGene_tx=XM_023242918.1;refGene_gn=DDX31;refGene_tx=XM_023242919.1;refGene_gn=DDX31;refGene_tx=XM_023242920.1;refGene_gn=DDX31;refGene_tx=XM_023242921.1
D4 92459144 92459215
refGene_gn=DDX31;refGene_tx=XM_023242915.1;refGene_gn=DDX31;refGene_tx=XM_023242916.1;refGene_gn=DDX31;refGene_tx=XM_023242917.1;refGene_gn=DDX31;refGene_tx=XM_023242918.1;refGene_gn=DDX31;refGene_tx=XM_023242919.1;refGene_gn=DDX31;refGene_tx=XM_023242920.1;refGene_gn=DDX31;refGene_tx=XM_023242921.1
D4 92460327 92460371
refGene_gn=DDX31;refGene_tx=XM_023242915.1;refGene_gn=DDX31;refGene_tx=XM_023242916.1;refGene_gn=DDX31;refGene_tx=XM_023242917.1;refGene_gn=

DDX31;refGene_tx=XM_023242918.1;refGene_gn=DDX31;refGene_tx=XM_0232429
19.1;refGene_gn=DDX31;refGene_tx=XM_023242920.1;refGene_gn=DDX31;refGe
ne_tx=XM_023242921.1
D4 92461310 92461386
refGene_gn=DDX31;refGene_tx=XM_023242915.1;refGene_gn=DDX31;refGene_tx
=XM_023242916.1;refGene_gn=DDX31;refGene_tx=XM_023242917.1;refGene_gn=
DDX31;refGene_tx=XM_023242918.1;refGene_gn=DDX31;refGene_tx=XM_0232429
19.1;refGene_gn=DDX31;refGene_tx=XM_023242920.1;refGene_gn=DDX31;refGe
ne_tx=XM_023242921.1
D4 92461492 92461749
refGene_gn=DDX31;refGene_tx=XM_023242915.1;refGene_gn=DDX31;refGene_tx
=XM_023242916.1;refGene_gn=DDX31;refGene_tx=XM_023242917.1;refGene_gn=
DDX31;refGene_tx=XM_023242918.1;refGene_gn=DDX31;refGene_tx=XM_0232429
19.1;refGene_gn=DDX31;refGene_tx=XM_023242920.1;refGene_gn=DDX31;refGe
ne_tx=XM_023242921.1
D4 92466929 92468039
refGene_gn=DDX31;refGene_tx=XM_023242915.1;refGene_gn=DDX31;refGene_tx
=XM_023242916.1;refGene_gn=DDX31;refGene_tx=XM_023242917.1;refGene_gn=
DDX31;refGene_tx=XM_023242918.1;refGene_gn=DDX31;refGene_tx=XM_0232429
19.1;refGene_gn=DDX31;refGene_tx=XM_023242920.1;refGene_gn=DDX31;refGe
ne_tx=XM_023242921.1
D4 92475128 92476748 refGene_gn=GTF3C4;refGene_tx=XM_023242319.1
D4 92477108 92477180 refGene_gn=GTF3C4;refGene_tx=XM_023242319.1
D4 92502389 92502627
refGene_gn=AK8;refGene_tx=XM_011288706.3;refGene_gn=AK8;refGene_tx=XM_
019816695.2;refGene_gn=AK8;refGene_tx=XM_019816697.1;refGene_gn=AK8;re
fGene_tx=XM_019816698.2;refGene_gn=AK8;refGene_tx=XM_019816702.1
D4 92503748 92503829
refGene_gn=AK8;refGene_tx=XM_011288706.3;refGene_gn=AK8;refGene_tx=XM_
019816695.2;refGene_gn=AK8;refGene_tx=XM_019816697.1;refGene_gn=AK8;re
fGene_tx=XM_019816698.2;refGene_gn=AK8;refGene_tx=XM_019816702.1
D4 92553780 92553922
refGene_gn=AK8;refGene_tx=XM_011288706.3;refGene_gn=AK8;refGene_tx=XM_
019816695.2;refGene_gn=AK8;refGene_tx=XM_019816697.1;refGene_gn=AK8;re
fGene_tx=XM_019816698.2;refGene_gn=AK8;refGene_tx=XM_019816702.1
D4 92574448 92574538
refGene_gn=AK8;refGene_tx=XM_011288706.3;refGene_gn=AK8;refGene_tx=XM_
019816695.2;refGene_gn=AK8;refGene_tx=XM_019816697.1;refGene_gn=AK8;re
fGene_tx=XM_019816698.2;refGene_gn=AK8;refGene_tx=XM_019816702.1
D4 92582092 92582224
refGene_gn=AK8;refGene_tx=XM_011288706.3;refGene_gn=AK8;refGene_tx=XM_
019816695.2;refGene_gn=AK8;refGene_tx=XM_019816697.1;refGene_gn=AK8;re
fGene_tx=XM_019816698.2;refGene_gn=AK8;refGene_tx=XM_019816702.1
D4 92585330 92585531
refGene_gn=AK8;refGene_tx=XM_011288706.3;refGene_gn=AK8;refGene_tx=XM_
019816695.2;refGene_gn=AK8;refGene_tx=XM_019816697.1;refGene_gn=AK8;re
fGene_tx=XM_019816698.2;refGene_gn=AK8;refGene_tx=XM_019816702.1
D4 92586130 92586202
refGene_gn=AK8;refGene_tx=XM_011288706.3;refGene_gn=AK8;refGene_tx=XM_
019816695.2;refGene_gn=AK8;refGene_tx=XM_019816697.1;refGene_gn=AK8;re

fGene_tx=XM_019816698.2;refGene_gn=AK8;refGene_tx=XM_019816702.1
D4 92586955 92587037
refGene_gn=AK8;refGene_tx=XM_011288706.3;refGene_gn=AK8;refGene_tx=XM_019816695.2;refGene_gn=AK8;refGene_tx=XM_019816697.1;refGene_gn=AK8;refGene_tx=XM_019816698.2;refGene_gn=AK8;refGene_tx=XM_019816702.1
D4 92609960 92610029
refGene_gn=AK8;refGene_tx=XM_011288706.3;refGene_gn=AK8;refGene_tx=XM_019816697.1;refGene_gn=AK8;refGene_tx=XM_019816698.2;refGene_gn=AK8;refGene_tx=XM_019816702.1
D4 92619332 92619446
refGene_gn=AK8;refGene_tx=XM_011288706.3;refGene_gn=AK8;refGene_tx=XM_019816695.2;refGene_gn=AK8;refGene_tx=XM_019816697.1;refGene_gn=AK8;refGene_tx=XM_019816698.2;refGene_gn=AK8;refGene_tx=XM_019816702.1
D4 92622113 92622163
refGene_gn=AK8;refGene_tx=XM_011288706.3;refGene_gn=AK8;refGene_tx=XM_019816695.2;refGene_gn=AK8;refGene_tx=XM_019816697.1;refGene_gn=AK8;refGene_tx=XM_019816698.2
D4 92627280 92627365
refGene_gn=AK8;refGene_tx=XM_011288706.3;refGene_gn=AK8;refGene_tx=XM_019816695.2;refGene_gn=AK8;refGene_tx=XM_019816697.1;refGene_gn=AK8;refGene_tx=XM_019816698.2;refGene_gn=AK8;refGene_tx=XM_019816702.1
D4 92630045 92630195
refGene_gn=AK8;refGene_tx=XM_011288706.3;refGene_gn=AK8;refGene_tx=XM_019816695.2
D4 92631843 92631885
refGene_gn=AK8;refGene_tx=XM_019816697.1;refGene_gn=AK8;refGene_tx=XM_019816698.2
D4 92635492 92635636
refGene_gn=SPACA9;refGene_tx=XM_006939580.4;refGene_gn=SPACA9;refGene_tx=XM_006939581.4;refGene_gn=SPACA9;refGene_tx=XM_006939582.4;refGene_gn=SPACA9;refGene_tx=XM_006939583.4;refGene_gn=SPACA9;refGene_tx=XM_006939584.4;refGene_gn=SPACA9;refGene_tx=XM_011288703.3;refGene_gn=SPACA9;refGene_tx=XM_011288704.3;refGene_gn=SPACA9;refGene_tx=XM_019816704.2;refGene_gn=SPACA9;refGene_tx=XM_019816705.2;refGene_gn=SPACA9;refGene_tx=XM_019816707.1
D4 92637271 92637474
refGene_gn=SPACA9;refGene_tx=XM_006939580.4;refGene_gn=SPACA9;refGene_tx=XM_006939581.4;refGene_gn=SPACA9;refGene_tx=XM_006939582.4;refGene_gn=SPACA9;refGene_tx=XM_006939583.4;refGene_gn=SPACA9;refGene_tx=XM_006939584.4;refGene_gn=SPACA9;refGene_tx=XM_011288703.3;refGene_gn=SPACA9;refGene_tx=XM_011288704.3;refGene_gn=SPACA9;refGene_tx=XM_019816704.2;refGene_gn=SPACA9;refGene_tx=XM_019816705.2;refGene_gn=SPACA9;refGene_tx=XM_019816706.2;refGene_gn=SPACA9;refGene_tx=XM_019816707.1
D4 92638197 92638531
refGene_gn=SPACA9;refGene_tx=XM_006939580.4;refGene_gn=SPACA9;refGene_tx=XM_006939581.4;refGene_gn=SPACA9;refGene_tx=XM_006939582.4;refGene_gn=SPACA9;refGene_tx=XM_006939583.4;refGene_gn=SPACA9;refGene_tx=XM_006939584.4;refGene_gn=SPACA9;refGene_tx=XM_011288703.3;refGene_gn=SPACA9;refGene_tx=XM_011288704.3;refGene_gn=SPACA9;refGene_tx=XM_019816704.2;refGene_gn=SPACA9;refGene_tx=XM_019816705.2;refGene_gn=SPACA9;refGen

e_tx=XM_019816706.2;refGene_gn=SPACA9;refGene_tx=XM_019816707.1
D4 92638946 92639012
refGene_gn=SPACA9;refGene_tx=XM_019816704.2;refGene_gn=SPACA9;refGene_tx=XM_019816705.2
D4 92639156 92639168 refGene_gn=SPACA9;refGene_tx=XM_019816707.1
D4 92644911 92645425
refGene_gn=TSC1;refGene_tx=XM_019816708.2;refGene_gn=TSC1;refGene_tx=XM_023242924.1;refGene_gn=TSC1;refGene_tx=XM_023242925.1;refGene_gn=TSC1;refGene_tx=XM_023242926.1;refGene_gn=TSC1;refGene_tx=XM_023242927.1;refGene_gn=TSC1;refGene_tx=XM_023242928.1;refGene_gn=TSC1;refGene_tx=XM_023242929.1;refGene_gn=TSC1;refGene_tx=XM_023242930.1;refGene_gn=TSC1;refGene_tx=XM_023242931.1;refGene_gn=TSC1;refGene_tx=XM_023242933.1;refGene_gn=TSC1;refGene_tx=XM_023242934.1
D4 92645796 92645958
refGene_gn=TSC1;refGene_tx=XM_019816708.2;refGene_gn=TSC1;refGene_tx=XM_023242924.1;refGene_gn=TSC1;refGene_tx=XM_023242925.1;refGene_gn=TSC1;refGene_tx=XM_023242926.1;refGene_gn=TSC1;refGene_tx=XM_023242927.1;refGene_gn=TSC1;refGene_tx=XM_023242928.1;refGene_gn=TSC1;refGene_tx=XM_023242929.1;refGene_gn=TSC1;refGene_tx=XM_023242930.1;refGene_gn=TSC1;refGene_tx=XM_023242931.1;refGene_gn=TSC1;refGene_tx=XM_023242933.1;refGene_gn=TSC1;refGene_tx=XM_023242934.1
D4 92646040 92646228
refGene_gn=TSC1;refGene_tx=XM_019816708.2;refGene_gn=TSC1;refGene_tx=XM_023242924.1;refGene_gn=TSC1;refGene_tx=XM_023242925.1;refGene_gn=TSC1;refGene_tx=XM_023242926.1;refGene_gn=TSC1;refGene_tx=XM_023242927.1;refGene_gn=TSC1;refGene_tx=XM_023242928.1;refGene_gn=TSC1;refGene_tx=XM_023242929.1;refGene_gn=TSC1;refGene_tx=XM_023242930.1;refGene_gn=TSC1;refGene_tx=XM_023242931.1;refGene_gn=TSC1;refGene_tx=XM_023242933.1;refGene_gn=TSC1;refGene_tx=XM_023242934.1
D4 92649016 92649139
refGene_gn=TSC1;refGene_tx=XM_019816708.2;refGene_gn=TSC1;refGene_tx=XM_023242924.1;refGene_gn=TSC1;refGene_tx=XM_023242925.1;refGene_gn=TSC1;refGene_tx=XM_023242926.1;refGene_gn=TSC1;refGene_tx=XM_023242927.1;refGene_gn=TSC1;refGene_tx=XM_023242928.1;refGene_gn=TSC1;refGene_tx=XM_023242929.1;refGene_gn=TSC1;refGene_tx=XM_023242930.1;refGene_gn=TSC1;refGene_tx=XM_023242931.1;refGene_gn=TSC1;refGene_tx=XM_023242933.1;refGene_gn=TSC1;refGene_tx=XM_023242934.1
D4 92650043 92650154
refGene_gn=TSC1;refGene_tx=XM_019816708.2;refGene_gn=TSC1;refGene_tx=XM_023242924.1;refGene_gn=TSC1;refGene_tx=XM_023242925.1;refGene_gn=TSC1;refGene_tx=XM_023242926.1;refGene_gn=TSC1;refGene_tx=XM_023242927.1;refGene_gn=TSC1;refGene_tx=XM_023242928.1;refGene_gn=TSC1;refGene_tx=XM_023242929.1;refGene_gn=TSC1;refGene_tx=XM_023242930.1;refGene_gn=TSC1;refGene_tx=XM_023242931.1;refGene_gn=TSC1;refGene_tx=XM_023242933.1;refGene_gn=TSC1;refGene_tx=XM_023242934.1
D4 92651016 92651199
refGene_gn=TSC1;refGene_tx=XM_019816708.2;refGene_gn=TSC1;refGene_tx=XM_023242924.1;refGene_gn=TSC1;refGene_tx=XM_023242925.1;refGene_gn=TSC1;refGene_tx=XM_023242926.1;refGene_gn=TSC1;refGene_tx=XM_023242927.1;refGene_gn=TSC1;refGene_tx=XM_023242928.1;refGene_gn=TSC1;refGene_tx=X

M_023242929.1;refGene_gn=TSC1;refGene_tx=XM_023242930.1;refGene_gn=TSC1;refGene_tx=XM_023242931.1;refGene_gn=TSC1;refGene_tx=XM_023242933.1;refGene_gn=TSC1;refGene_tx=XM_023242934.1
D4 92652080 92652247
refGene_gn=TSC1;refGene_tx=XM_019816708.2;refGene_gn=TSC1;refGene_tx=XM_023242924.1;refGene_gn=TSC1;refGene_tx=XM_023242925.1;refGene_gn=TSC1;refGene_tx=XM_023242926.1;refGene_gn=TSC1;refGene_tx=XM_023242927.1;refGene_gn=TSC1;refGene_tx=XM_023242928.1;refGene_gn=TSC1;refGene_tx=XM_023242929.1;refGene_gn=TSC1;refGene_tx=XM_023242930.1;refGene_gn=TSC1;refGene_tx=XM_023242931.1;refGene_gn=TSC1;refGene_tx=XM_023242933.1;refGene_gn=TSC1;refGene_tx=XM_023242934.1
D4 92652834 92652878
refGene_gn=TSC1;refGene_tx=XM_019816708.2;refGene_gn=TSC1;refGene_tx=XM_023242924.1;refGene_gn=TSC1;refGene_tx=XM_023242925.1;refGene_gn=TSC1;refGene_tx=XM_023242926.1;refGene_gn=TSC1;refGene_tx=XM_023242927.1;refGene_gn=TSC1;refGene_tx=XM_023242928.1;refGene_gn=TSC1;refGene_tx=XM_023242929.1;refGene_gn=TSC1;refGene_tx=XM_023242930.1;refGene_gn=TSC1;refGene_tx=XM_023242931.1;refGene_gn=TSC1;refGene_tx=XM_023242933.1;refGene_gn=TSC1;refGene_tx=XM_023242934.1
D4 92653904 92654463
refGene_gn=TSC1;refGene_tx=XM_019816708.2;refGene_gn=TSC1;refGene_tx=XM_023242924.1;refGene_gn=TSC1;refGene_tx=XM_023242925.1;refGene_gn=TSC1;refGene_tx=XM_023242926.1;refGene_gn=TSC1;refGene_tx=XM_023242927.1;refGene_gn=TSC1;refGene_tx=XM_023242928.1;refGene_gn=TSC1;refGene_tx=XM_023242929.1;refGene_gn=TSC1;refGene_tx=XM_023242930.1;refGene_gn=TSC1;refGene_tx=XM_023242931.1;refGene_gn=TSC1;refGene_tx=XM_023242933.1;refGene_gn=TSC1;refGene_tx=XM_023242934.1
D4 92654721 92654826
refGene_gn=TSC1;refGene_tx=XM_019816708.2;refGene_gn=TSC1;refGene_tx=XM_023242924.1;refGene_gn=TSC1;refGene_tx=XM_023242925.1;refGene_gn=TSC1;refGene_tx=XM_023242926.1;refGene_gn=TSC1;refGene_tx=XM_023242927.1;refGene_gn=TSC1;refGene_tx=XM_023242928.1;refGene_gn=TSC1;refGene_tx=XM_023242929.1;refGene_gn=TSC1;refGene_tx=XM_023242930.1;refGene_gn=TSC1;refGene_tx=XM_023242931.1;refGene_gn=TSC1;refGene_tx=XM_023242933.1;refGene_gn=TSC1;refGene_tx=XM_023242934.1
D4 92655478 92655548
refGene_gn=TSC1;refGene_tx=XM_019816708.2;refGene_gn=TSC1;refGene_tx=XM_023242924.1;refGene_gn=TSC1;refGene_tx=XM_023242925.1;refGene_gn=TSC1;refGene_tx=XM_023242926.1;refGene_gn=TSC1;refGene_tx=XM_023242927.1;refGene_gn=TSC1;refGene_tx=XM_023242928.1;refGene_gn=TSC1;refGene_tx=XM_023242929.1;refGene_gn=TSC1;refGene_tx=XM_023242930.1;refGene_gn=TSC1;refGene_tx=XM_023242931.1;refGene_gn=TSC1;refGene_tx=XM_023242933.1;refGene_gn=TSC1;refGene_tx=XM_023242934.1
D4 92657126 92657245
refGene_gn=TSC1;refGene_tx=XM_019816708.2;refGene_gn=TSC1;refGene_tx=XM_023242924.1;refGene_gn=TSC1;refGene_tx=XM_023242925.1;refGene_gn=TSC1;refGene_tx=XM_023242926.1;refGene_gn=TSC1;refGene_tx=XM_023242927.1;refGene_gn=TSC1;refGene_tx=XM_023242928.1;refGene_gn=TSC1;refGene_tx=XM_023242929.1;refGene_gn=TSC1;refGene_tx=XM_023242930.1;refGene_gn=TSC1;refGene_tx=XM_023242931.1;refGene_gn=TSC1;refGene_tx=XM_023242933.1;

refGene_gn=TSC1;refGene_tx=XM_023242934.1
D4 92657726 92657838
refGene_gn=TSC1;refGene_tx=XM_019816708.2;refGene_gn=TSC1;refGene_tx=XM_023242924.1;refGene_gn=TSC1;refGene_tx=XM_023242925.1;refGene_gn=TSC1;refGene_tx=XM_023242926.1;refGene_gn=TSC1;refGene_tx=XM_023242927.1;refGene_gn=TSC1;refGene_tx=XM_023242928.1;refGene_gn=TSC1;refGene_tx=XM_023242929.1;refGene_gn=TSC1;refGene_tx=XM_023242930.1;refGene_gn=TSC1;refGene_tx=XM_023242931.1;refGene_gn=TSC1;refGene_tx=XM_023242933.1;refGene_gn=TSC1;refGene_tx=XM_023242934.1
D4 92658154 92658270
refGene_gn=TSC1;refGene_tx=XM_019816708.2;refGene_gn=TSC1;refGene_tx=XM_023242924.1;refGene_gn=TSC1;refGene_tx=XM_023242925.1;refGene_gn=TSC1;refGene_tx=XM_023242926.1;refGene_gn=TSC1;refGene_tx=XM_023242927.1;refGene_gn=TSC1;refGene_tx=XM_023242928.1;refGene_gn=TSC1;refGene_tx=XM_023242929.1;refGene_gn=TSC1;refGene_tx=XM_023242930.1;refGene_gn=TSC1;refGene_tx=XM_023242931.1;refGene_gn=TSC1;refGene_tx=XM_023242933.1;refGene_gn=TSC1;refGene_tx=XM_023242934.1
D4 92658973 92659149
refGene_gn=TSC1;refGene_tx=XM_019816708.2;refGene_gn=TSC1;refGene_tx=XM_023242924.1;refGene_gn=TSC1;refGene_tx=XM_023242925.1;refGene_gn=TSC1;refGene_tx=XM_023242926.1;refGene_gn=TSC1;refGene_tx=XM_023242927.1;refGene_gn=TSC1;refGene_tx=XM_023242928.1;refGene_gn=TSC1;refGene_tx=XM_023242929.1;refGene_gn=TSC1;refGene_tx=XM_023242930.1;refGene_gn=TSC1;refGene_tx=XM_023242931.1;refGene_gn=TSC1;refGene_tx=XM_023242933.1;refGene_gn=TSC1;refGene_tx=XM_023242934.1
D4 92665330 92665404
refGene_gn=TSC1;refGene_tx=XM_019816708.2;refGene_gn=TSC1;refGene_tx=XM_023242924.1;refGene_gn=TSC1;refGene_tx=XM_023242925.1;refGene_gn=TSC1;refGene_tx=XM_023242926.1;refGene_gn=TSC1;refGene_tx=XM_023242927.1;refGene_gn=TSC1;refGene_tx=XM_023242928.1;refGene_gn=TSC1;refGene_tx=XM_023242929.1;refGene_gn=TSC1;refGene_tx=XM_023242930.1;refGene_gn=TSC1;refGene_tx=XM_023242931.1;refGene_gn=TSC1;refGene_tx=XM_023242933.1;refGene_gn=TSC1;refGene_tx=XM_023242934.1
D4 92665779 92665934
refGene_gn=TSC1;refGene_tx=XM_019816708.2;refGene_gn=TSC1;refGene_tx=XM_023242924.1;refGene_gn=TSC1;refGene_tx=XM_023242925.1;refGene_gn=TSC1;refGene_tx=XM_023242926.1;refGene_gn=TSC1;refGene_tx=XM_023242927.1;refGene_gn=TSC1;refGene_tx=XM_023242928.1;refGene_gn=TSC1;refGene_tx=XM_023242929.1;refGene_gn=TSC1;refGene_tx=XM_023242930.1;refGene_gn=TSC1;refGene_tx=XM_023242931.1;refGene_gn=TSC1;refGene_tx=XM_023242933.1;refGene_gn=TSC1;refGene_tx=XM_023242934.1
D4 92667413 92667558
refGene_gn=TSC1;refGene_tx=XM_019816708.2;refGene_gn=TSC1;refGene_tx=XM_023242924.1;refGene_gn=TSC1;refGene_tx=XM_023242925.1;refGene_gn=TSC1;refGene_tx=XM_023242926.1;refGene_gn=TSC1;refGene_tx=XM_023242927.1;refGene_gn=TSC1;refGene_tx=XM_023242928.1;refGene_gn=TSC1;refGene_tx=XM_023242929.1;refGene_gn=TSC1;refGene_tx=XM_023242930.1;refGene_gn=TSC1;refGene_tx=XM_023242931.1;refGene_gn=TSC1;refGene_tx=XM_023242933.1;refGene_gn=TSC1;refGene_tx=XM_023242934.1
D4 92669972 92670125

refGene_gn=TSC1;refGene_tx=XM_019816708.2;refGene_gn=TSC1;refGene_tx=XM_023242924.1;refGene_gn=TSC1;refGene_tx=XM_023242925.1;refGene_gn=TSC1;refGene_tx=XM_023242926.1;refGene_gn=TSC1;refGene_tx=XM_023242927.1;refGene_gn=TSC1;refGene_tx=XM_023242928.1;refGene_gn=TSC1;refGene_tx=XM_023242929.1;refGene_gn=TSC1;refGene_tx=XM_023242930.1;refGene_gn=TSC1;refGene_tx=XM_023242931.1;refGene_gn=TSC1;refGene_tx=XM_023242933.1;refGene_gn=TSC1;refGene_tx=XM_023242934.1
D4 92671113 92671217
refGene_gn=TSC1;refGene_tx=XM_019816708.2;refGene_gn=TSC1;refGene_tx=XM_023242924.1;refGene_gn=TSC1;refGene_tx=XM_023242925.1;refGene_gn=TSC1;refGene_tx=XM_023242926.1;refGene_gn=TSC1;refGene_tx=XM_023242927.1;refGene_gn=TSC1;refGene_tx=XM_023242928.1;refGene_gn=TSC1;refGene_tx=XM_023242929.1;refGene_gn=TSC1;refGene_tx=XM_023242930.1;refGene_gn=TSC1;refGene_tx=XM_023242931.1;refGene_gn=TSC1;refGene_tx=XM_023242933.1;refGene_gn=TSC1;refGene_tx=XM_023242934.1
D4 92673047 92673153
refGene_gn=TSC1;refGene_tx=XM_019816708.2;refGene_gn=TSC1;refGene_tx=XM_023242924.1;refGene_gn=TSC1;refGene_tx=XM_023242925.1;refGene_gn=TSC1;refGene_tx=XM_023242926.1;refGene_gn=TSC1;refGene_tx=XM_023242927.1;refGene_gn=TSC1;refGene_tx=XM_023242928.1;refGene_gn=TSC1;refGene_tx=XM_023242929.1;refGene_gn=TSC1;refGene_tx=XM_023242930.1;refGene_gn=TSC1;refGene_tx=XM_023242931.1;refGene_gn=TSC1;refGene_tx=XM_023242933.1;refGene_gn=TSC1;refGene_tx=XM_023242934.1
D4 92713979 92714079
refGene_gn=GFI1B;refGene_tx=XM_003995985.3;refGene_gn=GFI1B;refGene_tx=XM_019816711.1;refGene_gn=GFI1B;refGene_tx=XM_023242935.1
D4 92714536 92714674
refGene_gn=GFI1B;refGene_tx=XM_003995985.3;refGene_gn=GFI1B;refGene_tx=XM_019816711.1;refGene_gn=GFI1B;refGene_tx=XM_023242935.1
D4 92715115 92715378
refGene_gn=GFI1B;refGene_tx=XM_003995985.3;refGene_gn=GFI1B;refGene_tx=XM_019816711.1;refGene_gn=GFI1B;refGene_tx=XM_019816713.1;refGene_gn=GFI1B;refGene_tx=XM_023242935.1
D4 92715978 92716116
refGene_gn=GFI1B;refGene_tx=XM_003995985.3;refGene_gn=GFI1B;refGene_tx=XM_019816711.1;refGene_gn=GFI1B;refGene_tx=XM_019816713.1;refGene_gn=GFI1B;refGene_tx=XM_023242935.1
D4 92717120 92717385
refGene_gn=GFI1B;refGene_tx=XM_003995985.3;refGene_gn=GFI1B;refGene_tx=XM_019816711.1;refGene_gn=GFI1B;refGene_tx=XM_019816713.1;refGene_gn=GFI1B;refGene_tx=XM_023242935.1
D4 92718185 92718364
refGene_gn=GFI1B;refGene_tx=XM_003995985.3;refGene_gn=GFI1B;refGene_tx=XM_019816711.1;refGene_gn=GFI1B;refGene_tx=XM_019816713.1;refGene_gn=GFI1B;refGene_tx=XM_023242935.1
D4 92738425 92738572 refGene_gn=GTF3C5;refGene_tx=XM_003996028.5
D4 92740713 92740933 refGene_gn=GTF3C5;refGene_tx=XM_003996028.5
D4 92742205 92742404 refGene_gn=GTF3C5;refGene_tx=XM_003996028.5
D4 92745388 92745584 refGene_gn=GTF3C5;refGene_tx=XM_003996028.5
D4 92746736 92746841 refGene_gn=GTF3C5;refGene_tx=XM_003996028.5

D4 92747935 92748050 refGene_gn=GTF3C5;refGene_tx=XM_003996028.5
D4 92748542 92748623 refGene_gn=GTF3C5;refGene_tx=XM_003996028.5
D4 92748813 92748911 refGene_gn=GTF3C5;refGene_tx=XM_003996028.5
D4 92749799 92749882 refGene_gn=GTF3C5;refGene_tx=XM_003996028.5
D4 92750544 92750687 refGene_gn=GTF3C5;refGene_tx=XM_003996028.5
D4 92751536 92751691 refGene_gn=GTF3C5;refGene_tx=XM_003996028.5
D4 92754289 92754355 refGene_gn=CEL;refGene_tx=XM_011288708.3
D4 92756403 92756554 refGene_gn=CEL;refGene_tx=XM_011288708.3
D4 92756646 92756769 refGene_gn=CEL;refGene_tx=XM_011288708.3
D4 92756881 92757079 refGene_gn=CEL;refGene_tx=XM_011288708.3
D4 92757680 92757811 refGene_gn=CEL;refGene_tx=XM_011288708.3
D4 92757974 92758082 refGene_gn=CEL;refGene_tx=XM_011288708.3
D4 92758203 92758321 refGene_gn=CEL;refGene_tx=XM_011288708.3
D4 92759454 92759641 refGene_gn=CEL;refGene_tx=XM_011288708.3
D4 92759820 92760024 refGene_gn=CEL;refGene_tx=XM_011288708.3
D4 92760932 92761130 refGene_gn=CEL;refGene_tx=XM_011288708.3
D4 92761458 92762038 refGene_gn=CEL;refGene_tx=XM_011288708.3
D4 92768557 92768733
refGene_gn=RALGDS;refGene_tx=XM_006939589.3;refGene_gn=RALGDS;refGene_
tx=XM_006939590.2;refGene_gn=RALGDS;refGene_tx=XM_019816714.1;refGene_
gn=RALGDS;refGene_tx=XM_019816715.1;refGene_gn=RALGDS;refGene_tx=XM_01
9816716.1;refGene_gn=RALGDS;refGene_tx=XM_019816718.2
D4 92770737 92770852
refGene_gn=RALGDS;refGene_tx=XM_006939589.3;refGene_gn=RALGDS;refGene_
tx=XM_006939590.2;refGene_gn=RALGDS;refGene_tx=XM_019816714.1;refGene_
gn=RALGDS;refGene_tx=XM_019816715.1;refGene_gn=RALGDS;refGene_tx=XM_01
9816716.1;refGene_gn=RALGDS;refGene_tx=XM_019816718.2
D4 92772272 92772515
refGene_gn=RALGDS;refGene_tx=XM_006939589.3;refGene_gn=RALGDS;refGene_
tx=XM_006939590.2;refGene_gn=RALGDS;refGene_tx=XM_019816714.1;refGene_
gn=RALGDS;refGene_tx=XM_019816715.1;refGene_gn=RALGDS;refGene_tx=XM_01
9816716.1;refGene_gn=RALGDS;refGene_tx=XM_019816718.2
D4 92772680 92772882
refGene_gn=RALGDS;refGene_tx=XM_006939589.3;refGene_gn=RALGDS;refGene_
tx=XM_006939590.2;refGene_gn=RALGDS;refGene_tx=XM_019816714.1;refGene_
gn=RALGDS;refGene_tx=XM_019816715.1;refGene_gn=RALGDS;refGene_tx=XM_01
9816716.1;refGene_gn=RALGDS;refGene_tx=XM_019816718.2
D4 92773161 92773257
refGene_gn=RALGDS;refGene_tx=XM_006939589.3;refGene_gn=RALGDS;refGene_
tx=XM_006939590.2;refGene_gn=RALGDS;refGene_tx=XM_019816714.1;refGene_
gn=RALGDS;refGene_tx=XM_019816715.1;refGene_gn=RALGDS;refGene_tx=XM_01
9816716.1;refGene_gn=RALGDS;refGene_tx=XM_019816718.2
D4 92773444 92773566
refGene_gn=RALGDS;refGene_tx=XM_006939589.3;refGene_gn=RALGDS;refGene_
tx=XM_006939590.2;refGene_gn=RALGDS;refGene_tx=XM_019816714.1;refGene_
gn=RALGDS;refGene_tx=XM_019816715.1;refGene_gn=RALGDS;refGene_tx=XM_01
9816716.1;refGene_gn=RALGDS;refGene_tx=XM_019816718.2
D4 92773927 92773960
refGene_gn=RALGDS;refGene_tx=XM_006939589.3;refGene_gn=RALGDS;refGene_
tx=XM_006939590.2;refGene_gn=RALGDS;refGene_tx=XM_019816714.1;refGene_

gn=RALGDS;refGene_tx=XM_019816715.1;refGene_gn=RALGDS;refGene_tx=XM_01
9816716.1;refGene_gn=RALGDS;refGene_tx=XM_019816718.2
D4 92774533 92774620
refGene_gn=RALGDS;refGene_tx=XM_006939589.3;refGene_gn=RALGDS;refGene_
tx=XM_006939590.2;refGene_gn=RALGDS;refGene_tx=XM_019816714.1;refGene_
gn=RALGDS;refGene_tx=XM_019816715.1;refGene_gn=RALGDS;refGene_tx=XM_01
9816716.1;refGene_gn=RALGDS;refGene_tx=XM_019816718.2
D4 92774967 92775036
refGene_gn=RALGDS;refGene_tx=XM_006939589.3;refGene_gn=RALGDS;refGene_
tx=XM_006939590.2;refGene_gn=RALGDS;refGene_tx=XM_019816714.1;refGene_
gn=RALGDS;refGene_tx=XM_019816715.1;refGene_gn=RALGDS;refGene_tx=XM_01
9816716.1;refGene_gn=RALGDS;refGene_tx=XM_019816718.2
D4 92776012 92776097
refGene_gn=RALGDS;refGene_tx=XM_006939589.3;refGene_gn=RALGDS;refGene_
tx=XM_006939590.2;refGene_gn=RALGDS;refGene_tx=XM_019816714.1;refGene_
gn=RALGDS;refGene_tx=XM_019816715.1;refGene_gn=RALGDS;refGene_tx=XM_01
9816716.1;refGene_gn=RALGDS;refGene_tx=XM_019816718.2
D4 92776562 92776666
refGene_gn=RALGDS;refGene_tx=XM_006939589.3;refGene_gn=RALGDS;refGene_
tx=XM_006939590.2;refGene_gn=RALGDS;refGene_tx=XM_019816714.1;refGene_
gn=RALGDS;refGene_tx=XM_019816715.1;refGene_gn=RALGDS;refGene_tx=XM_01
9816716.1;refGene_gn=RALGDS;refGene_tx=XM_019816718.2
D4 92776992 92777208
refGene_gn=RALGDS;refGene_tx=XM_006939589.3;refGene_gn=RALGDS;refGene_
tx=XM_006939590.2;refGene_gn=RALGDS;refGene_tx=XM_019816714.1;refGene_
gn=RALGDS;refGene_tx=XM_019816715.1;refGene_gn=RALGDS;refGene_tx=XM_01
9816716.1;refGene_gn=RALGDS;refGene_tx=XM_019816718.2
D4 92777930 92778262
refGene_gn=RALGDS;refGene_tx=XM_006939589.3;refGene_gn=RALGDS;refGene_
tx=XM_006939590.2;refGene_gn=RALGDS;refGene_tx=XM_019816714.1;refGene_
gn=RALGDS;refGene_tx=XM_019816715.1;refGene_gn=RALGDS;refGene_tx=XM_01
9816716.1;refGene_gn=RALGDS;refGene_tx=XM_019816718.2
D4 92778491 92778685
refGene_gn=RALGDS;refGene_tx=XM_006939589.3;refGene_gn=RALGDS;refGene_
tx=XM_006939590.2;refGene_gn=RALGDS;refGene_tx=XM_019816714.1;refGene_
gn=RALGDS;refGene_tx=XM_019816715.1;refGene_gn=RALGDS;refGene_tx=XM_01
9816716.1;refGene_gn=RALGDS;refGene_tx=XM_019816718.2
D4 92779573 92779672
refGene_gn=RALGDS;refGene_tx=XM_006939589.3;refGene_gn=RALGDS;refGene_
tx=XM_006939590.2;refGene_gn=RALGDS;refGene_tx=XM_019816714.1;refGene_
gn=RALGDS;refGene_tx=XM_019816715.1;refGene_gn=RALGDS;refGene_tx=XM_01
9816716.1;refGene_gn=RALGDS;refGene_tx=XM_019816718.2
D4 92780122 92780316
refGene_gn=RALGDS;refGene_tx=XM_006939589.3;refGene_gn=RALGDS;refGene_
tx=XM_006939590.2;refGene_gn=RALGDS;refGene_tx=XM_019816714.1;refGene_
gn=RALGDS;refGene_tx=XM_019816715.1;refGene_gn=RALGDS;refGene_tx=XM_01
9816716.1;refGene_gn=RALGDS;refGene_tx=XM_019816718.2
D4 92781747 92781858
refGene_gn=RALGDS;refGene_tx=XM_006939589.3;refGene_gn=RALGDS;refGene_
tx=XM_006939590.2;refGene_gn=RALGDS;refGene_tx=XM_019816714.1;refGene_

gn=RALGDS;refGene_tx=XM_019816715.1;refGene_gn=RALGDS;refGene_tx=XM_01
9816716.1;refGene_gn=RALGDS;refGene_tx=XM_019816718.2
D4 92792058 92792241
refGene_gn=RALGDS;refGene_tx=XM_006939589.3;refGene_gn=RALGDS;refGene_
tx=XM_006939590.2;refGene_gn=RALGDS;refGene_tx=XM_019816714.1
D4 92798247 92798409
refGene_gn=RALGDS;refGene_tx=XM_019816715.1;refGene_gn=RALGDS;refGene_
tx=XM_019816716.1
D4 92799560 92799671 refGene_gn=RALGDS;refGene_tx=XM_019816718.2
D4 92819612 92820297
refGene_gn=GBGT1;refGene_tx=XM_023242939.1;refGene_gn=GBGT1;refGene_tx
=XM_023242940.1
D4 92821693 92821828
refGene_gn=GBGT1;refGene_tx=XM_023242939.1;refGene_gn=GBGT1;refGene_tx
=XM_023242940.1
D4 92822367 92822403
refGene_gn=GBGT1;refGene_tx=XM_023242939.1;refGene_gn=GBGT1;refGene_tx
=XM_023242940.1
D4 92822486 92822537
refGene_gn=GBGT1;refGene_tx=XM_023242939.1;refGene_gn=GBGT1;refGene_tx
=XM_023242940.1
D4 92827131 92827197 refGene_gn=GBGT1;refGene_tx=XM_023242939.1
D4 92827735 92827821 refGene_gn=GBGT1;refGene_tx=XM_023242939.1
D4 92837885 92838123
refGene_gn=LOC101081682;refGene_tx=XM_023242320.1
D4 92838997 92839093
refGene_gn=LOC101081682;refGene_tx=XM_023242320.1
D4 92839546 92839731
refGene_gn=LOC101081682;refGene_tx=XM_023242320.1
D4 92841273 92841465
refGene_gn=LOC101081682;refGene_tx=XM_023242320.1
D4 92841591 92841690
refGene_gn=LOC101081682;refGene_tx=XM_023242320.1
D4 92842246 92842832 refGene_gn=MRPS2;refGene_tx=XM_023242941.1
D4 92843419 92843549 refGene_gn=MRPS2;refGene_tx=XM_023242941.1
D4 92844194 92844320 refGene_gn=MRPS2;refGene_tx=XM_023242941.1
D4 92844564 92844604 refGene_gn=MRPS2;refGene_tx=XM_023242941.1
D4 92845295 92845453
refGene_gn=CD4H9orf116;refGene_tx=XM_003996030.5
D4 92846278 92846357
refGene_gn=CD4H9orf116;refGene_tx=XM_003996030.5
D4 92849198 92849387
refGene_gn=CD4H9orf116;refGene_tx=XM_003996030.5
D4 92854337 92857835
refGene_gn=PPP1R26;refGene_tx=XM_011288710.3;refGene_gn=PPP1R26;refGen
e_tx=XM_019816723.2;refGene_gn=PPP1R26;refGene_tx=XM_019816724.2;refGe
ne_gn=PPP1R26;refGene_tx=XM_019816725.2;refGene_gn=PPP1R26;refGene_tx=
XM_019816726.2;refGene_gn=PPP1R26;refGene_tx=XM_023242942.1;refGene_gn
=PPP1R26;refGene_tx=XM_023242944.1;refGene_gn=PPP1R26;refGene_tx=XM_02
3242945.1

D4 93032074 93033034
refGene_gn=LOC111557566;refGene_tx=XM_023242947.1;refGene_gn=LOC111557
566;refGene_tx=XM_023242948.1
D4 93048724 93049399
refGene_gn=OLFM1;refGene_tx=XM_003996031.5;refGene_gn=OLFM1;refGene_tx
=XM_006939613.3;refGene_gn=OLFM1;refGene_tx=XM_023242949.1
D4 93056300 93056407
refGene_gn=OLFM1;refGene_tx=XM_003996031.5;refGene_gn=OLFM1;refGene_tx
=XM_006939613.3;refGene_gn=OLFM1;refGene_tx=XM_023242949.1
D4 93063513 93063733
refGene_gn=OLFM1;refGene_tx=XM_003996031.5;refGene_gn=OLFM1;refGene_tx
=XM_006939613.3;refGene_gn=OLFM1;refGene_tx=XM_023242949.1
D4 93065690 93065846
refGene_gn=OLFM1;refGene_tx=XM_003996031.5;refGene_gn=OLFM1;refGene_tx
=XM_006939613.3;refGene_gn=OLFM1;refGene_tx=XM_023242949.1
D4 93070609 93070759
refGene_gn=OLFM1;refGene_tx=XM_003996031.5;refGene_gn=OLFM1;refGene_tx
=XM_023242949.1
D4 93072671 93072821 refGene_gn=OLFM1;refGene_tx=XM_023242949.1
D4 93084436 93084502 refGene_gn=OLFM1;refGene_tx=XM_003996031.5
D4 93185153 93185300
refGene_gn=COL5A1;refGene_tx=XM_023242950.1;refGene_gn=COL5A1;refGene_
tx=XM_023242951.1
D4 93189791 93190025
refGene_gn=COL5A1;refGene_tx=XM_023242950.1;refGene_gn=COL5A1;refGene_
tx=XM_023242951.1
D4 93193052 93193121 refGene_gn=COL5A1;refGene_tx=XM_023242951.1
D4 93193183 93193252 refGene_gn=COL5A1;refGene_tx=XM_023242950.1
D4 93195165 93195278
refGene_gn=COL5A1;refGene_tx=XM_023242950.1;refGene_gn=COL5A1;refGene_
tx=XM_023242951.1
D4 93196158 93196414
refGene_gn=COL5A1;refGene_tx=XM_023242950.1;refGene_gn=COL5A1;refGene_
tx=XM_023242951.1
D4 93197046 93197100
refGene_gn=COL5A1;refGene_tx=XM_023242950.1;refGene_gn=COL5A1;refGene_
tx=XM_023242951.1
D4 93197502 93197538
refGene_gn=COL5A1;refGene_tx=XM_023242950.1;refGene_gn=COL5A1;refGene_
tx=XM_023242951.1
D4 93198296 93198350
refGene_gn=COL5A1;refGene_tx=XM_023242950.1;refGene_gn=COL5A1;refGene_
tx=XM_023242951.1
D4 93199820 93199928
refGene_gn=COL5A1;refGene_tx=XM_023242950.1;refGene_gn=COL5A1;refGene_
tx=XM_023242951.1
D4 93201057 93201111
refGene_gn=COL5A1;refGene_tx=XM_023242950.1;refGene_gn=COL5A1;refGene_
tx=XM_023242951.1
D4 93201199 93201253

refGene_gn=COL5A1;refGene_tx=XM_023242950.1;refGene_gn=COL5A1;refGene_tx=XM_023242951.1
D4 93201354 93201462
refGene_gn=COL5A1;refGene_tx=XM_023242950.1;refGene_gn=COL5A1;refGene_tx=XM_023242951.1
D4 93202121 93202175
refGene_gn=COL5A1;refGene_tx=XM_023242950.1;refGene_gn=COL5A1;refGene_tx=XM_023242951.1
D4 93202802 93202856
refGene_gn=COL5A1;refGene_tx=XM_023242950.1;refGene_gn=COL5A1;refGene_tx=XM_023242951.1
D4 93203769 93203823
refGene_gn=COL5A1;refGene_tx=XM_023242950.1;refGene_gn=COL5A1;refGene_tx=XM_023242951.1
D4 93204081 93204135
refGene_gn=COL5A1;refGene_tx=XM_023242950.1;refGene_gn=COL5A1;refGene_tx=XM_023242951.1
D4 93204746 93204854
refGene_gn=COL5A1;refGene_tx=XM_023242950.1;refGene_gn=COL5A1;refGene_tx=XM_023242951.1
D4 93205615 93205669
refGene_gn=COL5A1;refGene_tx=XM_023242950.1;refGene_gn=COL5A1;refGene_tx=XM_023242951.1
D4 93206751 93206859
refGene_gn=COL5A1;refGene_tx=XM_023242950.1;refGene_gn=COL5A1;refGene_tx=XM_023242951.1
D4 93206963 93207017
refGene_gn=COL5A1;refGene_tx=XM_023242950.1;refGene_gn=COL5A1;refGene_tx=XM_023242951.1
D4 93208182 93208290
refGene_gn=COL5A1;refGene_tx=XM_023242950.1;refGene_gn=COL5A1;refGene_tx=XM_023242951.1
D4 93208380 93208434
refGene_gn=COL5A1;refGene_tx=XM_023242950.1;refGene_gn=COL5A1;refGene_tx=XM_023242951.1
D4 93209333 93209387
refGene_gn=COL5A1;refGene_tx=XM_023242950.1;refGene_gn=COL5A1;refGene_tx=XM_023242951.1
D4 93210289 93210397
refGene_gn=COL5A1;refGene_tx=XM_023242950.1;refGene_gn=COL5A1;refGene_tx=XM_023242951.1
D4 93212451 93212559
refGene_gn=COL5A1;refGene_tx=XM_023242950.1;refGene_gn=COL5A1;refGene_tx=XM_023242951.1
D4 93213848 93213902
refGene_gn=COL5A1;refGene_tx=XM_023242950.1;refGene_gn=COL5A1;refGene_tx=XM_023242951.1
D4 93213998 93214088
refGene_gn=COL5A1;refGene_tx=XM_023242950.1;refGene_gn=COL5A1;refGene_tx=XM_023242951.1

D4 93215444 93215552
refGene_gn=COL5A1;refGene_tx=XM_023242950.1;refGene_gn=COL5A1;refGene_tx=XM_023242951.1
D4 93216239 93216293
refGene_gn=COL5A1;refGene_tx=XM_023242950.1;refGene_gn=COL5A1;refGene_tx=XM_023242951.1
D4 93219223 93219277
refGene_gn=COL5A1;refGene_tx=XM_023242950.1;refGene_gn=COL5A1;refGene_tx=XM_023242951.1
D4 93220233 93220287
refGene_gn=COL5A1;refGene_tx=XM_023242950.1;refGene_gn=COL5A1;refGene_tx=XM_023242951.1
D4 93220727 93220772
refGene_gn=COL5A1;refGene_tx=XM_023242950.1;refGene_gn=COL5A1;refGene_tx=XM_023242951.1
D4 93221757 93221811
refGene_gn=COL5A1;refGene_tx=XM_023242950.1;refGene_gn=COL5A1;refGene_tx=XM_023242951.1
D4 93221936 93221981
refGene_gn=COL5A1;refGene_tx=XM_023242950.1;refGene_gn=COL5A1;refGene_tx=XM_023242951.1
D4 93226368 93226422
refGene_gn=COL5A1;refGene_tx=XM_023242950.1;refGene_gn=COL5A1;refGene_tx=XM_023242951.1
D4 93228815 93228869
refGene_gn=COL5A1;refGene_tx=XM_023242950.1;refGene_gn=COL5A1;refGene_tx=XM_023242951.1
D4 93229556 93229664
refGene_gn=COL5A1;refGene_tx=XM_023242950.1;refGene_gn=COL5A1;refGene_tx=XM_023242951.1
D4 93231850 93231904
refGene_gn=COL5A1;refGene_tx=XM_023242950.1;refGene_gn=COL5A1;refGene_tx=XM_023242951.1
D4 93233614 93233659
refGene_gn=COL5A1;refGene_tx=XM_023242950.1;refGene_gn=COL5A1;refGene_tx=XM_023242951.1
D4 93236677 93236731
refGene_gn=COL5A1;refGene_tx=XM_023242950.1;refGene_gn=COL5A1;refGene_tx=XM_023242951.1
D4 93237551 93237596
refGene_gn=COL5A1;refGene_tx=XM_023242950.1;refGene_gn=COL5A1;refGene_tx=XM_023242951.1
D4 93241868 93241922
refGene_gn=COL5A1;refGene_tx=XM_023242950.1;refGene_gn=COL5A1;refGene_tx=XM_023242951.1
D4 93242734 93242779
refGene_gn=COL5A1;refGene_tx=XM_023242950.1;refGene_gn=COL5A1;refGene_tx=XM_023242951.1
D4 93242984 93243038
refGene_gn=COL5A1;refGene_tx=XM_023242950.1;refGene_gn=COL5A1;refGene_

tx=XM_023242951.1
D4	93243581 93243626
refGene_gn=COL5A1;refGene_tx=XM_023242950.1;refGene_gn=COL5A1;refGene_tx=XM_023242951.1
D4	93244146 93244200
refGene_gn=COL5A1;refGene_tx=XM_023242950.1;refGene_gn=COL5A1;refGene_tx=XM_023242951.1
D4	93246581 93246626
refGene_gn=COL5A1;refGene_tx=XM_023242950.1;refGene_gn=COL5A1;refGene_tx=XM_023242951.1
D4	93248184 93248238
refGene_gn=COL5A1;refGene_tx=XM_023242950.1;refGene_gn=COL5A1;refGene_tx=XM_023242951.1
D4	93249413 93249467
refGene_gn=COL5A1;refGene_tx=XM_023242950.1;refGene_gn=COL5A1;refGene_tx=XM_023242951.1
D4	93250779 93250833
refGene_gn=COL5A1;refGene_tx=XM_023242950.1;refGene_gn=COL5A1;refGene_tx=XM_023242951.1
D4	93253295 93253349
refGene_gn=COL5A1;refGene_tx=XM_023242950.1;refGene_gn=COL5A1;refGene_tx=XM_023242951.1
D4	93253696 93253750
refGene_gn=COL5A1;refGene_tx=XM_023242950.1;refGene_gn=COL5A1;refGene_tx=XM_023242951.1
D4	93254668 93254725
refGene_gn=COL5A1;refGene_tx=XM_023242950.1;refGene_gn=COL5A1;refGene_tx=XM_023242951.1
D4	93255966 93256059
refGene_gn=COL5A1;refGene_tx=XM_023242950.1;refGene_gn=COL5A1;refGene_tx=XM_023242951.1
D4	93256803 93256878
refGene_gn=COL5A1;refGene_tx=XM_023242950.1;refGene_gn=COL5A1;refGene_tx=XM_023242951.1
D4	93264046 93264109
refGene_gn=COL5A1;refGene_tx=XM_023242950.1;refGene_gn=COL5A1;refGene_tx=XM_023242951.1
D4	93264331 93264373
refGene_gn=COL5A1;refGene_tx=XM_023242950.1;refGene_gn=COL5A1;refGene_tx=XM_023242951.1
D4	93268192 93268249
refGene_gn=COL5A1;refGene_tx=XM_023242950.1;refGene_gn=COL5A1;refGene_tx=XM_023242951.1
D4	93268397 93268556
refGene_gn=COL5A1;refGene_tx=XM_023242950.1;refGene_gn=COL5A1;refGene_tx=XM_023242951.1
D4	93269120 93269357
refGene_gn=COL5A1;refGene_tx=XM_023242950.1;refGene_gn=COL5A1;refGene_tx=XM_023242951.1
D4	93269945 93270083

refGene_gn=COL5A1;refGene_tx=XM_023242950.1;refGene_gn=COL5A1;refGene_tx=XM_023242951.1
D4 93270900 93271032
refGene_gn=COL5A1;refGene_tx=XM_023242950.1;refGene_gn=COL5A1;refGene_tx=XM_023242951.1
D4 93289851 93290014
refGene_gn=COL5A1;refGene_tx=XM_023242950.1;refGene_gn=COL5A1;refGene_tx=XM_023242951.1
D4 93291018 93291232
refGene_gn=COL5A1;refGene_tx=XM_023242950.1;refGene_gn=COL5A1;refGene_tx=XM_023242951.1
D4 93295796 93295964
refGene_gn=COL5A1;refGene_tx=XM_023242950.1;refGene_gn=COL5A1;refGene_tx=XM_023242951.1
D4 93321316 93322282
refGene_gn=LOC109494034;refGene_tx=XM_019816727.2;refGene_gn=LOC109494034;refGene_tx=XM_019816728.2
D4 93331589 93331707
refGene_gn=COL5A1;refGene_tx=XM_023242950.1;refGene_gn=COL5A1;refGene_tx=XM_023242951.1
D4 93469993 93470141
refGene_gn=RXRA;refGene_tx=XM_006939617.4;refGene_gn=RXRA;refGene_tx=XM_019816733.2;refGene_gn=RXRA;refGene_tx=XM_019816734.2;refGene_gn=RXRA;refGene_tx=XM_019816735.2;refGene_gn=RXRA;refGene_tx=XM_023242953.1
D4 93471715 93471821
refGene_gn=RXRA;refGene_tx=XM_006939617.4;refGene_gn=RXRA;refGene_tx=XM_019816733.2;refGene_gn=RXRA;refGene_tx=XM_019816734.2;refGene_gn=RXRA;refGene_tx=XM_019816735.2;refGene_gn=RXRA;refGene_tx=XM_023242953.1
D4 93473521 93473613
refGene_gn=RXRA;refGene_tx=XM_006939617.4;refGene_gn=RXRA;refGene_tx=XM_019816733.2;refGene_gn=RXRA;refGene_tx=XM_019816734.2;refGene_gn=RXRA;refGene_tx=XM_019816735.2;refGene_gn=RXRA;refGene_tx=XM_023242953.1
D4 93476335 93476468
refGene_gn=RXRA;refGene_tx=XM_006939617.4;refGene_gn=RXRA;refGene_tx=XM_019816733.2;refGene_gn=RXRA;refGene_tx=XM_019816734.2;refGene_gn=RXRA;refGene_tx=XM_019816735.2;refGene_gn=RXRA;refGene_tx=XM_023242953.1
D4 93478162 93478292
refGene_gn=RXRA;refGene_tx=XM_006939617.4;refGene_gn=RXRA;refGene_tx=XM_019816733.2;refGene_gn=RXRA;refGene_tx=XM_019816734.2;refGene_gn=RXRA;refGene_tx=XM_019816735.2;refGene_gn=RXRA;refGene_tx=XM_023242953.1
D4 93480915 93481085
refGene_gn=RXRA;refGene_tx=XM_006939617.4;refGene_gn=RXRA;refGene_tx=XM_019816733.2;refGene_gn=RXRA;refGene_tx=XM_019816734.2;refGene_gn=RXRA;refGene_tx=XM_019816735.2;refGene_gn=RXRA;refGene_tx=XM_023242953.1
D4 93488910 93489090
refGene_gn=RXRA;refGene_tx=XM_006939617.4;refGene_gn=RXRA;refGene_tx=XM_019816733.2;refGene_gn=RXRA;refGene_tx=XM_019816734.2;refGene_gn=RXRA;refGene_tx=XM_019816735.2;refGene_gn=RXRA;refGene_tx=XM_023242953.1
D4 93489413 93489564
refGene_gn=RXRA;refGene_tx=XM_006939617.4;refGene_gn=RXRA;refGene_tx=X

M_019816733.2;refGene_gn=RXRA;refGene_tx=XM_019816734.2;refGene_gn=RXRA;refGene_tx=XM_019816735.2;refGene_gn=RXRA;refGene_tx=XM_023242953.1
D4 93494862 93495128 refGene_gn=RXRA;refGene_tx=XM_006939617.4;refGene_gn=RXRA;refGene_tx=XM_019816733.2;refGene_gn=RXRA;refGene_tx=XM_019816734.2;refGene_gn=RXRA;refGene_tx=XM_019816735.2;refGene_gn=RXRA;refGene_tx=XM_023242953.1
D4 93552670 93552725 refGene_gn=RXRA;refGene_tx=XM_006939617.4
D4 93562459 93562487 refGene_gn=RXRA;refGene_tx=XM_023242953.1
D4 93610450 93610496 refGene_gn=LOC109493590;refGene_tx=XM_023242321.1
D4 93612129 93612627 refGene_gn=LOC109493590;refGene_tx=XM_023242321.1
D4 93612845 93613219 refGene_gn=LOC109493590;refGene_tx=XM_023242321.1
D4 93644561 93645473 refGene_gn=LOC109494041;refGene_tx=XM_023242955.1;refGene_gn=LOC109494041;refGene_tx=XM_023242956.1
D4 93680454 93680555 refGene_gn=WDR5;refGene_tx=XM_006939619.4;refGene_gn=WDR5;refGene_tx=XM_006939620.4
D4 93681590 93681678 refGene_gn=WDR5;refGene_tx=XM_006939619.4;refGene_gn=WDR5;refGene_tx=XM_006939620.4
D4 93682212 93682287 refGene_gn=WDR5;refGene_tx=XM_006939619.4;refGene_gn=WDR5;refGene_tx=XM_006939620.4
D4 93682606 93682640 refGene_gn=WDR5;refGene_tx=XM_006939619.4;refGene_gn=WDR5;refGene_tx=XM_006939620.4
D4 93683390 93683466 refGene_gn=WDR5;refGene_tx=XM_006939619.4;refGene_gn=WDR5;refGene_tx=XM_006939620.4
D4 93685806 93685853 refGene_gn=WDR5;refGene_tx=XM_006939619.4;refGene_gn=WDR5;refGene_tx=XM_006939620.4
D4 93689774 93689830 refGene_gn=WDR5;refGene_tx=XM_006939619.4;refGene_gn=WDR5;refGene_tx=XM_006939620.4
D4 93693184 93693268 refGene_gn=WDR5;refGene_tx=XM_006939619.4;refGene_gn=WDR5;refGene_tx=XM_006939620.4
D4 93693467 93693557 refGene_gn=WDR5;refGene_tx=XM_006939619.4;refGene_gn=WDR5;refGene_tx=XM_006939620.4
D4 93693792 93693882 refGene_gn=WDR5;refGene_tx=XM_006939619.4;refGene_gn=WDR5;refGene_tx=XM_006939620.4
D4 93694253 93694327 refGene_gn=WDR5;refGene_tx=XM_006939619.4;refGene_gn=WDR5;refGene_tx=XM_006939620.4

D4 93694653 93694762
refGene_gn=WDR5;refGene_tx=XM_006939619.4;refGene_gn=WDR5;refGene_tx=XM_006939620.4
D4 93695368 93695449
refGene_gn=WDR5;refGene_tx=XM_006939619.4;refGene_gn=WDR5;refGene_tx=XM_006939620.4
D4 93740472 93740586
refGene_gn=LOC109494043;refGene_tx=XM_019816738.2
D4 93740894 93741040
refGene_gn=LOC109494043;refGene_tx=XM_019816738.2
D4 93741719 93742125
refGene_gn=LOC109494043;refGene_tx=XM_019816738.2
D4 93743927 93744140
refGene_gn=BRD3;refGene_tx=XM_011288711.3;refGene_gn=BRD3;refGene_tx=XM_019816736.2;refGene_gn=BRD3;refGene_tx=XM_023242981.1;refGene_gn=BRD3;refGene_tx=XM_023242982.1
D4 93744878 93745016
refGene_gn=BRD3;refGene_tx=XM_011288711.3;refGene_gn=BRD3;refGene_tx=XM_019816736.2;refGene_gn=BRD3;refGene_tx=XM_023242981.1;refGene_gn=BRD3;refGene_tx=XM_023242982.1
D4 93745655 93745803
refGene_gn=BRD3;refGene_tx=XM_011288711.3;refGene_gn=BRD3;refGene_tx=XM_019816736.2;refGene_gn=BRD3;refGene_tx=XM_023242981.1;refGene_gn=BRD3;refGene_tx=XM_023242982.1
D4 93746616 93746831
refGene_gn=BRD3;refGene_tx=XM_011288711.3;refGene_gn=BRD3;refGene_tx=XM_019816736.2;refGene_gn=BRD3;refGene_tx=XM_023242981.1;refGene_gn=BRD3;refGene_tx=XM_023242982.1
D4 93748718 93749090
refGene_gn=BRD3;refGene_tx=XM_011288711.3;refGene_gn=BRD3;refGene_tx=XM_019816736.2;refGene_gn=BRD3;refGene_tx=XM_023242981.1;refGene_gn=BRD3;refGene_tx=XM_023242982.1
D4 93751626 93751755
refGene_gn=BRD3;refGene_tx=XM_011288711.3;refGene_gn=BRD3;refGene_tx=XM_019816736.2;refGene_gn=BRD3;refGene_tx=XM_023242981.1;refGene_gn=BRD3;refGene_tx=XM_023242982.1
D4 93753817 93754009
refGene_gn=BRD3;refGene_tx=XM_011288711.3;refGene_gn=BRD3;refGene_tx=XM_019816736.2;refGene_gn=BRD3;refGene_tx=XM_023242981.1;refGene_gn=BRD3;refGene_tx=XM_023242982.1
D4 93755148 93755390
refGene_gn=BRD3;refGene_tx=XM_011288711.3;refGene_gn=BRD3;refGene_tx=XM_019816736.2;refGene_gn=BRD3;refGene_tx=XM_023242981.1;refGene_gn=BRD3;refGene_tx=XM_023242982.1
D4 93758721 93759014
refGene_gn=BRD3;refGene_tx=XM_011288711.3;refGene_gn=BRD3;refGene_tx=XM_019816736.2;refGene_gn=BRD3;refGene_tx=XM_023242981.1;refGene_gn=BRD3;refGene_tx=XM_023242982.1
D4 93760137 93760263
refGene_gn=BRD3;refGene_tx=XM_011288711.3;refGene_gn=BRD3;refGene_tx=X

M_019816736.2;refGene_gn=BRD3;refGene_tx=XM_023242981.1;refGene_gn=BRD3;refGene_tx=XM_023242982.1
D4 93761963 93762079
refGene_gn=BRD3;refGene_tx=XM_011288711.3;refGene_gn=BRD3;refGene_tx=XM_019816736.2;refGene_gn=BRD3;refGene_tx=XM_023242981.1;refGene_gn=BRD3;refGene_tx=XM_023242982.1
D4 93791107 93791311
refGene_gn=VAV2;refGene_tx=XM_023242972.1;refGene_gn=VAV2;refGene_tx=XM_023242974.1;refGene_gn=VAV2;refGene_tx=XM_023242975.1;refGene_gn=VAV2;refGene_tx=XM_023242976.1;refGene_gn=VAV2;refGene_tx=XM_023242977.1;refGene_gn=VAV2;refGene_tx=XM_023242978.1;refGene_gn=VAV2;refGene_tx=XM_023242979.1;refGene_gn=VAV2;refGene_tx=XM_023242980.1
D4 93823686 93823803
refGene_gn=VAV2;refGene_tx=XM_023242972.1;refGene_gn=VAV2;refGene_tx=XM_023242974.1;refGene_gn=VAV2;refGene_tx=XM_023242975.1;refGene_gn=VAV2;refGene_tx=XM_023242976.1;refGene_gn=VAV2;refGene_tx=XM_023242977.1;refGene_gn=VAV2;refGene_tx=XM_023242978.1;refGene_gn=VAV2;refGene_tx=XM_023242979.1;refGene_gn=VAV2;refGene_tx=XM_023242980.1
D4 93883833 93883892
refGene_gn=VAV2;refGene_tx=XM_023242972.1;refGene_gn=VAV2;refGene_tx=XM_023242974.1;refGene_gn=VAV2;refGene_tx=XM_023242975.1;refGene_gn=VAV2;refGene_tx=XM_023242976.1;refGene_gn=VAV2;refGene_tx=XM_023242977.1;refGene_gn=VAV2;refGene_tx=XM_023242978.1;refGene_gn=VAV2;refGene_tx=XM_023242979.1;refGene_gn=VAV2;refGene_tx=XM_023242980.1
D4 93902490 93902559
refGene_gn=VAV2;refGene_tx=XM_023242972.1;refGene_gn=VAV2;refGene_tx=XM_023242974.1;refGene_gn=VAV2;refGene_tx=XM_023242975.1;refGene_gn=VAV2;refGene_tx=XM_023242976.1;refGene_gn=VAV2;refGene_tx=XM_023242977.1;refGene_gn=VAV2;refGene_tx=XM_023242978.1;refGene_gn=VAV2;refGene_tx=XM_023242979.1;refGene_gn=VAV2;refGene_tx=XM_023242980.1
D4 93918066 93918169
refGene_gn=VAV2;refGene_tx=XM_023242972.1;refGene_gn=VAV2;refGene_tx=XM_023242974.1;refGene_gn=VAV2;refGene_tx=XM_023242975.1;refGene_gn=VAV2;refGene_tx=XM_023242976.1;refGene_gn=VAV2;refGene_tx=XM_023242977.1;refGene_gn=VAV2;refGene_tx=XM_023242978.1;refGene_gn=VAV2;refGene_tx=XM_023242979.1;refGene_gn=VAV2;refGene_tx=XM_023242980.1
D4 93919919 93919934
refGene_gn=VAV2;refGene_tx=XM_023242972.1;refGene_gn=VAV2;refGene_tx=XM_023242974.1;refGene_gn=VAV2;refGene_tx=XM_023242976.1;refGene_gn=VAV2;refGene_tx=XM_023242979.1
D4 93920970 93921069
refGene_gn=VAV2;refGene_tx=XM_023242972.1;refGene_gn=VAV2;refGene_tx=XM_023242974.1;refGene_gn=VAV2;refGene_tx=XM_023242975.1;refGene_gn=VAV2;refGene_tx=XM_023242976.1;refGene_gn=VAV2;refGene_tx=XM_023242977.1;refGene_gn=VAV2;refGene_tx=XM_023242978.1;refGene_gn=VAV2;refGene_tx=XM_023242979.1;refGene_gn=VAV2;refGene_tx=XM_023242980.1
D4 93923031 93923100
refGene_gn=VAV2;refGene_tx=XM_023242972.1;refGene_gn=VAV2;refGene_tx=XM_023242974.1;refGene_gn=VAV2;refGene_tx=XM_023242975.1;refGene_gn=VAV2;refGene_tx=XM_023242976.1;refGene_gn=VAV2;refGene_tx=XM_023242977.1;

refGene_gn=VAV2;refGene_tx=XM_023242978.1;refGene_gn=VAV2;refGene_tx=XM_023242979.1;refGene_gn=VAV2;refGene_tx=XM_023242980.1
D4 93924696 93924797
refGene_gn=VAV2;refGene_tx=XM_023242972.1;refGene_gn=VAV2;refGene_tx=XM_023242974.1;refGene_gn=VAV2;refGene_tx=XM_023242975.1;refGene_gn=VAV2;refGene_tx=XM_023242976.1;refGene_gn=VAV2;refGene_tx=XM_023242977.1;refGene_gn=VAV2;refGene_tx=XM_023242978.1;refGene_gn=VAV2;refGene_tx=XM_023242979.1;refGene_gn=VAV2;refGene_tx=XM_023242980.1
D4 93932199 93932299
refGene_gn=VAV2;refGene_tx=XM_023242972.1;refGene_gn=VAV2;refGene_tx=XM_023242974.1;refGene_gn=VAV2;refGene_tx=XM_023242975.1;refGene_gn=VAV2;refGene_tx=XM_023242976.1;refGene_gn=VAV2;refGene_tx=XM_023242977.1;refGene_gn=VAV2;refGene_tx=XM_023242978.1;refGene_gn=VAV2;refGene_tx=XM_023242979.1;refGene_gn=VAV2;refGene_tx=XM_023242980.1
D4 93933268 93933364
refGene_gn=VAV2;refGene_tx=XM_023242972.1;refGene_gn=VAV2;refGene_tx=XM_023242974.1;refGene_gn=VAV2;refGene_tx=XM_023242975.1;refGene_gn=VAV2;refGene_tx=XM_023242976.1;refGene_gn=VAV2;refGene_tx=XM_023242977.1;refGene_gn=VAV2;refGene_tx=XM_023242978.1;refGene_gn=VAV2;refGene_tx=XM_023242979.1;refGene_gn=VAV2;refGene_tx=XM_023242980.1
D4 93934026 93934095
refGene_gn=VAV2;refGene_tx=XM_023242972.1;refGene_gn=VAV2;refGene_tx=XM_023242974.1;refGene_gn=VAV2;refGene_tx=XM_023242975.1;refGene_gn=VAV2;refGene_tx=XM_023242976.1;refGene_gn=VAV2;refGene_tx=XM_023242977.1;refGene_gn=VAV2;refGene_tx=XM_023242978.1;refGene_gn=VAV2;refGene_tx=XM_023242979.1;refGene_gn=VAV2;refGene_tx=XM_023242980.1
D4 93935770 93935857
refGene_gn=VAV2;refGene_tx=XM_023242972.1;refGene_gn=VAV2;refGene_tx=XM_023242974.1;refGene_gn=VAV2;refGene_tx=XM_023242975.1;refGene_gn=VAV2;refGene_tx=XM_023242976.1;refGene_gn=VAV2;refGene_tx=XM_023242977.1;refGene_gn=VAV2;refGene_tx=XM_023242978.1;refGene_gn=VAV2;refGene_tx=XM_023242979.1;refGene_gn=VAV2;refGene_tx=XM_023242980.1
D4 93937659 93937745
refGene_gn=VAV2;refGene_tx=XM_023242972.1;refGene_gn=VAV2;refGene_tx=XM_023242974.1;refGene_gn=VAV2;refGene_tx=XM_023242975.1;refGene_gn=VAV2;refGene_tx=XM_023242976.1;refGene_gn=VAV2;refGene_tx=XM_023242977.1;refGene_gn=VAV2;refGene_tx=XM_023242978.1;refGene_gn=VAV2;refGene_tx=XM_023242979.1;refGene_gn=VAV2;refGene_tx=XM_023242980.1
D4 93938450 93938583
refGene_gn=VAV2;refGene_tx=XM_023242972.1;refGene_gn=VAV2;refGene_tx=XM_023242974.1;refGene_gn=VAV2;refGene_tx=XM_023242975.1;refGene_gn=VAV2;refGene_tx=XM_023242976.1;refGene_gn=VAV2;refGene_tx=XM_023242977.1;refGene_gn=VAV2;refGene_tx=XM_023242978.1;refGene_gn=VAV2;refGene_tx=XM_023242979.1;refGene_gn=VAV2;refGene_tx=XM_023242980.1
D4 93939822 93939837
refGene_gn=VAV2;refGene_tx=XM_023242972.1;refGene_gn=VAV2;refGene_tx=XM_023242974.1;refGene_gn=VAV2;refGene_tx=XM_023242975.1;refGene_gn=VAV2;refGene_tx=XM_023242977.1;refGene_gn=VAV2;refGene_tx=XM_023242979.1
D4 93940820 93940930
refGene_gn=VAV2;refGene_tx=XM_023242972.1;refGene_gn=VAV2;refGene_tx=X

M_023242974.1;refGene_gn=VAV2;refGene_tx=XM_023242975.1;refGene_gn=VAV2;refGene_tx=XM_023242976.1;refGene_gn=VAV2;refGene_tx=XM_023242977.1;refGene_gn=VAV2;refGene_tx=XM_023242978.1;refGene_gn=VAV2;refGene_tx=XM_023242979.1;refGene_gn=VAV2;refGene_tx=XM_023242980.1
D4 93941826 93941928
refGene_gn=VAV2;refGene_tx=XM_023242972.1;refGene_gn=VAV2;refGene_tx=XM_023242974.1;refGene_gn=VAV2;refGene_tx=XM_023242975.1;refGene_gn=VAV2;refGene_tx=XM_023242976.1;refGene_gn=VAV2;refGene_tx=XM_023242977.1;refGene_gn=VAV2;refGene_tx=XM_023242978.1;refGene_gn=VAV2;refGene_tx=XM_023242979.1;refGene_gn=VAV2;refGene_tx=XM_023242980.1
D4 93942381 93942470
refGene_gn=VAV2;refGene_tx=XM_023242972.1;refGene_gn=VAV2;refGene_tx=XM_023242974.1;refGene_gn=VAV2;refGene_tx=XM_023242975.1;refGene_gn=VAV2;refGene_tx=XM_023242976.1;refGene_gn=VAV2;refGene_tx=XM_023242977.1;refGene_gn=VAV2;refGene_tx=XM_023242978.1;refGene_gn=VAV2;refGene_tx=XM_023242979.1;refGene_gn=VAV2;refGene_tx=XM_023242980.1
D4 93944689 93944706
refGene_gn=VAV2;refGene_tx=XM_023242972.1;refGene_gn=VAV2;refGene_tx=XM_023242974.1;refGene_gn=VAV2;refGene_tx=XM_023242975.1;refGene_gn=VAV2;refGene_tx=XM_023242976.1;refGene_gn=VAV2;refGene_tx=XM_023242977.1;refGene_gn=VAV2;refGene_tx=XM_023242978.1;refGene_gn=VAV2;refGene_tx=XM_023242979.1;refGene_gn=VAV2;refGene_tx=XM_023242980.1
D4 93945558 93945580
refGene_gn=VAV2;refGene_tx=XM_023242972.1;refGene_gn=VAV2;refGene_tx=XM_023242974.1;refGene_gn=VAV2;refGene_tx=XM_023242975.1;refGene_gn=VAV2;refGene_tx=XM_023242976.1;refGene_gn=VAV2;refGene_tx=XM_023242977.1;refGene_gn=VAV2;refGene_tx=XM_023242978.1;refGene_gn=VAV2;refGene_tx=XM_023242979.1;refGene_gn=VAV2;refGene_tx=XM_023242980.1
D4 93946078 93946206
refGene_gn=VAV2;refGene_tx=XM_023242972.1;refGene_gn=VAV2;refGene_tx=XM_023242974.1;refGene_gn=VAV2;refGene_tx=XM_023242975.1;refGene_gn=VAV2;refGene_tx=XM_023242976.1;refGene_gn=VAV2;refGene_tx=XM_023242977.1;refGene_gn=VAV2;refGene_tx=XM_023242978.1;refGene_gn=VAV2;refGene_tx=XM_023242979.1;refGene_gn=VAV2;refGene_tx=XM_023242980.1
D4 93946974 93947049
refGene_gn=VAV2;refGene_tx=XM_023242972.1;refGene_gn=VAV2;refGene_tx=XM_023242974.1;refGene_gn=VAV2;refGene_tx=XM_023242975.1;refGene_gn=VAV2;refGene_tx=XM_023242976.1;refGene_gn=VAV2;refGene_tx=XM_023242977.1;refGene_gn=VAV2;refGene_tx=XM_023242978.1;refGene_gn=VAV2;refGene_tx=XM_023242979.1;refGene_gn=VAV2;refGene_tx=XM_023242980.1
D4 93948371 93948424
refGene_gn=VAV2;refGene_tx=XM_023242972.1;refGene_gn=VAV2;refGene_tx=XM_023242974.1;refGene_gn=VAV2;refGene_tx=XM_023242975.1;refGene_gn=VAV2;refGene_tx=XM_023242976.1;refGene_gn=VAV2;refGene_tx=XM_023242977.1;refGene_gn=VAV2;refGene_tx=XM_023242978.1;refGene_gn=VAV2;refGene_tx=XM_023242979.1;refGene_gn=VAV2;refGene_tx=XM_023242980.1
D4 93949337 93949454
refGene_gn=VAV2;refGene_tx=XM_023242972.1;refGene_gn=VAV2;refGene_tx=XM_023242974.1;refGene_gn=VAV2;refGene_tx=XM_023242975.1;refGene_gn=VAV2;refGene_tx=XM_023242976.1;refGene_gn=VAV2;refGene_tx=XM_023242977.1;

refGene_gn=VAV2;refGene_tx=XM_023242978.1;refGene_gn=VAV2;refGene_tx=XM_023242979.1;refGene_gn=VAV2;refGene_tx=XM_023242980.1
D4 93951601 93951689
refGene_gn=VAV2;refGene_tx=XM_023242972.1;refGene_gn=VAV2;refGene_tx=XM_023242974.1;refGene_gn=VAV2;refGene_tx=XM_023242975.1;refGene_gn=VAV2;refGene_tx=XM_023242976.1;refGene_gn=VAV2;refGene_tx=XM_023242977.1;refGene_gn=VAV2;refGene_tx=XM_023242978.1;refGene_gn=VAV2;refGene_tx=XM_023242979.1;refGene_gn=VAV2;refGene_tx=XM_023242980.1
D4 93952619 93952743
refGene_gn=VAV2;refGene_tx=XM_023242972.1;refGene_gn=VAV2;refGene_tx=XM_023242974.1;refGene_gn=VAV2;refGene_tx=XM_023242975.1;refGene_gn=VAV2;refGene_tx=XM_023242976.1;refGene_gn=VAV2;refGene_tx=XM_023242977.1;refGene_gn=VAV2;refGene_tx=XM_023242978.1;refGene_gn=VAV2;refGene_tx=XM_023242979.1;refGene_gn=VAV2;refGene_tx=XM_023242980.1
D4 93953418 93953505
refGene_gn=VAV2;refGene_tx=XM_023242972.1;refGene_gn=VAV2;refGene_tx=XM_023242974.1;refGene_gn=VAV2;refGene_tx=XM_023242975.1;refGene_gn=VAV2;refGene_tx=XM_023242976.1;refGene_gn=VAV2;refGene_tx=XM_023242977.1;refGene_gn=VAV2;refGene_tx=XM_023242978.1
D4 93954432 93954587
refGene_gn=VAV2;refGene_tx=XM_023242972.1;refGene_gn=VAV2;refGene_tx=XM_023242974.1;refGene_gn=VAV2;refGene_tx=XM_023242975.1;refGene_gn=VAV2;refGene_tx=XM_023242976.1;refGene_gn=VAV2;refGene_tx=XM_023242977.1;refGene_gn=VAV2;refGene_tx=XM_023242978.1;refGene_gn=VAV2;refGene_tx=XM_023242979.1;refGene_gn=VAV2;refGene_tx=XM_023242980.1
D4 93957902 93957950
refGene_gn=VAV2;refGene_tx=XM_023242972.1;refGene_gn=VAV2;refGene_tx=XM_023242974.1;refGene_gn=VAV2;refGene_tx=XM_023242975.1;refGene_gn=VAV2;refGene_tx=XM_023242976.1;refGene_gn=VAV2;refGene_tx=XM_023242977.1;refGene_gn=VAV2;refGene_tx=XM_023242978.1;refGene_gn=VAV2;refGene_tx=XM_023242979.1;refGene_gn=VAV2;refGene_tx=XM_023242980.1
D4 93971581 93971912
refGene_gn=SARDH;refGene_tx=XM_019816743.2;refGene_gn=SARDH;refGene_tx=XM_023242968.1;refGene_gn=SARDH;refGene_tx=XM_023242969.1;refGene_gn=SARDH;refGene_tx=XM_023242970.1;refGene_gn=SARDH;refGene_tx=XM_023242971.1
D4 93973223 93973402
refGene_gn=SARDH;refGene_tx=XM_019816743.2;refGene_gn=SARDH;refGene_tx=XM_023242968.1;refGene_gn=SARDH;refGene_tx=XM_023242969.1;refGene_gn=SARDH;refGene_tx=XM_023242970.1;refGene_gn=SARDH;refGene_tx=XM_023242971.1
D4 93974463 93974643
refGene_gn=SARDH;refGene_tx=XM_019816743.2;refGene_gn=SARDH;refGene_tx=XM_023242968.1;refGene_gn=SARDH;refGene_tx=XM_023242969.1;refGene_gn=SARDH;refGene_tx=XM_023242970.1;refGene_gn=SARDH;refGene_tx=XM_023242971.1
D4 93975334 93975458
refGene_gn=SARDH;refGene_tx=XM_019816743.2;refGene_gn=SARDH;refGene_tx=XM_023242968.1;refGene_gn=SARDH;refGene_tx=XM_023242969.1;refGene_gn=SARDH;refGene_tx=XM_0232429

71.1
D4 93975662 93975763
refGene_gn=SARDH;refGene_tx=XM_019816743.2;refGene_gn=SARDH;refGene_tx=XM_023242968.1;refGene_gn=SARDH;refGene_tx=XM_023242969.1;refGene_gn=SARDH;refGene_tx=XM_023242970.1;refGene_gn=SARDH;refGene_tx=XM_023242971.1
D4 93982767 93982872
refGene_gn=SARDH;refGene_tx=XM_019816743.2;refGene_gn=SARDH;refGene_tx=XM_023242968.1;refGene_gn=SARDH;refGene_tx=XM_023242969.1;refGene_gn=SARDH;refGene_tx=XM_023242970.1;refGene_gn=SARDH;refGene_tx=XM_023242971.1
D4 93984132 93984262
refGene_gn=SARDH;refGene_tx=XM_019816743.2;refGene_gn=SARDH;refGene_tx=XM_023242968.1;refGene_gn=SARDH;refGene_tx=XM_023242969.1;refGene_gn=SARDH;refGene_tx=XM_023242970.1;refGene_gn=SARDH;refGene_tx=XM_023242971.1
D4 93984643 93984727
refGene_gn=SARDH;refGene_tx=XM_019816743.2;refGene_gn=SARDH;refGene_tx=XM_023242968.1;refGene_gn=SARDH;refGene_tx=XM_023242969.1;refGene_gn=SARDH;refGene_tx=XM_023242971.1
D4 93986732 93986819
refGene_gn=SARDH;refGene_tx=XM_019816743.2;refGene_gn=SARDH;refGene_tx=XM_023242968.1;refGene_gn=SARDH;refGene_tx=XM_023242969.1;refGene_gn=SARDH;refGene_tx=XM_023242970.1;refGene_gn=SARDH;refGene_tx=XM_023242971.1
D4 93987124 93987215
refGene_gn=SARDH;refGene_tx=XM_019816743.2;refGene_gn=SARDH;refGene_tx=XM_023242968.1;refGene_gn=SARDH;refGene_tx=XM_023242969.1;refGene_gn=SARDH;refGene_tx=XM_023242970.1;refGene_gn=SARDH;refGene_tx=XM_023242971.1
D4 93990398 93990540
refGene_gn=SARDH;refGene_tx=XM_019816743.2;refGene_gn=SARDH;refGene_tx=XM_023242968.1;refGene_gn=SARDH;refGene_tx=XM_023242969.1;refGene_gn=SARDH;refGene_tx=XM_023242970.1;refGene_gn=SARDH;refGene_tx=XM_023242971.1
D4 93993558 93993642
refGene_gn=SARDH;refGene_tx=XM_019816743.2;refGene_gn=SARDH;refGene_tx=XM_023242968.1;refGene_gn=SARDH;refGene_tx=XM_023242969.1;refGene_gn=SARDH;refGene_tx=XM_023242970.1
D4 93995360 93995474
refGene_gn=SARDH;refGene_tx=XM_019816743.2;refGene_gn=SARDH;refGene_tx=XM_023242968.1;refGene_gn=SARDH;refGene_tx=XM_023242969.1;refGene_gn=SARDH;refGene_tx=XM_023242970.1
D4 94000327 94000466
refGene_gn=SARDH;refGene_tx=XM_019816743.2;refGene_gn=SARDH;refGene_tx=XM_023242968.1;refGene_gn=SARDH;refGene_tx=XM_023242969.1;refGene_gn=SARDH;refGene_tx=XM_023242970.1;refGene_gn=SARDH;refGene_tx=XM_023242971.1
D4 94002068 94002182
refGene_gn=SARDH;refGene_tx=XM_019816743.2;refGene_gn=SARDH;refGene_tx

=XM_023242968.1;refGene_gn=SARDH;refGene_tx=XM_023242969.1;refGene_gn=SARDH;refGene_tx=XM_023242970.1;refGene_gn=SARDH;refGene_tx=XM_023242971.1
D4 94007391 94007539
refGene_gn=SARDH;refGene_tx=XM_019816743.2;refGene_gn=SARDH;refGene_tx=XM_023242968.1;refGene_gn=SARDH;refGene_tx=XM_023242969.1;refGene_gn=SARDH;refGene_tx=XM_023242970.1;refGene_gn=SARDH;refGene_tx=XM_023242971.1
D4 94011685 94011779
refGene_gn=SARDH;refGene_tx=XM_019816743.2;refGene_gn=SARDH;refGene_tx=XM_023242968.1;refGene_gn=SARDH;refGene_tx=XM_023242969.1;refGene_gn=SARDH;refGene_tx=XM_023242970.1;refGene_gn=SARDH;refGene_tx=XM_023242971.1
D4 94024358 94024521
refGene_gn=SARDH;refGene_tx=XM_019816743.2;refGene_gn=SARDH;refGene_tx=XM_023242968.1;refGene_gn=SARDH;refGene_tx=XM_023242969.1;refGene_gn=SARDH;refGene_tx=XM_023242970.1;refGene_gn=SARDH;refGene_tx=XM_023242971.1
D4 94025012 94025181
refGene_gn=SARDH;refGene_tx=XM_019816743.2;refGene_gn=SARDH;refGene_tx=XM_023242968.1;refGene_gn=SARDH;refGene_tx=XM_023242969.1;refGene_gn=SARDH;refGene_tx=XM_023242970.1;refGene_gn=SARDH;refGene_tx=XM_023242971.1
D4 94026816 94026952
refGene_gn=SARDH;refGene_tx=XM_019816743.2;refGene_gn=SARDH;refGene_tx=XM_023242968.1;refGene_gn=SARDH;refGene_tx=XM_023242969.1;refGene_gn=SARDH;refGene_tx=XM_023242970.1;refGene_gn=SARDH;refGene_tx=XM_023242971.1
D4 94028385 94028511
refGene_gn=SARDH;refGene_tx=XM_019816743.2;refGene_gn=SARDH;refGene_tx=XM_023242968.1;refGene_gn=SARDH;refGene_tx=XM_023242969.1;refGene_gn=SARDH;refGene_tx=XM_023242970.1;refGene_gn=SARDH;refGene_tx=XM_023242971.1
D4 94035414 94035546 refGene_gn=DBH;refGene_tx=XM_003996036.5
D4 94036491 94036651 refGene_gn=DBH;refGene_tx=XM_003996036.5
D4 94037025 94037153 refGene_gn=DBH;refGene_tx=XM_003996036.5
D4 94040658 94040718 refGene_gn=DBH;refGene_tx=XM_003996036.5
D4 94041381 94041420 refGene_gn=DBH;refGene_tx=XM_003996036.5
D4 94041815 94041959 refGene_gn=DBH;refGene_tx=XM_003996036.5
D4 94044843 94045010 refGene_gn=DBH;refGene_tx=XM_003996036.5
D4 94048320 94048423 refGene_gn=DBH;refGene_tx=XM_003996036.5
D4 94049076 94049253 refGene_gn=DBH;refGene_tx=XM_003996036.5
D4 94050152 94050410 refGene_gn=DBH;refGene_tx=XM_003996036.5
D4 94052024 94052171 refGene_gn=DBH;refGene_tx=XM_003996036.5
D4 94054961 94055279 refGene_gn=DBH;refGene_tx=XM_003996036.5
D4 94098086 94098179 refGene_gn=FAM163B;refGene_tx=XM_023242986.1
D4 94098696 94099095 refGene_gn=FAM163B;refGene_tx=XM_023242986.1
D4 94102769 94102888
refGene_gn=ADAMTSL2;refGene_tx=XM_006939621.4;refGene_gn=ADAMTSL2;refGene_tx=XM_023242967.1

D4	94103582 94103727
refGene_gn=ADAMTSL2;refGene_tx=XM_006939621.4;refGene_gn=ADAMTSL2;refGene_tx=XM_023242967.1
D4	94105770 94105947
refGene_gn=ADAMTSL2;refGene_tx=XM_006939621.4;refGene_gn=ADAMTSL2;refGene_tx=XM_023242967.1
D4	94106657 94106828
refGene_gn=ADAMTSL2;refGene_tx=XM_006939621.4;refGene_gn=ADAMTSL2;refGene_tx=XM_023242967.1
D4	94107354 94107510
refGene_gn=ADAMTSL2;refGene_tx=XM_006939621.4;refGene_gn=ADAMTSL2;refGene_tx=XM_023242967.1
D4	94107593 94107807
refGene_gn=ADAMTSL2;refGene_tx=XM_006939621.4;refGene_gn=ADAMTSL2;refGene_tx=XM_023242967.1
D4	94108890 94109017
refGene_gn=ADAMTSL2;refGene_tx=XM_006939621.4;refGene_gn=ADAMTSL2;refGene_tx=XM_023242967.1
D4	94113421 94113519
refGene_gn=ADAMTSL2;refGene_tx=XM_006939621.4;refGene_gn=ADAMTSL2;refGene_tx=XM_023242967.1
D4	94118094 94118467
refGene_gn=ADAMTSL2;refGene_tx=XM_006939621.4;refGene_gn=ADAMTSL2;refGene_tx=XM_023242967.1
D4	94121212 94121561
refGene_gn=ADAMTSL2;refGene_tx=XM_006939621.4;refGene_gn=ADAMTSL2;refGene_tx=XM_023242967.1
D4	94127265 94127441
refGene_gn=ADAMTSL2;refGene_tx=XM_006939621.4;refGene_gn=ADAMTSL2;refGene_tx=XM_023242967.1
D4	94129915 94129996
refGene_gn=ADAMTSL2;refGene_tx=XM_006939621.4;refGene_gn=ADAMTSL2;refGene_tx=XM_023242967.1
D4	94132464 94132588
refGene_gn=ADAMTSL2;refGene_tx=XM_006939621.4;refGene_gn=ADAMTSL2;refGene_tx=XM_023242967.1
D4	94132725 94132871
refGene_gn=ADAMTSL2;refGene_tx=XM_006939621.4;refGene_gn=ADAMTSL2;refGene_tx=XM_023242967.1
D4	94133249 94133352
refGene_gn=ADAMTSL2;refGene_tx=XM_006939621.4;refGene_gn=ADAMTSL2;refGene_tx=XM_023242967.1
D4	94134401 94134477
refGene_gn=ADAMTSL2;refGene_tx=XM_006939621.4;refGene_gn=ADAMTSL2;refGene_tx=XM_023242967.1
D4	94135138 94135281
refGene_gn=ADAMTSL2;refGene_tx=XM_006939621.4;refGene_gn=ADAMTSL2;refGene_tx=XM_023242967.1
D4	94135814 94135904
refGene_gn=ADAMTSL2;refGene_tx=XM_006939621.4;refGene_gn=ADAMTSL2;refG

ene_tx=XM_023242967.1
D4 94138469 94139069
refGene_gn=LOC111557567;refGene_tx=XM_023242983.1
D4 94139598 94139694
refGene_gn=LOC111557567;refGene_tx=XM_023242983.1
D4 94146651 94146786 refGene_gn=MYMK;refGene_tx=XM_006939622.4
D4 94151044 94151159 refGene_gn=MYMK;refGene_tx=XM_006939622.4
D4 94151963 94152112 refGene_gn=MYMK;refGene_tx=XM_006939622.4
D4 94154907 94155024 refGene_gn=MYMK;refGene_tx=XM_006939622.4
D4 94155506 94155656 refGene_gn=MYMK;refGene_tx=XM_006939622.4
D4 94177245 94177337 refGene_gn=SLC2A6;refGene_tx=XM_003996039.5
D4 94177844 94178007 refGene_gn=SLC2A6;refGene_tx=XM_003996039.5
D4 94179087 94179294 refGene_gn=SLC2A6;refGene_tx=XM_003996039.5
D4 94179890 94179990 refGene_gn=SLC2A6;refGene_tx=XM_003996039.5
D4 94180521 94180733 refGene_gn=SLC2A6;refGene_tx=XM_003996039.5
D4 94181110 94181263 refGene_gn=SLC2A6;refGene_tx=XM_003996039.5
D4 94181923 94182032 refGene_gn=SLC2A6;refGene_tx=XM_003996039.5
D4 94182444 94182630 refGene_gn=SLC2A6;refGene_tx=XM_003996039.5
D4 94182841 94182987 refGene_gn=SLC2A6;refGene_tx=XM_003996039.5
D4 94183688 94183844 refGene_gn=SLC2A6;refGene_tx=XM_003996039.5
D4 94187082 94187173
refGene_gn=CACFD1;refGene_tx=XM_023242984.1;refGene_gn=CACFD1;refGene_
tx=XM_023242985.1
D4 94187680 94187788
refGene_gn=CACFD1;refGene_tx=XM_023242984.1;refGene_gn=CACFD1;refGene_
tx=XM_023242985.1
D4 94190622 94190748
refGene_gn=CACFD1;refGene_tx=XM_023242984.1;refGene_gn=CACFD1;refGene_
tx=XM_023242985.1
D4 94192175 94192248
refGene_gn=CACFD1;refGene_tx=XM_023242984.1;refGene_gn=CACFD1;refGene_
tx=XM_023242985.1
D4 94195202 94195323
refGene_gn=CACFD1;refGene_tx=XM_023242984.1;refGene_gn=CACFD1;refGene_
tx=XM_023242985.1
D4 94196075 94196261
refGene_gn=ADAMTS13;refGene_tx=XM_023242957.1;refGene_gn=ADAMTS13;refG
ene_tx=XM_023242958.1;refGene_gn=ADAMTS13;refGene_tx=XM_023242959.1;re
fGene_gn=ADAMTS13;refGene_tx=XM_023242960.1;refGene_gn=ADAMTS13;refGen
e_tx=XM_023242961.1;refGene_gn=ADAMTS13;refGene_tx=XM_023242962.1;refG
ene_gn=ADAMTS13;refGene_tx=XM_023242963.1
D4 94196870 94197046
refGene_gn=ADAMTS13;refGene_tx=XM_023242957.1;refGene_gn=ADAMTS13;refG
ene_tx=XM_023242958.1;refGene_gn=ADAMTS13;refGene_tx=XM_023242959.1;re
fGene_gn=ADAMTS13;refGene_tx=XM_023242960.1;refGene_gn=ADAMTS13;refGen
e_tx=XM_023242961.1;refGene_gn=ADAMTS13;refGene_tx=XM_023242962.1;refG
ene_gn=ADAMTS13;refGene_tx=XM_023242963.1
D4 94198053 94198230
refGene_gn=ADAMTS13;refGene_tx=XM_023242957.1;refGene_gn=ADAMTS13;refG
ene_tx=XM_023242958.1;refGene_gn=ADAMTS13;refGene_tx=XM_023242959.1;re

fGene_gn=ADAMTS13;refGene_tx=XM_023242960.1;refGene_gn=ADAMTS13;refGene_tx=XM_023242961.1;refGene_gn=ADAMTS13;refGene_tx=XM_023242962.1;refGene_gn=ADAMTS13;refGene_tx=XM_023242963.1
D4 94198563 94198710
refGene_gn=ADAMTS13;refGene_tx=XM_023242957.1;refGene_gn=ADAMTS13;refGene_tx=XM_023242958.1;refGene_gn=ADAMTS13;refGene_tx=XM_023242959.1;refGene_gn=ADAMTS13;refGene_tx=XM_023242960.1;refGene_gn=ADAMTS13;refGene_tx=XM_023242961.1;refGene_gn=ADAMTS13;refGene_tx=XM_023242962.1;refGene_gn=ADAMTS13;refGene_tx=XM_023242963.1
D4 94199227 94199519
refGene_gn=ADAMTS13;refGene_tx=XM_023242957.1;refGene_gn=ADAMTS13;refGene_tx=XM_023242958.1;refGene_gn=ADAMTS13;refGene_tx=XM_023242959.1;refGene_gn=ADAMTS13;refGene_tx=XM_023242960.1;refGene_gn=ADAMTS13;refGene_tx=XM_023242961.1;refGene_gn=ADAMTS13;refGene_tx=XM_023242962.1;refGene_gn=ADAMTS13;refGene_tx=XM_023242963.1
D4 94200151 94200356
refGene_gn=ADAMTS13;refGene_tx=XM_023242957.1;refGene_gn=ADAMTS13;refGene_tx=XM_023242958.1;refGene_gn=ADAMTS13;refGene_tx=XM_023242959.1;refGene_gn=ADAMTS13;refGene_tx=XM_023242960.1;refGene_gn=ADAMTS13;refGene_tx=XM_023242961.1;refGene_gn=ADAMTS13;refGene_tx=XM_023242962.1;refGene_gn=ADAMTS13;refGene_tx=XM_023242963.1;refGene_gn=ADAMTS13;refGene_tx=XM_023242965.1
D4 94202508 94203117
refGene_gn=ADAMTS13;refGene_tx=XM_023242957.1;refGene_gn=ADAMTS13;refGene_tx=XM_023242958.1;refGene_gn=ADAMTS13;refGene_tx=XM_023242959.1;refGene_gn=ADAMTS13;refGene_tx=XM_023242960.1;refGene_gn=ADAMTS13;refGene_tx=XM_023242961.1;refGene_gn=ADAMTS13;refGene_tx=XM_023242962.1;refGene_gn=ADAMTS13;refGene_tx=XM_023242963.1;refGene_gn=ADAMTS13;refGene_tx=XM_023242964.1;refGene_gn=ADAMTS13;refGene_tx=XM_023242965.1
D4 94203783 94203994
refGene_gn=ADAMTS13;refGene_tx=XM_023242957.1;refGene_gn=ADAMTS13;refGene_tx=XM_023242958.1;refGene_gn=ADAMTS13;refGene_tx=XM_023242959.1;refGene_gn=ADAMTS13;refGene_tx=XM_023242960.1;refGene_gn=ADAMTS13;refGene_tx=XM_023242962.1;refGene_gn=ADAMTS13;refGene_tx=XM_023242964.1;refGene_gn=ADAMTS13;refGene_tx=XM_023242965.1
D4 94204072 94204202
refGene_gn=ADAMTS13;refGene_tx=XM_023242957.1;refGene_gn=ADAMTS13;refGene_tx=XM_023242958.1;refGene_gn=ADAMTS13;refGene_tx=XM_023242959.1;refGene_gn=ADAMTS13;refGene_tx=XM_023242960.1;refGene_gn=ADAMTS13;refGene_tx=XM_023242961.1;refGene_gn=ADAMTS13;refGene_tx=XM_023242962.1;refGene_gn=ADAMTS13;refGene_tx=XM_023242963.1;refGene_gn=ADAMTS13;refGene_tx=XM_023242964.1;refGene_gn=ADAMTS13;refGene_tx=XM_023242965.1
D4 94205641 94205762
refGene_gn=ADAMTS13;refGene_tx=XM_023242957.1;refGene_gn=ADAMTS13;refGene_tx=XM_023242958.1;refGene_gn=ADAMTS13;refGene_tx=XM_023242959.1;refGene_gn=ADAMTS13;refGene_tx=XM_023242960.1;refGene_gn=ADAMTS13;refGene_tx=XM_023242961.1;refGene_gn=ADAMTS13;refGene_tx=XM_023242962.1;refGene_gn=ADAMTS13;refGene_tx=XM_023242963.1;refGene_gn=ADAMTS13;refGene_tx=XM_023242964.1;refGene_gn=ADAMTS13;refGene_tx=XM_023242965.1
D4 94206369 94206553

refGene_gn=ADAMTS13;refGene_tx=XM_023242957.1;refGene_gn=ADAMTS13;refGene_tx=XM_023242958.1;refGene_gn=ADAMTS13;refGene_tx=XM_023242959.1;refGene_gn=ADAMTS13;refGene_tx=XM_023242960.1;refGene_gn=ADAMTS13;refGene_tx=XM_023242961.1;refGene_gn=ADAMTS13;refGene_tx=XM_023242962.1;refGene_gn=ADAMTS13;refGene_tx=XM_023242963.1;refGene_gn=ADAMTS13;refGene_tx=XM_023242964.1;refGene_gn=ADAMTS13;refGene_tx=XM_023242965.1
D4 94206758 94206962
refGene_gn=ADAMTS13;refGene_tx=XM_023242957.1;refGene_gn=ADAMTS13;refGene_tx=XM_023242958.1;refGene_gn=ADAMTS13;refGene_tx=XM_023242959.1;refGene_gn=ADAMTS13;refGene_tx=XM_023242960.1;refGene_gn=ADAMTS13;refGene_tx=XM_023242961.1;refGene_gn=ADAMTS13;refGene_tx=XM_023242964.1;refGene_gn=ADAMTS13;refGene_tx=XM_023242965.1
D4 94208193 94208323
refGene_gn=ADAMTS13;refGene_tx=XM_023242957.1;refGene_gn=ADAMTS13;refGene_tx=XM_023242958.1;refGene_gn=ADAMTS13;refGene_tx=XM_023242959.1;refGene_gn=ADAMTS13;refGene_tx=XM_023242960.1;refGene_gn=ADAMTS13;refGene_tx=XM_023242961.1;refGene_gn=ADAMTS13;refGene_tx=XM_023242962.1;refGene_gn=ADAMTS13;refGene_tx=XM_023242963.1;refGene_gn=ADAMTS13;refGene_tx=XM_023242964.1;refGene_gn=ADAMTS13;refGene_tx=XM_023242965.1
D4 94208432 94208568
refGene_gn=ADAMTS13;refGene_tx=XM_023242957.1;refGene_gn=ADAMTS13;refGene_tx=XM_023242958.1;refGene_gn=ADAMTS13;refGene_tx=XM_023242959.1;refGene_gn=ADAMTS13;refGene_tx=XM_023242960.1;refGene_gn=ADAMTS13;refGene_tx=XM_023242961.1;refGene_gn=ADAMTS13;refGene_tx=XM_023242962.1;refGene_gn=ADAMTS13;refGene_tx=XM_023242963.1;refGene_gn=ADAMTS13;refGene_tx=XM_023242964.1;refGene_gn=ADAMTS13;refGene_tx=XM_023242965.1
D4 94210463 94210645
refGene_gn=ADAMTS13;refGene_tx=XM_023242957.1;refGene_gn=ADAMTS13;refGene_tx=XM_023242958.1;refGene_gn=ADAMTS13;refGene_tx=XM_023242959.1;refGene_gn=ADAMTS13;refGene_tx=XM_023242960.1;refGene_gn=ADAMTS13;refGene_tx=XM_023242961.1;refGene_gn=ADAMTS13;refGene_tx=XM_023242962.1;refGene_gn=ADAMTS13;refGene_tx=XM_023242963.1;refGene_gn=ADAMTS13;refGene_tx=XM_023242964.1;refGene_gn=ADAMTS13;refGene_tx=XM_023242965.1
D4 94211804 94211885
refGene_gn=ADAMTS13;refGene_tx=XM_023242957.1;refGene_gn=ADAMTS13;refGene_tx=XM_023242958.1;refGene_gn=ADAMTS13;refGene_tx=XM_023242959.1;refGene_gn=ADAMTS13;refGene_tx=XM_023242960.1;refGene_gn=ADAMTS13;refGene_tx=XM_023242961.1;refGene_gn=ADAMTS13;refGene_tx=XM_023242962.1;refGene_gn=ADAMTS13;refGene_tx=XM_023242963.1;refGene_gn=ADAMTS13;refGene_tx=XM_023242964.1;refGene_gn=ADAMTS13;refGene_tx=XM_023242965.1
D4 94212609 94212730
refGene_gn=ADAMTS13;refGene_tx=XM_023242957.1;refGene_gn=ADAMTS13;refGene_tx=XM_023242958.1;refGene_gn=ADAMTS13;refGene_tx=XM_023242959.1;refGene_gn=ADAMTS13;refGene_tx=XM_023242960.1;refGene_gn=ADAMTS13;refGene_tx=XM_023242961.1;refGene_gn=ADAMTS13;refGene_tx=XM_023242962.1;refGene_gn=ADAMTS13;refGene_tx=XM_023242963.1;refGene_gn=ADAMTS13;refGene_tx=XM_023242964.1;refGene_gn=ADAMTS13;refGene_tx=XM_023242965.1
D4 94213075 94213224
refGene_gn=ADAMTS13;refGene_tx=XM_023242957.1;refGene_gn=ADAMTS13;refGene_tx=XM_023242958.1;refGene_gn=ADAMTS13;refGene_tx=XM_023242959.1;re

fGene_gn=ADAMTS13;refGene_tx=XM_023242960.1;refGene_gn=ADAMTS13;refGen
e_tx=XM_023242961.1;refGene_gn=ADAMTS13;refGene_tx=XM_023242962.1;refG
ene_gn=ADAMTS13;refGene_tx=XM_023242963.1;refGene_gn=ADAMTS13;refGene_
tx=XM_023242964.1;refGene_gn=ADAMTS13;refGene_tx=XM_023242965.1
D4 94213625 94213752
refGene_gn=ADAMTS13;refGene_tx=XM_023242957.1;refGene_gn=ADAMTS13;refG
ene_tx=XM_023242958.1;refGene_gn=ADAMTS13;refGene_tx=XM_023242959.1;re
fGene_gn=ADAMTS13;refGene_tx=XM_023242960.1;refGene_gn=ADAMTS13;refGen
e_tx=XM_023242961.1;refGene_gn=ADAMTS13;refGene_tx=XM_023242962.1;refG
ene_gn=ADAMTS13;refGene_tx=XM_023242963.1;refGene_gn=ADAMTS13;refGene_
tx=XM_023242964.1;refGene_gn=ADAMTS13;refGene_tx=XM_023242965.1
D4 94215933 94215997
refGene_gn=ADAMTS13;refGene_tx=XM_023242957.1;refGene_gn=ADAMTS13;refG
ene_tx=XM_023242958.1;refGene_gn=ADAMTS13;refGene_tx=XM_023242959.1;re
fGene_gn=ADAMTS13;refGene_tx=XM_023242960.1;refGene_gn=ADAMTS13;refGen
e_tx=XM_023242961.1;refGene_gn=ADAMTS13;refGene_tx=XM_023242962.1;refG
ene_gn=ADAMTS13;refGene_tx=XM_023242963.1;refGene_gn=ADAMTS13;refGene_
tx=XM_023242964.1;refGene_gn=ADAMTS13;refGene_tx=XM_023242965.1
D4 94216094 94216246
refGene_gn=ADAMTS13;refGene_tx=XM_023242957.1;refGene_gn=ADAMTS13;refG
ene_tx=XM_023242959.1;refGene_gn=ADAMTS13;refGene_tx=XM_023242960.1;re
fGene_gn=ADAMTS13;refGene_tx=XM_023242961.1;refGene_gn=ADAMTS13;refGen
e_tx=XM_023242962.1;refGene_gn=ADAMTS13;refGene_tx=XM_023242963.1;refG
ene_gn=ADAMTS13;refGene_tx=XM_023242964.1;refGene_gn=ADAMTS13;refGene_
tx=XM_023242965.1
D4 94216399 94216644
refGene_gn=ADAMTS13;refGene_tx=XM_023242957.1;refGene_gn=ADAMTS13;refG
ene_tx=XM_023242958.1;refGene_gn=ADAMTS13;refGene_tx=XM_023242959.1;re
fGene_gn=ADAMTS13;refGene_tx=XM_023242960.1;refGene_gn=ADAMTS13;refGen
e_tx=XM_023242961.1;refGene_gn=ADAMTS13;refGene_tx=XM_023242962.1;refG
ene_gn=ADAMTS13;refGene_tx=XM_023242963.1;refGene_gn=ADAMTS13;refGene_
tx=XM_023242964.1;refGene_gn=ADAMTS13;refGene_tx=XM_023242965.1
D4 94217376 94217539
refGene_gn=ADAMTS13;refGene_tx=XM_023242957.1;refGene_gn=ADAMTS13;refG
ene_tx=XM_023242958.1;refGene_gn=ADAMTS13;refGene_tx=XM_023242959.1;re
fGene_gn=ADAMTS13;refGene_tx=XM_023242960.1;refGene_gn=ADAMTS13;refGen
e_tx=XM_023242961.1;refGene_gn=ADAMTS13;refGene_tx=XM_023242962.1;refG
ene_gn=ADAMTS13;refGene_tx=XM_023242963.1;refGene_gn=ADAMTS13;refGene_
tx=XM_023242964.1;refGene_gn=ADAMTS13;refGene_tx=XM_023242965.1
D4 94218002 94218143
refGene_gn=ADAMTS13;refGene_tx=XM_023242957.1;refGene_gn=ADAMTS13;refG
ene_tx=XM_023242958.1;refGene_gn=ADAMTS13;refGene_tx=XM_023242959.1;re
fGene_gn=ADAMTS13;refGene_tx=XM_023242960.1;refGene_gn=ADAMTS13;refGen
e_tx=XM_023242961.1;refGene_gn=ADAMTS13;refGene_tx=XM_023242962.1;refG
ene_gn=ADAMTS13;refGene_tx=XM_023242963.1;refGene_gn=ADAMTS13;refGene_
tx=XM_023242964.1;refGene_gn=ADAMTS13;refGene_tx=XM_023242965.1
D4 94220515 94220662
refGene_gn=ADAMTS13;refGene_tx=XM_023242957.1;refGene_gn=ADAMTS13;refG
ene_tx=XM_023242958.1;refGene_gn=ADAMTS13;refGene_tx=XM_023242959.1;re
fGene_gn=ADAMTS13;refGene_tx=XM_023242960.1;refGene_gn=ADAMTS13;refGen

e_tx=XM_023242961.1;refGene_gn=ADAMTS13;refGene_tx=XM_023242962.1;refGene_gn=ADAMTS13;refGene_tx=XM_023242963.1;refGene_gn=ADAMTS13;refGene_tx=XM_023242964.1;refGene_gn=ADAMTS13;refGene_tx=XM_023242965.1
D4 94220780 94220905
refGene_gn=ADAMTS13;refGene_tx=XM_023242957.1;refGene_gn=ADAMTS13;refGene_tx=XM_023242958.1;refGene_gn=ADAMTS13;refGene_tx=XM_023242959.1;refGene_gn=ADAMTS13;refGene_tx=XM_023242960.1;refGene_gn=ADAMTS13;refGene_tx=XM_023242961.1;refGene_gn=ADAMTS13;refGene_tx=XM_023242962.1;refGene_gn=ADAMTS13;refGene_tx=XM_023242963.1;refGene_gn=ADAMTS13;refGene_tx=XM_023242964.1;refGene_gn=ADAMTS13;refGene_tx=XM_023242965.1
D4 94221062 94221146
refGene_gn=ADAMTS13;refGene_tx=XM_023242957.1;refGene_gn=ADAMTS13;refGene_tx=XM_023242958.1;refGene_gn=ADAMTS13;refGene_tx=XM_023242959.1;refGene_gn=ADAMTS13;refGene_tx=XM_023242960.1;refGene_gn=ADAMTS13;refGene_tx=XM_023242961.1;refGene_gn=ADAMTS13;refGene_tx=XM_023242962.1;refGene_gn=ADAMTS13;refGene_tx=XM_023242963.1;refGene_gn=ADAMTS13;refGene_tx=XM_023242964.1;refGene_gn=ADAMTS13;refGene_tx=XM_023242965.1
D4 94223091 94223237
refGene_gn=ADAMTS13;refGene_tx=XM_023242957.1;refGene_gn=ADAMTS13;refGene_tx=XM_023242958.1;refGene_gn=ADAMTS13;refGene_tx=XM_023242959.1;refGene_gn=ADAMTS13;refGene_tx=XM_023242960.1;refGene_gn=ADAMTS13;refGene_tx=XM_023242961.1;refGene_gn=ADAMTS13;refGene_tx=XM_023242962.1;refGene_gn=ADAMTS13;refGene_tx=XM_023242963.1;refGene_gn=ADAMTS13;refGene_tx=XM_023242964.1;refGene_gn=ADAMTS13;refGene_tx=XM_023242965.1
D4 94224269 94224411
refGene_gn=ADAMTS13;refGene_tx=XM_023242957.1;refGene_gn=ADAMTS13;refGene_tx=XM_023242958.1;refGene_gn=ADAMTS13;refGene_tx=XM_023242959.1;refGene_gn=ADAMTS13;refGene_tx=XM_023242960.1;refGene_gn=ADAMTS13;refGene_tx=XM_023242961.1;refGene_gn=ADAMTS13;refGene_tx=XM_023242962.1;refGene_gn=ADAMTS13;refGene_tx=XM_023242963.1;refGene_gn=ADAMTS13;refGene_tx=XM_023242964.1;refGene_gn=ADAMTS13;refGene_tx=XM_023242965.1
D4 94224490 94224771
refGene_gn=ADAMTS13;refGene_tx=XM_023242957.1;refGene_gn=ADAMTS13;refGene_tx=XM_023242958.1;refGene_gn=ADAMTS13;refGene_tx=XM_023242959.1;refGene_gn=ADAMTS13;refGene_tx=XM_023242960.1;refGene_gn=ADAMTS13;refGene_tx=XM_023242961.1;refGene_gn=ADAMTS13;refGene_tx=XM_023242962.1;refGene_gn=ADAMTS13;refGene_tx=XM_023242963.1;refGene_gn=ADAMTS13;refGene_tx=XM_023242964.1;refGene_gn=ADAMTS13;refGene_tx=XM_023242965.1
D4 94225782 94225849
refGene_gn=ADAMTS13;refGene_tx=XM_023242957.1;refGene_gn=ADAMTS13;refGene_tx=XM_023242958.1;refGene_gn=ADAMTS13;refGene_tx=XM_023242959.1;refGene_gn=ADAMTS13;refGene_tx=XM_023242960.1;refGene_gn=ADAMTS13;refGene_tx=XM_023242961.1;refGene_gn=ADAMTS13;refGene_tx=XM_023242962.1;refGene_gn=ADAMTS13;refGene_tx=XM_023242963.1;refGene_gn=ADAMTS13;refGene_tx=XM_023242964.1;refGene_gn=ADAMTS13;refGene_tx=XM_023242965.1
D4 94226194 94226366
refGene_gn=ADAMTS13;refGene_tx=XM_023242957.1;refGene_gn=ADAMTS13;refGene_tx=XM_023242958.1;refGene_gn=ADAMTS13;refGene_tx=XM_023242959.1;refGene_gn=ADAMTS13;refGene_tx=XM_023242960.1;refGene_gn=ADAMTS13;refGene_tx=XM_023242961.1;refGene_gn=ADAMTS13;refGene_tx=XM_023242962.1;refG

ene_gn=ADAMTS13;refGene_tx=XM_023242963.1;refGene_gn=ADAMTS13;refGene_tx=XM_023242964.1;refGene_gn=ADAMTS13;refGene_tx=XM_023242965.1
D4 94226608 94226646 refGene_gn=ADAMTS13;refGene_tx=XM_023242957.1;refGene_gn=ADAMTS13;refGene_tx=XM_023242958.1;refGene_gn=ADAMTS13;refGene_tx=XM_023242959.1;refGene_gn=ADAMTS13;refGene_tx=XM_023242960.1;refGene_gn=ADAMTS13;refGene_tx=XM_023242961.1;refGene_gn=ADAMTS13;refGene_tx=XM_023242962.1;refGene_gn=ADAMTS13;refGene_tx=XM_023242963.1;refGene_gn=ADAMTS13;refGene_tx=XM_023242964.1;refGene_gn=ADAMTS13;refGene_tx=XM_023242965.1
D4 94228848 94229304 refGene_gn=REXO4;refGene_tx=XM_023242989.1;refGene_gn=REXO4;refGene_tx=XM_023242990.1
D4 94231136 94231459 refGene_gn=REXO4;refGene_tx=XM_023242989.1;refGene_gn=REXO4;refGene_tx=XM_023242990.1
D4 94232672 94232810 refGene_gn=REXO4;refGene_tx=XM_023242989.1;refGene_gn=REXO4;refGene_tx=XM_023242990.1
D4 94232988 94233182 refGene_gn=REXO4;refGene_tx=XM_023242989.1;refGene_gn=REXO4;refGene_tx=XM_023242990.1
D4 94234554 94234643 refGene_gn=REXO4;refGene_tx=XM_023242989.1;refGene_gn=REXO4;refGene_tx=XM_023242990.1
D4 94236490 94236565 refGene_gn=REXO4;refGene_tx=XM_023242989.1;refGene_gn=REXO4;refGene_tx=XM_023242990.1
D4 94236978 94237053 refGene_gn=REXO4;refGene_tx=XM_023242989.1
D4 94237723 94237888 refGene_gn=REXO4;refGene_tx=XM_023242989.1;refGene_gn=REXO4;refGene_tx=XM_023242990.1
D4 94238964 94239054 refGene_gn=STKLD1;refGene_tx=XM_023242988.1
D4 94239533 94239613 refGene_gn=STKLD1;refGene_tx=XM_023242988.1
D4 94239928 94240069 refGene_gn=STKLD1;refGene_tx=XM_023242988.1
D4 94241335 94241464 refGene_gn=STKLD1;refGene_tx=XM_023242988.1
D4 94241534 94241663 refGene_gn=STKLD1;refGene_tx=XM_023242988.1
D4 94242358 94242487 refGene_gn=STKLD1;refGene_tx=XM_023242988.1
D4 94243309 94243450 refGene_gn=STKLD1;refGene_tx=XM_023242988.1
D4 94243735 94243852 refGene_gn=STKLD1;refGene_tx=XM_023242988.1
D4 94245913 94245997 refGene_gn=STKLD1;refGene_tx=XM_023242988.1
D4 94247227 94247358 refGene_gn=STKLD1;refGene_tx=XM_023242988.1
D4 94247964 94248128 refGene_gn=STKLD1;refGene_tx=XM_023242988.1
D4 94248686 94248805 refGene_gn=STKLD1;refGene_tx=XM_023242988.1
D4 94249927 94250043 refGene_gn=STKLD1;refGene_tx=XM_023242988.1
D4 94250898 94250969 refGene_gn=STKLD1;refGene_tx=XM_023242988.1
D4 94253597 94253699 refGene_gn=STKLD1;refGene_tx=XM_023242988.1
D4 94255157 94255232 refGene_gn=STKLD1;refGene_tx=XM_023242988.1
D4 94256747 94256792 refGene_gn=STKLD1;refGene_tx=XM_023242988.1
D4 94259650 94259737 refGene_gn=STKLD1;refGene_tx=XM_023242988.1
D4 94261399 94261414 refGene_gn=STKLD1;refGene_tx=XM_023242988.1

D4	94262067 94262115 refGene_gn=SURF4;refGene_tx=XM_023242992.1
D4	94269162 94269349 refGene_gn=SURF4;refGene_tx=XM_023242992.1
D4	94270037 94270114 refGene_gn=SURF4;refGene_tx=XM_023242992.1
D4	94270714 94270758 refGene_gn=SURF4;refGene_tx=XM_023242992.1
D4	94271267 94271454 refGene_gn=SURF4;refGene_tx=XM_023242992.1
D4	94272138 94272405 refGene_gn=SURF4;refGene_tx=XM_023242992.1
D4	94274890 94274971 refGene_gn=SURF2;refGene_tx=XM_006939624.4
D4	94275383 94275550 refGene_gn=SURF2;refGene_tx=XM_006939624.4
D4	94275731 94275920 refGene_gn=SURF2;refGene_tx=XM_006939624.4
D4	94277846 94277950 refGene_gn=SURF2;refGene_tx=XM_006939624.4
D4	94278623 94278778 refGene_gn=SURF2;refGene_tx=XM_006939624.4
D4	94278863 94279119
refGene_gn=LOC101085249;refGene_tx=XM_006939625.3;refGene_gn=LOC101085
249;refGene_tx=XM_019816755.2;refGene_gn=SURF2;refGene_tx=XM_006939624
.4
D4	94279210 94279262
refGene_gn=LOC101085249;refGene_tx=XM_006939625.3;refGene_gn=LOC101085
249;refGene_tx=XM_019816755.2
D4	94280071 94280223
refGene_gn=LOC101085249;refGene_tx=XM_006939625.3;refGene_gn=LOC101085
249;refGene_tx=XM_019816755.2
D4	94280548 94280631
refGene_gn=LOC101085249;refGene_tx=XM_006939625.3;refGene_gn=LOC101085
249;refGene_tx=XM_019816755.2
D4	94281120 94281312
refGene_gn=LOC101085249;refGene_tx=XM_006939625.3;refGene_gn=LOC101085
249;refGene_tx=XM_019816755.2
D4	94281585 94281658
refGene_gn=LOC101085249;refGene_tx=XM_006939625.3;refGene_gn=LOC101085
249;refGene_tx=XM_019816755.2
D4	94281743 94281906
refGene_gn=LOC101085249;refGene_tx=XM_006939625.3;refGene_gn=LOC101085
249;refGene_tx=XM_019816755.2
D4	94282220 94282302
refGene_gn=LOC101085249;refGene_tx=XM_006939625.3;refGene_gn=LOC101085
249;refGene_tx=XM_019816755.2
D4	94282397 94282467
refGene_gn=LOC101085249;refGene_tx=XM_006939625.3;refGene_gn=LOC101085
249;refGene_tx=XM_019816755.2
D4	94282871 94282976 refGene_gn=RPL7A;refGene_tx=XM_003996041.5
D4	94283164 94283234 refGene_gn=RPL7A;refGene_tx=XM_003996041.5
D4	94283518 94283649 refGene_gn=RPL7A;refGene_tx=XM_003996041.5
D4	94283948 94284028 refGene_gn=RPL7A;refGene_tx=XM_003996041.5
D4	94284238 94284379 refGene_gn=RPL7A;refGene_tx=XM_003996041.5
D4	94284873 94285023 refGene_gn=RPL7A;refGene_tx=XM_003996041.5
D4	94285422 94285543 refGene_gn=RPL7A;refGene_tx=XM_003996041.5
D4	94286138 94286141 refGene_gn=RPL7A;refGene_tx=XM_003996041.5
D4	94286875 94286998
refGene_gn=MED22;refGene_tx=XM_003996043.5;refGene_gn=MED22;refGene_tx
=XM_019816756.2

D4 94288497 94288578
refGene_gn=MED22;refGene_tx=XM_003996043.5;refGene_gn=MED22;refGene_tx=XM_019816756.2
D4 94289335 94289544
refGene_gn=MED22;refGene_tx=XM_003996043.5;refGene_gn=MED22;refGene_tx=XM_019816756.2
D4 94291991 94292181
refGene_gn=MED22;refGene_tx=XM_003996043.5;refGene_gn=MED22;refGene_tx=XM_019816756.2
D4 94296352 94296446 refGene_gn=SURF6;refGene_tx=XM_003996044.5
D4 94296543 94296679 refGene_gn=SURF6;refGene_tx=XM_019816754.2
D4 94297451 94297655
refGene_gn=SURF6;refGene_tx=XM_003996044.5;refGene_gn=SURF6;refGene_tx=XM_019816754.2
D4 94298222 94298311
refGene_gn=SURF6;refGene_tx=XM_003996044.5;refGene_gn=SURF6;refGene_tx=XM_019816754.2
D4 94299403 94299604
refGene_gn=SURF6;refGene_tx=XM_003996044.5;refGene_gn=SURF6;refGene_tx=XM_019816754.2
D4 94299795 94300275
refGene_gn=SURF6;refGene_tx=XM_003996044.5;refGene_gn=SURF6;refGene_tx=XM_019816754.2
D4 94309812 94310134
refGene_gn=LOC101087791;refGene_tx=XM_019816747.2
D4 94315998 94316071
refGene_gn=LOC101087791;refGene_tx=XM_019816747.2
D4 94317437 94317488
refGene_gn=LOC101087791;refGene_tx=XM_019816747.2
D4 94318070 94318121
refGene_gn=LOC101087791;refGene_tx=XM_019816747.2
D4 94318366 94318423
refGene_gn=LOC101087791;refGene_tx=XM_019816747.2
D4 94318833 94318881
refGene_gn=LOC101087791;refGene_tx=XM_019816747.2
D4 94319724 94319760
refGene_gn=LOC101087791;refGene_tx=XM_019816747.2
D4 94321966 94322095
refGene_gn=LOC101087791;refGene_tx=XM_019816747.2
D4 94329094 94329782
refGene_gn=LOC101087791;refGene_tx=XM_019816747.2
D4 94348432 94348457
refGene_gn=LOC105260069;refGene_tx=XM_023242322.1
D4 94348530 94348600
refGene_gn=LOC105260069;refGene_tx=XM_023242322.1
D4 94349180 94349234
refGene_gn=LOC105260069;refGene_tx=XM_023242322.1
D4 94351369 94351417
refGene_gn=LOC105260069;refGene_tx=XM_023242322.1
D4 94353201 94353237

refGene_gn=LOC105260069;refGene_tx=XM_023242322.1
D4	94353765	94353901
refGene_gn=LOC105260069;refGene_tx=XM_023242322.1
D4	94448525	94448707
refGene_gn=AGPAT2;refGene_tx=XM_023242995.1;refGene_gn=AGPAT2;refGene_tx=XM_023242996.1
D4	94456323	94456457
refGene_gn=AGPAT2;refGene_tx=XM_023242995.1;refGene_gn=AGPAT2;refGene_tx=XM_023242996.1
D4	94456869	94457045
refGene_gn=AGPAT2;refGene_tx=XM_023242995.1;refGene_gn=AGPAT2;refGene_tx=XM_023242996.1
D4	94458088	94458184
refGene_gn=AGPAT2;refGene_tx=XM_023242995.1;refGene_gn=AGPAT2;refGene_tx=XM_023242996.1
D4	94459325	94459472	refGene_gn=AGPAT2;refGene_tx=XM_023242995.1
D4	94459661	94459734
refGene_gn=AGPAT2;refGene_tx=XM_023242995.1;refGene_gn=AGPAT2;refGene_tx=XM_023242996.1
D4	94460566	94460742
refGene_gn=AGPAT2;refGene_tx=XM_023242995.1;refGene_gn=AGPAT2;refGene_tx=XM_023242996.1
D4	94461956	94461979
refGene_gn=EGFL7;refGene_tx=XM_019816758.2;refGene_gn=EGFL7;refGene_tx=XM_023242997.1;refGene_gn=EGFL7;refGene_tx=XM_023242998.1
D4	94462126	94462327
refGene_gn=EGFL7;refGene_tx=XM_006939646.3;refGene_gn=EGFL7;refGene_tx=XM_019816758.2;refGene_gn=EGFL7;refGene_tx=XM_023242997.1;refGene_gn=EGFL7;refGene_tx=XM_023242998.1
D4	94462819	94462884
refGene_gn=EGFL7;refGene_tx=XM_006939646.3;refGene_gn=EGFL7;refGene_tx=XM_019816758.2;refGene_gn=EGFL7;refGene_tx=XM_023242997.1;refGene_gn=EGFL7;refGene_tx=XM_023242998.1
D4	94463598	94463763
refGene_gn=EGFL7;refGene_tx=XM_006939646.3;refGene_gn=EGFL7;refGene_tx=XM_019816758.2;refGene_gn=EGFL7;refGene_tx=XM_023242997.1;refGene_gn=EGFL7;refGene_tx=XM_023242998.1
D4	94463902	94463998
refGene_gn=EGFL7;refGene_tx=XM_006939646.3;refGene_gn=EGFL7;refGene_tx=XM_019816758.2;refGene_gn=EGFL7;refGene_tx=XM_023242997.1;refGene_gn=EGFL7;refGene_tx=XM_023242998.1
D4	94464234	94464350
refGene_gn=EGFL7;refGene_tx=XM_006939646.3;refGene_gn=EGFL7;refGene_tx=XM_019816758.2;refGene_gn=EGFL7;refGene_tx=XM_023242997.1;refGene_gn=EGFL7;refGene_tx=XM_023242998.1
D4	94465253	94465370
refGene_gn=EGFL7;refGene_tx=XM_006939646.3;refGene_gn=EGFL7;refGene_tx=XM_019816758.2;refGene_gn=EGFL7;refGene_tx=XM_023242997.1;refGene_gn=EGFL7;refGene_tx=XM_023242998.1
D4	94465494	94465574

refGene_gn=EGFL7;refGene_tx=XM_006939646.3;refGene_gn=EGFL7;refGene_tx=XM_019816758.2;refGene_gn=EGFL7;refGene_tx=XM_023242997.1;refGene_gn=EGFL7;refGene_tx=XM_023242998.1
D4 94474729 94475094
refGene_gn=LOC111557484;refGene_tx=XM_023242323.1
D4 94476396 94476491
refGene_gn=LOC111557484;refGene_tx=XM_023242323.1
D4 94477507 94477938
refGene_gn=LOC111557484;refGene_tx=XM_023242323.1
D4 94479624 94480239
refGene_gn=LOC111557484;refGene_tx=XM_023242323.1
D4 94531080 94531147
refGene_gn=LOC109494050;refGene_tx=XM_023243000.1
D4 94531482 94531877
refGene_gn=LOC109494050;refGene_tx=XM_023243000.1
D4 94532202 94532331
refGene_gn=LOC109494050;refGene_tx=XM_023243000.1
D4 94533519 94533637
refGene_gn=LOC109494050;refGene_tx=XM_023243000.1
D4 94535466 94535578
refGene_gn=LOC109494050;refGene_tx=XM_023243000.1
D4 94535952 94536181
refGene_gn=LOC109494050;refGene_tx=XM_023243000.1
D4 94556608 94556669 refGene_gn=NOTCH1;refGene_tx=XM_023243001.1
D4 94558357 94558439 refGene_gn=NOTCH1;refGene_tx=XM_023243001.1
D4 94574749 94575012 refGene_gn=NOTCH1;refGene_tx=XM_023243001.1
D4 94575503 94575842 refGene_gn=NOTCH1;refGene_tx=XM_023243001.1
D4 94577892 94578015 refGene_gn=NOTCH1;refGene_tx=XM_023243001.1
D4 94578429 94578663 refGene_gn=NOTCH1;refGene_tx=XM_023243001.1
D4 94579018 94579174 refGene_gn=NOTCH1;refGene_tx=XM_023243001.1
D4 94579401 94579587 refGene_gn=NOTCH1;refGene_tx=XM_023243001.1
D4 94579898 94580012 refGene_gn=NOTCH1;refGene_tx=XM_023243001.1
D4 94580792 94580906 refGene_gn=NOTCH1;refGene_tx=XM_023243001.1
D4 94581071 94581305 refGene_gn=NOTCH1;refGene_tx=XM_023243001.1
D4 94581402 94581513 refGene_gn=NOTCH1;refGene_tx=XM_023243001.1
D4 94582057 94582250 refGene_gn=NOTCH1;refGene_tx=XM_023243001.1
D4 94583428 94583574 refGene_gn=NOTCH1;refGene_tx=XM_023243001.1
D4 94583789 94583903 refGene_gn=NOTCH1;refGene_tx=XM_023243001.1
D4 94585566 94585686 refGene_gn=NOTCH1;refGene_tx=XM_023243001.1
D4 94585987 94586140 refGene_gn=NOTCH1;refGene_tx=XM_023243001.1
D4 94586704 94586933 refGene_gn=NOTCH1;refGene_tx=XM_023243001.1
D4 94587554 94587756 refGene_gn=NOTCH1;refGene_tx=XM_023243001.1
D4 94588390 94588544 refGene_gn=NOTCH1;refGene_tx=XM_023243001.1
D4 94588653 94588838 refGene_gn=NOTCH1;refGene_tx=XM_023243001.1
D4 94589356 94589489 refGene_gn=NOTCH1;refGene_tx=XM_023243001.1
D4 94589817 94590075 refGene_gn=NOTCH1;refGene_tx=XM_023243001.1
D4 94590148 94590261 refGene_gn=NOTCH1;refGene_tx=XM_023243001.1
D4 94590775 94591347 refGene_gn=NOTCH1;refGene_tx=XM_023243001.1
D4 94591579 94592008 refGene_gn=NOTCH1;refGene_tx=XM_023243001.1
D4 94592528 94592677 refGene_gn=NOTCH1;refGene_tx=XM_023243001.1

D4 94593406 94593623 refGene_gn=NOTCH1;refGene_tx=XM_023243001.1
D4 94593791 94593879 refGene_gn=NOTCH1;refGene_tx=XM_023243001.1
D4 94593976 94594142 refGene_gn=NOTCH1;refGene_tx=XM_023243001.1
D4 94594654 94594950 refGene_gn=NOTCH1;refGene_tx=XM_023243001.1
D4 94596028 94596176 refGene_gn=NOTCH1;refGene_tx=XM_023243001.1
D4 94596262 94596360 refGene_gn=NOTCH1;refGene_tx=XM_023243001.1
D4 94597500 94598952 refGene_gn=NOTCH1;refGene_tx=XM_023243001.1
D4 94614600 94618158
refGene_gn=SEC16A;refGene_tx=XM_011288765.3;refGene_gn=SEC16A;refGene_
tx=XM_011288769.3;refGene_gn=SEC16A;refGene_tx=XM_011288770.3;refGene_
gn=SEC16A;refGene_tx=XM_019816769.2;refGene_gn=SEC16A;refGene_tx=XM_01
9816770.2;refGene_gn=SEC16A;refGene_tx=XM_019816771.2;refGene_gn=SEC16
A;refGene_tx=XM_019816772.2;refGene_gn=SEC16A;refGene_tx=XM_019816774.
2;refGene_gn=SEC16A;refGene_tx=XM_019816775.2
D4 94619931 94620071
refGene_gn=SEC16A;refGene_tx=XM_011288765.3;refGene_gn=SEC16A;refGene_
tx=XM_011288769.3;refGene_gn=SEC16A;refGene_tx=XM_011288770.3;refGene_
gn=SEC16A;refGene_tx=XM_019816769.2;refGene_gn=SEC16A;refGene_tx=XM_01
9816770.2;refGene_gn=SEC16A;refGene_tx=XM_019816771.2;refGene_gn=SEC16
A;refGene_tx=XM_019816772.2;refGene_gn=SEC16A;refGene_tx=XM_019816774.
2;refGene_gn=SEC16A;refGene_tx=XM_019816775.2
D4 94622473 94622571
refGene_gn=SEC16A;refGene_tx=XM_011288765.3;refGene_gn=SEC16A;refGene_
tx=XM_011288769.3;refGene_gn=SEC16A;refGene_tx=XM_011288770.3;refGene_
gn=SEC16A;refGene_tx=XM_019816769.2;refGene_gn=SEC16A;refGene_tx=XM_01
9816770.2;refGene_gn=SEC16A;refGene_tx=XM_019816771.2;refGene_gn=SEC16
A;refGene_tx=XM_019816772.2;refGene_gn=SEC16A;refGene_tx=XM_019816774.
2;refGene_gn=SEC16A;refGene_tx=XM_019816775.2
D4 94623927 94624054
refGene_gn=SEC16A;refGene_tx=XM_011288765.3;refGene_gn=SEC16A;refGene_
tx=XM_011288769.3;refGene_gn=SEC16A;refGene_tx=XM_011288770.3;refGene_
gn=SEC16A;refGene_tx=XM_019816769.2;refGene_gn=SEC16A;refGene_tx=XM_01
9816770.2;refGene_gn=SEC16A;refGene_tx=XM_019816771.2;refGene_gn=SEC16
A;refGene_tx=XM_019816772.2;refGene_gn=SEC16A;refGene_tx=XM_019816774.
2;refGene_gn=SEC16A;refGene_tx=XM_019816775.2
D4 94624570 94624742
refGene_gn=SEC16A;refGene_tx=XM_011288765.3;refGene_gn=SEC16A;refGene_
tx=XM_011288769.3;refGene_gn=SEC16A;refGene_tx=XM_011288770.3;refGene_
gn=SEC16A;refGene_tx=XM_019816769.2;refGene_gn=SEC16A;refGene_tx=XM_01
9816770.2;refGene_gn=SEC16A;refGene_tx=XM_019816771.2;refGene_gn=SEC16
A;refGene_tx=XM_019816772.2;refGene_gn=SEC16A;refGene_tx=XM_019816774.
2;refGene_gn=SEC16A;refGene_tx=XM_019816775.2
D4 94624831 94624997
refGene_gn=SEC16A;refGene_tx=XM_011288765.3;refGene_gn=SEC16A;refGene_
tx=XM_011288769.3;refGene_gn=SEC16A;refGene_tx=XM_011288770.3;refGene_
gn=SEC16A;refGene_tx=XM_019816769.2;refGene_gn=SEC16A;refGene_tx=XM_01
9816770.2;refGene_gn=SEC16A;refGene_tx=XM_019816771.2;refGene_gn=SEC16
A;refGene_tx=XM_019816772.2;refGene_gn=SEC16A;refGene_tx=XM_019816774.
2;refGene_gn=SEC16A;refGene_tx=XM_019816775.2
D4 94625871 94626014

refGene_gn=SEC16A;refGene_tx=XM_011288765.3;refGene_gn=SEC16A;refGene_
tx=XM_011288769.3;refGene_gn=SEC16A;refGene_tx=XM_011288770.3;refGene_
gn=SEC16A;refGene_tx=XM_019816769.2;refGene_gn=SEC16A;refGene_tx=XM_01
9816770.2;refGene_gn=SEC16A;refGene_tx=XM_019816771.2;refGene_gn=SEC16
A;refGene_tx=XM_019816772.2;refGene_gn=SEC16A;refGene_tx=XM_019816774.
2;refGene_gn=SEC16A;refGene_tx=XM_019816775.2
D4 94626586 94626648
refGene_gn=SEC16A;refGene_tx=XM_011288765.3;refGene_gn=SEC16A;refGene_
tx=XM_011288769.3;refGene_gn=SEC16A;refGene_tx=XM_011288770.3;refGene_
gn=SEC16A;refGene_tx=XM_019816769.2;refGene_gn=SEC16A;refGene_tx=XM_01
9816770.2;refGene_gn=SEC16A;refGene_tx=XM_019816771.2;refGene_gn=SEC16
A;refGene_tx=XM_019816772.2;refGene_gn=SEC16A;refGene_tx=XM_019816774.
2;refGene_gn=SEC16A;refGene_tx=XM_019816775.2
D4 94626756 94626895
refGene_gn=SEC16A;refGene_tx=XM_011288765.3;refGene_gn=SEC16A;refGene_
tx=XM_011288769.3;refGene_gn=SEC16A;refGene_tx=XM_011288770.3;refGene_
gn=SEC16A;refGene_tx=XM_019816769.2;refGene_gn=SEC16A;refGene_tx=XM_01
9816770.2;refGene_gn=SEC16A;refGene_tx=XM_019816771.2;refGene_gn=SEC16
A;refGene_tx=XM_019816772.2;refGene_gn=SEC16A;refGene_tx=XM_019816774.
2;refGene_gn=SEC16A;refGene_tx=XM_019816775.2
D4 94627301 94627547
refGene_gn=SEC16A;refGene_tx=XM_011288765.3;refGene_gn=SEC16A;refGene_
tx=XM_011288769.3;refGene_gn=SEC16A;refGene_tx=XM_011288770.3;refGene_
gn=SEC16A;refGene_tx=XM_019816769.2;refGene_gn=SEC16A;refGene_tx=XM_01
9816770.2;refGene_gn=SEC16A;refGene_tx=XM_019816771.2;refGene_gn=SEC16
A;refGene_tx=XM_019816772.2;refGene_gn=SEC16A;refGene_tx=XM_019816774.
2;refGene_gn=SEC16A;refGene_tx=XM_019816775.2
D4 94628593 94628691
refGene_gn=SEC16A;refGene_tx=XM_011288765.3;refGene_gn=SEC16A;refGene_
tx=XM_011288769.3;refGene_gn=SEC16A;refGene_tx=XM_011288770.3;refGene_
gn=SEC16A;refGene_tx=XM_019816769.2;refGene_gn=SEC16A;refGene_tx=XM_01
9816770.2;refGene_gn=SEC16A;refGene_tx=XM_019816771.2;refGene_gn=SEC16
A;refGene_tx=XM_019816772.2;refGene_gn=SEC16A;refGene_tx=XM_019816774.
2;refGene_gn=SEC16A;refGene_tx=XM_019816775.2
D4 94629068 94629150
refGene_gn=SEC16A;refGene_tx=XM_011288765.3;refGene_gn=SEC16A;refGene_
tx=XM_011288769.3;refGene_gn=SEC16A;refGene_tx=XM_011288770.3;refGene_
gn=SEC16A;refGene_tx=XM_019816769.2;refGene_gn=SEC16A;refGene_tx=XM_01
9816770.2;refGene_gn=SEC16A;refGene_tx=XM_019816771.2;refGene_gn=SEC16
A;refGene_tx=XM_019816772.2;refGene_gn=SEC16A;refGene_tx=XM_019816774.
2;refGene_gn=SEC16A;refGene_tx=XM_019816775.2
D4 94629361 94629479
refGene_gn=SEC16A;refGene_tx=XM_011288765.3;refGene_gn=SEC16A;refGene_
tx=XM_011288769.3;refGene_gn=SEC16A;refGene_tx=XM_011288770.3;refGene_
gn=SEC16A;refGene_tx=XM_019816769.2;refGene_gn=SEC16A;refGene_tx=XM_01
9816770.2;refGene_gn=SEC16A;refGene_tx=XM_019816771.2;refGene_gn=SEC16
A;refGene_tx=XM_019816772.2;refGene_gn=SEC16A;refGene_tx=XM_019816774.
2;refGene_gn=SEC16A;refGene_tx=XM_019816775.2
D4 94629657 94629769
refGene_gn=SEC16A;refGene_tx=XM_011288765.3;refGene_gn=SEC16A;refGene_

tx=XM_011288769.3;refGene_gn=SEC16A;refGene_tx=XM_011288770.3;refGene_gn=SEC16A;refGene_tx=XM_019816769.2;refGene_gn=SEC16A;refGene_tx=XM_019816770.2;refGene_gn=SEC16A;refGene_tx=XM_019816771.2;refGene_gn=SEC16A;refGene_tx=XM_019816772.2;refGene_gn=SEC16A;refGene_tx=XM_019816774.2;refGene_gn=SEC16A;refGene_tx=XM_019816775.2
D4 94629933 94630039
refGene_gn=SEC16A;refGene_tx=XM_011288765.3;refGene_gn=SEC16A;refGene_tx=XM_011288769.3;refGene_gn=SEC16A;refGene_tx=XM_011288770.3;refGene_gn=SEC16A;refGene_tx=XM_019816769.2;refGene_gn=SEC16A;refGene_tx=XM_019816770.2;refGene_gn=SEC16A;refGene_tx=XM_019816771.2;refGene_gn=SEC16A;refGene_tx=XM_019816772.2;refGene_gn=SEC16A;refGene_tx=XM_019816774.2;refGene_gn=SEC16A;refGene_tx=XM_019816775.2
D4 94630682 94630823
refGene_gn=SEC16A;refGene_tx=XM_011288765.3;refGene_gn=SEC16A;refGene_tx=XM_011288769.3;refGene_gn=SEC16A;refGene_tx=XM_011288770.3;refGene_gn=SEC16A;refGene_tx=XM_019816769.2;refGene_gn=SEC16A;refGene_tx=XM_019816770.2;refGene_gn=SEC16A;refGene_tx=XM_019816771.2;refGene_gn=SEC16A;refGene_tx=XM_019816772.2;refGene_gn=SEC16A;refGene_tx=XM_019816774.2;refGene_gn=SEC16A;refGene_tx=XM_019816775.2
D4 94631800 94631914
refGene_gn=SEC16A;refGene_tx=XM_011288765.3;refGene_gn=SEC16A;refGene_tx=XM_011288769.3;refGene_gn=SEC16A;refGene_tx=XM_011288770.3;refGene_gn=SEC16A;refGene_tx=XM_019816769.2;refGene_gn=SEC16A;refGene_tx=XM_019816770.2;refGene_gn=SEC16A;refGene_tx=XM_019816771.2;refGene_gn=SEC16A;refGene_tx=XM_019816772.2;refGene_gn=SEC16A;refGene_tx=XM_019816774.2;refGene_gn=SEC16A;refGene_tx=XM_019816775.2
D4 94632157 94632335
refGene_gn=SEC16A;refGene_tx=XM_011288765.3;refGene_gn=SEC16A;refGene_tx=XM_011288769.3;refGene_gn=SEC16A;refGene_tx=XM_011288770.3;refGene_gn=SEC16A;refGene_tx=XM_019816769.2;refGene_gn=SEC16A;refGene_tx=XM_019816770.2;refGene_gn=SEC16A;refGene_tx=XM_019816771.2;refGene_gn=SEC16A;refGene_tx=XM_019816772.2;refGene_gn=SEC16A;refGene_tx=XM_019816774.2;refGene_gn=SEC16A;refGene_tx=XM_019816775.2
D4 94632824 94633070
refGene_gn=SEC16A;refGene_tx=XM_011288765.3;refGene_gn=SEC16A;refGene_tx=XM_011288769.3;refGene_gn=SEC16A;refGene_tx=XM_011288770.3;refGene_gn=SEC16A;refGene_tx=XM_019816769.2;refGene_gn=SEC16A;refGene_tx=XM_019816770.2;refGene_gn=SEC16A;refGene_tx=XM_019816771.2;refGene_gn=SEC16A;refGene_tx=XM_019816772.2;refGene_gn=SEC16A;refGene_tx=XM_019816774.2;refGene_gn=SEC16A;refGene_tx=XM_019816775.2
D4 94633491 94633574
refGene_gn=SEC16A;refGene_tx=XM_011288765.3;refGene_gn=SEC16A;refGene_tx=XM_011288769.3;refGene_gn=SEC16A;refGene_tx=XM_011288770.3;refGene_gn=SEC16A;refGene_tx=XM_019816769.2;refGene_gn=SEC16A;refGene_tx=XM_019816770.2;refGene_gn=SEC16A;refGene_tx=XM_019816771.2;refGene_gn=SEC16A;refGene_tx=XM_019816772.2;refGene_gn=SEC16A;refGene_tx=XM_019816774.2;refGene_gn=SEC16A;refGene_tx=XM_019816775.2
D4 94634478 94634625
refGene_gn=SEC16A;refGene_tx=XM_011288765.3;refGene_gn=SEC16A;refGene_tx=XM_011288769.3;refGene_gn=SEC16A;refGene_tx=XM_011288770.3;refGene_

gn=SEC16A;refGene_tx=XM_019816769.2;refGene_gn=SEC16A;refGene_tx=XM_01
9816770.2;refGene_gn=SEC16A;refGene_tx=XM_019816771.2;refGene_gn=SEC16
A;refGene_tx=XM_019816772.2;refGene_gn=SEC16A;refGene_tx=XM_019816774.
2;refGene_gn=SEC16A;refGene_tx=XM_019816775.2
D4 94636371 94636452
refGene_gn=SEC16A;refGene_tx=XM_011288765.3;refGene_gn=SEC16A;refGene_
tx=XM_011288769.3;refGene_gn=SEC16A;refGene_tx=XM_011288770.3;refGene_
gn=SEC16A;refGene_tx=XM_019816769.2;refGene_gn=SEC16A;refGene_tx=XM_01
9816770.2;refGene_gn=SEC16A;refGene_tx=XM_019816771.2;refGene_gn=SEC16
A;refGene_tx=XM_019816772.2;refGene_gn=SEC16A;refGene_tx=XM_019816774.
2;refGene_gn=SEC16A;refGene_tx=XM_019816775.2
D4 94636649 94636706
refGene_gn=SEC16A;refGene_tx=XM_011288765.3;refGene_gn=SEC16A;refGene_
tx=XM_011288769.3;refGene_gn=SEC16A;refGene_tx=XM_011288770.3;refGene_
gn=SEC16A;refGene_tx=XM_019816769.2;refGene_gn=SEC16A;refGene_tx=XM_01
9816770.2;refGene_gn=SEC16A;refGene_tx=XM_019816771.2;refGene_gn=SEC16
A;refGene_tx=XM_019816772.2;refGene_gn=SEC16A;refGene_tx=XM_019816774.
2;refGene_gn=SEC16A;refGene_tx=XM_019816775.2
D4 94636843 94636955
refGene_gn=SEC16A;refGene_tx=XM_011288765.3;refGene_gn=SEC16A;refGene_
tx=XM_011288769.3;refGene_gn=SEC16A;refGene_tx=XM_011288770.3;refGene_
gn=SEC16A;refGene_tx=XM_019816769.2;refGene_gn=SEC16A;refGene_tx=XM_01
9816770.2;refGene_gn=SEC16A;refGene_tx=XM_019816771.2;refGene_gn=SEC16
A;refGene_tx=XM_019816772.2;refGene_gn=SEC16A;refGene_tx=XM_019816774.
2;refGene_gn=SEC16A;refGene_tx=XM_019816775.2
D4 94637279 94637417
refGene_gn=SEC16A;refGene_tx=XM_011288765.3;refGene_gn=SEC16A;refGene_
tx=XM_011288769.3;refGene_gn=SEC16A;refGene_tx=XM_011288770.3;refGene_
gn=SEC16A;refGene_tx=XM_019816769.2;refGene_gn=SEC16A;refGene_tx=XM_01
9816770.2;refGene_gn=SEC16A;refGene_tx=XM_019816771.2;refGene_gn=SEC16
A;refGene_tx=XM_019816772.2;refGene_gn=SEC16A;refGene_tx=XM_019816774.
2;refGene_gn=SEC16A;refGene_tx=XM_019816775.2
D4 94637664 94637759
refGene_gn=SEC16A;refGene_tx=XM_011288765.3;refGene_gn=SEC16A;refGene_
tx=XM_011288769.3;refGene_gn=SEC16A;refGene_tx=XM_011288770.3;refGene_
gn=SEC16A;refGene_tx=XM_019816769.2;refGene_gn=SEC16A;refGene_tx=XM_01
9816770.2;refGene_gn=SEC16A;refGene_tx=XM_019816771.2;refGene_gn=SEC16
A;refGene_tx=XM_019816772.2;refGene_gn=SEC16A;refGene_tx=XM_019816774.
2;refGene_gn=SEC16A;refGene_tx=XM_019816775.2
D4 94638616 94638691
refGene_gn=SEC16A;refGene_tx=XM_011288765.3;refGene_gn=SEC16A;refGene_
tx=XM_011288769.3;refGene_gn=SEC16A;refGene_tx=XM_011288770.3;refGene_
gn=SEC16A;refGene_tx=XM_019816769.2;refGene_gn=SEC16A;refGene_tx=XM_01
9816770.2;refGene_gn=SEC16A;refGene_tx=XM_019816771.2;refGene_gn=SEC16
A;refGene_tx=XM_019816772.2;refGene_gn=SEC16A;refGene_tx=XM_019816774.
2;refGene_gn=SEC16A;refGene_tx=XM_019816775.2
D4 94639638 94639698
refGene_gn=SEC16A;refGene_tx=XM_011288765.3;refGene_gn=SEC16A;refGene_
tx=XM_011288769.3;refGene_gn=SEC16A;refGene_tx=XM_019816769.2;refGene_
gn=SEC16A;refGene_tx=XM_019816770.2;refGene_gn=SEC16A;refGene_tx=XM_01

9816771.2;refGene_gn=SEC16A;refGene_tx=XM_019816772.2;refGene_gn=SEC16A;refGene_tx=XM_019816774.2
D4 94640571 94640820
refGene_gn=SEC16A;refGene_tx=XM_011288765.3;refGene_gn=SEC16A;refGene_tx=XM_011288769.3;refGene_gn=SEC16A;refGene_tx=XM_011288770.3;refGene_gn=SEC16A;refGene_tx=XM_019816769.2;refGene_gn=SEC16A;refGene_tx=XM_019816770.2;refGene_gn=SEC16A;refGene_tx=XM_019816771.2;refGene_gn=SEC16A;refGene_tx=XM_019816772.2;refGene_gn=SEC16A;refGene_tx=XM_019816774.2;refGene_gn=SEC16A;refGene_tx=XM_019816775.2
D4 94641648 94641717
refGene_gn=SEC16A;refGene_tx=XM_011288765.3;refGene_gn=SEC16A;refGene_tx=XM_011288769.3;refGene_gn=SEC16A;refGene_tx=XM_011288770.3
D4 94644482 94645270
refGene_gn=INPP5E;refGene_tx=XM_006939643.4;refGene_gn=INPP5E;refGene_tx=XM_023243017.1
D4 94649461 94649585
refGene_gn=INPP5E;refGene_tx=XM_006939643.4;refGene_gn=INPP5E;refGene_tx=XM_023243017.1
D4 94650784 94650882
refGene_gn=INPP5E;refGene_tx=XM_006939643.4;refGene_gn=INPP5E;refGene_tx=XM_023243017.1
D4 94651391 94651516
refGene_gn=INPP5E;refGene_tx=XM_006939643.4;refGene_gn=INPP5E;refGene_tx=XM_023243017.1
D4 94651636 94651756
refGene_gn=INPP5E;refGene_tx=XM_006939643.4;refGene_gn=INPP5E;refGene_tx=XM_023243017.1
D4 94651969 94652077
refGene_gn=INPP5E;refGene_tx=XM_006939643.4;refGene_gn=INPP5E;refGene_tx=XM_023243017.1
D4 94652641 94652803
refGene_gn=INPP5E;refGene_tx=XM_006939643.4;refGene_gn=INPP5E;refGene_tx=XM_023243017.1
D4 94652895 94653011
refGene_gn=INPP5E;refGene_tx=XM_006939643.4;refGene_gn=INPP5E;refGene_tx=XM_023243017.1
D4 94653558 94653695
refGene_gn=INPP5E;refGene_tx=XM_006939643.4;refGene_gn=INPP5E;refGene_tx=XM_023243017.1
D4 94654003 94654136
refGene_gn=INPP5E;refGene_tx=XM_006939643.4;refGene_gn=INPP5E;refGene_tx=XM_023243017.1
D4 94657004 94657174 refGene_gn=PMPCA;refGene_tx=XM_023243018.1
D4 94658135 94658280 refGene_gn=PMPCA;refGene_tx=XM_023243018.1
D4 94659939 94660002 refGene_gn=PMPCA;refGene_tx=XM_023243018.1
D4 94660165 94660256 refGene_gn=PMPCA;refGene_tx=XM_023243018.1
D4 94660368 94660487 refGene_gn=PMPCA;refGene_tx=XM_023243018.1
D4 94660566 94660659 refGene_gn=PMPCA;refGene_tx=XM_023243018.1
D4 94661000 94661264 refGene_gn=PMPCA;refGene_tx=XM_023243018.1
D4 94661903 94662004 refGene_gn=PMPCA;refGene_tx=XM_023243018.1

D4 94662989 94663084 refGene_gn=PMPCA;refGene_tx=XM_023243018.1
D4 94663990 94664073 refGene_gn=PMPCA;refGene_tx=XM_023243018.1
D4 94664309 94664389 refGene_gn=PMPCA;refGene_tx=XM_023243018.1
D4 94664772 94664975 refGene_gn=PMPCA;refGene_tx=XM_023243018.1
D4 94666127 94666204 refGene_gn=PMPCA;refGene_tx=XM_023243018.1
D4 94666457 94666605
refGene_gn=SDCCAG3;refGene_tx=XM_023243019.1;refGene_gn=SDCCAG3;refGen
e_tx=XM_023243020.1;refGene_gn=SDCCAG3;refGene_tx=XM_023243021.1
D4 94666660 94666810 refGene_gn=SDCCAG3;refGene_tx=XM_023243019.1
D4 94667471 94667540
refGene_gn=SDCCAG3;refGene_tx=XM_023243019.1;refGene_gn=SDCCAG3;refGen
e_tx=XM_023243020.1;refGene_gn=SDCCAG3;refGene_tx=XM_023243021.1
D4 94668326 94668442
refGene_gn=SDCCAG3;refGene_tx=XM_023243019.1;refGene_gn=SDCCAG3;refGen
e_tx=XM_023243020.1;refGene_gn=SDCCAG3;refGene_tx=XM_023243021.1
D4 94668703 94669111
refGene_gn=SDCCAG3;refGene_tx=XM_023243019.1;refGene_gn=SDCCAG3;refGen
e_tx=XM_023243020.1;refGene_gn=SDCCAG3;refGene_tx=XM_023243021.1
D4 94669678 94669752
refGene_gn=SDCCAG3;refGene_tx=XM_023243019.1;refGene_gn=SDCCAG3;refGen
e_tx=XM_023243020.1;refGene_gn=SDCCAG3;refGene_tx=XM_023243021.1
D4 94670118 94670230
refGene_gn=SDCCAG3;refGene_tx=XM_023243019.1;refGene_gn=SDCCAG3;refGen
e_tx=XM_023243020.1;refGene_gn=SDCCAG3;refGene_tx=XM_023243021.1
D4 94670448 94670511
refGene_gn=SDCCAG3;refGene_tx=XM_023243019.1;refGene_gn=SDCCAG3;refGen
e_tx=XM_023243020.1;refGene_gn=SDCCAG3;refGene_tx=XM_023243021.1
D4 94670947 94671087
refGene_gn=SDCCAG3;refGene_tx=XM_023243019.1;refGene_gn=SDCCAG3;refGen
e_tx=XM_023243020.1;refGene_gn=SDCCAG3;refGene_tx=XM_023243021.1
D4 94671662 94671762
refGene_gn=SDCCAG3;refGene_tx=XM_023243019.1;refGene_gn=SDCCAG3;refGen
e_tx=XM_023243020.1;refGene_gn=SDCCAG3;refGene_tx=XM_023243021.1
D4 94675218 94675348
refGene_gn=SNAPC4;refGene_tx=XM_023243004.1;refGene_gn=SNAPC4;refGene_
tx=XM_023243005.1;refGene_gn=SNAPC4;refGene_tx=XM_023243007.1;refGene_
gn=SNAPC4;refGene_tx=XM_023243008.1;refGene_gn=SNAPC4;refGene_tx=XM_02
3243010.1
D4 94676134 94676181
refGene_gn=SNAPC4;refGene_tx=XM_023243004.1;refGene_gn=SNAPC4;refGene_
tx=XM_023243005.1;refGene_gn=SNAPC4;refGene_tx=XM_023243007.1;refGene_
gn=SNAPC4;refGene_tx=XM_023243008.1;refGene_gn=SNAPC4;refGene_tx=XM_02
3243010.1
D4 94677626 94677791
refGene_gn=SNAPC4;refGene_tx=XM_023243004.1;refGene_gn=SNAPC4;refGene_
tx=XM_023243005.1;refGene_gn=SNAPC4;refGene_tx=XM_023243007.1;refGene_
gn=SNAPC4;refGene_tx=XM_023243008.1;refGene_gn=SNAPC4;refGene_tx=XM_02
3243009.1;refGene_gn=SNAPC4;refGene_tx=XM_023243010.1
D4 94677937 94678063
refGene_gn=SNAPC4;refGene_tx=XM_023243004.1;refGene_gn=SNAPC4;refGene_

tx=XM_023243005.1;refGene_gn=SNAPC4;refGene_tx=XM_023243007.1;refGene_gn=SNAPC4;refGene_tx=XM_023243008.1;refGene_gn=SNAPC4;refGene_tx=XM_023243009.1;refGene_gn=SNAPC4;refGene_tx=XM_023243010.1
D4 94678604 94678683
refGene_gn=SNAPC4;refGene_tx=XM_023243004.1;refGene_gn=SNAPC4;refGene_tx=XM_023243005.1;refGene_gn=SNAPC4;refGene_tx=XM_023243007.1;refGene_gn=SNAPC4;refGene_tx=XM_023243008.1;refGene_gn=SNAPC4;refGene_tx=XM_023243009.1;refGene_gn=SNAPC4;refGene_tx=XM_023243010.1
D4 94679040 94679122
refGene_gn=SNAPC4;refGene_tx=XM_023243004.1;refGene_gn=SNAPC4;refGene_tx=XM_023243005.1;refGene_gn=SNAPC4;refGene_tx=XM_023243007.1;refGene_gn=SNAPC4;refGene_tx=XM_023243008.1;refGene_gn=SNAPC4;refGene_tx=XM_023243009.1;refGene_gn=SNAPC4;refGene_tx=XM_023243010.1
D4 94679957 94680062
refGene_gn=SNAPC4;refGene_tx=XM_023243004.1;refGene_gn=SNAPC4;refGene_tx=XM_023243005.1;refGene_gn=SNAPC4;refGene_tx=XM_023243007.1;refGene_gn=SNAPC4;refGene_tx=XM_023243008.1;refGene_gn=SNAPC4;refGene_tx=XM_023243009.1;refGene_gn=SNAPC4;refGene_tx=XM_023243010.1
D4 94680148 94680221
refGene_gn=SNAPC4;refGene_tx=XM_023243004.1;refGene_gn=SNAPC4;refGene_tx=XM_023243005.1;refGene_gn=SNAPC4;refGene_tx=XM_023243007.1;refGene_gn=SNAPC4;refGene_tx=XM_023243008.1;refGene_gn=SNAPC4;refGene_tx=XM_023243009.1;refGene_gn=SNAPC4;refGene_tx=XM_023243010.1
D4 94680587 94680752
refGene_gn=SNAPC4;refGene_tx=XM_023243004.1;refGene_gn=SNAPC4;refGene_tx=XM_023243005.1;refGene_gn=SNAPC4;refGene_tx=XM_023243007.1;refGene_gn=SNAPC4;refGene_tx=XM_023243008.1;refGene_gn=SNAPC4;refGene_tx=XM_023243009.1;refGene_gn=SNAPC4;refGene_tx=XM_023243010.1
D4 94683161 94683309
refGene_gn=SNAPC4;refGene_tx=XM_023243004.1;refGene_gn=SNAPC4;refGene_tx=XM_023243005.1;refGene_gn=SNAPC4;refGene_tx=XM_023243007.1;refGene_gn=SNAPC4;refGene_tx=XM_023243008.1;refGene_gn=SNAPC4;refGene_tx=XM_023243009.1;refGene_gn=SNAPC4;refGene_tx=XM_023243010.1
D4 94683437 94683544
refGene_gn=SNAPC4;refGene_tx=XM_023243004.1;refGene_gn=SNAPC4;refGene_tx=XM_023243005.1;refGene_gn=SNAPC4;refGene_tx=XM_023243007.1;refGene_gn=SNAPC4;refGene_tx=XM_023243008.1;refGene_gn=SNAPC4;refGene_tx=XM_023243009.1;refGene_gn=SNAPC4;refGene_tx=XM_023243010.1
D4 94683678 94683773
refGene_gn=SNAPC4;refGene_tx=XM_023243004.1;refGene_gn=SNAPC4;refGene_tx=XM_023243005.1;refGene_gn=SNAPC4;refGene_tx=XM_023243007.1;refGene_gn=SNAPC4;refGene_tx=XM_023243008.1;refGene_gn=SNAPC4;refGene_tx=XM_023243009.1;refGene_gn=SNAPC4;refGene_tx=XM_023243010.1
D4 94686097 94686192
refGene_gn=SNAPC4;refGene_tx=XM_023243004.1;refGene_gn=SNAPC4;refGene_tx=XM_023243005.1;refGene_gn=SNAPC4;refGene_tx=XM_023243007.1;refGene_gn=SNAPC4;refGene_tx=XM_023243008.1;refGene_gn=SNAPC4;refGene_tx=XM_023243009.1;refGene_gn=SNAPC4;refGene_tx=XM_023243010.1
D4 94687117 94687197
refGene_gn=SNAPC4;refGene_tx=XM_023243004.1;refGene_gn=SNAPC4;refGene_

tx=XM_023243005.1;refGene_gn=SNAPC4;refGene_tx=XM_023243007.1;refGene_gn=SNAPC4;refGene_tx=XM_023243008.1;refGene_gn=SNAPC4;refGene_tx=XM_023243009.1;refGene_gn=SNAPC4;refGene_tx=XM_023243010.1
D4 94687502 94687985
refGene_gn=SNAPC4;refGene_tx=XM_023243004.1;refGene_gn=SNAPC4;refGene_tx=XM_023243005.1;refGene_gn=SNAPC4;refGene_tx=XM_023243007.1;refGene_gn=SNAPC4;refGene_tx=XM_023243008.1;refGene_gn=SNAPC4;refGene_tx=XM_023243009.1;refGene_gn=SNAPC4;refGene_tx=XM_023243010.1
D4 94688755 94688839
refGene_gn=SNAPC4;refGene_tx=XM_023243004.1;refGene_gn=SNAPC4;refGene_tx=XM_023243005.1;refGene_gn=SNAPC4;refGene_tx=XM_023243007.1;refGene_gn=SNAPC4;refGene_tx=XM_023243008.1;refGene_gn=SNAPC4;refGene_tx=XM_023243009.1;refGene_gn=SNAPC4;refGene_tx=XM_023243010.1
D4 94689406 94689653
refGene_gn=SNAPC4;refGene_tx=XM_023243004.1;refGene_gn=SNAPC4;refGene_tx=XM_023243005.1;refGene_gn=SNAPC4;refGene_tx=XM_023243007.1;refGene_gn=SNAPC4;refGene_tx=XM_023243008.1;refGene_gn=SNAPC4;refGene_tx=XM_023243009.1;refGene_gn=SNAPC4;refGene_tx=XM_023243010.1
D4 94690085 94690156
refGene_gn=SNAPC4;refGene_tx=XM_023243004.1;refGene_gn=SNAPC4;refGene_tx=XM_023243005.1;refGene_gn=SNAPC4;refGene_tx=XM_023243007.1;refGene_gn=SNAPC4;refGene_tx=XM_023243008.1;refGene_gn=SNAPC4;refGene_tx=XM_023243009.1;refGene_gn=SNAPC4;refGene_tx=XM_023243010.1
D4 94690610 94690721
refGene_gn=SNAPC4;refGene_tx=XM_023243004.1;refGene_gn=SNAPC4;refGene_tx=XM_023243005.1;refGene_gn=SNAPC4;refGene_tx=XM_023243007.1;refGene_gn=SNAPC4;refGene_tx=XM_023243008.1;refGene_gn=SNAPC4;refGene_tx=XM_023243009.1;refGene_gn=SNAPC4;refGene_tx=XM_023243010.1
D4 94691394 94691422
refGene_gn=SNAPC4;refGene_tx=XM_023243004.1;refGene_gn=SNAPC4;refGene_tx=XM_023243005.1;refGene_gn=SNAPC4;refGene_tx=XM_023243007.1;refGene_gn=SNAPC4;refGene_tx=XM_023243008.1;refGene_gn=SNAPC4;refGene_tx=XM_023243009.1;refGene_gn=SNAPC4;refGene_tx=XM_023243010.1
D4 94692016 94693734
refGene_gn=SNAPC4;refGene_tx=XM_023243004.1;refGene_gn=SNAPC4;refGene_tx=XM_023243005.1;refGene_gn=SNAPC4;refGene_tx=XM_023243007.1;refGene_gn=SNAPC4;refGene_tx=XM_023243008.1;refGene_gn=SNAPC4;refGene_tx=XM_023243009.1;refGene_gn=SNAPC4;refGene_tx=XM_023243010.1
D4 94694704 94694821
refGene_gn=SNAPC4;refGene_tx=XM_023243004.1;refGene_gn=SNAPC4;refGene_tx=XM_023243005.1;refGene_gn=SNAPC4;refGene_tx=XM_023243007.1;refGene_gn=SNAPC4;refGene_tx=XM_023243008.1;refGene_gn=SNAPC4;refGene_tx=XM_023243009.1
D4 94697196 94697854
refGene_gn=CARD9;refGene_tx=XM_023243011.1;refGene_gn=CARD9;refGene_tx=XM_023243012.1;refGene_gn=CARD9;refGene_tx=XM_023243013.1
D4 94698893 94699109
refGene_gn=CARD9;refGene_tx=XM_023243011.1;refGene_gn=CARD9;refGene_tx=XM_023243012.1;refGene_gn=CARD9;refGene_tx=XM_023243013.1;refGene_gn=CARD9;refGene_tx=XM_023243014.1;refGene_gn=CARD9;refGene_tx=XM_0232430

15.1
D4	94699288	94699426
refGene_gn=CARD9;refGene_tx=XM_023243011.1;refGene_gn=CARD9;refGene_tx
=XM_023243012.1;refGene_gn=CARD9;refGene_tx=XM_023243013.1;refGene_gn=
CARD9;refGene_tx=XM_023243014.1;refGene_gn=CARD9;refGene_tx=XM_0232430
15.1
D4	94699783	94700088
refGene_gn=CARD9;refGene_tx=XM_023243011.1;refGene_gn=CARD9;refGene_tx
=XM_023243012.1;refGene_gn=CARD9;refGene_tx=XM_023243013.1;refGene_gn=
CARD9;refGene_tx=XM_023243014.1;refGene_gn=CARD9;refGene_tx=XM_0232430
15.1
D4	94700185	94700365
refGene_gn=CARD9;refGene_tx=XM_023243011.1;refGene_gn=CARD9;refGene_tx
=XM_023243012.1;refGene_gn=CARD9;refGene_tx=XM_023243013.1;refGene_gn=
CARD9;refGene_tx=XM_023243014.1;refGene_gn=CARD9;refGene_tx=XM_0232430
15.1
D4	94700777	94700921
refGene_gn=CARD9;refGene_tx=XM_023243011.1;refGene_gn=CARD9;refGene_tx
=XM_023243012.1;refGene_gn=CARD9;refGene_tx=XM_023243013.1;refGene_gn=
CARD9;refGene_tx=XM_023243014.1;refGene_gn=CARD9;refGene_tx=XM_0232430
15.1
D4	94701153	94701279
refGene_gn=CARD9;refGene_tx=XM_023243011.1;refGene_gn=CARD9;refGene_tx
=XM_023243012.1;refGene_gn=CARD9;refGene_tx=XM_023243013.1;refGene_gn=
CARD9;refGene_tx=XM_023243014.1;refGene_gn=CARD9;refGene_tx=XM_0232430
15.1
D4	94702910	94703102
refGene_gn=CARD9;refGene_tx=XM_023243011.1;refGene_gn=CARD9;refGene_tx
=XM_023243012.1;refGene_gn=CARD9;refGene_tx=XM_023243013.1;refGene_gn=
CARD9;refGene_tx=XM_023243014.1;refGene_gn=CARD9;refGene_tx=XM_0232430
15.1
D4	94703575	94703635
refGene_gn=CARD9;refGene_tx=XM_023243011.1;refGene_gn=CARD9;refGene_tx
=XM_023243012.1;refGene_gn=CARD9;refGene_tx=XM_023243014.1;refGene_gn=
CARD9;refGene_tx=XM_023243015.1
D4	94703832	94703878
refGene_gn=CARD9;refGene_tx=XM_023243011.1;refGene_gn=CARD9;refGene_tx
=XM_023243012.1;refGene_gn=CARD9;refGene_tx=XM_023243013.1;refGene_gn=
CARD9;refGene_tx=XM_023243014.1;refGene_gn=CARD9;refGene_tx=XM_0232430
15.1
D4	94705114	94705191
refGene_gn=CARD9;refGene_tx=XM_023243011.1;refGene_gn=CARD9;refGene_tx
=XM_023243012.1;refGene_gn=CARD9;refGene_tx=XM_023243013.1;refGene_gn=
CARD9;refGene_tx=XM_023243014.1;refGene_gn=CARD9;refGene_tx=XM_0232430
15.1
D4	94705574	94705651
refGene_gn=CARD9;refGene_tx=XM_023243011.1;refGene_gn=CARD9;refGene_tx
=XM_023243012.1;refGene_gn=CARD9;refGene_tx=XM_023243013.1;refGene_gn=
CARD9;refGene_tx=XM_023243014.1;refGene_gn=CARD9;refGene_tx=XM_0232430
15.1

D4	94705748	94705848
refGene_gn=CARD9;refGene_tx=XM_023243011.1;refGene_gn=CARD9;refGene_tx=XM_023243012.1;refGene_gn=CARD9;refGene_tx=XM_023243013.1;refGene_gn=CARD9;refGene_tx=XM_023243014.1;refGene_gn=CARD9;refGene_tx=XM_023243015.1
D4	94706224	94706425	refGene_gn=DNLZ;refGene_tx=XM_023243022.1
D4	94706809	94706949	refGene_gn=DNLZ;refGene_tx=XM_023243022.1
D4	94707755	94707924	refGene_gn=DNLZ;refGene_tx=XM_023243022.1
D4	94711868	94712075	refGene_gn=GPSM1;refGene_tx=XM_023243023.1
D4	94713103	94713312	refGene_gn=GPSM1;refGene_tx=XM_023243023.1
D4	94713773	94713930	refGene_gn=GPSM1;refGene_tx=XM_023243023.1
D4	94718434	94718611	refGene_gn=GPSM1;refGene_tx=XM_023243023.1
D4	94719355	94719426	refGene_gn=GPSM1;refGene_tx=XM_023243023.1
D4	94724438	94724562	refGene_gn=GPSM1;refGene_tx=XM_023243023.1
D4	94725586	94725695	refGene_gn=GPSM1;refGene_tx=XM_023243023.1
D4	94726024	94726180	refGene_gn=GPSM1;refGene_tx=XM_023243023.1
D4	94726880	94726996	refGene_gn=GPSM1;refGene_tx=XM_023243023.1
D4	94727190	94727314	refGene_gn=GPSM1;refGene_tx=XM_023243023.1
D4	94727913	94728056	refGene_gn=GPSM1;refGene_tx=XM_023243023.1
D4	94728642	94728710	refGene_gn=GPSM1;refGene_tx=XM_023243023.1
D4	94728712	94728777	refGene_gn=GPSM1;refGene_tx=XM_023243023.1
D4	94729548	94729770	refGene_gn=GPSM1;refGene_tx=XM_023243023.1
D4	94736701	94736766	refGene_gn=GPSM1;refGene_tx=XM_023243023.1
D4	94750405	94750522
refGene_gn=LOC102899124;refGene_tx=XM_023243037.1
D4	94750597	94751106
refGene_gn=LOC102899124;refGene_tx=XM_023243037.1
D4	94751226	94751284
refGene_gn=LOC102899124;refGene_tx=XM_023243037.1
D4	94751535	94751768
refGene_gn=CCDC187;refGene_tx=XM_023243024.1;refGene_gn=CCDC187;refGene_tx=XM_023243025.1;refGene_gn=CCDC187;refGene_tx=XM_023243026.1;refGene_gn=CCDC187;refGene_tx=XM_023243027.1;refGene_gn=CCDC187;refGene_tx=XM_023243029.1;refGene_gn=CCDC187;refGene_tx=XM_023243030.1;refGene_gn=CCDC187;refGene_tx=XM_023243031.1;refGene_gn=CCDC187;refGene_tx=XM_023243032.1;refGene_gn=CCDC187;refGene_tx=XM_023243033.1;refGene_gn=CCDC187;refGene_tx=XM_023243035.1;refGene_gn=CCDC187;refGene_tx=XM_023243036.1
D4	94752209	94752739
refGene_gn=CCDC187;refGene_tx=XM_023243024.1;refGene_gn=CCDC187;refGene_tx=XM_023243025.1;refGene_gn=CCDC187;refGene_tx=XM_023243026.1;refGene_gn=CCDC187;refGene_tx=XM_023243027.1;refGene_gn=CCDC187;refGene_tx=XM_023243029.1;refGene_gn=CCDC187;refGene_tx=XM_023243030.1;refGene_gn=CCDC187;refGene_tx=XM_023243031.1;refGene_gn=CCDC187;refGene_tx=XM_023243032.1;refGene_gn=CCDC187;refGene_tx=XM_023243033.1;refGene_gn=CCDC187;refGene_tx=XM_023243034.1;refGene_gn=CCDC187;refGene_tx=XM_023243035.1;refGene_gn=CCDC187;refGene_tx=XM_023243036.1
D4	94752837	94752936
refGene_gn=CCDC187;refGene_tx=XM_023243024.1;refGene_gn=CCDC187;refGene_tx=XM_023243025.1;refGene_gn=CCDC187;refGene_tx=XM_023243026.1;refGe

ne_gn=CCDC187;refGene_tx=XM_023243027.1;refGene_gn=CCDC187;refGene_tx=XM_023243029.1;refGene_gn=CCDC187;refGene_tx=XM_023243030.1;refGene_gn=CCDC187;refGene_tx=XM_023243031.1;refGene_gn=CCDC187;refGene_tx=XM_023243032.1;refGene_gn=CCDC187;refGene_tx=XM_023243033.1;refGene_gn=CCDC187;refGene_tx=XM_023243034.1;refGene_gn=CCDC187;refGene_tx=XM_023243035.1;refGene_gn=CCDC187;refGene_tx=XM_023243036.1
D4 94754875 94754971
refGene_gn=CCDC187;refGene_tx=XM_023243024.1;refGene_gn=CCDC187;refGene_tx=XM_023243025.1;refGene_gn=CCDC187;refGene_tx=XM_023243026.1;refGene_gn=CCDC187;refGene_tx=XM_023243027.1;refGene_gn=CCDC187;refGene_tx=XM_023243029.1;refGene_gn=CCDC187;refGene_tx=XM_023243030.1;refGene_gn=CCDC187;refGene_tx=XM_023243031.1;refGene_gn=CCDC187;refGene_tx=XM_023243032.1;refGene_gn=CCDC187;refGene_tx=XM_023243033.1;refGene_gn=CCDC187;refGene_tx=XM_023243034.1;refGene_gn=CCDC187;refGene_tx=XM_023243035.1;refGene_gn=CCDC187;refGene_tx=XM_023243036.1
D4 94758297 94758432
refGene_gn=CCDC187;refGene_tx=XM_023243024.1;refGene_gn=CCDC187;refGene_tx=XM_023243025.1;refGene_gn=CCDC187;refGene_tx=XM_023243026.1;refGene_gn=CCDC187;refGene_tx=XM_023243027.1;refGene_gn=CCDC187;refGene_tx=XM_023243029.1;refGene_gn=CCDC187;refGene_tx=XM_023243030.1;refGene_gn=CCDC187;refGene_tx=XM_023243031.1;refGene_gn=CCDC187;refGene_tx=XM_023243032.1;refGene_gn=CCDC187;refGene_tx=XM_023243034.1;refGene_gn=CCDC187;refGene_tx=XM_023243035.1
D4 94758957 94759976
refGene_gn=CCDC187;refGene_tx=XM_023243024.1;refGene_gn=CCDC187;refGene_tx=XM_023243025.1;refGene_gn=CCDC187;refGene_tx=XM_023243026.1;refGene_gn=CCDC187;refGene_tx=XM_023243027.1;refGene_gn=CCDC187;refGene_tx=XM_023243029.1;refGene_gn=CCDC187;refGene_tx=XM_023243030.1;refGene_gn=CCDC187;refGene_tx=XM_023243031.1;refGene_gn=CCDC187;refGene_tx=XM_023243032.1;refGene_gn=CCDC187;refGene_tx=XM_023243033.1;refGene_gn=CCDC187;refGene_tx=XM_023243034.1;refGene_gn=CCDC187;refGene_tx=XM_023243035.1;refGene_gn=CCDC187;refGene_tx=XM_023243036.1
D4 94760204 94760299
refGene_gn=CCDC187;refGene_tx=XM_023243024.1;refGene_gn=CCDC187;refGene_tx=XM_023243025.1;refGene_gn=CCDC187;refGene_tx=XM_023243026.1;refGene_gn=CCDC187;refGene_tx=XM_023243027.1;refGene_gn=CCDC187;refGene_tx=XM_023243029.1;refGene_gn=CCDC187;refGene_tx=XM_023243030.1;refGene_gn=CCDC187;refGene_tx=XM_023243031.1;refGene_gn=CCDC187;refGene_tx=XM_023243032.1;refGene_gn=CCDC187;refGene_tx=XM_023243033.1;refGene_gn=CCDC187;refGene_tx=XM_023243034.1;refGene_gn=CCDC187;refGene_tx=XM_023243035.1;refGene_gn=CCDC187;refGene_tx=XM_023243036.1
D4 94762524 94763090
refGene_gn=CCDC187;refGene_tx=XM_023243024.1;refGene_gn=CCDC187;refGene_tx=XM_023243025.1;refGene_gn=CCDC187;refGene_tx=XM_023243026.1;refGene_gn=CCDC187;refGene_tx=XM_023243027.1;refGene_gn=CCDC187;refGene_tx=XM_023243029.1;refGene_gn=CCDC187;refGene_tx=XM_023243030.1;refGene_gn=CCDC187;refGene_tx=XM_023243031.1;refGene_gn=CCDC187;refGene_tx=XM_023243032.1;refGene_gn=CCDC187;refGene_tx=XM_023243033.1;refGene_gn=CCDC187;refGene_tx=XM_023243034.1;refGene_gn=CCDC187;refGene_tx=XM_023243035.1;refGene_gn=CCDC187;refGene_tx=XM_023243036.1

D4 94763449 94763537
refGene_gn=CCDC187;refGene_tx=XM_023243024.1;refGene_gn=CCDC187;refGene_tx=XM_023243025.1;refGene_gn=CCDC187;refGene_tx=XM_023243026.1;refGene_gn=CCDC187;refGene_tx=XM_023243027.1;refGene_gn=CCDC187;refGene_tx=XM_023243029.1;refGene_gn=CCDC187;refGene_tx=XM_023243030.1;refGene_gn=CCDC187;refGene_tx=XM_023243031.1;refGene_gn=CCDC187;refGene_tx=XM_023243032.1;refGene_gn=CCDC187;refGene_tx=XM_023243033.1;refGene_gn=CCDC187;refGene_tx=XM_023243034.1;refGene_gn=CCDC187;refGene_tx=XM_023243035.1;refGene_gn=CCDC187;refGene_tx=XM_023243036.1
D4 94766423 94766533
refGene_gn=CCDC187;refGene_tx=XM_023243024.1;refGene_gn=CCDC187;refGene_tx=XM_023243025.1;refGene_gn=CCDC187;refGene_tx=XM_023243026.1;refGene_gn=CCDC187;refGene_tx=XM_023243027.1;refGene_gn=CCDC187;refGene_tx=XM_023243029.1;refGene_gn=CCDC187;refGene_tx=XM_023243030.1;refGene_gn=CCDC187;refGene_tx=XM_023243031.1;refGene_gn=CCDC187;refGene_tx=XM_023243032.1;refGene_gn=CCDC187;refGene_tx=XM_023243033.1;refGene_gn=CCDC187;refGene_tx=XM_023243034.1;refGene_gn=CCDC187;refGene_tx=XM_023243035.1;refGene_gn=CCDC187;refGene_tx=XM_023243036.1
D4 94771110 94771175
refGene_gn=CCDC187;refGene_tx=XM_023243024.1;refGene_gn=CCDC187;refGene_tx=XM_023243025.1;refGene_gn=CCDC187;refGene_tx=XM_023243026.1;refGene_gn=CCDC187;refGene_tx=XM_023243027.1;refGene_gn=CCDC187;refGene_tx=XM_023243029.1;refGene_gn=CCDC187;refGene_tx=XM_023243030.1;refGene_gn=CCDC187;refGene_tx=XM_023243031.1;refGene_gn=CCDC187;refGene_tx=XM_023243032.1;refGene_gn=CCDC187;refGene_tx=XM_023243033.1;refGene_gn=CCDC187;refGene_tx=XM_023243034.1;refGene_gn=CCDC187;refGene_tx=XM_023243035.1;refGene_gn=CCDC187;refGene_tx=XM_023243036.1
D4 94771581 94771695
refGene_gn=CCDC187;refGene_tx=XM_023243024.1;refGene_gn=CCDC187;refGene_tx=XM_023243025.1;refGene_gn=CCDC187;refGene_tx=XM_023243026.1;refGene_gn=CCDC187;refGene_tx=XM_023243027.1;refGene_gn=CCDC187;refGene_tx=XM_023243029.1;refGene_gn=CCDC187;refGene_tx=XM_023243030.1;refGene_gn=CCDC187;refGene_tx=XM_023243031.1;refGene_gn=CCDC187;refGene_tx=XM_023243032.1;refGene_gn=CCDC187;refGene_tx=XM_023243033.1;refGene_gn=CCDC187;refGene_tx=XM_023243034.1;refGene_gn=CCDC187;refGene_tx=XM_023243035.1;refGene_gn=CCDC187;refGene_tx=XM_023243036.1
D4 94772634 94772756
refGene_gn=CCDC187;refGene_tx=XM_023243024.1;refGene_gn=CCDC187;refGene_tx=XM_023243025.1;refGene_gn=CCDC187;refGene_tx=XM_023243026.1;refGene_gn=CCDC187;refGene_tx=XM_023243027.1;refGene_gn=CCDC187;refGene_tx=XM_023243029.1;refGene_gn=CCDC187;refGene_tx=XM_023243030.1;refGene_gn=CCDC187;refGene_tx=XM_023243031.1;refGene_gn=CCDC187;refGene_tx=XM_023243032.1;refGene_gn=CCDC187;refGene_tx=XM_023243033.1;refGene_gn=CCDC187;refGene_tx=XM_023243034.1;refGene_gn=CCDC187;refGene_tx=XM_023243035.1;refGene_gn=CCDC187;refGene_tx=XM_023243036.1
D4 94772871 94772963
refGene_gn=CCDC187;refGene_tx=XM_023243024.1;refGene_gn=CCDC187;refGene_tx=XM_023243025.1;refGene_gn=CCDC187;refGene_tx=XM_023243026.1;refGene_gn=CCDC187;refGene_tx=XM_023243027.1;refGene_gn=CCDC187;refGene_tx=XM_023243029.1;refGene_gn=CCDC187;refGene_tx=XM_023243030.1;refGene_gn

=CCDC187;refGene_tx=XM_023243031.1;refGene_gn=CCDC187;refGene_tx=XM_023243032.1;refGene_gn=CCDC187;refGene_tx=XM_023243033.1;refGene_gn=CCDC187;refGene_tx=XM_023243034.1;refGene_gn=CCDC187;refGene_tx=XM_023243035.1;refGene_gn=CCDC187;refGene_tx=XM_023243036.1
D4 94775992 94776078
refGene_gn=CCDC187;refGene_tx=XM_023243024.1;refGene_gn=CCDC187;refGene_tx=XM_023243025.1;refGene_gn=CCDC187;refGene_tx=XM_023243026.1;refGene_gn=CCDC187;refGene_tx=XM_023243027.1;refGene_gn=CCDC187;refGene_tx=XM_023243029.1;refGene_gn=CCDC187;refGene_tx=XM_023243030.1;refGene_gn=CCDC187;refGene_tx=XM_023243031.1;refGene_gn=CCDC187;refGene_tx=XM_023243032.1;refGene_gn=CCDC187;refGene_tx=XM_023243033.1;refGene_gn=CCDC187;refGene_tx=XM_023243034.1;refGene_gn=CCDC187;refGene_tx=XM_023243035.1;refGene_gn=CCDC187;refGene_tx=XM_023243036.1
D4 94776619 94776873
refGene_gn=CCDC187;refGene_tx=XM_023243024.1;refGene_gn=CCDC187;refGene_tx=XM_023243025.1;refGene_gn=CCDC187;refGene_tx=XM_023243026.1;refGene_gn=CCDC187;refGene_tx=XM_023243027.1;refGene_gn=CCDC187;refGene_tx=XM_023243029.1;refGene_gn=CCDC187;refGene_tx=XM_023243030.1;refGene_gn=CCDC187;refGene_tx=XM_023243031.1;refGene_gn=CCDC187;refGene_tx=XM_023243032.1;refGene_gn=CCDC187;refGene_tx=XM_023243033.1;refGene_gn=CCDC187;refGene_tx=XM_023243034.1;refGene_gn=CCDC187;refGene_tx=XM_023243035.1;refGene_gn=CCDC187;refGene_tx=XM_023243036.1
D4 94777551 94777639
refGene_gn=CCDC187;refGene_tx=XM_023243024.1;refGene_gn=CCDC187;refGene_tx=XM_023243025.1;refGene_gn=CCDC187;refGene_tx=XM_023243026.1;refGene_gn=CCDC187;refGene_tx=XM_023243027.1;refGene_gn=CCDC187;refGene_tx=XM_023243029.1;refGene_gn=CCDC187;refGene_tx=XM_023243030.1;refGene_gn=CCDC187;refGene_tx=XM_023243031.1;refGene_gn=CCDC187;refGene_tx=XM_023243032.1;refGene_gn=CCDC187;refGene_tx=XM_023243033.1;refGene_gn=CCDC187;refGene_tx=XM_023243034.1;refGene_gn=CCDC187;refGene_tx=XM_023243035.1;refGene_gn=CCDC187;refGene_tx=XM_023243036.1
D4 94778964 94779111
refGene_gn=CCDC187;refGene_tx=XM_023243024.1;refGene_gn=CCDC187;refGene_tx=XM_023243025.1;refGene_gn=CCDC187;refGene_tx=XM_023243026.1;refGene_gn=CCDC187;refGene_tx=XM_023243027.1;refGene_gn=CCDC187;refGene_tx=XM_023243029.1;refGene_gn=CCDC187;refGene_tx=XM_023243030.1;refGene_gn=CCDC187;refGene_tx=XM_023243031.1;refGene_gn=CCDC187;refGene_tx=XM_023243032.1;refGene_gn=CCDC187;refGene_tx=XM_023243033.1;refGene_gn=CCDC187;refGene_tx=XM_023243034.1;refGene_gn=CCDC187;refGene_tx=XM_023243035.1;refGene_gn=CCDC187;refGene_tx=XM_023243036.1
D4 94780022 94780186
refGene_gn=CCDC187;refGene_tx=XM_023243024.1;refGene_gn=CCDC187;refGene_tx=XM_023243025.1;refGene_gn=CCDC187;refGene_tx=XM_023243026.1;refGene_gn=CCDC187;refGene_tx=XM_023243027.1;refGene_gn=CCDC187;refGene_tx=XM_023243029.1;refGene_gn=CCDC187;refGene_tx=XM_023243030.1;refGene_gn=CCDC187;refGene_tx=XM_023243031.1;refGene_gn=CCDC187;refGene_tx=XM_023243032.1;refGene_gn=CCDC187;refGene_tx=XM_023243033.1;refGene_gn=CCDC187;refGene_tx=XM_023243034.1;refGene_gn=CCDC187;refGene_tx=XM_023243035.1;refGene_gn=CCDC187;refGene_tx=XM_023243036.1
D4 94782388 94782551

refGene_gn=CCDC187;refGene_tx=XM_023243024.1;refGene_gn=CCDC187;refGene_tx=XM_023243025.1;refGene_gn=CCDC187;refGene_tx=XM_023243026.1;refGene_gn=CCDC187;refGene_tx=XM_023243027.1;refGene_gn=CCDC187;refGene_tx=XM_023243030.1;refGene_gn=CCDC187;refGene_tx=XM_023243031.1;refGene_gn=CCDC187;refGene_tx=XM_023243032.1;refGene_gn=CCDC187;refGene_tx=XM_023243033.1;refGene_gn=CCDC187;refGene_tx=XM_023243034.1;refGene_gn=CCDC187;refGene_tx=XM_023243035.1;refGene_gn=CCDC187;refGene_tx=XM_023243036.1
D4 94782958 94783092
refGene_gn=CCDC187;refGene_tx=XM_023243024.1;refGene_gn=CCDC187;refGene_tx=XM_023243025.1;refGene_gn=CCDC187;refGene_tx=XM_023243026.1;refGene_gn=CCDC187;refGene_tx=XM_023243027.1;refGene_gn=CCDC187;refGene_tx=XM_023243029.1;refGene_gn=CCDC187;refGene_tx=XM_023243030.1;refGene_gn=CCDC187;refGene_tx=XM_023243031.1;refGene_gn=CCDC187;refGene_tx=XM_023243032.1;refGene_gn=CCDC187;refGene_tx=XM_023243033.1;refGene_gn=CCDC187;refGene_tx=XM_023243034.1;refGene_gn=CCDC187;refGene_tx=XM_023243035.1;refGene_gn=CCDC187;refGene_tx=XM_023243036.1
D4 94783451 94783459 refGene_gn=CCDC187;refGene_tx=XM_023243029.1
D4 94784090 94784176
refGene_gn=LOC111557485;refGene_tx=XM_023242324.1
D4 94784372 94784451
refGene_gn=LOC111557485;refGene_tx=XM_023242324.1
D4 94784653 94786399
refGene_gn=LOC111557485;refGene_tx=XM_023242324.1
D4 94793172 94793470 refGene_gn=QSOX2;refGene_tx=XM_023242325.1
D4 94806941 94807042 refGene_gn=QSOX2;refGene_tx=XM_023242325.1
D4 94808957 94809006 refGene_gn=QSOX2;refGene_tx=XM_023242325.1
D4 94809661 94809767 refGene_gn=QSOX2;refGene_tx=XM_023242325.1
D4 94809911 94810002 refGene_gn=QSOX2;refGene_tx=XM_023242325.1
D4 94811533 94811679 refGene_gn=QSOX2;refGene_tx=XM_023242325.1
D4 94812950 94813085 refGene_gn=QSOX2;refGene_tx=XM_023242325.1
D4 94813341 94813471 refGene_gn=QSOX2;refGene_tx=XM_023242325.1
D4 94814989 94815112 refGene_gn=QSOX2;refGene_tx=XM_023242325.1
D4 94816218 94816369 refGene_gn=QSOX2;refGene_tx=XM_023242325.1
D4 94818858 94819047 refGene_gn=QSOX2;refGene_tx=XM_023242325.1
D4 94820900 94821433 refGene_gn=QSOX2;refGene_tx=XM_023242325.1
D4 94825182 94825270 refGene_gn=LHX3;refGene_tx=XM_023243040.1
D4 94826970 94827064 refGene_gn=LHX3;refGene_tx=XM_023243039.1
D4 94829253 94829425
refGene_gn=LHX3;refGene_tx=XM_023243039.1;refGene_gn=LHX3;refGene_tx=XM_023243040.1
D4 94829962 94830165
refGene_gn=LHX3;refGene_tx=XM_023243039.1;refGene_gn=LHX3;refGene_tx=XM_023243040.1
D4 94830728 94830880
refGene_gn=LHX3;refGene_tx=XM_023243039.1;refGene_gn=LHX3;refGene_tx=XM_023243040.1
D4 94830969 94831141
refGene_gn=LHX3;refGene_tx=XM_023243039.1;refGene_gn=LHX3;refGene_tx=XM_023243040.1

D4 94831943 94832362
refGene_gn=LHX3;refGene_tx=XM_023243039.1;refGene_gn=LHX3;refGene_tx=XM_023243040.1
D4 94888210 94888699
refGene_gn=TMEM250;refGene_tx=XM_023243042.1;refGene_gn=TMEM250;refGene_tx=XM_023243043.1;refGene_gn=TMEM250;refGene_tx=XM_023243044.1;refGene_gn=TMEM250;refGene_tx=XM_023243046.1
D4 94892142 94893333
refGene_gn=LOC109493594;refGene_tx=XM_023243041.1
D4 94944416 94945287
refGene_gn=NACC2;refGene_tx=XM_023243047.1;refGene_gn=NACC2;refGene_tx=XM_023243048.1;refGene_gn=NACC2;refGene_tx=XM_023243049.1;refGene_gn=NACC2;refGene_tx=XM_023243050.1;refGene_gn=NACC2;refGene_tx=XM_023243051.1
D4 94968106 94968271
refGene_gn=NACC2;refGene_tx=XM_023243047.1;refGene_gn=NACC2;refGene_tx=XM_023243048.1;refGene_gn=NACC2;refGene_tx=XM_023243049.1;refGene_gn=NACC2;refGene_tx=XM_023243050.1;refGene_gn=NACC2;refGene_tx=XM_023243051.1
D4 94970267 94970373
refGene_gn=NACC2;refGene_tx=XM_023243047.1;refGene_gn=NACC2;refGene_tx=XM_023243048.1;refGene_gn=NACC2;refGene_tx=XM_023243049.1;refGene_gn=NACC2;refGene_tx=XM_023243050.1;refGene_gn=NACC2;refGene_tx=XM_023243051.1
D4 94970724 94970822
refGene_gn=NACC2;refGene_tx=XM_023243047.1;refGene_gn=NACC2;refGene_tx=XM_023243048.1;refGene_gn=NACC2;refGene_tx=XM_023243049.1;refGene_gn=NACC2;refGene_tx=XM_023243050.1;refGene_gn=NACC2;refGene_tx=XM_023243051.1
D4 94972159 94972665
refGene_gn=NACC2;refGene_tx=XM_023243047.1;refGene_gn=NACC2;refGene_tx=XM_023243048.1;refGene_gn=NACC2;refGene_tx=XM_023243049.1;refGene_gn=NACC2;refGene_tx=XM_023243050.1;refGene_gn=NACC2;refGene_tx=XM_023243051.1
D4 95006302 95006440
refGene_gn=UBAC1;refGene_tx=XM_011288761.3;refGene_gn=UBAC1;refGene_tx=XM_023243052.1;refGene_gn=UBAC1;refGene_tx=XM_023243053.1
D4 95010007 95010128
refGene_gn=UBAC1;refGene_tx=XM_011288761.3;refGene_gn=UBAC1;refGene_tx=XM_023243052.1;refGene_gn=UBAC1;refGene_tx=XM_023243053.1
D4 95011035 95011109
refGene_gn=UBAC1;refGene_tx=XM_011288761.3;refGene_gn=UBAC1;refGene_tx=XM_023243052.1;refGene_gn=UBAC1;refGene_tx=XM_023243053.1
D4 95015369 95015477
refGene_gn=UBAC1;refGene_tx=XM_011288761.3;refGene_gn=UBAC1;refGene_tx=XM_023243052.1;refGene_gn=UBAC1;refGene_tx=XM_023243053.1
D4 95016356 95016459
refGene_gn=UBAC1;refGene_tx=XM_011288761.3;refGene_gn=UBAC1;refGene_tx=XM_023243052.1;refGene_gn=UBAC1;refGene_tx=XM_023243053.1
D4 95016731 95016840

refGene_gn=UBAC1;refGene_tx=XM_011288761.3;refGene_gn=UBAC1;refGene_tx=XM_023243052.1;refGene_gn=UBAC1;refGene_tx=XM_023243053.1
D4 95016938 95016980
refGene_gn=UBAC1;refGene_tx=XM_023243052.1;refGene_gn=UBAC1;refGene_tx=XM_023243053.1
D4 95017495 95017718
refGene_gn=UBAC1;refGene_tx=XM_011288761.3;refGene_gn=UBAC1;refGene_tx=XM_023243052.1;refGene_gn=UBAC1;refGene_tx=XM_023243053.1
D4 95019463 95019559 refGene_gn=UBAC1;refGene_tx=XM_023243053.1
D4 95020364 95020451
refGene_gn=UBAC1;refGene_tx=XM_011288761.3;refGene_gn=UBAC1;refGene_tx=XM_023243052.1
D4 95021215 95021354
refGene_gn=UBAC1;refGene_tx=XM_011288761.3;refGene_gn=UBAC1;refGene_tx=XM_023243052.1
D4 95023844 95023960
refGene_gn=UBAC1;refGene_tx=XM_011288761.3;refGene_gn=UBAC1;refGene_tx=XM_023243052.1
D4 95042421 95042581 refGene_gn=CAMSAP1;refGene_tx=XM_023243054.1
D4 95062424 95062645 refGene_gn=CAMSAP1;refGene_tx=XM_023243054.1
D4 95062646 95062688 refGene_gn=CAMSAP1;refGene_tx=XM_023243054.1
D4 95063503 95063665 refGene_gn=CAMSAP1;refGene_tx=XM_023243054.1
D4 95073704 95073785 refGene_gn=CAMSAP1;refGene_tx=XM_023243054.1
D4 95077309 95077451 refGene_gn=CAMSAP1;refGene_tx=XM_023243054.1
D4 95084605 95084745 refGene_gn=CAMSAP1;refGene_tx=XM_023243054.1
D4 95084834 95084931 refGene_gn=CAMSAP1;refGene_tx=XM_023243054.1
D4 95087333 95087511 refGene_gn=CAMSAP1;refGene_tx=XM_023243054.1
D4 95087910 95088002 refGene_gn=CAMSAP1;refGene_tx=XM_023243054.1
D4 95088702 95088787 refGene_gn=CAMSAP1;refGene_tx=XM_023243054.1
D4 95089467 95091811 refGene_gn=CAMSAP1;refGene_tx=XM_023243054.1
D4 95092748 95092885 refGene_gn=CAMSAP1;refGene_tx=XM_023243054.1
D4 95093724 95093933 refGene_gn=CAMSAP1;refGene_tx=XM_023243054.1
D4 95094016 95094119 refGene_gn=CAMSAP1;refGene_tx=XM_023243054.1
D4 95095472 95095588 refGene_gn=CAMSAP1;refGene_tx=XM_023243054.1
D4 95095987 95096106 refGene_gn=CAMSAP1;refGene_tx=XM_023243054.1
D4 95099164 95099467 refGene_gn=CAMSAP1;refGene_tx=XM_023243054.1
D4 95109556 95109830
refGene_gn=KCNT1;refGene_tx=XM_023243055.1;refGene_gn=KCNT1;refGene_tx=XM_023243056.1;refGene_gn=KCNT1;refGene_tx=XM_023243057.1
D4 95110001 95110122
refGene_gn=KCNT1;refGene_tx=XM_023243059.1;refGene_gn=KCNT1;refGene_tx=XM_023243060.1;refGene_gn=KCNT1;refGene_tx=XM_023243061.1;refGene_gn=KCNT1;refGene_tx=XM_023243062.1;refGene_gn=KCNT1;refGene_tx=XM_023243065.1;refGene_gn=KCNT1;refGene_tx=XM_023243066.1
D4 95110203 95110288
refGene_gn=KCNT1;refGene_tx=XM_023243055.1;refGene_gn=KCNT1;refGene_tx=XM_023243056.1;refGene_gn=KCNT1;refGene_tx=XM_023243057.1;refGene_gn=KCNT1;refGene_tx=XM_023243059.1;refGene_gn=KCNT1;refGene_tx=XM_023243060.1;refGene_gn=KCNT1;refGene_tx=XM_023243061.1;refGene_gn=KCNT1;refGene_tx=XM_023243062.1;refGene_gn=KCNT1;refGene_tx=XM_023243065.1;refGen

e_gn=KCNT1;refGene_tx=XM_023243066.1
D4 95113099 95113162
refGene_gn=KCNT1;refGene_tx=XM_023243055.1;refGene_gn=KCNT1;refGene_tx=XM_023243056.1;refGene_gn=KCNT1;refGene_tx=XM_023243059.1;refGene_gn=KCNT1;refGene_tx=XM_023243060.1;refGene_gn=KCNT1;refGene_tx=XM_023243062.1;refGene_gn=KCNT1;refGene_tx=XM_023243066.1
D4 95114312 95114640
refGene_gn=KCNT1;refGene_tx=XM_023243055.1;refGene_gn=KCNT1;refGene_tx=XM_023243056.1;refGene_gn=KCNT1;refGene_tx=XM_023243057.1;refGene_gn=KCNT1;refGene_tx=XM_023243059.1;refGene_gn=KCNT1;refGene_tx=XM_023243060.1;refGene_gn=KCNT1;refGene_tx=XM_023243061.1;refGene_gn=KCNT1;refGene_tx=XM_023243062.1;refGene_gn=KCNT1;refGene_tx=XM_023243065.1;refGene_gn=KCNT1;refGene_tx=XM_023243066.1
D4 95115260 95115281
refGene_gn=KCNT1;refGene_tx=XM_023243055.1;refGene_gn=KCNT1;refGene_tx=XM_023243056.1;refGene_gn=KCNT1;refGene_tx=XM_023243057.1;refGene_gn=KCNT1;refGene_tx=XM_023243059.1;refGene_gn=KCNT1;refGene_tx=XM_023243060.1;refGene_gn=KCNT1;refGene_tx=XM_023243061.1;refGene_gn=KCNT1;refGene_tx=XM_023243062.1;refGene_gn=KCNT1;refGene_tx=XM_023243063.1;refGene_gn=KCNT1;refGene_tx=XM_023243064.1;refGene_gn=KCNT1;refGene_tx=XM_023243065.1;refGene_gn=KCNT1;refGene_tx=XM_023243066.1
D4 95115589 95115718
refGene_gn=KCNT1;refGene_tx=XM_023243055.1;refGene_gn=KCNT1;refGene_tx=XM_023243056.1;refGene_gn=KCNT1;refGene_tx=XM_023243057.1;refGene_gn=KCNT1;refGene_tx=XM_023243059.1;refGene_gn=KCNT1;refGene_tx=XM_023243060.1;refGene_gn=KCNT1;refGene_tx=XM_023243061.1;refGene_gn=KCNT1;refGene_tx=XM_023243062.1;refGene_gn=KCNT1;refGene_tx=XM_023243063.1;refGene_gn=KCNT1;refGene_tx=XM_023243064.1;refGene_gn=KCNT1;refGene_tx=XM_023243065.1;refGene_gn=KCNT1;refGene_tx=XM_023243066.1
D4 95115801 95115885
refGene_gn=KCNT1;refGene_tx=XM_023243055.1;refGene_gn=KCNT1;refGene_tx=XM_023243056.1;refGene_gn=KCNT1;refGene_tx=XM_023243057.1;refGene_gn=KCNT1;refGene_tx=XM_023243059.1;refGene_gn=KCNT1;refGene_tx=XM_023243060.1;refGene_gn=KCNT1;refGene_tx=XM_023243061.1;refGene_gn=KCNT1;refGene_tx=XM_023243062.1;refGene_gn=KCNT1;refGene_tx=XM_023243063.1;refGene_gn=KCNT1;refGene_tx=XM_023243064.1;refGene_gn=KCNT1;refGene_tx=XM_023243065.1;refGene_gn=KCNT1;refGene_tx=XM_023243066.1
D4 95116169 95116271
refGene_gn=KCNT1;refGene_tx=XM_023243055.1;refGene_gn=KCNT1;refGene_tx=XM_023243056.1;refGene_gn=KCNT1;refGene_tx=XM_023243057.1;refGene_gn=KCNT1;refGene_tx=XM_023243059.1;refGene_gn=KCNT1;refGene_tx=XM_023243060.1;refGene_gn=KCNT1;refGene_tx=XM_023243061.1;refGene_gn=KCNT1;refGene_tx=XM_023243062.1;refGene_gn=KCNT1;refGene_tx=XM_023243063.1;refGene_gn=KCNT1;refGene_tx=XM_023243064.1;refGene_gn=KCNT1;refGene_tx=XM_023243065.1;refGene_gn=KCNT1;refGene_tx=XM_023243066.1
D4 95118690 95118802
refGene_gn=KCNT1;refGene_tx=XM_023243055.1;refGene_gn=KCNT1;refGene_tx=XM_023243056.1;refGene_gn=KCNT1;refGene_tx=XM_023243057.1;refGene_gn=KCNT1;refGene_tx=XM_023243059.1;refGene_gn=KCNT1;refGene_tx=XM_023243060.1;refGene_gn=KCNT1;refGene_tx=XM_023243061.1;refGene_gn=KCNT1;refGe

ne_tx=XM_023243062.1;refGene_gn=KCNT1;refGene_tx=XM_023243063.1;refGene_gn=KCNT1;refGene_tx=XM_023243064.1;refGene_gn=KCNT1;refGene_tx=XM_023243065.1;refGene_gn=KCNT1;refGene_tx=XM_023243066.1
D4 95118946 95119081
refGene_gn=KCNT1;refGene_tx=XM_023243055.1;refGene_gn=KCNT1;refGene_tx=XM_023243056.1;refGene_gn=KCNT1;refGene_tx=XM_023243057.1;refGene_gn=KCNT1;refGene_tx=XM_023243059.1;refGene_gn=KCNT1;refGene_tx=XM_023243060.1;refGene_gn=KCNT1;refGene_tx=XM_023243061.1;refGene_gn=KCNT1;refGene_tx=XM_023243062.1;refGene_gn=KCNT1;refGene_tx=XM_023243063.1;refGene_gn=KCNT1;refGene_tx=XM_023243064.1;refGene_gn=KCNT1;refGene_tx=XM_023243065.1;refGene_gn=KCNT1;refGene_tx=XM_023243066.1
D4 95119331 95119403
refGene_gn=KCNT1;refGene_tx=XM_023243055.1;refGene_gn=KCNT1;refGene_tx=XM_023243056.1;refGene_gn=KCNT1;refGene_tx=XM_023243057.1;refGene_gn=KCNT1;refGene_tx=XM_023243059.1;refGene_gn=KCNT1;refGene_tx=XM_023243060.1;refGene_gn=KCNT1;refGene_tx=XM_023243061.1;refGene_gn=KCNT1;refGene_tx=XM_023243062.1;refGene_gn=KCNT1;refGene_tx=XM_023243063.1;refGene_gn=KCNT1;refGene_tx=XM_023243064.1;refGene_gn=KCNT1;refGene_tx=XM_023243065.1;refGene_gn=KCNT1;refGene_tx=XM_023243066.1
D4 95119770 95119943
refGene_gn=KCNT1;refGene_tx=XM_023243055.1;refGene_gn=KCNT1;refGene_tx=XM_023243056.1;refGene_gn=KCNT1;refGene_tx=XM_023243057.1;refGene_gn=KCNT1;refGene_tx=XM_023243059.1;refGene_gn=KCNT1;refGene_tx=XM_023243060.1;refGene_gn=KCNT1;refGene_tx=XM_023243061.1;refGene_gn=KCNT1;refGene_tx=XM_023243062.1;refGene_gn=KCNT1;refGene_tx=XM_023243063.1;refGene_gn=KCNT1;refGene_tx=XM_023243064.1;refGene_gn=KCNT1;refGene_tx=XM_023243065.1;refGene_gn=KCNT1;refGene_tx=XM_023243066.1
D4 95120853 95120959
refGene_gn=KCNT1;refGene_tx=XM_023243055.1;refGene_gn=KCNT1;refGene_tx=XM_023243056.1;refGene_gn=KCNT1;refGene_tx=XM_023243057.1;refGene_gn=KCNT1;refGene_tx=XM_023243059.1;refGene_gn=KCNT1;refGene_tx=XM_023243060.1;refGene_gn=KCNT1;refGene_tx=XM_023243061.1;refGene_gn=KCNT1;refGene_tx=XM_023243062.1;refGene_gn=KCNT1;refGene_tx=XM_023243063.1;refGene_gn=KCNT1;refGene_tx=XM_023243064.1;refGene_gn=KCNT1;refGene_tx=XM_023243065.1;refGene_gn=KCNT1;refGene_tx=XM_023243066.1
D4 95121842 95122077
refGene_gn=KCNT1;refGene_tx=XM_023243055.1;refGene_gn=KCNT1;refGene_tx=XM_023243056.1;refGene_gn=KCNT1;refGene_tx=XM_023243057.1;refGene_gn=KCNT1;refGene_tx=XM_023243059.1;refGene_gn=KCNT1;refGene_tx=XM_023243060.1;refGene_gn=KCNT1;refGene_tx=XM_023243061.1;refGene_gn=KCNT1;refGene_tx=XM_023243062.1;refGene_gn=KCNT1;refGene_tx=XM_023243063.1;refGene_gn=KCNT1;refGene_tx=XM_023243064.1;refGene_gn=KCNT1;refGene_tx=XM_023243065.1;refGene_gn=KCNT1;refGene_tx=XM_023243066.1
D4 95122238 95122474
refGene_gn=KCNT1;refGene_tx=XM_023243055.1;refGene_gn=KCNT1;refGene_tx=XM_023243056.1;refGene_gn=KCNT1;refGene_tx=XM_023243057.1;refGene_gn=KCNT1;refGene_tx=XM_023243059.1;refGene_gn=KCNT1;refGene_tx=XM_023243060.1;refGene_gn=KCNT1;refGene_tx=XM_023243061.1;refGene_gn=KCNT1;refGene_tx=XM_023243062.1;refGene_gn=KCNT1;refGene_tx=XM_023243063.1;refGene_gn=KCNT1;refGene_tx=XM_023243064.1;refGene_gn=KCNT1;refGene_tx=XM_02

3243065.1;refGene_gn=KCNT1;refGene_tx=XM_023243066.1
D4 95122611 95122761
refGene_gn=KCNT1;refGene_tx=XM_023243055.1;refGene_gn=KCNT1;refGene_tx=XM_023243056.1;refGene_gn=KCNT1;refGene_tx=XM_023243057.1;refGene_gn=KCNT1;refGene_tx=XM_023243059.1;refGene_gn=KCNT1;refGene_tx=XM_023243060.1;refGene_gn=KCNT1;refGene_tx=XM_023243061.1;refGene_gn=KCNT1;refGene_tx=XM_023243062.1;refGene_gn=KCNT1;refGene_tx=XM_023243063.1;refGene_gn=KCNT1;refGene_tx=XM_023243064.1;refGene_gn=KCNT1;refGene_tx=XM_023243065.1;refGene_gn=KCNT1;refGene_tx=XM_023243066.1
D4 95123092 95123201
refGene_gn=KCNT1;refGene_tx=XM_023243055.1;refGene_gn=KCNT1;refGene_tx=XM_023243056.1;refGene_gn=KCNT1;refGene_tx=XM_023243057.1;refGene_gn=KCNT1;refGene_tx=XM_023243059.1;refGene_gn=KCNT1;refGene_tx=XM_023243060.1;refGene_gn=KCNT1;refGene_tx=XM_023243061.1;refGene_gn=KCNT1;refGene_tx=XM_023243062.1;refGene_gn=KCNT1;refGene_tx=XM_023243063.1;refGene_gn=KCNT1;refGene_tx=XM_023243064.1;refGene_gn=KCNT1;refGene_tx=XM_023243065.1;refGene_gn=KCNT1;refGene_tx=XM_023243066.1
D4 95123681 95123790
refGene_gn=KCNT1;refGene_tx=XM_023243055.1;refGene_gn=KCNT1;refGene_tx=XM_023243056.1;refGene_gn=KCNT1;refGene_tx=XM_023243057.1;refGene_gn=KCNT1;refGene_tx=XM_023243059.1;refGene_gn=KCNT1;refGene_tx=XM_023243060.1;refGene_gn=KCNT1;refGene_tx=XM_023243061.1;refGene_gn=KCNT1;refGene_tx=XM_023243062.1;refGene_gn=KCNT1;refGene_tx=XM_023243063.1;refGene_gn=KCNT1;refGene_tx=XM_023243064.1;refGene_gn=KCNT1;refGene_tx=XM_023243065.1;refGene_gn=KCNT1;refGene_tx=XM_023243066.1
D4 95123903 95123967
refGene_gn=KCNT1;refGene_tx=XM_023243055.1;refGene_gn=KCNT1;refGene_tx=XM_023243056.1;refGene_gn=KCNT1;refGene_tx=XM_023243057.1;refGene_gn=KCNT1;refGene_tx=XM_023243059.1;refGene_gn=KCNT1;refGene_tx=XM_023243060.1;refGene_gn=KCNT1;refGene_tx=XM_023243061.1;refGene_gn=KCNT1;refGene_tx=XM_023243062.1;refGene_gn=KCNT1;refGene_tx=XM_023243063.1;refGene_gn=KCNT1;refGene_tx=XM_023243064.1;refGene_gn=KCNT1;refGene_tx=XM_023243065.1;refGene_gn=KCNT1;refGene_tx=XM_023243066.1
D4 95124954 95125091
refGene_gn=KCNT1;refGene_tx=XM_023243055.1;refGene_gn=KCNT1;refGene_tx=XM_023243056.1;refGene_gn=KCNT1;refGene_tx=XM_023243057.1;refGene_gn=KCNT1;refGene_tx=XM_023243059.1;refGene_gn=KCNT1;refGene_tx=XM_023243060.1;refGene_gn=KCNT1;refGene_tx=XM_023243061.1;refGene_gn=KCNT1;refGene_tx=XM_023243062.1;refGene_gn=KCNT1;refGene_tx=XM_023243063.1;refGene_gn=KCNT1;refGene_tx=XM_023243064.1;refGene_gn=KCNT1;refGene_tx=XM_023243065.1;refGene_gn=KCNT1;refGene_tx=XM_023243066.1
D4 95125228 95125393
refGene_gn=KCNT1;refGene_tx=XM_023243055.1;refGene_gn=KCNT1;refGene_tx=XM_023243056.1;refGene_gn=KCNT1;refGene_tx=XM_023243057.1;refGene_gn=KCNT1;refGene_tx=XM_023243059.1;refGene_gn=KCNT1;refGene_tx=XM_023243060.1;refGene_gn=KCNT1;refGene_tx=XM_023243061.1;refGene_gn=KCNT1;refGene_tx=XM_023243062.1;refGene_gn=KCNT1;refGene_tx=XM_023243063.1;refGene_gn=KCNT1;refGene_tx=XM_023243064.1;refGene_gn=KCNT1;refGene_tx=XM_023243065.1;refGene_gn=KCNT1;refGene_tx=XM_023243066.1
D4 95126983 95127209

refGene_gn=KCNT1;refGene_tx=XM_023243055.1;refGene_gn=KCNT1;refGene_tx=XM_023243056.1;refGene_gn=KCNT1;refGene_tx=XM_023243057.1;refGene_gn=KCNT1;refGene_tx=XM_023243059.1;refGene_gn=KCNT1;refGene_tx=XM_023243060.1;refGene_gn=KCNT1;refGene_tx=XM_023243061.1;refGene_gn=KCNT1;refGene_tx=XM_023243062.1;refGene_gn=KCNT1;refGene_tx=XM_023243063.1;refGene_gn=KCNT1;refGene_tx=XM_023243064.1;refGene_gn=KCNT1;refGene_tx=XM_023243065.1;refGene_gn=KCNT1;refGene_tx=XM_023243066.1
D4 95128325 95128420
refGene_gn=KCNT1;refGene_tx=XM_023243055.1;refGene_gn=KCNT1;refGene_tx=XM_023243056.1;refGene_gn=KCNT1;refGene_tx=XM_023243057.1;refGene_gn=KCNT1;refGene_tx=XM_023243059.1;refGene_gn=KCNT1;refGene_tx=XM_023243060.1;refGene_gn=KCNT1;refGene_tx=XM_023243061.1;refGene_gn=KCNT1;refGene_tx=XM_023243062.1;refGene_gn=KCNT1;refGene_tx=XM_023243063.1;refGene_gn=KCNT1;refGene_tx=XM_023243064.1;refGene_gn=KCNT1;refGene_tx=XM_023243065.1;refGene_gn=KCNT1;refGene_tx=XM_023243066.1
D4 95128710 95128794
refGene_gn=KCNT1;refGene_tx=XM_023243055.1;refGene_gn=KCNT1;refGene_tx=XM_023243056.1;refGene_gn=KCNT1;refGene_tx=XM_023243057.1;refGene_gn=KCNT1;refGene_tx=XM_023243059.1;refGene_gn=KCNT1;refGene_tx=XM_023243060.1;refGene_gn=KCNT1;refGene_tx=XM_023243061.1;refGene_gn=KCNT1;refGene_tx=XM_023243062.1;refGene_gn=KCNT1;refGene_tx=XM_023243063.1;refGene_gn=KCNT1;refGene_tx=XM_023243064.1;refGene_gn=KCNT1;refGene_tx=XM_023243065.1;refGene_gn=KCNT1;refGene_tx=XM_023243066.1
D4 95128861 95128936
refGene_gn=KCNT1;refGene_tx=XM_023243055.1;refGene_gn=KCNT1;refGene_tx=XM_023243056.1;refGene_gn=KCNT1;refGene_tx=XM_023243057.1;refGene_gn=KCNT1;refGene_tx=XM_023243059.1;refGene_gn=KCNT1;refGene_tx=XM_023243060.1;refGene_gn=KCNT1;refGene_tx=XM_023243061.1;refGene_gn=KCNT1;refGene_tx=XM_023243062.1;refGene_gn=KCNT1;refGene_tx=XM_023243063.1;refGene_gn=KCNT1;refGene_tx=XM_023243064.1;refGene_gn=KCNT1;refGene_tx=XM_023243065.1;refGene_gn=KCNT1;refGene_tx=XM_023243066.1
D4 95129048 95129108
refGene_gn=KCNT1;refGene_tx=XM_023243055.1;refGene_gn=KCNT1;refGene_tx=XM_023243056.1;refGene_gn=KCNT1;refGene_tx=XM_023243057.1;refGene_gn=KCNT1;refGene_tx=XM_023243059.1;refGene_gn=KCNT1;refGene_tx=XM_023243060.1;refGene_gn=KCNT1;refGene_tx=XM_023243061.1;refGene_gn=KCNT1;refGene_tx=XM_023243062.1;refGene_gn=KCNT1;refGene_tx=XM_023243063.1;refGene_gn=KCNT1;refGene_tx=XM_023243064.1;refGene_gn=KCNT1;refGene_tx=XM_023243065.1;refGene_gn=KCNT1;refGene_tx=XM_023243066.1
D4 95129178 95129227
refGene_gn=KCNT1;refGene_tx=XM_023243055.1;refGene_gn=KCNT1;refGene_tx=XM_023243056.1;refGene_gn=KCNT1;refGene_tx=XM_023243057.1;refGene_gn=KCNT1;refGene_tx=XM_023243059.1;refGene_gn=KCNT1;refGene_tx=XM_023243060.1;refGene_gn=KCNT1;refGene_tx=XM_023243061.1;refGene_gn=KCNT1;refGene_tx=XM_023243062.1;refGene_gn=KCNT1;refGene_tx=XM_023243063.1;refGene_gn=KCNT1;refGene_tx=XM_023243064.1;refGene_gn=KCNT1;refGene_tx=XM_023243065.1;refGene_gn=KCNT1;refGene_tx=XM_023243066.1
D4 95130021 95130078
refGene_gn=KCNT1;refGene_tx=XM_023243055.1;refGene_gn=KCNT1;refGene_tx=XM_023243056.1;refGene_gn=KCNT1;refGene_tx=XM_023243057.1;refGene_gn=

KCNT1;refGene_tx=XM_023243059.1;refGene_gn=KCNT1;refGene_tx=XM_023243060.1;refGene_gn=KCNT1;refGene_tx=XM_023243061.1;refGene_gn=KCNT1;refGene_tx=XM_023243062.1;refGene_gn=KCNT1;refGene_tx=XM_023243063.1;refGene_gn=KCNT1;refGene_tx=XM_023243064.1;refGene_gn=KCNT1;refGene_tx=XM_023243065.1;refGene_gn=KCNT1;refGene_tx=XM_023243066.1
D4 95132198 95132298
refGene_gn=KCNT1;refGene_tx=XM_023243055.1;refGene_gn=KCNT1;refGene_tx=XM_023243056.1;refGene_gn=KCNT1;refGene_tx=XM_023243057.1;refGene_gn=KCNT1;refGene_tx=XM_023243059.1;refGene_gn=KCNT1;refGene_tx=XM_023243060.1;refGene_gn=KCNT1;refGene_tx=XM_023243061.1;refGene_gn=KCNT1;refGene_tx=XM_023243062.1;refGene_gn=KCNT1;refGene_tx=XM_023243063.1;refGene_gn=KCNT1;refGene_tx=XM_023243064.1;refGene_gn=KCNT1;refGene_tx=XM_023243065.1;refGene_gn=KCNT1;refGene_tx=XM_023243066.1
D4 95132823 95132903
refGene_gn=KCNT1;refGene_tx=XM_023243055.1;refGene_gn=KCNT1;refGene_tx=XM_023243056.1;refGene_gn=KCNT1;refGene_tx=XM_023243057.1;refGene_gn=KCNT1;refGene_tx=XM_023243059.1;refGene_gn=KCNT1;refGene_tx=XM_023243060.1;refGene_gn=KCNT1;refGene_tx=XM_023243061.1;refGene_gn=KCNT1;refGene_tx=XM_023243062.1;refGene_gn=KCNT1;refGene_tx=XM_023243063.1;refGene_gn=KCNT1;refGene_tx=XM_023243064.1;refGene_gn=KCNT1;refGene_tx=XM_023243065.1;refGene_gn=KCNT1;refGene_tx=XM_023243066.1
D4 95138939 95138981
refGene_gn=KCNT1;refGene_tx=XM_023243055.1;refGene_gn=KCNT1;refGene_tx=XM_023243057.1;refGene_gn=KCNT1;refGene_tx=XM_023243059.1;refGene_gn=KCNT1;refGene_tx=XM_023243061.1;refGene_gn=KCNT1;refGene_tx=XM_023243066.1
D4 95144051 95144206
refGene_gn=KCNT1;refGene_tx=XM_023243055.1;refGene_gn=KCNT1;refGene_tx=XM_023243057.1;refGene_gn=KCNT1;refGene_tx=XM_023243059.1;refGene_gn=KCNT1;refGene_tx=XM_023243060.1;refGene_gn=KCNT1;refGene_tx=XM_023243061.1
D4 95158805 95158900
refGene_gn=KCNT1;refGene_tx=XM_023243056.1;refGene_gn=KCNT1;refGene_tx=XM_023243062.1;refGene_gn=KCNT1;refGene_tx=XM_023243063.1;refGene_gn=KCNT1;refGene_tx=XM_023243064.1;refGene_gn=KCNT1;refGene_tx=XM_023243065.1;refGene_gn=KCNT1;refGene_tx=XM_023243066.1
D4 95171738 95171779 refGene_gn=SOHLH1;refGene_tx=XM_023243388.1
D4 95171891 95172079
refGene_gn=SOHLH1;refGene_tx=XM_023243387.1;refGene_gn=SOHLH1;refGene_tx=XM_023243388.1
D4 95172488 95172639
refGene_gn=SOHLH1;refGene_tx=XM_023243387.1;refGene_gn=SOHLH1;refGene_tx=XM_023243388.1
D4 95173004 95173126
refGene_gn=SOHLH1;refGene_tx=XM_023243387.1;refGene_gn=SOHLH1;refGene_tx=XM_023243388.1
D4 95173211 95173297
refGene_gn=SOHLH1;refGene_tx=XM_023243387.1;refGene_gn=SOHLH1;refGene_tx=XM_023243388.1
D4 95174582 95174769

refGene_gn=SOHLH1;refGene_tx=XM_023243387.1;refGene_gn=SOHLH1;refGene_tx=XM_023243388.1
D4 95175259 95175330
refGene_gn=SOHLH1;refGene_tx=XM_023243387.1;refGene_gn=SOHLH1;refGene_tx=XM_023243388.1
D4 95175877 95176095
refGene_gn=SOHLH1;refGene_tx=XM_023243387.1;refGene_gn=SOHLH1;refGene_tx=XM_023243388.1
D4 95180750 95180776
refGene_gn=LOC109493595;refGene_tx=XM_019815586.2
D4 95180863 95180965
refGene_gn=LOC109493595;refGene_tx=XM_019815586.2
D4 95182515 95182614
refGene_gn=LOC109493595;refGene_tx=XM_019815586.2
D4 95182978 95183052
refGene_gn=LOC109493595;refGene_tx=XM_019815586.2
D4 95183298 95183432
refGene_gn=LOC109493595;refGene_tx=XM_019815586.2
D4 95183574 95183805
refGene_gn=LOC109493595;refGene_tx=XM_019815586.2
D4 95183902 95183956
refGene_gn=LOC109493595;refGene_tx=XM_019815586.2
D4 95210892 95211017 refGene_gn=LCN9;refGene_tx=XM_023243389.1
D4 95211319 95211393 refGene_gn=LCN9;refGene_tx=XM_023243389.1
D4 95211502 95211639 refGene_gn=LCN9;refGene_tx=XM_023243389.1
D4 95212119 95212178 refGene_gn=LCN9;refGene_tx=XM_023243389.1
D4 95212180 95212214 refGene_gn=LCN9;refGene_tx=XM_023243389.1
D4 95216354 95216428
refGene_gn=GLT6D1;refGene_tx=XM_003996064.4;refGene_gn=GLT6D1;refGene_tx=XM_019816804.2
D4 95217486 95217521
refGene_gn=GLT6D1;refGene_tx=XM_019816803.2;refGene_gn=GLT6D1;refGene_tx=XM_019816805.2
D4 95219249 95219294
refGene_gn=GLT6D1;refGene_tx=XM_003996064.4;refGene_gn=GLT6D1;refGene_tx=XM_019816803.2;refGene_gn=GLT6D1;refGene_tx=XM_019816804.2;refGene_gn=GLT6D1;refGene_tx=XM_019816805.2
D4 95223609 95223638 refGene_gn=GLT6D1;refGene_tx=XM_023243068.1
D4 95223698 95223713
refGene_gn=LOC109494055;refGene_tx=XM_023243069.1
D4 95223723 95223891
refGene_gn=GLT6D1;refGene_tx=XM_003996064.4;refGene_gn=GLT6D1;refGene_tx=XM_019816803.2;refGene_gn=GLT6D1;refGene_tx=XM_023243068.1;refGene_gn=LOC109494055;refGene_tx=XM_023243069.1
D4 95224305 95224969
refGene_gn=GLT6D1;refGene_tx=XM_003996064.4;refGene_gn=GLT6D1;refGene_tx=XM_019816803.2;refGene_gn=GLT6D1;refGene_tx=XM_019816804.2;refGene_gn=GLT6D1;refGene_tx=XM_019816805.2;refGene_gn=GLT6D1;refGene_tx=XM_023243068.1
D4 95227902 95228196

refGene_gn=LOC109494055;refGene_tx=XM_023243069.1
D4	95228463 95228489
refGene_gn=LOC111556219;refGene_tx=XM_019816800.2
D4	95229138 95229234
refGene_gn=LOC111556219;refGene_tx=XM_019816800.2
D4	95229657 95230034
refGene_gn=LOC111556219;refGene_tx=XM_019816798.2;refGene_gn=LOC111556219;refGene_tx=XM_019816800.2
D4	95230232 95230306
refGene_gn=LOC111556219;refGene_tx=XM_019816798.2;refGene_gn=LOC111556219;refGene_tx=XM_019816800.2
D4	95231153 95231287
refGene_gn=LOC111556219;refGene_tx=XM_019816798.2;refGene_gn=LOC111556219;refGene_tx=XM_019816800.2
D4	95231767 95231920
refGene_gn=LOC111556219;refGene_tx=XM_019816798.2;refGene_gn=LOC111556219;refGene_tx=XM_019816800.2
D4	95243052 95243069
refGene_gn=LOC109494056;refGene_tx=XM_023243070.1
D4	95243296 95243401
refGene_gn=LOC109494056;refGene_tx=XM_023243070.1
D4	95243978 95244089
refGene_gn=LOC109494056;refGene_tx=XM_023243070.1
D4	95245068 95245142
refGene_gn=LOC109494056;refGene_tx=XM_023243070.1
D4	95245696 95245836
refGene_gn=LOC109494056;refGene_tx=XM_023243070.1
D4	95246229 95246325
refGene_gn=LOC109494056;refGene_tx=XM_023243070.1
D4	95254658 95254675 refGene_gn=BLGIII;refGene_tx=XM_011288758.3
D4	95254904 95255009 refGene_gn=BLGIII;refGene_tx=XM_011288758.3
D4	95255546 95255660 refGene_gn=BLGIII;refGene_tx=XM_011288758.3
D4	95256981 95257055 refGene_gn=BLGIII;refGene_tx=XM_011288758.3
D4	95257663 95257803 refGene_gn=BLGIII;refGene_tx=XM_011288758.3
D4	95258244 95258340 refGene_gn=BLGIII;refGene_tx=XM_011288758.3
D4	95267210 95267227 refGene_gn=BLGI;refGene_tx=XM_011288759.3
D4	95267454 95267559 refGene_gn=BLGI;refGene_tx=XM_011288759.3
D4	95268136 95268247 refGene_gn=BLGI;refGene_tx=XM_011288759.3
D4	95269124 95269198 refGene_gn=BLGI;refGene_tx=XM_011288759.3
D4	95269490 95269630 refGene_gn=BLGI;refGene_tx=XM_011288759.3
D4	95270021 95270117 refGene_gn=BLGI;refGene_tx=XM_011288759.3
D4	95274309 95274323
refGene_gn=LOC100533977;refGene_tx=XM_019815490.2
D4	95285456 95285470
refGene_gn=LOC100533977;refGene_tx=NM_001204777.1
D4	95286194 95286221
refGene_gn=LOC100533977;refGene_tx=NM_001204777.1;refGene_gn=LOC100533977;refGene_tx=XM_019815490.2
D4	95286402 95286501
refGene_gn=LOC100533977;refGene_tx=NM_001204777.1;refGene_gn=LOC100533

977;refGene_tx=XM_019815490.2
D4 95287234 95287345
refGene_gn=LOC100533977;refGene_tx=NM_001204777.1;refGene_gn=LOC100533
977;refGene_tx=XM_019815490.2
D4 95287822 95287893
refGene_gn=LOC100533977;refGene_tx=NM_001204777.1;refGene_gn=LOC100533
977;refGene_tx=XM_019815490.2
D4 95288327 95288458
refGene_gn=LOC100533977;refGene_tx=NM_001204777.1;refGene_gn=LOC100533
977;refGene_tx=XM_019815490.2
D4 95288743 95288833
refGene_gn=LOC100533977;refGene_tx=NM_001204777.1;refGene_gn=LOC100533
977;refGene_tx=XM_019815490.2
D4 95410284 95410474 refGene_gn=LCN6;refGene_tx=XM_023243390.1
D4 95412827 95412967 refGene_gn=LCN6;refGene_tx=XM_023243390.1
D4 95413200 95413290 refGene_gn=LCN6;refGene_tx=XM_023243390.1
D4 95423364 95423429 refGene_gn=LCN8;refGene_tx=XM_023243073.1
D4 95423527 95423617 refGene_gn=LCN8;refGene_tx=XM_023243073.1
D4 95423698 95423808 refGene_gn=LCN8;refGene_tx=XM_023243073.1
D4 95424094 95424160 refGene_gn=LCN8;refGene_tx=XM_023243073.1
D4 95424647 95424778 refGene_gn=LCN8;refGene_tx=XM_023243073.1
D4 95425619 95425703 refGene_gn=LCN8;refGene_tx=XM_023243073.1
D4 95429494 95429529 refGene_gn=LCN15;refGene_tx=XM_023243072.1
D4 95429768 95429870 refGene_gn=LCN15;refGene_tx=XM_023243072.1
D4 95430563 95430674 refGene_gn=LCN15;refGene_tx=XM_023243072.1
D4 95430899 95430970 refGene_gn=LCN15;refGene_tx=XM_023243072.1
D4 95431052 95431192 refGene_gn=LCN15;refGene_tx=XM_023243072.1
D4 95431570 95431672 refGene_gn=LCN15;refGene_tx=XM_023243072.1
D4 95439774 95439828 refGene_gn=TMEM141;refGene_tx=XM_003996082.5
D4 95440160 95440227 refGene_gn=TMEM141;refGene_tx=XM_003996082.5
D4 95440388 95440472 refGene_gn=TMEM141;refGene_tx=XM_003996082.5
D4 95440695 95440803 refGene_gn=TMEM141;refGene_tx=XM_003996082.5
D4 95441171 95441185 refGene_gn=TMEM141;refGene_tx=XM_003996082.5
D4 95444354 95444424
refGene_gn=CCDC183;refGene_tx=XM_003996083.5;refGene_gn=CCDC183;refGen
e_tx=XM_019816815.2
D4 95444831 95444953
refGene_gn=CCDC183;refGene_tx=XM_003996083.5;refGene_gn=CCDC183;refGen
e_tx=XM_019816815.2
D4 95445419 95445497
refGene_gn=CCDC183;refGene_tx=XM_003996083.5;refGene_gn=CCDC183;refGen
e_tx=XM_019816815.2
D4 95445704 95445872
refGene_gn=CCDC183;refGene_tx=XM_003996083.5;refGene_gn=CCDC183;refGen
e_tx=XM_019816815.2
D4 95446089 95446194
refGene_gn=CCDC183;refGene_tx=XM_003996083.5;refGene_gn=CCDC183;refGen
e_tx=XM_019816815.2
D4 95448072 95448195
refGene_gn=CCDC183;refGene_tx=XM_003996083.5;refGene_gn=CCDC183;refGen

e_tx=XM_019816815.2
D4 95449463 95449589
refGene_gn=CCDC183;refGene_tx=XM_003996083.5;refGene_gn=CCDC183;refGene_tx=XM_019816815.2
D4 95449681 95449748
refGene_gn=CCDC183;refGene_tx=XM_003996083.5;refGene_gn=CCDC183;refGene_tx=XM_019816815.2
D4 95450301 95450402
refGene_gn=CCDC183;refGene_tx=XM_003996083.5;refGene_gn=CCDC183;refGene_tx=XM_019816815.2
D4 95451001 95451162
refGene_gn=CCDC183;refGene_tx=XM_003996083.5;refGene_gn=CCDC183;refGene_tx=XM_019816815.2
D4 95451433 95451602
refGene_gn=CCDC183;refGene_tx=XM_003996083.5;refGene_gn=CCDC183;refGene_tx=XM_019816815.2
D4 95451690 95451798
refGene_gn=CCDC183;refGene_tx=XM_003996083.5;refGene_gn=CCDC183;refGene_tx=XM_019816815.2
D4 95451868 95451965
refGene_gn=CCDC183;refGene_tx=XM_003996083.5;refGene_gn=CCDC183;refGene_tx=XM_019816815.2
D4 95452687 95452806
refGene_gn=CCDC183;refGene_tx=XM_003996083.5;refGene_gn=CCDC183;refGene_tx=XM_019816815.2
D4 95453528 95453658 refGene_gn=RABL6;refGene_tx=XM_003996084.5
D4 95461516 95461567 refGene_gn=RABL6;refGene_tx=XM_003996084.5
D4 95461569 95461647 refGene_gn=RABL6;refGene_tx=XM_003996084.5
D4 95463354 95463402 refGene_gn=RABL6;refGene_tx=XM_003996084.5
D4 95465387 95465440 refGene_gn=RABL6;refGene_tx=XM_003996084.5
D4 95466359 95466451 refGene_gn=RABL6;refGene_tx=XM_003996084.5
D4 95468211 95468352 refGene_gn=RABL6;refGene_tx=XM_003996084.5
D4 95468704 95468810 refGene_gn=RABL6;refGene_tx=XM_003996084.5
D4 95472023 95472127 refGene_gn=RABL6;refGene_tx=XM_003996084.5
D4 95474053 95474358 refGene_gn=RABL6;refGene_tx=XM_003996084.5
D4 95474570 95474721 refGene_gn=RABL6;refGene_tx=XM_003996084.5
D4 95475633 95476126 refGene_gn=RABL6;refGene_tx=XM_003996084.5
D4 95476430 95476602 refGene_gn=RABL6;refGene_tx=XM_003996084.5
D4 95476960 95477019 refGene_gn=RABL6;refGene_tx=XM_003996084.5
D4 95477106 95477298 refGene_gn=RABL6;refGene_tx=XM_003996084.5
D4 95481276 95484216
refGene_gn=AJM1;refGene_tx=XM_011288757.3;refGene_gn=AJM1;refGene_tx=XM_023243071.1
D4 95487317 95487477
refGene_gn=PHPT1;refGene_tx=XM_003996092.5;refGene_gn=PHPT1;refGene_tx=XM_011288756.3
D4 95487808 95487933
refGene_gn=PHPT1;refGene_tx=XM_003996092.5;refGene_gn=PHPT1;refGene_tx=XM_011288756.3
D4 95488274 95488394 refGene_gn=PHPT1;refGene_tx=XM_011288756.3

D4 95488468 95488561 refGene_gn=PHPT1;refGene_tx=XM_003996092.5
D4 95489956 95489996
refGene_gn=MAMDC4;refGene_tx=XM_023243074.1;refGene_gn=MAMDC4;refGene_tx=XM_023243075.1;refGene_gn=MAMDC4;refGene_tx=XM_023243076.1;refGene_gn=MAMDC4;refGene_tx=XM_023243077.1;refGene_gn=MAMDC4;refGene_tx=XM_023243078.1;refGene_gn=MAMDC4;refGene_tx=XM_023243079.1
D4 95490510 95490618
refGene_gn=MAMDC4;refGene_tx=XM_023243074.1;refGene_gn=MAMDC4;refGene_tx=XM_023243075.1;refGene_gn=MAMDC4;refGene_tx=XM_023243076.1;refGene_gn=MAMDC4;refGene_tx=XM_023243077.1;refGene_gn=MAMDC4;refGene_tx=XM_023243078.1;refGene_gn=MAMDC4;refGene_tx=XM_023243079.1
D4 95490698 95490872
refGene_gn=MAMDC4;refGene_tx=XM_023243074.1;refGene_gn=MAMDC4;refGene_tx=XM_023243075.1;refGene_gn=MAMDC4;refGene_tx=XM_023243076.1;refGene_gn=MAMDC4;refGene_tx=XM_023243077.1;refGene_gn=MAMDC4;refGene_tx=XM_023243078.1;refGene_gn=MAMDC4;refGene_tx=XM_023243079.1
D4 95490951 95491077
refGene_gn=MAMDC4;refGene_tx=XM_023243074.1;refGene_gn=MAMDC4;refGene_tx=XM_023243075.1;refGene_gn=MAMDC4;refGene_tx=XM_023243076.1;refGene_gn=MAMDC4;refGene_tx=XM_023243077.1;refGene_gn=MAMDC4;refGene_tx=XM_023243078.1;refGene_gn=MAMDC4;refGene_tx=XM_023243079.1
D4 95491271 95491402
refGene_gn=MAMDC4;refGene_tx=XM_023243074.1;refGene_gn=MAMDC4;refGene_tx=XM_023243075.1;refGene_gn=MAMDC4;refGene_tx=XM_023243076.1;refGene_gn=MAMDC4;refGene_tx=XM_023243077.1;refGene_gn=MAMDC4;refGene_tx=XM_023243078.1;refGene_gn=MAMDC4;refGene_tx=XM_023243079.1
D4 95491485 95491570
refGene_gn=MAMDC4;refGene_tx=XM_023243074.1;refGene_gn=MAMDC4;refGene_tx=XM_023243075.1;refGene_gn=MAMDC4;refGene_tx=XM_023243076.1;refGene_gn=MAMDC4;refGene_tx=XM_023243077.1;refGene_gn=MAMDC4;refGene_tx=XM_023243078.1;refGene_gn=MAMDC4;refGene_tx=XM_023243079.1
D4 95491712 95492072
refGene_gn=MAMDC4;refGene_tx=XM_023243074.1;refGene_gn=MAMDC4;refGene_tx=XM_023243075.1;refGene_gn=MAMDC4;refGene_tx=XM_023243076.1;refGene_gn=MAMDC4;refGene_tx=XM_023243077.1;refGene_gn=MAMDC4;refGene_tx=XM_023243078.1;refGene_gn=MAMDC4;refGene_tx=XM_023243079.1
D4 95492153 95492242
refGene_gn=MAMDC4;refGene_tx=XM_023243074.1;refGene_gn=MAMDC4;refGene_tx=XM_023243075.1;refGene_gn=MAMDC4;refGene_tx=XM_023243076.1;refGene_gn=MAMDC4;refGene_tx=XM_023243077.1;refGene_gn=MAMDC4;refGene_tx=XM_023243078.1;refGene_gn=MAMDC4;refGene_tx=XM_023243079.1
D4 95492325 95492499
refGene_gn=MAMDC4;refGene_tx=XM_023243074.1;refGene_gn=MAMDC4;refGene_tx=XM_023243075.1;refGene_gn=MAMDC4;refGene_tx=XM_023243076.1;refGene_gn=MAMDC4;refGene_tx=XM_023243077.1;refGene_gn=MAMDC4;refGene_tx=XM_023243078.1;refGene_gn=MAMDC4;refGene_tx=XM_023243079.1
D4 95492810 95492895
refGene_gn=MAMDC4;refGene_tx=XM_023243074.1;refGene_gn=MAMDC4;refGene_tx=XM_023243075.1;refGene_gn=MAMDC4;refGene_tx=XM_023243076.1;refGene_gn=MAMDC4;refGene_tx=XM_023243077.1;refGene_gn=MAMDC4;refGene_tx=XM_02

3243078.1;refGene_gn=MAMDC4;refGene_tx=XM_023243079.1
D4 95492968 95493148
refGene_gn=MAMDC4;refGene_tx=XM_023243074.1;refGene_gn=MAMDC4;refGene_tx=XM_023243075.1;refGene_gn=MAMDC4;refGene_tx=XM_023243076.1;refGene_gn=MAMDC4;refGene_tx=XM_023243077.1;refGene_gn=MAMDC4;refGene_tx=XM_023243078.1;refGene_gn=MAMDC4;refGene_tx=XM_023243079.1
D4 95493235 95493364
refGene_gn=MAMDC4;refGene_tx=XM_023243074.1;refGene_gn=MAMDC4;refGene_tx=XM_023243075.1;refGene_gn=MAMDC4;refGene_tx=XM_023243076.1;refGene_gn=MAMDC4;refGene_tx=XM_023243077.1;refGene_gn=MAMDC4;refGene_tx=XM_023243078.1;refGene_gn=MAMDC4;refGene_tx=XM_023243079.1
D4 95493523 95493655
refGene_gn=MAMDC4;refGene_tx=XM_023243074.1;refGene_gn=MAMDC4;refGene_tx=XM_023243075.1;refGene_gn=MAMDC4;refGene_tx=XM_023243076.1;refGene_gn=MAMDC4;refGene_tx=XM_023243077.1;refGene_gn=MAMDC4;refGene_tx=XM_023243078.1;refGene_gn=MAMDC4;refGene_tx=XM_023243079.1
D4 95493765 95493887
refGene_gn=MAMDC4;refGene_tx=XM_023243074.1;refGene_gn=MAMDC4;refGene_tx=XM_023243075.1;refGene_gn=MAMDC4;refGene_tx=XM_023243076.1;refGene_gn=MAMDC4;refGene_tx=XM_023243077.1;refGene_gn=MAMDC4;refGene_tx=XM_023243078.1;refGene_gn=MAMDC4;refGene_tx=XM_023243079.1
D4 95493967 95494079
refGene_gn=MAMDC4;refGene_tx=XM_023243074.1;refGene_gn=MAMDC4;refGene_tx=XM_023243075.1;refGene_gn=MAMDC4;refGene_tx=XM_023243076.1;refGene_gn=MAMDC4;refGene_tx=XM_023243077.1;refGene_gn=MAMDC4;refGene_tx=XM_023243078.1;refGene_gn=MAMDC4;refGene_tx=XM_023243079.1
D4 95494255 95494372
refGene_gn=MAMDC4;refGene_tx=XM_023243074.1;refGene_gn=MAMDC4;refGene_tx=XM_023243075.1;refGene_gn=MAMDC4;refGene_tx=XM_023243076.1;refGene_gn=MAMDC4;refGene_tx=XM_023243077.1;refGene_gn=MAMDC4;refGene_tx=XM_023243078.1;refGene_gn=MAMDC4;refGene_tx=XM_023243079.1
D4 95494450 95494609
refGene_gn=MAMDC4;refGene_tx=XM_023243074.1;refGene_gn=MAMDC4;refGene_tx=XM_023243075.1;refGene_gn=MAMDC4;refGene_tx=XM_023243076.1;refGene_gn=MAMDC4;refGene_tx=XM_023243077.1;refGene_gn=MAMDC4;refGene_tx=XM_023243078.1;refGene_gn=MAMDC4;refGene_tx=XM_023243079.1
D4 95494718 95494855
refGene_gn=MAMDC4;refGene_tx=XM_023243074.1;refGene_gn=MAMDC4;refGene_tx=XM_023243075.1;refGene_gn=MAMDC4;refGene_tx=XM_023243076.1;refGene_gn=MAMDC4;refGene_tx=XM_023243077.1;refGene_gn=MAMDC4;refGene_tx=XM_023243078.1;refGene_gn=MAMDC4;refGene_tx=XM_023243079.1
D4 95494923 95495143
refGene_gn=MAMDC4;refGene_tx=XM_023243074.1;refGene_gn=MAMDC4;refGene_tx=XM_023243075.1;refGene_gn=MAMDC4;refGene_tx=XM_023243076.1;refGene_gn=MAMDC4;refGene_tx=XM_023243077.1;refGene_gn=MAMDC4;refGene_tx=XM_023243078.1;refGene_gn=MAMDC4;refGene_tx=XM_023243079.1
D4 95495212 95495350
refGene_gn=MAMDC4;refGene_tx=XM_023243074.1;refGene_gn=MAMDC4;refGene_tx=XM_023243075.1;refGene_gn=MAMDC4;refGene_tx=XM_023243076.1;refGene_gn=MAMDC4;refGene_tx=XM_023243077.1;refGene_gn=MAMDC4;refGene_tx=XM_02

3243078.1;refGene_gn=MAMDC4;refGene_tx=XM_023243079.1
D4	95495440	95495559
refGene_gn=MAMDC4;refGene_tx=XM_023243074.1;refGene_gn=MAMDC4;refGene_tx=XM_023243075.1;refGene_gn=MAMDC4;refGene_tx=XM_023243076.1;refGene_gn=MAMDC4;refGene_tx=XM_023243077.1;refGene_gn=MAMDC4;refGene_tx=XM_023243078.1;refGene_gn=MAMDC4;refGene_tx=XM_023243079.1
D4	95495642	95495733
refGene_gn=MAMDC4;refGene_tx=XM_023243074.1;refGene_gn=MAMDC4;refGene_tx=XM_023243075.1;refGene_gn=MAMDC4;refGene_tx=XM_023243076.1;refGene_gn=MAMDC4;refGene_tx=XM_023243077.1;refGene_gn=MAMDC4;refGene_tx=XM_023243078.1;refGene_gn=MAMDC4;refGene_tx=XM_023243079.1
D4	95495934	95496081
refGene_gn=MAMDC4;refGene_tx=XM_023243074.1;refGene_gn=MAMDC4;refGene_tx=XM_023243075.1;refGene_gn=MAMDC4;refGene_tx=XM_023243076.1;refGene_gn=MAMDC4;refGene_tx=XM_023243077.1;refGene_gn=MAMDC4;refGene_tx=XM_023243078.1;refGene_gn=MAMDC4;refGene_tx=XM_023243079.1
D4	95496245	95496380
refGene_gn=MAMDC4;refGene_tx=XM_023243074.1;refGene_gn=MAMDC4;refGene_tx=XM_023243075.1;refGene_gn=MAMDC4;refGene_tx=XM_023243076.1;refGene_gn=MAMDC4;refGene_tx=XM_023243077.1;refGene_gn=MAMDC4;refGene_tx=XM_023243078.1;refGene_gn=MAMDC4;refGene_tx=XM_023243079.1
D4	95496491	95496619
refGene_gn=MAMDC4;refGene_tx=XM_023243074.1;refGene_gn=MAMDC4;refGene_tx=XM_023243075.1;refGene_gn=MAMDC4;refGene_tx=XM_023243076.1;refGene_gn=MAMDC4;refGene_tx=XM_023243077.1;refGene_gn=MAMDC4;refGene_tx=XM_023243078.1;refGene_gn=MAMDC4;refGene_tx=XM_023243079.1
D4	95496716	95497160
refGene_gn=MAMDC4;refGene_tx=XM_023243074.1;refGene_gn=MAMDC4;refGene_tx=XM_023243075.1;refGene_gn=MAMDC4;refGene_tx=XM_023243076.1;refGene_gn=MAMDC4;refGene_tx=XM_023243077.1;refGene_gn=MAMDC4;refGene_tx=XM_023243078.1;refGene_gn=MAMDC4;refGene_tx=XM_023243079.1
D4	95497310	95497486
refGene_gn=MAMDC4;refGene_tx=XM_023243077.1;refGene_gn=MAMDC4;refGene_tx=XM_023243078.1;refGene_gn=MAMDC4;refGene_tx=XM_023243079.1
D4	95497619	95497661
refGene_gn=MAMDC4;refGene_tx=XM_023243077.1;refGene_gn=MAMDC4;refGene_tx=XM_023243078.1;refGene_gn=MAMDC4;refGene_tx=XM_023243079.1
D4	95499667	95499729	refGene_gn=EDF1;refGene_tx=XM_011288754.2
D4	95500368	95500462	refGene_gn=EDF1;refGene_tx=XM_011288754.2
D4	95500762	95500923	refGene_gn=EDF1;refGene_tx=XM_011288754.2
D4	95501279	95501331	refGene_gn=EDF1;refGene_tx=XM_011288754.2
D4	95502886	95502964	refGene_gn=EDF1;refGene_tx=XM_011288754.2
D4	95522133	95522321
refGene_gn=TRAF2;refGene_tx=XM_006939666.4;refGene_gn=TRAF2;refGene_tx=XM_023243084.1;refGene_gn=TRAF2;refGene_tx=XM_023243085.1
D4	95523160	95523239
refGene_gn=TRAF2;refGene_tx=XM_006939666.4;refGene_gn=TRAF2;refGene_tx=XM_023243084.1;refGene_gn=TRAF2;refGene_tx=XM_023243085.1
D4	95523473	95523572
refGene_gn=TRAF2;refGene_tx=XM_006939666.4;refGene_gn=TRAF2;refGene_tx

=XM_023243084.1;refGene_gn=TRAF2;refGene_tx=XM_023243085.1
D4 95528261 95528423
refGene_gn=TRAF2;refGene_tx=XM_006939666.4;refGene_gn=TRAF2;refGene_tx
=XM_023243084.1;refGene_gn=TRAF2;refGene_tx=XM_023243085.1
D4 95529415 95529490
refGene_gn=TRAF2;refGene_tx=XM_006939666.4;refGene_gn=TRAF2;refGene_tx
=XM_023243084.1;refGene_gn=TRAF2;refGene_tx=XM_023243085.1
D4 95530958 95531033
refGene_gn=TRAF2;refGene_tx=XM_006939666.4;refGene_gn=TRAF2;refGene_tx
=XM_023243084.1;refGene_gn=TRAF2;refGene_tx=XM_023243085.1
D4 95532453 95532801
refGene_gn=TRAF2;refGene_tx=XM_006939666.4;refGene_gn=TRAF2;refGene_tx
=XM_023243084.1;refGene_gn=TRAF2;refGene_tx=XM_023243085.1
D4 95533285 95533463
refGene_gn=TRAF2;refGene_tx=XM_006939666.4;refGene_gn=TRAF2;refGene_tx
=XM_023243084.1;refGene_gn=TRAF2;refGene_tx=XM_023243085.1
D4 95535150 95535299
refGene_gn=TRAF2;refGene_tx=XM_006939666.4;refGene_gn=TRAF2;refGene_tx
=XM_023243084.1;refGene_gn=TRAF2;refGene_tx=XM_023243085.1
D4 95535888 95536107
refGene_gn=TRAF2;refGene_tx=XM_006939666.4;refGene_gn=TRAF2;refGene_tx
=XM_023243084.1;refGene_gn=TRAF2;refGene_tx=XM_023243085.1
D4 95545436 95545680
refGene_gn=FBXW5;refGene_tx=XM_006939669.3;refGene_gn=FBXW5;refGene_tx
=XM_023243080.1;refGene_gn=FBXW5;refGene_tx=XM_023243081.1;refGene_gn=
FBXW5;refGene_tx=XM_023243082.1
D4 95545763 95545976
refGene_gn=FBXW5;refGene_tx=XM_006939669.3;refGene_gn=FBXW5;refGene_tx
=XM_023243080.1;refGene_gn=FBXW5;refGene_tx=XM_023243081.1;refGene_gn=
FBXW5;refGene_tx=XM_023243082.1
D4 95546052 95546200
refGene_gn=FBXW5;refGene_tx=XM_006939669.3;refGene_gn=FBXW5;refGene_tx
=XM_023243080.1;refGene_gn=FBXW5;refGene_tx=XM_023243081.1;refGene_gn=
FBXW5;refGene_tx=XM_023243082.1
D4 95546437 95547073
refGene_gn=FBXW5;refGene_tx=XM_006939669.3;refGene_gn=FBXW5;refGene_tx
=XM_023243080.1;refGene_gn=FBXW5;refGene_tx=XM_023243081.1;refGene_gn=
FBXW5;refGene_tx=XM_023243082.1
D4 95547191 95547366
refGene_gn=FBXW5;refGene_tx=XM_006939669.3;refGene_gn=FBXW5;refGene_tx
=XM_023243080.1;refGene_gn=FBXW5;refGene_tx=XM_023243081.1;refGene_gn=
FBXW5;refGene_tx=XM_023243082.1
D4 95547640 95547798
refGene_gn=FBXW5;refGene_tx=XM_006939669.3;refGene_gn=FBXW5;refGene_tx
=XM_023243080.1;refGene_gn=FBXW5;refGene_tx=XM_023243081.1;refGene_gn=
FBXW5;refGene_tx=XM_023243082.1
D4 95548168 95548384
refGene_gn=FBXW5;refGene_tx=XM_006939669.3;refGene_gn=FBXW5;refGene_tx
=XM_023243080.1;refGene_gn=FBXW5;refGene_tx=XM_023243081.1
D4 95549355 95549494

refGene_gn=FBXW5;refGene_tx=XM_023243080.1;refGene_gn=FBXW5;refGene_tx=XM_023243081.1;refGene_gn=FBXW5;refGene_tx=XM_023243082.1
D4 95549972 95550110 refGene_gn=C8G;refGene_tx=XM_006939670.4;refGene_gn=C8G;refGene_tx=XM_019816824.2
D4 95550192 95550329 refGene_gn=C8G;refGene_tx=XM_006939670.4;refGene_gn=C8G;refGene_tx=XM_019816824.2
D4 95550465 95550536 refGene_gn=C8G;refGene_tx=XM_006939670.4;refGene_gn=C8G;refGene_tx=XM_019816824.2
D4 95550609 95550717 refGene_gn=C8G;refGene_tx=XM_006939670.4;refGene_gn=C8G;refGene_tx=XM_019816824.2
D4 95550993 95551095 refGene_gn=C8G;refGene_tx=XM_006939670.4;refGene_gn=C8G;refGene_tx=XM_019816824.2
D4 95551186 95551225 refGene_gn=C8G;refGene_tx=XM_006939670.4;refGene_gn=C8G;refGene_tx=XM_019816824.2
D4 95551303 95551317 refGene_gn=C8G;refGene_tx=XM_006939670.4;refGene_gn=C8G;refGene_tx=XM_019816824.2
D4 95552004 95552065 refGene_gn=LCN12;refGene_tx=XM_019816826.2
D4 95553514 95553568 refGene_gn=LCN12;refGene_tx=XM_019816826.2
D4 95555698 95555786 refGene_gn=LCN12;refGene_tx=XM_019816826.2
D4 95556188 95556485 refGene_gn=LCN12;refGene_tx=XM_019816826.2
D4 95558080 95558160 refGene_gn=LCN12;refGene_tx=XM_019816826.2
D4 95558233 95558350 refGene_gn=LCN12;refGene_tx=XM_019816826.2
D4 95558465 95558611 refGene_gn=LCN12;refGene_tx=XM_019816826.2
D4 95575244 95575358 refGene_gn=PTGDS;refGene_tx=NM_001009300.1
D4 95576091 95576231 refGene_gn=PTGDS;refGene_tx=NM_001009300.1
D4 95576347 95576424 refGene_gn=PTGDS;refGene_tx=NM_001009300.1
D4 95577123 95577240 refGene_gn=PTGDS;refGene_tx=NM_001009300.1
D4 95577390 95577492 refGene_gn=PTGDS;refGene_tx=NM_001009300.1
D4 95578144 95578170 refGene_gn=PTGDS;refGene_tx=NM_001009300.1
D4 95579632 95579731 refGene_gn=LCNL1;refGene_tx=XM_006939671.4
D4 95580070 95580210 refGene_gn=LCNL1;refGene_tx=XM_006939671.4
D4 95580594 95580668 refGene_gn=LCNL1;refGene_tx=XM_006939671.4
D4 95580755 95580866 refGene_gn=LCNL1;refGene_tx=XM_006939671.4
D4 95581209 95581311 refGene_gn=LCNL1;refGene_tx=XM_006939671.4
D4 95581551 95581577 refGene_gn=LCNL1;refGene_tx=XM_006939671.4
D4 95586194 95586793 refGene_gn=PAXX;refGene_tx=XM_006939677.3;refGene_gn=PAXX;refGene_tx=XM_023243087.1;refGene_gn=PAXX;refGene_tx=XM_023243088.1
D4 95586877 95586992 refGene_gn=PAXX;refGene_tx=XM_006939677.3;refGene_gn=PAXX;refGene_tx=XM_023243087.1;refGene_gn=PAXX;refGene_tx=XM_023243088.1
D4 95587192 95587242 refGene_gn=PAXX;refGene_tx=XM_006939677.3;refGene_gn=PAXX;refGene_tx=X

M_023243087.1;refGene_gn=PAXX;refGene_tx=XM_023243088.1
D4	95587322	95587513
refGene_gn=PAXX;refGene_tx=XM_006939677.3;refGene_gn=PAXX;refGene_tx=XM_023243087.1
D4	95587591	95587660
refGene_gn=PAXX;refGene_tx=XM_006939677.3;refGene_gn=PAXX;refGene_tx=XM_023243087.1;refGene_gn=PAXX;refGene_tx=XM_023243088.1
D4	95587830	95587915
refGene_gn=PAXX;refGene_tx=XM_006939677.3;refGene_gn=PAXX;refGene_tx=XM_023243087.1;refGene_gn=PAXX;refGene_tx=XM_023243088.1
D4	95588015	95588055
refGene_gn=PAXX;refGene_tx=XM_006939677.3;refGene_gn=PAXX;refGene_tx=XM_023243087.1;refGene_gn=PAXX;refGene_tx=XM_023243088.1
D4	95588938	95589097
refGene_gn=CLIC3;refGene_tx=XM_023243089.1;refGene_gn=CLIC3;refGene_tx=XM_023243090.1
D4	95589177	95589353
refGene_gn=CLIC3;refGene_tx=XM_023243089.1;refGene_gn=CLIC3;refGene_tx=XM_023243090.1
D4	95589434	95589541
refGene_gn=CLIC3;refGene_tx=XM_023243089.1;refGene_gn=CLIC3;refGene_tx=XM_023243090.1
D4	95589688	95589814
refGene_gn=CLIC3;refGene_tx=XM_023243089.1;refGene_gn=CLIC3;refGene_tx=XM_023243090.1
D4	95589897	95590007
refGene_gn=CLIC3;refGene_tx=XM_023243089.1;refGene_gn=CLIC3;refGene_tx=XM_023243090.1
D4	95590806	95590842
refGene_gn=CLIC3;refGene_tx=XM_023243089.1;refGene_gn=CLIC3;refGene_tx=XM_023243090.1
D4	95595136	95595172	refGene_gn=ABCA2;refGene_tx=XM_023243091.1
D4	95595571	95595778	refGene_gn=ABCA2;refGene_tx=XM_023243091.1
D4	95595854	95595992	refGene_gn=ABCA2;refGene_tx=XM_023243091.1
D4	95596202	95596305	refGene_gn=ABCA2;refGene_tx=XM_023243091.1
D4	95596395	95596488	refGene_gn=ABCA2;refGene_tx=XM_023243091.1
D4	95596581	95596685	refGene_gn=ABCA2;refGene_tx=XM_023243091.1
D4	95596768	95596903	refGene_gn=ABCA2;refGene_tx=XM_023243091.1
D4	95596976	95597118	refGene_gn=ABCA2;refGene_tx=XM_023243091.1
D4	95597189	95597368	refGene_gn=ABCA2;refGene_tx=XM_023243091.1
D4	95597895	95598013	refGene_gn=ABCA2;refGene_tx=XM_023243091.1
D4	95598136	95598269	refGene_gn=ABCA2;refGene_tx=XM_023243091.1
D4	95598390	95598514	refGene_gn=ABCA2;refGene_tx=XM_023243091.1
D4	95598589	95598737	refGene_gn=ABCA2;refGene_tx=XM_023243091.1
D4	95598816	95598932	refGene_gn=ABCA2;refGene_tx=XM_023243091.1
D4	95599105	95599280	refGene_gn=ABCA2;refGene_tx=XM_023243091.1
D4	95599351	95599412	refGene_gn=ABCA2;refGene_tx=XM_023243091.1
D4	95599498	95599610	refGene_gn=ABCA2;refGene_tx=XM_023243091.1
D4	95599706	95599812	refGene_gn=ABCA2;refGene_tx=XM_023243091.1
D4	95599904	95600118	refGene_gn=ABCA2;refGene_tx=XM_023243091.1

D4 95600190 95600510 refGene_gn=ABCA2;refGene_tx=XM_023243091.1
D4 95600629 95600732 refGene_gn=ABCA2;refGene_tx=XM_023243091.1
D4 95600984 95601191 refGene_gn=ABCA2;refGene_tx=XM_023243091.1
D4 95601273 95601507 refGene_gn=ABCA2;refGene_tx=XM_023243091.1
D4 95601787 95601908 refGene_gn=ABCA2;refGene_tx=XM_023243091.1
D4 95602057 95602209 refGene_gn=ABCA2;refGene_tx=XM_023243091.1
D4 95602211 95602241 refGene_gn=ABCA2;refGene_tx=XM_023243091.1
D4 95602473 95602656 refGene_gn=ABCA2;refGene_tx=XM_023243091.1
D4 95602743 95602940 refGene_gn=ABCA2;refGene_tx=XM_023243091.1
D4 95603065 95603278 refGene_gn=ABCA2;refGene_tx=XM_023243091.1
D4 95603374 95603555 refGene_gn=ABCA2;refGene_tx=XM_023243091.1
D4 95603644 95603809 refGene_gn=ABCA2;refGene_tx=XM_023243091.1
D4 95603942 95604147 refGene_gn=ABCA2;refGene_tx=XM_023243091.1
D4 95604220 95604371 refGene_gn=ABCA2;refGene_tx=XM_023243091.1
D4 95604566 95604757 refGene_gn=ABCA2;refGene_tx=XM_023243091.1
D4 95604984 95605099 refGene_gn=ABCA2;refGene_tx=XM_023243091.1
D4 95605185 95605288 refGene_gn=ABCA2;refGene_tx=XM_023243091.1
D4 95605363 95605537 refGene_gn=ABCA2;refGene_tx=XM_023243091.1
D4 95605996 95606093 refGene_gn=ABCA2;refGene_tx=XM_023243091.1
D4 95606161 95606329 refGene_gn=ABCA2;refGene_tx=XM_023243091.1
D4 95606452 95606581 refGene_gn=ABCA2;refGene_tx=XM_023243091.1
D4 95607360 95607520 refGene_gn=ABCA2;refGene_tx=XM_023243091.1
D4 95607760 95608131 refGene_gn=ABCA2;refGene_tx=XM_023243091.1
D4 95608467 95608686 refGene_gn=ABCA2;refGene_tx=XM_023243091.1
D4 95609038 95609137 refGene_gn=ABCA2;refGene_tx=XM_023243091.1
D4 95609342 95609470 refGene_gn=ABCA2;refGene_tx=XM_023243091.1
D4 95609780 95609944 refGene_gn=ABCA2;refGene_tx=XM_023243091.1
D4 95610020 95610132 refGene_gn=ABCA2;refGene_tx=XM_023243091.1
D4 95610816 95610819 refGene_gn=ABCA2;refGene_tx=XM_023243091.1
D4 95611090 95611184 refGene_gn=ABCA2;refGene_tx=XM_023243091.1
D4 95614938 95615004 refGene_gn=ABCA2;refGene_tx=XM_023243091.1
D4 95617620 95618637 refGene_gn=FUT7;refGene_tx=XM_011288752.3
D4 95618741 95618747 refGene_gn=FUT7;refGene_tx=XM_011288752.3
D4 95624588 95624657
refGene_gn=NPDC1;refGene_tx=XM_023243094.1;refGene_gn=NPDC1;refGene_tx
=XM_023243095.1
D4 95624747 95624844
refGene_gn=NPDC1;refGene_tx=XM_023243094.1;refGene_gn=NPDC1;refGene_tx
=XM_023243095.1
D4 95625076 95625156
refGene_gn=NPDC1;refGene_tx=XM_023243094.1;refGene_gn=NPDC1;refGene_tx
=XM_023243095.1
D4 95625241 95625326
refGene_gn=NPDC1;refGene_tx=XM_023243094.1;refGene_gn=NPDC1;refGene_tx
=XM_023243095.1
D4 95625402 95625469
refGene_gn=NPDC1;refGene_tx=XM_023243094.1;refGene_gn=NPDC1;refGene_tx
=XM_023243095.1
D4 95625557 95625731
refGene_gn=NPDC1;refGene_tx=XM_023243094.1;refGene_gn=NPDC1;refGene_tx

=XM_023243095.1
D4 95625824 95625959
refGene_gn=NPDC1;refGene_tx=XM_023243094.1;refGene_gn=NPDC1;refGene_tx=XM_023243095.1
D4 95627065 95627212
refGene_gn=NPDC1;refGene_tx=XM_023243094.1;refGene_gn=NPDC1;refGene_tx=XM_023243095.1
D4 95629587 95629669 refGene_gn=NPDC1;refGene_tx=XM_023243094.1
D4 95629840 95629952 refGene_gn=NPDC1;refGene_tx=XM_023243095.1
D4 95632119 95632323
refGene_gn=ENTPD2;refGene_tx=XM_023243092.1;refGene_gn=ENTPD2;refGene_tx=XM_023243093.1
D4 95632397 95632532
refGene_gn=ENTPD2;refGene_tx=XM_023243092.1;refGene_gn=ENTPD2;refGene_tx=XM_023243093.1
D4 95633309 95633429
refGene_gn=ENTPD2;refGene_tx=XM_023243092.1;refGene_gn=ENTPD2;refGene_tx=XM_023243093.1
D4 95633511 95633766
refGene_gn=ENTPD2;refGene_tx=XM_023243092.1;refGene_gn=ENTPD2;refGene_tx=XM_023243093.1
D4 95634221 95634449 refGene_gn=ENTPD2;refGene_tx=XM_023243092.1
D4 95634529 95634689
refGene_gn=ENTPD2;refGene_tx=XM_023243092.1;refGene_gn=ENTPD2;refGene_tx=XM_023243093.1
D4 95634872 95635023
refGene_gn=ENTPD2;refGene_tx=XM_023243092.1;refGene_gn=ENTPD2;refGene_tx=XM_023243093.1
D4 95635542 95635660
refGene_gn=ENTPD2;refGene_tx=XM_023243092.1;refGene_gn=ENTPD2;refGene_tx=XM_023243093.1
D4 95636918 95637035
refGene_gn=ENTPD2;refGene_tx=XM_023243092.1;refGene_gn=ENTPD2;refGene_tx=XM_023243093.1
D4 95650452 95650581 refGene_gn=SAPCD2;refGene_tx=XM_023243096.1
D4 95650654 95650777 refGene_gn=SAPCD2;refGene_tx=XM_023243096.1
D4 95650889 95650997 refGene_gn=SAPCD2;refGene_tx=XM_023243096.1
D4 95651333 95651480 refGene_gn=SAPCD2;refGene_tx=XM_023243096.1
D4 95652119 95652232 refGene_gn=SAPCD2;refGene_tx=XM_023243096.1
D4 95654610 95655196 refGene_gn=SAPCD2;refGene_tx=XM_023243096.1
D4 95659949 95660238 refGene_gn=UAP1L1;refGene_tx=XM_023243097.1
D4 95660915 95661120 refGene_gn=UAP1L1;refGene_tx=XM_023243097.1
D4 95661313 95661489 refGene_gn=UAP1L1;refGene_tx=XM_023243097.1
D4 95661791 95661964 refGene_gn=UAP1L1;refGene_tx=XM_023243097.1
D4 95662062 95662256 refGene_gn=UAP1L1;refGene_tx=XM_023243097.1
D4 95662727 95662868 refGene_gn=UAP1L1;refGene_tx=XM_023243097.1
D4 95663440 95663626 refGene_gn=UAP1L1;refGene_tx=XM_023243097.1
D4 95664199 95664266 refGene_gn=UAP1L1;refGene_tx=XM_023243097.1
D4 95664721 95664734 refGene_gn=UAP1L1;refGene_tx=XM_023243097.1
D4 95664736 95664804 refGene_gn=UAP1L1;refGene_tx=XM_023243097.1

D4 95667757 95667868
refGene_gn=LOC105261146;refGene_tx=XM_023243098.1
D4 95668372 95668423
refGene_gn=LOC105261146;refGene_tx=XM_023243098.1
D4 95668425 95668483
refGene_gn=LOC105261146;refGene_tx=XM_023243098.1
D4 95669169 95669285
refGene_gn=LOC105261146;refGene_tx=XM_023243098.1
D4 95670783 95670938
refGene_gn=LOC105261146;refGene_tx=XM_023243098.1
D4 95671228 95671341
refGene_gn=LOC105261146;refGene_tx=XM_023243098.1
D4 95672660 95672846
refGene_gn=LOC105261146;refGene_tx=XM_023243098.1
D4 95673541 95673690
refGene_gn=LOC105261146;refGene_tx=XM_023243098.1
D4 95674044 95674233
refGene_gn=LOC105261146;refGene_tx=XM_023243098.1
D4 95676137 95676328
refGene_gn=LOC105261146;refGene_tx=XM_023243098.1
D4 95677197 95677318
refGene_gn=LOC105261146;refGene_tx=XM_023243098.1
D4 95678040 95678238
refGene_gn=LOC105261146;refGene_tx=XM_023243098.1
D4 95678358 95678490
refGene_gn=LOC105261146;refGene_tx=XM_023243098.1
D4 95679187 95679400
refGene_gn=LOC105261146;refGene_tx=XM_023243098.1
D4 95688565 95688647
refGene_gn=DPP7;refGene_tx=XM_006944308.4;refGene_gn=DPP7;refGene_tx=XM_019816832.2
D4 95688721 95688835
refGene_gn=DPP7;refGene_tx=XM_006944308.4;refGene_gn=DPP7;refGene_tx=XM_019816832.2
D4 95688914 95689054
refGene_gn=DPP7;refGene_tx=XM_006944308.4;refGene_gn=DPP7;refGene_tx=XM_019816832.2
D4 95689135 95689299
refGene_gn=DPP7;refGene_tx=XM_006944308.4;refGene_gn=DPP7;refGene_tx=XM_019816832.2
D4 95689480 95689616
refGene_gn=DPP7;refGene_tx=XM_006944308.4;refGene_gn=DPP7;refGene_tx=XM_019816832.2
D4 95689689 95689771
refGene_gn=DPP7;refGene_tx=XM_006944308.4;refGene_gn=DPP7;refGene_tx=XM_019816832.2
D4 95689880 95690047
refGene_gn=DPP7;refGene_tx=XM_006944308.4;refGene_gn=DPP7;refGene_tx=XM_019816832.2
D4 95690220 95690281

refGene_gn=DPP7;refGene_tx=XM_006944308.4;refGene_gn=DPP7;refGene_tx=XM_019816832.2
D4 95690486 95690722
refGene_gn=DPP7;refGene_tx=XM_006944308.4;refGene_gn=DPP7;refGene_tx=XM_019816832.2
D4 95690800 95690957
refGene_gn=DPP7;refGene_tx=XM_006944308.4;refGene_gn=DPP7;refGene_tx=XM_019816832.2
D4 95691076 95691141
refGene_gn=DPP7;refGene_tx=XM_006944308.4;refGene_gn=DPP7;refGene_tx=XM_019816832.2
D4 95691501 95691572
refGene_gn=DPP7;refGene_tx=XM_006944308.4;refGene_gn=DPP7;refGene_tx=XM_019816832.2
D4 95691695 95691897
refGene_gn=DPP7;refGene_tx=XM_006944308.4;refGene_gn=DPP7;refGene_tx=XM_019816832.2
D4 95693459 95693672
refGene_gn=LOC111557571;refGene_tx=XM_023243099.1
D4 95694373 95694505
refGene_gn=LOC111557571;refGene_tx=XM_023243099.1
D4 95694639 95694704
refGene_gn=LOC111557571;refGene_tx=XM_023243099.1
D4 95694706 95694836
refGene_gn=LOC111557571;refGene_tx=XM_023243099.1
D4 95695694 95695815
refGene_gn=LOC111557571;refGene_tx=XM_023243099.1
D4 95696686 95696877
refGene_gn=LOC111557571;refGene_tx=XM_023243099.1
D4 95698776 95698965
refGene_gn=LOC111557571;refGene_tx=XM_023243099.1
D4 95699320 95699469
refGene_gn=LOC111557571;refGene_tx=XM_023243099.1
D4 95700165 95700351
refGene_gn=LOC111557571;refGene_tx=XM_023243099.1
D4 95701671 95701784
refGene_gn=LOC111557571;refGene_tx=XM_023243099.1
D4 95702074 95702229
refGene_gn=LOC111557571;refGene_tx=XM_023243099.1
D4 95703725 95703841
refGene_gn=LOC111557571;refGene_tx=XM_023243099.1
D4 95704529 95704638
refGene_gn=LOC111557571;refGene_tx=XM_023243099.1
D4 95705140 95705251
refGene_gn=LOC111557571;refGene_tx=XM_023243099.1
D4 95717424 95717682
refGene_gn=GRIN1;refGene_tx=XM_011288748.3;refGene_gn=GRIN1;refGene_tx=XM_011288749.3;refGene_gn=GRIN1;refGene_tx=XM_011288750.3;refGene_gn=GRIN1;refGene_tx=XM_019816834.2;refGene_gn=GRIN1;refGene_tx=XM_019816837.2;refGene_gn=GRIN1;refGene_tx=XM_019816838.2;refGene_gn=GRIN1;refGe

ne_tx=XM_019816839.2;refGene_gn=GRIN1;refGene_tx=XM_019816840.2;refGene_gn=GRIN1;refGene_tx=XM_019816843.2
D4 95720214 95720349
refGene_gn=GRIN1;refGene_tx=XM_011288748.3;refGene_gn=GRIN1;refGene_tx=XM_011288749.3;refGene_gn=GRIN1;refGene_tx=XM_011288750.3;refGene_gn=GRIN1;refGene_tx=XM_019816834.2;refGene_gn=GRIN1;refGene_tx=XM_019816837.2;refGene_gn=GRIN1;refGene_tx=XM_019816838.2;refGene_gn=GRIN1;refGene_tx=XM_019816839.2;refGene_gn=GRIN1;refGene_tx=XM_019816840.2;refGene_gn=GRIN1;refGene_tx=XM_019816842.2;refGene_gn=GRIN1;refGene_tx=XM_019816843.2
D4 95722854 95723031
refGene_gn=GRIN1;refGene_tx=XM_011288748.3;refGene_gn=GRIN1;refGene_tx=XM_011288749.3;refGene_gn=GRIN1;refGene_tx=XM_011288750.3;refGene_gn=GRIN1;refGene_tx=XM_019816834.2;refGene_gn=GRIN1;refGene_tx=XM_019816837.2;refGene_gn=GRIN1;refGene_tx=XM_019816838.2;refGene_gn=GRIN1;refGene_tx=XM_019816839.2;refGene_gn=GRIN1;refGene_tx=XM_019816840.2;refGene_gn=GRIN1;refGene_tx=XM_019816842.2;refGene_gn=GRIN1;refGene_tx=XM_019816843.2
D4 95725189 95725252
refGene_gn=GRIN1;refGene_tx=XM_011288748.3;refGene_gn=GRIN1;refGene_tx=XM_011288749.3;refGene_gn=GRIN1;refGene_tx=XM_011288750.3;refGene_gn=GRIN1;refGene_tx=XM_019816834.2;refGene_gn=GRIN1;refGene_tx=XM_019816838.2;refGene_gn=GRIN1;refGene_tx=XM_019816842.2
D4 95726000 95726101
refGene_gn=GRIN1;refGene_tx=XM_011288748.3;refGene_gn=GRIN1;refGene_tx=XM_011288749.3;refGene_gn=GRIN1;refGene_tx=XM_011288750.3;refGene_gn=GRIN1;refGene_tx=XM_019816834.2;refGene_gn=GRIN1;refGene_tx=XM_019816837.2;refGene_gn=GRIN1;refGene_tx=XM_019816838.2;refGene_gn=GRIN1;refGene_tx=XM_019816839.2;refGene_gn=GRIN1;refGene_tx=XM_019816840.2;refGene_gn=GRIN1;refGene_tx=XM_019816842.2;refGene_gn=GRIN1;refGene_tx=XM_019816843.2
D4 95730155 95730277
refGene_gn=GRIN1;refGene_tx=XM_011288748.3;refGene_gn=GRIN1;refGene_tx=XM_011288749.3;refGene_gn=GRIN1;refGene_tx=XM_011288750.3;refGene_gn=GRIN1;refGene_tx=XM_019816834.2;refGene_gn=GRIN1;refGene_tx=XM_019816837.2;refGene_gn=GRIN1;refGene_tx=XM_019816838.2;refGene_gn=GRIN1;refGene_tx=XM_019816839.2;refGene_gn=GRIN1;refGene_tx=XM_019816840.2;refGene_gn=GRIN1;refGene_tx=XM_019816842.2;refGene_gn=GRIN1;refGene_tx=XM_019816843.2
D4 95730352 95730527
refGene_gn=GRIN1;refGene_tx=XM_011288748.3;refGene_gn=GRIN1;refGene_tx=XM_011288749.3;refGene_gn=GRIN1;refGene_tx=XM_011288750.3;refGene_gn=GRIN1;refGene_tx=XM_019816834.2;refGene_gn=GRIN1;refGene_tx=XM_019816837.2;refGene_gn=GRIN1;refGene_tx=XM_019816838.2;refGene_gn=GRIN1;refGene_tx=XM_019816839.2;refGene_gn=GRIN1;refGene_tx=XM_019816840.2;refGene_gn=GRIN1;refGene_tx=XM_019816842.2;refGene_gn=GRIN1;refGene_tx=XM_019816843.2
D4 95731825 95731970
refGene_gn=GRIN1;refGene_tx=XM_011288748.3;refGene_gn=GRIN1;refGene_tx=XM_011288749.3;refGene_gn=GRIN1;refGene_tx=XM_011288750.3;refGene_gn=

GRIN1;refGene_tx=XM_019816834.2;refGene_gn=GRIN1;refGene_tx=XM_0198168
37.2;refGene_gn=GRIN1;refGene_tx=XM_019816838.2;refGene_gn=GRIN1;refGe
ne_tx=XM_019816839.2;refGene_gn=GRIN1;refGene_tx=XM_019816840.2;refGen
e_gn=GRIN1;refGene_tx=XM_019816842.2;refGene_gn=GRIN1;refGene_tx=XM_01
9816843.2
D4 95732067 95732151
refGene_gn=GRIN1;refGene_tx=XM_011288748.3;refGene_gn=GRIN1;refGene_tx
=XM_011288749.3;refGene_gn=GRIN1;refGene_tx=XM_011288750.3;refGene_gn=
GRIN1;refGene_tx=XM_019816834.2;refGene_gn=GRIN1;refGene_tx=XM_0198168
37.2;refGene_gn=GRIN1;refGene_tx=XM_019816838.2;refGene_gn=GRIN1;refGe
ne_tx=XM_019816839.2;refGene_gn=GRIN1;refGene_tx=XM_019816840.2;refGen
e_gn=GRIN1;refGene_tx=XM_019816842.2;refGene_gn=GRIN1;refGene_tx=XM_01
9816843.2
D4 95734086 95734228
refGene_gn=GRIN1;refGene_tx=XM_011288748.3;refGene_gn=GRIN1;refGene_tx
=XM_011288749.3;refGene_gn=GRIN1;refGene_tx=XM_011288750.3;refGene_gn=
GRIN1;refGene_tx=XM_019816834.2;refGene_gn=GRIN1;refGene_tx=XM_0198168
37.2;refGene_gn=GRIN1;refGene_tx=XM_019816838.2;refGene_gn=GRIN1;refGe
ne_tx=XM_019816839.2;refGene_gn=GRIN1;refGene_tx=XM_019816840.2;refGen
e_gn=GRIN1;refGene_tx=XM_019816842.2;refGene_gn=GRIN1;refGene_tx=XM_01
9816843.2
D4 95734312 95734440
refGene_gn=GRIN1;refGene_tx=XM_011288748.3;refGene_gn=GRIN1;refGene_tx
=XM_011288749.3;refGene_gn=GRIN1;refGene_tx=XM_011288750.3;refGene_gn=
GRIN1;refGene_tx=XM_019816834.2;refGene_gn=GRIN1;refGene_tx=XM_0198168
37.2;refGene_gn=GRIN1;refGene_tx=XM_019816838.2;refGene_gn=GRIN1;refGe
ne_tx=XM_019816839.2;refGene_gn=GRIN1;refGene_tx=XM_019816840.2;refGen
e_gn=GRIN1;refGene_tx=XM_019816842.2;refGene_gn=GRIN1;refGene_tx=XM_01
9816843.2
D4 95734713 95734878
refGene_gn=GRIN1;refGene_tx=XM_011288748.3;refGene_gn=GRIN1;refGene_tx
=XM_011288749.3;refGene_gn=GRIN1;refGene_tx=XM_011288750.3;refGene_gn=
GRIN1;refGene_tx=XM_019816834.2;refGene_gn=GRIN1;refGene_tx=XM_0198168
37.2;refGene_gn=GRIN1;refGene_tx=XM_019816838.2;refGene_gn=GRIN1;refGe
ne_tx=XM_019816839.2;refGene_gn=GRIN1;refGene_tx=XM_019816840.2;refGen
e_gn=GRIN1;refGene_tx=XM_019816842.2;refGene_gn=GRIN1;refGene_tx=XM_01
9816843.2
D4 95734953 95735072
refGene_gn=GRIN1;refGene_tx=XM_011288748.3;refGene_gn=GRIN1;refGene_tx
=XM_011288749.3;refGene_gn=GRIN1;refGene_tx=XM_011288750.3;refGene_gn=
GRIN1;refGene_tx=XM_019816834.2;refGene_gn=GRIN1;refGene_tx=XM_0198168
37.2;refGene_gn=GRIN1;refGene_tx=XM_019816838.2;refGene_gn=GRIN1;refGe
ne_tx=XM_019816839.2;refGene_gn=GRIN1;refGene_tx=XM_019816840.2;refGen
e_gn=GRIN1;refGene_tx=XM_019816842.2;refGene_gn=GRIN1;refGene_tx=XM_01
9816843.2
D4 95735202 95735315
refGene_gn=GRIN1;refGene_tx=XM_011288748.3;refGene_gn=GRIN1;refGene_tx
=XM_011288749.3;refGene_gn=GRIN1;refGene_tx=XM_011288750.3;refGene_gn=
GRIN1;refGene_tx=XM_019816834.2;refGene_gn=GRIN1;refGene_tx=XM_0198168
37.2;refGene_gn=GRIN1;refGene_tx=XM_019816838.2;refGene_gn=GRIN1;refGe

ne_tx=XM_019816839.2;refGene_gn=GRIN1;refGene_tx=XM_019816840.2;refGene_gn=GRIN1;refGene_tx=XM_019816842.2;refGene_gn=GRIN1;refGene_tx=XM_019816843.2
D4 95735390 95735539 refGene_gn=GRIN1;refGene_tx=XM_011288748.3;refGene_gn=GRIN1;refGene_tx=XM_011288749.3;refGene_gn=GRIN1;refGene_tx=XM_011288750.3;refGene_gn=GRIN1;refGene_tx=XM_019816834.2;refGene_gn=GRIN1;refGene_tx=XM_019816837.2;refGene_gn=GRIN1;refGene_tx=XM_019816838.2;refGene_gn=GRIN1;refGene_tx=XM_019816839.2;refGene_gn=GRIN1;refGene_tx=XM_019816840.2;refGene_gn=GRIN1;refGene_tx=XM_019816842.2;refGene_gn=GRIN1;refGene_tx=XM_019816843.2
D4 95735639 95735797 refGene_gn=GRIN1;refGene_tx=XM_011288748.3;refGene_gn=GRIN1;refGene_tx=XM_011288749.3;refGene_gn=GRIN1;refGene_tx=XM_011288750.3;refGene_gn=GRIN1;refGene_tx=XM_019816834.2;refGene_gn=GRIN1;refGene_tx=XM_019816837.2;refGene_gn=GRIN1;refGene_tx=XM_019816838.2;refGene_gn=GRIN1;refGene_tx=XM_019816839.2;refGene_gn=GRIN1;refGene_tx=XM_019816840.2;refGene_gn=GRIN1;refGene_tx=XM_019816842.2;refGene_gn=GRIN1;refGene_tx=XM_019816843.2
D4 95736041 95736203 refGene_gn=GRIN1;refGene_tx=XM_011288748.3;refGene_gn=GRIN1;refGene_tx=XM_011288749.3;refGene_gn=GRIN1;refGene_tx=XM_011288750.3;refGene_gn=GRIN1;refGene_tx=XM_019816834.2;refGene_gn=GRIN1;refGene_tx=XM_019816837.2;refGene_gn=GRIN1;refGene_tx=XM_019816838.2;refGene_gn=GRIN1;refGene_tx=XM_019816839.2;refGene_gn=GRIN1;refGene_tx=XM_019816840.2;refGene_gn=GRIN1;refGene_tx=XM_019816842.2;refGene_gn=GRIN1;refGene_tx=XM_019816843.2
D4 95736385 95736495 refGene_gn=GRIN1;refGene_tx=XM_011288748.3;refGene_gn=GRIN1;refGene_tx=XM_011288749.3;refGene_gn=GRIN1;refGene_tx=XM_011288750.3;refGene_gn=GRIN1;refGene_tx=XM_019816834.2;refGene_gn=GRIN1;refGene_tx=XM_019816837.2;refGene_gn=GRIN1;refGene_tx=XM_019816838.2;refGene_gn=GRIN1;refGene_tx=XM_019816839.2;refGene_gn=GRIN1;refGene_tx=XM_019816840.2;refGene_gn=GRIN1;refGene_tx=XM_019816842.2;refGene_gn=GRIN1;refGene_tx=XM_019816843.2
D4 95736601 95736747 refGene_gn=GRIN1;refGene_tx=XM_011288748.3;refGene_gn=GRIN1;refGene_tx=XM_011288749.3;refGene_gn=GRIN1;refGene_tx=XM_011288750.3;refGene_gn=GRIN1;refGene_tx=XM_019816834.2;refGene_gn=GRIN1;refGene_tx=XM_019816837.2;refGene_gn=GRIN1;refGene_tx=XM_019816838.2;refGene_gn=GRIN1;refGene_tx=XM_019816839.2;refGene_gn=GRIN1;refGene_tx=XM_019816840.2;refGene_gn=GRIN1;refGene_tx=XM_019816842.2;refGene_gn=GRIN1;refGene_tx=XM_019816843.2
D4 95738015 95738126 refGene_gn=GRIN1;refGene_tx=XM_011288748.3;refGene_gn=GRIN1;refGene_tx=XM_011288749.3;refGene_gn=GRIN1;refGene_tx=XM_019816834.2;refGene_gn=GRIN1;refGene_tx=XM_019816837.2;refGene_gn=GRIN1;refGene_tx=XM_019816842.2;refGene_gn=GRIN1;refGene_tx=XM_019816843.2
D4 95738989 95739058 refGene_gn=GRIN1;refGene_tx=XM_019816834.2
D4 95740051 95740168

refGene_gn=GRIN1;refGene_tx=XM_011288748.3;refGene_gn=GRIN1;refGene_tx=XM_019816838.2;refGene_gn=GRIN1;refGene_tx=XM_019816839.2;refGene_gn=GRIN1;refGene_tx=XM_019816842.2;refGene_gn=GRIN1;refGene_tx=XM_019816843.2
D4 95740419 95740488
refGene_gn=GRIN1;refGene_tx=XM_011288749.3;refGene_gn=GRIN1;refGene_tx=XM_011288750.3;refGene_gn=GRIN1;refGene_tx=XM_019816837.2;refGene_gn=GRIN1;refGene_tx=XM_019816840.2
D4 95741412 95741702 refGene_gn=LRRC26;refGene_tx=XM_023243102.1
D4 95741790 95742463 refGene_gn=LRRC26;refGene_tx=XM_023243102.1
D4 95743531 95743733 refGene_gn=TMEM210;refGene_tx=XM_023243103.1
D4 95743922 95743958 refGene_gn=TMEM210;refGene_tx=XM_023243103.1
D4 95744167 95744303 refGene_gn=TMEM210;refGene_tx=XM_023243103.1
D4 95744438 95744526 refGene_gn=TMEM210;refGene_tx=XM_023243103.1
D4 95746632 95746845 refGene_gn=ANAPC2;refGene_tx=XM_011288746.3
D4 95746950 95747186 refGene_gn=ANAPC2;refGene_tx=XM_011288746.3
D4 95747382 95747512 refGene_gn=ANAPC2;refGene_tx=XM_011288746.3
D4 95750509 95750713 refGene_gn=ANAPC2;refGene_tx=XM_011288746.3
D4 95750802 95750878 refGene_gn=ANAPC2;refGene_tx=XM_011288746.3
D4 95751186 95751328 refGene_gn=ANAPC2;refGene_tx=XM_011288746.3
D4 95751679 95751861 refGene_gn=ANAPC2;refGene_tx=XM_011288746.3
D4 95752578 95752696 refGene_gn=ANAPC2;refGene_tx=XM_011288746.3
D4 95753050 95753170 refGene_gn=ANAPC2;refGene_tx=XM_011288746.3
D4 95753547 95753722 refGene_gn=ANAPC2;refGene_tx=XM_011288746.3
D4 95754727 95754860 refGene_gn=ANAPC2;refGene_tx=XM_011288746.3
D4 95755856 95756479 refGene_gn=ANAPC2;refGene_tx=XM_011288746.3
D4 95756742 95756847 refGene_gn=ANAPC2;refGene_tx=XM_011288746.3
D4 95757014 95757066 refGene_gn=SSNA1;refGene_tx=XM_023243104.1
D4 95757538 95757738 refGene_gn=SSNA1;refGene_tx=XM_023243104.1
D4 95758148 95758256 refGene_gn=SSNA1;refGene_tx=XM_023243104.1
D4 95760421 95760484 refGene_gn=TPRN;refGene_tx=XM_023243100.1
D4 95760593 95760700 refGene_gn=TPRN;refGene_tx=XM_023243100.1
D4 95760779 95761023 refGene_gn=TPRN;refGene_tx=XM_023243100.1
D4 95766488 95768321 refGene_gn=TPRN;refGene_tx=XM_023243100.1
D4 95771661 95772072 refGene_gn=TMEM203;refGene_tx=XM_003996118.3
D4 95772444 95772579
refGene_gn=NDOR1;refGene_tx=XM_003996105.5;refGene_gn=NDOR1;refGene_tx=XM_006939684.4;refGene_gn=NDOR1;refGene_tx=XM_019816846.1
D4 95772921 95772999
refGene_gn=NDOR1;refGene_tx=XM_003996105.5;refGene_gn=NDOR1;refGene_tx=XM_006939684.4;refGene_gn=NDOR1;refGene_tx=XM_019816846.1;refGene_gn=NDOR1;refGene_tx=XM_019816847.2
D4 95776886 95776984
refGene_gn=NDOR1;refGene_tx=XM_003996105.5;refGene_gn=NDOR1;refGene_tx=XM_006939684.4;refGene_gn=NDOR1;refGene_tx=XM_019816846.1;refGene_gn=NDOR1;refGene_tx=XM_019816847.2;refGene_gn=NDOR1;refGene_tx=XM_019816848.1;refGene_gn=NDOR1;refGene_tx=XM_019816849.1
D4 95777900 95777999
refGene_gn=NDOR1;refGene_tx=XM_003996105.5;refGene_gn=NDOR1;refGene_tx=XM_006939684.4;refGene_gn=NDOR1;refGene_tx=XM_019816846.1;refGene_gn=

NDOR1;refGene_tx=XM_019816847.2;refGene_gn=NDOR1;refGene_tx=XM_0198168
48.1;refGene_gn=NDOR1;refGene_tx=XM_019816849.1
D4 95778083 95778185
refGene_gn=NDOR1;refGene_tx=XM_003996105.5;refGene_gn=NDOR1;refGene_tx
=XM_006939684.4;refGene_gn=NDOR1;refGene_tx=XM_019816846.1;refGene_gn=
NDOR1;refGene_tx=XM_019816847.2;refGene_gn=NDOR1;refGene_tx=XM_0198168
48.1;refGene_gn=NDOR1;refGene_tx=XM_019816849.1
D4 95778295 95778505
refGene_gn=NDOR1;refGene_tx=XM_003996105.5;refGene_gn=NDOR1;refGene_tx
=XM_006939684.4;refGene_gn=NDOR1;refGene_tx=XM_019816846.1;refGene_gn=
NDOR1;refGene_tx=XM_019816847.2;refGene_gn=NDOR1;refGene_tx=XM_0198168
48.1;refGene_gn=NDOR1;refGene_tx=XM_019816849.1
D4 95778674 95778796
refGene_gn=NDOR1;refGene_tx=XM_003996105.5;refGene_gn=NDOR1;refGene_tx
=XM_006939684.4;refGene_gn=NDOR1;refGene_tx=XM_019816846.1;refGene_gn=
NDOR1;refGene_tx=XM_019816847.2;refGene_gn=NDOR1;refGene_tx=XM_0198168
48.1;refGene_gn=NDOR1;refGene_tx=XM_019816849.1
D4 95778906 95779127
refGene_gn=NDOR1;refGene_tx=XM_003996105.5;refGene_gn=NDOR1;refGene_tx
=XM_006939684.4;refGene_gn=NDOR1;refGene_tx=XM_019816846.1;refGene_gn=
NDOR1;refGene_tx=XM_019816847.2;refGene_gn=NDOR1;refGene_tx=XM_0198168
48.1;refGene_gn=NDOR1;refGene_tx=XM_019816849.1
D4 95779236 95779344
refGene_gn=NDOR1;refGene_tx=XM_003996105.5;refGene_gn=NDOR1;refGene_tx
=XM_006939684.4;refGene_gn=NDOR1;refGene_tx=XM_019816846.1;refGene_gn=
NDOR1;refGene_tx=XM_019816847.2;refGene_gn=NDOR1;refGene_tx=XM_0198168
48.1;refGene_gn=NDOR1;refGene_tx=XM_019816849.1
D4 95779540 95779655
refGene_gn=NDOR1;refGene_tx=XM_003996105.5;refGene_gn=NDOR1;refGene_tx
=XM_006939684.4;refGene_gn=NDOR1;refGene_tx=XM_019816846.1;refGene_gn=
NDOR1;refGene_tx=XM_019816847.2;refGene_gn=NDOR1;refGene_tx=XM_0198168
48.1;refGene_gn=NDOR1;refGene_tx=XM_019816849.1
D4 95779790 95779937
refGene_gn=NDOR1;refGene_tx=XM_003996105.5;refGene_gn=NDOR1;refGene_tx
=XM_006939684.4;refGene_gn=NDOR1;refGene_tx=XM_019816846.1;refGene_gn=
NDOR1;refGene_tx=XM_019816847.2;refGene_gn=NDOR1;refGene_tx=XM_0198168
48.1;refGene_gn=NDOR1;refGene_tx=XM_019816849.1
D4 95780035 95780154
refGene_gn=NDOR1;refGene_tx=XM_003996105.5;refGene_gn=NDOR1;refGene_tx
=XM_006939684.4;refGene_gn=NDOR1;refGene_tx=XM_019816846.1;refGene_gn=
NDOR1;refGene_tx=XM_019816847.2;refGene_gn=NDOR1;refGene_tx=XM_0198168
48.1;refGene_gn=NDOR1;refGene_tx=XM_019816849.1
D4 95780267 95780362
refGene_gn=NDOR1;refGene_tx=XM_003996105.5;refGene_gn=NDOR1;refGene_tx
=XM_019816846.1;refGene_gn=NDOR1;refGene_tx=XM_019816847.2;refGene_gn=
NDOR1;refGene_tx=XM_019816848.1;refGene_gn=NDOR1;refGene_tx=XM_0198168
49.1
D4 95780449 95780594
refGene_gn=NDOR1;refGene_tx=XM_003996105.5;refGene_gn=NDOR1;refGene_tx
=XM_019816846.1;refGene_gn=NDOR1;refGene_tx=XM_019816847.2;refGene_gn=

NDOR1;refGene_tx=XM_019816848.1;refGene_gn=NDOR1;refGene_tx=XM_019816849.1
D4	95781298	95781382	refGene_gn=LOC102901825;refGene_tx=XM_006939680.3
D4	95781613	95782666	refGene_gn=LOC102901825;refGene_tx=XM_006939680.3
D4	95784197	95784995	refGene_gn=RNF208;refGene_tx=XM_023243111.1
D4	95788169	95788300	refGene_gn=CYSRT1;refGene_tx=XM_011288744.3
D4	95788800	95788856	refGene_gn=CYSRT1;refGene_tx=XM_011288744.3
D4	95789142	95789567	refGene_gn=CYSRT1;refGene_tx=XM_011288744.3
D4	95791703	95791801	refGene_gn=RNF224;refGene_tx=XM_003996119.5
D4	95792078	95792544	refGene_gn=RNF224;refGene_tx=XM_003996119.5
D4	95794234	95794343	refGene_gn=SLC34A3;refGene_tx=XM_023243105.1;refGene_gn=SLC34A3;refGene_tx=XM_023243106.1;refGene_gn=SLC34A3;refGene_tx=XM_023243107.1;refGene_gn=SLC34A3;refGene_tx=XM_023243109.1
D4	95795040	95795166	refGene_gn=SLC34A3;refGene_tx=XM_023243105.1;refGene_gn=SLC34A3;refGene_tx=XM_023243106.1;refGene_gn=SLC34A3;refGene_tx=XM_023243107.1;refGene_gn=SLC34A3;refGene_tx=XM_023243108.1;refGene_gn=SLC34A3;refGene_tx=XM_023243109.1
D4	95795435	95795525	refGene_gn=SLC34A3;refGene_tx=XM_023243105.1;refGene_gn=SLC34A3;refGene_tx=XM_023243106.1;refGene_gn=SLC34A3;refGene_tx=XM_023243107.1;refGene_gn=SLC34A3;refGene_tx=XM_023243108.1;refGene_gn=SLC34A3;refGene_tx=XM_023243109.1
D4	95796003	95796132	refGene_gn=SLC34A3;refGene_tx=XM_023243105.1;refGene_gn=SLC34A3;refGene_tx=XM_023243106.1;refGene_gn=SLC34A3;refGene_tx=XM_023243107.1;refGene_gn=SLC34A3;refGene_tx=XM_023243108.1;refGene_gn=SLC34A3;refGene_tx=XM_023243109.1
D4	95796222	95796366	refGene_gn=SLC34A3;refGene_tx=XM_023243105.1;refGene_gn=SLC34A3;refGene_tx=XM_023243106.1;refGene_gn=SLC34A3;refGene_tx=XM_023243107.1;refGene_gn=SLC34A3;refGene_tx=XM_023243108.1;refGene_gn=SLC34A3;refGene_tx=XM_023243109.1
D4	95796441	95796553	refGene_gn=SLC34A3;refGene_tx=XM_023243105.1;refGene_gn=SLC34A3;refGene_tx=XM_023243106.1;refGene_gn=SLC34A3;refGene_tx=XM_023243107.1;refGene_gn=SLC34A3;refGene_tx=XM_023243108.1;refGene_gn=SLC34A3;refGene_tx=XM_023243109.1
D4	95796638	95796834	refGene_gn=SLC34A3;refGene_tx=XM_023243105.1;refGene_gn=SLC34A3;refGene_tx=XM_023243106.1;refGene_gn=SLC34A3;refGene_tx=XM_023243107.1;refGene_gn=SLC34A3;refGene_tx=XM_023243108.1;refGene_gn=SLC34A3;refGene_tx=XM_023243109.1
D4	95797020	95797110	refGene_gn=SLC34A3;refGene_tx=XM_023243105.1;refGene_gn=SLC34A3;refGene_tx=XM_023243106.1;refGene_gn=SLC34A3;refGene_tx=XM_023243107.1;refGene_gn=SLC34A3;refGene_tx=XM_023243108.1;refGene_gn=SLC34A3;refGene_tx=

XM_023243109.1
D4	95797206	95797294
refGene_gn=SLC34A3;refGene_tx=XM_023243105.1;refGene_gn=SLC34A3;refGene_tx=XM_023243106.1;refGene_gn=SLC34A3;refGene_tx=XM_023243107.1;refGene_gn=SLC34A3;refGene_tx=XM_023243108.1;refGene_gn=SLC34A3;refGene_tx=XM_023243109.1
D4	95797375	95797543
refGene_gn=SLC34A3;refGene_tx=XM_023243105.1;refGene_gn=SLC34A3;refGene_tx=XM_023243106.1;refGene_gn=SLC34A3;refGene_tx=XM_023243107.1;refGene_gn=SLC34A3;refGene_tx=XM_023243108.1;refGene_gn=SLC34A3;refGene_tx=XM_023243109.1
D4	95797624	95797741
refGene_gn=SLC34A3;refGene_tx=XM_023243105.1;refGene_gn=SLC34A3;refGene_tx=XM_023243106.1;refGene_gn=SLC34A3;refGene_tx=XM_023243107.1;refGene_gn=SLC34A3;refGene_tx=XM_023243108.1;refGene_gn=SLC34A3;refGene_tx=XM_023243109.1
D4	95797839	95797964
refGene_gn=SLC34A3;refGene_tx=XM_023243105.1;refGene_gn=SLC34A3;refGene_tx=XM_023243106.1;refGene_gn=SLC34A3;refGene_tx=XM_023243107.1;refGene_gn=SLC34A3;refGene_tx=XM_023243108.1
D4	95799188	95799653
refGene_gn=SLC34A3;refGene_tx=XM_023243105.1;refGene_gn=SLC34A3;refGene_tx=XM_023243106.1;refGene_gn=SLC34A3;refGene_tx=XM_023243107.1;refGene_gn=SLC34A3;refGene_tx=XM_023243108.1;refGene_gn=SLC34A3;refGene_tx=XM_023243109.1
D4	95803833	95803890	refGene_gn=TUBB4B;refGene_tx=XM_011288741.3
D4	95804168	95804277	refGene_gn=TUBB4B;refGene_tx=XM_011288741.3
D4	95804379	95804490	refGene_gn=TUBB4B;refGene_tx=XM_011288741.3
D4	95805019	95805045	refGene_gn=TUBB4B;refGene_tx=XM_011288741.3
D4	95805046	95806081	refGene_gn=TUBB4B;refGene_tx=XM_011288741.3
D4	95806227	95806316	refGene_gn=FAM166A;refGene_tx=XM_023242326.1
D4	95806662	95806780	refGene_gn=FAM166A;refGene_tx=XM_023242326.1
D4	95807144	95807239	refGene_gn=FAM166A;refGene_tx=XM_023242326.1
D4	95807575	95807702	refGene_gn=FAM166A;refGene_tx=XM_023242326.1
D4	95807789	95808055	refGene_gn=FAM166A;refGene_tx=XM_023242326.1
D4	95808136	95808463	refGene_gn=FAM166A;refGene_tx=XM_023242326.1
D4	95811033	95811079	refGene_gn=FAM166A;refGene_tx=XM_023242326.1
D4	95817449	95817551	refGene_gn=NELFB;refGene_tx=XM_023243114.1
D4	95817986	95818150	refGene_gn=NELFB;refGene_tx=XM_023243114.1
D4	95818370	95818470	refGene_gn=NELFB;refGene_tx=XM_023243114.1
D4	95818892	95819123	refGene_gn=NELFB;refGene_tx=XM_023243114.1
D4	95821483	95821669	refGene_gn=NELFB;refGene_tx=XM_023243114.1
D4	95822385	95822498	refGene_gn=NELFB;refGene_tx=XM_023243114.1
D4	95823044	95823147	refGene_gn=NELFB;refGene_tx=XM_023243114.1
D4	95823366	95823462	refGene_gn=NELFB;refGene_tx=XM_023243114.1
D4	95823694	95823837	refGene_gn=NELFB;refGene_tx=XM_023243114.1
D4	95823944	95824051	refGene_gn=NELFB;refGene_tx=XM_023243114.1
D4	95826596	95826738	refGene_gn=NELFB;refGene_tx=XM_023243114.1
D4	95826848	95826957	refGene_gn=NELFB;refGene_tx=XM_023243114.1
D4	95827058	95827205	refGene_gn=NELFB;refGene_tx=XM_023243114.1

D4 95830528 95831779 refGene_gn=TOR4A;refGene_tx=XM_023243115.1
D4 95856432 95856777 refGene_gn=NRARP;refGene_tx=XM_023243116.1
D4 95862417 95862701
refGene_gn=EXD3;refGene_tx=XM_023243117.1;refGene_gn=EXD3;refGene_tx=XM_023243120.1;refGene_gn=EXD3;refGene_tx=XM_023243121.1;refGene_gn=EXD3;refGene_tx=XM_023243122.1;refGene_gn=EXD3;refGene_tx=XM_023243123.1;refGene_gn=EXD3;refGene_tx=XM_023243124.1
D4 95863023 95863062
refGene_gn=EXD3;refGene_tx=XM_023243117.1;refGene_gn=EXD3;refGene_tx=XM_023243120.1;refGene_gn=EXD3;refGene_tx=XM_023243121.1;refGene_gn=EXD3;refGene_tx=XM_023243122.1;refGene_gn=EXD3;refGene_tx=XM_023243123.1;refGene_gn=EXD3;refGene_tx=XM_023243124.1
D4 95864778 95864872
refGene_gn=EXD3;refGene_tx=XM_023243117.1;refGene_gn=EXD3;refGene_tx=XM_023243120.1;refGene_gn=EXD3;refGene_tx=XM_023243121.1;refGene_gn=EXD3;refGene_tx=XM_023243122.1;refGene_gn=EXD3;refGene_tx=XM_023243123.1;refGene_gn=EXD3;refGene_tx=XM_023243124.1
D4 95878806 95878938
refGene_gn=EXD3;refGene_tx=XM_023243117.1;refGene_gn=EXD3;refGene_tx=XM_023243120.1;refGene_gn=EXD3;refGene_tx=XM_023243121.1;refGene_gn=EXD3;refGene_tx=XM_023243122.1;refGene_gn=EXD3;refGene_tx=XM_023243123.1;refGene_gn=EXD3;refGene_tx=XM_023243124.1
D4 95879174 95879228
refGene_gn=EXD3;refGene_tx=XM_023243117.1;refGene_gn=EXD3;refGene_tx=XM_023243120.1;refGene_gn=EXD3;refGene_tx=XM_023243121.1;refGene_gn=EXD3;refGene_tx=XM_023243122.1;refGene_gn=EXD3;refGene_tx=XM_023243123.1;refGene_gn=EXD3;refGene_tx=XM_023243124.1
D4 95888576 95889026 refGene_gn=EXD3;refGene_tx=XM_023243118.1
D4 95889458 95889620
refGene_gn=EXD3;refGene_tx=XM_023243117.1;refGene_gn=EXD3;refGene_tx=XM_023243118.1;refGene_gn=EXD3;refGene_tx=XM_023243120.1;refGene_gn=EXD3;refGene_tx=XM_023243121.1;refGene_gn=EXD3;refGene_tx=XM_023243122.1;refGene_gn=EXD3;refGene_tx=XM_023243123.1;refGene_gn=EXD3;refGene_tx=XM_023243124.1
D4 95890077 95890244
refGene_gn=EXD3;refGene_tx=XM_023243117.1;refGene_gn=EXD3;refGene_tx=XM_023243118.1;refGene_gn=EXD3;refGene_tx=XM_023243120.1;refGene_gn=EXD3;refGene_tx=XM_023243121.1;refGene_gn=EXD3;refGene_tx=XM_023243122.1;refGene_gn=EXD3;refGene_tx=XM_023243123.1;refGene_gn=EXD3;refGene_tx=XM_023243124.1
D4 95890325 95890488
refGene_gn=EXD3;refGene_tx=XM_023243117.1;refGene_gn=EXD3;refGene_tx=XM_023243118.1;refGene_gn=EXD3;refGene_tx=XM_023243120.1;refGene_gn=EXD3;refGene_tx=XM_023243121.1;refGene_gn=EXD3;refGene_tx=XM_023243122.1;refGene_gn=EXD3;refGene_tx=XM_023243123.1;refGene_gn=EXD3;refGene_tx=XM_023243124.1
D4 95890940 95891050
refGene_gn=EXD3;refGene_tx=XM_023243117.1;refGene_gn=EXD3;refGene_tx=XM_023243118.1;refGene_gn=EXD3;refGene_tx=XM_023243120.1;refGene_gn=EXD3;refGene_tx=XM_023243121.1;refGene_gn=EXD3;refGene_tx=XM_023243122.1;

refGene_gn=EXD3;refGene_tx=XM_023243123.1;refGene_gn=EXD3;refGene_tx=XM_023243124.1
D4 95891145 95891356
refGene_gn=EXD3;refGene_tx=XM_023243117.1;refGene_gn=EXD3;refGene_tx=XM_023243118.1;refGene_gn=EXD3;refGene_tx=XM_023243120.1;refGene_gn=EXD3;refGene_tx=XM_023243121.1;refGene_gn=EXD3;refGene_tx=XM_023243122.1;refGene_gn=EXD3;refGene_tx=XM_023243123.1;refGene_gn=EXD3;refGene_tx=XM_023243124.1
D4 95891787 95891911
refGene_gn=EXD3;refGene_tx=XM_023243117.1;refGene_gn=EXD3;refGene_tx=XM_023243118.1;refGene_gn=EXD3;refGene_tx=XM_023243120.1;refGene_gn=EXD3;refGene_tx=XM_023243121.1;refGene_gn=EXD3;refGene_tx=XM_023243122.1;refGene_gn=EXD3;refGene_tx=XM_023243123.1;refGene_gn=EXD3;refGene_tx=XM_023243124.1
D4 95892422 95892589
refGene_gn=EXD3;refGene_tx=XM_023243117.1;refGene_gn=EXD3;refGene_tx=XM_023243118.1;refGene_gn=EXD3;refGene_tx=XM_023243120.1;refGene_gn=EXD3;refGene_tx=XM_023243121.1;refGene_gn=EXD3;refGene_tx=XM_023243122.1;refGene_gn=EXD3;refGene_tx=XM_023243123.1;refGene_gn=EXD3;refGene_tx=XM_023243124.1
D4 95894383 95894422
refGene_gn=EXD3;refGene_tx=XM_023243117.1;refGene_gn=EXD3;refGene_tx=XM_023243118.1;refGene_gn=EXD3;refGene_tx=XM_023243120.1;refGene_gn=EXD3;refGene_tx=XM_023243121.1;refGene_gn=EXD3;refGene_tx=XM_023243122.1;refGene_gn=EXD3;refGene_tx=XM_023243123.1;refGene_gn=EXD3;refGene_tx=XM_023243124.1
D4 95894542 95894616
refGene_gn=EXD3;refGene_tx=XM_023243117.1;refGene_gn=EXD3;refGene_tx=XM_023243118.1;refGene_gn=EXD3;refGene_tx=XM_023243120.1;refGene_gn=EXD3;refGene_tx=XM_023243121.1;refGene_gn=EXD3;refGene_tx=XM_023243122.1;refGene_gn=EXD3;refGene_tx=XM_023243123.1;refGene_gn=EXD3;refGene_tx=XM_023243124.1
D4 95899215 95899313
refGene_gn=EXD3;refGene_tx=XM_023243117.1;refGene_gn=EXD3;refGene_tx=XM_023243118.1;refGene_gn=EXD3;refGene_tx=XM_023243120.1;refGene_gn=EXD3;refGene_tx=XM_023243121.1;refGene_gn=EXD3;refGene_tx=XM_023243122.1;refGene_gn=EXD3;refGene_tx=XM_023243123.1;refGene_gn=EXD3;refGene_tx=XM_023243124.1
D4 95901998 95902138
refGene_gn=EXD3;refGene_tx=XM_023243117.1;refGene_gn=EXD3;refGene_tx=XM_023243118.1;refGene_gn=EXD3;refGene_tx=XM_023243120.1;refGene_gn=EXD3;refGene_tx=XM_023243121.1;refGene_gn=EXD3;refGene_tx=XM_023243122.1;refGene_gn=EXD3;refGene_tx=XM_023243123.1;refGene_gn=EXD3;refGene_tx=XM_023243124.1
D4 95902216 95902555 refGene_gn=EXD3;refGene_tx=XM_023243122.1
D4 95903089 95903143
refGene_gn=EXD3;refGene_tx=XM_023243117.1;refGene_gn=EXD3;refGene_tx=XM_023243118.1;refGene_gn=EXD3;refGene_tx=XM_023243120.1;refGene_gn=EXD3;refGene_tx=XM_023243121.1;refGene_gn=EXD3;refGene_tx=XM_023243123.1;refGene_gn=EXD3;refGene_tx=XM_023243124.1

D4 95903389 95903542 refGene_gn=EXD3;refGene_tx=XM_023243123.1
D4 95905548 95905662 refGene_gn=EXD3;refGene_tx=XM_023243124.1
D4 95908644 95908812
refGene_gn=EXD3;refGene_tx=XM_023243117.1;refGene_gn=EXD3;refGene_tx=XM_023243118.1;refGene_gn=EXD3;refGene_tx=XM_023243120.1;refGene_gn=EXD3;refGene_tx=XM_023243121.1
D4 95909155 95909329
refGene_gn=EXD3;refGene_tx=XM_023243117.1;refGene_gn=EXD3;refGene_tx=XM_023243118.1;refGene_gn=EXD3;refGene_tx=XM_023243120.1;refGene_gn=EXD3;refGene_tx=XM_023243121.1
D4 95911856 95912354
refGene_gn=EXD3;refGene_tx=XM_023243117.1;refGene_gn=EXD3;refGene_tx=XM_023243118.1
D4 95915059 95915124
refGene_gn=EXD3;refGene_tx=XM_023243120.1;refGene_gn=EXD3;refGene_tx=XM_023243121.1
D4 95922536 95922639
refGene_gn=EXD3;refGene_tx=XM_023243120.1;refGene_gn=EXD3;refGene_tx=XM_023243121.1
D4 95926962 95927022
refGene_gn=LOC101085079;refGene_tx=XM_011288778.2
D4 95927824 95927901
refGene_gn=LOC101085079;refGene_tx=XM_011288778.2
D4 95934479 95935920
refGene_gn=LOC101085079;refGene_tx=XM_011288778.2
D4 95941889 95942066 refGene_gn=NOXA1;refGene_tx=XM_023242327.1
D4 95945007 95945090 refGene_gn=NOXA1;refGene_tx=XM_023242327.1
D4 95946054 95946163 refGene_gn=NOXA1;refGene_tx=XM_023242327.1
D4 95947767 95947902 refGene_gn=NOXA1;refGene_tx=XM_023242327.1
D4 95948148 95948256 refGene_gn=NOXA1;refGene_tx=XM_023242327.1
D4 95949204 95949264 refGene_gn=NOXA1;refGene_tx=XM_023242327.1
D4 95949541 95949585 refGene_gn=NOXA1;refGene_tx=XM_023242327.1
D4 95950077 95950198 refGene_gn=NOXA1;refGene_tx=XM_023242327.1
D4 95951003 95951048 refGene_gn=NOXA1;refGene_tx=XM_023242327.1
D4 95951192 95951304 refGene_gn=NOXA1;refGene_tx=XM_023242327.1
D4 95951499 95951558 refGene_gn=NOXA1;refGene_tx=XM_023242327.1
D4 95951798 95951953 refGene_gn=NOXA1;refGene_tx=XM_023242327.1
D4 95952112 95952227 refGene_gn=NOXA1;refGene_tx=XM_023242327.1
D4 95952331 95952446 refGene_gn=NOXA1;refGene_tx=XM_023242327.1
D4 95952534 95952623 refGene_gn=NOXA1;refGene_tx=XM_023242327.1
D4 95953215 95953407 refGene_gn=ENTPD8;refGene_tx=XM_003996126.3
D4 95953490 95953625 refGene_gn=ENTPD8;refGene_tx=XM_003996126.3
D4 95953770 95953881 refGene_gn=ENTPD8;refGene_tx=XM_003996126.3
D4 95954163 95954427 refGene_gn=ENTPD8;refGene_tx=XM_003996126.3
D4 95954516 95954747 refGene_gn=ENTPD8;refGene_tx=XM_003996126.3
D4 95954842 95955002 refGene_gn=ENTPD8;refGene_tx=XM_003996126.3
D4 95955147 95955298 refGene_gn=ENTPD8;refGene_tx=XM_003996126.3
D4 95956181 95956299 refGene_gn=ENTPD8;refGene_tx=XM_003996126.3
D4 95956390 95956516 refGene_gn=ENTPD8;refGene_tx=XM_003996126.3
D4 95971942 95972040

refGene_gn=NSMF;refGene_tx=XM_023243128.1;refGene_gn=NSMF;refGene_tx=XM_023243129.1;refGene_gn=NSMF;refGene_tx=XM_023243130.1;refGene_gn=NSMF;refGene_tx=XM_023243131.1;refGene_gn=NSMF;refGene_tx=XM_023243132.1;refGene_gn=NSMF;refGene_tx=XM_023243133.1;refGene_gn=NSMF;refGene_tx=XM_023243135.1;refGene_gn=NSMF;refGene_tx=XM_023243137.1;refGene_gn=NSMF;refGene_tx=XM_023243139.1;refGene_gn=NSMF;refGene_tx=XM_023243140.1;refGene_gn=NSMF;refGene_tx=XM_023243141.1
D4 95972128 95972204
refGene_gn=NSMF;refGene_tx=XM_023243128.1;refGene_gn=NSMF;refGene_tx=XM_023243129.1;refGene_gn=NSMF;refGene_tx=XM_023243130.1;refGene_gn=NSMF;refGene_tx=XM_023243131.1;refGene_gn=NSMF;refGene_tx=XM_023243132.1;refGene_gn=NSMF;refGene_tx=XM_023243133.1;refGene_gn=NSMF;refGene_tx=XM_023243135.1;refGene_gn=NSMF;refGene_tx=XM_023243136.1;refGene_gn=NSMF;refGene_tx=XM_023243137.1;refGene_gn=NSMF;refGene_tx=XM_023243138.1;refGene_gn=NSMF;refGene_tx=XM_023243139.1;refGene_gn=NSMF;refGene_tx=XM_023243140.1;refGene_gn=NSMF;refGene_tx=XM_023243141.1
D4 95972433 95972536
refGene_gn=NSMF;refGene_tx=XM_023243128.1;refGene_gn=NSMF;refGene_tx=XM_023243129.1;refGene_gn=NSMF;refGene_tx=XM_023243130.1;refGene_gn=NSMF;refGene_tx=XM_023243131.1;refGene_gn=NSMF;refGene_tx=XM_023243132.1;refGene_gn=NSMF;refGene_tx=XM_023243133.1;refGene_gn=NSMF;refGene_tx=XM_023243135.1;refGene_gn=NSMF;refGene_tx=XM_023243136.1;refGene_gn=NSMF;refGene_tx=XM_023243137.1;refGene_gn=NSMF;refGene_tx=XM_023243138.1;refGene_gn=NSMF;refGene_tx=XM_023243139.1;refGene_gn=NSMF;refGene_tx=XM_023243140.1;refGene_gn=NSMF;refGene_tx=XM_023243141.1
D4 95972727 95972817
refGene_gn=NSMF;refGene_tx=XM_023243128.1;refGene_gn=NSMF;refGene_tx=XM_023243129.1;refGene_gn=NSMF;refGene_tx=XM_023243130.1;refGene_gn=NSMF;refGene_tx=XM_023243131.1;refGene_gn=NSMF;refGene_tx=XM_023243132.1;refGene_gn=NSMF;refGene_tx=XM_023243133.1;refGene_gn=NSMF;refGene_tx=XM_023243135.1;refGene_gn=NSMF;refGene_tx=XM_023243136.1;refGene_gn=NSMF;refGene_tx=XM_023243137.1;refGene_gn=NSMF;refGene_tx=XM_023243138.1;refGene_gn=NSMF;refGene_tx=XM_023243139.1;refGene_gn=NSMF;refGene_tx=XM_023243140.1;refGene_gn=NSMF;refGene_tx=XM_023243141.1
D4 95972952 95973023
refGene_gn=NSMF;refGene_tx=XM_023243128.1;refGene_gn=NSMF;refGene_tx=XM_023243129.1;refGene_gn=NSMF;refGene_tx=XM_023243130.1;refGene_gn=NSMF;refGene_tx=XM_023243131.1;refGene_gn=NSMF;refGene_tx=XM_023243132.1;refGene_gn=NSMF;refGene_tx=XM_023243133.1;refGene_gn=NSMF;refGene_tx=XM_023243135.1;refGene_gn=NSMF;refGene_tx=XM_023243136.1;refGene_gn=NSMF;refGene_tx=XM_023243137.1;refGene_gn=NSMF;refGene_tx=XM_023243138.1;refGene_gn=NSMF;refGene_tx=XM_023243139.1;refGene_gn=NSMF;refGene_tx=XM_023243140.1;refGene_gn=NSMF;refGene_tx=XM_023243141.1
D4 95973126 95973160
refGene_gn=NSMF;refGene_tx=XM_023243128.1;refGene_gn=NSMF;refGene_tx=XM_023243129.1;refGene_gn=NSMF;refGene_tx=XM_023243130.1;refGene_gn=NSMF;refGene_tx=XM_023243131.1;refGene_gn=NSMF;refGene_tx=XM_023243132.1;refGene_gn=NSMF;refGene_tx=XM_023243133.1;refGene_gn=NSMF;refGene_tx=XM_023243135.1;refGene_gn=NSMF;refGene_tx=XM_023243136.1;refGene_gn=NSMF;refGene_tx=XM_023243137.1;refGene_gn=NSMF;refGene_tx=XM_023243138.1;

refGene_gn=NSMF;refGene_tx=XM_023243139.1;refGene_gn=NSMF;refGene_tx=XM_023243140.1;refGene_gn=NSMF;refGene_tx=XM_023243141.1
D4 95973245 95973329
refGene_gn=NSMF;refGene_tx=XM_023243128.1;refGene_gn=NSMF;refGene_tx=XM_023243129.1;refGene_gn=NSMF;refGene_tx=XM_023243130.1;refGene_gn=NSMF;refGene_tx=XM_023243131.1;refGene_gn=NSMF;refGene_tx=XM_023243132.1;refGene_gn=NSMF;refGene_tx=XM_023243133.1;refGene_gn=NSMF;refGene_tx=XM_023243135.1;refGene_gn=NSMF;refGene_tx=XM_023243136.1;refGene_gn=NSMF;refGene_tx=XM_023243137.1;refGene_gn=NSMF;refGene_tx=XM_023243138.1;refGene_gn=NSMF;refGene_tx=XM_023243139.1;refGene_gn=NSMF;refGene_tx=XM_023243140.1;refGene_gn=NSMF;refGene_tx=XM_023243141.1
D4 95973542 95973667
refGene_gn=NSMF;refGene_tx=XM_023243128.1;refGene_gn=NSMF;refGene_tx=XM_023243129.1;refGene_gn=NSMF;refGene_tx=XM_023243130.1;refGene_gn=NSMF;refGene_tx=XM_023243131.1;refGene_gn=NSMF;refGene_tx=XM_023243132.1;refGene_gn=NSMF;refGene_tx=XM_023243133.1;refGene_gn=NSMF;refGene_tx=XM_023243135.1;refGene_gn=NSMF;refGene_tx=XM_023243136.1;refGene_gn=NSMF;refGene_tx=XM_023243137.1;refGene_gn=NSMF;refGene_tx=XM_023243138.1;refGene_gn=NSMF;refGene_tx=XM_023243139.1;refGene_gn=NSMF;refGene_tx=XM_023243140.1;refGene_gn=NSMF;refGene_tx=XM_023243141.1
D4 95974165 95974255
refGene_gn=NSMF;refGene_tx=XM_023243128.1;refGene_gn=NSMF;refGene_tx=XM_023243129.1;refGene_gn=NSMF;refGene_tx=XM_023243130.1;refGene_gn=NSMF;refGene_tx=XM_023243131.1;refGene_gn=NSMF;refGene_tx=XM_023243133.1;refGene_gn=NSMF;refGene_tx=XM_023243135.1;refGene_gn=NSMF;refGene_tx=XM_023243136.1;refGene_gn=NSMF;refGene_tx=XM_023243138.1
D4 95975039 95975092
refGene_gn=NSMF;refGene_tx=XM_023243128.1;refGene_gn=NSMF;refGene_tx=XM_023243129.1;refGene_gn=NSMF;refGene_tx=XM_023243130.1;refGene_gn=NSMF;refGene_tx=XM_023243131.1;refGene_gn=NSMF;refGene_tx=XM_023243132.1;refGene_gn=NSMF;refGene_tx=XM_023243133.1;refGene_gn=NSMF;refGene_tx=XM_023243135.1;refGene_gn=NSMF;refGene_tx=XM_023243136.1;refGene_gn=NSMF;refGene_tx=XM_023243137.1;refGene_gn=NSMF;refGene_tx=XM_023243138.1;refGene_gn=NSMF;refGene_tx=XM_023243139.1;refGene_gn=NSMF;refGene_tx=XM_023243140.1;refGene_gn=NSMF;refGene_tx=XM_023243141.1
D4 95976030 95976123
refGene_gn=NSMF;refGene_tx=XM_023243128.1;refGene_gn=NSMF;refGene_tx=XM_023243129.1;refGene_gn=NSMF;refGene_tx=XM_023243130.1;refGene_gn=NSMF;refGene_tx=XM_023243131.1;refGene_gn=NSMF;refGene_tx=XM_023243132.1;refGene_gn=NSMF;refGene_tx=XM_023243136.1;refGene_gn=NSMF;refGene_tx=XM_023243137.1;refGene_gn=NSMF;refGene_tx=XM_023243138.1;refGene_gn=NSMF;refGene_tx=XM_023243139.1
D4 95976419 95976425
refGene_gn=NSMF;refGene_tx=XM_023243128.1;refGene_gn=NSMF;refGene_tx=XM_023243130.1;refGene_gn=NSMF;refGene_tx=XM_023243132.1;refGene_gn=NSMF;refGene_tx=XM_023243133.1;refGene_gn=NSMF;refGene_tx=XM_023243136.1;refGene_gn=NSMF;refGene_tx=XM_023243137.1;refGene_gn=NSMF;refGene_tx=XM_023243138.1;refGene_gn=NSMF;refGene_tx=XM_023243140.1
D4 95977307 95977383
refGene_gn=NSMF;refGene_tx=XM_023243128.1;refGene_gn=NSMF;refGene_tx=X

M_023243129.1;refGene_gn=NSMF;refGene_tx=XM_023243130.1;refGene_gn=NSMF;refGene_tx=XM_023243131.1;refGene_gn=NSMF;refGene_tx=XM_023243132.1;refGene_gn=NSMF;refGene_tx=XM_023243133.1;refGene_gn=NSMF;refGene_tx=XM_023243135.1;refGene_gn=NSMF;refGene_tx=XM_023243136.1;refGene_gn=NSMF;refGene_tx=XM_023243137.1;refGene_gn=NSMF;refGene_tx=XM_023243138.1;refGene_gn=NSMF;refGene_tx=XM_023243139.1;refGene_gn=NSMF;refGene_tx=XM_023243140.1;refGene_gn=NSMF;refGene_tx=XM_023243141.1
D4 95978567 95979074
refGene_gn=NSMF;refGene_tx=XM_023243128.1;refGene_gn=NSMF;refGene_tx=XM_023243129.1;refGene_gn=NSMF;refGene_tx=XM_023243130.1;refGene_gn=NSMF;refGene_tx=XM_023243131.1;refGene_gn=NSMF;refGene_tx=XM_023243132.1;refGene_gn=NSMF;refGene_tx=XM_023243133.1;refGene_gn=NSMF;refGene_tx=XM_023243135.1;refGene_gn=NSMF;refGene_tx=XM_023243136.1;refGene_gn=NSMF;refGene_tx=XM_023243137.1;refGene_gn=NSMF;refGene_tx=XM_023243138.1;refGene_gn=NSMF;refGene_tx=XM_023243139.1;refGene_gn=NSMF;refGene_tx=XM_023243140.1;refGene_gn=NSMF;refGene_tx=XM_023243141.1
D4 95979768 95979830
refGene_gn=NSMF;refGene_tx=XM_023243128.1;refGene_gn=NSMF;refGene_tx=XM_023243129.1;refGene_gn=NSMF;refGene_tx=XM_023243130.1;refGene_gn=NSMF;refGene_tx=XM_023243131.1;refGene_gn=NSMF;refGene_tx=XM_023243132.1;refGene_gn=NSMF;refGene_tx=XM_023243133.1;refGene_gn=NSMF;refGene_tx=XM_023243135.1;refGene_gn=NSMF;refGene_tx=XM_023243136.1;refGene_gn=NSMF;refGene_tx=XM_023243137.1;refGene_gn=NSMF;refGene_tx=XM_023243138.1;refGene_gn=NSMF;refGene_tx=XM_023243139.1;refGene_gn=NSMF;refGene_tx=XM_023243140.1;refGene_gn=NSMF;refGene_tx=XM_023243141.1
D4 95980273 95980344
refGene_gn=NSMF;refGene_tx=XM_023243128.1;refGene_gn=NSMF;refGene_tx=XM_023243129.1;refGene_gn=NSMF;refGene_tx=XM_023243130.1;refGene_gn=NSMF;refGene_tx=XM_023243131.1;refGene_gn=NSMF;refGene_tx=XM_023243132.1;refGene_gn=NSMF;refGene_tx=XM_023243133.1;refGene_gn=NSMF;refGene_tx=XM_023243135.1;refGene_gn=NSMF;refGene_tx=XM_023243136.1;refGene_gn=NSMF;refGene_tx=XM_023243137.1;refGene_gn=NSMF;refGene_tx=XM_023243138.1;refGene_gn=NSMF;refGene_tx=XM_023243139.1;refGene_gn=NSMF;refGene_tx=XM_023243140.1;refGene_gn=NSMF;refGene_tx=XM_023243141.1
D4 95981521 95981632
refGene_gn=PNPLA7;refGene_tx=XM_003996128.5;refGene_gn=PNPLA7;refGene_tx=XM_019816858.2;refGene_gn=PNPLA7;refGene_tx=XM_023243125.1;refGene_gn=PNPLA7;refGene_tx=XM_023243126.1
D4 95981803 95981907
refGene_gn=PNPLA7;refGene_tx=XM_003996128.5;refGene_gn=PNPLA7;refGene_tx=XM_019816858.2;refGene_gn=PNPLA7;refGene_tx=XM_023243125.1;refGene_gn=PNPLA7;refGene_tx=XM_023243126.1
D4 95982113 95982198
refGene_gn=PNPLA7;refGene_tx=XM_003996128.5;refGene_gn=PNPLA7;refGene_tx=XM_019816858.2;refGene_gn=PNPLA7;refGene_tx=XM_023243125.1;refGene_gn=PNPLA7;refGene_tx=XM_023243126.1
D4 95982556 95982667
refGene_gn=PNPLA7;refGene_tx=XM_003996128.5;refGene_gn=PNPLA7;refGene_tx=XM_019816858.2;refGene_gn=PNPLA7;refGene_tx=XM_019816859.2;refGene_gn=PNPLA7;refGene_tx=XM_023243125.1;refGene_gn=PNPLA7;refGene_tx=XM_02

3243126.1
D4	95982784	95982937
refGene_gn=PNPLA7;refGene_tx=XM_003996128.5;refGene_gn=PNPLA7;refGene_tx=XM_019816858.2;refGene_gn=PNPLA7;refGene_tx=XM_019816859.2;refGene_gn=PNPLA7;refGene_tx=XM_023243125.1;refGene_gn=PNPLA7;refGene_tx=XM_023243126.1
D4	95983368	95983517
refGene_gn=PNPLA7;refGene_tx=XM_003996128.5;refGene_gn=PNPLA7;refGene_tx=XM_019816858.2;refGene_gn=PNPLA7;refGene_tx=XM_019816859.2;refGene_gn=PNPLA7;refGene_tx=XM_023243125.1;refGene_gn=PNPLA7;refGene_tx=XM_023243126.1
D4	95984051	95984168
refGene_gn=PNPLA7;refGene_tx=XM_003996128.5;refGene_gn=PNPLA7;refGene_tx=XM_019816858.2;refGene_gn=PNPLA7;refGene_tx=XM_019816859.2;refGene_gn=PNPLA7;refGene_tx=XM_023243125.1;refGene_gn=PNPLA7;refGene_tx=XM_023243126.1
D4	95984276	95984434	refGene_gn=PNPLA7;refGene_tx=XM_011288725.3
D4	95984969	95985039
refGene_gn=PNPLA7;refGene_tx=XM_003996128.5;refGene_gn=PNPLA7;refGene_tx=XM_011288725.3;refGene_gn=PNPLA7;refGene_tx=XM_019816858.2;refGene_gn=PNPLA7;refGene_tx=XM_019816859.2;refGene_gn=PNPLA7;refGene_tx=XM_023243125.1;refGene_gn=PNPLA7;refGene_tx=XM_023243126.1
D4	95985217	95985334
refGene_gn=PNPLA7;refGene_tx=XM_003996128.5;refGene_gn=PNPLA7;refGene_tx=XM_011288725.3;refGene_gn=PNPLA7;refGene_tx=XM_019816858.2;refGene_gn=PNPLA7;refGene_tx=XM_019816859.2;refGene_gn=PNPLA7;refGene_tx=XM_023243125.1;refGene_gn=PNPLA7;refGene_tx=XM_023243126.1
D4	95987703	95987860
refGene_gn=PNPLA7;refGene_tx=XM_003996128.5;refGene_gn=PNPLA7;refGene_tx=XM_011288725.3;refGene_gn=PNPLA7;refGene_tx=XM_019816858.2;refGene_gn=PNPLA7;refGene_tx=XM_019816859.2;refGene_gn=PNPLA7;refGene_tx=XM_023243125.1;refGene_gn=PNPLA7;refGene_tx=XM_023243126.1
D4	95994467	95994586
refGene_gn=PNPLA7;refGene_tx=XM_003996128.5;refGene_gn=PNPLA7;refGene_tx=XM_011288725.3;refGene_gn=PNPLA7;refGene_tx=XM_019816858.2;refGene_gn=PNPLA7;refGene_tx=XM_019816859.2;refGene_gn=PNPLA7;refGene_tx=XM_023243125.1;refGene_gn=PNPLA7;refGene_tx=XM_023243126.1
D4	95995423	95995606
refGene_gn=PNPLA7;refGene_tx=XM_003996128.5;refGene_gn=PNPLA7;refGene_tx=XM_011288725.3;refGene_gn=PNPLA7;refGene_tx=XM_019816858.2;refGene_gn=PNPLA7;refGene_tx=XM_019816859.2;refGene_gn=PNPLA7;refGene_tx=XM_023243125.1
D4	95996634	95996803
refGene_gn=PNPLA7;refGene_tx=XM_003996128.5;refGene_gn=PNPLA7;refGene_tx=XM_011288725.3;refGene_gn=PNPLA7;refGene_tx=XM_019816858.2;refGene_gn=PNPLA7;refGene_tx=XM_019816859.2;refGene_gn=PNPLA7;refGene_tx=XM_023243125.1;refGene_gn=PNPLA7;refGene_tx=XM_023243126.1
D4	95997000	95997064
refGene_gn=PNPLA7;refGene_tx=XM_003996128.5;refGene_gn=PNPLA7;refGene_tx=XM_011288725.3;refGene_gn=PNPLA7;refGene_tx=XM_019816858.2;refGene_

gn=PNPLA7;refGene_tx=XM_019816859.2;refGene_gn=PNPLA7;refGene_tx=XM_023243125.1;refGene_gn=PNPLA7;refGene_tx=XM_023243126.1
D4 96005920 96006070
refGene_gn=PNPLA7;refGene_tx=XM_003996128.5;refGene_gn=PNPLA7;refGene_tx=XM_011288725.3;refGene_gn=PNPLA7;refGene_tx=XM_019816858.2;refGene_gn=PNPLA7;refGene_tx=XM_019816859.2;refGene_gn=PNPLA7;refGene_tx=XM_023243125.1;refGene_gn=PNPLA7;refGene_tx=XM_023243126.1
D4 96010085 96010155
refGene_gn=PNPLA7;refGene_tx=XM_003996128.5;refGene_gn=PNPLA7;refGene_tx=XM_011288725.3;refGene_gn=PNPLA7;refGene_tx=XM_019816858.2;refGene_gn=PNPLA7;refGene_tx=XM_019816859.2;refGene_gn=PNPLA7;refGene_tx=XM_023243125.1;refGene_gn=PNPLA7;refGene_tx=XM_023243126.1
D4 96014175 96014289
refGene_gn=PNPLA7;refGene_tx=XM_003996128.5;refGene_gn=PNPLA7;refGene_tx=XM_011288725.3;refGene_gn=PNPLA7;refGene_tx=XM_019816858.2;refGene_gn=PNPLA7;refGene_tx=XM_019816859.2;refGene_gn=PNPLA7;refGene_tx=XM_023243125.1;refGene_gn=PNPLA7;refGene_tx=XM_023243126.1
D4 96014793 96014917
refGene_gn=PNPLA7;refGene_tx=XM_003996128.5;refGene_gn=PNPLA7;refGene_tx=XM_011288725.3;refGene_gn=PNPLA7;refGene_tx=XM_019816858.2;refGene_gn=PNPLA7;refGene_tx=XM_019816859.2;refGene_gn=PNPLA7;refGene_tx=XM_023243125.1;refGene_gn=PNPLA7;refGene_tx=XM_023243126.1
D4 96015728 96015860
refGene_gn=PNPLA7;refGene_tx=XM_003996128.5;refGene_gn=PNPLA7;refGene_tx=XM_011288725.3;refGene_gn=PNPLA7;refGene_tx=XM_019816858.2;refGene_gn=PNPLA7;refGene_tx=XM_019816859.2;refGene_gn=PNPLA7;refGene_tx=XM_023243125.1;refGene_gn=PNPLA7;refGene_tx=XM_023243126.1
D4 96017210 96017416
refGene_gn=PNPLA7;refGene_tx=XM_003996128.5;refGene_gn=PNPLA7;refGene_tx=XM_011288725.3;refGene_gn=PNPLA7;refGene_tx=XM_019816858.2;refGene_gn=PNPLA7;refGene_tx=XM_019816859.2;refGene_gn=PNPLA7;refGene_tx=XM_023243125.1;refGene_gn=PNPLA7;refGene_tx=XM_023243126.1
D4 96018253 96018331
refGene_gn=PNPLA7;refGene_tx=XM_003996128.5;refGene_gn=PNPLA7;refGene_tx=XM_011288725.3;refGene_gn=PNPLA7;refGene_tx=XM_019816858.2;refGene_gn=PNPLA7;refGene_tx=XM_019816859.2;refGene_gn=PNPLA7;refGene_tx=XM_023243125.1;refGene_gn=PNPLA7;refGene_tx=XM_023243126.1
D4 96027588 96027735
refGene_gn=PNPLA7;refGene_tx=XM_003996128.5;refGene_gn=PNPLA7;refGene_tx=XM_011288725.3;refGene_gn=PNPLA7;refGene_tx=XM_019816858.2;refGene_gn=PNPLA7;refGene_tx=XM_019816859.2;refGene_gn=PNPLA7;refGene_tx=XM_023243125.1;refGene_gn=PNPLA7;refGene_tx=XM_023243126.1
D4 96027852 96027953
refGene_gn=PNPLA7;refGene_tx=XM_003996128.5;refGene_gn=PNPLA7;refGene_tx=XM_011288725.3;refGene_gn=PNPLA7;refGene_tx=XM_019816858.2;refGene_gn=PNPLA7;refGene_tx=XM_019816859.2;refGene_gn=PNPLA7;refGene_tx=XM_023243125.1;refGene_gn=PNPLA7;refGene_tx=XM_023243126.1
D4 96032210 96032351
refGene_gn=PNPLA7;refGene_tx=XM_003996128.5;refGene_gn=PNPLA7;refGene_tx=XM_011288725.3;refGene_gn=PNPLA7;refGene_tx=XM_019816858.2;refGene_

gn=PNPLA7;refGene_tx=XM_019816859.2;refGene_gn=PNPLA7;refGene_tx=XM_02
3243125.1;refGene_gn=PNPLA7;refGene_tx=XM_023243126.1
D4 96032889 96033016
refGene_gn=PNPLA7;refGene_tx=XM_003996128.5;refGene_gn=PNPLA7;refGene_
tx=XM_011288725.3;refGene_gn=PNPLA7;refGene_tx=XM_019816858.2;refGene_
gn=PNPLA7;refGene_tx=XM_019816859.2;refGene_gn=PNPLA7;refGene_tx=XM_02
3243125.1;refGene_gn=PNPLA7;refGene_tx=XM_023243126.1
D4 96034234 96034315
refGene_gn=PNPLA7;refGene_tx=XM_003996128.5;refGene_gn=PNPLA7;refGene_
tx=XM_011288725.3;refGene_gn=PNPLA7;refGene_tx=XM_019816858.2;refGene_
gn=PNPLA7;refGene_tx=XM_019816859.2;refGene_gn=PNPLA7;refGene_tx=XM_02
3243125.1;refGene_gn=PNPLA7;refGene_tx=XM_023243126.1
D4 96034970 96035099
refGene_gn=PNPLA7;refGene_tx=XM_003996128.5;refGene_gn=PNPLA7;refGene_
tx=XM_011288725.3;refGene_gn=PNPLA7;refGene_tx=XM_019816858.2;refGene_
gn=PNPLA7;refGene_tx=XM_019816859.2;refGene_gn=PNPLA7;refGene_tx=XM_02
3243125.1;refGene_gn=PNPLA7;refGene_tx=XM_023243126.1
D4 96037554 96037635
refGene_gn=PNPLA7;refGene_tx=XM_003996128.5;refGene_gn=PNPLA7;refGene_
tx=XM_011288725.3;refGene_gn=PNPLA7;refGene_tx=XM_019816858.2;refGene_
gn=PNPLA7;refGene_tx=XM_019816859.2;refGene_gn=PNPLA7;refGene_tx=XM_02
3243125.1;refGene_gn=PNPLA7;refGene_tx=XM_023243126.1
D4 96039648 96039808
refGene_gn=PNPLA7;refGene_tx=XM_003996128.5;refGene_gn=PNPLA7;refGene_
tx=XM_011288725.3;refGene_gn=PNPLA7;refGene_tx=XM_019816858.2;refGene_
gn=PNPLA7;refGene_tx=XM_019816859.2;refGene_gn=PNPLA7;refGene_tx=XM_02
3243125.1;refGene_gn=PNPLA7;refGene_tx=XM_023243126.1
D4 96040383 96040524
refGene_gn=PNPLA7;refGene_tx=XM_003996128.5;refGene_gn=PNPLA7;refGene_
tx=XM_011288725.3;refGene_gn=PNPLA7;refGene_tx=XM_019816858.2;refGene_
gn=PNPLA7;refGene_tx=XM_019816859.2;refGene_gn=PNPLA7;refGene_tx=XM_02
3243125.1;refGene_gn=PNPLA7;refGene_tx=XM_023243126.1
D4 96040704 96040796
refGene_gn=PNPLA7;refGene_tx=XM_003996128.5;refGene_gn=PNPLA7;refGene_
tx=XM_011288725.3;refGene_gn=PNPLA7;refGene_tx=XM_019816859.2;refGene_
gn=PNPLA7;refGene_tx=XM_023243125.1;refGene_gn=PNPLA7;refGene_tx=XM_02
3243126.1
D4 96041763 96041843
refGene_gn=PNPLA7;refGene_tx=XM_003996128.5;refGene_gn=PNPLA7;refGene_
tx=XM_011288725.3;refGene_gn=PNPLA7;refGene_tx=XM_019816859.2;refGene_
gn=PNPLA7;refGene_tx=XM_023243125.1;refGene_gn=PNPLA7;refGene_tx=XM_02
3243126.1
D4 96042332 96042420
refGene_gn=PNPLA7;refGene_tx=XM_003996128.5;refGene_gn=PNPLA7;refGene_
tx=XM_011288725.3;refGene_gn=PNPLA7;refGene_tx=XM_019816859.2;refGene_
gn=PNPLA7;refGene_tx=XM_023243125.1;refGene_gn=PNPLA7;refGene_tx=XM_02
3243126.1
D4 96042586 96042661
refGene_gn=PNPLA7;refGene_tx=XM_003996128.5;refGene_gn=PNPLA7;refGene_
tx=XM_011288725.3;refGene_gn=PNPLA7;refGene_tx=XM_019816859.2;refGene_

gn=PNPLA7;refGene_tx=XM_023243125.1;refGene_gn=PNPLA7;refGene_tx=XM_023243126.1
D4 96044311 96044341 refGene_gn=PNPLA7;refGene_tx=XM_003996128.5;refGene_gn=PNPLA7;refGene_tx=XM_011288725.3;refGene_gn=PNPLA7;refGene_tx=XM_019816858.2;refGene_gn=PNPLA7;refGene_tx=XM_019816859.2;refGene_gn=PNPLA7;refGene_tx=XM_023243125.1;refGene_gn=PNPLA7;refGene_tx=XM_023243126.1
D4 96046177 96046585 refGene_gn=MRPL41;refGene_tx=XM_003996129.5
D4 96047996 96048451 refGene_gn=DPH7;refGene_tx=XM_023242328.1
D4 96054338 96054511 refGene_gn=DPH7;refGene_tx=XM_023242328.1
D4 96054990 96055056 refGene_gn=DPH7;refGene_tx=XM_023242328.1
D4 96055181 96055251 refGene_gn=DPH7;refGene_tx=XM_023242328.1
D4 96055354 96055459 refGene_gn=DPH7;refGene_tx=XM_023242328.1
D4 96060161 96060334 refGene_gn=DPH7;refGene_tx=XM_023242328.1
D4 96060741 96060833 refGene_gn=DPH7;refGene_tx=XM_023242328.1
D4 96062561 96062649 refGene_gn=DPH7;refGene_tx=XM_023242328.1
D4 96063028 96063162 refGene_gn=DPH7;refGene_tx=XM_023242328.1
D4 96064277 96064613 refGene_gn=DPH7;refGene_tx=XM_023242328.1
D4 96064662 96064831 refGene_gn=DPH7;refGene_tx=XM_023242328.1
D4 96066396 96066491 refGene_gn=DPH7;refGene_tx=XM_023242328.1
D4 96068348 96068556 refGene_gn=ZMYND19;refGene_tx=XM_023243142.1;refGene_gn=ZMYND19;refGene_tx=XM_023243144.1
D4 96068773 96068954 refGene_gn=ZMYND19;refGene_tx=XM_023243142.1
D4 96072190 96072331 refGene_gn=ZMYND19;refGene_tx=XM_023243142.1;refGene_gn=ZMYND19;refGene_tx=XM_023243144.1
D4 96072930 96073037 refGene_gn=ZMYND19;refGene_tx=XM_023243142.1;refGene_gn=ZMYND19;refGene_tx=XM_023243144.1
D4 96074810 96074870 refGene_gn=ZMYND19;refGene_tx=XM_023243142.1;refGene_gn=ZMYND19;refGene_tx=XM_023243144.1
D4 96075966 96076017 refGene_gn=ZMYND19;refGene_tx=XM_023243142.1;refGene_gn=ZMYND19;refGene_tx=XM_023243144.1
D4 96084443 96084561 refGene_gn=ARRDC1;refGene_tx=XM_003996111.4
D4 96090546 96090657 refGene_gn=ARRDC1;refGene_tx=XM_003996111.4;refGene_gn=ARRDC1;refGene_tx=XM_006939696.4;refGene_gn=ARRDC1;refGene_tx=XM_023243145.1
D4 96091141 96091192 refGene_gn=ARRDC1;refGene_tx=XM_003996111.4;refGene_gn=ARRDC1;refGene_tx=XM_006939696.4;refGene_gn=ARRDC1;refGene_tx=XM_023243145.1
D4 96091292 96091447 refGene_gn=ARRDC1;refGene_tx=XM_003996111.4;refGene_gn=ARRDC1;refGene_tx=XM_006939696.4;refGene_gn=ARRDC1;refGene_tx=XM_023243145.1
D4 96091682 96091865 refGene_gn=ARRDC1;refGene_tx=XM_003996111.4;refGene_gn=ARRDC1;refGene_tx=XM_006939696.4;refGene_gn=ARRDC1;refGene_tx=XM_023243145.1
D4 96091962 96092139

refGene_gn=ARRDC1;refGene_tx=XM_003996111.4;refGene_gn=ARRDC1;refGene_tx=XM_006939696.4;refGene_gn=ARRDC1;refGene_tx=XM_023243145.1
D4 96092212 96092648
refGene_gn=ARRDC1;refGene_tx=XM_003996111.4;refGene_gn=ARRDC1;refGene_tx=XM_006939696.4;refGene_gn=ARRDC1;refGene_tx=XM_023243145.1
D4 96092724 96092789
refGene_gn=ARRDC1;refGene_tx=XM_003996111.4;refGene_gn=ARRDC1;refGene_tx=XM_006939696.4;refGene_gn=ARRDC1;refGene_tx=XM_023243145.1
D4 96096444 96096465
refGene_gn=EHMT1;refGene_tx=XM_023243147.1;refGene_gn=EHMT1;refGene_tx=XM_023243148.1
D4 96163571 96163638
refGene_gn=EHMT1;refGene_tx=XM_023243146.1;refGene_gn=EHMT1;refGene_tx=XM_023243147.1;refGene_gn=EHMT1;refGene_tx=XM_023243148.1;refGene_gn=EHMT1;refGene_tx=XM_023243149.1;refGene_gn=EHMT1;refGene_tx=XM_023243152.1
D4 96171251 96171808
refGene_gn=EHMT1;refGene_tx=XM_023243146.1;refGene_gn=EHMT1;refGene_tx=XM_023243147.1;refGene_gn=EHMT1;refGene_tx=XM_023243148.1;refGene_gn=EHMT1;refGene_tx=XM_023243149.1;refGene_gn=EHMT1;refGene_tx=XM_023243150.1;refGene_gn=EHMT1;refGene_tx=XM_023243152.1
D4 96178814 96178995
refGene_gn=EHMT1;refGene_tx=XM_023243146.1;refGene_gn=EHMT1;refGene_tx=XM_023243147.1;refGene_gn=EHMT1;refGene_tx=XM_023243148.1;refGene_gn=EHMT1;refGene_tx=XM_023243149.1;refGene_gn=EHMT1;refGene_tx=XM_023243150.1;refGene_gn=EHMT1;refGene_tx=XM_023243152.1
D4 96181554 96181712
refGene_gn=EHMT1;refGene_tx=XM_023243146.1;refGene_gn=EHMT1;refGene_tx=XM_023243147.1;refGene_gn=EHMT1;refGene_tx=XM_023243148.1;refGene_gn=EHMT1;refGene_tx=XM_023243149.1;refGene_gn=EHMT1;refGene_tx=XM_023243150.1;refGene_gn=EHMT1;refGene_tx=XM_023243152.1
D4 96182039 96182228
refGene_gn=EHMT1;refGene_tx=XM_023243146.1;refGene_gn=EHMT1;refGene_tx=XM_023243147.1;refGene_gn=EHMT1;refGene_tx=XM_023243148.1;refGene_gn=EHMT1;refGene_tx=XM_023243149.1;refGene_gn=EHMT1;refGene_tx=XM_023243150.1;refGene_gn=EHMT1;refGene_tx=XM_023243152.1
D4 96192657 96192735
refGene_gn=EHMT1;refGene_tx=XM_023243146.1;refGene_gn=EHMT1;refGene_tx=XM_023243147.1;refGene_gn=EHMT1;refGene_tx=XM_023243148.1;refGene_gn=EHMT1;refGene_tx=XM_023243149.1;refGene_gn=EHMT1;refGene_tx=XM_023243150.1;refGene_gn=EHMT1;refGene_tx=XM_023243152.1
D4 96194271 96194392
refGene_gn=EHMT1;refGene_tx=XM_023243146.1;refGene_gn=EHMT1;refGene_tx=XM_023243147.1;refGene_gn=EHMT1;refGene_tx=XM_023243148.1;refGene_gn=EHMT1;refGene_tx=XM_023243149.1;refGene_gn=EHMT1;refGene_tx=XM_023243150.1;refGene_gn=EHMT1;refGene_tx=XM_023243152.1
D4 96198012 96198144
refGene_gn=EHMT1;refGene_tx=XM_023243146.1;refGene_gn=EHMT1;refGene_tx=XM_023243147.1;refGene_gn=EHMT1;refGene_tx=XM_023243148.1;refGene_gn=EHMT1;refGene_tx=XM_023243149.1;refGene_gn=EHMT1;refGene_tx=XM_0232431

50.1;refGene_gn=EHMT1;refGene_tx=XM_023243152.1
D4 96201117 96201263
refGene_gn=EHMT1;refGene_tx=XM_023243146.1;refGene_gn=EHMT1;refGene_tx=XM_023243147.1;refGene_gn=EHMT1;refGene_tx=XM_023243148.1;refGene_gn=EHMT1;refGene_tx=XM_023243149.1;refGene_gn=EHMT1;refGene_tx=XM_023243150.1;refGene_gn=EHMT1;refGene_tx=XM_023243152.1
D4 96207372 96207516
refGene_gn=EHMT1;refGene_tx=XM_023243146.1;refGene_gn=EHMT1;refGene_tx=XM_023243147.1;refGene_gn=EHMT1;refGene_tx=XM_023243148.1;refGene_gn=EHMT1;refGene_tx=XM_023243150.1
D4 96208529 96208756
refGene_gn=EHMT1;refGene_tx=XM_023243146.1;refGene_gn=EHMT1;refGene_tx=XM_023243147.1;refGene_gn=EHMT1;refGene_tx=XM_023243148.1;refGene_gn=EHMT1;refGene_tx=XM_023243149.1;refGene_gn=EHMT1;refGene_tx=XM_023243150.1;refGene_gn=EHMT1;refGene_tx=XM_023243152.1
D4 96209830 96210004
refGene_gn=EHMT1;refGene_tx=XM_023243146.1;refGene_gn=EHMT1;refGene_tx=XM_023243147.1;refGene_gn=EHMT1;refGene_tx=XM_023243148.1;refGene_gn=EHMT1;refGene_tx=XM_023243149.1;refGene_gn=EHMT1;refGene_tx=XM_023243150.1;refGene_gn=EHMT1;refGene_tx=XM_023243152.1
D4 96211054 96211137
refGene_gn=EHMT1;refGene_tx=XM_023243146.1;refGene_gn=EHMT1;refGene_tx=XM_023243147.1;refGene_gn=EHMT1;refGene_tx=XM_023243148.1;refGene_gn=EHMT1;refGene_tx=XM_023243149.1;refGene_gn=EHMT1;refGene_tx=XM_023243150.1;refGene_gn=EHMT1;refGene_tx=XM_023243152.1
D4 96212713 96212820
refGene_gn=EHMT1;refGene_tx=XM_023243146.1;refGene_gn=EHMT1;refGene_tx=XM_023243147.1;refGene_gn=EHMT1;refGene_tx=XM_023243148.1;refGene_gn=EHMT1;refGene_tx=XM_023243149.1;refGene_gn=EHMT1;refGene_tx=XM_023243150.1;refGene_gn=EHMT1;refGene_tx=XM_023243152.1
D4 96221414 96221537
refGene_gn=EHMT1;refGene_tx=XM_023243146.1;refGene_gn=EHMT1;refGene_tx=XM_023243147.1;refGene_gn=EHMT1;refGene_tx=XM_023243148.1;refGene_gn=EHMT1;refGene_tx=XM_023243149.1;refGene_gn=EHMT1;refGene_tx=XM_023243150.1;refGene_gn=EHMT1;refGene_tx=XM_023243152.1
D4 96228887 96228989
refGene_gn=EHMT1;refGene_tx=XM_023243146.1;refGene_gn=EHMT1;refGene_tx=XM_023243147.1;refGene_gn=EHMT1;refGene_tx=XM_023243148.1;refGene_gn=EHMT1;refGene_tx=XM_023243149.1;refGene_gn=EHMT1;refGene_tx=XM_023243150.1;refGene_gn=EHMT1;refGene_tx=XM_023243152.1
D4 96230624 96230729
refGene_gn=EHMT1;refGene_tx=XM_023243146.1;refGene_gn=EHMT1;refGene_tx=XM_023243147.1;refGene_gn=EHMT1;refGene_tx=XM_023243148.1;refGene_gn=EHMT1;refGene_tx=XM_023243149.1;refGene_gn=EHMT1;refGene_tx=XM_023243150.1;refGene_gn=EHMT1;refGene_tx=XM_023243152.1
D4 96236534 96236689
refGene_gn=EHMT1;refGene_tx=XM_023243146.1;refGene_gn=EHMT1;refGene_tx=XM_023243147.1;refGene_gn=EHMT1;refGene_tx=XM_023243148.1;refGene_gn=EHMT1;refGene_tx=XM_023243149.1;refGene_gn=EHMT1;refGene_tx=XM_023243150.1;refGene_gn=EHMT1;refGene_tx=XM_023243152.1

D4 96237470 96237638
refGene_gn=EHMT1;refGene_tx=XM_023243146.1;refGene_gn=EHMT1;refGene_tx=XM_023243147.1;refGene_gn=EHMT1;refGene_tx=XM_023243148.1;refGene_gn=EHMT1;refGene_tx=XM_023243149.1;refGene_gn=EHMT1;refGene_tx=XM_023243150.1;refGene_gn=EHMT1;refGene_tx=XM_023243152.1
D4 96237823 96237968
refGene_gn=EHMT1;refGene_tx=XM_023243146.1;refGene_gn=EHMT1;refGene_tx=XM_023243147.1;refGene_gn=EHMT1;refGene_tx=XM_023243148.1;refGene_gn=EHMT1;refGene_tx=XM_023243149.1;refGene_gn=EHMT1;refGene_tx=XM_023243150.1;refGene_gn=EHMT1;refGene_tx=XM_023243152.1
D4 96238657 96238735
refGene_gn=EHMT1;refGene_tx=XM_023243146.1;refGene_gn=EHMT1;refGene_tx=XM_023243147.1;refGene_gn=EHMT1;refGene_tx=XM_023243148.1;refGene_gn=EHMT1;refGene_tx=XM_023243149.1;refGene_gn=EHMT1;refGene_tx=XM_023243150.1;refGene_gn=EHMT1;refGene_tx=XM_023243152.1
D4 96239490 96239606
refGene_gn=EHMT1;refGene_tx=XM_023243146.1;refGene_gn=EHMT1;refGene_tx=XM_023243147.1;refGene_gn=EHMT1;refGene_tx=XM_023243148.1;refGene_gn=EHMT1;refGene_tx=XM_023243149.1;refGene_gn=EHMT1;refGene_tx=XM_023243150.1;refGene_gn=EHMT1;refGene_tx=XM_023243152.1
D4 96240805 96240892
refGene_gn=EHMT1;refGene_tx=XM_023243146.1;refGene_gn=EHMT1;refGene_tx=XM_023243147.1;refGene_gn=EHMT1;refGene_tx=XM_023243148.1;refGene_gn=EHMT1;refGene_tx=XM_023243149.1;refGene_gn=EHMT1;refGene_tx=XM_023243150.1;refGene_gn=EHMT1;refGene_tx=XM_023243152.1
D4 96241313 96241392
refGene_gn=EHMT1;refGene_tx=XM_023243146.1;refGene_gn=EHMT1;refGene_tx=XM_023243147.1;refGene_gn=EHMT1;refGene_tx=XM_023243148.1;refGene_gn=EHMT1;refGene_tx=XM_023243149.1;refGene_gn=EHMT1;refGene_tx=XM_023243150.1;refGene_gn=EHMT1;refGene_tx=XM_023243152.1
D4 96248512 96248688
refGene_gn=EHMT1;refGene_tx=XM_023243146.1;refGene_gn=EHMT1;refGene_tx=XM_023243147.1;refGene_gn=EHMT1;refGene_tx=XM_023243148.1;refGene_gn=EHMT1;refGene_tx=XM_023243149.1;refGene_gn=EHMT1;refGene_tx=XM_023243150.1;refGene_gn=EHMT1;refGene_tx=XM_023243152.1
D4 96248868 96249046
refGene_gn=EHMT1;refGene_tx=XM_023243146.1;refGene_gn=EHMT1;refGene_tx=XM_023243147.1;refGene_gn=EHMT1;refGene_tx=XM_023243148.1;refGene_gn=EHMT1;refGene_tx=XM_023243149.1;refGene_gn=EHMT1;refGene_tx=XM_023243150.1;refGene_gn=EHMT1;refGene_tx=XM_023243152.1
D4 96272088 96272369
refGene_gn=CACNA1B;refGene_tx=XM_023243153.1;refGene_gn=CACNA1B;refGene_tx=XM_023243154.1;refGene_gn=CACNA1B;refGene_tx=XM_023243155.1;refGene_gn=CACNA1B;refGene_tx=XM_023243156.1;refGene_gn=CACNA1B;refGene_tx=XM_023243157.1;refGene_gn=CACNA1B;refGene_tx=XM_023243158.1;refGene_gn=CACNA1B;refGene_tx=XM_023243159.1;refGene_gn=CACNA1B;refGene_tx=XM_023243161.1;refGene_gn=CACNA1B;refGene_tx=XM_023243162.1;refGene_gn=CACNA1B;refGene_tx=XM_023243163.1;refGene_gn=CACNA1B;refGene_tx=XM_023243164.1;refGene_gn=CACNA1B;refGene_tx=XM_023243165.1;refGene_gn=CACNA1B;refGene_tx=XM_023243166.1;refGene_gn=CACNA1B;refGene_tx=XM_023243167.1;

refGene_gn=CACNA1B;refGene_tx=XM_023243168.1;refGene_gn=CACNA1B;refGene_tx=XM_023243169.1;refGene_gn=CACNA1B;refGene_tx=XM_023243170.1;refGene_gn=CACNA1B;refGene_tx=XM_023243172.1;refGene_gn=CACNA1B;refGene_tx=XM_023243173.1
D4 96273398 96273504
refGene_gn=CACNA1B;refGene_tx=XM_023243153.1;refGene_gn=CACNA1B;refGene_tx=XM_023243154.1;refGene_gn=CACNA1B;refGene_tx=XM_023243155.1;refGene_gn=CACNA1B;refGene_tx=XM_023243156.1;refGene_gn=CACNA1B;refGene_tx=XM_023243157.1;refGene_gn=CACNA1B;refGene_tx=XM_023243158.1;refGene_gn=CACNA1B;refGene_tx=XM_023243159.1;refGene_gn=CACNA1B;refGene_tx=XM_023243161.1;refGene_gn=CACNA1B;refGene_tx=XM_023243162.1;refGene_gn=CACNA1B;refGene_tx=XM_023243163.1;refGene_gn=CACNA1B;refGene_tx=XM_023243164.1;refGene_gn=CACNA1B;refGene_tx=XM_023243165.1;refGene_gn=CACNA1B;refGene_tx=XM_023243166.1;refGene_gn=CACNA1B;refGene_tx=XM_023243167.1;refGene_gn=CACNA1B;refGene_tx=XM_023243168.1;refGene_gn=CACNA1B;refGene_tx=XM_023243169.1;refGene_gn=CACNA1B;refGene_tx=XM_023243170.1;refGene_gn=CACNA1B;refGene_tx=XM_023243172.1;refGene_gn=CACNA1B;refGene_tx=XM_023243173.1
D4 96277317 96277457
refGene_gn=CACNA1B;refGene_tx=XM_023243153.1;refGene_gn=CACNA1B;refGene_tx=XM_023243154.1;refGene_gn=CACNA1B;refGene_tx=XM_023243155.1;refGene_gn=CACNA1B;refGene_tx=XM_023243156.1;refGene_gn=CACNA1B;refGene_tx=XM_023243157.1;refGene_gn=CACNA1B;refGene_tx=XM_023243158.1;refGene_gn=CACNA1B;refGene_tx=XM_023243159.1;refGene_gn=CACNA1B;refGene_tx=XM_023243161.1;refGene_gn=CACNA1B;refGene_tx=XM_023243162.1;refGene_gn=CACNA1B;refGene_tx=XM_023243163.1;refGene_gn=CACNA1B;refGene_tx=XM_023243164.1;refGene_gn=CACNA1B;refGene_tx=XM_023243165.1;refGene_gn=CACNA1B;refGene_tx=XM_023243166.1;refGene_gn=CACNA1B;refGene_tx=XM_023243167.1;refGene_gn=CACNA1B;refGene_tx=XM_023243168.1;refGene_gn=CACNA1B;refGene_tx=XM_023243169.1;refGene_gn=CACNA1B;refGene_tx=XM_023243170.1;refGene_gn=CACNA1B;refGene_tx=XM_023243172.1;refGene_gn=CACNA1B;refGene_tx=XM_023243173.1
D4 96309586 96309678
refGene_gn=CACNA1B;refGene_tx=XM_023243153.1;refGene_gn=CACNA1B;refGene_tx=XM_023243154.1;refGene_gn=CACNA1B;refGene_tx=XM_023243155.1;refGene_gn=CACNA1B;refGene_tx=XM_023243156.1;refGene_gn=CACNA1B;refGene_tx=XM_023243157.1;refGene_gn=CACNA1B;refGene_tx=XM_023243158.1;refGene_gn=CACNA1B;refGene_tx=XM_023243159.1;refGene_gn=CACNA1B;refGene_tx=XM_023243161.1;refGene_gn=CACNA1B;refGene_tx=XM_023243162.1;refGene_gn=CACNA1B;refGene_tx=XM_023243163.1;refGene_gn=CACNA1B;refGene_tx=XM_023243164.1;refGene_gn=CACNA1B;refGene_tx=XM_023243165.1;refGene_gn=CACNA1B;refGene_tx=XM_023243166.1;refGene_gn=CACNA1B;refGene_tx=XM_023243167.1;refGene_gn=CACNA1B;refGene_tx=XM_023243168.1;refGene_gn=CACNA1B;refGene_tx=XM_023243169.1;refGene_gn=CACNA1B;refGene_tx=XM_023243170.1;refGene_gn=CACNA1B;refGene_tx=XM_023243172.1;refGene_gn=CACNA1B;refGene_tx=XM_023243173.1
D4 96310902 96311055
refGene_gn=CACNA1B;refGene_tx=XM_023243153.1;refGene_gn=CACNA1B;refGene_tx=XM_023243154.1;refGene_gn=CACNA1B;refGene_tx=XM_023243155.1;refGene_gn=CACNA1B;refGene_tx=XM_023243156.1;refGene_gn=CACNA1B;refGene_tx=

XM_023243157.1;refGene_gn=CACNA1B;refGene_tx=XM_023243158.1;refGene_gn=CACNA1B;refGene_tx=XM_023243159.1;refGene_gn=CACNA1B;refGene_tx=XM_023243161.1;refGene_gn=CACNA1B;refGene_tx=XM_023243162.1;refGene_gn=CACNA1B;refGene_tx=XM_023243163.1;refGene_gn=CACNA1B;refGene_tx=XM_023243164.1;refGene_gn=CACNA1B;refGene_tx=XM_023243165.1;refGene_gn=CACNA1B;refGene_tx=XM_023243166.1;refGene_gn=CACNA1B;refGene_tx=XM_023243167.1;refGene_gn=CACNA1B;refGene_tx=XM_023243168.1;refGene_gn=CACNA1B;refGene_tx=XM_023243169.1;refGene_gn=CACNA1B;refGene_tx=XM_023243170.1;refGene_gn=CACNA1B;refGene_tx=XM_023243172.1;refGene_gn=CACNA1B;refGene_tx=XM_023243173.1
D4 96312269 96312460
refGene_gn=CACNA1B;refGene_tx=XM_023243153.1;refGene_gn=CACNA1B;refGene_tx=XM_023243154.1;refGene_gn=CACNA1B;refGene_tx=XM_023243155.1;refGene_gn=CACNA1B;refGene_tx=XM_023243156.1;refGene_gn=CACNA1B;refGene_tx=XM_023243157.1;refGene_gn=CACNA1B;refGene_tx=XM_023243158.1;refGene_gn=CACNA1B;refGene_tx=XM_023243159.1;refGene_gn=CACNA1B;refGene_tx=XM_023243161.1;refGene_gn=CACNA1B;refGene_tx=XM_023243162.1;refGene_gn=CACNA1B;refGene_tx=XM_023243163.1;refGene_gn=CACNA1B;refGene_tx=XM_023243164.1;refGene_gn=CACNA1B;refGene_tx=XM_023243165.1;refGene_gn=CACNA1B;refGene_tx=XM_023243166.1;refGene_gn=CACNA1B;refGene_tx=XM_023243167.1;refGene_gn=CACNA1B;refGene_tx=XM_023243168.1;refGene_gn=CACNA1B;refGene_tx=XM_023243169.1;refGene_gn=CACNA1B;refGene_tx=XM_023243170.1;refGene_gn=CACNA1B;refGene_tx=XM_023243172.1;refGene_gn=CACNA1B;refGene_tx=XM_023243173.1
D4 96323825 96323929
refGene_gn=CACNA1B;refGene_tx=XM_023243153.1;refGene_gn=CACNA1B;refGene_tx=XM_023243154.1;refGene_gn=CACNA1B;refGene_tx=XM_023243155.1;refGene_gn=CACNA1B;refGene_tx=XM_023243156.1;refGene_gn=CACNA1B;refGene_tx=XM_023243157.1;refGene_gn=CACNA1B;refGene_tx=XM_023243158.1;refGene_gn=CACNA1B;refGene_tx=XM_023243159.1;refGene_gn=CACNA1B;refGene_tx=XM_023243161.1;refGene_gn=CACNA1B;refGene_tx=XM_023243162.1;refGene_gn=CACNA1B;refGene_tx=XM_023243163.1;refGene_gn=CACNA1B;refGene_tx=XM_023243164.1;refGene_gn=CACNA1B;refGene_tx=XM_023243165.1;refGene_gn=CACNA1B;refGene_tx=XM_023243166.1;refGene_gn=CACNA1B;refGene_tx=XM_023243167.1;refGene_gn=CACNA1B;refGene_tx=XM_023243168.1;refGene_gn=CACNA1B;refGene_tx=XM_023243169.1;refGene_gn=CACNA1B;refGene_tx=XM_023243170.1;refGene_gn=CACNA1B;refGene_tx=XM_023243172.1;refGene_gn=CACNA1B;refGene_tx=XM_023243173.1
D4 96326959 96327075
refGene_gn=CACNA1B;refGene_tx=XM_023243153.1;refGene_gn=CACNA1B;refGene_tx=XM_023243154.1;refGene_gn=CACNA1B;refGene_tx=XM_023243155.1;refGene_gn=CACNA1B;refGene_tx=XM_023243156.1;refGene_gn=CACNA1B;refGene_tx=XM_023243157.1;refGene_gn=CACNA1B;refGene_tx=XM_023243158.1;refGene_gn=CACNA1B;refGene_tx=XM_023243159.1;refGene_gn=CACNA1B;refGene_tx=XM_023243161.1;refGene_gn=CACNA1B;refGene_tx=XM_023243162.1;refGene_gn=CACNA1B;refGene_tx=XM_023243163.1;refGene_gn=CACNA1B;refGene_tx=XM_023243164.1;refGene_gn=CACNA1B;refGene_tx=XM_023243165.1;refGene_gn=CACNA1B;refGene_tx=XM_023243166.1;refGene_gn=CACNA1B;refGene_tx=XM_023243167.1;refGene_gn=CACNA1B;refGene_tx=XM_023243168.1;refGene_gn=CACNA1B;refGene_tx=XM_023243169.1;refGene_gn=CACNA1B;refGene_tx=XM_023243170.1;refGe

ne_gn=CACNA1B;refGene_tx=XM_023243172.1;refGene_gn=CACNA1B;refGene_tx=XM_023243173.1
D4 96327757 96327814
refGene_gn=CACNA1B;refGene_tx=XM_023243153.1;refGene_gn=CACNA1B;refGene_tx=XM_023243154.1;refGene_gn=CACNA1B;refGene_tx=XM_023243155.1;refGene_gn=CACNA1B;refGene_tx=XM_023243156.1;refGene_gn=CACNA1B;refGene_tx=XM_023243157.1;refGene_gn=CACNA1B;refGene_tx=XM_023243158.1;refGene_gn=CACNA1B;refGene_tx=XM_023243159.1;refGene_gn=CACNA1B;refGene_tx=XM_023243161.1;refGene_gn=CACNA1B;refGene_tx=XM_023243162.1;refGene_gn=CACNA1B;refGene_tx=XM_023243163.1;refGene_gn=CACNA1B;refGene_tx=XM_023243164.1;refGene_gn=CACNA1B;refGene_tx=XM_023243165.1;refGene_gn=CACNA1B;refGene_tx=XM_023243166.1;refGene_gn=CACNA1B;refGene_tx=XM_023243167.1;refGene_gn=CACNA1B;refGene_tx=XM_023243168.1;refGene_gn=CACNA1B;refGene_tx=XM_023243169.1;refGene_gn=CACNA1B;refGene_tx=XM_023243170.1;refGene_gn=CACNA1B;refGene_tx=XM_023243172.1;refGene_gn=CACNA1B;refGene_tx=XM_023243173.1
D4 96328890 96328983
refGene_gn=CACNA1B;refGene_tx=XM_023243153.1;refGene_gn=CACNA1B;refGene_tx=XM_023243154.1;refGene_gn=CACNA1B;refGene_tx=XM_023243155.1;refGene_gn=CACNA1B;refGene_tx=XM_023243156.1;refGene_gn=CACNA1B;refGene_tx=XM_023243157.1;refGene_gn=CACNA1B;refGene_tx=XM_023243158.1;refGene_gn=CACNA1B;refGene_tx=XM_023243159.1;refGene_gn=CACNA1B;refGene_tx=XM_023243161.1;refGene_gn=CACNA1B;refGene_tx=XM_023243162.1;refGene_gn=CACNA1B;refGene_tx=XM_023243163.1;refGene_gn=CACNA1B;refGene_tx=XM_023243164.1;refGene_gn=CACNA1B;refGene_tx=XM_023243165.1;refGene_gn=CACNA1B;refGene_tx=XM_023243166.1;refGene_gn=CACNA1B;refGene_tx=XM_023243167.1;refGene_gn=CACNA1B;refGene_tx=XM_023243168.1;refGene_gn=CACNA1B;refGene_tx=XM_023243169.1;refGene_gn=CACNA1B;refGene_tx=XM_023243170.1;refGene_gn=CACNA1B;refGene_tx=XM_023243172.1;refGene_gn=CACNA1B;refGene_tx=XM_023243173.1
D4 96337238 96337448
refGene_gn=CACNA1B;refGene_tx=XM_023243153.1;refGene_gn=CACNA1B;refGene_tx=XM_023243154.1;refGene_gn=CACNA1B;refGene_tx=XM_023243155.1;refGene_gn=CACNA1B;refGene_tx=XM_023243156.1;refGene_gn=CACNA1B;refGene_tx=XM_023243157.1;refGene_gn=CACNA1B;refGene_tx=XM_023243158.1;refGene_gn=CACNA1B;refGene_tx=XM_023243159.1;refGene_gn=CACNA1B;refGene_tx=XM_023243161.1;refGene_gn=CACNA1B;refGene_tx=XM_023243162.1;refGene_gn=CACNA1B;refGene_tx=XM_023243163.1;refGene_gn=CACNA1B;refGene_tx=XM_023243164.1;refGene_gn=CACNA1B;refGene_tx=XM_023243165.1;refGene_gn=CACNA1B;refGene_tx=XM_023243166.1;refGene_gn=CACNA1B;refGene_tx=XM_023243167.1;refGene_gn=CACNA1B;refGene_tx=XM_023243168.1;refGene_gn=CACNA1B;refGene_tx=XM_023243169.1;refGene_gn=CACNA1B;refGene_tx=XM_023243170.1;refGene_gn=CACNA1B;refGene_tx=XM_023243172.1;refGene_gn=CACNA1B;refGene_tx=XM_023243173.1
D4 96341922 96342035
refGene_gn=CACNA1B;refGene_tx=XM_023243153.1;refGene_gn=CACNA1B;refGene_tx=XM_023243154.1;refGene_gn=CACNA1B;refGene_tx=XM_023243155.1;refGene_gn=CACNA1B;refGene_tx=XM_023243156.1;refGene_gn=CACNA1B;refGene_tx=XM_023243157.1;refGene_gn=CACNA1B;refGene_tx=XM_023243158.1;refGene_gn=CACNA1B;refGene_tx=XM_023243159.1;refGene_gn=CACNA1B;refGene_tx=XM_02

3243161.1;refGene_gn=CACNA1B;refGene_tx=XM_023243162.1;refGene_gn=CACNA1B;refGene_tx=XM_023243163.1;refGene_gn=CACNA1B;refGene_tx=XM_023243164.1;refGene_gn=CACNA1B;refGene_tx=XM_023243165.1;refGene_gn=CACNA1B;refGene_tx=XM_023243166.1;refGene_gn=CACNA1B;refGene_tx=XM_023243167.1;refGene_gn=CACNA1B;refGene_tx=XM_023243168.1;refGene_gn=CACNA1B;refGene_tx=XM_023243169.1;refGene_gn=CACNA1B;refGene_tx=XM_023243170.1;refGene_gn=CACNA1B;refGene_tx=XM_023243172.1;refGene_gn=CACNA1B;refGene_tx=XM_023243173.1
D4 96346389 96346502
refGene_gn=CACNA1B;refGene_tx=XM_023243153.1;refGene_gn=CACNA1B;refGene_tx=XM_023243154.1;refGene_gn=CACNA1B;refGene_tx=XM_023243155.1;refGene_gn=CACNA1B;refGene_tx=XM_023243156.1;refGene_gn=CACNA1B;refGene_tx=XM_023243157.1;refGene_gn=CACNA1B;refGene_tx=XM_023243158.1;refGene_gn=CACNA1B;refGene_tx=XM_023243159.1;refGene_gn=CACNA1B;refGene_tx=XM_023243161.1;refGene_gn=CACNA1B;refGene_tx=XM_023243162.1;refGene_gn=CACNA1B;refGene_tx=XM_023243163.1;refGene_gn=CACNA1B;refGene_tx=XM_023243164.1;refGene_gn=CACNA1B;refGene_tx=XM_023243165.1;refGene_gn=CACNA1B;refGene_tx=XM_023243166.1;refGene_gn=CACNA1B;refGene_tx=XM_023243167.1;refGene_gn=CACNA1B;refGene_tx=XM_023243168.1;refGene_gn=CACNA1B;refGene_tx=XM_023243169.1;refGene_gn=CACNA1B;refGene_tx=XM_023243170.1;refGene_gn=CACNA1B;refGene_tx=XM_023243172.1;refGene_gn=CACNA1B;refGene_tx=XM_023243173.1
D4 96348734 96348866
refGene_gn=CACNA1B;refGene_tx=XM_023243153.1;refGene_gn=CACNA1B;refGene_tx=XM_023243154.1;refGene_gn=CACNA1B;refGene_tx=XM_023243155.1;refGene_gn=CACNA1B;refGene_tx=XM_023243156.1;refGene_gn=CACNA1B;refGene_tx=XM_023243157.1;refGene_gn=CACNA1B;refGene_tx=XM_023243158.1;refGene_gn=CACNA1B;refGene_tx=XM_023243159.1;refGene_gn=CACNA1B;refGene_tx=XM_023243161.1;refGene_gn=CACNA1B;refGene_tx=XM_023243162.1;refGene_gn=CACNA1B;refGene_tx=XM_023243163.1;refGene_gn=CACNA1B;refGene_tx=XM_023243164.1;refGene_gn=CACNA1B;refGene_tx=XM_023243165.1;refGene_gn=CACNA1B;refGene_tx=XM_023243166.1;refGene_gn=CACNA1B;refGene_tx=XM_023243167.1;refGene_gn=CACNA1B;refGene_tx=XM_023243168.1;refGene_gn=CACNA1B;refGene_tx=XM_023243169.1;refGene_gn=CACNA1B;refGene_tx=XM_023243170.1;refGene_gn=CACNA1B;refGene_tx=XM_023243172.1;refGene_gn=CACNA1B;refGene_tx=XM_023243173.1
D4 96349095 96349168
refGene_gn=CACNA1B;refGene_tx=XM_023243153.1;refGene_gn=CACNA1B;refGene_tx=XM_023243154.1;refGene_gn=CACNA1B;refGene_tx=XM_023243155.1;refGene_gn=CACNA1B;refGene_tx=XM_023243156.1;refGene_gn=CACNA1B;refGene_tx=XM_023243157.1;refGene_gn=CACNA1B;refGene_tx=XM_023243158.1;refGene_gn=CACNA1B;refGene_tx=XM_023243159.1;refGene_gn=CACNA1B;refGene_tx=XM_023243161.1;refGene_gn=CACNA1B;refGene_tx=XM_023243162.1;refGene_gn=CACNA1B;refGene_tx=XM_023243163.1;refGene_gn=CACNA1B;refGene_tx=XM_023243164.1;refGene_gn=CACNA1B;refGene_tx=XM_023243165.1;refGene_gn=CACNA1B;refGene_tx=XM_023243166.1;refGene_gn=CACNA1B;refGene_tx=XM_023243167.1;refGene_gn=CACNA1B;refGene_tx=XM_023243168.1;refGene_gn=CACNA1B;refGene_tx=XM_023243169.1;refGene_gn=CACNA1B;refGene_tx=XM_023243170.1;refGene_gn=CACNA1B;refGene_tx=XM_023243172.1;refGene_gn=CACNA1B;refGene_tx=XM_023243173.1

D4 96358488 96358606
refGene_gn=CACNA1B;refGene_tx=XM_023243153.1;refGene_gn=CACNA1B;refGene_tx=XM_023243154.1;refGene_gn=CACNA1B;refGene_tx=XM_023243155.1;refGene_gn=CACNA1B;refGene_tx=XM_023243156.1;refGene_gn=CACNA1B;refGene_tx=XM_023243157.1;refGene_gn=CACNA1B;refGene_tx=XM_023243158.1;refGene_gn=CACNA1B;refGene_tx=XM_023243159.1;refGene_gn=CACNA1B;refGene_tx=XM_023243161.1;refGene_gn=CACNA1B;refGene_tx=XM_023243162.1;refGene_gn=CACNA1B;refGene_tx=XM_023243163.1;refGene_gn=CACNA1B;refGene_tx=XM_023243164.1;refGene_gn=CACNA1B;refGene_tx=XM_023243165.1;refGene_gn=CACNA1B;refGene_tx=XM_023243166.1;refGene_gn=CACNA1B;refGene_tx=XM_023243167.1;refGene_gn=CACNA1B;refGene_tx=XM_023243168.1;refGene_gn=CACNA1B;refGene_tx=XM_023243169.1;refGene_gn=CACNA1B;refGene_tx=XM_023243170.1;refGene_gn=CACNA1B;refGene_tx=XM_023243172.1;refGene_gn=CACNA1B;refGene_tx=XM_023243173.1
D4 96361643 96361711
refGene_gn=CACNA1B;refGene_tx=XM_023243153.1;refGene_gn=CACNA1B;refGene_tx=XM_023243154.1;refGene_gn=CACNA1B;refGene_tx=XM_023243155.1;refGene_gn=CACNA1B;refGene_tx=XM_023243156.1;refGene_gn=CACNA1B;refGene_tx=XM_023243157.1;refGene_gn=CACNA1B;refGene_tx=XM_023243158.1;refGene_gn=CACNA1B;refGene_tx=XM_023243159.1;refGene_gn=CACNA1B;refGene_tx=XM_023243161.1;refGene_gn=CACNA1B;refGene_tx=XM_023243162.1;refGene_gn=CACNA1B;refGene_tx=XM_023243163.1;refGene_gn=CACNA1B;refGene_tx=XM_023243164.1;refGene_gn=CACNA1B;refGene_tx=XM_023243165.1;refGene_gn=CACNA1B;refGene_tx=XM_023243166.1;refGene_gn=CACNA1B;refGene_tx=XM_023243167.1;refGene_gn=CACNA1B;refGene_tx=XM_023243168.1;refGene_gn=CACNA1B;refGene_tx=XM_023243169.1;refGene_gn=CACNA1B;refGene_tx=XM_023243170.1;refGene_gn=CACNA1B;refGene_tx=XM_023243172.1;refGene_gn=CACNA1B;refGene_tx=XM_023243173.1
D4 96364745 96364855
refGene_gn=CACNA1B;refGene_tx=XM_023243153.1;refGene_gn=CACNA1B;refGene_tx=XM_023243154.1;refGene_gn=CACNA1B;refGene_tx=XM_023243155.1;refGene_gn=CACNA1B;refGene_tx=XM_023243156.1;refGene_gn=CACNA1B;refGene_tx=XM_023243157.1;refGene_gn=CACNA1B;refGene_tx=XM_023243158.1;refGene_gn=CACNA1B;refGene_tx=XM_023243159.1;refGene_gn=CACNA1B;refGene_tx=XM_023243161.1;refGene_gn=CACNA1B;refGene_tx=XM_023243162.1;refGene_gn=CACNA1B;refGene_tx=XM_023243163.1;refGene_gn=CACNA1B;refGene_tx=XM_023243164.1;refGene_gn=CACNA1B;refGene_tx=XM_023243165.1;refGene_gn=CACNA1B;refGene_tx=XM_023243166.1;refGene_gn=CACNA1B;refGene_tx=XM_023243167.1;refGene_gn=CACNA1B;refGene_tx=XM_023243168.1;refGene_gn=CACNA1B;refGene_tx=XM_023243169.1;refGene_gn=CACNA1B;refGene_tx=XM_023243170.1;refGene_gn=CACNA1B;refGene_tx=XM_023243172.1;refGene_gn=CACNA1B;refGene_tx=XM_023243173.1
D4 96368663 96368726
refGene_gn=CACNA1B;refGene_tx=XM_023243153.1;refGene_gn=CACNA1B;refGene_tx=XM_023243154.1;refGene_gn=CACNA1B;refGene_tx=XM_023243155.1;refGene_gn=CACNA1B;refGene_tx=XM_023243156.1;refGene_gn=CACNA1B;refGene_tx=XM_023243157.1;refGene_gn=CACNA1B;refGene_tx=XM_023243158.1;refGene_gn=CACNA1B;refGene_tx=XM_023243159.1;refGene_gn=CACNA1B;refGene_tx=XM_023243161.1;refGene_gn=CACNA1B;refGene_tx=XM_023243162.1;refGene_gn=CACNA1B;refGene_tx=XM_023243170.1;refGene_gn=CACNA1B;refGene_tx=XM_0232431

72.1
D4 96372312 96373083
refGene_gn=CACNA1B;refGene_tx=XM_023243153.1;refGene_gn=CACNA1B;refGen
e_tx=XM_023243154.1;refGene_gn=CACNA1B;refGene_tx=XM_023243155.1;refGe
ne_gn=CACNA1B;refGene_tx=XM_023243156.1;refGene_gn=CACNA1B;refGene_tx=
XM_023243157.1;refGene_gn=CACNA1B;refGene_tx=XM_023243158.1;refGene_gn
=CACNA1B;refGene_tx=XM_023243159.1;refGene_gn=CACNA1B;refGene_tx=XM_02
3243161.1;refGene_gn=CACNA1B;refGene_tx=XM_023243162.1;refGene_gn=CACN
A1B;refGene_tx=XM_023243163.1;refGene_gn=CACNA1B;refGene_tx=XM_0232431
64.1;refGene_gn=CACNA1B;refGene_tx=XM_023243165.1;refGene_gn=CACNA1B;r
efGene_tx=XM_023243166.1;refGene_gn=CACNA1B;refGene_tx=XM_023243167.1;
refGene_gn=CACNA1B;refGene_tx=XM_023243168.1;refGene_gn=CACNA1B;refGen
e_tx=XM_023243169.1;refGene_gn=CACNA1B;refGene_tx=XM_023243170.1;refGe
ne_gn=CACNA1B;refGene_tx=XM_023243172.1;refGene_gn=CACNA1B;refGene_tx=
XM_023243173.1
D4 96374008 96374226
refGene_gn=CACNA1B;refGene_tx=XM_023243153.1;refGene_gn=CACNA1B;refGen
e_tx=XM_023243154.1;refGene_gn=CACNA1B;refGene_tx=XM_023243155.1;refGe
ne_gn=CACNA1B;refGene_tx=XM_023243156.1;refGene_gn=CACNA1B;refGene_tx=
XM_023243157.1;refGene_gn=CACNA1B;refGene_tx=XM_023243158.1;refGene_gn
=CACNA1B;refGene_tx=XM_023243159.1;refGene_gn=CACNA1B;refGene_tx=XM_02
3243161.1;refGene_gn=CACNA1B;refGene_tx=XM_023243162.1;refGene_gn=CACN
A1B;refGene_tx=XM_023243163.1;refGene_gn=CACNA1B;refGene_tx=XM_0232431
64.1;refGene_gn=CACNA1B;refGene_tx=XM_023243165.1;refGene_gn=CACNA1B;r
efGene_tx=XM_023243166.1;refGene_gn=CACNA1B;refGene_tx=XM_023243167.1;
refGene_gn=CACNA1B;refGene_tx=XM_023243168.1;refGene_gn=CACNA1B;refGen
e_tx=XM_023243169.1;refGene_gn=CACNA1B;refGene_tx=XM_023243170.1;refGe
ne_gn=CACNA1B;refGene_tx=XM_023243172.1;refGene_gn=CACNA1B;refGene_tx=
XM_023243173.1
D4 96402771 96402898
refGene_gn=CACNA1B;refGene_tx=XM_023243153.1;refGene_gn=CACNA1B;refGen
e_tx=XM_023243154.1;refGene_gn=CACNA1B;refGene_tx=XM_023243155.1;refGe
ne_gn=CACNA1B;refGene_tx=XM_023243156.1;refGene_gn=CACNA1B;refGene_tx=
XM_023243157.1;refGene_gn=CACNA1B;refGene_tx=XM_023243158.1;refGene_gn
=CACNA1B;refGene_tx=XM_023243159.1;refGene_gn=CACNA1B;refGene_tx=XM_02
3243161.1;refGene_gn=CACNA1B;refGene_tx=XM_023243162.1;refGene_gn=CACN
A1B;refGene_tx=XM_023243163.1;refGene_gn=CACNA1B;refGene_tx=XM_0232431
64.1;refGene_gn=CACNA1B;refGene_tx=XM_023243165.1;refGene_gn=CACNA1B;r
efGene_tx=XM_023243166.1;refGene_gn=CACNA1B;refGene_tx=XM_023243167.1;
refGene_gn=CACNA1B;refGene_tx=XM_023243168.1;refGene_gn=CACNA1B;refGen
e_tx=XM_023243169.1;refGene_gn=CACNA1B;refGene_tx=XM_023243170.1;refGe
ne_gn=CACNA1B;refGene_tx=XM_023243172.1;refGene_gn=CACNA1B;refGene_tx=
XM_023243173.1
D4 96405550 96405680
refGene_gn=CACNA1B;refGene_tx=XM_023243153.1;refGene_gn=CACNA1B;refGen
e_tx=XM_023243154.1;refGene_gn=CACNA1B;refGene_tx=XM_023243155.1;refGe
ne_gn=CACNA1B;refGene_tx=XM_023243156.1;refGene_gn=CACNA1B;refGene_tx=
XM_023243157.1;refGene_gn=CACNA1B;refGene_tx=XM_023243158.1;refGene_gn
=CACNA1B;refGene_tx=XM_023243159.1;refGene_gn=CACNA1B;refGene_tx=XM_02
3243161.1;refGene_gn=CACNA1B;refGene_tx=XM_023243162.1;refGene_gn=CACN

A1B;refGene_tx=XM_023243163.1;refGene_gn=CACNA1B;refGene_tx=XM_0232431
64.1;refGene_gn=CACNA1B;refGene_tx=XM_023243165.1;refGene_gn=CACNA1B;r
efGene_tx=XM_023243166.1;refGene_gn=CACNA1B;refGene_tx=XM_023243167.1;
refGene_gn=CACNA1B;refGene_tx=XM_023243168.1;refGene_gn=CACNA1B;refGen
e_tx=XM_023243169.1;refGene_gn=CACNA1B;refGene_tx=XM_023243170.1;refGe
ne_gn=CACNA1B;refGene_tx=XM_023243172.1;refGene_gn=CACNA1B;refGene_tx=
XM_023243173.1
D4 96405989 96406049
refGene_gn=CACNA1B;refGene_tx=XM_023243153.1;refGene_gn=CACNA1B;refGen
e_tx=XM_023243154.1;refGene_gn=CACNA1B;refGene_tx=XM_023243155.1;refGe
ne_gn=CACNA1B;refGene_tx=XM_023243156.1;refGene_gn=CACNA1B;refGene_tx=
XM_023243157.1;refGene_gn=CACNA1B;refGene_tx=XM_023243158.1;refGene_gn
=CACNA1B;refGene_tx=XM_023243159.1;refGene_gn=CACNA1B;refGene_tx=XM_02
3243161.1;refGene_gn=CACNA1B;refGene_tx=XM_023243162.1;refGene_gn=CACN
A1B;refGene_tx=XM_023243163.1;refGene_gn=CACNA1B;refGene_tx=XM_0232431
64.1;refGene_gn=CACNA1B;refGene_tx=XM_023243165.1;refGene_gn=CACNA1B;r
efGene_tx=XM_023243166.1;refGene_gn=CACNA1B;refGene_tx=XM_023243167.1;
refGene_gn=CACNA1B;refGene_tx=XM_023243168.1;refGene_gn=CACNA1B;refGen
e_tx=XM_023243169.1;refGene_gn=CACNA1B;refGene_tx=XM_023243170.1;refGe
ne_gn=CACNA1B;refGene_tx=XM_023243172.1;refGene_gn=CACNA1B;refGene_tx=
XM_023243173.1
D4 96407423 96407530
refGene_gn=CACNA1B;refGene_tx=XM_023243153.1;refGene_gn=CACNA1B;refGen
e_tx=XM_023243154.1;refGene_gn=CACNA1B;refGene_tx=XM_023243155.1;refGe
ne_gn=CACNA1B;refGene_tx=XM_023243156.1;refGene_gn=CACNA1B;refGene_tx=
XM_023243157.1;refGene_gn=CACNA1B;refGene_tx=XM_023243158.1;refGene_gn
=CACNA1B;refGene_tx=XM_023243159.1;refGene_gn=CACNA1B;refGene_tx=XM_02
3243161.1;refGene_gn=CACNA1B;refGene_tx=XM_023243162.1;refGene_gn=CACN
A1B;refGene_tx=XM_023243163.1;refGene_gn=CACNA1B;refGene_tx=XM_0232431
64.1;refGene_gn=CACNA1B;refGene_tx=XM_023243165.1;refGene_gn=CACNA1B;r
efGene_tx=XM_023243166.1;refGene_gn=CACNA1B;refGene_tx=XM_023243167.1;
refGene_gn=CACNA1B;refGene_tx=XM_023243168.1;refGene_gn=CACNA1B;refGen
e_tx=XM_023243169.1;refGene_gn=CACNA1B;refGene_tx=XM_023243170.1;refGe
ne_gn=CACNA1B;refGene_tx=XM_023243172.1;refGene_gn=CACNA1B;refGene_tx=
XM_023243173.1
D4 96408131 96408143
refGene_gn=CACNA1B;refGene_tx=XM_023243153.1;refGene_gn=CACNA1B;refGen
e_tx=XM_023243154.1;refGene_gn=CACNA1B;refGene_tx=XM_023243155.1;refGe
ne_gn=CACNA1B;refGene_tx=XM_023243156.1;refGene_gn=CACNA1B;refGene_tx=
XM_023243157.1;refGene_gn=CACNA1B;refGene_tx=XM_023243163.1;refGene_gn
=CACNA1B;refGene_tx=XM_023243164.1;refGene_gn=CACNA1B;refGene_tx=XM_02
3243165.1;refGene_gn=CACNA1B;refGene_tx=XM_023243166.1;refGene_gn=CACN
A1B;refGene_tx=XM_023243170.1;refGene_gn=CACNA1B;refGene_tx=XM_0232431
72.1
D4 96410296 96410393
refGene_gn=CACNA1B;refGene_tx=XM_023243153.1;refGene_gn=CACNA1B;refGen
e_tx=XM_023243154.1;refGene_gn=CACNA1B;refGene_tx=XM_023243155.1;refGe
ne_gn=CACNA1B;refGene_tx=XM_023243156.1;refGene_gn=CACNA1B;refGene_tx=
XM_023243157.1;refGene_gn=CACNA1B;refGene_tx=XM_023243158.1;refGene_gn
=CACNA1B;refGene_tx=XM_023243159.1;refGene_gn=CACNA1B;refGene_tx=XM_02

3243161.1;refGene_gn=CACNA1B;refGene_tx=XM_023243162.1;refGene_gn=CACNA1B;refGene_tx=XM_023243163.1;refGene_gn=CACNA1B;refGene_tx=XM_023243164.1;refGene_gn=CACNA1B;refGene_tx=XM_023243165.1;refGene_gn=CACNA1B;refGene_tx=XM_023243166.1;refGene_gn=CACNA1B;refGene_tx=XM_023243167.1;refGene_gn=CACNA1B;refGene_tx=XM_023243168.1;refGene_gn=CACNA1B;refGene_tx=XM_023243169.1;refGene_gn=CACNA1B;refGene_tx=XM_023243170.1;refGene_gn=CACNA1B;refGene_tx=XM_023243172.1;refGene_gn=CACNA1B;refGene_tx=XM_023243173.1

D4 96411995 96412156
refGene_gn=CACNA1B;refGene_tx=XM_023243153.1;refGene_gn=CACNA1B;refGene_tx=XM_023243154.1;refGene_gn=CACNA1B;refGene_tx=XM_023243155.1;refGene_gn=CACNA1B;refGene_tx=XM_023243156.1;refGene_gn=CACNA1B;refGene_tx=XM_023243157.1;refGene_gn=CACNA1B;refGene_tx=XM_023243158.1;refGene_gn=CACNA1B;refGene_tx=XM_023243159.1;refGene_gn=CACNA1B;refGene_tx=XM_023243161.1;refGene_gn=CACNA1B;refGene_tx=XM_023243162.1;refGene_gn=CACNA1B;refGene_tx=XM_023243163.1;refGene_gn=CACNA1B;refGene_tx=XM_023243164.1;refGene_gn=CACNA1B;refGene_tx=XM_023243165.1;refGene_gn=CACNA1B;refGene_tx=XM_023243166.1;refGene_gn=CACNA1B;refGene_tx=XM_023243167.1;refGene_gn=CACNA1B;refGene_tx=XM_023243168.1;refGene_gn=CACNA1B;refGene_tx=XM_023243169.1;refGene_gn=CACNA1B;refGene_tx=XM_023243170.1;refGene_gn=CACNA1B;refGene_tx=XM_023243172.1;refGene_gn=CACNA1B;refGene_tx=XM_023243173.1

D4 96415308 96415446
refGene_gn=CACNA1B;refGene_tx=XM_023243153.1;refGene_gn=CACNA1B;refGene_tx=XM_023243154.1;refGene_gn=CACNA1B;refGene_tx=XM_023243155.1;refGene_gn=CACNA1B;refGene_tx=XM_023243156.1;refGene_gn=CACNA1B;refGene_tx=XM_023243157.1;refGene_gn=CACNA1B;refGene_tx=XM_023243158.1;refGene_gn=CACNA1B;refGene_tx=XM_023243159.1;refGene_gn=CACNA1B;refGene_tx=XM_023243161.1;refGene_gn=CACNA1B;refGene_tx=XM_023243162.1;refGene_gn=CACNA1B;refGene_tx=XM_023243163.1;refGene_gn=CACNA1B;refGene_tx=XM_023243164.1;refGene_gn=CACNA1B;refGene_tx=XM_023243165.1;refGene_gn=CACNA1B;refGene_tx=XM_023243166.1;refGene_gn=CACNA1B;refGene_tx=XM_023243167.1;refGene_gn=CACNA1B;refGene_tx=XM_023243168.1;refGene_gn=CACNA1B;refGene_tx=XM_023243169.1;refGene_gn=CACNA1B;refGene_tx=XM_023243170.1;refGene_gn=CACNA1B;refGene_tx=XM_023243172.1;refGene_gn=CACNA1B;refGene_tx=XM_023243173.1

D4 96416226 96416428
refGene_gn=CACNA1B;refGene_tx=XM_023243153.1;refGene_gn=CACNA1B;refGene_tx=XM_023243154.1;refGene_gn=CACNA1B;refGene_tx=XM_023243155.1;refGene_gn=CACNA1B;refGene_tx=XM_023243156.1;refGene_gn=CACNA1B;refGene_tx=XM_023243157.1;refGene_gn=CACNA1B;refGene_tx=XM_023243158.1;refGene_gn=CACNA1B;refGene_tx=XM_023243159.1;refGene_gn=CACNA1B;refGene_tx=XM_023243161.1;refGene_gn=CACNA1B;refGene_tx=XM_023243162.1;refGene_gn=CACNA1B;refGene_tx=XM_023243163.1;refGene_gn=CACNA1B;refGene_tx=XM_023243164.1;refGene_gn=CACNA1B;refGene_tx=XM_023243165.1;refGene_gn=CACNA1B;refGene_tx=XM_023243166.1;refGene_gn=CACNA1B;refGene_tx=XM_023243167.1;refGene_gn=CACNA1B;refGene_tx=XM_023243168.1;refGene_gn=CACNA1B;refGene_tx=XM_023243169.1;refGene_gn=CACNA1B;refGene_tx=XM_023243170.1;refGene_gn=CACNA1B;refGene_tx=XM_023243172.1;refGene_gn=CACNA1B;refGene_tx=XM_023243173.1

D4 96416661 96416826
refGene_gn=CACNA1B;refGene_tx=XM_023243153.1;refGene_gn=CACNA1B;refGene_tx=XM_023243154.1;refGene_gn=CACNA1B;refGene_tx=XM_023243155.1;refGene_gn=CACNA1B;refGene_tx=XM_023243156.1;refGene_gn=CACNA1B;refGene_tx=XM_023243157.1;refGene_gn=CACNA1B;refGene_tx=XM_023243158.1;refGene_gn=CACNA1B;refGene_tx=XM_023243159.1;refGene_gn=CACNA1B;refGene_tx=XM_023243161.1;refGene_gn=CACNA1B;refGene_tx=XM_023243162.1;refGene_gn=CACNA1B;refGene_tx=XM_023243163.1;refGene_gn=CACNA1B;refGene_tx=XM_023243164.1;refGene_gn=CACNA1B;refGene_tx=XM_023243165.1;refGene_gn=CACNA1B;refGene_tx=XM_023243166.1;refGene_gn=CACNA1B;refGene_tx=XM_023243167.1;refGene_gn=CACNA1B;refGene_tx=XM_023243168.1;refGene_gn=CACNA1B;refGene_tx=XM_023243169.1;refGene_gn=CACNA1B;refGene_tx=XM_023243170.1;refGene_gn=CACNA1B;refGene_tx=XM_023243172.1;refGene_gn=CACNA1B;refGene_tx=XM_023243173.1
D4 96417128 96417239
refGene_gn=CACNA1B;refGene_tx=XM_023243153.1;refGene_gn=CACNA1B;refGene_tx=XM_023243154.1;refGene_gn=CACNA1B;refGene_tx=XM_023243155.1;refGene_gn=CACNA1B;refGene_tx=XM_023243156.1;refGene_gn=CACNA1B;refGene_tx=XM_023243157.1;refGene_gn=CACNA1B;refGene_tx=XM_023243158.1;refGene_gn=CACNA1B;refGene_tx=XM_023243159.1;refGene_gn=CACNA1B;refGene_tx=XM_023243161.1;refGene_gn=CACNA1B;refGene_tx=XM_023243162.1;refGene_gn=CACNA1B;refGene_tx=XM_023243163.1;refGene_gn=CACNA1B;refGene_tx=XM_023243164.1;refGene_gn=CACNA1B;refGene_tx=XM_023243165.1;refGene_gn=CACNA1B;refGene_tx=XM_023243166.1;refGene_gn=CACNA1B;refGene_tx=XM_023243167.1;refGene_gn=CACNA1B;refGene_tx=XM_023243168.1;refGene_gn=CACNA1B;refGene_tx=XM_023243169.1;refGene_gn=CACNA1B;refGene_tx=XM_023243170.1;refGene_gn=CACNA1B;refGene_tx=XM_023243172.1;refGene_gn=CACNA1B;refGene_tx=XM_023243173.1
D4 96417660 96417744
refGene_gn=CACNA1B;refGene_tx=XM_023243153.1;refGene_gn=CACNA1B;refGene_tx=XM_023243154.1;refGene_gn=CACNA1B;refGene_tx=XM_023243155.1;refGene_gn=CACNA1B;refGene_tx=XM_023243156.1;refGene_gn=CACNA1B;refGene_tx=XM_023243157.1;refGene_gn=CACNA1B;refGene_tx=XM_023243158.1;refGene_gn=CACNA1B;refGene_tx=XM_023243159.1;refGene_gn=CACNA1B;refGene_tx=XM_023243161.1;refGene_gn=CACNA1B;refGene_tx=XM_023243162.1;refGene_gn=CACNA1B;refGene_tx=XM_023243163.1;refGene_gn=CACNA1B;refGene_tx=XM_023243164.1;refGene_gn=CACNA1B;refGene_tx=XM_023243165.1;refGene_gn=CACNA1B;refGene_tx=XM_023243166.1;refGene_gn=CACNA1B;refGene_tx=XM_023243167.1;refGene_gn=CACNA1B;refGene_tx=XM_023243168.1;refGene_gn=CACNA1B;refGene_tx=XM_023243169.1;refGene_gn=CACNA1B;refGene_tx=XM_023243170.1;refGene_gn=CACNA1B;refGene_tx=XM_023243172.1;refGene_gn=CACNA1B;refGene_tx=XM_023243173.1
D4 96426611 96426617
refGene_gn=CACNA1B;refGene_tx=XM_023243153.1;refGene_gn=CACNA1B;refGene_tx=XM_023243154.1;refGene_gn=CACNA1B;refGene_tx=XM_023243155.1;refGene_gn=CACNA1B;refGene_tx=XM_023243156.1;refGene_gn=CACNA1B;refGene_tx=XM_023243158.1;refGene_gn=CACNA1B;refGene_tx=XM_023243159.1;refGene_gn=CACNA1B;refGene_tx=XM_023243163.1;refGene_gn=CACNA1B;refGene_tx=XM_023243164.1;refGene_gn=CACNA1B;refGene_tx=XM_023243167.1;refGene_gn=CACNA1B;refGene_tx=XM_023243170.1;refGene_gn=CACNA1B;refGene_tx=XM_0232431

72.1;refGene_gn=CACNA1B;refGene_tx=XM_023243173.1
D4 96430318 96430435
refGene_gn=CACNA1B;refGene_tx=XM_023243153.1;refGene_gn=CACNA1B;refGene_tx=XM_023243154.1;refGene_gn=CACNA1B;refGene_tx=XM_023243155.1;refGene_gn=CACNA1B;refGene_tx=XM_023243156.1;refGene_gn=CACNA1B;refGene_tx=XM_023243157.1;refGene_gn=CACNA1B;refGene_tx=XM_023243158.1;refGene_gn=CACNA1B;refGene_tx=XM_023243159.1;refGene_gn=CACNA1B;refGene_tx=XM_023243161.1;refGene_gn=CACNA1B;refGene_tx=XM_023243162.1;refGene_gn=CACNA1B;refGene_tx=XM_023243163.1;refGene_gn=CACNA1B;refGene_tx=XM_023243164.1;refGene_gn=CACNA1B;refGene_tx=XM_023243165.1;refGene_gn=CACNA1B;refGene_tx=XM_023243166.1;refGene_gn=CACNA1B;refGene_tx=XM_023243167.1;refGene_gn=CACNA1B;refGene_tx=XM_023243168.1;refGene_gn=CACNA1B;refGene_tx=XM_023243169.1;refGene_gn=CACNA1B;refGene_tx=XM_023243170.1;refGene_gn=CACNA1B;refGene_tx=XM_023243172.1;refGene_gn=CACNA1B;refGene_tx=XM_023243173.1
D4 96430831 96430897
refGene_gn=CACNA1B;refGene_tx=XM_023243153.1;refGene_gn=CACNA1B;refGene_tx=XM_023243154.1;refGene_gn=CACNA1B;refGene_tx=XM_023243155.1;refGene_gn=CACNA1B;refGene_tx=XM_023243156.1;refGene_gn=CACNA1B;refGene_tx=XM_023243157.1;refGene_gn=CACNA1B;refGene_tx=XM_023243158.1;refGene_gn=CACNA1B;refGene_tx=XM_023243159.1;refGene_gn=CACNA1B;refGene_tx=XM_023243161.1;refGene_gn=CACNA1B;refGene_tx=XM_023243162.1;refGene_gn=CACNA1B;refGene_tx=XM_023243163.1;refGene_gn=CACNA1B;refGene_tx=XM_023243164.1;refGene_gn=CACNA1B;refGene_tx=XM_023243165.1;refGene_gn=CACNA1B;refGene_tx=XM_023243166.1;refGene_gn=CACNA1B;refGene_tx=XM_023243167.1;refGene_gn=CACNA1B;refGene_tx=XM_023243168.1;refGene_gn=CACNA1B;refGene_tx=XM_023243169.1;refGene_gn=CACNA1B;refGene_tx=XM_023243170.1;refGene_gn=CACNA1B;refGene_tx=XM_023243172.1;refGene_gn=CACNA1B;refGene_tx=XM_023243173.1
D4 96432249 96432341
refGene_gn=CACNA1B;refGene_tx=XM_023243153.1;refGene_gn=CACNA1B;refGene_tx=XM_023243154.1;refGene_gn=CACNA1B;refGene_tx=XM_023243155.1;refGene_gn=CACNA1B;refGene_tx=XM_023243156.1;refGene_gn=CACNA1B;refGene_tx=XM_023243157.1;refGene_gn=CACNA1B;refGene_tx=XM_023243158.1;refGene_gn=CACNA1B;refGene_tx=XM_023243159.1;refGene_gn=CACNA1B;refGene_tx=XM_023243161.1;refGene_gn=CACNA1B;refGene_tx=XM_023243162.1;refGene_gn=CACNA1B;refGene_tx=XM_023243163.1;refGene_gn=CACNA1B;refGene_tx=XM_023243164.1;refGene_gn=CACNA1B;refGene_tx=XM_023243165.1;refGene_gn=CACNA1B;refGene_tx=XM_023243166.1;refGene_gn=CACNA1B;refGene_tx=XM_023243167.1;refGene_gn=CACNA1B;refGene_tx=XM_023243168.1;refGene_gn=CACNA1B;refGene_tx=XM_023243169.1;refGene_gn=CACNA1B;refGene_tx=XM_023243170.1;refGene_gn=CACNA1B;refGene_tx=XM_023243172.1;refGene_gn=CACNA1B;refGene_tx=XM_023243173.1
D4 96434772 96434917
refGene_gn=CACNA1B;refGene_tx=XM_023243153.1;refGene_gn=CACNA1B;refGene_tx=XM_023243154.1;refGene_gn=CACNA1B;refGene_tx=XM_023243155.1;refGene_gn=CACNA1B;refGene_tx=XM_023243156.1;refGene_gn=CACNA1B;refGene_tx=XM_023243157.1;refGene_gn=CACNA1B;refGene_tx=XM_023243158.1;refGene_gn=CACNA1B;refGene_tx=XM_023243159.1;refGene_gn=CACNA1B;refGene_tx=XM_023243161.1;refGene_gn=CACNA1B;refGene_tx=XM_023243162.1;refGene_gn=CACN

A1B;refGene_tx=XM_023243163.1;refGene_gn=CACNA1B;refGene_tx=XM_0232431
64.1;refGene_gn=CACNA1B;refGene_tx=XM_023243165.1;refGene_gn=CACNA1B;r
efGene_tx=XM_023243166.1;refGene_gn=CACNA1B;refGene_tx=XM_023243167.1;
refGene_gn=CACNA1B;refGene_tx=XM_023243168.1;refGene_gn=CACNA1B;refGen
e_tx=XM_023243169.1;refGene_gn=CACNA1B;refGene_tx=XM_023243170.1;refGe
ne_gn=CACNA1B;refGene_tx=XM_023243172.1;refGene_gn=CACNA1B;refGene_tx=
XM_023243173.1
D4 96440411 96440539
refGene_gn=CACNA1B;refGene_tx=XM_023243153.1;refGene_gn=CACNA1B;refGen
e_tx=XM_023243154.1;refGene_gn=CACNA1B;refGene_tx=XM_023243155.1;refGe
ne_gn=CACNA1B;refGene_tx=XM_023243156.1;refGene_gn=CACNA1B;refGene_tx=
XM_023243157.1;refGene_gn=CACNA1B;refGene_tx=XM_023243158.1;refGene_gn
=CACNA1B;refGene_tx=XM_023243159.1;refGene_gn=CACNA1B;refGene_tx=XM_02
3243161.1;refGene_gn=CACNA1B;refGene_tx=XM_023243162.1;refGene_gn=CACN
A1B;refGene_tx=XM_023243163.1;refGene_gn=CACNA1B;refGene_tx=XM_0232431
64.1;refGene_gn=CACNA1B;refGene_tx=XM_023243165.1;refGene_gn=CACNA1B;r
efGene_tx=XM_023243166.1;refGene_gn=CACNA1B;refGene_tx=XM_023243167.1;
refGene_gn=CACNA1B;refGene_tx=XM_023243168.1;refGene_gn=CACNA1B;refGen
e_tx=XM_023243169.1;refGene_gn=CACNA1B;refGene_tx=XM_023243170.1;refGe
ne_gn=CACNA1B;refGene_tx=XM_023243172.1;refGene_gn=CACNA1B;refGene_tx=
XM_023243173.1
D4 96444701 96444798
refGene_gn=CACNA1B;refGene_tx=XM_023243154.1;refGene_gn=CACNA1B;refGen
e_tx=XM_023243162.1
D4 96446359 96446456
refGene_gn=CACNA1B;refGene_tx=XM_023243153.1;refGene_gn=CACNA1B;refGen
e_tx=XM_023243155.1;refGene_gn=CACNA1B;refGene_tx=XM_023243156.1;refGe
ne_gn=CACNA1B;refGene_tx=XM_023243157.1;refGene_gn=CACNA1B;refGene_tx=
XM_023243158.1;refGene_gn=CACNA1B;refGene_tx=XM_023243159.1;refGene_gn
=CACNA1B;refGene_tx=XM_023243161.1;refGene_gn=CACNA1B;refGene_tx=XM_02
3243163.1;refGene_gn=CACNA1B;refGene_tx=XM_023243164.1;refGene_gn=CACN
A1B;refGene_tx=XM_023243165.1;refGene_gn=CACNA1B;refGene_tx=XM_0232431
66.1;refGene_gn=CACNA1B;refGene_tx=XM_023243167.1;refGene_gn=CACNA1B;r
efGene_tx=XM_023243168.1;refGene_gn=CACNA1B;refGene_tx=XM_023243169.1;
refGene_gn=CACNA1B;refGene_tx=XM_023243170.1;refGene_gn=CACNA1B;refGen
e_tx=XM_023243172.1;refGene_gn=CACNA1B;refGene_tx=XM_023243173.1
D4 96449356 96449465
refGene_gn=CACNA1B;refGene_tx=XM_023243153.1;refGene_gn=CACNA1B;refGen
e_tx=XM_023243154.1;refGene_gn=CACNA1B;refGene_tx=XM_023243155.1;refGe
ne_gn=CACNA1B;refGene_tx=XM_023243156.1;refGene_gn=CACNA1B;refGene_tx=
XM_023243157.1;refGene_gn=CACNA1B;refGene_tx=XM_023243158.1;refGene_gn
=CACNA1B;refGene_tx=XM_023243159.1;refGene_gn=CACNA1B;refGene_tx=XM_02
3243161.1;refGene_gn=CACNA1B;refGene_tx=XM_023243162.1;refGene_gn=CACN
A1B;refGene_tx=XM_023243163.1;refGene_gn=CACNA1B;refGene_tx=XM_0232431
64.1;refGene_gn=CACNA1B;refGene_tx=XM_023243165.1;refGene_gn=CACNA1B;r
efGene_tx=XM_023243166.1;refGene_gn=CACNA1B;refGene_tx=XM_023243167.1;
refGene_gn=CACNA1B;refGene_tx=XM_023243168.1;refGene_gn=CACNA1B;refGen
e_tx=XM_023243169.1;refGene_gn=CACNA1B;refGene_tx=XM_023243170.1;refGe
ne_gn=CACNA1B;refGene_tx=XM_023243172.1;refGene_gn=CACNA1B;refGene_tx=
XM_023243173.1

D4 96451712 96451738 refGene_gn=CACNA1B;refGene_tx=XM_023243172.1
D4 96454373 96454481
refGene_gn=CACNA1B;refGene_tx=XM_023243153.1;refGene_gn=CACNA1B;refGene_tx=XM_023243154.1;refGene_gn=CACNA1B;refGene_tx=XM_023243155.1;refGene_gn=CACNA1B;refGene_tx=XM_023243156.1;refGene_gn=CACNA1B;refGene_tx=XM_023243157.1;refGene_gn=CACNA1B;refGene_tx=XM_023243158.1;refGene_gn=CACNA1B;refGene_tx=XM_023243159.1;refGene_gn=CACNA1B;refGene_tx=XM_023243161.1;refGene_gn=CACNA1B;refGene_tx=XM_023243162.1;refGene_gn=CACNA1B;refGene_tx=XM_023243163.1;refGene_gn=CACNA1B;refGene_tx=XM_023243164.1;refGene_gn=CACNA1B;refGene_tx=XM_023243165.1;refGene_gn=CACNA1B;refGene_tx=XM_023243166.1;refGene_gn=CACNA1B;refGene_tx=XM_023243167.1;refGene_gn=CACNA1B;refGene_tx=XM_023243168.1;refGene_gn=CACNA1B;refGene_tx=XM_023243169.1;refGene_gn=CACNA1B;refGene_tx=XM_023243170.1;refGene_gn=CACNA1B;refGene_tx=XM_023243173.1
D4 96455013 96455126
refGene_gn=CACNA1B;refGene_tx=XM_023243153.1;refGene_gn=CACNA1B;refGene_tx=XM_023243154.1;refGene_gn=CACNA1B;refGene_tx=XM_023243155.1;refGene_gn=CACNA1B;refGene_tx=XM_023243156.1;refGene_gn=CACNA1B;refGene_tx=XM_023243157.1;refGene_gn=CACNA1B;refGene_tx=XM_023243158.1;refGene_gn=CACNA1B;refGene_tx=XM_023243159.1;refGene_gn=CACNA1B;refGene_tx=XM_023243161.1;refGene_gn=CACNA1B;refGene_tx=XM_023243162.1;refGene_gn=CACNA1B;refGene_tx=XM_023243163.1;refGene_gn=CACNA1B;refGene_tx=XM_023243164.1;refGene_gn=CACNA1B;refGene_tx=XM_023243165.1;refGene_gn=CACNA1B;refGene_tx=XM_023243166.1;refGene_gn=CACNA1B;refGene_tx=XM_023243167.1;refGene_gn=CACNA1B;refGene_tx=XM_023243168.1;refGene_gn=CACNA1B;refGene_tx=XM_023243169.1;refGene_gn=CACNA1B;refGene_tx=XM_023243170.1;refGene_gn=CACNA1B;refGene_tx=XM_023243173.1
D4 96456009 96456137
refGene_gn=CACNA1B;refGene_tx=XM_023243153.1;refGene_gn=CACNA1B;refGene_tx=XM_023243154.1;refGene_gn=CACNA1B;refGene_tx=XM_023243155.1;refGene_gn=CACNA1B;refGene_tx=XM_023243156.1;refGene_gn=CACNA1B;refGene_tx=XM_023243157.1;refGene_gn=CACNA1B;refGene_tx=XM_023243158.1;refGene_gn=CACNA1B;refGene_tx=XM_023243159.1;refGene_gn=CACNA1B;refGene_tx=XM_023243161.1;refGene_gn=CACNA1B;refGene_tx=XM_023243162.1;refGene_gn=CACNA1B;refGene_tx=XM_023243163.1;refGene_gn=CACNA1B;refGene_tx=XM_023243164.1;refGene_gn=CACNA1B;refGene_tx=XM_023243165.1;refGene_gn=CACNA1B;refGene_tx=XM_023243166.1;refGene_gn=CACNA1B;refGene_tx=XM_023243167.1;refGene_gn=CACNA1B;refGene_tx=XM_023243168.1;refGene_gn=CACNA1B;refGene_tx=XM_023243169.1;refGene_gn=CACNA1B;refGene_tx=XM_023243170.1;refGene_gn=CACNA1B;refGene_tx=XM_023243173.1
D4 96458190 96458326
refGene_gn=CACNA1B;refGene_tx=XM_023243153.1;refGene_gn=CACNA1B;refGene_tx=XM_023243154.1;refGene_gn=CACNA1B;refGene_tx=XM_023243155.1;refGene_gn=CACNA1B;refGene_tx=XM_023243156.1;refGene_gn=CACNA1B;refGene_tx=XM_023243157.1;refGene_gn=CACNA1B;refGene_tx=XM_023243158.1;refGene_gn=CACNA1B;refGene_tx=XM_023243159.1;refGene_gn=CACNA1B;refGene_tx=XM_023243161.1;refGene_gn=CACNA1B;refGene_tx=XM_023243162.1;refGene_gn=CACNA1B;refGene_tx=XM_023243163.1;refGene_gn=CACNA1B;refGene_tx=XM_023243164.1;refGene_gn=CACNA1B;refGene_tx=XM_023243165.1;refGene_gn=CACNA1B;refGene_tx=XM_023243166.1;refGene_gn=CACNA1B;refGene_tx=XM_023243167.1;

refGene_gn=CACNA1B;refGene_tx=XM_023243168.1;refGene_gn=CACNA1B;refGen
e_tx=XM_023243169.1;refGene_gn=CACNA1B;refGene_tx=XM_023243170.1;refGe
ne_gn=CACNA1B;refGene_tx=XM_023243173.1
D4 96458464 96458581
refGene_gn=CACNA1B;refGene_tx=XM_023243153.1;refGene_gn=CACNA1B;refGen
e_tx=XM_023243154.1;refGene_gn=CACNA1B;refGene_tx=XM_023243155.1;refGe
ne_gn=CACNA1B;refGene_tx=XM_023243156.1;refGene_gn=CACNA1B;refGene_tx=
XM_023243157.1;refGene_gn=CACNA1B;refGene_tx=XM_023243158.1;refGene_gn
=CACNA1B;refGene_tx=XM_023243159.1;refGene_gn=CACNA1B;refGene_tx=XM_02
3243161.1;refGene_gn=CACNA1B;refGene_tx=XM_023243162.1;refGene_gn=CACN
A1B;refGene_tx=XM_023243163.1;refGene_gn=CACNA1B;refGene_tx=XM_0232431
64.1;refGene_gn=CACNA1B;refGene_tx=XM_023243165.1;refGene_gn=CACNA1B;r
efGene_tx=XM_023243166.1;refGene_gn=CACNA1B;refGene_tx=XM_023243167.1;
refGene_gn=CACNA1B;refGene_tx=XM_023243168.1;refGene_gn=CACNA1B;refGen
e_tx=XM_023243169.1;refGene_gn=CACNA1B;refGene_tx=XM_023243173.1
D4 96459751 96459959
refGene_gn=CACNA1B;refGene_tx=XM_023243153.1;refGene_gn=CACNA1B;refGen
e_tx=XM_023243154.1;refGene_gn=CACNA1B;refGene_tx=XM_023243155.1;refGe
ne_gn=CACNA1B;refGene_tx=XM_023243156.1;refGene_gn=CACNA1B;refGene_tx=
XM_023243157.1;refGene_gn=CACNA1B;refGene_tx=XM_023243158.1;refGene_gn
=CACNA1B;refGene_tx=XM_023243159.1;refGene_gn=CACNA1B;refGene_tx=XM_02
3243161.1;refGene_gn=CACNA1B;refGene_tx=XM_023243162.1;refGene_gn=CACN
A1B;refGene_tx=XM_023243163.1;refGene_gn=CACNA1B;refGene_tx=XM_0232431
64.1;refGene_gn=CACNA1B;refGene_tx=XM_023243165.1;refGene_gn=CACNA1B;r
efGene_tx=XM_023243166.1;refGene_gn=CACNA1B;refGene_tx=XM_023243167.1;
refGene_gn=CACNA1B;refGene_tx=XM_023243168.1;refGene_gn=CACNA1B;refGen
e_tx=XM_023243169.1;refGene_gn=CACNA1B;refGene_tx=XM_023243170.1;refGe
ne_gn=CACNA1B;refGene_tx=XM_023243173.1
D4 96460215 96460460
refGene_gn=CACNA1B;refGene_tx=XM_023243153.1;refGene_gn=CACNA1B;refGen
e_tx=XM_023243154.1;refGene_gn=CACNA1B;refGene_tx=XM_023243155.1;refGe
ne_gn=CACNA1B;refGene_tx=XM_023243156.1;refGene_gn=CACNA1B;refGene_tx=
XM_023243157.1;refGene_gn=CACNA1B;refGene_tx=XM_023243158.1;refGene_gn
=CACNA1B;refGene_tx=XM_023243159.1;refGene_gn=CACNA1B;refGene_tx=XM_02
3243161.1;refGene_gn=CACNA1B;refGene_tx=XM_023243162.1;refGene_gn=CACN
A1B;refGene_tx=XM_023243163.1;refGene_gn=CACNA1B;refGene_tx=XM_0232431
64.1;refGene_gn=CACNA1B;refGene_tx=XM_023243165.1;refGene_gn=CACNA1B;r
efGene_tx=XM_023243166.1;refGene_gn=CACNA1B;refGene_tx=XM_023243167.1;
refGene_gn=CACNA1B;refGene_tx=XM_023243168.1;refGene_gn=CACNA1B;refGen
e_tx=XM_023243169.1;refGene_gn=CACNA1B;refGene_tx=XM_023243170.1;refGe
ne_gn=CACNA1B;refGene_tx=XM_023243173.1
D4 96460891 96461422
refGene_gn=CACNA1B;refGene_tx=XM_023243153.1;refGene_gn=CACNA1B;refGen
e_tx=XM_023243154.1;refGene_gn=CACNA1B;refGene_tx=XM_023243155.1;refGe
ne_gn=CACNA1B;refGene_tx=XM_023243156.1;refGene_gn=CACNA1B;refGene_tx=
XM_023243157.1;refGene_gn=CACNA1B;refGene_tx=XM_023243158.1;refGene_gn
=CACNA1B;refGene_tx=XM_023243159.1;refGene_gn=CACNA1B;refGene_tx=XM_02
3243161.1;refGene_gn=CACNA1B;refGene_tx=XM_023243162.1;refGene_gn=CACN
A1B;refGene_tx=XM_023243163.1;refGene_gn=CACNA1B;refGene_tx=XM_0232431
64.1;refGene_gn=CACNA1B;refGene_tx=XM_023243165.1;refGene_gn=CACNA1B;r

efGene_tx=XM_023243166.1;refGene_gn=CACNA1B;refGene_tx=XM_023243167.1;refGene_gn=CACNA1B;refGene_tx=XM_023243168.1;refGene_gn=CACNA1B;refGene_tx=XM_023243169.1;refGene_gn=CACNA1B;refGene_tx=XM_023243170.1;refGene_gn=CACNA1B;refGene_tx=XM_023243173.1
E1 203649 203730
refGene_gn=ZZEF1;refGene_tx=XM_011288812.2;refGene_gn=ZZEF1;refGene_tx=XM_023243632.1;refGene_gn=ZZEF1;refGene_tx=XM_023243633.1;refGene_gn=ZZEF1;refGene_tx=XM_023243635.1
E1 205427 205499
refGene_gn=ZZEF1;refGene_tx=XM_011288812.2;refGene_gn=ZZEF1;refGene_tx=XM_023243632.1;refGene_gn=ZZEF1;refGene_tx=XM_023243633.1;refGene_gn=ZZEF1;refGene_tx=XM_023243635.1
E1 206103 206257
refGene_gn=ZZEF1;refGene_tx=XM_011288812.2;refGene_gn=ZZEF1;refGene_tx=XM_023243632.1;refGene_gn=ZZEF1;refGene_tx=XM_023243633.1;refGene_gn=ZZEF1;refGene_tx=XM_023243635.1
E1 210940 211106
refGene_gn=ZZEF1;refGene_tx=XM_011288812.2;refGene_gn=ZZEF1;refGene_tx=XM_023243632.1;refGene_gn=ZZEF1;refGene_tx=XM_023243633.1;refGene_gn=ZZEF1;refGene_tx=XM_023243635.1
E1 211507 211606
refGene_gn=ZZEF1;refGene_tx=XM_011288812.2;refGene_gn=ZZEF1;refGene_tx=XM_023243632.1;refGene_gn=ZZEF1;refGene_tx=XM_023243633.1;refGene_gn=ZZEF1;refGene_tx=XM_023243635.1
E1 211974 212143
refGene_gn=ZZEF1;refGene_tx=XM_011288812.2;refGene_gn=ZZEF1;refGene_tx=XM_023243632.1;refGene_gn=ZZEF1;refGene_tx=XM_023243633.1;refGene_gn=ZZEF1;refGene_tx=XM_023243635.1
E1 212643 212787
refGene_gn=ZZEF1;refGene_tx=XM_011288812.2;refGene_gn=ZZEF1;refGene_tx=XM_023243632.1;refGene_gn=ZZEF1;refGene_tx=XM_023243633.1;refGene_gn=ZZEF1;refGene_tx=XM_023243635.1
E1 214608 214968
refGene_gn=ZZEF1;refGene_tx=XM_011288812.2;refGene_gn=ZZEF1;refGene_tx=XM_023243632.1;refGene_gn=ZZEF1;refGene_tx=XM_023243633.1;refGene_gn=ZZEF1;refGene_tx=XM_023243635.1
E1 215070 215206
refGene_gn=ZZEF1;refGene_tx=XM_011288812.2;refGene_gn=ZZEF1;refGene_tx=XM_023243632.1;refGene_gn=ZZEF1;refGene_tx=XM_023243633.1;refGene_gn=ZZEF1;refGene_tx=XM_023243635.1
E1 216314 216415
refGene_gn=ZZEF1;refGene_tx=XM_011288812.2;refGene_gn=ZZEF1;refGene_tx=XM_023243632.1;refGene_gn=ZZEF1;refGene_tx=XM_023243633.1;refGene_gn=ZZEF1;refGene_tx=XM_023243635.1
E1 217184 217376
refGene_gn=ZZEF1;refGene_tx=XM_011288812.2;refGene_gn=ZZEF1;refGene_tx=XM_023243632.1;refGene_gn=ZZEF1;refGene_tx=XM_023243633.1;refGene_gn=ZZEF1;refGene_tx=XM_023243635.1
E1 218941 219061
refGene_gn=ZZEF1;refGene_tx=XM_011288812.2;refGene_gn=ZZEF1;refGene_tx

=XM_023243632.1;refGene_gn=ZZEF1;refGene_tx=XM_023243633.1;refGene_gn=ZZEF1;refGene_tx=XM_023243635.1
E1 220767 220961
refGene_gn=ZZEF1;refGene_tx=XM_011288812.2;refGene_gn=ZZEF1;refGene_tx=XM_023243632.1;refGene_gn=ZZEF1;refGene_tx=XM_023243633.1;refGene_gn=ZZEF1;refGene_tx=XM_023243635.1
E1 225407 225540
refGene_gn=ZZEF1;refGene_tx=XM_011288812.2;refGene_gn=ZZEF1;refGene_tx=XM_023243632.1;refGene_gn=ZZEF1;refGene_tx=XM_023243633.1;refGene_gn=ZZEF1;refGene_tx=XM_023243635.1
E1 226100 226275
refGene_gn=ZZEF1;refGene_tx=XM_011288812.2;refGene_gn=ZZEF1;refGene_tx=XM_023243632.1;refGene_gn=ZZEF1;refGene_tx=XM_023243633.1;refGene_gn=ZZEF1;refGene_tx=XM_023243635.1
E1 226926 227204
refGene_gn=ZZEF1;refGene_tx=XM_011288812.2;refGene_gn=ZZEF1;refGene_tx=XM_023243632.1;refGene_gn=ZZEF1;refGene_tx=XM_023243633.1;refGene_gn=ZZEF1;refGene_tx=XM_023243635.1
E1 227732 227872
refGene_gn=ZZEF1;refGene_tx=XM_011288812.2;refGene_gn=ZZEF1;refGene_tx=XM_023243632.1;refGene_gn=ZZEF1;refGene_tx=XM_023243633.1;refGene_gn=ZZEF1;refGene_tx=XM_023243635.1
E1 229158 229288
refGene_gn=ZZEF1;refGene_tx=XM_011288812.2;refGene_gn=ZZEF1;refGene_tx=XM_023243632.1;refGene_gn=ZZEF1;refGene_tx=XM_023243633.1;refGene_gn=ZZEF1;refGene_tx=XM_023243635.1
E1 232204 232365
refGene_gn=ZZEF1;refGene_tx=XM_011288812.2;refGene_gn=ZZEF1;refGene_tx=XM_023243632.1;refGene_gn=ZZEF1;refGene_tx=XM_023243633.1;refGene_gn=ZZEF1;refGene_tx=XM_023243635.1
E1 233248 233511
refGene_gn=ZZEF1;refGene_tx=XM_011288812.2;refGene_gn=ZZEF1;refGene_tx=XM_023243632.1;refGene_gn=ZZEF1;refGene_tx=XM_023243633.1;refGene_gn=ZZEF1;refGene_tx=XM_023243635.1
E1 234051 234217
refGene_gn=ZZEF1;refGene_tx=XM_011288812.2;refGene_gn=ZZEF1;refGene_tx=XM_023243632.1;refGene_gn=ZZEF1;refGene_tx=XM_023243633.1;refGene_gn=ZZEF1;refGene_tx=XM_023243635.1
E1 235707 235846
refGene_gn=ZZEF1;refGene_tx=XM_011288812.2;refGene_gn=ZZEF1;refGene_tx=XM_023243632.1;refGene_gn=ZZEF1;refGene_tx=XM_023243633.1;refGene_gn=ZZEF1;refGene_tx=XM_023243635.1
E1 237394 237524
refGene_gn=ZZEF1;refGene_tx=XM_011288812.2;refGene_gn=ZZEF1;refGene_tx=XM_023243632.1;refGene_gn=ZZEF1;refGene_tx=XM_023243633.1;refGene_gn=ZZEF1;refGene_tx=XM_023243635.1
E1 239254 239449
refGene_gn=ZZEF1;refGene_tx=XM_011288812.2;refGene_gn=ZZEF1;refGene_tx=XM_023243632.1;refGene_gn=ZZEF1;refGene_tx=XM_023243633.1;refGene_gn=ZZEF1;refGene_tx=XM_023243635.1

E1	240092	240212
refGene_gn=ZZEF1;refGene_tx=XM_011288812.2;refGene_gn=ZZEF1;refGene_tx=XM_023243632.1;refGene_gn=ZZEF1;refGene_tx=XM_023243633.1;refGene_gn=ZZEF1;refGene_tx=XM_023243635.1
E1	242720	242885
refGene_gn=ZZEF1;refGene_tx=XM_011288812.2;refGene_gn=ZZEF1;refGene_tx=XM_023243632.1;refGene_gn=ZZEF1;refGene_tx=XM_023243633.1;refGene_gn=ZZEF1;refGene_tx=XM_023243635.1
E1	243548	244017
refGene_gn=ZZEF1;refGene_tx=XM_011288812.2;refGene_gn=ZZEF1;refGene_tx=XM_023243632.1;refGene_gn=ZZEF1;refGene_tx=XM_023243633.1;refGene_gn=ZZEF1;refGene_tx=XM_023243635.1
E1	245441	245535
refGene_gn=ZZEF1;refGene_tx=XM_011288812.2;refGene_gn=ZZEF1;refGene_tx=XM_023243632.1;refGene_gn=ZZEF1;refGene_tx=XM_023243633.1;refGene_gn=ZZEF1;refGene_tx=XM_023243635.1
E1	247635	247715
refGene_gn=ZZEF1;refGene_tx=XM_011288812.2;refGene_gn=ZZEF1;refGene_tx=XM_023243632.1;refGene_gn=ZZEF1;refGene_tx=XM_023243633.1;refGene_gn=ZZEF1;refGene_tx=XM_023243635.1
E1	250097	250338
refGene_gn=ZZEF1;refGene_tx=XM_011288812.2;refGene_gn=ZZEF1;refGene_tx=XM_023243632.1;refGene_gn=ZZEF1;refGene_tx=XM_023243633.1;refGene_gn=ZZEF1;refGene_tx=XM_023243635.1
E1	251428	251577
refGene_gn=ZZEF1;refGene_tx=XM_011288812.2;refGene_gn=ZZEF1;refGene_tx=XM_023243632.1;refGene_gn=ZZEF1;refGene_tx=XM_023243633.1;refGene_gn=ZZEF1;refGene_tx=XM_023243635.1
E1	252396	252598
refGene_gn=ZZEF1;refGene_tx=XM_011288812.2;refGene_gn=ZZEF1;refGene_tx=XM_023243632.1;refGene_gn=ZZEF1;refGene_tx=XM_023243633.1;refGene_gn=ZZEF1;refGene_tx=XM_023243635.1
E1	253360	253442
refGene_gn=ZZEF1;refGene_tx=XM_011288812.2;refGene_gn=ZZEF1;refGene_tx=XM_023243632.1;refGene_gn=ZZEF1;refGene_tx=XM_023243633.1;refGene_gn=ZZEF1;refGene_tx=XM_023243635.1
E1	253519	253686
refGene_gn=ZZEF1;refGene_tx=XM_011288812.2;refGene_gn=ZZEF1;refGene_tx=XM_023243632.1;refGene_gn=ZZEF1;refGene_tx=XM_023243633.1;refGene_gn=ZZEF1;refGene_tx=XM_023243635.1
E1	254504	254627
refGene_gn=ZZEF1;refGene_tx=XM_011288812.2;refGene_gn=ZZEF1;refGene_tx=XM_023243632.1;refGene_gn=ZZEF1;refGene_tx=XM_023243633.1;refGene_gn=ZZEF1;refGene_tx=XM_023243635.1
E1	254731	254853
refGene_gn=ZZEF1;refGene_tx=XM_011288812.2;refGene_gn=ZZEF1;refGene_tx=XM_023243632.1;refGene_gn=ZZEF1;refGene_tx=XM_023243633.1;refGene_gn=ZZEF1;refGene_tx=XM_023243635.1
E1	255969	256129
refGene_gn=ZZEF1;refGene_tx=XM_011288812.2;refGene_gn=ZZEF1;refGene_tx

=XM_023243632.1;refGene_gn=ZZEF1;refGene_tx=XM_023243633.1;refGene_gn=ZZEF1;refGene_tx=XM_023243635.1
E1 258690 258805
refGene_gn=ZZEF1;refGene_tx=XM_011288812.2;refGene_gn=ZZEF1;refGene_tx=XM_023243632.1;refGene_gn=ZZEF1;refGene_tx=XM_023243633.1;refGene_gn=ZZEF1;refGene_tx=XM_023243635.1
E1 259653 259724
refGene_gn=ZZEF1;refGene_tx=XM_011288812.2;refGene_gn=ZZEF1;refGene_tx=XM_023243632.1;refGene_gn=ZZEF1;refGene_tx=XM_023243633.1;refGene_gn=ZZEF1;refGene_tx=XM_023243635.1
E1 262299 262433
refGene_gn=ZZEF1;refGene_tx=XM_011288812.2;refGene_gn=ZZEF1;refGene_tx=XM_023243632.1;refGene_gn=ZZEF1;refGene_tx=XM_023243633.1;refGene_gn=ZZEF1;refGene_tx=XM_023243635.1
E1 262970 263140
refGene_gn=ZZEF1;refGene_tx=XM_011288812.2;refGene_gn=ZZEF1;refGene_tx=XM_023243632.1;refGene_gn=ZZEF1;refGene_tx=XM_023243633.1;refGene_gn=ZZEF1;refGene_tx=XM_023243635.1
E1 263834 263935
refGene_gn=ZZEF1;refGene_tx=XM_011288812.2;refGene_gn=ZZEF1;refGene_tx=XM_023243632.1;refGene_gn=ZZEF1;refGene_tx=XM_023243633.1;refGene_gn=ZZEF1;refGene_tx=XM_023243635.1
E1 264925 265141
refGene_gn=ZZEF1;refGene_tx=XM_011288812.2;refGene_gn=ZZEF1;refGene_tx=XM_023243632.1;refGene_gn=ZZEF1;refGene_tx=XM_023243633.1;refGene_gn=ZZEF1;refGene_tx=XM_023243635.1
E1 266098 266210
refGene_gn=ZZEF1;refGene_tx=XM_011288812.2;refGene_gn=ZZEF1;refGene_tx=XM_023243632.1;refGene_gn=ZZEF1;refGene_tx=XM_023243633.1;refGene_gn=ZZEF1;refGene_tx=XM_023243635.1
E1 270456 270605
refGene_gn=ZZEF1;refGene_tx=XM_011288812.2;refGene_gn=ZZEF1;refGene_tx=XM_023243632.1;refGene_gn=ZZEF1;refGene_tx=XM_023243633.1;refGene_gn=ZZEF1;refGene_tx=XM_023243635.1
E1 271448 271540
refGene_gn=ZZEF1;refGene_tx=XM_011288812.2;refGene_gn=ZZEF1;refGene_tx=XM_023243632.1;refGene_gn=ZZEF1;refGene_tx=XM_023243633.1;refGene_gn=ZZEF1;refGene_tx=XM_023243635.1
E1 272220 272319
refGene_gn=ZZEF1;refGene_tx=XM_011288812.2;refGene_gn=ZZEF1;refGene_tx=XM_023243632.1;refGene_gn=ZZEF1;refGene_tx=XM_023243633.1;refGene_gn=ZZEF1;refGene_tx=XM_023243635.1
E1 274004 274183
refGene_gn=ZZEF1;refGene_tx=XM_011288812.2;refGene_gn=ZZEF1;refGene_tx=XM_023243632.1;refGene_gn=ZZEF1;refGene_tx=XM_023243633.1;refGene_gn=ZZEF1;refGene_tx=XM_023243635.1
E1 275041 275158
refGene_gn=ZZEF1;refGene_tx=XM_011288812.2;refGene_gn=ZZEF1;refGene_tx=XM_023243632.1;refGene_gn=ZZEF1;refGene_tx=XM_023243633.1;refGene_gn=ZZEF1;refGene_tx=XM_023243635.1

E1	279333	279544
refGene_gn=ZZEF1;refGene_tx=XM_011288812.2;refGene_gn=ZZEF1;refGene_tx=XM_023243632.1;refGene_gn=ZZEF1;refGene_tx=XM_023243633.1;refGene_gn=ZZEF1;refGene_tx=XM_023243635.1
E1	280919	281119
refGene_gn=ZZEF1;refGene_tx=XM_011288812.2;refGene_gn=ZZEF1;refGene_tx=XM_023243632.1;refGene_gn=ZZEF1;refGene_tx=XM_023243633.1;refGene_gn=ZZEF1;refGene_tx=XM_023243635.1
E1	281960	282132
refGene_gn=ZZEF1;refGene_tx=XM_011288812.2;refGene_gn=ZZEF1;refGene_tx=XM_023243632.1;refGene_gn=ZZEF1;refGene_tx=XM_023243633.1;refGene_gn=ZZEF1;refGene_tx=XM_023243635.1
E1	283934	284129
refGene_gn=ZZEF1;refGene_tx=XM_011288812.2;refGene_gn=ZZEF1;refGene_tx=XM_023243632.1;refGene_gn=ZZEF1;refGene_tx=XM_023243633.1;refGene_gn=ZZEF1;refGene_tx=XM_023243635.1
E1	287449	287594
refGene_gn=ZZEF1;refGene_tx=XM_011288812.2;refGene_gn=ZZEF1;refGene_tx=XM_023243632.1;refGene_gn=ZZEF1;refGene_tx=XM_023243633.1;refGene_gn=ZZEF1;refGene_tx=XM_023243635.1
E1	303784	304138
refGene_gn=ZZEF1;refGene_tx=XM_011288812.2;refGene_gn=ZZEF1;refGene_tx=XM_023243632.1;refGene_gn=ZZEF1;refGene_tx=XM_023243633.1;refGene_gn=ZZEF1;refGene_tx=XM_023243635.1
E1	304664	304965
refGene_gn=LOC101086514;refGene_tx=XM_003996217.5
E1	309550	309691
refGene_gn=LOC101086514;refGene_tx=XM_003996217.5
E1	312969	313156
refGene_gn=LOC101086514;refGene_tx=XM_003996217.5
E1	314831	315051
refGene_gn=LOC101086514;refGene_tx=XM_003996217.5
E1	322150	322283	refGene_gn=ANKFY1;refGene_tx=XM_019817008.2
E1	322660	322751	refGene_gn=ANKFY1;refGene_tx=XM_019817008.2
E1	323789	323936	refGene_gn=ANKFY1;refGene_tx=XM_019817008.2
E1	325254	325379	refGene_gn=ANKFY1;refGene_tx=XM_019817008.2
E1	325683	325774	refGene_gn=ANKFY1;refGene_tx=XM_019817008.2
E1	326232	326380	refGene_gn=ANKFY1;refGene_tx=XM_019817008.2
E1	330969	331146	refGene_gn=ANKFY1;refGene_tx=XM_019817008.2
E1	332540	332741	refGene_gn=ANKFY1;refGene_tx=XM_019817008.2
E1	333398	333555	refGene_gn=ANKFY1;refGene_tx=XM_019817008.2
E1	334407	334526	refGene_gn=ANKFY1;refGene_tx=XM_019817008.2
E1	335271	335440	refGene_gn=ANKFY1;refGene_tx=XM_019817008.2
E1	336028	336182	refGene_gn=ANKFY1;refGene_tx=XM_019817008.2
E1	336366	336465	refGene_gn=ANKFY1;refGene_tx=XM_019817008.2
E1	337359	337567	refGene_gn=ANKFY1;refGene_tx=XM_019817008.2
E1	339778	339876	refGene_gn=ANKFY1;refGene_tx=XM_019817008.2
E1	343821	344021	refGene_gn=ANKFY1;refGene_tx=XM_019817008.2
E1	344237	344306	refGene_gn=ANKFY1;refGene_tx=XM_019817008.2
E1	345353	345558	refGene_gn=ANKFY1;refGene_tx=XM_019817008.2

E1 351222 351388 refGene_gn=ANKFY1;refGene_tx=XM_019817008.2
E1 352549 352699 refGene_gn=ANKFY1;refGene_tx=XM_019817008.2
E1 352813 352937 refGene_gn=ANKFY1;refGene_tx=XM_019817008.2
E1 357421 357557 refGene_gn=ANKFY1;refGene_tx=XM_019817008.2
E1 368214 368333 refGene_gn=ANKFY1;refGene_tx=XM_019817008.2
E1 372449 372642 refGene_gn=ANKFY1;refGene_tx=XM_019817008.2
E1 393423 393493 refGene_gn=ANKFY1;refGene_tx=XM_019817008.2
E1 414258 414345 refGene_gn=UBE2G1;refGene_tx=XM_023243636.1
E1 419040 419219 refGene_gn=UBE2G1;refGene_tx=XM_023243636.1
E1 425268 425366 refGene_gn=UBE2G1;refGene_tx=XM_023243636.1
E1 439204 439307 refGene_gn=UBE2G1;refGene_tx=XM_023243636.1
E1 510024 510070 refGene_gn=UBE2G1;refGene_tx=XM_023243636.1
E1 548856 549067
refGene_gn=SPNS3;refGene_tx=XM_006939708.2;refGene_gn=SPNS3;refGene_tx=XM_006939709.2;refGene_gn=SPNS3;refGene_tx=XM_006939710.3;refGene_gn=SPNS3;refGene_tx=XM_006939711.3;refGene_gn=SPNS3;refGene_tx=XM_011288807.2;refGene_gn=SPNS3;refGene_tx=XM_019817011.1;refGene_gn=SPNS3;refGene_tx=XM_019817012.1;refGene_gn=SPNS3;refGene_tx=XM_019817013.1
E1 552134 552200
refGene_gn=SPNS3;refGene_tx=XM_006939708.2;refGene_gn=SPNS3;refGene_tx=XM_006939709.2;refGene_gn=SPNS3;refGene_tx=XM_006939710.3;refGene_gn=SPNS3;refGene_tx=XM_006939711.3;refGene_gn=SPNS3;refGene_tx=XM_011288807.2;refGene_gn=SPNS3;refGene_tx=XM_019817011.1;refGene_gn=SPNS3;refGene_tx=XM_019817012.1;refGene_gn=SPNS3;refGene_tx=XM_019817013.1
E1 555510 555640 refGene_gn=SPNS3;refGene_tx=XM_019817010.1
E1 556248 556414
refGene_gn=SPNS3;refGene_tx=XM_006939708.2;refGene_gn=SPNS3;refGene_tx=XM_006939709.2;refGene_gn=SPNS3;refGene_tx=XM_006939710.3;refGene_gn=SPNS3;refGene_tx=XM_006939711.3;refGene_gn=SPNS3;refGene_tx=XM_011288807.2;refGene_gn=SPNS3;refGene_tx=XM_019817010.1;refGene_gn=SPNS3;refGene_tx=XM_019817011.1;refGene_gn=SPNS3;refGene_tx=XM_019817012.1;refGene_gn=SPNS3;refGene_tx=XM_019817013.1;refGene_gn=SPNS3;refGene_tx=XM_019817014.1
E1 556919 557071
refGene_gn=SPNS3;refGene_tx=XM_006939708.2;refGene_gn=SPNS3;refGene_tx=XM_006939709.2;refGene_gn=SPNS3;refGene_tx=XM_006939710.3;refGene_gn=SPNS3;refGene_tx=XM_006939711.3;refGene_gn=SPNS3;refGene_tx=XM_019817010.1;refGene_gn=SPNS3;refGene_tx=XM_019817011.1;refGene_gn=SPNS3;refGene_tx=XM_019817013.1;refGene_gn=SPNS3;refGene_tx=XM_019817014.1
E1 557791 557858
refGene_gn=SPNS3;refGene_tx=XM_006939708.2;refGene_gn=SPNS3;refGene_tx=XM_006939710.3;refGene_gn=SPNS3;refGene_tx=XM_006939711.3;refGene_gn=SPNS3;refGene_tx=XM_019817010.1;refGene_gn=SPNS3;refGene_tx=XM_019817011.1;refGene_gn=SPNS3;refGene_tx=XM_019817013.1
E1 558470 558619
refGene_gn=SPNS3;refGene_tx=XM_006939708.2;refGene_gn=SPNS3;refGene_tx=XM_006939710.3;refGene_gn=SPNS3;refGene_tx=XM_006939711.3;refGene_gn=SPNS3;refGene_tx=XM_019817010.1;refGene_gn=SPNS3;refGene_tx=XM_019817012.1;refGene_gn=SPNS3;refGene_tx=XM_019817014.1
E1 559393 559546

refGene_gn=SPNS3;refGene_tx=XM_006939708.2;refGene_gn=SPNS3;refGene_tx=XM_006939709.2;refGene_gn=SPNS3;refGene_tx=XM_006939710.3;refGene_gn=SPNS3;refGene_tx=XM_006939711.3;refGene_gn=SPNS3;refGene_tx=XM_011288807.2;refGene_gn=SPNS3;refGene_tx=XM_019817010.1;refGene_gn=SPNS3;refGene_tx=XM_019817011.1;refGene_gn=SPNS3;refGene_tx=XM_019817012.1;refGene_gn=SPNS3;refGene_tx=XM_019817013.1;refGene_gn=SPNS3;refGene_tx=XM_019817014.1
E1 562852 563046
refGene_gn=SPNS3;refGene_tx=XM_006939708.2;refGene_gn=SPNS3;refGene_tx=XM_006939709.2;refGene_gn=SPNS3;refGene_tx=XM_006939710.3;refGene_gn=SPNS3;refGene_tx=XM_006939711.3;refGene_gn=SPNS3;refGene_tx=XM_011288807.2;refGene_gn=SPNS3;refGene_tx=XM_019817010.1;refGene_gn=SPNS3;refGene_tx=XM_019817011.1;refGene_gn=SPNS3;refGene_tx=XM_019817012.1;refGene_gn=SPNS3;refGene_tx=XM_019817013.1;refGene_gn=SPNS3;refGene_tx=XM_019817014.1
E1 577186 577252
refGene_gn=SPNS3;refGene_tx=XM_006939708.2;refGene_gn=SPNS3;refGene_tx=XM_006939709.2;refGene_gn=SPNS3;refGene_tx=XM_006939710.3;refGene_gn=SPNS3;refGene_tx=XM_011288807.2;refGene_gn=SPNS3;refGene_tx=XM_019817010.1;refGene_gn=SPNS3;refGene_tx=XM_019817011.1;refGene_gn=SPNS3;refGene_tx=XM_019817012.1;refGene_gn=SPNS3;refGene_tx=XM_019817014.1
E1 582535 582634
refGene_gn=SPNS3;refGene_tx=XM_006939708.2;refGene_gn=SPNS3;refGene_tx=XM_006939709.2;refGene_gn=SPNS3;refGene_tx=XM_006939710.3;refGene_gn=SPNS3;refGene_tx=XM_006939711.3;refGene_gn=SPNS3;refGene_tx=XM_011288807.2;refGene_gn=SPNS3;refGene_tx=XM_019817010.1;refGene_gn=SPNS3;refGene_tx=XM_019817011.1;refGene_gn=SPNS3;refGene_tx=XM_019817012.1;refGene_gn=SPNS3;refGene_tx=XM_019817013.1;refGene_gn=SPNS3;refGene_tx=XM_019817014.1
E1 582733 582905
refGene_gn=SPNS3;refGene_tx=XM_006939708.2;refGene_gn=SPNS3;refGene_tx=XM_006939709.2;refGene_gn=SPNS3;refGene_tx=XM_006939710.3;refGene_gn=SPNS3;refGene_tx=XM_006939711.3;refGene_gn=SPNS3;refGene_tx=XM_011288807.2;refGene_gn=SPNS3;refGene_tx=XM_019817010.1;refGene_gn=SPNS3;refGene_tx=XM_019817011.1;refGene_gn=SPNS3;refGene_tx=XM_019817012.1;refGene_gn=SPNS3;refGene_tx=XM_019817013.1;refGene_gn=SPNS3;refGene_tx=XM_019817014.1
E1 583731 584056
refGene_gn=SPNS3;refGene_tx=XM_006939708.2;refGene_gn=SPNS3;refGene_tx=XM_006939709.2;refGene_gn=SPNS3;refGene_tx=XM_006939710.3;refGene_gn=SPNS3;refGene_tx=XM_006939711.3;refGene_gn=SPNS3;refGene_tx=XM_011288807.2;refGene_gn=SPNS3;refGene_tx=XM_019817010.1;refGene_gn=SPNS3;refGene_tx=XM_019817011.1;refGene_gn=SPNS3;refGene_tx=XM_019817012.1;refGene_gn=SPNS3;refGene_tx=XM_019817013.1;refGene_gn=SPNS3;refGene_tx=XM_019817014.1
E1 591506 591876
refGene_gn=SPNS2;refGene_tx=XM_023244677.1;refGene_gn=SPNS2;refGene_tx=XM_023244678.1;refGene_gn=SPNS2;refGene_tx=XM_023244679.1;refGene_gn=SPNS2;refGene_tx=XM_023244680.1;refGene_gn=SPNS2;refGene_tx=XM_023244682.1;refGene_gn=SPNS2;refGene_tx=XM_023244683.1

E1 599520 599670
refGene_gn=SPNS2;refGene_tx=XM_023244677.1;refGene_gn=SPNS2;refGene_tx=XM_023244680.1;refGene_gn=SPNS2;refGene_tx=XM_023244682.1
E1 603166 603232
refGene_gn=SPNS2;refGene_tx=XM_023244677.1;refGene_gn=SPNS2;refGene_tx=XM_023244678.1;refGene_gn=SPNS2;refGene_tx=XM_023244680.1;refGene_gn=SPNS2;refGene_tx=XM_023244682.1;refGene_gn=SPNS2;refGene_tx=XM_023244683.1
E1 611150 611287
refGene_gn=SPNS2;refGene_tx=XM_023244677.1;refGene_gn=SPNS2;refGene_tx=XM_023244678.1;refGene_gn=SPNS2;refGene_tx=XM_023244679.1;refGene_gn=SPNS2;refGene_tx=XM_023244680.1;refGene_gn=SPNS2;refGene_tx=XM_023244682.1;refGene_gn=SPNS2;refGene_tx=XM_023244683.1
E1 614971 615123
refGene_gn=SPNS2;refGene_tx=XM_023244677.1;refGene_gn=SPNS2;refGene_tx=XM_023244678.1;refGene_gn=SPNS2;refGene_tx=XM_023244679.1;refGene_gn=SPNS2;refGene_tx=XM_023244680.1;refGene_gn=SPNS2;refGene_tx=XM_023244682.1;refGene_gn=SPNS2;refGene_tx=XM_023244683.1
E1 615272 615339
refGene_gn=SPNS2;refGene_tx=XM_023244677.1;refGene_gn=SPNS2;refGene_tx=XM_023244678.1;refGene_gn=SPNS2;refGene_tx=XM_023244679.1;refGene_gn=SPNS2;refGene_tx=XM_023244680.1;refGene_gn=SPNS2;refGene_tx=XM_023244682.1;refGene_gn=SPNS2;refGene_tx=XM_023244683.1
E1 616134 616277
refGene_gn=SPNS2;refGene_tx=XM_023244677.1;refGene_gn=SPNS2;refGene_tx=XM_023244678.1;refGene_gn=SPNS2;refGene_tx=XM_023244679.1;refGene_gn=SPNS2;refGene_tx=XM_023244680.1;refGene_gn=SPNS2;refGene_tx=XM_023244682.1;refGene_gn=SPNS2;refGene_tx=XM_023244683.1
E1 616481 616634
refGene_gn=SPNS2;refGene_tx=XM_023244677.1;refGene_gn=SPNS2;refGene_tx=XM_023244678.1;refGene_gn=SPNS2;refGene_tx=XM_023244679.1;refGene_gn=SPNS2;refGene_tx=XM_023244680.1;refGene_gn=SPNS2;refGene_tx=XM_023244682.1;refGene_gn=SPNS2;refGene_tx=XM_023244683.1
E1 616740 616930
refGene_gn=SPNS2;refGene_tx=XM_023244677.1;refGene_gn=SPNS2;refGene_tx=XM_023244678.1;refGene_gn=SPNS2;refGene_tx=XM_023244679.1;refGene_gn=SPNS2;refGene_tx=XM_023244680.1;refGene_gn=SPNS2;refGene_tx=XM_023244682.1;refGene_gn=SPNS2;refGene_tx=XM_023244683.1
E1 617807 617873
refGene_gn=SPNS2;refGene_tx=XM_023244677.1;refGene_gn=SPNS2;refGene_tx=XM_023244678.1;refGene_gn=SPNS2;refGene_tx=XM_023244679.1;refGene_gn=SPNS2;refGene_tx=XM_023244680.1;refGene_gn=SPNS2;refGene_tx=XM_023244682.1;refGene_gn=SPNS2;refGene_tx=XM_023244683.1
E1 619311 619410
refGene_gn=SPNS2;refGene_tx=XM_023244677.1;refGene_gn=SPNS2;refGene_tx=XM_023244678.1;refGene_gn=SPNS2;refGene_tx=XM_023244679.1;refGene_gn=SPNS2;refGene_tx=XM_023244680.1;refGene_gn=SPNS2;refGene_tx=XM_023244682.1;refGene_gn=SPNS2;refGene_tx=XM_023244683.1
E1 619479 619643
refGene_gn=SPNS2;refGene_tx=XM_023244677.1;refGene_gn=SPNS2;refGene_tx

=XM_023244678.1;refGene_gn=SPNS2;refGene_tx=XM_023244679.1;refGene_gn=SPNS2;refGene_tx=XM_023244680.1;refGene_gn=SPNS2;refGene_tx=XM_023244682.1;refGene_gn=SPNS2;refGene_tx=XM_023244683.1
E1 619765 619957
refGene_gn=SPNS2;refGene_tx=XM_023244677.1;refGene_gn=SPNS2;refGene_tx=XM_023244678.1;refGene_gn=SPNS2;refGene_tx=XM_023244679.1;refGene_gn=SPNS2;refGene_tx=XM_023244680.1;refGene_gn=SPNS2;refGene_tx=XM_023244682.1;refGene_gn=SPNS2;refGene_tx=XM_023244683.1
E1 620379 620776
refGene_gn=SPNS2;refGene_tx=XM_023244677.1;refGene_gn=SPNS2;refGene_tx=XM_023244678.1;refGene_gn=SPNS2;refGene_tx=XM_023244679.1;refGene_gn=SPNS2;refGene_tx=XM_023244680.1
E1 621513 622078 refGene_gn=MYBBP1A;refGene_tx=XM_011288806.3
E1 622282 622419 refGene_gn=MYBBP1A;refGene_tx=XM_011288806.3
E1 622959 623061 refGene_gn=MYBBP1A;refGene_tx=XM_011288806.3
E1 623250 623364 refGene_gn=MYBBP1A;refGene_tx=XM_011288806.3
E1 623878 623947 refGene_gn=MYBBP1A;refGene_tx=XM_011288806.3
E1 624027 624162 refGene_gn=MYBBP1A;refGene_tx=XM_011288806.3
E1 624355 624596 refGene_gn=MYBBP1A;refGene_tx=XM_011288806.3
E1 625492 625650 refGene_gn=MYBBP1A;refGene_tx=XM_011288806.3
E1 625736 625919 refGene_gn=MYBBP1A;refGene_tx=XM_011288806.3
E1 625996 626152 refGene_gn=MYBBP1A;refGene_tx=XM_011288806.3
E1 626238 626319 refGene_gn=MYBBP1A;refGene_tx=XM_011288806.3
E1 626627 626779 refGene_gn=MYBBP1A;refGene_tx=XM_011288806.3
E1 626875 626972 refGene_gn=MYBBP1A;refGene_tx=XM_011288806.3
E1 628253 628359 refGene_gn=MYBBP1A;refGene_tx=XM_011288806.3
E1 628446 628614 refGene_gn=MYBBP1A;refGene_tx=XM_011288806.3
E1 628701 628827 refGene_gn=MYBBP1A;refGene_tx=XM_011288806.3
E1 629292 629403 refGene_gn=MYBBP1A;refGene_tx=XM_011288806.3
E1 629622 629918 refGene_gn=MYBBP1A;refGene_tx=XM_011288806.3
E1 630876 630994 refGene_gn=MYBBP1A;refGene_tx=XM_011288806.3
E1 631107 631275 refGene_gn=MYBBP1A;refGene_tx=XM_011288806.3
E1 631391 631567 refGene_gn=MYBBP1A;refGene_tx=XM_011288806.3
E1 632428 632536 refGene_gn=MYBBP1A;refGene_tx=XM_011288806.3
E1 632635 632710 refGene_gn=MYBBP1A;refGene_tx=XM_011288806.3
E1 632795 632879 refGene_gn=MYBBP1A;refGene_tx=XM_011288806.3
E1 633573 633669 refGene_gn=MYBBP1A;refGene_tx=XM_011288806.3
E1 633839 634046 refGene_gn=MYBBP1A;refGene_tx=XM_011288806.3
E1 635418 636449
refGene_gn=GGT6;refGene_tx=XM_011288805.3;refGene_gn=GGT6;refGene_tx=XM_019817018.2
E1 636704 636818
refGene_gn=GGT6;refGene_tx=XM_011288805.3;refGene_gn=GGT6;refGene_tx=XM_019817018.2
E1 636908 637093
refGene_gn=GGT6;refGene_tx=XM_011288805.3;refGene_gn=GGT6;refGene_tx=XM_019817018.2
E1 637654 637800
refGene_gn=GGT6;refGene_tx=XM_011288805.3;refGene_gn=GGT6;refGene_tx=XM_019817018.2

E1 638773 639040 refGene_gn=GGT6;refGene_tx=XM_019817018.2
E1 639179 639233 refGene_gn=GGT6;refGene_tx=XM_019817018.2
E1 661516 661924 refGene_gn=SMTNL2;refGene_tx=XM_023243730.1
E1 668248 668339 refGene_gn=SMTNL2;refGene_tx=XM_023243730.1
E1 668794 669037 refGene_gn=SMTNL2;refGene_tx=XM_023243730.1
E1 669592 669659 refGene_gn=SMTNL2;refGene_tx=XM_023243730.1
E1 670354 670501 refGene_gn=SMTNL2;refGene_tx=XM_023243730.1
E1 671231 671349 refGene_gn=SMTNL2;refGene_tx=XM_023243730.1
E1 671555 671707 refGene_gn=SMTNL2;refGene_tx=XM_023243730.1
E1 677555 677682 refGene_gn=SMTNL2;refGene_tx=XM_023243730.1
E1 687017 687208
refGene_gn=LOC102900493;refGene_tx=XM_019817021.1
E1 688214 688369
refGene_gn=LOC102900493;refGene_tx=XM_019817021.1
E1 688977 689507
refGene_gn=LOC102900493;refGene_tx=XM_019817021.1
E1 700993 701057 refGene_gn=TEKT1;refGene_tx=XM_019817023.2
E1 702490 702697
refGene_gn=TEKT1;refGene_tx=XM_011288804.3;refGene_gn=TEKT1;refGene_tx
=XM_019817023.2;refGene_gn=TEKT1;refGene_tx=XM_023243958.1
E1 706604 706770
refGene_gn=TEKT1;refGene_tx=XM_011288804.3;refGene_gn=TEKT1;refGene_tx
=XM_019817023.2;refGene_gn=TEKT1;refGene_tx=XM_023243958.1
E1 708232 708361
refGene_gn=TEKT1;refGene_tx=XM_011288804.3;refGene_gn=TEKT1;refGene_tx
=XM_019817023.2;refGene_gn=TEKT1;refGene_tx=XM_023243958.1
E1 708783 708927
refGene_gn=TEKT1;refGene_tx=XM_011288804.3;refGene_gn=TEKT1;refGene_tx
=XM_019817023.2;refGene_gn=TEKT1;refGene_tx=XM_023243958.1
E1 709326 709549
refGene_gn=TEKT1;refGene_tx=XM_011288804.3;refGene_gn=TEKT1;refGene_tx
=XM_019817023.2;refGene_gn=TEKT1;refGene_tx=XM_023243958.1
E1 715860 716057
refGene_gn=TEKT1;refGene_tx=XM_011288804.3;refGene_gn=TEKT1;refGene_tx
=XM_019817023.2;refGene_gn=TEKT1;refGene_tx=XM_023243958.1
E1 716681 716889
refGene_gn=TEKT1;refGene_tx=XM_011288804.3;refGene_gn=TEKT1;refGene_tx
=XM_019817023.2;refGene_gn=TEKT1;refGene_tx=XM_023243958.1
E1 722828 722957
refGene_gn=FBXO39;refGene_tx=XM_003996137.5;refGene_gn=FBXO39;refGene_
tx=XM_006939713.3;refGene_gn=FBXO39;refGene_tx=XM_011288803.3;refGene_
gn=FBXO39;refGene_tx=XM_019817024.2;refGene_gn=FBXO39;refGene_tx=XM_01
9817025.2;refGene_gn=FBXO39;refGene_tx=XM_019817026.2
E1 723138 723315
refGene_gn=FBXO39;refGene_tx=XM_003996137.5;refGene_gn=FBXO39;refGene_
tx=XM_006939713.3;refGene_gn=FBXO39;refGene_tx=XM_011288803.3;refGene_
gn=FBXO39;refGene_tx=XM_019817024.2;refGene_gn=FBXO39;refGene_tx=XM_01
9817025.2;refGene_gn=FBXO39;refGene_tx=XM_019817026.2
E1 724138 725164
refGene_gn=FBXO39;refGene_tx=XM_003996137.5;refGene_gn=FBXO39;refGene_

tx=XM_006939713.3;refGene_gn=FBXO39;refGene_tx=XM_011288803.3;refGene_gn=FBXO39;refGene_tx=XM_019817024.2;refGene_gn=FBXO39;refGene_tx=XM_019817025.2;refGene_gn=FBXO39;refGene_tx=XM_019817026.2
E1 748304 748409 refGene_gn=SLC13A5;refGene_tx=XM_003996138.4
E1 753513 753642
refGene_gn=SLC13A5;refGene_tx=XM_003996138.4;refGene_gn=SLC13A5;refGene_tx=XM_019817022.1
E1 753852 753989
refGene_gn=SLC13A5;refGene_tx=XM_003996138.4;refGene_gn=SLC13A5;refGene_tx=XM_019817022.1
E1 755836 756021
refGene_gn=SLC13A5;refGene_tx=XM_003996138.4;refGene_gn=SLC13A5;refGene_tx=XM_019817022.1
E1 756579 756748
refGene_gn=SLC13A5;refGene_tx=XM_003996138.4;refGene_gn=SLC13A5;refGene_tx=XM_019817022.1
E1 757744 757867
refGene_gn=SLC13A5;refGene_tx=XM_003996138.4;refGene_gn=SLC13A5;refGene_tx=XM_019817022.1
E1 759629 759845
refGene_gn=SLC13A5;refGene_tx=XM_003996138.4;refGene_gn=SLC13A5;refGene_tx=XM_019817022.1
E1 760545 760646
refGene_gn=SLC13A5;refGene_tx=XM_003996138.4;refGene_gn=SLC13A5;refGene_tx=XM_019817022.1
E1 761583 761702
refGene_gn=SLC13A5;refGene_tx=XM_003996138.4;refGene_gn=SLC13A5;refGene_tx=XM_019817022.1
E1 763899 764061
refGene_gn=SLC13A5;refGene_tx=XM_003996138.4;refGene_gn=SLC13A5;refGene_tx=XM_019817022.1
E1 766623 766761
refGene_gn=SLC13A5;refGene_tx=XM_003996138.4;refGene_gn=SLC13A5;refGene_tx=XM_019817022.1
E1 767427 767559
refGene_gn=SLC13A5;refGene_tx=XM_003996138.4;refGene_gn=SLC13A5;refGene_tx=XM_019817022.1
E1 787303 787331 refGene_gn=MED31;refGene_tx=XM_003996140.4
E1 788024 788102 refGene_gn=MED31;refGene_tx=XM_003996140.4
E1 788423 788520 refGene_gn=MED31;refGene_tx=XM_003996140.4
E1 794269 794462 refGene_gn=MED31;refGene_tx=XM_003996140.4
E1 796064 796133 refGene_gn=TXNDC17;refGene_tx=XM_011288802.2
E1 796766 796842 refGene_gn=TXNDC17;refGene_tx=XM_011288802.2
E1 797107 797189 refGene_gn=TXNDC17;refGene_tx=XM_011288802.2
E1 797580 798119
refGene_gn=LOC105261147;refGene_tx=XM_023244499.1;refGene_gn=TXNDC17;refGene_tx=XM_011288802.2
E1 798211 800940
refGene_gn=LOC105261147;refGene_tx=XM_023244499.1
E1 801645 801793

refGene_gn=LOC105261147;refGene_tx=XM_023244499.1
E1 802360 802753
refGene_gn=LOC105261147;refGene_tx=XM_023244499.1
E1 806354 806366
refGene_gn=KIAA0753;refGene_tx=XM_006939714.4;refGene_gn=KIAA0753;refGene_tx=XM_006939716.4;refGene_gn=KIAA0753;refGene_tx=XM_006939718.4;refGene_gn=KIAA0753;refGene_tx=XM_011288801.3;refGene_gn=KIAA0753;refGene_tx=XM_019817028.2;refGene_gn=KIAA0753;refGene_tx=XM_019817029.2;refGene_gn=KIAA0753;refGene_tx=XM_019817030.2;refGene_gn=KIAA0753;refGene_tx=XM_023243757.1
E1 808034 808617
refGene_gn=KIAA0753;refGene_tx=XM_006939714.4;refGene_gn=KIAA0753;refGene_tx=XM_006939716.4;refGene_gn=KIAA0753;refGene_tx=XM_006939718.4;refGene_gn=KIAA0753;refGene_tx=XM_011288801.3;refGene_gn=KIAA0753;refGene_tx=XM_019817028.2;refGene_gn=KIAA0753;refGene_tx=XM_019817029.2;refGene_gn=KIAA0753;refGene_tx=XM_019817030.2;refGene_gn=KIAA0753;refGene_tx=XM_023243757.1
E1 811078 811185
refGene_gn=KIAA0753;refGene_tx=XM_006939714.4;refGene_gn=KIAA0753;refGene_tx=XM_006939716.4;refGene_gn=KIAA0753;refGene_tx=XM_006939718.4;refGene_gn=KIAA0753;refGene_tx=XM_011288801.3;refGene_gn=KIAA0753;refGene_tx=XM_019817028.2;refGene_gn=KIAA0753;refGene_tx=XM_019817029.2;refGene_gn=KIAA0753;refGene_tx=XM_019817030.2;refGene_gn=KIAA0753;refGene_tx=XM_023243757.1
E1 812638 812701
refGene_gn=KIAA0753;refGene_tx=XM_006939714.4;refGene_gn=KIAA0753;refGene_tx=XM_006939716.4;refGene_gn=KIAA0753;refGene_tx=XM_006939718.4;refGene_gn=KIAA0753;refGene_tx=XM_011288801.3;refGene_gn=KIAA0753;refGene_tx=XM_019817028.2;refGene_gn=KIAA0753;refGene_tx=XM_019817029.2;refGene_gn=KIAA0753;refGene_tx=XM_019817030.2;refGene_gn=KIAA0753;refGene_tx=XM_023243757.1
E1 813101 813317
refGene_gn=KIAA0753;refGene_tx=XM_006939714.4;refGene_gn=KIAA0753;refGene_tx=XM_006939716.4;refGene_gn=KIAA0753;refGene_tx=XM_006939718.4;refGene_gn=KIAA0753;refGene_tx=XM_011288801.3;refGene_gn=KIAA0753;refGene_tx=XM_019817028.2;refGene_gn=KIAA0753;refGene_tx=XM_019817029.2;refGene_gn=KIAA0753;refGene_tx=XM_023243757.1
E1 815051 815262
refGene_gn=KIAA0753;refGene_tx=XM_006939714.4;refGene_gn=KIAA0753;refGene_tx=XM_006939716.4;refGene_gn=KIAA0753;refGene_tx=XM_006939718.4;refGene_gn=KIAA0753;refGene_tx=XM_011288801.3;refGene_gn=KIAA0753;refGene_tx=XM_019817028.2;refGene_gn=KIAA0753;refGene_tx=XM_019817029.2;refGene_gn=KIAA0753;refGene_tx=XM_019817030.2;refGene_gn=KIAA0753;refGene_tx=XM_023243757.1
E1 816557 816784
refGene_gn=KIAA0753;refGene_tx=XM_006939714.4;refGene_gn=KIAA0753;refGene_tx=XM_006939716.4;refGene_gn=KIAA0753;refGene_tx=XM_006939718.4;refGene_gn=KIAA0753;refGene_tx=XM_011288801.3;refGene_gn=KIAA0753;refGene_tx=XM_019817028.2;refGene_gn=KIAA0753;refGene_tx=XM_019817029.2;refGene_gn=KIAA0753;refGene_tx=XM_019817030.2;refGene_gn=KIAA0753;refGene_

tx=XM_023243757.1
E1 818224 818391
refGene_gn=KIAA0753;refGene_tx=XM_006939714.4;refGene_gn=KIAA0753;refGene_tx=XM_006939716.4;refGene_gn=KIAA0753;refGene_tx=XM_006939718.4;refGene_gn=KIAA0753;refGene_tx=XM_011288801.3;refGene_gn=KIAA0753;refGene_tx=XM_019817028.2;refGene_gn=KIAA0753;refGene_tx=XM_019817029.2;refGene_gn=KIAA0753;refGene_tx=XM_019817030.2;refGene_gn=KIAA0753;refGene_tx=XM_023243757.1
E1 819700 819820
refGene_gn=KIAA0753;refGene_tx=XM_006939714.4;refGene_gn=KIAA0753;refGene_tx=XM_006939716.4;refGene_gn=KIAA0753;refGene_tx=XM_006939718.4;refGene_gn=KIAA0753;refGene_tx=XM_011288801.3;refGene_gn=KIAA0753;refGene_tx=XM_019817028.2;refGene_gn=KIAA0753;refGene_tx=XM_019817029.2;refGene_gn=KIAA0753;refGene_tx=XM_019817030.2;refGene_gn=KIAA0753;refGene_tx=XM_023243757.1
E1 821221 821311
refGene_gn=KIAA0753;refGene_tx=XM_006939714.4;refGene_gn=KIAA0753;refGene_tx=XM_006939716.4;refGene_gn=KIAA0753;refGene_tx=XM_006939718.4;refGene_gn=KIAA0753;refGene_tx=XM_011288801.3;refGene_gn=KIAA0753;refGene_tx=XM_019817028.2;refGene_gn=KIAA0753;refGene_tx=XM_019817029.2;refGene_gn=KIAA0753;refGene_tx=XM_019817030.2;refGene_gn=KIAA0753;refGene_tx=XM_023243757.1
E1 823683 823762
refGene_gn=KIAA0753;refGene_tx=XM_006939714.4;refGene_gn=KIAA0753;refGene_tx=XM_006939716.4;refGene_gn=KIAA0753;refGene_tx=XM_006939718.4;refGene_gn=KIAA0753;refGene_tx=XM_011288801.3;refGene_gn=KIAA0753;refGene_tx=XM_019817028.2;refGene_gn=KIAA0753;refGene_tx=XM_019817029.2;refGene_gn=KIAA0753;refGene_tx=XM_019817030.2;refGene_gn=KIAA0753;refGene_tx=XM_023243757.1
E1 824667 824748
refGene_gn=KIAA0753;refGene_tx=XM_006939714.4;refGene_gn=KIAA0753;refGene_tx=XM_006939716.4;refGene_gn=KIAA0753;refGene_tx=XM_006939718.4;refGene_gn=KIAA0753;refGene_tx=XM_011288801.3;refGene_gn=KIAA0753;refGene_tx=XM_019817028.2;refGene_gn=KIAA0753;refGene_tx=XM_019817029.2;refGene_gn=KIAA0753;refGene_tx=XM_019817030.2;refGene_gn=KIAA0753;refGene_tx=XM_023243757.1
E1 827310 827478
refGene_gn=KIAA0753;refGene_tx=XM_006939714.4;refGene_gn=KIAA0753;refGene_tx=XM_006939716.4;refGene_gn=KIAA0753;refGene_tx=XM_006939718.4;refGene_gn=KIAA0753;refGene_tx=XM_011288801.3;refGene_gn=KIAA0753;refGene_tx=XM_019817028.2;refGene_gn=KIAA0753;refGene_tx=XM_019817029.2;refGene_gn=KIAA0753;refGene_tx=XM_019817030.2;refGene_gn=KIAA0753;refGene_tx=XM_023243757.1
E1 828318 828400
refGene_gn=KIAA0753;refGene_tx=XM_006939714.4;refGene_gn=KIAA0753;refGene_tx=XM_006939716.4;refGene_gn=KIAA0753;refGene_tx=XM_006939718.4;refGene_gn=KIAA0753;refGene_tx=XM_011288801.3;refGene_gn=KIAA0753;refGene_tx=XM_019817028.2;refGene_gn=KIAA0753;refGene_tx=XM_019817029.2;refGene_gn=KIAA0753;refGene_tx=XM_019817030.2;refGene_gn=KIAA0753;refGene_tx=XM_023243757.1

E1 831420 831538
refGene_gn=KIAA0753;refGene_tx=XM_006939714.4;refGene_gn=KIAA0753;refGene_tx=XM_006939716.4;refGene_gn=KIAA0753;refGene_tx=XM_006939718.4;refGene_gn=KIAA0753;refGene_tx=XM_011288801.3;refGene_gn=KIAA0753;refGene_tx=XM_019817028.2;refGene_gn=KIAA0753;refGene_tx=XM_019817029.2;refGene_gn=KIAA0753;refGene_tx=XM_019817030.2;refGene_gn=KIAA0753;refGene_tx=XM_023243757.1
E1 831943 832168
refGene_gn=KIAA0753;refGene_tx=XM_006939714.4;refGene_gn=KIAA0753;refGene_tx=XM_006939716.4;refGene_gn=KIAA0753;refGene_tx=XM_006939718.4;refGene_gn=KIAA0753;refGene_tx=XM_011288801.3;refGene_gn=KIAA0753;refGene_tx=XM_019817028.2;refGene_gn=KIAA0753;refGene_tx=XM_019817029.2;refGene_gn=KIAA0753;refGene_tx=XM_019817030.2;refGene_gn=KIAA0753;refGene_tx=XM_023243757.1
E1 836563 836681
refGene_gn=KIAA0753;refGene_tx=XM_006939714.4;refGene_gn=KIAA0753;refGene_tx=XM_006939716.4;refGene_gn=KIAA0753;refGene_tx=XM_006939718.4;refGene_gn=KIAA0753;refGene_tx=XM_011288801.3;refGene_gn=KIAA0753;refGene_tx=XM_019817028.2;refGene_gn=KIAA0753;refGene_tx=XM_019817029.2;refGene_gn=KIAA0753;refGene_tx=XM_019817030.2;refGene_gn=KIAA0753;refGene_tx=XM_023243757.1
E1 849683 849705
refGene_gn=PITPNM3;refGene_tx=XM_023243754.1;refGene_gn=PITPNM3;refGene_tx=XM_023243755.1;refGene_gn=PITPNM3;refGene_tx=XM_023243756.1
E1 866940 867036
refGene_gn=PITPNM3;refGene_tx=XM_023243754.1;refGene_gn=PITPNM3;refGene_tx=XM_023243755.1;refGene_gn=PITPNM3;refGene_tx=XM_023243756.1
E1 872727 872835
refGene_gn=PITPNM3;refGene_tx=XM_023243754.1;refGene_gn=PITPNM3;refGene_tx=XM_023243756.1
E1 888442 888490
refGene_gn=PITPNM3;refGene_tx=XM_023243754.1;refGene_gn=PITPNM3;refGene_tx=XM_023243755.1;refGene_gn=PITPNM3;refGene_tx=XM_023243756.1
E1 898247 898324
refGene_gn=PITPNM3;refGene_tx=XM_023243754.1;refGene_gn=PITPNM3;refGene_tx=XM_023243755.1;refGene_gn=PITPNM3;refGene_tx=XM_023243756.1
E1 898649 898885
refGene_gn=PITPNM3;refGene_tx=XM_023243754.1;refGene_gn=PITPNM3;refGene_tx=XM_023243755.1;refGene_gn=PITPNM3;refGene_tx=XM_023243756.1
E1 900310 900500
refGene_gn=PITPNM3;refGene_tx=XM_023243754.1;refGene_gn=PITPNM3;refGene_tx=XM_023243755.1;refGene_gn=PITPNM3;refGene_tx=XM_023243756.1
E1 900935 901058
refGene_gn=PITPNM3;refGene_tx=XM_023243754.1;refGene_gn=PITPNM3;refGene_tx=XM_023243755.1;refGene_gn=PITPNM3;refGene_tx=XM_023243756.1
E1 901384 901575
refGene_gn=PITPNM3;refGene_tx=XM_023243754.1;refGene_gn=PITPNM3;refGene_tx=XM_023243755.1;refGene_gn=PITPNM3;refGene_tx=XM_023243756.1
E1 903683 903856
refGene_gn=PITPNM3;refGene_tx=XM_023243754.1;refGene_gn=PITPNM3;refGen

e_tx=XM_023243755.1;refGene_gn=PITPNM3;refGene_tx=XM_023243756.1
E1 905032 905203
refGene_gn=PITPNM3;refGene_tx=XM_023243754.1;refGene_gn=PITPNM3;refGen
e_tx=XM_023243755.1;refGene_gn=PITPNM3;refGene_tx=XM_023243756.1
E1 906404 906599
refGene_gn=PITPNM3;refGene_tx=XM_023243754.1;refGene_gn=PITPNM3;refGen
e_tx=XM_023243755.1;refGene_gn=PITPNM3;refGene_tx=XM_023243756.1
E1 907001 907150
refGene_gn=PITPNM3;refGene_tx=XM_023243754.1;refGene_gn=PITPNM3;refGen
e_tx=XM_023243755.1;refGene_gn=PITPNM3;refGene_tx=XM_023243756.1
E1 908960 909077
refGene_gn=PITPNM3;refGene_tx=XM_023243754.1;refGene_gn=PITPNM3;refGen
e_tx=XM_023243755.1;refGene_gn=PITPNM3;refGene_tx=XM_023243756.1
E1 911284 911401
refGene_gn=PITPNM3;refGene_tx=XM_023243754.1;refGene_gn=PITPNM3;refGen
e_tx=XM_023243755.1;refGene_gn=PITPNM3;refGene_tx=XM_023243756.1
E1 911582 911731
refGene_gn=PITPNM3;refGene_tx=XM_023243754.1;refGene_gn=PITPNM3;refGen
e_tx=XM_023243755.1;refGene_gn=PITPNM3;refGene_tx=XM_023243756.1
E1 911967 912117
refGene_gn=PITPNM3;refGene_tx=XM_023243754.1;refGene_gn=PITPNM3;refGen
e_tx=XM_023243755.1;refGene_gn=PITPNM3;refGene_tx=XM_023243756.1
E1 915889 916073
refGene_gn=PITPNM3;refGene_tx=XM_023243754.1;refGene_gn=PITPNM3;refGen
e_tx=XM_023243755.1
E1 918246 918375
refGene_gn=PITPNM3;refGene_tx=XM_023243754.1;refGene_gn=PITPNM3;refGen
e_tx=XM_023243755.1;refGene_gn=PITPNM3;refGene_tx=XM_023243756.1
E1 919631 919934
refGene_gn=PITPNM3;refGene_tx=XM_023243754.1;refGene_gn=PITPNM3;refGen
e_tx=XM_023243755.1;refGene_gn=PITPNM3;refGene_tx=XM_023243756.1
E1 924867 924955
refGene_gn=PIMREG;refGene_tx=XM_023243759.1;refGene_gn=PIMREG;refGene_
tx=XM_023243760.1
E1 925457 925553
refGene_gn=PIMREG;refGene_tx=XM_023243759.1;refGene_gn=PIMREG;refGene_
tx=XM_023243760.1
E1 925917 926213
refGene_gn=PIMREG;refGene_tx=XM_023243759.1;refGene_gn=PIMREG;refGene_
tx=XM_023243760.1
E1 927705 927999
refGene_gn=PIMREG;refGene_tx=XM_023243759.1;refGene_gn=PIMREG;refGene_
tx=XM_023243760.1
E1 933336 933432
refGene_gn=AIPL1;refGene_tx=XM_023243858.1;refGene_gn=AIPL1;refGene_tx
=XM_023243859.1
E1 933955 934135
refGene_gn=AIPL1;refGene_tx=XM_023243858.1;refGene_gn=AIPL1;refGene_tx
=XM_023243859.1
E1 939057 939246

refGene_gn=AIPL1;refGene_tx=XM_023243858.1;refGene_gn=AIPL1;refGene_tx
=XM_023243859.1
E1 940333 940510
refGene_gn=AIPL1;refGene_tx=XM_023243858.1;refGene_gn=AIPL1;refGene_tx
=XM_023243859.1
E1 940643 940785
refGene_gn=AIPL1;refGene_tx=XM_023243858.1;refGene_gn=AIPL1;refGene_tx
=XM_023243859.1
E1 941209 941412 refGene_gn=AIPL1;refGene_tx=XM_023243858.1
E1 961474 961659 refGene_gn=AIPL1;refGene_tx=XM_023243859.1
E1 1094484 1094837
refGene_gn=WSCD1;refGene_tx=XM_023244810.1;refGene_gn=WSCD1;refGene_tx
=XM_023244811.1;refGene_gn=WSCD1;refGene_tx=XM_023244812.1;refGene_gn=
WSCD1;refGene_tx=XM_023244813.1;refGene_gn=WSCD1;refGene_tx=XM_0232448
15.1
E1 1095438 1095660
refGene_gn=WSCD1;refGene_tx=XM_023244810.1;refGene_gn=WSCD1;refGene_tx
=XM_023244811.1;refGene_gn=WSCD1;refGene_tx=XM_023244812.1;refGene_gn=
WSCD1;refGene_tx=XM_023244813.1;refGene_gn=WSCD1;refGene_tx=XM_0232448
15.1
E1 1096931 1097096
refGene_gn=WSCD1;refGene_tx=XM_023244810.1;refGene_gn=WSCD1;refGene_tx
=XM_023244811.1;refGene_gn=WSCD1;refGene_tx=XM_023244812.1;refGene_gn=
WSCD1;refGene_tx=XM_023244813.1;refGene_gn=WSCD1;refGene_tx=XM_0232448
15.1
E1 1097429 1097589
refGene_gn=WSCD1;refGene_tx=XM_023244810.1;refGene_gn=WSCD1;refGene_tx
=XM_023244811.1;refGene_gn=WSCD1;refGene_tx=XM_023244812.1;refGene_gn=
WSCD1;refGene_tx=XM_023244813.1;refGene_gn=WSCD1;refGene_tx=XM_0232448
15.1
E1 1104249 1104371
refGene_gn=WSCD1;refGene_tx=XM_023244810.1;refGene_gn=WSCD1;refGene_tx
=XM_023244811.1;refGene_gn=WSCD1;refGene_tx=XM_023244812.1;refGene_gn=
WSCD1;refGene_tx=XM_023244813.1;refGene_gn=WSCD1;refGene_tx=XM_0232448
15.1
E1 1107683 1107868
refGene_gn=WSCD1;refGene_tx=XM_023244810.1;refGene_gn=WSCD1;refGene_tx
=XM_023244811.1;refGene_gn=WSCD1;refGene_tx=XM_023244812.1;refGene_gn=
WSCD1;refGene_tx=XM_023244813.1;refGene_gn=WSCD1;refGene_tx=XM_0232448
15.1
E1 1109187 1109302
refGene_gn=WSCD1;refGene_tx=XM_023244810.1;refGene_gn=WSCD1;refGene_tx
=XM_023244811.1;refGene_gn=WSCD1;refGene_tx=XM_023244812.1;refGene_gn=
WSCD1;refGene_tx=XM_023244813.1;refGene_gn=WSCD1;refGene_tx=XM_0232448
15.1
E1 1113755 1114182
refGene_gn=WSCD1;refGene_tx=XM_023244810.1;refGene_gn=WSCD1;refGene_tx
=XM_023244811.1;refGene_gn=WSCD1;refGene_tx=XM_023244812.1;refGene_gn=
WSCD1;refGene_tx=XM_023244813.1;refGene_gn=WSCD1;refGene_tx=XM_0232448
15.1

E1 1389555 1389864
refGene_gn=LOC109494148;refGene_tx=XM_023244500.1
E1 1389865 1389977
refGene_gn=LOC109494148;refGene_tx=XM_023244500.1
E1 1389979 1390209
refGene_gn=LOC109494148;refGene_tx=XM_023244500.1
E1 1398092 1398713
refGene_gn=MIS12;refGene_tx=XM_006939725.4;refGene_gn=MIS12;refGene_tx=XM_011288798.2;refGene_gn=MIS12;refGene_tx=XM_023244565.1
E1 1403239 1403332
refGene_gn=DERL2;refGene_tx=XM_003996142.4;refGene_gn=DERL2;refGene_tx=XM_019817036.2
E1 1404162 1404228
refGene_gn=DERL2;refGene_tx=XM_003996142.4;refGene_gn=DERL2;refGene_tx=XM_019817036.2
E1 1406170 1406244
refGene_gn=DERL2;refGene_tx=XM_003996142.4;refGene_gn=DERL2;refGene_tx=XM_019817036.2
E1 1407290 1407384
refGene_gn=DERL2;refGene_tx=XM_003996142.4;refGene_gn=DERL2;refGene_tx=XM_019817036.2
E1 1408836 1409032
refGene_gn=DERL2;refGene_tx=XM_003996142.4;refGene_gn=DERL2;refGene_tx=XM_019817036.2
E1 1409272 1409363
refGene_gn=DERL2;refGene_tx=XM_003996142.4;refGene_gn=DERL2;refGene_tx=XM_019817036.2
E1 1413152 1413258
refGene_gn=DERL2;refGene_tx=XM_003996142.4;refGene_gn=DERL2;refGene_tx=XM_019817036.2
E1 1419315 1419580
refGene_gn=DHX33;refGene_tx=XM_023244562.1;refGene_gn=DHX33;refGene_tx=XM_023244563.1;refGene_gn=DHX33;refGene_tx=XM_023244564.1
E1 1421645 1421806
refGene_gn=DHX33;refGene_tx=XM_023244562.1;refGene_gn=DHX33;refGene_tx=XM_023244563.1;refGene_gn=DHX33;refGene_tx=XM_023244564.1
E1 1422443 1422671
refGene_gn=DHX33;refGene_tx=XM_023244562.1;refGene_gn=DHX33;refGene_tx=XM_023244563.1;refGene_gn=DHX33;refGene_tx=XM_023244564.1
E1 1422814 1422985
refGene_gn=DHX33;refGene_tx=XM_023244562.1;refGene_gn=DHX33;refGene_tx=XM_023244563.1;refGene_gn=DHX33;refGene_tx=XM_023244564.1
E1 1425027 1425213
refGene_gn=DHX33;refGene_tx=XM_023244562.1;refGene_gn=DHX33;refGene_tx=XM_023244563.1;refGene_gn=DHX33;refGene_tx=XM_023244564.1
E1 1425790 1425902
refGene_gn=DHX33;refGene_tx=XM_023244562.1;refGene_gn=DHX33;refGene_tx=XM_023244563.1;refGene_gn=DHX33;refGene_tx=XM_023244564.1
E1 1426874 1427034
refGene_gn=DHX33;refGene_tx=XM_023244562.1;refGene_gn=DHX33;refGene_tx

=XM_023244563.1;refGene_gn=DHX33;refGene_tx=XM_023244564.1
E1 1427132 1427221
refGene_gn=DHX33;refGene_tx=XM_023244562.1;refGene_gn=DHX33;refGene_tx=XM_023244563.1;refGene_gn=DHX33;refGene_tx=XM_023244564.1
E1 1428101 1428229
refGene_gn=DHX33;refGene_tx=XM_023244562.1;refGene_gn=DHX33;refGene_tx=XM_023244563.1;refGene_gn=DHX33;refGene_tx=XM_023244564.1
E1 1428621 1428825
refGene_gn=DHX33;refGene_tx=XM_023244562.1;refGene_gn=DHX33;refGene_tx=XM_023244563.1;refGene_gn=DHX33;refGene_tx=XM_023244564.1
E1 1428901 1429039 refGene_gn=DHX33;refGene_tx=XM_023244564.1
E1 1430488 1430575
refGene_gn=DHX33;refGene_tx=XM_023244562.1;refGene_gn=DHX33;refGene_tx=XM_023244563.1
E1 1431053 1431110 refGene_gn=DHX33;refGene_tx=XM_023244563.1
E1 1433548 1433857 refGene_gn=DHX33;refGene_tx=XM_023244562.1
E1 1438298 1438504 refGene_gn=C1QBP;refGene_tx=XM_023244638.1
E1 1438605 1438607 refGene_gn=C1QBP;refGene_tx=XM_023244638.1
E1 1439277 1439425 refGene_gn=C1QBP;refGene_tx=XM_023244638.1
E1 1440855 1440949 refGene_gn=C1QBP;refGene_tx=XM_023244638.1
E1 1442090 1442186 refGene_gn=C1QBP;refGene_tx=XM_023244638.1
E1 1442383 1442506 refGene_gn=C1QBP;refGene_tx=XM_023244638.1
E1 1442612 1442762 refGene_gn=C1QBP;refGene_tx=XM_023244638.1
E1 1443171 1443201
refGene_gn=RPAIN;refGene_tx=XM_023244634.1;refGene_gn=RPAIN;refGene_tx=XM_023244635.1;refGene_gn=RPAIN;refGene_tx=XM_023244636.1
E1 1445149 1445290
refGene_gn=RPAIN;refGene_tx=XM_023244634.1;refGene_gn=RPAIN;refGene_tx=XM_023244635.1;refGene_gn=RPAIN;refGene_tx=XM_023244636.1
E1 1445641 1445705
refGene_gn=RPAIN;refGene_tx=XM_023244634.1;refGene_gn=RPAIN;refGene_tx=XM_023244635.1;refGene_gn=RPAIN;refGene_tx=XM_023244636.1
E1 1445870 1445982
refGene_gn=RPAIN;refGene_tx=XM_023244634.1;refGene_gn=RPAIN;refGene_tx=XM_023244635.1;refGene_gn=RPAIN;refGene_tx=XM_023244636.1
E1 1446726 1446734 refGene_gn=RPAIN;refGene_tx=XM_023244637.1
E1 1447265 1447326
refGene_gn=RPAIN;refGene_tx=XM_023244634.1;refGene_gn=RPAIN;refGene_tx=XM_023244635.1;refGene_gn=RPAIN;refGene_tx=XM_023244636.1;refGene_gn=RPAIN;refGene_tx=XM_023244637.1
E1 1448322 1448493
refGene_gn=RPAIN;refGene_tx=XM_023244634.1;refGene_gn=RPAIN;refGene_tx=XM_023244635.1;refGene_gn=RPAIN;refGene_tx=XM_023244636.1;refGene_gn=RPAIN;refGene_tx=XM_023244637.1
E1 1449699 1449780
refGene_gn=RPAIN;refGene_tx=XM_023244634.1;refGene_gn=RPAIN;refGene_tx=XM_023244637.1
E1 1450278 1450575
refGene_gn=NUP88;refGene_tx=XM_023244632.1;refGene_gn=NUP88;refGene_tx=XM_023244633.1

E1 1452460 1452630
refGene_gn=NUP88;refGene_tx=XM_023244632.1;refGene_gn=NUP88;refGene_tx
=XM_023244633.1
E1 1453383 1453509
refGene_gn=NUP88;refGene_tx=XM_023244632.1;refGene_gn=NUP88;refGene_tx
=XM_023244633.1
E1 1456564 1456651
refGene_gn=NUP88;refGene_tx=XM_023244632.1;refGene_gn=NUP88;refGene_tx
=XM_023244633.1
E1 1459228 1459405
refGene_gn=NUP88;refGene_tx=XM_023244632.1;refGene_gn=NUP88;refGene_tx
=XM_023244633.1
E1 1462691 1462878
refGene_gn=NUP88;refGene_tx=XM_023244632.1;refGene_gn=NUP88;refGene_tx
=XM_023244633.1
E1 1463314 1463462
refGene_gn=NUP88;refGene_tx=XM_023244632.1;refGene_gn=NUP88;refGene_tx
=XM_023244633.1
E1 1466367 1466466
refGene_gn=NUP88;refGene_tx=XM_023244632.1;refGene_gn=NUP88;refGene_tx
=XM_023244633.1
E1 1470758 1470849
refGene_gn=NUP88;refGene_tx=XM_023244632.1;refGene_gn=NUP88;refGene_tx
=XM_023244633.1
E1 1472212 1472314 refGene_gn=NUP88;refGene_tx=XM_023244632.1
E1 1473804 1473963
refGene_gn=NUP88;refGene_tx=XM_023244632.1;refGene_gn=NUP88;refGene_tx
=XM_023244633.1
E1 1474173 1474299
refGene_gn=NUP88;refGene_tx=XM_023244632.1;refGene_gn=NUP88;refGene_tx
=XM_023244633.1
E1 1474381 1474447
refGene_gn=NUP88;refGene_tx=XM_023244632.1;refGene_gn=NUP88;refGene_tx
=XM_023244633.1
E1 1474548 1474629
refGene_gn=NUP88;refGene_tx=XM_023244632.1;refGene_gn=NUP88;refGene_tx
=XM_023244633.1
E1 1474855 1474982
refGene_gn=NUP88;refGene_tx=XM_023244632.1;refGene_gn=NUP88;refGene_tx
=XM_023244633.1
E1 1475088 1475207
refGene_gn=NUP88;refGene_tx=XM_023244632.1;refGene_gn=NUP88;refGene_tx
=XM_023244633.1
E1 1475457 1475521
refGene_gn=NUP88;refGene_tx=XM_023244632.1;refGene_gn=NUP88;refGene_tx
=XM_023244633.1
E1 1478113 1478215
refGene_gn=RABEP1;refGene_tx=XM_023244630.1;refGene_gn=RABEP1;refGene_
tx=XM_023244631.1
E1 1478982 1479099

refGene_gn=RABEP1;refGene_tx=XM_023244630.1;refGene_gn=RABEP1;refGene_tx=XM_023244631.1
E1 1479797 1479896
refGene_gn=RABEP1;refGene_tx=XM_023244630.1;refGene_gn=RABEP1;refGene_tx=XM_023244631.1
E1 1480088 1480144
refGene_gn=RABEP1;refGene_tx=XM_023244630.1;refGene_gn=RABEP1;refGene_tx=XM_023244631.1
E1 1480605 1480795
refGene_gn=RABEP1;refGene_tx=XM_023244630.1;refGene_gn=RABEP1;refGene_tx=XM_023244631.1
E1 1483513 1483654
refGene_gn=RABEP1;refGene_tx=XM_023244630.1;refGene_gn=RABEP1;refGene_tx=XM_023244631.1
E1 1486817 1486916
refGene_gn=RABEP1;refGene_tx=XM_023244630.1;refGene_gn=RABEP1;refGene_tx=XM_023244631.1
E1 1492352 1492469
refGene_gn=RABEP1;refGene_tx=XM_023244630.1;refGene_gn=RABEP1;refGene_tx=XM_023244631.1
E1 1493550 1493655
refGene_gn=RABEP1;refGene_tx=XM_023244630.1;refGene_gn=RABEP1;refGene_tx=XM_023244631.1
E1 1494800 1495274
refGene_gn=RABEP1;refGene_tx=XM_023244630.1;refGene_gn=RABEP1;refGene_tx=XM_023244631.1
E1 1499111 1499243
refGene_gn=RABEP1;refGene_tx=XM_023244630.1;refGene_gn=RABEP1;refGene_tx=XM_023244631.1
E1 1501090 1501269
refGene_gn=RABEP1;refGene_tx=XM_023244630.1;refGene_gn=RABEP1;refGene_tx=XM_023244631.1
E1 1504109 1504245
refGene_gn=RABEP1;refGene_tx=XM_023244630.1;refGene_gn=RABEP1;refGene_tx=XM_023244631.1
E1 1509002 1509122
refGene_gn=RABEP1;refGene_tx=XM_023244630.1;refGene_gn=RABEP1;refGene_tx=XM_023244631.1
E1 1513793 1513954
refGene_gn=RABEP1;refGene_tx=XM_023244630.1;refGene_gn=RABEP1;refGene_tx=XM_023244631.1
E1 1515931 1516135
refGene_gn=RABEP1;refGene_tx=XM_023244630.1;refGene_gn=RABEP1;refGene_tx=XM_023244631.1
E1 1538497 1538626 refGene_gn=RABEP1;refGene_tx=XM_023244630.1
E1 1579580 1579614
refGene_gn=RABEP1;refGene_tx=XM_023244630.1;refGene_gn=RABEP1;refGene_tx=XM_023244631.1
E1 1606296 1606317 refGene_gn=SCIMP;refGene_tx=XM_006939729.4
E1 1617735 1617859 refGene_gn=SCIMP;refGene_tx=XM_006939729.4

E1	1619677	1619741	refGene_gn=SCIMP;refGene_tx=XM_006939729.4
E1	1622869	1622943	refGene_gn=SCIMP;refGene_tx=XM_006939729.4
E1	1624738	1624890	refGene_gn=SCIMP;refGene_tx=XM_006939729.4
E1	1651866	1653264
refGene_gn=ZFP3;refGene_tx=XM_023243455.1;refGene_gn=ZFP3;refGene_tx=XM_023243456.1;refGene_gn=ZFP3;refGene_tx=XM_023243457.1
E1	1678984	1679662
refGene_gn=KIF1C;refGene_tx=XM_023243538.1;refGene_gn=KIF1C;refGene_tx=XM_023243539.1;refGene_gn=KIF1C;refGene_tx=XM_023243540.1
E1	1681045	1681654
refGene_gn=KIF1C;refGene_tx=XM_023243538.1;refGene_gn=KIF1C;refGene_tx=XM_023243539.1;refGene_gn=KIF1C;refGene_tx=XM_023243540.1
E1	1682342	1682415
refGene_gn=KIF1C;refGene_tx=XM_023243538.1;refGene_gn=KIF1C;refGene_tx=XM_023243539.1;refGene_gn=KIF1C;refGene_tx=XM_023243540.1
E1	1682521	1682708
refGene_gn=KIF1C;refGene_tx=XM_023243538.1;refGene_gn=KIF1C;refGene_tx=XM_023243539.1;refGene_gn=KIF1C;refGene_tx=XM_023243540.1
E1	1683061	1683145
refGene_gn=KIF1C;refGene_tx=XM_023243538.1;refGene_gn=KIF1C;refGene_tx=XM_023243539.1;refGene_gn=KIF1C;refGene_tx=XM_023243540.1
E1	1686279	1686374
refGene_gn=KIF1C;refGene_tx=XM_023243538.1;refGene_gn=KIF1C;refGene_tx=XM_023243539.1;refGene_gn=KIF1C;refGene_tx=XM_023243540.1
E1	1687231	1687311
refGene_gn=KIF1C;refGene_tx=XM_023243538.1;refGene_gn=KIF1C;refGene_tx=XM_023243539.1;refGene_gn=KIF1C;refGene_tx=XM_023243540.1
E1	1691432	1691508
refGene_gn=KIF1C;refGene_tx=XM_023243538.1;refGene_gn=KIF1C;refGene_tx=XM_023243539.1;refGene_gn=KIF1C;refGene_tx=XM_023243540.1
E1	1691620	1691700
refGene_gn=KIF1C;refGene_tx=XM_023243538.1;refGene_gn=KIF1C;refGene_tx=XM_023243539.1;refGene_gn=KIF1C;refGene_tx=XM_023243540.1
E1	1691892	1692062
refGene_gn=KIF1C;refGene_tx=XM_023243538.1;refGene_gn=KIF1C;refGene_tx=XM_023243539.1;refGene_gn=KIF1C;refGene_tx=XM_023243540.1
E1	1692985	1693131
refGene_gn=KIF1C;refGene_tx=XM_023243538.1;refGene_gn=KIF1C;refGene_tx=XM_023243539.1;refGene_gn=KIF1C;refGene_tx=XM_023243540.1
E1	1693282	1693361
refGene_gn=KIF1C;refGene_tx=XM_023243538.1;refGene_gn=KIF1C;refGene_tx=XM_023243539.1;refGene_gn=KIF1C;refGene_tx=XM_023243540.1
E1	1693821	1693897
refGene_gn=KIF1C;refGene_tx=XM_023243538.1;refGene_gn=KIF1C;refGene_tx=XM_023243539.1;refGene_gn=KIF1C;refGene_tx=XM_023243540.1
E1	1693980	1694046
refGene_gn=KIF1C;refGene_tx=XM_023243538.1;refGene_gn=KIF1C;refGene_tx=XM_023243539.1;refGene_gn=KIF1C;refGene_tx=XM_023243540.1
E1	1694156	1694234
refGene_gn=KIF1C;refGene_tx=XM_023243538.1;refGene_gn=KIF1C;refGene_tx

=XM_023243539.1;refGene_gn=KIF1C;refGene_tx=XM_023243540.1
E1 1694360 1694472
refGene_gn=KIF1C;refGene_tx=XM_023243538.1;refGene_gn=KIF1C;refGene_tx
=XM_023243539.1;refGene_gn=KIF1C;refGene_tx=XM_023243540.1
E1 1694560 1694739
refGene_gn=KIF1C;refGene_tx=XM_023243538.1;refGene_gn=KIF1C;refGene_tx
=XM_023243539.1;refGene_gn=KIF1C;refGene_tx=XM_023243540.1
E1 1695122 1695188
refGene_gn=KIF1C;refGene_tx=XM_023243538.1;refGene_gn=KIF1C;refGene_tx
=XM_023243539.1;refGene_gn=KIF1C;refGene_tx=XM_023243540.1
E1 1695875 1696055
refGene_gn=KIF1C;refGene_tx=XM_023243538.1;refGene_gn=KIF1C;refGene_tx
=XM_023243539.1;refGene_gn=KIF1C;refGene_tx=XM_023243540.1
E1 1696439 1696516
refGene_gn=KIF1C;refGene_tx=XM_023243538.1;refGene_gn=KIF1C;refGene_tx
=XM_023243539.1;refGene_gn=KIF1C;refGene_tx=XM_023243540.1
E1 1696869 1696975
refGene_gn=KIF1C;refGene_tx=XM_023243538.1;refGene_gn=KIF1C;refGene_tx
=XM_023243539.1;refGene_gn=KIF1C;refGene_tx=XM_023243540.1
E1 1700650 1700733 refGene_gn=INCA1;refGene_tx=XM_023244560.1
E1 1705523 1705637 refGene_gn=INCA1;refGene_tx=XM_023244560.1
E1 1705856 1705896 refGene_gn=INCA1;refGene_tx=XM_023244560.1
E1 1706178 1706372 refGene_gn=INCA1;refGene_tx=XM_023244560.1
E1 1706828 1706994 refGene_gn=INCA1;refGene_tx=XM_023244560.1
E1 1707185 1707215 refGene_gn=INCA1;refGene_tx=XM_023244560.1
E1 1707216 1707333 refGene_gn=INCA1;refGene_tx=XM_023244560.1
E1 1708016 1708053
refGene_gn=CAMTA2;refGene_tx=XM_023244549.1;refGene_gn=CAMTA2;refGene_
tx=XM_023244550.1;refGene_gn=CAMTA2;refGene_tx=XM_023244551.1;refGene_
gn=CAMTA2;refGene_tx=XM_023244552.1;refGene_gn=CAMTA2;refGene_tx=XM_02
3244553.1;refGene_gn=CAMTA2;refGene_tx=XM_023244554.1;refGene_gn=CAMTA
2;refGene_tx=XM_023244556.1;refGene_gn=CAMTA2;refGene_tx=XM_023244557.
1;refGene_gn=CAMTA2;refGene_tx=XM_023244558.1
E1 1708582 1708587 refGene_gn=CAMTA2;refGene_tx=XM_023244555.1
E1 1709350 1709445
refGene_gn=CAMTA2;refGene_tx=XM_023244549.1;refGene_gn=CAMTA2;refGene_
tx=XM_023244550.1;refGene_gn=CAMTA2;refGene_tx=XM_023244551.1;refGene_
gn=CAMTA2;refGene_tx=XM_023244552.1;refGene_gn=CAMTA2;refGene_tx=XM_02
3244553.1;refGene_gn=CAMTA2;refGene_tx=XM_023244554.1;refGene_gn=CAMTA
2;refGene_tx=XM_023244555.1;refGene_gn=CAMTA2;refGene_tx=XM_023244556.
1;refGene_gn=CAMTA2;refGene_tx=XM_023244557.1;refGene_gn=CAMTA2;refGen
e_tx=XM_023244558.1;refGene_gn=CAMTA2;refGene_tx=XM_023244559.1
E1 1709607 1709711
refGene_gn=CAMTA2;refGene_tx=XM_023244549.1;refGene_gn=CAMTA2;refGene_
tx=XM_023244550.1;refGene_gn=CAMTA2;refGene_tx=XM_023244551.1;refGene_
gn=CAMTA2;refGene_tx=XM_023244552.1;refGene_gn=CAMTA2;refGene_tx=XM_02
3244553.1;refGene_gn=CAMTA2;refGene_tx=XM_023244554.1;refGene_gn=CAMTA
2;refGene_tx=XM_023244555.1;refGene_gn=CAMTA2;refGene_tx=XM_023244556.
1;refGene_gn=CAMTA2;refGene_tx=XM_023244557.1;refGene_gn=CAMTA2;refGen
e_tx=XM_023244558.1;refGene_gn=CAMTA2;refGene_tx=XM_023244559.1

E1 1710843 1710911
refGene_gn=CAMTA2;refGene_tx=XM_023244549.1;refGene_gn=CAMTA2;refGene_tx=XM_023244550.1;refGene_gn=CAMTA2;refGene_tx=XM_023244551.1;refGene_gn=CAMTA2;refGene_tx=XM_023244552.1;refGene_gn=CAMTA2;refGene_tx=XM_023244553.1;refGene_gn=CAMTA2;refGene_tx=XM_023244554.1;refGene_gn=CAMTA2;refGene_tx=XM_023244555.1;refGene_gn=CAMTA2;refGene_tx=XM_023244556.1;refGene_gn=CAMTA2;refGene_tx=XM_023244557.1;refGene_gn=CAMTA2;refGene_tx=XM_023244558.1;refGene_gn=CAMTA2;refGene_tx=XM_023244559.1
E1 1710990 1711126
refGene_gn=CAMTA2;refGene_tx=XM_023244549.1;refGene_gn=CAMTA2;refGene_tx=XM_023244550.1;refGene_gn=CAMTA2;refGene_tx=XM_023244551.1;refGene_gn=CAMTA2;refGene_tx=XM_023244552.1;refGene_gn=CAMTA2;refGene_tx=XM_023244553.1;refGene_gn=CAMTA2;refGene_tx=XM_023244554.1;refGene_gn=CAMTA2;refGene_tx=XM_023244555.1;refGene_gn=CAMTA2;refGene_tx=XM_023244556.1;refGene_gn=CAMTA2;refGene_tx=XM_023244557.1;refGene_gn=CAMTA2;refGene_tx=XM_023244558.1;refGene_gn=CAMTA2;refGene_tx=XM_023244559.1
E1 1711596 1711683
refGene_gn=CAMTA2;refGene_tx=XM_023244549.1;refGene_gn=CAMTA2;refGene_tx=XM_023244550.1;refGene_gn=CAMTA2;refGene_tx=XM_023244551.1;refGene_gn=CAMTA2;refGene_tx=XM_023244552.1;refGene_gn=CAMTA2;refGene_tx=XM_023244553.1;refGene_gn=CAMTA2;refGene_tx=XM_023244554.1;refGene_gn=CAMTA2;refGene_tx=XM_023244555.1;refGene_gn=CAMTA2;refGene_tx=XM_023244556.1;refGene_gn=CAMTA2;refGene_tx=XM_023244559.1
E1 1711867 1712021
refGene_gn=CAMTA2;refGene_tx=XM_023244549.1;refGene_gn=CAMTA2;refGene_tx=XM_023244550.1;refGene_gn=CAMTA2;refGene_tx=XM_023244551.1;refGene_gn=CAMTA2;refGene_tx=XM_023244552.1;refGene_gn=CAMTA2;refGene_tx=XM_023244553.1;refGene_gn=CAMTA2;refGene_tx=XM_023244554.1;refGene_gn=CAMTA2;refGene_tx=XM_023244555.1;refGene_gn=CAMTA2;refGene_tx=XM_023244556.1;refGene_gn=CAMTA2;refGene_tx=XM_023244557.1;refGene_gn=CAMTA2;refGene_tx=XM_023244558.1;refGene_gn=CAMTA2;refGene_tx=XM_023244559.1
E1 1712343 1712478
refGene_gn=CAMTA2;refGene_tx=XM_023244549.1;refGene_gn=CAMTA2;refGene_tx=XM_023244550.1;refGene_gn=CAMTA2;refGene_tx=XM_023244551.1;refGene_gn=CAMTA2;refGene_tx=XM_023244552.1;refGene_gn=CAMTA2;refGene_tx=XM_023244553.1;refGene_gn=CAMTA2;refGene_tx=XM_023244554.1;refGene_gn=CAMTA2;refGene_tx=XM_023244555.1;refGene_gn=CAMTA2;refGene_tx=XM_023244556.1;refGene_gn=CAMTA2;refGene_tx=XM_023244557.1;refGene_gn=CAMTA2;refGene_tx=XM_023244558.1;refGene_gn=CAMTA2;refGene_tx=XM_023244559.1
E1 1713006 1713950
refGene_gn=CAMTA2;refGene_tx=XM_023244549.1;refGene_gn=CAMTA2;refGene_tx=XM_023244550.1;refGene_gn=CAMTA2;refGene_tx=XM_023244551.1;refGene_gn=CAMTA2;refGene_tx=XM_023244552.1;refGene_gn=CAMTA2;refGene_tx=XM_023244553.1;refGene_gn=CAMTA2;refGene_tx=XM_023244554.1;refGene_gn=CAMTA2;refGene_tx=XM_023244555.1;refGene_gn=CAMTA2;refGene_tx=XM_023244556.1;refGene_gn=CAMTA2;refGene_tx=XM_023244557.1;refGene_gn=CAMTA2;refGene_tx=XM_023244558.1;refGene_gn=CAMTA2;refGene_tx=XM_023244559.1
E1 1714501 1714628
refGene_gn=CAMTA2;refGene_tx=XM_023244549.1;refGene_gn=CAMTA2;refGene_tx=XM_023244550.1;refGene_gn=CAMTA2;refGene_tx=XM_023244551.1;refGene_

gn=CAMTA2;refGene_tx=XM_023244552.1;refGene_gn=CAMTA2;refGene_tx=XM_023244553.1;refGene_gn=CAMTA2;refGene_tx=XM_023244554.1;refGene_gn=CAMTA2;refGene_tx=XM_023244555.1;refGene_gn=CAMTA2;refGene_tx=XM_023244556.1;refGene_gn=CAMTA2;refGene_tx=XM_023244557.1;refGene_gn=CAMTA2;refGene_tx=XM_023244558.1;refGene_gn=CAMTA2;refGene_tx=XM_023244559.1
E1 1715854 1715989
refGene_gn=CAMTA2;refGene_tx=XM_023244549.1;refGene_gn=CAMTA2;refGene_tx=XM_023244550.1;refGene_gn=CAMTA2;refGene_tx=XM_023244551.1;refGene_gn=CAMTA2;refGene_tx=XM_023244552.1;refGene_gn=CAMTA2;refGene_tx=XM_023244553.1;refGene_gn=CAMTA2;refGene_tx=XM_023244554.1;refGene_gn=CAMTA2;refGene_tx=XM_023244555.1;refGene_gn=CAMTA2;refGene_tx=XM_023244556.1;refGene_gn=CAMTA2;refGene_tx=XM_023244557.1;refGene_gn=CAMTA2;refGene_tx=XM_023244558.1;refGene_gn=CAMTA2;refGene_tx=XM_023244559.1
E1 1717697 1717813
refGene_gn=CAMTA2;refGene_tx=XM_023244549.1;refGene_gn=CAMTA2;refGene_tx=XM_023244550.1;refGene_gn=CAMTA2;refGene_tx=XM_023244551.1;refGene_gn=CAMTA2;refGene_tx=XM_023244552.1;refGene_gn=CAMTA2;refGene_tx=XM_023244553.1;refGene_gn=CAMTA2;refGene_tx=XM_023244554.1;refGene_gn=CAMTA2;refGene_tx=XM_023244555.1;refGene_gn=CAMTA2;refGene_tx=XM_023244556.1;refGene_gn=CAMTA2;refGene_tx=XM_023244557.1;refGene_gn=CAMTA2;refGene_tx=XM_023244558.1;refGene_gn=CAMTA2;refGene_tx=XM_023244559.1
E1 1718304 1718489
refGene_gn=CAMTA2;refGene_tx=XM_023244549.1;refGene_gn=CAMTA2;refGene_tx=XM_023244550.1;refGene_gn=CAMTA2;refGene_tx=XM_023244551.1;refGene_gn=CAMTA2;refGene_tx=XM_023244552.1;refGene_gn=CAMTA2;refGene_tx=XM_023244553.1;refGene_gn=CAMTA2;refGene_tx=XM_023244554.1;refGene_gn=CAMTA2;refGene_tx=XM_023244555.1;refGene_gn=CAMTA2;refGene_tx=XM_023244556.1;refGene_gn=CAMTA2;refGene_tx=XM_023244557.1;refGene_gn=CAMTA2;refGene_tx=XM_023244558.1;refGene_gn=CAMTA2;refGene_tx=XM_023244559.1
E1 1718735 1718814
refGene_gn=CAMTA2;refGene_tx=XM_023244549.1;refGene_gn=CAMTA2;refGene_tx=XM_023244550.1;refGene_gn=CAMTA2;refGene_tx=XM_023244551.1;refGene_gn=CAMTA2;refGene_tx=XM_023244552.1;refGene_gn=CAMTA2;refGene_tx=XM_023244553.1;refGene_gn=CAMTA2;refGene_tx=XM_023244554.1;refGene_gn=CAMTA2;refGene_tx=XM_023244555.1;refGene_gn=CAMTA2;refGene_tx=XM_023244556.1;refGene_gn=CAMTA2;refGene_tx=XM_023244557.1;refGene_gn=CAMTA2;refGene_tx=XM_023244558.1;refGene_gn=CAMTA2;refGene_tx=XM_023244559.1
E1 1718962 1719185
refGene_gn=CAMTA2;refGene_tx=XM_023244549.1;refGene_gn=CAMTA2;refGene_tx=XM_023244550.1;refGene_gn=CAMTA2;refGene_tx=XM_023244551.1;refGene_gn=CAMTA2;refGene_tx=XM_023244552.1;refGene_gn=CAMTA2;refGene_tx=XM_023244553.1;refGene_gn=CAMTA2;refGene_tx=XM_023244554.1;refGene_gn=CAMTA2;refGene_tx=XM_023244555.1;refGene_gn=CAMTA2;refGene_tx=XM_023244556.1;refGene_gn=CAMTA2;refGene_tx=XM_023244557.1;refGene_gn=CAMTA2;refGene_tx=XM_023244558.1;refGene_gn=CAMTA2;refGene_tx=XM_023244559.1
E1 1719324 1719629
refGene_gn=CAMTA2;refGene_tx=XM_023244549.1;refGene_gn=CAMTA2;refGene_tx=XM_023244550.1;refGene_gn=CAMTA2;refGene_tx=XM_023244551.1;refGene_gn=CAMTA2;refGene_tx=XM_023244552.1;refGene_gn=CAMTA2;refGene_tx=XM_023244553.1;refGene_gn=CAMTA2;refGene_tx=XM_023244554.1;refGene_gn=CAMTA

2;refGene_tx=XM_023244555.1;refGene_gn=CAMTA2;refGene_tx=XM_023244556.1;refGene_gn=CAMTA2;refGene_tx=XM_023244557.1;refGene_gn=CAMTA2;refGene_tx=XM_023244558.1;refGene_gn=CAMTA2;refGene_tx=XM_023244559.1
E1 1720652 1720849
refGene_gn=CAMTA2;refGene_tx=XM_023244549.1;refGene_gn=CAMTA2;refGene_tx=XM_023244550.1;refGene_gn=CAMTA2;refGene_tx=XM_023244551.1;refGene_gn=CAMTA2;refGene_tx=XM_023244552.1;refGene_gn=CAMTA2;refGene_tx=XM_023244553.1;refGene_gn=CAMTA2;refGene_tx=XM_023244554.1;refGene_gn=CAMTA2;refGene_tx=XM_023244555.1;refGene_gn=CAMTA2;refGene_tx=XM_023244556.1;refGene_gn=CAMTA2;refGene_tx=XM_023244557.1;refGene_gn=CAMTA2;refGene_tx=XM_023244558.1;refGene_gn=CAMTA2;refGene_tx=XM_023244559.1
E1 1721063 1721247
refGene_gn=CAMTA2;refGene_tx=XM_023244549.1;refGene_gn=CAMTA2;refGene_tx=XM_023244550.1;refGene_gn=CAMTA2;refGene_tx=XM_023244551.1;refGene_gn=CAMTA2;refGene_tx=XM_023244552.1;refGene_gn=CAMTA2;refGene_tx=XM_023244553.1;refGene_gn=CAMTA2;refGene_tx=XM_023244554.1;refGene_gn=CAMTA2;refGene_tx=XM_023244555.1;refGene_gn=CAMTA2;refGene_tx=XM_023244556.1;refGene_gn=CAMTA2;refGene_tx=XM_023244557.1;refGene_gn=CAMTA2;refGene_tx=XM_023244558.1;refGene_gn=CAMTA2;refGene_tx=XM_023244559.1
E1 1721344 1721416
refGene_gn=CAMTA2;refGene_tx=XM_023244549.1;refGene_gn=CAMTA2;refGene_tx=XM_023244550.1;refGene_gn=CAMTA2;refGene_tx=XM_023244551.1;refGene_gn=CAMTA2;refGene_tx=XM_023244552.1;refGene_gn=CAMTA2;refGene_tx=XM_023244553.1;refGene_gn=CAMTA2;refGene_tx=XM_023244554.1;refGene_gn=CAMTA2;refGene_tx=XM_023244555.1;refGene_gn=CAMTA2;refGene_tx=XM_023244556.1;refGene_gn=CAMTA2;refGene_tx=XM_023244557.1;refGene_gn=CAMTA2;refGene_tx=XM_023244558.1;refGene_gn=CAMTA2;refGene_tx=XM_023244559.1
E1 1721502 1721523
refGene_gn=CAMTA2;refGene_tx=XM_023244549.1;refGene_gn=CAMTA2;refGene_tx=XM_023244550.1;refGene_gn=CAMTA2;refGene_tx=XM_023244551.1;refGene_gn=CAMTA2;refGene_tx=XM_023244552.1;refGene_gn=CAMTA2;refGene_tx=XM_023244555.1;refGene_gn=CAMTA2;refGene_tx=XM_023244557.1;refGene_gn=CAMTA2;refGene_tx=XM_023244558.1;refGene_gn=CAMTA2;refGene_tx=XM_023244559.1
E1 1721687 1721857
refGene_gn=CAMTA2;refGene_tx=XM_023244549.1;refGene_gn=CAMTA2;refGene_tx=XM_023244550.1;refGene_gn=CAMTA2;refGene_tx=XM_023244551.1;refGene_gn=CAMTA2;refGene_tx=XM_023244552.1;refGene_gn=CAMTA2;refGene_tx=XM_023244553.1;refGene_gn=CAMTA2;refGene_tx=XM_023244554.1;refGene_gn=CAMTA2;refGene_tx=XM_023244555.1;refGene_gn=CAMTA2;refGene_tx=XM_023244556.1;refGene_gn=CAMTA2;refGene_tx=XM_023244557.1;refGene_gn=CAMTA2;refGene_tx=XM_023244558.1;refGene_gn=CAMTA2;refGene_tx=XM_023244559.1
E1 1721968 1722046
refGene_gn=CAMTA2;refGene_tx=XM_023244549.1;refGene_gn=CAMTA2;refGene_tx=XM_023244550.1;refGene_gn=CAMTA2;refGene_tx=XM_023244551.1;refGene_gn=CAMTA2;refGene_tx=XM_023244552.1;refGene_gn=CAMTA2;refGene_tx=XM_023244553.1;refGene_gn=CAMTA2;refGene_tx=XM_023244554.1;refGene_gn=CAMTA2;refGene_tx=XM_023244555.1;refGene_gn=CAMTA2;refGene_tx=XM_023244556.1;refGene_gn=CAMTA2;refGene_tx=XM_023244557.1;refGene_gn=CAMTA2;refGene_tx=XM_023244558.1;refGene_gn=CAMTA2;refGene_tx=XM_023244559.1

E1	1722143	1722207
refGene_gn=CAMTA2;refGene_tx=XM_023244549.1;refGene_gn=CAMTA2;refGene_tx=XM_023244550.1;refGene_gn=CAMTA2;refGene_tx=XM_023244551.1;refGene_gn=CAMTA2;refGene_tx=XM_023244552.1;refGene_gn=CAMTA2;refGene_tx=XM_023244553.1;refGene_gn=CAMTA2;refGene_tx=XM_023244554.1;refGene_gn=CAMTA2;refGene_tx=XM_023244555.1;refGene_gn=CAMTA2;refGene_tx=XM_023244556.1;refGene_gn=CAMTA2;refGene_tx=XM_023244557.1;refGene_gn=CAMTA2;refGene_tx=XM_023244558.1;refGene_gn=CAMTA2;refGene_tx=XM_023244559.1
E1	1722963	1723048	refGene_gn=SPAG7;refGene_tx=XM_003996144.5
E1	1725340	1725408	refGene_gn=SPAG7;refGene_tx=XM_003996144.5
E1	1725636	1725725	refGene_gn=SPAG7;refGene_tx=XM_003996144.5
E1	1725858	1725943	refGene_gn=SPAG7;refGene_tx=XM_003996144.5
E1	1726062	1726152	refGene_gn=SPAG7;refGene_tx=XM_003996144.5
E1	1726239	1726396	refGene_gn=SPAG7;refGene_tx=XM_003996144.5
E1	1726502	1726612	refGene_gn=SPAG7;refGene_tx=XM_003996144.5
E1	1727461	1727531	refGene_gn=ENO3;refGene_tx=XM_003996145.4
E1	1727624	1727683	refGene_gn=ENO3;refGene_tx=XM_003996145.4
E1	1727839	1727948	refGene_gn=ENO3;refGene_tx=XM_003996145.4
E1	1728317	1728519	refGene_gn=ENO3;refGene_tx=XM_003996145.4
E1	1728638	1728836	refGene_gn=ENO3;refGene_tx=XM_003996145.4
E1	1728940	1729163	refGene_gn=ENO3;refGene_tx=XM_003996145.4
E1	1730053	1730187	refGene_gn=ENO3;refGene_tx=XM_003996145.4
E1	1730550	1730620	refGene_gn=ENO3;refGene_tx=XM_003996145.4
E1	1730799	1730858	refGene_gn=ENO3;refGene_tx=XM_003996145.4
E1	1731025	1731121	refGene_gn=ENO3;refGene_tx=XM_003996145.4
E1	1731354	1731439	refGene_gn=ENO3;refGene_tx=XM_003996145.4
E1	1734603	1734735	refGene_gn=PFN1;refGene_tx=XM_023243426.1
E1	1736197	1736390	refGene_gn=PFN1;refGene_tx=XM_023243426.1
E1	1736990	1737088	refGene_gn=PFN1;refGene_tx=XM_023243426.1
E1	1738060	1738362	refGene_gn=RNF167;refGene_tx=XM_003996238.5
E1	1738460	1738541	refGene_gn=RNF167;refGene_tx=XM_003996238.5
E1	1739061	1739155	refGene_gn=RNF167;refGene_tx=XM_003996238.5
E1	1739297	1739403	refGene_gn=RNF167;refGene_tx=XM_003996238.5
E1	1739592	1739683	refGene_gn=RNF167;refGene_tx=XM_003996238.5
E1	1739861	1739949	refGene_gn=RNF167;refGene_tx=XM_003996238.5
E1	1740066	1740192	refGene_gn=RNF167;refGene_tx=XM_003996238.5
E1	1741309	1741390	refGene_gn=RNF167;refGene_tx=XM_003996238.5
E1	1741475	1741559	refGene_gn=RNF167;refGene_tx=XM_003996238.5
E1	1742582	1742677	refGene_gn=SLC25A11;refGene_tx=XM_003996147.5
E1	1743244	1743397	refGene_gn=SLC25A11;refGene_tx=XM_003996147.5
E1	1743482	1743689	refGene_gn=SLC25A11;refGene_tx=XM_003996147.5
E1	1743873	1743964	refGene_gn=SLC25A11;refGene_tx=XM_003996147.5
E1	1744046	1744137	refGene_gn=SLC25A11;refGene_tx=XM_003996147.5
E1	1744221	1744321	refGene_gn=SLC25A11;refGene_tx=XM_003996147.5
E1	1744401	1744453	refGene_gn=SLC25A11;refGene_tx=XM_003996147.5
E1	1744559	1744715	refGene_gn=SLC25A11;refGene_tx=XM_003996147.5
E1	1747490	1748228	refGene_gn=GP1BA;refGene_tx=XM_023243425.1
E1	1748230	1749352	refGene_gn=GP1BA;refGene_tx=XM_023243425.1
E1	1767553	1767788
refGene_gn=CHRNE;refGene_tx=XM_023243749.1;refGene_gn=CHRNE;refGene_tx

=XM_023243750.1
E1	1768003	1768146
refGene_gn=CHRNE;refGene_tx=XM_023243749.1;refGene_gn=CHRNE;refGene_tx
=XM_023243750.1
E1	1768228	1768273
refGene_gn=CHRNE;refGene_tx=XM_023243749.1;refGene_gn=CHRNE;refGene_tx
=XM_023243750.1
E1	1768375	1768485
refGene_gn=CHRNE;refGene_tx=XM_023243749.1;refGene_gn=CHRNE;refGene_tx
=XM_023243750.1
E1	1768570	1768726
refGene_gn=CHRNE;refGene_tx=XM_023243749.1;refGene_gn=CHRNE;refGene_tx
=XM_023243750.1
E1	1768856	1768957
refGene_gn=CHRNE;refGene_tx=XM_023243749.1;refGene_gn=CHRNE;refGene_tx
=XM_023243750.1
E1	1769356	1769554
refGene_gn=CHRNE;refGene_tx=XM_023243749.1;refGene_gn=CHRNE;refGene_tx
=XM_023243750.1
E1	1769630	1769745
refGene_gn=CHRNE;refGene_tx=XM_023243749.1;refGene_gn=CHRNE;refGene_tx
=XM_023243750.1
E1	1769868	1770039
refGene_gn=CE1H17orf107;refGene_tx=XM_023243751.1;refGene_gn=CE1H17orf
107;refGene_tx=XM_023243752.1
E1	1770127	1770406
refGene_gn=CE1H17orf107;refGene_tx=XM_023243751.1;refGene_gn=CE1H17orf
107;refGene_tx=XM_023243752.1
E1	1770513	1770705
refGene_gn=CE1H17orf107;refGene_tx=XM_023243751.1;refGene_gn=CE1H17orf
107;refGene_tx=XM_023243752.1
E1	1770804	1770870
refGene_gn=CE1H17orf107;refGene_tx=XM_023243751.1;refGene_gn=CE1H17orf
107;refGene_tx=XM_023243752.1
E1	1770998	1771113
refGene_gn=CHRNE;refGene_tx=XM_023243749.1;refGene_gn=CHRNE;refGene_tx
=XM_023243750.1
E1	1771211	1771398
refGene_gn=CHRNE;refGene_tx=XM_023243749.1;refGene_gn=CHRNE;refGene_tx
=XM_023243750.1
E1	1771498	1771605
refGene_gn=CHRNE;refGene_tx=XM_023243749.1;refGene_gn=CHRNE;refGene_tx
=XM_023243750.1
E1	1771690	1771840
refGene_gn=CHRNE;refGene_tx=XM_023243749.1;refGene_gn=CHRNE;refGene_tx
=XM_023243750.1
E1	1773257	1773341
refGene_gn=MINK1;refGene_tx=XM_023243731.1;refGene_gn=MINK1;refGene_tx
=XM_023243733.1;refGene_gn=MINK1;refGene_tx=XM_023243734.1;refGene_gn=
MINK1;refGene_tx=XM_023243735.1;refGene_gn=MINK1;refGene_tx=XM_0232437

36.1;refGene_gn=MINK1;refGene_tx=XM_023243737.1;refGene_gn=MINK1;refGene_tx=XM_023243738.1;refGene_gn=MINK1;refGene_tx=XM_023243739.1;refGene_gn=MINK1;refGene_tx=XM_023243740.1;refGene_gn=MINK1;refGene_tx=XM_023243741.1;refGene_gn=MINK1;refGene_tx=XM_023243742.1;refGene_gn=MINK1;refGene_tx=XM_023243743.1;refGene_gn=MINK1;refGene_tx=XM_023243744.1;refGene_gn=MINK1;refGene_tx=XM_023243745.1
E1 1773662 1773802
refGene_gn=MINK1;refGene_tx=XM_023243731.1;refGene_gn=MINK1;refGene_tx=XM_023243733.1;refGene_gn=MINK1;refGene_tx=XM_023243734.1;refGene_gn=MINK1;refGene_tx=XM_023243735.1;refGene_gn=MINK1;refGene_tx=XM_023243736.1;refGene_gn=MINK1;refGene_tx=XM_023243737.1;refGene_gn=MINK1;refGene_tx=XM_023243738.1;refGene_gn=MINK1;refGene_tx=XM_023243739.1;refGene_gn=MINK1;refGene_tx=XM_023243740.1;refGene_gn=MINK1;refGene_tx=XM_023243741.1;refGene_gn=MINK1;refGene_tx=XM_023243742.1;refGene_gn=MINK1;refGene_tx=XM_023243743.1;refGene_gn=MINK1;refGene_tx=XM_023243744.1;refGene_gn=MINK1;refGene_tx=XM_023243745.1
E1 1773887 1774047
refGene_gn=MINK1;refGene_tx=XM_023243731.1;refGene_gn=MINK1;refGene_tx=XM_023243733.1;refGene_gn=MINK1;refGene_tx=XM_023243734.1;refGene_gn=MINK1;refGene_tx=XM_023243735.1;refGene_gn=MINK1;refGene_tx=XM_023243736.1;refGene_gn=MINK1;refGene_tx=XM_023243737.1;refGene_gn=MINK1;refGene_tx=XM_023243738.1;refGene_gn=MINK1;refGene_tx=XM_023243739.1;refGene_gn=MINK1;refGene_tx=XM_023243740.1;refGene_gn=MINK1;refGene_tx=XM_023243741.1;refGene_gn=MINK1;refGene_tx=XM_023243742.1;refGene_gn=MINK1;refGene_tx=XM_023243743.1;refGene_gn=MINK1;refGene_tx=XM_023243744.1;refGene_gn=MINK1;refGene_tx=XM_023243745.1
E1 1774129 1774279
refGene_gn=MINK1;refGene_tx=XM_023243731.1;refGene_gn=MINK1;refGene_tx=XM_023243733.1;refGene_gn=MINK1;refGene_tx=XM_023243734.1;refGene_gn=MINK1;refGene_tx=XM_023243735.1;refGene_gn=MINK1;refGene_tx=XM_023243736.1;refGene_gn=MINK1;refGene_tx=XM_023243737.1;refGene_gn=MINK1;refGene_tx=XM_023243738.1;refGene_gn=MINK1;refGene_tx=XM_023243739.1;refGene_gn=MINK1;refGene_tx=XM_023243740.1;refGene_gn=MINK1;refGene_tx=XM_023243741.1;refGene_gn=MINK1;refGene_tx=XM_023243742.1;refGene_gn=MINK1;refGene_tx=XM_023243743.1;refGene_gn=MINK1;refGene_tx=XM_023243744.1;refGene_gn=MINK1;refGene_tx=XM_023243745.1
E1 1774351 1774452
refGene_gn=MINK1;refGene_tx=XM_023243731.1;refGene_gn=MINK1;refGene_tx=XM_023243733.1;refGene_gn=MINK1;refGene_tx=XM_023243734.1;refGene_gn=MINK1;refGene_tx=XM_023243735.1;refGene_gn=MINK1;refGene_tx=XM_023243736.1;refGene_gn=MINK1;refGene_tx=XM_023243737.1;refGene_gn=MINK1;refGene_tx=XM_023243738.1;refGene_gn=MINK1;refGene_tx=XM_023243739.1;refGene_gn=MINK1;refGene_tx=XM_023243740.1;refGene_gn=MINK1;refGene_tx=XM_023243741.1;refGene_gn=MINK1;refGene_tx=XM_023243742.1;refGene_gn=MINK1;refGene_tx=XM_023243743.1;refGene_gn=MINK1;refGene_tx=XM_023243744.1;refGene_gn=MINK1;refGene_tx=XM_023243745.1
E1 1774608 1774743
refGene_gn=MINK1;refGene_tx=XM_023243731.1;refGene_gn=MINK1;refGene_tx=XM_023243733.1;refGene_gn=MINK1;refGene_tx=XM_023243734.1;refGene_gn=MINK1;refGene_tx=XM_023243735.1;refGene_gn=MINK1;refGene_tx=XM_0232437

36.1;refGene_gn=MINK1;refGene_tx=XM_023243737.1;refGene_gn=MINK1;refGene_tx=XM_023243738.1;refGene_gn=MINK1;refGene_tx=XM_023243739.1;refGene_gn=MINK1;refGene_tx=XM_023243740.1;refGene_gn=MINK1;refGene_tx=XM_023243741.1;refGene_gn=MINK1;refGene_tx=XM_023243742.1;refGene_gn=MINK1;refGene_tx=XM_023243743.1;refGene_gn=MINK1;refGene_tx=XM_023243744.1;refGene_gn=MINK1;refGene_tx=XM_023243745.1
E1 1774924 1775082
refGene_gn=MINK1;refGene_tx=XM_023243731.1;refGene_gn=MINK1;refGene_tx=XM_023243733.1;refGene_gn=MINK1;refGene_tx=XM_023243734.1;refGene_gn=MINK1;refGene_tx=XM_023243735.1;refGene_gn=MINK1;refGene_tx=XM_023243736.1;refGene_gn=MINK1;refGene_tx=XM_023243737.1;refGene_gn=MINK1;refGene_tx=XM_023243738.1;refGene_gn=MINK1;refGene_tx=XM_023243739.1;refGene_gn=MINK1;refGene_tx=XM_023243740.1;refGene_gn=MINK1;refGene_tx=XM_023243741.1;refGene_gn=MINK1;refGene_tx=XM_023243742.1;refGene_gn=MINK1;refGene_tx=XM_023243743.1;refGene_gn=MINK1;refGene_tx=XM_023243744.1;refGene_gn=MINK1;refGene_tx=XM_023243745.1;refGene_gn=MINK1;refGene_tx=XM_023243746.1
E1 1775172 1775340
refGene_gn=MINK1;refGene_tx=XM_023243731.1;refGene_gn=MINK1;refGene_tx=XM_023243733.1;refGene_gn=MINK1;refGene_tx=XM_023243734.1;refGene_gn=MINK1;refGene_tx=XM_023243735.1;refGene_gn=MINK1;refGene_tx=XM_023243736.1;refGene_gn=MINK1;refGene_tx=XM_023243737.1;refGene_gn=MINK1;refGene_tx=XM_023243738.1;refGene_gn=MINK1;refGene_tx=XM_023243739.1;refGene_gn=MINK1;refGene_tx=XM_023243740.1;refGene_gn=MINK1;refGene_tx=XM_023243741.1;refGene_gn=MINK1;refGene_tx=XM_023243742.1;refGene_gn=MINK1;refGene_tx=XM_023243743.1;refGene_gn=MINK1;refGene_tx=XM_023243744.1;refGene_gn=MINK1;refGene_tx=XM_023243745.1;refGene_gn=MINK1;refGene_tx=XM_023243746.1
E1 1775702 1775811
refGene_gn=MINK1;refGene_tx=XM_023243731.1;refGene_gn=MINK1;refGene_tx=XM_023243733.1;refGene_gn=MINK1;refGene_tx=XM_023243734.1;refGene_gn=MINK1;refGene_tx=XM_023243735.1;refGene_gn=MINK1;refGene_tx=XM_023243736.1;refGene_gn=MINK1;refGene_tx=XM_023243737.1;refGene_gn=MINK1;refGene_tx=XM_023243738.1;refGene_gn=MINK1;refGene_tx=XM_023243739.1;refGene_gn=MINK1;refGene_tx=XM_023243740.1;refGene_gn=MINK1;refGene_tx=XM_023243741.1;refGene_gn=MINK1;refGene_tx=XM_023243742.1;refGene_gn=MINK1;refGene_tx=XM_023243743.1;refGene_gn=MINK1;refGene_tx=XM_023243744.1;refGene_gn=MINK1;refGene_tx=XM_023243745.1;refGene_gn=MINK1;refGene_tx=XM_023243746.1
E1 1775970 1776108
refGene_gn=MINK1;refGene_tx=XM_023243731.1;refGene_gn=MINK1;refGene_tx=XM_023243733.1;refGene_gn=MINK1;refGene_tx=XM_023243734.1;refGene_gn=MINK1;refGene_tx=XM_023243735.1;refGene_gn=MINK1;refGene_tx=XM_023243736.1;refGene_gn=MINK1;refGene_tx=XM_023243737.1;refGene_gn=MINK1;refGene_tx=XM_023243738.1;refGene_gn=MINK1;refGene_tx=XM_023243739.1;refGene_gn=MINK1;refGene_tx=XM_023243740.1;refGene_gn=MINK1;refGene_tx=XM_023243741.1;refGene_gn=MINK1;refGene_tx=XM_023243742.1;refGene_gn=MINK1;refGene_tx=XM_023243743.1;refGene_gn=MINK1;refGene_tx=XM_023243744.1;refGene_gn=MINK1;refGene_tx=XM_023243745.1;refGene_gn=MINK1;refGene_tx=XM_023243746.1

E1	1776190	1776335
refGene_gn=MINK1;refGene_tx=XM_023243731.1;refGene_gn=MINK1;refGene_tx=XM_023243733.1;refGene_gn=MINK1;refGene_tx=XM_023243734.1;refGene_gn=MINK1;refGene_tx=XM_023243735.1;refGene_gn=MINK1;refGene_tx=XM_023243736.1;refGene_gn=MINK1;refGene_tx=XM_023243737.1;refGene_gn=MINK1;refGene_tx=XM_023243738.1;refGene_gn=MINK1;refGene_tx=XM_023243739.1;refGene_gn=MINK1;refGene_tx=XM_023243740.1;refGene_gn=MINK1;refGene_tx=XM_023243741.1;refGene_gn=MINK1;refGene_tx=XM_023243742.1;refGene_gn=MINK1;refGene_tx=XM_023243743.1;refGene_gn=MINK1;refGene_tx=XM_023243744.1;refGene_gn=MINK1;refGene_tx=XM_023243745.1;refGene_gn=MINK1;refGene_tx=XM_023243746.1
E1	1776511	1776672
refGene_gn=MINK1;refGene_tx=XM_023243731.1;refGene_gn=MINK1;refGene_tx=XM_023243733.1;refGene_gn=MINK1;refGene_tx=XM_023243734.1;refGene_gn=MINK1;refGene_tx=XM_023243735.1;refGene_gn=MINK1;refGene_tx=XM_023243736.1;refGene_gn=MINK1;refGene_tx=XM_023243737.1;refGene_gn=MINK1;refGene_tx=XM_023243738.1;refGene_gn=MINK1;refGene_tx=XM_023243739.1;refGene_gn=MINK1;refGene_tx=XM_023243740.1;refGene_gn=MINK1;refGene_tx=XM_023243741.1;refGene_gn=MINK1;refGene_tx=XM_023243742.1;refGene_gn=MINK1;refGene_tx=XM_023243743.1;refGene_gn=MINK1;refGene_tx=XM_023243744.1;refGene_gn=MINK1;refGene_tx=XM_023243745.1;refGene_gn=MINK1;refGene_tx=XM_023243746.1
E1	1776763	1776787
refGene_gn=MINK1;refGene_tx=XM_023243731.1;refGene_gn=MINK1;refGene_tx=XM_023243734.1;refGene_gn=MINK1;refGene_tx=XM_023243735.1;refGene_gn=MINK1;refGene_tx=XM_023243736.1;refGene_gn=MINK1;refGene_tx=XM_023243738.1;refGene_gn=MINK1;refGene_tx=XM_023243739.1;refGene_gn=MINK1;refGene_tx=XM_023243740.1;refGene_gn=MINK1;refGene_tx=XM_023243741.1;refGene_gn=MINK1;refGene_tx=XM_023243744.1;refGene_gn=MINK1;refGene_tx=XM_023243746.1
E1	1776987	1777076
refGene_gn=MINK1;refGene_tx=XM_023243731.1;refGene_gn=MINK1;refGene_tx=XM_023243733.1;refGene_gn=MINK1;refGene_tx=XM_023243734.1;refGene_gn=MINK1;refGene_tx=XM_023243735.1;refGene_gn=MINK1;refGene_tx=XM_023243736.1;refGene_gn=MINK1;refGene_tx=XM_023243737.1;refGene_gn=MINK1;refGene_tx=XM_023243738.1;refGene_gn=MINK1;refGene_tx=XM_023243739.1;refGene_gn=MINK1;refGene_tx=XM_023243740.1;refGene_gn=MINK1;refGene_tx=XM_023243741.1;refGene_gn=MINK1;refGene_tx=XM_023243742.1;refGene_gn=MINK1;refGene_tx=XM_023243743.1;refGene_gn=MINK1;refGene_tx=XM_023243744.1;refGene_gn=MINK1;refGene_tx=XM_023243745.1;refGene_gn=MINK1;refGene_tx=XM_023243746.1
E1	1777276	1777386
refGene_gn=MINK1;refGene_tx=XM_023243731.1;refGene_gn=MINK1;refGene_tx=XM_023243733.1;refGene_gn=MINK1;refGene_tx=XM_023243734.1;refGene_gn=MINK1;refGene_tx=XM_023243735.1;refGene_gn=MINK1;refGene_tx=XM_023243736.1;refGene_gn=MINK1;refGene_tx=XM_023243737.1;refGene_gn=MINK1;refGene_tx=XM_023243738.1;refGene_gn=MINK1;refGene_tx=XM_023243739.1;refGene_gn=MINK1;refGene_tx=XM_023243740.1;refGene_gn=MINK1;refGene_tx=XM_023243741.1;refGene_gn=MINK1;refGene_tx=XM_023243742.1;refGene_gn=MINK1;refGene_tx=XM_023243743.1;refGene_gn=MINK1;refGene_tx=XM_023243744.1;r

efGene_gn=MINK1;refGene_tx=XM_023243745.1;refGene_gn=MINK1;refGene_tx=
XM_023243746.1
E1 1777533 1777644
refGene_gn=MINK1;refGene_tx=XM_023243731.1;refGene_gn=MINK1;refGene_tx
=XM_023243733.1;refGene_gn=MINK1;refGene_tx=XM_023243734.1;refGene_gn=
MINK1;refGene_tx=XM_023243735.1;refGene_gn=MINK1;refGene_tx=XM_0232437
39.1;refGene_gn=MINK1;refGene_tx=XM_023243740.1;refGene_gn=MINK1;refGe
ne_tx=XM_023243741.1;refGene_gn=MINK1;refGene_tx=XM_023243742.1;refGen
e_gn=MINK1;refGene_tx=XM_023243743.1;refGene_gn=MINK1;refGene_tx=XM_02
3243746.1
E1 1777799 1777885
refGene_gn=MINK1;refGene_tx=XM_023243731.1;refGene_gn=MINK1;refGene_tx
=XM_023243733.1;refGene_gn=MINK1;refGene_tx=XM_023243734.1;refGene_gn=
MINK1;refGene_tx=XM_023243735.1;refGene_gn=MINK1;refGene_tx=XM_0232437
36.1;refGene_gn=MINK1;refGene_tx=XM_023243737.1;refGene_gn=MINK1;refGe
ne_tx=XM_023243738.1;refGene_gn=MINK1;refGene_tx=XM_023243739.1;refGen
e_gn=MINK1;refGene_tx=XM_023243740.1;refGene_gn=MINK1;refGene_tx=XM_02
3243741.1;refGene_gn=MINK1;refGene_tx=XM_023243742.1;refGene_gn=MINK1;
refGene_tx=XM_023243743.1;refGene_gn=MINK1;refGene_tx=XM_023243744.1;r
efGene_gn=MINK1;refGene_tx=XM_023243745.1;refGene_gn=MINK1;refGene_tx=
XM_023243746.1
E1 1778314 1778575
refGene_gn=MINK1;refGene_tx=XM_023243731.1;refGene_gn=MINK1;refGene_tx
=XM_023243733.1;refGene_gn=MINK1;refGene_tx=XM_023243734.1;refGene_gn=
MINK1;refGene_tx=XM_023243735.1;refGene_gn=MINK1;refGene_tx=XM_0232437
36.1;refGene_gn=MINK1;refGene_tx=XM_023243737.1;refGene_gn=MINK1;refGe
ne_tx=XM_023243738.1;refGene_gn=MINK1;refGene_tx=XM_023243739.1;refGen
e_gn=MINK1;refGene_tx=XM_023243740.1;refGene_gn=MINK1;refGene_tx=XM_02
3243741.1;refGene_gn=MINK1;refGene_tx=XM_023243742.1;refGene_gn=MINK1;
refGene_tx=XM_023243743.1;refGene_gn=MINK1;refGene_tx=XM_023243744.1;r
efGene_gn=MINK1;refGene_tx=XM_023243745.1;refGene_gn=MINK1;refGene_tx=
XM_023243746.1
E1 1778863 1779037
refGene_gn=MINK1;refGene_tx=XM_023243731.1;refGene_gn=MINK1;refGene_tx
=XM_023243733.1;refGene_gn=MINK1;refGene_tx=XM_023243734.1;refGene_gn=
MINK1;refGene_tx=XM_023243735.1;refGene_gn=MINK1;refGene_tx=XM_0232437
36.1;refGene_gn=MINK1;refGene_tx=XM_023243737.1;refGene_gn=MINK1;refGe
ne_tx=XM_023243738.1;refGene_gn=MINK1;refGene_tx=XM_023243739.1;refGen
e_gn=MINK1;refGene_tx=XM_023243740.1;refGene_gn=MINK1;refGene_tx=XM_02
3243741.1;refGene_gn=MINK1;refGene_tx=XM_023243742.1;refGene_gn=MINK1;
refGene_tx=XM_023243743.1;refGene_gn=MINK1;refGene_tx=XM_023243744.1;r
efGene_gn=MINK1;refGene_tx=XM_023243745.1;refGene_gn=MINK1;refGene_tx=
XM_023243746.1
E1 1779196 1779724
refGene_gn=MINK1;refGene_tx=XM_023243731.1;refGene_gn=MINK1;refGene_tx
=XM_023243733.1;refGene_gn=MINK1;refGene_tx=XM_023243734.1;refGene_gn=
MINK1;refGene_tx=XM_023243735.1;refGene_gn=MINK1;refGene_tx=XM_0232437
36.1;refGene_gn=MINK1;refGene_tx=XM_023243737.1;refGene_gn=MINK1;refGe
ne_tx=XM_023243738.1;refGene_gn=MINK1;refGene_tx=XM_023243739.1;refGen
e_gn=MINK1;refGene_tx=XM_023243740.1;refGene_gn=MINK1;refGene_tx=XM_02

3243741.1;refGene_gn=MINK1;refGene_tx=XM_023243742.1;refGene_gn=MINK1;
refGene_tx=XM_023243743.1;refGene_gn=MINK1;refGene_tx=XM_023243744.1;r
efGene_gn=MINK1;refGene_tx=XM_023243745.1;refGene_gn=MINK1;refGene_tx=
XM_023243746.1
E1 1780173 1780287
refGene_gn=MINK1;refGene_tx=XM_023243731.1;refGene_gn=MINK1;refGene_tx
=XM_023243733.1;refGene_gn=MINK1;refGene_tx=XM_023243734.1;refGene_gn=
MINK1;refGene_tx=XM_023243735.1;refGene_gn=MINK1;refGene_tx=XM_0232437
36.1;refGene_gn=MINK1;refGene_tx=XM_023243737.1;refGene_gn=MINK1;refGe
ne_tx=XM_023243738.1;refGene_gn=MINK1;refGene_tx=XM_023243739.1;refGen
e_gn=MINK1;refGene_tx=XM_023243740.1;refGene_gn=MINK1;refGene_tx=XM_02
3243741.1;refGene_gn=MINK1;refGene_tx=XM_023243742.1;refGene_gn=MINK1;
refGene_tx=XM_023243743.1;refGene_gn=MINK1;refGene_tx=XM_023243744.1;r
efGene_gn=MINK1;refGene_tx=XM_023243745.1;refGene_gn=MINK1;refGene_tx=
XM_023243746.1
E1 1781755 1781975
refGene_gn=MINK1;refGene_tx=XM_023243731.1;refGene_gn=MINK1;refGene_tx
=XM_023243733.1;refGene_gn=MINK1;refGene_tx=XM_023243734.1;refGene_gn=
MINK1;refGene_tx=XM_023243735.1;refGene_gn=MINK1;refGene_tx=XM_0232437
36.1;refGene_gn=MINK1;refGene_tx=XM_023243737.1;refGene_gn=MINK1;refGe
ne_tx=XM_023243738.1;refGene_gn=MINK1;refGene_tx=XM_023243739.1;refGen
e_gn=MINK1;refGene_tx=XM_023243740.1;refGene_gn=MINK1;refGene_tx=XM_02
3243741.1;refGene_gn=MINK1;refGene_tx=XM_023243742.1;refGene_gn=MINK1;
refGene_tx=XM_023243743.1;refGene_gn=MINK1;refGene_tx=XM_023243744.1;r
efGene_gn=MINK1;refGene_tx=XM_023243745.1;refGene_gn=MINK1;refGene_tx=
XM_023243746.1
E1 1782265 1782335
refGene_gn=MINK1;refGene_tx=XM_023243731.1;refGene_gn=MINK1;refGene_tx
=XM_023243733.1;refGene_gn=MINK1;refGene_tx=XM_023243734.1;refGene_gn=
MINK1;refGene_tx=XM_023243735.1;refGene_gn=MINK1;refGene_tx=XM_0232437
36.1;refGene_gn=MINK1;refGene_tx=XM_023243737.1;refGene_gn=MINK1;refGe
ne_tx=XM_023243738.1;refGene_gn=MINK1;refGene_tx=XM_023243739.1;refGen
e_gn=MINK1;refGene_tx=XM_023243740.1;refGene_gn=MINK1;refGene_tx=XM_02
3243741.1;refGene_gn=MINK1;refGene_tx=XM_023243742.1;refGene_gn=MINK1;
refGene_tx=XM_023243743.1;refGene_gn=MINK1;refGene_tx=XM_023243744.1;r
efGene_gn=MINK1;refGene_tx=XM_023243745.1;refGene_gn=MINK1;refGene_tx=
XM_023243746.1
E1 1782689 1782865
refGene_gn=MINK1;refGene_tx=XM_023243731.1;refGene_gn=MINK1;refGene_tx
=XM_023243733.1;refGene_gn=MINK1;refGene_tx=XM_023243734.1;refGene_gn=
MINK1;refGene_tx=XM_023243735.1;refGene_gn=MINK1;refGene_tx=XM_0232437
36.1;refGene_gn=MINK1;refGene_tx=XM_023243737.1;refGene_gn=MINK1;refGe
ne_tx=XM_023243738.1;refGene_gn=MINK1;refGene_tx=XM_023243739.1;refGen
e_gn=MINK1;refGene_tx=XM_023243740.1;refGene_gn=MINK1;refGene_tx=XM_02
3243741.1;refGene_gn=MINK1;refGene_tx=XM_023243742.1;refGene_gn=MINK1;
refGene_tx=XM_023243743.1;refGene_gn=MINK1;refGene_tx=XM_023243744.1;r
efGene_gn=MINK1;refGene_tx=XM_023243745.1;refGene_gn=MINK1;refGene_tx=
XM_023243746.1
E1 1783280 1783359
refGene_gn=MINK1;refGene_tx=XM_023243731.1;refGene_gn=MINK1;refGene_tx

=XM_023243733.1;refGene_gn=MINK1;refGene_tx=XM_023243734.1;refGene_gn=MINK1;refGene_tx=XM_023243735.1;refGene_gn=MINK1;refGene_tx=XM_023243736.1;refGene_gn=MINK1;refGene_tx=XM_023243737.1;refGene_gn=MINK1;refGene_tx=XM_023243738.1;refGene_gn=MINK1;refGene_tx=XM_023243739.1;refGene_gn=MINK1;refGene_tx=XM_023243740.1;refGene_gn=MINK1;refGene_tx=XM_023243741.1;refGene_gn=MINK1;refGene_tx=XM_023243742.1;refGene_gn=MINK1;refGene_tx=XM_023243743.1;refGene_gn=MINK1;refGene_tx=XM_023243744.1;refGene_gn=MINK1;refGene_tx=XM_023243745.1;refGene_gn=MINK1;refGene_tx=XM_023243746.1
E1 1783463 1783518
refGene_gn=MINK1;refGene_tx=XM_023243731.1;refGene_gn=MINK1;refGene_tx=XM_023243733.1;refGene_gn=MINK1;refGene_tx=XM_023243734.1;refGene_gn=MINK1;refGene_tx=XM_023243735.1;refGene_gn=MINK1;refGene_tx=XM_023243736.1;refGene_gn=MINK1;refGene_tx=XM_023243737.1;refGene_gn=MINK1;refGene_tx=XM_023243738.1;refGene_gn=MINK1;refGene_tx=XM_023243739.1;refGene_gn=MINK1;refGene_tx=XM_023243740.1;refGene_gn=MINK1;refGene_tx=XM_023243741.1;refGene_gn=MINK1;refGene_tx=XM_023243742.1;refGene_gn=MINK1;refGene_tx=XM_023243743.1;refGene_gn=MINK1;refGene_tx=XM_023243744.1;refGene_gn=MINK1;refGene_tx=XM_023243745.1;refGene_gn=MINK1;refGene_tx=XM_023243746.1
E1 1783744 1783875
refGene_gn=MINK1;refGene_tx=XM_023243731.1;refGene_gn=MINK1;refGene_tx=XM_023243733.1;refGene_gn=MINK1;refGene_tx=XM_023243734.1;refGene_gn=MINK1;refGene_tx=XM_023243735.1;refGene_gn=MINK1;refGene_tx=XM_023243736.1;refGene_gn=MINK1;refGene_tx=XM_023243737.1;refGene_gn=MINK1;refGene_tx=XM_023243738.1;refGene_gn=MINK1;refGene_tx=XM_023243739.1;refGene_gn=MINK1;refGene_tx=XM_023243740.1;refGene_gn=MINK1;refGene_tx=XM_023243741.1;refGene_gn=MINK1;refGene_tx=XM_023243742.1;refGene_gn=MINK1;refGene_tx=XM_023243743.1;refGene_gn=MINK1;refGene_tx=XM_023243744.1;refGene_gn=MINK1;refGene_tx=XM_023243745.1;refGene_gn=MINK1;refGene_tx=XM_023243746.1
E1 1784279 1784370
refGene_gn=MINK1;refGene_tx=XM_023243731.1;refGene_gn=MINK1;refGene_tx=XM_023243733.1;refGene_gn=MINK1;refGene_tx=XM_023243734.1;refGene_gn=MINK1;refGene_tx=XM_023243735.1;refGene_gn=MINK1;refGene_tx=XM_023243736.1;refGene_gn=MINK1;refGene_tx=XM_023243737.1;refGene_gn=MINK1;refGene_tx=XM_023243738.1;refGene_gn=MINK1;refGene_tx=XM_023243739.1;refGene_gn=MINK1;refGene_tx=XM_023243740.1;refGene_gn=MINK1;refGene_tx=XM_023243741.1;refGene_gn=MINK1;refGene_tx=XM_023243742.1;refGene_gn=MINK1;refGene_tx=XM_023243743.1;refGene_gn=MINK1;refGene_tx=XM_023243744.1;refGene_gn=MINK1;refGene_tx=XM_023243745.1;refGene_gn=MINK1;refGene_tx=XM_023243746.1
E1 1784738 1784849
refGene_gn=MINK1;refGene_tx=XM_023243731.1;refGene_gn=MINK1;refGene_tx=XM_023243733.1;refGene_gn=MINK1;refGene_tx=XM_023243734.1;refGene_gn=MINK1;refGene_tx=XM_023243735.1;refGene_gn=MINK1;refGene_tx=XM_023243736.1;refGene_gn=MINK1;refGene_tx=XM_023243737.1;refGene_gn=MINK1;refGene_tx=XM_023243738.1;refGene_gn=MINK1;refGene_tx=XM_023243739.1;refGene_gn=MINK1;refGene_tx=XM_023243740.1;refGene_gn=MINK1;refGene_tx=XM_023243741.1;refGene_gn=MINK1;refGene_tx=XM_023243742.1;refGene_gn=MINK1;

refGene_tx=XM_023243743.1;refGene_gn=MINK1;refGene_tx=XM_023243744.1;refGene_gn=MINK1;refGene_tx=XM_023243745.1;refGene_gn=MINK1;refGene_tx=XM_023243746.1
E1 1786673 1786799 refGene_gn=MINK1;refGene_tx=XM_023243731.1;refGene_gn=MINK1;refGene_tx=XM_023243733.1;refGene_gn=MINK1;refGene_tx=XM_023243734.1;refGene_gn=MINK1;refGene_tx=XM_023243735.1;refGene_gn=MINK1;refGene_tx=XM_023243736.1;refGene_gn=MINK1;refGene_tx=XM_023243737.1;refGene_gn=MINK1;refGene_tx=XM_023243738.1;refGene_gn=MINK1;refGene_tx=XM_023243739.1;refGene_gn=MINK1;refGene_tx=XM_023243740.1;refGene_gn=MINK1;refGene_tx=XM_023243741.1;refGene_gn=MINK1;refGene_tx=XM_023243742.1;refGene_gn=MINK1;refGene_tx=XM_023243743.1;refGene_gn=MINK1;refGene_tx=XM_023243744.1;refGene_gn=MINK1;refGene_tx=XM_023243745.1;refGene_gn=MINK1;refGene_tx=XM_023243746.1
E1 1786880 1786937 refGene_gn=MINK1;refGene_tx=XM_023243731.1;refGene_gn=MINK1;refGene_tx=XM_023243733.1;refGene_gn=MINK1;refGene_tx=XM_023243734.1;refGene_gn=MINK1;refGene_tx=XM_023243735.1;refGene_gn=MINK1;refGene_tx=XM_023243736.1;refGene_gn=MINK1;refGene_tx=XM_023243737.1;refGene_gn=MINK1;refGene_tx=XM_023243738.1;refGene_gn=MINK1;refGene_tx=XM_023243739.1;refGene_gn=MINK1;refGene_tx=XM_023243740.1;refGene_gn=MINK1;refGene_tx=XM_023243741.1;refGene_gn=MINK1;refGene_tx=XM_023243742.1;refGene_gn=MINK1;refGene_tx=XM_023243743.1;refGene_gn=MINK1;refGene_tx=XM_023243744.1;refGene_gn=MINK1;refGene_tx=XM_023243745.1;refGene_gn=MINK1;refGene_tx=XM_023243746.1
E1 1788397 1788463 refGene_gn=MINK1;refGene_tx=XM_023243731.1;refGene_gn=MINK1;refGene_tx=XM_023243733.1;refGene_gn=MINK1;refGene_tx=XM_023243734.1;refGene_gn=MINK1;refGene_tx=XM_023243735.1;refGene_gn=MINK1;refGene_tx=XM_023243736.1;refGene_gn=MINK1;refGene_tx=XM_023243737.1;refGene_gn=MINK1;refGene_tx=XM_023243738.1;refGene_gn=MINK1;refGene_tx=XM_023243739.1;refGene_gn=MINK1;refGene_tx=XM_023243740.1;refGene_gn=MINK1;refGene_tx=XM_023243741.1;refGene_gn=MINK1;refGene_tx=XM_023243742.1;refGene_gn=MINK1;refGene_tx=XM_023243743.1;refGene_gn=MINK1;refGene_tx=XM_023243744.1;refGene_gn=MINK1;refGene_tx=XM_023243745.1;refGene_gn=MINK1;refGene_tx=XM_023243746.1
E1 1816654 1816711 refGene_gn=MINK1;refGene_tx=XM_023243731.1;refGene_gn=MINK1;refGene_tx=XM_023243733.1;refGene_gn=MINK1;refGene_tx=XM_023243734.1;refGene_gn=MINK1;refGene_tx=XM_023243735.1;refGene_gn=MINK1;refGene_tx=XM_023243736.1;refGene_gn=MINK1;refGene_tx=XM_023243737.1;refGene_gn=MINK1;refGene_tx=XM_023243738.1;refGene_gn=MINK1;refGene_tx=XM_023243739.1;refGene_gn=MINK1;refGene_tx=XM_023243740.1;refGene_gn=MINK1;refGene_tx=XM_023243741.1;refGene_gn=MINK1;refGene_tx=XM_023243742.1;refGene_gn=MINK1;refGene_tx=XM_023243743.1;refGene_gn=MINK1;refGene_tx=XM_023243744.1;refGene_gn=MINK1;refGene_tx=XM_023243745.1;refGene_gn=MINK1;refGene_tx=XM_023243746.1
E1 1823278 1823503 refGene_gn=PLD2;refGene_tx=XM_023243748.1
E1 1823738 1823853 refGene_gn=PLD2;refGene_tx=XM_023243748.1
E1 1824598 1824752 refGene_gn=PLD2;refGene_tx=XM_023243748.1

E1	1824935	1825070	refGene_gn=PLD2;refGene_tx=XM_023243748.1
E1	1825261	1825311	refGene_gn=PLD2;refGene_tx=XM_023243748.1
E1	1825486	1825600	refGene_gn=PLD2;refGene_tx=XM_023243748.1
E1	1825711	1825800	refGene_gn=PLD2;refGene_tx=XM_023243748.1
E1	1826006	1826111	refGene_gn=PLD2;refGene_tx=XM_023243748.1
E1	1826228	1826342	refGene_gn=PLD2;refGene_tx=XM_023243748.1
E1	1826425	1826544	refGene_gn=PLD2;refGene_tx=XM_023243748.1
E1	1826703	1826830	refGene_gn=PLD2;refGene_tx=XM_023243748.1
E1	1827180	1827351	refGene_gn=PLD2;refGene_tx=XM_023243748.1
E1	1827781	1827892	refGene_gn=PLD2;refGene_tx=XM_023243748.1
E1	1828588	1828667	refGene_gn=PLD2;refGene_tx=XM_023243748.1
E1	1828799	1828883	refGene_gn=PLD2;refGene_tx=XM_023243748.1
E1	1831724	1831874	refGene_gn=PLD2;refGene_tx=XM_023243748.1
E1	1832369	1832522	refGene_gn=PLD2;refGene_tx=XM_023243748.1
E1	1832613	1832706	refGene_gn=PLD2;refGene_tx=XM_023243748.1
E1	1832877	1832936	refGene_gn=PLD2;refGene_tx=XM_023243748.1
E1	1833066	1833132	refGene_gn=PLD2;refGene_tx=XM_023243748.1
E1	1833221	1833327	refGene_gn=PLD2;refGene_tx=XM_023243748.1
E1	1833712	1833855	refGene_gn=PLD2;refGene_tx=XM_023243748.1
E1	1833969	1834100	refGene_gn=PLD2;refGene_tx=XM_023243748.1
E1	1834191	1834300	refGene_gn=PLD2;refGene_tx=XM_023243748.1
E1	1843723	1843866	refGene_gn=PSMB6;refGene_tx=XM_003996148.4
E1	1843991	1844136	refGene_gn=PSMB6;refGene_tx=XM_003996148.4
E1	1844334	1844464	refGene_gn=PSMB6;refGene_tx=XM_003996148.4
E1	1844579	1844711	refGene_gn=PSMB6;refGene_tx=XM_003996148.4
E1	1845043	1845111	refGene_gn=PSMB6;refGene_tx=XM_003996148.4
E1	1845531	1845633	refGene_gn=PSMB6;refGene_tx=XM_003996148.4
E1	1851546	1852084
refGene_gn=GLTPD2;refGene_tx=XM_023243798.1;refGene_gn=GLTPD2;refGene_
tx=XM_023243799.1
E1	1852172	1852339
refGene_gn=GLTPD2;refGene_tx=XM_023243798.1;refGene_gn=GLTPD2;refGene_
tx=XM_023243799.1
E1	1852597	1852876
refGene_gn=GLTPD2;refGene_tx=XM_023243798.1;refGene_gn=GLTPD2;refGene_
tx=XM_023243799.1
E1	1854696	1854888	refGene_gn=VMO1;refGene_tx=XM_003996152.5
E1	1855017	1855133	refGene_gn=VMO1;refGene_tx=XM_003996152.5
E1	1855402	1855700	refGene_gn=VMO1;refGene_tx=XM_003996152.5
E1	1856481	1856962
refGene_gn=TM4SF5;refGene_tx=XM_023243800.1;refGene_gn=TM4SF5;refGene_
tx=XM_023243801.1;refGene_gn=TM4SF5;refGene_tx=XM_023243802.1
E1	1857169	1857306
refGene_gn=TM4SF5;refGene_tx=XM_023243800.1;refGene_gn=TM4SF5;refGene_
tx=XM_023243801.1;refGene_gn=TM4SF5;refGene_tx=XM_023243802.1
E1	1857644	1857725
refGene_gn=TM4SF5;refGene_tx=XM_023243800.1;refGene_gn=TM4SF5;refGene_
tx=XM_023243801.1;refGene_gn=TM4SF5;refGene_tx=XM_023243802.1
E1	1862775	1862949
refGene_gn=TM4SF5;refGene_tx=XM_023243800.1;refGene_gn=TM4SF5;refGene_

tx=XM_023243801.1;refGene_gn=TM4SF5;refGene_tx=XM_023243802.1
E1 1879754 1879926 refGene_gn=ZMYND15;refGene_tx=XM_023243618.1
E1 1880123 1880284 refGene_gn=ZMYND15;refGene_tx=XM_023243618.1
E1 1880545 1880604 refGene_gn=ZMYND15;refGene_tx=XM_023243618.1
E1 1880685 1880839 refGene_gn=ZMYND15;refGene_tx=XM_023243618.1
E1 1880971 1881070 refGene_gn=ZMYND15;refGene_tx=XM_023243618.1
E1 1881151 1881240 refGene_gn=ZMYND15;refGene_tx=XM_023243618.1
E1 1881351 1881468 refGene_gn=ZMYND15;refGene_tx=XM_023243618.1
E1 1881613 1881694 refGene_gn=ZMYND15;refGene_tx=XM_023243618.1
E1 1882009 1882162 refGene_gn=ZMYND15;refGene_tx=XM_023243618.1
E1 1882610 1882771 refGene_gn=ZMYND15;refGene_tx=XM_023243618.1
E1 1883024 1883180 refGene_gn=ZMYND15;refGene_tx=XM_023243618.1
E1 1883278 1883513 refGene_gn=ZMYND15;refGene_tx=XM_023243618.1
E1 1883939 1884528 refGene_gn=ZMYND15;refGene_tx=XM_023243618.1
E1 1885568 1885635 refGene_gn=CXCL16;refGene_tx=XM_003996243.5
E1 1885993 1886132 refGene_gn=CXCL16;refGene_tx=XM_003996243.5
E1 1886398 1886481 refGene_gn=CXCL16;refGene_tx=XM_003996243.5
E1 1887350 1887770 refGene_gn=CXCL16;refGene_tx=XM_003996243.5
E1 1888394 1888441 refGene_gn=CXCL16;refGene_tx=XM_003996243.5
E1 1889361 1889499 refGene_gn=MED11;refGene_tx=XM_003996154.5
E1 1890144 1890275 refGene_gn=MED11;refGene_tx=XM_003996154.5
E1 1890485 1890570 refGene_gn=MED11;refGene_tx=XM_003996154.5
E1 1899385 1899479
refGene_gn=ARRB2;refGene_tx=XM_023243688.1;refGene_gn=ARRB2;refGene_tx
=XM_023243690.1
E1 1899771 1899826
refGene_gn=ARRB2;refGene_tx=XM_023243688.1;refGene_gn=ARRB2;refGene_tx
=XM_023243689.1;refGene_gn=ARRB2;refGene_tx=XM_023243690.1
E1 1899936 1900016
refGene_gn=ARRB2;refGene_tx=XM_023243688.1;refGene_gn=ARRB2;refGene_tx
=XM_023243689.1;refGene_gn=ARRB2;refGene_tx=XM_023243690.1
E1 1900116 1900200
refGene_gn=ARRB2;refGene_tx=XM_023243688.1;refGene_gn=ARRB2;refGene_tx
=XM_023243689.1;refGene_gn=ARRB2;refGene_tx=XM_023243690.1
E1 1901009 1901147
refGene_gn=ARRB2;refGene_tx=XM_023243688.1;refGene_gn=ARRB2;refGene_tx
=XM_023243689.1;refGene_gn=ARRB2;refGene_tx=XM_023243690.1
E1 1901423 1901496
refGene_gn=ARRB2;refGene_tx=XM_023243688.1;refGene_gn=ARRB2;refGene_tx
=XM_023243689.1;refGene_gn=ARRB2;refGene_tx=XM_023243690.1
E1 1901667 1901752
refGene_gn=ARRB2;refGene_tx=XM_023243688.1;refGene_gn=ARRB2;refGene_tx
=XM_023243689.1;refGene_gn=ARRB2;refGene_tx=XM_023243690.1
E1 1901948 1902084
refGene_gn=ARRB2;refGene_tx=XM_023243688.1;refGene_gn=ARRB2;refGene_tx
=XM_023243689.1;refGene_gn=ARRB2;refGene_tx=XM_023243690.1
E1 1902230 1902298
refGene_gn=ARRB2;refGene_tx=XM_023243688.1;refGene_gn=ARRB2;refGene_tx
=XM_023243689.1;refGene_gn=ARRB2;refGene_tx=XM_023243690.1
E1 1902678 1902738

refGene_gn=ARRB2;refGene_tx=XM_023243688.1;refGene_gn=ARRB2;refGene_tx
=XM_023243689.1;refGene_gn=ARRB2;refGene_tx=XM_023243690.1
E1 1903027 1903224
refGene_gn=ARRB2;refGene_tx=XM_023243688.1;refGene_gn=ARRB2;refGene_tx
=XM_023243689.1;refGene_gn=ARRB2;refGene_tx=XM_023243690.1
E1 1903407 1903452
refGene_gn=ARRB2;refGene_tx=XM_023243688.1;refGene_gn=ARRB2;refGene_tx
=XM_023243689.1;refGene_gn=ARRB2;refGene_tx=XM_023243690.1
E1 1903557 1903618
refGene_gn=ARRB2;refGene_tx=XM_023243688.1;refGene_gn=ARRB2;refGene_tx
=XM_023243689.1;refGene_gn=ARRB2;refGene_tx=XM_023243690.1
E1 1904297 1904328
refGene_gn=ARRB2;refGene_tx=XM_023243688.1;refGene_gn=ARRB2;refGene_tx
=XM_023243689.1;refGene_gn=ARRB2;refGene_tx=XM_023243690.1
E1 1906352 1906633
refGene_gn=ARRB2;refGene_tx=XM_023243688.1;refGene_gn=ARRB2;refGene_tx
=XM_023243689.1
E1 1906917 1906940 refGene_gn=ARRB2;refGene_tx=XM_023243690.1
E1 1913075 1913324 refGene_gn=PELP1;refGene_tx=XM_023243559.1
E1 1922875 1922940 refGene_gn=PELP1;refGene_tx=XM_023243559.1
E1 1923322 1923428 refGene_gn=PELP1;refGene_tx=XM_023243559.1
E1 1927318 1927468
refGene_gn=PELP1;refGene_tx=XM_019817066.2;refGene_gn=PELP1;refGene_tx
=XM_023243559.1
E1 1927639 1927711
refGene_gn=PELP1;refGene_tx=XM_019817066.2;refGene_gn=PELP1;refGene_tx
=XM_023243559.1
E1 1931420 1931480
refGene_gn=PELP1;refGene_tx=XM_019817066.2;refGene_gn=PELP1;refGene_tx
=XM_023243559.1
E1 1931736 1931887
refGene_gn=PELP1;refGene_tx=XM_019817066.2;refGene_gn=PELP1;refGene_tx
=XM_023243559.1
E1 1932032 1932162
refGene_gn=PELP1;refGene_tx=XM_019817066.2;refGene_gn=PELP1;refGene_tx
=XM_023243559.1
E1 1932282 1932370
refGene_gn=PELP1;refGene_tx=XM_019817066.2;refGene_gn=PELP1;refGene_tx
=XM_023243559.1
E1 1932755 1932844
refGene_gn=PELP1;refGene_tx=XM_019817066.2;refGene_gn=PELP1;refGene_tx
=XM_023243559.1
E1 1932935 1933052
refGene_gn=PELP1;refGene_tx=XM_019817066.2;refGene_gn=PELP1;refGene_tx
=XM_023243559.1
E1 1933153 1933301
refGene_gn=PELP1;refGene_tx=XM_019817066.2;refGene_gn=PELP1;refGene_tx
=XM_023243559.1
E1 1933405 1933565
refGene_gn=PELP1;refGene_tx=XM_019817066.2;refGene_gn=PELP1;refGene_tx

=XM_023243559.1
E1	1934160	1934216
refGene_gn=PELP1;refGene_tx=XM_019817066.2;refGene_gn=PELP1;refGene_tx
=XM_023243559.1
E1	1934334	1934541
refGene_gn=PELP1;refGene_tx=XM_019817066.2;refGene_gn=PELP1;refGene_tx
=XM_023243559.1
E1	1934672	1936124
refGene_gn=PELP1;refGene_tx=XM_019817066.2;refGene_gn=PELP1;refGene_tx
=XM_023243559.1
E1	1936223	1936316
refGene_gn=PELP1;refGene_tx=XM_019817066.2;refGene_gn=PELP1;refGene_tx
=XM_023243559.1
E1	1962474	1962561
refGene_gn=LOC102901906;refGene_tx=XM_023243567.1
E1	1962649	1963378
refGene_gn=LOC102901906;refGene_tx=XM_023243567.1
E1	2031087	2031222
refGene_gn=LOC101094304;refGene_tx=XM_023243560.1;refGene_gn=LOC101094
304;refGene_tx=XM_023243561.1;refGene_gn=LOC101094304;refGene_tx=XM_02
3243562.1;refGene_gn=LOC101094304;refGene_tx=XM_023243563.1;refGene_gn
=LOC101094304;refGene_tx=XM_023243564.1;refGene_gn=LOC101094304;refGen
e_tx=XM_023243565.1;refGene_gn=LOC101094304;refGene_tx=XM_023243566.1
E1	2031723	2031928
refGene_gn=LOC101094304;refGene_tx=XM_023243560.1;refGene_gn=LOC101094
304;refGene_tx=XM_023243561.1;refGene_gn=LOC101094304;refGene_tx=XM_02
3243562.1;refGene_gn=LOC101094304;refGene_tx=XM_023243563.1;refGene_gn
=LOC101094304;refGene_tx=XM_023243564.1;refGene_gn=LOC101094304;refGen
e_tx=XM_023243565.1;refGene_gn=LOC101094304;refGene_tx=XM_023243566.1
E1	2032971	2033053
refGene_gn=LOC101094304;refGene_tx=XM_023243560.1;refGene_gn=LOC101094
304;refGene_tx=XM_023243561.1;refGene_gn=LOC101094304;refGene_tx=XM_02
3243562.1;refGene_gn=LOC101094304;refGene_tx=XM_023243563.1;refGene_gn
=LOC101094304;refGene_tx=XM_023243564.1;refGene_gn=LOC101094304;refGen
e_tx=XM_023243565.1;refGene_gn=LOC101094304;refGene_tx=XM_023243566.1
E1	2033134	2033257
refGene_gn=LOC101094304;refGene_tx=XM_023243560.1;refGene_gn=LOC101094
304;refGene_tx=XM_023243561.1;refGene_gn=LOC101094304;refGene_tx=XM_02
3243562.1;refGene_gn=LOC101094304;refGene_tx=XM_023243563.1;refGene_gn
=LOC101094304;refGene_tx=XM_023243564.1;refGene_gn=LOC101094304;refGen
e_tx=XM_023243565.1;refGene_gn=LOC101094304;refGene_tx=XM_023243566.1
E1	2033355	2033459
refGene_gn=LOC101094304;refGene_tx=XM_023243560.1;refGene_gn=LOC101094
304;refGene_tx=XM_023243561.1;refGene_gn=LOC101094304;refGene_tx=XM_02
3243562.1;refGene_gn=LOC101094304;refGene_tx=XM_023243563.1;refGene_gn
=LOC101094304;refGene_tx=XM_023243564.1;refGene_gn=LOC101094304;refGen
e_tx=XM_023243565.1;refGene_gn=LOC101094304;refGene_tx=XM_023243566.1
E1	2033706	2033867
refGene_gn=LOC101094304;refGene_tx=XM_023243560.1;refGene_gn=LOC101094
304;refGene_tx=XM_023243561.1;refGene_gn=LOC101094304;refGene_tx=XM_02

3243562.1;refGene_gn=LOC101094304;refGene_tx=XM_023243563.1;refGene_gn
=LOC101094304;refGene_tx=XM_023243564.1;refGene_gn=LOC101094304;refGen
e_tx=XM_023243565.1;refGene_gn=LOC101094304;refGene_tx=XM_023243566.1
E1 2034684 2034828
refGene_gn=LOC101094304;refGene_tx=XM_023243560.1;refGene_gn=LOC101094
304;refGene_tx=XM_023243561.1;refGene_gn=LOC101094304;refGene_tx=XM_02
3243562.1;refGene_gn=LOC101094304;refGene_tx=XM_023243563.1;refGene_gn
=LOC101094304;refGene_tx=XM_023243564.1;refGene_gn=LOC101094304;refGen
e_tx=XM_023243565.1;refGene_gn=LOC101094304;refGene_tx=XM_023243566.1
E1 2035578 2035788
refGene_gn=LOC101094304;refGene_tx=XM_023243560.1;refGene_gn=LOC101094
304;refGene_tx=XM_023243561.1;refGene_gn=LOC101094304;refGene_tx=XM_02
3243562.1;refGene_gn=LOC101094304;refGene_tx=XM_023243563.1;refGene_gn
=LOC101094304;refGene_tx=XM_023243564.1;refGene_gn=LOC101094304;refGen
e_tx=XM_023243565.1;refGene_gn=LOC101094304;refGene_tx=XM_023243566.1
E1 2037989 2038076
refGene_gn=LOC101094304;refGene_tx=XM_023243560.1;refGene_gn=LOC101094
304;refGene_tx=XM_023243561.1;refGene_gn=LOC101094304;refGene_tx=XM_02
3243562.1;refGene_gn=LOC101094304;refGene_tx=XM_023243563.1;refGene_gn
=LOC101094304;refGene_tx=XM_023243564.1;refGene_gn=LOC101094304;refGen
e_tx=XM_023243565.1;refGene_gn=LOC101094304;refGene_tx=XM_023243566.1
E1 2038290 2038460
refGene_gn=LOC101094304;refGene_tx=XM_023243560.1;refGene_gn=LOC101094
304;refGene_tx=XM_023243561.1;refGene_gn=LOC101094304;refGene_tx=XM_02
3243562.1;refGene_gn=LOC101094304;refGene_tx=XM_023243563.1;refGene_gn
=LOC101094304;refGene_tx=XM_023243564.1;refGene_gn=LOC101094304;refGen
e_tx=XM_023243565.1;refGene_gn=LOC101094304;refGene_tx=XM_023243566.1
E1 2039229 2039351
refGene_gn=LOC101094304;refGene_tx=XM_023243560.1;refGene_gn=LOC101094
304;refGene_tx=XM_023243561.1;refGene_gn=LOC101094304;refGene_tx=XM_02
3243562.1;refGene_gn=LOC101094304;refGene_tx=XM_023243563.1;refGene_gn
=LOC101094304;refGene_tx=XM_023243564.1;refGene_gn=LOC101094304;refGen
e_tx=XM_023243565.1;refGene_gn=LOC101094304;refGene_tx=XM_023243566.1
E1 2039439 2039702
refGene_gn=LOC101094304;refGene_tx=XM_023243562.1;refGene_gn=LOC101094
304;refGene_tx=XM_023243563.1;refGene_gn=LOC101094304;refGene_tx=XM_02
3243564.1
E1 2040549 2040554
refGene_gn=LOC101094304;refGene_tx=XM_023243566.1
E1 2041423 2041530
refGene_gn=LOC101094304;refGene_tx=XM_023243560.1;refGene_gn=LOC101094
304;refGene_tx=XM_023243561.1;refGene_gn=LOC101094304;refGene_tx=XM_02
3243565.1
E1 2041644 2041815
refGene_gn=LOC101094304;refGene_tx=XM_023243560.1;refGene_gn=LOC101094
304;refGene_tx=XM_023243561.1
E1 2041919 2042099
refGene_gn=LOC101094304;refGene_tx=XM_023243560.1;refGene_gn=LOC101094
304;refGene_tx=XM_023243561.1
E1 2044136 2044379 refGene_gn=RNASEK;refGene_tx=XM_003996247.3

E1 2045040 2045117 refGene_gn=RNASEK;refGene_tx=XM_003996247.3
E1 2045537 2045679 refGene_gn=RNASEK;refGene_tx=XM_003996247.3
E1 2046209 2046230
refGene_gn=CE1H17orf49;refGene_tx=XM_023243568.1;refGene_gn=CE1H17orf4
9;refGene_tx=XM_023243569.1;refGene_gn=CE1H17orf49;refGene_tx=XM_02324
3570.1
E1 2046425 2046520
refGene_gn=CE1H17orf49;refGene_tx=XM_023243568.1;refGene_gn=CE1H17orf4
9;refGene_tx=XM_023243569.1;refGene_gn=CE1H17orf49;refGene_tx=XM_02324
3570.1
E1 2046960 2047062
refGene_gn=CE1H17orf49;refGene_tx=XM_023243568.1;refGene_gn=CE1H17orf4
9;refGene_tx=XM_023243569.1
E1 2047687 2047878
refGene_gn=CE1H17orf49;refGene_tx=XM_023243568.1;refGene_gn=CE1H17orf4
9;refGene_tx=XM_023243569.1;refGene_gn=CE1H17orf49;refGene_tx=XM_02324
3570.1
E1 2048077 2048182
refGene_gn=CE1H17orf49;refGene_tx=XM_023243568.1;refGene_gn=CE1H17orf4
9;refGene_tx=XM_023243569.1;refGene_gn=CE1H17orf49;refGene_tx=XM_02324
3570.1
E1 2048441 2048506
refGene_gn=CE1H17orf49;refGene_tx=XM_023243568.1;refGene_gn=CE1H17orf4
9;refGene_tx=XM_023243569.1;refGene_gn=CE1H17orf49;refGene_tx=XM_02324
3570.1
E1 2054382 2054385 refGene_gn=BCL6B;refGene_tx=XM_011288886.3
E1 2054876 2055067
refGene_gn=BCL6B;refGene_tx=XM_011288886.3;refGene_gn=BCL6B;refGene_tx
=XM_011288887.3
E1 2055260 2055482
refGene_gn=BCL6B;refGene_tx=XM_011288886.3;refGene_gn=BCL6B;refGene_tx
=XM_011288887.3
E1 2055583 2055916
refGene_gn=BCL6B;refGene_tx=XM_011288886.3;refGene_gn=BCL6B;refGene_tx
=XM_011288887.3
E1 2056174 2056299
refGene_gn=BCL6B;refGene_tx=XM_011288886.3;refGene_gn=BCL6B;refGene_tx
=XM_011288887.3
E1 2056863 2057028
refGene_gn=BCL6B;refGene_tx=XM_011288886.3;refGene_gn=BCL6B;refGene_tx
=XM_011288887.3
E1 2057126 2057257
refGene_gn=BCL6B;refGene_tx=XM_011288886.3;refGene_gn=BCL6B;refGene_tx
=XM_011288887.3
E1 2057368 2057506
refGene_gn=BCL6B;refGene_tx=XM_011288886.3;refGene_gn=BCL6B;refGene_tx
=XM_011288887.3
E1 2057898 2058015
refGene_gn=BCL6B;refGene_tx=XM_011288886.3;refGene_gn=BCL6B;refGene_tx
=XM_011288887.3

E1 2063019 2063218 refGene_gn=SLC16A13;refGene_tx=XM_003996156.3
E1 2063367 2063511 refGene_gn=SLC16A13;refGene_tx=XM_003996156.3
E1 2064125 2064863 refGene_gn=SLC16A13;refGene_tx=XM_003996156.3
E1 2065522 2065722 refGene_gn=SLC16A13;refGene_tx=XM_003996156.3
E1 2066721 2066957
refGene_gn=SLC16A11;refGene_tx=XM_023243674.1;refGene_gn=SLC16A11;refGene_tx=XM_023243675.1
E1 2067067 2068094
refGene_gn=SLC16A11;refGene_tx=XM_023243674.1;refGene_gn=SLC16A11;refGene_tx=XM_023243675.1
E1 2087838 2087992
refGene_gn=CLEC10A;refGene_tx=XM_023243843.1;refGene_gn=CLEC10A;refGene_tx=XM_023243844.1
E1 2088119 2088226
refGene_gn=CLEC10A;refGene_tx=XM_023243843.1;refGene_gn=CLEC10A;refGene_tx=XM_023243844.1
E1 2088483 2088635
refGene_gn=CLEC10A;refGene_tx=XM_023243843.1;refGene_gn=CLEC10A;refGene_tx=XM_023243844.1
E1 2088725 2088812
refGene_gn=CLEC10A;refGene_tx=XM_023243843.1;refGene_gn=CLEC10A;refGene_tx=XM_023243844.1
E1 2088966 2089038
refGene_gn=CLEC10A;refGene_tx=XM_023243843.1;refGene_gn=CLEC10A;refGene_tx=XM_023243844.1
E1 2089137 2089209
refGene_gn=CLEC10A;refGene_tx=XM_023243843.1;refGene_gn=CLEC10A;refGene_tx=XM_023243844.1
E1 2089307 2089403
refGene_gn=CLEC10A;refGene_tx=XM_023243843.1;refGene_gn=CLEC10A;refGene_tx=XM_023243844.1
E1 2089689 2089806
refGene_gn=CLEC10A;refGene_tx=XM_023243843.1;refGene_gn=CLEC10A;refGene_tx=XM_023243844.1
E1 2090372 2090439
refGene_gn=CLEC10A;refGene_tx=XM_023243843.1;refGene_gn=CLEC10A;refGene_tx=XM_023243844.1
E1 2103000 2103157
refGene_gn=ASGR2;refGene_tx=XM_023243845.1;refGene_gn=ASGR2;refGene_tx=XM_023243846.1
E1 2103396 2103503
refGene_gn=ASGR2;refGene_tx=XM_023243845.1;refGene_gn=ASGR2;refGene_tx=XM_023243846.1
E1 2107837 2107983
refGene_gn=ASGR2;refGene_tx=XM_023243845.1;refGene_gn=ASGR2;refGene_tx=XM_023243846.1
E1 2108069 2108156
refGene_gn=ASGR2;refGene_tx=XM_023243845.1;refGene_gn=ASGR2;refGene_tx=XM_023243846.1
E1 2108323 2108395

refGene_gn=ASGR2;refGene_tx=XM_023243845.1;refGene_gn=ASGR2;refGene_tx=XM_023243846.1
E1 2108789 2108885
refGene_gn=ASGR2;refGene_tx=XM_023243845.1;refGene_gn=ASGR2;refGene_tx=XM_023243846.1
E1 2108991 2109111
refGene_gn=ASGR2;refGene_tx=XM_023243845.1;refGene_gn=ASGR2;refGene_tx=XM_023243846.1
E1 2110517 2110617 refGene_gn=ASGR2;refGene_tx=XM_023243846.1
E1 2112820 2112944 refGene_gn=ASGR2;refGene_tx=XM_023243845.1
E1 2143874 2144031
refGene_gn=LOC102902281;refGene_tx=XM_023244113.1;refGene_gn=LOC102902281;refGene_tx=XM_023244114.1;refGene_gn=LOC102902281;refGene_tx=XM_023244115.1;refGene_gn=LOC102902281;refGene_tx=XM_023244116.1
E1 2144123 2144230
refGene_gn=LOC102902281;refGene_tx=XM_023244113.1;refGene_gn=LOC102902281;refGene_tx=XM_023244114.1;refGene_gn=LOC102902281;refGene_tx=XM_023244115.1;refGene_gn=LOC102902281;refGene_tx=XM_023244116.1
E1 2144353 2144505
refGene_gn=LOC102902281;refGene_tx=XM_023244113.1;refGene_gn=LOC102902281;refGene_tx=XM_023244114.1;refGene_gn=LOC102902281;refGene_tx=XM_023244115.1;refGene_gn=LOC102902281;refGene_tx=XM_023244116.1
E1 2144602 2144689
refGene_gn=LOC102902281;refGene_tx=XM_023244113.1;refGene_gn=LOC102902281;refGene_tx=XM_023244114.1;refGene_gn=LOC102902281;refGene_tx=XM_023244115.1;refGene_gn=LOC102902281;refGene_tx=XM_023244116.1
E1 2144874 2144946
refGene_gn=LOC102902281;refGene_tx=XM_023244113.1;refGene_gn=LOC102902281;refGene_tx=XM_023244114.1;refGene_gn=LOC102902281;refGene_tx=XM_023244115.1;refGene_gn=LOC102902281;refGene_tx=XM_023244116.1
E1 2145031 2145127
refGene_gn=LOC102902281;refGene_tx=XM_023244113.1;refGene_gn=LOC102902281;refGene_tx=XM_023244114.1;refGene_gn=LOC102902281;refGene_tx=XM_023244115.1;refGene_gn=LOC102902281;refGene_tx=XM_023244116.1
E1 2145246 2145363
refGene_gn=LOC102902281;refGene_tx=XM_023244113.1;refGene_gn=LOC102902281;refGene_tx=XM_023244114.1;refGene_gn=LOC102902281;refGene_tx=XM_023244115.1;refGene_gn=LOC102902281;refGene_tx=XM_023244116.1
E1 2146471 2146697
refGene_gn=LOC102902281;refGene_tx=XM_023244113.1;refGene_gn=LOC102902281;refGene_tx=XM_023244114.1;refGene_gn=LOC102902281;refGene_tx=XM_023244115.1;refGene_gn=LOC102902281;refGene_tx=XM_023244116.1
E1 2158716 2158823
refGene_gn=DLG4;refGene_tx=XM_011288883.3;refGene_gn=DLG4;refGene_tx=XM_011288884.3;refGene_gn=DLG4;refGene_tx=XM_011288885.3;refGene_gn=DLG4;refGene_tx=XM_019817072.2;refGene_gn=DLG4;refGene_tx=XM_019817076.2;refGene_gn=DLG4;refGene_tx=XM_019817078.2;refGene_gn=DLG4;refGene_tx=XM_019817080.2;refGene_gn=DLG4;refGene_tx=XM_019817081.2;refGene_gn=DLG4;refGene_tx=XM_019817082.2;refGene_gn=DLG4;refGene_tx=XM_019817083.2;refGene_gn=DLG4;refGene_tx=XM_019817084.2;refGene_gn=DLG4;refGene_tx=X

M_019817086.2;refGene_gn=DLG4;refGene_tx=XM_023243650.1;refGene_gn=DLG4;refGene_tx=XM_023243651.1;refGene_gn=DLG4;refGene_tx=XM_023243652.1;refGene_gn=DLG4;refGene_tx=XM_023243653.1;refGene_gn=DLG4;refGene_tx=XM_023243655.1
E1 2158993 2159085
refGene_gn=DLG4;refGene_tx=XM_011288883.3;refGene_gn=DLG4;refGene_tx=XM_011288884.3;refGene_gn=DLG4;refGene_tx=XM_011288885.3;refGene_gn=DLG4;refGene_tx=XM_019817072.2;refGene_gn=DLG4;refGene_tx=XM_019817076.2;refGene_gn=DLG4;refGene_tx=XM_019817078.2;refGene_gn=DLG4;refGene_tx=XM_019817080.2;refGene_gn=DLG4;refGene_tx=XM_019817081.2;refGene_gn=DLG4;refGene_tx=XM_019817082.2;refGene_gn=DLG4;refGene_tx=XM_019817083.2;refGene_gn=DLG4;refGene_tx=XM_019817084.2;refGene_gn=DLG4;refGene_tx=XM_019817086.2;refGene_gn=DLG4;refGene_tx=XM_023243650.1;refGene_gn=DLG4;refGene_tx=XM_023243651.1;refGene_gn=DLG4;refGene_tx=XM_023243652.1;refGene_gn=DLG4;refGene_tx=XM_023243653.1;refGene_gn=DLG4;refGene_tx=XM_023243655.1
E1 2159582 2159692
refGene_gn=DLG4;refGene_tx=XM_011288883.3;refGene_gn=DLG4;refGene_tx=XM_011288884.3;refGene_gn=DLG4;refGene_tx=XM_011288885.3;refGene_gn=DLG4;refGene_tx=XM_019817072.2;refGene_gn=DLG4;refGene_tx=XM_019817076.2;refGene_gn=DLG4;refGene_tx=XM_019817078.2;refGene_gn=DLG4;refGene_tx=XM_019817080.2;refGene_gn=DLG4;refGene_tx=XM_019817081.2;refGene_gn=DLG4;refGene_tx=XM_019817082.2;refGene_gn=DLG4;refGene_tx=XM_019817083.2;refGene_gn=DLG4;refGene_tx=XM_019817084.2;refGene_gn=DLG4;refGene_tx=XM_019817086.2;refGene_gn=DLG4;refGene_tx=XM_023243650.1;refGene_gn=DLG4;refGene_tx=XM_023243651.1;refGene_gn=DLG4;refGene_tx=XM_023243652.1;refGene_gn=DLG4;refGene_tx=XM_023243653.1;refGene_gn=DLG4;refGene_tx=XM_023243655.1
E1 2160605 2160778
refGene_gn=DLG4;refGene_tx=XM_011288883.3;refGene_gn=DLG4;refGene_tx=XM_011288884.3;refGene_gn=DLG4;refGene_tx=XM_011288885.3;refGene_gn=DLG4;refGene_tx=XM_019817072.2;refGene_gn=DLG4;refGene_tx=XM_019817076.2;refGene_gn=DLG4;refGene_tx=XM_019817078.2;refGene_gn=DLG4;refGene_tx=XM_019817080.2;refGene_gn=DLG4;refGene_tx=XM_019817081.2;refGene_gn=DLG4;refGene_tx=XM_019817082.2;refGene_gn=DLG4;refGene_tx=XM_019817083.2;refGene_gn=DLG4;refGene_tx=XM_019817084.2;refGene_gn=DLG4;refGene_tx=XM_019817086.2;refGene_gn=DLG4;refGene_tx=XM_023243650.1;refGene_gn=DLG4;refGene_tx=XM_023243651.1;refGene_gn=DLG4;refGene_tx=XM_023243652.1;refGene_gn=DLG4;refGene_tx=XM_023243653.1;refGene_gn=DLG4;refGene_tx=XM_023243655.1
E1 2161058 2161160
refGene_gn=DLG4;refGene_tx=XM_011288883.3;refGene_gn=DLG4;refGene_tx=XM_011288884.3;refGene_gn=DLG4;refGene_tx=XM_011288885.3;refGene_gn=DLG4;refGene_tx=XM_019817072.2;refGene_gn=DLG4;refGene_tx=XM_019817076.2;refGene_gn=DLG4;refGene_tx=XM_019817078.2;refGene_gn=DLG4;refGene_tx=XM_019817080.2;refGene_gn=DLG4;refGene_tx=XM_019817081.2;refGene_gn=DLG4;refGene_tx=XM_019817082.2;refGene_gn=DLG4;refGene_tx=XM_019817083.2;refGene_gn=DLG4;refGene_tx=XM_019817084.2;refGene_gn=DLG4;refGene_tx=XM_019817086.2;refGene_gn=DLG4;refGene_tx=XM_023243650.1;refGene_gn=DLG4;refGene_tx=XM_023243651.1;refGene_gn=DLG4;refGene_tx=XM_023243652.1;

refGene_gn=DLG4;refGene_tx=XM_023243653.1;refGene_gn=DLG4;refGene_tx=XM_023243655.1
E1 2161953 2162001
refGene_gn=DLG4;refGene_tx=XM_011288883.3;refGene_gn=DLG4;refGene_tx=XM_011288884.3;refGene_gn=DLG4;refGene_tx=XM_011288885.3;refGene_gn=DLG4;refGene_tx=XM_019817072.2;refGene_gn=DLG4;refGene_tx=XM_019817076.2;refGene_gn=DLG4;refGene_tx=XM_019817078.2;refGene_gn=DLG4;refGene_tx=XM_019817080.2;refGene_gn=DLG4;refGene_tx=XM_019817081.2;refGene_gn=DLG4;refGene_tx=XM_019817082.2;refGene_gn=DLG4;refGene_tx=XM_019817083.2;refGene_gn=DLG4;refGene_tx=XM_019817084.2;refGene_gn=DLG4;refGene_tx=XM_019817086.2;refGene_gn=DLG4;refGene_tx=XM_023243650.1;refGene_gn=DLG4;refGene_tx=XM_023243651.1;refGene_gn=DLG4;refGene_tx=XM_023243652.1;refGene_gn=DLG4;refGene_tx=XM_023243653.1;refGene_gn=DLG4;refGene_tx=XM_023243655.1
E1 2162129 2162157
refGene_gn=DLG4;refGene_tx=XM_011288883.3;refGene_gn=DLG4;refGene_tx=XM_011288884.3;refGene_gn=DLG4;refGene_tx=XM_011288885.3;refGene_gn=DLG4;refGene_tx=XM_019817072.2;refGene_gn=DLG4;refGene_tx=XM_019817076.2;refGene_gn=DLG4;refGene_tx=XM_019817078.2;refGene_gn=DLG4;refGene_tx=XM_019817080.2;refGene_gn=DLG4;refGene_tx=XM_019817081.2;refGene_gn=DLG4;refGene_tx=XM_019817082.2;refGene_gn=DLG4;refGene_tx=XM_019817083.2;refGene_gn=DLG4;refGene_tx=XM_019817084.2;refGene_gn=DLG4;refGene_tx=XM_019817086.2;refGene_gn=DLG4;refGene_tx=XM_023243650.1;refGene_gn=DLG4;refGene_tx=XM_023243651.1;refGene_gn=DLG4;refGene_tx=XM_023243652.1;refGene_gn=DLG4;refGene_tx=XM_023243653.1;refGene_gn=DLG4;refGene_tx=XM_023243655.1
E1 2162251 2162288
refGene_gn=DLG4;refGene_tx=XM_011288883.3;refGene_gn=DLG4;refGene_tx=XM_011288884.3;refGene_gn=DLG4;refGene_tx=XM_011288885.3;refGene_gn=DLG4;refGene_tx=XM_019817072.2;refGene_gn=DLG4;refGene_tx=XM_019817076.2;refGene_gn=DLG4;refGene_tx=XM_019817078.2;refGene_gn=DLG4;refGene_tx=XM_019817080.2;refGene_gn=DLG4;refGene_tx=XM_019817081.2;refGene_gn=DLG4;refGene_tx=XM_019817082.2;refGene_gn=DLG4;refGene_tx=XM_019817083.2;refGene_gn=DLG4;refGene_tx=XM_019817084.2;refGene_gn=DLG4;refGene_tx=XM_019817086.2;refGene_gn=DLG4;refGene_tx=XM_023243650.1;refGene_gn=DLG4;refGene_tx=XM_023243651.1;refGene_gn=DLG4;refGene_tx=XM_023243652.1;refGene_gn=DLG4;refGene_tx=XM_023243653.1;refGene_gn=DLG4;refGene_tx=XM_023243655.1
E1 2162570 2162747
refGene_gn=DLG4;refGene_tx=XM_011288883.3;refGene_gn=DLG4;refGene_tx=XM_011288884.3;refGene_gn=DLG4;refGene_tx=XM_011288885.3;refGene_gn=DLG4;refGene_tx=XM_019817072.2;refGene_gn=DLG4;refGene_tx=XM_019817076.2;refGene_gn=DLG4;refGene_tx=XM_019817078.2;refGene_gn=DLG4;refGene_tx=XM_019817080.2;refGene_gn=DLG4;refGene_tx=XM_019817081.2;refGene_gn=DLG4;refGene_tx=XM_019817082.2;refGene_gn=DLG4;refGene_tx=XM_019817083.2;refGene_gn=DLG4;refGene_tx=XM_019817084.2;refGene_gn=DLG4;refGene_tx=XM_019817086.2;refGene_gn=DLG4;refGene_tx=XM_023243650.1;refGene_gn=DLG4;refGene_tx=XM_023243651.1;refGene_gn=DLG4;refGene_tx=XM_023243652.1;refGene_gn=DLG4;refGene_tx=XM_023243653.1;refGene_gn=DLG4;refGene_tx=XM_023243655.1

E1 2163953 2164068
refGene_gn=DLG4;refGene_tx=XM_011288883.3;refGene_gn=DLG4;refGene_tx=XM_011288884.3;refGene_gn=DLG4;refGene_tx=XM_011288885.3;refGene_gn=DLG4;refGene_tx=XM_019817072.2;refGene_gn=DLG4;refGene_tx=XM_019817076.2;refGene_gn=DLG4;refGene_tx=XM_019817078.2;refGene_gn=DLG4;refGene_tx=XM_019817080.2;refGene_gn=DLG4;refGene_tx=XM_019817081.2;refGene_gn=DLG4;refGene_tx=XM_019817082.2;refGene_gn=DLG4;refGene_tx=XM_019817083.2;refGene_gn=DLG4;refGene_tx=XM_019817084.2;refGene_gn=DLG4;refGene_tx=XM_019817086.2;refGene_gn=DLG4;refGene_tx=XM_023243650.1;refGene_gn=DLG4;refGene_tx=XM_023243651.1;refGene_gn=DLG4;refGene_tx=XM_023243652.1;refGene_gn=DLG4;refGene_tx=XM_023243653.1;refGene_gn=DLG4;refGene_tx=XM_023243655.1
E1 2164216 2164319
refGene_gn=DLG4;refGene_tx=XM_011288883.3;refGene_gn=DLG4;refGene_tx=XM_011288884.3;refGene_gn=DLG4;refGene_tx=XM_011288885.3;refGene_gn=DLG4;refGene_tx=XM_019817072.2;refGene_gn=DLG4;refGene_tx=XM_019817076.2;refGene_gn=DLG4;refGene_tx=XM_019817078.2;refGene_gn=DLG4;refGene_tx=XM_019817080.2;refGene_gn=DLG4;refGene_tx=XM_019817081.2;refGene_gn=DLG4;refGene_tx=XM_019817082.2;refGene_gn=DLG4;refGene_tx=XM_019817083.2;refGene_gn=DLG4;refGene_tx=XM_019817084.2;refGene_gn=DLG4;refGene_tx=XM_019817086.2;refGene_gn=DLG4;refGene_tx=XM_023243650.1;refGene_gn=DLG4;refGene_tx=XM_023243651.1;refGene_gn=DLG4;refGene_tx=XM_023243652.1;refGene_gn=DLG4;refGene_tx=XM_023243653.1;refGene_gn=DLG4;refGene_tx=XM_023243655.1
E1 2164415 2164711
refGene_gn=DLG4;refGene_tx=XM_011288883.3;refGene_gn=DLG4;refGene_tx=XM_011288884.3;refGene_gn=DLG4;refGene_tx=XM_011288885.3;refGene_gn=DLG4;refGene_tx=XM_019817072.2;refGene_gn=DLG4;refGene_tx=XM_019817076.2;refGene_gn=DLG4;refGene_tx=XM_019817078.2;refGene_gn=DLG4;refGene_tx=XM_019817080.2;refGene_gn=DLG4;refGene_tx=XM_019817081.2;refGene_gn=DLG4;refGene_tx=XM_019817082.2;refGene_gn=DLG4;refGene_tx=XM_019817083.2;refGene_gn=DLG4;refGene_tx=XM_019817084.2;refGene_gn=DLG4;refGene_tx=XM_019817086.2;refGene_gn=DLG4;refGene_tx=XM_023243650.1;refGene_gn=DLG4;refGene_tx=XM_023243651.1;refGene_gn=DLG4;refGene_tx=XM_023243652.1;refGene_gn=DLG4;refGene_tx=XM_023243653.1;refGene_gn=DLG4;refGene_tx=XM_023243655.1
E1 2167041 2167186
refGene_gn=DLG4;refGene_tx=XM_011288883.3;refGene_gn=DLG4;refGene_tx=XM_011288884.3;refGene_gn=DLG4;refGene_tx=XM_011288885.3;refGene_gn=DLG4;refGene_tx=XM_019817072.2;refGene_gn=DLG4;refGene_tx=XM_019817076.2;refGene_gn=DLG4;refGene_tx=XM_019817078.2;refGene_gn=DLG4;refGene_tx=XM_019817080.2;refGene_gn=DLG4;refGene_tx=XM_019817081.2;refGene_gn=DLG4;refGene_tx=XM_019817082.2;refGene_gn=DLG4;refGene_tx=XM_019817083.2;refGene_gn=DLG4;refGene_tx=XM_019817084.2;refGene_gn=DLG4;refGene_tx=XM_019817086.2;refGene_gn=DLG4;refGene_tx=XM_023243650.1;refGene_gn=DLG4;refGene_tx=XM_023243651.1;refGene_gn=DLG4;refGene_tx=XM_023243652.1;refGene_gn=DLG4;refGene_tx=XM_023243653.1;refGene_gn=DLG4;refGene_tx=XM_023243655.1
E1 2167336 2167473
refGene_gn=DLG4;refGene_tx=XM_011288883.3;refGene_gn=DLG4;refGene_tx=X

M_011288884.3;refGene_gn=DLG4;refGene_tx=XM_011288885.3;refGene_gn=DLG4;refGene_tx=XM_019817072.2;refGene_gn=DLG4;refGene_tx=XM_019817076.2;refGene_gn=DLG4;refGene_tx=XM_019817078.2;refGene_gn=DLG4;refGene_tx=XM_019817080.2;refGene_gn=DLG4;refGene_tx=XM_019817081.2;refGene_gn=DLG4;refGene_tx=XM_019817082.2;refGene_gn=DLG4;refGene_tx=XM_019817083.2;refGene_gn=DLG4;refGene_tx=XM_019817084.2;refGene_gn=DLG4;refGene_tx=XM_019817086.2;refGene_gn=DLG4;refGene_tx=XM_023243650.1;refGene_gn=DLG4;refGene_tx=XM_023243651.1;refGene_gn=DLG4;refGene_tx=XM_023243652.1;refGene_gn=DLG4;refGene_tx=XM_023243653.1;refGene_gn=DLG4;refGene_tx=XM_023243655.1
E1 2167575 2167745
refGene_gn=DLG4;refGene_tx=XM_011288883.3;refGene_gn=DLG4;refGene_tx=XM_011288884.3;refGene_gn=DLG4;refGene_tx=XM_011288885.3;refGene_gn=DLG4;refGene_tx=XM_019817072.2;refGene_gn=DLG4;refGene_tx=XM_019817076.2;refGene_gn=DLG4;refGene_tx=XM_019817078.2;refGene_gn=DLG4;refGene_tx=XM_019817080.2;refGene_gn=DLG4;refGene_tx=XM_019817081.2;refGene_gn=DLG4;refGene_tx=XM_019817082.2;refGene_gn=DLG4;refGene_tx=XM_019817083.2;refGene_gn=DLG4;refGene_tx=XM_019817084.2;refGene_gn=DLG4;refGene_tx=XM_019817086.2;refGene_gn=DLG4;refGene_tx=XM_023243650.1;refGene_gn=DLG4;refGene_tx=XM_023243651.1;refGene_gn=DLG4;refGene_tx=XM_023243652.1;refGene_gn=DLG4;refGene_tx=XM_023243653.1;refGene_gn=DLG4;refGene_tx=XM_023243655.1
E1 2167843 2167968
refGene_gn=DLG4;refGene_tx=XM_011288883.3;refGene_gn=DLG4;refGene_tx=XM_011288884.3;refGene_gn=DLG4;refGene_tx=XM_011288885.3;refGene_gn=DLG4;refGene_tx=XM_019817072.2;refGene_gn=DLG4;refGene_tx=XM_019817076.2;refGene_gn=DLG4;refGene_tx=XM_019817078.2;refGene_gn=DLG4;refGene_tx=XM_019817080.2;refGene_gn=DLG4;refGene_tx=XM_019817081.2;refGene_gn=DLG4;refGene_tx=XM_019817082.2;refGene_gn=DLG4;refGene_tx=XM_019817083.2;refGene_gn=DLG4;refGene_tx=XM_019817084.2;refGene_gn=DLG4;refGene_tx=XM_019817086.2;refGene_gn=DLG4;refGene_tx=XM_023243650.1;refGene_gn=DLG4;refGene_tx=XM_023243651.1;refGene_gn=DLG4;refGene_tx=XM_023243652.1;refGene_gn=DLG4;refGene_tx=XM_023243653.1;refGene_gn=DLG4;refGene_tx=XM_023243655.1
E1 2168188 2168248
refGene_gn=DLG4;refGene_tx=XM_011288883.3;refGene_gn=DLG4;refGene_tx=XM_011288884.3;refGene_gn=DLG4;refGene_tx=XM_011288885.3;refGene_gn=DLG4;refGene_tx=XM_019817072.2;refGene_gn=DLG4;refGene_tx=XM_019817076.2;refGene_gn=DLG4;refGene_tx=XM_019817078.2;refGene_gn=DLG4;refGene_tx=XM_019817080.2;refGene_gn=DLG4;refGene_tx=XM_019817081.2;refGene_gn=DLG4;refGene_tx=XM_019817082.2;refGene_gn=DLG4;refGene_tx=XM_019817083.2;refGene_gn=DLG4;refGene_tx=XM_019817084.2;refGene_gn=DLG4;refGene_tx=XM_019817086.2;refGene_gn=DLG4;refGene_tx=XM_023243650.1;refGene_gn=DLG4;refGene_tx=XM_023243651.1;refGene_gn=DLG4;refGene_tx=XM_023243652.1;refGene_gn=DLG4;refGene_tx=XM_023243653.1;refGene_gn=DLG4;refGene_tx=XM_023243655.1
E1 2168385 2168439
refGene_gn=DLG4;refGene_tx=XM_011288883.3;refGene_gn=DLG4;refGene_tx=XM_011288884.3;refGene_gn=DLG4;refGene_tx=XM_011288885.3;refGene_gn=DLG4;refGene_tx=XM_019817072.2;refGene_gn=DLG4;refGene_tx=XM_019817076.2;

refGene_gn=DLG4;refGene_tx=XM_019817078.2;refGene_gn=DLG4;refGene_tx=XM_019817080.2;refGene_gn=DLG4;refGene_tx=XM_019817081.2;refGene_gn=DLG4;refGene_tx=XM_019817082.2;refGene_gn=DLG4;refGene_tx=XM_019817083.2;refGene_gn=DLG4;refGene_tx=XM_019817084.2;refGene_gn=DLG4;refGene_tx=XM_019817086.2;refGene_gn=DLG4;refGene_tx=XM_023243652.1;refGene_gn=DLG4;refGene_tx=XM_023243653.1
E1 2171783 2171948
refGene_gn=DLG4;refGene_tx=XM_011288883.3;refGene_gn=DLG4;refGene_tx=XM_011288884.3;refGene_gn=DLG4;refGene_tx=XM_011288885.3;refGene_gn=DLG4;refGene_tx=XM_019817072.2;refGene_gn=DLG4;refGene_tx=XM_019817076.2;refGene_gn=DLG4;refGene_tx=XM_019817078.2;refGene_gn=DLG4;refGene_tx=XM_019817080.2;refGene_gn=DLG4;refGene_tx=XM_019817081.2;refGene_gn=DLG4;refGene_tx=XM_019817082.2;refGene_gn=DLG4;refGene_tx=XM_023243652.1;refGene_gn=DLG4;refGene_tx=XM_023243653.1
E1 2174766 2174808
refGene_gn=DLG4;refGene_tx=XM_019817082.2;refGene_gn=DLG4;refGene_tx=XM_023243652.1;refGene_gn=DLG4;refGene_tx=XM_023243653.1
E1 2179693 2179723
refGene_gn=DLG4;refGene_tx=XM_011288883.3;refGene_gn=DLG4;refGene_tx=XM_011288884.3;refGene_gn=DLG4;refGene_tx=XM_011288885.3;refGene_gn=DLG4;refGene_tx=XM_019817081.2;refGene_gn=DLG4;refGene_tx=XM_019817083.2;refGene_gn=DLG4;refGene_tx=XM_019817084.2;refGene_gn=DLG4;refGene_tx=XM_019817086.2
E1 2180737 2180778
refGene_gn=DLG4;refGene_tx=XM_019817072.2;refGene_gn=DLG4;refGene_tx=XM_019817076.2;refGene_gn=DLG4;refGene_tx=XM_019817078.2;refGene_gn=DLG4;refGene_tx=XM_019817080.2
E1 2181049 2181148
refGene_gn=DLG4;refGene_tx=XM_019817072.2;refGene_gn=DLG4;refGene_tx=XM_019817076.2;refGene_gn=DLG4;refGene_tx=XM_019817078.2;refGene_gn=DLG4;refGene_tx=XM_019817080.2
E1 2181323 2181342
refGene_gn=DLG4;refGene_tx=XM_019817072.2;refGene_gn=DLG4;refGene_tx=XM_019817076.2;refGene_gn=DLG4;refGene_tx=XM_019817078.2;refGene_gn=DLG4;refGene_tx=XM_019817080.2
E1 2182455 2182517 refGene_gn=ACADVL;refGene_tx=XM_006939745.4
E1 2182601 2182677 refGene_gn=ACADVL;refGene_tx=XM_006939745.4
E1 2182941 2183007 refGene_gn=ACADVL;refGene_tx=XM_006939745.4
E1 2183092 2183165 refGene_gn=ACADVL;refGene_tx=XM_006939745.4
E1 2183254 2183319 refGene_gn=ACADVL;refGene_tx=XM_006939745.4
E1 2183647 2183782 refGene_gn=ACADVL;refGene_tx=XM_006939745.4
E1 2184192 2184337 refGene_gn=ACADVL;refGene_tx=XM_006939745.4
E1 2184619 2184749 refGene_gn=ACADVL;refGene_tx=XM_006939745.4
E1 2184852 2184978 refGene_gn=ACADVL;refGene_tx=XM_006939745.4
E1 2185349 2185548 refGene_gn=ACADVL;refGene_tx=XM_006939745.4
E1 2185743 2185848 refGene_gn=ACADVL;refGene_tx=XM_006939745.4
E1 2186284 2186371 refGene_gn=ACADVL;refGene_tx=XM_006939745.4
E1 2186455 2186518 refGene_gn=ACADVL;refGene_tx=XM_006939745.4
E1 2186619 2186721 refGene_gn=ACADVL;refGene_tx=XM_006939745.4
E1 2186803 2186901 refGene_gn=ACADVL;refGene_tx=XM_006939745.4

E1	2186976	2187049	refGene_gn=ACADVL;refGene_tx=XM_006939745.4
E1	2187134	2187207	refGene_gn=ACADVL;refGene_tx=XM_006939745.4
E1	2187331	2187404	refGene_gn=ACADVL;refGene_tx=XM_006939745.4
E1	2187494	2187570	refGene_gn=ACADVL;refGene_tx=XM_006939745.4
E1	2187644	2187785	refGene_gn=ACADVL;refGene_tx=XM_006939745.4
E1	2188560	2189009
refGene_gn=DVL2;refGene_tx=XM_003996159.4;refGene_gn=DVL2;refGene_tx=XM_023243657.1
E1	2189137	2189353
refGene_gn=DVL2;refGene_tx=XM_003996159.4;refGene_gn=DVL2;refGene_tx=XM_023243657.1
E1	2189755	2189935
refGene_gn=DVL2;refGene_tx=XM_003996159.4;refGene_gn=DVL2;refGene_tx=XM_023243657.1
E1	2190071	2190203
refGene_gn=DVL2;refGene_tx=XM_003996159.4;refGene_gn=DVL2;refGene_tx=XM_023243657.1
E1	2190291	2190420
refGene_gn=DVL2;refGene_tx=XM_003996159.4;refGene_gn=DVL2;refGene_tx=XM_023243657.1
E1	2190591	2190659
refGene_gn=DVL2;refGene_tx=XM_003996159.4;refGene_gn=DVL2;refGene_tx=XM_023243657.1
E1	2191143	2191220
refGene_gn=DVL2;refGene_tx=XM_003996159.4;refGene_gn=DVL2;refGene_tx=XM_023243657.1
E1	2191299	2191439
refGene_gn=DVL2;refGene_tx=XM_003996159.4;refGene_gn=DVL2;refGene_tx=XM_023243657.1
E1	2191528	2191598
refGene_gn=DVL2;refGene_tx=XM_003996159.4;refGene_gn=DVL2;refGene_tx=XM_023243657.1
E1	2191743	2191834
refGene_gn=DVL2;refGene_tx=XM_003996159.4;refGene_gn=DVL2;refGene_tx=XM_023243657.1
E1	2191958	2192094
refGene_gn=DVL2;refGene_tx=XM_003996159.4;refGene_gn=DVL2;refGene_tx=XM_023243657.1
E1	2192198	2192308
refGene_gn=DVL2;refGene_tx=XM_003996159.4;refGene_gn=DVL2;refGene_tx=XM_023243657.1
E1	2192402	2192548
refGene_gn=DVL2;refGene_tx=XM_003996159.4;refGene_gn=DVL2;refGene_tx=XM_023243657.1
E1	2192816	2192886
refGene_gn=DVL2;refGene_tx=XM_003996159.4;refGene_gn=DVL2;refGene_tx=XM_023243657.1
E1	2195272	2195466
refGene_gn=DVL2;refGene_tx=XM_003996159.4;refGene_gn=DVL2;refGene_tx=XM_023243657.1

E1 2196793 2197008
refGene_gn=PHF23;refGene_tx=XM_011288880.3;refGene_gn=PHF23;refGene_tx
=XM_011288881.2;refGene_gn=PHF23;refGene_tx=XM_023243658.1
E1 2197111 2197949
refGene_gn=PHF23;refGene_tx=XM_011288880.3;refGene_gn=PHF23;refGene_tx
=XM_011288881.2;refGene_gn=PHF23;refGene_tx=XM_023243658.1
E1 2198493 2198586
refGene_gn=PHF23;refGene_tx=XM_011288880.3;refGene_gn=PHF23;refGene_tx
=XM_011288881.2;refGene_gn=PHF23;refGene_tx=XM_023243658.1
E1 2198776 2198808
refGene_gn=PHF23;refGene_tx=XM_011288880.3;refGene_gn=PHF23;refGene_tx
=XM_011288881.2;refGene_gn=PHF23;refGene_tx=XM_023243658.1
E1 2199357 2199481
refGene_gn=PHF23;refGene_tx=XM_011288881.2;refGene_gn=PHF23;refGene_tx
=XM_023243658.1
E1 2200394 2200428 refGene_gn=PHF23;refGene_tx=XM_011288880.3
E1 2201974 2202040 refGene_gn=GABARAP;refGene_tx=XM_003996160.5
E1 2202515 2202634 refGene_gn=GABARAP;refGene_tx=XM_003996160.5
E1 2202777 2202856 refGene_gn=GABARAP;refGene_tx=XM_003996160.5
E1 2203407 2203497 refGene_gn=GABARAP;refGene_tx=XM_003996160.5
E1 2205281 2205342
refGene_gn=CTDNEP1;refGene_tx=XM_003996161.4;refGene_gn=CTDNEP1;refGen
e_tx=XM_011288877.3
E1 2205585 2205670
refGene_gn=CTDNEP1;refGene_tx=XM_003996161.4;refGene_gn=CTDNEP1;refGen
e_tx=XM_011288877.3
E1 2206324 2206454
refGene_gn=CTDNEP1;refGene_tx=XM_003996161.4;refGene_gn=CTDNEP1;refGen
e_tx=XM_011288877.3
E1 2206642 2206759
refGene_gn=CTDNEP1;refGene_tx=XM_003996161.4;refGene_gn=CTDNEP1;refGen
e_tx=XM_011288877.3
E1 2207426 2207498
refGene_gn=CTDNEP1;refGene_tx=XM_003996161.4;refGene_gn=CTDNEP1;refGen
e_tx=XM_011288877.3
E1 2207729 2207848
refGene_gn=CTDNEP1;refGene_tx=XM_003996161.4;refGene_gn=CTDNEP1;refGen
e_tx=XM_011288877.3
E1 2207945 2208012
refGene_gn=CTDNEP1;refGene_tx=XM_003996161.4;refGene_gn=CTDNEP1;refGen
e_tx=XM_011288877.3
E1 2210545 2210647
refGene_gn=CTDNEP1;refGene_tx=XM_003996161.4;refGene_gn=CTDNEP1;refGen
e_tx=XM_011288877.3
E1 2211904 2211947 refGene_gn=ELP5;refGene_tx=XM_006939746.4
E1 2212104 2212165 refGene_gn=ELP5;refGene_tx=XM_006939746.4
E1 2212252 2212333 refGene_gn=ELP5;refGene_tx=XM_006939746.4
E1 2213882 2214103 refGene_gn=ELP5;refGene_tx=XM_006939746.4
E1 2215444 2215626 refGene_gn=ELP5;refGene_tx=XM_006939746.4
E1 2216152 2216248 refGene_gn=ELP5;refGene_tx=XM_006939746.4

```
E1	2216402	2216503	refGene_gn=ELP5;refGene_tx=XM_006939746.4
E1	2217121	2217236	refGene_gn=ELP5;refGene_tx=XM_006939746.4
E1	2217976	2218139	refGene_gn=CLDN7;refGene_tx=XM_011288873.2
E1	2218234	2218319	refGene_gn=CLDN7;refGene_tx=XM_011288873.2
E1	2218415	2218580	refGene_gn=CLDN7;refGene_tx=XM_011288873.2
E1	2219323	2219546	refGene_gn=CLDN7;refGene_tx=XM_011288873.2
E1	2228993	2229346
refGene_gn=LOC109494081;refGene_tx=XM_019816899.1
E1	2230002	2230251
refGene_gn=LOC109494081;refGene_tx=XM_019816899.1
E1	2230377	2230414
refGene_gn=LOC109494081;refGene_tx=XM_019816899.1
E1	2230572	2230605	refGene_gn=SLC2A4;refGene_tx=XM_023244732.1
E1	2231880	2231997	refGene_gn=SLC2A4;refGene_tx=XM_023244732.1
E1	2232113	2232286	refGene_gn=SLC2A4;refGene_tx=XM_023244732.1
E1	2232373	2232498	refGene_gn=SLC2A4;refGene_tx=XM_023244732.1
E1	2232616	2232732	refGene_gn=SLC2A4;refGene_tx=XM_023244732.1
E1	2232833	2232996	refGene_gn=SLC2A4;refGene_tx=XM_023244732.1
E1	2233126	2233314	refGene_gn=SLC2A4;refGene_tx=XM_023244732.1
E1	2233485	2233590	refGene_gn=SLC2A4;refGene_tx=XM_023244732.1
E1	2233703	2233805	refGene_gn=SLC2A4;refGene_tx=XM_023244732.1
E1	2234512	2234716	refGene_gn=SLC2A4;refGene_tx=XM_023244732.1
E1	2235199	2235406	refGene_gn=SLC2A4;refGene_tx=XM_023244732.1
E1	2237201	2237252	refGene_gn=YBX2;refGene_tx=XM_023244733.1
E1	2237576	2237754	refGene_gn=YBX2;refGene_tx=XM_023244733.1
E1	2238044	2238148	refGene_gn=YBX2;refGene_tx=XM_023244733.1
E1	2238291	2238576	refGene_gn=YBX2;refGene_tx=XM_023244733.1
E1	2238844	2238934	refGene_gn=YBX2;refGene_tx=XM_023244733.1
E1	2239794	2239828	refGene_gn=YBX2;refGene_tx=XM_023244733.1
E1	2241224	2241288	refGene_gn=YBX2;refGene_tx=XM_023244733.1
E1	2241886	2242151	refGene_gn=YBX2;refGene_tx=XM_023244733.1
E1	2259983	2260148
refGene_gn=EIF5A;refGene_tx=XM_003996163.5;refGene_gn=EIF5A;refGene_tx
=XM_006939748.4;refGene_gn=EIF5A;refGene_tx=XM_019817087.1;refGene_gn=
EIF5A;refGene_tx=XM_019817088.1
E1	2261254	2261359
refGene_gn=EIF5A;refGene_tx=XM_003996163.5;refGene_gn=EIF5A;refGene_tx
=XM_006939748.4;refGene_gn=EIF5A;refGene_tx=XM_019817087.1;refGene_gn=
EIF5A;refGene_tx=XM_019817088.1
E1	2261527	2261659
refGene_gn=EIF5A;refGene_tx=XM_003996163.5;refGene_gn=EIF5A;refGene_tx
=XM_006939748.4;refGene_gn=EIF5A;refGene_tx=XM_019817087.1;refGene_gn=
EIF5A;refGene_tx=XM_019817088.1
E1	2261754	2261817
refGene_gn=EIF5A;refGene_tx=XM_003996163.5;refGene_gn=EIF5A;refGene_tx
=XM_006939748.4;refGene_gn=EIF5A;refGene_tx=XM_019817087.1;refGene_gn=
EIF5A;refGene_tx=XM_019817088.1
E1	2262946	2263030	refGene_gn=GPS2;refGene_tx=XM_023244734.1
E1	2263162	2263258	refGene_gn=GPS2;refGene_tx=XM_023244734.1
E1	2263354	2263434	refGene_gn=GPS2;refGene_tx=XM_023244734.1
```

```
E1	2263521	2263611	refGene_gn=GPS2;refGene_tx=XM_023244734.1
E1	2263705	2263859	refGene_gn=GPS2;refGene_tx=XM_023244734.1
E1	2264061	2264144	refGene_gn=GPS2;refGene_tx=XM_023244734.1
E1	2264235	2264315	refGene_gn=GPS2;refGene_tx=XM_023244734.1
E1	2264444	2264557	refGene_gn=GPS2;refGene_tx=XM_023244734.1
E1	2264635	2264745	refGene_gn=GPS2;refGene_tx=XM_023244734.1
E1	2265102	2265196	refGene_gn=GPS2;refGene_tx=XM_023244734.1
E1	2266112	2266317	refGene_gn=NEURL4;refGene_tx=XM_011288868.2
E1	2266694	2266860	refGene_gn=NEURL4;refGene_tx=XM_011288868.2
E1	2266943	2267056	refGene_gn=NEURL4;refGene_tx=XM_011288868.2
E1	2267218	2267363	refGene_gn=NEURL4;refGene_tx=XM_011288868.2
E1	2267451	2267559	refGene_gn=NEURL4;refGene_tx=XM_011288868.2
E1	2267644	2267732	refGene_gn=NEURL4;refGene_tx=XM_011288868.2
E1	2267837	2267942	refGene_gn=NEURL4;refGene_tx=XM_011288868.2
E1	2267943	2267964	refGene_gn=NEURL4;refGene_tx=XM_011288868.2
E1	2267965	2268019	refGene_gn=NEURL4;refGene_tx=XM_011288868.2
E1	2268422	2268581	refGene_gn=NEURL4;refGene_tx=XM_011288868.2
E1	2269135	2269300	refGene_gn=NEURL4;refGene_tx=XM_011288868.2
E1	2269494	2269656	refGene_gn=NEURL4;refGene_tx=XM_011288868.2
E1	2269739	2269838	refGene_gn=NEURL4;refGene_tx=XM_011288868.2
E1	2269934	2270159	refGene_gn=NEURL4;refGene_tx=XM_011288868.2
E1	2270259	2270405	refGene_gn=NEURL4;refGene_tx=XM_011288868.2
E1	2270793	2270925	refGene_gn=NEURL4;refGene_tx=XM_011288868.2
E1	2271012	2271188	refGene_gn=NEURL4;refGene_tx=XM_011288868.2
E1	2271991	2272070	refGene_gn=NEURL4;refGene_tx=XM_011288868.2
E1	2272169	2272246	refGene_gn=NEURL4;refGene_tx=XM_011288868.2
E1	2272336	2272535	refGene_gn=NEURL4;refGene_tx=XM_011288868.2
E1	2272630	2272793	refGene_gn=NEURL4;refGene_tx=XM_011288868.2
E1	2272914	2273000	refGene_gn=NEURL4;refGene_tx=XM_011288868.2
E1	2273321	2273503	refGene_gn=NEURL4;refGene_tx=XM_011288868.2
E1	2273751	2274016	refGene_gn=NEURL4;refGene_tx=XM_011288868.2
E1	2274132	2274205	refGene_gn=NEURL4;refGene_tx=XM_011288868.2
E1	2274986	2275075	refGene_gn=NEURL4;refGene_tx=XM_011288868.2
E1	2275159	2275271	refGene_gn=NEURL4;refGene_tx=XM_011288868.2
E1	2275438	2275737	refGene_gn=NEURL4;refGene_tx=XM_011288868.2
E1	2275888	2275954	refGene_gn=NEURL4;refGene_tx=XM_011288868.2
E1	2276189	2276634	refGene_gn=NEURL4;refGene_tx=XM_011288868.2
E1	2277561	2277843	refGene_gn=NEURL4;refGene_tx=XM_011288868.2
E1	2279022	2279513
refGene_gn=LOC102899307;refGene_tx=XM_006939750.2;refGene_gn=LOC102899
307;refGene_tx=XM_006939751.3
E1	2279876	2279984
refGene_gn=LOC102899307;refGene_tx=XM_006939750.2;refGene_gn=LOC102899
307;refGene_tx=XM_006939751.3
E1	2280119	2280191
refGene_gn=LOC102899307;refGene_tx=XM_006939750.2;refGene_gn=LOC102899
307;refGene_tx=XM_006939751.3
E1	2280419	2280508
refGene_gn=LOC102899307;refGene_tx=XM_006939750.2;refGene_gn=LOC102899
307;refGene_tx=XM_006939751.3
```

E1 2280605 2280860
refGene_gn=LOC102899307;refGene_tx=XM_006939750.2;refGene_gn=LOC102899
307;refGene_tx=XM_006939751.3
E1 2281190 2281397
refGene_gn=LOC102899307;refGene_tx=XM_006939750.2;refGene_gn=LOC102899
307;refGene_tx=XM_006939751.3
E1 2281488 2281626
refGene_gn=LOC102899307;refGene_tx=XM_006939750.2;refGene_gn=LOC102899
307;refGene_tx=XM_006939751.3
E1 2281871 2282028
refGene_gn=LOC102899307;refGene_tx=XM_006939750.2
E1 2282179 2282238
refGene_gn=LOC102899307;refGene_tx=XM_006939750.2;refGene_gn=LOC102899
307;refGene_tx=XM_006939751.3
E1 2283036 2283089 refGene_gn=ACAP1;refGene_tx=XM_023244544.1
E1 2283642 2283700 refGene_gn=ACAP1;refGene_tx=XM_023244544.1
E1 2287745 2287865 refGene_gn=ACAP1;refGene_tx=XM_023244544.1
E1 2288051 2288105 refGene_gn=ACAP1;refGene_tx=XM_023244544.1
E1 2288196 2288255 refGene_gn=ACAP1;refGene_tx=XM_023244544.1
E1 2288518 2288702 refGene_gn=ACAP1;refGene_tx=XM_023244544.1
E1 2288815 2288860 refGene_gn=ACAP1;refGene_tx=XM_023244544.1
E1 2288985 2289081 refGene_gn=ACAP1;refGene_tx=XM_023244544.1
E1 2289196 2289271 refGene_gn=ACAP1;refGene_tx=XM_023244544.1
E1 2289361 2289471 refGene_gn=ACAP1;refGene_tx=XM_023244544.1
E1 2291003 2291055 refGene_gn=ACAP1;refGene_tx=XM_023244544.1
E1 2291146 2291247 refGene_gn=ACAP1;refGene_tx=XM_023244544.1
E1 2291604 2291728 refGene_gn=ACAP1;refGene_tx=XM_023244544.1
E1 2291827 2292039 refGene_gn=ACAP1;refGene_tx=XM_023244544.1
E1 2292557 2292627 refGene_gn=ACAP1;refGene_tx=XM_023244544.1
E1 2292771 2292866 refGene_gn=ACAP1;refGene_tx=XM_023244544.1
E1 2292950 2293120 refGene_gn=ACAP1;refGene_tx=XM_023244544.1
E1 2293558 2293731 refGene_gn=ACAP1;refGene_tx=XM_023244544.1
E1 2294172 2294282 refGene_gn=ACAP1;refGene_tx=XM_023244544.1
E1 2294364 2294475 refGene_gn=ACAP1;refGene_tx=XM_023244544.1
E1 2294798 2294848 refGene_gn=ACAP1;refGene_tx=XM_023244544.1
E1 2295140 2295241 refGene_gn=ACAP1;refGene_tx=XM_023244544.1
E1 2296821 2297520 refGene_gn=KCTD11;refGene_tx=XM_006939754.4
E1 2299098 2299306
refGene_gn=TMEM95;refGene_tx=XM_019816989.2;refGene_gn=TMEM95;refGene_
tx=XM_019816991.2
E1 2299434 2299491
refGene_gn=TMEM95;refGene_tx=XM_019816989.2;refGene_gn=TMEM95;refGene_
tx=XM_019816991.2
E1 2299696 2299777
refGene_gn=TMEM95;refGene_tx=XM_019816989.2;refGene_gn=TMEM95;refGene_
tx=XM_019816991.2
E1 2299879 2299900
refGene_gn=TMEM95;refGene_tx=XM_019816989.2;refGene_gn=TMEM95;refGene_
tx=XM_019816991.2
E1 2300039 2300120 refGene_gn=TMEM95;refGene_tx=XM_019816989.2

E1	2300251	2300339
refGene_gn=TMEM95;refGene_tx=XM_019816989.2;refGene_gn=TMEM95;refGene_tx=XM_019816991.2
E1	2300472	2300506
refGene_gn=TMEM95;refGene_tx=XM_019816989.2;refGene_gn=TMEM95;refGene_tx=XM_019816991.2
E1	2310557	2310720
refGene_gn=TNK1;refGene_tx=XM_003996167.4;refGene_gn=TNK1;refGene_tx=XM_011288863.2;refGene_gn=TNK1;refGene_tx=XM_011288864.2;refGene_gn=TNK1;refGene_tx=XM_011288865.2
E1	2310897	2310968
refGene_gn=TNK1;refGene_tx=XM_003996167.4;refGene_gn=TNK1;refGene_tx=XM_011288863.2;refGene_gn=TNK1;refGene_tx=XM_011288864.2;refGene_gn=TNK1;refGene_tx=XM_011288865.2
E1	2311082	2311277
refGene_gn=TNK1;refGene_tx=XM_003996167.4;refGene_gn=TNK1;refGene_tx=XM_011288863.2;refGene_gn=TNK1;refGene_tx=XM_011288864.2;refGene_gn=TNK1;refGene_tx=XM_011288865.2
E1	2311413	2311569
refGene_gn=TNK1;refGene_tx=XM_003996167.4;refGene_gn=TNK1;refGene_tx=XM_011288863.2;refGene_gn=TNK1;refGene_tx=XM_011288864.2;refGene_gn=TNK1;refGene_tx=XM_011288865.2
E1	2311661	2311945
refGene_gn=TNK1;refGene_tx=XM_003996167.4;refGene_gn=TNK1;refGene_tx=XM_011288863.2;refGene_gn=TNK1;refGene_tx=XM_011288864.2;refGene_gn=TNK1;refGene_tx=XM_011288865.2
E1	2312127	2312401
refGene_gn=TNK1;refGene_tx=XM_003996167.4;refGene_gn=TNK1;refGene_tx=XM_011288863.2;refGene_gn=TNK1;refGene_tx=XM_011288864.2;refGene_gn=TNK1;refGene_tx=XM_011288865.2
E1	2313501	2313596
refGene_gn=TNK1;refGene_tx=XM_003996167.4;refGene_gn=TNK1;refGene_tx=XM_011288863.2;refGene_gn=TNK1;refGene_tx=XM_011288864.2;refGene_gn=TNK1;refGene_tx=XM_011288865.2
E1	2313926	2314091
refGene_gn=TNK1;refGene_tx=XM_003996167.4;refGene_gn=TNK1;refGene_tx=XM_011288863.2;refGene_gn=TNK1;refGene_tx=XM_011288864.2;refGene_gn=TNK1;refGene_tx=XM_011288865.2
E1	2314325	2314405
refGene_gn=TNK1;refGene_tx=XM_003996167.4;refGene_gn=TNK1;refGene_tx=XM_011288863.2;refGene_gn=TNK1;refGene_tx=XM_011288864.2;refGene_gn=TNK1;refGene_tx=XM_011288865.2
E1	2314708	2315007
refGene_gn=TNK1;refGene_tx=XM_003996167.4;refGene_gn=TNK1;refGene_tx=XM_011288863.2;refGene_gn=TNK1;refGene_tx=XM_011288864.2;refGene_gn=TNK1;refGene_tx=XM_011288865.2
E1	2315088	2315184
refGene_gn=TNK1;refGene_tx=XM_003996167.4;refGene_gn=TNK1;refGene_tx=XM_011288863.2;refGene_gn=TNK1;refGene_tx=XM_011288864.2;refGene_gn=TNK1;refGene_tx=XM_011288865.2

E1	2315267	2315381
refGene_gn=TNK1;refGene_tx=XM_003996167.4;refGene_gn=TNK1;refGene_tx=XM_011288863.2;refGene_gn=TNK1;refGene_tx=XM_011288864.2;refGene_gn=TNK1;refGene_tx=XM_011288865.2
E1	2316615	2316672
refGene_gn=PLSCR3;refGene_tx=XM_003996168.3;refGene_gn=PLSCR3;refGene_tx=XM_011288867.2;refGene_gn=PLSCR3;refGene_tx=XM_019817090.2
E1	2316861	2317023
refGene_gn=PLSCR3;refGene_tx=XM_003996168.3;refGene_gn=PLSCR3;refGene_tx=XM_011288867.2;refGene_gn=PLSCR3;refGene_tx=XM_019817090.2
E1	2318564	2318726
refGene_gn=PLSCR3;refGene_tx=XM_003996168.3;refGene_gn=PLSCR3;refGene_tx=XM_011288867.2;refGene_gn=PLSCR3;refGene_tx=XM_019817090.2
E1	2318971	2319192
refGene_gn=PLSCR3;refGene_tx=XM_003996168.3;refGene_gn=PLSCR3;refGene_tx=XM_011288867.2;refGene_gn=PLSCR3;refGene_tx=XM_019817090.2
E1	2319291	2319334
refGene_gn=PLSCR3;refGene_tx=XM_003996168.3;refGene_gn=PLSCR3;refGene_tx=XM_011288867.2;refGene_gn=PLSCR3;refGene_tx=XM_019817090.2
E1	2319446	2319682
refGene_gn=PLSCR3;refGene_tx=XM_003996168.3;refGene_gn=PLSCR3;refGene_tx=XM_011288867.2;refGene_gn=PLSCR3;refGene_tx=XM_019817090.2
E1	2319940	2319947
refGene_gn=PLSCR3;refGene_tx=XM_003996168.3;refGene_gn=PLSCR3;refGene_tx=XM_011288867.2;refGene_gn=PLSCR3;refGene_tx=XM_019817090.2
E1	2327877	2328021	refGene_gn=TMEM256;refGene_tx=XM_003996169.3
E1	2328115	2328196	refGene_gn=TMEM256;refGene_tx=XM_003996169.3
E1	2328446	2328478	refGene_gn=TMEM256;refGene_tx=XM_003996169.3
E1	2328719	2328804	refGene_gn=TMEM256;refGene_tx=XM_003996169.3
E1	2333088	2333542
refGene_gn=NLGN2;refGene_tx=XM_023244702.1;refGene_gn=NLGN2;refGene_tx=XM_023244703.1
E1	2336395	2336446	refGene_gn=NLGN2;refGene_tx=XM_023244702.1
E1	2338578	2338728
refGene_gn=NLGN2;refGene_tx=XM_023244702.1;refGene_gn=NLGN2;refGene_tx=XM_023244703.1
E1	2338901	2339087
refGene_gn=NLGN2;refGene_tx=XM_023244702.1;refGene_gn=NLGN2;refGene_tx=XM_023244703.1
E1	2339221	2339414
refGene_gn=NLGN2;refGene_tx=XM_023244702.1;refGene_gn=NLGN2;refGene_tx=XM_023244703.1
E1	2339765	2340362
refGene_gn=NLGN2;refGene_tx=XM_023244702.1;refGene_gn=NLGN2;refGene_tx=XM_023244703.1
E1	2341266	2342140
refGene_gn=NLGN2;refGene_tx=XM_023244702.1;refGene_gn=NLGN2;refGene_tx=XM_023244703.1
E1	2344816	2344960
refGene_gn=SPEM1;refGene_tx=XM_011288861.3;refGene_gn=SPEM1;refGene_tx

=XM_019817095.2
E1 2345057 2345115
refGene_gn=SPEM1;refGene_tx=XM_011288861.3;refGene_gn=SPEM1;refGene_tx=XM_019817095.2;refGene_gn=SPEM1;refGene_tx=XM_023244704.1
E1 2345296 2346040
refGene_gn=SPEM1;refGene_tx=XM_011288861.3;refGene_gn=SPEM1;refGene_tx=XM_019817095.2;refGene_gn=SPEM1;refGene_tx=XM_023244704.1
E1 2347794 2347830
refGene_gn=SPEM2;refGene_tx=XM_019817092.1;refGene_gn=SPEM2;refGene_tx=XM_019817093.1;refGene_gn=SPEM2;refGene_tx=XM_019817094.1
E1 2347918 2348056
refGene_gn=SPEM2;refGene_tx=XM_011288862.3;refGene_gn=SPEM2;refGene_tx=XM_019817092.1;refGene_gn=SPEM2;refGene_tx=XM_019817093.1;refGene_gn=SPEM2;refGene_tx=XM_019817094.1
E1 2348152 2348210
refGene_gn=SPEM2;refGene_tx=XM_011288862.3;refGene_gn=SPEM2;refGene_tx=XM_019817093.1;refGene_gn=SPEM2;refGene_tx=XM_019817094.1
E1 2348351 2349649
refGene_gn=SPEM2;refGene_tx=XM_011288862.3;refGene_gn=SPEM2;refGene_tx=XM_019817092.1;refGene_gn=SPEM2;refGene_tx=XM_019817093.1;refGene_gn=SPEM2;refGene_tx=XM_019817094.1
E1 2350416 2350554
refGene_gn=LOC102900997;refGene_tx=XM_019817096.2
E1 2350676 2350734
refGene_gn=LOC102900997;refGene_tx=XM_019817096.2
E1 2350834 2354418
refGene_gn=LOC102900997;refGene_tx=XM_019817096.2
E1 2356899 2357113 refGene_gn=TMEM102;refGene_tx=XM_003996170.3
E1 2357239 2358558 refGene_gn=TMEM102;refGene_tx=XM_003996170.3
E1 2360641 2360834
refGene_gn=FGF11;refGene_tx=XM_003996171.5;refGene_gn=FGF11;refGene_tx=XM_006939756.4;refGene_gn=FGF11;refGene_tx=XM_019817097.2
E1 2362489 2362600
refGene_gn=FGF11;refGene_tx=XM_003996171.5;refGene_gn=FGF11;refGene_tx=XM_006939756.4
E1 2362806 2362910
refGene_gn=FGF11;refGene_tx=XM_003996171.5;refGene_gn=FGF11;refGene_tx=XM_006939756.4;refGene_gn=FGF11;refGene_tx=XM_019817097.2
E1 2363700 2363899
refGene_gn=FGF11;refGene_tx=XM_003996171.5;refGene_gn=FGF11;refGene_tx=XM_006939756.4;refGene_gn=FGF11;refGene_tx=XM_019817097.2
E1 2364182 2364253
refGene_gn=FGF11;refGene_tx=XM_003996171.5;refGene_gn=FGF11;refGene_tx=XM_006939756.4;refGene_gn=FGF11;refGene_tx=XM_019817097.2
E1 2366207 2366268 refGene_gn=CHRNB1;refGene_tx=XM_003996172.4
E1 2366353 2366493 refGene_gn=CHRNB1;refGene_tx=XM_003996172.4
E1 2367086 2367131 refGene_gn=CHRNB1;refGene_tx=XM_003996172.4
E1 2367810 2367920 refGene_gn=CHRNB1;refGene_tx=XM_003996172.4
E1 2368009 2368118 refGene_gn=CHRNB1;refGene_tx=XM_003996172.4
E1 2368686 2368834 refGene_gn=CHRNB1;refGene_tx=XM_003996172.4

E1	2369163	2369373	refGene_gn=CHRNB1;refGene_tx=XM_003996172.4
E1	2371776	2372000	refGene_gn=CHRNB1;refGene_tx=XM_003996172.4
E1	2372496	2372678	refGene_gn=CHRNB1;refGene_tx=XM_003996172.4
E1	2373000	2373148	refGene_gn=CHRNB1;refGene_tx=XM_003996172.4
E1	2373438	2373579	refGene_gn=CHRNB1;refGene_tx=XM_003996172.4
E1	2377795	2379776
refGene_gn=ZBTB4;refGene_tx=XM_023243602.1;refGene_gn=ZBTB4;refGene_tx=XM_023243603.1;refGene_gn=ZBTB4;refGene_tx=XM_023243604.1;refGene_gn=ZBTB4;refGene_tx=XM_023243605.1;refGene_gn=ZBTB4;refGene_tx=XM_023243606.1
E1	2380429	2381499
refGene_gn=ZBTB4;refGene_tx=XM_023243602.1;refGene_gn=ZBTB4;refGene_tx=XM_023243603.1;refGene_gn=ZBTB4;refGene_tx=XM_023243604.1;refGene_gn=ZBTB4;refGene_tx=XM_023243605.1;refGene_gn=ZBTB4;refGene_tx=XM_023243606.1
E1	2389520	2389647	refGene_gn=SLC35G6;refGene_tx=XM_023244501.1
E1	2390270	2390497	refGene_gn=SLC35G6;refGene_tx=XM_023244501.1
E1	2391189	2391384	refGene_gn=SLC35G6;refGene_tx=XM_023244501.1
E1	2391872	2392886	refGene_gn=SLC35G6;refGene_tx=XM_023244501.1
E1	2394765	2394858	refGene_gn=POLR2A;refGene_tx=XM_023243607.1
E1	2404859	2404992	refGene_gn=POLR2A;refGene_tx=XM_023243607.1
E1	2405107	2405244	refGene_gn=POLR2A;refGene_tx=XM_023243607.1
E1	2405356	2405530	refGene_gn=POLR2A;refGene_tx=XM_023243607.1
E1	2405642	2405924	refGene_gn=POLR2A;refGene_tx=XM_023243607.1
E1	2406179	2406341	refGene_gn=POLR2A;refGene_tx=XM_023243607.1
E1	2406438	2406659	refGene_gn=POLR2A;refGene_tx=XM_023243607.1
E1	2406795	2406928	refGene_gn=POLR2A;refGene_tx=XM_023243607.1
E1	2407409	2407534	refGene_gn=POLR2A;refGene_tx=XM_023243607.1
E1	2407631	2407842	refGene_gn=POLR2A;refGene_tx=XM_023243607.1
E1	2408438	2408648	refGene_gn=POLR2A;refGene_tx=XM_023243607.1
E1	2408784	2408953	refGene_gn=POLR2A;refGene_tx=XM_023243607.1
E1	2409122	2409217	refGene_gn=POLR2A;refGene_tx=XM_023243607.1
E1	2409326	2409530	refGene_gn=POLR2A;refGene_tx=XM_023243607.1
E1	2409655	2409878	refGene_gn=POLR2A;refGene_tx=XM_023243607.1
E1	2410357	2410541	refGene_gn=POLR2A;refGene_tx=XM_023243607.1
E1	2410920	2411092	refGene_gn=POLR2A;refGene_tx=XM_023243607.1
E1	2411184	2411290	refGene_gn=POLR2A;refGene_tx=XM_023243607.1
E1	2411385	2411591	refGene_gn=POLR2A;refGene_tx=XM_023243607.1
E1	2413265	2413490	refGene_gn=POLR2A;refGene_tx=XM_023243607.1
E1	2413627	2413874	refGene_gn=POLR2A;refGene_tx=XM_023243607.1
E1	2414121	2414222	refGene_gn=POLR2A;refGene_tx=XM_023243607.1
E1	2415105	2415210	refGene_gn=POLR2A;refGene_tx=XM_023243607.1
E1	2415297	2415480	refGene_gn=POLR2A;refGene_tx=XM_023243607.1
E1	2415675	2415879	refGene_gn=POLR2A;refGene_tx=XM_023243607.1
E1	2416058	2416245	refGene_gn=POLR2A;refGene_tx=XM_023243607.1
E1	2416336	2416450	refGene_gn=POLR2A;refGene_tx=XM_023243607.1
E1	2416596	2416746	refGene_gn=POLR2A;refGene_tx=XM_023243607.1
E1	2416851	2417053	refGene_gn=POLR2A;refGene_tx=XM_023243607.1
E1	2417144	2418111	refGene_gn=POLR2A;refGene_tx=XM_023243607.1
E1	2454601	2454760

refGene_gn=LOC101098755;refGene_tx=XM_023243851.1;refGene_gn=LOC101098755;refGene_tx=XM_023243852.1
E1 2454967 2455018
refGene_gn=LOC101098755;refGene_tx=XM_023243851.1;refGene_gn=LOC101098755;refGene_tx=XM_023243852.1
E1 2455403 2455479
refGene_gn=LOC101098755;refGene_tx=XM_023243851.1;refGene_gn=LOC101098755;refGene_tx=XM_023243852.1
E1 2456177 2456231
refGene_gn=LOC101098755;refGene_tx=XM_023243851.1;refGene_gn=LOC101098755;refGene_tx=XM_023243852.1
E1 2456316 2456352
refGene_gn=LOC101098755;refGene_tx=XM_023243851.1;refGene_gn=LOC101098755;refGene_tx=XM_023243852.1
E1 2461365 2461490
refGene_gn=LOC101098755;refGene_tx=XM_023243851.1;refGene_gn=LOC101098755;refGene_tx=XM_023243852.1
E1 2461673 2461925
refGene_gn=LOC101098755;refGene_tx=XM_023243851.1;refGene_gn=LOC101098755;refGene_tx=XM_023243852.1
E1 2464021 2464279
refGene_gn=TNFSF13;refGene_tx=NM_001171065.1;refGene_gn=TNFSF13;refGene_tx=XM_019816876.1;refGene_gn=TNFSF13;refGene_tx=XM_019816877.2
E1 2464628 2464707
refGene_gn=TNFSF13;refGene_tx=NM_001171065.1;refGene_gn=TNFSF13;refGene_tx=XM_019816876.1;refGene_gn=TNFSF13;refGene_tx=XM_019816877.2
E1 2464857 2464905
refGene_gn=TNFSF13;refGene_tx=NM_001171065.1;refGene_gn=TNFSF13;refGene_tx=XM_019816876.1;refGene_gn=TNFSF13;refGene_tx=XM_019816877.2
E1 2465050 2465169
refGene_gn=TNFSF13;refGene_tx=NM_001171065.1;refGene_gn=TNFSF13;refGene_tx=XM_019816876.1;refGene_gn=TNFSF13;refGene_tx=XM_019816877.2
E1 2465374 2465513
refGene_gn=TNFSF13;refGene_tx=NM_001171065.1;refGene_gn=TNFSF13;refGene_tx=XM_019816876.1;refGene_gn=TNFSF13;refGene_tx=XM_019816877.2
E1 2465750 2465860
refGene_gn=TNFSF13;refGene_tx=NM_001171065.1;refGene_gn=TNFSF13;refGene_tx=XM_019816876.1;refGene_gn=TNFSF13;refGene_tx=XM_019816877.2
E1 2468133 2468848 refGene_gn=SENP3;refGene_tx=XM_003996174.4
E1 2469646 2469886 refGene_gn=SENP3;refGene_tx=XM_003996174.4
E1 2469981 2470093 refGene_gn=SENP3;refGene_tx=XM_003996174.4
E1 2470471 2470619 refGene_gn=SENP3;refGene_tx=XM_003996174.4
E1 2470747 2470795 refGene_gn=SENP3;refGene_tx=XM_003996174.4
E1 2471193 2471271 refGene_gn=SENP3;refGene_tx=XM_003996174.4
E1 2472837 2472975 refGene_gn=SENP3;refGene_tx=XM_003996174.4
E1 2473157 2473241 refGene_gn=SENP3;refGene_tx=XM_003996174.4
E1 2473358 2473409 refGene_gn=SENP3;refGene_tx=XM_003996174.4
E1 2473848 2473959 refGene_gn=SENP3;refGene_tx=XM_003996174.4
E1 2475232 2475255 refGene_gn=EIF4A1;refGene_tx=XM_003996175.5
E1 2476534 2476583 refGene_gn=EIF4A1;refGene_tx=XM_003996175.5

| | | | |
|---|---|---|---|
| E1 | 2476968 | 2477101 | refGene_gn=EIF4A1;refGene_tx=XM_003996175.5 |
| E1 | 2477560 | 2477700 | refGene_gn=EIF4A1;refGene_tx=XM_003996175.5 |
| E1 | 2478672 | 2478841 | refGene_gn=EIF4A1;refGene_tx=XM_003996175.5 |
| E1 | 2479167 | 2479277 | refGene_gn=EIF4A1;refGene_tx=XM_003996175.5 |
| E1 | 2479469 | 2479613 | refGene_gn=EIF4A1;refGene_tx=XM_003996175.5 |
| E1 | 2479703 | 2479841 | refGene_gn=EIF4A1;refGene_tx=XM_003996175.5 |
| E1 | 2479933 | 2480023 | refGene_gn=EIF4A1;refGene_tx=XM_003996175.5 |
| E1 | 2480303 | 2480383 | refGene_gn=EIF4A1;refGene_tx=XM_003996175.5 |
| E1 | 2480489 | 2480634 | refGene_gn=EIF4A1;refGene_tx=XM_003996175.5 |
| E1 | 2481761 | 2481810 | refGene_gn=CD68;refGene_tx=XM_019817098.2 |
| E1 | 2481892 | 2482305 | refGene_gn=CD68;refGene_tx=XM_019817098.2 |
| E1 | 2482454 | 2482574 | refGene_gn=CD68;refGene_tx=XM_019817098.2 |
| E1 | 2482701 | 2482774 | refGene_gn=CD68;refGene_tx=XM_019817098.2 |
| E1 | 2482895 | 2483066 | refGene_gn=CD68;refGene_tx=XM_019817098.2 |
| E1 | 2483383 | 2483517 | refGene_gn=CD68;refGene_tx=XM_019817098.2 |
| E1 | 2485361 | 2485464 | refGene_gn=MPDU1;refGene_tx=XM_003996178.4 |
| E1 | 2486423 | 2486489 | refGene_gn=MPDU1;refGene_tx=XM_003996178.4 |
| E1 | 2486673 | 2486806 | refGene_gn=MPDU1;refGene_tx=XM_003996178.4 |
| E1 | 2487362 | 2487448 | refGene_gn=MPDU1;refGene_tx=XM_003996178.4 |
| E1 | 2487573 | 2487692 | refGene_gn=MPDU1;refGene_tx=XM_003996178.4 |
| E1 | 2487781 | 2487892 | refGene_gn=MPDU1;refGene_tx=XM_003996178.4 |
| E1 | 2487988 | 2488111 | refGene_gn=MPDU1;refGene_tx=XM_003996178.4 |
| E1 | 2488886 | 2489055 | refGene_gn=SOX15;refGene_tx=XM_003996179.4 |
| E1 | 2489688 | 2490221 | refGene_gn=SOX15;refGene_tx=XM_003996179.4 |
| E1 | 2492345 | 2492441 | refGene_gn=FXR2;refGene_tx=XM_023243850.1 |
| E1 | 2492752 | 2492853 | refGene_gn=FXR2;refGene_tx=XM_023243850.1 |
| E1 | 2493004 | 2493100 | refGene_gn=FXR2;refGene_tx=XM_023243850.1 |
| E1 | 2493196 | 2493397 | refGene_gn=FXR2;refGene_tx=XM_023243850.1 |
| E1 | 2493489 | 2493687 | refGene_gn=FXR2;refGene_tx=XM_023243850.1 |
| E1 | 2493855 | 2494078 | refGene_gn=FXR2;refGene_tx=XM_023243850.1 |
| E1 | 2494388 | 2494475 | refGene_gn=FXR2;refGene_tx=XM_023243850.1 |
| E1 | 2494700 | 2494810 | refGene_gn=FXR2;refGene_tx=XM_023243850.1 |
| E1 | 2495138 | 2495217 | refGene_gn=FXR2;refGene_tx=XM_023243850.1 |
| E1 | 2495414 | 2495585 | refGene_gn=FXR2;refGene_tx=XM_023243850.1 |
| E1 | 2497988 | 2498105 | refGene_gn=FXR2;refGene_tx=XM_023243850.1 |
| E1 | 2498589 | 2498683 | refGene_gn=FXR2;refGene_tx=XM_023243850.1 |
| E1 | 2498919 | 2499068 | refGene_gn=FXR2;refGene_tx=XM_023243850.1 |
| E1 | 2499186 | 2499258 | refGene_gn=FXR2;refGene_tx=XM_023243850.1 |
| E1 | 2499664 | 2499758 | refGene_gn=FXR2;refGene_tx=XM_023243850.1 |
| E1 | 2500101 | 2500154 | refGene_gn=FXR2;refGene_tx=XM_023243850.1 |
| E1 | 2505961 | 2506042 | refGene_gn=FXR2;refGene_tx=XM_023243850.1 |
| E1 | 2515824 | 2515992 | refGene_gn=SAT2;refGene_tx=XM_003996180.5 |
| E1 | 2516127 | 2516168 | refGene_gn=SAT2;refGene_tx=XM_003996180.5 |
| E1 | 2516351 | 2516456 | refGene_gn=SAT2;refGene_tx=XM_003996180.5 |
| E1 | 2516557 | 2516641 | refGene_gn=SAT2;refGene_tx=XM_003996180.5 |
| E1 | 2516761 | 2516813 | refGene_gn=SAT2;refGene_tx=XM_003996180.5 |
| E1 | 2517073 | 2517145 | refGene_gn=SAT2;refGene_tx=XM_003996180.5 |
| E1 | 2518599 | 2518686 | refGene_gn=SHBG;refGene_tx=XM_003996181.4;refGene_gn=SHBG;refGene_tx=XM_006939759.3;refGene_gn=SHBG;refGene_tx=XM_006939760.3 |

E1 2518887 2518979
refGene_gn=SHBG;refGene_tx=XM_003996181.4;refGene_gn=SHBG;refGene_tx=XM_006939759.3;refGene_gn=SHBG;refGene_tx=XM_006939760.3
E1 2518981 2518986 refGene_gn=SHBG;refGene_tx=XM_006939761.2
E1 2519145 2519335
refGene_gn=SHBG;refGene_tx=XM_003996181.4;refGene_gn=SHBG;refGene_tx=XM_006939759.3;refGene_gn=SHBG;refGene_tx=XM_006939760.3;refGene_gn=SHBG;refGene_tx=XM_006939761.2;refGene_gn=SHBG;refGene_tx=XM_011288859.2
E1 2519628 2519790
refGene_gn=SHBG;refGene_tx=XM_003996181.4;refGene_gn=SHBG;refGene_tx=XM_006939759.3;refGene_gn=SHBG;refGene_tx=XM_006939760.3;refGene_gn=SHBG;refGene_tx=XM_006939761.2;refGene_gn=SHBG;refGene_tx=XM_011288859.2
E1 2520028 2520191
refGene_gn=SHBG;refGene_tx=XM_003996181.4;refGene_gn=SHBG;refGene_tx=XM_006939759.3;refGene_gn=SHBG;refGene_tx=XM_006939760.3;refGene_gn=SHBG;refGene_tx=XM_006939761.2;refGene_gn=SHBG;refGene_tx=XM_011288859.2
E1 2520324 2520461
refGene_gn=SHBG;refGene_tx=XM_003996181.4;refGene_gn=SHBG;refGene_tx=XM_006939759.3;refGene_gn=SHBG;refGene_tx=XM_006939760.3;refGene_gn=SHBG;refGene_tx=XM_006939761.2;refGene_gn=SHBG;refGene_tx=XM_011288859.2
E1 2521410 2521618
refGene_gn=SHBG;refGene_tx=XM_003996181.4;refGene_gn=SHBG;refGene_tx=XM_006939759.3;refGene_gn=SHBG;refGene_tx=XM_006939760.3;refGene_gn=SHBG;refGene_tx=XM_006939761.2;refGene_gn=SHBG;refGene_tx=XM_011288859.2
E1 2521870 2522019
refGene_gn=SHBG;refGene_tx=XM_003996181.4;refGene_gn=SHBG;refGene_tx=XM_006939759.3;refGene_gn=SHBG;refGene_tx=XM_006939760.3;refGene_gn=SHBG;refGene_tx=XM_006939761.2;refGene_gn=SHBG;refGene_tx=XM_011288859.2
E1 2528276 2528388
refGene_gn=ATP1B2;refGene_tx=XM_003996185.5;refGene_gn=ATP1B2;refGene_tx=XM_019817099.2;refGene_gn=ATP1B2;refGene_tx=XM_019817100.2
E1 2530163 2530292
refGene_gn=ATP1B2;refGene_tx=XM_003996185.5;refGene_gn=ATP1B2;refGene_tx=XM_019817099.2;refGene_gn=ATP1B2;refGene_tx=XM_019817100.2;refGene_gn=ATP1B2;refGene_tx=XM_019817101.2
E1 2530537 2530642
refGene_gn=ATP1B2;refGene_tx=XM_003996185.5;refGene_gn=ATP1B2;refGene_tx=XM_019817099.2;refGene_gn=ATP1B2;refGene_tx=XM_019817100.2;refGene_gn=ATP1B2;refGene_tx=XM_019817101.2
E1 2530757 2530963
refGene_gn=ATP1B2;refGene_tx=XM_003996185.5;refGene_gn=ATP1B2;refGene_tx=XM_019817099.2;refGene_gn=ATP1B2;refGene_tx=XM_019817100.2;refGene_gn=ATP1B2;refGene_tx=XM_019817101.2
E1 2531184 2531241
refGene_gn=ATP1B2;refGene_tx=XM_003996185.5;refGene_gn=ATP1B2;refGene_tx=XM_019817099.2;refGene_gn=ATP1B2;refGene_tx=XM_019817100.2;refGene_gn=ATP1B2;refGene_tx=XM_019817101.2
E1 2531335 2531362
refGene_gn=ATP1B2;refGene_tx=XM_019817099.2;refGene_gn=ATP1B2;refGene_tx=XM_019817100.2;refGene_gn=ATP1B2;refGene_tx=XM_019817101.2

E1	2531745	2531766
refGene_gn=ATP1B2;refGene_tx=XM_019817099.2;refGene_gn=ATP1B2;refGene_tx=XM_019817101.2
E1	2532015	2532114
refGene_gn=ATP1B2;refGene_tx=XM_003996185.5;refGene_gn=ATP1B2;refGene_tx=XM_019817099.2;refGene_gn=ATP1B2;refGene_tx=XM_019817100.2;refGene_gn=ATP1B2;refGene_tx=XM_019817101.2
E1	2532225	2532390
refGene_gn=ATP1B2;refGene_tx=XM_003996185.5;refGene_gn=ATP1B2;refGene_tx=XM_019817099.2;refGene_gn=ATP1B2;refGene_tx=XM_019817100.2;refGene_gn=ATP1B2;refGene_tx=XM_019817101.2
E1	2540122	2540204	refGene_gn=TP53;refGene_tx=NM_001009294.1
E1	2540739	2540846	refGene_gn=TP53;refGene_tx=NM_001009294.1
E1	2541119	2541193	refGene_gn=TP53;refGene_tx=NM_001009294.1
E1	2541280	2541417	refGene_gn=TP53;refGene_tx=NM_001009294.1
E1	2541684	2541794	refGene_gn=TP53;refGene_tx=NM_001009294.1
E1	2542025	2542138	refGene_gn=TP53;refGene_tx=NM_001009294.1
E1	2542219	2542406	refGene_gn=TP53;refGene_tx=NM_001009294.1
E1	2542842	2543097	refGene_gn=TP53;refGene_tx=NM_001009294.1
E1	2543190	2543212	refGene_gn=TP53;refGene_tx=NM_001009294.1
E1	2543353	2543427	refGene_gn=TP53;refGene_tx=NM_001009294.1
E1	2552395	2552790	refGene_gn=WRAP53;refGene_tx=XM_003996266.5
E1	2552925	2553024	refGene_gn=WRAP53;refGene_tx=XM_003996266.5
E1	2553288	2553400	refGene_gn=WRAP53;refGene_tx=XM_003996266.5
E1	2561630	2561719	refGene_gn=WRAP53;refGene_tx=XM_003996266.5
E1	2561949	2562040	refGene_gn=WRAP53;refGene_tx=XM_003996266.5
E1	2562133	2562266	refGene_gn=WRAP53;refGene_tx=XM_003996266.5
E1	2562832	2563041	refGene_gn=WRAP53;refGene_tx=XM_003996266.5
E1	2563234	2563338	refGene_gn=WRAP53;refGene_tx=XM_003996266.5
E1	2563481	2563616	refGene_gn=WRAP53;refGene_tx=XM_003996266.5
E1	2563730	2563977	refGene_gn=WRAP53;refGene_tx=XM_003996266.5
E1	2566152	2566274
refGene_gn=EFNB3;refGene_tx=NM_001114343.1;refGene_gn=EFNB3;refGene_tx=XM_023243402.1
E1	2568458	2568751
refGene_gn=EFNB3;refGene_tx=NM_001114343.1;refGene_gn=EFNB3;refGene_tx=XM_023243402.1
E1	2568940	2569033
refGene_gn=EFNB3;refGene_tx=NM_001114343.1;refGene_gn=EFNB3;refGene_tx=XM_023243402.1
E1	2569148	2569253
refGene_gn=EFNB3;refGene_tx=NM_001114343.1;refGene_gn=EFNB3;refGene_tx=XM_023243402.1
E1	2569711	2570121	refGene_gn=EFNB3;refGene_tx=NM_001114343.1
E1	2570561	2570977	refGene_gn=EFNB3;refGene_tx=XM_023243402.1
E1	2579270	2579424	refGene_gn=DNAH2;refGene_tx=XM_023244573.1
E1	2585475	2585534	refGene_gn=DNAH2;refGene_tx=XM_023244573.1
E1	2587252	2587423	refGene_gn=DNAH2;refGene_tx=XM_023244573.1
E1	2595706	2595935	refGene_gn=DNAH2;refGene_tx=XM_023244573.1
E1	2596496	2596607	refGene_gn=DNAH2;refGene_tx=XM_023244573.1

| | | | |
|---|---|---|---|
| E1 | 2596776 | 2597015 | refGene_gn=DNAH2;refGene_tx=XM_023244573.1 |
| E1 | 2598220 | 2598412 | refGene_gn=DNAH2;refGene_tx=XM_023244573.1 |
| E1 | 2598877 | 2599035 | refGene_gn=DNAH2;refGene_tx=XM_023244573.1 |
| E1 | 2599448 | 2599536 | refGene_gn=DNAH2;refGene_tx=XM_023244573.1 |
| E1 | 2599870 | 2600076 | refGene_gn=DNAH2;refGene_tx=XM_023244573.1 |
| E1 | 2600438 | 2600568 | refGene_gn=DNAH2;refGene_tx=XM_023244573.1 |
| E1 | 2600836 | 2601019 | refGene_gn=DNAH2;refGene_tx=XM_023244573.1 |
| E1 | 2601823 | 2602038 | refGene_gn=DNAH2;refGene_tx=XM_023244573.1 |
| E1 | 2612131 | 2612278 | refGene_gn=DNAH2;refGene_tx=XM_023244573.1 |
| E1 | 2613289 | 2613446 | refGene_gn=DNAH2;refGene_tx=XM_023244573.1 |
| E1 | 2613730 | 2613970 | refGene_gn=DNAH2;refGene_tx=XM_023244573.1 |
| E1 | 2614272 | 2614461 | refGene_gn=DNAH2;refGene_tx=XM_023244573.1 |
| E1 | 2614668 | 2614816 | refGene_gn=DNAH2;refGene_tx=XM_023244573.1 |
| E1 | 2615054 | 2615247 | refGene_gn=DNAH2;refGene_tx=XM_023244573.1 |
| E1 | 2616180 | 2616381 | refGene_gn=DNAH2;refGene_tx=XM_023244573.1 |
| E1 | 2616468 | 2616625 | refGene_gn=DNAH2;refGene_tx=XM_023244573.1 |
| E1 | 2616984 | 2617159 | refGene_gn=DNAH2;refGene_tx=XM_023244573.1 |
| E1 | 2617590 | 2617754 | refGene_gn=DNAH2;refGene_tx=XM_023244573.1 |
| E1 | 2618129 | 2618291 | refGene_gn=DNAH2;refGene_tx=XM_023244573.1 |
| E1 | 2618379 | 2618483 | refGene_gn=DNAH2;refGene_tx=XM_023244573.1 |
| E1 | 2620205 | 2620362 | refGene_gn=DNAH2;refGene_tx=XM_023244573.1 |
| E1 | 2620515 | 2620598 | refGene_gn=DNAH2;refGene_tx=XM_023244573.1 |
| E1 | 2620728 | 2620909 | refGene_gn=DNAH2;refGene_tx=XM_023244573.1 |
| E1 | 2621767 | 2621906 | refGene_gn=DNAH2;refGene_tx=XM_023244573.1 |
| E1 | 2624179 | 2624397 | refGene_gn=DNAH2;refGene_tx=XM_023244573.1 |
| E1 | 2624664 | 2624766 | refGene_gn=DNAH2;refGene_tx=XM_023244573.1 |
| E1 | 2625103 | 2625229 | refGene_gn=DNAH2;refGene_tx=XM_023244573.1 |
| E1 | 2625497 | 2625608 | refGene_gn=DNAH2;refGene_tx=XM_023244573.1 |
| E1 | 2625842 | 2626031 | refGene_gn=DNAH2;refGene_tx=XM_023244573.1 |
| E1 | 2626408 | 2626512 | refGene_gn=DNAH2;refGene_tx=XM_023244573.1 |
| E1 | 2626599 | 2626789 | refGene_gn=DNAH2;refGene_tx=XM_023244573.1 |
| E1 | 2627317 | 2627498 | refGene_gn=DNAH2;refGene_tx=XM_023244573.1 |
| E1 | 2628174 | 2628302 | refGene_gn=DNAH2;refGene_tx=XM_023244573.1 |
| E1 | 2628649 | 2628802 | refGene_gn=DNAH2;refGene_tx=XM_023244573.1 |
| E1 | 2629077 | 2629203 | refGene_gn=DNAH2;refGene_tx=XM_023244573.1 |
| E1 | 2629996 | 2630215 | refGene_gn=DNAH2;refGene_tx=XM_023244573.1 |
| E1 | 2630450 | 2630568 | refGene_gn=DNAH2;refGene_tx=XM_023244573.1 |
| E1 | 2630789 | 2630926 | refGene_gn=DNAH2;refGene_tx=XM_023244573.1 |
| E1 | 2631393 | 2631531 | refGene_gn=DNAH2;refGene_tx=XM_023244573.1 |
| E1 | 2631616 | 2631775 | refGene_gn=DNAH2;refGene_tx=XM_023244573.1 |
| E1 | 2634908 | 2635061 | refGene_gn=DNAH2;refGene_tx=XM_023244573.1 |
| E1 | 2635258 | 2635350 | refGene_gn=DNAH2;refGene_tx=XM_023244573.1 |
| E1 | 2635630 | 2635829 | refGene_gn=DNAH2;refGene_tx=XM_023244573.1 |
| E1 | 2635961 | 2636186 | refGene_gn=DNAH2;refGene_tx=XM_023244573.1 |
| E1 | 2637176 | 2637281 | refGene_gn=DNAH2;refGene_tx=XM_023244573.1 |
| E1 | 2638425 | 2638614 | refGene_gn=DNAH2;refGene_tx=XM_023244573.1 |
| E1 | 2638715 | 2638801 | refGene_gn=DNAH2;refGene_tx=XM_023244573.1 |
| E1 | 2638926 | 2639057 | refGene_gn=DNAH2;refGene_tx=XM_023244573.1 |
| E1 | 2639206 | 2639356 | refGene_gn=DNAH2;refGene_tx=XM_023244573.1 |
| E1 | 2639688 | 2639856 | refGene_gn=DNAH2;refGene_tx=XM_023244573.1 |

E1	2640058	2640219	refGene_gn=DNAH2;refGene_tx=XM_023244573.1
E1	2640459	2640599	refGene_gn=DNAH2;refGene_tx=XM_023244573.1
E1	2642166	2642299	refGene_gn=DNAH2;refGene_tx=XM_023244573.1
E1	2642397	2642537	refGene_gn=DNAH2;refGene_tx=XM_023244573.1
E1	2644323	2644534	refGene_gn=DNAH2;refGene_tx=XM_023244573.1
E1	2644763	2644880	refGene_gn=DNAH2;refGene_tx=XM_023244573.1
E1	2645078	2645220	refGene_gn=DNAH2;refGene_tx=XM_023244573.1
E1	2646000	2646170	refGene_gn=DNAH2;refGene_tx=XM_023244573.1
E1	2646281	2646398	refGene_gn=DNAH2;refGene_tx=XM_023244573.1
E1	2655357	2655522	refGene_gn=DNAH2;refGene_tx=XM_023244573.1
E1	2656075	2656201	refGene_gn=DNAH2;refGene_tx=XM_023244573.1
E1	2656376	2656525	refGene_gn=DNAH2;refGene_tx=XM_023244573.1
E1	2656596	2656651	refGene_gn=DNAH2;refGene_tx=XM_023244573.1
E1	2656652	2656664	refGene_gn=DNAH2;refGene_tx=XM_023244573.1
E1	2656813	2656964	refGene_gn=DNAH2;refGene_tx=XM_023244573.1
E1	2657196	2657345	refGene_gn=DNAH2;refGene_tx=XM_023244573.1
E1	2657541	2657675	refGene_gn=DNAH2;refGene_tx=XM_023244573.1
E1	2657792	2657937	refGene_gn=DNAH2;refGene_tx=XM_023244573.1
E1	2658654	2658854	refGene_gn=DNAH2;refGene_tx=XM_023244573.1
E1	2659772	2659899	refGene_gn=DNAH2;refGene_tx=XM_023244573.1
E1	2661459	2661646	refGene_gn=DNAH2;refGene_tx=XM_023244573.1
E1	2661741	2661902	refGene_gn=DNAH2;refGene_tx=XM_023244573.1
E1	2662046	2662221	refGene_gn=DNAH2;refGene_tx=XM_023244573.1
E1	2662442	2662582	refGene_gn=DNAH2;refGene_tx=XM_023244573.1
E1	2662583	2662634	refGene_gn=DNAH2;refGene_tx=XM_023244573.1
E1	2667657	2667849	refGene_gn=DNAH2;refGene_tx=XM_023244573.1
E1	2668009	2668194	refGene_gn=DNAH2;refGene_tx=XM_023244573.1
E1	2668461	2668690	refGene_gn=DNAH2;refGene_tx=XM_023244573.1
E1	2668778	2668930	refGene_gn=DNAH2;refGene_tx=XM_023244573.1
E1	2669044	2669159	refGene_gn=DNAH2;refGene_tx=XM_023244573.1
E1	2669638	2669815	refGene_gn=DNAH2;refGene_tx=XM_023244573.1
E1	2669917	2669992	refGene_gn=DNAH2;refGene_tx=XM_023244573.1
E1	2670152	2670303	refGene_gn=DNAH2;refGene_tx=XM_023244573.1
E1	2670425	2670580	refGene_gn=DNAH2;refGene_tx=XM_023244573.1
E1	2683859	2683996
refGene_gn=KDM6B;refGene_tx=XM_023243442.1;refGene_gn=KDM6B;refGene_tx=XM_023243443.1;refGene_gn=KDM6B;refGene_tx=XM_023243444.1;refGene_gn=KDM6B;refGene_tx=XM_023243445.1;refGene_gn=KDM6B;refGene_tx=XM_023243446.1;refGene_gn=KDM6B;refGene_tx=XM_023243447.1;refGene_gn=KDM6B;refGene_tx=XM_023243448.1;refGene_gn=KDM6B;refGene_tx=XM_023243449.1;refGene_gn=KDM6B;refGene_tx=XM_023243450.1
E1	2684179	2684278
refGene_gn=KDM6B;refGene_tx=XM_023243442.1;refGene_gn=KDM6B;refGene_tx=XM_023243443.1;refGene_gn=KDM6B;refGene_tx=XM_023243444.1;refGene_gn=KDM6B;refGene_tx=XM_023243445.1;refGene_gn=KDM6B;refGene_tx=XM_023243446.1;refGene_gn=KDM6B;refGene_tx=XM_023243447.1;refGene_gn=KDM6B;refGene_tx=XM_023243448.1;refGene_gn=KDM6B;refGene_tx=XM_023243449.1;refGene_gn=KDM6B;refGene_tx=XM_023243450.1
E1	2684380	2684600
refGene_gn=KDM6B;refGene_tx=XM_023243442.1;refGene_gn=KDM6B;refGene_tx

=XM_023243443.1;refGene_gn=KDM6B;refGene_tx=XM_023243444.1;refGene_gn=KDM6B;refGene_tx=XM_023243445.1;refGene_gn=KDM6B;refGene_tx=XM_023243446.1;refGene_gn=KDM6B;refGene_tx=XM_023243447.1;refGene_gn=KDM6B;refGene_tx=XM_023243448.1;refGene_gn=KDM6B;refGene_tx=XM_023243449.1;refGene_gn=KDM6B;refGene_tx=XM_023243450.1
E1 2684682 2684775
refGene_gn=KDM6B;refGene_tx=XM_023243442.1;refGene_gn=KDM6B;refGene_tx=XM_023243443.1;refGene_gn=KDM6B;refGene_tx=XM_023243444.1;refGene_gn=KDM6B;refGene_tx=XM_023243445.1;refGene_gn=KDM6B;refGene_tx=XM_023243446.1;refGene_gn=KDM6B;refGene_tx=XM_023243447.1;refGene_gn=KDM6B;refGene_tx=XM_023243448.1;refGene_gn=KDM6B;refGene_tx=XM_023243449.1;refGene_gn=KDM6B;refGene_tx=XM_023243450.1
E1 2684861 2685023
refGene_gn=KDM6B;refGene_tx=XM_023243442.1;refGene_gn=KDM6B;refGene_tx=XM_023243443.1;refGene_gn=KDM6B;refGene_tx=XM_023243444.1;refGene_gn=KDM6B;refGene_tx=XM_023243445.1;refGene_gn=KDM6B;refGene_tx=XM_023243446.1;refGene_gn=KDM6B;refGene_tx=XM_023243447.1;refGene_gn=KDM6B;refGene_tx=XM_023243448.1;refGene_gn=KDM6B;refGene_tx=XM_023243449.1;refGene_gn=KDM6B;refGene_tx=XM_023243450.1
E1 2685102 2685303
refGene_gn=KDM6B;refGene_tx=XM_023243442.1;refGene_gn=KDM6B;refGene_tx=XM_023243443.1;refGene_gn=KDM6B;refGene_tx=XM_023243444.1;refGene_gn=KDM6B;refGene_tx=XM_023243445.1;refGene_gn=KDM6B;refGene_tx=XM_023243446.1;refGene_gn=KDM6B;refGene_tx=XM_023243447.1;refGene_gn=KDM6B;refGene_tx=XM_023243448.1;refGene_gn=KDM6B;refGene_tx=XM_023243449.1;refGene_gn=KDM6B;refGene_tx=XM_023243450.1
E1 2685390 2685738
refGene_gn=KDM6B;refGene_tx=XM_023243442.1;refGene_gn=KDM6B;refGene_tx=XM_023243443.1;refGene_gn=KDM6B;refGene_tx=XM_023243444.1;refGene_gn=KDM6B;refGene_tx=XM_023243445.1;refGene_gn=KDM6B;refGene_tx=XM_023243446.1;refGene_gn=KDM6B;refGene_tx=XM_023243447.1;refGene_gn=KDM6B;refGene_tx=XM_023243448.1;refGene_gn=KDM6B;refGene_tx=XM_023243449.1;refGene_gn=KDM6B;refGene_tx=XM_023243450.1
E1 2685859 2688015
refGene_gn=KDM6B;refGene_tx=XM_023243442.1;refGene_gn=KDM6B;refGene_tx=XM_023243443.1;refGene_gn=KDM6B;refGene_tx=XM_023243444.1;refGene_gn=KDM6B;refGene_tx=XM_023243445.1;refGene_gn=KDM6B;refGene_tx=XM_023243446.1;refGene_gn=KDM6B;refGene_tx=XM_023243447.1;refGene_gn=KDM6B;refGene_tx=XM_023243448.1;refGene_gn=KDM6B;refGene_tx=XM_023243449.1;refGene_gn=KDM6B;refGene_tx=XM_023243450.1
E1 2688131 2688258
refGene_gn=KDM6B;refGene_tx=XM_023243442.1;refGene_gn=KDM6B;refGene_tx=XM_023243443.1;refGene_gn=KDM6B;refGene_tx=XM_023243444.1;refGene_gn=KDM6B;refGene_tx=XM_023243445.1;refGene_gn=KDM6B;refGene_tx=XM_023243446.1;refGene_gn=KDM6B;refGene_tx=XM_023243447.1;refGene_gn=KDM6B;refGene_tx=XM_023243448.1;refGene_gn=KDM6B;refGene_tx=XM_023243449.1;refGene_gn=KDM6B;refGene_tx=XM_023243450.1
E1 2688382 2688488
refGene_gn=KDM6B;refGene_tx=XM_023243442.1;refGene_gn=KDM6B;refGene_tx=XM_023243443.1;refGene_gn=KDM6B;refGene_tx=XM_023243444.1;refGene_gn=

KDM6B;refGene_tx=XM_023243445.1;refGene_gn=KDM6B;refGene_tx=XM_0232434
46.1;refGene_gn=KDM6B;refGene_tx=XM_023243447.1;refGene_gn=KDM6B;refGe
ne_tx=XM_023243448.1;refGene_gn=KDM6B;refGene_tx=XM_023243449.1;refGen
e_gn=KDM6B;refGene_tx=XM_023243450.1
E1 2689511 2689717
refGene_gn=KDM6B;refGene_tx=XM_023243442.1;refGene_gn=KDM6B;refGene_tx
=XM_023243443.1;refGene_gn=KDM6B;refGene_tx=XM_023243444.1;refGene_gn=
KDM6B;refGene_tx=XM_023243445.1;refGene_gn=KDM6B;refGene_tx=XM_0232434
46.1;refGene_gn=KDM6B;refGene_tx=XM_023243447.1;refGene_gn=KDM6B;refGe
ne_tx=XM_023243448.1;refGene_gn=KDM6B;refGene_tx=XM_023243449.1;refGen
e_gn=KDM6B;refGene_tx=XM_023243450.1
E1 2689818 2689883
refGene_gn=KDM6B;refGene_tx=XM_023243442.1;refGene_gn=KDM6B;refGene_tx
=XM_023243443.1;refGene_gn=KDM6B;refGene_tx=XM_023243444.1;refGene_gn=
KDM6B;refGene_tx=XM_023243445.1;refGene_gn=KDM6B;refGene_tx=XM_0232434
46.1;refGene_gn=KDM6B;refGene_tx=XM_023243447.1;refGene_gn=KDM6B;refGe
ne_tx=XM_023243448.1;refGene_gn=KDM6B;refGene_tx=XM_023243449.1;refGen
e_gn=KDM6B;refGene_tx=XM_023243450.1
E1 2689970 2690042
refGene_gn=KDM6B;refGene_tx=XM_023243442.1;refGene_gn=KDM6B;refGene_tx
=XM_023243443.1;refGene_gn=KDM6B;refGene_tx=XM_023243444.1;refGene_gn=
KDM6B;refGene_tx=XM_023243445.1;refGene_gn=KDM6B;refGene_tx=XM_0232434
46.1;refGene_gn=KDM6B;refGene_tx=XM_023243447.1;refGene_gn=KDM6B;refGe
ne_tx=XM_023243448.1;refGene_gn=KDM6B;refGene_tx=XM_023243449.1;refGen
e_gn=KDM6B;refGene_tx=XM_023243450.1
E1 2690128 2690277
refGene_gn=KDM6B;refGene_tx=XM_023243442.1;refGene_gn=KDM6B;refGene_tx
=XM_023243443.1;refGene_gn=KDM6B;refGene_tx=XM_023243444.1;refGene_gn=
KDM6B;refGene_tx=XM_023243445.1;refGene_gn=KDM6B;refGene_tx=XM_0232434
46.1;refGene_gn=KDM6B;refGene_tx=XM_023243447.1;refGene_gn=KDM6B;refGe
ne_tx=XM_023243448.1;refGene_gn=KDM6B;refGene_tx=XM_023243449.1;refGen
e_gn=KDM6B;refGene_tx=XM_023243450.1
E1 2690412 2690527
refGene_gn=KDM6B;refGene_tx=XM_023243442.1;refGene_gn=KDM6B;refGene_tx
=XM_023243443.1;refGene_gn=KDM6B;refGene_tx=XM_023243444.1;refGene_gn=
KDM6B;refGene_tx=XM_023243445.1;refGene_gn=KDM6B;refGene_tx=XM_0232434
46.1;refGene_gn=KDM6B;refGene_tx=XM_023243447.1;refGene_gn=KDM6B;refGe
ne_tx=XM_023243448.1;refGene_gn=KDM6B;refGene_tx=XM_023243449.1;refGen
e_gn=KDM6B;refGene_tx=XM_023243450.1
E1 2690625 2690813
refGene_gn=KDM6B;refGene_tx=XM_023243442.1;refGene_gn=KDM6B;refGene_tx
=XM_023243443.1;refGene_gn=KDM6B;refGene_tx=XM_023243444.1;refGene_gn=
KDM6B;refGene_tx=XM_023243445.1;refGene_gn=KDM6B;refGene_tx=XM_0232434
46.1;refGene_gn=KDM6B;refGene_tx=XM_023243447.1;refGene_gn=KDM6B;refGe
ne_tx=XM_023243448.1;refGene_gn=KDM6B;refGene_tx=XM_023243449.1;refGen
e_gn=KDM6B;refGene_tx=XM_023243450.1
E1 2690952 2691094
refGene_gn=KDM6B;refGene_tx=XM_023243442.1;refGene_gn=KDM6B;refGene_tx
=XM_023243443.1;refGene_gn=KDM6B;refGene_tx=XM_023243444.1;refGene_gn=
KDM6B;refGene_tx=XM_023243445.1;refGene_gn=KDM6B;refGene_tx=XM_0232434

46.1;refGene_gn=KDM6B;refGene_tx=XM_023243447.1;refGene_gn=KDM6B;refGene_tx=XM_023243448.1;refGene_gn=KDM6B;refGene_tx=XM_023243449.1;refGene_gn=KDM6B;refGene_tx=XM_023243450.1
E1	2691393	2691520
refGene_gn=KDM6B;refGene_tx=XM_023243442.1;refGene_gn=KDM6B;refGene_tx=XM_023243443.1;refGene_gn=KDM6B;refGene_tx=XM_023243444.1;refGene_gn=KDM6B;refGene_tx=XM_023243445.1;refGene_gn=KDM6B;refGene_tx=XM_023243446.1;refGene_gn=KDM6B;refGene_tx=XM_023243447.1;refGene_gn=KDM6B;refGene_tx=XM_023243448.1;refGene_gn=KDM6B;refGene_tx=XM_023243449.1;refGene_gn=KDM6B;refGene_tx=XM_023243450.1
E1	2691608	2691779
refGene_gn=KDM6B;refGene_tx=XM_023243442.1;refGene_gn=KDM6B;refGene_tx=XM_023243443.1;refGene_gn=KDM6B;refGene_tx=XM_023243444.1;refGene_gn=KDM6B;refGene_tx=XM_023243445.1;refGene_gn=KDM6B;refGene_tx=XM_023243446.1;refGene_gn=KDM6B;refGene_tx=XM_023243447.1;refGene_gn=KDM6B;refGene_tx=XM_023243448.1;refGene_gn=KDM6B;refGene_tx=XM_023243449.1;refGene_gn=KDM6B;refGene_tx=XM_023243450.1
E1	2691917	2691941
refGene_gn=KDM6B;refGene_tx=XM_023243442.1;refGene_gn=KDM6B;refGene_tx=XM_023243443.1;refGene_gn=KDM6B;refGene_tx=XM_023243444.1;refGene_gn=KDM6B;refGene_tx=XM_023243445.1;refGene_gn=KDM6B;refGene_tx=XM_023243446.1;refGene_gn=KDM6B;refGene_tx=XM_023243447.1;refGene_gn=KDM6B;refGene_tx=XM_023243448.1;refGene_gn=KDM6B;refGene_tx=XM_023243449.1;refGene_gn=KDM6B;refGene_tx=XM_023243450.1
E1	2693579	2693789	refGene_gn=TMEM88;refGene_tx=XM_011288897.2
E1	2693905	2694175	refGene_gn=TMEM88;refGene_tx=XM_011288897.2
E1	2695161	2695274	refGene_gn=NAA38;refGene_tx=XM_003996186.5
E1	2695417	2695601	refGene_gn=NAA38;refGene_tx=XM_003996186.5
E1	2695780	2695861	refGene_gn=NAA38;refGene_tx=XM_003996186.5
E1	2696484	2696668
refGene_gn=LOC101100259;refGene_tx=XM_023243936.1;refGene_gn=LOC101100259;refGene_tx=XM_023243937.1;refGene_gn=LOC101100259;refGene_tx=XM_023243938.1
E1	2696775	2696852
refGene_gn=LOC101100259;refGene_tx=XM_023243936.1;refGene_gn=LOC101100259;refGene_tx=XM_023243937.1;refGene_gn=LOC101100259;refGene_tx=XM_023243938.1
E1	2696988	2697207
refGene_gn=LOC101100259;refGene_tx=XM_023243936.1;refGene_gn=LOC101100259;refGene_tx=XM_023243937.1;refGene_gn=LOC101100259;refGene_tx=XM_023243938.1
E1	2697769	2698000
refGene_gn=LOC101100259;refGene_tx=XM_023243936.1;refGene_gn=LOC101100259;refGene_tx=XM_023243937.1;refGene_gn=LOC101100259;refGene_tx=XM_023243938.1
E1	2722193	2722470
refGene_gn=CHD3;refGene_tx=XM_023243920.1;refGene_gn=CHD3;refGene_tx=XM_023243921.1;refGene_gn=CHD3;refGene_tx=XM_023243922.1;refGene_gn=CHD3;refGene_tx=XM_023243923.1;refGene_gn=CHD3;refGene_tx=XM_023243924.1;refGene_gn=CHD3;refGene_tx=XM_023243925.1;refGene_gn=CHD3;refGene_tx=X

M_023243926.1;refGene_gn=CHD3;refGene_tx=XM_023243927.1;refGene_gn=CHD3;refGene_tx=XM_023243928.1;refGene_gn=CHD3;refGene_tx=XM_023243929.1;refGene_gn=CHD3;refGene_tx=XM_023243930.1;refGene_gn=CHD3;refGene_tx=XM_023243931.1;refGene_gn=CHD3;refGene_tx=XM_023243932.1;refGene_gn=CHD3;refGene_tx=XM_023243933.1;refGene_gn=CHD3;refGene_tx=XM_023243934.1;refGene_gn=CHD3;refGene_tx=XM_023243935.1
E1 2727115 2727228
refGene_gn=CHD3;refGene_tx=XM_023243920.1;refGene_gn=CHD3;refGene_tx=XM_023243921.1;refGene_gn=CHD3;refGene_tx=XM_023243922.1;refGene_gn=CHD3;refGene_tx=XM_023243923.1;refGene_gn=CHD3;refGene_tx=XM_023243924.1;refGene_gn=CHD3;refGene_tx=XM_023243925.1;refGene_gn=CHD3;refGene_tx=XM_023243926.1;refGene_gn=CHD3;refGene_tx=XM_023243927.1;refGene_gn=CHD3;refGene_tx=XM_023243928.1;refGene_gn=CHD3;refGene_tx=XM_023243929.1;refGene_gn=CHD3;refGene_tx=XM_023243930.1;refGene_gn=CHD3;refGene_tx=XM_023243931.1;refGene_gn=CHD3;refGene_tx=XM_023243932.1;refGene_gn=CHD3;refGene_tx=XM_023243933.1;refGene_gn=CHD3;refGene_tx=XM_023243934.1;refGene_gn=CHD3;refGene_tx=XM_023243935.1
E1 2727747 2727918
refGene_gn=CHD3;refGene_tx=XM_023243920.1;refGene_gn=CHD3;refGene_tx=XM_023243921.1;refGene_gn=CHD3;refGene_tx=XM_023243922.1;refGene_gn=CHD3;refGene_tx=XM_023243923.1;refGene_gn=CHD3;refGene_tx=XM_023243924.1;refGene_gn=CHD3;refGene_tx=XM_023243925.1;refGene_gn=CHD3;refGene_tx=XM_023243926.1;refGene_gn=CHD3;refGene_tx=XM_023243927.1;refGene_gn=CHD3;refGene_tx=XM_023243928.1;refGene_gn=CHD3;refGene_tx=XM_023243929.1;refGene_gn=CHD3;refGene_tx=XM_023243930.1;refGene_gn=CHD3;refGene_tx=XM_023243931.1;refGene_gn=CHD3;refGene_tx=XM_023243932.1;refGene_gn=CHD3;refGene_tx=XM_023243933.1;refGene_gn=CHD3;refGene_tx=XM_023243934.1;refGene_gn=CHD3;refGene_tx=XM_023243935.1
E1 2728093 2728236
refGene_gn=CHD3;refGene_tx=XM_023243920.1;refGene_gn=CHD3;refGene_tx=XM_023243921.1;refGene_gn=CHD3;refGene_tx=XM_023243922.1;refGene_gn=CHD3;refGene_tx=XM_023243923.1;refGene_gn=CHD3;refGene_tx=XM_023243924.1;refGene_gn=CHD3;refGene_tx=XM_023243925.1;refGene_gn=CHD3;refGene_tx=XM_023243926.1;refGene_gn=CHD3;refGene_tx=XM_023243927.1;refGene_gn=CHD3;refGene_tx=XM_023243928.1;refGene_gn=CHD3;refGene_tx=XM_023243929.1;refGene_gn=CHD3;refGene_tx=XM_023243930.1;refGene_gn=CHD3;refGene_tx=XM_023243931.1;refGene_gn=CHD3;refGene_tx=XM_023243932.1;refGene_gn=CHD3;refGene_tx=XM_023243933.1;refGene_gn=CHD3;refGene_tx=XM_023243934.1;refGene_gn=CHD3;refGene_tx=XM_023243935.1
E1 2729948 2730232
refGene_gn=CHD3;refGene_tx=XM_023243920.1;refGene_gn=CHD3;refGene_tx=XM_023243921.1;refGene_gn=CHD3;refGene_tx=XM_023243922.1;refGene_gn=CHD3;refGene_tx=XM_023243923.1;refGene_gn=CHD3;refGene_tx=XM_023243924.1;refGene_gn=CHD3;refGene_tx=XM_023243925.1;refGene_gn=CHD3;refGene_tx=XM_023243926.1;refGene_gn=CHD3;refGene_tx=XM_023243927.1;refGene_gn=CHD3;refGene_tx=XM_023243928.1;refGene_gn=CHD3;refGene_tx=XM_023243929.1;refGene_gn=CHD3;refGene_tx=XM_023243930.1;refGene_gn=CHD3;refGene_tx=XM_023243931.1;refGene_gn=CHD3;refGene_tx=XM_023243932.1;refGene_gn=CHD3;refGene_tx=XM_023243933.1;refGene_gn=CHD3;refGene_tx=XM_023243934.1;refGene_gn=CHD3;refGene_tx=XM_023243935.1

E1 2730449 2730580
refGene_gn=CHD3;refGene_tx=XM_023243920.1;refGene_gn=CHD3;refGene_tx=X
M_023243921.1;refGene_gn=CHD3;refGene_tx=XM_023243922.1;refGene_gn=CHD
3;refGene_tx=XM_023243923.1;refGene_gn=CHD3;refGene_tx=XM_023243924.1;
refGene_gn=CHD3;refGene_tx=XM_023243925.1;refGene_gn=CHD3;refGene_tx=X
M_023243926.1;refGene_gn=CHD3;refGene_tx=XM_023243927.1;refGene_gn=CHD
3;refGene_tx=XM_023243928.1;refGene_gn=CHD3;refGene_tx=XM_023243929.1;
refGene_gn=CHD3;refGene_tx=XM_023243930.1;refGene_gn=CHD3;refGene_tx=X
M_023243931.1;refGene_gn=CHD3;refGene_tx=XM_023243932.1;refGene_gn=CHD
3;refGene_tx=XM_023243933.1;refGene_gn=CHD3;refGene_tx=XM_023243934.1;
refGene_gn=CHD3;refGene_tx=XM_023243935.1
E1 2730751 2730902
refGene_gn=CHD3;refGene_tx=XM_023243920.1;refGene_gn=CHD3;refGene_tx=X
M_023243921.1;refGene_gn=CHD3;refGene_tx=XM_023243922.1;refGene_gn=CHD
3;refGene_tx=XM_023243923.1;refGene_gn=CHD3;refGene_tx=XM_023243924.1;
refGene_gn=CHD3;refGene_tx=XM_023243925.1;refGene_gn=CHD3;refGene_tx=X
M_023243926.1;refGene_gn=CHD3;refGene_tx=XM_023243927.1;refGene_gn=CHD
3;refGene_tx=XM_023243928.1;refGene_gn=CHD3;refGene_tx=XM_023243929.1;
refGene_gn=CHD3;refGene_tx=XM_023243930.1;refGene_gn=CHD3;refGene_tx=X
M_023243931.1;refGene_gn=CHD3;refGene_tx=XM_023243932.1;refGene_gn=CHD
3;refGene_tx=XM_023243933.1;refGene_gn=CHD3;refGene_tx=XM_023243934.1;
refGene_gn=CHD3;refGene_tx=XM_023243935.1
E1 2731063 2731260
refGene_gn=CHD3;refGene_tx=XM_023243920.1;refGene_gn=CHD3;refGene_tx=X
M_023243921.1;refGene_gn=CHD3;refGene_tx=XM_023243922.1;refGene_gn=CHD
3;refGene_tx=XM_023243923.1;refGene_gn=CHD3;refGene_tx=XM_023243924.1;
refGene_gn=CHD3;refGene_tx=XM_023243925.1;refGene_gn=CHD3;refGene_tx=X
M_023243926.1;refGene_gn=CHD3;refGene_tx=XM_023243927.1;refGene_gn=CHD
3;refGene_tx=XM_023243928.1;refGene_gn=CHD3;refGene_tx=XM_023243929.1;
refGene_gn=CHD3;refGene_tx=XM_023243930.1;refGene_gn=CHD3;refGene_tx=X
M_023243931.1;refGene_gn=CHD3;refGene_tx=XM_023243932.1;refGene_gn=CHD
3;refGene_tx=XM_023243933.1;refGene_gn=CHD3;refGene_tx=XM_023243934.1;
refGene_gn=CHD3;refGene_tx=XM_023243935.1
E1 2731568 2731802
refGene_gn=CHD3;refGene_tx=XM_023243920.1;refGene_gn=CHD3;refGene_tx=X
M_023243921.1;refGene_gn=CHD3;refGene_tx=XM_023243922.1;refGene_gn=CHD
3;refGene_tx=XM_023243923.1;refGene_gn=CHD3;refGene_tx=XM_023243924.1;
refGene_gn=CHD3;refGene_tx=XM_023243925.1;refGene_gn=CHD3;refGene_tx=X
M_023243926.1;refGene_gn=CHD3;refGene_tx=XM_023243927.1;refGene_gn=CHD
3;refGene_tx=XM_023243928.1;refGene_gn=CHD3;refGene_tx=XM_023243929.1;
refGene_gn=CHD3;refGene_tx=XM_023243930.1;refGene_gn=CHD3;refGene_tx=X
M_023243931.1;refGene_gn=CHD3;refGene_tx=XM_023243932.1;refGene_gn=CHD
3;refGene_tx=XM_023243933.1;refGene_gn=CHD3;refGene_tx=XM_023243934.1;
refGene_gn=CHD3;refGene_tx=XM_023243935.1
E1 2732000 2732204
refGene_gn=CHD3;refGene_tx=XM_023243920.1;refGene_gn=CHD3;refGene_tx=X
M_023243921.1;refGene_gn=CHD3;refGene_tx=XM_023243922.1;refGene_gn=CHD
3;refGene_tx=XM_023243923.1;refGene_gn=CHD3;refGene_tx=XM_023243924.1;
refGene_gn=CHD3;refGene_tx=XM_023243925.1;refGene_gn=CHD3;refGene_tx=X
M_023243926.1;refGene_gn=CHD3;refGene_tx=XM_023243927.1;refGene_gn=CHD

3;refGene_tx=XM_023243928.1;refGene_gn=CHD3;refGene_tx=XM_023243929.1;
refGene_gn=CHD3;refGene_tx=XM_023243930.1;refGene_gn=CHD3;refGene_tx=X
M_023243931.1;refGene_gn=CHD3;refGene_tx=XM_023243932.1;refGene_gn=CHD
3;refGene_tx=XM_023243933.1;refGene_gn=CHD3;refGene_tx=XM_023243934.1;
refGene_gn=CHD3;refGene_tx=XM_023243935.1
E1 2732695 2732907
refGene_gn=CHD3;refGene_tx=XM_023243920.1;refGene_gn=CHD3;refGene_tx=X
M_023243921.1;refGene_gn=CHD3;refGene_tx=XM_023243922.1;refGene_gn=CHD
3;refGene_tx=XM_023243923.1;refGene_gn=CHD3;refGene_tx=XM_023243924.1;
refGene_gn=CHD3;refGene_tx=XM_023243925.1;refGene_gn=CHD3;refGene_tx=X
M_023243926.1;refGene_gn=CHD3;refGene_tx=XM_023243927.1;refGene_gn=CHD
3;refGene_tx=XM_023243928.1;refGene_gn=CHD3;refGene_tx=XM_023243929.1;
refGene_gn=CHD3;refGene_tx=XM_023243930.1;refGene_gn=CHD3;refGene_tx=X
M_023243931.1;refGene_gn=CHD3;refGene_tx=XM_023243932.1;refGene_gn=CHD
3;refGene_tx=XM_023243933.1;refGene_gn=CHD3;refGene_tx=XM_023243934.1;
refGene_gn=CHD3;refGene_tx=XM_023243935.1
E1 2733836 2733968
refGene_gn=CHD3;refGene_tx=XM_023243920.1;refGene_gn=CHD3;refGene_tx=X
M_023243921.1;refGene_gn=CHD3;refGene_tx=XM_023243922.1;refGene_gn=CHD
3;refGene_tx=XM_023243923.1;refGene_gn=CHD3;refGene_tx=XM_023243924.1;
refGene_gn=CHD3;refGene_tx=XM_023243925.1;refGene_gn=CHD3;refGene_tx=X
M_023243926.1;refGene_gn=CHD3;refGene_tx=XM_023243927.1;refGene_gn=CHD
3;refGene_tx=XM_023243928.1;refGene_gn=CHD3;refGene_tx=XM_023243929.1;
refGene_gn=CHD3;refGene_tx=XM_023243930.1;refGene_gn=CHD3;refGene_tx=X
M_023243931.1;refGene_gn=CHD3;refGene_tx=XM_023243932.1;refGene_gn=CHD
3;refGene_tx=XM_023243933.1;refGene_gn=CHD3;refGene_tx=XM_023243934.1;
refGene_gn=CHD3;refGene_tx=XM_023243935.1
E1 2734350 2734450
refGene_gn=CHD3;refGene_tx=XM_023243920.1;refGene_gn=CHD3;refGene_tx=X
M_023243921.1;refGene_gn=CHD3;refGene_tx=XM_023243922.1;refGene_gn=CHD
3;refGene_tx=XM_023243923.1;refGene_gn=CHD3;refGene_tx=XM_023243924.1;
refGene_gn=CHD3;refGene_tx=XM_023243925.1;refGene_gn=CHD3;refGene_tx=X
M_023243926.1;refGene_gn=CHD3;refGene_tx=XM_023243927.1;refGene_gn=CHD
3;refGene_tx=XM_023243928.1;refGene_gn=CHD3;refGene_tx=XM_023243929.1;
refGene_gn=CHD3;refGene_tx=XM_023243930.1;refGene_gn=CHD3;refGene_tx=X
M_023243931.1;refGene_gn=CHD3;refGene_tx=XM_023243932.1;refGene_gn=CHD
3;refGene_tx=XM_023243933.1;refGene_gn=CHD3;refGene_tx=XM_023243934.1;
refGene_gn=CHD3;refGene_tx=XM_023243935.1
E1 2734862 2735054
refGene_gn=CHD3;refGene_tx=XM_023243920.1;refGene_gn=CHD3;refGene_tx=X
M_023243921.1;refGene_gn=CHD3;refGene_tx=XM_023243922.1;refGene_gn=CHD
3;refGene_tx=XM_023243923.1;refGene_gn=CHD3;refGene_tx=XM_023243924.1;
refGene_gn=CHD3;refGene_tx=XM_023243925.1;refGene_gn=CHD3;refGene_tx=X
M_023243926.1;refGene_gn=CHD3;refGene_tx=XM_023243927.1;refGene_gn=CHD
3;refGene_tx=XM_023243928.1;refGene_gn=CHD3;refGene_tx=XM_023243929.1;
refGene_gn=CHD3;refGene_tx=XM_023243930.1;refGene_gn=CHD3;refGene_tx=X
M_023243931.1;refGene_gn=CHD3;refGene_tx=XM_023243932.1;refGene_gn=CHD
3;refGene_tx=XM_023243933.1;refGene_gn=CHD3;refGene_tx=XM_023243934.1;
refGene_gn=CHD3;refGene_tx=XM_023243935.1
E1 2735195 2735396

refGene_gn=CHD3;refGene_tx=XM_023243920.1;refGene_gn=CHD3;refGene_tx=XM_023243921.1;refGene_gn=CHD3;refGene_tx=XM_023243922.1;refGene_gn=CHD3;refGene_tx=XM_023243923.1;refGene_gn=CHD3;refGene_tx=XM_023243924.1;refGene_gn=CHD3;refGene_tx=XM_023243925.1;refGene_gn=CHD3;refGene_tx=XM_023243926.1;refGene_gn=CHD3;refGene_tx=XM_023243927.1;refGene_gn=CHD3;refGene_tx=XM_023243928.1;refGene_gn=CHD3;refGene_tx=XM_023243929.1;refGene_gn=CHD3;refGene_tx=XM_023243930.1;refGene_gn=CHD3;refGene_tx=XM_023243931.1;refGene_gn=CHD3;refGene_tx=XM_023243932.1;refGene_gn=CHD3;refGene_tx=XM_023243933.1;refGene_gn=CHD3;refGene_tx=XM_023243934.1;refGene_gn=CHD3;refGene_tx=XM_023243935.1
E1 2735744 2735882
refGene_gn=CHD3;refGene_tx=XM_023243920.1;refGene_gn=CHD3;refGene_tx=XM_023243921.1;refGene_gn=CHD3;refGene_tx=XM_023243922.1;refGene_gn=CHD3;refGene_tx=XM_023243923.1;refGene_gn=CHD3;refGene_tx=XM_023243924.1;refGene_gn=CHD3;refGene_tx=XM_023243925.1;refGene_gn=CHD3;refGene_tx=XM_023243926.1;refGene_gn=CHD3;refGene_tx=XM_023243927.1;refGene_gn=CHD3;refGene_tx=XM_023243928.1;refGene_gn=CHD3;refGene_tx=XM_023243929.1;refGene_gn=CHD3;refGene_tx=XM_023243930.1;refGene_gn=CHD3;refGene_tx=XM_023243931.1;refGene_gn=CHD3;refGene_tx=XM_023243932.1;refGene_gn=CHD3;refGene_tx=XM_023243933.1;refGene_gn=CHD3;refGene_tx=XM_023243934.1;refGene_gn=CHD3;refGene_tx=XM_023243935.1
E1 2736112 2736234
refGene_gn=CHD3;refGene_tx=XM_023243920.1;refGene_gn=CHD3;refGene_tx=XM_023243921.1;refGene_gn=CHD3;refGene_tx=XM_023243922.1;refGene_gn=CHD3;refGene_tx=XM_023243923.1;refGene_gn=CHD3;refGene_tx=XM_023243924.1;refGene_gn=CHD3;refGene_tx=XM_023243925.1;refGene_gn=CHD3;refGene_tx=XM_023243926.1;refGene_gn=CHD3;refGene_tx=XM_023243927.1;refGene_gn=CHD3;refGene_tx=XM_023243928.1;refGene_gn=CHD3;refGene_tx=XM_023243929.1;refGene_gn=CHD3;refGene_tx=XM_023243930.1;refGene_gn=CHD3;refGene_tx=XM_023243931.1;refGene_gn=CHD3;refGene_tx=XM_023243932.1;refGene_gn=CHD3;refGene_tx=XM_023243933.1;refGene_gn=CHD3;refGene_tx=XM_023243934.1;refGene_gn=CHD3;refGene_tx=XM_023243935.1
E1 2736389 2736563
refGene_gn=CHD3;refGene_tx=XM_023243920.1;refGene_gn=CHD3;refGene_tx=XM_023243921.1;refGene_gn=CHD3;refGene_tx=XM_023243922.1;refGene_gn=CHD3;refGene_tx=XM_023243923.1;refGene_gn=CHD3;refGene_tx=XM_023243924.1;refGene_gn=CHD3;refGene_tx=XM_023243925.1;refGene_gn=CHD3;refGene_tx=XM_023243926.1;refGene_gn=CHD3;refGene_tx=XM_023243927.1;refGene_gn=CHD3;refGene_tx=XM_023243928.1;refGene_gn=CHD3;refGene_tx=XM_023243929.1;refGene_gn=CHD3;refGene_tx=XM_023243930.1;refGene_gn=CHD3;refGene_tx=XM_023243931.1;refGene_gn=CHD3;refGene_tx=XM_023243932.1;refGene_gn=CHD3;refGene_tx=XM_023243933.1;refGene_gn=CHD3;refGene_tx=XM_023243934.1;refGene_gn=CHD3;refGene_tx=XM_023243935.1
E1 2736690 2736832
refGene_gn=CHD3;refGene_tx=XM_023243920.1;refGene_gn=CHD3;refGene_tx=XM_023243921.1;refGene_gn=CHD3;refGene_tx=XM_023243922.1;refGene_gn=CHD3;refGene_tx=XM_023243923.1;refGene_gn=CHD3;refGene_tx=XM_023243924.1;refGene_gn=CHD3;refGene_tx=XM_023243925.1;refGene_gn=CHD3;refGene_tx=XM_023243926.1;refGene_gn=CHD3;refGene_tx=XM_023243927.1;refGene_gn=CHD3;refGene_tx=XM_023243928.1;refGene_gn=CHD3;refGene_tx=XM_023243929.1;

refGene_gn=CHD3;refGene_tx=XM_023243930.1;refGene_gn=CHD3;refGene_tx=XM_023243931.1;refGene_gn=CHD3;refGene_tx=XM_023243932.1;refGene_gn=CHD3;refGene_tx=XM_023243933.1;refGene_gn=CHD3;refGene_tx=XM_023243934.1;refGene_gn=CHD3;refGene_tx=XM_023243935.1
E1 2737055 2737187
refGene_gn=CHD3;refGene_tx=XM_023243920.1;refGene_gn=CHD3;refGene_tx=XM_023243921.1;refGene_gn=CHD3;refGene_tx=XM_023243922.1;refGene_gn=CHD3;refGene_tx=XM_023243923.1;refGene_gn=CHD3;refGene_tx=XM_023243924.1;refGene_gn=CHD3;refGene_tx=XM_023243925.1;refGene_gn=CHD3;refGene_tx=XM_023243926.1;refGene_gn=CHD3;refGene_tx=XM_023243927.1;refGene_gn=CHD3;refGene_tx=XM_023243928.1;refGene_gn=CHD3;refGene_tx=XM_023243929.1;refGene_gn=CHD3;refGene_tx=XM_023243930.1;refGene_gn=CHD3;refGene_tx=XM_023243931.1;refGene_gn=CHD3;refGene_tx=XM_023243932.1;refGene_gn=CHD3;refGene_tx=XM_023243933.1;refGene_gn=CHD3;refGene_tx=XM_023243934.1;refGene_gn=CHD3;refGene_tx=XM_023243935.1
E1 2737804 2737922
refGene_gn=CHD3;refGene_tx=XM_023243920.1;refGene_gn=CHD3;refGene_tx=XM_023243921.1;refGene_gn=CHD3;refGene_tx=XM_023243922.1;refGene_gn=CHD3;refGene_tx=XM_023243923.1;refGene_gn=CHD3;refGene_tx=XM_023243924.1;refGene_gn=CHD3;refGene_tx=XM_023243925.1;refGene_gn=CHD3;refGene_tx=XM_023243926.1;refGene_gn=CHD3;refGene_tx=XM_023243927.1;refGene_gn=CHD3;refGene_tx=XM_023243928.1;refGene_gn=CHD3;refGene_tx=XM_023243929.1;refGene_gn=CHD3;refGene_tx=XM_023243930.1;refGene_gn=CHD3;refGene_tx=XM_023243931.1;refGene_gn=CHD3;refGene_tx=XM_023243932.1;refGene_gn=CHD3;refGene_tx=XM_023243933.1;refGene_gn=CHD3;refGene_tx=XM_023243934.1;refGene_gn=CHD3;refGene_tx=XM_023243935.1
E1 2738095 2738220
refGene_gn=CHD3;refGene_tx=XM_023243920.1;refGene_gn=CHD3;refGene_tx=XM_023243921.1;refGene_gn=CHD3;refGene_tx=XM_023243922.1;refGene_gn=CHD3;refGene_tx=XM_023243923.1;refGene_gn=CHD3;refGene_tx=XM_023243924.1;refGene_gn=CHD3;refGene_tx=XM_023243925.1;refGene_gn=CHD3;refGene_tx=XM_023243926.1;refGene_gn=CHD3;refGene_tx=XM_023243927.1;refGene_gn=CHD3;refGene_tx=XM_023243928.1;refGene_gn=CHD3;refGene_tx=XM_023243929.1;refGene_gn=CHD3;refGene_tx=XM_023243930.1;refGene_gn=CHD3;refGene_tx=XM_023243931.1;refGene_gn=CHD3;refGene_tx=XM_023243932.1;refGene_gn=CHD3;refGene_tx=XM_023243933.1;refGene_gn=CHD3;refGene_tx=XM_023243934.1;refGene_gn=CHD3;refGene_tx=XM_023243935.1
E1 2738416 2738648
refGene_gn=CHD3;refGene_tx=XM_023243920.1;refGene_gn=CHD3;refGene_tx=XM_023243921.1;refGene_gn=CHD3;refGene_tx=XM_023243922.1;refGene_gn=CHD3;refGene_tx=XM_023243923.1;refGene_gn=CHD3;refGene_tx=XM_023243924.1;refGene_gn=CHD3;refGene_tx=XM_023243925.1;refGene_gn=CHD3;refGene_tx=XM_023243926.1;refGene_gn=CHD3;refGene_tx=XM_023243927.1;refGene_gn=CHD3;refGene_tx=XM_023243928.1;refGene_gn=CHD3;refGene_tx=XM_023243929.1;refGene_gn=CHD3;refGene_tx=XM_023243930.1;refGene_gn=CHD3;refGene_tx=XM_023243931.1;refGene_gn=CHD3;refGene_tx=XM_023243932.1;refGene_gn=CHD3;refGene_tx=XM_023243933.1;refGene_gn=CHD3;refGene_tx=XM_023243934.1;refGene_gn=CHD3;refGene_tx=XM_023243935.1
E1 2738950 2739117
refGene_gn=CHD3;refGene_tx=XM_023243920.1;refGene_gn=CHD3;refGene_tx=X

M_023243921.1;refGene_gn=CHD3;refGene_tx=XM_023243922.1;refGene_gn=CHD
3;refGene_tx=XM_023243923.1;refGene_gn=CHD3;refGene_tx=XM_023243924.1;
refGene_gn=CHD3;refGene_tx=XM_023243925.1;refGene_gn=CHD3;refGene_tx=X
M_023243926.1;refGene_gn=CHD3;refGene_tx=XM_023243927.1;refGene_gn=CHD
3;refGene_tx=XM_023243928.1;refGene_gn=CHD3;refGene_tx=XM_023243929.1;
refGene_gn=CHD3;refGene_tx=XM_023243930.1;refGene_gn=CHD3;refGene_tx=X
M_023243931.1;refGene_gn=CHD3;refGene_tx=XM_023243932.1;refGene_gn=CHD
3;refGene_tx=XM_023243933.1;refGene_gn=CHD3;refGene_tx=XM_023243934.1;
refGene_gn=CHD3;refGene_tx=XM_023243935.1
E1 2739597 2739775
refGene_gn=CHD3;refGene_tx=XM_023243920.1;refGene_gn=CHD3;refGene_tx=X
M_023243921.1;refGene_gn=CHD3;refGene_tx=XM_023243922.1;refGene_gn=CHD
3;refGene_tx=XM_023243923.1;refGene_gn=CHD3;refGene_tx=XM_023243924.1;
refGene_gn=CHD3;refGene_tx=XM_023243925.1;refGene_gn=CHD3;refGene_tx=X
M_023243926.1;refGene_gn=CHD3;refGene_tx=XM_023243927.1;refGene_gn=CHD
3;refGene_tx=XM_023243928.1;refGene_gn=CHD3;refGene_tx=XM_023243929.1;
refGene_gn=CHD3;refGene_tx=XM_023243930.1;refGene_gn=CHD3;refGene_tx=X
M_023243931.1;refGene_gn=CHD3;refGene_tx=XM_023243932.1;refGene_gn=CHD
3;refGene_tx=XM_023243933.1;refGene_gn=CHD3;refGene_tx=XM_023243934.1;
refGene_gn=CHD3;refGene_tx=XM_023243935.1
E1 2740177 2740243
refGene_gn=CHD3;refGene_tx=XM_023243920.1;refGene_gn=CHD3;refGene_tx=X
M_023243921.1;refGene_gn=CHD3;refGene_tx=XM_023243922.1;refGene_gn=CHD
3;refGene_tx=XM_023243923.1;refGene_gn=CHD3;refGene_tx=XM_023243924.1;
refGene_gn=CHD3;refGene_tx=XM_023243925.1;refGene_gn=CHD3;refGene_tx=X
M_023243926.1;refGene_gn=CHD3;refGene_tx=XM_023243927.1;refGene_gn=CHD
3;refGene_tx=XM_023243928.1;refGene_gn=CHD3;refGene_tx=XM_023243929.1;
refGene_gn=CHD3;refGene_tx=XM_023243930.1;refGene_gn=CHD3;refGene_tx=X
M_023243931.1;refGene_gn=CHD3;refGene_tx=XM_023243932.1;refGene_gn=CHD
3;refGene_tx=XM_023243933.1;refGene_gn=CHD3;refGene_tx=XM_023243934.1;
refGene_gn=CHD3;refGene_tx=XM_023243935.1
E1 2740654 2740740
refGene_gn=CHD3;refGene_tx=XM_023243920.1;refGene_gn=CHD3;refGene_tx=X
M_023243921.1;refGene_gn=CHD3;refGene_tx=XM_023243922.1;refGene_gn=CHD
3;refGene_tx=XM_023243923.1;refGene_gn=CHD3;refGene_tx=XM_023243924.1;
refGene_gn=CHD3;refGene_tx=XM_023243925.1;refGene_gn=CHD3;refGene_tx=X
M_023243926.1;refGene_gn=CHD3;refGene_tx=XM_023243927.1;refGene_gn=CHD
3;refGene_tx=XM_023243928.1;refGene_gn=CHD3;refGene_tx=XM_023243929.1;
refGene_gn=CHD3;refGene_tx=XM_023243930.1;refGene_gn=CHD3;refGene_tx=X
M_023243931.1;refGene_gn=CHD3;refGene_tx=XM_023243932.1;refGene_gn=CHD
3;refGene_tx=XM_023243933.1;refGene_gn=CHD3;refGene_tx=XM_023243934.1;
refGene_gn=CHD3;refGene_tx=XM_023243935.1
E1 2740888 2741022
refGene_gn=CHD3;refGene_tx=XM_023243920.1;refGene_gn=CHD3;refGene_tx=X
M_023243921.1;refGene_gn=CHD3;refGene_tx=XM_023243922.1;refGene_gn=CHD
3;refGene_tx=XM_023243923.1;refGene_gn=CHD3;refGene_tx=XM_023243924.1;
refGene_gn=CHD3;refGene_tx=XM_023243925.1;refGene_gn=CHD3;refGene_tx=X
M_023243926.1;refGene_gn=CHD3;refGene_tx=XM_023243927.1;refGene_gn=CHD
3;refGene_tx=XM_023243928.1;refGene_gn=CHD3;refGene_tx=XM_023243929.1;
refGene_gn=CHD3;refGene_tx=XM_023243930.1;refGene_gn=CHD3;refGene_tx=X

M_023243931.1;refGene_gn=CHD3;refGene_tx=XM_023243932.1;refGene_gn=CHD3;refGene_tx=XM_023243933.1;refGene_gn=CHD3;refGene_tx=XM_023243934.1;refGene_gn=CHD3;refGene_tx=XM_023243935.1
E1 2741581 2741726
refGene_gn=CHD3;refGene_tx=XM_023243920.1;refGene_gn=CHD3;refGene_tx=XM_023243921.1;refGene_gn=CHD3;refGene_tx=XM_023243922.1;refGene_gn=CHD3;refGene_tx=XM_023243923.1;refGene_gn=CHD3;refGene_tx=XM_023243924.1;refGene_gn=CHD3;refGene_tx=XM_023243925.1;refGene_gn=CHD3;refGene_tx=XM_023243926.1;refGene_gn=CHD3;refGene_tx=XM_023243927.1;refGene_gn=CHD3;refGene_tx=XM_023243928.1;refGene_gn=CHD3;refGene_tx=XM_023243929.1;refGene_gn=CHD3;refGene_tx=XM_023243930.1;refGene_gn=CHD3;refGene_tx=XM_023243931.1;refGene_gn=CHD3;refGene_tx=XM_023243932.1;refGene_gn=CHD3;refGene_tx=XM_023243933.1;refGene_gn=CHD3;refGene_tx=XM_023243934.1;refGene_gn=CHD3;refGene_tx=XM_023243935.1
E1 2741906 2742069
refGene_gn=CHD3;refGene_tx=XM_023243920.1;refGene_gn=CHD3;refGene_tx=XM_023243921.1;refGene_gn=CHD3;refGene_tx=XM_023243922.1;refGene_gn=CHD3;refGene_tx=XM_023243923.1;refGene_gn=CHD3;refGene_tx=XM_023243924.1;refGene_gn=CHD3;refGene_tx=XM_023243925.1;refGene_gn=CHD3;refGene_tx=XM_023243926.1;refGene_gn=CHD3;refGene_tx=XM_023243927.1;refGene_gn=CHD3;refGene_tx=XM_023243928.1;refGene_gn=CHD3;refGene_tx=XM_023243929.1;refGene_gn=CHD3;refGene_tx=XM_023243930.1;refGene_gn=CHD3;refGene_tx=XM_023243931.1;refGene_gn=CHD3;refGene_tx=XM_023243932.1;refGene_gn=CHD3;refGene_tx=XM_023243933.1;refGene_gn=CHD3;refGene_tx=XM_023243934.1;refGene_gn=CHD3;refGene_tx=XM_023243935.1
E1 2742167 2742280
refGene_gn=CHD3;refGene_tx=XM_023243920.1;refGene_gn=CHD3;refGene_tx=XM_023243921.1;refGene_gn=CHD3;refGene_tx=XM_023243922.1;refGene_gn=CHD3;refGene_tx=XM_023243923.1;refGene_gn=CHD3;refGene_tx=XM_023243924.1;refGene_gn=CHD3;refGene_tx=XM_023243925.1;refGene_gn=CHD3;refGene_tx=XM_023243926.1;refGene_gn=CHD3;refGene_tx=XM_023243927.1;refGene_gn=CHD3;refGene_tx=XM_023243928.1;refGene_gn=CHD3;refGene_tx=XM_023243929.1;refGene_gn=CHD3;refGene_tx=XM_023243930.1;refGene_gn=CHD3;refGene_tx=XM_023243931.1;refGene_gn=CHD3;refGene_tx=XM_023243932.1;refGene_gn=CHD3;refGene_tx=XM_023243933.1;refGene_gn=CHD3;refGene_tx=XM_023243934.1;refGene_gn=CHD3;refGene_tx=XM_023243935.1
E1 2742368 2742504
refGene_gn=CHD3;refGene_tx=XM_023243920.1;refGene_gn=CHD3;refGene_tx=XM_023243921.1;refGene_gn=CHD3;refGene_tx=XM_023243922.1;refGene_gn=CHD3;refGene_tx=XM_023243923.1;refGene_gn=CHD3;refGene_tx=XM_023243924.1;refGene_gn=CHD3;refGene_tx=XM_023243925.1;refGene_gn=CHD3;refGene_tx=XM_023243926.1;refGene_gn=CHD3;refGene_tx=XM_023243927.1;refGene_gn=CHD3;refGene_tx=XM_023243928.1;refGene_gn=CHD3;refGene_tx=XM_023243929.1;refGene_gn=CHD3;refGene_tx=XM_023243930.1;refGene_gn=CHD3;refGene_tx=XM_023243931.1;refGene_gn=CHD3;refGene_tx=XM_023243932.1;refGene_gn=CHD3;refGene_tx=XM_023243933.1;refGene_gn=CHD3;refGene_tx=XM_023243934.1;refGene_gn=CHD3;refGene_tx=XM_023243935.1
E1 2742617 2742719
refGene_gn=CHD3;refGene_tx=XM_023243920.1;refGene_gn=CHD3;refGene_tx=XM_023243921.1;refGene_gn=CHD3;refGene_tx=XM_023243922.1;refGene_gn=CHD

3;refGene_tx=XM_023243923.1;refGene_gn=CHD3;refGene_tx=XM_023243924.1;
refGene_gn=CHD3;refGene_tx=XM_023243925.1;refGene_gn=CHD3;refGene_tx=X
M_023243928.1;refGene_gn=CHD3;refGene_tx=XM_023243929.1;refGene_gn=CHD
3;refGene_tx=XM_023243930.1;refGene_gn=CHD3;refGene_tx=XM_023243931.1;
refGene_gn=CHD3;refGene_tx=XM_023243932.1;refGene_gn=CHD3;refGene_tx=X
M_023243934.1;refGene_gn=CHD3;refGene_tx=XM_023243935.1
E1 2742899 2743031
refGene_gn=CHD3;refGene_tx=XM_023243920.1;refGene_gn=CHD3;refGene_tx=X
M_023243921.1;refGene_gn=CHD3;refGene_tx=XM_023243922.1;refGene_gn=CHD
3;refGene_tx=XM_023243923.1;refGene_gn=CHD3;refGene_tx=XM_023243924.1;
refGene_gn=CHD3;refGene_tx=XM_023243925.1;refGene_gn=CHD3;refGene_tx=X
M_023243926.1;refGene_gn=CHD3;refGene_tx=XM_023243927.1;refGene_gn=CHD
3;refGene_tx=XM_023243928.1;refGene_gn=CHD3;refGene_tx=XM_023243929.1;
refGene_gn=CHD3;refGene_tx=XM_023243930.1;refGene_gn=CHD3;refGene_tx=X
M_023243931.1;refGene_gn=CHD3;refGene_tx=XM_023243932.1;refGene_gn=CHD
3;refGene_tx=XM_023243933.1;refGene_gn=CHD3;refGene_tx=XM_023243934.1;
refGene_gn=CHD3;refGene_tx=XM_023243935.1
E1 2743181 2743290
refGene_gn=CHD3;refGene_tx=XM_023243920.1;refGene_gn=CHD3;refGene_tx=X
M_023243921.1;refGene_gn=CHD3;refGene_tx=XM_023243922.1;refGene_gn=CHD
3;refGene_tx=XM_023243923.1;refGene_gn=CHD3;refGene_tx=XM_023243924.1;
refGene_gn=CHD3;refGene_tx=XM_023243925.1;refGene_gn=CHD3;refGene_tx=X
M_023243926.1;refGene_gn=CHD3;refGene_tx=XM_023243927.1;refGene_gn=CHD
3;refGene_tx=XM_023243928.1;refGene_gn=CHD3;refGene_tx=XM_023243929.1;
refGene_gn=CHD3;refGene_tx=XM_023243930.1;refGene_gn=CHD3;refGene_tx=X
M_023243931.1;refGene_gn=CHD3;refGene_tx=XM_023243932.1;refGene_gn=CHD
3;refGene_tx=XM_023243933.1;refGene_gn=CHD3;refGene_tx=XM_023243934.1;
refGene_gn=CHD3;refGene_tx=XM_023243935.1
E1 2743727 2743860
refGene_gn=CHD3;refGene_tx=XM_023243920.1;refGene_gn=CHD3;refGene_tx=X
M_023243921.1;refGene_gn=CHD3;refGene_tx=XM_023243922.1;refGene_gn=CHD
3;refGene_tx=XM_023243923.1;refGene_gn=CHD3;refGene_tx=XM_023243924.1;
refGene_gn=CHD3;refGene_tx=XM_023243925.1;refGene_gn=CHD3;refGene_tx=X
M_023243926.1;refGene_gn=CHD3;refGene_tx=XM_023243927.1;refGene_gn=CHD
3;refGene_tx=XM_023243928.1;refGene_gn=CHD3;refGene_tx=XM_023243929.1;
refGene_gn=CHD3;refGene_tx=XM_023243930.1;refGene_gn=CHD3;refGene_tx=X
M_023243931.1;refGene_gn=CHD3;refGene_tx=XM_023243932.1;refGene_gn=CHD
3;refGene_tx=XM_023243933.1;refGene_gn=CHD3;refGene_tx=XM_023243934.1;
refGene_gn=CHD3;refGene_tx=XM_023243935.1
E1 2744120 2744362
refGene_gn=CHD3;refGene_tx=XM_023243920.1;refGene_gn=CHD3;refGene_tx=X
M_023243921.1;refGene_gn=CHD3;refGene_tx=XM_023243922.1;refGene_gn=CHD
3;refGene_tx=XM_023243923.1;refGene_gn=CHD3;refGene_tx=XM_023243924.1;
refGene_gn=CHD3;refGene_tx=XM_023243925.1;refGene_gn=CHD3;refGene_tx=X
M_023243926.1;refGene_gn=CHD3;refGene_tx=XM_023243927.1;refGene_gn=CHD
3;refGene_tx=XM_023243928.1;refGene_gn=CHD3;refGene_tx=XM_023243929.1;
refGene_gn=CHD3;refGene_tx=XM_023243930.1;refGene_gn=CHD3;refGene_tx=X
M_023243931.1;refGene_gn=CHD3;refGene_tx=XM_023243932.1;refGene_gn=CHD
3;refGene_tx=XM_023243933.1;refGene_gn=CHD3;refGene_tx=XM_023243934.1;
refGene_gn=CHD3;refGene_tx=XM_023243935.1

E1	2744604	2744681	refGene_gn=CHD3;refGene_tx=XM_023243935.1
E1	2745079	2745587
refGene_gn=CHD3;refGene_tx=XM_023243920.1;refGene_gn=CHD3;refGene_tx=XM_023243921.1;refGene_gn=CHD3;refGene_tx=XM_023243922.1;refGene_gn=CHD3;refGene_tx=XM_023243923.1;refGene_gn=CHD3;refGene_tx=XM_023243924.1;refGene_gn=CHD3;refGene_tx=XM_023243925.1;refGene_gn=CHD3;refGene_tx=XM_023243926.1;refGene_gn=CHD3;refGene_tx=XM_023243927.1;refGene_gn=CHD3;refGene_tx=XM_023243928.1;refGene_gn=CHD3;refGene_tx=XM_023243929.1;refGene_gn=CHD3;refGene_tx=XM_023243930.1;refGene_gn=CHD3;refGene_tx=XM_023243931.1;refGene_gn=CHD3;refGene_tx=XM_023243932.1;refGene_gn=CHD3;refGene_tx=XM_023243933.1;refGene_gn=CHD3;refGene_tx=XM_023243934.1
E1	2745820	2745947
refGene_gn=CHD3;refGene_tx=XM_023243920.1;refGene_gn=CHD3;refGene_tx=XM_023243921.1;refGene_gn=CHD3;refGene_tx=XM_023243922.1;refGene_gn=CHD3;refGene_tx=XM_023243923.1;refGene_gn=CHD3;refGene_tx=XM_023243924.1;refGene_gn=CHD3;refGene_tx=XM_023243925.1;refGene_gn=CHD3;refGene_tx=XM_023243926.1;refGene_gn=CHD3;refGene_tx=XM_023243927.1;refGene_gn=CHD3;refGene_tx=XM_023243928.1;refGene_gn=CHD3;refGene_tx=XM_023243929.1;refGene_gn=CHD3;refGene_tx=XM_023243933.1
E1	2746396	2746637
refGene_gn=CHD3;refGene_tx=XM_023243920.1;refGene_gn=CHD3;refGene_tx=XM_023243921.1;refGene_gn=CHD3;refGene_tx=XM_023243922.1;refGene_gn=CHD3;refGene_tx=XM_023243923.1;refGene_gn=CHD3;refGene_tx=XM_023243924.1;refGene_gn=CHD3;refGene_tx=XM_023243925.1;refGene_gn=CHD3;refGene_tx=XM_023243926.1;refGene_gn=CHD3;refGene_tx=XM_023243927.1;refGene_gn=CHD3;refGene_tx=XM_023243928.1;refGene_gn=CHD3;refGene_tx=XM_023243929.1;refGene_gn=CHD3;refGene_tx=XM_023243931.1;refGene_gn=CHD3;refGene_tx=XM_023243933.1;refGene_gn=CHD3;refGene_tx=XM_023243934.1
E1	2750207	2750263
refGene_gn=LOC111557863;refGene_tx=XM_023244600.1
E1	2750389	2750879
refGene_gn=LOC111557863;refGene_tx=XM_023244600.1
E1	2756122	2756200
refGene_gn=KCNAB3;refGene_tx=XM_003996187.4;refGene_gn=KCNAB3;refGene_tx=XM_019817106.2;refGene_gn=KCNAB3;refGene_tx=XM_023244599.1
E1	2756464	2756553
refGene_gn=KCNAB3;refGene_tx=XM_003996187.4;refGene_gn=KCNAB3;refGene_tx=XM_019817106.2;refGene_gn=KCNAB3;refGene_tx=XM_019817107.2;refGene_gn=KCNAB3;refGene_tx=XM_023244599.1
E1	2756609	2756621	refGene_gn=KCNAB3;refGene_tx=XM_019817108.2
E1	2756722	2756843
refGene_gn=KCNAB3;refGene_tx=XM_003996187.4;refGene_gn=KCNAB3;refGene_tx=XM_019817106.2;refGene_gn=KCNAB3;refGene_tx=XM_019817107.2;refGene_gn=KCNAB3;refGene_tx=XM_019817108.2;refGene_gn=KCNAB3;refGene_tx=XM_023244599.1
E1	2756938	2757030
refGene_gn=KCNAB3;refGene_tx=XM_003996187.4;refGene_gn=KCNAB3;refGene_tx=XM_019817106.2;refGene_gn=KCNAB3;refGene_tx=XM_019817107.2;refGene_gn=KCNAB3;refGene_tx=XM_019817108.2;refGene_gn=KCNAB3;refGene_tx=XM_023244599.1

E1 2757114 2757235
refGene_gn=KCNAB3;refGene_tx=XM_003996187.4;refGene_gn=KCNAB3;refGene_tx=XM_019817106.2;refGene_gn=KCNAB3;refGene_tx=XM_019817107.2;refGene_gn=KCNAB3;refGene_tx=XM_019817108.2;refGene_gn=KCNAB3;refGene_tx=XM_023244599.1
E1 2757390 2757476
refGene_gn=KCNAB3;refGene_tx=XM_003996187.4;refGene_gn=KCNAB3;refGene_tx=XM_019817106.2;refGene_gn=KCNAB3;refGene_tx=XM_019817107.2;refGene_gn=KCNAB3;refGene_tx=XM_019817108.2;refGene_gn=KCNAB3;refGene_tx=XM_023244599.1
E1 2757750 2757837
refGene_gn=KCNAB3;refGene_tx=XM_003996187.4;refGene_gn=KCNAB3;refGene_tx=XM_019817106.2;refGene_gn=KCNAB3;refGene_tx=XM_019817107.2;refGene_gn=KCNAB3;refGene_tx=XM_019817108.2;refGene_gn=KCNAB3;refGene_tx=XM_023244599.1
E1 2758035 2758079
refGene_gn=KCNAB3;refGene_tx=XM_003996187.4;refGene_gn=KCNAB3;refGene_tx=XM_019817106.2;refGene_gn=KCNAB3;refGene_tx=XM_019817107.2;refGene_gn=KCNAB3;refGene_tx=XM_019817108.2;refGene_gn=KCNAB3;refGene_tx=XM_023244599.1
E1 2758278 2758323
refGene_gn=KCNAB3;refGene_tx=XM_003996187.4;refGene_gn=KCNAB3;refGene_tx=XM_019817106.2;refGene_gn=KCNAB3;refGene_tx=XM_019817107.2;refGene_gn=KCNAB3;refGene_tx=XM_019817108.2;refGene_gn=KCNAB3;refGene_tx=XM_023244599.1
E1 2758406 2758451
refGene_gn=KCNAB3;refGene_tx=XM_003996187.4;refGene_gn=KCNAB3;refGene_tx=XM_019817107.2;refGene_gn=KCNAB3;refGene_tx=XM_019817108.2;refGene_gn=KCNAB3;refGene_tx=XM_023244599.1
E1 2759773 2759853
refGene_gn=KCNAB3;refGene_tx=XM_003996187.4;refGene_gn=KCNAB3;refGene_tx=XM_019817106.2;refGene_gn=KCNAB3;refGene_tx=XM_019817107.2;refGene_gn=KCNAB3;refGene_tx=XM_019817108.2;refGene_gn=KCNAB3;refGene_tx=XM_023244599.1
E1 2760095 2760133
refGene_gn=KCNAB3;refGene_tx=XM_003996187.4;refGene_gn=KCNAB3;refGene_tx=XM_019817106.2;refGene_gn=KCNAB3;refGene_tx=XM_019817107.2;refGene_gn=KCNAB3;refGene_tx=XM_019817108.2;refGene_gn=KCNAB3;refGene_tx=XM_023244599.1
E1 2760206 2760300
refGene_gn=KCNAB3;refGene_tx=XM_003996187.4;refGene_gn=KCNAB3;refGene_tx=XM_019817106.2;refGene_gn=KCNAB3;refGene_tx=XM_019817107.2;refGene_gn=KCNAB3;refGene_tx=XM_019817108.2;refGene_gn=KCNAB3;refGene_tx=XM_023244599.1
E1 2761553 2761795
refGene_gn=KCNAB3;refGene_tx=XM_003996187.4;refGene_gn=KCNAB3;refGene_tx=XM_019817106.2;refGene_gn=KCNAB3;refGene_tx=XM_019817107.2;refGene_gn=KCNAB3;refGene_tx=XM_019817108.2
E1 2763651 2763780 refGene_gn=TRAPPC1;refGene_tx=XM_003996188.4
E1 2764080 2764219 refGene_gn=TRAPPC1;refGene_tx=XM_003996188.4

E1 2764578 2764649 refGene_gn=TRAPPC1;refGene_tx=XM_003996188.4
E1 2764804 2764903 refGene_gn=TRAPPC1;refGene_tx=XM_003996188.4
E1 2766134 2766416
refGene_gn=CNTROB;refGene_tx=XM_023244592.1;refGene_gn=CNTROB;refGene_tx=XM_023244593.1;refGene_gn=CNTROB;refGene_tx=XM_023244594.1;refGene_gn=CNTROB;refGene_tx=XM_023244595.1;refGene_gn=CNTROB;refGene_tx=XM_023244596.1;refGene_gn=CNTROB;refGene_tx=XM_023244597.1
E1 2767382 2767471
refGene_gn=CNTROB;refGene_tx=XM_023244592.1;refGene_gn=CNTROB;refGene_tx=XM_023244593.1;refGene_gn=CNTROB;refGene_tx=XM_023244594.1;refGene_gn=CNTROB;refGene_tx=XM_023244595.1;refGene_gn=CNTROB;refGene_tx=XM_023244596.1;refGene_gn=CNTROB;refGene_tx=XM_023244597.1;refGene_gn=CNTROB;refGene_tx=XM_023244598.1
E1 2767690 2767772
refGene_gn=CNTROB;refGene_tx=XM_023244592.1;refGene_gn=CNTROB;refGene_tx=XM_023244593.1;refGene_gn=CNTROB;refGene_tx=XM_023244594.1;refGene_gn=CNTROB;refGene_tx=XM_023244595.1;refGene_gn=CNTROB;refGene_tx=XM_023244596.1;refGene_gn=CNTROB;refGene_tx=XM_023244597.1;refGene_gn=CNTROB;refGene_tx=XM_023244598.1
E1 2768239 2768396
refGene_gn=CNTROB;refGene_tx=XM_023244592.1;refGene_gn=CNTROB;refGene_tx=XM_023244593.1;refGene_gn=CNTROB;refGene_tx=XM_023244594.1;refGene_gn=CNTROB;refGene_tx=XM_023244595.1;refGene_gn=CNTROB;refGene_tx=XM_023244596.1;refGene_gn=CNTROB;refGene_tx=XM_023244597.1;refGene_gn=CNTROB;refGene_tx=XM_023244598.1
E1 2769286 2769403
refGene_gn=CNTROB;refGene_tx=XM_023244592.1;refGene_gn=CNTROB;refGene_tx=XM_023244593.1;refGene_gn=CNTROB;refGene_tx=XM_023244594.1;refGene_gn=CNTROB;refGene_tx=XM_023244595.1;refGene_gn=CNTROB;refGene_tx=XM_023244596.1;refGene_gn=CNTROB;refGene_tx=XM_023244597.1;refGene_gn=CNTROB;refGene_tx=XM_023244598.1
E1 2769611 2769728
refGene_gn=CNTROB;refGene_tx=XM_023244592.1;refGene_gn=CNTROB;refGene_tx=XM_023244593.1;refGene_gn=CNTROB;refGene_tx=XM_023244594.1;refGene_gn=CNTROB;refGene_tx=XM_023244595.1;refGene_gn=CNTROB;refGene_tx=XM_023244596.1;refGene_gn=CNTROB;refGene_tx=XM_023244597.1;refGene_gn=CNTROB;refGene_tx=XM_023244598.1
E1 2770315 2770414
refGene_gn=CNTROB;refGene_tx=XM_023244592.1;refGene_gn=CNTROB;refGene_tx=XM_023244593.1;refGene_gn=CNTROB;refGene_tx=XM_023244594.1;refGene_gn=CNTROB;refGene_tx=XM_023244595.1;refGene_gn=CNTROB;refGene_tx=XM_023244596.1;refGene_gn=CNTROB;refGene_tx=XM_023244597.1;refGene_gn=CNTROB;refGene_tx=XM_023244598.1
E1 2771604 2771841
refGene_gn=CNTROB;refGene_tx=XM_023244592.1;refGene_gn=CNTROB;refGene_tx=XM_023244593.1;refGene_gn=CNTROB;refGene_tx=XM_023244594.1;refGene_gn=CNTROB;refGene_tx=XM_023244595.1;refGene_gn=CNTROB;refGene_tx=XM_023244596.1;refGene_gn=CNTROB;refGene_tx=XM_023244597.1;refGene_gn=CNTROB;refGene_tx=XM_023244598.1
E1 2772149 2772296

refGene_gn=CNTROB;refGene_tx=XM_023244592.1;refGene_gn=CNTROB;refGene_
tx=XM_023244593.1;refGene_gn=CNTROB;refGene_tx=XM_023244594.1;refGene_
gn=CNTROB;refGene_tx=XM_023244595.1;refGene_gn=CNTROB;refGene_tx=XM_02
3244596.1;refGene_gn=CNTROB;refGene_tx=XM_023244597.1;refGene_gn=CNTRO
B;refGene_tx=XM_023244598.1
E1 2784255 2784389
refGene_gn=CNTROB;refGene_tx=XM_023244592.1;refGene_gn=CNTROB;refGene_
tx=XM_023244593.1;refGene_gn=CNTROB;refGene_tx=XM_023244594.1;refGene_
gn=CNTROB;refGene_tx=XM_023244595.1;refGene_gn=CNTROB;refGene_tx=XM_02
3244596.1;refGene_gn=CNTROB;refGene_tx=XM_023244597.1;refGene_gn=CNTRO
B;refGene_tx=XM_023244598.1
E1 2784919 2785045
refGene_gn=CNTROB;refGene_tx=XM_023244592.1;refGene_gn=CNTROB;refGene_
tx=XM_023244593.1;refGene_gn=CNTROB;refGene_tx=XM_023244594.1;refGene_
gn=CNTROB;refGene_tx=XM_023244595.1;refGene_gn=CNTROB;refGene_tx=XM_02
3244596.1;refGene_gn=CNTROB;refGene_tx=XM_023244597.1;refGene_gn=CNTRO
B;refGene_tx=XM_023244598.1
E1 2785188 2785351
refGene_gn=CNTROB;refGene_tx=XM_023244592.1;refGene_gn=CNTROB;refGene_
tx=XM_023244593.1;refGene_gn=CNTROB;refGene_tx=XM_023244594.1;refGene_
gn=CNTROB;refGene_tx=XM_023244595.1;refGene_gn=CNTROB;refGene_tx=XM_02
3244596.1;refGene_gn=CNTROB;refGene_tx=XM_023244597.1;refGene_gn=CNTRO
B;refGene_tx=XM_023244598.1
E1 2786470 2786729
refGene_gn=CNTROB;refGene_tx=XM_023244592.1;refGene_gn=CNTROB;refGene_
tx=XM_023244593.1;refGene_gn=CNTROB;refGene_tx=XM_023244594.1;refGene_
gn=CNTROB;refGene_tx=XM_023244595.1;refGene_gn=CNTROB;refGene_tx=XM_02
3244596.1;refGene_gn=CNTROB;refGene_tx=XM_023244597.1;refGene_gn=CNTRO
B;refGene_tx=XM_023244598.1
E1 2787948 2788103
refGene_gn=CNTROB;refGene_tx=XM_023244592.1;refGene_gn=CNTROB;refGene_
tx=XM_023244593.1;refGene_gn=CNTROB;refGene_tx=XM_023244594.1;refGene_
gn=CNTROB;refGene_tx=XM_023244595.1;refGene_gn=CNTROB;refGene_tx=XM_02
3244596.1;refGene_gn=CNTROB;refGene_tx=XM_023244597.1;refGene_gn=CNTRO
B;refGene_tx=XM_023244598.1
E1 2788355 2788419
refGene_gn=CNTROB;refGene_tx=XM_023244592.1;refGene_gn=CNTROB;refGene_
tx=XM_023244593.1;refGene_gn=CNTROB;refGene_tx=XM_023244594.1;refGene_
gn=CNTROB;refGene_tx=XM_023244595.1;refGene_gn=CNTROB;refGene_tx=XM_02
3244596.1;refGene_gn=CNTROB;refGene_tx=XM_023244597.1;refGene_gn=CNTRO
B;refGene_tx=XM_023244598.1
E1 2788534 2788720
refGene_gn=CNTROB;refGene_tx=XM_023244592.1;refGene_gn=CNTROB;refGene_
tx=XM_023244593.1;refGene_gn=CNTROB;refGene_tx=XM_023244594.1;refGene_
gn=CNTROB;refGene_tx=XM_023244595.1;refGene_gn=CNTROB;refGene_tx=XM_02
3244596.1;refGene_gn=CNTROB;refGene_tx=XM_023244598.1
E1 2788803 2788936
refGene_gn=CNTROB;refGene_tx=XM_023244592.1;refGene_gn=CNTROB;refGene_
tx=XM_023244593.1;refGene_gn=CNTROB;refGene_tx=XM_023244594.1;refGene_
gn=CNTROB;refGene_tx=XM_023244595.1;refGene_gn=CNTROB;refGene_tx=XM_02

3244596.1;refGene_gn=CNTROB;refGene_tx=XM_023244597.1;refGene_gn=CNTRO
B;refGene_tx=XM_023244598.1
E1	2789214	2789287
refGene_gn=CNTROB;refGene_tx=XM_023244592.1;refGene_gn=CNTROB;refGene_
tx=XM_023244593.1;refGene_gn=CNTROB;refGene_tx=XM_023244594.1;refGene_
gn=CNTROB;refGene_tx=XM_023244595.1;refGene_gn=CNTROB;refGene_tx=XM_02
3244596.1;refGene_gn=CNTROB;refGene_tx=XM_023244597.1;refGene_gn=CNTRO
B;refGene_tx=XM_023244598.1
E1	2789504	2789633
refGene_gn=CNTROB;refGene_tx=XM_023244592.1;refGene_gn=CNTROB;refGene_
tx=XM_023244593.1;refGene_gn=CNTROB;refGene_tx=XM_023244594.1;refGene_
gn=CNTROB;refGene_tx=XM_023244595.1;refGene_gn=CNTROB;refGene_tx=XM_02
3244596.1;refGene_gn=CNTROB;refGene_tx=XM_023244597.1;refGene_gn=CNTRO
B;refGene_tx=XM_023244598.1
E1	2823071	2823777	refGene_gn=GUCY2D;refGene_tx=XM_023243814.1
E1	2823872	2824177	refGene_gn=GUCY2D;refGene_tx=XM_023243814.1
E1	2827421	2827773	refGene_gn=GUCY2D;refGene_tx=XM_023243814.1
E1	2828120	2828205	refGene_gn=GUCY2D;refGene_tx=XM_023243814.1
E1	2828485	2828588	refGene_gn=GUCY2D;refGene_tx=XM_023243814.1
E1	2829003	2829105	refGene_gn=GUCY2D;refGene_tx=XM_023243814.1
E1	2830870	2830951	refGene_gn=GUCY2D;refGene_tx=XM_023243814.1
E1	2832036	2832243	refGene_gn=GUCY2D;refGene_tx=XM_023243814.1
E1	2832336	2832493	refGene_gn=GUCY2D;refGene_tx=XM_023243814.1
E1	2832967	2833117	refGene_gn=GUCY2D;refGene_tx=XM_023243814.1
E1	2833718	2833867	refGene_gn=GUCY2D;refGene_tx=XM_023243814.1
E1	2834404	2834568	refGene_gn=GUCY2D;refGene_tx=XM_023243814.1
E1	2834671	2834864	refGene_gn=GUCY2D;refGene_tx=XM_023243814.1
E1	2835076	2835251	refGene_gn=GUCY2D;refGene_tx=XM_023243814.1
E1	2835512	2835611	refGene_gn=GUCY2D;refGene_tx=XM_023243814.1
E1	2835738	2835833	refGene_gn=GUCY2D;refGene_tx=XM_023243814.1
E1	2835975	2836061	refGene_gn=GUCY2D;refGene_tx=XM_023243814.1
E1	2836149	2836243	refGene_gn=GUCY2D;refGene_tx=XM_023243814.1
E1	2855986	2856133	refGene_gn=ALOX15B;refGene_tx=XM_023243919.1
E1	2856215	2856438	refGene_gn=ALOX15B;refGene_tx=XM_023243919.1
E1	2856840	2856922	refGene_gn=ALOX15B;refGene_tx=XM_023243919.1
E1	2859703	2859826	refGene_gn=ALOX15B;refGene_tx=XM_023243919.1
E1	2860107	2860211	refGene_gn=ALOX15B;refGene_tx=XM_023243919.1
E1	2861349	2861522	refGene_gn=ALOX15B;refGene_tx=XM_023243919.1
E1	2861712	2861859	refGene_gn=ALOX15B;refGene_tx=XM_023243919.1
E1	2861951	2862155	refGene_gn=ALOX15B;refGene_tx=XM_023243919.1
E1	2863070	2863157	refGene_gn=ALOX15B;refGene_tx=XM_023243919.1
E1	2863315	2863485	refGene_gn=ALOX15B;refGene_tx=XM_023243919.1
E1	2863634	2863756	refGene_gn=ALOX15B;refGene_tx=XM_023243919.1
E1	2863852	2863953	refGene_gn=ALOX15B;refGene_tx=XM_023243919.1
E1	2864031	2864202	refGene_gn=ALOX15B;refGene_tx=XM_023243919.1
E1	2864646	2864826	refGene_gn=ALOX15B;refGene_tx=XM_023243919.1
E1	2875377	2875557	refGene_gn=ALOX12B;refGene_tx=XM_003996189.2
E1	2875712	2875883	refGene_gn=ALOX12B;refGene_tx=XM_003996189.2
E1	2876055	2876156	refGene_gn=ALOX12B;refGene_tx=XM_003996189.2
E1	2877627	2877749	refGene_gn=ALOX12B;refGene_tx=XM_003996189.2

E1 2877972 2878142 refGene_gn=ALOX12B;refGene_tx=XM_003996189.2
E1 2878408 2878495 refGene_gn=ALOX12B;refGene_tx=XM_003996189.2
E1 2878709 2878913 refGene_gn=ALOX12B;refGene_tx=XM_003996189.2
E1 2880032 2880176 refGene_gn=ALOX12B;refGene_tx=XM_003996189.2
E1 2880387 2880560 refGene_gn=ALOX12B;refGene_tx=XM_003996189.2
E1 2880873 2880977 refGene_gn=ALOX12B;refGene_tx=XM_003996189.2
E1 2881236 2881359 refGene_gn=ALOX12B;refGene_tx=XM_003996189.2
E1 2881476 2881563 refGene_gn=ALOX12B;refGene_tx=XM_003996189.2
E1 2881703 2881785 refGene_gn=ALOX12B;refGene_tx=XM_003996189.2
E1 2884135 2884340 refGene_gn=ALOX12B;refGene_tx=XM_003996189.2
E1 2885392 2885539 refGene_gn=ALOX12B;refGene_tx=XM_003996189.2
E1 2893300 2893480
refGene_gn=ALOXE3;refGene_tx=XM_003996190.5;refGene_gn=ALOXE3;refGene_tx=XM_019817110.2;refGene_gn=ALOXE3;refGene_tx=XM_019817111.1;refGene_gn=ALOXE3;refGene_tx=XM_019817112.1;refGene_gn=ALOXE3;refGene_tx=XM_019817113.2
E1 2897875 2898046
refGene_gn=ALOXE3;refGene_tx=XM_003996190.5;refGene_gn=ALOXE3;refGene_tx=XM_019817110.2;refGene_gn=ALOXE3;refGene_tx=XM_019817111.1;refGene_gn=ALOXE3;refGene_tx=XM_019817112.1;refGene_gn=ALOXE3;refGene_tx=XM_019817113.2
E1 2898586 2898687
refGene_gn=ALOXE3;refGene_tx=XM_003996190.5;refGene_gn=ALOXE3;refGene_tx=XM_019817110.2;refGene_gn=ALOXE3;refGene_tx=XM_019817111.1;refGene_gn=ALOXE3;refGene_tx=XM_019817112.1;refGene_gn=ALOXE3;refGene_tx=XM_019817113.2
E1 2902489 2902611
refGene_gn=ALOXE3;refGene_tx=XM_003996190.5;refGene_gn=ALOXE3;refGene_tx=XM_019817110.2;refGene_gn=ALOXE3;refGene_tx=XM_019817111.1;refGene_gn=ALOXE3;refGene_tx=XM_019817112.1;refGene_gn=ALOXE3;refGene_tx=XM_019817113.2
E1 2903163 2903333
refGene_gn=ALOXE3;refGene_tx=XM_003996190.5;refGene_gn=ALOXE3;refGene_tx=XM_019817110.2;refGene_gn=ALOXE3;refGene_tx=XM_019817111.1;refGene_gn=ALOXE3;refGene_tx=XM_019817112.1;refGene_gn=ALOXE3;refGene_tx=XM_019817113.2
E1 2903950 2904037
refGene_gn=ALOXE3;refGene_tx=XM_003996190.5;refGene_gn=ALOXE3;refGene_tx=XM_019817110.2;refGene_gn=ALOXE3;refGene_tx=XM_019817111.1;refGene_gn=ALOXE3;refGene_tx=XM_019817112.1;refGene_gn=ALOXE3;refGene_tx=XM_019817113.2
E1 2904129 2904333
refGene_gn=ALOXE3;refGene_tx=XM_003996190.5;refGene_gn=ALOXE3;refGene_tx=XM_019817110.2;refGene_gn=ALOXE3;refGene_tx=XM_019817111.1;refGene_gn=ALOXE3;refGene_tx=XM_019817112.1;refGene_gn=ALOXE3;refGene_tx=XM_019817113.2
E1 2904420 2904564
refGene_gn=ALOXE3;refGene_tx=XM_003996190.5;refGene_gn=ALOXE3;refGene_tx=XM_019817110.2;refGene_gn=ALOXE3;refGene_tx=XM_019817111.1;refGene_gn=ALOXE3;refGene_tx=XM_019817112.1;refGene_gn=ALOXE3;refGene_tx=XM_01

9817113.2
E1	2905300	2905473
refGene_gn=ALOXE3;refGene_tx=XM_003996190.5;refGene_gn=ALOXE3;refGene_tx=XM_019817110.2;refGene_gn=ALOXE3;refGene_tx=XM_019817111.1;refGene_gn=ALOXE3;refGene_tx=XM_019817112.1;refGene_gn=ALOXE3;refGene_tx=XM_019817113.2
E1	2906027	2906131
refGene_gn=ALOXE3;refGene_tx=XM_003996190.5;refGene_gn=ALOXE3;refGene_tx=XM_019817110.2;refGene_gn=ALOXE3;refGene_tx=XM_019817111.1;refGene_gn=ALOXE3;refGene_tx=XM_019817112.1;refGene_gn=ALOXE3;refGene_tx=XM_019817113.2
E1	2909210	2909336
refGene_gn=ALOXE3;refGene_tx=XM_003996190.5;refGene_gn=ALOXE3;refGene_tx=XM_019817110.2;refGene_gn=ALOXE3;refGene_tx=XM_019817111.1;refGene_gn=ALOXE3;refGene_tx=XM_019817112.1;refGene_gn=ALOXE3;refGene_tx=XM_019817113.2
E1	2909624	2909744
refGene_gn=ALOXE3;refGene_tx=XM_003996190.5;refGene_gn=ALOXE3;refGene_tx=XM_019817110.2;refGene_gn=ALOXE3;refGene_tx=XM_019817111.1;refGene_gn=ALOXE3;refGene_tx=XM_019817112.1;refGene_gn=ALOXE3;refGene_tx=XM_019817113.2
E1	2910261	2910343
refGene_gn=ALOXE3;refGene_tx=XM_003996190.5;refGene_gn=ALOXE3;refGene_tx=XM_019817110.2;refGene_gn=ALOXE3;refGene_tx=XM_019817111.1;refGene_gn=ALOXE3;refGene_tx=XM_019817112.1;refGene_gn=ALOXE3;refGene_tx=XM_019817113.2
E1	2911587	2911792
refGene_gn=ALOXE3;refGene_tx=XM_003996190.5;refGene_gn=ALOXE3;refGene_tx=XM_019817110.2;refGene_gn=ALOXE3;refGene_tx=XM_019817111.1;refGene_gn=ALOXE3;refGene_tx=XM_019817112.1;refGene_gn=ALOXE3;refGene_tx=XM_019817113.2
E1	2912413	2912560
refGene_gn=ALOXE3;refGene_tx=XM_003996190.5;refGene_gn=ALOXE3;refGene_tx=XM_019817110.2;refGene_gn=ALOXE3;refGene_tx=XM_019817111.1;refGene_gn=ALOXE3;refGene_tx=XM_019817112.1;refGene_gn=ALOXE3;refGene_tx=XM_019817113.2
E1	2916156	2916617
refGene_gn=HES7;refGene_tx=XM_003996191.4;refGene_gn=HES7;refGene_tx=XM_019817116.2
E1	2916929	2917017
refGene_gn=HES7;refGene_tx=XM_003996191.4;refGene_gn=HES7;refGene_tx=XM_019817116.2
E1	2917608	2917704
refGene_gn=HES7;refGene_tx=XM_003996191.4;refGene_gn=HES7;refGene_tx=XM_019817116.2
E1	2918697	2918739	refGene_gn=HES7;refGene_tx=XM_003996191.4
E1	2929486	2929765
refGene_gn=PER1;refGene_tx=XM_003996192.5;refGene_gn=PER1;refGene_tx=XM_006939763.4;refGene_gn=PER1;refGene_tx=XM_019817122.2;refGene_gn=PER1;refGene_tx=XM_019817123.2;refGene_gn=PER1;refGene_tx=XM_023244700.1

E1 2930165 2930316
refGene_gn=PER1;refGene_tx=XM_003996192.5;refGene_gn=PER1;refGene_tx=XM_006939763.4;refGene_gn=PER1;refGene_tx=XM_019817122.2;refGene_gn=PER1;refGene_tx=XM_019817123.2;refGene_gn=PER1;refGene_tx=XM_023244700.1
E1 2930652 2930842
refGene_gn=PER1;refGene_tx=XM_003996192.5;refGene_gn=PER1;refGene_tx=XM_006939763.4;refGene_gn=PER1;refGene_tx=XM_019817122.2;refGene_gn=PER1;refGene_tx=XM_019817123.2;refGene_gn=PER1;refGene_tx=XM_023244700.1
E1 2931005 2931183
refGene_gn=PER1;refGene_tx=XM_003996192.5;refGene_gn=PER1;refGene_tx=XM_006939763.4;refGene_gn=PER1;refGene_tx=XM_019817122.2;refGene_gn=PER1;refGene_tx=XM_019817123.2;refGene_gn=PER1;refGene_tx=XM_023244700.1
E1 2931627 2932235
refGene_gn=PER1;refGene_tx=XM_003996192.5;refGene_gn=PER1;refGene_tx=XM_006939763.4;refGene_gn=PER1;refGene_tx=XM_019817122.2;refGene_gn=PER1;refGene_tx=XM_019817123.2;refGene_gn=PER1;refGene_tx=XM_023244700.1
E1 2933105 2933348
refGene_gn=PER1;refGene_tx=XM_003996192.5;refGene_gn=PER1;refGene_tx=XM_006939763.4;refGene_gn=PER1;refGene_tx=XM_019817122.2;refGene_gn=PER1;refGene_tx=XM_019817123.2;refGene_gn=PER1;refGene_tx=XM_023244700.1
E1 2933956 2934133
refGene_gn=PER1;refGene_tx=XM_003996192.5;refGene_gn=PER1;refGene_tx=XM_006939763.4;refGene_gn=PER1;refGene_tx=XM_019817123.2;refGene_gn=PER1;refGene_tx=XM_023244700.1
E1 2934348 2934476
refGene_gn=PER1;refGene_tx=XM_003996192.5;refGene_gn=PER1;refGene_tx=XM_006939763.4;refGene_gn=PER1;refGene_tx=XM_019817122.2;refGene_gn=PER1;refGene_tx=XM_019817123.2;refGene_gn=PER1;refGene_tx=XM_023244700.1
E1 2934569 2934741
refGene_gn=PER1;refGene_tx=XM_003996192.5;refGene_gn=PER1;refGene_tx=XM_006939763.4;refGene_gn=PER1;refGene_tx=XM_019817122.2;refGene_gn=PER1;refGene_tx=XM_019817123.2;refGene_gn=PER1;refGene_tx=XM_023244700.1
E1 2934844 2934950
refGene_gn=PER1;refGene_tx=XM_003996192.5;refGene_gn=PER1;refGene_tx=XM_006939763.4;refGene_gn=PER1;refGene_tx=XM_019817122.2;refGene_gn=PER1;refGene_tx=XM_019817123.2;refGene_gn=PER1;refGene_tx=XM_023244700.1
E1 2935121 2935253
refGene_gn=PER1;refGene_tx=XM_003996192.5;refGene_gn=PER1;refGene_tx=XM_006939763.4;refGene_gn=PER1;refGene_tx=XM_019817122.2;refGene_gn=PER1;refGene_tx=XM_019817123.2;refGene_gn=PER1;refGene_tx=XM_023244700.1
E1 2935347 2935456
refGene_gn=PER1;refGene_tx=XM_003996192.5;refGene_gn=PER1;refGene_tx=XM_006939763.4;refGene_gn=PER1;refGene_tx=XM_019817122.2;refGene_gn=PER1;refGene_tx=XM_019817123.2;refGene_gn=PER1;refGene_tx=XM_023244700.1
E1 2935539 2935693
refGene_gn=PER1;refGene_tx=XM_003996192.5;refGene_gn=PER1;refGene_tx=XM_006939763.4;refGene_gn=PER1;refGene_tx=XM_019817122.2;refGene_gn=PER1;refGene_tx=XM_019817123.2;refGene_gn=PER1;refGene_tx=XM_023244700.1
E1 2935871 2935978
refGene_gn=PER1;refGene_tx=XM_003996192.5;refGene_gn=PER1;refGene_tx=X

M_006939763.4;refGene_gn=PER1;refGene_tx=XM_019817122.2;refGene_gn=PER1;refGene_tx=XM_019817123.2;refGene_gn=PER1;refGene_tx=XM_023244700.1
E1 2936056 2936135
refGene_gn=PER1;refGene_tx=XM_003996192.5;refGene_gn=PER1;refGene_tx=XM_006939763.4;refGene_gn=PER1;refGene_tx=XM_019817122.2;refGene_gn=PER1;refGene_tx=XM_019817123.2;refGene_gn=PER1;refGene_tx=XM_023244700.1
E1 2936521 2936664
refGene_gn=PER1;refGene_tx=XM_003996192.5;refGene_gn=PER1;refGene_tx=XM_006939763.4;refGene_gn=PER1;refGene_tx=XM_019817122.2;refGene_gn=PER1;refGene_tx=XM_019817123.2;refGene_gn=PER1;refGene_tx=XM_023244700.1
E1 2936802 2936854
refGene_gn=PER1;refGene_tx=XM_003996192.5;refGene_gn=PER1;refGene_tx=XM_006939763.4;refGene_gn=PER1;refGene_tx=XM_019817122.2;refGene_gn=PER1;refGene_tx=XM_019817123.2;refGene_gn=PER1;refGene_tx=XM_023244700.1
E1 2936967 2937169
refGene_gn=PER1;refGene_tx=XM_003996192.5;refGene_gn=PER1;refGene_tx=XM_006939763.4;refGene_gn=PER1;refGene_tx=XM_019817122.2;refGene_gn=PER1;refGene_tx=XM_019817123.2;refGene_gn=PER1;refGene_tx=XM_023244700.1
E1 2937262 2937384
refGene_gn=PER1;refGene_tx=XM_003996192.5;refGene_gn=PER1;refGene_tx=XM_006939763.4;refGene_gn=PER1;refGene_tx=XM_019817122.2;refGene_gn=PER1;refGene_tx=XM_019817123.2;refGene_gn=PER1;refGene_tx=XM_023244700.1
E1 2937494 2937649
refGene_gn=PER1;refGene_tx=XM_003996192.5;refGene_gn=PER1;refGene_tx=XM_006939763.4;refGene_gn=PER1;refGene_tx=XM_019817122.2;refGene_gn=PER1;refGene_tx=XM_019817123.2;refGene_gn=PER1;refGene_tx=XM_023244700.1
E1 2937744 2937843
refGene_gn=PER1;refGene_tx=XM_003996192.5;refGene_gn=PER1;refGene_tx=XM_006939763.4;refGene_gn=PER1;refGene_tx=XM_019817122.2;refGene_gn=PER1;refGene_tx=XM_019817123.2;refGene_gn=PER1;refGene_tx=XM_023244700.1
E1 2937956 2938378
refGene_gn=PER1;refGene_tx=XM_003996192.5;refGene_gn=PER1;refGene_tx=XM_006939763.4;refGene_gn=PER1;refGene_tx=XM_019817122.2;refGene_gn=PER1;refGene_tx=XM_019817123.2;refGene_gn=PER1;refGene_tx=XM_023244700.1
E1 2939905 2940019 refGene_gn=PER1;refGene_tx=XM_003996192.5
E1 2947246 2947263 refGene_gn=VAMP2;refGene_tx=XM_023244701.1
E1 2947859 2947911 refGene_gn=VAMP2;refGene_tx=XM_023244701.1
E1 2948013 2948172 refGene_gn=VAMP2;refGene_tx=XM_023244701.1
E1 2948633 2948754 refGene_gn=VAMP2;refGene_tx=XM_023244701.1
E1 2949257 2949259 refGene_gn=VAMP2;refGene_tx=XM_023244701.1
E1 2962146 2962216 refGene_gn=TMEM107;refGene_tx=XM_003996193.4
E1 2962447 2962544 refGene_gn=TMEM107;refGene_tx=XM_003996193.4
E1 2963488 2963589 refGene_gn=TMEM107;refGene_tx=XM_003996193.4
E1 2963691 2963759 refGene_gn=TMEM107;refGene_tx=XM_003996193.4
E1 2963942 2964029 refGene_gn=TMEM107;refGene_tx=XM_003996193.4
E1 2974350 2975415 refGene_gn=BORCS6;refGene_tx=XM_019816906.2
E1 2983280 2983454
refGene_gn=AURKB;refGene_tx=XM_019817124.1;refGene_gn=AURKB;refGene_tx=XM_019817126.2;refGene_gn=AURKB;refGene_tx=XM_019817127.2;refGene_gn=AURKB;refGene_tx=XM_019817128.1;refGene_gn=AURKB;refGene_tx=XM_0198171

29.1;refGene_gn=AURKB;refGene_tx=XM_019817130.1;refGene_gn=AURKB;refGene_tx=XM_019817131.2;refGene_gn=AURKB;refGene_tx=XM_019817132.2;refGene_gn=AURKB;refGene_tx=XM_019817133.1;refGene_gn=AURKB;refGene_tx=XM_019817135.1;refGene_gn=AURKB;refGene_tx=XM_023243857.1
E1 2983595 2983770
refGene_gn=AURKB;refGene_tx=XM_019817124.1;refGene_gn=AURKB;refGene_tx=XM_019817126.2;refGene_gn=AURKB;refGene_tx=XM_019817127.2;refGene_gn=AURKB;refGene_tx=XM_019817128.1;refGene_gn=AURKB;refGene_tx=XM_019817129.1;refGene_gn=AURKB;refGene_tx=XM_019817130.1;refGene_gn=AURKB;refGene_tx=XM_019817131.2;refGene_gn=AURKB;refGene_tx=XM_019817132.2;refGene_gn=AURKB;refGene_tx=XM_019817133.1;refGene_gn=AURKB;refGene_tx=XM_019817134.1;refGene_gn=AURKB;refGene_tx=XM_019817135.1;refGene_gn=AURKB;refGene_tx=XM_023243857.1
E1 2984827 2984976
refGene_gn=AURKB;refGene_tx=XM_019817124.1;refGene_gn=AURKB;refGene_tx=XM_019817126.2;refGene_gn=AURKB;refGene_tx=XM_019817127.2;refGene_gn=AURKB;refGene_tx=XM_019817128.1;refGene_gn=AURKB;refGene_tx=XM_019817129.1;refGene_gn=AURKB;refGene_tx=XM_019817130.1;refGene_gn=AURKB;refGene_tx=XM_019817131.2;refGene_gn=AURKB;refGene_tx=XM_019817132.2;refGene_gn=AURKB;refGene_tx=XM_019817133.1;refGene_gn=AURKB;refGene_tx=XM_019817134.1;refGene_gn=AURKB;refGene_tx=XM_019817135.1;refGene_gn=AURKB;refGene_tx=XM_023243857.1
E1 2985084 2985233
refGene_gn=AURKB;refGene_tx=XM_019817124.1;refGene_gn=AURKB;refGene_tx=XM_019817126.2;refGene_gn=AURKB;refGene_tx=XM_019817127.2;refGene_gn=AURKB;refGene_tx=XM_019817128.1;refGene_gn=AURKB;refGene_tx=XM_019817129.1;refGene_gn=AURKB;refGene_tx=XM_019817130.1;refGene_gn=AURKB;refGene_tx=XM_019817131.2;refGene_gn=AURKB;refGene_tx=XM_019817132.2;refGene_gn=AURKB;refGene_tx=XM_019817133.1;refGene_gn=AURKB;refGene_tx=XM_019817134.1;refGene_gn=AURKB;refGene_tx=XM_019817135.1;refGene_gn=AURKB;refGene_tx=XM_023243857.1
E1 2985455 2985650
refGene_gn=AURKB;refGene_tx=XM_019817124.1;refGene_gn=AURKB;refGene_tx=XM_019817126.2;refGene_gn=AURKB;refGene_tx=XM_019817127.2;refGene_gn=AURKB;refGene_tx=XM_019817128.1;refGene_gn=AURKB;refGene_tx=XM_019817129.1;refGene_gn=AURKB;refGene_tx=XM_019817130.1;refGene_gn=AURKB;refGene_tx=XM_019817131.2;refGene_gn=AURKB;refGene_tx=XM_019817132.2;refGene_gn=AURKB;refGene_tx=XM_019817133.1;refGene_gn=AURKB;refGene_tx=XM_019817134.1;refGene_gn=AURKB;refGene_tx=XM_023243857.1
E1 2985843 2985898
refGene_gn=AURKB;refGene_tx=XM_019817126.2;refGene_gn=AURKB;refGene_tx=XM_019817127.2;refGene_gn=AURKB;refGene_tx=XM_019817130.1;refGene_gn=AURKB;refGene_tx=XM_019817131.2;refGene_gn=AURKB;refGene_tx=XM_019817132.2;refGene_gn=AURKB;refGene_tx=XM_019817133.1;refGene_gn=AURKB;refGene_tx=XM_023243857.1
E1 2986006 2986109
refGene_gn=AURKB;refGene_tx=XM_019817124.1;refGene_gn=AURKB;refGene_tx=XM_019817126.2;refGene_gn=AURKB;refGene_tx=XM_019817127.2;refGene_gn=AURKB;refGene_tx=XM_019817128.1;refGene_gn=AURKB;refGene_tx=XM_019817129.1;refGene_gn=AURKB;refGene_tx=XM_019817130.1;refGene_gn=AURKB;refGe

ne_tx=XM_019817131.2;refGene_gn=AURKB;refGene_tx=XM_019817132.2;refGene_gn=AURKB;refGene_tx=XM_019817133.1;refGene_gn=AURKB;refGene_tx=XM_019817134.1;refGene_gn=AURKB;refGene_tx=XM_023243857.1
E1	2988181	2988278
refGene_gn=AURKB;refGene_tx=XM_019817124.1;refGene_gn=AURKB;refGene_tx=XM_019817128.1;refGene_gn=AURKB;refGene_tx=XM_019817130.1;refGene_gn=AURKB;refGene_tx=XM_019817131.2;refGene_gn=AURKB;refGene_tx=XM_019817132.2;refGene_gn=AURKB;refGene_tx=XM_019817134.1;refGene_gn=AURKB;refGene_tx=XM_023243857.1
E1	2988579	2988702
refGene_gn=AURKB;refGene_tx=XM_019817124.1;refGene_gn=AURKB;refGene_tx=XM_019817128.1;refGene_gn=AURKB;refGene_tx=XM_019817134.1
E1	2997766	2997932	refGene_gn=AURKB;refGene_tx=XM_019817129.1
E1	3003106	3003246
refGene_gn=CTC1;refGene_tx=XM_006939768.3;refGene_gn=CTC1;refGene_tx=XM_019817137.1;refGene_gn=CTC1;refGene_tx=XM_019817138.1
E1	3003401	3003528
refGene_gn=CTC1;refGene_tx=XM_006939768.3;refGene_gn=CTC1;refGene_tx=XM_019817137.1;refGene_gn=CTC1;refGene_tx=XM_019817138.1
E1	3003678	3003844
refGene_gn=CTC1;refGene_tx=XM_006939768.3;refGene_gn=CTC1;refGene_tx=XM_019817137.1;refGene_gn=CTC1;refGene_tx=XM_019817138.1
E1	3004104	3004173
refGene_gn=CTC1;refGene_tx=XM_006939768.3;refGene_gn=CTC1;refGene_tx=XM_006939769.3;refGene_gn=CTC1;refGene_tx=XM_011288854.2;refGene_gn=CTC1;refGene_tx=XM_011288855.2;refGene_gn=CTC1;refGene_tx=XM_019817137.1;refGene_gn=CTC1;refGene_tx=XM_019817138.1;refGene_gn=CTC1;refGene_tx=XM_019817139.1;refGene_gn=CTC1;refGene_tx=XM_019817140.1
E1	3004263	3004408
refGene_gn=CTC1;refGene_tx=XM_006939768.3;refGene_gn=CTC1;refGene_tx=XM_006939769.3;refGene_gn=CTC1;refGene_tx=XM_019817137.1;refGene_gn=CTC1;refGene_tx=XM_019817138.1
E1	3004839	3004917
refGene_gn=CTC1;refGene_tx=XM_006939768.3;refGene_gn=CTC1;refGene_tx=XM_006939769.3;refGene_gn=CTC1;refGene_tx=XM_019817137.1;refGene_gn=CTC1;refGene_tx=XM_019817138.1;refGene_gn=CTC1;refGene_tx=XM_019817139.1;refGene_gn=CTC1;refGene_tx=XM_019817140.1
E1	3005185	3005360
refGene_gn=CTC1;refGene_tx=XM_006939768.3;refGene_gn=CTC1;refGene_tx=XM_006939769.3;refGene_gn=CTC1;refGene_tx=XM_011288854.2;refGene_gn=CTC1;refGene_tx=XM_011288855.2;refGene_gn=CTC1;refGene_tx=XM_019817137.1;refGene_gn=CTC1;refGene_tx=XM_019817138.1;refGene_gn=CTC1;refGene_tx=XM_019817139.1;refGene_gn=CTC1;refGene_tx=XM_019817140.1
E1	3005452	3005541
refGene_gn=CTC1;refGene_tx=XM_006939768.3;refGene_gn=CTC1;refGene_tx=XM_006939769.3;refGene_gn=CTC1;refGene_tx=XM_011288854.2;refGene_gn=CTC1;refGene_tx=XM_011288855.2;refGene_gn=CTC1;refGene_tx=XM_019817137.1;refGene_gn=CTC1;refGene_tx=XM_019817138.1;refGene_gn=CTC1;refGene_tx=XM_019817139.1;refGene_gn=CTC1;refGene_tx=XM_019817140.1
E1	3005916	3006110

refGene_gn=CTC1;refGene_tx=XM_006939768.3;refGene_gn=CTC1;refGene_tx=XM_006939769.3;refGene_gn=CTC1;refGene_tx=XM_011288854.2;refGene_gn=CTC1;refGene_tx=XM_011288855.2;refGene_gn=CTC1;refGene_tx=XM_019817137.1;refGene_gn=CTC1;refGene_tx=XM_019817138.1;refGene_gn=CTC1;refGene_tx=XM_019817139.1;refGene_gn=CTC1;refGene_tx=XM_019817140.1
E1 3006354 3006444
refGene_gn=CTC1;refGene_tx=XM_006939768.3;refGene_gn=CTC1;refGene_tx=XM_006939769.3;refGene_gn=CTC1;refGene_tx=XM_011288854.2;refGene_gn=CTC1;refGene_tx=XM_011288855.2;refGene_gn=CTC1;refGene_tx=XM_019817139.1;refGene_gn=CTC1;refGene_tx=XM_019817140.1
E1 3006540 3006868
refGene_gn=CTC1;refGene_tx=XM_006939768.3;refGene_gn=CTC1;refGene_tx=XM_006939769.3;refGene_gn=CTC1;refGene_tx=XM_011288854.2;refGene_gn=CTC1;refGene_tx=XM_011288855.2;refGene_gn=CTC1;refGene_tx=XM_019817137.1;refGene_gn=CTC1;refGene_tx=XM_019817138.1;refGene_gn=CTC1;refGene_tx=XM_019817139.1;refGene_gn=CTC1;refGene_tx=XM_019817140.1
E1 3007018 3007133
refGene_gn=CTC1;refGene_tx=XM_006939768.3;refGene_gn=CTC1;refGene_tx=XM_006939769.3;refGene_gn=CTC1;refGene_tx=XM_011288854.2;refGene_gn=CTC1;refGene_tx=XM_011288855.2;refGene_gn=CTC1;refGene_tx=XM_019817137.1;refGene_gn=CTC1;refGene_tx=XM_019817138.1;refGene_gn=CTC1;refGene_tx=XM_019817139.1;refGene_gn=CTC1;refGene_tx=XM_019817140.1
E1 3007722 3007849
refGene_gn=CTC1;refGene_tx=XM_006939768.3;refGene_gn=CTC1;refGene_tx=XM_006939769.3;refGene_gn=CTC1;refGene_tx=XM_011288854.2;refGene_gn=CTC1;refGene_tx=XM_011288855.2;refGene_gn=CTC1;refGene_tx=XM_019817137.1;refGene_gn=CTC1;refGene_tx=XM_019817138.1;refGene_gn=CTC1;refGene_tx=XM_019817139.1;refGene_gn=CTC1;refGene_tx=XM_019817140.1
E1 3008356 3008557
refGene_gn=CTC1;refGene_tx=XM_006939768.3;refGene_gn=CTC1;refGene_tx=XM_006939769.3;refGene_gn=CTC1;refGene_tx=XM_011288854.2;refGene_gn=CTC1;refGene_tx=XM_011288855.2;refGene_gn=CTC1;refGene_tx=XM_019817137.1;refGene_gn=CTC1;refGene_tx=XM_019817138.1;refGene_gn=CTC1;refGene_tx=XM_019817139.1;refGene_gn=CTC1;refGene_tx=XM_019817140.1
E1 3008645 3008823
refGene_gn=CTC1;refGene_tx=XM_006939768.3;refGene_gn=CTC1;refGene_tx=XM_006939769.3;refGene_gn=CTC1;refGene_tx=XM_011288854.2;refGene_gn=CTC1;refGene_tx=XM_011288855.2;refGene_gn=CTC1;refGene_tx=XM_019817137.1;refGene_gn=CTC1;refGene_tx=XM_019817138.1;refGene_gn=CTC1;refGene_tx=XM_019817139.1;refGene_gn=CTC1;refGene_tx=XM_019817140.1
E1 3008946 3009179
refGene_gn=CTC1;refGene_tx=XM_006939768.3;refGene_gn=CTC1;refGene_tx=XM_006939769.3;refGene_gn=CTC1;refGene_tx=XM_011288854.2;refGene_gn=CTC1;refGene_tx=XM_011288855.2;refGene_gn=CTC1;refGene_tx=XM_019817137.1;refGene_gn=CTC1;refGene_tx=XM_019817138.1;refGene_gn=CTC1;refGene_tx=XM_019817139.1;refGene_gn=CTC1;refGene_tx=XM_019817140.1
E1 3009694 3009823
refGene_gn=CTC1;refGene_tx=XM_006939768.3;refGene_gn=CTC1;refGene_tx=XM_006939769.3;refGene_gn=CTC1;refGene_tx=XM_011288854.2;refGene_gn=CTC1;refGene_tx=XM_011288855.2;refGene_gn=CTC1;refGene_tx=XM_019817137.1;

refGene_gn=CTC1;refGene_tx=XM_019817138.1;refGene_gn=CTC1;refGene_tx=XM_019817139.1;refGene_gn=CTC1;refGene_tx=XM_019817140.1
E1 3009925 3010210
refGene_gn=CTC1;refGene_tx=XM_006939768.3;refGene_gn=CTC1;refGene_tx=XM_006939769.3;refGene_gn=CTC1;refGene_tx=XM_011288854.2;refGene_gn=CTC1;refGene_tx=XM_011288855.2;refGene_gn=CTC1;refGene_tx=XM_019817137.1;refGene_gn=CTC1;refGene_tx=XM_019817138.1;refGene_gn=CTC1;refGene_tx=XM_019817139.1;refGene_gn=CTC1;refGene_tx=XM_019817140.1
E1 3012849 3012994
refGene_gn=CTC1;refGene_tx=XM_006939768.3;refGene_gn=CTC1;refGene_tx=XM_006939769.3;refGene_gn=CTC1;refGene_tx=XM_011288854.2;refGene_gn=CTC1;refGene_tx=XM_011288855.2;refGene_gn=CTC1;refGene_tx=XM_019817137.1;refGene_gn=CTC1;refGene_tx=XM_019817138.1;refGene_gn=CTC1;refGene_tx=XM_019817139.1;refGene_gn=CTC1;refGene_tx=XM_019817140.1
E1 3013324 3013536
refGene_gn=CTC1;refGene_tx=XM_006939768.3;refGene_gn=CTC1;refGene_tx=XM_006939769.3;refGene_gn=CTC1;refGene_tx=XM_011288854.2;refGene_gn=CTC1;refGene_tx=XM_011288855.2;refGene_gn=CTC1;refGene_tx=XM_019817137.1;refGene_gn=CTC1;refGene_tx=XM_019817138.1;refGene_gn=CTC1;refGene_tx=XM_019817139.1;refGene_gn=CTC1;refGene_tx=XM_019817140.1
E1 3013685 3013923
refGene_gn=CTC1;refGene_tx=XM_006939768.3;refGene_gn=CTC1;refGene_tx=XM_006939769.3;refGene_gn=CTC1;refGene_tx=XM_011288854.2;refGene_gn=CTC1;refGene_tx=XM_011288855.2;refGene_gn=CTC1;refGene_tx=XM_019817137.1;refGene_gn=CTC1;refGene_tx=XM_019817138.1;refGene_gn=CTC1;refGene_tx=XM_019817139.1;refGene_gn=CTC1;refGene_tx=XM_019817140.1
E1 3018433 3018597
refGene_gn=CTC1;refGene_tx=XM_006939768.3;refGene_gn=CTC1;refGene_tx=XM_006939769.3;refGene_gn=CTC1;refGene_tx=XM_011288854.2;refGene_gn=CTC1;refGene_tx=XM_011288855.2;refGene_gn=CTC1;refGene_tx=XM_019817137.1;refGene_gn=CTC1;refGene_tx=XM_019817138.1;refGene_gn=CTC1;refGene_tx=XM_019817139.1;refGene_gn=CTC1;refGene_tx=XM_019817140.1
E1 3023250 3023283
refGene_gn=CTC1;refGene_tx=XM_006939768.3;refGene_gn=CTC1;refGene_tx=XM_006939769.3;refGene_gn=CTC1;refGene_tx=XM_011288854.2;refGene_gn=CTC1;refGene_tx=XM_011288855.2;refGene_gn=CTC1;refGene_tx=XM_019817137.1;refGene_gn=CTC1;refGene_tx=XM_019817138.1;refGene_gn=CTC1;refGene_tx=XM_019817139.1;refGene_gn=CTC1;refGene_tx=XM_019817140.1
E1 3031800 3031942
refGene_gn=PFAS;refGene_tx=XM_003996196.5;refGene_gn=PFAS;refGene_tx=XM_006939771.2;refGene_gn=PFAS;refGene_tx=XM_006939772.3
E1 3032028 3032164
refGene_gn=PFAS;refGene_tx=XM_003996196.5;refGene_gn=PFAS;refGene_tx=XM_006939771.2;refGene_gn=PFAS;refGene_tx=XM_006939772.3
E1 3032394 3032500
refGene_gn=PFAS;refGene_tx=XM_003996196.5;refGene_gn=PFAS;refGene_tx=XM_006939771.2;refGene_gn=PFAS;refGene_tx=XM_006939772.3
E1 3032840 3033030
refGene_gn=PFAS;refGene_tx=XM_003996196.5;refGene_gn=PFAS;refGene_tx=XM_006939771.2;refGene_gn=PFAS;refGene_tx=XM_006939772.3

E1 3033142 3033248
refGene_gn=PFAS;refGene_tx=XM_003996196.5;refGene_gn=PFAS;refGene_tx=XM_006939771.2;refGene_gn=PFAS;refGene_tx=XM_006939772.3
E1 3033615 3033756
refGene_gn=PFAS;refGene_tx=XM_003996196.5;refGene_gn=PFAS;refGene_tx=XM_006939771.2;refGene_gn=PFAS;refGene_tx=XM_006939772.3
E1 3033874 3033999
refGene_gn=PFAS;refGene_tx=XM_003996196.5;refGene_gn=PFAS;refGene_tx=XM_006939771.2;refGene_gn=PFAS;refGene_tx=XM_006939772.3
E1 3034168 3034297
refGene_gn=PFAS;refGene_tx=XM_003996196.5;refGene_gn=PFAS;refGene_tx=XM_006939771.2;refGene_gn=PFAS;refGene_tx=XM_006939772.3
E1 3035265 3035397
refGene_gn=PFAS;refGene_tx=XM_003996196.5;refGene_gn=PFAS;refGene_tx=XM_006939771.2;refGene_gn=PFAS;refGene_tx=XM_006939772.3
E1 3035534 3035663
refGene_gn=PFAS;refGene_tx=XM_003996196.5;refGene_gn=PFAS;refGene_tx=XM_006939771.2;refGene_gn=PFAS;refGene_tx=XM_006939772.3
E1 3039527 3039601
refGene_gn=PFAS;refGene_tx=XM_003996196.5;refGene_gn=PFAS;refGene_tx=XM_006939771.2;refGene_gn=PFAS;refGene_tx=XM_006939772.3
E1 3039711 3039868
refGene_gn=PFAS;refGene_tx=XM_003996196.5;refGene_gn=PFAS;refGene_tx=XM_006939771.2;refGene_gn=PFAS;refGene_tx=XM_006939772.3
E1 3040182 3040244
refGene_gn=PFAS;refGene_tx=XM_003996196.5;refGene_gn=PFAS;refGene_tx=XM_006939771.2;refGene_gn=PFAS;refGene_tx=XM_006939772.3
E1 3040372 3040534
refGene_gn=PFAS;refGene_tx=XM_003996196.5;refGene_gn=PFAS;refGene_tx=XM_006939771.2;refGene_gn=PFAS;refGene_tx=XM_006939772.3
E1 3040783 3040903
refGene_gn=PFAS;refGene_tx=XM_003996196.5;refGene_gn=PFAS;refGene_tx=XM_006939771.2;refGene_gn=PFAS;refGene_tx=XM_006939772.3
E1 3041036 3041168
refGene_gn=PFAS;refGene_tx=XM_003996196.5;refGene_gn=PFAS;refGene_tx=XM_006939771.2;refGene_gn=PFAS;refGene_tx=XM_006939772.3
E1 3041376 3041607
refGene_gn=PFAS;refGene_tx=XM_003996196.5;refGene_gn=PFAS;refGene_tx=XM_006939771.2;refGene_gn=PFAS;refGene_tx=XM_006939772.3
E1 3041778 3041959
refGene_gn=PFAS;refGene_tx=XM_003996196.5;refGene_gn=PFAS;refGene_tx=XM_006939771.2;refGene_gn=PFAS;refGene_tx=XM_006939772.3
E1 3042044 3042128
refGene_gn=PFAS;refGene_tx=XM_003996196.5;refGene_gn=PFAS;refGene_tx=XM_006939771.2;refGene_gn=PFAS;refGene_tx=XM_006939772.3
E1 3042273 3042429
refGene_gn=PFAS;refGene_tx=XM_003996196.5;refGene_gn=PFAS;refGene_tx=XM_006939771.2;refGene_gn=PFAS;refGene_tx=XM_006939772.3
E1 3042625 3042745
refGene_gn=PFAS;refGene_tx=XM_003996196.5;refGene_gn=PFAS;refGene_tx=X

M_006939771.2;refGene_gn=PFAS;refGene_tx=XM_006939772.3
E1 3043085 3043231
refGene_gn=PFAS;refGene_tx=XM_003996196.5;refGene_gn=PFAS;refGene_tx=X
M_006939771.2;refGene_gn=PFAS;refGene_tx=XM_006939772.3
E1 3043333 3043541
refGene_gn=PFAS;refGene_tx=XM_003996196.5;refGene_gn=PFAS;refGene_tx=X
M_006939771.2;refGene_gn=PFAS;refGene_tx=XM_006939772.3
E1 3043692 3043784
refGene_gn=PFAS;refGene_tx=XM_003996196.5;refGene_gn=PFAS;refGene_tx=X
M_006939771.2;refGene_gn=PFAS;refGene_tx=XM_006939772.3
E1 3043863 3043978
refGene_gn=PFAS;refGene_tx=XM_003996196.5;refGene_gn=PFAS;refGene_tx=X
M_006939771.2;refGene_gn=PFAS;refGene_tx=XM_006939772.3
E1 3045233 3045557
refGene_gn=PFAS;refGene_tx=XM_003996196.5;refGene_gn=PFAS;refGene_tx=X
M_006939771.2;refGene_gn=PFAS;refGene_tx=XM_006939772.3
E1 3045660 3045971
refGene_gn=PFAS;refGene_tx=XM_003996196.5;refGene_gn=PFAS;refGene_tx=X
M_006939771.2;refGene_gn=PFAS;refGene_tx=XM_006939772.3
E1 3058094 3058279
refGene_gn=RANGRF;refGene_tx=XM_003996197.5;refGene_gn=RANGRF;refGene_
tx=XM_011288852.3;refGene_gn=RANGRF;refGene_tx=XM_019817142.2
E1 3058367 3058484
refGene_gn=RANGRF;refGene_tx=XM_003996197.5;refGene_gn=RANGRF;refGene_
tx=XM_011288852.3;refGene_gn=RANGRF;refGene_tx=XM_019817142.2
E1 3058630 3058639 refGene_gn=SLC25A35;refGene_tx=XM_019817141.2
E1 3058701 3058890
refGene_gn=RANGRF;refGene_tx=XM_003996197.5;refGene_gn=RANGRF;refGene_
tx=XM_011288852.3;refGene_gn=RANGRF;refGene_tx=XM_019817142.2
E1 3058924 3059051
refGene_gn=RANGRF;refGene_tx=XM_003996197.5;refGene_gn=RANGRF;refGene_
tx=XM_011288852.3
E1 3059223 3059347
refGene_gn=RANGRF;refGene_tx=XM_003996197.5;refGene_gn=RANGRF;refGene_
tx=XM_019817142.2
E1 3059913 3060087 refGene_gn=SLC25A35;refGene_tx=XM_006939773.4
E1 3060259 3060394
refGene_gn=SLC25A35;refGene_tx=XM_006939773.4;refGene_gn=SLC25A35;refG
ene_tx=XM_019817141.2
E1 3060779 3060932
refGene_gn=SLC25A35;refGene_tx=XM_006939773.4;refGene_gn=SLC25A35;refG
ene_tx=XM_019817141.2
E1 3061373 3061439
refGene_gn=SLC25A35;refGene_tx=XM_006939773.4;refGene_gn=SLC25A35;refG
ene_tx=XM_019817141.2
E1 3062331 3062706
refGene_gn=SLC25A35;refGene_tx=XM_006939773.4;refGene_gn=SLC25A35;refG
ene_tx=XM_019817141.2
E1 3068979 3069085
refGene_gn=LOC102901445;refGene_tx=XM_006939774.2

E1	3071919	3071980
refGene_gn=LOC102901445;refGene_tx=XM_023243597.1
E1	3072610	3072714
refGene_gn=LOC102901445;refGene_tx=XM_006939774.2;refGene_gn=LOC102901
445;refGene_tx=XM_023243597.1
E1	3074370	3074841
refGene_gn=LOC102901445;refGene_tx=XM_006939774.2;refGene_gn=LOC102901
445;refGene_tx=XM_023243597.1
E1	3075238	3075845
refGene_gn=ARHGEF15;refGene_tx=XM_023243590.1;refGene_gn=ARHGEF15;refG
ene_tx=XM_023243591.1;refGene_gn=ARHGEF15;refGene_tx=XM_023243592.1;re
fGene_gn=ARHGEF15;refGene_tx=XM_023243593.1;refGene_gn=ARHGEF15;refGen
e_tx=XM_023243594.1;refGene_gn=ARHGEF15;refGene_tx=XM_023243596.1
E1	3076863	3077196
refGene_gn=ARHGEF15;refGene_tx=XM_023243590.1;refGene_gn=ARHGEF15;refG
ene_tx=XM_023243591.1;refGene_gn=ARHGEF15;refGene_tx=XM_023243592.1;re
fGene_gn=ARHGEF15;refGene_tx=XM_023243593.1;refGene_gn=ARHGEF15;refGen
e_tx=XM_023243594.1;refGene_gn=ARHGEF15;refGene_tx=XM_023243596.1
E1	3077459	3077514
refGene_gn=ARHGEF15;refGene_tx=XM_023243590.1;refGene_gn=ARHGEF15;refG
ene_tx=XM_023243591.1;refGene_gn=ARHGEF15;refGene_tx=XM_023243592.1;re
fGene_gn=ARHGEF15;refGene_tx=XM_023243593.1;refGene_gn=ARHGEF15;refGen
e_tx=XM_023243594.1;refGene_gn=ARHGEF15;refGene_tx=XM_023243596.1
E1	3078742	3078801
refGene_gn=ARHGEF15;refGene_tx=XM_023243590.1;refGene_gn=ARHGEF15;refG
ene_tx=XM_023243591.1;refGene_gn=ARHGEF15;refGene_tx=XM_023243592.1;re
fGene_gn=ARHGEF15;refGene_tx=XM_023243593.1;refGene_gn=ARHGEF15;refGen
e_tx=XM_023243594.1;refGene_gn=ARHGEF15;refGene_tx=XM_023243596.1
E1	3078909	3079121
refGene_gn=ARHGEF15;refGene_tx=XM_023243590.1;refGene_gn=ARHGEF15;refG
ene_tx=XM_023243591.1;refGene_gn=ARHGEF15;refGene_tx=XM_023243592.1;re
fGene_gn=ARHGEF15;refGene_tx=XM_023243593.1;refGene_gn=ARHGEF15;refGen
e_tx=XM_023243594.1;refGene_gn=ARHGEF15;refGene_tx=XM_023243596.1
E1	3079239	3079400
refGene_gn=ARHGEF15;refGene_tx=XM_023243590.1;refGene_gn=ARHGEF15;refG
ene_tx=XM_023243591.1;refGene_gn=ARHGEF15;refGene_tx=XM_023243592.1;re
fGene_gn=ARHGEF15;refGene_tx=XM_023243593.1;refGene_gn=ARHGEF15;refGen
e_tx=XM_023243594.1;refGene_gn=ARHGEF15;refGene_tx=XM_023243596.1
E1	3079581	3079734
refGene_gn=ARHGEF15;refGene_tx=XM_023243590.1;refGene_gn=ARHGEF15;refG
ene_tx=XM_023243591.1;refGene_gn=ARHGEF15;refGene_tx=XM_023243592.1;re
fGene_gn=ARHGEF15;refGene_tx=XM_023243593.1;refGene_gn=ARHGEF15;refGen
e_tx=XM_023243594.1;refGene_gn=ARHGEF15;refGene_tx=XM_023243596.1
E1	3079834	3079964
refGene_gn=ARHGEF15;refGene_tx=XM_023243590.1;refGene_gn=ARHGEF15;refG
ene_tx=XM_023243591.1;refGene_gn=ARHGEF15;refGene_tx=XM_023243592.1;re
fGene_gn=ARHGEF15;refGene_tx=XM_023243593.1;refGene_gn=ARHGEF15;refGen
e_tx=XM_023243594.1;refGene_gn=ARHGEF15;refGene_tx=XM_023243596.1
E1	3081228	3081303
refGene_gn=ARHGEF15;refGene_tx=XM_023243590.1;refGene_gn=ARHGEF15;refG

ene_tx=XM_023243591.1;refGene_gn=ARHGEF15;refGene_tx=XM_023243592.1;refGene_gn=ARHGEF15;refGene_tx=XM_023243593.1;refGene_gn=ARHGEF15;refGene_tx=XM_023243594.1;refGene_gn=ARHGEF15;refGene_tx=XM_023243596.1
E1 3081420 3081513
refGene_gn=ARHGEF15;refGene_tx=XM_023243590.1;refGene_gn=ARHGEF15;refGene_tx=XM_023243591.1;refGene_gn=ARHGEF15;refGene_tx=XM_023243592.1;refGene_gn=ARHGEF15;refGene_tx=XM_023243593.1;refGene_gn=ARHGEF15;refGene_tx=XM_023243594.1;refGene_gn=ARHGEF15;refGene_tx=XM_023243596.1
E1 3081603 3081764
refGene_gn=ARHGEF15;refGene_tx=XM_023243590.1;refGene_gn=ARHGEF15;refGene_tx=XM_023243591.1;refGene_gn=ARHGEF15;refGene_tx=XM_023243592.1;refGene_gn=ARHGEF15;refGene_tx=XM_023243593.1;refGene_gn=ARHGEF15;refGene_tx=XM_023243594.1;refGene_gn=ARHGEF15;refGene_tx=XM_023243596.1
E1 3081870 3082023
refGene_gn=ARHGEF15;refGene_tx=XM_023243590.1;refGene_gn=ARHGEF15;refGene_tx=XM_023243591.1;refGene_gn=ARHGEF15;refGene_tx=XM_023243592.1;refGene_gn=ARHGEF15;refGene_tx=XM_023243593.1;refGene_gn=ARHGEF15;refGene_tx=XM_023243594.1;refGene_gn=ARHGEF15;refGene_tx=XM_023243596.1
E1 3082175 3082258
refGene_gn=ARHGEF15;refGene_tx=XM_023243590.1;refGene_gn=ARHGEF15;refGene_tx=XM_023243591.1;refGene_gn=ARHGEF15;refGene_tx=XM_023243592.1;refGene_gn=ARHGEF15;refGene_tx=XM_023243593.1;refGene_gn=ARHGEF15;refGene_tx=XM_023243594.1;refGene_gn=ARHGEF15;refGene_tx=XM_023243596.1
E1 3082368 3082473
refGene_gn=ARHGEF15;refGene_tx=XM_023243590.1;refGene_gn=ARHGEF15;refGene_tx=XM_023243591.1;refGene_gn=ARHGEF15;refGene_tx=XM_023243592.1;refGene_gn=ARHGEF15;refGene_tx=XM_023243593.1;refGene_gn=ARHGEF15;refGene_tx=XM_023243594.1;refGene_gn=ARHGEF15;refGene_tx=XM_023243596.1
E1 3083852 3084004
refGene_gn=ARHGEF15;refGene_tx=XM_023243590.1;refGene_gn=ARHGEF15;refGene_tx=XM_023243591.1;refGene_gn=ARHGEF15;refGene_tx=XM_023243592.1;refGene_gn=ARHGEF15;refGene_tx=XM_023243593.1;refGene_gn=ARHGEF15;refGene_tx=XM_023243594.1;refGene_gn=ARHGEF15;refGene_tx=XM_023243596.1
E1 3093486 3093931 refGene_gn=ODF4;refGene_tx=XM_003996279.5
E1 3096540 3096714 refGene_gn=ODF4;refGene_tx=XM_003996279.5
E1 3096941 3097076 refGene_gn=ODF4;refGene_tx=XM_003996279.5
E1 3097273 3097458 refGene_gn=ODF4;refGene_tx=XM_003996279.5
E1 3116728 3117984
refGene_gn=KRBA2;refGene_tx=XM_006939775.4;refGene_gn=KRBA2;refGene_tx=XM_006939776.4;refGene_gn=KRBA2;refGene_tx=XM_006939777.3;refGene_gn=KRBA2;refGene_tx=XM_006939778.4;refGene_gn=KRBA2;refGene_tx=XM_006939780.3;refGene_gn=KRBA2;refGene_tx=XM_019817146.2;refGene_gn=KRBA2;refGene_tx=XM_019817147.1;refGene_gn=KRBA2;refGene_tx=XM_019817148.2;refGene_gn=KRBA2;refGene_tx=XM_019817149.1;refGene_gn=KRBA2;refGene_tx=XM_019817150.2;refGene_gn=KRBA2;refGene_tx=XM_023243427.1
E1 3119514 3119548
refGene_gn=KRBA2;refGene_tx=XM_006939775.4;refGene_gn=KRBA2;refGene_tx=XM_006939776.4;refGene_gn=KRBA2;refGene_tx=XM_006939777.3;refGene_gn=KRBA2;refGene_tx=XM_006939778.4;refGene_gn=KRBA2;refGene_tx=XM_006939780.3;refGene_gn=KRBA2;refGene_tx=XM_019817146.2;refGene_gn=KRBA2;refGe

ne_tx=XM_019817147.1;refGene_gn=KRBA2;refGene_tx=XM_019817148.2;refGen
e_gn=KRBA2;refGene_tx=XM_023243427.1
E1 3123401 3123530 refGene_gn=RPL26;refGene_tx=XM_006939783.2
E1 3126844 3126985 refGene_gn=RPL26;refGene_tx=XM_006939783.2
E1 3128102 3128270 refGene_gn=RPL26;refGene_tx=XM_006939783.2
E1 3138237 3138831
refGene_gn=RNF222;refGene_tx=XM_023244272.1;refGene_gn=RNF222;refGene_
tx=XM_023244273.1;refGene_gn=RNF222;refGene_tx=XM_023244274.1
E1 3186113 3186199
refGene_gn=NDEL1;refGene_tx=XM_019817151.2;refGene_gn=NDEL1;refGene_tx
=XM_019817152.2;refGene_gn=NDEL1;refGene_tx=XM_019817153.2;refGene_gn=
NDEL1;refGene_tx=XM_023244691.1;refGene_gn=NDEL1;refGene_tx=XM_0232446
93.1;refGene_gn=NDEL1;refGene_tx=XM_023244694.1;refGene_gn=NDEL1;refGe
ne_tx=XM_023244695.1;refGene_gn=NDEL1;refGene_tx=XM_023244696.1;refGen
e_gn=NDEL1;refGene_tx=XM_023244697.1
E1 3187901 3188055
refGene_gn=NDEL1;refGene_tx=XM_019817151.2;refGene_gn=NDEL1;refGene_tx
=XM_019817152.2;refGene_gn=NDEL1;refGene_tx=XM_019817153.2;refGene_gn=
NDEL1;refGene_tx=XM_023244691.1;refGene_gn=NDEL1;refGene_tx=XM_0232446
93.1;refGene_gn=NDEL1;refGene_tx=XM_023244694.1;refGene_gn=NDEL1;refGe
ne_tx=XM_023244695.1;refGene_gn=NDEL1;refGene_tx=XM_023244696.1;refGen
e_gn=NDEL1;refGene_tx=XM_023244697.1
E1 3188988 3189137
refGene_gn=NDEL1;refGene_tx=XM_019817151.2;refGene_gn=NDEL1;refGene_tx
=XM_019817152.2;refGene_gn=NDEL1;refGene_tx=XM_019817153.2;refGene_gn=
NDEL1;refGene_tx=XM_023244691.1;refGene_gn=NDEL1;refGene_tx=XM_0232446
93.1;refGene_gn=NDEL1;refGene_tx=XM_023244694.1;refGene_gn=NDEL1;refGe
ne_tx=XM_023244695.1;refGene_gn=NDEL1;refGene_tx=XM_023244696.1;refGen
e_gn=NDEL1;refGene_tx=XM_023244697.1
E1 3191125 3191262
refGene_gn=NDEL1;refGene_tx=XM_019817151.2;refGene_gn=NDEL1;refGene_tx
=XM_019817152.2;refGene_gn=NDEL1;refGene_tx=XM_019817153.2;refGene_gn=
NDEL1;refGene_tx=XM_023244691.1;refGene_gn=NDEL1;refGene_tx=XM_0232446
93.1;refGene_gn=NDEL1;refGene_tx=XM_023244694.1;refGene_gn=NDEL1;refGe
ne_tx=XM_023244695.1;refGene_gn=NDEL1;refGene_tx=XM_023244696.1;refGen
e_gn=NDEL1;refGene_tx=XM_023244697.1
E1 3193094 3193268
refGene_gn=NDEL1;refGene_tx=XM_019817151.2;refGene_gn=NDEL1;refGene_tx
=XM_019817152.2;refGene_gn=NDEL1;refGene_tx=XM_019817153.2;refGene_gn=
NDEL1;refGene_tx=XM_023244691.1;refGene_gn=NDEL1;refGene_tx=XM_0232446
93.1;refGene_gn=NDEL1;refGene_tx=XM_023244694.1;refGene_gn=NDEL1;refGe
ne_tx=XM_023244695.1;refGene_gn=NDEL1;refGene_tx=XM_023244696.1;refGen
e_gn=NDEL1;refGene_tx=XM_023244697.1
E1 3197458 3197550
refGene_gn=NDEL1;refGene_tx=XM_019817151.2;refGene_gn=NDEL1;refGene_tx
=XM_019817152.2;refGene_gn=NDEL1;refGene_tx=XM_019817153.2;refGene_gn=
NDEL1;refGene_tx=XM_023244691.1;refGene_gn=NDEL1;refGene_tx=XM_0232446
93.1;refGene_gn=NDEL1;refGene_tx=XM_023244694.1;refGene_gn=NDEL1;refGe
ne_tx=XM_023244695.1;refGene_gn=NDEL1;refGene_tx=XM_023244696.1;refGen
e_gn=NDEL1;refGene_tx=XM_023244697.1

E1 3201095 3201247
refGene_gn=NDEL1;refGene_tx=XM_019817151.2;refGene_gn=NDEL1;refGene_tx
=XM_019817152.2;refGene_gn=NDEL1;refGene_tx=XM_019817153.2;refGene_gn=
NDEL1;refGene_tx=XM_023244691.1;refGene_gn=NDEL1;refGene_tx=XM_0232446
93.1;refGene_gn=NDEL1;refGene_tx=XM_023244694.1;refGene_gn=NDEL1;refGe
ne_tx=XM_023244695.1;refGene_gn=NDEL1;refGene_tx=XM_023244696.1;refGen
e_gn=NDEL1;refGene_tx=XM_023244697.1
E1 3203625 3203752 refGene_gn=NDEL1;refGene_tx=XM_023244695.1
E1 3204295 3204330 refGene_gn=NDEL1;refGene_tx=XM_023244697.1
E1 3207814 3207908
refGene_gn=NDEL1;refGene_tx=XM_023244696.1;refGene_gn=NDEL1;refGene_tx
=XM_023244697.1
E1 3217542 3217687
refGene_gn=MYH10;refGene_tx=XM_023244684.1;refGene_gn=MYH10;refGene_tx
=XM_023244685.1;refGene_gn=MYH10;refGene_tx=XM_023244686.1;refGene_gn=
MYH10;refGene_tx=XM_023244687.1;refGene_gn=MYH10;refGene_tx=XM_0232446
88.1;refGene_gn=MYH10;refGene_tx=XM_023244689.1;refGene_gn=MYH10;refGe
ne_tx=XM_023244690.1
E1 3218482 3218655
refGene_gn=MYH10;refGene_tx=XM_023244684.1;refGene_gn=MYH10;refGene_tx
=XM_023244685.1;refGene_gn=MYH10;refGene_tx=XM_023244686.1;refGene_gn=
MYH10;refGene_tx=XM_023244687.1;refGene_gn=MYH10;refGene_tx=XM_0232446
88.1;refGene_gn=MYH10;refGene_tx=XM_023244689.1;refGene_gn=MYH10;refGe
ne_tx=XM_023244690.1
E1 3219576 3219685
refGene_gn=MYH10;refGene_tx=XM_023244684.1;refGene_gn=MYH10;refGene_tx
=XM_023244685.1;refGene_gn=MYH10;refGene_tx=XM_023244686.1;refGene_gn=
MYH10;refGene_tx=XM_023244687.1;refGene_gn=MYH10;refGene_tx=XM_0232446
88.1;refGene_gn=MYH10;refGene_tx=XM_023244689.1;refGene_gn=MYH10;refGe
ne_tx=XM_023244690.1
E1 3220591 3220800
refGene_gn=MYH10;refGene_tx=XM_023244684.1;refGene_gn=MYH10;refGene_tx
=XM_023244685.1;refGene_gn=MYH10;refGene_tx=XM_023244686.1;refGene_gn=
MYH10;refGene_tx=XM_023244687.1;refGene_gn=MYH10;refGene_tx=XM_0232446
88.1;refGene_gn=MYH10;refGene_tx=XM_023244689.1;refGene_gn=MYH10;refGe
ne_tx=XM_023244690.1
E1 3220883 3221007
refGene_gn=MYH10;refGene_tx=XM_023244684.1;refGene_gn=MYH10;refGene_tx
=XM_023244685.1;refGene_gn=MYH10;refGene_tx=XM_023244686.1;refGene_gn=
MYH10;refGene_tx=XM_023244687.1;refGene_gn=MYH10;refGene_tx=XM_0232446
88.1;refGene_gn=MYH10;refGene_tx=XM_023244689.1;refGene_gn=MYH10;refGe
ne_tx=XM_023244690.1
E1 3221762 3221851
refGene_gn=MYH10;refGene_tx=XM_023244684.1;refGene_gn=MYH10;refGene_tx
=XM_023244685.1;refGene_gn=MYH10;refGene_tx=XM_023244686.1;refGene_gn=
MYH10;refGene_tx=XM_023244687.1;refGene_gn=MYH10;refGene_tx=XM_0232446
88.1;refGene_gn=MYH10;refGene_tx=XM_023244689.1;refGene_gn=MYH10;refGe
ne_tx=XM_023244690.1
E1 3224433 3224562
refGene_gn=MYH10;refGene_tx=XM_023244684.1;refGene_gn=MYH10;refGene_tx

=XM_023244685.1;refGene_gn=MYH10;refGene_tx=XM_023244686.1;refGene_gn=MYH10;refGene_tx=XM_023244687.1;refGene_gn=MYH10;refGene_tx=XM_023244688.1;refGene_gn=MYH10;refGene_tx=XM_023244689.1;refGene_gn=MYH10;refGene_tx=XM_023244690.1
E1 3228505 3228667 refGene_gn=MYH10;refGene_tx=XM_023244684.1;refGene_gn=MYH10;refGene_tx=XM_023244685.1;refGene_gn=MYH10;refGene_tx=XM_023244686.1;refGene_gn=MYH10;refGene_tx=XM_023244687.1;refGene_gn=MYH10;refGene_tx=XM_023244688.1;refGene_gn=MYH10;refGene_tx=XM_023244689.1;refGene_gn=MYH10;refGene_tx=XM_023244690.1
E1 3229766 3229979 refGene_gn=MYH10;refGene_tx=XM_023244684.1;refGene_gn=MYH10;refGene_tx=XM_023244685.1;refGene_gn=MYH10;refGene_tx=XM_023244686.1;refGene_gn=MYH10;refGene_tx=XM_023244687.1;refGene_gn=MYH10;refGene_tx=XM_023244688.1;refGene_gn=MYH10;refGene_tx=XM_023244689.1;refGene_gn=MYH10;refGene_tx=XM_023244690.1
E1 3231608 3231821 refGene_gn=MYH10;refGene_tx=XM_023244684.1;refGene_gn=MYH10;refGene_tx=XM_023244685.1;refGene_gn=MYH10;refGene_tx=XM_023244686.1;refGene_gn=MYH10;refGene_tx=XM_023244687.1;refGene_gn=MYH10;refGene_tx=XM_023244688.1;refGene_gn=MYH10;refGene_tx=XM_023244689.1;refGene_gn=MYH10;refGene_tx=XM_023244690.1
E1 3232078 3232327 refGene_gn=MYH10;refGene_tx=XM_023244684.1;refGene_gn=MYH10;refGene_tx=XM_023244685.1;refGene_gn=MYH10;refGene_tx=XM_023244686.1;refGene_gn=MYH10;refGene_tx=XM_023244687.1;refGene_gn=MYH10;refGene_tx=XM_023244688.1;refGene_gn=MYH10;refGene_tx=XM_023244689.1;refGene_gn=MYH10;refGene_tx=XM_023244690.1
E1 3232715 3232868 refGene_gn=MYH10;refGene_tx=XM_023244684.1;refGene_gn=MYH10;refGene_tx=XM_023244685.1;refGene_gn=MYH10;refGene_tx=XM_023244686.1;refGene_gn=MYH10;refGene_tx=XM_023244687.1;refGene_gn=MYH10;refGene_tx=XM_023244688.1;refGene_gn=MYH10;refGene_tx=XM_023244689.1;refGene_gn=MYH10;refGene_tx=XM_023244690.1
E1 3234115 3234220 refGene_gn=MYH10;refGene_tx=XM_023244684.1;refGene_gn=MYH10;refGene_tx=XM_023244685.1;refGene_gn=MYH10;refGene_tx=XM_023244686.1;refGene_gn=MYH10;refGene_tx=XM_023244687.1;refGene_gn=MYH10;refGene_tx=XM_023244688.1;refGene_gn=MYH10;refGene_tx=XM_023244689.1;refGene_gn=MYH10;refGene_tx=XM_023244690.1
E1 3235078 3235274 refGene_gn=NDEL1;refGene_tx=XM_023244694.1
E1 3236885 3237092 refGene_gn=MYH10;refGene_tx=XM_023244684.1;refGene_gn=MYH10;refGene_tx=XM_023244685.1;refGene_gn=MYH10;refGene_tx=XM_023244686.1;refGene_gn=MYH10;refGene_tx=XM_023244687.1;refGene_gn=MYH10;refGene_tx=XM_023244688.1;refGene_gn=MYH10;refGene_tx=XM_023244689.1;refGene_gn=MYH10;refGene_tx=XM_023244690.1
E1 3238406 3238551 refGene_gn=MYH10;refGene_tx=XM_023244684.1;refGene_gn=MYH10;refGene_tx=XM_023244685.1;refGene_gn=MYH10;refGene_tx=XM_023244686.1;refGene_gn=

MYH10;refGene_tx=XM_023244687.1;refGene_gn=MYH10;refGene_tx=XM_0232446
88.1;refGene_gn=MYH10;refGene_tx=XM_023244689.1;refGene_gn=MYH10;refGe
ne_tx=XM_023244690.1
E1 3239349 3239593 refGene_gn=NDEL1;refGene_tx=XM_023244693.1
E1 3241795 3242008
refGene_gn=MYH10;refGene_tx=XM_023244684.1;refGene_gn=MYH10;refGene_tx
=XM_023244685.1;refGene_gn=MYH10;refGene_tx=XM_023244686.1;refGene_gn=
MYH10;refGene_tx=XM_023244687.1;refGene_gn=MYH10;refGene_tx=XM_0232446
88.1;refGene_gn=MYH10;refGene_tx=XM_023244689.1;refGene_gn=MYH10;refGe
ne_tx=XM_023244690.1
E1 3243312 3243484
refGene_gn=MYH10;refGene_tx=XM_023244684.1;refGene_gn=MYH10;refGene_tx
=XM_023244685.1;refGene_gn=MYH10;refGene_tx=XM_023244686.1;refGene_gn=
MYH10;refGene_tx=XM_023244687.1;refGene_gn=MYH10;refGene_tx=XM_0232446
88.1;refGene_gn=MYH10;refGene_tx=XM_023244689.1;refGene_gn=MYH10;refGe
ne_tx=XM_023244690.1
E1 3244828 3244952
refGene_gn=MYH10;refGene_tx=XM_023244684.1;refGene_gn=MYH10;refGene_tx
=XM_023244685.1;refGene_gn=MYH10;refGene_tx=XM_023244686.1;refGene_gn=
MYH10;refGene_tx=XM_023244687.1;refGene_gn=MYH10;refGene_tx=XM_0232446
88.1;refGene_gn=MYH10;refGene_tx=XM_023244689.1;refGene_gn=MYH10;refGe
ne_tx=XM_023244690.1
E1 3245741 3245879
refGene_gn=MYH10;refGene_tx=XM_023244684.1;refGene_gn=MYH10;refGene_tx
=XM_023244685.1;refGene_gn=MYH10;refGene_tx=XM_023244686.1;refGene_gn=
MYH10;refGene_tx=XM_023244687.1;refGene_gn=MYH10;refGene_tx=XM_0232446
88.1;refGene_gn=MYH10;refGene_tx=XM_023244689.1;refGene_gn=MYH10;refGe
ne_tx=XM_023244690.1
E1 3249247 3249454
refGene_gn=MYH10;refGene_tx=XM_023244684.1;refGene_gn=MYH10;refGene_tx
=XM_023244685.1;refGene_gn=MYH10;refGene_tx=XM_023244686.1;refGene_gn=
MYH10;refGene_tx=XM_023244687.1;refGene_gn=MYH10;refGene_tx=XM_0232446
88.1;refGene_gn=MYH10;refGene_tx=XM_023244689.1;refGene_gn=MYH10;refGe
ne_tx=XM_023244690.1
E1 3250154 3250286
refGene_gn=MYH10;refGene_tx=XM_023244684.1;refGene_gn=MYH10;refGene_tx
=XM_023244685.1;refGene_gn=MYH10;refGene_tx=XM_023244686.1;refGene_gn=
MYH10;refGene_tx=XM_023244687.1;refGene_gn=MYH10;refGene_tx=XM_0232446
88.1;refGene_gn=MYH10;refGene_tx=XM_023244689.1;refGene_gn=MYH10;refGe
ne_tx=XM_023244690.1
E1 3250403 3250512
refGene_gn=MYH10;refGene_tx=XM_023244684.1;refGene_gn=MYH10;refGene_tx
=XM_023244685.1;refGene_gn=MYH10;refGene_tx=XM_023244686.1;refGene_gn=
MYH10;refGene_tx=XM_023244687.1;refGene_gn=MYH10;refGene_tx=XM_0232446
88.1;refGene_gn=MYH10;refGene_tx=XM_023244689.1;refGene_gn=MYH10;refGe
ne_tx=XM_023244690.1
E1 3252021 3252520
refGene_gn=NDEL1;refGene_tx=XM_019817151.2;refGene_gn=NDEL1;refGene_tx
=XM_019817152.2;refGene_gn=NDEL1;refGene_tx=XM_019817153.2;refGene_gn=
NDEL1;refGene_tx=XM_023244691.1

E1 3253835 3253996
refGene_gn=MYH10;refGene_tx=XM_023244684.1;refGene_gn=MYH10;refGene_tx=XM_023244685.1;refGene_gn=MYH10;refGene_tx=XM_023244686.1;refGene_gn=MYH10;refGene_tx=XM_023244687.1;refGene_gn=MYH10;refGene_tx=XM_023244688.1;refGene_gn=MYH10;refGene_tx=XM_023244689.1;refGene_gn=MYH10;refGene_tx=XM_023244690.1
E1 3254090 3254160
refGene_gn=MYH10;refGene_tx=XM_023244684.1;refGene_gn=MYH10;refGene_tx=XM_023244685.1;refGene_gn=MYH10;refGene_tx=XM_023244686.1;refGene_gn=MYH10;refGene_tx=XM_023244687.1;refGene_gn=MYH10;refGene_tx=XM_023244688.1;refGene_gn=MYH10;refGene_tx=XM_023244689.1;refGene_gn=MYH10;refGene_tx=XM_023244690.1
E1 3256147 3256269
refGene_gn=MYH10;refGene_tx=XM_023244684.1;refGene_gn=MYH10;refGene_tx=XM_023244685.1;refGene_gn=MYH10;refGene_tx=XM_023244686.1;refGene_gn=MYH10;refGene_tx=XM_023244687.1;refGene_gn=MYH10;refGene_tx=XM_023244688.1;refGene_gn=MYH10;refGene_tx=XM_023244689.1;refGene_gn=MYH10;refGene_tx=XM_023244690.1
E1 3256361 3256555
refGene_gn=MYH10;refGene_tx=XM_023244684.1;refGene_gn=MYH10;refGene_tx=XM_023244685.1;refGene_gn=MYH10;refGene_tx=XM_023244686.1;refGene_gn=MYH10;refGene_tx=XM_023244687.1;refGene_gn=MYH10;refGene_tx=XM_023244688.1;refGene_gn=MYH10;refGene_tx=XM_023244689.1;refGene_gn=MYH10;refGene_tx=XM_023244690.1
E1 3265435 3265498
refGene_gn=MYH10;refGene_tx=XM_023244684.1;refGene_gn=MYH10;refGene_tx=XM_023244685.1;refGene_gn=MYH10;refGene_tx=XM_023244686.1;refGene_gn=MYH10;refGene_tx=XM_023244689.1
E1 3269700 3269815
refGene_gn=MYH10;refGene_tx=XM_023244684.1;refGene_gn=MYH10;refGene_tx=XM_023244685.1;refGene_gn=MYH10;refGene_tx=XM_023244686.1;refGene_gn=MYH10;refGene_tx=XM_023244687.1;refGene_gn=MYH10;refGene_tx=XM_023244688.1;refGene_gn=MYH10;refGene_tx=XM_023244689.1;refGene_gn=MYH10;refGene_tx=XM_023244690.1
E1 3270053 3270227
refGene_gn=MYH10;refGene_tx=XM_023244684.1;refGene_gn=MYH10;refGene_tx=XM_023244685.1;refGene_gn=MYH10;refGene_tx=XM_023244686.1;refGene_gn=MYH10;refGene_tx=XM_023244687.1;refGene_gn=MYH10;refGene_tx=XM_023244688.1;refGene_gn=MYH10;refGene_tx=XM_023244689.1;refGene_gn=MYH10;refGene_tx=XM_023244690.1
E1 3272289 3272463
refGene_gn=MYH10;refGene_tx=XM_023244684.1;refGene_gn=MYH10;refGene_tx=XM_023244685.1;refGene_gn=MYH10;refGene_tx=XM_023244686.1;refGene_gn=MYH10;refGene_tx=XM_023244687.1;refGene_gn=MYH10;refGene_tx=XM_023244688.1;refGene_gn=MYH10;refGene_tx=XM_023244689.1;refGene_gn=MYH10;refGene_tx=XM_023244690.1
E1 3275881 3276034
refGene_gn=MYH10;refGene_tx=XM_023244684.1;refGene_gn=MYH10;refGene_tx=XM_023244685.1;refGene_gn=MYH10;refGene_tx=XM_023244686.1;refGene_gn=MYH10;refGene_tx=XM_023244687.1;refGene_gn=MYH10;refGene_tx=XM_0232446

88.1;refGene_gn=MYH10;refGene_tx=XM_023244689.1;refGene_gn=MYH10;refGene_tx=XM_023244690.1
E1 3276658 3276777
refGene_gn=MYH10;refGene_tx=XM_023244684.1;refGene_gn=MYH10;refGene_tx=XM_023244685.1;refGene_gn=MYH10;refGene_tx=XM_023244686.1;refGene_gn=MYH10;refGene_tx=XM_023244687.1;refGene_gn=MYH10;refGene_tx=XM_023244688.1;refGene_gn=MYH10;refGene_tx=XM_023244689.1;refGene_gn=MYH10;refGene_tx=XM_023244690.1
E1 3278288 3278384
refGene_gn=MYH10;refGene_tx=XM_023244684.1;refGene_gn=MYH10;refGene_tx=XM_023244685.1;refGene_gn=MYH10;refGene_tx=XM_023244686.1;refGene_gn=MYH10;refGene_tx=XM_023244687.1;refGene_gn=MYH10;refGene_tx=XM_023244688.1;refGene_gn=MYH10;refGene_tx=XM_023244689.1;refGene_gn=MYH10;refGene_tx=XM_023244690.1
E1 3278632 3278776
refGene_gn=MYH10;refGene_tx=XM_023244684.1;refGene_gn=MYH10;refGene_tx=XM_023244685.1;refGene_gn=MYH10;refGene_tx=XM_023244686.1;refGene_gn=MYH10;refGene_tx=XM_023244687.1;refGene_gn=MYH10;refGene_tx=XM_023244688.1;refGene_gn=MYH10;refGene_tx=XM_023244689.1;refGene_gn=MYH10;refGene_tx=XM_023244690.1
E1 3281453 3281552
refGene_gn=MYH10;refGene_tx=XM_023244684.1;refGene_gn=MYH10;refGene_tx=XM_023244685.1;refGene_gn=MYH10;refGene_tx=XM_023244686.1;refGene_gn=MYH10;refGene_tx=XM_023244687.1;refGene_gn=MYH10;refGene_tx=XM_023244688.1;refGene_gn=MYH10;refGene_tx=XM_023244689.1;refGene_gn=MYH10;refGene_tx=XM_023244690.1
E1 3283444 3283508
refGene_gn=MYH10;refGene_tx=XM_023244684.1;refGene_gn=MYH10;refGene_tx=XM_023244685.1;refGene_gn=MYH10;refGene_tx=XM_023244686.1;refGene_gn=MYH10;refGene_tx=XM_023244687.1;refGene_gn=MYH10;refGene_tx=XM_023244688.1;refGene_gn=MYH10;refGene_tx=XM_023244689.1;refGene_gn=MYH10;refGene_tx=XM_023244690.1
E1 3294023 3294116
refGene_gn=MYH10;refGene_tx=XM_023244684.1;refGene_gn=MYH10;refGene_tx=XM_023244685.1;refGene_gn=MYH10;refGene_tx=XM_023244686.1;refGene_gn=MYH10;refGene_tx=XM_023244687.1;refGene_gn=MYH10;refGene_tx=XM_023244688.1;refGene_gn=MYH10;refGene_tx=XM_023244689.1;refGene_gn=MYH10;refGene_tx=XM_023244690.1
E1 3299922 3299952
refGene_gn=MYH10;refGene_tx=XM_023244684.1;refGene_gn=MYH10;refGene_tx=XM_023244685.1;refGene_gn=MYH10;refGene_tx=XM_023244687.1;refGene_gn=MYH10;refGene_tx=XM_023244688.1;refGene_gn=MYH10;refGene_tx=XM_023244689.1
E1 3300547 3300650
refGene_gn=MYH10;refGene_tx=XM_023244684.1;refGene_gn=MYH10;refGene_tx=XM_023244685.1;refGene_gn=MYH10;refGene_tx=XM_023244686.1;refGene_gn=MYH10;refGene_tx=XM_023244687.1;refGene_gn=MYH10;refGene_tx=XM_023244688.1;refGene_gn=MYH10;refGene_tx=XM_023244689.1;refGene_gn=MYH10;refGene_tx=XM_023244690.1
E1 3316468 3316496

refGene_gn=MYH10;refGene_tx=XM_023244684.1;refGene_gn=MYH10;refGene_tx
=XM_023244685.1;refGene_gn=MYH10;refGene_tx=XM_023244686.1;refGene_gn=
MYH10;refGene_tx=XM_023244687.1;refGene_gn=MYH10;refGene_tx=XM_0232446
88.1;refGene_gn=MYH10;refGene_tx=XM_023244689.1;refGene_gn=MYH10;refGe
ne_tx=XM_023244690.1
E1 3323741 3323898
refGene_gn=MYH10;refGene_tx=XM_023244684.1;refGene_gn=MYH10;refGene_tx
=XM_023244685.1;refGene_gn=MYH10;refGene_tx=XM_023244686.1;refGene_gn=
MYH10;refGene_tx=XM_023244687.1;refGene_gn=MYH10;refGene_tx=XM_0232446
88.1;refGene_gn=MYH10;refGene_tx=XM_023244689.1;refGene_gn=MYH10;refGe
ne_tx=XM_023244690.1
E1 3340142 3340518
refGene_gn=MYH10;refGene_tx=XM_023244684.1;refGene_gn=MYH10;refGene_tx
=XM_023244685.1;refGene_gn=MYH10;refGene_tx=XM_023244686.1;refGene_gn=
MYH10;refGene_tx=XM_023244687.1;refGene_gn=MYH10;refGene_tx=XM_0232446
88.1;refGene_gn=MYH10;refGene_tx=XM_023244689.1;refGene_gn=MYH10;refGe
ne_tx=XM_023244690.1
E1 3347236 3347295
refGene_gn=MYH10;refGene_tx=XM_023244684.1;refGene_gn=MYH10;refGene_tx
=XM_023244685.1;refGene_gn=MYH10;refGene_tx=XM_023244686.1;refGene_gn=
MYH10;refGene_tx=XM_023244687.1;refGene_gn=MYH10;refGene_tx=XM_0232446
88.1;refGene_gn=MYH10;refGene_tx=XM_023244690.1
E1 3425706 3425784 refGene_gn=CCDC42;refGene_tx=XM_023244099.1
E1 3428626 3428638 refGene_gn=CCDC42;refGene_tx=XM_023244100.1
E1 3428854 3429013
refGene_gn=CCDC42;refGene_tx=XM_023244099.1;refGene_gn=CCDC42;refGene_
tx=XM_023244100.1
E1 3429106 3429328
refGene_gn=CCDC42;refGene_tx=XM_023244099.1;refGene_gn=CCDC42;refGene_
tx=XM_023244100.1
E1 3434525 3434723
refGene_gn=CCDC42;refGene_tx=XM_023244099.1;refGene_gn=CCDC42;refGene_
tx=XM_023244100.1
E1 3436217 3436322
refGene_gn=CCDC42;refGene_tx=XM_023244099.1;refGene_gn=CCDC42;refGene_
tx=XM_023244100.1
E1 3436577 3436683
refGene_gn=CCDC42;refGene_tx=XM_023244099.1;refGene_gn=CCDC42;refGene_
tx=XM_023244100.1
E1 3436964 3437225
refGene_gn=CCDC42;refGene_tx=XM_023244099.1;refGene_gn=CCDC42;refGene_
tx=XM_023244100.1
E1 3437800 3437871
refGene_gn=CCDC42;refGene_tx=XM_023244099.1;refGene_gn=CCDC42;refGene_
tx=XM_023244100.1
E1 3461684 3463448 refGene_gn=MFSD6L;refGene_tx=XM_003996281.5
E1 3466402 3466556
refGene_gn=PIK3R6;refGene_tx=XM_023243588.1;refGene_gn=PIK3R6;refGene_
tx=XM_023243589.1
E1 3467130 3467243

refGene_gn=PIK3R6;refGene_tx=XM_023243588.1;refGene_gn=PIK3R6;refGene_tx=XM_023243589.1
E1 3479240 3479356
refGene_gn=PIK3R6;refGene_tx=XM_023243588.1;refGene_gn=PIK3R6;refGene_tx=XM_023243589.1
E1 3480949 3481040
refGene_gn=PIK3R6;refGene_tx=XM_023243588.1;refGene_gn=PIK3R6;refGene_tx=XM_023243589.1
E1 3481527 3481598
refGene_gn=PIK3R6;refGene_tx=XM_023243588.1;refGene_gn=PIK3R6;refGene_tx=XM_023243589.1
E1 3481878 3481966
refGene_gn=PIK3R6;refGene_tx=XM_023243588.1;refGene_gn=PIK3R6;refGene_tx=XM_023243589.1
E1 3483783 3483894
refGene_gn=PIK3R6;refGene_tx=XM_023243588.1;refGene_gn=PIK3R6;refGene_tx=XM_023243589.1
E1 3487522 3487645
refGene_gn=PIK3R6;refGene_tx=XM_023243588.1;refGene_gn=PIK3R6;refGene_tx=XM_023243589.1
E1 3488351 3488430
refGene_gn=PIK3R6;refGene_tx=XM_023243588.1;refGene_gn=PIK3R6;refGene_tx=XM_023243589.1
E1 3488720 3489144
refGene_gn=PIK3R6;refGene_tx=XM_023243588.1;refGene_gn=PIK3R6;refGene_tx=XM_023243589.1
E1 3490447 3490534
refGene_gn=PIK3R6;refGene_tx=XM_023243588.1;refGene_gn=PIK3R6;refGene_tx=XM_023243589.1
E1 3490812 3490978
refGene_gn=PIK3R6;refGene_tx=XM_023243588.1;refGene_gn=PIK3R6;refGene_tx=XM_023243589.1
E1 3493528 3493712
refGene_gn=PIK3R6;refGene_tx=XM_023243588.1;refGene_gn=PIK3R6;refGene_tx=XM_023243589.1
E1 3494911 3494981
refGene_gn=PIK3R6;refGene_tx=XM_023243588.1;refGene_gn=PIK3R6;refGene_tx=XM_023243589.1
E1 3497077 3497244
refGene_gn=PIK3R6;refGene_tx=XM_023243588.1;refGene_gn=PIK3R6;refGene_tx=XM_023243589.1
E1 3498071 3498140
refGene_gn=PIK3R6;refGene_tx=XM_023243588.1;refGene_gn=PIK3R6;refGene_tx=XM_023243589.1
E1 3498741 3498833 refGene_gn=PIK3R6;refGene_tx=XM_023243588.1
E1 3499269 3499353 refGene_gn=PIK3R6;refGene_tx=XM_023243588.1
E1 3508228 3508241 refGene_gn=PIK3R6;refGene_tx=XM_023243588.1
E1 3535529 3535677
refGene_gn=PIK3R5;refGene_tx=XM_023243497.1;refGene_gn=PIK3R5;refGene_tx=XM_023243498.1;refGene_gn=PIK3R5;refGene_tx=XM_023243499.1;refGene_

gn=PIK3R5;refGene_tx=XM_023243500.1;refGene_gn=PIK3R5;refGene_tx=XM_02
3243501.1
E1 3535792 3535905
refGene_gn=PIK3R5;refGene_tx=XM_023243497.1;refGene_gn=PIK3R5;refGene_
tx=XM_023243498.1;refGene_gn=PIK3R5;refGene_tx=XM_023243499.1;refGene_
gn=PIK3R5;refGene_tx=XM_023243500.1;refGene_gn=PIK3R5;refGene_tx=XM_02
3243501.1
E1 3536340 3536423
refGene_gn=PIK3R5;refGene_tx=XM_023243497.1;refGene_gn=PIK3R5;refGene_
tx=XM_023243498.1;refGene_gn=PIK3R5;refGene_tx=XM_023243499.1;refGene_
gn=PIK3R5;refGene_tx=XM_023243500.1;refGene_gn=PIK3R5;refGene_tx=XM_02
3243501.1
E1 3536497 3536591
refGene_gn=PIK3R5;refGene_tx=XM_023243497.1;refGene_gn=PIK3R5;refGene_
tx=XM_023243498.1;refGene_gn=PIK3R5;refGene_tx=XM_023243499.1;refGene_
gn=PIK3R5;refGene_tx=XM_023243500.1;refGene_gn=PIK3R5;refGene_tx=XM_02
3243501.1
E1 3538764 3538841
refGene_gn=PIK3R5;refGene_tx=XM_023243497.1;refGene_gn=PIK3R5;refGene_
tx=XM_023243498.1;refGene_gn=PIK3R5;refGene_tx=XM_023243499.1;refGene_
gn=PIK3R5;refGene_tx=XM_023243500.1;refGene_gn=PIK3R5;refGene_tx=XM_02
3243501.1
E1 3538932 3539026
refGene_gn=PIK3R5;refGene_tx=XM_023243497.1;refGene_gn=PIK3R5;refGene_
tx=XM_023243498.1;refGene_gn=PIK3R5;refGene_tx=XM_023243499.1;refGene_
gn=PIK3R5;refGene_tx=XM_023243500.1;refGene_gn=PIK3R5;refGene_tx=XM_02
3243501.1
E1 3539141 3539270
refGene_gn=PIK3R5;refGene_tx=XM_023243497.1;refGene_gn=PIK3R5;refGene_
tx=XM_023243498.1;refGene_gn=PIK3R5;refGene_tx=XM_023243499.1;refGene_
gn=PIK3R5;refGene_tx=XM_023243500.1;refGene_gn=PIK3R5;refGene_tx=XM_02
3243501.1
E1 3539494 3539602
refGene_gn=PIK3R5;refGene_tx=XM_023243497.1;refGene_gn=PIK3R5;refGene_
tx=XM_023243498.1;refGene_gn=PIK3R5;refGene_tx=XM_023243499.1;refGene_
gn=PIK3R5;refGene_tx=XM_023243500.1;refGene_gn=PIK3R5;refGene_tx=XM_02
3243501.1
E1 3539834 3539985
refGene_gn=PIK3R5;refGene_tx=XM_023243497.1;refGene_gn=PIK3R5;refGene_
tx=XM_023243498.1;refGene_gn=PIK3R5;refGene_tx=XM_023243499.1;refGene_
gn=PIK3R5;refGene_tx=XM_023243500.1;refGene_gn=PIK3R5;refGene_tx=XM_02
3243501.1
E1 3540112 3540836
refGene_gn=PIK3R5;refGene_tx=XM_023243497.1;refGene_gn=PIK3R5;refGene_
tx=XM_023243498.1;refGene_gn=PIK3R5;refGene_tx=XM_023243499.1;refGene_
gn=PIK3R5;refGene_tx=XM_023243500.1;refGene_gn=PIK3R5;refGene_tx=XM_02
3243501.1
E1 3541105 3541189
refGene_gn=PIK3R5;refGene_tx=XM_023243497.1;refGene_gn=PIK3R5;refGene_
tx=XM_023243498.1;refGene_gn=PIK3R5;refGene_tx=XM_023243499.1;refGene_

gn=PIK3R5;refGene_tx=XM_023243500.1;refGene_gn=PIK3R5;refGene_tx=XM_02
3243501.1
E1 3541743 3541897
refGene_gn=PIK3R5;refGene_tx=XM_023243497.1;refGene_gn=PIK3R5;refGene_
tx=XM_023243498.1;refGene_gn=PIK3R5;refGene_tx=XM_023243499.1;refGene_
gn=PIK3R5;refGene_tx=XM_023243500.1;refGene_gn=PIK3R5;refGene_tx=XM_02
3243501.1
E1 3542319 3542514
refGene_gn=PIK3R5;refGene_tx=XM_023243497.1;refGene_gn=PIK3R5;refGene_
tx=XM_023243498.1;refGene_gn=PIK3R5;refGene_tx=XM_023243499.1;refGene_
gn=PIK3R5;refGene_tx=XM_023243500.1;refGene_gn=PIK3R5;refGene_tx=XM_02
3243501.1
E1 3544921 3545179
refGene_gn=PIK3R5;refGene_tx=XM_023243497.1;refGene_gn=PIK3R5;refGene_
tx=XM_023243498.1;refGene_gn=PIK3R5;refGene_tx=XM_023243499.1;refGene_
gn=PIK3R5;refGene_tx=XM_023243500.1;refGene_gn=PIK3R5;refGene_tx=XM_02
3243501.1
E1 3552897 3553036
refGene_gn=PIK3R5;refGene_tx=XM_023243497.1;refGene_gn=PIK3R5;refGene_
tx=XM_023243498.1;refGene_gn=PIK3R5;refGene_tx=XM_023243499.1;refGene_
gn=PIK3R5;refGene_tx=XM_023243500.1
E1 3553887 3553956
refGene_gn=PIK3R5;refGene_tx=XM_023243497.1;refGene_gn=PIK3R5;refGene_
tx=XM_023243498.1;refGene_gn=PIK3R5;refGene_tx=XM_023243499.1;refGene_
gn=PIK3R5;refGene_tx=XM_023243500.1
E1 3556183 3556284
refGene_gn=PIK3R5;refGene_tx=XM_023243497.1;refGene_gn=PIK3R5;refGene_
tx=XM_023243498.1;refGene_gn=PIK3R5;refGene_tx=XM_023243499.1;refGene_
gn=PIK3R5;refGene_tx=XM_023243500.1
E1 3558295 3558398
refGene_gn=PIK3R5;refGene_tx=XM_023243497.1;refGene_gn=PIK3R5;refGene_
tx=XM_023243498.1;refGene_gn=PIK3R5;refGene_tx=XM_023243499.1;refGene_
gn=PIK3R5;refGene_tx=XM_023243500.1
E1 3646789 3647807
refGene_gn=NTN1;refGene_tx=XM_023244470.1;refGene_gn=NTN1;refGene_tx=X
M_023244471.1
E1 3752311 3752500
refGene_gn=NTN1;refGene_tx=XM_023244470.1;refGene_gn=NTN1;refGene_tx=X
M_023244471.1
E1 3765422 3765572
refGene_gn=NTN1;refGene_tx=XM_023244470.1;refGene_gn=NTN1;refGene_tx=X
M_023244471.1
E1 3768056 3768110
refGene_gn=NTN1;refGene_tx=XM_023244470.1;refGene_gn=NTN1;refGene_tx=X
M_023244471.1
E1 3799979 3800054
refGene_gn=NTN1;refGene_tx=XM_023244470.1;refGene_gn=NTN1;refGene_tx=X
M_023244471.1
E1 3815384 3815713
refGene_gn=NTN1;refGene_tx=XM_023244470.1;refGene_gn=NTN1;refGene_tx=X

M_023244471.1
E1	3824530	3824627
refGene_gn=STX8;refGene_tx=XM_011288848.3;refGene_gn=STX8;refGene_tx=XM_019817162.2;refGene_gn=STX8;refGene_tx=XM_019817164.2
E1	3857509	3857697	refGene_gn=STX8;refGene_tx=XM_019817163.2
E1	3906705	3906817
refGene_gn=STX8;refGene_tx=XM_019817162.2;refGene_gn=STX8;refGene_tx=XM_019817164.2
E1	3925287	3925389
refGene_gn=STX8;refGene_tx=XM_011288848.3;refGene_gn=STX8;refGene_tx=XM_019817162.2;refGene_gn=STX8;refGene_tx=XM_019817163.2;refGene_gn=STX8;refGene_tx=XM_019817164.2
E1	4013739	4013832
refGene_gn=STX8;refGene_tx=XM_011288848.3;refGene_gn=STX8;refGene_tx=XM_019817162.2;refGene_gn=STX8;refGene_tx=XM_019817163.2;refGene_gn=STX8;refGene_tx=XM_019817164.2
E1	4027943	4028068
refGene_gn=STX8;refGene_tx=XM_011288848.3;refGene_gn=STX8;refGene_tx=XM_019817162.2;refGene_gn=STX8;refGene_tx=XM_019817163.2;refGene_gn=STX8;refGene_tx=XM_019817164.2
E1	4054322	4054433
refGene_gn=STX8;refGene_tx=XM_011288848.3;refGene_gn=STX8;refGene_tx=XM_019817162.2;refGene_gn=STX8;refGene_tx=XM_019817163.2
E1	4066792	4066887
refGene_gn=STX8;refGene_tx=XM_011288848.3;refGene_gn=STX8;refGene_tx=XM_019817162.2;refGene_gn=STX8;refGene_tx=XM_019817163.2
E1	4074822	4074922
refGene_gn=STX8;refGene_tx=XM_011288848.3;refGene_gn=STX8;refGene_tx=XM_019817162.2;refGene_gn=STX8;refGene_tx=XM_019817163.2;refGene_gn=STX8;refGene_tx=XM_019817164.2
E1	4080813	4080830
refGene_gn=STX8;refGene_tx=XM_011288848.3;refGene_gn=STX8;refGene_tx=XM_019817162.2;refGene_gn=STX8;refGene_tx=XM_019817163.2
E1	4081570	4081658	refGene_gn=CFAP52;refGene_tx=XM_003996206.5
E1	4087828	4088028	refGene_gn=CFAP52;refGene_tx=XM_003996206.5
E1	4088896	4089033	refGene_gn=CFAP52;refGene_tx=XM_003996206.5
E1	4093322	4093451	refGene_gn=CFAP52;refGene_tx=XM_003996206.5
E1	4097159	4097259	refGene_gn=CFAP52;refGene_tx=XM_003996206.5
E1	4098189	4098306	refGene_gn=CFAP52;refGene_tx=XM_003996206.5
E1	4101704	4101805	refGene_gn=CFAP52;refGene_tx=XM_003996206.5
E1	4104636	4104807	refGene_gn=CFAP52;refGene_tx=XM_003996206.5
E1	4111604	4111753	refGene_gn=CFAP52;refGene_tx=XM_003996206.5
E1	4114143	4114289	refGene_gn=CFAP52;refGene_tx=XM_003996206.5
E1	4117144	4117296	refGene_gn=CFAP52;refGene_tx=XM_003996206.5
E1	4119849	4119952	refGene_gn=CFAP52;refGene_tx=XM_003996206.5
E1	4123278	4123390	refGene_gn=CFAP52;refGene_tx=XM_003996206.5
E1	4124913	4125089	refGene_gn=CFAP52;refGene_tx=XM_003996206.5
E1	4127719	4128017	refGene_gn=USP43;refGene_tx=XM_023243763.1
E1	4128019	4128216	refGene_gn=USP43;refGene_tx=XM_023243763.1
E1	4139520	4139652	refGene_gn=USP43;refGene_tx=XM_023243763.1

E1 4146727 4146831 refGene_gn=USP43;refGene_tx=XM_023243763.1
E1 4154586 4154679 refGene_gn=USP43;refGene_tx=XM_023243763.1
E1 4156162 4156298 refGene_gn=USP43;refGene_tx=XM_023243763.1
E1 4159715 4159851 refGene_gn=USP43;refGene_tx=XM_023243763.1
E1 4161155 4161291 refGene_gn=USP43;refGene_tx=XM_023243763.1
E1 4163768 4163880 refGene_gn=USP43;refGene_tx=XM_023243763.1
E1 4167046 4167150 refGene_gn=USP43;refGene_tx=XM_023243763.1
E1 4170996 4171074 refGene_gn=USP43;refGene_tx=XM_023243763.1
E1 4172344 4172471 refGene_gn=USP43;refGene_tx=XM_023243763.1
E1 4172574 4172923 refGene_gn=USP43;refGene_tx=XM_023243763.1
E1 4175878 4176037 refGene_gn=USP43;refGene_tx=XM_023243763.1
E1 4177948 4178114 refGene_gn=USP43;refGene_tx=XM_023243763.1
E1 4193318 4194360 refGene_gn=USP43;refGene_tx=XM_023243763.1
E1 4224271 4224480
refGene_gn=DHRS7C;refGene_tx=XM_003996208.4;refGene_gn=DHRS7C;refGene_
tx=XM_006939786.3
E1 4226008 4226164
refGene_gn=DHRS7C;refGene_tx=XM_003996208.4;refGene_gn=DHRS7C;refGene_
tx=XM_006939786.3
E1 4230766 4230859
refGene_gn=DHRS7C;refGene_tx=XM_003996208.4;refGene_gn=DHRS7C;refGene_
tx=XM_006939786.3
E1 4232855 4233069
refGene_gn=DHRS7C;refGene_tx=XM_003996208.4;refGene_gn=DHRS7C;refGene_
tx=XM_006939786.3
E1 4235052 4235165
refGene_gn=DHRS7C;refGene_tx=XM_003996208.4;refGene_gn=DHRS7C;refGene_
tx=XM_006939786.3
E1 4240267 4240421
refGene_gn=DHRS7C;refGene_tx=XM_003996208.4;refGene_gn=DHRS7C;refGene_
tx=XM_006939786.3
E1 4251535 4251794 refGene_gn=GSG1L2;refGene_tx=XM_003996287.5
E1 4255020 4255132 refGene_gn=GSG1L2;refGene_tx=XM_003996287.5
E1 4257152 4257305 refGene_gn=GSG1L2;refGene_tx=XM_003996287.5
E1 4258937 4258985 refGene_gn=GSG1L2;refGene_tx=XM_003996287.5
E1 4267313 4267620 refGene_gn=GSG1L2;refGene_tx=XM_003996287.5
E1 4270869 4270968
refGene_gn=GLP2R;refGene_tx=XM_019817165.2;refGene_gn=GLP2R;refGene_tx
=XM_019817166.1
E1 4276900 4276988
refGene_gn=GLP2R;refGene_tx=XM_019817165.2;refGene_gn=GLP2R;refGene_tx
=XM_019817166.1
E1 4279216 4279321
refGene_gn=GLP2R;refGene_tx=XM_019817165.2;refGene_gn=GLP2R;refGene_tx
=XM_019817166.1
E1 4282732 4282854
refGene_gn=GLP2R;refGene_tx=XM_019817165.2;refGene_gn=GLP2R;refGene_tx
=XM_019817166.1
E1 4290154 4290261
refGene_gn=GLP2R;refGene_tx=XM_019817165.2;refGene_gn=GLP2R;refGene_tx

=XM_019817166.1
E1 4292140 4292294
refGene_gn=GLP2R;refGene_tx=XM_019817165.2;refGene_gn=GLP2R;refGene_tx
=XM_019817166.1
E1 4294209 4294369
refGene_gn=GLP2R;refGene_tx=XM_019817165.2;refGene_gn=GLP2R;refGene_tx
=XM_019817166.1
E1 4295781 4295842
refGene_gn=GLP2R;refGene_tx=XM_019817165.2;refGene_gn=GLP2R;refGene_tx
=XM_019817166.1
E1 4297214 4297284
refGene_gn=GLP2R;refGene_tx=XM_019817165.2;refGene_gn=GLP2R;refGene_tx
=XM_019817166.1
E1 4305909 4305998
refGene_gn=GLP2R;refGene_tx=XM_019817165.2;refGene_gn=GLP2R;refGene_tx
=XM_019817166.1
E1 4312996 4313135
refGene_gn=GLP2R;refGene_tx=XM_019817165.2;refGene_gn=GLP2R;refGene_tx
=XM_019817166.1
E1 4322188 4322230
refGene_gn=GLP2R;refGene_tx=XM_019817165.2;refGene_gn=GLP2R;refGene_tx
=XM_019817166.1
E1 4323095 4323431
refGene_gn=GLP2R;refGene_tx=XM_019817165.2;refGene_gn=GLP2R;refGene_tx
=XM_019817166.1
E1 4327363 4327479 refGene_gn=RCVRN;refGene_tx=XM_003996209.4
E1 4330572 4330684 refGene_gn=RCVRN;refGene_tx=XM_003996209.4
E1 4334111 4334492 refGene_gn=RCVRN;refGene_tx=XM_003996209.4
E1 4344112 4344226
refGene_gn=GAS7;refGene_tx=XM_003996210.5;refGene_gn=GAS7;refGene_tx=X
M_006939787.4;refGene_gn=GAS7;refGene_tx=XM_006939788.2;refGene_gn=GAS
7;refGene_tx=XM_006939789.4;refGene_gn=GAS7;refGene_tx=XM_011288845.3;
refGene_gn=GAS7;refGene_tx=XM_019817167.2;refGene_gn=GAS7;refGene_tx=X
M_019817168.1;refGene_gn=GAS7;refGene_tx=XM_019817169.1;refGene_gn=GAS
7;refGene_tx=XM_023243853.1;refGene_gn=GAS7;refGene_tx=XM_023243855.1
E1 4345290 4345389
refGene_gn=GAS7;refGene_tx=XM_003996210.5;refGene_gn=GAS7;refGene_tx=X
M_006939787.4;refGene_gn=GAS7;refGene_tx=XM_006939788.2;refGene_gn=GAS
7;refGene_tx=XM_006939789.4;refGene_gn=GAS7;refGene_tx=XM_011288845.3;
refGene_gn=GAS7;refGene_tx=XM_019817167.2;refGene_gn=GAS7;refGene_tx=X
M_019817168.1;refGene_gn=GAS7;refGene_tx=XM_019817169.1;refGene_gn=GAS
7;refGene_tx=XM_023243853.1;refGene_gn=GAS7;refGene_tx=XM_023243855.1
E1 4346704 4346867
refGene_gn=GAS7;refGene_tx=XM_003996210.5;refGene_gn=GAS7;refGene_tx=X
M_006939787.4;refGene_gn=GAS7;refGene_tx=XM_006939788.2;refGene_gn=GAS
7;refGene_tx=XM_006939789.4;refGene_gn=GAS7;refGene_tx=XM_011288845.3;
refGene_gn=GAS7;refGene_tx=XM_019817167.2;refGene_gn=GAS7;refGene_tx=X
M_019817168.1;refGene_gn=GAS7;refGene_tx=XM_019817169.1;refGene_gn=GAS
7;refGene_tx=XM_023243853.1;refGene_gn=GAS7;refGene_tx=XM_023243854.1;
refGene_gn=GAS7;refGene_tx=XM_023243855.1

E1 4351871 4351995
refGene_gn=GAS7;refGene_tx=XM_003996210.5;refGene_gn=GAS7;refGene_tx=XM_006939787.4;refGene_gn=GAS7;refGene_tx=XM_006939788.2;refGene_gn=GAS7;refGene_tx=XM_006939789.4;refGene_gn=GAS7;refGene_tx=XM_011288845.3;refGene_gn=GAS7;refGene_tx=XM_019817167.2;refGene_gn=GAS7;refGene_tx=XM_019817168.1;refGene_gn=GAS7;refGene_tx=XM_019817169.1;refGene_gn=GAS7;refGene_tx=XM_023243853.1;refGene_gn=GAS7;refGene_tx=XM_023243854.1;refGene_gn=GAS7;refGene_tx=XM_023243855.1
E1 4352594 4352723
refGene_gn=GAS7;refGene_tx=XM_003996210.5;refGene_gn=GAS7;refGene_tx=XM_006939787.4;refGene_gn=GAS7;refGene_tx=XM_006939788.2;refGene_gn=GAS7;refGene_tx=XM_006939789.4;refGene_gn=GAS7;refGene_tx=XM_011288845.3;refGene_gn=GAS7;refGene_tx=XM_019817167.2;refGene_gn=GAS7;refGene_tx=XM_019817168.1;refGene_gn=GAS7;refGene_tx=XM_019817169.1;refGene_gn=GAS7;refGene_tx=XM_023243853.1;refGene_gn=GAS7;refGene_tx=XM_023243854.1;refGene_gn=GAS7;refGene_tx=XM_023243855.1
E1 4360131 4360210
refGene_gn=GAS7;refGene_tx=XM_003996210.5;refGene_gn=GAS7;refGene_tx=XM_006939787.4;refGene_gn=GAS7;refGene_tx=XM_006939788.2;refGene_gn=GAS7;refGene_tx=XM_006939789.4;refGene_gn=GAS7;refGene_tx=XM_011288845.3;refGene_gn=GAS7;refGene_tx=XM_019817167.2;refGene_gn=GAS7;refGene_tx=XM_019817168.1;refGene_gn=GAS7;refGene_tx=XM_019817169.1;refGene_gn=GAS7;refGene_tx=XM_023243853.1;refGene_gn=GAS7;refGene_tx=XM_023243854.1;refGene_gn=GAS7;refGene_tx=XM_023243855.1
E1 4364859 4364934
refGene_gn=GAS7;refGene_tx=XM_003996210.5;refGene_gn=GAS7;refGene_tx=XM_006939787.4;refGene_gn=GAS7;refGene_tx=XM_006939788.2;refGene_gn=GAS7;refGene_tx=XM_006939789.4;refGene_gn=GAS7;refGene_tx=XM_011288845.3;refGene_gn=GAS7;refGene_tx=XM_019817167.2;refGene_gn=GAS7;refGene_tx=XM_019817168.1;refGene_gn=GAS7;refGene_tx=XM_019817169.1;refGene_gn=GAS7;refGene_tx=XM_023243853.1;refGene_gn=GAS7;refGene_tx=XM_023243854.1;refGene_gn=GAS7;refGene_tx=XM_023243855.1
E1 4367618 4367734
refGene_gn=GAS7;refGene_tx=XM_003996210.5;refGene_gn=GAS7;refGene_tx=XM_006939787.4;refGene_gn=GAS7;refGene_tx=XM_006939788.2;refGene_gn=GAS7;refGene_tx=XM_006939789.4;refGene_gn=GAS7;refGene_tx=XM_011288845.3;refGene_gn=GAS7;refGene_tx=XM_019817167.2;refGene_gn=GAS7;refGene_tx=XM_019817168.1;refGene_gn=GAS7;refGene_tx=XM_019817169.1;refGene_gn=GAS7;refGene_tx=XM_023243853.1;refGene_gn=GAS7;refGene_tx=XM_023243854.1;refGene_gn=GAS7;refGene_tx=XM_023243855.1
E1 4370714 4370804
refGene_gn=GAS7;refGene_tx=XM_003996210.5;refGene_gn=GAS7;refGene_tx=XM_006939787.4;refGene_gn=GAS7;refGene_tx=XM_006939788.2;refGene_gn=GAS7;refGene_tx=XM_006939789.4;refGene_gn=GAS7;refGene_tx=XM_011288845.3;refGene_gn=GAS7;refGene_tx=XM_019817167.2;refGene_gn=GAS7;refGene_tx=XM_019817168.1;refGene_gn=GAS7;refGene_tx=XM_019817169.1;refGene_gn=GAS7;refGene_tx=XM_023243853.1;refGene_gn=GAS7;refGene_tx=XM_023243854.1;refGene_gn=GAS7;refGene_tx=XM_023243855.1
E1 4377949 4378003
refGene_gn=GAS7;refGene_tx=XM_003996210.5;refGene_gn=GAS7;refGene_tx=X

M_006939787.4;refGene_gn=GAS7;refGene_tx=XM_006939788.2;refGene_gn=GAS
7;refGene_tx=XM_006939789.4;refGene_gn=GAS7;refGene_tx=XM_011288845.3;
refGene_gn=GAS7;refGene_tx=XM_019817167.2;refGene_gn=GAS7;refGene_tx=X
M_019817168.1;refGene_gn=GAS7;refGene_tx=XM_019817169.1;refGene_gn=GAS
7;refGene_tx=XM_023243853.1;refGene_gn=GAS7;refGene_tx=XM_023243854.1;
refGene_gn=GAS7;refGene_tx=XM_023243855.1
E1 4386583 4386669
refGene_gn=GAS7;refGene_tx=XM_003996210.5;refGene_gn=GAS7;refGene_tx=X
M_006939787.4;refGene_gn=GAS7;refGene_tx=XM_006939788.2;refGene_gn=GAS
7;refGene_tx=XM_006939789.4;refGene_gn=GAS7;refGene_tx=XM_011288845.3;
refGene_gn=GAS7;refGene_tx=XM_019817167.2;refGene_gn=GAS7;refGene_tx=X
M_023243853.1;refGene_gn=GAS7;refGene_tx=XM_023243854.1
E1 4399649 4399730
refGene_gn=GAS7;refGene_tx=XM_003996210.5;refGene_gn=GAS7;refGene_tx=X
M_006939787.4;refGene_gn=GAS7;refGene_tx=XM_006939788.2;refGene_gn=GAS
7;refGene_tx=XM_006939789.4;refGene_gn=GAS7;refGene_tx=XM_011288845.3;
refGene_gn=GAS7;refGene_tx=XM_019817167.2;refGene_gn=GAS7;refGene_tx=X
M_019817168.1;refGene_gn=GAS7;refGene_tx=XM_019817169.1;refGene_gn=GAS
7;refGene_tx=XM_023243853.1;refGene_gn=GAS7;refGene_tx=XM_023243854.1;
refGene_gn=GAS7;refGene_tx=XM_023243855.1
E1 4426188 4426309
refGene_gn=GAS7;refGene_tx=XM_003996210.5;refGene_gn=GAS7;refGene_tx=X
M_006939787.4;refGene_gn=GAS7;refGene_tx=XM_006939788.2;refGene_gn=GAS
7;refGene_tx=XM_006939789.4;refGene_gn=GAS7;refGene_tx=XM_011288845.3;
refGene_gn=GAS7;refGene_tx=XM_019817167.2;refGene_gn=GAS7;refGene_tx=X
M_019817168.1;refGene_gn=GAS7;refGene_tx=XM_019817169.1;refGene_gn=GAS
7;refGene_tx=XM_023243853.1;refGene_gn=GAS7;refGene_tx=XM_023243854.1;
refGene_gn=GAS7;refGene_tx=XM_023243855.1
E1 4430331 4430381 refGene_gn=GAS7;refGene_tx=XM_019817169.1
E1 4438404 4438506 refGene_gn=GAS7;refGene_tx=XM_019817167.2
E1 4440685 4440688 refGene_gn=GAS7;refGene_tx=XM_003996210.5
E1 4454367 4454414 refGene_gn=GAS7;refGene_tx=XM_023243855.1
E1 4459404 4459490 refGene_gn=GAS7;refGene_tx=XM_019817168.1
E1 4557711 4557894
refGene_gn=GAS7;refGene_tx=XM_023243853.1;refGene_gn=GAS7;refGene_tx=X
M_023243854.1
E1 4627999 4628014
refGene_gn=LOC101083886;refGene_tx=XM_023243778.1
E1 4628417 4628552
refGene_gn=LOC101083886;refGene_tx=XM_023243778.1
E1 4629950 4630046
refGene_gn=LOC101083886;refGene_tx=XM_023243778.1
E1 4630134 4630239
refGene_gn=LOC101083886;refGene_tx=XM_023243778.1
E1 4632883 4633054
refGene_gn=LOC101083886;refGene_tx=XM_023243778.1
E1 4633337 4633463
refGene_gn=LOC101083886;refGene_tx=XM_023243778.1
E1 4634779 4634983
refGene_gn=LOC101083886;refGene_tx=XM_023243778.1

E1 4635058 4635367
refGene_gn=LOC101083886;refGene_tx=XM_023243778.1
E1 4636139 4636264
refGene_gn=LOC101083886;refGene_tx=XM_023243778.1
E1 4637159 4637325
refGene_gn=LOC101083886;refGene_tx=XM_023243778.1
E1 4637705 4637889
refGene_gn=LOC101083886;refGene_tx=XM_023243778.1
E1 4638276 4638473
refGene_gn=LOC101083886;refGene_tx=XM_023243778.1
E1 4639910 4640029
refGene_gn=LOC101083886;refGene_tx=XM_023243778.1
E1 4640108 4640235
refGene_gn=LOC101083886;refGene_tx=XM_023243778.1
E1 4642983 4643373
refGene_gn=LOC101083886;refGene_tx=XM_023243778.1
E1 4644879 4644970
refGene_gn=LOC101083886;refGene_tx=XM_023243778.1
E1 4645058 4645204
refGene_gn=LOC101083886;refGene_tx=XM_023243778.1
E1 4645800 4645977
refGene_gn=LOC101083886;refGene_tx=XM_023243778.1
E1 4648030 4648274
refGene_gn=LOC101083886;refGene_tx=XM_023243778.1
E1 4650513 4650765
refGene_gn=LOC101083886;refGene_tx=XM_023243778.1
E1 4651482 4651619
refGene_gn=LOC101083886;refGene_tx=XM_023243778.1
E1 4652858 4652982
refGene_gn=LOC101083886;refGene_tx=XM_023243778.1
E1 4654055 4654173
refGene_gn=LOC101083886;refGene_tx=XM_023243778.1
E1 4658420 4658508
refGene_gn=LOC101083886;refGene_tx=XM_023243778.1
E1 4658603 4658677
refGene_gn=LOC101083886;refGene_tx=XM_023243778.1
E1 4662045 4662355
refGene_gn=LOC101083886;refGene_tx=XM_023243778.1
E1 4663818 4663989
refGene_gn=LOC101083886;refGene_tx=XM_023243778.1
E1 4664105 4664255
refGene_gn=LOC101083886;refGene_tx=XM_023243778.1
E1 4665499 4665618
refGene_gn=LOC101083886;refGene_tx=XM_023243778.1
E1 4668198 4668337
refGene_gn=LOC101083886;refGene_tx=XM_023243778.1
E1 4671656 4671760
refGene_gn=LOC101083886;refGene_tx=XM_023243778.1
E1 4671869 4671968
refGene_gn=LOC101083886;refGene_tx=XM_023243778.1

E1 4672059 4672123
refGene_gn=LOC101083886;refGene_tx=XM_023243778.1
E1 4674357 4674450
refGene_gn=LOC101083886;refGene_tx=XM_023243778.1
E1 4675574 4675686
refGene_gn=LOC101083886;refGene_tx=XM_023243778.1
E1 4675773 4675801
refGene_gn=LOC101083886;refGene_tx=XM_023243778.1
E1 4678147 4678304
refGene_gn=LOC101083886;refGene_tx=XM_023243778.1
E1 4678391 4678535
refGene_gn=LOC101083886;refGene_tx=XM_023243778.1
E1 4680105 4680322
refGene_gn=LOC101083886;refGene_tx=XM_023243778.1
E1 4688900 4688951
refGene_gn=LOC101083886;refGene_tx=XM_023243778.1
E1 4692826 4692963
refGene_gn=LOC101083886;refGene_tx=XM_023243778.1
E1 4708459 4708609
refGene_gn=LOC105261153;refGene_tx=XM_011288842.2
E1 4709500 4709596
refGene_gn=LOC105261153;refGene_tx=XM_011288842.2
E1 4710165 4710270
refGene_gn=LOC105261153;refGene_tx=XM_011288842.2
E1 4710452 4710623
refGene_gn=LOC105261153;refGene_tx=XM_011288842.2
E1 4710710 4710836
refGene_gn=LOC105261153;refGene_tx=XM_011288842.2
E1 4711891 4712095
refGene_gn=LOC105261153;refGene_tx=XM_011288842.2
E1 4712798 4713125
refGene_gn=LOC105261153;refGene_tx=XM_011288842.2;refGene_gn=LOC105261
153;refGene_tx=XM_019817172.2
E1 4714425 4714550
refGene_gn=LOC105261153;refGene_tx=XM_011288842.2;refGene_gn=LOC105261
153;refGene_tx=XM_019817172.2
E1 4714647 4714813
refGene_gn=LOC105261153;refGene_tx=XM_011288842.2;refGene_gn=LOC105261
153;refGene_tx=XM_019817172.2
E1 4714897 4715081
refGene_gn=LOC105261153;refGene_tx=XM_011288842.2;refGene_gn=LOC105261
153;refGene_tx=XM_019817172.2
E1 4716195 4716392
refGene_gn=LOC105261153;refGene_tx=XM_011288842.2;refGene_gn=LOC105261
153;refGene_tx=XM_019817172.2
E1 4716519 4716638
refGene_gn=LOC105261153;refGene_tx=XM_011288842.2;refGene_gn=LOC105261
153;refGene_tx=XM_019817172.2
E1 4717208 4717335
refGene_gn=LOC105261153;refGene_tx=XM_011288842.2;refGene_gn=LOC105261

153;refGene_tx=XM_019817172.2
E1 4717984 4718374
refGene_gn=LOC105261153;refGene_tx=XM_011288842.2;refGene_gn=LOC105261
153;refGene_tx=XM_019817172.2
E1 4718460 4718551
refGene_gn=LOC105261153;refGene_tx=XM_011288842.2;refGene_gn=LOC105261
153;refGene_tx=XM_019817172.2
E1 4718634 4718780
refGene_gn=LOC105261153;refGene_tx=XM_011288842.2;refGene_gn=LOC105261
153;refGene_tx=XM_019817172.2
E1 4718860 4719037
refGene_gn=LOC105261153;refGene_tx=XM_011288842.2;refGene_gn=LOC105261
153;refGene_tx=XM_019817172.2
E1 4719130 4719373
refGene_gn=LOC105261153;refGene_tx=XM_011288842.2;refGene_gn=LOC105261
153;refGene_tx=XM_019817172.2
E1 4721270 4721526
refGene_gn=LOC105261153;refGene_tx=XM_011288842.2;refGene_gn=LOC105261
153;refGene_tx=XM_019817172.2
E1 4723402 4723539
refGene_gn=LOC105261153;refGene_tx=XM_011288842.2;refGene_gn=LOC105261
153;refGene_tx=XM_019817172.2
E1 4723634 4723758
refGene_gn=LOC105261153;refGene_tx=XM_011288842.2;refGene_gn=LOC105261
153;refGene_tx=XM_019817172.2
E1 4724133 4724251
refGene_gn=LOC105261153;refGene_tx=XM_011288842.2;refGene_gn=LOC105261
153;refGene_tx=XM_019817172.2
E1 4724334 4724422
refGene_gn=LOC105261153;refGene_tx=XM_011288842.2;refGene_gn=LOC105261
153;refGene_tx=XM_019817172.2
E1 4726251 4726319
refGene_gn=LOC105261153;refGene_tx=XM_011288842.2;refGene_gn=LOC105261
153;refGene_tx=XM_019817172.2
E1 4726429 4726739
refGene_gn=LOC105261153;refGene_tx=XM_011288842.2;refGene_gn=LOC105261
153;refGene_tx=XM_019817172.2
E1 4727616 4727787
refGene_gn=LOC105261153;refGene_tx=XM_011288842.2;refGene_gn=LOC105261
153;refGene_tx=XM_019817172.2
E1 4727903 4728053
refGene_gn=LOC105261153;refGene_tx=XM_011288842.2;refGene_gn=LOC105261
153;refGene_tx=XM_019817172.2
E1 4728143 4728262
refGene_gn=LOC105261153;refGene_tx=XM_011288842.2;refGene_gn=LOC105261
153;refGene_tx=XM_019817172.2
E1 4729427 4729566
refGene_gn=LOC105261153;refGene_tx=XM_011288842.2;refGene_gn=LOC105261
153;refGene_tx=XM_019817172.2
E1 4729800 4729904

refGene_gn=LOC105261153;refGene_tx=XM_011288842.2;refGene_gn=LOC105261153;refGene_tx=XM_019817172.2
E1 4729995 4730094
refGene_gn=LOC105261153;refGene_tx=XM_011288842.2;refGene_gn=LOC105261153;refGene_tx=XM_019817172.2
E1 4730732 4730796
refGene_gn=LOC105261153;refGene_tx=XM_011288842.2;refGene_gn=LOC105261153;refGene_tx=XM_019817172.2
E1 4730928 4731021
refGene_gn=LOC105261153;refGene_tx=XM_011288842.2;refGene_gn=LOC105261153;refGene_tx=XM_019817172.2
E1 4731104 4731213
refGene_gn=LOC105261153;refGene_tx=XM_011288842.2;refGene_gn=LOC105261153;refGene_tx=XM_019817172.2
E1 4731314 4731342
refGene_gn=LOC105261153;refGene_tx=XM_011288842.2;refGene_gn=LOC105261153;refGene_tx=XM_019817172.2
E1 4734039 4734196
refGene_gn=LOC105261153;refGene_tx=XM_011288842.2;refGene_gn=LOC105261153;refGene_tx=XM_019817172.2
E1 4734281 4734425
refGene_gn=LOC105261153;refGene_tx=XM_011288842.2;refGene_gn=LOC105261153;refGene_tx=XM_019817172.2
E1 4735381 4735591
refGene_gn=LOC105261153;refGene_tx=XM_011288842.2;refGene_gn=LOC105261153;refGene_tx=XM_019817172.2
E1 4753914 4754067
refGene_gn=LOC101085080;refGene_tx=XM_019817180.2
E1 4755045 4755141
refGene_gn=LOC101085080;refGene_tx=XM_019817180.2
E1 4755245 4755602
refGene_gn=LOC101085080;refGene_tx=XM_019817180.2
E1 4755689 4755815
refGene_gn=LOC101085080;refGene_tx=XM_019817180.2
E1 4757407 4757611
refGene_gn=LOC101085080;refGene_tx=XM_019817180.2
E1 4758175 4758484
refGene_gn=LOC101085080;refGene_tx=XM_019817180.2
E1 4759134 4759259
refGene_gn=LOC101085080;refGene_tx=XM_019817180.2
E1 4759360 4759526
refGene_gn=LOC101085080;refGene_tx=XM_019817180.2
E1 4759605 4759789
refGene_gn=LOC101085080;refGene_tx=XM_019817180.2
E1 4761012 4761209
refGene_gn=LOC101085080;refGene_tx=XM_019817180.2
E1 4761336 4761455
refGene_gn=LOC101085080;refGene_tx=XM_019817180.2
E1 4761948 4762075
refGene_gn=LOC101085080;refGene_tx=XM_019817180.2

E1 4762547 4762937
refGene_gn=LOC101085080;refGene_tx=XM_019817180.2
E1 4763023 4763114
refGene_gn=LOC101085080;refGene_tx=XM_019817180.2
E1 4763401 4763547
refGene_gn=LOC101085080;refGene_tx=XM_019817180.2
E1 4763751 4763928
refGene_gn=LOC101085080;refGene_tx=XM_019817180.2
E1 4764228 4764471
refGene_gn=LOC101085080;refGene_tx=XM_019817180.2
E1 4765157 4765413
refGene_gn=LOC101085080;refGene_tx=XM_019817180.2
E1 4765528 4765665
refGene_gn=LOC101085080;refGene_tx=XM_019817180.2
E1 4765756 4765880
refGene_gn=LOC101085080;refGene_tx=XM_019817180.2
E1 4766411 4766529
refGene_gn=LOC101085080;refGene_tx=XM_019817180.2
E1 4766636 4766724
refGene_gn=LOC101085080;refGene_tx=XM_019817180.2
E1 4767133 4767204
refGene_gn=LOC101085080;refGene_tx=XM_019817180.2
E1 4768038 4768348
refGene_gn=LOC101085080;refGene_tx=XM_019817180.2
E1 4769840 4770011
refGene_gn=LOC101085080;refGene_tx=XM_019817180.2
E1 4770127 4770277
refGene_gn=LOC101085080;refGene_tx=XM_019817180.2
E1 4770367 4770486
refGene_gn=LOC101085080;refGene_tx=XM_019817180.2
E1 4771113 4771252
refGene_gn=LOC101085080;refGene_tx=XM_019817180.2
E1 4772935 4773039
refGene_gn=LOC101085080;refGene_tx=XM_019817180.2
E1 4773165 4773264
refGene_gn=LOC101085080;refGene_tx=XM_019817180.2
E1 4773406 4773470
refGene_gn=LOC101085080;refGene_tx=XM_019817180.2
E1 4773624 4773717
refGene_gn=LOC101085080;refGene_tx=XM_019817180.2
E1 4774135 4774250
refGene_gn=LOC101085080;refGene_tx=XM_019817180.2
E1 4774339 4774367
refGene_gn=LOC101085080;refGene_tx=XM_019817180.2
E1 4775091 4775248
refGene_gn=LOC101085080;refGene_tx=XM_019817180.2
E1 4775587 4775731
refGene_gn=LOC101085080;refGene_tx=XM_019817180.2
E1 4775816 4776020
refGene_gn=LOC101085080;refGene_tx=XM_019817180.2

E1	4790931	4791084
refGene_gn=LOC101084401;refGene_tx=XM_023243774.1
E1	4792915	4793011
refGene_gn=LOC101084401;refGene_tx=XM_023243774.1
E1	4793133	4793238
refGene_gn=LOC101084401;refGene_tx=XM_023243774.1
E1	4793499	4793670
refGene_gn=LOC101084401;refGene_tx=XM_023243774.1
E1	4793757	4793883
refGene_gn=LOC101084401;refGene_tx=XM_023243774.1
E1	4794205	4794409
refGene_gn=LOC101084401;refGene_tx=XM_023243774.1
E1	4794496	4794805
refGene_gn=LOC101084401;refGene_tx=XM_023243774.1
E1	4795208	4795333
refGene_gn=LOC101084401;refGene_tx=XM_023243774.1
E1	4795422	4795588
refGene_gn=LOC101084401;refGene_tx=XM_023243774.1
E1	4795872	4796056
refGene_gn=LOC101084401;refGene_tx=XM_023243774.1
E1	4797327	4797524
refGene_gn=LOC101084401;refGene_tx=XM_023243774.1
E1	4797678	4797797
refGene_gn=LOC101084401;refGene_tx=XM_023243774.1
E1	4799378	4799505
refGene_gn=LOC101084401;refGene_tx=XM_023243774.1
E1	4800078	4800468
refGene_gn=LOC101084401;refGene_tx=XM_023243774.1
E1	4800553	4800644
refGene_gn=LOC101084401;refGene_tx=XM_023243774.1
E1	4800721	4800867
refGene_gn=LOC101084401;refGene_tx=XM_023243774.1
E1	4801695	4801872
refGene_gn=LOC101084401;refGene_tx=XM_023243774.1
E1	4801965	4802208
refGene_gn=LOC101084401;refGene_tx=XM_023243774.1
E1	4804292	4804548
refGene_gn=LOC101084401;refGene_tx=XM_023243774.1
E1	4804646	4804783
refGene_gn=LOC101084401;refGene_tx=XM_023243774.1
E1	4804871	4804995
refGene_gn=LOC101084401;refGene_tx=XM_023243774.1
E1	4805311	4805429
refGene_gn=LOC101084401;refGene_tx=XM_023243774.1
E1	4805511	4805599
refGene_gn=LOC101084401;refGene_tx=XM_023243774.1
E1	4806756	4806827
refGene_gn=LOC101084401;refGene_tx=XM_023243774.1
E1	4806924	4807234
refGene_gn=LOC101084401;refGene_tx=XM_023243774.1

E1 4807999 4808170
refGene_gn=LOC101084401;refGene_tx=XM_023243774.1
E1 4808286 4808436
refGene_gn=LOC101084401;refGene_tx=XM_023243774.1
E1 4808526 4808645
refGene_gn=LOC101084401;refGene_tx=XM_023243774.1
E1 4808825 4808964
refGene_gn=LOC101084401;refGene_tx=XM_023243774.1
E1 4809121 4809225
refGene_gn=LOC101084401;refGene_tx=XM_023243774.1
E1 4809316 4809415
refGene_gn=LOC101084401;refGene_tx=XM_023243774.1
E1 4810046 4810110
refGene_gn=LOC101084401;refGene_tx=XM_023243774.1
E1 4810244 4810337
refGene_gn=LOC101084401;refGene_tx=XM_023243774.1
E1 4810420 4810535
refGene_gn=LOC101084401;refGene_tx=XM_023243774.1
E1 4811270 4811298
refGene_gn=LOC101084401;refGene_tx=XM_023243774.1
E1 4812366 4812523
refGene_gn=LOC101084401;refGene_tx=XM_023243774.1
E1 4812641 4812785
refGene_gn=LOC101084401;refGene_tx=XM_023243774.1
E1 4812879 4813083
refGene_gn=LOC101084401;refGene_tx=XM_023243774.1
E1 4817990 4818143
refGene_gn=LOC101085328;refGene_tx=XM_006939791.4
E1 4820222 4820318
refGene_gn=LOC101085328;refGene_tx=XM_006939791.4
E1 4820437 4820542
refGene_gn=LOC101085328;refGene_tx=XM_006939791.4
E1 4820625 4820796
refGene_gn=LOC101085328;refGene_tx=XM_006939791.4
E1 4820883 4820900
refGene_gn=LOC101085328;refGene_tx=XM_006939791.4
E1 4820902 4821008
refGene_gn=LOC101085328;refGene_tx=XM_006939791.4
E1 4821325 4821529
refGene_gn=LOC101085328;refGene_tx=XM_006939791.4
E1 4821616 4821925
refGene_gn=LOC101085328;refGene_tx=XM_006939791.4
E1 4822087 4822212
refGene_gn=LOC101085328;refGene_tx=XM_006939791.4
E1 4822313 4822479
refGene_gn=LOC101085328;refGene_tx=XM_006939791.4
E1 4822556 4822740
refGene_gn=LOC101085328;refGene_tx=XM_006939791.4
E1 4823662 4823859
refGene_gn=LOC101085328;refGene_tx=XM_006939791.4

E1 4824001 4824120
refGene_gn=LOC101085328;refGene_tx=XM_006939791.4
E1 4825301 4825428
refGene_gn=LOC101085328;refGene_tx=XM_006939791.4
E1 4826354 4826744
refGene_gn=LOC101085328;refGene_tx=XM_006939791.4
E1 4826829 4826920
refGene_gn=LOC101085328;refGene_tx=XM_006939791.4
E1 4827003 4827149
refGene_gn=LOC101085328;refGene_tx=XM_006939791.4
E1 4827231 4827408
refGene_gn=LOC101085328;refGene_tx=XM_006939791.4
E1 4827501 4827744
refGene_gn=LOC101085328;refGene_tx=XM_006939791.4
E1 4829569 4829825
refGene_gn=LOC101085328;refGene_tx=XM_006939791.4
E1 4831452 4831589
refGene_gn=LOC101085328;refGene_tx=XM_006939791.4
E1 4831677 4831801
refGene_gn=LOC101085328;refGene_tx=XM_006939791.4
E1 4832729 4832847
refGene_gn=LOC101085328;refGene_tx=XM_006939791.4
E1 4832939 4833027
refGene_gn=LOC101085328;refGene_tx=XM_006939791.4
E1 4833866 4833940
refGene_gn=LOC101085328;refGene_tx=XM_006939791.4
E1 4834554 4834864
refGene_gn=LOC101085328;refGene_tx=XM_006939791.4
E1 4834966 4835137
refGene_gn=LOC101085328;refGene_tx=XM_006939791.4
E1 4835253 4835403
refGene_gn=LOC101085328;refGene_tx=XM_006939791.4
E1 4835491 4835610
refGene_gn=LOC101085328;refGene_tx=XM_006939791.4
E1 4835969 4836108
refGene_gn=LOC101085328;refGene_tx=XM_006939791.4
E1 4836603 4836707
refGene_gn=LOC101085328;refGene_tx=XM_006939791.4
E1 4837348 4837447
refGene_gn=LOC101085328;refGene_tx=XM_006939791.4
E1 4837573 4837637
refGene_gn=LOC101085328;refGene_tx=XM_006939791.4
E1 4838169 4838262
refGene_gn=LOC101085328;refGene_tx=XM_006939791.4
E1 4838344 4838453
refGene_gn=LOC101085328;refGene_tx=XM_006939791.4
E1 4838536 4838564
refGene_gn=LOC101085328;refGene_tx=XM_006939791.4
E1 4839632 4839789
refGene_gn=LOC101085328;refGene_tx=XM_006939791.4

E1 4841506 4841650
refGene_gn=LOC101085328;refGene_tx=XM_006939791.4
E1 4841741 4841945
refGene_gn=LOC101085328;refGene_tx=XM_006939791.4
E1 4918468 4918495
refGene_gn=LOC101084662;refGene_tx=XM_011288844.3
E1 4919052 4919190
refGene_gn=LOC101084662;refGene_tx=XM_011288844.3
E1 4919300 4919396
refGene_gn=LOC101084662;refGene_tx=XM_011288844.3
E1 4919564 4919669
refGene_gn=LOC101084662;refGene_tx=XM_011288844.3
E1 4919756 4919927
refGene_gn=LOC101084662;refGene_tx=XM_011288844.3
E1 4920811 4920937
refGene_gn=LOC101084662;refGene_tx=XM_011288844.3
E1 4921019 4921223
refGene_gn=LOC101084662;refGene_tx=XM_011288844.3
E1 4921343 4921652
refGene_gn=LOC101084662;refGene_tx=XM_011288844.3
E1 4922633 4922758
refGene_gn=LOC101084662;refGene_tx=XM_011288844.3
E1 4923030 4923196
refGene_gn=LOC101084662;refGene_tx=XM_011288844.3
E1 4923870 4924054
refGene_gn=LOC101084662;refGene_tx=XM_011288844.3
E1 4924675 4924872
refGene_gn=LOC101084662;refGene_tx=XM_011288844.3
E1 4925030 4925149
refGene_gn=LOC101084662;refGene_tx=XM_011288844.3
E1 4926516 4926643
refGene_gn=LOC101084662;refGene_tx=XM_011288844.3
E1 4926745 4927135
refGene_gn=LOC101084662;refGene_tx=XM_011288844.3
E1 4927562 4927653
refGene_gn=LOC101084662;refGene_tx=XM_011288844.3
E1 4927732 4927878
refGene_gn=LOC101084662;refGene_tx=XM_011288844.3
E1 4927987 4928164
refGene_gn=LOC101084662;refGene_tx=XM_011288844.3
E1 4928255 4928498
refGene_gn=LOC101084662;refGene_tx=XM_011288844.3
E1 4928689 4928945
refGene_gn=LOC101084662;refGene_tx=XM_011288844.3
E1 4929041 4929178
refGene_gn=LOC101084662;refGene_tx=XM_011288844.3
E1 4929267 4929391
refGene_gn=LOC101084662;refGene_tx=XM_011288844.3
E1 4929790 4929908
refGene_gn=LOC101084662;refGene_tx=XM_011288844.3

E1 4929992 4930080
refGene_gn=LOC101084662;refGene_tx=XM_011288844.3
E1 4930682 4930753
refGene_gn=LOC101084662;refGene_tx=XM_011288844.3
E1 4930868 4931175
refGene_gn=LOC101084662;refGene_tx=XM_011288844.3
E1 4931277 4931448
refGene_gn=LOC101084662;refGene_tx=XM_011288844.3
E1 4932200 4932350
refGene_gn=LOC101084662;refGene_tx=XM_011288844.3
E1 4932433 4932552
refGene_gn=LOC101084662;refGene_tx=XM_011288844.3
E1 4933201 4933340
refGene_gn=LOC101084662;refGene_tx=XM_011288844.3
E1 4933426 4933530
refGene_gn=LOC101084662;refGene_tx=XM_011288844.3
E1 4934321 4934420
refGene_gn=LOC101084662;refGene_tx=XM_011288844.3
E1 4934498 4934562
refGene_gn=LOC101084662;refGene_tx=XM_011288844.3
E1 4935330 4935423
refGene_gn=LOC101084662;refGene_tx=XM_011288844.3
E1 4936564 4936673
refGene_gn=LOC101084662;refGene_tx=XM_011288844.3
E1 4937252 4937280
refGene_gn=LOC101084662;refGene_tx=XM_011288844.3
E1 4937717 4937874
refGene_gn=LOC101084662;refGene_tx=XM_011288844.3
E1 4938316 4938460
refGene_gn=LOC101084662;refGene_tx=XM_011288844.3
E1 4939529 4939733
refGene_gn=LOC101084662;refGene_tx=XM_011288844.3
E1 4964323 4964458
refGene_gn=LOC101084915;refGene_tx=XM_003996293.5;refGene_gn=LOC101084
915;refGene_tx=XM_019817179.2
E1 4969572 4969688
refGene_gn=LOC101084915;refGene_tx=XM_003996293.5;refGene_gn=LOC101084
915;refGene_tx=XM_019817179.2
E1 4973796 4973889
refGene_gn=LOC101084915;refGene_tx=XM_003996293.5;refGene_gn=LOC101084
915;refGene_tx=XM_019817179.2
E1 4974616 4974814
refGene_gn=LOC101084915;refGene_tx=XM_003996293.5;refGene_gn=LOC101084
915;refGene_tx=XM_019817179.2
E1 4978145 4978236
refGene_gn=LOC101084915;refGene_tx=XM_003996293.5;refGene_gn=LOC101084
915;refGene_tx=XM_019817179.2
E1 4979536 4979821
refGene_gn=LOC101084915;refGene_tx=XM_003996293.5;refGene_gn=LOC101084
915;refGene_tx=XM_019817179.2

E1 4980731 4980819
refGene_gn=ADPRM;refGene_tx=XM_011288836.3;refGene_gn=ADPRM;refGene_tx
=XM_011288837.3;refGene_gn=ADPRM;refGene_tx=XM_023243775.1
E1 4988047 4988665
refGene_gn=ADPRM;refGene_tx=XM_011288836.3;refGene_gn=ADPRM;refGene_tx
=XM_011288837.3;refGene_gn=ADPRM;refGene_tx=XM_011288838.2;refGene_gn=
ADPRM;refGene_tx=XM_023243775.1
E1 4989457 4989574
refGene_gn=ADPRM;refGene_tx=XM_011288836.3;refGene_gn=ADPRM;refGene_tx
=XM_011288837.3;refGene_gn=ADPRM;refGene_tx=XM_011288838.2;refGene_gn=
ADPRM;refGene_tx=XM_023243775.1
E1 4992710 4993021
refGene_gn=ADPRM;refGene_tx=XM_011288836.3;refGene_gn=ADPRM;refGene_tx
=XM_011288837.3;refGene_gn=ADPRM;refGene_tx=XM_011288838.2
E1 4996818 4996954 refGene_gn=TMEM220;refGene_tx=XM_023243777.1
E1 4999031 4999091 refGene_gn=TMEM220;refGene_tx=XM_023243777.1
E1 5000395 5000519 refGene_gn=TMEM220;refGene_tx=XM_023243777.1
E1 5002472 5002533 refGene_gn=TMEM220;refGene_tx=XM_023243777.1
E1 5005184 5005214 refGene_gn=TMEM220;refGene_tx=XM_023243777.1
E1 5005689 5005782 refGene_gn=TMEM220;refGene_tx=XM_023243777.1
E1 5014920 5014952 refGene_gn=ADPRM;refGene_tx=XM_023243775.1
E1 5094577 5094985 refGene_gn=PIRT;refGene_tx=XM_006939842.4
E1 5421391 5422032
refGene_gn=SHISA6;refGene_tx=XM_003996297.5;refGene_gn=SHISA6;refGene_
tx=XM_023244668.1;refGene_gn=SHISA6;refGene_tx=XM_023244669.1;refGene_
gn=SHISA6;refGene_tx=XM_023244670.1
E1 5445906 5446067
refGene_gn=SHISA6;refGene_tx=XM_003996297.5;refGene_gn=SHISA6;refGene_
tx=XM_023244668.1;refGene_gn=SHISA6;refGene_tx=XM_023244669.1;refGene_
gn=SHISA6;refGene_tx=XM_023244670.1
E1 5555466 5555562
refGene_gn=SHISA6;refGene_tx=XM_003996297.5;refGene_gn=SHISA6;refGene_
tx=XM_023244668.1
E1 5695926 5695983
refGene_gn=SHISA6;refGene_tx=XM_003996297.5;refGene_gn=SHISA6;refGene_
tx=XM_023244669.1
E1 5699468 5699621
refGene_gn=SHISA6;refGene_tx=XM_003996297.5;refGene_gn=SHISA6;refGene_
tx=XM_023244668.1;refGene_gn=SHISA6;refGene_tx=XM_023244669.1;refGene_
gn=SHISA6;refGene_tx=XM_023244670.1
E1 5701350 5701898
refGene_gn=SHISA6;refGene_tx=XM_003996297.5;refGene_gn=SHISA6;refGene_
tx=XM_023244668.1;refGene_gn=SHISA6;refGene_tx=XM_023244669.1;refGene_
gn=SHISA6;refGene_tx=XM_023244670.1
E1 5721785 5722178 refGene_gn=DNAH9;refGene_tx=XM_023244531.1
E1 5731008 5731205 refGene_gn=DNAH9;refGene_tx=XM_023244531.1
E1 5733179 5733338 refGene_gn=DNAH9;refGene_tx=XM_023244531.1
E1 5734377 5734508 refGene_gn=DNAH9;refGene_tx=XM_023244531.1
E1 5739589 5739801 refGene_gn=DNAH9;refGene_tx=XM_023244531.1
E1 5741359 5741593 refGene_gn=DNAH9;refGene_tx=XM_023244531.1

E1 5749212 5749380 refGene_gn=DNAH9;refGene_tx=XM_023244531.1
E1 5750864 5750981 refGene_gn=DNAH9;refGene_tx=XM_023244531.1
E1 5752578 5752729 refGene_gn=DNAH9;refGene_tx=XM_023244531.1
E1 5756192 5756307 refGene_gn=DNAH9;refGene_tx=XM_023244531.1
E1 5761339 5761408 refGene_gn=DNAH9;refGene_tx=XM_023244531.1
E1 5762726 5762853 refGene_gn=DNAH9;refGene_tx=XM_023244531.1
E1 5766329 5766585 refGene_gn=DNAH9;refGene_tx=XM_023244531.1
E1 5767786 5768028 refGene_gn=DNAH9;refGene_tx=XM_023244531.1
E1 5773911 5774047 refGene_gn=DNAH9;refGene_tx=XM_023244531.1
E1 5778036 5778233 refGene_gn=DNAH9;refGene_tx=XM_023244531.1
E1 5778323 5778748 refGene_gn=DNAH9;refGene_tx=XM_023244531.1
E1 5799235 5799458 refGene_gn=DNAH9;refGene_tx=XM_023244531.1
E1 5800106 5800273 refGene_gn=DNAH9;refGene_tx=XM_023244531.1
E1 5809175 5810046 refGene_gn=DNAH9;refGene_tx=XM_023244531.1
E1 5813241 5813372 refGene_gn=DNAH9;refGene_tx=XM_023244531.1
E1 5813712 5813839 refGene_gn=DNAH9;refGene_tx=XM_023244531.1
E1 5818308 5818461 refGene_gn=DNAH9;refGene_tx=XM_023244531.1
E1 5819453 5819579 refGene_gn=DNAH9;refGene_tx=XM_023244531.1
E1 5821770 5822010 refGene_gn=DNAH9;refGene_tx=XM_023244531.1
E1 5822324 5822485 refGene_gn=DNAH9;refGene_tx=XM_023244531.1
E1 5835327 5835484 refGene_gn=DNAH9;refGene_tx=XM_023244531.1
E1 5841092 5841197 refGene_gn=DNAH9;refGene_tx=XM_023244531.1
E1 5851830 5851988 refGene_gn=DNAH9;refGene_tx=XM_023244531.1
E1 5854480 5854619 refGene_gn=DNAH9;refGene_tx=XM_023244531.1
E1 5856349 5856637 refGene_gn=DNAH9;refGene_tx=XM_023244531.1
E1 5867708 5867919 refGene_gn=DNAH9;refGene_tx=XM_023244531.1
E1 5872807 5872935 refGene_gn=DNAH9;refGene_tx=XM_023244531.1
E1 5874266 5874377 refGene_gn=DNAH9;refGene_tx=XM_023244529.1
E1 5876156 5876265
refGene_gn=DNAH9;refGene_tx=XM_023244529.1;refGene_gn=DNAH9;refGene_tx
=XM_023244531.1
E1 5877330 5877478
refGene_gn=DNAH9;refGene_tx=XM_023244529.1;refGene_gn=DNAH9;refGene_tx
=XM_023244531.1
E1 5879592 5879767
refGene_gn=DNAH9;refGene_tx=XM_023244529.1;refGene_gn=DNAH9;refGene_tx
=XM_023244531.1
E1 5885924 5886098
refGene_gn=DNAH9;refGene_tx=XM_023244529.1;refGene_gn=DNAH9;refGene_tx
=XM_023244531.1
E1 5886568 5886776
refGene_gn=DNAH9;refGene_tx=XM_023244529.1;refGene_gn=DNAH9;refGene_tx
=XM_023244531.1
E1 5900182 5900348
refGene_gn=DNAH9;refGene_tx=XM_023244529.1;refGene_gn=DNAH9;refGene_tx
=XM_023244531.1
E1 5901992 5902095
refGene_gn=DNAH9;refGene_tx=XM_023244529.1;refGene_gn=DNAH9;refGene_tx
=XM_023244531.1
E1 5902665 5902905

refGene_gn=DNAH9;refGene_tx=XM_023244529.1;refGene_gn=DNAH9;refGene_tx=XM_023244531.1
E1	5913224	5913386
refGene_gn=DNAH9;refGene_tx=XM_023244529.1;refGene_gn=DNAH9;refGene_tx=XM_023244531.1
E1	5917712	5917909
refGene_gn=DNAH9;refGene_tx=XM_023244529.1;refGene_gn=DNAH9;refGene_tx=XM_023244531.1
E1	5927585	5927748
refGene_gn=DNAH9;refGene_tx=XM_023244529.1;refGene_gn=DNAH9;refGene_tx=XM_023244531.1
E1	5929560	5929684
refGene_gn=DNAH9;refGene_tx=XM_023244529.1;refGene_gn=DNAH9;refGene_tx=XM_023244531.1
E1	5942023	5942166
refGene_gn=DNAH9;refGene_tx=XM_023244529.1;refGene_gn=DNAH9;refGene_tx=XM_023244531.1
E1	5942549	5942711
refGene_gn=DNAH9;refGene_tx=XM_023244529.1;refGene_gn=DNAH9;refGene_tx=XM_023244531.1
E1	5942918	5943152
refGene_gn=DNAH9;refGene_tx=XM_023244529.1;refGene_gn=DNAH9;refGene_tx=XM_023244531.1
E1	5952270	5952531
refGene_gn=DNAH9;refGene_tx=XM_023244529.1;refGene_gn=DNAH9;refGene_tx=XM_023244531.1
E1	5961319	5961745
refGene_gn=DNAH9;refGene_tx=XM_023244529.1;refGene_gn=DNAH9;refGene_tx=XM_023244531.1
E1	5969545	5969665
refGene_gn=DNAH9;refGene_tx=XM_023244529.1;refGene_gn=DNAH9;refGene_tx=XM_023244531.1
E1	5972335	5972524
refGene_gn=DNAH9;refGene_tx=XM_023244529.1;refGene_gn=DNAH9;refGene_tx=XM_023244531.1
E1	5973593	5973829
refGene_gn=DNAH9;refGene_tx=XM_023244529.1;refGene_gn=DNAH9;refGene_tx=XM_023244531.1
E1	5979060	5979183
refGene_gn=DNAH9;refGene_tx=XM_023244529.1;refGene_gn=DNAH9;refGene_tx=XM_023244531.1
E1	5980163	5980368
refGene_gn=DNAH9;refGene_tx=XM_023244529.1;refGene_gn=DNAH9;refGene_tx=XM_023244531.1
E1	5981362	5981527
refGene_gn=DNAH9;refGene_tx=XM_023244529.1;refGene_gn=DNAH9;refGene_tx=XM_023244531.1
E1	5986737	5986878
refGene_gn=DNAH9;refGene_tx=XM_023244529.1;refGene_gn=DNAH9;refGene_tx=XM_023244531.1

E1 5991417 5991588
refGene_gn=DNAH9;refGene_tx=XM_023244529.1;refGene_gn=DNAH9;refGene_tx
=XM_023244531.1
E1 5993577 5993700
refGene_gn=DNAH9;refGene_tx=XM_023244529.1;refGene_gn=DNAH9;refGene_tx
=XM_023244531.1
E1 6000982 6001176
refGene_gn=DNAH9;refGene_tx=XM_023244529.1;refGene_gn=DNAH9;refGene_tx
=XM_023244531.1
E1 6003441 6003590
refGene_gn=DNAH9;refGene_tx=XM_023244529.1;refGene_gn=DNAH9;refGene_tx
=XM_023244531.1
E1 6017356 6017484
refGene_gn=DNAH9;refGene_tx=XM_023244529.1;refGene_gn=DNAH9;refGene_tx
=XM_023244531.1
E1 6019933 6020161
refGene_gn=DNAH9;refGene_tx=XM_023244529.1;refGene_gn=DNAH9;refGene_tx
=XM_023244531.1
E1 6023415 6023607
refGene_gn=DNAH9;refGene_tx=XM_023244529.1;refGene_gn=DNAH9;refGene_tx
=XM_023244531.1
E1 6024612 6024804
refGene_gn=DNAH9;refGene_tx=XM_023244529.1;refGene_gn=DNAH9;refGene_tx
=XM_023244531.1
E1 6027377 6027548
refGene_gn=DNAH9;refGene_tx=XM_023244529.1;refGene_gn=DNAH9;refGene_tx
=XM_023244531.1
E1 6032097 6032280
refGene_gn=DNAH9;refGene_tx=XM_023244529.1;refGene_gn=DNAH9;refGene_tx
=XM_023244531.1
E1 6050110 6050500
refGene_gn=DNAH9;refGene_tx=XM_023244529.1;refGene_gn=DNAH9;refGene_tx
=XM_023244531.1
E1 6057416 6057644
refGene_gn=DNAH9;refGene_tx=XM_023244529.1;refGene_gn=DNAH9;refGene_tx
=XM_023244531.1
E1 6060244 6061023
refGene_gn=ZNF18;refGene_tx=XM_003996215.4;refGene_gn=ZNF18;refGene_tx
=XM_019817184.2;refGene_gn=ZNF18;refGene_tx=XM_023244532.1
E1 6066474 6066582
refGene_gn=ZNF18;refGene_tx=XM_003996215.4;refGene_gn=ZNF18;refGene_tx
=XM_019817184.2;refGene_gn=ZNF18;refGene_tx=XM_023244532.1
E1 6067386 6067471
refGene_gn=ZNF18;refGene_tx=XM_003996215.4;refGene_gn=ZNF18;refGene_tx
=XM_019817184.2;refGene_gn=ZNF18;refGene_tx=XM_023244532.1
E1 6074628 6074708
refGene_gn=ZNF18;refGene_tx=XM_003996215.4;refGene_gn=ZNF18;refGene_tx
=XM_019817184.2;refGene_gn=ZNF18;refGene_tx=XM_023244532.1
E1 6075108 6075298
refGene_gn=ZNF18;refGene_tx=XM_003996215.4;refGene_gn=ZNF18;refGene_tx

=XM_019817184.2;refGene_gn=ZNF18;refGene_tx=XM_023244532.1
E1 6077073 6077463
refGene_gn=ZNF18;refGene_tx=XM_003996215.4;refGene_gn=ZNF18;refGene_tx
=XM_019817184.2;refGene_gn=ZNF18;refGene_tx=XM_023244532.1
E1 6111751 6111869
refGene_gn=MAP2K4;refGene_tx=XM_023244457.1;refGene_gn=MAP2K4;refGene_
tx=XM_023244458.1
E1 6125100 6125133 refGene_gn=MAP2K4;refGene_tx=XM_023244457.1
E1 6150083 6150186
refGene_gn=MAP2K4;refGene_tx=XM_023244457.1;refGene_gn=MAP2K4;refGene_
tx=XM_023244458.1
E1 6180827 6181002
refGene_gn=MAP2K4;refGene_tx=XM_023244457.1;refGene_gn=MAP2K4;refGene_
tx=XM_023244458.1
E1 6195451 6195571
refGene_gn=MAP2K4;refGene_tx=XM_023244457.1;refGene_gn=MAP2K4;refGene_
tx=XM_023244458.1
E1 6211395 6211515
refGene_gn=MAP2K4;refGene_tx=XM_023244457.1;refGene_gn=MAP2K4;refGene_
tx=XM_023244458.1
E1 6213845 6213897
refGene_gn=MAP2K4;refGene_tx=XM_023244457.1;refGene_gn=MAP2K4;refGene_
tx=XM_023244458.1
E1 6217426 6217554
refGene_gn=MAP2K4;refGene_tx=XM_023244457.1;refGene_gn=MAP2K4;refGene_
tx=XM_023244458.1
E1 6235923 6236001
refGene_gn=MAP2K4;refGene_tx=XM_023244457.1;refGene_gn=MAP2K4;refGene_
tx=XM_023244458.1
E1 6237710 6237859
refGene_gn=MAP2K4;refGene_tx=XM_023244457.1;refGene_gn=MAP2K4;refGene_
tx=XM_023244458.1
E1 6249593 6249639
refGene_gn=MAP2K4;refGene_tx=XM_023244457.1;refGene_gn=MAP2K4;refGene_
tx=XM_023244458.1
E1 6251337 6251451
refGene_gn=MAP2K4;refGene_tx=XM_023244457.1;refGene_gn=MAP2K4;refGene_
tx=XM_023244458.1
E1 6712783 6712838
refGene_gn=MYOCD;refGene_tx=XM_006939797.4;refGene_gn=MYOCD;refGene_tx
=XM_011288833.3;refGene_gn=MYOCD;refGene_tx=XM_019817187.2
E1 6758450 6758516
refGene_gn=MYOCD;refGene_tx=XM_006939797.4;refGene_gn=MYOCD;refGene_tx
=XM_011288833.3
E1 6767961 6768017
refGene_gn=MYOCD;refGene_tx=XM_006939797.4;refGene_gn=MYOCD;refGene_tx
=XM_011288833.3;refGene_gn=MYOCD;refGene_tx=XM_019817187.2
E1 6769479 6769555
refGene_gn=MYOCD;refGene_tx=XM_006939797.4;refGene_gn=MYOCD;refGene_tx
=XM_011288833.3;refGene_gn=MYOCD;refGene_tx=XM_019817187.2;refGene_gn=

MYOCD;refGene_tx=XM_019817188.2;refGene_gn=MYOCD;refGene_tx=XM_0198171
89.2
E1 6772677 6772839
refGene_gn=MYOCD;refGene_tx=XM_006939797.4;refGene_gn=MYOCD;refGene_tx
=XM_011288833.3;refGene_gn=MYOCD;refGene_tx=XM_019817187.2;refGene_gn=
MYOCD;refGene_tx=XM_019817188.2;refGene_gn=MYOCD;refGene_tx=XM_0198171
89.2
E1 6791078 6791257
refGene_gn=MYOCD;refGene_tx=XM_006939797.4;refGene_gn=MYOCD;refGene_tx
=XM_011288833.3;refGene_gn=MYOCD;refGene_tx=XM_019817187.2;refGene_gn=
MYOCD;refGene_tx=XM_019817188.2;refGene_gn=MYOCD;refGene_tx=XM_0198171
89.2
E1 6793543 6793669
refGene_gn=MYOCD;refGene_tx=XM_006939797.4;refGene_gn=MYOCD;refGene_tx
=XM_011288833.3;refGene_gn=MYOCD;refGene_tx=XM_019817187.2;refGene_gn=
MYOCD;refGene_tx=XM_019817188.2;refGene_gn=MYOCD;refGene_tx=XM_0198171
89.2
E1 6799329 6799574
refGene_gn=MYOCD;refGene_tx=XM_006939797.4;refGene_gn=MYOCD;refGene_tx
=XM_011288833.3;refGene_gn=MYOCD;refGene_tx=XM_019817187.2;refGene_gn=
MYOCD;refGene_tx=XM_019817188.2;refGene_gn=MYOCD;refGene_tx=XM_0198171
89.2
E1 6801496 6801650
refGene_gn=MYOCD;refGene_tx=XM_006939797.4;refGene_gn=MYOCD;refGene_tx
=XM_011288833.3;refGene_gn=MYOCD;refGene_tx=XM_019817187.2;refGene_gn=
MYOCD;refGene_tx=XM_019817188.2;refGene_gn=MYOCD;refGene_tx=XM_0198171
89.2
E1 6813937 6814822
refGene_gn=MYOCD;refGene_tx=XM_006939797.4;refGene_gn=MYOCD;refGene_tx
=XM_011288833.3;refGene_gn=MYOCD;refGene_tx=XM_019817187.2;refGene_gn=
MYOCD;refGene_tx=XM_019817188.2;refGene_gn=MYOCD;refGene_tx=XM_0198171
89.2
E1 6816765 6816909
refGene_gn=MYOCD;refGene_tx=XM_006939797.4;refGene_gn=MYOCD;refGene_tx
=XM_011288833.3;refGene_gn=MYOCD;refGene_tx=XM_019817187.2;refGene_gn=
MYOCD;refGene_tx=XM_019817188.2;refGene_gn=MYOCD;refGene_tx=XM_0198171
89.2
E1 6819395 6819524
refGene_gn=MYOCD;refGene_tx=XM_006939797.4;refGene_gn=MYOCD;refGene_tx
=XM_011288833.3;refGene_gn=MYOCD;refGene_tx=XM_019817187.2;refGene_gn=
MYOCD;refGene_tx=XM_019817188.2;refGene_gn=MYOCD;refGene_tx=XM_0198171
89.2
E1 6822734 6822792
refGene_gn=MYOCD;refGene_tx=XM_006939797.4;refGene_gn=MYOCD;refGene_tx
=XM_011288833.3;refGene_gn=MYOCD;refGene_tx=XM_019817187.2;refGene_gn=
MYOCD;refGene_tx=XM_019817188.2;refGene_gn=MYOCD;refGene_tx=XM_0198171
89.2
E1 6825041 6825595
refGene_gn=MYOCD;refGene_tx=XM_006939797.4;refGene_gn=MYOCD;refGene_tx
=XM_011288833.3;refGene_gn=MYOCD;refGene_tx=XM_019817187.2;refGene_gn=

MYOCD;refGene_tx=XM_019817188.2;refGene_gn=MYOCD;refGene_tx=XM_0198171
89.2
E1	6848896	6848949
refGene_gn=ARHGAP44;refGene_tx=XM_023244131.1;refGene_gn=ARHGAP44;refG
ene_tx=XM_023244132.1;refGene_gn=ARHGAP44;refGene_tx=XM_023244134.1;re
fGene_gn=ARHGAP44;refGene_tx=XM_023244136.1
E1	6944132	6944172
refGene_gn=ARHGAP44;refGene_tx=XM_023244131.1;refGene_gn=ARHGAP44;refG
ene_tx=XM_023244132.1;refGene_gn=ARHGAP44;refGene_tx=XM_023244134.1;re
fGene_gn=ARHGAP44;refGene_tx=XM_023244136.1
E1	6945065	6945170
refGene_gn=ARHGAP44;refGene_tx=XM_023244131.1;refGene_gn=ARHGAP44;refG
ene_tx=XM_023244132.1;refGene_gn=ARHGAP44;refGene_tx=XM_023244134.1;re
fGene_gn=ARHGAP44;refGene_tx=XM_023244136.1
E1	6959200	6959277
refGene_gn=ARHGAP44;refGene_tx=XM_023244131.1;refGene_gn=ARHGAP44;refG
ene_tx=XM_023244132.1;refGene_gn=ARHGAP44;refGene_tx=XM_023244134.1;re
fGene_gn=ARHGAP44;refGene_tx=XM_023244136.1
E1	6966925	6967037
refGene_gn=ARHGAP44;refGene_tx=XM_023244131.1;refGene_gn=ARHGAP44;refG
ene_tx=XM_023244132.1;refGene_gn=ARHGAP44;refGene_tx=XM_023244134.1;re
fGene_gn=ARHGAP44;refGene_tx=XM_023244135.1;refGene_gn=ARHGAP44;refGen
e_tx=XM_023244136.1
E1	6971405	6971482
refGene_gn=ARHGAP44;refGene_tx=XM_023244131.1;refGene_gn=ARHGAP44;refG
ene_tx=XM_023244132.1;refGene_gn=ARHGAP44;refGene_tx=XM_023244134.1;re
fGene_gn=ARHGAP44;refGene_tx=XM_023244135.1;refGene_gn=ARHGAP44;refGen
e_tx=XM_023244136.1
E1	6979893	6980011
refGene_gn=ARHGAP44;refGene_tx=XM_023244131.1;refGene_gn=ARHGAP44;refG
ene_tx=XM_023244132.1;refGene_gn=ARHGAP44;refGene_tx=XM_023244134.1;re
fGene_gn=ARHGAP44;refGene_tx=XM_023244135.1;refGene_gn=ARHGAP44;refGen
e_tx=XM_023244136.1
E1	6987701	6987770
refGene_gn=ARHGAP44;refGene_tx=XM_023244131.1;refGene_gn=ARHGAP44;refG
ene_tx=XM_023244132.1;refGene_gn=ARHGAP44;refGene_tx=XM_023244134.1;re
fGene_gn=ARHGAP44;refGene_tx=XM_023244135.1;refGene_gn=ARHGAP44;refGen
e_tx=XM_023244136.1
E1	6990446	6990528
refGene_gn=ARHGAP44;refGene_tx=XM_023244131.1;refGene_gn=ARHGAP44;refG
ene_tx=XM_023244132.1;refGene_gn=ARHGAP44;refGene_tx=XM_023244134.1;re
fGene_gn=ARHGAP44;refGene_tx=XM_023244135.1;refGene_gn=ARHGAP44;refGen
e_tx=XM_023244136.1
E1	6991155	6991283
refGene_gn=ARHGAP44;refGene_tx=XM_023244131.1;refGene_gn=ARHGAP44;refG
ene_tx=XM_023244132.1;refGene_gn=ARHGAP44;refGene_tx=XM_023244134.1;re
fGene_gn=ARHGAP44;refGene_tx=XM_023244135.1;refGene_gn=ARHGAP44;refGen
e_tx=XM_023244136.1
E1	6995970	6996082
refGene_gn=ARHGAP44;refGene_tx=XM_023244131.1;refGene_gn=ARHGAP44;refG

ene_tx=XM_023244132.1;refGene_gn=ARHGAP44;refGene_tx=XM_023244134.1;refGene_gn=ARHGAP44;refGene_tx=XM_023244135.1;refGene_gn=ARHGAP44;refGene_tx=XM_023244136.1
E1 6996492 6996574
refGene_gn=ARHGAP44;refGene_tx=XM_023244131.1;refGene_gn=ARHGAP44;refGene_tx=XM_023244132.1;refGene_gn=ARHGAP44;refGene_tx=XM_023244134.1;refGene_gn=ARHGAP44;refGene_tx=XM_023244135.1;refGene_gn=ARHGAP44;refGene_tx=XM_023244136.1
E1 6999057 6999138
refGene_gn=ARHGAP44;refGene_tx=XM_023244131.1;refGene_gn=ARHGAP44;refGene_tx=XM_023244132.1;refGene_gn=ARHGAP44;refGene_tx=XM_023244134.1;refGene_gn=ARHGAP44;refGene_tx=XM_023244135.1;refGene_gn=ARHGAP44;refGene_tx=XM_023244136.1
E1 7002982 7003096
refGene_gn=ARHGAP44;refGene_tx=XM_023244131.1;refGene_gn=ARHGAP44;refGene_tx=XM_023244132.1;refGene_gn=ARHGAP44;refGene_tx=XM_023244134.1;refGene_gn=ARHGAP44;refGene_tx=XM_023244135.1;refGene_gn=ARHGAP44;refGene_tx=XM_023244136.1
E1 7003680 7003772
refGene_gn=ARHGAP44;refGene_tx=XM_023244131.1;refGene_gn=ARHGAP44;refGene_tx=XM_023244132.1;refGene_gn=ARHGAP44;refGene_tx=XM_023244134.1;refGene_gn=ARHGAP44;refGene_tx=XM_023244135.1;refGene_gn=ARHGAP44;refGene_tx=XM_023244136.1
E1 7006059 7006240
refGene_gn=ARHGAP44;refGene_tx=XM_023244131.1;refGene_gn=ARHGAP44;refGene_tx=XM_023244132.1;refGene_gn=ARHGAP44;refGene_tx=XM_023244134.1;refGene_gn=ARHGAP44;refGene_tx=XM_023244135.1;refGene_gn=ARHGAP44;refGene_tx=XM_023244136.1
E1 7016689 7016707
refGene_gn=ARHGAP44;refGene_tx=XM_023244132.1;refGene_gn=ARHGAP44;refGene_tx=XM_023244135.1;refGene_gn=ARHGAP44;refGene_tx=XM_023244136.1
E1 7017446 7017689
refGene_gn=ARHGAP44;refGene_tx=XM_023244131.1;refGene_gn=ARHGAP44;refGene_tx=XM_023244132.1;refGene_gn=ARHGAP44;refGene_tx=XM_023244135.1;refGene_gn=ARHGAP44;refGene_tx=XM_023244136.1
E1 7021264 7021422
refGene_gn=ARHGAP44;refGene_tx=XM_023244131.1;refGene_gn=ARHGAP44;refGene_tx=XM_023244132.1;refGene_gn=ARHGAP44;refGene_tx=XM_023244134.1;refGene_gn=ARHGAP44;refGene_tx=XM_023244135.1;refGene_gn=ARHGAP44;refGene_tx=XM_023244136.1
E1 7023646 7024027
refGene_gn=ARHGAP44;refGene_tx=XM_023244131.1;refGene_gn=ARHGAP44;refGene_tx=XM_023244132.1;refGene_gn=ARHGAP44;refGene_tx=XM_023244134.1;refGene_gn=ARHGAP44;refGene_tx=XM_023244135.1;refGene_gn=ARHGAP44;refGene_tx=XM_023244136.1
E1 7025875 7025883 refGene_gn=ARHGAP44;refGene_tx=XM_023244132.1
E1 7027814 7027954
refGene_gn=ARHGAP44;refGene_tx=XM_023244131.1;refGene_gn=ARHGAP44;refGene_tx=XM_023244134.1;refGene_gn=ARHGAP44;refGene_tx=XM_023244135.1;refGene_gn=ARHGAP44;refGene_tx=XM_023244136.1

E1	7030098	7030332
refGene_gn=ELAC2;refGene_tx=XM_011288910.3;refGene_gn=ELAC2;refGene_tx=XM_023244130.1
E1	7030826	7030971
refGene_gn=ELAC2;refGene_tx=XM_011288910.3;refGene_gn=ELAC2;refGene_tx=XM_023244130.1
E1	7031505	7031584
refGene_gn=ELAC2;refGene_tx=XM_011288910.3;refGene_gn=ELAC2;refGene_tx=XM_023244130.1
E1	7031691	7031812
refGene_gn=ELAC2;refGene_tx=XM_011288910.3;refGene_gn=ELAC2;refGene_tx=XM_023244130.1
E1	7031880	7031980
refGene_gn=ELAC2;refGene_tx=XM_011288910.3;refGene_gn=ELAC2;refGene_tx=XM_023244130.1
E1	7032899	7033009
refGene_gn=ELAC2;refGene_tx=XM_011288910.3;refGene_gn=ELAC2;refGene_tx=XM_023244130.1
E1	7033130	7033169
refGene_gn=ELAC2;refGene_tx=XM_011288910.3;refGene_gn=ELAC2;refGene_tx=XM_023244130.1
E1	7033606	7033745
refGene_gn=ELAC2;refGene_tx=XM_011288910.3;refGene_gn=ELAC2;refGene_tx=XM_023244130.1
E1	7035313	7035410
refGene_gn=ELAC2;refGene_tx=XM_011288910.3;refGene_gn=ELAC2;refGene_tx=XM_023244130.1
E1	7036738	7036857
refGene_gn=ELAC2;refGene_tx=XM_011288910.3;refGene_gn=ELAC2;refGene_tx=XM_023244130.1
E1	7038734	7038820
refGene_gn=ELAC2;refGene_tx=XM_011288910.3;refGene_gn=ELAC2;refGene_tx=XM_023244130.1
E1	7038901	7039040
refGene_gn=ELAC2;refGene_tx=XM_011288910.3;refGene_gn=ELAC2;refGene_tx=XM_023244130.1
E1	7039824	7039920
refGene_gn=ELAC2;refGene_tx=XM_011288910.3;refGene_gn=ELAC2;refGene_tx=XM_023244130.1
E1	7041539	7041652
refGene_gn=ELAC2;refGene_tx=XM_011288910.3;refGene_gn=ELAC2;refGene_tx=XM_023244130.1
E1	7042205	7042278
refGene_gn=ELAC2;refGene_tx=XM_011288910.3;refGene_gn=ELAC2;refGene_tx=XM_023244130.1
E1	7042366	7042425
refGene_gn=ELAC2;refGene_tx=XM_011288910.3;refGene_gn=ELAC2;refGene_tx=XM_023244130.1
E1	7045634	7045693
refGene_gn=ELAC2;refGene_tx=XM_011288910.3;refGene_gn=ELAC2;refGene_tx

=XM_023244130.1
E1 7046588 7046696 refGene_gn=ELAC2;refGene_tx=XM_011288910.3
E1 7048055 7048124
refGene_gn=ELAC2;refGene_tx=XM_011288910.3;refGene_gn=ELAC2;refGene_tx
=XM_023244130.1
E1 7049864 7049922
refGene_gn=ELAC2;refGene_tx=XM_011288910.3;refGene_gn=ELAC2;refGene_tx
=XM_023244130.1
E1 7050988 7051053
refGene_gn=ELAC2;refGene_tx=XM_011288910.3;refGene_gn=ELAC2;refGene_tx
=XM_023244130.1
E1 7051734 7051805
refGene_gn=ELAC2;refGene_tx=XM_011288910.3;refGene_gn=ELAC2;refGene_tx
=XM_023244130.1
E1 7051931 7051982
refGene_gn=ELAC2;refGene_tx=XM_011288910.3;refGene_gn=ELAC2;refGene_tx
=XM_023244130.1
E1 7052568 7052801
refGene_gn=ELAC2;refGene_tx=XM_011288910.3;refGene_gn=ELAC2;refGene_tx
=XM_023244130.1
E1 7481953 7482575 refGene_gn=HS3ST3A1;refGene_tx=XM_023243537.1
E1 7581533 7582132 refGene_gn=HS3ST3A1;refGene_tx=XM_023243537.1
E1 7955427 7955470
refGene_gn=LOC101091737;refGene_tx=XM_011288832.3
E1 7960838 7960972
refGene_gn=LOC101091737;refGene_tx=XM_011288832.3
E1 7963177 7963499
refGene_gn=LOC101091737;refGene_tx=XM_011288832.3
E1 7989753 7989878
refGene_gn=LOC101091737;refGene_tx=XM_011288832.3
E1 8051872 8051943
refGene_gn=LOC101091737;refGene_tx=XM_011288832.3
E1 8078879 8079112
refGene_gn=LOC101091737;refGene_tx=XM_011288832.3
E1 8093632 8094039
refGene_gn=LOC101091737;refGene_tx=XM_011288832.3
E1 8135779 8136333
refGene_gn=LOC101091978;refGene_tx=XM_011288909.3
E1 8172022 8172641
refGene_gn=LOC101091978;refGene_tx=XM_011288909.3
E1 8881230 8881394
refGene_gn=PMP22;refGene_tx=XM_023243571.1;refGene_gn=PMP22;refGene_tx
=XM_023243572.1
E1 8889605 8889746
refGene_gn=PMP22;refGene_tx=XM_023243571.1;refGene_gn=PMP22;refGene_tx
=XM_023243572.1
E1 8908048 8908148
refGene_gn=PMP22;refGene_tx=XM_023243571.1;refGene_gn=PMP22;refGene_tx
=XM_023243572.1
E1 8909505 8909583

refGene_gn=PMP22;refGene_tx=XM_023243571.1;refGene_gn=PMP22;refGene_tx=XM_023243572.1
E1 8946916 8947133 refGene_gn=TEKT3;refGene_tx=XM_003996304.5;refGene_gn=TEKT3;refGene_tx=XM_006939849.4;refGene_gn=TEKT3;refGene_tx=XM_023244103.1
E1 8953845 8954000 refGene_gn=TEKT3;refGene_tx=XM_003996304.5;refGene_gn=TEKT3;refGene_tx=XM_006939849.4;refGene_gn=TEKT3;refGene_tx=XM_023244103.1
E1 8956558 8956781 refGene_gn=TEKT3;refGene_tx=XM_003996304.5;refGene_gn=TEKT3;refGene_tx=XM_006939849.4;refGene_gn=TEKT3;refGene_tx=XM_023244103.1
E1 8958304 8958448 refGene_gn=TEKT3;refGene_tx=XM_003996304.5;refGene_gn=TEKT3;refGene_tx=XM_006939849.4;refGene_gn=TEKT3;refGene_tx=XM_023244103.1
E1 8963037 8963108 refGene_gn=TEKT3;refGene_tx=XM_003996304.5;refGene_gn=TEKT3;refGene_tx=XM_006939849.4;refGene_gn=TEKT3;refGene_tx=XM_023244103.1
E1 8968228 8968312 refGene_gn=TEKT3;refGene_tx=XM_003996304.5;refGene_gn=TEKT3;refGene_tx=XM_006939849.4;refGene_gn=TEKT3;refGene_tx=XM_023244103.1
E1 8971679 8972285 refGene_gn=TEKT3;refGene_tx=XM_003996304.5;refGene_gn=TEKT3;refGene_tx=XM_006939849.4;refGene_gn=TEKT3;refGene_tx=XM_023244103.1
E1 8981579 8981612 refGene_gn=TEKT3;refGene_tx=XM_003996304.5
E1 9050686 9051144 refGene_gn=CDRT4;refGene_tx=XM_019817196.2;refGene_gn=CDRT4;refGene_tx=XM_019817197.2;refGene_gn=CDRT4;refGene_tx=XM_023243992.1;refGene_gn=CDRT4;refGene_tx=XM_023243993.1
E1 9052634 9052712 refGene_gn=CDRT4;refGene_tx=XM_019817196.2;refGene_gn=CDRT4;refGene_tx=XM_019817197.2;refGene_gn=CDRT4;refGene_tx=XM_023243992.1;refGene_gn=CDRT4;refGene_tx=XM_023243993.1
E1 9081060 9081499 refGene_gn=CDRT4;refGene_tx=XM_019817196.2;refGene_gn=CDRT4;refGene_tx=XM_019817197.2;refGene_gn=CDRT4;refGene_tx=XM_023243992.1
E1 9147260 9147287 refGene_gn=TVP23B;refGene_tx=NM_001281857.1
E1 9147720 9147849 refGene_gn=TVP23B;refGene_tx=NM_001281857.1
E1 9154133 9154265 refGene_gn=TVP23B;refGene_tx=NM_001281857.1
E1 9155457 9155547 refGene_gn=TVP23B;refGene_tx=NM_001281857.1
E1 9162430 9162575 refGene_gn=TVP23B;refGene_tx=NM_001281857.1
E1 9164331 9164414 refGene_gn=TVP23B;refGene_tx=NM_001281857.1
E1 9170119 9170131 refGene_gn=TVP23B;refGene_tx=NM_001281857.1
E1 9173420 9174229 refGene_gn=CDRT1;refGene_tx=XM_023243648.1
E1 9182399 9182668 refGene_gn=CDRT1;refGene_tx=XM_023243648.1
E1 9184872 9185031 refGene_gn=CDRT1;refGene_tx=XM_023243648.1
E1 9187834 9187977 refGene_gn=CDRT1;refGene_tx=XM_023243648.1
E1 9188943 9189092 refGene_gn=CDRT1;refGene_tx=XM_023243648.1
E1 9193480 9193602 refGene_gn=CDRT1;refGene_tx=XM_023243648.1
E1 9198309 9198510 refGene_gn=CDRT1;refGene_tx=XM_023243648.1
E1 9199814 9199924 refGene_gn=CDRT1;refGene_tx=XM_023243648.1

E1 9202887 9202998 refGene_gn=CDRT1;refGene_tx=XM_023243648.1
E1 9203790 9203918 refGene_gn=CDRT1;refGene_tx=XM_023243648.1
E1 9204017 9204215 refGene_gn=CDRT1;refGene_tx=XM_023243648.1
E1 9204724 9204898 refGene_gn=CDRT1;refGene_tx=XM_023243648.1
E1 9206868 9207373 refGene_gn=CDRT1;refGene_tx=XM_023243648.1
E1 9219307 9219891
refGene_gn=TRIM16;refGene_tx=XM_003996307.5;refGene_gn=TRIM16;refGene_
tx=XM_006939855.2;refGene_gn=TRIM16;refGene_tx=XM_023244652.1
E1 9222466 9222562
refGene_gn=TRIM16;refGene_tx=XM_003996307.5;refGene_gn=TRIM16;refGene_
tx=XM_006939855.2;refGene_gn=TRIM16;refGene_tx=XM_023244652.1
E1 9223032 9223198
refGene_gn=TRIM16;refGene_tx=XM_003996307.5;refGene_gn=TRIM16;refGene_
tx=XM_006939855.2;refGene_gn=TRIM16;refGene_tx=XM_023244652.1
E1 9226670 9226904
refGene_gn=TRIM16;refGene_tx=XM_003996307.5;refGene_gn=TRIM16;refGene_
tx=XM_006939855.2;refGene_gn=TRIM16;refGene_tx=XM_023244652.1
E1 9231805 9231901
refGene_gn=TRIM16;refGene_tx=XM_003996307.5;refGene_gn=TRIM16;refGene_
tx=XM_006939855.2;refGene_gn=TRIM16;refGene_tx=XM_023244652.1
E1 9238109 9238628
refGene_gn=TRIM16;refGene_tx=XM_003996307.5;refGene_gn=TRIM16;refGene_
tx=XM_006939855.2;refGene_gn=TRIM16;refGene_tx=XM_023244652.1
E1 9245012 9245026 refGene_gn=TRIM16;refGene_tx=XM_006939855.2
E1 9264273 9264296 refGene_gn=TRIM16;refGene_tx=XM_023244652.1
E1 9284323 9285861
refGene_gn=LOC101092714;refGene_tx=XM_019817200.1;refGene_gn=LOC101092
714;refGene_tx=XM_019817202.2;refGene_gn=LOC101092714;refGene_tx=XM_01
9817203.2;refGene_gn=LOC101092714;refGene_tx=XM_019817207.2;refGene_gn
=LOC101092714;refGene_tx=XM_019817208.2
E1 9288474 9288511
refGene_gn=LOC101092714;refGene_tx=XM_019817200.1;refGene_gn=LOC101092
714;refGene_tx=XM_019817202.2;refGene_gn=LOC101092714;refGene_tx=XM_01
9817203.2
E1 9312035 9312063 refGene_gn=ZNF624;refGene_tx=XM_006939859.3
E1 9312994 9313083 refGene_gn=ZNF624;refGene_tx=XM_006939859.3
E1 9316406 9316521 refGene_gn=ZNF624;refGene_tx=XM_006939859.3
E1 9317883 9317976 refGene_gn=ZNF624;refGene_tx=XM_006939859.3
E1 9326562 9327790 refGene_gn=ZNF624;refGene_tx=XM_006939859.3
E1 9346574 9346681 refGene_gn=ZNF624;refGene_tx=XM_006939859.3
E1 9357955 9358044 refGene_gn=ZNF624;refGene_tx=XM_006939859.3
E1 9360263 9360329 refGene_gn=ZNF624;refGene_tx=XM_006939859.3
E1 9368002 9368129 refGene_gn=ZNF624;refGene_tx=XM_006939859.3
E1 9368888 9368984 refGene_gn=ZNF624;refGene_tx=XM_006939859.3
E1 9378946 9381165 refGene_gn=ZNF624;refGene_tx=XM_006939859.3
E1 9420280 9420683
refGene_gn=ZNF287;refGene_tx=XM_006939865.4;refGene_gn=ZNF287;refGene_
tx=XM_019817209.2;refGene_gn=ZNF287;refGene_tx=XM_019817211.2;refGene_
gn=ZNF287;refGene_tx=XM_023244404.1
E1 9421331 9421432

refGene_gn=ZNF287;refGene_tx=XM_006939865.4;refGene_gn=ZNF287;refGene_tx=XM_019817209.2;refGene_gn=ZNF287;refGene_tx=XM_019817211.2;refGene_gn=ZNF287;refGene_tx=XM_023244404.1
E1 9424035 9424162
refGene_gn=ZNF287;refGene_tx=XM_006939865.4;refGene_gn=ZNF287;refGene_tx=XM_019817209.2;refGene_gn=ZNF287;refGene_tx=XM_019817211.2;refGene_gn=ZNF287;refGene_tx=XM_023244404.1
E1 9424646 9424733
refGene_gn=ZNF287;refGene_tx=XM_006939865.4;refGene_gn=ZNF287;refGene_tx=XM_019817209.2;refGene_gn=ZNF287;refGene_tx=XM_019817211.2;refGene_gn=ZNF287;refGene_tx=XM_023244404.1
E1 9433368 9434936
refGene_gn=ZNF287;refGene_tx=XM_006939865.4;refGene_gn=ZNF287;refGene_tx=XM_019817209.2;refGene_gn=ZNF287;refGene_tx=XM_019817211.2;refGene_gn=ZNF287;refGene_tx=XM_023244404.1
E1 9472380 9472632 refGene_gn=LRRC75A;refGene_tx=XM_023244643.1
E1 9503118 9503247 refGene_gn=LRRC75A;refGene_tx=XM_023244643.1
E1 9513625 9513741 refGene_gn=LRRC75A;refGene_tx=XM_023244643.1
E1 9516803 9517344 refGene_gn=LRRC75A;refGene_tx=XM_023244643.1
E1 9522909 9523004
refGene_gn=TRPV2;refGene_tx=XM_011288828.3;refGene_gn=TRPV2;refGene_tx=XM_019817212.2
E1 9524844 9524924
refGene_gn=TRPV2;refGene_tx=XM_011288828.3;refGene_gn=TRPV2;refGene_tx=XM_019817212.2
E1 9525688 9525816
refGene_gn=TRPV2;refGene_tx=XM_011288828.3;refGene_gn=TRPV2;refGene_tx=XM_019817212.2
E1 9526761 9527099
refGene_gn=TRPV2;refGene_tx=XM_011288828.3;refGene_gn=TRPV2;refGene_tx=XM_019817212.2
E1 9527210 9527277
refGene_gn=TRPV2;refGene_tx=XM_011288828.3;refGene_gn=TRPV2;refGene_tx=XM_019817212.2
E1 9529277 9529443
refGene_gn=TRPV2;refGene_tx=XM_011288828.3;refGene_gn=TRPV2;refGene_tx=XM_019817212.2
E1 9529883 9529954
refGene_gn=TRPV2;refGene_tx=XM_011288828.3;refGene_gn=TRPV2;refGene_tx=XM_019817212.2
E1 9530840 9530936
refGene_gn=TRPV2;refGene_tx=XM_011288828.3;refGene_gn=TRPV2;refGene_tx=XM_019817212.2
E1 9531544 9531700
refGene_gn=TRPV2;refGene_tx=XM_011288828.3;refGene_gn=TRPV2;refGene_tx=XM_019817212.2
E1 9532072 9532246
refGene_gn=TRPV2;refGene_tx=XM_011288828.3;refGene_gn=TRPV2;refGene_tx=XM_019817212.2
E1 9533807 9534106

refGene_gn=TRPV2;refGene_tx=XM_011288828.3;refGene_gn=TRPV2;refGene_tx=XM_019817212.2
E1 9534725 9535019
refGene_gn=TRPV2;refGene_tx=XM_011288828.3;refGene_gn=TRPV2;refGene_tx=XM_019817212.2
E1 9536224 9536358
refGene_gn=TRPV2;refGene_tx=XM_011288828.3;refGene_gn=TRPV2;refGene_tx=XM_019817212.2
E1 9538713 9538913
refGene_gn=TRPV2;refGene_tx=XM_011288828.3;refGene_gn=TRPV2;refGene_tx=XM_019817212.2
E1 9561107 9562025
refGene_gn=UBB;refGene_tx=XM_003996309.5;refGene_gn=UBB;refGene_tx=XM_023243723.1
E1 9597762 9598157
refGene_gn=CENPV;refGene_tx=XM_023243704.1;refGene_gn=CENPV;refGene_tx=XM_023243705.1;refGene_gn=CENPV;refGene_tx=XM_023243706.1;refGene_gn=CENPV;refGene_tx=XM_023243707.1;refGene_gn=CENPV;refGene_tx=XM_023243708.1
E1 9600816 9600915
refGene_gn=CENPV;refGene_tx=XM_023243704.1;refGene_gn=CENPV;refGene_tx=XM_023243705.1;refGene_gn=CENPV;refGene_tx=XM_023243706.1;refGene_gn=CENPV;refGene_tx=XM_023243707.1;refGene_gn=CENPV;refGene_tx=XM_023243708.1
E1 9602508 9602578
refGene_gn=CENPV;refGene_tx=XM_023243704.1;refGene_gn=CENPV;refGene_tx=XM_023243705.1;refGene_gn=CENPV;refGene_tx=XM_023243706.1;refGene_gn=CENPV;refGene_tx=XM_023243707.1;refGene_gn=CENPV;refGene_tx=XM_023243708.1
E1 9605792 9605882 refGene_gn=CENPV;refGene_tx=XM_023243708.1
E1 9608918 9609033
refGene_gn=CENPV;refGene_tx=XM_023243704.1;refGene_gn=CENPV;refGene_tx=XM_023243705.1;refGene_gn=CENPV;refGene_tx=XM_023243706.1;refGene_gn=CENPV;refGene_tx=XM_023243707.1
E1 9609932 9610022
refGene_gn=CENPV;refGene_tx=XM_023243704.1;refGene_gn=CENPV;refGene_tx=XM_023243705.1
E1 9610349 9610474
refGene_gn=CENPV;refGene_tx=XM_023243704.1;refGene_gn=CENPV;refGene_tx=XM_023243705.1;refGene_gn=CENPV;refGene_tx=XM_023243706.1
E1 9612475 9612495 refGene_gn=CENPV;refGene_tx=XM_023243707.1
E1 9618836 9618851
refGene_gn=PIGL;refGene_tx=XM_011288826.3;refGene_gn=PIGL;refGene_tx=XM_011288827.3
E1 9624517 9624616
refGene_gn=PIGL;refGene_tx=XM_003996321.5;refGene_gn=PIGL;refGene_tx=XM_011288826.3
E1 9635356 9635490
refGene_gn=PIGL;refGene_tx=XM_003996321.5;refGene_gn=PIGL;refGene_tx=XM_011288826.3;refGene_gn=PIGL;refGene_tx=XM_011288827.3

E1 9637042 9637074
refGene_gn=PIGL;refGene_tx=XM_003996321.5;refGene_gn=PIGL;refGene_tx=XM_011288826.3;refGene_gn=PIGL;refGene_tx=XM_011288827.3
E1 9638844 9638912
refGene_gn=PIGL;refGene_tx=XM_003996321.5;refGene_gn=PIGL;refGene_tx=XM_011288826.3;refGene_gn=PIGL;refGene_tx=XM_011288827.3
E1 9645020 9645111
refGene_gn=PIGL;refGene_tx=XM_003996321.5;refGene_gn=PIGL;refGene_tx=XM_011288826.3;refGene_gn=PIGL;refGene_tx=XM_011288827.3
E1 9694089 9694189
refGene_gn=PIGL;refGene_tx=XM_003996321.5;refGene_gn=PIGL;refGene_tx=XM_011288826.3;refGene_gn=PIGL;refGene_tx=XM_011288827.3
E1 9706573 9706808
refGene_gn=PIGL;refGene_tx=XM_003996321.5;refGene_gn=PIGL;refGene_tx=XM_011288826.3;refGene_gn=PIGL;refGene_tx=XM_011288827.3
E1 9730677 9730785
refGene_gn=NCOR1;refGene_tx=XM_023244071.1;refGene_gn=NCOR1;refGene_tx=XM_023244072.1;refGene_gn=NCOR1;refGene_tx=XM_023244073.1;refGene_gn=NCOR1;refGene_tx=XM_023244074.1;refGene_gn=NCOR1;refGene_tx=XM_023244075.1;refGene_gn=NCOR1;refGene_tx=XM_023244076.1;refGene_gn=NCOR1;refGene_tx=XM_023244077.1;refGene_gn=NCOR1;refGene_tx=XM_023244078.1;refGene_gn=NCOR1;refGene_tx=XM_023244079.1;refGene_gn=NCOR1;refGene_tx=XM_023244080.1;refGene_gn=NCOR1;refGene_tx=XM_023244081.1;refGene_gn=NCOR1;refGene_tx=XM_023244082.1;refGene_gn=NCOR1;refGene_tx=XM_023244083.1
E1 9740260 9740394
refGene_gn=NCOR1;refGene_tx=XM_023244071.1;refGene_gn=NCOR1;refGene_tx=XM_023244072.1;refGene_gn=NCOR1;refGene_tx=XM_023244073.1;refGene_gn=NCOR1;refGene_tx=XM_023244074.1;refGene_gn=NCOR1;refGene_tx=XM_023244075.1;refGene_gn=NCOR1;refGene_tx=XM_023244076.1;refGene_gn=NCOR1;refGene_tx=XM_023244077.1;refGene_gn=NCOR1;refGene_tx=XM_023244078.1;refGene_gn=NCOR1;refGene_tx=XM_023244079.1;refGene_gn=NCOR1;refGene_tx=XM_023244080.1;refGene_gn=NCOR1;refGene_tx=XM_023244081.1;refGene_gn=NCOR1;refGene_tx=XM_023244082.1;refGene_gn=NCOR1;refGene_tx=XM_023244083.1
E1 9751170 9751363
refGene_gn=NCOR1;refGene_tx=XM_023244071.1;refGene_gn=NCOR1;refGene_tx=XM_023244072.1;refGene_gn=NCOR1;refGene_tx=XM_023244073.1;refGene_gn=NCOR1;refGene_tx=XM_023244074.1;refGene_gn=NCOR1;refGene_tx=XM_023244075.1;refGene_gn=NCOR1;refGene_tx=XM_023244076.1;refGene_gn=NCOR1;refGene_tx=XM_023244077.1;refGene_gn=NCOR1;refGene_tx=XM_023244078.1;refGene_gn=NCOR1;refGene_tx=XM_023244079.1;refGene_gn=NCOR1;refGene_tx=XM_023244080.1;refGene_gn=NCOR1;refGene_tx=XM_023244081.1;refGene_gn=NCOR1;refGene_tx=XM_023244082.1;refGene_gn=NCOR1;refGene_tx=XM_023244083.1
E1 9757009 9757192
refGene_gn=NCOR1;refGene_tx=XM_023244071.1;refGene_gn=NCOR1;refGene_tx=XM_023244072.1;refGene_gn=NCOR1;refGene_tx=XM_023244073.1;refGene_gn=NCOR1;refGene_tx=XM_023244074.1;refGene_gn=NCOR1;refGene_tx=XM_023244075.1;refGene_gn=NCOR1;refGene_tx=XM_023244076.1;refGene_gn=NCOR1;refGene_tx=XM_023244077.1;refGene_gn=NCOR1;refGene_tx=XM_023244078.1;refGene_gn=NCOR1;refGene_tx=XM_023244079.1;refGene_gn=NCOR1;refGene_tx=XM_023244080.1;refGene_gn=NCOR1;refGene_tx=XM_023244081.1;refGene_gn=NCOR1;

refGene_tx=XM_023244082.1;refGene_gn=NCOR1;refGene_tx=XM_023244083.1
E1 9766288 9766402
refGene_gn=NCOR1;refGene_tx=XM_023244071.1;refGene_gn=NCOR1;refGene_tx=XM_023244072.1;refGene_gn=NCOR1;refGene_tx=XM_023244073.1;refGene_gn=NCOR1;refGene_tx=XM_023244074.1;refGene_gn=NCOR1;refGene_tx=XM_023244075.1;refGene_gn=NCOR1;refGene_tx=XM_023244076.1;refGene_gn=NCOR1;refGene_tx=XM_023244077.1;refGene_gn=NCOR1;refGene_tx=XM_023244078.1;refGene_gn=NCOR1;refGene_tx=XM_023244079.1;refGene_gn=NCOR1;refGene_tx=XM_023244080.1;refGene_gn=NCOR1;refGene_tx=XM_023244081.1;refGene_gn=NCOR1;refGene_tx=XM_023244082.1;refGene_gn=NCOR1;refGene_tx=XM_023244083.1
E1 9772299 9772356
refGene_gn=NCOR1;refGene_tx=XM_023244071.1;refGene_gn=NCOR1;refGene_tx=XM_023244072.1;refGene_gn=NCOR1;refGene_tx=XM_023244073.1;refGene_gn=NCOR1;refGene_tx=XM_023244074.1;refGene_gn=NCOR1;refGene_tx=XM_023244075.1;refGene_gn=NCOR1;refGene_tx=XM_023244076.1;refGene_gn=NCOR1;refGene_tx=XM_023244077.1;refGene_gn=NCOR1;refGene_tx=XM_023244078.1;refGene_gn=NCOR1;refGene_tx=XM_023244079.1;refGene_gn=NCOR1;refGene_tx=XM_023244080.1;refGene_gn=NCOR1;refGene_tx=XM_023244081.1;refGene_gn=NCOR1;refGene_tx=XM_023244082.1;refGene_gn=NCOR1;refGene_tx=XM_023244083.1
E1 9773156 9773209
refGene_gn=NCOR1;refGene_tx=XM_023244071.1;refGene_gn=NCOR1;refGene_tx=XM_023244072.1;refGene_gn=NCOR1;refGene_tx=XM_023244073.1;refGene_gn=NCOR1;refGene_tx=XM_023244074.1;refGene_gn=NCOR1;refGene_tx=XM_023244075.1;refGene_gn=NCOR1;refGene_tx=XM_023244076.1;refGene_gn=NCOR1;refGene_tx=XM_023244077.1;refGene_gn=NCOR1;refGene_tx=XM_023244078.1;refGene_gn=NCOR1;refGene_tx=XM_023244079.1;refGene_gn=NCOR1;refGene_tx=XM_023244080.1;refGene_gn=NCOR1;refGene_tx=XM_023244081.1;refGene_gn=NCOR1;refGene_tx=XM_023244082.1;refGene_gn=NCOR1;refGene_tx=XM_023244083.1
E1 9773541 9773568
refGene_gn=NCOR1;refGene_tx=XM_023244071.1;refGene_gn=NCOR1;refGene_tx=XM_023244072.1;refGene_gn=NCOR1;refGene_tx=XM_023244073.1;refGene_gn=NCOR1;refGene_tx=XM_023244076.1;refGene_gn=NCOR1;refGene_tx=XM_023244077.1;refGene_gn=NCOR1;refGene_tx=XM_023244078.1;refGene_gn=NCOR1;refGene_tx=XM_023244079.1;refGene_gn=NCOR1;refGene_tx=XM_023244082.1
E1 9775657 9775724
refGene_gn=NCOR1;refGene_tx=XM_023244071.1;refGene_gn=NCOR1;refGene_tx=XM_023244072.1;refGene_gn=NCOR1;refGene_tx=XM_023244073.1;refGene_gn=NCOR1;refGene_tx=XM_023244074.1;refGene_gn=NCOR1;refGene_tx=XM_023244075.1;refGene_gn=NCOR1;refGene_tx=XM_023244076.1;refGene_gn=NCOR1;refGene_tx=XM_023244077.1;refGene_gn=NCOR1;refGene_tx=XM_023244078.1;refGene_gn=NCOR1;refGene_tx=XM_023244079.1;refGene_gn=NCOR1;refGene_tx=XM_023244080.1;refGene_gn=NCOR1;refGene_tx=XM_023244081.1;refGene_gn=NCOR1;refGene_tx=XM_023244082.1;refGene_gn=NCOR1;refGene_tx=XM_023244083.1
E1 9778108 9778281
refGene_gn=NCOR1;refGene_tx=XM_023244071.1;refGene_gn=NCOR1;refGene_tx=XM_023244072.1;refGene_gn=NCOR1;refGene_tx=XM_023244073.1;refGene_gn=NCOR1;refGene_tx=XM_023244074.1;refGene_gn=NCOR1;refGene_tx=XM_023244075.1;refGene_gn=NCOR1;refGene_tx=XM_023244076.1;refGene_gn=NCOR1;refGene_tx=XM_023244077.1;refGene_gn=NCOR1;refGene_tx=XM_023244078.1;refGene_gn=NCOR1;refGene_tx=XM_023244079.1;refGene_gn=NCOR1;refGene_tx=XM_02

3244080.1;refGene_gn=NCOR1;refGene_tx=XM_023244081.1;refGene_gn=NCOR1;
refGene_tx=XM_023244082.1;refGene_gn=NCOR1;refGene_tx=XM_023244083.1
E1 9779677 9779768
refGene_gn=NCOR1;refGene_tx=XM_023244071.1;refGene_gn=NCOR1;refGene_tx
=XM_023244072.1;refGene_gn=NCOR1;refGene_tx=XM_023244073.1;refGene_gn=
NCOR1;refGene_tx=XM_023244074.1;refGene_gn=NCOR1;refGene_tx=XM_0232440
75.1;refGene_gn=NCOR1;refGene_tx=XM_023244076.1;refGene_gn=NCOR1;refGe
ne_tx=XM_023244077.1;refGene_gn=NCOR1;refGene_tx=XM_023244078.1;refGen
e_gn=NCOR1;refGene_tx=XM_023244079.1;refGene_gn=NCOR1;refGene_tx=XM_02
3244080.1;refGene_gn=NCOR1;refGene_tx=XM_023244081.1;refGene_gn=NCOR1;
refGene_tx=XM_023244082.1;refGene_gn=NCOR1;refGene_tx=XM_023244083.1
E1 9782341 9782520
refGene_gn=NCOR1;refGene_tx=XM_023244071.1;refGene_gn=NCOR1;refGene_tx
=XM_023244072.1;refGene_gn=NCOR1;refGene_tx=XM_023244073.1;refGene_gn=
NCOR1;refGene_tx=XM_023244074.1;refGene_gn=NCOR1;refGene_tx=XM_0232440
75.1;refGene_gn=NCOR1;refGene_tx=XM_023244076.1;refGene_gn=NCOR1;refGe
ne_tx=XM_023244077.1;refGene_gn=NCOR1;refGene_tx=XM_023244078.1;refGen
e_gn=NCOR1;refGene_tx=XM_023244079.1;refGene_gn=NCOR1;refGene_tx=XM_02
3244080.1;refGene_gn=NCOR1;refGene_tx=XM_023244081.1;refGene_gn=NCOR1;
refGene_tx=XM_023244082.1;refGene_gn=NCOR1;refGene_tx=XM_023244083.1
E1 9783060 9783115
refGene_gn=NCOR1;refGene_tx=XM_023244071.1;refGene_gn=NCOR1;refGene_tx
=XM_023244072.1;refGene_gn=NCOR1;refGene_tx=XM_023244073.1;refGene_gn=
NCOR1;refGene_tx=XM_023244074.1;refGene_gn=NCOR1;refGene_tx=XM_0232440
75.1;refGene_gn=NCOR1;refGene_tx=XM_023244076.1;refGene_gn=NCOR1;refGe
ne_tx=XM_023244077.1;refGene_gn=NCOR1;refGene_tx=XM_023244078.1;refGen
e_gn=NCOR1;refGene_tx=XM_023244079.1;refGene_gn=NCOR1;refGene_tx=XM_02
3244080.1;refGene_gn=NCOR1;refGene_tx=XM_023244081.1;refGene_gn=NCOR1;
refGene_tx=XM_023244082.1;refGene_gn=NCOR1;refGene_tx=XM_023244083.1
E1 9783774 9783876
refGene_gn=NCOR1;refGene_tx=XM_023244071.1;refGene_gn=NCOR1;refGene_tx
=XM_023244072.1;refGene_gn=NCOR1;refGene_tx=XM_023244073.1;refGene_gn=
NCOR1;refGene_tx=XM_023244074.1;refGene_gn=NCOR1;refGene_tx=XM_0232440
75.1;refGene_gn=NCOR1;refGene_tx=XM_023244076.1;refGene_gn=NCOR1;refGe
ne_tx=XM_023244077.1;refGene_gn=NCOR1;refGene_tx=XM_023244078.1;refGen
e_gn=NCOR1;refGene_tx=XM_023244079.1;refGene_gn=NCOR1;refGene_tx=XM_02
3244080.1;refGene_gn=NCOR1;refGene_tx=XM_023244081.1;refGene_gn=NCOR1;
refGene_tx=XM_023244082.1;refGene_gn=NCOR1;refGene_tx=XM_023244083.1
E1 9792453 9792581
refGene_gn=NCOR1;refGene_tx=XM_023244071.1;refGene_gn=NCOR1;refGene_tx
=XM_023244072.1;refGene_gn=NCOR1;refGene_tx=XM_023244073.1;refGene_gn=
NCOR1;refGene_tx=XM_023244074.1;refGene_gn=NCOR1;refGene_tx=XM_0232440
75.1;refGene_gn=NCOR1;refGene_tx=XM_023244076.1;refGene_gn=NCOR1;refGe
ne_tx=XM_023244077.1;refGene_gn=NCOR1;refGene_tx=XM_023244078.1;refGen
e_gn=NCOR1;refGene_tx=XM_023244079.1;refGene_gn=NCOR1;refGene_tx=XM_02
3244080.1;refGene_gn=NCOR1;refGene_tx=XM_023244081.1;refGene_gn=NCOR1;
refGene_tx=XM_023244082.1;refGene_gn=NCOR1;refGene_tx=XM_023244083.1
E1 9796932 9797150
refGene_gn=NCOR1;refGene_tx=XM_023244071.1;refGene_gn=NCOR1;refGene_tx
=XM_023244072.1;refGene_gn=NCOR1;refGene_tx=XM_023244073.1;refGene_gn=

NCOR1;refGene_tx=XM_023244074.1;refGene_gn=NCOR1;refGene_tx=XM_0232440
75.1;refGene_gn=NCOR1;refGene_tx=XM_023244076.1;refGene_gn=NCOR1;refGe
ne_tx=XM_023244077.1;refGene_gn=NCOR1;refGene_tx=XM_023244078.1;refGen
e_gn=NCOR1;refGene_tx=XM_023244079.1;refGene_gn=NCOR1;refGene_tx=XM_02
3244080.1;refGene_gn=NCOR1;refGene_tx=XM_023244081.1;refGene_gn=NCOR1;
refGene_tx=XM_023244082.1;refGene_gn=NCOR1;refGene_tx=XM_023244083.1
E1 9800736 9800799
refGene_gn=NCOR1;refGene_tx=XM_023244071.1;refGene_gn=NCOR1;refGene_tx
=XM_023244072.1;refGene_gn=NCOR1;refGene_tx=XM_023244073.1;refGene_gn=
NCOR1;refGene_tx=XM_023244074.1;refGene_gn=NCOR1;refGene_tx=XM_0232440
75.1;refGene_gn=NCOR1;refGene_tx=XM_023244076.1;refGene_gn=NCOR1;refGe
ne_tx=XM_023244077.1;refGene_gn=NCOR1;refGene_tx=XM_023244078.1;refGen
e_gn=NCOR1;refGene_tx=XM_023244079.1;refGene_gn=NCOR1;refGene_tx=XM_02
3244080.1;refGene_gn=NCOR1;refGene_tx=XM_023244081.1;refGene_gn=NCOR1;
refGene_tx=XM_023244082.1;refGene_gn=NCOR1;refGene_tx=XM_023244083.1
E1 9803681 9803821
refGene_gn=NCOR1;refGene_tx=XM_023244071.1;refGene_gn=NCOR1;refGene_tx
=XM_023244072.1;refGene_gn=NCOR1;refGene_tx=XM_023244073.1;refGene_gn=
NCOR1;refGene_tx=XM_023244074.1;refGene_gn=NCOR1;refGene_tx=XM_0232440
75.1;refGene_gn=NCOR1;refGene_tx=XM_023244076.1;refGene_gn=NCOR1;refGe
ne_tx=XM_023244077.1;refGene_gn=NCOR1;refGene_tx=XM_023244078.1;refGen
e_gn=NCOR1;refGene_tx=XM_023244079.1;refGene_gn=NCOR1;refGene_tx=XM_02
3244080.1;refGene_gn=NCOR1;refGene_tx=XM_023244081.1;refGene_gn=NCOR1;
refGene_tx=XM_023244082.1;refGene_gn=NCOR1;refGene_tx=XM_023244083.1
E1 9809084 9809211
refGene_gn=NCOR1;refGene_tx=XM_023244071.1;refGene_gn=NCOR1;refGene_tx
=XM_023244072.1;refGene_gn=NCOR1;refGene_tx=XM_023244073.1;refGene_gn=
NCOR1;refGene_tx=XM_023244074.1;refGene_gn=NCOR1;refGene_tx=XM_0232440
75.1;refGene_gn=NCOR1;refGene_tx=XM_023244076.1;refGene_gn=NCOR1;refGe
ne_tx=XM_023244077.1;refGene_gn=NCOR1;refGene_tx=XM_023244078.1;refGen
e_gn=NCOR1;refGene_tx=XM_023244079.1;refGene_gn=NCOR1;refGene_tx=XM_02
3244080.1;refGene_gn=NCOR1;refGene_tx=XM_023244081.1;refGene_gn=NCOR1;
refGene_tx=XM_023244082.1;refGene_gn=NCOR1;refGene_tx=XM_023244083.1
E1 9816731 9817284
refGene_gn=NCOR1;refGene_tx=XM_023244071.1;refGene_gn=NCOR1;refGene_tx
=XM_023244072.1;refGene_gn=NCOR1;refGene_tx=XM_023244073.1;refGene_gn=
NCOR1;refGene_tx=XM_023244074.1;refGene_gn=NCOR1;refGene_tx=XM_0232440
75.1;refGene_gn=NCOR1;refGene_tx=XM_023244076.1;refGene_gn=NCOR1;refGe
ne_tx=XM_023244077.1;refGene_gn=NCOR1;refGene_tx=XM_023244078.1;refGen
e_gn=NCOR1;refGene_tx=XM_023244079.1;refGene_gn=NCOR1;refGene_tx=XM_02
3244080.1;refGene_gn=NCOR1;refGene_tx=XM_023244081.1;refGene_gn=NCOR1;
refGene_tx=XM_023244082.1;refGene_gn=NCOR1;refGene_tx=XM_023244083.1
E1 9819468 9819598
refGene_gn=NCOR1;refGene_tx=XM_023244071.1;refGene_gn=NCOR1;refGene_tx
=XM_023244072.1;refGene_gn=NCOR1;refGene_tx=XM_023244073.1;refGene_gn=
NCOR1;refGene_tx=XM_023244074.1;refGene_gn=NCOR1;refGene_tx=XM_0232440
75.1;refGene_gn=NCOR1;refGene_tx=XM_023244076.1;refGene_gn=NCOR1;refGe
ne_tx=XM_023244077.1;refGene_gn=NCOR1;refGene_tx=XM_023244078.1;refGen
e_gn=NCOR1;refGene_tx=XM_023244079.1;refGene_gn=NCOR1;refGene_tx=XM_02
3244080.1;refGene_gn=NCOR1;refGene_tx=XM_023244081.1;refGene_gn=NCOR1;

refGene_tx=XM_023244082.1;refGene_gn=NCOR1;refGene_tx=XM_023244083.1
E1 9822652 9822848
refGene_gn=NCOR1;refGene_tx=XM_023244071.1;refGene_gn=NCOR1;refGene_tx=XM_023244072.1;refGene_gn=NCOR1;refGene_tx=XM_023244073.1;refGene_gn=NCOR1;refGene_tx=XM_023244074.1;refGene_gn=NCOR1;refGene_tx=XM_023244075.1;refGene_gn=NCOR1;refGene_tx=XM_023244076.1;refGene_gn=NCOR1;refGene_tx=XM_023244077.1;refGene_gn=NCOR1;refGene_tx=XM_023244078.1;refGene_gn=NCOR1;refGene_tx=XM_023244079.1;refGene_gn=NCOR1;refGene_tx=XM_023244080.1;refGene_gn=NCOR1;refGene_tx=XM_023244081.1;refGene_gn=NCOR1;refGene_tx=XM_023244082.1;refGene_gn=NCOR1;refGene_tx=XM_023244083.1
E1 9829003 9829174
refGene_gn=NCOR1;refGene_tx=XM_023244071.1;refGene_gn=NCOR1;refGene_tx=XM_023244072.1;refGene_gn=NCOR1;refGene_tx=XM_023244073.1;refGene_gn=NCOR1;refGene_tx=XM_023244074.1;refGene_gn=NCOR1;refGene_tx=XM_023244075.1;refGene_gn=NCOR1;refGene_tx=XM_023244076.1;refGene_gn=NCOR1;refGene_tx=XM_023244077.1;refGene_gn=NCOR1;refGene_tx=XM_023244079.1;refGene_gn=NCOR1;refGene_tx=XM_023244082.1
E1 9829913 9830074
refGene_gn=NCOR1;refGene_tx=XM_023244071.1;refGene_gn=NCOR1;refGene_tx=XM_023244072.1;refGene_gn=NCOR1;refGene_tx=XM_023244073.1;refGene_gn=NCOR1;refGene_tx=XM_023244074.1;refGene_gn=NCOR1;refGene_tx=XM_023244075.1;refGene_gn=NCOR1;refGene_tx=XM_023244076.1;refGene_gn=NCOR1;refGene_tx=XM_023244077.1;refGene_gn=NCOR1;refGene_tx=XM_023244078.1;refGene_gn=NCOR1;refGene_tx=XM_023244079.1;refGene_gn=NCOR1;refGene_tx=XM_023244080.1;refGene_gn=NCOR1;refGene_tx=XM_023244081.1;refGene_gn=NCOR1;refGene_tx=XM_023244082.1;refGene_gn=NCOR1;refGene_tx=XM_023244083.1
E1 9833340 9833461
refGene_gn=NCOR1;refGene_tx=XM_023244071.1;refGene_gn=NCOR1;refGene_tx=XM_023244072.1;refGene_gn=NCOR1;refGene_tx=XM_023244073.1;refGene_gn=NCOR1;refGene_tx=XM_023244074.1;refGene_gn=NCOR1;refGene_tx=XM_023244075.1;refGene_gn=NCOR1;refGene_tx=XM_023244076.1;refGene_gn=NCOR1;refGene_tx=XM_023244077.1;refGene_gn=NCOR1;refGene_tx=XM_023244078.1;refGene_gn=NCOR1;refGene_tx=XM_023244079.1;refGene_gn=NCOR1;refGene_tx=XM_023244080.1;refGene_gn=NCOR1;refGene_tx=XM_023244081.1;refGene_gn=NCOR1;refGene_tx=XM_023244082.1;refGene_gn=NCOR1;refGene_tx=XM_023244083.1
E1 9833561 9833663
refGene_gn=NCOR1;refGene_tx=XM_023244071.1;refGene_gn=NCOR1;refGene_tx=XM_023244072.1;refGene_gn=NCOR1;refGene_tx=XM_023244073.1;refGene_gn=NCOR1;refGene_tx=XM_023244074.1;refGene_gn=NCOR1;refGene_tx=XM_023244075.1;refGene_gn=NCOR1;refGene_tx=XM_023244076.1;refGene_gn=NCOR1;refGene_tx=XM_023244077.1;refGene_gn=NCOR1;refGene_tx=XM_023244078.1;refGene_gn=NCOR1;refGene_tx=XM_023244079.1;refGene_gn=NCOR1;refGene_tx=XM_023244080.1;refGene_gn=NCOR1;refGene_tx=XM_023244081.1;refGene_gn=NCOR1;refGene_tx=XM_023244082.1;refGene_gn=NCOR1;refGene_tx=XM_023244083.1
E1 9833997 9834098
refGene_gn=NCOR1;refGene_tx=XM_023244071.1;refGene_gn=NCOR1;refGene_tx=XM_023244072.1;refGene_gn=NCOR1;refGene_tx=XM_023244073.1;refGene_gn=NCOR1;refGene_tx=XM_023244074.1;refGene_gn=NCOR1;refGene_tx=XM_023244075.1;refGene_gn=NCOR1;refGene_tx=XM_023244076.1;refGene_gn=NCOR1;refGene_tx=XM_023244077.1;refGene_gn=NCOR1;refGene_tx=XM_023244078.1;refGen

e_gn=NCOR1;refGene_tx=XM_023244079.1;refGene_gn=NCOR1;refGene_tx=XM_023244080.1;refGene_gn=NCOR1;refGene_tx=XM_023244081.1;refGene_gn=NCOR1;refGene_tx=XM_023244082.1;refGene_gn=NCOR1;refGene_tx=XM_023244083.1
E1 9836292 9836461
refGene_gn=NCOR1;refGene_tx=XM_023244071.1;refGene_gn=NCOR1;refGene_tx=XM_023244072.1;refGene_gn=NCOR1;refGene_tx=XM_023244073.1;refGene_gn=NCOR1;refGene_tx=XM_023244074.1;refGene_gn=NCOR1;refGene_tx=XM_023244075.1;refGene_gn=NCOR1;refGene_tx=XM_023244076.1;refGene_gn=NCOR1;refGene_tx=XM_023244077.1;refGene_gn=NCOR1;refGene_tx=XM_023244078.1;refGene_gn=NCOR1;refGene_tx=XM_023244079.1;refGene_gn=NCOR1;refGene_tx=XM_023244080.1;refGene_gn=NCOR1;refGene_tx=XM_023244081.1;refGene_gn=NCOR1;refGene_tx=XM_023244082.1;refGene_gn=NCOR1;refGene_tx=XM_023244083.1
E1 9838608 9838749
refGene_gn=NCOR1;refGene_tx=XM_023244071.1;refGene_gn=NCOR1;refGene_tx=XM_023244072.1;refGene_gn=NCOR1;refGene_tx=XM_023244073.1;refGene_gn=NCOR1;refGene_tx=XM_023244074.1;refGene_gn=NCOR1;refGene_tx=XM_023244075.1;refGene_gn=NCOR1;refGene_tx=XM_023244076.1;refGene_gn=NCOR1;refGene_tx=XM_023244077.1;refGene_gn=NCOR1;refGene_tx=XM_023244078.1;refGene_gn=NCOR1;refGene_tx=XM_023244079.1;refGene_gn=NCOR1;refGene_tx=XM_023244080.1;refGene_gn=NCOR1;refGene_tx=XM_023244081.1;refGene_gn=NCOR1;refGene_tx=XM_023244082.1;refGene_gn=NCOR1;refGene_tx=XM_023244083.1
E1 9840050 9840134
refGene_gn=NCOR1;refGene_tx=XM_023244071.1;refGene_gn=NCOR1;refGene_tx=XM_023244072.1;refGene_gn=NCOR1;refGene_tx=XM_023244073.1;refGene_gn=NCOR1;refGene_tx=XM_023244074.1;refGene_gn=NCOR1;refGene_tx=XM_023244075.1;refGene_gn=NCOR1;refGene_tx=XM_023244076.1;refGene_gn=NCOR1;refGene_tx=XM_023244077.1;refGene_gn=NCOR1;refGene_tx=XM_023244078.1;refGene_gn=NCOR1;refGene_tx=XM_023244079.1;refGene_gn=NCOR1;refGene_tx=XM_023244080.1;refGene_gn=NCOR1;refGene_tx=XM_023244081.1;refGene_gn=NCOR1;refGene_tx=XM_023244082.1;refGene_gn=NCOR1;refGene_tx=XM_023244083.1
E1 9840609 9840866
refGene_gn=NCOR1;refGene_tx=XM_023244071.1;refGene_gn=NCOR1;refGene_tx=XM_023244072.1;refGene_gn=NCOR1;refGene_tx=XM_023244073.1;refGene_gn=NCOR1;refGene_tx=XM_023244074.1;refGene_gn=NCOR1;refGene_tx=XM_023244075.1;refGene_gn=NCOR1;refGene_tx=XM_023244076.1;refGene_gn=NCOR1;refGene_tx=XM_023244077.1;refGene_gn=NCOR1;refGene_tx=XM_023244078.1;refGene_gn=NCOR1;refGene_tx=XM_023244079.1;refGene_gn=NCOR1;refGene_tx=XM_023244080.1;refGene_gn=NCOR1;refGene_tx=XM_023244081.1;refGene_gn=NCOR1;refGene_tx=XM_023244082.1;refGene_gn=NCOR1;refGene_tx=XM_023244083.1
E1 9841215 9841576
refGene_gn=NCOR1;refGene_tx=XM_023244071.1;refGene_gn=NCOR1;refGene_tx=XM_023244072.1;refGene_gn=NCOR1;refGene_tx=XM_023244073.1;refGene_gn=NCOR1;refGene_tx=XM_023244074.1;refGene_gn=NCOR1;refGene_tx=XM_023244075.1;refGene_gn=NCOR1;refGene_tx=XM_023244076.1;refGene_gn=NCOR1;refGene_tx=XM_023244077.1;refGene_gn=NCOR1;refGene_tx=XM_023244078.1;refGene_gn=NCOR1;refGene_tx=XM_023244079.1;refGene_gn=NCOR1;refGene_tx=XM_023244080.1;refGene_gn=NCOR1;refGene_tx=XM_023244081.1;refGene_gn=NCOR1;refGene_tx=XM_023244082.1;refGene_gn=NCOR1;refGene_tx=XM_023244083.1
E1 9843614 9843842
refGene_gn=NCOR1;refGene_tx=XM_023244071.1;refGene_gn=NCOR1;refGene_tx

=XM_023244072.1;refGene_gn=NCOR1;refGene_tx=XM_023244073.1;refGene_gn=NCOR1;refGene_tx=XM_023244074.1;refGene_gn=NCOR1;refGene_tx=XM_023244075.1;refGene_gn=NCOR1;refGene_tx=XM_023244076.1;refGene_gn=NCOR1;refGene_tx=XM_023244077.1;refGene_gn=NCOR1;refGene_tx=XM_023244078.1;refGene_gn=NCOR1;refGene_tx=XM_023244079.1;refGene_gn=NCOR1;refGene_tx=XM_023244080.1;refGene_gn=NCOR1;refGene_tx=XM_023244081.1;refGene_gn=NCOR1;refGene_tx=XM_023244082.1;refGene_gn=NCOR1;refGene_tx=XM_023244083.1
E1 9846309 9846519
refGene_gn=NCOR1;refGene_tx=XM_023244071.1;refGene_gn=NCOR1;refGene_tx=XM_023244072.1;refGene_gn=NCOR1;refGene_tx=XM_023244073.1;refGene_gn=NCOR1;refGene_tx=XM_023244074.1;refGene_gn=NCOR1;refGene_tx=XM_023244075.1;refGene_gn=NCOR1;refGene_tx=XM_023244076.1;refGene_gn=NCOR1;refGene_tx=XM_023244077.1;refGene_gn=NCOR1;refGene_tx=XM_023244078.1;refGene_gn=NCOR1;refGene_tx=XM_023244079.1;refGene_gn=NCOR1;refGene_tx=XM_023244080.1;refGene_gn=NCOR1;refGene_tx=XM_023244081.1;refGene_gn=NCOR1;refGene_tx=XM_023244082.1;refGene_gn=NCOR1;refGene_tx=XM_023244083.1
E1 9846997 9847147
refGene_gn=NCOR1;refGene_tx=XM_023244071.1;refGene_gn=NCOR1;refGene_tx=XM_023244072.1;refGene_gn=NCOR1;refGene_tx=XM_023244073.1;refGene_gn=NCOR1;refGene_tx=XM_023244074.1;refGene_gn=NCOR1;refGene_tx=XM_023244075.1;refGene_gn=NCOR1;refGene_tx=XM_023244076.1;refGene_gn=NCOR1;refGene_tx=XM_023244077.1;refGene_gn=NCOR1;refGene_tx=XM_023244078.1;refGene_gn=NCOR1;refGene_tx=XM_023244079.1;refGene_gn=NCOR1;refGene_tx=XM_023244080.1;refGene_gn=NCOR1;refGene_tx=XM_023244081.1;refGene_gn=NCOR1;refGene_tx=XM_023244082.1;refGene_gn=NCOR1;refGene_tx=XM_023244083.1
E1 9847475 9847577
refGene_gn=NCOR1;refGene_tx=XM_023244071.1;refGene_gn=NCOR1;refGene_tx=XM_023244072.1;refGene_gn=NCOR1;refGene_tx=XM_023244073.1;refGene_gn=NCOR1;refGene_tx=XM_023244074.1;refGene_gn=NCOR1;refGene_tx=XM_023244075.1;refGene_gn=NCOR1;refGene_tx=XM_023244076.1;refGene_gn=NCOR1;refGene_tx=XM_023244077.1;refGene_gn=NCOR1;refGene_tx=XM_023244078.1;refGene_gn=NCOR1;refGene_tx=XM_023244079.1;refGene_gn=NCOR1;refGene_tx=XM_023244080.1;refGene_gn=NCOR1;refGene_tx=XM_023244081.1;refGene_gn=NCOR1;refGene_tx=XM_023244082.1;refGene_gn=NCOR1;refGene_tx=XM_023244083.1
E1 9850700 9850866
refGene_gn=NCOR1;refGene_tx=XM_023244071.1;refGene_gn=NCOR1;refGene_tx=XM_023244072.1;refGene_gn=NCOR1;refGene_tx=XM_023244073.1;refGene_gn=NCOR1;refGene_tx=XM_023244074.1;refGene_gn=NCOR1;refGene_tx=XM_023244075.1;refGene_gn=NCOR1;refGene_tx=XM_023244076.1;refGene_gn=NCOR1;refGene_tx=XM_023244077.1;refGene_gn=NCOR1;refGene_tx=XM_023244078.1;refGene_gn=NCOR1;refGene_tx=XM_023244079.1;refGene_gn=NCOR1;refGene_tx=XM_023244080.1;refGene_gn=NCOR1;refGene_tx=XM_023244081.1;refGene_gn=NCOR1;refGene_tx=XM_023244082.1;refGene_gn=NCOR1;refGene_tx=XM_023244083.1
E1 9851046 9851540
refGene_gn=NCOR1;refGene_tx=XM_023244071.1;refGene_gn=NCOR1;refGene_tx=XM_023244072.1;refGene_gn=NCOR1;refGene_tx=XM_023244073.1;refGene_gn=NCOR1;refGene_tx=XM_023244074.1;refGene_gn=NCOR1;refGene_tx=XM_023244075.1;refGene_gn=NCOR1;refGene_tx=XM_023244076.1;refGene_gn=NCOR1;refGene_tx=XM_023244077.1;refGene_gn=NCOR1;refGene_tx=XM_023244078.1;refGene_gn=NCOR1;refGene_tx=XM_023244079.1;refGene_gn=NCOR1;refGene_tx=XM_02

3244080.1;refGene_gn=NCOR1;refGene_tx=XM_023244081.1;refGene_gn=NCOR1;
refGene_tx=XM_023244082.1;refGene_gn=NCOR1;refGene_tx=XM_023244083.1
E1 9854285 9854414
refGene_gn=NCOR1;refGene_tx=XM_023244071.1;refGene_gn=NCOR1;refGene_tx
=XM_023244072.1;refGene_gn=NCOR1;refGene_tx=XM_023244073.1;refGene_gn=
NCOR1;refGene_tx=XM_023244074.1;refGene_gn=NCOR1;refGene_tx=XM_0232440
75.1;refGene_gn=NCOR1;refGene_tx=XM_023244076.1;refGene_gn=NCOR1;refGe
ne_tx=XM_023244077.1;refGene_gn=NCOR1;refGene_tx=XM_023244078.1;refGen
e_gn=NCOR1;refGene_tx=XM_023244079.1;refGene_gn=NCOR1;refGene_tx=XM_02
3244080.1;refGene_gn=NCOR1;refGene_tx=XM_023244081.1;refGene_gn=NCOR1;
refGene_tx=XM_023244082.1;refGene_gn=NCOR1;refGene_tx=XM_023244083.1
E1 9854835 9854993
refGene_gn=NCOR1;refGene_tx=XM_023244071.1;refGene_gn=NCOR1;refGene_tx
=XM_023244072.1;refGene_gn=NCOR1;refGene_tx=XM_023244073.1;refGene_gn=
NCOR1;refGene_tx=XM_023244074.1;refGene_gn=NCOR1;refGene_tx=XM_0232440
75.1;refGene_gn=NCOR1;refGene_tx=XM_023244076.1;refGene_gn=NCOR1;refGe
ne_tx=XM_023244077.1;refGene_gn=NCOR1;refGene_tx=XM_023244078.1;refGen
e_gn=NCOR1;refGene_tx=XM_023244079.1;refGene_gn=NCOR1;refGene_tx=XM_02
3244080.1;refGene_gn=NCOR1;refGene_tx=XM_023244081.1;refGene_gn=NCOR1;
refGene_tx=XM_023244082.1;refGene_gn=NCOR1;refGene_tx=XM_023244083.1
E1 9855139 9855363
refGene_gn=NCOR1;refGene_tx=XM_023244071.1;refGene_gn=NCOR1;refGene_tx
=XM_023244072.1;refGene_gn=NCOR1;refGene_tx=XM_023244073.1;refGene_gn=
NCOR1;refGene_tx=XM_023244074.1;refGene_gn=NCOR1;refGene_tx=XM_0232440
75.1;refGene_gn=NCOR1;refGene_tx=XM_023244076.1;refGene_gn=NCOR1;refGe
ne_tx=XM_023244077.1;refGene_gn=NCOR1;refGene_tx=XM_023244078.1;refGen
e_gn=NCOR1;refGene_tx=XM_023244079.1;refGene_gn=NCOR1;refGene_tx=XM_02
3244080.1;refGene_gn=NCOR1;refGene_tx=XM_023244081.1;refGene_gn=NCOR1;
refGene_tx=XM_023244082.1;refGene_gn=NCOR1;refGene_tx=XM_023244083.1
E1 9856608 9856749
refGene_gn=NCOR1;refGene_tx=XM_023244071.1;refGene_gn=NCOR1;refGene_tx
=XM_023244072.1;refGene_gn=NCOR1;refGene_tx=XM_023244073.1;refGene_gn=
NCOR1;refGene_tx=XM_023244074.1;refGene_gn=NCOR1;refGene_tx=XM_0232440
75.1;refGene_gn=NCOR1;refGene_tx=XM_023244076.1;refGene_gn=NCOR1;refGe
ne_tx=XM_023244077.1;refGene_gn=NCOR1;refGene_tx=XM_023244078.1;refGen
e_gn=NCOR1;refGene_tx=XM_023244079.1;refGene_gn=NCOR1;refGene_tx=XM_02
3244080.1;refGene_gn=NCOR1;refGene_tx=XM_023244081.1;refGene_gn=NCOR1;
refGene_tx=XM_023244082.1;refGene_gn=NCOR1;refGene_tx=XM_023244083.1
E1 9858436 9858579
refGene_gn=NCOR1;refGene_tx=XM_023244071.1;refGene_gn=NCOR1;refGene_tx
=XM_023244072.1;refGene_gn=NCOR1;refGene_tx=XM_023244073.1;refGene_gn=
NCOR1;refGene_tx=XM_023244074.1;refGene_gn=NCOR1;refGene_tx=XM_0232440
75.1;refGene_gn=NCOR1;refGene_tx=XM_023244076.1;refGene_gn=NCOR1;refGe
ne_tx=XM_023244077.1;refGene_gn=NCOR1;refGene_tx=XM_023244078.1;refGen
e_gn=NCOR1;refGene_tx=XM_023244079.1;refGene_gn=NCOR1;refGene_tx=XM_02
3244080.1;refGene_gn=NCOR1;refGene_tx=XM_023244081.1;refGene_gn=NCOR1;
refGene_tx=XM_023244082.1;refGene_gn=NCOR1;refGene_tx=XM_023244083.1
E1 9859617 9859797 refGene_gn=TTC19;refGene_tx=XM_023244085.1
E1 9862515 9862569
refGene_gn=NCOR1;refGene_tx=XM_023244071.1;refGene_gn=NCOR1;refGene_tx

=XM_023244072.1;refGene_gn=NCOR1;refGene_tx=XM_023244073.1;refGene_gn=NCOR1;refGene_tx=XM_023244074.1;refGene_gn=NCOR1;refGene_tx=XM_023244075.1;refGene_gn=NCOR1;refGene_tx=XM_023244076.1;refGene_gn=NCOR1;refGene_tx=XM_023244077.1;refGene_gn=NCOR1;refGene_tx=XM_023244078.1;refGene_gn=NCOR1;refGene_tx=XM_023244079.1;refGene_gn=NCOR1;refGene_tx=XM_023244080.1;refGene_gn=NCOR1;refGene_tx=XM_023244081.1;refGene_gn=NCOR1;refGene_tx=XM_023244082.1;refGene_gn=NCOR1;refGene_tx=XM_023244083.1
E1 9863012 9863234
refGene_gn=NCOR1;refGene_tx=XM_023244071.1;refGene_gn=NCOR1;refGene_tx=XM_023244072.1;refGene_gn=NCOR1;refGene_tx=XM_023244073.1;refGene_gn=NCOR1;refGene_tx=XM_023244074.1;refGene_gn=NCOR1;refGene_tx=XM_023244075.1;refGene_gn=NCOR1;refGene_tx=XM_023244076.1;refGene_gn=NCOR1;refGene_tx=XM_023244077.1;refGene_gn=NCOR1;refGene_tx=XM_023244078.1;refGene_gn=NCOR1;refGene_tx=XM_023244079.1;refGene_gn=NCOR1;refGene_tx=XM_023244080.1;refGene_gn=NCOR1;refGene_tx=XM_023244081.1;refGene_gn=NCOR1;refGene_tx=XM_023244082.1;refGene_gn=NCOR1;refGene_tx=XM_023244083.1
E1 9865608 9865788
refGene_gn=NCOR1;refGene_tx=XM_023244071.1;refGene_gn=NCOR1;refGene_tx=XM_023244072.1;refGene_gn=NCOR1;refGene_tx=XM_023244073.1;refGene_gn=NCOR1;refGene_tx=XM_023244074.1;refGene_gn=NCOR1;refGene_tx=XM_023244075.1;refGene_gn=NCOR1;refGene_tx=XM_023244076.1;refGene_gn=NCOR1;refGene_tx=XM_023244077.1;refGene_gn=NCOR1;refGene_tx=XM_023244078.1;refGene_gn=NCOR1;refGene_tx=XM_023244080.1;refGene_gn=NCOR1;refGene_tx=XM_023244081.1;refGene_gn=NCOR1;refGene_tx=XM_023244082.1;refGene_gn=NCOR1;refGene_tx=XM_023244083.1
E1 9867280 9867468
refGene_gn=NCOR1;refGene_tx=XM_023244071.1;refGene_gn=NCOR1;refGene_tx=XM_023244072.1;refGene_gn=NCOR1;refGene_tx=XM_023244073.1;refGene_gn=NCOR1;refGene_tx=XM_023244074.1;refGene_gn=NCOR1;refGene_tx=XM_023244075.1;refGene_gn=NCOR1;refGene_tx=XM_023244076.1;refGene_gn=NCOR1;refGene_tx=XM_023244077.1;refGene_gn=NCOR1;refGene_tx=XM_023244078.1;refGene_gn=NCOR1;refGene_tx=XM_023244079.1;refGene_gn=NCOR1;refGene_tx=XM_023244080.1;refGene_gn=NCOR1;refGene_tx=XM_023244081.1;refGene_gn=NCOR1;refGene_tx=XM_023244082.1;refGene_gn=NCOR1;refGene_tx=XM_023244083.1
E1 9869535 9869625 refGene_gn=TTC19;refGene_tx=XM_023244086.1
E1 9871716 9871850
refGene_gn=TTC19;refGene_tx=XM_023244084.1;refGene_gn=TTC19;refGene_tx=XM_023244087.1
E1 9872362 9872525
refGene_gn=TTC19;refGene_tx=XM_023244084.1;refGene_gn=TTC19;refGene_tx=XM_023244087.1
E1 9873966 9874121
refGene_gn=TTC19;refGene_tx=XM_023244084.1;refGene_gn=TTC19;refGene_tx=XM_023244085.1;refGene_gn=TTC19;refGene_tx=XM_023244086.1;refGene_gn=TTC19;refGene_tx=XM_023244087.1
E1 9894123 9894218
refGene_gn=TTC19;refGene_tx=XM_023244084.1;refGene_gn=TTC19;refGene_tx=XM_023244085.1;refGene_gn=TTC19;refGene_tx=XM_023244086.1;refGene_gn=TTC19;refGene_tx=XM_023244087.1
E1 9896312 9896374

refGene_gn=TTC19;refGene_tx=XM_023244084.1;refGene_gn=TTC19;refGene_tx=XM_023244085.1;refGene_gn=TTC19;refGene_tx=XM_023244086.1;refGene_gn=TTC19;refGene_tx=XM_023244087.1
E1 9896693 9896750 refGene_gn=TTC19;refGene_tx=XM_023244084.1;refGene_gn=TTC19;refGene_tx=XM_023244085.1;refGene_gn=TTC19;refGene_tx=XM_023244086.1;refGene_gn=TTC19;refGene_tx=XM_023244087.1
E1 9897691 9897730 refGene_gn=TTC19;refGene_tx=XM_023244084.1;refGene_gn=TTC19;refGene_tx=XM_023244085.1;refGene_gn=TTC19;refGene_tx=XM_023244086.1;refGene_gn=TTC19;refGene_tx=XM_023244087.1
E1 9898181 9898292 refGene_gn=TTC19;refGene_tx=XM_023244085.1;refGene_gn=TTC19;refGene_tx=XM_023244086.1;refGene_gn=TTC19;refGene_tx=XM_023244087.1
E1 9899998 9900132 refGene_gn=TTC19;refGene_tx=XM_023244084.1;refGene_gn=TTC19;refGene_tx=XM_023244085.1;refGene_gn=TTC19;refGene_tx=XM_023244086.1;refGene_gn=TTC19;refGene_tx=XM_023244087.1
E1 9900217 9900401 refGene_gn=TTC19;refGene_tx=XM_023244084.1;refGene_gn=TTC19;refGene_tx=XM_023244085.1;refGene_gn=TTC19;refGene_tx=XM_023244086.1;refGene_gn=TTC19;refGene_tx=XM_023244087.1
E1 9900659 9900750 refGene_gn=ZSWIM7;refGene_tx=XM_023244089.1;refGene_gn=ZSWIM7;refGene_tx=XM_023244090.1
E1 9903227 9903249 refGene_gn=ZSWIM7;refGene_tx=XM_023244089.1;refGene_gn=ZSWIM7;refGene_tx=XM_023244090.1
E1 9907378 9907481 refGene_gn=ZSWIM7;refGene_tx=XM_023244089.1;refGene_gn=ZSWIM7;refGene_tx=XM_023244090.1
E1 9911665 9911770 refGene_gn=ZSWIM7;refGene_tx=XM_023244089.1;refGene_gn=ZSWIM7;refGene_tx=XM_023244090.1
E1 9912221 9912299 refGene_gn=ZSWIM7;refGene_tx=XM_023244090.1
E1 9913060 9913177 refGene_gn=ZSWIM7;refGene_tx=XM_023244089.1
E1 9915104 9915771 refGene_gn=ADORA2B;refGene_tx=XM_023244088.1
E1 9940270 9940605 refGene_gn=ADORA2B;refGene_tx=XM_023244088.1
E1 9975581 9975671 refGene_gn=SPECC1;refGene_tx=XM_006939885.4;refGene_gn=SPECC1;refGene_tx=XM_019817229.2;refGene_gn=SPECC1;refGene_tx=XM_019817232.2;refGene_gn=SPECC1;refGene_tx=XM_019817233.2;refGene_gn=SPECC1;refGene_tx=XM_023244466.1
E1 9986649 9986709 refGene_gn=SPECC1;refGene_tx=XM_006939885.4;refGene_gn=SPECC1;refGene_tx=XM_019817229.2;refGene_gn=SPECC1;refGene_tx=XM_019817232.2;refGene_gn=SPECC1;refGene_tx=XM_019817233.2;refGene_gn=SPECC1;refGene_tx=XM_023244466.1
E1 9993189 9993306 refGene_gn=SPECC1;refGene_tx=XM_006939885.4;refGene_gn=SPECC1;refGene_

tx=XM_019817229.2;refGene_gn=SPECC1;refGene_tx=XM_019817232.2;refGene_gn=SPECC1;refGene_tx=XM_019817233.2;refGene_gn=SPECC1;refGene_tx=XM_023244466.1
E1 10008759 10008873 refGene_gn=SPECC1;refGene_tx=XM_019817231.2
E1 10034049 10034152
refGene_gn=SPECC1;refGene_tx=XM_006939885.4;refGene_gn=SPECC1;refGene_tx=XM_019817229.2;refGene_gn=SPECC1;refGene_tx=XM_019817231.2;refGene_gn=SPECC1;refGene_tx=XM_019817232.2;refGene_gn=SPECC1;refGene_tx=XM_019817233.2;refGene_gn=SPECC1;refGene_tx=XM_023244466.1
E1 10038644 10038801
refGene_gn=SPECC1;refGene_tx=XM_006939885.4;refGene_gn=SPECC1;refGene_tx=XM_019817229.2;refGene_gn=SPECC1;refGene_tx=XM_019817231.2;refGene_gn=SPECC1;refGene_tx=XM_019817232.2;refGene_gn=SPECC1;refGene_tx=XM_019817233.2;refGene_gn=SPECC1;refGene_tx=XM_023244466.1
E1 10041836 10041918
refGene_gn=SPECC1;refGene_tx=XM_006939885.4;refGene_gn=SPECC1;refGene_tx=XM_019817229.2;refGene_gn=SPECC1;refGene_tx=XM_019817231.2;refGene_gn=SPECC1;refGene_tx=XM_019817232.2;refGene_gn=SPECC1;refGene_tx=XM_019817233.2;refGene_gn=SPECC1;refGene_tx=XM_023244466.1
E1 10044933 10045034
refGene_gn=SPECC1;refGene_tx=XM_006939885.4;refGene_gn=SPECC1;refGene_tx=XM_019817229.2;refGene_gn=SPECC1;refGene_tx=XM_019817231.2;refGene_gn=SPECC1;refGene_tx=XM_019817232.2;refGene_gn=SPECC1;refGene_tx=XM_019817233.2;refGene_gn=SPECC1;refGene_tx=XM_023244466.1
E1 10046205 10046351
refGene_gn=SPECC1;refGene_tx=XM_006939885.4;refGene_gn=SPECC1;refGene_tx=XM_019817229.2;refGene_gn=SPECC1;refGene_tx=XM_019817231.2;refGene_gn=SPECC1;refGene_tx=XM_019817232.2;refGene_gn=SPECC1;refGene_tx=XM_019817233.2;refGene_gn=SPECC1;refGene_tx=XM_023244466.1
E1 10053892 10053914 refGene_gn=SPECC1;refGene_tx=XM_019817234.2
E1 10058910 10059089
refGene_gn=SPECC1;refGene_tx=XM_006939885.4;refGene_gn=SPECC1;refGene_tx=XM_019817229.2;refGene_gn=SPECC1;refGene_tx=XM_019817231.2;refGene_gn=SPECC1;refGene_tx=XM_019817232.2;refGene_gn=SPECC1;refGene_tx=XM_019817233.2;refGene_gn=SPECC1;refGene_tx=XM_019817234.2;refGene_gn=SPECC1;refGene_tx=XM_023244466.1
E1 10059500 10059574
refGene_gn=SPECC1;refGene_tx=XM_006939885.4;refGene_gn=SPECC1;refGene_tx=XM_019817229.2;refGene_gn=SPECC1;refGene_tx=XM_019817231.2;refGene_gn=SPECC1;refGene_tx=XM_019817232.2;refGene_gn=SPECC1;refGene_tx=XM_019817233.2;refGene_gn=SPECC1;refGene_tx=XM_019817234.2;refGene_gn=SPECC1;refGene_tx=XM_023244466.1
E1 10063293 10063501
refGene_gn=SPECC1;refGene_tx=XM_006939885.4;refGene_gn=SPECC1;refGene_tx=XM_019817229.2;refGene_gn=SPECC1;refGene_tx=XM_019817231.2;refGene_gn=SPECC1;refGene_tx=XM_019817232.2;refGene_gn=SPECC1;refGene_tx=XM_019817233.2;refGene_gn=SPECC1;refGene_tx=XM_019817234.2;refGene_gn=SPECC1;refGene_tx=XM_023244466.1
E1 10076606 10078189
refGene_gn=SPECC1;refGene_tx=XM_006939885.4;refGene_gn=SPECC1;refGene_

tx=XM_019817229.2;refGene_gn=SPECC1;refGene_tx=XM_019817231.2;refGene_gn=SPECC1;refGene_tx=XM_019817232.2;refGene_gn=SPECC1;refGene_tx=XM_019817233.2;refGene_gn=SPECC1;refGene_tx=XM_019817234.2;refGene_gn=SPECC1;refGene_tx=XM_023244466.1
E1 10126674 10126714 refGene_gn=SPECC1;refGene_tx=XM_019817233.2
E1 10166482 10166618
refGene_gn=SPECC1;refGene_tx=XM_006939885.4;refGene_gn=SPECC1;refGene_tx=XM_019817229.2;refGene_gn=SPECC1;refGene_tx=XM_019817231.2;refGene_gn=SPECC1;refGene_tx=XM_019817232.2;refGene_gn=SPECC1;refGene_tx=XM_019817234.2;refGene_gn=SPECC1;refGene_tx=XM_023244466.1
E1 10179590 10179737
refGene_gn=SPECC1;refGene_tx=XM_006939885.4;refGene_gn=SPECC1;refGene_tx=XM_019817229.2;refGene_gn=SPECC1;refGene_tx=XM_019817231.2;refGene_gn=SPECC1;refGene_tx=XM_019817232.2;refGene_gn=SPECC1;refGene_tx=XM_019817234.2;refGene_gn=SPECC1;refGene_tx=XM_023244466.1
E1 10300459 10300547 refGene_gn=AKAP10;refGene_tx=XM_003996325.5
E1 10310420 10310468 refGene_gn=AKAP10;refGene_tx=XM_003996325.5
E1 10315773 10315956 refGene_gn=AKAP10;refGene_tx=XM_003996325.5
E1 10320001 10320559 refGene_gn=AKAP10;refGene_tx=XM_003996325.5
E1 10344770 10344869 refGene_gn=AKAP10;refGene_tx=XM_003996325.5
E1 10349901 10349986 refGene_gn=AKAP10;refGene_tx=XM_003996325.5
E1 10350792 10350916 refGene_gn=AKAP10;refGene_tx=XM_003996325.5
E1 10352024 10352161 refGene_gn=AKAP10;refGene_tx=XM_003996325.5
E1 10354639 10354784 refGene_gn=AKAP10;refGene_tx=XM_003996325.5
E1 10356819 10356993 refGene_gn=AKAP10;refGene_tx=XM_003996325.5
E1 10361978 10362088 refGene_gn=AKAP10;refGene_tx=XM_003996325.5
E1 10365429 10365512 refGene_gn=AKAP10;refGene_tx=XM_003996325.5
E1 10369418 10369471 refGene_gn=AKAP10;refGene_tx=XM_003996325.5
E1 10370063 10370159 refGene_gn=AKAP10;refGene_tx=XM_003996325.5
E1 10379447 10379453 refGene_gn=AKAP10;refGene_tx=XM_003996325.5
E1 10401658 10401748 refGene_gn=ULK2;refGene_tx=XM_023244419.1
E1 10402999 10403092 refGene_gn=ULK2;refGene_tx=XM_023244419.1
E1 10404180 10404222 refGene_gn=ULK2;refGene_tx=XM_023244419.1
E1 10414521 10414554 refGene_gn=ULK2;refGene_tx=XM_023244419.1
E1 10414843 10414880 refGene_gn=ULK2;refGene_tx=XM_023244419.1
E1 10419433 10419607 refGene_gn=ULK2;refGene_tx=XM_023244419.1
E1 10420528 10420602 refGene_gn=ULK2;refGene_tx=XM_023244419.1
E1 10422675 10422777 refGene_gn=ULK2;refGene_tx=XM_023244419.1
E1 10423920 10423979 refGene_gn=ULK2;refGene_tx=XM_023244419.1
E1 10424525 10424608 refGene_gn=ULK2;refGene_tx=XM_023244419.1
E1 10428873 10428921 refGene_gn=ULK2;refGene_tx=XM_023244419.1
E1 10429745 10429834 refGene_gn=ULK2;refGene_tx=XM_023244419.1
E1 10435594 10435766 refGene_gn=ULK2;refGene_tx=XM_023244419.1
E1 10450531 10450592 refGene_gn=ULK2;refGene_tx=XM_023244419.1
E1 10457425 10457563 refGene_gn=ULK2;refGene_tx=XM_023244419.1
E1 10461053 10461199 refGene_gn=ULK2;refGene_tx=XM_023244419.1
E1 10463109 10463190 refGene_gn=ULK2;refGene_tx=XM_023244419.1
E1 10465015 10465302 refGene_gn=ULK2;refGene_tx=XM_023244419.1
E1 10466676 10466864 refGene_gn=ULK2;refGene_tx=XM_023244419.1
E1 10467269 10467373 refGene_gn=ULK2;refGene_tx=XM_023244419.1

E1 10475570 10475720 refGene_gn=ULK2;refGene_tx=XM_023244419.1
E1 10477043 10477252 refGene_gn=ULK2;refGene_tx=XM_023244419.1
E1 10479242 10479433 refGene_gn=ULK2;refGene_tx=XM_023244419.1
E1 10479884 10479976 refGene_gn=ULK2;refGene_tx=XM_023244419.1
E1 10479978 10480002 refGene_gn=ULK2;refGene_tx=XM_023244419.1
E1 10480845 10481003 refGene_gn=ULK2;refGene_tx=XM_023244419.1
E1 10483321 10483457 refGene_gn=ULK2;refGene_tx=XM_023244419.1
E1 10484374 10484433 refGene_gn=ULK2;refGene_tx=XM_023244419.1
E1 10516925 10517087 refGene_gn=ALDH3A1;refGene_tx=XM_006939887.2
E1 10518489 10518721 refGene_gn=ALDH3A1;refGene_tx=XM_006939887.2
E1 10519194 10519280 refGene_gn=ALDH3A1;refGene_tx=XM_006939887.2
E1 10519498 10519707 refGene_gn=ALDH3A1;refGene_tx=XM_006939887.2
E1 10520289 10520407 refGene_gn=ALDH3A1;refGene_tx=XM_006939887.2
E1 10521119 10521261 refGene_gn=ALDH3A1;refGene_tx=XM_006939887.2
E1 10521866 10522033 refGene_gn=ALDH3A1;refGene_tx=XM_006939887.2
E1 10522329 10522429 refGene_gn=ALDH3A1;refGene_tx=XM_006939887.2
E1 10522866 10522997 refGene_gn=ALDH3A1;refGene_tx=XM_006939887.2
E1 10523099 10523114 refGene_gn=ALDH3A1;refGene_tx=XM_006939887.2
E1 10544487 10544682
refGene_gn=LOC101084495;refGene_tx=XM_003996361.4;refGene_gn=LOC101084495;refGene_tx=XM_019817235.1;refGene_gn=LOC101084495;refGene_tx=XM_019817236.2;refGene_gn=LOC101084495;refGene_tx=XM_019817237.2
E1 10545904 10546006
refGene_gn=LOC101084495;refGene_tx=XM_003996361.4;refGene_gn=LOC101084495;refGene_tx=XM_019817235.1;refGene_gn=LOC101084495;refGene_tx=XM_019817236.2;refGene_gn=LOC101084495;refGene_tx=XM_019817237.2;refGene_gn=LOC101084495;refGene_tx=XM_019817238.2
E1 10546268 10546337
refGene_gn=LOC101084495;refGene_tx=XM_003996361.4;refGene_gn=LOC101084495;refGene_tx=XM_019817235.1;refGene_gn=LOC101084495;refGene_tx=XM_019817236.2;refGene_gn=LOC101084495;refGene_tx=XM_019817237.2;refGene_gn=LOC101084495;refGene_tx=XM_019817238.2;refGene_gn=LOC101084495;refGene_tx=XM_019817239.1
E1 10547086 10547235
refGene_gn=LOC101084495;refGene_tx=XM_003996361.4;refGene_gn=LOC101084495;refGene_tx=XM_019817235.1;refGene_gn=LOC101084495;refGene_tx=XM_019817236.2;refGene_gn=LOC101084495;refGene_tx=XM_019817237.2;refGene_gn=LOC101084495;refGene_tx=XM_019817238.2;refGene_gn=LOC101084495;refGene_tx=XM_019817239.1
E1 10547951 10547994
refGene_gn=LOC101084495;refGene_tx=XM_003996361.4;refGene_gn=LOC101084495;refGene_tx=XM_019817235.1;refGene_gn=LOC101084495;refGene_tx=XM_019817236.2;refGene_gn=LOC101084495;refGene_tx=XM_019817237.2;refGene_gn=LOC101084495;refGene_tx=XM_019817238.2;refGene_gn=LOC101084495;refGene_tx=XM_019817239.1
E1 10551657 10551702
refGene_gn=LOC101084495;refGene_tx=XM_003996361.4;refGene_gn=LOC101084495;refGene_tx=XM_019817235.1;refGene_gn=LOC101084495;refGene_tx=XM_019817236.2;refGene_gn=LOC101084495;refGene_tx=XM_019817237.2;refGene_gn=LOC101084495;refGene_tx=XM_019817238.2;refGene_gn=LOC101084495;refGen

e_tx=XM_019817239.1
E1 10551935 10552033
refGene_gn=LOC101084495;refGene_tx=XM_003996361.4;refGene_gn=LOC101084495;refGene_tx=XM_019817235.1;refGene_gn=LOC101084495;refGene_tx=XM_019817236.2;refGene_gn=LOC101084495;refGene_tx=XM_019817237.2;refGene_gn=LOC101084495;refGene_tx=XM_019817238.2;refGene_gn=LOC101084495;refGene_tx=XM_019817239.1
E1 10552464 10552562
refGene_gn=LOC101084495;refGene_tx=XM_003996361.4;refGene_gn=LOC101084495;refGene_tx=XM_019817235.1;refGene_gn=LOC101084495;refGene_tx=XM_019817236.2;refGene_gn=LOC101084495;refGene_tx=XM_019817237.2;refGene_gn=LOC101084495;refGene_tx=XM_019817238.2;refGene_gn=LOC101084495;refGene_tx=XM_019817239.1
E1 10553529 10553661
refGene_gn=LOC101084495;refGene_tx=XM_003996361.4;refGene_gn=LOC101084495;refGene_tx=XM_019817235.1;refGene_gn=LOC101084495;refGene_tx=XM_019817236.2;refGene_gn=LOC101084495;refGene_tx=XM_019817237.2;refGene_gn=LOC101084495;refGene_tx=XM_019817238.2;refGene_gn=LOC101084495;refGene_tx=XM_019817239.1
E1 10554389 10554457
refGene_gn=LOC101084495;refGene_tx=XM_003996361.4;refGene_gn=LOC101084495;refGene_tx=XM_019817235.1;refGene_gn=LOC101084495;refGene_tx=XM_019817236.2;refGene_gn=LOC101084495;refGene_tx=XM_019817237.2;refGene_gn=LOC101084495;refGene_tx=XM_019817238.2;refGene_gn=LOC101084495;refGene_tx=XM_019817239.1
E1 10555691 10555800
refGene_gn=LOC101084495;refGene_tx=XM_003996361.4;refGene_gn=LOC101084495;refGene_tx=XM_019817235.1;refGene_gn=LOC101084495;refGene_tx=XM_019817236.2;refGene_gn=LOC101084495;refGene_tx=XM_019817237.2;refGene_gn=LOC101084495;refGene_tx=XM_019817238.2;refGene_gn=LOC101084495;refGene_tx=XM_019817239.1
E1 10556665 10556741
refGene_gn=LOC101084495;refGene_tx=XM_003996361.4;refGene_gn=LOC101084495;refGene_tx=XM_019817235.1;refGene_gn=LOC101084495;refGene_tx=XM_019817236.2;refGene_gn=LOC101084495;refGene_tx=XM_019817237.2;refGene_gn=LOC101084495;refGene_tx=XM_019817238.2;refGene_gn=LOC101084495;refGene_tx=XM_019817239.1
E1 10557163 10557233
refGene_gn=LOC101084495;refGene_tx=XM_003996361.4;refGene_gn=LOC101084495;refGene_tx=XM_019817235.1;refGene_gn=LOC101084495;refGene_tx=XM_019817236.2;refGene_gn=LOC101084495;refGene_tx=XM_019817237.2;refGene_gn=LOC101084495;refGene_tx=XM_019817238.2;refGene_gn=LOC101084495;refGene_tx=XM_019817239.1
E1 10561500 10561633
refGene_gn=LOC101084495;refGene_tx=XM_003996361.4;refGene_gn=LOC101084495;refGene_tx=XM_019817235.1;refGene_gn=LOC101084495;refGene_tx=XM_019817236.2;refGene_gn=LOC101084495;refGene_tx=XM_019817237.2;refGene_gn=LOC101084495;refGene_tx=XM_019817238.2;refGene_gn=LOC101084495;refGene_tx=XM_019817239.1
E1 10561705 10561800

refGene_gn=LOC101084495;refGene_tx=XM_003996361.4;refGene_gn=LOC101084495;refGene_tx=XM_019817235.1;refGene_gn=LOC101084495;refGene_tx=XM_019817236.2;refGene_gn=LOC101084495;refGene_tx=XM_019817237.2;refGene_gn=LOC101084495;refGene_tx=XM_019817238.2;refGene_gn=LOC101084495;refGene_tx=XM_019817239.1
E1 10562688 10562791
refGene_gn=LOC101084495;refGene_tx=XM_003996361.4;refGene_gn=LOC101084495;refGene_tx=XM_019817235.1;refGene_gn=LOC101084495;refGene_tx=XM_019817236.2;refGene_gn=LOC101084495;refGene_tx=XM_019817237.2;refGene_gn=LOC101084495;refGene_tx=XM_019817238.2;refGene_gn=LOC101084495;refGene_tx=XM_019817239.1
E1 10562910 10562918
refGene_gn=LOC101084495;refGene_tx=XM_019817237.2
E1 10562919 10562948
refGene_gn=LOC101084495;refGene_tx=XM_019817236.2
E1 10563767 10563988
refGene_gn=LOC101084495;refGene_tx=XM_003996361.4;refGene_gn=LOC101084495;refGene_tx=XM_019817235.1;refGene_gn=LOC101084495;refGene_tx=XM_019817238.2;refGene_gn=LOC101084495;refGene_tx=XM_019817239.1
E1 10567272 10567325
refGene_gn=ALDH3A2;refGene_tx=XM_003996327.3;refGene_gn=ALDH3A2;refGene_tx=XM_019817242.1
E1 10569394 10569472
refGene_gn=ALDH3A2;refGene_tx=XM_006939888.2;refGene_gn=ALDH3A2;refGene_tx=XM_006939889.3
E1 10570592 10570828
refGene_gn=ALDH3A2;refGene_tx=XM_003996327.3;refGene_gn=ALDH3A2;refGene_tx=XM_006939888.2;refGene_gn=ALDH3A2;refGene_tx=XM_006939889.3
E1 10574368 10574468
refGene_gn=ALDH3A2;refGene_tx=XM_003996327.3;refGene_gn=ALDH3A2;refGene_tx=XM_006939888.2;refGene_gn=ALDH3A2;refGene_tx=XM_006939889.3;refGene_gn=ALDH3A2;refGene_tx=XM_019817242.1
E1 10576121 10576288
refGene_gn=ALDH3A2;refGene_tx=XM_003996327.3;refGene_gn=ALDH3A2;refGene_tx=XM_006939888.2;refGene_gn=ALDH3A2;refGene_tx=XM_006939889.3;refGene_gn=ALDH3A2;refGene_tx=XM_019817242.1
E1 10578278 10578420
refGene_gn=ALDH3A2;refGene_tx=XM_003996327.3;refGene_gn=ALDH3A2;refGene_tx=XM_006939888.2;refGene_gn=ALDH3A2;refGene_tx=XM_006939889.3;refGene_gn=ALDH3A2;refGene_tx=XM_019817242.1
E1 10580250 10580368
refGene_gn=ALDH3A2;refGene_tx=XM_003996327.3;refGene_gn=ALDH3A2;refGene_tx=XM_006939888.2;refGene_gn=ALDH3A2;refGene_tx=XM_006939889.3;refGene_gn=ALDH3A2;refGene_tx=XM_019817242.1
E1 10581978 10582187
refGene_gn=ALDH3A2;refGene_tx=XM_003996327.3;refGene_gn=ALDH3A2;refGene_tx=XM_006939888.2;refGene_gn=ALDH3A2;refGene_tx=XM_006939889.3;refGene_gn=ALDH3A2;refGene_tx=XM_019817242.1
E1 10583798 10583884
refGene_gn=ALDH3A2;refGene_tx=XM_003996327.3;refGene_gn=ALDH3A2;refGen

e_tx=XM_006939888.2;refGene_gn=ALDH3A2;refGene_tx=XM_006939889.3;refGene_gn=ALDH3A2;refGene_tx=XM_019817242.1
E1 10585430 10585662
refGene_gn=ALDH3A2;refGene_tx=XM_003996327.3;refGene_gn=ALDH3A2;refGene_tx=XM_006939888.2;refGene_gn=ALDH3A2;refGene_tx=XM_006939889.3;refGene_gn=ALDH3A2;refGene_tx=XM_019817242.1
E1 10587994 10588147
refGene_gn=ALDH3A2;refGene_tx=XM_003996327.3;refGene_gn=ALDH3A2;refGene_tx=XM_006939888.2;refGene_gn=ALDH3A2;refGene_tx=XM_006939889.3;refGene_gn=ALDH3A2;refGene_tx=XM_019817242.1
E1 10627082 10627312
refGene_gn=LOC101084751;refGene_tx=XM_023243779.1;refGene_gn=LOC101084751;refGene_tx=XM_023243781.1;refGene_gn=LOC101084751;refGene_tx=XM_023243782.1
E1 10627614 10627669
refGene_gn=LOC101084751;refGene_tx=XM_023243779.1;refGene_gn=LOC101084751;refGene_tx=XM_023243781.1;refGene_gn=LOC101084751;refGene_tx=XM_023243782.1
E1 10628054 10628093
refGene_gn=LOC101084751;refGene_tx=XM_023243783.1;refGene_gn=LOC101084751;refGene_tx=XM_023243784.1
E1 10628685 10628780
refGene_gn=LOC101084751;refGene_tx=XM_023243779.1;refGene_gn=LOC101084751;refGene_tx=XM_023243781.1;refGene_gn=LOC101084751;refGene_tx=XM_023243782.1;refGene_gn=LOC101084751;refGene_tx=XM_023243783.1;refGene_gn=LOC101084751;refGene_tx=XM_023243784.1
E1 10628941 10629134
refGene_gn=LOC101084751;refGene_tx=XM_023243781.1;refGene_gn=LOC101084751;refGene_tx=XM_023243782.1;refGene_gn=LOC101084751;refGene_tx=XM_023243783.1
E1 10635429 10635562
refGene_gn=LOC101084751;refGene_tx=XM_023243779.1;refGene_gn=LOC101084751;refGene_tx=XM_023243781.1;refGene_gn=LOC101084751;refGene_tx=XM_023243782.1;refGene_gn=LOC101084751;refGene_tx=XM_023243783.1;refGene_gn=LOC101084751;refGene_tx=XM_023243784.1
E1 10639115 10639185
refGene_gn=LOC101084751;refGene_tx=XM_023243779.1;refGene_gn=LOC101084751;refGene_tx=XM_023243781.1;refGene_gn=LOC101084751;refGene_tx=XM_023243782.1;refGene_gn=LOC101084751;refGene_tx=XM_023243783.1;refGene_gn=LOC101084751;refGene_tx=XM_023243784.1
E1 10644191 10644267
refGene_gn=LOC101084751;refGene_tx=XM_023243779.1;refGene_gn=LOC101084751;refGene_tx=XM_023243781.1;refGene_gn=LOC101084751;refGene_tx=XM_023243782.1;refGene_gn=LOC101084751;refGene_tx=XM_023243783.1;refGene_gn=LOC101084751;refGene_tx=XM_023243784.1
E1 10644828 10644937
refGene_gn=LOC101084751;refGene_tx=XM_023243779.1;refGene_gn=LOC101084751;refGene_tx=XM_023243781.1;refGene_gn=LOC101084751;refGene_tx=XM_023243782.1;refGene_gn=LOC101084751;refGene_tx=XM_023243783.1;refGene_gn=LOC101084751;refGene_tx=XM_023243784.1

E1 10653490 10653558
refGene_gn=LOC101084751;refGene_tx=XM_023243779.1;refGene_gn=LOC101084751;refGene_tx=XM_023243781.1;refGene_gn=LOC101084751;refGene_tx=XM_023243782.1;refGene_gn=LOC101084751;refGene_tx=XM_023243783.1;refGene_gn=LOC101084751;refGene_tx=XM_023243784.1
E1 10654460 10654574
refGene_gn=LOC101084751;refGene_tx=XM_023243779.1;refGene_gn=LOC101084751;refGene_tx=XM_023243781.1;refGene_gn=LOC101084751;refGene_tx=XM_023243782.1;refGene_gn=LOC101084751;refGene_tx=XM_023243783.1;refGene_gn=LOC101084751;refGene_tx=XM_023243784.1
E1 10671490 10671588
refGene_gn=LOC101084751;refGene_tx=XM_023243779.1;refGene_gn=LOC101084751;refGene_tx=XM_023243781.1;refGene_gn=LOC101084751;refGene_tx=XM_023243782.1;refGene_gn=LOC101084751;refGene_tx=XM_023243783.1;refGene_gn=LOC101084751;refGene_tx=XM_023243784.1
E1 10673185 10673283
refGene_gn=LOC101084751;refGene_tx=XM_023243779.1;refGene_gn=LOC101084751;refGene_tx=XM_023243781.1;refGene_gn=LOC101084751;refGene_tx=XM_023243782.1;refGene_gn=LOC101084751;refGene_tx=XM_023243783.1;refGene_gn=LOC101084751;refGene_tx=XM_023243784.1
E1 10674918 10674963
refGene_gn=LOC101084751;refGene_tx=XM_023243779.1;refGene_gn=LOC101084751;refGene_tx=XM_023243781.1;refGene_gn=LOC101084751;refGene_tx=XM_023243782.1;refGene_gn=LOC101084751;refGene_tx=XM_023243783.1;refGene_gn=LOC101084751;refGene_tx=XM_023243784.1
E1 10675524 10675567
refGene_gn=LOC101084751;refGene_tx=XM_023243779.1;refGene_gn=LOC101084751;refGene_tx=XM_023243781.1;refGene_gn=LOC101084751;refGene_tx=XM_023243782.1;refGene_gn=LOC101084751;refGene_tx=XM_023243783.1;refGene_gn=LOC101084751;refGene_tx=XM_023243784.1
E1 10676354 10676503
refGene_gn=LOC101084751;refGene_tx=XM_023243779.1;refGene_gn=LOC101084751;refGene_tx=XM_023243781.1;refGene_gn=LOC101084751;refGene_tx=XM_023243782.1;refGene_gn=LOC101084751;refGene_tx=XM_023243783.1;refGene_gn=LOC101084751;refGene_tx=XM_023243784.1
E1 10679706 10679775
refGene_gn=LOC101084751;refGene_tx=XM_023243779.1;refGene_gn=LOC101084751;refGene_tx=XM_023243781.1;refGene_gn=LOC101084751;refGene_tx=XM_023243782.1;refGene_gn=LOC101084751;refGene_tx=XM_023243783.1;refGene_gn=LOC101084751;refGene_tx=XM_023243784.1
E1 10686248 10686350
refGene_gn=LOC101084751;refGene_tx=XM_023243779.1;refGene_gn=LOC101084751;refGene_tx=XM_023243781.1;refGene_gn=LOC101084751;refGene_tx=XM_023243782.1;refGene_gn=LOC101084751;refGene_tx=XM_023243783.1;refGene_gn=LOC101084751;refGene_tx=XM_023243784.1
E1 10686493 10686652
refGene_gn=LOC101084751;refGene_tx=XM_023243779.1;refGene_gn=LOC101084751;refGene_tx=XM_023243781.1;refGene_gn=LOC101084751;refGene_tx=XM_023243782.1;refGene_gn=LOC101084751;refGene_tx=XM_023243783.1;refGene_gn=LOC101084751;refGene_tx=XM_023243784.1

E1 10689002 10689205
refGene_gn=LOC101085002;refGene_tx=XM_003996362.5
E1 10691078 10691163
refGene_gn=LOC101085002;refGene_tx=XM_003996362.5
E1 10691865 10691960
refGene_gn=LOC101085002;refGene_tx=XM_003996362.5
E1 10703378 10703511
refGene_gn=LOC101085002;refGene_tx=XM_003996362.5
E1 10704039 10704109
refGene_gn=LOC101085002;refGene_tx=XM_003996362.5
E1 10707346 10707422
refGene_gn=LOC101085002;refGene_tx=XM_003996362.5
E1 10707541 10707650
refGene_gn=LOC101085002;refGene_tx=XM_003996362.5
E1 10708920 10708988
refGene_gn=LOC101085002;refGene_tx=XM_003996362.5
E1 10709071 10709185
refGene_gn=LOC101085002;refGene_tx=XM_003996362.5
E1 10709291 10709389
refGene_gn=LOC101085002;refGene_tx=XM_003996362.5
E1 10709599 10709697
refGene_gn=LOC101085002;refGene_tx=XM_003996362.5
E1 10712880 10712925
refGene_gn=LOC101085002;refGene_tx=XM_003996362.5
E1 10714655 10714698
refGene_gn=LOC101085002;refGene_tx=XM_003996362.5
E1 10715545 10715694
refGene_gn=LOC101085002;refGene_tx=XM_003996362.5
E1 10718057 10718126
refGene_gn=LOC101085002;refGene_tx=XM_003996362.5
E1 10719886 10719988
refGene_gn=LOC101085002;refGene_tx=XM_003996362.5
E1 10727089 10727218
refGene_gn=LOC101085002;refGene_tx=XM_003996362.5
E1 10833580 10834051
refGene_gn=RNF112;refGene_tx=XM_023243895.1;refGene_gn=RNF112;refGene_tx=XM_023243896.1;refGene_gn=RNF112;refGene_tx=XM_023243897.1;refGene_gn=RNF112;refGene_tx=XM_023243898.1
E1 10834163 10834238
refGene_gn=RNF112;refGene_tx=XM_023243895.1;refGene_gn=RNF112;refGene_tx=XM_023243896.1;refGene_gn=RNF112;refGene_tx=XM_023243897.1;refGene_gn=RNF112;refGene_tx=XM_023243898.1
E1 10834379 10834433
refGene_gn=RNF112;refGene_tx=XM_023243895.1;refGene_gn=RNF112;refGene_tx=XM_023243896.1;refGene_gn=RNF112;refGene_tx=XM_023243897.1;refGene_gn=RNF112;refGene_tx=XM_023243898.1
E1 10834532 10834699
refGene_gn=RNF112;refGene_tx=XM_023243895.1;refGene_gn=RNF112;refGene_tx=XM_023243896.1;refGene_gn=RNF112;refGene_tx=XM_023243897.1;refGene_gn=RNF112;refGene_tx=XM_023243898.1

E1 10834856 10834974
refGene_gn=RNF112;refGene_tx=XM_023243895.1;refGene_gn=RNF112;refGene_tx=XM_023243896.1;refGene_gn=RNF112;refGene_tx=XM_023243897.1;refGene_gn=RNF112;refGene_tx=XM_023243898.1
E1 10835114 10835189
refGene_gn=RNF112;refGene_tx=XM_023243895.1;refGene_gn=RNF112;refGene_tx=XM_023243896.1;refGene_gn=RNF112;refGene_tx=XM_023243897.1;refGene_gn=RNF112;refGene_tx=XM_023243898.1
E1 10835269 10835326
refGene_gn=RNF112;refGene_tx=XM_023243895.1;refGene_gn=RNF112;refGene_tx=XM_023243896.1;refGene_gn=RNF112;refGene_tx=XM_023243897.1;refGene_gn=RNF112;refGene_tx=XM_023243898.1
E1 10835632 10835683
refGene_gn=RNF112;refGene_tx=XM_023243895.1;refGene_gn=RNF112;refGene_tx=XM_023243896.1;refGene_gn=RNF112;refGene_tx=XM_023243897.1;refGene_gn=RNF112;refGene_tx=XM_023243898.1
E1 10836118 10836223
refGene_gn=RNF112;refGene_tx=XM_023243895.1;refGene_gn=RNF112;refGene_tx=XM_023243896.1;refGene_gn=RNF112;refGene_tx=XM_023243897.1;refGene_gn=RNF112;refGene_tx=XM_023243898.1
E1 10836463 10836592
refGene_gn=RNF112;refGene_tx=XM_023243895.1;refGene_gn=RNF112;refGene_tx=XM_023243896.1;refGene_gn=RNF112;refGene_tx=XM_023243897.1
E1 10836680 10836887
refGene_gn=RNF112;refGene_tx=XM_023243895.1;refGene_gn=RNF112;refGene_tx=XM_023243896.1;refGene_gn=RNF112;refGene_tx=XM_023243897.1;refGene_gn=RNF112;refGene_tx=XM_023243898.1
E1 10837080 10837366
refGene_gn=RNF112;refGene_tx=XM_023243895.1;refGene_gn=RNF112;refGene_tx=XM_023243896.1;refGene_gn=RNF112;refGene_tx=XM_023243897.1;refGene_gn=RNF112;refGene_tx=XM_023243898.1
E1 10838111 10838152
refGene_gn=RNF112;refGene_tx=XM_023243895.1;refGene_gn=RNF112;refGene_tx=XM_023243896.1;refGene_gn=RNF112;refGene_tx=XM_023243897.1;refGene_gn=RNF112;refGene_tx=XM_023243898.1
E1 10838350 10838404
refGene_gn=RNF112;refGene_tx=XM_023243895.1;refGene_gn=RNF112;refGene_tx=XM_023243896.1;refGene_gn=RNF112;refGene_tx=XM_023243897.1;refGene_gn=RNF112;refGene_tx=XM_023243898.1
E1 10862382 10862388 refGene_gn=MFAP4;refGene_tx=XM_006939892.3
E1 10862711 10862790 refGene_gn=MFAP4;refGene_tx=XM_006939892.3
E1 10863030 10863185 refGene_gn=MFAP4;refGene_tx=XM_006939892.3
E1 10863677 10863774 refGene_gn=MFAP4;refGene_tx=XM_006939892.3
E1 10863872 10864055 refGene_gn=MFAP4;refGene_tx=XM_006939892.3
E1 10864443 10864691 refGene_gn=MFAP4;refGene_tx=XM_006939892.3
E1 10865983 10866137
refGene_gn=MAPK7;refGene_tx=XM_006939891.4;refGene_gn=MAPK7;refGene_tx=XM_023243598.1;refGene_gn=MAPK7;refGene_tx=XM_023243599.1;refGene_gn=MAPK7;refGene_tx=XM_023243600.1
E1 10866284 10866418

refGene_gn=MAPK7;refGene_tx=XM_006939891.4;refGene_gn=MAPK7;refGene_tx=XM_023243598.1;refGene_gn=MAPK7;refGene_tx=XM_023243599.1;refGene_gn=MAPK7;refGene_tx=XM_023243600.1
E1 10866833 10867483 refGene_gn=MAPK7;refGene_tx=XM_006939891.4;refGene_gn=MAPK7;refGene_tx=XM_023243598.1;refGene_gn=MAPK7;refGene_tx=XM_023243599.1;refGene_gn=MAPK7;refGene_tx=XM_023243600.1
E1 10867590 10868669 refGene_gn=MAPK7;refGene_tx=XM_006939891.4;refGene_gn=MAPK7;refGene_tx=XM_023243598.1;refGene_gn=MAPK7;refGene_tx=XM_023243599.1;refGene_gn=MAPK7;refGene_tx=XM_023243600.1
E1 10869287 10869453 refGene_gn=MAPK7;refGene_tx=XM_006939891.4;refGene_gn=MAPK7;refGene_tx=XM_023243598.1;refGene_gn=MAPK7;refGene_tx=XM_023243599.1;refGene_gn=MAPK7;refGene_tx=XM_023243600.1
E1 10870356 10870588 refGene_gn=MAPK7;refGene_tx=XM_006939891.4;refGene_gn=MAPK7;refGene_tx=XM_023243598.1;refGene_gn=MAPK7;refGene_tx=XM_023243599.1;refGene_gn=MAPK7;refGene_tx=XM_023243600.1
E1 10892601 10892635 refGene_gn=LOC102901427;refGene_tx=XM_006939893.3
E1 10893425 10893576 refGene_gn=LOC102901427;refGene_tx=XM_006939893.3
E1 10893657 10893917 refGene_gn=LOC102901427;refGene_tx=XM_006939893.3
E1 10894443 10894610 refGene_gn=LOC102901427;refGene_tx=XM_006939893.3
E1 10895235 10895457 refGene_gn=LOC102901427;refGene_tx=XM_006939893.3
E1 10896656 10896719 refGene_gn=B9D1;refGene_tx=XM_003996329.5
E1 10898371 10898440 refGene_gn=B9D1;refGene_tx=XM_003996329.5
E1 10900791 10900903 refGene_gn=B9D1;refGene_tx=XM_003996329.5
E1 10905578 10905675 refGene_gn=B9D1;refGene_tx=XM_003996329.5
E1 10906818 10906881 refGene_gn=B9D1;refGene_tx=XM_003996329.5
E1 10910685 10910753 refGene_gn=B9D1;refGene_tx=XM_003996329.5
E1 10911022 10911165 refGene_gn=B9D1;refGene_tx=XM_003996329.5
E1 10918728 10919210 refGene_gn=EPN2;refGene_tx=XM_023244545.1;refGene_gn=EPN2;refGene_tx=XM_023244546.1;refGene_gn=EPN2;refGene_tx=XM_023244547.1
E1 10921135 10921351 refGene_gn=EPN2;refGene_tx=XM_023244545.1;refGene_gn=EPN2;refGene_tx=XM_023244546.1;refGene_gn=EPN2;refGene_tx=XM_023244547.1
E1 10923370 10923457 refGene_gn=EPN2;refGene_tx=XM_023244545.1;refGene_gn=EPN2;refGene_tx=XM_023244546.1;refGene_gn=EPN2;refGene_tx=XM_023244547.1
E1 10924028 10924202 refGene_gn=EPN2;refGene_tx=XM_023244545.1;refGene_gn=EPN2;refGene_tx=XM_023244546.1;refGene_gn=EPN2;refGene_tx=XM_023244547.1
E1 10936794 10936969 refGene_gn=EPN2;refGene_tx=XM_023244545.1;refGene_gn=EPN2;refGene_tx=X

M_023244546.1;refGene_gn=EPN2;refGene_tx=XM_023244547.1
E1 10937942 10938035
refGene_gn=EPN2;refGene_tx=XM_023244545.1;refGene_gn=EPN2;refGene_tx=XM_023244546.1;refGene_gn=EPN2;refGene_tx=XM_023244547.1
E1 10940120 10940233
refGene_gn=EPN2;refGene_tx=XM_023244545.1;refGene_gn=EPN2;refGene_tx=XM_023244546.1;refGene_gn=EPN2;refGene_tx=XM_023244547.1
E1 10959303 10959474
refGene_gn=EPN2;refGene_tx=XM_023244545.1;refGene_gn=EPN2;refGene_tx=XM_023244546.1
E1 10961627 10962222
refGene_gn=EPN2;refGene_tx=XM_023244545.1;refGene_gn=EPN2;refGene_tx=XM_023244546.1;refGene_gn=EPN2;refGene_tx=XM_023244547.1
E1 11062139 11062217
refGene_gn=GRAP;refGene_tx=XM_003996365.4;refGene_gn=GRAP;refGene_tx=XM_006939895.3
E1 11071469 11071567
refGene_gn=GRAP;refGene_tx=XM_003996365.4;refGene_gn=GRAP;refGene_tx=XM_006939895.3
E1 11075550 11075673
refGene_gn=GRAP;refGene_tx=XM_003996365.4;refGene_gn=GRAP;refGene_tx=XM_006939895.3
E1 11084129 11084298
refGene_gn=GRAP;refGene_tx=XM_003996365.4;refGene_gn=GRAP;refGene_tx=XM_006939895.3
E1 11086105 11086279
refGene_gn=GRAP;refGene_tx=XM_003996365.4;refGene_gn=GRAP;refGene_tx=XM_006939895.3
E1 11086977 11087061
refGene_gn=GRAP;refGene_tx=XM_003996365.4;refGene_gn=GRAP;refGene_tx=XM_006939895.3
E1 11087753 11087860 refGene_gn=SLC5A10;refGene_tx=XM_003996333.2
E1 11087981 11088039 refGene_gn=SLC5A10;refGene_tx=XM_003996333.2
E1 11088426 11088642 refGene_gn=SLC5A10;refGene_tx=XM_003996333.2
E1 11088814 11088983 refGene_gn=SLC5A10;refGene_tx=XM_003996333.2
E1 11092872 11093023 refGene_gn=SLC5A10;refGene_tx=XM_003996333.2
E1 11095332 11095440 refGene_gn=SLC5A10;refGene_tx=XM_003996333.2
E1 11105110 11105632 refGene_gn=FAM83G;refGene_tx=XM_003996334.5
E1 11118369 11118537 refGene_gn=FAM83G;refGene_tx=XM_003996334.5
E1 11125762 11125887 refGene_gn=FAM83G;refGene_tx=XM_003996334.5
E1 11126628 11127886 refGene_gn=FAM83G;refGene_tx=XM_003996334.5
E1 11128499 11128635 refGene_gn=SLC5A10;refGene_tx=XM_003996333.2
E1 11131579 11131990 refGene_gn=FAM83G;refGene_tx=XM_003996334.5
E1 11132162 11132368 refGene_gn=SLC5A10;refGene_tx=XM_003996333.2
E1 11133863 11133944 refGene_gn=SLC5A10;refGene_tx=XM_003996333.2
E1 11134237 11134343 refGene_gn=SLC5A10;refGene_tx=XM_003996333.2
E1 11139072 11139177 refGene_gn=SLC5A10;refGene_tx=XM_003996333.2
E1 11139739 11139799 refGene_gn=SLC5A10;refGene_tx=XM_003996333.2
E1 11140122 11140227 refGene_gn=SLC5A10;refGene_tx=XM_003996333.2
E1 11140485 11140557 refGene_gn=SLC5A10;refGene_tx=XM_003996333.2

E1	11145274	11145388	refGene_gn=SLC5A10;refGene_tx=XM_003996333.2
E1	11170306	11170323	refGene_gn=PRPSAP2;refGene_tx=XM_006939899.4
E1	11171029	11171188
refGene_gn=PRPSAP2;refGene_tx=XM_003996335.5;refGene_gn=PRPSAP2;refGen
e_tx=XM_006939898.4;refGene_gn=PRPSAP2;refGene_tx=XM_006939899.4
E1	11174668	11174815
refGene_gn=PRPSAP2;refGene_tx=XM_003996335.5;refGene_gn=PRPSAP2;refGen
e_tx=XM_006939898.4;refGene_gn=PRPSAP2;refGene_tx=XM_006939899.4
E1	11180089	11180160
refGene_gn=PRPSAP2;refGene_tx=XM_003996335.5;refGene_gn=PRPSAP2;refGen
e_tx=XM_006939898.4;refGene_gn=PRPSAP2;refGene_tx=XM_006939899.4
E1	11186083	11186232
refGene_gn=PRPSAP2;refGene_tx=XM_003996335.5;refGene_gn=PRPSAP2;refGen
e_tx=XM_006939898.4;refGene_gn=PRPSAP2;refGene_tx=XM_006939899.4
E1	11193563	11193619
refGene_gn=PRPSAP2;refGene_tx=XM_003996335.5;refGene_gn=PRPSAP2;refGen
e_tx=XM_006939898.4;refGene_gn=PRPSAP2;refGene_tx=XM_006939899.4
E1	11200723	11200839
refGene_gn=PRPSAP2;refGene_tx=XM_003996335.5;refGene_gn=PRPSAP2;refGen
e_tx=XM_006939898.4;refGene_gn=PRPSAP2;refGene_tx=XM_006939899.4
E1	11204259	11204432
refGene_gn=PRPSAP2;refGene_tx=XM_003996335.5;refGene_gn=PRPSAP2;refGen
e_tx=XM_006939898.4;refGene_gn=PRPSAP2;refGene_tx=XM_006939899.4
E1	11207560	11207627
refGene_gn=PRPSAP2;refGene_tx=XM_003996335.5;refGene_gn=PRPSAP2;refGen
e_tx=XM_006939898.4;refGene_gn=PRPSAP2;refGene_tx=XM_006939899.4
E1	11210175	11210228
refGene_gn=PRPSAP2;refGene_tx=XM_003996335.5;refGene_gn=PRPSAP2;refGen
e_tx=XM_006939898.4;refGene_gn=PRPSAP2;refGene_tx=XM_006939899.4
E1	11211648	11211767
refGene_gn=PRPSAP2;refGene_tx=XM_003996335.5;refGene_gn=PRPSAP2;refGen
e_tx=XM_006939898.4;refGene_gn=PRPSAP2;refGene_tx=XM_006939899.4
E1	11262037	11262133	refGene_gn=SHMT1;refGene_tx=XM_003996366.5
E1	11263456	11263602	refGene_gn=SHMT1;refGene_tx=XM_003996366.5
E1	11266779	11266895	refGene_gn=SHMT1;refGene_tx=XM_003996366.5
E1	11267262	11267423	refGene_gn=SHMT1;refGene_tx=XM_003996366.5
E1	11271741	11271823	refGene_gn=SHMT1;refGene_tx=XM_003996366.5
E1	11272667	11272880	refGene_gn=SHMT1;refGene_tx=XM_003996366.5
E1	11274883	11275000	refGene_gn=SHMT1;refGene_tx=XM_003996366.5
E1	11276494	11276617	refGene_gn=SHMT1;refGene_tx=XM_003996366.5
E1	11278683	11278800	refGene_gn=SHMT1;refGene_tx=XM_003996366.5
E1	11279395	11279506	refGene_gn=SHMT1;refGene_tx=XM_003996366.5
E1	11279849	11280022	refGene_gn=SHMT1;refGene_tx=XM_003996366.5
E1	11285483	11285937	refGene_gn=SMCR8;refGene_tx=XM_003996337.4
E1	11291837	11294200	refGene_gn=SMCR8;refGene_tx=XM_003996337.4
E1	11295187	11295367
refGene_gn=TOP3A;refGene_tx=XM_003996367.5;refGene_gn=TOP3A;refGene_tx
=XM_019817252.2
E1	11297470	11297530
refGene_gn=TOP3A;refGene_tx=XM_003996367.5;refGene_gn=TOP3A;refGene_tx

=XM_019817252.2
E1 11298709 11298783
refGene_gn=TOP3A;refGene_tx=XM_003996367.5;refGene_gn=TOP3A;refGene_tx
=XM_019817252.2
E1 11299595 11299671
refGene_gn=TOP3A;refGene_tx=XM_003996367.5;refGene_gn=TOP3A;refGene_tx
=XM_019817252.2
E1 11300812 11300921
refGene_gn=TOP3A;refGene_tx=XM_003996367.5;refGene_gn=TOP3A;refGene_tx
=XM_019817252.2
E1 11301935 11302079
refGene_gn=TOP3A;refGene_tx=XM_003996367.5;refGene_gn=TOP3A;refGene_tx
=XM_019817252.2
E1 11302232 11302403
refGene_gn=TOP3A;refGene_tx=XM_003996367.5;refGene_gn=TOP3A;refGene_tx
=XM_019817252.2
E1 11302678 11302779
refGene_gn=TOP3A;refGene_tx=XM_003996367.5;refGene_gn=TOP3A;refGene_tx
=XM_019817252.2
E1 11304401 11304476 refGene_gn=TOP3A;refGene_tx=XM_003996367.5
E1 11305788 11305871
refGene_gn=TOP3A;refGene_tx=XM_003996367.5;refGene_gn=TOP3A;refGene_tx
=XM_019817252.2
E1 11306947 11307155
refGene_gn=TOP3A;refGene_tx=XM_003996367.5;refGene_gn=TOP3A;refGene_tx
=XM_019817252.2
E1 11309000 11309186
refGene_gn=TOP3A;refGene_tx=XM_003996367.5;refGene_gn=TOP3A;refGene_tx
=XM_019817252.2
E1 11309370 11309500
refGene_gn=TOP3A;refGene_tx=XM_003996367.5;refGene_gn=TOP3A;refGene_tx
=XM_019817252.2
E1 11312195 11312309
refGene_gn=TOP3A;refGene_tx=XM_003996367.5;refGene_gn=TOP3A;refGene_tx
=XM_019817252.2
E1 11312410 11312576
refGene_gn=TOP3A;refGene_tx=XM_003996367.5;refGene_gn=TOP3A;refGene_tx
=XM_019817252.2
E1 11313760 11313904
refGene_gn=TOP3A;refGene_tx=XM_003996367.5;refGene_gn=TOP3A;refGene_tx
=XM_019817252.2
E1 11316418 11316541
refGene_gn=TOP3A;refGene_tx=XM_003996367.5;refGene_gn=TOP3A;refGene_tx
=XM_019817252.2
E1 11317570 11318214
refGene_gn=TOP3A;refGene_tx=XM_003996367.5;refGene_gn=TOP3A;refGene_tx
=XM_019817252.2
E1 11319572 11319748
refGene_gn=TOP3A;refGene_tx=XM_003996367.5;refGene_gn=TOP3A;refGene_tx
=XM_019817252.2

E1 11327456 11328511
refGene_gn=MIEF2;refGene_tx=XM_006939900.4;refGene_gn=MIEF2;refGene_tx=XM_019817254.2;refGene_gn=MIEF2;refGene_tx=XM_019817257.2;refGene_gn=MIEF2;refGene_tx=XM_019817259.2;refGene_gn=MIEF2;refGene_tx=XM_023243808.1;refGene_gn=MIEF2;refGene_tx=XM_023243809.1;refGene_gn=MIEF2;refGene_tx=XM_023243811.1
E1 11328617 11328722
refGene_gn=MIEF2;refGene_tx=XM_006939900.4;refGene_gn=MIEF2;refGene_tx=XM_019817254.2;refGene_gn=MIEF2;refGene_tx=XM_019817257.2;refGene_gn=MIEF2;refGene_tx=XM_023243808.1;refGene_gn=MIEF2;refGene_tx=XM_023243809.1;refGene_gn=MIEF2;refGene_tx=XM_023243811.1
E1 11329037 11329218
refGene_gn=MIEF2;refGene_tx=XM_006939900.4;refGene_gn=MIEF2;refGene_tx=XM_019817254.2;refGene_gn=MIEF2;refGene_tx=XM_019817257.2;refGene_gn=MIEF2;refGene_tx=XM_019817259.2;refGene_gn=MIEF2;refGene_tx=XM_023243808.1;refGene_gn=MIEF2;refGene_tx=XM_023243809.1;refGene_gn=MIEF2;refGene_tx=XM_023243811.1
E1 11329407 11329761
refGene_gn=MIEF2;refGene_tx=XM_006939900.4;refGene_gn=MIEF2;refGene_tx=XM_019817254.2;refGene_gn=MIEF2;refGene_tx=XM_019817257.2;refGene_gn=MIEF2;refGene_tx=XM_019817259.2;refGene_gn=MIEF2;refGene_tx=XM_023243808.1;refGene_gn=MIEF2;refGene_tx=XM_023243809.1;refGene_gn=MIEF2;refGene_tx=XM_023243811.1
E1 11330709 11330782 refGene_gn=MIEF2;refGene_tx=XM_006939900.4
E1 11333051 11333114
refGene_gn=FLII;refGene_tx=XM_003996339.4;refGene_gn=FLII;refGene_tx=XM_006939906.3
E1 11334793 11334904
refGene_gn=FLII;refGene_tx=XM_003996339.4;refGene_gn=FLII;refGene_tx=XM_006939906.3
E1 11335212 11335284
refGene_gn=FLII;refGene_tx=XM_003996339.4;refGene_gn=FLII;refGene_tx=XM_006939906.3
E1 11336481 11336562
refGene_gn=FLII;refGene_tx=XM_003996339.4;refGene_gn=FLII;refGene_tx=XM_006939906.3
E1 11336860 11336946
refGene_gn=FLII;refGene_tx=XM_003996339.4;refGene_gn=FLII;refGene_tx=XM_006939906.3
E1 11337071 11337233
refGene_gn=FLII;refGene_tx=XM_003996339.4;refGene_gn=FLII;refGene_tx=XM_006939906.3
E1 11337527 11337631
refGene_gn=FLII;refGene_tx=XM_003996339.4;refGene_gn=FLII;refGene_tx=XM_006939906.3
E1 11339424 11339600
refGene_gn=FLII;refGene_tx=XM_003996339.4;refGene_gn=FLII;refGene_tx=XM_006939906.3
E1 11339694 11339852
refGene_gn=FLII;refGene_tx=XM_003996339.4;refGene_gn=FLII;refGene_tx=X

M_006939906.3
E1 11340919 11341004
refGene_gn=FLII;refGene_tx=XM_003996339.4;refGene_gn=FLII;refGene_tx=X
M_006939906.3
E1 11341412 11341560
refGene_gn=FLII;refGene_tx=XM_003996339.4;refGene_gn=FLII;refGene_tx=X
M_006939906.3
E1 11341730 11341867
refGene_gn=FLII;refGene_tx=XM_003996339.4;refGene_gn=FLII;refGene_tx=X
M_006939906.3
E1 11342065 11342278
refGene_gn=FLII;refGene_tx=XM_003996339.4;refGene_gn=FLII;refGene_tx=X
M_006939906.3
E1 11342564 11342744
refGene_gn=FLII;refGene_tx=XM_003996339.4;refGene_gn=FLII;refGene_tx=X
M_006939906.3
E1 11343387 11343470
refGene_gn=FLII;refGene_tx=XM_003996339.4;refGene_gn=FLII;refGene_tx=X
M_006939906.3
E1 11343603 11343678
refGene_gn=FLII;refGene_tx=XM_003996339.4;refGene_gn=FLII;refGene_tx=X
M_006939906.3
E1 11343918 11344002
refGene_gn=FLII;refGene_tx=XM_003996339.4;refGene_gn=FLII;refGene_tx=X
M_006939906.3
E1 11344118 11344290
refGene_gn=FLII;refGene_tx=XM_003996339.4;refGene_gn=FLII;refGene_tx=X
M_006939906.3
E1 11344836 11344941
refGene_gn=FLII;refGene_tx=XM_003996339.4;refGene_gn=FLII;refGene_tx=X
M_006939906.3
E1 11345031 11345223
refGene_gn=FLII;refGene_tx=XM_003996339.4;refGene_gn=FLII;refGene_tx=X
M_006939906.3
E1 11345386 11345575
refGene_gn=FLII;refGene_tx=XM_003996339.4;refGene_gn=FLII;refGene_tx=X
M_006939906.3
E1 11345668 11345808
refGene_gn=FLII;refGene_tx=XM_003996339.4;refGene_gn=FLII;refGene_tx=X
M_006939906.3
E1 11345903 11346150
refGene_gn=FLII;refGene_tx=XM_003996339.4;refGene_gn=FLII;refGene_tx=X
M_006939906.3
E1 11346294 11346449
refGene_gn=FLII;refGene_tx=XM_003996339.4;refGene_gn=FLII;refGene_tx=X
M_006939906.3
E1 11346564 11346625
refGene_gn=FLII;refGene_tx=XM_003996339.4;refGene_gn=FLII;refGene_tx=X
M_006939906.3
E1 11346710 11346839

refGene_gn=FLII;refGene_tx=XM_003996339.4;refGene_gn=FLII;refGene_tx=XM_006939906.3
E1 11346922 11347029
refGene_gn=FLII;refGene_tx=XM_003996339.4;refGene_gn=FLII;refGene_tx=XM_006939906.3
E1 11347122 11347228
refGene_gn=FLII;refGene_tx=XM_003996339.4;refGene_gn=FLII;refGene_tx=XM_006939906.3
E1 11347363 11347429
refGene_gn=FLII;refGene_tx=XM_003996339.4;refGene_gn=FLII;refGene_tx=XM_006939906.3
E1 11347513 11347648
refGene_gn=FLII;refGene_tx=XM_003996339.4;refGene_gn=FLII;refGene_tx=XM_006939906.3
E1 11349196 11349275
refGene_gn=LLGL1;refGene_tx=XM_011288818.3;refGene_gn=LLGL1;refGene_tx=XM_011288819.2;refGene_gn=LLGL1;refGene_tx=XM_019817260.2
E1 11349429 11349550
refGene_gn=LLGL1;refGene_tx=XM_011288818.3;refGene_gn=LLGL1;refGene_tx=XM_011288819.2;refGene_gn=LLGL1;refGene_tx=XM_019817260.2
E1 11349834 11349947
refGene_gn=LLGL1;refGene_tx=XM_011288818.3;refGene_gn=LLGL1;refGene_tx=XM_011288819.2;refGene_gn=LLGL1;refGene_tx=XM_019817260.2
E1 11350144 11350259
refGene_gn=LLGL1;refGene_tx=XM_011288818.3;refGene_gn=LLGL1;refGene_tx=XM_011288819.2;refGene_gn=LLGL1;refGene_tx=XM_019817260.2
E1 11350445 11350710
refGene_gn=LLGL1;refGene_tx=XM_011288818.3;refGene_gn=LLGL1;refGene_tx=XM_011288819.2;refGene_gn=LLGL1;refGene_tx=XM_019817260.2
E1 11351223 11351519
refGene_gn=LLGL1;refGene_tx=XM_011288818.3;refGene_gn=LLGL1;refGene_tx=XM_011288819.2;refGene_gn=LLGL1;refGene_tx=XM_019817260.2
E1 11353465 11353619
refGene_gn=LLGL1;refGene_tx=XM_011288818.3;refGene_gn=LLGL1;refGene_tx=XM_011288819.2;refGene_gn=LLGL1;refGene_tx=XM_019817260.2
E1 11353879 11354027
refGene_gn=LLGL1;refGene_tx=XM_011288818.3;refGene_gn=LLGL1;refGene_tx=XM_011288819.2;refGene_gn=LLGL1;refGene_tx=XM_019817260.2
E1 11354364 11354657
refGene_gn=LLGL1;refGene_tx=XM_011288818.3;refGene_gn=LLGL1;refGene_tx=XM_011288819.2;refGene_gn=LLGL1;refGene_tx=XM_019817260.2
E1 11355365 11355470
refGene_gn=LLGL1;refGene_tx=XM_011288818.3;refGene_gn=LLGL1;refGene_tx=XM_011288819.2;refGene_gn=LLGL1;refGene_tx=XM_019817260.2
E1 11355543 11355697
refGene_gn=LLGL1;refGene_tx=XM_011288818.3;refGene_gn=LLGL1;refGene_tx=XM_011288819.2;refGene_gn=LLGL1;refGene_tx=XM_019817260.2
E1 11356752 11356820
refGene_gn=LLGL1;refGene_tx=XM_011288818.3;refGene_gn=LLGL1;refGene_tx=XM_011288819.2;refGene_gn=LLGL1;refGene_tx=XM_019817260.2

E1 11357015 11357239
refGene_gn=LLGL1;refGene_tx=XM_011288818.3;refGene_gn=LLGL1;refGene_tx
=XM_011288819.2;refGene_gn=LLGL1;refGene_tx=XM_019817260.2
E1 11357324 11357479
refGene_gn=LLGL1;refGene_tx=XM_011288818.3;refGene_gn=LLGL1;refGene_tx
=XM_011288819.2;refGene_gn=LLGL1;refGene_tx=XM_019817260.2
E1 11357633 11357688
refGene_gn=LLGL1;refGene_tx=XM_011288818.3;refGene_gn=LLGL1;refGene_tx
=XM_011288819.2;refGene_gn=LLGL1;refGene_tx=XM_019817260.2
E1 11358012 11358148
refGene_gn=LLGL1;refGene_tx=XM_011288818.3;refGene_gn=LLGL1;refGene_tx
=XM_011288819.2;refGene_gn=LLGL1;refGene_tx=XM_019817260.2
E1 11358233 11358396
refGene_gn=LLGL1;refGene_tx=XM_011288818.3;refGene_gn=LLGL1;refGene_tx
=XM_011288819.2;refGene_gn=LLGL1;refGene_tx=XM_019817260.2
E1 11358475 11358634
refGene_gn=LLGL1;refGene_tx=XM_011288818.3;refGene_gn=LLGL1;refGene_tx
=XM_011288819.2;refGene_gn=LLGL1;refGene_tx=XM_019817260.2
E1 11359589 11359720
refGene_gn=LLGL1;refGene_tx=XM_011288818.3;refGene_gn=LLGL1;refGene_tx
=XM_011288819.2;refGene_gn=LLGL1;refGene_tx=XM_019817260.2
E1 11359807 11359889
refGene_gn=LLGL1;refGene_tx=XM_011288818.3;refGene_gn=LLGL1;refGene_tx
=XM_011288819.2;refGene_gn=LLGL1;refGene_tx=XM_019817260.2
E1 11362043 11362141
refGene_gn=LLGL1;refGene_tx=XM_011288818.3;refGene_gn=LLGL1;refGene_tx
=XM_011288819.2;refGene_gn=LLGL1;refGene_tx=XM_019817260.2
E1 11365611 11365692
refGene_gn=LLGL1;refGene_tx=XM_011288818.3;refGene_gn=LLGL1;refGene_tx
=XM_019817260.2
E1 11376233 11376411 refGene_gn=ALKBH5;refGene_tx=XM_003996341.5
E1 11377310 11377466 refGene_gn=ALKBH5;refGene_tx=XM_003996341.5
E1 11386510 11386591 refGene_gn=ALKBH5;refGene_tx=XM_003996341.5
E1 11394838 11395608 refGene_gn=ALKBH5;refGene_tx=XM_003996341.5
E1 11402095 11402197 refGene_gn=MYO15A;refGene_tx=XM_023243764.1
E1 11405292 11405433 refGene_gn=MYO15A;refGene_tx=XM_023243764.1
E1 11406603 11406737 refGene_gn=MYO15A;refGene_tx=XM_023243764.1
E1 11407064 11407198 refGene_gn=MYO15A;refGene_tx=XM_023243764.1
E1 11409176 11409310 refGene_gn=MYO15A;refGene_tx=XM_023243764.1
E1 11410360 11410392 refGene_gn=MYO15A;refGene_tx=XM_023243764.1
E1 11410394 11410520 refGene_gn=MYO15A;refGene_tx=XM_023243764.1
E1 11411533 11411630 refGene_gn=MYO15A;refGene_tx=XM_023243764.1
E1 11411921 11411999 refGene_gn=MYO15A;refGene_tx=XM_023243764.1
E1 11412252 11412347 refGene_gn=MYO15A;refGene_tx=XM_023243764.1
E1 11413748 11413879 refGene_gn=MYO15A;refGene_tx=XM_023243764.1
E1 11415352 11415435 refGene_gn=MYO15A;refGene_tx=XM_023243764.1
E1 11415683 11415757 refGene_gn=MYO15A;refGene_tx=XM_023243764.1
E1 11416012 11416085 refGene_gn=MYO15A;refGene_tx=XM_023243764.1
E1 11416352 11416425 refGene_gn=MYO15A;refGene_tx=XM_023243764.1
E1 11416699 11416815 refGene_gn=MYO15A;refGene_tx=XM_023243764.1

| | | | |
|---|---|---|---|
| E1 | 11417455 | 11417634 | refGene_gn=MYO15A;refGene_tx=XM_023243764.1 |
| E1 | 11418137 | 11418212 | refGene_gn=MYO15A;refGene_tx=XM_023243764.1 |
| E1 | 11418300 | 11418412 | refGene_gn=MYO15A;refGene_tx=XM_023243764.1 |
| E1 | 11418732 | 11418874 | refGene_gn=MYO15A;refGene_tx=XM_023243764.1 |
| E1 | 11419675 | 11419794 | refGene_gn=MYO15A;refGene_tx=XM_023243764.1 |
| E1 | 11419881 | 11419997 | refGene_gn=MYO15A;refGene_tx=XM_023243764.1 |
| E1 | 11420321 | 11420397 | refGene_gn=MYO15A;refGene_tx=XM_023243764.1 |
| E1 | 11420866 | 11420926 | refGene_gn=MYO15A;refGene_tx=XM_023243764.1 |
| E1 | 11421166 | 11421288 | refGene_gn=MYO15A;refGene_tx=XM_023243764.1 |
| E1 | 11422431 | 11422504 | refGene_gn=MYO15A;refGene_tx=XM_023243764.1 |
| E1 | 11422636 | 11422742 | refGene_gn=MYO15A;refGene_tx=XM_023243764.1 |
| E1 | 11423060 | 11423193 | refGene_gn=MYO15A;refGene_tx=XM_023243764.1 |
| E1 | 11423292 | 11423473 | refGene_gn=MYO15A;refGene_tx=XM_023243764.1 |
| E1 | 11423615 | 11423693 | refGene_gn=MYO15A;refGene_tx=XM_023243764.1 |
| E1 | 11423766 | 11423834 | refGene_gn=MYO15A;refGene_tx=XM_023243764.1 |
| E1 | 11424046 | 11424173 | refGene_gn=MYO15A;refGene_tx=XM_023243764.1 |
| E1 | 11424717 | 11424812 | refGene_gn=MYO15A;refGene_tx=XM_023243764.1 |
| E1 | 11424967 | 11425080 | refGene_gn=MYO15A;refGene_tx=XM_023243764.1 |
| E1 | 11425398 | 11425590 | refGene_gn=MYO15A;refGene_tx=XM_023243764.1 |
| E1 | 11425782 | 11425855 | refGene_gn=MYO15A;refGene_tx=XM_023243764.1 |
| E1 | 11426180 | 11426362 | refGene_gn=MYO15A;refGene_tx=XM_023243764.1 |
| E1 | 11427844 | 11428080 | refGene_gn=MYO15A;refGene_tx=XM_023243764.1 |
| E1 | 11429112 | 11429208 | refGene_gn=MYO15A;refGene_tx=XM_023243764.1 |
| E1 | 11429714 | 11429845 | refGene_gn=MYO15A;refGene_tx=XM_023243764.1 |
| E1 | 11429928 | 11430010 | refGene_gn=MYO15A;refGene_tx=XM_023243764.1 |
| E1 | 11430105 | 11430159 | refGene_gn=MYO15A;refGene_tx=XM_023243764.1 |
| E1 | 11430885 | 11430970 | refGene_gn=MYO15A;refGene_tx=XM_023243764.1 |
| E1 | 11431479 | 11431655 | refGene_gn=MYO15A;refGene_tx=XM_023243764.1 |
| E1 | 11431981 | 11432099 | refGene_gn=MYO15A;refGene_tx=XM_023243764.1 |
| E1 | 11432540 | 11432665 | refGene_gn=MYO15A;refGene_tx=XM_023243764.1 |
| E1 | 11432856 | 11432902 | refGene_gn=MYO15A;refGene_tx=XM_023243764.1 |
| E1 | 11433016 | 11433165 | refGene_gn=MYO15A;refGene_tx=XM_023243764.1 |
| E1 | 11433951 | 11434029 | refGene_gn=MYO15A;refGene_tx=XM_023243764.1 |
| E1 | 11434729 | 11434855 | refGene_gn=MYO15A;refGene_tx=XM_023243764.1 |
| E1 | 11435402 | 11435534 | refGene_gn=MYO15A;refGene_tx=XM_023243764.1 |
| E1 | 11435952 | 11436048 | refGene_gn=MYO15A;refGene_tx=XM_023243764.1 |
| E1 | 11436303 | 11436427 | refGene_gn=MYO15A;refGene_tx=XM_023243764.1 |
| E1 | 11436513 | 11436572 | refGene_gn=MYO15A;refGene_tx=XM_023243764.1 |
| E1 | 11437041 | 11437155 | refGene_gn=MYO15A;refGene_tx=XM_023243764.1 |
| E1 | 11438524 | 11438686 | refGene_gn=MYO15A;refGene_tx=XM_023243764.1 |
| E1 | 11439227 | 11439341 | refGene_gn=MYO15A;refGene_tx=XM_023243764.1 |
| E1 | 11440221 | 11440285 | refGene_gn=MYO15A;refGene_tx=XM_023243764.1 |
| E1 | 11440420 | 11440524 | refGene_gn=MYO15A;refGene_tx=XM_023243764.1 |
| E1 | 11444585 | 11444682 | refGene_gn=MYO15A;refGene_tx=XM_023243764.1 |
| E1 | 11444932 | 11445007 | refGene_gn=MYO15A;refGene_tx=XM_023243764.1 |
| E1 | 11445291 | 11445401 | refGene_gn=MYO15A;refGene_tx=XM_023243764.1 |
| E1 | 11446221 | 11446285 | refGene_gn=MYO15A;refGene_tx=XM_023243764.1 |
| E1 | 11446877 | 11446960 | refGene_gn=MYO15A;refGene_tx=XM_023243764.1 |
| E1 | 11449064 | 11452655 | refGene_gn=MYO15A;refGene_tx=XM_023243764.1 |
| E1 | 11461435 | 11461522 | refGene_gn=DRG2;refGene_tx=XM_003996342.5 |

E1 11461884 11461938 refGene_gn=DRG2;refGene_tx=XM_003996342.5
E1 11465114 11465173 refGene_gn=DRG2;refGene_tx=XM_003996342.5
E1 11465965 11466054 refGene_gn=DRG2;refGene_tx=XM_003996342.5
E1 11467746 11467823 refGene_gn=DRG2;refGene_tx=XM_003996342.5
E1 11468232 11468330 refGene_gn=DRG2;refGene_tx=XM_003996342.5
E1 11469187 11469278 refGene_gn=DRG2;refGene_tx=XM_003996342.5
E1 11469374 11469447 refGene_gn=DRG2;refGene_tx=XM_003996342.5
E1 11470101 11470192 refGene_gn=DRG2;refGene_tx=XM_003996342.5
E1 11472124 11472185 refGene_gn=DRG2;refGene_tx=XM_003996342.5
E1 11472834 11472924 refGene_gn=DRG2;refGene_tx=XM_003996342.5
E1 11477735 11477896 refGene_gn=DRG2;refGene_tx=XM_003996342.5
E1 11481519 11481583 refGene_gn=DRG2;refGene_tx=XM_003996342.5
E1 11493434 11493498 refGene_gn=GID4;refGene_tx=XM_023244619.1
E1 11496337 11496468 refGene_gn=GID4;refGene_tx=XM_023244619.1
E1 11498443 11498545 refGene_gn=GID4;refGene_tx=XM_023244619.1
E1 11508053 11508161 refGene_gn=GID4;refGene_tx=XM_023244619.1
E1 11513732 11513792 refGene_gn=GID4;refGene_tx=XM_023244619.1
E1 11518125 11518222 refGene_gn=ATPAF2;refGene_tx=XM_023244618.1
E1 11519101 11519290 refGene_gn=GID4;refGene_tx=XM_023244619.1
E1 11519924 11520057
refGene_gn=ATPAF2;refGene_tx=XM_003996343.4;refGene_gn=ATPAF2;refGene_tx=XM_019817272.2
E1 11530587 11530632
refGene_gn=ATPAF2;refGene_tx=XM_003996343.4;refGene_gn=ATPAF2;refGene_tx=XM_019817272.2;refGene_gn=ATPAF2;refGene_tx=XM_023244618.1
E1 11530890 11531036
refGene_gn=ATPAF2;refGene_tx=XM_003996343.4;refGene_gn=ATPAF2;refGene_tx=XM_019817272.2;refGene_gn=ATPAF2;refGene_tx=XM_023244618.1
E1 11531548 11531646
refGene_gn=ATPAF2;refGene_tx=XM_003996343.4;refGene_gn=ATPAF2;refGene_tx=XM_019817272.2;refGene_gn=ATPAF2;refGene_tx=XM_023244618.1
E1 11533265 11533346
refGene_gn=ATPAF2;refGene_tx=XM_003996343.4;refGene_gn=ATPAF2;refGene_tx=XM_019817272.2;refGene_gn=ATPAF2;refGene_tx=XM_023244618.1
E1 11534555 11534668
refGene_gn=ATPAF2;refGene_tx=XM_003996343.4;refGene_gn=ATPAF2;refGene_tx=XM_019817272.2;refGene_gn=ATPAF2;refGene_tx=XM_023244618.1
E1 11535192 11535308
refGene_gn=ATPAF2;refGene_tx=XM_003996343.4;refGene_gn=ATPAF2;refGene_tx=XM_019817272.2;refGene_gn=ATPAF2;refGene_tx=XM_023244618.1
E1 11537002 11537212
refGene_gn=ATPAF2;refGene_tx=XM_003996343.4;refGene_gn=ATPAF2;refGene_tx=XM_019817272.2;refGene_gn=ATPAF2;refGene_tx=XM_023244618.1
E1 11538667 11538781
refGene_gn=DRC3;refGene_tx=XM_006939907.4;refGene_gn=DRC3;refGene_tx=XM_019817265.2;refGene_gn=DRC3;refGene_tx=XM_019817267.2;refGene_gn=DRC3;refGene_tx=XM_023244617.1
E1 11539380 11539512
refGene_gn=DRC3;refGene_tx=XM_006939907.4;refGene_gn=DRC3;refGene_tx=XM_019817265.2;refGene_gn=DRC3;refGene_tx=XM_023244617.1

E1 11544274 11544398
refGene_gn=DRC3;refGene_tx=XM_006939907.4;refGene_gn=DRC3;refGene_tx=X
M_019817265.2;refGene_gn=DRC3;refGene_tx=XM_019817267.2;refGene_gn=DRC
3;refGene_tx=XM_023244617.1
E1 11546237 11546308
refGene_gn=DRC3;refGene_tx=XM_006939907.4;refGene_gn=DRC3;refGene_tx=X
M_019817265.2;refGene_gn=DRC3;refGene_tx=XM_019817267.2;refGene_gn=DRC
3;refGene_tx=XM_023244617.1
E1 11548564 11548696
refGene_gn=DRC3;refGene_tx=XM_006939907.4;refGene_gn=DRC3;refGene_tx=X
M_019817265.2;refGene_gn=DRC3;refGene_tx=XM_019817267.2;refGene_gn=DRC
3;refGene_tx=XM_023244617.1
E1 11553541 11553716
refGene_gn=DRC3;refGene_tx=XM_006939907.4;refGene_gn=DRC3;refGene_tx=X
M_019817265.2;refGene_gn=DRC3;refGene_tx=XM_019817267.2;refGene_gn=DRC
3;refGene_tx=XM_023244617.1
E1 11555359 11555472
refGene_gn=DRC3;refGene_tx=XM_006939907.4;refGene_gn=DRC3;refGene_tx=X
M_019817265.2;refGene_gn=DRC3;refGene_tx=XM_019817267.2;refGene_gn=DRC
3;refGene_tx=XM_023244617.1
E1 11556144 11556264
refGene_gn=DRC3;refGene_tx=XM_006939907.4;refGene_gn=DRC3;refGene_tx=X
M_019817265.2;refGene_gn=DRC3;refGene_tx=XM_019817267.2;refGene_gn=DRC
3;refGene_tx=XM_023244617.1
E1 11559448 11559508
refGene_gn=DRC3;refGene_tx=XM_019817265.2;refGene_gn=DRC3;refGene_tx=X
M_019817267.2;refGene_gn=DRC3;refGene_tx=XM_023244617.1
E1 11560637 11560784
refGene_gn=DRC3;refGene_tx=XM_006939907.4;refGene_gn=DRC3;refGene_tx=X
M_019817265.2;refGene_gn=DRC3;refGene_tx=XM_019817267.2;refGene_gn=DRC
3;refGene_tx=XM_023244617.1
E1 11562297 11562464
refGene_gn=DRC3;refGene_tx=XM_006939907.4;refGene_gn=DRC3;refGene_tx=X
M_019817265.2;refGene_gn=DRC3;refGene_tx=XM_019817267.2;refGene_gn=DRC
3;refGene_tx=XM_023244617.1
E1 11565649 11565766
refGene_gn=DRC3;refGene_tx=XM_006939907.4;refGene_gn=DRC3;refGene_tx=X
M_019817265.2;refGene_gn=DRC3;refGene_tx=XM_019817267.2;refGene_gn=DRC
3;refGene_tx=XM_023244617.1
E1 11569217 11569377
refGene_gn=DRC3;refGene_tx=XM_006939907.4;refGene_gn=DRC3;refGene_tx=X
M_019817265.2;refGene_gn=DRC3;refGene_tx=XM_019817267.2;refGene_gn=DRC
3;refGene_tx=XM_023244617.1
E1 11574374 11574426
refGene_gn=TOM1L2;refGene_tx=XM_003996345.4;refGene_gn=TOM1L2;refGene_
tx=XM_006939908.3;refGene_gn=TOM1L2;refGene_tx=XM_006939909.3;refGene_
gn=TOM1L2;refGene_tx=XM_006939912.3;refGene_gn=TOM1L2;refGene_tx=XM_01
9817271.1
E1 11636212 11636297
refGene_gn=TOM1L2;refGene_tx=XM_003996345.4;refGene_gn=TOM1L2;refGene_

tx=XM_006939908.3;refGene_gn=TOM1L2;refGene_tx=XM_006939909.3;refGene_gn=TOM1L2;refGene_tx=XM_006939912.3;refGene_gn=TOM1L2;refGene_tx=XM_019817271.1
E1 11641815 11641826 refGene_gn=TOM1L2;refGene_tx=XM_019817270.2
E1 11643635 11643714
refGene_gn=TOM1L2;refGene_tx=XM_003996345.4;refGene_gn=TOM1L2;refGene_tx=XM_006939908.3;refGene_gn=TOM1L2;refGene_tx=XM_006939909.3;refGene_gn=TOM1L2;refGene_tx=XM_006939912.3;refGene_gn=TOM1L2;refGene_tx=XM_019817270.2;refGene_gn=TOM1L2;refGene_tx=XM_019817271.1
E1 11650135 11650285
refGene_gn=TOM1L2;refGene_tx=XM_003996345.4;refGene_gn=TOM1L2;refGene_tx=XM_006939908.3;refGene_gn=TOM1L2;refGene_tx=XM_006939909.3;refGene_gn=TOM1L2;refGene_tx=XM_006939912.3;refGene_gn=TOM1L2;refGene_tx=XM_019817270.2;refGene_gn=TOM1L2;refGene_tx=XM_019817271.1
E1 11660279 11660414
refGene_gn=TOM1L2;refGene_tx=XM_003996345.4;refGene_gn=TOM1L2;refGene_tx=XM_006939908.3;refGene_gn=TOM1L2;refGene_tx=XM_006939909.3;refGene_gn=TOM1L2;refGene_tx=XM_006939912.3;refGene_gn=TOM1L2;refGene_tx=XM_019817270.2;refGene_gn=TOM1L2;refGene_tx=XM_019817271.1
E1 11661887 11662046
refGene_gn=TOM1L2;refGene_tx=XM_003996345.4;refGene_gn=TOM1L2;refGene_tx=XM_006939908.3;refGene_gn=TOM1L2;refGene_tx=XM_006939909.3;refGene_gn=TOM1L2;refGene_tx=XM_006939912.3;refGene_gn=TOM1L2;refGene_tx=XM_019817270.2
E1 11664740 11664857
refGene_gn=TOM1L2;refGene_tx=XM_003996345.4;refGene_gn=TOM1L2;refGene_tx=XM_006939908.3;refGene_gn=TOM1L2;refGene_tx=XM_006939909.3;refGene_gn=TOM1L2;refGene_tx=XM_006939912.3;refGene_gn=TOM1L2;refGene_tx=XM_019817270.2;refGene_gn=TOM1L2;refGene_tx=XM_019817271.1
E1 11674156 11674290
refGene_gn=TOM1L2;refGene_tx=XM_003996345.4;refGene_gn=TOM1L2;refGene_tx=XM_006939908.3;refGene_gn=TOM1L2;refGene_tx=XM_006939909.3;refGene_gn=TOM1L2;refGene_tx=XM_006939912.3;refGene_gn=TOM1L2;refGene_tx=XM_019817270.2;refGene_gn=TOM1L2;refGene_tx=XM_019817271.1
E1 11676580 11676629
refGene_gn=TOM1L2;refGene_tx=XM_003996345.4;refGene_gn=TOM1L2;refGene_tx=XM_006939908.3;refGene_gn=TOM1L2;refGene_tx=XM_006939909.3;refGene_gn=TOM1L2;refGene_tx=XM_006939912.3;refGene_gn=TOM1L2;refGene_tx=XM_019817270.2;refGene_gn=TOM1L2;refGene_tx=XM_019817271.1
E1 11677422 11677546
refGene_gn=TOM1L2;refGene_tx=XM_003996345.4;refGene_gn=TOM1L2;refGene_tx=XM_006939908.3;refGene_gn=TOM1L2;refGene_tx=XM_006939909.3;refGene_gn=TOM1L2;refGene_tx=XM_006939912.3;refGene_gn=TOM1L2;refGene_tx=XM_019817270.2;refGene_gn=TOM1L2;refGene_tx=XM_019817271.1
E1 11679354 11679472
refGene_gn=TOM1L2;refGene_tx=XM_003996345.4;refGene_gn=TOM1L2;refGene_tx=XM_006939908.3;refGene_gn=TOM1L2;refGene_tx=XM_006939909.3;refGene_gn=TOM1L2;refGene_tx=XM_006939912.3;refGene_gn=TOM1L2;refGene_tx=XM_019817270.2;refGene_gn=TOM1L2;refGene_tx=XM_019817271.1
E1 11680459 11680535

refGene_gn=TOM1L2;refGene_tx=XM_003996345.4;refGene_gn=TOM1L2;refGene_tx=XM_006939908.3;refGene_gn=TOM1L2;refGene_tx=XM_006939909.3;refGene_gn=TOM1L2;refGene_tx=XM_006939912.3;refGene_gn=TOM1L2;refGene_tx=XM_019817270.2;refGene_gn=TOM1L2;refGene_tx=XM_019817271.1
E1 11682929 11682995
refGene_gn=TOM1L2;refGene_tx=XM_006939908.3;refGene_gn=TOM1L2;refGene_tx=XM_006939909.3;refGene_gn=TOM1L2;refGene_tx=XM_019817270.2;refGene_gn=TOM1L2;refGene_tx=XM_019817271.1
E1 11688619 11688679
refGene_gn=TOM1L2;refGene_tx=XM_003996345.4;refGene_gn=TOM1L2;refGene_tx=XM_006939908.3;refGene_gn=TOM1L2;refGene_tx=XM_006939909.3;refGene_gn=TOM1L2;refGene_tx=XM_019817270.2;refGene_gn=TOM1L2;refGene_tx=XM_019817271.1
E1 11690553 11690590
refGene_gn=TOM1L2;refGene_tx=XM_003996345.4;refGene_gn=TOM1L2;refGene_tx=XM_006939908.3;refGene_gn=TOM1L2;refGene_tx=XM_006939909.3;refGene_gn=TOM1L2;refGene_tx=XM_006939912.3;refGene_gn=TOM1L2;refGene_tx=XM_019817270.2;refGene_gn=TOM1L2;refGene_tx=XM_019817271.1
E1 11691651 11691806
refGene_gn=TOM1L2;refGene_tx=XM_003996345.4;refGene_gn=TOM1L2;refGene_tx=XM_006939908.3;refGene_gn=TOM1L2;refGene_tx=XM_006939909.3;refGene_gn=TOM1L2;refGene_tx=XM_006939912.3;refGene_gn=TOM1L2;refGene_tx=XM_019817270.2;refGene_gn=TOM1L2;refGene_tx=XM_019817271.1
E1 11703744 11703835
refGene_gn=SREBF1;refGene_tx=XM_023243911.1;refGene_gn=SREBF1;refGene_tx=XM_023243912.1
E1 11715256 11715275 refGene_gn=SREBF1;refGene_tx=XM_023243913.1
E1 11716786 11717218
refGene_gn=SREBF1;refGene_tx=XM_023243911.1;refGene_gn=SREBF1;refGene_tx=XM_023243912.1;refGene_gn=SREBF1;refGene_tx=XM_023243913.1
E1 11717626 11717814
refGene_gn=SREBF1;refGene_tx=XM_023243911.1;refGene_gn=SREBF1;refGene_tx=XM_023243912.1;refGene_gn=SREBF1;refGene_tx=XM_023243913.1
E1 11717899 11718037
refGene_gn=SREBF1;refGene_tx=XM_023243911.1;refGene_gn=SREBF1;refGene_tx=XM_023243912.1;refGene_gn=SREBF1;refGene_tx=XM_023243913.1
E1 11718117 11718339
refGene_gn=SREBF1;refGene_tx=XM_023243911.1;refGene_gn=SREBF1;refGene_tx=XM_023243912.1;refGene_gn=SREBF1;refGene_tx=XM_023243913.1
E1 11719010 11719125
refGene_gn=SREBF1;refGene_tx=XM_023243911.1;refGene_gn=SREBF1;refGene_tx=XM_023243912.1;refGene_gn=SREBF1;refGene_tx=XM_023243913.1
E1 11719346 11719567
refGene_gn=SREBF1;refGene_tx=XM_023243911.1;refGene_gn=SREBF1;refGene_tx=XM_023243912.1;refGene_gn=SREBF1;refGene_tx=XM_023243913.1
E1 11719821 11720023
refGene_gn=SREBF1;refGene_tx=XM_023243911.1;refGene_gn=SREBF1;refGene_tx=XM_023243912.1;refGene_gn=SREBF1;refGene_tx=XM_023243913.1
E1 11720153 11720332
refGene_gn=SREBF1;refGene_tx=XM_023243911.1;refGene_gn=SREBF1;refGene_

tx=XM_023243912.1;refGene_gn=SREBF1;refGene_tx=XM_023243913.1
E1 11720614 11720876
refGene_gn=SREBF1;refGene_tx=XM_023243911.1;refGene_gn=SREBF1;refGene_
tx=XM_023243912.1;refGene_gn=SREBF1;refGene_tx=XM_023243913.1
E1 11720959 11721126
refGene_gn=SREBF1;refGene_tx=XM_023243911.1;refGene_gn=SREBF1;refGene_
tx=XM_023243912.1;refGene_gn=SREBF1;refGene_tx=XM_023243913.1
E1 11721313 11721482
refGene_gn=SREBF1;refGene_tx=XM_023243911.1;refGene_gn=SREBF1;refGene_
tx=XM_023243912.1;refGene_gn=SREBF1;refGene_tx=XM_023243913.1
E1 11721903 11722012
refGene_gn=SREBF1;refGene_tx=XM_023243911.1;refGene_gn=SREBF1;refGene_
tx=XM_023243912.1;refGene_gn=SREBF1;refGene_tx=XM_023243913.1
E1 11722307 11722417
refGene_gn=SREBF1;refGene_tx=XM_023243911.1;refGene_gn=SREBF1;refGene_
tx=XM_023243912.1;refGene_gn=SREBF1;refGene_tx=XM_023243913.1
E1 11722518 11722651
refGene_gn=SREBF1;refGene_tx=XM_023243911.1;refGene_gn=SREBF1;refGene_
tx=XM_023243912.1;refGene_gn=SREBF1;refGene_tx=XM_023243913.1
E1 11722834 11723000
refGene_gn=SREBF1;refGene_tx=XM_023243911.1;refGene_gn=SREBF1;refGene_
tx=XM_023243912.1;refGene_gn=SREBF1;refGene_tx=XM_023243913.1
E1 11723457 11723658
refGene_gn=SREBF1;refGene_tx=XM_023243911.1;refGene_gn=SREBF1;refGene_
tx=XM_023243912.1;refGene_gn=SREBF1;refGene_tx=XM_023243913.1
E1 11723748 11723857
refGene_gn=SREBF1;refGene_tx=XM_023243911.1;refGene_gn=SREBF1;refGene_
tx=XM_023243912.1;refGene_gn=SREBF1;refGene_tx=XM_023243913.1
E1 11724221 11724451
refGene_gn=SREBF1;refGene_tx=XM_023243911.1;refGene_gn=SREBF1;refGene_
tx=XM_023243912.1;refGene_gn=SREBF1;refGene_tx=XM_023243913.1
E1 11727171 11727186
refGene_gn=RAI1;refGene_tx=XM_003996372.4;refGene_gn=RAI1;refGene_tx=X
M_023243910.1
E1 11727664 11727714
refGene_gn=RAI1;refGene_tx=XM_003996372.4;refGene_gn=RAI1;refGene_tx=X
M_023243910.1
E1 11732340 11732434
refGene_gn=RAI1;refGene_tx=XM_003996372.4;refGene_gn=RAI1;refGene_tx=X
M_023243910.1
E1 11736600 11742123
refGene_gn=RAI1;refGene_tx=XM_003996372.4;refGene_gn=RAI1;refGene_tx=X
M_023243910.1
E1 11834049 11834317
refGene_gn=LOC111557840;refGene_tx=XM_023244507.1
E1 11844618 11846117
refGene_gn=LOC111557840;refGene_tx=XM_023244507.1
E1 11906564 11906627 refGene_gn=PEMT;refGene_tx=XM_006939913.4
E1 11921857 11921965
refGene_gn=PEMT;refGene_tx=XM_006939913.4;refGene_gn=PEMT;refGene_tx=X

M_019817277.2;refGene_gn=PEMT;refGene_tx=XM_019817278.2;refGene_gn=PEMT;refGene_tx=XM_019817279.2
E1 11971863 11971979 refGene_gn=PEMT;refGene_tx=XM_006939913.4;refGene_gn=PEMT;refGene_tx=XM_019817277.2;refGene_gn=PEMT;refGene_tx=XM_019817278.2;refGene_gn=PEMT;refGene_tx=XM_019817279.2
E1 11985522 11985668 refGene_gn=PEMT;refGene_tx=XM_006939913.4;refGene_gn=PEMT;refGene_tx=XM_019817277.2;refGene_gn=PEMT;refGene_tx=XM_019817278.2;refGene_gn=PEMT;refGene_tx=XM_019817279.2
E1 11988209 11988321 refGene_gn=PEMT;refGene_tx=XM_006939913.4;refGene_gn=PEMT;refGene_tx=XM_019817277.2;refGene_gn=PEMT;refGene_tx=XM_019817278.2;refGene_gn=PEMT;refGene_tx=XM_019817279.2
E1 11990783 11990858 refGene_gn=PEMT;refGene_tx=XM_006939913.4;refGene_gn=PEMT;refGene_tx=XM_019817277.2;refGene_gn=PEMT;refGene_tx=XM_019817278.2;refGene_gn=PEMT;refGene_tx=XM_019817279.2
E1 11991066 11991124 refGene_gn=PEMT;refGene_tx=XM_006939913.4;refGene_gn=PEMT;refGene_tx=XM_019817277.2;refGene_gn=PEMT;refGene_tx=XM_019817278.2;refGene_gn=PEMT;refGene_tx=XM_019817279.2
E1 11994307 11994331 refGene_gn=LOC109494219;refGene_tx=XM_023244566.1
E1 11995693 11996395 refGene_gn=LOC109494219;refGene_tx=XM_023244566.1
E1 11996922 11997208 refGene_gn=RASD1;refGene_tx=XM_003996349.5;refGene_gn=RASD1;refGene_tx=XM_011288815.3
E1 11997438 11997992 refGene_gn=RASD1;refGene_tx=XM_003996349.5;refGene_gn=RASD1;refGene_tx=XM_011288815.3
E1 12001608 12001825 refGene_gn=MED9;refGene_tx=XM_011288814.3
E1 12014048 12014275 refGene_gn=MED9;refGene_tx=XM_011288814.3
E1 12119449 12119592 refGene_gn=NT5M;refGene_tx=XM_003996374.4;refGene_gn=NT5M;refGene_tx=XM_023244726.1
E1 12120945 12121078 refGene_gn=NT5M;refGene_tx=XM_003996374.4;refGene_gn=NT5M;refGene_tx=XM_023244726.1
E1 12132011 12132170 refGene_gn=NT5M;refGene_tx=XM_023244728.1
E1 12133820 12134072 refGene_gn=NT5M;refGene_tx=XM_023244727.1
E1 12135551 12135612 refGene_gn=NT5M;refGene_tx=XM_003996374.4;refGene_gn=NT5M;refGene_tx=XM_023244726.1;refGene_gn=NT5M;refGene_tx=XM_023244727.1;refGene_gn=NT5M;refGene_tx=XM_023244728.1
E1 12148241 12148342 refGene_gn=NT5M;refGene_tx=XM_003996374.4;refGene_gn=NT5M;refGene_tx=XM_023244726.1;refGene_gn=NT5M;refGene_tx=XM_023244727.1;refGene_gn=NT5M;refGene_tx=XM_023244728.1

E1 12151821 12152091
refGene_gn=NT5M;refGene_tx=XM_003996374.4;refGene_gn=NT5M;refGene_tx=XM_023244726.1;refGene_gn=NT5M;refGene_tx=XM_023244727.1;refGene_gn=NT5M;refGene_tx=XM_023244728.1
E1 12169637 12169692 refGene_gn=COPS3;refGene_tx=XM_003996352.4
E1 12173275 12173405 refGene_gn=COPS3;refGene_tx=XM_003996352.4
E1 12176465 12176578 refGene_gn=COPS3;refGene_tx=XM_003996352.4
E1 12176670 12176720 refGene_gn=COPS3;refGene_tx=XM_003996352.4
E1 12178485 12178578 refGene_gn=COPS3;refGene_tx=XM_003996352.4
E1 12179999 12180179 refGene_gn=COPS3;refGene_tx=XM_003996352.4
E1 12182959 12183100 refGene_gn=COPS3;refGene_tx=XM_003996352.4
E1 12184707 12184881 refGene_gn=COPS3;refGene_tx=XM_003996352.4
E1 12187779 12187866 refGene_gn=COPS3;refGene_tx=XM_003996352.4
E1 12194753 12194867 refGene_gn=COPS3;refGene_tx=XM_003996352.4
E1 12196021 12196102 refGene_gn=COPS3;refGene_tx=XM_003996352.4
E1 12196402 12196456 refGene_gn=COPS3;refGene_tx=XM_003996352.4
E1 12208239 12208488
refGene_gn=FLCN;refGene_tx=XM_006939916.3;refGene_gn=FLCN;refGene_tx=XM_006939917.3;refGene_gn=FLCN;refGene_tx=XM_006939918.3;refGene_gn=FLCN;refGene_tx=XM_019817281.2;refGene_gn=FLCN;refGene_tx=XM_019817282.2
E1 12209777 12209924
refGene_gn=FLCN;refGene_tx=XM_006939916.3;refGene_gn=FLCN;refGene_tx=XM_006939917.3;refGene_gn=FLCN;refGene_tx=XM_006939918.3;refGene_gn=FLCN;refGene_tx=XM_019817281.2;refGene_gn=FLCN;refGene_tx=XM_019817282.2
E1 12211018 12211240
refGene_gn=FLCN;refGene_tx=XM_006939916.3;refGene_gn=FLCN;refGene_tx=XM_006939917.3;refGene_gn=FLCN;refGene_tx=XM_006939918.3;refGene_gn=FLCN;refGene_tx=XM_019817281.2;refGene_gn=FLCN;refGene_tx=XM_019817282.2
E1 12212442 12212603
refGene_gn=FLCN;refGene_tx=XM_006939916.3;refGene_gn=FLCN;refGene_tx=XM_006939917.3;refGene_gn=FLCN;refGene_tx=XM_006939918.3;refGene_gn=FLCN;refGene_tx=XM_019817281.2;refGene_gn=FLCN;refGene_tx=XM_019817282.2
E1 12214551 12214643
refGene_gn=FLCN;refGene_tx=XM_006939916.3;refGene_gn=FLCN;refGene_tx=XM_006939917.3;refGene_gn=FLCN;refGene_tx=XM_006939918.3;refGene_gn=FLCN;refGene_tx=XM_019817281.2;refGene_gn=FLCN;refGene_tx=XM_019817282.2
E1 12216259 12216450
refGene_gn=FLCN;refGene_tx=XM_006939916.3;refGene_gn=FLCN;refGene_tx=XM_006939917.3;refGene_gn=FLCN;refGene_tx=XM_006939918.3;refGene_gn=FLCN;refGene_tx=XM_019817281.2;refGene_gn=FLCN;refGene_tx=XM_019817282.2
E1 12217940 12218054
refGene_gn=FLCN;refGene_tx=XM_006939916.3;refGene_gn=FLCN;refGene_tx=XM_006939917.3;refGene_gn=FLCN;refGene_tx=XM_006939918.3;refGene_gn=FLCN;refGene_tx=XM_019817281.2;refGene_gn=FLCN;refGene_tx=XM_019817282.2
E1 12218857 12218981
refGene_gn=FLCN;refGene_tx=XM_006939916.3;refGene_gn=FLCN;refGene_tx=XM_006939917.3;refGene_gn=FLCN;refGene_tx=XM_006939918.3;refGene_gn=FLCN;refGene_tx=XM_019817281.2;refGene_gn=FLCN;refGene_tx=XM_019817282.2
E1 12219994 12220126
refGene_gn=FLCN;refGene_tx=XM_006939916.3;refGene_gn=FLCN;refGene_tx=X

M_006939917.3;refGene_gn=FLCN;refGene_tx=XM_006939918.3;refGene_gn=FLCN;refGene_tx=XM_019817281.2;refGene_gn=FLCN;refGene_tx=XM_019817282.2
E1 12220229 12220335
refGene_gn=FLCN;refGene_tx=XM_006939916.3;refGene_gn=FLCN;refGene_tx=XM_006939917.3;refGene_gn=FLCN;refGene_tx=XM_006939918.3;refGene_gn=FLCN;refGene_tx=XM_019817281.2;refGene_gn=FLCN;refGene_tx=XM_019817282.2
E1 12221002 12221204
refGene_gn=FLCN;refGene_tx=XM_006939916.3;refGene_gn=FLCN;refGene_tx=XM_006939917.3;refGene_gn=FLCN;refGene_tx=XM_006939918.3;refGene_gn=FLCN;refGene_tx=XM_019817281.2;refGene_gn=FLCN;refGene_tx=XM_019817282.2
E1 12234847 12235289 refGene_gn=PLD6;refGene_tx=XM_006939920.4
E1 12237778 12238026 refGene_gn=PLD6;refGene_tx=XM_006939920.4
E1 12252088 12252160
refGene_gn=MPRIP;refGene_tx=XM_019817284.2;refGene_gn=MPRIP;refGene_tx=XM_023244645.1;refGene_gn=MPRIP;refGene_tx=XM_023244648.1;refGene_gn=MPRIP;refGene_tx=XM_023244649.1;refGene_gn=MPRIP;refGene_tx=XM_023244650.1
E1 12256070 12256103
refGene_gn=MPRIP;refGene_tx=XM_023244646.1;refGene_gn=MPRIP;refGene_tx=XM_023244647.1;refGene_gn=MPRIP;refGene_tx=XM_023244651.1
E1 12256642 12256728
refGene_gn=MPRIP;refGene_tx=XM_019817284.2;refGene_gn=MPRIP;refGene_tx=XM_023244645.1;refGene_gn=MPRIP;refGene_tx=XM_023244646.1;refGene_gn=MPRIP;refGene_tx=XM_023244647.1;refGene_gn=MPRIP;refGene_tx=XM_023244648.1;refGene_gn=MPRIP;refGene_tx=XM_023244649.1;refGene_gn=MPRIP;refGene_tx=XM_023244650.1;refGene_gn=MPRIP;refGene_tx=XM_023244651.1
E1 12258076 12258239
refGene_gn=MPRIP;refGene_tx=XM_019817284.2;refGene_gn=MPRIP;refGene_tx=XM_023244645.1;refGene_gn=MPRIP;refGene_tx=XM_023244646.1;refGene_gn=MPRIP;refGene_tx=XM_023244647.1;refGene_gn=MPRIP;refGene_tx=XM_023244648.1;refGene_gn=MPRIP;refGene_tx=XM_023244649.1;refGene_gn=MPRIP;refGene_tx=XM_023244650.1;refGene_gn=MPRIP;refGene_tx=XM_023244651.1
E1 12259058 12259145
refGene_gn=MPRIP;refGene_tx=XM_019817284.2;refGene_gn=MPRIP;refGene_tx=XM_023244645.1;refGene_gn=MPRIP;refGene_tx=XM_023244646.1;refGene_gn=MPRIP;refGene_tx=XM_023244647.1;refGene_gn=MPRIP;refGene_tx=XM_023244648.1;refGene_gn=MPRIP;refGene_tx=XM_023244649.1;refGene_gn=MPRIP;refGene_tx=XM_023244650.1;refGene_gn=MPRIP;refGene_tx=XM_023244651.1
E1 12259897 12260017
refGene_gn=MPRIP;refGene_tx=XM_019817284.2;refGene_gn=MPRIP;refGene_tx=XM_023244645.1;refGene_gn=MPRIP;refGene_tx=XM_023244646.1;refGene_gn=MPRIP;refGene_tx=XM_023244647.1;refGene_gn=MPRIP;refGene_tx=XM_023244648.1;refGene_gn=MPRIP;refGene_tx=XM_023244649.1;refGene_gn=MPRIP;refGene_tx=XM_023244650.1;refGene_gn=MPRIP;refGene_tx=XM_023244651.1
E1 12261080 12261240
refGene_gn=MPRIP;refGene_tx=XM_019817284.2;refGene_gn=MPRIP;refGene_tx=XM_023244645.1;refGene_gn=MPRIP;refGene_tx=XM_023244646.1;refGene_gn=MPRIP;refGene_tx=XM_023244647.1;refGene_gn=MPRIP;refGene_tx=XM_023244648.1;refGene_gn=MPRIP;refGene_tx=XM_023244649.1;refGene_gn=MPRIP;refGene_tx=XM_023244650.1;refGene_gn=MPRIP;refGene_tx=XM_023244651.1

E1 12262097 12262215
refGene_gn=MPRIP;refGene_tx=XM_019817284.2;refGene_gn=MPRIP;refGene_tx=XM_023244645.1;refGene_gn=MPRIP;refGene_tx=XM_023244646.1;refGene_gn=MPRIP;refGene_tx=XM_023244647.1;refGene_gn=MPRIP;refGene_tx=XM_023244648.1;refGene_gn=MPRIP;refGene_tx=XM_023244649.1;refGene_gn=MPRIP;refGene_tx=XM_023244650.1;refGene_gn=MPRIP;refGene_tx=XM_023244651.1
E1 12262923 12263071
refGene_gn=MPRIP;refGene_tx=XM_019817284.2;refGene_gn=MPRIP;refGene_tx=XM_023244645.1;refGene_gn=MPRIP;refGene_tx=XM_023244646.1;refGene_gn=MPRIP;refGene_tx=XM_023244647.1;refGene_gn=MPRIP;refGene_tx=XM_023244648.1;refGene_gn=MPRIP;refGene_tx=XM_023244649.1;refGene_gn=MPRIP;refGene_tx=XM_023244650.1;refGene_gn=MPRIP;refGene_tx=XM_023244651.1
E1 12266544 12270414
refGene_gn=MPRIP;refGene_tx=XM_019817284.2;refGene_gn=MPRIP;refGene_tx=XM_023244645.1;refGene_gn=MPRIP;refGene_tx=XM_023244646.1;refGene_gn=MPRIP;refGene_tx=XM_023244647.1;refGene_gn=MPRIP;refGene_tx=XM_023244648.1;refGene_gn=MPRIP;refGene_tx=XM_023244649.1
E1 12273080 12273197
refGene_gn=MPRIP;refGene_tx=XM_019817284.2;refGene_gn=MPRIP;refGene_tx=XM_023244645.1;refGene_gn=MPRIP;refGene_tx=XM_023244646.1;refGene_gn=MPRIP;refGene_tx=XM_023244647.1;refGene_gn=MPRIP;refGene_tx=XM_023244648.1;refGene_gn=MPRIP;refGene_tx=XM_023244649.1;refGene_gn=MPRIP;refGene_tx=XM_023244650.1;refGene_gn=MPRIP;refGene_tx=XM_023244651.1
E1 12275458 12276023
refGene_gn=MPRIP;refGene_tx=XM_019817284.2;refGene_gn=MPRIP;refGene_tx=XM_023244645.1;refGene_gn=MPRIP;refGene_tx=XM_023244646.1;refGene_gn=MPRIP;refGene_tx=XM_023244647.1;refGene_gn=MPRIP;refGene_tx=XM_023244648.1;refGene_gn=MPRIP;refGene_tx=XM_023244649.1;refGene_gn=MPRIP;refGene_tx=XM_023244650.1;refGene_gn=MPRIP;refGene_tx=XM_023244651.1
E1 12279145 12279255
refGene_gn=MPRIP;refGene_tx=XM_019817284.2;refGene_gn=MPRIP;refGene_tx=XM_023244645.1;refGene_gn=MPRIP;refGene_tx=XM_023244646.1;refGene_gn=MPRIP;refGene_tx=XM_023244647.1;refGene_gn=MPRIP;refGene_tx=XM_023244648.1;refGene_gn=MPRIP;refGene_tx=XM_023244649.1;refGene_gn=MPRIP;refGene_tx=XM_023244650.1;refGene_gn=MPRIP;refGene_tx=XM_023244651.1
E1 12283341 12283431
refGene_gn=MPRIP;refGene_tx=XM_019817284.2;refGene_gn=MPRIP;refGene_tx=XM_023244645.1;refGene_gn=MPRIP;refGene_tx=XM_023244646.1;refGene_gn=MPRIP;refGene_tx=XM_023244647.1;refGene_gn=MPRIP;refGene_tx=XM_023244649.1;refGene_gn=MPRIP;refGene_tx=XM_023244650.1;refGene_gn=MPRIP;refGene_tx=XM_023244651.1
E1 12286187 12286256
refGene_gn=MPRIP;refGene_tx=XM_019817284.2;refGene_gn=MPRIP;refGene_tx=XM_023244645.1;refGene_gn=MPRIP;refGene_tx=XM_023244646.1;refGene_gn=MPRIP;refGene_tx=XM_023244647.1;refGene_gn=MPRIP;refGene_tx=XM_023244648.1;refGene_gn=MPRIP;refGene_tx=XM_023244649.1;refGene_gn=MPRIP;refGene_tx=XM_023244650.1;refGene_gn=MPRIP;refGene_tx=XM_023244651.1
E1 12287512 12287569
refGene_gn=MPRIP;refGene_tx=XM_019817284.2;refGene_gn=MPRIP;refGene_tx=XM_023244645.1;refGene_gn=MPRIP;refGene_tx=XM_023244646.1;refGene_gn=

MPRIP;refGene_tx=XM_023244647.1;refGene_gn=MPRIP;refGene_tx=XM_0232446
48.1;refGene_gn=MPRIP;refGene_tx=XM_023244649.1;refGene_gn=MPRIP;refGe
ne_tx=XM_023244650.1;refGene_gn=MPRIP;refGene_tx=XM_023244651.1
E1 12289140 12289254
refGene_gn=MPRIP;refGene_tx=XM_019817284.2;refGene_gn=MPRIP;refGene_tx
=XM_023244645.1;refGene_gn=MPRIP;refGene_tx=XM_023244646.1;refGene_gn=
MPRIP;refGene_tx=XM_023244647.1;refGene_gn=MPRIP;refGene_tx=XM_0232446
48.1;refGene_gn=MPRIP;refGene_tx=XM_023244650.1;refGene_gn=MPRIP;refGe
ne_tx=XM_023244651.1
E1 12290151 12290293
refGene_gn=MPRIP;refGene_tx=XM_019817284.2;refGene_gn=MPRIP;refGene_tx
=XM_023244645.1;refGene_gn=MPRIP;refGene_tx=XM_023244646.1;refGene_gn=
MPRIP;refGene_tx=XM_023244647.1;refGene_gn=MPRIP;refGene_tx=XM_0232446
48.1;refGene_gn=MPRIP;refGene_tx=XM_023244649.1;refGene_gn=MPRIP;refGe
ne_tx=XM_023244650.1;refGene_gn=MPRIP;refGene_tx=XM_023244651.1
E1 12302859 12303019
refGene_gn=MPRIP;refGene_tx=XM_019817284.2;refGene_gn=MPRIP;refGene_tx
=XM_023244645.1;refGene_gn=MPRIP;refGene_tx=XM_023244646.1;refGene_gn=
MPRIP;refGene_tx=XM_023244647.1;refGene_gn=MPRIP;refGene_tx=XM_0232446
48.1;refGene_gn=MPRIP;refGene_tx=XM_023244649.1;refGene_gn=MPRIP;refGe
ne_tx=XM_023244650.1;refGene_gn=MPRIP;refGene_tx=XM_023244651.1
E1 12304104 12304324
refGene_gn=MPRIP;refGene_tx=XM_019817284.2;refGene_gn=MPRIP;refGene_tx
=XM_023244645.1;refGene_gn=MPRIP;refGene_tx=XM_023244646.1;refGene_gn=
MPRIP;refGene_tx=XM_023244647.1;refGene_gn=MPRIP;refGene_tx=XM_0232446
48.1;refGene_gn=MPRIP;refGene_tx=XM_023244649.1;refGene_gn=MPRIP;refGe
ne_tx=XM_023244650.1;refGene_gn=MPRIP;refGene_tx=XM_023244651.1
E1 12310073 12310158
refGene_gn=MPRIP;refGene_tx=XM_019817284.2;refGene_gn=MPRIP;refGene_tx
=XM_023244645.1;refGene_gn=MPRIP;refGene_tx=XM_023244646.1;refGene_gn=
MPRIP;refGene_tx=XM_023244647.1;refGene_gn=MPRIP;refGene_tx=XM_0232446
48.1;refGene_gn=MPRIP;refGene_tx=XM_023244649.1;refGene_gn=MPRIP;refGe
ne_tx=XM_023244650.1;refGene_gn=MPRIP;refGene_tx=XM_023244651.1
E1 12315361 12315513
refGene_gn=MPRIP;refGene_tx=XM_019817284.2;refGene_gn=MPRIP;refGene_tx
=XM_023244645.1;refGene_gn=MPRIP;refGene_tx=XM_023244646.1;refGene_gn=
MPRIP;refGene_tx=XM_023244647.1;refGene_gn=MPRIP;refGene_tx=XM_0232446
48.1;refGene_gn=MPRIP;refGene_tx=XM_023244649.1;refGene_gn=MPRIP;refGe
ne_tx=XM_023244650.1;refGene_gn=MPRIP;refGene_tx=XM_023244651.1
E1 12323842 12324085 refGene_gn=MPRIP;refGene_tx=XM_019817284.2
E1 12364190 12364256
refGene_gn=MPRIP;refGene_tx=XM_023244645.1;refGene_gn=MPRIP;refGene_tx
=XM_023244646.1;refGene_gn=MPRIP;refGene_tx=XM_023244647.1;refGene_gn=
MPRIP;refGene_tx=XM_023244648.1;refGene_gn=MPRIP;refGene_tx=XM_0232446
49.1;refGene_gn=MPRIP;refGene_tx=XM_023244650.1;refGene_gn=MPRIP;refGe
ne_tx=XM_023244651.1
E1 12366096 12366174
refGene_gn=MPRIP;refGene_tx=XM_023244645.1;refGene_gn=MPRIP;refGene_tx
=XM_023244646.1;refGene_gn=MPRIP;refGene_tx=XM_023244647.1;refGene_gn=
MPRIP;refGene_tx=XM_023244648.1;refGene_gn=MPRIP;refGene_tx=XM_0232446

49.1;refGene_gn=MPRIP;refGene_tx=XM_023244650.1;refGene_gn=MPRIP;refGene_tx=XM_023244651.1
E1	12399706	12399829	refGene_gn=MPRIP;refGene_tx=XM_023244645.1;refGene_gn=MPRIP;refGene_tx=XM_023244646.1;refGene_gn=MPRIP;refGene_tx=XM_023244647.1;refGene_gn=MPRIP;refGene_tx=XM_023244648.1;refGene_gn=MPRIP;refGene_tx=XM_023244649.1;refGene_gn=MPRIP;refGene_tx=XM_023244650.1;refGene_gn=MPRIP;refGene_tx=XM_023244651.1
E1	12471837	12471895	refGene_gn=TNFRSF13B;refGene_tx=XM_023244016.1;refGene_gn=TNFRSF13B;refGene_tx=XM_023244017.1
E1	12491563	12491701	refGene_gn=TNFRSF13B;refGene_tx=XM_023244016.1;refGene_gn=TNFRSF13B;refGene_tx=XM_023244017.1
E1	12495721	12495967	refGene_gn=TNFRSF13B;refGene_tx=XM_023244016.1;refGene_gn=TNFRSF13B;refGene_tx=XM_023244017.1
E1	12506124	12506381	refGene_gn=TNFRSF13B;refGene_tx=XM_023244017.1
E1	12506829	12507006	refGene_gn=TNFRSF13B;refGene_tx=XM_023244016.1
E1	12507382	12507633	refGene_gn=TNFRSF13B;refGene_tx=XM_023244016.1
E1	12521603	12521646	refGene_gn=USP22;refGene_tx=XM_023244640.1
E1	12522403	12522553	refGene_gn=USP22;refGene_tx=XM_023244640.1
E1	12523468	12523531	refGene_gn=USP22;refGene_tx=XM_023244640.1
E1	12524342	12524434	refGene_gn=USP22;refGene_tx=XM_023244640.1
E1	12525564	12525691	refGene_gn=USP22;refGene_tx=XM_023244640.1
E1	12530177	12530336	refGene_gn=USP22;refGene_tx=XM_023244640.1
E1	12533439	12533545	refGene_gn=USP22;refGene_tx=XM_023244640.1
E1	12535716	12535864	refGene_gn=USP22;refGene_tx=XM_023244640.1
E1	12536884	12537054	refGene_gn=USP22;refGene_tx=XM_023244640.1
E1	12537755	12537857	refGene_gn=USP22;refGene_tx=XM_023244640.1
E1	12539651	12539765	refGene_gn=USP22;refGene_tx=XM_023244640.1
E1	12545728	12545861	refGene_gn=USP22;refGene_tx=XM_023244640.1
E1	12563557	12563728	refGene_gn=USP22;refGene_tx=XM_023244640.1
E1	12606591	12606611	refGene_gn=DHRS7B;refGene_tx=XM_006939923.3
E1	12649291	12649470	refGene_gn=DHRS7B;refGene_tx=XM_006939923.3;refGene_gn=DHRS7B;refGene_tx=XM_006939924.4;refGene_gn=DHRS7B;refGene_tx=XM_019817294.2;refGene_gn=DHRS7B;refGene_tx=XM_019817295.2;refGene_gn=DHRS7B;refGene_tx=XM_023244805.1
E1	12654250	12654360	refGene_gn=DHRS7B;refGene_tx=XM_006939923.3;refGene_gn=DHRS7B;refGene_tx=XM_006939924.4;refGene_gn=DHRS7B;refGene_tx=XM_019817294.2;refGene_gn=DHRS7B;refGene_tx=XM_019817295.2;refGene_gn=DHRS7B;refGene_tx=XM_023244805.1
E1	12662832	12663049	refGene_gn=DHRS7B;refGene_tx=XM_006939923.3;refGene_gn=DHRS7B;refGene_tx=XM_006939924.4;refGene_gn=DHRS7B;refGene_tx=XM_019817294.2;refGene_

gn=DHRS7B;refGene_tx=XM_019817295.2;refGene_gn=DHRS7B;refGene_tx=XM_02
3244805.1
E1	12663683	12663776
refGene_gn=DHRS7B;refGene_tx=XM_006939923.3;refGene_gn=DHRS7B;refGene_
tx=XM_006939924.4;refGene_gn=DHRS7B;refGene_tx=XM_019817294.2;refGene_
gn=DHRS7B;refGene_tx=XM_019817295.2;refGene_gn=DHRS7B;refGene_tx=XM_02
3244805.1
E1	12666988	12667141
refGene_gn=DHRS7B;refGene_tx=XM_006939923.3;refGene_gn=DHRS7B;refGene_
tx=XM_006939924.4;refGene_gn=DHRS7B;refGene_tx=XM_019817294.2;refGene_
gn=DHRS7B;refGene_tx=XM_019817295.2;refGene_gn=DHRS7B;refGene_tx=XM_02
3244805.1
E1	12669790	12669996
refGene_gn=DHRS7B;refGene_tx=XM_006939923.3;refGene_gn=DHRS7B;refGene_
tx=XM_006939924.4;refGene_gn=DHRS7B;refGene_tx=XM_019817294.2;refGene_
gn=DHRS7B;refGene_tx=XM_019817295.2;refGene_gn=DHRS7B;refGene_tx=XM_02
3244805.1
E1	12675635	12676152	refGene_gn=TMEM11;refGene_tx=XM_003996354.3
E1	12689491	12689553	refGene_gn=TMEM11;refGene_tx=XM_003996354.3
E1	12706451	12706568
refGene_gn=NATD1;refGene_tx=XM_006939925.2;refGene_gn=NATD1;refGene_tx
=XM_023243430.1
E1	12708493	12708787
refGene_gn=NATD1;refGene_tx=XM_006939925.2;refGene_gn=NATD1;refGene_tx
=XM_023243430.1
E1	12716103	12716209	refGene_gn=NATD1;refGene_tx=XM_023243430.1
E1	12739450	12739499	refGene_gn=MAP2K3;refGene_tx=XM_003996355.3
E1	12754495	12754562	refGene_gn=MAP2K3;refGene_tx=XM_003996355.3
E1	12754948	12754997	refGene_gn=MAP2K3;refGene_tx=XM_003996355.3
E1	12756036	12756150	refGene_gn=MAP2K3;refGene_tx=XM_003996355.3
E1	12756326	12756446	refGene_gn=MAP2K3;refGene_tx=XM_003996355.3
E1	12757769	12757886	refGene_gn=MAP2K3;refGene_tx=XM_003996355.3
E1	12759148	12759200	refGene_gn=MAP2K3;refGene_tx=XM_003996355.3
E1	12760626	12760754	refGene_gn=MAP2K3;refGene_tx=XM_003996355.3
E1	12761218	12761296	refGene_gn=MAP2K3;refGene_tx=XM_003996355.3
E1	12765475	12765615	refGene_gn=MAP2K3;refGene_tx=XM_003996355.3
E1	12766755	12766801	refGene_gn=MAP2K3;refGene_tx=XM_003996355.3
E1	12767429	12767513	refGene_gn=MAP2K3;refGene_tx=XM_003996355.3
E1	12867547	12867797
refGene_gn=KCNJ12;refGene_tx=XM_019817298.2;refGene_gn=KCNJ12;refGene_
tx=XM_023243493.1
E1	12869865	12869884	refGene_gn=KCNJ12;refGene_tx=XM_023243494.1
E1	12869991	12871349
refGene_gn=KCNJ12;refGene_tx=XM_019817298.2;refGene_gn=KCNJ12;refGene_
tx=XM_023243493.1;refGene_gn=KCNJ12;refGene_tx=XM_023243494.1;refGene_
gn=KCNJ12;refGene_tx=XM_023243495.1;refGene_gn=KCNJ12;refGene_tx=XM_02
3243496.1
E1	13073791	13074284
refGene_gn=LOC111557844;refGene_tx=XM_023244512.1
E1	13075499	13076258

refGene_gn=LOC111557844;refGene_tx=XM_023244512.1
E1 13076738 13077183
refGene_gn=LOC111557844;refGene_tx=XM_023244512.1
E1 13078460 13078593
refGene_gn=LOC111557844;refGene_tx=XM_023244512.1
E1 13218091 13218209
refGene_gn=ATP2A3;refGene_tx=XM_011289053.2;refGene_gn=ATP2A3;refGene_tx=XM_011289054.2;refGene_gn=ATP2A3;refGene_tx=XM_011289055.2;refGene_gn=ATP2A3;refGene_tx=XM_019817299.2;refGene_gn=ATP2A3;refGene_tx=XM_019817300.2
E1 13227867 13227885
refGene_gn=ATP2A3;refGene_tx=XM_011289053.2;refGene_gn=ATP2A3;refGene_tx=XM_011289054.2;refGene_gn=ATP2A3;refGene_tx=XM_011289055.2;refGene_gn=ATP2A3;refGene_tx=XM_019817299.2;refGene_gn=ATP2A3;refGene_tx=XM_019817300.2
E1 13228168 13228251
refGene_gn=ATP2A3;refGene_tx=XM_011289053.2;refGene_gn=ATP2A3;refGene_tx=XM_011289054.2;refGene_gn=ATP2A3;refGene_tx=XM_011289055.2;refGene_gn=ATP2A3;refGene_tx=XM_019817299.2
E1 13229758 13229863
refGene_gn=ATP2A3;refGene_tx=XM_011289053.2;refGene_gn=ATP2A3;refGene_tx=XM_011289054.2;refGene_gn=ATP2A3;refGene_tx=XM_011289055.2;refGene_gn=ATP2A3;refGene_tx=XM_019817300.2;refGene_gn=ATP2A3;refGene_tx=XM_019817301.1
E1 13230051 13230190
refGene_gn=ATP2A3;refGene_tx=XM_011289053.2;refGene_gn=ATP2A3;refGene_tx=XM_011289054.2;refGene_gn=ATP2A3;refGene_tx=XM_011289055.2;refGene_gn=ATP2A3;refGene_tx=XM_019817299.2;refGene_gn=ATP2A3;refGene_tx=XM_019817300.2;refGene_gn=ATP2A3;refGene_tx=XM_019817301.1
E1 13230481 13230562
refGene_gn=ATP2A3;refGene_tx=XM_011289053.2;refGene_gn=ATP2A3;refGene_tx=XM_011289054.2;refGene_gn=ATP2A3;refGene_tx=XM_011289055.2;refGene_gn=ATP2A3;refGene_tx=XM_019817299.2;refGene_gn=ATP2A3;refGene_tx=XM_019817300.2;refGene_gn=ATP2A3;refGene_tx=XM_019817301.1
E1 13230653 13230739
refGene_gn=ATP2A3;refGene_tx=XM_011289053.2;refGene_gn=ATP2A3;refGene_tx=XM_011289054.2;refGene_gn=ATP2A3;refGene_tx=XM_011289055.2;refGene_gn=ATP2A3;refGene_tx=XM_019817299.2;refGene_gn=ATP2A3;refGene_tx=XM_019817300.2;refGene_gn=ATP2A3;refGene_tx=XM_019817301.1
E1 13232320 13232785
refGene_gn=ATP2A3;refGene_tx=XM_011289053.2;refGene_gn=ATP2A3;refGene_tx=XM_011289054.2;refGene_gn=ATP2A3;refGene_tx=XM_011289055.2;refGene_gn=ATP2A3;refGene_tx=XM_019817299.2;refGene_gn=ATP2A3;refGene_tx=XM_019817300.2;refGene_gn=ATP2A3;refGene_tx=XM_019817301.1
E1 13234336 13234425
refGene_gn=ATP2A3;refGene_tx=XM_011289053.2;refGene_gn=ATP2A3;refGene_tx=XM_011289054.2;refGene_gn=ATP2A3;refGene_tx=XM_011289055.2;refGene_gn=ATP2A3;refGene_tx=XM_019817299.2;refGene_gn=ATP2A3;refGene_tx=XM_019817300.2;refGene_gn=ATP2A3;refGene_tx=XM_019817301.1
E1 13234599 13234702

refGene_gn=ATP2A3;refGene_tx=XM_011289053.2;refGene_gn=ATP2A3;refGene_tx=XM_011289054.2;refGene_gn=ATP2A3;refGene_tx=XM_011289055.2;refGene_gn=ATP2A3;refGene_tx=XM_019817299.2;refGene_gn=ATP2A3;refGene_tx=XM_019817300.2;refGene_gn=ATP2A3;refGene_tx=XM_019817301.1
E1 13235966 13236098
refGene_gn=ATP2A3;refGene_tx=XM_011289053.2;refGene_gn=ATP2A3;refGene_tx=XM_011289054.2;refGene_gn=ATP2A3;refGene_tx=XM_011289055.2;refGene_gn=ATP2A3;refGene_tx=XM_019817299.2;refGene_gn=ATP2A3;refGene_tx=XM_019817300.2;refGene_gn=ATP2A3;refGene_tx=XM_019817301.1
E1 13236491 13236617
refGene_gn=ATP2A3;refGene_tx=XM_011289053.2;refGene_gn=ATP2A3;refGene_tx=XM_011289054.2;refGene_gn=ATP2A3;refGene_tx=XM_011289055.2;refGene_gn=ATP2A3;refGene_tx=XM_019817299.2;refGene_gn=ATP2A3;refGene_tx=XM_019817300.2;refGene_gn=ATP2A3;refGene_tx=XM_019817301.1
E1 13237808 13238027
refGene_gn=ATP2A3;refGene_tx=XM_011289053.2;refGene_gn=ATP2A3;refGene_tx=XM_011289054.2;refGene_gn=ATP2A3;refGene_tx=XM_011289055.2;refGene_gn=ATP2A3;refGene_tx=XM_019817299.2;refGene_gn=ATP2A3;refGene_tx=XM_019817300.2;refGene_gn=ATP2A3;refGene_tx=XM_019817301.1
E1 13238142 13238478
refGene_gn=ATP2A3;refGene_tx=XM_011289053.2;refGene_gn=ATP2A3;refGene_tx=XM_011289054.2;refGene_gn=ATP2A3;refGene_tx=XM_011289055.2;refGene_gn=ATP2A3;refGene_tx=XM_019817299.2;refGene_gn=ATP2A3;refGene_tx=XM_019817300.2;refGene_gn=ATP2A3;refGene_tx=XM_019817301.1
E1 13240260 13240481
refGene_gn=ATP2A3;refGene_tx=XM_011289053.2;refGene_gn=ATP2A3;refGene_tx=XM_011289054.2;refGene_gn=ATP2A3;refGene_tx=XM_011289055.2;refGene_gn=ATP2A3;refGene_tx=XM_019817299.2;refGene_gn=ATP2A3;refGene_tx=XM_019817300.2;refGene_gn=ATP2A3;refGene_tx=XM_019817301.1
E1 13241400 13241603
refGene_gn=ATP2A3;refGene_tx=XM_011289053.2;refGene_gn=ATP2A3;refGene_tx=XM_011289054.2;refGene_gn=ATP2A3;refGene_tx=XM_011289055.2;refGene_gn=ATP2A3;refGene_tx=XM_019817299.2;refGene_gn=ATP2A3;refGene_tx=XM_019817300.2;refGene_gn=ATP2A3;refGene_tx=XM_019817301.1
E1 13242196 13242282
refGene_gn=ATP2A3;refGene_tx=XM_011289053.2;refGene_gn=ATP2A3;refGene_tx=XM_011289054.2;refGene_gn=ATP2A3;refGene_tx=XM_011289055.2;refGene_gn=ATP2A3;refGene_tx=XM_019817299.2;refGene_gn=ATP2A3;refGene_tx=XM_019817300.2;refGene_gn=ATP2A3;refGene_tx=XM_019817301.1
E1 13246151 13246285
refGene_gn=ATP2A3;refGene_tx=XM_011289053.2;refGene_gn=ATP2A3;refGene_tx=XM_011289054.2;refGene_gn=ATP2A3;refGene_tx=XM_011289055.2;refGene_gn=ATP2A3;refGene_tx=XM_019817299.2;refGene_gn=ATP2A3;refGene_tx=XM_019817300.2;refGene_gn=ATP2A3;refGene_tx=XM_019817301.1
E1 13246705 13246823
refGene_gn=ATP2A3;refGene_tx=XM_011289053.2;refGene_gn=ATP2A3;refGene_tx=XM_011289054.2;refGene_gn=ATP2A3;refGene_tx=XM_011289055.2;refGene_gn=ATP2A3;refGene_tx=XM_019817299.2;refGene_gn=ATP2A3;refGene_tx=XM_019817300.2;refGene_gn=ATP2A3;refGene_tx=XM_019817301.1
E1 13247328 13247446

refGene_gn=ATP2A3;refGene_tx=XM_011289053.2;refGene_gn=ATP2A3;refGene_tx=XM_011289054.2;refGene_gn=ATP2A3;refGene_tx=XM_011289055.2;refGene_gn=ATP2A3;refGene_tx=XM_019817299.2;refGene_gn=ATP2A3;refGene_tx=XM_019817300.2;refGene_gn=ATP2A3;refGene_tx=XM_019817301.1
E1 13247796 13247883 refGene_gn=ATP2A3;refGene_tx=XM_011289055.2
E1 13250496 13250561
refGene_gn=ATP2A3;refGene_tx=XM_011289053.2;refGene_gn=ATP2A3;refGene_tx=XM_011289054.2;refGene_gn=ATP2A3;refGene_tx=XM_011289055.2;refGene_gn=ATP2A3;refGene_tx=XM_019817299.2;refGene_gn=ATP2A3;refGene_tx=XM_019817300.2;refGene_gn=ATP2A3;refGene_tx=XM_019817301.1
E1 13257752 13257889
refGene_gn=P2RX1;refGene_tx=XM_003996380.4;refGene_gn=P2RX1;refGene_tx=XM_006939939.3
E1 13266728 13266876
refGene_gn=P2RX1;refGene_tx=XM_003996380.4;refGene_gn=P2RX1;refGene_tx=XM_006939939.3
E1 13267167 13267239
refGene_gn=P2RX1;refGene_tx=XM_003996380.4;refGene_gn=P2RX1;refGene_tx=XM_006939939.3
E1 13267682 13267752
refGene_gn=P2RX1;refGene_tx=XM_003996380.4;refGene_gn=P2RX1;refGene_tx=XM_006939939.3
E1 13268000 13268097
refGene_gn=P2RX1;refGene_tx=XM_003996380.4;refGene_gn=P2RX1;refGene_tx=XM_006939939.3
E1 13268397 13268478
refGene_gn=P2RX1;refGene_tx=XM_003996380.4;refGene_gn=P2RX1;refGene_tx=XM_006939939.3
E1 13268737 13268879
refGene_gn=P2RX1;refGene_tx=XM_003996380.4;refGene_gn=P2RX1;refGene_tx=XM_006939939.3
E1 13270613 13270741
refGene_gn=P2RX1;refGene_tx=XM_003996380.4;refGene_gn=P2RX1;refGene_tx=XM_006939939.3
E1 13271150 13271241
refGene_gn=P2RX1;refGene_tx=XM_003996380.4;refGene_gn=P2RX1;refGene_tx=XM_006939939.3
E1 13271668 13271734
refGene_gn=P2RX1;refGene_tx=XM_003996380.4;refGene_gn=P2RX1;refGene_tx=XM_006939939.3
E1 13272081 13272183
refGene_gn=P2RX1;refGene_tx=XM_003996380.4;refGene_gn=P2RX1;refGene_tx=XM_006939939.3
E1 13272294 13272360
refGene_gn=P2RX1;refGene_tx=XM_003996380.4;refGene_gn=P2RX1;refGene_tx=XM_006939939.3
E1 13284746 13285106
refGene_gn=CAMKK1;refGene_tx=XM_006939941.3;refGene_gn=CAMKK1;refGene_tx=XM_006939942.3;refGene_gn=CAMKK1;refGene_tx=XM_011289051.3
E1 13286048 13286096

refGene_gn=CAMKK1;refGene_tx=XM_006939941.3;refGene_gn=CAMKK1;refGene_tx=XM_006939942.3;refGene_gn=CAMKK1;refGene_tx=XM_011289051.3
E1 13286544 13286598
refGene_gn=CAMKK1;refGene_tx=XM_006939941.3;refGene_gn=CAMKK1;refGene_tx=XM_006939942.3;refGene_gn=CAMKK1;refGene_tx=XM_011289051.3
E1 13286995 13287047
refGene_gn=CAMKK1;refGene_tx=XM_006939941.3;refGene_gn=CAMKK1;refGene_tx=XM_006939942.3;refGene_gn=CAMKK1;refGene_tx=XM_011289051.3
E1 13287302 13287436
refGene_gn=CAMKK1;refGene_tx=XM_006939941.3;refGene_gn=CAMKK1;refGene_tx=XM_006939942.3;refGene_gn=CAMKK1;refGene_tx=XM_011289051.3
E1 13287871 13287908
refGene_gn=CAMKK1;refGene_tx=XM_006939941.3;refGene_gn=CAMKK1;refGene_tx=XM_006939942.3;refGene_gn=CAMKK1;refGene_tx=XM_011289051.3
E1 13288709 13288731
refGene_gn=CAMKK1;refGene_tx=XM_006939941.3;refGene_gn=CAMKK1;refGene_tx=XM_006939942.3
E1 13289931 13290020
refGene_gn=CAMKK1;refGene_tx=XM_006939941.3;refGene_gn=CAMKK1;refGene_tx=XM_006939942.3;refGene_gn=CAMKK1;refGene_tx=XM_011289051.3
E1 13293231 13293431
refGene_gn=CAMKK1;refGene_tx=XM_006939941.3;refGene_gn=CAMKK1;refGene_tx=XM_006939942.3
E1 13295376 13295430
refGene_gn=CAMKK1;refGene_tx=XM_006939941.3;refGene_gn=CAMKK1;refGene_tx=XM_006939942.3;refGene_gn=CAMKK1;refGene_tx=XM_011289051.3
E1 13296211 13296285
refGene_gn=CAMKK1;refGene_tx=XM_006939941.3;refGene_gn=CAMKK1;refGene_tx=XM_006939942.3;refGene_gn=CAMKK1;refGene_tx=XM_011289051.3
E1 13298024 13298112
refGene_gn=CAMKK1;refGene_tx=XM_006939941.3;refGene_gn=CAMKK1;refGene_tx=XM_006939942.3;refGene_gn=CAMKK1;refGene_tx=XM_011289051.3
E1 13298333 13298462
refGene_gn=CAMKK1;refGene_tx=XM_006939941.3;refGene_gn=CAMKK1;refGene_tx=XM_006939942.3;refGene_gn=CAMKK1;refGene_tx=XM_011289051.3
E1 13300635 13300739
refGene_gn=CAMKK1;refGene_tx=XM_006939941.3;refGene_gn=CAMKK1;refGene_tx=XM_006939942.3;refGene_gn=CAMKK1;refGene_tx=XM_011289051.3
E1 13301406 13301491
refGene_gn=CAMKK1;refGene_tx=XM_006939941.3;refGene_gn=CAMKK1;refGene_tx=XM_011289051.3
E1 13303568 13303641 refGene_gn=CAMKK1;refGene_tx=XM_006939942.3
E1 13314115 13314298
refGene_gn=NCBP3;refGene_tx=XM_003996382.5;refGene_gn=NCBP3;refGene_tx=XM_006939943.3
E1 13316405 13316471
refGene_gn=NCBP3;refGene_tx=XM_003996382.5;refGene_gn=NCBP3;refGene_tx=XM_006939943.3
E1 13318334 13318440
refGene_gn=NCBP3;refGene_tx=XM_003996382.5;refGene_gn=NCBP3;refGene_tx

=XM_006939943.3
E1	13331457 13331583
refGene_gn=NCBP3;refGene_tx=XM_003996382.5;refGene_gn=NCBP3;refGene_tx
=XM_006939943.3
E1	13333680 13333809
refGene_gn=NCBP3;refGene_tx=XM_003996382.5;refGene_gn=NCBP3;refGene_tx
=XM_006939943.3
E1	13334020 13334097
refGene_gn=NCBP3;refGene_tx=XM_003996382.5;refGene_gn=NCBP3;refGene_tx
=XM_006939943.3
E1	13334841 13334950
refGene_gn=NCBP3;refGene_tx=XM_003996382.5;refGene_gn=NCBP3;refGene_tx
=XM_006939943.3
E1	13338462 13338562
refGene_gn=NCBP3;refGene_tx=XM_003996382.5;refGene_gn=NCBP3;refGene_tx
=XM_006939943.3
E1	13338852 13338956
refGene_gn=NCBP3;refGene_tx=XM_003996382.5;refGene_gn=NCBP3;refGene_tx
=XM_006939943.3
E1	13340741 13341036
refGene_gn=NCBP3;refGene_tx=XM_003996382.5;refGene_gn=NCBP3;refGene_tx
=XM_006939943.3
E1	13342560 13342715
refGene_gn=NCBP3;refGene_tx=XM_003996382.5;refGene_gn=NCBP3;refGene_tx
=XM_006939943.3
E1	13344231 13344393
refGene_gn=NCBP3;refGene_tx=XM_003996382.5;refGene_gn=NCBP3;refGene_tx
=XM_006939943.3
E1	13345415 13345651
refGene_gn=NCBP3;refGene_tx=XM_003996382.5;refGene_gn=NCBP3;refGene_tx
=XM_006939943.3
E1	13346044 13346206
refGene_gn=ITGAE;refGene_tx=XM_019816881.2;refGene_gn=NCBP3;refGene_tx
=XM_006939943.3
E1	13356265 13356299 refGene_gn=ITGAE;refGene_tx=NM_001009219.1
E1	13370262 13370287 refGene_gn=ITGAE;refGene_tx=XM_023243405.1
E1	13371253 13371374
refGene_gn=ITGAE;refGene_tx=NM_001009219.1;refGene_gn=ITGAE;refGene_tx
=XM_019816881.2;refGene_gn=ITGAE;refGene_tx=XM_023243405.1
E1	13381245 13381337
refGene_gn=ITGAE;refGene_tx=NM_001009219.1;refGene_gn=ITGAE;refGene_tx
=XM_019816881.2;refGene_gn=ITGAE;refGene_tx=XM_023243405.1
E1	13382631 13382699
refGene_gn=ITGAE;refGene_tx=NM_001009219.1;refGene_gn=ITGAE;refGene_tx
=XM_019816881.2;refGene_gn=ITGAE;refGene_tx=XM_023243405.1
E1	13383097 13383215
refGene_gn=ITGAE;refGene_tx=NM_001009219.1;refGene_gn=ITGAE;refGene_tx
=XM_019816881.2;refGene_gn=ITGAE;refGene_tx=XM_023243405.1;refGene_gn=
ITGAE;refGene_tx=XM_023243406.1
E1	13383431 13383563

refGene_gn=ITGAE;refGene_tx=NM_001009219.1;refGene_gn=ITGAE;refGene_tx=XM_019816881.2;refGene_gn=ITGAE;refGene_tx=XM_023243405.1;refGene_gn=ITGAE;refGene_tx=XM_023243406.1
E1 13383978 13384094
refGene_gn=ITGAE;refGene_tx=NM_001009219.1;refGene_gn=ITGAE;refGene_tx=XM_019816881.2;refGene_gn=ITGAE;refGene_tx=XM_023243405.1;refGene_gn=ITGAE;refGene_tx=XM_023243406.1
E1 13384736 13384888
refGene_gn=ITGAE;refGene_tx=NM_001009219.1;refGene_gn=ITGAE;refGene_tx=XM_019816881.2;refGene_gn=ITGAE;refGene_tx=XM_023243405.1;refGene_gn=ITGAE;refGene_tx=XM_023243406.1
E1 13385864 13386018
refGene_gn=ITGAE;refGene_tx=NM_001009219.1;refGene_gn=ITGAE;refGene_tx=XM_019816881.2;refGene_gn=ITGAE;refGene_tx=XM_023243405.1;refGene_gn=ITGAE;refGene_tx=XM_023243406.1
E1 13386589 13386734
refGene_gn=ITGAE;refGene_tx=NM_001009219.1;refGene_gn=ITGAE;refGene_tx=XM_019816881.2;refGene_gn=ITGAE;refGene_tx=XM_023243405.1;refGene_gn=ITGAE;refGene_tx=XM_023243406.1
E1 13387212 13387280
refGene_gn=ITGAE;refGene_tx=NM_001009219.1;refGene_gn=ITGAE;refGene_tx=XM_019816881.2;refGene_gn=ITGAE;refGene_tx=XM_023243405.1;refGene_gn=ITGAE;refGene_tx=XM_023243406.1
E1 13388066 13388202
refGene_gn=ITGAE;refGene_tx=NM_001009219.1;refGene_gn=ITGAE;refGene_tx=XM_019816881.2;refGene_gn=ITGAE;refGene_tx=XM_023243405.1;refGene_gn=ITGAE;refGene_tx=XM_023243406.1
E1 13388786 13388929
refGene_gn=ITGAE;refGene_tx=NM_001009219.1;refGene_gn=ITGAE;refGene_tx=XM_019816881.2;refGene_gn=ITGAE;refGene_tx=XM_023243405.1;refGene_gn=ITGAE;refGene_tx=XM_023243406.1
E1 13389283 13389424
refGene_gn=ITGAE;refGene_tx=NM_001009219.1;refGene_gn=ITGAE;refGene_tx=XM_019816881.2;refGene_gn=ITGAE;refGene_tx=XM_023243405.1;refGene_gn=ITGAE;refGene_tx=XM_023243406.1
E1 13390091 13390316
refGene_gn=ITGAE;refGene_tx=NM_001009219.1;refGene_gn=ITGAE;refGene_tx=XM_019816881.2;refGene_gn=ITGAE;refGene_tx=XM_023243405.1;refGene_gn=ITGAE;refGene_tx=XM_023243406.1
E1 13391326 13391457
refGene_gn=ITGAE;refGene_tx=NM_001009219.1;refGene_gn=ITGAE;refGene_tx=XM_019816881.2;refGene_gn=ITGAE;refGene_tx=XM_023243405.1;refGene_gn=ITGAE;refGene_tx=XM_023243406.1
E1 13393028 13393159
refGene_gn=ITGAE;refGene_tx=NM_001009219.1;refGene_gn=ITGAE;refGene_tx=XM_019816881.2;refGene_gn=ITGAE;refGene_tx=XM_023243405.1;refGene_gn=ITGAE;refGene_tx=XM_023243406.1
E1 13394417 13394584
refGene_gn=ITGAE;refGene_tx=NM_001009219.1;refGene_gn=ITGAE;refGene_tx=XM_019816881.2;refGene_gn=ITGAE;refGene_tx=XM_023243405.1;refGene_gn=

ITGAE;refGene_tx=XM_023243406.1
E1 13395956 13396085
refGene_gn=ITGAE;refGene_tx=NM_001009219.1;refGene_gn=ITGAE;refGene_tx=XM_019816881.2;refGene_gn=ITGAE;refGene_tx=XM_023243405.1;refGene_gn=ITGAE;refGene_tx=XM_023243406.1
E1 13399851 13399925
refGene_gn=ITGAE;refGene_tx=NM_001009219.1;refGene_gn=ITGAE;refGene_tx=XM_019816881.2;refGene_gn=ITGAE;refGene_tx=XM_023243405.1;refGene_gn=ITGAE;refGene_tx=XM_023243406.1
E1 13403300 13403433
refGene_gn=ITGAE;refGene_tx=NM_001009219.1;refGene_gn=ITGAE;refGene_tx=XM_019816881.2;refGene_gn=ITGAE;refGene_tx=XM_023243405.1;refGene_gn=ITGAE;refGene_tx=XM_023243406.1
E1 13404824 13404923
refGene_gn=ITGAE;refGene_tx=NM_001009219.1;refGene_gn=ITGAE;refGene_tx=XM_019816881.2;refGene_gn=ITGAE;refGene_tx=XM_023243405.1;refGene_gn=ITGAE;refGene_tx=XM_023243406.1
E1 13405444 13405524
refGene_gn=ITGAE;refGene_tx=NM_001009219.1;refGene_gn=ITGAE;refGene_tx=XM_019816881.2;refGene_gn=ITGAE;refGene_tx=XM_023243405.1;refGene_gn=ITGAE;refGene_tx=XM_023243406.1
E1 13406264 13406342
refGene_gn=ITGAE;refGene_tx=NM_001009219.1;refGene_gn=ITGAE;refGene_tx=XM_019816881.2;refGene_gn=ITGAE;refGene_tx=XM_023243405.1;refGene_gn=ITGAE;refGene_tx=XM_023243406.1
E1 13406849 13406913
refGene_gn=ITGAE;refGene_tx=NM_001009219.1;refGene_gn=ITGAE;refGene_tx=XM_019816881.2;refGene_gn=ITGAE;refGene_tx=XM_023243405.1;refGene_gn=ITGAE;refGene_tx=XM_023243406.1
E1 13406996 13407104
refGene_gn=ITGAE;refGene_tx=NM_001009219.1;refGene_gn=ITGAE;refGene_tx=XM_019816881.2;refGene_gn=ITGAE;refGene_tx=XM_023243405.1;refGene_gn=ITGAE;refGene_tx=XM_023243406.1
E1 13407929 13410278 refGene_gn=HASPIN;refGene_tx=XM_003996383.5
E1 13410476 13410533
refGene_gn=ITGAE;refGene_tx=NM_001009219.1;refGene_gn=ITGAE;refGene_tx=XM_019816881.2;refGene_gn=ITGAE;refGene_tx=XM_023243405.1;refGene_gn=ITGAE;refGene_tx=XM_023243406.1
E1 13410845 13410941
refGene_gn=ITGAE;refGene_tx=NM_001009219.1;refGene_gn=ITGAE;refGene_tx=XM_019816881.2;refGene_gn=ITGAE;refGene_tx=XM_023243405.1;refGene_gn=ITGAE;refGene_tx=XM_023243406.1
E1 13412700 13412796
refGene_gn=ITGAE;refGene_tx=NM_001009219.1;refGene_gn=ITGAE;refGene_tx=XM_019816881.2;refGene_gn=ITGAE;refGene_tx=XM_023243405.1;refGene_gn=ITGAE;refGene_tx=XM_023243406.1
E1 13417668 13417773
refGene_gn=ITGAE;refGene_tx=NM_001009219.1;refGene_gn=ITGAE;refGene_tx=XM_019816881.2;refGene_gn=ITGAE;refGene_tx=XM_023243405.1;refGene_gn=ITGAE;refGene_tx=XM_023243406.1

E1 13419135 13419231
refGene_gn=ITGAE;refGene_tx=NM_001009219.1;refGene_gn=ITGAE;refGene_tx=XM_019816881.2;refGene_gn=ITGAE;refGene_tx=XM_023243405.1;refGene_gn=ITGAE;refGene_tx=XM_023243406.1
E1 13436150 13436287
refGene_gn=P2RX5;refGene_tx=XM_011289042.2;refGene_gn=P2RX5;refGene_tx=XM_011289049.3;refGene_gn=P2RX5;refGene_tx=XM_019817310.2;refGene_gn=P2RX5;refGene_tx=XM_019817311.2;refGene_gn=P2RX5;refGene_tx=XM_019817314.2;refGene_gn=P2RX5;refGene_tx=XM_019817315.2;refGene_gn=P2RX5;refGene_tx=XM_023244107.1;refGene_gn=P2RX5;refGene_tx=XM_023244108.1;refGene_gn=P2RX5;refGene_tx=XM_023244111.1;refGene_gn=P2RX5;refGene_tx=XM_023244112.1
E1 13437319 13437360
refGene_gn=P2RX5;refGene_tx=XM_023244109.1;refGene_gn=P2RX5;refGene_tx=XM_023244110.1
E1 13440505 13440656
refGene_gn=P2RX5;refGene_tx=XM_011289042.2;refGene_gn=P2RX5;refGene_tx=XM_011289049.3;refGene_gn=P2RX5;refGene_tx=XM_019817310.2;refGene_gn=P2RX5;refGene_tx=XM_019817311.2;refGene_gn=P2RX5;refGene_tx=XM_019817314.2;refGene_gn=P2RX5;refGene_tx=XM_019817315.2;refGene_gn=P2RX5;refGene_tx=XM_023244107.1;refGene_gn=P2RX5;refGene_tx=XM_023244108.1;refGene_gn=P2RX5;refGene_tx=XM_023244109.1;refGene_gn=P2RX5;refGene_tx=XM_023244110.1;refGene_gn=P2RX5;refGene_tx=XM_023244111.1;refGene_gn=P2RX5;refGene_tx=XM_023244112.1
E1 13441305 13441377
refGene_gn=P2RX5;refGene_tx=XM_011289042.2;refGene_gn=P2RX5;refGene_tx=XM_011289049.3;refGene_gn=P2RX5;refGene_tx=XM_019817310.2;refGene_gn=P2RX5;refGene_tx=XM_019817311.2;refGene_gn=P2RX5;refGene_tx=XM_019817314.2;refGene_gn=P2RX5;refGene_tx=XM_019817315.2;refGene_gn=P2RX5;refGene_tx=XM_023244107.1;refGene_gn=P2RX5;refGene_tx=XM_023244109.1;refGene_gn=P2RX5;refGene_tx=XM_023244110.1;refGene_gn=P2RX5;refGene_tx=XM_023244111.1;refGene_gn=P2RX5;refGene_tx=XM_023244112.1
E1 13441663 13441739
refGene_gn=P2RX5;refGene_tx=XM_011289042.2;refGene_gn=P2RX5;refGene_tx=XM_011289049.3;refGene_gn=P2RX5;refGene_tx=XM_019817310.2;refGene_gn=P2RX5;refGene_tx=XM_019817311.2;refGene_gn=P2RX5;refGene_tx=XM_019817314.2;refGene_gn=P2RX5;refGene_tx=XM_019817315.2;refGene_gn=P2RX5;refGene_tx=XM_023244107.1;refGene_gn=P2RX5;refGene_tx=XM_023244108.1;refGene_gn=P2RX5;refGene_tx=XM_023244109.1;refGene_gn=P2RX5;refGene_tx=XM_023244110.1;refGene_gn=P2RX5;refGene_tx=XM_023244111.1;refGene_gn=P2RX5;refGene_tx=XM_023244112.1
E1 13441826 13441923
refGene_gn=P2RX5;refGene_tx=XM_011289042.2;refGene_gn=P2RX5;refGene_tx=XM_011289049.3;refGene_gn=P2RX5;refGene_tx=XM_019817310.2;refGene_gn=P2RX5;refGene_tx=XM_019817311.2;refGene_gn=P2RX5;refGene_tx=XM_019817314.2;refGene_gn=P2RX5;refGene_tx=XM_019817315.2;refGene_gn=P2RX5;refGene_tx=XM_023244107.1;refGene_gn=P2RX5;refGene_tx=XM_023244108.1;refGene_gn=P2RX5;refGene_tx=XM_023244109.1;refGene_gn=P2RX5;refGene_tx=XM_023244110.1;refGene_gn=P2RX5;refGene_tx=XM_023244111.1;refGene_gn=P2RX5;refGene_tx=XM_023244112.1

E1 13442199 13442280
refGene_gn=P2RX5;refGene_tx=XM_011289042.2;refGene_gn=P2RX5;refGene_tx
=XM_011289049.3;refGene_gn=P2RX5;refGene_tx=XM_019817310.2;refGene_gn=
P2RX5;refGene_tx=XM_019817311.2;refGene_gn=P2RX5;refGene_tx=XM_0198173
14.2;refGene_gn=P2RX5;refGene_tx=XM_019817315.2;refGene_gn=P2RX5;refGe
ne_tx=XM_023244107.1;refGene_gn=P2RX5;refGene_tx=XM_023244108.1;refGen
e_gn=P2RX5;refGene_tx=XM_023244109.1;refGene_gn=P2RX5;refGene_tx=XM_02
3244110.1;refGene_gn=P2RX5;refGene_tx=XM_023244111.1;refGene_gn=P2RX5;
refGene_tx=XM_023244112.1
E1 13442611 13442750
refGene_gn=P2RX5;refGene_tx=XM_011289042.2;refGene_gn=P2RX5;refGene_tx
=XM_011289049.3;refGene_gn=P2RX5;refGene_tx=XM_019817310.2;refGene_gn=
P2RX5;refGene_tx=XM_019817311.2;refGene_gn=P2RX5;refGene_tx=XM_0198173
14.2;refGene_gn=P2RX5;refGene_tx=XM_019817315.2;refGene_gn=P2RX5;refGe
ne_tx=XM_023244107.1;refGene_gn=P2RX5;refGene_tx=XM_023244108.1;refGen
e_gn=P2RX5;refGene_tx=XM_023244109.1;refGene_gn=P2RX5;refGene_tx=XM_02
3244110.1;refGene_gn=P2RX5;refGene_tx=XM_023244111.1;refGene_gn=P2RX5;
refGene_tx=XM_023244112.1
E1 13443309 13443446
refGene_gn=P2RX5;refGene_tx=XM_011289042.2;refGene_gn=P2RX5;refGene_tx
=XM_011289049.3;refGene_gn=P2RX5;refGene_tx=XM_019817310.2;refGene_gn=
P2RX5;refGene_tx=XM_019817311.2;refGene_gn=P2RX5;refGene_tx=XM_0198173
14.2;refGene_gn=P2RX5;refGene_tx=XM_019817315.2;refGene_gn=P2RX5;refGe
ne_tx=XM_023244107.1;refGene_gn=P2RX5;refGene_tx=XM_023244108.1;refGen
e_gn=P2RX5;refGene_tx=XM_023244109.1;refGene_gn=P2RX5;refGene_tx=XM_02
3244110.1;refGene_gn=P2RX5;refGene_tx=XM_023244111.1;refGene_gn=P2RX5;
refGene_tx=XM_023244112.1
E1 13443901 13443995
refGene_gn=P2RX5;refGene_tx=XM_011289042.2;refGene_gn=P2RX5;refGene_tx
=XM_011289049.3;refGene_gn=P2RX5;refGene_tx=XM_019817310.2;refGene_gn=
P2RX5;refGene_tx=XM_019817311.2;refGene_gn=P2RX5;refGene_tx=XM_0198173
14.2;refGene_gn=P2RX5;refGene_tx=XM_019817315.2;refGene_gn=P2RX5;refGe
ne_tx=XM_023244107.1;refGene_gn=P2RX5;refGene_tx=XM_023244108.1;refGen
e_gn=P2RX5;refGene_tx=XM_023244109.1;refGene_gn=P2RX5;refGene_tx=XM_02
3244110.1;refGene_gn=P2RX5;refGene_tx=XM_023244111.1;refGene_gn=P2RX5;
refGene_tx=XM_023244112.1
E1 13444117 13444183
refGene_gn=P2RX5;refGene_tx=XM_011289042.2;refGene_gn=P2RX5;refGene_tx
=XM_011289049.3;refGene_gn=P2RX5;refGene_tx=XM_019817310.2;refGene_gn=
P2RX5;refGene_tx=XM_019817311.2;refGene_gn=P2RX5;refGene_tx=XM_0198173
14.2;refGene_gn=P2RX5;refGene_tx=XM_019817315.2;refGene_gn=P2RX5;refGe
ne_tx=XM_023244107.1;refGene_gn=P2RX5;refGene_tx=XM_023244108.1;refGen
e_gn=P2RX5;refGene_tx=XM_023244109.1;refGene_gn=P2RX5;refGene_tx=XM_02
3244110.1;refGene_gn=P2RX5;refGene_tx=XM_023244111.1;refGene_gn=P2RX5;
refGene_tx=XM_023244112.1
E1 13445435 13445534
refGene_gn=P2RX5;refGene_tx=XM_019817310.2;refGene_gn=P2RX5;refGene_tx
=XM_019817311.2;refGene_gn=P2RX5;refGene_tx=XM_019817314.2;refGene_gn=
P2RX5;refGene_tx=XM_019817315.2;refGene_gn=P2RX5;refGene_tx=XM_0232441
07.1;refGene_gn=P2RX5;refGene_tx=XM_023244108.1;refGene_gn=P2RX5;refGe

ne_tx=XM_023244109.1;refGene_gn=P2RX5;refGene_tx=XM_023244110.1;refGene_gn=P2RX5;refGene_tx=XM_023244112.1
E1 13446175 13446258
refGene_gn=P2RX5;refGene_tx=XM_011289042.2;refGene_gn=P2RX5;refGene_tx=XM_011289049.3;refGene_gn=P2RX5;refGene_tx=XM_019817310.2;refGene_gn=P2RX5;refGene_tx=XM_019817311.2;refGene_gn=P2RX5;refGene_tx=XM_019817314.2;refGene_gn=P2RX5;refGene_tx=XM_019817315.2;refGene_gn=P2RX5;refGene_tx=XM_023244107.1;refGene_gn=P2RX5;refGene_tx=XM_023244108.1;refGene_gn=P2RX5;refGene_tx=XM_023244109.1;refGene_gn=P2RX5;refGene_tx=XM_023244110.1;refGene_gn=P2RX5;refGene_tx=XM_023244111.1;refGene_gn=P2RX5;refGene_tx=XM_023244112.1
E1 13447202 13447342
refGene_gn=P2RX5;refGene_tx=XM_019817314.2;refGene_gn=P2RX5;refGene_tx=XM_023244107.1;refGene_gn=P2RX5;refGene_tx=XM_023244108.1;refGene_gn=P2RX5;refGene_tx=XM_023244109.1;refGene_gn=P2RX5;refGene_tx=XM_023244110.1;refGene_gn=P2RX5;refGene_tx=XM_023244111.1;refGene_gn=P2RX5;refGene_tx=XM_023244112.1
E1 13448744 13448814
refGene_gn=P2RX5;refGene_tx=XM_011289049.3;refGene_gn=P2RX5;refGene_tx=XM_019817314.2;refGene_gn=P2RX5;refGene_tx=XM_019817315.2
E1 13448909 13448995
refGene_gn=P2RX5;refGene_tx=XM_023244107.1;refGene_gn=P2RX5;refGene_tx=XM_023244108.1;refGene_gn=P2RX5;refGene_tx=XM_023244109.1;refGene_gn=P2RX5;refGene_tx=XM_023244110.1;refGene_gn=P2RX5;refGene_tx=XM_023244111.1;refGene_gn=P2RX5;refGene_tx=XM_023244112.1
E1 13449805 13450013
refGene_gn=P2RX5;refGene_tx=XM_011289042.2;refGene_gn=P2RX5;refGene_tx=XM_019817310.2;refGene_gn=P2RX5;refGene_tx=XM_019817311.2
E1 13454561 13454894
refGene_gn=EMC6;refGene_tx=XM_006939950.3;refGene_gn=EMC6;refGene_tx=XM_006939951.4
E1 13455480 13455519 refGene_gn=TAX1BP3;refGene_tx=XM_003996387.4
E1 13458039 13458057 refGene_gn=TAX1BP3;refGene_tx=XM_006939953.4
E1 13458472 13458592
refGene_gn=TAX1BP3;refGene_tx=XM_003996387.4;refGene_gn=TAX1BP3;refGene_tx=XM_006939953.4
E1 13458981 13459059
refGene_gn=TAX1BP3;refGene_tx=XM_003996387.4;refGene_gn=TAX1BP3;refGene_tx=XM_006939953.4
E1 13459368 13459506
refGene_gn=TAX1BP3;refGene_tx=XM_003996387.4;refGene_gn=TAX1BP3;refGene_tx=XM_006939953.4
E1 13462269 13462351
refGene_gn=CTNS;refGene_tx=XM_011289039.3;refGene_gn=CTNS;refGene_tx=XM_011289040.2;refGene_gn=CTNS;refGene_tx=XM_011289041.3
E1 13462689 13462804
refGene_gn=CTNS;refGene_tx=XM_011289039.3;refGene_gn=CTNS;refGene_tx=XM_011289040.2
E1 13463090 13463208
refGene_gn=CTNS;refGene_tx=XM_011289039.3;refGene_gn=CTNS;refGene_tx=X

M_011289040.2;refGene_gn=CTNS;refGene_tx=XM_011289041.3
E1 13465058 13465229
refGene_gn=CTNS;refGene_tx=XM_011289039.3;refGene_gn=CTNS;refGene_tx=XM_011289040.2;refGene_gn=CTNS;refGene_tx=XM_011289041.3
E1 13466236 13466356
refGene_gn=CTNS;refGene_tx=XM_011289039.3;refGene_gn=CTNS;refGene_tx=XM_011289040.2;refGene_gn=CTNS;refGene_tx=XM_011289041.3
E1 13466439 13466539
refGene_gn=CTNS;refGene_tx=XM_011289039.3;refGene_gn=CTNS;refGene_tx=XM_011289040.2;refGene_gn=CTNS;refGene_tx=XM_011289041.3
E1 13467308 13467440
refGene_gn=CTNS;refGene_tx=XM_011289039.3;refGene_gn=CTNS;refGene_tx=XM_011289040.2;refGene_gn=CTNS;refGene_tx=XM_011289041.3
E1 13467554 13467658
refGene_gn=CTNS;refGene_tx=XM_011289039.3;refGene_gn=CTNS;refGene_tx=XM_011289040.2;refGene_gn=CTNS;refGene_tx=XM_011289041.3
E1 13470522 13470607
refGene_gn=CTNS;refGene_tx=XM_011289039.3;refGene_gn=CTNS;refGene_tx=XM_011289040.2;refGene_gn=CTNS;refGene_tx=XM_011289041.3
E1 13472134 13472213
refGene_gn=CTNS;refGene_tx=XM_011289039.3;refGene_gn=CTNS;refGene_tx=XM_011289040.2;refGene_gn=CTNS;refGene_tx=XM_011289041.3
E1 13476359 13476420
refGene_gn=CTNS;refGene_tx=XM_011289039.3;refGene_gn=CTNS;refGene_tx=XM_011289040.2;refGene_gn=CTNS;refGene_tx=XM_011289041.3
E1 13479608 13479776 refGene_gn=SHPK;refGene_tx=XM_003996389.4
E1 13485285 13485427
refGene_gn=SHPK;refGene_tx=XM_003996389.4;refGene_gn=SHPK;refGene_tx=XM_019817316.1
E1 13493453 13493637
refGene_gn=SHPK;refGene_tx=XM_003996389.4;refGene_gn=SHPK;refGene_tx=XM_019817316.1
E1 13494177 13494330
refGene_gn=SHPK;refGene_tx=XM_003996389.4;refGene_gn=SHPK;refGene_tx=XM_019817316.1
E1 13495313 13495489
refGene_gn=SHPK;refGene_tx=XM_003996389.4;refGene_gn=SHPK;refGene_tx=XM_019817316.1
E1 13499247 13499448
refGene_gn=SHPK;refGene_tx=XM_003996389.4;refGene_gn=SHPK;refGene_tx=XM_019817316.1
E1 13503663 13504076
refGene_gn=SHPK;refGene_tx=XM_003996389.4;refGene_gn=SHPK;refGene_tx=XM_019817316.1
E1 13517253 13517540
refGene_gn=TRPV1;refGene_tx=XM_003996390.5;refGene_gn=TRPV1;refGene_tx=XM_019817317.2;refGene_gn=TRPV1;refGene_tx=XM_019817318.2;refGene_gn=TRPV1;refGene_tx=XM_019817321.2;refGene_gn=TRPV1;refGene_tx=XM_019817322.2;refGene_gn=TRPV1;refGene_tx=XM_019817323.2;refGene_gn=TRPV1;refGene_tx=XM_023244720.1;refGene_gn=TRPV1;refGene_tx=XM_023244721.1;refGen

e_gn=TRPV1;refGene_tx=XM_023244722.1
E1 13518343 13518510
refGene_gn=TRPV1;refGene_tx=XM_003996390.5;refGene_gn=TRPV1;refGene_tx=XM_019817317.2;refGene_gn=TRPV1;refGene_tx=XM_019817318.2;refGene_gn=TRPV1;refGene_tx=XM_019817321.2;refGene_gn=TRPV1;refGene_tx=XM_019817322.2;refGene_gn=TRPV1;refGene_tx=XM_019817323.2;refGene_gn=TRPV1;refGene_tx=XM_023244720.1;refGene_gn=TRPV1;refGene_tx=XM_023244721.1;refGene_gn=TRPV1;refGene_tx=XM_023244722.1
E1 13518587 13518740
refGene_gn=TRPV1;refGene_tx=XM_003996390.5;refGene_gn=TRPV1;refGene_tx=XM_019817317.2;refGene_gn=TRPV1;refGene_tx=XM_019817318.2;refGene_gn=TRPV1;refGene_tx=XM_019817321.2;refGene_gn=TRPV1;refGene_tx=XM_019817322.2;refGene_gn=TRPV1;refGene_tx=XM_019817323.2;refGene_gn=TRPV1;refGene_tx=XM_023244720.1;refGene_gn=TRPV1;refGene_tx=XM_023244721.1;refGene_gn=TRPV1;refGene_tx=XM_023244722.1
E1 13519220 13519361
refGene_gn=TRPV1;refGene_tx=XM_003996390.5;refGene_gn=TRPV1;refGene_tx=XM_019817317.2;refGene_gn=TRPV1;refGene_tx=XM_019817318.2;refGene_gn=TRPV1;refGene_tx=XM_019817322.2;refGene_gn=TRPV1;refGene_tx=XM_019817323.2;refGene_gn=TRPV1;refGene_tx=XM_023244720.1;refGene_gn=TRPV1;refGene_tx=XM_023244721.1;refGene_gn=TRPV1;refGene_tx=XM_023244722.1
E1 13519508 13519807
refGene_gn=TRPV1;refGene_tx=XM_003996390.5;refGene_gn=TRPV1;refGene_tx=XM_019817317.2;refGene_gn=TRPV1;refGene_tx=XM_019817318.2;refGene_gn=TRPV1;refGene_tx=XM_019817321.2;refGene_gn=TRPV1;refGene_tx=XM_019817322.2;refGene_gn=TRPV1;refGene_tx=XM_019817323.2;refGene_gn=TRPV1;refGene_tx=XM_023244720.1;refGene_gn=TRPV1;refGene_tx=XM_023244721.1;refGene_gn=TRPV1;refGene_tx=XM_023244722.1
E1 13520384 13520564
refGene_gn=TRPV1;refGene_tx=XM_003996390.5;refGene_gn=TRPV1;refGene_tx=XM_019817317.2;refGene_gn=TRPV1;refGene_tx=XM_019817318.2;refGene_gn=TRPV1;refGene_tx=XM_019817321.2;refGene_gn=TRPV1;refGene_tx=XM_019817322.2;refGene_gn=TRPV1;refGene_tx=XM_019817323.2;refGene_gn=TRPV1;refGene_tx=XM_023244720.1;refGene_gn=TRPV1;refGene_tx=XM_023244721.1;refGene_gn=TRPV1;refGene_tx=XM_023244722.1
E1 13522555 13522714
refGene_gn=TRPV1;refGene_tx=XM_003996390.5;refGene_gn=TRPV1;refGene_tx=XM_019817317.2;refGene_gn=TRPV1;refGene_tx=XM_019817318.2;refGene_gn=TRPV1;refGene_tx=XM_019817321.2;refGene_gn=TRPV1;refGene_tx=XM_023244720.1;refGene_gn=TRPV1;refGene_tx=XM_023244721.1;refGene_gn=TRPV1;refGene_tx=XM_023244722.1
E1 13524159 13524255
refGene_gn=TRPV1;refGene_tx=XM_003996390.5;refGene_gn=TRPV1;refGene_tx=XM_019817317.2;refGene_gn=TRPV1;refGene_tx=XM_019817318.2;refGene_gn=TRPV1;refGene_tx=XM_019817321.2;refGene_gn=TRPV1;refGene_tx=XM_019817322.2;refGene_gn=TRPV1;refGene_tx=XM_023244720.1;refGene_gn=TRPV1;refGene_tx=XM_023244721.1;refGene_gn=TRPV1;refGene_tx=XM_023244722.1
E1 13525955 13526026
refGene_gn=TRPV1;refGene_tx=XM_003996390.5;refGene_gn=TRPV1;refGene_tx=XM_019817317.2;refGene_gn=TRPV1;refGene_tx=XM_019817318.2;refGene_gn=

TRPV1;refGene_tx=XM_019817321.2;refGene_gn=TRPV1;refGene_tx=XM_0198173
22.2;refGene_gn=TRPV1;refGene_tx=XM_019817323.2;refGene_gn=TRPV1;refGe
ne_tx=XM_023244720.1;refGene_gn=TRPV1;refGene_tx=XM_023244721.1;refGen
e_gn=TRPV1;refGene_tx=XM_023244722.1
E1 13530148 13530314
refGene_gn=TRPV1;refGene_tx=XM_003996390.5;refGene_gn=TRPV1;refGene_tx
=XM_019817317.2;refGene_gn=TRPV1;refGene_tx=XM_019817318.2;refGene_gn=
TRPV1;refGene_tx=XM_019817321.2;refGene_gn=TRPV1;refGene_tx=XM_0198173
22.2;refGene_gn=TRPV1;refGene_tx=XM_019817323.2;refGene_gn=TRPV1;refGe
ne_tx=XM_023244720.1;refGene_gn=TRPV1;refGene_tx=XM_023244721.1;refGen
e_gn=TRPV1;refGene_tx=XM_023244722.1
E1 13530743 13530810
refGene_gn=TRPV1;refGene_tx=XM_003996390.5;refGene_gn=TRPV1;refGene_tx
=XM_019817317.2;refGene_gn=TRPV1;refGene_tx=XM_019817318.2;refGene_gn=
TRPV1;refGene_tx=XM_019817321.2;refGene_gn=TRPV1;refGene_tx=XM_0198173
22.2;refGene_gn=TRPV1;refGene_tx=XM_019817323.2;refGene_gn=TRPV1;refGe
ne_tx=XM_023244720.1;refGene_gn=TRPV1;refGene_tx=XM_023244721.1;refGen
e_gn=TRPV1;refGene_tx=XM_023244722.1
E1 13533269 13533592
refGene_gn=TRPV1;refGene_tx=XM_003996390.5;refGene_gn=TRPV1;refGene_tx
=XM_019817317.2;refGene_gn=TRPV1;refGene_tx=XM_019817318.2;refGene_gn=
TRPV1;refGene_tx=XM_019817321.2;refGene_gn=TRPV1;refGene_tx=XM_0198173
22.2;refGene_gn=TRPV1;refGene_tx=XM_019817323.2;refGene_gn=TRPV1;refGe
ne_tx=XM_023244720.1;refGene_gn=TRPV1;refGene_tx=XM_023244721.1;refGen
e_gn=TRPV1;refGene_tx=XM_023244722.1
E1 13534999 13535127
refGene_gn=TRPV1;refGene_tx=XM_003996390.5;refGene_gn=TRPV1;refGene_tx
=XM_019817317.2;refGene_gn=TRPV1;refGene_tx=XM_019817318.2;refGene_gn=
TRPV1;refGene_tx=XM_019817321.2;refGene_gn=TRPV1;refGene_tx=XM_0198173
22.2;refGene_gn=TRPV1;refGene_tx=XM_019817323.2;refGene_gn=TRPV1;refGe
ne_tx=XM_023244720.1;refGene_gn=TRPV1;refGene_tx=XM_023244721.1;refGen
e_gn=TRPV1;refGene_tx=XM_023244722.1
E1 13535672 13535788
refGene_gn=TRPV1;refGene_tx=XM_003996390.5;refGene_gn=TRPV1;refGene_tx
=XM_019817317.2;refGene_gn=TRPV1;refGene_tx=XM_019817318.2;refGene_gn=
TRPV1;refGene_tx=XM_019817321.2;refGene_gn=TRPV1;refGene_tx=XM_0198173
22.2;refGene_gn=TRPV1;refGene_tx=XM_019817323.2;refGene_gn=TRPV1;refGe
ne_tx=XM_023244720.1;refGene_gn=TRPV1;refGene_tx=XM_023244721.1;refGen
e_gn=TRPV1;refGene_tx=XM_023244722.1
E1 13540321 13540494
refGene_gn=TRPV1;refGene_tx=XM_003996390.5;refGene_gn=TRPV1;refGene_tx
=XM_019817317.2;refGene_gn=TRPV1;refGene_tx=XM_019817318.2;refGene_gn=
TRPV1;refGene_tx=XM_019817321.2;refGene_gn=TRPV1;refGene_tx=XM_0198173
22.2;refGene_gn=TRPV1;refGene_tx=XM_019817323.2;refGene_gn=TRPV1;refGe
ne_tx=XM_023244720.1;refGene_gn=TRPV1;refGene_tx=XM_023244721.1;refGen
e_gn=TRPV1;refGene_tx=XM_023244722.1
E1 13547772 13547891
refGene_gn=TRPV3;refGene_tx=XM_006939955.3;refGene_gn=TRPV3;refGene_tx
=XM_019817324.2;refGene_gn=TRPV3;refGene_tx=XM_019817325.2;refGene_gn=
TRPV3;refGene_tx=XM_019817326.2

E1 13554038 13554143
refGene_gn=TRPV3;refGene_tx=XM_006939955.3;refGene_gn=TRPV3;refGene_tx=XM_019817324.2;refGene_gn=TRPV3;refGene_tx=XM_019817325.2;refGene_gn=TRPV3;refGene_tx=XM_019817326.2
E1 13554316 13554403
refGene_gn=TRPV3;refGene_tx=XM_006939955.3;refGene_gn=TRPV3;refGene_tx=XM_019817324.2;refGene_gn=TRPV3;refGene_tx=XM_019817325.2;refGene_gn=TRPV3;refGene_tx=XM_019817326.2
E1 13555262 13555417
refGene_gn=TRPV3;refGene_tx=XM_006939955.3;refGene_gn=TRPV3;refGene_tx=XM_019817324.2;refGene_gn=TRPV3;refGene_tx=XM_019817325.2;refGene_gn=TRPV3;refGene_tx=XM_019817326.2
E1 13556158 13556335
refGene_gn=TRPV3;refGene_tx=XM_006939955.3;refGene_gn=TRPV3;refGene_tx=XM_019817324.2;refGene_gn=TRPV3;refGene_tx=XM_019817325.2;refGene_gn=TRPV3;refGene_tx=XM_019817326.2
E1 13561541 13561682
refGene_gn=TRPV3;refGene_tx=XM_006939955.3;refGene_gn=TRPV3;refGene_tx=XM_019817324.2;refGene_gn=TRPV3;refGene_tx=XM_019817325.2;refGene_gn=TRPV3;refGene_tx=XM_019817326.2
E1 13563452 13563733
refGene_gn=TRPV3;refGene_tx=XM_006939955.3;refGene_gn=TRPV3;refGene_tx=XM_019817324.2;refGene_gn=TRPV3;refGene_tx=XM_019817325.2;refGene_gn=TRPV3;refGene_tx=XM_019817326.2
E1 13565868 13566045
refGene_gn=TRPV3;refGene_tx=XM_006939955.3;refGene_gn=TRPV3;refGene_tx=XM_019817324.2;refGene_gn=TRPV3;refGene_tx=XM_019817325.2;refGene_gn=TRPV3;refGene_tx=XM_019817326.2
E1 13566995 13567154
refGene_gn=TRPV3;refGene_tx=XM_006939955.3;refGene_gn=TRPV3;refGene_tx=XM_019817324.2;refGene_gn=TRPV3;refGene_tx=XM_019817325.2;refGene_gn=TRPV3;refGene_tx=XM_019817326.2
E1 13567829 13567931
refGene_gn=TRPV3;refGene_tx=XM_006939955.3;refGene_gn=TRPV3;refGene_tx=XM_019817324.2;refGene_gn=TRPV3;refGene_tx=XM_019817325.2;refGene_gn=TRPV3;refGene_tx=XM_019817326.2
E1 13569869 13569943
refGene_gn=TRPV3;refGene_tx=XM_006939955.3;refGene_gn=TRPV3;refGene_tx=XM_019817324.2;refGene_gn=TRPV3;refGene_tx=XM_019817325.2;refGene_gn=TRPV3;refGene_tx=XM_019817326.2
E1 13572120 13572525
refGene_gn=TRPV3;refGene_tx=XM_006939955.3;refGene_gn=TRPV3;refGene_tx=XM_019817324.2;refGene_gn=TRPV3;refGene_tx=XM_019817325.2;refGene_gn=TRPV3;refGene_tx=XM_019817326.2
E1 13573798 13573865
refGene_gn=TRPV3;refGene_tx=XM_006939955.3;refGene_gn=TRPV3;refGene_tx=XM_019817324.2;refGene_gn=TRPV3;refGene_tx=XM_019817325.2
E1 13575199 13575474
refGene_gn=TRPV3;refGene_tx=XM_006939955.3;refGene_gn=TRPV3;refGene_tx=XM_019817324.2;refGene_gn=TRPV3;refGene_tx=XM_019817325.2

E1 13576492 13576623
refGene_gn=TRPV3;refGene_tx=XM_006939955.3;refGene_gn=TRPV3;refGene_tx=XM_019817324.2;refGene_gn=TRPV3;refGene_tx=XM_019817325.2
E1 13577751 13577831
refGene_gn=TRPV3;refGene_tx=XM_006939955.3;refGene_gn=TRPV3;refGene_tx=XM_019817324.2;refGene_gn=TRPV3;refGene_tx=XM_019817325.2
E1 13578334 13578429
refGene_gn=TRPV3;refGene_tx=XM_006939955.3;refGene_gn=TRPV3;refGene_tx=XM_019817324.2
E1 13581751 13581765 refGene_gn=TRPV3;refGene_tx=XM_019817325.2
E1 13585526 13585724
refGene_gn=ASPA;refGene_tx=XM_006939957.4;refGene_gn=ASPA;refGene_tx=XM_006939958.3
E1 13589406 13589516
refGene_gn=ASPA;refGene_tx=XM_006939957.4;refGene_gn=ASPA;refGene_tx=XM_006939958.3
E1 13593763 13593871
refGene_gn=ASPA;refGene_tx=XM_006939957.4;refGene_gn=ASPA;refGene_tx=XM_006939958.3
E1 13596965 13597059
refGene_gn=ASPA;refGene_tx=XM_006939957.4;refGene_gn=ASPA;refGene_tx=XM_006939958.3
E1 13598753 13598949
refGene_gn=ASPA;refGene_tx=XM_006939957.4;refGene_gn=ASPA;refGene_tx=XM_006939958.3
E1 13602590 13602826
refGene_gn=ASPA;refGene_tx=XM_006939957.4;refGene_gn=ASPA;refGene_tx=XM_006939958.3
E1 13609771 13609814 refGene_gn=SPATA22;refGene_tx=XM_023243768.1
E1 13611531 13611564 refGene_gn=SPATA22;refGene_tx=XM_023243768.1
E1 13611566 13611656 refGene_gn=SPATA22;refGene_tx=XM_023243768.1
E1 13613842 13613903 refGene_gn=SPATA22;refGene_tx=XM_023243768.1
E1 13614033 13614129 refGene_gn=SPATA22;refGene_tx=XM_023243768.1
E1 13615783 13616126 refGene_gn=SPATA22;refGene_tx=XM_023243768.1
E1 13618272 13618402 refGene_gn=SPATA22;refGene_tx=XM_023243768.1
E1 13620475 13620573 refGene_gn=SPATA22;refGene_tx=XM_023243768.1
E1 13629762 13630707
refGene_gn=LOC111557757;refGene_tx=XM_023243770.1
E1 13654823 13655768
refGene_gn=LOC101095449;refGene_tx=XM_003996400.4
E1 13672693 13673650
refGene_gn=LOC111557756;refGene_tx=XM_023243769.1
E1 13676327 13676382 refGene_gn=SPATA22;refGene_tx=XM_023243768.1
E1 13679591 13680536
refGene_gn=LOC101081011;refGene_tx=XM_023243941.1
E1 13717428 13718376
refGene_gn=LOC102900819;refGene_tx=XM_023243940.1
E1 13743247 13744195
refGene_gn=LOC101082023;refGene_tx=XM_003996427.2
E1 13765630 13766578

refGene_gn=LOC101095695;refGene_tx=XM_003996401.1
E1 13779172 13780120
refGene_gn=LOC101095929;refGene_tx=XM_003996402.1
E1 13790236 13791184
refGene_gn=LOC101082275;refGene_tx=XM_006939962.3
E1 13799511 13800453
refGene_gn=LOC101096182;refGene_tx=XM_023244383.1
E1 13815057 13816011
refGene_gn=LOC101096427;refGene_tx=XM_003996403.3
E1 13847095 13848025
refGene_gn=LOC101096683;refGene_tx=XM_003996404.1
E1 13869612 13870542
refGene_gn=LOC101096930;refGene_tx=XM_003996405.1
E1 13882242 13883172
refGene_gn=LOC101097188;refGene_tx=XM_003996406.1
E1 13896114 13897170
refGene_gn=LOC101097434;refGene_tx=XM_003996407.4
E1 13908515 13909457
refGene_gn=LOC101082530;refGene_tx=XM_006939963.3
E1 13934266 13935208
refGene_gn=LOC102900923;refGene_tx=XM_006939992.4
E1 13943852 13944794
refGene_gn=LOC102899079;refGene_tx=XM_006939964.4
E1 13987388 13987397
refGene_gn=RAP1GAP2;refGene_tx=XM_006939965.4;refGene_gn=RAP1GAP2;refGene_tx=XM_011289035.3;refGene_gn=RAP1GAP2;refGene_tx=XM_019817338.2;refGene_gn=RAP1GAP2;refGene_tx=XM_019817340.2;refGene_gn=RAP1GAP2;refGene_tx=XM_019817342.2;refGene_gn=RAP1GAP2;refGene_tx=XM_023244673.1;refGene_gn=RAP1GAP2;refGene_tx=XM_023244674.1
E1 13987795 13987872
refGene_gn=RAP1GAP2;refGene_tx=XM_006939965.4;refGene_gn=RAP1GAP2;refGene_tx=XM_011289035.3;refGene_gn=RAP1GAP2;refGene_tx=XM_019817338.2;refGene_gn=RAP1GAP2;refGene_tx=XM_019817340.2;refGene_gn=RAP1GAP2;refGene_tx=XM_019817342.2;refGene_gn=RAP1GAP2;refGene_tx=XM_023244673.1;refGene_gn=RAP1GAP2;refGene_tx=XM_023244674.1
E1 13990885 13991012
refGene_gn=RAP1GAP2;refGene_tx=XM_006939965.4;refGene_gn=RAP1GAP2;refGene_tx=XM_011289035.3;refGene_gn=RAP1GAP2;refGene_tx=XM_019817338.2;refGene_gn=RAP1GAP2;refGene_tx=XM_019817340.2;refGene_gn=RAP1GAP2;refGene_tx=XM_019817342.2;refGene_gn=RAP1GAP2;refGene_tx=XM_023244673.1;refGene_gn=RAP1GAP2;refGene_tx=XM_023244674.1
E1 13991503 13991621
refGene_gn=RAP1GAP2;refGene_tx=XM_006939965.4;refGene_gn=RAP1GAP2;refGene_tx=XM_011289035.3;refGene_gn=RAP1GAP2;refGene_tx=XM_019817338.2;refGene_gn=RAP1GAP2;refGene_tx=XM_019817340.2;refGene_gn=RAP1GAP2;refGene_tx=XM_019817342.2;refGene_gn=RAP1GAP2;refGene_tx=XM_023244673.1;refGene_gn=RAP1GAP2;refGene_tx=XM_023244674.1
E1 13991852 13991966
refGene_gn=RAP1GAP2;refGene_tx=XM_006939965.4;refGene_gn=RAP1GAP2;refGene_tx=XM_011289035.3;refGene_gn=RAP1GAP2;refGene_tx=XM_019817338.2;re

fGene_gn=RAP1GAP2;refGene_tx=XM_019817340.2;refGene_gn=RAP1GAP2;refGen
e_tx=XM_019817342.2;refGene_gn=RAP1GAP2;refGene_tx=XM_023244673.1;refG
ene_gn=RAP1GAP2;refGene_tx=XM_023244674.1
E1 13994330 13994452
refGene_gn=RAP1GAP2;refGene_tx=XM_006939965.4;refGene_gn=RAP1GAP2;refG
ene_tx=XM_011289035.3;refGene_gn=RAP1GAP2;refGene_tx=XM_019817338.2;re
fGene_gn=RAP1GAP2;refGene_tx=XM_019817340.2;refGene_gn=RAP1GAP2;refGen
e_tx=XM_019817342.2;refGene_gn=RAP1GAP2;refGene_tx=XM_023244673.1;refG
ene_gn=RAP1GAP2;refGene_tx=XM_023244674.1
E1 13996089 13996227
refGene_gn=RAP1GAP2;refGene_tx=XM_006939965.4;refGene_gn=RAP1GAP2;refG
ene_tx=XM_011289035.3;refGene_gn=RAP1GAP2;refGene_tx=XM_019817338.2;re
fGene_gn=RAP1GAP2;refGene_tx=XM_019817340.2;refGene_gn=RAP1GAP2;refGen
e_tx=XM_019817342.2;refGene_gn=RAP1GAP2;refGene_tx=XM_023244673.1;refG
ene_gn=RAP1GAP2;refGene_tx=XM_023244674.1
E1 14003752 14003887
refGene_gn=RAP1GAP2;refGene_tx=XM_006939965.4;refGene_gn=RAP1GAP2;refG
ene_tx=XM_011289035.3;refGene_gn=RAP1GAP2;refGene_tx=XM_019817338.2;re
fGene_gn=RAP1GAP2;refGene_tx=XM_019817340.2;refGene_gn=RAP1GAP2;refGen
e_tx=XM_019817342.2;refGene_gn=RAP1GAP2;refGene_tx=XM_023244673.1;refG
ene_gn=RAP1GAP2;refGene_tx=XM_023244674.1
E1 14005262 14005349
refGene_gn=RAP1GAP2;refGene_tx=XM_006939965.4;refGene_gn=RAP1GAP2;refG
ene_tx=XM_011289035.3;refGene_gn=RAP1GAP2;refGene_tx=XM_019817338.2;re
fGene_gn=RAP1GAP2;refGene_tx=XM_019817340.2;refGene_gn=RAP1GAP2;refGen
e_tx=XM_019817342.2;refGene_gn=RAP1GAP2;refGene_tx=XM_023244673.1;refG
ene_gn=RAP1GAP2;refGene_tx=XM_023244674.1
E1 14005869 14005941
refGene_gn=RAP1GAP2;refGene_tx=XM_006939965.4;refGene_gn=RAP1GAP2;refG
ene_tx=XM_011289035.3;refGene_gn=RAP1GAP2;refGene_tx=XM_019817338.2;re
fGene_gn=RAP1GAP2;refGene_tx=XM_019817340.2;refGene_gn=RAP1GAP2;refGen
e_tx=XM_019817342.2;refGene_gn=RAP1GAP2;refGene_tx=XM_023244673.1;refG
ene_gn=RAP1GAP2;refGene_tx=XM_023244674.1
E1 14011039 14011195
refGene_gn=RAP1GAP2;refGene_tx=XM_006939965.4;refGene_gn=RAP1GAP2;refG
ene_tx=XM_011289035.3;refGene_gn=RAP1GAP2;refGene_tx=XM_019817338.2;re
fGene_gn=RAP1GAP2;refGene_tx=XM_019817340.2;refGene_gn=RAP1GAP2;refGen
e_tx=XM_019817342.2;refGene_gn=RAP1GAP2;refGene_tx=XM_023244673.1;refG
ene_gn=RAP1GAP2;refGene_tx=XM_023244674.1
E1 14015476 14015606
refGene_gn=RAP1GAP2;refGene_tx=XM_006939965.4;refGene_gn=RAP1GAP2;refG
ene_tx=XM_011289035.3;refGene_gn=RAP1GAP2;refGene_tx=XM_019817338.2;re
fGene_gn=RAP1GAP2;refGene_tx=XM_019817340.2;refGene_gn=RAP1GAP2;refGen
e_tx=XM_019817342.2;refGene_gn=RAP1GAP2;refGene_tx=XM_023244673.1;refG
ene_gn=RAP1GAP2;refGene_tx=XM_023244674.1
E1 14017889 14017990
refGene_gn=RAP1GAP2;refGene_tx=XM_006939965.4;refGene_gn=RAP1GAP2;refG
ene_tx=XM_011289035.3;refGene_gn=RAP1GAP2;refGene_tx=XM_019817338.2;re
fGene_gn=RAP1GAP2;refGene_tx=XM_019817340.2;refGene_gn=RAP1GAP2;refGen
e_tx=XM_019817342.2;refGene_gn=RAP1GAP2;refGene_tx=XM_023244673.1;refG

ene_gn=RAP1GAP2;refGene_tx=XM_023244674.1
E1 14023555 14023639
refGene_gn=RAP1GAP2;refGene_tx=XM_006939965.4;refGene_gn=RAP1GAP2;refGene_tx=XM_011289035.3;refGene_gn=RAP1GAP2;refGene_tx=XM_019817338.2;refGene_gn=RAP1GAP2;refGene_tx=XM_019817340.2;refGene_gn=RAP1GAP2;refGene_tx=XM_019817342.2;refGene_gn=RAP1GAP2;refGene_tx=XM_023244673.1;refGene_gn=RAP1GAP2;refGene_tx=XM_023244674.1
E1 14026088 14026142
refGene_gn=RAP1GAP2;refGene_tx=XM_006939965.4;refGene_gn=RAP1GAP2;refGene_tx=XM_011289035.3;refGene_gn=RAP1GAP2;refGene_tx=XM_019817338.2;refGene_gn=RAP1GAP2;refGene_tx=XM_019817340.2;refGene_gn=RAP1GAP2;refGene_tx=XM_019817342.2;refGene_gn=RAP1GAP2;refGene_tx=XM_023244673.1;refGene_gn=RAP1GAP2;refGene_tx=XM_023244674.1
E1 14026921 14027000
refGene_gn=RAP1GAP2;refGene_tx=XM_006939965.4;refGene_gn=RAP1GAP2;refGene_tx=XM_011289035.3;refGene_gn=RAP1GAP2;refGene_tx=XM_019817338.2;refGene_gn=RAP1GAP2;refGene_tx=XM_019817340.2;refGene_gn=RAP1GAP2;refGene_tx=XM_019817342.2;refGene_gn=RAP1GAP2;refGene_tx=XM_023244673.1;refGene_gn=RAP1GAP2;refGene_tx=XM_023244674.1
E1 14036979 14037083
refGene_gn=RAP1GAP2;refGene_tx=XM_006939965.4;refGene_gn=RAP1GAP2;refGene_tx=XM_011289035.3;refGene_gn=RAP1GAP2;refGene_tx=XM_019817338.2;refGene_gn=RAP1GAP2;refGene_tx=XM_019817340.2;refGene_gn=RAP1GAP2;refGene_tx=XM_019817342.2;refGene_gn=RAP1GAP2;refGene_tx=XM_023244673.1;refGene_gn=RAP1GAP2;refGene_tx=XM_023244674.1
E1 14038265 14038478
refGene_gn=RAP1GAP2;refGene_tx=XM_006939965.4;refGene_gn=RAP1GAP2;refGene_tx=XM_011289035.3;refGene_gn=RAP1GAP2;refGene_tx=XM_019817338.2;refGene_gn=RAP1GAP2;refGene_tx=XM_019817340.2;refGene_gn=RAP1GAP2;refGene_tx=XM_019817342.2;refGene_gn=RAP1GAP2;refGene_tx=XM_023244673.1;refGene_gn=RAP1GAP2;refGene_tx=XM_023244674.1
E1 14038870 14038903
refGene_gn=RAP1GAP2;refGene_tx=XM_006939965.4;refGene_gn=RAP1GAP2;refGene_tx=XM_011289035.3;refGene_gn=RAP1GAP2;refGene_tx=XM_019817338.2;refGene_gn=RAP1GAP2;refGene_tx=XM_019817340.2;refGene_gn=RAP1GAP2;refGene_tx=XM_019817342.2;refGene_gn=RAP1GAP2;refGene_tx=XM_023244673.1;refGene_gn=RAP1GAP2;refGene_tx=XM_023244674.1
E1 14039617 14039662
refGene_gn=RAP1GAP2;refGene_tx=XM_006939965.4;refGene_gn=RAP1GAP2;refGene_tx=XM_011289035.3;refGene_gn=RAP1GAP2;refGene_tx=XM_019817338.2;refGene_gn=RAP1GAP2;refGene_tx=XM_019817340.2;refGene_gn=RAP1GAP2;refGene_tx=XM_019817342.2;refGene_gn=RAP1GAP2;refGene_tx=XM_023244673.1;refGene_gn=RAP1GAP2;refGene_tx=XM_023244674.1
E1 14044700 14044736
refGene_gn=RAP1GAP2;refGene_tx=XM_006939965.4;refGene_gn=RAP1GAP2;refGene_tx=XM_011289035.3;refGene_gn=RAP1GAP2;refGene_tx=XM_019817338.2;refGene_gn=RAP1GAP2;refGene_tx=XM_019817340.2;refGene_gn=RAP1GAP2;refGene_tx=XM_019817342.2;refGene_gn=RAP1GAP2;refGene_tx=XM_023244673.1;refGene_gn=RAP1GAP2;refGene_tx=XM_023244674.1
E1 14052680 14052923 refGene_gn=RAP1GAP2;refGene_tx=XM_023244674.1

E1 14084182 14084267
refGene_gn=RAP1GAP2;refGene_tx=XM_006939965.4;refGene_gn=RAP1GAP2;refGene_tx=XM_011289035.3;refGene_gn=RAP1GAP2;refGene_tx=XM_019817338.2;refGene_gn=RAP1GAP2;refGene_tx=XM_023244673.1
E1 14121273 14121380 refGene_gn=RAP1GAP2;refGene_tx=XM_006939965.4
E1 14161332 14161349 refGene_gn=RAP1GAP2;refGene_tx=XM_011289035.3
E1 14170175 14170211
refGene_gn=RAP1GAP2;refGene_tx=XM_019817338.2;refGene_gn=RAP1GAP2;refGene_tx=XM_019817340.2;refGene_gn=RAP1GAP2;refGene_tx=XM_023244673.1
E1 14189636 14189693 refGene_gn=RAP1GAP2;refGene_tx=XM_019817340.2
E1 14195420 14195537 refGene_gn=RAP1GAP2;refGene_tx=XM_023244673.1
E1 14208576 14208626 refGene_gn=RAP1GAP2;refGene_tx=XM_023244673.1
E1 14229904 14230088 refGene_gn=CCDC92B;refGene_tx=XM_023244011.1
E1 14234738 14234786 refGene_gn=CCDC92B;refGene_tx=XM_023244011.1
E1 14237356 14238018 refGene_gn=CCDC92B;refGene_tx=XM_023244011.1
E1 14249442 14249542 refGene_gn=CLUH;refGene_tx=XM_023243506.1
E1 14256320 14256523
refGene_gn=CLUH;refGene_tx=XM_011289031.3;refGene_gn=CLUH;refGene_tx=XM_011289032.3;refGene_gn=CLUH;refGene_tx=XM_023243506.1
E1 14256670 14256842
refGene_gn=CLUH;refGene_tx=XM_011289031.3;refGene_gn=CLUH;refGene_tx=XM_011289032.3;refGene_gn=CLUH;refGene_tx=XM_023243506.1
E1 14257875 14258019
refGene_gn=CLUH;refGene_tx=XM_011289031.3;refGene_gn=CLUH;refGene_tx=XM_011289032.3;refGene_gn=CLUH;refGene_tx=XM_023243506.1
E1 14258217 14258342
refGene_gn=CLUH;refGene_tx=XM_011289031.3;refGene_gn=CLUH;refGene_tx=XM_011289032.3;refGene_gn=CLUH;refGene_tx=XM_011289034.2;refGene_gn=CLUH;refGene_tx=XM_023243506.1
E1 14258441 14258596
refGene_gn=CLUH;refGene_tx=XM_011289031.3;refGene_gn=CLUH;refGene_tx=XM_011289032.3;refGene_gn=CLUH;refGene_tx=XM_011289034.2;refGene_gn=CLUH;refGene_tx=XM_023243506.1
E1 14258677 14258803
refGene_gn=CLUH;refGene_tx=XM_011289031.3;refGene_gn=CLUH;refGene_tx=XM_011289032.3;refGene_gn=CLUH;refGene_tx=XM_011289034.2;refGene_gn=CLUH;refGene_tx=XM_023243506.1
E1 14259123 14259271
refGene_gn=CLUH;refGene_tx=XM_011289031.3;refGene_gn=CLUH;refGene_tx=XM_011289032.3;refGene_gn=CLUH;refGene_tx=XM_011289034.2;refGene_gn=CLUH;refGene_tx=XM_023243506.1
E1 14259485 14259578
refGene_gn=CLUH;refGene_tx=XM_011289031.3;refGene_gn=CLUH;refGene_tx=XM_011289032.3;refGene_gn=CLUH;refGene_tx=XM_011289034.2;refGene_gn=CLUH;refGene_tx=XM_023243506.1
E1 14260379 14261074
refGene_gn=CLUH;refGene_tx=XM_011289031.3;refGene_gn=CLUH;refGene_tx=XM_011289032.3;refGene_gn=CLUH;refGene_tx=XM_011289034.2;refGene_gn=CLUH;refGene_tx=XM_023243506.1
E1 14261939 14262163

refGene_gn=CLUH;refGene_tx=XM_011289031.3;refGene_gn=CLUH;refGene_tx=XM_011289032.3;refGene_gn=CLUH;refGene_tx=XM_011289034.2;refGene_gn=CLUH;refGene_tx=XM_023243506.1
E1 14262358 14262463
refGene_gn=CLUH;refGene_tx=XM_011289031.3;refGene_gn=CLUH;refGene_tx=XM_011289032.3;refGene_gn=CLUH;refGene_tx=XM_011289034.2;refGene_gn=CLUH;refGene_tx=XM_023243506.1
E1 14262597 14262698
refGene_gn=CLUH;refGene_tx=XM_011289031.3;refGene_gn=CLUH;refGene_tx=XM_011289032.3;refGene_gn=CLUH;refGene_tx=XM_011289034.2;refGene_gn=CLUH;refGene_tx=XM_023243506.1
E1 14263580 14263733
refGene_gn=CLUH;refGene_tx=XM_011289031.3;refGene_gn=CLUH;refGene_tx=XM_011289032.3;refGene_gn=CLUH;refGene_tx=XM_011289034.2;refGene_gn=CLUH;refGene_tx=XM_023243506.1
E1 14263825 14263888
refGene_gn=CLUH;refGene_tx=XM_011289031.3;refGene_gn=CLUH;refGene_tx=XM_011289032.3;refGene_gn=CLUH;refGene_tx=XM_011289034.2;refGene_gn=CLUH;refGene_tx=XM_023243506.1
E1 14264005 14264250
refGene_gn=CLUH;refGene_tx=XM_011289031.3;refGene_gn=CLUH;refGene_tx=XM_011289032.3;refGene_gn=CLUH;refGene_tx=XM_011289034.2;refGene_gn=CLUH;refGene_tx=XM_023243506.1
E1 14264481 14264566
refGene_gn=CLUH;refGene_tx=XM_011289031.3;refGene_gn=CLUH;refGene_tx=XM_011289032.3;refGene_gn=CLUH;refGene_tx=XM_011289034.2;refGene_gn=CLUH;refGene_tx=XM_023243506.1
E1 14264723 14264877
refGene_gn=CLUH;refGene_tx=XM_011289031.3;refGene_gn=CLUH;refGene_tx=XM_011289032.3;refGene_gn=CLUH;refGene_tx=XM_011289034.2;refGene_gn=CLUH;refGene_tx=XM_023243506.1
E1 14264956 14265096
refGene_gn=CLUH;refGene_tx=XM_011289031.3;refGene_gn=CLUH;refGene_tx=XM_011289032.3;refGene_gn=CLUH;refGene_tx=XM_011289034.2;refGene_gn=CLUH;refGene_tx=XM_023243506.1
E1 14265801 14265882
refGene_gn=CLUH;refGene_tx=XM_011289031.3;refGene_gn=CLUH;refGene_tx=XM_011289032.3;refGene_gn=CLUH;refGene_tx=XM_011289034.2;refGene_gn=CLUH;refGene_tx=XM_023243506.1
E1 14265956 14266082
refGene_gn=CLUH;refGene_tx=XM_011289031.3;refGene_gn=CLUH;refGene_tx=XM_011289032.3;refGene_gn=CLUH;refGene_tx=XM_011289034.2;refGene_gn=CLUH;refGene_tx=XM_023243506.1
E1 14266163 14266285
refGene_gn=CLUH;refGene_tx=XM_011289031.3;refGene_gn=CLUH;refGene_tx=XM_011289032.3;refGene_gn=CLUH;refGene_tx=XM_011289034.2;refGene_gn=CLUH;refGene_tx=XM_023243506.1
E1 14266554 14266648
refGene_gn=CLUH;refGene_tx=XM_011289031.3;refGene_gn=CLUH;refGene_tx=XM_011289032.3;refGene_gn=CLUH;refGene_tx=XM_011289034.2;refGene_gn=CLU

H;refGene_tx=XM_023243506.1
E1 14266738 14266873
refGene_gn=CLUH;refGene_tx=XM_011289031.3;refGene_gn=CLUH;refGene_tx=XM_011289032.3;refGene_gn=CLUH;refGene_tx=XM_011289034.2;refGene_gn=CLUH;refGene_tx=XM_023243506.1
E1 14266971 14267045
refGene_gn=CLUH;refGene_tx=XM_011289031.3;refGene_gn=CLUH;refGene_tx=XM_011289032.3;refGene_gn=CLUH;refGene_tx=XM_011289034.2;refGene_gn=CLUH;refGene_tx=XM_023243506.1
E1 14267855 14268042
refGene_gn=CLUH;refGene_tx=XM_011289031.3;refGene_gn=CLUH;refGene_tx=XM_011289032.3;refGene_gn=CLUH;refGene_tx=XM_011289034.2;refGene_gn=CLUH;refGene_tx=XM_023243506.1
E1 14278039 14278113
refGene_gn=PAFAH1B1;refGene_tx=NM_001114340.1;refGene_gn=PAFAH1B1;refGene_tx=XM_006939934.4;refGene_gn=PAFAH1B1;refGene_tx=XM_019816874.1;refGene_gn=PAFAH1B1;refGene_tx=XM_023243400.1
E1 14278992 14279149
refGene_gn=PAFAH1B1;refGene_tx=NM_001114340.1;refGene_gn=PAFAH1B1;refGene_tx=XM_006939934.4;refGene_gn=PAFAH1B1;refGene_tx=XM_019816874.1;refGene_gn=PAFAH1B1;refGene_tx=XM_023243400.1
E1 14283235 14283337
refGene_gn=PAFAH1B1;refGene_tx=NM_001114340.1;refGene_gn=PAFAH1B1;refGene_tx=XM_006939934.4;refGene_gn=PAFAH1B1;refGene_tx=XM_019816874.1;refGene_gn=PAFAH1B1;refGene_tx=XM_023243400.1
E1 14286656 14286885
refGene_gn=PAFAH1B1;refGene_tx=NM_001114340.1;refGene_gn=PAFAH1B1;refGene_tx=XM_006939934.4;refGene_gn=PAFAH1B1;refGene_tx=XM_019816874.1;refGene_gn=PAFAH1B1;refGene_tx=XM_023243400.1
E1 14287640 14287743
refGene_gn=PAFAH1B1;refGene_tx=NM_001114340.1;refGene_gn=PAFAH1B1;refGene_tx=XM_006939934.4;refGene_gn=PAFAH1B1;refGene_tx=XM_019816874.1;refGene_gn=PAFAH1B1;refGene_tx=XM_023243400.1
E1 14290361 14290530
refGene_gn=PAFAH1B1;refGene_tx=NM_001114340.1;refGene_gn=PAFAH1B1;refGene_tx=XM_006939934.4;refGene_gn=PAFAH1B1;refGene_tx=XM_019816874.1;refGene_gn=PAFAH1B1;refGene_tx=XM_023243400.1
E1 14296779 14296986
refGene_gn=PAFAH1B1;refGene_tx=NM_001114340.1;refGene_gn=PAFAH1B1;refGene_tx=XM_006939934.4;refGene_gn=PAFAH1B1;refGene_tx=XM_019816874.1;refGene_gn=PAFAH1B1;refGene_tx=XM_023243400.1
E1 14297983 14298058
refGene_gn=PAFAH1B1;refGene_tx=NM_001114340.1;refGene_gn=PAFAH1B1;refGene_tx=XM_006939934.4;refGene_gn=PAFAH1B1;refGene_tx=XM_019816874.1;refGene_gn=PAFAH1B1;refGene_tx=XM_023243400.1
E1 14298308 14298393
refGene_gn=PAFAH1B1;refGene_tx=NM_001114340.1;refGene_gn=PAFAH1B1;refGene_tx=XM_006939934.4;refGene_gn=PAFAH1B1;refGene_tx=XM_019816874.1;refGene_gn=PAFAH1B1;refGene_tx=XM_023243400.1
E1 14313703 14313735

refGene_gn=PAFAH1B1;refGene_tx=NM_001114340.1;refGene_gn=PAFAH1B1;refGene_tx=XM_006939934.4;refGene_gn=PAFAH1B1;refGene_tx=XM_023243400.1
E1 14444984 14445112
refGene_gn=METTL16;refGene_tx=XM_003996410.5;refGene_gn=METTL16;refGene_tx=XM_019817344.2;refGene_gn=METTL16;refGene_tx=XM_019817345.2
E1 14467999 14468199
refGene_gn=METTL16;refGene_tx=XM_003996410.5;refGene_gn=METTL16;refGene_tx=XM_019817344.2;refGene_gn=METTL16;refGene_tx=XM_019817345.2
E1 14473696 14473837
refGene_gn=METTL16;refGene_tx=XM_003996410.5;refGene_gn=METTL16;refGene_tx=XM_019817344.2;refGene_gn=METTL16;refGene_tx=XM_019817345.2
E1 14475708 14475824
refGene_gn=METTL16;refGene_tx=XM_003996410.5;refGene_gn=METTL16;refGene_tx=XM_019817344.2;refGene_gn=METTL16;refGene_tx=XM_019817345.2
E1 14477735 14477878
refGene_gn=METTL16;refGene_tx=XM_003996410.5;refGene_gn=METTL16;refGene_tx=XM_019817344.2;refGene_gn=METTL16;refGene_tx=XM_019817345.2
E1 14496156 14496226
refGene_gn=METTL16;refGene_tx=XM_003996410.5;refGene_gn=METTL16;refGene_tx=XM_019817344.2;refGene_gn=METTL16;refGene_tx=XM_019817345.2
E1 14497300 14497390
refGene_gn=METTL16;refGene_tx=XM_003996410.5;refGene_gn=METTL16;refGene_tx=XM_019817344.2;refGene_gn=METTL16;refGene_tx=XM_019817345.2
E1 14506967 14507141
refGene_gn=METTL16;refGene_tx=XM_003996410.5;refGene_gn=METTL16;refGene_tx=XM_019817344.2;refGene_gn=METTL16;refGene_tx=XM_019817345.2
E1 14507262 14507886
refGene_gn=METTL16;refGene_tx=XM_003996410.5;refGene_gn=METTL16;refGene_tx=XM_019817344.2;refGene_gn=METTL16;refGene_tx=XM_019817345.2
E1 14523337 14523410 refGene_gn=MNT;refGene_tx=XM_003996411.5
E1 14528740 14529320 refGene_gn=MNT;refGene_tx=XM_003996411.5
E1 14529928 14529970 refGene_gn=MNT;refGene_tx=XM_003996411.5
E1 14530111 14530223 refGene_gn=MNT;refGene_tx=XM_003996411.5
E1 14535535 14535728 refGene_gn=MNT;refGene_tx=XM_003996411.5
E1 14535864 14536622 refGene_gn=MNT;refGene_tx=XM_003996411.5
E1 14540003 14540017 refGene_gn=SGSM2;refGene_tx=XM_023244239.1
E1 14543471 14543575
refGene_gn=SGSM2;refGene_tx=XM_023244237.1;refGene_gn=SGSM2;refGene_tx=XM_023244238.1;refGene_gn=SGSM2;refGene_tx=XM_023244240.1;refGene_gn=SGSM2;refGene_tx=XM_023244241.1;refGene_gn=SGSM2;refGene_tx=XM_023244242.1;refGene_gn=SGSM2;refGene_tx=XM_023244243.1;refGene_gn=SGSM2;refGene_tx=XM_023244244.1
E1 14543780 14543948
refGene_gn=SGSM2;refGene_tx=XM_023244237.1;refGene_gn=SGSM2;refGene_tx=XM_023244238.1;refGene_gn=SGSM2;refGene_tx=XM_023244239.1;refGene_gn=SGSM2;refGene_tx=XM_023244240.1;refGene_gn=SGSM2;refGene_tx=XM_023244241.1;refGene_gn=SGSM2;refGene_tx=XM_023244242.1;refGene_gn=SGSM2;refGene_tx=XM_023244243.1;refGene_gn=SGSM2;refGene_tx=XM_023244244.1
E1 14544469 14544566
refGene_gn=SGSM2;refGene_tx=XM_023244237.1;refGene_gn=SGSM2;refGene_tx

=XM_023244238.1;refGene_gn=SGSM2;refGene_tx=XM_023244239.1;refGene_gn=SGSM2;refGene_tx=XM_023244240.1;refGene_gn=SGSM2;refGene_tx=XM_023244241.1;refGene_gn=SGSM2;refGene_tx=XM_023244242.1;refGene_gn=SGSM2;refGene_tx=XM_023244243.1;refGene_gn=SGSM2;refGene_tx=XM_023244244.1
E1 14544978 14545088
refGene_gn=SGSM2;refGene_tx=XM_023244237.1;refGene_gn=SGSM2;refGene_tx=XM_023244238.1;refGene_gn=SGSM2;refGene_tx=XM_023244239.1;refGene_gn=SGSM2;refGene_tx=XM_023244240.1;refGene_gn=SGSM2;refGene_tx=XM_023244241.1;refGene_gn=SGSM2;refGene_tx=XM_023244242.1;refGene_gn=SGSM2;refGene_tx=XM_023244243.1;refGene_gn=SGSM2;refGene_tx=XM_023244244.1
E1 14545299 14545382
refGene_gn=SGSM2;refGene_tx=XM_023244237.1;refGene_gn=SGSM2;refGene_tx=XM_023244238.1;refGene_gn=SGSM2;refGene_tx=XM_023244239.1;refGene_gn=SGSM2;refGene_tx=XM_023244240.1;refGene_gn=SGSM2;refGene_tx=XM_023244241.1;refGene_gn=SGSM2;refGene_tx=XM_023244242.1;refGene_gn=SGSM2;refGene_tx=XM_023244243.1;refGene_gn=SGSM2;refGene_tx=XM_023244244.1
E1 14545831 14545956
refGene_gn=SGSM2;refGene_tx=XM_023244237.1;refGene_gn=SGSM2;refGene_tx=XM_023244238.1;refGene_gn=SGSM2;refGene_tx=XM_023244239.1;refGene_gn=SGSM2;refGene_tx=XM_023244240.1;refGene_gn=SGSM2;refGene_tx=XM_023244241.1;refGene_gn=SGSM2;refGene_tx=XM_023244242.1;refGene_gn=SGSM2;refGene_tx=XM_023244243.1;refGene_gn=SGSM2;refGene_tx=XM_023244244.1
E1 14546210 14546600
refGene_gn=SGSM2;refGene_tx=XM_023244237.1;refGene_gn=SGSM2;refGene_tx=XM_023244238.1;refGene_gn=SGSM2;refGene_tx=XM_023244239.1;refGene_gn=SGSM2;refGene_tx=XM_023244240.1;refGene_gn=SGSM2;refGene_tx=XM_023244241.1;refGene_gn=SGSM2;refGene_tx=XM_023244242.1;refGene_gn=SGSM2;refGene_tx=XM_023244243.1;refGene_gn=SGSM2;refGene_tx=XM_023244244.1
E1 14548366 14548549
refGene_gn=SGSM2;refGene_tx=XM_023244237.1;refGene_gn=SGSM2;refGene_tx=XM_023244238.1;refGene_gn=SGSM2;refGene_tx=XM_023244239.1;refGene_gn=SGSM2;refGene_tx=XM_023244240.1;refGene_gn=SGSM2;refGene_tx=XM_023244241.1;refGene_gn=SGSM2;refGene_tx=XM_023244242.1;refGene_gn=SGSM2;refGene_tx=XM_023244243.1;refGene_gn=SGSM2;refGene_tx=XM_023244244.1
E1 14548753 14548882
refGene_gn=SGSM2;refGene_tx=XM_023244237.1;refGene_gn=SGSM2;refGene_tx=XM_023244238.1;refGene_gn=SGSM2;refGene_tx=XM_023244239.1;refGene_gn=SGSM2;refGene_tx=XM_023244240.1;refGene_gn=SGSM2;refGene_tx=XM_023244241.1;refGene_gn=SGSM2;refGene_tx=XM_023244242.1;refGene_gn=SGSM2;refGene_tx=XM_023244243.1;refGene_gn=SGSM2;refGene_tx=XM_023244244.1
E1 14549368 14549514
refGene_gn=SGSM2;refGene_tx=XM_023244237.1;refGene_gn=SGSM2;refGene_tx=XM_023244238.1;refGene_gn=SGSM2;refGene_tx=XM_023244239.1;refGene_gn=SGSM2;refGene_tx=XM_023244240.1;refGene_gn=SGSM2;refGene_tx=XM_023244241.1;refGene_gn=SGSM2;refGene_tx=XM_023244242.1;refGene_gn=SGSM2;refGene_tx=XM_023244243.1;refGene_gn=SGSM2;refGene_tx=XM_023244244.1
E1 14549600 14549665
refGene_gn=SGSM2;refGene_tx=XM_023244237.1;refGene_gn=SGSM2;refGene_tx=XM_023244238.1;refGene_gn=SGSM2;refGene_tx=XM_023244239.1;refGene_gn=SGSM2;refGene_tx=XM_023244240.1;refGene_gn=SGSM2;refGene_tx=XM_0232442

41.1;refGene_gn=SGSM2;refGene_tx=XM_023244242.1;refGene_gn=SGSM2;refGene_tx=XM_023244243.1;refGene_gn=SGSM2;refGene_tx=XM_023244244.1
E1 14550466 14550620
refGene_gn=SGSM2;refGene_tx=XM_023244237.1;refGene_gn=SGSM2;refGene_tx=XM_023244238.1;refGene_gn=SGSM2;refGene_tx=XM_023244239.1;refGene_gn=SGSM2;refGene_tx=XM_023244240.1;refGene_gn=SGSM2;refGene_tx=XM_023244241.1;refGene_gn=SGSM2;refGene_tx=XM_023244242.1;refGene_gn=SGSM2;refGene_tx=XM_023244243.1;refGene_gn=SGSM2;refGene_tx=XM_023244244.1
E1 14554222 14554357
refGene_gn=SGSM2;refGene_tx=XM_023244237.1;refGene_gn=SGSM2;refGene_tx=XM_023244238.1;refGene_gn=SGSM2;refGene_tx=XM_023244239.1;refGene_gn=SGSM2;refGene_tx=XM_023244242.1;refGene_gn=SGSM2;refGene_tx=XM_023244243.1;refGene_gn=SGSM2;refGene_tx=XM_023244244.1
E1 14555948 14556075
refGene_gn=SGSM2;refGene_tx=XM_023244237.1;refGene_gn=SGSM2;refGene_tx=XM_023244238.1;refGene_gn=SGSM2;refGene_tx=XM_023244239.1;refGene_gn=SGSM2;refGene_tx=XM_023244240.1;refGene_gn=SGSM2;refGene_tx=XM_023244241.1;refGene_gn=SGSM2;refGene_tx=XM_023244242.1;refGene_gn=SGSM2;refGene_tx=XM_023244243.1;refGene_gn=SGSM2;refGene_tx=XM_023244244.1
E1 14556249 14556410
refGene_gn=SGSM2;refGene_tx=XM_023244237.1;refGene_gn=SGSM2;refGene_tx=XM_023244238.1;refGene_gn=SGSM2;refGene_tx=XM_023244239.1;refGene_gn=SGSM2;refGene_tx=XM_023244240.1;refGene_gn=SGSM2;refGene_tx=XM_023244241.1;refGene_gn=SGSM2;refGene_tx=XM_023244242.1;refGene_gn=SGSM2;refGene_tx=XM_023244243.1;refGene_gn=SGSM2;refGene_tx=XM_023244244.1
E1 14556617 14556685
refGene_gn=SGSM2;refGene_tx=XM_023244237.1;refGene_gn=SGSM2;refGene_tx=XM_023244238.1;refGene_gn=SGSM2;refGene_tx=XM_023244239.1;refGene_gn=SGSM2;refGene_tx=XM_023244240.1;refGene_gn=SGSM2;refGene_tx=XM_023244241.1;refGene_gn=SGSM2;refGene_tx=XM_023244242.1;refGene_gn=SGSM2;refGene_tx=XM_023244243.1;refGene_gn=SGSM2;refGene_tx=XM_023244244.1
E1 14557081 14557206
refGene_gn=SGSM2;refGene_tx=XM_023244237.1;refGene_gn=SGSM2;refGene_tx=XM_023244238.1;refGene_gn=SGSM2;refGene_tx=XM_023244239.1;refGene_gn=SGSM2;refGene_tx=XM_023244240.1;refGene_gn=SGSM2;refGene_tx=XM_023244241.1;refGene_gn=SGSM2;refGene_tx=XM_023244242.1;refGene_gn=SGSM2;refGene_tx=XM_023244243.1;refGene_gn=SGSM2;refGene_tx=XM_023244244.1
E1 14557730 14557865
refGene_gn=SGSM2;refGene_tx=XM_023244237.1;refGene_gn=SGSM2;refGene_tx=XM_023244238.1;refGene_gn=SGSM2;refGene_tx=XM_023244239.1;refGene_gn=SGSM2;refGene_tx=XM_023244240.1;refGene_gn=SGSM2;refGene_tx=XM_023244241.1;refGene_gn=SGSM2;refGene_tx=XM_023244242.1;refGene_gn=SGSM2;refGene_tx=XM_023244243.1;refGene_gn=SGSM2;refGene_tx=XM_023244244.1
E1 14558203 14558349
refGene_gn=SGSM2;refGene_tx=XM_023244237.1;refGene_gn=SGSM2;refGene_tx=XM_023244238.1;refGene_gn=SGSM2;refGene_tx=XM_023244239.1;refGene_gn=SGSM2;refGene_tx=XM_023244240.1;refGene_gn=SGSM2;refGene_tx=XM_023244241.1;refGene_gn=SGSM2;refGene_tx=XM_023244242.1;refGene_gn=SGSM2;refGene_tx=XM_023244243.1;refGene_gn=SGSM2;refGene_tx=XM_023244244.1
E1 14558447 14558515

refGene_gn=SGSM2;refGene_tx=XM_023244237.1;refGene_gn=SGSM2;refGene_tx=XM_023244238.1;refGene_gn=SGSM2;refGene_tx=XM_023244239.1;refGene_gn=SGSM2;refGene_tx=XM_023244240.1;refGene_gn=SGSM2;refGene_tx=XM_023244241.1;refGene_gn=SGSM2;refGene_tx=XM_023244242.1;refGene_gn=SGSM2;refGene_tx=XM_023244243.1;refGene_gn=SGSM2;refGene_tx=XM_023244244.1
E1 14558828 14558990
refGene_gn=SGSM2;refGene_tx=XM_023244237.1;refGene_gn=SGSM2;refGene_tx=XM_023244238.1;refGene_gn=SGSM2;refGene_tx=XM_023244239.1;refGene_gn=SGSM2;refGene_tx=XM_023244240.1;refGene_gn=SGSM2;refGene_tx=XM_023244241.1;refGene_gn=SGSM2;refGene_tx=XM_023244242.1;refGene_gn=SGSM2;refGene_tx=XM_023244243.1;refGene_gn=SGSM2;refGene_tx=XM_023244244.1
E1 14559261 14559424
refGene_gn=SGSM2;refGene_tx=XM_023244237.1;refGene_gn=SGSM2;refGene_tx=XM_023244238.1;refGene_gn=SGSM2;refGene_tx=XM_023244239.1;refGene_gn=SGSM2;refGene_tx=XM_023244240.1;refGene_gn=SGSM2;refGene_tx=XM_023244241.1;refGene_gn=SGSM2;refGene_tx=XM_023244242.1;refGene_gn=SGSM2;refGene_tx=XM_023244243.1;refGene_gn=SGSM2;refGene_tx=XM_023244244.1
E1 14573487 14573563
refGene_gn=SGSM2;refGene_tx=XM_023244237.1;refGene_gn=SGSM2;refGene_tx=XM_023244238.1;refGene_gn=SGSM2;refGene_tx=XM_023244239.1;refGene_gn=SGSM2;refGene_tx=XM_023244240.1;refGene_gn=SGSM2;refGene_tx=XM_023244241.1
E1 14577016 14577073
refGene_gn=SGSM2;refGene_tx=XM_023244237.1;refGene_gn=SGSM2;refGene_tx=XM_023244238.1;refGene_gn=SGSM2;refGene_tx=XM_023244239.1;refGene_gn=SGSM2;refGene_tx=XM_023244240.1;refGene_gn=SGSM2;refGene_tx=XM_023244241.1
E1 14578391 14578488
refGene_gn=TSR1;refGene_tx=XM_003996412.5;refGene_gn=TSR1;refGene_tx=XM_023244246.1
E1 14578665 14578769
refGene_gn=TSR1;refGene_tx=XM_003996412.5;refGene_gn=TSR1;refGene_tx=XM_023244246.1
E1 14579093 14579313
refGene_gn=TSR1;refGene_tx=XM_003996412.5;refGene_gn=TSR1;refGene_tx=XM_023244246.1
E1 14579429 14579564
refGene_gn=TSR1;refGene_tx=XM_003996412.5;refGene_gn=TSR1;refGene_tx=XM_023244246.1
E1 14579957 14580379
refGene_gn=TSR1;refGene_tx=XM_003996412.5;refGene_gn=TSR1;refGene_tx=XM_023244246.1
E1 14581389 14581549
refGene_gn=TSR1;refGene_tx=XM_003996412.5;refGene_gn=TSR1;refGene_tx=XM_023244246.1
E1 14582220 14582375
refGene_gn=TSR1;refGene_tx=XM_003996412.5;refGene_gn=TSR1;refGene_tx=XM_023244246.1
E1 14582633 14582830
refGene_gn=TSR1;refGene_tx=XM_003996412.5;refGene_gn=TSR1;refGene_tx=X

M_023244246.1
E1 14583508 14583671
refGene_gn=TSR1;refGene_tx=XM_003996412.5;refGene_gn=TSR1;refGene_tx=XM_023244246.1
E1 14584013 14584124
refGene_gn=TSR1;refGene_tx=XM_003996412.5;refGene_gn=TSR1;refGene_tx=XM_023244246.1
E1 14584801 14584964
refGene_gn=TSR1;refGene_tx=XM_003996412.5;refGene_gn=TSR1;refGene_tx=XM_023244246.1
E1 14586294 14586411
refGene_gn=TSR1;refGene_tx=XM_003996412.5;refGene_gn=TSR1;refGene_tx=XM_023244246.1
E1 14586828 14586969
refGene_gn=TSR1;refGene_tx=XM_003996412.5;refGene_gn=TSR1;refGene_tx=XM_023244246.1
E1 14587073 14587148
refGene_gn=TSR1;refGene_tx=XM_003996412.5;refGene_gn=TSR1;refGene_tx=XM_023244246.1
E1 14587277 14587456
refGene_gn=TSR1;refGene_tx=XM_003996412.5;refGene_gn=TSR1;refGene_tx=XM_023244246.1
E1 14587800 14588016
refGene_gn=SRR;refGene_tx=XM_006939966.4;refGene_gn=SRR;refGene_tx=XM_019817353.2;refGene_gn=SRR;refGene_tx=XM_019817354.2
E1 14588360 14588570
refGene_gn=SRR;refGene_tx=XM_006939966.4;refGene_gn=SRR;refGene_tx=XM_019817353.2;refGene_gn=SRR;refGene_tx=XM_019817354.2
E1 14589324 14589399
refGene_gn=SRR;refGene_tx=XM_006939966.4;refGene_gn=SRR;refGene_tx=XM_019817353.2;refGene_gn=SRR;refGene_tx=XM_019817354.2
E1 14589522 14589642
refGene_gn=SRR;refGene_tx=XM_006939966.4;refGene_gn=SRR;refGene_tx=XM_019817353.2;refGene_gn=SRR;refGene_tx=XM_019817354.2
E1 14591034 14591138
refGene_gn=SRR;refGene_tx=XM_006939966.4;refGene_gn=SRR;refGene_tx=XM_019817353.2
E1 14591332 14591459
refGene_gn=SRR;refGene_tx=XM_006939966.4;refGene_gn=SRR;refGene_tx=XM_019817353.2
E1 14594647 14594819
refGene_gn=SRR;refGene_tx=XM_006939966.4;refGene_gn=SRR;refGene_tx=XM_019817353.2;refGene_gn=SRR;refGene_tx=XM_019817354.2
E1 14605340 14605428 refGene_gn=SMG6;refGene_tx=XM_023244236.1
E1 14606633 14606644
refGene_gn=SRR;refGene_tx=XM_019817353.2;refGene_gn=SRR;refGene_tx=XM_019817354.2
E1 14607657 14609398 refGene_gn=SMG6;refGene_tx=XM_023244236.1
E1 14609902 14610095 refGene_gn=SMG6;refGene_tx=XM_023244236.1
E1 14610585 14610696 refGene_gn=SMG6;refGene_tx=XM_023244236.1

E1 14613632 14613739 refGene_gn=SMG6;refGene_tx=XM_023244236.1
E1 14613993 14614072 refGene_gn=SMG6;refGene_tx=XM_023244236.1
E1 14621721 14621832 refGene_gn=SMG6;refGene_tx=XM_023244236.1
E1 14622203 14622416 refGene_gn=SMG6;refGene_tx=XM_023244236.1
E1 14654181 14654243 refGene_gn=SMG6;refGene_tx=XM_023244236.1
E1 14660934 14661079 refGene_gn=SMG6;refGene_tx=XM_023244236.1
E1 14707188 14707306 refGene_gn=SMG6;refGene_tx=XM_023244236.1
E1 14708906 14709075 refGene_gn=SMG6;refGene_tx=XM_023244236.1
E1 14720665 14720867 refGene_gn=SMG6;refGene_tx=XM_023244236.1
E1 14811547 14811724 refGene_gn=SMG6;refGene_tx=XM_023244236.1
E1 14815515 14815662 refGene_gn=SMG6;refGene_tx=XM_023244236.1
E1 14825407 14825561 refGene_gn=SMG6;refGene_tx=XM_023244236.1
E1 14828503 14828715 refGene_gn=SMG6;refGene_tx=XM_023244236.1
E1 14829024 14829106 refGene_gn=SMG6;refGene_tx=XM_023244236.1
E1 14832329 14832460 refGene_gn=SMG6;refGene_tx=XM_023244236.1
E1 14835171 14837333
refGene_gn=HIC1;refGene_tx=XM_023244247.1;refGene_gn=HIC1;refGene_tx=XM_023244248.1
E1 14837689 14837726 refGene_gn=HIC1;refGene_tx=XM_023244247.1
E1 14847810 14848310 refGene_gn=OVCA2;refGene_tx=XM_006939976.3
E1 14848656 14848963 refGene_gn=OVCA2;refGene_tx=XM_006939976.3
E1 14849045 14849141
refGene_gn=DPH1;refGene_tx=XM_003996430.5;refGene_gn=DPH1;refGene_tx=XM_019817356.2;refGene_gn=DPH1;refGene_tx=XM_019817357.2
E1 14849312 14849453
refGene_gn=DPH1;refGene_tx=XM_003996430.5;refGene_gn=DPH1;refGene_tx=XM_019817356.2;refGene_gn=DPH1;refGene_tx=XM_019817357.2
E1 14849731 14849810
refGene_gn=DPH1;refGene_tx=XM_003996430.5;refGene_gn=DPH1;refGene_tx=XM_019817356.2;refGene_gn=DPH1;refGene_tx=XM_019817357.2
E1 14850314 14850415
refGene_gn=DPH1;refGene_tx=XM_003996430.5;refGene_gn=DPH1;refGene_tx=XM_019817356.2;refGene_gn=DPH1;refGene_tx=XM_019817357.2
E1 14850547 14850704
refGene_gn=DPH1;refGene_tx=XM_003996430.5;refGene_gn=DPH1;refGene_tx=XM_019817356.2;refGene_gn=DPH1;refGene_tx=XM_019817357.2
E1 14851107 14851176
refGene_gn=DPH1;refGene_tx=XM_003996430.5;refGene_gn=DPH1;refGene_tx=XM_019817356.2;refGene_gn=DPH1;refGene_tx=XM_019817357.2
E1 14852816 14852938
refGene_gn=DPH1;refGene_tx=XM_003996430.5;refGene_gn=DPH1;refGene_tx=XM_019817356.2;refGene_gn=DPH1;refGene_tx=XM_019817357.2
E1 14853066 14853224
refGene_gn=DPH1;refGene_tx=XM_003996430.5;refGene_gn=DPH1;refGene_tx=XM_019817356.2;refGene_gn=DPH1;refGene_tx=XM_019817357.2
E1 14853639 14853761
refGene_gn=DPH1;refGene_tx=XM_003996430.5;refGene_gn=DPH1;refGene_tx=XM_019817356.2;refGene_gn=DPH1;refGene_tx=XM_019817357.2
E1 14855420 14855484
refGene_gn=DPH1;refGene_tx=XM_003996430.5;refGene_gn=DPH1;refGene_tx=X

M_019817356.2;refGene_gn=DPH1;refGene_tx=XM_019817357.2
E1 14855600 14855753
refGene_gn=DPH1;refGene_tx=XM_003996430.5;refGene_gn=DPH1;refGene_tx=XM_019817356.2
E1 14857858 14857919
refGene_gn=DPH1;refGene_tx=XM_003996430.5;refGene_gn=DPH1;refGene_tx=XM_019817356.2
E1 14862378 14862391 refGene_gn=RTN4RL1;refGene_tx=XM_006939977.4
E1 14931413 14932738 refGene_gn=RTN4RL1;refGene_tx=XM_006939977.4
E1 14961901 14962009 refGene_gn=RPA1;refGene_tx=XM_006939978.4
E1 14963522 14963609 refGene_gn=RPA1;refGene_tx=XM_006939978.4
E1 14966873 14966981 refGene_gn=RPA1;refGene_tx=XM_006939978.4
E1 14969183 14969360 refGene_gn=RPA1;refGene_tx=XM_006939978.4
E1 14972289 14972422 refGene_gn=RPA1;refGene_tx=XM_006939978.4
E1 14974604 14974753 refGene_gn=RPA1;refGene_tx=XM_006939978.4
E1 14975461 14975601 refGene_gn=RPA1;refGene_tx=XM_006939978.4
E1 14975759 14975952 refGene_gn=RPA1;refGene_tx=XM_006939978.4
E1 14976091 14976160 refGene_gn=RPA1;refGene_tx=XM_006939978.4
E1 14978247 14978350 refGene_gn=RPA1;refGene_tx=XM_006939978.4
E1 14980010 14980146 refGene_gn=RPA1;refGene_tx=XM_006939978.4
E1 14983219 14983312 refGene_gn=RPA1;refGene_tx=XM_006939978.4
E1 15004612 15004701 refGene_gn=RPA1;refGene_tx=XM_006939978.4
E1 15014527 15014636 refGene_gn=RPA1;refGene_tx=XM_006939978.4
E1 15015297 15015376 refGene_gn=RPA1;refGene_tx=XM_006939978.4
E1 15016875 15016926 refGene_gn=RPA1;refGene_tx=XM_006939978.4
E1 15034307 15034340 refGene_gn=RPA1;refGene_tx=XM_006939978.4
E1 15035979 15036113
refGene_gn=SMYD4;refGene_tx=XM_003996433.4;refGene_gn=SMYD4;refGene_tx=XM_006939979.3;refGene_gn=SMYD4;refGene_tx=XM_019817359.2;refGene_gn=SMYD4;refGene_tx=XM_019817360.2
E1 15053127 15053272
refGene_gn=SMYD4;refGene_tx=XM_003996433.4;refGene_gn=SMYD4;refGene_tx=XM_006939979.3;refGene_gn=SMYD4;refGene_tx=XM_019817359.2;refGene_gn=SMYD4;refGene_tx=XM_019817360.2
E1 15059857 15059947
refGene_gn=SMYD4;refGene_tx=XM_003996433.4;refGene_gn=SMYD4;refGene_tx=XM_006939979.3;refGene_gn=SMYD4;refGene_tx=XM_019817359.2;refGene_gn=SMYD4;refGene_tx=XM_019817360.2
E1 15061448 15062583
refGene_gn=SMYD4;refGene_tx=XM_003996433.4;refGene_gn=SMYD4;refGene_tx=XM_006939979.3;refGene_gn=SMYD4;refGene_tx=XM_019817359.2;refGene_gn=SMYD4;refGene_tx=XM_019817360.2
E1 15075668 15075851
refGene_gn=SMYD4;refGene_tx=XM_003996433.4;refGene_gn=SMYD4;refGene_tx=XM_006939979.3;refGene_gn=SMYD4;refGene_tx=XM_019817359.2;refGene_gn=SMYD4;refGene_tx=XM_019817360.2
E1 15076763 15076927
refGene_gn=SMYD4;refGene_tx=XM_003996433.4;refGene_gn=SMYD4;refGene_tx=XM_006939979.3;refGene_gn=SMYD4;refGene_tx=XM_019817359.2;refGene_gn=SMYD4;refGene_tx=XM_019817360.2

E1 15078874 15079007
refGene_gn=SMYD4;refGene_tx=XM_003996433.4;refGene_gn=SMYD4;refGene_tx=XM_006939979.3;refGene_gn=SMYD4;refGene_tx=XM_019817359.2;refGene_gn=SMYD4;refGene_tx=XM_019817360.2
E1 15079843 15079960
refGene_gn=SMYD4;refGene_tx=XM_003996433.4;refGene_gn=SMYD4;refGene_tx=XM_006939979.3;refGene_gn=SMYD4;refGene_tx=XM_019817359.2;refGene_gn=SMYD4;refGene_tx=XM_019817360.2
E1 15080150 15080274
refGene_gn=SMYD4;refGene_tx=XM_003996433.4;refGene_gn=SMYD4;refGene_tx=XM_006939979.3;refGene_gn=SMYD4;refGene_tx=XM_019817359.2;refGene_gn=SMYD4;refGene_tx=XM_019817360.2
E1 15081194 15081219 refGene_gn=SMYD4;refGene_tx=XM_019817360.2
E1 15081371 15081549 refGene_gn=SMYD4;refGene_tx=XM_003996433.4
E1 15083805 15084065 refGene_gn=SERPINF1;refGene_tx=XM_003996416.4
E1 15084507 15084718 refGene_gn=SERPINF1;refGene_tx=XM_003996416.4
E1 15086078 15086221 refGene_gn=SERPINF1;refGene_tx=XM_003996416.4
E1 15087293 15087497 refGene_gn=SERPINF1;refGene_tx=XM_003996416.4
E1 15087905 15088061 refGene_gn=SERPINF1;refGene_tx=XM_003996416.4
E1 15088722 15088921 refGene_gn=SERPINF1;refGene_tx=XM_003996416.4
E1 15090476 15090557 refGene_gn=SERPINF1;refGene_tx=XM_003996416.4
E1 15095306 15095316 refGene_gn=SMYD4;refGene_tx=XM_006939979.3
E1 15101291 15101704
refGene_gn=SERPINF2;refGene_tx=XM_006939980.3;refGene_gn=SERPINF2;refGene_tx=XM_011289028.2;refGene_gn=SERPINF2;refGene_tx=XM_019817361.1;refGene_gn=SERPINF2;refGene_tx=XM_019817362.1
E1 15102371 15102576
refGene_gn=SERPINF2;refGene_tx=XM_006939980.3;refGene_gn=SERPINF2;refGene_tx=XM_011289028.2;refGene_gn=SERPINF2;refGene_tx=XM_019817361.1;refGene_gn=SERPINF2;refGene_tx=XM_019817362.1
E1 15103722 15103882 refGene_gn=SMYD4;refGene_tx=XM_019817359.2
E1 15104471 15104614
refGene_gn=SERPINF2;refGene_tx=XM_006939980.3;refGene_gn=SERPINF2;refGene_tx=XM_011289028.2;refGene_gn=SERPINF2;refGene_tx=XM_019817361.1;refGene_gn=SERPINF2;refGene_tx=XM_019817362.1
E1 15105106 15105310
refGene_gn=SERPINF2;refGene_tx=XM_006939980.3;refGene_gn=SERPINF2;refGene_tx=XM_011289028.2;refGene_gn=SERPINF2;refGene_tx=XM_019817361.1;refGene_gn=SERPINF2;refGene_tx=XM_019817362.1
E1 15105424 15105568
refGene_gn=SERPINF2;refGene_tx=XM_006939980.3;refGene_gn=SERPINF2;refGene_tx=XM_011289028.2;refGene_gn=SERPINF2;refGene_tx=XM_019817361.1;refGene_gn=SERPINF2;refGene_tx=XM_019817362.1
E1 15106054 15106256
refGene_gn=SERPINF2;refGene_tx=XM_006939980.3;refGene_gn=SERPINF2;refGene_tx=XM_011289028.2;refGene_gn=SERPINF2;refGene_tx=XM_019817361.1;refGene_gn=SERPINF2;refGene_tx=XM_019817362.1
E1 15106552 15106615
refGene_gn=SERPINF2;refGene_tx=XM_006939980.3;refGene_gn=SERPINF2;refGene_tx=XM_011289028.2;refGene_gn=SERPINF2;refGene_tx=XM_019817361.1;re

fGene_gn=SERPINF2;refGene_tx=XM_019817362.1
E1	15106734	15106773
refGene_gn=SERPINF2;refGene_tx=XM_006939980.3;refGene_gn=SERPINF2;refGene_tx=XM_011289028.2;refGene_gn=SERPINF2;refGene_tx=XM_019817361.1;refGene_gn=SERPINF2;refGene_tx=XM_019817362.1
E1	15106952	15107022
refGene_gn=SERPINF2;refGene_tx=XM_006939980.3;refGene_gn=SERPINF2;refGene_tx=XM_011289028.2;refGene_gn=SERPINF2;refGene_tx=XM_019817361.1;refGene_gn=SERPINF2;refGene_tx=XM_019817362.1
E1	15107232	15107249	refGene_gn=SERPINF2;refGene_tx=XM_006939980.3
E1	15110007	15110328	refGene_gn=WDR81;refGene_tx=XM_003996418.4
E1	15111368	15111548	refGene_gn=WDR81;refGene_tx=XM_003996418.4
E1	15111828	15111974
refGene_gn=WDR81;refGene_tx=XM_003996418.4;refGene_gn=WDR81;refGene_tx=XM_011289026.3
E1	15112630	15113331
refGene_gn=WDR81;refGene_tx=XM_003996418.4;refGene_gn=WDR81;refGene_tx=XM_011289026.3;refGene_gn=WDR81;refGene_tx=XM_011289027.3
E1	15113998	15114164
refGene_gn=WDR81;refGene_tx=XM_003996418.4;refGene_gn=WDR81;refGene_tx=XM_011289026.3;refGene_gn=WDR81;refGene_tx=XM_011289027.3
E1	15114252	15114418
refGene_gn=WDR81;refGene_tx=XM_003996418.4;refGene_gn=WDR81;refGene_tx=XM_011289026.3;refGene_gn=WDR81;refGene_tx=XM_011289027.3
E1	15114729	15114920
refGene_gn=WDR81;refGene_tx=XM_003996418.4;refGene_gn=WDR81;refGene_tx=XM_011289026.3;refGene_gn=WDR81;refGene_tx=XM_011289027.3
E1	15115038	15115229
refGene_gn=WDR81;refGene_tx=XM_003996418.4;refGene_gn=WDR81;refGene_tx=XM_011289026.3;refGene_gn=WDR81;refGene_tx=XM_011289027.3
E1	15115489	15115597
refGene_gn=WDR81;refGene_tx=XM_003996418.4;refGene_gn=WDR81;refGene_tx=XM_011289026.3;refGene_gn=WDR81;refGene_tx=XM_011289027.3
E1	15119415	15123091
refGene_gn=WDR81;refGene_tx=XM_003996418.4;refGene_gn=WDR81;refGene_tx=XM_011289026.3;refGene_gn=WDR81;refGene_tx=XM_011289027.3
E1	15134768	15134938	refGene_gn=TLCD2;refGene_tx=XM_006939981.4
E1	15135476	15135559	refGene_gn=TLCD2;refGene_tx=XM_006939981.4
E1	15135762	15135845	refGene_gn=TLCD2;refGene_tx=XM_006939981.4
E1	15136409	15136862	refGene_gn=TLCD2;refGene_tx=XM_006939981.4
E1	15155839	15155939	refGene_gn=PRPF8;refGene_tx=XM_003996419.3
E1	15156436	15156605	refGene_gn=PRPF8;refGene_tx=XM_003996419.3
E1	15157980	15158145	refGene_gn=PRPF8;refGene_tx=XM_003996419.3
E1	15158223	15158442	refGene_gn=PRPF8;refGene_tx=XM_003996419.3
E1	15158567	15158780	refGene_gn=PRPF8;refGene_tx=XM_003996419.3
E1	15158906	15159032	refGene_gn=PRPF8;refGene_tx=XM_003996419.3
E1	15159112	15159218	refGene_gn=PRPF8;refGene_tx=XM_003996419.3
E1	15160120	15160311	refGene_gn=PRPF8;refGene_tx=XM_003996419.3
E1	15160440	15160560	refGene_gn=PRPF8;refGene_tx=XM_003996419.3
E1	15160650	15160840	refGene_gn=PRPF8;refGene_tx=XM_003996419.3

E1 15160975 15161095 refGene_gn=PRPF8;refGene_tx=XM_003996419.3
E1 15161204 15161339 refGene_gn=PRPF8;refGene_tx=XM_003996419.3
E1 15161691 15161821 refGene_gn=PRPF8;refGene_tx=XM_003996419.3
E1 15162229 15162426 refGene_gn=PRPF8;refGene_tx=XM_003996419.3
E1 15162894 15163101 refGene_gn=PRPF8;refGene_tx=XM_003996419.3
E1 15163225 15163389 refGene_gn=PRPF8;refGene_tx=XM_003996419.3
E1 15163541 15163668 refGene_gn=PRPF8;refGene_tx=XM_003996419.3
E1 15163798 15163991 refGene_gn=PRPF8;refGene_tx=XM_003996419.3
E1 15164266 15164454 refGene_gn=PRPF8;refGene_tx=XM_003996419.3
E1 15164619 15164858 refGene_gn=PRPF8;refGene_tx=XM_003996419.3
E1 15165092 15165239 refGene_gn=PRPF8;refGene_tx=XM_003996419.3
E1 15165429 15165640 refGene_gn=PRPF8;refGene_tx=XM_003996419.3
E1 15165842 15165959 refGene_gn=PRPF8;refGene_tx=XM_003996419.3
E1 15177439 15177687 refGene_gn=PRPF8;refGene_tx=XM_003996419.3
E1 15177788 15177968 refGene_gn=PRPF8;refGene_tx=XM_003996419.3
E1 15178163 15178299 refGene_gn=PRPF8;refGene_tx=XM_003996419.3
E1 15178418 15178588 refGene_gn=PRPF8;refGene_tx=XM_003996419.3
E1 15178733 15178863 refGene_gn=PRPF8;refGene_tx=XM_003996419.3
E1 15178988 15179135 refGene_gn=PRPF8;refGene_tx=XM_003996419.3
E1 15179618 15179779 refGene_gn=PRPF8;refGene_tx=XM_003996419.3
E1 15179967 15180159 refGene_gn=PRPF8;refGene_tx=XM_003996419.3
E1 15180473 15180711 refGene_gn=PRPF8;refGene_tx=XM_003996419.3
E1 15180842 15180971 refGene_gn=PRPF8;refGene_tx=XM_003996419.3
E1 15182080 15182194 refGene_gn=PRPF8;refGene_tx=XM_003996419.3
E1 15182273 15182447 refGene_gn=PRPF8;refGene_tx=XM_003996419.3
E1 15182741 15182935 refGene_gn=PRPF8;refGene_tx=XM_003996419.3
E1 15183651 15183891 refGene_gn=PRPF8;refGene_tx=XM_003996419.3
E1 15183985 15184127 refGene_gn=PRPF8;refGene_tx=XM_003996419.3
E1 15186042 15186183 refGene_gn=PRPF8;refGene_tx=XM_003996419.3
E1 15186318 15186458 refGene_gn=PRPF8;refGene_tx=XM_003996419.3
E1 15186562 15186765 refGene_gn=PRPF8;refGene_tx=XM_003996419.3
E1 15186909 15187064 refGene_gn=PRPF8;refGene_tx=XM_003996419.3
E1 15188267 15188495 refGene_gn=RILP;refGene_tx=XM_006939983.3
E1 15188569 15188663 refGene_gn=RILP;refGene_tx=XM_006939983.3
E1 15188747 15188854 refGene_gn=RILP;refGene_tx=XM_006939983.3
E1 15188962 15189208 refGene_gn=RILP;refGene_tx=XM_006939983.3
E1 15189509 15189637 refGene_gn=RILP;refGene_tx=XM_006939983.3
E1 15190019 15190142 refGene_gn=RILP;refGene_tx=XM_006939983.3
E1 15190822 15190906 refGene_gn=RILP;refGene_tx=XM_006939983.3
E1 15191185 15191272 refGene_gn=RILP;refGene_tx=XM_006939983.3
E1 15191303 15191409 refGene_gn=RILP;refGene_tx=XM_006939983.3
E1 15192039 15192257
refGene_gn=SCARF1;refGene_tx=XM_011289025.3;refGene_gn=SCARF1;refGene_
tx=XM_019817363.2
E1 15192486 15192646
refGene_gn=SCARF1;refGene_tx=XM_011289025.3;refGene_gn=SCARF1;refGene_
tx=XM_019817363.2;refGene_gn=SCARF1;refGene_tx=XM_023243575.1
E1 15192904 15193006
refGene_gn=SCARF1;refGene_tx=XM_011289025.3;refGene_gn=SCARF1;refGene_
tx=XM_019817363.2;refGene_gn=SCARF1;refGene_tx=XM_023243575.1

E1 15193880 15194406
refGene_gn=SCARF1;refGene_tx=XM_011289025.3;refGene_gn=SCARF1;refGene_tx=XM_019817363.2;refGene_gn=SCARF1;refGene_tx=XM_023243575.1
E1 15196432 15196651
refGene_gn=SCARF1;refGene_tx=XM_011289025.3;refGene_gn=SCARF1;refGene_tx=XM_019817363.2;refGene_gn=SCARF1;refGene_tx=XM_023243575.1
E1 15197059 15197188
refGene_gn=SCARF1;refGene_tx=XM_011289025.3;refGene_gn=SCARF1;refGene_tx=XM_019817363.2;refGene_gn=SCARF1;refGene_tx=XM_023243575.1
E1 15198001 15198105
refGene_gn=SCARF1;refGene_tx=XM_011289025.3;refGene_gn=SCARF1;refGene_tx=XM_019817363.2;refGene_gn=SCARF1;refGene_tx=XM_023243575.1
E1 15198591 15198712
refGene_gn=SCARF1;refGene_tx=XM_011289025.3;refGene_gn=SCARF1;refGene_tx=XM_019817363.2;refGene_gn=SCARF1;refGene_tx=XM_023243575.1
E1 15199955 15200077
refGene_gn=SCARF1;refGene_tx=XM_011289025.3;refGene_gn=SCARF1;refGene_tx=XM_019817363.2;refGene_gn=SCARF1;refGene_tx=XM_023243575.1
E1 15200161 15200308
refGene_gn=SCARF1;refGene_tx=XM_011289025.3;refGene_gn=SCARF1;refGene_tx=XM_019817363.2;refGene_gn=SCARF1;refGene_tx=XM_023243575.1
E1 15200965 15201825
refGene_gn=SCARF1;refGene_tx=XM_011289025.3;refGene_gn=SCARF1;refGene_tx=XM_019817363.2;refGene_gn=SCARF1;refGene_tx=XM_023243575.1
E1 15210927 15211087
refGene_gn=SLC43A2;refGene_tx=XM_003996420.5;refGene_gn=SLC43A2;refGene_tx=XM_019817364.2;refGene_gn=SLC43A2;refGene_tx=XM_019817365.2;refGene_gn=SLC43A2;refGene_tx=XM_019817366.1;refGene_gn=SLC43A2;refGene_tx=XM_019817367.1
E1 15219771 15219979
refGene_gn=SLC43A2;refGene_tx=XM_003996420.5;refGene_gn=SLC43A2;refGene_tx=XM_019817364.2;refGene_gn=SLC43A2;refGene_tx=XM_019817365.2;refGene_gn=SLC43A2;refGene_tx=XM_019817366.1;refGene_gn=SLC43A2;refGene_tx=XM_019817367.1
E1 15221270 15221326
refGene_gn=SLC43A2;refGene_tx=XM_003996420.5;refGene_gn=SLC43A2;refGene_tx=XM_019817364.2;refGene_gn=SLC43A2;refGene_tx=XM_019817365.2;refGene_gn=SLC43A2;refGene_tx=XM_019817366.1;refGene_gn=SLC43A2;refGene_tx=XM_019817367.1
E1 15222476 15222553
refGene_gn=SLC43A2;refGene_tx=XM_003996420.5;refGene_gn=SLC43A2;refGene_tx=XM_019817364.2;refGene_gn=SLC43A2;refGene_tx=XM_019817365.2;refGene_gn=SLC43A2;refGene_tx=XM_019817366.1;refGene_gn=SLC43A2;refGene_tx=XM_019817367.1
E1 15234356 15234449
refGene_gn=SLC43A2;refGene_tx=XM_003996420.5;refGene_gn=SLC43A2;refGene_tx=XM_019817364.2;refGene_gn=SLC43A2;refGene_tx=XM_019817365.2;refGene_gn=SLC43A2;refGene_tx=XM_019817366.1;refGene_gn=SLC43A2;refGene_tx=XM_019817367.1
E1 15235770 15235904

refGene_gn=SLC43A2;refGene_tx=XM_003996420.5;refGene_gn=SLC43A2;refGene_tx=XM_019817364.2;refGene_gn=SLC43A2;refGene_tx=XM_019817365.2;refGene_gn=SLC43A2;refGene_tx=XM_019817366.1;refGene_gn=SLC43A2;refGene_tx=XM_019817367.1
E1 15235987 15236190
refGene_gn=SLC43A2;refGene_tx=XM_003996420.5;refGene_gn=SLC43A2;refGene_tx=XM_019817364.2;refGene_gn=SLC43A2;refGene_tx=XM_019817365.2;refGene_gn=SLC43A2;refGene_tx=XM_019817366.1;refGene_gn=SLC43A2;refGene_tx=XM_019817367.1
E1 15236644 15236791
refGene_gn=SLC43A2;refGene_tx=XM_003996420.5;refGene_gn=SLC43A2;refGene_tx=XM_019817364.2;refGene_gn=SLC43A2;refGene_tx=XM_019817365.2;refGene_gn=SLC43A2;refGene_tx=XM_019817366.1;refGene_gn=SLC43A2;refGene_tx=XM_019817367.1
E1 15239302 15239314
refGene_gn=SLC43A2;refGene_tx=XM_019817364.2;refGene_gn=SLC43A2;refGene_tx=XM_019817365.2;refGene_gn=SLC43A2;refGene_tx=XM_019817366.1;refGene_gn=SLC43A2;refGene_tx=XM_019817367.1
E1 15239900 15240039
refGene_gn=SLC43A2;refGene_tx=XM_003996420.5;refGene_gn=SLC43A2;refGene_tx=XM_019817364.2;refGene_gn=SLC43A2;refGene_tx=XM_019817365.2;refGene_gn=SLC43A2;refGene_tx=XM_019817366.1;refGene_gn=SLC43A2;refGene_tx=XM_019817367.1
E1 15241338 15241468
refGene_gn=SLC43A2;refGene_tx=XM_003996420.5;refGene_gn=SLC43A2;refGene_tx=XM_019817364.2;refGene_gn=SLC43A2;refGene_tx=XM_019817365.2;refGene_gn=SLC43A2;refGene_tx=XM_019817366.1;refGene_gn=SLC43A2;refGene_tx=XM_019817367.1
E1 15243513 15243587
refGene_gn=SLC43A2;refGene_tx=XM_003996420.5;refGene_gn=SLC43A2;refGene_tx=XM_019817364.2;refGene_gn=SLC43A2;refGene_tx=XM_019817365.2;refGene_gn=SLC43A2;refGene_tx=XM_019817366.1;refGene_gn=SLC43A2;refGene_tx=XM_019817367.1
E1 15244881 15245005
refGene_gn=SLC43A2;refGene_tx=XM_003996420.5;refGene_gn=SLC43A2;refGene_tx=XM_019817364.2;refGene_gn=SLC43A2;refGene_tx=XM_019817365.2;refGene_gn=SLC43A2;refGene_tx=XM_019817366.1;refGene_gn=SLC43A2;refGene_tx=XM_019817367.1
E1 15245519 15245681
refGene_gn=SLC43A2;refGene_tx=XM_003996420.5;refGene_gn=SLC43A2;refGene_tx=XM_019817364.2;refGene_gn=SLC43A2;refGene_tx=XM_019817365.2;refGene_gn=SLC43A2;refGene_tx=XM_019817366.1;refGene_gn=SLC43A2;refGene_tx=XM_019817367.1
E1 15258085 15258105 refGene_gn=PITPNA;refGene_tx=XM_011289023.3
E1 15261802 15261833 refGene_gn=PITPNA;refGene_tx=XM_011289023.3
E1 15266650 15266796 refGene_gn=PITPNA;refGene_tx=XM_011289023.3
E1 15270832 15270924 refGene_gn=PITPNA;refGene_tx=XM_011289023.3
E1 15274968 15274976 refGene_gn=PITPNA;refGene_tx=XM_011289023.3
E1 15276032 15276107 refGene_gn=PITPNA;refGene_tx=XM_011289023.3
E1 15278253 15278337 refGene_gn=PITPNA;refGene_tx=XM_011289023.3

E1 15280229 15280307 refGene_gn=PITPNA;refGene_tx=XM_011289023.3
E1 15280457 15280568 refGene_gn=PITPNA;refGene_tx=XM_011289023.3
E1 15281340 15281463 refGene_gn=PITPNA;refGene_tx=XM_011289023.3
E1 15295161 15295206 refGene_gn=PITPNA;refGene_tx=XM_011289023.3
E1 15300187 15300240 refGene_gn=INPP5K;refGene_tx=XM_011289021.2
E1 15301795 15301920
refGene_gn=INPP5K;refGene_tx=XM_019817368.2;refGene_gn=INPP5K;refGene_tx=XM_019817369.2
E1 15302052 15302160
refGene_gn=INPP5K;refGene_tx=XM_011289021.2;refGene_gn=INPP5K;refGene_tx=XM_019817368.2;refGene_gn=INPP5K;refGene_tx=XM_019817369.2
E1 15302475 15302584
refGene_gn=INPP5K;refGene_tx=XM_011289021.2;refGene_gn=INPP5K;refGene_tx=XM_011289022.3;refGene_gn=INPP5K;refGene_tx=XM_019817368.2;refGene_gn=INPP5K;refGene_tx=XM_019817369.2
E1 15306484 15306601
refGene_gn=INPP5K;refGene_tx=XM_011289021.2;refGene_gn=INPP5K;refGene_tx=XM_011289022.3;refGene_gn=INPP5K;refGene_tx=XM_019817368.2
E1 15306916 15307092
refGene_gn=INPP5K;refGene_tx=XM_011289021.2;refGene_gn=INPP5K;refGene_tx=XM_011289022.3;refGene_gn=INPP5K;refGene_tx=XM_019817368.2;refGene_gn=INPP5K;refGene_tx=XM_019817369.2
E1 15307607 15307719
refGene_gn=INPP5K;refGene_tx=XM_011289021.2;refGene_gn=INPP5K;refGene_tx=XM_011289022.3;refGene_gn=INPP5K;refGene_tx=XM_019817368.2;refGene_gn=INPP5K;refGene_tx=XM_019817369.2
E1 15308376 15308486
refGene_gn=INPP5K;refGene_tx=XM_011289021.2;refGene_gn=INPP5K;refGene_tx=XM_011289022.3;refGene_gn=INPP5K;refGene_tx=XM_019817368.2;refGene_gn=INPP5K;refGene_tx=XM_019817369.2
E1 15315182 15315369
refGene_gn=INPP5K;refGene_tx=XM_011289021.2;refGene_gn=INPP5K;refGene_tx=XM_011289022.3;refGene_gn=INPP5K;refGene_tx=XM_019817368.2;refGene_gn=INPP5K;refGene_tx=XM_019817369.2
E1 15316237 15316375
refGene_gn=INPP5K;refGene_tx=XM_011289021.2;refGene_gn=INPP5K;refGene_tx=XM_011289022.3;refGene_gn=INPP5K;refGene_tx=XM_019817368.2;refGene_gn=INPP5K;refGene_tx=XM_019817369.2
E1 15316638 15316722
refGene_gn=INPP5K;refGene_tx=XM_011289021.2;refGene_gn=INPP5K;refGene_tx=XM_011289022.3;refGene_gn=INPP5K;refGene_tx=XM_019817368.2;refGene_gn=INPP5K;refGene_tx=XM_019817369.2
E1 15316890 15316995
refGene_gn=INPP5K;refGene_tx=XM_011289021.2;refGene_gn=INPP5K;refGene_tx=XM_011289022.3;refGene_gn=INPP5K;refGene_tx=XM_019817368.2;refGene_gn=INPP5K;refGene_tx=XM_019817369.2
E1 15317192 15317252
refGene_gn=INPP5K;refGene_tx=XM_011289021.2;refGene_gn=INPP5K;refGene_tx=XM_011289022.3;refGene_gn=INPP5K;refGene_tx=XM_019817368.2;refGene_gn=INPP5K;refGene_tx=XM_019817369.2

E1 15320648 15320723 refGene_gn=MYO1C;refGene_tx=XM_003996421.4
E1 15326280 15326298 refGene_gn=MYO1C;refGene_tx=XM_011289020.2
E1 15327668 15327824
refGene_gn=MYO1C;refGene_tx=XM_003996421.4;refGene_gn=MYO1C;refGene_tx=XM_006939987.3;refGene_gn=MYO1C;refGene_tx=XM_011289020.2;refGene_gn=MYO1C;refGene_tx=XM_023243434.1
E1 15327940 15328056
refGene_gn=MYO1C;refGene_tx=XM_003996421.4;refGene_gn=MYO1C;refGene_tx=XM_006939987.3;refGene_gn=MYO1C;refGene_tx=XM_011289020.2;refGene_gn=MYO1C;refGene_tx=XM_023243434.1
E1 15328603 15328802
refGene_gn=MYO1C;refGene_tx=XM_003996421.4;refGene_gn=MYO1C;refGene_tx=XM_006939987.3;refGene_gn=MYO1C;refGene_tx=XM_011289020.2;refGene_gn=MYO1C;refGene_tx=XM_023243434.1
E1 15328882 15328963
refGene_gn=MYO1C;refGene_tx=XM_003996421.4;refGene_gn=MYO1C;refGene_tx=XM_006939987.3;refGene_gn=MYO1C;refGene_tx=XM_011289020.2;refGene_gn=MYO1C;refGene_tx=XM_023243434.1
E1 15330344 15330524
refGene_gn=MYO1C;refGene_tx=XM_003996421.4;refGene_gn=MYO1C;refGene_tx=XM_006939987.3;refGene_gn=MYO1C;refGene_tx=XM_011289020.2;refGene_gn=MYO1C;refGene_tx=XM_023243434.1
E1 15330605 15330704
refGene_gn=MYO1C;refGene_tx=XM_003996421.4;refGene_gn=MYO1C;refGene_tx=XM_006939987.3;refGene_gn=MYO1C;refGene_tx=XM_011289020.2;refGene_gn=MYO1C;refGene_tx=XM_023243434.1
E1 15330902 15331016
refGene_gn=MYO1C;refGene_tx=XM_003996421.4;refGene_gn=MYO1C;refGene_tx=XM_006939987.3;refGene_gn=MYO1C;refGene_tx=XM_011289020.2;refGene_gn=MYO1C;refGene_tx=XM_023243434.1
E1 15331102 15331174
refGene_gn=MYO1C;refGene_tx=XM_003996421.4;refGene_gn=MYO1C;refGene_tx=XM_006939987.3;refGene_gn=MYO1C;refGene_tx=XM_011289020.2;refGene_gn=MYO1C;refGene_tx=XM_023243434.1
E1 15331458 15331578
refGene_gn=MYO1C;refGene_tx=XM_003996421.4;refGene_gn=MYO1C;refGene_tx=XM_006939987.3;refGene_gn=MYO1C;refGene_tx=XM_011289020.2;refGene_gn=MYO1C;refGene_tx=XM_023243434.1
E1 15331701 15331784
refGene_gn=MYO1C;refGene_tx=XM_003996421.4;refGene_gn=MYO1C;refGene_tx=XM_006939987.3;refGene_gn=MYO1C;refGene_tx=XM_011289020.2;refGene_gn=MYO1C;refGene_tx=XM_023243434.1
E1 15332005 15332111
refGene_gn=MYO1C;refGene_tx=XM_003996421.4;refGene_gn=MYO1C;refGene_tx=XM_006939987.3;refGene_gn=MYO1C;refGene_tx=XM_011289020.2;refGene_gn=MYO1C;refGene_tx=XM_023243434.1
E1 15332228 15332309
refGene_gn=MYO1C;refGene_tx=XM_003996421.4;refGene_gn=MYO1C;refGene_tx=XM_006939987.3;refGene_gn=MYO1C;refGene_tx=XM_011289020.2;refGene_gn=MYO1C;refGene_tx=XM_023243434.1

E1 15332611 15332703
refGene_gn=MYO1C;refGene_tx=XM_003996421.4;refGene_gn=MYO1C;refGene_tx=XM_006939987.3;refGene_gn=MYO1C;refGene_tx=XM_011289020.2;refGene_gn=MYO1C;refGene_tx=XM_023243434.1
E1 15335326 15335421
refGene_gn=MYO1C;refGene_tx=XM_003996421.4;refGene_gn=MYO1C;refGene_tx=XM_006939987.3;refGene_gn=MYO1C;refGene_tx=XM_011289020.2;refGene_gn=MYO1C;refGene_tx=XM_023243434.1
E1 15335501 15335548
refGene_gn=MYO1C;refGene_tx=XM_003996421.4;refGene_gn=MYO1C;refGene_tx=XM_006939987.3;refGene_gn=MYO1C;refGene_tx=XM_011289020.2;refGene_gn=MYO1C;refGene_tx=XM_023243434.1
E1 15335656 15335737
refGene_gn=MYO1C;refGene_tx=XM_003996421.4;refGene_gn=MYO1C;refGene_tx=XM_006939987.3;refGene_gn=MYO1C;refGene_tx=XM_011289020.2;refGene_gn=MYO1C;refGene_tx=XM_023243434.1
E1 15337973 15338079
refGene_gn=MYO1C;refGene_tx=XM_003996421.4;refGene_gn=MYO1C;refGene_tx=XM_006939987.3;refGene_gn=MYO1C;refGene_tx=XM_011289020.2;refGene_gn=MYO1C;refGene_tx=XM_023243434.1
E1 15338188 15338306
refGene_gn=MYO1C;refGene_tx=XM_003996421.4;refGene_gn=MYO1C;refGene_tx=XM_006939987.3;refGene_gn=MYO1C;refGene_tx=XM_011289020.2;refGene_gn=MYO1C;refGene_tx=XM_023243434.1
E1 15339150 15339264
refGene_gn=MYO1C;refGene_tx=XM_003996421.4;refGene_gn=MYO1C;refGene_tx=XM_006939987.3;refGene_gn=MYO1C;refGene_tx=XM_011289020.2;refGene_gn=MYO1C;refGene_tx=XM_023243434.1
E1 15339343 15339420
refGene_gn=MYO1C;refGene_tx=XM_003996421.4;refGene_gn=MYO1C;refGene_tx=XM_006939987.3;refGene_gn=MYO1C;refGene_tx=XM_011289020.2;refGene_gn=MYO1C;refGene_tx=XM_023243434.1
E1 15339771 15339840
refGene_gn=MYO1C;refGene_tx=XM_003996421.4;refGene_gn=MYO1C;refGene_tx=XM_006939987.3;refGene_gn=MYO1C;refGene_tx=XM_011289020.2;refGene_gn=MYO1C;refGene_tx=XM_023243434.1
E1 15339938 15340023
refGene_gn=MYO1C;refGene_tx=XM_003996421.4;refGene_gn=MYO1C;refGene_tx=XM_006939987.3;refGene_gn=MYO1C;refGene_tx=XM_011289020.2;refGene_gn=MYO1C;refGene_tx=XM_023243434.1
E1 15340118 15340278
refGene_gn=MYO1C;refGene_tx=XM_003996421.4;refGene_gn=MYO1C;refGene_tx=XM_006939987.3;refGene_gn=MYO1C;refGene_tx=XM_011289020.2;refGene_gn=MYO1C;refGene_tx=XM_023243434.1
E1 15340567 15340651
refGene_gn=MYO1C;refGene_tx=XM_003996421.4;refGene_gn=MYO1C;refGene_tx=XM_006939987.3;refGene_gn=MYO1C;refGene_tx=XM_011289020.2;refGene_gn=MYO1C;refGene_tx=XM_023243434.1
E1 15341389 15341483
refGene_gn=MYO1C;refGene_tx=XM_003996421.4;refGene_gn=MYO1C;refGene_tx

=XM_006939987.3;refGene_gn=MYO1C;refGene_tx=XM_011289020.2;refGene_gn=MYO1C;refGene_tx=XM_023243434.1
E1 15341577 15341633
refGene_gn=MYO1C;refGene_tx=XM_003996421.4;refGene_gn=MYO1C;refGene_tx=XM_006939987.3;refGene_gn=MYO1C;refGene_tx=XM_011289020.2;refGene_gn=MYO1C;refGene_tx=XM_023243434.1
E1 15341759 15341895
refGene_gn=MYO1C;refGene_tx=XM_003996421.4;refGene_gn=MYO1C;refGene_tx=XM_006939987.3;refGene_gn=MYO1C;refGene_tx=XM_011289020.2;refGene_gn=MYO1C;refGene_tx=XM_023243434.1
E1 15341972 15342043
refGene_gn=MYO1C;refGene_tx=XM_003996421.4;refGene_gn=MYO1C;refGene_tx=XM_006939987.3;refGene_gn=MYO1C;refGene_tx=XM_011289020.2;refGene_gn=MYO1C;refGene_tx=XM_023243434.1
E1 15342132 15342230
refGene_gn=MYO1C;refGene_tx=XM_003996421.4;refGene_gn=MYO1C;refGene_tx=XM_006939987.3;refGene_gn=MYO1C;refGene_tx=XM_011289020.2;refGene_gn=MYO1C;refGene_tx=XM_023243434.1
E1 15342304 15342404
refGene_gn=MYO1C;refGene_tx=XM_003996421.4;refGene_gn=MYO1C;refGene_tx=XM_006939987.3;refGene_gn=MYO1C;refGene_tx=XM_011289020.2;refGene_gn=MYO1C;refGene_tx=XM_023243434.1
E1 15342859 15342886
refGene_gn=MYO1C;refGene_tx=XM_003996421.4;refGene_gn=MYO1C;refGene_tx=XM_006939987.3;refGene_gn=MYO1C;refGene_tx=XM_011289020.2;refGene_gn=MYO1C;refGene_tx=XM_023243434.1
E1 15350290 15350531
refGene_gn=CRK;refGene_tx=XM_023244818.1;refGene_gn=CRK;refGene_tx=XM_023244819.1
E1 15360974 15361510
refGene_gn=CRK;refGene_tx=XM_023244818.1;refGene_gn=CRK;refGene_tx=XM_023244819.1
E1 15372947 15373085
refGene_gn=CRK;refGene_tx=XM_023244818.1;refGene_gn=CRK;refGene_tx=XM_023244819.1
E1 15407361 15407425
refGene_gn=YWHAE;refGene_tx=XM_003996422.5;refGene_gn=YWHAE;refGene_tx=XM_006939989.4
E1 15432294 15432358 refGene_gn=YWHAE;refGene_tx=XM_019817370.2
E1 15449405 15449605
refGene_gn=YWHAE;refGene_tx=XM_003996422.5;refGene_gn=YWHAE;refGene_tx=XM_006939989.4;refGene_gn=YWHAE;refGene_tx=XM_019817370.2
E1 15453281 15453388
refGene_gn=YWHAE;refGene_tx=XM_003996422.5;refGene_gn=YWHAE;refGene_tx=XM_006939989.4;refGene_gn=YWHAE;refGene_tx=XM_019817370.2
E1 15454042 15454249
refGene_gn=YWHAE;refGene_tx=XM_003996422.5;refGene_gn=YWHAE;refGene_tx=XM_006939989.4;refGene_gn=YWHAE;refGene_tx=XM_019817370.2
E1 15456385 15456522
refGene_gn=YWHAE;refGene_tx=XM_003996422.5;refGene_gn=YWHAE;refGene_tx

=XM_006939989.4;refGene_gn=YWHAE;refGene_tx=XM_019817370.2
E1 15459419 15459427 refGene_gn=YWHAE;refGene_tx=XM_006939989.4
E1 15461129 15461182
refGene_gn=YWHAE;refGene_tx=XM_003996422.5;refGene_gn=YWHAE;refGene_tx
=XM_019817370.2
E1 15479434 15479571
refGene_gn=DOC2B;refGene_tx=XM_023244655.1;refGene_gn=DOC2B;refGene_tx
=XM_023244656.1
E1 15480240 15480337
refGene_gn=DOC2B;refGene_tx=XM_023244655.1;refGene_gn=DOC2B;refGene_tx
=XM_023244656.1
E1 15481251 15481333
refGene_gn=DOC2B;refGene_tx=XM_023244655.1;refGene_gn=DOC2B;refGene_tx
=XM_023244656.1
E1 15492622 15492780
refGene_gn=DOC2B;refGene_tx=XM_023244655.1;refGene_gn=DOC2B;refGene_tx
=XM_023244656.1
E1 15495424 15495551
refGene_gn=DOC2B;refGene_tx=XM_023244655.1;refGene_gn=DOC2B;refGene_tx
=XM_023244656.1
E1 15495733 15495843
refGene_gn=DOC2B;refGene_tx=XM_023244655.1;refGene_gn=DOC2B;refGene_tx
=XM_023244656.1
E1 15498159 15498234
refGene_gn=DOC2B;refGene_tx=XM_023244655.1;refGene_gn=DOC2B;refGene_tx
=XM_023244656.1
E1 15499721 15499748 refGene_gn=DOC2B;refGene_tx=XM_023244655.1
E1 15500286 15500366
refGene_gn=DOC2B;refGene_tx=XM_023244655.1;refGene_gn=DOC2B;refGene_tx
=XM_023244656.1
E1 15508451 15508824
refGene_gn=DOC2B;refGene_tx=XM_023244655.1;refGene_gn=DOC2B;refGene_tx
=XM_023244656.1
E1 15538192 15538534 refGene_gn=RPH3AL;refGene_tx=XM_023244657.1
E1 15541591 15541621 refGene_gn=RPH3AL;refGene_tx=XM_023244657.1
E1 15541622 15541738 refGene_gn=RPH3AL;refGene_tx=XM_023244657.1
E1 15543490 15543604 refGene_gn=RPH3AL;refGene_tx=XM_023244657.1
E1 15564157 15564332 refGene_gn=RPH3AL;refGene_tx=XM_023244657.1
E1 15593805 15593892 refGene_gn=RPH3AL;refGene_tx=XM_023244657.1
E1 15621637 15621767 refGene_gn=RPH3AL;refGene_tx=XM_023244657.1
E1 15623964 15624108 refGene_gn=RPH3AL;refGene_tx=XM_023244657.1
E1 15629504 15629581 refGene_gn=RPH3AL;refGene_tx=XM_023244657.1
E1 15711819 15712036
refGene_gn=CE1H17orf97;refGene_tx=XM_011289014.2;refGene_gn=CE1H17orf9
7;refGene_tx=XM_019817374.1
E1 15714740 15715204
refGene_gn=CE1H17orf97;refGene_tx=XM_011289014.2;refGene_gn=CE1H17orf9
7;refGene_tx=XM_019817374.1
E1 15724571 15724902 refGene_gn=RFLNB;refGene_tx=XM_023243672.1
E1 15726745 15726855 refGene_gn=RFLNB;refGene_tx=XM_023243672.1

E1	15729608 15729824 refGene_gn=RFLNB;refGene_tx=XM_023243672.1
E1	15749289 15749460 refGene_gn=VPS53;refGene_tx=XM_011289012.3
E1	15749956 15750061 refGene_gn=VPS53;refGene_tx=XM_011289012.3
E1	15751575 15751713 refGene_gn=VPS53;refGene_tx=XM_011289012.3
E1	15763995 15764065 refGene_gn=VPS53;refGene_tx=XM_011289012.3
E1	15768190 15768339 refGene_gn=VPS53;refGene_tx=XM_011289012.3
E1	15781582 15781661 refGene_gn=VPS53;refGene_tx=XM_011289012.3
E1	15782612 15782695 refGene_gn=VPS53;refGene_tx=XM_011289012.3
E1	15788953 15789101 refGene_gn=VPS53;refGene_tx=XM_011289012.3
E1	15791874 15792117 refGene_gn=VPS53;refGene_tx=XM_011289012.3
E1	15801687 15801782 refGene_gn=VPS53;refGene_tx=XM_011289012.3
E1	15808768 15808870 refGene_gn=VPS53;refGene_tx=XM_011289012.3
E1	15822782 15822924 refGene_gn=VPS53;refGene_tx=XM_011289012.3
E1	15825768 15825911 refGene_gn=VPS53;refGene_tx=XM_011289012.3
E1	15826365 15826509 refGene_gn=VPS53;refGene_tx=XM_011289012.3
E1	15829341 15829420 refGene_gn=VPS53;refGene_tx=XM_011289012.3
E1	15840037 15840157 refGene_gn=VPS53;refGene_tx=XM_011289012.3
E1	15842325 15842441 refGene_gn=VPS53;refGene_tx=XM_011289012.3
E1	15846609 15846696 refGene_gn=VPS53;refGene_tx=XM_011289012.3
E1	15869740 15869807 refGene_gn=VPS53;refGene_tx=XM_011289012.3
E1	15871487 15871537 refGene_gn=VPS53;refGene_tx=XM_011289012.3
E1	15882025 15882106 refGene_gn=VPS53;refGene_tx=XM_011289012.3
E1	15883892 15883979 refGene_gn=VPS53;refGene_tx=XM_011289012.3
E1	15898714 15898836
refGene_gn=FAM57A;refGene_tx=XM_003996445.5;refGene_gn=FAM57A;refGene_tx=XM_019817378.2;refGene_gn=FAM57A;refGene_tx=XM_019817379.2
E1	15899267 15899351
refGene_gn=FAM57A;refGene_tx=XM_003996445.5;refGene_gn=FAM57A;refGene_tx=XM_019817379.2
E1	15902297 15902502
refGene_gn=FAM57A;refGene_tx=XM_003996445.5;refGene_gn=FAM57A;refGene_tx=XM_019817378.2
E1	15903663 15903759
refGene_gn=FAM57A;refGene_tx=XM_003996445.5;refGene_gn=FAM57A;refGene_tx=XM_019817378.2;refGene_gn=FAM57A;refGene_tx=XM_019817379.2
E1	15904206 15904476
refGene_gn=FAM57A;refGene_tx=XM_003996445.5;refGene_gn=FAM57A;refGene_tx=XM_019817378.2
E1	15907115 15910282
refGene_gn=GEMIN4;refGene_tx=XM_003996446.4;refGene_gn=GEMIN4;refGene_tx=XM_019817375.2
E1	15913161 15913444
refGene_gn=GEMIN4;refGene_tx=XM_003996446.4;refGene_gn=GEMIN4;refGene_tx=XM_019817375.2;refGene_gn=LOC101088630;refGene_tx=XM_023244675.1
E1	15914072 15914103
refGene_gn=LOC101088630;refGene_tx=XM_023244675.1
E1	15914175 15914439
refGene_gn=LOC101088630;refGene_tx=XM_019817380.2
E1	15919527 15919593 refGene_gn=GLOD4;refGene_tx=XM_003996448.4
E1	15927255 15927342 refGene_gn=GLOD4;refGene_tx=XM_003996448.4

E1 15927431 15927545 refGene_gn=GLOD4;refGene_tx=XM_003996448.4
E1 15927823 15927910 refGene_gn=GLOD4;refGene_tx=XM_003996448.4
E1 15928661 15928798 refGene_gn=GLOD4;refGene_tx=XM_003996448.4
E1 15931442 15931587 refGene_gn=GLOD4;refGene_tx=XM_003996448.4
E1 15936174 15936295 refGene_gn=GLOD4;refGene_tx=XM_003996448.4
E1 15937100 15937150 refGene_gn=GLOD4;refGene_tx=XM_003996448.4
E1 15939018 15939129 refGene_gn=GLOD4;refGene_tx=XM_003996448.4
E1 15939204 15939545 refGene_gn=MRM3;refGene_tx=XM_003996449.4
E1 15939803 15940048
refGene_gn=MRM3;refGene_tx=XM_003996449.4;refGene_gn=MRM3;refGene_tx=XM_019817376.1
E1 15942601 15942769
refGene_gn=MRM3;refGene_tx=XM_003996449.4;refGene_gn=MRM3;refGene_tx=XM_019817376.1;refGene_gn=MRM3;refGene_tx=XM_019817377.1
E1 15945102 15945641
refGene_gn=MRM3;refGene_tx=XM_003996449.4;refGene_gn=MRM3;refGene_tx=XM_019817376.1;refGene_gn=MRM3;refGene_tx=XM_019817377.1
E1 15952695 15952878 refGene_gn=NXN;refGene_tx=XM_023244495.1
E1 15955730 15955855 refGene_gn=NXN;refGene_tx=XM_023244495.1
E1 15957406 15957586 refGene_gn=NXN;refGene_tx=XM_023244495.1
E1 15976902 15977009 refGene_gn=NXN;refGene_tx=XM_023244495.1
E1 15978913 15979014 refGene_gn=NXN;refGene_tx=XM_023244495.1
E1 15979494 15979628 refGene_gn=NXN;refGene_tx=XM_023244495.1
E1 15981238 15981356 refGene_gn=NXN;refGene_tx=XM_023244495.1
E1 16110442 16110814 refGene_gn=NXN;refGene_tx=XM_023244495.1
E1 16125914 16126152 refGene_gn=TIMM22;refGene_tx=XM_003996450.5
E1 16128287 16128484 refGene_gn=TIMM22;refGene_tx=XM_003996450.5
E1 16130236 16130309 refGene_gn=TIMM22;refGene_tx=XM_003996450.5
E1 16131687 16131764 refGene_gn=TIMM22;refGene_tx=XM_003996450.5
E1 16136874 16136964
refGene_gn=ABR;refGene_tx=XM_003996452.5;refGene_gn=ABR;refGene_tx=XM_003996453.5;refGene_gn=ABR;refGene_tx=XM_019817386.2;refGene_gn=ABR;refGene_tx=XM_023243676.1;refGene_gn=ABR;refGene_tx=XM_023243677.1;refGene_gn=ABR;refGene_tx=XM_023243678.1;refGene_gn=ABR;refGene_tx=XM_023243679.1;refGene_gn=ABR;refGene_tx=XM_023243681.1
E1 16137552 16137700
refGene_gn=ABR;refGene_tx=XM_003996452.5;refGene_gn=ABR;refGene_tx=XM_003996453.5;refGene_gn=ABR;refGene_tx=XM_019817386.2;refGene_gn=ABR;refGene_tx=XM_023243676.1;refGene_gn=ABR;refGene_tx=XM_023243677.1;refGene_gn=ABR;refGene_tx=XM_023243678.1;refGene_gn=ABR;refGene_tx=XM_023243679.1;refGene_gn=ABR;refGene_tx=XM_023243681.1
E1 16139768 16139874
refGene_gn=ABR;refGene_tx=XM_003996452.5;refGene_gn=ABR;refGene_tx=XM_003996453.5;refGene_gn=ABR;refGene_tx=XM_019817386.2;refGene_gn=ABR;refGene_tx=XM_023243676.1;refGene_gn=ABR;refGene_tx=XM_023243677.1;refGene_gn=ABR;refGene_tx=XM_023243678.1;refGene_gn=ABR;refGene_tx=XM_023243679.1;refGene_gn=ABR;refGene_tx=XM_023243681.1
E1 16140939 16141074
refGene_gn=ABR;refGene_tx=XM_003996452.5;refGene_gn=ABR;refGene_tx=XM_003996453.5;refGene_gn=ABR;refGene_tx=XM_019817386.2;refGene_gn=ABR;re

fGene_tx=XM_023243676.1;refGene_gn=ABR;refGene_tx=XM_023243677.1;refGene_gn=ABR;refGene_tx=XM_023243678.1;refGene_gn=ABR;refGene_tx=XM_023243679.1;refGene_gn=ABR;refGene_tx=XM_023243681.1
E1 16142139 16142279
refGene_gn=ABR;refGene_tx=XM_003996452.5;refGene_gn=ABR;refGene_tx=XM_003996453.5;refGene_gn=ABR;refGene_tx=XM_019817386.2;refGene_gn=ABR;refGene_tx=XM_023243676.1;refGene_gn=ABR;refGene_tx=XM_023243677.1;refGene_gn=ABR;refGene_tx=XM_023243678.1;refGene_gn=ABR;refGene_tx=XM_023243679.1;refGene_gn=ABR;refGene_tx=XM_023243681.1
E1 16142967 16143077
refGene_gn=ABR;refGene_tx=XM_003996452.5;refGene_gn=ABR;refGene_tx=XM_003996453.5;refGene_gn=ABR;refGene_tx=XM_019817386.2;refGene_gn=ABR;refGene_tx=XM_023243676.1;refGene_gn=ABR;refGene_tx=XM_023243677.1;refGene_gn=ABR;refGene_tx=XM_023243678.1;refGene_gn=ABR;refGene_tx=XM_023243679.1;refGene_gn=ABR;refGene_tx=XM_023243681.1
E1 16143291 16143387
refGene_gn=ABR;refGene_tx=XM_003996452.5;refGene_gn=ABR;refGene_tx=XM_003996453.5;refGene_gn=ABR;refGene_tx=XM_019817386.2;refGene_gn=ABR;refGene_tx=XM_023243676.1;refGene_gn=ABR;refGene_tx=XM_023243677.1;refGene_gn=ABR;refGene_tx=XM_023243678.1;refGene_gn=ABR;refGene_tx=XM_023243679.1;refGene_gn=ABR;refGene_tx=XM_023243681.1
E1 16157258 16157420 refGene_gn=ABR;refGene_tx=XM_003996453.5
E1 16160586 16160697 refGene_gn=ABR;refGene_tx=XM_023243680.1
E1 16172234 16172366
refGene_gn=ABR;refGene_tx=XM_003996452.5;refGene_gn=ABR;refGene_tx=XM_019817386.2;refGene_gn=ABR;refGene_tx=XM_023243676.1;refGene_gn=ABR;refGene_tx=XM_023243677.1;refGene_gn=ABR;refGene_tx=XM_023243678.1;refGene_gn=ABR;refGene_tx=XM_023243679.1;refGene_gn=ABR;refGene_tx=XM_023243680.1;refGene_gn=ABR;refGene_tx=XM_023243681.1
E1 16172775 16172873
refGene_gn=ABR;refGene_tx=XM_003996452.5;refGene_gn=ABR;refGene_tx=XM_019817386.2;refGene_gn=ABR;refGene_tx=XM_023243676.1;refGene_gn=ABR;refGene_tx=XM_023243677.1;refGene_gn=ABR;refGene_tx=XM_023243678.1;refGene_gn=ABR;refGene_tx=XM_023243679.1;refGene_gn=ABR;refGene_tx=XM_023243680.1;refGene_gn=ABR;refGene_tx=XM_023243681.1
E1 16175608 16175683
refGene_gn=ABR;refGene_tx=XM_003996452.5;refGene_gn=ABR;refGene_tx=XM_019817386.2;refGene_gn=ABR;refGene_tx=XM_023243676.1;refGene_gn=ABR;refGene_tx=XM_023243677.1;refGene_gn=ABR;refGene_tx=XM_023243678.1;refGene_gn=ABR;refGene_tx=XM_023243679.1;refGene_gn=ABR;refGene_tx=XM_023243680.1;refGene_gn=ABR;refGene_tx=XM_023243681.1
E1 16176623 16176728
refGene_gn=ABR;refGene_tx=XM_003996452.5;refGene_gn=ABR;refGene_tx=XM_019817386.2;refGene_gn=ABR;refGene_tx=XM_023243676.1;refGene_gn=ABR;refGene_tx=XM_023243677.1;refGene_gn=ABR;refGene_tx=XM_023243678.1;refGene_gn=ABR;refGene_tx=XM_023243679.1;refGene_gn=ABR;refGene_tx=XM_023243680.1;refGene_gn=ABR;refGene_tx=XM_023243681.1
E1 16177219 16177295
refGene_gn=ABR;refGene_tx=XM_003996452.5;refGene_gn=ABR;refGene_tx=XM_019817386.2;refGene_gn=ABR;refGene_tx=XM_023243676.1;refGene_gn=ABR;re

fGene_tx=XM_023243677.1;refGene_gn=ABR;refGene_tx=XM_023243678.1;refGene_gn=ABR;refGene_tx=XM_023243679.1;refGene_gn=ABR;refGene_tx=XM_023243680.1;refGene_gn=ABR;refGene_tx=XM_023243681.1
E1 16177624 16177747
refGene_gn=ABR;refGene_tx=XM_003996452.5;refGene_gn=ABR;refGene_tx=XM_019817386.2;refGene_gn=ABR;refGene_tx=XM_023243676.1;refGene_gn=ABR;refGene_tx=XM_023243677.1;refGene_gn=ABR;refGene_tx=XM_023243678.1;refGene_gn=ABR;refGene_tx=XM_023243679.1;refGene_gn=ABR;refGene_tx=XM_023243680.1;refGene_gn=ABR;refGene_tx=XM_023243681.1
E1 16182104 16182270
refGene_gn=ABR;refGene_tx=XM_003996452.5;refGene_gn=ABR;refGene_tx=XM_019817386.2;refGene_gn=ABR;refGene_tx=XM_023243676.1;refGene_gn=ABR;refGene_tx=XM_023243677.1;refGene_gn=ABR;refGene_tx=XM_023243678.1;refGene_gn=ABR;refGene_tx=XM_023243679.1;refGene_gn=ABR;refGene_tx=XM_023243680.1;refGene_gn=ABR;refGene_tx=XM_023243681.1
E1 16183986 16184108
refGene_gn=ABR;refGene_tx=XM_003996452.5;refGene_gn=ABR;refGene_tx=XM_019817386.2;refGene_gn=ABR;refGene_tx=XM_023243676.1;refGene_gn=ABR;refGene_tx=XM_023243677.1;refGene_gn=ABR;refGene_tx=XM_023243678.1;refGene_gn=ABR;refGene_tx=XM_023243679.1;refGene_gn=ABR;refGene_tx=XM_023243680.1;refGene_gn=ABR;refGene_tx=XM_023243681.1
E1 16185853 16185994
refGene_gn=ABR;refGene_tx=XM_003996452.5;refGene_gn=ABR;refGene_tx=XM_019817386.2;refGene_gn=ABR;refGene_tx=XM_023243676.1;refGene_gn=ABR;refGene_tx=XM_023243677.1;refGene_gn=ABR;refGene_tx=XM_023243678.1;refGene_gn=ABR;refGene_tx=XM_023243679.1;refGene_gn=ABR;refGene_tx=XM_023243680.1;refGene_gn=ABR;refGene_tx=XM_023243681.1
E1 16186561 16186614
refGene_gn=ABR;refGene_tx=XM_003996452.5;refGene_gn=ABR;refGene_tx=XM_019817386.2;refGene_gn=ABR;refGene_tx=XM_023243676.1;refGene_gn=ABR;refGene_tx=XM_023243677.1;refGene_gn=ABR;refGene_tx=XM_023243678.1;refGene_gn=ABR;refGene_tx=XM_023243679.1;refGene_gn=ABR;refGene_tx=XM_023243680.1;refGene_gn=ABR;refGene_tx=XM_023243681.1
E1 16190934 16190980 refGene_gn=ABR;refGene_tx=XM_019817386.2
E1 16191360 16191442
refGene_gn=ABR;refGene_tx=XM_003996452.5;refGene_gn=ABR;refGene_tx=XM_023243676.1;refGene_gn=ABR;refGene_tx=XM_023243677.1;refGene_gn=ABR;refGene_tx=XM_023243678.1;refGene_gn=ABR;refGene_tx=XM_023243679.1;refGene_gn=ABR;refGene_tx=XM_023243680.1;refGene_gn=ABR;refGene_tx=XM_023243681.1
E1 16194431 16194539
refGene_gn=ABR;refGene_tx=XM_003996452.5;refGene_gn=ABR;refGene_tx=XM_023243676.1;refGene_gn=ABR;refGene_tx=XM_023243677.1;refGene_gn=ABR;refGene_tx=XM_023243678.1;refGene_gn=ABR;refGene_tx=XM_023243679.1;refGene_gn=ABR;refGene_tx=XM_023243680.1;refGene_gn=ABR;refGene_tx=XM_023243681.1
E1 16199126 16199312
refGene_gn=ABR;refGene_tx=XM_003996452.5;refGene_gn=ABR;refGene_tx=XM_023243676.1;refGene_gn=ABR;refGene_tx=XM_023243677.1;refGene_gn=ABR;refGene_tx=XM_023243678.1;refGene_gn=ABR;refGene_tx=XM_023243679.1;refGe

ne_gn=ABR;refGene_tx=XM_023243680.1;refGene_gn=ABR;refGene_tx=XM_023243681.1
E1 16206030 16206129
refGene_gn=ABR;refGene_tx=XM_003996452.5;refGene_gn=ABR;refGene_tx=XM_023243676.1;refGene_gn=ABR;refGene_tx=XM_023243677.1;refGene_gn=ABR;refGene_tx=XM_023243678.1;refGene_gn=ABR;refGene_tx=XM_023243679.1;refGene_gn=ABR;refGene_tx=XM_023243680.1;refGene_gn=ABR;refGene_tx=XM_023243681.1
E1 16214281 16214416 refGene_gn=ABR;refGene_tx=XM_003996452.5
E1 16228696 16228881
refGene_gn=ABR;refGene_tx=XM_023243676.1;refGene_gn=ABR;refGene_tx=XM_023243677.1;refGene_gn=ABR;refGene_tx=XM_023243678.1;refGene_gn=ABR;refGene_tx=XM_023243679.1;refGene_gn=ABR;refGene_tx=XM_023243680.1;refGene_gn=ABR;refGene_tx=XM_023243681.1
E1 16272425 16272486 refGene_gn=ABR;refGene_tx=XM_023243681.1
E1 16316295 16317340
refGene_gn=ABR;refGene_tx=XM_023243676.1;refGene_gn=ABR;refGene_tx=XM_023243677.1;refGene_gn=ABR;refGene_tx=XM_023243678.1;refGene_gn=ABR;refGene_tx=XM_023243679.1;refGene_gn=ABR;refGene_tx=XM_023243680.1
E1 16364536 16365193 refGene_gn=BHLHA9;refGene_tx=XM_003996505.2
E1 16373639 16374026 refGene_gn=TUSC5;refGene_tx=XM_006940001.3
E1 16386103 16386236 refGene_gn=TUSC5;refGene_tx=XM_006940001.3
E1 16387807 16387821 refGene_gn=TUSC5;refGene_tx=XM_006940001.3
E1 16422641 16422766
refGene_gn=GOSR1;refGene_tx=XM_003996454.3;refGene_gn=GOSR1;refGene_tx=XM_006940002.2
E1 16426571 16426654
refGene_gn=GOSR1;refGene_tx=XM_003996454.3;refGene_gn=GOSR1;refGene_tx=XM_006940002.2
E1 16433638 16433668
refGene_gn=GOSR1;refGene_tx=XM_003996454.3;refGene_gn=GOSR1;refGene_tx=XM_006940002.2
E1 16452773 16452848
refGene_gn=GOSR1;refGene_tx=XM_003996454.3;refGene_gn=GOSR1;refGene_tx=XM_006940002.2
E1 16455660 16455752
refGene_gn=GOSR1;refGene_tx=XM_003996454.3;refGene_gn=GOSR1;refGene_tx=XM_006940002.2
E1 16459959 16460067
refGene_gn=GOSR1;refGene_tx=XM_003996454.3;refGene_gn=GOSR1;refGene_tx=XM_006940002.2
E1 16460450 16460538
refGene_gn=GOSR1;refGene_tx=XM_003996454.3;refGene_gn=GOSR1;refGene_tx=XM_006940002.2
E1 16464811 16464932
refGene_gn=GOSR1;refGene_tx=XM_003996454.3;refGene_gn=GOSR1;refGene_tx=XM_006940002.2
E1 16467880 16467911
refGene_gn=GOSR1;refGene_tx=XM_003996454.3;refGene_gn=GOSR1;refGene_tx=XM_006940002.2

E1 16481843 16482070 refGene_gn=CPD;refGene_tx=XM_011289068.3
E1 16483636 16483662 refGene_gn=CPD;refGene_tx=XM_019817392.2
E1 16484195 16484295
refGene_gn=CPD;refGene_tx=XM_011289068.3;refGene_gn=CPD;refGene_tx=XM_019817392.2
E1 16484589 16484775
refGene_gn=CPD;refGene_tx=XM_011289068.3;refGene_gn=CPD;refGene_tx=XM_019817392.2
E1 16485332 16485464
refGene_gn=CPD;refGene_tx=XM_011289068.3;refGene_gn=CPD;refGene_tx=XM_019817392.2
E1 16489221 16489286
refGene_gn=CPD;refGene_tx=XM_011289068.3;refGene_gn=CPD;refGene_tx=XM_019817392.2
E1 16489388 16489484
refGene_gn=CPD;refGene_tx=XM_011289068.3;refGene_gn=CPD;refGene_tx=XM_019817392.2
E1 16491227 16491359
refGene_gn=CPD;refGene_tx=XM_011289068.3;refGene_gn=CPD;refGene_tx=XM_019817392.2
E1 16492708 16492844
refGene_gn=CPD;refGene_tx=XM_011289068.3;refGene_gn=CPD;refGene_tx=XM_019817392.2
E1 16495670 16495866
refGene_gn=CPD;refGene_tx=XM_011289068.3;refGene_gn=CPD;refGene_tx=XM_019817392.2
E1 16498564 16498888
refGene_gn=CPD;refGene_tx=XM_011289068.3;refGene_gn=CPD;refGene_tx=XM_019817392.2
E1 16503225 16503395
refGene_gn=CPD;refGene_tx=XM_011289068.3;refGene_gn=CPD;refGene_tx=XM_019817392.2
E1 16504713 16504856
refGene_gn=CPD;refGene_tx=XM_011289068.3;refGene_gn=CPD;refGene_tx=XM_019817392.2
E1 16505785 16505888
refGene_gn=CPD;refGene_tx=XM_011289068.3;refGene_gn=CPD;refGene_tx=XM_019817392.2
E1 16514456 16514566
refGene_gn=CPD;refGene_tx=XM_011289068.3;refGene_gn=CPD;refGene_tx=XM_019817392.2
E1 16523332 16523500
refGene_gn=CPD;refGene_tx=XM_011289068.3;refGene_gn=CPD;refGene_tx=XM_019817392.2
E1 16524607 16524799
refGene_gn=CPD;refGene_tx=XM_011289068.3;refGene_gn=CPD;refGene_tx=XM_019817392.2
E1 16526198 16526548
refGene_gn=CPD;refGene_tx=XM_011289068.3;refGene_gn=CPD;refGene_tx=XM_019817392.2

E1	16527567	16527737	refGene_gn=CPD;refGene_tx=XM_011289068.3;refGene_gn=CPD;refGene_tx=XM_019817392.2
E1	16528536	16528679	refGene_gn=CPD;refGene_tx=XM_011289068.3;refGene_gn=CPD;refGene_tx=XM_019817392.2
E1	16553015	16553263	refGene_gn=CPD;refGene_tx=XM_011289068.3;refGene_gn=CPD;refGene_tx=XM_019817392.2
E1	16559243	16559980	refGene_gn=CPD;refGene_tx=XM_011289068.3;refGene_gn=CPD;refGene_tx=XM_019817392.2
E1	16599338	16599420	refGene_gn=TMIGD1;refGene_tx=XM_023244795.1
E1	16601299	16601578	refGene_gn=TMIGD1;refGene_tx=XM_023244795.1
E1	16606518	16606794	refGene_gn=TMIGD1;refGene_tx=XM_023244795.1
E1	16612814	16612918	refGene_gn=TMIGD1;refGene_tx=XM_023244795.1
E1	16614161	16614202	refGene_gn=TMIGD1;refGene_tx=XM_023244795.1
E1	16614736	16614740	refGene_gn=TMIGD1;refGene_tx=XM_023244795.1
E1	16636383	16636396	refGene_gn=BLMH;refGene_tx=XM_006940004.4
E1	16636708	16636906	refGene_gn=BLMH;refGene_tx=XM_006940004.4
E1	16642802	16642912	refGene_gn=BLMH;refGene_tx=XM_006940004.4
E1	16643599	16643741	refGene_gn=BLMH;refGene_tx=XM_006940004.4
E1	16644559	16644648	refGene_gn=BLMH;refGene_tx=XM_006940004.4
E1	16646074	16646167	refGene_gn=BLMH;refGene_tx=XM_006940004.4
E1	16662385	16662541	refGene_gn=BLMH;refGene_tx=XM_006940004.4
E1	16663432	16663591	refGene_gn=BLMH;refGene_tx=XM_006940004.4
E1	16663704	16663772	refGene_gn=BLMH;refGene_tx=XM_006940004.4
E1	16665117	16665235	refGene_gn=BLMH;refGene_tx=XM_006940004.4
E1	16667949	16668019	refGene_gn=BLMH;refGene_tx=XM_006940004.4
E1	16682792	16682944	refGene_gn=BLMH;refGene_tx=XM_006940004.4
E1	16708934	16709277	refGene_gn=SLC6A4;refGene_tx=XM_019817396.2;refGene_gn=SLC6A4;refGene_tx=XM_019817397.2
E1	16711623	16711758	refGene_gn=SLC6A4;refGene_tx=XM_019817396.2
E1	16713171	16713391	refGene_gn=SLC6A4;refGene_tx=XM_019817396.2;refGene_gn=SLC6A4;refGene_tx=XM_019817397.2
E1	16714686	16714825	refGene_gn=SLC6A4;refGene_tx=XM_019817396.2;refGene_gn=SLC6A4;refGene_tx=XM_019817397.2
E1	16715383	16715518	refGene_gn=SLC6A4;refGene_tx=XM_019817396.2;refGene_gn=SLC6A4;refGene_tx=XM_019817397.2
E1	16715851	16715955	refGene_gn=SLC6A4;refGene_tx=XM_019817396.2;refGene_gn=SLC6A4;refGene_tx=XM_019817397.2
E1	16717711	16717839	refGene_gn=SLC6A4;refGene_tx=XM_019817396.2;refGene_gn=SLC6A4;refGene_tx=XM_019817397.2
E1	16719079	16719192

refGene_gn=SLC6A4;refGene_tx=XM_019817396.2;refGene_gn=SLC6A4;refGene_tx=XM_019817397.2
E1 16719923 16720055
refGene_gn=SLC6A4;refGene_tx=XM_019817396.2;refGene_gn=SLC6A4;refGene_tx=XM_019817397.2
E1 16721314 16721414
refGene_gn=SLC6A4;refGene_tx=XM_019817396.2;refGene_gn=SLC6A4;refGene_tx=XM_019817397.2
E1 16723070 16723171
refGene_gn=SLC6A4;refGene_tx=XM_019817396.2;refGene_gn=SLC6A4;refGene_tx=XM_019817397.2
E1 16727316 16727484
refGene_gn=SLC6A4;refGene_tx=XM_019817396.2;refGene_gn=SLC6A4;refGene_tx=XM_019817397.2
E1 16730986 16731061
refGene_gn=SLC6A4;refGene_tx=XM_019817396.2;refGene_gn=SLC6A4;refGene_tx=XM_019817397.2
E1 16752991 16754045
refGene_gn=NSRP1;refGene_tx=XM_003996459.4;refGene_gn=NSRP1;refGene_tx=XM_023244452.1;refGene_gn=NSRP1;refGene_tx=XM_023244453.1;refGene_gn=NSRP1;refGene_tx=XM_023244454.1
E1 16755899 16756008
refGene_gn=NSRP1;refGene_tx=XM_003996459.4;refGene_gn=NSRP1;refGene_tx=XM_023244452.1;refGene_gn=NSRP1;refGene_tx=XM_023244453.1;refGene_gn=NSRP1;refGene_tx=XM_023244454.1
E1 16756967 16757175
refGene_gn=NSRP1;refGene_tx=XM_003996459.4;refGene_gn=NSRP1;refGene_tx=XM_023244452.1;refGene_gn=NSRP1;refGene_tx=XM_023244453.1;refGene_gn=NSRP1;refGene_tx=XM_023244454.1
E1 16758055 16758184
refGene_gn=NSRP1;refGene_tx=XM_003996459.4;refGene_gn=NSRP1;refGene_tx=XM_023244452.1;refGene_gn=NSRP1;refGene_tx=XM_023244453.1;refGene_gn=NSRP1;refGene_tx=XM_023244454.1
E1 16766985 16767042
refGene_gn=NSRP1;refGene_tx=XM_003996459.4;refGene_gn=NSRP1;refGene_tx=XM_023244452.1;refGene_gn=NSRP1;refGene_tx=XM_023244453.1;refGene_gn=NSRP1;refGene_tx=XM_023244454.1
E1 16812331 16812425 refGene_gn=NSRP1;refGene_tx=XM_003996459.4
E1 16813629 16813649 refGene_gn=NSRP1;refGene_tx=XM_003996459.4
E1 16823974 16824168
refGene_gn=EFCAB5;refGene_tx=XM_011289005.3;refGene_gn=EFCAB5;refGene_tx=XM_019817398.2;refGene_gn=EFCAB5;refGene_tx=XM_019817399.2;refGene_gn=EFCAB5;refGene_tx=XM_019817400.2;refGene_gn=EFCAB5;refGene_tx=XM_019817404.2;refGene_gn=EFCAB5;refGene_tx=XM_023244432.1;refGene_gn=EFCAB5;refGene_tx=XM_023244433.1;refGene_gn=EFCAB5;refGene_tx=XM_023244434.1;refGene_gn=EFCAB5;refGene_tx=XM_023244435.1;refGene_gn=EFCAB5;refGene_tx=XM_023244436.1;refGene_gn=EFCAB5;refGene_tx=XM_023244446.1
E1 16826218 16826379 refGene_gn=EFCAB5;refGene_tx=XM_023244437.1
E1 16835668 16835682 refGene_gn=EFCAB5;refGene_tx=XM_023244439.1
E1 16836679 16836776

refGene_gn=EFCAB5;refGene_tx=XM_011289005.3;refGene_gn=EFCAB5;refGene_
tx=XM_019817398.2;refGene_gn=EFCAB5;refGene_tx=XM_019817399.2;refGene_
gn=EFCAB5;refGene_tx=XM_019817400.2;refGene_gn=EFCAB5;refGene_tx=XM_01
9817404.2;refGene_gn=EFCAB5;refGene_tx=XM_023244432.1;refGene_gn=EFCAB
5;refGene_tx=XM_023244433.1;refGene_gn=EFCAB5;refGene_tx=XM_023244434.
1;refGene_gn=EFCAB5;refGene_tx=XM_023244435.1;refGene_gn=EFCAB5;refGen
e_tx=XM_023244436.1;refGene_gn=EFCAB5;refGene_tx=XM_023244437.1;refGen
e_gn=EFCAB5;refGene_tx=XM_023244439.1;refGene_gn=EFCAB5;refGene_tx=XM_
023244442.1;refGene_gn=EFCAB5;refGene_tx=XM_023244443.1;refGene_gn=EFC
AB5;refGene_tx=XM_023244444.1;refGene_gn=EFCAB5;refGene_tx=XM_02324444
5.1;refGene_gn=EFCAB5;refGene_tx=XM_023244446.1
E1 16836830 16836854 refGene_gn=EFCAB5;refGene_tx=XM_023244440.1
E1 16837381 16837668
refGene_gn=EFCAB5;refGene_tx=XM_011289005.3;refGene_gn=EFCAB5;refGene_
tx=XM_019817398.2;refGene_gn=EFCAB5;refGene_tx=XM_019817399.2;refGene_
gn=EFCAB5;refGene_tx=XM_019817400.2;refGene_gn=EFCAB5;refGene_tx=XM_01
9817404.2;refGene_gn=EFCAB5;refGene_tx=XM_023244432.1;refGene_gn=EFCAB
5;refGene_tx=XM_023244433.1;refGene_gn=EFCAB5;refGene_tx=XM_023244434.
1;refGene_gn=EFCAB5;refGene_tx=XM_023244435.1;refGene_gn=EFCAB5;refGen
e_tx=XM_023244436.1;refGene_gn=EFCAB5;refGene_tx=XM_023244437.1;refGen
e_gn=EFCAB5;refGene_tx=XM_023244439.1;refGene_gn=EFCAB5;refGene_tx=XM_
023244440.1;refGene_gn=EFCAB5;refGene_tx=XM_023244446.1;refGene_gn=EFC
AB5;refGene_tx=XM_023244447.1
E1 16838848 16839102
refGene_gn=EFCAB5;refGene_tx=XM_011289005.3;refGene_gn=EFCAB5;refGene_
tx=XM_019817398.2;refGene_gn=EFCAB5;refGene_tx=XM_019817399.2;refGene_
gn=EFCAB5;refGene_tx=XM_019817400.2;refGene_gn=EFCAB5;refGene_tx=XM_01
9817404.2;refGene_gn=EFCAB5;refGene_tx=XM_023244432.1;refGene_gn=EFCAB
5;refGene_tx=XM_023244433.1;refGene_gn=EFCAB5;refGene_tx=XM_023244434.
1;refGene_gn=EFCAB5;refGene_tx=XM_023244435.1;refGene_gn=EFCAB5;refGen
e_tx=XM_023244436.1;refGene_gn=EFCAB5;refGene_tx=XM_023244437.1;refGen
e_gn=EFCAB5;refGene_tx=XM_023244439.1;refGene_gn=EFCAB5;refGene_tx=XM_
023244440.1;refGene_gn=EFCAB5;refGene_tx=XM_023244446.1
E1 16840450 16840556 refGene_gn=EFCAB5;refGene_tx=XM_023244441.1
E1 16841253 16841357
refGene_gn=EFCAB5;refGene_tx=XM_011289005.3;refGene_gn=EFCAB5;refGene_
tx=XM_019817398.2;refGene_gn=EFCAB5;refGene_tx=XM_019817399.2;refGene_
gn=EFCAB5;refGene_tx=XM_019817400.2;refGene_gn=EFCAB5;refGene_tx=XM_01
9817404.2;refGene_gn=EFCAB5;refGene_tx=XM_023244432.1;refGene_gn=EFCAB
5;refGene_tx=XM_023244433.1;refGene_gn=EFCAB5;refGene_tx=XM_023244434.
1;refGene_gn=EFCAB5;refGene_tx=XM_023244435.1;refGene_gn=EFCAB5;refGen
e_tx=XM_023244436.1;refGene_gn=EFCAB5;refGene_tx=XM_023244437.1;refGen
e_gn=EFCAB5;refGene_tx=XM_023244439.1;refGene_gn=EFCAB5;refGene_tx=XM_
023244440.1;refGene_gn=EFCAB5;refGene_tx=XM_023244441.1;refGene_gn=EFC
AB5;refGene_tx=XM_023244442.1;refGene_gn=EFCAB5;refGene_tx=XM_02324444
3.1;refGene_gn=EFCAB5;refGene_tx=XM_023244444.1;refGene_gn=EFCAB5;refG
ene_tx=XM_023244445.1;refGene_gn=EFCAB5;refGene_tx=XM_023244446.1;refG
ene_gn=EFCAB5;refGene_tx=XM_023244447.1
E1 16852837 16852990
refGene_gn=EFCAB5;refGene_tx=XM_011289005.3;refGene_gn=EFCAB5;refGene_

tx=XM_019817398.2;refGene_gn=EFCAB5;refGene_tx=XM_019817399.2;refGene_gn=EFCAB5;refGene_tx=XM_019817400.2;refGene_gn=EFCAB5;refGene_tx=XM_019817404.2;refGene_gn=EFCAB5;refGene_tx=XM_023244432.1;refGene_gn=EFCAB5;refGene_tx=XM_023244433.1;refGene_gn=EFCAB5;refGene_tx=XM_023244434.1;refGene_gn=EFCAB5;refGene_tx=XM_023244435.1;refGene_gn=EFCAB5;refGene_tx=XM_023244436.1;refGene_gn=EFCAB5;refGene_tx=XM_023244437.1;refGene_gn=EFCAB5;refGene_tx=XM_023244439.1;refGene_gn=EFCAB5;refGene_tx=XM_023244440.1;refGene_gn=EFCAB5;refGene_tx=XM_023244441.1;refGene_gn=EFCAB5;refGene_tx=XM_023244442.1;refGene_gn=EFCAB5;refGene_tx=XM_023244443.1;refGene_gn=EFCAB5;refGene_tx=XM_023244444.1;refGene_gn=EFCAB5;refGene_tx=XM_023244445.1;refGene_gn=EFCAB5;refGene_tx=XM_023244446.1;refGene_gn=EFCAB5;refGene_tx=XM_023244447.1
E1 16855793 16856022
refGene_gn=EFCAB5;refGene_tx=XM_011289005.3;refGene_gn=EFCAB5;refGene_tx=XM_019817398.2;refGene_gn=EFCAB5;refGene_tx=XM_019817399.2;refGene_gn=EFCAB5;refGene_tx=XM_019817400.2;refGene_gn=EFCAB5;refGene_tx=XM_019817404.2;refGene_gn=EFCAB5;refGene_tx=XM_023244432.1;refGene_gn=EFCAB5;refGene_tx=XM_023244433.1;refGene_gn=EFCAB5;refGene_tx=XM_023244434.1;refGene_gn=EFCAB5;refGene_tx=XM_023244435.1;refGene_gn=EFCAB5;refGene_tx=XM_023244436.1;refGene_gn=EFCAB5;refGene_tx=XM_023244437.1;refGene_gn=EFCAB5;refGene_tx=XM_023244439.1;refGene_gn=EFCAB5;refGene_tx=XM_023244440.1;refGene_gn=EFCAB5;refGene_tx=XM_023244441.1;refGene_gn=EFCAB5;refGene_tx=XM_023244442.1;refGene_gn=EFCAB5;refGene_tx=XM_023244443.1;refGene_gn=EFCAB5;refGene_tx=XM_023244444.1;refGene_gn=EFCAB5;refGene_tx=XM_023244445.1;refGene_gn=EFCAB5;refGene_tx=XM_023244446.1;refGene_gn=EFCAB5;refGene_tx=XM_023244447.1
E1 16856512 16856682
refGene_gn=EFCAB5;refGene_tx=XM_011289005.3;refGene_gn=EFCAB5;refGene_tx=XM_019817398.2;refGene_gn=EFCAB5;refGene_tx=XM_019817399.2;refGene_gn=EFCAB5;refGene_tx=XM_019817400.2;refGene_gn=EFCAB5;refGene_tx=XM_019817404.2;refGene_gn=EFCAB5;refGene_tx=XM_023244432.1;refGene_gn=EFCAB5;refGene_tx=XM_023244433.1;refGene_gn=EFCAB5;refGene_tx=XM_023244434.1;refGene_gn=EFCAB5;refGene_tx=XM_023244435.1;refGene_gn=EFCAB5;refGene_tx=XM_023244436.1;refGene_gn=EFCAB5;refGene_tx=XM_023244437.1;refGene_gn=EFCAB5;refGene_tx=XM_023244439.1;refGene_gn=EFCAB5;refGene_tx=XM_023244440.1;refGene_gn=EFCAB5;refGene_tx=XM_023244441.1;refGene_gn=EFCAB5;refGene_tx=XM_023244442.1;refGene_gn=EFCAB5;refGene_tx=XM_023244443.1;refGene_gn=EFCAB5;refGene_tx=XM_023244444.1;refGene_gn=EFCAB5;refGene_tx=XM_023244445.1;refGene_gn=EFCAB5;refGene_tx=XM_023244446.1;refGene_gn=EFCAB5;refGene_tx=XM_023244447.1
E1 16859449 16859739
refGene_gn=EFCAB5;refGene_tx=XM_011289005.3;refGene_gn=EFCAB5;refGene_tx=XM_019817398.2;refGene_gn=EFCAB5;refGene_tx=XM_019817399.2;refGene_gn=EFCAB5;refGene_tx=XM_019817400.2;refGene_gn=EFCAB5;refGene_tx=XM_019817404.2;refGene_gn=EFCAB5;refGene_tx=XM_023244432.1;refGene_gn=EFCAB5;refGene_tx=XM_023244433.1;refGene_gn=EFCAB5;refGene_tx=XM_023244434.1;refGene_gn=EFCAB5;refGene_tx=XM_023244435.1;refGene_gn=EFCAB5;refGene_tx=XM_023244436.1;refGene_gn=EFCAB5;refGene_tx=XM_023244437.1;refGene_gn=EFCAB5;refGene_tx=XM_023244439.1;refGene_gn=EFCAB5;refGene_tx=XM_023244440.1;refGene_gn=EFCAB5;refGene_tx=XM_023244441.1;refGene_gn=EFC

AB5;refGene_tx=XM_023244442.1;refGene_gn=EFCAB5;refGene_tx=XM_02324444
3.1;refGene_gn=EFCAB5;refGene_tx=XM_023244444.1;refGene_gn=EFCAB5;refG
ene_tx=XM_023244445.1;refGene_gn=EFCAB5;refGene_tx=XM_023244446.1;refG
ene_gn=EFCAB5;refGene_tx=XM_023244447.1
E1 16876737 16876894
refGene_gn=EFCAB5;refGene_tx=XM_011289005.3;refGene_gn=EFCAB5;refGene_
tx=XM_019817398.2;refGene_gn=EFCAB5;refGene_tx=XM_019817399.2;refGene_
gn=EFCAB5;refGene_tx=XM_019817400.2;refGene_gn=EFCAB5;refGene_tx=XM_01
9817404.2;refGene_gn=EFCAB5;refGene_tx=XM_023244432.1;refGene_gn=EFCAB
5;refGene_tx=XM_023244433.1;refGene_gn=EFCAB5;refGene_tx=XM_023244434.
1;refGene_gn=EFCAB5;refGene_tx=XM_023244435.1;refGene_gn=EFCAB5;refGen
e_tx=XM_023244436.1;refGene_gn=EFCAB5;refGene_tx=XM_023244437.1;refGen
e_gn=EFCAB5;refGene_tx=XM_023244439.1;refGene_gn=EFCAB5;refGene_tx=XM_
023244440.1;refGene_gn=EFCAB5;refGene_tx=XM_023244441.1;refGene_gn=EFC
AB5;refGene_tx=XM_023244442.1;refGene_gn=EFCAB5;refGene_tx=XM_02324444
3.1;refGene_gn=EFCAB5;refGene_tx=XM_023244444.1;refGene_gn=EFCAB5;refG
ene_tx=XM_023244445.1;refGene_gn=EFCAB5;refGene_tx=XM_023244446.1;refG
ene_gn=EFCAB5;refGene_tx=XM_023244447.1
E1 16883517 16883732
refGene_gn=EFCAB5;refGene_tx=XM_011289005.3;refGene_gn=EFCAB5;refGene_
tx=XM_019817398.2;refGene_gn=EFCAB5;refGene_tx=XM_019817399.2;refGene_
gn=EFCAB5;refGene_tx=XM_019817400.2;refGene_gn=EFCAB5;refGene_tx=XM_01
9817404.2;refGene_gn=EFCAB5;refGene_tx=XM_023244432.1;refGene_gn=EFCAB
5;refGene_tx=XM_023244433.1;refGene_gn=EFCAB5;refGene_tx=XM_023244434.
1;refGene_gn=EFCAB5;refGene_tx=XM_023244435.1;refGene_gn=EFCAB5;refGen
e_tx=XM_023244436.1;refGene_gn=EFCAB5;refGene_tx=XM_023244437.1;refGen
e_gn=EFCAB5;refGene_tx=XM_023244439.1;refGene_gn=EFCAB5;refGene_tx=XM_
023244440.1;refGene_gn=EFCAB5;refGene_tx=XM_023244441.1;refGene_gn=EFC
AB5;refGene_tx=XM_023244442.1;refGene_gn=EFCAB5;refGene_tx=XM_02324444
3.1;refGene_gn=EFCAB5;refGene_tx=XM_023244444.1;refGene_gn=EFCAB5;refG
ene_tx=XM_023244445.1;refGene_gn=EFCAB5;refGene_tx=XM_023244446.1;refG
ene_gn=EFCAB5;refGene_tx=XM_023244447.1
E1 16885235 16885328
refGene_gn=EFCAB5;refGene_tx=XM_011289005.3;refGene_gn=EFCAB5;refGene_
tx=XM_019817398.2;refGene_gn=EFCAB5;refGene_tx=XM_019817399.2;refGene_
gn=EFCAB5;refGene_tx=XM_019817400.2;refGene_gn=EFCAB5;refGene_tx=XM_01
9817404.2;refGene_gn=EFCAB5;refGene_tx=XM_023244432.1;refGene_gn=EFCAB
5;refGene_tx=XM_023244433.1;refGene_gn=EFCAB5;refGene_tx=XM_023244434.
1;refGene_gn=EFCAB5;refGene_tx=XM_023244435.1;refGene_gn=EFCAB5;refGen
e_tx=XM_023244436.1;refGene_gn=EFCAB5;refGene_tx=XM_023244437.1;refGen
e_gn=EFCAB5;refGene_tx=XM_023244439.1;refGene_gn=EFCAB5;refGene_tx=XM_
023244440.1;refGene_gn=EFCAB5;refGene_tx=XM_023244441.1;refGene_gn=EFC
AB5;refGene_tx=XM_023244442.1;refGene_gn=EFCAB5;refGene_tx=XM_02324444
3.1;refGene_gn=EFCAB5;refGene_tx=XM_023244444.1;refGene_gn=EFCAB5;refG
ene_tx=XM_023244445.1;refGene_gn=EFCAB5;refGene_tx=XM_023244446.1;refG
ene_gn=EFCAB5;refGene_tx=XM_023244447.1
E1 16885427 16885502
refGene_gn=EFCAB5;refGene_tx=XM_011289005.3;refGene_gn=EFCAB5;refGene_
tx=XM_019817398.2;refGene_gn=EFCAB5;refGene_tx=XM_019817399.2;refGene_
gn=EFCAB5;refGene_tx=XM_019817400.2;refGene_gn=EFCAB5;refGene_tx=XM_01

9817404.2;refGene_gn=EFCAB5;refGene_tx=XM_023244432.1;refGene_gn=EFCAB5;refGene_tx=XM_023244433.1;refGene_gn=EFCAB5;refGene_tx=XM_023244434.1;refGene_gn=EFCAB5;refGene_tx=XM_023244435.1;refGene_gn=EFCAB5;refGene_tx=XM_023244436.1;refGene_gn=EFCAB5;refGene_tx=XM_023244437.1;refGene_gn=EFCAB5;refGene_tx=XM_023244439.1;refGene_gn=EFCAB5;refGene_tx=XM_023244440.1;refGene_gn=EFCAB5;refGene_tx=XM_023244441.1;refGene_gn=EFCAB5;refGene_tx=XM_023244442.1;refGene_gn=EFCAB5;refGene_tx=XM_023244443.1;refGene_gn=EFCAB5;refGene_tx=XM_023244444.1;refGene_gn=EFCAB5;refGene_tx=XM_023244445.1;refGene_gn=EFCAB5;refGene_tx=XM_023244446.1;refGene_gn=EFCAB5;refGene_tx=XM_023244447.1
E1 16887184 16888015
refGene_gn=EFCAB5;refGene_tx=XM_011289005.3;refGene_gn=EFCAB5;refGene_tx=XM_019817398.2;refGene_gn=EFCAB5;refGene_tx=XM_019817399.2;refGene_gn=EFCAB5;refGene_tx=XM_019817400.2;refGene_gn=EFCAB5;refGene_tx=XM_019817404.2;refGene_gn=EFCAB5;refGene_tx=XM_023244432.1;refGene_gn=EFCAB5;refGene_tx=XM_023244433.1;refGene_gn=EFCAB5;refGene_tx=XM_023244434.1;refGene_gn=EFCAB5;refGene_tx=XM_023244435.1;refGene_gn=EFCAB5;refGene_tx=XM_023244436.1;refGene_gn=EFCAB5;refGene_tx=XM_023244437.1;refGene_gn=EFCAB5;refGene_tx=XM_023244439.1;refGene_gn=EFCAB5;refGene_tx=XM_023244440.1;refGene_gn=EFCAB5;refGene_tx=XM_023244441.1;refGene_gn=EFCAB5;refGene_tx=XM_023244442.1;refGene_gn=EFCAB5;refGene_tx=XM_023244443.1;refGene_gn=EFCAB5;refGene_tx=XM_023244444.1;refGene_gn=EFCAB5;refGene_tx=XM_023244445.1;refGene_gn=EFCAB5;refGene_tx=XM_023244446.1;refGene_gn=EFCAB5;refGene_tx=XM_023244447.1
E1 16891084 16891184
refGene_gn=EFCAB5;refGene_tx=XM_011289005.3;refGene_gn=EFCAB5;refGene_tx=XM_019817398.2;refGene_gn=EFCAB5;refGene_tx=XM_019817399.2;refGene_gn=EFCAB5;refGene_tx=XM_019817400.2;refGene_gn=EFCAB5;refGene_tx=XM_019817404.2;refGene_gn=EFCAB5;refGene_tx=XM_023244432.1;refGene_gn=EFCAB5;refGene_tx=XM_023244433.1;refGene_gn=EFCAB5;refGene_tx=XM_023244434.1;refGene_gn=EFCAB5;refGene_tx=XM_023244435.1;refGene_gn=EFCAB5;refGene_tx=XM_023244436.1;refGene_gn=EFCAB5;refGene_tx=XM_023244437.1;refGene_gn=EFCAB5;refGene_tx=XM_023244439.1;refGene_gn=EFCAB5;refGene_tx=XM_023244440.1;refGene_gn=EFCAB5;refGene_tx=XM_023244441.1;refGene_gn=EFCAB5;refGene_tx=XM_023244442.1;refGene_gn=EFCAB5;refGene_tx=XM_023244443.1;refGene_gn=EFCAB5;refGene_tx=XM_023244444.1;refGene_gn=EFCAB5;refGene_tx=XM_023244445.1;refGene_gn=EFCAB5;refGene_tx=XM_023244446.1;refGene_gn=EFCAB5;refGene_tx=XM_023244447.1
E1 16912153 16912309
refGene_gn=EFCAB5;refGene_tx=XM_011289005.3;refGene_gn=EFCAB5;refGene_tx=XM_019817398.2;refGene_gn=EFCAB5;refGene_tx=XM_019817399.2;refGene_gn=EFCAB5;refGene_tx=XM_019817400.2;refGene_gn=EFCAB5;refGene_tx=XM_019817404.2;refGene_gn=EFCAB5;refGene_tx=XM_023244432.1;refGene_gn=EFCAB5;refGene_tx=XM_023244433.1;refGene_gn=EFCAB5;refGene_tx=XM_023244434.1;refGene_gn=EFCAB5;refGene_tx=XM_023244435.1;refGene_gn=EFCAB5;refGene_tx=XM_023244436.1;refGene_gn=EFCAB5;refGene_tx=XM_023244437.1;refGene_gn=EFCAB5;refGene_tx=XM_023244439.1;refGene_gn=EFCAB5;refGene_tx=XM_023244440.1;refGene_gn=EFCAB5;refGene_tx=XM_023244441.1;refGene_gn=EFCAB5;refGene_tx=XM_023244442.1;refGene_gn=EFCAB5;refGene_tx=XM_023244443.1;refGene_gn=EFCAB5;refGene_tx=XM_023244444.1;refGene_gn=EFCAB5;refG

ene_tx=XM_023244445.1;refGene_gn=EFCAB5;refGene_tx=XM_023244446.1;refGene_gn=EFCAB5;refGene_tx=XM_023244447.1
E1 16930221 16930292
refGene_gn=EFCAB5;refGene_tx=XM_011289005.3;refGene_gn=EFCAB5;refGene_tx=XM_019817398.2;refGene_gn=EFCAB5;refGene_tx=XM_019817399.2;refGene_gn=EFCAB5;refGene_tx=XM_019817400.2;refGene_gn=EFCAB5;refGene_tx=XM_019817404.2;refGene_gn=EFCAB5;refGene_tx=XM_023244432.1;refGene_gn=EFCAB5;refGene_tx=XM_023244433.1;refGene_gn=EFCAB5;refGene_tx=XM_023244434.1;refGene_gn=EFCAB5;refGene_tx=XM_023244435.1;refGene_gn=EFCAB5;refGene_tx=XM_023244436.1;refGene_gn=EFCAB5;refGene_tx=XM_023244437.1;refGene_gn=EFCAB5;refGene_tx=XM_023244439.1;refGene_gn=EFCAB5;refGene_tx=XM_023244440.1;refGene_gn=EFCAB5;refGene_tx=XM_023244441.1;refGene_gn=EFCAB5;refGene_tx=XM_023244442.1;refGene_gn=EFCAB5;refGene_tx=XM_023244443.1;refGene_gn=EFCAB5;refGene_tx=XM_023244444.1;refGene_gn=EFCAB5;refGene_tx=XM_023244445.1;refGene_gn=EFCAB5;refGene_tx=XM_023244446.1;refGene_gn=EFCAB5;refGene_tx=XM_023244447.1
E1 16935434 16935483
refGene_gn=EFCAB5;refGene_tx=XM_011289005.3;refGene_gn=EFCAB5;refGene_tx=XM_019817398.2;refGene_gn=EFCAB5;refGene_tx=XM_019817399.2;refGene_gn=EFCAB5;refGene_tx=XM_019817400.2;refGene_gn=EFCAB5;refGene_tx=XM_019817404.2;refGene_gn=EFCAB5;refGene_tx=XM_023244432.1;refGene_gn=EFCAB5;refGene_tx=XM_023244433.1;refGene_gn=EFCAB5;refGene_tx=XM_023244434.1;refGene_gn=EFCAB5;refGene_tx=XM_023244435.1;refGene_gn=EFCAB5;refGene_tx=XM_023244436.1;refGene_gn=EFCAB5;refGene_tx=XM_023244437.1;refGene_gn=EFCAB5;refGene_tx=XM_023244439.1;refGene_gn=EFCAB5;refGene_tx=XM_023244440.1;refGene_gn=EFCAB5;refGene_tx=XM_023244441.1;refGene_gn=EFCAB5;refGene_tx=XM_023244442.1;refGene_gn=EFCAB5;refGene_tx=XM_023244443.1;refGene_gn=EFCAB5;refGene_tx=XM_023244444.1;refGene_gn=EFCAB5;refGene_tx=XM_023244445.1;refGene_gn=EFCAB5;refGene_tx=XM_023244446.1;refGene_gn=EFCAB5;refGene_tx=XM_023244447.1
E1 16938882 16939039
refGene_gn=EFCAB5;refGene_tx=XM_011289005.3;refGene_gn=EFCAB5;refGene_tx=XM_019817398.2;refGene_gn=EFCAB5;refGene_tx=XM_019817399.2;refGene_gn=EFCAB5;refGene_tx=XM_019817400.2;refGene_gn=EFCAB5;refGene_tx=XM_019817404.2;refGene_gn=EFCAB5;refGene_tx=XM_023244432.1;refGene_gn=EFCAB5;refGene_tx=XM_023244433.1;refGene_gn=EFCAB5;refGene_tx=XM_023244434.1;refGene_gn=EFCAB5;refGene_tx=XM_023244435.1;refGene_gn=EFCAB5;refGene_tx=XM_023244436.1;refGene_gn=EFCAB5;refGene_tx=XM_023244437.1;refGene_gn=EFCAB5;refGene_tx=XM_023244439.1;refGene_gn=EFCAB5;refGene_tx=XM_023244440.1;refGene_gn=EFCAB5;refGene_tx=XM_023244441.1;refGene_gn=EFCAB5;refGene_tx=XM_023244442.1;refGene_gn=EFCAB5;refGene_tx=XM_023244443.1;refGene_gn=EFCAB5;refGene_tx=XM_023244444.1;refGene_gn=EFCAB5;refGene_tx=XM_023244445.1;refGene_gn=EFCAB5;refGene_tx=XM_023244446.1;refGene_gn=EFCAB5;refGene_tx=XM_023244447.1
E1 16968487 16969064
refGene_gn=EFCAB5;refGene_tx=XM_011289005.3;refGene_gn=EFCAB5;refGene_tx=XM_019817398.2;refGene_gn=EFCAB5;refGene_tx=XM_019817399.2;refGene_gn=EFCAB5;refGene_tx=XM_019817400.2;refGene_gn=EFCAB5;refGene_tx=XM_019817404.2;refGene_gn=EFCAB5;refGene_tx=XM_023244432.1;refGene_gn=EFCAB5;refGene_tx=XM_023244433.1;refGene_gn=EFCAB5;refGene_tx=XM_023244434.

1;refGene_gn=EFCAB5;refGene_tx=XM_023244435.1;refGene_gn=EFCAB5;refGene_tx=XM_023244436.1;refGene_gn=EFCAB5;refGene_tx=XM_023244437.1;refGene_gn=EFCAB5;refGene_tx=XM_023244439.1;refGene_gn=EFCAB5;refGene_tx=XM_023244440.1;refGene_gn=EFCAB5;refGene_tx=XM_023244441.1;refGene_gn=EFCAB5;refGene_tx=XM_023244442.1;refGene_gn=EFCAB5;refGene_tx=XM_023244443.1;refGene_gn=EFCAB5;refGene_tx=XM_023244444.1;refGene_gn=EFCAB5;refGene_tx=XM_023244445.1;refGene_gn=EFCAB5;refGene_tx=XM_023244447.1
E1 16997286 16997371
refGene_gn=EFCAB5;refGene_tx=XM_011289005.3;refGene_gn=EFCAB5;refGene_tx=XM_019817398.2;refGene_gn=EFCAB5;refGene_tx=XM_019817399.2;refGene_gn=EFCAB5;refGene_tx=XM_019817400.2;refGene_gn=EFCAB5;refGene_tx=XM_019817404.2;refGene_gn=EFCAB5;refGene_tx=XM_023244432.1;refGene_gn=EFCAB5;refGene_tx=XM_023244433.1;refGene_gn=EFCAB5;refGene_tx=XM_023244434.1;refGene_gn=EFCAB5;refGene_tx=XM_023244435.1;refGene_gn=EFCAB5;refGene_tx=XM_023244436.1;refGene_gn=EFCAB5;refGene_tx=XM_023244437.1;refGene_gn=EFCAB5;refGene_tx=XM_023244439.1;refGene_gn=EFCAB5;refGene_tx=XM_023244440.1;refGene_gn=EFCAB5;refGene_tx=XM_023244441.1;refGene_gn=EFCAB5;refGene_tx=XM_023244442.1;refGene_gn=EFCAB5;refGene_tx=XM_023244443.1;refGene_gn=EFCAB5;refGene_tx=XM_023244444.1;refGene_gn=EFCAB5;refGene_tx=XM_023244445.1;refGene_gn=EFCAB5;refGene_tx=XM_023244447.1
E1 16998783 16998846
refGene_gn=EFCAB5;refGene_tx=XM_011289005.3;refGene_gn=EFCAB5;refGene_tx=XM_019817398.2;refGene_gn=EFCAB5;refGene_tx=XM_019817399.2;refGene_gn=EFCAB5;refGene_tx=XM_019817400.2;refGene_gn=EFCAB5;refGene_tx=XM_019817404.2;refGene_gn=EFCAB5;refGene_tx=XM_023244432.1;refGene_gn=EFCAB5;refGene_tx=XM_023244433.1;refGene_gn=EFCAB5;refGene_tx=XM_023244434.1;refGene_gn=EFCAB5;refGene_tx=XM_023244435.1;refGene_gn=EFCAB5;refGene_tx=XM_023244436.1;refGene_gn=EFCAB5;refGene_tx=XM_023244437.1;refGene_gn=EFCAB5;refGene_tx=XM_023244439.1;refGene_gn=EFCAB5;refGene_tx=XM_023244440.1;refGene_gn=EFCAB5;refGene_tx=XM_023244441.1;refGene_gn=EFCAB5;refGene_tx=XM_023244442.1;refGene_gn=EFCAB5;refGene_tx=XM_023244443.1;refGene_gn=EFCAB5;refGene_tx=XM_023244444.1;refGene_gn=EFCAB5;refGene_tx=XM_023244445.1;refGene_gn=EFCAB5;refGene_tx=XM_023244447.1
E1 16999468 16999510
refGene_gn=EFCAB5;refGene_tx=XM_011289005.3;refGene_gn=EFCAB5;refGene_tx=XM_019817398.2;refGene_gn=EFCAB5;refGene_tx=XM_019817399.2;refGene_gn=EFCAB5;refGene_tx=XM_019817400.2;refGene_gn=EFCAB5;refGene_tx=XM_019817404.2;refGene_gn=EFCAB5;refGene_tx=XM_023244432.1;refGene_gn=EFCAB5;refGene_tx=XM_023244433.1;refGene_gn=EFCAB5;refGene_tx=XM_023244434.1;refGene_gn=EFCAB5;refGene_tx=XM_023244435.1;refGene_gn=EFCAB5;refGene_tx=XM_023244436.1;refGene_gn=EFCAB5;refGene_tx=XM_023244437.1;refGene_gn=EFCAB5;refGene_tx=XM_023244439.1;refGene_gn=EFCAB5;refGene_tx=XM_023244440.1;refGene_gn=EFCAB5;refGene_tx=XM_023244441.1;refGene_gn=EFCAB5;refGene_tx=XM_023244442.1;refGene_gn=EFCAB5;refGene_tx=XM_023244443.1;refGene_gn=EFCAB5;refGene_tx=XM_023244444.1;refGene_gn=EFCAB5;refGene_tx=XM_023244445.1;refGene_gn=EFCAB5;refGene_tx=XM_023244447.1
E1 17025166 17025229
refGene_gn=SSH2;refGene_tx=XM_023244448.1;refGene_gn=SSH2;refGene_tx=XM_023244449.1;refGene_gn=SSH2;refGene_tx=XM_023244450.1
E1 17083863 17083944

refGene_gn=SSH2;refGene_tx=XM_023244448.1;refGene_gn=SSH2;refGene_tx=XM_023244449.1
E1 17122503 17122547
refGene_gn=SSH2;refGene_tx=XM_023244448.1;refGene_gn=SSH2;refGene_tx=XM_023244449.1;refGene_gn=SSH2;refGene_tx=XM_023244450.1
E1 17169799 17169927
refGene_gn=SSH2;refGene_tx=XM_006940006.4;refGene_gn=SSH2;refGene_tx=XM_019817417.2
E1 17212088 17212192
refGene_gn=SSH2;refGene_tx=XM_006940006.4;refGene_gn=SSH2;refGene_tx=XM_019817417.2;refGene_gn=SSH2;refGene_tx=XM_023244448.1;refGene_gn=SSH2;refGene_tx=XM_023244449.1;refGene_gn=SSH2;refGene_tx=XM_023244450.1
E1 17215315 17215380
refGene_gn=SSH2;refGene_tx=XM_006940006.4;refGene_gn=SSH2;refGene_tx=XM_019817417.2;refGene_gn=SSH2;refGene_tx=XM_023244448.1;refGene_gn=SSH2;refGene_tx=XM_023244449.1;refGene_gn=SSH2;refGene_tx=XM_023244450.1
E1 17225742 17225864
refGene_gn=SSH2;refGene_tx=XM_006940006.4;refGene_gn=SSH2;refGene_tx=XM_019817417.2;refGene_gn=SSH2;refGene_tx=XM_023244448.1;refGene_gn=SSH2;refGene_tx=XM_023244449.1;refGene_gn=SSH2;refGene_tx=XM_023244450.1
E1 17231027 17231096
refGene_gn=SSH2;refGene_tx=XM_006940006.4;refGene_gn=SSH2;refGene_tx=XM_019817417.2;refGene_gn=SSH2;refGene_tx=XM_023244448.1;refGene_gn=SSH2;refGene_tx=XM_023244449.1;refGene_gn=SSH2;refGene_tx=XM_023244450.1
E1 17231890 17231956
refGene_gn=SSH2;refGene_tx=XM_006940006.4;refGene_gn=SSH2;refGene_tx=XM_019817417.2;refGene_gn=SSH2;refGene_tx=XM_023244448.1;refGene_gn=SSH2;refGene_tx=XM_023244449.1;refGene_gn=SSH2;refGene_tx=XM_023244450.1
E1 17233791 17233847 refGene_gn=SSH2;refGene_tx=XM_019817417.2
E1 17236518 17236713
refGene_gn=SSH2;refGene_tx=XM_006940006.4;refGene_gn=SSH2;refGene_tx=XM_023244448.1;refGene_gn=SSH2;refGene_tx=XM_023244449.1;refGene_gn=SSH2;refGene_tx=XM_023244450.1
E1 17240352 17240446
refGene_gn=SSH2;refGene_tx=XM_006940006.4;refGene_gn=SSH2;refGene_tx=XM_023244448.1;refGene_gn=SSH2;refGene_tx=XM_023244449.1;refGene_gn=SSH2;refGene_tx=XM_023244450.1
E1 17240797 17240926
refGene_gn=SSH2;refGene_tx=XM_006940006.4;refGene_gn=SSH2;refGene_tx=XM_023244448.1;refGene_gn=SSH2;refGene_tx=XM_023244449.1;refGene_gn=SSH2;refGene_tx=XM_023244450.1
E1 17251031 17251078
refGene_gn=SSH2;refGene_tx=XM_006940006.4;refGene_gn=SSH2;refGene_tx=XM_023244448.1;refGene_gn=SSH2;refGene_tx=XM_023244449.1;refGene_gn=SSH2;refGene_tx=XM_023244450.1
E1 17255093 17255240
refGene_gn=SSH2;refGene_tx=XM_006940006.4;refGene_gn=SSH2;refGene_tx=XM_023244448.1;refGene_gn=SSH2;refGene_tx=XM_023244449.1;refGene_gn=SSH2;refGene_tx=XM_023244450.1
E1 17258501 17258702

refGene_gn=SSH2;refGene_tx=XM_006940006.4;refGene_gn=SSH2;refGene_tx=XM_023244448.1;refGene_gn=SSH2;refGene_tx=XM_023244449.1;refGene_gn=SSH2;refGene_tx=XM_023244450.1
E1 17267536 17268371
refGene_gn=SSH2;refGene_tx=XM_006940006.4;refGene_gn=SSH2;refGene_tx=XM_023244448.1;refGene_gn=SSH2;refGene_tx=XM_023244449.1;refGene_gn=SSH2;refGene_tx=XM_023244450.1
E1 17270744 17272730
refGene_gn=SSH2;refGene_tx=XM_006940006.4;refGene_gn=SSH2;refGene_tx=XM_023244448.1;refGene_gn=SSH2;refGene_tx=XM_023244449.1;refGene_gn=SSH2;refGene_tx=XM_023244450.1
E1 17282865 17283063
refGene_gn=CORO6;refGene_tx=XM_003996460.5;refGene_gn=CORO6;refGene_tx=XM_011289001.3;refGene_gn=CORO6;refGene_tx=XM_019817418.2;refGene_gn=CORO6;refGene_tx=XM_019817419.2;refGene_gn=CORO6;refGene_tx=XM_019817420.2;refGene_gn=CORO6;refGene_tx=XM_019817421.2;refGene_gn=CORO6;refGene_tx=XM_019817422.2
E1 17284458 17284581
refGene_gn=CORO6;refGene_tx=XM_003996460.5;refGene_gn=CORO6;refGene_tx=XM_011289001.3;refGene_gn=CORO6;refGene_tx=XM_019817418.2;refGene_gn=CORO6;refGene_tx=XM_019817419.2;refGene_gn=CORO6;refGene_tx=XM_019817420.2;refGene_gn=CORO6;refGene_tx=XM_019817421.2;refGene_gn=CORO6;refGene_tx=XM_019817422.2
E1 17285042 17285172
refGene_gn=CORO6;refGene_tx=XM_003996460.5;refGene_gn=CORO6;refGene_tx=XM_011289001.3;refGene_gn=CORO6;refGene_tx=XM_019817418.2;refGene_gn=CORO6;refGene_tx=XM_019817419.2;refGene_gn=CORO6;refGene_tx=XM_019817420.2;refGene_gn=CORO6;refGene_tx=XM_019817421.2;refGene_gn=CORO6;refGene_tx=XM_019817422.2
E1 17285264 17285446
refGene_gn=CORO6;refGene_tx=XM_003996460.5;refGene_gn=CORO6;refGene_tx=XM_011289001.3;refGene_gn=CORO6;refGene_tx=XM_019817418.2;refGene_gn=CORO6;refGene_tx=XM_019817419.2;refGene_gn=CORO6;refGene_tx=XM_019817420.2;refGene_gn=CORO6;refGene_tx=XM_019817421.2;refGene_gn=CORO6;refGene_tx=XM_019817422.2
E1 17286053 17286173
refGene_gn=CORO6;refGene_tx=XM_019817420.2;refGene_gn=CORO6;refGene_tx=XM_019817422.2
E1 17286606 17286726
refGene_gn=CORO6;refGene_tx=XM_003996460.5;refGene_gn=CORO6;refGene_tx=XM_011289001.3;refGene_gn=CORO6;refGene_tx=XM_019817418.2;refGene_gn=CORO6;refGene_tx=XM_019817419.2;refGene_gn=CORO6;refGene_tx=XM_019817421.2
E1 17287149 17287254
refGene_gn=CORO6;refGene_tx=XM_003996460.5;refGene_gn=CORO6;refGene_tx=XM_011289001.3;refGene_gn=CORO6;refGene_tx=XM_019817418.2;refGene_gn=CORO6;refGene_tx=XM_019817419.2;refGene_gn=CORO6;refGene_tx=XM_019817420.2;refGene_gn=CORO6;refGene_tx=XM_019817421.2;refGene_gn=CORO6;refGene_tx=XM_019817422.2
E1 17287342 17287488

refGene_gn=CORO6;refGene_tx=XM_003996460.5;refGene_gn=CORO6;refGene_tx
=XM_011289001.3;refGene_gn=CORO6;refGene_tx=XM_019817418.2;refGene_gn=
CORO6;refGene_tx=XM_019817419.2;refGene_gn=CORO6;refGene_tx=XM_0198174
20.2;refGene_gn=CORO6;refGene_tx=XM_019817421.2;refGene_gn=CORO6;refGe
ne_tx=XM_019817422.2
E1 17287808 17287866
refGene_gn=CORO6;refGene_tx=XM_003996460.5;refGene_gn=CORO6;refGene_tx
=XM_011289001.3;refGene_gn=CORO6;refGene_tx=XM_019817418.2;refGene_gn=
CORO6;refGene_tx=XM_019817419.2;refGene_gn=CORO6;refGene_tx=XM_0198174
20.2;refGene_gn=CORO6;refGene_tx=XM_019817421.2;refGene_gn=CORO6;refGe
ne_tx=XM_019817422.2
E1 17287974 17288205
refGene_gn=CORO6;refGene_tx=XM_003996460.5;refGene_gn=CORO6;refGene_tx
=XM_011289001.3;refGene_gn=CORO6;refGene_tx=XM_019817418.2;refGene_gn=
CORO6;refGene_tx=XM_019817419.2;refGene_gn=CORO6;refGene_tx=XM_0198174
20.2;refGene_gn=CORO6;refGene_tx=XM_019817421.2;refGene_gn=CORO6;refGe
ne_tx=XM_019817422.2
E1 17288299 17288425
refGene_gn=CORO6;refGene_tx=XM_003996460.5;refGene_gn=CORO6;refGene_tx
=XM_011289001.3;refGene_gn=CORO6;refGene_tx=XM_019817418.2;refGene_gn=
CORO6;refGene_tx=XM_019817419.2;refGene_gn=CORO6;refGene_tx=XM_0198174
20.2;refGene_gn=CORO6;refGene_tx=XM_019817421.2;refGene_gn=CORO6;refGe
ne_tx=XM_019817422.2
E1 17289033 17289076
refGene_gn=CORO6;refGene_tx=XM_019817418.2;refGene_gn=CORO6;refGene_tx
=XM_019817419.2;refGene_gn=CORO6;refGene_tx=XM_019817420.2;refGene_gn=
CORO6;refGene_tx=XM_019817421.2
E1 17290624 17290901
refGene_gn=ANKRD13B;refGene_tx=XM_006940036.4;refGene_gn=ANKRD13B;refG
ene_tx=XM_011289000.3
E1 17291372 17291449
refGene_gn=ANKRD13B;refGene_tx=XM_006940036.4;refGene_gn=ANKRD13B;refG
ene_tx=XM_011289000.3;refGene_gn=ANKRD13B;refGene_tx=XM_019817423.2
E1 17291526 17291690
refGene_gn=ANKRD13B;refGene_tx=XM_006940036.4;refGene_gn=ANKRD13B;refG
ene_tx=XM_011289000.3;refGene_gn=ANKRD13B;refGene_tx=XM_019817423.2
E1 17291813 17291966
refGene_gn=ANKRD13B;refGene_tx=XM_006940036.4;refGene_gn=ANKRD13B;refG
ene_tx=XM_011289000.3;refGene_gn=ANKRD13B;refGene_tx=XM_019817423.2
E1 17292106 17292264
refGene_gn=ANKRD13B;refGene_tx=XM_006940036.4;refGene_gn=ANKRD13B;refG
ene_tx=XM_011289000.3;refGene_gn=ANKRD13B;refGene_tx=XM_019817423.2
E1 17292391 17292522
refGene_gn=ANKRD13B;refGene_tx=XM_006940036.4;refGene_gn=ANKRD13B;refG
ene_tx=XM_011289000.3;refGene_gn=ANKRD13B;refGene_tx=XM_019817423.2
E1 17292748 17292813
refGene_gn=ANKRD13B;refGene_tx=XM_006940036.4;refGene_gn=ANKRD13B;refG
ene_tx=XM_011289000.3;refGene_gn=ANKRD13B;refGene_tx=XM_019817423.2
E1 17293595 17293677
refGene_gn=ANKRD13B;refGene_tx=XM_006940036.4;refGene_gn=ANKRD13B;refG

ene_tx=XM_011289000.3;refGene_gn=ANKRD13B;refGene_tx=XM_019817423.2
E1 17294712 17294779
refGene_gn=ANKRD13B;refGene_tx=XM_006940036.4;refGene_gn=ANKRD13B;refGene_tx=XM_011289000.3;refGene_gn=ANKRD13B;refGene_tx=XM_019817423.2
E1 17294855 17295045
refGene_gn=ANKRD13B;refGene_tx=XM_006940036.4;refGene_gn=ANKRD13B;refGene_tx=XM_011289000.3;refGene_gn=ANKRD13B;refGene_tx=XM_019817423.2
E1 17295137 17295281
refGene_gn=ANKRD13B;refGene_tx=XM_006940036.4;refGene_gn=ANKRD13B;refGene_tx=XM_011289000.3;refGene_gn=ANKRD13B;refGene_tx=XM_019817423.2
E1 17295896 17295942
refGene_gn=ANKRD13B;refGene_tx=XM_006940036.4;refGene_gn=ANKRD13B;refGene_tx=XM_011289000.3;refGene_gn=ANKRD13B;refGene_tx=XM_019817423.2
E1 17296028 17296153
refGene_gn=ANKRD13B;refGene_tx=XM_006940036.4;refGene_gn=ANKRD13B;refGene_tx=XM_011289000.3;refGene_gn=ANKRD13B;refGene_tx=XM_019817423.2
E1 17296283 17296419
refGene_gn=ANKRD13B;refGene_tx=XM_006940036.4;refGene_gn=ANKRD13B;refGene_tx=XM_011289000.3;refGene_gn=ANKRD13B;refGene_tx=XM_019817423.2
E1 17307056 17307170
refGene_gn=ANKRD13B;refGene_tx=XM_006940036.4;refGene_gn=ANKRD13B;refGene_tx=XM_011289000.3;refGene_gn=ANKRD13B;refGene_tx=XM_019817423.2
E1 17311391 17311443
refGene_gn=GIT1;refGene_tx=XM_023244606.1;refGene_gn=GIT1;refGene_tx=XM_023244607.1
E1 17317029 17317163
refGene_gn=GIT1;refGene_tx=XM_023244606.1;refGene_gn=GIT1;refGene_tx=XM_023244607.1
E1 17317621 17317734
refGene_gn=GIT1;refGene_tx=XM_023244606.1;refGene_gn=GIT1;refGene_tx=XM_023244607.1
E1 17317859 17317965
refGene_gn=GIT1;refGene_tx=XM_023244606.1;refGene_gn=GIT1;refGene_tx=XM_023244607.1
E1 17318492 17318710
refGene_gn=GIT1;refGene_tx=XM_023244606.1;refGene_gn=GIT1;refGene_tx=XM_023244607.1
E1 17318795 17318890
refGene_gn=GIT1;refGene_tx=XM_023244606.1;refGene_gn=GIT1;refGene_tx=XM_023244607.1
E1 17319228 17319271
refGene_gn=GIT1;refGene_tx=XM_023244606.1;refGene_gn=GIT1;refGene_tx=XM_023244607.1
E1 17321044 17321071 refGene_gn=GIT1;refGene_tx=XM_023244607.1
E1 17321250 17321299
refGene_gn=GIT1;refGene_tx=XM_023244606.1;refGene_gn=GIT1;refGene_tx=XM_023244607.1
E1 17321747 17321820
refGene_gn=GIT1;refGene_tx=XM_023244606.1;refGene_gn=GIT1;refGene_tx=XM_023244607.1

E1 17322459 17322557
refGene_gn=GIT1;refGene_tx=XM_023244606.1;refGene_gn=GIT1;refGene_tx=X
M_023244607.1
E1 17322916 17323028
refGene_gn=GIT1;refGene_tx=XM_023244606.1;refGene_gn=GIT1;refGene_tx=X
M_023244607.1
E1 17323158 17323292
refGene_gn=GIT1;refGene_tx=XM_023244606.1;refGene_gn=GIT1;refGene_tx=X
M_023244607.1
E1 17323456 17323609
refGene_gn=GIT1;refGene_tx=XM_023244606.1;refGene_gn=GIT1;refGene_tx=X
M_023244607.1
E1 17323721 17323952
refGene_gn=GIT1;refGene_tx=XM_023244606.1;refGene_gn=GIT1;refGene_tx=X
M_023244607.1
E1 17324040 17324094
refGene_gn=GIT1;refGene_tx=XM_023244606.1;refGene_gn=GIT1;refGene_tx=X
M_023244607.1
E1 17324281 17324368
refGene_gn=GIT1;refGene_tx=XM_023244606.1;refGene_gn=GIT1;refGene_tx=X
M_023244607.1
E1 17324466 17324540
refGene_gn=GIT1;refGene_tx=XM_023244606.1;refGene_gn=GIT1;refGene_tx=X
M_023244607.1
E1 17324643 17324826
refGene_gn=GIT1;refGene_tx=XM_023244606.1;refGene_gn=GIT1;refGene_tx=X
M_023244607.1
E1 17324938 17325002
refGene_gn=GIT1;refGene_tx=XM_023244606.1;refGene_gn=GIT1;refGene_tx=X
M_023244607.1
E1 17325165 17325378
refGene_gn=GIT1;refGene_tx=XM_023244606.1;refGene_gn=GIT1;refGene_tx=X
M_023244607.1
E1 17327191 17327304
refGene_gn=TP53I13;refGene_tx=XM_003996464.4;refGene_gn=TP53I13;refGen
e_tx=XM_019817424.1;refGene_gn=TP53I13;refGene_tx=XM_019817425.1;refGe
ne_gn=TP53I13;refGene_tx=XM_019817426.2;refGene_gn=TP53I13;refGene_tx=
XM_019817427.1
E1 17327396 17327952
refGene_gn=TP53I13;refGene_tx=XM_003996464.4;refGene_gn=TP53I13;refGen
e_tx=XM_019817424.1;refGene_gn=TP53I13;refGene_tx=XM_019817425.1;refGe
ne_gn=TP53I13;refGene_tx=XM_019817426.2;refGene_gn=TP53I13;refGene_tx=
XM_019817427.1
E1 17328036 17328240
refGene_gn=TP53I13;refGene_tx=XM_003996464.4;refGene_gn=TP53I13;refGen
e_tx=XM_019817424.1;refGene_gn=TP53I13;refGene_tx=XM_019817425.1;refGe
ne_gn=TP53I13;refGene_tx=XM_019817426.2;refGene_gn=TP53I13;refGene_tx=
XM_019817427.1
E1 17328365 17328494
refGene_gn=TP53I13;refGene_tx=XM_003996464.4;refGene_gn=TP53I13;refGen

e_tx=XM_019817424.1;refGene_gn=TP53I13;refGene_tx=XM_019817426.2
E1 17330359 17330401
refGene_gn=TP53I13;refGene_tx=XM_003996464.4;refGene_gn=TP53I13;refGene_tx=XM_019817424.1;refGene_gn=TP53I13;refGene_tx=XM_019817425.1;refGene_gn=TP53I13;refGene_tx=XM_019817426.2;refGene_gn=TP53I13;refGene_tx=XM_019817427.1
E1 17330629 17330698
refGene_gn=TP53I13;refGene_tx=XM_003996464.4;refGene_gn=TP53I13;refGene_tx=XM_019817425.1
E1 17330886 17330958
refGene_gn=TP53I13;refGene_tx=XM_003996464.4;refGene_gn=TP53I13;refGene_tx=XM_019817424.1;refGene_gn=TP53I13;refGene_tx=XM_019817425.1;refGene_gn=TP53I13;refGene_tx=XM_019817427.1
E1 17333047 17333916 refGene_gn=ABHD15;refGene_tx=XM_023244608.1
E1 17336972 17337498 refGene_gn=ABHD15;refGene_tx=XM_023244608.1
E1 17353432 17353894
refGene_gn=TAOK1;refGene_tx=XM_003996465.5;refGene_gn=TAOK1;refGene_tx=XM_019817428.2
E1 17362355 17362538
refGene_gn=TAOK1;refGene_tx=XM_003996465.5;refGene_gn=TAOK1;refGene_tx=XM_019817428.2
E1 17364439 17364652
refGene_gn=TAOK1;refGene_tx=XM_003996465.5;refGene_gn=TAOK1;refGene_tx=XM_019817428.2
E1 17368793 17369033
refGene_gn=TAOK1;refGene_tx=XM_003996465.5;refGene_gn=TAOK1;refGene_tx=XM_019817428.2
E1 17371332 17371536
refGene_gn=TAOK1;refGene_tx=XM_003996465.5;refGene_gn=TAOK1;refGene_tx=XM_019817428.2
E1 17373916 17374045
refGene_gn=TAOK1;refGene_tx=XM_003996465.5;refGene_gn=TAOK1;refGene_tx=XM_019817428.2
E1 17376173 17376410
refGene_gn=TAOK1;refGene_tx=XM_003996465.5;refGene_gn=TAOK1;refGene_tx=XM_019817428.2
E1 17378953 17379088
refGene_gn=TAOK1;refGene_tx=XM_003996465.5;refGene_gn=TAOK1;refGene_tx=XM_019817428.2
E1 17381210 17381414
refGene_gn=TAOK1;refGene_tx=XM_003996465.5;refGene_gn=TAOK1;refGene_tx=XM_019817428.2
E1 17383104 17383272
refGene_gn=TAOK1;refGene_tx=XM_003996465.5;refGene_gn=TAOK1;refGene_tx=XM_019817428.2
E1 17388335 17388417
refGene_gn=TAOK1;refGene_tx=XM_003996465.5;refGene_gn=TAOK1;refGene_tx=XM_019817428.2
E1 17391142 17391236
refGene_gn=TAOK1;refGene_tx=XM_003996465.5;refGene_gn=TAOK1;refGene_tx

=XM_019817428.2
E1	17393167	17393259	refGene_gn=TAOK1;refGene_tx=XM_003996465.5
E1	17396145	17396259	refGene_gn=TAOK1;refGene_tx=XM_003996465.5
E1	17398569	17398666	refGene_gn=TAOK1;refGene_tx=XM_003996465.5
E1	17399211	17399257	refGene_gn=TAOK1;refGene_tx=XM_003996465.5
E1	17400311	17400413	refGene_gn=TAOK1;refGene_tx=XM_003996465.5
E1	17403073	17403145	refGene_gn=TAOK1;refGene_tx=XM_003996465.5
E1	17432364	17432496	refGene_gn=TAOK1;refGene_tx=XM_003996465.5
E1	17578580	17578887	refGene_gn=NUFIP2;refGene_tx=XM_003996466.5
E1	17584081	17585806	refGene_gn=NUFIP2;refGene_tx=XM_003996466.5
E1	17599520	17599553	refGene_gn=NUFIP2;refGene_tx=XM_003996466.5
E1	17601708	17601761	refGene_gn=NUFIP2;refGene_tx=XM_003996466.5
E1	17611031	17611179	refGene_gn=CRYBA1;refGene_tx=XM_003996467.3
E1	17611598	17611741	refGene_gn=CRYBA1;refGene_tx=XM_003996467.3
E1	17612658	17612800	refGene_gn=CRYBA1;refGene_tx=XM_003996467.3
E1	17614012	17614131	refGene_gn=CRYBA1;refGene_tx=XM_003996467.3
E1	17615101	17615166	refGene_gn=CRYBA1;refGene_tx=XM_003996467.3
E1	17616765	17616796	refGene_gn=CRYBA1;refGene_tx=XM_003996467.3
E1	17685885	17686884
refGene_gn=MYO18A;refGene_tx=XM_019817430.2;refGene_gn=MYO18A;refGene_tx=XM_019817431.2;refGene_gn=MYO18A;refGene_tx=XM_023243860.1;refGene_gn=MYO18A;refGene_tx=XM_023243862.1;refGene_gn=MYO18A;refGene_tx=XM_023243863.1;refGene_gn=MYO18A;refGene_tx=XM_023243864.1;refGene_gn=MYO18A;refGene_tx=XM_023243865.1;refGene_gn=MYO18A;refGene_tx=XM_023243866.1;refGene_gn=MYO18A;refGene_tx=XM_023243867.1;refGene_gn=MYO18A;refGene_tx=XM_023243868.1;refGene_gn=MYO18A;refGene_tx=XM_023243869.1;refGene_gn=MYO18A;refGene_tx=XM_023243871.1;refGene_gn=MYO18A;refGene_tx=XM_023243872.1;refGene_gn=MYO18A;refGene_tx=XM_023243873.1;refGene_gn=MYO18A;refGene_tx=XM_023243874.1;refGene_gn=MYO18A;refGene_tx=XM_023243875.1;refGene_gn=MYO18A;refGene_tx=XM_023243876.1;refGene_gn=MYO18A;refGene_tx=XM_023243877.1
E1	17709745	17709751	refGene_gn=MYO18A;refGene_tx=XM_019817439.1
E1	17715696	17715735	refGene_gn=MYO18A;refGene_tx=XM_019817438.1
E1	17717904	17717964	refGene_gn=MYO18A;refGene_tx=XM_019817438.1
E1	17720374	17721040	refGene_gn=MYO18A;refGene_tx=XM_019817438.1
E1	17723166	17723244
refGene_gn=MYO18A;refGene_tx=XM_019817430.2;refGene_gn=MYO18A;refGene_tx=XM_019817431.2;refGene_gn=MYO18A;refGene_tx=XM_019817438.1;refGene_gn=MYO18A;refGene_tx=XM_023243860.1;refGene_gn=MYO18A;refGene_tx=XM_023243862.1;refGene_gn=MYO18A;refGene_tx=XM_023243863.1;refGene_gn=MYO18A;refGene_tx=XM_023243864.1;refGene_gn=MYO18A;refGene_tx=XM_023243865.1;refGene_gn=MYO18A;refGene_tx=XM_023243871.1;refGene_gn=MYO18A;refGene_tx=XM_023243872.1;refGene_gn=MYO18A;refGene_tx=XM_023243873.1
E1	17724619	17724655
refGene_gn=MYO18A;refGene_tx=XM_019817430.2;refGene_gn=MYO18A;refGene_tx=XM_019817431.2;refGene_gn=MYO18A;refGene_tx=XM_019817438.1;refGene_gn=MYO18A;refGene_tx=XM_023243860.1;refGene_gn=MYO18A;refGene_tx=XM_023243862.1;refGene_gn=MYO18A;refGene_tx=XM_023243863.1;refGene_gn=MYO18A;refGene_tx=XM_023243864.1;refGene_gn=MYO18A;refGene_tx=XM_023243865.1;refGene_gn=MYO18A;refGene_tx=XM_023243866.1;refGene_gn=MYO18A;refGen

e_tx=XM_023243867.1;refGene_gn=MYO18A;refGene_tx=XM_023243871.1;refGene_gn=MYO18A;refGene_tx=XM_023243872.1;refGene_gn=MYO18A;refGene_tx=XM_023243873.1;refGene_gn=MYO18A;refGene_tx=XM_023243874.1;refGene_gn=MYO18A;refGene_tx=XM_023243876.1
E1 17726397 17726485
refGene_gn=MYO18A;refGene_tx=XM_019817430.2;refGene_gn=MYO18A;refGene_tx=XM_019817431.2;refGene_gn=MYO18A;refGene_tx=XM_019817438.1;refGene_gn=MYO18A;refGene_tx=XM_019817439.1;refGene_gn=MYO18A;refGene_tx=XM_023243860.1;refGene_gn=MYO18A;refGene_tx=XM_023243862.1;refGene_gn=MYO18A;refGene_tx=XM_023243863.1;refGene_gn=MYO18A;refGene_tx=XM_023243864.1;refGene_gn=MYO18A;refGene_tx=XM_023243865.1;refGene_gn=MYO18A;refGene_tx=XM_023243866.1;refGene_gn=MYO18A;refGene_tx=XM_023243867.1;refGene_gn=MYO18A;refGene_tx=XM_023243868.1;refGene_gn=MYO18A;refGene_tx=XM_023243869.1;refGene_gn=MYO18A;refGene_tx=XM_023243871.1;refGene_gn=MYO18A;refGene_tx=XM_023243872.1;refGene_gn=MYO18A;refGene_tx=XM_023243873.1;refGene_gn=MYO18A;refGene_tx=XM_023243874.1;refGene_gn=MYO18A;refGene_tx=XM_023243875.1;refGene_gn=MYO18A;refGene_tx=XM_023243876.1;refGene_gn=MYO18A;refGene_tx=XM_023243877.1
E1 17726688 17726795
refGene_gn=MYO18A;refGene_tx=XM_019817430.2;refGene_gn=MYO18A;refGene_tx=XM_019817431.2;refGene_gn=MYO18A;refGene_tx=XM_019817438.1;refGene_gn=MYO18A;refGene_tx=XM_019817439.1;refGene_gn=MYO18A;refGene_tx=XM_023243860.1;refGene_gn=MYO18A;refGene_tx=XM_023243862.1;refGene_gn=MYO18A;refGene_tx=XM_023243863.1;refGene_gn=MYO18A;refGene_tx=XM_023243864.1;refGene_gn=MYO18A;refGene_tx=XM_023243865.1;refGene_gn=MYO18A;refGene_tx=XM_023243866.1;refGene_gn=MYO18A;refGene_tx=XM_023243867.1;refGene_gn=MYO18A;refGene_tx=XM_023243868.1;refGene_gn=MYO18A;refGene_tx=XM_023243869.1;refGene_gn=MYO18A;refGene_tx=XM_023243871.1;refGene_gn=MYO18A;refGene_tx=XM_023243872.1;refGene_gn=MYO18A;refGene_tx=XM_023243873.1;refGene_gn=MYO18A;refGene_tx=XM_023243874.1;refGene_gn=MYO18A;refGene_tx=XM_023243875.1;refGene_gn=MYO18A;refGene_tx=XM_023243876.1;refGene_gn=MYO18A;refGene_tx=XM_023243877.1
E1 17726904 17727081
refGene_gn=MYO18A;refGene_tx=XM_019817430.2;refGene_gn=MYO18A;refGene_tx=XM_019817431.2;refGene_gn=MYO18A;refGene_tx=XM_019817438.1;refGene_gn=MYO18A;refGene_tx=XM_019817439.1;refGene_gn=MYO18A;refGene_tx=XM_023243860.1;refGene_gn=MYO18A;refGene_tx=XM_023243862.1;refGene_gn=MYO18A;refGene_tx=XM_023243863.1;refGene_gn=MYO18A;refGene_tx=XM_023243864.1;refGene_gn=MYO18A;refGene_tx=XM_023243865.1;refGene_gn=MYO18A;refGene_tx=XM_023243866.1;refGene_gn=MYO18A;refGene_tx=XM_023243867.1;refGene_gn=MYO18A;refGene_tx=XM_023243868.1;refGene_gn=MYO18A;refGene_tx=XM_023243869.1;refGene_gn=MYO18A;refGene_tx=XM_023243871.1;refGene_gn=MYO18A;refGene_tx=XM_023243872.1;refGene_gn=MYO18A;refGene_tx=XM_023243873.1;refGene_gn=MYO18A;refGene_tx=XM_023243874.1;refGene_gn=MYO18A;refGene_tx=XM_023243875.1;refGene_gn=MYO18A;refGene_tx=XM_023243876.1;refGene_gn=MYO18A;refGene_tx=XM_023243877.1
E1 17727438 17727652
refGene_gn=MYO18A;refGene_tx=XM_019817430.2;refGene_gn=MYO18A;refGene_tx=XM_019817431.2;refGene_gn=MYO18A;refGene_tx=XM_019817438.1;refGene_gn=MYO18A;refGene_tx=XM_019817439.1;refGene_gn=MYO18A;refGene_tx=XM_02

3243860.1;refGene_gn=MYO18A;refGene_tx=XM_023243862.1;refGene_gn=MYO18A;refGene_tx=XM_023243863.1;refGene_gn=MYO18A;refGene_tx=XM_023243864.1;refGene_gn=MYO18A;refGene_tx=XM_023243865.1;refGene_gn=MYO18A;refGene_tx=XM_023243866.1;refGene_gn=MYO18A;refGene_tx=XM_023243867.1;refGene_gn=MYO18A;refGene_tx=XM_023243868.1;refGene_gn=MYO18A;refGene_tx=XM_023243869.1;refGene_gn=MYO18A;refGene_tx=XM_023243871.1;refGene_gn=MYO18A;refGene_tx=XM_023243872.1;refGene_gn=MYO18A;refGene_tx=XM_023243873.1;refGene_gn=MYO18A;refGene_tx=XM_023243874.1;refGene_gn=MYO18A;refGene_tx=XM_023243875.1;refGene_gn=MYO18A;refGene_tx=XM_023243876.1;refGene_gn=MYO18A;refGene_tx=XM_023243877.1
E1 17728010 17728153
refGene_gn=MYO18A;refGene_tx=XM_019817430.2;refGene_gn=MYO18A;refGene_tx=XM_019817431.2;refGene_gn=MYO18A;refGene_tx=XM_019817438.1;refGene_gn=MYO18A;refGene_tx=XM_019817439.1;refGene_gn=MYO18A;refGene_tx=XM_023243860.1;refGene_gn=MYO18A;refGene_tx=XM_023243862.1;refGene_gn=MYO18A;refGene_tx=XM_023243863.1;refGene_gn=MYO18A;refGene_tx=XM_023243864.1;refGene_gn=MYO18A;refGene_tx=XM_023243865.1;refGene_gn=MYO18A;refGene_tx=XM_023243866.1;refGene_gn=MYO18A;refGene_tx=XM_023243867.1;refGene_gn=MYO18A;refGene_tx=XM_023243868.1;refGene_gn=MYO18A;refGene_tx=XM_023243869.1;refGene_gn=MYO18A;refGene_tx=XM_023243871.1;refGene_gn=MYO18A;refGene_tx=XM_023243872.1;refGene_gn=MYO18A;refGene_tx=XM_023243873.1;refGene_gn=MYO18A;refGene_tx=XM_023243874.1;refGene_gn=MYO18A;refGene_tx=XM_023243875.1;refGene_gn=MYO18A;refGene_tx=XM_023243876.1;refGene_gn=MYO18A;refGene_tx=XM_023243877.1
E1 17728764 17728865
refGene_gn=MYO18A;refGene_tx=XM_019817430.2;refGene_gn=MYO18A;refGene_tx=XM_019817431.2;refGene_gn=MYO18A;refGene_tx=XM_019817438.1;refGene_gn=MYO18A;refGene_tx=XM_019817439.1;refGene_gn=MYO18A;refGene_tx=XM_023243860.1;refGene_gn=MYO18A;refGene_tx=XM_023243862.1;refGene_gn=MYO18A;refGene_tx=XM_023243863.1;refGene_gn=MYO18A;refGene_tx=XM_023243864.1;refGene_gn=MYO18A;refGene_tx=XM_023243865.1;refGene_gn=MYO18A;refGene_tx=XM_023243866.1;refGene_gn=MYO18A;refGene_tx=XM_023243867.1;refGene_gn=MYO18A;refGene_tx=XM_023243868.1;refGene_gn=MYO18A;refGene_tx=XM_023243869.1;refGene_gn=MYO18A;refGene_tx=XM_023243871.1;refGene_gn=MYO18A;refGene_tx=XM_023243872.1;refGene_gn=MYO18A;refGene_tx=XM_023243873.1;refGene_gn=MYO18A;refGene_tx=XM_023243874.1;refGene_gn=MYO18A;refGene_tx=XM_023243875.1;refGene_gn=MYO18A;refGene_tx=XM_023243876.1;refGene_gn=MYO18A;refGene_tx=XM_023243877.1
E1 17729690 17729754
refGene_gn=MYO18A;refGene_tx=XM_019817430.2;refGene_gn=MYO18A;refGene_tx=XM_019817431.2;refGene_gn=MYO18A;refGene_tx=XM_019817438.1;refGene_gn=MYO18A;refGene_tx=XM_019817439.1;refGene_gn=MYO18A;refGene_tx=XM_023243860.1;refGene_gn=MYO18A;refGene_tx=XM_023243862.1;refGene_gn=MYO18A;refGene_tx=XM_023243863.1;refGene_gn=MYO18A;refGene_tx=XM_023243864.1;refGene_gn=MYO18A;refGene_tx=XM_023243865.1;refGene_gn=MYO18A;refGene_tx=XM_023243866.1;refGene_gn=MYO18A;refGene_tx=XM_023243867.1;refGene_gn=MYO18A;refGene_tx=XM_023243868.1;refGene_gn=MYO18A;refGene_tx=XM_023243869.1;refGene_gn=MYO18A;refGene_tx=XM_023243871.1;refGene_gn=MYO18A;refGene_tx=XM_023243872.1;refGene_gn=MYO18A;refGene_tx=XM_023243873.1;refGene_gn=MYO18A;refGene_tx=XM_023243874.1;refGene_gn=MYO18A;refG

ene_tx=XM_023243875.1;refGene_gn=MYO18A;refGene_tx=XM_023243876.1;refGene_gn=MYO18A;refGene_tx=XM_023243877.1
E1 17729978 17730123
refGene_gn=MYO18A;refGene_tx=XM_019817430.2;refGene_gn=MYO18A;refGene_tx=XM_019817431.2;refGene_gn=MYO18A;refGene_tx=XM_019817438.1;refGene_gn=MYO18A;refGene_tx=XM_019817439.1;refGene_gn=MYO18A;refGene_tx=XM_023243860.1;refGene_gn=MYO18A;refGene_tx=XM_023243862.1;refGene_gn=MYO18A;refGene_tx=XM_023243863.1;refGene_gn=MYO18A;refGene_tx=XM_023243864.1;refGene_gn=MYO18A;refGene_tx=XM_023243865.1;refGene_gn=MYO18A;refGene_tx=XM_023243866.1;refGene_gn=MYO18A;refGene_tx=XM_023243867.1;refGene_gn=MYO18A;refGene_tx=XM_023243868.1;refGene_gn=MYO18A;refGene_tx=XM_023243869.1;refGene_gn=MYO18A;refGene_tx=XM_023243871.1;refGene_gn=MYO18A;refGene_tx=XM_023243872.1;refGene_gn=MYO18A;refGene_tx=XM_023243873.1;refGene_gn=MYO18A;refGene_tx=XM_023243874.1;refGene_gn=MYO18A;refGene_tx=XM_023243875.1;refGene_gn=MYO18A;refGene_tx=XM_023243876.1;refGene_gn=MYO18A;refGene_tx=XM_023243877.1
E1 17730447 17730468
refGene_gn=MYO18A;refGene_tx=XM_019817430.2;refGene_gn=MYO18A;refGene_tx=XM_019817431.2;refGene_gn=MYO18A;refGene_tx=XM_019817438.1;refGene_gn=MYO18A;refGene_tx=XM_019817439.1;refGene_gn=MYO18A;refGene_tx=XM_023243860.1;refGene_gn=MYO18A;refGene_tx=XM_023243862.1;refGene_gn=MYO18A;refGene_tx=XM_023243865.1;refGene_gn=MYO18A;refGene_tx=XM_023243866.1;refGene_gn=MYO18A;refGene_tx=XM_023243868.1;refGene_gn=MYO18A;refGene_tx=XM_023243871.1;refGene_gn=MYO18A;refGene_tx=XM_023243872.1;refGene_gn=MYO18A;refGene_tx=XM_023243874.1
E1 17731706 17731718
refGene_gn=MYO18A;refGene_tx=XM_019817430.2;refGene_gn=MYO18A;refGene_tx=XM_019817431.2;refGene_gn=MYO18A;refGene_tx=XM_019817438.1;refGene_gn=MYO18A;refGene_tx=XM_019817439.1;refGene_gn=MYO18A;refGene_tx=XM_023243860.1;refGene_gn=MYO18A;refGene_tx=XM_023243863.1;refGene_gn=MYO18A;refGene_tx=XM_023243865.1;refGene_gn=MYO18A;refGene_tx=XM_023243866.1;refGene_gn=MYO18A;refGene_tx=XM_023243867.1;refGene_gn=MYO18A;refGene_tx=XM_023243868.1;refGene_gn=MYO18A;refGene_tx=XM_023243869.1;refGene_gn=MYO18A;refGene_tx=XM_023243871.1;refGene_gn=MYO18A;refGene_tx=XM_023243872.1;refGene_gn=MYO18A;refGene_tx=XM_023243873.1;refGene_gn=MYO18A;refGene_tx=XM_023243874.1;refGene_gn=MYO18A;refGene_tx=XM_023243875.1;refGene_gn=MYO18A;refGene_tx=XM_023243876.1;refGene_gn=MYO18A;refGene_tx=XM_023243877.1
E1 17732333 17732510
refGene_gn=MYO18A;refGene_tx=XM_019817430.2;refGene_gn=MYO18A;refGene_tx=XM_019817431.2;refGene_gn=MYO18A;refGene_tx=XM_019817438.1;refGene_gn=MYO18A;refGene_tx=XM_019817439.1;refGene_gn=MYO18A;refGene_tx=XM_023243860.1;refGene_gn=MYO18A;refGene_tx=XM_023243862.1;refGene_gn=MYO18A;refGene_tx=XM_023243863.1;refGene_gn=MYO18A;refGene_tx=XM_023243864.1;refGene_gn=MYO18A;refGene_tx=XM_023243865.1;refGene_gn=MYO18A;refGene_tx=XM_023243866.1;refGene_gn=MYO18A;refGene_tx=XM_023243867.1;refGene_gn=MYO18A;refGene_tx=XM_023243868.1;refGene_gn=MYO18A;refGene_tx=XM_023243869.1;refGene_gn=MYO18A;refGene_tx=XM_023243871.1;refGene_gn=MYO18A;refGene_tx=XM_023243872.1;refGene_gn=MYO18A;refGene_tx=XM_023243873.1;refGene_gn=MYO18A;refGene_tx=XM_023243874.1;refGene_gn=MYO18A;refG

ene_tx=XM_023243875.1;refGene_gn=MYO18A;refGene_tx=XM_023243876.1;refGene_gn=MYO18A;refGene_tx=XM_023243877.1
E1 17732734 17732825
refGene_gn=MYO18A;refGene_tx=XM_019817430.2;refGene_gn=MYO18A;refGene_tx=XM_019817431.2;refGene_gn=MYO18A;refGene_tx=XM_019817438.1;refGene_gn=MYO18A;refGene_tx=XM_019817439.1;refGene_gn=MYO18A;refGene_tx=XM_023243860.1;refGene_gn=MYO18A;refGene_tx=XM_023243862.1;refGene_gn=MYO18A;refGene_tx=XM_023243863.1;refGene_gn=MYO18A;refGene_tx=XM_023243864.1;refGene_gn=MYO18A;refGene_tx=XM_023243865.1;refGene_gn=MYO18A;refGene_tx=XM_023243866.1;refGene_gn=MYO18A;refGene_tx=XM_023243867.1;refGene_gn=MYO18A;refGene_tx=XM_023243868.1;refGene_gn=MYO18A;refGene_tx=XM_023243869.1;refGene_gn=MYO18A;refGene_tx=XM_023243871.1;refGene_gn=MYO18A;refGene_tx=XM_023243872.1;refGene_gn=MYO18A;refGene_tx=XM_023243873.1;refGene_gn=MYO18A;refGene_tx=XM_023243874.1;refGene_gn=MYO18A;refGene_tx=XM_023243875.1;refGene_gn=MYO18A;refGene_tx=XM_023243876.1;refGene_gn=MYO18A;refGene_tx=XM_023243877.1
E1 17733123 17733316
refGene_gn=MYO18A;refGene_tx=XM_019817430.2;refGene_gn=MYO18A;refGene_tx=XM_019817431.2;refGene_gn=MYO18A;refGene_tx=XM_019817438.1;refGene_gn=MYO18A;refGene_tx=XM_019817439.1;refGene_gn=MYO18A;refGene_tx=XM_023243860.1;refGene_gn=MYO18A;refGene_tx=XM_023243862.1;refGene_gn=MYO18A;refGene_tx=XM_023243863.1;refGene_gn=MYO18A;refGene_tx=XM_023243864.1;refGene_gn=MYO18A;refGene_tx=XM_023243865.1;refGene_gn=MYO18A;refGene_tx=XM_023243866.1;refGene_gn=MYO18A;refGene_tx=XM_023243867.1;refGene_gn=MYO18A;refGene_tx=XM_023243868.1;refGene_gn=MYO18A;refGene_tx=XM_023243869.1;refGene_gn=MYO18A;refGene_tx=XM_023243871.1;refGene_gn=MYO18A;refGene_tx=XM_023243872.1;refGene_gn=MYO18A;refGene_tx=XM_023243873.1;refGene_gn=MYO18A;refGene_tx=XM_023243874.1;refGene_gn=MYO18A;refGene_tx=XM_023243875.1;refGene_gn=MYO18A;refGene_tx=XM_023243876.1;refGene_gn=MYO18A;refGene_tx=XM_023243877.1
E1 17734056 17734143
refGene_gn=MYO18A;refGene_tx=XM_019817430.2;refGene_gn=MYO18A;refGene_tx=XM_019817431.2;refGene_gn=MYO18A;refGene_tx=XM_019817438.1;refGene_gn=MYO18A;refGene_tx=XM_019817439.1;refGene_gn=MYO18A;refGene_tx=XM_023243860.1;refGene_gn=MYO18A;refGene_tx=XM_023243862.1;refGene_gn=MYO18A;refGene_tx=XM_023243863.1;refGene_gn=MYO18A;refGene_tx=XM_023243864.1;refGene_gn=MYO18A;refGene_tx=XM_023243865.1;refGene_gn=MYO18A;refGene_tx=XM_023243866.1;refGene_gn=MYO18A;refGene_tx=XM_023243867.1;refGene_gn=MYO18A;refGene_tx=XM_023243868.1;refGene_gn=MYO18A;refGene_tx=XM_023243869.1;refGene_gn=MYO18A;refGene_tx=XM_023243871.1;refGene_gn=MYO18A;refGene_tx=XM_023243872.1;refGene_gn=MYO18A;refGene_tx=XM_023243873.1;refGene_gn=MYO18A;refGene_tx=XM_023243874.1;refGene_gn=MYO18A;refGene_tx=XM_023243875.1;refGene_gn=MYO18A;refGene_tx=XM_023243876.1;refGene_gn=MYO18A;refGene_tx=XM_023243877.1
E1 17736108 17736250
refGene_gn=MYO18A;refGene_tx=XM_019817430.2;refGene_gn=MYO18A;refGene_tx=XM_019817431.2;refGene_gn=MYO18A;refGene_tx=XM_019817438.1;refGene_gn=MYO18A;refGene_tx=XM_019817439.1;refGene_gn=MYO18A;refGene_tx=XM_023243860.1;refGene_gn=MYO18A;refGene_tx=XM_023243862.1;refGene_gn=MYO18A;refGene_tx=XM_023243863.1;refGene_gn=MYO18A;refGene_tx=XM_023243864.

1;refGene_gn=MYO18A;refGene_tx=XM_023243865.1;refGene_gn=MYO18A;refGene_tx=XM_023243866.1;refGene_gn=MYO18A;refGene_tx=XM_023243867.1;refGene_gn=MYO18A;refGene_tx=XM_023243868.1;refGene_gn=MYO18A;refGene_tx=XM_023243869.1;refGene_gn=MYO18A;refGene_tx=XM_023243871.1;refGene_gn=MYO18A;refGene_tx=XM_023243872.1;refGene_gn=MYO18A;refGene_tx=XM_023243873.1;refGene_gn=MYO18A;refGene_tx=XM_023243874.1;refGene_gn=MYO18A;refGene_tx=XM_023243875.1;refGene_gn=MYO18A;refGene_tx=XM_023243876.1;refGene_gn=MYO18A;refGene_tx=XM_023243877.1

E1 17736405 17736565

refGene_gn=MYO18A;refGene_tx=XM_019817430.2;refGene_gn=MYO18A;refGene_tx=XM_019817431.2;refGene_gn=MYO18A;refGene_tx=XM_019817438.1;refGene_gn=MYO18A;refGene_tx=XM_019817439.1;refGene_gn=MYO18A;refGene_tx=XM_023243860.1;refGene_gn=MYO18A;refGene_tx=XM_023243862.1;refGene_gn=MYO18A;refGene_tx=XM_023243863.1;refGene_gn=MYO18A;refGene_tx=XM_023243864.1;refGene_gn=MYO18A;refGene_tx=XM_023243865.1;refGene_gn=MYO18A;refGene_tx=XM_023243866.1;refGene_gn=MYO18A;refGene_tx=XM_023243867.1;refGene_gn=MYO18A;refGene_tx=XM_023243868.1;refGene_gn=MYO18A;refGene_tx=XM_023243869.1;refGene_gn=MYO18A;refGene_tx=XM_023243871.1;refGene_gn=MYO18A;refGene_tx=XM_023243872.1;refGene_gn=MYO18A;refGene_tx=XM_023243873.1;refGene_gn=MYO18A;refGene_tx=XM_023243874.1;refGene_gn=MYO18A;refGene_tx=XM_023243875.1;refGene_gn=MYO18A;refGene_tx=XM_023243876.1;refGene_gn=MYO18A;refGene_tx=XM_023243877.1

E1 17737566 17737753

refGene_gn=MYO18A;refGene_tx=XM_019817430.2;refGene_gn=MYO18A;refGene_tx=XM_019817431.2;refGene_gn=MYO18A;refGene_tx=XM_019817438.1;refGene_gn=MYO18A;refGene_tx=XM_019817439.1;refGene_gn=MYO18A;refGene_tx=XM_023243860.1;refGene_gn=MYO18A;refGene_tx=XM_023243862.1;refGene_gn=MYO18A;refGene_tx=XM_023243863.1;refGene_gn=MYO18A;refGene_tx=XM_023243864.1;refGene_gn=MYO18A;refGene_tx=XM_023243865.1;refGene_gn=MYO18A;refGene_tx=XM_023243866.1;refGene_gn=MYO18A;refGene_tx=XM_023243867.1;refGene_gn=MYO18A;refGene_tx=XM_023243868.1;refGene_gn=MYO18A;refGene_tx=XM_023243869.1;refGene_gn=MYO18A;refGene_tx=XM_023243871.1;refGene_gn=MYO18A;refGene_tx=XM_023243872.1;refGene_gn=MYO18A;refGene_tx=XM_023243873.1;refGene_gn=MYO18A;refGene_tx=XM_023243874.1;refGene_gn=MYO18A;refGene_tx=XM_023243875.1;refGene_gn=MYO18A;refGene_tx=XM_023243876.1;refGene_gn=MYO18A;refGene_tx=XM_023243877.1

E1 17738052 17738296

refGene_gn=MYO18A;refGene_tx=XM_019817430.2;refGene_gn=MYO18A;refGene_tx=XM_019817431.2;refGene_gn=MYO18A;refGene_tx=XM_019817438.1;refGene_gn=MYO18A;refGene_tx=XM_019817439.1;refGene_gn=MYO18A;refGene_tx=XM_023243860.1;refGene_gn=MYO18A;refGene_tx=XM_023243862.1;refGene_gn=MYO18A;refGene_tx=XM_023243863.1;refGene_gn=MYO18A;refGene_tx=XM_023243864.1;refGene_gn=MYO18A;refGene_tx=XM_023243865.1;refGene_gn=MYO18A;refGene_tx=XM_023243866.1;refGene_gn=MYO18A;refGene_tx=XM_023243867.1;refGene_gn=MYO18A;refGene_tx=XM_023243868.1;refGene_gn=MYO18A;refGene_tx=XM_023243869.1;refGene_gn=MYO18A;refGene_tx=XM_023243871.1;refGene_gn=MYO18A;refGene_tx=XM_023243872.1;refGene_gn=MYO18A;refGene_tx=XM_023243873.1;refGene_gn=MYO18A;refGene_tx=XM_023243874.1;refGene_gn=MYO18A;refGene_tx=XM_023243875.1;refGene_gn=MYO18A;refGene_tx=XM_023243876.1;refGene_gn=MYO18A;refGene_tx=XM_023243877.1

E1 17742609 17742719
refGene_gn=MYO18A;refGene_tx=XM_019817430.2;refGene_gn=MYO18A;refGene_tx=XM_019817431.2;refGene_gn=MYO18A;refGene_tx=XM_019817438.1;refGene_gn=MYO18A;refGene_tx=XM_019817439.1;refGene_gn=MYO18A;refGene_tx=XM_023243860.1;refGene_gn=MYO18A;refGene_tx=XM_023243862.1;refGene_gn=MYO18A;refGene_tx=XM_023243863.1;refGene_gn=MYO18A;refGene_tx=XM_023243864.1;refGene_gn=MYO18A;refGene_tx=XM_023243865.1;refGene_gn=MYO18A;refGene_tx=XM_023243866.1;refGene_gn=MYO18A;refGene_tx=XM_023243867.1;refGene_gn=MYO18A;refGene_tx=XM_023243868.1;refGene_gn=MYO18A;refGene_tx=XM_023243869.1;refGene_gn=MYO18A;refGene_tx=XM_023243871.1;refGene_gn=MYO18A;refGene_tx=XM_023243872.1;refGene_gn=MYO18A;refGene_tx=XM_023243873.1;refGene_gn=MYO18A;refGene_tx=XM_023243874.1;refGene_gn=MYO18A;refGene_tx=XM_023243875.1;refGene_gn=MYO18A;refGene_tx=XM_023243876.1;refGene_gn=MYO18A;refGene_tx=XM_023243877.1
E1 17745813 17745879
refGene_gn=MYO18A;refGene_tx=XM_019817430.2;refGene_gn=MYO18A;refGene_tx=XM_019817431.2;refGene_gn=MYO18A;refGene_tx=XM_019817438.1;refGene_gn=MYO18A;refGene_tx=XM_019817439.1;refGene_gn=MYO18A;refGene_tx=XM_023243860.1;refGene_gn=MYO18A;refGene_tx=XM_023243862.1;refGene_gn=MYO18A;refGene_tx=XM_023243863.1;refGene_gn=MYO18A;refGene_tx=XM_023243864.1;refGene_gn=MYO18A;refGene_tx=XM_023243865.1;refGene_gn=MYO18A;refGene_tx=XM_023243866.1;refGene_gn=MYO18A;refGene_tx=XM_023243867.1;refGene_gn=MYO18A;refGene_tx=XM_023243868.1;refGene_gn=MYO18A;refGene_tx=XM_023243869.1;refGene_gn=MYO18A;refGene_tx=XM_023243871.1;refGene_gn=MYO18A;refGene_tx=XM_023243872.1;refGene_gn=MYO18A;refGene_tx=XM_023243873.1;refGene_gn=MYO18A;refGene_tx=XM_023243874.1;refGene_gn=MYO18A;refGene_tx=XM_023243875.1;refGene_gn=MYO18A;refGene_tx=XM_023243876.1;refGene_gn=MYO18A;refGene_tx=XM_023243877.1
E1 17748859 17748988
refGene_gn=MYO18A;refGene_tx=XM_019817430.2;refGene_gn=MYO18A;refGene_tx=XM_019817431.2;refGene_gn=MYO18A;refGene_tx=XM_019817438.1;refGene_gn=MYO18A;refGene_tx=XM_019817439.1;refGene_gn=MYO18A;refGene_tx=XM_023243860.1;refGene_gn=MYO18A;refGene_tx=XM_023243862.1;refGene_gn=MYO18A;refGene_tx=XM_023243863.1;refGene_gn=MYO18A;refGene_tx=XM_023243864.1;refGene_gn=MYO18A;refGene_tx=XM_023243865.1;refGene_gn=MYO18A;refGene_tx=XM_023243866.1;refGene_gn=MYO18A;refGene_tx=XM_023243867.1;refGene_gn=MYO18A;refGene_tx=XM_023243868.1;refGene_gn=MYO18A;refGene_tx=XM_023243869.1;refGene_gn=MYO18A;refGene_tx=XM_023243871.1;refGene_gn=MYO18A;refGene_tx=XM_023243872.1;refGene_gn=MYO18A;refGene_tx=XM_023243873.1;refGene_gn=MYO18A;refGene_tx=XM_023243874.1;refGene_gn=MYO18A;refGene_tx=XM_023243875.1;refGene_gn=MYO18A;refGene_tx=XM_023243876.1;refGene_gn=MYO18A;refGene_tx=XM_023243877.1
E1 17749693 17749837
refGene_gn=MYO18A;refGene_tx=XM_019817430.2;refGene_gn=MYO18A;refGene_tx=XM_019817431.2;refGene_gn=MYO18A;refGene_tx=XM_019817438.1;refGene_gn=MYO18A;refGene_tx=XM_019817439.1;refGene_gn=MYO18A;refGene_tx=XM_023243860.1;refGene_gn=MYO18A;refGene_tx=XM_023243862.1;refGene_gn=MYO18A;refGene_tx=XM_023243863.1;refGene_gn=MYO18A;refGene_tx=XM_023243864.1;refGene_gn=MYO18A;refGene_tx=XM_023243865.1;refGene_gn=MYO18A;refGene_tx=XM_023243866.1;refGene_gn=MYO18A;refGene_tx=XM_023243867.1;refGen

e_gn=MYO18A;refGene_tx=XM_023243868.1;refGene_gn=MYO18A;refGene_tx=XM_023243869.1;refGene_gn=MYO18A;refGene_tx=XM_023243871.1;refGene_gn=MYO18A;refGene_tx=XM_023243872.1;refGene_gn=MYO18A;refGene_tx=XM_023243873.1;refGene_gn=MYO18A;refGene_tx=XM_023243874.1;refGene_gn=MYO18A;refGene_tx=XM_023243875.1;refGene_gn=MYO18A;refGene_tx=XM_023243876.1;refGene_gn=MYO18A;refGene_tx=XM_023243877.1
E1 17750221 17750311
refGene_gn=MYO18A;refGene_tx=XM_019817430.2;refGene_gn=MYO18A;refGene_tx=XM_019817431.2;refGene_gn=MYO18A;refGene_tx=XM_019817438.1;refGene_gn=MYO18A;refGene_tx=XM_019817439.1;refGene_gn=MYO18A;refGene_tx=XM_023243860.1;refGene_gn=MYO18A;refGene_tx=XM_023243862.1;refGene_gn=MYO18A;refGene_tx=XM_023243863.1;refGene_gn=MYO18A;refGene_tx=XM_023243864.1;refGene_gn=MYO18A;refGene_tx=XM_023243865.1;refGene_gn=MYO18A;refGene_tx=XM_023243866.1;refGene_gn=MYO18A;refGene_tx=XM_023243867.1;refGene_gn=MYO18A;refGene_tx=XM_023243868.1;refGene_gn=MYO18A;refGene_tx=XM_023243869.1;refGene_gn=MYO18A;refGene_tx=XM_023243871.1;refGene_gn=MYO18A;refGene_tx=XM_023243872.1;refGene_gn=MYO18A;refGene_tx=XM_023243873.1;refGene_gn=MYO18A;refGene_tx=XM_023243874.1;refGene_gn=MYO18A;refGene_tx=XM_023243875.1;refGene_gn=MYO18A;refGene_tx=XM_023243876.1;refGene_gn=MYO18A;refGene_tx=XM_023243877.1
E1 17750458 17750578
refGene_gn=MYO18A;refGene_tx=XM_019817430.2;refGene_gn=MYO18A;refGene_tx=XM_019817431.2;refGene_gn=MYO18A;refGene_tx=XM_019817438.1;refGene_gn=MYO18A;refGene_tx=XM_019817439.1;refGene_gn=MYO18A;refGene_tx=XM_023243860.1;refGene_gn=MYO18A;refGene_tx=XM_023243862.1;refGene_gn=MYO18A;refGene_tx=XM_023243863.1;refGene_gn=MYO18A;refGene_tx=XM_023243864.1;refGene_gn=MYO18A;refGene_tx=XM_023243865.1;refGene_gn=MYO18A;refGene_tx=XM_023243866.1;refGene_gn=MYO18A;refGene_tx=XM_023243867.1;refGene_gn=MYO18A;refGene_tx=XM_023243868.1;refGene_gn=MYO18A;refGene_tx=XM_023243869.1;refGene_gn=MYO18A;refGene_tx=XM_023243871.1;refGene_gn=MYO18A;refGene_tx=XM_023243872.1;refGene_gn=MYO18A;refGene_tx=XM_023243873.1;refGene_gn=MYO18A;refGene_tx=XM_023243874.1;refGene_gn=MYO18A;refGene_tx=XM_023243875.1;refGene_gn=MYO18A;refGene_tx=XM_023243876.1;refGene_gn=MYO18A;refGene_tx=XM_023243877.1
E1 17750780 17750892
refGene_gn=MYO18A;refGene_tx=XM_019817430.2;refGene_gn=MYO18A;refGene_tx=XM_019817431.2;refGene_gn=MYO18A;refGene_tx=XM_019817438.1;refGene_gn=MYO18A;refGene_tx=XM_019817439.1;refGene_gn=MYO18A;refGene_tx=XM_023243860.1;refGene_gn=MYO18A;refGene_tx=XM_023243862.1;refGene_gn=MYO18A;refGene_tx=XM_023243863.1;refGene_gn=MYO18A;refGene_tx=XM_023243864.1;refGene_gn=MYO18A;refGene_tx=XM_023243865.1;refGene_gn=MYO18A;refGene_tx=XM_023243866.1;refGene_gn=MYO18A;refGene_tx=XM_023243867.1;refGene_gn=MYO18A;refGene_tx=XM_023243868.1;refGene_gn=MYO18A;refGene_tx=XM_023243869.1;refGene_gn=MYO18A;refGene_tx=XM_023243871.1;refGene_gn=MYO18A;refGene_tx=XM_023243872.1;refGene_gn=MYO18A;refGene_tx=XM_023243873.1;refGene_gn=MYO18A;refGene_tx=XM_023243874.1;refGene_gn=MYO18A;refGene_tx=XM_023243875.1;refGene_gn=MYO18A;refGene_tx=XM_023243876.1;refGene_gn=MYO18A;refGene_tx=XM_023243877.1
E1 17751306 17751434
refGene_gn=MYO18A;refGene_tx=XM_019817430.2;refGene_gn=MYO18A;refGene_

tx=XM_019817431.2;refGene_gn=MYO18A;refGene_tx=XM_019817438.1;refGene_gn=MYO18A;refGene_tx=XM_019817439.1;refGene_gn=MYO18A;refGene_tx=XM_023243860.1;refGene_gn=MYO18A;refGene_tx=XM_023243862.1;refGene_gn=MYO18A;refGene_tx=XM_023243863.1;refGene_gn=MYO18A;refGene_tx=XM_023243864.1;refGene_gn=MYO18A;refGene_tx=XM_023243865.1;refGene_gn=MYO18A;refGene_tx=XM_023243866.1;refGene_gn=MYO18A;refGene_tx=XM_023243867.1;refGene_gn=MYO18A;refGene_tx=XM_023243868.1;refGene_gn=MYO18A;refGene_tx=XM_023243869.1;refGene_gn=MYO18A;refGene_tx=XM_023243871.1;refGene_gn=MYO18A;refGene_tx=XM_023243872.1;refGene_gn=MYO18A;refGene_tx=XM_023243873.1;refGene_gn=MYO18A;refGene_tx=XM_023243874.1;refGene_gn=MYO18A;refGene_tx=XM_023243875.1;refGene_gn=MYO18A;refGene_tx=XM_023243876.1;refGene_gn=MYO18A;refGene_tx=XM_023243877.1
E1 17751696 17751851
refGene_gn=MYO18A;refGene_tx=XM_019817430.2;refGene_gn=MYO18A;refGene_tx=XM_019817431.2;refGene_gn=MYO18A;refGene_tx=XM_019817438.1;refGene_gn=MYO18A;refGene_tx=XM_019817439.1;refGene_gn=MYO18A;refGene_tx=XM_023243860.1;refGene_gn=MYO18A;refGene_tx=XM_023243862.1;refGene_gn=MYO18A;refGene_tx=XM_023243863.1;refGene_gn=MYO18A;refGene_tx=XM_023243864.1;refGene_gn=MYO18A;refGene_tx=XM_023243865.1;refGene_gn=MYO18A;refGene_tx=XM_023243866.1;refGene_gn=MYO18A;refGene_tx=XM_023243867.1;refGene_gn=MYO18A;refGene_tx=XM_023243868.1;refGene_gn=MYO18A;refGene_tx=XM_023243869.1;refGene_gn=MYO18A;refGene_tx=XM_023243871.1;refGene_gn=MYO18A;refGene_tx=XM_023243872.1;refGene_gn=MYO18A;refGene_tx=XM_023243873.1;refGene_gn=MYO18A;refGene_tx=XM_023243874.1;refGene_gn=MYO18A;refGene_tx=XM_023243875.1;refGene_gn=MYO18A;refGene_tx=XM_023243876.1;refGene_gn=MYO18A;refGene_tx=XM_023243877.1
E1 17753622 17753746
refGene_gn=MYO18A;refGene_tx=XM_019817430.2;refGene_gn=MYO18A;refGene_tx=XM_019817431.2;refGene_gn=MYO18A;refGene_tx=XM_019817438.1;refGene_gn=MYO18A;refGene_tx=XM_019817439.1;refGene_gn=MYO18A;refGene_tx=XM_023243860.1;refGene_gn=MYO18A;refGene_tx=XM_023243862.1;refGene_gn=MYO18A;refGene_tx=XM_023243863.1;refGene_gn=MYO18A;refGene_tx=XM_023243864.1;refGene_gn=MYO18A;refGene_tx=XM_023243865.1;refGene_gn=MYO18A;refGene_tx=XM_023243866.1;refGene_gn=MYO18A;refGene_tx=XM_023243867.1;refGene_gn=MYO18A;refGene_tx=XM_023243868.1;refGene_gn=MYO18A;refGene_tx=XM_023243869.1;refGene_gn=MYO18A;refGene_tx=XM_023243871.1;refGene_gn=MYO18A;refGene_tx=XM_023243872.1;refGene_gn=MYO18A;refGene_tx=XM_023243873.1;refGene_gn=MYO18A;refGene_tx=XM_023243874.1;refGene_gn=MYO18A;refGene_tx=XM_023243875.1;refGene_gn=MYO18A;refGene_tx=XM_023243876.1;refGene_gn=MYO18A;refGene_tx=XM_023243877.1
E1 17753832 17754033
refGene_gn=MYO18A;refGene_tx=XM_019817430.2;refGene_gn=MYO18A;refGene_tx=XM_019817431.2;refGene_gn=MYO18A;refGene_tx=XM_019817438.1;refGene_gn=MYO18A;refGene_tx=XM_019817439.1;refGene_gn=MYO18A;refGene_tx=XM_023243860.1;refGene_gn=MYO18A;refGene_tx=XM_023243862.1;refGene_gn=MYO18A;refGene_tx=XM_023243863.1;refGene_gn=MYO18A;refGene_tx=XM_023243864.1;refGene_gn=MYO18A;refGene_tx=XM_023243865.1;refGene_gn=MYO18A;refGene_tx=XM_023243866.1;refGene_gn=MYO18A;refGene_tx=XM_023243867.1;refGene_gn=MYO18A;refGene_tx=XM_023243868.1;refGene_gn=MYO18A;refGene_tx=XM_023243869.1;refGene_gn=MYO18A;refGene_tx=XM_023243871.1;refGene_gn=MYO

18A;refGene_tx=XM_023243872.1;refGene_gn=MYO18A;refGene_tx=XM_02324387
3.1;refGene_gn=MYO18A;refGene_tx=XM_023243874.1;refGene_gn=MYO18A;refG
ene_tx=XM_023243875.1;refGene_gn=MYO18A;refGene_tx=XM_023243876.1;refG
ene_gn=MYO18A;refGene_tx=XM_023243877.1
E1 17754594 17754705
refGene_gn=MYO18A;refGene_tx=XM_019817430.2;refGene_gn=MYO18A;refGene_
tx=XM_019817431.2;refGene_gn=MYO18A;refGene_tx=XM_019817438.1;refGene_
gn=MYO18A;refGene_tx=XM_019817439.1;refGene_gn=MYO18A;refGene_tx=XM_02
3243860.1;refGene_gn=MYO18A;refGene_tx=XM_023243862.1;refGene_gn=MYO18
A;refGene_tx=XM_023243863.1;refGene_gn=MYO18A;refGene_tx=XM_023243864.
1;refGene_gn=MYO18A;refGene_tx=XM_023243865.1;refGene_gn=MYO18A;refGen
e_tx=XM_023243866.1;refGene_gn=MYO18A;refGene_tx=XM_023243867.1;refGen
e_gn=MYO18A;refGene_tx=XM_023243868.1;refGene_gn=MYO18A;refGene_tx=XM_
023243869.1;refGene_gn=MYO18A;refGene_tx=XM_023243871.1;refGene_gn=MYO
18A;refGene_tx=XM_023243872.1;refGene_gn=MYO18A;refGene_tx=XM_02324387
3.1;refGene_gn=MYO18A;refGene_tx=XM_023243874.1;refGene_gn=MYO18A;refG
ene_tx=XM_023243875.1;refGene_gn=MYO18A;refGene_tx=XM_023243876.1;refG
ene_gn=MYO18A;refGene_tx=XM_023243877.1
E1 17755452 17755557
refGene_gn=MYO18A;refGene_tx=XM_019817430.2;refGene_gn=MYO18A;refGene_
tx=XM_019817431.2;refGene_gn=MYO18A;refGene_tx=XM_019817438.1;refGene_
gn=MYO18A;refGene_tx=XM_019817439.1;refGene_gn=MYO18A;refGene_tx=XM_02
3243860.1;refGene_gn=MYO18A;refGene_tx=XM_023243862.1;refGene_gn=MYO18
A;refGene_tx=XM_023243863.1;refGene_gn=MYO18A;refGene_tx=XM_023243864.
1;refGene_gn=MYO18A;refGene_tx=XM_023243865.1;refGene_gn=MYO18A;refGen
e_tx=XM_023243866.1;refGene_gn=MYO18A;refGene_tx=XM_023243867.1;refGen
e_gn=MYO18A;refGene_tx=XM_023243868.1;refGene_gn=MYO18A;refGene_tx=XM_
023243869.1;refGene_gn=MYO18A;refGene_tx=XM_023243871.1;refGene_gn=MYO
18A;refGene_tx=XM_023243872.1;refGene_gn=MYO18A;refGene_tx=XM_02324387
3.1;refGene_gn=MYO18A;refGene_tx=XM_023243874.1;refGene_gn=MYO18A;refG
ene_tx=XM_023243875.1;refGene_gn=MYO18A;refGene_tx=XM_023243876.1;refG
ene_gn=MYO18A;refGene_tx=XM_023243877.1
E1 17755879 17756026
refGene_gn=MYO18A;refGene_tx=XM_019817430.2;refGene_gn=MYO18A;refGene_
tx=XM_019817431.2;refGene_gn=MYO18A;refGene_tx=XM_019817438.1;refGene_
gn=MYO18A;refGene_tx=XM_019817439.1;refGene_gn=MYO18A;refGene_tx=XM_02
3243860.1;refGene_gn=MYO18A;refGene_tx=XM_023243862.1;refGene_gn=MYO18
A;refGene_tx=XM_023243863.1;refGene_gn=MYO18A;refGene_tx=XM_023243864.
1;refGene_gn=MYO18A;refGene_tx=XM_023243865.1;refGene_gn=MYO18A;refGen
e_tx=XM_023243866.1;refGene_gn=MYO18A;refGene_tx=XM_023243867.1;refGen
e_gn=MYO18A;refGene_tx=XM_023243868.1;refGene_gn=MYO18A;refGene_tx=XM_
023243869.1;refGene_gn=MYO18A;refGene_tx=XM_023243871.1;refGene_gn=MYO
18A;refGene_tx=XM_023243872.1;refGene_gn=MYO18A;refGene_tx=XM_02324387
3.1;refGene_gn=MYO18A;refGene_tx=XM_023243874.1;refGene_gn=MYO18A;refG
ene_tx=XM_023243875.1;refGene_gn=MYO18A;refGene_tx=XM_023243876.1;refG
ene_gn=MYO18A;refGene_tx=XM_023243877.1
E1 17756404 17756518
refGene_gn=MYO18A;refGene_tx=XM_019817430.2;refGene_gn=MYO18A;refGene_
tx=XM_019817431.2;refGene_gn=MYO18A;refGene_tx=XM_019817438.1;refGene_
gn=MYO18A;refGene_tx=XM_019817439.1;refGene_gn=MYO18A;refGene_tx=XM_02

3243860.1;refGene_gn=MYO18A;refGene_tx=XM_023243862.1;refGene_gn=MYO18A;refGene_tx=XM_023243863.1;refGene_gn=MYO18A;refGene_tx=XM_023243864.1;refGene_gn=MYO18A;refGene_tx=XM_023243865.1;refGene_gn=MYO18A;refGene_tx=XM_023243866.1;refGene_gn=MYO18A;refGene_tx=XM_023243867.1;refGene_gn=MYO18A;refGene_tx=XM_023243868.1;refGene_gn=MYO18A;refGene_tx=XM_023243869.1;refGene_gn=MYO18A;refGene_tx=XM_023243871.1;refGene_gn=MYO18A;refGene_tx=XM_023243872.1;refGene_gn=MYO18A;refGene_tx=XM_023243873.1;refGene_gn=MYO18A;refGene_tx=XM_023243874.1;refGene_gn=MYO18A;refGene_tx=XM_023243875.1;refGene_gn=MYO18A;refGene_tx=XM_023243876.1;refGene_gn=MYO18A;refGene_tx=XM_023243877.1

E1 17757870 17757987

refGene_gn=MYO18A;refGene_tx=XM_019817430.2;refGene_gn=MYO18A;refGene_tx=XM_019817431.2;refGene_gn=MYO18A;refGene_tx=XM_019817438.1;refGene_gn=MYO18A;refGene_tx=XM_019817439.1;refGene_gn=MYO18A;refGene_tx=XM_023243860.1;refGene_gn=MYO18A;refGene_tx=XM_023243862.1;refGene_gn=MYO18A;refGene_tx=XM_023243863.1;refGene_gn=MYO18A;refGene_tx=XM_023243864.1;refGene_gn=MYO18A;refGene_tx=XM_023243865.1;refGene_gn=MYO18A;refGene_tx=XM_023243866.1;refGene_gn=MYO18A;refGene_tx=XM_023243867.1;refGene_gn=MYO18A;refGene_tx=XM_023243868.1;refGene_gn=MYO18A;refGene_tx=XM_023243869.1;refGene_gn=MYO18A;refGene_tx=XM_023243871.1;refGene_gn=MYO18A;refGene_tx=XM_023243872.1;refGene_gn=MYO18A;refGene_tx=XM_023243873.1;refGene_gn=MYO18A;refGene_tx=XM_023243874.1;refGene_gn=MYO18A;refGene_tx=XM_023243875.1;refGene_gn=MYO18A;refGene_tx=XM_023243876.1;refGene_gn=MYO18A;refGene_tx=XM_023243877.1

E1 17758164 17758248

refGene_gn=MYO18A;refGene_tx=XM_019817430.2;refGene_gn=MYO18A;refGene_tx=XM_019817431.2;refGene_gn=MYO18A;refGene_tx=XM_019817438.1;refGene_gn=MYO18A;refGene_tx=XM_019817439.1;refGene_gn=MYO18A;refGene_tx=XM_023243860.1;refGene_gn=MYO18A;refGene_tx=XM_023243862.1;refGene_gn=MYO18A;refGene_tx=XM_023243863.1;refGene_gn=MYO18A;refGene_tx=XM_023243864.1;refGene_gn=MYO18A;refGene_tx=XM_023243865.1;refGene_gn=MYO18A;refGene_tx=XM_023243866.1;refGene_gn=MYO18A;refGene_tx=XM_023243867.1;refGene_gn=MYO18A;refGene_tx=XM_023243868.1;refGene_gn=MYO18A;refGene_tx=XM_023243869.1;refGene_gn=MYO18A;refGene_tx=XM_023243871.1;refGene_gn=MYO18A;refGene_tx=XM_023243872.1;refGene_gn=MYO18A;refGene_tx=XM_023243873.1;refGene_gn=MYO18A;refGene_tx=XM_023243874.1;refGene_gn=MYO18A;refGene_tx=XM_023243875.1;refGene_gn=MYO18A;refGene_tx=XM_023243876.1;refGene_gn=MYO18A;refGene_tx=XM_023243877.1

E1 17758665 17758803

refGene_gn=MYO18A;refGene_tx=XM_019817430.2;refGene_gn=MYO18A;refGene_tx=XM_019817431.2;refGene_gn=MYO18A;refGene_tx=XM_019817438.1;refGene_gn=MYO18A;refGene_tx=XM_019817439.1;refGene_gn=MYO18A;refGene_tx=XM_023243860.1;refGene_gn=MYO18A;refGene_tx=XM_023243862.1;refGene_gn=MYO18A;refGene_tx=XM_023243863.1;refGene_gn=MYO18A;refGene_tx=XM_023243864.1;refGene_gn=MYO18A;refGene_tx=XM_023243865.1;refGene_gn=MYO18A;refGene_tx=XM_023243866.1;refGene_gn=MYO18A;refGene_tx=XM_023243867.1;refGene_gn=MYO18A;refGene_tx=XM_023243868.1;refGene_gn=MYO18A;refGene_tx=XM_023243869.1;refGene_gn=MYO18A;refGene_tx=XM_023243871.1;refGene_gn=MYO18A;refGene_tx=XM_023243872.1;refGene_gn=MYO18A;refGene_tx=XM_023243873.1;refGene_gn=MYO18A;refGene_tx=XM_023243874.1;refGene_gn=MYO18A;refG

ene_tx=XM_023243875.1;refGene_gn=MYO18A;refGene_tx=XM_023243876.1;refGene_gn=MYO18A;refGene_tx=XM_023243877.1
E1 17761683 17761869
refGene_gn=MYO18A;refGene_tx=XM_019817430.2;refGene_gn=MYO18A;refGene_tx=XM_019817431.2;refGene_gn=MYO18A;refGene_tx=XM_019817438.1;refGene_gn=MYO18A;refGene_tx=XM_019817439.1;refGene_gn=MYO18A;refGene_tx=XM_023243860.1;refGene_gn=MYO18A;refGene_tx=XM_023243862.1;refGene_gn=MYO18A;refGene_tx=XM_023243863.1;refGene_gn=MYO18A;refGene_tx=XM_023243864.1;refGene_gn=MYO18A;refGene_tx=XM_023243865.1;refGene_gn=MYO18A;refGene_tx=XM_023243866.1;refGene_gn=MYO18A;refGene_tx=XM_023243867.1;refGene_gn=MYO18A;refGene_tx=XM_023243868.1;refGene_gn=MYO18A;refGene_tx=XM_023243869.1;refGene_gn=MYO18A;refGene_tx=XM_023243871.1;refGene_gn=MYO18A;refGene_tx=XM_023243872.1;refGene_gn=MYO18A;refGene_tx=XM_023243873.1;refGene_gn=MYO18A;refGene_tx=XM_023243874.1;refGene_gn=MYO18A;refGene_tx=XM_023243875.1;refGene_gn=MYO18A;refGene_tx=XM_023243876.1;refGene_gn=MYO18A;refGene_tx=XM_023243877.1
E1 17762167 17762307
refGene_gn=MYO18A;refGene_tx=XM_019817430.2;refGene_gn=MYO18A;refGene_tx=XM_019817431.2;refGene_gn=MYO18A;refGene_tx=XM_019817438.1;refGene_gn=MYO18A;refGene_tx=XM_019817439.1;refGene_gn=MYO18A;refGene_tx=XM_023243860.1;refGene_gn=MYO18A;refGene_tx=XM_023243862.1;refGene_gn=MYO18A;refGene_tx=XM_023243863.1;refGene_gn=MYO18A;refGene_tx=XM_023243864.1;refGene_gn=MYO18A;refGene_tx=XM_023243865.1;refGene_gn=MYO18A;refGene_tx=XM_023243866.1;refGene_gn=MYO18A;refGene_tx=XM_023243867.1;refGene_gn=MYO18A;refGene_tx=XM_023243868.1;refGene_gn=MYO18A;refGene_tx=XM_023243869.1;refGene_gn=MYO18A;refGene_tx=XM_023243871.1;refGene_gn=MYO18A;refGene_tx=XM_023243872.1;refGene_gn=MYO18A;refGene_tx=XM_023243873.1;refGene_gn=MYO18A;refGene_tx=XM_023243874.1;refGene_gn=MYO18A;refGene_tx=XM_023243875.1;refGene_gn=MYO18A;refGene_tx=XM_023243876.1;refGene_gn=MYO18A;refGene_tx=XM_023243877.1
E1 17763133 17763178
refGene_gn=MYO18A;refGene_tx=XM_019817430.2;refGene_gn=MYO18A;refGene_tx=XM_019817431.2;refGene_gn=MYO18A;refGene_tx=XM_019817438.1;refGene_gn=MYO18A;refGene_tx=XM_019817439.1;refGene_gn=MYO18A;refGene_tx=XM_023243860.1;refGene_gn=MYO18A;refGene_tx=XM_023243862.1;refGene_gn=MYO18A;refGene_tx=XM_023243863.1;refGene_gn=MYO18A;refGene_tx=XM_023243864.1;refGene_gn=MYO18A;refGene_tx=XM_023243866.1;refGene_gn=MYO18A;refGene_tx=XM_023243867.1;refGene_gn=MYO18A;refGene_tx=XM_023243868.1;refGene_gn=MYO18A;refGene_tx=XM_023243869.1;refGene_gn=MYO18A;refGene_tx=XM_023243871.1;refGene_gn=MYO18A;refGene_tx=XM_023243875.1
E1 17766444 17766567
refGene_gn=MYO18A;refGene_tx=XM_019817430.2;refGene_gn=MYO18A;refGene_tx=XM_019817431.2;refGene_gn=MYO18A;refGene_tx=XM_019817438.1;refGene_gn=MYO18A;refGene_tx=XM_019817439.1;refGene_gn=MYO18A;refGene_tx=XM_023243860.1;refGene_gn=MYO18A;refGene_tx=XM_023243862.1;refGene_gn=MYO18A;refGene_tx=XM_023243863.1;refGene_gn=MYO18A;refGene_tx=XM_023243864.1;refGene_gn=MYO18A;refGene_tx=XM_023243865.1;refGene_gn=MYO18A;refGene_tx=XM_023243866.1;refGene_gn=MYO18A;refGene_tx=XM_023243867.1;refGene_gn=MYO18A;refGene_tx=XM_023243868.1;refGene_gn=MYO18A;refGene_tx=XM_023243869.1;refGene_gn=MYO18A;refGene_tx=XM_023243871.1;refGene_gn=MYO

18A;refGene_tx=XM_023243872.1;refGene_gn=MYO18A;refGene_tx=XM_02324387
3.1;refGene_gn=MYO18A;refGene_tx=XM_023243874.1;refGene_gn=MYO18A;refG
ene_tx=XM_023243875.1;refGene_gn=MYO18A;refGene_tx=XM_023243876.1;refG
ene_gn=MYO18A;refGene_tx=XM_023243877.1
E1 17767951 17769206
refGene_gn=MYO18A;refGene_tx=XM_019817430.2;refGene_gn=MYO18A;refGene_
tx=XM_019817431.2;refGene_gn=MYO18A;refGene_tx=XM_019817438.1;refGene_
gn=MYO18A;refGene_tx=XM_019817439.1;refGene_gn=MYO18A;refGene_tx=XM_02
3243860.1;refGene_gn=MYO18A;refGene_tx=XM_023243862.1;refGene_gn=MYO18
A;refGene_tx=XM_023243863.1;refGene_gn=MYO18A;refGene_tx=XM_023243864.
1;refGene_gn=MYO18A;refGene_tx=XM_023243865.1;refGene_gn=MYO18A;refGen
e_tx=XM_023243866.1;refGene_gn=MYO18A;refGene_tx=XM_023243867.1;refGen
e_gn=MYO18A;refGene_tx=XM_023243868.1;refGene_gn=MYO18A;refGene_tx=XM_
023243869.1
E1 17773764 17773909
refGene_gn=MYO18A;refGene_tx=XM_023243871.1;refGene_gn=MYO18A;refGene_
tx=XM_023243872.1;refGene_gn=MYO18A;refGene_tx=XM_023243873.1;refGene_
gn=MYO18A;refGene_tx=XM_023243874.1;refGene_gn=MYO18A;refGene_tx=XM_02
3243875.1;refGene_gn=MYO18A;refGene_tx=XM_023243876.1;refGene_gn=MYO18
A;refGene_tx=XM_023243877.1
E1 17789733 17789864
refGene_gn=PIPOX;refGene_tx=XM_003996472.3;refGene_gn=PIPOX;refGene_tx
=XM_023243771.1
E1 17790246 17790322
refGene_gn=PIPOX;refGene_tx=XM_003996472.3;refGene_gn=PIPOX;refGene_tx
=XM_023243771.1
E1 17790912 17791071
refGene_gn=PIPOX;refGene_tx=XM_003996472.3;refGene_gn=PIPOX;refGene_tx
=XM_023243771.1
E1 17791460 17791613
refGene_gn=PIPOX;refGene_tx=XM_003996472.3;refGene_gn=PIPOX;refGene_tx
=XM_023243771.1
E1 17792630 17792813
refGene_gn=PIPOX;refGene_tx=XM_003996472.3;refGene_gn=PIPOX;refGene_tx
=XM_023243771.1
E1 17793090 17793304
refGene_gn=PIPOX;refGene_tx=XM_003996472.3;refGene_gn=PIPOX;refGene_tx
=XM_023243771.1
E1 17800603 17800752
refGene_gn=PIPOX;refGene_tx=XM_003996472.3;refGene_gn=PIPOX;refGene_tx
=XM_023243771.1
E1 17801963 17802077 refGene_gn=PIPOX;refGene_tx=XM_003996472.3
E1 17840013 17840068
refGene_gn=SEZ6;refGene_tx=XM_023244601.1;refGene_gn=SEZ6;refGene_tx=X
M_023244602.1;refGene_gn=SEZ6;refGene_tx=XM_023244603.1;refGene_gn=SEZ
6;refGene_tx=XM_023244604.1
E1 17861176 17861824
refGene_gn=SEZ6;refGene_tx=XM_023244601.1;refGene_gn=SEZ6;refGene_tx=X
M_023244602.1;refGene_gn=SEZ6;refGene_tx=XM_023244603.1;refGene_gn=SEZ
6;refGene_tx=XM_023244604.1

E1 17862359 17862493
refGene_gn=SEZ6;refGene_tx=XM_023244601.1;refGene_gn=SEZ6;refGene_tx=XM_023244602.1;refGene_gn=SEZ6;refGene_tx=XM_023244603.1;refGene_gn=SEZ6;refGene_tx=XM_023244604.1
E1 17872607 17872803
refGene_gn=SEZ6;refGene_tx=XM_023244601.1;refGene_gn=SEZ6;refGene_tx=XM_023244602.1;refGene_gn=SEZ6;refGene_tx=XM_023244603.1;refGene_gn=SEZ6;refGene_tx=XM_023244604.1
E1 17877066 17877252
refGene_gn=SEZ6;refGene_tx=XM_023244601.1;refGene_gn=SEZ6;refGene_tx=XM_023244602.1;refGene_gn=SEZ6;refGene_tx=XM_023244603.1;refGene_gn=SEZ6;refGene_tx=XM_023244604.1
E1 17880099 17880268
refGene_gn=SEZ6;refGene_tx=XM_023244601.1;refGene_gn=SEZ6;refGene_tx=XM_023244602.1;refGene_gn=SEZ6;refGene_tx=XM_023244603.1;refGene_gn=SEZ6;refGene_tx=XM_023244604.1
E1 17880410 17880577
refGene_gn=SEZ6;refGene_tx=XM_023244601.1;refGene_gn=SEZ6;refGene_tx=XM_023244602.1;refGene_gn=SEZ6;refGene_tx=XM_023244603.1;refGene_gn=SEZ6;refGene_tx=XM_023244604.1
E1 17881154 17881349
refGene_gn=SEZ6;refGene_tx=XM_023244601.1;refGene_gn=SEZ6;refGene_tx=XM_023244602.1;refGene_gn=SEZ6;refGene_tx=XM_023244603.1;refGene_gn=SEZ6;refGene_tx=XM_023244604.1
E1 17881596 17881735
refGene_gn=SEZ6;refGene_tx=XM_023244601.1;refGene_gn=SEZ6;refGene_tx=XM_023244602.1;refGene_gn=SEZ6;refGene_tx=XM_023244603.1;refGene_gn=SEZ6;refGene_tx=XM_023244604.1
E1 17881845 17882042
refGene_gn=SEZ6;refGene_tx=XM_023244601.1;refGene_gn=SEZ6;refGene_tx=XM_023244602.1;refGene_gn=SEZ6;refGene_tx=XM_023244603.1;refGene_gn=SEZ6;refGene_tx=XM_023244604.1
E1 17882634 17882829
refGene_gn=SEZ6;refGene_tx=XM_023244601.1;refGene_gn=SEZ6;refGene_tx=XM_023244602.1;refGene_gn=SEZ6;refGene_tx=XM_023244603.1;refGene_gn=SEZ6;refGene_tx=XM_023244604.1
E1 17883235 17883427
refGene_gn=SEZ6;refGene_tx=XM_023244601.1;refGene_gn=SEZ6;refGene_tx=XM_023244602.1;refGene_gn=SEZ6;refGene_tx=XM_023244603.1;refGene_gn=SEZ6;refGene_tx=XM_023244604.1
E1 17883533 17883731
refGene_gn=SEZ6;refGene_tx=XM_023244601.1;refGene_gn=SEZ6;refGene_tx=XM_023244602.1;refGene_gn=SEZ6;refGene_tx=XM_023244603.1;refGene_gn=SEZ6;refGene_tx=XM_023244604.1
E1 17883999 17884038
refGene_gn=SEZ6;refGene_tx=XM_023244601.1;refGene_gn=SEZ6;refGene_tx=XM_023244602.1
E1 17884289 17884407
refGene_gn=SEZ6;refGene_tx=XM_023244601.1;refGene_gn=SEZ6;refGene_tx=XM_023244602.1;refGene_gn=SEZ6;refGene_tx=XM_023244603.1;refGene_gn=SEZ

6;refGene_tx=XM_023244604.1
E1	17884491	17884594
refGene_gn=SEZ6;refGene_tx=XM_023244601.1;refGene_gn=SEZ6;refGene_tx=XM_023244602.1;refGene_gn=SEZ6;refGene_tx=XM_023244603.1;refGene_gn=SEZ6;refGene_tx=XM_023244604.1
E1	17884757	17884803
refGene_gn=SEZ6;refGene_tx=XM_023244601.1;refGene_gn=SEZ6;refGene_tx=XM_023244602.1;refGene_gn=SEZ6;refGene_tx=XM_023244603.1;refGene_gn=SEZ6;refGene_tx=XM_023244604.1
E1	17889556	17889622	refGene_gn=PHF12;refGene_tx=XM_003996473.5
E1	17890266	17890448	refGene_gn=PHF12;refGene_tx=XM_003996473.5
E1	17911474	17911547	refGene_gn=PHF12;refGene_tx=XM_003996473.5
E1	17914098	17914492	refGene_gn=PHF12;refGene_tx=XM_003996473.5
E1	17915785	17915906	refGene_gn=PHF12;refGene_tx=XM_003996473.5
E1	17917160	17917293	refGene_gn=PHF12;refGene_tx=XM_003996473.5
E1	17919175	17919340	refGene_gn=PHF12;refGene_tx=XM_003996473.5
E1	17922318	17922477	refGene_gn=PHF12;refGene_tx=XM_003996473.5
E1	17923056	17923852	refGene_gn=PHF12;refGene_tx=XM_003996473.5
E1	17925103	17925229	refGene_gn=PHF12;refGene_tx=XM_003996473.5
E1	17925883	17926027	refGene_gn=PHF12;refGene_tx=XM_003996473.5
E1	17927106	17927205	refGene_gn=PHF12;refGene_tx=XM_003996473.5
E1	17927952	17928035	refGene_gn=PHF12;refGene_tx=XM_003996473.5
E1	17928606	17928745	refGene_gn=PHF12;refGene_tx=XM_003996473.5
E1	17929091	17929426	refGene_gn=PHF12;refGene_tx=XM_003996473.5
E1	17932836	17932963
refGene_gn=DHRS13;refGene_tx=XM_011288996.3;refGene_gn=DHRS13;refGene_tx=XM_019817451.2
E1	17933060	17933179
refGene_gn=DHRS13;refGene_tx=XM_011288996.3;refGene_gn=DHRS13;refGene_tx=XM_019817451.2
E1	17934192	17934316
refGene_gn=DHRS13;refGene_tx=XM_011288996.3;refGene_gn=DHRS13;refGene_tx=XM_019817451.2
E1	17934485	17934809
refGene_gn=DHRS13;refGene_tx=XM_011288996.3;refGene_gn=DHRS13;refGene_tx=XM_019817451.2
E1	17936416	17936859
refGene_gn=DHRS13;refGene_tx=XM_011288996.3;refGene_gn=DHRS13;refGene_tx=XM_019817451.2
E1	17937705	17937754
refGene_gn=FLOT2;refGene_tx=XM_011288992.2;refGene_gn=FLOT2;refGene_tx=XM_011288993.2
E1	17944313	17944395
refGene_gn=FLOT2;refGene_tx=XM_011288992.2;refGene_gn=FLOT2;refGene_tx=XM_011288993.2
E1	17947339	17947430	refGene_gn=FLOT2;refGene_tx=XM_011288993.2
E1	17949949	17950040
refGene_gn=FLOT2;refGene_tx=XM_011288992.2;refGene_gn=FLOT2;refGene_tx=XM_011288994.2;refGene_gn=FLOT2;refGene_tx=XM_019817452.2
E1	17951037	17951161

refGene_gn=FLOT2;refGene_tx=XM_011288992.2;refGene_gn=FLOT2;refGene_tx
=XM_011288993.2;refGene_gn=FLOT2;refGene_tx=XM_011288994.2;refGene_gn=
FLOT2;refGene_tx=XM_011288995.2;refGene_gn=FLOT2;refGene_tx=XM_0198174
52.2
E1 17951573 17951692
refGene_gn=FLOT2;refGene_tx=XM_011288992.2;refGene_gn=FLOT2;refGene_tx
=XM_011288993.2;refGene_gn=FLOT2;refGene_tx=XM_011288994.2;refGene_gn=
FLOT2;refGene_tx=XM_011288995.2;refGene_gn=FLOT2;refGene_tx=XM_0198174
52.2
E1 17951809 17951923
refGene_gn=FLOT2;refGene_tx=XM_011288992.2;refGene_gn=FLOT2;refGene_tx
=XM_011288993.2;refGene_gn=FLOT2;refGene_tx=XM_011288994.2;refGene_gn=
FLOT2;refGene_tx=XM_011288995.2;refGene_gn=FLOT2;refGene_tx=XM_0198174
52.2
E1 17952024 17952144
refGene_gn=FLOT2;refGene_tx=XM_011288992.2;refGene_gn=FLOT2;refGene_tx
=XM_011288993.2;refGene_gn=FLOT2;refGene_tx=XM_011288994.2;refGene_gn=
FLOT2;refGene_tx=XM_011288995.2;refGene_gn=FLOT2;refGene_tx=XM_0198174
52.2
E1 17952233 17952448
refGene_gn=FLOT2;refGene_tx=XM_011288992.2;refGene_gn=FLOT2;refGene_tx
=XM_011288993.2;refGene_gn=FLOT2;refGene_tx=XM_011288994.2;refGene_gn=
FLOT2;refGene_tx=XM_011288995.2;refGene_gn=FLOT2;refGene_tx=XM_0198174
52.2
E1 17952857 17953041
refGene_gn=FLOT2;refGene_tx=XM_011288992.2;refGene_gn=FLOT2;refGene_tx
=XM_011288993.2;refGene_gn=FLOT2;refGene_tx=XM_011288994.2;refGene_gn=
FLOT2;refGene_tx=XM_011288995.2;refGene_gn=FLOT2;refGene_tx=XM_0198174
52.2
E1 17953341 17953491
refGene_gn=FLOT2;refGene_tx=XM_011288992.2;refGene_gn=FLOT2;refGene_tx
=XM_011288993.2;refGene_gn=FLOT2;refGene_tx=XM_011288994.2;refGene_gn=
FLOT2;refGene_tx=XM_011288995.2;refGene_gn=FLOT2;refGene_tx=XM_0198174
52.2
E1 17953568 17953607
refGene_gn=FLOT2;refGene_tx=XM_011288992.2;refGene_gn=FLOT2;refGene_tx
=XM_011288993.2;refGene_gn=FLOT2;refGene_tx=XM_011288994.2;refGene_gn=
FLOT2;refGene_tx=XM_011288995.2;refGene_gn=FLOT2;refGene_tx=XM_0198174
52.2
E1 17973304 17973427 refGene_gn=ERAL1;refGene_tx=XM_003996475.4
E1 17973541 17973622 refGene_gn=ERAL1;refGene_tx=XM_003996475.4
E1 17973745 17973895 refGene_gn=ERAL1;refGene_tx=XM_003996475.4
E1 17974039 17974288 refGene_gn=ERAL1;refGene_tx=XM_003996475.4
E1 17974379 17974492 refGene_gn=ERAL1;refGene_tx=XM_003996475.4
E1 17974661 17974723 refGene_gn=ERAL1;refGene_tx=XM_003996475.4
E1 17974876 17974923 refGene_gn=ERAL1;refGene_tx=XM_003996475.4
E1 17975319 17975397 refGene_gn=ERAL1;refGene_tx=XM_003996475.4
E1 17975502 17975630 refGene_gn=ERAL1;refGene_tx=XM_003996475.4
E1 17976930 17977213 refGene_gn=ERAL1;refGene_tx=XM_003996475.4
E1 18047626 18047708

refGene_gn=FAM222B;refGene_tx=XM_003996476.5;refGene_gn=FAM222B;refGene_tx=XM_003996477.5
E1 18058294 18059901
refGene_gn=FAM222B;refGene_tx=XM_003996476.5;refGene_gn=FAM222B;refGene_tx=XM_003996477.5;refGene_gn=FAM222B;refGene_tx=XM_011288991.3
E1 18064856 18065489 refGene_gn=TRAF4;refGene_tx=XM_003996478.4
E1 18065765 18065921 refGene_gn=TRAF4;refGene_tx=XM_003996478.4
E1 18066018 18066180 refGene_gn=TRAF4;refGene_tx=XM_003996478.4
E1 18066267 18066429 refGene_gn=TRAF4;refGene_tx=XM_003996478.4
E1 18066499 18066604 refGene_gn=TRAF4;refGene_tx=XM_003996478.4
E1 18067210 18067262 refGene_gn=TRAF4;refGene_tx=XM_003996478.4
E1 18070220 18070363 refGene_gn=TRAF4;refGene_tx=XM_003996478.4
E1 18071157 18071285
refGene_gn=NEK8;refGene_tx=XM_019817454.1;refGene_gn=NEK8;refGene_tx=XM_019817455.2;refGene_gn=NEK8;refGene_tx=XM_019817456.1;refGene_gn=NEK8;refGene_tx=XM_019817457.1;refGene_gn=NEK8;refGene_tx=XM_019817458.1
E1 18071780 18071809
refGene_gn=NEK8;refGene_tx=XM_011288989.2;refGene_gn=NEK8;refGene_tx=XM_019817459.1
E1 18072186 18072345
refGene_gn=NEK8;refGene_tx=XM_011288989.2;refGene_gn=NEK8;refGene_tx=XM_019817454.1;refGene_gn=NEK8;refGene_tx=XM_019817455.2;refGene_gn=NEK8;refGene_tx=XM_019817456.1;refGene_gn=NEK8;refGene_tx=XM_019817457.1;refGene_gn=NEK8;refGene_tx=XM_019817458.1;refGene_gn=NEK8;refGene_tx=XM_019817459.1
E1 18072510 18072669
refGene_gn=NEK8;refGene_tx=XM_011288989.2;refGene_gn=NEK8;refGene_tx=XM_019817454.1;refGene_gn=NEK8;refGene_tx=XM_019817455.2;refGene_gn=NEK8;refGene_tx=XM_019817456.1;refGene_gn=NEK8;refGene_tx=XM_019817457.1;refGene_gn=NEK8;refGene_tx=XM_019817458.1;refGene_gn=NEK8;refGene_tx=XM_019817459.1
E1 18072774 18072938
refGene_gn=NEK8;refGene_tx=XM_011288989.2;refGene_gn=NEK8;refGene_tx=XM_019817454.1;refGene_gn=NEK8;refGene_tx=XM_019817455.2;refGene_gn=NEK8;refGene_tx=XM_019817456.1;refGene_gn=NEK8;refGene_tx=XM_019817457.1;refGene_gn=NEK8;refGene_tx=XM_019817458.1;refGene_gn=NEK8;refGene_tx=XM_019817459.1
E1 18073172 18073323
refGene_gn=NEK8;refGene_tx=XM_011288989.2;refGene_gn=NEK8;refGene_tx=XM_019817454.1;refGene_gn=NEK8;refGene_tx=XM_019817455.2;refGene_gn=NEK8;refGene_tx=XM_019817456.1;refGene_gn=NEK8;refGene_tx=XM_019817457.1;refGene_gn=NEK8;refGene_tx=XM_019817458.1;refGene_gn=NEK8;refGene_tx=XM_019817459.1
E1 18074202 18074320
refGene_gn=NEK8;refGene_tx=XM_011288989.2;refGene_gn=NEK8;refGene_tx=XM_019817454.1;refGene_gn=NEK8;refGene_tx=XM_019817455.2;refGene_gn=NEK8;refGene_tx=XM_019817456.1;refGene_gn=NEK8;refGene_tx=XM_019817457.1;refGene_gn=NEK8;refGene_tx=XM_019817458.1;refGene_gn=NEK8;refGene_tx=XM_019817459.1
E1 18074654 18074827

refGene_gn=NEK8;refGene_tx=XM_011288989.2;refGene_gn=NEK8;refGene_tx=XM_019817454.1;refGene_gn=NEK8;refGene_tx=XM_019817455.2;refGene_gn=NEK8;refGene_tx=XM_019817456.1;refGene_gn=NEK8;refGene_tx=XM_019817457.1;refGene_gn=NEK8;refGene_tx=XM_019817458.1;refGene_gn=NEK8;refGene_tx=XM_019817459.1
E1 18075414 18075565
refGene_gn=NEK8;refGene_tx=XM_011288989.2;refGene_gn=NEK8;refGene_tx=XM_019817454.1;refGene_gn=NEK8;refGene_tx=XM_019817455.2;refGene_gn=NEK8;refGene_tx=XM_019817456.1;refGene_gn=NEK8;refGene_tx=XM_019817457.1;refGene_gn=NEK8;refGene_tx=XM_019817458.1;refGene_gn=NEK8;refGene_tx=XM_019817459.1
E1 18075669 18075851
refGene_gn=NEK8;refGene_tx=XM_011288989.2;refGene_gn=NEK8;refGene_tx=XM_019817454.1;refGene_gn=NEK8;refGene_tx=XM_019817455.2;refGene_gn=NEK8;refGene_tx=XM_019817456.1;refGene_gn=NEK8;refGene_tx=XM_019817457.1;refGene_gn=NEK8;refGene_tx=XM_019817458.1;refGene_gn=NEK8;refGene_tx=XM_019817459.1
E1 18075933 18076013
refGene_gn=NEK8;refGene_tx=XM_011288989.2;refGene_gn=NEK8;refGene_tx=XM_019817454.1;refGene_gn=NEK8;refGene_tx=XM_019817455.2;refGene_gn=NEK8;refGene_tx=XM_019817456.1;refGene_gn=NEK8;refGene_tx=XM_019817457.1;refGene_gn=NEK8;refGene_tx=XM_019817458.1;refGene_gn=NEK8;refGene_tx=XM_019817459.1
E1 18076176 18076385
refGene_gn=NEK8;refGene_tx=XM_011288989.2;refGene_gn=NEK8;refGene_tx=XM_019817454.1;refGene_gn=NEK8;refGene_tx=XM_019817455.2;refGene_gn=NEK8;refGene_tx=XM_019817456.1;refGene_gn=NEK8;refGene_tx=XM_019817457.1;refGene_gn=NEK8;refGene_tx=XM_019817458.1;refGene_gn=NEK8;refGene_tx=XM_019817459.1
E1 18076855 18076987
refGene_gn=NEK8;refGene_tx=XM_011288989.2;refGene_gn=NEK8;refGene_tx=XM_019817454.1;refGene_gn=NEK8;refGene_tx=XM_019817455.2;refGene_gn=NEK8;refGene_tx=XM_019817456.1;refGene_gn=NEK8;refGene_tx=XM_019817457.1;refGene_gn=NEK8;refGene_tx=XM_019817458.1;refGene_gn=NEK8;refGene_tx=XM_019817459.1
E1 18077217 18077450
refGene_gn=NEK8;refGene_tx=XM_011288989.2;refGene_gn=NEK8;refGene_tx=XM_019817454.1;refGene_gn=NEK8;refGene_tx=XM_019817455.2;refGene_gn=NEK8;refGene_tx=XM_019817456.1;refGene_gn=NEK8;refGene_tx=XM_019817457.1;refGene_gn=NEK8;refGene_tx=XM_019817458.1;refGene_gn=NEK8;refGene_tx=XM_019817459.1
E1 18077687 18077893
refGene_gn=NEK8;refGene_tx=XM_011288989.2;refGene_gn=NEK8;refGene_tx=XM_019817454.1;refGene_gn=NEK8;refGene_tx=XM_019817455.2;refGene_gn=NEK8;refGene_tx=XM_019817456.1;refGene_gn=NEK8;refGene_tx=XM_019817457.1;refGene_gn=NEK8;refGene_tx=XM_019817458.1;refGene_gn=NEK8;refGene_tx=XM_019817459.1
E1 18079156 18079158 refGene_gn=NEK8;refGene_tx=XM_019817455.2
E1 18079711 18079758
refGene_gn=NEK8;refGene_tx=XM_011288989.2;refGene_gn=NEK8;refGene_tx=X

M_019817454.1;refGene_gn=NEK8;refGene_tx=XM_019817456.1;refGene_gn=NEK8;refGene_tx=XM_019817457.1;refGene_gn=NEK8;refGene_tx=XM_019817458.1;refGene_gn=NEK8;refGene_tx=XM_019817459.1
E1 18082140 18082334 refGene_gn=TLCD1;refGene_tx=XM_011288988.3
E1 18082664 18082747 refGene_gn=TLCD1;refGene_tx=XM_011288988.3
E1 18082835 18082918 refGene_gn=TLCD1;refGene_tx=XM_011288988.3
E1 18083182 18083566 refGene_gn=TLCD1;refGene_tx=XM_011288988.3
E1 18084216 18084231 refGene_gn=RPL23A;refGene_tx=XM_003996479.3
E1 18084442 18084512 refGene_gn=RPL23A;refGene_tx=XM_003996479.3
E1 18084947 18085124 refGene_gn=RPL23A;refGene_tx=XM_003996479.3
E1 18086628 18086812 refGene_gn=RPL23A;refGene_tx=XM_003996479.3
E1 18087414 18087439 refGene_gn=RPL23A;refGene_tx=XM_003996479.3
E1 18088924 18089059 refGene_gn=RAB34;refGene_tx=XM_019817461.1
E1 18089858 18089948
refGene_gn=RAB34;refGene_tx=XM_006940024.4;refGene_gn=RAB34;refGene_tx=XM_006940025.4;refGene_gn=RAB34;refGene_tx=XM_019817460.2;refGene_gn=RAB34;refGene_tx=XM_019817461.1;refGene_gn=RAB34;refGene_tx=XM_023243646.1
E1 18090155 18090247
refGene_gn=RAB34;refGene_tx=XM_006940024.4;refGene_gn=RAB34;refGene_tx=XM_006940025.4;refGene_gn=RAB34;refGene_tx=XM_019817460.2;refGene_gn=RAB34;refGene_tx=XM_019817461.1;refGene_gn=RAB34;refGene_tx=XM_023243646.1
E1 18091127 18091193
refGene_gn=RAB34;refGene_tx=XM_006940024.4;refGene_gn=RAB34;refGene_tx=XM_006940025.4;refGene_gn=RAB34;refGene_tx=XM_019817460.2;refGene_gn=RAB34;refGene_tx=XM_019817461.1;refGene_gn=RAB34;refGene_tx=XM_023243646.1
E1 18091296 18091398
refGene_gn=RAB34;refGene_tx=XM_006940024.4;refGene_gn=RAB34;refGene_tx=XM_006940025.4;refGene_gn=RAB34;refGene_tx=XM_019817460.2;refGene_gn=RAB34;refGene_tx=XM_019817461.1;refGene_gn=RAB34;refGene_tx=XM_023243646.1
E1 18091504 18091569
refGene_gn=RAB34;refGene_tx=XM_006940024.4;refGene_gn=RAB34;refGene_tx=XM_006940025.4;refGene_gn=RAB34;refGene_tx=XM_019817460.2;refGene_gn=RAB34;refGene_tx=XM_019817461.1;refGene_gn=RAB34;refGene_tx=XM_023243646.1
E1 18091680 18091732
refGene_gn=RAB34;refGene_tx=XM_006940024.4;refGene_gn=RAB34;refGene_tx=XM_006940025.4;refGene_gn=RAB34;refGene_tx=XM_019817460.2;refGene_gn=RAB34;refGene_tx=XM_019817461.1;refGene_gn=RAB34;refGene_tx=XM_023243646.1
E1 18091919 18092001
refGene_gn=RAB34;refGene_tx=XM_006940024.4;refGene_gn=RAB34;refGene_tx=XM_006940025.4;refGene_gn=RAB34;refGene_tx=XM_019817460.2;refGene_gn=RAB34;refGene_tx=XM_019817461.1;refGene_gn=RAB34;refGene_tx=XM_023243646.1
E1 18092085 18092176
refGene_gn=RAB34;refGene_tx=XM_006940024.4;refGene_gn=RAB34;refGene_tx

=XM_006940025.4;refGene_gn=RAB34;refGene_tx=XM_019817461.1;refGene_gn=RAB34;refGene_tx=XM_023243646.1
E1 18092247 18092416
refGene_gn=RAB34;refGene_tx=XM_006940024.4;refGene_gn=RAB34;refGene_tx=XM_006940025.4;refGene_gn=RAB34;refGene_tx=XM_019817460.2;refGene_gn=RAB34;refGene_tx=XM_019817461.1;refGene_gn=RAB34;refGene_tx=XM_023243646.1
E1 18092498 18092566
refGene_gn=RAB34;refGene_tx=XM_006940024.4;refGene_gn=RAB34;refGene_tx=XM_006940025.4;refGene_gn=RAB34;refGene_tx=XM_019817460.2;refGene_gn=RAB34;refGene_tx=XM_019817461.1;refGene_gn=RAB34;refGene_tx=XM_023243646.1
E1 18095733 18095821
refGene_gn=PROCA1;refGene_tx=XM_006940026.4;refGene_gn=PROCA1;refGene_tx=XM_006940028.4;refGene_gn=PROCA1;refGene_tx=XM_019817463.2
E1 18096466 18096479
refGene_gn=PROCA1;refGene_tx=XM_006940027.4;refGene_gn=PROCA1;refGene_tx=XM_019817465.2
E1 18098657 18098796
refGene_gn=PROCA1;refGene_tx=XM_006940026.4;refGene_gn=PROCA1;refGene_tx=XM_006940027.4;refGene_gn=PROCA1;refGene_tx=XM_019817463.2;refGene_gn=PROCA1;refGene_tx=XM_019817464.2
E1 18101855 18101991
refGene_gn=PROCA1;refGene_tx=XM_006940026.4;refGene_gn=PROCA1;refGene_tx=XM_006940027.4;refGene_gn=PROCA1;refGene_tx=XM_006940028.4;refGene_gn=PROCA1;refGene_tx=XM_019817463.2;refGene_gn=PROCA1;refGene_tx=XM_019817464.2;refGene_gn=PROCA1;refGene_tx=XM_019817465.2
E1 18102268 18102427
refGene_gn=PROCA1;refGene_tx=XM_006940026.4;refGene_gn=PROCA1;refGene_tx=XM_006940027.4;refGene_gn=PROCA1;refGene_tx=XM_006940028.4;refGene_gn=PROCA1;refGene_tx=XM_019817463.2;refGene_gn=PROCA1;refGene_tx=XM_019817464.2;refGene_gn=PROCA1;refGene_tx=XM_019817465.2
E1 18102535 18103214
refGene_gn=PROCA1;refGene_tx=XM_006940026.4;refGene_gn=PROCA1;refGene_tx=XM_006940027.4;refGene_gn=PROCA1;refGene_tx=XM_006940028.4;refGene_gn=PROCA1;refGene_tx=XM_019817463.2;refGene_gn=PROCA1;refGene_tx=XM_019817464.2;refGene_gn=PROCA1;refGene_tx=XM_019817465.2
E1 18105329 18105516
refGene_gn=SUPT6H;refGene_tx=XM_003996485.5;refGene_gn=SUPT6H;refGene_tx=XM_006940029.4;refGene_gn=SUPT6H;refGene_tx=XM_006940030.4
E1 18105795 18105983
refGene_gn=SUPT6H;refGene_tx=XM_003996485.5;refGene_gn=SUPT6H;refGene_tx=XM_006940029.4;refGene_gn=SUPT6H;refGene_tx=XM_006940030.4
E1 18106439 18106606
refGene_gn=SUPT6H;refGene_tx=XM_003996485.5;refGene_gn=SUPT6H;refGene_tx=XM_006940029.4;refGene_gn=SUPT6H;refGene_tx=XM_006940030.4
E1 18106698 18106776
refGene_gn=SUPT6H;refGene_tx=XM_003996485.5;refGene_gn=SUPT6H;refGene_tx=XM_006940029.4;refGene_gn=SUPT6H;refGene_tx=XM_006940030.4
E1 18107054 18107167

refGene_gn=SUPT6H;refGene_tx=XM_003996485.5;refGene_gn=SUPT6H;refGene_tx=XM_006940029.4;refGene_gn=SUPT6H;refGene_tx=XM_006940030.4
E1 18108842 18108967
refGene_gn=SUPT6H;refGene_tx=XM_003996485.5;refGene_gn=SUPT6H;refGene_tx=XM_006940029.4;refGene_gn=SUPT6H;refGene_tx=XM_006940030.4
E1 18109138 18109252
refGene_gn=SUPT6H;refGene_tx=XM_003996485.5;refGene_gn=SUPT6H;refGene_tx=XM_006940029.4;refGene_gn=SUPT6H;refGene_tx=XM_006940030.4
E1 18109793 18110032
refGene_gn=SUPT6H;refGene_tx=XM_003996485.5;refGene_gn=SUPT6H;refGene_tx=XM_006940029.4;refGene_gn=SUPT6H;refGene_tx=XM_006940030.4
E1 18110656 18110852
refGene_gn=SUPT6H;refGene_tx=XM_003996485.5;refGene_gn=SUPT6H;refGene_tx=XM_006940029.4;refGene_gn=SUPT6H;refGene_tx=XM_006940030.4
E1 18111897 18112038
refGene_gn=SUPT6H;refGene_tx=XM_003996485.5;refGene_gn=SUPT6H;refGene_tx=XM_006940029.4;refGene_gn=SUPT6H;refGene_tx=XM_006940030.4
E1 18112793 18112936
refGene_gn=SUPT6H;refGene_tx=XM_003996485.5;refGene_gn=SUPT6H;refGene_tx=XM_006940029.4;refGene_gn=SUPT6H;refGene_tx=XM_006940030.4
E1 18113372 18113520
refGene_gn=SUPT6H;refGene_tx=XM_003996485.5;refGene_gn=SUPT6H;refGene_tx=XM_006940029.4;refGene_gn=SUPT6H;refGene_tx=XM_006940030.4
E1 18114111 18114319
refGene_gn=SUPT6H;refGene_tx=XM_003996485.5;refGene_gn=SUPT6H;refGene_tx=XM_006940029.4;refGene_gn=SUPT6H;refGene_tx=XM_006940030.4
E1 18115453 18115581
refGene_gn=SUPT6H;refGene_tx=XM_003996485.5;refGene_gn=SUPT6H;refGene_tx=XM_006940029.4;refGene_gn=SUPT6H;refGene_tx=XM_006940030.4
E1 18116760 18116928
refGene_gn=SUPT6H;refGene_tx=XM_003996485.5;refGene_gn=SUPT6H;refGene_tx=XM_006940029.4;refGene_gn=SUPT6H;refGene_tx=XM_006940030.4
E1 18117005 18117143
refGene_gn=SUPT6H;refGene_tx=XM_003996485.5;refGene_gn=SUPT6H;refGene_tx=XM_006940029.4;refGene_gn=SUPT6H;refGene_tx=XM_006940030.4
E1 18117444 18117580
refGene_gn=SUPT6H;refGene_tx=XM_003996485.5;refGene_gn=SUPT6H;refGene_tx=XM_006940029.4;refGene_gn=SUPT6H;refGene_tx=XM_006940030.4
E1 18117857 18117934
refGene_gn=SUPT6H;refGene_tx=XM_003996485.5;refGene_gn=SUPT6H;refGene_tx=XM_006940029.4;refGene_gn=SUPT6H;refGene_tx=XM_006940030.4
E1 18119060 18119178
refGene_gn=SUPT6H;refGene_tx=XM_003996485.5;refGene_gn=SUPT6H;refGene_tx=XM_006940029.4;refGene_gn=SUPT6H;refGene_tx=XM_006940030.4
E1 18119294 18119434
refGene_gn=SUPT6H;refGene_tx=XM_003996485.5;refGene_gn=SUPT6H;refGene_tx=XM_006940029.4;refGene_gn=SUPT6H;refGene_tx=XM_006940030.4
E1 18119791 18119987
refGene_gn=SUPT6H;refGene_tx=XM_003996485.5;refGene_gn=SUPT6H;refGene_tx=XM_006940029.4;refGene_gn=SUPT6H;refGene_tx=XM_006940030.4

E1 18120180 18120335
refGene_gn=SUPT6H;refGene_tx=XM_003996485.5;refGene_gn=SUPT6H;refGene_tx=XM_006940029.4;refGene_gn=SUPT6H;refGene_tx=XM_006940030.4
E1 18120511 18120662
refGene_gn=SUPT6H;refGene_tx=XM_003996485.5;refGene_gn=SUPT6H;refGene_tx=XM_006940029.4;refGene_gn=SUPT6H;refGene_tx=XM_006940030.4
E1 18120748 18120878
refGene_gn=SUPT6H;refGene_tx=XM_003996485.5;refGene_gn=SUPT6H;refGene_tx=XM_006940029.4;refGene_gn=SUPT6H;refGene_tx=XM_006940030.4
E1 18121407 18121506
refGene_gn=SUPT6H;refGene_tx=XM_003996485.5;refGene_gn=SUPT6H;refGene_tx=XM_006940029.4;refGene_gn=SUPT6H;refGene_tx=XM_006940030.4
E1 18121656 18121805
refGene_gn=SUPT6H;refGene_tx=XM_003996485.5;refGene_gn=SUPT6H;refGene_tx=XM_006940029.4;refGene_gn=SUPT6H;refGene_tx=XM_006940030.4
E1 18122448 18122591
refGene_gn=SUPT6H;refGene_tx=XM_003996485.5;refGene_gn=SUPT6H;refGene_tx=XM_006940029.4;refGene_gn=SUPT6H;refGene_tx=XM_006940030.4
E1 18122772 18122862
refGene_gn=SUPT6H;refGene_tx=XM_003996485.5;refGene_gn=SUPT6H;refGene_tx=XM_006940029.4;refGene_gn=SUPT6H;refGene_tx=XM_006940030.4
E1 18123204 18123321
refGene_gn=SUPT6H;refGene_tx=XM_003996485.5;refGene_gn=SUPT6H;refGene_tx=XM_006940029.4;refGene_gn=SUPT6H;refGene_tx=XM_006940030.4
E1 18123571 18123673
refGene_gn=SUPT6H;refGene_tx=XM_003996485.5;refGene_gn=SUPT6H;refGene_tx=XM_006940029.4;refGene_gn=SUPT6H;refGene_tx=XM_006940030.4
E1 18124056 18124330
refGene_gn=SUPT6H;refGene_tx=XM_003996485.5;refGene_gn=SUPT6H;refGene_tx=XM_006940029.4;refGene_gn=SUPT6H;refGene_tx=XM_006940030.4
E1 18125211 18125296
refGene_gn=SUPT6H;refGene_tx=XM_003996485.5;refGene_gn=SUPT6H;refGene_tx=XM_006940029.4;refGene_gn=SUPT6H;refGene_tx=XM_006940030.4
E1 18125529 18125722
refGene_gn=SUPT6H;refGene_tx=XM_003996485.5;refGene_gn=SUPT6H;refGene_tx=XM_006940029.4;refGene_gn=SUPT6H;refGene_tx=XM_006940030.4
E1 18126078 18126155
refGene_gn=SUPT6H;refGene_tx=XM_003996485.5;refGene_gn=SUPT6H;refGene_tx=XM_006940029.4;refGene_gn=SUPT6H;refGene_tx=XM_006940030.4
E1 18126251 18126410
refGene_gn=SUPT6H;refGene_tx=XM_003996485.5;refGene_gn=SUPT6H;refGene_tx=XM_006940029.4;refGene_gn=SUPT6H;refGene_tx=XM_006940030.4
E1 18127192 18127301
refGene_gn=SUPT6H;refGene_tx=XM_003996485.5;refGene_gn=SUPT6H;refGene_tx=XM_006940029.4;refGene_gn=SUPT6H;refGene_tx=XM_006940030.4
E1 18137331 18137482 refGene_gn=SDF2;refGene_tx=XM_003996486.4
E1 18137804 18137913 refGene_gn=SDF2;refGene_tx=XM_019817466.1
E1 18140843 18141040
refGene_gn=SDF2;refGene_tx=XM_003996486.4;refGene_gn=SDF2;refGene_tx=XM_019817466.1

E1 18148095 18148383
refGene_gn=SDF2;refGene_tx=XM_003996486.4;refGene_gn=SDF2;refGene_tx=XM_019817466.1
E1 18152120 18152185 refGene_gn=KIAA0100;refGene_tx=XM_011288983.3
E1 18153050 18153174 refGene_gn=KIAA0100;refGene_tx=XM_011288983.3
E1 18153683 18153733 refGene_gn=KIAA0100;refGene_tx=XM_011288983.3
E1 18154030 18154178 refGene_gn=KIAA0100;refGene_tx=XM_011288983.3
E1 18154355 18154474 refGene_gn=KIAA0100;refGene_tx=XM_011288983.3
E1 18154706 18154767 refGene_gn=KIAA0100;refGene_tx=XM_011288983.3
E1 18154933 18155123 refGene_gn=KIAA0100;refGene_tx=XM_011288983.3
E1 18155260 18155384 refGene_gn=KIAA0100;refGene_tx=XM_011288983.3
E1 18155796 18155921 refGene_gn=KIAA0100;refGene_tx=XM_011288983.3
E1 18156166 18156234 refGene_gn=KIAA0100;refGene_tx=XM_011288983.3
E1 18156347 18156493 refGene_gn=KIAA0100;refGene_tx=XM_011288983.3
E1 18157110 18157229 refGene_gn=KIAA0100;refGene_tx=XM_011288983.3
E1 18157340 18157504 refGene_gn=KIAA0100;refGene_tx=XM_011288983.3
E1 18157663 18157922 refGene_gn=KIAA0100;refGene_tx=XM_011288983.3
E1 18159058 18159264 refGene_gn=KIAA0100;refGene_tx=XM_011288983.3
E1 18160594 18161687 refGene_gn=KIAA0100;refGene_tx=XM_011288983.3
E1 18162121 18162313 refGene_gn=KIAA0100;refGene_tx=XM_011288983.3
E1 18162443 18162646 refGene_gn=KIAA0100;refGene_tx=XM_011288983.3
E1 18162787 18162906 refGene_gn=KIAA0100;refGene_tx=XM_011288983.3
E1 18163020 18163241 refGene_gn=KIAA0100;refGene_tx=XM_011288983.3
E1 18163533 18163724 refGene_gn=KIAA0100;refGene_tx=XM_011288983.3
E1 18163846 18163946 refGene_gn=KIAA0100;refGene_tx=XM_011288983.3
E1 18164071 18164274 refGene_gn=KIAA0100;refGene_tx=XM_011288983.3
E1 18165302 18165597 refGene_gn=KIAA0100;refGene_tx=XM_011288983.3
E1 18168006 18168164 refGene_gn=KIAA0100;refGene_tx=XM_011288983.3
E1 18168432 18168639 refGene_gn=KIAA0100;refGene_tx=XM_011288983.3
E1 18172703 18172803 refGene_gn=KIAA0100;refGene_tx=XM_011288983.3
E1 18173014 18173197 refGene_gn=KIAA0100;refGene_tx=XM_011288983.3
E1 18173533 18173655 refGene_gn=KIAA0100;refGene_tx=XM_011288983.3
E1 18174137 18174293 refGene_gn=KIAA0100;refGene_tx=XM_011288983.3
E1 18174378 18174500 refGene_gn=KIAA0100;refGene_tx=XM_011288983.3
E1 18175788 18175969 refGene_gn=KIAA0100;refGene_tx=XM_011288983.3
E1 18176607 18176694 refGene_gn=KIAA0100;refGene_tx=XM_011288983.3
E1 18176894 18177005 refGene_gn=KIAA0100;refGene_tx=XM_011288983.3
E1 18177142 18177278 refGene_gn=KIAA0100;refGene_tx=XM_011288983.3
E1 18177382 18177513 refGene_gn=KIAA0100;refGene_tx=XM_011288983.3
E1 18177693 18177795 refGene_gn=KIAA0100;refGene_tx=XM_011288983.3
E1 18178144 18178279 refGene_gn=KIAA0100;refGene_tx=XM_011288983.3
E1 18178701 18178884 refGene_gn=KIAA0100;refGene_tx=XM_011288983.3
E1 18179886 18179943
refGene_gn=LOC101085585;refGene_tx=XM_011288982.3;refGene_gn=LOC101085585;refGene_tx=XM_019817469.2;refGene_gn=LOC101085585;refGene_tx=XM_019817470.2
E1 18180338 18180587
refGene_gn=LOC101085585;refGene_tx=XM_011288982.3;refGene_gn=LOC101085585;refGene_tx=XM_019817469.2;refGene_gn=LOC101085585;refGene_tx=XM_019817470.2

E1 18180679 18180763
refGene_gn=LOC101085585;refGene_tx=XM_011288982.3;refGene_gn=LOC101085585;refGene_tx=XM_019817469.2;refGene_gn=LOC101085585;refGene_tx=XM_019817470.2
E1 18180876 18180945
refGene_gn=LOC101085585;refGene_tx=XM_011288982.3;refGene_gn=LOC101085585;refGene_tx=XM_019817469.2;refGene_gn=LOC101085585;refGene_tx=XM_019817470.2
E1 18181325 18181395
refGene_gn=LOC101085585;refGene_tx=XM_011288982.3;refGene_gn=LOC101085585;refGene_tx=XM_019817469.2;refGene_gn=LOC101085585;refGene_tx=XM_019817470.2
E1 18181621 18181726
refGene_gn=LOC101085585;refGene_tx=XM_011288982.3;refGene_gn=LOC101085585;refGene_tx=XM_019817469.2;refGene_gn=LOC101085585;refGene_tx=XM_019817470.2
E1 18181902 18181943
refGene_gn=LOC101085585;refGene_tx=XM_011288982.3;refGene_gn=LOC101085585;refGene_tx=XM_019817469.2;refGene_gn=LOC101085585;refGene_tx=XM_019817470.2
E1 18182187 18182215
refGene_gn=LOC101085585;refGene_tx=XM_011288982.3;refGene_gn=LOC101085585;refGene_tx=XM_019817469.2;refGene_gn=LOC101085585;refGene_tx=XM_019817470.2
E1 18182326 18182416
refGene_gn=LOC101085585;refGene_tx=XM_011288982.3;refGene_gn=LOC101085585;refGene_tx=XM_019817469.2;refGene_gn=LOC101085585;refGene_tx=XM_019817470.2
E1 18182499 18182588
refGene_gn=LOC101085585;refGene_tx=XM_011288982.3;refGene_gn=LOC101085585;refGene_tx=XM_019817469.2;refGene_gn=LOC101085585;refGene_tx=XM_019817470.2
E1 18182724 18182835
refGene_gn=LOC101085585;refGene_tx=XM_011288982.3;refGene_gn=LOC101085585;refGene_tx=XM_019817469.2;refGene_gn=LOC101085585;refGene_tx=XM_019817470.2
E1 18183215 18183437
refGene_gn=LOC101085585;refGene_tx=XM_011288982.3;refGene_gn=LOC101085585;refGene_tx=XM_019817469.2;refGene_gn=LOC101085585;refGene_tx=XM_019817470.2
E1 18199295 18199346
refGene_gn=SPAG5;refGene_tx=XM_003996487.4;refGene_gn=SPAG5;refGene_tx=XM_019817467.2;refGene_gn=SPAG5;refGene_tx=XM_019817468.2
E1 18199587 18199713
refGene_gn=SPAG5;refGene_tx=XM_003996487.4;refGene_gn=SPAG5;refGene_tx=XM_019817467.2;refGene_gn=SPAG5;refGene_tx=XM_019817468.2
E1 18203976 18205064
refGene_gn=SPAG5;refGene_tx=XM_003996487.4;refGene_gn=SPAG5;refGene_tx=XM_019817467.2;refGene_gn=SPAG5;refGene_tx=XM_019817468.2
E1 18205166 18205344

refGene_gn=SPAG5;refGene_tx=XM_003996487.4;refGene_gn=SPAG5;refGene_tx=XM_019817467.2;refGene_gn=SPAG5;refGene_tx=XM_019817468.2
E1 18210326 18210401
refGene_gn=SPAG5;refGene_tx=XM_003996487.4;refGene_gn=SPAG5;refGene_tx=XM_019817467.2;refGene_gn=SPAG5;refGene_tx=XM_019817468.2
E1 18210640 18210733
refGene_gn=SPAG5;refGene_tx=XM_003996487.4;refGene_gn=SPAG5;refGene_tx=XM_019817467.2;refGene_gn=SPAG5;refGene_tx=XM_019817468.2
E1 18210824 18210959
refGene_gn=SPAG5;refGene_tx=XM_003996487.4;refGene_gn=SPAG5;refGene_tx=XM_019817467.2;refGene_gn=SPAG5;refGene_tx=XM_019817468.2
E1 18211177 18211297
refGene_gn=SPAG5;refGene_tx=XM_003996487.4;refGene_gn=SPAG5;refGene_tx=XM_019817467.2;refGene_gn=SPAG5;refGene_tx=XM_019817468.2
E1 18211417 18211510
refGene_gn=SPAG5;refGene_tx=XM_003996487.4;refGene_gn=SPAG5;refGene_tx=XM_019817467.2;refGene_gn=SPAG5;refGene_tx=XM_019817468.2
E1 18211612 18211726
refGene_gn=SPAG5;refGene_tx=XM_003996487.4;refGene_gn=SPAG5;refGene_tx=XM_019817467.2;refGene_gn=SPAG5;refGene_tx=XM_019817468.2
E1 18212099 18212193
refGene_gn=SPAG5;refGene_tx=XM_003996487.4;refGene_gn=SPAG5;refGene_tx=XM_019817467.2;refGene_gn=SPAG5;refGene_tx=XM_019817468.2
E1 18212408 18212556
refGene_gn=SPAG5;refGene_tx=XM_003996487.4;refGene_gn=SPAG5;refGene_tx=XM_019817467.2;refGene_gn=SPAG5;refGene_tx=XM_019817468.2
E1 18212635 18212738
refGene_gn=SPAG5;refGene_tx=XM_003996487.4;refGene_gn=SPAG5;refGene_tx=XM_019817467.2;refGene_gn=SPAG5;refGene_tx=XM_019817468.2
E1 18212931 18213065
refGene_gn=SPAG5;refGene_tx=XM_003996487.4;refGene_gn=SPAG5;refGene_tx=XM_019817467.2;refGene_gn=SPAG5;refGene_tx=XM_019817468.2
E1 18213264 18213403
refGene_gn=SPAG5;refGene_tx=XM_003996487.4;refGene_gn=SPAG5;refGene_tx=XM_019817467.2;refGene_gn=SPAG5;refGene_tx=XM_019817468.2
E1 18216887 18216999
refGene_gn=SPAG5;refGene_tx=XM_003996487.4;refGene_gn=SPAG5;refGene_tx=XM_019817467.2;refGene_gn=SPAG5;refGene_tx=XM_019817468.2
E1 18217166 18217253
refGene_gn=SPAG5;refGene_tx=XM_003996487.4;refGene_gn=SPAG5;refGene_tx=XM_019817467.2;refGene_gn=SPAG5;refGene_tx=XM_019817468.2
E1 18217516 18217637
refGene_gn=SPAG5;refGene_tx=XM_003996487.4;refGene_gn=SPAG5;refGene_tx=XM_019817467.2;refGene_gn=SPAG5;refGene_tx=XM_019817468.2
E1 18217746 18217858
refGene_gn=SPAG5;refGene_tx=XM_003996487.4;refGene_gn=SPAG5;refGene_tx=XM_019817467.2;refGene_gn=SPAG5;refGene_tx=XM_019817468.2
E1 18218234 18218315
refGene_gn=SPAG5;refGene_tx=XM_003996487.4;refGene_gn=SPAG5;refGene_tx=XM_019817467.2;refGene_gn=SPAG5;refGene_tx=XM_019817468.2

E1 18218479 18218635
refGene_gn=SPAG5;refGene_tx=XM_003996487.4;refGene_gn=SPAG5;refGene_tx=XM_019817467.2;refGene_gn=SPAG5;refGene_tx=XM_019817468.2
E1 18218722 18218797
refGene_gn=SPAG5;refGene_tx=XM_003996487.4;refGene_gn=SPAG5;refGene_tx=XM_019817467.2;refGene_gn=SPAG5;refGene_tx=XM_019817468.2
E1 18218928 18219009
refGene_gn=SPAG5;refGene_tx=XM_003996487.4;refGene_gn=SPAG5;refGene_tx=XM_019817467.2;refGene_gn=SPAG5;refGene_tx=XM_019817468.2
E1 18219242 18219314
refGene_gn=SPAG5;refGene_tx=XM_003996487.4;refGene_gn=SPAG5;refGene_tx=XM_019817467.2;refGene_gn=SPAG5;refGene_tx=XM_019817468.2
E1 18221271 18221383 refGene_gn=ALDOC;refGene_tx=XM_003996488.5
E1 18221471 18221683 refGene_gn=ALDOC;refGene_tx=XM_003996488.5
E1 18221794 18221849 refGene_gn=ALDOC;refGene_tx=XM_003996488.5
E1 18221943 18222104 refGene_gn=ALDOC;refGene_tx=XM_003996488.5
E1 18222224 18222308 refGene_gn=ALDOC;refGene_tx=XM_003996488.5
E1 18222537 18222712 refGene_gn=ALDOC;refGene_tx=XM_003996488.5
E1 18222838 18223038 refGene_gn=ALDOC;refGene_tx=XM_003996488.5
E1 18223187 18223283 refGene_gn=ALDOC;refGene_tx=XM_003996488.5
E1 18225497 18225531 refGene_gn=PIGS;refGene_tx=XM_003996489.4
E1 18225808 18225948 refGene_gn=PIGS;refGene_tx=XM_003996489.4
E1 18226033 18226145 refGene_gn=PIGS;refGene_tx=XM_003996489.4
E1 18232221 18232311 refGene_gn=PIGS;refGene_tx=XM_003996489.4
E1 18232579 18232671 refGene_gn=PIGS;refGene_tx=XM_003996489.4
E1 18234407 18234615 refGene_gn=PIGS;refGene_tx=XM_003996489.4
E1 18234895 18235038 refGene_gn=PIGS;refGene_tx=XM_003996489.4
E1 18235743 18235858 refGene_gn=PIGS;refGene_tx=XM_003996489.4
E1 18236950 18237096 refGene_gn=PIGS;refGene_tx=XM_003996489.4
E1 18237670 18237771 refGene_gn=PIGS;refGene_tx=XM_003996489.4
E1 18238789 18239000 refGene_gn=PIGS;refGene_tx=XM_003996489.4
E1 18239359 18239635 refGene_gn=PIGS;refGene_tx=XM_003996489.4
E1 18241368 18241594 refGene_gn=UNC119;refGene_tx=XM_023243435.1
E1 18245116 18245230 refGene_gn=UNC119;refGene_tx=XM_023243435.1
E1 18245695 18245798 refGene_gn=UNC119;refGene_tx=XM_023243435.1
E1 18245991 18246164 refGene_gn=UNC119;refGene_tx=XM_023243435.1
E1 18246452 18246565 refGene_gn=UNC119;refGene_tx=XM_023243435.1
E1 18253164 18253484
refGene_gn=FOXN1;refGene_tx=XM_003996492.5;refGene_gn=FOXN1;refGene_tx=XM_019817472.2;refGene_gn=FOXN1;refGene_tx=XM_019817473.2;refGene_gn=FOXN1;refGene_tx=XM_019817474.2;refGene_gn=FOXN1;refGene_tx=XM_019817475.2;refGene_gn=FOXN1;refGene_tx=XM_019817476.2;refGene_gn=FOXN1;refGene_tx=XM_019817477.2
E1 18255087 18255579
refGene_gn=FOXN1;refGene_tx=XM_003996492.5;refGene_gn=FOXN1;refGene_tx=XM_019817472.2;refGene_gn=FOXN1;refGene_tx=XM_019817473.2;refGene_gn=FOXN1;refGene_tx=XM_019817474.2;refGene_gn=FOXN1;refGene_tx=XM_019817475.2;refGene_gn=FOXN1;refGene_tx=XM_019817476.2;refGene_gn=FOXN1;refGene_tx=XM_019817477.2
E1 18255780 18255988

refGene_gn=FOXN1;refGene_tx=XM_003996492.5;refGene_gn=FOXN1;refGene_tx=XM_019817472.2;refGene_gn=FOXN1;refGene_tx=XM_019817473.2;refGene_gn=FOXN1;refGene_tx=XM_019817474.2;refGene_gn=FOXN1;refGene_tx=XM_019817475.2;refGene_gn=FOXN1;refGene_tx=XM_019817476.2;refGene_gn=FOXN1;refGene_tx=XM_019817477.2
E1 18259418 18259515
refGene_gn=FOXN1;refGene_tx=XM_003996492.5;refGene_gn=FOXN1;refGene_tx=XM_019817472.2;refGene_gn=FOXN1;refGene_tx=XM_019817473.2;refGene_gn=FOXN1;refGene_tx=XM_019817474.2;refGene_gn=FOXN1;refGene_tx=XM_019817475.2;refGene_gn=FOXN1;refGene_tx=XM_019817476.2;refGene_gn=FOXN1;refGene_tx=XM_019817477.2
E1 18260265 18260396
refGene_gn=FOXN1;refGene_tx=XM_003996492.5;refGene_gn=FOXN1;refGene_tx=XM_019817472.2;refGene_gn=FOXN1;refGene_tx=XM_019817473.2;refGene_gn=FOXN1;refGene_tx=XM_019817474.2;refGene_gn=FOXN1;refGene_tx=XM_019817475.2;refGene_gn=FOXN1;refGene_tx=XM_019817476.2;refGene_gn=FOXN1;refGene_tx=XM_019817477.2
E1 18262173 18262287
refGene_gn=FOXN1;refGene_tx=XM_003996492.5;refGene_gn=FOXN1;refGene_tx=XM_019817472.2;refGene_gn=FOXN1;refGene_tx=XM_019817473.2;refGene_gn=FOXN1;refGene_tx=XM_019817474.2;refGene_gn=FOXN1;refGene_tx=XM_019817475.2;refGene_gn=FOXN1;refGene_tx=XM_019817476.2;refGene_gn=FOXN1;refGene_tx=XM_019817477.2
E1 18264594 18265053
refGene_gn=FOXN1;refGene_tx=XM_003996492.5;refGene_gn=FOXN1;refGene_tx=XM_019817472.2;refGene_gn=FOXN1;refGene_tx=XM_019817473.2;refGene_gn=FOXN1;refGene_tx=XM_019817474.2;refGene_gn=FOXN1;refGene_tx=XM_019817475.2;refGene_gn=FOXN1;refGene_tx=XM_019817476.2;refGene_gn=FOXN1;refGene_tx=XM_019817477.2
E1 18265408 18265540
refGene_gn=FOXN1;refGene_tx=XM_003996492.5;refGene_gn=FOXN1;refGene_tx=XM_019817472.2;refGene_gn=FOXN1;refGene_tx=XM_019817473.2;refGene_gn=FOXN1;refGene_tx=XM_019817474.2;refGene_gn=FOXN1;refGene_tx=XM_019817475.2;refGene_gn=FOXN1;refGene_tx=XM_019817476.2;refGene_gn=FOXN1;refGene_tx=XM_019817477.2
E1 18273195 18273240
refGene_gn=FOXN1;refGene_tx=XM_003996492.5;refGene_gn=FOXN1;refGene_tx=XM_019817472.2;refGene_gn=FOXN1;refGene_tx=XM_019817477.2
E1 18291111 18291282
refGene_gn=SLC13A2;refGene_tx=XM_003996493.4;refGene_gn=SLC13A2;refGene_tx=XM_019817479.1;refGene_gn=SLC13A2;refGene_tx=XM_019817480.2
E1 18291782 18291920
refGene_gn=SLC13A2;refGene_tx=XM_003996493.4;refGene_gn=SLC13A2;refGene_tx=XM_019817479.1;refGene_gn=SLC13A2;refGene_tx=XM_019817480.2
E1 18292565 18292727
refGene_gn=SLC13A2;refGene_tx=XM_003996493.4;refGene_gn=SLC13A2;refGene_tx=XM_019817479.1;refGene_gn=SLC13A2;refGene_tx=XM_019817480.2
E1 18293809 18293931
refGene_gn=SLC13A2;refGene_tx=XM_003996493.4;refGene_gn=SLC13A2;refGene_tx=XM_019817479.1;refGene_gn=SLC13A2;refGene_tx=XM_019817480.2

E1 18294214 18294303
refGene_gn=SLC13A2;refGene_tx=XM_003996493.4;refGene_gn=SLC13A2;refGene_tx=XM_019817479.1;refGene_gn=SLC13A2;refGene_tx=XM_019817480.2
E1 18294498 18294717
refGene_gn=SLC13A2;refGene_tx=XM_003996493.4;refGene_gn=SLC13A2;refGene_tx=XM_019817479.1;refGene_gn=SLC13A2;refGene_tx=XM_019817480.2
E1 18296959 18297082
refGene_gn=SLC13A2;refGene_tx=XM_003996493.4;refGene_gn=SLC13A2;refGene_tx=XM_019817479.1;refGene_gn=SLC13A2;refGene_tx=XM_019817480.2
E1 18297176 18297357
refGene_gn=SLC13A2;refGene_tx=XM_003996493.4;refGene_gn=SLC13A2;refGene_tx=XM_019817479.1;refGene_gn=SLC13A2;refGene_tx=XM_019817480.2
E1 18298627 18298833
refGene_gn=SLC13A2;refGene_tx=XM_003996493.4;refGene_gn=SLC13A2;refGene_tx=XM_019817479.1;refGene_gn=SLC13A2;refGene_tx=XM_019817480.2
E1 18298945 18299082
refGene_gn=SLC13A2;refGene_tx=XM_003996493.4;refGene_gn=SLC13A2;refGene_tx=XM_019817479.1;refGene_gn=SLC13A2;refGene_tx=XM_019817480.2
E1 18299892 18300021
refGene_gn=SLC13A2;refGene_tx=XM_003996493.4;refGene_gn=SLC13A2;refGene_tx=XM_019817479.1;refGene_gn=SLC13A2;refGene_tx=XM_019817480.2
E1 18305856 18306090 refGene_gn=SLC13A2;refGene_tx=XM_019817479.1
E1 18316675 18316777 refGene_gn=SLC13A2;refGene_tx=XM_003996493.4
E1 18358503 18358731 refGene_gn=SLC46A1;refGene_tx=XM_003996494.5
E1 18359147 18360000 refGene_gn=SLC46A1;refGene_tx=XM_003996494.5
E1 18362229 18362313 refGene_gn=SLC46A1;refGene_tx=XM_003996494.5
E1 18363761 18363918 refGene_gn=SLC46A1;refGene_tx=XM_003996494.5
E1 18364461 18364519 refGene_gn=SLC46A1;refGene_tx=XM_003996494.5
E1 18368367 18368497
refGene_gn=SARM1;refGene_tx=XM_003996496.5;refGene_gn=SARM1;refGene_tx=XM_019817478.2
E1 18368580 18368702
refGene_gn=SARM1;refGene_tx=XM_003996496.5;refGene_gn=SARM1;refGene_tx=XM_019817478.2
E1 18373651 18373841
refGene_gn=SARM1;refGene_tx=XM_003996496.5;refGene_gn=SARM1;refGene_tx=XM_019817478.2
E1 18373917 18374020
refGene_gn=SARM1;refGene_tx=XM_003996496.5;refGene_gn=SARM1;refGene_tx=XM_019817478.2
E1 18375981 18376217
refGene_gn=SARM1;refGene_tx=XM_003996496.5;refGene_gn=SARM1;refGene_tx=XM_019817478.2
E1 18376309 18376401
refGene_gn=SARM1;refGene_tx=XM_003996496.5;refGene_gn=SARM1;refGene_tx=XM_019817478.2
E1 18376665 18376878
refGene_gn=SARM1;refGene_tx=XM_003996496.5;refGene_gn=SARM1;refGene_tx=XM_019817478.2
E1 18378862 18379481

refGene_gn=SARM1;refGene_tx=XM_003996496.5;refGene_gn=SARM1;refGene_tx=XM_019817478.2
E1 18384872 18385342 refGene_gn=SARM1;refGene_tx=XM_003996496.5
E1 18387109 18387173 refGene_gn=VTN;refGene_tx=XM_003996497.5
E1 18387276 18387396 refGene_gn=VTN;refGene_tx=XM_003996497.5
E1 18387472 18387790 refGene_gn=VTN;refGene_tx=XM_003996497.5
E1 18387870 18388010 refGene_gn=VTN;refGene_tx=XM_003996497.5
E1 18388267 18388424 refGene_gn=VTN;refGene_tx=XM_003996497.5
E1 18388618 18388771 refGene_gn=VTN;refGene_tx=XM_003996497.5
E1 18389198 18389540 refGene_gn=VTN;refGene_tx=XM_003996497.5
E1 18389780 18389893 refGene_gn=VTN;refGene_tx=XM_003996497.5
E1 18391158 18391334 refGene_gn=SARM1;refGene_tx=XM_019817478.2
E1 18391348 18391384
refGene_gn=SEBOX;refGene_tx=XM_019817485.2;refGene_gn=TMEM199;refGene_tx=XM_019817484.1
E1 18391548 18391709 refGene_gn=SEBOX;refGene_tx=XM_019817485.2
E1 18391848 18392229 refGene_gn=SEBOX;refGene_tx=XM_019817485.2
E1 18395818 18395914 refGene_gn=TMEM199;refGene_tx=XM_003996498.4
E1 18395936 18395981 refGene_gn=TMEM199;refGene_tx=XM_019817483.1
E1 18396187 18396300
refGene_gn=TMEM199;refGene_tx=XM_003996498.4;refGene_gn=TMEM199;refGene_tx=XM_019817483.1;refGene_gn=TMEM199;refGene_tx=XM_019817484.1
E1 18396525 18396568
refGene_gn=TMEM199;refGene_tx=XM_003996498.4;refGene_gn=TMEM199;refGene_tx=XM_019817483.1;refGene_gn=TMEM199;refGene_tx=XM_019817484.1
E1 18397083 18397173
refGene_gn=TMEM199;refGene_tx=XM_003996498.4;refGene_gn=TMEM199;refGene_tx=XM_019817483.1;refGene_gn=TMEM199;refGene_tx=XM_019817484.1
E1 18397527 18397601
refGene_gn=TMEM199;refGene_tx=XM_003996498.4;refGene_gn=TMEM199;refGene_tx=XM_019817483.1;refGene_gn=TMEM199;refGene_tx=XM_019817484.1
E1 18398621 18398832
refGene_gn=TMEM199;refGene_tx=XM_003996498.4;refGene_gn=TMEM199;refGene_tx=XM_019817483.1;refGene_gn=TMEM199;refGene_tx=XM_019817484.1
E1 18399043 18399204
refGene_gn=POLDIP2;refGene_tx=XM_011288980.2;refGene_gn=POLDIP2;refGene_tx=XM_019817482.1
E1 18400546 18400628
refGene_gn=POLDIP2;refGene_tx=XM_011288980.2;refGene_gn=POLDIP2;refGene_tx=XM_019817482.1
E1 18401491 18401589
refGene_gn=POLDIP2;refGene_tx=XM_011288980.2;refGene_gn=POLDIP2;refGene_tx=XM_019817482.1
E1 18402279 18402376
refGene_gn=POLDIP2;refGene_tx=XM_011288980.2;refGene_gn=POLDIP2;refGene_tx=XM_019817482.1
E1 18402546 18402622
refGene_gn=POLDIP2;refGene_tx=XM_011288980.2;refGene_gn=POLDIP2;refGene_tx=XM_019817482.1
E1 18402857 18402965

refGene_gn=POLDIP2;refGene_tx=XM_011288980.2;refGene_gn=POLDIP2;refGene_tx=XM_019817482.1
E1 18403428 18403565
refGene_gn=POLDIP2;refGene_tx=XM_011288980.2;refGene_gn=POLDIP2;refGene_tx=XM_019817482.1
E1 18404335 18404362 refGene_gn=POLDIP2;refGene_tx=XM_011288980.2
E1 18404567 18404693
refGene_gn=POLDIP2;refGene_tx=XM_011288980.2;refGene_gn=POLDIP2;refGene_tx=XM_019817482.1
E1 18405729 18405809
refGene_gn=POLDIP2;refGene_tx=XM_011288980.2;refGene_gn=POLDIP2;refGene_tx=XM_019817482.1
E1 18406623 18406738
refGene_gn=POLDIP2;refGene_tx=XM_011288980.2;refGene_gn=POLDIP2;refGene_tx=XM_019817482.1
E1 18410261 18410498 refGene_gn=TNFAIP1;refGene_tx=XM_003996499.5
E1 18412137 18412333 refGene_gn=TNFAIP1;refGene_tx=XM_003996499.5
E1 18413307 18413360 refGene_gn=TNFAIP1;refGene_tx=XM_003996499.5
E1 18413436 18413526 refGene_gn=TNFAIP1;refGene_tx=XM_003996499.5
E1 18413944 18414114 refGene_gn=TNFAIP1;refGene_tx=XM_003996499.5
E1 18414725 18414930 refGene_gn=TNFAIP1;refGene_tx=XM_003996499.5
E1 18422596 18422723
refGene_gn=IFT20;refGene_tx=XM_003996500.4;refGene_gn=IFT20;refGene_tx=XM_003996501.4
E1 18423884 18423970
refGene_gn=IFT20;refGene_tx=XM_003996500.4;refGene_gn=IFT20;refGene_tx=XM_003996501.4
E1 18424497 18424601
refGene_gn=IFT20;refGene_tx=XM_003996500.4;refGene_gn=IFT20;refGene_tx=XM_003996501.4
E1 18425067 18425149
refGene_gn=IFT20;refGene_tx=XM_003996500.4;refGene_gn=IFT20;refGene_tx=XM_003996501.4
E1 18426625 18426885 refGene_gn=TMEM97;refGene_tx=XM_003996502.3
E1 18427933 18428078 refGene_gn=TMEM97;refGene_tx=XM_003996502.3
E1 18433964 18434090 refGene_gn=TMEM97;refGene_tx=XM_003996502.3
E1 18454603 18454658 refGene_gn=NLK;refGene_tx=XM_003996503.5
E1 18456720 18456814 refGene_gn=NLK;refGene_tx=XM_003996503.5
E1 18457732 18457931 refGene_gn=NLK;refGene_tx=XM_003996503.5
E1 18465313 18465400 refGene_gn=NLK;refGene_tx=XM_003996503.5
E1 18474037 18474139 refGene_gn=NLK;refGene_tx=XM_003996503.5
E1 18477413 18477623 refGene_gn=NLK;refGene_tx=XM_003996503.5
E1 18481748 18481834 refGene_gn=NLK;refGene_tx=XM_003996503.5
E1 18483306 18483413 refGene_gn=NLK;refGene_tx=XM_003996503.5
E1 18514137 18514193 refGene_gn=NLK;refGene_tx=XM_003996503.5
E1 18523841 18523971 refGene_gn=NLK;refGene_tx=XM_003996503.5
E1 18616449 18616907 refGene_gn=NLK;refGene_tx=XM_003996503.5
E1 18617003 18617035
refGene_gn=LOC111557845;refGene_tx=XM_023244515.1
E1 18617755 18618207

refGene_gn=LOC111557845;refGene_tx=XM_023244515.1
E1 18618386 18618529
refGene_gn=LOC111557845;refGene_tx=XM_023244515.1
E1 18725915 18725927 refGene_gn=LYRM9;refGene_tx=XM_003996520.5
E1 18745269 18745413
refGene_gn=LYRM9;refGene_tx=XM_003996520.5;refGene_gn=LYRM9;refGene_tx=XM_023243794.1
E1 18747086 18747179
refGene_gn=LYRM9;refGene_tx=XM_003996520.5;refGene_gn=LYRM9;refGene_tx=XM_023243794.1
E1 18748427 18748445
refGene_gn=LYRM9;refGene_tx=XM_003996520.5;refGene_gn=LYRM9;refGene_tx=XM_023243794.1
E1 18824523 18824633 refGene_gn=NOS2;refGene_tx=XM_023243518.1
E1 18829274 18829381 refGene_gn=NOS2;refGene_tx=XM_019817487.2
E1 18832439 18832524
refGene_gn=NOS2;refGene_tx=XM_019817487.2;refGene_gn=NOS2;refGene_tx=XM_023243518.1
E1 18833304 18833427
refGene_gn=NOS2;refGene_tx=XM_019817487.2;refGene_gn=NOS2;refGene_tx=XM_023243518.1
E1 18835001 18835150
refGene_gn=NOS2;refGene_tx=XM_019817487.2;refGene_gn=NOS2;refGene_tx=XM_023243518.1
E1 18838531 18838694
refGene_gn=NOS2;refGene_tx=XM_019817487.2;refGene_gn=NOS2;refGene_tx=XM_023243518.1
E1 18839645 18839737
refGene_gn=NOS2;refGene_tx=XM_019817487.2;refGene_gn=NOS2;refGene_tx=XM_023243518.1
E1 18840692 18840834
refGene_gn=NOS2;refGene_tx=XM_019817487.2;refGene_gn=NOS2;refGene_tx=XM_023243518.1
E1 18840926 18841066
refGene_gn=NOS2;refGene_tx=XM_019817487.2;refGene_gn=NOS2;refGene_tx=XM_023243518.1
E1 18842434 18842609
refGene_gn=NOS2;refGene_tx=XM_019817487.2;refGene_gn=NOS2;refGene_tx=XM_023243518.1
E1 18842703 18842805
refGene_gn=NOS2;refGene_tx=XM_019817487.2;refGene_gn=NOS2;refGene_tx=XM_023243518.1
E1 18846187 18846382
refGene_gn=NOS2;refGene_tx=XM_019817487.2;refGene_gn=NOS2;refGene_tx=XM_023243518.1
E1 18847393 18847476
refGene_gn=NOS2;refGene_tx=XM_019817487.2;refGene_gn=NOS2;refGene_tx=XM_023243518.1
E1 18847862 18848007
refGene_gn=NOS2;refGene_tx=XM_019817487.2;refGene_gn=NOS2;refGene_tx=X

M_023243518.1
E1 18848980 18849085
refGene_gn=NOS2;refGene_tx=XM_019817487.2;refGene_gn=NOS2;refGene_tx=XM_023243518.1
E1 18850255 18850305
refGene_gn=NOS2;refGene_tx=XM_019817487.2;refGene_gn=NOS2;refGene_tx=XM_023243518.1
E1 18850728 18850903
refGene_gn=NOS2;refGene_tx=XM_019817487.2;refGene_gn=NOS2;refGene_tx=XM_023243518.1
E1 18851680 18851813
refGene_gn=NOS2;refGene_tx=XM_019817487.2;refGene_gn=NOS2;refGene_tx=XM_023243518.1
E1 18852806 18852885
refGene_gn=NOS2;refGene_tx=XM_019817487.2;refGene_gn=NOS2;refGene_tx=XM_023243518.1
E1 18853657 18853839
refGene_gn=NOS2;refGene_tx=XM_019817487.2;refGene_gn=NOS2;refGene_tx=XM_023243518.1
E1 18855938 18856102
refGene_gn=NOS2;refGene_tx=XM_019817487.2;refGene_gn=NOS2;refGene_tx=XM_023243518.1
E1 18857114 18857322
refGene_gn=NOS2;refGene_tx=XM_019817487.2;refGene_gn=NOS2;refGene_tx=XM_023243518.1
E1 18859114 18859202
refGene_gn=NOS2;refGene_tx=XM_019817487.2;refGene_gn=NOS2;refGene_tx=XM_023243518.1
E1 18859655 18859777
refGene_gn=NOS2;refGene_tx=XM_019817487.2;refGene_gn=NOS2;refGene_tx=XM_023243518.1
E1 18860086 18860235
refGene_gn=NOS2;refGene_tx=XM_019817487.2;refGene_gn=NOS2;refGene_tx=XM_023243518.1
E1 18861146 18861341
refGene_gn=NOS2;refGene_tx=XM_019817487.2;refGene_gn=NOS2;refGene_tx=XM_023243518.1
E1 18861913 18862033
refGene_gn=NOS2;refGene_tx=XM_019817487.2;refGene_gn=NOS2;refGene_tx=XM_023243518.1
E1 18864735 18864882
refGene_gn=LGALS9;refGene_tx=XM_003996522.3;refGene_gn=LGALS9;refGene_tx=XM_003996523.5;refGene_gn=LGALS9;refGene_tx=XM_019817494.2
E1 18866644 18866807
refGene_gn=LGALS9;refGene_tx=XM_003996522.3;refGene_gn=LGALS9;refGene_tx=XM_003996523.5
E1 18866937 18867026
refGene_gn=LGALS9;refGene_tx=XM_003996522.3;refGene_gn=LGALS9;refGene_tx=XM_003996523.5;refGene_gn=LGALS9;refGene_tx=XM_019817494.2
E1 18867467 18867509

refGene_gn=LGALS9;refGene_tx=XM_003996522.3;refGene_gn=LGALS9;refGene_tx=XM_003996523.5;refGene_gn=LGALS9;refGene_tx=XM_019817494.2
E1 18868046 18868097
refGene_gn=LGALS9;refGene_tx=XM_003996522.3;refGene_gn=LGALS9;refGene_tx=XM_003996523.5;refGene_gn=LGALS9;refGene_tx=XM_019817494.2
E1 18868667 18868703
refGene_gn=LGALS9;refGene_tx=XM_003996522.3;refGene_gn=LGALS9;refGene_tx=XM_003996523.5;refGene_gn=LGALS9;refGene_tx=XM_019817494.2
E1 18870395 18870491 refGene_gn=LGALS9;refGene_tx=XM_003996522.3
E1 18871673 18871784
refGene_gn=LGALS9;refGene_tx=XM_003996522.3;refGene_gn=LGALS9;refGene_tx=XM_003996523.5;refGene_gn=LGALS9;refGene_tx=XM_019817494.2
E1 18872880 18873082
refGene_gn=LGALS9;refGene_tx=XM_003996522.3;refGene_gn=LGALS9;refGene_tx=XM_003996523.5;refGene_gn=LGALS9;refGene_tx=XM_019817494.2
E1 18875502 18875594
refGene_gn=LGALS9;refGene_tx=XM_003996522.3;refGene_gn=LGALS9;refGene_tx=XM_003996523.5;refGene_gn=LGALS9;refGene_tx=XM_019817494.2
E1 18878628 18878667
refGene_gn=LGALS9;refGene_tx=XM_003996522.3;refGene_gn=LGALS9;refGene_tx=XM_003996523.5;refGene_gn=LGALS9;refGene_tx=XM_019817494.2
E1 18896206 18896285
refGene_gn=KSR1;refGene_tx=XM_006940043.4;refGene_gn=KSR1;refGene_tx=XM_006940047.4;refGene_gn=KSR1;refGene_tx=XM_006940048.4;refGene_gn=KSR1;refGene_tx=XM_011288974.3;refGene_gn=KSR1;refGene_tx=XM_011288976.2;refGene_gn=KSR1;refGene_tx=XM_011288978.2;refGene_gn=KSR1;refGene_tx=XM_019817492.2;refGene_gn=KSR1;refGene_tx=XM_019817493.2;refGene_gn=KSR1;refGene_tx=XM_023244117.1;refGene_gn=KSR1;refGene_tx=XM_023244118.1;refGene_gn=KSR1;refGene_tx=XM_023244121.1
E1 18897564 18897571 refGene_gn=KSR1;refGene_tx=XM_023244119.1
E1 18898183 18898264
refGene_gn=KSR1;refGene_tx=XM_006940043.4;refGene_gn=KSR1;refGene_tx=XM_006940047.4;refGene_gn=KSR1;refGene_tx=XM_006940048.4;refGene_gn=KSR1;refGene_tx=XM_011288974.3;refGene_gn=KSR1;refGene_tx=XM_011288976.2;refGene_gn=KSR1;refGene_tx=XM_019817492.2;refGene_gn=KSR1;refGene_tx=XM_019817493.2;refGene_gn=KSR1;refGene_tx=XM_023244117.1;refGene_gn=KSR1;refGene_tx=XM_023244118.1;refGene_gn=KSR1;refGene_tx=XM_023244121.1
E1 18900836 18900974
refGene_gn=KSR1;refGene_tx=XM_006940043.4;refGene_gn=KSR1;refGene_tx=XM_006940047.4;refGene_gn=KSR1;refGene_tx=XM_006940048.4;refGene_gn=KSR1;refGene_tx=XM_011288974.3;refGene_gn=KSR1;refGene_tx=XM_011288976.2;refGene_gn=KSR1;refGene_tx=XM_011288978.2;refGene_gn=KSR1;refGene_tx=XM_019817492.2;refGene_gn=KSR1;refGene_tx=XM_019817493.2;refGene_gn=KSR1;refGene_tx=XM_023244117.1;refGene_gn=KSR1;refGene_tx=XM_023244118.1;refGene_gn=KSR1;refGene_tx=XM_023244119.1;refGene_gn=KSR1;refGene_tx=XM_023244120.1;refGene_gn=KSR1;refGene_tx=XM_023244121.1
E1 18905536 18905672
refGene_gn=KSR1;refGene_tx=XM_006940043.4;refGene_gn=KSR1;refGene_tx=XM_006940047.4;refGene_gn=KSR1;refGene_tx=XM_006940048.4;refGene_gn=KSR1;refGene_tx=XM_011288974.3;refGene_gn=KSR1;refGene_tx=XM_011288976.2;

refGene_gn=KSR1;refGene_tx=XM_011288978.2;refGene_gn=KSR1;refGene_tx=XM_019817492.2;refGene_gn=KSR1;refGene_tx=XM_019817493.2;refGene_gn=KSR1;refGene_tx=XM_023244117.1;refGene_gn=KSR1;refGene_tx=XM_023244118.1;refGene_gn=KSR1;refGene_tx=XM_023244119.1;refGene_gn=KSR1;refGene_tx=XM_023244120.1;refGene_gn=KSR1;refGene_tx=XM_023244121.1
E1 18906847 18906979
refGene_gn=KSR1;refGene_tx=XM_006940043.4;refGene_gn=KSR1;refGene_tx=XM_006940047.4;refGene_gn=KSR1;refGene_tx=XM_006940048.4;refGene_gn=KSR1;refGene_tx=XM_011288974.3;refGene_gn=KSR1;refGene_tx=XM_011288976.2;refGene_gn=KSR1;refGene_tx=XM_011288978.2;refGene_gn=KSR1;refGene_tx=XM_019817492.2;refGene_gn=KSR1;refGene_tx=XM_019817493.2;refGene_gn=KSR1;refGene_tx=XM_023244117.1;refGene_gn=KSR1;refGene_tx=XM_023244118.1;refGene_gn=KSR1;refGene_tx=XM_023244119.1;refGene_gn=KSR1;refGene_tx=XM_023244120.1;refGene_gn=KSR1;refGene_tx=XM_023244121.1
E1 18907714 18907848
refGene_gn=KSR1;refGene_tx=XM_006940043.4;refGene_gn=KSR1;refGene_tx=XM_006940047.4;refGene_gn=KSR1;refGene_tx=XM_006940048.4;refGene_gn=KSR1;refGene_tx=XM_011288974.3;refGene_gn=KSR1;refGene_tx=XM_011288976.2;refGene_gn=KSR1;refGene_tx=XM_011288978.2;refGene_gn=KSR1;refGene_tx=XM_019817492.2;refGene_gn=KSR1;refGene_tx=XM_019817493.2;refGene_gn=KSR1;refGene_tx=XM_023244117.1;refGene_gn=KSR1;refGene_tx=XM_023244118.1;refGene_gn=KSR1;refGene_tx=XM_023244119.1;refGene_gn=KSR1;refGene_tx=XM_023244120.1;refGene_gn=KSR1;refGene_tx=XM_023244121.1
E1 18910803 18910900
refGene_gn=KSR1;refGene_tx=XM_006940043.4;refGene_gn=KSR1;refGene_tx=XM_006940047.4;refGene_gn=KSR1;refGene_tx=XM_006940048.4;refGene_gn=KSR1;refGene_tx=XM_011288974.3;refGene_gn=KSR1;refGene_tx=XM_011288976.2;refGene_gn=KSR1;refGene_tx=XM_011288978.2;refGene_gn=KSR1;refGene_tx=XM_019817492.2;refGene_gn=KSR1;refGene_tx=XM_019817493.2;refGene_gn=KSR1;refGene_tx=XM_023244117.1;refGene_gn=KSR1;refGene_tx=XM_023244118.1;refGene_gn=KSR1;refGene_tx=XM_023244119.1;refGene_gn=KSR1;refGene_tx=XM_023244120.1;refGene_gn=KSR1;refGene_tx=XM_023244121.1
E1 18913553 18913933
refGene_gn=KSR1;refGene_tx=XM_006940043.4;refGene_gn=KSR1;refGene_tx=XM_006940047.4;refGene_gn=KSR1;refGene_tx=XM_006940048.4;refGene_gn=KSR1;refGene_tx=XM_011288974.3;refGene_gn=KSR1;refGene_tx=XM_011288976.2;refGene_gn=KSR1;refGene_tx=XM_011288978.2;refGene_gn=KSR1;refGene_tx=XM_019817492.2;refGene_gn=KSR1;refGene_tx=XM_019817493.2;refGene_gn=KSR1;refGene_tx=XM_023244117.1;refGene_gn=KSR1;refGene_tx=XM_023244118.1;refGene_gn=KSR1;refGene_tx=XM_023244119.1;refGene_gn=KSR1;refGene_tx=XM_023244120.1;refGene_gn=KSR1;refGene_tx=XM_023244121.1
E1 18914773 18914822
refGene_gn=KSR1;refGene_tx=XM_006940043.4;refGene_gn=KSR1;refGene_tx=XM_006940047.4;refGene_gn=KSR1;refGene_tx=XM_006940048.4;refGene_gn=KSR1;refGene_tx=XM_011288974.3;refGene_gn=KSR1;refGene_tx=XM_011288976.2;refGene_gn=KSR1;refGene_tx=XM_011288978.2;refGene_gn=KSR1;refGene_tx=XM_019817492.2;refGene_gn=KSR1;refGene_tx=XM_019817493.2;refGene_gn=KSR1;refGene_tx=XM_023244117.1;refGene_gn=KSR1;refGene_tx=XM_023244118.1;refGene_gn=KSR1;refGene_tx=XM_023244119.1;refGene_gn=KSR1;refGene_tx=XM_023244120.1;refGene_gn=KSR1;refGene_tx=XM_023244121.1

E1 18916032 18916087
refGene_gn=KSR1;refGene_tx=XM_006940043.4;refGene_gn=KSR1;refGene_tx=XM_006940047.4;refGene_gn=KSR1;refGene_tx=XM_006940048.4;refGene_gn=KSR1;refGene_tx=XM_011288974.3;refGene_gn=KSR1;refGene_tx=XM_011288976.2;refGene_gn=KSR1;refGene_tx=XM_011288978.2;refGene_gn=KSR1;refGene_tx=XM_019817492.2;refGene_gn=KSR1;refGene_tx=XM_019817493.2;refGene_gn=KSR1;refGene_tx=XM_023244117.1;refGene_gn=KSR1;refGene_tx=XM_023244118.1;refGene_gn=KSR1;refGene_tx=XM_023244119.1;refGene_gn=KSR1;refGene_tx=XM_023244120.1;refGene_gn=KSR1;refGene_tx=XM_023244121.1
E1 18918450 18918492
refGene_gn=KSR1;refGene_tx=XM_006940043.4;refGene_gn=KSR1;refGene_tx=XM_006940047.4;refGene_gn=KSR1;refGene_tx=XM_006940048.4;refGene_gn=KSR1;refGene_tx=XM_011288974.3;refGene_gn=KSR1;refGene_tx=XM_011288976.2;refGene_gn=KSR1;refGene_tx=XM_011288978.2;refGene_gn=KSR1;refGene_tx=XM_019817492.2;refGene_gn=KSR1;refGene_tx=XM_019817493.2;refGene_gn=KSR1;refGene_tx=XM_023244118.1;refGene_gn=KSR1;refGene_tx=XM_023244119.1;refGene_gn=KSR1;refGene_tx=XM_023244120.1;refGene_gn=KSR1;refGene_tx=XM_023244121.1
E1 18920313 18920482
refGene_gn=KSR1;refGene_tx=XM_006940043.4;refGene_gn=KSR1;refGene_tx=XM_006940047.4;refGene_gn=KSR1;refGene_tx=XM_006940048.4;refGene_gn=KSR1;refGene_tx=XM_011288974.3;refGene_gn=KSR1;refGene_tx=XM_011288976.2;refGene_gn=KSR1;refGene_tx=XM_011288978.2;refGene_gn=KSR1;refGene_tx=XM_019817492.2;refGene_gn=KSR1;refGene_tx=XM_019817493.2;refGene_gn=KSR1;refGene_tx=XM_023244117.1;refGene_gn=KSR1;refGene_tx=XM_023244118.1;refGene_gn=KSR1;refGene_tx=XM_023244119.1;refGene_gn=KSR1;refGene_tx=XM_023244120.1;refGene_gn=KSR1;refGene_tx=XM_023244121.1
E1 18923847 18923954
refGene_gn=KSR1;refGene_tx=XM_006940043.4;refGene_gn=KSR1;refGene_tx=XM_006940047.4;refGene_gn=KSR1;refGene_tx=XM_006940048.4;refGene_gn=KSR1;refGene_tx=XM_011288974.3;refGene_gn=KSR1;refGene_tx=XM_011288976.2;refGene_gn=KSR1;refGene_tx=XM_011288978.2;refGene_gn=KSR1;refGene_tx=XM_019817492.2;refGene_gn=KSR1;refGene_tx=XM_019817493.2;refGene_gn=KSR1;refGene_tx=XM_023244117.1;refGene_gn=KSR1;refGene_tx=XM_023244118.1;refGene_gn=KSR1;refGene_tx=XM_023244119.1;refGene_gn=KSR1;refGene_tx=XM_023244120.1;refGene_gn=KSR1;refGene_tx=XM_023244121.1
E1 18924055 18924117
refGene_gn=KSR1;refGene_tx=XM_006940043.4;refGene_gn=KSR1;refGene_tx=XM_006940047.4;refGene_gn=KSR1;refGene_tx=XM_006940048.4;refGene_gn=KSR1;refGene_tx=XM_011288974.3;refGene_gn=KSR1;refGene_tx=XM_011288976.2;refGene_gn=KSR1;refGene_tx=XM_011288978.2;refGene_gn=KSR1;refGene_tx=XM_019817492.2;refGene_gn=KSR1;refGene_tx=XM_019817493.2;refGene_gn=KSR1;refGene_tx=XM_023244117.1;refGene_gn=KSR1;refGene_tx=XM_023244118.1;refGene_gn=KSR1;refGene_tx=XM_023244119.1;refGene_gn=KSR1;refGene_tx=XM_023244120.1;refGene_gn=KSR1;refGene_tx=XM_023244121.1
E1 18925675 18925759
refGene_gn=KSR1;refGene_tx=XM_006940043.4;refGene_gn=KSR1;refGene_tx=XM_006940047.4;refGene_gn=KSR1;refGene_tx=XM_006940048.4;refGene_gn=KSR1;refGene_tx=XM_011288974.3;refGene_gn=KSR1;refGene_tx=XM_011288976.2;refGene_gn=KSR1;refGene_tx=XM_011288978.2;refGene_gn=KSR1;refGene_tx=X

M_019817492.2;refGene_gn=KSR1;refGene_tx=XM_019817493.2;refGene_gn=KSR1;refGene_tx=XM_023244117.1;refGene_gn=KSR1;refGene_tx=XM_023244118.1;refGene_gn=KSR1;refGene_tx=XM_023244119.1;refGene_gn=KSR1;refGene_tx=XM_023244120.1;refGene_gn=KSR1;refGene_tx=XM_023244121.1
E1 18930111 18930172 refGene_gn=KSR1;refGene_tx=XM_006940043.4;refGene_gn=KSR1;refGene_tx=XM_006940047.4;refGene_gn=KSR1;refGene_tx=XM_006940048.4;refGene_gn=KSR1;refGene_tx=XM_011288974.3;refGene_gn=KSR1;refGene_tx=XM_011288976.2;refGene_gn=KSR1;refGene_tx=XM_011288978.2;refGene_gn=KSR1;refGene_tx=XM_019817492.2;refGene_gn=KSR1;refGene_tx=XM_019817493.2;refGene_gn=KSR1;refGene_tx=XM_023244117.1;refGene_gn=KSR1;refGene_tx=XM_023244118.1;refGene_gn=KSR1;refGene_tx=XM_023244119.1;refGene_gn=KSR1;refGene_tx=XM_023244120.1;refGene_gn=KSR1;refGene_tx=XM_023244121.1
E1 18932830 18932835 refGene_gn=KSR1;refGene_tx=XM_006940043.4;refGene_gn=KSR1;refGene_tx=XM_006940047.4;refGene_gn=KSR1;refGene_tx=XM_006940048.4;refGene_gn=KSR1;refGene_tx=XM_011288974.3;refGene_gn=KSR1;refGene_tx=XM_011288976.2;refGene_gn=KSR1;refGene_tx=XM_011288978.2;refGene_gn=KSR1;refGene_tx=XM_019817492.2;refGene_gn=KSR1;refGene_tx=XM_019817493.2;refGene_gn=KSR1;refGene_tx=XM_023244117.1;refGene_gn=KSR1;refGene_tx=XM_023244118.1;refGene_gn=KSR1;refGene_tx=XM_023244119.1;refGene_gn=KSR1;refGene_tx=XM_023244120.1;refGene_gn=KSR1;refGene_tx=XM_023244121.1
E1 18935022 18935482 refGene_gn=KSR1;refGene_tx=XM_006940043.4;refGene_gn=KSR1;refGene_tx=XM_006940047.4;refGene_gn=KSR1;refGene_tx=XM_006940048.4;refGene_gn=KSR1;refGene_tx=XM_011288974.3;refGene_gn=KSR1;refGene_tx=XM_011288976.2;refGene_gn=KSR1;refGene_tx=XM_011288978.2;refGene_gn=KSR1;refGene_tx=XM_019817492.2;refGene_gn=KSR1;refGene_tx=XM_019817493.2;refGene_gn=KSR1;refGene_tx=XM_023244117.1;refGene_gn=KSR1;refGene_tx=XM_023244118.1;refGene_gn=KSR1;refGene_tx=XM_023244119.1;refGene_gn=KSR1;refGene_tx=XM_023244120.1;refGene_gn=KSR1;refGene_tx=XM_023244121.1
E1 18940247 18940395 refGene_gn=KSR1;refGene_tx=XM_006940043.4;refGene_gn=KSR1;refGene_tx=XM_006940047.4;refGene_gn=KSR1;refGene_tx=XM_006940048.4;refGene_gn=KSR1;refGene_tx=XM_011288974.3;refGene_gn=KSR1;refGene_tx=XM_011288976.2;refGene_gn=KSR1;refGene_tx=XM_011288978.2;refGene_gn=KSR1;refGene_tx=XM_019817492.2;refGene_gn=KSR1;refGene_tx=XM_019817493.2;refGene_gn=KSR1;refGene_tx=XM_023244117.1;refGene_gn=KSR1;refGene_tx=XM_023244118.1;refGene_gn=KSR1;refGene_tx=XM_023244119.1;refGene_gn=KSR1;refGene_tx=XM_023244120.1;refGene_gn=KSR1;refGene_tx=XM_023244121.1
E1 18960395 18960410 refGene_gn=KSR1;refGene_tx=XM_006940047.4
E1 18969356 18969497 refGene_gn=KSR1;refGene_tx=XM_011288976.2;refGene_gn=KSR1;refGene_tx=XM_011288978.2;refGene_gn=KSR1;refGene_tx=XM_023244117.1;refGene_gn=KSR1;refGene_tx=XM_023244118.1;refGene_gn=KSR1;refGene_tx=XM_023244119.1;refGene_gn=KSR1;refGene_tx=XM_023244120.1;refGene_gn=KSR1;refGene_tx=XM_023244121.1
E1 19022491 19022494 refGene_gn=KSR1;refGene_tx=XM_023244121.1
E1 19057016 19057247 refGene_gn=KSR1;refGene_tx=XM_011288976.2;refGene_gn=KSR1;refGene_tx=X

M_011288978.2;refGene_gn=KSR1;refGene_tx=XM_023244117.1;refGene_gn=KSR1;refGene_tx=XM_023244118.1;refGene_gn=KSR1;refGene_tx=XM_023244119.1;refGene_gn=KSR1;refGene_tx=XM_023244120.1
E1 19149206 19149366 refGene_gn=WSB1;refGene_tx=XM_003996524.5
E1 19149841 19149949 refGene_gn=WSB1;refGene_tx=XM_003996524.5
E1 19151199 19151313 refGene_gn=WSB1;refGene_tx=XM_003996524.5
E1 19151839 19152012 refGene_gn=WSB1;refGene_tx=XM_003996524.5
E1 19153371 19153395 refGene_gn=WSB1;refGene_tx=XM_023243441.1
E1 19154258 19154359
refGene_gn=WSB1;refGene_tx=XM_003996524.5;refGene_gn=WSB1;refGene_tx=XM_023243441.1
E1 19155985 19156117
refGene_gn=WSB1;refGene_tx=XM_003996524.5;refGene_gn=WSB1;refGene_tx=XM_023243441.1
E1 19157256 19157525
refGene_gn=WSB1;refGene_tx=XM_003996524.5;refGene_gn=WSB1;refGene_tx=XM_023243441.1
E1 19158444 19158613
refGene_gn=WSB1;refGene_tx=XM_003996524.5;refGene_gn=WSB1;refGene_tx=XM_023243441.1
E1 19164781 19164821
refGene_gn=WSB1;refGene_tx=XM_003996524.5;refGene_gn=WSB1;refGene_tx=XM_023243441.1
E1 19278924 19278984
refGene_gn=NF1;refGene_tx=XM_023243526.1;refGene_gn=NF1;refGene_tx=XM_023243527.1;refGene_gn=NF1;refGene_tx=XM_023243528.1;refGene_gn=NF1;refGene_tx=XM_023243529.1;refGene_gn=NF1;refGene_tx=XM_023243530.1;refGene_gn=NF1;refGene_tx=XM_023243531.1;refGene_gn=NF1;refGene_tx=XM_023243532.1;refGene_gn=NF1;refGene_tx=XM_023243533.1;refGene_gn=NF1;refGene_tx=XM_023243534.1
E1 19335013 19335157
refGene_gn=NF1;refGene_tx=XM_023243526.1;refGene_gn=NF1;refGene_tx=XM_023243527.1;refGene_gn=NF1;refGene_tx=XM_023243528.1;refGene_gn=NF1;refGene_tx=XM_023243529.1;refGene_gn=NF1;refGene_tx=XM_023243530.1;refGene_gn=NF1;refGene_tx=XM_023243531.1;refGene_gn=NF1;refGene_tx=XM_023243532.1;refGene_gn=NF1;refGene_tx=XM_023243533.1;refGene_gn=NF1;refGene_tx=XM_023243534.1
E1 19336842 19336926
refGene_gn=NF1;refGene_tx=XM_023243526.1;refGene_gn=NF1;refGene_tx=XM_023243527.1;refGene_gn=NF1;refGene_tx=XM_023243528.1;refGene_gn=NF1;refGene_tx=XM_023243529.1;refGene_gn=NF1;refGene_tx=XM_023243530.1;refGene_gn=NF1;refGene_tx=XM_023243531.1;refGene_gn=NF1;refGene_tx=XM_023243532.1;refGene_gn=NF1;refGene_tx=XM_023243533.1;refGene_gn=NF1;refGene_tx=XM_023243534.1
E1 19340459 19340650
refGene_gn=NF1;refGene_tx=XM_023243526.1;refGene_gn=NF1;refGene_tx=XM_023243527.1;refGene_gn=NF1;refGene_tx=XM_023243528.1;refGene_gn=NF1;refGene_tx=XM_023243529.1;refGene_gn=NF1;refGene_tx=XM_023243530.1;refGene_gn=NF1;refGene_tx=XM_023243531.1;refGene_gn=NF1;refGene_tx=XM_023243532.1;refGene_gn=NF1;refGene_tx=XM_023243533.1;refGene_gn=NF1;refGene

_tx=XM_023243534.1
E1 19346962 19347069
refGene_gn=NF1;refGene_tx=XM_023243526.1;refGene_gn=NF1;refGene_tx=XM_023243527.1;refGene_gn=NF1;refGene_tx=XM_023243528.1;refGene_gn=NF1;refGene_tx=XM_023243529.1;refGene_gn=NF1;refGene_tx=XM_023243530.1;refGene_gn=NF1;refGene_tx=XM_023243531.1;refGene_gn=NF1;refGene_tx=XM_023243532.1;refGene_gn=NF1;refGene_tx=XM_023243533.1;refGene_gn=NF1;refGene_tx=XM_023243534.1
E1 19354555 19354623
refGene_gn=NF1;refGene_tx=XM_023243526.1;refGene_gn=NF1;refGene_tx=XM_023243527.1;refGene_gn=NF1;refGene_tx=XM_023243528.1;refGene_gn=NF1;refGene_tx=XM_023243529.1;refGene_gn=NF1;refGene_tx=XM_023243530.1;refGene_gn=NF1;refGene_tx=XM_023243531.1;refGene_gn=NF1;refGene_tx=XM_023243532.1;refGene_gn=NF1;refGene_tx=XM_023243533.1;refGene_gn=NF1;refGene_tx=XM_023243534.1
E1 19354874 19354950
refGene_gn=NF1;refGene_tx=XM_023243526.1;refGene_gn=NF1;refGene_tx=XM_023243527.1;refGene_gn=NF1;refGene_tx=XM_023243528.1;refGene_gn=NF1;refGene_tx=XM_023243529.1;refGene_gn=NF1;refGene_tx=XM_023243530.1;refGene_gn=NF1;refGene_tx=XM_023243531.1;refGene_gn=NF1;refGene_tx=XM_023243532.1;refGene_gn=NF1;refGene_tx=XM_023243533.1;refGene_gn=NF1;refGene_tx=XM_023243534.1
E1 19355654 19355812
refGene_gn=NF1;refGene_tx=XM_023243526.1;refGene_gn=NF1;refGene_tx=XM_023243527.1;refGene_gn=NF1;refGene_tx=XM_023243528.1;refGene_gn=NF1;refGene_tx=XM_023243529.1;refGene_gn=NF1;refGene_tx=XM_023243530.1;refGene_gn=NF1;refGene_tx=XM_023243531.1;refGene_gn=NF1;refGene_tx=XM_023243532.1;refGene_gn=NF1;refGene_tx=XM_023243533.1;refGene_gn=NF1;refGene_tx=XM_023243534.1
E1 19363144 19363318
refGene_gn=NF1;refGene_tx=XM_023243526.1;refGene_gn=NF1;refGene_tx=XM_023243527.1;refGene_gn=NF1;refGene_tx=XM_023243528.1;refGene_gn=NF1;refGene_tx=XM_023243529.1;refGene_gn=NF1;refGene_tx=XM_023243530.1;refGene_gn=NF1;refGene_tx=XM_023243531.1;refGene_gn=NF1;refGene_tx=XM_023243532.1;refGene_gn=NF1;refGene_tx=XM_023243533.1;refGene_gn=NF1;refGene_tx=XM_023243534.1
E1 19363801 19363924
refGene_gn=NF1;refGene_tx=XM_023243526.1;refGene_gn=NF1;refGene_tx=XM_023243527.1;refGene_gn=NF1;refGene_tx=XM_023243528.1;refGene_gn=NF1;refGene_tx=XM_023243529.1;refGene_gn=NF1;refGene_tx=XM_023243530.1;refGene_gn=NF1;refGene_tx=XM_023243531.1;refGene_gn=NF1;refGene_tx=XM_023243532.1;refGene_gn=NF1;refGene_tx=XM_023243533.1;refGene_gn=NF1;refGene_tx=XM_023243534.1
E1 19364165 19364240
refGene_gn=NF1;refGene_tx=XM_023243526.1;refGene_gn=NF1;refGene_tx=XM_023243527.1;refGene_gn=NF1;refGene_tx=XM_023243528.1;refGene_gn=NF1;refGene_tx=XM_023243529.1;refGene_gn=NF1;refGene_tx=XM_023243530.1;refGene_gn=NF1;refGene_tx=XM_023243531.1;refGene_gn=NF1;refGene_tx=XM_023243532.1;refGene_gn=NF1;refGene_tx=XM_023243533.1;refGene_gn=NF1;refGene_tx=XM_023243534.1

E1 19366064 19366094
refGene_gn=NF1;refGene_tx=XM_023243526.1;refGene_gn=NF1;refGene_tx=XM_023243527.1;refGene_gn=NF1;refGene_tx=XM_023243529.1;refGene_gn=NF1;refGene_tx=XM_023243530.1;refGene_gn=NF1;refGene_tx=XM_023243531.1;refGene_gn=NF1;refGene_tx=XM_023243534.1
E1 19369546 19369678
refGene_gn=NF1;refGene_tx=XM_023243526.1;refGene_gn=NF1;refGene_tx=XM_023243527.1;refGene_gn=NF1;refGene_tx=XM_023243528.1;refGene_gn=NF1;refGene_tx=XM_023243529.1;refGene_gn=NF1;refGene_tx=XM_023243530.1;refGene_gn=NF1;refGene_tx=XM_023243531.1;refGene_gn=NF1;refGene_tx=XM_023243532.1;refGene_gn=NF1;refGene_tx=XM_023243533.1;refGene_gn=NF1;refGene_tx=XM_023243534.1;refGene_gn=NF1;refGene_tx=XM_023243535.1
E1 19376379 19376514
refGene_gn=NF1;refGene_tx=XM_023243526.1;refGene_gn=NF1;refGene_tx=XM_023243527.1;refGene_gn=NF1;refGene_tx=XM_023243528.1;refGene_gn=NF1;refGene_tx=XM_023243529.1;refGene_gn=NF1;refGene_tx=XM_023243530.1;refGene_gn=NF1;refGene_tx=XM_023243531.1;refGene_gn=NF1;refGene_tx=XM_023243532.1;refGene_gn=NF1;refGene_tx=XM_023243533.1;refGene_gn=NF1;refGene_tx=XM_023243534.1;refGene_gn=NF1;refGene_tx=XM_023243535.1
E1 19379986 19380100
refGene_gn=NF1;refGene_tx=XM_023243526.1;refGene_gn=NF1;refGene_tx=XM_023243527.1;refGene_gn=NF1;refGene_tx=XM_023243528.1;refGene_gn=NF1;refGene_tx=XM_023243529.1;refGene_gn=NF1;refGene_tx=XM_023243530.1;refGene_gn=NF1;refGene_tx=XM_023243531.1;refGene_gn=NF1;refGene_tx=XM_023243532.1;refGene_gn=NF1;refGene_tx=XM_023243533.1;refGene_gn=NF1;refGene_tx=XM_023243534.1;refGene_gn=NF1;refGene_tx=XM_023243535.1
E1 19384816 19384896
refGene_gn=NF1;refGene_tx=XM_023243526.1;refGene_gn=NF1;refGene_tx=XM_023243527.1;refGene_gn=NF1;refGene_tx=XM_023243528.1;refGene_gn=NF1;refGene_tx=XM_023243529.1;refGene_gn=NF1;refGene_tx=XM_023243530.1;refGene_gn=NF1;refGene_tx=XM_023243531.1;refGene_gn=NF1;refGene_tx=XM_023243532.1;refGene_gn=NF1;refGene_tx=XM_023243533.1;refGene_gn=NF1;refGene_tx=XM_023243534.1;refGene_gn=NF1;refGene_tx=XM_023243535.1
E1 19386907 19387031
refGene_gn=NF1;refGene_tx=XM_023243526.1;refGene_gn=NF1;refGene_tx=XM_023243527.1;refGene_gn=NF1;refGene_tx=XM_023243528.1;refGene_gn=NF1;refGene_tx=XM_023243529.1;refGene_gn=NF1;refGene_tx=XM_023243530.1;refGene_gn=NF1;refGene_tx=XM_023243531.1;refGene_gn=NF1;refGene_tx=XM_023243532.1;refGene_gn=NF1;refGene_tx=XM_023243533.1;refGene_gn=NF1;refGene_tx=XM_023243534.1;refGene_gn=NF1;refGene_tx=XM_023243535.1
E1 19388401 19388563
refGene_gn=NF1;refGene_tx=XM_023243526.1;refGene_gn=NF1;refGene_tx=XM_023243527.1;refGene_gn=NF1;refGene_tx=XM_023243528.1;refGene_gn=NF1;refGene_tx=XM_023243529.1;refGene_gn=NF1;refGene_tx=XM_023243530.1;refGene_gn=NF1;refGene_tx=XM_023243531.1;refGene_gn=NF1;refGene_tx=XM_023243532.1;refGene_gn=NF1;refGene_tx=XM_023243533.1;refGene_gn=NF1;refGene_tx=XM_023243534.1;refGene_gn=NF1;refGene_tx=XM_023243535.1
E1 19390284 19390534
refGene_gn=NF1;refGene_tx=XM_023243526.1;refGene_gn=NF1;refGene_tx=XM_023243527.1;refGene_gn=NF1;refGene_tx=XM_023243528.1;refGene_gn=NF1;re

fGene_tx=XM_023243529.1;refGene_gn=NF1;refGene_tx=XM_023243530.1;refGene_gn=NF1;refGene_tx=XM_023243531.1;refGene_gn=NF1;refGene_tx=XM_023243532.1;refGene_gn=NF1;refGene_tx=XM_023243533.1;refGene_gn=NF1;refGene_tx=XM_023243534.1;refGene_gn=NF1;refGene_tx=XM_023243535.1
E1 19391079 19391153
refGene_gn=NF1;refGene_tx=XM_023243526.1;refGene_gn=NF1;refGene_tx=XM_023243527.1;refGene_gn=NF1;refGene_tx=XM_023243528.1;refGene_gn=NF1;refGene_tx=XM_023243529.1;refGene_gn=NF1;refGene_tx=XM_023243530.1;refGene_gn=NF1;refGene_tx=XM_023243531.1;refGene_gn=NF1;refGene_tx=XM_023243532.1;refGene_gn=NF1;refGene_tx=XM_023243533.1;refGene_gn=NF1;refGene_tx=XM_023243534.1;refGene_gn=NF1;refGene_tx=XM_023243535.1
E1 19391494 19391578
refGene_gn=NF1;refGene_tx=XM_023243526.1;refGene_gn=NF1;refGene_tx=XM_023243527.1;refGene_gn=NF1;refGene_tx=XM_023243528.1;refGene_gn=NF1;refGene_tx=XM_023243529.1;refGene_gn=NF1;refGene_tx=XM_023243530.1;refGene_gn=NF1;refGene_tx=XM_023243531.1;refGene_gn=NF1;refGene_tx=XM_023243532.1;refGene_gn=NF1;refGene_tx=XM_023243533.1;refGene_gn=NF1;refGene_tx=XM_023243534.1;refGene_gn=NF1;refGene_tx=XM_023243535.1
E1 19399127 19399568
refGene_gn=NF1;refGene_tx=XM_023243526.1;refGene_gn=NF1;refGene_tx=XM_023243527.1;refGene_gn=NF1;refGene_tx=XM_023243528.1;refGene_gn=NF1;refGene_tx=XM_023243529.1;refGene_gn=NF1;refGene_tx=XM_023243530.1;refGene_gn=NF1;refGene_tx=XM_023243531.1;refGene_gn=NF1;refGene_tx=XM_023243532.1;refGene_gn=NF1;refGene_tx=XM_023243533.1;refGene_gn=NF1;refGene_tx=XM_023243534.1;refGene_gn=NF1;refGene_tx=XM_023243535.1
E1 19399924 19400064
refGene_gn=NF1;refGene_tx=XM_023243526.1;refGene_gn=NF1;refGene_tx=XM_023243527.1;refGene_gn=NF1;refGene_tx=XM_023243528.1;refGene_gn=NF1;refGene_tx=XM_023243529.1;refGene_gn=NF1;refGene_tx=XM_023243530.1;refGene_gn=NF1;refGene_tx=XM_023243531.1;refGene_gn=NF1;refGene_tx=XM_023243532.1;refGene_gn=NF1;refGene_tx=XM_023243533.1;refGene_gn=NF1;refGene_tx=XM_023243534.1;refGene_gn=NF1;refGene_tx=XM_023243535.1
E1 19400336 19400459
refGene_gn=NF1;refGene_tx=XM_023243526.1;refGene_gn=NF1;refGene_tx=XM_023243527.1;refGene_gn=NF1;refGene_tx=XM_023243528.1;refGene_gn=NF1;refGene_tx=XM_023243529.1;refGene_gn=NF1;refGene_tx=XM_023243530.1;refGene_gn=NF1;refGene_tx=XM_023243531.1;refGene_gn=NF1;refGene_tx=XM_023243532.1;refGene_gn=NF1;refGene_tx=XM_023243533.1;refGene_gn=NF1;refGene_tx=XM_023243534.1;refGene_gn=NF1;refGene_tx=XM_023243535.1
E1 19400962 19401046
refGene_gn=NF1;refGene_tx=XM_023243526.1;refGene_gn=NF1;refGene_tx=XM_023243527.1;refGene_gn=NF1;refGene_tx=XM_023243528.1;refGene_gn=NF1;refGene_tx=XM_023243529.1;refGene_gn=NF1;refGene_tx=XM_023243530.1;refGene_gn=NF1;refGene_tx=XM_023243531.1;refGene_gn=NF1;refGene_tx=XM_023243532.1;refGene_gn=NF1;refGene_tx=XM_023243533.1;refGene_gn=NF1;refGene_tx=XM_023243534.1;refGene_gn=NF1;refGene_tx=XM_023243535.1
E1 19401603 19401720
refGene_gn=NF1;refGene_tx=XM_023243526.1;refGene_gn=NF1;refGene_tx=XM_023243527.1;refGene_gn=NF1;refGene_tx=XM_023243528.1;refGene_gn=NF1;refGene_tx=XM_023243529.1;refGene_gn=NF1;refGene_tx=XM_023243530.1;refGe

ne_gn=NF1;refGene_tx=XM_023243531.1;refGene_gn=NF1;refGene_tx=XM_02324
3532.1;refGene_gn=NF1;refGene_tx=XM_023243533.1;refGene_gn=NF1;refGene
_tx=XM_023243534.1;refGene_gn=NF1;refGene_tx=XM_023243535.1
E1 19402316 19402498
refGene_gn=NF1;refGene_tx=XM_023243526.1;refGene_gn=NF1;refGene_tx=XM_
023243527.1;refGene_gn=NF1;refGene_tx=XM_023243528.1;refGene_gn=NF1;re
fGene_tx=XM_023243529.1;refGene_gn=NF1;refGene_tx=XM_023243530.1;refGe
ne_gn=NF1;refGene_tx=XM_023243531.1;refGene_gn=NF1;refGene_tx=XM_02324
3532.1;refGene_gn=NF1;refGene_tx=XM_023243533.1;refGene_gn=NF1;refGene
_tx=XM_023243534.1;refGene_gn=NF1;refGene_tx=XM_023243535.1
E1 19402613 19402825
refGene_gn=NF1;refGene_tx=XM_023243526.1;refGene_gn=NF1;refGene_tx=XM_
023243527.1;refGene_gn=NF1;refGene_tx=XM_023243528.1;refGene_gn=NF1;re
fGene_tx=XM_023243529.1;refGene_gn=NF1;refGene_tx=XM_023243530.1;refGe
ne_gn=NF1;refGene_tx=XM_023243531.1;refGene_gn=NF1;refGene_tx=XM_02324
3532.1;refGene_gn=NF1;refGene_tx=XM_023243533.1;refGene_gn=NF1;refGene
_tx=XM_023243534.1;refGene_gn=NF1;refGene_tx=XM_023243535.1
E1 19405931 19406093
refGene_gn=NF1;refGene_tx=XM_023243526.1;refGene_gn=NF1;refGene_tx=XM_
023243527.1;refGene_gn=NF1;refGene_tx=XM_023243528.1;refGene_gn=NF1;re
fGene_tx=XM_023243529.1;refGene_gn=NF1;refGene_tx=XM_023243530.1;refGe
ne_gn=NF1;refGene_tx=XM_023243531.1;refGene_gn=NF1;refGene_tx=XM_02324
3532.1;refGene_gn=NF1;refGene_tx=XM_023243533.1;refGene_gn=NF1;refGene
_tx=XM_023243534.1;refGene_gn=NF1;refGene_tx=XM_023243535.1
E1 19406230 19406334
refGene_gn=NF1;refGene_tx=XM_023243526.1;refGene_gn=NF1;refGene_tx=XM_
023243527.1;refGene_gn=NF1;refGene_tx=XM_023243528.1;refGene_gn=NF1;re
fGene_tx=XM_023243529.1;refGene_gn=NF1;refGene_tx=XM_023243530.1;refGe
ne_gn=NF1;refGene_tx=XM_023243531.1;refGene_gn=NF1;refGene_tx=XM_02324
3532.1;refGene_gn=NF1;refGene_tx=XM_023243533.1;refGene_gn=NF1;refGene
_tx=XM_023243534.1;refGene_gn=NF1;refGene_tx=XM_023243535.1
E1 19409736 19409872
refGene_gn=NF1;refGene_tx=XM_023243526.1;refGene_gn=NF1;refGene_tx=XM_
023243527.1;refGene_gn=NF1;refGene_tx=XM_023243528.1;refGene_gn=NF1;re
fGene_tx=XM_023243529.1;refGene_gn=NF1;refGene_tx=XM_023243530.1;refGe
ne_gn=NF1;refGene_tx=XM_023243531.1;refGene_gn=NF1;refGene_tx=XM_02324
3532.1;refGene_gn=NF1;refGene_tx=XM_023243533.1;refGene_gn=NF1;refGene
_tx=XM_023243534.1;refGene_gn=NF1;refGene_tx=XM_023243535.1
E1 19420458 19420521
refGene_gn=NF1;refGene_tx=XM_023243526.1;refGene_gn=NF1;refGene_tx=XM_
023243527.1;refGene_gn=NF1;refGene_tx=XM_023243528.1;refGene_gn=NF1;re
fGene_tx=XM_023243530.1;refGene_gn=NF1;refGene_tx=XM_023243531.1;refGe
ne_gn=NF1;refGene_tx=XM_023243532.1;refGene_gn=NF1;refGene_tx=XM_02324
3534.1;refGene_gn=NF1;refGene_tx=XM_023243535.1
E1 19425144 19425303
refGene_gn=NF1;refGene_tx=XM_023243526.1;refGene_gn=NF1;refGene_tx=XM_
023243527.1;refGene_gn=NF1;refGene_tx=XM_023243528.1;refGene_gn=NF1;re
fGene_tx=XM_023243529.1;refGene_gn=NF1;refGene_tx=XM_023243530.1;refGe
ne_gn=NF1;refGene_tx=XM_023243531.1;refGene_gn=NF1;refGene_tx=XM_02324
3532.1;refGene_gn=NF1;refGene_tx=XM_023243533.1;refGene_gn=NF1;refGene

_tx=XM_023243534.1;refGene_gn=NF1;refGene_tx=XM_023243535.1
E1 19425914 19426012
refGene_gn=NF1;refGene_tx=XM_023243526.1;refGene_gn=NF1;refGene_tx=XM_023243527.1;refGene_gn=NF1;refGene_tx=XM_023243528.1;refGene_gn=NF1;refGene_tx=XM_023243529.1;refGene_gn=NF1;refGene_tx=XM_023243530.1;refGene_gn=NF1;refGene_tx=XM_023243531.1;refGene_gn=NF1;refGene_tx=XM_023243532.1;refGene_gn=NF1;refGene_tx=XM_023243533.1;refGene_gn=NF1;refGene_tx=XM_023243534.1;refGene_gn=NF1;refGene_tx=XM_023243535.1
E1 19427221 19427368
refGene_gn=NF1;refGene_tx=XM_023243526.1;refGene_gn=NF1;refGene_tx=XM_023243527.1;refGene_gn=NF1;refGene_tx=XM_023243528.1;refGene_gn=NF1;refGene_tx=XM_023243529.1;refGene_gn=NF1;refGene_tx=XM_023243530.1;refGene_gn=NF1;refGene_tx=XM_023243531.1;refGene_gn=NF1;refGene_tx=XM_023243532.1;refGene_gn=NF1;refGene_tx=XM_023243533.1;refGene_gn=NF1;refGene_tx=XM_023243534.1;refGene_gn=NF1;refGene_tx=XM_023243535.1
E1 19429560 19429707
refGene_gn=NF1;refGene_tx=XM_023243526.1;refGene_gn=NF1;refGene_tx=XM_023243527.1;refGene_gn=NF1;refGene_tx=XM_023243528.1;refGene_gn=NF1;refGene_tx=XM_023243529.1;refGene_gn=NF1;refGene_tx=XM_023243530.1;refGene_gn=NF1;refGene_tx=XM_023243531.1;refGene_gn=NF1;refGene_tx=XM_023243532.1;refGene_gn=NF1;refGene_tx=XM_023243533.1;refGene_gn=NF1;refGene_tx=XM_023243534.1;refGene_gn=NF1;refGene_tx=XM_023243535.1
E1 19433252 19433363
refGene_gn=NF1;refGene_tx=XM_023243526.1;refGene_gn=NF1;refGene_tx=XM_023243527.1;refGene_gn=NF1;refGene_tx=XM_023243528.1;refGene_gn=NF1;refGene_tx=XM_023243529.1;refGene_gn=NF1;refGene_tx=XM_023243530.1;refGene_gn=NF1;refGene_tx=XM_023243531.1;refGene_gn=NF1;refGene_tx=XM_023243532.1;refGene_gn=NF1;refGene_tx=XM_023243533.1;refGene_gn=NF1;refGene_tx=XM_023243534.1;refGene_gn=NF1;refGene_tx=XM_023243535.1
E1 19457798 19459127 refGene_gn=OMG;refGene_tx=XM_006940053.3
E1 19459966 19459969 refGene_gn=OMG;refGene_tx=XM_006940053.3
E1 19468690 19470049
refGene_gn=EVI2B;refGene_tx=XM_003996528.3;refGene_gn=EVI2B;refGene_tx=XM_023243536.1
E1 19479557 19480262 refGene_gn=EVI2A;refGene_tx=XM_003996529.5
E1 19486540 19486973
refGene_gn=NF1;refGene_tx=XM_023243526.1;refGene_gn=NF1;refGene_tx=XM_023243527.1;refGene_gn=NF1;refGene_tx=XM_023243528.1;refGene_gn=NF1;refGene_tx=XM_023243529.1;refGene_gn=NF1;refGene_tx=XM_023243530.1;refGene_gn=NF1;refGene_tx=XM_023243531.1;refGene_gn=NF1;refGene_tx=XM_023243532.1;refGene_gn=NF1;refGene_tx=XM_023243533.1;refGene_gn=NF1;refGene_tx=XM_023243534.1;refGene_gn=NF1;refGene_tx=XM_023243535.1
E1 19487570 19487911
refGene_gn=NF1;refGene_tx=XM_023243526.1;refGene_gn=NF1;refGene_tx=XM_023243527.1;refGene_gn=NF1;refGene_tx=XM_023243528.1;refGene_gn=NF1;refGene_tx=XM_023243529.1;refGene_gn=NF1;refGene_tx=XM_023243530.1;refGene_gn=NF1;refGene_tx=XM_023243531.1;refGene_gn=NF1;refGene_tx=XM_023243532.1;refGene_gn=NF1;refGene_tx=XM_023243533.1;refGene_gn=NF1;refGene_tx=XM_023243534.1;refGene_gn=NF1;refGene_tx=XM_023243535.1
E1 19490249 19490452

refGene_gn=NF1;refGene_tx=XM_023243526.1;refGene_gn=NF1;refGene_tx=XM_023243527.1;refGene_gn=NF1;refGene_tx=XM_023243528.1;refGene_gn=NF1;refGene_tx=XM_023243529.1;refGene_gn=NF1;refGene_tx=XM_023243530.1;refGene_gn=NF1;refGene_tx=XM_023243531.1;refGene_gn=NF1;refGene_tx=XM_023243532.1;refGene_gn=NF1;refGene_tx=XM_023243533.1;refGene_gn=NF1;refGene_tx=XM_023243534.1;refGene_gn=NF1;refGene_tx=XM_023243535.1
E1 19494128 19494322
refGene_gn=NF1;refGene_tx=XM_023243526.1;refGene_gn=NF1;refGene_tx=XM_023243527.1;refGene_gn=NF1;refGene_tx=XM_023243528.1;refGene_gn=NF1;refGene_tx=XM_023243529.1;refGene_gn=NF1;refGene_tx=XM_023243530.1;refGene_gn=NF1;refGene_tx=XM_023243531.1;refGene_gn=NF1;refGene_tx=XM_023243532.1;refGene_gn=NF1;refGene_tx=XM_023243533.1;refGene_gn=NF1;refGene_tx=XM_023243534.1;refGene_gn=NF1;refGene_tx=XM_023243535.1
E1 19495275 19495416
refGene_gn=NF1;refGene_tx=XM_023243526.1;refGene_gn=NF1;refGene_tx=XM_023243527.1;refGene_gn=NF1;refGene_tx=XM_023243528.1;refGene_gn=NF1;refGene_tx=XM_023243529.1;refGene_gn=NF1;refGene_tx=XM_023243530.1;refGene_gn=NF1;refGene_tx=XM_023243531.1;refGene_gn=NF1;refGene_tx=XM_023243532.1;refGene_gn=NF1;refGene_tx=XM_023243533.1;refGene_gn=NF1;refGene_tx=XM_023243534.1;refGene_gn=NF1;refGene_tx=XM_023243535.1
E1 19495582 19495862
refGene_gn=NF1;refGene_tx=XM_023243526.1;refGene_gn=NF1;refGene_tx=XM_023243527.1;refGene_gn=NF1;refGene_tx=XM_023243528.1;refGene_gn=NF1;refGene_tx=XM_023243529.1;refGene_gn=NF1;refGene_tx=XM_023243530.1;refGene_gn=NF1;refGene_tx=XM_023243531.1;refGene_gn=NF1;refGene_tx=XM_023243532.1;refGene_gn=NF1;refGene_tx=XM_023243533.1;refGene_gn=NF1;refGene_tx=XM_023243534.1;refGene_gn=NF1;refGene_tx=XM_023243535.1
E1 19496328 19496543
refGene_gn=NF1;refGene_tx=XM_023243526.1;refGene_gn=NF1;refGene_tx=XM_023243527.1;refGene_gn=NF1;refGene_tx=XM_023243528.1;refGene_gn=NF1;refGene_tx=XM_023243529.1;refGene_gn=NF1;refGene_tx=XM_023243530.1;refGene_gn=NF1;refGene_tx=XM_023243531.1;refGene_gn=NF1;refGene_tx=XM_023243532.1;refGene_gn=NF1;refGene_tx=XM_023243533.1;refGene_gn=NF1;refGene_tx=XM_023243534.1;refGene_gn=NF1;refGene_tx=XM_023243535.1
E1 19496776 19496838
refGene_gn=NF1;refGene_tx=XM_023243526.1;refGene_gn=NF1;refGene_tx=XM_023243527.1;refGene_gn=NF1;refGene_tx=XM_023243528.1;refGene_gn=NF1;refGene_tx=XM_023243529.1;refGene_gn=NF1;refGene_tx=XM_023243530.1;refGene_gn=NF1;refGene_tx=XM_023243531.1;refGene_gn=NF1;refGene_tx=XM_023243532.1;refGene_gn=NF1;refGene_tx=XM_023243533.1;refGene_gn=NF1;refGene_tx=XM_023243534.1;refGene_gn=NF1;refGene_tx=XM_023243535.1
E1 19496981 19497096
refGene_gn=NF1;refGene_tx=XM_023243526.1;refGene_gn=NF1;refGene_tx=XM_023243527.1;refGene_gn=NF1;refGene_tx=XM_023243528.1;refGene_gn=NF1;refGene_tx=XM_023243529.1;refGene_gn=NF1;refGene_tx=XM_023243530.1;refGene_gn=NF1;refGene_tx=XM_023243531.1;refGene_gn=NF1;refGene_tx=XM_023243532.1;refGene_gn=NF1;refGene_tx=XM_023243533.1;refGene_gn=NF1;refGene_tx=XM_023243534.1;refGene_gn=NF1;refGene_tx=XM_023243535.1
E1 19497628 19497730
refGene_gn=NF1;refGene_tx=XM_023243526.1;refGene_gn=NF1;refGene_tx=XM_

023243527.1;refGene_gn=NF1;refGene_tx=XM_023243528.1;refGene_gn=NF1;re
fGene_tx=XM_023243529.1;refGene_gn=NF1;refGene_tx=XM_023243530.1;refGe
ne_gn=NF1;refGene_tx=XM_023243531.1;refGene_gn=NF1;refGene_tx=XM_02324
3532.1;refGene_gn=NF1;refGene_tx=XM_023243533.1;refGene_gn=NF1;refGene
_tx=XM_023243534.1;refGene_gn=NF1;refGene_tx=XM_023243535.1
E1 19499464 19499605
refGene_gn=NF1;refGene_tx=XM_023243526.1;refGene_gn=NF1;refGene_tx=XM_
023243527.1;refGene_gn=NF1;refGene_tx=XM_023243528.1;refGene_gn=NF1;re
fGene_tx=XM_023243529.1;refGene_gn=NF1;refGene_tx=XM_023243530.1;refGe
ne_gn=NF1;refGene_tx=XM_023243531.1;refGene_gn=NF1;refGene_tx=XM_02324
3532.1;refGene_gn=NF1;refGene_tx=XM_023243533.1;refGene_gn=NF1;refGene
_tx=XM_023243534.1;refGene_gn=NF1;refGene_tx=XM_023243535.1
E1 19504195 19504322
refGene_gn=NF1;refGene_tx=XM_023243526.1;refGene_gn=NF1;refGene_tx=XM_
023243527.1;refGene_gn=NF1;refGene_tx=XM_023243528.1;refGene_gn=NF1;re
fGene_tx=XM_023243529.1;refGene_gn=NF1;refGene_tx=XM_023243530.1;refGe
ne_gn=NF1;refGene_tx=XM_023243531.1;refGene_gn=NF1;refGene_tx=XM_02324
3532.1;refGene_gn=NF1;refGene_tx=XM_023243533.1;refGene_gn=NF1;refGene
_tx=XM_023243534.1;refGene_gn=NF1;refGene_tx=XM_023243535.1
E1 19505885 19506017
refGene_gn=NF1;refGene_tx=XM_023243526.1;refGene_gn=NF1;refGene_tx=XM_
023243527.1;refGene_gn=NF1;refGene_tx=XM_023243528.1;refGene_gn=NF1;re
fGene_tx=XM_023243529.1;refGene_gn=NF1;refGene_tx=XM_023243530.1;refGe
ne_gn=NF1;refGene_tx=XM_023243531.1;refGene_gn=NF1;refGene_tx=XM_02324
3532.1;refGene_gn=NF1;refGene_tx=XM_023243533.1;refGene_gn=NF1;refGene
_tx=XM_023243534.1;refGene_gn=NF1;refGene_tx=XM_023243535.1
E1 19507003 19507139
refGene_gn=NF1;refGene_tx=XM_023243526.1;refGene_gn=NF1;refGene_tx=XM_
023243527.1;refGene_gn=NF1;refGene_tx=XM_023243528.1;refGene_gn=NF1;re
fGene_tx=XM_023243529.1;refGene_gn=NF1;refGene_tx=XM_023243530.1;refGe
ne_gn=NF1;refGene_tx=XM_023243531.1;refGene_gn=NF1;refGene_tx=XM_02324
3532.1;refGene_gn=NF1;refGene_tx=XM_023243533.1;refGene_gn=NF1;refGene
_tx=XM_023243534.1;refGene_gn=NF1;refGene_tx=XM_023243535.1
E1 19508608 19508766
refGene_gn=NF1;refGene_tx=XM_023243526.1;refGene_gn=NF1;refGene_tx=XM_
023243527.1;refGene_gn=NF1;refGene_tx=XM_023243528.1;refGene_gn=NF1;re
fGene_tx=XM_023243529.1;refGene_gn=NF1;refGene_tx=XM_023243530.1;refGe
ne_gn=NF1;refGene_tx=XM_023243531.1;refGene_gn=NF1;refGene_tx=XM_02324
3532.1;refGene_gn=NF1;refGene_tx=XM_023243533.1;refGene_gn=NF1;refGene
_tx=XM_023243534.1;refGene_gn=NF1;refGene_tx=XM_023243535.1
E1 19511254 19511377
refGene_gn=NF1;refGene_tx=XM_023243526.1;refGene_gn=NF1;refGene_tx=XM_
023243527.1;refGene_gn=NF1;refGene_tx=XM_023243528.1;refGene_gn=NF1;re
fGene_tx=XM_023243529.1;refGene_gn=NF1;refGene_tx=XM_023243530.1;refGe
ne_gn=NF1;refGene_tx=XM_023243532.1;refGene_gn=NF1;refGene_tx=XM_02324
3533.1;refGene_gn=NF1;refGene_tx=XM_023243534.1;refGene_gn=NF1;refGene
_tx=XM_023243535.1
E1 19511767 19511898
refGene_gn=NF1;refGene_tx=XM_023243526.1;refGene_gn=NF1;refGene_tx=XM_
023243527.1;refGene_gn=NF1;refGene_tx=XM_023243528.1;refGene_gn=NF1;re

fGene_tx=XM_023243529.1;refGene_gn=NF1;refGene_tx=XM_023243530.1;refGene_gn=NF1;refGene_tx=XM_023243531.1;refGene_gn=NF1;refGene_tx=XM_023243532.1;refGene_gn=NF1;refGene_tx=XM_023243533.1;refGene_gn=NF1;refGene_tx=XM_023243534.1;refGene_gn=NF1;refGene_tx=XM_023243535.1
E1 19512085 19512186
refGene_gn=NF1;refGene_tx=XM_023243526.1;refGene_gn=NF1;refGene_tx=XM_023243527.1;refGene_gn=NF1;refGene_tx=XM_023243528.1;refGene_gn=NF1;refGene_tx=XM_023243529.1;refGene_gn=NF1;refGene_tx=XM_023243530.1;refGene_gn=NF1;refGene_tx=XM_023243531.1;refGene_gn=NF1;refGene_tx=XM_023243532.1;refGene_gn=NF1;refGene_tx=XM_023243533.1;refGene_gn=NF1;refGene_tx=XM_023243534.1;refGene_gn=NF1;refGene_tx=XM_023243535.1
E1 19513462 19513605
refGene_gn=NF1;refGene_tx=XM_023243526.1;refGene_gn=NF1;refGene_tx=XM_023243527.1;refGene_gn=NF1;refGene_tx=XM_023243528.1;refGene_gn=NF1;refGene_tx=XM_023243529.1;refGene_gn=NF1;refGene_tx=XM_023243530.1;refGene_gn=NF1;refGene_tx=XM_023243531.1;refGene_gn=NF1;refGene_tx=XM_023243532.1;refGene_gn=NF1;refGene_tx=XM_023243533.1;refGene_gn=NF1;refGene_tx=XM_023243534.1;refGene_gn=NF1;refGene_tx=XM_023243535.1
E1 19513988 19514035
refGene_gn=NF1;refGene_tx=XM_023243526.1;refGene_gn=NF1;refGene_tx=XM_023243527.1;refGene_gn=NF1;refGene_tx=XM_023243528.1;refGene_gn=NF1;refGene_tx=XM_023243529.1;refGene_gn=NF1;refGene_tx=XM_023243530.1;refGene_gn=NF1;refGene_tx=XM_023243531.1;refGene_gn=NF1;refGene_tx=XM_023243532.1;refGene_gn=NF1;refGene_tx=XM_023243533.1;refGene_gn=NF1;refGene_tx=XM_023243534.1;refGene_gn=NF1;refGene_tx=XM_023243535.1
E1 19515245 19515462
refGene_gn=NF1;refGene_tx=XM_023243526.1;refGene_gn=NF1;refGene_tx=XM_023243527.1;refGene_gn=NF1;refGene_tx=XM_023243528.1;refGene_gn=NF1;refGene_tx=XM_023243529.1;refGene_gn=NF1;refGene_tx=XM_023243530.1;refGene_gn=NF1;refGene_tx=XM_023243531.1;refGene_gn=NF1;refGene_tx=XM_023243532.1;refGene_gn=NF1;refGene_tx=XM_023243533.1;refGene_gn=NF1;refGene_tx=XM_023243535.1
E1 19516810 19516909
refGene_gn=NF1;refGene_tx=XM_023243526.1;refGene_gn=NF1;refGene_tx=XM_023243527.1;refGene_gn=NF1;refGene_tx=XM_023243528.1;refGene_gn=NF1;refGene_tx=XM_023243529.1;refGene_gn=NF1;refGene_tx=XM_023243531.1;refGene_gn=NF1;refGene_tx=XM_023243535.1
E1 19526910 19527087
refGene_gn=NF1;refGene_tx=XM_023243526.1;refGene_gn=NF1;refGene_tx=XM_023243527.1;refGene_gn=NF1;refGene_tx=XM_023243528.1;refGene_gn=NF1;refGene_tx=XM_023243529.1;refGene_gn=NF1;refGene_tx=XM_023243530.1;refGene_gn=NF1;refGene_tx=XM_023243531.1;refGene_gn=NF1;refGene_tx=XM_023243532.1;refGene_gn=NF1;refGene_tx=XM_023243533.1;refGene_gn=NF1;refGene_tx=XM_023243534.1;refGene_gn=NF1;refGene_tx=XM_023243535.1
E1 19543693 19543852
refGene_gn=RAB11FIP4;refGene_tx=XM_003996530.5
E1 19587176 19587264
refGene_gn=RAB11FIP4;refGene_tx=XM_003996530.5
E1 19588629 19588718
refGene_gn=RAB11FIP4;refGene_tx=XM_003996530.5

E1 19630614 19630644
refGene_gn=RAB11FIP4;refGene_tx=XM_011288972.3
E1 19652187 19652417
refGene_gn=RAB11FIP4;refGene_tx=XM_003996530.5;refGene_gn=RAB11FIP4;refGene_tx=XM_011288972.3
E1 19654707 19654902
refGene_gn=RAB11FIP4;refGene_tx=XM_003996530.5;refGene_gn=RAB11FIP4;refGene_tx=XM_011288972.3
E1 19655449 19655584
refGene_gn=RAB11FIP4;refGene_tx=XM_003996530.5;refGene_gn=RAB11FIP4;refGene_tx=XM_011288972.3
E1 19655930 19655966
refGene_gn=RAB11FIP4;refGene_tx=XM_003996530.5;refGene_gn=RAB11FIP4;refGene_tx=XM_011288972.3
E1 19657155 19657255
refGene_gn=RAB11FIP4;refGene_tx=XM_003996530.5;refGene_gn=RAB11FIP4;refGene_tx=XM_011288972.3
E1 19657521 19657625
refGene_gn=RAB11FIP4;refGene_tx=XM_003996530.5;refGene_gn=RAB11FIP4;refGene_tx=XM_011288972.3
E1 19658806 19658947
refGene_gn=RAB11FIP4;refGene_tx=XM_003996530.5;refGene_gn=RAB11FIP4;refGene_tx=XM_011288972.3
E1 19660367 19660449
refGene_gn=RAB11FIP4;refGene_tx=XM_003996530.5;refGene_gn=RAB11FIP4;refGene_tx=XM_011288972.3
E1 19660934 19661072
refGene_gn=RAB11FIP4;refGene_tx=XM_003996530.5;refGene_gn=RAB11FIP4;refGene_tx=XM_011288972.3
E1 19661155 19661314
refGene_gn=RAB11FIP4;refGene_tx=XM_003996530.5;refGene_gn=RAB11FIP4;refGene_tx=XM_011288972.3
E1 19662352 19662496
refGene_gn=RAB11FIP4;refGene_tx=XM_003996530.5;refGene_gn=RAB11FIP4;refGene_tx=XM_011288972.3
E1 19663673 19663790
refGene_gn=RAB11FIP4;refGene_tx=XM_003996530.5;refGene_gn=RAB11FIP4;refGene_tx=XM_011288972.3
E1 19932464 19932634
refGene_gn=COPRS;refGene_tx=XM_006940073.4;refGene_gn=COPRS;refGene_tx=XM_023243545.1
E1 19933544 19933766
refGene_gn=COPRS;refGene_tx=XM_006940073.4;refGene_gn=COPRS;refGene_tx=XM_023243545.1
E1 19935411 19935478
refGene_gn=COPRS;refGene_tx=XM_006940073.4;refGene_gn=COPRS;refGene_tx=XM_023243545.1
E1 19937544 19937607 refGene_gn=COPRS;refGene_tx=XM_006940073.4
E1 19937859 19937958 refGene_gn=COPRS;refGene_tx=XM_023243545.1
E1 19941659 19941817 refGene_gn=UTP6;refGene_tx=XM_023243546.1

E1	19942743	19942816	refGene_gn=UTP6;refGene_tx=XM_023243546.1
E1	19946441	19946508	refGene_gn=UTP6;refGene_tx=XM_023243546.1
E1	19947623	19947733	refGene_gn=UTP6;refGene_tx=XM_023243546.1
E1	19947833	19947913	refGene_gn=UTP6;refGene_tx=XM_023243546.1
E1	19950041	19950234	refGene_gn=UTP6;refGene_tx=XM_023243546.1
E1	19955669	19955747	refGene_gn=UTP6;refGene_tx=XM_023243546.1
E1	19956102	19956182	refGene_gn=UTP6;refGene_tx=XM_023243546.1
E1	19956864	19957049	refGene_gn=UTP6;refGene_tx=XM_023243546.1
E1	19958141	19958223	refGene_gn=UTP6;refGene_tx=XM_023243546.1
E1	19959264	19959346	refGene_gn=UTP6;refGene_tx=XM_023243546.1
E1	19960102	19960180	refGene_gn=UTP6;refGene_tx=XM_023243546.1
E1	19962118	19962237	refGene_gn=UTP6;refGene_tx=XM_023243546.1
E1	19964255	19964319	refGene_gn=UTP6;refGene_tx=XM_023243546.1
E1	19964760	19964808	refGene_gn=UTP6;refGene_tx=XM_023243546.1
E1	19965902	19965995	refGene_gn=UTP6;refGene_tx=XM_023243546.1
E1	19966445	19966487	refGene_gn=UTP6;refGene_tx=XM_023243546.1
E1	19969359	19969444	refGene_gn=UTP6;refGene_tx=XM_023243546.1
E1	19970914	19971006	refGene_gn=UTP6;refGene_tx=XM_023243546.1
E1	19996466	19996737
refGene_gn=SUZ12;refGene_tx=XM_011289064.3;refGene_gn=SUZ12;refGene_tx=XM_023244424.1;refGene_gn=SUZ12;refGene_tx=XM_023244425.1
E1	19999109	19999156
refGene_gn=SUZ12;refGene_tx=XM_011289064.3;refGene_gn=SUZ12;refGene_tx=XM_023244424.1;refGene_gn=SUZ12;refGene_tx=XM_023244425.1
E1	19999243	19999308
refGene_gn=SUZ12;refGene_tx=XM_011289064.3;refGene_gn=SUZ12;refGene_tx=XM_023244424.1;refGene_gn=SUZ12;refGene_tx=XM_023244425.1
E1	20003562	20003631
refGene_gn=SUZ12;refGene_tx=XM_011289064.3;refGene_gn=SUZ12;refGene_tx=XM_023244425.1
E1	20014644	20014694
refGene_gn=SUZ12;refGene_tx=XM_011289064.3;refGene_gn=SUZ12;refGene_tx=XM_023244424.1;refGene_gn=SUZ12;refGene_tx=XM_023244425.1
E1	20019033	20019119
refGene_gn=SUZ12;refGene_tx=XM_011289064.3;refGene_gn=SUZ12;refGene_tx=XM_023244424.1;refGene_gn=SUZ12;refGene_tx=XM_023244425.1
E1	20020731	20020963
refGene_gn=SUZ12;refGene_tx=XM_011289064.3;refGene_gn=SUZ12;refGene_tx=XM_023244424.1;refGene_gn=SUZ12;refGene_tx=XM_023244425.1
E1	20021875	20021969
refGene_gn=SUZ12;refGene_tx=XM_011289064.3;refGene_gn=SUZ12;refGene_tx=XM_023244424.1;refGene_gn=SUZ12;refGene_tx=XM_023244425.1
E1	20028125	20028231
refGene_gn=SUZ12;refGene_tx=XM_011289064.3;refGene_gn=SUZ12;refGene_tx=XM_023244424.1;refGene_gn=SUZ12;refGene_tx=XM_023244425.1
E1	20031541	20031719
refGene_gn=SUZ12;refGene_tx=XM_011289064.3;refGene_gn=SUZ12;refGene_tx=XM_023244424.1;refGene_gn=SUZ12;refGene_tx=XM_023244425.1
E1	20033548	20033640
refGene_gn=SUZ12;refGene_tx=XM_011289064.3;refGene_gn=SUZ12;refGene_tx

=XM_023244424.1;refGene_gn=SUZ12;refGene_tx=XM_023244425.1
E1 20033845 20033989
refGene_gn=SUZ12;refGene_tx=XM_011289064.3;refGene_gn=SUZ12;refGene_tx=XM_023244424.1;refGene_gn=SUZ12;refGene_tx=XM_023244425.1
E1 20034787 20034945
refGene_gn=SUZ12;refGene_tx=XM_011289064.3;refGene_gn=SUZ12;refGene_tx=XM_023244424.1;refGene_gn=SUZ12;refGene_tx=XM_023244425.1
E1 20035328 20035527
refGene_gn=SUZ12;refGene_tx=XM_011289064.3;refGene_gn=SUZ12;refGene_tx=XM_023244424.1;refGene_gn=SUZ12;refGene_tx=XM_023244425.1
E1 20037426 20037506
refGene_gn=SUZ12;refGene_tx=XM_011289064.3;refGene_gn=SUZ12;refGene_tx=XM_023244424.1;refGene_gn=SUZ12;refGene_tx=XM_023244425.1
E1 20040003 20040349
refGene_gn=SUZ12;refGene_tx=XM_011289064.3;refGene_gn=SUZ12;refGene_tx=XM_023244424.1
E1 20045439 20045503 refGene_gn=SUZ12;refGene_tx=XM_023244425.1
E1 20055476 20055733 refGene_gn=CRLF3;refGene_tx=XM_003996532.4
E1 20056594 20056707 refGene_gn=CRLF3;refGene_tx=XM_003996532.4
E1 20075060 20075193 refGene_gn=CRLF3;refGene_tx=XM_003996532.4
E1 20075872 20076095 refGene_gn=CRLF3;refGene_tx=XM_003996532.4
E1 20078650 20078828 refGene_gn=CRLF3;refGene_tx=XM_003996532.4
E1 20079319 20079407 refGene_gn=CRLF3;refGene_tx=XM_003996532.4
E1 20082357 20082565 refGene_gn=CRLF3;refGene_tx=XM_003996532.4
E1 20097544 20097661 refGene_gn=CRLF3;refGene_tx=XM_003996532.4
E1 20107973 20108039
refGene_gn=ATAD5;refGene_tx=XM_023243726.1;refGene_gn=ATAD5;refGene_tx=XM_023243727.1
E1 20108349 20108352 refGene_gn=ATAD5;refGene_tx=XM_023243728.1
E1 20112293 20114188
refGene_gn=ATAD5;refGene_tx=XM_023243726.1;refGene_gn=ATAD5;refGene_tx=XM_023243727.1;refGene_gn=ATAD5;refGene_tx=XM_023243728.1
E1 20114983 20115092
refGene_gn=ATAD5;refGene_tx=XM_023243726.1;refGene_gn=ATAD5;refGene_tx=XM_023243727.1;refGene_gn=ATAD5;refGene_tx=XM_023243728.1
E1 20118396 20118561
refGene_gn=ATAD5;refGene_tx=XM_023243726.1;refGene_gn=ATAD5;refGene_tx=XM_023243727.1;refGene_gn=ATAD5;refGene_tx=XM_023243728.1
E1 20120309 20120436
refGene_gn=ATAD5;refGene_tx=XM_023243726.1;refGene_gn=ATAD5;refGene_tx=XM_023243727.1;refGene_gn=ATAD5;refGene_tx=XM_023243728.1
E1 20121491 20121573
refGene_gn=ATAD5;refGene_tx=XM_023243726.1;refGene_gn=ATAD5;refGene_tx=XM_023243727.1;refGene_gn=ATAD5;refGene_tx=XM_023243728.1
E1 20129847 20130032
refGene_gn=ATAD5;refGene_tx=XM_023243726.1;refGene_gn=ATAD5;refGene_tx=XM_023243727.1;refGene_gn=ATAD5;refGene_tx=XM_023243728.1
E1 20131975 20132133
refGene_gn=ATAD5;refGene_tx=XM_023243726.1;refGene_gn=ATAD5;refGene_tx=XM_023243727.1;refGene_gn=ATAD5;refGene_tx=XM_023243728.1

E1 20133189 20133349
refGene_gn=ATAD5;refGene_tx=XM_023243726.1;refGene_gn=ATAD5;refGene_tx
=XM_023243727.1;refGene_gn=ATAD5;refGene_tx=XM_023243728.1
E1 20133812 20134142
refGene_gn=ATAD5;refGene_tx=XM_023243726.1;refGene_gn=ATAD5;refGene_tx
=XM_023243727.1;refGene_gn=ATAD5;refGene_tx=XM_023243728.1
E1 20135699 20135796
refGene_gn=ATAD5;refGene_tx=XM_023243726.1;refGene_gn=ATAD5;refGene_tx
=XM_023243727.1;refGene_gn=ATAD5;refGene_tx=XM_023243728.1
E1 20137197 20137277
refGene_gn=ATAD5;refGene_tx=XM_023243726.1;refGene_gn=ATAD5;refGene_tx
=XM_023243727.1;refGene_gn=ATAD5;refGene_tx=XM_023243728.1
E1 20137457 20137600
refGene_gn=ATAD5;refGene_tx=XM_023243726.1;refGene_gn=ATAD5;refGene_tx
=XM_023243727.1;refGene_gn=ATAD5;refGene_tx=XM_023243728.1
E1 20137697 20137848
refGene_gn=ATAD5;refGene_tx=XM_023243726.1;refGene_gn=ATAD5;refGene_tx
=XM_023243727.1;refGene_gn=ATAD5;refGene_tx=XM_023243728.1
E1 20143430 20143604
refGene_gn=ATAD5;refGene_tx=XM_023243726.1;refGene_gn=ATAD5;refGene_tx
=XM_023243727.1;refGene_gn=ATAD5;refGene_tx=XM_023243728.1
E1 20144346 20144480
refGene_gn=ATAD5;refGene_tx=XM_023243726.1;refGene_gn=ATAD5;refGene_tx
=XM_023243727.1;refGene_gn=ATAD5;refGene_tx=XM_023243728.1
E1 20145188 20145282
refGene_gn=ATAD5;refGene_tx=XM_023243726.1;refGene_gn=ATAD5;refGene_tx
=XM_023243727.1;refGene_gn=ATAD5;refGene_tx=XM_023243728.1
E1 20146541 20146606
refGene_gn=ATAD5;refGene_tx=XM_023243726.1;refGene_gn=ATAD5;refGene_tx
=XM_023243727.1;refGene_gn=ATAD5;refGene_tx=XM_023243728.1
E1 20148618 20148799
refGene_gn=ATAD5;refGene_tx=XM_023243726.1;refGene_gn=ATAD5;refGene_tx
=XM_023243727.1;refGene_gn=ATAD5;refGene_tx=XM_023243728.1
E1 20150676 20150858
refGene_gn=ATAD5;refGene_tx=XM_023243726.1;refGene_gn=ATAD5;refGene_tx
=XM_023243727.1;refGene_gn=ATAD5;refGene_tx=XM_023243728.1
E1 20152157 20153023
refGene_gn=ATAD5;refGene_tx=XM_023243726.1;refGene_gn=ATAD5;refGene_tx
=XM_023243727.1;refGene_gn=ATAD5;refGene_tx=XM_023243728.1
E1 20153706 20153865
refGene_gn=ATAD5;refGene_tx=XM_023243726.1;refGene_gn=ATAD5;refGene_tx
=XM_023243727.1;refGene_gn=ATAD5;refGene_tx=XM_023243728.1
E1 20153969 20154048
refGene_gn=ATAD5;refGene_tx=XM_023243726.1;refGene_gn=ATAD5;refGene_tx
=XM_023243727.1;refGene_gn=ATAD5;refGene_tx=XM_023243728.1
E1 20157389 20157827
refGene_gn=TEFM;refGene_tx=XM_003996533.4;refGene_gn=TEFM;refGene_tx=X
M_006940055.3;refGene_gn=TEFM;refGene_tx=XM_006940056.3
E1 20159643 20159793
refGene_gn=TEFM;refGene_tx=XM_003996533.4;refGene_gn=TEFM;refGene_tx=X

M_006940055.3
E1	20162627	20163091
refGene_gn=TEFM;refGene_tx=XM_003996533.4;refGene_gn=TEFM;refGene_tx=XM_006940055.3;refGene_gn=TEFM;refGene_tx=XM_006940056.3
E1	20163807	20163859
refGene_gn=TEFM;refGene_tx=XM_003996533.4;refGene_gn=TEFM;refGene_tx=XM_006940055.3;refGene_gn=TEFM;refGene_tx=XM_006940056.3
E1	20180038	20180132
refGene_gn=ADAP2;refGene_tx=XM_023244215.1;refGene_gn=ADAP2;refGene_tx=XM_023244216.1;refGene_gn=ADAP2;refGene_tx=XM_023244218.1;refGene_gn=ADAP2;refGene_tx=XM_023244219.1;refGene_gn=ADAP2;refGene_tx=XM_023244220.1;refGene_gn=ADAP2;refGene_tx=XM_023244221.1
E1	20181024	20181155
refGene_gn=ADAP2;refGene_tx=XM_023244215.1;refGene_gn=ADAP2;refGene_tx=XM_023244216.1;refGene_gn=ADAP2;refGene_tx=XM_023244218.1;refGene_gn=ADAP2;refGene_tx=XM_023244219.1;refGene_gn=ADAP2;refGene_tx=XM_023244220.1;refGene_gn=ADAP2;refGene_tx=XM_023244221.1
E1	20182778	20182870
refGene_gn=ADAP2;refGene_tx=XM_023244215.1;refGene_gn=ADAP2;refGene_tx=XM_023244216.1;refGene_gn=ADAP2;refGene_tx=XM_023244218.1;refGene_gn=ADAP2;refGene_tx=XM_023244219.1;refGene_gn=ADAP2;refGene_tx=XM_023244220.1;refGene_gn=ADAP2;refGene_tx=XM_023244221.1;refGene_gn=ADAP2;refGene_tx=XM_023244222.1
E1	20185968	20186048
refGene_gn=ADAP2;refGene_tx=XM_023244215.1;refGene_gn=ADAP2;refGene_tx=XM_023244216.1;refGene_gn=ADAP2;refGene_tx=XM_023244218.1;refGene_gn=ADAP2;refGene_tx=XM_023244219.1;refGene_gn=ADAP2;refGene_tx=XM_023244220.1;refGene_gn=ADAP2;refGene_tx=XM_023244221.1;refGene_gn=ADAP2;refGene_tx=XM_023244222.1
E1	20186959	20187072
refGene_gn=ADAP2;refGene_tx=XM_023244215.1;refGene_gn=ADAP2;refGene_tx=XM_023244216.1;refGene_gn=ADAP2;refGene_tx=XM_023244218.1;refGene_gn=ADAP2;refGene_tx=XM_023244219.1;refGene_gn=ADAP2;refGene_tx=XM_023244220.1;refGene_gn=ADAP2;refGene_tx=XM_023244221.1;refGene_gn=ADAP2;refGene_tx=XM_023244222.1
E1	20199521	20199668
refGene_gn=ADAP2;refGene_tx=XM_023244215.1;refGene_gn=ADAP2;refGene_tx=XM_023244216.1;refGene_gn=ADAP2;refGene_tx=XM_023244218.1;refGene_gn=ADAP2;refGene_tx=XM_023244219.1;refGene_gn=ADAP2;refGene_tx=XM_023244220.1;refGene_gn=ADAP2;refGene_tx=XM_023244222.1
E1	20204792	20204876
refGene_gn=ADAP2;refGene_tx=XM_023244215.1;refGene_gn=ADAP2;refGene_tx=XM_023244216.1;refGene_gn=ADAP2;refGene_tx=XM_023244218.1;refGene_gn=ADAP2;refGene_tx=XM_023244219.1;refGene_gn=ADAP2;refGene_tx=XM_023244221.1;refGene_gn=ADAP2;refGene_tx=XM_023244222.1
E1	20208124	20208187
refGene_gn=ADAP2;refGene_tx=XM_023244215.1;refGene_gn=ADAP2;refGene_tx=XM_023244216.1;refGene_gn=ADAP2;refGene_tx=XM_023244218.1;refGene_gn=ADAP2;refGene_tx=XM_023244219.1;refGene_gn=ADAP2;refGene_tx=XM_023244220.1;refGene_gn=ADAP2;refGene_tx=XM_023244221.1;refGene_gn=ADAP2;refGe

ne_tx=XM_023244222.1
E1 20209498 20209576
refGene_gn=ADAP2;refGene_tx=XM_023244215.1;refGene_gn=ADAP2;refGene_tx=XM_023244216.1;refGene_gn=ADAP2;refGene_tx=XM_023244218.1;refGene_gn=ADAP2;refGene_tx=XM_023244219.1;refGene_gn=ADAP2;refGene_tx=XM_023244220.1;refGene_gn=ADAP2;refGene_tx=XM_023244221.1;refGene_gn=ADAP2;refGene_tx=XM_023244222.1
E1 20210222 20210451
refGene_gn=ADAP2;refGene_tx=XM_023244215.1;refGene_gn=ADAP2;refGene_tx=XM_023244216.1;refGene_gn=ADAP2;refGene_tx=XM_023244218.1;refGene_gn=ADAP2;refGene_tx=XM_023244219.1;refGene_gn=ADAP2;refGene_tx=XM_023244220.1;refGene_gn=ADAP2;refGene_tx=XM_023244221.1;refGene_gn=ADAP2;refGene_tx=XM_023244222.1
E1 20211624 20211713
refGene_gn=ADAP2;refGene_tx=XM_023244215.1;refGene_gn=ADAP2;refGene_tx=XM_023244218.1;refGene_gn=ADAP2;refGene_tx=XM_023244220.1;refGene_gn=ADAP2;refGene_tx=XM_023244221.1;refGene_gn=ADAP2;refGene_tx=XM_023244222.1
E1 20216255 20216269 refGene_gn=ADAP2;refGene_tx=XM_023244219.1
E1 20219242 20219605 refGene_gn=RNF135;refGene_tx=XM_023244214.1
E1 20225180 20225324
refGene_gn=ADAP2;refGene_tx=XM_023244216.1;refGene_gn=RNF135;refGene_tx=XM_023244214.1
E1 20228676 20228797 refGene_gn=RNF135;refGene_tx=XM_023244214.1
E1 20232711 20232801 refGene_gn=RNF135;refGene_tx=XM_023244214.1
E1 20234666 20235196 refGene_gn=RNF135;refGene_tx=XM_023244214.1
E1 20247517 20247593
refGene_gn=RHOT1;refGene_tx=XM_023244210.1;refGene_gn=RHOT1;refGene_tx=XM_023244211.1;refGene_gn=RHOT1;refGene_tx=XM_023244212.1;refGene_gn=RHOT1;refGene_tx=XM_023244213.1
E1 20266139 20266198
refGene_gn=RHOT1;refGene_tx=XM_006940065.4;refGene_gn=RHOT1;refGene_tx=XM_019817498.2;refGene_gn=RHOT1;refGene_tx=XM_019817499.2;refGene_gn=RHOT1;refGene_tx=XM_023244210.1;refGene_gn=RHOT1;refGene_tx=XM_023244211.1;refGene_gn=RHOT1;refGene_tx=XM_023244212.1;refGene_gn=RHOT1;refGene_tx=XM_023244213.1
E1 20268837 20268919
refGene_gn=RHOT1;refGene_tx=XM_006940065.4;refGene_gn=RHOT1;refGene_tx=XM_019817498.2;refGene_gn=RHOT1;refGene_tx=XM_019817499.2;refGene_gn=RHOT1;refGene_tx=XM_023244210.1;refGene_gn=RHOT1;refGene_tx=XM_023244211.1;refGene_gn=RHOT1;refGene_tx=XM_023244212.1;refGene_gn=RHOT1;refGene_tx=XM_023244213.1
E1 20270820 20270864
refGene_gn=RHOT1;refGene_tx=XM_006940065.4;refGene_gn=RHOT1;refGene_tx=XM_019817498.2;refGene_gn=RHOT1;refGene_tx=XM_019817499.2;refGene_gn=RHOT1;refGene_tx=XM_023244210.1;refGene_gn=RHOT1;refGene_tx=XM_023244211.1;refGene_gn=RHOT1;refGene_tx=XM_023244212.1;refGene_gn=RHOT1;refGene_tx=XM_023244213.1
E1 20271167 20271221
refGene_gn=RHOT1;refGene_tx=XM_006940065.4;refGene_gn=RHOT1;refGene_tx

=XM_019817498.2;refGene_gn=RHOT1;refGene_tx=XM_019817499.2;refGene_gn=RHOT1;refGene_tx=XM_023244210.1;refGene_gn=RHOT1;refGene_tx=XM_023244211.1;refGene_gn=RHOT1;refGene_tx=XM_023244212.1;refGene_gn=RHOT1;refGene_tx=XM_023244213.1
E1 20271533 20271586
refGene_gn=RHOT1;refGene_tx=XM_006940065.4;refGene_gn=RHOT1;refGene_tx=XM_019817498.2;refGene_gn=RHOT1;refGene_tx=XM_019817499.2;refGene_gn=RHOT1;refGene_tx=XM_023244210.1;refGene_gn=RHOT1;refGene_tx=XM_023244211.1;refGene_gn=RHOT1;refGene_tx=XM_023244212.1;refGene_gn=RHOT1;refGene_tx=XM_023244213.1
E1 20274909 20275018
refGene_gn=RHOT1;refGene_tx=XM_006940065.4;refGene_gn=RHOT1;refGene_tx=XM_019817498.2;refGene_gn=RHOT1;refGene_tx=XM_019817499.2;refGene_gn=RHOT1;refGene_tx=XM_023244210.1;refGene_gn=RHOT1;refGene_tx=XM_023244211.1;refGene_gn=RHOT1;refGene_tx=XM_023244212.1;refGene_gn=RHOT1;refGene_tx=XM_023244213.1
E1 20275326 20275428
refGene_gn=RHOT1;refGene_tx=XM_006940065.4;refGene_gn=RHOT1;refGene_tx=XM_019817498.2;refGene_gn=RHOT1;refGene_tx=XM_019817499.2;refGene_gn=RHOT1;refGene_tx=XM_023244210.1;refGene_gn=RHOT1;refGene_tx=XM_023244211.1;refGene_gn=RHOT1;refGene_tx=XM_023244212.1;refGene_gn=RHOT1;refGene_tx=XM_023244213.1
E1 20283825 20283924
refGene_gn=RHOT1;refGene_tx=XM_006940065.4;refGene_gn=RHOT1;refGene_tx=XM_019817498.2;refGene_gn=RHOT1;refGene_tx=XM_019817499.2;refGene_gn=RHOT1;refGene_tx=XM_023244210.1;refGene_gn=RHOT1;refGene_tx=XM_023244211.1;refGene_gn=RHOT1;refGene_tx=XM_023244212.1;refGene_gn=RHOT1;refGene_tx=XM_023244213.1
E1 20284449 20284558
refGene_gn=RHOT1;refGene_tx=XM_006940065.4;refGene_gn=RHOT1;refGene_tx=XM_019817498.2;refGene_gn=RHOT1;refGene_tx=XM_019817499.2;refGene_gn=RHOT1;refGene_tx=XM_023244210.1;refGene_gn=RHOT1;refGene_tx=XM_023244211.1;refGene_gn=RHOT1;refGene_tx=XM_023244212.1;refGene_gn=RHOT1;refGene_tx=XM_023244213.1
E1 20285314 20285435
refGene_gn=RHOT1;refGene_tx=XM_006940065.4;refGene_gn=RHOT1;refGene_tx=XM_019817498.2;refGene_gn=RHOT1;refGene_tx=XM_019817499.2;refGene_gn=RHOT1;refGene_tx=XM_023244210.1;refGene_gn=RHOT1;refGene_tx=XM_023244211.1;refGene_gn=RHOT1;refGene_tx=XM_023244212.1;refGene_gn=RHOT1;refGene_tx=XM_023244213.1
E1 20287623 20287708
refGene_gn=RHOT1;refGene_tx=XM_006940065.4;refGene_gn=RHOT1;refGene_tx=XM_019817498.2;refGene_gn=RHOT1;refGene_tx=XM_019817499.2;refGene_gn=RHOT1;refGene_tx=XM_023244210.1;refGene_gn=RHOT1;refGene_tx=XM_023244211.1;refGene_gn=RHOT1;refGene_tx=XM_023244212.1;refGene_gn=RHOT1;refGene_tx=XM_023244213.1
E1 20288077 20288223
refGene_gn=RHOT1;refGene_tx=XM_006940065.4;refGene_gn=RHOT1;refGene_tx=XM_019817498.2;refGene_gn=RHOT1;refGene_tx=XM_019817499.2;refGene_gn=RHOT1;refGene_tx=XM_023244210.1;refGene_gn=RHOT1;refGene_tx=XM_0232442

11.1;refGene_gn=RHOT1;refGene_tx=XM_023244212.1;refGene_gn=RHOT1;refGene_tx=XM_023244213.1
E1 20289637 20289738
refGene_gn=RHOT1;refGene_tx=XM_006940065.4;refGene_gn=RHOT1;refGene_tx=XM_019817498.2;refGene_gn=RHOT1;refGene_tx=XM_019817499.2;refGene_gn=RHOT1;refGene_tx=XM_023244210.1;refGene_gn=RHOT1;refGene_tx=XM_023244211.1;refGene_gn=RHOT1;refGene_tx=XM_023244212.1;refGene_gn=RHOT1;refGene_tx=XM_023244213.1
E1 20291779 20291910
refGene_gn=RHOT1;refGene_tx=XM_006940065.4;refGene_gn=RHOT1;refGene_tx=XM_019817498.2;refGene_gn=RHOT1;refGene_tx=XM_019817499.2;refGene_gn=RHOT1;refGene_tx=XM_023244210.1;refGene_gn=RHOT1;refGene_tx=XM_023244211.1;refGene_gn=RHOT1;refGene_tx=XM_023244212.1;refGene_gn=RHOT1;refGene_tx=XM_023244213.1
E1 20292629 20292713
refGene_gn=RHOT1;refGene_tx=XM_006940065.4;refGene_gn=RHOT1;refGene_tx=XM_019817498.2;refGene_gn=RHOT1;refGene_tx=XM_019817499.2;refGene_gn=RHOT1;refGene_tx=XM_023244210.1;refGene_gn=RHOT1;refGene_tx=XM_023244211.1;refGene_gn=RHOT1;refGene_tx=XM_023244212.1;refGene_gn=RHOT1;refGene_tx=XM_023244213.1
E1 20294284 20294404
refGene_gn=RHOT1;refGene_tx=XM_006940065.4;refGene_gn=RHOT1;refGene_tx=XM_019817498.2;refGene_gn=RHOT1;refGene_tx=XM_019817499.2;refGene_gn=RHOT1;refGene_tx=XM_023244210.1;refGene_gn=RHOT1;refGene_tx=XM_023244211.1;refGene_gn=RHOT1;refGene_tx=XM_023244212.1;refGene_gn=RHOT1;refGene_tx=XM_023244213.1
E1 20296791 20296994
refGene_gn=RHOT1;refGene_tx=XM_006940065.4;refGene_gn=RHOT1;refGene_tx=XM_019817498.2;refGene_gn=RHOT1;refGene_tx=XM_019817499.2;refGene_gn=RHOT1;refGene_tx=XM_023244210.1;refGene_gn=RHOT1;refGene_tx=XM_023244211.1;refGene_gn=RHOT1;refGene_tx=XM_023244212.1;refGene_gn=RHOT1;refGene_tx=XM_023244213.1
E1 20297978 20298074
refGene_gn=RHOT1;refGene_tx=XM_006940065.4;refGene_gn=RHOT1;refGene_tx=XM_019817498.2;refGene_gn=RHOT1;refGene_tx=XM_019817499.2;refGene_gn=RHOT1;refGene_tx=XM_023244210.1;refGene_gn=RHOT1;refGene_tx=XM_023244212.1
E1 20299736 20299859
refGene_gn=RHOT1;refGene_tx=XM_006940065.4;refGene_gn=RHOT1;refGene_tx=XM_019817498.2;refGene_gn=RHOT1;refGene_tx=XM_019817499.2;refGene_gn=RHOT1;refGene_tx=XM_023244210.1;refGene_gn=RHOT1;refGene_tx=XM_023244211.1
E1 20311060 20311178
refGene_gn=RHOT1;refGene_tx=XM_006940065.4;refGene_gn=RHOT1;refGene_tx=XM_019817498.2;refGene_gn=RHOT1;refGene_tx=XM_019817499.2;refGene_gn=RHOT1;refGene_tx=XM_023244210.1;refGene_gn=RHOT1;refGene_tx=XM_023244211.1;refGene_gn=RHOT1;refGene_tx=XM_023244212.1;refGene_gn=RHOT1;refGene_tx=XM_023244213.1
E1 20350599 20350710
refGene_gn=RHBDL3;refGene_tx=XM_023243711.1;refGene_gn=RHBDL3;refGene_

tx=XM_023243713.1;refGene_gn=RHBDL3;refGene_tx=XM_023243714.1
E1 20352249 20352273
refGene_gn=RHBDL3;refGene_tx=XM_023243711.1;refGene_gn=RHBDL3;refGene_tx=XM_023243714.1
E1 20361621 20361738 refGene_gn=RHBDL3;refGene_tx=XM_023243712.1
E1 20366476 20366635
refGene_gn=RHBDL3;refGene_tx=XM_023243711.1;refGene_gn=RHBDL3;refGene_tx=XM_023243712.1;refGene_gn=RHBDL3;refGene_tx=XM_023243713.1
E1 20370538 20370763
refGene_gn=RHBDL3;refGene_tx=XM_023243711.1;refGene_gn=RHBDL3;refGene_tx=XM_023243712.1;refGene_gn=RHBDL3;refGene_tx=XM_023243713.1;refGene_gn=RHBDL3;refGene_tx=XM_023243714.1
E1 20373922 20374071
refGene_gn=RHBDL3;refGene_tx=XM_023243711.1;refGene_gn=RHBDL3;refGene_tx=XM_023243712.1;refGene_gn=RHBDL3;refGene_tx=XM_023243713.1;refGene_gn=RHBDL3;refGene_tx=XM_023243714.1
E1 20376416 20376529
refGene_gn=RHBDL3;refGene_tx=XM_023243711.1;refGene_gn=RHBDL3;refGene_tx=XM_023243712.1;refGene_gn=RHBDL3;refGene_tx=XM_023243713.1;refGene_gn=RHBDL3;refGene_tx=XM_023243714.1
E1 20382220 20382321
refGene_gn=RHBDL3;refGene_tx=XM_023243711.1;refGene_gn=RHBDL3;refGene_tx=XM_023243712.1;refGene_gn=RHBDL3;refGene_tx=XM_023243713.1;refGene_gn=RHBDL3;refGene_tx=XM_023243714.1
E1 20391872 20391933
refGene_gn=RHBDL3;refGene_tx=XM_023243711.1;refGene_gn=RHBDL3;refGene_tx=XM_023243712.1;refGene_gn=RHBDL3;refGene_tx=XM_023243713.1;refGene_gn=RHBDL3;refGene_tx=XM_023243714.1
E1 20395126 20395398
refGene_gn=RHBDL3;refGene_tx=XM_023243711.1;refGene_gn=RHBDL3;refGene_tx=XM_023243712.1;refGene_gn=RHBDL3;refGene_tx=XM_023243713.1;refGene_gn=RHBDL3;refGene_tx=XM_023243714.1
E1 20403028 20403244
refGene_gn=CE1H17orf75;refGene_tx=XM_003996534.5
E1 20403865 20403969
refGene_gn=CE1H17orf75;refGene_tx=XM_003996534.5
E1 20405090 20405227
refGene_gn=CE1H17orf75;refGene_tx=XM_003996534.5
E1 20405386 20405451
refGene_gn=CE1H17orf75;refGene_tx=XM_003996534.5
E1 20406173 20406293
refGene_gn=CE1H17orf75;refGene_tx=XM_003996534.5
E1 20408038 20408096
refGene_gn=CE1H17orf75;refGene_tx=XM_003996534.5
E1 20408334 20408478
refGene_gn=CE1H17orf75;refGene_tx=XM_003996534.5
E1 20409365 20409491
refGene_gn=CE1H17orf75;refGene_tx=XM_003996534.5
E1 20410561 20410642
refGene_gn=CE1H17orf75;refGene_tx=XM_003996534.5

E1 20411340 20411480
refGene_gn=CE1H17orf75;refGene_tx=XM_003996534.5
E1 20415874 20415915
refGene_gn=ZNF207;refGene_tx=XM_006940067.4;refGene_gn=ZNF207;refGene_tx=XM_006940068.4;refGene_gn=ZNF207;refGene_tx=XM_006940069.4;refGene_gn=ZNF207;refGene_tx=XM_006940070.4;refGene_gn=ZNF207;refGene_tx=XM_006940071.4;refGene_gn=ZNF207;refGene_tx=XM_006940072.4;refGene_gn=ZNF207;refGene_tx=XM_019817503.2
E1 20417292 20417419
refGene_gn=ZNF207;refGene_tx=XM_006940067.4;refGene_gn=ZNF207;refGene_tx=XM_006940068.4;refGene_gn=ZNF207;refGene_tx=XM_006940069.4;refGene_gn=ZNF207;refGene_tx=XM_006940070.4;refGene_gn=ZNF207;refGene_tx=XM_006940071.4;refGene_gn=ZNF207;refGene_tx=XM_006940072.4;refGene_gn=ZNF207;refGene_tx=XM_019817503.2
E1 20421685 20421824
refGene_gn=ZNF207;refGene_tx=XM_006940067.4;refGene_gn=ZNF207;refGene_tx=XM_006940068.4;refGene_gn=ZNF207;refGene_tx=XM_006940069.4;refGene_gn=ZNF207;refGene_tx=XM_006940070.4;refGene_gn=ZNF207;refGene_tx=XM_006940071.4;refGene_gn=ZNF207;refGene_tx=XM_006940072.4;refGene_gn=ZNF207;refGene_tx=XM_019817503.2
E1 20423196 20423367
refGene_gn=ZNF207;refGene_tx=XM_006940067.4;refGene_gn=ZNF207;refGene_tx=XM_006940068.4;refGene_gn=ZNF207;refGene_tx=XM_006940069.4;refGene_gn=ZNF207;refGene_tx=XM_006940070.4;refGene_gn=ZNF207;refGene_tx=XM_006940071.4;refGene_gn=ZNF207;refGene_tx=XM_006940072.4;refGene_gn=ZNF207;refGene_tx=XM_019817503.2
E1 20423492 20423568
refGene_gn=ZNF207;refGene_tx=XM_006940067.4;refGene_gn=ZNF207;refGene_tx=XM_006940068.4;refGene_gn=ZNF207;refGene_tx=XM_006940069.4;refGene_gn=ZNF207;refGene_tx=XM_006940070.4;refGene_gn=ZNF207;refGene_tx=XM_006940071.4;refGene_gn=ZNF207;refGene_tx=XM_006940072.4;refGene_gn=ZNF207;refGene_tx=XM_019817503.2
E1 20424080 20424128
refGene_gn=ZNF207;refGene_tx=XM_006940067.4;refGene_gn=ZNF207;refGene_tx=XM_006940068.4;refGene_gn=ZNF207;refGene_tx=XM_006940071.4;refGene_gn=ZNF207;refGene_tx=XM_006940072.4;refGene_gn=ZNF207;refGene_tx=XM_019817503.2
E1 20425193 20425264
refGene_gn=ZNF207;refGene_tx=XM_006940067.4;refGene_gn=ZNF207;refGene_tx=XM_006940068.4;refGene_gn=ZNF207;refGene_tx=XM_006940069.4;refGene_gn=ZNF207;refGene_tx=XM_006940070.4;refGene_gn=ZNF207;refGene_tx=XM_006940071.4;refGene_gn=ZNF207;refGene_tx=XM_006940072.4;refGene_gn=ZNF207;refGene_tx=XM_019817503.2
E1 20427069 20427227
refGene_gn=ZNF207;refGene_tx=XM_006940067.4;refGene_gn=ZNF207;refGene_tx=XM_006940068.4;refGene_gn=ZNF207;refGene_tx=XM_006940069.4;refGene_gn=ZNF207;refGene_tx=XM_006940070.4;refGene_gn=ZNF207;refGene_tx=XM_006940071.4;refGene_gn=ZNF207;refGene_tx=XM_006940072.4;refGene_gn=ZNF207;refGene_tx=XM_019817503.2
E1 20428384 20428477

refGene_gn=ZNF207;refGene_tx=XM_006940067.4;refGene_gn=ZNF207;refGene_tx=XM_006940068.4;refGene_gn=ZNF207;refGene_tx=XM_006940069.4;refGene_gn=ZNF207;refGene_tx=XM_006940070.4;refGene_gn=ZNF207;refGene_tx=XM_019817503.2
E1 20429494 20429737
refGene_gn=ZNF207;refGene_tx=XM_006940067.4;refGene_gn=ZNF207;refGene_tx=XM_006940068.4;refGene_gn=ZNF207;refGene_tx=XM_006940069.4;refGene_gn=ZNF207;refGene_tx=XM_006940070.4;refGene_gn=ZNF207;refGene_tx=XM_006940071.4;refGene_gn=ZNF207;refGene_tx=XM_006940072.4;refGene_gn=ZNF207;refGene_tx=XM_019817503.2
E1 20430334 20430494
refGene_gn=ZNF207;refGene_tx=XM_006940067.4;refGene_gn=ZNF207;refGene_tx=XM_006940068.4;refGene_gn=ZNF207;refGene_tx=XM_006940069.4;refGene_gn=ZNF207;refGene_tx=XM_006940070.4;refGene_gn=ZNF207;refGene_tx=XM_006940071.4;refGene_gn=ZNF207;refGene_tx=XM_006940072.4;refGene_gn=ZNF207;refGene_tx=XM_019817503.2
E1 20430627 20430788
refGene_gn=ZNF207;refGene_tx=XM_006940067.4;refGene_gn=ZNF207;refGene_tx=XM_006940068.4;refGene_gn=ZNF207;refGene_tx=XM_006940069.4;refGene_gn=ZNF207;refGene_tx=XM_006940070.4;refGene_gn=ZNF207;refGene_tx=XM_006940071.4;refGene_gn=ZNF207;refGene_tx=XM_006940072.4;refGene_gn=ZNF207;refGene_tx=XM_019817503.2
E1 20497246 20497337 refGene_gn=PSMD11;refGene_tx=XM_023243436.1
E1 20499667 20499769 refGene_gn=PSMD11;refGene_tx=XM_023243436.1
E1 20505544 20505669 refGene_gn=PSMD11;refGene_tx=XM_023243436.1
E1 20511763 20511835 refGene_gn=PSMD11;refGene_tx=XM_023243436.1
E1 20512208 20512266 refGene_gn=PSMD11;refGene_tx=XM_023243436.1
E1 20514666 20514861 refGene_gn=PSMD11;refGene_tx=XM_023243436.1
E1 20520148 20520293 refGene_gn=PSMD11;refGene_tx=XM_023243436.1
E1 20521037 20521098 refGene_gn=PSMD11;refGene_tx=XM_023243436.1
E1 20522719 20522782 refGene_gn=PSMD11;refGene_tx=XM_023243436.1
E1 20524250 20524376 refGene_gn=PSMD11;refGene_tx=XM_023243436.1
E1 20524828 20524864 refGene_gn=PSMD11;refGene_tx=XM_023243436.1
E1 20525119 20525171 refGene_gn=PSMD11;refGene_tx=XM_023243436.1
E1 20525469 20525612 refGene_gn=PSMD11;refGene_tx=XM_023243436.1
E1 20531875 20532799 refGene_gn=CDK5R1;refGene_tx=XM_011288969.3
E1 20539276 20539433
refGene_gn=MYO1D;refGene_tx=XM_019817507.2;refGene_gn=MYO1D;refGene_tx=XM_023244611.1;refGene_gn=MYO1D;refGene_tx=XM_023244612.1
E1 20624493 20624648
refGene_gn=MYO1D;refGene_tx=XM_019817507.2;refGene_gn=MYO1D;refGene_tx=XM_023244611.1;refGene_gn=MYO1D;refGene_tx=XM_023244612.1
E1 20679906 20680020
refGene_gn=MYO1D;refGene_tx=XM_019817507.2;refGene_gn=MYO1D;refGene_tx=XM_023244611.1;refGene_gn=MYO1D;refGene_tx=XM_023244612.1
E1 20693649 20693754
refGene_gn=MYO1D;refGene_tx=XM_019817507.2;refGene_gn=MYO1D;refGene_tx=XM_023244611.1;refGene_gn=MYO1D;refGene_tx=XM_023244612.1
E1 20694300 20694445
refGene_gn=MYO1D;refGene_tx=XM_019817507.2;refGene_gn=MYO1D;refGene_tx

=XM_023244611.1;refGene_gn=MYO1D;refGene_tx=XM_023244612.1
E1	20700179	20700403
refGene_gn=MYO1D;refGene_tx=XM_019817507.2;refGene_gn=MYO1D;refGene_tx
=XM_023244611.1;refGene_gn=MYO1D;refGene_tx=XM_023244612.1
E1	20742221	20742429
refGene_gn=MYO1D;refGene_tx=XM_019817507.2;refGene_gn=MYO1D;refGene_tx
=XM_023244611.1;refGene_gn=MYO1D;refGene_tx=XM_023244612.1
E1	20752597	20752764
refGene_gn=MYO1D;refGene_tx=XM_019817507.2;refGene_gn=MYO1D;refGene_tx
=XM_023244611.1;refGene_gn=MYO1D;refGene_tx=XM_023244612.1
E1	20767333	20767466
refGene_gn=MYO1D;refGene_tx=XM_019817507.2;refGene_gn=MYO1D;refGene_tx
=XM_023244611.1;refGene_gn=MYO1D;refGene_tx=XM_023244612.1
E1	20773354	20773429
refGene_gn=MYO1D;refGene_tx=XM_019817507.2;refGene_gn=MYO1D;refGene_tx
=XM_023244611.1;refGene_gn=MYO1D;refGene_tx=XM_023244612.1
E1	20776802	20776873
refGene_gn=MYO1D;refGene_tx=XM_019817507.2;refGene_gn=MYO1D;refGene_tx
=XM_023244611.1;refGene_gn=MYO1D;refGene_tx=XM_023244612.1
E1	20780601	20780772
refGene_gn=MYO1D;refGene_tx=XM_019817507.2;refGene_gn=MYO1D;refGene_tx
=XM_023244611.1;refGene_gn=MYO1D;refGene_tx=XM_023244612.1
E1	20785288	20785403
refGene_gn=MYO1D;refGene_tx=XM_019817507.2;refGene_gn=MYO1D;refGene_tx
=XM_023244611.1;refGene_gn=MYO1D;refGene_tx=XM_023244612.1
E1	20785480	20785626
refGene_gn=MYO1D;refGene_tx=XM_019817507.2;refGene_gn=MYO1D;refGene_tx
=XM_023244611.1;refGene_gn=MYO1D;refGene_tx=XM_023244612.1
E1	20789198	20789402
refGene_gn=MYO1D;refGene_tx=XM_019817507.2;refGene_gn=MYO1D;refGene_tx
=XM_023244611.1;refGene_gn=MYO1D;refGene_tx=XM_023244612.1
E1	20791072	20791189
refGene_gn=MYO1D;refGene_tx=XM_019817507.2;refGene_gn=MYO1D;refGene_tx
=XM_023244611.1;refGene_gn=MYO1D;refGene_tx=XM_023244612.1
E1	20795256	20795352
refGene_gn=MYO1D;refGene_tx=XM_019817507.2;refGene_gn=MYO1D;refGene_tx
=XM_023244611.1;refGene_gn=MYO1D;refGene_tx=XM_023244612.1
E1	20796816	20796870
refGene_gn=MYO1D;refGene_tx=XM_019817507.2;refGene_gn=MYO1D;refGene_tx
=XM_023244611.1;refGene_gn=MYO1D;refGene_tx=XM_023244612.1
E1	20798531	20798697
refGene_gn=MYO1D;refGene_tx=XM_019817507.2;refGene_gn=MYO1D;refGene_tx
=XM_023244611.1;refGene_gn=MYO1D;refGene_tx=XM_023244612.1
E1	20800977	20801071
refGene_gn=MYO1D;refGene_tx=XM_019817507.2;refGene_gn=MYO1D;refGene_tx
=XM_023244611.1
E1	20803036	20803245
refGene_gn=MYO1D;refGene_tx=XM_019817507.2;refGene_gn=MYO1D;refGene_tx
=XM_023244611.1
E1	20855745	20855759	refGene_gn=MYO1D;refGene_tx=XM_019817507.2

E1 20902544 20902639
refGene_gn=MYO1D;refGene_tx=XM_023244611.1;refGene_gn=MYO1D;refGene_tx
=XM_023244612.1
E1 20915636 20915768
refGene_gn=LOC111557864;refGene_tx=XM_023244616.1
E1 20918724 20918925
refGene_gn=LOC111557864;refGene_tx=XM_023244616.1
E1 20932917 20933048
refGene_gn=TMEM98;refGene_tx=XM_011288968.3;refGene_gn=TMEM98;refGene_
tx=XM_023244614.1
E1 20934177 20934309
refGene_gn=TMEM98;refGene_tx=XM_011288968.3;refGene_gn=TMEM98;refGene_
tx=XM_023244614.1
E1 20935275 20935309
refGene_gn=TMEM98;refGene_tx=XM_011288968.3;refGene_gn=TMEM98;refGene_
tx=XM_023244614.1
E1 20937782 20937898
refGene_gn=TMEM98;refGene_tx=XM_011288968.3;refGene_gn=TMEM98;refGene_
tx=XM_023244614.1
E1 20940675 20940735
refGene_gn=TMEM98;refGene_tx=XM_011288968.3;refGene_gn=TMEM98;refGene_
tx=XM_023244614.1
E1 20941988 20942196
refGene_gn=TMEM98;refGene_tx=XM_011288968.3;refGene_gn=TMEM98;refGene_
tx=XM_023244614.1
E1 20962823 20963014 refGene_gn=SPACA3;refGene_tx=XM_023244613.1
E1 20978277 20978552
refGene_gn=SPACA3;refGene_tx=XM_006940080.3;refGene_gn=SPACA3;refGene_
tx=XM_019817510.2;refGene_gn=SPACA3;refGene_tx=XM_019817511.2;refGene_
gn=SPACA3;refGene_tx=XM_019817512.2;refGene_gn=SPACA3;refGene_tx=XM_02
3244613.1
E1 20979381 20979540
refGene_gn=SPACA3;refGene_tx=XM_006940080.3;refGene_gn=SPACA3;refGene_
tx=XM_019817510.2;refGene_gn=SPACA3;refGene_tx=XM_019817511.2;refGene_
gn=SPACA3;refGene_tx=XM_019817512.2;refGene_gn=SPACA3;refGene_tx=XM_02
3244613.1
E1 20979931 20980010
refGene_gn=SPACA3;refGene_tx=XM_006940080.3;refGene_gn=SPACA3;refGene_
tx=XM_019817510.2;refGene_gn=SPACA3;refGene_tx=XM_019817511.2;refGene_
gn=SPACA3;refGene_tx=XM_019817512.2;refGene_gn=SPACA3;refGene_tx=XM_02
3244613.1
E1 20982255 20982322
refGene_gn=SPACA3;refGene_tx=XM_006940080.3;refGene_gn=SPACA3;refGene_
tx=XM_019817510.2;refGene_gn=SPACA3;refGene_tx=XM_019817511.2;refGene_
gn=SPACA3;refGene_tx=XM_019817512.2;refGene_gn=SPACA3;refGene_tx=XM_02
3244613.1
E1 20994572 20994674
refGene_gn=ASIC2;refGene_tx=XM_003996555.5;refGene_gn=ASIC2;refGene_tx
=XM_023243509.1
E1 20996480 20996549

refGene_gn=ASIC2;refGene_tx=XM_003996555.5;refGene_gn=ASIC2;refGene_tx=XM_023243509.1
E1 20997990 20998070
refGene_gn=ASIC2;refGene_tx=XM_003996555.5;refGene_gn=ASIC2;refGene_tx=XM_023243509.1
E1 21001620 21001712
refGene_gn=ASIC2;refGene_tx=XM_003996555.5;refGene_gn=ASIC2;refGene_tx=XM_023243509.1
E1 21004387 21004541
refGene_gn=ASIC2;refGene_tx=XM_003996555.5;refGene_gn=ASIC2;refGene_tx=XM_023243509.1
E1 21005922 21005979
refGene_gn=ASIC2;refGene_tx=XM_003996555.5;refGene_gn=ASIC2;refGene_tx=XM_023243509.1
E1 21008216 21008367
refGene_gn=ASIC2;refGene_tx=XM_003996555.5;refGene_gn=ASIC2;refGene_tx=XM_023243509.1
E1 21058923 21059051
refGene_gn=ASIC2;refGene_tx=XM_003996555.5;refGene_gn=ASIC2;refGene_tx=XM_023243509.1
E1 21081786 21081937
refGene_gn=ASIC2;refGene_tx=XM_003996555.5;refGene_gn=ASIC2;refGene_tx=XM_023243509.1
E1 21263156 21263870 refGene_gn=ASIC2;refGene_tx=XM_023243509.1
E1 22008373 22008928 refGene_gn=ASIC2;refGene_tx=XM_003996555.5
E1 22009376 22009657
refGene_gn=LOC111557726;refGene_tx=XM_023243510.1;refGene_gn=LOC111557726;refGene_tx=XM_023243511.1;refGene_gn=LOC111557726;refGene_tx=XM_023243512.1;refGene_gn=LOC111557726;refGene_tx=XM_023243513.1
E1 22011471 22011567
refGene_gn=LOC111557726;refGene_tx=XM_023243510.1;refGene_gn=LOC111557726;refGene_tx=XM_023243511.1;refGene_gn=LOC111557726;refGene_tx=XM_023243512.1
E1 22012394 22012505
refGene_gn=LOC111557726;refGene_tx=XM_023243510.1;refGene_gn=LOC111557726;refGene_tx=XM_023243512.1
E1 22012646 22012743
refGene_gn=LOC111557726;refGene_tx=XM_023243510.1;refGene_gn=LOC111557726;refGene_tx=XM_023243511.1;refGene_gn=LOC111557726;refGene_tx=XM_023243512.1;refGene_gn=LOC111557726;refGene_tx=XM_023243513.1
E1 22017085 22017230
refGene_gn=LOC111557726;refGene_tx=XM_023243510.1;refGene_gn=LOC111557726;refGene_tx=XM_023243511.1;refGene_gn=LOC111557726;refGene_tx=XM_023243513.1
E1 22086118 22086194 refGene_gn=MCP-1;refGene_tx=XM_003996556.4
E1 22086937 22087055 refGene_gn=MCP-1;refGene_tx=XM_003996556.4
E1 22087399 22087505 refGene_gn=MCP-1;refGene_tx=XM_003996556.4
E1 22096994 22097070
refGene_gn=LOC101096931;refGene_tx=XM_003996557.4
E1 22097803 22097921

refGene_gn=LOC101096931;refGene_tx=XM_003996557.4
E1 22098311 22098417
refGene_gn=LOC101096931;refGene_tx=XM_003996557.4
E1 22134214 22134290
refGene_gn=LOC101097189;refGene_tx=XM_003996558.3
E1 22135075 22135193
refGene_gn=LOC101097189;refGene_tx=XM_003996558.3
E1 22135601 22135707
refGene_gn=LOC101097189;refGene_tx=XM_003996558.3
E1 22145179 22145255
refGene_gn=LOC101099760;refGene_tx=XM_003996569.4
E1 22145801 22145916
refGene_gn=LOC101099760;refGene_tx=XM_003996569.4
E1 22146356 22146462
refGene_gn=LOC101099760;refGene_tx=XM_003996569.4
E1 22148620 22148717 refGene_gn=CCL1;refGene_tx=XM_003996570.2
E1 22150097 22150209 refGene_gn=CCL1;refGene_tx=XM_003996570.2
E1 22151413 22151489 refGene_gn=CCL1;refGene_tx=XM_003996570.2
E1 22395153 22395386 refGene_gn=TMEM132E;refGene_tx=XM_023244516.1
E1 22395939 22395999 refGene_gn=TMEM132E;refGene_tx=XM_023244516.1
E1 22409050 22410097 refGene_gn=TMEM132E;refGene_tx=XM_023244516.1
E1 22411539 22411686 refGene_gn=TMEM132E;refGene_tx=XM_023244516.1
E1 22411962 22412155 refGene_gn=TMEM132E;refGene_tx=XM_023244516.1
E1 22412960 22413104 refGene_gn=TMEM132E;refGene_tx=XM_023244516.1
E1 22415411 22415617 refGene_gn=TMEM132E;refGene_tx=XM_023244516.1
E1 22417197 22417486 refGene_gn=TMEM132E;refGene_tx=XM_023244516.1
E1 22418401 22418593 refGene_gn=TMEM132E;refGene_tx=XM_023244516.1
E1 22419585 22420641 refGene_gn=TMEM132E;refGene_tx=XM_023244516.1
E1 22689341 22689414
refGene_gn=CCT6B;refGene_tx=XM_006940082.4;refGene_gn=CCT6B;refGene_tx
=XM_019817513.2;refGene_gn=CCT6B;refGene_tx=XM_019817515.2;refGene_gn=
CCT6B;refGene_tx=XM_019817516.2
E1 22689930 22690003
refGene_gn=CCT6B;refGene_tx=XM_006940082.4;refGene_gn=CCT6B;refGene_tx
=XM_019817513.2;refGene_gn=CCT6B;refGene_tx=XM_019817515.2;refGene_gn=
CCT6B;refGene_tx=XM_019817516.2
E1 22691286 22691389
refGene_gn=CCT6B;refGene_tx=XM_006940082.4;refGene_gn=CCT6B;refGene_tx
=XM_019817513.2;refGene_gn=CCT6B;refGene_tx=XM_019817515.2;refGene_gn=
CCT6B;refGene_tx=XM_019817516.2
E1 22693268 22693402
refGene_gn=CCT6B;refGene_tx=XM_006940082.4;refGene_gn=CCT6B;refGene_tx
=XM_019817513.2;refGene_gn=CCT6B;refGene_tx=XM_019817515.2;refGene_gn=
CCT6B;refGene_tx=XM_019817516.2
E1 22702336 22702484
refGene_gn=CCT6B;refGene_tx=XM_006940082.4;refGene_gn=CCT6B;refGene_tx
=XM_019817513.2;refGene_gn=CCT6B;refGene_tx=XM_019817515.2;refGene_gn=
CCT6B;refGene_tx=XM_019817516.2
E1 22702788 22702909
refGene_gn=CCT6B;refGene_tx=XM_006940082.4;refGene_gn=CCT6B;refGene_tx

=XM_019817513.2;refGene_gn=CCT6B;refGene_tx=XM_019817515.2;refGene_gn=CCT6B;refGene_tx=XM_019817516.2
E1 22703468 22703551
refGene_gn=CCT6B;refGene_tx=XM_006940082.4;refGene_gn=CCT6B;refGene_tx=XM_019817513.2;refGene_gn=CCT6B;refGene_tx=XM_019817515.2;refGene_gn=CCT6B;refGene_tx=XM_019817516.2
E1 22705780 22705940
refGene_gn=CCT6B;refGene_tx=XM_006940082.4;refGene_gn=CCT6B;refGene_tx=XM_019817513.2;refGene_gn=CCT6B;refGene_tx=XM_019817515.2;refGene_gn=CCT6B;refGene_tx=XM_019817516.2
E1 22706327 22706438
refGene_gn=CCT6B;refGene_tx=XM_006940082.4;refGene_gn=CCT6B;refGene_tx=XM_019817513.2;refGene_gn=CCT6B;refGene_tx=XM_019817516.2
E1 22712285 22712389
refGene_gn=CCT6B;refGene_tx=XM_006940082.4;refGene_gn=CCT6B;refGene_tx=XM_019817513.2;refGene_gn=CCT6B;refGene_tx=XM_019817515.2;refGene_gn=CCT6B;refGene_tx=XM_019817516.2
E1 22714493 22714667
refGene_gn=CCT6B;refGene_tx=XM_006940082.4;refGene_gn=CCT6B;refGene_tx=XM_019817513.2;refGene_gn=CCT6B;refGene_tx=XM_019817515.2;refGene_gn=CCT6B;refGene_tx=XM_019817516.2
E1 22720598 22720733
refGene_gn=CCT6B;refGene_tx=XM_006940082.4;refGene_gn=CCT6B;refGene_tx=XM_019817513.2;refGene_gn=CCT6B;refGene_tx=XM_019817515.2;refGene_gn=CCT6B;refGene_tx=XM_019817516.2
E1 22721530 22721594
refGene_gn=CCT6B;refGene_tx=XM_006940082.4;refGene_gn=CCT6B;refGene_tx=XM_019817513.2;refGene_gn=CCT6B;refGene_tx=XM_019817515.2;refGene_gn=CCT6B;refGene_tx=XM_019817516.2
E1 22723223 22723360
refGene_gn=CCT6B;refGene_tx=XM_006940082.4;refGene_gn=CCT6B;refGene_tx=XM_019817513.2;refGene_gn=CCT6B;refGene_tx=XM_019817515.2
E1 22723538 22724651 refGene_gn=ZNF830;refGene_tx=XM_003996560.5
E1 22744329 22744358
refGene_gn=LIG3;refGene_tx=XM_003996561.5;refGene_gn=LIG3;refGene_tx=XM_006940083.4;refGene_gn=LIG3;refGene_tx=XM_023243786.1;refGene_gn=LIG3;refGene_tx=XM_023243788.1
E1 22745571 22746125
refGene_gn=LIG3;refGene_tx=XM_003996561.5;refGene_gn=LIG3;refGene_tx=XM_006940083.4;refGene_gn=LIG3;refGene_tx=XM_023243785.1;refGene_gn=LIG3;refGene_tx=XM_023243786.1;refGene_gn=LIG3;refGene_tx=XM_023243788.1
E1 22751174 22751318
refGene_gn=LIG3;refGene_tx=XM_003996561.5;refGene_gn=LIG3;refGene_tx=XM_006940083.4;refGene_gn=LIG3;refGene_tx=XM_023243785.1;refGene_gn=LIG3;refGene_tx=XM_023243786.1;refGene_gn=LIG3;refGene_tx=XM_023243788.1
E1 22753139 22753337
refGene_gn=LIG3;refGene_tx=XM_003996561.5;refGene_gn=LIG3;refGene_tx=XM_006940083.4;refGene_gn=LIG3;refGene_tx=XM_023243785.1;refGene_gn=LIG3;refGene_tx=XM_023243786.1;refGene_gn=LIG3;refGene_tx=XM_023243788.1
E1 22754315 22754467

refGene_gn=LIG3;refGene_tx=XM_003996561.5;refGene_gn=LIG3;refGene_tx=XM_006940083.4;refGene_gn=LIG3;refGene_tx=XM_023243785.1;refGene_gn=LIG3;refGene_tx=XM_023243786.1;refGene_gn=LIG3;refGene_tx=XM_023243788.1
E1 22755160 22755327
refGene_gn=LIG3;refGene_tx=XM_003996561.5;refGene_gn=LIG3;refGene_tx=XM_006940083.4;refGene_gn=LIG3;refGene_tx=XM_023243785.1;refGene_gn=LIG3;refGene_tx=XM_023243786.1;refGene_gn=LIG3;refGene_tx=XM_023243788.1
E1 22755439 22755517
refGene_gn=LIG3;refGene_tx=XM_003996561.5;refGene_gn=LIG3;refGene_tx=XM_006940083.4;refGene_gn=LIG3;refGene_tx=XM_023243785.1;refGene_gn=LIG3;refGene_tx=XM_023243786.1;refGene_gn=LIG3;refGene_tx=XM_023243788.1
E1 22755981 22756150
refGene_gn=LIG3;refGene_tx=XM_003996561.5;refGene_gn=LIG3;refGene_tx=XM_006940083.4;refGene_gn=LIG3;refGene_tx=XM_023243785.1;refGene_gn=LIG3;refGene_tx=XM_023243786.1;refGene_gn=LIG3;refGene_tx=XM_023243788.1
E1 22757618 22757774
refGene_gn=LIG3;refGene_tx=XM_003996561.5;refGene_gn=LIG3;refGene_tx=XM_006940083.4;refGene_gn=LIG3;refGene_tx=XM_023243785.1;refGene_gn=LIG3;refGene_tx=XM_023243786.1;refGene_gn=LIG3;refGene_tx=XM_023243788.1
E1 22759513 22759645
refGene_gn=LIG3;refGene_tx=XM_003996561.5;refGene_gn=LIG3;refGene_tx=XM_006940083.4;refGene_gn=LIG3;refGene_tx=XM_023243785.1;refGene_gn=LIG3;refGene_tx=XM_023243788.1
E1 22760038 22760118
refGene_gn=LIG3;refGene_tx=XM_003996561.5;refGene_gn=LIG3;refGene_tx=XM_006940083.4;refGene_gn=LIG3;refGene_tx=XM_023243785.1;refGene_gn=LIG3;refGene_tx=XM_023243786.1;refGene_gn=LIG3;refGene_tx=XM_023243788.1
E1 22760456 22760544
refGene_gn=LIG3;refGene_tx=XM_003996561.5;refGene_gn=LIG3;refGene_tx=XM_006940083.4;refGene_gn=LIG3;refGene_tx=XM_023243785.1;refGene_gn=LIG3;refGene_tx=XM_023243786.1;refGene_gn=LIG3;refGene_tx=XM_023243788.1
E1 22760916 22760994
refGene_gn=LIG3;refGene_tx=XM_003996561.5;refGene_gn=LIG3;refGene_tx=XM_006940083.4;refGene_gn=LIG3;refGene_tx=XM_023243785.1;refGene_gn=LIG3;refGene_tx=XM_023243786.1;refGene_gn=LIG3;refGene_tx=XM_023243788.1
E1 22761324 22761448
refGene_gn=LIG3;refGene_tx=XM_003996561.5;refGene_gn=LIG3;refGene_tx=XM_006940083.4;refGene_gn=LIG3;refGene_tx=XM_023243785.1;refGene_gn=LIG3;refGene_tx=XM_023243786.1;refGene_gn=LIG3;refGene_tx=XM_023243788.1
E1 22762055 22762198
refGene_gn=LIG3;refGene_tx=XM_003996561.5;refGene_gn=LIG3;refGene_tx=XM_006940083.4;refGene_gn=LIG3;refGene_tx=XM_023243785.1;refGene_gn=LIG3;refGene_tx=XM_023243786.1;refGene_gn=LIG3;refGene_tx=XM_023243788.1
E1 22762541 22762616
refGene_gn=LIG3;refGene_tx=XM_003996561.5;refGene_gn=LIG3;refGene_tx=XM_006940083.4;refGene_gn=LIG3;refGene_tx=XM_023243785.1;refGene_gn=LIG3;refGene_tx=XM_023243786.1;refGene_gn=LIG3;refGene_tx=XM_023243788.1
E1 22762971 22763019 refGene_gn=LIG3;refGene_tx=XM_023243788.1
E1 22763029 22763176
refGene_gn=LIG3;refGene_tx=XM_003996561.5;refGene_gn=LIG3;refGene_tx=X

M_006940083.4;refGene_gn=LIG3;refGene_tx=XM_023243785.1;refGene_gn=LIG3;refGene_tx=XM_023243786.1
E1 22763818 22764011 refGene_gn=LIG3;refGene_tx=XM_003996561.5;refGene_gn=LIG3;refGene_tx=XM_006940083.4;refGene_gn=LIG3;refGene_tx=XM_023243785.1;refGene_gn=LIG3;refGene_tx=XM_023243786.1
E1 22764396 22764479 refGene_gn=LIG3;refGene_tx=XM_003996561.5;refGene_gn=LIG3;refGene_tx=XM_006940083.4;refGene_gn=LIG3;refGene_tx=XM_023243785.1;refGene_gn=LIG3;refGene_tx=XM_023243786.1
E1 22764659 22764716 refGene_gn=LIG3;refGene_tx=XM_006940083.4
E1 22765597 22765831 refGene_gn=LIG3;refGene_tx=XM_003996561.5;refGene_gn=LIG3;refGene_tx=XM_023243785.1;refGene_gn=LIG3;refGene_tx=XM_023243786.1
E1 22771295 22771477 refGene_gn=RFFL;refGene_tx=XM_006940088.4;refGene_gn=RFFL;refGene_tx=XM_011288966.3;refGene_gn=RFFL;refGene_tx=XM_023243789.1;refGene_gn=RFFL;refGene_tx=XM_023243790.1;refGene_gn=RFFL;refGene_tx=XM_023243791.1;refGene_gn=RFFL;refGene_tx=XM_023243793.1
E1 22773921 22773945 refGene_gn=RFFL;refGene_tx=XM_006940088.4;refGene_gn=RFFL;refGene_tx=XM_011288966.3;refGene_gn=RFFL;refGene_tx=XM_023243789.1;refGene_gn=RFFL;refGene_tx=XM_023243790.1;refGene_gn=RFFL;refGene_tx=XM_023243791.1;refGene_gn=RFFL;refGene_tx=XM_023243793.1
E1 22775188 22775399 refGene_gn=RFFL;refGene_tx=XM_006940088.4;refGene_gn=RFFL;refGene_tx=XM_011288966.3;refGene_gn=RFFL;refGene_tx=XM_023243789.1;refGene_gn=RFFL;refGene_tx=XM_023243790.1;refGene_gn=RFFL;refGene_tx=XM_023243791.1;refGene_gn=RFFL;refGene_tx=XM_023243793.1
E1 22776363 22776447 refGene_gn=RFFL;refGene_tx=XM_006940088.4;refGene_gn=RFFL;refGene_tx=XM_011288966.3;refGene_gn=RFFL;refGene_tx=XM_023243789.1;refGene_gn=RFFL;refGene_tx=XM_023243790.1;refGene_gn=RFFL;refGene_tx=XM_023243793.1
E1 22779548 22779962 refGene_gn=RFFL;refGene_tx=XM_006940088.4;refGene_gn=RFFL;refGene_tx=XM_011288966.3;refGene_gn=RFFL;refGene_tx=XM_023243789.1;refGene_gn=RFFL;refGene_tx=XM_023243790.1;refGene_gn=RFFL;refGene_tx=XM_023243791.1;refGene_gn=RFFL;refGene_tx=XM_023243793.1
E1 22786739 22786927 refGene_gn=RFFL;refGene_tx=XM_006940088.4;refGene_gn=RFFL;refGene_tx=XM_011288966.3;refGene_gn=RFFL;refGene_tx=XM_023243789.1;refGene_gn=RFFL;refGene_tx=XM_023243790.1;refGene_gn=RFFL;refGene_tx=XM_023243791.1;refGene_gn=RFFL;refGene_tx=XM_023243793.1
E1 22787030 22787115 refGene_gn=RFFL;refGene_tx=XM_011288966.3
E1 22835280 22835470 refGene_gn=RFFL;refGene_tx=XM_023243789.1;refGene_gn=RFFL;refGene_tx=XM_023243790.1;refGene_gn=RFFL;refGene_tx=XM_023243791.1
E1 22842052 22842136 refGene_gn=RAD51D;refGene_tx=XM_003996572.5
E1 22842278 22842443 refGene_gn=RAD51D;refGene_tx=XM_003996572.5
E1 22844638 22844709 refGene_gn=RAD51D;refGene_tx=XM_003996572.5

E1 22844830 22844921 refGene_gn=RAD51D;refGene_tx=XM_003996572.5
E1 22848295 22848391 refGene_gn=RAD51D;refGene_tx=XM_003996572.5
E1 22848897 22849032 refGene_gn=RAD51D;refGene_tx=XM_003996572.5
E1 22849262 22849344 refGene_gn=RAD51D;refGene_tx=XM_003996572.5
E1 22857381 22857500 refGene_gn=RAD51D;refGene_tx=XM_003996572.5
E1 22857715 22857777 refGene_gn=RAD51D;refGene_tx=XM_003996572.5
E1 22858112 22858194 refGene_gn=RAD51D;refGene_tx=XM_003996572.5
E1 22859831 22860040
refGene_gn=FNDC8;refGene_tx=XM_003996565.5;refGene_gn=FNDC8;refGene_tx
=XM_019817519.2
E1 22863473 22863843
refGene_gn=FNDC8;refGene_tx=XM_003996565.5;refGene_gn=FNDC8;refGene_tx
=XM_019817519.2
E1 22865314 22865551
refGene_gn=FNDC8;refGene_tx=XM_003996565.5;refGene_gn=FNDC8;refGene_tx
=XM_019817519.2
E1 22866081 22866234
refGene_gn=FNDC8;refGene_tx=XM_003996565.5;refGene_gn=FNDC8;refGene_tx
=XM_019817519.2
E1 22868141 22868154
refGene_gn=NLE1;refGene_tx=XM_006940089.3;refGene_gn=NLE1;refGene_tx=X
M_006940090.3
E1 22868823 22868894
refGene_gn=NLE1;refGene_tx=XM_006940089.3;refGene_gn=NLE1;refGene_tx=X
M_006940090.3
E1 22869010 22869170
refGene_gn=NLE1;refGene_tx=XM_006940089.3;refGene_gn=NLE1;refGene_tx=X
M_006940090.3
E1 22870075 22870278
refGene_gn=NLE1;refGene_tx=XM_006940089.3;refGene_gn=NLE1;refGene_tx=X
M_006940090.3
E1 22871007 22871054
refGene_gn=NLE1;refGene_tx=XM_006940089.3;refGene_gn=NLE1;refGene_tx=X
M_006940090.3
E1 22871202 22871338
refGene_gn=NLE1;refGene_tx=XM_006940089.3;refGene_gn=NLE1;refGene_tx=X
M_006940090.3
E1 22871849 22872042
refGene_gn=NLE1;refGene_tx=XM_006940089.3;refGene_gn=NLE1;refGene_tx=X
M_006940090.3
E1 22872381 22872479
refGene_gn=NLE1;refGene_tx=XM_006940089.3;refGene_gn=NLE1;refGene_tx=X
M_006940090.3
E1 22872651 22872728
refGene_gn=NLE1;refGene_tx=XM_006940089.3;refGene_gn=NLE1;refGene_tx=X
M_006940090.3
E1 22873641 22873721
refGene_gn=NLE1;refGene_tx=XM_006940089.3;refGene_gn=NLE1;refGene_tx=X
M_006940090.3
E1 22874314 22874532

refGene_gn=NLE1;refGene_tx=XM_006940089.3;refGene_gn=NLE1;refGene_tx=XM_006940090.3
E1 22875497 22875647
refGene_gn=NLE1;refGene_tx=XM_006940089.3;refGene_gn=NLE1;refGene_tx=XM_006940090.3
E1 22875758 22875782
refGene_gn=NLE1;refGene_tx=XM_006940089.3;refGene_gn=NLE1;refGene_tx=XM_006940090.3
E1 22880089 22880257
refGene_gn=UNC45B;refGene_tx=XM_003996567.5;refGene_gn=UNC45B;refGene_tx=XM_006940091.4;refGene_gn=UNC45B;refGene_tx=XM_019817521.2
E1 22880818 22880855
refGene_gn=UNC45B;refGene_tx=XM_003996567.5;refGene_gn=UNC45B;refGene_tx=XM_006940091.4;refGene_gn=UNC45B;refGene_tx=XM_019817521.2
E1 22882470 22882646
refGene_gn=UNC45B;refGene_tx=XM_003996567.5;refGene_gn=UNC45B;refGene_tx=XM_006940091.4;refGene_gn=UNC45B;refGene_tx=XM_019817521.2
E1 22884355 22884445
refGene_gn=UNC45B;refGene_tx=XM_003996567.5;refGene_gn=UNC45B;refGene_tx=XM_006940091.4;refGene_gn=UNC45B;refGene_tx=XM_019817521.2
E1 22886742 22886910
refGene_gn=UNC45B;refGene_tx=XM_003996567.5;refGene_gn=UNC45B;refGene_tx=XM_006940091.4;refGene_gn=UNC45B;refGene_tx=XM_019817521.2
E1 22887495 22887664
refGene_gn=UNC45B;refGene_tx=XM_003996567.5;refGene_gn=UNC45B;refGene_tx=XM_006940091.4;refGene_gn=UNC45B;refGene_tx=XM_019817521.2
E1 22889614 22889785
refGene_gn=UNC45B;refGene_tx=XM_003996567.5;refGene_gn=UNC45B;refGene_tx=XM_006940091.4;refGene_gn=UNC45B;refGene_tx=XM_019817521.2
E1 22892381 22892553
refGene_gn=UNC45B;refGene_tx=XM_003996567.5;refGene_gn=UNC45B;refGene_tx=XM_006940091.4;refGene_gn=UNC45B;refGene_tx=XM_019817521.2
E1 22895192 22895493
refGene_gn=UNC45B;refGene_tx=XM_003996567.5;refGene_gn=UNC45B;refGene_tx=XM_006940091.4;refGene_gn=UNC45B;refGene_tx=XM_019817521.2
E1 22896387 22896482
refGene_gn=UNC45B;refGene_tx=XM_003996567.5;refGene_gn=UNC45B;refGene_tx=XM_006940091.4;refGene_gn=UNC45B;refGene_tx=XM_019817521.2
E1 22896651 22896793
refGene_gn=UNC45B;refGene_tx=XM_003996567.5;refGene_gn=UNC45B;refGene_tx=XM_006940091.4;refGene_gn=UNC45B;refGene_tx=XM_019817521.2
E1 22897480 22897621
refGene_gn=UNC45B;refGene_tx=XM_003996567.5;refGene_gn=UNC45B;refGene_tx=XM_006940091.4;refGene_gn=UNC45B;refGene_tx=XM_019817521.2
E1 22898794 22898922
refGene_gn=UNC45B;refGene_tx=XM_003996567.5;refGene_gn=UNC45B;refGene_tx=XM_006940091.4;refGene_gn=UNC45B;refGene_tx=XM_019817521.2
E1 22899799 22899866
refGene_gn=UNC45B;refGene_tx=XM_003996567.5;refGene_gn=UNC45B;refGene_tx=XM_006940091.4;refGene_gn=UNC45B;refGene_tx=XM_019817521.2

E1 22900892 22901006
refGene_gn=UNC45B;refGene_tx=XM_003996567.5;refGene_gn=UNC45B;refGene_tx=XM_006940091.4;refGene_gn=UNC45B;refGene_tx=XM_019817521.2
E1 22901366 22901482
refGene_gn=UNC45B;refGene_tx=XM_003996567.5;refGene_gn=UNC45B;refGene_tx=XM_006940091.4;refGene_gn=UNC45B;refGene_tx=XM_019817521.2
E1 22902360 22902478
refGene_gn=UNC45B;refGene_tx=XM_003996567.5;refGene_gn=UNC45B;refGene_tx=XM_006940091.4;refGene_gn=UNC45B;refGene_tx=XM_019817521.2
E1 22910657 22910813
refGene_gn=UNC45B;refGene_tx=XM_003996567.5;refGene_gn=UNC45B;refGene_tx=XM_006940091.4;refGene_gn=UNC45B;refGene_tx=XM_019817521.2
E1 22913812 22914073
refGene_gn=UNC45B;refGene_tx=XM_003996567.5;refGene_gn=UNC45B;refGene_tx=XM_006940091.4;refGene_gn=UNC45B;refGene_tx=XM_019817521.2
E1 22950610 22951622
refGene_gn=LOC101100766;refGene_tx=XM_006940092.2;refGene_gn=LOC101100766;refGene_tx=XM_019817522.1;refGene_gn=LOC101100766;refGene_tx=XM_019817525.1;refGene_gn=LOC101100766;refGene_tx=XM_019817526.1;refGene_gn=LOC101100766;refGene_tx=XM_019817529.2;refGene_gn=LOC101100766;refGene_tx=XM_019817530.1
E1 22952895 22953021
refGene_gn=LOC101100766;refGene_tx=XM_006940092.2;refGene_gn=LOC101100766;refGene_tx=XM_019817522.1;refGene_gn=LOC101100766;refGene_tx=XM_019817525.1;refGene_gn=LOC101100766;refGene_tx=XM_019817526.1;refGene_gn=LOC101100766;refGene_tx=XM_019817529.2;refGene_gn=LOC101100766;refGene_tx=XM_019817530.1
E1 22954818 22954892
refGene_gn=LOC101100766;refGene_tx=XM_019817530.1
E1 22957637 22958358
refGene_gn=LOC101100766;refGene_tx=XM_006940092.2;refGene_gn=LOC101100766;refGene_tx=XM_019817522.1;refGene_gn=LOC101100766;refGene_tx=XM_019817525.1;refGene_gn=LOC101100766;refGene_tx=XM_019817526.1;refGene_gn=LOC101100766;refGene_tx=XM_019817529.2
E1 22958527 22959335
refGene_gn=LOC101100766;refGene_tx=XM_006940092.2;refGene_gn=LOC101100766;refGene_tx=XM_019817522.1;refGene_gn=LOC101100766;refGene_tx=XM_019817525.1;refGene_gn=LOC101100766;refGene_tx=XM_019817526.1;refGene_gn=LOC101100766;refGene_tx=XM_019817529.2
E1 22997072 22997229 refGene_gn=SLFN14;refGene_tx=XM_023243943.1
E1 22997230 22997526 refGene_gn=SLFN14;refGene_tx=XM_023243943.1
E1 22997527 22997867 refGene_gn=SLFN14;refGene_tx=XM_023243943.1
E1 22999884 23000599 refGene_gn=SLFN14;refGene_tx=XM_023243943.1
E1 23003453 23003582 refGene_gn=SLFN14;refGene_tx=XM_023243943.1
E1 23005113 23005589 refGene_gn=SLFN14;refGene_tx=XM_023243943.1
E1 23005829 23005944 refGene_gn=SLFN14;refGene_tx=XM_023243943.1
E1 23005946 23006553 refGene_gn=SLFN14;refGene_tx=XM_023243943.1
E1 23016602 23017002 refGene_gn=PEX12;refGene_tx=XM_003996568.5
E1 23017857 23018411 refGene_gn=PEX12;refGene_tx=XM_003996568.5
E1 23018693 23018819 refGene_gn=PEX12;refGene_tx=XM_003996568.5

E1 23031906 23031943
refGene_gn=AP2B1;refGene_tx=XM_023244427.1;refGene_gn=AP2B1;refGene_tx=XM_023244428.1;refGene_gn=AP2B1;refGene_tx=XM_023244429.1
E1 23038399 23038505
refGene_gn=AP2B1;refGene_tx=XM_023244427.1;refGene_gn=AP2B1;refGene_tx=XM_023244428.1;refGene_gn=AP2B1;refGene_tx=XM_023244429.1;refGene_gn=AP2B1;refGene_tx=XM_023244430.1
E1 23043323 23043459
refGene_gn=AP2B1;refGene_tx=XM_023244427.1;refGene_gn=AP2B1;refGene_tx=XM_023244428.1;refGene_gn=AP2B1;refGene_tx=XM_023244429.1;refGene_gn=AP2B1;refGene_tx=XM_023244430.1
E1 23045333 23045579
refGene_gn=AP2B1;refGene_tx=XM_023244427.1;refGene_gn=AP2B1;refGene_tx=XM_023244428.1;refGene_gn=AP2B1;refGene_tx=XM_023244429.1;refGene_gn=AP2B1;refGene_tx=XM_023244430.1
E1 23060469 23060660
refGene_gn=AP2B1;refGene_tx=XM_023244427.1;refGene_gn=AP2B1;refGene_tx=XM_023244428.1;refGene_gn=AP2B1;refGene_tx=XM_023244429.1;refGene_gn=AP2B1;refGene_tx=XM_023244430.1
E1 23063144 23063366
refGene_gn=AP2B1;refGene_tx=XM_023244427.1;refGene_gn=AP2B1;refGene_tx=XM_023244428.1;refGene_gn=AP2B1;refGene_tx=XM_023244429.1;refGene_gn=AP2B1;refGene_tx=XM_023244430.1
E1 23064570 23064691
refGene_gn=AP2B1;refGene_tx=XM_023244427.1;refGene_gn=AP2B1;refGene_tx=XM_023244428.1;refGene_gn=AP2B1;refGene_tx=XM_023244429.1;refGene_gn=AP2B1;refGene_tx=XM_023244430.1
E1 23064820 23064916
refGene_gn=AP2B1;refGene_tx=XM_023244427.1;refGene_gn=AP2B1;refGene_tx=XM_023244428.1;refGene_gn=AP2B1;refGene_tx=XM_023244429.1;refGene_gn=AP2B1;refGene_tx=XM_023244430.1
E1 23072181 23072297
refGene_gn=AP2B1;refGene_tx=XM_023244427.1;refGene_gn=AP2B1;refGene_tx=XM_023244428.1;refGene_gn=AP2B1;refGene_tx=XM_023244429.1;refGene_gn=AP2B1;refGene_tx=XM_023244430.1
E1 23072788 23072954
refGene_gn=AP2B1;refGene_tx=XM_023244427.1;refGene_gn=AP2B1;refGene_tx=XM_023244428.1;refGene_gn=AP2B1;refGene_tx=XM_023244429.1;refGene_gn=AP2B1;refGene_tx=XM_023244430.1
E1 23074408 23074507
refGene_gn=AP2B1;refGene_tx=XM_023244427.1;refGene_gn=AP2B1;refGene_tx=XM_023244428.1;refGene_gn=AP2B1;refGene_tx=XM_023244429.1;refGene_gn=AP2B1;refGene_tx=XM_023244430.1
E1 23083327 23083587
refGene_gn=AP2B1;refGene_tx=XM_023244427.1;refGene_gn=AP2B1;refGene_tx=XM_023244428.1;refGene_gn=AP2B1;refGene_tx=XM_023244429.1;refGene_gn=AP2B1;refGene_tx=XM_023244430.1
E1 23094998 23095191
refGene_gn=AP2B1;refGene_tx=XM_023244427.1;refGene_gn=AP2B1;refGene_tx=XM_023244428.1;refGene_gn=AP2B1;refGene_tx=XM_023244429.1;refGene_gn=

AP2B1;refGene_tx=XM_023244430.1
E1 23107105 23107147
refGene_gn=AP2B1;refGene_tx=XM_023244427.1;refGene_gn=AP2B1;refGene_tx=XM_023244428.1;refGene_gn=AP2B1;refGene_tx=XM_023244430.1
E1 23108003 23108150
refGene_gn=AP2B1;refGene_tx=XM_023244427.1;refGene_gn=AP2B1;refGene_tx=XM_023244428.1;refGene_gn=AP2B1;refGene_tx=XM_023244429.1;refGene_gn=AP2B1;refGene_tx=XM_023244430.1
E1 23110612 23110758
refGene_gn=AP2B1;refGene_tx=XM_023244427.1;refGene_gn=AP2B1;refGene_tx=XM_023244428.1;refGene_gn=AP2B1;refGene_tx=XM_023244429.1;refGene_gn=AP2B1;refGene_tx=XM_023244430.1
E1 23116791 23116921
refGene_gn=AP2B1;refGene_tx=XM_023244427.1;refGene_gn=AP2B1;refGene_tx=XM_023244428.1;refGene_gn=AP2B1;refGene_tx=XM_023244429.1;refGene_gn=AP2B1;refGene_tx=XM_023244430.1
E1 23137515 23137600
refGene_gn=AP2B1;refGene_tx=XM_023244427.1;refGene_gn=AP2B1;refGene_tx=XM_023244428.1;refGene_gn=AP2B1;refGene_tx=XM_023244429.1;refGene_gn=AP2B1;refGene_tx=XM_023244430.1
E1 23138520 23138607
refGene_gn=AP2B1;refGene_tx=XM_023244427.1;refGene_gn=AP2B1;refGene_tx=XM_023244428.1;refGene_gn=AP2B1;refGene_tx=XM_023244429.1;refGene_gn=AP2B1;refGene_tx=XM_023244430.1
E1 23145537 23145692
refGene_gn=AP2B1;refGene_tx=XM_023244427.1;refGene_gn=AP2B1;refGene_tx=XM_023244428.1;refGene_gn=AP2B1;refGene_tx=XM_023244429.1;refGene_gn=AP2B1;refGene_tx=XM_023244430.1
E1 23152334 23152409
refGene_gn=AP2B1;refGene_tx=XM_023244427.1;refGene_gn=AP2B1;refGene_tx=XM_023244428.1;refGene_gn=AP2B1;refGene_tx=XM_023244429.1;refGene_gn=AP2B1;refGene_tx=XM_023244430.1
E1 23164601 23164817 refGene_gn=RASL10B;refGene_tx=XM_023243686.1
E1 23169380 23169505 refGene_gn=RASL10B;refGene_tx=XM_023243686.1
E1 23170060 23170331 refGene_gn=RASL10B;refGene_tx=XM_023243686.1
E1 23173530 23175085 refGene_gn=GAS2L2;refGene_tx=XM_003996576.3
E1 23175620 23175873 refGene_gn=GAS2L2;refGene_tx=XM_003996576.3
E1 23176662 23176759 refGene_gn=GAS2L2;refGene_tx=XM_003996576.3
E1 23177858 23177966 refGene_gn=GAS2L2;refGene_tx=XM_003996576.3
E1 23178825 23179067 refGene_gn=GAS2L2;refGene_tx=XM_003996576.3
E1 23180495 23180880 refGene_gn=GAS2L2;refGene_tx=XM_003996576.3
E1 23185097 23185110
refGene_gn=CE1H17orf50;refGene_tx=XM_003996592.4
E1 23187230 23187564
refGene_gn=CE1H17orf50;refGene_tx=XM_003996592.4
E1 23187656 23187867
refGene_gn=CE1H17orf50;refGene_tx=XM_003996592.4
E1 23189671 23190066
refGene_gn=MMP28;refGene_tx=XM_023243582.1;refGene_gn=MMP28;refGene_tx=XM_023243583.1

E1 23190902 23191070
refGene_gn=MMP28;refGene_tx=XM_023243582.1;refGene_gn=MMP28;refGene_tx
=XM_023243583.1
E1 23191285 23191308 refGene_gn=MMP28;refGene_tx=XM_023243585.1
E1 23191365 23191515 refGene_gn=MMP28;refGene_tx=XM_023243582.1
E1 23192760 23193006
refGene_gn=MMP28;refGene_tx=XM_023243582.1;refGene_gn=MMP28;refGene_tx
=XM_023243583.1;refGene_gn=MMP28;refGene_tx=XM_023243585.1
E1 23194972 23195197
refGene_gn=MMP28;refGene_tx=XM_023243582.1;refGene_gn=MMP28;refGene_tx
=XM_023243583.1;refGene_gn=MMP28;refGene_tx=XM_023243585.1
E1 23200490 23200678
refGene_gn=MMP28;refGene_tx=XM_023243582.1;refGene_gn=MMP28;refGene_tx
=XM_023243583.1;refGene_gn=MMP28;refGene_tx=XM_023243585.1
E1 23200888 23200968
refGene_gn=MMP28;refGene_tx=XM_023243582.1;refGene_gn=MMP28;refGene_tx
=XM_023243583.1;refGene_gn=MMP28;refGene_tx=XM_023243585.1
E1 23212586 23212697
refGene_gn=MMP28;refGene_tx=XM_023243582.1;refGene_gn=MMP28;refGene_tx
=XM_023243583.1;refGene_gn=MMP28;refGene_tx=XM_023243585.1
E1 23225734 23225741
refGene_gn=TAF15;refGene_tx=XM_023244806.1;refGene_gn=TAF15;refGene_tx
=XM_023244807.1;refGene_gn=TAF15;refGene_tx=XM_023244808.1
E1 23231937 23231977
refGene_gn=TAF15;refGene_tx=XM_023244806.1;refGene_gn=TAF15;refGene_tx
=XM_023244807.1;refGene_gn=TAF15;refGene_tx=XM_023244808.1
E1 23233452 23233504
refGene_gn=TAF15;refGene_tx=XM_023244806.1;refGene_gn=TAF15;refGene_tx
=XM_023244807.1;refGene_gn=TAF15;refGene_tx=XM_023244808.1
E1 23233591 23233676
refGene_gn=TAF15;refGene_tx=XM_023244806.1;refGene_gn=TAF15;refGene_tx
=XM_023244807.1;refGene_gn=TAF15;refGene_tx=XM_023244808.1
E1 23233758 23233861
refGene_gn=TAF15;refGene_tx=XM_023244806.1;refGene_gn=TAF15;refGene_tx
=XM_023244807.1;refGene_gn=TAF15;refGene_tx=XM_023244808.1
E1 23236013 23236207
refGene_gn=TAF15;refGene_tx=XM_023244806.1;refGene_gn=TAF15;refGene_tx
=XM_023244807.1;refGene_gn=TAF15;refGene_tx=XM_023244808.1
E1 23237258 23237379
refGene_gn=TAF15;refGene_tx=XM_023244806.1;refGene_gn=TAF15;refGene_tx
=XM_023244807.1;refGene_gn=TAF15;refGene_tx=XM_023244808.1
E1 23238619 23238664 refGene_gn=TAF15;refGene_tx=XM_023244806.1
E1 23244431 23244466
refGene_gn=TAF15;refGene_tx=XM_023244806.1;refGene_gn=TAF15;refGene_tx
=XM_023244807.1;refGene_gn=TAF15;refGene_tx=XM_023244808.1
E1 23245085 23245118
refGene_gn=TAF15;refGene_tx=XM_023244806.1;refGene_gn=TAF15;refGene_tx
=XM_023244807.1;refGene_gn=TAF15;refGene_tx=XM_023244808.1
E1 23246332 23246442
refGene_gn=TAF15;refGene_tx=XM_023244806.1;refGene_gn=TAF15;refGene_tx

=XM_023244807.1;refGene_gn=TAF15;refGene_tx=XM_023244808.1
E1	23247444	23247574
refGene_gn=TAF15;refGene_tx=XM_023244806.1;refGene_gn=TAF15;refGene_tx=XM_023244807.1;refGene_gn=TAF15;refGene_tx=XM_023244808.1
E1	23250568	23250661
refGene_gn=TAF15;refGene_tx=XM_023244806.1;refGene_gn=TAF15;refGene_tx=XM_023244807.1;refGene_gn=TAF15;refGene_tx=XM_023244808.1
E1	23251791	23251873
refGene_gn=TAF15;refGene_tx=XM_023244806.1;refGene_gn=TAF15;refGene_tx=XM_023244807.1;refGene_gn=TAF15;refGene_tx=XM_023244808.1
E1	23251989	23252078
refGene_gn=TAF15;refGene_tx=XM_023244806.1;refGene_gn=TAF15;refGene_tx=XM_023244807.1;refGene_gn=TAF15;refGene_tx=XM_023244808.1
E1	23252186	23252703
refGene_gn=TAF15;refGene_tx=XM_023244806.1;refGene_gn=TAF15;refGene_tx=XM_023244807.1;refGene_gn=TAF15;refGene_tx=XM_023244808.1
E1	23253987	23254027
refGene_gn=TAF15;refGene_tx=XM_023244806.1;refGene_gn=TAF15;refGene_tx=XM_023244807.1;refGene_gn=TAF15;refGene_tx=XM_023244808.1
E1	23261739	23262120
refGene_gn=HEATR9;refGene_tx=XM_003996595.5;refGene_gn=HEATR9;refGene_tx=XM_023243908.1;refGene_gn=HEATR9;refGene_tx=XM_023243909.1
E1	23262350	23262437
refGene_gn=HEATR9;refGene_tx=XM_003996595.5;refGene_gn=HEATR9;refGene_tx=XM_023243908.1;refGene_gn=HEATR9;refGene_tx=XM_023243909.1
E1	23262879	23262931
refGene_gn=HEATR9;refGene_tx=XM_003996595.5;refGene_gn=HEATR9;refGene_tx=XM_023243908.1;refGene_gn=HEATR9;refGene_tx=XM_023243909.1
E1	23263620	23263694
refGene_gn=HEATR9;refGene_tx=XM_003996595.5;refGene_gn=HEATR9;refGene_tx=XM_023243908.1;refGene_gn=HEATR9;refGene_tx=XM_023243909.1
E1	23264765	23264885
refGene_gn=HEATR9;refGene_tx=XM_003996595.5;refGene_gn=HEATR9;refGene_tx=XM_023243908.1;refGene_gn=HEATR9;refGene_tx=XM_023243909.1
E1	23264994	23265087
refGene_gn=HEATR9;refGene_tx=XM_003996595.5;refGene_gn=HEATR9;refGene_tx=XM_023243908.1;refGene_gn=HEATR9;refGene_tx=XM_023243909.1
E1	23265699	23265882
refGene_gn=HEATR9;refGene_tx=XM_003996595.5;refGene_gn=HEATR9;refGene_tx=XM_023243908.1;refGene_gn=HEATR9;refGene_tx=XM_023243909.1
E1	23269523	23269687
refGene_gn=HEATR9;refGene_tx=XM_003996595.5;refGene_gn=HEATR9;refGene_tx=XM_023243908.1;refGene_gn=HEATR9;refGene_tx=XM_023243909.1
E1	23270031	23270089
refGene_gn=HEATR9;refGene_tx=XM_003996595.5;refGene_gn=HEATR9;refGene_tx=XM_023243908.1;refGene_gn=HEATR9;refGene_tx=XM_023243909.1
E1	23270754	23270811
refGene_gn=HEATR9;refGene_tx=XM_003996595.5;refGene_gn=HEATR9;refGene_tx=XM_023243908.1;refGene_gn=HEATR9;refGene_tx=XM_023243909.1
E1	23271003	23271060

refGene_gn=HEATR9;refGene_tx=XM_003996595.5;refGene_gn=HEATR9;refGene_tx=XM_023243908.1;refGene_gn=HEATR9;refGene_tx=XM_023243909.1
E1 23271267 23271394 refGene_gn=HEATR9;refGene_tx=XM_003996595.5;refGene_gn=HEATR9;refGene_tx=XM_023243908.1;refGene_gn=HEATR9;refGene_tx=XM_023243909.1
E1 23271653 23271835 refGene_gn=HEATR9;refGene_tx=XM_003996595.5;refGene_gn=HEATR9;refGene_tx=XM_023243908.1;refGene_gn=HEATR9;refGene_tx=XM_023243909.1
E1 23273547 23273597 refGene_gn=HEATR9;refGene_tx=XM_003996595.5;refGene_gn=HEATR9;refGene_tx=XM_023243908.1;refGene_gn=HEATR9;refGene_tx=XM_023243909.1
E1 23274750 23274838 refGene_gn=HEATR9;refGene_tx=XM_003996595.5;refGene_gn=HEATR9;refGene_tx=XM_023243908.1
E1 23276460 23276548 refGene_gn=CCL5;refGene_tx=NM_001009827.1
E1 23281555 23281667 refGene_gn=CCL5;refGene_tx=NM_001009827.1
E1 23282867 23282943 refGene_gn=CCL5;refGene_tx=NM_001009827.1
E1 23316060 23316193 refGene_gn=CCL16;refGene_tx=XM_006940098.3
E1 23316846 23316967 refGene_gn=CCL16;refGene_tx=XM_006940098.3
E1 23319226 23319296 refGene_gn=CCL16;refGene_tx=XM_006940098.3
E1 23321632 23321720 refGene_gn=CCL14;refGene_tx=XM_003996596.5
E1 23322154 23322269 refGene_gn=CCL14;refGene_tx=XM_003996596.5
E1 23324022 23324107 refGene_gn=CCL14;refGene_tx=XM_003996596.5
E1 23334367 23334455 refGene_gn=LOC101086516;refGene_tx=XM_003996597.5
E1 23334855 23334967 refGene_gn=LOC101086516;refGene_tx=XM_003996597.5
E1 23335451 23335499 refGene_gn=LOC101086516;refGene_tx=XM_003996597.5
E1 23338107 23338183 refGene_gn=LOC101086516;refGene_tx=XM_003996597.5
E1 23388969 23389045 refGene_gn=LOC101081091;refGene_tx=XM_003996577.4
E1 23389640 23389755 refGene_gn=LOC101081091;refGene_tx=XM_003996577.4
E1 23390158 23390249 refGene_gn=LOC101081091;refGene_tx=XM_003996577.4
E1 23406672 23406763 refGene_gn=LOC100302540;refGene_tx=NM_001163657.1
E1 23407189 23407304 refGene_gn=LOC100302540;refGene_tx=NM_001163657.1
E1 23407994 23408070 refGene_gn=LOC100302540;refGene_tx=NM_001163657.1
E1 23417953 23418029 refGene_gn=LOC100302541;refGene_tx=NM_001163658.1
E1 23418539 23418654 refGene_gn=LOC100302541;refGene_tx=NM_001163658.1
E1 23419201 23419289 refGene_gn=LOC100302541;refGene_tx=NM_001163658.1
E1 23492745 23492964

refGene_gn=HEATR6;refGene_tx=XM_003996579.4;refGene_gn=HEATR6;refGene_
tx=XM_006940099.4;refGene_gn=HEATR6;refGene_tx=XM_019817540.2
E1 23493206 23493221 refGene_gn=HEATR6;refGene_tx=XM_019817539.2
E1 23496322 23496430
refGene_gn=HEATR6;refGene_tx=XM_003996579.4;refGene_gn=HEATR6;refGene_
tx=XM_006940099.4;refGene_gn=HEATR6;refGene_tx=XM_019817539.2;refGene_
gn=HEATR6;refGene_tx=XM_019817540.2
E1 23499484 23499625
refGene_gn=HEATR6;refGene_tx=XM_003996579.4;refGene_gn=HEATR6;refGene_
tx=XM_006940099.4;refGene_gn=HEATR6;refGene_tx=XM_019817539.2;refGene_
gn=HEATR6;refGene_tx=XM_019817541.1;refGene_gn=HEATR6;refGene_tx=XM_02
3243517.1
E1 23500097 23500213
refGene_gn=HEATR6;refGene_tx=XM_003996579.4;refGene_gn=HEATR6;refGene_
tx=XM_006940099.4;refGene_gn=HEATR6;refGene_tx=XM_019817539.2;refGene_
gn=HEATR6;refGene_tx=XM_019817541.1;refGene_gn=HEATR6;refGene_tx=XM_02
3243517.1
E1 23501025 23501140
refGene_gn=HEATR6;refGene_tx=XM_003996579.4;refGene_gn=HEATR6;refGene_
tx=XM_006940099.4;refGene_gn=HEATR6;refGene_tx=XM_019817539.2;refGene_
gn=HEATR6;refGene_tx=XM_019817541.1;refGene_gn=HEATR6;refGene_tx=XM_02
3243517.1
E1 23502670 23502772
refGene_gn=HEATR6;refGene_tx=XM_003996579.4;refGene_gn=HEATR6;refGene_
tx=XM_006940099.4;refGene_gn=HEATR6;refGene_tx=XM_019817539.2;refGene_
gn=HEATR6;refGene_tx=XM_019817540.2;refGene_gn=HEATR6;refGene_tx=XM_01
9817541.1;refGene_gn=HEATR6;refGene_tx=XM_023243517.1
E1 23503643 23503781
refGene_gn=HEATR6;refGene_tx=XM_003996579.4;refGene_gn=HEATR6;refGene_
tx=XM_006940099.4;refGene_gn=HEATR6;refGene_tx=XM_019817539.2;refGene_
gn=HEATR6;refGene_tx=XM_019817540.2;refGene_gn=HEATR6;refGene_tx=XM_01
9817541.1;refGene_gn=HEATR6;refGene_tx=XM_023243517.1
E1 23505459 23505755
refGene_gn=HEATR6;refGene_tx=XM_003996579.4;refGene_gn=HEATR6;refGene_
tx=XM_006940099.4;refGene_gn=HEATR6;refGene_tx=XM_019817539.2;refGene_
gn=HEATR6;refGene_tx=XM_019817540.2;refGene_gn=HEATR6;refGene_tx=XM_01
9817541.1;refGene_gn=HEATR6;refGene_tx=XM_023243517.1
E1 23506495 23506673
refGene_gn=HEATR6;refGene_tx=XM_003996579.4;refGene_gn=HEATR6;refGene_
tx=XM_006940099.4;refGene_gn=HEATR6;refGene_tx=XM_019817539.2;refGene_
gn=HEATR6;refGene_tx=XM_019817540.2;refGene_gn=HEATR6;refGene_tx=XM_01
9817541.1;refGene_gn=HEATR6;refGene_tx=XM_023243517.1
E1 23511901 23512108
refGene_gn=HEATR6;refGene_tx=XM_003996579.4;refGene_gn=HEATR6;refGene_
tx=XM_006940099.4;refGene_gn=HEATR6;refGene_tx=XM_019817539.2;refGene_
gn=HEATR6;refGene_tx=XM_019817540.2;refGene_gn=HEATR6;refGene_tx=XM_01
9817541.1;refGene_gn=HEATR6;refGene_tx=XM_023243517.1
E1 23512546 23512646
refGene_gn=HEATR6;refGene_tx=XM_003996579.4;refGene_gn=HEATR6;refGene_
tx=XM_006940099.4;refGene_gn=HEATR6;refGene_tx=XM_019817539.2;refGene_

gn=HEATR6;refGene_tx=XM_019817540.2;refGene_gn=HEATR6;refGene_tx=XM_019817541.1;refGene_gn=HEATR6;refGene_tx=XM_023243517.1
E1 23514955 23515311
refGene_gn=HEATR6;refGene_tx=XM_003996579.4;refGene_gn=HEATR6;refGene_tx=XM_006940099.4;refGene_gn=HEATR6;refGene_tx=XM_019817539.2;refGene_gn=HEATR6;refGene_tx=XM_019817540.2;refGene_gn=HEATR6;refGene_tx=XM_019817541.1;refGene_gn=HEATR6;refGene_tx=XM_023243517.1
E1 23516129 23516252
refGene_gn=HEATR6;refGene_tx=XM_003996579.4;refGene_gn=HEATR6;refGene_tx=XM_006940099.4;refGene_gn=HEATR6;refGene_tx=XM_019817539.2;refGene_gn=HEATR6;refGene_tx=XM_019817540.2;refGene_gn=HEATR6;refGene_tx=XM_019817541.1;refGene_gn=HEATR6;refGene_tx=XM_023243517.1
E1 23516621 23516708
refGene_gn=HEATR6;refGene_tx=XM_003996579.4;refGene_gn=HEATR6;refGene_tx=XM_006940099.4;refGene_gn=HEATR6;refGene_tx=XM_019817539.2;refGene_gn=HEATR6;refGene_tx=XM_019817540.2;refGene_gn=HEATR6;refGene_tx=XM_019817541.1;refGene_gn=HEATR6;refGene_tx=XM_023243517.1
E1 23520260 23520395
refGene_gn=HEATR6;refGene_tx=XM_003996579.4;refGene_gn=HEATR6;refGene_tx=XM_006940099.4;refGene_gn=HEATR6;refGene_tx=XM_019817539.2;refGene_gn=HEATR6;refGene_tx=XM_019817540.2;refGene_gn=HEATR6;refGene_tx=XM_019817541.1;refGene_gn=HEATR6;refGene_tx=XM_023243517.1
E1 23521107 23521230
refGene_gn=HEATR6;refGene_tx=XM_003996579.4;refGene_gn=HEATR6;refGene_tx=XM_006940099.4;refGene_gn=HEATR6;refGene_tx=XM_019817539.2;refGene_gn=HEATR6;refGene_tx=XM_019817540.2;refGene_gn=HEATR6;refGene_tx=XM_019817541.1;refGene_gn=HEATR6;refGene_tx=XM_023243517.1
E1 23521939 23522064
refGene_gn=HEATR6;refGene_tx=XM_003996579.4;refGene_gn=HEATR6;refGene_tx=XM_006940099.4;refGene_gn=HEATR6;refGene_tx=XM_019817539.2;refGene_gn=HEATR6;refGene_tx=XM_019817540.2;refGene_gn=HEATR6;refGene_tx=XM_019817541.1;refGene_gn=HEATR6;refGene_tx=XM_023243517.1
E1 23522884 23522981
refGene_gn=HEATR6;refGene_tx=XM_003996579.4;refGene_gn=HEATR6;refGene_tx=XM_006940099.4;refGene_gn=HEATR6;refGene_tx=XM_019817539.2;refGene_gn=HEATR6;refGene_tx=XM_019817540.2;refGene_gn=HEATR6;refGene_tx=XM_019817541.1;refGene_gn=HEATR6;refGene_tx=XM_023243517.1
E1 23524055 23524260
refGene_gn=HEATR6;refGene_tx=XM_003996579.4;refGene_gn=HEATR6;refGene_tx=XM_006940099.4;refGene_gn=HEATR6;refGene_tx=XM_019817539.2;refGene_gn=HEATR6;refGene_tx=XM_019817540.2;refGene_gn=HEATR6;refGene_tx=XM_019817541.1;refGene_gn=HEATR6;refGene_tx=XM_023243517.1
E1 23526479 23527051
refGene_gn=HEATR6;refGene_tx=XM_003996579.4;refGene_gn=HEATR6;refGene_tx=XM_006940099.4;refGene_gn=HEATR6;refGene_tx=XM_019817539.2;refGene_gn=HEATR6;refGene_tx=XM_019817540.2;refGene_gn=HEATR6;refGene_tx=XM_019817541.1;refGene_gn=HEATR6;refGene_tx=XM_023243517.1
E1 23527471 23527489 refGene_gn=HEATR6;refGene_tx=XM_006940099.4
E1 23617542 23617886
refGene_gn=HNF1B;refGene_tx=XM_003996580.5;refGene_gn=HNF1B;refGene_tx

=XM_003996581.5;refGene_gn=HNF1B;refGene_tx=XM_019817542.2;refGene_gn=HNF1B;refGene_tx=XM_019817543.2;refGene_gn=HNF1B;refGene_tx=XM_019817544.2;refGene_gn=HNF1B;refGene_tx=XM_019817545.2
E1 23622615 23622815
refGene_gn=HNF1B;refGene_tx=XM_003996580.5;refGene_gn=HNF1B;refGene_tx=XM_003996581.5;refGene_gn=HNF1B;refGene_tx=XM_019817542.2;refGene_gn=HNF1B;refGene_tx=XM_019817543.2;refGene_gn=HNF1B;refGene_tx=XM_019817544.2;refGene_gn=HNF1B;refGene_tx=XM_019817545.2
E1 23627700 23627965
refGene_gn=HNF1B;refGene_tx=XM_003996580.5;refGene_gn=HNF1B;refGene_tx=XM_003996581.5;refGene_gn=HNF1B;refGene_tx=XM_019817542.2;refGene_gn=HNF1B;refGene_tx=XM_019817543.2;refGene_gn=HNF1B;refGene_tx=XM_019817544.2;refGene_gn=HNF1B;refGene_tx=XM_019817545.2
E1 23629626 23629862
refGene_gn=HNF1B;refGene_tx=XM_003996580.5;refGene_gn=HNF1B;refGene_tx=XM_003996581.5;refGene_gn=HNF1B;refGene_tx=XM_019817542.2;refGene_gn=HNF1B;refGene_tx=XM_019817543.2;refGene_gn=HNF1B;refGene_tx=XM_019817544.2;refGene_gn=HNF1B;refGene_tx=XM_019817545.2
E1 23649229 23649390
refGene_gn=HNF1B;refGene_tx=XM_003996580.5;refGene_gn=HNF1B;refGene_tx=XM_003996581.5;refGene_gn=HNF1B;refGene_tx=XM_019817542.2;refGene_gn=HNF1B;refGene_tx=XM_019817543.2;refGene_gn=HNF1B;refGene_tx=XM_019817544.2;refGene_gn=HNF1B;refGene_tx=XM_019817545.2
E1 23655080 23655216
refGene_gn=HNF1B;refGene_tx=XM_003996580.5;refGene_gn=HNF1B;refGene_tx=XM_003996581.5;refGene_gn=HNF1B;refGene_tx=XM_019817542.2;refGene_gn=HNF1B;refGene_tx=XM_019817543.2;refGene_gn=HNF1B;refGene_tx=XM_019817544.2;refGene_gn=HNF1B;refGene_tx=XM_019817545.2
E1 23659235 23659430
refGene_gn=HNF1B;refGene_tx=XM_003996580.5;refGene_gn=HNF1B;refGene_tx=XM_003996581.5;refGene_gn=HNF1B;refGene_tx=XM_019817542.2;refGene_gn=HNF1B;refGene_tx=XM_019817543.2;refGene_gn=HNF1B;refGene_tx=XM_019817544.2;refGene_gn=HNF1B;refGene_tx=XM_019817545.2
E1 23662083 23662202
refGene_gn=HNF1B;refGene_tx=XM_003996580.5;refGene_gn=HNF1B;refGene_tx=XM_003996581.5;refGene_gn=HNF1B;refGene_tx=XM_019817542.2;refGene_gn=HNF1B;refGene_tx=XM_019817544.2
E1 23670139 23670252
refGene_gn=HNF1B;refGene_tx=XM_003996580.5;refGene_gn=HNF1B;refGene_tx=XM_003996581.5;refGene_gn=HNF1B;refGene_tx=XM_019817542.2;refGene_gn=HNF1B;refGene_tx=XM_019817543.2;refGene_gn=HNF1B;refGene_tx=XM_019817544.2;refGene_gn=HNF1B;refGene_tx=XM_019817545.2
E1 23720191 23720278 refGene_gn=DDX52;refGene_tx=XM_003996582.4
E1 23721573 23721769
refGene_gn=DDX52;refGene_tx=XM_003996582.4;refGene_gn=DDX52;refGene_tx=XM_011288961.2
E1 23726276 23726404
refGene_gn=DDX52;refGene_tx=XM_003996582.4;refGene_gn=DDX52;refGene_tx=XM_011288961.2
E1 23727342 23727528

refGene_gn=DDX52;refGene_tx=XM_003996582.4;refGene_gn=DDX52;refGene_tx=XM_011288961.2
E1 23729319 23729463
refGene_gn=DDX52;refGene_tx=XM_003996582.4;refGene_gn=DDX52;refGene_tx=XM_011288961.2
E1 23730124 23730236
refGene_gn=DDX52;refGene_tx=XM_003996582.4;refGene_gn=DDX52;refGene_tx=XM_011288961.2
E1 23731087 23731160
refGene_gn=DDX52;refGene_tx=XM_003996582.4;refGene_gn=DDX52;refGene_tx=XM_011288961.2
E1 23731639 23731843
refGene_gn=DDX52;refGene_tx=XM_003996582.4;refGene_gn=DDX52;refGene_tx=XM_011288961.2
E1 23733404 23733495
refGene_gn=DDX52;refGene_tx=XM_003996582.4;refGene_gn=DDX52;refGene_tx=XM_011288961.2
E1 23736232 23736355
refGene_gn=DDX52;refGene_tx=XM_003996582.4;refGene_gn=DDX52;refGene_tx=XM_011288961.2
E1 23736488 23736639
refGene_gn=DDX52;refGene_tx=XM_003996582.4;refGene_gn=DDX52;refGene_tx=XM_011288961.2
E1 23736832 23736908
refGene_gn=DDX52;refGene_tx=XM_003996582.4;refGene_gn=DDX52;refGene_tx=XM_011288961.2
E1 23738186 23738258
refGene_gn=DDX52;refGene_tx=XM_003996582.4;refGene_gn=DDX52;refGene_tx=XM_011288961.2
E1 23738835 23738928
refGene_gn=DDX52;refGene_tx=XM_003996582.4;refGene_gn=DDX52;refGene_tx=XM_011288961.2
E1 23742181 23742239
refGene_gn=DDX52;refGene_tx=XM_003996582.4;refGene_gn=DDX52;refGene_tx=XM_011288961.2
E1 23745830 23745907
refGene_gn=SYNRG;refGene_tx=XM_003996583.5;refGene_gn=SYNRG;refGene_tx=XM_006940101.4;refGene_gn=SYNRG;refGene_tx=XM_006940102.4;refGene_gn=SYNRG;refGene_tx=XM_006940103.4;refGene_gn=SYNRG;refGene_tx=XM_006940104.4;refGene_gn=SYNRG;refGene_tx=XM_006940105.4;refGene_gn=SYNRG;refGene_tx=XM_006940106.4;refGene_gn=SYNRG;refGene_tx=XM_006940107.4;refGene_gn=SYNRG;refGene_tx=XM_006940108.4;refGene_gn=SYNRG;refGene_tx=XM_006940109.4;refGene_gn=SYNRG;refGene_tx=XM_006940110.4;refGene_gn=SYNRG;refGene_tx=XM_006940111.4;refGene_gn=SYNRG;refGene_tx=XM_006940112.4;refGene_gn=SYNRG;refGene_tx=XM_011288960.3;refGene_gn=SYNRG;refGene_tx=XM_019817546.2;refGene_gn=SYNRG;refGene_tx=XM_019817547.2
E1 23752695 23752736
refGene_gn=SYNRG;refGene_tx=XM_003996583.5;refGene_gn=SYNRG;refGene_tx=XM_006940101.4;refGene_gn=SYNRG;refGene_tx=XM_006940102.4;refGene_gn=SYNRG;refGene_tx=XM_006940103.4;refGene_gn=SYNRG;refGene_tx=XM_0069401

04.4;refGene_gn=SYNRG;refGene_tx=XM_006940105.4;refGene_gn=SYNRG;refGene_tx=XM_006940106.4;refGene_gn=SYNRG;refGene_tx=XM_006940107.4;refGene_gn=SYNRG;refGene_tx=XM_006940108.4;refGene_gn=SYNRG;refGene_tx=XM_006940109.4;refGene_gn=SYNRG;refGene_tx=XM_006940110.4;refGene_gn=SYNRG;refGene_tx=XM_006940111.4;refGene_gn=SYNRG;refGene_tx=XM_006940112.4;refGene_gn=SYNRG;refGene_tx=XM_011288960.3;refGene_gn=SYNRG;refGene_tx=XM_019817546.2;refGene_gn=SYNRG;refGene_tx=XM_019817547.2
E1 23756528 23756650
refGene_gn=SYNRG;refGene_tx=XM_003996583.5;refGene_gn=SYNRG;refGene_tx=XM_006940101.4;refGene_gn=SYNRG;refGene_tx=XM_006940102.4;refGene_gn=SYNRG;refGene_tx=XM_006940103.4;refGene_gn=SYNRG;refGene_tx=XM_006940104.4;refGene_gn=SYNRG;refGene_tx=XM_006940105.4;refGene_gn=SYNRG;refGene_tx=XM_006940106.4;refGene_gn=SYNRG;refGene_tx=XM_006940107.4;refGene_gn=SYNRG;refGene_tx=XM_006940108.4;refGene_gn=SYNRG;refGene_tx=XM_006940109.4;refGene_gn=SYNRG;refGene_tx=XM_006940110.4;refGene_gn=SYNRG;refGene_tx=XM_006940111.4;refGene_gn=SYNRG;refGene_tx=XM_006940112.4;refGene_gn=SYNRG;refGene_tx=XM_011288960.3;refGene_gn=SYNRG;refGene_tx=XM_019817546.2;refGene_gn=SYNRG;refGene_tx=XM_019817547.2
E1 23770630 23770761
refGene_gn=SYNRG;refGene_tx=XM_003996583.5;refGene_gn=SYNRG;refGene_tx=XM_006940101.4;refGene_gn=SYNRG;refGene_tx=XM_006940102.4;refGene_gn=SYNRG;refGene_tx=XM_006940103.4;refGene_gn=SYNRG;refGene_tx=XM_006940104.4;refGene_gn=SYNRG;refGene_tx=XM_006940105.4;refGene_gn=SYNRG;refGene_tx=XM_006940106.4;refGene_gn=SYNRG;refGene_tx=XM_006940107.4;refGene_gn=SYNRG;refGene_tx=XM_006940108.4;refGene_gn=SYNRG;refGene_tx=XM_006940109.4;refGene_gn=SYNRG;refGene_tx=XM_006940110.4;refGene_gn=SYNRG;refGene_tx=XM_006940111.4;refGene_gn=SYNRG;refGene_tx=XM_006940112.4;refGene_gn=SYNRG;refGene_tx=XM_011288960.3;refGene_gn=SYNRG;refGene_tx=XM_019817546.2;refGene_gn=SYNRG;refGene_tx=XM_019817547.2
E1 23771405 23771511
refGene_gn=SYNRG;refGene_tx=XM_003996583.5;refGene_gn=SYNRG;refGene_tx=XM_006940101.4;refGene_gn=SYNRG;refGene_tx=XM_006940102.4;refGene_gn=SYNRG;refGene_tx=XM_006940103.4;refGene_gn=SYNRG;refGene_tx=XM_006940104.4;refGene_gn=SYNRG;refGene_tx=XM_006940105.4;refGene_gn=SYNRG;refGene_tx=XM_006940106.4;refGene_gn=SYNRG;refGene_tx=XM_006940107.4;refGene_gn=SYNRG;refGene_tx=XM_006940108.4;refGene_gn=SYNRG;refGene_tx=XM_006940109.4;refGene_gn=SYNRG;refGene_tx=XM_006940110.4;refGene_gn=SYNRG;refGene_tx=XM_006940111.4;refGene_gn=SYNRG;refGene_tx=XM_006940112.4;refGene_gn=SYNRG;refGene_tx=XM_011288960.3;refGene_gn=SYNRG;refGene_tx=XM_019817546.2;refGene_gn=SYNRG;refGene_tx=XM_019817547.2
E1 23772047 23772159
refGene_gn=SYNRG;refGene_tx=XM_003996583.5;refGene_gn=SYNRG;refGene_tx=XM_006940101.4;refGene_gn=SYNRG;refGene_tx=XM_006940102.4;refGene_gn=SYNRG;refGene_tx=XM_006940103.4;refGene_gn=SYNRG;refGene_tx=XM_006940104.4;refGene_gn=SYNRG;refGene_tx=XM_006940105.4;refGene_gn=SYNRG;refGene_tx=XM_006940106.4;refGene_gn=SYNRG;refGene_tx=XM_006940107.4;refGene_gn=SYNRG;refGene_tx=XM_006940108.4;refGene_gn=SYNRG;refGene_tx=XM_006940109.4;refGene_gn=SYNRG;refGene_tx=XM_006940110.4;refGene_gn=SYNRG;refGene_tx=XM_006940111.4;refGene_gn=SYNRG;refGene_tx=XM_006940112.4;refGene_gn=SYNRG;refGene_tx=XM_011288960.3;refGene_gn=SYNRG;refGene_tx=

XM_019817546.2;refGene_gn=SYNRG;refGene_tx=XM_019817547.2
E1 23779697 23780000
refGene_gn=SYNRG;refGene_tx=XM_006940101.4;refGene_gn=SYNRG;refGene_tx=XM_006940102.4;refGene_gn=SYNRG;refGene_tx=XM_006940103.4;refGene_gn=SYNRG;refGene_tx=XM_006940104.4;refGene_gn=SYNRG;refGene_tx=XM_006940105.4;refGene_gn=SYNRG;refGene_tx=XM_006940106.4;refGene_gn=SYNRG;refGene_tx=XM_006940107.4;refGene_gn=SYNRG;refGene_tx=XM_006940108.4;refGene_gn=SYNRG;refGene_tx=XM_019817546.2;refGene_gn=SYNRG;refGene_tx=XM_019817547.2
E1 23780909 23781143
refGene_gn=SYNRG;refGene_tx=XM_006940101.4;refGene_gn=SYNRG;refGene_tx=XM_006940102.4;refGene_gn=SYNRG;refGene_tx=XM_006940103.4;refGene_gn=SYNRG;refGene_tx=XM_006940104.4;refGene_gn=SYNRG;refGene_tx=XM_006940105.4;refGene_gn=SYNRG;refGene_tx=XM_006940106.4;refGene_gn=SYNRG;refGene_tx=XM_006940107.4;refGene_gn=SYNRG;refGene_tx=XM_011288960.3;refGene_gn=SYNRG;refGene_tx=XM_019817546.2
E1 23782052 23782130
refGene_gn=SYNRG;refGene_tx=XM_003996583.5;refGene_gn=SYNRG;refGene_tx=XM_006940101.4;refGene_gn=SYNRG;refGene_tx=XM_006940102.4;refGene_gn=SYNRG;refGene_tx=XM_006940103.4;refGene_gn=SYNRG;refGene_tx=XM_006940104.4;refGene_gn=SYNRG;refGene_tx=XM_006940105.4;refGene_gn=SYNRG;refGene_tx=XM_006940106.4;refGene_gn=SYNRG;refGene_tx=XM_006940107.4;refGene_gn=SYNRG;refGene_tx=XM_006940108.4;refGene_gn=SYNRG;refGene_tx=XM_006940109.4;refGene_gn=SYNRG;refGene_tx=XM_006940110.4;refGene_gn=SYNRG;refGene_tx=XM_006940111.4;refGene_gn=SYNRG;refGene_tx=XM_006940112.4;refGene_gn=SYNRG;refGene_tx=XM_011288960.3;refGene_gn=SYNRG;refGene_tx=XM_019817546.2;refGene_gn=SYNRG;refGene_tx=XM_019817547.2
E1 23784818 23785015
refGene_gn=SYNRG;refGene_tx=XM_003996583.5;refGene_gn=SYNRG;refGene_tx=XM_006940101.4;refGene_gn=SYNRG;refGene_tx=XM_006940102.4;refGene_gn=SYNRG;refGene_tx=XM_006940103.4;refGene_gn=SYNRG;refGene_tx=XM_006940104.4;refGene_gn=SYNRG;refGene_tx=XM_006940105.4;refGene_gn=SYNRG;refGene_tx=XM_006940106.4;refGene_gn=SYNRG;refGene_tx=XM_006940107.4;refGene_gn=SYNRG;refGene_tx=XM_006940108.4;refGene_gn=SYNRG;refGene_tx=XM_006940109.4;refGene_gn=SYNRG;refGene_tx=XM_006940110.4;refGene_gn=SYNRG;refGene_tx=XM_006940111.4;refGene_gn=SYNRG;refGene_tx=XM_006940112.4;refGene_gn=SYNRG;refGene_tx=XM_011288960.3;refGene_gn=SYNRG;refGene_tx=XM_019817546.2;refGene_gn=SYNRG;refGene_tx=XM_019817547.2
E1 23785615 23785864
refGene_gn=SYNRG;refGene_tx=XM_003996583.5;refGene_gn=SYNRG;refGene_tx=XM_006940101.4;refGene_gn=SYNRG;refGene_tx=XM_006940102.4;refGene_gn=SYNRG;refGene_tx=XM_006940103.4;refGene_gn=SYNRG;refGene_tx=XM_006940104.4;refGene_gn=SYNRG;refGene_tx=XM_006940105.4;refGene_gn=SYNRG;refGene_tx=XM_006940106.4;refGene_gn=SYNRG;refGene_tx=XM_006940107.4;refGene_gn=SYNRG;refGene_tx=XM_006940108.4;refGene_gn=SYNRG;refGene_tx=XM_006940109.4;refGene_gn=SYNRG;refGene_tx=XM_006940110.4;refGene_gn=SYNRG;refGene_tx=XM_006940111.4;refGene_gn=SYNRG;refGene_tx=XM_006940112.4;refGene_gn=SYNRG;refGene_tx=XM_011288960.3;refGene_gn=SYNRG;refGene_tx=XM_019817546.2;refGene_gn=SYNRG;refGene_tx=XM_019817547.2
E1 23787734 23787868

refGene_gn=SYNRG;refGene_tx=XM_003996583.5;refGene_gn=SYNRG;refGene_tx=XM_006940101.4;refGene_gn=SYNRG;refGene_tx=XM_006940102.4;refGene_gn=SYNRG;refGene_tx=XM_006940103.4;refGene_gn=SYNRG;refGene_tx=XM_006940104.4;refGene_gn=SYNRG;refGene_tx=XM_006940105.4;refGene_gn=SYNRG;refGene_tx=XM_006940106.4;refGene_gn=SYNRG;refGene_tx=XM_006940107.4;refGene_gn=SYNRG;refGene_tx=XM_006940108.4;refGene_gn=SYNRG;refGene_tx=XM_006940109.4;refGene_gn=SYNRG;refGene_tx=XM_006940110.4;refGene_gn=SYNRG;refGene_tx=XM_006940111.4;refGene_gn=SYNRG;refGene_tx=XM_006940112.4;refGene_gn=SYNRG;refGene_tx=XM_011288960.3;refGene_gn=SYNRG;refGene_tx=XM_019817546.2;refGene_gn=SYNRG;refGene_tx=XM_019817547.2
E1 23792708 23792827
refGene_gn=SYNRG;refGene_tx=XM_003996583.5;refGene_gn=SYNRG;refGene_tx=XM_006940101.4;refGene_gn=SYNRG;refGene_tx=XM_006940102.4;refGene_gn=SYNRG;refGene_tx=XM_006940103.4;refGene_gn=SYNRG;refGene_tx=XM_006940104.4;refGene_gn=SYNRG;refGene_tx=XM_006940105.4;refGene_gn=SYNRG;refGene_tx=XM_006940106.4;refGene_gn=SYNRG;refGene_tx=XM_006940107.4;refGene_gn=SYNRG;refGene_tx=XM_006940108.4;refGene_gn=SYNRG;refGene_tx=XM_006940109.4;refGene_gn=SYNRG;refGene_tx=XM_006940110.4;refGene_gn=SYNRG;refGene_tx=XM_006940111.4;refGene_gn=SYNRG;refGene_tx=XM_006940112.4;refGene_gn=SYNRG;refGene_tx=XM_011288960.3;refGene_gn=SYNRG;refGene_tx=XM_019817546.2;refGene_gn=SYNRG;refGene_tx=XM_019817547.2
E1 23793029 23793092
refGene_gn=SYNRG;refGene_tx=XM_003996583.5;refGene_gn=SYNRG;refGene_tx=XM_006940101.4;refGene_gn=SYNRG;refGene_tx=XM_006940102.4;refGene_gn=SYNRG;refGene_tx=XM_006940103.4;refGene_gn=SYNRG;refGene_tx=XM_006940104.4;refGene_gn=SYNRG;refGene_tx=XM_006940105.4;refGene_gn=SYNRG;refGene_tx=XM_006940106.4;refGene_gn=SYNRG;refGene_tx=XM_006940107.4;refGene_gn=SYNRG;refGene_tx=XM_006940108.4;refGene_gn=SYNRG;refGene_tx=XM_006940109.4;refGene_gn=SYNRG;refGene_tx=XM_006940110.4;refGene_gn=SYNRG;refGene_tx=XM_006940111.4;refGene_gn=SYNRG;refGene_tx=XM_006940112.4;refGene_gn=SYNRG;refGene_tx=XM_011288960.3;refGene_gn=SYNRG;refGene_tx=XM_019817546.2;refGene_gn=SYNRG;refGene_tx=XM_019817547.2
E1 23796927 23797875
refGene_gn=SYNRG;refGene_tx=XM_003996583.5;refGene_gn=SYNRG;refGene_tx=XM_006940101.4;refGene_gn=SYNRG;refGene_tx=XM_006940102.4;refGene_gn=SYNRG;refGene_tx=XM_006940103.4;refGene_gn=SYNRG;refGene_tx=XM_006940104.4;refGene_gn=SYNRG;refGene_tx=XM_006940105.4;refGene_gn=SYNRG;refGene_tx=XM_006940106.4;refGene_gn=SYNRG;refGene_tx=XM_006940107.4;refGene_gn=SYNRG;refGene_tx=XM_006940108.4;refGene_gn=SYNRG;refGene_tx=XM_006940109.4;refGene_gn=SYNRG;refGene_tx=XM_006940110.4;refGene_gn=SYNRG;refGene_tx=XM_006940111.4;refGene_gn=SYNRG;refGene_tx=XM_006940112.4;refGene_gn=SYNRG;refGene_tx=XM_011288960.3;refGene_gn=SYNRG;refGene_tx=XM_019817546.2;refGene_gn=SYNRG;refGene_tx=XM_019817547.2
E1 23807432 23808026
refGene_gn=SYNRG;refGene_tx=XM_003996583.5;refGene_gn=SYNRG;refGene_tx=XM_006940101.4;refGene_gn=SYNRG;refGene_tx=XM_006940102.4;refGene_gn=SYNRG;refGene_tx=XM_006940103.4;refGene_gn=SYNRG;refGene_tx=XM_006940104.4;refGene_gn=SYNRG;refGene_tx=XM_006940105.4;refGene_gn=SYNRG;refGene_tx=XM_006940106.4;refGene_gn=SYNRG;refGene_tx=XM_006940107.4;refGene_gn=SYNRG;refGene_tx=XM_006940108.4;refGene_gn=SYNRG;refGene_tx=XM_00

6940109.4;refGene_gn=SYNRG;refGene_tx=XM_006940110.4;refGene_gn=SYNRG;
refGene_tx=XM_006940111.4;refGene_gn=SYNRG;refGene_tx=XM_006940112.4;r
efGene_gn=SYNRG;refGene_tx=XM_011288960.3;refGene_gn=SYNRG;refGene_tx=
XM_019817546.2;refGene_gn=SYNRG;refGene_tx=XM_019817547.2
E1 23809802 23809966
refGene_gn=SYNRG;refGene_tx=XM_003996583.5;refGene_gn=SYNRG;refGene_tx
=XM_006940101.4;refGene_gn=SYNRG;refGene_tx=XM_006940102.4;refGene_gn=
SYNRG;refGene_tx=XM_006940103.4;refGene_gn=SYNRG;refGene_tx=XM_0069401
04.4;refGene_gn=SYNRG;refGene_tx=XM_006940105.4;refGene_gn=SYNRG;refGe
ne_tx=XM_006940106.4;refGene_gn=SYNRG;refGene_tx=XM_006940107.4;refGen
e_gn=SYNRG;refGene_tx=XM_006940108.4;refGene_gn=SYNRG;refGene_tx=XM_00
6940109.4;refGene_gn=SYNRG;refGene_tx=XM_006940110.4;refGene_gn=SYNRG;
refGene_tx=XM_006940111.4;refGene_gn=SYNRG;refGene_tx=XM_006940112.4;r
efGene_gn=SYNRG;refGene_tx=XM_011288960.3;refGene_gn=SYNRG;refGene_tx=
XM_019817546.2;refGene_gn=SYNRG;refGene_tx=XM_019817547.2
E1 23810766 23810820
refGene_gn=SYNRG;refGene_tx=XM_003996583.5;refGene_gn=SYNRG;refGene_tx
=XM_006940101.4;refGene_gn=SYNRG;refGene_tx=XM_006940102.4;refGene_gn=
SYNRG;refGene_tx=XM_006940103.4;refGene_gn=SYNRG;refGene_tx=XM_0069401
04.4;refGene_gn=SYNRG;refGene_tx=XM_006940105.4;refGene_gn=SYNRG;refGe
ne_tx=XM_006940106.4;refGene_gn=SYNRG;refGene_tx=XM_006940107.4;refGen
e_gn=SYNRG;refGene_tx=XM_006940108.4;refGene_gn=SYNRG;refGene_tx=XM_00
6940109.4;refGene_gn=SYNRG;refGene_tx=XM_006940110.4;refGene_gn=SYNRG;
refGene_tx=XM_006940111.4;refGene_gn=SYNRG;refGene_tx=XM_006940112.4;r
efGene_gn=SYNRG;refGene_tx=XM_011288960.3;refGene_gn=SYNRG;refGene_tx=
XM_019817546.2;refGene_gn=SYNRG;refGene_tx=XM_019817547.2
E1 23811712 23811809
refGene_gn=SYNRG;refGene_tx=XM_003996583.5;refGene_gn=SYNRG;refGene_tx
=XM_006940101.4;refGene_gn=SYNRG;refGene_tx=XM_006940102.4;refGene_gn=
SYNRG;refGene_tx=XM_006940103.4;refGene_gn=SYNRG;refGene_tx=XM_0069401
04.4;refGene_gn=SYNRG;refGene_tx=XM_006940105.4;refGene_gn=SYNRG;refGe
ne_tx=XM_006940106.4;refGene_gn=SYNRG;refGene_tx=XM_006940107.4;refGen
e_gn=SYNRG;refGene_tx=XM_006940108.4;refGene_gn=SYNRG;refGene_tx=XM_00
6940109.4;refGene_gn=SYNRG;refGene_tx=XM_006940110.4;refGene_gn=SYNRG;
refGene_tx=XM_006940111.4;refGene_gn=SYNRG;refGene_tx=XM_006940112.4;r
efGene_gn=SYNRG;refGene_tx=XM_011288960.3;refGene_gn=SYNRG;refGene_tx=
XM_019817546.2;refGene_gn=SYNRG;refGene_tx=XM_019817547.2
E1 23813629 23813778
refGene_gn=SYNRG;refGene_tx=XM_003996583.5;refGene_gn=SYNRG;refGene_tx
=XM_006940101.4;refGene_gn=SYNRG;refGene_tx=XM_006940102.4;refGene_gn=
SYNRG;refGene_tx=XM_006940103.4;refGene_gn=SYNRG;refGene_tx=XM_0069401
04.4;refGene_gn=SYNRG;refGene_tx=XM_006940105.4;refGene_gn=SYNRG;refGe
ne_tx=XM_006940106.4;refGene_gn=SYNRG;refGene_tx=XM_006940107.4;refGen
e_gn=SYNRG;refGene_tx=XM_006940108.4;refGene_gn=SYNRG;refGene_tx=XM_00
6940109.4;refGene_gn=SYNRG;refGene_tx=XM_006940110.4;refGene_gn=SYNRG;
refGene_tx=XM_006940111.4;refGene_gn=SYNRG;refGene_tx=XM_006940112.4;r
efGene_gn=SYNRG;refGene_tx=XM_011288960.3;refGene_gn=SYNRG;refGene_tx=
XM_019817546.2;refGene_gn=SYNRG;refGene_tx=XM_019817547.2
E1 23828135 23828246
refGene_gn=SYNRG;refGene_tx=XM_003996583.5;refGene_gn=SYNRG;refGene_tx

=XM_006940101.4;refGene_gn=SYNRG;refGene_tx=XM_006940102.4;refGene_gn=SYNRG;refGene_tx=XM_006940103.4;refGene_gn=SYNRG;refGene_tx=XM_006940104.4;refGene_gn=SYNRG;refGene_tx=XM_006940105.4;refGene_gn=SYNRG;refGene_tx=XM_006940106.4;refGene_gn=SYNRG;refGene_tx=XM_006940107.4;refGene_gn=SYNRG;refGene_tx=XM_006940108.4;refGene_gn=SYNRG;refGene_tx=XM_006940109.4;refGene_gn=SYNRG;refGene_tx=XM_006940110.4;refGene_gn=SYNRG;refGene_tx=XM_006940111.4;refGene_gn=SYNRG;refGene_tx=XM_006940112.4;refGene_gn=SYNRG;refGene_tx=XM_011288960.3;refGene_gn=SYNRG;refGene_tx=XM_019817546.2;refGene_gn=SYNRG;refGene_tx=XM_019817547.2
E1 23828598 23828634
refGene_gn=SYNRG;refGene_tx=XM_006940101.4;refGene_gn=SYNRG;refGene_tx=XM_006940102.4;refGene_gn=SYNRG;refGene_tx=XM_006940104.4;refGene_gn=SYNRG;refGene_tx=XM_006940105.4;refGene_gn=SYNRG;refGene_tx=XM_006940106.4;refGene_gn=SYNRG;refGene_tx=XM_006940107.4;refGene_gn=SYNRG;refGene_tx=XM_006940108.4;refGene_gn=SYNRG;refGene_tx=XM_006940109.4;refGene_gn=SYNRG;refGene_tx=XM_006940110.4;refGene_gn=SYNRG;refGene_tx=XM_006940112.4;refGene_gn=SYNRG;refGene_tx=XM_011288960.3;refGene_gn=SYNRG;refGene_tx=XM_019817546.2;refGene_gn=SYNRG;refGene_tx=XM_019817547.2
E1 23829639 23829771
refGene_gn=SYNRG;refGene_tx=XM_003996583.5;refGene_gn=SYNRG;refGene_tx=XM_006940101.4;refGene_gn=SYNRG;refGene_tx=XM_006940102.4;refGene_gn=SYNRG;refGene_tx=XM_006940103.4;refGene_gn=SYNRG;refGene_tx=XM_006940104.4;refGene_gn=SYNRG;refGene_tx=XM_006940105.4;refGene_gn=SYNRG;refGene_tx=XM_006940106.4;refGene_gn=SYNRG;refGene_tx=XM_006940107.4;refGene_gn=SYNRG;refGene_tx=XM_006940108.4;refGene_gn=SYNRG;refGene_tx=XM_006940109.4;refGene_gn=SYNRG;refGene_tx=XM_006940110.4;refGene_gn=SYNRG;refGene_tx=XM_006940111.4;refGene_gn=SYNRG;refGene_tx=XM_006940112.4;refGene_gn=SYNRG;refGene_tx=XM_011288960.3;refGene_gn=SYNRG;refGene_tx=XM_019817546.2;refGene_gn=SYNRG;refGene_tx=XM_019817547.2
E1 23834263 23834860 refGene_gn=DUSP14;refGene_tx=XM_006940100.4
E1 23868444 23868630
refGene_gn=TADA2A;refGene_tx=XM_019817548.1;refGene_gn=TADA2A;refGene_tx=XM_023243547.1;refGene_gn=TADA2A;refGene_tx=XM_023243548.1;refGene_gn=TADA2A;refGene_tx=XM_023243549.1;refGene_gn=TADA2A;refGene_tx=XM_023243550.1
E1 23870148 23870222
refGene_gn=TADA2A;refGene_tx=XM_019817548.1;refGene_gn=TADA2A;refGene_tx=XM_023243547.1;refGene_gn=TADA2A;refGene_tx=XM_023243548.1;refGene_gn=TADA2A;refGene_tx=XM_023243549.1;refGene_gn=TADA2A;refGene_tx=XM_023243550.1
E1 23872285 23872329
refGene_gn=TADA2A;refGene_tx=XM_019817548.1;refGene_gn=TADA2A;refGene_tx=XM_023243547.1;refGene_gn=TADA2A;refGene_tx=XM_023243548.1;refGene_gn=TADA2A;refGene_tx=XM_023243549.1;refGene_gn=TADA2A;refGene_tx=XM_023243550.1
E1 23873126 23873259
refGene_gn=TADA2A;refGene_tx=XM_019817548.1;refGene_gn=TADA2A;refGene_tx=XM_023243547.1;refGene_gn=TADA2A;refGene_tx=XM_023243548.1;refGene_gn=TADA2A;refGene_tx=XM_023243549.1;refGene_gn=TADA2A;refGene_tx=XM_023243550.1

E1 23876292 23876364
refGene_gn=TADA2A;refGene_tx=XM_019817548.1;refGene_gn=TADA2A;refGene_tx=XM_023243547.1;refGene_gn=TADA2A;refGene_tx=XM_023243549.1
E1 23878507 23878618
refGene_gn=TADA2A;refGene_tx=XM_019817548.1;refGene_gn=TADA2A;refGene_tx=XM_023243547.1;refGene_gn=TADA2A;refGene_tx=XM_023243548.1;refGene_gn=TADA2A;refGene_tx=XM_023243549.1
E1 23881894 23881938
refGene_gn=TADA2A;refGene_tx=XM_019817548.1;refGene_gn=TADA2A;refGene_tx=XM_023243547.1;refGene_gn=TADA2A;refGene_tx=XM_023243548.1;refGene_gn=TADA2A;refGene_tx=XM_023243549.1;refGene_gn=TADA2A;refGene_tx=XM_023243550.1
E1 23884042 23884106
refGene_gn=TADA2A;refGene_tx=XM_019817548.1;refGene_gn=TADA2A;refGene_tx=XM_023243547.1;refGene_gn=TADA2A;refGene_tx=XM_023243548.1;refGene_gn=TADA2A;refGene_tx=XM_023243549.1;refGene_gn=TADA2A;refGene_tx=XM_023243550.1
E1 23893040 23893113
refGene_gn=TADA2A;refGene_tx=XM_019817548.1;refGene_gn=TADA2A;refGene_tx=XM_023243547.1;refGene_gn=TADA2A;refGene_tx=XM_023243548.1;refGene_gn=TADA2A;refGene_tx=XM_023243550.1
E1 23894539 23894628
refGene_gn=TADA2A;refGene_tx=XM_019817548.1;refGene_gn=TADA2A;refGene_tx=XM_023243547.1;refGene_gn=TADA2A;refGene_tx=XM_023243548.1;refGene_gn=TADA2A;refGene_tx=XM_023243550.1
E1 23895884 23896042
refGene_gn=TADA2A;refGene_tx=XM_019817548.1;refGene_gn=TADA2A;refGene_tx=XM_023243547.1;refGene_gn=TADA2A;refGene_tx=XM_023243548.1;refGene_gn=TADA2A;refGene_tx=XM_023243549.1;refGene_gn=TADA2A;refGene_tx=XM_023243550.1
E1 23897763 23897855
refGene_gn=TADA2A;refGene_tx=XM_019817548.1;refGene_gn=TADA2A;refGene_tx=XM_023243547.1;refGene_gn=TADA2A;refGene_tx=XM_023243548.1;refGene_gn=TADA2A;refGene_tx=XM_023243549.1;refGene_gn=TADA2A;refGene_tx=XM_023243550.1
E1 23907230 23907290
refGene_gn=TADA2A;refGene_tx=XM_019817548.1;refGene_gn=TADA2A;refGene_tx=XM_023243547.1;refGene_gn=TADA2A;refGene_tx=XM_023243548.1;refGene_gn=TADA2A;refGene_tx=XM_023243549.1;refGene_gn=TADA2A;refGene_tx=XM_023243550.1
E1 23908307 23908414
refGene_gn=TADA2A;refGene_tx=XM_019817548.1;refGene_gn=TADA2A;refGene_tx=XM_023243547.1;refGene_gn=TADA2A;refGene_tx=XM_023243548.1;refGene_gn=TADA2A;refGene_tx=XM_023243549.1;refGene_gn=TADA2A;refGene_tx=XM_023243550.1
E1 23917708 23917733
refGene_gn=TADA2A;refGene_tx=XM_023243547.1;refGene_gn=TADA2A;refGene_tx=XM_023243548.1;refGene_gn=TADA2A;refGene_tx=XM_023243549.1;refGene_gn=TADA2A;refGene_tx=XM_023243550.1
E1 23938898 23938976

refGene_gn=CE1H17orf78;refGene_tx=XM_019816928.2
E1 23942185 23942320
refGene_gn=CE1H17orf78;refGene_tx=XM_019816928.2
E1 23942763 23942885
refGene_gn=CE1H17orf78;refGene_tx=XM_019816928.2
E1 23943894 23944020
refGene_gn=CE1H17orf78;refGene_tx=XM_019816928.2
E1 23944276 23944382
refGene_gn=CE1H17orf78;refGene_tx=XM_019816928.2
E1 23948855 23949029
refGene_gn=CE1H17orf78;refGene_tx=XM_019816928.2
E1 23951029 23951086
refGene_gn=CE1H17orf78;refGene_tx=XM_019816928.2
E1 23975951 23976178
refGene_gn=ACACA;refGene_tx=XM_006940115.4;refGene_gn=ACACA;refGene_tx=XM_006940119.4;refGene_gn=ACACA;refGene_tx=XM_011288954.1;refGene_gn=ACACA;refGene_tx=XM_011288955.1;refGene_gn=ACACA;refGene_tx=XM_019817551.1;refGene_gn=ACACA;refGene_tx=XM_019817552.1;refGene_gn=ACACA;refGene_tx=XM_019817554.1;refGene_gn=ACACA;refGene_tx=XM_019817555.2;refGene_gn=ACACA;refGene_tx=XM_019817560.1;refGene_gn=ACACA;refGene_tx=XM_023244661.1;refGene_gn=ACACA;refGene_tx=XM_023244663.1;refGene_gn=ACACA;refGene_tx=XM_023244664.1;refGene_gn=ACACA;refGene_tx=XM_023244665.1
E1 23997381 23997431 refGene_gn=ACACA;refGene_tx=XM_019817558.2
E1 23999759 23999812
refGene_gn=ACACA;refGene_tx=XM_019817556.2;refGene_gn=ACACA;refGene_tx=XM_019817557.2
E1 24008512 24008645
refGene_gn=ACACA;refGene_tx=XM_006940115.4;refGene_gn=ACACA;refGene_tx=XM_006940119.4;refGene_gn=ACACA;refGene_tx=XM_011288954.1;refGene_gn=ACACA;refGene_tx=XM_011288955.1;refGene_gn=ACACA;refGene_tx=XM_019817551.1;refGene_gn=ACACA;refGene_tx=XM_019817552.1;refGene_gn=ACACA;refGene_tx=XM_019817554.1;refGene_gn=ACACA;refGene_tx=XM_019817555.2;refGene_gn=ACACA;refGene_tx=XM_019817556.2;refGene_gn=ACACA;refGene_tx=XM_019817557.2;refGene_gn=ACACA;refGene_tx=XM_019817558.2;refGene_gn=ACACA;refGene_tx=XM_019817560.1;refGene_gn=ACACA;refGene_tx=XM_023244661.1;refGene_gn=ACACA;refGene_tx=XM_023244663.1;refGene_gn=ACACA;refGene_tx=XM_023244664.1;refGene_gn=ACACA;refGene_tx=XM_023244665.1
E1 24010179 24010318
refGene_gn=ACACA;refGene_tx=XM_006940115.4;refGene_gn=ACACA;refGene_tx=XM_006940119.4;refGene_gn=ACACA;refGene_tx=XM_011288954.1;refGene_gn=ACACA;refGene_tx=XM_011288955.1;refGene_gn=ACACA;refGene_tx=XM_019817551.1;refGene_gn=ACACA;refGene_tx=XM_019817552.1;refGene_gn=ACACA;refGene_tx=XM_019817554.1;refGene_gn=ACACA;refGene_tx=XM_019817555.2;refGene_gn=ACACA;refGene_tx=XM_019817556.2;refGene_gn=ACACA;refGene_tx=XM_019817557.2;refGene_gn=ACACA;refGene_tx=XM_019817558.2;refGene_gn=ACACA;refGene_tx=XM_019817560.1;refGene_gn=ACACA;refGene_tx=XM_023244661.1;refGene_gn=ACACA;refGene_tx=XM_023244663.1;refGene_gn=ACACA;refGene_tx=XM_023244664.1;refGene_gn=ACACA;refGene_tx=XM_023244665.1
E1 24016653 24016763
refGene_gn=ACACA;refGene_tx=XM_006940115.4;refGene_gn=ACACA;refGene_tx

=XM_006940119.4;refGene_gn=ACACA;refGene_tx=XM_011288954.1;refGene_gn=ACACA;refGene_tx=XM_011288955.1;refGene_gn=ACACA;refGene_tx=XM_019817551.1;refGene_gn=ACACA;refGene_tx=XM_019817552.1;refGene_gn=ACACA;refGene_tx=XM_019817554.1;refGene_gn=ACACA;refGene_tx=XM_019817555.2;refGene_gn=ACACA;refGene_tx=XM_019817556.2;refGene_gn=ACACA;refGene_tx=XM_019817557.2;refGene_gn=ACACA;refGene_tx=XM_019817558.2;refGene_gn=ACACA;refGene_tx=XM_019817560.1;refGene_gn=ACACA;refGene_tx=XM_023244661.1;refGene_gn=ACACA;refGene_tx=XM_023244663.1;refGene_gn=ACACA;refGene_tx=XM_023244664.1;refGene_gn=ACACA;refGene_tx=XM_023244665.1
E1 24017584 24017666
refGene_gn=ACACA;refGene_tx=XM_006940115.4;refGene_gn=ACACA;refGene_tx=XM_006940119.4;refGene_gn=ACACA;refGene_tx=XM_011288954.1;refGene_gn=ACACA;refGene_tx=XM_011288955.1;refGene_gn=ACACA;refGene_tx=XM_019817551.1;refGene_gn=ACACA;refGene_tx=XM_019817552.1;refGene_gn=ACACA;refGene_tx=XM_019817554.1;refGene_gn=ACACA;refGene_tx=XM_019817555.2;refGene_gn=ACACA;refGene_tx=XM_019817556.2;refGene_gn=ACACA;refGene_tx=XM_019817557.2;refGene_gn=ACACA;refGene_tx=XM_019817558.2;refGene_gn=ACACA;refGene_tx=XM_019817560.1;refGene_gn=ACACA;refGene_tx=XM_023244661.1;refGene_gn=ACACA;refGene_tx=XM_023244663.1;refGene_gn=ACACA;refGene_tx=XM_023244664.1;refGene_gn=ACACA;refGene_tx=XM_023244665.1
E1 24018891 24018990
refGene_gn=ACACA;refGene_tx=XM_006940115.4;refGene_gn=ACACA;refGene_tx=XM_006940119.4;refGene_gn=ACACA;refGene_tx=XM_011288954.1;refGene_gn=ACACA;refGene_tx=XM_011288955.1;refGene_gn=ACACA;refGene_tx=XM_019817551.1;refGene_gn=ACACA;refGene_tx=XM_019817552.1;refGene_gn=ACACA;refGene_tx=XM_019817554.1;refGene_gn=ACACA;refGene_tx=XM_019817555.2;refGene_gn=ACACA;refGene_tx=XM_019817556.2;refGene_gn=ACACA;refGene_tx=XM_019817557.2;refGene_gn=ACACA;refGene_tx=XM_019817558.2;refGene_gn=ACACA;refGene_tx=XM_019817560.1;refGene_gn=ACACA;refGene_tx=XM_023244661.1;refGene_gn=ACACA;refGene_tx=XM_023244663.1;refGene_gn=ACACA;refGene_tx=XM_023244664.1;refGene_gn=ACACA;refGene_tx=XM_023244665.1
E1 24020305 24020412
refGene_gn=ACACA;refGene_tx=XM_006940115.4;refGene_gn=ACACA;refGene_tx=XM_006940119.4;refGene_gn=ACACA;refGene_tx=XM_011288954.1;refGene_gn=ACACA;refGene_tx=XM_011288955.1;refGene_gn=ACACA;refGene_tx=XM_019817551.1;refGene_gn=ACACA;refGene_tx=XM_019817552.1;refGene_gn=ACACA;refGene_tx=XM_019817554.1;refGene_gn=ACACA;refGene_tx=XM_019817555.2;refGene_gn=ACACA;refGene_tx=XM_019817556.2;refGene_gn=ACACA;refGene_tx=XM_019817557.2;refGene_gn=ACACA;refGene_tx=XM_019817558.2;refGene_gn=ACACA;refGene_tx=XM_019817560.1;refGene_gn=ACACA;refGene_tx=XM_023244661.1;refGene_gn=ACACA;refGene_tx=XM_023244663.1;refGene_gn=ACACA;refGene_tx=XM_023244664.1;refGene_gn=ACACA;refGene_tx=XM_023244665.1
E1 24024634 24024745
refGene_gn=ACACA;refGene_tx=XM_006940115.4;refGene_gn=ACACA;refGene_tx=XM_006940119.4;refGene_gn=ACACA;refGene_tx=XM_011288954.1;refGene_gn=ACACA;refGene_tx=XM_011288955.1;refGene_gn=ACACA;refGene_tx=XM_019817551.1;refGene_gn=ACACA;refGene_tx=XM_019817552.1;refGene_gn=ACACA;refGene_tx=XM_019817554.1;refGene_gn=ACACA;refGene_tx=XM_019817555.2;refGene_gn=ACACA;refGene_tx=XM_019817556.2;refGene_gn=ACACA;refGene_tx=XM_019817557.2;refGene_gn=ACACA;refGene_tx=XM_019817558.2;refGene_gn=ACACA;

refGene_tx=XM_019817560.1;refGene_gn=ACACA;refGene_tx=XM_023244661.1;refGene_gn=ACACA;refGene_tx=XM_023244663.1;refGene_gn=ACACA;refGene_tx=XM_023244664.1;refGene_gn=ACACA;refGene_tx=XM_023244665.1
E1 24030276 24030486
refGene_gn=ACACA;refGene_tx=XM_006940115.4;refGene_gn=ACACA;refGene_tx=XM_006940119.4;refGene_gn=ACACA;refGene_tx=XM_011288954.1;refGene_gn=ACACA;refGene_tx=XM_011288955.1;refGene_gn=ACACA;refGene_tx=XM_011288956.2;refGene_gn=ACACA;refGene_tx=XM_019817551.1;refGene_gn=ACACA;refGene_tx=XM_019817552.1;refGene_gn=ACACA;refGene_tx=XM_019817554.1;refGene_gn=ACACA;refGene_tx=XM_019817555.2;refGene_gn=ACACA;refGene_tx=XM_019817556.2;refGene_gn=ACACA;refGene_tx=XM_019817557.2;refGene_gn=ACACA;refGene_tx=XM_019817558.2;refGene_gn=ACACA;refGene_tx=XM_019817560.1;refGene_gn=ACACA;refGene_tx=XM_023244661.1;refGene_gn=ACACA;refGene_tx=XM_023244663.1;refGene_gn=ACACA;refGene_tx=XM_023244664.1;refGene_gn=ACACA;refGene_tx=XM_023244665.1
E1 24033906 24034077
refGene_gn=ACACA;refGene_tx=XM_006940115.4;refGene_gn=ACACA;refGene_tx=XM_006940119.4;refGene_gn=ACACA;refGene_tx=XM_011288954.1;refGene_gn=ACACA;refGene_tx=XM_011288955.1;refGene_gn=ACACA;refGene_tx=XM_011288956.2;refGene_gn=ACACA;refGene_tx=XM_019817551.1;refGene_gn=ACACA;refGene_tx=XM_019817552.1;refGene_gn=ACACA;refGene_tx=XM_019817554.1;refGene_gn=ACACA;refGene_tx=XM_019817555.2;refGene_gn=ACACA;refGene_tx=XM_019817556.2;refGene_gn=ACACA;refGene_tx=XM_019817557.2;refGene_gn=ACACA;refGene_tx=XM_019817558.2;refGene_gn=ACACA;refGene_tx=XM_019817560.1;refGene_gn=ACACA;refGene_tx=XM_023244661.1;refGene_gn=ACACA;refGene_tx=XM_023244663.1;refGene_gn=ACACA;refGene_tx=XM_023244664.1;refGene_gn=ACACA;refGene_tx=XM_023244665.1
E1 24034970 24035132
refGene_gn=ACACA;refGene_tx=XM_006940115.4;refGene_gn=ACACA;refGene_tx=XM_006940119.4;refGene_gn=ACACA;refGene_tx=XM_011288954.1;refGene_gn=ACACA;refGene_tx=XM_011288955.1;refGene_gn=ACACA;refGene_tx=XM_011288956.2;refGene_gn=ACACA;refGene_tx=XM_019817551.1;refGene_gn=ACACA;refGene_tx=XM_019817552.1;refGene_gn=ACACA;refGene_tx=XM_019817554.1;refGene_gn=ACACA;refGene_tx=XM_019817555.2;refGene_gn=ACACA;refGene_tx=XM_019817556.2;refGene_gn=ACACA;refGene_tx=XM_019817557.2;refGene_gn=ACACA;refGene_tx=XM_019817558.2;refGene_gn=ACACA;refGene_tx=XM_019817560.1;refGene_gn=ACACA;refGene_tx=XM_023244661.1;refGene_gn=ACACA;refGene_tx=XM_023244663.1;refGene_gn=ACACA;refGene_tx=XM_023244664.1;refGene_gn=ACACA;refGene_tx=XM_023244665.1
E1 24035458 24035622
refGene_gn=ACACA;refGene_tx=XM_006940115.4;refGene_gn=ACACA;refGene_tx=XM_006940119.4;refGene_gn=ACACA;refGene_tx=XM_011288954.1;refGene_gn=ACACA;refGene_tx=XM_011288955.1;refGene_gn=ACACA;refGene_tx=XM_011288956.2;refGene_gn=ACACA;refGene_tx=XM_019817551.1;refGene_gn=ACACA;refGene_tx=XM_019817552.1;refGene_gn=ACACA;refGene_tx=XM_019817554.1;refGene_gn=ACACA;refGene_tx=XM_019817555.2;refGene_gn=ACACA;refGene_tx=XM_019817556.2;refGene_gn=ACACA;refGene_tx=XM_019817557.2;refGene_gn=ACACA;refGene_tx=XM_019817558.2;refGene_gn=ACACA;refGene_tx=XM_019817560.1;refGene_gn=ACACA;refGene_tx=XM_023244661.1;refGene_gn=ACACA;refGene_tx=XM_023244663.1;refGene_gn=ACACA;refGene_tx=XM_023244664.1;refGene_gn=A

CACA;refGene_tx=XM_023244665.1
E1 24039678 24039829
refGene_gn=ACACA;refGene_tx=XM_006940115.4;refGene_gn=ACACA;refGene_tx
=XM_006940119.4;refGene_gn=ACACA;refGene_tx=XM_011288954.1;refGene_gn=
ACACA;refGene_tx=XM_011288955.1;refGene_gn=ACACA;refGene_tx=XM_0112889
56.2;refGene_gn=ACACA;refGene_tx=XM_019817551.1;refGene_gn=ACACA;refGe
ne_tx=XM_019817552.1;refGene_gn=ACACA;refGene_tx=XM_019817554.1;refGen
e_gn=ACACA;refGene_tx=XM_019817555.2;refGene_gn=ACACA;refGene_tx=XM_01
9817556.2;refGene_gn=ACACA;refGene_tx=XM_019817557.2;refGene_gn=ACACA;
refGene_tx=XM_019817558.2;refGene_gn=ACACA;refGene_tx=XM_019817560.1;r
efGene_gn=ACACA;refGene_tx=XM_023244661.1;refGene_gn=ACACA;refGene_tx=
XM_023244663.1;refGene_gn=ACACA;refGene_tx=XM_023244664.1;refGene_gn=A
CACA;refGene_tx=XM_023244665.1
E1 24040793 24040897
refGene_gn=ACACA;refGene_tx=XM_006940115.4;refGene_gn=ACACA;refGene_tx
=XM_006940119.4;refGene_gn=ACACA;refGene_tx=XM_011288954.1;refGene_gn=
ACACA;refGene_tx=XM_011288955.1;refGene_gn=ACACA;refGene_tx=XM_0112889
56.2;refGene_gn=ACACA;refGene_tx=XM_019817551.1;refGene_gn=ACACA;refGe
ne_tx=XM_019817552.1;refGene_gn=ACACA;refGene_tx=XM_019817554.1;refGen
e_gn=ACACA;refGene_tx=XM_019817555.2;refGene_gn=ACACA;refGene_tx=XM_01
9817556.2;refGene_gn=ACACA;refGene_tx=XM_019817557.2;refGene_gn=ACACA;
refGene_tx=XM_019817558.2;refGene_gn=ACACA;refGene_tx=XM_019817560.1;r
efGene_gn=ACACA;refGene_tx=XM_023244661.1;refGene_gn=ACACA;refGene_tx=
XM_023244663.1;refGene_gn=ACACA;refGene_tx=XM_023244664.1;refGene_gn=A
CACA;refGene_tx=XM_023244665.1
E1 24042892 24042974
refGene_gn=ACACA;refGene_tx=XM_006940115.4;refGene_gn=ACACA;refGene_tx
=XM_006940119.4;refGene_gn=ACACA;refGene_tx=XM_011288954.1;refGene_gn=
ACACA;refGene_tx=XM_011288955.1;refGene_gn=ACACA;refGene_tx=XM_0112889
56.2;refGene_gn=ACACA;refGene_tx=XM_019817551.1;refGene_gn=ACACA;refGe
ne_tx=XM_019817552.1;refGene_gn=ACACA;refGene_tx=XM_019817554.1;refGen
e_gn=ACACA;refGene_tx=XM_019817555.2;refGene_gn=ACACA;refGene_tx=XM_01
9817556.2;refGene_gn=ACACA;refGene_tx=XM_019817557.2;refGene_gn=ACACA;
refGene_tx=XM_019817558.2;refGene_gn=ACACA;refGene_tx=XM_019817560.1;r
efGene_gn=ACACA;refGene_tx=XM_023244661.1;refGene_gn=ACACA;refGene_tx=
XM_023244663.1;refGene_gn=ACACA;refGene_tx=XM_023244664.1;refGene_gn=A
CACA;refGene_tx=XM_023244665.1
E1 24043372 24043518
refGene_gn=ACACA;refGene_tx=XM_006940115.4;refGene_gn=ACACA;refGene_tx
=XM_006940119.4;refGene_gn=ACACA;refGene_tx=XM_011288954.1;refGene_gn=
ACACA;refGene_tx=XM_011288955.1;refGene_gn=ACACA;refGene_tx=XM_0112889
56.2;refGene_gn=ACACA;refGene_tx=XM_019817551.1;refGene_gn=ACACA;refGe
ne_tx=XM_019817552.1;refGene_gn=ACACA;refGene_tx=XM_019817554.1;refGen
e_gn=ACACA;refGene_tx=XM_019817555.2;refGene_gn=ACACA;refGene_tx=XM_01
9817556.2;refGene_gn=ACACA;refGene_tx=XM_019817557.2;refGene_gn=ACACA;
refGene_tx=XM_019817558.2;refGene_gn=ACACA;refGene_tx=XM_019817560.1;r
efGene_gn=ACACA;refGene_tx=XM_023244661.1;refGene_gn=ACACA;refGene_tx=
XM_023244663.1;refGene_gn=ACACA;refGene_tx=XM_023244664.1;refGene_gn=A
CACA;refGene_tx=XM_023244665.1
E1 24044283 24044434

refGene_gn=ACACA;refGene_tx=XM_006940115.4;refGene_gn=ACACA;refGene_tx
=XM_006940119.4;refGene_gn=ACACA;refGene_tx=XM_011288954.1;refGene_gn=
ACACA;refGene_tx=XM_011288955.1;refGene_gn=ACACA;refGene_tx=XM_0112889
56.2;refGene_gn=ACACA;refGene_tx=XM_019817551.1;refGene_gn=ACACA;refGe
ne_tx=XM_019817552.1;refGene_gn=ACACA;refGene_tx=XM_019817554.1;refGen
e_gn=ACACA;refGene_tx=XM_019817555.2;refGene_gn=ACACA;refGene_tx=XM_01
9817556.2;refGene_gn=ACACA;refGene_tx=XM_019817557.2;refGene_gn=ACACA;
refGene_tx=XM_019817558.2;refGene_gn=ACACA;refGene_tx=XM_019817560.1;r
efGene_gn=ACACA;refGene_tx=XM_023244661.1;refGene_gn=ACACA;refGene_tx=
XM_023244663.1;refGene_gn=ACACA;refGene_tx=XM_023244664.1;refGene_gn=A
CACA;refGene_tx=XM_023244665.1
E1 24046140 24046275
refGene_gn=ACACA;refGene_tx=XM_006940115.4;refGene_gn=ACACA;refGene_tx
=XM_006940119.4;refGene_gn=ACACA;refGene_tx=XM_011288954.1;refGene_gn=
ACACA;refGene_tx=XM_011288955.1;refGene_gn=ACACA;refGene_tx=XM_0112889
56.2;refGene_gn=ACACA;refGene_tx=XM_019817551.1;refGene_gn=ACACA;refGe
ne_tx=XM_019817552.1;refGene_gn=ACACA;refGene_tx=XM_019817554.1;refGen
e_gn=ACACA;refGene_tx=XM_019817555.2;refGene_gn=ACACA;refGene_tx=XM_01
9817556.2;refGene_gn=ACACA;refGene_tx=XM_019817557.2;refGene_gn=ACACA;
refGene_tx=XM_019817558.2;refGene_gn=ACACA;refGene_tx=XM_019817560.1;r
efGene_gn=ACACA;refGene_tx=XM_023244661.1;refGene_gn=ACACA;refGene_tx=
XM_023244663.1;refGene_gn=ACACA;refGene_tx=XM_023244664.1;refGene_gn=A
CACA;refGene_tx=XM_023244665.1
E1 24046629 24046776
refGene_gn=ACACA;refGene_tx=XM_006940115.4;refGene_gn=ACACA;refGene_tx
=XM_006940119.4;refGene_gn=ACACA;refGene_tx=XM_011288954.1;refGene_gn=
ACACA;refGene_tx=XM_011288955.1;refGene_gn=ACACA;refGene_tx=XM_0112889
56.2;refGene_gn=ACACA;refGene_tx=XM_019817551.1;refGene_gn=ACACA;refGe
ne_tx=XM_019817552.1;refGene_gn=ACACA;refGene_tx=XM_019817554.1;refGen
e_gn=ACACA;refGene_tx=XM_019817555.2;refGene_gn=ACACA;refGene_tx=XM_01
9817556.2;refGene_gn=ACACA;refGene_tx=XM_019817557.2;refGene_gn=ACACA;
refGene_tx=XM_019817558.2;refGene_gn=ACACA;refGene_tx=XM_019817560.1;r
efGene_gn=ACACA;refGene_tx=XM_023244661.1;refGene_gn=ACACA;refGene_tx=
XM_023244663.1;refGene_gn=ACACA;refGene_tx=XM_023244664.1;refGene_gn=A
CACA;refGene_tx=XM_023244665.1
E1 24047113 24047302
refGene_gn=ACACA;refGene_tx=XM_006940115.4;refGene_gn=ACACA;refGene_tx
=XM_006940119.4;refGene_gn=ACACA;refGene_tx=XM_011288954.1;refGene_gn=
ACACA;refGene_tx=XM_011288955.1;refGene_gn=ACACA;refGene_tx=XM_0112889
56.2;refGene_gn=ACACA;refGene_tx=XM_019817551.1;refGene_gn=ACACA;refGe
ne_tx=XM_019817552.1;refGene_gn=ACACA;refGene_tx=XM_019817554.1;refGen
e_gn=ACACA;refGene_tx=XM_019817555.2;refGene_gn=ACACA;refGene_tx=XM_01
9817556.2;refGene_gn=ACACA;refGene_tx=XM_019817557.2;refGene_gn=ACACA;
refGene_tx=XM_019817558.2;refGene_gn=ACACA;refGene_tx=XM_019817560.1;r
efGene_gn=ACACA;refGene_tx=XM_023244661.1;refGene_gn=ACACA;refGene_tx=
XM_023244663.1;refGene_gn=ACACA;refGene_tx=XM_023244664.1;refGene_gn=A
CACA;refGene_tx=XM_023244665.1
E1 24047979 24048080
refGene_gn=ACACA;refGene_tx=XM_006940115.4;refGene_gn=ACACA;refGene_tx
=XM_006940119.4;refGene_gn=ACACA;refGene_tx=XM_011288954.1;refGene_gn=

ACACA;refGene_tx=XM_011288955.1;refGene_gn=ACACA;refGene_tx=XM_0112889
56.2;refGene_gn=ACACA;refGene_tx=XM_019817551.1;refGene_gn=ACACA;refGe
ne_tx=XM_019817552.1;refGene_gn=ACACA;refGene_tx=XM_019817554.1;refGen
e_gn=ACACA;refGene_tx=XM_019817555.2;refGene_gn=ACACA;refGene_tx=XM_01
9817556.2;refGene_gn=ACACA;refGene_tx=XM_019817557.2;refGene_gn=ACACA;
refGene_tx=XM_019817558.2;refGene_gn=ACACA;refGene_tx=XM_019817560.1;r
efGene_gn=ACACA;refGene_tx=XM_023244661.1;refGene_gn=ACACA;refGene_tx=
XM_023244663.1;refGene_gn=ACACA;refGene_tx=XM_023244664.1;refGene_gn=A
CACA;refGene_tx=XM_023244665.1
E1 24048772 24048861
refGene_gn=ACACA;refGene_tx=XM_006940115.4;refGene_gn=ACACA;refGene_tx
=XM_006940119.4;refGene_gn=ACACA;refGene_tx=XM_011288954.1;refGene_gn=
ACACA;refGene_tx=XM_011288955.1;refGene_gn=ACACA;refGene_tx=XM_0112889
56.2;refGene_gn=ACACA;refGene_tx=XM_019817551.1;refGene_gn=ACACA;refGe
ne_tx=XM_019817552.1;refGene_gn=ACACA;refGene_tx=XM_019817554.1;refGen
e_gn=ACACA;refGene_tx=XM_019817555.2;refGene_gn=ACACA;refGene_tx=XM_01
9817556.2;refGene_gn=ACACA;refGene_tx=XM_019817557.2;refGene_gn=ACACA;
refGene_tx=XM_019817558.2;refGene_gn=ACACA;refGene_tx=XM_019817560.1;r
efGene_gn=ACACA;refGene_tx=XM_023244661.1;refGene_gn=ACACA;refGene_tx=
XM_023244663.1;refGene_gn=ACACA;refGene_tx=XM_023244664.1;refGene_gn=A
CACA;refGene_tx=XM_023244665.1
E1 24053984 24054109
refGene_gn=ACACA;refGene_tx=XM_006940115.4;refGene_gn=ACACA;refGene_tx
=XM_006940119.4;refGene_gn=ACACA;refGene_tx=XM_011288954.1;refGene_gn=
ACACA;refGene_tx=XM_011288955.1;refGene_gn=ACACA;refGene_tx=XM_0112889
56.2;refGene_gn=ACACA;refGene_tx=XM_019817551.1;refGene_gn=ACACA;refGe
ne_tx=XM_019817552.1;refGene_gn=ACACA;refGene_tx=XM_019817554.1;refGen
e_gn=ACACA;refGene_tx=XM_019817555.2;refGene_gn=ACACA;refGene_tx=XM_01
9817556.2;refGene_gn=ACACA;refGene_tx=XM_019817557.2;refGene_gn=ACACA;
refGene_tx=XM_019817558.2;refGene_gn=ACACA;refGene_tx=XM_019817560.1;r
efGene_gn=ACACA;refGene_tx=XM_023244661.1;refGene_gn=ACACA;refGene_tx=
XM_023244663.1;refGene_gn=ACACA;refGene_tx=XM_023244664.1;refGene_gn=A
CACA;refGene_tx=XM_023244665.1
E1 24061609 24061723
refGene_gn=ACACA;refGene_tx=XM_006940115.4;refGene_gn=ACACA;refGene_tx
=XM_006940119.4;refGene_gn=ACACA;refGene_tx=XM_011288954.1;refGene_gn=
ACACA;refGene_tx=XM_011288955.1;refGene_gn=ACACA;refGene_tx=XM_0112889
56.2;refGene_gn=ACACA;refGene_tx=XM_019817551.1;refGene_gn=ACACA;refGe
ne_tx=XM_019817552.1;refGene_gn=ACACA;refGene_tx=XM_019817554.1;refGen
e_gn=ACACA;refGene_tx=XM_019817555.2;refGene_gn=ACACA;refGene_tx=XM_01
9817556.2;refGene_gn=ACACA;refGene_tx=XM_019817557.2;refGene_gn=ACACA;
refGene_tx=XM_019817558.2;refGene_gn=ACACA;refGene_tx=XM_019817560.1;r
efGene_gn=ACACA;refGene_tx=XM_023244661.1;refGene_gn=ACACA;refGene_tx=
XM_023244663.1;refGene_gn=ACACA;refGene_tx=XM_023244664.1;refGene_gn=A
CACA;refGene_tx=XM_023244665.1
E1 24063224 24063338
refGene_gn=ACACA;refGene_tx=XM_006940115.4;refGene_gn=ACACA;refGene_tx
=XM_006940119.4;refGene_gn=ACACA;refGene_tx=XM_011288954.1;refGene_gn=
ACACA;refGene_tx=XM_011288955.1;refGene_gn=ACACA;refGene_tx=XM_0112889
56.2;refGene_gn=ACACA;refGene_tx=XM_019817551.1;refGene_gn=ACACA;refGe

ne_tx=XM_019817552.1;refGene_gn=ACACA;refGene_tx=XM_019817554.1;refGene_gn=ACACA;refGene_tx=XM_019817555.2;refGene_gn=ACACA;refGene_tx=XM_019817556.2;refGene_gn=ACACA;refGene_tx=XM_019817557.2;refGene_gn=ACACA;refGene_tx=XM_019817558.2;refGene_gn=ACACA;refGene_tx=XM_019817560.1;refGene_gn=ACACA;refGene_tx=XM_023244661.1;refGene_gn=ACACA;refGene_tx=XM_023244663.1;refGene_gn=ACACA;refGene_tx=XM_023244664.1;refGene_gn=ACACA;refGene_tx=XM_023244665.1
E1 24064323 24064413
refGene_gn=ACACA;refGene_tx=XM_006940115.4;refGene_gn=ACACA;refGene_tx=XM_006940119.4;refGene_gn=ACACA;refGene_tx=XM_011288954.1;refGene_gn=ACACA;refGene_tx=XM_011288955.1;refGene_gn=ACACA;refGene_tx=XM_011288956.2;refGene_gn=ACACA;refGene_tx=XM_019817551.1;refGene_gn=ACACA;refGene_tx=XM_019817552.1;refGene_gn=ACACA;refGene_tx=XM_019817554.1;refGene_gn=ACACA;refGene_tx=XM_019817555.2;refGene_gn=ACACA;refGene_tx=XM_019817556.2;refGene_gn=ACACA;refGene_tx=XM_019817557.2;refGene_gn=ACACA;refGene_tx=XM_019817558.2;refGene_gn=ACACA;refGene_tx=XM_019817560.1;refGene_gn=ACACA;refGene_tx=XM_023244661.1;refGene_gn=ACACA;refGene_tx=XM_023244663.1;refGene_gn=ACACA;refGene_tx=XM_023244664.1;refGene_gn=ACACA;refGene_tx=XM_023244665.1
E1 24066191 24066310
refGene_gn=ACACA;refGene_tx=XM_006940115.4;refGene_gn=ACACA;refGene_tx=XM_006940119.4;refGene_gn=ACACA;refGene_tx=XM_011288954.1;refGene_gn=ACACA;refGene_tx=XM_011288955.1;refGene_gn=ACACA;refGene_tx=XM_011288956.2;refGene_gn=ACACA;refGene_tx=XM_019817551.1;refGene_gn=ACACA;refGene_tx=XM_019817552.1;refGene_gn=ACACA;refGene_tx=XM_019817554.1;refGene_gn=ACACA;refGene_tx=XM_019817555.2;refGene_gn=ACACA;refGene_tx=XM_019817556.2;refGene_gn=ACACA;refGene_tx=XM_019817557.2;refGene_gn=ACACA;refGene_tx=XM_019817558.2;refGene_gn=ACACA;refGene_tx=XM_019817560.1;refGene_gn=ACACA;refGene_tx=XM_023244661.1;refGene_gn=ACACA;refGene_tx=XM_023244663.1;refGene_gn=ACACA;refGene_tx=XM_023244664.1;refGene_gn=ACACA;refGene_tx=XM_023244665.1
E1 24078821 24078845
refGene_gn=ACACA;refGene_tx=XM_006940115.4;refGene_gn=ACACA;refGene_tx=XM_006940119.4;refGene_gn=ACACA;refGene_tx=XM_011288954.1;refGene_gn=ACACA;refGene_tx=XM_011288956.2;refGene_gn=ACACA;refGene_tx=XM_019817551.1;refGene_gn=ACACA;refGene_tx=XM_019817555.2;refGene_gn=ACACA;refGene_tx=XM_019817556.2;refGene_gn=ACACA;refGene_tx=XM_019817557.2;refGene_gn=ACACA;refGene_tx=XM_019817558.2;refGene_gn=ACACA;refGene_tx=XM_019817560.1;refGene_gn=ACACA;refGene_tx=XM_023244661.1;refGene_gn=ACACA;refGene_tx=XM_023244663.1;refGene_gn=ACACA;refGene_tx=XM_023244664.1;refGene_gn=ACACA;refGene_tx=XM_023244665.1
E1 24081488 24081632
refGene_gn=ACACA;refGene_tx=XM_006940115.4;refGene_gn=ACACA;refGene_tx=XM_006940119.4;refGene_gn=ACACA;refGene_tx=XM_011288954.1;refGene_gn=ACACA;refGene_tx=XM_011288955.1;refGene_gn=ACACA;refGene_tx=XM_011288956.2;refGene_gn=ACACA;refGene_tx=XM_019817551.1;refGene_gn=ACACA;refGene_tx=XM_019817552.1;refGene_gn=ACACA;refGene_tx=XM_019817554.1;refGene_gn=ACACA;refGene_tx=XM_019817555.2;refGene_gn=ACACA;refGene_tx=XM_019817556.2;refGene_gn=ACACA;refGene_tx=XM_019817557.2;refGene_gn=ACACA;refGene_tx=XM_019817558.2;refGene_gn=ACACA;refGene_tx=XM_019817560.1;r

efGene_gn=ACACA;refGene_tx=XM_023244661.1;refGene_gn=ACACA;refGene_tx=XM_023244663.1;refGene_gn=ACACA;refGene_tx=XM_023244664.1;refGene_gn=ACACA;refGene_tx=XM_023244665.1
E1 24082417 24082514
refGene_gn=ACACA;refGene_tx=XM_006940115.4;refGene_gn=ACACA;refGene_tx=XM_006940119.4;refGene_gn=ACACA;refGene_tx=XM_011288954.1;refGene_gn=ACACA;refGene_tx=XM_011288955.1;refGene_gn=ACACA;refGene_tx=XM_011288956.2;refGene_gn=ACACA;refGene_tx=XM_019817551.1;refGene_gn=ACACA;refGene_tx=XM_019817552.1;refGene_gn=ACACA;refGene_tx=XM_019817554.1;refGene_gn=ACACA;refGene_tx=XM_019817555.2;refGene_gn=ACACA;refGene_tx=XM_019817556.2;refGene_gn=ACACA;refGene_tx=XM_019817557.2;refGene_gn=ACACA;refGene_tx=XM_019817558.2;refGene_gn=ACACA;refGene_tx=XM_019817560.1;refGene_gn=ACACA;refGene_tx=XM_023244661.1;refGene_gn=ACACA;refGene_tx=XM_023244663.1;refGene_gn=ACACA;refGene_tx=XM_023244664.1;refGene_gn=ACACA;refGene_tx=XM_023244665.1
E1 24083655 24083769
refGene_gn=ACACA;refGene_tx=XM_006940115.4;refGene_gn=ACACA;refGene_tx=XM_006940119.4;refGene_gn=ACACA;refGene_tx=XM_011288954.1;refGene_gn=ACACA;refGene_tx=XM_011288955.1;refGene_gn=ACACA;refGene_tx=XM_011288956.2;refGene_gn=ACACA;refGene_tx=XM_019817551.1;refGene_gn=ACACA;refGene_tx=XM_019817552.1;refGene_gn=ACACA;refGene_tx=XM_019817554.1;refGene_gn=ACACA;refGene_tx=XM_019817555.2;refGene_gn=ACACA;refGene_tx=XM_019817556.2;refGene_gn=ACACA;refGene_tx=XM_019817557.2;refGene_gn=ACACA;refGene_tx=XM_019817558.2;refGene_gn=ACACA;refGene_tx=XM_019817560.1;refGene_gn=ACACA;refGene_tx=XM_023244661.1;refGene_gn=ACACA;refGene_tx=XM_023244663.1;refGene_gn=ACACA;refGene_tx=XM_023244664.1;refGene_gn=ACACA;refGene_tx=XM_023244665.1
E1 24087909 24087966
refGene_gn=ACACA;refGene_tx=XM_006940115.4;refGene_gn=ACACA;refGene_tx=XM_006940119.4;refGene_gn=ACACA;refGene_tx=XM_011288954.1;refGene_gn=ACACA;refGene_tx=XM_011288955.1;refGene_gn=ACACA;refGene_tx=XM_011288956.2;refGene_gn=ACACA;refGene_tx=XM_019817551.1;refGene_gn=ACACA;refGene_tx=XM_019817552.1;refGene_gn=ACACA;refGene_tx=XM_019817554.1;refGene_gn=ACACA;refGene_tx=XM_019817555.2;refGene_gn=ACACA;refGene_tx=XM_019817556.2;refGene_gn=ACACA;refGene_tx=XM_019817557.2;refGene_gn=ACACA;refGene_tx=XM_019817558.2;refGene_gn=ACACA;refGene_tx=XM_019817560.1;refGene_gn=ACACA;refGene_tx=XM_023244661.1;refGene_gn=ACACA;refGene_tx=XM_023244663.1;refGene_gn=ACACA;refGene_tx=XM_023244664.1;refGene_gn=ACACA;refGene_tx=XM_023244665.1
E1 24088198 24088243
refGene_gn=ACACA;refGene_tx=XM_006940115.4;refGene_gn=ACACA;refGene_tx=XM_006940119.4;refGene_gn=ACACA;refGene_tx=XM_011288954.1;refGene_gn=ACACA;refGene_tx=XM_011288955.1;refGene_gn=ACACA;refGene_tx=XM_011288956.2;refGene_gn=ACACA;refGene_tx=XM_019817555.2;refGene_gn=ACACA;refGene_tx=XM_019817556.2;refGene_gn=ACACA;refGene_tx=XM_019817557.2;refGene_gn=ACACA;refGene_tx=XM_019817558.2;refGene_gn=ACACA;refGene_tx=XM_019817560.1;refGene_gn=ACACA;refGene_tx=XM_023244661.1;refGene_gn=ACACA;refGene_tx=XM_023244663.1;refGene_gn=ACACA;refGene_tx=XM_023244664.1;refGene_gn=ACACA;refGene_tx=XM_023244665.1
E1 24095582 24095624

refGene_gn=ACACA;refGene_tx=XM_006940115.4;refGene_gn=ACACA;refGene_tx=XM_006940119.4;refGene_gn=ACACA;refGene_tx=XM_011288954.1;refGene_gn=ACACA;refGene_tx=XM_011288955.1;refGene_gn=ACACA;refGene_tx=XM_011288956.2;refGene_gn=ACACA;refGene_tx=XM_019817551.1;refGene_gn=ACACA;refGene_tx=XM_019817552.1;refGene_gn=ACACA;refGene_tx=XM_019817554.1;refGene_gn=ACACA;refGene_tx=XM_019817555.2;refGene_gn=ACACA;refGene_tx=XM_019817556.2;refGene_gn=ACACA;refGene_tx=XM_019817557.2;refGene_gn=ACACA;refGene_tx=XM_019817558.2;refGene_gn=ACACA;refGene_tx=XM_019817560.1;refGene_gn=ACACA;refGene_tx=XM_023244661.1;refGene_gn=ACACA;refGene_tx=XM_023244663.1;refGene_gn=ACACA;refGene_tx=XM_023244664.1;refGene_gn=ACACA;refGene_tx=XM_023244665.1
E1 24096449 24096665
refGene_gn=ACACA;refGene_tx=XM_006940115.4;refGene_gn=ACACA;refGene_tx=XM_006940119.4;refGene_gn=ACACA;refGene_tx=XM_011288954.1;refGene_gn=ACACA;refGene_tx=XM_011288955.1;refGene_gn=ACACA;refGene_tx=XM_011288956.2;refGene_gn=ACACA;refGene_tx=XM_019817551.1;refGene_gn=ACACA;refGene_tx=XM_019817552.1;refGene_gn=ACACA;refGene_tx=XM_019817554.1;refGene_gn=ACACA;refGene_tx=XM_019817555.2;refGene_gn=ACACA;refGene_tx=XM_019817556.2;refGene_gn=ACACA;refGene_tx=XM_019817557.2;refGene_gn=ACACA;refGene_tx=XM_019817558.2;refGene_gn=ACACA;refGene_tx=XM_019817560.1;refGene_gn=ACACA;refGene_tx=XM_023244661.1;refGene_gn=ACACA;refGene_tx=XM_023244663.1;refGene_gn=ACACA;refGene_tx=XM_023244664.1;refGene_gn=ACACA;refGene_tx=XM_023244665.1
E1 24097787 24097943
refGene_gn=ACACA;refGene_tx=XM_006940115.4;refGene_gn=ACACA;refGene_tx=XM_006940119.4;refGene_gn=ACACA;refGene_tx=XM_011288954.1;refGene_gn=ACACA;refGene_tx=XM_011288955.1;refGene_gn=ACACA;refGene_tx=XM_011288956.2;refGene_gn=ACACA;refGene_tx=XM_019817551.1;refGene_gn=ACACA;refGene_tx=XM_019817552.1;refGene_gn=ACACA;refGene_tx=XM_019817554.1;refGene_gn=ACACA;refGene_tx=XM_019817555.2;refGene_gn=ACACA;refGene_tx=XM_019817556.2;refGene_gn=ACACA;refGene_tx=XM_019817557.2;refGene_gn=ACACA;refGene_tx=XM_019817558.2;refGene_gn=ACACA;refGene_tx=XM_019817560.1;refGene_gn=ACACA;refGene_tx=XM_023244661.1;refGene_gn=ACACA;refGene_tx=XM_023244663.1;refGene_gn=ACACA;refGene_tx=XM_023244664.1;refGene_gn=ACACA;refGene_tx=XM_023244665.1
E1 24100084 24100288
refGene_gn=ACACA;refGene_tx=XM_006940115.4;refGene_gn=ACACA;refGene_tx=XM_006940119.4;refGene_gn=ACACA;refGene_tx=XM_011288954.1;refGene_gn=ACACA;refGene_tx=XM_011288955.1;refGene_gn=ACACA;refGene_tx=XM_011288956.2;refGene_gn=ACACA;refGene_tx=XM_019817551.1;refGene_gn=ACACA;refGene_tx=XM_019817552.1;refGene_gn=ACACA;refGene_tx=XM_019817554.1;refGene_gn=ACACA;refGene_tx=XM_019817555.2;refGene_gn=ACACA;refGene_tx=XM_019817556.2;refGene_gn=ACACA;refGene_tx=XM_019817557.2;refGene_gn=ACACA;refGene_tx=XM_019817558.2;refGene_gn=ACACA;refGene_tx=XM_019817560.1;refGene_gn=ACACA;refGene_tx=XM_023244661.1;refGene_gn=ACACA;refGene_tx=XM_023244663.1;refGene_gn=ACACA;refGene_tx=XM_023244664.1;refGene_gn=ACACA;refGene_tx=XM_023244665.1
E1 24104907 24105063
refGene_gn=ACACA;refGene_tx=XM_006940115.4;refGene_gn=ACACA;refGene_tx=XM_006940119.4;refGene_gn=ACACA;refGene_tx=XM_011288954.1;refGene_gn=

ACACA;refGene_tx=XM_011288955.1;refGene_gn=ACACA;refGene_tx=XM_011288956.2;refGene_gn=ACACA;refGene_tx=XM_019817551.1;refGene_gn=ACACA;refGene_tx=XM_019817552.1;refGene_gn=ACACA;refGene_tx=XM_019817554.1;refGene_gn=ACACA;refGene_tx=XM_019817555.2;refGene_gn=ACACA;refGene_tx=XM_019817556.2;refGene_gn=ACACA;refGene_tx=XM_019817557.2;refGene_gn=ACACA;refGene_tx=XM_019817558.2;refGene_gn=ACACA;refGene_tx=XM_019817560.1;refGene_gn=ACACA;refGene_tx=XM_023244661.1;refGene_gn=ACACA;refGene_tx=XM_023244663.1;refGene_gn=ACACA;refGene_tx=XM_023244664.1;refGene_gn=ACACA;refGene_tx=XM_023244665.1

E1 24106830 24106977

refGene_gn=ACACA;refGene_tx=XM_006940115.4;refGene_gn=ACACA;refGene_tx=XM_006940119.4;refGene_gn=ACACA;refGene_tx=XM_011288954.1;refGene_gn=ACACA;refGene_tx=XM_011288955.1;refGene_gn=ACACA;refGene_tx=XM_011288956.2;refGene_gn=ACACA;refGene_tx=XM_019817551.1;refGene_gn=ACACA;refGene_tx=XM_019817552.1;refGene_gn=ACACA;refGene_tx=XM_019817554.1;refGene_gn=ACACA;refGene_tx=XM_019817555.2;refGene_gn=ACACA;refGene_tx=XM_019817556.2;refGene_gn=ACACA;refGene_tx=XM_019817557.2;refGene_gn=ACACA;refGene_tx=XM_019817558.2;refGene_gn=ACACA;refGene_tx=XM_019817560.1;refGene_gn=ACACA;refGene_tx=XM_023244661.1;refGene_gn=ACACA;refGene_tx=XM_023244663.1;refGene_gn=ACACA;refGene_tx=XM_023244664.1;refGene_gn=ACACA;refGene_tx=XM_023244665.1

E1 24122294 24122564

refGene_gn=ACACA;refGene_tx=XM_006940115.4;refGene_gn=ACACA;refGene_tx=XM_006940119.4;refGene_gn=ACACA;refGene_tx=XM_011288954.1;refGene_gn=ACACA;refGene_tx=XM_011288955.1;refGene_gn=ACACA;refGene_tx=XM_011288956.2;refGene_gn=ACACA;refGene_tx=XM_019817551.1;refGene_gn=ACACA;refGene_tx=XM_019817552.1;refGene_gn=ACACA;refGene_tx=XM_019817554.1;refGene_gn=ACACA;refGene_tx=XM_019817555.2;refGene_gn=ACACA;refGene_tx=XM_019817556.2;refGene_gn=ACACA;refGene_tx=XM_019817557.2;refGene_gn=ACACA;refGene_tx=XM_019817558.2;refGene_gn=ACACA;refGene_tx=XM_019817560.1;refGene_gn=ACACA;refGene_tx=XM_023244661.1;refGene_gn=ACACA;refGene_tx=XM_023244663.1;refGene_gn=ACACA;refGene_tx=XM_023244664.1;refGene_gn=ACACA;refGene_tx=XM_023244665.1

E1 24126965 24127063

refGene_gn=ACACA;refGene_tx=XM_006940115.4;refGene_gn=ACACA;refGene_tx=XM_006940119.4;refGene_gn=ACACA;refGene_tx=XM_011288954.1;refGene_gn=ACACA;refGene_tx=XM_011288955.1;refGene_gn=ACACA;refGene_tx=XM_011288956.2;refGene_gn=ACACA;refGene_tx=XM_019817551.1;refGene_gn=ACACA;refGene_tx=XM_019817552.1;refGene_gn=ACACA;refGene_tx=XM_019817554.1;refGene_gn=ACACA;refGene_tx=XM_019817555.2;refGene_gn=ACACA;refGene_tx=XM_019817556.2;refGene_gn=ACACA;refGene_tx=XM_019817557.2;refGene_gn=ACACA;refGene_tx=XM_019817558.2;refGene_gn=ACACA;refGene_tx=XM_019817560.1;refGene_gn=ACACA;refGene_tx=XM_023244661.1;refGene_gn=ACACA;refGene_tx=XM_023244663.1;refGene_gn=ACACA;refGene_tx=XM_023244664.1;refGene_gn=ACACA;refGene_tx=XM_023244665.1

E1 24131085 24131206

refGene_gn=ACACA;refGene_tx=XM_006940115.4;refGene_gn=ACACA;refGene_tx=XM_006940119.4;refGene_gn=ACACA;refGene_tx=XM_011288954.1;refGene_gn=ACACA;refGene_tx=XM_011288955.1;refGene_gn=ACACA;refGene_tx=XM_011288956.2;refGene_gn=ACACA;refGene_tx=XM_019817551.1;refGene_gn=ACACA;refGe

ne_tx=XM_019817552.1;refGene_gn=ACACA;refGene_tx=XM_019817554.1;refGene_gn=ACACA;refGene_tx=XM_019817555.2;refGene_gn=ACACA;refGene_tx=XM_019817556.2;refGene_gn=ACACA;refGene_tx=XM_019817557.2;refGene_gn=ACACA;refGene_tx=XM_019817558.2;refGene_gn=ACACA;refGene_tx=XM_019817560.1;refGene_gn=ACACA;refGene_tx=XM_023244661.1;refGene_gn=ACACA;refGene_tx=XM_023244663.1;refGene_gn=ACACA;refGene_tx=XM_023244664.1;refGene_gn=ACACA;refGene_tx=XM_023244665.1
E1 24131812 24131923
refGene_gn=ACACA;refGene_tx=XM_006940115.4;refGene_gn=ACACA;refGene_tx=XM_006940119.4;refGene_gn=ACACA;refGene_tx=XM_011288954.1;refGene_gn=ACACA;refGene_tx=XM_011288955.1;refGene_gn=ACACA;refGene_tx=XM_011288956.2;refGene_gn=ACACA;refGene_tx=XM_019817551.1;refGene_gn=ACACA;refGene_tx=XM_019817552.1;refGene_gn=ACACA;refGene_tx=XM_019817554.1;refGene_gn=ACACA;refGene_tx=XM_019817555.2;refGene_gn=ACACA;refGene_tx=XM_019817556.2;refGene_gn=ACACA;refGene_tx=XM_019817557.2;refGene_gn=ACACA;refGene_tx=XM_019817558.2;refGene_gn=ACACA;refGene_tx=XM_019817560.1;refGene_gn=ACACA;refGene_tx=XM_023244661.1;refGene_gn=ACACA;refGene_tx=XM_023244663.1;refGene_gn=ACACA;refGene_tx=XM_023244664.1;refGene_gn=ACACA;refGene_tx=XM_023244665.1
E1 24133672 24133741 refGene_gn=ACACA;refGene_tx=XM_019817560.1
E1 24159588 24159732
refGene_gn=ACACA;refGene_tx=XM_006940115.4;refGene_gn=ACACA;refGene_tx=XM_006940119.4;refGene_gn=ACACA;refGene_tx=XM_011288954.1;refGene_gn=ACACA;refGene_tx=XM_011288955.1;refGene_gn=ACACA;refGene_tx=XM_011288956.2;refGene_gn=ACACA;refGene_tx=XM_019817551.1;refGene_gn=ACACA;refGene_tx=XM_019817552.1;refGene_gn=ACACA;refGene_tx=XM_019817554.1;refGene_gn=ACACA;refGene_tx=XM_019817555.2;refGene_gn=ACACA;refGene_tx=XM_019817556.2;refGene_gn=ACACA;refGene_tx=XM_019817557.2;refGene_gn=ACACA;refGene_tx=XM_019817558.2;refGene_gn=ACACA;refGene_tx=XM_023244661.1;refGene_gn=ACACA;refGene_tx=XM_023244663.1;refGene_gn=ACACA;refGene_tx=XM_023244664.1;refGene_gn=ACACA;refGene_tx=XM_023244665.1
E1 24160322 24160443
refGene_gn=ACACA;refGene_tx=XM_006940115.4;refGene_gn=ACACA;refGene_tx=XM_006940119.4;refGene_gn=ACACA;refGene_tx=XM_011288954.1;refGene_gn=ACACA;refGene_tx=XM_011288955.1;refGene_gn=ACACA;refGene_tx=XM_011288956.2;refGene_gn=ACACA;refGene_tx=XM_019817551.1;refGene_gn=ACACA;refGene_tx=XM_019817552.1;refGene_gn=ACACA;refGene_tx=XM_019817554.1;refGene_gn=ACACA;refGene_tx=XM_019817555.2;refGene_gn=ACACA;refGene_tx=XM_019817556.2;refGene_gn=ACACA;refGene_tx=XM_019817557.2;refGene_gn=ACACA;refGene_tx=XM_019817558.2;refGene_gn=ACACA;refGene_tx=XM_023244661.1;refGene_gn=ACACA;refGene_tx=XM_023244663.1;refGene_gn=ACACA;refGene_tx=XM_023244664.1;refGene_gn=ACACA;refGene_tx=XM_023244665.1
E1 24163644 24163741
refGene_gn=ACACA;refGene_tx=XM_006940115.4;refGene_gn=ACACA;refGene_tx=XM_006940119.4;refGene_gn=ACACA;refGene_tx=XM_011288954.1;refGene_gn=ACACA;refGene_tx=XM_011288955.1;refGene_gn=ACACA;refGene_tx=XM_011288956.2;refGene_gn=ACACA;refGene_tx=XM_019817551.1;refGene_gn=ACACA;refGene_tx=XM_019817552.1;refGene_gn=ACACA;refGene_tx=XM_019817554.1;refGene_gn=ACACA;refGene_tx=XM_019817555.2;refGene_gn=ACACA;refGene_tx=XM_019817556.2;refGene_gn=ACACA;refGene_tx=XM_019817557.2;refGene_gn=ACACA;

refGene_tx=XM_019817558.2;refGene_gn=ACACA;refGene_tx=XM_023244661.1;refGene_gn=ACACA;refGene_tx=XM_023244663.1;refGene_gn=ACACA;refGene_tx=XM_023244664.1;refGene_gn=ACACA;refGene_tx=XM_023244665.1
E1 24166858 24166955
refGene_gn=ACACA;refGene_tx=XM_006940115.4;refGene_gn=ACACA;refGene_tx=XM_006940119.4;refGene_gn=ACACA;refGene_tx=XM_011288954.1;refGene_gn=ACACA;refGene_tx=XM_011288955.1;refGene_gn=ACACA;refGene_tx=XM_011288956.2;refGene_gn=ACACA;refGene_tx=XM_019817551.1;refGene_gn=ACACA;refGene_tx=XM_019817552.1;refGene_gn=ACACA;refGene_tx=XM_019817554.1;refGene_gn=ACACA;refGene_tx=XM_019817555.2;refGene_gn=ACACA;refGene_tx=XM_019817556.2;refGene_gn=ACACA;refGene_tx=XM_019817557.2;refGene_gn=ACACA;refGene_tx=XM_019817558.2;refGene_gn=ACACA;refGene_tx=XM_023244661.1;refGene_gn=ACACA;refGene_tx=XM_023244663.1;refGene_gn=ACACA;refGene_tx=XM_023244664.1;refGene_gn=ACACA;refGene_tx=XM_023244665.1
E1 24168064 24168200
refGene_gn=ACACA;refGene_tx=XM_006940115.4;refGene_gn=ACACA;refGene_tx=XM_006940119.4;refGene_gn=ACACA;refGene_tx=XM_011288954.1;refGene_gn=ACACA;refGene_tx=XM_011288955.1;refGene_gn=ACACA;refGene_tx=XM_011288956.2;refGene_gn=ACACA;refGene_tx=XM_019817551.1;refGene_gn=ACACA;refGene_tx=XM_019817552.1;refGene_gn=ACACA;refGene_tx=XM_019817554.1;refGene_gn=ACACA;refGene_tx=XM_019817555.2;refGene_gn=ACACA;refGene_tx=XM_019817556.2;refGene_gn=ACACA;refGene_tx=XM_019817557.2;refGene_gn=ACACA;refGene_tx=XM_019817558.2;refGene_gn=ACACA;refGene_tx=XM_023244661.1;refGene_gn=ACACA;refGene_tx=XM_023244663.1;refGene_gn=ACACA;refGene_tx=XM_023244664.1;refGene_gn=ACACA;refGene_tx=XM_023244665.1
E1 24175270 24175448
refGene_gn=ACACA;refGene_tx=XM_006940115.4;refGene_gn=ACACA;refGene_tx=XM_006940119.4;refGene_gn=ACACA;refGene_tx=XM_011288954.1;refGene_gn=ACACA;refGene_tx=XM_011288955.1;refGene_gn=ACACA;refGene_tx=XM_011288956.2;refGene_gn=ACACA;refGene_tx=XM_019817551.1;refGene_gn=ACACA;refGene_tx=XM_019817552.1;refGene_gn=ACACA;refGene_tx=XM_019817554.1;refGene_gn=ACACA;refGene_tx=XM_019817555.2;refGene_gn=ACACA;refGene_tx=XM_019817556.2;refGene_gn=ACACA;refGene_tx=XM_019817557.2;refGene_gn=ACACA;refGene_tx=XM_019817558.2;refGene_gn=ACACA;refGene_tx=XM_023244661.1;refGene_gn=ACACA;refGene_tx=XM_023244663.1;refGene_gn=ACACA;refGene_tx=XM_023244664.1;refGene_gn=ACACA;refGene_tx=XM_023244665.1
E1 24177198 24177311
refGene_gn=ACACA;refGene_tx=XM_006940115.4;refGene_gn=ACACA;refGene_tx=XM_006940119.4;refGene_gn=ACACA;refGene_tx=XM_011288954.1;refGene_gn=ACACA;refGene_tx=XM_011288955.1;refGene_gn=ACACA;refGene_tx=XM_011288956.2;refGene_gn=ACACA;refGene_tx=XM_019817551.1;refGene_gn=ACACA;refGene_tx=XM_019817552.1;refGene_gn=ACACA;refGene_tx=XM_019817554.1;refGene_gn=ACACA;refGene_tx=XM_019817555.2;refGene_gn=ACACA;refGene_tx=XM_019817556.2;refGene_gn=ACACA;refGene_tx=XM_019817557.2;refGene_gn=ACACA;refGene_tx=XM_019817558.2;refGene_gn=ACACA;refGene_tx=XM_023244661.1;refGene_gn=ACACA;refGene_tx=XM_023244663.1;refGene_gn=ACACA;refGene_tx=XM_023244664.1;refGene_gn=ACACA;refGene_tx=XM_023244665.1
E1 24189976 24190131
refGene_gn=ACACA;refGene_tx=XM_006940115.4;refGene_gn=ACACA;refGene_tx=XM_006940119.4;refGene_gn=ACACA;refGene_tx=XM_011288954.1;refGene_gn=

ACACA;refGene_tx=XM_011288955.1;refGene_gn=ACACA;refGene_tx=XM_0112889
56.2;refGene_gn=ACACA;refGene_tx=XM_019817551.1;refGene_gn=ACACA;refGe
ne_tx=XM_019817552.1;refGene_gn=ACACA;refGene_tx=XM_019817554.1;refGen
e_gn=ACACA;refGene_tx=XM_019817555.2;refGene_gn=ACACA;refGene_tx=XM_01
9817556.2;refGene_gn=ACACA;refGene_tx=XM_019817557.2;refGene_gn=ACACA;
refGene_tx=XM_019817558.2;refGene_gn=ACACA;refGene_tx=XM_023244661.1;r
efGene_gn=ACACA;refGene_tx=XM_023244663.1;refGene_gn=ACACA;refGene_tx=
XM_023244664.1;refGene_gn=ACACA;refGene_tx=XM_023244665.1
E1 24190800 24190971
refGene_gn=ACACA;refGene_tx=XM_006940115.4;refGene_gn=ACACA;refGene_tx
=XM_006940119.4;refGene_gn=ACACA;refGene_tx=XM_011288954.1;refGene_gn=
ACACA;refGene_tx=XM_011288955.1;refGene_gn=ACACA;refGene_tx=XM_0112889
56.2;refGene_gn=ACACA;refGene_tx=XM_019817551.1;refGene_gn=ACACA;refGe
ne_tx=XM_019817552.1;refGene_gn=ACACA;refGene_tx=XM_019817554.1;refGen
e_gn=ACACA;refGene_tx=XM_019817555.2;refGene_gn=ACACA;refGene_tx=XM_01
9817556.2;refGene_gn=ACACA;refGene_tx=XM_019817557.2;refGene_gn=ACACA;
refGene_tx=XM_019817558.2;refGene_gn=ACACA;refGene_tx=XM_023244661.1;r
efGene_gn=ACACA;refGene_tx=XM_023244663.1;refGene_gn=ACACA;refGene_tx=
XM_023244664.1;refGene_gn=ACACA;refGene_tx=XM_023244665.1
E1 24198058 24198195
refGene_gn=ACACA;refGene_tx=XM_006940115.4;refGene_gn=ACACA;refGene_tx
=XM_006940119.4;refGene_gn=ACACA;refGene_tx=XM_011288954.1;refGene_gn=
ACACA;refGene_tx=XM_011288955.1;refGene_gn=ACACA;refGene_tx=XM_0112889
56.2;refGene_gn=ACACA;refGene_tx=XM_019817551.1;refGene_gn=ACACA;refGe
ne_tx=XM_019817552.1;refGene_gn=ACACA;refGene_tx=XM_019817554.1;refGen
e_gn=ACACA;refGene_tx=XM_019817555.2;refGene_gn=ACACA;refGene_tx=XM_01
9817556.2;refGene_gn=ACACA;refGene_tx=XM_019817557.2;refGene_gn=ACACA;
refGene_tx=XM_019817558.2;refGene_gn=ACACA;refGene_tx=XM_023244661.1;r
efGene_gn=ACACA;refGene_tx=XM_023244663.1;refGene_gn=ACACA;refGene_tx=
XM_023244664.1;refGene_gn=ACACA;refGene_tx=XM_023244665.1
E1 24199745 24199869
refGene_gn=ACACA;refGene_tx=XM_006940115.4;refGene_gn=ACACA;refGene_tx
=XM_006940119.4;refGene_gn=ACACA;refGene_tx=XM_011288954.1;refGene_gn=
ACACA;refGene_tx=XM_011288955.1;refGene_gn=ACACA;refGene_tx=XM_0112889
56.2;refGene_gn=ACACA;refGene_tx=XM_019817551.1;refGene_gn=ACACA;refGe
ne_tx=XM_019817552.1;refGene_gn=ACACA;refGene_tx=XM_019817554.1;refGen
e_gn=ACACA;refGene_tx=XM_019817555.2;refGene_gn=ACACA;refGene_tx=XM_01
9817556.2;refGene_gn=ACACA;refGene_tx=XM_019817557.2;refGene_gn=ACACA;
refGene_tx=XM_019817558.2;refGene_gn=ACACA;refGene_tx=XM_023244661.1;r
efGene_gn=ACACA;refGene_tx=XM_023244663.1;refGene_gn=ACACA;refGene_tx=
XM_023244664.1;refGene_gn=ACACA;refGene_tx=XM_023244665.1
E1 24228765 24228829 refGene_gn=AATF;refGene_tx=XM_006940121.4
E1 24248874 24248914 refGene_gn=AATF;refGene_tx=XM_023244757.1
E1 24249983 24250055
refGene_gn=AATF;refGene_tx=XM_006940121.4;refGene_gn=AATF;refGene_tx=X
M_023244757.1
E1 24257971 24258052
refGene_gn=AATF;refGene_tx=XM_006940121.4;refGene_gn=AATF;refGene_tx=X
M_023244757.1
E1 24259903 24259971

refGene_gn=AATF;refGene_tx=XM_006940121.4;refGene_gn=AATF;refGene_tx=XM_023244757.1
E1 24287087 24287171
refGene_gn=AATF;refGene_tx=XM_006940121.4;refGene_gn=AATF;refGene_tx=XM_023244757.1
E1 24288667 24288832
refGene_gn=AATF;refGene_tx=XM_006940121.4;refGene_gn=AATF;refGene_tx=XM_023244757.1
E1 24289421 24289623
refGene_gn=AATF;refGene_tx=XM_006940121.4;refGene_gn=AATF;refGene_tx=XM_023244757.1
E1 24291027 24291142
refGene_gn=AATF;refGene_tx=XM_006940121.4;refGene_gn=AATF;refGene_tx=XM_023244757.1
E1 24324284 24324422
refGene_gn=AATF;refGene_tx=XM_006940121.4;refGene_gn=AATF;refGene_tx=XM_023244757.1
E1 24324878 24325313
refGene_gn=AATF;refGene_tx=XM_006940121.4;refGene_gn=AATF;refGene_tx=XM_023244757.1
E1 24328330 24328522
refGene_gn=AATF;refGene_tx=XM_006940121.4;refGene_gn=AATF;refGene_tx=XM_023244757.1
E1 24329473 24329564
refGene_gn=AATF;refGene_tx=XM_006940121.4;refGene_gn=AATF;refGene_tx=XM_023244757.1
E1 24335636 24336016 refGene_gn=LHX1;refGene_tx=XM_003996587.4
E1 24336333 24336499 refGene_gn=LHX1;refGene_tx=XM_003996587.4
E1 24337863 24338141 refGene_gn=LHX1;refGene_tx=XM_003996587.4
E1 24338235 24338462 refGene_gn=LHX1;refGene_tx=XM_003996587.4
E1 24340403 24340573 refGene_gn=LHX1;refGene_tx=XM_003996587.4
E1 24609123 24609143 refGene_gn=MRM1;refGene_tx=XM_019817562.1
E1 24623978 24624049 refGene_gn=MRM1;refGene_tx=XM_003996601.5
E1 24624481 24624601
refGene_gn=MRM1;refGene_tx=XM_003996601.5;refGene_gn=MRM1;refGene_tx=XM_019817562.1
E1 24624702 24624835
refGene_gn=MRM1;refGene_tx=XM_003996601.5;refGene_gn=MRM1;refGene_tx=XM_019817562.1
E1 24626520 24626614
refGene_gn=MRM1;refGene_tx=XM_003996601.5;refGene_gn=MRM1;refGene_tx=XM_019817562.1
E1 24626831 24627394
refGene_gn=MRM1;refGene_tx=XM_003996601.5;refGene_gn=MRM1;refGene_tx=XM_019817562.1
E1 24628823 24628981 refGene_gn=DHRS11;refGene_tx=XM_019817564.2
E1 24629212 24629279 refGene_gn=DHRS11;refGene_tx=XM_019817564.2
E1 24629480 24629573 refGene_gn=DHRS11;refGene_tx=XM_019817564.2
E1 24630194 24630324 refGene_gn=DHRS11;refGene_tx=XM_019817564.2
E1 24630724 24630819 refGene_gn=DHRS11;refGene_tx=XM_019817564.2

E1 24634089 24634299 refGene_gn=DHRS11;refGene_tx=XM_019817564.2
E1 24636967 24637056 refGene_gn=DHRS11;refGene_tx=XM_019817564.2
E1 24637058 24637323 refGene_gn=DHRS11;refGene_tx=XM_019817564.2
E1 24639754 24639958
refGene_gn=GGNBP2;refGene_tx=XM_011288946.3;refGene_gn=GGNBP2;refGene_tx=XM_011288947.3;refGene_gn=GGNBP2;refGene_tx=XM_011288948.3;refGene_gn=GGNBP2;refGene_tx=XM_011288949.3;refGene_gn=GGNBP2;refGene_tx=XM_011288950.3;refGene_gn=GGNBP2;refGene_tx=XM_011288951.3;refGene_gn=GGNBP2;refGene_tx=XM_011288952.3
E1 24641268 24641517
refGene_gn=GGNBP2;refGene_tx=XM_011288946.3;refGene_gn=GGNBP2;refGene_tx=XM_011288947.3;refGene_gn=GGNBP2;refGene_tx=XM_011288948.3;refGene_gn=GGNBP2;refGene_tx=XM_011288949.3;refGene_gn=GGNBP2;refGene_tx=XM_011288950.3;refGene_gn=GGNBP2;refGene_tx=XM_011288951.3;refGene_gn=GGNBP2;refGene_tx=XM_011288952.3
E1 24642054 24642203
refGene_gn=GGNBP2;refGene_tx=XM_011288946.3;refGene_gn=GGNBP2;refGene_tx=XM_011288947.3;refGene_gn=GGNBP2;refGene_tx=XM_011288948.3;refGene_gn=GGNBP2;refGene_tx=XM_011288949.3;refGene_gn=GGNBP2;refGene_tx=XM_011288950.3;refGene_gn=GGNBP2;refGene_tx=XM_011288951.3;refGene_gn=GGNBP2;refGene_tx=XM_011288952.3
E1 24642285 24642411
refGene_gn=GGNBP2;refGene_tx=XM_011288946.3;refGene_gn=GGNBP2;refGene_tx=XM_011288947.3;refGene_gn=GGNBP2;refGene_tx=XM_011288948.3;refGene_gn=GGNBP2;refGene_tx=XM_011288949.3;refGene_gn=GGNBP2;refGene_tx=XM_011288950.3;refGene_gn=GGNBP2;refGene_tx=XM_011288951.3;refGene_gn=GGNBP2;refGene_tx=XM_011288952.3
E1 24642704 24642852
refGene_gn=GGNBP2;refGene_tx=XM_011288946.3;refGene_gn=GGNBP2;refGene_tx=XM_011288947.3;refGene_gn=GGNBP2;refGene_tx=XM_011288948.3;refGene_gn=GGNBP2;refGene_tx=XM_011288949.3;refGene_gn=GGNBP2;refGene_tx=XM_011288950.3;refGene_gn=GGNBP2;refGene_tx=XM_011288951.3;refGene_gn=GGNBP2;refGene_tx=XM_011288952.3
E1 24644649 24644844
refGene_gn=GGNBP2;refGene_tx=XM_011288946.3;refGene_gn=GGNBP2;refGene_tx=XM_011288947.3;refGene_gn=GGNBP2;refGene_tx=XM_011288948.3;refGene_gn=GGNBP2;refGene_tx=XM_011288949.3;refGene_gn=GGNBP2;refGene_tx=XM_011288950.3;refGene_gn=GGNBP2;refGene_tx=XM_011288951.3;refGene_gn=GGNBP2;refGene_tx=XM_011288952.3
E1 24646532 24646707
refGene_gn=GGNBP2;refGene_tx=XM_011288946.3;refGene_gn=GGNBP2;refGene_tx=XM_011288947.3;refGene_gn=GGNBP2;refGene_tx=XM_011288948.3;refGene_gn=GGNBP2;refGene_tx=XM_011288949.3;refGene_gn=GGNBP2;refGene_tx=XM_011288950.3;refGene_gn=GGNBP2;refGene_tx=XM_011288951.3;refGene_gn=GGNBP2;refGene_tx=XM_011288952.3
E1 24647770 24647980
refGene_gn=GGNBP2;refGene_tx=XM_011288946.3;refGene_gn=GGNBP2;refGene_tx=XM_011288947.3;refGene_gn=GGNBP2;refGene_tx=XM_011288948.3;refGene_gn=GGNBP2;refGene_tx=XM_011288949.3;refGene_gn=GGNBP2;refGene_tx=XM_011288950.3;refGene_gn=GGNBP2;refGene_tx=XM_011288951.3;refGene_gn=GGNBP

2;refGene_tx=XM_011288952.3
E1 24655508 24655622
refGene_gn=GGNBP2;refGene_tx=XM_011288946.3;refGene_gn=GGNBP2;refGene_tx=XM_011288947.3;refGene_gn=GGNBP2;refGene_tx=XM_011288948.3;refGene_gn=GGNBP2;refGene_tx=XM_011288949.3;refGene_gn=GGNBP2;refGene_tx=XM_011288950.3;refGene_gn=GGNBP2;refGene_tx=XM_011288951.3;refGene_gn=GGNBP2;refGene_tx=XM_011288952.3
E1 24655715 24655841
refGene_gn=GGNBP2;refGene_tx=XM_011288946.3;refGene_gn=GGNBP2;refGene_tx=XM_011288947.3;refGene_gn=GGNBP2;refGene_tx=XM_011288948.3;refGene_gn=GGNBP2;refGene_tx=XM_011288949.3;refGene_gn=GGNBP2;refGene_tx=XM_011288950.3
E1 24659621 24659720
refGene_gn=GGNBP2;refGene_tx=XM_011288946.3;refGene_gn=GGNBP2;refGene_tx=XM_011288947.3;refGene_gn=GGNBP2;refGene_tx=XM_011288948.3;refGene_gn=GGNBP2;refGene_tx=XM_011288949.3;refGene_gn=GGNBP2;refGene_tx=XM_011288950.3;refGene_gn=GGNBP2;refGene_tx=XM_011288951.3;refGene_gn=GGNBP2;refGene_tx=XM_011288952.3
E1 24662611 24662865
refGene_gn=GGNBP2;refGene_tx=XM_011288946.3;refGene_gn=GGNBP2;refGene_tx=XM_011288947.3;refGene_gn=GGNBP2;refGene_tx=XM_011288948.3;refGene_gn=GGNBP2;refGene_tx=XM_011288949.3;refGene_gn=GGNBP2;refGene_tx=XM_011288950.3;refGene_gn=GGNBP2;refGene_tx=XM_011288951.3;refGene_gn=GGNBP2;refGene_tx=XM_011288952.3
E1 24664120 24664201
refGene_gn=GGNBP2;refGene_tx=XM_011288946.3;refGene_gn=GGNBP2;refGene_tx=XM_011288947.3;refGene_gn=GGNBP2;refGene_tx=XM_011288948.3;refGene_gn=GGNBP2;refGene_tx=XM_011288949.3;refGene_gn=GGNBP2;refGene_tx=XM_011288950.3;refGene_gn=GGNBP2;refGene_tx=XM_011288951.3;refGene_gn=GGNBP2;refGene_tx=XM_011288952.3
E1 24673376 24673469
refGene_gn=GGNBP2;refGene_tx=XM_011288946.3;refGene_gn=GGNBP2;refGene_tx=XM_011288947.3;refGene_gn=GGNBP2;refGene_tx=XM_011288948.3;refGene_gn=GGNBP2;refGene_tx=XM_011288949.3;refGene_gn=GGNBP2;refGene_tx=XM_011288950.3;refGene_gn=GGNBP2;refGene_tx=XM_011288951.3;refGene_gn=GGNBP2;refGene_tx=XM_011288952.3
E1 24680705 24682211
refGene_gn=PIGW;refGene_tx=XM_006940127.4;refGene_gn=PIGW;refGene_tx=XM_006940128.4;refGene_gn=PIGW;refGene_tx=XM_023243580.1;refGene_gn=PIGW;refGene_tx=XM_023243581.1
E1 24686414 24686426
refGene_gn=MYO19;refGene_tx=XM_006940126.4;refGene_gn=MYO19;refGene_tx=XM_019817565.2;refGene_gn=MYO19;refGene_tx=XM_019817566.2
E1 24689980 24690131
refGene_gn=MYO19;refGene_tx=XM_006940124.2;refGene_gn=MYO19;refGene_tx=XM_006940125.2;refGene_gn=MYO19;refGene_tx=XM_006940126.4;refGene_gn=MYO19;refGene_tx=XM_019817565.2;refGene_gn=MYO19;refGene_tx=XM_019817566.2
E1 24690582 24690731
refGene_gn=MYO19;refGene_tx=XM_006940124.2;refGene_gn=MYO19;refGene_tx

=XM_006940125.2;refGene_gn=MYO19;refGene_tx=XM_006940126.4;refGene_gn=MYO19;refGene_tx=XM_019817565.2;refGene_gn=MYO19;refGene_tx=XM_019817566.2
E1 24692479 24692593
refGene_gn=MYO19;refGene_tx=XM_006940124.2;refGene_gn=MYO19;refGene_tx=XM_006940125.2;refGene_gn=MYO19;refGene_tx=XM_006940126.4;refGene_gn=MYO19;refGene_tx=XM_019817565.2;refGene_gn=MYO19;refGene_tx=XM_019817566.2
E1 24696921 24697054
refGene_gn=MYO19;refGene_tx=XM_006940124.2;refGene_gn=MYO19;refGene_tx=XM_006940125.2;refGene_gn=MYO19;refGene_tx=XM_006940126.4;refGene_gn=MYO19;refGene_tx=XM_011288945.2;refGene_gn=MYO19;refGene_tx=XM_019817565.2;refGene_gn=MYO19;refGene_tx=XM_019817566.2
E1 24697731 24697801
refGene_gn=MYO19;refGene_tx=XM_006940124.2;refGene_gn=MYO19;refGene_tx=XM_006940125.2;refGene_gn=MYO19;refGene_tx=XM_006940126.4;refGene_gn=MYO19;refGene_tx=XM_011288945.2;refGene_gn=MYO19;refGene_tx=XM_019817565.2;refGene_gn=MYO19;refGene_tx=XM_019817566.2
E1 24698355 24698458
refGene_gn=MYO19;refGene_tx=XM_006940124.2;refGene_gn=MYO19;refGene_tx=XM_006940125.2;refGene_gn=MYO19;refGene_tx=XM_006940126.4;refGene_gn=MYO19;refGene_tx=XM_011288945.2;refGene_gn=MYO19;refGene_tx=XM_019817565.2;refGene_gn=MYO19;refGene_tx=XM_019817566.2
E1 24699101 24699198
refGene_gn=MYO19;refGene_tx=XM_006940124.2;refGene_gn=MYO19;refGene_tx=XM_006940125.2;refGene_gn=MYO19;refGene_tx=XM_006940126.4;refGene_gn=MYO19;refGene_tx=XM_011288945.2;refGene_gn=MYO19;refGene_tx=XM_019817565.2;refGene_gn=MYO19;refGene_tx=XM_019817566.2
E1 24699339 24699416
refGene_gn=MYO19;refGene_tx=XM_006940124.2;refGene_gn=MYO19;refGene_tx=XM_006940125.2;refGene_gn=MYO19;refGene_tx=XM_006940126.4;refGene_gn=MYO19;refGene_tx=XM_011288945.2;refGene_gn=MYO19;refGene_tx=XM_019817565.2;refGene_gn=MYO19;refGene_tx=XM_019817566.2
E1 24701102 24701250
refGene_gn=MYO19;refGene_tx=XM_006940124.2;refGene_gn=MYO19;refGene_tx=XM_006940125.2;refGene_gn=MYO19;refGene_tx=XM_006940126.4;refGene_gn=MYO19;refGene_tx=XM_011288945.2;refGene_gn=MYO19;refGene_tx=XM_019817565.2;refGene_gn=MYO19;refGene_tx=XM_019817566.2
E1 24701611 24701783
refGene_gn=MYO19;refGene_tx=XM_006940124.2;refGene_gn=MYO19;refGene_tx=XM_006940125.2;refGene_gn=MYO19;refGene_tx=XM_006940126.4;refGene_gn=MYO19;refGene_tx=XM_011288945.2;refGene_gn=MYO19;refGene_tx=XM_019817565.2;refGene_gn=MYO19;refGene_tx=XM_019817566.2
E1 24703513 24703587
refGene_gn=MYO19;refGene_tx=XM_006940124.2;refGene_gn=MYO19;refGene_tx=XM_006940125.2;refGene_gn=MYO19;refGene_tx=XM_006940126.4;refGene_gn=MYO19;refGene_tx=XM_011288945.2;refGene_gn=MYO19;refGene_tx=XM_019817565.2;refGene_gn=MYO19;refGene_tx=XM_019817566.2
E1 24704271 24704393
refGene_gn=MYO19;refGene_tx=XM_006940124.2;refGene_gn=MYO19;refGene_tx

=XM_006940125.2;refGene_gn=MYO19;refGene_tx=XM_006940126.4;refGene_gn=MYO19;refGene_tx=XM_011288945.2;refGene_gn=MYO19;refGene_tx=XM_019817565.2;refGene_gn=MYO19;refGene_tx=XM_019817566.2
E1 24704561 24704675
refGene_gn=MYO19;refGene_tx=XM_006940124.2;refGene_gn=MYO19;refGene_tx=XM_006940125.2;refGene_gn=MYO19;refGene_tx=XM_006940126.4;refGene_gn=MYO19;refGene_tx=XM_011288945.2;refGene_gn=MYO19;refGene_tx=XM_019817565.2;refGene_gn=MYO19;refGene_tx=XM_019817566.2
E1 24704934 24705111
refGene_gn=MYO19;refGene_tx=XM_006940124.2;refGene_gn=MYO19;refGene_tx=XM_006940125.2;refGene_gn=MYO19;refGene_tx=XM_006940126.4;refGene_gn=MYO19;refGene_tx=XM_011288945.2;refGene_gn=MYO19;refGene_tx=XM_019817565.2;refGene_gn=MYO19;refGene_tx=XM_019817566.2
E1 24705426 24705579
refGene_gn=MYO19;refGene_tx=XM_006940124.2;refGene_gn=MYO19;refGene_tx=XM_006940125.2;refGene_gn=MYO19;refGene_tx=XM_006940126.4;refGene_gn=MYO19;refGene_tx=XM_011288945.2;refGene_gn=MYO19;refGene_tx=XM_019817565.2;refGene_gn=MYO19;refGene_tx=XM_019817566.2
E1 24706609 24706717
refGene_gn=MYO19;refGene_tx=XM_006940124.2;refGene_gn=MYO19;refGene_tx=XM_006940125.2;refGene_gn=MYO19;refGene_tx=XM_006940126.4;refGene_gn=MYO19;refGene_tx=XM_011288945.2;refGene_gn=MYO19;refGene_tx=XM_019817565.2;refGene_gn=MYO19;refGene_tx=XM_019817566.2
E1 24707548 24707619
refGene_gn=MYO19;refGene_tx=XM_006940124.2;refGene_gn=MYO19;refGene_tx=XM_006940125.2;refGene_gn=MYO19;refGene_tx=XM_006940126.4;refGene_gn=MYO19;refGene_tx=XM_011288945.2;refGene_gn=MYO19;refGene_tx=XM_019817565.2;refGene_gn=MYO19;refGene_tx=XM_019817566.2
E1 24709402 24709506
refGene_gn=MYO19;refGene_tx=XM_006940124.2;refGene_gn=MYO19;refGene_tx=XM_006940125.2;refGene_gn=MYO19;refGene_tx=XM_006940126.4;refGene_gn=MYO19;refGene_tx=XM_011288945.2;refGene_gn=MYO19;refGene_tx=XM_019817565.2;refGene_gn=MYO19;refGene_tx=XM_019817566.2
E1 24711074 24711241
refGene_gn=MYO19;refGene_tx=XM_006940124.2;refGene_gn=MYO19;refGene_tx=XM_006940125.2;refGene_gn=MYO19;refGene_tx=XM_006940126.4;refGene_gn=MYO19;refGene_tx=XM_011288945.2;refGene_gn=MYO19;refGene_tx=XM_019817565.2;refGene_gn=MYO19;refGene_tx=XM_019817566.2
E1 24711354 24711484
refGene_gn=MYO19;refGene_tx=XM_006940124.2;refGene_gn=MYO19;refGene_tx=XM_006940125.2;refGene_gn=MYO19;refGene_tx=XM_006940126.4;refGene_gn=MYO19;refGene_tx=XM_011288945.2;refGene_gn=MYO19;refGene_tx=XM_019817565.2;refGene_gn=MYO19;refGene_tx=XM_019817566.2
E1 24713290 24713376
refGene_gn=MYO19;refGene_tx=XM_006940124.2;refGene_gn=MYO19;refGene_tx=XM_006940125.2;refGene_gn=MYO19;refGene_tx=XM_006940126.4;refGene_gn=MYO19;refGene_tx=XM_011288945.2;refGene_gn=MYO19;refGene_tx=XM_019817565.2;refGene_gn=MYO19;refGene_tx=XM_019817566.2
E1 24713834 24714137
refGene_gn=MYO19;refGene_tx=XM_006940124.2;refGene_gn=MYO19;refGene_tx

=XM_006940125.2;refGene_gn=MYO19;refGene_tx=XM_006940126.4;refGene_gn=MYO19;refGene_tx=XM_011288945.2;refGene_gn=MYO19;refGene_tx=XM_019817565.2;refGene_gn=MYO19;refGene_tx=XM_019817566.2
E1 24715064 24715193
refGene_gn=MYO19;refGene_tx=XM_006940124.2;refGene_gn=MYO19;refGene_tx=XM_006940125.2;refGene_gn=MYO19;refGene_tx=XM_006940126.4;refGene_gn=MYO19;refGene_tx=XM_011288945.2;refGene_gn=MYO19;refGene_tx=XM_019817565.2;refGene_gn=MYO19;refGene_tx=XM_019817566.2
E1 26990789 26990971 refGene_gn=ZNHIT3;refGene_tx=XM_003996603.4
E1 26992442 26992523 refGene_gn=ZNHIT3;refGene_tx=XM_003996603.4
E1 26993764 26993848 refGene_gn=ZNHIT3;refGene_tx=XM_003996603.4
E1 26995663 26995695 refGene_gn=ZNHIT3;refGene_tx=XM_003996603.4
E1 26995823 26995909 refGene_gn=ZNHIT3;refGene_tx=XM_003996603.4
E1 27045200 27045258 refGene_gn=CA4;refGene_tx=XM_003996604.5
E1 27049818 27049872 refGene_gn=CA4;refGene_tx=XM_003996604.5
E1 27050689 27050845 refGene_gn=CA4;refGene_tx=XM_003996604.5
E1 27051550 27051696 refGene_gn=CA4;refGene_tx=XM_003996604.5
E1 27051814 27051913 refGene_gn=CA4;refGene_tx=XM_003996604.5
E1 27052171 27052238 refGene_gn=CA4;refGene_tx=XM_003996604.5
E1 27052379 27052546 refGene_gn=CA4;refGene_tx=XM_003996604.5
E1 27053622 27053817 refGene_gn=CA4;refGene_tx=XM_003996604.5
E1 27067368 27067576
refGene_gn=LOC111557847;refGene_tx=XM_023244519.1
E1 27067578 27067790
refGene_gn=LOC111557847;refGene_tx=XM_023244519.1
E1 27067791 27067881
refGene_gn=LOC111557847;refGene_tx=XM_023244519.1
E1 27072517 27072691
refGene_gn=USP32;refGene_tx=XM_003996605.5;refGene_gn=USP32;refGene_tx=XM_011289074.3
E1 27073350 27073443
refGene_gn=USP32;refGene_tx=XM_003996605.5;refGene_gn=USP32;refGene_tx=XM_011289074.3
E1 27073943 27074368
refGene_gn=USP32;refGene_tx=XM_003996605.5;refGene_gn=USP32;refGene_tx=XM_011289074.3
E1 27076075 27076364
refGene_gn=USP32;refGene_tx=XM_003996605.5;refGene_gn=USP32;refGene_tx=XM_011289074.3
E1 27078353 27078545
refGene_gn=USP32;refGene_tx=XM_003996605.5;refGene_gn=USP32;refGene_tx=XM_011289074.3
E1 27086701 27086822
refGene_gn=USP32;refGene_tx=XM_003996605.5;refGene_gn=USP32;refGene_tx=XM_011289074.3
E1 27087751 27087838
refGene_gn=USP32;refGene_tx=XM_003996605.5;refGene_gn=USP32;refGene_tx=XM_011289074.3
E1 27091886 27092071
refGene_gn=USP32;refGene_tx=XM_003996605.5;refGene_gn=USP32;refGene_tx

=XM_011289074.3
E1	27098590	27098802
refGene_gn=USP32;refGene_tx=XM_003996605.5;refGene_gn=USP32;refGene_tx=XM_011289074.3
E1	27100180	27100292
refGene_gn=USP32;refGene_tx=XM_003996605.5;refGene_gn=USP32;refGene_tx=XM_011289074.3
E1	27101517	27101669
refGene_gn=USP32;refGene_tx=XM_003996605.5;refGene_gn=USP32;refGene_tx=XM_011289074.3
E1	27102349	27102524
refGene_gn=USP32;refGene_tx=XM_003996605.5;refGene_gn=USP32;refGene_tx=XM_011289074.3
E1	27103242	27103416
refGene_gn=USP32;refGene_tx=XM_003996605.5;refGene_gn=USP32;refGene_tx=XM_011289074.3
E1	27105629	27105735
refGene_gn=USP32;refGene_tx=XM_003996605.5;refGene_gn=USP32;refGene_tx=XM_011289074.3
E1	27105940	27106079
refGene_gn=USP32;refGene_tx=XM_003996605.5;refGene_gn=USP32;refGene_tx=XM_011289074.3
E1	27106892	27106967
refGene_gn=USP32;refGene_tx=XM_003996605.5;refGene_gn=USP32;refGene_tx=XM_011289074.3
E1	27107773	27107855
refGene_gn=USP32;refGene_tx=XM_003996605.5;refGene_gn=USP32;refGene_tx=XM_011289074.3
E1	27109102	27109257
refGene_gn=USP32;refGene_tx=XM_003996605.5;refGene_gn=USP32;refGene_tx=XM_011289074.3
E1	27113061	27113221
refGene_gn=USP32;refGene_tx=XM_003996605.5;refGene_gn=USP32;refGene_tx=XM_011289074.3
E1	27118646	27118787
refGene_gn=USP32;refGene_tx=XM_003996605.5;refGene_gn=USP32;refGene_tx=XM_011289074.3
E1	27119567	27119743
refGene_gn=USP32;refGene_tx=XM_003996605.5;refGene_gn=USP32;refGene_tx=XM_011289074.3
E1	27124558	27124754
refGene_gn=USP32;refGene_tx=XM_003996605.5;refGene_gn=USP32;refGene_tx=XM_011289074.3
E1	27135011	27135114
refGene_gn=USP32;refGene_tx=XM_003996605.5;refGene_gn=USP32;refGene_tx=XM_011289074.3
E1	27146569	27146631
refGene_gn=USP32;refGene_tx=XM_003996605.5;refGene_gn=USP32;refGene_tx=XM_011289074.3
E1	27155003	27155087

refGene_gn=USP32;refGene_tx=XM_003996605.5;refGene_gn=USP32;refGene_tx=XM_011289074.3
E1 27157480 27157543
refGene_gn=USP32;refGene_tx=XM_003996605.5;refGene_gn=USP32;refGene_tx=XM_011289074.3
E1 27158748 27158864
refGene_gn=USP32;refGene_tx=XM_003996605.5;refGene_gn=USP32;refGene_tx=XM_011289074.3
E1 27160142 27160250
refGene_gn=USP32;refGene_tx=XM_003996605.5;refGene_gn=USP32;refGene_tx=XM_011289074.3
E1 27161797 27161929
refGene_gn=USP32;refGene_tx=XM_003996605.5;refGene_gn=USP32;refGene_tx=XM_011289074.3
E1 27184508 27184668
refGene_gn=USP32;refGene_tx=XM_003996605.5;refGene_gn=USP32;refGene_tx=XM_011289074.3
E1 27190603 27190722
refGene_gn=USP32;refGene_tx=XM_003996605.5;refGene_gn=USP32;refGene_tx=XM_011289074.3
E1 27195123 27195229
refGene_gn=USP32;refGene_tx=XM_003996605.5;refGene_gn=USP32;refGene_tx=XM_011289074.3
E1 27240524 27240652
refGene_gn=USP32;refGene_tx=XM_003996605.5;refGene_gn=USP32;refGene_tx=XM_011289074.3
E1 27312075 27312133
refGene_gn=USP32;refGene_tx=XM_003996605.5;refGene_gn=USP32;refGene_tx=XM_011289074.3
E1 27333079 27333130
refGene_gn=CE1H17orf64;refGene_tx=XM_003996606.3;refGene_gn=CE1H17orf64;refGene_tx=XM_019817571.2
E1 27334274 27334376
refGene_gn=CE1H17orf64;refGene_tx=XM_003996606.3;refGene_gn=CE1H17orf64;refGene_tx=XM_019817571.2
E1 27334633 27334809
refGene_gn=CE1H17orf64;refGene_tx=XM_003996606.3;refGene_gn=CE1H17orf64;refGene_tx=XM_019817571.2
E1 27335280 27335380
refGene_gn=CE1H17orf64;refGene_tx=XM_003996606.3;refGene_gn=CE1H17orf64;refGene_tx=XM_019817571.2
E1 27339730 27339926
refGene_gn=CE1H17orf64;refGene_tx=XM_003996606.3;refGene_gn=CE1H17orf64;refGene_tx=XM_019817571.2
E1 27342844 27342930
refGene_gn=CE1H17orf64;refGene_tx=XM_003996606.3
E1 27358516 27358770 refGene_gn=APPBP2;refGene_tx=XM_023244758.1
E1 27359807 27359973 refGene_gn=APPBP2;refGene_tx=XM_023244758.1
E1 27361009 27361200 refGene_gn=APPBP2;refGene_tx=XM_023244758.1
E1 27363329 27363415 refGene_gn=APPBP2;refGene_tx=XM_023244758.1

E1 27366324 27366449 refGene_gn=APPBP2;refGene_tx=XM_023244758.1
E1 27367143 27367249 refGene_gn=APPBP2;refGene_tx=XM_023244758.1
E1 27367329 27367397 refGene_gn=APPBP2;refGene_tx=XM_023244758.1
E1 27369817 27369907 refGene_gn=APPBP2;refGene_tx=XM_023244758.1
E1 27374170 27374339 refGene_gn=APPBP2;refGene_tx=XM_023244758.1
E1 27380317 27380441 refGene_gn=APPBP2;refGene_tx=XM_023244758.1
E1 27383164 27383316 refGene_gn=APPBP2;refGene_tx=XM_023244758.1
E1 27385827 27385916 refGene_gn=APPBP2;refGene_tx=XM_023244758.1
E1 27406347 27406413
refGene_gn=CE1H17orf64;refGene_tx=XM_019817571.2
E1 27406840 27406978 refGene_gn=APPBP2;refGene_tx=XM_023244758.1
E1 27409774 27409893
refGene_gn=CE1H17orf64;refGene_tx=XM_019817571.2
E1 27481036 27481508 refGene_gn=PPM1D;refGene_tx=XM_003996608.5
E1 27497910 27498139 refGene_gn=PPM1D;refGene_tx=XM_003996608.5
E1 27505468 27505593 refGene_gn=PPM1D;refGene_tx=XM_003996608.5
E1 27518019 27518210 refGene_gn=PPM1D;refGene_tx=XM_003996608.5
E1 27520025 27520268 refGene_gn=PPM1D;refGene_tx=XM_003996608.5
E1 27526568 27527126 refGene_gn=PPM1D;refGene_tx=XM_003996608.5
E1 27540412 27540422 refGene_gn=BCAS3;refGene_tx=XM_011289076.3
E1 27540518 27540602 refGene_gn=BCAS3;refGene_tx=XM_011289076.3
E1 27541137 27541215 refGene_gn=BCAS3;refGene_tx=XM_011289076.3
E1 27542559 27542647 refGene_gn=BCAS3;refGene_tx=XM_011289076.3
E1 27545650 27545705 refGene_gn=BCAS3;refGene_tx=XM_011289076.3
E1 27553237 27553313 refGene_gn=BCAS3;refGene_tx=XM_011289076.3
E1 27570544 27570651 refGene_gn=BCAS3;refGene_tx=XM_011289076.3
E1 27595960 27596042 refGene_gn=BCAS3;refGene_tx=XM_011289076.3
E1 27639766 27639839 refGene_gn=BCAS3;refGene_tx=XM_011289076.3
E1 27688761 27688869 refGene_gn=BCAS3;refGene_tx=XM_011289076.3
E1 27692606 27692683 refGene_gn=BCAS3;refGene_tx=XM_011289076.3
E1 27708559 27708636 refGene_gn=BCAS3;refGene_tx=XM_011289076.3
E1 27715623 27715707 refGene_gn=BCAS3;refGene_tx=XM_011289076.3
E1 27728946 27729117 refGene_gn=BCAS3;refGene_tx=XM_011289076.3
E1 27743725 27743819 refGene_gn=BCAS3;refGene_tx=XM_011289076.3
E1 27773715 27773849 refGene_gn=BCAS3;refGene_tx=XM_011289076.3
E1 27794582 27794847 refGene_gn=BCAS3;refGene_tx=XM_011289076.3
E1 27827615 27827766 refGene_gn=BCAS3;refGene_tx=XM_011289076.3
E1 27843755 27843800 refGene_gn=BCAS3;refGene_tx=XM_011289076.3
E1 27851445 27851570 refGene_gn=BCAS3;refGene_tx=XM_011289076.3
E1 27853553 27853719 refGene_gn=BCAS3;refGene_tx=XM_011289076.3
E1 27855653 27855754 refGene_gn=BCAS3;refGene_tx=XM_011289076.3
E1 27881310 27881411 refGene_gn=BCAS3;refGene_tx=XM_011289076.3
E1 27884112 27884309 refGene_gn=BCAS3;refGene_tx=XM_011289076.3
E1 27890210 27890308 refGene_gn=BCAS3;refGene_tx=XM_011289076.3
E1 28131832 28132000 refGene_gn=BCAS3;refGene_tx=XM_011289076.3
E1 28143464 28143530 refGene_gn=BCAS3;refGene_tx=XM_011289076.3
E1 28155515 28155664 refGene_gn=BCAS3;refGene_tx=XM_011289076.3
E1 28163679 28164074 refGene_gn=TBX2;refGene_tx=XM_023244584.1
E1 28165187 28165455 refGene_gn=TBX2;refGene_tx=XM_023244584.1
E1 28166426 28166573 refGene_gn=TBX2;refGene_tx=XM_023244584.1

E1 28167778 28167855 refGene_gn=TBX2;refGene_tx=XM_023244584.1
E1 28167955 28168119 refGene_gn=TBX2;refGene_tx=XM_023244584.1
E1 28168578 28169216 refGene_gn=TBX2;refGene_tx=XM_023244584.1
E1 28171343 28171796 refGene_gn=TBX2;refGene_tx=XM_023244584.1
E1 28214437 28214638
refGene_gn=TBX4;refGene_tx=XM_023243831.1;refGene_gn=TBX4;refGene_tx=XM_023243832.1;refGene_gn=TBX4;refGene_tx=XM_023243833.1
E1 28215454 28215549
refGene_gn=TBX4;refGene_tx=XM_023243831.1;refGene_gn=TBX4;refGene_tx=XM_023243832.1;refGene_gn=TBX4;refGene_tx=XM_023243833.1
E1 28223611 28223731
refGene_gn=TBX4;refGene_tx=XM_023243831.1;refGene_gn=TBX4;refGene_tx=XM_023243832.1;refGene_gn=TBX4;refGene_tx=XM_023243833.1
E1 28225018 28225166
refGene_gn=TBX4;refGene_tx=XM_023243831.1;refGene_gn=TBX4;refGene_tx=XM_023243832.1;refGene_gn=TBX4;refGene_tx=XM_023243833.1
E1 28235863 28236016
refGene_gn=TBX4;refGene_tx=XM_023243831.1;refGene_gn=TBX4;refGene_tx=XM_023243832.1;refGene_gn=TBX4;refGene_tx=XM_023243833.1
E1 28236978 28237067
refGene_gn=TBX4;refGene_tx=XM_023243831.1;refGene_gn=TBX4;refGene_tx=XM_023243832.1;refGene_gn=TBX4;refGene_tx=XM_023243833.1
E1 28237192 28237428
refGene_gn=TBX4;refGene_tx=XM_023243831.1;refGene_gn=TBX4;refGene_tx=XM_023243832.1;refGene_gn=TBX4;refGene_tx=XM_023243833.1
E1 28239937 28240557
refGene_gn=TBX4;refGene_tx=XM_023243831.1;refGene_gn=TBX4;refGene_tx=XM_023243832.1;refGene_gn=TBX4;refGene_tx=XM_023243833.1
E1 28240576 28240643 refGene_gn=TBX4;refGene_tx=XM_023243833.1
E1 28251346 28251447 refGene_gn=TBX4;refGene_tx=XM_023243833.1
E1 28418363 28418461 refGene_gn=BRIP1;refGene_tx=XM_019817574.2
E1 28418463 28418513 refGene_gn=BRIP1;refGene_tx=XM_019817574.2
E1 28418514 28419151 refGene_gn=BRIP1;refGene_tx=XM_019817574.2
E1 28425716 28426043 refGene_gn=BRIP1;refGene_tx=XM_019817574.2
E1 28436153 28436236 refGene_gn=BRIP1;refGene_tx=XM_019817574.2
E1 28456176 28456289 refGene_gn=BRIP1;refGene_tx=XM_019817574.2
E1 28488509 28488631 refGene_gn=BRIP1;refGene_tx=XM_019817574.2
E1 28489622 28489782 refGene_gn=BRIP1;refGene_tx=XM_019817574.2
E1 28537564 28537726 refGene_gn=BRIP1;refGene_tx=XM_019817574.2
E1 28542751 28542892 refGene_gn=BRIP1;refGene_tx=XM_019817574.2
E1 28543009 28543175 refGene_gn=BRIP1;refGene_tx=XM_019817574.2
E1 28546097 28546252 refGene_gn=BRIP1;refGene_tx=XM_019817574.2
E1 28555757 28555890 refGene_gn=BRIP1;refGene_tx=XM_019817574.2
E1 28561560 28561760 refGene_gn=BRIP1;refGene_tx=XM_019817574.2
E1 28564279 28564501 refGene_gn=BRIP1;refGene_tx=XM_019817574.2
E1 28569234 28569525 refGene_gn=BRIP1;refGene_tx=XM_019817574.2
E1 28606243 28606363 refGene_gn=BRIP1;refGene_tx=XM_019817574.2
E1 28607271 28607402 refGene_gn=BRIP1;refGene_tx=XM_019817574.2
E1 28618703 28618877 refGene_gn=BRIP1;refGene_tx=XM_019817574.2
E1 28620472 28620584 refGene_gn=BRIP1;refGene_tx=XM_019817574.2

E1 28622128 28622224 refGene_gn=BRIP1;refGene_tx=XM_019817574.2
E1 28627373 28627543
refGene_gn=INTS2;refGene_tx=XM_011289083.3;refGene_gn=INTS2;refGene_tx=XM_011289084.3
E1 28627654 28627831
refGene_gn=INTS2;refGene_tx=XM_011289083.3;refGene_gn=INTS2;refGene_tx=XM_011289084.3
E1 28629195 28629301
refGene_gn=INTS2;refGene_tx=XM_011289083.3;refGene_gn=INTS2;refGene_tx=XM_011289084.3
E1 28629685 28629793
refGene_gn=INTS2;refGene_tx=XM_011289083.3;refGene_gn=INTS2;refGene_tx=XM_011289084.3
E1 28630393 28630645
refGene_gn=INTS2;refGene_tx=XM_011289083.3;refGene_gn=INTS2;refGene_tx=XM_011289084.3
E1 28634515 28634714
refGene_gn=INTS2;refGene_tx=XM_011289083.3;refGene_gn=INTS2;refGene_tx=XM_011289084.3
E1 28638750 28638876
refGene_gn=INTS2;refGene_tx=XM_011289083.3;refGene_gn=INTS2;refGene_tx=XM_011289084.3
E1 28642707 28642909
refGene_gn=INTS2;refGene_tx=XM_011289083.3;refGene_gn=INTS2;refGene_tx=XM_011289084.3
E1 28643983 28644148
refGene_gn=INTS2;refGene_tx=XM_011289083.3;refGene_gn=INTS2;refGene_tx=XM_011289084.3
E1 28646613 28646718
refGene_gn=INTS2;refGene_tx=XM_011289083.3;refGene_gn=INTS2;refGene_tx=XM_011289084.3
E1 28658475 28658584
refGene_gn=INTS2;refGene_tx=XM_011289083.3;refGene_gn=INTS2;refGene_tx=XM_011289084.3
E1 28659367 28659544
refGene_gn=INTS2;refGene_tx=XM_011289083.3;refGene_gn=INTS2;refGene_tx=XM_011289084.3
E1 28661058 28661193
refGene_gn=INTS2;refGene_tx=XM_011289083.3;refGene_gn=INTS2;refGene_tx=XM_011289084.3
E1 28663146 28663215
refGene_gn=INTS2;refGene_tx=XM_011289083.3;refGene_gn=INTS2;refGene_tx=XM_011289084.3
E1 28664807 28664922
refGene_gn=INTS2;refGene_tx=XM_011289083.3;refGene_gn=INTS2;refGene_tx=XM_011289084.3
E1 28665005 28665077
refGene_gn=INTS2;refGene_tx=XM_011289083.3;refGene_gn=INTS2;refGene_tx=XM_011289084.3
E1 28669842 28669968

refGene_gn=INTS2;refGene_tx=XM_011289083.3;refGene_gn=INTS2;refGene_tx=XM_011289084.3
E1 28672085 28672312
refGene_gn=INTS2;refGene_tx=XM_011289083.3;refGene_gn=INTS2;refGene_tx=XM_011289084.3
E1 28675415 28675589
refGene_gn=INTS2;refGene_tx=XM_011289083.3;refGene_gn=INTS2;refGene_tx=XM_011289084.3
E1 28675819 28675950
refGene_gn=INTS2;refGene_tx=XM_011289083.3;refGene_gn=INTS2;refGene_tx=XM_011289084.3
E1 28679406 28679520
refGene_gn=INTS2;refGene_tx=XM_011289083.3;refGene_gn=INTS2;refGene_tx=XM_011289084.3
E1 28680833 28680936
refGene_gn=INTS2;refGene_tx=XM_011289083.3;refGene_gn=INTS2;refGene_tx=XM_011289084.3
E1 28684494 28684633
refGene_gn=INTS2;refGene_tx=XM_011289083.3;refGene_gn=INTS2;refGene_tx=XM_011289084.3
E1 28685258 28685565
refGene_gn=INTS2;refGene_tx=XM_011289083.3;refGene_gn=INTS2;refGene_tx=XM_011289084.3
E1 28686843 28686866
refGene_gn=INTS2;refGene_tx=XM_011289083.3;refGene_gn=INTS2;refGene_tx=XM_011289084.3
E1 28697834 28697967 refGene_gn=MED13;refGene_tx=XM_011289087.3
E1 28698558 28698659 refGene_gn=MED13;refGene_tx=XM_011289087.3
E1 28701524 28701698 refGene_gn=MED13;refGene_tx=XM_011289087.3
E1 28704031 28704180 refGene_gn=MED13;refGene_tx=XM_011289087.3
E1 28706080 28706266 refGene_gn=MED13;refGene_tx=XM_011289087.3
E1 28706376 28706535 refGene_gn=MED13;refGene_tx=XM_011289087.3
E1 28706993 28707136 refGene_gn=MED13;refGene_tx=XM_011289087.3
E1 28709839 28710063 refGene_gn=MED13;refGene_tx=XM_011289087.3
E1 28710254 28710446 refGene_gn=MED13;refGene_tx=XM_011289087.3
E1 28711291 28711511 refGene_gn=MED13;refGene_tx=XM_011289087.3
E1 28713612 28714075 refGene_gn=MED13;refGene_tx=XM_011289087.3
E1 28715023 28715213 refGene_gn=MED13;refGene_tx=XM_011289087.3
E1 28716630 28716854 refGene_gn=MED13;refGene_tx=XM_011289087.3
E1 28719931 28720093 refGene_gn=MED13;refGene_tx=XM_011289087.3
E1 28732255 28733169 refGene_gn=MED13;refGene_tx=XM_011289087.3
E1 28733919 28734116 refGene_gn=MED13;refGene_tx=XM_011289087.3
E1 28735498 28735713 refGene_gn=MED13;refGene_tx=XM_011289087.3
E1 28735852 28735943 refGene_gn=MED13;refGene_tx=XM_011289087.3
E1 28737630 28737752 refGene_gn=MED13;refGene_tx=XM_011289087.3
E1 28739757 28739839 refGene_gn=MED13;refGene_tx=XM_011289087.3
E1 28742244 28742458 refGene_gn=MED13;refGene_tx=XM_011289087.3
E1 28754141 28754825 refGene_gn=MED13;refGene_tx=XM_011289087.3
E1 28767859 28767970 refGene_gn=MED13;refGene_tx=XM_011289087.3
E1 28768162 28768325 refGene_gn=MED13;refGene_tx=XM_011289087.3

E1 28768906 28769101 refGene_gn=MED13;refGene_tx=XM_011289087.3
E1 28770729 28770927 refGene_gn=MED13;refGene_tx=XM_011289087.3
E1 28771952 28772098 refGene_gn=MED13;refGene_tx=XM_011289087.3
E1 28794430 28794599 refGene_gn=MED13;refGene_tx=XM_011289087.3
E1 28803267 28803502 refGene_gn=MED13;refGene_tx=XM_011289087.3
E1 28805289 28805355 refGene_gn=MED13;refGene_tx=XM_011289087.3
E1 28980664 28980680 refGene_gn=RPS6KB1;refGene_tx=XM_019817580.2
E1 28982041 28982063 refGene_gn=RPS6KB1;refGene_tx=XM_019817579.2
E1 28993874 28993925
refGene_gn=RPS6KB1;refGene_tx=XM_019817579.2;refGene_gn=RPS6KB1;refGene_tx=XM_019817580.2;refGene_gn=RPS6KB1;refGene_tx=XM_019817582.2
E1 28997390 28997793
refGene_gn=RNFT1;refGene_tx=XM_011289088.3;refGene_gn=RNFT1;refGene_tx=XM_011289089.3;refGene_gn=RNFT1;refGene_tx=XM_019817588.1
E1 28997982 28998059
refGene_gn=RNFT1;refGene_tx=XM_011289088.3;refGene_gn=RNFT1;refGene_tx=XM_011289089.3;refGene_gn=RNFT1;refGene_tx=XM_019817588.1
E1 28999108 28999209
refGene_gn=RNFT1;refGene_tx=XM_011289088.3;refGene_gn=RNFT1;refGene_tx=XM_011289089.3;refGene_gn=RNFT1;refGene_tx=XM_019817588.1
E1 29000683 29000837
refGene_gn=RNFT1;refGene_tx=XM_011289088.3;refGene_gn=RNFT1;refGene_tx=XM_011289089.3;refGene_gn=RNFT1;refGene_tx=XM_019817588.1
E1 29001291 29001450
refGene_gn=RNFT1;refGene_tx=XM_011289088.3;refGene_gn=RNFT1;refGene_tx=XM_011289089.3;refGene_gn=RNFT1;refGene_tx=XM_019817588.1
E1 29002559 29002625
refGene_gn=RNFT1;refGene_tx=XM_011289088.3;refGene_gn=RNFT1;refGene_tx=XM_011289089.3;refGene_gn=RNFT1;refGene_tx=XM_019817588.1
E1 29005850 29005952
refGene_gn=RNFT1;refGene_tx=XM_011289088.3;refGene_gn=RNFT1;refGene_tx=XM_011289089.3;refGene_gn=RNFT1;refGene_tx=XM_019817588.1
E1 29006586 29006721
refGene_gn=RNFT1;refGene_tx=XM_011289088.3;refGene_gn=RNFT1;refGene_tx=XM_011289089.3;refGene_gn=RNFT1;refGene_tx=XM_019817588.1
E1 29013222 29013460
refGene_gn=RPS6KB1;refGene_tx=XM_003996616.4;refGene_gn=RPS6KB1;refGene_tx=XM_003996617.4;refGene_gn=RPS6KB1;refGene_tx=XM_023244405.1;refGene_gn=RPS6KB1;refGene_tx=XM_023244406.1
E1 29013462 29013478 refGene_gn=RPS6KB1;refGene_tx=XM_019817584.2
E1 29014079 29014155 refGene_gn=RPS6KB1;refGene_tx=XM_023244407.1
E1 29014410 29014523
refGene_gn=RPS6KB1;refGene_tx=XM_003996616.4;refGene_gn=RPS6KB1;refGene_tx=XM_003996617.4;refGene_gn=RPS6KB1;refGene_tx=XM_019817579.2;refGene_gn=RPS6KB1;refGene_tx=XM_019817580.2;refGene_gn=RPS6KB1;refGene_tx=XM_019817582.2;refGene_gn=RPS6KB1;refGene_tx=XM_019817584.2;refGene_gn=RPS6KB1;refGene_tx=XM_023244405.1;refGene_gn=RPS6KB1;refGene_tx=XM_023244406.1;refGene_gn=RPS6KB1;refGene_tx=XM_023244407.1
E1 29017341 29017449
refGene_gn=RPS6KB1;refGene_tx=XM_003996616.4;refGene_gn=RPS6KB1;refGen

e_tx=XM_003996617.4;refGene_gn=RPS6KB1;refGene_tx=XM_019817579.2;refGene_gn=RPS6KB1;refGene_tx=XM_019817580.2;refGene_gn=RPS6KB1;refGene_tx=XM_019817582.2;refGene_gn=RPS6KB1;refGene_tx=XM_019817584.2;refGene_gn=RPS6KB1;refGene_tx=XM_023244405.1;refGene_gn=RPS6KB1;refGene_tx=XM_023244406.1;refGene_gn=RPS6KB1;refGene_tx=XM_023244407.1
E1 29023290 29023368
refGene_gn=RPS6KB1;refGene_tx=XM_003996616.4;refGene_gn=RPS6KB1;refGene_tx=XM_003996617.4;refGene_gn=RPS6KB1;refGene_tx=XM_019817579.2;refGene_gn=RPS6KB1;refGene_tx=XM_019817580.2;refGene_gn=RPS6KB1;refGene_tx=XM_019817582.2;refGene_gn=RPS6KB1;refGene_tx=XM_019817584.2;refGene_gn=RPS6KB1;refGene_tx=XM_023244405.1;refGene_gn=RPS6KB1;refGene_tx=XM_023244406.1;refGene_gn=RPS6KB1;refGene_tx=XM_023244407.1
E1 29023554 29023617
refGene_gn=RPS6KB1;refGene_tx=XM_003996616.4;refGene_gn=RPS6KB1;refGene_tx=XM_003996617.4;refGene_gn=RPS6KB1;refGene_tx=XM_019817579.2;refGene_gn=RPS6KB1;refGene_tx=XM_019817580.2;refGene_gn=RPS6KB1;refGene_tx=XM_019817582.2;refGene_gn=RPS6KB1;refGene_tx=XM_019817584.2;refGene_gn=RPS6KB1;refGene_tx=XM_023244405.1;refGene_gn=RPS6KB1;refGene_tx=XM_023244406.1;refGene_gn=RPS6KB1;refGene_tx=XM_023244407.1
E1 29023911 29024019
refGene_gn=RPS6KB1;refGene_tx=XM_003996616.4;refGene_gn=RPS6KB1;refGene_tx=XM_003996617.4;refGene_gn=RPS6KB1;refGene_tx=XM_019817579.2;refGene_gn=RPS6KB1;refGene_tx=XM_019817580.2;refGene_gn=RPS6KB1;refGene_tx=XM_019817582.2;refGene_gn=RPS6KB1;refGene_tx=XM_019817584.2;refGene_gn=RPS6KB1;refGene_tx=XM_023244405.1;refGene_gn=RPS6KB1;refGene_tx=XM_023244406.1;refGene_gn=RPS6KB1;refGene_tx=XM_023244407.1
E1 29024505 29024596
refGene_gn=RPS6KB1;refGene_tx=XM_003996616.4;refGene_gn=RPS6KB1;refGene_tx=XM_003996617.4;refGene_gn=RPS6KB1;refGene_tx=XM_019817579.2;refGene_gn=RPS6KB1;refGene_tx=XM_019817580.2;refGene_gn=RPS6KB1;refGene_tx=XM_019817582.2;refGene_gn=RPS6KB1;refGene_tx=XM_019817584.2;refGene_gn=RPS6KB1;refGene_tx=XM_023244405.1;refGene_gn=RPS6KB1;refGene_tx=XM_023244406.1;refGene_gn=RPS6KB1;refGene_tx=XM_023244407.1
E1 29024782 29024873
refGene_gn=RPS6KB1;refGene_tx=XM_003996616.4;refGene_gn=RPS6KB1;refGene_tx=XM_003996617.4;refGene_gn=RPS6KB1;refGene_tx=XM_019817579.2;refGene_gn=RPS6KB1;refGene_tx=XM_019817580.2;refGene_gn=RPS6KB1;refGene_tx=XM_019817582.2;refGene_gn=RPS6KB1;refGene_tx=XM_019817584.2;refGene_gn=RPS6KB1;refGene_tx=XM_023244405.1;refGene_gn=RPS6KB1;refGene_tx=XM_023244406.1;refGene_gn=RPS6KB1;refGene_tx=XM_023244407.1
E1 29026855 29026956
refGene_gn=RPS6KB1;refGene_tx=XM_003996616.4;refGene_gn=RPS6KB1;refGene_tx=XM_003996617.4;refGene_gn=RPS6KB1;refGene_tx=XM_019817579.2;refGene_gn=RPS6KB1;refGene_tx=XM_019817580.2;refGene_gn=RPS6KB1;refGene_tx=XM_019817582.2;refGene_gn=RPS6KB1;refGene_tx=XM_019817584.2;refGene_gn=RPS6KB1;refGene_tx=XM_023244405.1;refGene_gn=RPS6KB1;refGene_tx=XM_023244406.1;refGene_gn=RPS6KB1;refGene_tx=XM_023244407.1
E1 29028397 29028455
refGene_gn=RPS6KB1;refGene_tx=XM_003996616.4;refGene_gn=RPS6KB1;refGene_tx=XM_003996617.4;refGene_gn=RPS6KB1;refGene_tx=XM_019817579.2;refGe

ne_gn=RPS6KB1;refGene_tx=XM_019817580.2;refGene_gn=RPS6KB1;refGene_tx=XM_019817582.2;refGene_gn=RPS6KB1;refGene_tx=XM_019817584.2;refGene_gn=RPS6KB1;refGene_tx=XM_023244405.1;refGene_gn=RPS6KB1;refGene_tx=XM_023244406.1;refGene_gn=RPS6KB1;refGene_tx=XM_023244407.1
E1 29030568 29030716
refGene_gn=RPS6KB1;refGene_tx=XM_003996616.4;refGene_gn=RPS6KB1;refGene_tx=XM_003996617.4;refGene_gn=RPS6KB1;refGene_tx=XM_019817579.2;refGene_gn=RPS6KB1;refGene_tx=XM_019817580.2;refGene_gn=RPS6KB1;refGene_tx=XM_019817582.2;refGene_gn=RPS6KB1;refGene_tx=XM_019817584.2;refGene_gn=RPS6KB1;refGene_tx=XM_023244405.1;refGene_gn=RPS6KB1;refGene_tx=XM_023244406.1;refGene_gn=RPS6KB1;refGene_tx=XM_023244407.1
E1 29040574 29040643
refGene_gn=RPS6KB1;refGene_tx=XM_003996616.4;refGene_gn=RPS6KB1;refGene_tx=XM_019817579.2;refGene_gn=RPS6KB1;refGene_tx=XM_019817580.2;refGene_gn=RPS6KB1;refGene_tx=XM_019817582.2;refGene_gn=RPS6KB1;refGene_tx=XM_019817584.2;refGene_gn=RPS6KB1;refGene_tx=XM_023244405.1;refGene_gn=RPS6KB1;refGene_tx=XM_023244406.1;refGene_gn=RPS6KB1;refGene_tx=XM_023244407.1
E1 29041847 29041968
refGene_gn=RPS6KB1;refGene_tx=XM_003996616.4;refGene_gn=RPS6KB1;refGene_tx=XM_003996617.4;refGene_gn=RPS6KB1;refGene_tx=XM_019817579.2;refGene_gn=RPS6KB1;refGene_tx=XM_019817580.2;refGene_gn=RPS6KB1;refGene_tx=XM_019817582.2;refGene_gn=RPS6KB1;refGene_tx=XM_019817584.2;refGene_gn=RPS6KB1;refGene_tx=XM_023244405.1;refGene_gn=RPS6KB1;refGene_tx=XM_023244406.1;refGene_gn=RPS6KB1;refGene_tx=XM_023244407.1
E1 29043015 29043065
refGene_gn=RPS6KB1;refGene_tx=XM_003996616.4;refGene_gn=RPS6KB1;refGene_tx=XM_003996617.4;refGene_gn=RPS6KB1;refGene_tx=XM_019817579.2;refGene_gn=RPS6KB1;refGene_tx=XM_019817580.2;refGene_gn=RPS6KB1;refGene_tx=XM_019817582.2;refGene_gn=RPS6KB1;refGene_tx=XM_019817584.2;refGene_gn=RPS6KB1;refGene_tx=XM_023244405.1;refGene_gn=RPS6KB1;refGene_tx=XM_023244406.1;refGene_gn=RPS6KB1;refGene_tx=XM_023244407.1
E1 29052165 29052306
refGene_gn=RPS6KB1;refGene_tx=XM_003996616.4;refGene_gn=RPS6KB1;refGene_tx=XM_003996617.4;refGene_gn=RPS6KB1;refGene_tx=XM_019817579.2;refGene_gn=RPS6KB1;refGene_tx=XM_019817580.2;refGene_gn=RPS6KB1;refGene_tx=XM_019817582.2;refGene_gn=RPS6KB1;refGene_tx=XM_019817584.2;refGene_gn=RPS6KB1;refGene_tx=XM_023244407.1
E1 29053652 29053824 refGene_gn=TUBD1;refGene_tx=XM_011289092.2
E1 29058499 29058647 refGene_gn=TUBD1;refGene_tx=XM_011289092.2
E1 29059889 29060106 refGene_gn=TUBD1;refGene_tx=XM_011289092.2
E1 29060847 29061079 refGene_gn=TUBD1;refGene_tx=XM_011289092.2
E1 29062631 29062752 refGene_gn=TUBD1;refGene_tx=XM_023244408.1
E1 29070642 29070807
refGene_gn=TUBD1;refGene_tx=XM_011289092.2;refGene_gn=TUBD1;refGene_tx=XM_023244408.1
E1 29084738 29084879
refGene_gn=TUBD1;refGene_tx=XM_011289092.2;refGene_gn=TUBD1;refGene_tx=XM_023244408.1
E1 29086907 29087091

refGene_gn=TUBD1;refGene_tx=XM_011289092.2;refGene_gn=TUBD1;refGene_tx=XM_023244408.1
E1 29088882 29088985
refGene_gn=TUBD1;refGene_tx=XM_011289092.2;refGene_gn=TUBD1;refGene_tx=XM_023244408.1
E1 29106541 29106685
refGene_gn=VMP1;refGene_tx=XM_023243665.1;refGene_gn=VMP1;refGene_tx=XM_023243666.1;refGene_gn=VMP1;refGene_tx=XM_023243667.1
E1 29107767 29107870
refGene_gn=VMP1;refGene_tx=XM_023243665.1;refGene_gn=VMP1;refGene_tx=XM_023243666.1;refGene_gn=VMP1;refGene_tx=XM_023243667.1
E1 29125504 29125566
refGene_gn=VMP1;refGene_tx=XM_023243665.1;refGene_gn=VMP1;refGene_tx=XM_023243666.1;refGene_gn=VMP1;refGene_tx=XM_023243667.1
E1 29130745 29130862
refGene_gn=VMP1;refGene_tx=XM_023243665.1;refGene_gn=VMP1;refGene_tx=XM_023243666.1;refGene_gn=VMP1;refGene_tx=XM_023243667.1
E1 29136183 29136264
refGene_gn=VMP1;refGene_tx=XM_023243665.1;refGene_gn=VMP1;refGene_tx=XM_023243666.1;refGene_gn=VMP1;refGene_tx=XM_023243667.1
E1 29169864 29169996
refGene_gn=VMP1;refGene_tx=XM_023243665.1;refGene_gn=VMP1;refGene_tx=XM_023243666.1
E1 29178108 29178276
refGene_gn=VMP1;refGene_tx=XM_023243665.1;refGene_gn=VMP1;refGene_tx=XM_023243666.1;refGene_gn=VMP1;refGene_tx=XM_023243667.1
E1 29203595 29203706
refGene_gn=VMP1;refGene_tx=XM_023243665.1;refGene_gn=VMP1;refGene_tx=XM_023243666.1;refGene_gn=VMP1;refGene_tx=XM_023243667.1
E1 29204961 29205052
refGene_gn=VMP1;refGene_tx=XM_023243665.1;refGene_gn=VMP1;refGene_tx=XM_023243666.1;refGene_gn=VMP1;refGene_tx=XM_023243667.1
E1 29209693 29209829
refGene_gn=VMP1;refGene_tx=XM_023243665.1;refGene_gn=VMP1;refGene_tx=XM_023243666.1;refGene_gn=VMP1;refGene_tx=XM_023243667.1
E1 29212381 29212457
refGene_gn=VMP1;refGene_tx=XM_023243665.1;refGene_gn=VMP1;refGene_tx=XM_023243666.1;refGene_gn=VMP1;refGene_tx=XM_023243667.1
E1 29229010 29229469
refGene_gn=LOC101081092;refGene_tx=XM_011289101.2
E1 29237507 29237528 refGene_gn=PTRH2;refGene_tx=XM_023243668.1
E1 29243246 29243786
refGene_gn=PTRH2;refGene_tx=XM_019817590.2;refGene_gn=PTRH2;refGene_tx=XM_023243668.1;refGene_gn=PTRH2;refGene_tx=XM_023243669.1
E1 29248634 29248759 refGene_gn=CLTC;refGene_tx=XM_019817592.2
E1 29251243 29251319 refGene_gn=CLTC;refGene_tx=XM_019817592.2
E1 29256256 29256304 refGene_gn=CLTC;refGene_tx=XM_019817592.2
E1 29256305 29256479 refGene_gn=CLTC;refGene_tx=XM_019817592.2
E1 29257066 29257237 refGene_gn=CLTC;refGene_tx=XM_019817592.2
E1 29258151 29258262 refGene_gn=CLTC;refGene_tx=XM_019817592.2

E1 29258376 29258508 refGene_gn=CLTC;refGene_tx=XM_019817592.2
E1 29258594 29258744 refGene_gn=CLTC;refGene_tx=XM_019817592.2
E1 29259102 29259270 refGene_gn=CLTC;refGene_tx=XM_019817592.2
E1 29259352 29259460 refGene_gn=CLTC;refGene_tx=XM_019817592.2
E1 29259580 29259745 refGene_gn=CLTC;refGene_tx=XM_019817592.2
E1 29259948 29260106 refGene_gn=CLTC;refGene_tx=XM_019817592.2
E1 29260575 29260768 refGene_gn=CLTC;refGene_tx=XM_019817592.2
E1 29260969 29261153 refGene_gn=CLTC;refGene_tx=XM_019817592.2
E1 29261622 29261768 refGene_gn=CLTC;refGene_tx=XM_019817592.2
E1 29262704 29262827 refGene_gn=CLTC;refGene_tx=XM_019817592.2
E1 29263507 29263742 refGene_gn=CLTC;refGene_tx=XM_019817592.2
E1 29265535 29265678 refGene_gn=CLTC;refGene_tx=XM_019817592.2
E1 29266315 29266441 refGene_gn=CLTC;refGene_tx=XM_019817592.2
E1 29273502 29273666 refGene_gn=CLTC;refGene_tx=XM_019817592.2
E1 29275253 29275434 refGene_gn=CLTC;refGene_tx=XM_019817592.2
E1 29275663 29275828 refGene_gn=CLTC;refGene_tx=XM_019817592.2
E1 29276039 29276177 refGene_gn=CLTC;refGene_tx=XM_019817592.2
E1 29277040 29277163 refGene_gn=CLTC;refGene_tx=XM_019817592.2
E1 29277781 29277934 refGene_gn=CLTC;refGene_tx=XM_019817592.2
E1 29280052 29280253 refGene_gn=CLTC;refGene_tx=XM_019817592.2
E1 29280973 29281171 refGene_gn=CLTC;refGene_tx=XM_019817592.2
E1 29284125 29284299 refGene_gn=CLTC;refGene_tx=XM_019817592.2
E1 29284715 29284829 refGene_gn=CLTC;refGene_tx=XM_019817592.2
E1 29287900 29288062 refGene_gn=CLTC;refGene_tx=XM_019817592.2
E1 29288618 29288887 refGene_gn=CLTC;refGene_tx=XM_019817592.2
E1 29290281 29290284 refGene_gn=CLTC;refGene_tx=XM_019817592.2
E1 29292086 29292294 refGene_gn=CLTC;refGene_tx=XM_019817592.2
E1 29315480 29315522 refGene_gn=CLTC;refGene_tx=XM_019817592.2
E1 29327526 29327666 refGene_gn=DHX40;refGene_tx=XM_011289115.3
E1 29329128 29329357 refGene_gn=DHX40;refGene_tx=XM_011289115.3
E1 29330290 29330360 refGene_gn=DHX40;refGene_tx=XM_011289115.3
E1 29334556 29334651 refGene_gn=DHX40;refGene_tx=XM_011289115.3
E1 29335363 29335472 refGene_gn=DHX40;refGene_tx=XM_011289115.3
E1 29336549 29336664 refGene_gn=DHX40;refGene_tx=XM_011289115.3
E1 29350435 29350593 refGene_gn=DHX40;refGene_tx=XM_011289115.3
E1 29351768 29351849 refGene_gn=DHX40;refGene_tx=XM_011289115.3
E1 29356090 29356273 refGene_gn=DHX40;refGene_tx=XM_011289115.3
E1 29356414 29356501 refGene_gn=DHX40;refGene_tx=XM_011289115.3
E1 29358710 29358810 refGene_gn=DHX40;refGene_tx=XM_011289115.3
E1 29368930 29369062 refGene_gn=DHX40;refGene_tx=XM_011289115.3
E1 29370026 29370093 refGene_gn=DHX40;refGene_tx=XM_011289115.3
E1 29370325 29370553 refGene_gn=DHX40;refGene_tx=XM_011289115.3
E1 29370742 29370862 refGene_gn=DHX40;refGene_tx=XM_011289115.3
E1 29372715 29372861 refGene_gn=DHX40;refGene_tx=XM_011289115.3
E1 29374135 29374303 refGene_gn=DHX40;refGene_tx=XM_011289115.3
E1 29375142 29375254 refGene_gn=DHX40;refGene_tx=XM_011289115.3
E1 29507533 29507623
refGene_gn=YPEL2;refGene_tx=XM_023244589.1;refGene_gn=YPEL2;refGene_tx
=XM_023244590.1
E1 29511948 29512057

refGene_gn=YPEL2;refGene_tx=XM_023244589.1;refGene_gn=YPEL2;refGene_tx=XM_023244590.1
E1 29513159 29513203
refGene_gn=YPEL2;refGene_tx=XM_023244589.1;refGene_gn=YPEL2;refGene_tx=XM_023244590.1
E1 29545622 29545739
refGene_gn=YPEL2;refGene_tx=XM_023244589.1;refGene_gn=YPEL2;refGene_tx=XM_023244590.1
E1 29617672 29617795 refGene_gn=GDPD1;refGene_tx=XM_011289103.3
E1 29618491 29618543 refGene_gn=GDPD1;refGene_tx=XM_011289103.3
E1 29619743 29619803 refGene_gn=GDPD1;refGene_tx=XM_011289103.3
E1 29625725 29625859 refGene_gn=GDPD1;refGene_tx=XM_011289103.3
E1 29631348 29631438 refGene_gn=GDPD1;refGene_tx=XM_011289103.3
E1 29631937 29632056 refGene_gn=GDPD1;refGene_tx=XM_011289103.3
E1 29640619 29640665 refGene_gn=GDPD1;refGene_tx=XM_011289103.3
E1 29643241 29643377 refGene_gn=GDPD1;refGene_tx=XM_011289103.3
E1 29655636 29655679 refGene_gn=GDPD1;refGene_tx=XM_011289103.3
E1 29668300 29668442 refGene_gn=GDPD1;refGene_tx=XM_011289103.3
E1 29677949 29678147 refGene_gn=SMG8;refGene_tx=XM_003996623.5
E1 29678574 29679447 refGene_gn=SMG8;refGene_tx=XM_003996623.5
E1 29680477 29680623 refGene_gn=SMG8;refGene_tx=XM_003996623.5
E1 29681383 29683148 refGene_gn=SMG8;refGene_tx=XM_003996623.5
E1 29692389 29692458
refGene_gn=PRR11;refGene_tx=XM_003996624.4;refGene_gn=PRR11;refGene_tx=XM_019817595.2;refGene_gn=PRR11;refGene_tx=XM_019817596.2;refGene_gn=PRR11;refGene_tx=XM_023243670.1
E1 29693289 29693386
refGene_gn=PRR11;refGene_tx=XM_003996624.4;refGene_gn=PRR11;refGene_tx=XM_019817595.2;refGene_gn=PRR11;refGene_tx=XM_019817596.2;refGene_gn=PRR11;refGene_tx=XM_023243670.1
E1 29693474 29693534
refGene_gn=PRR11;refGene_tx=XM_003996624.4;refGene_gn=PRR11;refGene_tx=XM_019817595.2;refGene_gn=PRR11;refGene_tx=XM_019817596.2;refGene_gn=PRR11;refGene_tx=XM_023243670.1
E1 29695415 29695528
refGene_gn=PRR11;refGene_tx=XM_003996624.4;refGene_gn=PRR11;refGene_tx=XM_019817595.2;refGene_gn=PRR11;refGene_tx=XM_019817596.2;refGene_gn=PRR11;refGene_tx=XM_023243670.1
E1 29696258 29696357
refGene_gn=PRR11;refGene_tx=XM_003996624.4;refGene_gn=PRR11;refGene_tx=XM_019817595.2;refGene_gn=PRR11;refGene_tx=XM_019817596.2;refGene_gn=PRR11;refGene_tx=XM_023243670.1
E1 29698051 29698312
refGene_gn=PRR11;refGene_tx=XM_003996624.4;refGene_gn=PRR11;refGene_tx=XM_019817595.2;refGene_gn=PRR11;refGene_tx=XM_019817596.2;refGene_gn=PRR11;refGene_tx=XM_023243670.1
E1 29701535 29701658
refGene_gn=PRR11;refGene_tx=XM_003996624.4;refGene_gn=PRR11;refGene_tx=XM_019817595.2;refGene_gn=PRR11;refGene_tx=XM_019817596.2;refGene_gn=PRR11;refGene_tx=XM_023243670.1

E1 29701900 29702051
refGene_gn=PRR11;refGene_tx=XM_003996624.4;refGene_gn=PRR11;refGene_tx
=XM_019817595.2;refGene_gn=PRR11;refGene_tx=XM_019817596.2;refGene_gn=
PRR11;refGene_tx=XM_023243670.1
E1 29704972 29705105
refGene_gn=PRR11;refGene_tx=XM_003996624.4;refGene_gn=PRR11;refGene_tx
=XM_019817595.2;refGene_gn=PRR11;refGene_tx=XM_019817596.2;refGene_gn=
PRR11;refGene_tx=XM_023243670.1
E1 29708987 29709012 refGene_gn=PRR11;refGene_tx=XM_019817595.2
E1 29717073 29717106
refGene_gn=SKA2;refGene_tx=XM_019817598.1;refGene_gn=SKA2;refGene_tx=X
M_019817599.2
E1 29717153 29717497 refGene_gn=SKA2;refGene_tx=XM_019817597.1
E1 29741372 29741459
refGene_gn=SKA2;refGene_tx=XM_019817597.1;refGene_gn=SKA2;refGene_tx=X
M_019817598.1
E1 29759043 29759220
refGene_gn=SKA2;refGene_tx=XM_019817598.1;refGene_gn=SKA2;refGene_tx=X
M_019817599.2
E1 29765539 29765683
refGene_gn=SKA2;refGene_tx=XM_019817597.1;refGene_gn=SKA2;refGene_tx=X
M_019817598.1;refGene_gn=SKA2;refGene_tx=XM_019817599.2
E1 29771501 29771522
refGene_gn=TRIM37;refGene_tx=XM_003996626.5;refGene_gn=TRIM37;refGene_
tx=XM_011289104.3;refGene_gn=TRIM37;refGene_tx=XM_011289105.3;refGene_
gn=TRIM37;refGene_tx=XM_019817600.2;refGene_gn=TRIM37;refGene_tx=XM_01
9817601.2
E1 29773223 29773325
refGene_gn=TRIM37;refGene_tx=XM_003996626.5;refGene_gn=TRIM37;refGene_
tx=XM_011289104.3;refGene_gn=TRIM37;refGene_tx=XM_011289105.3;refGene_
gn=TRIM37;refGene_tx=XM_019817600.2;refGene_gn=TRIM37;refGene_tx=XM_01
9817601.2
E1 29782317 29782358
refGene_gn=TRIM37;refGene_tx=XM_003996626.5;refGene_gn=TRIM37;refGene_
tx=XM_011289104.3;refGene_gn=TRIM37;refGene_tx=XM_011289105.3;refGene_
gn=TRIM37;refGene_tx=XM_019817600.2;refGene_gn=TRIM37;refGene_tx=XM_01
9817601.2
E1 29786782 29786899
refGene_gn=TRIM37;refGene_tx=XM_003996626.5;refGene_gn=TRIM37;refGene_
tx=XM_011289104.3;refGene_gn=TRIM37;refGene_tx=XM_011289105.3;refGene_
gn=TRIM37;refGene_tx=XM_019817600.2;refGene_gn=TRIM37;refGene_tx=XM_01
9817601.2
E1 29790118 29790206
refGene_gn=TRIM37;refGene_tx=XM_003996626.5;refGene_gn=TRIM37;refGene_
tx=XM_011289104.3;refGene_gn=TRIM37;refGene_tx=XM_011289105.3;refGene_
gn=TRIM37;refGene_tx=XM_019817600.2;refGene_gn=TRIM37;refGene_tx=XM_01
9817601.2
E1 29793111 29793234
refGene_gn=TRIM37;refGene_tx=XM_003996626.5;refGene_gn=TRIM37;refGene_
tx=XM_011289104.3;refGene_gn=TRIM37;refGene_tx=XM_011289105.3;refGene_

gn=TRIM37;refGene_tx=XM_019817600.2;refGene_gn=TRIM37;refGene_tx=XM_019817601.2
E1 29797885 29798009
refGene_gn=TRIM37;refGene_tx=XM_003996626.5;refGene_gn=TRIM37;refGene_tx=XM_011289104.3;refGene_gn=TRIM37;refGene_tx=XM_011289105.3;refGene_gn=TRIM37;refGene_tx=XM_019817600.2;refGene_gn=TRIM37;refGene_tx=XM_019817601.2
E1 29799089 29799157
refGene_gn=TRIM37;refGene_tx=XM_003996626.5;refGene_gn=TRIM37;refGene_tx=XM_011289104.3;refGene_gn=TRIM37;refGene_tx=XM_011289105.3;refGene_gn=TRIM37;refGene_tx=XM_019817600.2;refGene_gn=TRIM37;refGene_tx=XM_019817601.2
E1 29815430 29815555
refGene_gn=TRIM37;refGene_tx=XM_003996626.5;refGene_gn=TRIM37;refGene_tx=XM_011289104.3;refGene_gn=TRIM37;refGene_tx=XM_011289105.3;refGene_gn=TRIM37;refGene_tx=XM_019817600.2;refGene_gn=TRIM37;refGene_tx=XM_019817601.2
E1 29822073 29822124
refGene_gn=TRIM37;refGene_tx=XM_003996626.5;refGene_gn=TRIM37;refGene_tx=XM_011289104.3;refGene_gn=TRIM37;refGene_tx=XM_011289105.3;refGene_gn=TRIM37;refGene_tx=XM_019817600.2;refGene_gn=TRIM37;refGene_tx=XM_019817601.2
E1 29823749 29823831
refGene_gn=TRIM37;refGene_tx=XM_003996626.5;refGene_gn=TRIM37;refGene_tx=XM_011289104.3;refGene_gn=TRIM37;refGene_tx=XM_011289105.3;refGene_gn=TRIM37;refGene_tx=XM_019817600.2;refGene_gn=TRIM37;refGene_tx=XM_019817601.2
E1 29832344 29832421
refGene_gn=TRIM37;refGene_tx=XM_003996626.5;refGene_gn=TRIM37;refGene_tx=XM_011289104.3;refGene_gn=TRIM37;refGene_tx=XM_011289105.3;refGene_gn=TRIM37;refGene_tx=XM_019817600.2;refGene_gn=TRIM37;refGene_tx=XM_019817601.2
E1 29844678 29844858
refGene_gn=TRIM37;refGene_tx=XM_003996626.5;refGene_gn=TRIM37;refGene_tx=XM_011289104.3;refGene_gn=TRIM37;refGene_tx=XM_011289105.3;refGene_gn=TRIM37;refGene_tx=XM_019817600.2;refGene_gn=TRIM37;refGene_tx=XM_019817601.2
E1 29848507 29848622
refGene_gn=TRIM37;refGene_tx=XM_003996626.5;refGene_gn=TRIM37;refGene_tx=XM_011289104.3;refGene_gn=TRIM37;refGene_tx=XM_011289105.3;refGene_gn=TRIM37;refGene_tx=XM_019817600.2;refGene_gn=TRIM37;refGene_tx=XM_019817601.2
E1 29849749 29849965
refGene_gn=TRIM37;refGene_tx=XM_003996626.5;refGene_gn=TRIM37;refGene_tx=XM_011289104.3;refGene_gn=TRIM37;refGene_tx=XM_011289105.3;refGene_gn=TRIM37;refGene_tx=XM_019817600.2;refGene_gn=TRIM37;refGene_tx=XM_019817601.2
E1 29851205 29851342
refGene_gn=TRIM37;refGene_tx=XM_003996626.5;refGene_gn=TRIM37;refGene_tx=XM_011289104.3;refGene_gn=TRIM37;refGene_tx=XM_011289105.3;refGene_

gn=TRIM37;refGene_tx=XM_019817600.2;refGene_gn=TRIM37;refGene_tx=XM_01
9817601.2
E1 29854767 29854853
refGene_gn=TRIM37;refGene_tx=XM_003996626.5;refGene_gn=TRIM37;refGene_
tx=XM_011289104.3;refGene_gn=TRIM37;refGene_tx=XM_011289105.3;refGene_
gn=TRIM37;refGene_tx=XM_019817600.2;refGene_gn=TRIM37;refGene_tx=XM_01
9817601.2
E1 29865466 29865661
refGene_gn=TRIM37;refGene_tx=XM_003996626.5;refGene_gn=TRIM37;refGene_
tx=XM_011289104.3;refGene_gn=TRIM37;refGene_tx=XM_011289105.3;refGene_
gn=TRIM37;refGene_tx=XM_019817600.2;refGene_gn=TRIM37;refGene_tx=XM_01
9817601.2
E1 29874528 29874837
refGene_gn=TRIM37;refGene_tx=XM_003996626.5;refGene_gn=TRIM37;refGene_
tx=XM_011289104.3;refGene_gn=TRIM37;refGene_tx=XM_011289105.3;refGene_
gn=TRIM37;refGene_tx=XM_019817600.2;refGene_gn=TRIM37;refGene_tx=XM_01
9817601.2
E1 29891185 29891314
refGene_gn=TRIM37;refGene_tx=XM_003996626.5;refGene_gn=TRIM37;refGene_
tx=XM_011289104.3;refGene_gn=TRIM37;refGene_tx=XM_011289105.3
E1 29891670 29891860
refGene_gn=TRIM37;refGene_tx=XM_003996626.5;refGene_gn=TRIM37;refGene_
tx=XM_011289104.3;refGene_gn=TRIM37;refGene_tx=XM_011289105.3;refGene_
gn=TRIM37;refGene_tx=XM_019817600.2;refGene_gn=TRIM37;refGene_tx=XM_01
9817601.2
E1 29894687 29894827
refGene_gn=TRIM37;refGene_tx=XM_003996626.5;refGene_gn=TRIM37;refGene_
tx=XM_011289104.3;refGene_gn=TRIM37;refGene_tx=XM_011289105.3;refGene_
gn=TRIM37;refGene_tx=XM_019817600.2;refGene_gn=TRIM37;refGene_tx=XM_01
9817601.2
E1 29905375 29905486
refGene_gn=TRIM37;refGene_tx=XM_003996626.5;refGene_gn=TRIM37;refGene_
tx=XM_011289104.3;refGene_gn=TRIM37;refGene_tx=XM_011289105.3;refGene_
gn=TRIM37;refGene_tx=XM_019817600.2;refGene_gn=TRIM37;refGene_tx=XM_01
9817601.2
E1 29906840 29906923
refGene_gn=TRIM37;refGene_tx=XM_003996626.5;refGene_gn=TRIM37;refGene_
tx=XM_011289104.3;refGene_gn=TRIM37;refGene_tx=XM_011289105.3;refGene_
gn=TRIM37;refGene_tx=XM_019817600.2;refGene_gn=TRIM37;refGene_tx=XM_01
9817601.2
E1 29923319 29923323 refGene_gn=TRIM37;refGene_tx=XM_011289105.3
E1 29925158 29926216 refGene_gn=PPM1E;refGene_tx=XM_023243797.1
E1 29932170 29932264 refGene_gn=PPM1E;refGene_tx=XM_023243797.1
E1 29932498 29932642 refGene_gn=PPM1E;refGene_tx=XM_023243797.1
E1 29935000 29935189 refGene_gn=PPM1E;refGene_tx=XM_023243797.1
E1 29938515 29938715 refGene_gn=PPM1E;refGene_tx=XM_023243797.1
E1 29946232 29946351 refGene_gn=PPM1E;refGene_tx=XM_023243797.1
E1 30151351 30151809 refGene_gn=PPM1E;refGene_tx=XM_023243797.1
E1 30175150 30175240 refGene_gn=RAD51C;refGene_tx=XM_011289106.3
E1 30176701 30176762 refGene_gn=RAD51C;refGene_tx=XM_011289106.3

E1 30185849 30185910 refGene_gn=RAD51C;refGene_tx=XM_011289106.3
E1 30186843 30186910 refGene_gn=RAD51C;refGene_tx=XM_011289106.3
E1 30195647 30195779 refGene_gn=RAD51C;refGene_tx=XM_011289106.3
E1 30207119 30207253 refGene_gn=RAD51C;refGene_tx=XM_011289106.3
E1 30208696 30208863 refGene_gn=RAD51C;refGene_tx=XM_011289106.3
E1 30209481 30209740 refGene_gn=RAD51C;refGene_tx=XM_011289106.3
E1 30212751 30212869 refGene_gn=RAD51C;refGene_tx=XM_011289106.3
E1 30243516 30243652
refGene_gn=TEX14;refGene_tx=XM_006940144.4;refGene_gn=TEX14;refGene_tx=XM_019817604.2;refGene_gn=TEX14;refGene_tx=XM_023244159.1;refGene_gn=TEX14;refGene_tx=XM_023244160.1;refGene_gn=TEX14;refGene_tx=XM_023244161.1
E1 30257904 30257917 refGene_gn=TEX14;refGene_tx=XM_006940145.4
E1 30258555 30258670
refGene_gn=TEX14;refGene_tx=XM_006940144.4;refGene_gn=TEX14;refGene_tx=XM_006940145.4;refGene_gn=TEX14;refGene_tx=XM_019817604.2;refGene_gn=TEX14;refGene_tx=XM_023244159.1;refGene_gn=TEX14;refGene_tx=XM_023244160.1;refGene_gn=TEX14;refGene_tx=XM_023244161.1
E1 30265071 30265237
refGene_gn=TEX14;refGene_tx=XM_006940144.4;refGene_gn=TEX14;refGene_tx=XM_006940145.4;refGene_gn=TEX14;refGene_tx=XM_019817604.2;refGene_gn=TEX14;refGene_tx=XM_023244159.1;refGene_gn=TEX14;refGene_tx=XM_023244160.1;refGene_gn=TEX14;refGene_tx=XM_023244161.1
E1 30266261 30266398
refGene_gn=TEX14;refGene_tx=XM_006940144.4;refGene_gn=TEX14;refGene_tx=XM_006940145.4;refGene_gn=TEX14;refGene_tx=XM_019817604.2;refGene_gn=TEX14;refGene_tx=XM_023244159.1;refGene_gn=TEX14;refGene_tx=XM_023244160.1;refGene_gn=TEX14;refGene_tx=XM_023244161.1
E1 30270366 30270448
refGene_gn=TEX14;refGene_tx=XM_006940144.4;refGene_gn=TEX14;refGene_tx=XM_006940145.4;refGene_gn=TEX14;refGene_tx=XM_019817604.2;refGene_gn=TEX14;refGene_tx=XM_023244159.1;refGene_gn=TEX14;refGene_tx=XM_023244160.1;refGene_gn=TEX14;refGene_tx=XM_023244161.1
E1 30271337 30271468
refGene_gn=TEX14;refGene_tx=XM_006940144.4;refGene_gn=TEX14;refGene_tx=XM_006940145.4;refGene_gn=TEX14;refGene_tx=XM_019817604.2;refGene_gn=TEX14;refGene_tx=XM_023244159.1;refGene_gn=TEX14;refGene_tx=XM_023244160.1;refGene_gn=TEX14;refGene_tx=XM_023244161.1
E1 30273237 30273351
refGene_gn=TEX14;refGene_tx=XM_006940144.4;refGene_gn=TEX14;refGene_tx=XM_006940145.4;refGene_gn=TEX14;refGene_tx=XM_019817604.2;refGene_gn=TEX14;refGene_tx=XM_023244159.1;refGene_gn=TEX14;refGene_tx=XM_023244160.1
E1 30274407 30274531
refGene_gn=TEX14;refGene_tx=XM_006940144.4;refGene_gn=TEX14;refGene_tx=XM_006940145.4;refGene_gn=TEX14;refGene_tx=XM_019817604.2;refGene_gn=TEX14;refGene_tx=XM_023244159.1;refGene_gn=TEX14;refGene_tx=XM_023244160.1;refGene_gn=TEX14;refGene_tx=XM_023244161.1
E1 30277649 30277828
refGene_gn=TEX14;refGene_tx=XM_006940144.4;refGene_gn=TEX14;refGene_tx

=XM_006940145.4;refGene_gn=TEX14;refGene_tx=XM_019817604.2;refGene_gn=TEX14;refGene_tx=XM_023244159.1;refGene_gn=TEX14;refGene_tx=XM_023244160.1;refGene_gn=TEX14;refGene_tx=XM_023244161.1
E1 30282580 30282732
refGene_gn=TEX14;refGene_tx=XM_006940144.4;refGene_gn=TEX14;refGene_tx=XM_006940145.4;refGene_gn=TEX14;refGene_tx=XM_019817604.2;refGene_gn=TEX14;refGene_tx=XM_023244159.1;refGene_gn=TEX14;refGene_tx=XM_023244160.1;refGene_gn=TEX14;refGene_tx=XM_023244161.1
E1 30283876 30284067
refGene_gn=TEX14;refGene_tx=XM_006940144.4;refGene_gn=TEX14;refGene_tx=XM_006940145.4;refGene_gn=TEX14;refGene_tx=XM_019817604.2;refGene_gn=TEX14;refGene_tx=XM_023244159.1;refGene_gn=TEX14;refGene_tx=XM_023244160.1;refGene_gn=TEX14;refGene_tx=XM_023244161.1
E1 30284686 30284837
refGene_gn=TEX14;refGene_tx=XM_006940144.4;refGene_gn=TEX14;refGene_tx=XM_006940145.4;refGene_gn=TEX14;refGene_tx=XM_019817604.2;refGene_gn=TEX14;refGene_tx=XM_023244159.1;refGene_gn=TEX14;refGene_tx=XM_023244160.1;refGene_gn=TEX14;refGene_tx=XM_023244161.1
E1 30286410 30287312
refGene_gn=TEX14;refGene_tx=XM_006940144.4;refGene_gn=TEX14;refGene_tx=XM_006940145.4;refGene_gn=TEX14;refGene_tx=XM_019817604.2;refGene_gn=TEX14;refGene_tx=XM_023244159.1;refGene_gn=TEX14;refGene_tx=XM_023244160.1;refGene_gn=TEX14;refGene_tx=XM_023244161.1
E1 30290659 30290766
refGene_gn=TEX14;refGene_tx=XM_006940144.4;refGene_gn=TEX14;refGene_tx=XM_006940145.4;refGene_gn=TEX14;refGene_tx=XM_019817604.2;refGene_gn=TEX14;refGene_tx=XM_023244159.1;refGene_gn=TEX14;refGene_tx=XM_023244160.1;refGene_gn=TEX14;refGene_tx=XM_023244161.1
E1 30293035 30293124
refGene_gn=TEX14;refGene_tx=XM_006940144.4;refGene_gn=TEX14;refGene_tx=XM_006940145.4;refGene_gn=TEX14;refGene_tx=XM_019817604.2;refGene_gn=TEX14;refGene_tx=XM_023244159.1;refGene_gn=TEX14;refGene_tx=XM_023244160.1;refGene_gn=TEX14;refGene_tx=XM_023244161.1
E1 30294465 30294747
refGene_gn=TEX14;refGene_tx=XM_006940144.4;refGene_gn=TEX14;refGene_tx=XM_006940145.4;refGene_gn=TEX14;refGene_tx=XM_019817604.2;refGene_gn=TEX14;refGene_tx=XM_023244159.1;refGene_gn=TEX14;refGene_tx=XM_023244160.1;refGene_gn=TEX14;refGene_tx=XM_023244161.1
E1 30295629 30295730
refGene_gn=TEX14;refGene_tx=XM_006940144.4;refGene_gn=TEX14;refGene_tx=XM_006940145.4;refGene_gn=TEX14;refGene_tx=XM_019817604.2;refGene_gn=TEX14;refGene_tx=XM_023244159.1;refGene_gn=TEX14;refGene_tx=XM_023244160.1;refGene_gn=TEX14;refGene_tx=XM_023244161.1
E1 30297305 30297425
refGene_gn=TEX14;refGene_tx=XM_006940144.4;refGene_gn=TEX14;refGene_tx=XM_006940145.4;refGene_gn=TEX14;refGene_tx=XM_019817604.2;refGene_gn=TEX14;refGene_tx=XM_023244159.1;refGene_gn=TEX14;refGene_tx=XM_023244160.1;refGene_gn=TEX14;refGene_tx=XM_023244161.1
E1 30298828 30298895
refGene_gn=TEX14;refGene_tx=XM_006940144.4;refGene_gn=TEX14;refGene_tx

=XM_006940145.4;refGene_gn=TEX14;refGene_tx=XM_019817604.2;refGene_gn=TEX14;refGene_tx=XM_023244159.1;refGene_gn=TEX14;refGene_tx=XM_023244160.1;refGene_gn=TEX14;refGene_tx=XM_023244161.1
E1 30301827 30301909
refGene_gn=TEX14;refGene_tx=XM_006940144.4;refGene_gn=TEX14;refGene_tx=XM_006940145.4;refGene_gn=TEX14;refGene_tx=XM_019817604.2;refGene_gn=TEX14;refGene_tx=XM_023244159.1;refGene_gn=TEX14;refGene_tx=XM_023244160.1;refGene_gn=TEX14;refGene_tx=XM_023244161.1
E1 30305167 30305230
refGene_gn=TEX14;refGene_tx=XM_006940144.4;refGene_gn=TEX14;refGene_tx=XM_006940145.4;refGene_gn=TEX14;refGene_tx=XM_019817604.2;refGene_gn=TEX14;refGene_tx=XM_023244160.1;refGene_gn=TEX14;refGene_tx=XM_023244161.1
E1 30305743 30305871
refGene_gn=TEX14;refGene_tx=XM_006940144.4;refGene_gn=TEX14;refGene_tx=XM_006940145.4;refGene_gn=TEX14;refGene_tx=XM_019817604.2;refGene_gn=TEX14;refGene_tx=XM_023244159.1;refGene_gn=TEX14;refGene_tx=XM_023244160.1;refGene_gn=TEX14;refGene_tx=XM_023244161.1
E1 30306784 30306990
refGene_gn=TEX14;refGene_tx=XM_006940144.4;refGene_gn=TEX14;refGene_tx=XM_006940145.4;refGene_gn=TEX14;refGene_tx=XM_019817604.2;refGene_gn=TEX14;refGene_tx=XM_023244159.1;refGene_gn=TEX14;refGene_tx=XM_023244160.1;refGene_gn=TEX14;refGene_tx=XM_023244161.1
E1 30308033 30308130
refGene_gn=TEX14;refGene_tx=XM_006940144.4;refGene_gn=TEX14;refGene_tx=XM_006940145.4;refGene_gn=TEX14;refGene_tx=XM_019817604.2;refGene_gn=TEX14;refGene_tx=XM_023244159.1;refGene_gn=TEX14;refGene_tx=XM_023244160.1;refGene_gn=TEX14;refGene_tx=XM_023244161.1
E1 30309536 30309611
refGene_gn=TEX14;refGene_tx=XM_006940144.4;refGene_gn=TEX14;refGene_tx=XM_006940145.4;refGene_gn=TEX14;refGene_tx=XM_019817604.2;refGene_gn=TEX14;refGene_tx=XM_023244159.1;refGene_gn=TEX14;refGene_tx=XM_023244160.1;refGene_gn=TEX14;refGene_tx=XM_023244161.1
E1 30312317 30312395
refGene_gn=TEX14;refGene_tx=XM_006940144.4;refGene_gn=TEX14;refGene_tx=XM_006940145.4;refGene_gn=TEX14;refGene_tx=XM_019817604.2;refGene_gn=TEX14;refGene_tx=XM_023244159.1;refGene_gn=TEX14;refGene_tx=XM_023244160.1;refGene_gn=TEX14;refGene_tx=XM_023244161.1
E1 30313117 30313217
refGene_gn=TEX14;refGene_tx=XM_006940144.4;refGene_gn=TEX14;refGene_tx=XM_006940145.4;refGene_gn=TEX14;refGene_tx=XM_019817604.2;refGene_gn=TEX14;refGene_tx=XM_023244159.1;refGene_gn=TEX14;refGene_tx=XM_023244160.1;refGene_gn=TEX14;refGene_tx=XM_023244161.1
E1 30318372 30318465
refGene_gn=TEX14;refGene_tx=XM_006940144.4;refGene_gn=TEX14;refGene_tx=XM_006940145.4;refGene_gn=TEX14;refGene_tx=XM_019817604.2;refGene_gn=TEX14;refGene_tx=XM_023244159.1;refGene_gn=TEX14;refGene_tx=XM_023244161.1
E1 30320740 30320850
refGene_gn=TEX14;refGene_tx=XM_006940144.4;refGene_gn=TEX14;refGene_tx

=XM_006940145.4;refGene_gn=TEX14;refGene_tx=XM_019817604.2;refGene_gn=TEX14;refGene_tx=XM_023244159.1;refGene_gn=TEX14;refGene_tx=XM_023244160.1;refGene_gn=TEX14;refGene_tx=XM_023244161.1
E1 30322097 30322149
refGene_gn=TEX14;refGene_tx=XM_006940144.4;refGene_gn=TEX14;refGene_tx=XM_006940145.4;refGene_gn=TEX14;refGene_tx=XM_019817604.2;refGene_gn=TEX14;refGene_tx=XM_023244159.1;refGene_gn=TEX14;refGene_tx=XM_023244160.1;refGene_gn=TEX14;refGene_tx=XM_023244161.1
E1 30322606 30322643
refGene_gn=TEX14;refGene_tx=XM_006940144.4;refGene_gn=TEX14;refGene_tx=XM_006940145.4;refGene_gn=TEX14;refGene_tx=XM_019817604.2;refGene_gn=TEX14;refGene_tx=XM_023244159.1;refGene_gn=TEX14;refGene_tx=XM_023244160.1;refGene_gn=TEX14;refGene_tx=XM_023244161.1
E1 30333437 30335037
refGene_gn=LOC101096522;refGene_tx=XM_006940146.4;refGene_gn=LOC101096522;refGene_tx=XM_019817610.2
E1 30335681 30335726
refGene_gn=LOC101096522;refGene_tx=XM_006940146.4;refGene_gn=LOC101096522;refGene_tx=XM_019817610.2
E1 30336978 30336986
refGene_gn=LOC101096522;refGene_tx=XM_006940146.4;refGene_gn=LOC101096522;refGene_tx=XM_019817610.2
E1 30349968 30350262
refGene_gn=LOC101096522;refGene_tx=XM_006940146.4;refGene_gn=LOC101096522;refGene_tx=XM_019817610.2
E1 30350641 30350735
refGene_gn=LOC101096522;refGene_tx=XM_006940146.4;refGene_gn=LOC101096522;refGene_tx=XM_019817610.2
E1 30351178 30351265
refGene_gn=LOC101096522;refGene_tx=XM_006940146.4;refGene_gn=LOC101096522;refGene_tx=XM_019817610.2
E1 30351769 30351893
refGene_gn=LOC101096522;refGene_tx=XM_006940146.4;refGene_gn=LOC101096522;refGene_tx=XM_019817610.2
E1 30354817 30354952
refGene_gn=LOC101096522;refGene_tx=XM_006940146.4;refGene_gn=LOC101096522;refGene_tx=XM_019817610.2
E1 30355064 30355182
refGene_gn=LOC101096522;refGene_tx=XM_006940146.4;refGene_gn=LOC101096522;refGene_tx=XM_019817610.2
E1 30355270 30355372
refGene_gn=LOC101096522;refGene_tx=XM_006940146.4;refGene_gn=LOC101096522;refGene_tx=XM_019817610.2
E1 30355558 30355655
refGene_gn=LOC101096522;refGene_tx=XM_006940146.4;refGene_gn=LOC101096522;refGene_tx=XM_019817610.2
E1 30355743 30355906
refGene_gn=LOC101096522;refGene_tx=XM_006940146.4;refGene_gn=LOC101096522;refGene_tx=XM_019817610.2
E1 30356069 30356169

refGene_gn=LOC101096522;refGene_tx=XM_006940146.4;refGene_gn=LOC101096522;refGene_tx=XM_019817610.2
E1 30356380 30356857
refGene_gn=LOC101096522;refGene_tx=XM_006940146.4;refGene_gn=LOC101096522;refGene_tx=XM_019817610.2
E1 30359574 30359577 refGene_gn=MTMR4;refGene_tx=XM_011289108.3
E1 30361803 30361848 refGene_gn=MTMR4;refGene_tx=XM_011289107.3
E1 30363278 30363368
refGene_gn=MTMR4;refGene_tx=XM_011289107.3;refGene_gn=MTMR4;refGene_tx=XM_011289108.3
E1 30363725 30363842
refGene_gn=MTMR4;refGene_tx=XM_011289107.3;refGene_gn=MTMR4;refGene_tx=XM_011289108.3;refGene_gn=MTMR4;refGene_tx=XM_023243437.1
E1 30364381 30364464
refGene_gn=MTMR4;refGene_tx=XM_011289107.3;refGene_gn=MTMR4;refGene_tx=XM_011289108.3
E1 30366489 30366650
refGene_gn=MTMR4;refGene_tx=XM_011289107.3;refGene_gn=MTMR4;refGene_tx=XM_011289108.3;refGene_gn=MTMR4;refGene_tx=XM_023243437.1
E1 30366761 30366858
refGene_gn=MTMR4;refGene_tx=XM_011289107.3;refGene_gn=MTMR4;refGene_tx=XM_011289108.3;refGene_gn=MTMR4;refGene_tx=XM_023243437.1
E1 30367029 30367143
refGene_gn=MTMR4;refGene_tx=XM_011289107.3;refGene_gn=MTMR4;refGene_tx=XM_011289108.3;refGene_gn=MTMR4;refGene_tx=XM_023243437.1
E1 30367992 30368189
refGene_gn=MTMR4;refGene_tx=XM_011289107.3;refGene_gn=MTMR4;refGene_tx=XM_011289108.3;refGene_gn=MTMR4;refGene_tx=XM_023243437.1
E1 30368437 30368566
refGene_gn=MTMR4;refGene_tx=XM_011289107.3;refGene_gn=MTMR4;refGene_tx=XM_011289108.3;refGene_gn=MTMR4;refGene_tx=XM_023243437.1
E1 30368994 30369106
refGene_gn=MTMR4;refGene_tx=XM_011289107.3;refGene_gn=MTMR4;refGene_tx=XM_011289108.3;refGene_gn=MTMR4;refGene_tx=XM_023243437.1
E1 30369593 30369789
refGene_gn=MTMR4;refGene_tx=XM_011289107.3;refGene_gn=MTMR4;refGene_tx=XM_011289108.3;refGene_gn=MTMR4;refGene_tx=XM_023243437.1
E1 30370025 30370211
refGene_gn=MTMR4;refGene_tx=XM_011289107.3;refGene_gn=MTMR4;refGene_tx=XM_011289108.3;refGene_gn=MTMR4;refGene_tx=XM_023243437.1
E1 30370287 30370458
refGene_gn=MTMR4;refGene_tx=XM_011289107.3;refGene_gn=MTMR4;refGene_tx=XM_011289108.3;refGene_gn=MTMR4;refGene_tx=XM_023243437.1
E1 30370608 30370763
refGene_gn=MTMR4;refGene_tx=XM_011289107.3;refGene_gn=MTMR4;refGene_tx=XM_011289108.3;refGene_gn=MTMR4;refGene_tx=XM_023243437.1
E1 30376877 30378276
refGene_gn=MTMR4;refGene_tx=XM_011289107.3;refGene_gn=MTMR4;refGene_tx=XM_011289108.3;refGene_gn=MTMR4;refGene_tx=XM_023243437.1
E1 30380722 30380833

refGene_gn=MTMR4;refGene_tx=XM_011289107.3;refGene_gn=MTMR4;refGene_tx=XM_011289108.3;refGene_gn=MTMR4;refGene_tx=XM_023243437.1
E1 30381068 30381157
refGene_gn=MTMR4;refGene_tx=XM_011289107.3;refGene_gn=MTMR4;refGene_tx=XM_011289108.3;refGene_gn=MTMR4;refGene_tx=XM_023243437.1
E1 30381610 30381788
refGene_gn=MTMR4;refGene_tx=XM_011289107.3;refGene_gn=MTMR4;refGene_tx=XM_011289108.3;refGene_gn=MTMR4;refGene_tx=XM_023243437.1
E1 30385001 30386052 refGene_gn=HSF5;refGene_tx=XM_011289117.3
E1 30391109 30391484 refGene_gn=HSF5;refGene_tx=XM_011289117.3
E1 30404210 30404308 refGene_gn=HSF5;refGene_tx=XM_011289117.3
E1 30406892 30407414 refGene_gn=HSF5;refGene_tx=XM_011289117.3
E1 30409719 30409897 refGene_gn=HSF5;refGene_tx=XM_011289117.3
E1 30436041 30436112 refGene_gn=HSF5;refGene_tx=XM_011289117.3
E1 30443183 30443435
refGene_gn=RNF43;refGene_tx=XM_006940148.3;refGene_gn=RNF43;refGene_tx=XM_019817611.2
E1 30489959 30490082
refGene_gn=RNF43;refGene_tx=XM_006940148.3;refGene_gn=RNF43;refGene_tx=XM_019817611.2
E1 30496973 30497048
refGene_gn=RNF43;refGene_tx=XM_006940148.3;refGene_gn=RNF43;refGene_tx=XM_019817611.2
E1 30497159 30497291
refGene_gn=RNF43;refGene_tx=XM_006940148.3;refGene_gn=RNF43;refGene_tx=XM_019817611.2
E1 30497924 30498029
refGene_gn=RNF43;refGene_tx=XM_006940148.3;refGene_gn=RNF43;refGene_tx=XM_019817611.2
E1 30499397 30499559
refGene_gn=RNF43;refGene_tx=XM_006940148.3;refGene_gn=RNF43;refGene_tx=XM_019817611.2
E1 30500100 30500203
refGene_gn=RNF43;refGene_tx=XM_006940148.3;refGene_gn=RNF43;refGene_tx=XM_019817611.2
E1 30500866 30502216
refGene_gn=RNF43;refGene_tx=XM_006940148.3;refGene_gn=RNF43;refGene_tx=XM_019817611.2
E1 30504585 30504632
refGene_gn=RNF43;refGene_tx=XM_006940148.3;refGene_gn=RNF43;refGene_tx=XM_019817611.2
E1 30507563 30507632 refGene_gn=SUPT4H1;refGene_tx=XM_011289109.3
E1 30508175 30508282 refGene_gn=SUPT4H1;refGene_tx=XM_011289109.3
E1 30511609 30511665 refGene_gn=SUPT4H1;refGene_tx=XM_011289109.3
E1 30511929 30511983 refGene_gn=SUPT4H1;refGene_tx=XM_011289109.3
E1 30512417 30512485 refGene_gn=SUPT4H1;refGene_tx=XM_011289109.3
E1 30529566 30529902
refGene_gn=TSPOAP1;refGene_tx=XM_006940154.4;refGene_gn=TSPOAP1;refGene_tx=XM_019817613.2;refGene_gn=TSPOAP1;refGene_tx=XM_019817614.2;refGene_gn=TSPOAP1;refGene_tx=XM_019817615.2;refGene_gn=TSPOAP1;refGene_tx=

XM_019817616.2;refGene_gn=TSPOAP1;refGene_tx=XM_019817617.2;refGene_gn=TSPOAP1;refGene_tx=XM_019817618.2
E1 30530705 30530813
refGene_gn=TSPOAP1;refGene_tx=XM_006940154.4;refGene_gn=TSPOAP1;refGene_tx=XM_019817613.2;refGene_gn=TSPOAP1;refGene_tx=XM_019817614.2;refGene_gn=TSPOAP1;refGene_tx=XM_019817615.2;refGene_gn=TSPOAP1;refGene_tx=XM_019817616.2;refGene_gn=TSPOAP1;refGene_tx=XM_019817617.2;refGene_gn=TSPOAP1;refGene_tx=XM_019817618.2
E1 30531098 30531227
refGene_gn=TSPOAP1;refGene_tx=XM_006940154.4;refGene_gn=TSPOAP1;refGene_tx=XM_019817613.2;refGene_gn=TSPOAP1;refGene_tx=XM_019817614.2;refGene_gn=TSPOAP1;refGene_tx=XM_019817615.2;refGene_gn=TSPOAP1;refGene_tx=XM_019817616.2;refGene_gn=TSPOAP1;refGene_tx=XM_019817617.2;refGene_gn=TSPOAP1;refGene_tx=XM_019817618.2
E1 30531802 30531982
refGene_gn=TSPOAP1;refGene_tx=XM_006940154.4;refGene_gn=TSPOAP1;refGene_tx=XM_019817613.2;refGene_gn=TSPOAP1;refGene_tx=XM_019817614.2;refGene_gn=TSPOAP1;refGene_tx=XM_019817615.2;refGene_gn=TSPOAP1;refGene_tx=XM_019817616.2;refGene_gn=TSPOAP1;refGene_tx=XM_019817617.2;refGene_gn=TSPOAP1;refGene_tx=XM_019817618.2
E1 30532534 30532720
refGene_gn=TSPOAP1;refGene_tx=XM_006940154.4;refGene_gn=TSPOAP1;refGene_tx=XM_019817613.2;refGene_gn=TSPOAP1;refGene_tx=XM_019817614.2;refGene_gn=TSPOAP1;refGene_tx=XM_019817615.2;refGene_gn=TSPOAP1;refGene_tx=XM_019817616.2;refGene_gn=TSPOAP1;refGene_tx=XM_019817617.2;refGene_gn=TSPOAP1;refGene_tx=XM_019817618.2
E1 30533977 30534055
refGene_gn=TSPOAP1;refGene_tx=XM_006940154.4;refGene_gn=TSPOAP1;refGene_tx=XM_019817613.2;refGene_gn=TSPOAP1;refGene_tx=XM_019817614.2;refGene_gn=TSPOAP1;refGene_tx=XM_019817615.2;refGene_gn=TSPOAP1;refGene_tx=XM_019817616.2;refGene_gn=TSPOAP1;refGene_tx=XM_019817617.2;refGene_gn=TSPOAP1;refGene_tx=XM_019817618.2
E1 30534141 30534225
refGene_gn=TSPOAP1;refGene_tx=XM_006940154.4;refGene_gn=TSPOAP1;refGene_tx=XM_019817613.2;refGene_gn=TSPOAP1;refGene_tx=XM_019817614.2;refGene_gn=TSPOAP1;refGene_tx=XM_019817615.2;refGene_gn=TSPOAP1;refGene_tx=XM_019817616.2;refGene_gn=TSPOAP1;refGene_tx=XM_019817617.2;refGene_gn=TSPOAP1;refGene_tx=XM_019817618.2
E1 30534439 30534529
refGene_gn=TSPOAP1;refGene_tx=XM_006940154.4;refGene_gn=TSPOAP1;refGene_tx=XM_019817613.2;refGene_gn=TSPOAP1;refGene_tx=XM_019817614.2;refGene_gn=TSPOAP1;refGene_tx=XM_019817615.2;refGene_gn=TSPOAP1;refGene_tx=XM_019817616.2;refGene_gn=TSPOAP1;refGene_tx=XM_019817617.2;refGene_gn=TSPOAP1;refGene_tx=XM_019817618.2
E1 30534710 30534833
refGene_gn=TSPOAP1;refGene_tx=XM_006940154.4;refGene_gn=TSPOAP1;refGene_tx=XM_019817613.2;refGene_gn=TSPOAP1;refGene_tx=XM_019817614.2;refGene_gn=TSPOAP1;refGene_tx=XM_019817615.2;refGene_gn=TSPOAP1;refGene_tx=XM_019817616.2;refGene_gn=TSPOAP1;refGene_tx=XM_019817617.2;refGene_gn=TSPOAP1;refGene_tx=XM_019817618.2

E1 30535088 30535193
refGene_gn=TSPOAP1;refGene_tx=XM_006940154.4;refGene_gn=TSPOAP1;refGene_tx=XM_019817613.2;refGene_gn=TSPOAP1;refGene_tx=XM_019817614.2;refGene_gn=TSPOAP1;refGene_tx=XM_019817615.2;refGene_gn=TSPOAP1;refGene_tx=XM_019817616.2;refGene_gn=TSPOAP1;refGene_tx=XM_019817617.2;refGene_gn=TSPOAP1;refGene_tx=XM_019817618.2;refGene_gn=TSPOAP1;refGene_tx=XM_019817620.2
E1 30536605 30536656
refGene_gn=TSPOAP1;refGene_tx=XM_006940154.4;refGene_gn=TSPOAP1;refGene_tx=XM_019817613.2;refGene_gn=TSPOAP1;refGene_tx=XM_019817614.2;refGene_gn=TSPOAP1;refGene_tx=XM_019817615.2;refGene_gn=TSPOAP1;refGene_tx=XM_019817616.2;refGene_gn=TSPOAP1;refGene_tx=XM_019817617.2;refGene_gn=TSPOAP1;refGene_tx=XM_019817618.2;refGene_gn=TSPOAP1;refGene_tx=XM_019817620.2
E1 30537061 30537082
refGene_gn=TSPOAP1;refGene_tx=XM_006940154.4;refGene_gn=TSPOAP1;refGene_tx=XM_019817613.2;refGene_gn=TSPOAP1;refGene_tx=XM_019817614.2;refGene_gn=TSPOAP1;refGene_tx=XM_019817615.2;refGene_gn=TSPOAP1;refGene_tx=XM_019817616.2;refGene_gn=TSPOAP1;refGene_tx=XM_019817617.2;refGene_gn=TSPOAP1;refGene_tx=XM_019817618.2;refGene_gn=TSPOAP1;refGene_tx=XM_019817620.2
E1 30537888 30538093
refGene_gn=TSPOAP1;refGene_tx=XM_006940154.4;refGene_gn=TSPOAP1;refGene_tx=XM_019817613.2;refGene_gn=TSPOAP1;refGene_tx=XM_019817614.2;refGene_gn=TSPOAP1;refGene_tx=XM_019817615.2;refGene_gn=TSPOAP1;refGene_tx=XM_019817616.2;refGene_gn=TSPOAP1;refGene_tx=XM_019817617.2;refGene_gn=TSPOAP1;refGene_tx=XM_019817618.2;refGene_gn=TSPOAP1;refGene_tx=XM_019817620.2
E1 30538703 30538876
refGene_gn=TSPOAP1;refGene_tx=XM_006940154.4;refGene_gn=TSPOAP1;refGene_tx=XM_019817613.2;refGene_gn=TSPOAP1;refGene_tx=XM_019817614.2;refGene_gn=TSPOAP1;refGene_tx=XM_019817615.2;refGene_gn=TSPOAP1;refGene_tx=XM_019817616.2;refGene_gn=TSPOAP1;refGene_tx=XM_019817617.2;refGene_gn=TSPOAP1;refGene_tx=XM_019817618.2;refGene_gn=TSPOAP1;refGene_tx=XM_019817620.2
E1 30540076 30540192
refGene_gn=TSPOAP1;refGene_tx=XM_006940154.4;refGene_gn=TSPOAP1;refGene_tx=XM_019817613.2;refGene_gn=TSPOAP1;refGene_tx=XM_019817614.2;refGene_gn=TSPOAP1;refGene_tx=XM_019817615.2;refGene_gn=TSPOAP1;refGene_tx=XM_019817616.2;refGene_gn=TSPOAP1;refGene_tx=XM_019817617.2;refGene_gn=TSPOAP1;refGene_tx=XM_019817618.2;refGene_gn=TSPOAP1;refGene_tx=XM_019817620.2
E1 30540477 30540587
refGene_gn=TSPOAP1;refGene_tx=XM_006940154.4;refGene_gn=TSPOAP1;refGene_tx=XM_019817613.2;refGene_gn=TSPOAP1;refGene_tx=XM_019817614.2;refGene_gn=TSPOAP1;refGene_tx=XM_019817615.2;refGene_gn=TSPOAP1;refGene_tx=XM_019817616.2;refGene_gn=TSPOAP1;refGene_tx=XM_019817617.2;refGene_gn=TSPOAP1;refGene_tx=XM_019817618.2;refGene_gn=TSPOAP1;refGene_tx=XM_019817620.2
E1 30543389 30544220

refGene_gn=TSPOAP1;refGene_tx=XM_006940154.4;refGene_gn=TSPOAP1;refGene_tx=XM_019817613.2;refGene_gn=TSPOAP1;refGene_tx=XM_019817614.2;refGene_gn=TSPOAP1;refGene_tx=XM_019817615.2;refGene_gn=TSPOAP1;refGene_tx=XM_019817616.2;refGene_gn=TSPOAP1;refGene_tx=XM_019817617.2;refGene_gn=TSPOAP1;refGene_tx=XM_019817618.2;refGene_gn=TSPOAP1;refGene_tx=XM_019817620.2
E1 30544368 30544520
refGene_gn=TSPOAP1;refGene_tx=XM_006940154.4;refGene_gn=TSPOAP1;refGene_tx=XM_019817613.2;refGene_gn=TSPOAP1;refGene_tx=XM_019817614.2;refGene_gn=TSPOAP1;refGene_tx=XM_019817615.2;refGene_gn=TSPOAP1;refGene_tx=XM_019817616.2;refGene_gn=TSPOAP1;refGene_tx=XM_019817617.2;refGene_gn=TSPOAP1;refGene_tx=XM_019817618.2;refGene_gn=TSPOAP1;refGene_tx=XM_019817620.2
E1 30544859 30545237
refGene_gn=TSPOAP1;refGene_tx=XM_006940154.4;refGene_gn=TSPOAP1;refGene_tx=XM_019817613.2;refGene_gn=TSPOAP1;refGene_tx=XM_019817614.2;refGene_gn=TSPOAP1;refGene_tx=XM_019817615.2;refGene_gn=TSPOAP1;refGene_tx=XM_019817616.2;refGene_gn=TSPOAP1;refGene_tx=XM_019817617.2;refGene_gn=TSPOAP1;refGene_tx=XM_019817618.2;refGene_gn=TSPOAP1;refGene_tx=XM_019817620.2
E1 30545322 30545541
refGene_gn=TSPOAP1;refGene_tx=XM_006940154.4;refGene_gn=TSPOAP1;refGene_tx=XM_019817613.2;refGene_gn=TSPOAP1;refGene_tx=XM_019817614.2;refGene_gn=TSPOAP1;refGene_tx=XM_019817615.2;refGene_gn=TSPOAP1;refGene_tx=XM_019817616.2;refGene_gn=TSPOAP1;refGene_tx=XM_019817617.2;refGene_gn=TSPOAP1;refGene_tx=XM_019817618.2;refGene_gn=TSPOAP1;refGene_tx=XM_019817620.2
E1 30545876 30546065
refGene_gn=TSPOAP1;refGene_tx=XM_006940154.4;refGene_gn=TSPOAP1;refGene_tx=XM_019817613.2;refGene_gn=TSPOAP1;refGene_tx=XM_019817614.2;refGene_gn=TSPOAP1;refGene_tx=XM_019817615.2;refGene_gn=TSPOAP1;refGene_tx=XM_019817616.2;refGene_gn=TSPOAP1;refGene_tx=XM_019817617.2;refGene_gn=TSPOAP1;refGene_tx=XM_019817618.2;refGene_gn=TSPOAP1;refGene_tx=XM_019817620.2
E1 30546635 30547499
refGene_gn=TSPOAP1;refGene_tx=XM_006940154.4;refGene_gn=TSPOAP1;refGene_tx=XM_019817613.2;refGene_gn=TSPOAP1;refGene_tx=XM_019817614.2;refGene_gn=TSPOAP1;refGene_tx=XM_019817615.2;refGene_gn=TSPOAP1;refGene_tx=XM_019817616.2;refGene_gn=TSPOAP1;refGene_tx=XM_019817617.2;refGene_gn=TSPOAP1;refGene_tx=XM_019817618.2;refGene_gn=TSPOAP1;refGene_tx=XM_019817620.2
E1 30548133 30548233
refGene_gn=TSPOAP1;refGene_tx=XM_006940154.4;refGene_gn=TSPOAP1;refGene_tx=XM_019817613.2;refGene_gn=TSPOAP1;refGene_tx=XM_019817614.2;refGene_gn=TSPOAP1;refGene_tx=XM_019817615.2;refGene_gn=TSPOAP1;refGene_tx=XM_019817616.2;refGene_gn=TSPOAP1;refGene_tx=XM_019817617.2;refGene_gn=TSPOAP1;refGene_tx=XM_019817618.2;refGene_gn=TSPOAP1;refGene_tx=XM_019817620.2
E1 30548324 30548479
refGene_gn=TSPOAP1;refGene_tx=XM_006940154.4;refGene_gn=TSPOAP1;refGen

e_tx=XM_019817613.2;refGene_gn=TSPOAP1;refGene_tx=XM_019817614.2;refGene_gn=TSPOAP1;refGene_tx=XM_019817615.2;refGene_gn=TSPOAP1;refGene_tx=XM_019817616.2;refGene_gn=TSPOAP1;refGene_tx=XM_019817617.2;refGene_gn=TSPOAP1;refGene_tx=XM_019817618.2;refGene_gn=TSPOAP1;refGene_tx=XM_019817620.2
E1 30549052 30549221
refGene_gn=TSPOAP1;refGene_tx=XM_006940154.4;refGene_gn=TSPOAP1;refGene_tx=XM_019817613.2;refGene_gn=TSPOAP1;refGene_tx=XM_019817614.2;refGene_gn=TSPOAP1;refGene_tx=XM_019817615.2;refGene_gn=TSPOAP1;refGene_tx=XM_019817616.2;refGene_gn=TSPOAP1;refGene_tx=XM_019817617.2;refGene_gn=TSPOAP1;refGene_tx=XM_019817618.2;refGene_gn=TSPOAP1;refGene_tx=XM_019817620.2
E1 30549588 30549660
refGene_gn=TSPOAP1;refGene_tx=XM_006940154.4;refGene_gn=TSPOAP1;refGene_tx=XM_019817613.2;refGene_gn=TSPOAP1;refGene_tx=XM_019817614.2;refGene_gn=TSPOAP1;refGene_tx=XM_019817615.2;refGene_gn=TSPOAP1;refGene_tx=XM_019817616.2;refGene_gn=TSPOAP1;refGene_tx=XM_019817617.2;refGene_gn=TSPOAP1;refGene_tx=XM_019817618.2;refGene_gn=TSPOAP1;refGene_tx=XM_019817620.2
E1 30550082 30550115
refGene_gn=TSPOAP1;refGene_tx=XM_006940154.4;refGene_gn=TSPOAP1;refGene_tx=XM_019817613.2;refGene_gn=TSPOAP1;refGene_tx=XM_019817614.2;refGene_gn=TSPOAP1;refGene_tx=XM_019817616.2;refGene_gn=TSPOAP1;refGene_tx=XM_019817617.2;refGene_gn=TSPOAP1;refGene_tx=XM_019817618.2;refGene_gn=TSPOAP1;refGene_tx=XM_019817620.2
E1 30550292 30550396
refGene_gn=TSPOAP1;refGene_tx=XM_006940154.4;refGene_gn=TSPOAP1;refGene_tx=XM_019817613.2;refGene_gn=TSPOAP1;refGene_tx=XM_019817614.2;refGene_gn=TSPOAP1;refGene_tx=XM_019817615.2;refGene_gn=TSPOAP1;refGene_tx=XM_019817616.2;refGene_gn=TSPOAP1;refGene_tx=XM_019817617.2;refGene_gn=TSPOAP1;refGene_tx=XM_019817618.2;refGene_gn=TSPOAP1;refGene_tx=XM_019817620.2
E1 30550480 30550552
refGene_gn=TSPOAP1;refGene_tx=XM_006940154.4;refGene_gn=TSPOAP1;refGene_tx=XM_019817613.2;refGene_gn=TSPOAP1;refGene_tx=XM_019817614.2;refGene_gn=TSPOAP1;refGene_tx=XM_019817615.2;refGene_gn=TSPOAP1;refGene_tx=XM_019817616.2;refGene_gn=TSPOAP1;refGene_tx=XM_019817620.2
E1 30550810 30550921
refGene_gn=TSPOAP1;refGene_tx=XM_006940154.4;refGene_gn=TSPOAP1;refGene_tx=XM_019817613.2;refGene_gn=TSPOAP1;refGene_tx=XM_019817614.2;refGene_gn=TSPOAP1;refGene_tx=XM_019817615.2;refGene_gn=TSPOAP1;refGene_tx=XM_019817616.2;refGene_gn=TSPOAP1;refGene_tx=XM_019817620.2
E1 30551572 30551602
refGene_gn=TSPOAP1;refGene_tx=XM_006940154.4;refGene_gn=TSPOAP1;refGene_tx=XM_019817618.2
E1 30553340 30553520
refGene_gn=TSPOAP1;refGene_tx=XM_019817613.2;refGene_gn=TSPOAP1;refGene_tx=XM_019817614.2;refGene_gn=TSPOAP1;refGene_tx=XM_019817615.2;refGene_gn=TSPOAP1;refGene_tx=XM_019817616.2;refGene_gn=TSPOAP1;refGene_tx=XM_019817617.2;refGene_gn=TSPOAP1;refGene_tx=XM_019817620.2

E1 30564997 30565022 refGene_gn=MPO;refGene_tx=XM_019816875.2
E1 30565119 30565200
refGene_gn=MPO;refGene_tx=NM_001122746.2;refGene_gn=MPO;refGene_tx=XM_006940138.4;refGene_gn=MPO;refGene_tx=XM_019816875.2
E1 30565329 30565423
refGene_gn=MPO;refGene_tx=NM_001122746.2;refGene_gn=MPO;refGene_tx=XM_006940138.4;refGene_gn=MPO;refGene_tx=XM_019816875.2
E1 30565725 30565901
refGene_gn=MPO;refGene_tx=NM_001122746.2;refGene_gn=MPO;refGene_tx=XM_006940138.4;refGene_gn=MPO;refGene_tx=XM_019816875.2
E1 30566114 30566238
refGene_gn=MPO;refGene_tx=NM_001122746.2;refGene_gn=MPO;refGene_tx=XM_006940138.4;refGene_gn=MPO;refGene_tx=XM_019816875.2
E1 30566350 30566480
refGene_gn=MPO;refGene_tx=NM_001122746.2;refGene_gn=MPO;refGene_tx=XM_006940138.4;refGene_gn=MPO;refGene_tx=XM_019816875.2
E1 30566565 30566772
refGene_gn=MPO;refGene_tx=NM_001122746.2;refGene_gn=MPO;refGene_tx=XM_006940138.4;refGene_gn=MPO;refGene_tx=XM_019816875.2
E1 30567626 30567945
refGene_gn=MPO;refGene_tx=NM_001122746.2;refGene_gn=MPO;refGene_tx=XM_006940138.4;refGene_gn=MPO;refGene_tx=XM_019816875.2
E1 30569785 30569946
refGene_gn=MPO;refGene_tx=NM_001122746.2;refGene_gn=MPO;refGene_tx=XM_006940138.4;refGene_gn=MPO;refGene_tx=XM_019816875.2
E1 30571139 30571395
refGene_gn=MPO;refGene_tx=NM_001122746.2;refGene_gn=MPO;refGene_tx=XM_006940138.4;refGene_gn=MPO;refGene_tx=XM_019816875.2
E1 30571840 30572011
refGene_gn=MPO;refGene_tx=NM_001122746.2;refGene_gn=MPO;refGene_tx=XM_006940138.4;refGene_gn=MPO;refGene_tx=XM_019816875.2
E1 30572613 30572851
refGene_gn=MPO;refGene_tx=NM_001122746.2;refGene_gn=MPO;refGene_tx=XM_006940138.4;refGene_gn=MPO;refGene_tx=XM_019816875.2
E1 30573611 30573819
refGene_gn=MPO;refGene_tx=NM_001122746.2;refGene_gn=MPO;refGene_tx=XM_006940138.4;refGene_gn=MPO;refGene_tx=XM_019816875.2
E1 30576247 30576594
refGene_gn=LPO;refGene_tx=XM_023244021.1;refGene_gn=LPO;refGene_tx=XM_023244022.1
E1 30576780 30577018 refGene_gn=LPO;refGene_tx=XM_023244022.1
E1 30578018 30578192
refGene_gn=LPO;refGene_tx=XM_023244021.1;refGene_gn=LPO;refGene_tx=XM_023244022.1
E1 30579054 30579307
refGene_gn=LPO;refGene_tx=XM_023244021.1;refGene_gn=LPO;refGene_tx=XM_023244022.1
E1 30584898 30585059
refGene_gn=LPO;refGene_tx=XM_023244021.1;refGene_gn=LPO;refGene_tx=XM_023244022.1

E1 30586949 30587274
refGene_gn=LPO;refGene_tx=XM_023244021.1;refGene_gn=LPO;refGene_tx=XM_
023244022.1
E1 30588780 30588987
refGene_gn=LPO;refGene_tx=XM_023244021.1;refGene_gn=LPO;refGene_tx=XM_
023244022.1
E1 30589756 30589886
refGene_gn=LPO;refGene_tx=XM_023244021.1;refGene_gn=LPO;refGene_tx=XM_
023244022.1
E1 30590234 30590395
refGene_gn=LPO;refGene_tx=XM_023244021.1;refGene_gn=LPO;refGene_tx=XM_
023244022.1
E1 30591988 30592136 refGene_gn=LPO;refGene_tx=XM_023244022.1
E1 30626511 30626591
refGene_gn=MKS1;refGene_tx=XM_003996637.3;refGene_gn=MKS1;refGene_tx=X
M_019817622.1
E1 30627010 30627120
refGene_gn=MKS1;refGene_tx=XM_003996637.3;refGene_gn=MKS1;refGene_tx=X
M_019817622.1
E1 30628551 30628622
refGene_gn=MKS1;refGene_tx=XM_003996637.3;refGene_gn=MKS1;refGene_tx=X
M_019817622.1
E1 30629292 30629448
refGene_gn=MKS1;refGene_tx=XM_003996637.3;refGene_gn=MKS1;refGene_tx=X
M_019817622.1
E1 30630532 30630630
refGene_gn=MKS1;refGene_tx=XM_003996637.3;refGene_gn=MKS1;refGene_tx=X
M_019817622.1
E1 30630822 30630951
refGene_gn=MKS1;refGene_tx=XM_003996637.3;refGene_gn=MKS1;refGene_tx=X
M_019817622.1
E1 30631214 30631319
refGene_gn=MKS1;refGene_tx=XM_003996637.3;refGene_gn=MKS1;refGene_tx=X
M_019817622.1
E1 30631907 30632016
refGene_gn=MKS1;refGene_tx=XM_003996637.3;refGene_gn=MKS1;refGene_tx=X
M_019817622.1
E1 30632820 30632877
refGene_gn=MKS1;refGene_tx=XM_003996637.3;refGene_gn=MKS1;refGene_tx=X
M_019817622.1
E1 30633928 30633971
refGene_gn=MKS1;refGene_tx=XM_003996637.3;refGene_gn=MKS1;refGene_tx=X
M_019817622.1
E1 30634228 30634294
refGene_gn=MKS1;refGene_tx=XM_003996637.3;refGene_gn=MKS1;refGene_tx=X
M_019817622.1
E1 30635739 30635810
refGene_gn=MKS1;refGene_tx=XM_003996637.3;refGene_gn=MKS1;refGene_tx=X
M_019817622.1
E1 30636128 30636198

refGene_gn=MKS1;refGene_tx=XM_003996637.3;refGene_gn=MKS1;refGene_tx=X
M_019817622.1
E1 30636297 30636405
refGene_gn=MKS1;refGene_tx=XM_003996637.3;refGene_gn=MKS1;refGene_tx=X
M_019817622.1
E1 30637082 30637279
refGene_gn=MKS1;refGene_tx=XM_003996637.3;refGene_gn=MKS1;refGene_tx=X
M_019817622.1
E1 30637745 30637828
refGene_gn=MKS1;refGene_tx=XM_003996637.3;refGene_gn=MKS1;refGene_tx=X
M_019817622.1
E1 30637913 30638011
refGene_gn=MKS1;refGene_tx=XM_003996637.3;refGene_gn=MKS1;refGene_tx=X
M_019817622.1
E1 30638112 30638204
refGene_gn=MKS1;refGene_tx=XM_003996637.3;refGene_gn=MKS1;refGene_tx=X
M_019817622.1
E1 30656243 30657179
refGene_gn=LOC101098177;refGene_tx=XM_003996638.3
E1 30667702 30668638
refGene_gn=LOC101098416;refGene_tx=XM_023244234.1
E1 30682366 30683302
refGene_gn=LOC101098667;refGene_tx=XM_003996640.3
E1 30754907 30755045 refGene_gn=DYNLL2;refGene_tx=XM_023243619.1
E1 30756955 30757087 refGene_gn=DYNLL2;refGene_tx=XM_023243619.1
E1 30826407 30826601
refGene_gn=SRSF1;refGene_tx=XM_019817632.2;refGene_gn=SRSF1;refGene_tx
=XM_023244278.1
E1 30827010 30827195
refGene_gn=SRSF1;refGene_tx=XM_019817632.2;refGene_gn=SRSF1;refGene_tx
=XM_023244278.1
E1 30827577 30827750
refGene_gn=SRSF1;refGene_tx=XM_019817632.2;refGene_gn=SRSF1;refGene_tx
=XM_023244278.1
E1 30827951 30828138 refGene_gn=SRSF1;refGene_tx=XM_019817632.2
E1 30828515 30828554
refGene_gn=SRSF1;refGene_tx=XM_019817632.2;refGene_gn=SRSF1;refGene_tx
=XM_023244278.1
E1 30844181 30844235
refGene_gn=VEZF1;refGene_tx=XM_019817629.2;refGene_gn=VEZF1;refGene_tx
=XM_019817630.2;refGene_gn=VEZF1;refGene_tx=XM_023244275.1
E1 30845788 30845821
refGene_gn=VEZF1;refGene_tx=XM_023244276.1;refGene_gn=VEZF1;refGene_tx
=XM_023244277.1
E1 30850528 30851223
refGene_gn=VEZF1;refGene_tx=XM_019817629.2;refGene_gn=VEZF1;refGene_tx
=XM_019817630.2;refGene_gn=VEZF1;refGene_tx=XM_023244275.1;refGene_gn=
VEZF1;refGene_tx=XM_023244276.1;refGene_gn=VEZF1;refGene_tx=XM_0232442
77.1
E1 30852002 30852066

refGene_gn=VEZF1;refGene_tx=XM_019817629.2;refGene_gn=VEZF1;refGene_tx=XM_019817630.2;refGene_gn=VEZF1;refGene_tx=XM_023244275.1;refGene_gn=VEZF1;refGene_tx=XM_023244276.1;refGene_gn=VEZF1;refGene_tx=XM_023244277.1
E1 30853145 30853347
refGene_gn=VEZF1;refGene_tx=XM_019817629.2;refGene_gn=VEZF1;refGene_tx=XM_019817630.2;refGene_gn=VEZF1;refGene_tx=XM_023244275.1;refGene_gn=VEZF1;refGene_tx=XM_023244276.1;refGene_gn=VEZF1;refGene_tx=XM_023244277.1
E1 30854609 30854747
refGene_gn=VEZF1;refGene_tx=XM_019817629.2;refGene_gn=VEZF1;refGene_tx=XM_019817630.2;refGene_gn=VEZF1;refGene_tx=XM_023244275.1;refGene_gn=VEZF1;refGene_tx=XM_023244276.1;refGene_gn=VEZF1;refGene_tx=XM_023244277.1
E1 30857600 30858028
refGene_gn=VEZF1;refGene_tx=XM_019817629.2;refGene_gn=VEZF1;refGene_tx=XM_019817630.2;refGene_gn=VEZF1;refGene_tx=XM_023244275.1;refGene_gn=VEZF1;refGene_tx=XM_023244276.1;refGene_gn=VEZF1;refGene_tx=XM_023244277.1
E1 30939802 30940138
refGene_gn=CUEDC1;refGene_tx=XM_023244509.1;refGene_gn=CUEDC1;refGene_tx=XM_023244510.1;refGene_gn=CUEDC1;refGene_tx=XM_023244511.1
E1 30944262 30944390
refGene_gn=CUEDC1;refGene_tx=XM_023244509.1;refGene_gn=CUEDC1;refGene_tx=XM_023244510.1;refGene_gn=CUEDC1;refGene_tx=XM_023244511.1
E1 30949259 30949386
refGene_gn=CUEDC1;refGene_tx=XM_023244509.1;refGene_gn=CUEDC1;refGene_tx=XM_023244510.1;refGene_gn=CUEDC1;refGene_tx=XM_023244511.1
E1 30950114 30950307
refGene_gn=CUEDC1;refGene_tx=XM_023244509.1;refGene_gn=CUEDC1;refGene_tx=XM_023244510.1;refGene_gn=CUEDC1;refGene_tx=XM_023244511.1
E1 30951406 30951508
refGene_gn=CUEDC1;refGene_tx=XM_023244509.1;refGene_gn=CUEDC1;refGene_tx=XM_023244510.1;refGene_gn=CUEDC1;refGene_tx=XM_023244511.1
E1 30954068 30954140
refGene_gn=CUEDC1;refGene_tx=XM_023244509.1;refGene_gn=CUEDC1;refGene_tx=XM_023244510.1;refGene_gn=CUEDC1;refGene_tx=XM_023244511.1
E1 30954966 30955060
refGene_gn=CUEDC1;refGene_tx=XM_023244509.1;refGene_gn=CUEDC1;refGene_tx=XM_023244510.1
E1 30955761 30955820
refGene_gn=CUEDC1;refGene_tx=XM_023244509.1;refGene_gn=CUEDC1;refGene_tx=XM_023244510.1
E1 30956654 30956722
refGene_gn=CUEDC1;refGene_tx=XM_023244509.1;refGene_gn=CUEDC1;refGene_tx=XM_023244510.1;refGene_gn=CUEDC1;refGene_tx=XM_023244511.1
E1 30970313 30970357 refGene_gn=MRPS23;refGene_tx=XM_003996649.5
E1 30970915 30971086 refGene_gn=MRPS23;refGene_tx=XM_003996649.5
E1 30973108 30973186 refGene_gn=MRPS23;refGene_tx=XM_003996649.5
E1 30973402 30973475 refGene_gn=MRPS23;refGene_tx=XM_003996649.5

E1 30973476 30973530 refGene_gn=MRPS23;refGene_tx=XM_003996649.5
E1 30973941 30974094 refGene_gn=MRPS23;refGene_tx=XM_003996649.5
E1 31056320 31056782 refGene_gn=CCDC182;refGene_tx=XM_003996666.4
E1 31116714 31116776
refGene_gn=MSI2;refGene_tx=XM_023244138.1;refGene_gn=MSI2;refGene_tx=XM_023244146.1;refGene_gn=MSI2;refGene_tx=XM_023244147.1
E1 31119228 31119270
refGene_gn=MSI2;refGene_tx=XM_023244137.1;refGene_gn=MSI2;refGene_tx=XM_023244140.1;refGene_gn=MSI2;refGene_tx=XM_023244141.1;refGene_gn=MSI2;refGene_tx=XM_023244142.1
E1 31121511 31121720
refGene_gn=MSI2;refGene_tx=XM_023244137.1;refGene_gn=MSI2;refGene_tx=XM_023244138.1;refGene_gn=MSI2;refGene_tx=XM_023244140.1;refGene_gn=MSI2;refGene_tx=XM_023244141.1;refGene_gn=MSI2;refGene_tx=XM_023244142.1;refGene_gn=MSI2;refGene_tx=XM_023244143.1;refGene_gn=MSI2;refGene_tx=XM_023244144.1;refGene_gn=MSI2;refGene_tx=XM_023244145.1
E1 31139511 31139561
refGene_gn=MSI2;refGene_tx=XM_023244143.1;refGene_gn=MSI2;refGene_tx=XM_023244144.1;refGene_gn=MSI2;refGene_tx=XM_023244145.1
E1 31143153 31143216
refGene_gn=MSI2;refGene_tx=XM_023244137.1;refGene_gn=MSI2;refGene_tx=XM_023244138.1;refGene_gn=MSI2;refGene_tx=XM_023244140.1;refGene_gn=MSI2;refGene_tx=XM_023244141.1;refGene_gn=MSI2;refGene_tx=XM_023244142.1;refGene_gn=MSI2;refGene_tx=XM_023244143.1;refGene_gn=MSI2;refGene_tx=XM_023244144.1;refGene_gn=MSI2;refGene_tx=XM_023244145.1;refGene_gn=MSI2;refGene_tx=XM_023244146.1;refGene_gn=MSI2;refGene_tx=XM_023244147.1
E1 31144562 31144586
refGene_gn=MSI2;refGene_tx=XM_023244137.1;refGene_gn=MSI2;refGene_tx=XM_023244138.1;refGene_gn=MSI2;refGene_tx=XM_023244141.1;refGene_gn=MSI2;refGene_tx=XM_023244143.1;refGene_gn=MSI2;refGene_tx=XM_023244144.1;refGene_gn=MSI2;refGene_tx=XM_023244146.1
E1 31160338 31160379 refGene_gn=MSI2;refGene_tx=XM_023244148.1
E1 31164293 31164368
refGene_gn=MSI2;refGene_tx=XM_023244137.1;refGene_gn=MSI2;refGene_tx=XM_023244138.1;refGene_gn=MSI2;refGene_tx=XM_023244140.1;refGene_gn=MSI2;refGene_tx=XM_023244141.1;refGene_gn=MSI2;refGene_tx=XM_023244142.1;refGene_gn=MSI2;refGene_tx=XM_023244143.1;refGene_gn=MSI2;refGene_tx=XM_023244144.1;refGene_gn=MSI2;refGene_tx=XM_023244145.1;refGene_gn=MSI2;refGene_tx=XM_023244146.1;refGene_gn=MSI2;refGene_tx=XM_023244147.1;refGene_gn=MSI2;refGene_tx=XM_023244148.1
E1 31174244 31174359
refGene_gn=MSI2;refGene_tx=XM_023244137.1;refGene_gn=MSI2;refGene_tx=XM_023244138.1;refGene_gn=MSI2;refGene_tx=XM_023244140.1;refGene_gn=MSI2;refGene_tx=XM_023244141.1;refGene_gn=MSI2;refGene_tx=XM_023244142.1;refGene_gn=MSI2;refGene_tx=XM_023244143.1;refGene_gn=MSI2;refGene_tx=XM_023244144.1;refGene_gn=MSI2;refGene_tx=XM_023244145.1;refGene_gn=MSI2;refGene_tx=XM_023244146.1;refGene_gn=MSI2;refGene_tx=XM_023244147.1;refGene_gn=MSI2;refGene_tx=XM_023244148.1
E1 31191492 31191575
refGene_gn=MSI2;refGene_tx=XM_023244137.1;refGene_gn=MSI2;refGene_tx=X

M_023244138.1;refGene_gn=MSI2;refGene_tx=XM_023244140.1;refGene_gn=MSI2;refGene_tx=XM_023244141.1;refGene_gn=MSI2;refGene_tx=XM_023244142.1;refGene_gn=MSI2;refGene_tx=XM_023244143.1;refGene_gn=MSI2;refGene_tx=XM_023244144.1;refGene_gn=MSI2;refGene_tx=XM_023244145.1;refGene_gn=MSI2;refGene_tx=XM_023244146.1;refGene_gn=MSI2;refGene_tx=XM_023244147.1;refGene_gn=MSI2;refGene_tx=XM_023244148.1
E1 31255161 31255210
refGene_gn=MSI2;refGene_tx=XM_023244137.1;refGene_gn=MSI2;refGene_tx=XM_023244138.1;refGene_gn=MSI2;refGene_tx=XM_023244140.1;refGene_gn=MSI2;refGene_tx=XM_023244141.1;refGene_gn=MSI2;refGene_tx=XM_023244142.1;refGene_gn=MSI2;refGene_tx=XM_023244143.1;refGene_gn=MSI2;refGene_tx=XM_023244144.1;refGene_gn=MSI2;refGene_tx=XM_023244145.1;refGene_gn=MSI2;refGene_tx=XM_023244146.1;refGene_gn=MSI2;refGene_tx=XM_023244147.1;refGene_gn=MSI2;refGene_tx=XM_023244148.1
E1 31372218 31372311
refGene_gn=MSI2;refGene_tx=XM_023244137.1;refGene_gn=MSI2;refGene_tx=XM_023244138.1;refGene_gn=MSI2;refGene_tx=XM_023244140.1;refGene_gn=MSI2;refGene_tx=XM_023244141.1;refGene_gn=MSI2;refGene_tx=XM_023244142.1;refGene_gn=MSI2;refGene_tx=XM_023244143.1;refGene_gn=MSI2;refGene_tx=XM_023244144.1;refGene_gn=MSI2;refGene_tx=XM_023244145.1;refGene_gn=MSI2;refGene_tx=XM_023244146.1;refGene_gn=MSI2;refGene_tx=XM_023244147.1;refGene_gn=MSI2;refGene_tx=XM_023244148.1
E1 31500872 31500914
refGene_gn=MSI2;refGene_tx=XM_023244137.1;refGene_gn=MSI2;refGene_tx=XM_023244138.1;refGene_gn=MSI2;refGene_tx=XM_023244140.1;refGene_gn=MSI2;refGene_tx=XM_023244141.1;refGene_gn=MSI2;refGene_tx=XM_023244142.1;refGene_gn=MSI2;refGene_tx=XM_023244143.1;refGene_gn=MSI2;refGene_tx=XM_023244144.1;refGene_gn=MSI2;refGene_tx=XM_023244145.1;refGene_gn=MSI2;refGene_tx=XM_023244146.1;refGene_gn=MSI2;refGene_tx=XM_023244147.1;refGene_gn=MSI2;refGene_tx=XM_023244148.1
E1 31504769 31504854
refGene_gn=MSI2;refGene_tx=XM_023244137.1;refGene_gn=MSI2;refGene_tx=XM_023244138.1;refGene_gn=MSI2;refGene_tx=XM_023244140.1;refGene_gn=MSI2;refGene_tx=XM_023244141.1;refGene_gn=MSI2;refGene_tx=XM_023244142.1;refGene_gn=MSI2;refGene_tx=XM_023244143.1;refGene_gn=MSI2;refGene_tx=XM_023244144.1;refGene_gn=MSI2;refGene_tx=XM_023244145.1;refGene_gn=MSI2;refGene_tx=XM_023244146.1;refGene_gn=MSI2;refGene_tx=XM_023244147.1;refGene_gn=MSI2;refGene_tx=XM_023244148.1
E1 31505562 31505644
refGene_gn=MSI2;refGene_tx=XM_023244137.1;refGene_gn=MSI2;refGene_tx=XM_023244138.1;refGene_gn=MSI2;refGene_tx=XM_023244140.1;refGene_gn=MSI2;refGene_tx=XM_023244141.1;refGene_gn=MSI2;refGene_tx=XM_023244142.1;refGene_gn=MSI2;refGene_tx=XM_023244143.1;refGene_gn=MSI2;refGene_tx=XM_023244144.1;refGene_gn=MSI2;refGene_tx=XM_023244145.1;refGene_gn=MSI2;refGene_tx=XM_023244146.1;refGene_gn=MSI2;refGene_tx=XM_023244147.1;refGene_gn=MSI2;refGene_tx=XM_023244148.1
E1 31505978 31506019
refGene_gn=MSI2;refGene_tx=XM_023244137.1;refGene_gn=MSI2;refGene_tx=XM_023244138.1;refGene_gn=MSI2;refGene_tx=XM_023244140.1;refGene_gn=MSI2;refGene_tx=XM_023244141.1;refGene_gn=MSI2;refGene_tx=XM_023244142.1;

refGene_gn=MSI2;refGene_tx=XM_023244143.1;refGene_gn=MSI2;refGene_tx=XM_023244144.1;refGene_gn=MSI2;refGene_tx=XM_023244145.1;refGene_gn=MSI2;refGene_tx=XM_023244146.1;refGene_gn=MSI2;refGene_tx=XM_023244147.1;refGene_gn=MSI2;refGene_tx=XM_023244148.1
E1 31506312 31506374
refGene_gn=MSI2;refGene_tx=XM_023244137.1;refGene_gn=MSI2;refGene_tx=XM_023244138.1;refGene_gn=MSI2;refGene_tx=XM_023244140.1;refGene_gn=MSI2;refGene_tx=XM_023244141.1;refGene_gn=MSI2;refGene_tx=XM_023244142.1;refGene_gn=MSI2;refGene_tx=XM_023244143.1;refGene_gn=MSI2;refGene_tx=XM_023244144.1;refGene_gn=MSI2;refGene_tx=XM_023244145.1;refGene_gn=MSI2;refGene_tx=XM_023244146.1;refGene_gn=MSI2;refGene_tx=XM_023244147.1;refGene_gn=MSI2;refGene_tx=XM_023244148.1
E1 31612911 31612986
refGene_gn=AKAP1;refGene_tx=XM_006940163.4;refGene_gn=AKAP1;refGene_tx=XM_023244708.1;refGene_gn=AKAP1;refGene_tx=XM_023244709.1;refGene_gn=AKAP1;refGene_tx=XM_023244710.1;refGene_gn=AKAP1;refGene_tx=XM_023244711.1;refGene_gn=AKAP1;refGene_tx=XM_023244713.1;refGene_gn=AKAP1;refGene_tx=XM_023244714.1;refGene_gn=AKAP1;refGene_tx=XM_023244715.1
E1 31613533 31613596
refGene_gn=AKAP1;refGene_tx=XM_006940163.4;refGene_gn=AKAP1;refGene_tx=XM_023244708.1;refGene_gn=AKAP1;refGene_tx=XM_023244709.1;refGene_gn=AKAP1;refGene_tx=XM_023244710.1;refGene_gn=AKAP1;refGene_tx=XM_023244711.1;refGene_gn=AKAP1;refGene_tx=XM_023244713.1;refGene_gn=AKAP1;refGene_tx=XM_023244714.1;refGene_gn=AKAP1;refGene_tx=XM_023244715.1
E1 31613800 31613874
refGene_gn=AKAP1;refGene_tx=XM_006940163.4;refGene_gn=AKAP1;refGene_tx=XM_023244708.1;refGene_gn=AKAP1;refGene_tx=XM_023244709.1;refGene_gn=AKAP1;refGene_tx=XM_023244710.1;refGene_gn=AKAP1;refGene_tx=XM_023244711.1;refGene_gn=AKAP1;refGene_tx=XM_023244713.1;refGene_gn=AKAP1;refGene_tx=XM_023244714.1;refGene_gn=AKAP1;refGene_tx=XM_023244715.1
E1 31615282 31615350
refGene_gn=AKAP1;refGene_tx=XM_006940163.4;refGene_gn=AKAP1;refGene_tx=XM_023244708.1;refGene_gn=AKAP1;refGene_tx=XM_023244709.1;refGene_gn=AKAP1;refGene_tx=XM_023244710.1;refGene_gn=AKAP1;refGene_tx=XM_023244711.1;refGene_gn=AKAP1;refGene_tx=XM_023244713.1;refGene_gn=AKAP1;refGene_tx=XM_023244714.1;refGene_gn=AKAP1;refGene_tx=XM_023244715.1
E1 31615804 31615955
refGene_gn=AKAP1;refGene_tx=XM_006940163.4;refGene_gn=AKAP1;refGene_tx=XM_023244708.1;refGene_gn=AKAP1;refGene_tx=XM_023244709.1;refGene_gn=AKAP1;refGene_tx=XM_023244710.1;refGene_gn=AKAP1;refGene_tx=XM_023244711.1;refGene_gn=AKAP1;refGene_tx=XM_023244713.1;refGene_gn=AKAP1;refGene_tx=XM_023244714.1;refGene_gn=AKAP1;refGene_tx=XM_023244715.1
E1 31617363 31617541
refGene_gn=AKAP1;refGene_tx=XM_006940163.4;refGene_gn=AKAP1;refGene_tx=XM_023244708.1;refGene_gn=AKAP1;refGene_tx=XM_023244709.1;refGene_gn=AKAP1;refGene_tx=XM_023244710.1;refGene_gn=AKAP1;refGene_tx=XM_023244711.1;refGene_gn=AKAP1;refGene_tx=XM_023244713.1;refGene_gn=AKAP1;refGene_tx=XM_023244714.1;refGene_gn=AKAP1;refGene_tx=XM_023244715.1
E1 31619880 31620008
refGene_gn=AKAP1;refGene_tx=XM_006940163.4;refGene_gn=AKAP1;refGene_tx

=XM_023244708.1;refGene_gn=AKAP1;refGene_tx=XM_023244709.1;refGene_gn=AKAP1;refGene_tx=XM_023244710.1;refGene_gn=AKAP1;refGene_tx=XM_023244711.1;refGene_gn=AKAP1;refGene_tx=XM_023244713.1;refGene_gn=AKAP1;refGene_tx=XM_023244714.1;refGene_gn=AKAP1;refGene_tx=XM_023244715.1
E1 31620554 31620681
refGene_gn=AKAP1;refGene_tx=XM_006940163.4;refGene_gn=AKAP1;refGene_tx=XM_023244708.1;refGene_gn=AKAP1;refGene_tx=XM_023244709.1;refGene_gn=AKAP1;refGene_tx=XM_023244710.1;refGene_gn=AKAP1;refGene_tx=XM_023244711.1;refGene_gn=AKAP1;refGene_tx=XM_023244713.1;refGene_gn=AKAP1;refGene_tx=XM_023244714.1;refGene_gn=AKAP1;refGene_tx=XM_023244715.1
E1 31622268 31622444
refGene_gn=AKAP1;refGene_tx=XM_006940163.4;refGene_gn=AKAP1;refGene_tx=XM_023244708.1;refGene_gn=AKAP1;refGene_tx=XM_023244709.1;refGene_gn=AKAP1;refGene_tx=XM_023244710.1;refGene_gn=AKAP1;refGene_tx=XM_023244711.1;refGene_gn=AKAP1;refGene_tx=XM_023244713.1;refGene_gn=AKAP1;refGene_tx=XM_023244714.1;refGene_gn=AKAP1;refGene_tx=XM_023244715.1
E1 31624960 31626575
refGene_gn=AKAP1;refGene_tx=XM_006940163.4;refGene_gn=AKAP1;refGene_tx=XM_023244708.1;refGene_gn=AKAP1;refGene_tx=XM_023244709.1;refGene_gn=AKAP1;refGene_tx=XM_023244710.1;refGene_gn=AKAP1;refGene_tx=XM_023244711.1;refGene_gn=AKAP1;refGene_tx=XM_023244713.1;refGene_gn=AKAP1;refGene_tx=XM_023244714.1;refGene_gn=AKAP1;refGene_tx=XM_023244715.1
E1 31718465 31718528 refGene_gn=SCPEP1;refGene_tx=XM_011289112.2
E1 31722163 31722327 refGene_gn=SCPEP1;refGene_tx=XM_011289112.2
E1 31723250 31723388 refGene_gn=SCPEP1;refGene_tx=XM_011289112.2
E1 31725640 31725754 refGene_gn=SCPEP1;refGene_tx=XM_011289112.2
E1 31726466 31726560 refGene_gn=SCPEP1;refGene_tx=XM_011289112.2
E1 31727150 31727279 refGene_gn=SCPEP1;refGene_tx=XM_011289112.2
E1 31728036 31728074 refGene_gn=SCPEP1;refGene_tx=XM_011289112.2
E1 31734077 31734150 refGene_gn=SCPEP1;refGene_tx=XM_011289112.2
E1 31738964 31739039 refGene_gn=SCPEP1;refGene_tx=XM_011289112.2
E1 31739419 31739575 refGene_gn=SCPEP1;refGene_tx=XM_011289112.2
E1 31743349 31743439 refGene_gn=SCPEP1;refGene_tx=XM_011289112.2
E1 31746367 31746516 refGene_gn=SCPEP1;refGene_tx=XM_011289112.2
E1 31749812 31749879 refGene_gn=SCPEP1;refGene_tx=XM_011289112.2
E1 31759569 31759814
refGene_gn=COIL;refGene_tx=XM_003996651.5;refGene_gn=COIL;refGene_tx=XM_006940170.3
E1 31768149 31769263
refGene_gn=COIL;refGene_tx=XM_003996651.5;refGene_gn=COIL;refGene_tx=XM_006940170.3
E1 31769385 31769469
refGene_gn=COIL;refGene_tx=XM_003996651.5;refGene_gn=COIL;refGene_tx=XM_006940170.3
E1 31769881 31769929
refGene_gn=COIL;refGene_tx=XM_003996651.5;refGene_gn=COIL;refGene_tx=XM_006940170.3
E1 31772479 31772549
refGene_gn=COIL;refGene_tx=XM_003996651.5;refGene_gn=COIL;refGene_tx=XM_006940170.3

E1 31775172 31775261 refGene_gn=COIL;refGene_tx=XM_003996651.5
E1 31776828 31776918
refGene_gn=COIL;refGene_tx=XM_003996651.5;refGene_gn=COIL;refGene_tx=XM_006940170.3
E1 31784791 31785394
refGene_gn=TRIM25;refGene_tx=NM_001290251.1;refGene_gn=TRIM25;refGene_tx=XM_003996669.4
E1 31789439 31789535
refGene_gn=TRIM25;refGene_tx=NM_001290251.1;refGene_gn=TRIM25;refGene_tx=XM_003996669.4
E1 31792307 31792541
refGene_gn=TRIM25;refGene_tx=NM_001290251.1;refGene_gn=TRIM25;refGene_tx=XM_003996669.4
E1 31795648 31795820
refGene_gn=TRIM25;refGene_tx=NM_001290251.1;refGene_gn=TRIM25;refGene_tx=XM_003996669.4
E1 31797600 31797666
refGene_gn=TRIM25;refGene_tx=NM_001290251.1;refGene_gn=TRIM25;refGene_tx=XM_003996669.4
E1 31801128 31801155
refGene_gn=TRIM25;refGene_tx=NM_001290251.1;refGene_gn=TRIM25;refGene_tx=XM_003996669.4
E1 31801498 31801580 refGene_gn=TRIM25;refGene_tx=XM_003996669.4
E1 31801659 31801760
refGene_gn=TRIM25;refGene_tx=NM_001290251.1;refGene_gn=TRIM25;refGene_tx=XM_003996669.4
E1 31804499 31805029
refGene_gn=TRIM25;refGene_tx=NM_001290251.1;refGene_gn=TRIM25;refGene_tx=XM_003996669.4
E1 31825300 31825480 refGene_gn=DGKE;refGene_tx=XM_023244705.1
E1 31826088 31826200 refGene_gn=DGKE;refGene_tx=XM_023244705.1
E1 31826867 31826995 refGene_gn=DGKE;refGene_tx=XM_023244705.1
E1 31829254 31829326 refGene_gn=DGKE;refGene_tx=XM_023244705.1
E1 31830332 31830446 refGene_gn=DGKE;refGene_tx=XM_023244705.1
E1 31831677 31831729 refGene_gn=DGKE;refGene_tx=XM_023244705.1
E1 31832060 31832218 refGene_gn=DGKE;refGene_tx=XM_023244705.1
E1 31832492 31832636 refGene_gn=DGKE;refGene_tx=XM_023244705.1
E1 31834418 31834538 refGene_gn=DGKE;refGene_tx=XM_023244705.1
E1 31836337 31836497 refGene_gn=DGKE;refGene_tx=XM_023244705.1
E1 31846044 31846499 refGene_gn=DGKE;refGene_tx=XM_023244705.1
E1 31848794 31848832
refGene_gn=CE1H17orf67;refGene_tx=XM_006940172.4;refGene_gn=CE1H17orf67;refGene_tx=XM_006940173.4;refGene_gn=CE1H17orf67;refGene_tx=XM_023244706.1
E1 31851904 31852107
refGene_gn=CE1H17orf67;refGene_tx=XM_006940172.4;refGene_gn=CE1H17orf67;refGene_tx=XM_006940173.4;refGene_gn=CE1H17orf67;refGene_tx=XM_023244706.1
E1 31857972 31858027
refGene_gn=CE1H17orf67;refGene_tx=XM_003996653.5

E1 31859353 31859454
refGene_gn=CE1H17orf67;refGene_tx=XM_003996653.5;refGene_gn=CE1H17orf67;refGene_tx=XM_006940172.4;refGene_gn=CE1H17orf67;refGene_tx=XM_006940173.4;refGene_gn=CE1H17orf67;refGene_tx=XM_023244706.1
E1 31861191 31861350
refGene_gn=CE1H17orf67;refGene_tx=XM_023244706.1
E1 31880323 31880440
refGene_gn=CE1H17orf67;refGene_tx=XM_003996653.5;refGene_gn=CE1H17orf67;refGene_tx=XM_006940172.4;refGene_gn=CE1H17orf67;refGene_tx=XM_006940173.4
E1 32063589 32064288 refGene_gn=NOG;refGene_tx=XM_011279931.3
E1 32139724 32140509
refGene_gn=ANKFN1;refGene_tx=XM_023243438.1;refGene_gn=ANKFN1;refGene_tx=XM_023243439.1
E1 32154677 32154891
refGene_gn=ANKFN1;refGene_tx=XM_023243438.1;refGene_gn=ANKFN1;refGene_tx=XM_023243439.1
E1 32161028 32161195
refGene_gn=ANKFN1;refGene_tx=XM_023243438.1;refGene_gn=ANKFN1;refGene_tx=XM_023243439.1
E1 32172396 32172588
refGene_gn=ANKFN1;refGene_tx=XM_023243438.1;refGene_gn=ANKFN1;refGene_tx=XM_023243439.1;refGene_gn=ANKFN1;refGene_tx=XM_023243440.1
E1 32174119 32174270
refGene_gn=ANKFN1;refGene_tx=XM_023243438.1;refGene_gn=ANKFN1;refGene_tx=XM_023243439.1;refGene_gn=ANKFN1;refGene_tx=XM_023243440.1
E1 32176989 32177156
refGene_gn=ANKFN1;refGene_tx=XM_023243438.1;refGene_gn=ANKFN1;refGene_tx=XM_023243439.1;refGene_gn=ANKFN1;refGene_tx=XM_023243440.1
E1 32186621 32186837
refGene_gn=ANKFN1;refGene_tx=XM_023243438.1;refGene_gn=ANKFN1;refGene_tx=XM_023243439.1;refGene_gn=ANKFN1;refGene_tx=XM_023243440.1
E1 32195382 32195499
refGene_gn=ANKFN1;refGene_tx=XM_023243438.1;refGene_gn=ANKFN1;refGene_tx=XM_023243439.1;refGene_gn=ANKFN1;refGene_tx=XM_023243440.1
E1 32195962 32196095
refGene_gn=ANKFN1;refGene_tx=XM_023243438.1;refGene_gn=ANKFN1;refGene_tx=XM_023243439.1;refGene_gn=ANKFN1;refGene_tx=XM_023243440.1
E1 32196403 32196503
refGene_gn=ANKFN1;refGene_tx=XM_023243438.1;refGene_gn=ANKFN1;refGene_tx=XM_023243439.1;refGene_gn=ANKFN1;refGene_tx=XM_023243440.1
E1 32204573 32204681
refGene_gn=ANKFN1;refGene_tx=XM_023243438.1;refGene_gn=ANKFN1;refGene_tx=XM_023243439.1;refGene_gn=ANKFN1;refGene_tx=XM_023243440.1
E1 32210040 32210131
refGene_gn=ANKFN1;refGene_tx=XM_023243438.1;refGene_gn=ANKFN1;refGene_tx=XM_023243439.1;refGene_gn=ANKFN1;refGene_tx=XM_023243440.1
E1 32212454 32212552
refGene_gn=ANKFN1;refGene_tx=XM_023243438.1;refGene_gn=ANKFN1;refGene_tx=XM_023243439.1;refGene_gn=ANKFN1;refGene_tx=XM_023243440.1

E1 32258522 32258636
refGene_gn=ANKFN1;refGene_tx=XM_023243438.1;refGene_gn=ANKFN1;refGene_
tx=XM_023243440.1
E1 32258911 32258978 refGene_gn=ANKFN1;refGene_tx=XM_023243438.1
E1 32260587 32260782 refGene_gn=ANKFN1;refGene_tx=XM_023243440.1
E1 32274645 32274856 refGene_gn=ANKFN1;refGene_tx=XM_023243440.1
E1 32278086 32278288 refGene_gn=ANKFN1;refGene_tx=XM_023243440.1
E1 32292739 32292874 refGene_gn=ANKFN1;refGene_tx=XM_023243440.1
E1 32383503 32383544 refGene_gn=ANKFN1;refGene_tx=XM_023243440.1
E1 32440682 32440703 refGene_gn=ANKFN1;refGene_tx=XM_023243440.1
E1 32539358 32539687
refGene_gn=LOC111557720;refGene_tx=XM_023243459.1
E1 32601363 32601461
refGene_gn=LOC111557720;refGene_tx=XM_023243459.1
E1 32604796 32604916
refGene_gn=LOC111557720;refGene_tx=XM_023243459.1
E1 32606901 32606957
refGene_gn=LOC111557720;refGene_tx=XM_023243459.1
E1 32744992 32745058
refGene_gn=PCTP;refGene_tx=XM_003996654.4;refGene_gn=PCTP;refGene_tx=X
M_006940174.3;refGene_gn=PCTP;refGene_tx=XM_019817647.2
E1 32746190 32746258
refGene_gn=PCTP;refGene_tx=XM_003996654.4;refGene_gn=PCTP;refGene_tx=X
M_006940174.3;refGene_gn=PCTP;refGene_tx=XM_019817647.2
E1 32747122 32747294
refGene_gn=PCTP;refGene_tx=XM_003996654.4;refGene_gn=PCTP;refGene_tx=X
M_006940174.3;refGene_gn=PCTP;refGene_tx=XM_019817647.2
E1 32750450 32750530
refGene_gn=PCTP;refGene_tx=XM_003996654.4;refGene_gn=PCTP;refGene_tx=X
M_006940174.3;refGene_gn=PCTP;refGene_tx=XM_019817647.2
E1 32753074 32753192
refGene_gn=PCTP;refGene_tx=XM_003996654.4;refGene_gn=PCTP;refGene_tx=X
M_006940174.3;refGene_gn=PCTP;refGene_tx=XM_019817647.2
E1 32771262 32771406
refGene_gn=PCTP;refGene_tx=XM_003996654.4;refGene_gn=PCTP;refGene_tx=X
M_006940174.3
E1 32801422 32801827
refGene_gn=TMEM100;refGene_tx=XM_003996655.5;refGene_gn=TMEM100;refGen
e_tx=XM_019817648.2;refGene_gn=TMEM100;refGene_tx=XM_019817649.2
E1 33059663 33059689 refGene_gn=MMD;refGene_tx=XM_023243505.1
E1 33064415 33064570
refGene_gn=MMD;refGene_tx=XM_023243502.1;refGene_gn=MMD;refGene_tx=XM_
023243503.1;refGene_gn=MMD;refGene_tx=XM_023243504.1
E1 33066118 33066148
refGene_gn=MMD;refGene_tx=XM_023243502.1;refGene_gn=MMD;refGene_tx=XM_
023243504.1
E1 33067100 33067182
refGene_gn=MMD;refGene_tx=XM_023243502.1;refGene_gn=MMD;refGene_tx=XM_
023243503.1;refGene_gn=MMD;refGene_tx=XM_023243504.1;refGene_gn=MMD;re
fGene_tx=XM_023243505.1

E1 33071015 33071176
refGene_gn=MMD;refGene_tx=XM_023243502.1;refGene_gn=MMD;refGene_tx=XM_
023243503.1;refGene_gn=MMD;refGene_tx=XM_023243504.1;refGene_gn=MMD;re
fGene_tx=XM_023243505.1
E1 33074082 33074157
refGene_gn=MMD;refGene_tx=XM_023243502.1;refGene_gn=MMD;refGene_tx=XM_
023243503.1;refGene_gn=MMD;refGene_tx=XM_023243504.1;refGene_gn=MMD;re
fGene_tx=XM_023243505.1
E1 33075919 33076021
refGene_gn=MMD;refGene_tx=XM_023243502.1;refGene_gn=MMD;refGene_tx=XM_
023243503.1;refGene_gn=MMD;refGene_tx=XM_023243504.1;refGene_gn=MMD;re
fGene_tx=XM_023243505.1
E1 33077050 33077151 refGene_gn=MMD;refGene_tx=XM_023243504.1
E1 33077855 33077925
refGene_gn=MMD;refGene_tx=XM_023243502.1;refGene_gn=MMD;refGene_tx=XM_
023243503.1;refGene_gn=MMD;refGene_tx=XM_023243504.1;refGene_gn=MMD;re
fGene_tx=XM_023243505.1
E1 33083884 33084085
refGene_gn=MMD;refGene_tx=XM_023243502.1;refGene_gn=MMD;refGene_tx=XM_
023243503.1;refGene_gn=MMD;refGene_tx=XM_023243505.1
E1 33139345 33139564
refGene_gn=HLF;refGene_tx=XM_003996672.4;refGene_gn=HLF;refGene_tx=XM_
006940189.3
E1 33144532 33144753
refGene_gn=HLF;refGene_tx=XM_003996672.4;refGene_gn=HLF;refGene_tx=XM_
006940189.3
E1 33185931 33186267
refGene_gn=HLF;refGene_tx=XM_003996672.4;refGene_gn=HLF;refGene_tx=XM_
006940189.3
E1 33188428 33188543
refGene_gn=HLF;refGene_tx=XM_003996672.4;refGene_gn=HLF;refGene_tx=XM_
006940189.3
E1 33296725 33296852
refGene_gn=STXBP4;refGene_tx=XM_006940190.4;refGene_gn=STXBP4;refGene_
tx=XM_019817652.2;refGene_gn=STXBP4;refGene_tx=XM_019817653.2
E1 33316918 33316976
refGene_gn=STXBP4;refGene_tx=XM_006940190.4;refGene_gn=STXBP4;refGene_
tx=XM_019817652.2;refGene_gn=STXBP4;refGene_tx=XM_019817653.2
E1 33363293 33363427
refGene_gn=STXBP4;refGene_tx=XM_006940190.4;refGene_gn=STXBP4;refGene_
tx=XM_019817652.2;refGene_gn=STXBP4;refGene_tx=XM_019817653.2
E1 33365473 33365523
refGene_gn=STXBP4;refGene_tx=XM_006940190.4;refGene_gn=STXBP4;refGene_
tx=XM_019817652.2;refGene_gn=STXBP4;refGene_tx=XM_019817653.2
E1 33366044 33366161
refGene_gn=STXBP4;refGene_tx=XM_006940190.4;refGene_gn=STXBP4;refGene_
tx=XM_019817652.2;refGene_gn=STXBP4;refGene_tx=XM_019817653.2
E1 33373300 33373477
refGene_gn=STXBP4;refGene_tx=XM_006940190.4;refGene_gn=STXBP4;refGene_
tx=XM_019817652.2;refGene_gn=STXBP4;refGene_tx=XM_019817653.2

E1 33400550 33400616
refGene_gn=STXBP4;refGene_tx=XM_006940190.4;refGene_gn=STXBP4;refGene_tx=XM_019817652.2;refGene_gn=STXBP4;refGene_tx=XM_019817653.2
E1 33404305 33404395
refGene_gn=STXBP4;refGene_tx=XM_006940190.4;refGene_gn=STXBP4;refGene_tx=XM_019817652.2;refGene_gn=STXBP4;refGene_tx=XM_019817653.2
E1 33415761 33415853
refGene_gn=STXBP4;refGene_tx=XM_006940190.4;refGene_gn=STXBP4;refGene_tx=XM_019817652.2;refGene_gn=STXBP4;refGene_tx=XM_019817653.2
E1 33418180 33418277
refGene_gn=STXBP4;refGene_tx=XM_006940190.4;refGene_gn=STXBP4;refGene_tx=XM_019817652.2;refGene_gn=STXBP4;refGene_tx=XM_019817653.2
E1 33428730 33428825
refGene_gn=STXBP4;refGene_tx=XM_006940190.4;refGene_gn=STXBP4;refGene_tx=XM_019817652.2;refGene_gn=STXBP4;refGene_tx=XM_019817653.2
E1 33435261 33435340
refGene_gn=STXBP4;refGene_tx=XM_006940190.4;refGene_gn=STXBP4;refGene_tx=XM_019817652.2;refGene_gn=STXBP4;refGene_tx=XM_019817653.2
E1 33436328 33436548
refGene_gn=STXBP4;refGene_tx=XM_006940190.4;refGene_gn=STXBP4;refGene_tx=XM_019817652.2;refGene_gn=STXBP4;refGene_tx=XM_019817653.2
E1 33436734 33436841
refGene_gn=STXBP4;refGene_tx=XM_006940190.4;refGene_gn=STXBP4;refGene_tx=XM_019817652.2;refGene_gn=STXBP4;refGene_tx=XM_019817653.2
E1 33446305 33446438
refGene_gn=STXBP4;refGene_tx=XM_006940190.4;refGene_gn=STXBP4;refGene_tx=XM_019817652.2;refGene_gn=STXBP4;refGene_tx=XM_019817653.2
E1 33450401 33450448
refGene_gn=STXBP4;refGene_tx=XM_006940190.4;refGene_gn=STXBP4;refGene_tx=XM_019817652.2
E1 33465729 33466092
refGene_gn=LOC101085843;refGene_tx=XM_003996673.5
E1 33469251 33469407
refGene_gn=LOC101085843;refGene_tx=XM_003996673.5
E1 33472135 33472261
refGene_gn=LOC101085843;refGene_tx=XM_003996673.5
E1 33472646 33472829
refGene_gn=LOC101085843;refGene_tx=XM_003996673.5
E1 33474838 33474902
refGene_gn=TOM1L1;refGene_tx=XM_019817657.2;refGene_gn=TOM1L1;refGene_tx=XM_019817658.2;refGene_gn=TOM1L1;refGene_tx=XM_019817659.2;refGene_gn=TOM1L1;refGene_tx=XM_019817661.2;refGene_gn=TOM1L1;refGene_tx=XM_023244403.1
E1 33477314 33477396
refGene_gn=TOM1L1;refGene_tx=XM_019817657.2;refGene_gn=TOM1L1;refGene_tx=XM_019817658.2;refGene_gn=TOM1L1;refGene_tx=XM_019817659.2;refGene_gn=TOM1L1;refGene_tx=XM_019817661.2;refGene_gn=TOM1L1;refGene_tx=XM_023244403.1
E1 33477843 33477949
refGene_gn=TOM1L1;refGene_tx=XM_019817657.2;refGene_gn=TOM1L1;refGene_

tx=XM_019817658.2;refGene_gn=TOM1L1;refGene_tx=XM_019817659.2;refGene_gn=TOM1L1;refGene_tx=XM_019817661.2;refGene_gn=TOM1L1;refGene_tx=XM_023244403.1
E1 33479816 33479868
refGene_gn=TOM1L1;refGene_tx=XM_019817657.2;refGene_gn=TOM1L1;refGene_tx=XM_019817658.2;refGene_gn=TOM1L1;refGene_tx=XM_019817659.2;refGene_gn=TOM1L1;refGene_tx=XM_019817661.2;refGene_gn=TOM1L1;refGene_tx=XM_023244403.1
E1 33489476 33489573
refGene_gn=TOM1L1;refGene_tx=XM_019817657.2;refGene_gn=TOM1L1;refGene_tx=XM_019817658.2;refGene_gn=TOM1L1;refGene_tx=XM_019817659.2;refGene_gn=TOM1L1;refGene_tx=XM_019817661.2;refGene_gn=TOM1L1;refGene_tx=XM_023244403.1
E1 33490468 33490586
refGene_gn=TOM1L1;refGene_tx=XM_019817657.2;refGene_gn=TOM1L1;refGene_tx=XM_019817658.2;refGene_gn=TOM1L1;refGene_tx=XM_019817659.2;refGene_gn=TOM1L1;refGene_tx=XM_019817661.2;refGene_gn=TOM1L1;refGene_tx=XM_023244403.1
E1 33490971 33491032
refGene_gn=TOM1L1;refGene_tx=XM_019817657.2;refGene_gn=TOM1L1;refGene_tx=XM_019817658.2;refGene_gn=TOM1L1;refGene_tx=XM_019817659.2;refGene_gn=TOM1L1;refGene_tx=XM_019817661.2;refGene_gn=TOM1L1;refGene_tx=XM_023244403.1
E1 33496485 33496619
refGene_gn=TOM1L1;refGene_tx=XM_019817657.2;refGene_gn=TOM1L1;refGene_tx=XM_019817658.2;refGene_gn=TOM1L1;refGene_tx=XM_019817659.2;refGene_gn=TOM1L1;refGene_tx=XM_019817661.2;refGene_gn=TOM1L1;refGene_tx=XM_023244403.1
E1 33507653 33507770
refGene_gn=TOM1L1;refGene_tx=XM_019817657.2;refGene_gn=TOM1L1;refGene_tx=XM_019817658.2;refGene_gn=TOM1L1;refGene_tx=XM_019817659.2;refGene_gn=TOM1L1;refGene_tx=XM_019817661.2;refGene_gn=TOM1L1;refGene_tx=XM_023244403.1
E1 33509208 33509310
refGene_gn=TOM1L1;refGene_tx=XM_019817657.2;refGene_gn=TOM1L1;refGene_tx=XM_019817658.2;refGene_gn=TOM1L1;refGene_tx=XM_019817659.2;refGene_gn=TOM1L1;refGene_tx=XM_019817661.2;refGene_gn=TOM1L1;refGene_tx=XM_023244403.1
E1 33510788 33510914
refGene_gn=TOM1L1;refGene_tx=XM_019817657.2;refGene_gn=TOM1L1;refGene_tx=XM_019817658.2;refGene_gn=TOM1L1;refGene_tx=XM_019817659.2;refGene_gn=TOM1L1;refGene_tx=XM_023244403.1
E1 33511521 33511671
refGene_gn=TOM1L1;refGene_tx=XM_019817657.2;refGene_gn=TOM1L1;refGene_tx=XM_019817658.2;refGene_gn=TOM1L1;refGene_tx=XM_019817659.2;refGene_gn=TOM1L1;refGene_tx=XM_023244403.1
E1 33517583 33517662
refGene_gn=TOM1L1;refGene_tx=XM_019817657.2;refGene_gn=TOM1L1;refGene_tx=XM_019817658.2;refGene_gn=TOM1L1;refGene_tx=XM_019817659.2;refGene_gn=TOM1L1;refGene_tx=XM_019817661.2;refGene_gn=TOM1L1;refGene_tx=XM_02

3244403.1
E1 33519738 33519823
refGene_gn=TOM1L1;refGene_tx=XM_019817657.2;refGene_gn=TOM1L1;refGene_tx=XM_019817658.2;refGene_gn=TOM1L1;refGene_tx=XM_019817659.2;refGene_gn=TOM1L1;refGene_tx=XM_019817661.2;refGene_gn=TOM1L1;refGene_tx=XM_023244403.1
E1 33522187 33522245
refGene_gn=TOM1L1;refGene_tx=XM_019817657.2;refGene_gn=TOM1L1;refGene_tx=XM_019817661.2;refGene_gn=TOM1L1;refGene_tx=XM_023244403.1
E1 34333033 34335073 refGene_gn=KIF2B;refGene_tx=XM_003996679.5
E1 35735832 35736117
refGene_gn=LOC109494088;refGene_tx=XM_023243460.1
E1 35736236 35736923
refGene_gn=LOC109494088;refGene_tx=XM_023243460.1
E1 35737158 35737219 refGene_gn=CA10;refGene_tx=XM_003996677.5
E1 35817886 35817961 refGene_gn=CA10;refGene_tx=XM_003996677.5
E1 35943523 35943666 refGene_gn=CA10;refGene_tx=XM_003996677.5
E1 36125114 36125300 refGene_gn=CA10;refGene_tx=XM_003996677.5
E1 36203848 36203944 refGene_gn=CA10;refGene_tx=XM_003996677.5
E1 36208466 36208539 refGene_gn=CA10;refGene_tx=XM_003996677.5
E1 36222774 36222929 refGene_gn=CA10;refGene_tx=XM_003996677.5
E1 36224833 36224957 refGene_gn=CA10;refGene_tx=XM_003996677.5
E1 36224959 36225007 refGene_gn=CA10;refGene_tx=XM_003996677.5
E1 36227598 36227621 refGene_gn=CA10;refGene_tx=XM_003996677.5
E1 36499180 36499190 refGene_gn=UTP18;refGene_tx=XM_023244494.1
E1 36499975 36500000 refGene_gn=UTP18;refGene_tx=XM_023244493.1
E1 36501582 36501725
refGene_gn=UTP18;refGene_tx=XM_023244493.1;refGene_gn=UTP18;refGene_tx=XM_023244494.1
E1 36506633 36506808
refGene_gn=UTP18;refGene_tx=XM_023244493.1;refGene_gn=UTP18;refGene_tx=XM_023244494.1
E1 36516650 36516774
refGene_gn=UTP18;refGene_tx=XM_023244493.1;refGene_gn=UTP18;refGene_tx=XM_023244494.1
E1 36521735 36521826
refGene_gn=UTP18;refGene_tx=XM_023244493.1;refGene_gn=UTP18;refGene_tx=XM_023244494.1
E1 36522058 36522159
refGene_gn=UTP18;refGene_tx=XM_023244493.1;refGene_gn=UTP18;refGene_tx=XM_023244494.1
E1 36527371 36527546
refGene_gn=UTP18;refGene_tx=XM_023244493.1;refGene_gn=UTP18;refGene_tx=XM_023244494.1
E1 36527645 36527771
refGene_gn=UTP18;refGene_tx=XM_023244493.1;refGene_gn=UTP18;refGene_tx=XM_023244494.1
E1 36529628 36529717
refGene_gn=UTP18;refGene_tx=XM_023244493.1;refGene_gn=UTP18;refGene_tx=XM_023244494.1

E1 36533609 36533677
refGene_gn=UTP18;refGene_tx=XM_023244493.1;refGene_gn=UTP18;refGene_tx
=XM_023244494.1
E1 36537541 36537640
refGene_gn=UTP18;refGene_tx=XM_023244493.1;refGene_gn=UTP18;refGene_tx
=XM_023244494.1
E1 36538983 36539096
refGene_gn=UTP18;refGene_tx=XM_023244493.1;refGene_gn=UTP18;refGene_tx
=XM_023244494.1
E1 36541057 36541396
refGene_gn=UTP18;refGene_tx=XM_023244493.1;refGene_gn=UTP18;refGene_tx
=XM_023244494.1
E1 36542131 36542152
refGene_gn=MBTD1;refGene_tx=XM_019817663.2;refGene_gn=MBTD1;refGene_tx
=XM_019817664.2;refGene_gn=MBTD1;refGene_tx=XM_019817665.2;refGene_gn=
MBTD1;refGene_tx=XM_019817666.2;refGene_gn=MBTD1;refGene_tx=XM_0198176
74.2;refGene_gn=MBTD1;refGene_tx=XM_019817675.2;refGene_gn=MBTD1;refGe
ne_tx=XM_019817678.2
E1 36569288 36569490
refGene_gn=MBTD1;refGene_tx=XM_006940197.3;refGene_gn=MBTD1;refGene_tx
=XM_006940199.4;refGene_gn=MBTD1;refGene_tx=XM_006940201.3;refGene_gn=
MBTD1;refGene_tx=XM_019817663.2;refGene_gn=MBTD1;refGene_tx=XM_0198176
64.2;refGene_gn=MBTD1;refGene_tx=XM_019817665.2;refGene_gn=MBTD1;refGe
ne_tx=XM_019817666.2;refGene_gn=MBTD1;refGene_tx=XM_019817667.2;refGen
e_gn=MBTD1;refGene_tx=XM_019817668.2;refGene_gn=MBTD1;refGene_tx=XM_01
9817669.2;refGene_gn=MBTD1;refGene_tx=XM_019817670.2;refGene_gn=MBTD1;
refGene_tx=XM_019817671.2;refGene_gn=MBTD1;refGene_tx=XM_019817672.2;r
efGene_gn=MBTD1;refGene_tx=XM_019817673.2;refGene_gn=MBTD1;refGene_tx=
XM_019817674.2;refGene_gn=MBTD1;refGene_tx=XM_019817675.2;refGene_gn=M
BTD1;refGene_tx=XM_019817678.2
E1 36576624 36576764
refGene_gn=MBTD1;refGene_tx=XM_006940197.3;refGene_gn=MBTD1;refGene_tx
=XM_006940199.4;refGene_gn=MBTD1;refGene_tx=XM_006940201.3;refGene_gn=
MBTD1;refGene_tx=XM_019817663.2;refGene_gn=MBTD1;refGene_tx=XM_0198176
64.2;refGene_gn=MBTD1;refGene_tx=XM_019817665.2;refGene_gn=MBTD1;refGe
ne_tx=XM_019817666.2;refGene_gn=MBTD1;refGene_tx=XM_019817667.2;refGen
e_gn=MBTD1;refGene_tx=XM_019817668.2;refGene_gn=MBTD1;refGene_tx=XM_01
9817669.2;refGene_gn=MBTD1;refGene_tx=XM_019817670.2;refGene_gn=MBTD1;
refGene_tx=XM_019817671.2;refGene_gn=MBTD1;refGene_tx=XM_019817672.2;r
efGene_gn=MBTD1;refGene_tx=XM_019817673.2;refGene_gn=MBTD1;refGene_tx=
XM_019817674.2;refGene_gn=MBTD1;refGene_tx=XM_019817675.2;refGene_gn=M
BTD1;refGene_tx=XM_019817678.2
E1 36578072 36578187
refGene_gn=MBTD1;refGene_tx=XM_006940197.3;refGene_gn=MBTD1;refGene_tx
=XM_006940199.4;refGene_gn=MBTD1;refGene_tx=XM_006940201.3;refGene_gn=
MBTD1;refGene_tx=XM_019817663.2;refGene_gn=MBTD1;refGene_tx=XM_0198176
64.2;refGene_gn=MBTD1;refGene_tx=XM_019817665.2;refGene_gn=MBTD1;refGe
ne_tx=XM_019817666.2;refGene_gn=MBTD1;refGene_tx=XM_019817667.2;refGen
e_gn=MBTD1;refGene_tx=XM_019817668.2;refGene_gn=MBTD1;refGene_tx=XM_01
9817669.2;refGene_gn=MBTD1;refGene_tx=XM_019817670.2;refGene_gn=MBTD1;

refGene_tx=XM_019817671.2;refGene_gn=MBTD1;refGene_tx=XM_019817672.2;refGene_gn=MBTD1;refGene_tx=XM_019817673.2;refGene_gn=MBTD1;refGene_tx=XM_019817674.2;refGene_gn=MBTD1;refGene_tx=XM_019817675.2;refGene_gn=MBTD1;refGene_tx=XM_019817678.2
E1 36580595 36580681
refGene_gn=MBTD1;refGene_tx=XM_006940197.3;refGene_gn=MBTD1;refGene_tx=XM_006940199.4;refGene_gn=MBTD1;refGene_tx=XM_006940201.3;refGene_gn=MBTD1;refGene_tx=XM_019817663.2;refGene_gn=MBTD1;refGene_tx=XM_019817664.2;refGene_gn=MBTD1;refGene_tx=XM_019817665.2;refGene_gn=MBTD1;refGene_tx=XM_019817666.2;refGene_gn=MBTD1;refGene_tx=XM_019817667.2;refGene_gn=MBTD1;refGene_tx=XM_019817668.2;refGene_gn=MBTD1;refGene_tx=XM_019817669.2;refGene_gn=MBTD1;refGene_tx=XM_019817670.2;refGene_gn=MBTD1;refGene_tx=XM_019817671.2;refGene_gn=MBTD1;refGene_tx=XM_019817672.2;refGene_gn=MBTD1;refGene_tx=XM_019817673.2;refGene_gn=MBTD1;refGene_tx=XM_019817674.2;refGene_gn=MBTD1;refGene_tx=XM_019817675.2;refGene_gn=MBTD1;refGene_tx=XM_019817676.2;refGene_gn=MBTD1;refGene_tx=XM_019817678.2
E1 36589760 36589878
refGene_gn=MBTD1;refGene_tx=XM_006940197.3;refGene_gn=MBTD1;refGene_tx=XM_006940199.4;refGene_gn=MBTD1;refGene_tx=XM_006940201.3;refGene_gn=MBTD1;refGene_tx=XM_019817663.2;refGene_gn=MBTD1;refGene_tx=XM_019817664.2;refGene_gn=MBTD1;refGene_tx=XM_019817665.2;refGene_gn=MBTD1;refGene_tx=XM_019817666.2;refGene_gn=MBTD1;refGene_tx=XM_019817667.2;refGene_gn=MBTD1;refGene_tx=XM_019817668.2;refGene_gn=MBTD1;refGene_tx=XM_019817669.2;refGene_gn=MBTD1;refGene_tx=XM_019817670.2;refGene_gn=MBTD1;refGene_tx=XM_019817671.2;refGene_gn=MBTD1;refGene_tx=XM_019817672.2;refGene_gn=MBTD1;refGene_tx=XM_019817673.2;refGene_gn=MBTD1;refGene_tx=XM_019817674.2;refGene_gn=MBTD1;refGene_tx=XM_019817675.2;refGene_gn=MBTD1;refGene_tx=XM_019817676.2;refGene_gn=MBTD1;refGene_tx=XM_019817678.2;refGene_gn=MBTD1;refGene_tx=XM_019817679.2
E1 36592215 36592350
refGene_gn=MBTD1;refGene_tx=XM_006940197.3;refGene_gn=MBTD1;refGene_tx=XM_006940199.4;refGene_gn=MBTD1;refGene_tx=XM_006940201.3;refGene_gn=MBTD1;refGene_tx=XM_019817663.2;refGene_gn=MBTD1;refGene_tx=XM_019817664.2;refGene_gn=MBTD1;refGene_tx=XM_019817665.2;refGene_gn=MBTD1;refGene_tx=XM_019817666.2;refGene_gn=MBTD1;refGene_tx=XM_019817667.2;refGene_gn=MBTD1;refGene_tx=XM_019817668.2;refGene_gn=MBTD1;refGene_tx=XM_019817669.2;refGene_gn=MBTD1;refGene_tx=XM_019817670.2;refGene_gn=MBTD1;refGene_tx=XM_019817671.2;refGene_gn=MBTD1;refGene_tx=XM_019817672.2;refGene_gn=MBTD1;refGene_tx=XM_019817673.2;refGene_gn=MBTD1;refGene_tx=XM_019817674.2;refGene_gn=MBTD1;refGene_tx=XM_019817675.2;refGene_gn=MBTD1;refGene_tx=XM_019817676.2;refGene_gn=MBTD1;refGene_tx=XM_019817678.2;refGene_gn=MBTD1;refGene_tx=XM_019817679.2
E1 36592628 36592717
refGene_gn=MBTD1;refGene_tx=XM_006940197.3;refGene_gn=MBTD1;refGene_tx=XM_006940199.4;refGene_gn=MBTD1;refGene_tx=XM_006940201.3;refGene_gn=MBTD1;refGene_tx=XM_019817663.2;refGene_gn=MBTD1;refGene_tx=XM_019817664.2;refGene_gn=MBTD1;refGene_tx=XM_019817665.2;refGene_gn=MBTD1;refGene_tx=XM_019817666.2;refGene_gn=MBTD1;refGene_tx=XM_019817667.2;refGene_gn=MBTD1;refGene_tx=XM_019817668.2;refGene_gn=MBTD1;refGene_tx=XM_01

9817669.2;refGene_gn=MBTD1;refGene_tx=XM_019817670.2;refGene_gn=MBTD1;refGene_tx=XM_019817671.2;refGene_gn=MBTD1;refGene_tx=XM_019817672.2;refGene_gn=MBTD1;refGene_tx=XM_019817673.2;refGene_gn=MBTD1;refGene_tx=XM_019817674.2;refGene_gn=MBTD1;refGene_tx=XM_019817675.2;refGene_gn=MBTD1;refGene_tx=XM_019817676.2;refGene_gn=MBTD1;refGene_tx=XM_019817678.2;refGene_gn=MBTD1;refGene_tx=XM_019817679.2
E1 36592929 36593164
refGene_gn=MBTD1;refGene_tx=XM_006940197.3;refGene_gn=MBTD1;refGene_tx=XM_006940199.4;refGene_gn=MBTD1;refGene_tx=XM_006940201.3;refGene_gn=MBTD1;refGene_tx=XM_019817663.2;refGene_gn=MBTD1;refGene_tx=XM_019817664.2;refGene_gn=MBTD1;refGene_tx=XM_019817665.2;refGene_gn=MBTD1;refGene_tx=XM_019817666.2;refGene_gn=MBTD1;refGene_tx=XM_019817667.2;refGene_gn=MBTD1;refGene_tx=XM_019817668.2;refGene_gn=MBTD1;refGene_tx=XM_019817669.2;refGene_gn=MBTD1;refGene_tx=XM_019817670.2;refGene_gn=MBTD1;refGene_tx=XM_019817671.2;refGene_gn=MBTD1;refGene_tx=XM_019817672.2;refGene_gn=MBTD1;refGene_tx=XM_019817673.2;refGene_gn=MBTD1;refGene_tx=XM_019817674.2;refGene_gn=MBTD1;refGene_tx=XM_019817675.2;refGene_gn=MBTD1;refGene_tx=XM_019817676.2;refGene_gn=MBTD1;refGene_tx=XM_019817678.2;refGene_gn=MBTD1;refGene_tx=XM_019817679.2
E1 36593782 36593838
refGene_gn=MBTD1;refGene_tx=XM_006940197.3;refGene_gn=MBTD1;refGene_tx=XM_006940199.4;refGene_gn=MBTD1;refGene_tx=XM_006940201.3;refGene_gn=MBTD1;refGene_tx=XM_019817663.2;refGene_gn=MBTD1;refGene_tx=XM_019817664.2;refGene_gn=MBTD1;refGene_tx=XM_019817665.2;refGene_gn=MBTD1;refGene_tx=XM_019817666.2;refGene_gn=MBTD1;refGene_tx=XM_019817667.2;refGene_gn=MBTD1;refGene_tx=XM_019817668.2;refGene_gn=MBTD1;refGene_tx=XM_019817669.2;refGene_gn=MBTD1;refGene_tx=XM_019817670.2;refGene_gn=MBTD1;refGene_tx=XM_019817671.2;refGene_gn=MBTD1;refGene_tx=XM_019817672.2;refGene_gn=MBTD1;refGene_tx=XM_019817673.2;refGene_gn=MBTD1;refGene_tx=XM_019817674.2;refGene_gn=MBTD1;refGene_tx=XM_019817675.2;refGene_gn=MBTD1;refGene_tx=XM_019817676.2;refGene_gn=MBTD1;refGene_tx=XM_019817678.2;refGene_gn=MBTD1;refGene_tx=XM_019817679.2
E1 36594177 36594282
refGene_gn=MBTD1;refGene_tx=XM_006940197.3;refGene_gn=MBTD1;refGene_tx=XM_006940199.4;refGene_gn=MBTD1;refGene_tx=XM_006940201.3;refGene_gn=MBTD1;refGene_tx=XM_019817663.2;refGene_gn=MBTD1;refGene_tx=XM_019817664.2;refGene_gn=MBTD1;refGene_tx=XM_019817665.2;refGene_gn=MBTD1;refGene_tx=XM_019817666.2;refGene_gn=MBTD1;refGene_tx=XM_019817667.2;refGene_gn=MBTD1;refGene_tx=XM_019817668.2;refGene_gn=MBTD1;refGene_tx=XM_019817669.2;refGene_gn=MBTD1;refGene_tx=XM_019817670.2;refGene_gn=MBTD1;refGene_tx=XM_019817671.2;refGene_gn=MBTD1;refGene_tx=XM_019817672.2;refGene_gn=MBTD1;refGene_tx=XM_019817673.2;refGene_gn=MBTD1;refGene_tx=XM_019817674.2;refGene_gn=MBTD1;refGene_tx=XM_019817675.2;refGene_gn=MBTD1;refGene_tx=XM_019817676.2;refGene_gn=MBTD1;refGene_tx=XM_019817678.2;refGene_gn=MBTD1;refGene_tx=XM_019817679.2
E1 36597736 36597884
refGene_gn=MBTD1;refGene_tx=XM_006940197.3;refGene_gn=MBTD1;refGene_tx=XM_006940199.4;refGene_gn=MBTD1;refGene_tx=XM_006940201.3;refGene_gn=MBTD1;refGene_tx=XM_019817663.2;refGene_gn=MBTD1;refGene_tx=XM_019817664.2;refGene_gn=MBTD1;refGene_tx=XM_019817665.2;refGene_gn=MBTD1;refGe

ne_tx=XM_019817666.2;refGene_gn=MBTD1;refGene_tx=XM_019817667.2;refGen
e_gn=MBTD1;refGene_tx=XM_019817668.2;refGene_gn=MBTD1;refGene_tx=XM_01
9817669.2;refGene_gn=MBTD1;refGene_tx=XM_019817670.2;refGene_gn=MBTD1;
refGene_tx=XM_019817671.2;refGene_gn=MBTD1;refGene_tx=XM_019817672.2;r
efGene_gn=MBTD1;refGene_tx=XM_019817673.2;refGene_gn=MBTD1;refGene_tx=
XM_019817674.2;refGene_gn=MBTD1;refGene_tx=XM_019817675.2;refGene_gn=M
BTD1;refGene_tx=XM_019817676.2;refGene_gn=MBTD1;refGene_tx=XM_01981767
8.2;refGene_gn=MBTD1;refGene_tx=XM_019817679.2
E1 36598754 36598837
refGene_gn=MBTD1;refGene_tx=XM_006940197.3;refGene_gn=MBTD1;refGene_tx
=XM_006940199.4;refGene_gn=MBTD1;refGene_tx=XM_006940201.3;refGene_gn=
MBTD1;refGene_tx=XM_019817663.2;refGene_gn=MBTD1;refGene_tx=XM_0198176
64.2;refGene_gn=MBTD1;refGene_tx=XM_019817665.2;refGene_gn=MBTD1;refGe
ne_tx=XM_019817666.2;refGene_gn=MBTD1;refGene_tx=XM_019817667.2;refGen
e_gn=MBTD1;refGene_tx=XM_019817668.2;refGene_gn=MBTD1;refGene_tx=XM_01
9817669.2;refGene_gn=MBTD1;refGene_tx=XM_019817670.2;refGene_gn=MBTD1;
refGene_tx=XM_019817671.2;refGene_gn=MBTD1;refGene_tx=XM_019817672.2;r
efGene_gn=MBTD1;refGene_tx=XM_019817673.2;refGene_gn=MBTD1;refGene_tx=
XM_019817674.2;refGene_gn=MBTD1;refGene_tx=XM_019817675.2;refGene_gn=M
BTD1;refGene_tx=XM_019817676.2;refGene_gn=MBTD1;refGene_tx=XM_01981767
9.2
E1 36599224 36599459
refGene_gn=MBTD1;refGene_tx=XM_006940197.3;refGene_gn=MBTD1;refGene_tx
=XM_006940199.4;refGene_gn=MBTD1;refGene_tx=XM_006940201.3;refGene_gn=
MBTD1;refGene_tx=XM_019817663.2;refGene_gn=MBTD1;refGene_tx=XM_0198176
64.2;refGene_gn=MBTD1;refGene_tx=XM_019817665.2;refGene_gn=MBTD1;refGe
ne_tx=XM_019817666.2;refGene_gn=MBTD1;refGene_tx=XM_019817667.2;refGen
e_gn=MBTD1;refGene_tx=XM_019817668.2;refGene_gn=MBTD1;refGene_tx=XM_01
9817669.2;refGene_gn=MBTD1;refGene_tx=XM_019817670.2;refGene_gn=MBTD1;
refGene_tx=XM_019817671.2;refGene_gn=MBTD1;refGene_tx=XM_019817672.2;r
efGene_gn=MBTD1;refGene_tx=XM_019817673.2;refGene_gn=MBTD1;refGene_tx=
XM_019817674.2;refGene_gn=MBTD1;refGene_tx=XM_019817675.2;refGene_gn=M
BTD1;refGene_tx=XM_019817676.2;refGene_gn=MBTD1;refGene_tx=XM_01981767
8.2;refGene_gn=MBTD1;refGene_tx=XM_019817679.2
E1 36599995 36600073
refGene_gn=MBTD1;refGene_tx=XM_006940197.3;refGene_gn=MBTD1;refGene_tx
=XM_006940199.4;refGene_gn=MBTD1;refGene_tx=XM_006940201.3;refGene_gn=
MBTD1;refGene_tx=XM_019817663.2;refGene_gn=MBTD1;refGene_tx=XM_0198176
64.2;refGene_gn=MBTD1;refGene_tx=XM_019817665.2;refGene_gn=MBTD1;refGe
ne_tx=XM_019817666.2;refGene_gn=MBTD1;refGene_tx=XM_019817667.2;refGen
e_gn=MBTD1;refGene_tx=XM_019817668.2;refGene_gn=MBTD1;refGene_tx=XM_01
9817669.2;refGene_gn=MBTD1;refGene_tx=XM_019817670.2;refGene_gn=MBTD1;
refGene_tx=XM_019817671.2;refGene_gn=MBTD1;refGene_tx=XM_019817672.2;r
efGene_gn=MBTD1;refGene_tx=XM_019817673.2;refGene_gn=MBTD1;refGene_tx=
XM_019817674.2;refGene_gn=MBTD1;refGene_tx=XM_019817675.2;refGene_gn=M
BTD1;refGene_tx=XM_019817676.2;refGene_gn=MBTD1;refGene_tx=XM_01981767
9.2
E1 36604162 36604339
refGene_gn=MBTD1;refGene_tx=XM_006940197.3;refGene_gn=MBTD1;refGene_tx
=XM_006940199.4;refGene_gn=MBTD1;refGene_tx=XM_006940201.3;refGene_gn=

MBTD1;refGene_tx=XM_019817663.2;refGene_gn=MBTD1;refGene_tx=XM_019817664.2;refGene_gn=MBTD1;refGene_tx=XM_019817665.2;refGene_gn=MBTD1;refGene_tx=XM_019817666.2;refGene_gn=MBTD1;refGene_tx=XM_019817667.2;refGene_gn=MBTD1;refGene_tx=XM_019817668.2;refGene_gn=MBTD1;refGene_tx=XM_019817669.2;refGene_gn=MBTD1;refGene_tx=XM_019817670.2;refGene_gn=MBTD1;refGene_tx=XM_019817671.2;refGene_gn=MBTD1;refGene_tx=XM_019817672.2;refGene_gn=MBTD1;refGene_tx=XM_019817673.2;refGene_gn=MBTD1;refGene_tx=XM_019817676.2;refGene_gn=MBTD1;refGene_tx=XM_019817679.2
E1 36609145 36609264
refGene_gn=MBTD1;refGene_tx=XM_006940197.3;refGene_gn=MBTD1;refGene_tx=XM_006940199.4;refGene_gn=MBTD1;refGene_tx=XM_006940201.3;refGene_gn=MBTD1;refGene_tx=XM_019817663.2;refGene_gn=MBTD1;refGene_tx=XM_019817664.2;refGene_gn=MBTD1;refGene_tx=XM_019817665.2;refGene_gn=MBTD1;refGene_tx=XM_019817666.2;refGene_gn=MBTD1;refGene_tx=XM_019817667.2;refGene_gn=MBTD1;refGene_tx=XM_019817668.2;refGene_gn=MBTD1;refGene_tx=XM_019817669.2;refGene_gn=MBTD1;refGene_tx=XM_019817670.2;refGene_gn=MBTD1;refGene_tx=XM_019817671.2;refGene_gn=MBTD1;refGene_tx=XM_019817672.2;refGene_gn=MBTD1;refGene_tx=XM_019817673.2;refGene_gn=MBTD1;refGene_tx=XM_019817674.2;refGene_gn=MBTD1;refGene_tx=XM_019817675.2;refGene_gn=MBTD1;refGene_tx=XM_019817676.2;refGene_gn=MBTD1;refGene_tx=XM_019817679.2
E1 36619351 36619469 refGene_gn=NME2;refGene_tx=XM_023243660.1
E1 36620225 36620338 refGene_gn=NME2;refGene_tx=XM_023243660.1
E1 36620758 36620860 refGene_gn=NME2;refGene_tx=XM_023243660.1
E1 36621738 36621864 refGene_gn=NME2;refGene_tx=XM_023243660.1
E1 36626147 36626265
refGene_gn=LOC101088196;refGene_tx=XM_006940203.3;refGene_gn=LOC101088196;refGene_tx=XM_019817680.1
E1 36627014 36627127
refGene_gn=LOC101088196;refGene_tx=XM_006940203.3;refGene_gn=LOC101088196;refGene_tx=XM_019817680.1
E1 36628199 36628301
refGene_gn=LOC101088196;refGene_tx=XM_006940203.3;refGene_gn=LOC101088196;refGene_tx=XM_019817680.1
E1 36630768 36630894
refGene_gn=LOC101088196;refGene_tx=XM_006940203.3;refGene_gn=LOC101088196;refGene_tx=XM_019817680.1
E1 36664059 36664362
refGene_gn=SPAG9;refGene_tx=XM_023244410.1;refGene_gn=SPAG9;refGene_tx=XM_023244411.1;refGene_gn=SPAG9;refGene_tx=XM_023244412.1;refGene_gn=SPAG9;refGene_tx=XM_023244413.1;refGene_gn=SPAG9;refGene_tx=XM_023244414.1;refGene_gn=SPAG9;refGene_tx=XM_023244416.1
E1 36697653 36697774
refGene_gn=SPAG9;refGene_tx=XM_023244410.1;refGene_gn=SPAG9;refGene_tx=XM_023244411.1;refGene_gn=SPAG9;refGene_tx=XM_023244412.1;refGene_gn=SPAG9;refGene_tx=XM_023244413.1;refGene_gn=SPAG9;refGene_tx=XM_023244414.1;refGene_gn=SPAG9;refGene_tx=XM_023244416.1
E1 36715666 36715737
refGene_gn=SPAG9;refGene_tx=XM_019817688.2;refGene_gn=SPAG9;refGene_tx=XM_023244410.1;refGene_gn=SPAG9;refGene_tx=XM_023244411.1;refGene_gn=

SPAG9;refGene_tx=XM_023244412.1;refGene_gn=SPAG9;refGene_tx=XM_0232444
13.1;refGene_gn=SPAG9;refGene_tx=XM_023244414.1;refGene_gn=SPAG9;refGe
ne_tx=XM_023244416.1;refGene_gn=SPAG9;refGene_tx=XM_023244417.1
E1 36722876 36722971
refGene_gn=SPAG9;refGene_tx=XM_023244410.1;refGene_gn=SPAG9;refGene_tx
=XM_023244411.1;refGene_gn=SPAG9;refGene_tx=XM_023244412.1;refGene_gn=
SPAG9;refGene_tx=XM_023244413.1;refGene_gn=SPAG9;refGene_tx=XM_0232444
14.1;refGene_gn=SPAG9;refGene_tx=XM_023244416.1
E1 36723689 36723850
refGene_gn=SPAG9;refGene_tx=XM_003996685.5;refGene_gn=SPAG9;refGene_tx
=XM_011289199.3;refGene_gn=SPAG9;refGene_tx=XM_019817681.2;refGene_gn=
SPAG9;refGene_tx=XM_019817685.2;refGene_gn=SPAG9;refGene_tx=XM_0198176
86.2
E1 36728517 36728668
refGene_gn=SPAG9;refGene_tx=XM_003996685.5;refGene_gn=SPAG9;refGene_tx
=XM_011289199.3;refGene_gn=SPAG9;refGene_tx=XM_019817681.2;refGene_gn=
SPAG9;refGene_tx=XM_019817685.2;refGene_gn=SPAG9;refGene_tx=XM_0198176
86.2;refGene_gn=SPAG9;refGene_tx=XM_019817688.2;refGene_gn=SPAG9;refGe
ne_tx=XM_023244410.1;refGene_gn=SPAG9;refGene_tx=XM_023244411.1;refGen
e_gn=SPAG9;refGene_tx=XM_023244412.1;refGene_gn=SPAG9;refGene_tx=XM_02
3244413.1;refGene_gn=SPAG9;refGene_tx=XM_023244414.1;refGene_gn=SPAG9;
refGene_tx=XM_023244416.1;refGene_gn=SPAG9;refGene_tx=XM_023244417.1
E1 36747745 36747953
refGene_gn=SPAG9;refGene_tx=XM_003996685.5;refGene_gn=SPAG9;refGene_tx
=XM_011289199.3;refGene_gn=SPAG9;refGene_tx=XM_019817681.2;refGene_gn=
SPAG9;refGene_tx=XM_019817685.2;refGene_gn=SPAG9;refGene_tx=XM_0198176
86.2;refGene_gn=SPAG9;refGene_tx=XM_019817688.2;refGene_gn=SPAG9;refGe
ne_tx=XM_023244410.1;refGene_gn=SPAG9;refGene_tx=XM_023244411.1;refGen
e_gn=SPAG9;refGene_tx=XM_023244412.1;refGene_gn=SPAG9;refGene_tx=XM_02
3244413.1;refGene_gn=SPAG9;refGene_tx=XM_023244414.1;refGene_gn=SPAG9;
refGene_tx=XM_023244416.1;refGene_gn=SPAG9;refGene_tx=XM_023244417.1
E1 36748472 36748572
refGene_gn=SPAG9;refGene_tx=XM_003996685.5;refGene_gn=SPAG9;refGene_tx
=XM_011289199.3;refGene_gn=SPAG9;refGene_tx=XM_019817681.2;refGene_gn=
SPAG9;refGene_tx=XM_019817685.2;refGene_gn=SPAG9;refGene_tx=XM_0198176
86.2;refGene_gn=SPAG9;refGene_tx=XM_019817688.2;refGene_gn=SPAG9;refGe
ne_tx=XM_023244410.1;refGene_gn=SPAG9;refGene_tx=XM_023244411.1;refGen
e_gn=SPAG9;refGene_tx=XM_023244412.1;refGene_gn=SPAG9;refGene_tx=XM_02
3244413.1;refGene_gn=SPAG9;refGene_tx=XM_023244414.1;refGene_gn=SPAG9;
refGene_tx=XM_023244416.1;refGene_gn=SPAG9;refGene_tx=XM_023244417.1
E1 36754394 36754516
refGene_gn=SPAG9;refGene_tx=XM_003996685.5;refGene_gn=SPAG9;refGene_tx
=XM_011289199.3;refGene_gn=SPAG9;refGene_tx=XM_019817681.2;refGene_gn=
SPAG9;refGene_tx=XM_019817685.2;refGene_gn=SPAG9;refGene_tx=XM_0198176
86.2;refGene_gn=SPAG9;refGene_tx=XM_019817688.2;refGene_gn=SPAG9;refGe
ne_tx=XM_023244410.1;refGene_gn=SPAG9;refGene_tx=XM_023244411.1;refGen
e_gn=SPAG9;refGene_tx=XM_023244412.1;refGene_gn=SPAG9;refGene_tx=XM_02
3244413.1;refGene_gn=SPAG9;refGene_tx=XM_023244414.1;refGene_gn=SPAG9;
refGene_tx=XM_023244416.1;refGene_gn=SPAG9;refGene_tx=XM_023244417.1
E1 36759310 36759322

refGene_gn=SPAG9;refGene_tx=XM_019817681.2;refGene_gn=SPAG9;refGene_tx
=XM_019817686.2;refGene_gn=SPAG9;refGene_tx=XM_019817688.2;refGene_gn=
SPAG9;refGene_tx=XM_023244410.1;refGene_gn=SPAG9;refGene_tx=XM_0232444
12.1;refGene_gn=SPAG9;refGene_tx=XM_023244414.1;refGene_gn=SPAG9;refGe
ne_tx=XM_023244417.1
E1 36760566 36760581
refGene_gn=SPAG9;refGene_tx=XM_019817681.2;refGene_gn=SPAG9;refGene_tx
=XM_019817685.2;refGene_gn=SPAG9;refGene_tx=XM_019817688.2;refGene_gn=
SPAG9;refGene_tx=XM_023244410.1;refGene_gn=SPAG9;refGene_tx=XM_0232444
11.1;refGene_gn=SPAG9;refGene_tx=XM_023244414.1;refGene_gn=SPAG9;refGe
ne_tx=XM_023244417.1
E1 36761053 36761111
refGene_gn=SPAG9;refGene_tx=XM_003996685.5;refGene_gn=SPAG9;refGene_tx
=XM_011289199.3;refGene_gn=SPAG9;refGene_tx=XM_019817681.2;refGene_gn=
SPAG9;refGene_tx=XM_019817685.2;refGene_gn=SPAG9;refGene_tx=XM_0198176
86.2;refGene_gn=SPAG9;refGene_tx=XM_019817688.2;refGene_gn=SPAG9;refGe
ne_tx=XM_023244410.1;refGene_gn=SPAG9;refGene_tx=XM_023244411.1;refGen
e_gn=SPAG9;refGene_tx=XM_023244412.1;refGene_gn=SPAG9;refGene_tx=XM_02
3244413.1;refGene_gn=SPAG9;refGene_tx=XM_023244414.1;refGene_gn=SPAG9;
refGene_tx=XM_023244416.1;refGene_gn=SPAG9;refGene_tx=XM_023244417.1
E1 36762116 36762269
refGene_gn=SPAG9;refGene_tx=XM_003996685.5;refGene_gn=SPAG9;refGene_tx
=XM_011289199.3;refGene_gn=SPAG9;refGene_tx=XM_019817681.2;refGene_gn=
SPAG9;refGene_tx=XM_019817685.2;refGene_gn=SPAG9;refGene_tx=XM_0198176
86.2;refGene_gn=SPAG9;refGene_tx=XM_019817688.2;refGene_gn=SPAG9;refGe
ne_tx=XM_023244410.1;refGene_gn=SPAG9;refGene_tx=XM_023244411.1;refGen
e_gn=SPAG9;refGene_tx=XM_023244412.1;refGene_gn=SPAG9;refGene_tx=XM_02
3244413.1;refGene_gn=SPAG9;refGene_tx=XM_023244414.1;refGene_gn=SPAG9;
refGene_tx=XM_023244416.1;refGene_gn=SPAG9;refGene_tx=XM_023244417.1
E1 36762705 36762757
refGene_gn=SPAG9;refGene_tx=XM_003996685.5;refGene_gn=SPAG9;refGene_tx
=XM_011289199.3;refGene_gn=SPAG9;refGene_tx=XM_019817681.2;refGene_gn=
SPAG9;refGene_tx=XM_019817685.2;refGene_gn=SPAG9;refGene_tx=XM_0198176
86.2;refGene_gn=SPAG9;refGene_tx=XM_019817688.2;refGene_gn=SPAG9;refGe
ne_tx=XM_023244410.1;refGene_gn=SPAG9;refGene_tx=XM_023244411.1;refGen
e_gn=SPAG9;refGene_tx=XM_023244412.1;refGene_gn=SPAG9;refGene_tx=XM_02
3244413.1;refGene_gn=SPAG9;refGene_tx=XM_023244414.1;refGene_gn=SPAG9;
refGene_tx=XM_023244416.1;refGene_gn=SPAG9;refGene_tx=XM_023244417.1
E1 36765469 36765600
refGene_gn=SPAG9;refGene_tx=XM_003996685.5;refGene_gn=SPAG9;refGene_tx
=XM_011289199.3;refGene_gn=SPAG9;refGene_tx=XM_019817681.2;refGene_gn=
SPAG9;refGene_tx=XM_019817685.2;refGene_gn=SPAG9;refGene_tx=XM_0198176
86.2;refGene_gn=SPAG9;refGene_tx=XM_019817688.2;refGene_gn=SPAG9;refGe
ne_tx=XM_023244410.1;refGene_gn=SPAG9;refGene_tx=XM_023244411.1;refGen
e_gn=SPAG9;refGene_tx=XM_023244412.1;refGene_gn=SPAG9;refGene_tx=XM_02
3244413.1;refGene_gn=SPAG9;refGene_tx=XM_023244414.1;refGene_gn=SPAG9;
refGene_tx=XM_023244416.1;refGene_gn=SPAG9;refGene_tx=XM_023244417.1
E1 36767883 36767940
refGene_gn=SPAG9;refGene_tx=XM_003996685.5;refGene_gn=SPAG9;refGene_tx
=XM_011289199.3;refGene_gn=SPAG9;refGene_tx=XM_019817681.2;refGene_gn=

SPAG9;refGene_tx=XM_019817685.2;refGene_gn=SPAG9;refGene_tx=XM_019817686.2;refGene_gn=SPAG9;refGene_tx=XM_019817688.2;refGene_gn=SPAG9;refGene_tx=XM_023244410.1;refGene_gn=SPAG9;refGene_tx=XM_023244411.1;refGene_gn=SPAG9;refGene_tx=XM_023244412.1;refGene_gn=SPAG9;refGene_tx=XM_023244413.1;refGene_gn=SPAG9;refGene_tx=XM_023244414.1;refGene_gn=SPAG9;refGene_tx=XM_023244416.1;refGene_gn=SPAG9;refGene_tx=XM_023244417.1
E1 36768965 36769139
refGene_gn=SPAG9;refGene_tx=XM_003996685.5;refGene_gn=SPAG9;refGene_tx=XM_011289199.3;refGene_gn=SPAG9;refGene_tx=XM_019817681.2;refGene_gn=SPAG9;refGene_tx=XM_019817685.2;refGene_gn=SPAG9;refGene_tx=XM_019817686.2;refGene_gn=SPAG9;refGene_tx=XM_019817688.2;refGene_gn=SPAG9;refGene_tx=XM_023244410.1;refGene_gn=SPAG9;refGene_tx=XM_023244411.1;refGene_gn=SPAG9;refGene_tx=XM_023244412.1;refGene_gn=SPAG9;refGene_tx=XM_023244413.1;refGene_gn=SPAG9;refGene_tx=XM_023244414.1;refGene_gn=SPAG9;refGene_tx=XM_023244416.1;refGene_gn=SPAG9;refGene_tx=XM_023244417.1
E1 36770232 36770362
refGene_gn=SPAG9;refGene_tx=XM_003996685.5;refGene_gn=SPAG9;refGene_tx=XM_011289199.3;refGene_gn=SPAG9;refGene_tx=XM_019817681.2;refGene_gn=SPAG9;refGene_tx=XM_019817685.2;refGene_gn=SPAG9;refGene_tx=XM_019817686.2;refGene_gn=SPAG9;refGene_tx=XM_019817688.2;refGene_gn=SPAG9;refGene_tx=XM_023244410.1;refGene_gn=SPAG9;refGene_tx=XM_023244411.1;refGene_gn=SPAG9;refGene_tx=XM_023244412.1;refGene_gn=SPAG9;refGene_tx=XM_023244413.1;refGene_gn=SPAG9;refGene_tx=XM_023244414.1;refGene_gn=SPAG9;refGene_tx=XM_023244416.1;refGene_gn=SPAG9;refGene_tx=XM_023244417.1
E1 36772254 36772344
refGene_gn=SPAG9;refGene_tx=XM_003996685.5;refGene_gn=SPAG9;refGene_tx=XM_011289199.3;refGene_gn=SPAG9;refGene_tx=XM_019817681.2;refGene_gn=SPAG9;refGene_tx=XM_019817685.2;refGene_gn=SPAG9;refGene_tx=XM_019817686.2;refGene_gn=SPAG9;refGene_tx=XM_019817688.2;refGene_gn=SPAG9;refGene_tx=XM_023244410.1;refGene_gn=SPAG9;refGene_tx=XM_023244411.1;refGene_gn=SPAG9;refGene_tx=XM_023244412.1;refGene_gn=SPAG9;refGene_tx=XM_023244413.1;refGene_gn=SPAG9;refGene_tx=XM_023244414.1;refGene_gn=SPAG9;refGene_tx=XM_023244416.1;refGene_gn=SPAG9;refGene_tx=XM_023244417.1
E1 36772828 36772996
refGene_gn=SPAG9;refGene_tx=XM_003996685.5;refGene_gn=SPAG9;refGene_tx=XM_011289199.3;refGene_gn=SPAG9;refGene_tx=XM_019817681.2;refGene_gn=SPAG9;refGene_tx=XM_019817685.2;refGene_gn=SPAG9;refGene_tx=XM_019817686.2;refGene_gn=SPAG9;refGene_tx=XM_019817688.2;refGene_gn=SPAG9;refGene_tx=XM_023244410.1;refGene_gn=SPAG9;refGene_tx=XM_023244411.1;refGene_gn=SPAG9;refGene_tx=XM_023244412.1;refGene_gn=SPAG9;refGene_tx=XM_023244413.1;refGene_gn=SPAG9;refGene_tx=XM_023244414.1;refGene_gn=SPAG9;refGene_tx=XM_023244416.1;refGene_gn=SPAG9;refGene_tx=XM_023244417.1
E1 36773413 36773585
refGene_gn=SPAG9;refGene_tx=XM_003996685.5;refGene_gn=SPAG9;refGene_tx=XM_011289199.3;refGene_gn=SPAG9;refGene_tx=XM_019817681.2;refGene_gn=SPAG9;refGene_tx=XM_019817685.2;refGene_gn=SPAG9;refGene_tx=XM_019817686.2;refGene_gn=SPAG9;refGene_tx=XM_019817688.2;refGene_gn=SPAG9;refGene_tx=XM_023244410.1;refGene_gn=SPAG9;refGene_tx=XM_023244411.1;refGene_gn=SPAG9;refGene_tx=XM_023244412.1;refGene_gn=SPAG9;refGene_tx=XM_023244413.1;refGene_gn=SPAG9;refGene_tx=XM_023244414.1;refGene_gn=SPAG9;

refGene_tx=XM_023244416.1;refGene_gn=SPAG9;refGene_tx=XM_023244417.1
E1 36775596 36775815
refGene_gn=SPAG9;refGene_tx=XM_003996685.5;refGene_gn=SPAG9;refGene_tx=XM_011289199.3;refGene_gn=SPAG9;refGene_tx=XM_019817681.2;refGene_gn=SPAG9;refGene_tx=XM_019817685.2;refGene_gn=SPAG9;refGene_tx=XM_019817686.2;refGene_gn=SPAG9;refGene_tx=XM_019817688.2;refGene_gn=SPAG9;refGene_tx=XM_023244410.1;refGene_gn=SPAG9;refGene_tx=XM_023244411.1;refGene_gn=SPAG9;refGene_tx=XM_023244412.1;refGene_gn=SPAG9;refGene_tx=XM_023244413.1;refGene_gn=SPAG9;refGene_tx=XM_023244414.1;refGene_gn=SPAG9;refGene_tx=XM_023244416.1;refGene_gn=SPAG9;refGene_tx=XM_023244417.1
E1 36776095 36776291
refGene_gn=SPAG9;refGene_tx=XM_003996685.5;refGene_gn=SPAG9;refGene_tx=XM_011289199.3;refGene_gn=SPAG9;refGene_tx=XM_019817681.2;refGene_gn=SPAG9;refGene_tx=XM_019817685.2;refGene_gn=SPAG9;refGene_tx=XM_019817686.2;refGene_gn=SPAG9;refGene_tx=XM_019817688.2;refGene_gn=SPAG9;refGene_tx=XM_023244410.1;refGene_gn=SPAG9;refGene_tx=XM_023244411.1;refGene_gn=SPAG9;refGene_tx=XM_023244412.1;refGene_gn=SPAG9;refGene_tx=XM_023244413.1;refGene_gn=SPAG9;refGene_tx=XM_023244414.1;refGene_gn=SPAG9;refGene_tx=XM_023244416.1;refGene_gn=SPAG9;refGene_tx=XM_023244417.1
E1 36777686 36777812
refGene_gn=SPAG9;refGene_tx=XM_003996685.5;refGene_gn=SPAG9;refGene_tx=XM_011289199.3;refGene_gn=SPAG9;refGene_tx=XM_019817681.2;refGene_gn=SPAG9;refGene_tx=XM_019817685.2;refGene_gn=SPAG9;refGene_tx=XM_019817686.2;refGene_gn=SPAG9;refGene_tx=XM_019817688.2;refGene_gn=SPAG9;refGene_tx=XM_023244410.1;refGene_gn=SPAG9;refGene_tx=XM_023244411.1;refGene_gn=SPAG9;refGene_tx=XM_023244412.1;refGene_gn=SPAG9;refGene_tx=XM_023244413.1;refGene_gn=SPAG9;refGene_tx=XM_023244414.1;refGene_gn=SPAG9;refGene_tx=XM_023244416.1;refGene_gn=SPAG9;refGene_tx=XM_023244417.1
E1 36779077 36779158
refGene_gn=SPAG9;refGene_tx=XM_003996685.5;refGene_gn=SPAG9;refGene_tx=XM_011289199.3;refGene_gn=SPAG9;refGene_tx=XM_019817681.2;refGene_gn=SPAG9;refGene_tx=XM_019817685.2;refGene_gn=SPAG9;refGene_tx=XM_019817686.2;refGene_gn=SPAG9;refGene_tx=XM_019817688.2;refGene_gn=SPAG9;refGene_tx=XM_023244410.1;refGene_gn=SPAG9;refGene_tx=XM_023244411.1;refGene_gn=SPAG9;refGene_tx=XM_023244412.1;refGene_gn=SPAG9;refGene_tx=XM_023244413.1;refGene_gn=SPAG9;refGene_tx=XM_023244414.1;refGene_gn=SPAG9;refGene_tx=XM_023244416.1;refGene_gn=SPAG9;refGene_tx=XM_023244417.1
E1 36780115 36780183
refGene_gn=SPAG9;refGene_tx=XM_003996685.5;refGene_gn=SPAG9;refGene_tx=XM_011289199.3;refGene_gn=SPAG9;refGene_tx=XM_019817681.2;refGene_gn=SPAG9;refGene_tx=XM_019817685.2;refGene_gn=SPAG9;refGene_tx=XM_019817686.2;refGene_gn=SPAG9;refGene_tx=XM_019817688.2;refGene_gn=SPAG9;refGene_tx=XM_023244410.1;refGene_gn=SPAG9;refGene_tx=XM_023244411.1;refGene_gn=SPAG9;refGene_tx=XM_023244412.1;refGene_gn=SPAG9;refGene_tx=XM_023244413.1;refGene_gn=SPAG9;refGene_tx=XM_023244414.1;refGene_gn=SPAG9;refGene_tx=XM_023244416.1;refGene_gn=SPAG9;refGene_tx=XM_023244417.1
E1 36782379 36782528
refGene_gn=SPAG9;refGene_tx=XM_003996685.5;refGene_gn=SPAG9;refGene_tx=XM_011289199.3;refGene_gn=SPAG9;refGene_tx=XM_019817681.2;refGene_gn=SPAG9;refGene_tx=XM_019817685.2;refGene_gn=SPAG9;refGene_tx=XM_0198176

86.2;refGene_gn=SPAG9;refGene_tx=XM_019817688.2;refGene_gn=SPAG9;refGene_tx=XM_023244410.1;refGene_gn=SPAG9;refGene_tx=XM_023244411.1;refGene_gn=SPAG9;refGene_tx=XM_023244412.1;refGene_gn=SPAG9;refGene_tx=XM_023244413.1;refGene_gn=SPAG9;refGene_tx=XM_023244414.1;refGene_gn=SPAG9;refGene_tx=XM_023244416.1;refGene_gn=SPAG9;refGene_tx=XM_023244417.1
E1 36783827 36783999
refGene_gn=SPAG9;refGene_tx=XM_003996685.5;refGene_gn=SPAG9;refGene_tx=XM_011289199.3;refGene_gn=SPAG9;refGene_tx=XM_019817681.2;refGene_gn=SPAG9;refGene_tx=XM_019817685.2;refGene_gn=SPAG9;refGene_tx=XM_019817686.2;refGene_gn=SPAG9;refGene_tx=XM_019817688.2;refGene_gn=SPAG9;refGene_tx=XM_023244410.1;refGene_gn=SPAG9;refGene_tx=XM_023244411.1;refGene_gn=SPAG9;refGene_tx=XM_023244412.1;refGene_gn=SPAG9;refGene_tx=XM_023244413.1;refGene_gn=SPAG9;refGene_tx=XM_023244414.1;refGene_gn=SPAG9;refGene_tx=XM_023244416.1;refGene_gn=SPAG9;refGene_tx=XM_023244417.1
E1 36785870 36785984
refGene_gn=SPAG9;refGene_tx=XM_003996685.5;refGene_gn=SPAG9;refGene_tx=XM_011289199.3;refGene_gn=SPAG9;refGene_tx=XM_019817681.2;refGene_gn=SPAG9;refGene_tx=XM_019817685.2;refGene_gn=SPAG9;refGene_tx=XM_019817686.2;refGene_gn=SPAG9;refGene_tx=XM_019817688.2;refGene_gn=SPAG9;refGene_tx=XM_023244410.1;refGene_gn=SPAG9;refGene_tx=XM_023244411.1;refGene_gn=SPAG9;refGene_tx=XM_023244412.1;refGene_gn=SPAG9;refGene_tx=XM_023244413.1;refGene_gn=SPAG9;refGene_tx=XM_023244414.1;refGene_gn=SPAG9;refGene_tx=XM_023244416.1;refGene_gn=SPAG9;refGene_tx=XM_023244417.1
E1 36787560 36787599
refGene_gn=SPAG9;refGene_tx=XM_003996685.5;refGene_gn=SPAG9;refGene_tx=XM_019817681.2;refGene_gn=SPAG9;refGene_tx=XM_019817688.2;refGene_gn=SPAG9;refGene_tx=XM_023244410.1;refGene_gn=SPAG9;refGene_tx=XM_023244411.1;refGene_gn=SPAG9;refGene_tx=XM_023244412.1;refGene_gn=SPAG9;refGene_tx=XM_023244413.1;refGene_gn=SPAG9;refGene_tx=XM_023244417.1
E1 36788548 36788725
refGene_gn=SPAG9;refGene_tx=XM_003996685.5;refGene_gn=SPAG9;refGene_tx=XM_011289199.3;refGene_gn=SPAG9;refGene_tx=XM_019817681.2;refGene_gn=SPAG9;refGene_tx=XM_019817685.2;refGene_gn=SPAG9;refGene_tx=XM_019817686.2;refGene_gn=SPAG9;refGene_tx=XM_019817688.2;refGene_gn=SPAG9;refGene_tx=XM_023244410.1;refGene_gn=SPAG9;refGene_tx=XM_023244411.1;refGene_gn=SPAG9;refGene_tx=XM_023244412.1;refGene_gn=SPAG9;refGene_tx=XM_023244413.1;refGene_gn=SPAG9;refGene_tx=XM_023244414.1;refGene_gn=SPAG9;refGene_tx=XM_023244416.1;refGene_gn=SPAG9;refGene_tx=XM_023244417.1
E1 36790440 36790590
refGene_gn=SPAG9;refGene_tx=XM_003996685.5;refGene_gn=SPAG9;refGene_tx=XM_011289199.3;refGene_gn=SPAG9;refGene_tx=XM_019817681.2;refGene_gn=SPAG9;refGene_tx=XM_019817685.2;refGene_gn=SPAG9;refGene_tx=XM_019817686.2;refGene_gn=SPAG9;refGene_tx=XM_019817688.2;refGene_gn=SPAG9;refGene_tx=XM_023244410.1;refGene_gn=SPAG9;refGene_tx=XM_023244411.1;refGene_gn=SPAG9;refGene_tx=XM_023244412.1;refGene_gn=SPAG9;refGene_tx=XM_023244413.1;refGene_gn=SPAG9;refGene_tx=XM_023244414.1;refGene_gn=SPAG9;refGene_tx=XM_023244416.1;refGene_gn=SPAG9;refGene_tx=XM_023244417.1
E1 36792263 36792379
refGene_gn=SPAG9;refGene_tx=XM_003996685.5;refGene_gn=SPAG9;refGene_tx=XM_011289199.3;refGene_gn=SPAG9;refGene_tx=XM_019817681.2;refGene_gn=

SPAG9;refGene_tx=XM_019817685.2;refGene_gn=SPAG9;refGene_tx=XM_0198176
86.2;refGene_gn=SPAG9;refGene_tx=XM_019817688.2;refGene_gn=SPAG9;refGe
ne_tx=XM_023244410.1;refGene_gn=SPAG9;refGene_tx=XM_023244411.1;refGen
e_gn=SPAG9;refGene_tx=XM_023244412.1;refGene_gn=SPAG9;refGene_tx=XM_02
3244413.1;refGene_gn=SPAG9;refGene_tx=XM_023244414.1;refGene_gn=SPAG9;
refGene_tx=XM_023244416.1;refGene_gn=SPAG9;refGene_tx=XM_023244417.1
E1 36885605 36886643
refGene_gn=TOB1;refGene_tx=XM_011289196.3;refGene_gn=TOB1;refGene_tx=X
M_023243725.1
E1 36904815 36906336
refGene_gn=WFIKKN2;refGene_tx=XM_003996686.5;refGene_gn=WFIKKN2;refGen
e_tx=XM_006940205.4
E1 36909006 36909207 refGene_gn=WFIKKN2;refGene_tx=XM_003996686.5
E1 36969817 36969851 refGene_gn=LUC7L3;refGene_tx=XM_006940206.4
E1 36970722 36970747
refGene_gn=LUC7L3;refGene_tx=XM_019817690.2;refGene_gn=LUC7L3;refGene_
tx=XM_019817691.2
E1 36971112 36971303
refGene_gn=LUC7L3;refGene_tx=XM_006940206.4;refGene_gn=LUC7L3;refGene_
tx=XM_011289194.3;refGene_gn=LUC7L3;refGene_tx=XM_011289195.3;refGene_
gn=LUC7L3;refGene_tx=XM_019817690.2;refGene_gn=LUC7L3;refGene_tx=XM_01
9817691.2;refGene_gn=LUC7L3;refGene_tx=XM_019817692.2
E1 36971388 36971549
refGene_gn=LUC7L3;refGene_tx=XM_006940206.4;refGene_gn=LUC7L3;refGene_
tx=XM_006940207.4;refGene_gn=LUC7L3;refGene_tx=XM_011289194.3;refGene_
gn=LUC7L3;refGene_tx=XM_019817690.2;refGene_gn=LUC7L3;refGene_tx=XM_01
9817691.2;refGene_gn=LUC7L3;refGene_tx=XM_019817692.2
E1 36972811 36972837 refGene_gn=LUC7L3;refGene_tx=XM_006940208.4
E1 36974964 36975125
refGene_gn=LUC7L3;refGene_tx=XM_006940206.4;refGene_gn=LUC7L3;refGene_
tx=XM_006940207.4;refGene_gn=LUC7L3;refGene_tx=XM_006940208.4;refGene_
gn=LUC7L3;refGene_tx=XM_011289194.3;refGene_gn=LUC7L3;refGene_tx=XM_01
1289195.3;refGene_gn=LUC7L3;refGene_tx=XM_019817690.2;refGene_gn=LUC7L
3;refGene_tx=XM_019817691.2;refGene_gn=LUC7L3;refGene_tx=XM_019817692.
2
E1 36976174 36976458
refGene_gn=LUC7L3;refGene_tx=XM_006940206.4;refGene_gn=LUC7L3;refGene_
tx=XM_006940207.4;refGene_gn=LUC7L3;refGene_tx=XM_006940208.4;refGene_
gn=LUC7L3;refGene_tx=XM_011289194.3;refGene_gn=LUC7L3;refGene_tx=XM_01
1289195.3;refGene_gn=LUC7L3;refGene_tx=XM_019817690.2;refGene_gn=LUC7L
3;refGene_tx=XM_019817691.2;refGene_gn=LUC7L3;refGene_tx=XM_019817692.
2
E1 36976850 36977012
refGene_gn=LUC7L3;refGene_tx=XM_006940206.4;refGene_gn=LUC7L3;refGene_
tx=XM_006940207.4;refGene_gn=LUC7L3;refGene_tx=XM_006940208.4;refGene_
gn=LUC7L3;refGene_tx=XM_011289194.3;refGene_gn=LUC7L3;refGene_tx=XM_01
1289195.3;refGene_gn=LUC7L3;refGene_tx=XM_019817690.2;refGene_gn=LUC7L
3;refGene_tx=XM_019817691.2;refGene_gn=LUC7L3;refGene_tx=XM_019817692.
2
E1 36979144 36979249

refGene_gn=LUC7L3;refGene_tx=XM_006940206.4;refGene_gn=LUC7L3;refGene_tx=XM_006940207.4;refGene_gn=LUC7L3;refGene_tx=XM_006940208.4;refGene_gn=LUC7L3;refGene_tx=XM_011289194.3;refGene_gn=LUC7L3;refGene_tx=XM_011289195.3;refGene_gn=LUC7L3;refGene_tx=XM_019817690.2;refGene_gn=LUC7L3;refGene_tx=XM_019817691.2;refGene_gn=LUC7L3;refGene_tx=XM_019817692.2
E1 36980221 36980296
refGene_gn=LUC7L3;refGene_tx=XM_006940206.4;refGene_gn=LUC7L3;refGene_tx=XM_006940207.4;refGene_gn=LUC7L3;refGene_tx=XM_006940208.4;refGene_gn=LUC7L3;refGene_tx=XM_011289194.3;refGene_gn=LUC7L3;refGene_tx=XM_011289195.3;refGene_gn=LUC7L3;refGene_tx=XM_019817690.2;refGene_gn=LUC7L3;refGene_tx=XM_019817691.2;refGene_gn=LUC7L3;refGene_tx=XM_019817692.2
E1 36980660 36980805
refGene_gn=LUC7L3;refGene_tx=XM_006940206.4;refGene_gn=LUC7L3;refGene_tx=XM_006940207.4;refGene_gn=LUC7L3;refGene_tx=XM_006940208.4;refGene_gn=LUC7L3;refGene_tx=XM_011289194.3;refGene_gn=LUC7L3;refGene_tx=XM_011289195.3;refGene_gn=LUC7L3;refGene_tx=XM_019817690.2;refGene_gn=LUC7L3;refGene_tx=XM_019817691.2;refGene_gn=LUC7L3;refGene_tx=XM_019817692.2
E1 36981782 36981822
refGene_gn=LUC7L3;refGene_tx=XM_006940206.4;refGene_gn=LUC7L3;refGene_tx=XM_006940207.4;refGene_gn=LUC7L3;refGene_tx=XM_006940208.4;refGene_gn=LUC7L3;refGene_tx=XM_011289194.3;refGene_gn=LUC7L3;refGene_tx=XM_011289195.3;refGene_gn=LUC7L3;refGene_tx=XM_019817690.2;refGene_gn=LUC7L3;refGene_tx=XM_019817691.2
E1 36984838 36984905
refGene_gn=LUC7L3;refGene_tx=XM_006940206.4;refGene_gn=LUC7L3;refGene_tx=XM_006940207.4;refGene_gn=LUC7L3;refGene_tx=XM_006940208.4;refGene_gn=LUC7L3;refGene_tx=XM_011289194.3;refGene_gn=LUC7L3;refGene_tx=XM_011289195.3;refGene_gn=LUC7L3;refGene_tx=XM_019817690.2;refGene_gn=LUC7L3;refGene_tx=XM_019817691.2
E1 36995761 36995860
refGene_gn=LUC7L3;refGene_tx=XM_006940206.4;refGene_gn=LUC7L3;refGene_tx=XM_006940207.4;refGene_gn=LUC7L3;refGene_tx=XM_006940208.4;refGene_gn=LUC7L3;refGene_tx=XM_011289194.3;refGene_gn=LUC7L3;refGene_tx=XM_011289195.3;refGene_gn=LUC7L3;refGene_tx=XM_019817690.2;refGene_gn=LUC7L3;refGene_tx=XM_019817691.2
E1 37005766 37005900 refGene_gn=ANKRD40;refGene_tx=XM_023243551.1
E1 37011122 37011271 refGene_gn=ANKRD40;refGene_tx=XM_023243551.1
E1 37012429 37012915 refGene_gn=ANKRD40;refGene_tx=XM_023243551.1
E1 37015681 37015863 refGene_gn=ANKRD40;refGene_tx=XM_023243551.1
E1 37017431 37017578 refGene_gn=ANKRD40;refGene_tx=XM_023243551.1
E1 37022166 37022275 refGene_gn=ABCC3;refGene_tx=XM_023243461.1
E1 37024779 37024974 refGene_gn=ABCC3;refGene_tx=XM_023243461.1
E1 37027501 37027668 refGene_gn=ABCC3;refGene_tx=XM_023243461.1
E1 37028227 37028386 refGene_gn=ABCC3;refGene_tx=XM_023243461.1
E1 37028525 37028672 refGene_gn=ABCC3;refGene_tx=XM_023243461.1
E1 37031876 37031978 refGene_gn=ABCC3;refGene_tx=XM_023243461.1
E1 37033275 37033402 refGene_gn=ABCC3;refGene_tx=XM_023243461.1

E1	37033560	37033760	refGene_gn=ABCC3;refGene_tx=XM_023243461.1
E1	37035085	37035396	refGene_gn=ABCC3;refGene_tx=XM_023243461.1
E1	37035685	37035897	refGene_gn=ABCC3;refGene_tx=XM_023243461.1
E1	37035982	37036126	refGene_gn=ABCC3;refGene_tx=XM_023243461.1
E1	37036302	37036417	refGene_gn=ABCC3;refGene_tx=XM_023243461.1
E1	37037470	37037663	refGene_gn=ABCC3;refGene_tx=XM_023243461.1
E1	37037956	37038124	refGene_gn=ABCC3;refGene_tx=XM_023243461.1
E1	37043285	37043462	refGene_gn=ABCC3;refGene_tx=XM_023243461.1
E1	37043542	37043736	refGene_gn=ABCC3;refGene_tx=XM_023243461.1
E1	37044105	37044193	refGene_gn=ABCC3;refGene_tx=XM_023243461.1
E1	37044568	37044716	refGene_gn=ABCC3;refGene_tx=XM_023243461.1
E1	37044824	37045021	refGene_gn=ABCC3;refGene_tx=XM_023243461.1
E1	37046495	37046604	refGene_gn=ABCC3;refGene_tx=XM_023243461.1
E1	37046797	37047094	refGene_gn=ABCC3;refGene_tx=XM_023243461.1
E1	37050115	37050307	refGene_gn=ABCC3;refGene_tx=XM_023243461.1
E1	37051528	37051577	refGene_gn=ABCC3;refGene_tx=XM_023243461.1
E1	37051579	37051659	refGene_gn=ABCC3;refGene_tx=XM_023243461.1
E1	37052370	37052432	refGene_gn=ABCC3;refGene_tx=XM_023243461.1
E1	37052644	37052696	refGene_gn=ABCC3;refGene_tx=XM_023243461.1
E1	37052697	37052750	refGene_gn=ABCC3;refGene_tx=XM_023243461.1
E1	37053579	37053717	refGene_gn=ABCC3;refGene_tx=XM_023243461.1
E1	37053934	37054060	refGene_gn=ABCC3;refGene_tx=XM_023243461.1
E1	37054674	37054851	refGene_gn=ABCC3;refGene_tx=XM_023243461.1
E1	37065627	37065657	refGene_gn=ABCC3;refGene_tx=XM_023243461.1
E1	37073066	37073801
refGene_gn=CACNA1G;refGene_tx=XM_023244164.1;refGene_gn=CACNA1G;refGene_tx=XM_023244165.1;refGene_gn=CACNA1G;refGene_tx=XM_023244166.1;refGene_gn=CACNA1G;refGene_tx=XM_023244167.1;refGene_gn=CACNA1G;refGene_tx=XM_023244168.1;refGene_gn=CACNA1G;refGene_tx=XM_023244169.1;refGene_gn=CACNA1G;refGene_tx=XM_023244170.1;refGene_gn=CACNA1G;refGene_tx=XM_023244171.1;refGene_gn=CACNA1G;refGene_tx=XM_023244172.1;refGene_gn=CACNA1G;refGene_tx=XM_023244173.1;refGene_gn=CACNA1G;refGene_tx=XM_023244174.1;refGene_gn=CACNA1G;refGene_tx=XM_023244175.1;refGene_gn=CACNA1G;refGene_tx=XM_023244176.1;refGene_gn=CACNA1G;refGene_tx=XM_023244177.1;refGene_gn=CACNA1G;refGene_tx=XM_023244178.1;refGene_gn=CACNA1G;refGene_tx=XM_023244179.1;refGene_gn=CACNA1G;refGene_tx=XM_023244180.1;refGene_gn=CACNA1G;refGene_tx=XM_023244181.1;refGene_gn=CACNA1G;refGene_tx=XM_023244182.1;refGene_gn=CACNA1G;refGene_tx=XM_023244183.1;refGene_gn=CACNA1G;refGene_tx=XM_023244184.1;refGene_gn=CACNA1G;refGene_tx=XM_023244185.1;refGene_gn=CACNA1G;refGene_tx=XM_023244186.1;refGene_gn=CACNA1G;refGene_tx=XM_023244187.1
E1	37074980	37075150
refGene_gn=CACNA1G;refGene_tx=XM_023244164.1;refGene_gn=CACNA1G;refGene_tx=XM_023244165.1;refGene_gn=CACNA1G;refGene_tx=XM_023244166.1;refGene_gn=CACNA1G;refGene_tx=XM_023244167.1;refGene_gn=CACNA1G;refGene_tx=XM_023244168.1;refGene_gn=CACNA1G;refGene_tx=XM_023244169.1;refGene_gn=CACNA1G;refGene_tx=XM_023244170.1;refGene_gn=CACNA1G;refGene_tx=XM_023244171.1;refGene_gn=CACNA1G;refGene_tx=XM_023244172.1;refGene_gn=CACNA1G;refGene_tx=XM_023244173.1;refGene_gn=CACNA1G;refGene_tx=XM_023244174.1;refGene_gn=CACNA1G;refGene_tx=XM_023244175.1;refGene_gn=CACNA1G;r

efGene_tx=XM_023244176.1;refGene_gn=CACNA1G;refGene_tx=XM_023244177.1;refGene_gn=CACNA1G;refGene_tx=XM_023244178.1;refGene_gn=CACNA1G;refGene_tx=XM_023244179.1;refGene_gn=CACNA1G;refGene_tx=XM_023244180.1;refGene_gn=CACNA1G;refGene_tx=XM_023244181.1;refGene_gn=CACNA1G;refGene_tx=XM_023244182.1;refGene_gn=CACNA1G;refGene_tx=XM_023244183.1;refGene_gn=CACNA1G;refGene_tx=XM_023244184.1;refGene_gn=CACNA1G;refGene_tx=XM_023244185.1;refGene_gn=CACNA1G;refGene_tx=XM_023244186.1;refGene_gn=CACNA1G;refGene_tx=XM_023244187.1

E1 37075447 37075625
refGene_gn=CACNA1G;refGene_tx=XM_023244164.1;refGene_gn=CACNA1G;refGene_tx=XM_023244165.1;refGene_gn=CACNA1G;refGene_tx=XM_023244166.1;refGene_gn=CACNA1G;refGene_tx=XM_023244167.1;refGene_gn=CACNA1G;refGene_tx=XM_023244168.1;refGene_gn=CACNA1G;refGene_tx=XM_023244169.1;refGene_gn=CACNA1G;refGene_tx=XM_023244170.1;refGene_gn=CACNA1G;refGene_tx=XM_023244171.1;refGene_gn=CACNA1G;refGene_tx=XM_023244172.1;refGene_gn=CACNA1G;refGene_tx=XM_023244173.1;refGene_gn=CACNA1G;refGene_tx=XM_023244174.1;refGene_gn=CACNA1G;refGene_tx=XM_023244175.1;refGene_gn=CACNA1G;refGene_tx=XM_023244176.1;refGene_gn=CACNA1G;refGene_tx=XM_023244177.1;refGene_gn=CACNA1G;refGene_tx=XM_023244178.1;refGene_gn=CACNA1G;refGene_tx=XM_023244179.1;refGene_gn=CACNA1G;refGene_tx=XM_023244180.1;refGene_gn=CACNA1G;refGene_tx=XM_023244181.1;refGene_gn=CACNA1G;refGene_tx=XM_023244182.1;refGene_gn=CACNA1G;refGene_tx=XM_023244183.1;refGene_gn=CACNA1G;refGene_tx=XM_023244184.1;refGene_gn=CACNA1G;refGene_tx=XM_023244185.1;refGene_gn=CACNA1G;refGene_tx=XM_023244186.1;refGene_gn=CACNA1G;refGene_tx=XM_023244187.1

E1 37077352 37077487
refGene_gn=CACNA1G;refGene_tx=XM_023244164.1;refGene_gn=CACNA1G;refGene_tx=XM_023244165.1;refGene_gn=CACNA1G;refGene_tx=XM_023244166.1;refGene_gn=CACNA1G;refGene_tx=XM_023244167.1;refGene_gn=CACNA1G;refGene_tx=XM_023244168.1;refGene_gn=CACNA1G;refGene_tx=XM_023244169.1;refGene_gn=CACNA1G;refGene_tx=XM_023244170.1;refGene_gn=CACNA1G;refGene_tx=XM_023244172.1;refGene_gn=CACNA1G;refGene_tx=XM_023244176.1;refGene_gn=CACNA1G;refGene_tx=XM_023244187.1

E1 37079445 37079589
refGene_gn=CACNA1G;refGene_tx=XM_023244164.1;refGene_gn=CACNA1G;refGene_tx=XM_023244165.1;refGene_gn=CACNA1G;refGene_tx=XM_023244166.1;refGene_gn=CACNA1G;refGene_tx=XM_023244167.1;refGene_gn=CACNA1G;refGene_tx=XM_023244168.1;refGene_gn=CACNA1G;refGene_tx=XM_023244169.1;refGene_gn=CACNA1G;refGene_tx=XM_023244170.1;refGene_gn=CACNA1G;refGene_tx=XM_023244171.1;refGene_gn=CACNA1G;refGene_tx=XM_023244173.1;refGene_gn=CACNA1G;refGene_tx=XM_023244174.1;refGene_gn=CACNA1G;refGene_tx=XM_023244175.1;refGene_gn=CACNA1G;refGene_tx=XM_023244177.1;refGene_gn=CACNA1G;refGene_tx=XM_023244187.1

E1 37080234 37080573
refGene_gn=CACNA1G;refGene_tx=XM_023244164.1;refGene_gn=CACNA1G;refGene_tx=XM_023244165.1;refGene_gn=CACNA1G;refGene_tx=XM_023244166.1;refGene_gn=CACNA1G;refGene_tx=XM_023244167.1;refGene_gn=CACNA1G;refGene_tx=XM_023244168.1;refGene_gn=CACNA1G;refGene_tx=XM_023244169.1;refGene_gn=CACNA1G;refGene_tx=XM_023244170.1;refGene_gn=CACNA1G;refGene_tx=XM_023244171.1;refGene_gn=CACNA1G;refGene_tx=XM_023244172.1;refGene_gn=CACN

A1G;refGene_tx=XM_023244173.1;refGene_gn=CACNA1G;refGene_tx=XM_0232441
74.1;refGene_gn=CACNA1G;refGene_tx=XM_023244175.1;refGene_gn=CACNA1G;r
efGene_tx=XM_023244176.1;refGene_gn=CACNA1G;refGene_tx=XM_023244177.1;
refGene_gn=CACNA1G;refGene_tx=XM_023244178.1;refGene_gn=CACNA1G;refGen
e_tx=XM_023244179.1;refGene_gn=CACNA1G;refGene_tx=XM_023244180.1;refGe
ne_gn=CACNA1G;refGene_tx=XM_023244181.1;refGene_gn=CACNA1G;refGene_tx=
XM_023244182.1;refGene_gn=CACNA1G;refGene_tx=XM_023244183.1;refGene_gn
=CACNA1G;refGene_tx=XM_023244184.1;refGene_gn=CACNA1G;refGene_tx=XM_02
3244185.1;refGene_gn=CACNA1G;refGene_tx=XM_023244186.1;refGene_gn=CACN
A1G;refGene_tx=XM_023244187.1
E1 37080776 37080898
refGene_gn=CACNA1G;refGene_tx=XM_023244164.1;refGene_gn=CACNA1G;refGen
e_tx=XM_023244165.1;refGene_gn=CACNA1G;refGene_tx=XM_023244166.1;refGe
ne_gn=CACNA1G;refGene_tx=XM_023244167.1;refGene_gn=CACNA1G;refGene_tx=
XM_023244168.1;refGene_gn=CACNA1G;refGene_tx=XM_023244169.1;refGene_gn
=CACNA1G;refGene_tx=XM_023244170.1;refGene_gn=CACNA1G;refGene_tx=XM_02
3244171.1;refGene_gn=CACNA1G;refGene_tx=XM_023244172.1;refGene_gn=CACN
A1G;refGene_tx=XM_023244173.1;refGene_gn=CACNA1G;refGene_tx=XM_0232441
74.1;refGene_gn=CACNA1G;refGene_tx=XM_023244175.1;refGene_gn=CACNA1G;r
efGene_tx=XM_023244176.1;refGene_gn=CACNA1G;refGene_tx=XM_023244177.1;
refGene_gn=CACNA1G;refGene_tx=XM_023244178.1;refGene_gn=CACNA1G;refGen
e_tx=XM_023244179.1;refGene_gn=CACNA1G;refGene_tx=XM_023244180.1;refGe
ne_gn=CACNA1G;refGene_tx=XM_023244181.1;refGene_gn=CACNA1G;refGene_tx=
XM_023244182.1;refGene_gn=CACNA1G;refGene_tx=XM_023244183.1;refGene_gn
=CACNA1G;refGene_tx=XM_023244184.1;refGene_gn=CACNA1G;refGene_tx=XM_02
3244185.1;refGene_gn=CACNA1G;refGene_tx=XM_023244186.1;refGene_gn=CACN
A1G;refGene_tx=XM_023244187.1;refGene_gn=CACNA1G;refGene_tx=XM_0232441
88.1
E1 37080992 37081071
refGene_gn=CACNA1G;refGene_tx=XM_023244164.1;refGene_gn=CACNA1G;refGen
e_tx=XM_023244165.1;refGene_gn=CACNA1G;refGene_tx=XM_023244166.1;refGe
ne_gn=CACNA1G;refGene_tx=XM_023244167.1;refGene_gn=CACNA1G;refGene_tx=
XM_023244168.1;refGene_gn=CACNA1G;refGene_tx=XM_023244169.1;refGene_gn
=CACNA1G;refGene_tx=XM_023244170.1;refGene_gn=CACNA1G;refGene_tx=XM_02
3244171.1;refGene_gn=CACNA1G;refGene_tx=XM_023244172.1;refGene_gn=CACN
A1G;refGene_tx=XM_023244173.1;refGene_gn=CACNA1G;refGene_tx=XM_0232441
74.1;refGene_gn=CACNA1G;refGene_tx=XM_023244175.1;refGene_gn=CACNA1G;r
efGene_tx=XM_023244176.1;refGene_gn=CACNA1G;refGene_tx=XM_023244177.1;
refGene_gn=CACNA1G;refGene_tx=XM_023244178.1;refGene_gn=CACNA1G;refGen
e_tx=XM_023244179.1;refGene_gn=CACNA1G;refGene_tx=XM_023244180.1;refGe
ne_gn=CACNA1G;refGene_tx=XM_023244181.1;refGene_gn=CACNA1G;refGene_tx=
XM_023244182.1;refGene_gn=CACNA1G;refGene_tx=XM_023244183.1;refGene_gn
=CACNA1G;refGene_tx=XM_023244184.1;refGene_gn=CACNA1G;refGene_tx=XM_02
3244185.1;refGene_gn=CACNA1G;refGene_tx=XM_023244186.1;refGene_gn=CACN
A1G;refGene_tx=XM_023244187.1;refGene_gn=CACNA1G;refGene_tx=XM_0232441
88.1
E1 37081198 37081269
refGene_gn=CACNA1G;refGene_tx=XM_023244164.1;refGene_gn=CACNA1G;refGen
e_tx=XM_023244165.1;refGene_gn=CACNA1G;refGene_tx=XM_023244166.1;refGe
ne_gn=CACNA1G;refGene_tx=XM_023244167.1;refGene_gn=CACNA1G;refGene_tx=

XM_023244168.1;refGene_gn=CACNA1G;refGene_tx=XM_023244169.1;refGene_gn
=CACNA1G;refGene_tx=XM_023244170.1;refGene_gn=CACNA1G;refGene_tx=XM_02
3244171.1;refGene_gn=CACNA1G;refGene_tx=XM_023244172.1;refGene_gn=CACN
A1G;refGene_tx=XM_023244173.1;refGene_gn=CACNA1G;refGene_tx=XM_0232441
74.1;refGene_gn=CACNA1G;refGene_tx=XM_023244175.1;refGene_gn=CACNA1G;r
efGene_tx=XM_023244176.1;refGene_gn=CACNA1G;refGene_tx=XM_023244177.1;
refGene_gn=CACNA1G;refGene_tx=XM_023244178.1;refGene_gn=CACNA1G;refGen
e_tx=XM_023244179.1;refGene_gn=CACNA1G;refGene_tx=XM_023244180.1;refGe
ne_gn=CACNA1G;refGene_tx=XM_023244181.1;refGene_gn=CACNA1G;refGene_tx=
XM_023244182.1;refGene_gn=CACNA1G;refGene_tx=XM_023244183.1;refGene_gn
=CACNA1G;refGene_tx=XM_023244184.1;refGene_gn=CACNA1G;refGene_tx=XM_02
3244185.1;refGene_gn=CACNA1G;refGene_tx=XM_023244186.1;refGene_gn=CACN
A1G;refGene_tx=XM_023244187.1;refGene_gn=CACNA1G;refGene_tx=XM_0232441
88.1
E1 37081558 37081692
refGene_gn=CACNA1G;refGene_tx=XM_023244164.1;refGene_gn=CACNA1G;refGen
e_tx=XM_023244165.1;refGene_gn=CACNA1G;refGene_tx=XM_023244166.1;refGe
ne_gn=CACNA1G;refGene_tx=XM_023244167.1;refGene_gn=CACNA1G;refGene_tx=
XM_023244168.1;refGene_gn=CACNA1G;refGene_tx=XM_023244169.1;refGene_gn
=CACNA1G;refGene_tx=XM_023244170.1;refGene_gn=CACNA1G;refGene_tx=XM_02
3244171.1;refGene_gn=CACNA1G;refGene_tx=XM_023244172.1;refGene_gn=CACN
A1G;refGene_tx=XM_023244173.1;refGene_gn=CACNA1G;refGene_tx=XM_0232441
74.1;refGene_gn=CACNA1G;refGene_tx=XM_023244175.1;refGene_gn=CACNA1G;r
efGene_tx=XM_023244176.1;refGene_gn=CACNA1G;refGene_tx=XM_023244177.1;
refGene_gn=CACNA1G;refGene_tx=XM_023244178.1;refGene_gn=CACNA1G;refGen
e_tx=XM_023244179.1;refGene_gn=CACNA1G;refGene_tx=XM_023244180.1;refGe
ne_gn=CACNA1G;refGene_tx=XM_023244181.1;refGene_gn=CACNA1G;refGene_tx=
XM_023244182.1;refGene_gn=CACNA1G;refGene_tx=XM_023244183.1;refGene_gn
=CACNA1G;refGene_tx=XM_023244184.1;refGene_gn=CACNA1G;refGene_tx=XM_02
3244185.1;refGene_gn=CACNA1G;refGene_tx=XM_023244186.1;refGene_gn=CACN
A1G;refGene_tx=XM_023244187.1;refGene_gn=CACNA1G;refGene_tx=XM_0232441
88.1
E1 37082431 37082541
refGene_gn=CACNA1G;refGene_tx=XM_023244164.1;refGene_gn=CACNA1G;refGen
e_tx=XM_023244165.1;refGene_gn=CACNA1G;refGene_tx=XM_023244166.1;refGe
ne_gn=CACNA1G;refGene_tx=XM_023244167.1;refGene_gn=CACNA1G;refGene_tx=
XM_023244168.1;refGene_gn=CACNA1G;refGene_tx=XM_023244169.1;refGene_gn
=CACNA1G;refGene_tx=XM_023244170.1;refGene_gn=CACNA1G;refGene_tx=XM_02
3244171.1;refGene_gn=CACNA1G;refGene_tx=XM_023244172.1;refGene_gn=CACN
A1G;refGene_tx=XM_023244173.1;refGene_gn=CACNA1G;refGene_tx=XM_0232441
74.1;refGene_gn=CACNA1G;refGene_tx=XM_023244175.1;refGene_gn=CACNA1G;r
efGene_tx=XM_023244176.1;refGene_gn=CACNA1G;refGene_tx=XM_023244177.1;
refGene_gn=CACNA1G;refGene_tx=XM_023244178.1;refGene_gn=CACNA1G;refGen
e_tx=XM_023244179.1;refGene_gn=CACNA1G;refGene_tx=XM_023244180.1;refGe
ne_gn=CACNA1G;refGene_tx=XM_023244181.1;refGene_gn=CACNA1G;refGene_tx=
XM_023244182.1;refGene_gn=CACNA1G;refGene_tx=XM_023244183.1;refGene_gn
=CACNA1G;refGene_tx=XM_023244184.1;refGene_gn=CACNA1G;refGene_tx=XM_02
3244185.1;refGene_gn=CACNA1G;refGene_tx=XM_023244186.1;refGene_gn=CACN
A1G;refGene_tx=XM_023244187.1;refGene_gn=CACNA1G;refGene_tx=XM_0232441
88.1

E1 37083266 37083418
refGene_gn=CACNA1G;refGene_tx=XM_023244164.1;refGene_gn=CACNA1G;refGene_tx=XM_023244165.1;refGene_gn=CACNA1G;refGene_tx=XM_023244166.1;refGene_gn=CACNA1G;refGene_tx=XM_023244167.1;refGene_gn=CACNA1G;refGene_tx=XM_023244168.1;refGene_gn=CACNA1G;refGene_tx=XM_023244169.1;refGene_gn=CACNA1G;refGene_tx=XM_023244170.1;refGene_gn=CACNA1G;refGene_tx=XM_023244171.1;refGene_gn=CACNA1G;refGene_tx=XM_023244172.1;refGene_gn=CACNA1G;refGene_tx=XM_023244173.1;refGene_gn=CACNA1G;refGene_tx=XM_023244174.1;refGene_gn=CACNA1G;refGene_tx=XM_023244175.1;refGene_gn=CACNA1G;refGene_tx=XM_023244176.1;refGene_gn=CACNA1G;refGene_tx=XM_023244177.1;refGene_gn=CACNA1G;refGene_tx=XM_023244178.1;refGene_gn=CACNA1G;refGene_tx=XM_023244179.1;refGene_gn=CACNA1G;refGene_tx=XM_023244180.1;refGene_gn=CACNA1G;refGene_tx=XM_023244181.1;refGene_gn=CACNA1G;refGene_tx=XM_023244182.1;refGene_gn=CACNA1G;refGene_tx=XM_023244183.1;refGene_gn=CACNA1G;refGene_tx=XM_023244184.1;refGene_gn=CACNA1G;refGene_tx=XM_023244185.1;refGene_gn=CACNA1G;refGene_tx=XM_023244186.1;refGene_gn=CACNA1G;refGene_tx=XM_023244187.1;refGene_gn=CACNA1G;refGene_tx=XM_023244188.1
E1 37088424 37088478
refGene_gn=CACNA1G;refGene_tx=XM_023244164.1;refGene_gn=CACNA1G;refGene_tx=XM_023244165.1;refGene_gn=CACNA1G;refGene_tx=XM_023244166.1;refGene_gn=CACNA1G;refGene_tx=XM_023244168.1;refGene_gn=CACNA1G;refGene_tx=XM_023244171.1;refGene_gn=CACNA1G;refGene_tx=XM_023244172.1;refGene_gn=CACNA1G;refGene_tx=XM_023244173.1;refGene_gn=CACNA1G;refGene_tx=XM_023244176.1;refGene_gn=CACNA1G;refGene_tx=XM_023244178.1;refGene_gn=CACNA1G;refGene_tx=XM_023244179.1;refGene_gn=CACNA1G;refGene_tx=XM_023244181.1;refGene_gn=CACNA1G;refGene_tx=XM_023244183.1;refGene_gn=CACNA1G;refGene_tx=XM_023244185.1;refGene_gn=CACNA1G;refGene_tx=XM_023244187.1;refGene_gn=CACNA1G;refGene_tx=XM_023244188.1
E1 37090348 37090562
refGene_gn=CACNA1G;refGene_tx=XM_023244164.1;refGene_gn=CACNA1G;refGene_tx=XM_023244165.1;refGene_gn=CACNA1G;refGene_tx=XM_023244166.1;refGene_gn=CACNA1G;refGene_tx=XM_023244167.1;refGene_gn=CACNA1G;refGene_tx=XM_023244168.1;refGene_gn=CACNA1G;refGene_tx=XM_023244169.1;refGene_gn=CACNA1G;refGene_tx=XM_023244170.1;refGene_gn=CACNA1G;refGene_tx=XM_023244171.1;refGene_gn=CACNA1G;refGene_tx=XM_023244172.1;refGene_gn=CACNA1G;refGene_tx=XM_023244173.1;refGene_gn=CACNA1G;refGene_tx=XM_023244174.1;refGene_gn=CACNA1G;refGene_tx=XM_023244175.1;refGene_gn=CACNA1G;refGene_tx=XM_023244176.1;refGene_gn=CACNA1G;refGene_tx=XM_023244177.1;refGene_gn=CACNA1G;refGene_tx=XM_023244178.1;refGene_gn=CACNA1G;refGene_tx=XM_023244179.1;refGene_gn=CACNA1G;refGene_tx=XM_023244180.1;refGene_gn=CACNA1G;refGene_tx=XM_023244181.1;refGene_gn=CACNA1G;refGene_tx=XM_023244182.1;refGene_gn=CACNA1G;refGene_tx=XM_023244183.1;refGene_gn=CACNA1G;refGene_tx=XM_023244184.1;refGene_gn=CACNA1G;refGene_tx=XM_023244185.1;refGene_gn=CACNA1G;refGene_tx=XM_023244186.1;refGene_gn=CACNA1G;refGene_tx=XM_023244187.1;refGene_gn=CACNA1G;refGene_tx=XM_023244188.1
E1 37091112 37091202
refGene_gn=CACNA1G;refGene_tx=XM_023244164.1;refGene_gn=CACNA1G;refGene_tx=XM_023244165.1;refGene_gn=CACNA1G;refGene_tx=XM_023244166.1;refGe

ne_gn=CACNA1G;refGene_tx=XM_023244167.1;refGene_gn=CACNA1G;refGene_tx=
XM_023244168.1;refGene_gn=CACNA1G;refGene_tx=XM_023244169.1;refGene_gn
=CACNA1G;refGene_tx=XM_023244170.1;refGene_gn=CACNA1G;refGene_tx=XM_02
3244171.1;refGene_gn=CACNA1G;refGene_tx=XM_023244172.1;refGene_gn=CACN
A1G;refGene_tx=XM_023244173.1;refGene_gn=CACNA1G;refGene_tx=XM_0232441
74.1;refGene_gn=CACNA1G;refGene_tx=XM_023244175.1;refGene_gn=CACNA1G;r
efGene_tx=XM_023244176.1;refGene_gn=CACNA1G;refGene_tx=XM_023244177.1;
refGene_gn=CACNA1G;refGene_tx=XM_023244178.1;refGene_gn=CACNA1G;refGen
e_tx=XM_023244179.1;refGene_gn=CACNA1G;refGene_tx=XM_023244180.1;refGe
ne_gn=CACNA1G;refGene_tx=XM_023244181.1;refGene_gn=CACNA1G;refGene_tx=
XM_023244182.1;refGene_gn=CACNA1G;refGene_tx=XM_023244183.1;refGene_gn
=CACNA1G;refGene_tx=XM_023244184.1;refGene_gn=CACNA1G;refGene_tx=XM_02
3244185.1;refGene_gn=CACNA1G;refGene_tx=XM_023244186.1;refGene_gn=CACN
A1G;refGene_tx=XM_023244187.1;refGene_gn=CACNA1G;refGene_tx=XM_0232441
88.1
E1 37091830 37091956
refGene_gn=CACNA1G;refGene_tx=XM_023244164.1;refGene_gn=CACNA1G;refGen
e_tx=XM_023244165.1;refGene_gn=CACNA1G;refGene_tx=XM_023244166.1;refGe
ne_gn=CACNA1G;refGene_tx=XM_023244167.1;refGene_gn=CACNA1G;refGene_tx=
XM_023244168.1;refGene_gn=CACNA1G;refGene_tx=XM_023244169.1;refGene_gn
=CACNA1G;refGene_tx=XM_023244170.1;refGene_gn=CACNA1G;refGene_tx=XM_02
3244171.1;refGene_gn=CACNA1G;refGene_tx=XM_023244172.1;refGene_gn=CACN
A1G;refGene_tx=XM_023244173.1;refGene_gn=CACNA1G;refGene_tx=XM_0232441
74.1;refGene_gn=CACNA1G;refGene_tx=XM_023244175.1;refGene_gn=CACNA1G;r
efGene_tx=XM_023244176.1;refGene_gn=CACNA1G;refGene_tx=XM_023244177.1;
refGene_gn=CACNA1G;refGene_tx=XM_023244178.1;refGene_gn=CACNA1G;refGen
e_tx=XM_023244179.1;refGene_gn=CACNA1G;refGene_tx=XM_023244180.1;refGe
ne_gn=CACNA1G;refGene_tx=XM_023244181.1;refGene_gn=CACNA1G;refGene_tx=
XM_023244182.1;refGene_gn=CACNA1G;refGene_tx=XM_023244183.1;refGene_gn
=CACNA1G;refGene_tx=XM_023244184.1;refGene_gn=CACNA1G;refGene_tx=XM_02
3244185.1;refGene_gn=CACNA1G;refGene_tx=XM_023244186.1;refGene_gn=CACN
A1G;refGene_tx=XM_023244187.1;refGene_gn=CACNA1G;refGene_tx=XM_0232441
88.1
E1 37093487 37093614
refGene_gn=CACNA1G;refGene_tx=XM_023244164.1;refGene_gn=CACNA1G;refGen
e_tx=XM_023244165.1;refGene_gn=CACNA1G;refGene_tx=XM_023244166.1;refGe
ne_gn=CACNA1G;refGene_tx=XM_023244167.1;refGene_gn=CACNA1G;refGene_tx=
XM_023244168.1;refGene_gn=CACNA1G;refGene_tx=XM_023244169.1;refGene_gn
=CACNA1G;refGene_tx=XM_023244170.1;refGene_gn=CACNA1G;refGene_tx=XM_02
3244171.1;refGene_gn=CACNA1G;refGene_tx=XM_023244172.1;refGene_gn=CACN
A1G;refGene_tx=XM_023244173.1;refGene_gn=CACNA1G;refGene_tx=XM_0232441
74.1;refGene_gn=CACNA1G;refGene_tx=XM_023244175.1;refGene_gn=CACNA1G;r
efGene_tx=XM_023244176.1;refGene_gn=CACNA1G;refGene_tx=XM_023244177.1;
refGene_gn=CACNA1G;refGene_tx=XM_023244178.1;refGene_gn=CACNA1G;refGen
e_tx=XM_023244179.1;refGene_gn=CACNA1G;refGene_tx=XM_023244180.1;refGe
ne_gn=CACNA1G;refGene_tx=XM_023244181.1;refGene_gn=CACNA1G;refGene_tx=
XM_023244182.1;refGene_gn=CACNA1G;refGene_tx=XM_023244183.1;refGene_gn
=CACNA1G;refGene_tx=XM_023244184.1;refGene_gn=CACNA1G;refGene_tx=XM_02
3244185.1;refGene_gn=CACNA1G;refGene_tx=XM_023244186.1;refGene_gn=CACN
A1G;refGene_tx=XM_023244187.1;refGene_gn=CACNA1G;refGene_tx=XM_0232441

88.1
E1 37094523 37094708
refGene_gn=CACNA1G;refGene_tx=XM_023244164.1;refGene_gn=CACNA1G;refGene_tx=XM_023244165.1;refGene_gn=CACNA1G;refGene_tx=XM_023244166.1;refGene_gn=CACNA1G;refGene_tx=XM_023244167.1;refGene_gn=CACNA1G;refGene_tx=XM_023244168.1;refGene_gn=CACNA1G;refGene_tx=XM_023244169.1;refGene_gn=CACNA1G;refGene_tx=XM_023244170.1;refGene_gn=CACNA1G;refGene_tx=XM_023244171.1;refGene_gn=CACNA1G;refGene_tx=XM_023244172.1;refGene_gn=CACNA1G;refGene_tx=XM_023244173.1;refGene_gn=CACNA1G;refGene_tx=XM_023244174.1;refGene_gn=CACNA1G;refGene_tx=XM_023244175.1;refGene_gn=CACNA1G;refGene_tx=XM_023244176.1;refGene_gn=CACNA1G;refGene_tx=XM_023244177.1;refGene_gn=CACNA1G;refGene_tx=XM_023244178.1;refGene_gn=CACNA1G;refGene_tx=XM_023244179.1;refGene_gn=CACNA1G;refGene_tx=XM_023244180.1;refGene_gn=CACNA1G;refGene_tx=XM_023244181.1;refGene_gn=CACNA1G;refGene_tx=XM_023244182.1;refGene_gn=CACNA1G;refGene_tx=XM_023244183.1;refGene_gn=CACNA1G;refGene_tx=XM_023244184.1;refGene_gn=CACNA1G;refGene_tx=XM_023244185.1;refGene_gn=CACNA1G;refGene_tx=XM_023244186.1;refGene_gn=CACNA1G;refGene_tx=XM_023244187.1;refGene_gn=CACNA1G;refGene_tx=XM_023244188.1
E1 37094858 37094927
refGene_gn=CACNA1G;refGene_tx=XM_023244164.1;refGene_gn=CACNA1G;refGene_tx=XM_023244165.1;refGene_gn=CACNA1G;refGene_tx=XM_023244166.1;refGene_gn=CACNA1G;refGene_tx=XM_023244167.1;refGene_gn=CACNA1G;refGene_tx=XM_023244168.1;refGene_gn=CACNA1G;refGene_tx=XM_023244169.1;refGene_gn=CACNA1G;refGene_tx=XM_023244170.1;refGene_gn=CACNA1G;refGene_tx=XM_023244171.1;refGene_gn=CACNA1G;refGene_tx=XM_023244172.1;refGene_gn=CACNA1G;refGene_tx=XM_023244173.1;refGene_gn=CACNA1G;refGene_tx=XM_023244174.1;refGene_gn=CACNA1G;refGene_tx=XM_023244175.1;refGene_gn=CACNA1G;refGene_tx=XM_023244176.1;refGene_gn=CACNA1G;refGene_tx=XM_023244177.1;refGene_gn=CACNA1G;refGene_tx=XM_023244178.1;refGene_gn=CACNA1G;refGene_tx=XM_023244179.1;refGene_gn=CACNA1G;refGene_tx=XM_023244180.1;refGene_gn=CACNA1G;refGene_tx=XM_023244181.1;refGene_gn=CACNA1G;refGene_tx=XM_023244182.1;refGene_gn=CACNA1G;refGene_tx=XM_023244183.1;refGene_gn=CACNA1G;refGene_tx=XM_023244184.1;refGene_gn=CACNA1G;refGene_tx=XM_023244185.1;refGene_gn=CACNA1G;refGene_tx=XM_023244186.1;refGene_gn=CACNA1G;refGene_tx=XM_023244187.1;refGene_gn=CACNA1G;refGene_tx=XM_023244188.1
E1 37096558 37096682
refGene_gn=CACNA1G;refGene_tx=XM_023244164.1;refGene_gn=CACNA1G;refGene_tx=XM_023244165.1;refGene_gn=CACNA1G;refGene_tx=XM_023244166.1;refGene_gn=CACNA1G;refGene_tx=XM_023244167.1;refGene_gn=CACNA1G;refGene_tx=XM_023244168.1;refGene_gn=CACNA1G;refGene_tx=XM_023244169.1;refGene_gn=CACNA1G;refGene_tx=XM_023244170.1;refGene_gn=CACNA1G;refGene_tx=XM_023244171.1;refGene_gn=CACNA1G;refGene_tx=XM_023244172.1;refGene_gn=CACNA1G;refGene_tx=XM_023244173.1;refGene_gn=CACNA1G;refGene_tx=XM_023244174.1;refGene_gn=CACNA1G;refGene_tx=XM_023244175.1;refGene_gn=CACNA1G;refGene_tx=XM_023244176.1;refGene_gn=CACNA1G;refGene_tx=XM_023244177.1;refGene_gn=CACNA1G;refGene_tx=XM_023244178.1;refGene_gn=CACNA1G;refGene_tx=XM_023244179.1;refGene_gn=CACNA1G;refGene_tx=XM_023244180.1;refGene_gn=CACNA1G;refGene_tx=XM_023244181.1;refGene_gn=CACNA1G;refGene_tx=

XM_023244182.1;refGene_gn=CACNA1G;refGene_tx=XM_023244183.1;refGene_gn=CACNA1G;refGene_tx=XM_023244184.1;refGene_gn=CACNA1G;refGene_tx=XM_023244185.1;refGene_gn=CACNA1G;refGene_tx=XM_023244186.1;refGene_gn=CACNA1G;refGene_tx=XM_023244187.1;refGene_gn=CACNA1G;refGene_tx=XM_023244188.1
E1 37096871 37096972
refGene_gn=CACNA1G;refGene_tx=XM_023244164.1;refGene_gn=CACNA1G;refGene_tx=XM_023244165.1;refGene_gn=CACNA1G;refGene_tx=XM_023244166.1;refGene_gn=CACNA1G;refGene_tx=XM_023244167.1;refGene_gn=CACNA1G;refGene_tx=XM_023244168.1;refGene_gn=CACNA1G;refGene_tx=XM_023244169.1;refGene_gn=CACNA1G;refGene_tx=XM_023244170.1;refGene_gn=CACNA1G;refGene_tx=XM_023244171.1;refGene_gn=CACNA1G;refGene_tx=XM_023244172.1;refGene_gn=CACNA1G;refGene_tx=XM_023244173.1;refGene_gn=CACNA1G;refGene_tx=XM_023244174.1;refGene_gn=CACNA1G;refGene_tx=XM_023244175.1;refGene_gn=CACNA1G;refGene_tx=XM_023244176.1;refGene_gn=CACNA1G;refGene_tx=XM_023244177.1;refGene_gn=CACNA1G;refGene_tx=XM_023244178.1;refGene_gn=CACNA1G;refGene_tx=XM_023244179.1;refGene_gn=CACNA1G;refGene_tx=XM_023244180.1;refGene_gn=CACNA1G;refGene_tx=XM_023244181.1;refGene_gn=CACNA1G;refGene_tx=XM_023244182.1;refGene_gn=CACNA1G;refGene_tx=XM_023244183.1;refGene_gn=CACNA1G;refGene_tx=XM_023244184.1;refGene_gn=CACNA1G;refGene_tx=XM_023244185.1;refGene_gn=CACNA1G;refGene_tx=XM_023244186.1;refGene_gn=CACNA1G;refGene_tx=XM_023244187.1;refGene_gn=CACNA1G;refGene_tx=XM_023244188.1
E1 37097895 37098327
refGene_gn=CACNA1G;refGene_tx=XM_023244164.1;refGene_gn=CACNA1G;refGene_tx=XM_023244165.1;refGene_gn=CACNA1G;refGene_tx=XM_023244166.1;refGene_gn=CACNA1G;refGene_tx=XM_023244167.1;refGene_gn=CACNA1G;refGene_tx=XM_023244168.1;refGene_gn=CACNA1G;refGene_tx=XM_023244169.1;refGene_gn=CACNA1G;refGene_tx=XM_023244170.1;refGene_gn=CACNA1G;refGene_tx=XM_023244171.1;refGene_gn=CACNA1G;refGene_tx=XM_023244172.1;refGene_gn=CACNA1G;refGene_tx=XM_023244173.1;refGene_gn=CACNA1G;refGene_tx=XM_023244174.1;refGene_gn=CACNA1G;refGene_tx=XM_023244175.1;refGene_gn=CACNA1G;refGene_tx=XM_023244176.1;refGene_gn=CACNA1G;refGene_tx=XM_023244177.1;refGene_gn=CACNA1G;refGene_tx=XM_023244178.1;refGene_gn=CACNA1G;refGene_tx=XM_023244179.1;refGene_gn=CACNA1G;refGene_tx=XM_023244180.1;refGene_gn=CACNA1G;refGene_tx=XM_023244181.1;refGene_gn=CACNA1G;refGene_tx=XM_023244182.1;refGene_gn=CACNA1G;refGene_tx=XM_023244183.1;refGene_gn=CACNA1G;refGene_tx=XM_023244184.1;refGene_gn=CACNA1G;refGene_tx=XM_023244185.1;refGene_gn=CACNA1G;refGene_tx=XM_023244186.1;refGene_gn=CACNA1G;refGene_tx=XM_023244187.1;refGene_gn=CACNA1G;refGene_tx=XM_023244188.1
E1 37099098 37099292
refGene_gn=CACNA1G;refGene_tx=XM_023244164.1;refGene_gn=CACNA1G;refGene_tx=XM_023244165.1;refGene_gn=CACNA1G;refGene_tx=XM_023244166.1;refGene_gn=CACNA1G;refGene_tx=XM_023244167.1;refGene_gn=CACNA1G;refGene_tx=XM_023244168.1;refGene_gn=CACNA1G;refGene_tx=XM_023244169.1;refGene_gn=CACNA1G;refGene_tx=XM_023244170.1;refGene_gn=CACNA1G;refGene_tx=XM_023244171.1;refGene_gn=CACNA1G;refGene_tx=XM_023244172.1;refGene_gn=CACNA1G;refGene_tx=XM_023244173.1;refGene_gn=CACNA1G;refGene_tx=XM_023244174.1;refGene_gn=CACNA1G;refGene_tx=XM_023244175.1;refGene_gn=CACNA1G;r

efGene_tx=XM_023244176.1;refGene_gn=CACNA1G;refGene_tx=XM_023244177.1;refGene_gn=CACNA1G;refGene_tx=XM_023244178.1;refGene_gn=CACNA1G;refGene_tx=XM_023244179.1;refGene_gn=CACNA1G;refGene_tx=XM_023244180.1;refGene_gn=CACNA1G;refGene_tx=XM_023244181.1;refGene_gn=CACNA1G;refGene_tx=XM_023244182.1;refGene_gn=CACNA1G;refGene_tx=XM_023244183.1;refGene_gn=CACNA1G;refGene_tx=XM_023244184.1;refGene_gn=CACNA1G;refGene_tx=XM_023244185.1;refGene_gn=CACNA1G;refGene_tx=XM_023244186.1;refGene_gn=CACNA1G;refGene_tx=XM_023244187.1;refGene_gn=CACNA1G;refGene_tx=XM_023244188.1
E1 37099380 37099465
refGene_gn=CACNA1G;refGene_tx=XM_023244164.1;refGene_gn=CACNA1G;refGene_tx=XM_023244165.1;refGene_gn=CACNA1G;refGene_tx=XM_023244166.1;refGene_gn=CACNA1G;refGene_tx=XM_023244167.1;refGene_gn=CACNA1G;refGene_tx=XM_023244168.1;refGene_gn=CACNA1G;refGene_tx=XM_023244169.1;refGene_gn=CACNA1G;refGene_tx=XM_023244170.1;refGene_gn=CACNA1G;refGene_tx=XM_023244171.1;refGene_gn=CACNA1G;refGene_tx=XM_023244172.1;refGene_gn=CACNA1G;refGene_tx=XM_023244173.1;refGene_gn=CACNA1G;refGene_tx=XM_023244174.1;refGene_gn=CACNA1G;refGene_tx=XM_023244175.1;refGene_gn=CACNA1G;refGene_tx=XM_023244176.1;refGene_gn=CACNA1G;refGene_tx=XM_023244177.1;refGene_gn=CACNA1G;refGene_tx=XM_023244178.1;refGene_gn=CACNA1G;refGene_tx=XM_023244179.1;refGene_gn=CACNA1G;refGene_tx=XM_023244180.1;refGene_gn=CACNA1G;refGene_tx=XM_023244181.1;refGene_gn=CACNA1G;refGene_tx=XM_023244182.1;refGene_gn=CACNA1G;refGene_tx=XM_023244183.1;refGene_gn=CACNA1G;refGene_tx=XM_023244184.1;refGene_gn=CACNA1G;refGene_tx=XM_023244185.1;refGene_gn=CACNA1G;refGene_tx=XM_023244186.1;refGene_gn=CACNA1G;refGene_tx=XM_023244187.1;refGene_gn=CACNA1G;refGene_tx=XM_023244188.1
E1 37100958 37101027
refGene_gn=CACNA1G;refGene_tx=XM_023244164.1;refGene_gn=CACNA1G;refGene_tx=XM_023244165.1;refGene_gn=CACNA1G;refGene_tx=XM_023244166.1;refGene_gn=CACNA1G;refGene_tx=XM_023244167.1;refGene_gn=CACNA1G;refGene_tx=XM_023244169.1;refGene_gn=CACNA1G;refGene_tx=XM_023244171.1;refGene_gn=CACNA1G;refGene_tx=XM_023244172.1;refGene_gn=CACNA1G;refGene_tx=XM_023244173.1;refGene_gn=CACNA1G;refGene_tx=XM_023244174.1;refGene_gn=CACNA1G;refGene_tx=XM_023244175.1;refGene_gn=CACNA1G;refGene_tx=XM_023244178.1;refGene_gn=CACNA1G;refGene_tx=XM_023244179.1;refGene_gn=CACNA1G;refGene_tx=XM_023244180.1;refGene_gn=CACNA1G;refGene_tx=XM_023244182.1;refGene_gn=CACNA1G;refGene_tx=XM_023244187.1
E1 37103880 37104036
refGene_gn=CACNA1G;refGene_tx=XM_023244164.1;refGene_gn=CACNA1G;refGene_tx=XM_023244165.1;refGene_gn=CACNA1G;refGene_tx=XM_023244166.1;refGene_gn=CACNA1G;refGene_tx=XM_023244167.1;refGene_gn=CACNA1G;refGene_tx=XM_023244168.1;refGene_gn=CACNA1G;refGene_tx=XM_023244169.1;refGene_gn=CACNA1G;refGene_tx=XM_023244170.1;refGene_gn=CACNA1G;refGene_tx=XM_023244171.1;refGene_gn=CACNA1G;refGene_tx=XM_023244172.1;refGene_gn=CACNA1G;refGene_tx=XM_023244173.1;refGene_gn=CACNA1G;refGene_tx=XM_023244174.1;refGene_gn=CACNA1G;refGene_tx=XM_023244175.1;refGene_gn=CACNA1G;refGene_tx=XM_023244176.1;refGene_gn=CACNA1G;refGene_tx=XM_023244177.1;refGene_gn=CACNA1G;refGene_tx=XM_023244178.1;refGene_gn=CACNA1G;refGene_tx=XM_023244179.1;refGene_gn=CACNA1G;refGene_tx=XM_023244180.1;refGe

ne_gn=CACNA1G;refGene_tx=XM_023244181.1;refGene_gn=CACNA1G;refGene_tx=
XM_023244182.1;refGene_gn=CACNA1G;refGene_tx=XM_023244183.1;refGene_gn
=CACNA1G;refGene_tx=XM_023244184.1;refGene_gn=CACNA1G;refGene_tx=XM_02
3244185.1;refGene_gn=CACNA1G;refGene_tx=XM_023244186.1;refGene_gn=CACN
A1G;refGene_tx=XM_023244187.1;refGene_gn=CACNA1G;refGene_tx=XM_0232441
88.1
E1 37104131 37104246
refGene_gn=CACNA1G;refGene_tx=XM_023244164.1;refGene_gn=CACNA1G;refGen
e_tx=XM_023244165.1;refGene_gn=CACNA1G;refGene_tx=XM_023244166.1;refGe
ne_gn=CACNA1G;refGene_tx=XM_023244167.1;refGene_gn=CACNA1G;refGene_tx=
XM_023244168.1;refGene_gn=CACNA1G;refGene_tx=XM_023244169.1;refGene_gn
=CACNA1G;refGene_tx=XM_023244170.1;refGene_gn=CACNA1G;refGene_tx=XM_02
3244171.1;refGene_gn=CACNA1G;refGene_tx=XM_023244172.1;refGene_gn=CACN
A1G;refGene_tx=XM_023244173.1;refGene_gn=CACNA1G;refGene_tx=XM_0232441
74.1;refGene_gn=CACNA1G;refGene_tx=XM_023244175.1;refGene_gn=CACNA1G;r
efGene_tx=XM_023244176.1;refGene_gn=CACNA1G;refGene_tx=XM_023244177.1;
refGene_gn=CACNA1G;refGene_tx=XM_023244178.1;refGene_gn=CACNA1G;refGen
e_tx=XM_023244179.1;refGene_gn=CACNA1G;refGene_tx=XM_023244180.1;refGe
ne_gn=CACNA1G;refGene_tx=XM_023244181.1;refGene_gn=CACNA1G;refGene_tx=
XM_023244182.1;refGene_gn=CACNA1G;refGene_tx=XM_023244183.1;refGene_gn
=CACNA1G;refGene_tx=XM_023244184.1;refGene_gn=CACNA1G;refGene_tx=XM_02
3244185.1;refGene_gn=CACNA1G;refGene_tx=XM_023244186.1;refGene_gn=CACN
A1G;refGene_tx=XM_023244187.1;refGene_gn=CACNA1G;refGene_tx=XM_0232441
88.1
E1 37104346 37104532
refGene_gn=CACNA1G;refGene_tx=XM_023244164.1;refGene_gn=CACNA1G;refGen
e_tx=XM_023244165.1;refGene_gn=CACNA1G;refGene_tx=XM_023244166.1;refGe
ne_gn=CACNA1G;refGene_tx=XM_023244167.1;refGene_gn=CACNA1G;refGene_tx=
XM_023244168.1;refGene_gn=CACNA1G;refGene_tx=XM_023244169.1;refGene_gn
=CACNA1G;refGene_tx=XM_023244170.1;refGene_gn=CACNA1G;refGene_tx=XM_02
3244171.1;refGene_gn=CACNA1G;refGene_tx=XM_023244172.1;refGene_gn=CACN
A1G;refGene_tx=XM_023244173.1;refGene_gn=CACNA1G;refGene_tx=XM_0232441
74.1;refGene_gn=CACNA1G;refGene_tx=XM_023244175.1;refGene_gn=CACNA1G;r
efGene_tx=XM_023244176.1;refGene_gn=CACNA1G;refGene_tx=XM_023244177.1;
refGene_gn=CACNA1G;refGene_tx=XM_023244178.1;refGene_gn=CACNA1G;refGen
e_tx=XM_023244179.1;refGene_gn=CACNA1G;refGene_tx=XM_023244180.1;refGe
ne_gn=CACNA1G;refGene_tx=XM_023244181.1;refGene_gn=CACNA1G;refGene_tx=
XM_023244182.1;refGene_gn=CACNA1G;refGene_tx=XM_023244183.1;refGene_gn
=CACNA1G;refGene_tx=XM_023244184.1;refGene_gn=CACNA1G;refGene_tx=XM_02
3244185.1;refGene_gn=CACNA1G;refGene_tx=XM_023244186.1;refGene_gn=CACN
A1G;refGene_tx=XM_023244187.1;refGene_gn=CACNA1G;refGene_tx=XM_0232441
88.1
E1 37105213 37105365
refGene_gn=CACNA1G;refGene_tx=XM_023244164.1;refGene_gn=CACNA1G;refGen
e_tx=XM_023244165.1;refGene_gn=CACNA1G;refGene_tx=XM_023244166.1;refGe
ne_gn=CACNA1G;refGene_tx=XM_023244167.1;refGene_gn=CACNA1G;refGene_tx=
XM_023244168.1;refGene_gn=CACNA1G;refGene_tx=XM_023244169.1;refGene_gn
=CACNA1G;refGene_tx=XM_023244170.1;refGene_gn=CACNA1G;refGene_tx=XM_02
3244171.1;refGene_gn=CACNA1G;refGene_tx=XM_023244172.1;refGene_gn=CACN
A1G;refGene_tx=XM_023244173.1;refGene_gn=CACNA1G;refGene_tx=XM_0232441

74.1;refGene_gn=CACNA1G;refGene_tx=XM_023244175.1;refGene_gn=CACNA1G;r
efGene_tx=XM_023244176.1;refGene_gn=CACNA1G;refGene_tx=XM_023244177.1;
refGene_gn=CACNA1G;refGene_tx=XM_023244178.1;refGene_gn=CACNA1G;refGen
e_tx=XM_023244179.1;refGene_gn=CACNA1G;refGene_tx=XM_023244180.1;refGe
ne_gn=CACNA1G;refGene_tx=XM_023244181.1;refGene_gn=CACNA1G;refGene_tx=
XM_023244182.1;refGene_gn=CACNA1G;refGene_tx=XM_023244183.1;refGene_gn
=CACNA1G;refGene_tx=XM_023244184.1;refGene_gn=CACNA1G;refGene_tx=XM_02
3244185.1;refGene_gn=CACNA1G;refGene_tx=XM_023244186.1;refGene_gn=CACN
A1G;refGene_tx=XM_023244187.1;refGene_gn=CACNA1G;refGene_tx=XM_0232441
88.1
E1 37117099 37117476
refGene_gn=CACNA1G;refGene_tx=XM_023244164.1;refGene_gn=CACNA1G;refGen
e_tx=XM_023244165.1;refGene_gn=CACNA1G;refGene_tx=XM_023244166.1;refGe
ne_gn=CACNA1G;refGene_tx=XM_023244167.1;refGene_gn=CACNA1G;refGene_tx=
XM_023244168.1;refGene_gn=CACNA1G;refGene_tx=XM_023244169.1;refGene_gn
=CACNA1G;refGene_tx=XM_023244170.1;refGene_gn=CACNA1G;refGene_tx=XM_02
3244171.1;refGene_gn=CACNA1G;refGene_tx=XM_023244172.1;refGene_gn=CACN
A1G;refGene_tx=XM_023244173.1;refGene_gn=CACNA1G;refGene_tx=XM_0232441
74.1;refGene_gn=CACNA1G;refGene_tx=XM_023244175.1;refGene_gn=CACNA1G;r
efGene_tx=XM_023244176.1;refGene_gn=CACNA1G;refGene_tx=XM_023244177.1;
refGene_gn=CACNA1G;refGene_tx=XM_023244178.1;refGene_gn=CACNA1G;refGen
e_tx=XM_023244179.1;refGene_gn=CACNA1G;refGene_tx=XM_023244180.1;refGe
ne_gn=CACNA1G;refGene_tx=XM_023244181.1;refGene_gn=CACNA1G;refGene_tx=
XM_023244182.1;refGene_gn=CACNA1G;refGene_tx=XM_023244183.1;refGene_gn
=CACNA1G;refGene_tx=XM_023244184.1;refGene_gn=CACNA1G;refGene_tx=XM_02
3244185.1;refGene_gn=CACNA1G;refGene_tx=XM_023244186.1;refGene_gn=CACN
A1G;refGene_tx=XM_023244187.1;refGene_gn=CACNA1G;refGene_tx=XM_0232441
88.1
E1 37119651 37120435
refGene_gn=CACNA1G;refGene_tx=XM_023244164.1;refGene_gn=CACNA1G;refGen
e_tx=XM_023244165.1;refGene_gn=CACNA1G;refGene_tx=XM_023244166.1;refGe
ne_gn=CACNA1G;refGene_tx=XM_023244167.1;refGene_gn=CACNA1G;refGene_tx=
XM_023244168.1;refGene_gn=CACNA1G;refGene_tx=XM_023244169.1;refGene_gn
=CACNA1G;refGene_tx=XM_023244170.1;refGene_gn=CACNA1G;refGene_tx=XM_02
3244171.1;refGene_gn=CACNA1G;refGene_tx=XM_023244172.1;refGene_gn=CACN
A1G;refGene_tx=XM_023244173.1;refGene_gn=CACNA1G;refGene_tx=XM_0232441
74.1;refGene_gn=CACNA1G;refGene_tx=XM_023244175.1;refGene_gn=CACNA1G;r
efGene_tx=XM_023244176.1;refGene_gn=CACNA1G;refGene_tx=XM_023244177.1;
refGene_gn=CACNA1G;refGene_tx=XM_023244178.1;refGene_gn=CACNA1G;refGen
e_tx=XM_023244179.1;refGene_gn=CACNA1G;refGene_tx=XM_023244180.1;refGe
ne_gn=CACNA1G;refGene_tx=XM_023244181.1;refGene_gn=CACNA1G;refGene_tx=
XM_023244182.1;refGene_gn=CACNA1G;refGene_tx=XM_023244183.1;refGene_gn
=CACNA1G;refGene_tx=XM_023244184.1;refGene_gn=CACNA1G;refGene_tx=XM_02
3244185.1;refGene_gn=CACNA1G;refGene_tx=XM_023244186.1;refGene_gn=CACN
A1G;refGene_tx=XM_023244187.1;refGene_gn=CACNA1G;refGene_tx=XM_0232441
88.1
E1 37122370 37122463
refGene_gn=CACNA1G;refGene_tx=XM_023244164.1;refGene_gn=CACNA1G;refGen
e_tx=XM_023244165.1;refGene_gn=CACNA1G;refGene_tx=XM_023244166.1;refGe
ne_gn=CACNA1G;refGene_tx=XM_023244167.1;refGene_gn=CACNA1G;refGene_tx=

XM_023244168.1;refGene_gn=CACNA1G;refGene_tx=XM_023244169.1;refGene_gn
=CACNA1G;refGene_tx=XM_023244170.1;refGene_gn=CACNA1G;refGene_tx=XM_02
3244171.1;refGene_gn=CACNA1G;refGene_tx=XM_023244172.1;refGene_gn=CACN
A1G;refGene_tx=XM_023244173.1;refGene_gn=CACNA1G;refGene_tx=XM_0232441
74.1;refGene_gn=CACNA1G;refGene_tx=XM_023244175.1;refGene_gn=CACNA1G;r
efGene_tx=XM_023244176.1;refGene_gn=CACNA1G;refGene_tx=XM_023244177.1;
refGene_gn=CACNA1G;refGene_tx=XM_023244178.1;refGene_gn=CACNA1G;refGen
e_tx=XM_023244179.1;refGene_gn=CACNA1G;refGene_tx=XM_023244180.1;refGe
ne_gn=CACNA1G;refGene_tx=XM_023244181.1;refGene_gn=CACNA1G;refGene_tx=
XM_023244182.1;refGene_gn=CACNA1G;refGene_tx=XM_023244183.1;refGene_gn
=CACNA1G;refGene_tx=XM_023244184.1;refGene_gn=CACNA1G;refGene_tx=XM_02
3244185.1;refGene_gn=CACNA1G;refGene_tx=XM_023244186.1;refGene_gn=CACN
A1G;refGene_tx=XM_023244187.1;refGene_gn=CACNA1G;refGene_tx=XM_0232441
88.1
E1 37122629 37122930
refGene_gn=CACNA1G;refGene_tx=XM_023244164.1;refGene_gn=CACNA1G;refGen
e_tx=XM_023244165.1;refGene_gn=CACNA1G;refGene_tx=XM_023244166.1;refGe
ne_gn=CACNA1G;refGene_tx=XM_023244167.1;refGene_gn=CACNA1G;refGene_tx=
XM_023244168.1;refGene_gn=CACNA1G;refGene_tx=XM_023244169.1;refGene_gn
=CACNA1G;refGene_tx=XM_023244170.1;refGene_gn=CACNA1G;refGene_tx=XM_02
3244171.1;refGene_gn=CACNA1G;refGene_tx=XM_023244172.1;refGene_gn=CACN
A1G;refGene_tx=XM_023244173.1;refGene_gn=CACNA1G;refGene_tx=XM_0232441
74.1;refGene_gn=CACNA1G;refGene_tx=XM_023244175.1;refGene_gn=CACNA1G;r
efGene_tx=XM_023244176.1;refGene_gn=CACNA1G;refGene_tx=XM_023244177.1;
refGene_gn=CACNA1G;refGene_tx=XM_023244178.1;refGene_gn=CACNA1G;refGen
e_tx=XM_023244179.1;refGene_gn=CACNA1G;refGene_tx=XM_023244180.1;refGe
ne_gn=CACNA1G;refGene_tx=XM_023244181.1;refGene_gn=CACNA1G;refGene_tx=
XM_023244182.1;refGene_gn=CACNA1G;refGene_tx=XM_023244183.1;refGene_gn
=CACNA1G;refGene_tx=XM_023244184.1;refGene_gn=CACNA1G;refGene_tx=XM_02
3244185.1;refGene_gn=CACNA1G;refGene_tx=XM_023244186.1;refGene_gn=CACN
A1G;refGene_tx=XM_023244187.1;refGene_gn=CACNA1G;refGene_tx=XM_0232441
88.1
E1 37123427 37123587
refGene_gn=CACNA1G;refGene_tx=XM_023244164.1;refGene_gn=CACNA1G;refGen
e_tx=XM_023244165.1;refGene_gn=CACNA1G;refGene_tx=XM_023244166.1;refGe
ne_gn=CACNA1G;refGene_tx=XM_023244167.1;refGene_gn=CACNA1G;refGene_tx=
XM_023244168.1;refGene_gn=CACNA1G;refGene_tx=XM_023244169.1;refGene_gn
=CACNA1G;refGene_tx=XM_023244170.1;refGene_gn=CACNA1G;refGene_tx=XM_02
3244171.1;refGene_gn=CACNA1G;refGene_tx=XM_023244172.1;refGene_gn=CACN
A1G;refGene_tx=XM_023244173.1;refGene_gn=CACNA1G;refGene_tx=XM_0232441
74.1;refGene_gn=CACNA1G;refGene_tx=XM_023244175.1;refGene_gn=CACNA1G;r
efGene_tx=XM_023244176.1;refGene_gn=CACNA1G;refGene_tx=XM_023244177.1;
refGene_gn=CACNA1G;refGene_tx=XM_023244178.1;refGene_gn=CACNA1G;refGen
e_tx=XM_023244179.1;refGene_gn=CACNA1G;refGene_tx=XM_023244180.1;refGe
ne_gn=CACNA1G;refGene_tx=XM_023244181.1;refGene_gn=CACNA1G;refGene_tx=
XM_023244182.1;refGene_gn=CACNA1G;refGene_tx=XM_023244183.1;refGene_gn
=CACNA1G;refGene_tx=XM_023244184.1;refGene_gn=CACNA1G;refGene_tx=XM_02
3244185.1;refGene_gn=CACNA1G;refGene_tx=XM_023244186.1;refGene_gn=CACN
A1G;refGene_tx=XM_023244187.1;refGene_gn=CACNA1G;refGene_tx=XM_0232441
88.1

E1 37125720 37125818
refGene_gn=CACNA1G;refGene_tx=XM_023244164.1;refGene_gn=CACNA1G;refGene_tx=XM_023244165.1;refGene_gn=CACNA1G;refGene_tx=XM_023244166.1;refGene_gn=CACNA1G;refGene_tx=XM_023244167.1;refGene_gn=CACNA1G;refGene_tx=XM_023244168.1;refGene_gn=CACNA1G;refGene_tx=XM_023244169.1;refGene_gn=CACNA1G;refGene_tx=XM_023244170.1;refGene_gn=CACNA1G;refGene_tx=XM_023244171.1;refGene_gn=CACNA1G;refGene_tx=XM_023244172.1;refGene_gn=CACNA1G;refGene_tx=XM_023244173.1;refGene_gn=CACNA1G;refGene_tx=XM_023244174.1;refGene_gn=CACNA1G;refGene_tx=XM_023244175.1;refGene_gn=CACNA1G;refGene_tx=XM_023244176.1;refGene_gn=CACNA1G;refGene_tx=XM_023244177.1;refGene_gn=CACNA1G;refGene_tx=XM_023244178.1;refGene_gn=CACNA1G;refGene_tx=XM_023244179.1;refGene_gn=CACNA1G;refGene_tx=XM_023244180.1;refGene_gn=CACNA1G;refGene_tx=XM_023244181.1;refGene_gn=CACNA1G;refGene_tx=XM_023244182.1;refGene_gn=CACNA1G;refGene_tx=XM_023244183.1;refGene_gn=CACNA1G;refGene_tx=XM_023244184.1;refGene_gn=CACNA1G;refGene_tx=XM_023244185.1;refGene_gn=CACNA1G;refGene_tx=XM_023244186.1;refGene_gn=CACNA1G;refGene_tx=XM_023244187.1;refGene_gn=CACNA1G;refGene_tx=XM_023244188.1
E1 37126283 37126417
refGene_gn=CACNA1G;refGene_tx=XM_023244164.1;refGene_gn=CACNA1G;refGene_tx=XM_023244165.1;refGene_gn=CACNA1G;refGene_tx=XM_023244166.1;refGene_gn=CACNA1G;refGene_tx=XM_023244167.1;refGene_gn=CACNA1G;refGene_tx=XM_023244168.1;refGene_gn=CACNA1G;refGene_tx=XM_023244169.1;refGene_gn=CACNA1G;refGene_tx=XM_023244170.1;refGene_gn=CACNA1G;refGene_tx=XM_023244171.1;refGene_gn=CACNA1G;refGene_tx=XM_023244172.1;refGene_gn=CACNA1G;refGene_tx=XM_023244173.1;refGene_gn=CACNA1G;refGene_tx=XM_023244174.1;refGene_gn=CACNA1G;refGene_tx=XM_023244175.1;refGene_gn=CACNA1G;refGene_tx=XM_023244176.1;refGene_gn=CACNA1G;refGene_tx=XM_023244177.1;refGene_gn=CACNA1G;refGene_tx=XM_023244178.1;refGene_gn=CACNA1G;refGene_tx=XM_023244179.1;refGene_gn=CACNA1G;refGene_tx=XM_023244180.1;refGene_gn=CACNA1G;refGene_tx=XM_023244181.1;refGene_gn=CACNA1G;refGene_tx=XM_023244182.1;refGene_gn=CACNA1G;refGene_tx=XM_023244183.1;refGene_gn=CACNA1G;refGene_tx=XM_023244184.1;refGene_gn=CACNA1G;refGene_tx=XM_023244185.1;refGene_gn=CACNA1G;refGene_tx=XM_023244186.1;refGene_gn=CACNA1G;refGene_tx=XM_023244188.1
E1 37126523 37126635
refGene_gn=CACNA1G;refGene_tx=XM_023244164.1;refGene_gn=CACNA1G;refGene_tx=XM_023244165.1;refGene_gn=CACNA1G;refGene_tx=XM_023244166.1;refGene_gn=CACNA1G;refGene_tx=XM_023244167.1;refGene_gn=CACNA1G;refGene_tx=XM_023244168.1;refGene_gn=CACNA1G;refGene_tx=XM_023244169.1;refGene_gn=CACNA1G;refGene_tx=XM_023244170.1;refGene_gn=CACNA1G;refGene_tx=XM_023244171.1;refGene_gn=CACNA1G;refGene_tx=XM_023244172.1;refGene_gn=CACNA1G;refGene_tx=XM_023244173.1;refGene_gn=CACNA1G;refGene_tx=XM_023244174.1;refGene_gn=CACNA1G;refGene_tx=XM_023244175.1;refGene_gn=CACNA1G;refGene_tx=XM_023244176.1;refGene_gn=CACNA1G;refGene_tx=XM_023244177.1;refGene_gn=CACNA1G;refGene_tx=XM_023244178.1;refGene_gn=CACNA1G;refGene_tx=XM_023244179.1;refGene_gn=CACNA1G;refGene_tx=XM_023244180.1;refGene_gn=CACNA1G;refGene_tx=XM_023244181.1;refGene_gn=CACNA1G;refGene_tx=XM_023244182.1;refGene_gn=CACNA1G;refGene_tx=XM_023244183.1;refGene_gn=CACNA1G;refGene_tx=XM_023244184.1;refGene_gn=CACNA1G;refGene_tx=XM_02

3244185.1;refGene_gn=CACNA1G;refGene_tx=XM_023244186.1;refGene_gn=CACNA1G;refGene_tx=XM_023244188.1
E1 37133680 37133922
refGene_gn=CACNA1G;refGene_tx=XM_023244164.1;refGene_gn=CACNA1G;refGene_tx=XM_023244165.1;refGene_gn=CACNA1G;refGene_tx=XM_023244166.1;refGene_gn=CACNA1G;refGene_tx=XM_023244167.1;refGene_gn=CACNA1G;refGene_tx=XM_023244168.1;refGene_gn=CACNA1G;refGene_tx=XM_023244169.1;refGene_gn=CACNA1G;refGene_tx=XM_023244170.1;refGene_gn=CACNA1G;refGene_tx=XM_023244171.1;refGene_gn=CACNA1G;refGene_tx=XM_023244172.1;refGene_gn=CACNA1G;refGene_tx=XM_023244173.1;refGene_gn=CACNA1G;refGene_tx=XM_023244174.1;refGene_gn=CACNA1G;refGene_tx=XM_023244175.1;refGene_gn=CACNA1G;refGene_tx=XM_023244176.1;refGene_gn=CACNA1G;refGene_tx=XM_023244177.1;refGene_gn=CACNA1G;refGene_tx=XM_023244178.1;refGene_gn=CACNA1G;refGene_tx=XM_023244179.1;refGene_gn=CACNA1G;refGene_tx=XM_023244180.1;refGene_gn=CACNA1G;refGene_tx=XM_023244181.1;refGene_gn=CACNA1G;refGene_tx=XM_023244182.1;refGene_gn=CACNA1G;refGene_tx=XM_023244183.1;refGene_gn=CACNA1G;refGene_tx=XM_023244184.1;refGene_gn=CACNA1G;refGene_tx=XM_023244185.1;refGene_gn=CACNA1G;refGene_tx=XM_023244186.1;refGene_gn=CACNA1G;refGene_tx=XM_023244188.1
E1 37134544 37134582
refGene_gn=LOC109494089;refGene_tx=XM_023243462.1
E1 37134721 37134791
refGene_gn=LOC109494089;refGene_tx=XM_023243462.1
E1 37134905 37135143
refGene_gn=LOC109494089;refGene_tx=XM_023243462.1
E1 37135461 37136066
refGene_gn=LOC109494089;refGene_tx=XM_023243462.1
E1 37137824 37137890
refGene_gn=LOC109494089;refGene_tx=XM_023243462.1
E1 37139572 37139743
refGene_gn=SPATA20;refGene_tx=XM_011289191.3;refGene_gn=SPATA20;refGene_tx=XM_023244189.1;refGene_gn=SPATA20;refGene_tx=XM_023244190.1;refGene_gn=SPATA20;refGene_tx=XM_023244191.1
E1 37139911 37139992
refGene_gn=SPATA20;refGene_tx=XM_011289191.3;refGene_gn=SPATA20;refGene_tx=XM_023244189.1;refGene_gn=SPATA20;refGene_tx=XM_023244190.1;refGene_gn=SPATA20;refGene_tx=XM_023244191.1
E1 37140658 37140858
refGene_gn=SPATA20;refGene_tx=XM_011289191.3;refGene_gn=SPATA20;refGene_tx=XM_023244189.1;refGene_gn=SPATA20;refGene_tx=XM_023244190.1;refGene_gn=SPATA20;refGene_tx=XM_023244191.1
E1 37142112 37142324
refGene_gn=SPATA20;refGene_tx=XM_011289191.3;refGene_gn=SPATA20;refGene_tx=XM_023244189.1;refGene_gn=SPATA20;refGene_tx=XM_023244190.1;refGene_gn=SPATA20;refGene_tx=XM_023244191.1
E1 37142624 37142793
refGene_gn=SPATA20;refGene_tx=XM_011289191.3;refGene_gn=SPATA20;refGene_tx=XM_023244189.1;refGene_gn=SPATA20;refGene_tx=XM_023244190.1;refGene_gn=SPATA20;refGene_tx=XM_023244191.1
E1 37143116 37143309

refGene_gn=SPATA20;refGene_tx=XM_011289191.3;refGene_gn=SPATA20;refGene_tx=XM_023244189.1;refGene_gn=SPATA20;refGene_tx=XM_023244190.1;refGene_gn=SPATA20;refGene_tx=XM_023244191.1
E1 37143392 37143602
refGene_gn=SPATA20;refGene_tx=XM_011289191.3;refGene_gn=SPATA20;refGene_tx=XM_023244189.1;refGene_gn=SPATA20;refGene_tx=XM_023244190.1;refGene_gn=SPATA20;refGene_tx=XM_023244191.1
E1 37143701 37143776
refGene_gn=SPATA20;refGene_tx=XM_011289191.3;refGene_gn=SPATA20;refGene_tx=XM_023244189.1;refGene_gn=SPATA20;refGene_tx=XM_023244190.1;refGene_gn=SPATA20;refGene_tx=XM_023244191.1
E1 37144015 37144120
refGene_gn=SPATA20;refGene_tx=XM_011289191.3;refGene_gn=SPATA20;refGene_tx=XM_023244189.1;refGene_gn=SPATA20;refGene_tx=XM_023244190.1;refGene_gn=SPATA20;refGene_tx=XM_023244191.1
E1 37144210 37144341
refGene_gn=SPATA20;refGene_tx=XM_011289191.3;refGene_gn=SPATA20;refGene_tx=XM_023244189.1;refGene_gn=SPATA20;refGene_tx=XM_023244190.1;refGene_gn=SPATA20;refGene_tx=XM_023244191.1
E1 37144698 37144900
refGene_gn=SPATA20;refGene_tx=XM_011289191.3;refGene_gn=SPATA20;refGene_tx=XM_023244189.1;refGene_gn=SPATA20;refGene_tx=XM_023244190.1;refGene_gn=SPATA20;refGene_tx=XM_023244191.1
E1 37145025 37145169
refGene_gn=SPATA20;refGene_tx=XM_011289191.3;refGene_gn=SPATA20;refGene_tx=XM_023244189.1;refGene_gn=SPATA20;refGene_tx=XM_023244190.1;refGene_gn=SPATA20;refGene_tx=XM_023244191.1
E1 37145241 37145396
refGene_gn=SPATA20;refGene_tx=XM_011289191.3;refGene_gn=SPATA20;refGene_tx=XM_023244189.1;refGene_gn=SPATA20;refGene_tx=XM_023244190.1;refGene_gn=SPATA20;refGene_tx=XM_023244191.1
E1 37145560 37145625
refGene_gn=SPATA20;refGene_tx=XM_011289191.3;refGene_gn=SPATA20;refGene_tx=XM_023244189.1;refGene_gn=SPATA20;refGene_tx=XM_023244190.1;refGene_gn=SPATA20;refGene_tx=XM_023244191.1
E1 37145735 37145915
refGene_gn=SPATA20;refGene_tx=XM_011289191.3;refGene_gn=SPATA20;refGene_tx=XM_023244189.1;refGene_gn=SPATA20;refGene_tx=XM_023244190.1;refGene_gn=SPATA20;refGene_tx=XM_023244191.1
E1 37146053 37146130 refGene_gn=SPATA20;refGene_tx=XM_011289191.3
E1 37146919 37146996 refGene_gn=SPATA20;refGene_tx=XM_023244191.1
E1 37151369 37151704
refGene_gn=EPN3;refGene_tx=XM_023244192.1;refGene_gn=EPN3;refGene_tx=XM_023244193.1
E1 37151910 37152141
refGene_gn=EPN3;refGene_tx=XM_023244192.1;refGene_gn=EPN3;refGene_tx=XM_023244193.1
E1 37152270 37152387 refGene_gn=EPN3;refGene_tx=XM_023244192.1
E1 37152566 37152836
refGene_gn=EPN3;refGene_tx=XM_023244192.1;refGene_gn=EPN3;refGene_tx=X

M_023244193.1
E1 37153294 37153382
refGene_gn=EPN3;refGene_tx=XM_023244192.1;refGene_gn=EPN3;refGene_tx=X
M_023244193.1
E1 37154325 37154451
refGene_gn=EPN3;refGene_tx=XM_023244192.1;refGene_gn=EPN3;refGene_tx=X
M_023244193.1
E1 37154676 37154757
refGene_gn=EPN3;refGene_tx=XM_023244192.1;refGene_gn=EPN3;refGene_tx=X
M_023244193.1
E1 37155403 37155522
refGene_gn=EPN3;refGene_tx=XM_023244192.1;refGene_gn=EPN3;refGene_tx=X
M_023244193.1
E1 37156688 37157226
refGene_gn=EPN3;refGene_tx=XM_023244192.1;refGene_gn=EPN3;refGene_tx=X
M_023244193.1
E1 37161916 37162018 refGene_gn=MYCBPAP;refGene_tx=XM_023243463.1
E1 37163892 37164063 refGene_gn=MYCBPAP;refGene_tx=XM_023243463.1
E1 37164242 37164388 refGene_gn=MYCBPAP;refGene_tx=XM_023243463.1
E1 37164759 37164875 refGene_gn=MYCBPAP;refGene_tx=XM_023243463.1
E1 37165517 37165639 refGene_gn=MYCBPAP;refGene_tx=XM_023243463.1
E1 37166023 37166434 refGene_gn=MYCBPAP;refGene_tx=XM_023243463.1
E1 37166781 37166928 refGene_gn=MYCBPAP;refGene_tx=XM_023243463.1
E1 37167600 37167788 refGene_gn=MYCBPAP;refGene_tx=XM_023243463.1
E1 37168268 37168458 refGene_gn=MYCBPAP;refGene_tx=XM_023243463.1
E1 37168946 37169058 refGene_gn=MYCBPAP;refGene_tx=XM_023243463.1
E1 37169507 37169623 refGene_gn=MYCBPAP;refGene_tx=XM_023243463.1
E1 37169707 37169823 refGene_gn=MYCBPAP;refGene_tx=XM_023243463.1
E1 37170761 37170912 refGene_gn=MYCBPAP;refGene_tx=XM_023243463.1
E1 37171723 37171839 refGene_gn=MYCBPAP;refGene_tx=XM_023243463.1
E1 37172090 37172280 refGene_gn=MYCBPAP;refGene_tx=XM_023243463.1
E1 37173733 37173837 refGene_gn=MYCBPAP;refGene_tx=XM_023243463.1
E1 37173988 37174148 refGene_gn=MYCBPAP;refGene_tx=XM_023243463.1
E1 37175119 37175247 refGene_gn=MYCBPAP;refGene_tx=XM_023243463.1
E1 37181209 37181417 refGene_gn=MYCBPAP;refGene_tx=XM_023243463.1
E1 37202673 37202791
refGene_gn=RSAD1;refGene_tx=XM_003996691.4;refGene_gn=RSAD1;refGene_tx
=XM_011289190.3
E1 37202979 37203083
refGene_gn=RSAD1;refGene_tx=XM_003996691.4;refGene_gn=RSAD1;refGene_tx
=XM_011289190.3
E1 37203459 37203514
refGene_gn=RSAD1;refGene_tx=XM_003996691.4;refGene_gn=RSAD1;refGene_tx
=XM_011289190.3
E1 37203732 37203880
refGene_gn=RSAD1;refGene_tx=XM_003996691.4;refGene_gn=RSAD1;refGene_tx
=XM_011289190.3
E1 37204289 37204353
refGene_gn=RSAD1;refGene_tx=XM_003996691.4;refGene_gn=RSAD1;refGene_tx
=XM_011289190.3

E1 37204548 37204914
refGene_gn=RSAD1;refGene_tx=XM_003996691.4;refGene_gn=RSAD1;refGene_tx
=XM_011289190.3
E1 37207838 37208043
refGene_gn=RSAD1;refGene_tx=XM_003996691.4;refGene_gn=RSAD1;refGene_tx
=XM_011289190.3
E1 37208156 37208290
refGene_gn=RSAD1;refGene_tx=XM_003996691.4;refGene_gn=RSAD1;refGene_tx
=XM_011289190.3
E1 37208698 37208833
refGene_gn=RSAD1;refGene_tx=XM_003996691.4;refGene_gn=RSAD1;refGene_tx
=XM_011289190.3
E1 37211057 37211108 refGene_gn=ACSF2;refGene_tx=XM_003996692.5
E1 37211337 37211406 refGene_gn=ACSF2;refGene_tx=XM_003996692.5
E1 37211544 37211655 refGene_gn=ACSF2;refGene_tx=XM_003996692.5
E1 37211744 37211886 refGene_gn=ACSF2;refGene_tx=XM_003996692.5
E1 37212890 37213042 refGene_gn=ACSF2;refGene_tx=XM_003996692.5
E1 37213904 37214012 refGene_gn=ACSF2;refGene_tx=XM_003996692.5
E1 37216183 37216957
refGene_gn=CHAD;refGene_tx=XM_023243514.1;refGene_gn=CHAD;refGene_tx=X
M_023243515.1
E1 37217635 37217933 refGene_gn=ACSF2;refGene_tx=XM_019817700.2
E1 37218055 37218114 refGene_gn=ACSF2;refGene_tx=XM_019817700.2
E1 37218812 37218976
refGene_gn=CHAD;refGene_tx=XM_023243514.1;refGene_gn=CHAD;refGene_tx=X
M_023243515.1
E1 37219252 37219394
refGene_gn=CHAD;refGene_tx=XM_023243514.1;refGene_gn=CHAD;refGene_tx=X
M_023243515.1
E1 37220341 37220418
refGene_gn=ACSF2;refGene_tx=XM_003996692.5;refGene_gn=ACSF2;refGene_tx
=XM_019817700.2
E1 37220699 37220791
refGene_gn=ACSF2;refGene_tx=XM_003996692.5;refGene_gn=ACSF2;refGene_tx
=XM_019817700.2
E1 37221088 37221246
refGene_gn=ACSF2;refGene_tx=XM_003996692.5;refGene_gn=ACSF2;refGene_tx
=XM_019817700.2
E1 37221386 37221482
refGene_gn=ACSF2;refGene_tx=XM_003996692.5;refGene_gn=ACSF2;refGene_tx
=XM_019817700.2
E1 37222054 37222220
refGene_gn=ACSF2;refGene_tx=XM_003996692.5;refGene_gn=ACSF2;refGene_tx
=XM_019817700.2
E1 37222379 37222498
refGene_gn=ACSF2;refGene_tx=XM_003996692.5;refGene_gn=ACSF2;refGene_tx
=XM_019817700.2
E1 37222860 37222914
refGene_gn=ACSF2;refGene_tx=XM_003996692.5;refGene_gn=ACSF2;refGene_tx
=XM_019817700.2

E1 37223165 37223294
refGene_gn=ACSF2;refGene_tx=XM_003996692.5;refGene_gn=ACSF2;refGene_tx
=XM_019817700.2
E1 37223668 37223864
refGene_gn=ACSF2;refGene_tx=XM_003996692.5;refGene_gn=ACSF2;refGene_tx
=XM_019817700.2
E1 37244076 37244204
refGene_gn=ACSF2;refGene_tx=XM_003996692.5;refGene_gn=ACSF2;refGene_tx
=XM_019817700.2
E1 37256133 37256238 refGene_gn=LRRC59;refGene_tx=XM_003996693.5
E1 37257569 37257629 refGene_gn=LRRC59;refGene_tx=XM_003996693.5
E1 37259418 37259577 refGene_gn=LRRC59;refGene_tx=XM_003996693.5
E1 37259805 37259910 refGene_gn=LRRC59;refGene_tx=XM_003996693.5
E1 37262998 37263071 refGene_gn=LRRC59;refGene_tx=XM_003996693.5
E1 37265109 37265283 refGene_gn=LRRC59;refGene_tx=XM_003996693.5
E1 37266977 37267222 refGene_gn=LRRC59;refGene_tx=XM_003996693.5
E1 37269238 37269415
refGene_gn=EME1;refGene_tx=XM_003996695.4;refGene_gn=EME1;refGene_tx=X
M_019817701.2
E1 37269645 37269835
refGene_gn=EME1;refGene_tx=XM_003996695.4;refGene_gn=EME1;refGene_tx=X
M_019817701.2
E1 37270607 37270723
refGene_gn=EME1;refGene_tx=XM_003996695.4;refGene_gn=EME1;refGene_tx=X
M_019817701.2
E1 37270963 37271087
refGene_gn=EME1;refGene_tx=XM_003996695.4;refGene_gn=EME1;refGene_tx=X
M_019817701.2
E1 37271283 37271405
refGene_gn=EME1;refGene_tx=XM_003996695.4;refGene_gn=EME1;refGene_tx=X
M_019817701.2
E1 37271492 37271582 refGene_gn=EME1;refGene_tx=XM_003996695.4
E1 37273850 37273978
refGene_gn=EME1;refGene_tx=XM_003996695.4;refGene_gn=EME1;refGene_tx=X
M_019817701.2
E1 37274067 37274851
refGene_gn=EME1;refGene_tx=XM_003996695.4;refGene_gn=EME1;refGene_tx=X
M_019817701.2
E1 37275900 37275940 refGene_gn=MRPL27;refGene_tx=XM_003996697.5
E1 37278028 37278160 refGene_gn=MRPL27;refGene_tx=XM_003996697.5
E1 37278576 37278644 refGene_gn=MRPL27;refGene_tx=XM_003996697.5
E1 37280372 37280579 refGene_gn=MRPL27;refGene_tx=XM_003996697.5
E1 37287551 37287874 refGene_gn=XYLT2;refGene_tx=XM_023243625.1
E1 37289390 37289724 refGene_gn=XYLT2;refGene_tx=XM_023243625.1
E1 37290360 37290556 refGene_gn=XYLT2;refGene_tx=XM_023243625.1
E1 37290818 37291081 refGene_gn=XYLT2;refGene_tx=XM_023243625.1
E1 37291321 37291498 refGene_gn=XYLT2;refGene_tx=XM_023243625.1
E1 37291603 37291820 refGene_gn=XYLT2;refGene_tx=XM_023243625.1
E1 37291973 37292054 refGene_gn=XYLT2;refGene_tx=XM_023243625.1
E1 37292470 37292673 refGene_gn=XYLT2;refGene_tx=XM_023243625.1

E1	37292966	37293142	refGene_gn=XYLT2;refGene_tx=XM_023243625.1
E1	37293376	37293869	refGene_gn=XYLT2;refGene_tx=XM_023243625.1
E1	37300190	37300325	refGene_gn=XYLT2;refGene_tx=XM_023243625.1
E1	37355764	37355872
refGene_gn=TMEM92;refGene_tx=XM_003996698.3;refGene_gn=TMEM92;refGene_tx=XM_011289189.2
E1	37356099	37356295
refGene_gn=TMEM92;refGene_tx=XM_003996698.3;refGene_gn=TMEM92;refGene_tx=XM_011289189.2
E1	37356431	37356506
refGene_gn=TMEM92;refGene_tx=XM_003996698.3;refGene_gn=TMEM92;refGene_tx=XM_011289189.2
E1	37357333	37357359
refGene_gn=TMEM92;refGene_tx=XM_003996698.3;refGene_gn=TMEM92;refGene_tx=XM_011289189.2
E1	37360583	37360667
refGene_gn=TMEM92;refGene_tx=XM_003996698.3;refGene_gn=TMEM92;refGene_tx=XM_011289189.2
E1	37421157	37421251
refGene_gn=COL1A1;refGene_tx=XM_003996699.4;refGene_gn=COL1A1;refGene_tx=XM_006940215.3
E1	37422791	37422986
refGene_gn=COL1A1;refGene_tx=XM_003996699.4;refGene_gn=COL1A1;refGene_tx=XM_006940215.3
E1	37423077	37423112
refGene_gn=COL1A1;refGene_tx=XM_003996699.4;refGene_gn=COL1A1;refGene_tx=XM_006940215.3
E1	37423222	37423258
refGene_gn=COL1A1;refGene_tx=XM_003996699.4;refGene_gn=COL1A1;refGene_tx=XM_006940215.3
E1	37423345	37423447
refGene_gn=COL1A1;refGene_tx=XM_003996699.4;refGene_gn=COL1A1;refGene_tx=XM_006940215.3
E1	37424252	37424324
refGene_gn=COL1A1;refGene_tx=XM_003996699.4;refGene_gn=COL1A1;refGene_tx=XM_006940215.3
E1	37424574	37424619
refGene_gn=COL1A1;refGene_tx=XM_003996699.4;refGene_gn=COL1A1;refGene_tx=XM_006940215.3
E1	37424780	37424834
refGene_gn=COL1A1;refGene_tx=XM_003996699.4;refGene_gn=COL1A1;refGene_tx=XM_006940215.3
E1	37425033	37425087
refGene_gn=COL1A1;refGene_tx=XM_003996699.4;refGene_gn=COL1A1;refGene_tx=XM_006940215.3
E1	37425598	37425652
refGene_gn=COL1A1;refGene_tx=XM_003996699.4;refGene_gn=COL1A1;refGene_tx=XM_006940215.3
E1	37425769	37425823
refGene_gn=COL1A1;refGene_tx=XM_003996699.4;refGene_gn=COL1A1;refGene_

tx=XM_006940215.3
E1 37426134 37426188
refGene_gn=COL1A1;refGene_tx=XM_003996699.4;refGene_gn=COL1A1;refGene_
tx=XM_006940215.3
E1 37426271 37426316
refGene_gn=COL1A1;refGene_tx=XM_003996699.4;refGene_gn=COL1A1;refGene_
tx=XM_006940215.3
E1 37426431 37426485
refGene_gn=COL1A1;refGene_tx=XM_003996699.4;refGene_gn=COL1A1;refGene_
tx=XM_006940215.3
E1 37426603 37426648
refGene_gn=COL1A1;refGene_tx=XM_003996699.4;refGene_gn=COL1A1;refGene_
tx=XM_006940215.3
E1 37426811 37426865
refGene_gn=COL1A1;refGene_tx=XM_003996699.4;refGene_gn=COL1A1;refGene_
tx=XM_006940215.3
E1 37427091 37427190
refGene_gn=COL1A1;refGene_tx=XM_003996699.4;refGene_gn=COL1A1;refGene_
tx=XM_006940215.3
E1 37427272 37427317
refGene_gn=COL1A1;refGene_tx=XM_003996699.4;refGene_gn=COL1A1;refGene_
tx=XM_006940215.3
E1 37427414 37427513
refGene_gn=COL1A1;refGene_tx=XM_003996699.4;refGene_gn=COL1A1;refGene_
tx=XM_006940215.3
E1 37427630 37427684
refGene_gn=COL1A1;refGene_tx=XM_003996699.4;refGene_gn=COL1A1;refGene_
tx=XM_006940215.3
E1 37427811 37427919
refGene_gn=COL1A1;refGene_tx=XM_003996699.4;refGene_gn=COL1A1;refGene_
tx=XM_006940215.3
E1 37427999 37428053
refGene_gn=COL1A1;refGene_tx=XM_003996699.4;refGene_gn=COL1A1;refGene_
tx=XM_006940215.3
E1 37428326 37428425
refGene_gn=COL1A1;refGene_tx=XM_003996699.4;refGene_gn=COL1A1;refGene_
tx=XM_006940215.3
E1 37428584 37428638
refGene_gn=COL1A1;refGene_tx=XM_003996699.4;refGene_gn=COL1A1;refGene_
tx=XM_006940215.3
E1 37428728 37428827
refGene_gn=COL1A1;refGene_tx=XM_003996699.4;refGene_gn=COL1A1;refGene_
tx=XM_006940215.3
E1 37429278 37429332
refGene_gn=COL1A1;refGene_tx=XM_003996699.4;refGene_gn=COL1A1;refGene_
tx=XM_006940215.3
E1 37429555 37429609
refGene_gn=COL1A1;refGene_tx=XM_003996699.4;refGene_gn=COL1A1;refGene_
tx=XM_006940215.3
E1 37429721 37429775

refGene_gn=COL1A1;refGene_tx=XM_003996699.4;refGene_gn=COL1A1;refGene_tx=XM_006940215.3
E1 37429876 37429930
refGene_gn=COL1A1;refGene_tx=XM_003996699.4;refGene_gn=COL1A1;refGene_tx=XM_006940215.3
E1 37430364 37430409
refGene_gn=COL1A1;refGene_tx=XM_003996699.4;refGene_gn=COL1A1;refGene_tx=XM_006940215.3
E1 37430494 37430593
refGene_gn=COL1A1;refGene_tx=XM_003996699.4;refGene_gn=COL1A1;refGene_tx=XM_006940215.3
E1 37430868 37430976
refGene_gn=COL1A1;refGene_tx=XM_003996699.4;refGene_gn=COL1A1;refGene_tx=XM_006940215.3
E1 37431412 37431520
refGene_gn=COL1A1;refGene_tx=XM_003996699.4;refGene_gn=COL1A1;refGene_tx=XM_006940215.3
E1 37431727 37431781
refGene_gn=COL1A1;refGene_tx=XM_003996699.4;refGene_gn=COL1A1;refGene_tx=XM_006940215.3
E1 37431982 37432036
refGene_gn=COL1A1;refGene_tx=XM_003996699.4;refGene_gn=COL1A1;refGene_tx=XM_006940215.3
E1 37432248 37432356
refGene_gn=COL1A1;refGene_tx=XM_003996699.4;refGene_gn=COL1A1;refGene_tx=XM_006940215.3
E1 37432450 37432504
refGene_gn=COL1A1;refGene_tx=XM_003996699.4;refGene_gn=COL1A1;refGene_tx=XM_006940215.3
E1 37432624 37432678
refGene_gn=COL1A1;refGene_tx=XM_003996699.4;refGene_gn=COL1A1;refGene_tx=XM_006940215.3
E1 37432813 37432975 refGene_gn=COL1A1;refGene_tx=XM_003996699.4
E1 37433058 37433166 refGene_gn=COL1A1;refGene_tx=XM_003996699.4
E1 37433346 37433454
refGene_gn=COL1A1;refGene_tx=XM_003996699.4;refGene_gn=COL1A1;refGene_tx=XM_006940215.3
E1 37433561 37433615
refGene_gn=COL1A1;refGene_tx=XM_003996699.4;refGene_gn=COL1A1;refGene_tx=XM_006940215.3
E1 37433722 37433830
refGene_gn=COL1A1;refGene_tx=XM_003996699.4;refGene_gn=COL1A1;refGene_tx=XM_006940215.3
E1 37434211 37434265
refGene_gn=COL1A1;refGene_tx=XM_003996699.4;refGene_gn=COL1A1;refGene_tx=XM_006940215.3
E1 37434379 37434487
refGene_gn=COL1A1;refGene_tx=XM_003996699.4;refGene_gn=COL1A1;refGene_tx=XM_006940215.3
E1 37434798 37434852

refGene_gn=COL1A1;refGene_tx=XM_003996699.4;refGene_gn=COL1A1;refGene_tx=XM_006940215.3
E1 37435136 37435244
refGene_gn=COL1A1;refGene_tx=XM_003996699.4;refGene_gn=COL1A1;refGene_tx=XM_006940215.3
E1 37435336 37435619
refGene_gn=COL1A1;refGene_tx=XM_003996699.4;refGene_gn=COL1A1;refGene_tx=XM_006940215.3
E1 37435800 37435991
refGene_gn=COL1A1;refGene_tx=XM_003996699.4;refGene_gn=COL1A1;refGene_tx=XM_006940215.3
E1 37436253 37436496
refGene_gn=COL1A1;refGene_tx=XM_003996699.4;refGene_gn=COL1A1;refGene_tx=XM_006940215.3
E1 37436594 37436741
refGene_gn=COL1A1;refGene_tx=XM_003996699.4;refGene_gn=COL1A1;refGene_tx=XM_006940215.3
E1 37445981 37446162 refGene_gn=SGCA;refGene_tx=XM_003996700.4
E1 37451118 37451145 refGene_gn=SGCA;refGene_tx=XM_003996700.4
E1 37451312 37451521 refGene_gn=SGCA;refGene_tx=XM_003996700.4
E1 37452428 37452591 refGene_gn=SGCA;refGene_tx=XM_003996700.4
E1 37452884 37453083 refGene_gn=SGCA;refGene_tx=XM_003996700.4
E1 37453433 37453506 refGene_gn=SGCA;refGene_tx=XM_003996700.4
E1 37453666 37453821 refGene_gn=SGCA;refGene_tx=XM_003996700.4
E1 37453918 37454038 refGene_gn=SGCA;refGene_tx=XM_003996700.4
E1 37454883 37454920 refGene_gn=SGCA;refGene_tx=XM_003996700.4
E1 37468756 37470139 refGene_gn=PPP1R9B;refGene_tx=XM_003996702.4
E1 37473922 37474055 refGene_gn=PPP1R9B;refGene_tx=XM_003996702.4
E1 37475440 37475561 refGene_gn=PPP1R9B;refGene_tx=XM_003996702.4
E1 37477481 37477586 refGene_gn=PPP1R9B;refGene_tx=XM_003996702.4
E1 37478873 37479009 refGene_gn=PPP1R9B;refGene_tx=XM_003996702.4
E1 37479634 37479787 refGene_gn=PPP1R9B;refGene_tx=XM_003996702.4
E1 37479904 37479958 refGene_gn=PPP1R9B;refGene_tx=XM_003996702.4
E1 37481434 37481664 refGene_gn=PPP1R9B;refGene_tx=XM_003996702.4
E1 37482036 37482133 refGene_gn=PPP1R9B;refGene_tx=XM_003996702.4
E1 37482320 37482374 refGene_gn=PPP1R9B;refGene_tx=XM_003996702.4
E1 37492574 37492617 refGene_gn=SAMD14;refGene_tx=XM_023244659.1
E1 37498642 37498809 refGene_gn=SAMD14;refGene_tx=XM_023244659.1
E1 37499227 37499516 refGene_gn=SAMD14;refGene_tx=XM_023244659.1
E1 37500302 37500390 refGene_gn=SAMD14;refGene_tx=XM_023244659.1
E1 37500489 37500564 refGene_gn=SAMD14;refGene_tx=XM_023244659.1
E1 37500669 37500829 refGene_gn=SAMD14;refGene_tx=XM_023244659.1
E1 37502052 37502172 refGene_gn=SAMD14;refGene_tx=XM_023244659.1
E1 37502257 37502413 refGene_gn=SAMD14;refGene_tx=XM_023244659.1
E1 37503245 37503401 refGene_gn=SAMD14;refGene_tx=XM_023244659.1
E1 37506252 37506393 refGene_gn=PDK2;refGene_tx=XM_003996734.3
E1 37506971 37507085 refGene_gn=PDK2;refGene_tx=XM_003996734.3
E1 37507705 37507813 refGene_gn=PDK2;refGene_tx=XM_003996734.3
E1 37507977 37508076 refGene_gn=PDK2;refGene_tx=XM_003996734.3
E1 37508167 37508244 refGene_gn=PDK2;refGene_tx=XM_003996734.3

E1	37510865	37510943	refGene_gn=PDK2;refGene_tx=XM_003996734.3
E1	37511153	37511243	refGene_gn=PDK2;refGene_tx=XM_003996734.3
E1	37511757	37511942	refGene_gn=PDK2;refGene_tx=XM_003996734.3
E1	37512335	37512407	refGene_gn=PDK2;refGene_tx=XM_003996734.3
E1	37518918	37519060	refGene_gn=PDK2;refGene_tx=XM_003996734.3
E1	37520940	37521058	refGene_gn=PDK2;refGene_tx=XM_003996734.3
E1	37527058	37527214	refGene_gn=ITGA3;refGene_tx=XM_019817709.2
E1	37528002	37528113	refGene_gn=ITGA3;refGene_tx=XM_011289188.3
E1	37528499	37528625
refGene_gn=ITGA3;refGene_tx=XM_011289188.3;refGene_gn=ITGA3;refGene_tx
=XM_019817709.2
E1	37532876	37532975
refGene_gn=ITGA3;refGene_tx=XM_011289188.3;refGene_gn=ITGA3;refGene_tx
=XM_019817709.2
E1	37534625	37534739
refGene_gn=ITGA3;refGene_tx=XM_011289188.3;refGene_gn=ITGA3;refGene_tx
=XM_019817709.2
E1	37535351	37535474
refGene_gn=ITGA3;refGene_tx=XM_011289188.3;refGene_gn=ITGA3;refGene_tx
=XM_019817709.2
E1	37535673	37535856
refGene_gn=ITGA3;refGene_tx=XM_011289188.3;refGene_gn=ITGA3;refGene_tx
=XM_019817709.2
E1	37536008	37536111
refGene_gn=ITGA3;refGene_tx=XM_011289188.3;refGene_gn=ITGA3;refGene_tx
=XM_019817709.2
E1	37536675	37536753
refGene_gn=ITGA3;refGene_tx=XM_011289188.3;refGene_gn=ITGA3;refGene_tx
=XM_019817709.2
E1	37536861	37536941
refGene_gn=ITGA3;refGene_tx=XM_011289188.3;refGene_gn=ITGA3;refGene_tx
=XM_019817709.2
E1	37537565	37537634
refGene_gn=ITGA3;refGene_tx=XM_011289188.3;refGene_gn=ITGA3;refGene_tx
=XM_019817709.2
E1	37537874	37538022
refGene_gn=ITGA3;refGene_tx=XM_011289188.3;refGene_gn=ITGA3;refGene_tx
=XM_019817709.2
E1	37538355	37538453
refGene_gn=ITGA3;refGene_tx=XM_011289188.3;refGene_gn=ITGA3;refGene_tx
=XM_019817709.2
E1	37538550	37538700
refGene_gn=ITGA3;refGene_tx=XM_011289188.3;refGene_gn=ITGA3;refGene_tx
=XM_019817709.2
E1	37539476	37539613
refGene_gn=ITGA3;refGene_tx=XM_011289188.3;refGene_gn=ITGA3;refGene_tx
=XM_019817709.2
E1	37539686	37539754
refGene_gn=ITGA3;refGene_tx=XM_011289188.3;refGene_gn=ITGA3;refGene_tx
=XM_019817709.2

E1	37540740	37540827
refGene_gn=ITGA3;refGene_tx=XM_011289188.3;refGene_gn=ITGA3;refGene_tx=XM_019817709.2
E1	37540998	37541135
refGene_gn=ITGA3;refGene_tx=XM_011289188.3;refGene_gn=ITGA3;refGene_tx=XM_019817709.2
E1	37541262	37541351
refGene_gn=ITGA3;refGene_tx=XM_011289188.3;refGene_gn=ITGA3;refGene_tx=XM_019817709.2
E1	37543769	37543966
refGene_gn=ITGA3;refGene_tx=XM_011289188.3;refGene_gn=ITGA3;refGene_tx=XM_019817709.2
E1	37544423	37544631
refGene_gn=ITGA3;refGene_tx=XM_011289188.3;refGene_gn=ITGA3;refGene_tx=XM_019817709.2
E1	37545024	37545111
refGene_gn=ITGA3;refGene_tx=XM_011289188.3;refGene_gn=ITGA3;refGene_tx=XM_019817709.2
E1	37546975	37547225
refGene_gn=ITGA3;refGene_tx=XM_011289188.3;refGene_gn=ITGA3;refGene_tx=XM_019817709.2
E1	37548763	37548843
refGene_gn=ITGA3;refGene_tx=XM_011289188.3;refGene_gn=ITGA3;refGene_tx=XM_019817709.2
E1	37549180	37549308
refGene_gn=ITGA3;refGene_tx=XM_011289188.3;refGene_gn=ITGA3;refGene_tx=XM_019817709.2
E1	37556569	37556775
refGene_gn=ITGA3;refGene_tx=XM_011289188.3;refGene_gn=ITGA3;refGene_tx=XM_019817709.2
E1	37595968	37596293	refGene_gn=DLX3;refGene_tx=XM_023244224.1
E1	37597306	37597497	refGene_gn=DLX3;refGene_tx=XM_023244224.1
E1	37599130	37599478	refGene_gn=DLX3;refGene_tx=XM_023244224.1
E1	37616658	37616907	refGene_gn=DLX4;refGene_tx=XM_019817715.2
E1	37617148	37617345	refGene_gn=DLX4;refGene_tx=XM_019817715.2
E1	37620821	37621077	refGene_gn=DLX4;refGene_tx=XM_019817715.2
E1	37729412	37729532
refGene_gn=TAC4;refGene_tx=XM_006940216.4;refGene_gn=TAC4;refGene_tx=XM_011289187.3
E1	37732719	37732813
refGene_gn=TAC4;refGene_tx=XM_006940216.4;refGene_gn=TAC4;refGene_tx=XM_011289187.3
E1	37733441	37733474
refGene_gn=TAC4;refGene_tx=XM_006940216.4;refGene_gn=TAC4;refGene_tx=XM_011289187.3
E1	37734448	37734519	refGene_gn=TAC4;refGene_tx=XM_006940216.4
E1	37735288	37735332	refGene_gn=TAC4;refGene_tx=XM_011289187.3
E1	37745486	37745588
refGene_gn=KAT7;refGene_tx=XM_003996705.5;refGene_gn=KAT7;refGene_tx=XM_003996706.5;refGene_gn=KAT7;refGene_tx=XM_019817717.2

E1 37746501 37746608
refGene_gn=KAT7;refGene_tx=XM_003996705.5;refGene_gn=KAT7;refGene_tx=XM_003996706.5;refGene_gn=KAT7;refGene_tx=XM_019817717.2
E1 37747225 37747372
refGene_gn=KAT7;refGene_tx=XM_003996705.5;refGene_gn=KAT7;refGene_tx=XM_003996706.5;refGene_gn=KAT7;refGene_tx=XM_019817717.2
E1 37749819 37749913
refGene_gn=KAT7;refGene_tx=XM_003996705.5;refGene_gn=KAT7;refGene_tx=XM_003996706.5;refGene_gn=KAT7;refGene_tx=XM_019817717.2
E1 37751006 37751147
refGene_gn=KAT7;refGene_tx=XM_003996705.5;refGene_gn=KAT7;refGene_tx=XM_003996706.5;refGene_gn=KAT7;refGene_tx=XM_019817717.2
E1 37751388 37751478
refGene_gn=KAT7;refGene_tx=XM_003996705.5;refGene_gn=KAT7;refGene_tx=XM_003996706.5;refGene_gn=KAT7;refGene_tx=XM_019817717.2
E1 37754334 37754526
refGene_gn=KAT7;refGene_tx=XM_003996705.5;refGene_gn=KAT7;refGene_tx=XM_003996706.5;refGene_gn=KAT7;refGene_tx=XM_019817717.2
E1 37756038 37756149
refGene_gn=KAT7;refGene_tx=XM_003996705.5;refGene_gn=KAT7;refGene_tx=XM_003996706.5;refGene_gn=KAT7;refGene_tx=XM_019817717.2
E1 37758828 37758927
refGene_gn=KAT7;refGene_tx=XM_003996705.5;refGene_gn=KAT7;refGene_tx=XM_003996706.5;refGene_gn=KAT7;refGene_tx=XM_019817717.2
E1 37760399 37760489
refGene_gn=KAT7;refGene_tx=XM_003996705.5;refGene_gn=KAT7;refGene_tx=XM_019817717.2
E1 37763775 37763858
refGene_gn=KAT7;refGene_tx=XM_003996705.5;refGene_gn=KAT7;refGene_tx=XM_003996706.5;refGene_gn=KAT7;refGene_tx=XM_019817717.2
E1 37768945 37769185
refGene_gn=KAT7;refGene_tx=XM_003996705.5;refGene_gn=KAT7;refGene_tx=XM_003996706.5;refGene_gn=KAT7;refGene_tx=XM_019817717.2
E1 37770819 37770996
refGene_gn=KAT7;refGene_tx=XM_003996705.5;refGene_gn=KAT7;refGene_tx=XM_003996706.5
E1 37773354 37773502
refGene_gn=KAT7;refGene_tx=XM_003996705.5;refGene_gn=KAT7;refGene_tx=XM_003996706.5;refGene_gn=KAT7;refGene_tx=XM_019817717.2
E1 37775947 37775962
refGene_gn=KAT7;refGene_tx=XM_003996705.5;refGene_gn=KAT7;refGene_tx=XM_003996706.5;refGene_gn=KAT7;refGene_tx=XM_019817717.2
E1 37796729 37796925 refGene_gn=FAM117A;refGene_tx=XM_023244642.1
E1 37823526 37823696 refGene_gn=FAM117A;refGene_tx=XM_023244642.1
E1 37832119 37832215 refGene_gn=FAM117A;refGene_tx=XM_023244642.1
E1 37834176 37834287 refGene_gn=FAM117A;refGene_tx=XM_023244642.1
E1 37834781 37834916 refGene_gn=FAM117A;refGene_tx=XM_023244642.1
E1 37836073 37836275 refGene_gn=FAM117A;refGene_tx=XM_023244642.1
E1 37837728 37837879 refGene_gn=FAM117A;refGene_tx=XM_023244642.1
E1 37841217 37841518 refGene_gn=FAM117A;refGene_tx=XM_023244642.1

E1 37844833 37845039 refGene_gn=SLC35B1;refGene_tx=XM_003996710.5
E1 37845717 37845821 refGene_gn=SLC35B1;refGene_tx=XM_003996710.5
E1 37846379 37846510 refGene_gn=SLC35B1;refGene_tx=XM_003996710.5
E1 37846779 37846810 refGene_gn=SLC35B1;refGene_tx=XM_003996710.5
E1 37847851 37848009 refGene_gn=SLC35B1;refGene_tx=XM_003996710.5
E1 37849002 37849129 refGene_gn=SLC35B1;refGene_tx=XM_003996710.5
E1 37849832 37849939 refGene_gn=SLC35B1;refGene_tx=XM_003996710.5
E1 37850101 37850255 refGene_gn=SLC35B1;refGene_tx=XM_003996710.5
E1 37850797 37850850 refGene_gn=SLC35B1;refGene_tx=XM_003996710.5
E1 37914195 37914273
refGene_gn=SPOP;refGene_tx=XM_023244729.1;refGene_gn=SPOP;refGene_tx=XM_023244730.1;refGene_gn=SPOP;refGene_tx=XM_023244731.1
E1 37914939 37915061
refGene_gn=SPOP;refGene_tx=XM_019817719.2;refGene_gn=SPOP;refGene_tx=XM_023244729.1;refGene_gn=SPOP;refGene_tx=XM_023244730.1;refGene_gn=SPOP;refGene_tx=XM_023244731.1
E1 37916998 37917150
refGene_gn=SPOP;refGene_tx=XM_019817719.2;refGene_gn=SPOP;refGene_tx=XM_023244729.1;refGene_gn=SPOP;refGene_tx=XM_023244730.1;refGene_gn=SPOP;refGene_tx=XM_023244731.1
E1 37917279 37917407
refGene_gn=SPOP;refGene_tx=XM_019817719.2;refGene_gn=SPOP;refGene_tx=XM_023244729.1;refGene_gn=SPOP;refGene_tx=XM_023244730.1;refGene_gn=SPOP;refGene_tx=XM_023244731.1
E1 37923973 37924151
refGene_gn=SPOP;refGene_tx=XM_019817719.2;refGene_gn=SPOP;refGene_tx=XM_023244729.1;refGene_gn=SPOP;refGene_tx=XM_023244730.1;refGene_gn=SPOP;refGene_tx=XM_023244731.1
E1 37927178 37927234
refGene_gn=SPOP;refGene_tx=XM_019817719.2;refGene_gn=SPOP;refGene_tx=XM_023244729.1;refGene_gn=SPOP;refGene_tx=XM_023244730.1;refGene_gn=SPOP;refGene_tx=XM_023244731.1
E1 37927704 37927827
refGene_gn=SPOP;refGene_tx=XM_019817719.2;refGene_gn=SPOP;refGene_tx=XM_023244729.1;refGene_gn=SPOP;refGene_tx=XM_023244730.1;refGene_gn=SPOP;refGene_tx=XM_023244731.1
E1 37931851 37931994
refGene_gn=SPOP;refGene_tx=XM_019817719.2;refGene_gn=SPOP;refGene_tx=XM_023244729.1;refGene_gn=SPOP;refGene_tx=XM_023244730.1;refGene_gn=SPOP;refGene_tx=XM_023244731.1
E1 37932141 37932181 refGene_gn=SPOP;refGene_tx=XM_023244730.1
E1 37933313 37933458
refGene_gn=SPOP;refGene_tx=XM_019817719.2;refGene_gn=SPOP;refGene_tx=XM_023244729.1
E1 37934880 37934881 refGene_gn=SPOP;refGene_tx=XM_023244731.1
E1 37952245 37952950
refGene_gn=NXPH3;refGene_tx=XM_019817722.2;refGene_gn=NXPH3;refGene_tx=XM_023243848.1
E1 37954195 37954342 refGene_gn=NXPH3;refGene_tx=XM_019817722.2
E1 37955420 37955474 refGene_gn=NXPH3;refGene_tx=XM_023243848.1

E1 37997338 37997640 refGene_gn=NGFR;refGene_tx=XM_023243766.1
E1 37998295 37998456 refGene_gn=NGFR;refGene_tx=XM_023243766.1
E1 38000255 38000508 refGene_gn=NGFR;refGene_tx=XM_023243766.1
E1 38004139 38004499 refGene_gn=NGFR;refGene_tx=XM_023243766.1
E1 38007963 38008105 refGene_gn=NGFR;refGene_tx=XM_023243766.1
E1 38014963 38015011 refGene_gn=NGFR;refGene_tx=XM_023243766.1
E1 38015012 38015039 refGene_gn=NGFR;refGene_tx=XM_023243766.1
E1 38086912 38087000 refGene_gn=PHB;refGene_tx=NM_001110448.1
E1 38088337 38088499 refGene_gn=PHB;refGene_tx=NM_001110448.1
E1 38090739 38090882 refGene_gn=PHB;refGene_tx=NM_001110448.1
E1 38091027 38091144 refGene_gn=PHB;refGene_tx=NM_001110448.1
E1 38094485 38094581 refGene_gn=PHB;refGene_tx=NM_001110448.1
E1 38095955 38096150 refGene_gn=PHB;refGene_tx=NM_001110448.1
E1 38168425 38169328
refGene_gn=ZNF652;refGene_tx=XM_019817724.2;refGene_gn=ZNF652;refGene_tx=XM_023243408.1;refGene_gn=ZNF652;refGene_tx=XM_023243410.1
E1 38171778 38171926
refGene_gn=ZNF652;refGene_tx=XM_019817724.2;refGene_gn=ZNF652;refGene_tx=XM_023243408.1;refGene_gn=ZNF652;refGene_tx=XM_023243410.1
E1 38172305 38172421
refGene_gn=ZNF652;refGene_tx=XM_019817724.2;refGene_gn=ZNF652;refGene_tx=XM_023243408.1;refGene_gn=ZNF652;refGene_tx=XM_023243410.1
E1 38172779 38172924
refGene_gn=ZNF652;refGene_tx=XM_019817724.2;refGene_gn=ZNF652;refGene_tx=XM_023243408.1;refGene_gn=ZNF652;refGene_tx=XM_023243410.1
E1 38181878 38182390
refGene_gn=ZNF652;refGene_tx=XM_019817724.2;refGene_gn=ZNF652;refGene_tx=XM_023243408.1;refGene_gn=ZNF652;refGene_tx=XM_023243410.1
E1 38224359 38224404
refGene_gn=PHOSPHO1;refGene_tx=XM_006940221.4;refGene_gn=PHOSPHO1;refGene_tx=XM_019817740.2;refGene_gn=PHOSPHO1;refGene_tx=XM_023243413.1;refGene_gn=PHOSPHO1;refGene_tx=XM_023243414.1;refGene_gn=PHOSPHO1;refGene_tx=XM_023243415.1;refGene_gn=PHOSPHO1;refGene_tx=XM_023243416.1;refGene_gn=PHOSPHO1;refGene_tx=XM_023243417.1;refGene_gn=PHOSPHO1;refGene_tx=XM_023243418.1;refGene_gn=PHOSPHO1;refGene_tx=XM_023243419.1;refGene_gn=PHOSPHO1;refGene_tx=XM_023243420.1;refGene_gn=PHOSPHO1;refGene_tx=XM_023243421.1
E1 38225241 38225363
refGene_gn=PHOSPHO1;refGene_tx=XM_006940221.4;refGene_gn=PHOSPHO1;refGene_tx=XM_023243413.1;refGene_gn=PHOSPHO1;refGene_tx=XM_023243414.1;refGene_gn=PHOSPHO1;refGene_tx=XM_023243415.1;refGene_gn=PHOSPHO1;refGene_tx=XM_023243416.1;refGene_gn=PHOSPHO1;refGene_tx=XM_023243417.1;refGene_gn=PHOSPHO1;refGene_tx=XM_023243418.1;refGene_gn=PHOSPHO1;refGene_tx=XM_023243419.1;refGene_gn=PHOSPHO1;refGene_tx=XM_023243420.1;refGene_gn=PHOSPHO1;refGene_tx=XM_023243421.1
E1 38226055 38226941
refGene_gn=PHOSPHO1;refGene_tx=XM_006940221.4;refGene_gn=PHOSPHO1;refGene_tx=XM_019817740.2;refGene_gn=PHOSPHO1;refGene_tx=XM_023243413.1;refGene_gn=PHOSPHO1;refGene_tx=XM_023243414.1;refGene_gn=PHOSPHO1;refGene_tx=XM_023243415.1;refGene_gn=PHOSPHO1;refGene_tx=XM_023243416.1;refG

ene_gn=PHOSPHO1;refGene_tx=XM_023243417.1;refGene_gn=PHOSPHO1;refGene_tx=XM_023243418.1;refGene_gn=PHOSPHO1;refGene_tx=XM_023243419.1;refGene_gn=PHOSPHO1;refGene_tx=XM_023243420.1;refGene_gn=PHOSPHO1;refGene_tx=XM_023243421.1;refGene_gn=PHOSPHO1;refGene_tx=XM_023243422.1;refGene_gn=PHOSPHO1;refGene_tx=XM_023243423.1
E1 38228499 38228663
refGene_gn=ABI3;refGene_tx=XM_023243411.1;refGene_gn=ABI3;refGene_tx=XM_023243412.1
E1 38229031 38229163
refGene_gn=ABI3;refGene_tx=XM_023243411.1;refGene_gn=ABI3;refGene_tx=XM_023243412.1
E1 38229669 38229827
refGene_gn=ABI3;refGene_tx=XM_023243411.1;refGene_gn=ABI3;refGene_tx=XM_023243412.1
E1 38230014 38230110
refGene_gn=ABI3;refGene_tx=XM_023243411.1;refGene_gn=ABI3;refGene_tx=XM_023243412.1
E1 38230239 38230325
refGene_gn=ABI3;refGene_tx=XM_023243411.1;refGene_gn=ABI3;refGene_tx=XM_023243412.1
E1 38231689 38231866
refGene_gn=ABI3;refGene_tx=XM_023243411.1;refGene_gn=ABI3;refGene_tx=XM_023243412.1
E1 38233178 38233346
refGene_gn=ABI3;refGene_tx=XM_023243411.1;refGene_gn=ABI3;refGene_tx=XM_023243412.1
E1 38237882 38237999
refGene_gn=ABI3;refGene_tx=XM_023243411.1;refGene_gn=ABI3;refGene_tx=XM_023243412.1
E1 38240231 38240315
refGene_gn=GNGT2;refGene_tx=XM_006940231.4;refGene_gn=GNGT2;refGene_tx=XM_023243424.1
E1 38240695 38240821
refGene_gn=GNGT2;refGene_tx=XM_006940231.4;refGene_gn=GNGT2;refGene_tx=XM_023243424.1
E1 38264767 38264958 refGene_gn=B4GALNT2;refGene_tx=XM_023243464.1
E1 38265496 38265716 refGene_gn=B4GALNT2;refGene_tx=XM_023243464.1
E1 38268360 38268501 refGene_gn=B4GALNT2;refGene_tx=XM_023243464.1
E1 38269440 38269628 refGene_gn=B4GALNT2;refGene_tx=XM_023243464.1
E1 38280098 38280185 refGene_gn=B4GALNT2;refGene_tx=XM_023243464.1
E1 38281009 38281190 refGene_gn=B4GALNT2;refGene_tx=XM_023243464.1
E1 38283987 38284025 refGene_gn=B4GALNT2;refGene_tx=XM_023243464.1
E1 38285046 38285153 refGene_gn=B4GALNT2;refGene_tx=XM_023243464.1
E1 38289934 38290072 refGene_gn=B4GALNT2;refGene_tx=XM_023243464.1
E1 38290602 38290791 refGene_gn=B4GALNT2;refGene_tx=XM_023243464.1
E1 38341201 38341311 refGene_gn=B4GALNT2;refGene_tx=XM_023243464.1
E1 38357822 38357915
refGene_gn=IGF2BP1;refGene_tx=XM_003996719.4;refGene_gn=IGF2BP1;refGene_tx=XM_019817742.2
E1 38359760 38359874

refGene_gn=IGF2BP1;refGene_tx=XM_003996719.4;refGene_gn=IGF2BP1;refGene_tx=XM_019817742.2
E1 38360122 38360254
refGene_gn=IGF2BP1;refGene_tx=XM_003996719.4;refGene_gn=IGF2BP1;refGene_tx=XM_019817742.2
E1 38361308 38361383
refGene_gn=IGF2BP1;refGene_tx=XM_003996719.4;refGene_gn=IGF2BP1;refGene_tx=XM_019817742.2
E1 38361957 38362077
refGene_gn=IGF2BP1;refGene_tx=XM_003996719.4;refGene_gn=IGF2BP1;refGene_tx=XM_019817742.2
E1 38362408 38362531
refGene_gn=IGF2BP1;refGene_tx=XM_003996719.4;refGene_gn=IGF2BP1;refGene_tx=XM_019817742.2
E1 38365632 38365768
refGene_gn=IGF2BP1;refGene_tx=XM_003996719.4;refGene_gn=IGF2BP1;refGene_tx=XM_019817742.2
E1 38367046 38367169
refGene_gn=IGF2BP1;refGene_tx=XM_003996719.4;refGene_gn=IGF2BP1;refGene_tx=XM_019817742.2
E1 38368058 38368193
refGene_gn=IGF2BP1;refGene_tx=XM_003996719.4;refGene_gn=IGF2BP1;refGene_tx=XM_019817742.2
E1 38369112 38369394
refGene_gn=IGF2BP1;refGene_tx=XM_003996719.4;refGene_gn=IGF2BP1;refGene_tx=XM_019817742.2
E1 38370970 38371034
refGene_gn=IGF2BP1;refGene_tx=XM_003996719.4;refGene_gn=IGF2BP1;refGene_tx=XM_019817742.2
E1 38373518 38373570
refGene_gn=IGF2BP1;refGene_tx=XM_003996719.4;refGene_gn=IGF2BP1;refGene_tx=XM_019817742.2
E1 38374102 38374151
refGene_gn=IGF2BP1;refGene_tx=XM_003996719.4;refGene_gn=IGF2BP1;refGene_tx=XM_019817742.2
E1 38398703 38398764
refGene_gn=IGF2BP1;refGene_tx=XM_003996719.4;refGene_gn=IGF2BP1;refGene_tx=XM_019817742.2
E1 38399994 38400169
refGene_gn=IGF2BP1;refGene_tx=XM_003996719.4;refGene_gn=IGF2BP1;refGene_tx=XM_019817742.2
E1 38422090 38422176 refGene_gn=GIP;refGene_tx=XM_023244264.1
E1 38423789 38423936 refGene_gn=GIP;refGene_tx=XM_023244264.1
E1 38424546 38424643 refGene_gn=GIP;refGene_tx=XM_023244264.1
E1 38431161 38431215 refGene_gn=SNF8;refGene_tx=XM_011289185.3
E1 38431581 38431632 refGene_gn=SNF8;refGene_tx=XM_011289185.3
E1 38434273 38434412
refGene_gn=SNF8;refGene_tx=XM_011289185.3;refGene_gn=SNF8;refGene_tx=XM_023244699.1
E1 38435729 38435834

refGene_gn=SNF8;refGene_tx=XM_011289185.3;refGene_gn=SNF8;refGene_tx=XM_023244699.1
E1 38436504 38436577
refGene_gn=SNF8;refGene_tx=XM_011289185.3;refGene_gn=SNF8;refGene_tx=XM_023244699.1
E1 38437523 38437665
refGene_gn=SNF8;refGene_tx=XM_011289185.3;refGene_gn=SNF8;refGene_tx=XM_023244699.1
E1 38438533 38438608
refGene_gn=SNF8;refGene_tx=XM_011289185.3;refGene_gn=SNF8;refGene_tx=XM_023244699.1
E1 38439384 38439522
refGene_gn=SNF8;refGene_tx=XM_011289185.3;refGene_gn=SNF8;refGene_tx=XM_023244699.1
E1 38442915 38443086 refGene_gn=UBE2Z;refGene_tx=XM_023244698.1
E1 38445488 38445579 refGene_gn=UBE2Z;refGene_tx=XM_023244698.1
E1 38446016 38446129 refGene_gn=UBE2Z;refGene_tx=XM_023244698.1
E1 38449260 38449372 refGene_gn=UBE2Z;refGene_tx=XM_023244698.1
E1 38451436 38451624 refGene_gn=UBE2Z;refGene_tx=XM_023244698.1
E1 38453573 38453646 refGene_gn=UBE2Z;refGene_tx=XM_023244698.1
E1 38455657 38455974 refGene_gn=UBE2Z;refGene_tx=XM_023244698.1
E1 38462872 38462987
refGene_gn=ATP5MC1;refGene_tx=XM_003996721.5;refGene_gn=ATP5MC1;refGene_tx=XM_003996722.4
E1 38463317 38463496
refGene_gn=ATP5MC1;refGene_tx=XM_003996721.5;refGene_gn=ATP5MC1;refGene_tx=XM_003996722.4
E1 38464215 38464293
refGene_gn=ATP5MC1;refGene_tx=XM_003996721.5;refGene_gn=ATP5MC1;refGene_tx=XM_003996722.4
E1 38465182 38465221
refGene_gn=ATP5MC1;refGene_tx=XM_003996721.5;refGene_gn=ATP5MC1;refGene_tx=XM_003996722.4
E1 38508611 38508788
refGene_gn=CALCOCO2;refGene_tx=XM_003996724.5;refGene_gn=CALCOCO2;refGene_tx=XM_006940233.4;refGene_gn=CALCOCO2;refGene_tx=XM_019817746.2;refGene_gn=CALCOCO2;refGene_tx=XM_023243507.1
E1 38509316 38509345
refGene_gn=CALCOCO2;refGene_tx=XM_003996724.5;refGene_gn=CALCOCO2;refGene_tx=XM_006940233.4;refGene_gn=CALCOCO2;refGene_tx=XM_019817746.2;refGene_gn=CALCOCO2;refGene_tx=XM_023243507.1
E1 38510401 38510537
refGene_gn=CALCOCO2;refGene_tx=XM_003996724.5;refGene_gn=CALCOCO2;refGene_tx=XM_006940233.4;refGene_gn=CALCOCO2;refGene_tx=XM_019817746.2;refGene_gn=CALCOCO2;refGene_tx=XM_023243507.1
E1 38512603 38512699
refGene_gn=CALCOCO2;refGene_tx=XM_003996724.5;refGene_gn=CALCOCO2;refGene_tx=XM_006940233.4;refGene_gn=CALCOCO2;refGene_tx=XM_019817746.2;refGene_gn=CALCOCO2;refGene_tx=XM_023243507.1
E1 38513470 38513554 refGene_gn=CALCOCO2;refGene_tx=XM_023243508.1

E1 38513855 38514014 refGene_gn=CALCOCO2;refGene_tx=XM_023243508.1
E1 38515478 38515568
refGene_gn=CALCOCO2;refGene_tx=XM_003996724.5;refGene_gn=CALCOCO2;refGene_tx=XM_006940233.4;refGene_gn=CALCOCO2;refGene_tx=XM_019817746.2;refGene_gn=CALCOCO2;refGene_tx=XM_023243507.1;refGene_gn=CALCOCO2;refGene_tx=XM_023243508.1
E1 38515851 38515974
refGene_gn=CALCOCO2;refGene_tx=XM_003996724.5;refGene_gn=CALCOCO2;refGene_tx=XM_006940233.4;refGene_gn=CALCOCO2;refGene_tx=XM_019817746.2;refGene_gn=CALCOCO2;refGene_tx=XM_023243507.1;refGene_gn=CALCOCO2;refGene_tx=XM_023243508.1
E1 38516921 38516991
refGene_gn=CALCOCO2;refGene_tx=XM_003996724.5;refGene_gn=CALCOCO2;refGene_tx=XM_006940233.4;refGene_gn=CALCOCO2;refGene_tx=XM_019817746.2;refGene_gn=CALCOCO2;refGene_tx=XM_023243507.1;refGene_gn=CALCOCO2;refGene_tx=XM_023243508.1
E1 38517322 38517411
refGene_gn=CALCOCO2;refGene_tx=XM_003996724.5;refGene_gn=CALCOCO2;refGene_tx=XM_006940233.4;refGene_gn=CALCOCO2;refGene_tx=XM_019817746.2;refGene_gn=CALCOCO2;refGene_tx=XM_023243507.1;refGene_gn=CALCOCO2;refGene_tx=XM_023243508.1
E1 38519680 38519806
refGene_gn=CALCOCO2;refGene_tx=XM_003996724.5;refGene_gn=CALCOCO2;refGene_tx=XM_006940233.4;refGene_gn=CALCOCO2;refGene_tx=XM_019817746.2;refGene_gn=CALCOCO2;refGene_tx=XM_023243508.1
E1 38521771 38521905
refGene_gn=CALCOCO2;refGene_tx=XM_003996724.5;refGene_gn=CALCOCO2;refGene_tx=XM_006940233.4;refGene_gn=CALCOCO2;refGene_tx=XM_019817746.2;refGene_gn=CALCOCO2;refGene_tx=XM_023243507.1;refGene_gn=CALCOCO2;refGene_tx=XM_023243508.1
E1 38522064 38522167
refGene_gn=CALCOCO2;refGene_tx=XM_003996724.5;refGene_gn=CALCOCO2;refGene_tx=XM_006940233.4;refGene_gn=CALCOCO2;refGene_tx=XM_019817746.2;refGene_gn=CALCOCO2;refGene_tx=XM_023243507.1;refGene_gn=CALCOCO2;refGene_tx=XM_023243508.1
E1 38527274 38527454
refGene_gn=CALCOCO2;refGene_tx=XM_003996724.5;refGene_gn=CALCOCO2;refGene_tx=XM_006940233.4;refGene_gn=CALCOCO2;refGene_tx=XM_019817746.2;refGene_gn=CALCOCO2;refGene_tx=XM_023243507.1;refGene_gn=CALCOCO2;refGene_tx=XM_023243508.1
E1 38553957 38554034 refGene_gn=TTLL6;refGene_tx=XM_023243879.1
E1 38556243 38556462 refGene_gn=TTLL6;refGene_tx=XM_023243879.1
E1 38557537 38557578 refGene_gn=TTLL6;refGene_tx=XM_023243879.1
E1 38558722 38558841 refGene_gn=TTLL6;refGene_tx=XM_023243879.1
E1 38558980 38559111 refGene_gn=TTLL6;refGene_tx=XM_023243879.1
E1 38560880 38561037 refGene_gn=TTLL6;refGene_tx=XM_023243879.1
E1 38561780 38561924 refGene_gn=TTLL6;refGene_tx=XM_023243879.1
E1 38562521 38562607 refGene_gn=TTLL6;refGene_tx=XM_023243879.1
E1 38567871 38568097 refGene_gn=TTLL6;refGene_tx=XM_023243879.1
E1 38569335 38569511 refGene_gn=TTLL6;refGene_tx=XM_023243879.1

E1	38571871	38572060	refGene_gn=TTLL6;refGene_tx=XM_023243879.1
E1	38573612	38573787	refGene_gn=TTLL6;refGene_tx=XM_023243879.1
E1	38574977	38575079	refGene_gn=TTLL6;refGene_tx=XM_023243879.1
E1	38575080	38575239	refGene_gn=TTLL6;refGene_tx=XM_023243879.1
E1	38585428	38585789	refGene_gn=TTLL6;refGene_tx=XM_023243879.1
E1	38586670	38586897	refGene_gn=TTLL6;refGene_tx=XM_023243879.1
E1	38619627	38620231	refGene_gn=HOXB13;refGene_tx=XM_003996743.4
E1	38621215	38621469	refGene_gn=HOXB13;refGene_tx=XM_003996743.4
E1	38696589	38696596	refGene_gn=HOXB9;refGene_tx=XM_003996744.5
E1	38697890	38698409	refGene_gn=HOXB9;refGene_tx=XM_003996744.5
E1	38701152	38701388	refGene_gn=HOXB9;refGene_tx=XM_003996744.5
E1	38709728	38709984	refGene_gn=HOXB8;refGene_tx=XM_011289183.3
E1	38709986	38710151	refGene_gn=HOXB8;refGene_tx=XM_011289183.3
E1	38711043	38711351	refGene_gn=HOXB8;refGene_tx=XM_011289183.3
E1	38713637	38714037	refGene_gn=HOXB7;refGene_tx=XM_003996745.4
E1	38716099	38716353	refGene_gn=HOXB7;refGene_tx=XM_003996745.4
E1	38726090	38726505	refGene_gn=HOXB6;refGene_tx=XM_006940250.3
E1	38727565	38727825	refGene_gn=HOXB6;refGene_tx=XM_006940250.3
E1	38730478	38731040	refGene_gn=HOXB5;refGene_tx=XM_003996746.4
E1	38731730	38731978	refGene_gn=HOXB5;refGene_tx=XM_003996746.4
E1	38746209	38746657	refGene_gn=HOXB4;refGene_tx=XM_023244423.1
E1	38747521	38747820	refGene_gn=HOXB4;refGene_tx=XM_023244423.1
E1	38772539	38772987
refGene_gn=HOXB3;refGene_tx=XM_011289180.3;refGene_gn=HOXB3;refGene_tx
=XM_019817750.2;refGene_gn=HOXB3;refGene_tx=XM_023244420.1
E1	38773850	38774701
refGene_gn=HOXB3;refGene_tx=XM_011289180.3;refGene_gn=HOXB3;refGene_tx
=XM_019817750.2;refGene_gn=HOXB3;refGene_tx=XM_023244420.1
E1	38780035	38780417	refGene_gn=HOXB2;refGene_tx=XM_023244422.1
E1	38781177	38781836	refGene_gn=HOXB2;refGene_tx=XM_023244422.1
E1	38791646	38792232	refGene_gn=HOXB1;refGene_tx=XM_003996749.4
E1	38792715	38793044	refGene_gn=HOXB1;refGene_tx=XM_003996749.4
E1	38885273	38885319	refGene_gn=SKAP1;refGene_tx=XM_006940253.4
E1	38917386	38917492	refGene_gn=SKAP1;refGene_tx=XM_006940253.4
E1	38944370	38944396	refGene_gn=SKAP1;refGene_tx=XM_006940253.4
E1	38962725	38962842	refGene_gn=SKAP1;refGene_tx=XM_006940253.4
E1	39118975	39119053	refGene_gn=SKAP1;refGene_tx=XM_006940253.4
E1	39120160	39120244	refGene_gn=SKAP1;refGene_tx=XM_006940253.4
E1	39122897	39123022	refGene_gn=SKAP1;refGene_tx=XM_006940253.4
E1	39126046	39126110	refGene_gn=SKAP1;refGene_tx=XM_006940253.4
E1	39128266	39128458	refGene_gn=SKAP1;refGene_tx=XM_006940253.4
E1	39139308	39139359	refGene_gn=SKAP1;refGene_tx=XM_006940253.4
E1	39147103	39147204	refGene_gn=SKAP1;refGene_tx=XM_006940253.4
E1	39167769	39167871	refGene_gn=SKAP1;refGene_tx=XM_006940253.4
E1	39181460	39181734
refGene_gn=SNX11;refGene_tx=XM_003996750.4;refGene_gn=SNX11;refGene_tx
=XM_006940254.4;refGene_gn=SNX11;refGene_tx=XM_006940255.3;refGene_gn=
SNX11;refGene_tx=XM_019817754.2
E1	39182858	39183071
refGene_gn=SNX11;refGene_tx=XM_003996750.4;refGene_gn=SNX11;refGene_tx

=XM_006940254.4;refGene_gn=SNX11;refGene_tx=XM_006940255.3;refGene_gn=SNX11;refGene_tx=XM_019817754.2
E1 39183302 39183398
refGene_gn=SNX11;refGene_tx=XM_003996750.4;refGene_gn=SNX11;refGene_tx=XM_006940254.4;refGene_gn=SNX11;refGene_tx=XM_006940255.3;refGene_gn=SNX11;refGene_tx=XM_019817754.2
E1 39185234 39185335
refGene_gn=SNX11;refGene_tx=XM_003996750.4;refGene_gn=SNX11;refGene_tx=XM_006940254.4;refGene_gn=SNX11;refGene_tx=XM_006940255.3;refGene_gn=SNX11;refGene_tx=XM_019817754.2
E1 39185639 39185726
refGene_gn=SNX11;refGene_tx=XM_003996750.4;refGene_gn=SNX11;refGene_tx=XM_006940254.4;refGene_gn=SNX11;refGene_tx=XM_006940255.3
E1 39186163 39186218
refGene_gn=SNX11;refGene_tx=XM_003996750.4;refGene_gn=SNX11;refGene_tx=XM_006940254.4;refGene_gn=SNX11;refGene_tx=XM_006940255.3;refGene_gn=SNX11;refGene_tx=XM_019817754.2
E1 39187190 39187250
refGene_gn=SNX11;refGene_tx=XM_006940254.4;refGene_gn=SNX11;refGene_tx=XM_019817754.2
E1 39187621 39187752
refGene_gn=SNX11;refGene_tx=XM_006940254.4;refGene_gn=SNX11;refGene_tx=XM_019817754.2
E1 39214544 39214684
refGene_gn=CBX1;refGene_tx=XM_003996751.4;refGene_gn=CBX1;refGene_tx=XM_019817755.1
E1 39215013 39215191
refGene_gn=CBX1;refGene_tx=XM_003996751.4;refGene_gn=CBX1;refGene_tx=XM_019817755.1
E1 39215755 39215850
refGene_gn=CBX1;refGene_tx=XM_003996751.4;refGene_gn=CBX1;refGene_tx=XM_019817755.1
E1 39218325 39218470
refGene_gn=CBX1;refGene_tx=XM_003996751.4;refGene_gn=CBX1;refGene_tx=XM_019817755.1
E1 39227791 39229186
refGene_gn=NFE2L1;refGene_tx=XM_003996752.5;refGene_gn=NFE2L1;refGene_tx=XM_006940256.3;refGene_gn=NFE2L1;refGene_tx=XM_006940257.4;refGene_gn=NFE2L1;refGene_tx=XM_019817756.2;refGene_gn=NFE2L1;refGene_tx=XM_019817757.2;refGene_gn=NFE2L1;refGene_tx=XM_019817758.2;refGene_gn=NFE2L1;refGene_tx=XM_019817759.2
E1 39229990 39230149
refGene_gn=NFE2L1;refGene_tx=XM_003996752.5;refGene_gn=NFE2L1;refGene_tx=XM_006940256.3;refGene_gn=NFE2L1;refGene_tx=XM_006940257.4;refGene_gn=NFE2L1;refGene_tx=XM_019817756.2;refGene_gn=NFE2L1;refGene_tx=XM_019817757.2;refGene_gn=NFE2L1;refGene_tx=XM_019817758.2;refGene_gn=NFE2L1;refGene_tx=XM_019817759.2
E1 39230374 39230464
refGene_gn=NFE2L1;refGene_tx=XM_003996752.5;refGene_gn=NFE2L1;refGene_tx=XM_006940256.3;refGene_gn=NFE2L1;refGene_tx=XM_006940257.4;refGene_

gn=NFE2L1;refGene_tx=XM_019817756.2;refGene_gn=NFE2L1;refGene_tx=XM_01
9817758.2;refGene_gn=NFE2L1;refGene_tx=XM_019817759.2
E1 39230876 39231092
refGene_gn=NFE2L1;refGene_tx=XM_003996752.5;refGene_gn=NFE2L1;refGene_
tx=XM_006940256.3;refGene_gn=NFE2L1;refGene_tx=XM_006940257.4;refGene_
gn=NFE2L1;refGene_tx=XM_019817756.2;refGene_gn=NFE2L1;refGene_tx=XM_01
9817757.2;refGene_gn=NFE2L1;refGene_tx=XM_019817758.2;refGene_gn=NFE2L
1;refGene_tx=XM_019817759.2
E1 39232296 39232428
refGene_gn=NFE2L1;refGene_tx=XM_019817758.2;refGene_gn=NFE2L1;refGene_
tx=XM_019817759.2
E1 39235819 39236329
refGene_gn=NFE2L1;refGene_tx=XM_003996752.5;refGene_gn=NFE2L1;refGene_
tx=XM_006940256.3;refGene_gn=NFE2L1;refGene_tx=XM_006940257.4;refGene_
gn=NFE2L1;refGene_tx=XM_019817756.2;refGene_gn=NFE2L1;refGene_tx=XM_01
9817757.2
E1 39248520 39248631
refGene_gn=COPZ2;refGene_tx=XM_023244226.1;refGene_gn=COPZ2;refGene_tx
=XM_023244227.1;refGene_gn=COPZ2;refGene_tx=XM_023244228.1;refGene_gn=
COPZ2;refGene_tx=XM_023244229.1;refGene_gn=COPZ2;refGene_tx=XM_0232442
30.1;refGene_gn=COPZ2;refGene_tx=XM_023244231.1
E1 39249396 39249471
refGene_gn=COPZ2;refGene_tx=XM_023244226.1;refGene_gn=COPZ2;refGene_tx
=XM_023244227.1;refGene_gn=COPZ2;refGene_tx=XM_023244228.1;refGene_gn=
COPZ2;refGene_tx=XM_023244229.1;refGene_gn=COPZ2;refGene_tx=XM_0232442
30.1;refGene_gn=COPZ2;refGene_tx=XM_023244231.1
E1 39251062 39251144
refGene_gn=COPZ2;refGene_tx=XM_023244226.1;refGene_gn=COPZ2;refGene_tx
=XM_023244227.1;refGene_gn=COPZ2;refGene_tx=XM_023244228.1;refGene_gn=
COPZ2;refGene_tx=XM_023244229.1;refGene_gn=COPZ2;refGene_tx=XM_0232442
30.1;refGene_gn=COPZ2;refGene_tx=XM_023244231.1
E1 39251683 39251775
refGene_gn=COPZ2;refGene_tx=XM_023244226.1;refGene_gn=COPZ2;refGene_tx
=XM_023244227.1;refGene_gn=COPZ2;refGene_tx=XM_023244228.1;refGene_gn=
COPZ2;refGene_tx=XM_023244229.1;refGene_gn=COPZ2;refGene_tx=XM_0232442
30.1;refGene_gn=COPZ2;refGene_tx=XM_023244231.1
E1 39252231 39252287
refGene_gn=COPZ2;refGene_tx=XM_023244226.1;refGene_gn=COPZ2;refGene_tx
=XM_023244227.1;refGene_gn=COPZ2;refGene_tx=XM_023244228.1;refGene_gn=
COPZ2;refGene_tx=XM_023244229.1;refGene_gn=COPZ2;refGene_tx=XM_0232442
30.1;refGene_gn=COPZ2;refGene_tx=XM_023244231.1
E1 39252378 39252522
refGene_gn=COPZ2;refGene_tx=XM_023244226.1;refGene_gn=COPZ2;refGene_tx
=XM_023244227.1;refGene_gn=COPZ2;refGene_tx=XM_023244228.1
E1 39252997 39253075
refGene_gn=COPZ2;refGene_tx=XM_023244226.1;refGene_gn=COPZ2;refGene_tx
=XM_023244227.1;refGene_gn=COPZ2;refGene_tx=XM_023244228.1;refGene_gn=
COPZ2;refGene_tx=XM_023244229.1;refGene_gn=COPZ2;refGene_tx=XM_0232442
30.1;refGene_gn=COPZ2;refGene_tx=XM_023244231.1
E1 39255374 39255426

refGene_gn=COPZ2;refGene_tx=XM_023244226.1;refGene_gn=COPZ2;refGene_tx=XM_023244227.1;refGene_gn=COPZ2;refGene_tx=XM_023244228.1;refGene_gn=COPZ2;refGene_tx=XM_023244229.1;refGene_gn=COPZ2;refGene_tx=XM_023244230.1;refGene_gn=COPZ2;refGene_tx=XM_023244231.1
E1 39256035 39256074
refGene_gn=COPZ2;refGene_tx=XM_023244226.1;refGene_gn=COPZ2;refGene_tx=XM_023244227.1;refGene_gn=COPZ2;refGene_tx=XM_023244229.1;refGene_gn=COPZ2;refGene_tx=XM_023244230.1
E1 39256715 39256787
refGene_gn=COPZ2;refGene_tx=XM_023244226.1;refGene_gn=COPZ2;refGene_tx=XM_023244229.1
E1 39258028 39258076
refGene_gn=COPZ2;refGene_tx=XM_023244227.1;refGene_gn=COPZ2;refGene_tx=XM_023244228.1;refGene_gn=COPZ2;refGene_tx=XM_023244230.1;refGene_gn=COPZ2;refGene_tx=XM_023244231.1
E1 39295173 39295239
refGene_gn=CDK5RAP3;refGene_tx=XM_003996753.5;refGene_gn=CDK5RAP3;refGene_tx=XM_019817760.1;refGene_gn=CDK5RAP3;refGene_tx=XM_019817761.2
E1 39295349 39295521
refGene_gn=CDK5RAP3;refGene_tx=XM_003996753.5;refGene_gn=CDK5RAP3;refGene_tx=XM_019817760.1;refGene_gn=CDK5RAP3;refGene_tx=XM_019817761.2
E1 39295876 39296082
refGene_gn=CDK5RAP3;refGene_tx=XM_003996753.5;refGene_gn=CDK5RAP3;refGene_tx=XM_019817760.1;refGene_gn=CDK5RAP3;refGene_tx=XM_019817761.2
E1 39296937 39297026
refGene_gn=CDK5RAP3;refGene_tx=XM_003996753.5;refGene_gn=CDK5RAP3;refGene_tx=XM_019817760.1;refGene_gn=CDK5RAP3;refGene_tx=XM_019817761.2
E1 39297648 39297724
refGene_gn=CDK5RAP3;refGene_tx=XM_003996753.5;refGene_gn=CDK5RAP3;refGene_tx=XM_019817760.1;refGene_gn=CDK5RAP3;refGene_tx=XM_019817761.2
E1 39298132 39298243
refGene_gn=CDK5RAP3;refGene_tx=XM_003996753.5;refGene_gn=CDK5RAP3;refGene_tx=XM_019817760.1;refGene_gn=CDK5RAP3;refGene_tx=XM_019817761.2
E1 39298634 39298779
refGene_gn=CDK5RAP3;refGene_tx=XM_003996753.5;refGene_gn=CDK5RAP3;refGene_tx=XM_019817760.1;refGene_gn=CDK5RAP3;refGene_tx=XM_019817761.2
E1 39298996 39299133
refGene_gn=CDK5RAP3;refGene_tx=XM_003996753.5;refGene_gn=CDK5RAP3;refGene_tx=XM_019817760.1;refGene_gn=CDK5RAP3;refGene_tx=XM_019817761.2
E1 39299329 39299508
refGene_gn=CDK5RAP3;refGene_tx=XM_003996753.5;refGene_gn=CDK5RAP3;refGene_tx=XM_019817760.1;refGene_gn=CDK5RAP3;refGene_tx=XM_019817761.2
E1 39300162 39300211 refGene_gn=CDK5RAP3;refGene_tx=XM_003996753.5
E1 39300585 39300686
refGene_gn=CDK5RAP3;refGene_tx=XM_003996753.5;refGene_gn=CDK5RAP3;refGene_tx=XM_019817760.1;refGene_gn=CDK5RAP3;refGene_tx=XM_019817761.2
E1 39300960 39301092
refGene_gn=CDK5RAP3;refGene_tx=XM_003996753.5;refGene_gn=CDK5RAP3;refGene_tx=XM_019817760.1;refGene_gn=CDK5RAP3;refGene_tx=XM_019817761.2
E1 39303129 39303175

refGene_gn=CDK5RAP3;refGene_tx=XM_003996753.5;refGene_gn=CDK5RAP3;refGene_tx=XM_019817760.1
E1 39303262 39303377 refGene_gn=CDK5RAP3;refGene_tx=XM_019817760.1
E1 39303400 39303406 refGene_gn=CDK5RAP3;refGene_tx=XM_003996753.5
E1 39315660 39315963 refGene_gn=PRR15L;refGene_tx=XM_006940260.3;refGene_gn=PRR15L;refGene_tx=XM_019817763.2
E1 39321155 39321324 refGene_gn=PNPO;refGene_tx=XM_003996754.5
E1 39321554 39321625 refGene_gn=PNPO;refGene_tx=XM_003996754.5
E1 39321924 39322053 refGene_gn=PNPO;refGene_tx=XM_003996754.5
E1 39322280 39322334 refGene_gn=PNPO;refGene_tx=XM_003996754.5
E1 39323201 39323301 refGene_gn=PNPO;refGene_tx=XM_003996754.5
E1 39324691 39324816 refGene_gn=PNPO;refGene_tx=XM_003996754.5
E1 39325691 39325770 refGene_gn=PNPO;refGene_tx=XM_003996754.5
E1 39325771 39325827 refGene_gn=PNPO;refGene_tx=XM_003996754.5
E1 39333533 39333634 refGene_gn=SP2;refGene_tx=XM_006940261.4;refGene_gn=SP2;refGene_tx=XM_006940262.3;refGene_gn=SP2;refGene_tx=XM_006940263.3;refGene_gn=SP2;refGene_tx=XM_019817764.1;refGene_gn=SP2;refGene_tx=XM_019817765.1;refGene_gn=SP2;refGene_tx=XM_019817767.2;refGene_gn=SP2;refGene_tx=XM_023244725.1
E1 39335015 39335209 refGene_gn=SP2;refGene_tx=XM_006940261.4;refGene_gn=SP2;refGene_tx=XM_006940262.3;refGene_gn=SP2;refGene_tx=XM_006940263.3;refGene_gn=SP2;refGene_tx=XM_019817764.1;refGene_gn=SP2;refGene_tx=XM_019817765.1;refGene_gn=SP2;refGene_tx=XM_019817767.2;refGene_gn=SP2;refGene_tx=XM_023244725.1
E1 39335452 39335627 refGene_gn=SP2;refGene_tx=XM_006940261.4;refGene_gn=SP2;refGene_tx=XM_006940262.3;refGene_gn=SP2;refGene_tx=XM_006940263.3;refGene_gn=SP2;refGene_tx=XM_019817764.1;refGene_gn=SP2;refGene_tx=XM_019817765.1;refGene_gn=SP2;refGene_tx=XM_019817767.2;refGene_gn=SP2;refGene_tx=XM_023244725.1
E1 39337350 39337663 refGene_gn=SP2;refGene_tx=XM_006940261.4;refGene_gn=SP2;refGene_tx=XM_006940262.3;refGene_gn=SP2;refGene_tx=XM_006940263.3;refGene_gn=SP2;refGene_tx=XM_019817764.1;refGene_gn=SP2;refGene_tx=XM_019817765.1;refGene_gn=SP2;refGene_tx=XM_019817767.2;refGene_gn=SP2;refGene_tx=XM_023244725.1
E1 39342666 39343641 refGene_gn=SP2;refGene_tx=XM_006940261.4;refGene_gn=SP2;refGene_tx=XM_006940262.3;refGene_gn=SP2;refGene_tx=XM_006940263.3;refGene_gn=SP2;refGene_tx=XM_019817764.1;refGene_gn=SP2;refGene_tx=XM_019817765.1;refGene_gn=SP2;refGene_tx=XM_019817767.2;refGene_gn=SP2;refGene_tx=XM_023244725.1
E1 39344343 39344423 refGene_gn=SP2;refGene_tx=XM_006940261.4;refGene_gn=SP2;refGene_tx=XM_006940262.3;refGene_gn=SP2;refGene_tx=XM_006940263.3;refGene_gn=SP2;refGene_tx=XM_019817764.1;refGene_gn=SP2;refGene_tx=XM_019817765.1;refGene_gn=SP2;refGene_tx=XM_019817767.2;refGene_gn=SP2;refGene_tx=XM_02324

4725.1
E1 39358148 39358155
refGene_gn=SP2;refGene_tx=XM_006940261.4;refGene_gn=SP2;refGene_tx=XM_
019817764.1
E1 39399976 39401107 refGene_gn=SP6;refGene_tx=XM_006940264.4
E1 39408139 39408313
refGene_gn=SCRN2;refGene_tx=XM_006940265.2;refGene_gn=SCRN2;refGene_tx
=XM_006940266.2
E1 39408537 39408719
refGene_gn=SCRN2;refGene_tx=XM_006940265.2;refGene_gn=SCRN2;refGene_tx
=XM_006940266.2
E1 39409201 39409506
refGene_gn=SCRN2;refGene_tx=XM_006940265.2;refGene_gn=SCRN2;refGene_tx
=XM_006940266.2
E1 39409956 39410172
refGene_gn=SCRN2;refGene_tx=XM_006940265.2;refGene_gn=SCRN2;refGene_tx
=XM_006940266.2
E1 39410310 39410476
refGene_gn=SCRN2;refGene_tx=XM_006940265.2;refGene_gn=SCRN2;refGene_tx
=XM_006940266.2
E1 39410577 39410758
refGene_gn=SCRN2;refGene_tx=XM_006940265.2;refGene_gn=SCRN2;refGene_tx
=XM_006940266.2
E1 39411033 39411192
refGene_gn=SCRN2;refGene_tx=XM_006940265.2;refGene_gn=SCRN2;refGene_tx
=XM_006940266.2
E1 39411934 39412302
refGene_gn=LRRC46;refGene_tx=XM_003996806.5;refGene_gn=LRRC46;refGene_
tx=XM_006940267.4
E1 39412600 39412749
refGene_gn=LRRC46;refGene_tx=XM_003996806.5;refGene_gn=LRRC46;refGene_
tx=XM_006940267.4
E1 39413103 39413173
refGene_gn=LRRC46;refGene_tx=XM_003996806.5;refGene_gn=LRRC46;refGene_
tx=XM_006940267.4
E1 39413366 39413476
refGene_gn=LRRC46;refGene_tx=XM_003996806.5;refGene_gn=LRRC46;refGene_
tx=XM_006940267.4
E1 39413753 39413800
refGene_gn=LRRC46;refGene_tx=XM_003996806.5;refGene_gn=LRRC46;refGene_
tx=XM_006940267.4
E1 39414958 39415067
refGene_gn=LRRC46;refGene_tx=XM_003996806.5;refGene_gn=LRRC46;refGene_
tx=XM_006940267.4
E1 39416383 39416489
refGene_gn=LRRC46;refGene_tx=XM_003996806.5;refGene_gn=LRRC46;refGene_
tx=XM_006940267.4
E1 39416580 39416590
refGene_gn=LRRC46;refGene_tx=XM_003996806.5;refGene_gn=LRRC46;refGene_
tx=XM_006940267.4

E1 39417076 39417128 refGene_gn=MRPL10;refGene_tx=XM_003996758.4
E1 39419017 39419187
refGene_gn=MRPL10;refGene_tx=XM_003996758.4;refGene_gn=MRPL10;refGene_tx=XM_006940268.2
E1 39420257 39420422
refGene_gn=MRPL10;refGene_tx=XM_003996758.4;refGene_gn=MRPL10;refGene_tx=XM_006940268.2
E1 39420636 39420781
refGene_gn=MRPL10;refGene_tx=XM_003996758.4;refGene_gn=MRPL10;refGene_tx=XM_006940268.2
E1 39421563 39421820
refGene_gn=MRPL10;refGene_tx=XM_003996758.4;refGene_gn=MRPL10;refGene_tx=XM_006940268.2
E1 39426035 39426110 refGene_gn=OSBPL7;refGene_tx=XM_011289176.2
E1 39426225 39426351 refGene_gn=OSBPL7;refGene_tx=XM_011289176.2
E1 39426534 39426588 refGene_gn=OSBPL7;refGene_tx=XM_011289176.2
E1 39427222 39427336 refGene_gn=OSBPL7;refGene_tx=XM_011289176.2
E1 39427693 39427804 refGene_gn=OSBPL7;refGene_tx=XM_011289176.2
E1 39427924 39428042 refGene_gn=OSBPL7;refGene_tx=XM_011289176.2
E1 39428690 39428794 refGene_gn=OSBPL7;refGene_tx=XM_011289176.2
E1 39429164 39429257 refGene_gn=OSBPL7;refGene_tx=XM_011289176.2
E1 39429342 39429475 refGene_gn=OSBPL7;refGene_tx=XM_011289176.2
E1 39429557 39429652 refGene_gn=OSBPL7;refGene_tx=XM_011289176.2
E1 39429841 39429937 refGene_gn=OSBPL7;refGene_tx=XM_011289176.2
E1 39431514 39431652 refGene_gn=OSBPL7;refGene_tx=XM_011289176.2
E1 39432201 39432295 refGene_gn=OSBPL7;refGene_tx=XM_011289176.2
E1 39432958 39433206 refGene_gn=OSBPL7;refGene_tx=XM_011289176.2
E1 39434450 39434588 refGene_gn=OSBPL7;refGene_tx=XM_011289176.2
E1 39437808 39437872 refGene_gn=OSBPL7;refGene_tx=XM_011289176.2
E1 39437999 39438078 refGene_gn=OSBPL7;refGene_tx=XM_011289176.2
E1 39439488 39439633 refGene_gn=OSBPL7;refGene_tx=XM_011289176.2
E1 39439736 39439881 refGene_gn=OSBPL7;refGene_tx=XM_011289176.2
E1 39439969 39440096 refGene_gn=OSBPL7;refGene_tx=XM_011289176.2
E1 39440306 39440429 refGene_gn=OSBPL7;refGene_tx=XM_011289176.2
E1 39440580 39440689 refGene_gn=OSBPL7;refGene_tx=XM_011289176.2
E1 39488428 39489047 refGene_gn=TBX21;refGene_tx=XM_023244122.1
E1 39489286 39489348 refGene_gn=TBX21;refGene_tx=XM_023244122.1
E1 39489485 39489644 refGene_gn=TBX21;refGene_tx=XM_023244122.1
E1 39490188 39490310 refGene_gn=TBX21;refGene_tx=XM_023244122.1
E1 39490662 39490817 refGene_gn=TBX21;refGene_tx=XM_023244122.1
E1 39498131 39498622 refGene_gn=TBX21;refGene_tx=XM_023244122.1
E1 39521119 39521248
refGene_gn=TBKBP1;refGene_tx=XM_019817773.2;refGene_gn=TBKBP1;refGene_tx=XM_023243522.1;refGene_gn=TBKBP1;refGene_tx=XM_023243523.1;refGene_gn=TBKBP1;refGene_tx=XM_023243524.1;refGene_gn=TBKBP1;refGene_tx=XM_023243525.1
E1 39522187 39522915
refGene_gn=TBKBP1;refGene_tx=XM_019817773.2;refGene_gn=TBKBP1;refGene_tx=XM_023243522.1;refGene_gn=TBKBP1;refGene_tx=XM_023243523.1;refGene_gn=TBKBP1;refGene_tx=XM_023243524.1;refGene_gn=TBKBP1;refGene_tx=XM_02

3243525.1
E1 39523163 39523282
refGene_gn=TBKBP1;refGene_tx=XM_019817773.2;refGene_gn=TBKBP1;refGene_tx=XM_023243522.1;refGene_gn=TBKBP1;refGene_tx=XM_023243523.1;refGene_gn=TBKBP1;refGene_tx=XM_023243524.1;refGene_gn=TBKBP1;refGene_tx=XM_023243525.1
E1 39531918 39531980
refGene_gn=TBKBP1;refGene_tx=XM_019817773.2;refGene_gn=TBKBP1;refGene_tx=XM_023243522.1;refGene_gn=TBKBP1;refGene_tx=XM_023243523.1;refGene_gn=TBKBP1;refGene_tx=XM_023243524.1;refGene_gn=TBKBP1;refGene_tx=XM_023243525.1
E1 39532144 39532320
refGene_gn=TBKBP1;refGene_tx=XM_019817773.2;refGene_gn=TBKBP1;refGene_tx=XM_023243522.1;refGene_gn=TBKBP1;refGene_tx=XM_023243523.1;refGene_gn=TBKBP1;refGene_tx=XM_023243524.1;refGene_gn=TBKBP1;refGene_tx=XM_023243525.1
E1 39532901 39533082
refGene_gn=TBKBP1;refGene_tx=XM_019817773.2;refGene_gn=TBKBP1;refGene_tx=XM_023243522.1;refGene_gn=TBKBP1;refGene_tx=XM_023243523.1;refGene_gn=TBKBP1;refGene_tx=XM_023243524.1;refGene_gn=TBKBP1;refGene_tx=XM_023243525.1
E1 39534122 39534227
refGene_gn=TBKBP1;refGene_tx=XM_019817773.2;refGene_gn=TBKBP1;refGene_tx=XM_023243522.1;refGene_gn=TBKBP1;refGene_tx=XM_023243523.1;refGene_gn=TBKBP1;refGene_tx=XM_023243524.1;refGene_gn=TBKBP1;refGene_tx=XM_023243525.1
E1 39534507 39534630
refGene_gn=TBKBP1;refGene_tx=XM_019817773.2;refGene_gn=TBKBP1;refGene_tx=XM_023243522.1;refGene_gn=TBKBP1;refGene_tx=XM_023243523.1;refGene_gn=TBKBP1;refGene_tx=XM_023243524.1;refGene_gn=TBKBP1;refGene_tx=XM_023243525.1
E1 39534987 39535299
refGene_gn=TBKBP1;refGene_tx=XM_019817773.2;refGene_gn=TBKBP1;refGene_tx=XM_023243522.1;refGene_gn=TBKBP1;refGene_tx=XM_023243523.1;refGene_gn=TBKBP1;refGene_tx=XM_023243524.1;refGene_gn=TBKBP1;refGene_tx=XM_023243525.1
E1 39547663 39547664 refGene_gn=KPNB1;refGene_tx=XM_023243587.1
E1 39548583 39548745 refGene_gn=KPNB1;refGene_tx=XM_023243587.1
E1 39549748 39549863 refGene_gn=KPNB1;refGene_tx=XM_023243587.1
E1 39550506 39550612 refGene_gn=KPNB1;refGene_tx=XM_023243587.1
E1 39550723 39550867 refGene_gn=KPNB1;refGene_tx=XM_023243587.1
E1 39551237 39551345 refGene_gn=KPNB1;refGene_tx=XM_023243587.1
E1 39551841 39551924 refGene_gn=KPNB1;refGene_tx=XM_023243587.1
E1 39553116 39553261 refGene_gn=KPNB1;refGene_tx=XM_023243587.1
E1 39554450 39554522 refGene_gn=KPNB1;refGene_tx=XM_023243587.1
E1 39554847 39554995 refGene_gn=KPNB1;refGene_tx=XM_023243587.1
E1 39556881 39557012 refGene_gn=KPNB1;refGene_tx=XM_023243587.1
E1 39557788 39557980 refGene_gn=KPNB1;refGene_tx=XM_023243587.1
E1 39559095 39559320 refGene_gn=KPNB1;refGene_tx=XM_023243587.1
E1 39561138 39561240 refGene_gn=KPNB1;refGene_tx=XM_023243587.1

E1 39562042 39562153 refGene_gn=KPNB1;refGene_tx=XM_023243587.1
E1 39562507 39562597 refGene_gn=KPNB1;refGene_tx=XM_023243587.1
E1 39563415 39563475 refGene_gn=KPNB1;refGene_tx=XM_023243587.1
E1 39565575 39565728 refGene_gn=KPNB1;refGene_tx=XM_023243587.1
E1 39566666 39566867 refGene_gn=KPNB1;refGene_tx=XM_023243587.1
E1 39570004 39570187 refGene_gn=KPNB1;refGene_tx=XM_023243587.1
E1 39572367 39572426 refGene_gn=KPNB1;refGene_tx=XM_023243587.1
E1 39572529 39572569 refGene_gn=KPNB1;refGene_tx=XM_023243587.1
E1 39604448 39604601 refGene_gn=NPEPPS;refGene_tx=XM_023243586.1
E1 39606708 39606756 refGene_gn=NPEPPS;refGene_tx=XM_023243586.1
E1 39607045 39607201 refGene_gn=NPEPPS;refGene_tx=XM_023243586.1
E1 39607684 39607792 refGene_gn=NPEPPS;refGene_tx=XM_023243586.1
E1 39610706 39610763 refGene_gn=NPEPPS;refGene_tx=XM_023243586.1
E1 39611190 39611333 refGene_gn=NPEPPS;refGene_tx=XM_023243586.1
E1 39615824 39616044 refGene_gn=NPEPPS;refGene_tx=XM_023243586.1
E1 39617509 39617644 refGene_gn=NPEPPS;refGene_tx=XM_023243586.1
E1 39620343 39620483 refGene_gn=NPEPPS;refGene_tx=XM_023243586.1
E1 39622206 39622270 refGene_gn=NPEPPS;refGene_tx=XM_023243586.1
E1 39624108 39624218 refGene_gn=NPEPPS;refGene_tx=XM_023243586.1
E1 39626807 39626868 refGene_gn=NPEPPS;refGene_tx=XM_023243586.1
E1 39627285 39627390 refGene_gn=NPEPPS;refGene_tx=XM_023243586.1
E1 39628218 39628383 refGene_gn=NPEPPS;refGene_tx=XM_023243586.1
E1 39630421 39630536 refGene_gn=NPEPPS;refGene_tx=XM_023243586.1
E1 39631378 39631412 refGene_gn=NPEPPS;refGene_tx=XM_023243586.1
E1 39631523 39631620 refGene_gn=NPEPPS;refGene_tx=XM_023243586.1
E1 39632065 39632266 refGene_gn=NPEPPS;refGene_tx=XM_023243586.1
E1 39633738 39633846 refGene_gn=NPEPPS;refGene_tx=XM_023243586.1
E1 39635277 39635399 refGene_gn=NPEPPS;refGene_tx=XM_023243586.1
E1 39644062 39644140 refGene_gn=NPEPPS;refGene_tx=XM_023243586.1
E1 39688697 39688782 refGene_gn=NPEPPS;refGene_tx=XM_023243586.1
E1 39698287 39698542 refGene_gn=NPEPPS;refGene_tx=XM_023243586.1
E1 39730878 39730944
refGene_gn=MRPL45;refGene_tx=XM_011289173.3;refGene_gn=MRPL45;refGene_tx=XM_011289174.3;refGene_gn=MRPL45;refGene_tx=XM_019817781.2
E1 39731833 39732011
refGene_gn=MRPL45;refGene_tx=XM_011289173.3;refGene_gn=MRPL45;refGene_tx=XM_011289174.3;refGene_gn=MRPL45;refGene_tx=XM_019817781.2
E1 39732851 39732969
refGene_gn=MRPL45;refGene_tx=XM_011289173.3;refGene_gn=MRPL45;refGene_tx=XM_011289174.3;refGene_gn=MRPL45;refGene_tx=XM_019817781.2
E1 39736139 39736238
refGene_gn=MRPL45;refGene_tx=XM_011289173.3;refGene_gn=MRPL45;refGene_tx=XM_019817781.2
E1 39739816 39739865
refGene_gn=MRPL45;refGene_tx=XM_011289173.3;refGene_gn=MRPL45;refGene_tx=XM_011289174.3;refGene_gn=MRPL45;refGene_tx=XM_019817781.2
E1 39741496 39741646
refGene_gn=MRPL45;refGene_tx=XM_011289173.3;refGene_gn=MRPL45;refGene_tx=XM_011289174.3;refGene_gn=MRPL45;refGene_tx=XM_019817781.2
E1 39742089 39742263

refGene_gn=MRPL45;refGene_tx=XM_011289173.3;refGene_gn=MRPL45;refGene_tx=XM_011289174.3
E1	39742445 39742532
refGene_gn=MRPL45;refGene_tx=XM_011289173.3;refGene_gn=MRPL45;refGene_tx=XM_011289174.3;refGene_gn=MRPL45;refGene_tx=XM_019817781.2
E1	39746410 39751519
refGene_gn=GPR179;refGene_tx=XM_003996811.2;refGene_gn=GPR179;refGene_tx=XM_019817778.1
E1	39753347 39753494
refGene_gn=GPR179;refGene_tx=XM_003996811.2;refGene_gn=GPR179;refGene_tx=XM_019817778.1
E1	39754042 39754148
refGene_gn=GPR179;refGene_tx=XM_003996811.2;refGene_gn=GPR179;refGene_tx=XM_019817778.1
E1	39754830 39754969
refGene_gn=GPR179;refGene_tx=XM_003996811.2;refGene_gn=GPR179;refGene_tx=XM_019817778.1
E1	39755163 39755399
refGene_gn=GPR179;refGene_tx=XM_003996811.2;refGene_gn=GPR179;refGene_tx=XM_019817778.1
E1	39755695 39755805
refGene_gn=GPR179;refGene_tx=XM_003996811.2;refGene_gn=GPR179;refGene_tx=XM_019817778.1
E1	39756430 39756499
refGene_gn=GPR179;refGene_tx=XM_003996811.2;refGene_gn=GPR179;refGene_tx=XM_019817778.1
E1	39757346 39757582
refGene_gn=GPR179;refGene_tx=XM_003996811.2;refGene_gn=GPR179;refGene_tx=XM_019817778.1
E1	39758003 39758091
refGene_gn=GPR179;refGene_tx=XM_003996811.2;refGene_gn=GPR179;refGene_tx=XM_019817778.1
E1	39760309 39760418
refGene_gn=GPR179;refGene_tx=XM_003996811.2;refGene_gn=GPR179;refGene_tx=XM_019817778.1
E1	39761748 39762542 refGene_gn=GPR179;refGene_tx=XM_003996811.2
E1	39768856 39769821
refGene_gn=SOCS7;refGene_tx=XM_023244796.1;refGene_gn=SOCS7;refGene_tx=XM_023244798.1;refGene_gn=SOCS7;refGene_tx=XM_023244799.1
E1	39776541 39776606
refGene_gn=SOCS7;refGene_tx=XM_023244796.1;refGene_gn=SOCS7;refGene_tx=XM_023244798.1;refGene_gn=SOCS7;refGene_tx=XM_023244799.1
E1	39780262 39780367
refGene_gn=SOCS7;refGene_tx=XM_023244796.1;refGene_gn=SOCS7;refGene_tx=XM_023244798.1;refGene_gn=SOCS7;refGene_tx=XM_023244799.1
E1	39780815 39780917
refGene_gn=SOCS7;refGene_tx=XM_023244796.1;refGene_gn=SOCS7;refGene_tx=XM_023244799.1
E1	39781782 39781913
refGene_gn=SOCS7;refGene_tx=XM_023244796.1;refGene_gn=SOCS7;refGene_tx

=XM_023244798.1;refGene_gn=SOCS7;refGene_tx=XM_023244799.1
E1 39782422 39782591
refGene_gn=SOCS7;refGene_tx=XM_023244796.1;refGene_gn=SOCS7;refGene_tx
=XM_023244798.1;refGene_gn=SOCS7;refGene_tx=XM_023244799.1
E1 39784620 39784749
refGene_gn=SOCS7;refGene_tx=XM_023244796.1;refGene_gn=SOCS7;refGene_tx
=XM_023244798.1;refGene_gn=SOCS7;refGene_tx=XM_023244799.1
E1 39787725 39787868 refGene_gn=SOCS7;refGene_tx=XM_023244799.1
E1 39794714 39794850
refGene_gn=SOCS7;refGene_tx=XM_023244796.1;refGene_gn=SOCS7;refGene_tx
=XM_023244798.1
E1 39795264 39795385
refGene_gn=SOCS7;refGene_tx=XM_023244796.1;refGene_gn=SOCS7;refGene_tx
=XM_023244798.1
E1 39814238 39814247 refGene_gn=ARHGAP23;refGene_tx=XM_023244568.1
E1 39822020 39822083 refGene_gn=ARHGAP23;refGene_tx=XM_023244567.1
E1 39847016 39847178
refGene_gn=ARHGAP23;refGene_tx=XM_023244567.1;refGene_gn=ARHGAP23;refG
ene_tx=XM_023244568.1
E1 39849751 39849779
refGene_gn=ARHGAP23;refGene_tx=XM_023244567.1;refGene_gn=ARHGAP23;refG
ene_tx=XM_023244568.1
E1 39851427 39851526
refGene_gn=ARHGAP23;refGene_tx=XM_023244567.1;refGene_gn=ARHGAP23;refG
ene_tx=XM_023244568.1;refGene_gn=ARHGAP23;refGene_tx=XM_023244569.1;re
fGene_gn=ARHGAP23;refGene_tx=XM_023244570.1
E1 39851712 39851791
refGene_gn=ARHGAP23;refGene_tx=XM_023244567.1;refGene_gn=ARHGAP23;refG
ene_tx=XM_023244568.1;refGene_gn=ARHGAP23;refGene_tx=XM_023244569.1;re
fGene_gn=ARHGAP23;refGene_tx=XM_023244570.1
E1 39851954 39852009
refGene_gn=ARHGAP23;refGene_tx=XM_023244567.1;refGene_gn=ARHGAP23;refG
ene_tx=XM_023244568.1;refGene_gn=ARHGAP23;refGene_tx=XM_023244569.1;re
fGene_gn=ARHGAP23;refGene_tx=XM_023244570.1;refGene_gn=ARHGAP23;refGen
e_tx=XM_023244571.1
E1 39854579 39855744
refGene_gn=ARHGAP23;refGene_tx=XM_023244567.1;refGene_gn=ARHGAP23;refG
ene_tx=XM_023244568.1;refGene_gn=ARHGAP23;refGene_tx=XM_023244569.1;re
fGene_gn=ARHGAP23;refGene_tx=XM_023244570.1;refGene_gn=ARHGAP23;refGen
e_tx=XM_023244571.1;refGene_gn=ARHGAP23;refGene_tx=XM_023244572.1
E1 39857346 39857502
refGene_gn=ARHGAP23;refGene_tx=XM_023244567.1;refGene_gn=ARHGAP23;refG
ene_tx=XM_023244568.1;refGene_gn=ARHGAP23;refGene_tx=XM_023244569.1;re
fGene_gn=ARHGAP23;refGene_tx=XM_023244570.1;refGene_gn=ARHGAP23;refGen
e_tx=XM_023244571.1;refGene_gn=ARHGAP23;refGene_tx=XM_023244572.1
E1 39857713 39857825
refGene_gn=ARHGAP23;refGene_tx=XM_023244567.1;refGene_gn=ARHGAP23;refG
ene_tx=XM_023244568.1;refGene_gn=ARHGAP23;refGene_tx=XM_023244569.1;re
fGene_gn=ARHGAP23;refGene_tx=XM_023244570.1;refGene_gn=ARHGAP23;refGen
e_tx=XM_023244571.1;refGene_gn=ARHGAP23;refGene_tx=XM_023244572.1

E1 39858008 39858066
refGene_gn=ARHGAP23;refGene_tx=XM_023244567.1;refGene_gn=ARHGAP23;refGene_tx=XM_023244568.1;refGene_gn=ARHGAP23;refGene_tx=XM_023244569.1;refGene_gn=ARHGAP23;refGene_tx=XM_023244570.1;refGene_gn=ARHGAP23;refGene_tx=XM_023244571.1;refGene_gn=ARHGAP23;refGene_tx=XM_023244572.1
E1 39859583 39859727
refGene_gn=ARHGAP23;refGene_tx=XM_023244567.1;refGene_gn=ARHGAP23;refGene_tx=XM_023244568.1;refGene_gn=ARHGAP23;refGene_tx=XM_023244569.1;refGene_gn=ARHGAP23;refGene_tx=XM_023244570.1;refGene_gn=ARHGAP23;refGene_tx=XM_023244571.1;refGene_gn=ARHGAP23;refGene_tx=XM_023244572.1
E1 39866392 39866713
refGene_gn=ARHGAP23;refGene_tx=XM_023244567.1;refGene_gn=ARHGAP23;refGene_tx=XM_023244568.1;refGene_gn=ARHGAP23;refGene_tx=XM_023244569.1;refGene_gn=ARHGAP23;refGene_tx=XM_023244570.1;refGene_gn=ARHGAP23;refGene_tx=XM_023244571.1;refGene_gn=ARHGAP23;refGene_tx=XM_023244572.1
E1 39868016 39868078
refGene_gn=ARHGAP23;refGene_tx=XM_023244567.1;refGene_gn=ARHGAP23;refGene_tx=XM_023244568.1;refGene_gn=ARHGAP23;refGene_tx=XM_023244569.1;refGene_gn=ARHGAP23;refGene_tx=XM_023244570.1;refGene_gn=ARHGAP23;refGene_tx=XM_023244571.1;refGene_gn=ARHGAP23;refGene_tx=XM_023244572.1
E1 39868357 39868488
refGene_gn=ARHGAP23;refGene_tx=XM_023244567.1;refGene_gn=ARHGAP23;refGene_tx=XM_023244568.1;refGene_gn=ARHGAP23;refGene_tx=XM_023244569.1;refGene_gn=ARHGAP23;refGene_tx=XM_023244570.1;refGene_gn=ARHGAP23;refGene_tx=XM_023244571.1;refGene_gn=ARHGAP23;refGene_tx=XM_023244572.1
E1 39869130 39869252
refGene_gn=ARHGAP23;refGene_tx=XM_023244567.1;refGene_gn=ARHGAP23;refGene_tx=XM_023244568.1;refGene_gn=ARHGAP23;refGene_tx=XM_023244569.1;refGene_gn=ARHGAP23;refGene_tx=XM_023244570.1;refGene_gn=ARHGAP23;refGene_tx=XM_023244571.1;refGene_gn=ARHGAP23;refGene_tx=XM_023244572.1
E1 39869641 39869797
refGene_gn=ARHGAP23;refGene_tx=XM_023244567.1;refGene_gn=ARHGAP23;refGene_tx=XM_023244568.1;refGene_gn=ARHGAP23;refGene_tx=XM_023244569.1;refGene_gn=ARHGAP23;refGene_tx=XM_023244570.1;refGene_gn=ARHGAP23;refGene_tx=XM_023244571.1;refGene_gn=ARHGAP23;refGene_tx=XM_023244572.1
E1 39875003 39875082
refGene_gn=ARHGAP23;refGene_tx=XM_023244567.1;refGene_gn=ARHGAP23;refGene_tx=XM_023244568.1;refGene_gn=ARHGAP23;refGene_tx=XM_023244569.1;refGene_gn=ARHGAP23;refGene_tx=XM_023244570.1;refGene_gn=ARHGAP23;refGene_tx=XM_023244571.1;refGene_gn=ARHGAP23;refGene_tx=XM_023244572.1
E1 39877832 39877906
refGene_gn=ARHGAP23;refGene_tx=XM_023244567.1;refGene_gn=ARHGAP23;refGene_tx=XM_023244568.1;refGene_gn=ARHGAP23;refGene_tx=XM_023244569.1;refGene_gn=ARHGAP23;refGene_tx=XM_023244570.1;refGene_gn=ARHGAP23;refGene_tx=XM_023244571.1;refGene_gn=ARHGAP23;refGene_tx=XM_023244572.1
E1 39878198 39878288
refGene_gn=ARHGAP23;refGene_tx=XM_023244567.1;refGene_gn=ARHGAP23;refGene_tx=XM_023244568.1;refGene_gn=ARHGAP23;refGene_tx=XM_023244569.1;refGene_gn=ARHGAP23;refGene_tx=XM_023244570.1;refGene_gn=ARHGAP23;refGene_tx=XM_023244571.1;refGene_gn=ARHGAP23;refGene_tx=XM_023244572.1

E1 39879082 39879208
refGene_gn=ARHGAP23;refGene_tx=XM_023244567.1;refGene_gn=ARHGAP23;refGene_tx=XM_023244568.1;refGene_gn=ARHGAP23;refGene_tx=XM_023244569.1;refGene_gn=ARHGAP23;refGene_tx=XM_023244570.1;refGene_gn=ARHGAP23;refGene_tx=XM_023244571.1;refGene_gn=ARHGAP23;refGene_tx=XM_023244572.1
E1 39884944 39884986
refGene_gn=ARHGAP23;refGene_tx=XM_023244567.1;refGene_gn=ARHGAP23;refGene_tx=XM_023244568.1;refGene_gn=ARHGAP23;refGene_tx=XM_023244569.1;refGene_gn=ARHGAP23;refGene_tx=XM_023244570.1;refGene_gn=ARHGAP23;refGene_tx=XM_023244571.1;refGene_gn=ARHGAP23;refGene_tx=XM_023244572.1
E1 39885544 39885641
refGene_gn=ARHGAP23;refGene_tx=XM_023244567.1;refGene_gn=ARHGAP23;refGene_tx=XM_023244568.1;refGene_gn=ARHGAP23;refGene_tx=XM_023244569.1;refGene_gn=ARHGAP23;refGene_tx=XM_023244570.1;refGene_gn=ARHGAP23;refGene_tx=XM_023244571.1;refGene_gn=ARHGAP23;refGene_tx=XM_023244572.1
E1 39887763 39887795
refGene_gn=ARHGAP23;refGene_tx=XM_023244567.1;refGene_gn=ARHGAP23;refGene_tx=XM_023244568.1;refGene_gn=ARHGAP23;refGene_tx=XM_023244569.1;refGene_gn=ARHGAP23;refGene_tx=XM_023244570.1;refGene_gn=ARHGAP23;refGene_tx=XM_023244571.1;refGene_gn=ARHGAP23;refGene_tx=XM_023244572.1
E1 39891873 39892908
refGene_gn=ARHGAP23;refGene_tx=XM_023244567.1;refGene_gn=ARHGAP23;refGene_tx=XM_023244568.1;refGene_gn=ARHGAP23;refGene_tx=XM_023244569.1;refGene_gn=ARHGAP23;refGene_tx=XM_023244570.1;refGene_gn=ARHGAP23;refGene_tx=XM_023244571.1;refGene_gn=ARHGAP23;refGene_tx=XM_023244572.1
E1 39906050 39906185 refGene_gn=SRCIN1;refGene_tx=XM_023244791.1
E1 39912180 39912327 refGene_gn=SRCIN1;refGene_tx=XM_023244791.1
E1 39916633 39916786 refGene_gn=SRCIN1;refGene_tx=XM_023244791.1
E1 39917049 39917204 refGene_gn=SRCIN1;refGene_tx=XM_023244791.1
E1 39918819 39919054 refGene_gn=SRCIN1;refGene_tx=XM_023244791.1
E1 39919513 39919757 refGene_gn=SRCIN1;refGene_tx=XM_023244791.1
E1 39920097 39920245 refGene_gn=SRCIN1;refGene_tx=XM_023244791.1
E1 39920398 39920529 refGene_gn=SRCIN1;refGene_tx=XM_023244791.1
E1 39925953 39926129 refGene_gn=SRCIN1;refGene_tx=XM_023244791.1
E1 39927178 39927262 refGene_gn=SRCIN1;refGene_tx=XM_023244791.1
E1 39927263 39927367 refGene_gn=SRCIN1;refGene_tx=XM_023244791.1
E1 39927685 39927729 refGene_gn=SRCIN1;refGene_tx=XM_023244791.1
E1 39927974 39928067 refGene_gn=SRCIN1;refGene_tx=XM_023244791.1
E1 39929199 39930074 refGene_gn=SRCIN1;refGene_tx=XM_023244791.1
E1 39930583 39930677 refGene_gn=SRCIN1;refGene_tx=XM_023244791.1
E1 39931084 39931283 refGene_gn=SRCIN1;refGene_tx=XM_023244791.1
E1 39931951 39932147 refGene_gn=SRCIN1;refGene_tx=XM_023244791.1
E1 39935781 39935802 refGene_gn=SRCIN1;refGene_tx=XM_023244791.1
E1 39945554 39945856 refGene_gn=SRCIN1;refGene_tx=XM_023244791.1
E1 39960647 39960710 refGene_gn=SRCIN1;refGene_tx=XM_023244791.1
E1 39969054 39969178 refGene_gn=SRCIN1;refGene_tx=XM_023244791.1
E1 40020365 40021505 refGene_gn=EPOP;refGene_tx=XM_011289216.3
E1 40044119 40044228
refGene_gn=MLLT6;refGene_tx=XM_023244251.1;refGene_gn=MLLT6;refGene_tx=XM_023244252.1;refGene_gn=MLLT6;refGene_tx=XM_023244253.1;refGene_gn=

MLLT6;refGene_tx=XM_023244254.1;refGene_gn=MLLT6;refGene_tx=XM_0232442
55.1
E1 40045445 40045525
refGene_gn=MLLT6;refGene_tx=XM_023244251.1;refGene_gn=MLLT6;refGene_tx
=XM_023244252.1;refGene_gn=MLLT6;refGene_tx=XM_023244253.1;refGene_gn=
MLLT6;refGene_tx=XM_023244254.1;refGene_gn=MLLT6;refGene_tx=XM_0232442
55.1
E1 40045991 40046046
refGene_gn=MLLT6;refGene_tx=XM_023244251.1;refGene_gn=MLLT6;refGene_tx
=XM_023244252.1;refGene_gn=MLLT6;refGene_tx=XM_023244253.1;refGene_gn=
MLLT6;refGene_tx=XM_023244254.1;refGene_gn=MLLT6;refGene_tx=XM_0232442
55.1
E1 40046276 40046386
refGene_gn=MLLT6;refGene_tx=XM_023244251.1;refGene_gn=MLLT6;refGene_tx
=XM_023244252.1;refGene_gn=MLLT6;refGene_tx=XM_023244253.1;refGene_gn=
MLLT6;refGene_tx=XM_023244254.1;refGene_gn=MLLT6;refGene_tx=XM_0232442
55.1
E1 40047365 40047469
refGene_gn=MLLT6;refGene_tx=XM_023244251.1;refGene_gn=MLLT6;refGene_tx
=XM_023244252.1;refGene_gn=MLLT6;refGene_tx=XM_023244253.1;refGene_gn=
MLLT6;refGene_tx=XM_023244254.1;refGene_gn=MLLT6;refGene_tx=XM_0232442
55.1
E1 40047698 40047792
refGene_gn=MLLT6;refGene_tx=XM_023244251.1;refGene_gn=MLLT6;refGene_tx
=XM_023244252.1;refGene_gn=MLLT6;refGene_tx=XM_023244253.1;refGene_gn=
MLLT6;refGene_tx=XM_023244254.1;refGene_gn=MLLT6;refGene_tx=XM_0232442
55.1
E1 40050085 40050244
refGene_gn=MLLT6;refGene_tx=XM_023244251.1;refGene_gn=MLLT6;refGene_tx
=XM_023244252.1;refGene_gn=MLLT6;refGene_tx=XM_023244253.1;refGene_gn=
MLLT6;refGene_tx=XM_023244254.1;refGene_gn=MLLT6;refGene_tx=XM_0232442
55.1
E1 40050911 40051010
refGene_gn=MLLT6;refGene_tx=XM_023244251.1;refGene_gn=MLLT6;refGene_tx
=XM_023244252.1;refGene_gn=MLLT6;refGene_tx=XM_023244253.1;refGene_gn=
MLLT6;refGene_tx=XM_023244254.1;refGene_gn=MLLT6;refGene_tx=XM_0232442
55.1
E1 40053453 40053679
refGene_gn=MLLT6;refGene_tx=XM_023244251.1;refGene_gn=MLLT6;refGene_tx
=XM_023244252.1;refGene_gn=MLLT6;refGene_tx=XM_023244253.1;refGene_gn=
MLLT6;refGene_tx=XM_023244254.1;refGene_gn=MLLT6;refGene_tx=XM_0232442
55.1
E1 40054192 40054807
refGene_gn=MLLT6;refGene_tx=XM_023244251.1;refGene_gn=MLLT6;refGene_tx
=XM_023244252.1;refGene_gn=MLLT6;refGene_tx=XM_023244253.1;refGene_gn=
MLLT6;refGene_tx=XM_023244254.1;refGene_gn=MLLT6;refGene_tx=XM_0232442
55.1
E1 40055250 40055438
refGene_gn=MLLT6;refGene_tx=XM_023244251.1;refGene_gn=MLLT6;refGene_tx
=XM_023244252.1;refGene_gn=MLLT6;refGene_tx=XM_023244253.1;refGene_gn=

MLLT6;refGene_tx=XM_023244254.1;refGene_gn=MLLT6;refGene_tx=XM_0232442
55.1
E1 40055651 40055760
refGene_gn=MLLT6;refGene_tx=XM_023244251.1;refGene_gn=MLLT6;refGene_tx
=XM_023244252.1;refGene_gn=MLLT6;refGene_tx=XM_023244253.1;refGene_gn=
MLLT6;refGene_tx=XM_023244254.1;refGene_gn=MLLT6;refGene_tx=XM_0232442
55.1
E1 40056957 40057024
refGene_gn=MLLT6;refGene_tx=XM_023244251.1;refGene_gn=MLLT6;refGene_tx
=XM_023244252.1;refGene_gn=MLLT6;refGene_tx=XM_023244253.1;refGene_gn=
MLLT6;refGene_tx=XM_023244254.1;refGene_gn=MLLT6;refGene_tx=XM_0232442
55.1
E1 40057223 40057369
refGene_gn=MLLT6;refGene_tx=XM_023244251.1;refGene_gn=MLLT6;refGene_tx
=XM_023244252.1;refGene_gn=MLLT6;refGene_tx=XM_023244253.1;refGene_gn=
MLLT6;refGene_tx=XM_023244254.1;refGene_gn=MLLT6;refGene_tx=XM_0232442
55.1
E1 40057886 40058084
refGene_gn=MLLT6;refGene_tx=XM_023244251.1;refGene_gn=MLLT6;refGene_tx
=XM_023244252.1;refGene_gn=MLLT6;refGene_tx=XM_023244253.1;refGene_gn=
MLLT6;refGene_tx=XM_023244254.1;refGene_gn=MLLT6;refGene_tx=XM_0232442
55.1
E1 40058187 40058276
refGene_gn=MLLT6;refGene_tx=XM_023244251.1;refGene_gn=MLLT6;refGene_tx
=XM_023244252.1;refGene_gn=MLLT6;refGene_tx=XM_023244253.1;refGene_gn=
MLLT6;refGene_tx=XM_023244254.1;refGene_gn=MLLT6;refGene_tx=XM_0232442
55.1
E1 40059397 40059747
refGene_gn=MLLT6;refGene_tx=XM_023244251.1;refGene_gn=MLLT6;refGene_tx
=XM_023244252.1;refGene_gn=MLLT6;refGene_tx=XM_023244253.1;refGene_gn=
MLLT6;refGene_tx=XM_023244254.1
E1 40060185 40060276
refGene_gn=MLLT6;refGene_tx=XM_023244251.1;refGene_gn=MLLT6;refGene_tx
=XM_023244252.1;refGene_gn=MLLT6;refGene_tx=XM_023244253.1;refGene_gn=
MLLT6;refGene_tx=XM_023244254.1;refGene_gn=MLLT6;refGene_tx=XM_0232442
55.1
E1 40061297 40061648
refGene_gn=MLLT6;refGene_tx=XM_023244251.1;refGene_gn=MLLT6;refGene_tx
=XM_023244252.1;refGene_gn=MLLT6;refGene_tx=XM_023244253.1;refGene_gn=
MLLT6;refGene_tx=XM_023244254.1;refGene_gn=MLLT6;refGene_tx=XM_0232442
55.1
E1 40062268 40062310
refGene_gn=MLLT6;refGene_tx=XM_023244254.1;refGene_gn=MLLT6;refGene_tx
=XM_023244255.1
E1 40067209 40067266
refGene_gn=MLLT6;refGene_tx=XM_023244251.1;refGene_gn=MLLT6;refGene_tx
=XM_023244252.1;refGene_gn=MLLT6;refGene_tx=XM_023244253.1
E1 40067831 40067951
refGene_gn=MLLT6;refGene_tx=XM_023244251.1;refGene_gn=MLLT6;refGene_tx
=XM_023244252.1;refGene_gn=MLLT6;refGene_tx=XM_023244253.1

E1 40069868 40070048
refGene_gn=MLLT6;refGene_tx=XM_023244251.1;refGene_gn=MLLT6;refGene_tx=XM_023244252.1;refGene_gn=MLLT6;refGene_tx=XM_023244253.1
E1 40071798 40072176
refGene_gn=PCGF2;refGene_tx=XM_019817797.2;refGene_gn=PCGF2;refGene_tx=XM_019817798.2;refGene_gn=PCGF2;refGene_tx=XM_019817800.2;refGene_gn=PCGF2;refGene_tx=XM_023244256.1
E1 40072719 40072800
refGene_gn=PCGF2;refGene_tx=XM_019817797.2;refGene_gn=PCGF2;refGene_tx=XM_019817798.2;refGene_gn=PCGF2;refGene_tx=XM_019817800.2;refGene_gn=PCGF2;refGene_tx=XM_023244256.1
E1 40074074 40074173
refGene_gn=PCGF2;refGene_tx=XM_019817797.2;refGene_gn=PCGF2;refGene_tx=XM_019817798.2;refGene_gn=PCGF2;refGene_tx=XM_019817800.2;refGene_gn=PCGF2;refGene_tx=XM_023244256.1
E1 40074262 40074317
refGene_gn=PCGF2;refGene_tx=XM_019817797.2;refGene_gn=PCGF2;refGene_tx=XM_019817798.2;refGene_gn=PCGF2;refGene_tx=XM_019817800.2;refGene_gn=PCGF2;refGene_tx=XM_023244256.1
E1 40074493 40074602
refGene_gn=PCGF2;refGene_tx=XM_019817797.2;refGene_gn=PCGF2;refGene_tx=XM_019817798.2;refGene_gn=PCGF2;refGene_tx=XM_019817800.2;refGene_gn=PCGF2;refGene_tx=XM_023244256.1
E1 40074803 40074854
refGene_gn=PCGF2;refGene_tx=XM_019817797.2;refGene_gn=PCGF2;refGene_tx=XM_019817798.2;refGene_gn=PCGF2;refGene_tx=XM_019817800.2;refGene_gn=PCGF2;refGene_tx=XM_023244256.1
E1 40074936 40074992
refGene_gn=PCGF2;refGene_tx=XM_019817797.2;refGene_gn=PCGF2;refGene_tx=XM_019817798.2;refGene_gn=PCGF2;refGene_tx=XM_019817800.2;refGene_gn=PCGF2;refGene_tx=XM_023244256.1
E1 40075291 40075388
refGene_gn=PCGF2;refGene_tx=XM_019817797.2;refGene_gn=PCGF2;refGene_tx=XM_019817798.2;refGene_gn=PCGF2;refGene_tx=XM_019817800.2;refGene_gn=PCGF2;refGene_tx=XM_023244256.1
E1 40076009 40076383
refGene_gn=PCGF2;refGene_tx=XM_019817797.2;refGene_gn=PCGF2;refGene_tx=XM_019817798.2;refGene_gn=PCGF2;refGene_tx=XM_019817800.2;refGene_gn=PCGF2;refGene_tx=XM_023244256.1
E1 40081328 40081487 refGene_gn=PCGF2;refGene_tx=XM_019817800.2
E1 40086773 40086776 refGene_gn=PSMB3;refGene_tx=XM_003996762.5
E1 40087084 40087269 refGene_gn=PSMB3;refGene_tx=XM_003996762.5
E1 40089019 40089127 refGene_gn=PSMB3;refGene_tx=XM_003996762.5
E1 40093660 40093838 refGene_gn=PSMB3;refGene_tx=XM_003996762.5
E1 40095300 40095395 refGene_gn=PSMB3;refGene_tx=XM_003996762.5
E1 40096308 40096357 refGene_gn=PSMB3;refGene_tx=XM_003996762.5
E1 40102355 40102436
refGene_gn=PIP4K2B;refGene_tx=XM_019817801.2;refGene_gn=PIP4K2B;refGene_tx=XM_019817802.2;refGene_gn=PIP4K2B;refGene_tx=XM_019817803.2
E1 40103019 40103123

refGene_gn=PIP4K2B;refGene_tx=XM_019817801.2;refGene_gn=PIP4K2B;refGene_tx=XM_019817802.2;refGene_gn=PIP4K2B;refGene_tx=XM_019817803.2
E1 40103219 40103478
refGene_gn=PIP4K2B;refGene_tx=XM_019817801.2;refGene_gn=PIP4K2B;refGene_tx=XM_019817802.2;refGene_gn=PIP4K2B;refGene_tx=XM_019817803.2
E1 40105794 40105908
refGene_gn=PIP4K2B;refGene_tx=XM_019817801.2;refGene_gn=PIP4K2B;refGene_tx=XM_019817802.2;refGene_gn=PIP4K2B;refGene_tx=XM_019817803.2
E1 40106423 40106462
refGene_gn=PIP4K2B;refGene_tx=XM_019817801.2;refGene_gn=PIP4K2B;refGene_tx=XM_019817802.2;refGene_gn=PIP4K2B;refGene_tx=XM_019817803.2
E1 40107445 40107592
refGene_gn=PIP4K2B;refGene_tx=XM_019817801.2;refGene_gn=PIP4K2B;refGene_tx=XM_019817802.2;refGene_gn=PIP4K2B;refGene_tx=XM_019817803.2
E1 40108489 40108642
refGene_gn=PIP4K2B;refGene_tx=XM_019817801.2;refGene_gn=PIP4K2B;refGene_tx=XM_019817802.2;refGene_gn=PIP4K2B;refGene_tx=XM_019817803.2
E1 40110885 40110982
refGene_gn=PIP4K2B;refGene_tx=XM_019817801.2;refGene_gn=PIP4K2B;refGene_tx=XM_019817802.2;refGene_gn=PIP4K2B;refGene_tx=XM_019817803.2
E1 40113794 40113892
refGene_gn=PIP4K2B;refGene_tx=XM_019817801.2;refGene_gn=PIP4K2B;refGene_tx=XM_019817802.2;refGene_gn=PIP4K2B;refGene_tx=XM_019817803.2
E1 40123964 40124123 refGene_gn=PIP4K2B;refGene_tx=XM_019817801.2
E1 40126549 40126664 refGene_gn=CWC25;refGene_tx=XM_003996763.5
E1 40127391 40127553 refGene_gn=CWC25;refGene_tx=XM_003996763.5
E1 40135197 40135296 refGene_gn=CWC25;refGene_tx=XM_003996763.5
E1 40135663 40135875 refGene_gn=CWC25;refGene_tx=XM_003996763.5
E1 40136478 40136542 refGene_gn=CWC25;refGene_tx=XM_003996763.5
E1 40137214 40137342 refGene_gn=CWC25;refGene_tx=XM_003996763.5
E1 40138308 40138378 refGene_gn=CWC25;refGene_tx=XM_003996763.5
E1 40141131 40141368 refGene_gn=CWC25;refGene_tx=XM_003996763.5
E1 40146369 40146542 refGene_gn=CWC25;refGene_tx=XM_003996763.5
E1 40148840 40148858 refGene_gn=CWC25;refGene_tx=XM_003996763.5
E1 40158810 40158968
refGene_gn=CE1H17orf98;refGene_tx=XM_003996764.5
E1 40159861 40159961
refGene_gn=CE1H17orf98;refGene_tx=XM_003996764.5
E1 40161896 40162103
refGene_gn=CE1H17orf98;refGene_tx=XM_003996764.5
E1 40164785 40164868 refGene_gn=RPL23;refGene_tx=XM_003996765.5
E1 40164967 40165081 refGene_gn=RPL23;refGene_tx=XM_003996765.5
E1 40169171 40169300 refGene_gn=RPL23;refGene_tx=XM_003996765.5
E1 40169632 40169716 refGene_gn=RPL23;refGene_tx=XM_003996765.5
E1 40170295 40170308 refGene_gn=RPL23;refGene_tx=XM_003996765.5
E1 40182301 40182370 refGene_gn=LASP1;refGene_tx=XM_006940285.4
E1 40189041 40189136 refGene_gn=LASP1;refGene_tx=XM_006940285.4
E1 40199289 40199374 refGene_gn=LASP1;refGene_tx=XM_006940285.4
E1 40206672 40206780 refGene_gn=LASP1;refGene_tx=XM_006940285.4
E1 40215007 40215158 refGene_gn=LASP1;refGene_tx=XM_006940285.4

E1 40215516 40215620 refGene_gn=LASP1;refGene_tx=XM_006940285.4
E1 40218929 40219103 refGene_gn=LASP1;refGene_tx=XM_006940285.4
E1 40233301 40233414
refGene_gn=FBXO47;refGene_tx=XM_003996767.5;refGene_gn=FBXO47;refGene_tx=XM_023243915.1
E1 40233965 40234223
refGene_gn=FBXO47;refGene_tx=XM_003996767.5;refGene_gn=FBXO47;refGene_tx=XM_023243915.1;refGene_gn=FBXO47;refGene_tx=XM_023243916.1
E1 40237018 40237161
refGene_gn=FBXO47;refGene_tx=XM_003996767.5;refGene_gn=FBXO47;refGene_tx=XM_023243915.1;refGene_gn=FBXO47;refGene_tx=XM_023243916.1
E1 40237828 40237975
refGene_gn=FBXO47;refGene_tx=XM_003996767.5;refGene_gn=FBXO47;refGene_tx=XM_023243915.1;refGene_gn=FBXO47;refGene_tx=XM_023243916.1
E1 40238941 40239118
refGene_gn=FBXO47;refGene_tx=XM_003996767.5;refGene_gn=FBXO47;refGene_tx=XM_023243915.1;refGene_gn=FBXO47;refGene_tx=XM_023243916.1
E1 40243907 40244016
refGene_gn=FBXO47;refGene_tx=XM_003996767.5;refGene_gn=FBXO47;refGene_tx=XM_023243915.1;refGene_gn=FBXO47;refGene_tx=XM_023243916.1
E1 40246639 40246717
refGene_gn=FBXO47;refGene_tx=XM_003996767.5;refGene_gn=FBXO47;refGene_tx=XM_023243915.1;refGene_gn=FBXO47;refGene_tx=XM_023243916.1
E1 40248485 40248562
refGene_gn=FBXO47;refGene_tx=XM_003996767.5;refGene_gn=FBXO47;refGene_tx=XM_023243915.1;refGene_gn=FBXO47;refGene_tx=XM_023243916.1
E1 40251654 40251825
refGene_gn=FBXO47;refGene_tx=XM_003996767.5;refGene_gn=FBXO47;refGene_tx=XM_023243915.1;refGene_gn=FBXO47;refGene_tx=XM_023243916.1
E1 40252175 40252359
refGene_gn=FBXO47;refGene_tx=XM_003996767.5;refGene_gn=FBXO47;refGene_tx=XM_023243915.1;refGene_gn=FBXO47;refGene_tx=XM_023243916.1
E1 40303809 40303929 refGene_gn=PLXDC1;refGene_tx=XM_011289170.2
E1 40305182 40305343 refGene_gn=PLXDC1;refGene_tx=XM_011289170.2
E1 40307594 40307630 refGene_gn=PLXDC1;refGene_tx=XM_011289170.2
E1 40313840 40313976 refGene_gn=PLXDC1;refGene_tx=XM_011289170.2
E1 40315912 40315973 refGene_gn=PLXDC1;refGene_tx=XM_011289170.2
E1 40319107 40319189 refGene_gn=PLXDC1;refGene_tx=XM_011289170.2
E1 40323125 40323159 refGene_gn=PLXDC1;refGene_tx=XM_011289170.2
E1 40323160 40323222 refGene_gn=PLXDC1;refGene_tx=XM_011289170.2
E1 40333269 40333369 refGene_gn=PLXDC1;refGene_tx=XM_011289170.2
E1 40334916 40335035 refGene_gn=PLXDC1;refGene_tx=XM_011289170.2
E1 40335415 40335538 refGene_gn=PLXDC1;refGene_tx=XM_011289170.2
E1 40336169 40336239 refGene_gn=PLXDC1;refGene_tx=XM_011289170.2
E1 40336532 40336676 refGene_gn=PLXDC1;refGene_tx=XM_011289170.2
E1 40358409 40358511 refGene_gn=PLXDC1;refGene_tx=XM_011289170.2
E1 40358512 40358589 refGene_gn=PLXDC1;refGene_tx=XM_011289170.2
E1 40366454 40366530 refGene_gn=PLXDC1;refGene_tx=XM_011289170.2
E1 40370865 40370914 refGene_gn=ARL5C;refGene_tx=XM_023244155.1
E1 40372142 40372294 refGene_gn=ARL5C;refGene_tx=XM_023244155.1

E1 40372751 40372835 refGene_gn=ARL5C;refGene_tx=XM_023244155.1
E1 40373495 40373643 refGene_gn=ARL5C;refGene_tx=XM_023244155.1
E1 40374795 40374856 refGene_gn=ARL5C;refGene_tx=XM_023244155.1
E1 40375468 40375514 refGene_gn=ARL5C;refGene_tx=XM_023244155.1
E1 40381602 40382067
refGene_gn=CACNB1;refGene_tx=XM_006940287.4;refGene_gn=CACNB1;refGene_tx=XM_019817811.2;refGene_gn=CACNB1;refGene_tx=XM_019817812.2;refGene_gn=CACNB1;refGene_tx=XM_023243671.1
E1 40383868 40384159
refGene_gn=CACNB1;refGene_tx=XM_003996770.5;refGene_gn=CACNB1;refGene_tx=XM_006940287.4;refGene_gn=CACNB1;refGene_tx=XM_019817811.2;refGene_gn=CACNB1;refGene_tx=XM_019817812.2;refGene_gn=CACNB1;refGene_tx=XM_023243671.1
E1 40384586 40384682
refGene_gn=CACNB1;refGene_tx=XM_003996770.5;refGene_gn=CACNB1;refGene_tx=XM_006940287.4;refGene_gn=CACNB1;refGene_tx=XM_019817811.2;refGene_gn=CACNB1;refGene_tx=XM_019817812.2;refGene_gn=CACNB1;refGene_tx=XM_023243671.1
E1 40387932 40388084
refGene_gn=CACNB1;refGene_tx=XM_003996770.5;refGene_gn=CACNB1;refGene_tx=XM_006940287.4;refGene_gn=CACNB1;refGene_tx=XM_019817811.2;refGene_gn=CACNB1;refGene_tx=XM_019817812.2;refGene_gn=CACNB1;refGene_tx=XM_023243671.1
E1 40388230 40388340
refGene_gn=CACNB1;refGene_tx=XM_003996770.5;refGene_gn=CACNB1;refGene_tx=XM_006940287.4;refGene_gn=CACNB1;refGene_tx=XM_019817811.2;refGene_gn=CACNB1;refGene_tx=XM_019817812.2;refGene_gn=CACNB1;refGene_tx=XM_023243671.1
E1 40388505 40388564
refGene_gn=CACNB1;refGene_tx=XM_003996770.5;refGene_gn=CACNB1;refGene_tx=XM_006940287.4;refGene_gn=CACNB1;refGene_tx=XM_019817811.2;refGene_gn=CACNB1;refGene_tx=XM_019817812.2;refGene_gn=CACNB1;refGene_tx=XM_023243671.1
E1 40388989 40389070
refGene_gn=CACNB1;refGene_tx=XM_003996770.5;refGene_gn=CACNB1;refGene_tx=XM_006940287.4;refGene_gn=CACNB1;refGene_tx=XM_019817811.2;refGene_gn=CACNB1;refGene_tx=XM_019817812.2;refGene_gn=CACNB1;refGene_tx=XM_023243671.1
E1 40389350 40389370 refGene_gn=CACNB1;refGene_tx=XM_019817812.2
E1 40390768 40390788 refGene_gn=CACNB1;refGene_tx=XM_023243671.1
E1 40391617 40391772
refGene_gn=CACNB1;refGene_tx=XM_003996770.5;refGene_gn=CACNB1;refGene_tx=XM_006940287.4;refGene_gn=CACNB1;refGene_tx=XM_019817811.2
E1 40392185 40392262
refGene_gn=CACNB1;refGene_tx=XM_003996770.5;refGene_gn=CACNB1;refGene_tx=XM_006940287.4;refGene_gn=CACNB1;refGene_tx=XM_019817811.2;refGene_gn=CACNB1;refGene_tx=XM_019817812.2;refGene_gn=CACNB1;refGene_tx=XM_023243671.1
E1 40392479 40392616
refGene_gn=CACNB1;refGene_tx=XM_003996770.5;refGene_gn=CACNB1;refGene_

tx=XM_006940287.4;refGene_gn=CACNB1;refGene_tx=XM_019817811.2;refGene_gn=CACNB1;refGene_tx=XM_019817812.2;refGene_gn=CACNB1;refGene_tx=XM_023243671.1
E1 40393079 40393202
refGene_gn=CACNB1;refGene_tx=XM_003996770.5;refGene_gn=CACNB1;refGene_tx=XM_006940287.4;refGene_gn=CACNB1;refGene_tx=XM_019817811.2;refGene_gn=CACNB1;refGene_tx=XM_019817812.2;refGene_gn=CACNB1;refGene_tx=XM_023243671.1
E1 40394069 40394189
refGene_gn=CACNB1;refGene_tx=XM_003996770.5;refGene_gn=CACNB1;refGene_tx=XM_006940287.4;refGene_gn=CACNB1;refGene_tx=XM_019817811.2;refGene_gn=CACNB1;refGene_tx=XM_019817812.2;refGene_gn=CACNB1;refGene_tx=XM_023243671.1
E1 40396296 40396326 refGene_gn=CACNB1;refGene_tx=XM_006940287.4
E1 40397671 40397758
refGene_gn=CACNB1;refGene_tx=XM_003996770.5;refGene_gn=CACNB1;refGene_tx=XM_019817811.2;refGene_gn=CACNB1;refGene_tx=XM_019817812.2;refGene_gn=CACNB1;refGene_tx=XM_023243671.1
E1 40400216 40400300
refGene_gn=CACNB1;refGene_tx=XM_003996770.5;refGene_gn=CACNB1;refGene_tx=XM_019817811.2;refGene_gn=CACNB1;refGene_tx=XM_019817812.2;refGene_gn=CACNB1;refGene_tx=XM_023243671.1
E1 40403129 40403134 refGene_gn=RPL19;refGene_tx=XM_011289169.2
E1 40404065 40404172 refGene_gn=RPL19;refGene_tx=XM_011289169.2
E1 40405464 40405587 refGene_gn=RPL19;refGene_tx=XM_011289169.2
E1 40406452 40406573 refGene_gn=RPL19;refGene_tx=XM_011289169.2
E1 40407119 40407230 refGene_gn=RPL19;refGene_tx=XM_011289169.2
E1 40407598 40407722 refGene_gn=RPL19;refGene_tx=XM_011289169.2
E1 40413885 40413990
refGene_gn=STAC2;refGene_tx=XM_003996771.5;refGene_gn=STAC2;refGene_tx=XM_019817813.2
E1 40414576 40414714
refGene_gn=STAC2;refGene_tx=XM_003996771.5;refGene_gn=STAC2;refGene_tx=XM_019817813.2
E1 40415062 40415114
refGene_gn=STAC2;refGene_tx=XM_003996771.5;refGene_gn=STAC2;refGene_tx=XM_019817813.2
E1 40415499 40415597
refGene_gn=STAC2;refGene_tx=XM_003996771.5;refGene_gn=STAC2;refGene_tx=XM_019817813.2
E1 40416084 40416146
refGene_gn=STAC2;refGene_tx=XM_003996771.5;refGene_gn=STAC2;refGene_tx=XM_019817813.2
E1 40416238 40416311
refGene_gn=STAC2;refGene_tx=XM_003996771.5;refGene_gn=STAC2;refGene_tx=XM_019817813.2
E1 40416400 40416513
refGene_gn=STAC2;refGene_tx=XM_003996771.5;refGene_gn=STAC2;refGene_tx=XM_019817813.2
E1 40417130 40417221

refGene_gn=STAC2;refGene_tx=XM_003996771.5;refGene_gn=STAC2;refGene_tx=XM_019817813.2
E1 40417402 40417500
refGene_gn=STAC2;refGene_tx=XM_003996771.5;refGene_gn=STAC2;refGene_tx=XM_019817813.2
E1 40417844 40418151
refGene_gn=STAC2;refGene_tx=XM_003996771.5;refGene_gn=STAC2;refGene_tx=XM_019817813.2
E1 40426642 40426732
refGene_gn=STAC2;refGene_tx=XM_003996771.5;refGene_gn=STAC2;refGene_tx=XM_019817813.2
E1 40437229 40437312
refGene_gn=LOC101100850;refGene_tx=XM_006940321.3
E1 40438113 40438399
refGene_gn=LOC101100850;refGene_tx=XM_006940321.3;refGene_gn=LOC101100850;refGene_tx=XM_019817814.2
E1 40438534 40438608
refGene_gn=LOC101100850;refGene_tx=XM_006940321.3;refGene_gn=LOC101100850;refGene_tx=XM_019817814.2;refGene_gn=LOC101100850;refGene_tx=XM_019817815.2
E1 40439366 40439472
refGene_gn=LOC101100850;refGene_tx=XM_006940321.3;refGene_gn=LOC101100850;refGene_tx=XM_019817814.2;refGene_gn=LOC101100850;refGene_tx=XM_019817815.2
E1 40439580 40439687
refGene_gn=LOC101100850;refGene_tx=XM_006940321.3;refGene_gn=LOC101100850;refGene_tx=XM_019817814.2;refGene_gn=LOC101100850;refGene_tx=XM_019817815.2
E1 40441956 40442109
refGene_gn=LOC101100850;refGene_tx=XM_006940321.3;refGene_gn=LOC101100850;refGene_tx=XM_019817814.2;refGene_gn=LOC101100850;refGene_tx=XM_019817815.2
E1 40442334 40442445
refGene_gn=LOC101100850;refGene_tx=XM_006940321.3;refGene_gn=LOC101100850;refGene_tx=XM_019817814.2;refGene_gn=LOC101100850;refGene_tx=XM_019817815.2
E1 40442935 40443033
refGene_gn=LOC101100850;refGene_tx=XM_006940321.3;refGene_gn=LOC101100850;refGene_tx=XM_019817814.2;refGene_gn=LOC101100850;refGene_tx=XM_019817815.2
E1 40443249 40443304
refGene_gn=LOC101100850;refGene_tx=XM_006940321.3;refGene_gn=LOC101100850;refGene_tx=XM_019817814.2;refGene_gn=LOC101100850;refGene_tx=XM_019817815.2
E1 40443579 40443673
refGene_gn=LOC101100850;refGene_tx=XM_006940321.3;refGene_gn=LOC101100850;refGene_tx=XM_019817814.2;refGene_gn=LOC101100850;refGene_tx=XM_019817815.2
E1 40444068 40444159
refGene_gn=LOC101100850;refGene_tx=XM_006940321.3;refGene_gn=LOC101100

850;refGene_tx=XM_019817814.2;refGene_gn=LOC101100850;refGene_tx=XM_019817815.2
E1 40444617 40444700
refGene_gn=LOC101100850;refGene_tx=XM_006940321.3;refGene_gn=LOC101100850;refGene_tx=XM_019817814.2;refGene_gn=LOC101100850;refGene_tx=XM_019817815.2
E1 40445349 40445495
refGene_gn=LOC101100850;refGene_tx=XM_006940321.3;refGene_gn=LOC101100850;refGene_tx=XM_019817814.2;refGene_gn=LOC101100850;refGene_tx=XM_019817815.2
E1 40445636 40445688
refGene_gn=LOC101100850;refGene_tx=XM_006940321.3;refGene_gn=LOC101100850;refGene_tx=XM_019817814.2;refGene_gn=LOC101100850;refGene_tx=XM_019817815.2
E1 40446293 40446331
refGene_gn=LOC101100850;refGene_tx=XM_006940321.3;refGene_gn=LOC101100850;refGene_tx=XM_019817814.2;refGene_gn=LOC101100850;refGene_tx=XM_019817815.2
E1 40455901 40456009
refGene_gn=FBXL20;refGene_tx=XM_003996772.5;refGene_gn=FBXL20;refGene_tx=XM_006940293.4;refGene_gn=FBXL20;refGene_tx=XM_023243645.1
E1 40457989 40458202
refGene_gn=FBXL20;refGene_tx=XM_003996772.5;refGene_gn=FBXL20;refGene_tx=XM_006940293.4;refGene_gn=FBXL20;refGene_tx=XM_023243645.1
E1 40459154 40459211
refGene_gn=FBXL20;refGene_tx=XM_003996772.5;refGene_gn=FBXL20;refGene_tx=XM_006940293.4;refGene_gn=FBXL20;refGene_tx=XM_023243645.1
E1 40460809 40460854
refGene_gn=FBXL20;refGene_tx=XM_003996772.5;refGene_gn=FBXL20;refGene_tx=XM_006940293.4;refGene_gn=FBXL20;refGene_tx=XM_023243645.1
E1 40461794 40461855
refGene_gn=FBXL20;refGene_tx=XM_003996772.5;refGene_gn=FBXL20;refGene_tx=XM_006940293.4;refGene_gn=FBXL20;refGene_tx=XM_023243645.1
E1 40463846 40463977
refGene_gn=FBXL20;refGene_tx=XM_003996772.5;refGene_gn=FBXL20;refGene_tx=XM_006940293.4;refGene_gn=FBXL20;refGene_tx=XM_023243645.1
E1 40466651 40466726
refGene_gn=FBXL20;refGene_tx=XM_003996772.5;refGene_gn=FBXL20;refGene_tx=XM_006940293.4;refGene_gn=FBXL20;refGene_tx=XM_023243645.1
E1 40468211 40468338
refGene_gn=FBXL20;refGene_tx=XM_003996772.5;refGene_gn=FBXL20;refGene_tx=XM_006940293.4;refGene_gn=FBXL20;refGene_tx=XM_023243645.1
E1 40470303 40470399
refGene_gn=FBXL20;refGene_tx=XM_003996772.5;refGene_gn=FBXL20;refGene_tx=XM_006940293.4;refGene_gn=FBXL20;refGene_tx=XM_023243645.1
E1 40479266 40479335
refGene_gn=FBXL20;refGene_tx=XM_003996772.5;refGene_gn=FBXL20;refGene_tx=XM_006940293.4;refGene_gn=FBXL20;refGene_tx=XM_023243645.1
E1 40480702 40480797
refGene_gn=FBXL20;refGene_tx=XM_003996772.5;refGene_gn=FBXL20;refGene_

tx=XM_006940293.4;refGene_gn=FBXL20;refGene_tx=XM_023243645.1
E1 40482828 40482903
refGene_gn=FBXL20;refGene_tx=XM_003996772.5;refGene_gn=FBXL20;refGene_tx=XM_006940293.4;refGene_gn=FBXL20;refGene_tx=XM_023243645.1
E1 40485476 40485531
refGene_gn=FBXL20;refGene_tx=XM_003996772.5;refGene_gn=FBXL20;refGene_tx=XM_006940293.4;refGene_gn=FBXL20;refGene_tx=XM_023243645.1
E1 40495230 40495292
refGene_gn=FBXL20;refGene_tx=XM_003996772.5;refGene_gn=FBXL20;refGene_tx=XM_006940293.4;refGene_gn=FBXL20;refGene_tx=XM_023243645.1
E1 40552357 40552399 refGene_gn=FBXL20;refGene_tx=XM_023243645.1
E1 40553201 40553249 refGene_gn=FBXL20;refGene_tx=XM_006940293.4
E1 40557978 40561231
refGene_gn=MED1;refGene_tx=XM_003996775.4;refGene_gn=MED1;refGene_tx=XM_023243643.1;refGene_gn=MED1;refGene_tx=XM_023243644.1
E1 40562569 40562675
refGene_gn=MED1;refGene_tx=XM_003996775.4;refGene_gn=MED1;refGene_tx=XM_023243643.1;refGene_gn=MED1;refGene_tx=XM_023243644.1
E1 40562791 40562887
refGene_gn=MED1;refGene_tx=XM_003996775.4;refGene_gn=MED1;refGene_tx=XM_023243643.1;refGene_gn=MED1;refGene_tx=XM_023243644.1
E1 40564424 40564626
refGene_gn=MED1;refGene_tx=XM_003996775.4;refGene_gn=MED1;refGene_tx=XM_023243643.1;refGene_gn=MED1;refGene_tx=XM_023243644.1
E1 40565616 40565735
refGene_gn=MED1;refGene_tx=XM_003996775.4;refGene_gn=MED1;refGene_tx=XM_023243643.1;refGene_gn=MED1;refGene_tx=XM_023243644.1
E1 40565996 40566121
refGene_gn=MED1;refGene_tx=XM_003996775.4;refGene_gn=MED1;refGene_tx=XM_023243643.1;refGene_gn=MED1;refGene_tx=XM_023243644.1
E1 40568699 40568811
refGene_gn=MED1;refGene_tx=XM_003996775.4;refGene_gn=MED1;refGene_tx=XM_023243643.1;refGene_gn=MED1;refGene_tx=XM_023243644.1
E1 40570185 40570275
refGene_gn=MED1;refGene_tx=XM_003996775.4;refGene_gn=MED1;refGene_tx=XM_023243643.1;refGene_gn=MED1;refGene_tx=XM_023243644.1
E1 40570637 40570711
refGene_gn=MED1;refGene_tx=XM_003996775.4;refGene_gn=MED1;refGene_tx=XM_023243643.1;refGene_gn=MED1;refGene_tx=XM_023243644.1
E1 40571157 40571232
refGene_gn=MED1;refGene_tx=XM_003996775.4;refGene_gn=MED1;refGene_tx=XM_023243643.1;refGene_gn=MED1;refGene_tx=XM_023243644.1
E1 40571885 40571957 refGene_gn=MED1;refGene_tx=XM_003996775.4
E1 40574370 40574399 refGene_gn=MED1;refGene_tx=XM_003996775.4
E1 40575119 40575252 refGene_gn=MED1;refGene_tx=XM_003996775.4
E1 40575355 40575410 refGene_gn=MED1;refGene_tx=XM_003996775.4
E1 40578861 40578940 refGene_gn=MED1;refGene_tx=XM_003996775.4
E1 40581074 40581181 refGene_gn=MED1;refGene_tx=XM_003996775.4
E1 40583600 40583625 refGene_gn=MED1;refGene_tx=XM_003996775.4
E1 40590677 40591726

refGene_gn=CDK12;refGene_tx=XM_003996776.5;refGene_gn=CDK12;refGene_tx=XM_003996777.5;refGene_gn=CDK12;refGene_tx=XM_006940294.4
E1 40596679 40597567
refGene_gn=CDK12;refGene_tx=XM_003996776.5;refGene_gn=CDK12;refGene_tx=XM_003996777.5;refGene_gn=CDK12;refGene_tx=XM_006940294.4
E1 40606597 40606774
refGene_gn=CDK12;refGene_tx=XM_003996776.5;refGene_gn=CDK12;refGene_tx=XM_003996777.5;refGene_gn=CDK12;refGene_tx=XM_006940294.4
E1 40608422 40608562
refGene_gn=CDK12;refGene_tx=XM_003996776.5;refGene_gn=CDK12;refGene_tx=XM_003996777.5;refGene_gn=CDK12;refGene_tx=XM_006940294.4
E1 40609067 40609238
refGene_gn=CDK12;refGene_tx=XM_003996776.5;refGene_gn=CDK12;refGene_tx=XM_003996777.5;refGene_gn=CDK12;refGene_tx=XM_006940294.4
E1 40613297 40613487
refGene_gn=CDK12;refGene_tx=XM_003996776.5;refGene_gn=CDK12;refGene_tx=XM_003996777.5;refGene_gn=CDK12;refGene_tx=XM_006940294.4
E1 40619047 40619104
refGene_gn=CDK12;refGene_tx=XM_003996776.5;refGene_gn=CDK12;refGene_tx=XM_003996777.5;refGene_gn=CDK12;refGene_tx=XM_006940294.4
E1 40619428 40619530
refGene_gn=CDK12;refGene_tx=XM_003996776.5;refGene_gn=CDK12;refGene_tx=XM_003996777.5;refGene_gn=CDK12;refGene_tx=XM_006940294.4
E1 40625441 40625519
refGene_gn=CDK12;refGene_tx=XM_003996776.5;refGene_gn=CDK12;refGene_tx=XM_003996777.5;refGene_gn=CDK12;refGene_tx=XM_006940294.4
E1 40626554 40626671
refGene_gn=CDK12;refGene_tx=XM_003996776.5;refGene_gn=CDK12;refGene_tx=XM_003996777.5;refGene_gn=CDK12;refGene_tx=XM_006940294.4
E1 40627145 40627277
refGene_gn=CDK12;refGene_tx=XM_003996776.5;refGene_gn=CDK12;refGene_tx=XM_003996777.5;refGene_gn=CDK12;refGene_tx=XM_006940294.4
E1 40630354 40630566
refGene_gn=CDK12;refGene_tx=XM_003996776.5;refGene_gn=CDK12;refGene_tx=XM_003996777.5;refGene_gn=CDK12;refGene_tx=XM_006940294.4
E1 40631785 40632238
refGene_gn=CDK12;refGene_tx=XM_003996776.5;refGene_gn=CDK12;refGene_tx=XM_003996777.5;refGene_gn=CDK12;refGene_tx=XM_006940294.4
E1 40635975 40636688
refGene_gn=CDK12;refGene_tx=XM_003996776.5;refGene_gn=CDK12;refGene_tx=XM_003996777.5;refGene_gn=CDK12;refGene_tx=XM_006940294.4
E1 40699453 40700602 refGene_gn=NEUROD2;refGene_tx=XM_023244092.1
E1 40720110 40720191 refGene_gn=PPP1R1B;refGene_tx=XM_023244759.1
E1 40721854 40721915
refGene_gn=PPP1R1B;refGene_tx=XM_003996778.5;refGene_gn=PPP1R1B;refGene_tx=XM_011289167.3;refGene_gn=PPP1R1B;refGene_tx=XM_023244759.1
E1 40722230 40722253
refGene_gn=PPP1R1B;refGene_tx=XM_003996778.5;refGene_gn=PPP1R1B;refGene_tx=XM_011289167.3;refGene_gn=PPP1R1B;refGene_tx=XM_023244759.1
E1 40722659 40722735

refGene_gn=PPP1R1B;refGene_tx=XM_003996778.5;refGene_gn=PPP1R1B;refGene_tx=XM_011289167.3;refGene_gn=PPP1R1B;refGene_tx=XM_023244759.1
E1 40726517 40726721
refGene_gn=PPP1R1B;refGene_tx=XM_003996778.5;refGene_gn=PPP1R1B;refGene_tx=XM_011289167.3;refGene_gn=PPP1R1B;refGene_tx=XM_023244759.1
E1 40728003 40728123
refGene_gn=PPP1R1B;refGene_tx=XM_003996778.5;refGene_gn=PPP1R1B;refGene_tx=XM_011289167.3;refGene_gn=PPP1R1B;refGene_tx=XM_023244759.1
E1 40728226 40728276
refGene_gn=PPP1R1B;refGene_tx=XM_003996778.5;refGene_gn=PPP1R1B;refGene_tx=XM_011289167.3;refGene_gn=PPP1R1B;refGene_tx=XM_023244759.1
E1 40741364 40741586
refGene_gn=STARD3;refGene_tx=XM_011289165.3;refGene_gn=STARD3;refGene_tx=XM_011289166.3;refGene_gn=STARD3;refGene_tx=XM_019817817.2;refGene_gn=STARD3;refGene_tx=XM_019817818.2
E1 40744393 40744471
refGene_gn=STARD3;refGene_tx=XM_011289165.3;refGene_gn=STARD3;refGene_tx=XM_019817817.2;refGene_gn=STARD3;refGene_tx=XM_019817818.2
E1 40744857 40744935
refGene_gn=STARD3;refGene_tx=XM_011289165.3;refGene_gn=STARD3;refGene_tx=XM_011289166.3;refGene_gn=STARD3;refGene_tx=XM_019817817.2;refGene_gn=STARD3;refGene_tx=XM_019817818.2
E1 40745055 40745109
refGene_gn=STARD3;refGene_tx=XM_011289165.3;refGene_gn=STARD3;refGene_tx=XM_011289166.3;refGene_gn=STARD3;refGene_tx=XM_019817817.2;refGene_gn=STARD3;refGene_tx=XM_019817818.2
E1 40745504 40745622
refGene_gn=STARD3;refGene_tx=XM_011289165.3;refGene_gn=STARD3;refGene_tx=XM_011289166.3;refGene_gn=STARD3;refGene_tx=XM_019817817.2;refGene_gn=STARD3;refGene_tx=XM_019817818.2
E1 40745830 40745929
refGene_gn=STARD3;refGene_tx=XM_011289165.3;refGene_gn=STARD3;refGene_tx=XM_011289166.3;refGene_gn=STARD3;refGene_tx=XM_019817817.2;refGene_gn=STARD3;refGene_tx=XM_019817818.2
E1 40746153 40746206
refGene_gn=STARD3;refGene_tx=XM_011289165.3;refGene_gn=STARD3;refGene_tx=XM_011289166.3;refGene_gn=STARD3;refGene_tx=XM_019817817.2;refGene_gn=STARD3;refGene_tx=XM_019817818.2
E1 40746554 40746647
refGene_gn=STARD3;refGene_tx=XM_011289165.3;refGene_gn=STARD3;refGene_tx=XM_011289166.3;refGene_gn=STARD3;refGene_tx=XM_019817817.2;refGene_gn=STARD3;refGene_tx=XM_019817818.2
E1 40747302 40747365
refGene_gn=STARD3;refGene_tx=XM_011289165.3;refGene_gn=STARD3;refGene_tx=XM_011289166.3;refGene_gn=STARD3;refGene_tx=XM_019817817.2;refGene_gn=STARD3;refGene_tx=XM_019817818.2
E1 40747529 40747625
refGene_gn=STARD3;refGene_tx=XM_011289165.3;refGene_gn=STARD3;refGene_tx=XM_011289166.3;refGene_gn=STARD3;refGene_tx=XM_019817817.2;refGene_gn=STARD3;refGene_tx=XM_019817818.2

E1 40747985 40748065
refGene_gn=STARD3;refGene_tx=XM_011289165.3;refGene_gn=STARD3;refGene_tx=XM_011289166.3;refGene_gn=STARD3;refGene_tx=XM_019817817.2;refGene_gn=STARD3;refGene_tx=XM_019817818.2
E1 40748160 40748265
refGene_gn=STARD3;refGene_tx=XM_011289165.3;refGene_gn=STARD3;refGene_tx=XM_011289166.3;refGene_gn=STARD3;refGene_tx=XM_019817817.2;refGene_gn=STARD3;refGene_tx=XM_019817818.2
E1 40749378 40749472
refGene_gn=STARD3;refGene_tx=XM_011289165.3;refGene_gn=STARD3;refGene_tx=XM_011289166.3;refGene_gn=STARD3;refGene_tx=XM_019817817.2;refGene_gn=STARD3;refGene_tx=XM_019817818.2
E1 40749928 40750033
refGene_gn=STARD3;refGene_tx=XM_011289165.3;refGene_gn=STARD3;refGene_tx=XM_011289166.3;refGene_gn=STARD3;refGene_tx=XM_019817817.2;refGene_gn=STARD3;refGene_tx=XM_019817818.2
E1 40752571 40752681 refGene_gn=TCAP;refGene_tx=XM_003996780.4
E1 40752928 40753322 refGene_gn=TCAP;refGene_tx=XM_003996780.4
E1 40755625 40755833 refGene_gn=PNMT;refGene_tx=XM_003996821.5
E1 40756443 40756595 refGene_gn=PNMT;refGene_tx=XM_003996821.5
E1 40756596 40756652 refGene_gn=PNMT;refGene_tx=XM_003996821.5
E1 40756755 40757197 refGene_gn=PNMT;refGene_tx=XM_003996821.5
E1 40759555 40759619 refGene_gn=PGAP3;refGene_tx=XM_003996781.5
E1 40759783 40759988 refGene_gn=PGAP3;refGene_tx=XM_003996781.5
E1 40760287 40760424 refGene_gn=PGAP3;refGene_tx=XM_003996781.5
E1 40760778 40760840 refGene_gn=PGAP3;refGene_tx=XM_003996781.5
E1 40761449 40761512 refGene_gn=PGAP3;refGene_tx=XM_003996781.5
E1 40769475 40769628 refGene_gn=PGAP3;refGene_tx=XM_003996781.5
E1 40770683 40770781 refGene_gn=PGAP3;refGene_tx=XM_003996781.5
E1 40772016 40772197 refGene_gn=PGAP3;refGene_tx=XM_003996781.5
E1 40780249 40780322 refGene_gn=ERBB2;refGene_tx=NM_001048163.1
E1 40786772 40786924
refGene_gn=ERBB2;refGene_tx=NM_001048163.1;refGene_gn=ERBB2;refGene_tx=XM_006940248.4
E1 40787797 40788011
refGene_gn=ERBB2;refGene_tx=NM_001048163.1;refGene_gn=ERBB2;refGene_tx=XM_006940248.4
E1 40788825 40788960
refGene_gn=ERBB2;refGene_tx=NM_001048163.1;refGene_gn=ERBB2;refGene_tx=XM_006940248.4
E1 40789251 40789320
refGene_gn=ERBB2;refGene_tx=NM_001048163.1;refGene_gn=ERBB2;refGene_tx=XM_006940248.4
E1 40789479 40789595
refGene_gn=ERBB2;refGene_tx=NM_001048163.1;refGene_gn=ERBB2;refGene_tx=XM_006940248.4
E1 40789754 40789896
refGene_gn=ERBB2;refGene_tx=NM_001048163.1;refGene_gn=ERBB2;refGene_tx=XM_006940248.4
E1 40790963 40791083

refGene_gn=ERBB2;refGene_tx=NM_001048163.1;refGene_gn=ERBB2;refGene_tx=XM_006940248.4
E1 40791372 40791499
refGene_gn=ERBB2;refGene_tx=NM_001048163.1;refGene_gn=ERBB2;refGene_tx=XM_006940248.4
E1 40793526 40793600
refGene_gn=ERBB2;refGene_tx=NM_001048163.1;refGene_gn=ERBB2;refGene_tx=XM_006940248.4
E1 40793688 40793779
refGene_gn=ERBB2;refGene_tx=NM_001048163.1;refGene_gn=ERBB2;refGene_tx=XM_006940248.4
E1 40793972 40794172
refGene_gn=ERBB2;refGene_tx=NM_001048163.1;refGene_gn=ERBB2;refGene_tx=XM_006940248.4
E1 40794528 40794661
refGene_gn=ERBB2;refGene_tx=NM_001048163.1;refGene_gn=ERBB2;refGene_tx=XM_006940248.4
E1 40794746 40794837
refGene_gn=ERBB2;refGene_tx=NM_001048163.1;refGene_gn=ERBB2;refGene_tx=XM_006940248.4
E1 40795537 40795698
refGene_gn=ERBB2;refGene_tx=NM_001048163.1;refGene_gn=ERBB2;refGene_tx=XM_006940248.4
E1 40798158 40798206
refGene_gn=ERBB2;refGene_tx=NM_001048163.1;refGene_gn=ERBB2;refGene_tx=XM_006940248.4
E1 40799531 40799670
refGene_gn=ERBB2;refGene_tx=NM_001048163.1;refGene_gn=ERBB2;refGene_tx=XM_006940248.4
E1 40799753 40799876
refGene_gn=ERBB2;refGene_tx=NM_001048163.1;refGene_gn=ERBB2;refGene_tx=XM_006940248.4
E1 40800158 40800257
refGene_gn=ERBB2;refGene_tx=NM_001048163.1;refGene_gn=ERBB2;refGene_tx=XM_006940248.4
E1 40800532 40800718
refGene_gn=ERBB2;refGene_tx=NM_001048163.1;refGene_gn=ERBB2;refGene_tx=XM_006940248.4
E1 40800853 40801009
refGene_gn=ERBB2;refGene_tx=NM_001048163.1;refGene_gn=ERBB2;refGene_tx=XM_006940248.4
E1 40801122 40801198
refGene_gn=ERBB2;refGene_tx=NM_001048163.1;refGene_gn=ERBB2;refGene_tx=XM_006940248.4
E1 40801508 40801655
refGene_gn=ERBB2;refGene_tx=NM_001048163.1;refGene_gn=ERBB2;refGene_tx=XM_006940248.4
E1 40802073 40802171
refGene_gn=ERBB2;refGene_tx=NM_001048163.1;refGene_gn=ERBB2;refGene_tx=XM_006940248.4

E1 40802364 40802553
refGene_gn=ERBB2;refGene_tx=NM_001048163.1;refGene_gn=ERBB2;refGene_tx
=XM_006940248.4
E1 40802830 40803083
refGene_gn=ERBB2;refGene_tx=NM_001048163.1;refGene_gn=ERBB2;refGene_tx
=XM_006940248.4
E1 40803227 40803598
refGene_gn=ERBB2;refGene_tx=NM_001048163.1;refGene_gn=ERBB2;refGene_tx
=XM_006940248.4
E1 40805044 40805128 refGene_gn=MIEN1;refGene_tx=XM_003996783.5
E1 40805213 40805290 refGene_gn=MIEN1;refGene_tx=XM_003996783.5
E1 40805733 40805831 refGene_gn=MIEN1;refGene_tx=XM_003996783.5
E1 40805934 40806023 refGene_gn=MIEN1;refGene_tx=XM_003996783.5
E1 40815376 40815528
refGene_gn=GRB7;refGene_tx=XM_019817821.2;refGene_gn=GRB7;refGene_tx=X
M_019817822.2
E1 40815640 40815794
refGene_gn=GRB7;refGene_tx=XM_019817821.2;refGene_gn=GRB7;refGene_tx=X
M_019817822.2
E1 40815977 40816134
refGene_gn=GRB7;refGene_tx=XM_019817821.2;refGene_gn=GRB7;refGene_tx=X
M_019817822.2
E1 40816261 40816383
refGene_gn=GRB7;refGene_tx=XM_019817821.2;refGene_gn=GRB7;refGene_tx=X
M_019817822.2
E1 40816474 40816552
refGene_gn=GRB7;refGene_tx=XM_019817821.2;refGene_gn=GRB7;refGene_tx=X
M_019817822.2
E1 40816820 40816958
refGene_gn=GRB7;refGene_tx=XM_019817821.2;refGene_gn=GRB7;refGene_tx=X
M_019817822.2
E1 40817299 40817410
refGene_gn=GRB7;refGene_tx=XM_019817821.2;refGene_gn=GRB7;refGene_tx=X
M_019817822.2
E1 40817608 40817707
refGene_gn=GRB7;refGene_tx=XM_019817821.2;refGene_gn=GRB7;refGene_tx=X
M_019817822.2
E1 40817957 40818038
refGene_gn=GRB7;refGene_tx=XM_019817821.2;refGene_gn=GRB7;refGene_tx=X
M_019817822.2
E1 40818117 40818234
refGene_gn=GRB7;refGene_tx=XM_019817821.2;refGene_gn=GRB7;refGene_tx=X
M_019817822.2
E1 40818419 40818474
refGene_gn=GRB7;refGene_tx=XM_019817821.2;refGene_gn=GRB7;refGene_tx=X
M_019817822.2
E1 40818601 40818689 refGene_gn=GRB7;refGene_tx=XM_019817821.2
E1 40818794 40818888
refGene_gn=GRB7;refGene_tx=XM_019817821.2;refGene_gn=GRB7;refGene_tx=X
M_019817822.2

E1 40819072 40819219
refGene_gn=GRB7;refGene_tx=XM_019817821.2;refGene_gn=GRB7;refGene_tx=X
M_019817822.2
E1 40835009 40835713
refGene_gn=IKZF3;refGene_tx=XM_006940304.4;refGene_gn=IKZF3;refGene_tx
=XM_019817823.2;refGene_gn=IKZF3;refGene_tx=XM_019817824.2
E1 40847385 40847502
refGene_gn=IKZF3;refGene_tx=XM_006940304.4;refGene_gn=IKZF3;refGene_tx
=XM_019817823.2;refGene_gn=IKZF3;refGene_tx=XM_019817824.2
E1 40857109 40857226
refGene_gn=IKZF3;refGene_tx=XM_019817823.2;refGene_gn=IKZF3;refGene_tx
=XM_019817824.2
E1 40859192 40859360
refGene_gn=IKZF3;refGene_tx=XM_019817823.2;refGene_gn=IKZF3;refGene_tx
=XM_019817824.2
E1 40860275 40860536
refGene_gn=IKZF3;refGene_tx=XM_019817823.2;refGene_gn=IKZF3;refGene_tx
=XM_019817824.2
E1 40895133 40895235
refGene_gn=IKZF3;refGene_tx=XM_019817823.2;refGene_gn=IKZF3;refGene_tx
=XM_019817824.2
E1 40897396 40897450
refGene_gn=IKZF3;refGene_tx=XM_019817823.2;refGene_gn=IKZF3;refGene_tx
=XM_019817824.2
E1 40921675 40921682
refGene_gn=IKZF3;refGene_tx=XM_019817823.2;refGene_gn=IKZF3;refGene_tx
=XM_019817824.2
E1 40925651 40925700 refGene_gn=ZPBP2;refGene_tx=XM_003996822.5
E1 40925843 40925909 refGene_gn=ZPBP2;refGene_tx=XM_003996822.5
E1 40927296 40927422
refGene_gn=ZPBP2;refGene_tx=XM_003996822.5;refGene_gn=ZPBP2;refGene_tx
=XM_023244102.1
E1 40928855 40929017
refGene_gn=ZPBP2;refGene_tx=XM_003996822.5;refGene_gn=ZPBP2;refGene_tx
=XM_023244102.1
E1 40929596 40929815
refGene_gn=ZPBP2;refGene_tx=XM_003996822.5;refGene_gn=ZPBP2;refGene_tx
=XM_023244102.1
E1 40930456 40930539
refGene_gn=ZPBP2;refGene_tx=XM_003996822.5;refGene_gn=ZPBP2;refGene_tx
=XM_023244102.1
E1 40932206 40932387
refGene_gn=ZPBP2;refGene_tx=XM_003996822.5;refGene_gn=ZPBP2;refGene_tx
=XM_023244102.1
E1 40932940 40933035
refGene_gn=ZPBP2;refGene_tx=XM_003996822.5;refGene_gn=ZPBP2;refGene_tx
=XM_023244102.1
E1 40951974 40952114 refGene_gn=GSDMB;refGene_tx=XM_019816946.2
E1 40952552 40952627 refGene_gn=GSDMB;refGene_tx=XM_019816946.2
E1 40953006 40953079 refGene_gn=GSDMB;refGene_tx=XM_019816946.2

E1 40953080 40953150 refGene_gn=GSDMB;refGene_tx=XM_019816946.2
E1 40953242 40953420 refGene_gn=GSDMB;refGene_tx=XM_019816946.2
E1 40956244 40956321 refGene_gn=GSDMB;refGene_tx=XM_019816946.2
E1 40956529 40956597 refGene_gn=GSDMB;refGene_tx=XM_019816946.2
E1 40956848 40956989 refGene_gn=GSDMB;refGene_tx=XM_019816946.2
E1 40956990 40957033 refGene_gn=GSDMB;refGene_tx=XM_019816946.2
E1 40958662 40958693 refGene_gn=GSDMB;refGene_tx=XM_019816946.2
E1 40958695 40958833 refGene_gn=GSDMB;refGene_tx=XM_019816946.2
E1 40961684 40961919 refGene_gn=GSDMB;refGene_tx=XM_019816946.2
E1 40966473 40966609
refGene_gn=ORMDL3;refGene_tx=XM_003996789.5;refGene_gn=ORMDL3;refGene_
tx=XM_003996790.5;refGene_gn=ORMDL3;refGene_tx=XM_006940305.4;refGene_
gn=ORMDL3;refGene_tx=XM_006940306.4
E1 40967029 40967181
refGene_gn=ORMDL3;refGene_tx=XM_003996789.5;refGene_gn=ORMDL3;refGene_
tx=XM_003996790.5;refGene_gn=ORMDL3;refGene_tx=XM_006940305.4;refGene_
gn=ORMDL3;refGene_tx=XM_006940306.4
E1 40967900 40968074
refGene_gn=ORMDL3;refGene_tx=XM_003996789.5;refGene_gn=ORMDL3;refGene_
tx=XM_003996790.5;refGene_gn=ORMDL3;refGene_tx=XM_006940305.4;refGene_
gn=ORMDL3;refGene_tx=XM_006940306.4
E1 40984279 40985101 refGene_gn=LRRC3C;refGene_tx=XM_019817830.2
E1 40998520 40998734
refGene_gn=GSDMA;refGene_tx=XM_003996792.4;refGene_gn=GSDMA;refGene_tx
=XM_019817829.2
E1 40998914 40999095
refGene_gn=GSDMA;refGene_tx=XM_003996792.4;refGene_gn=GSDMA;refGene_tx
=XM_019817829.2
E1 41002401 41002567
refGene_gn=GSDMA;refGene_tx=XM_003996792.4;refGene_gn=GSDMA;refGene_tx
=XM_019817829.2
E1 41003358 41003455
refGene_gn=GSDMA;refGene_tx=XM_003996792.4;refGene_gn=GSDMA;refGene_tx
=XM_019817829.2
E1 41003980 41004028
refGene_gn=GSDMA;refGene_tx=XM_003996792.4;refGene_gn=GSDMA;refGene_tx
=XM_019817829.2
E1 41004437 41004461
refGene_gn=GSDMA;refGene_tx=XM_003996792.4;refGene_gn=GSDMA;refGene_tx
=XM_019817829.2
E1 41004972 41005005
refGene_gn=GSDMA;refGene_tx=XM_003996792.4;refGene_gn=GSDMA;refGene_tx
=XM_019817829.2
E1 41005629 41005784
refGene_gn=GSDMA;refGene_tx=XM_003996792.4;refGene_gn=GSDMA;refGene_tx
=XM_019817829.2
E1 41006211 41006326
refGene_gn=GSDMA;refGene_tx=XM_003996792.4;refGene_gn=GSDMA;refGene_tx
=XM_019817829.2
E1 41007406 41007480

refGene_gn=GSDMA;refGene_tx=XM_003996792.4;refGene_gn=GSDMA;refGene_tx=XM_019817829.2
E1 41008447 41008690
refGene_gn=GSDMA;refGene_tx=XM_003996792.4;refGene_gn=GSDMA;refGene_tx=XM_019817829.2
E1 41012352 41012566
refGene_gn=PSMD3;refGene_tx=XM_003996793.5;refGene_gn=PSMD3;refGene_tx=XM_019817828.2
E1 41014773 41014964
refGene_gn=PSMD3;refGene_tx=XM_003996793.5;refGene_gn=PSMD3;refGene_tx=XM_019817828.2
E1 41016497 41016635
refGene_gn=PSMD3;refGene_tx=XM_003996793.5;refGene_gn=PSMD3;refGene_tx=XM_019817828.2
E1 41018661 41018798
refGene_gn=PSMD3;refGene_tx=XM_003996793.5;refGene_gn=PSMD3;refGene_tx=XM_019817828.2
E1 41019533 41019724
refGene_gn=PSMD3;refGene_tx=XM_003996793.5;refGene_gn=PSMD3;refGene_tx=XM_019817828.2
E1 41019864 41019968
refGene_gn=PSMD3;refGene_tx=XM_003996793.5;refGene_gn=PSMD3;refGene_tx=XM_019817828.2
E1 41024095 41024210
refGene_gn=PSMD3;refGene_tx=XM_003996793.5;refGene_gn=PSMD3;refGene_tx=XM_019817828.2
E1 41024313 41024433
refGene_gn=PSMD3;refGene_tx=XM_003996793.5;refGene_gn=PSMD3;refGene_tx=XM_019817828.2
E1 41024577 41024681
refGene_gn=PSMD3;refGene_tx=XM_003996793.5;refGene_gn=PSMD3;refGene_tx=XM_019817828.2
E1 41024949 41025105
refGene_gn=PSMD3;refGene_tx=XM_003996793.5;refGene_gn=PSMD3;refGene_tx=XM_019817828.2
E1 41026319 41026370
refGene_gn=PSMD3;refGene_tx=XM_003996793.5;refGene_gn=PSMD3;refGene_tx=XM_019817828.2
E1 41026546 41026684
refGene_gn=PSMD3;refGene_tx=XM_003996793.5;refGene_gn=PSMD3;refGene_tx=XM_019817828.2
E1 41033796 41033809
refGene_gn=CSF3;refGene_tx=NM_001009227.1;refGene_gn=CSF3;refGene_tx=XM_006940247.2
E1 41033931 41034086
refGene_gn=CSF3;refGene_tx=NM_001009227.1;refGene_gn=CSF3;refGene_tx=XM_006940247.2
E1 41034496 41034604
refGene_gn=CSF3;refGene_tx=NM_001009227.1;refGene_gn=CSF3;refGene_tx=XM_006940247.2

E1 41034743 41034890
refGene_gn=CSF3;refGene_tx=NM_001009227.1;refGene_gn=CSF3;refGene_tx=X
M_006940247.2
E1 41035247 41035412
refGene_gn=CSF3;refGene_tx=NM_001009227.1;refGene_gn=CSF3;refGene_tx=X
M_006940247.2
E1 41038370 41038487
refGene_gn=MED24;refGene_tx=XM_003996794.5;refGene_gn=MED24;refGene_tx
=XM_006940307.4;refGene_gn=MED24;refGene_tx=XM_006940308.4;refGene_gn=
MED24;refGene_tx=XM_011289159.3;refGene_gn=MED24;refGene_tx=XM_0112891
60.3
E1 41038669 41038818
refGene_gn=MED24;refGene_tx=XM_003996794.5;refGene_gn=MED24;refGene_tx
=XM_006940307.4;refGene_gn=MED24;refGene_tx=XM_006940308.4;refGene_gn=
MED24;refGene_tx=XM_011289159.3;refGene_gn=MED24;refGene_tx=XM_0112891
60.3
E1 41039097 41039178
refGene_gn=MED24;refGene_tx=XM_003996794.5;refGene_gn=MED24;refGene_tx
=XM_006940307.4;refGene_gn=MED24;refGene_tx=XM_006940308.4;refGene_gn=
MED24;refGene_tx=XM_011289159.3;refGene_gn=MED24;refGene_tx=XM_0112891
60.3
E1 41040063 41040163
refGene_gn=MED24;refGene_tx=XM_003996794.5;refGene_gn=MED24;refGene_tx
=XM_006940307.4;refGene_gn=MED24;refGene_tx=XM_006940308.4;refGene_gn=
MED24;refGene_tx=XM_011289159.3;refGene_gn=MED24;refGene_tx=XM_0112891
60.3
E1 41040456 41040547
refGene_gn=MED24;refGene_tx=XM_003996794.5;refGene_gn=MED24;refGene_tx
=XM_006940307.4;refGene_gn=MED24;refGene_tx=XM_006940308.4;refGene_gn=
MED24;refGene_tx=XM_011289159.3;refGene_gn=MED24;refGene_tx=XM_0112891
60.3
E1 41040703 41040885
refGene_gn=MED24;refGene_tx=XM_003996794.5;refGene_gn=MED24;refGene_tx
=XM_006940307.4;refGene_gn=MED24;refGene_tx=XM_006940308.4;refGene_gn=
MED24;refGene_tx=XM_011289159.3;refGene_gn=MED24;refGene_tx=XM_0112891
60.3
E1 41041276 41041541
refGene_gn=MED24;refGene_tx=XM_003996794.5;refGene_gn=MED24;refGene_tx
=XM_006940307.4;refGene_gn=MED24;refGene_tx=XM_006940308.4;refGene_gn=
MED24;refGene_tx=XM_011289159.3;refGene_gn=MED24;refGene_tx=XM_0112891
60.3
E1 41043420 41043596
refGene_gn=MED24;refGene_tx=XM_003996794.5;refGene_gn=MED24;refGene_tx
=XM_006940307.4;refGene_gn=MED24;refGene_tx=XM_006940308.4;refGene_gn=
MED24;refGene_tx=XM_011289159.3;refGene_gn=MED24;refGene_tx=XM_0112891
60.3
E1 41043912 41044012
refGene_gn=MED24;refGene_tx=XM_003996794.5;refGene_gn=MED24;refGene_tx
=XM_006940307.4;refGene_gn=MED24;refGene_tx=XM_006940308.4;refGene_gn=
MED24;refGene_tx=XM_011289159.3;refGene_gn=MED24;refGene_tx=XM_0112891

60.3
E1 41044127 41044306
refGene_gn=MED24;refGene_tx=XM_003996794.5;refGene_gn=MED24;refGene_tx=XM_006940307.4;refGene_gn=MED24;refGene_tx=XM_006940308.4;refGene_gn=MED24;refGene_tx=XM_011289159.3;refGene_gn=MED24;refGene_tx=XM_011289160.3
E1 41044656 41044739
refGene_gn=MED24;refGene_tx=XM_003996794.5;refGene_gn=MED24;refGene_tx=XM_006940307.4;refGene_gn=MED24;refGene_tx=XM_006940308.4;refGene_gn=MED24;refGene_tx=XM_011289159.3;refGene_gn=MED24;refGene_tx=XM_011289160.3
E1 41045181 41045219
refGene_gn=MED24;refGene_tx=XM_003996794.5;refGene_gn=MED24;refGene_tx=XM_006940307.4;refGene_gn=MED24;refGene_tx=XM_006940308.4;refGene_gn=MED24;refGene_tx=XM_011289159.3;refGene_gn=MED24;refGene_tx=XM_011289160.3
E1 41045851 41045994
refGene_gn=MED24;refGene_tx=XM_003996794.5;refGene_gn=MED24;refGene_tx=XM_006940307.4;refGene_gn=MED24;refGene_tx=XM_006940308.4;refGene_gn=MED24;refGene_tx=XM_011289159.3;refGene_gn=MED24;refGene_tx=XM_011289160.3
E1 41046495 41046607
refGene_gn=MED24;refGene_tx=XM_003996794.5;refGene_gn=MED24;refGene_tx=XM_006940307.4;refGene_gn=MED24;refGene_tx=XM_006940308.4;refGene_gn=MED24;refGene_tx=XM_011289159.3;refGene_gn=MED24;refGene_tx=XM_011289160.3
E1 41047037 41047124
refGene_gn=MED24;refGene_tx=XM_003996794.5;refGene_gn=MED24;refGene_tx=XM_006940307.4;refGene_gn=MED24;refGene_tx=XM_006940308.4;refGene_gn=MED24;refGene_tx=XM_011289159.3;refGene_gn=MED24;refGene_tx=XM_011289160.3
E1 41047391 41047474
refGene_gn=MED24;refGene_tx=XM_003996794.5;refGene_gn=MED24;refGene_tx=XM_006940307.4;refGene_gn=MED24;refGene_tx=XM_006940308.4;refGene_gn=MED24;refGene_tx=XM_011289159.3;refGene_gn=MED24;refGene_tx=XM_011289160.3
E1 41047859 41047907
refGene_gn=MED24;refGene_tx=XM_003996794.5;refGene_gn=MED24;refGene_tx=XM_006940307.4;refGene_gn=MED24;refGene_tx=XM_006940308.4;refGene_gn=MED24;refGene_tx=XM_011289159.3;refGene_gn=MED24;refGene_tx=XM_011289160.3
E1 41048453 41048567
refGene_gn=MED24;refGene_tx=XM_003996794.5;refGene_gn=MED24;refGene_tx=XM_006940307.4;refGene_gn=MED24;refGene_tx=XM_006940308.4;refGene_gn=MED24;refGene_tx=XM_011289159.3;refGene_gn=MED24;refGene_tx=XM_011289160.3
E1 41048870 41049021
refGene_gn=MED24;refGene_tx=XM_003996794.5;refGene_gn=MED24;refGene_tx=XM_006940307.4;refGene_gn=MED24;refGene_tx=XM_006940308.4;refGene_gn=MED24;refGene_tx=XM_011289159.3;refGene_gn=MED24;refGene_tx=XM_0112891

60.3
E1 41049163 41049275
refGene_gn=MED24;refGene_tx=XM_003996794.5;refGene_gn=MED24;refGene_tx=XM_006940307.4;refGene_gn=MED24;refGene_tx=XM_006940308.4;refGene_gn=MED24;refGene_tx=XM_011289159.3;refGene_gn=MED24;refGene_tx=XM_011289160.3
E1 41050715 41050772
refGene_gn=MED24;refGene_tx=XM_006940307.4;refGene_gn=MED24;refGene_tx=XM_006940308.4;refGene_gn=MED24;refGene_tx=XM_011289159.3;refGene_gn=MED24;refGene_tx=XM_011289160.3
E1 41050916 41051149
refGene_gn=MED24;refGene_tx=XM_003996794.5;refGene_gn=MED24;refGene_tx=XM_006940307.4;refGene_gn=MED24;refGene_tx=XM_006940308.4;refGene_gn=MED24;refGene_tx=XM_011289159.3;refGene_gn=MED24;refGene_tx=XM_011289160.3
E1 41051500 41051574
refGene_gn=MED24;refGene_tx=XM_003996794.5;refGene_gn=MED24;refGene_tx=XM_006940307.4;refGene_gn=MED24;refGene_tx=XM_006940308.4;refGene_gn=MED24;refGene_tx=XM_011289159.3;refGene_gn=MED24;refGene_tx=XM_011289160.3
E1 41051922 41051961
refGene_gn=MED24;refGene_tx=XM_003996794.5;refGene_gn=MED24;refGene_tx=XM_006940307.4;refGene_gn=MED24;refGene_tx=XM_006940308.4;refGene_gn=MED24;refGene_tx=XM_011289159.3;refGene_gn=MED24;refGene_tx=XM_011289160.3
E1 41061141 41061224
refGene_gn=MED24;refGene_tx=XM_003996794.5;refGene_gn=MED24;refGene_tx=XM_006940307.4;refGene_gn=MED24;refGene_tx=XM_006940308.4;refGene_gn=MED24;refGene_tx=XM_011289159.3;refGene_gn=MED24;refGene_tx=XM_011289160.3
E1 41061313 41061443
refGene_gn=MED24;refGene_tx=XM_003996794.5;refGene_gn=MED24;refGene_tx=XM_006940307.4;refGene_gn=MED24;refGene_tx=XM_006940308.4;refGene_gn=MED24;refGene_tx=XM_011289159.3;refGene_gn=MED24;refGene_tx=XM_011289160.3
E1 41080491 41080544 refGene_gn=THRA;refGene_tx=XM_011289162.3
E1 41082918 41082986 refGene_gn=THRA;refGene_tx=XM_011289162.3
E1 41083579 41083680 refGene_gn=THRA;refGene_tx=XM_011289162.3
E1 41085670 41085818 refGene_gn=THRA;refGene_tx=XM_011289162.3
E1 41086430 41086636 refGene_gn=THRA;refGene_tx=XM_011289162.3
E1 41087632 41087779 refGene_gn=THRA;refGene_tx=XM_011289162.3
E1 41088691 41088950 refGene_gn=THRA;refGene_tx=XM_011289162.3
E1 41089597 41089848 refGene_gn=THRA;refGene_tx=XM_011289162.3
E1 41093510 41093710 refGene_gn=NR1D1;refGene_tx=XM_003996795.5
E1 41094395 41094606 refGene_gn=NR1D1;refGene_tx=XM_003996795.5
E1 41095160 41095346 refGene_gn=NR1D1;refGene_tx=XM_003996795.5
E1 41095657 41096298 refGene_gn=NR1D1;refGene_tx=XM_003996795.5
E1 41096652 41096797 refGene_gn=NR1D1;refGene_tx=XM_003996795.5
E1 41096901 41096990 refGene_gn=NR1D1;refGene_tx=XM_003996795.5
E1 41097245 41097578 refGene_gn=NR1D1;refGene_tx=XM_003996795.5

E1 41100315 41100346 refGene_gn=NR1D1;refGene_tx=XM_003996795.5
E1 41120231 41120999
refGene_gn=MSL1;refGene_tx=XM_003996799.4;refGene_gn=MSL1;refGene_tx=XM_019817831.2
E1 41123493 41123717
refGene_gn=MSL1;refGene_tx=XM_003996799.4;refGene_gn=MSL1;refGene_tx=XM_019817831.2
E1 41125008 41125391
refGene_gn=MSL1;refGene_tx=XM_003996799.4;refGene_gn=MSL1;refGene_tx=XM_019817831.2
E1 41127298 41127346 refGene_gn=MSL1;refGene_tx=XM_003996799.4
E1 41127805 41127870
refGene_gn=MSL1;refGene_tx=XM_003996799.4;refGene_gn=MSL1;refGene_tx=XM_019817831.2
E1 41128501 41128569
refGene_gn=MSL1;refGene_tx=XM_003996799.4;refGene_gn=MSL1;refGene_tx=XM_019817831.2
E1 41129007 41129132
refGene_gn=MSL1;refGene_tx=XM_003996799.4;refGene_gn=MSL1;refGene_tx=XM_019817831.2
E1 41129300 41129373
refGene_gn=MSL1;refGene_tx=XM_003996799.4;refGene_gn=MSL1;refGene_tx=XM_019817831.2
E1 41129734 41129825
refGene_gn=MSL1;refGene_tx=XM_003996799.4;refGene_gn=MSL1;refGene_tx=XM_019817831.2
E1 41134375 41134606 refGene_gn=CASC3;refGene_tx=XM_006940311.4
E1 41134974 41135002 refGene_gn=CASC3;refGene_tx=XM_006940311.4
E1 41135343 41135381 refGene_gn=CASC3;refGene_tx=XM_006940311.4
E1 41147579 41147738 refGene_gn=CASC3;refGene_tx=XM_006940311.4
E1 41147828 41147980 refGene_gn=CASC3;refGene_tx=XM_006940311.4
E1 41148462 41148639 refGene_gn=CASC3;refGene_tx=XM_006940311.4
E1 41148825 41149511 refGene_gn=CASC3;refGene_tx=XM_006940311.4
E1 41152159 41152224 refGene_gn=CASC3;refGene_tx=XM_006940311.4
E1 41152663 41152778 refGene_gn=CASC3;refGene_tx=XM_006940311.4
E1 41153048 41153147 refGene_gn=CASC3;refGene_tx=XM_006940311.4
E1 41153402 41153617 refGene_gn=CASC3;refGene_tx=XM_006940311.4
E1 41154241 41154364 refGene_gn=CASC3;refGene_tx=XM_006940311.4
E1 41154505 41154529 refGene_gn=CASC3;refGene_tx=XM_006940311.4
E1 41160759 41161279 refGene_gn=RAPGEFL1;refGene_tx=XM_023243729.1
E1 41164091 41164170 refGene_gn=RAPGEFL1;refGene_tx=XM_023243729.1
E1 41165093 41165229 refGene_gn=RAPGEFL1;refGene_tx=XM_023243729.1
E1 41165457 41165555 refGene_gn=RAPGEFL1;refGene_tx=XM_023243729.1
E1 41167944 41168057 refGene_gn=RAPGEFL1;refGene_tx=XM_023243729.1
E1 41168301 41168469 refGene_gn=RAPGEFL1;refGene_tx=XM_023243729.1
E1 41169183 41169281 refGene_gn=RAPGEFL1;refGene_tx=XM_023243729.1
E1 41169393 41169516 refGene_gn=RAPGEFL1;refGene_tx=XM_023243729.1
E1 41170030 41170209 refGene_gn=RAPGEFL1;refGene_tx=XM_023243729.1
E1 41170295 41170386 refGene_gn=RAPGEFL1;refGene_tx=XM_023243729.1
E1 41170939 41170990 refGene_gn=RAPGEFL1;refGene_tx=XM_023243729.1

E1 41171321 41171409 refGene_gn=RAPGEFL1;refGene_tx=XM_023243729.1
E1 41171594 41171659 refGene_gn=RAPGEFL1;refGene_tx=XM_023243729.1
E1 41172043 41172098 refGene_gn=RAPGEFL1;refGene_tx=XM_023243729.1
E1 41172334 41172459 refGene_gn=RAPGEFL1;refGene_tx=XM_023243729.1
E1 41211788 41211851
refGene_gn=WIPF2;refGene_tx=XM_011289154.3;refGene_gn=WIPF2;refGene_tx=XM_011289155.3;refGene_gn=WIPF2;refGene_tx=XM_011289156.3;refGene_gn=WIPF2;refGene_tx=XM_011289157.3
E1 41214971 41214974 refGene_gn=WIPF2;refGene_tx=XM_011289158.3
E1 41218513 41218646
refGene_gn=WIPF2;refGene_tx=XM_011289154.3;refGene_gn=WIPF2;refGene_tx=XM_011289155.3;refGene_gn=WIPF2;refGene_tx=XM_011289156.3;refGene_gn=WIPF2;refGene_tx=XM_011289157.3;refGene_gn=WIPF2;refGene_tx=XM_011289158.3
E1 41219364 41219481
refGene_gn=WIPF2;refGene_tx=XM_011289154.3;refGene_gn=WIPF2;refGene_tx=XM_011289155.3;refGene_gn=WIPF2;refGene_tx=XM_011289156.3;refGene_gn=WIPF2;refGene_tx=XM_011289157.3;refGene_gn=WIPF2;refGene_tx=XM_011289158.3
E1 41219813 41220470
refGene_gn=WIPF2;refGene_tx=XM_011289154.3;refGene_gn=WIPF2;refGene_tx=XM_011289155.3;refGene_gn=WIPF2;refGene_tx=XM_011289156.3;refGene_gn=WIPF2;refGene_tx=XM_011289157.3;refGene_gn=WIPF2;refGene_tx=XM_011289158.3
E1 41229910 41230120
refGene_gn=WIPF2;refGene_tx=XM_011289154.3;refGene_gn=WIPF2;refGene_tx=XM_011289155.3;refGene_gn=WIPF2;refGene_tx=XM_011289156.3;refGene_gn=WIPF2;refGene_tx=XM_011289157.3;refGene_gn=WIPF2;refGene_tx=XM_011289158.3
E1 41231493 41231595
refGene_gn=WIPF2;refGene_tx=XM_011289154.3;refGene_gn=WIPF2;refGene_tx=XM_011289155.3;refGene_gn=WIPF2;refGene_tx=XM_011289156.3;refGene_gn=WIPF2;refGene_tx=XM_011289157.3;refGene_gn=WIPF2;refGene_tx=XM_011289158.3
E1 41231841 41231882
refGene_gn=WIPF2;refGene_tx=XM_011289154.3;refGene_gn=WIPF2;refGene_tx=XM_011289155.3;refGene_gn=WIPF2;refGene_tx=XM_011289156.3;refGene_gn=WIPF2;refGene_tx=XM_011289157.3;refGene_gn=WIPF2;refGene_tx=XM_011289158.3
E1 41240579 41240754
refGene_gn=CDC6;refGene_tx=XM_003996826.5;refGene_gn=CDC6;refGene_tx=XM_019817832.2;refGene_gn=CDC6;refGene_tx=XM_019817833.1;refGene_gn=CDC6;refGene_tx=XM_023243620.1
E1 41241828 41242110
refGene_gn=CDC6;refGene_tx=XM_003996826.5;refGene_gn=CDC6;refGene_tx=XM_019817832.2;refGene_gn=CDC6;refGene_tx=XM_019817833.1;refGene_gn=CDC6;refGene_tx=XM_023243620.1
E1 41242233 41242433
refGene_gn=CDC6;refGene_tx=XM_003996826.5;refGene_gn=CDC6;refGene_tx=XM_019817832.2;refGene_gn=CDC6;refGene_tx=XM_019817833.1;refGene_gn=CDC

6;refGene_tx=XM_023243620.1
E1 41242926 41243102
refGene_gn=CDC6;refGene_tx=XM_003996826.5;refGene_gn=CDC6;refGene_tx=XM_019817832.2;refGene_gn=CDC6;refGene_tx=XM_019817833.1;refGene_gn=CDC6;refGene_tx=XM_023243620.1
E1 41243507 41243614
refGene_gn=CDC6;refGene_tx=XM_003996826.5;refGene_gn=CDC6;refGene_tx=XM_019817832.2;refGene_gn=CDC6;refGene_tx=XM_019817833.1;refGene_gn=CDC6;refGene_tx=XM_023243620.1
E1 41244047 41244187
refGene_gn=CDC6;refGene_tx=XM_003996826.5;refGene_gn=CDC6;refGene_tx=XM_019817832.2;refGene_gn=CDC6;refGene_tx=XM_019817833.1;refGene_gn=CDC6;refGene_tx=XM_023243620.1
E1 41245083 41245184
refGene_gn=CDC6;refGene_tx=XM_003996826.5;refGene_gn=CDC6;refGene_tx=XM_019817832.2;refGene_gn=CDC6;refGene_tx=XM_019817833.1;refGene_gn=CDC6;refGene_tx=XM_023243620.1
E1 41246332 41246397
refGene_gn=CDC6;refGene_tx=XM_003996826.5;refGene_gn=CDC6;refGene_tx=XM_019817832.2;refGene_gn=CDC6;refGene_tx=XM_019817833.1;refGene_gn=CDC6;refGene_tx=XM_023243620.1
E1 41248673 41248876
refGene_gn=CDC6;refGene_tx=XM_003996826.5;refGene_gn=CDC6;refGene_tx=XM_019817832.2;refGene_gn=CDC6;refGene_tx=XM_019817833.1;refGene_gn=CDC6;refGene_tx=XM_023243620.1
E1 41249287 41249428
refGene_gn=CDC6;refGene_tx=XM_003996826.5;refGene_gn=CDC6;refGene_tx=XM_019817832.2;refGene_gn=CDC6;refGene_tx=XM_019817833.1;refGene_gn=CDC6;refGene_tx=XM_023243620.1
E1 41249962 41250052
refGene_gn=CDC6;refGene_tx=XM_003996826.5;refGene_gn=CDC6;refGene_tx=XM_019817832.2;refGene_gn=CDC6;refGene_tx=XM_019817833.1;refGene_gn=CDC6;refGene_tx=XM_023243620.1
E1 41277560 41277738 refGene_gn=RARA;refGene_tx=XM_019817836.2
E1 41286140 41286313
refGene_gn=RARA;refGene_tx=XM_003996829.5;refGene_gn=RARA;refGene_tx=XM_019817838.2
E1 41290196 41290329 refGene_gn=RARA;refGene_tx=XM_006940327.4
E1 41291620 41291769
refGene_gn=RARA;refGene_tx=XM_003996829.5;refGene_gn=RARA;refGene_tx=XM_006940327.4;refGene_gn=RARA;refGene_tx=XM_019817836.2;refGene_gn=RARA;refGene_tx=XM_019817838.2
E1 41293005 41293147
refGene_gn=RARA;refGene_tx=XM_003996829.5;refGene_gn=RARA;refGene_tx=XM_006940327.4;refGene_gn=RARA;refGene_tx=XM_019817836.2;refGene_gn=RARA;refGene_tx=XM_019817838.2
E1 41295169 41295330
refGene_gn=RARA;refGene_tx=XM_003996829.5;refGene_gn=RARA;refGene_tx=XM_006940327.4;refGene_gn=RARA;refGene_tx=XM_019817836.2;refGene_gn=RARA;refGene_tx=XM_019817838.2

E1 41295574 41295751
refGene_gn=RARA;refGene_tx=XM_003996829.5;refGene_gn=RARA;refGene_tx=XM_006940327.4;refGene_gn=RARA;refGene_tx=XM_019817836.2;refGene_gn=RARA;refGene_tx=XM_019817838.2
E1 41297105 41297310
refGene_gn=RARA;refGene_tx=XM_003996829.5;refGene_gn=RARA;refGene_tx=XM_006940327.4;refGene_gn=RARA;refGene_tx=XM_019817836.2;refGene_gn=RARA;refGene_tx=XM_019817838.2
E1 41298132 41298291
refGene_gn=RARA;refGene_tx=XM_003996829.5;refGene_gn=RARA;refGene_tx=XM_006940327.4;refGene_gn=RARA;refGene_tx=XM_019817836.2;refGene_gn=RARA;refGene_tx=XM_019817838.2
E1 41298910 41299128
refGene_gn=RARA;refGene_tx=XM_003996829.5;refGene_gn=RARA;refGene_tx=XM_006940327.4;refGene_gn=RARA;refGene_tx=XM_019817836.2;refGene_gn=RARA;refGene_tx=XM_019817838.2
E1 41306326 41307106 refGene_gn=GJD3;refGene_tx=XM_023243622.1
E1 41316453 41316582 refGene_gn=TOP2A;refGene_tx=XM_003996831.5
E1 41316893 41317093 refGene_gn=TOP2A;refGene_tx=XM_003996831.5
E1 41317896 41318037 refGene_gn=TOP2A;refGene_tx=XM_003996831.5
E1 41318421 41318465 refGene_gn=TOP2A;refGene_tx=XM_003996831.5
E1 41318584 41318711 refGene_gn=TOP2A;refGene_tx=XM_003996831.5
E1 41318945 41319098 refGene_gn=TOP2A;refGene_tx=XM_003996831.5
E1 41324625 41324716 refGene_gn=TOP2A;refGene_tx=XM_003996831.5
E1 41325050 41325233 refGene_gn=TOP2A;refGene_tx=XM_003996831.5
E1 41326437 41326521 refGene_gn=TOP2A;refGene_tx=XM_003996831.5
E1 41326652 41326817 refGene_gn=TOP2A;refGene_tx=XM_003996831.5
E1 41326918 41327010 refGene_gn=TOP2A;refGene_tx=XM_003996831.5
E1 41328289 41328485 refGene_gn=TOP2A;refGene_tx=XM_003996831.5
E1 41328647 41328848 refGene_gn=TOP2A;refGene_tx=XM_003996831.5
E1 41328972 41329107 refGene_gn=TOP2A;refGene_tx=XM_003996831.5
E1 41329299 41329531 refGene_gn=TOP2A;refGene_tx=XM_003996831.5
E1 41331326 41331475 refGene_gn=TOP2A;refGene_tx=XM_003996831.5
E1 41333224 41333346 refGene_gn=TOP2A;refGene_tx=XM_003996831.5
E1 41333447 41333562 refGene_gn=TOP2A;refGene_tx=XM_003996831.5
E1 41334386 41334479 refGene_gn=TOP2A;refGene_tx=XM_003996831.5
E1 41334694 41334804 refGene_gn=TOP2A;refGene_tx=XM_003996831.5
E1 41334895 41335001 refGene_gn=TOP2A;refGene_tx=XM_003996831.5
E1 41335139 41335250 refGene_gn=TOP2A;refGene_tx=XM_003996831.5
E1 41335508 41335634 refGene_gn=TOP2A;refGene_tx=XM_003996831.5
E1 41335927 41336085 refGene_gn=TOP2A;refGene_tx=XM_003996831.5
E1 41337290 41337429 refGene_gn=TOP2A;refGene_tx=XM_003996831.5
E1 41338705 41338843 refGene_gn=TOP2A;refGene_tx=XM_003996831.5
E1 41338951 41339053 refGene_gn=TOP2A;refGene_tx=XM_003996831.5
E1 41339238 41339412 refGene_gn=TOP2A;refGene_tx=XM_003996831.5
E1 41339780 41339993 refGene_gn=TOP2A;refGene_tx=XM_003996831.5
E1 41340431 41340529 refGene_gn=TOP2A;refGene_tx=XM_003996831.5
E1 41340678 41340824 refGene_gn=TOP2A;refGene_tx=XM_003996831.5
E1 41342307 41342371 refGene_gn=TOP2A;refGene_tx=XM_003996831.5
E1 41342630 41342721 refGene_gn=TOP2A;refGene_tx=XM_003996831.5

E1 41342941 41343097 refGene_gn=TOP2A;refGene_tx=XM_003996831.5
E1 41343949 41343970 refGene_gn=TOP2A;refGene_tx=XM_003996831.5
E1 41365367 41365716 refGene_gn=IGFBP4;refGene_tx=XM_003996921.4
E1 41373155 41373313 refGene_gn=IGFBP4;refGene_tx=XM_003996921.4
E1 41374069 41374204 refGene_gn=IGFBP4;refGene_tx=XM_003996921.4
E1 41376197 41376332 refGene_gn=IGFBP4;refGene_tx=XM_003996921.4
E1 41390267 41390409
refGene_gn=TNS4;refGene_tx=XM_006940328.3;refGene_gn=TNS4;refGene_tx=XM_019817840.2
E1 41390917 41390944 refGene_gn=TNS4;refGene_tx=XM_006940328.3
E1 41391201 41391270
refGene_gn=TNS4;refGene_tx=XM_006940328.3;refGene_gn=TNS4;refGene_tx=XM_019817840.2
E1 41391953 41392122
refGene_gn=TNS4;refGene_tx=XM_006940328.3;refGene_gn=TNS4;refGene_tx=XM_019817840.2
E1 41392839 41392908
refGene_gn=TNS4;refGene_tx=XM_006940328.3;refGene_gn=TNS4;refGene_tx=XM_019817840.2
E1 41393718 41393796
refGene_gn=TNS4;refGene_tx=XM_006940328.3;refGene_gn=TNS4;refGene_tx=XM_019817840.2
E1 41393901 41393994
refGene_gn=TNS4;refGene_tx=XM_006940328.3;refGene_gn=TNS4;refGene_tx=XM_019817840.2
E1 41395174 41395300
refGene_gn=TNS4;refGene_tx=XM_006940328.3;refGene_gn=TNS4;refGene_tx=XM_019817840.2
E1 41395634 41395721
refGene_gn=TNS4;refGene_tx=XM_006940328.3;refGene_gn=TNS4;refGene_tx=XM_019817840.2
E1 41397802 41398227
refGene_gn=TNS4;refGene_tx=XM_006940328.3;refGene_gn=TNS4;refGene_tx=XM_019817840.2
E1 41399299 41399723
refGene_gn=TNS4;refGene_tx=XM_006940328.3;refGene_gn=TNS4;refGene_tx=XM_019817840.2
E1 41405957 41406396
refGene_gn=TNS4;refGene_tx=XM_006940328.3;refGene_gn=TNS4;refGene_tx=XM_019817840.2
E1 41458843 41459923 refGene_gn=CCR7;refGene_tx=XM_003996833.4
E1 41462864 41462914 refGene_gn=CCR7;refGene_tx=XM_003996833.4
E1 41469144 41469154 refGene_gn=CCR7;refGene_tx=XM_003996833.4
E1 41512201 41512410
refGene_gn=SMARCE1;refGene_tx=XM_003996834.5;refGene_gn=SMARCE1;refGene_tx=XM_019817842.2
E1 41514368 41514579
refGene_gn=SMARCE1;refGene_tx=XM_003996834.5;refGene_gn=SMARCE1;refGene_tx=XM_019817842.2
E1 41515210 41515312

refGene_gn=SMARCE1;refGene_tx=XM_003996834.5;refGene_gn=SMARCE1;refGene_tx=XM_019817842.2
E1 41516072 41516245
refGene_gn=SMARCE1;refGene_tx=XM_003996834.5;refGene_gn=SMARCE1;refGene_tx=XM_019817842.2
E1 41519240 41519412
refGene_gn=SMARCE1;refGene_tx=XM_003996834.5;refGene_gn=SMARCE1;refGene_tx=XM_019817842.2
E1 41519672 41519804
refGene_gn=SMARCE1;refGene_tx=XM_003996834.5;refGene_gn=SMARCE1;refGene_tx=XM_019817842.2
E1 41520739 41520820
refGene_gn=SMARCE1;refGene_tx=XM_003996834.5;refGene_gn=SMARCE1;refGene_tx=XM_019817842.2
E1 41525779 41525884 refGene_gn=SMARCE1;refGene_tx=XM_003996834.5
E1 41528861 41528905 refGene_gn=SMARCE1;refGene_tx=XM_003996834.5
E1 41529078 41529085 refGene_gn=SMARCE1;refGene_tx=XM_003996834.5
E1 41545511 41545740
refGene_gn=KRT222;refGene_tx=XM_006940329.4;refGene_gn=KRT222;refGene_tx=XM_019817845.2;refGene_gn=KRT222;refGene_tx=XM_019817846.2;refGene_gn=KRT222;refGene_tx=XM_019817847.2
E1 41546173 41546309
refGene_gn=KRT222;refGene_tx=XM_006940329.4;refGene_gn=KRT222;refGene_tx=XM_019817845.2;refGene_gn=KRT222;refGene_tx=XM_019817846.2;refGene_gn=KRT222;refGene_tx=XM_019817847.2
E1 41546483 41546560
refGene_gn=KRT222;refGene_tx=XM_006940329.4;refGene_gn=KRT222;refGene_tx=XM_019817845.2;refGene_gn=KRT222;refGene_tx=XM_019817846.2;refGene_gn=KRT222;refGene_tx=XM_019817847.2
E1 41549108 41549329
refGene_gn=KRT222;refGene_tx=XM_006940329.4;refGene_gn=KRT222;refGene_tx=XM_019817845.2;refGene_gn=KRT222;refGene_tx=XM_019817846.2;refGene_gn=KRT222;refGene_tx=XM_019817847.2
E1 41550033 41550162
refGene_gn=KRT222;refGene_tx=XM_006940329.4;refGene_gn=KRT222;refGene_tx=XM_019817845.2;refGene_gn=KRT222;refGene_tx=XM_019817846.2;refGene_gn=KRT222;refGene_tx=XM_019817847.2
E1 41553261 41553357 refGene_gn=KRT222;refGene_tx=XM_006940329.4
E1 41585079 41585183
refGene_gn=KRT24;refGene_tx=XM_003996835.5;refGene_gn=KRT24;refGene_tx=XM_006940330.4
E1 41585823 41585954
refGene_gn=KRT24;refGene_tx=XM_003996835.5;refGene_gn=KRT24;refGene_tx=XM_006940330.4
E1 41586163 41586381
refGene_gn=KRT24;refGene_tx=XM_003996835.5;refGene_gn=KRT24;refGene_tx=XM_006940330.4
E1 41586721 41586847
refGene_gn=KRT24;refGene_tx=XM_003996835.5;refGene_gn=KRT24;refGene_tx=XM_006940330.4

E1 41586942 41587104
refGene_gn=KRT24;refGene_tx=XM_003996835.5;refGene_gn=KRT24;refGene_tx
=XM_006940330.4
E1 41587830 41587987
refGene_gn=KRT24;refGene_tx=XM_003996835.5;refGene_gn=KRT24;refGene_tx
=XM_006940330.4
E1 41588620 41588703
refGene_gn=KRT24;refGene_tx=XM_003996835.5;refGene_gn=KRT24;refGene_tx
=XM_006940330.4
E1 41589746 41590358
refGene_gn=KRT24;refGene_tx=XM_003996835.5;refGene_gn=KRT24;refGene_tx
=XM_006940330.4
E1 41615978 41616088 refGene_gn=KRT25;refGene_tx=XM_011289152.3
E1 41617125 41617193 refGene_gn=KRT25;refGene_tx=XM_011289152.3
E1 41617558 41617776 refGene_gn=KRT25;refGene_tx=XM_011289152.3
E1 41618376 41618502 refGene_gn=KRT25;refGene_tx=XM_011289152.3
E1 41618587 41618749 refGene_gn=KRT25;refGene_tx=XM_011289152.3
E1 41620655 41620812 refGene_gn=KRT25;refGene_tx=XM_011289152.3
E1 41621231 41621314 refGene_gn=KRT25;refGene_tx=XM_011289152.3
E1 41621604 41622033 refGene_gn=KRT25;refGene_tx=XM_011289152.3
E1 41632161 41632316
refGene_gn=LOC101083308;refGene_tx=XM_006940332.4
E1 41633357 41633425
refGene_gn=LOC101083308;refGene_tx=XM_006940332.4
E1 41634675 41634893
refGene_gn=LOC101083308;refGene_tx=XM_006940332.4
E1 41635485 41635611
refGene_gn=LOC101083308;refGene_tx=XM_006940332.4
E1 41635696 41635858
refGene_gn=LOC101083308;refGene_tx=XM_006940332.4
E1 41635980 41636137
refGene_gn=LOC101083308;refGene_tx=XM_006940332.4
E1 41636523 41636606
refGene_gn=LOC101083308;refGene_tx=XM_006940332.4
E1 41637067 41637496
refGene_gn=LOC101083308;refGene_tx=XM_006940332.4
E1 41645011 41645154
refGene_gn=LOC111556209;refGene_tx=XM_003996838.4
E1 41645269 41645319
refGene_gn=LOC111556209;refGene_tx=XM_003996838.4
E1 41645749 41645967
refGene_gn=LOC111556209;refGene_tx=XM_003996838.4
E1 41647329 41647455
refGene_gn=LOC111556209;refGene_tx=XM_003996838.4
E1 41647540 41647702
refGene_gn=LOC111556209;refGene_tx=XM_003996838.4
E1 41647980 41648137
refGene_gn=LOC111556209;refGene_tx=XM_003996838.4
E1 41648502 41648585
refGene_gn=LOC111556209;refGene_tx=XM_003996838.4

E1	41649296 41649710
refGene_gn=LOC111556209;refGene_tx=XM_003996838.4
E1	41666297 41666455
refGene_gn=KRT10;refGene_tx=XM_023244763.1;refGene_gn=KRT10;refGene_tx=XM_023244764.1;refGene_gn=KRT10;refGene_tx=XM_023244765.1
E1	41666968 41667024
refGene_gn=KRT10;refGene_tx=XM_023244763.1;refGene_gn=KRT10;refGene_tx=XM_023244764.1;refGene_gn=KRT10;refGene_tx=XM_023244765.1;refGene_gn=KRT10;refGene_tx=XM_023244766.1
E1	41667951 41668169
refGene_gn=KRT10;refGene_tx=XM_023244763.1;refGene_gn=KRT10;refGene_tx=XM_023244764.1;refGene_gn=KRT10;refGene_tx=XM_023244765.1;refGene_gn=KRT10;refGene_tx=XM_023244766.1
E1	41671105 41671231
refGene_gn=KRT10;refGene_tx=XM_023244763.1;refGene_gn=KRT10;refGene_tx=XM_023244764.1;refGene_gn=KRT10;refGene_tx=XM_023244765.1;refGene_gn=KRT10;refGene_tx=XM_023244766.1
E1	41671312 41671474
refGene_gn=KRT10;refGene_tx=XM_023244763.1;refGene_gn=KRT10;refGene_tx=XM_023244764.1;refGene_gn=KRT10;refGene_tx=XM_023244765.1;refGene_gn=KRT10;refGene_tx=XM_023244766.1
E1	41672255 41672412
refGene_gn=KRT10;refGene_tx=XM_023244763.1;refGene_gn=KRT10;refGene_tx=XM_023244764.1;refGene_gn=KRT10;refGene_tx=XM_023244765.1;refGene_gn=KRT10;refGene_tx=XM_023244766.1
E1	41673511 41673594
refGene_gn=KRT10;refGene_tx=XM_023244763.1;refGene_gn=KRT10;refGene_tx=XM_023244764.1;refGene_gn=KRT10;refGene_tx=XM_023244765.1;refGene_gn=KRT10;refGene_tx=XM_023244766.1
E1	41674483 41674517
refGene_gn=KRT10;refGene_tx=XM_023244763.1;refGene_gn=KRT10;refGene_tx=XM_023244764.1;refGene_gn=KRT10;refGene_tx=XM_023244765.1;refGene_gn=KRT10;refGene_tx=XM_023244766.1
E1	41684260 41684367
refGene_gn=KRT10;refGene_tx=XM_023244767.1;refGene_gn=KRT10;refGene_tx=XM_023244768.1
E1	41684706 41685032
refGene_gn=KRT10;refGene_tx=XM_023244763.1;refGene_gn=KRT10;refGene_tx=XM_023244764.1;refGene_gn=KRT10;refGene_tx=XM_023244765.1;refGene_gn=KRT10;refGene_tx=XM_023244766.1;refGene_gn=KRT10;refGene_tx=XM_023244767.1;refGene_gn=KRT10;refGene_tx=XM_023244768.1
E1	41685386 41685604
refGene_gn=KRT10;refGene_tx=XM_023244763.1;refGene_gn=KRT10;refGene_tx=XM_023244764.1;refGene_gn=KRT10;refGene_tx=XM_023244765.1;refGene_gn=KRT10;refGene_tx=XM_023244766.1;refGene_gn=KRT10;refGene_tx=XM_023244767.1;refGene_gn=KRT10;refGene_tx=XM_023244768.1
E1	41685952 41686078
refGene_gn=KRT10;refGene_tx=XM_023244763.1;refGene_gn=KRT10;refGene_tx=XM_023244764.1;refGene_gn=KRT10;refGene_tx=XM_023244765.1;refGene_gn=KRT10;refGene_tx=XM_023244766.1;refGene_gn=KRT10;refGene_tx=XM_0232447

67.1;refGene_gn=KRT10;refGene_tx=XM_023244768.1
E1 41686169 41686331
refGene_gn=KRT10;refGene_tx=XM_023244763.1;refGene_gn=KRT10;refGene_tx
=XM_023244764.1;refGene_gn=KRT10;refGene_tx=XM_023244765.1;refGene_gn=
KRT10;refGene_tx=XM_023244766.1;refGene_gn=KRT10;refGene_tx=XM_0232447
67.1;refGene_gn=KRT10;refGene_tx=XM_023244768.1
E1 41686430 41686587
refGene_gn=KRT10;refGene_tx=XM_023244763.1;refGene_gn=KRT10;refGene_tx
=XM_023244764.1;refGene_gn=KRT10;refGene_tx=XM_023244765.1;refGene_gn=
KRT10;refGene_tx=XM_023244766.1;refGene_gn=KRT10;refGene_tx=XM_0232447
67.1;refGene_gn=KRT10;refGene_tx=XM_023244768.1
E1 41686961 41687044
refGene_gn=KRT10;refGene_tx=XM_023244763.1;refGene_gn=KRT10;refGene_tx
=XM_023244764.1;refGene_gn=KRT10;refGene_tx=XM_023244765.1;refGene_gn=
KRT10;refGene_tx=XM_023244766.1;refGene_gn=KRT10;refGene_tx=XM_0232447
67.1;refGene_gn=KRT10;refGene_tx=XM_023244768.1
E1 41687883 41688465
refGene_gn=KRT10;refGene_tx=XM_023244763.1;refGene_gn=KRT10;refGene_tx
=XM_023244764.1;refGene_gn=KRT10;refGene_tx=XM_023244765.1;refGene_gn=
KRT10;refGene_tx=XM_023244766.1;refGene_gn=KRT10;refGene_tx=XM_0232447
67.1;refGene_gn=KRT10;refGene_tx=XM_023244768.1
E1 41724150 41724242 refGene_gn=KRT12;refGene_tx=XM_003996840.4
E1 41724768 41724839 refGene_gn=KRT12;refGene_tx=XM_003996840.4
E1 41725502 41725723 refGene_gn=KRT12;refGene_tx=XM_003996840.4
E1 41725864 41725990 refGene_gn=KRT12;refGene_tx=XM_003996840.4
E1 41726088 41726250 refGene_gn=KRT12;refGene_tx=XM_003996840.4
E1 41727365 41727522 refGene_gn=KRT12;refGene_tx=XM_003996840.4
E1 41728500 41728583 refGene_gn=KRT12;refGene_tx=XM_003996840.4
E1 41728967 41729212 refGene_gn=KRT12;refGene_tx=XM_003996840.4
E1 41729214 41729578 refGene_gn=KRT12;refGene_tx=XM_003996840.4
E1 41736076 41736174 refGene_gn=KRT20;refGene_tx=XM_019817854.2
E1 41736702 41736752 refGene_gn=KRT20;refGene_tx=XM_019817854.2
E1 41737549 41737770 refGene_gn=KRT20;refGene_tx=XM_019817854.2
E1 41738691 41738817 refGene_gn=KRT20;refGene_tx=XM_019817854.2
E1 41738942 41739104 refGene_gn=KRT20;refGene_tx=XM_019817854.2
E1 41739381 41739538 refGene_gn=KRT20;refGene_tx=XM_019817854.2
E1 41740825 41740908 refGene_gn=KRT20;refGene_tx=XM_019817854.2
E1 41742514 41742919 refGene_gn=KRT20;refGene_tx=XM_019817854.2
E1 41780036 41780131 refGene_gn=KRT23;refGene_tx=XM_011289149.3
E1 41781536 41781568 refGene_gn=KRT23;refGene_tx=XM_011289149.3
E1 41783022 41783243 refGene_gn=KRT23;refGene_tx=XM_011289149.3
E1 41786192 41786315 refGene_gn=KRT23;refGene_tx=XM_011289149.3
E1 41786402 41786564 refGene_gn=KRT23;refGene_tx=XM_011289149.3
E1 41787618 41787775 refGene_gn=KRT23;refGene_tx=XM_011289149.3
E1 41788902 41788985 refGene_gn=KRT23;refGene_tx=XM_011289149.3
E1 41795905 41796301 refGene_gn=KRT23;refGene_tx=XM_011289149.3
E1 41814903 41815156 refGene_gn=KRT39;refGene_tx=XM_019817851.2
E1 41816525 41816746 refGene_gn=KRT39;refGene_tx=XM_019817851.2
E1 41818281 41818407 refGene_gn=KRT39;refGene_tx=XM_019817851.2
E1 41818518 41818680 refGene_gn=KRT39;refGene_tx=XM_019817851.2

E1 41819344 41819501 refGene_gn=KRT39;refGene_tx=XM_019817851.2
E1 41820187 41820270 refGene_gn=KRT39;refGene_tx=XM_019817851.2
E1 41821521 41821944 refGene_gn=KRT39;refGene_tx=XM_019817851.2
E1 41831774 41831874 refGene_gn=KRT40;refGene_tx=XM_003996843.5
E1 41832382 41832603 refGene_gn=KRT40;refGene_tx=XM_003996843.5
E1 41833944 41834070 refGene_gn=KRT40;refGene_tx=XM_003996843.5
E1 41834149 41834311 refGene_gn=KRT40;refGene_tx=XM_003996843.5
E1 41835158 41835173 refGene_gn=KRT40;refGene_tx=XM_003996843.5
E1 41835174 41835319 refGene_gn=KRT40;refGene_tx=XM_003996843.5
E1 41835581 41835634 refGene_gn=KRT40;refGene_tx=XM_003996843.5
E1 41835636 41835666 refGene_gn=KRT40;refGene_tx=XM_003996843.5
E1 41836373 41836820 refGene_gn=KRT40;refGene_tx=XM_003996843.5
E1 41848906 41849192
refGene_gn=LOC101087113;refGene_tx=XM_006940334.4
E1 41855805 41855816
refGene_gn=LOC101087113;refGene_tx=XM_006940334.4
E1 41866500 41866794
refGene_gn=LOC101087623;refGene_tx=XM_003996846.5
E1 41886752 41887099
refGene_gn=LOC101087885;refGene_tx=XM_023243466.1
E1 41887101 41887333
refGene_gn=LOC101087885;refGene_tx=XM_023243466.1
E1 41888757 41888825
refGene_gn=LOC101087885;refGene_tx=XM_023243466.1
E1 41889650 41890149
refGene_gn=LOC101087885;refGene_tx=XM_023243466.1
E1 41896338 41896923
refGene_gn=LOC109494368;refGene_tx=XM_019817869.2
E1 41898740 41898748
refGene_gn=LOC109494368;refGene_tx=XM_019817869.2
E1 41901970 41902627
refGene_gn=LOC109494368;refGene_tx=XM_019817869.2
E1 41906145 41906533
refGene_gn=LOC109494368;refGene_tx=XM_019817869.2
E1 41912995 41913385
refGene_gn=LOC101085333;refGene_tx=XM_003996929.3
E1 41917642 41918029
refGene_gn=LOC101085588;refGene_tx=XM_023244788.1
E1 41922934 41923333
refGene_gn=LOC101085845;refGene_tx=XM_003996930.5
E1 41932136 41932273
refGene_gn=LOC105261176;refGene_tx=XM_023244786.1;refGene_gn=LOC105261
176;refGene_tx=XM_023244787.1
E1 41932609 41933126
refGene_gn=LOC105261176;refGene_tx=XM_023244786.1;refGene_gn=LOC105261
176;refGene_tx=XM_023244787.1
E1 41940626 41941253
refGene_gn=LOC101088126;refGene_tx=XM_023244785.1
E1 41947006 41947488
refGene_gn=LOC105261175;refGene_tx=XM_023244777.1;refGene_gn=LOC105261

175;refGene_tx=XM_023244778.1;refGene_gn=LOC105261175;refGene_tx=XM_023244779.1
E1 41952333 41952346
refGene_gn=LOC105261175;refGene_tx=XM_023244778.1
E1 41953466 41953923
refGene_gn=LOC105261175;refGene_tx=XM_023244777.1;refGene_gn=LOC105261175;refGene_tx=XM_023244778.1;refGene_gn=LOC105261175;refGene_tx=XM_023244779.1
E1 41965320 41966940
refGene_gn=LOC101088376;refGene_tx=XM_023244760.1
E1 41974919 41975867
refGene_gn=LOC101093843;refGene_tx=XM_004001257.5;refGene_gn=LOC101093843;refGene_tx=XM_023244771.1;refGene_gn=LOC101093843;refGene_tx=XM_023244772.1;refGene_gn=LOC101093843;refGene_tx=XM_023244773.1;refGene_gn=LOC101093843;refGene_tx=XM_023244774.1;refGene_gn=LOC101093843;refGene_tx=XM_023244775.1
E1 41987898 41988002
refGene_gn=LOC105260091;refGene_tx=XM_023244780.1
E1 41998756 41998901
refGene_gn=LOC105260091;refGene_tx=XM_023244780.1
E1 42003419 42003425
refGene_gn=LOC105260091;refGene_tx=XM_023244780.1
E1 42005585 42006176
refGene_gn=LOC105260091;refGene_tx=XM_023244780.1;refGene_gn=LOC105260091;refGene_tx=XM_023244781.1;refGene_gn=LOC105260091;refGene_tx=XM_023244782.1;refGene_gn=LOC105260091;refGene_tx=XM_023244783.1
E1 42011770 42011961
refGene_gn=LOC105260091;refGene_tx=XM_023244781.1;refGene_gn=LOC105260091;refGene_tx=XM_023244782.1
E1 42035412 42035557
refGene_gn=LOC102901573;refGene_tx=XM_023244769.1
E1 42051415 42052933
refGene_gn=LOC109497009;refGene_tx=XM_023244776.1
E1 42054302 42054394
refGene_gn=LOC102901573;refGene_tx=XM_023244769.1
E1 42071684 42072191
refGene_gn=LOC109494366;refGene_tx=XM_019817865.2
E1 42078239 42078331
refGene_gn=LOC101086104;refGene_tx=XM_023244409.1
E1 42082112 42082206
refGene_gn=LOC101086104;refGene_tx=XM_023244409.1
E1 42087412 42087710
refGene_gn=LOC101086104;refGene_tx=XM_023244409.1
E1 42109296 42110355
refGene_gn=LOC102902071;refGene_tx=XM_006940396.4
E1 42114941 42116392
refGene_gn=LOC101086358;refGene_tx=XM_019817870.2
E1 42118759 42118922
refGene_gn=LOC101086358;refGene_tx=XM_019817870.2
E1 42122804 42123143

refGene_gn=LOC111557764;refGene_tx=XM_023243847.1
E1 42151284 42151402
refGene_gn=LOC101089661;refGene_tx=XM_003996855.4
E1 42151613 42151834
refGene_gn=LOC101089661;refGene_tx=XM_003996855.4
E1 42151956 42152082
refGene_gn=LOC101089661;refGene_tx=XM_003996855.4
E1 42152176 42152338
refGene_gn=LOC101089661;refGene_tx=XM_003996855.4
E1 42153581 42153738
refGene_gn=LOC101089661;refGene_tx=XM_003996855.4
E1 42154287 42154370
refGene_gn=LOC101089661;refGene_tx=XM_003996855.4
E1 42155564 42155912
refGene_gn=LOC101089661;refGene_tx=XM_003996855.4
E1 42166891 42167018
refGene_gn=LOC101086611;refGene_tx=XM_006940344.3
E1 42167780 42167968
refGene_gn=LOC101086611;refGene_tx=XM_006940344.3
E1 42167969 42168002
refGene_gn=LOC101086611;refGene_tx=XM_006940344.3
E1 42168124 42168250
refGene_gn=LOC101086611;refGene_tx=XM_006940344.3
E1 42168344 42168506
refGene_gn=LOC101086611;refGene_tx=XM_006940344.3
E1 42169541 42169698
refGene_gn=LOC101086611;refGene_tx=XM_006940344.3
E1 42170662 42170745
refGene_gn=LOC101086611;refGene_tx=XM_006940344.3
E1 42172716 42173064
refGene_gn=LOC101086611;refGene_tx=XM_006940344.3
E1 42183957 42184042
refGene_gn=LOC101086861;refGene_tx=XM_006940343.4
E1 42184506 42184727
refGene_gn=LOC101086861;refGene_tx=XM_006940343.4
E1 42184849 42184975
refGene_gn=LOC101086861;refGene_tx=XM_006940343.4
E1 42185069 42185231
refGene_gn=LOC101086861;refGene_tx=XM_006940343.4
E1 42186108 42186265
refGene_gn=LOC101086861;refGene_tx=XM_006940343.4
E1 42186777 42186860
refGene_gn=LOC101086861;refGene_tx=XM_006940343.4
E1 42186962 42187310
refGene_gn=LOC101086861;refGene_tx=XM_006940343.4
E1 42196868 42197001
refGene_gn=LOC101087114;refGene_tx=XM_006940342.3;refGene_gn=LOC101087
114;refGene_tx=XM_019817880.2
E1 42197645 42197866
refGene_gn=LOC101087114;refGene_tx=XM_006940342.3;refGene_gn=LOC101087

114;refGene_tx=XM_019817880.2
E1 42197990 42198116
refGene_gn=LOC101087114;refGene_tx=XM_006940342.3;refGene_gn=LOC101087
114;refGene_tx=XM_019817880.2
E1 42198210 42198372
refGene_gn=LOC101087114;refGene_tx=XM_006940342.3;refGene_gn=LOC101087
114;refGene_tx=XM_019817880.2
E1 42199139 42199296
refGene_gn=LOC101087114;refGene_tx=XM_006940342.3;refGene_gn=LOC101087
114;refGene_tx=XM_019817880.2
E1 42199645 42199728
refGene_gn=LOC101087114;refGene_tx=XM_006940342.3;refGene_gn=LOC101087
114;refGene_tx=XM_019817880.2
E1 42199935 42200283
refGene_gn=LOC101087114;refGene_tx=XM_006940342.3
E1 42207997 42208127
refGene_gn=LOC101087114;refGene_tx=XM_006940397.3
E1 42208873 42209094
refGene_gn=LOC101087114;refGene_tx=XM_006940397.3
E1 42209452 42209578
refGene_gn=LOC101087114;refGene_tx=XM_006940397.3
E1 42209667 42209829
refGene_gn=LOC101087114;refGene_tx=XM_006940397.3
E1 42210185 42210342
refGene_gn=LOC101087114;refGene_tx=XM_006940397.3
E1 42211322 42211405
refGene_gn=LOC101087114;refGene_tx=XM_006940397.3
E1 42212806 42213235
refGene_gn=LOC101087114;refGene_tx=XM_006940397.3;refGene_gn=LOC101087
114;refGene_tx=XM_019817880.2
E1 42221514 42221593
refGene_gn=LOC102901728;refGene_tx=XM_011289139.3
E1 42222002 42222223
refGene_gn=LOC102901728;refGene_tx=XM_011289139.3
E1 42222670 42222796
refGene_gn=LOC102901728;refGene_tx=XM_011289139.3
E1 42222885 42222948
refGene_gn=LOC102901728;refGene_tx=XM_011289139.3
E1 42222949 42223048
refGene_gn=LOC102901728;refGene_tx=XM_011289139.3
E1 42223433 42223590
refGene_gn=LOC102901728;refGene_tx=XM_011289139.3
E1 42224381 42224464
refGene_gn=LOC102901728;refGene_tx=XM_011289139.3
E1 42224656 42225148
refGene_gn=LOC102901728;refGene_tx=XM_011289139.3
E1 42235300 42235433
refGene_gn=LOC101087365;refGene_tx=XM_006940341.4
E1 42236001 42236222
refGene_gn=LOC101087365;refGene_tx=XM_006940341.4

E1 42236670 42236796
refGene_gn=LOC101087365;refGene_tx=XM_006940341.4
E1 42236885 42237047
refGene_gn=LOC101087365;refGene_tx=XM_006940341.4
E1 42237432 42237589
refGene_gn=LOC101087365;refGene_tx=XM_006940341.4
E1 42238373 42238456
refGene_gn=LOC101087365;refGene_tx=XM_006940341.4
E1 42238648 42239140
refGene_gn=LOC101087365;refGene_tx=XM_006940341.4
E1 42251507 42251637 refGene_gn=KRT32;refGene_tx=XM_003996856.5
E1 42252466 42252584 refGene_gn=KRT32;refGene_tx=XM_019817879.2
E1 42253843 42254064
refGene_gn=KRT32;refGene_tx=XM_003996856.5;refGene_gn=KRT32;refGene_tx
=XM_019817879.2
E1 42254776 42254902
refGene_gn=KRT32;refGene_tx=XM_003996856.5;refGene_gn=KRT32;refGene_tx
=XM_019817879.2
E1 42254983 42255145
refGene_gn=KRT32;refGene_tx=XM_003996856.5;refGene_gn=KRT32;refGene_tx
=XM_019817879.2
E1 42256431 42256588
refGene_gn=KRT32;refGene_tx=XM_003996856.5;refGene_gn=KRT32;refGene_tx
=XM_019817879.2
E1 42256765 42256848
refGene_gn=KRT32;refGene_tx=XM_003996856.5;refGene_gn=KRT32;refGene_tx
=XM_019817879.2
E1 42257525 42257993
refGene_gn=KRT32;refGene_tx=XM_003996856.5;refGene_gn=KRT32;refGene_tx
=XM_019817879.2
E1 42263819 42263967
refGene_gn=KRT35;refGene_tx=XM_003996857.5;refGene_gn=KRT35;refGene_tx
=XM_023243710.1
E1 42264256 42264477
refGene_gn=KRT35;refGene_tx=XM_003996857.5;refGene_gn=KRT35;refGene_tx
=XM_023243710.1
E1 42264904 42265030
refGene_gn=KRT35;refGene_tx=XM_003996857.5;refGene_gn=KRT35;refGene_tx
=XM_023243710.1
E1 42265342 42265504
refGene_gn=KRT35;refGene_tx=XM_003996857.5;refGene_gn=KRT35;refGene_tx
=XM_023243710.1
E1 42265638 42265795
refGene_gn=KRT35;refGene_tx=XM_003996857.5;refGene_gn=KRT35;refGene_tx
=XM_023243710.1
E1 42265990 42266073
refGene_gn=KRT35;refGene_tx=XM_003996857.5;refGene_gn=KRT35;refGene_tx
=XM_023243710.1
E1 42266919 42267384
refGene_gn=KRT35;refGene_tx=XM_003996857.5;refGene_gn=KRT35;refGene_tx

=XM_023243710.1
E1 42273719 42273915
refGene_gn=KRT36;refGene_tx=XM_003996935.3;refGene_gn=KRT36;refGene_tx=XM_019817872.2;refGene_gn=KRT36;refGene_tx=XM_019817874.2;refGene_gn=KRT36;refGene_tx=XM_019817875.2
E1 42274301 42274522
refGene_gn=KRT36;refGene_tx=XM_003996935.3;refGene_gn=KRT36;refGene_tx=XM_019817872.2;refGene_gn=KRT36;refGene_tx=XM_019817874.2;refGene_gn=KRT36;refGene_tx=XM_019817875.2
E1 42274699 42274825
refGene_gn=KRT36;refGene_tx=XM_003996935.3;refGene_gn=KRT36;refGene_tx=XM_019817872.2;refGene_gn=KRT36;refGene_tx=XM_019817874.2;refGene_gn=KRT36;refGene_tx=XM_019817875.2
E1 42274950 42275112
refGene_gn=KRT36;refGene_tx=XM_003996935.3;refGene_gn=KRT36;refGene_tx=XM_019817872.2;refGene_gn=KRT36;refGene_tx=XM_019817874.2;refGene_gn=KRT36;refGene_tx=XM_019817875.2
E1 42275645 42275802
refGene_gn=KRT36;refGene_tx=XM_003996935.3;refGene_gn=KRT36;refGene_tx=XM_019817872.2;refGene_gn=KRT36;refGene_tx=XM_019817874.2;refGene_gn=KRT36;refGene_tx=XM_019817875.2
E1 42276065 42276148
refGene_gn=KRT36;refGene_tx=XM_003996935.3;refGene_gn=KRT36;refGene_tx=XM_019817872.2;refGene_gn=KRT36;refGene_tx=XM_019817874.2;refGene_gn=KRT36;refGene_tx=XM_019817875.2
E1 42276872 42277331
refGene_gn=KRT36;refGene_tx=XM_003996935.3;refGene_gn=KRT36;refGene_tx=XM_019817872.2;refGene_gn=KRT36;refGene_tx=XM_019817874.2;refGene_gn=KRT36;refGene_tx=XM_019817875.2
E1 42287280 42287351
refGene_gn=LOC101087886;refGene_tx=XM_019817881.2
E1 42287700 42287726
refGene_gn=LOC101087886;refGene_tx=XM_019817881.2
E1 42288378 42288599
refGene_gn=LOC101087886;refGene_tx=XM_019817881.2
E1 42288694 42288820
refGene_gn=LOC101087886;refGene_tx=XM_019817881.2
E1 42288908 42289070
refGene_gn=LOC101087886;refGene_tx=XM_019817881.2
E1 42289246 42289403
refGene_gn=LOC101087886;refGene_tx=XM_019817881.2
E1 42289576 42289659
refGene_gn=LOC101087886;refGene_tx=XM_019817881.2
E1 42291115 42291598
refGene_gn=LOC101087886;refGene_tx=XM_019817881.2
E1 42300105 42300197
refGene_gn=LOC101090406;refGene_tx=XM_006940345.3
E1 42300732 42300758
refGene_gn=LOC101090406;refGene_tx=XM_006940345.3
E1 42301067 42301288

refGene_gn=LOC101090406;refGene_tx=XM_006940345.3
E1 42301481 42301607
refGene_gn=LOC101090406;refGene_tx=XM_006940345.3
E1 42301693 42301855
refGene_gn=LOC101090406;refGene_tx=XM_006940345.3
E1 42302338 42302495
refGene_gn=LOC101090406;refGene_tx=XM_006940345.3
E1 42302635 42302718
refGene_gn=LOC101090406;refGene_tx=XM_006940345.3
E1 42303599 42304045
refGene_gn=LOC101090406;refGene_tx=XM_006940345.3
E1 42304047 42304141
refGene_gn=LOC101090406;refGene_tx=XM_006940345.3
E1 42309364 42309619 refGene_gn=KRT19;refGene_tx=XM_003996859.4
E1 42309749 42309875 refGene_gn=KRT19;refGene_tx=XM_003996859.4
E1 42309989 42310151 refGene_gn=KRT19;refGene_tx=XM_003996859.4
E1 42310455 42310612 refGene_gn=KRT19;refGene_tx=XM_003996859.4
E1 42310828 42310911 refGene_gn=KRT19;refGene_tx=XM_003996859.4
E1 42312614 42313031 refGene_gn=KRT19;refGene_tx=XM_003996859.4
E1 42344565 42344585 refGene_gn=KRT9;refGene_tx=XM_003996937.5
E1 42345678 42346310 refGene_gn=KRT9;refGene_tx=XM_003996937.5
E1 42346688 42346912 refGene_gn=KRT9;refGene_tx=XM_003996937.5
E1 42347037 42347163 refGene_gn=KRT9;refGene_tx=XM_003996937.5
E1 42347873 42348035 refGene_gn=KRT9;refGene_tx=XM_003996937.5
E1 42348309 42348466 refGene_gn=KRT9;refGene_tx=XM_003996937.5
E1 42348584 42348667 refGene_gn=KRT9;refGene_tx=XM_003996937.5
E1 42349770 42350403 refGene_gn=KRT9;refGene_tx=XM_003996937.5
E1 42360535 42360633 refGene_gn=KRT14;refGene_tx=XM_003996860.5
E1 42361193 42361240 refGene_gn=KRT14;refGene_tx=XM_003996860.5
E1 42361333 42361554 refGene_gn=KRT14;refGene_tx=XM_003996860.5
E1 42361645 42361771 refGene_gn=KRT14;refGene_tx=XM_003996860.5
E1 42361873 42362035 refGene_gn=KRT14;refGene_tx=XM_003996860.5
E1 42362201 42362358 refGene_gn=KRT14;refGene_tx=XM_003996860.5
E1 42363152 42363235 refGene_gn=KRT14;refGene_tx=XM_003996860.5
E1 42364441 42364972 refGene_gn=KRT14;refGene_tx=XM_003996860.5
E1 42385063 42385164 refGene_gn=KRT16;refGene_tx=XM_003996861.5
E1 42385275 42385322 refGene_gn=KRT16;refGene_tx=XM_003996861.5
E1 42385418 42385639 refGene_gn=KRT16;refGene_tx=XM_003996861.5
E1 42385947 42386073 refGene_gn=KRT16;refGene_tx=XM_003996861.5
E1 42386165 42386327 refGene_gn=KRT16;refGene_tx=XM_003996861.5
E1 42386420 42386577 refGene_gn=KRT16;refGene_tx=XM_003996861.5
E1 42386713 42386796 refGene_gn=KRT16;refGene_tx=XM_003996861.5
E1 42387271 42387817 refGene_gn=KRT16;refGene_tx=XM_003996861.5
E1 42395306 42395401 refGene_gn=KRT17;refGene_tx=XM_003996862.4
E1 42396234 42396260 refGene_gn=KRT17;refGene_tx=XM_003996862.4
E1 42396383 42396604 refGene_gn=KRT17;refGene_tx=XM_003996862.4
E1 42396696 42396822 refGene_gn=KRT17;refGene_tx=XM_003996862.4
E1 42397241 42397403 refGene_gn=KRT17;refGene_tx=XM_003996862.4
E1 42397997 42398154 refGene_gn=KRT17;refGene_tx=XM_003996862.4
E1 42398534 42398617 refGene_gn=KRT17;refGene_tx=XM_003996862.4

E1 42399699 42400131 refGene_gn=KRT17;refGene_tx=XM_003996862.4
E1 42402219 42402317
refGene_gn=LOC101088377;refGene_tx=XM_003996938.3
E1 42402439 42402489
refGene_gn=LOC101088377;refGene_tx=XM_003996938.3
E1 42403532 42403753
refGene_gn=LOC101088377;refGene_tx=XM_003996938.3
E1 42404007 42404133
refGene_gn=LOC101088377;refGene_tx=XM_003996938.3
E1 42404720 42404882
refGene_gn=LOC101088377;refGene_tx=XM_003996938.3
E1 42406015 42406172
refGene_gn=LOC101088377;refGene_tx=XM_003996938.3
E1 42406299 42406382
refGene_gn=LOC101088377;refGene_tx=XM_003996938.3
E1 42408190 42408709
refGene_gn=LOC101088377;refGene_tx=XM_003996938.3
E1 42452270 42452301 refGene_gn=EIF1;refGene_tx=XM_003996863.5
E1 42453009 42453173 refGene_gn=EIF1;refGene_tx=XM_003996863.5
E1 42453541 42453643 refGene_gn=EIF1;refGene_tx=XM_003996863.5
E1 42454225 42454270 refGene_gn=EIF1;refGene_tx=XM_003996863.5
E1 42475742 42475953
refGene_gn=GAST;refGene_tx=XM_003996865.4;refGene_gn=GAST;refGene_tx=XM_023244654.1
E1 42476075 42476179
refGene_gn=GAST;refGene_tx=XM_003996865.4;refGene_gn=GAST;refGene_tx=XM_023244654.1
E1 42481937 42482449 refGene_gn=HAP1;refGene_tx=XM_023243467.1
E1 42482701 42482987 refGene_gn=HAP1;refGene_tx=XM_023243467.1
E1 42483166 42483296 refGene_gn=HAP1;refGene_tx=XM_023243467.1
E1 42483471 42483560 refGene_gn=HAP1;refGene_tx=XM_023243467.1
E1 42484140 42484215 refGene_gn=HAP1;refGene_tx=XM_023243467.1
E1 42484595 42484876 refGene_gn=HAP1;refGene_tx=XM_023243467.1
E1 42485093 42485215 refGene_gn=HAP1;refGene_tx=XM_023243467.1
E1 42485604 42485729 refGene_gn=HAP1;refGene_tx=XM_023243467.1
E1 42486126 42486202 refGene_gn=HAP1;refGene_tx=XM_023243467.1
E1 42486404 42486829 refGene_gn=HAP1;refGene_tx=XM_023243467.1
E1 42487193 42487288 refGene_gn=HAP1;refGene_tx=XM_023243467.1
E1 42488330 42488811 refGene_gn=HAP1;refGene_tx=XM_023243467.1
E1 42506086 42506238
refGene_gn=JUP;refGene_tx=XM_011289138.3;refGene_gn=JUP;refGene_tx=XM_023243552.1;refGene_gn=JUP;refGene_tx=XM_023243553.1
E1 42506486 42506526
refGene_gn=JUP;refGene_tx=XM_011289138.3;refGene_gn=JUP;refGene_tx=XM_023243552.1;refGene_gn=JUP;refGene_tx=XM_023243553.1
E1 42507363 42507485
refGene_gn=JUP;refGene_tx=XM_011289138.3;refGene_gn=JUP;refGene_tx=XM_023243552.1;refGene_gn=JUP;refGene_tx=XM_023243553.1
E1 42507596 42507747
refGene_gn=JUP;refGene_tx=XM_011289138.3;refGene_gn=JUP;refGene_tx=XM_

023243552.1;refGene_gn=JUP;refGene_tx=XM_023243553.1
E1 42510343 42510463
refGene_gn=JUP;refGene_tx=XM_011289138.3;refGene_gn=JUP;refGene_tx=XM_
023243552.1;refGene_gn=JUP;refGene_tx=XM_023243553.1
E1 42510649 42510805
refGene_gn=JUP;refGene_tx=XM_011289138.3;refGene_gn=JUP;refGene_tx=XM_
023243552.1;refGene_gn=JUP;refGene_tx=XM_023243553.1
E1 42511989 42512328
refGene_gn=JUP;refGene_tx=XM_011289138.3;refGene_gn=JUP;refGene_tx=XM_
023243552.1;refGene_gn=JUP;refGene_tx=XM_023243553.1
E1 42513753 42513857
refGene_gn=JUP;refGene_tx=XM_011289138.3;refGene_gn=JUP;refGene_tx=XM_
023243552.1;refGene_gn=JUP;refGene_tx=XM_023243553.1
E1 42513967 42514112
refGene_gn=JUP;refGene_tx=XM_011289138.3;refGene_gn=JUP;refGene_tx=XM_
023243552.1;refGene_gn=JUP;refGene_tx=XM_023243553.1
E1 42514916 42515118
refGene_gn=JUP;refGene_tx=XM_011289138.3;refGene_gn=JUP;refGene_tx=XM_
023243552.1;refGene_gn=JUP;refGene_tx=XM_023243553.1
E1 42515872 42516111
refGene_gn=JUP;refGene_tx=XM_011289138.3;refGene_gn=JUP;refGene_tx=XM_
023243552.1;refGene_gn=JUP;refGene_tx=XM_023243553.1
E1 42516328 42516588
refGene_gn=JUP;refGene_tx=XM_011289138.3;refGene_gn=JUP;refGene_tx=XM_
023243552.1;refGene_gn=JUP;refGene_tx=XM_023243553.1
E1 42518249 42518454
refGene_gn=JUP;refGene_tx=XM_011289138.3;refGene_gn=JUP;refGene_tx=XM_
023243552.1;refGene_gn=JUP;refGene_tx=XM_023243553.1
E1 42540743 42540766 refGene_gn=P3H4;refGene_tx=XM_023243541.1
E1 42541080 42541225 refGene_gn=P3H4;refGene_tx=XM_023243541.1
E1 42542696 42542780 refGene_gn=P3H4;refGene_tx=XM_023243541.1
E1 42543423 42543569 refGene_gn=P3H4;refGene_tx=XM_023243541.1
E1 42544495 42544624 refGene_gn=P3H4;refGene_tx=XM_023243541.1
E1 42545297 42545469 refGene_gn=P3H4;refGene_tx=XM_023243541.1
E1 42545594 42545747 refGene_gn=P3H4;refGene_tx=XM_023243541.1
E1 42546101 42546581 refGene_gn=P3H4;refGene_tx=XM_023243541.1
E1 42547746 42547991 refGene_gn=FKBP10;refGene_tx=XM_011289137.3
E1 42551831 42551977 refGene_gn=FKBP10;refGene_tx=XM_011289137.3
E1 42552416 42552606 refGene_gn=FKBP10;refGene_tx=XM_011289137.3
E1 42552725 42552871 refGene_gn=FKBP10;refGene_tx=XM_011289137.3
E1 42553275 42553465 refGene_gn=FKBP10;refGene_tx=XM_011289137.3
E1 42553621 42553767 refGene_gn=FKBP10;refGene_tx=XM_011289137.3
E1 42554497 42554690 refGene_gn=FKBP10;refGene_tx=XM_011289137.3
E1 42555107 42555250 refGene_gn=FKBP10;refGene_tx=XM_011289137.3
E1 42555450 42555614 refGene_gn=FKBP10;refGene_tx=XM_011289137.3
E1 42555938 42556124 refGene_gn=FKBP10;refGene_tx=XM_011289137.3
E1 42559131 42559266 refGene_gn=NT5C3B;refGene_tx=XM_023244064.1
E1 42560371 42560572 refGene_gn=NT5C3B;refGene_tx=XM_023244064.1
E1 42561670 42561833 refGene_gn=NT5C3B;refGene_tx=XM_023244064.1
E1 42562766 42562856 refGene_gn=NT5C3B;refGene_tx=XM_023244064.1

E1 42563345 42563431 refGene_gn=NT5C3B;refGene_tx=XM_023244064.1
E1 42565489 42565536 refGene_gn=NT5C3B;refGene_tx=XM_023244064.1
E1 42565622 42565692 refGene_gn=NT5C3B;refGene_tx=XM_023244064.1
E1 42566266 42566365 refGene_gn=NT5C3B;refGene_tx=XM_023244064.1
E1 42566613 42566625 refGene_gn=NT5C3B;refGene_tx=XM_023244064.1
E1 42567535 42567729 refGene_gn=KLHL10;refGene_tx=XM_003996867.5
E1 42569874 42570364 refGene_gn=KLHL10;refGene_tx=XM_003996867.5
E1 42571077 42571695 refGene_gn=KLHL10;refGene_tx=XM_003996867.5
E1 42572212 42572362 refGene_gn=KLHL10;refGene_tx=XM_003996867.5
E1 42572600 42572975 refGene_gn=KLHL10;refGene_tx=XM_003996867.5
E1 42577085 42578667 refGene_gn=KLHL11;refGene_tx=XM_023244063.1
E1 42583959 42584504 refGene_gn=KLHL11;refGene_tx=XM_023244063.1
E1 42586539 42586634
refGene_gn=ACLY;refGene_tx=XM_003996869.5;refGene_gn=ACLY;refGene_tx=XM_019817885.2;refGene_gn=ACLY;refGene_tx=XM_019817886.2
E1 42587287 42587364
refGene_gn=ACLY;refGene_tx=XM_003996869.5;refGene_gn=ACLY;refGene_tx=XM_019817885.2;refGene_gn=ACLY;refGene_tx=XM_019817886.2
E1 42587940 42588023
refGene_gn=ACLY;refGene_tx=XM_003996869.5;refGene_gn=ACLY;refGene_tx=XM_019817885.2;refGene_gn=ACLY;refGene_tx=XM_019817886.2
E1 42588381 42588495
refGene_gn=ACLY;refGene_tx=XM_003996869.5;refGene_gn=ACLY;refGene_tx=XM_019817885.2;refGene_gn=ACLY;refGene_tx=XM_019817886.2
E1 42589624 42589768
refGene_gn=ACLY;refGene_tx=XM_003996869.5;refGene_gn=ACLY;refGene_tx=XM_019817885.2;refGene_gn=ACLY;refGene_tx=XM_019817886.2
E1 42589967 42590118
refGene_gn=ACLY;refGene_tx=XM_003996869.5;refGene_gn=ACLY;refGene_tx=XM_019817885.2;refGene_gn=ACLY;refGene_tx=XM_019817886.2
E1 42591392 42591547
refGene_gn=ACLY;refGene_tx=XM_003996869.5;refGene_gn=ACLY;refGene_tx=XM_019817885.2;refGene_gn=ACLY;refGene_tx=XM_019817886.2
E1 42593765 42593859
refGene_gn=ACLY;refGene_tx=XM_003996869.5;refGene_gn=ACLY;refGene_tx=XM_019817885.2;refGene_gn=ACLY;refGene_tx=XM_019817886.2
E1 42594513 42594641
refGene_gn=ACLY;refGene_tx=XM_003996869.5;refGene_gn=ACLY;refGene_tx=XM_019817885.2;refGene_gn=ACLY;refGene_tx=XM_019817886.2
E1 42599194 42599305
refGene_gn=ACLY;refGene_tx=XM_003996869.5;refGene_gn=ACLY;refGene_tx=XM_019817885.2;refGene_gn=ACLY;refGene_tx=XM_019817886.2
E1 42600698 42600780
refGene_gn=ACLY;refGene_tx=XM_003996869.5;refGene_gn=ACLY;refGene_tx=XM_019817885.2;refGene_gn=ACLY;refGene_tx=XM_019817886.2
E1 42604922 42605119
refGene_gn=ACLY;refGene_tx=XM_003996869.5;refGene_gn=ACLY;refGene_tx=XM_019817885.2;refGene_gn=ACLY;refGene_tx=XM_019817886.2
E1 42606170 42606275
refGene_gn=ACLY;refGene_tx=XM_003996869.5;refGene_gn=ACLY;refGene_tx=X

M_019817885.2;refGene_gn=ACLY;refGene_tx=XM_019817886.2
E1 42608380 42608549
refGene_gn=ACLY;refGene_tx=XM_003996869.5;refGene_gn=ACLY;refGene_tx=XM_019817885.2;refGene_gn=ACLY;refGene_tx=XM_019817886.2
E1 42608846 42608988
refGene_gn=ACLY;refGene_tx=XM_003996869.5;refGene_gn=ACLY;refGene_tx=XM_019817885.2;refGene_gn=ACLY;refGene_tx=XM_019817886.2
E1 42611570 42611600
refGene_gn=ACLY;refGene_tx=XM_003996869.5;refGene_gn=ACLY;refGene_tx=XM_019817885.2
E1 42612937 42613028
refGene_gn=ACLY;refGene_tx=XM_003996869.5;refGene_gn=ACLY;refGene_tx=XM_019817885.2;refGene_gn=ACLY;refGene_tx=XM_019817886.2
E1 42613637 42613792
refGene_gn=ACLY;refGene_tx=XM_003996869.5;refGene_gn=ACLY;refGene_tx=XM_019817885.2;refGene_gn=ACLY;refGene_tx=XM_019817886.2
E1 42615310 42615428
refGene_gn=ACLY;refGene_tx=XM_003996869.5;refGene_gn=ACLY;refGene_tx=XM_019817885.2;refGene_gn=ACLY;refGene_tx=XM_019817886.2
E1 42616606 42616668
refGene_gn=ACLY;refGene_tx=XM_003996869.5;refGene_gn=ACLY;refGene_tx=XM_019817885.2;refGene_gn=ACLY;refGene_tx=XM_019817886.2
E1 42617366 42617503
refGene_gn=ACLY;refGene_tx=XM_003996869.5;refGene_gn=ACLY;refGene_tx=XM_019817885.2;refGene_gn=ACLY;refGene_tx=XM_019817886.2
E1 42618791 42618910
refGene_gn=ACLY;refGene_tx=XM_003996869.5;refGene_gn=ACLY;refGene_tx=XM_019817885.2;refGene_gn=ACLY;refGene_tx=XM_019817886.2
E1 42619685 42619816
refGene_gn=ACLY;refGene_tx=XM_003996869.5;refGene_gn=ACLY;refGene_tx=XM_019817885.2;refGene_gn=ACLY;refGene_tx=XM_019817886.2
E1 42620956 42621036
refGene_gn=ACLY;refGene_tx=XM_003996869.5;refGene_gn=ACLY;refGene_tx=XM_019817885.2;refGene_gn=ACLY;refGene_tx=XM_019817886.2
E1 42621515 42621706
refGene_gn=ACLY;refGene_tx=XM_003996869.5;refGene_gn=ACLY;refGene_tx=XM_019817885.2;refGene_gn=ACLY;refGene_tx=XM_019817886.2
E1 42622299 42622362
refGene_gn=ACLY;refGene_tx=XM_003996869.5;refGene_gn=ACLY;refGene_tx=XM_019817885.2;refGene_gn=ACLY;refGene_tx=XM_019817886.2
E1 42624311 42624434
refGene_gn=ACLY;refGene_tx=XM_003996869.5;refGene_gn=ACLY;refGene_tx=XM_019817885.2;refGene_gn=ACLY;refGene_tx=XM_019817886.2
E1 42625553 42625735
refGene_gn=ACLY;refGene_tx=XM_003996869.5;refGene_gn=ACLY;refGene_tx=XM_019817885.2;refGene_gn=ACLY;refGene_tx=XM_019817886.2
E1 42634414 42634433
refGene_gn=ACLY;refGene_tx=XM_019817885.2;refGene_gn=ACLY;refGene_tx=XM_019817886.2
E1 42640519 42640633 refGene_gn=TTC25;refGene_tx=XM_006940348.4

E1 42642244 42642376 refGene_gn=TTC25;refGene_tx=XM_006940348.4
E1 42642578 42642729 refGene_gn=TTC25;refGene_tx=XM_006940348.4
E1 42643421 42643483 refGene_gn=TTC25;refGene_tx=XM_006940348.4
E1 42643781 42643947 refGene_gn=TTC25;refGene_tx=XM_006940348.4
E1 42645268 42645493 refGene_gn=TTC25;refGene_tx=XM_006940348.4
E1 42645665 42645873 refGene_gn=TTC25;refGene_tx=XM_006940348.4
E1 42652513 42652600 refGene_gn=TTC25;refGene_tx=XM_006940348.4
E1 42654431 42654628 refGene_gn=TTC25;refGene_tx=XM_006940348.4
E1 42656916 42657017 refGene_gn=TTC25;refGene_tx=XM_006940348.4
E1 42660655 42660740 refGene_gn=TTC25;refGene_tx=XM_006940348.4
E1 42663139 42663675 refGene_gn=TTC25;refGene_tx=XM_006940348.4
E1 42665292 42665295 refGene_gn=CNP;refGene_tx=XM_003996870.5
E1 42666479 42667149
refGene_gn=CNP;refGene_tx=XM_003996870.5;refGene_gn=CNP;refGene_tx=XM_006940350.4;refGene_gn=CNP;refGene_tx=XM_019817887.2
E1 42668855 42668995
refGene_gn=CNP;refGene_tx=XM_003996870.5;refGene_gn=CNP;refGene_tx=XM_006940350.4;refGene_gn=CNP;refGene_tx=XM_019817887.2
E1 42670367 42670817
refGene_gn=CNP;refGene_tx=XM_003996870.5;refGene_gn=CNP;refGene_tx=XM_006940350.4;refGene_gn=CNP;refGene_tx=XM_019817887.2
E1 42673441 42673479
refGene_gn=DNAJC7;refGene_tx=XM_003996871.5;refGene_gn=DNAJC7;refGene_tx=XM_006940351.4;refGene_gn=DNAJC7;refGene_tx=XM_006940352.4
E1 42673919 42673982
refGene_gn=DNAJC7;refGene_tx=XM_003996871.5;refGene_gn=DNAJC7;refGene_tx=XM_006940351.4;refGene_gn=DNAJC7;refGene_tx=XM_006940352.4
E1 42675495 42675648
refGene_gn=DNAJC7;refGene_tx=XM_003996871.5;refGene_gn=DNAJC7;refGene_tx=XM_006940351.4;refGene_gn=DNAJC7;refGene_tx=XM_006940352.4
E1 42676124 42676271
refGene_gn=DNAJC7;refGene_tx=XM_003996871.5;refGene_gn=DNAJC7;refGene_tx=XM_006940351.4;refGene_gn=DNAJC7;refGene_tx=XM_006940352.4
E1 42676664 42676738
refGene_gn=DNAJC7;refGene_tx=XM_003996871.5;refGene_gn=DNAJC7;refGene_tx=XM_006940351.4;refGene_gn=DNAJC7;refGene_tx=XM_006940352.4
E1 42679808 42679900
refGene_gn=DNAJC7;refGene_tx=XM_003996871.5;refGene_gn=DNAJC7;refGene_tx=XM_006940351.4;refGene_gn=DNAJC7;refGene_tx=XM_006940352.4
E1 42680720 42680885
refGene_gn=DNAJC7;refGene_tx=XM_003996871.5;refGene_gn=DNAJC7;refGene_tx=XM_006940351.4;refGene_gn=DNAJC7;refGene_tx=XM_006940352.4
E1 42681421 42681575
refGene_gn=DNAJC7;refGene_tx=XM_003996871.5;refGene_gn=DNAJC7;refGene_tx=XM_006940351.4;refGene_gn=DNAJC7;refGene_tx=XM_006940352.4
E1 42682074 42682193
refGene_gn=DNAJC7;refGene_tx=XM_003996871.5;refGene_gn=DNAJC7;refGene_tx=XM_006940351.4;refGene_gn=DNAJC7;refGene_tx=XM_006940352.4
E1 42686433 42686508
refGene_gn=DNAJC7;refGene_tx=XM_003996871.5;refGene_gn=DNAJC7;refGene_

tx=XM_006940351.4;refGene_gn=DNAJC7;refGene_tx=XM_006940352.4
E1 42687156 42687270
refGene_gn=DNAJC7;refGene_tx=XM_003996871.5;refGene_gn=DNAJC7;refGene_tx=XM_006940351.4;refGene_gn=DNAJC7;refGene_tx=XM_006940352.4
E1 42687758 42687883
refGene_gn=DNAJC7;refGene_tx=XM_003996871.5;refGene_gn=DNAJC7;refGene_tx=XM_006940351.4;refGene_gn=DNAJC7;refGene_tx=XM_006940352.4
E1 42688633 42688722
refGene_gn=DNAJC7;refGene_tx=XM_003996871.5;refGene_gn=DNAJC7;refGene_tx=XM_006940351.4
E1 42695701 42695730 refGene_gn=DNAJC7;refGene_tx=XM_006940351.4
E1 42699748 42699825 refGene_gn=DNAJC7;refGene_tx=XM_003996871.5
E1 42703420 42703514
refGene_gn=NKIRAS2;refGene_tx=XM_003996873.5;refGene_gn=NKIRAS2;refGene_tx=XM_011289135.3
E1 42703914 42704156
refGene_gn=NKIRAS2;refGene_tx=XM_003996873.5;refGene_gn=NKIRAS2;refGene_tx=XM_011289135.3
E1 42704828 42705068
refGene_gn=NKIRAS2;refGene_tx=XM_003996873.5;refGene_gn=NKIRAS2;refGene_tx=XM_011289135.3
E1 42707664 42707901
refGene_gn=ZNF385C;refGene_tx=XM_019817890.2;refGene_gn=ZNF385C;refGene_tx=XM_019817891.2;refGene_gn=ZNF385C;refGene_tx=XM_019817892.2;refGene_gn=ZNF385C;refGene_tx=XM_019817897.2;refGene_gn=ZNF385C;refGene_tx=XM_023243487.1;refGene_gn=ZNF385C;refGene_tx=XM_023243488.1;refGene_gn=ZNF385C;refGene_tx=XM_023243489.1;refGene_gn=ZNF385C;refGene_tx=XM_023243490.1;refGene_gn=ZNF385C;refGene_tx=XM_023243492.1
E1 42708377 42708488
refGene_gn=ZNF385C;refGene_tx=XM_019817890.2;refGene_gn=ZNF385C;refGene_tx=XM_019817891.2;refGene_gn=ZNF385C;refGene_tx=XM_019817892.2;refGene_gn=ZNF385C;refGene_tx=XM_019817897.2;refGene_gn=ZNF385C;refGene_tx=XM_023243487.1;refGene_gn=ZNF385C;refGene_tx=XM_023243488.1;refGene_gn=ZNF385C;refGene_tx=XM_023243489.1;refGene_gn=ZNF385C;refGene_tx=XM_023243490.1;refGene_gn=ZNF385C;refGene_tx=XM_023243492.1
E1 42708756 42708950
refGene_gn=ZNF385C;refGene_tx=XM_019817890.2;refGene_gn=ZNF385C;refGene_tx=XM_019817891.2;refGene_gn=ZNF385C;refGene_tx=XM_019817892.2;refGene_gn=ZNF385C;refGene_tx=XM_019817897.2;refGene_gn=ZNF385C;refGene_tx=XM_023243487.1;refGene_gn=ZNF385C;refGene_tx=XM_023243488.1;refGene_gn=ZNF385C;refGene_tx=XM_023243489.1;refGene_gn=ZNF385C;refGene_tx=XM_023243490.1;refGene_gn=ZNF385C;refGene_tx=XM_023243492.1
E1 42709502 42709796
refGene_gn=ZNF385C;refGene_tx=XM_019817890.2;refGene_gn=ZNF385C;refGene_tx=XM_019817891.2;refGene_gn=ZNF385C;refGene_tx=XM_019817892.2;refGene_gn=ZNF385C;refGene_tx=XM_019817897.2;refGene_gn=ZNF385C;refGene_tx=XM_023243487.1;refGene_gn=ZNF385C;refGene_tx=XM_023243488.1;refGene_gn=ZNF385C;refGene_tx=XM_023243489.1;refGene_gn=ZNF385C;refGene_tx=XM_023243490.1;refGene_gn=ZNF385C;refGene_tx=XM_023243492.1
E1 42711889 42712055

refGene_gn=ZNF385C;refGene_tx=XM_019817890.2;refGene_gn=ZNF385C;refGene_tx=XM_019817891.2;refGene_gn=ZNF385C;refGene_tx=XM_019817892.2;refGene_gn=ZNF385C;refGene_tx=XM_019817897.2;refGene_gn=ZNF385C;refGene_tx=XM_023243487.1;refGene_gn=ZNF385C;refGene_tx=XM_023243488.1;refGene_gn=ZNF385C;refGene_tx=XM_023243489.1;refGene_gn=ZNF385C;refGene_tx=XM_023243490.1;refGene_gn=ZNF385C;refGene_tx=XM_023243492.1
E1 42713468 42713579
refGene_gn=ZNF385C;refGene_tx=XM_019817890.2;refGene_gn=ZNF385C;refGene_tx=XM_019817891.2;refGene_gn=ZNF385C;refGene_tx=XM_019817892.2;refGene_gn=ZNF385C;refGene_tx=XM_019817897.2;refGene_gn=ZNF385C;refGene_tx=XM_023243487.1;refGene_gn=ZNF385C;refGene_tx=XM_023243488.1;refGene_gn=ZNF385C;refGene_tx=XM_023243489.1;refGene_gn=ZNF385C;refGene_tx=XM_023243490.1;refGene_gn=ZNF385C;refGene_tx=XM_023243492.1
E1 42713824 42713898
refGene_gn=ZNF385C;refGene_tx=XM_023243487.1;refGene_gn=ZNF385C;refGene_tx=XM_023243488.1
E1 42715451 42715600
refGene_gn=ZNF385C;refGene_tx=XM_019817890.2;refGene_gn=ZNF385C;refGene_tx=XM_019817891.2;refGene_gn=ZNF385C;refGene_tx=XM_019817892.2;refGene_gn=ZNF385C;refGene_tx=XM_019817897.2;refGene_gn=ZNF385C;refGene_tx=XM_023243487.1;refGene_gn=ZNF385C;refGene_tx=XM_023243488.1;refGene_gn=ZNF385C;refGene_tx=XM_023243489.1;refGene_gn=ZNF385C;refGene_tx=XM_023243490.1
E1 42716994 42718479
refGene_gn=ZNF385C;refGene_tx=XM_019817888.2;refGene_gn=ZNF385C;refGene_tx=XM_023243485.1;refGene_gn=ZNF385C;refGene_tx=XM_023243486.1
E1 42719770 42720313
refGene_gn=ZNF385C;refGene_tx=XM_019817888.2;refGene_gn=ZNF385C;refGene_tx=XM_023243485.1;refGene_gn=ZNF385C;refGene_tx=XM_023243486.1
E1 42724913 42725194
refGene_gn=ZNF385C;refGene_tx=XM_023243487.1;refGene_gn=ZNF385C;refGene_tx=XM_023243488.1
E1 42728258 42728302
refGene_gn=ZNF385C;refGene_tx=XM_019817897.2;refGene_gn=ZNF385C;refGene_tx=XM_023243492.1
E1 42734896 42735148
refGene_gn=ZNF385C;refGene_tx=XM_019817890.2;refGene_gn=ZNF385C;refGene_tx=XM_019817891.2;refGene_gn=ZNF385C;refGene_tx=XM_019817892.2;refGene_gn=ZNF385C;refGene_tx=XM_023243489.1;refGene_gn=ZNF385C;refGene_tx=XM_023243490.1;refGene_gn=ZNF385C;refGene_tx=XM_023243492.1
E1 42741317 42741351
refGene_gn=ZNF385C;refGene_tx=XM_023243489.1;refGene_gn=ZNF385C;refGene_tx=XM_023243490.1;refGene_gn=ZNF385C;refGene_tx=XM_023243492.1
E1 42759615 42759801 refGene_gn=DHX58;refGene_tx=XM_006940356.4
E1 42760096 42760193 refGene_gn=DHX58;refGene_tx=XM_006940356.4
E1 42761255 42761446 refGene_gn=DHX58;refGene_tx=XM_006940356.4
E1 42762294 42762456 refGene_gn=DHX58;refGene_tx=XM_006940356.4
E1 42762543 42762693 refGene_gn=DHX58;refGene_tx=XM_006940356.4
E1 42763210 42763464 refGene_gn=DHX58;refGene_tx=XM_006940356.4
E1 42764142 42764334 refGene_gn=DHX58;refGene_tx=XM_006940356.4

E1 42764490 42764617 refGene_gn=DHX58;refGene_tx=XM_006940356.4
E1 42765230 42765347 refGene_gn=DHX58;refGene_tx=XM_006940356.4
E1 42766093 42766284 refGene_gn=DHX58;refGene_tx=XM_006940356.4
E1 42766669 42766871 refGene_gn=DHX58;refGene_tx=XM_006940356.4
E1 42767076 42767245 refGene_gn=DHX58;refGene_tx=XM_006940356.4
E1 42767376 42767396 refGene_gn=DHX58;refGene_tx=XM_006940356.4
E1 42768518 42768712 refGene_gn=KAT2A;refGene_tx=XM_023243429.1
E1 42768846 42768931 refGene_gn=KAT2A;refGene_tx=XM_023243429.1
E1 42769067 42769131 refGene_gn=KAT2A;refGene_tx=XM_023243429.1
E1 42769210 42769247 refGene_gn=KAT2A;refGene_tx=XM_023243429.1
E1 42769342 42769457 refGene_gn=KAT2A;refGene_tx=XM_023243429.1
E1 42769720 42769864 refGene_gn=KAT2A;refGene_tx=XM_023243429.1
E1 42770331 42770442 refGene_gn=KAT2A;refGene_tx=XM_023243429.1
E1 42771302 42771438 refGene_gn=KAT2A;refGene_tx=XM_023243429.1
E1 42771649 42771858 refGene_gn=KAT2A;refGene_tx=XM_023243429.1
E1 42771944 42772081 refGene_gn=KAT2A;refGene_tx=XM_023243429.1
E1 42772164 42772275 refGene_gn=KAT2A;refGene_tx=XM_023243429.1
E1 42772479 42772586 refGene_gn=KAT2A;refGene_tx=XM_023243429.1
E1 42773478 42773670 refGene_gn=KAT2A;refGene_tx=XM_023243429.1
E1 42773765 42773947 refGene_gn=KAT2A;refGene_tx=XM_023243429.1
E1 42774237 42774327 refGene_gn=KAT2A;refGene_tx=XM_023243429.1
E1 42774433 42774579 refGene_gn=KAT2A;refGene_tx=XM_023243429.1
E1 42774913 42775037 refGene_gn=KAT2A;refGene_tx=XM_023243429.1
E1 42775176 42775515 refGene_gn=KAT2A;refGene_tx=XM_023243429.1
E1 42776835 42777324 refGene_gn=HSPB9;refGene_tx=XM_006940357.3
E1 42778868 42778984 refGene_gn=RAB5C;refGene_tx=XM_003996878.5
E1 42779716 42779810 refGene_gn=RAB5C;refGene_tx=XM_003996878.5
E1 42780724 42780847 refGene_gn=RAB5C;refGene_tx=XM_003996878.5
E1 42781053 42781205 refGene_gn=RAB5C;refGene_tx=XM_003996878.5
E1 42782720 42782886 refGene_gn=RAB5C;refGene_tx=XM_003996878.5
E1 42804540 42804936 refGene_gn=KCNH4;refGene_tx=XM_019817899.2
E1 42806159 42806233 refGene_gn=KCNH4;refGene_tx=XM_019817899.2
E1 42806843 42806950 refGene_gn=KCNH4;refGene_tx=XM_019817899.2
E1 42807224 42807577 refGene_gn=KCNH4;refGene_tx=XM_019817899.2
E1 42807780 42807819 refGene_gn=KCNH4;refGene_tx=XM_019817899.2
E1 42807937 42807988 refGene_gn=KCNH4;refGene_tx=XM_019817900.2
E1 42808468 42808713
refGene_gn=KCNH4;refGene_tx=XM_019817899.2;refGene_gn=KCNH4;refGene_tx
=XM_019817900.2
E1 42809288 42809538
refGene_gn=KCNH4;refGene_tx=XM_019817899.2;refGene_gn=KCNH4;refGene_tx
=XM_019817900.2
E1 42811177 42811377
refGene_gn=KCNH4;refGene_tx=XM_019817899.2;refGene_gn=KCNH4;refGene_tx
=XM_019817900.2
E1 42811769 42811964
refGene_gn=KCNH4;refGene_tx=XM_019817899.2;refGene_gn=KCNH4;refGene_tx
=XM_019817900.2
E1 42812956 42813164
refGene_gn=KCNH4;refGene_tx=XM_019817899.2;refGene_gn=KCNH4;refGene_tx

=XM_019817900.2
E1 42815901 42816059
refGene_gn=KCNH4;refGene_tx=XM_019817899.2;refGene_gn=KCNH4;refGene_tx
=XM_019817900.2
E1 42816404 42816648
refGene_gn=KCNH4;refGene_tx=XM_019817899.2;refGene_gn=KCNH4;refGene_tx
=XM_019817900.2
E1 42817479 42817613
refGene_gn=KCNH4;refGene_tx=XM_019817899.2;refGene_gn=KCNH4;refGene_tx
=XM_019817900.2
E1 42817738 42817885
refGene_gn=KCNH4;refGene_tx=XM_019817899.2;refGene_gn=KCNH4;refGene_tx
=XM_019817900.2
E1 42818219 42818453
refGene_gn=KCNH4;refGene_tx=XM_019817899.2;refGene_gn=KCNH4;refGene_tx
=XM_019817900.2
E1 42820344 42820420
refGene_gn=KCNH4;refGene_tx=XM_019817899.2;refGene_gn=KCNH4;refGene_tx
=XM_019817900.2
E1 42823112 42823484 refGene_gn=HCRT;refGene_tx=XM_023243939.1
E1 42824326 42824347 refGene_gn=HCRT;refGene_tx=XM_023243939.1
E1 42824457 42824505 refGene_gn=GHDC;refGene_tx=XM_019817901.2
E1 42826723 42826942
refGene_gn=GHDC;refGene_tx=XM_006940358.4;refGene_gn=GHDC;refGene_tx=X
M_011289134.3;refGene_gn=GHDC;refGene_tx=XM_019817902.2
E1 42827209 42827295
refGene_gn=GHDC;refGene_tx=XM_006940358.4;refGene_gn=GHDC;refGene_tx=X
M_011289134.3;refGene_gn=GHDC;refGene_tx=XM_019817901.2;refGene_gn=GHD
C;refGene_tx=XM_019817902.2
E1 42827390 42827410 refGene_gn=GHDC;refGene_tx=XM_019817903.2
E1 42827664 42827798
refGene_gn=GHDC;refGene_tx=XM_006940358.4;refGene_gn=GHDC;refGene_tx=X
M_011289134.3;refGene_gn=GHDC;refGene_tx=XM_019817901.2;refGene_gn=GHD
C;refGene_tx=XM_019817902.2;refGene_gn=GHDC;refGene_tx=XM_019817903.2
E1 42827871 42827943
refGene_gn=GHDC;refGene_tx=XM_006940358.4;refGene_gn=GHDC;refGene_tx=X
M_011289134.3;refGene_gn=GHDC;refGene_tx=XM_019817901.2;refGene_gn=GHD
C;refGene_tx=XM_019817902.2;refGene_gn=GHDC;refGene_tx=XM_019817903.2
E1 42828043 42828236
refGene_gn=GHDC;refGene_tx=XM_006940358.4;refGene_gn=GHDC;refGene_tx=X
M_011289134.3;refGene_gn=GHDC;refGene_tx=XM_019817901.2;refGene_gn=GHD
C;refGene_tx=XM_019817902.2;refGene_gn=GHDC;refGene_tx=XM_019817903.2
E1 42828633 42829135
refGene_gn=GHDC;refGene_tx=XM_006940358.4;refGene_gn=GHDC;refGene_tx=X
M_011289134.3;refGene_gn=GHDC;refGene_tx=XM_019817901.2;refGene_gn=GHD
C;refGene_tx=XM_019817902.2;refGene_gn=GHDC;refGene_tx=XM_019817903.2
E1 42829282 42829404
refGene_gn=GHDC;refGene_tx=XM_006940358.4;refGene_gn=GHDC;refGene_tx=X
M_011289134.3;refGene_gn=GHDC;refGene_tx=XM_019817901.2;refGene_gn=GHD
C;refGene_tx=XM_019817902.2;refGene_gn=GHDC;refGene_tx=XM_019817903.2

E1 42829727 42830042
refGene_gn=GHDC;refGene_tx=XM_006940358.4;refGene_gn=GHDC;refGene_tx=XM_011289134.3;refGene_gn=GHDC;refGene_tx=XM_019817901.2;refGene_gn=GHDC;refGene_tx=XM_019817902.2;refGene_gn=GHDC;refGene_tx=XM_019817903.2
E1 42830193 42830365
refGene_gn=GHDC;refGene_tx=XM_011289134.3;refGene_gn=GHDC;refGene_tx=XM_019817901.2;refGene_gn=GHDC;refGene_tx=XM_019817903.2
E1 42838901 42839028 refGene_gn=STAT5B;refGene_tx=XM_023244609.1
E1 42839560 42839668 refGene_gn=STAT5B;refGene_tx=XM_023244609.1
E1 42840000 42840052 refGene_gn=STAT5B;refGene_tx=XM_023244609.1
E1 42844308 42844479 refGene_gn=STAT5B;refGene_tx=XM_023244609.1
E1 42845333 42845464 refGene_gn=STAT5B;refGene_tx=XM_023244609.1
E1 42845564 42845659 refGene_gn=STAT5B;refGene_tx=XM_023244609.1
E1 42846683 42846890 refGene_gn=STAT5B;refGene_tx=XM_023244609.1
E1 42849193 42849286 refGene_gn=STAT5B;refGene_tx=XM_023244609.1
E1 42850779 42850902 refGene_gn=STAT5B;refGene_tx=XM_023244609.1
E1 42850980 42851068 refGene_gn=STAT5B;refGene_tx=XM_023244609.1
E1 42851451 42851631 refGene_gn=STAT5B;refGene_tx=XM_023244609.1
E1 42852058 42852214 refGene_gn=STAT5B;refGene_tx=XM_023244609.1
E1 42852711 42852730 refGene_gn=STAT5B;refGene_tx=XM_023244609.1
E1 42852732 42852862 refGene_gn=STAT5B;refGene_tx=XM_023244609.1
E1 42853120 42853251 refGene_gn=STAT5B;refGene_tx=XM_023244609.1
E1 42858034 42858209 refGene_gn=STAT5B;refGene_tx=XM_023244609.1
E1 42859398 42859488 refGene_gn=STAT5B;refGene_tx=XM_023244609.1
E1 42860616 42860773 refGene_gn=STAT5B;refGene_tx=XM_023244609.1
E1 42864245 42864373 refGene_gn=STAT5B;refGene_tx=XM_023244609.1
E1 42914228 42914356
refGene_gn=LOC101090913;refGene_tx=XM_023244066.1;refGene_gn=LOC101090913;refGene_tx=XM_023244068.1;refGene_gn=LOC101090913;refGene_tx=XM_023244069.1;refGene_gn=LOC101090913;refGene_tx=XM_023244070.1
E1 42914699 42914856
refGene_gn=LOC101090913;refGene_tx=XM_023244066.1;refGene_gn=LOC101090913;refGene_tx=XM_023244068.1;refGene_gn=LOC101090913;refGene_tx=XM_023244069.1;refGene_gn=LOC101090913;refGene_tx=XM_023244070.1
E1 42916148 42916238
refGene_gn=LOC101090913;refGene_tx=XM_023244066.1;refGene_gn=LOC101090913;refGene_tx=XM_023244068.1;refGene_gn=LOC101090913;refGene_tx=XM_023244069.1;refGene_gn=LOC101090913;refGene_tx=XM_023244070.1
E1 42919252 42919427
refGene_gn=LOC101090913;refGene_tx=XM_023244066.1;refGene_gn=LOC101090913;refGene_tx=XM_023244068.1;refGene_gn=LOC101090913;refGene_tx=XM_023244069.1;refGene_gn=LOC101090913;refGene_tx=XM_023244070.1
E1 42924621 42924752
refGene_gn=LOC101090913;refGene_tx=XM_023244066.1;refGene_gn=LOC101090913;refGene_tx=XM_023244068.1;refGene_gn=LOC101090913;refGene_tx=XM_023244069.1;refGene_gn=LOC101090913;refGene_tx=XM_023244070.1
E1 42925010 42925162
refGene_gn=LOC101090913;refGene_tx=XM_023244066.1;refGene_gn=LOC101090913;refGene_tx=XM_023244068.1;refGene_gn=LOC101090913;refGene_tx=XM_023244069.1;refGene_gn=LOC101090913;refGene_tx=XM_023244070.1

E1 42925657 42925813
refGene_gn=LOC101090913;refGene_tx=XM_023244066.1;refGene_gn=LOC101090913;refGene_tx=XM_023244068.1;refGene_gn=LOC101090913;refGene_tx=XM_023244069.1;refGene_gn=LOC101090913;refGene_tx=XM_023244070.1
E1 42926240 42926420
refGene_gn=LOC101090913;refGene_tx=XM_023244066.1;refGene_gn=LOC101090913;refGene_tx=XM_023244068.1;refGene_gn=LOC101090913;refGene_tx=XM_023244069.1;refGene_gn=LOC101090913;refGene_tx=XM_023244070.1
E1 42928225 42928313
refGene_gn=LOC101090913;refGene_tx=XM_023244066.1;refGene_gn=LOC101090913;refGene_tx=XM_023244068.1;refGene_gn=LOC101090913;refGene_tx=XM_023244069.1;refGene_gn=LOC101090913;refGene_tx=XM_023244070.1
E1 42928414 42928537
refGene_gn=LOC101090913;refGene_tx=XM_023244066.1;refGene_gn=LOC101090913;refGene_tx=XM_023244068.1;refGene_gn=LOC101090913;refGene_tx=XM_023244069.1;refGene_gn=LOC101090913;refGene_tx=XM_023244070.1
E1 42928898 42928991
refGene_gn=LOC101090913;refGene_tx=XM_023244066.1;refGene_gn=LOC101090913;refGene_tx=XM_023244068.1;refGene_gn=LOC101090913;refGene_tx=XM_023244069.1;refGene_gn=LOC101090913;refGene_tx=XM_023244070.1
E1 42929467 42929674
refGene_gn=LOC101090913;refGene_tx=XM_023244066.1;refGene_gn=LOC101090913;refGene_tx=XM_023244068.1;refGene_gn=LOC101090913;refGene_tx=XM_023244069.1;refGene_gn=LOC101090913;refGene_tx=XM_023244070.1
E1 42930259 42930354
refGene_gn=LOC101090913;refGene_tx=XM_023244066.1;refGene_gn=LOC101090913;refGene_tx=XM_023244068.1;refGene_gn=LOC101090913;refGene_tx=XM_023244069.1;refGene_gn=LOC101090913;refGene_tx=XM_023244070.1
E1 42930449 42930580
refGene_gn=LOC101090913;refGene_tx=XM_023244066.1;refGene_gn=LOC101090913;refGene_tx=XM_023244068.1;refGene_gn=LOC101090913;refGene_tx=XM_023244069.1;refGene_gn=LOC101090913;refGene_tx=XM_023244070.1
E1 42931060 42931216
refGene_gn=LOC101090913;refGene_tx=XM_023244066.1;refGene_gn=LOC101090913;refGene_tx=XM_023244068.1;refGene_gn=LOC101090913;refGene_tx=XM_023244069.1;refGene_gn=LOC101090913;refGene_tx=XM_023244070.1
E1 42931932 42931984
refGene_gn=LOC101090913;refGene_tx=XM_023244066.1
E1 42932316 42932424
refGene_gn=LOC101090913;refGene_tx=XM_023244066.1
E1 42933457 42933555
refGene_gn=LOC101090913;refGene_tx=XM_023244066.1;refGene_gn=LOC101090913;refGene_tx=XM_023244069.1;refGene_gn=LOC101090913;refGene_tx=XM_023244070.1
E1 42936593 42936658
refGene_gn=LOC101090913;refGene_tx=XM_023244066.1;refGene_gn=LOC101090913;refGene_tx=XM_023244068.1
E1 42938172 42938228
refGene_gn=STAT3;refGene_tx=XM_003996881.5;refGene_gn=STAT3;refGene_tx=XM_006940359.4;refGene_gn=STAT3;refGene_tx=XM_019817905.2

E1 42938900 42939013
refGene_gn=STAT3;refGene_tx=XM_003996881.5;refGene_gn=STAT3;refGene_tx=XM_006940359.4;refGene_gn=STAT3;refGene_tx=XM_006940360.4;refGene_gn=STAT3;refGene_tx=XM_006940361.4;refGene_gn=STAT3;refGene_tx=XM_019817905.2
E1 42939296 42939339
refGene_gn=STAT3;refGene_tx=XM_003996881.5;refGene_gn=STAT3;refGene_tx=XM_006940359.4;refGene_gn=STAT3;refGene_tx=XM_006940360.4;refGene_gn=STAT3;refGene_tx=XM_006940361.4;refGene_gn=STAT3;refGene_tx=XM_019817905.2
E1 42942316 42942529
refGene_gn=STAT3;refGene_tx=XM_003996881.5;refGene_gn=STAT3;refGene_tx=XM_006940359.4;refGene_gn=STAT3;refGene_tx=XM_006940360.4;refGene_gn=STAT3;refGene_tx=XM_006940361.4;refGene_gn=STAT3;refGene_tx=XM_019817905.2
E1 42943034 42943174
refGene_gn=STAT3;refGene_tx=XM_003996881.5;refGene_gn=STAT3;refGene_tx=XM_006940359.4;refGene_gn=STAT3;refGene_tx=XM_006940360.4;refGene_gn=STAT3;refGene_tx=XM_006940361.4;refGene_gn=STAT3;refGene_tx=XM_019817905.2
E1 42943290 42943385
refGene_gn=STAT3;refGene_tx=XM_003996881.5;refGene_gn=STAT3;refGene_tx=XM_006940359.4;refGene_gn=STAT3;refGene_tx=XM_006940360.4;refGene_gn=STAT3;refGene_tx=XM_006940361.4;refGene_gn=STAT3;refGene_tx=XM_019817905.2
E1 42943613 42943666
refGene_gn=STAT3;refGene_tx=XM_003996881.5;refGene_gn=STAT3;refGene_tx=XM_006940359.4;refGene_gn=STAT3;refGene_tx=XM_006940360.4;refGene_gn=STAT3;refGene_tx=XM_006940361.4;refGene_gn=STAT3;refGene_tx=XM_019817905.2
E1 42944939 42945075
refGene_gn=STAT3;refGene_tx=XM_003996881.5;refGene_gn=STAT3;refGene_tx=XM_006940359.4;refGene_gn=STAT3;refGene_tx=XM_006940360.4;refGene_gn=STAT3;refGene_tx=XM_006940361.4;refGene_gn=STAT3;refGene_tx=XM_019817905.2
E1 42945418 42945517
refGene_gn=STAT3;refGene_tx=XM_003996881.5;refGene_gn=STAT3;refGene_tx=XM_006940359.4;refGene_gn=STAT3;refGene_tx=XM_006940360.4;refGene_gn=STAT3;refGene_tx=XM_006940361.4;refGene_gn=STAT3;refGene_tx=XM_019817905.2
E1 42946341 42946425
refGene_gn=STAT3;refGene_tx=XM_003996881.5;refGene_gn=STAT3;refGene_tx=XM_006940359.4;refGene_gn=STAT3;refGene_tx=XM_006940360.4;refGene_gn=STAT3;refGene_tx=XM_006940361.4;refGene_gn=STAT3;refGene_tx=XM_019817905.2
E1 42949063 42949111
refGene_gn=STAT3;refGene_tx=XM_003996881.5;refGene_gn=STAT3;refGene_tx=XM_006940359.4;refGene_gn=STAT3;refGene_tx=XM_006940360.4;refGene_gn=STAT3;refGene_tx=XM_006940361.4;refGene_gn=STAT3;refGene_tx=XM_019817905.2

E1 42949220 42949314
refGene_gn=STAT3;refGene_tx=XM_003996881.5;refGene_gn=STAT3;refGene_tx=XM_006940359.4;refGene_gn=STAT3;refGene_tx=XM_006940360.4;refGene_gn=STAT3;refGene_tx=XM_006940361.4;refGene_gn=STAT3;refGene_tx=XM_019817905.2
E1 42949413 42949443
refGene_gn=STAT3;refGene_tx=XM_003996881.5;refGene_gn=STAT3;refGene_tx=XM_006940359.4;refGene_gn=STAT3;refGene_tx=XM_006940360.4;refGene_gn=STAT3;refGene_tx=XM_006940361.4;refGene_gn=STAT3;refGene_tx=XM_019817905.2
E1 42953831 42953891
refGene_gn=STAT3;refGene_tx=XM_003996881.5;refGene_gn=STAT3;refGene_tx=XM_006940359.4;refGene_gn=STAT3;refGene_tx=XM_006940360.4;refGene_gn=STAT3;refGene_tx=XM_006940361.4;refGene_gn=STAT3;refGene_tx=XM_019817905.2
E1 42954751 42954844
refGene_gn=STAT3;refGene_tx=XM_003996881.5;refGene_gn=STAT3;refGene_tx=XM_006940359.4;refGene_gn=STAT3;refGene_tx=XM_006940360.4;refGene_gn=STAT3;refGene_tx=XM_006940361.4;refGene_gn=STAT3;refGene_tx=XM_019817905.2
E1 42954974 42955133
refGene_gn=STAT3;refGene_tx=XM_003996881.5;refGene_gn=STAT3;refGene_tx=XM_006940359.4;refGene_gn=STAT3;refGene_tx=XM_006940360.4;refGene_gn=STAT3;refGene_tx=XM_006940361.4;refGene_gn=STAT3;refGene_tx=XM_019817905.2
E1 42958410 42958562
refGene_gn=STAT3;refGene_tx=XM_003996881.5;refGene_gn=STAT3;refGene_tx=XM_006940359.4;refGene_gn=STAT3;refGene_tx=XM_006940360.4;refGene_gn=STAT3;refGene_tx=XM_006940361.4;refGene_gn=STAT3;refGene_tx=XM_019817905.2
E1 42958697 42958792
refGene_gn=STAT3;refGene_tx=XM_003996881.5;refGene_gn=STAT3;refGene_tx=XM_006940359.4;refGene_gn=STAT3;refGene_tx=XM_006940360.4;refGene_gn=STAT3;refGene_tx=XM_006940361.4;refGene_gn=STAT3;refGene_tx=XM_019817905.2
E1 42959015 42959097
refGene_gn=STAT3;refGene_tx=XM_003996881.5;refGene_gn=STAT3;refGene_tx=XM_006940359.4;refGene_gn=STAT3;refGene_tx=XM_006940360.4;refGene_gn=STAT3;refGene_tx=XM_006940361.4;refGene_gn=STAT3;refGene_tx=XM_019817905.2
E1 42959280 42959376
refGene_gn=STAT3;refGene_tx=XM_003996881.5;refGene_gn=STAT3;refGene_tx=XM_006940359.4;refGene_gn=STAT3;refGene_tx=XM_006940360.4;refGene_gn=STAT3;refGene_tx=XM_006940361.4;refGene_gn=STAT3;refGene_tx=XM_019817905.2
E1 42965899 42965998
refGene_gn=STAT3;refGene_tx=XM_003996881.5;refGene_gn=STAT3;refGene_tx=XM_006940359.4;refGene_gn=STAT3;refGene_tx=XM_006940360.4;refGene_gn=STAT3;refGene_tx=XM_006940361.4;refGene_gn=STAT3;refGene_tx=XM_019817905.2

E1 42966869 42967014
refGene_gn=STAT3;refGene_tx=XM_003996881.5;refGene_gn=STAT3;refGene_tx=XM_006940359.4;refGene_gn=STAT3;refGene_tx=XM_006940360.4;refGene_gn=STAT3;refGene_tx=XM_006940361.4;refGene_gn=STAT3;refGene_tx=XM_019817905.2
E1 42967930 42968058
refGene_gn=STAT3;refGene_tx=XM_003996881.5;refGene_gn=STAT3;refGene_tx=XM_006940359.4;refGene_gn=STAT3;refGene_tx=XM_006940360.4;refGene_gn=STAT3;refGene_tx=XM_006940361.4;refGene_gn=STAT3;refGene_tx=XM_019817905.2
E1 43019051 43019753 refGene_gn=CAVIN1;refGene_tx=XM_003996883.5
E1 43029485 43029956 refGene_gn=CAVIN1;refGene_tx=XM_003996883.5
E1 43050522 43050639
refGene_gn=ATP6V0A1;refGene_tx=XM_003996884.3;refGene_gn=ATP6V0A1;refGene_tx=XM_003996885.3;refGene_gn=ATP6V0A1;refGene_tx=XM_019817908.1;refGene_gn=ATP6V0A1;refGene_tx=XM_019817909.1;refGene_gn=ATP6V0A1;refGene_tx=XM_019817911.1;refGene_gn=ATP6V0A1;refGene_tx=XM_019817912.1;refGene_gn=ATP6V0A1;refGene_tx=XM_019817913.1;refGene_gn=ATP6V0A1;refGene_tx=XM_019817914.1
E1 43062498 43062577
refGene_gn=ATP6V0A1;refGene_tx=XM_003996884.3;refGene_gn=ATP6V0A1;refGene_tx=XM_003996885.3;refGene_gn=ATP6V0A1;refGene_tx=XM_019817908.1;refGene_gn=ATP6V0A1;refGene_tx=XM_019817909.1;refGene_gn=ATP6V0A1;refGene_tx=XM_019817911.1;refGene_gn=ATP6V0A1;refGene_tx=XM_019817912.1;refGene_gn=ATP6V0A1;refGene_tx=XM_019817913.1;refGene_gn=ATP6V0A1;refGene_tx=XM_019817914.1
E1 43064050 43064148
refGene_gn=ATP6V0A1;refGene_tx=XM_003996884.3;refGene_gn=ATP6V0A1;refGene_tx=XM_003996885.3;refGene_gn=ATP6V0A1;refGene_tx=XM_019817908.1;refGene_gn=ATP6V0A1;refGene_tx=XM_019817909.1;refGene_gn=ATP6V0A1;refGene_tx=XM_019817911.1;refGene_gn=ATP6V0A1;refGene_tx=XM_019817912.1;refGene_gn=ATP6V0A1;refGene_tx=XM_019817913.1;refGene_gn=ATP6V0A1;refGene_tx=XM_019817914.1
E1 43064710 43064839
refGene_gn=ATP6V0A1;refGene_tx=XM_003996884.3;refGene_gn=ATP6V0A1;refGene_tx=XM_003996885.3;refGene_gn=ATP6V0A1;refGene_tx=XM_019817908.1;refGene_gn=ATP6V0A1;refGene_tx=XM_019817909.1;refGene_gn=ATP6V0A1;refGene_tx=XM_019817911.1;refGene_gn=ATP6V0A1;refGene_tx=XM_019817912.1;refGene_gn=ATP6V0A1;refGene_tx=XM_019817913.1;refGene_gn=ATP6V0A1;refGene_tx=XM_019817914.1
E1 43070761 43070865
refGene_gn=ATP6V0A1;refGene_tx=XM_003996884.3;refGene_gn=ATP6V0A1;refGene_tx=XM_003996885.3;refGene_gn=ATP6V0A1;refGene_tx=XM_019817908.1;refGene_gn=ATP6V0A1;refGene_tx=XM_019817909.1;refGene_gn=ATP6V0A1;refGene_tx=XM_019817911.1;refGene_gn=ATP6V0A1;refGene_tx=XM_019817912.1;refGene_gn=ATP6V0A1;refGene_tx=XM_019817913.1;refGene_gn=ATP6V0A1;refGene_tx=XM_019817914.1
E1 43071235 43071362
refGene_gn=ATP6V0A1;refGene_tx=XM_003996884.3;refGene_gn=ATP6V0A1;refGene_tx=XM_003996885.3;refGene_gn=ATP6V0A1;refGene_tx=XM_019817908.1;re

fGene_gn=ATP6V0A1;refGene_tx=XM_019817909.1;refGene_gn=ATP6V0A1;refGene_tx=XM_019817911.1;refGene_gn=ATP6V0A1;refGene_tx=XM_019817912.1;refGene_gn=ATP6V0A1;refGene_tx=XM_019817913.1;refGene_gn=ATP6V0A1;refGene_tx=XM_019817914.1
E1 43073419 43073502
refGene_gn=ATP6V0A1;refGene_tx=XM_003996884.3;refGene_gn=ATP6V0A1;refGene_tx=XM_003996885.3;refGene_gn=ATP6V0A1;refGene_tx=XM_019817908.1;refGene_gn=ATP6V0A1;refGene_tx=XM_019817909.1;refGene_gn=ATP6V0A1;refGene_tx=XM_019817911.1;refGene_gn=ATP6V0A1;refGene_tx=XM_019817912.1;refGene_gn=ATP6V0A1;refGene_tx=XM_019817913.1;refGene_gn=ATP6V0A1;refGene_tx=XM_019817914.1
E1 43074117 43074211
refGene_gn=ATP6V0A1;refGene_tx=XM_003996884.3;refGene_gn=ATP6V0A1;refGene_tx=XM_003996885.3;refGene_gn=ATP6V0A1;refGene_tx=XM_019817908.1;refGene_gn=ATP6V0A1;refGene_tx=XM_019817909.1;refGene_gn=ATP6V0A1;refGene_tx=XM_019817911.1;refGene_gn=ATP6V0A1;refGene_tx=XM_019817912.1;refGene_gn=ATP6V0A1;refGene_tx=XM_019817913.1;refGene_gn=ATP6V0A1;refGene_tx=XM_019817914.1
E1 43077952 43078165
refGene_gn=ATP6V0A1;refGene_tx=XM_003996884.3;refGene_gn=ATP6V0A1;refGene_tx=XM_003996885.3;refGene_gn=ATP6V0A1;refGene_tx=XM_019817908.1;refGene_gn=ATP6V0A1;refGene_tx=XM_019817909.1;refGene_gn=ATP6V0A1;refGene_tx=XM_019817911.1;refGene_gn=ATP6V0A1;refGene_tx=XM_019817912.1;refGene_gn=ATP6V0A1;refGene_tx=XM_019817913.1;refGene_gn=ATP6V0A1;refGene_tx=XM_019817914.1
E1 43078835 43078986
refGene_gn=ATP6V0A1;refGene_tx=XM_003996884.3;refGene_gn=ATP6V0A1;refGene_tx=XM_003996885.3;refGene_gn=ATP6V0A1;refGene_tx=XM_019817908.1;refGene_gn=ATP6V0A1;refGene_tx=XM_019817909.1;refGene_gn=ATP6V0A1;refGene_tx=XM_019817911.1;refGene_gn=ATP6V0A1;refGene_tx=XM_019817912.1;refGene_gn=ATP6V0A1;refGene_tx=XM_019817913.1;refGene_gn=ATP6V0A1;refGene_tx=XM_019817914.1
E1 43079869 43080009
refGene_gn=ATP6V0A1;refGene_tx=XM_003996884.3;refGene_gn=ATP6V0A1;refGene_tx=XM_003996885.3;refGene_gn=ATP6V0A1;refGene_tx=XM_019817908.1;refGene_gn=ATP6V0A1;refGene_tx=XM_019817909.1;refGene_gn=ATP6V0A1;refGene_tx=XM_019817911.1;refGene_gn=ATP6V0A1;refGene_tx=XM_019817912.1;refGene_gn=ATP6V0A1;refGene_tx=XM_019817913.1;refGene_gn=ATP6V0A1;refGene_tx=XM_019817914.1
E1 43080343 43080501
refGene_gn=ATP6V0A1;refGene_tx=XM_003996884.3;refGene_gn=ATP6V0A1;refGene_tx=XM_003996885.3;refGene_gn=ATP6V0A1;refGene_tx=XM_019817908.1;refGene_gn=ATP6V0A1;refGene_tx=XM_019817909.1;refGene_gn=ATP6V0A1;refGene_tx=XM_019817911.1;refGene_gn=ATP6V0A1;refGene_tx=XM_019817912.1;refGene_gn=ATP6V0A1;refGene_tx=XM_019817913.1;refGene_gn=ATP6V0A1;refGene_tx=XM_019817914.1
E1 43080956 43081047
refGene_gn=ATP6V0A1;refGene_tx=XM_003996884.3;refGene_gn=ATP6V0A1;refGene_tx=XM_003996885.3;refGene_gn=ATP6V0A1;refGene_tx=XM_019817908.1;refGene_gn=ATP6V0A1;refGene_tx=XM_019817909.1;refGene_gn=ATP6V0A1;refGen

e_tx=XM_019817911.1;refGene_gn=ATP6V0A1;refGene_tx=XM_019817912.1;refGene_gn=ATP6V0A1;refGene_tx=XM_019817913.1;refGene_gn=ATP6V0A1;refGene_tx=XM_019817914.1
E1 43084644 43084763
refGene_gn=ATP6V0A1;refGene_tx=XM_003996884.3;refGene_gn=ATP6V0A1;refGene_tx=XM_003996885.3;refGene_gn=ATP6V0A1;refGene_tx=XM_019817908.1;refGene_gn=ATP6V0A1;refGene_tx=XM_019817909.1;refGene_gn=ATP6V0A1;refGene_tx=XM_019817911.1;refGene_gn=ATP6V0A1;refGene_tx=XM_019817912.1;refGene_gn=ATP6V0A1;refGene_tx=XM_019817913.1;refGene_gn=ATP6V0A1;refGene_tx=XM_019817914.1
E1 43085702 43085919
refGene_gn=ATP6V0A1;refGene_tx=XM_003996884.3;refGene_gn=ATP6V0A1;refGene_tx=XM_003996885.3;refGene_gn=ATP6V0A1;refGene_tx=XM_019817908.1;refGene_gn=ATP6V0A1;refGene_tx=XM_019817909.1;refGene_gn=ATP6V0A1;refGene_tx=XM_019817911.1;refGene_gn=ATP6V0A1;refGene_tx=XM_019817912.1;refGene_gn=ATP6V0A1;refGene_tx=XM_019817913.1;refGene_gn=ATP6V0A1;refGene_tx=XM_019817914.1
E1 43086456 43086564
refGene_gn=ATP6V0A1;refGene_tx=XM_003996884.3;refGene_gn=ATP6V0A1;refGene_tx=XM_003996885.3;refGene_gn=ATP6V0A1;refGene_tx=XM_019817908.1;refGene_gn=ATP6V0A1;refGene_tx=XM_019817909.1;refGene_gn=ATP6V0A1;refGene_tx=XM_019817911.1;refGene_gn=ATP6V0A1;refGene_tx=XM_019817912.1;refGene_gn=ATP6V0A1;refGene_tx=XM_019817913.1;refGene_gn=ATP6V0A1;refGene_tx=XM_019817914.1
E1 43087399 43087474
refGene_gn=ATP6V0A1;refGene_tx=XM_019817908.1;refGene_gn=ATP6V0A1;refGene_tx=XM_019817909.1;refGene_gn=ATP6V0A1;refGene_tx=XM_019817911.1
E1 43091288 43091396
refGene_gn=ATP6V0A1;refGene_tx=XM_003996884.3;refGene_gn=ATP6V0A1;refGene_tx=XM_003996885.3;refGene_gn=ATP6V0A1;refGene_tx=XM_019817908.1;refGene_gn=ATP6V0A1;refGene_tx=XM_019817909.1;refGene_gn=ATP6V0A1;refGene_tx=XM_019817911.1;refGene_gn=ATP6V0A1;refGene_tx=XM_019817912.1;refGene_gn=ATP6V0A1;refGene_tx=XM_019817913.1
E1 43092382 43092400
refGene_gn=ATP6V0A1;refGene_tx=XM_019817908.1;refGene_gn=ATP6V0A1;refGene_tx=XM_019817911.1;refGene_gn=ATP6V0A1;refGene_tx=XM_019817912.1;refGene_gn=ATP6V0A1;refGene_tx=XM_019817913.1
E1 43097328 43097446
refGene_gn=ATP6V0A1;refGene_tx=XM_003996884.3;refGene_gn=ATP6V0A1;refGene_tx=XM_003996885.3;refGene_gn=ATP6V0A1;refGene_tx=XM_019817908.1;refGene_gn=ATP6V0A1;refGene_tx=XM_019817909.1;refGene_gn=ATP6V0A1;refGene_tx=XM_019817911.1;refGene_gn=ATP6V0A1;refGene_tx=XM_019817912.1;refGene_gn=ATP6V0A1;refGene_tx=XM_019817913.1;refGene_gn=ATP6V0A1;refGene_tx=XM_019817914.1
E1 43097760 43097932
refGene_gn=ATP6V0A1;refGene_tx=XM_003996884.3;refGene_gn=ATP6V0A1;refGene_tx=XM_003996885.3;refGene_gn=ATP6V0A1;refGene_tx=XM_019817908.1;refGene_gn=ATP6V0A1;refGene_tx=XM_019817909.1;refGene_gn=ATP6V0A1;refGene_tx=XM_019817911.1;refGene_gn=ATP6V0A1;refGene_tx=XM_019817912.1;refGene_gn=ATP6V0A1;refGene_tx=XM_019817913.1;refGene_gn=ATP6V0A1;refGene_

tx=XM_019817914.1
E1 43103429 43103523
refGene_gn=ATP6V0A1;refGene_tx=XM_003996884.3;refGene_gn=ATP6V0A1;refGene_tx=XM_003996885.3;refGene_gn=ATP6V0A1;refGene_tx=XM_019817908.1;refGene_gn=ATP6V0A1;refGene_tx=XM_019817909.1;refGene_gn=ATP6V0A1;refGene_tx=XM_019817911.1;refGene_gn=ATP6V0A1;refGene_tx=XM_019817912.1;refGene_gn=ATP6V0A1;refGene_tx=XM_019817913.1;refGene_gn=ATP6V0A1;refGene_tx=XM_019817914.1
E1 43112207 43112630
refGene_gn=NAGLU;refGene_tx=XM_023243555.1;refGene_gn=NAGLU;refGene_tx=XM_023243556.1
E1 43113973 43114121 refGene_gn=NAGLU;refGene_tx=XM_023243555.1
E1 43114839 43114986
refGene_gn=NAGLU;refGene_tx=XM_023243555.1;refGene_gn=NAGLU;refGene_tx=XM_023243556.1
E1 43115173 43115259
refGene_gn=NAGLU;refGene_tx=XM_023243555.1;refGene_gn=NAGLU;refGene_tx=XM_023243556.1
E1 43116236 43116493
refGene_gn=NAGLU;refGene_tx=XM_023243555.1;refGene_gn=NAGLU;refGene_tx=XM_023243556.1
E1 43117434 43118645
refGene_gn=NAGLU;refGene_tx=XM_023243555.1;refGene_gn=NAGLU;refGene_tx=XM_023243556.1
E1 43119407 43119504 refGene_gn=HSD17B1;refGene_tx=XM_006940403.4
E1 43119611 43119779 refGene_gn=HSD17B1;refGene_tx=XM_006940403.4
E1 43119920 43120100 refGene_gn=HSD17B1;refGene_tx=XM_006940403.4
E1 43120205 43120299 refGene_gn=HSD17B1;refGene_tx=XM_006940403.4
E1 43120708 43120886 refGene_gn=HSD17B1;refGene_tx=XM_006940403.4
E1 43120974 43121190 refGene_gn=HSD17B1;refGene_tx=XM_006940403.4
E1 43126905 43127605
refGene_gn=COASY;refGene_tx=XM_006940365.2;refGene_gn=COASY;refGene_tx=XM_006940366.2;refGene_gn=COASY;refGene_tx=XM_019817915.1
E1 43128239 43128454
refGene_gn=COASY;refGene_tx=XM_006940365.2;refGene_gn=COASY;refGene_tx=XM_006940366.2;refGene_gn=COASY;refGene_tx=XM_019817915.1
E1 43128706 43128838
refGene_gn=COASY;refGene_tx=XM_006940365.2;refGene_gn=COASY;refGene_tx=XM_006940366.2;refGene_gn=COASY;refGene_tx=XM_019817915.1
E1 43128964 43129154
refGene_gn=COASY;refGene_tx=XM_006940365.2;refGene_gn=COASY;refGene_tx=XM_006940366.2;refGene_gn=COASY;refGene_tx=XM_019817915.1
E1 43129238 43129303
refGene_gn=COASY;refGene_tx=XM_006940365.2;refGene_gn=COASY;refGene_tx=XM_006940366.2;refGene_gn=COASY;refGene_tx=XM_019817915.1
E1 43129487 43129572
refGene_gn=COASY;refGene_tx=XM_006940365.2;refGene_gn=COASY;refGene_tx=XM_006940366.2;refGene_gn=COASY;refGene_tx=XM_019817915.1
E1 43129738 43129836
refGene_gn=COASY;refGene_tx=XM_006940365.2;refGene_gn=COASY;refGene_tx

=XM_006940366.2;refGene_gn=COASY;refGene_tx=XM_019817915.1
E1 43129925 43130072
refGene_gn=COASY;refGene_tx=XM_006940365.2;refGene_gn=COASY;refGene_tx
=XM_006940366.2;refGene_gn=COASY;refGene_tx=XM_019817915.1
E1 43130173 43130224
refGene_gn=COASY;refGene_tx=XM_006940365.2;refGene_gn=COASY;refGene_tx
=XM_006940366.2;refGene_gn=COASY;refGene_tx=XM_019817915.1
E1 43131163 43131499
refGene_gn=MLX;refGene_tx=XM_003996892.4;refGene_gn=MLX;refGene_tx=XM_
003996893.4;refGene_gn=MLX;refGene_tx=XM_003996894.4;refGene_gn=MLX;re
fGene_tx=XM_019817916.1
E1 43131883 43131920
refGene_gn=MLX;refGene_tx=XM_003996892.4;refGene_gn=MLX;refGene_tx=XM_
003996893.4;refGene_gn=MLX;refGene_tx=XM_003996894.4;refGene_gn=MLX;re
fGene_tx=XM_019817916.1
E1 43132385 43132475
refGene_gn=MLX;refGene_tx=XM_003996892.4;refGene_gn=MLX;refGene_tx=XM_
003996893.4;refGene_gn=MLX;refGene_tx=XM_019817916.1
E1 43132751 43132858
refGene_gn=MLX;refGene_tx=XM_003996892.4;refGene_gn=MLX;refGene_tx=XM_
003996893.4;refGene_gn=MLX;refGene_tx=XM_003996894.4;refGene_gn=MLX;re
fGene_tx=XM_019817916.1
E1 43133109 43133209
refGene_gn=MLX;refGene_tx=XM_003996892.4;refGene_gn=MLX;refGene_tx=XM_
003996893.4;refGene_gn=MLX;refGene_tx=XM_003996894.4;refGene_gn=MLX;re
fGene_tx=XM_019817916.1
E1 43133400 43133500
refGene_gn=MLX;refGene_tx=XM_003996892.4;refGene_gn=MLX;refGene_tx=XM_
003996893.4;refGene_gn=MLX;refGene_tx=XM_003996894.4;refGene_gn=MLX;re
fGene_tx=XM_019817916.1
E1 43133842 43134044
refGene_gn=MLX;refGene_tx=XM_003996892.4;refGene_gn=MLX;refGene_tx=XM_
003996893.4;refGene_gn=MLX;refGene_tx=XM_003996894.4;refGene_gn=MLX;re
fGene_tx=XM_019817916.1
E1 43134682 43134694 refGene_gn=MLX;refGene_tx=XM_019817916.1
E1 43134747 43134804
refGene_gn=MLX;refGene_tx=XM_003996892.4;refGene_gn=MLX;refGene_tx=XM_
003996893.4;refGene_gn=MLX;refGene_tx=XM_003996894.4
E1 43136096 43136153 refGene_gn=PSMC3IP;refGene_tx=XM_003996895.5
E1 43136239 43136299 refGene_gn=PSMC3IP;refGene_tx=XM_003996895.5
E1 43136424 43136478 refGene_gn=PSMC3IP;refGene_tx=XM_003996895.5
E1 43136593 43136739 refGene_gn=PSMC3IP;refGene_tx=XM_003996895.5
E1 43137261 43137373 refGene_gn=PSMC3IP;refGene_tx=XM_003996895.5
E1 43139415 43139505 refGene_gn=PSMC3IP;refGene_tx=XM_003996895.5
E1 43139685 43139786 refGene_gn=PSMC3IP;refGene_tx=XM_003996895.5
E1 43139877 43139911 refGene_gn=PSMC3IP;refGene_tx=XM_003996895.5
E1 43143024 43143482 refGene_gn=RETREG3;refGene_tx=XM_003996897.5
E1 43143897 43144030 refGene_gn=RETREG3;refGene_tx=XM_003996897.5
E1 43144645 43144728 refGene_gn=RETREG3;refGene_tx=XM_003996897.5
E1 43145732 43145870 refGene_gn=RETREG3;refGene_tx=XM_003996897.5

E1 43146683 43146768 refGene_gn=RETREG3;refGene_tx=XM_003996897.5
E1 43147434 43147561 refGene_gn=RETREG3;refGene_tx=XM_003996897.5
E1 43148497 43148528 refGene_gn=RETREG3;refGene_tx=XM_003996897.5
E1 43150290 43150397 refGene_gn=RETREG3;refGene_tx=XM_003996897.5
E1 43164609 43164848 refGene_gn=RETREG3;refGene_tx=XM_003996897.5
E1 43165246 43165295 refGene_gn=TUBG1;refGene_tx=XM_011289129.3
E1 43165591 43165704 refGene_gn=TUBG1;refGene_tx=XM_011289129.3
E1 43165926 43166094 refGene_gn=TUBG1;refGene_tx=XM_011289129.3
E1 43167072 43167141 refGene_gn=TUBG1;refGene_tx=XM_011289129.3
E1 43167417 43167497 refGene_gn=TUBG1;refGene_tx=XM_011289129.3
E1 43167943 43168070 refGene_gn=TUBG1;refGene_tx=XM_011289129.3
E1 43168304 43168391 refGene_gn=TUBG1;refGene_tx=XM_011289129.3
E1 43168497 43168647 refGene_gn=TUBG1;refGene_tx=XM_011289129.3
E1 43169035 43169188 refGene_gn=TUBG1;refGene_tx=XM_011289129.3
E1 43169279 43169441 refGene_gn=TUBG1;refGene_tx=XM_011289129.3
E1 43169630 43169828 refGene_gn=TUBG1;refGene_tx=XM_011289129.3
E1 43202219 43202268
refGene_gn=LOC101099090;refGene_tx=XM_003996898.4;refGene_gn=LOC101099
090;refGene_tx=XM_019817917.2
E1 43202546 43202659
refGene_gn=LOC101099090;refGene_tx=XM_003996898.4;refGene_gn=LOC101099
090;refGene_tx=XM_019817917.2
E1 43202882 43203050
refGene_gn=LOC101099090;refGene_tx=XM_003996898.4;refGene_gn=LOC101099
090;refGene_tx=XM_019817917.2
E1 43203363 43203432
refGene_gn=LOC101099090;refGene_tx=XM_003996898.4;refGene_gn=LOC101099
090;refGene_tx=XM_019817917.2
E1 43204199 43204279
refGene_gn=LOC101099090;refGene_tx=XM_003996898.4
E1 43204690 43204817
refGene_gn=LOC101099090;refGene_tx=XM_003996898.4;refGene_gn=LOC101099
090;refGene_tx=XM_019817917.2
E1 43205421 43205508
refGene_gn=LOC101099090;refGene_tx=XM_003996898.4;refGene_gn=LOC101099
090;refGene_tx=XM_019817917.2
E1 43205614 43205764
refGene_gn=LOC101099090;refGene_tx=XM_003996898.4;refGene_gn=LOC101099
090;refGene_tx=XM_019817917.2
E1 43206151 43206304
refGene_gn=LOC101099090;refGene_tx=XM_003996898.4;refGene_gn=LOC101099
090;refGene_tx=XM_019817917.2
E1 43206395 43206557
refGene_gn=LOC101099090;refGene_tx=XM_003996898.4;refGene_gn=LOC101099
090;refGene_tx=XM_019817917.2
E1 43206684 43206882
refGene_gn=LOC101099090;refGene_tx=XM_003996898.4;refGene_gn=LOC101099
090;refGene_tx=XM_019817917.2
E1 43208212 43208389
refGene_gn=PLEKHH3;refGene_tx=XM_011289127.3;refGene_gn=PLEKHH3;refGen

e_tx=XM_023243576.1;refGene_gn=PLEKHH3;refGene_tx=XM_023243577.1
E1 43208962 43209154
refGene_gn=PLEKHH3;refGene_tx=XM_011289127.3;refGene_gn=PLEKHH3;refGene_tx=XM_023243576.1;refGene_gn=PLEKHH3;refGene_tx=XM_023243577.1
E1 43209451 43209913
refGene_gn=PLEKHH3;refGene_tx=XM_011289127.3;refGene_gn=PLEKHH3;refGene_tx=XM_023243576.1;refGene_gn=PLEKHH3;refGene_tx=XM_023243577.1
E1 43210104 43210237
refGene_gn=PLEKHH3;refGene_tx=XM_011289127.3;refGene_gn=PLEKHH3;refGene_tx=XM_023243576.1;refGene_gn=PLEKHH3;refGene_tx=XM_023243577.1
E1 43210509 43210646
refGene_gn=PLEKHH3;refGene_tx=XM_011289127.3;refGene_gn=PLEKHH3;refGene_tx=XM_023243576.1;refGene_gn=PLEKHH3;refGene_tx=XM_023243577.1
E1 43210865 43211073
refGene_gn=PLEKHH3;refGene_tx=XM_011289127.3;refGene_gn=PLEKHH3;refGene_tx=XM_023243576.1;refGene_gn=PLEKHH3;refGene_tx=XM_023243577.1
E1 43211627 43211934
refGene_gn=PLEKHH3;refGene_tx=XM_011289127.3;refGene_gn=PLEKHH3;refGene_tx=XM_023243576.1;refGene_gn=PLEKHH3;refGene_tx=XM_023243577.1
E1 43212716 43212837
refGene_gn=PLEKHH3;refGene_tx=XM_011289127.3;refGene_gn=PLEKHH3;refGene_tx=XM_023243576.1;refGene_gn=PLEKHH3;refGene_tx=XM_023243577.1
E1 43212944 43213102
refGene_gn=PLEKHH3;refGene_tx=XM_011289127.3;refGene_gn=PLEKHH3;refGene_tx=XM_023243576.1;refGene_gn=PLEKHH3;refGene_tx=XM_023243577.1
E1 43213186 43213378
refGene_gn=PLEKHH3;refGene_tx=XM_011289127.3;refGene_gn=PLEKHH3;refGene_tx=XM_023243576.1;refGene_gn=PLEKHH3;refGene_tx=XM_023243577.1
E1 43213477 43213557
refGene_gn=PLEKHH3;refGene_tx=XM_011289127.3;refGene_gn=PLEKHH3;refGene_tx=XM_023243576.1;refGene_gn=PLEKHH3;refGene_tx=XM_023243577.1
E1 43213899 43213955
refGene_gn=PLEKHH3;refGene_tx=XM_011289127.3;refGene_gn=PLEKHH3;refGene_tx=XM_023243576.1;refGene_gn=PLEKHH3;refGene_tx=XM_023243577.1
E1 43215327 43215489 refGene_gn=PLEKHH3;refGene_tx=XM_011289127.3
E1 43215915 43216077
refGene_gn=PLEKHH3;refGene_tx=XM_023243576.1;refGene_gn=PLEKHH3;refGene_tx=XM_023243577.1
E1 43218734 43219802 refGene_gn=CCR10;refGene_tx=XM_023244588.1
E1 43220627 43220651 refGene_gn=CCR10;refGene_tx=XM_023244588.1
E1 43221516 43221580 refGene_gn=CNTNAP1;refGene_tx=XM_003996899.5
E1 43222433 43222535 refGene_gn=CNTNAP1;refGene_tx=XM_003996899.5
E1 43222629 43222823
refGene_gn=CNTNAP1;refGene_tx=XM_003996899.5;refGene_gn=CNTNAP1;refGene_tx=XM_019817918.2
E1 43223281 43223429
refGene_gn=CNTNAP1;refGene_tx=XM_003996899.5;refGene_gn=CNTNAP1;refGene_tx=XM_019817918.2
E1 43223506 43223710
refGene_gn=CNTNAP1;refGene_tx=XM_003996899.5;refGene_gn=CNTNAP1;refGen

e_tx=XM_019817918.2
E1 43224336 43224521
refGene_gn=CNTNAP1;refGene_tx=XM_003996899.5;refGene_gn=CNTNAP1;refGen
e_tx=XM_019817918.2
E1 43225087 43225231
refGene_gn=CNTNAP1;refGene_tx=XM_003996899.5;refGene_gn=CNTNAP1;refGen
e_tx=XM_019817918.2
E1 43225903 43226165
refGene_gn=CNTNAP1;refGene_tx=XM_003996899.5;refGene_gn=CNTNAP1;refGen
e_tx=XM_019817918.2
E1 43226620 43226770
refGene_gn=CNTNAP1;refGene_tx=XM_003996899.5;refGene_gn=CNTNAP1;refGen
e_tx=XM_019817918.2
E1 43227028 43227200
refGene_gn=CNTNAP1;refGene_tx=XM_003996899.5;refGene_gn=CNTNAP1;refGen
e_tx=XM_019817918.2
E1 43227649 43227756
refGene_gn=CNTNAP1;refGene_tx=XM_003996899.5;refGene_gn=CNTNAP1;refGen
e_tx=XM_019817918.2
E1 43227902 43228022
refGene_gn=CNTNAP1;refGene_tx=XM_003996899.5;refGene_gn=CNTNAP1;refGen
e_tx=XM_019817918.2
E1 43228375 43228579
refGene_gn=CNTNAP1;refGene_tx=XM_003996899.5;refGene_gn=CNTNAP1;refGen
e_tx=XM_019817918.2
E1 43228776 43228933
refGene_gn=CNTNAP1;refGene_tx=XM_003996899.5;refGene_gn=CNTNAP1;refGen
e_tx=XM_019817918.2
E1 43229016 43229144
refGene_gn=CNTNAP1;refGene_tx=XM_003996899.5;refGene_gn=CNTNAP1;refGen
e_tx=XM_019817918.2
E1 43229453 43229639
refGene_gn=CNTNAP1;refGene_tx=XM_003996899.5;refGene_gn=CNTNAP1;refGen
e_tx=XM_019817918.2
E1 43230111 43230333
refGene_gn=CNTNAP1;refGene_tx=XM_003996899.5;refGene_gn=CNTNAP1;refGen
e_tx=XM_019817918.2
E1 43230589 43230829
refGene_gn=CNTNAP1;refGene_tx=XM_003996899.5;refGene_gn=CNTNAP1;refGen
e_tx=XM_019817918.2
E1 43232064 43232418
refGene_gn=CNTNAP1;refGene_tx=XM_003996899.5;refGene_gn=CNTNAP1;refGen
e_tx=XM_019817918.2
E1 43233035 43233163
refGene_gn=CNTNAP1;refGene_tx=XM_003996899.5;refGene_gn=CNTNAP1;refGen
e_tx=XM_019817918.2
E1 43233613 43233707
refGene_gn=CNTNAP1;refGene_tx=XM_003996899.5;refGene_gn=CNTNAP1;refGen
e_tx=XM_019817918.2
E1 43233905 43234151

refGene_gn=CNTNAP1;refGene_tx=XM_003996899.5;refGene_gn=CNTNAP1;refGene_tx=XM_019817918.2
E1 43234259 43234307
refGene_gn=CNTNAP1;refGene_tx=XM_003996899.5;refGene_gn=CNTNAP1;refGene_tx=XM_019817918.2
E1 43234930 43235217
refGene_gn=CNTNAP1;refGene_tx=XM_003996899.5;refGene_gn=CNTNAP1;refGene_tx=XM_019817918.2
E1 43238884 43238945
refGene_gn=EZH1;refGene_tx=XM_006940369.4;refGene_gn=EZH1;refGene_tx=XM_023244586.1;refGene_gn=EZH1;refGene_tx=XM_023244587.1
E1 43239220 43239305
refGene_gn=EZH1;refGene_tx=XM_006940369.4;refGene_gn=EZH1;refGene_tx=XM_023244586.1;refGene_gn=EZH1;refGene_tx=XM_023244587.1
E1 43239718 43239799
refGene_gn=EZH1;refGene_tx=XM_006940369.4;refGene_gn=EZH1;refGene_tx=XM_023244586.1;refGene_gn=EZH1;refGene_tx=XM_023244587.1
E1 43240253 43240335
refGene_gn=EZH1;refGene_tx=XM_006940369.4;refGene_gn=EZH1;refGene_tx=XM_023244586.1;refGene_gn=EZH1;refGene_tx=XM_023244587.1
E1 43240747 43240843
refGene_gn=EZH1;refGene_tx=XM_006940369.4;refGene_gn=EZH1;refGene_tx=XM_023244586.1;refGene_gn=EZH1;refGene_tx=XM_023244587.1
E1 43241363 43241542
refGene_gn=EZH1;refGene_tx=XM_006940369.4;refGene_gn=EZH1;refGene_tx=XM_023244586.1;refGene_gn=EZH1;refGene_tx=XM_023244587.1
E1 43242596 43242722
refGene_gn=EZH1;refGene_tx=XM_006940369.4;refGene_gn=EZH1;refGene_tx=XM_023244586.1;refGene_gn=EZH1;refGene_tx=XM_023244587.1
E1 43243218 43243259
refGene_gn=EZH1;refGene_tx=XM_006940369.4;refGene_gn=EZH1;refGene_tx=XM_023244586.1;refGene_gn=EZH1;refGene_tx=XM_023244587.1
E1 43244152 43244244
refGene_gn=EZH1;refGene_tx=XM_006940369.4;refGene_gn=EZH1;refGene_tx=XM_023244586.1;refGene_gn=EZH1;refGene_tx=XM_023244587.1
E1 43245764 43245961
refGene_gn=EZH1;refGene_tx=XM_006940369.4;refGene_gn=EZH1;refGene_tx=XM_023244586.1;refGene_gn=EZH1;refGene_tx=XM_023244587.1
E1 43246464 43246645
refGene_gn=EZH1;refGene_tx=XM_006940369.4;refGene_gn=EZH1;refGene_tx=XM_023244586.1;refGene_gn=EZH1;refGene_tx=XM_023244587.1
E1 43249801 43249893
refGene_gn=EZH1;refGene_tx=XM_006940369.4;refGene_gn=EZH1;refGene_tx=XM_023244586.1;refGene_gn=EZH1;refGene_tx=XM_023244587.1
E1 43250277 43250441
refGene_gn=EZH1;refGene_tx=XM_006940369.4;refGene_gn=EZH1;refGene_tx=XM_023244586.1;refGene_gn=EZH1;refGene_tx=XM_023244587.1
E1 43250928 43251031
refGene_gn=EZH1;refGene_tx=XM_006940369.4;refGene_gn=EZH1;refGene_tx=XM_023244586.1;refGene_gn=EZH1;refGene_tx=XM_023244587.1

E1 43251949 43252126
refGene_gn=EZH1;refGene_tx=XM_006940369.4;refGene_gn=EZH1;refGene_tx=XM_023244586.1;refGene_gn=EZH1;refGene_tx=XM_023244587.1
E1 43253143 43253264
refGene_gn=EZH1;refGene_tx=XM_006940369.4;refGene_gn=EZH1;refGene_tx=XM_023244586.1;refGene_gn=EZH1;refGene_tx=XM_023244587.1
E1 43254891 43255011
refGene_gn=EZH1;refGene_tx=XM_006940369.4;refGene_gn=EZH1;refGene_tx=XM_023244586.1;refGene_gn=EZH1;refGene_tx=XM_023244587.1
E1 43257020 43257149
refGene_gn=EZH1;refGene_tx=XM_006940369.4;refGene_gn=EZH1;refGene_tx=XM_023244586.1;refGene_gn=EZH1;refGene_tx=XM_023244587.1
E1 43257491 43257612
refGene_gn=EZH1;refGene_tx=XM_006940369.4;refGene_gn=EZH1;refGene_tx=XM_023244586.1;refGene_gn=EZH1;refGene_tx=XM_023244587.1
E1 43258216 43258221
refGene_gn=EZH1;refGene_tx=XM_006940369.4;refGene_gn=EZH1;refGene_tx=XM_023244586.1;refGene_gn=EZH1;refGene_tx=XM_023244587.1
E1 43278407 43278507
refGene_gn=RAMP2;refGene_tx=XM_023243804.1;refGene_gn=RAMP2;refGene_tx=XM_023243805.1
E1 43278855 43279011
refGene_gn=RAMP2;refGene_tx=XM_023243804.1;refGene_gn=RAMP2;refGene_tx=XM_023243805.1
E1 43279467 43279576
refGene_gn=RAMP2;refGene_tx=XM_023243804.1;refGene_gn=RAMP2;refGene_tx=XM_023243805.1
E1 43279703 43279959
refGene_gn=RAMP2;refGene_tx=XM_023243804.1;refGene_gn=RAMP2;refGene_tx=XM_023243805.1
E1 43284372 43284425 refGene_gn=VPS25;refGene_tx=XM_003996902.3
E1 43284638 43284784 refGene_gn=VPS25;refGene_tx=XM_003996902.3
E1 43285705 43285759 refGene_gn=VPS25;refGene_tx=XM_003996902.3
E1 43286388 43286477 refGene_gn=VPS25;refGene_tx=XM_003996902.3
E1 43287252 43287328 refGene_gn=VPS25;refGene_tx=XM_003996902.3
E1 43288727 43288840 refGene_gn=VPS25;refGene_tx=XM_003996902.3
E1 43290501 43291116
refGene_gn=WNK4;refGene_tx=XM_003996903.5;refGene_gn=WNK4;refGene_tx=XM_006940374.4;refGene_gn=WNK4;refGene_tx=XM_006940375.4;refGene_gn=WNK4;refGene_tx=XM_006940376.4;refGene_gn=WNK4;refGene_tx=XM_006940377.4;refGene_gn=WNK4;refGene_tx=XM_019817921.2;refGene_gn=WNK4;refGene_tx=XM_019817922.2;refGene_gn=WNK4;refGene_tx=XM_019817924.2
E1 43292525 43292698
refGene_gn=WNK4;refGene_tx=XM_003996903.5;refGene_gn=WNK4;refGene_tx=XM_006940374.4;refGene_gn=WNK4;refGene_tx=XM_006940375.4;refGene_gn=WNK4;refGene_tx=XM_006940376.4;refGene_gn=WNK4;refGene_tx=XM_006940377.4;refGene_gn=WNK4;refGene_tx=XM_019817921.2;refGene_gn=WNK4;refGene_tx=XM_019817922.2;refGene_gn=WNK4;refGene_tx=XM_019817924.2
E1 43293690 43293911
refGene_gn=WNK4;refGene_tx=XM_003996903.5;refGene_gn=WNK4;refGene_tx=X

M_006940374.4;refGene_gn=WNK4;refGene_tx=XM_006940375.4;refGene_gn=WNK4;refGene_tx=XM_006940376.4;refGene_gn=WNK4;refGene_tx=XM_006940377.4;refGene_gn=WNK4;refGene_tx=XM_019817921.2;refGene_gn=WNK4;refGene_tx=XM_019817922.2;refGene_gn=WNK4;refGene_tx=XM_019817924.2
E1 43294238 43294396
refGene_gn=WNK4;refGene_tx=XM_003996903.5;refGene_gn=WNK4;refGene_tx=XM_006940374.4;refGene_gn=WNK4;refGene_tx=XM_006940375.4;refGene_gn=WNK4;refGene_tx=XM_006940376.4;refGene_gn=WNK4;refGene_tx=XM_006940377.4;refGene_gn=WNK4;refGene_tx=XM_019817921.2;refGene_gn=WNK4;refGene_tx=XM_019817922.2;refGene_gn=WNK4;refGene_tx=XM_019817924.2
E1 43294870 43294959
refGene_gn=WNK4;refGene_tx=XM_003996903.5;refGene_gn=WNK4;refGene_tx=XM_006940374.4;refGene_gn=WNK4;refGene_tx=XM_006940375.4;refGene_gn=WNK4;refGene_tx=XM_006940376.4;refGene_gn=WNK4;refGene_tx=XM_006940377.4;refGene_gn=WNK4;refGene_tx=XM_019817921.2;refGene_gn=WNK4;refGene_tx=XM_019817922.2;refGene_gn=WNK4;refGene_tx=XM_019817924.2
E1 43295055 43295272
refGene_gn=WNK4;refGene_tx=XM_003996903.5;refGene_gn=WNK4;refGene_tx=XM_006940374.4;refGene_gn=WNK4;refGene_tx=XM_006940375.4;refGene_gn=WNK4;refGene_tx=XM_006940376.4;refGene_gn=WNK4;refGene_tx=XM_006940377.4;refGene_gn=WNK4;refGene_tx=XM_019817921.2;refGene_gn=WNK4;refGene_tx=XM_019817922.2;refGene_gn=WNK4;refGene_tx=XM_019817924.2
E1 43297240 43297505
refGene_gn=WNK4;refGene_tx=XM_003996903.5;refGene_gn=WNK4;refGene_tx=XM_006940374.4;refGene_gn=WNK4;refGene_tx=XM_006940375.4;refGene_gn=WNK4;refGene_tx=XM_006940376.4;refGene_gn=WNK4;refGene_tx=XM_006940377.4;refGene_gn=WNK4;refGene_tx=XM_019817921.2;refGene_gn=WNK4;refGene_tx=XM_019817922.2;refGene_gn=WNK4;refGene_tx=XM_019817924.2
E1 43297750 43297872
refGene_gn=WNK4;refGene_tx=XM_003996903.5;refGene_gn=WNK4;refGene_tx=XM_006940374.4;refGene_gn=WNK4;refGene_tx=XM_006940375.4;refGene_gn=WNK4;refGene_tx=XM_006940376.4;refGene_gn=WNK4;refGene_tx=XM_006940377.4;refGene_gn=WNK4;refGene_tx=XM_019817921.2;refGene_gn=WNK4;refGene_tx=XM_019817922.2;refGene_gn=WNK4;refGene_tx=XM_019817924.2
E1 43298099 43298158
refGene_gn=WNK4;refGene_tx=XM_003996903.5;refGene_gn=WNK4;refGene_tx=XM_006940374.4;refGene_gn=WNK4;refGene_tx=XM_006940375.4;refGene_gn=WNK4;refGene_tx=XM_006940376.4;refGene_gn=WNK4;refGene_tx=XM_006940377.4;refGene_gn=WNK4;refGene_tx=XM_019817921.2;refGene_gn=WNK4;refGene_tx=XM_019817922.2;refGene_gn=WNK4;refGene_tx=XM_019817924.2
E1 43298249 43298367
refGene_gn=WNK4;refGene_tx=XM_003996903.5;refGene_gn=WNK4;refGene_tx=XM_006940374.4;refGene_gn=WNK4;refGene_tx=XM_006940375.4;refGene_gn=WNK4;refGene_tx=XM_006940376.4;refGene_gn=WNK4;refGene_tx=XM_006940377.4;refGene_gn=WNK4;refGene_tx=XM_019817921.2;refGene_gn=WNK4;refGene_tx=XM_019817922.2;refGene_gn=WNK4;refGene_tx=XM_019817924.2
E1 43298641 43298758
refGene_gn=WNK4;refGene_tx=XM_003996903.5;refGene_gn=WNK4;refGene_tx=XM_006940374.4;refGene_gn=WNK4;refGene_tx=XM_006940375.4;refGene_gn=WNK4;refGene_tx=XM_006940376.4;refGene_gn=WNK4;refGene_tx=XM_006940377.4;

refGene_gn=WNK4;refGene_tx=XM_019817921.2;refGene_gn=WNK4;refGene_tx=XM_019817922.2;refGene_gn=WNK4;refGene_tx=XM_019817924.2
E1 43302990 43303128
refGene_gn=WNK4;refGene_tx=XM_003996903.5;refGene_gn=WNK4;refGene_tx=XM_006940374.4;refGene_gn=WNK4;refGene_tx=XM_006940375.4;refGene_gn=WNK4;refGene_tx=XM_006940376.4;refGene_gn=WNK4;refGene_tx=XM_006940377.4;refGene_gn=WNK4;refGene_tx=XM_019817921.2;refGene_gn=WNK4;refGene_tx=XM_019817922.2;refGene_gn=WNK4;refGene_tx=XM_019817924.2
E1 43303709 43303764
refGene_gn=WNK4;refGene_tx=XM_003996903.5;refGene_gn=WNK4;refGene_tx=XM_006940374.4;refGene_gn=WNK4;refGene_tx=XM_006940375.4;refGene_gn=WNK4;refGene_tx=XM_006940376.4;refGene_gn=WNK4;refGene_tx=XM_006940377.4;refGene_gn=WNK4;refGene_tx=XM_019817921.2;refGene_gn=WNK4;refGene_tx=XM_019817922.2;refGene_gn=WNK4;refGene_tx=XM_019817924.2
E1 43303867 43304463
refGene_gn=WNK4;refGene_tx=XM_003996903.5;refGene_gn=WNK4;refGene_tx=XM_006940374.4;refGene_gn=WNK4;refGene_tx=XM_006940375.4;refGene_gn=WNK4;refGene_tx=XM_006940376.4;refGene_gn=WNK4;refGene_tx=XM_006940377.4;refGene_gn=WNK4;refGene_tx=XM_019817921.2;refGene_gn=WNK4;refGene_tx=XM_019817922.2;refGene_gn=WNK4;refGene_tx=XM_019817924.2
E1 43304542 43304603
refGene_gn=WNK4;refGene_tx=XM_003996903.5;refGene_gn=WNK4;refGene_tx=XM_006940374.4;refGene_gn=WNK4;refGene_tx=XM_006940375.4;refGene_gn=WNK4;refGene_tx=XM_006940376.4;refGene_gn=WNK4;refGene_tx=XM_006940377.4;refGene_gn=WNK4;refGene_tx=XM_019817921.2;refGene_gn=WNK4;refGene_tx=XM_019817922.2;refGene_gn=WNK4;refGene_tx=XM_019817924.2
E1 43304702 43305111
refGene_gn=WNK4;refGene_tx=XM_003996903.5;refGene_gn=WNK4;refGene_tx=XM_006940374.4;refGene_gn=WNK4;refGene_tx=XM_006940375.4;refGene_gn=WNK4;refGene_tx=XM_006940376.4;refGene_gn=WNK4;refGene_tx=XM_006940377.4;refGene_gn=WNK4;refGene_tx=XM_019817921.2;refGene_gn=WNK4;refGene_tx=XM_019817922.2;refGene_gn=WNK4;refGene_tx=XM_019817924.2
E1 43305194 43305394
refGene_gn=WNK4;refGene_tx=XM_003996903.5;refGene_gn=WNK4;refGene_tx=XM_006940374.4;refGene_gn=WNK4;refGene_tx=XM_006940375.4;refGene_gn=WNK4;refGene_tx=XM_006940376.4;refGene_gn=WNK4;refGene_tx=XM_006940377.4;refGene_gn=WNK4;refGene_tx=XM_019817921.2;refGene_gn=WNK4;refGene_tx=XM_019817922.2;refGene_gn=WNK4;refGene_tx=XM_019817924.2
E1 43305560 43305658
refGene_gn=WNK4;refGene_tx=XM_003996903.5;refGene_gn=WNK4;refGene_tx=XM_006940374.4;refGene_gn=WNK4;refGene_tx=XM_006940376.4;refGene_gn=WNK4;refGene_tx=XM_006940377.4;refGene_gn=WNK4;refGene_tx=XM_019817924.2
E1 43305765 43305875
refGene_gn=WNK4;refGene_tx=XM_003996903.5;refGene_gn=WNK4;refGene_tx=XM_006940374.4;refGene_gn=WNK4;refGene_tx=XM_006940375.4;refGene_gn=WNK4;refGene_tx=XM_006940376.4;refGene_gn=WNK4;refGene_tx=XM_006940377.4;refGene_gn=WNK4;refGene_tx=XM_019817921.2;refGene_gn=WNK4;refGene_tx=XM_019817922.2;refGene_gn=WNK4;refGene_tx=XM_019817924.2
E1 43307118 43307225
refGene_gn=LOC101100523;refGene_tx=XM_003996904.5

E1 43307570 43307784
refGene_gn=LOC101100523;refGene_tx=XM_003996904.5
E1 43308177 43308346 refGene_gn=CNTD1;refGene_tx=XM_003996905.5
E1 43311311 43311387 refGene_gn=CNTD1;refGene_tx=XM_003996905.5
E1 43311634 43311803
refGene_gn=CNTD1;refGene_tx=XM_003996905.5;refGene_gn=CNTD1;refGene_tx
=XM_019817926.1
E1 43312771 43312934
refGene_gn=CNTD1;refGene_tx=XM_003996905.5;refGene_gn=CNTD1;refGene_tx
=XM_019817926.1
E1 43313844 43313989
refGene_gn=CNTD1;refGene_tx=XM_003996905.5;refGene_gn=CNTD1;refGene_tx
=XM_019817926.1
E1 43314601 43314698
refGene_gn=CNTD1;refGene_tx=XM_003996905.5;refGene_gn=CNTD1;refGene_tx
=XM_019817926.1
E1 43315275 43315443
refGene_gn=CNTD1;refGene_tx=XM_003996905.5;refGene_gn=CNTD1;refGene_tx
=XM_019817926.1
E1 43316710 43316879
refGene_gn=BECN1;refGene_tx=XM_003996906.5;refGene_gn=BECN1;refGene_tx
=XM_019817925.2
E1 43317594 43317737
refGene_gn=BECN1;refGene_tx=XM_003996906.5;refGene_gn=BECN1;refGene_tx
=XM_019817925.2
E1 43318180 43318241
refGene_gn=BECN1;refGene_tx=XM_003996906.5;refGene_gn=BECN1;refGene_tx
=XM_019817925.2
E1 43319014 43319164
refGene_gn=BECN1;refGene_tx=XM_003996906.5;refGene_gn=BECN1;refGene_tx
=XM_019817925.2
E1 43321250 43321397
refGene_gn=BECN1;refGene_tx=XM_003996906.5;refGene_gn=BECN1;refGene_tx
=XM_019817925.2
E1 43322309 43322504
refGene_gn=BECN1;refGene_tx=XM_003996906.5;refGene_gn=BECN1;refGene_tx
=XM_019817925.2
E1 43322664 43322801
refGene_gn=BECN1;refGene_tx=XM_003996906.5;refGene_gn=BECN1;refGene_tx
=XM_019817925.2
E1 43322914 43323005
refGene_gn=BECN1;refGene_tx=XM_003996906.5;refGene_gn=BECN1;refGene_tx
=XM_019817925.2
E1 43323160 43323229 refGene_gn=BECN1;refGene_tx=XM_019817925.2
E1 43323679 43323738
refGene_gn=BECN1;refGene_tx=XM_003996906.5;refGene_gn=BECN1;refGene_tx
=XM_019817925.2
E1 43324884 43324952
refGene_gn=BECN1;refGene_tx=XM_003996906.5;refGene_gn=BECN1;refGene_tx
=XM_019817925.2

E1 43328589 43328716
refGene_gn=BECN1;refGene_tx=XM_003996906.5;refGene_gn=BECN1;refGene_tx
=XM_019817925.2
E1 43338018 43338060 refGene_gn=PSME3;refGene_tx=XM_003996908.5
E1 43338576 43338651 refGene_gn=PSME3;refGene_tx=XM_003996909.5
E1 43338755 43338788
refGene_gn=PSME3;refGene_tx=XM_003996908.5;refGene_gn=PSME3;refGene_tx
=XM_003996909.5
E1 43338906 43338969
refGene_gn=PSME3;refGene_tx=XM_003996908.5;refGene_gn=PSME3;refGene_tx
=XM_003996909.5
E1 43339167 43339272
refGene_gn=PSME3;refGene_tx=XM_003996908.5;refGene_gn=PSME3;refGene_tx
=XM_003996909.5
E1 43340171 43340220
refGene_gn=PSME3;refGene_tx=XM_003996908.5;refGene_gn=PSME3;refGene_tx
=XM_003996909.5
E1 43340761 43340874
refGene_gn=PSME3;refGene_tx=XM_003996908.5;refGene_gn=PSME3;refGene_tx
=XM_003996909.5
E1 43341140 43341209
refGene_gn=PSME3;refGene_tx=XM_003996908.5;refGene_gn=PSME3;refGene_tx
=XM_003996909.5
E1 43341349 43341417
refGene_gn=PSME3;refGene_tx=XM_003996908.5;refGene_gn=PSME3;refGene_tx
=XM_003996909.5
E1 43341516 43341571
refGene_gn=PSME3;refGene_tx=XM_003996908.5;refGene_gn=PSME3;refGene_tx
=XM_003996909.5
E1 43341688 43341775
refGene_gn=PSME3;refGene_tx=XM_003996908.5;refGene_gn=PSME3;refGene_tx
=XM_003996909.5
E1 43342653 43342734
refGene_gn=PSME3;refGene_tx=XM_003996908.5;refGene_gn=PSME3;refGene_tx
=XM_003996909.5
E1 43345816 43347404
refGene_gn=LOC101080592;refGene_tx=XM_003996911.4;refGene_gn=LOC101080
592;refGene_tx=XM_023244639.1
E1 43349246 43349532
refGene_gn=LOC101080592;refGene_tx=XM_003996911.4
E1 43349766 43349896
refGene_gn=LOC101080592;refGene_tx=XM_003996911.4
E1 43350220 43350487
refGene_gn=LOC101080592;refGene_tx=XM_003996911.4
E1 43351622 43353222
refGene_gn=LOC101080592;refGene_tx=XM_011289125.3;refGene_gn=LOC101080
592;refGene_tx=XM_023244639.1
E1 43354693 43354979
refGene_gn=LOC101080592;refGene_tx=XM_011289125.3;refGene_gn=LOC101080
592;refGene_tx=XM_023244639.1

E1 43355376 43355506
refGene_gn=LOC101080592;refGene_tx=XM_011289125.3;refGene_gn=LOC101080592;refGene_tx=XM_023244639.1
E1 43356229 43356505
refGene_gn=LOC101080592;refGene_tx=XM_011289125.3;refGene_gn=LOC101080592;refGene_tx=XM_023244639.1
E1 43365719 43367310
refGene_gn=LOC101093157;refGene_tx=XM_003996955.5
E1 43371750 43371791
refGene_gn=LOC101093157;refGene_tx=XM_003996955.5
E1 43371792 43372037
refGene_gn=LOC101093157;refGene_tx=XM_003996955.5
E1 43372735 43372865
refGene_gn=LOC101093157;refGene_tx=XM_003996955.5
E1 43373108 43373384
refGene_gn=LOC101093157;refGene_tx=XM_003996955.5
E1 43389755 43389985
refGene_gn=G6PC;refGene_tx=NM_001042378.1;refGene_gn=G6PC;refGene_tx=XM_019816888.1
E1 43392338 43392448
refGene_gn=G6PC;refGene_tx=NM_001042378.1;refGene_gn=G6PC;refGene_tx=XM_019816888.1
E1 43393628 43393734
refGene_gn=G6PC;refGene_tx=NM_001042378.1;refGene_gn=G6PC;refGene_tx=XM_019816888.1
E1 43394377 43394493
refGene_gn=G6PC;refGene_tx=NM_001042378.1;refGene_gn=G6PC;refGene_tx=XM_019816888.1
E1 43396711 43397223
refGene_gn=G6PC;refGene_tx=NM_001042378.1;refGene_gn=G6PC;refGene_tx=XM_019816888.1
E1 43430346 43430482
refGene_gn=LOC101081013;refGene_tx=XM_003996914.4
E1 43432122 43432217
refGene_gn=LOC101081013;refGene_tx=XM_003996914.4
E1 43433225 43433280
refGene_gn=LOC101081013;refGene_tx=XM_003996914.4
E1 43433738 43433830
refGene_gn=LOC101081013;refGene_tx=XM_003996914.4
E1 43433972 43434039
refGene_gn=LOC101081013;refGene_tx=XM_003996914.4
E1 43434617 43434748
refGene_gn=LOC101081013;refGene_tx=XM_003996914.4
E1 43434971 43435088
refGene_gn=LOC101081013;refGene_tx=XM_003996914.4
E1 43435183 43435340
refGene_gn=LOC101081013;refGene_tx=XM_003996914.4
E1 43435765 43435823
refGene_gn=LOC101081013;refGene_tx=XM_003996914.4
E1 43436750 43436910

refGene_gn=LOC101081013;refGene_tx=XM_003996914.4
E1 43438924 43439056
refGene_gn=LOC101081013;refGene_tx=XM_003996914.4
E1 43439205 43439244
refGene_gn=LOC101081013;refGene_tx=XM_003996914.4
E1 43442253 43442261
refGene_gn=PTGES3L;refGene_tx=XM_006940380.4;refGene_gn=PTGES3L;refGene_tx=XM_019817929.2
E1 43442428 43442464
refGene_gn=PTGES3L;refGene_tx=XM_006940380.4;refGene_gn=PTGES3L;refGene_tx=XM_006940381.4;refGene_gn=PTGES3L;refGene_tx=XM_019817929.2;refGene_gn=PTGES3L;refGene_tx=XM_019817930.2
E1 43442968 43443022
refGene_gn=PTGES3L;refGene_tx=XM_006940380.4;refGene_gn=PTGES3L;refGene_tx=XM_006940381.4;refGene_gn=PTGES3L;refGene_tx=XM_019817929.2;refGene_gn=PTGES3L;refGene_tx=XM_019817930.2
E1 43443704 43443794
refGene_gn=PTGES3L;refGene_tx=XM_006940380.4;refGene_gn=PTGES3L;refGene_tx=XM_006940381.4;refGene_gn=PTGES3L;refGene_tx=XM_019817929.2;refGene_gn=PTGES3L;refGene_tx=XM_019817930.2
E1 43445800 43445899
refGene_gn=PTGES3L;refGene_tx=XM_006940380.4;refGene_gn=PTGES3L;refGene_tx=XM_006940381.4;refGene_gn=PTGES3L;refGene_tx=XM_019817929.2;refGene_gn=PTGES3L;refGene_tx=XM_019817930.2
E1 43446004 43446071
refGene_gn=PTGES3L;refGene_tx=XM_006940380.4;refGene_gn=PTGES3L;refGene_tx=XM_006940381.4;refGene_gn=PTGES3L;refGene_tx=XM_019817929.2;refGene_gn=PTGES3L;refGene_tx=XM_019817930.2
E1 43446177 43446291
refGene_gn=PTGES3L;refGene_tx=XM_006940380.4;refGene_gn=PTGES3L;refGene_tx=XM_006940381.4
E1 43446669 43446854
refGene_gn=PTGES3L;refGene_tx=XM_006940380.4;refGene_gn=PTGES3L;refGene_tx=XM_006940381.4;refGene_gn=PTGES3L;refGene_tx=XM_019817929.2;refGene_gn=PTGES3L;refGene_tx=XM_019817930.2
E1 43447211 43447709 refGene_gn=RUNDC1;refGene_tx=XM_023243431.1
E1 43451360 43451519 refGene_gn=RUNDC1;refGene_tx=XM_023243431.1
E1 43452749 43452948 refGene_gn=RUNDC1;refGene_tx=XM_023243431.1
E1 43453483 43453603 refGene_gn=RUNDC1;refGene_tx=XM_023243431.1
E1 43453993 43454859 refGene_gn=RUNDC1;refGene_tx=XM_023243431.1
E1 43461006 43461087 refGene_gn=RPL27;refGene_tx=XM_006940382.4
E1 43462023 43462193 refGene_gn=RPL27;refGene_tx=XM_006940382.4
E1 43463443 43463554 refGene_gn=RPL27;refGene_tx=XM_006940382.4
E1 43463657 43463706 refGene_gn=RPL27;refGene_tx=XM_006940382.4
E1 43465720 43465741
refGene_gn=IFI35;refGene_tx=XM_006940383.3;refGene_gn=IFI35;refGene_tx=XM_019817931.1
E1 43474298 43474400
refGene_gn=IFI35;refGene_tx=XM_006940383.3;refGene_gn=IFI35;refGene_tx=XM_019817931.1

E1	43474746	43474894
refGene_gn=IFI35;refGene_tx=XM_006940383.3;refGene_gn=IFI35;refGene_tx
=XM_019817931.1
E1	43474966	43475073
refGene_gn=IFI35;refGene_tx=XM_006940383.3;refGene_gn=IFI35;refGene_tx
=XM_019817931.1
E1	43475171	43475358
refGene_gn=IFI35;refGene_tx=XM_006940383.3;refGene_gn=IFI35;refGene_tx
=XM_019817931.1
E1	43475453	43475560
refGene_gn=IFI35;refGene_tx=XM_006940383.3;refGene_gn=IFI35;refGene_tx
=XM_019817931.1
E1	43475773	43475965	refGene_gn=IFI35;refGene_tx=XM_006940383.3
E1	43476145	43476154	refGene_gn=IFI35;refGene_tx=XM_019817931.1
E1	43477716	43477800	refGene_gn=VAT1;refGene_tx=XM_003996916.5
E1	43477950	43478192	refGene_gn=VAT1;refGene_tx=XM_003996916.5
E1	43478958	43479048	refGene_gn=VAT1;refGene_tx=XM_003996916.5
E1	43479159	43479330	refGene_gn=VAT1;refGene_tx=XM_003996916.5
E1	43479734	43479942	refGene_gn=VAT1;refGene_tx=XM_003996916.5
E1	43482910	43483318	refGene_gn=VAT1;refGene_tx=XM_003996916.5
E1	43484878	43484980	refGene_gn=RND2;refGene_tx=XM_023243773.1
E1	43485465	43485553	refGene_gn=RND2;refGene_tx=XM_023243773.1
E1	43486357	43486467	refGene_gn=RND2;refGene_tx=XM_023243773.1
E1	43487238	43487373	refGene_gn=RND2;refGene_tx=XM_023243773.1
E1	43487618	43487873	refGene_gn=RND2;refGene_tx=XM_023243773.1
E1	43502325	43502471
refGene_gn=BRCA1;refGene_tx=XM_019817933.2;refGene_gn=BRCA1;refGene_tx
=XM_019817934.1;refGene_gn=BRCA1;refGene_tx=XM_019817935.2;refGene_gn=
BRCA1;refGene_tx=XM_019817936.2;refGene_gn=BRCA1;refGene_tx=XM_0198179
37.2;refGene_gn=BRCA1;refGene_tx=XM_019817938.2;refGene_gn=BRCA1;refGe
ne_tx=XM_019817939.2
E1	43503330	43503391
refGene_gn=BRCA1;refGene_tx=XM_019817933.2;refGene_gn=BRCA1;refGene_tx
=XM_019817934.1;refGene_gn=BRCA1;refGene_tx=XM_019817935.2;refGene_gn=
BRCA1;refGene_tx=XM_019817936.2;refGene_gn=BRCA1;refGene_tx=XM_0198179
37.2;refGene_gn=BRCA1;refGene_tx=XM_019817938.2;refGene_gn=BRCA1;refGe
ne_tx=XM_019817939.2
E1	43503756	43503830
refGene_gn=BRCA1;refGene_tx=XM_019817933.2;refGene_gn=BRCA1;refGene_tx
=XM_019817934.1;refGene_gn=BRCA1;refGene_tx=XM_019817935.2;refGene_gn=
BRCA1;refGene_tx=XM_019817936.2;refGene_gn=BRCA1;refGene_tx=XM_0198179
37.2;refGene_gn=BRCA1;refGene_tx=XM_019817938.2;refGene_gn=BRCA1;refGe
ne_tx=XM_019817939.2
E1	43505656	43505711
refGene_gn=BRCA1;refGene_tx=XM_019817933.2;refGene_gn=BRCA1;refGene_tx
=XM_019817934.1;refGene_gn=BRCA1;refGene_tx=XM_019817935.2;refGene_gn=
BRCA1;refGene_tx=XM_019817936.2;refGene_gn=BRCA1;refGene_tx=XM_0198179
37.2;refGene_gn=BRCA1;refGene_tx=XM_019817938.2;refGene_gn=BRCA1;refGe
ne_tx=XM_019817939.2
E1	43509725	43509809

refGene_gn=BRCA1;refGene_tx=XM_019817933.2;refGene_gn=BRCA1;refGene_tx=XM_019817934.1;refGene_gn=BRCA1;refGene_tx=XM_019817935.2;refGene_gn=BRCA1;refGene_tx=XM_019817936.2;refGene_gn=BRCA1;refGene_tx=XM_019817937.2;refGene_gn=BRCA1;refGene_tx=XM_019817938.2;refGene_gn=BRCA1;refGene_tx=XM_019817939.2
E1 43512538 43512579
refGene_gn=BRCA1;refGene_tx=XM_019817933.2;refGene_gn=BRCA1;refGene_tx=XM_019817934.1;refGene_gn=BRCA1;refGene_tx=XM_019817935.2;refGene_gn=BRCA1;refGene_tx=XM_019817936.2;refGene_gn=BRCA1;refGene_tx=XM_019817937.2;refGene_gn=BRCA1;refGene_tx=XM_019817938.2;refGene_gn=BRCA1;refGene_tx=XM_019817939.2
E1 43513071 43513149
refGene_gn=BRCA1;refGene_tx=XM_019817933.2;refGene_gn=BRCA1;refGene_tx=XM_019817934.1;refGene_gn=BRCA1;refGene_tx=XM_019817935.2;refGene_gn=BRCA1;refGene_tx=XM_019817936.2;refGene_gn=BRCA1;refGene_tx=XM_019817937.2;refGene_gn=BRCA1;refGene_tx=XM_019817938.2;refGene_gn=BRCA1;refGene_tx=XM_019817939.2
E1 43515319 43515407
refGene_gn=BRCA1;refGene_tx=XM_019817933.2;refGene_gn=BRCA1;refGene_tx=XM_019817934.1;refGene_gn=BRCA1;refGene_tx=XM_019817935.2;refGene_gn=BRCA1;refGene_tx=XM_019817936.2;refGene_gn=BRCA1;refGene_tx=XM_019817937.2;refGene_gn=BRCA1;refGene_tx=XM_019817938.2;refGene_gn=BRCA1;refGene_tx=XM_019817939.2
E1 43517907 43518218
refGene_gn=BRCA1;refGene_tx=XM_019817933.2;refGene_gn=BRCA1;refGene_tx=XM_019817934.1;refGene_gn=BRCA1;refGene_tx=XM_019817935.2;refGene_gn=BRCA1;refGene_tx=XM_019817936.2;refGene_gn=BRCA1;refGene_tx=XM_019817937.2;refGene_gn=BRCA1;refGene_tx=XM_019817938.2;refGene_gn=BRCA1;refGene_tx=XM_019817939.2
E1 43519661 43519852
refGene_gn=BRCA1;refGene_tx=XM_019817933.2;refGene_gn=BRCA1;refGene_tx=XM_019817934.1;refGene_gn=BRCA1;refGene_tx=XM_019817935.2;refGene_gn=BRCA1;refGene_tx=XM_019817936.2;refGene_gn=BRCA1;refGene_tx=XM_019817937.2;refGene_gn=BRCA1;refGene_tx=XM_019817938.2;refGene_gn=BRCA1;refGene_tx=XM_019817939.2
E1 43522050 43522177
refGene_gn=BRCA1;refGene_tx=XM_019817933.2;refGene_gn=BRCA1;refGene_tx=XM_019817934.1;refGene_gn=BRCA1;refGene_tx=XM_019817935.2;refGene_gn=BRCA1;refGene_tx=XM_019817936.2;refGene_gn=BRCA1;refGene_tx=XM_019817937.2;refGene_gn=BRCA1;refGene_tx=XM_019817938.2;refGene_gn=BRCA1;refGene_tx=XM_019817939.2
E1 43527227 43527399
refGene_gn=BRCA1;refGene_tx=XM_019817933.2;refGene_gn=BRCA1;refGene_tx=XM_019817934.1;refGene_gn=BRCA1;refGene_tx=XM_019817935.2;refGene_gn=BRCA1;refGene_tx=XM_019817936.2;refGene_gn=BRCA1;refGene_tx=XM_019817937.2;refGene_gn=BRCA1;refGene_tx=XM_019817938.2;refGene_gn=BRCA1;refGene_tx=XM_019817939.2
E1 43533275 43533364
refGene_gn=BRCA1;refGene_tx=XM_019817933.2;refGene_gn=BRCA1;refGene_tx=XM_019817934.1;refGene_gn=BRCA1;refGene_tx=XM_019817935.2;refGene_gn=

BRCA1;refGene_tx=XM_019817936.2;refGene_gn=BRCA1;refGene_tx=XM_0198179
37.2;refGene_gn=BRCA1;refGene_tx=XM_019817938.2;refGene_gn=BRCA1;refGe
ne_tx=XM_019817939.2
E1 43533761 43537196
refGene_gn=BRCA1;refGene_tx=XM_019817933.2;refGene_gn=BRCA1;refGene_tx
=XM_019817934.1;refGene_gn=BRCA1;refGene_tx=XM_019817935.2;refGene_gn=
BRCA1;refGene_tx=XM_019817936.2;refGene_gn=BRCA1;refGene_tx=XM_0198179
37.2;refGene_gn=BRCA1;refGene_tx=XM_019817938.2;refGene_gn=BRCA1;refGe
ne_tx=XM_019817939.2
E1 43537810 43537887
refGene_gn=BRCA1;refGene_tx=XM_019817933.2;refGene_gn=BRCA1;refGene_tx
=XM_019817934.1;refGene_gn=BRCA1;refGene_tx=XM_019817935.2;refGene_gn=
BRCA1;refGene_tx=XM_019817936.2;refGene_gn=BRCA1;refGene_tx=XM_0198179
37.2;refGene_gn=BRCA1;refGene_tx=XM_019817938.2;refGene_gn=BRCA1;refGe
ne_tx=XM_019817939.2
E1 43539309 43539355
refGene_gn=BRCA1;refGene_tx=XM_019817933.2;refGene_gn=BRCA1;refGene_tx
=XM_019817934.1;refGene_gn=BRCA1;refGene_tx=XM_019817935.2;refGene_gn=
BRCA1;refGene_tx=XM_019817936.2;refGene_gn=BRCA1;refGene_tx=XM_0198179
37.2;refGene_gn=BRCA1;refGene_tx=XM_019817938.2;refGene_gn=BRCA1;refGe
ne_tx=XM_019817939.2
E1 43540327 43540433
refGene_gn=BRCA1;refGene_tx=XM_019817933.2;refGene_gn=BRCA1;refGene_tx
=XM_019817934.1;refGene_gn=BRCA1;refGene_tx=XM_019817935.2;refGene_gn=
BRCA1;refGene_tx=XM_019817936.2;refGene_gn=BRCA1;refGene_tx=XM_0198179
37.2;refGene_gn=BRCA1;refGene_tx=XM_019817938.2;refGene_gn=BRCA1;refGe
ne_tx=XM_019817939.2
E1 43541615 43541755
refGene_gn=BRCA1;refGene_tx=XM_019817933.2;refGene_gn=BRCA1;refGene_tx
=XM_019817934.1;refGene_gn=BRCA1;refGene_tx=XM_019817935.2;refGene_gn=
BRCA1;refGene_tx=XM_019817936.2;refGene_gn=BRCA1;refGene_tx=XM_0198179
37.2;refGene_gn=BRCA1;refGene_tx=XM_019817938.2;refGene_gn=BRCA1;refGe
ne_tx=XM_019817939.2
E1 43542324 43542413
refGene_gn=BRCA1;refGene_tx=XM_019817933.2;refGene_gn=BRCA1;refGene_tx
=XM_019817934.1;refGene_gn=BRCA1;refGene_tx=XM_019817935.2;refGene_gn=
BRCA1;refGene_tx=XM_019817936.2;refGene_gn=BRCA1;refGene_tx=XM_0198179
37.2;refGene_gn=BRCA1;refGene_tx=XM_019817938.2;refGene_gn=BRCA1;refGe
ne_tx=XM_019817939.2
E1 43543403 43543481
refGene_gn=BRCA1;refGene_tx=XM_019817933.2;refGene_gn=BRCA1;refGene_tx
=XM_019817934.1;refGene_gn=BRCA1;refGene_tx=XM_019817935.2;refGene_gn=
BRCA1;refGene_tx=XM_019817936.2;refGene_gn=BRCA1;refGene_tx=XM_0198179
37.2;refGene_gn=BRCA1;refGene_tx=XM_019817938.2;refGene_gn=BRCA1;refGe
ne_tx=XM_019817939.2
E1 43551263 43551317
refGene_gn=BRCA1;refGene_tx=XM_019817933.2;refGene_gn=BRCA1;refGene_tx
=XM_019817934.1;refGene_gn=BRCA1;refGene_tx=XM_019817935.2;refGene_gn=
BRCA1;refGene_tx=XM_019817936.2;refGene_gn=BRCA1;refGene_tx=XM_0198179
37.2;refGene_gn=BRCA1;refGene_tx=XM_019817938.2;refGene_gn=BRCA1;refGe

ne_tx=XM_019817939.2
E1 43558756 43558836
refGene_gn=BRCA1;refGene_tx=XM_019817933.2;refGene_gn=BRCA1;refGene_tx=XM_019817934.1;refGene_gn=BRCA1;refGene_tx=XM_019817935.2;refGene_gn=BRCA1;refGene_tx=XM_019817936.2;refGene_gn=BRCA1;refGene_tx=XM_019817937.2;refGene_gn=BRCA1;refGene_tx=XM_019817938.2;refGene_gn=BRCA1;refGene_tx=XM_019817939.2
E1 43564376 43564478
refGene_gn=NBR1;refGene_tx=XM_003996920.4;refGene_gn=NBR1;refGene_tx=XM_019817941.1;refGene_gn=NBR1;refGene_tx=XM_019817942.2;refGene_gn=NBR1;refGene_tx=XM_019817943.2;refGene_gn=NBR1;refGene_tx=XM_019817945.1;refGene_gn=NBR1;refGene_tx=XM_019817946.1
E1 43568748 43568811
refGene_gn=NBR1;refGene_tx=XM_003996920.4;refGene_gn=NBR1;refGene_tx=XM_019817941.1;refGene_gn=NBR1;refGene_tx=XM_019817942.2;refGene_gn=NBR1;refGene_tx=XM_019817943.2;refGene_gn=NBR1;refGene_tx=XM_019817945.1;refGene_gn=NBR1;refGene_tx=XM_019817946.1
E1 43569518 43569537
refGene_gn=NBR1;refGene_tx=XM_003996920.4;refGene_gn=NBR1;refGene_tx=XM_019817941.1;refGene_gn=NBR1;refGene_tx=XM_019817942.2;refGene_gn=NBR1;refGene_tx=XM_019817943.2;refGene_gn=NBR1;refGene_tx=XM_019817945.1;refGene_gn=NBR1;refGene_tx=XM_019817946.1
E1 43571489 43571512
refGene_gn=NBR1;refGene_tx=XM_003996920.4;refGene_gn=NBR1;refGene_tx=XM_019817941.1;refGene_gn=NBR1;refGene_tx=XM_019817942.2;refGene_gn=NBR1;refGene_tx=XM_019817943.2;refGene_gn=NBR1;refGene_tx=XM_019817945.1;refGene_gn=NBR1;refGene_tx=XM_019817946.1
E1 43574360 43574555
refGene_gn=NBR1;refGene_tx=XM_003996920.4;refGene_gn=NBR1;refGene_tx=XM_019817941.1;refGene_gn=NBR1;refGene_tx=XM_019817942.2;refGene_gn=NBR1;refGene_tx=XM_019817943.2;refGene_gn=NBR1;refGene_tx=XM_019817944.2;refGene_gn=NBR1;refGene_tx=XM_019817945.1;refGene_gn=NBR1;refGene_tx=XM_019817946.1
E1 43575098 43575176
refGene_gn=NBR1;refGene_tx=XM_003996920.4;refGene_gn=NBR1;refGene_tx=XM_019817941.1;refGene_gn=NBR1;refGene_tx=XM_019817942.2;refGene_gn=NBR1;refGene_tx=XM_019817943.2;refGene_gn=NBR1;refGene_tx=XM_019817944.2;refGene_gn=NBR1;refGene_tx=XM_019817945.1;refGene_gn=NBR1;refGene_tx=XM_019817946.1
E1 43575608 43575823
refGene_gn=NBR1;refGene_tx=XM_003996920.4;refGene_gn=NBR1;refGene_tx=XM_019817941.1;refGene_gn=NBR1;refGene_tx=XM_019817942.2;refGene_gn=NBR1;refGene_tx=XM_019817943.2;refGene_gn=NBR1;refGene_tx=XM_019817944.2;refGene_gn=NBR1;refGene_tx=XM_019817945.1;refGene_gn=NBR1;refGene_tx=XM_019817946.1
E1 43576359 43576527
refGene_gn=NBR1;refGene_tx=XM_003996920.4;refGene_gn=NBR1;refGene_tx=XM_019817941.1;refGene_gn=NBR1;refGene_tx=XM_019817942.2;refGene_gn=NBR1;refGene_tx=XM_019817943.2;refGene_gn=NBR1;refGene_tx=XM_019817944.2;refGene_gn=NBR1;refGene_tx=XM_019817945.1;refGene_gn=NBR1;refGene_tx=X

M_019817946.1
E1 43576797 43577007
refGene_gn=NBR1;refGene_tx=XM_003996920.4;refGene_gn=NBR1;refGene_tx=XM_019817941.1;refGene_gn=NBR1;refGene_tx=XM_019817942.2;refGene_gn=NBR1;refGene_tx=XM_019817943.2;refGene_gn=NBR1;refGene_tx=XM_019817944.2;refGene_gn=NBR1;refGene_tx=XM_019817945.1;refGene_gn=NBR1;refGene_tx=XM_019817946.1
E1 43578141 43578301
refGene_gn=NBR1;refGene_tx=XM_003996920.4;refGene_gn=NBR1;refGene_tx=XM_019817941.1;refGene_gn=NBR1;refGene_tx=XM_019817942.2;refGene_gn=NBR1;refGene_tx=XM_019817943.2;refGene_gn=NBR1;refGene_tx=XM_019817944.2;refGene_gn=NBR1;refGene_tx=XM_019817945.1;refGene_gn=NBR1;refGene_tx=XM_019817946.1
E1 43578395 43578686
refGene_gn=NBR1;refGene_tx=XM_003996920.4;refGene_gn=NBR1;refGene_tx=XM_019817941.1;refGene_gn=NBR1;refGene_tx=XM_019817942.2;refGene_gn=NBR1;refGene_tx=XM_019817943.2;refGene_gn=NBR1;refGene_tx=XM_019817944.2;refGene_gn=NBR1;refGene_tx=XM_019817945.1;refGene_gn=NBR1;refGene_tx=XM_019817946.1
E1 43579359 43579509
refGene_gn=NBR1;refGene_tx=XM_003996920.4;refGene_gn=NBR1;refGene_tx=XM_019817941.1;refGene_gn=NBR1;refGene_tx=XM_019817942.2;refGene_gn=NBR1;refGene_tx=XM_019817943.2;refGene_gn=NBR1;refGene_tx=XM_019817944.2;refGene_gn=NBR1;refGene_tx=XM_019817945.1;refGene_gn=NBR1;refGene_tx=XM_019817946.1
E1 43579991 43580067
refGene_gn=NBR1;refGene_tx=XM_003996920.4;refGene_gn=NBR1;refGene_tx=XM_019817941.1;refGene_gn=NBR1;refGene_tx=XM_019817942.2;refGene_gn=NBR1;refGene_tx=XM_019817943.2;refGene_gn=NBR1;refGene_tx=XM_019817944.2;refGene_gn=NBR1;refGene_tx=XM_019817945.1;refGene_gn=NBR1;refGene_tx=XM_019817946.1
E1 43580713 43580824
refGene_gn=NBR1;refGene_tx=XM_003996920.4;refGene_gn=NBR1;refGene_tx=XM_019817941.1;refGene_gn=NBR1;refGene_tx=XM_019817942.2;refGene_gn=NBR1;refGene_tx=XM_019817943.2;refGene_gn=NBR1;refGene_tx=XM_019817944.2;refGene_gn=NBR1;refGene_tx=XM_019817945.1;refGene_gn=NBR1;refGene_tx=XM_019817946.1
E1 43581012 43581177
refGene_gn=NBR1;refGene_tx=XM_003996920.4;refGene_gn=NBR1;refGene_tx=XM_019817941.1;refGene_gn=NBR1;refGene_tx=XM_019817942.2;refGene_gn=NBR1;refGene_tx=XM_019817943.2;refGene_gn=NBR1;refGene_tx=XM_019817944.2;refGene_gn=NBR1;refGene_tx=XM_019817945.1;refGene_gn=NBR1;refGene_tx=XM_019817946.1
E1 43582559 43583004
refGene_gn=NBR1;refGene_tx=XM_003996920.4;refGene_gn=NBR1;refGene_tx=XM_019817941.1;refGene_gn=NBR1;refGene_tx=XM_019817942.2;refGene_gn=NBR1;refGene_tx=XM_019817943.2;refGene_gn=NBR1;refGene_tx=XM_019817944.2;refGene_gn=NBR1;refGene_tx=XM_019817945.1;refGene_gn=NBR1;refGene_tx=XM_019817946.1
E1 43583756 43583851

refGene_gn=NBR1;refGene_tx=XM_003996920.4;refGene_gn=NBR1;refGene_tx=XM_019817941.1;refGene_gn=NBR1;refGene_tx=XM_019817942.2;refGene_gn=NBR1;refGene_tx=XM_019817943.2;refGene_gn=NBR1;refGene_tx=XM_019817944.2;refGene_gn=NBR1;refGene_tx=XM_019817945.1;refGene_gn=NBR1;refGene_tx=XM_019817946.1
E1 43584530 43584607 refGene_gn=NBR1;refGene_tx=XM_003996920.4;refGene_gn=NBR1;refGene_tx=XM_019817941.1;refGene_gn=NBR1;refGene_tx=XM_019817942.2;refGene_gn=NBR1;refGene_tx=XM_019817943.2;refGene_gn=NBR1;refGene_tx=XM_019817944.2;refGene_gn=NBR1;refGene_tx=XM_019817945.1;refGene_gn=NBR1;refGene_tx=XM_019817946.1
E1 43585315 43585424 refGene_gn=NBR1;refGene_tx=XM_003996920.4;refGene_gn=NBR1;refGene_tx=XM_019817941.1;refGene_gn=NBR1;refGene_tx=XM_019817942.2;refGene_gn=NBR1;refGene_tx=XM_019817943.2;refGene_gn=NBR1;refGene_tx=XM_019817944.2
E1 43588099 43588273 refGene_gn=NBR1;refGene_tx=XM_003996920.4;refGene_gn=NBR1;refGene_tx=XM_019817941.1;refGene_gn=NBR1;refGene_tx=XM_019817942.2;refGene_gn=NBR1;refGene_tx=XM_019817943.2;refGene_gn=NBR1;refGene_tx=XM_019817944.2
E1 43591347 43591558 refGene_gn=TMEM106A;refGene_tx=XM_003996959.5
E1 43592077 43592141 refGene_gn=TMEM106A;refGene_tx=XM_003996959.5
E1 43593178 43593332 refGene_gn=TMEM106A;refGene_tx=XM_003996959.5
E1 43593830 43593971 refGene_gn=TMEM106A;refGene_tx=XM_003996959.5
E1 43594075 43594119 refGene_gn=TMEM106A;refGene_tx=XM_003996959.5
E1 43594632 43594686 refGene_gn=TMEM106A;refGene_tx=XM_003996959.5
E1 43595019 43595140 refGene_gn=TMEM106A;refGene_tx=XM_003996959.5
E1 43597866 43597893 refGene_gn=LOC102901590;refGene_tx=XM_006940391.4;refGene_gn=LOC102901590;refGene_tx=XM_019817948.2
E1 43598627 43598686 refGene_gn=LOC102901590;refGene_tx=XM_006940391.4;refGene_gn=LOC102901590;refGene_tx=XM_019817948.2
E1 43599162 43599517 refGene_gn=LOC102901590;refGene_tx=XM_006940391.4;refGene_gn=LOC102901590;refGene_tx=XM_019817948.2
E1 43602060 43602183 refGene_gn=LOC102901590;refGene_tx=XM_006940391.4;refGene_gn=LOC102901590;refGene_tx=XM_019817948.2
E1 43661424 43662030 refGene_gn=ARL4D;refGene_tx=XM_006940407.3
E1 43723528 43723676 refGene_gn=DHX8;refGene_tx=XM_003996961.4
E1 43728218 43728304 refGene_gn=DHX8;refGene_tx=XM_003996961.4
E1 43728853 43728926 refGene_gn=DHX8;refGene_tx=XM_003996961.4
E1 43729311 43729397 refGene_gn=DHX8;refGene_tx=XM_003996961.4
E1 43731056 43731166 refGene_gn=DHX8;refGene_tx=XM_003996961.4
E1 43731546 43731912 refGene_gn=DHX8;refGene_tx=XM_003996961.4
E1 43732567 43732712 refGene_gn=DHX8;refGene_tx=XM_003996961.4
E1 43733055 43733259 refGene_gn=DHX8;refGene_tx=XM_003996961.4
E1 43733910 43733998 refGene_gn=DHX8;refGene_tx=XM_003996961.4
E1 43736193 43736291 refGene_gn=DHX8;refGene_tx=XM_003996961.4
E1 43737221 43737369 refGene_gn=DHX8;refGene_tx=XM_003996961.4

E1 43738630 43738812 refGene_gn=DHX8;refGene_tx=XM_003996961.4
E1 43740578 43740773 refGene_gn=DHX8;refGene_tx=XM_003996961.4
E1 43741077 43741263 refGene_gn=DHX8;refGene_tx=XM_003996961.4
E1 43741389 43741600 refGene_gn=DHX8;refGene_tx=XM_003996961.4
E1 43741950 43742132 refGene_gn=DHX8;refGene_tx=XM_003996961.4
E1 43744220 43744361 refGene_gn=DHX8;refGene_tx=XM_003996961.4
E1 43747628 43747784 refGene_gn=DHX8;refGene_tx=XM_003996961.4
E1 43750962 43750980 refGene_gn=DHX8;refGene_tx=XM_003996961.4
E1 43750981 43751101 refGene_gn=DHX8;refGene_tx=XM_003996961.4
E1 43751850 43751979 refGene_gn=DHX8;refGene_tx=XM_003996961.4
E1 43752159 43752356 refGene_gn=DHX8;refGene_tx=XM_003996961.4
E1 43752815 43752995 refGene_gn=DHX8;refGene_tx=XM_003996961.4
E1 43753728 43753948 refGene_gn=DHX8;refGene_tx=XM_003996961.4
E1 43758581 43758806
refGene_gn=ETV4;refGene_tx=XM_006940408.4;refGene_gn=ETV4;refGene_tx=XM_011289264.2;refGene_gn=ETV4;refGene_tx=XM_019817949.2;refGene_gn=ETV4;refGene_tx=XM_019817950.2;refGene_gn=ETV4;refGene_tx=XM_019817951.2;refGene_gn=ETV4;refGene_tx=XM_019817953.2;refGene_gn=ETV4;refGene_tx=XM_019817954.2;refGene_gn=ETV4;refGene_tx=XM_019817955.2;refGene_gn=ETV4;refGene_tx=XM_019817956.2;refGene_gn=ETV4;refGene_tx=XM_023243906.1
E1 43759173 43759275
refGene_gn=ETV4;refGene_tx=XM_006940408.4;refGene_gn=ETV4;refGene_tx=XM_011289264.2;refGene_gn=ETV4;refGene_tx=XM_019817949.2;refGene_gn=ETV4;refGene_tx=XM_019817950.2;refGene_gn=ETV4;refGene_tx=XM_019817951.2;refGene_gn=ETV4;refGene_tx=XM_019817953.2;refGene_gn=ETV4;refGene_tx=XM_019817954.2;refGene_gn=ETV4;refGene_tx=XM_019817955.2;refGene_gn=ETV4;refGene_tx=XM_019817956.2;refGene_gn=ETV4;refGene_tx=XM_023243906.1
E1 43759523 43759696
refGene_gn=ETV4;refGene_tx=XM_006940408.4;refGene_gn=ETV4;refGene_tx=XM_011289264.2;refGene_gn=ETV4;refGene_tx=XM_019817949.2;refGene_gn=ETV4;refGene_tx=XM_019817950.2;refGene_gn=ETV4;refGene_tx=XM_019817951.2;refGene_gn=ETV4;refGene_tx=XM_019817953.2;refGene_gn=ETV4;refGene_tx=XM_019817954.2;refGene_gn=ETV4;refGene_tx=XM_019817955.2;refGene_gn=ETV4;refGene_tx=XM_019817956.2;refGene_gn=ETV4;refGene_tx=XM_023243906.1
E1 43759847 43759916
refGene_gn=ETV4;refGene_tx=XM_006940408.4;refGene_gn=ETV4;refGene_tx=XM_011289264.2;refGene_gn=ETV4;refGene_tx=XM_019817949.2;refGene_gn=ETV4;refGene_tx=XM_019817950.2;refGene_gn=ETV4;refGene_tx=XM_019817951.2;refGene_gn=ETV4;refGene_tx=XM_019817953.2;refGene_gn=ETV4;refGene_tx=XM_019817954.2;refGene_gn=ETV4;refGene_tx=XM_019817955.2;refGene_gn=ETV4;refGene_tx=XM_019817956.2;refGene_gn=ETV4;refGene_tx=XM_023243906.1
E1 43760071 43760146
refGene_gn=ETV4;refGene_tx=XM_006940408.4;refGene_gn=ETV4;refGene_tx=XM_011289264.2;refGene_gn=ETV4;refGene_tx=XM_019817949.2;refGene_gn=ETV4;refGene_tx=XM_019817950.2;refGene_gn=ETV4;refGene_tx=XM_019817951.2;refGene_gn=ETV4;refGene_tx=XM_019817953.2;refGene_gn=ETV4;refGene_tx=XM_019817954.2;refGene_gn=ETV4;refGene_tx=XM_019817955.2;refGene_gn=ETV4;refGene_tx=XM_019817956.2;refGene_gn=ETV4;refGene_tx=XM_023243906.1
E1 43760322 43760353 refGene_gn=ETV4;refGene_tx=XM_023243906.1
E1 43762085 43762354

refGene_gn=ETV4;refGene_tx=XM_006940408.4;refGene_gn=ETV4;refGene_tx=XM_011289264.2;refGene_gn=ETV4;refGene_tx=XM_019817949.2;refGene_gn=ETV4;refGene_tx=XM_019817950.2;refGene_gn=ETV4;refGene_tx=XM_019817951.2;refGene_gn=ETV4;refGene_tx=XM_019817953.2;refGene_gn=ETV4;refGene_tx=XM_019817954.2;refGene_gn=ETV4;refGene_tx=XM_019817955.2;refGene_gn=ETV4;refGene_tx=XM_019817956.2
E1 43762638 43762800
refGene_gn=ETV4;refGene_tx=XM_006940408.4;refGene_gn=ETV4;refGene_tx=XM_011289264.2;refGene_gn=ETV4;refGene_tx=XM_019817949.2;refGene_gn=ETV4;refGene_tx=XM_019817950.2;refGene_gn=ETV4;refGene_tx=XM_019817951.2;refGene_gn=ETV4;refGene_tx=XM_019817953.2;refGene_gn=ETV4;refGene_tx=XM_019817954.2;refGene_gn=ETV4;refGene_tx=XM_019817955.2;refGene_gn=ETV4;refGene_tx=XM_019817956.2
E1 43763223 43763350
refGene_gn=ETV4;refGene_tx=XM_006940408.4;refGene_gn=ETV4;refGene_tx=XM_011289264.2;refGene_gn=ETV4;refGene_tx=XM_019817949.2;refGene_gn=ETV4;refGene_tx=XM_019817950.2;refGene_gn=ETV4;refGene_tx=XM_019817951.2;refGene_gn=ETV4;refGene_tx=XM_019817953.2;refGene_gn=ETV4;refGene_tx=XM_019817954.2;refGene_gn=ETV4;refGene_tx=XM_019817955.2;refGene_gn=ETV4;refGene_tx=XM_019817956.2
E1 43764559 43764613
refGene_gn=ETV4;refGene_tx=XM_006940408.4;refGene_gn=ETV4;refGene_tx=XM_011289264.2;refGene_gn=ETV4;refGene_tx=XM_019817949.2;refGene_gn=ETV4;refGene_tx=XM_019817950.2;refGene_gn=ETV4;refGene_tx=XM_019817951.2;refGene_gn=ETV4;refGene_tx=XM_019817953.2;refGene_gn=ETV4;refGene_tx=XM_019817954.2;refGene_gn=ETV4;refGene_tx=XM_019817955.2;refGene_gn=ETV4;refGene_tx=XM_019817956.2
E1 43769771 43769847
refGene_gn=ETV4;refGene_tx=XM_011289264.2;refGene_gn=ETV4;refGene_tx=XM_019817956.2
E1 43771682 43771730
refGene_gn=ETV4;refGene_tx=XM_006940408.4;refGene_gn=ETV4;refGene_tx=XM_019817949.2;refGene_gn=ETV4;refGene_tx=XM_019817950.2;refGene_gn=ETV4;refGene_tx=XM_019817951.2;refGene_gn=ETV4;refGene_tx=XM_019817953.2;refGene_gn=ETV4;refGene_tx=XM_019817954.2;refGene_gn=ETV4;refGene_tx=XM_019817955.2
E1 43771953 43772047
refGene_gn=ETV4;refGene_tx=XM_006940408.4;refGene_gn=ETV4;refGene_tx=XM_019817949.2;refGene_gn=ETV4;refGene_tx=XM_019817950.2;refGene_gn=ETV4;refGene_tx=XM_019817951.2;refGene_gn=ETV4;refGene_tx=XM_019817953.2;refGene_gn=ETV4;refGene_tx=XM_019817954.2;refGene_gn=ETV4;refGene_tx=XM_019817955.2
E1 43772228 43772288
refGene_gn=ETV4;refGene_tx=XM_006940408.4;refGene_gn=ETV4;refGene_tx=XM_019817949.2;refGene_gn=ETV4;refGene_tx=XM_019817950.2;refGene_gn=ETV4;refGene_tx=XM_019817951.2;refGene_gn=ETV4;refGene_tx=XM_019817953.2;refGene_gn=ETV4;refGene_tx=XM_019817954.2
E1 43848728 43848886
refGene_gn=MEOX1;refGene_tx=XM_003996963.4;refGene_gn=MEOX1;refGene_tx=XM_011289262.3;refGene_gn=MEOX1;refGene_tx=XM_023243806.1

E1 43850391 43850564
refGene_gn=MEOX1;refGene_tx=XM_003996963.4;refGene_gn=MEOX1;refGene_tx=XM_011289262.3
E1 43867106 43867575
refGene_gn=MEOX1;refGene_tx=XM_003996963.4;refGene_gn=MEOX1;refGene_tx=XM_011289262.3;refGene_gn=MEOX1;refGene_tx=XM_023243806.1
E1 43943089 43943511 refGene_gn=SOST;refGene_tx=XM_011289261.3
E1 43946241 43946458 refGene_gn=SOST;refGene_tx=XM_011289261.3
E1 43959763 43959969 refGene_gn=DUSP3;refGene_tx=XM_023243574.1
E1 43964865 43965092 refGene_gn=DUSP3;refGene_tx=XM_023243574.1
E1 43967943 43968068 refGene_gn=DUSP3;refGene_tx=XM_023243574.1
E1 43970527 43970651
refGene_gn=CE1H17orf105;refGene_tx=XM_006940410.4
E1 43971398 43971469
refGene_gn=CE1H17orf105;refGene_tx=XM_006940410.4
E1 43971738 43971857
refGene_gn=CE1H17orf105;refGene_tx=XM_006940410.4
E1 43972775 43972899
refGene_gn=CE1H17orf105;refGene_tx=XM_006940410.4
E1 43973640 43973697
refGene_gn=CE1H17orf105;refGene_tx=XM_006940410.4
E1 43982242 43982419
refGene_gn=MPP3;refGene_tx=XM_019817961.2;refGene_gn=MPP3;refGene_tx=XM_019817962.2;refGene_gn=MPP3;refGene_tx=XM_023243721.1;refGene_gn=MPP3;refGene_tx=XM_023243722.1
E1 43982744 43982753 refGene_gn=MPP3;refGene_tx=XM_019817963.2
E1 43985656 43985779
refGene_gn=MPP3;refGene_tx=XM_019817961.2;refGene_gn=MPP3;refGene_tx=XM_019817962.2;refGene_gn=MPP3;refGene_tx=XM_019817963.2;refGene_gn=MPP3;refGene_tx=XM_023243721.1;refGene_gn=MPP3;refGene_tx=XM_023243722.1
E1 43987221 43987330
refGene_gn=MPP3;refGene_tx=XM_019817961.2;refGene_gn=MPP3;refGene_tx=XM_019817962.2;refGene_gn=MPP3;refGene_tx=XM_019817963.2;refGene_gn=MPP3;refGene_tx=XM_023243721.1;refGene_gn=MPP3;refGene_tx=XM_023243722.1
E1 43987535 43987629
refGene_gn=MPP3;refGene_tx=XM_019817961.2;refGene_gn=MPP3;refGene_tx=XM_019817962.2;refGene_gn=MPP3;refGene_tx=XM_019817963.2;refGene_gn=MPP3;refGene_tx=XM_023243721.1;refGene_gn=MPP3;refGene_tx=XM_023243722.1
E1 43989743 43989824
refGene_gn=MPP3;refGene_tx=XM_019817961.2;refGene_gn=MPP3;refGene_tx=XM_019817962.2;refGene_gn=MPP3;refGene_tx=XM_019817963.2;refGene_gn=MPP3;refGene_tx=XM_023243721.1;refGene_gn=MPP3;refGene_tx=XM_023243722.1
E1 43989940 43990105
refGene_gn=MPP3;refGene_tx=XM_019817961.2;refGene_gn=MPP3;refGene_tx=XM_019817962.2;refGene_gn=MPP3;refGene_tx=XM_019817963.2;refGene_gn=MPP3;refGene_tx=XM_023243721.1;refGene_gn=MPP3;refGene_tx=XM_023243722.1
E1 43990424 43990466
refGene_gn=MPP3;refGene_tx=XM_019817961.2;refGene_gn=MPP3;refGene_tx=XM_019817962.2;refGene_gn=MPP3;refGene_tx=XM_019817963.2;refGene_gn=MPP3;refGene_tx=XM_023243721.1;refGene_gn=MPP3;refGene_tx=XM_023243722.1

E1 43990962 43990983
refGene_gn=MPP3;refGene_tx=XM_019817961.2;refGene_gn=MPP3;refGene_tx=XM_019817962.2;refGene_gn=MPP3;refGene_tx=XM_019817963.2;refGene_gn=MPP3;refGene_tx=XM_023243721.1;refGene_gn=MPP3;refGene_tx=XM_023243722.1
E1 43992296 43992361
refGene_gn=MPP3;refGene_tx=XM_019817961.2;refGene_gn=MPP3;refGene_tx=XM_019817962.2;refGene_gn=MPP3;refGene_tx=XM_019817963.2;refGene_gn=MPP3;refGene_tx=XM_023243721.1;refGene_gn=MPP3;refGene_tx=XM_023243722.1
E1 43994371 43994568
refGene_gn=MPP3;refGene_tx=XM_019817961.2;refGene_gn=MPP3;refGene_tx=XM_019817962.2;refGene_gn=MPP3;refGene_tx=XM_019817963.2;refGene_gn=MPP3;refGene_tx=XM_023243721.1;refGene_gn=MPP3;refGene_tx=XM_023243722.1
E1 43997267 43997342
refGene_gn=MPP3;refGene_tx=XM_019817961.2;refGene_gn=MPP3;refGene_tx=XM_019817962.2;refGene_gn=MPP3;refGene_tx=XM_019817963.2;refGene_gn=MPP3;refGene_tx=XM_023243721.1;refGene_gn=MPP3;refGene_tx=XM_023243722.1
E1 43998677 43998763
refGene_gn=MPP3;refGene_tx=XM_019817961.2;refGene_gn=MPP3;refGene_tx=XM_019817962.2;refGene_gn=MPP3;refGene_tx=XM_019817963.2;refGene_gn=MPP3;refGene_tx=XM_023243721.1;refGene_gn=MPP3;refGene_tx=XM_023243722.1
E1 43999920 44000002
refGene_gn=MPP3;refGene_tx=XM_019817961.2;refGene_gn=MPP3;refGene_tx=XM_019817962.2;refGene_gn=MPP3;refGene_tx=XM_019817963.2;refGene_gn=MPP3;refGene_tx=XM_023243721.1;refGene_gn=MPP3;refGene_tx=XM_023243722.1
E1 44004255 44004393
refGene_gn=MPP3;refGene_tx=XM_019817961.2;refGene_gn=MPP3;refGene_tx=XM_019817962.2;refGene_gn=MPP3;refGene_tx=XM_019817963.2;refGene_gn=MPP3;refGene_tx=XM_023243721.1;refGene_gn=MPP3;refGene_tx=XM_023243722.1
E1 44004645 44004726
refGene_gn=MPP3;refGene_tx=XM_019817961.2;refGene_gn=MPP3;refGene_tx=XM_019817962.2;refGene_gn=MPP3;refGene_tx=XM_019817963.2;refGene_gn=MPP3;refGene_tx=XM_023243721.1;refGene_gn=MPP3;refGene_tx=XM_023243722.1
E1 44006346 44006424
refGene_gn=MPP3;refGene_tx=XM_019817961.2;refGene_gn=MPP3;refGene_tx=XM_019817962.2;refGene_gn=MPP3;refGene_tx=XM_019817963.2;refGene_gn=MPP3;refGene_tx=XM_023243721.1;refGene_gn=MPP3;refGene_tx=XM_023243722.1
E1 44006689 44006808
refGene_gn=MPP3;refGene_tx=XM_019817961.2;refGene_gn=MPP3;refGene_tx=XM_019817962.2;refGene_gn=MPP3;refGene_tx=XM_019817963.2;refGene_gn=MPP3;refGene_tx=XM_023243721.1;refGene_gn=MPP3;refGene_tx=XM_023243722.1
E1 44007055 44007164
refGene_gn=MPP3;refGene_tx=XM_019817961.2;refGene_gn=MPP3;refGene_tx=XM_019817962.2;refGene_gn=MPP3;refGene_tx=XM_019817963.2;refGene_gn=MPP3;refGene_tx=XM_023243721.1;refGene_gn=MPP3;refGene_tx=XM_023243722.1
E1 44022040 44022089
refGene_gn=CD300LG;refGene_tx=XM_023243691.1;refGene_gn=CD300LG;refGene_tx=XM_023243693.1;refGene_gn=CD300LG;refGene_tx=XM_023243694.1;refGene_gn=CD300LG;refGene_tx=XM_023243695.1;refGene_gn=CD300LG;refGene_tx=XM_023243696.1;refGene_gn=CD300LG;refGene_tx=XM_023243697.1;refGene_gn=CD300LG;refGene_tx=XM_023243698.1;refGene_gn=CD300LG;refGene_tx=XM_02

3243699.1
E1 44023003 44023339
refGene_gn=CD300LG;refGene_tx=XM_023243691.1;refGene_gn=CD300LG;refGene_tx=XM_023243693.1;refGene_gn=CD300LG;refGene_tx=XM_023243694.1;refGene_gn=CD300LG;refGene_tx=XM_023243695.1;refGene_gn=CD300LG;refGene_tx=XM_023243696.1;refGene_gn=CD300LG;refGene_tx=XM_023243697.1;refGene_gn=CD300LG;refGene_tx=XM_023243698.1;refGene_gn=CD300LG;refGene_tx=XM_023243699.1
E1 44026025 44026127
refGene_gn=CD300LG;refGene_tx=XM_023243691.1;refGene_gn=CD300LG;refGene_tx=XM_023243693.1;refGene_gn=CD300LG;refGene_tx=XM_023243697.1;refGene_gn=CD300LG;refGene_tx=XM_023243698.1
E1 44026687 44026940
refGene_gn=CD300LG;refGene_tx=XM_023243691.1;refGene_gn=CD300LG;refGene_tx=XM_023243693.1;refGene_gn=CD300LG;refGene_tx=XM_023243694.1;refGene_gn=CD300LG;refGene_tx=XM_023243695.1;refGene_gn=CD300LG;refGene_tx=XM_023243696.1;refGene_gn=CD300LG;refGene_tx=XM_023243697.1;refGene_gn=CD300LG;refGene_tx=XM_023243698.1;refGene_gn=CD300LG;refGene_tx=XM_023243699.1
E1 44027621 44027734
refGene_gn=CD300LG;refGene_tx=XM_023243691.1;refGene_gn=CD300LG;refGene_tx=XM_023243693.1;refGene_gn=CD300LG;refGene_tx=XM_023243694.1;refGene_gn=CD300LG;refGene_tx=XM_023243697.1;refGene_gn=CD300LG;refGene_tx=XM_023243699.1
E1 44030119 44030211
refGene_gn=CD300LG;refGene_tx=XM_023243691.1;refGene_gn=CD300LG;refGene_tx=XM_023243693.1;refGene_gn=CD300LG;refGene_tx=XM_023243694.1;refGene_gn=CD300LG;refGene_tx=XM_023243695.1;refGene_gn=CD300LG;refGene_tx=XM_023243696.1;refGene_gn=CD300LG;refGene_tx=XM_023243697.1;refGene_gn=CD300LG;refGene_tx=XM_023243698.1;refGene_gn=CD300LG;refGene_tx=XM_023243699.1
E1 44032905 44033007
refGene_gn=CD300LG;refGene_tx=XM_023243691.1;refGene_gn=CD300LG;refGene_tx=XM_023243693.1;refGene_gn=CD300LG;refGene_tx=XM_023243694.1;refGene_gn=CD300LG;refGene_tx=XM_023243695.1;refGene_gn=CD300LG;refGene_tx=XM_023243696.1;refGene_gn=CD300LG;refGene_tx=XM_023243697.1;refGene_gn=CD300LG;refGene_tx=XM_023243698.1;refGene_gn=CD300LG;refGene_tx=XM_023243699.1
E1 44033497 44033721
refGene_gn=CD300LG;refGene_tx=XM_023243691.1;refGene_gn=CD300LG;refGene_tx=XM_023243693.1;refGene_gn=CD300LG;refGene_tx=XM_023243694.1;refGene_gn=CD300LG;refGene_tx=XM_023243695.1;refGene_gn=CD300LG;refGene_tx=XM_023243696.1;refGene_gn=CD300LG;refGene_tx=XM_023243697.1;refGene_gn=CD300LG;refGene_tx=XM_023243698.1;refGene_gn=CD300LG;refGene_tx=XM_023243699.1
E1 44044343 44044520
refGene_gn=MPP2;refGene_tx=XM_019817965.2;refGene_gn=MPP2;refGene_tx=XM_023243609.1;refGene_gn=MPP2;refGene_tx=XM_023243610.1;refGene_gn=MPP2;refGene_tx=XM_023243611.1;refGene_gn=MPP2;refGene_tx=XM_023243613.1;refGene_gn=MPP2;refGene_tx=XM_023243614.1;refGene_gn=MPP2;refGene_tx=X

M_023243615.1;refGene_gn=MPP2;refGene_tx=XM_023243616.1;refGene_gn=MPP2;refGene_tx=XM_023243617.1
E1 44045760 44045889
refGene_gn=MPP2;refGene_tx=XM_019817965.2;refGene_gn=MPP2;refGene_tx=XM_023243609.1;refGene_gn=MPP2;refGene_tx=XM_023243610.1;refGene_gn=MPP2;refGene_tx=XM_023243611.1;refGene_gn=MPP2;refGene_tx=XM_023243613.1;refGene_gn=MPP2;refGene_tx=XM_023243614.1;refGene_gn=MPP2;refGene_tx=XM_023243615.1;refGene_gn=MPP2;refGene_tx=XM_023243616.1;refGene_gn=MPP2;refGene_tx=XM_023243617.1
E1 44046232 44046435
refGene_gn=MPP2;refGene_tx=XM_019817965.2;refGene_gn=MPP2;refGene_tx=XM_023243609.1;refGene_gn=MPP2;refGene_tx=XM_023243610.1;refGene_gn=MPP2;refGene_tx=XM_023243611.1;refGene_gn=MPP2;refGene_tx=XM_023243613.1;refGene_gn=MPP2;refGene_tx=XM_023243614.1;refGene_gn=MPP2;refGene_tx=XM_023243615.1;refGene_gn=MPP2;refGene_tx=XM_023243616.1;refGene_gn=MPP2;refGene_tx=XM_023243617.1
E1 44047108 44047270
refGene_gn=MPP2;refGene_tx=XM_019817965.2;refGene_gn=MPP2;refGene_tx=XM_023243609.1;refGene_gn=MPP2;refGene_tx=XM_023243610.1;refGene_gn=MPP2;refGene_tx=XM_023243611.1;refGene_gn=MPP2;refGene_tx=XM_023243613.1;refGene_gn=MPP2;refGene_tx=XM_023243614.1;refGene_gn=MPP2;refGene_tx=XM_023243615.1;refGene_gn=MPP2;refGene_tx=XM_023243616.1;refGene_gn=MPP2;refGene_tx=XM_023243617.1
E1 44047502 44047571
refGene_gn=MPP2;refGene_tx=XM_019817965.2;refGene_gn=MPP2;refGene_tx=XM_023243609.1;refGene_gn=MPP2;refGene_tx=XM_023243610.1;refGene_gn=MPP2;refGene_tx=XM_023243611.1;refGene_gn=MPP2;refGene_tx=XM_023243613.1;refGene_gn=MPP2;refGene_tx=XM_023243614.1;refGene_gn=MPP2;refGene_tx=XM_023243615.1;refGene_gn=MPP2;refGene_tx=XM_023243616.1;refGene_gn=MPP2;refGene_tx=XM_023243617.1
E1 44047668 44047774
refGene_gn=MPP2;refGene_tx=XM_019817965.2;refGene_gn=MPP2;refGene_tx=XM_023243609.1;refGene_gn=MPP2;refGene_tx=XM_023243610.1;refGene_gn=MPP2;refGene_tx=XM_023243611.1;refGene_gn=MPP2;refGene_tx=XM_023243613.1;refGene_gn=MPP2;refGene_tx=XM_023243614.1;refGene_gn=MPP2;refGene_tx=XM_023243615.1;refGene_gn=MPP2;refGene_tx=XM_023243616.1;refGene_gn=MPP2;refGene_tx=XM_023243617.1
E1 44047880 44048012
refGene_gn=MPP2;refGene_tx=XM_019817965.2;refGene_gn=MPP2;refGene_tx=XM_023243609.1;refGene_gn=MPP2;refGene_tx=XM_023243610.1;refGene_gn=MPP2;refGene_tx=XM_023243611.1;refGene_gn=MPP2;refGene_tx=XM_023243613.1;refGene_gn=MPP2;refGene_tx=XM_023243614.1;refGene_gn=MPP2;refGene_tx=XM_023243615.1;refGene_gn=MPP2;refGene_tx=XM_023243616.1;refGene_gn=MPP2;refGene_tx=XM_023243617.1
E1 44048712 44048940
refGene_gn=MPP2;refGene_tx=XM_019817965.2;refGene_gn=MPP2;refGene_tx=XM_023243609.1;refGene_gn=MPP2;refGene_tx=XM_023243610.1;refGene_gn=MPP2;refGene_tx=XM_023243611.1;refGene_gn=MPP2;refGene_tx=XM_023243613.1;refGene_gn=MPP2;refGene_tx=XM_023243614.1;refGene_gn=MPP2;refGene_tx=XM_023243615.1;refGene_gn=MPP2;refGene_tx=XM_023243616.1;refGene_gn=MPP

2;refGene_tx=XM_023243617.1
E1 44049330 44049480
refGene_gn=MPP2;refGene_tx=XM_019817965.2;refGene_gn=MPP2;refGene_tx=XM_023243609.1;refGene_gn=MPP2;refGene_tx=XM_023243610.1;refGene_gn=MPP2;refGene_tx=XM_023243611.1;refGene_gn=MPP2;refGene_tx=XM_023243613.1;refGene_gn=MPP2;refGene_tx=XM_023243614.1;refGene_gn=MPP2;refGene_tx=XM_023243615.1;refGene_gn=MPP2;refGene_tx=XM_023243616.1;refGene_gn=MPP2;refGene_tx=XM_023243617.1
E1 44049619 44049772
refGene_gn=MPP2;refGene_tx=XM_019817965.2;refGene_gn=MPP2;refGene_tx=XM_023243609.1;refGene_gn=MPP2;refGene_tx=XM_023243610.1;refGene_gn=MPP2;refGene_tx=XM_023243611.1;refGene_gn=MPP2;refGene_tx=XM_023243613.1;refGene_gn=MPP2;refGene_tx=XM_023243614.1;refGene_gn=MPP2;refGene_tx=XM_023243615.1;refGene_gn=MPP2;refGene_tx=XM_023243616.1;refGene_gn=MPP2;refGene_tx=XM_023243617.1
E1 44062586 44062705
refGene_gn=MPP2;refGene_tx=XM_019817965.2;refGene_gn=MPP2;refGene_tx=XM_023243609.1;refGene_gn=MPP2;refGene_tx=XM_023243610.1;refGene_gn=MPP2;refGene_tx=XM_023243611.1;refGene_gn=MPP2;refGene_tx=XM_023243613.1;refGene_gn=MPP2;refGene_tx=XM_023243614.1;refGene_gn=MPP2;refGene_tx=XM_023243615.1;refGene_gn=MPP2;refGene_tx=XM_023243616.1;refGene_gn=MPP2;refGene_tx=XM_023243617.1
E1 44064268 44064350
refGene_gn=MPP2;refGene_tx=XM_019817965.2;refGene_gn=MPP2;refGene_tx=XM_023243609.1
E1 44067698 44067729
refGene_gn=MPP2;refGene_tx=XM_023243610.1;refGene_gn=MPP2;refGene_tx=XM_023243611.1;refGene_gn=MPP2;refGene_tx=XM_023243613.1;refGene_gn=MPP2;refGene_tx=XM_023243614.1;refGene_gn=MPP2;refGene_tx=XM_023243615.1;refGene_gn=MPP2;refGene_tx=XM_023243616.1;refGene_gn=MPP2;refGene_tx=XM_023243617.1
E1 44083942 44083967 refGene_gn=PPY;refGene_tx=XM_003996970.3
E1 44084119 44084191 refGene_gn=PPY;refGene_tx=XM_003996970.3
E1 44084404 44084595 refGene_gn=PPY;refGene_tx=XM_003996970.3
E1 44092946 44092971
refGene_gn=PYY;refGene_tx=XM_023243716.1;refGene_gn=PYY;refGene_tx=XM_023243717.1
E1 44093130 44093211
refGene_gn=PYY;refGene_tx=XM_023243716.1;refGene_gn=PYY;refGene_tx=XM_023243717.1
E1 44093324 44093512
refGene_gn=PYY;refGene_tx=XM_023243716.1;refGene_gn=PYY;refGene_tx=XM_023243717.1
E1 44093855 44094032 refGene_gn=PYY;refGene_tx=XM_023243716.1
E1 44094691 44094824 refGene_gn=TMEM101;refGene_tx=XM_023243624.1
E1 44098145 44098249 refGene_gn=TMEM101;refGene_tx=XM_023243624.1
E1 44128420 44128837 refGene_gn=NAGS;refGene_tx=XM_011289272.3
E1 44129403 44129678 refGene_gn=NAGS;refGene_tx=XM_011289272.3
E1 44129794 44130008 refGene_gn=NAGS;refGene_tx=XM_011289272.3
E1 44130278 44130459 refGene_gn=NAGS;refGene_tx=XM_011289272.3

E1 44131003 44131175 refGene_gn=NAGS;refGene_tx=XM_011289272.3
E1 44131266 44131449 refGene_gn=NAGS;refGene_tx=XM_011289272.3
E1 44132077 44132231 refGene_gn=NAGS;refGene_tx=XM_011289272.3
E1 44134413 44134722 refGene_gn=TMEM101;refGene_tx=XM_003996972.5
E1 44135500 44135647
refGene_gn=TMEM101;refGene_tx=XM_003996972.5;refGene_gn=TMEM101;refGene_tx=XM_023243624.1
E1 44136522 44136703
refGene_gn=TMEM101;refGene_tx=XM_003996972.5;refGene_gn=TMEM101;refGene_tx=XM_023243624.1
E1 44137002 44137139
refGene_gn=TMEM101;refGene_tx=XM_003996972.5;refGene_gn=TMEM101;refGene_tx=XM_023243624.1
E1 44150707 44150800 refGene_gn=LSM12;refGene_tx=XM_003996973.5
E1 44151585 44151712 refGene_gn=LSM12;refGene_tx=XM_003996973.5
E1 44153115 44153225 refGene_gn=LSM12;refGene_tx=XM_003996973.5
E1 44167179 44167313 refGene_gn=LSM12;refGene_tx=XM_003996973.5
E1 44169090 44169214 refGene_gn=LSM12;refGene_tx=XM_003996973.5
E1 44171965 44172183
refGene_gn=G6PC3;refGene_tx=XM_003996974.5;refGene_gn=G6PC3;refGene_tx=XM_019817969.2
E1 44174078 44174185
refGene_gn=G6PC3;refGene_tx=XM_003996974.5;refGene_gn=G6PC3;refGene_tx=XM_019817969.2
E1 44174565 44174656
refGene_gn=G6PC3;refGene_tx=XM_003996974.5;refGene_gn=G6PC3;refGene_tx=XM_019817969.2
E1 44174854 44174973 refGene_gn=G6PC3;refGene_tx=XM_003996974.5
E1 44175163 44175305
refGene_gn=G6PC3;refGene_tx=XM_003996974.5;refGene_gn=G6PC3;refGene_tx=XM_019817969.2
E1 44175538 44175902
refGene_gn=G6PC3;refGene_tx=XM_003996974.5;refGene_gn=G6PC3;refGene_tx=XM_019817969.2
E1 44178185 44178225
refGene_gn=HDAC5;refGene_tx=XM_019817968.2;refGene_gn=HDAC5;refGene_tx=XM_023244460.1;refGene_gn=HDAC5;refGene_tx=XM_023244461.1;refGene_gn=HDAC5;refGene_tx=XM_023244462.1;refGene_gn=HDAC5;refGene_tx=XM_023244463.1
E1 44178304 44178470
refGene_gn=HDAC5;refGene_tx=XM_019817968.2;refGene_gn=HDAC5;refGene_tx=XM_023244460.1;refGene_gn=HDAC5;refGene_tx=XM_023244461.1;refGene_gn=HDAC5;refGene_tx=XM_023244462.1;refGene_gn=HDAC5;refGene_tx=XM_023244463.1
E1 44178600 44178685
refGene_gn=HDAC5;refGene_tx=XM_019817968.2;refGene_gn=HDAC5;refGene_tx=XM_023244460.1;refGene_gn=HDAC5;refGene_tx=XM_023244461.1;refGene_gn=HDAC5;refGene_tx=XM_023244462.1;refGene_gn=HDAC5;refGene_tx=XM_023244463.1
E1 44179006 44179140

refGene_gn=HDAC5;refGene_tx=XM_019817968.2;refGene_gn=HDAC5;refGene_tx=XM_023244460.1;refGene_gn=HDAC5;refGene_tx=XM_023244461.1;refGene_gn=HDAC5;refGene_tx=XM_023244462.1;refGene_gn=HDAC5;refGene_tx=XM_023244463.1
E1 44179763 44179882
refGene_gn=HDAC5;refGene_tx=XM_019817968.2;refGene_gn=HDAC5;refGene_tx=XM_023244460.1;refGene_gn=HDAC5;refGene_tx=XM_023244461.1;refGene_gn=HDAC5;refGene_tx=XM_023244462.1;refGene_gn=HDAC5;refGene_tx=XM_023244463.1
E1 44180059 44180157
refGene_gn=HDAC5;refGene_tx=XM_019817968.2;refGene_gn=HDAC5;refGene_tx=XM_023244460.1;refGene_gn=HDAC5;refGene_tx=XM_023244461.1;refGene_gn=HDAC5;refGene_tx=XM_023244462.1;refGene_gn=HDAC5;refGene_tx=XM_023244463.1
E1 44180414 44180534
refGene_gn=HDAC5;refGene_tx=XM_019817968.2;refGene_gn=HDAC5;refGene_tx=XM_023244460.1;refGene_gn=HDAC5;refGene_tx=XM_023244461.1;refGene_gn=HDAC5;refGene_tx=XM_023244462.1;refGene_gn=HDAC5;refGene_tx=XM_023244463.1
E1 44181484 44181572
refGene_gn=HDAC5;refGene_tx=XM_019817968.2;refGene_gn=HDAC5;refGene_tx=XM_023244460.1;refGene_gn=HDAC5;refGene_tx=XM_023244461.1;refGene_gn=HDAC5;refGene_tx=XM_023244462.1;refGene_gn=HDAC5;refGene_tx=XM_023244463.1
E1 44181720 44181776
refGene_gn=HDAC5;refGene_tx=XM_019817968.2;refGene_gn=HDAC5;refGene_tx=XM_023244460.1;refGene_gn=HDAC5;refGene_tx=XM_023244461.1;refGene_gn=HDAC5;refGene_tx=XM_023244462.1;refGene_gn=HDAC5;refGene_tx=XM_023244463.1
E1 44182183 44182291
refGene_gn=HDAC5;refGene_tx=XM_019817968.2;refGene_gn=HDAC5;refGene_tx=XM_023244460.1;refGene_gn=HDAC5;refGene_tx=XM_023244461.1;refGene_gn=HDAC5;refGene_tx=XM_023244462.1;refGene_gn=HDAC5;refGene_tx=XM_023244463.1
E1 44182461 44182511
refGene_gn=HDAC5;refGene_tx=XM_019817968.2;refGene_gn=HDAC5;refGene_tx=XM_023244460.1;refGene_gn=HDAC5;refGene_tx=XM_023244461.1;refGene_gn=HDAC5;refGene_tx=XM_023244462.1;refGene_gn=HDAC5;refGene_tx=XM_023244463.1
E1 44182981 44183102
refGene_gn=HDAC5;refGene_tx=XM_019817968.2;refGene_gn=HDAC5;refGene_tx=XM_023244460.1;refGene_gn=HDAC5;refGene_tx=XM_023244461.1;refGene_gn=HDAC5;refGene_tx=XM_023244462.1;refGene_gn=HDAC5;refGene_tx=XM_023244463.1
E1 44183728 44183862
refGene_gn=HDAC5;refGene_tx=XM_019817968.2;refGene_gn=HDAC5;refGene_tx=XM_023244460.1;refGene_gn=HDAC5;refGene_tx=XM_023244461.1;refGene_gn=HDAC5;refGene_tx=XM_023244462.1;refGene_gn=HDAC5;refGene_tx=XM_023244463.1
E1 44184714 44184880

refGene_gn=HDAC5;refGene_tx=XM_019817968.2;refGene_gn=HDAC5;refGene_tx=XM_023244460.1;refGene_gn=HDAC5;refGene_tx=XM_023244461.1;refGene_gn=HDAC5;refGene_tx=XM_023244462.1;refGene_gn=HDAC5;refGene_tx=XM_023244463.1
E1 44185589 44185883
refGene_gn=HDAC5;refGene_tx=XM_019817968.2;refGene_gn=HDAC5;refGene_tx=XM_023244460.1;refGene_gn=HDAC5;refGene_tx=XM_023244461.1;refGene_gn=HDAC5;refGene_tx=XM_023244462.1;refGene_gn=HDAC5;refGene_tx=XM_023244463.1
E1 44186147 44186359
refGene_gn=HDAC5;refGene_tx=XM_019817968.2;refGene_gn=HDAC5;refGene_tx=XM_023244460.1;refGene_gn=HDAC5;refGene_tx=XM_023244461.1;refGene_gn=HDAC5;refGene_tx=XM_023244462.1;refGene_gn=HDAC5;refGene_tx=XM_023244463.1
E1 44188172 44188395
refGene_gn=HDAC5;refGene_tx=XM_019817968.2;refGene_gn=HDAC5;refGene_tx=XM_023244460.1;refGene_gn=HDAC5;refGene_tx=XM_023244461.1;refGene_gn=HDAC5;refGene_tx=XM_023244462.1;refGene_gn=HDAC5;refGene_tx=XM_023244463.1
E1 44188621 44188753
refGene_gn=HDAC5;refGene_tx=XM_019817968.2;refGene_gn=HDAC5;refGene_tx=XM_023244460.1;refGene_gn=HDAC5;refGene_tx=XM_023244461.1;refGene_gn=HDAC5;refGene_tx=XM_023244462.1;refGene_gn=HDAC5;refGene_tx=XM_023244463.1
E1 44189069 44189182
refGene_gn=HDAC5;refGene_tx=XM_019817968.2;refGene_gn=HDAC5;refGene_tx=XM_023244460.1;refGene_gn=HDAC5;refGene_tx=XM_023244461.1;refGene_gn=HDAC5;refGene_tx=XM_023244462.1;refGene_gn=HDAC5;refGene_tx=XM_023244463.1
E1 44189271 44189418
refGene_gn=HDAC5;refGene_tx=XM_019817968.2;refGene_gn=HDAC5;refGene_tx=XM_023244460.1;refGene_gn=HDAC5;refGene_tx=XM_023244461.1;refGene_gn=HDAC5;refGene_tx=XM_023244462.1;refGene_gn=HDAC5;refGene_tx=XM_023244463.1
E1 44189549 44189680
refGene_gn=HDAC5;refGene_tx=XM_019817968.2;refGene_gn=HDAC5;refGene_tx=XM_023244460.1;refGene_gn=HDAC5;refGene_tx=XM_023244461.1;refGene_gn=HDAC5;refGene_tx=XM_023244462.1;refGene_gn=HDAC5;refGene_tx=XM_023244463.1
E1 44189970 44190085
refGene_gn=HDAC5;refGene_tx=XM_019817968.2;refGene_gn=HDAC5;refGene_tx=XM_023244460.1;refGene_gn=HDAC5;refGene_tx=XM_023244461.1;refGene_gn=HDAC5;refGene_tx=XM_023244462.1;refGene_gn=HDAC5;refGene_tx=XM_023244463.1
E1 44190188 44190333
refGene_gn=HDAC5;refGene_tx=XM_019817968.2;refGene_gn=HDAC5;refGene_tx=XM_023244460.1;refGene_gn=HDAC5;refGene_tx=XM_023244461.1;refGene_gn=HDAC5;refGene_tx=XM_023244462.1;refGene_gn=HDAC5;refGene_tx=XM_023244463.1
E1 44190438 44190698

refGene_gn=HDAC5;refGene_tx=XM_019817968.2;refGene_gn=HDAC5;refGene_tx=XM_023244460.1;refGene_gn=HDAC5;refGene_tx=XM_023244461.1;refGene_gn=HDAC5;refGene_tx=XM_023244462.1;refGene_gn=HDAC5;refGene_tx=XM_023244463.1
E1 44201595 44201670
refGene_gn=HDAC5;refGene_tx=XM_019817968.2;refGene_gn=HDAC5;refGene_tx=XM_023244460.1;refGene_gn=HDAC5;refGene_tx=XM_023244461.1;refGene_gn=HDAC5;refGene_tx=XM_023244462.1
E1 44209286 44209308
refGene_gn=HDAC5;refGene_tx=XM_023244460.1;refGene_gn=HDAC5;refGene_tx=XM_023244461.1;refGene_gn=HDAC5;refGene_tx=XM_023244463.1
E1 44230242 44230245
refGene_gn=CE1H17orf53;refGene_tx=XM_006940416.4
E1 44231223 44231274
refGene_gn=CE1H17orf53;refGene_tx=XM_006940416.4
E1 44237075 44238023
refGene_gn=CE1H17orf53;refGene_tx=XM_006940416.4
E1 44238024 44238252
refGene_gn=CE1H17orf53;refGene_tx=XM_006940416.4
E1 44239673 44239757
refGene_gn=CE1H17orf53;refGene_tx=XM_006940416.4
E1 44241222 44241363
refGene_gn=CE1H17orf53;refGene_tx=XM_006940416.4
E1 44241955 44242064
refGene_gn=CE1H17orf53;refGene_tx=XM_006940416.4
E1 44242251 44242337
refGene_gn=CE1H17orf53;refGene_tx=XM_006940416.4
E1 44242685 44242811
refGene_gn=CE1H17orf53;refGene_tx=XM_006940416.4
E1 44244173 44244291
refGene_gn=CE1H17orf53;refGene_tx=XM_006940416.4
E1 44246468 44246560
refGene_gn=CE1H17orf53;refGene_tx=XM_006940416.4
E1 44251350 44251651 refGene_gn=ASB16;refGene_tx=XM_011289257.3
E1 44253427 44253695 refGene_gn=ASB16;refGene_tx=XM_011289257.3
E1 44255938 44256431 refGene_gn=ASB16;refGene_tx=XM_011289257.3
E1 44256813 44256927 refGene_gn=ASB16;refGene_tx=XM_011289257.3
E1 44257393 44257579 refGene_gn=ASB16;refGene_tx=XM_011289257.3
E1 44267571 44267606
refGene_gn=TMUB2;refGene_tx=XM_003996976.5;refGene_gn=TMUB2;refGene_tx=XM_006940418.4
E1 44268913 44269480
refGene_gn=TMUB2;refGene_tx=XM_003996976.5;refGene_gn=TMUB2;refGene_tx=XM_006940418.4;refGene_gn=TMUB2;refGene_tx=XM_019817974.2
E1 44269737 44270104
refGene_gn=TMUB2;refGene_tx=XM_003996976.5;refGene_gn=TMUB2;refGene_tx=XM_006940418.4;refGene_gn=TMUB2;refGene_tx=XM_019817974.2
E1 44273594 44273743
refGene_gn=ATXN7L3;refGene_tx=XM_011289255.3;refGene_gn=ATXN7L3;refGene_tx=XM_023244800.1;refGene_gn=ATXN7L3;refGene_tx=XM_023244801.1;refGe

ne_gn=ATXN7L3;refGene_tx=XM_023244802.1;refGene_gn=ATXN7L3;refGene_tx=XM_023244803.1
E1 44273856 44274014
refGene_gn=ATXN7L3;refGene_tx=XM_011289255.3;refGene_gn=ATXN7L3;refGene_tx=XM_023244800.1;refGene_gn=ATXN7L3;refGene_tx=XM_023244801.1;refGene_gn=ATXN7L3;refGene_tx=XM_023244802.1;refGene_gn=ATXN7L3;refGene_tx=XM_023244803.1
E1 44274107 44274179
refGene_gn=ATXN7L3;refGene_tx=XM_011289255.3;refGene_gn=ATXN7L3;refGene_tx=XM_023244800.1;refGene_gn=ATXN7L3;refGene_tx=XM_023244801.1;refGene_gn=ATXN7L3;refGene_tx=XM_023244802.1;refGene_gn=ATXN7L3;refGene_tx=XM_023244803.1
E1 44274457 44274501
refGene_gn=ATXN7L3;refGene_tx=XM_011289255.3;refGene_gn=ATXN7L3;refGene_tx=XM_023244800.1;refGene_gn=ATXN7L3;refGene_tx=XM_023244801.1;refGene_gn=ATXN7L3;refGene_tx=XM_023244802.1;refGene_gn=ATXN7L3;refGene_tx=XM_023244803.1
E1 44274774 44274843
refGene_gn=ATXN7L3;refGene_tx=XM_011289255.3;refGene_gn=ATXN7L3;refGene_tx=XM_023244800.1;refGene_gn=ATXN7L3;refGene_tx=XM_023244801.1;refGene_gn=ATXN7L3;refGene_tx=XM_023244802.1;refGene_gn=ATXN7L3;refGene_tx=XM_023244803.1
E1 44275151 44275237
refGene_gn=ATXN7L3;refGene_tx=XM_011289255.3;refGene_gn=ATXN7L3;refGene_tx=XM_023244800.1;refGene_gn=ATXN7L3;refGene_tx=XM_023244801.1;refGene_gn=ATXN7L3;refGene_tx=XM_023244802.1;refGene_gn=ATXN7L3;refGene_tx=XM_023244803.1
E1 44275391 44275458
refGene_gn=ATXN7L3;refGene_tx=XM_011289255.3;refGene_gn=ATXN7L3;refGene_tx=XM_023244800.1;refGene_gn=ATXN7L3;refGene_tx=XM_023244801.1;refGene_gn=ATXN7L3;refGene_tx=XM_023244802.1;refGene_gn=ATXN7L3;refGene_tx=XM_023244803.1
E1 44275765 44275788
refGene_gn=ATXN7L3;refGene_tx=XM_011289255.3;refGene_gn=ATXN7L3;refGene_tx=XM_023244800.1;refGene_gn=ATXN7L3;refGene_tx=XM_023244801.1;refGene_gn=ATXN7L3;refGene_tx=XM_023244802.1;refGene_gn=ATXN7L3;refGene_tx=XM_023244803.1
E1 44275961 44276059
refGene_gn=ATXN7L3;refGene_tx=XM_011289255.3;refGene_gn=ATXN7L3;refGene_tx=XM_023244800.1;refGene_gn=ATXN7L3;refGene_tx=XM_023244801.1;refGene_gn=ATXN7L3;refGene_tx=XM_023244802.1;refGene_gn=ATXN7L3;refGene_tx=XM_023244803.1
E1 44276250 44276422
refGene_gn=ATXN7L3;refGene_tx=XM_011289255.3;refGene_gn=ATXN7L3;refGene_tx=XM_023244800.1;refGene_gn=ATXN7L3;refGene_tx=XM_023244801.1;refGene_gn=ATXN7L3;refGene_tx=XM_023244802.1;refGene_gn=ATXN7L3;refGene_tx=XM_023244803.1
E1 44276626 44276759
refGene_gn=ATXN7L3;refGene_tx=XM_011289255.3;refGene_gn=ATXN7L3;refGene_tx=XM_023244800.1;refGene_gn=ATXN7L3;refGene_tx=XM_023244801.1;refGe

ne_gn=ATXN7L3;refGene_tx=XM_023244802.1;refGene_gn=ATXN7L3;refGene_tx=XM_023244803.1
E1 44277058 44277109
refGene_gn=ATXN7L3;refGene_tx=XM_011289255.3;refGene_gn=ATXN7L3;refGene_tx=XM_023244800.1;refGene_gn=ATXN7L3;refGene_tx=XM_023244801.1;refGene_gn=ATXN7L3;refGene_tx=XM_023244802.1;refGene_gn=ATXN7L3;refGene_tx=XM_023244803.1
E1 44286457 44286583
refGene_gn=UBTF;refGene_tx=XM_003996979.5;refGene_gn=UBTF;refGene_tx=XM_011289253.3;refGene_gn=UBTF;refGene_tx=XM_023243451.1;refGene_gn=UBTF;refGene_tx=XM_023243452.1;refGene_gn=UBTF;refGene_tx=XM_023243453.1;refGene_gn=UBTF;refGene_tx=XM_023243454.1
E1 44286666 44286810
refGene_gn=UBTF;refGene_tx=XM_003996979.5;refGene_gn=UBTF;refGene_tx=XM_011289253.3;refGene_gn=UBTF;refGene_tx=XM_023243451.1;refGene_gn=UBTF;refGene_tx=XM_023243452.1;refGene_gn=UBTF;refGene_tx=XM_023243453.1;refGene_gn=UBTF;refGene_tx=XM_023243454.1
E1 44286907 44286979
refGene_gn=UBTF;refGene_tx=XM_003996979.5;refGene_gn=UBTF;refGene_tx=XM_011289253.3;refGene_gn=UBTF;refGene_tx=XM_023243451.1;refGene_gn=UBTF;refGene_tx=XM_023243452.1;refGene_gn=UBTF;refGene_tx=XM_023243453.1;refGene_gn=UBTF;refGene_tx=XM_023243454.1
E1 44287082 44287130
refGene_gn=UBTF;refGene_tx=XM_003996979.5;refGene_gn=UBTF;refGene_tx=XM_011289253.3;refGene_gn=UBTF;refGene_tx=XM_023243451.1;refGene_gn=UBTF;refGene_tx=XM_023243452.1;refGene_gn=UBTF;refGene_tx=XM_023243453.1;refGene_gn=UBTF;refGene_tx=XM_023243454.1
E1 44287962 44288152
refGene_gn=UBTF;refGene_tx=XM_003996979.5;refGene_gn=UBTF;refGene_tx=XM_011289253.3;refGene_gn=UBTF;refGene_tx=XM_023243451.1;refGene_gn=UBTF;refGene_tx=XM_023243452.1;refGene_gn=UBTF;refGene_tx=XM_023243453.1;refGene_gn=UBTF;refGene_tx=XM_023243454.1
E1 44288254 44288343
refGene_gn=UBTF;refGene_tx=XM_003996979.5;refGene_gn=UBTF;refGene_tx=XM_011289253.3;refGene_gn=UBTF;refGene_tx=XM_023243451.1;refGene_gn=UBTF;refGene_tx=XM_023243452.1;refGene_gn=UBTF;refGene_tx=XM_023243453.1;refGene_gn=UBTF;refGene_tx=XM_023243454.1
E1 44288784 44288895
refGene_gn=UBTF;refGene_tx=XM_003996979.5;refGene_gn=UBTF;refGene_tx=XM_011289253.3;refGene_gn=UBTF;refGene_tx=XM_023243451.1;refGene_gn=UBTF;refGene_tx=XM_023243452.1;refGene_gn=UBTF;refGene_tx=XM_023243453.1;refGene_gn=UBTF;refGene_tx=XM_023243454.1
E1 44288997 44289153
refGene_gn=UBTF;refGene_tx=XM_003996979.5;refGene_gn=UBTF;refGene_tx=XM_011289253.3;refGene_gn=UBTF;refGene_tx=XM_023243451.1;refGene_gn=UBTF;refGene_tx=XM_023243452.1;refGene_gn=UBTF;refGene_tx=XM_023243453.1;refGene_gn=UBTF;refGene_tx=XM_023243454.1
E1 44289540 44289696
refGene_gn=UBTF;refGene_tx=XM_003996979.5;refGene_gn=UBTF;refGene_tx=XM_011289253.3;refGene_gn=UBTF;refGene_tx=XM_023243451.1;refGene_gn=UBT

F;refGene_tx=XM_023243452.1;refGene_gn=UBTF;refGene_tx=XM_023243453.1;
refGene_gn=UBTF;refGene_tx=XM_023243454.1
E1 44289786 44289900
refGene_gn=UBTF;refGene_tx=XM_003996979.5;refGene_gn=UBTF;refGene_tx=X
M_011289253.3;refGene_gn=UBTF;refGene_tx=XM_023243451.1;refGene_gn=UBT
F;refGene_tx=XM_023243452.1;refGene_gn=UBTF;refGene_tx=XM_023243453.1;
refGene_gn=UBTF;refGene_tx=XM_023243454.1
E1 44290027 44290069
refGene_gn=UBTF;refGene_tx=XM_003996979.5;refGene_gn=UBTF;refGene_tx=X
M_011289253.3;refGene_gn=UBTF;refGene_tx=XM_023243451.1;refGene_gn=UBT
F;refGene_tx=XM_023243452.1;refGene_gn=UBTF;refGene_tx=XM_023243453.1;
refGene_gn=UBTF;refGene_tx=XM_023243454.1
E1 44290330 44290472
refGene_gn=UBTF;refGene_tx=XM_003996979.5;refGene_gn=UBTF;refGene_tx=X
M_011289253.3;refGene_gn=UBTF;refGene_tx=XM_023243451.1;refGene_gn=UBT
F;refGene_tx=XM_023243452.1;refGene_gn=UBTF;refGene_tx=XM_023243453.1;
refGene_gn=UBTF;refGene_tx=XM_023243454.1
E1 44290631 44290765
refGene_gn=UBTF;refGene_tx=XM_003996979.5;refGene_gn=UBTF;refGene_tx=X
M_011289253.3;refGene_gn=UBTF;refGene_tx=XM_023243451.1;refGene_gn=UBT
F;refGene_tx=XM_023243452.1;refGene_gn=UBTF;refGene_tx=XM_023243453.1;
refGene_gn=UBTF;refGene_tx=XM_023243454.1
E1 44291107 44291218
refGene_gn=UBTF;refGene_tx=XM_003996979.5;refGene_gn=UBTF;refGene_tx=X
M_011289253.3;refGene_gn=UBTF;refGene_tx=XM_023243451.1;refGene_gn=UBT
F;refGene_tx=XM_023243452.1;refGene_gn=UBTF;refGene_tx=XM_023243453.1
E1 44291575 44291696
refGene_gn=UBTF;refGene_tx=XM_003996979.5;refGene_gn=UBTF;refGene_tx=X
M_011289253.3;refGene_gn=UBTF;refGene_tx=XM_023243451.1;refGene_gn=UBT
F;refGene_tx=XM_023243452.1;refGene_gn=UBTF;refGene_tx=XM_023243453.1;
refGene_gn=UBTF;refGene_tx=XM_023243454.1
E1 44291982 44292047
refGene_gn=UBTF;refGene_tx=XM_003996979.5;refGene_gn=UBTF;refGene_tx=X
M_011289253.3;refGene_gn=UBTF;refGene_tx=XM_023243451.1;refGene_gn=UBT
F;refGene_tx=XM_023243452.1;refGene_gn=UBTF;refGene_tx=XM_023243453.1;
refGene_gn=UBTF;refGene_tx=XM_023243454.1
E1 44294231 44294387
refGene_gn=UBTF;refGene_tx=XM_003996979.5;refGene_gn=UBTF;refGene_tx=X
M_011289253.3;refGene_gn=UBTF;refGene_tx=XM_023243451.1;refGene_gn=UBT
F;refGene_tx=XM_023243452.1;refGene_gn=UBTF;refGene_tx=XM_023243453.1;
refGene_gn=UBTF;refGene_tx=XM_023243454.1
E1 44294484 44294568
refGene_gn=UBTF;refGene_tx=XM_003996979.5;refGene_gn=UBTF;refGene_tx=X
M_011289253.3;refGene_gn=UBTF;refGene_tx=XM_023243451.1;refGene_gn=UBT
F;refGene_tx=XM_023243452.1;refGene_gn=UBTF;refGene_tx=XM_023243453.1;
refGene_gn=UBTF;refGene_tx=XM_023243454.1
E1 44295128 44295304
refGene_gn=UBTF;refGene_tx=XM_003996979.5;refGene_gn=UBTF;refGene_tx=X
M_011289253.3;refGene_gn=UBTF;refGene_tx=XM_023243451.1;refGene_gn=UBT
F;refGene_tx=XM_023243452.1;refGene_gn=UBTF;refGene_tx=XM_023243453.1;

refGene_gn=UBTF;refGene_tx=XM_023243454.1
E1 44296723 44296781
refGene_gn=UBTF;refGene_tx=XM_003996979.5;refGene_gn=UBTF;refGene_tx=XM_011289253.3;refGene_gn=UBTF;refGene_tx=XM_023243451.1;refGene_gn=UBTF;refGene_tx=XM_023243452.1;refGene_gn=UBTF;refGene_tx=XM_023243453.1;refGene_gn=UBTF;refGene_tx=XM_023243454.1
E1 44318532 44318613
refGene_gn=SLC4A1;refGene_tx=XM_019817978.2;refGene_gn=SLC4A1;refGene_tx=XM_019817979.2;refGene_gn=SLC4A1;refGene_tx=XM_019817980.2;refGene_gn=SLC4A1;refGene_tx=XM_019817981.2;refGene_gn=SLC4A1;refGene_tx=XM_019817982.2
E1 44319279 44319456
refGene_gn=SLC4A1;refGene_tx=XM_019817978.2;refGene_gn=SLC4A1;refGene_tx=XM_019817979.2;refGene_gn=SLC4A1;refGene_tx=XM_019817980.2;refGene_gn=SLC4A1;refGene_tx=XM_019817981.2;refGene_gn=SLC4A1;refGene_tx=XM_019817982.2
E1 44319550 44319720
refGene_gn=SLC4A1;refGene_tx=XM_019817978.2;refGene_gn=SLC4A1;refGene_tx=XM_019817979.2;refGene_gn=SLC4A1;refGene_tx=XM_019817980.2;refGene_gn=SLC4A1;refGene_tx=XM_019817981.2;refGene_gn=SLC4A1;refGene_tx=XM_019817982.2
E1 44320821 44321075
refGene_gn=SLC4A1;refGene_tx=XM_019817978.2;refGene_gn=SLC4A1;refGene_tx=XM_019817979.2;refGene_gn=SLC4A1;refGene_tx=XM_019817980.2;refGene_gn=SLC4A1;refGene_tx=XM_019817981.2;refGene_gn=SLC4A1;refGene_tx=XM_019817982.2
E1 44321567 44321734
refGene_gn=SLC4A1;refGene_tx=XM_019817978.2;refGene_gn=SLC4A1;refGene_tx=XM_019817979.2;refGene_gn=SLC4A1;refGene_tx=XM_019817980.2;refGene_gn=SLC4A1;refGene_tx=XM_019817981.2;refGene_gn=SLC4A1;refGene_tx=XM_019817982.2
E1 44322195 44322285
refGene_gn=SLC4A1;refGene_tx=XM_019817978.2;refGene_gn=SLC4A1;refGene_tx=XM_019817979.2;refGene_gn=SLC4A1;refGene_tx=XM_019817980.2;refGene_gn=SLC4A1;refGene_tx=XM_019817981.2;refGene_gn=SLC4A1;refGene_tx=XM_019817982.2
E1 44322729 44322903
refGene_gn=SLC4A1;refGene_tx=XM_019817978.2;refGene_gn=SLC4A1;refGene_tx=XM_019817979.2;refGene_gn=SLC4A1;refGene_tx=XM_019817980.2;refGene_gn=SLC4A1;refGene_tx=XM_019817981.2;refGene_gn=SLC4A1;refGene_tx=XM_019817982.2
E1 44323982 44324177
refGene_gn=SLC4A1;refGene_tx=XM_019817978.2;refGene_gn=SLC4A1;refGene_tx=XM_019817979.2;refGene_gn=SLC4A1;refGene_tx=XM_019817980.2;refGene_gn=SLC4A1;refGene_tx=XM_019817981.2;refGene_gn=SLC4A1;refGene_tx=XM_019817982.2
E1 44324265 44324414
refGene_gn=SLC4A1;refGene_tx=XM_019817978.2;refGene_gn=SLC4A1;refGene_tx=XM_019817979.2;refGene_gn=SLC4A1;refGene_tx=XM_019817980.2;refGene_gn=SLC4A1;refGene_tx=XM_019817981.2;refGene_gn=SLC4A1;refGene_tx=XM_01

9817982.2
E1 44324593 44324806
refGene_gn=SLC4A1;refGene_tx=XM_019817978.2;refGene_gn=SLC4A1;refGene_tx=XM_019817979.2;refGene_gn=SLC4A1;refGene_tx=XM_019817980.2;refGene_gn=SLC4A1;refGene_tx=XM_019817981.2;refGene_gn=SLC4A1;refGene_tx=XM_019817982.2
E1 44325454 44325659
refGene_gn=SLC4A1;refGene_tx=XM_019817978.2;refGene_gn=SLC4A1;refGene_tx=XM_019817979.2;refGene_gn=SLC4A1;refGene_tx=XM_019817980.2;refGene_gn=SLC4A1;refGene_tx=XM_019817981.2;refGene_gn=SLC4A1;refGene_tx=XM_019817982.2
E1 44326127 44326330
refGene_gn=SLC4A1;refGene_tx=XM_019817978.2;refGene_gn=SLC4A1;refGene_tx=XM_019817979.2;refGene_gn=SLC4A1;refGene_tx=XM_019817980.2;refGene_gn=SLC4A1;refGene_tx=XM_019817981.2;refGene_gn=SLC4A1;refGene_tx=XM_019817982.2
E1 44326484 44326569
refGene_gn=SLC4A1;refGene_tx=XM_019817978.2;refGene_gn=SLC4A1;refGene_tx=XM_019817979.2;refGene_gn=SLC4A1;refGene_tx=XM_019817980.2;refGene_gn=SLC4A1;refGene_tx=XM_019817981.2;refGene_gn=SLC4A1;refGene_tx=XM_019817982.2
E1 44326708 44326835
refGene_gn=SLC4A1;refGene_tx=XM_019817978.2;refGene_gn=SLC4A1;refGene_tx=XM_019817979.2;refGene_gn=SLC4A1;refGene_tx=XM_019817980.2;refGene_gn=SLC4A1;refGene_tx=XM_019817981.2;refGene_gn=SLC4A1;refGene_tx=XM_019817982.2
E1 44327283 44327419
refGene_gn=SLC4A1;refGene_tx=XM_019817978.2;refGene_gn=SLC4A1;refGene_tx=XM_019817979.2;refGene_gn=SLC4A1;refGene_tx=XM_019817980.2;refGene_gn=SLC4A1;refGene_tx=XM_019817981.2;refGene_gn=SLC4A1;refGene_tx=XM_019817982.2
E1 44327520 44327761
refGene_gn=SLC4A1;refGene_tx=XM_019817978.2;refGene_gn=SLC4A1;refGene_tx=XM_019817979.2;refGene_gn=SLC4A1;refGene_tx=XM_019817980.2;refGene_gn=SLC4A1;refGene_tx=XM_019817981.2;refGene_gn=SLC4A1;refGene_tx=XM_019817982.2
E1 44328567 44328629
refGene_gn=SLC4A1;refGene_tx=XM_019817978.2;refGene_gn=SLC4A1;refGene_tx=XM_019817979.2;refGene_gn=SLC4A1;refGene_tx=XM_019817980.2;refGene_gn=SLC4A1;refGene_tx=XM_019817981.2;refGene_gn=SLC4A1;refGene_tx=XM_019817982.2
E1 44329670 44329761
refGene_gn=SLC4A1;refGene_tx=XM_019817978.2;refGene_gn=SLC4A1;refGene_tx=XM_019817979.2;refGene_gn=SLC4A1;refGene_tx=XM_019817980.2;refGene_gn=SLC4A1;refGene_tx=XM_019817981.2;refGene_gn=SLC4A1;refGene_tx=XM_019817982.2
E1 44329886 44329901
refGene_gn=SLC4A1;refGene_tx=XM_019817978.2;refGene_gn=SLC4A1;refGene_tx=XM_019817979.2;refGene_gn=SLC4A1;refGene_tx=XM_019817980.2;refGene_gn=SLC4A1;refGene_tx=XM_019817981.2;refGene_gn=SLC4A1;refGene_tx=XM_01

9817982.2
E1 44358815 44358922
refGene_gn=RUNDC3A;refGene_tx=XM_003996982.5;refGene_gn=RUNDC3A;refGene_tx=XM_011289249.3;refGene_gn=RUNDC3A;refGene_tx=XM_011289250.2;refGene_gn=RUNDC3A;refGene_tx=XM_011289251.3;refGene_gn=RUNDC3A;refGene_tx=XM_019817983.2
E1 44362577 44362693
refGene_gn=RUNDC3A;refGene_tx=XM_003996982.5;refGene_gn=RUNDC3A;refGene_tx=XM_011289249.3;refGene_gn=RUNDC3A;refGene_tx=XM_011289250.2;refGene_gn=RUNDC3A;refGene_tx=XM_011289251.3;refGene_gn=RUNDC3A;refGene_tx=XM_019817983.2
E1 44363061 44363210
refGene_gn=RUNDC3A;refGene_tx=XM_003996982.5;refGene_gn=RUNDC3A;refGene_tx=XM_011289249.3;refGene_gn=RUNDC3A;refGene_tx=XM_011289250.2;refGene_gn=RUNDC3A;refGene_tx=XM_011289251.3;refGene_gn=RUNDC3A;refGene_tx=XM_019817983.2
E1 44363401 44363487
refGene_gn=RUNDC3A;refGene_tx=XM_003996982.5;refGene_gn=RUNDC3A;refGene_tx=XM_011289249.3;refGene_gn=RUNDC3A;refGene_tx=XM_011289250.2;refGene_gn=RUNDC3A;refGene_tx=XM_011289251.3;refGene_gn=RUNDC3A;refGene_tx=XM_019817983.2
E1 44364431 44364521
refGene_gn=RUNDC3A;refGene_tx=XM_003996982.5;refGene_gn=RUNDC3A;refGene_tx=XM_011289249.3;refGene_gn=RUNDC3A;refGene_tx=XM_011289250.2;refGene_gn=RUNDC3A;refGene_tx=XM_011289251.3;refGene_gn=RUNDC3A;refGene_tx=XM_019817983.2
E1 44364621 44364702
refGene_gn=RUNDC3A;refGene_tx=XM_003996982.5;refGene_gn=RUNDC3A;refGene_tx=XM_011289249.3;refGene_gn=RUNDC3A;refGene_tx=XM_011289250.2;refGene_gn=RUNDC3A;refGene_tx=XM_011289251.3;refGene_gn=RUNDC3A;refGene_tx=XM_019817983.2
E1 44364846 44365012
refGene_gn=RUNDC3A;refGene_tx=XM_003996982.5;refGene_gn=RUNDC3A;refGene_tx=XM_011289249.3;refGene_gn=RUNDC3A;refGene_tx=XM_011289250.2;refGene_gn=RUNDC3A;refGene_tx=XM_011289251.3;refGene_gn=RUNDC3A;refGene_tx=XM_019817983.2
E1 44365111 44365269
refGene_gn=RUNDC3A;refGene_tx=XM_003996982.5;refGene_gn=RUNDC3A;refGene_tx=XM_011289249.3;refGene_gn=RUNDC3A;refGene_tx=XM_011289250.2;refGene_gn=RUNDC3A;refGene_tx=XM_011289251.3;refGene_gn=RUNDC3A;refGene_tx=XM_019817983.2
E1 44366010 44366148
refGene_gn=RUNDC3A;refGene_tx=XM_003996982.5;refGene_gn=RUNDC3A;refGene_tx=XM_011289249.3;refGene_gn=RUNDC3A;refGene_tx=XM_011289250.2;refGene_gn=RUNDC3A;refGene_tx=XM_011289251.3;refGene_gn=RUNDC3A;refGene_tx=XM_019817983.2
E1 44366266 44366373
refGene_gn=RUNDC3A;refGene_tx=XM_003996982.5;refGene_gn=RUNDC3A;refGene_tx=XM_011289249.3;refGene_gn=RUNDC3A;refGene_tx=XM_011289250.2;refGene_gn=RUNDC3A;refGene_tx=XM_011289251.3;refGene_gn=RUNDC3A;refGene_tx=

XM_019817983.2
E1 44366798 44366818
refGene_gn=RUNDC3A;refGene_tx=XM_011289250.2;refGene_gn=RUNDC3A;refGene_tx=XM_011289251.3
E1 44367413 44367556
refGene_gn=RUNDC3A;refGene_tx=XM_003996982.5;refGene_gn=RUNDC3A;refGene_tx=XM_011289249.3;refGene_gn=RUNDC3A;refGene_tx=XM_019817983.2
E1 44369093 44369209
refGene_gn=SLC25A39;refGene_tx=XM_003996983.3;refGene_gn=SLC25A39;refGene_tx=XM_003996984.3;refGene_gn=SLC25A39;refGene_tx=XM_006940420.4
E1 44369301 44369382
refGene_gn=SLC25A39;refGene_tx=XM_003996983.3;refGene_gn=SLC25A39;refGene_tx=XM_003996984.3;refGene_gn=SLC25A39;refGene_tx=XM_006940420.4
E1 44369461 44369543
refGene_gn=SLC25A39;refGene_tx=XM_003996983.3;refGene_gn=SLC25A39;refGene_tx=XM_003996984.3;refGene_gn=SLC25A39;refGene_tx=XM_006940420.4
E1 44369769 44369879
refGene_gn=SLC25A39;refGene_tx=XM_003996983.3;refGene_gn=SLC25A39;refGene_tx=XM_003996984.3;refGene_gn=SLC25A39;refGene_tx=XM_006940420.4
E1 44370221 44370395
refGene_gn=SLC25A39;refGene_tx=XM_003996983.3;refGene_gn=SLC25A39;refGene_tx=XM_003996984.3;refGene_gn=SLC25A39;refGene_tx=XM_006940420.4
E1 44370576 44370701
refGene_gn=SLC25A39;refGene_tx=XM_003996983.3;refGene_gn=SLC25A39;refGene_tx=XM_003996984.3;refGene_gn=SLC25A39;refGene_tx=XM_006940420.4
E1 44370844 44370912
refGene_gn=SLC25A39;refGene_tx=XM_003996983.3;refGene_gn=SLC25A39;refGene_tx=XM_003996984.3;refGene_gn=SLC25A39;refGene_tx=XM_006940420.4
E1 44371255 44371389
refGene_gn=SLC25A39;refGene_tx=XM_003996983.3;refGene_gn=SLC25A39;refGene_tx=XM_003996984.3;refGene_gn=SLC25A39;refGene_tx=XM_006940420.4
E1 44371622 44371667
refGene_gn=SLC25A39;refGene_tx=XM_003996983.3;refGene_gn=SLC25A39;refGene_tx=XM_003996984.3;refGene_gn=SLC25A39;refGene_tx=XM_006940420.4
E1 44371791 44371851
refGene_gn=SLC25A39;refGene_tx=XM_003996983.3;refGene_gn=SLC25A39;refGene_tx=XM_003996984.3;refGene_gn=SLC25A39;refGene_tx=XM_006940420.4
E1 44371984 44372069
refGene_gn=SLC25A39;refGene_tx=XM_003996983.3;refGene_gn=SLC25A39;refGene_tx=XM_003996984.3;refGene_gn=SLC25A39;refGene_tx=XM_006940420.4
E1 44392593 44392731
refGene_gn=GRN;refGene_tx=XM_011289247.3;refGene_gn=GRN;refGene_tx=XM_011289248.3
E1 44392856 44392982
refGene_gn=GRN;refGene_tx=XM_011289247.3;refGene_gn=GRN;refGene_tx=XM_011289248.3
E1 44393090 44393175
refGene_gn=GRN;refGene_tx=XM_011289247.3;refGene_gn=GRN;refGene_tx=XM_011289248.3
E1 44393661 44393774

refGene_gn=GRN;refGene_tx=XM_011289247.3;refGene_gn=GRN;refGene_tx=XM_011289248.3
E1 44393870 44394006
refGene_gn=GRN;refGene_tx=XM_011289247.3;refGene_gn=GRN;refGene_tx=XM_011289248.3
E1 44394108 44394221
refGene_gn=GRN;refGene_tx=XM_011289247.3;refGene_gn=GRN;refGene_tx=XM_011289248.3
E1 44394431 44394555
refGene_gn=GRN;refGene_tx=XM_011289247.3;refGene_gn=GRN;refGene_tx=XM_011289248.3
E1 44394834 44394932
refGene_gn=GRN;refGene_tx=XM_011289247.3;refGene_gn=GRN;refGene_tx=XM_011289248.3
E1 44395022 44395268
refGene_gn=GRN;refGene_tx=XM_011289247.3;refGene_gn=GRN;refGene_tx=XM_011289248.3
E1 44395518 44395752
refGene_gn=GRN;refGene_tx=XM_011289247.3;refGene_gn=GRN;refGene_tx=XM_011289248.3
E1 44395842 44396073
refGene_gn=GRN;refGene_tx=XM_011289247.3;refGene_gn=GRN;refGene_tx=XM_011289248.3
E1 44396169 44396307
refGene_gn=GRN;refGene_tx=XM_011289247.3;refGene_gn=GRN;refGene_tx=XM_011289248.3
E1 44397284 44398764 refGene_gn=FAM171A2;refGene_tx=XM_023243573.1
E1 44399330 44399457 refGene_gn=FAM171A2;refGene_tx=XM_023243573.1
E1 44399578 44399695 refGene_gn=FAM171A2;refGene_tx=XM_023243573.1
E1 44399804 44399984 refGene_gn=FAM171A2;refGene_tx=XM_023243573.1
E1 44400068 44400227 refGene_gn=FAM171A2;refGene_tx=XM_023243573.1
E1 44402893 44402986 refGene_gn=FAM171A2;refGene_tx=XM_023243573.1
E1 44403219 44403447 refGene_gn=FAM171A2;refGene_tx=XM_023243573.1
E1 44406755 44406873 refGene_gn=FAM171A2;refGene_tx=XM_023243573.1
E1 44410454 44410511 refGene_gn=ITGA2B;refGene_tx=XM_003996987.5
E1 44411443 44411560 refGene_gn=ITGA2B;refGene_tx=XM_003996987.5
E1 44411774 44411876 refGene_gn=ITGA2B;refGene_tx=XM_003996987.5
E1 44412095 44412209 refGene_gn=ITGA2B;refGene_tx=XM_003996987.5
E1 44412684 44412810 refGene_gn=ITGA2B;refGene_tx=XM_003996987.5
E1 44412932 44413085 refGene_gn=ITGA2B;refGene_tx=XM_003996987.5
E1 44413188 44413285 refGene_gn=ITGA2B;refGene_tx=XM_003996987.5
E1 44413406 44413487 refGene_gn=ITGA2B;refGene_tx=XM_003996987.5
E1 44413807 44413887 refGene_gn=ITGA2B;refGene_tx=XM_003996987.5
E1 44414155 44414248 refGene_gn=ITGA2B;refGene_tx=XM_003996987.5
E1 44415267 44415415 refGene_gn=ITGA2B;refGene_tx=XM_003996987.5
E1 44415564 44415632 refGene_gn=ITGA2B;refGene_tx=XM_003996987.5
E1 44416024 44416150 refGene_gn=ITGA2B;refGene_tx=XM_003996987.5
E1 44416294 44416446 refGene_gn=ITGA2B;refGene_tx=XM_003996987.5
E1 44416534 44416590 refGene_gn=ITGA2B;refGene_tx=XM_003996987.5
E1 44416673 44416778 refGene_gn=ITGA2B;refGene_tx=XM_003996987.5

E1 44416877 44416923 refGene_gn=ITGA2B;refGene_tx=XM_003996987.5
E1 44417172 44417355 refGene_gn=ITGA2B;refGene_tx=XM_003996987.5
E1 44420093 44420305 refGene_gn=ITGA2B;refGene_tx=XM_003996987.5
E1 44420499 44420552 refGene_gn=ITGA2B;refGene_tx=XM_003996987.5
E1 44420689 44420743 refGene_gn=ITGA2B;refGene_tx=XM_003996987.5
E1 44420889 44420933 refGene_gn=ITGA2B;refGene_tx=XM_003996987.5
E1 44421087 44421135 refGene_gn=ITGA2B;refGene_tx=XM_003996987.5
E1 44421510 44421639 refGene_gn=ITGA2B;refGene_tx=XM_003996987.5
E1 44421732 44421778 refGene_gn=ITGA2B;refGene_tx=XM_003996987.5
E1 44421853 44421903 refGene_gn=ITGA2B;refGene_tx=XM_003996987.5
E1 44422178 44422344 refGene_gn=ITGA2B;refGene_tx=XM_003996987.5
E1 44422457 44422555 refGene_gn=ITGA2B;refGene_tx=XM_003996987.5
E1 44422643 44422765 refGene_gn=ITGA2B;refGene_tx=XM_003996987.5
E1 44424312 44424500 refGene_gn=ITGA2B;refGene_tx=XM_003996987.5
E1 44433383 44434901 refGene_gn=GPATCH8;refGene_tx=XM_003996988.5
E1 44434902 44435241 refGene_gn=GPATCH8;refGene_tx=XM_003996988.5
E1 44435242 44437274 refGene_gn=GPATCH8;refGene_tx=XM_003996988.5
E1 44440979 44441110 refGene_gn=GPATCH8;refGene_tx=XM_003996988.5
E1 44455451 44455595 refGene_gn=GPATCH8;refGene_tx=XM_003996988.5
E1 44474449 44474536 refGene_gn=GPATCH8;refGene_tx=XM_003996988.5
E1 44477198 44477266 refGene_gn=GPATCH8;refGene_tx=XM_003996988.5
E1 44499238 44499311 refGene_gn=GPATCH8;refGene_tx=XM_003996988.5
E1 44507674 44507749 refGene_gn=GPATCH8;refGene_tx=XM_003996988.5
E1 44535328 44535373 refGene_gn=GPATCH8;refGene_tx=XM_003996988.5
E1 44576143 44577841 refGene_gn=FZD2;refGene_tx=XM_011289270.2
E1 44646841 44646907 refGene_gn=MEIOC;refGene_tx=XM_023244517.1
E1 44647484 44647619 refGene_gn=MEIOC;refGene_tx=XM_023244517.1
E1 44650879 44651037 refGene_gn=MEIOC;refGene_tx=XM_023244517.1
E1 44653913 44654018 refGene_gn=MEIOC;refGene_tx=XM_023244517.1
E1 44655168 44657041 refGene_gn=MEIOC;refGene_tx=XM_023244517.1
E1 44657499 44657634 refGene_gn=MEIOC;refGene_tx=XM_023244517.1
E1 44661008 44661189 refGene_gn=MEIOC;refGene_tx=XM_023244517.1
E1 44661612 44661842 refGene_gn=MEIOC;refGene_tx=XM_023244517.1
E1 44666086 44666274
refGene_gn=CCDC43;refGene_tx=XM_003996989.5;refGene_gn=CCDC43;refGene_
tx=XM_006940423.4
E1 44668092 44668151 refGene_gn=CCDC43;refGene_tx=XM_003996989.5
E1 44669627 44669763
refGene_gn=CCDC43;refGene_tx=XM_003996989.5;refGene_gn=CCDC43;refGene_
tx=XM_006940423.4
E1 44671334 44671422
refGene_gn=CCDC43;refGene_tx=XM_003996989.5;refGene_gn=CCDC43;refGene_
tx=XM_006940423.4
E1 44675127 44675409
refGene_gn=CCDC43;refGene_tx=XM_003996989.5;refGene_gn=CCDC43;refGene_
tx=XM_006940423.4
E1 44700618 44700652 refGene_gn=DBF4B;refGene_tx=XM_023244518.1
E1 44713106 44713261 refGene_gn=DBF4B;refGene_tx=XM_023244518.1
E1 44716487 44716673 refGene_gn=DBF4B;refGene_tx=XM_023244518.1
E1 44717290 44717341 refGene_gn=DBF4B;refGene_tx=XM_023244518.1

E1 44718112 44718200 refGene_gn=DBF4B;refGene_tx=XM_023244518.1
E1 44720492 44720569 refGene_gn=DBF4B;refGene_tx=XM_023244518.1
E1 44722230 44722264 refGene_gn=DBF4B;refGene_tx=XM_023244518.1
E1 44723711 44723757 refGene_gn=DBF4B;refGene_tx=XM_023244518.1
E1 44726291 44726408 refGene_gn=DBF4B;refGene_tx=XM_023244518.1
E1 44729167 44729276 refGene_gn=DBF4B;refGene_tx=XM_023244518.1
E1 44729402 44729527 refGene_gn=DBF4B;refGene_tx=XM_023244518.1
E1 44730332 44731001 refGene_gn=DBF4B;refGene_tx=XM_023244518.1
E1 44731002 44731115 refGene_gn=DBF4B;refGene_tx=XM_023244518.1
E1 44731117 44731215 refGene_gn=DBF4B;refGene_tx=XM_023244518.1
E1 44737928 44737995
refGene_gn=ADAM11;refGene_tx=XM_023243682.1;refGene_gn=ADAM11;refGene_tx=XM_023243683.1;refGene_gn=ADAM11;refGene_tx=XM_023243684.1
E1 44738444 44738620
refGene_gn=ADAM11;refGene_tx=XM_023243682.1;refGene_gn=ADAM11;refGene_tx=XM_023243683.1;refGene_gn=ADAM11;refGene_tx=XM_023243684.1
E1 44747170 44747247
refGene_gn=ADAM11;refGene_tx=XM_023243682.1;refGene_gn=ADAM11;refGene_tx=XM_023243683.1;refGene_gn=ADAM11;refGene_tx=XM_023243684.1
E1 44747443 44747510
refGene_gn=ADAM11;refGene_tx=XM_023243682.1;refGene_gn=ADAM11;refGene_tx=XM_023243683.1;refGene_gn=ADAM11;refGene_tx=XM_023243684.1
E1 44748521 44748607
refGene_gn=ADAM11;refGene_tx=XM_023243682.1;refGene_gn=ADAM11;refGene_tx=XM_023243683.1;refGene_gn=ADAM11;refGene_tx=XM_023243684.1
E1 44748696 44748772
refGene_gn=ADAM11;refGene_tx=XM_023243682.1;refGene_gn=ADAM11;refGene_tx=XM_023243683.1;refGene_gn=ADAM11;refGene_tx=XM_023243684.1
E1 44749172 44749239
refGene_gn=ADAM11;refGene_tx=XM_019817998.2;refGene_gn=ADAM11;refGene_tx=XM_023243682.1;refGene_gn=ADAM11;refGene_tx=XM_023243683.1;refGene_gn=ADAM11;refGene_tx=XM_023243684.1
E1 44749307 44749375
refGene_gn=ADAM11;refGene_tx=XM_019817998.2;refGene_gn=ADAM11;refGene_tx=XM_023243682.1;refGene_gn=ADAM11;refGene_tx=XM_023243683.1;refGene_gn=ADAM11;refGene_tx=XM_023243684.1
E1 44749732 44749807
refGene_gn=ADAM11;refGene_tx=XM_019817998.2;refGene_gn=ADAM11;refGene_tx=XM_023243682.1;refGene_gn=ADAM11;refGene_tx=XM_023243683.1;refGene_gn=ADAM11;refGene_tx=XM_023243684.1
E1 44749887 44749959
refGene_gn=ADAM11;refGene_tx=XM_019817998.2;refGene_gn=ADAM11;refGene_tx=XM_023243682.1;refGene_gn=ADAM11;refGene_tx=XM_023243683.1;refGene_gn=ADAM11;refGene_tx=XM_023243684.1
E1 44750169 44750336
refGene_gn=ADAM11;refGene_tx=XM_019817998.2;refGene_gn=ADAM11;refGene_tx=XM_023243682.1;refGene_gn=ADAM11;refGene_tx=XM_023243683.1;refGene_gn=ADAM11;refGene_tx=XM_023243684.1
E1 44750872 44750957
refGene_gn=ADAM11;refGene_tx=XM_019817998.2;refGene_gn=ADAM11;refGene_

tx=XM_023243682.1;refGene_gn=ADAM11;refGene_tx=XM_023243683.1;refGene_gn=ADAM11;refGene_tx=XM_023243684.1
E1 44751075 44751166
refGene_gn=ADAM11;refGene_tx=XM_019817998.2;refGene_gn=ADAM11;refGene_tx=XM_023243682.1;refGene_gn=ADAM11;refGene_tx=XM_023243683.1;refGene_gn=ADAM11;refGene_tx=XM_023243684.1
E1 44751298 44751350
refGene_gn=ADAM11;refGene_tx=XM_019817998.2;refGene_gn=ADAM11;refGene_tx=XM_023243682.1;refGene_gn=ADAM11;refGene_tx=XM_023243683.1;refGene_gn=ADAM11;refGene_tx=XM_023243684.1
E1 44751852 44751952
refGene_gn=ADAM11;refGene_tx=XM_019817998.2;refGene_gn=ADAM11;refGene_tx=XM_023243682.1;refGene_gn=ADAM11;refGene_tx=XM_023243683.1;refGene_gn=ADAM11;refGene_tx=XM_023243684.1
E1 44752047 44752119
refGene_gn=ADAM11;refGene_tx=XM_019817998.2;refGene_gn=ADAM11;refGene_tx=XM_023243682.1;refGene_gn=ADAM11;refGene_tx=XM_023243683.1;refGene_gn=ADAM11;refGene_tx=XM_023243684.1
E1 44752232 44752325
refGene_gn=ADAM11;refGene_tx=XM_019817998.2;refGene_gn=ADAM11;refGene_tx=XM_023243682.1;refGene_gn=ADAM11;refGene_tx=XM_023243683.1;refGene_gn=ADAM11;refGene_tx=XM_023243684.1
E1 44752723 44752804
refGene_gn=ADAM11;refGene_tx=XM_019817998.2;refGene_gn=ADAM11;refGene_tx=XM_023243682.1;refGene_gn=ADAM11;refGene_tx=XM_023243683.1;refGene_gn=ADAM11;refGene_tx=XM_023243684.1
E1 44753324 44753375
refGene_gn=ADAM11;refGene_tx=XM_019817998.2;refGene_gn=ADAM11;refGene_tx=XM_023243682.1;refGene_gn=ADAM11;refGene_tx=XM_023243683.1;refGene_gn=ADAM11;refGene_tx=XM_023243684.1
E1 44753490 44753554
refGene_gn=ADAM11;refGene_tx=XM_019817998.2;refGene_gn=ADAM11;refGene_tx=XM_023243682.1;refGene_gn=ADAM11;refGene_tx=XM_023243683.1;refGene_gn=ADAM11;refGene_tx=XM_023243684.1
E1 44753769 44753869
refGene_gn=ADAM11;refGene_tx=XM_019817998.2;refGene_gn=ADAM11;refGene_tx=XM_023243682.1;refGene_gn=ADAM11;refGene_tx=XM_023243683.1;refGene_gn=ADAM11;refGene_tx=XM_023243684.1
E1 44754093 44754213
refGene_gn=ADAM11;refGene_tx=XM_019817998.2;refGene_gn=ADAM11;refGene_tx=XM_023243682.1;refGene_gn=ADAM11;refGene_tx=XM_023243683.1;refGene_gn=ADAM11;refGene_tx=XM_023243684.1
E1 44754306 44754475
refGene_gn=ADAM11;refGene_tx=XM_019817998.2;refGene_gn=ADAM11;refGene_tx=XM_023243683.1;refGene_gn=ADAM11;refGene_tx=XM_023243684.1
E1 44754566 44754681
refGene_gn=ADAM11;refGene_tx=XM_019817998.2;refGene_gn=ADAM11;refGene_tx=XM_023243682.1;refGene_gn=ADAM11;refGene_tx=XM_023243683.1;refGene_gn=ADAM11;refGene_tx=XM_023243684.1
E1 44754768 44754859

refGene_gn=ADAM11;refGene_tx=XM_019817998.2;refGene_gn=ADAM11;refGene_tx=XM_023243682.1;refGene_gn=ADAM11;refGene_tx=XM_023243683.1;refGene_gn=ADAM11;refGene_tx=XM_023243684.1
E1 44755486 44755535
refGene_gn=ADAM11;refGene_tx=XM_019817998.2;refGene_gn=ADAM11;refGene_tx=XM_023243682.1;refGene_gn=ADAM11;refGene_tx=XM_023243683.1;refGene_gn=ADAM11;refGene_tx=XM_023243684.1
E1 44756025 44756303
refGene_gn=ADAM11;refGene_tx=XM_023243682.1;refGene_gn=ADAM11;refGene_tx=XM_023243684.1
E1 44776601 44777792
refGene_gn=GJC1;refGene_tx=XM_011289239.3;refGene_gn=GJC1;refGene_tx=XM_011289240.3;refGene_gn=GJC1;refGene_tx=XM_011289241.3;refGene_gn=GJC1;refGene_tx=XM_011289242.3;refGene_gn=GJC1;refGene_tx=XM_019818000.2;refGene_gn=GJC1;refGene_tx=XM_023243558.1
E1 44819765 44819768 refGene_gn=HIGD1B;refGene_tx=XM_003996991.5
E1 44819857 44820008
refGene_gn=HIGD1B;refGene_tx=XM_003996991.5;refGene_gn=HIGD1B;refGene_tx=XM_019818006.2
E1 44820596 44820731
refGene_gn=HIGD1B;refGene_tx=XM_003996991.5;refGene_gn=HIGD1B;refGene_tx=XM_019818006.2
E1 44821333 44821398
refGene_gn=HIGD1B;refGene_tx=XM_003996991.5;refGene_gn=HIGD1B;refGene_tx=XM_019818006.2
E1 44822445 44822541 refGene_gn=EFTUD2;refGene_tx=XM_003996992.5
E1 44822883 44822991 refGene_gn=EFTUD2;refGene_tx=XM_003996992.5
E1 44823138 44823292 refGene_gn=EFTUD2;refGene_tx=XM_003996992.5
E1 44823792 44823887 refGene_gn=EFTUD2;refGene_tx=XM_003996992.5
E1 44824043 44824162 refGene_gn=EFTUD2;refGene_tx=XM_003996992.5
E1 44824827 44824915 refGene_gn=EFTUD2;refGene_tx=XM_003996992.5
E1 44825107 44825234 refGene_gn=EFTUD2;refGene_tx=XM_003996992.5
E1 44825416 44825503 refGene_gn=EFTUD2;refGene_tx=XM_003996992.5
E1 44826330 44826413 refGene_gn=EFTUD2;refGene_tx=XM_003996992.5
E1 44827390 44827492 refGene_gn=EFTUD2;refGene_tx=XM_003996992.5
E1 44828893 44829034 refGene_gn=EFTUD2;refGene_tx=XM_003996992.5
E1 44829398 44829510 refGene_gn=EFTUD2;refGene_tx=XM_003996992.5
E1 44830925 44831119 refGene_gn=EFTUD2;refGene_tx=XM_003996992.5
E1 44832226 44832354 refGene_gn=EFTUD2;refGene_tx=XM_003996992.5
E1 44833976 44834112 refGene_gn=EFTUD2;refGene_tx=XM_003996992.5
E1 44835545 44835636 refGene_gn=EFTUD2;refGene_tx=XM_003996992.5
E1 44836217 44836281 refGene_gn=EFTUD2;refGene_tx=XM_003996992.5
E1 44839643 44839768 refGene_gn=EFTUD2;refGene_tx=XM_003996992.5
E1 44842026 44842193 refGene_gn=EFTUD2;refGene_tx=XM_003996992.5
E1 44845274 44845357 refGene_gn=EFTUD2;refGene_tx=XM_003996992.5
E1 44846286 44846377 refGene_gn=EFTUD2;refGene_tx=XM_003996992.5
E1 44848813 44848849 refGene_gn=EFTUD2;refGene_tx=XM_003996992.5
E1 44849601 44849667 refGene_gn=EFTUD2;refGene_tx=XM_003996992.5
E1 44850149 44850225 refGene_gn=EFTUD2;refGene_tx=XM_003996992.5
E1 44851754 44851833 refGene_gn=EFTUD2;refGene_tx=XM_003996992.5

E1 44852498 44852664 refGene_gn=EFTUD2;refGene_tx=XM_003996992.5
E1 44857893 44857998 refGene_gn=EFTUD2;refGene_tx=XM_003996992.5
E1 44861646 44861676 refGene_gn=CCDC103;refGene_tx=XM_006940425.3
E1 44861958 44861961 refGene_gn=CCDC103;refGene_tx=XM_006940426.3
E1 44863378 44863530
refGene_gn=CCDC103;refGene_tx=XM_003996994.3;refGene_gn=CCDC103;refGen
e_tx=XM_006940425.3;refGene_gn=CCDC103;refGene_tx=XM_006940426.3;refGe
ne_gn=CCDC103;refGene_tx=XM_006940427.2;refGene_gn=CCDC103;refGene_tx=
XM_019818003.2;refGene_gn=CCDC103;refGene_tx=XM_019818004.2;refGene_gn
=CCDC103;refGene_tx=XM_019818005.2
E1 44863926 44864059
refGene_gn=CCDC103;refGene_tx=XM_003996994.3;refGene_gn=CCDC103;refGen
e_tx=XM_006940425.3;refGene_gn=CCDC103;refGene_tx=XM_006940426.3;refGe
ne_gn=CCDC103;refGene_tx=XM_006940427.2;refGene_gn=CCDC103;refGene_tx=
XM_019818003.2;refGene_gn=CCDC103;refGene_tx=XM_019818004.2;refGene_gn
=CCDC103;refGene_tx=XM_019818005.2
E1 44864758 44865220
refGene_gn=CCDC103;refGene_tx=XM_003996994.3;refGene_gn=CCDC103;refGen
e_tx=XM_006940425.3;refGene_gn=CCDC103;refGene_tx=XM_006940426.3;refGe
ne_gn=CCDC103;refGene_tx=XM_006940427.2;refGene_gn=CCDC103;refGene_tx=
XM_019818003.2;refGene_gn=CCDC103;refGene_tx=XM_019818004.2;refGene_gn
=CCDC103;refGene_tx=XM_019818005.2
E1 44866462 44867704
refGene_gn=LOC101086018;refGene_tx=XM_003997011.3
E1 44870013 44870055
refGene_gn=GFAP;refGene_tx=XM_003996995.4;refGene_gn=GFAP;refGene_tx=X
M_019818001.1
E1 44870697 44870783
refGene_gn=GFAP;refGene_tx=XM_003996995.4;refGene_gn=GFAP;refGene_tx=X
M_019818001.1
E1 44872434 44872554 refGene_gn=GFAP;refGene_tx=XM_019818001.1
E1 44872904 44872948
refGene_gn=GFAP;refGene_tx=XM_003996995.4;refGene_gn=GFAP;refGene_tx=X
M_019818001.1
E1 44873520 44873741
refGene_gn=GFAP;refGene_tx=XM_003996995.4;refGene_gn=GFAP;refGene_tx=X
M_019818001.1
E1 44873971 44874097
refGene_gn=GFAP;refGene_tx=XM_003996995.4;refGene_gn=GFAP;refGene_tx=X
M_019818001.1
E1 44875272 44875434
refGene_gn=GFAP;refGene_tx=XM_003996995.4;refGene_gn=GFAP;refGene_tx=X
M_019818001.1
E1 44876140 44876236
refGene_gn=GFAP;refGene_tx=XM_003996995.4;refGene_gn=GFAP;refGene_tx=X
M_019818001.1
E1 44876429 44876490
refGene_gn=GFAP;refGene_tx=XM_003996995.4;refGene_gn=GFAP;refGene_tx=X
M_019818001.1
E1 44877493 44877954

refGene_gn=GFAP;refGene_tx=XM_003996995.4;refGene_gn=GFAP;refGene_tx=XM_019818001.1
E1 44885995 44886102 refGene_gn=KIF18B;refGene_tx=XM_006940428.2;refGene_gn=KIF18B;refGene_tx=XM_019818007.2
E1 44886326 44886430 refGene_gn=KIF18B;refGene_tx=XM_006940428.2;refGene_gn=KIF18B;refGene_tx=XM_019818007.2
E1 44886937 44887027 refGene_gn=KIF18B;refGene_tx=XM_006940428.2;refGene_gn=KIF18B;refGene_tx=XM_019818007.2
E1 44887961 44888490 refGene_gn=KIF18B;refGene_tx=XM_006940428.2;refGene_gn=KIF18B;refGene_tx=XM_019818007.2
E1 44888711 44888911 refGene_gn=KIF18B;refGene_tx=XM_006940428.2;refGene_gn=KIF18B;refGene_tx=XM_019818007.2
E1 44891487 44891615 refGene_gn=KIF18B;refGene_tx=XM_006940428.2;refGene_gn=KIF18B;refGene_tx=XM_019818007.2
E1 44891977 44892155 refGene_gn=KIF18B;refGene_tx=XM_006940428.2;refGene_gn=KIF18B;refGene_tx=XM_019818007.2
E1 44892639 44892740 refGene_gn=KIF18B;refGene_tx=XM_006940428.2;refGene_gn=KIF18B;refGene_tx=XM_019818007.2
E1 44892879 44892954 refGene_gn=KIF18B;refGene_tx=XM_006940428.2;refGene_gn=KIF18B;refGene_tx=XM_019818007.2
E1 44893859 44894036 refGene_gn=KIF18B;refGene_tx=XM_006940428.2;refGene_gn=KIF18B;refGene_tx=XM_019818007.2
E1 44894177 44894375 refGene_gn=KIF18B;refGene_tx=XM_006940428.2;refGene_gn=KIF18B;refGene_tx=XM_019818007.2
E1 44894455 44894566 refGene_gn=KIF18B;refGene_tx=XM_006940428.2;refGene_gn=KIF18B;refGene_tx=XM_019818007.2
E1 44894757 44894862 refGene_gn=KIF18B;refGene_tx=XM_006940428.2;refGene_gn=KIF18B;refGene_tx=XM_019818007.2
E1 44895162 44895320 refGene_gn=KIF18B;refGene_tx=XM_006940428.2;refGene_gn=KIF18B;refGene_tx=XM_019818007.2
E1 44895960 44896287 refGene_gn=KIF18B;refGene_tx=XM_006940428.2;refGene_gn=KIF18B;refGene_tx=XM_019818007.2
E1 44897941 44897957 refGene_gn=KIF18B;refGene_tx=XM_019818007.2
E1 44915057 44915237 refGene_gn=C1QL1;refGene_tx=XM_003997012.5
E1 44919453 44919501 refGene_gn=C1QL1;refGene_tx=XM_023243762.1

E1	44920732	44921329
refGene_gn=C1QL1;refGene_tx=XM_003997012.5;refGene_gn=C1QL1;refGene_tx=XM_023243762.1
E1	44965473	44965765
refGene_gn=DCAKD;refGene_tx=XM_003996997.5;refGene_gn=DCAKD;refGene_tx=XM_006940429.4;refGene_gn=DCAKD;refGene_tx=XM_011289235.3;refGene_gn=DCAKD;refGene_tx=XM_011289236.3;refGene_gn=DCAKD;refGene_tx=XM_019818009.2
E1	44969114	44969202
refGene_gn=DCAKD;refGene_tx=XM_003996997.5;refGene_gn=DCAKD;refGene_tx=XM_006940429.4;refGene_gn=DCAKD;refGene_tx=XM_011289235.3;refGene_gn=DCAKD;refGene_tx=XM_011289236.3;refGene_gn=DCAKD;refGene_tx=XM_019818009.2
E1	44972639	44972843
refGene_gn=DCAKD;refGene_tx=XM_003996997.5;refGene_gn=DCAKD;refGene_tx=XM_006940429.4;refGene_gn=DCAKD;refGene_tx=XM_011289235.3;refGene_gn=DCAKD;refGene_tx=XM_011289236.3;refGene_gn=DCAKD;refGene_tx=XM_019818009.2
E1	44973481	44973593
refGene_gn=DCAKD;refGene_tx=XM_003996997.5;refGene_gn=DCAKD;refGene_tx=XM_006940429.4;refGene_gn=DCAKD;refGene_tx=XM_011289235.3;refGene_gn=DCAKD;refGene_tx=XM_019818009.2
E1	44991065	44991135	refGene_gn=DCAKD;refGene_tx=XM_011289236.3
E1	44991765	44991896	refGene_gn=NMT1;refGene_tx=XM_003996998.5
E1	45003356	45003465	refGene_gn=NMT1;refGene_tx=XM_003996998.5
E1	45005992	45006137	refGene_gn=NMT1;refGene_tx=XM_003996998.5
E1	45013078	45013197	refGene_gn=NMT1;refGene_tx=XM_003996998.5
E1	45013781	45013873	refGene_gn=NMT1;refGene_tx=XM_003996998.5
E1	45014744	45014861	refGene_gn=NMT1;refGene_tx=XM_003996998.5
E1	45015672	45015843	refGene_gn=NMT1;refGene_tx=XM_003996998.5
E1	45016647	45016756	refGene_gn=NMT1;refGene_tx=XM_003996998.5
E1	45019663	45019834	refGene_gn=NMT1;refGene_tx=XM_003996998.5
E1	45020458	45020626	refGene_gn=NMT1;refGene_tx=XM_003996998.5
E1	45021598	45021736	refGene_gn=NMT1;refGene_tx=XM_003996998.5
E1	45022393	45022414	refGene_gn=NMT1;refGene_tx=XM_003996998.5
E1	45028728	45028817
refGene_gn=PLCD3;refGene_tx=XM_023243637.1;refGene_gn=PLCD3;refGene_tx=XM_023243638.1;refGene_gn=PLCD3;refGene_tx=XM_023243640.1;refGene_gn=PLCD3;refGene_tx=XM_023243641.1;refGene_gn=PLCD3;refGene_tx=XM_023243642.1
E1	45028996	45029146
refGene_gn=PLCD3;refGene_tx=XM_023243637.1;refGene_gn=PLCD3;refGene_tx=XM_023243638.1;refGene_gn=PLCD3;refGene_tx=XM_023243640.1;refGene_gn=PLCD3;refGene_tx=XM_023243641.1;refGene_gn=PLCD3;refGene_tx=XM_023243642.1
E1	45029221	45029357
refGene_gn=PLCD3;refGene_tx=XM_023243637.1;refGene_gn=PLCD3;refGene_tx=XM_023243638.1;refGene_gn=PLCD3;refGene_tx=XM_023243640.1;refGene_gn=PLCD3;refGene_tx=XM_023243641.1;refGene_gn=PLCD3;refGene_tx=XM_023243642.1

E1 45029597 45029764
refGene_gn=PLCD3;refGene_tx=XM_023243637.1;refGene_gn=PLCD3;refGene_tx=XM_023243638.1;refGene_gn=PLCD3;refGene_tx=XM_023243640.1;refGene_gn=PLCD3;refGene_tx=XM_023243641.1;refGene_gn=PLCD3;refGene_tx=XM_023243642.1
E1 45030366 45030483
refGene_gn=PLCD3;refGene_tx=XM_023243637.1;refGene_gn=PLCD3;refGene_tx=XM_023243638.1;refGene_gn=PLCD3;refGene_tx=XM_023243640.1;refGene_gn=PLCD3;refGene_tx=XM_023243641.1;refGene_gn=PLCD3;refGene_tx=XM_023243642.1
E1 45031188 45031351
refGene_gn=PLCD3;refGene_tx=XM_023243637.1;refGene_gn=PLCD3;refGene_tx=XM_023243638.1;refGene_gn=PLCD3;refGene_tx=XM_023243640.1;refGene_gn=PLCD3;refGene_tx=XM_023243641.1;refGene_gn=PLCD3;refGene_tx=XM_023243642.1
E1 45031442 45031577
refGene_gn=PLCD3;refGene_tx=XM_023243637.1;refGene_gn=PLCD3;refGene_tx=XM_023243638.1;refGene_gn=PLCD3;refGene_tx=XM_023243640.1;refGene_gn=PLCD3;refGene_tx=XM_023243641.1;refGene_gn=PLCD3;refGene_tx=XM_023243642.1
E1 45033225 45033378
refGene_gn=PLCD3;refGene_tx=XM_023243637.1;refGene_gn=PLCD3;refGene_tx=XM_023243638.1;refGene_gn=PLCD3;refGene_tx=XM_023243640.1;refGene_gn=PLCD3;refGene_tx=XM_023243641.1;refGene_gn=PLCD3;refGene_tx=XM_023243642.1
E1 45034694 45034839
refGene_gn=PLCD3;refGene_tx=XM_023243637.1;refGene_gn=PLCD3;refGene_tx=XM_023243638.1;refGene_gn=PLCD3;refGene_tx=XM_023243640.1;refGene_gn=PLCD3;refGene_tx=XM_023243641.1;refGene_gn=PLCD3;refGene_tx=XM_023243642.1
E1 45035020 45035222
refGene_gn=PLCD3;refGene_tx=XM_023243637.1;refGene_gn=PLCD3;refGene_tx=XM_023243638.1;refGene_gn=PLCD3;refGene_tx=XM_023243640.1;refGene_gn=PLCD3;refGene_tx=XM_023243641.1;refGene_gn=PLCD3;refGene_tx=XM_023243642.1
E1 45035678 45035907
refGene_gn=PLCD3;refGene_tx=XM_023243637.1;refGene_gn=PLCD3;refGene_tx=XM_023243638.1;refGene_gn=PLCD3;refGene_tx=XM_023243640.1;refGene_gn=PLCD3;refGene_tx=XM_023243641.1;refGene_gn=PLCD3;refGene_tx=XM_023243642.1
E1 45036838 45036968
refGene_gn=PLCD3;refGene_tx=XM_023243637.1;refGene_gn=PLCD3;refGene_tx=XM_023243638.1;refGene_gn=PLCD3;refGene_tx=XM_023243640.1;refGene_gn=PLCD3;refGene_tx=XM_023243641.1;refGene_gn=PLCD3;refGene_tx=XM_023243642.1
E1 45037256 45037485
refGene_gn=PLCD3;refGene_tx=XM_023243637.1;refGene_gn=PLCD3;refGene_tx=XM_023243638.1;refGene_gn=PLCD3;refGene_tx=XM_023243640.1;refGene_gn=PLCD3;refGene_tx=XM_023243641.1;refGene_gn=PLCD3;refGene_tx=XM_023243642.1

E1 45037603 45037765
refGene_gn=PLCD3;refGene_tx=XM_023243637.1;refGene_gn=PLCD3;refGene_tx=XM_023243638.1;refGene_gn=PLCD3;refGene_tx=XM_023243640.1;refGene_gn=PLCD3;refGene_tx=XM_023243641.1;refGene_gn=PLCD3;refGene_tx=XM_023243642.1
E1 45043470 45043498
refGene_gn=PLCD3;refGene_tx=XM_023243637.1;refGene_gn=PLCD3;refGene_tx=XM_023243638.1
E1 45046990 45047129
refGene_gn=PLCD3;refGene_tx=XM_023243640.1;refGene_gn=PLCD3;refGene_tx=XM_023243641.1
E1 45049395 45049483
refGene_gn=ACBD4;refGene_tx=XM_003996999.4;refGene_gn=ACBD4;refGene_tx=XM_006940434.3;refGene_gn=ACBD4;refGene_tx=XM_006940435.3;refGene_gn=ACBD4;refGene_tx=XM_006940440.4;refGene_gn=ACBD4;refGene_tx=XM_011289234.2;refGene_gn=ACBD4;refGene_tx=XM_019818011.2;refGene_gn=ACBD4;refGene_tx=XM_019818012.2;refGene_gn=ACBD4;refGene_tx=XM_019818013.2;refGene_gn=ACBD4;refGene_tx=XM_019818014.1;refGene_gn=ACBD4;refGene_tx=XM_019818015.2
E1 45049748 45049869
refGene_gn=ACBD4;refGene_tx=XM_003996999.4;refGene_gn=ACBD4;refGene_tx=XM_006940434.3;refGene_gn=ACBD4;refGene_tx=XM_006940435.3;refGene_gn=ACBD4;refGene_tx=XM_006940440.4;refGene_gn=ACBD4;refGene_tx=XM_011289234.2;refGene_gn=ACBD4;refGene_tx=XM_019818011.2;refGene_gn=ACBD4;refGene_tx=XM_019818012.2;refGene_gn=ACBD4;refGene_tx=XM_019818013.2;refGene_gn=ACBD4;refGene_tx=XM_019818014.1;refGene_gn=ACBD4;refGene_tx=XM_019818015.2;refGene_gn=ACBD4;refGene_tx=XM_019818017.2
E1 45049939 45050024
refGene_gn=ACBD4;refGene_tx=XM_003996999.4;refGene_gn=ACBD4;refGene_tx=XM_006940434.3;refGene_gn=ACBD4;refGene_tx=XM_006940435.3;refGene_gn=ACBD4;refGene_tx=XM_006940440.4;refGene_gn=ACBD4;refGene_tx=XM_011289234.2;refGene_gn=ACBD4;refGene_tx=XM_019818011.2;refGene_gn=ACBD4;refGene_tx=XM_019818012.2;refGene_gn=ACBD4;refGene_tx=XM_019818013.2;refGene_gn=ACBD4;refGene_tx=XM_019818014.1;refGene_gn=ACBD4;refGene_tx=XM_019818015.2;refGene_gn=ACBD4;refGene_tx=XM_019818017.2
E1 45050284 45050405
refGene_gn=ACBD4;refGene_tx=XM_003996999.4;refGene_gn=ACBD4;refGene_tx=XM_006940434.3;refGene_gn=ACBD4;refGene_tx=XM_006940435.3;refGene_gn=ACBD4;refGene_tx=XM_006940440.4;refGene_gn=ACBD4;refGene_tx=XM_011289234.2;refGene_gn=ACBD4;refGene_tx=XM_019818011.2;refGene_gn=ACBD4;refGene_tx=XM_019818012.2;refGene_gn=ACBD4;refGene_tx=XM_019818013.2;refGene_gn=ACBD4;refGene_tx=XM_019818014.1;refGene_gn=ACBD4;refGene_tx=XM_019818015.2;refGene_gn=ACBD4;refGene_tx=XM_019818017.2
E1 45050653 45050743
refGene_gn=ACBD4;refGene_tx=XM_003996999.4;refGene_gn=ACBD4;refGene_tx=XM_006940434.3;refGene_gn=ACBD4;refGene_tx=XM_006940435.3;refGene_gn=ACBD4;refGene_tx=XM_006940440.4;refGene_gn=ACBD4;refGene_tx=XM_011289234.2;refGene_gn=ACBD4;refGene_tx=XM_019818011.2;refGene_gn=ACBD4;refGene_tx=XM_019818012.2;refGene_gn=ACBD4;refGene_tx=XM_019818013.2;refGene_gn=ACBD4;refGene_tx=XM_019818014.1;refGene_gn=ACBD4;refGene_tx=XM_01

9818015.2;refGene_gn=ACBD4;refGene_tx=XM_019818017.2
E1 45051018 45051128
refGene_gn=ACBD4;refGene_tx=XM_003996999.4;refGene_gn=ACBD4;refGene_tx=XM_006940434.3;refGene_gn=ACBD4;refGene_tx=XM_006940435.3;refGene_gn=ACBD4;refGene_tx=XM_006940440.4;refGene_gn=ACBD4;refGene_tx=XM_011289234.2;refGene_gn=ACBD4;refGene_tx=XM_019818011.2;refGene_gn=ACBD4;refGene_tx=XM_019818012.2;refGene_gn=ACBD4;refGene_tx=XM_019818013.2;refGene_gn=ACBD4;refGene_tx=XM_019818014.1;refGene_gn=ACBD4;refGene_tx=XM_019818015.2;refGene_gn=ACBD4;refGene_tx=XM_019818017.2
E1 45051216 45051295
refGene_gn=ACBD4;refGene_tx=XM_003996999.4;refGene_gn=ACBD4;refGene_tx=XM_006940434.3;refGene_gn=ACBD4;refGene_tx=XM_006940435.3;refGene_gn=ACBD4;refGene_tx=XM_006940440.4;refGene_gn=ACBD4;refGene_tx=XM_011289234.2;refGene_gn=ACBD4;refGene_tx=XM_019818011.2;refGene_gn=ACBD4;refGene_tx=XM_019818012.2;refGene_gn=ACBD4;refGene_tx=XM_019818013.2;refGene_gn=ACBD4;refGene_tx=XM_019818014.1;refGene_gn=ACBD4;refGene_tx=XM_019818015.2;refGene_gn=ACBD4;refGene_tx=XM_019818017.2
E1 45051887 45052018
refGene_gn=ACBD4;refGene_tx=XM_003996999.4;refGene_gn=ACBD4;refGene_tx=XM_006940434.3;refGene_gn=ACBD4;refGene_tx=XM_006940435.3;refGene_gn=ACBD4;refGene_tx=XM_006940440.4;refGene_gn=ACBD4;refGene_tx=XM_011289234.2;refGene_gn=ACBD4;refGene_tx=XM_019818011.2;refGene_gn=ACBD4;refGene_tx=XM_019818012.2;refGene_gn=ACBD4;refGene_tx=XM_019818013.2;refGene_gn=ACBD4;refGene_tx=XM_019818014.1;refGene_gn=ACBD4;refGene_tx=XM_019818015.2;refGene_gn=ACBD4;refGene_tx=XM_019818017.2
E1 45052395 45052473 refGene_gn=ACBD4;refGene_tx=XM_019818015.2
E1 45054438 45054564
refGene_gn=ACBD4;refGene_tx=XM_003996999.4;refGene_gn=ACBD4;refGene_tx=XM_006940434.3;refGene_gn=ACBD4;refGene_tx=XM_006940435.3;refGene_gn=ACBD4;refGene_tx=XM_006940440.4;refGene_gn=ACBD4;refGene_tx=XM_011289234.2;refGene_gn=ACBD4;refGene_tx=XM_019818011.2;refGene_gn=ACBD4;refGene_tx=XM_019818012.2;refGene_gn=ACBD4;refGene_tx=XM_019818013.2;refGene_gn=ACBD4;refGene_tx=XM_019818014.1;refGene_gn=ACBD4;refGene_tx=XM_019818017.2
E1 45059686 45060766 refGene_gn=HEXIM1;refGene_tx=XM_003997000.5
E1 45067936 45068080 refGene_gn=HEXIM2;refGene_tx=XM_011289267.1
E1 45069807 45069921 refGene_gn=HEXIM2;refGene_tx=XM_011289267.1
E1 45072570 45073365 refGene_gn=HEXIM2;refGene_tx=XM_011289267.1
E1 45100824 45100953
refGene_gn=FMNL1;refGene_tx=XM_006940443.4;refGene_gn=FMNL1;refGene_tx=XM_023244520.1;refGene_gn=FMNL1;refGene_tx=XM_023244521.1;refGene_gn=FMNL1;refGene_tx=XM_023244522.1;refGene_gn=FMNL1;refGene_tx=XM_023244523.1
E1 45109345 45109429
refGene_gn=FMNL1;refGene_tx=XM_006940443.4;refGene_gn=FMNL1;refGene_tx=XM_023244520.1;refGene_gn=FMNL1;refGene_tx=XM_023244521.1;refGene_gn=FMNL1;refGene_tx=XM_023244522.1;refGene_gn=FMNL1;refGene_tx=XM_023244523.1
E1 45110949 45111063
refGene_gn=FMNL1;refGene_tx=XM_006940443.4;refGene_gn=FMNL1;refGene_tx

=XM_023244520.1;refGene_gn=FMNL1;refGene_tx=XM_023244521.1;refGene_gn=FMNL1;refGene_tx=XM_023244522.1;refGene_gn=FMNL1;refGene_tx=XM_023244523.1
E1 45111729 45111803
refGene_gn=FMNL1;refGene_tx=XM_006940443.4;refGene_gn=FMNL1;refGene_tx=XM_023244520.1;refGene_gn=FMNL1;refGene_tx=XM_023244521.1;refGene_gn=FMNL1;refGene_tx=XM_023244522.1;refGene_gn=FMNL1;refGene_tx=XM_023244523.1
E1 45112187 45112271
refGene_gn=FMNL1;refGene_tx=XM_006940443.4;refGene_gn=FMNL1;refGene_tx=XM_023244520.1;refGene_gn=FMNL1;refGene_tx=XM_023244521.1;refGene_gn=FMNL1;refGene_tx=XM_023244522.1;refGene_gn=FMNL1;refGene_tx=XM_023244523.1
E1 45112670 45112808
refGene_gn=FMNL1;refGene_tx=XM_006940443.4;refGene_gn=FMNL1;refGene_tx=XM_023244520.1;refGene_gn=FMNL1;refGene_tx=XM_023244521.1;refGene_gn=FMNL1;refGene_tx=XM_023244522.1;refGene_gn=FMNL1;refGene_tx=XM_023244523.1
E1 45113103 45113121
refGene_gn=FMNL1;refGene_tx=XM_006940443.4;refGene_gn=FMNL1;refGene_tx=XM_023244520.1;refGene_gn=FMNL1;refGene_tx=XM_023244521.1
E1 45114413 45114522
refGene_gn=FMNL1;refGene_tx=XM_006940443.4;refGene_gn=FMNL1;refGene_tx=XM_023244520.1;refGene_gn=FMNL1;refGene_tx=XM_023244521.1;refGene_gn=FMNL1;refGene_tx=XM_023244522.1;refGene_gn=FMNL1;refGene_tx=XM_023244523.1
E1 45115146 45115223
refGene_gn=FMNL1;refGene_tx=XM_006940443.4;refGene_gn=FMNL1;refGene_tx=XM_023244520.1;refGene_gn=FMNL1;refGene_tx=XM_023244521.1;refGene_gn=FMNL1;refGene_tx=XM_023244522.1;refGene_gn=FMNL1;refGene_tx=XM_023244523.1
E1 45115433 45115527
refGene_gn=FMNL1;refGene_tx=XM_006940443.4;refGene_gn=FMNL1;refGene_tx=XM_023244520.1;refGene_gn=FMNL1;refGene_tx=XM_023244521.1;refGene_gn=FMNL1;refGene_tx=XM_023244522.1;refGene_gn=FMNL1;refGene_tx=XM_023244523.1
E1 45116450 45116525
refGene_gn=FMNL1;refGene_tx=XM_006940443.4;refGene_gn=FMNL1;refGene_tx=XM_023244520.1;refGene_gn=FMNL1;refGene_tx=XM_023244521.1;refGene_gn=FMNL1;refGene_tx=XM_023244522.1;refGene_gn=FMNL1;refGene_tx=XM_023244523.1
E1 45116836 45116947
refGene_gn=FMNL1;refGene_tx=XM_006940443.4;refGene_gn=FMNL1;refGene_tx=XM_023244520.1;refGene_gn=FMNL1;refGene_tx=XM_023244521.1;refGene_gn=FMNL1;refGene_tx=XM_023244522.1;refGene_gn=FMNL1;refGene_tx=XM_023244523.1
E1 45118280 45118430
refGene_gn=FMNL1;refGene_tx=XM_006940443.4;refGene_gn=FMNL1;refGene_tx=XM_023244520.1;refGene_gn=FMNL1;refGene_tx=XM_023244521.1;refGene_gn=FMNL1;refGene_tx=XM_023244522.1;refGene_gn=FMNL1;refGene_tx=XM_0232445

23.1
E1	45118955 45119057
refGene_gn=FMNL1;refGene_tx=XM_006940443.4;refGene_gn=FMNL1;refGene_tx=XM_023244520.1;refGene_gn=FMNL1;refGene_tx=XM_023244521.1;refGene_gn=FMNL1;refGene_tx=XM_023244522.1;refGene_gn=FMNL1;refGene_tx=XM_023244523.1
E1	45119220 45119491
refGene_gn=FMNL1;refGene_tx=XM_006940443.4;refGene_gn=FMNL1;refGene_tx=XM_023244520.1;refGene_gn=FMNL1;refGene_tx=XM_023244521.1;refGene_gn=FMNL1;refGene_tx=XM_023244522.1;refGene_gn=FMNL1;refGene_tx=XM_023244523.1
E1	45119739 45120045
refGene_gn=FMNL1;refGene_tx=XM_006940443.4;refGene_gn=FMNL1;refGene_tx=XM_023244520.1;refGene_gn=FMNL1;refGene_tx=XM_023244521.1;refGene_gn=FMNL1;refGene_tx=XM_023244522.1;refGene_gn=FMNL1;refGene_tx=XM_023244523.1
E1	45120235 45120360
refGene_gn=FMNL1;refGene_tx=XM_006940443.4;refGene_gn=FMNL1;refGene_tx=XM_023244520.1;refGene_gn=FMNL1;refGene_tx=XM_023244521.1;refGene_gn=FMNL1;refGene_tx=XM_023244522.1;refGene_gn=FMNL1;refGene_tx=XM_023244523.1
E1	45121256 45121459
refGene_gn=FMNL1;refGene_tx=XM_006940443.4;refGene_gn=FMNL1;refGene_tx=XM_023244520.1;refGene_gn=FMNL1;refGene_tx=XM_023244521.1;refGene_gn=FMNL1;refGene_tx=XM_023244522.1;refGene_gn=FMNL1;refGene_tx=XM_023244523.1
E1	45121931 45122166
refGene_gn=FMNL1;refGene_tx=XM_006940443.4;refGene_gn=FMNL1;refGene_tx=XM_023244520.1;refGene_gn=FMNL1;refGene_tx=XM_023244521.1;refGene_gn=FMNL1;refGene_tx=XM_023244522.1;refGene_gn=FMNL1;refGene_tx=XM_023244523.1
E1	45122411 45122480
refGene_gn=FMNL1;refGene_tx=XM_006940443.4;refGene_gn=FMNL1;refGene_tx=XM_023244520.1;refGene_gn=FMNL1;refGene_tx=XM_023244521.1;refGene_gn=FMNL1;refGene_tx=XM_023244522.1;refGene_gn=FMNL1;refGene_tx=XM_023244523.1
E1	45123510 45123591
refGene_gn=FMNL1;refGene_tx=XM_006940443.4;refGene_gn=FMNL1;refGene_tx=XM_023244520.1;refGene_gn=FMNL1;refGene_tx=XM_023244521.1;refGene_gn=FMNL1;refGene_tx=XM_023244522.1;refGene_gn=FMNL1;refGene_tx=XM_023244523.1
E1	45123674 45123804
refGene_gn=FMNL1;refGene_tx=XM_006940443.4;refGene_gn=FMNL1;refGene_tx=XM_023244520.1;refGene_gn=FMNL1;refGene_tx=XM_023244521.1;refGene_gn=FMNL1;refGene_tx=XM_023244522.1;refGene_gn=FMNL1;refGene_tx=XM_023244523.1
E1	45123957 45124121
refGene_gn=FMNL1;refGene_tx=XM_006940443.4;refGene_gn=FMNL1;refGene_tx=XM_023244520.1;refGene_gn=FMNL1;refGene_tx=XM_023244521.1;refGene_gn=FMNL1;refGene_tx=XM_023244522.1;refGene_gn=FMNL1;refGene_tx=XM_0232445

23.1
E1 45124319 45124421
refGene_gn=FMNL1;refGene_tx=XM_006940443.4;refGene_gn=FMNL1;refGene_tx=XM_023244520.1;refGene_gn=FMNL1;refGene_tx=XM_023244521.1;refGene_gn=FMNL1;refGene_tx=XM_023244522.1;refGene_gn=FMNL1;refGene_tx=XM_023244523.1
E1 45124565 45124661
refGene_gn=FMNL1;refGene_tx=XM_006940443.4;refGene_gn=FMNL1;refGene_tx=XM_023244520.1;refGene_gn=FMNL1;refGene_tx=XM_023244521.1;refGene_gn=FMNL1;refGene_tx=XM_023244522.1;refGene_gn=FMNL1;refGene_tx=XM_023244523.1
E1 45124942 45125063
refGene_gn=FMNL1;refGene_tx=XM_006940443.4;refGene_gn=FMNL1;refGene_tx=XM_023244520.1;refGene_gn=FMNL1;refGene_tx=XM_023244521.1;refGene_gn=FMNL1;refGene_tx=XM_023244522.1;refGene_gn=FMNL1;refGene_tx=XM_023244523.1
E1 45125225 45125317
refGene_gn=FMNL1;refGene_tx=XM_023244521.1;refGene_gn=FMNL1;refGene_tx=XM_023244523.1
E1 45125581 45125685
refGene_gn=FMNL1;refGene_tx=XM_006940443.4;refGene_gn=FMNL1;refGene_tx=XM_023244520.1;refGene_gn=FMNL1;refGene_tx=XM_023244522.1
E1 45126083 45126220
refGene_gn=SPATA32;refGene_tx=XM_011289230.2;refGene_gn=SPATA32;refGene_tx=XM_023244526.1
E1 45126548 45126707
refGene_gn=SPATA32;refGene_tx=XM_011289230.2;refGene_gn=SPATA32;refGene_tx=XM_023244526.1;refGene_gn=SPATA32;refGene_tx=XM_023244527.1
E1 45126828 45127007
refGene_gn=SPATA32;refGene_tx=XM_011289230.2;refGene_gn=SPATA32;refGene_tx=XM_023244526.1
E1 45127254 45127375
refGene_gn=SPATA32;refGene_tx=XM_011289230.2;refGene_gn=SPATA32;refGene_tx=XM_023244526.1;refGene_gn=SPATA32;refGene_tx=XM_023244527.1
E1 45127458 45127530
refGene_gn=SPATA32;refGene_tx=XM_011289230.2;refGene_gn=SPATA32;refGene_tx=XM_023244526.1;refGene_gn=SPATA32;refGene_tx=XM_023244527.1
E1 45127899 45128043
refGene_gn=SPATA32;refGene_tx=XM_011289230.2;refGene_gn=SPATA32;refGene_tx=XM_023244526.1;refGene_gn=SPATA32;refGene_tx=XM_023244527.1
E1 45128896 45128952
refGene_gn=SPATA32;refGene_tx=XM_011289230.2;refGene_gn=SPATA32;refGene_tx=XM_023244526.1;refGene_gn=SPATA32;refGene_tx=XM_023244527.1
E1 45129068 45129293
refGene_gn=SPATA32;refGene_tx=XM_011289230.2;refGene_gn=SPATA32;refGene_tx=XM_023244526.1;refGene_gn=SPATA32;refGene_tx=XM_023244527.1
E1 45131815 45132678
refGene_gn=SPATA32;refGene_tx=XM_011289230.2;refGene_gn=SPATA32;refGene_tx=XM_023244526.1;refGene_gn=SPATA32;refGene_tx=XM_023244527.1
E1 45132981 45133036

refGene_gn=SPATA32;refGene_tx=XM_011289230.2;refGene_gn=SPATA32;refGene_tx=XM_023244526.1;refGene_gn=SPATA32;refGene_tx=XM_023244527.1
E1 45133859 45133914
refGene_gn=SPATA32;refGene_tx=XM_011289230.2;refGene_gn=SPATA32;refGene_tx=XM_023244526.1;refGene_gn=SPATA32;refGene_tx=XM_023244527.1
E1 45140772 45140785
refGene_gn=SPATA32;refGene_tx=XM_011289230.2;refGene_gn=SPATA32;refGene_tx=XM_023244526.1;refGene_gn=SPATA32;refGene_tx=XM_023244527.1
E1 45143162 45143327
refGene_gn=MAP3K14;refGene_tx=XM_023244524.1;refGene_gn=MAP3K14;refGene_tx=XM_023244525.1
E1 45143662 45143763
refGene_gn=MAP3K14;refGene_tx=XM_023244524.1;refGene_gn=MAP3K14;refGene_tx=XM_023244525.1
E1 45144890 45145035
refGene_gn=MAP3K14;refGene_tx=XM_023244524.1;refGene_gn=MAP3K14;refGene_tx=XM_023244525.1
E1 45145465 45145572
refGene_gn=MAP3K14;refGene_tx=XM_023244524.1;refGene_gn=MAP3K14;refGene_tx=XM_023244525.1
E1 45145781 45146156
refGene_gn=MAP3K14;refGene_tx=XM_023244524.1;refGene_gn=MAP3K14;refGene_tx=XM_023244525.1
E1 45147107 45147258
refGene_gn=MAP3K14;refGene_tx=XM_023244524.1;refGene_gn=MAP3K14;refGene_tx=XM_023244525.1
E1 45147531 45147695
refGene_gn=MAP3K14;refGene_tx=XM_023244524.1;refGene_gn=MAP3K14;refGene_tx=XM_023244525.1
E1 45149351 45149456
refGene_gn=MAP3K14;refGene_tx=XM_023244524.1;refGene_gn=MAP3K14;refGene_tx=XM_023244525.1
E1 45149962 45150094
refGene_gn=MAP3K14;refGene_tx=XM_023244524.1;refGene_gn=MAP3K14;refGene_tx=XM_023244525.1
E1 45150280 45150410
refGene_gn=MAP3K14;refGene_tx=XM_023244524.1;refGene_gn=MAP3K14;refGene_tx=XM_023244525.1
E1 45156498 45156636
refGene_gn=MAP3K14;refGene_tx=XM_023244524.1;refGene_gn=MAP3K14;refGene_tx=XM_023244525.1
E1 45157826 45158444
refGene_gn=MAP3K14;refGene_tx=XM_023244524.1;refGene_gn=MAP3K14;refGene_tx=XM_023244525.1
E1 45158554 45158765
refGene_gn=MAP3K14;refGene_tx=XM_023244524.1;refGene_gn=MAP3K14;refGene_tx=XM_023244525.1
E1 45160371 45160441
refGene_gn=MAP3K14;refGene_tx=XM_023244524.1;refGene_gn=MAP3K14;refGene_tx=XM_023244525.1

E1 45161582 45161835
refGene_gn=MAP3K14;refGene_tx=XM_023244524.1;refGene_gn=MAP3K14;refGene_tx=XM_023244525.1
E1 45243479 45243657
refGene_gn=ARHGAP27;refGene_tx=XM_023243890.1;refGene_gn=ARHGAP27;refGene_tx=XM_023243891.1;refGene_gn=ARHGAP27;refGene_tx=XM_023243892.1;refGene_gn=ARHGAP27;refGene_tx=XM_023243893.1
E1 45243778 45243884
refGene_gn=ARHGAP27;refGene_tx=XM_023243890.1;refGene_gn=ARHGAP27;refGene_tx=XM_023243891.1;refGene_gn=ARHGAP27;refGene_tx=XM_023243892.1;refGene_gn=ARHGAP27;refGene_tx=XM_023243893.1
E1 45244003 45244138
refGene_gn=ARHGAP27;refGene_tx=XM_023243890.1;refGene_gn=ARHGAP27;refGene_tx=XM_023243891.1;refGene_gn=ARHGAP27;refGene_tx=XM_023243892.1;refGene_gn=ARHGAP27;refGene_tx=XM_023243893.1
E1 45244249 45244327
refGene_gn=ARHGAP27;refGene_tx=XM_023243890.1;refGene_gn=ARHGAP27;refGene_tx=XM_023243891.1;refGene_gn=ARHGAP27;refGene_tx=XM_023243892.1;refGene_gn=ARHGAP27;refGene_tx=XM_023243893.1
E1 45244489 45244588
refGene_gn=ARHGAP27;refGene_tx=XM_023243890.1;refGene_gn=ARHGAP27;refGene_tx=XM_023243891.1;refGene_gn=ARHGAP27;refGene_tx=XM_023243892.1;refGene_gn=ARHGAP27;refGene_tx=XM_023243893.1
E1 45244661 45244769
refGene_gn=ARHGAP27;refGene_tx=XM_023243890.1;refGene_gn=ARHGAP27;refGene_tx=XM_023243891.1;refGene_gn=ARHGAP27;refGene_tx=XM_023243892.1;refGene_gn=ARHGAP27;refGene_tx=XM_023243893.1
E1 45244898 45245004
refGene_gn=ARHGAP27;refGene_tx=XM_023243890.1;refGene_gn=ARHGAP27;refGene_tx=XM_023243891.1;refGene_gn=ARHGAP27;refGene_tx=XM_023243892.1;refGene_gn=ARHGAP27;refGene_tx=XM_023243893.1
E1 45245912 45246011
refGene_gn=ARHGAP27;refGene_tx=XM_023243890.1;refGene_gn=ARHGAP27;refGene_tx=XM_023243891.1;refGene_gn=ARHGAP27;refGene_tx=XM_023243892.1;refGene_gn=ARHGAP27;refGene_tx=XM_023243893.1
E1 45248341 45248446
refGene_gn=ARHGAP27;refGene_tx=XM_023243890.1;refGene_gn=ARHGAP27;refGene_tx=XM_023243891.1;refGene_gn=ARHGAP27;refGene_tx=XM_023243892.1;refGene_gn=ARHGAP27;refGene_tx=XM_023243893.1
E1 45248910 45249001
refGene_gn=ARHGAP27;refGene_tx=XM_023243890.1;refGene_gn=ARHGAP27;refGene_tx=XM_023243891.1;refGene_gn=ARHGAP27;refGene_tx=XM_023243892.1;refGene_gn=ARHGAP27;refGene_tx=XM_023243893.1
E1 45249439 45249507
refGene_gn=ARHGAP27;refGene_tx=XM_023243890.1;refGene_gn=ARHGAP27;refGene_tx=XM_023243891.1;refGene_gn=ARHGAP27;refGene_tx=XM_023243892.1;refGene_gn=ARHGAP27;refGene_tx=XM_023243893.1
E1 45249857 45249941
refGene_gn=ARHGAP27;refGene_tx=XM_023243890.1;refGene_gn=ARHGAP27;refGene_tx=XM_023243891.1;refGene_gn=ARHGAP27;refGene_tx=XM_023243892.1;re

fGene_gn=ARHGAP27;refGene_tx=XM_023243893.1
E1 45250013 45250094
refGene_gn=ARHGAP27;refGene_tx=XM_023243890.1;refGene_gn=ARHGAP27;refGene_tx=XM_023243891.1;refGene_gn=ARHGAP27;refGene_tx=XM_023243892.1;refGene_gn=ARHGAP27;refGene_tx=XM_023243893.1
E1 45250335 45250518
refGene_gn=ARHGAP27;refGene_tx=XM_023243890.1;refGene_gn=ARHGAP27;refGene_tx=XM_023243891.1;refGene_gn=ARHGAP27;refGene_tx=XM_023243892.1;refGene_gn=ARHGAP27;refGene_tx=XM_023243893.1
E1 45250840 45251245
refGene_gn=ARHGAP27;refGene_tx=XM_023243890.1;refGene_gn=ARHGAP27;refGene_tx=XM_023243891.1;refGene_gn=ARHGAP27;refGene_tx=XM_023243892.1;refGene_gn=ARHGAP27;refGene_tx=XM_023243893.1
E1 45254893 45254950 refGene_gn=ARHGAP27;refGene_tx=XM_023243892.1
E1 45270803 45271463
refGene_gn=ARHGAP27;refGene_tx=XM_023243890.1;refGene_gn=ARHGAP27;refGene_tx=XM_023243891.1
E1 45277712 45277824 refGene_gn=PLEKHM1;refGene_tx=XM_023243888.1
E1 45278526 45278846
refGene_gn=PLEKHM1;refGene_tx=XM_023243887.1;refGene_gn=PLEKHM1;refGene_tx=XM_023243888.1;refGene_gn=PLEKHM1;refGene_tx=XM_023243889.1
E1 45279300 45279364
refGene_gn=PLEKHM1;refGene_tx=XM_023243887.1;refGene_gn=PLEKHM1;refGene_tx=XM_023243888.1;refGene_gn=PLEKHM1;refGene_tx=XM_023243889.1
E1 45281088 45281524
refGene_gn=PLEKHM1;refGene_tx=XM_023243882.1;refGene_gn=PLEKHM1;refGene_tx=XM_023243884.1;refGene_gn=PLEKHM1;refGene_tx=XM_023243885.1;refGene_gn=PLEKHM1;refGene_tx=XM_023243886.1
E1 45284634 45284828
refGene_gn=PLEKHM1;refGene_tx=XM_023243882.1;refGene_gn=PLEKHM1;refGene_tx=XM_023243884.1;refGene_gn=PLEKHM1;refGene_tx=XM_023243885.1;refGene_gn=PLEKHM1;refGene_tx=XM_023243886.1;refGene_gn=PLEKHM1;refGene_tx=XM_023243887.1;refGene_gn=PLEKHM1;refGene_tx=XM_023243888.1;refGene_gn=PLEKHM1;refGene_tx=XM_023243889.1
E1 45288532 45288678
refGene_gn=PLEKHM1;refGene_tx=XM_023243882.1;refGene_gn=PLEKHM1;refGene_tx=XM_023243884.1;refGene_gn=PLEKHM1;refGene_tx=XM_023243885.1;refGene_gn=PLEKHM1;refGene_tx=XM_023243886.1;refGene_gn=PLEKHM1;refGene_tx=XM_023243887.1;refGene_gn=PLEKHM1;refGene_tx=XM_023243888.1;refGene_gn=PLEKHM1;refGene_tx=XM_023243889.1
E1 45290977 45291898
refGene_gn=PLEKHM1;refGene_tx=XM_023243882.1;refGene_gn=PLEKHM1;refGene_tx=XM_023243884.1;refGene_gn=PLEKHM1;refGene_tx=XM_023243885.1;refGene_gn=PLEKHM1;refGene_tx=XM_023243886.1;refGene_gn=PLEKHM1;refGene_tx=XM_023243887.1;refGene_gn=PLEKHM1;refGene_tx=XM_023243888.1;refGene_gn=PLEKHM1;refGene_tx=XM_023243889.1
E1 45294625 45294896
refGene_gn=PLEKHM1;refGene_tx=XM_023243882.1;refGene_gn=PLEKHM1;refGene_tx=XM_023243884.1;refGene_gn=PLEKHM1;refGene_tx=XM_023243885.1;refGene_gn=PLEKHM1;refGene_tx=XM_023243886.1;refGene_gn=PLEKHM1;refGene_tx=

XM_023243887.1;refGene_gn=PLEKHM1;refGene_tx=XM_023243888.1;refGene_gn=PLEKHM1;refGene_tx=XM_023243889.1
E1 45304471 45304883
refGene_gn=PLEKHM1;refGene_tx=XM_023243882.1;refGene_gn=PLEKHM1;refGene_tx=XM_023243884.1;refGene_gn=PLEKHM1;refGene_tx=XM_023243885.1;refGene_gn=PLEKHM1;refGene_tx=XM_023243886.1;refGene_gn=PLEKHM1;refGene_tx=XM_023243887.1;refGene_gn=PLEKHM1;refGene_tx=XM_023243888.1;refGene_gn=PLEKHM1;refGene_tx=XM_023243889.1
E1 45311343 45311970
refGene_gn=PLEKHM1;refGene_tx=XM_023243882.1;refGene_gn=PLEKHM1;refGene_tx=XM_023243884.1;refGene_gn=PLEKHM1;refGene_tx=XM_023243885.1;refGene_gn=PLEKHM1;refGene_tx=XM_023243886.1;refGene_gn=PLEKHM1;refGene_tx=XM_023243887.1;refGene_gn=PLEKHM1;refGene_tx=XM_023243888.1;refGene_gn=PLEKHM1;refGene_tx=XM_023243889.1
E1 45313708 45313956
refGene_gn=PLEKHM1;refGene_tx=XM_023243882.1;refGene_gn=PLEKHM1;refGene_tx=XM_023243884.1;refGene_gn=PLEKHM1;refGene_tx=XM_023243885.1;refGene_gn=PLEKHM1;refGene_tx=XM_023243886.1;refGene_gn=PLEKHM1;refGene_tx=XM_023243887.1;refGene_gn=PLEKHM1;refGene_tx=XM_023243888.1;refGene_gn=PLEKHM1;refGene_tx=XM_023243889.1
E1 45317974 45318022
refGene_gn=PLEKHM1;refGene_tx=XM_023243882.1;refGene_gn=PLEKHM1;refGene_tx=XM_023243884.1;refGene_gn=PLEKHM1;refGene_tx=XM_023243885.1;refGene_gn=PLEKHM1;refGene_tx=XM_023243886.1;refGene_gn=PLEKHM1;refGene_tx=XM_023243887.1;refGene_gn=PLEKHM1;refGene_tx=XM_023243888.1;refGene_gn=PLEKHM1;refGene_tx=XM_023243889.1
E1 45337672 45337769
refGene_gn=LOC105259662;refGene_tx=XM_023244472.1
E1 45337869 45338012
refGene_gn=LOC105259662;refGene_tx=XM_023244472.1
E1 45338091 45338146
refGene_gn=LOC105259662;refGene_tx=XM_023244472.1
E1 45339886 45339936
refGene_gn=LOC105259662;refGene_tx=XM_023244472.1
E1 45343019 45343058
refGene_gn=LOC105259662;refGene_tx=XM_023244472.1
E1 45343060 45343129
refGene_gn=LOC105259662;refGene_tx=XM_023244472.1
E1 45343507 45348324
refGene_gn=LOC105259662;refGene_tx=XM_023244472.1
E1 45350355 45350474
refGene_gn=LOC105259662;refGene_tx=XM_023244472.1
E1 45358517 45358592
refGene_gn=LOC105259662;refGene_tx=XM_023244472.1
E1 45359155 45359227
refGene_gn=LOC105259662;refGene_tx=XM_023244472.1
E1 45375616 45375688
refGene_gn=LOC105259662;refGene_tx=XM_023244472.1
E1 45377659 45377731
refGene_gn=LOC101100197;refGene_tx=XM_023244477.1;refGene_gn=LOC105259

662;refGene_tx=XM_023244472.1
E1 45379998 45383294
refGene_gn=LOC105259662;refGene_tx=XM_023244472.1
E1 45386199 45386212
refGene_gn=LOC101100197;refGene_tx=XM_023244473.1
E1 45386735 45389712
refGene_gn=LOC101100197;refGene_tx=XM_023244473.1;refGene_gn=LOC101100197;refGene_tx=XM_023244478.1;refGene_gn=LOC101100197;refGene_tx=XM_023244479.1
E1 45389816 45390011
refGene_gn=LOC101100197;refGene_tx=XM_023244474.1;refGene_gn=LOC101100197;refGene_tx=XM_023244475.1;refGene_gn=LOC101100197;refGene_tx=XM_023244476.1
E1 45391568 45391654
refGene_gn=LOC101100197;refGene_tx=XM_023244474.1;refGene_gn=LOC101100197;refGene_tx=XM_023244475.1;refGene_gn=LOC101100197;refGene_tx=XM_023244477.1;refGene_gn=LOC101100197;refGene_tx=XM_023244478.1;refGene_gn=LOC101100197;refGene_tx=XM_023244479.1;refGene_gn=LOC101100197;refGene_tx=XM_023244482.1
E1 45393987 45394086
refGene_gn=LOC101100197;refGene_tx=XM_023244473.1;refGene_gn=LOC101100197;refGene_tx=XM_023244474.1;refGene_gn=LOC101100197;refGene_tx=XM_023244476.1;refGene_gn=LOC101100197;refGene_tx=XM_023244477.1;refGene_gn=LOC101100197;refGene_tx=XM_023244478.1
E1 45394697 45394729
refGene_gn=LOC101100197;refGene_tx=XM_023244480.1;refGene_gn=LOC101100197;refGene_tx=XM_023244481.1
E1 45395039 45395208
refGene_gn=LOC101100197;refGene_tx=XM_023244473.1;refGene_gn=LOC101100197;refGene_tx=XM_023244474.1;refGene_gn=LOC101100197;refGene_tx=XM_023244475.1;refGene_gn=LOC101100197;refGene_tx=XM_023244476.1;refGene_gn=LOC101100197;refGene_tx=XM_023244477.1;refGene_gn=LOC101100197;refGene_tx=XM_023244478.1;refGene_gn=LOC101100197;refGene_tx=XM_023244479.1;refGene_gn=LOC101100197;refGene_tx=XM_023244480.1;refGene_gn=LOC101100197;refGene_tx=XM_023244481.1;refGene_gn=LOC101100197;refGene_tx=XM_023244482.1
E1 45396036 45396159
refGene_gn=LOC101100197;refGene_tx=XM_023244473.1;refGene_gn=LOC101100197;refGene_tx=XM_023244474.1;refGene_gn=LOC101100197;refGene_tx=XM_023244475.1;refGene_gn=LOC101100197;refGene_tx=XM_023244476.1;refGene_gn=LOC101100197;refGene_tx=XM_023244477.1;refGene_gn=LOC101100197;refGene_tx=XM_023244478.1;refGene_gn=LOC101100197;refGene_tx=XM_023244479.1;refGene_gn=LOC101100197;refGene_tx=XM_023244480.1;refGene_gn=LOC101100197;refGene_tx=XM_023244481.1;refGene_gn=LOC101100197;refGene_tx=XM_023244482.1
E1 45399478 45399658
refGene_gn=LOC101100197;refGene_tx=XM_023244473.1;refGene_gn=LOC101100197;refGene_tx=XM_023244474.1;refGene_gn=LOC101100197;refGene_tx=XM_023244475.1;refGene_gn=LOC101100197;refGene_tx=XM_023244476.1;refGene_gn=LOC101100197;refGene_tx=XM_023244477.1;refGene_gn=LOC101100197;refGen

e_tx=XM_023244478.1;refGene_gn=LOC101100197;refGene_tx=XM_023244479.1;refGene_gn=LOC101100197;refGene_tx=XM_023244480.1;refGene_gn=LOC101100197;refGene_tx=XM_023244481.1;refGene_gn=LOC101100197;refGene_tx=XM_023244482.1
E1 45400024 45400120
refGene_gn=LOC101100197;refGene_tx=XM_023244473.1;refGene_gn=LOC101100197;refGene_tx=XM_023244474.1;refGene_gn=LOC101100197;refGene_tx=XM_023244475.1;refGene_gn=LOC101100197;refGene_tx=XM_023244476.1;refGene_gn=LOC101100197;refGene_tx=XM_023244477.1;refGene_gn=LOC101100197;refGene_tx=XM_023244478.1;refGene_gn=LOC101100197;refGene_tx=XM_023244479.1;refGene_gn=LOC101100197;refGene_tx=XM_023244480.1;refGene_gn=LOC101100197;refGene_tx=XM_023244481.1;refGene_gn=LOC101100197;refGene_tx=XM_023244482.1
E1 45402788 45402854 refGene_gn=LYZL6;refGene_tx=XM_023244489.1
E1 45403707 45403786 refGene_gn=LYZL6;refGene_tx=XM_023244489.1
E1 45404651 45404838 refGene_gn=LYZL6;refGene_tx=XM_023244489.1
E1 45405777 45405916 refGene_gn=LYZL6;refGene_tx=XM_023244489.1
E1 45429443 45429806
refGene_gn=RPRML;refGene_tx=XM_023244491.1;refGene_gn=RPRML;refGene_tx=XM_023244492.1
E1 45456039 45456201
refGene_gn=GOSR2;refGene_tx=XM_023244483.1;refGene_gn=GOSR2;refGene_tx=XM_023244484.1;refGene_gn=GOSR2;refGene_tx=XM_023244485.1;refGene_gn=GOSR2;refGene_tx=XM_023244486.1;refGene_gn=GOSR2;refGene_tx=XM_023244487.1;refGene_gn=GOSR2;refGene_tx=XM_023244488.1
E1 45459404 45459545
refGene_gn=GOSR2;refGene_tx=XM_023244483.1;refGene_gn=GOSR2;refGene_tx=XM_023244484.1;refGene_gn=GOSR2;refGene_tx=XM_023244485.1;refGene_gn=GOSR2;refGene_tx=XM_023244486.1
E1 45461953 45462086
refGene_gn=GOSR2;refGene_tx=XM_023244483.1;refGene_gn=GOSR2;refGene_tx=XM_023244484.1;refGene_gn=GOSR2;refGene_tx=XM_023244485.1;refGene_gn=GOSR2;refGene_tx=XM_023244486.1;refGene_gn=GOSR2;refGene_tx=XM_023244487.1;refGene_gn=GOSR2;refGene_tx=XM_023244488.1
E1 45462906 45463015
refGene_gn=GOSR2;refGene_tx=XM_023244483.1;refGene_gn=GOSR2;refGene_tx=XM_023244484.1;refGene_gn=GOSR2;refGene_tx=XM_023244485.1;refGene_gn=GOSR2;refGene_tx=XM_023244486.1;refGene_gn=GOSR2;refGene_tx=XM_023244487.1;refGene_gn=GOSR2;refGene_tx=XM_023244488.1
E1 45464778 45464843
refGene_gn=GOSR2;refGene_tx=XM_023244483.1;refGene_gn=GOSR2;refGene_tx=XM_023244484.1;refGene_gn=GOSR2;refGene_tx=XM_023244485.1;refGene_gn=GOSR2;refGene_tx=XM_023244486.1;refGene_gn=GOSR2;refGene_tx=XM_023244487.1;refGene_gn=GOSR2;refGene_tx=XM_023244488.1
E1 45471332 45471361
refGene_gn=GOSR2;refGene_tx=XM_023244483.1;refGene_gn=GOSR2;refGene_tx=XM_023244484.1;refGene_gn=GOSR2;refGene_tx=XM_023244487.1;refGene_gn=GOSR2;refGene_tx=XM_023244488.1
E1 45505654 45506128 refGene_gn=WNT9B;refGene_tx=XM_023243469.1
E1 45506968 45507090 refGene_gn=WNT9B;refGene_tx=XM_023243469.1

E1 45507092 45507233 refGene_gn=WNT9B;refGene_tx=XM_023243469.1
E1 45509270 45509549 refGene_gn=WNT9B;refGene_tx=XM_023243469.1
E1 45516757 45516851 refGene_gn=WNT9B;refGene_tx=XM_023243469.1
E1 45553637 45553717
refGene_gn=WNT3;refGene_tx=XM_003997020.4;refGene_gn=WNT3;refGene_tx=XM_019818026.2
E1 45590265 45590507
refGene_gn=WNT3;refGene_tx=XM_003997020.4;refGene_gn=WNT3;refGene_tx=XM_019818026.2
E1 45594117 45594383
refGene_gn=WNT3;refGene_tx=XM_003997020.4;refGene_gn=WNT3;refGene_tx=XM_019818026.2
E1 45595093 45595573
refGene_gn=WNT3;refGene_tx=XM_003997020.4;refGene_gn=WNT3;refGene_tx=XM_019818026.2
E1 45606827 45606873 refGene_gn=NSF;refGene_tx=XM_023244667.1
E1 45607332 45607388 refGene_gn=NSF;refGene_tx=XM_023244667.1
E1 45610801 45610915 refGene_gn=NSF;refGene_tx=XM_023244667.1
E1 45612878 45613013 refGene_gn=NSF;refGene_tx=XM_023244667.1
E1 45634400 45634480 refGene_gn=NSF;refGene_tx=XM_023244667.1
E1 45636522 45636589 refGene_gn=NSF;refGene_tx=XM_023244667.1
E1 45649362 45649496 refGene_gn=NSF;refGene_tx=XM_023244667.1
E1 45652789 45652946 refGene_gn=NSF;refGene_tx=XM_023244667.1
E1 45655079 45655175 refGene_gn=NSF;refGene_tx=XM_023244667.1
E1 45662802 45662990 refGene_gn=NSF;refGene_tx=XM_023244667.1
E1 45663178 45663253 refGene_gn=NSF;refGene_tx=XM_023244667.1
E1 45664377 45664543 refGene_gn=NSF;refGene_tx=XM_023244667.1
E1 45679479 45679679 refGene_gn=NSF;refGene_tx=XM_023244667.1
E1 45710301 45710457 refGene_gn=NSF;refGene_tx=XM_023244667.1
E1 45713504 45713580 refGene_gn=NSF;refGene_tx=XM_023244667.1
E1 45714049 45714157 refGene_gn=NSF;refGene_tx=XM_023244667.1
E1 45717155 45717322 refGene_gn=NSF;refGene_tx=XM_023244667.1
E1 45726120 45726160 refGene_gn=NSF;refGene_tx=XM_023244667.1
E1 45729663 45729763 refGene_gn=NSF;refGene_tx=XM_023244667.1
E1 45732043 45732129 refGene_gn=NSF;refGene_tx=XM_023244667.1
E1 45755350 45755362 refGene_gn=NSF;refGene_tx=XM_023244667.1
E1 45766598 45766746
refGene_gn=LOC101089059;refGene_tx=XM_003997022.4
E1 45773956 45774067
refGene_gn=LOC101089059;refGene_tx=XM_003997022.4
E1 45779779 45779904
refGene_gn=LOC101089059;refGene_tx=XM_003997022.4
E1 45781843 45782005
refGene_gn=LOC101089059;refGene_tx=XM_003997022.4
E1 45920520 45920553
refGene_gn=CRHR1;refGene_tx=XM_004001535.5;refGene_gn=CRHR1;refGene_tx=XM_019818028.2;refGene_gn=CRHR1;refGene_tx=XM_019818029.2
E1 45941151 45941239
refGene_gn=CRHR1;refGene_tx=XM_004001535.5;refGene_gn=CRHR1;refGene_tx=XM_019818028.2;refGene_gn=CRHR1;refGene_tx=XM_019818029.2

E1 45949606 45949762
refGene_gn=CRHR1;refGene_tx=XM_004001535.5;refGene_gn=CRHR1;refGene_tx=XM_019818028.2;refGene_gn=CRHR1;refGene_tx=XM_019818029.2
E1 45953650 45953736
refGene_gn=CRHR1;refGene_tx=XM_004001535.5;refGene_gn=CRHR1;refGene_tx=XM_019818028.2;refGene_gn=CRHR1;refGene_tx=XM_019818029.2
E1 45959358 45959465
refGene_gn=CRHR1;refGene_tx=XM_004001535.5;refGene_gn=CRHR1;refGene_tx=XM_019818028.2;refGene_gn=CRHR1;refGene_tx=XM_019818029.2
E1 45960208 45960329
refGene_gn=CRHR1;refGene_tx=XM_004001535.5;refGene_gn=CRHR1;refGene_tx=XM_019818028.2;refGene_gn=CRHR1;refGene_tx=XM_019818029.2
E1 45960555 45960709
refGene_gn=CRHR1;refGene_tx=XM_004001535.5;refGene_gn=CRHR1;refGene_tx=XM_019818028.2;refGene_gn=CRHR1;refGene_tx=XM_019818029.2
E1 45961006 45961067
refGene_gn=CRHR1;refGene_tx=XM_004001535.5;refGene_gn=CRHR1;refGene_tx=XM_019818028.2;refGene_gn=CRHR1;refGene_tx=XM_019818029.2
E1 45963138 45963211
refGene_gn=CRHR1;refGene_tx=XM_004001535.5;refGene_gn=CRHR1;refGene_tx=XM_019818028.2;refGene_gn=CRHR1;refGene_tx=XM_019818029.2
E1 45963466 45963552
refGene_gn=CRHR1;refGene_tx=XM_004001535.5;refGene_gn=CRHR1;refGene_tx=XM_019818028.2;refGene_gn=CRHR1;refGene_tx=XM_019818029.2
E1 45963631 45963743 refGene_gn=CRHR1;refGene_tx=XM_019818029.2
E1 45963774 45963910
refGene_gn=CRHR1;refGene_tx=XM_004001535.5;refGene_gn=CRHR1;refGene_tx=XM_019818028.2
E1 45964090 45964132
refGene_gn=CRHR1;refGene_tx=XM_004001535.5;refGene_gn=CRHR1;refGene_tx=XM_019818028.2
E1 45964702 45964843
refGene_gn=CRHR1;refGene_tx=XM_004001535.5;refGene_gn=CRHR1;refGene_tx=XM_019818028.2
E1 45974134 45976258 refGene_gn=SPPL2C;refGene_tx=XM_011289275.3
E1 46056332 46056474
refGene_gn=MAPT;refGene_tx=XM_003997029.5;refGene_gn=MAPT;refGene_tx=XM_003997030.5;refGene_gn=MAPT;refGene_tx=XM_006940453.4;refGene_gn=MAPT;refGene_tx=XM_006940454.4;refGene_gn=MAPT;refGene_tx=XM_006940455.4;refGene_gn=MAPT;refGene_tx=XM_006940456.4;refGene_gn=MAPT;refGene_tx=XM_006940458.4;refGene_gn=MAPT;refGene_tx=XM_006940459.4;refGene_gn=MAPT;refGene_tx=XM_019818030.2;refGene_gn=MAPT;refGene_tx=XM_019818031.2;refGene_gn=MAPT;refGene_tx=XM_019818032.2;refGene_gn=MAPT;refGene_tx=XM_019818033.2;refGene_gn=MAPT;refGene_tx=XM_019818034.2;refGene_gn=MAPT;refGene_tx=XM_019818036.2;refGene_gn=MAPT;refGene_tx=XM_019818037.2;refGene_gn=MAPT;refGene_tx=XM_019818038.2;refGene_gn=MAPT;refGene_tx=XM_019818039.2
E1 46060825 46060912
refGene_gn=MAPT;refGene_tx=XM_003997029.5;refGene_gn=MAPT;refGene_tx=XM_003997030.5;refGene_gn=MAPT;refGene_tx=XM_006940453.4;refGene_gn=MAP

T;refGene_tx=XM_006940454.4;refGene_gn=MAPT;refGene_tx=XM_006940455.4;
refGene_gn=MAPT;refGene_tx=XM_006940456.4;refGene_gn=MAPT;refGene_tx=X
M_006940458.4;refGene_gn=MAPT;refGene_tx=XM_006940459.4;refGene_gn=MAP
T;refGene_tx=XM_019818030.2;refGene_gn=MAPT;refGene_tx=XM_019818032.2;
refGene_gn=MAPT;refGene_tx=XM_019818033.2;refGene_gn=MAPT;refGene_tx=X
M_019818036.2;refGene_gn=MAPT;refGene_tx=XM_019818038.2
E1 46063434 46063521
refGene_gn=MAPT;refGene_tx=XM_003997029.5;refGene_gn=MAPT;refGene_tx=X
M_003997030.5;refGene_gn=MAPT;refGene_tx=XM_006940453.4;refGene_gn=MAP
T;refGene_tx=XM_006940454.4;refGene_gn=MAPT;refGene_tx=XM_006940455.4;
refGene_gn=MAPT;refGene_tx=XM_006940456.4;refGene_gn=MAPT;refGene_tx=X
M_006940458.4;refGene_gn=MAPT;refGene_tx=XM_006940459.4;refGene_gn=MAP
T;refGene_tx=XM_019818032.2
E1 46067318 46067384
refGene_gn=MAPT;refGene_tx=XM_003997029.5;refGene_gn=MAPT;refGene_tx=X
M_003997030.5;refGene_gn=MAPT;refGene_tx=XM_006940453.4;refGene_gn=MAP
T;refGene_tx=XM_006940454.4;refGene_gn=MAPT;refGene_tx=XM_006940455.4;
refGene_gn=MAPT;refGene_tx=XM_006940456.4;refGene_gn=MAPT;refGene_tx=X
M_006940458.4;refGene_gn=MAPT;refGene_tx=XM_006940459.4;refGene_gn=MAP
T;refGene_tx=XM_019818030.2;refGene_gn=MAPT;refGene_tx=XM_019818031.2;
refGene_gn=MAPT;refGene_tx=XM_019818032.2;refGene_gn=MAPT;refGene_tx=X
M_019818033.2;refGene_gn=MAPT;refGene_tx=XM_019818034.2;refGene_gn=MAP
T;refGene_tx=XM_019818036.2;refGene_gn=MAPT;refGene_tx=XM_019818037.2;
refGene_gn=MAPT;refGene_tx=XM_019818038.2;refGene_gn=MAPT;refGene_tx=X
M_019818039.2
E1 46072624 46072930
refGene_gn=MAPT;refGene_tx=XM_006940453.4;refGene_gn=MAPT;refGene_tx=X
M_006940454.4;refGene_gn=MAPT;refGene_tx=XM_006940455.4;refGene_gn=MAP
T;refGene_tx=XM_006940456.4;refGene_gn=MAPT;refGene_tx=XM_019818030.2;
refGene_gn=MAPT;refGene_tx=XM_019818031.2;refGene_gn=MAPT;refGene_tx=X
M_019818032.2
E1 46075609 46075665
refGene_gn=MAPT;refGene_tx=XM_003997029.5;refGene_gn=MAPT;refGene_tx=X
M_003997030.5;refGene_gn=MAPT;refGene_tx=XM_006940453.4;refGene_gn=MAP
T;refGene_tx=XM_006940454.4;refGene_gn=MAPT;refGene_tx=XM_006940455.4;
refGene_gn=MAPT;refGene_tx=XM_006940456.4;refGene_gn=MAPT;refGene_tx=X
M_006940458.4;refGene_gn=MAPT;refGene_tx=XM_006940459.4;refGene_gn=MAP
T;refGene_tx=XM_019818030.2;refGene_gn=MAPT;refGene_tx=XM_019818031.2;
refGene_gn=MAPT;refGene_tx=XM_019818032.2;refGene_gn=MAPT;refGene_tx=X
M_019818033.2;refGene_gn=MAPT;refGene_tx=XM_019818034.2;refGene_gn=MAP
T;refGene_tx=XM_019818036.2;refGene_gn=MAPT;refGene_tx=XM_019818037.2;
refGene_gn=MAPT;refGene_tx=XM_019818038.2;refGene_gn=MAPT;refGene_tx=X
M_019818039.2
E1 46078250 46078448
refGene_gn=MAPT;refGene_tx=XM_006940453.4;refGene_gn=MAPT;refGene_tx=X
M_006940454.4;refGene_gn=MAPT;refGene_tx=XM_006940455.4;refGene_gn=MAP
T;refGene_tx=XM_006940458.4;refGene_gn=MAPT;refGene_tx=XM_019818030.2;
refGene_gn=MAPT;refGene_tx=XM_019818031.2;refGene_gn=MAPT;refGene_tx=X
M_019818033.2;refGene_gn=MAPT;refGene_tx=XM_019818034.2
E1 46079447 46079580

refGene_gn=MAPT;refGene_tx=XM_003997029.5;refGene_gn=MAPT;refGene_tx=XM_003997030.5;refGene_gn=MAPT;refGene_tx=XM_006940453.4;refGene_gn=MAPT;refGene_tx=XM_006940454.4;refGene_gn=MAPT;refGene_tx=XM_006940455.4;refGene_gn=MAPT;refGene_tx=XM_006940456.4;refGene_gn=MAPT;refGene_tx=XM_006940458.4;refGene_gn=MAPT;refGene_tx=XM_006940459.4;refGene_gn=MAPT;refGene_tx=XM_019818030.2;refGene_gn=MAPT;refGene_tx=XM_019818031.2;refGene_gn=MAPT;refGene_tx=XM_019818032.2;refGene_gn=MAPT;refGene_tx=XM_019818033.2;refGene_gn=MAPT;refGene_tx=XM_019818034.2;refGene_gn=MAPT;refGene_tx=XM_019818036.2;refGene_gn=MAPT;refGene_tx=XM_019818037.2;refGene_gn=MAPT;refGene_tx=XM_019818038.2;refGene_gn=MAPT;refGene_tx=XM_019818039.2
E1 46081079 46081133
refGene_gn=MAPT;refGene_tx=XM_006940453.4;refGene_gn=MAPT;refGene_tx=XM_006940455.4;refGene_gn=MAPT;refGene_tx=XM_006940456.4;refGene_gn=MAPT;refGene_tx=XM_006940458.4;refGene_gn=MAPT;refGene_tx=XM_006940459.4;refGene_gn=MAPT;refGene_tx=XM_019818030.2;refGene_gn=MAPT;refGene_tx=XM_019818031.2;refGene_gn=MAPT;refGene_tx=XM_019818033.2;refGene_gn=MAPT;refGene_tx=XM_019818034.2
E1 46083442 46083708
refGene_gn=MAPT;refGene_tx=XM_003997029.5;refGene_gn=MAPT;refGene_tx=XM_003997030.5;refGene_gn=MAPT;refGene_tx=XM_006940453.4;refGene_gn=MAPT;refGene_tx=XM_006940454.4;refGene_gn=MAPT;refGene_tx=XM_006940455.4;refGene_gn=MAPT;refGene_tx=XM_006940456.4;refGene_gn=MAPT;refGene_tx=XM_006940458.4;refGene_gn=MAPT;refGene_tx=XM_006940459.4;refGene_gn=MAPT;refGene_tx=XM_019818030.2;refGene_gn=MAPT;refGene_tx=XM_019818031.2;refGene_gn=MAPT;refGene_tx=XM_019818032.2;refGene_gn=MAPT;refGene_tx=XM_019818033.2;refGene_gn=MAPT;refGene_tx=XM_019818034.2;refGene_gn=MAPT;refGene_tx=XM_019818036.2;refGene_gn=MAPT;refGene_tx=XM_019818037.2;refGene_gn=MAPT;refGene_tx=XM_019818038.2;refGene_gn=MAPT;refGene_tx=XM_019818039.2
E1 46093925 46094018
refGene_gn=MAPT;refGene_tx=XM_003997030.5;refGene_gn=MAPT;refGene_tx=XM_006940453.4;refGene_gn=MAPT;refGene_tx=XM_006940454.4;refGene_gn=MAPT;refGene_tx=XM_006940456.4;refGene_gn=MAPT;refGene_tx=XM_006940458.4;refGene_gn=MAPT;refGene_tx=XM_006940459.4;refGene_gn=MAPT;refGene_tx=XM_019818030.2;refGene_gn=MAPT;refGene_tx=XM_019818031.2;refGene_gn=MAPT;refGene_tx=XM_019818032.2;refGene_gn=MAPT;refGene_tx=XM_019818033.2;refGene_gn=MAPT;refGene_tx=XM_019818034.2;refGene_gn=MAPT;refGene_tx=XM_019818036.2;refGene_gn=MAPT;refGene_tx=XM_019818037.2
E1 46097399 46097481
refGene_gn=MAPT;refGene_tx=XM_003997029.5;refGene_gn=MAPT;refGene_tx=XM_003997030.5;refGene_gn=MAPT;refGene_tx=XM_006940453.4;refGene_gn=MAPT;refGene_tx=XM_006940454.4;refGene_gn=MAPT;refGene_tx=XM_006940455.4;refGene_gn=MAPT;refGene_tx=XM_006940456.4;refGene_gn=MAPT;refGene_tx=XM_006940458.4;refGene_gn=MAPT;refGene_tx=XM_006940459.4;refGene_gn=MAPT;refGene_tx=XM_019818030.2;refGene_gn=MAPT;refGene_tx=XM_019818031.2;refGene_gn=MAPT;refGene_tx=XM_019818032.2;refGene_gn=MAPT;refGene_tx=XM_019818033.2;refGene_gn=MAPT;refGene_tx=XM_019818034.2;refGene_gn=MAPT;refGene_tx=XM_019818036.2;refGene_gn=MAPT;refGene_tx=XM_019818037.2;refGene_gn=MAPT;refGene_tx=XM_019818038.2;refGene_gn=MAPT;refGene_tx=X

M_019818039.2
E1 46098883 46098996
refGene_gn=MAPT;refGene_tx=XM_003997029.5;refGene_gn=MAPT;refGene_tx=XM_003997030.5;refGene_gn=MAPT;refGene_tx=XM_006940453.4;refGene_gn=MAPT;refGene_tx=XM_006940454.4;refGene_gn=MAPT;refGene_tx=XM_006940455.4;refGene_gn=MAPT;refGene_tx=XM_006940456.4;refGene_gn=MAPT;refGene_tx=XM_006940458.4;refGene_gn=MAPT;refGene_tx=XM_006940459.4;refGene_gn=MAPT;refGene_tx=XM_019818030.2;refGene_gn=MAPT;refGene_tx=XM_019818031.2;refGene_gn=MAPT;refGene_tx=XM_019818032.2;refGene_gn=MAPT;refGene_tx=XM_019818033.2;refGene_gn=MAPT;refGene_tx=XM_019818034.2;refGene_gn=MAPT;refGene_tx=XM_019818036.2;refGene_gn=MAPT;refGene_tx=XM_019818037.2;refGene_gn=MAPT;refGene_tx=XM_019818038.2;refGene_gn=MAPT;refGene_tx=XM_019818039.2
E1 46101885 46102101
refGene_gn=MAPT;refGene_tx=XM_003997029.5;refGene_gn=MAPT;refGene_tx=XM_003997030.5;refGene_gn=MAPT;refGene_tx=XM_006940453.4;refGene_gn=MAPT;refGene_tx=XM_006940454.4;refGene_gn=MAPT;refGene_tx=XM_006940455.4;refGene_gn=MAPT;refGene_tx=XM_006940456.4;refGene_gn=MAPT;refGene_tx=XM_006940458.4;refGene_gn=MAPT;refGene_tx=XM_006940459.4;refGene_gn=MAPT;refGene_tx=XM_019818030.2;refGene_gn=MAPT;refGene_tx=XM_019818031.2;refGene_gn=MAPT;refGene_tx=XM_019818032.2;refGene_gn=MAPT;refGene_tx=XM_019818033.2;refGene_gn=MAPT;refGene_tx=XM_019818034.2;refGene_gn=MAPT;refGene_tx=XM_019818036.2;refGene_gn=MAPT;refGene_tx=XM_019818037.2;refGene_gn=MAPT;refGene_tx=XM_019818038.2;refGene_gn=MAPT;refGene_tx=XM_019818039.2
E1 46108988 46109216
refGene_gn=KANSL1;refGene_tx=XM_011289277.3;refGene_gn=KANSL1;refGene_tx=XM_011289278.3;refGene_gn=KANSL1;refGene_tx=XM_011289279.3;refGene_gn=KANSL1;refGene_tx=XM_011289280.3;refGene_gn=KANSL1;refGene_tx=XM_019818040.2
E1 46109836 46110089
refGene_gn=KANSL1;refGene_tx=XM_011289277.3;refGene_gn=KANSL1;refGene_tx=XM_011289278.3;refGene_gn=KANSL1;refGene_tx=XM_011289279.3;refGene_gn=KANSL1;refGene_tx=XM_011289280.3;refGene_gn=KANSL1;refGene_tx=XM_019818040.2
E1 46110965 46111078
refGene_gn=KANSL1;refGene_tx=XM_011289277.3;refGene_gn=KANSL1;refGene_tx=XM_011289278.3;refGene_gn=KANSL1;refGene_tx=XM_011289279.3;refGene_gn=KANSL1;refGene_tx=XM_011289280.3;refGene_gn=KANSL1;refGene_tx=XM_019818040.2
E1 46111329 46111387
refGene_gn=KANSL1;refGene_tx=XM_011289277.3;refGene_gn=KANSL1;refGene_tx=XM_011289278.3;refGene_gn=KANSL1;refGene_tx=XM_011289279.3;refGene_gn=KANSL1;refGene_tx=XM_011289280.3;refGene_gn=KANSL1;refGene_tx=XM_019818040.2
E1 46111975 46112097
refGene_gn=KANSL1;refGene_tx=XM_011289277.3;refGene_gn=KANSL1;refGene_tx=XM_011289278.3;refGene_gn=KANSL1;refGene_tx=XM_011289279.3;refGene_gn=KANSL1;refGene_tx=XM_011289280.3;refGene_gn=KANSL1;refGene_tx=XM_019818040.2

E1 46115580 46115729
refGene_gn=KANSL1;refGene_tx=XM_011289277.3;refGene_gn=KANSL1;refGene_tx=XM_011289278.3;refGene_gn=KANSL1;refGene_tx=XM_011289279.3;refGene_gn=KANSL1;refGene_tx=XM_011289280.3;refGene_gn=KANSL1;refGene_tx=XM_019818040.2
E1 46116061 46116250
refGene_gn=KANSL1;refGene_tx=XM_011289277.3;refGene_gn=KANSL1;refGene_tx=XM_011289278.3;refGene_gn=KANSL1;refGene_tx=XM_011289279.3;refGene_gn=KANSL1;refGene_tx=XM_019818040.2
E1 46116762 46116945
refGene_gn=KANSL1;refGene_tx=XM_011289277.3;refGene_gn=KANSL1;refGene_tx=XM_011289278.3;refGene_gn=KANSL1;refGene_tx=XM_011289279.3;refGene_gn=KANSL1;refGene_tx=XM_011289280.3;refGene_gn=KANSL1;refGene_tx=XM_019818040.2
E1 46123211 46123383
refGene_gn=KANSL1;refGene_tx=XM_011289277.3;refGene_gn=KANSL1;refGene_tx=XM_011289278.3;refGene_gn=KANSL1;refGene_tx=XM_011289279.3;refGene_gn=KANSL1;refGene_tx=XM_011289280.3;refGene_gn=KANSL1;refGene_tx=XM_019818040.2
E1 46142798 46142994
refGene_gn=KANSL1;refGene_tx=XM_011289277.3;refGene_gn=KANSL1;refGene_tx=XM_011289278.3;refGene_gn=KANSL1;refGene_tx=XM_011289279.3;refGene_gn=KANSL1;refGene_tx=XM_011289280.3;refGene_gn=KANSL1;refGene_tx=XM_019818040.2
E1 46143831 46143941
refGene_gn=KANSL1;refGene_tx=XM_011289277.3;refGene_gn=KANSL1;refGene_tx=XM_011289278.3;refGene_gn=KANSL1;refGene_tx=XM_011289279.3;refGene_gn=KANSL1;refGene_tx=XM_011289280.3;refGene_gn=KANSL1;refGene_tx=XM_019818040.2
E1 46153949 46154054
refGene_gn=KANSL1;refGene_tx=XM_011289277.3;refGene_gn=KANSL1;refGene_tx=XM_011289278.3;refGene_gn=KANSL1;refGene_tx=XM_011289280.3;refGene_gn=KANSL1;refGene_tx=XM_019818040.2
E1 46162370 46162512
refGene_gn=KANSL1;refGene_tx=XM_011289277.3;refGene_gn=KANSL1;refGene_tx=XM_011289278.3;refGene_gn=KANSL1;refGene_tx=XM_011289279.3;refGene_gn=KANSL1;refGene_tx=XM_011289280.3;refGene_gn=KANSL1;refGene_tx=XM_019818040.2
E1 46243929 46245218
refGene_gn=KANSL1;refGene_tx=XM_011289277.3;refGene_gn=KANSL1;refGene_tx=XM_011289278.3;refGene_gn=KANSL1;refGene_tx=XM_011289279.3;refGene_gn=KANSL1;refGene_tx=XM_011289280.3;refGene_gn=KANSL1;refGene_tx=XM_019818040.2
E1 46318410 46318493
refGene_gn=CDC27;refGene_tx=XM_003997032.5;refGene_gn=CDC27;refGene_tx=XM_006940466.4;refGene_gn=CDC27;refGene_tx=XM_006940467.4
E1 46320357 46320514
refGene_gn=CDC27;refGene_tx=XM_003997032.5;refGene_gn=CDC27;refGene_tx=XM_006940466.4;refGene_gn=CDC27;refGene_tx=XM_006940467.4
E1 46321326 46321401

refGene_gn=CDC27;refGene_tx=XM_003997032.5;refGene_gn=CDC27;refGene_tx
=XM_006940466.4;refGene_gn=CDC27;refGene_tx=XM_006940467.4
E1 46327059 46327188
refGene_gn=CDC27;refGene_tx=XM_003997032.5;refGene_gn=CDC27;refGene_tx
=XM_006940466.4;refGene_gn=CDC27;refGene_tx=XM_006940467.4
E1 46331084 46331202
refGene_gn=CDC27;refGene_tx=XM_003997032.5;refGene_gn=CDC27;refGene_tx
=XM_006940466.4;refGene_gn=CDC27;refGene_tx=XM_006940467.4
E1 46335839 46336048
refGene_gn=CDC27;refGene_tx=XM_003997032.5;refGene_gn=CDC27;refGene_tx
=XM_006940466.4;refGene_gn=CDC27;refGene_tx=XM_006940467.4
E1 46337530 46337683
refGene_gn=CDC27;refGene_tx=XM_003997032.5;refGene_gn=CDC27;refGene_tx
=XM_006940466.4;refGene_gn=CDC27;refGene_tx=XM_006940467.4
E1 46340804 46340977
refGene_gn=CDC27;refGene_tx=XM_003997032.5;refGene_gn=CDC27;refGene_tx
=XM_006940466.4;refGene_gn=CDC27;refGene_tx=XM_006940467.4
E1 46341183 46341391
refGene_gn=CDC27;refGene_tx=XM_003997032.5;refGene_gn=CDC27;refGene_tx
=XM_006940466.4;refGene_gn=CDC27;refGene_tx=XM_006940467.4
E1 46342374 46342474
refGene_gn=CDC27;refGene_tx=XM_003997032.5;refGene_gn=CDC27;refGene_tx
=XM_006940466.4;refGene_gn=CDC27;refGene_tx=XM_006940467.4
E1 46349445 46349561
refGene_gn=CDC27;refGene_tx=XM_003997032.5;refGene_gn=CDC27;refGene_tx
=XM_006940466.4;refGene_gn=CDC27;refGene_tx=XM_006940467.4
E1 46352242 46352357
refGene_gn=CDC27;refGene_tx=XM_003997032.5;refGene_gn=CDC27;refGene_tx
=XM_006940466.4;refGene_gn=CDC27;refGene_tx=XM_006940467.4
E1 46353336 46353548
refGene_gn=CDC27;refGene_tx=XM_003997032.5;refGene_gn=CDC27;refGene_tx
=XM_006940466.4;refGene_gn=CDC27;refGene_tx=XM_006940467.4
E1 46353646 46353801
refGene_gn=CDC27;refGene_tx=XM_003997032.5;refGene_gn=CDC27;refGene_tx
=XM_006940466.4;refGene_gn=CDC27;refGene_tx=XM_006940467.4
E1 46354539 46354637
refGene_gn=CDC27;refGene_tx=XM_003997032.5;refGene_gn=CDC27;refGene_tx
=XM_006940466.4;refGene_gn=CDC27;refGene_tx=XM_006940467.4
E1 46358979 46359105
refGene_gn=CDC27;refGene_tx=XM_003997032.5;refGene_gn=CDC27;refGene_tx
=XM_006940466.4;refGene_gn=CDC27;refGene_tx=XM_006940467.4
E1 46360712 46360860
refGene_gn=CDC27;refGene_tx=XM_003997032.5;refGene_gn=CDC27;refGene_tx
=XM_006940466.4
E1 46370975 46371051
refGene_gn=CDC27;refGene_tx=XM_003997032.5;refGene_gn=CDC27;refGene_tx
=XM_006940466.4;refGene_gn=CDC27;refGene_tx=XM_006940467.4
E1 46386082 46386109
refGene_gn=CDC27;refGene_tx=XM_003997032.5;refGene_gn=CDC27;refGene_tx
=XM_006940466.4

E1 46402160 46402277
refGene_gn=MYL4;refGene_tx=XM_003997034.5;refGene_gn=MYL4;refGene_tx=XM_006940463.4;refGene_gn=MYL4;refGene_tx=XM_006940464.4;refGene_gn=MYL4;refGene_tx=XM_006940465.4;refGene_gn=MYL4;refGene_tx=XM_011289281.3;refGene_gn=MYL4;refGene_tx=XM_023244496.1
E1 46405767 46405795
refGene_gn=MYL4;refGene_tx=XM_003997034.5;refGene_gn=MYL4;refGene_tx=XM_006940463.4;refGene_gn=MYL4;refGene_tx=XM_006940464.4;refGene_gn=MYL4;refGene_tx=XM_006940465.4;refGene_gn=MYL4;refGene_tx=XM_011289281.3;refGene_gn=MYL4;refGene_tx=XM_023244496.1
E1 46412969 46413119
refGene_gn=MYL4;refGene_tx=XM_003997034.5;refGene_gn=MYL4;refGene_tx=XM_006940463.4;refGene_gn=MYL4;refGene_tx=XM_006940464.4;refGene_gn=MYL4;refGene_tx=XM_006940465.4;refGene_gn=MYL4;refGene_tx=XM_011289281.3;refGene_gn=MYL4;refGene_tx=XM_023244496.1
E1 46414360 46414534
refGene_gn=MYL4;refGene_tx=XM_003997034.5;refGene_gn=MYL4;refGene_tx=XM_006940463.4;refGene_gn=MYL4;refGene_tx=XM_006940464.4;refGene_gn=MYL4;refGene_tx=XM_006940465.4;refGene_gn=MYL4;refGene_tx=XM_011289281.3;refGene_gn=MYL4;refGene_tx=XM_023244496.1
E1 46415019 46415097
refGene_gn=MYL4;refGene_tx=XM_003997034.5;refGene_gn=MYL4;refGene_tx=XM_006940463.4;refGene_gn=MYL4;refGene_tx=XM_006940464.4;refGene_gn=MYL4;refGene_tx=XM_006940465.4;refGene_gn=MYL4;refGene_tx=XM_011289281.3;refGene_gn=MYL4;refGene_tx=XM_023244496.1
E1 46415653 46415682
refGene_gn=MYL4;refGene_tx=XM_003997034.5;refGene_gn=MYL4;refGene_tx=XM_006940463.4;refGene_gn=MYL4;refGene_tx=XM_006940464.4;refGene_gn=MYL4;refGene_tx=XM_006940465.4;refGene_gn=MYL4;refGene_tx=XM_011289281.3;refGene_gn=MYL4;refGene_tx=XM_023244496.1
E1 46436251 46436318 refGene_gn=ITGB3;refGene_tx=XM_003997035.5
E1 46453786 46453872 refGene_gn=ITGB3;refGene_tx=XM_003997035.5
E1 46455363 46455513 refGene_gn=ITGB3;refGene_tx=XM_019818041.1
E1 46461401 46461597
refGene_gn=ITGB3;refGene_tx=XM_003997035.5;refGene_gn=ITGB3;refGene_tx=XM_019818041.1
E1 46462740 46462993
refGene_gn=ITGB3;refGene_tx=XM_003997035.5;refGene_gn=ITGB3;refGene_tx=XM_019818041.1
E1 46464557 46464720
refGene_gn=ITGB3;refGene_tx=XM_003997035.5;refGene_gn=ITGB3;refGene_tx=XM_019818041.1
E1 46465353 46465515
refGene_gn=ITGB3;refGene_tx=XM_003997035.5;refGene_gn=ITGB3;refGene_tx=XM_019818041.1
E1 46468081 46468177
refGene_gn=ITGB3;refGene_tx=XM_003997035.5;refGene_gn=ITGB3;refGene_tx=XM_019818041.1
E1 46468602 46468692
refGene_gn=ITGB3;refGene_tx=XM_003997035.5;refGene_gn=ITGB3;refGene_tx

=XM_019818041.1
E1	46469393	46469528
refGene_gn=ITGB3;refGene_tx=XM_003997035.5;refGene_gn=ITGB3;refGene_tx=XM_019818041.1
E1	46470985	46471415
refGene_gn=ITGB3;refGene_tx=XM_003997035.5;refGene_gn=ITGB3;refGene_tx=XM_019818041.1
E1	46477370	46477593
refGene_gn=ITGB3;refGene_tx=XM_003997035.5;refGene_gn=ITGB3;refGene_tx=XM_019818041.1
E1	46478758	46478859
refGene_gn=ITGB3;refGene_tx=XM_003997035.5;refGene_gn=ITGB3;refGene_tx=XM_019818041.1
E1	46481031	46481151
refGene_gn=ITGB3;refGene_tx=XM_003997035.5;refGene_gn=ITGB3;refGene_tx=XM_019818041.1
E1	46487071	46487238
refGene_gn=ITGB3;refGene_tx=XM_003997035.5;refGene_gn=ITGB3;refGene_tx=XM_019818041.1
E1	46488567	46488633
refGene_gn=ITGB3;refGene_tx=XM_003997035.5;refGene_gn=ITGB3;refGene_tx=XM_019818041.1
E1	46505020	46505050
refGene_gn=LOC102899173;refGene_tx=XM_019818044.2;refGene_gn=LOC102899173;refGene_tx=XM_023244736.1;refGene_gn=LOC102899173;refGene_tx=XM_023244737.1;refGene_gn=LOC102899173;refGene_tx=XM_023244738.1;refGene_gn=LOC102899173;refGene_tx=XM_023244739.1;refGene_gn=LOC102899173;refGene_tx=XM_023244740.1;refGene_gn=LOC102899173;refGene_tx=XM_023244741.1;refGene_gn=LOC102899173;refGene_tx=XM_023244742.1;refGene_gn=LOC102899173;refGene_tx=XM_023244743.1;refGene_gn=LOC102899173;refGene_tx=XM_023244744.1;refGene_gn=LOC102899173;refGene_tx=XM_023244745.1;refGene_gn=LOC102899173;refGene_tx=XM_023244746.1;refGene_gn=LOC102899173;refGene_tx=XM_023244747.1;refGene_gn=LOC102899173;refGene_tx=XM_023244748.1;refGene_gn=LOC102899173;refGene_tx=XM_023244749.1;refGene_gn=LOC102899173;refGene_tx=XM_023244750.1;refGene_gn=LOC102899173;refGene_tx=XM_023244755.1;refGene_gn=LOC102899173;refGene_tx=XM_023244756.1
E1	46509649	46509810
refGene_gn=LOC102899173;refGene_tx=XM_019818044.2;refGene_gn=LOC102899173;refGene_tx=XM_023244736.1;refGene_gn=LOC102899173;refGene_tx=XM_023244737.1;refGene_gn=LOC102899173;refGene_tx=XM_023244738.1;refGene_gn=LOC102899173;refGene_tx=XM_023244739.1;refGene_gn=LOC102899173;refGene_tx=XM_023244740.1;refGene_gn=LOC102899173;refGene_tx=XM_023244741.1;refGene_gn=LOC102899173;refGene_tx=XM_023244742.1;refGene_gn=LOC102899173;refGene_tx=XM_023244743.1;refGene_gn=LOC102899173;refGene_tx=XM_023244744.1;refGene_gn=LOC102899173;refGene_tx=XM_023244745.1;refGene_gn=LOC102899173;refGene_tx=XM_023244746.1;refGene_gn=LOC102899173;refGene_tx=XM_023244747.1;refGene_gn=LOC102899173;refGene_tx=XM_023244748.1;refGene_gn=LOC102899173;refGene_tx=XM_023244749.1;refGene_gn=LOC102899173;refGene_tx=XM_023244750.1;refGene_gn=LOC102899173;refGene_tx=XM_023244755.1;refGene_gn=LOC102899173;refGene_tx=XM_023244756.1

E1 46518477 46518592
refGene_gn=LOC102899173;refGene_tx=XM_019818044.2;refGene_gn=LOC102899173;refGene_tx=XM_023244736.1;refGene_gn=LOC102899173;refGene_tx=XM_023244737.1;refGene_gn=LOC102899173;refGene_tx=XM_023244738.1;refGene_gn=LOC102899173;refGene_tx=XM_023244739.1;refGene_gn=LOC102899173;refGene_tx=XM_023244740.1;refGene_gn=LOC102899173;refGene_tx=XM_023244741.1;refGene_gn=LOC102899173;refGene_tx=XM_023244742.1;refGene_gn=LOC102899173;refGene_tx=XM_023244743.1;refGene_gn=LOC102899173;refGene_tx=XM_023244744.1;refGene_gn=LOC102899173;refGene_tx=XM_023244745.1;refGene_gn=LOC102899173;refGene_tx=XM_023244746.1;refGene_gn=LOC102899173;refGene_tx=XM_023244747.1;refGene_gn=LOC102899173;refGene_tx=XM_023244748.1;refGene_gn=LOC102899173;refGene_tx=XM_023244749.1;refGene_gn=LOC102899173;refGene_tx=XM_023244750.1;refGene_gn=LOC102899173;refGene_tx=XM_023244755.1;refGene_gn=LOC102899173;refGene_tx=XM_023244756.1
E1 46521531 46521662
refGene_gn=LOC102899173;refGene_tx=XM_019818044.2;refGene_gn=LOC102899173;refGene_tx=XM_023244736.1;refGene_gn=LOC102899173;refGene_tx=XM_023244737.1;refGene_gn=LOC102899173;refGene_tx=XM_023244738.1;refGene_gn=LOC102899173;refGene_tx=XM_023244739.1;refGene_gn=LOC102899173;refGene_tx=XM_023244740.1;refGene_gn=LOC102899173;refGene_tx=XM_023244741.1;refGene_gn=LOC102899173;refGene_tx=XM_023244742.1;refGene_gn=LOC102899173;refGene_tx=XM_023244743.1;refGene_gn=LOC102899173;refGene_tx=XM_023244744.1;refGene_gn=LOC102899173;refGene_tx=XM_023244745.1;refGene_gn=LOC102899173;refGene_tx=XM_023244746.1;refGene_gn=LOC102899173;refGene_tx=XM_023244747.1;refGene_gn=LOC102899173;refGene_tx=XM_023244748.1;refGene_gn=LOC102899173;refGene_tx=XM_023244749.1;refGene_gn=LOC102899173;refGene_tx=XM_023244750.1;refGene_gn=LOC102899173;refGene_tx=XM_023244755.1;refGene_gn=LOC102899173;refGene_tx=XM_023244756.1
E1 46522451 46522534
refGene_gn=LOC102899173;refGene_tx=XM_019818044.2;refGene_gn=LOC102899173;refGene_tx=XM_023244736.1;refGene_gn=LOC102899173;refGene_tx=XM_023244737.1;refGene_gn=LOC102899173;refGene_tx=XM_023244738.1;refGene_gn=LOC102899173;refGene_tx=XM_023244739.1;refGene_gn=LOC102899173;refGene_tx=XM_023244740.1;refGene_gn=LOC102899173;refGene_tx=XM_023244741.1;refGene_gn=LOC102899173;refGene_tx=XM_023244742.1;refGene_gn=LOC102899173;refGene_tx=XM_023244743.1;refGene_gn=LOC102899173;refGene_tx=XM_023244744.1;refGene_gn=LOC102899173;refGene_tx=XM_023244745.1;refGene_gn=LOC102899173;refGene_tx=XM_023244746.1;refGene_gn=LOC102899173;refGene_tx=XM_023244747.1;refGene_gn=LOC102899173;refGene_tx=XM_023244748.1;refGene_gn=LOC102899173;refGene_tx=XM_023244749.1;refGene_gn=LOC102899173;refGene_tx=XM_023244750.1;refGene_gn=LOC102899173;refGene_tx=XM_023244755.1;refGene_gn=LOC102899173;refGene_tx=XM_023244756.1
E1 46524837 46524981
refGene_gn=LOC102899173;refGene_tx=XM_019818044.2;refGene_gn=LOC102899173;refGene_tx=XM_023244736.1;refGene_gn=LOC102899173;refGene_tx=XM_023244737.1;refGene_gn=LOC102899173;refGene_tx=XM_023244738.1;refGene_gn=LOC102899173;refGene_tx=XM_023244739.1;refGene_gn=LOC102899173;refGene_tx=XM_023244740.1;refGene_gn=LOC102899173;refGene_tx=XM_023244741.1;refGene_gn=LOC102899173;refGene_tx=XM_023244742.1;refGene_gn=LOC102899173;refGene_tx=XM_023244743.1;refGene_gn=LOC102899173;refGene_tx=XM_02

3244744.1;refGene_gn=LOC102899173;refGene_tx=XM_023244745.1;refGene_gn=LOC102899173;refGene_tx=XM_023244746.1;refGene_gn=LOC102899173;refGene_tx=XM_023244747.1;refGene_gn=LOC102899173;refGene_tx=XM_023244748.1;refGene_gn=LOC102899173;refGene_tx=XM_023244749.1;refGene_gn=LOC102899173;refGene_tx=XM_023244750.1;refGene_gn=LOC102899173;refGene_tx=XM_023244755.1;refGene_gn=LOC102899173;refGene_tx=XM_023244756.1
E1 46533695 46533770
refGene_gn=LOC102899173;refGene_tx=XM_019818044.2;refGene_gn=LOC102899173;refGene_tx=XM_023244736.1;refGene_gn=LOC102899173;refGene_tx=XM_023244737.1;refGene_gn=LOC102899173;refGene_tx=XM_023244738.1;refGene_gn=LOC102899173;refGene_tx=XM_023244739.1;refGene_gn=LOC102899173;refGene_tx=XM_023244740.1;refGene_gn=LOC102899173;refGene_tx=XM_023244741.1;refGene_gn=LOC102899173;refGene_tx=XM_023244742.1;refGene_gn=LOC102899173;refGene_tx=XM_023244743.1;refGene_gn=LOC102899173;refGene_tx=XM_023244744.1;refGene_gn=LOC102899173;refGene_tx=XM_023244745.1;refGene_gn=LOC102899173;refGene_tx=XM_023244746.1;refGene_gn=LOC102899173;refGene_tx=XM_023244747.1;refGene_gn=LOC102899173;refGene_tx=XM_023244748.1;refGene_gn=LOC102899173;refGene_tx=XM_023244749.1;refGene_gn=LOC102899173;refGene_tx=XM_023244750.1;refGene_gn=LOC102899173;refGene_tx=XM_023244755.1;refGene_gn=LOC102899173;refGene_tx=XM_023244756.1
E1 46539006 46539150
refGene_gn=LOC102899173;refGene_tx=XM_019818044.2;refGene_gn=LOC102899173;refGene_tx=XM_023244736.1;refGene_gn=LOC102899173;refGene_tx=XM_023244737.1;refGene_gn=LOC102899173;refGene_tx=XM_023244738.1;refGene_gn=LOC102899173;refGene_tx=XM_023244739.1;refGene_gn=LOC102899173;refGene_tx=XM_023244740.1;refGene_gn=LOC102899173;refGene_tx=XM_023244741.1;refGene_gn=LOC102899173;refGene_tx=XM_023244742.1;refGene_gn=LOC102899173;refGene_tx=XM_023244743.1;refGene_gn=LOC102899173;refGene_tx=XM_023244744.1;refGene_gn=LOC102899173;refGene_tx=XM_023244745.1;refGene_gn=LOC102899173;refGene_tx=XM_023244746.1;refGene_gn=LOC102899173;refGene_tx=XM_023244747.1;refGene_gn=LOC102899173;refGene_tx=XM_023244748.1;refGene_gn=LOC102899173;refGene_tx=XM_023244749.1;refGene_gn=LOC102899173;refGene_tx=XM_023244750.1;refGene_gn=LOC102899173;refGene_tx=XM_023244755.1;refGene_gn=LOC102899173;refGene_tx=XM_023244756.1
E1 46545757 46545829
refGene_gn=LOC102899173;refGene_tx=XM_019818044.2;refGene_gn=LOC102899173;refGene_tx=XM_023244736.1;refGene_gn=LOC102899173;refGene_tx=XM_023244737.1;refGene_gn=LOC102899173;refGene_tx=XM_023244738.1;refGene_gn=LOC102899173;refGene_tx=XM_023244739.1;refGene_gn=LOC102899173;refGene_tx=XM_023244740.1;refGene_gn=LOC102899173;refGene_tx=XM_023244741.1;refGene_gn=LOC102899173;refGene_tx=XM_023244742.1;refGene_gn=LOC102899173;refGene_tx=XM_023244743.1;refGene_gn=LOC102899173;refGene_tx=XM_023244744.1;refGene_gn=LOC102899173;refGene_tx=XM_023244745.1;refGene_gn=LOC102899173;refGene_tx=XM_023244746.1;refGene_gn=LOC102899173;refGene_tx=XM_023244747.1;refGene_gn=LOC102899173;refGene_tx=XM_023244748.1;refGene_gn=LOC102899173;refGene_tx=XM_023244749.1;refGene_gn=LOC102899173;refGene_tx=XM_023244750.1;refGene_gn=LOC102899173;refGene_tx=XM_023244755.1;refGene_gn=LOC102899173;refGene_tx=XM_023244756.1
E1 46551568 46552063
refGene_gn=LOC102899173;refGene_tx=XM_019818044.2;refGene_gn=LOC102899

173;refGene_tx=XM_023244736.1;refGene_gn=LOC102899173;refGene_tx=XM_023244737.1;refGene_gn=LOC102899173;refGene_tx=XM_023244738.1;refGene_gn=LOC102899173;refGene_tx=XM_023244739.1;refGene_gn=LOC102899173;refGene_tx=XM_023244740.1;refGene_gn=LOC102899173;refGene_tx=XM_023244741.1;refGene_gn=LOC102899173;refGene_tx=XM_023244742.1;refGene_gn=LOC102899173;refGene_tx=XM_023244743.1;refGene_gn=LOC102899173;refGene_tx=XM_023244744.1;refGene_gn=LOC102899173;refGene_tx=XM_023244745.1;refGene_gn=LOC102899173;refGene_tx=XM_023244746.1;refGene_gn=LOC102899173;refGene_tx=XM_023244747.1;refGene_gn=LOC102899173;refGene_tx=XM_023244748.1;refGene_gn=LOC102899173;refGene_tx=XM_023244749.1;refGene_gn=LOC102899173;refGene_tx=XM_023244750.1;refGene_gn=LOC102899173;refGene_tx=XM_023244755.1;refGene_gn=LOC102899173;refGene_tx=XM_023244756.1
E1 46554095 46554233
refGene_gn=LOC102899173;refGene_tx=XM_019818044.2;refGene_gn=LOC102899173;refGene_tx=XM_023244736.1;refGene_gn=LOC102899173;refGene_tx=XM_023244737.1;refGene_gn=LOC102899173;refGene_tx=XM_023244738.1;refGene_gn=LOC102899173;refGene_tx=XM_023244739.1;refGene_gn=LOC102899173;refGene_tx=XM_023244740.1;refGene_gn=LOC102899173;refGene_tx=XM_023244741.1;refGene_gn=LOC102899173;refGene_tx=XM_023244742.1;refGene_gn=LOC102899173;refGene_tx=XM_023244743.1;refGene_gn=LOC102899173;refGene_tx=XM_023244744.1;refGene_gn=LOC102899173;refGene_tx=XM_023244745.1;refGene_gn=LOC102899173;refGene_tx=XM_023244746.1;refGene_gn=LOC102899173;refGene_tx=XM_023244747.1;refGene_gn=LOC102899173;refGene_tx=XM_023244748.1;refGene_gn=LOC102899173;refGene_tx=XM_023244749.1;refGene_gn=LOC102899173;refGene_tx=XM_023244750.1;refGene_gn=LOC102899173;refGene_tx=XM_023244755.1;refGene_gn=LOC102899173;refGene_tx=XM_023244756.1
E1 46555464 46555536
refGene_gn=LOC102899173;refGene_tx=XM_019818044.2;refGene_gn=LOC102899173;refGene_tx=XM_023244736.1;refGene_gn=LOC102899173;refGene_tx=XM_023244737.1;refGene_gn=LOC102899173;refGene_tx=XM_023244738.1;refGene_gn=LOC102899173;refGene_tx=XM_023244739.1;refGene_gn=LOC102899173;refGene_tx=XM_023244740.1;refGene_gn=LOC102899173;refGene_tx=XM_023244741.1;refGene_gn=LOC102899173;refGene_tx=XM_023244742.1;refGene_gn=LOC102899173;refGene_tx=XM_023244743.1;refGene_gn=LOC102899173;refGene_tx=XM_023244744.1;refGene_gn=LOC102899173;refGene_tx=XM_023244745.1;refGene_gn=LOC102899173;refGene_tx=XM_023244746.1;refGene_gn=LOC102899173;refGene_tx=XM_023244747.1;refGene_gn=LOC102899173;refGene_tx=XM_023244748.1;refGene_gn=LOC102899173;refGene_tx=XM_023244749.1;refGene_gn=LOC102899173;refGene_tx=XM_023244750.1;refGene_gn=LOC102899173;refGene_tx=XM_023244755.1;refGene_gn=LOC102899173;refGene_tx=XM_023244756.1
E1 46561189 46561333
refGene_gn=LOC102899173;refGene_tx=XM_019818044.2;refGene_gn=LOC102899173;refGene_tx=XM_023244736.1;refGene_gn=LOC102899173;refGene_tx=XM_023244737.1;refGene_gn=LOC102899173;refGene_tx=XM_023244738.1;refGene_gn=LOC102899173;refGene_tx=XM_023244739.1;refGene_gn=LOC102899173;refGene_tx=XM_023244740.1;refGene_gn=LOC102899173;refGene_tx=XM_023244741.1;refGene_gn=LOC102899173;refGene_tx=XM_023244742.1;refGene_gn=LOC102899173;refGene_tx=XM_023244743.1;refGene_gn=LOC102899173;refGene_tx=XM_023244744.1;refGene_gn=LOC102899173;refGene_tx=XM_023244745.1;refGene_gn=LOC102899173;refGene_tx=XM_023244746.1;refGene_gn=LOC102899173;refGen

e_tx=XM_023244747.1;refGene_gn=LOC102899173;refGene_tx=XM_023244748.1;refGene_gn=LOC102899173;refGene_tx=XM_023244749.1;refGene_gn=LOC102899173;refGene_tx=XM_023244750.1;refGene_gn=LOC102899173;refGene_tx=XM_023244755.1;refGene_gn=LOC102899173;refGene_tx=XM_023244756.1
E1 46564919 46564994
refGene_gn=LOC102899173;refGene_tx=XM_019818044.2;refGene_gn=LOC102899173;refGene_tx=XM_023244736.1;refGene_gn=LOC102899173;refGene_tx=XM_023244737.1;refGene_gn=LOC102899173;refGene_tx=XM_023244738.1;refGene_gn=LOC102899173;refGene_tx=XM_023244739.1;refGene_gn=LOC102899173;refGene_tx=XM_023244740.1;refGene_gn=LOC102899173;refGene_tx=XM_023244741.1;refGene_gn=LOC102899173;refGene_tx=XM_023244742.1;refGene_gn=LOC102899173;refGene_tx=XM_023244743.1;refGene_gn=LOC102899173;refGene_tx=XM_023244744.1;refGene_gn=LOC102899173;refGene_tx=XM_023244745.1;refGene_gn=LOC102899173;refGene_tx=XM_023244746.1;refGene_gn=LOC102899173;refGene_tx=XM_023244747.1;refGene_gn=LOC102899173;refGene_tx=XM_023244748.1;refGene_gn=LOC102899173;refGene_tx=XM_023244749.1;refGene_gn=LOC102899173;refGene_tx=XM_023244750.1;refGene_gn=LOC102899173;refGene_tx=XM_023244755.1;refGene_gn=LOC102899173;refGene_tx=XM_023244756.1
E1 46583363 46583507
refGene_gn=LOC102899173;refGene_tx=XM_019818044.2;refGene_gn=LOC102899173;refGene_tx=XM_023244736.1;refGene_gn=LOC102899173;refGene_tx=XM_023244737.1;refGene_gn=LOC102899173;refGene_tx=XM_023244738.1;refGene_gn=LOC102899173;refGene_tx=XM_023244739.1;refGene_gn=LOC102899173;refGene_tx=XM_023244740.1;refGene_gn=LOC102899173;refGene_tx=XM_023244741.1;refGene_gn=LOC102899173;refGene_tx=XM_023244742.1;refGene_gn=LOC102899173;refGene_tx=XM_023244743.1;refGene_gn=LOC102899173;refGene_tx=XM_023244744.1;refGene_gn=LOC102899173;refGene_tx=XM_023244745.1;refGene_gn=LOC102899173;refGene_tx=XM_023244746.1;refGene_gn=LOC102899173;refGene_tx=XM_023244747.1;refGene_gn=LOC102899173;refGene_tx=XM_023244748.1;refGene_gn=LOC102899173;refGene_tx=XM_023244749.1;refGene_gn=LOC102899173;refGene_tx=XM_023244750.1;refGene_gn=LOC102899173;refGene_tx=XM_023244755.1;refGene_gn=LOC102899173;refGene_tx=XM_023244756.1
E1 46585126 46585198
refGene_gn=LOC102899173;refGene_tx=XM_019818044.2;refGene_gn=LOC102899173;refGene_tx=XM_023244736.1;refGene_gn=LOC102899173;refGene_tx=XM_023244737.1;refGene_gn=LOC102899173;refGene_tx=XM_023244738.1;refGene_gn=LOC102899173;refGene_tx=XM_023244739.1;refGene_gn=LOC102899173;refGene_tx=XM_023244740.1;refGene_gn=LOC102899173;refGene_tx=XM_023244741.1;refGene_gn=LOC102899173;refGene_tx=XM_023244742.1;refGene_gn=LOC102899173;refGene_tx=XM_023244743.1;refGene_gn=LOC102899173;refGene_tx=XM_023244744.1;refGene_gn=LOC102899173;refGene_tx=XM_023244745.1;refGene_gn=LOC102899173;refGene_tx=XM_023244746.1;refGene_gn=LOC102899173;refGene_tx=XM_023244747.1;refGene_gn=LOC102899173;refGene_tx=XM_023244748.1;refGene_gn=LOC102899173;refGene_tx=XM_023244749.1;refGene_gn=LOC102899173;refGene_tx=XM_023244750.1;refGene_gn=LOC102899173;refGene_tx=XM_023244755.1;refGene_gn=LOC102899173;refGene_tx=XM_023244756.1
E1 46588222 46588366
refGene_gn=LOC102899173;refGene_tx=XM_019818044.2;refGene_gn=LOC102899173;refGene_tx=XM_023244736.1;refGene_gn=LOC102899173;refGene_tx=XM_023244737.1;refGene_gn=LOC102899173;refGene_tx=XM_023244738.1;refGene_gn

=LOC102899173;refGene_tx=XM_023244739.1;refGene_gn=LOC102899173;refGene_tx=XM_023244740.1;refGene_gn=LOC102899173;refGene_tx=XM_023244741.1;refGene_gn=LOC102899173;refGene_tx=XM_023244742.1;refGene_gn=LOC102899173;refGene_tx=XM_023244743.1;refGene_gn=LOC102899173;refGene_tx=XM_023244744.1;refGene_gn=LOC102899173;refGene_tx=XM_023244745.1;refGene_gn=LOC102899173;refGene_tx=XM_023244746.1;refGene_gn=LOC102899173;refGene_tx=XM_023244747.1;refGene_gn=LOC102899173;refGene_tx=XM_023244748.1;refGene_gn=LOC102899173;refGene_tx=XM_023244749.1;refGene_gn=LOC102899173;refGene_tx=XM_023244750.1;refGene_gn=LOC102899173;refGene_tx=XM_023244755.1;refGene_gn=LOC102899173;refGene_tx=XM_023244756.1
E1 46589177 46589249
refGene_gn=LOC102899173;refGene_tx=XM_019818044.2;refGene_gn=LOC102899173;refGene_tx=XM_023244736.1;refGene_gn=LOC102899173;refGene_tx=XM_023244737.1;refGene_gn=LOC102899173;refGene_tx=XM_023244738.1;refGene_gn=LOC102899173;refGene_tx=XM_023244739.1;refGene_gn=LOC102899173;refGene_tx=XM_023244740.1;refGene_gn=LOC102899173;refGene_tx=XM_023244741.1;refGene_gn=LOC102899173;refGene_tx=XM_023244742.1;refGene_gn=LOC102899173;refGene_tx=XM_023244743.1;refGene_gn=LOC102899173;refGene_tx=XM_023244744.1;refGene_gn=LOC102899173;refGene_tx=XM_023244745.1;refGene_gn=LOC102899173;refGene_tx=XM_023244746.1;refGene_gn=LOC102899173;refGene_tx=XM_023244747.1;refGene_gn=LOC102899173;refGene_tx=XM_023244748.1;refGene_gn=LOC102899173;refGene_tx=XM_023244749.1;refGene_gn=LOC102899173;refGene_tx=XM_023244750.1;refGene_gn=LOC102899173;refGene_tx=XM_023244755.1;refGene_gn=LOC102899173;refGene_tx=XM_023244756.1
E1 46592497 46592647
refGene_gn=LOC102899173;refGene_tx=XM_019818044.2;refGene_gn=LOC102899173;refGene_tx=XM_023244736.1;refGene_gn=LOC102899173;refGene_tx=XM_023244737.1;refGene_gn=LOC102899173;refGene_tx=XM_023244738.1;refGene_gn=LOC102899173;refGene_tx=XM_023244739.1;refGene_gn=LOC102899173;refGene_tx=XM_023244740.1;refGene_gn=LOC102899173;refGene_tx=XM_023244741.1;refGene_gn=LOC102899173;refGene_tx=XM_023244742.1;refGene_gn=LOC102899173;refGene_tx=XM_023244743.1;refGene_gn=LOC102899173;refGene_tx=XM_023244744.1;refGene_gn=LOC102899173;refGene_tx=XM_023244745.1;refGene_gn=LOC102899173;refGene_tx=XM_023244746.1;refGene_gn=LOC102899173;refGene_tx=XM_023244747.1;refGene_gn=LOC102899173;refGene_tx=XM_023244748.1;refGene_gn=LOC102899173;refGene_tx=XM_023244750.1;refGene_gn=LOC102899173;refGene_tx=XM_023244755.1;refGene_gn=LOC102899173;refGene_tx=XM_023244756.1
E1 46594160 46594211
refGene_gn=LOC102899173;refGene_tx=XM_019818044.2;refGene_gn=LOC102899173;refGene_tx=XM_023244736.1;refGene_gn=LOC102899173;refGene_tx=XM_023244737.1;refGene_gn=LOC102899173;refGene_tx=XM_023244738.1;refGene_gn=LOC102899173;refGene_tx=XM_023244739.1;refGene_gn=LOC102899173;refGene_tx=XM_023244740.1;refGene_gn=LOC102899173;refGene_tx=XM_023244741.1;refGene_gn=LOC102899173;refGene_tx=XM_023244742.1;refGene_gn=LOC102899173;refGene_tx=XM_023244743.1;refGene_gn=LOC102899173;refGene_tx=XM_023244744.1;refGene_gn=LOC102899173;refGene_tx=XM_023244745.1;refGene_gn=LOC102899173;refGene_tx=XM_023244746.1;refGene_gn=LOC102899173;refGene_tx=XM_023244747.1;refGene_gn=LOC102899173;refGene_tx=XM_023244748.1;refGene_gn=LOC102899173;refGene_tx=XM_023244750.1;refGene_gn=LOC102899

173;refGene_tx=XM_023244755.1;refGene_gn=LOC102899173;refGene_tx=XM_02
3244756.1
E1 46597836 46597980
refGene_gn=LOC102899173;refGene_tx=XM_019818044.2;refGene_gn=LOC102899
173;refGene_tx=XM_023244736.1;refGene_gn=LOC102899173;refGene_tx=XM_02
3244737.1;refGene_gn=LOC102899173;refGene_tx=XM_023244738.1;refGene_gn
=LOC102899173;refGene_tx=XM_023244739.1;refGene_gn=LOC102899173;refGen
e_tx=XM_023244740.1;refGene_gn=LOC102899173;refGene_tx=XM_023244741.1;
refGene_gn=LOC102899173;refGene_tx=XM_023244742.1;refGene_gn=LOC102899
173;refGene_tx=XM_023244743.1;refGene_gn=LOC102899173;refGene_tx=XM_02
3244744.1;refGene_gn=LOC102899173;refGene_tx=XM_023244745.1;refGene_gn
=LOC102899173;refGene_tx=XM_023244746.1;refGene_gn=LOC102899173;refGen
e_tx=XM_023244747.1;refGene_gn=LOC102899173;refGene_tx=XM_023244748.1;
refGene_gn=LOC102899173;refGene_tx=XM_023244749.1;refGene_gn=LOC102899
173;refGene_tx=XM_023244750.1;refGene_gn=LOC102899173;refGene_tx=XM_02
3244755.1;refGene_gn=LOC102899173;refGene_tx=XM_023244756.1
E1 46599772 46599844
refGene_gn=LOC102899173;refGene_tx=XM_019818044.2;refGene_gn=LOC102899
173;refGene_tx=XM_023244736.1;refGene_gn=LOC102899173;refGene_tx=XM_02
3244737.1;refGene_gn=LOC102899173;refGene_tx=XM_023244738.1;refGene_gn
=LOC102899173;refGene_tx=XM_023244739.1;refGene_gn=LOC102899173;refGen
e_tx=XM_023244740.1;refGene_gn=LOC102899173;refGene_tx=XM_023244741.1;
refGene_gn=LOC102899173;refGene_tx=XM_023244742.1;refGene_gn=LOC102899
173;refGene_tx=XM_023244743.1;refGene_gn=LOC102899173;refGene_tx=XM_02
3244744.1;refGene_gn=LOC102899173;refGene_tx=XM_023244745.1;refGene_gn
=LOC102899173;refGene_tx=XM_023244746.1;refGene_gn=LOC102899173;refGen
e_tx=XM_023244747.1;refGene_gn=LOC102899173;refGene_tx=XM_023244748.1;
refGene_gn=LOC102899173;refGene_tx=XM_023244749.1;refGene_gn=LOC102899
173;refGene_tx=XM_023244750.1;refGene_gn=LOC102899173;refGene_tx=XM_02
3244755.1;refGene_gn=LOC102899173;refGene_tx=XM_023244756.1
E1 46603492 46603636
refGene_gn=LOC102899173;refGene_tx=XM_019818044.2;refGene_gn=LOC102899
173;refGene_tx=XM_023244736.1;refGene_gn=LOC102899173;refGene_tx=XM_02
3244737.1;refGene_gn=LOC102899173;refGene_tx=XM_023244738.1;refGene_gn
=LOC102899173;refGene_tx=XM_023244739.1;refGene_gn=LOC102899173;refGen
e_tx=XM_023244740.1;refGene_gn=LOC102899173;refGene_tx=XM_023244741.1;
refGene_gn=LOC102899173;refGene_tx=XM_023244742.1;refGene_gn=LOC102899
173;refGene_tx=XM_023244743.1;refGene_gn=LOC102899173;refGene_tx=XM_02
3244744.1;refGene_gn=LOC102899173;refGene_tx=XM_023244745.1;refGene_gn
=LOC102899173;refGene_tx=XM_023244746.1;refGene_gn=LOC102899173;refGen
e_tx=XM_023244747.1;refGene_gn=LOC102899173;refGene_tx=XM_023244748.1;
refGene_gn=LOC102899173;refGene_tx=XM_023244749.1;refGene_gn=LOC102899
173;refGene_tx=XM_023244750.1;refGene_gn=LOC102899173;refGene_tx=XM_02
3244755.1;refGene_gn=LOC102899173;refGene_tx=XM_023244756.1
E1 46607765 46607837
refGene_gn=LOC102899173;refGene_tx=XM_019818044.2;refGene_gn=LOC102899
173;refGene_tx=XM_023244736.1;refGene_gn=LOC102899173;refGene_tx=XM_02
3244737.1;refGene_gn=LOC102899173;refGene_tx=XM_023244738.1;refGene_gn
=LOC102899173;refGene_tx=XM_023244739.1;refGene_gn=LOC102899173;refGen
e_tx=XM_023244740.1;refGene_gn=LOC102899173;refGene_tx=XM_023244741.1;

refGene_gn=LOC102899173;refGene_tx=XM_023244742.1;refGene_gn=LOC102899173;refGene_tx=XM_023244743.1;refGene_gn=LOC102899173;refGene_tx=XM_023244744.1;refGene_gn=LOC102899173;refGene_tx=XM_023244745.1;refGene_gn=LOC102899173;refGene_tx=XM_023244746.1;refGene_gn=LOC102899173;refGene_tx=XM_023244747.1;refGene_gn=LOC102899173;refGene_tx=XM_023244748.1;refGene_gn=LOC102899173;refGene_tx=XM_023244749.1;refGene_gn=LOC102899173;refGene_tx=XM_023244750.1;refGene_gn=LOC102899173;refGene_tx=XM_023244755.1;refGene_gn=LOC102899173;refGene_tx=XM_023244756.1
E1 46613819 46613963
refGene_gn=LOC102899173;refGene_tx=XM_019818044.2;refGene_gn=LOC102899173;refGene_tx=XM_023244736.1;refGene_gn=LOC102899173;refGene_tx=XM_023244737.1;refGene_gn=LOC102899173;refGene_tx=XM_023244738.1;refGene_gn=LOC102899173;refGene_tx=XM_023244739.1;refGene_gn=LOC102899173;refGene_tx=XM_023244740.1;refGene_gn=LOC102899173;refGene_tx=XM_023244741.1;refGene_gn=LOC102899173;refGene_tx=XM_023244742.1;refGene_gn=LOC102899173;refGene_tx=XM_023244743.1;refGene_gn=LOC102899173;refGene_tx=XM_023244744.1;refGene_gn=LOC102899173;refGene_tx=XM_023244745.1;refGene_gn=LOC102899173;refGene_tx=XM_023244746.1;refGene_gn=LOC102899173;refGene_tx=XM_023244747.1;refGene_gn=LOC102899173;refGene_tx=XM_023244748.1;refGene_gn=LOC102899173;refGene_tx=XM_023244749.1;refGene_gn=LOC102899173;refGene_tx=XM_023244750.1;refGene_gn=LOC102899173;refGene_tx=XM_023244755.1;refGene_gn=LOC102899173;refGene_tx=XM_023244756.1
E1 46614297 46614366
refGene_gn=LOC102899173;refGene_tx=XM_019818044.2;refGene_gn=LOC102899173;refGene_tx=XM_023244736.1;refGene_gn=LOC102899173;refGene_tx=XM_023244737.1;refGene_gn=LOC102899173;refGene_tx=XM_023244738.1;refGene_gn=LOC102899173;refGene_tx=XM_023244739.1;refGene_gn=LOC102899173;refGene_tx=XM_023244740.1;refGene_gn=LOC102899173;refGene_tx=XM_023244741.1;refGene_gn=LOC102899173;refGene_tx=XM_023244742.1;refGene_gn=LOC102899173;refGene_tx=XM_023244743.1;refGene_gn=LOC102899173;refGene_tx=XM_023244744.1;refGene_gn=LOC102899173;refGene_tx=XM_023244745.1;refGene_gn=LOC102899173;refGene_tx=XM_023244746.1;refGene_gn=LOC102899173;refGene_tx=XM_023244747.1;refGene_gn=LOC102899173;refGene_tx=XM_023244748.1;refGene_gn=LOC102899173;refGene_tx=XM_023244749.1;refGene_gn=LOC102899173;refGene_tx=XM_023244750.1;refGene_gn=LOC102899173;refGene_tx=XM_023244755.1;refGene_gn=LOC102899173;refGene_tx=XM_023244756.1
E1 46617804 46617945
refGene_gn=LOC102899173;refGene_tx=XM_019818044.2;refGene_gn=LOC102899173;refGene_tx=XM_023244736.1;refGene_gn=LOC102899173;refGene_tx=XM_023244737.1;refGene_gn=LOC102899173;refGene_tx=XM_023244738.1;refGene_gn=LOC102899173;refGene_tx=XM_023244739.1;refGene_gn=LOC102899173;refGene_tx=XM_023244740.1;refGene_gn=LOC102899173;refGene_tx=XM_023244741.1;refGene_gn=LOC102899173;refGene_tx=XM_023244742.1;refGene_gn=LOC102899173;refGene_tx=XM_023244743.1;refGene_gn=LOC102899173;refGene_tx=XM_023244744.1;refGene_gn=LOC102899173;refGene_tx=XM_023244745.1;refGene_gn=LOC102899173;refGene_tx=XM_023244746.1;refGene_gn=LOC102899173;refGene_tx=XM_023244747.1;refGene_gn=LOC102899173;refGene_tx=XM_023244748.1;refGene_gn=LOC102899173;refGene_tx=XM_023244749.1;refGene_gn=LOC102899173;refGene_tx=XM_023244750.1;refGene_gn=LOC102899173;refGene_tx=XM_023244755.1;refGene_gn=LOC102899173;refGene_tx=XM_023244756.1

E1 46622139 46622211
refGene_gn=LOC102899173;refGene_tx=XM_023244736.1;refGene_gn=LOC102899
173;refGene_tx=XM_023244737.1;refGene_gn=LOC102899173;refGene_tx=XM_02
3244738.1;refGene_gn=LOC102899173;refGene_tx=XM_023244739.1;refGene_gn
=LOC102899173;refGene_tx=XM_023244740.1;refGene_gn=LOC102899173;refGen
e_tx=XM_023244741.1;refGene_gn=LOC102899173;refGene_tx=XM_023244742.1;
refGene_gn=LOC102899173;refGene_tx=XM_023244743.1;refGene_gn=LOC102899
173;refGene_tx=XM_023244744.1;refGene_gn=LOC102899173;refGene_tx=XM_02
3244745.1;refGene_gn=LOC102899173;refGene_tx=XM_023244746.1;refGene_gn
=LOC102899173;refGene_tx=XM_023244747.1;refGene_gn=LOC102899173;refGen
e_tx=XM_023244748.1;refGene_gn=LOC102899173;refGene_tx=XM_023244749.1;
refGene_gn=LOC102899173;refGene_tx=XM_023244750.1;refGene_gn=LOC102899
173;refGene_tx=XM_023244755.1;refGene_gn=LOC102899173;refGene_tx=XM_02
3244756.1
E1 46623340 46623484
refGene_gn=LOC102899173;refGene_tx=XM_023244736.1;refGene_gn=LOC102899
173;refGene_tx=XM_023244737.1;refGene_gn=LOC102899173;refGene_tx=XM_02
3244738.1;refGene_gn=LOC102899173;refGene_tx=XM_023244739.1;refGene_gn
=LOC102899173;refGene_tx=XM_023244740.1;refGene_gn=LOC102899173;refGen
e_tx=XM_023244741.1;refGene_gn=LOC102899173;refGene_tx=XM_023244742.1;
refGene_gn=LOC102899173;refGene_tx=XM_023244743.1;refGene_gn=LOC102899
173;refGene_tx=XM_023244744.1;refGene_gn=LOC102899173;refGene_tx=XM_02
3244745.1;refGene_gn=LOC102899173;refGene_tx=XM_023244746.1;refGene_gn
=LOC102899173;refGene_tx=XM_023244747.1;refGene_gn=LOC102899173;refGen
e_tx=XM_023244748.1;refGene_gn=LOC102899173;refGene_tx=XM_023244749.1;
refGene_gn=LOC102899173;refGene_tx=XM_023244750.1;refGene_gn=LOC102899
173;refGene_tx=XM_023244755.1;refGene_gn=LOC102899173;refGene_tx=XM_02
3244756.1
E1 46623933 46624005
refGene_gn=LOC102899173;refGene_tx=XM_023244736.1;refGene_gn=LOC102899
173;refGene_tx=XM_023244737.1;refGene_gn=LOC102899173;refGene_tx=XM_02
3244738.1;refGene_gn=LOC102899173;refGene_tx=XM_023244739.1;refGene_gn
=LOC102899173;refGene_tx=XM_023244740.1;refGene_gn=LOC102899173;refGen
e_tx=XM_023244741.1;refGene_gn=LOC102899173;refGene_tx=XM_023244742.1;
refGene_gn=LOC102899173;refGene_tx=XM_023244743.1;refGene_gn=LOC102899
173;refGene_tx=XM_023244744.1;refGene_gn=LOC102899173;refGene_tx=XM_02
3244745.1;refGene_gn=LOC102899173;refGene_tx=XM_023244746.1;refGene_gn
=LOC102899173;refGene_tx=XM_023244747.1;refGene_gn=LOC102899173;refGen
e_tx=XM_023244748.1;refGene_gn=LOC102899173;refGene_tx=XM_023244749.1;
refGene_gn=LOC102899173;refGene_tx=XM_023244750.1;refGene_gn=LOC102899
173;refGene_tx=XM_023244755.1;refGene_gn=LOC102899173;refGene_tx=XM_02
3244756.1
E1 46625430 46625527
refGene_gn=LOC102899173;refGene_tx=XM_023244751.1
E1 46625975 46626047
refGene_gn=LOC102899173;refGene_tx=XM_023244751.1
E1 46630594 46630738
refGene_gn=LOC102899173;refGene_tx=XM_023244736.1;refGene_gn=LOC102899
173;refGene_tx=XM_023244737.1;refGene_gn=LOC102899173;refGene_tx=XM_02
3244738.1;refGene_gn=LOC102899173;refGene_tx=XM_023244739.1;refGene_gn

=LOC102899173;refGene_tx=XM_023244740.1;refGene_gn=LOC102899173;refGene_tx=XM_023244741.1;refGene_gn=LOC102899173;refGene_tx=XM_023244742.1;refGene_gn=LOC102899173;refGene_tx=XM_023244743.1;refGene_gn=LOC102899173;refGene_tx=XM_023244744.1;refGene_gn=LOC102899173;refGene_tx=XM_023244745.1;refGene_gn=LOC102899173;refGene_tx=XM_023244746.1;refGene_gn=LOC102899173;refGene_tx=XM_023244747.1;refGene_gn=LOC102899173;refGene_tx=XM_023244748.1;refGene_gn=LOC102899173;refGene_tx=XM_023244749.1;refGene_gn=LOC102899173;refGene_tx=XM_023244750.1;refGene_gn=LOC102899173;refGene_tx=XM_023244751.1;refGene_gn=LOC102899173;refGene_tx=XM_023244755.1;refGene_gn=LOC102899173;refGene_tx=XM_023244756.1
E1 46638776 46638848
refGene_gn=LOC102899173;refGene_tx=XM_023244736.1;refGene_gn=LOC102899173;refGene_tx=XM_023244737.1;refGene_gn=LOC102899173;refGene_tx=XM_023244738.1;refGene_gn=LOC102899173;refGene_tx=XM_023244739.1;refGene_gn=LOC102899173;refGene_tx=XM_023244740.1;refGene_gn=LOC102899173;refGene_tx=XM_023244741.1;refGene_gn=LOC102899173;refGene_tx=XM_023244742.1;refGene_gn=LOC102899173;refGene_tx=XM_023244743.1;refGene_gn=LOC102899173;refGene_tx=XM_023244744.1;refGene_gn=LOC102899173;refGene_tx=XM_023244745.1;refGene_gn=LOC102899173;refGene_tx=XM_023244746.1;refGene_gn=LOC102899173;refGene_tx=XM_023244747.1;refGene_gn=LOC102899173;refGene_tx=XM_023244748.1;refGene_gn=LOC102899173;refGene_tx=XM_023244749.1;refGene_gn=LOC102899173;refGene_tx=XM_023244750.1;refGene_gn=LOC102899173;refGene_tx=XM_023244751.1;refGene_gn=LOC102899173;refGene_tx=XM_023244755.1;refGene_gn=LOC102899173;refGene_tx=XM_023244756.1
E1 46646198 46646342
refGene_gn=LOC102899173;refGene_tx=XM_023244736.1;refGene_gn=LOC102899173;refGene_tx=XM_023244737.1;refGene_gn=LOC102899173;refGene_tx=XM_023244738.1;refGene_gn=LOC102899173;refGene_tx=XM_023244739.1;refGene_gn=LOC102899173;refGene_tx=XM_023244740.1;refGene_gn=LOC102899173;refGene_tx=XM_023244741.1;refGene_gn=LOC102899173;refGene_tx=XM_023244742.1;refGene_gn=LOC102899173;refGene_tx=XM_023244743.1;refGene_gn=LOC102899173;refGene_tx=XM_023244744.1;refGene_gn=LOC102899173;refGene_tx=XM_023244745.1;refGene_gn=LOC102899173;refGene_tx=XM_023244746.1;refGene_gn=LOC102899173;refGene_tx=XM_023244747.1;refGene_gn=LOC102899173;refGene_tx=XM_023244748.1;refGene_gn=LOC102899173;refGene_tx=XM_023244749.1;refGene_gn=LOC102899173;refGene_tx=XM_023244750.1;refGene_gn=LOC102899173;refGene_tx=XM_023244751.1;refGene_gn=LOC102899173;refGene_tx=XM_023244755.1;refGene_gn=LOC102899173;refGene_tx=XM_023244756.1
E1 46647242 46647314
refGene_gn=LOC102899173;refGene_tx=XM_023244736.1;refGene_gn=LOC102899173;refGene_tx=XM_023244737.1;refGene_gn=LOC102899173;refGene_tx=XM_023244738.1;refGene_gn=LOC102899173;refGene_tx=XM_023244739.1;refGene_gn=LOC102899173;refGene_tx=XM_023244740.1;refGene_gn=LOC102899173;refGene_tx=XM_023244741.1;refGene_gn=LOC102899173;refGene_tx=XM_023244742.1;refGene_gn=LOC102899173;refGene_tx=XM_023244743.1;refGene_gn=LOC102899173;refGene_tx=XM_023244744.1;refGene_gn=LOC102899173;refGene_tx=XM_023244745.1;refGene_gn=LOC102899173;refGene_tx=XM_023244746.1;refGene_gn=LOC102899173;refGene_tx=XM_023244747.1;refGene_gn=LOC102899173;refGene_tx=XM_023244748.1;refGene_gn=LOC102899173;refGene_tx=XM_023244749.1;refGene_gn=LOC102899173;refGene_tx=XM_023244750.1;refGene_gn=LOC102899

173;refGene_tx=XM_023244751.1;refGene_gn=LOC102899173;refGene_tx=XM_023244755.1;refGene_gn=LOC102899173;refGene_tx=XM_023244756.1
E1 46649654 46649798
refGene_gn=LOC102899173;refGene_tx=XM_023244736.1;refGene_gn=LOC102899173;refGene_tx=XM_023244737.1;refGene_gn=LOC102899173;refGene_tx=XM_023244738.1;refGene_gn=LOC102899173;refGene_tx=XM_023244739.1;refGene_gn=LOC102899173;refGene_tx=XM_023244740.1;refGene_gn=LOC102899173;refGene_tx=XM_023244741.1;refGene_gn=LOC102899173;refGene_tx=XM_023244742.1;refGene_gn=LOC102899173;refGene_tx=XM_023244743.1;refGene_gn=LOC102899173;refGene_tx=XM_023244744.1;refGene_gn=LOC102899173;refGene_tx=XM_023244745.1;refGene_gn=LOC102899173;refGene_tx=XM_023244746.1;refGene_gn=LOC102899173;refGene_tx=XM_023244747.1;refGene_gn=LOC102899173;refGene_tx=XM_023244748.1;refGene_gn=LOC102899173;refGene_tx=XM_023244749.1;refGene_gn=LOC102899173;refGene_tx=XM_023244750.1;refGene_gn=LOC102899173;refGene_tx=XM_023244751.1;refGene_gn=LOC102899173;refGene_tx=XM_023244755.1;refGene_gn=LOC102899173;refGene_tx=XM_023244756.1
E1 46652511 46652574
refGene_gn=LOC102899173;refGene_tx=XM_023244736.1;refGene_gn=LOC102899173;refGene_tx=XM_023244737.1;refGene_gn=LOC102899173;refGene_tx=XM_023244738.1;refGene_gn=LOC102899173;refGene_tx=XM_023244739.1;refGene_gn=LOC102899173;refGene_tx=XM_023244740.1;refGene_gn=LOC102899173;refGene_tx=XM_023244741.1;refGene_gn=LOC102899173;refGene_tx=XM_023244742.1;refGene_gn=LOC102899173;refGene_tx=XM_023244743.1;refGene_gn=LOC102899173;refGene_tx=XM_023244744.1;refGene_gn=LOC102899173;refGene_tx=XM_023244745.1;refGene_gn=LOC102899173;refGene_tx=XM_023244746.1;refGene_gn=LOC102899173;refGene_tx=XM_023244747.1;refGene_gn=LOC102899173;refGene_tx=XM_023244748.1;refGene_gn=LOC102899173;refGene_tx=XM_023244749.1;refGene_gn=LOC102899173;refGene_tx=XM_023244750.1;refGene_gn=LOC102899173;refGene_tx=XM_023244751.1;refGene_gn=LOC102899173;refGene_tx=XM_023244752.1;refGene_gn=LOC102899173;refGene_tx=XM_023244753.1;refGene_gn=LOC102899173;refGene_tx=XM_023244755.1;refGene_gn=LOC102899173;refGene_tx=XM_023244756.1
E1 46654346 46654400
refGene_gn=LOC102899173;refGene_tx=XM_023244736.1;refGene_gn=LOC102899173;refGene_tx=XM_023244737.1;refGene_gn=LOC102899173;refGene_tx=XM_023244738.1;refGene_gn=LOC102899173;refGene_tx=XM_023244739.1;refGene_gn=LOC102899173;refGene_tx=XM_023244740.1;refGene_gn=LOC102899173;refGene_tx=XM_023244742.1;refGene_gn=LOC102899173;refGene_tx=XM_023244743.1;refGene_gn=LOC102899173;refGene_tx=XM_023244744.1;refGene_gn=LOC102899173;refGene_tx=XM_023244745.1;refGene_gn=LOC102899173;refGene_tx=XM_023244747.1;refGene_gn=LOC102899173;refGene_tx=XM_023244748.1;refGene_gn=LOC102899173;refGene_tx=XM_023244749.1;refGene_gn=LOC102899173;refGene_tx=XM_023244750.1;refGene_gn=LOC102899173;refGene_tx=XM_023244751.1;refGene_gn=LOC102899173;refGene_tx=XM_023244752.1
E1 46657904 46658009
refGene_gn=LOC102899173;refGene_tx=XM_023244736.1;refGene_gn=LOC102899173;refGene_tx=XM_023244737.1;refGene_gn=LOC102899173;refGene_tx=XM_023244738.1;refGene_gn=LOC102899173;refGene_tx=XM_023244739.1;refGene_gn=LOC102899173;refGene_tx=XM_023244740.1;refGene_gn=LOC102899173;refGene_tx=XM_023244741.1;refGene_gn=LOC102899173;refGene_tx=XM_023244742.1;

refGene_gn=LOC102899173;refGene_tx=XM_023244743.1;refGene_gn=LOC102899173;refGene_tx=XM_023244745.1;refGene_gn=LOC102899173;refGene_tx=XM_023244747.1;refGene_gn=LOC102899173;refGene_tx=XM_023244748.1;refGene_gn=LOC102899173;refGene_tx=XM_023244749.1;refGene_gn=LOC102899173;refGene_tx=XM_023244750.1;refGene_gn=LOC102899173;refGene_tx=XM_023244751.1;refGene_gn=LOC102899173;refGene_tx=XM_023244752.1;refGene_gn=LOC102899173;refGene_tx=XM_023244753.1
E1 46670170 46670272
refGene_gn=LOC102899173;refGene_tx=XM_023244736.1;refGene_gn=LOC102899173;refGene_tx=XM_023244737.1;refGene_gn=LOC102899173;refGene_tx=XM_023244738.1;refGene_gn=LOC102899173;refGene_tx=XM_023244739.1;refGene_gn=LOC102899173;refGene_tx=XM_023244740.1;refGene_gn=LOC102899173;refGene_tx=XM_023244741.1;refGene_gn=LOC102899173;refGene_tx=XM_023244742.1;refGene_gn=LOC102899173;refGene_tx=XM_023244743.1;refGene_gn=LOC102899173;refGene_tx=XM_023244744.1;refGene_gn=LOC102899173;refGene_tx=XM_023244745.1;refGene_gn=LOC102899173;refGene_tx=XM_023244746.1;refGene_gn=LOC102899173;refGene_tx=XM_023244747.1;refGene_gn=LOC102899173;refGene_tx=XM_023244748.1;refGene_gn=LOC102899173;refGene_tx=XM_023244749.1;refGene_gn=LOC102899173;refGene_tx=XM_023244750.1;refGene_gn=LOC102899173;refGene_tx=XM_023244751.1;refGene_gn=LOC102899173;refGene_tx=XM_023244752.1;refGene_gn=LOC102899173;refGene_tx=XM_023244753.1
E1 46670940 46671000
refGene_gn=LOC102899173;refGene_tx=XM_023244736.1;refGene_gn=LOC102899173;refGene_tx=XM_023244737.1;refGene_gn=LOC102899173;refGene_tx=XM_023244738.1;refGene_gn=LOC102899173;refGene_tx=XM_023244739.1;refGene_gn=LOC102899173;refGene_tx=XM_023244740.1;refGene_gn=LOC102899173;refGene_tx=XM_023244741.1;refGene_gn=LOC102899173;refGene_tx=XM_023244742.1;refGene_gn=LOC102899173;refGene_tx=XM_023244743.1;refGene_gn=LOC102899173;refGene_tx=XM_023244744.1;refGene_gn=LOC102899173;refGene_tx=XM_023244745.1;refGene_gn=LOC102899173;refGene_tx=XM_023244746.1;refGene_gn=LOC102899173;refGene_tx=XM_023244747.1;refGene_gn=LOC102899173;refGene_tx=XM_023244748.1;refGene_gn=LOC102899173;refGene_tx=XM_023244749.1;refGene_gn=LOC102899173;refGene_tx=XM_023244750.1;refGene_gn=LOC102899173;refGene_tx=XM_023244751.1;refGene_gn=LOC102899173;refGene_tx=XM_023244752.1;refGene_gn=LOC102899173;refGene_tx=XM_023244753.1
E1 46671822 46671933
refGene_gn=LOC102899173;refGene_tx=XM_023244736.1;refGene_gn=LOC102899173;refGene_tx=XM_023244737.1;refGene_gn=LOC102899173;refGene_tx=XM_023244738.1;refGene_gn=LOC102899173;refGene_tx=XM_023244739.1;refGene_gn=LOC102899173;refGene_tx=XM_023244740.1;refGene_gn=LOC102899173;refGene_tx=XM_023244741.1;refGene_gn=LOC102899173;refGene_tx=XM_023244742.1;refGene_gn=LOC102899173;refGene_tx=XM_023244743.1;refGene_gn=LOC102899173;refGene_tx=XM_023244744.1;refGene_gn=LOC102899173;refGene_tx=XM_023244745.1;refGene_gn=LOC102899173;refGene_tx=XM_023244746.1;refGene_gn=LOC102899173;refGene_tx=XM_023244747.1;refGene_gn=LOC102899173;refGene_tx=XM_023244748.1;refGene_gn=LOC102899173;refGene_tx=XM_023244749.1;refGene_gn=LOC102899173;refGene_tx=XM_023244750.1;refGene_gn=LOC102899173;refGene_tx=XM_023244751.1;refGene_gn=LOC102899173;refGene_tx=XM_023244752.1;refGene_gn=LOC102899173;refGene_tx=XM_023244753.1
E1 46672313 46672355

refGene_gn=LOC102899173;refGene_tx=XM_023244736.1;refGene_gn=LOC102899173;refGene_tx=XM_023244737.1;refGene_gn=LOC102899173;refGene_tx=XM_023244738.1;refGene_gn=LOC102899173;refGene_tx=XM_023244739.1;refGene_gn=LOC102899173;refGene_tx=XM_023244740.1;refGene_gn=LOC102899173;refGene_tx=XM_023244741.1;refGene_gn=LOC102899173;refGene_tx=XM_023244742.1;refGene_gn=LOC102899173;refGene_tx=XM_023244743.1;refGene_gn=LOC102899173;refGene_tx=XM_023244744.1;refGene_gn=LOC102899173;refGene_tx=XM_023244745.1;refGene_gn=LOC102899173;refGene_tx=XM_023244746.1;refGene_gn=LOC102899173;refGene_tx=XM_023244747.1;refGene_gn=LOC102899173;refGene_tx=XM_023244748.1;refGene_gn=LOC102899173;refGene_tx=XM_023244749.1;refGene_gn=LOC102899173;refGene_tx=XM_023244750.1;refGene_gn=LOC102899173;refGene_tx=XM_023244751.1;refGene_gn=LOC102899173;refGene_tx=XM_023244752.1;refGene_gn=LOC102899173;refGene_tx=XM_023244753.1
E1 46678829 46678934
refGene_gn=LOC102899173;refGene_tx=XM_023244736.1;refGene_gn=LOC102899173;refGene_tx=XM_023244737.1;refGene_gn=LOC102899173;refGene_tx=XM_023244738.1;refGene_gn=LOC102899173;refGene_tx=XM_023244739.1;refGene_gn=LOC102899173;refGene_tx=XM_023244740.1;refGene_gn=LOC102899173;refGene_tx=XM_023244741.1;refGene_gn=LOC102899173;refGene_tx=XM_023244742.1;refGene_gn=LOC102899173;refGene_tx=XM_023244743.1;refGene_gn=LOC102899173;refGene_tx=XM_023244744.1;refGene_gn=LOC102899173;refGene_tx=XM_023244745.1;refGene_gn=LOC102899173;refGene_tx=XM_023244746.1;refGene_gn=LOC102899173;refGene_tx=XM_023244747.1;refGene_gn=LOC102899173;refGene_tx=XM_023244748.1;refGene_gn=LOC102899173;refGene_tx=XM_023244749.1;refGene_gn=LOC102899173;refGene_tx=XM_023244750.1;refGene_gn=LOC102899173;refGene_tx=XM_023244751.1;refGene_gn=LOC102899173;refGene_tx=XM_023244752.1;refGene_gn=LOC102899173;refGene_tx=XM_023244753.1
E1 46682318 46682372
refGene_gn=LOC102899173;refGene_tx=XM_023244736.1;refGene_gn=LOC102899173;refGene_tx=XM_023244737.1;refGene_gn=LOC102899173;refGene_tx=XM_023244738.1;refGene_gn=LOC102899173;refGene_tx=XM_023244739.1;refGene_gn=LOC102899173;refGene_tx=XM_023244740.1;refGene_gn=LOC102899173;refGene_tx=XM_023244741.1;refGene_gn=LOC102899173;refGene_tx=XM_023244742.1;refGene_gn=LOC102899173;refGene_tx=XM_023244743.1;refGene_gn=LOC102899173;refGene_tx=XM_023244744.1;refGene_gn=LOC102899173;refGene_tx=XM_023244745.1;refGene_gn=LOC102899173;refGene_tx=XM_023244746.1;refGene_gn=LOC102899173;refGene_tx=XM_023244747.1;refGene_gn=LOC102899173;refGene_tx=XM_023244748.1;refGene_gn=LOC102899173;refGene_tx=XM_023244749.1;refGene_gn=LOC102899173;refGene_tx=XM_023244750.1;refGene_gn=LOC102899173;refGene_tx=XM_023244751.1;refGene_gn=LOC102899173;refGene_tx=XM_023244752.1;refGene_gn=LOC102899173;refGene_tx=XM_023244753.1
E1 46683575 46683680
refGene_gn=LOC102899173;refGene_tx=XM_023244736.1;refGene_gn=LOC102899173;refGene_tx=XM_023244737.1;refGene_gn=LOC102899173;refGene_tx=XM_023244738.1;refGene_gn=LOC102899173;refGene_tx=XM_023244739.1;refGene_gn=LOC102899173;refGene_tx=XM_023244740.1;refGene_gn=LOC102899173;refGene_tx=XM_023244741.1;refGene_gn=LOC102899173;refGene_tx=XM_023244742.1;refGene_gn=LOC102899173;refGene_tx=XM_023244743.1;refGene_gn=LOC102899173;refGene_tx=XM_023244744.1;refGene_gn=LOC102899173;refGene_tx=XM_023244745.1;refGene_gn=LOC102899173;refGene_tx=XM_023244746.1;refGene_gn

=LOC102899173;refGene_tx=XM_023244747.1;refGene_gn=LOC102899173;refGen
e_tx=XM_023244749.1;refGene_gn=LOC102899173;refGene_tx=XM_023244750.1;
refGene_gn=LOC102899173;refGene_tx=XM_023244751.1;refGene_gn=LOC102899
173;refGene_tx=XM_023244752.1;refGene_gn=LOC102899173;refGene_tx=XM_02
3244753.1
E1 46684198 46684252
refGene_gn=LOC102899173;refGene_tx=XM_023244736.1;refGene_gn=LOC102899
173;refGene_tx=XM_023244737.1;refGene_gn=LOC102899173;refGene_tx=XM_02
3244738.1;refGene_gn=LOC102899173;refGene_tx=XM_023244739.1;refGene_gn
=LOC102899173;refGene_tx=XM_023244740.1;refGene_gn=LOC102899173;refGen
e_tx=XM_023244741.1;refGene_gn=LOC102899173;refGene_tx=XM_023244743.1;
refGene_gn=LOC102899173;refGene_tx=XM_023244744.1;refGene_gn=LOC102899
173;refGene_tx=XM_023244745.1;refGene_gn=LOC102899173;refGene_tx=XM_02
3244746.1;refGene_gn=LOC102899173;refGene_tx=XM_023244747.1;refGene_gn
=LOC102899173;refGene_tx=XM_023244749.1;refGene_gn=LOC102899173;refGen
e_tx=XM_023244750.1;refGene_gn=LOC102899173;refGene_tx=XM_023244751.1;
refGene_gn=LOC102899173;refGene_tx=XM_023244752.1;refGene_gn=LOC102899
173;refGene_tx=XM_023244753.1
E1 46685382 46685487
refGene_gn=LOC102899173;refGene_tx=XM_023244736.1;refGene_gn=LOC102899
173;refGene_tx=XM_023244737.1;refGene_gn=LOC102899173;refGene_tx=XM_02
3244738.1;refGene_gn=LOC102899173;refGene_tx=XM_023244739.1;refGene_gn
=LOC102899173;refGene_tx=XM_023244740.1;refGene_gn=LOC102899173;refGen
e_tx=XM_023244741.1;refGene_gn=LOC102899173;refGene_tx=XM_023244742.1;
refGene_gn=LOC102899173;refGene_tx=XM_023244743.1;refGene_gn=LOC102899
173;refGene_tx=XM_023244744.1;refGene_gn=LOC102899173;refGene_tx=XM_02
3244745.1;refGene_gn=LOC102899173;refGene_tx=XM_023244746.1;refGene_gn
=LOC102899173;refGene_tx=XM_023244747.1;refGene_gn=LOC102899173;refGen
e_tx=XM_023244748.1;refGene_gn=LOC102899173;refGene_tx=XM_023244749.1;
refGene_gn=LOC102899173;refGene_tx=XM_023244750.1;refGene_gn=LOC102899
173;refGene_tx=XM_023244751.1;refGene_gn=LOC102899173;refGene_tx=XM_02
3244752.1;refGene_gn=LOC102899173;refGene_tx=XM_023244753.1
E1 46688527 46688581
refGene_gn=LOC102899173;refGene_tx=XM_023244736.1;refGene_gn=LOC102899
173;refGene_tx=XM_023244737.1;refGene_gn=LOC102899173;refGene_tx=XM_02
3244738.1;refGene_gn=LOC102899173;refGene_tx=XM_023244739.1;refGene_gn
=LOC102899173;refGene_tx=XM_023244740.1;refGene_gn=LOC102899173;refGen
e_tx=XM_023244741.1;refGene_gn=LOC102899173;refGene_tx=XM_023244742.1;
refGene_gn=LOC102899173;refGene_tx=XM_023244743.1;refGene_gn=LOC102899
173;refGene_tx=XM_023244744.1;refGene_gn=LOC102899173;refGene_tx=XM_02
3244745.1;refGene_gn=LOC102899173;refGene_tx=XM_023244746.1;refGene_gn
=LOC102899173;refGene_tx=XM_023244747.1;refGene_gn=LOC102899173;refGen
e_tx=XM_023244748.1;refGene_gn=LOC102899173;refGene_tx=XM_023244749.1;
refGene_gn=LOC102899173;refGene_tx=XM_023244750.1;refGene_gn=LOC102899
173;refGene_tx=XM_023244751.1;refGene_gn=LOC102899173;refGene_tx=XM_02
3244752.1;refGene_gn=LOC102899173;refGene_tx=XM_023244753.1
E1 46690684 46690789
refGene_gn=LOC102899173;refGene_tx=XM_023244736.1;refGene_gn=LOC102899
173;refGene_tx=XM_023244737.1;refGene_gn=LOC102899173;refGene_tx=XM_02
3244738.1;refGene_gn=LOC102899173;refGene_tx=XM_023244739.1;refGene_gn

=LOC102899173;refGene_tx=XM_023244740.1;refGene_gn=LOC102899173;refGen
e_tx=XM_023244741.1;refGene_gn=LOC102899173;refGene_tx=XM_023244742.1;
refGene_gn=LOC102899173;refGene_tx=XM_023244743.1;refGene_gn=LOC102899
173;refGene_tx=XM_023244744.1;refGene_gn=LOC102899173;refGene_tx=XM_02
3244745.1;refGene_gn=LOC102899173;refGene_tx=XM_023244746.1;refGene_gn
=LOC102899173;refGene_tx=XM_023244747.1;refGene_gn=LOC102899173;refGen
e_tx=XM_023244748.1;refGene_gn=LOC102899173;refGene_tx=XM_023244749.1;
refGene_gn=LOC102899173;refGene_tx=XM_023244750.1;refGene_gn=LOC102899
173;refGene_tx=XM_023244751.1;refGene_gn=LOC102899173;refGene_tx=XM_02
3244752.1;refGene_gn=LOC102899173;refGene_tx=XM_023244753.1
E1 46691336 46691390
refGene_gn=LOC102899173;refGene_tx=XM_023244736.1;refGene_gn=LOC102899
173;refGene_tx=XM_023244737.1;refGene_gn=LOC102899173;refGene_tx=XM_02
3244738.1;refGene_gn=LOC102899173;refGene_tx=XM_023244739.1;refGene_gn
=LOC102899173;refGene_tx=XM_023244740.1;refGene_gn=LOC102899173;refGen
e_tx=XM_023244741.1;refGene_gn=LOC102899173;refGene_tx=XM_023244742.1;
refGene_gn=LOC102899173;refGene_tx=XM_023244743.1;refGene_gn=LOC102899
173;refGene_tx=XM_023244744.1;refGene_gn=LOC102899173;refGene_tx=XM_02
3244745.1;refGene_gn=LOC102899173;refGene_tx=XM_023244746.1;refGene_gn
=LOC102899173;refGene_tx=XM_023244747.1;refGene_gn=LOC102899173;refGen
e_tx=XM_023244748.1;refGene_gn=LOC102899173;refGene_tx=XM_023244749.1;
refGene_gn=LOC102899173;refGene_tx=XM_023244750.1;refGene_gn=LOC102899
173;refGene_tx=XM_023244751.1;refGene_gn=LOC102899173;refGene_tx=XM_02
3244752.1;refGene_gn=LOC102899173;refGene_tx=XM_023244753.1
E1 46696180 46696234
refGene_gn=LOC102899173;refGene_tx=XM_023244736.1;refGene_gn=LOC102899
173;refGene_tx=XM_023244737.1;refGene_gn=LOC102899173;refGene_tx=XM_02
3244738.1;refGene_gn=LOC102899173;refGene_tx=XM_023244739.1;refGene_gn
=LOC102899173;refGene_tx=XM_023244741.1;refGene_gn=LOC102899173;refGen
e_tx=XM_023244742.1;refGene_gn=LOC102899173;refGene_tx=XM_023244743.1;
refGene_gn=LOC102899173;refGene_tx=XM_023244744.1;refGene_gn=LOC102899
173;refGene_tx=XM_023244745.1;refGene_gn=LOC102899173;refGene_tx=XM_02
3244746.1;refGene_gn=LOC102899173;refGene_tx=XM_023244747.1;refGene_gn
=LOC102899173;refGene_tx=XM_023244748.1;refGene_gn=LOC102899173;refGen
e_tx=XM_023244749.1;refGene_gn=LOC102899173;refGene_tx=XM_023244751.1;
refGene_gn=LOC102899173;refGene_tx=XM_023244752.1;refGene_gn=LOC102899
173;refGene_tx=XM_023244753.1
E1 46696839 46696944
refGene_gn=LOC102899173;refGene_tx=XM_023244736.1;refGene_gn=LOC102899
173;refGene_tx=XM_023244737.1;refGene_gn=LOC102899173;refGene_tx=XM_02
3244738.1;refGene_gn=LOC102899173;refGene_tx=XM_023244739.1;refGene_gn
=LOC102899173;refGene_tx=XM_023244740.1;refGene_gn=LOC102899173;refGen
e_tx=XM_023244741.1;refGene_gn=LOC102899173;refGene_tx=XM_023244742.1;
refGene_gn=LOC102899173;refGene_tx=XM_023244743.1;refGene_gn=LOC102899
173;refGene_tx=XM_023244744.1;refGene_gn=LOC102899173;refGene_tx=XM_02
3244745.1;refGene_gn=LOC102899173;refGene_tx=XM_023244746.1;refGene_gn
=LOC102899173;refGene_tx=XM_023244747.1;refGene_gn=LOC102899173;refGen
e_tx=XM_023244748.1;refGene_gn=LOC102899173;refGene_tx=XM_023244749.1;
refGene_gn=LOC102899173;refGene_tx=XM_023244751.1;refGene_gn=LOC102899
173;refGene_tx=XM_023244752.1;refGene_gn=LOC102899173;refGene_tx=XM_02

3244753.1
E1 46697499 46697553
refGene_gn=LOC102899173;refGene_tx=XM_023244736.1;refGene_gn=LOC102899
173;refGene_tx=XM_023244737.1;refGene_gn=LOC102899173;refGene_tx=XM_02
3244738.1;refGene_gn=LOC102899173;refGene_tx=XM_023244739.1;refGene_gn
=LOC102899173;refGene_tx=XM_023244740.1;refGene_gn=LOC102899173;refGen
e_tx=XM_023244741.1;refGene_gn=LOC102899173;refGene_tx=XM_023244742.1;
refGene_gn=LOC102899173;refGene_tx=XM_023244743.1;refGene_gn=LOC102899
173;refGene_tx=XM_023244744.1;refGene_gn=LOC102899173;refGene_tx=XM_02
3244745.1;refGene_gn=LOC102899173;refGene_tx=XM_023244746.1;refGene_gn
=LOC102899173;refGene_tx=XM_023244747.1;refGene_gn=LOC102899173;refGen
e_tx=XM_023244748.1;refGene_gn=LOC102899173;refGene_tx=XM_023244749.1;
refGene_gn=LOC102899173;refGene_tx=XM_023244751.1;refGene_gn=LOC102899
173;refGene_tx=XM_023244752.1;refGene_gn=LOC102899173;refGene_tx=XM_02
3244753.1
E1 46699064 46699169
refGene_gn=LOC102899173;refGene_tx=XM_023244736.1;refGene_gn=LOC102899
173;refGene_tx=XM_023244737.1;refGene_gn=LOC102899173;refGene_tx=XM_02
3244738.1;refGene_gn=LOC102899173;refGene_tx=XM_023244739.1;refGene_gn
=LOC102899173;refGene_tx=XM_023244740.1;refGene_gn=LOC102899173;refGen
e_tx=XM_023244741.1;refGene_gn=LOC102899173;refGene_tx=XM_023244742.1;
refGene_gn=LOC102899173;refGene_tx=XM_023244743.1;refGene_gn=LOC102899
173;refGene_tx=XM_023244744.1;refGene_gn=LOC102899173;refGene_tx=XM_02
3244745.1;refGene_gn=LOC102899173;refGene_tx=XM_023244746.1;refGene_gn
=LOC102899173;refGene_tx=XM_023244747.1;refGene_gn=LOC102899173;refGen
e_tx=XM_023244748.1;refGene_gn=LOC102899173;refGene_tx=XM_023244749.1;
refGene_gn=LOC102899173;refGene_tx=XM_023244750.1;refGene_gn=LOC102899
173;refGene_tx=XM_023244751.1;refGene_gn=LOC102899173;refGene_tx=XM_02
3244752.1;refGene_gn=LOC102899173;refGene_tx=XM_023244753.1
E1 46702192 46702246
refGene_gn=LOC102899173;refGene_tx=XM_023244736.1;refGene_gn=LOC102899
173;refGene_tx=XM_023244737.1;refGene_gn=LOC102899173;refGene_tx=XM_02
3244738.1;refGene_gn=LOC102899173;refGene_tx=XM_023244739.1;refGene_gn
=LOC102899173;refGene_tx=XM_023244740.1;refGene_gn=LOC102899173;refGen
e_tx=XM_023244741.1;refGene_gn=LOC102899173;refGene_tx=XM_023244742.1;
refGene_gn=LOC102899173;refGene_tx=XM_023244743.1;refGene_gn=LOC102899
173;refGene_tx=XM_023244744.1;refGene_gn=LOC102899173;refGene_tx=XM_02
3244745.1;refGene_gn=LOC102899173;refGene_tx=XM_023244746.1;refGene_gn
=LOC102899173;refGene_tx=XM_023244747.1;refGene_gn=LOC102899173;refGen
e_tx=XM_023244748.1;refGene_gn=LOC102899173;refGene_tx=XM_023244749.1;
refGene_gn=LOC102899173;refGene_tx=XM_023244750.1;refGene_gn=LOC102899
173;refGene_tx=XM_023244751.1;refGene_gn=LOC102899173;refGene_tx=XM_02
3244752.1;refGene_gn=LOC102899173;refGene_tx=XM_023244753.1
E1 46702868 46702973
refGene_gn=LOC102899173;refGene_tx=XM_023244736.1;refGene_gn=LOC102899
173;refGene_tx=XM_023244737.1;refGene_gn=LOC102899173;refGene_tx=XM_02
3244738.1;refGene_gn=LOC102899173;refGene_tx=XM_023244739.1;refGene_gn
=LOC102899173;refGene_tx=XM_023244740.1;refGene_gn=LOC102899173;refGen
e_tx=XM_023244741.1;refGene_gn=LOC102899173;refGene_tx=XM_023244742.1;
refGene_gn=LOC102899173;refGene_tx=XM_023244743.1;refGene_gn=LOC102899

173;refGene_tx=XM_023244744.1;refGene_gn=LOC102899173;refGene_tx=XM_02
3244745.1;refGene_gn=LOC102899173;refGene_tx=XM_023244746.1;refGene_gn
=LOC102899173;refGene_tx=XM_023244747.1;refGene_gn=LOC102899173;refGen
e_tx=XM_023244748.1;refGene_gn=LOC102899173;refGene_tx=XM_023244749.1;
refGene_gn=LOC102899173;refGene_tx=XM_023244750.1;refGene_gn=LOC102899
173;refGene_tx=XM_023244751.1;refGene_gn=LOC102899173;refGene_tx=XM_02
3244752.1;refGene_gn=LOC102899173;refGene_tx=XM_023244753.1
E1 46703866 46703920
refGene_gn=LOC102899173;refGene_tx=XM_023244736.1;refGene_gn=LOC102899
173;refGene_tx=XM_023244737.1;refGene_gn=LOC102899173;refGene_tx=XM_02
3244738.1;refGene_gn=LOC102899173;refGene_tx=XM_023244739.1;refGene_gn
=LOC102899173;refGene_tx=XM_023244740.1;refGene_gn=LOC102899173;refGen
e_tx=XM_023244741.1;refGene_gn=LOC102899173;refGene_tx=XM_023244742.1;
refGene_gn=LOC102899173;refGene_tx=XM_023244743.1;refGene_gn=LOC102899
173;refGene_tx=XM_023244744.1;refGene_gn=LOC102899173;refGene_tx=XM_02
3244745.1;refGene_gn=LOC102899173;refGene_tx=XM_023244746.1;refGene_gn
=LOC102899173;refGene_tx=XM_023244747.1;refGene_gn=LOC102899173;refGen
e_tx=XM_023244748.1;refGene_gn=LOC102899173;refGene_tx=XM_023244749.1;
refGene_gn=LOC102899173;refGene_tx=XM_023244750.1;refGene_gn=LOC102899
173;refGene_tx=XM_023244751.1;refGene_gn=LOC102899173;refGene_tx=XM_02
3244752.1;refGene_gn=LOC102899173;refGene_tx=XM_023244753.1
E1 46704776 46704881
refGene_gn=LOC102899173;refGene_tx=XM_023244736.1;refGene_gn=LOC102899
173;refGene_tx=XM_023244737.1;refGene_gn=LOC102899173;refGene_tx=XM_02
3244738.1;refGene_gn=LOC102899173;refGene_tx=XM_023244739.1;refGene_gn
=LOC102899173;refGene_tx=XM_023244740.1;refGene_gn=LOC102899173;refGen
e_tx=XM_023244741.1;refGene_gn=LOC102899173;refGene_tx=XM_023244742.1;
refGene_gn=LOC102899173;refGene_tx=XM_023244743.1;refGene_gn=LOC102899
173;refGene_tx=XM_023244744.1;refGene_gn=LOC102899173;refGene_tx=XM_02
3244745.1;refGene_gn=LOC102899173;refGene_tx=XM_023244746.1;refGene_gn
=LOC102899173;refGene_tx=XM_023244747.1;refGene_gn=LOC102899173;refGen
e_tx=XM_023244748.1;refGene_gn=LOC102899173;refGene_tx=XM_023244749.1;
refGene_gn=LOC102899173;refGene_tx=XM_023244750.1;refGene_gn=LOC102899
173;refGene_tx=XM_023244751.1;refGene_gn=LOC102899173;refGene_tx=XM_02
3244752.1;refGene_gn=LOC102899173;refGene_tx=XM_023244753.1
E1 46709019 46709073
refGene_gn=LOC102899173;refGene_tx=XM_023244736.1;refGene_gn=LOC102899
173;refGene_tx=XM_023244737.1;refGene_gn=LOC102899173;refGene_tx=XM_02
3244738.1;refGene_gn=LOC102899173;refGene_tx=XM_023244739.1;refGene_gn
=LOC102899173;refGene_tx=XM_023244740.1;refGene_gn=LOC102899173;refGen
e_tx=XM_023244741.1;refGene_gn=LOC102899173;refGene_tx=XM_023244742.1;
refGene_gn=LOC102899173;refGene_tx=XM_023244743.1;refGene_gn=LOC102899
173;refGene_tx=XM_023244744.1;refGene_gn=LOC102899173;refGene_tx=XM_02
3244745.1;refGene_gn=LOC102899173;refGene_tx=XM_023244746.1;refGene_gn
=LOC102899173;refGene_tx=XM_023244748.1;refGene_gn=LOC102899173;refGen
e_tx=XM_023244749.1;refGene_gn=LOC102899173;refGene_tx=XM_023244750.1;
refGene_gn=LOC102899173;refGene_tx=XM_023244751.1;refGene_gn=LOC102899
173;refGene_tx=XM_023244752.1;refGene_gn=LOC102899173;refGene_tx=XM_02
3244753.1
E1 46710052 46710157

refGene_gn=LOC102899173;refGene_tx=XM_023244736.1;refGene_gn=LOC102899173;refGene_tx=XM_023244737.1;refGene_gn=LOC102899173;refGene_tx=XM_023244738.1;refGene_gn=LOC102899173;refGene_tx=XM_023244739.1;refGene_gn=LOC102899173;refGene_tx=XM_023244740.1;refGene_gn=LOC102899173;refGene_tx=XM_023244741.1;refGene_gn=LOC102899173;refGene_tx=XM_023244742.1;refGene_gn=LOC102899173;refGene_tx=XM_023244743.1;refGene_gn=LOC102899173;refGene_tx=XM_023244744.1;refGene_gn=LOC102899173;refGene_tx=XM_023244745.1;refGene_gn=LOC102899173;refGene_tx=XM_023244746.1;refGene_gn=LOC102899173;refGene_tx=XM_023244748.1;refGene_gn=LOC102899173;refGene_tx=XM_023244749.1;refGene_gn=LOC102899173;refGene_tx=XM_023244750.1;refGene_gn=LOC102899173;refGene_tx=XM_023244751.1;refGene_gn=LOC102899173;refGene_tx=XM_023244752.1;refGene_gn=LOC102899173;refGene_tx=XM_023244753.1

E1 46710916 46710970

refGene_gn=LOC102899173;refGene_tx=XM_023244736.1;refGene_gn=LOC102899173;refGene_tx=XM_023244737.1;refGene_gn=LOC102899173;refGene_tx=XM_023244738.1;refGene_gn=LOC102899173;refGene_tx=XM_023244739.1;refGene_gn=LOC102899173;refGene_tx=XM_023244740.1;refGene_gn=LOC102899173;refGene_tx=XM_023244741.1;refGene_gn=LOC102899173;refGene_tx=XM_023244742.1;refGene_gn=LOC102899173;refGene_tx=XM_023244743.1;refGene_gn=LOC102899173;refGene_tx=XM_023244744.1;refGene_gn=LOC102899173;refGene_tx=XM_023244745.1;refGene_gn=LOC102899173;refGene_tx=XM_023244746.1;refGene_gn=LOC102899173;refGene_tx=XM_023244747.1;refGene_gn=LOC102899173;refGene_tx=XM_023244748.1;refGene_gn=LOC102899173;refGene_tx=XM_023244749.1;refGene_gn=LOC102899173;refGene_tx=XM_023244750.1;refGene_gn=LOC102899173;refGene_tx=XM_023244751.1;refGene_gn=LOC102899173;refGene_tx=XM_023244752.1;refGene_gn=LOC102899173;refGene_tx=XM_023244753.1

E1 46712387 46712492

refGene_gn=LOC102899173;refGene_tx=XM_023244736.1;refGene_gn=LOC102899173;refGene_tx=XM_023244737.1;refGene_gn=LOC102899173;refGene_tx=XM_023244738.1;refGene_gn=LOC102899173;refGene_tx=XM_023244739.1;refGene_gn=LOC102899173;refGene_tx=XM_023244740.1;refGene_gn=LOC102899173;refGene_tx=XM_023244741.1;refGene_gn=LOC102899173;refGene_tx=XM_023244742.1;refGene_gn=LOC102899173;refGene_tx=XM_023244743.1;refGene_gn=LOC102899173;refGene_tx=XM_023244744.1;refGene_gn=LOC102899173;refGene_tx=XM_023244745.1;refGene_gn=LOC102899173;refGene_tx=XM_023244746.1;refGene_gn=LOC102899173;refGene_tx=XM_023244747.1;refGene_gn=LOC102899173;refGene_tx=XM_023244748.1;refGene_gn=LOC102899173;refGene_tx=XM_023244749.1;refGene_gn=LOC102899173;refGene_tx=XM_023244750.1;refGene_gn=LOC102899173;refGene_tx=XM_023244751.1;refGene_gn=LOC102899173;refGene_tx=XM_023244752.1;refGene_gn=LOC102899173;refGene_tx=XM_023244753.1

E1 46721792 46721846

refGene_gn=LOC102899173;refGene_tx=XM_023244736.1;refGene_gn=LOC102899173;refGene_tx=XM_023244737.1;refGene_gn=LOC102899173;refGene_tx=XM_023244738.1;refGene_gn=LOC102899173;refGene_tx=XM_023244739.1;refGene_gn=LOC102899173;refGene_tx=XM_023244740.1;refGene_gn=LOC102899173;refGene_tx=XM_023244741.1;refGene_gn=LOC102899173;refGene_tx=XM_023244742.1;refGene_gn=LOC102899173;refGene_tx=XM_023244743.1;refGene_gn=LOC102899173;refGene_tx=XM_023244744.1;refGene_gn=LOC102899173;refGene_tx=XM_023244745.1;refGene_gn=LOC102899173;refGene_tx=XM_023244746.1;refGene_gn

=LOC102899173;refGene_tx=XM_023244747.1;refGene_gn=LOC102899173;refGene_tx=XM_023244748.1;refGene_gn=LOC102899173;refGene_tx=XM_023244749.1;refGene_gn=LOC102899173;refGene_tx=XM_023244750.1;refGene_gn=LOC102899173;refGene_tx=XM_023244751.1;refGene_gn=LOC102899173;refGene_tx=XM_023244752.1;refGene_gn=LOC102899173;refGene_tx=XM_023244753.1
E1 46730013 46730067
refGene_gn=LOC102899173;refGene_tx=XM_023244736.1;refGene_gn=LOC102899173;refGene_tx=XM_023244737.1;refGene_gn=LOC102899173;refGene_tx=XM_023244738.1;refGene_gn=LOC102899173;refGene_tx=XM_023244739.1;refGene_gn=LOC102899173;refGene_tx=XM_023244740.1;refGene_gn=LOC102899173;refGene_tx=XM_023244741.1;refGene_gn=LOC102899173;refGene_tx=XM_023244742.1;refGene_gn=LOC102899173;refGene_tx=XM_023244743.1;refGene_gn=LOC102899173;refGene_tx=XM_023244744.1;refGene_gn=LOC102899173;refGene_tx=XM_023244745.1;refGene_gn=LOC102899173;refGene_tx=XM_023244746.1;refGene_gn=LOC102899173;refGene_tx=XM_023244747.1;refGene_gn=LOC102899173;refGene_tx=XM_023244748.1;refGene_gn=LOC102899173;refGene_tx=XM_023244749.1;refGene_gn=LOC102899173;refGene_tx=XM_023244750.1;refGene_gn=LOC102899173;refGene_tx=XM_023244751.1;refGene_gn=LOC102899173;refGene_tx=XM_023244752.1;refGene_gn=LOC102899173;refGene_tx=XM_023244753.1
E1 46731474 46731580
refGene_gn=LOC102899173;refGene_tx=XM_023244736.1;refGene_gn=LOC102899173;refGene_tx=XM_023244737.1;refGene_gn=LOC102899173;refGene_tx=XM_023244738.1;refGene_gn=LOC102899173;refGene_tx=XM_023244739.1;refGene_gn=LOC102899173;refGene_tx=XM_023244740.1;refGene_gn=LOC102899173;refGene_tx=XM_023244741.1;refGene_gn=LOC102899173;refGene_tx=XM_023244742.1;refGene_gn=LOC102899173;refGene_tx=XM_023244743.1;refGene_gn=LOC102899173;refGene_tx=XM_023244744.1;refGene_gn=LOC102899173;refGene_tx=XM_023244745.1;refGene_gn=LOC102899173;refGene_tx=XM_023244746.1;refGene_gn=LOC102899173;refGene_tx=XM_023244747.1;refGene_gn=LOC102899173;refGene_tx=XM_023244748.1;refGene_gn=LOC102899173;refGene_tx=XM_023244749.1;refGene_gn=LOC102899173;refGene_tx=XM_023244750.1;refGene_gn=LOC102899173;refGene_tx=XM_023244751.1;refGene_gn=LOC102899173;refGene_tx=XM_023244752.1;refGene_gn=LOC102899173;refGene_tx=XM_023244753.1;refGene_gn=LOC102899173;refGene_tx=XM_023244754.1
E1 46732251 46732305
refGene_gn=LOC102899173;refGene_tx=XM_023244736.1;refGene_gn=LOC102899173;refGene_tx=XM_023244737.1;refGene_gn=LOC102899173;refGene_tx=XM_023244738.1;refGene_gn=LOC102899173;refGene_tx=XM_023244739.1;refGene_gn=LOC102899173;refGene_tx=XM_023244740.1;refGene_gn=LOC102899173;refGene_tx=XM_023244741.1;refGene_gn=LOC102899173;refGene_tx=XM_023244742.1;refGene_gn=LOC102899173;refGene_tx=XM_023244743.1;refGene_gn=LOC102899173;refGene_tx=XM_023244744.1;refGene_gn=LOC102899173;refGene_tx=XM_023244745.1;refGene_gn=LOC102899173;refGene_tx=XM_023244746.1;refGene_gn=LOC102899173;refGene_tx=XM_023244747.1;refGene_gn=LOC102899173;refGene_tx=XM_023244748.1;refGene_gn=LOC102899173;refGene_tx=XM_023244749.1;refGene_gn=LOC102899173;refGene_tx=XM_023244750.1;refGene_gn=LOC102899173;refGene_tx=XM_023244751.1;refGene_gn=LOC102899173;refGene_tx=XM_023244752.1;refGene_gn=LOC102899173;refGene_tx=XM_023244753.1;refGene_gn=LOC102899173;refGene_tx=XM_023244754.1
E1 46733062 46733167

refGene_gn=LOC102899173;refGene_tx=XM_019818045.2;refGene_gn=LOC102899
173;refGene_tx=XM_023244736.1;refGene_gn=LOC102899173;refGene_tx=XM_02
3244737.1;refGene_gn=LOC102899173;refGene_tx=XM_023244738.1;refGene_gn
=LOC102899173;refGene_tx=XM_023244739.1;refGene_gn=LOC102899173;refGen
e_tx=XM_023244740.1;refGene_gn=LOC102899173;refGene_tx=XM_023244741.1;
refGene_gn=LOC102899173;refGene_tx=XM_023244742.1;refGene_gn=LOC102899
173;refGene_tx=XM_023244743.1;refGene_gn=LOC102899173;refGene_tx=XM_02
3244744.1;refGene_gn=LOC102899173;refGene_tx=XM_023244745.1;refGene_gn
=LOC102899173;refGene_tx=XM_023244746.1;refGene_gn=LOC102899173;refGen
e_tx=XM_023244747.1;refGene_gn=LOC102899173;refGene_tx=XM_023244748.1;
refGene_gn=LOC102899173;refGene_tx=XM_023244749.1;refGene_gn=LOC102899
173;refGene_tx=XM_023244750.1;refGene_gn=LOC102899173;refGene_tx=XM_02
3244751.1;refGene_gn=LOC102899173;refGene_tx=XM_023244752.1;refGene_gn
=LOC102899173;refGene_tx=XM_023244753.1;refGene_gn=LOC102899173;refGen
e_tx=XM_023244754.1
E1 46733767 46733821
refGene_gn=LOC102899173;refGene_tx=XM_019818045.2;refGene_gn=LOC102899
173;refGene_tx=XM_023244736.1;refGene_gn=LOC102899173;refGene_tx=XM_02
3244737.1;refGene_gn=LOC102899173;refGene_tx=XM_023244738.1;refGene_gn
=LOC102899173;refGene_tx=XM_023244739.1;refGene_gn=LOC102899173;refGen
e_tx=XM_023244740.1;refGene_gn=LOC102899173;refGene_tx=XM_023244741.1;
refGene_gn=LOC102899173;refGene_tx=XM_023244742.1;refGene_gn=LOC102899
173;refGene_tx=XM_023244743.1;refGene_gn=LOC102899173;refGene_tx=XM_02
3244744.1;refGene_gn=LOC102899173;refGene_tx=XM_023244745.1;refGene_gn
=LOC102899173;refGene_tx=XM_023244746.1;refGene_gn=LOC102899173;refGen
e_tx=XM_023244747.1;refGene_gn=LOC102899173;refGene_tx=XM_023244748.1;
refGene_gn=LOC102899173;refGene_tx=XM_023244749.1;refGene_gn=LOC102899
173;refGene_tx=XM_023244750.1;refGene_gn=LOC102899173;refGene_tx=XM_02
3244751.1;refGene_gn=LOC102899173;refGene_tx=XM_023244752.1;refGene_gn
=LOC102899173;refGene_tx=XM_023244753.1;refGene_gn=LOC102899173;refGen
e_tx=XM_023244754.1
E1 46734487 46734592
refGene_gn=LOC102899173;refGene_tx=XM_019818045.2;refGene_gn=LOC102899
173;refGene_tx=XM_023244736.1;refGene_gn=LOC102899173;refGene_tx=XM_02
3244737.1;refGene_gn=LOC102899173;refGene_tx=XM_023244738.1;refGene_gn
=LOC102899173;refGene_tx=XM_023244739.1;refGene_gn=LOC102899173;refGen
e_tx=XM_023244740.1;refGene_gn=LOC102899173;refGene_tx=XM_023244741.1;
refGene_gn=LOC102899173;refGene_tx=XM_023244742.1;refGene_gn=LOC102899
173;refGene_tx=XM_023244743.1;refGene_gn=LOC102899173;refGene_tx=XM_02
3244744.1;refGene_gn=LOC102899173;refGene_tx=XM_023244745.1;refGene_gn
=LOC102899173;refGene_tx=XM_023244746.1;refGene_gn=LOC102899173;refGen
e_tx=XM_023244747.1;refGene_gn=LOC102899173;refGene_tx=XM_023244748.1;
refGene_gn=LOC102899173;refGene_tx=XM_023244749.1;refGene_gn=LOC102899
173;refGene_tx=XM_023244750.1;refGene_gn=LOC102899173;refGene_tx=XM_02
3244751.1;refGene_gn=LOC102899173;refGene_tx=XM_023244752.1;refGene_gn
=LOC102899173;refGene_tx=XM_023244753.1;refGene_gn=LOC102899173;refGen
e_tx=XM_023244754.1
E1 46735164 46735218
refGene_gn=LOC102899173;refGene_tx=XM_019818045.2;refGene_gn=LOC102899
173;refGene_tx=XM_023244736.1;refGene_gn=LOC102899173;refGene_tx=XM_02

3244737.1;refGene_gn=LOC102899173;refGene_tx=XM_023244738.1;refGene_gn
=LOC102899173;refGene_tx=XM_023244739.1;refGene_gn=LOC102899173;refGen
e_tx=XM_023244740.1;refGene_gn=LOC102899173;refGene_tx=XM_023244741.1;
refGene_gn=LOC102899173;refGene_tx=XM_023244742.1;refGene_gn=LOC102899
173;refGene_tx=XM_023244743.1;refGene_gn=LOC102899173;refGene_tx=XM_02
3244744.1;refGene_gn=LOC102899173;refGene_tx=XM_023244745.1;refGene_gn
=LOC102899173;refGene_tx=XM_023244746.1;refGene_gn=LOC102899173;refGen
e_tx=XM_023244747.1;refGene_gn=LOC102899173;refGene_tx=XM_023244748.1;
refGene_gn=LOC102899173;refGene_tx=XM_023244749.1;refGene_gn=LOC102899
173;refGene_tx=XM_023244750.1;refGene_gn=LOC102899173;refGene_tx=XM_02
3244751.1;refGene_gn=LOC102899173;refGene_tx=XM_023244752.1;refGene_gn
=LOC102899173;refGene_tx=XM_023244753.1;refGene_gn=LOC102899173;refGen
e_tx=XM_023244754.1
E1 46735562 46735667
refGene_gn=LOC102899173;refGene_tx=XM_019818045.2;refGene_gn=LOC102899
173;refGene_tx=XM_023244736.1;refGene_gn=LOC102899173;refGene_tx=XM_02
3244737.1;refGene_gn=LOC102899173;refGene_tx=XM_023244738.1;refGene_gn
=LOC102899173;refGene_tx=XM_023244739.1;refGene_gn=LOC102899173;refGen
e_tx=XM_023244740.1;refGene_gn=LOC102899173;refGene_tx=XM_023244741.1;
refGene_gn=LOC102899173;refGene_tx=XM_023244742.1;refGene_gn=LOC102899
173;refGene_tx=XM_023244743.1;refGene_gn=LOC102899173;refGene_tx=XM_02
3244744.1;refGene_gn=LOC102899173;refGene_tx=XM_023244745.1;refGene_gn
=LOC102899173;refGene_tx=XM_023244746.1;refGene_gn=LOC102899173;refGen
e_tx=XM_023244747.1;refGene_gn=LOC102899173;refGene_tx=XM_023244748.1;
refGene_gn=LOC102899173;refGene_tx=XM_023244749.1;refGene_gn=LOC102899
173;refGene_tx=XM_023244750.1;refGene_gn=LOC102899173;refGene_tx=XM_02
3244751.1;refGene_gn=LOC102899173;refGene_tx=XM_023244752.1;refGene_gn
=LOC102899173;refGene_tx=XM_023244753.1;refGene_gn=LOC102899173;refGen
e_tx=XM_023244754.1
E1 46737317 46737371
refGene_gn=LOC102899173;refGene_tx=XM_019818045.2;refGene_gn=LOC102899
173;refGene_tx=XM_023244736.1;refGene_gn=LOC102899173;refGene_tx=XM_02
3244737.1;refGene_gn=LOC102899173;refGene_tx=XM_023244738.1;refGene_gn
=LOC102899173;refGene_tx=XM_023244739.1;refGene_gn=LOC102899173;refGen
e_tx=XM_023244740.1;refGene_gn=LOC102899173;refGene_tx=XM_023244741.1;
refGene_gn=LOC102899173;refGene_tx=XM_023244742.1;refGene_gn=LOC102899
173;refGene_tx=XM_023244743.1;refGene_gn=LOC102899173;refGene_tx=XM_02
3244744.1;refGene_gn=LOC102899173;refGene_tx=XM_023244745.1;refGene_gn
=LOC102899173;refGene_tx=XM_023244746.1;refGene_gn=LOC102899173;refGen
e_tx=XM_023244747.1;refGene_gn=LOC102899173;refGene_tx=XM_023244748.1;
refGene_gn=LOC102899173;refGene_tx=XM_023244749.1;refGene_gn=LOC102899
173;refGene_tx=XM_023244750.1;refGene_gn=LOC102899173;refGene_tx=XM_02
3244751.1;refGene_gn=LOC102899173;refGene_tx=XM_023244752.1;refGene_gn
=LOC102899173;refGene_tx=XM_023244753.1;refGene_gn=LOC102899173;refGen
e_tx=XM_023244754.1
E1 46737952 46738057
refGene_gn=LOC102899173;refGene_tx=XM_019818045.2;refGene_gn=LOC102899
173;refGene_tx=XM_023244736.1;refGene_gn=LOC102899173;refGene_tx=XM_02
3244737.1;refGene_gn=LOC102899173;refGene_tx=XM_023244738.1;refGene_gn
=LOC102899173;refGene_tx=XM_023244739.1;refGene_gn=LOC102899173;refGen

e_tx=XM_023244740.1;refGene_gn=LOC102899173;refGene_tx=XM_023244741.1;
refGene_gn=LOC102899173;refGene_tx=XM_023244742.1;refGene_gn=LOC102899
173;refGene_tx=XM_023244743.1;refGene_gn=LOC102899173;refGene_tx=XM_02
3244744.1;refGene_gn=LOC102899173;refGene_tx=XM_023244745.1;refGene_gn
=LOC102899173;refGene_tx=XM_023244746.1;refGene_gn=LOC102899173;refGen
e_tx=XM_023244747.1;refGene_gn=LOC102899173;refGene_tx=XM_023244748.1;
refGene_gn=LOC102899173;refGene_tx=XM_023244749.1;refGene_gn=LOC102899
173;refGene_tx=XM_023244750.1;refGene_gn=LOC102899173;refGene_tx=XM_02
3244751.1;refGene_gn=LOC102899173;refGene_tx=XM_023244752.1;refGene_gn
=LOC102899173;refGene_tx=XM_023244753.1;refGene_gn=LOC102899173;refGen
e_tx=XM_023244754.1
E1 46738874 46738928
refGene_gn=LOC102899173;refGene_tx=XM_019818045.2;refGene_gn=LOC102899
173;refGene_tx=XM_023244736.1;refGene_gn=LOC102899173;refGene_tx=XM_02
3244737.1;refGene_gn=LOC102899173;refGene_tx=XM_023244738.1;refGene_gn
=LOC102899173;refGene_tx=XM_023244739.1;refGene_gn=LOC102899173;refGen
e_tx=XM_023244740.1;refGene_gn=LOC102899173;refGene_tx=XM_023244741.1;
refGene_gn=LOC102899173;refGene_tx=XM_023244742.1;refGene_gn=LOC102899
173;refGene_tx=XM_023244743.1;refGene_gn=LOC102899173;refGene_tx=XM_02
3244744.1;refGene_gn=LOC102899173;refGene_tx=XM_023244745.1;refGene_gn
=LOC102899173;refGene_tx=XM_023244746.1;refGene_gn=LOC102899173;refGen
e_tx=XM_023244747.1;refGene_gn=LOC102899173;refGene_tx=XM_023244748.1;
refGene_gn=LOC102899173;refGene_tx=XM_023244749.1;refGene_gn=LOC102899
173;refGene_tx=XM_023244750.1;refGene_gn=LOC102899173;refGene_tx=XM_02
3244751.1;refGene_gn=LOC102899173;refGene_tx=XM_023244752.1;refGene_gn
=LOC102899173;refGene_tx=XM_023244753.1;refGene_gn=LOC102899173;refGen
e_tx=XM_023244754.1
E1 46740043 46740148
refGene_gn=LOC102899173;refGene_tx=XM_019818045.2;refGene_gn=LOC102899
173;refGene_tx=XM_023244736.1;refGene_gn=LOC102899173;refGene_tx=XM_02
3244737.1;refGene_gn=LOC102899173;refGene_tx=XM_023244738.1;refGene_gn
=LOC102899173;refGene_tx=XM_023244739.1;refGene_gn=LOC102899173;refGen
e_tx=XM_023244740.1;refGene_gn=LOC102899173;refGene_tx=XM_023244741.1;
refGene_gn=LOC102899173;refGene_tx=XM_023244742.1;refGene_gn=LOC102899
173;refGene_tx=XM_023244743.1;refGene_gn=LOC102899173;refGene_tx=XM_02
3244744.1;refGene_gn=LOC102899173;refGene_tx=XM_023244745.1;refGene_gn
=LOC102899173;refGene_tx=XM_023244746.1;refGene_gn=LOC102899173;refGen
e_tx=XM_023244747.1;refGene_gn=LOC102899173;refGene_tx=XM_023244748.1;
refGene_gn=LOC102899173;refGene_tx=XM_023244749.1;refGene_gn=LOC102899
173;refGene_tx=XM_023244750.1;refGene_gn=LOC102899173;refGene_tx=XM_02
3244751.1;refGene_gn=LOC102899173;refGene_tx=XM_023244752.1;refGene_gn
=LOC102899173;refGene_tx=XM_023244753.1;refGene_gn=LOC102899173;refGen
e_tx=XM_023244754.1
E1 46740819 46740873
refGene_gn=LOC102899173;refGene_tx=XM_019818045.2;refGene_gn=LOC102899
173;refGene_tx=XM_023244736.1;refGene_gn=LOC102899173;refGene_tx=XM_02
3244737.1;refGene_gn=LOC102899173;refGene_tx=XM_023244738.1;refGene_gn
=LOC102899173;refGene_tx=XM_023244739.1;refGene_gn=LOC102899173;refGen
e_tx=XM_023244740.1;refGene_gn=LOC102899173;refGene_tx=XM_023244741.1;
refGene_gn=LOC102899173;refGene_tx=XM_023244742.1;refGene_gn=LOC102899

173;refGene_tx=XM_023244743.1;refGene_gn=LOC102899173;refGene_tx=XM_023244744.1;refGene_gn=LOC102899173;refGene_tx=XM_023244745.1;refGene_gn=LOC102899173;refGene_tx=XM_023244746.1;refGene_gn=LOC102899173;refGene_tx=XM_023244747.1;refGene_gn=LOC102899173;refGene_tx=XM_023244748.1;refGene_gn=LOC102899173;refGene_tx=XM_023244749.1;refGene_gn=LOC102899173;refGene_tx=XM_023244750.1;refGene_gn=LOC102899173;refGene_tx=XM_023244751.1;refGene_gn=LOC102899173;refGene_tx=XM_023244752.1;refGene_gn=LOC102899173;refGene_tx=XM_023244753.1;refGene_gn=LOC102899173;refGene_tx=XM_023244754.1
E1 46741774 46741879
refGene_gn=LOC102899173;refGene_tx=XM_019818045.2;refGene_gn=LOC102899173;refGene_tx=XM_023244736.1;refGene_gn=LOC102899173;refGene_tx=XM_023244737.1;refGene_gn=LOC102899173;refGene_tx=XM_023244738.1;refGene_gn=LOC102899173;refGene_tx=XM_023244739.1;refGene_gn=LOC102899173;refGene_tx=XM_023244740.1;refGene_gn=LOC102899173;refGene_tx=XM_023244741.1;refGene_gn=LOC102899173;refGene_tx=XM_023244742.1;refGene_gn=LOC102899173;refGene_tx=XM_023244743.1;refGene_gn=LOC102899173;refGene_tx=XM_023244744.1;refGene_gn=LOC102899173;refGene_tx=XM_023244745.1;refGene_gn=LOC102899173;refGene_tx=XM_023244746.1;refGene_gn=LOC102899173;refGene_tx=XM_023244747.1;refGene_gn=LOC102899173;refGene_tx=XM_023244748.1;refGene_gn=LOC102899173;refGene_tx=XM_023244749.1;refGene_gn=LOC102899173;refGene_tx=XM_023244750.1;refGene_gn=LOC102899173;refGene_tx=XM_023244751.1;refGene_gn=LOC102899173;refGene_tx=XM_023244752.1;refGene_gn=LOC102899173;refGene_tx=XM_023244753.1;refGene_gn=LOC102899173;refGene_tx=XM_023244754.1
E1 46745002 46745056
refGene_gn=LOC102899173;refGene_tx=XM_019818045.2;refGene_gn=LOC102899173;refGene_tx=XM_023244736.1;refGene_gn=LOC102899173;refGene_tx=XM_023244737.1;refGene_gn=LOC102899173;refGene_tx=XM_023244738.1;refGene_gn=LOC102899173;refGene_tx=XM_023244739.1;refGene_gn=LOC102899173;refGene_tx=XM_023244740.1;refGene_gn=LOC102899173;refGene_tx=XM_023244741.1;refGene_gn=LOC102899173;refGene_tx=XM_023244742.1;refGene_gn=LOC102899173;refGene_tx=XM_023244743.1;refGene_gn=LOC102899173;refGene_tx=XM_023244744.1;refGene_gn=LOC102899173;refGene_tx=XM_023244745.1;refGene_gn=LOC102899173;refGene_tx=XM_023244746.1;refGene_gn=LOC102899173;refGene_tx=XM_023244747.1;refGene_gn=LOC102899173;refGene_tx=XM_023244748.1;refGene_gn=LOC102899173;refGene_tx=XM_023244749.1;refGene_gn=LOC102899173;refGene_tx=XM_023244750.1;refGene_gn=LOC102899173;refGene_tx=XM_023244751.1;refGene_gn=LOC102899173;refGene_tx=XM_023244752.1;refGene_gn=LOC102899173;refGene_tx=XM_023244753.1;refGene_gn=LOC102899173;refGene_tx=XM_023244754.1
E1 46745709 46745814
refGene_gn=LOC102899173;refGene_tx=XM_019818045.2;refGene_gn=LOC102899173;refGene_tx=XM_023244736.1;refGene_gn=LOC102899173;refGene_tx=XM_023244737.1;refGene_gn=LOC102899173;refGene_tx=XM_023244738.1;refGene_gn=LOC102899173;refGene_tx=XM_023244739.1;refGene_gn=LOC102899173;refGene_tx=XM_023244740.1;refGene_gn=LOC102899173;refGene_tx=XM_023244741.1;refGene_gn=LOC102899173;refGene_tx=XM_023244742.1;refGene_gn=LOC102899173;refGene_tx=XM_023244743.1;refGene_gn=LOC102899173;refGene_tx=XM_023244744.1;refGene_gn=LOC102899173;refGene_tx=XM_023244745.1;refGene_gn

=LOC102899173;refGene_tx=XM_023244746.1;refGene_gn=LOC102899173;refGene_tx=XM_023244747.1;refGene_gn=LOC102899173;refGene_tx=XM_023244748.1;refGene_gn=LOC102899173;refGene_tx=XM_023244749.1;refGene_gn=LOC102899173;refGene_tx=XM_023244750.1;refGene_gn=LOC102899173;refGene_tx=XM_023244751.1;refGene_gn=LOC102899173;refGene_tx=XM_023244752.1;refGene_gn=LOC102899173;refGene_tx=XM_023244753.1;refGene_gn=LOC102899173;refGene_tx=XM_023244754.1
E1 46745990 46746044
refGene_gn=LOC102899173;refGene_tx=XM_019818045.2;refGene_gn=LOC102899173;refGene_tx=XM_023244736.1;refGene_gn=LOC102899173;refGene_tx=XM_023244737.1;refGene_gn=LOC102899173;refGene_tx=XM_023244738.1;refGene_gn=LOC102899173;refGene_tx=XM_023244739.1;refGene_gn=LOC102899173;refGene_tx=XM_023244740.1;refGene_gn=LOC102899173;refGene_tx=XM_023244741.1;refGene_gn=LOC102899173;refGene_tx=XM_023244742.1;refGene_gn=LOC102899173;refGene_tx=XM_023244743.1;refGene_gn=LOC102899173;refGene_tx=XM_023244744.1;refGene_gn=LOC102899173;refGene_tx=XM_023244745.1;refGene_gn=LOC102899173;refGene_tx=XM_023244746.1;refGene_gn=LOC102899173;refGene_tx=XM_023244747.1;refGene_gn=LOC102899173;refGene_tx=XM_023244748.1;refGene_gn=LOC102899173;refGene_tx=XM_023244749.1;refGene_gn=LOC102899173;refGene_tx=XM_023244750.1;refGene_gn=LOC102899173;refGene_tx=XM_023244751.1;refGene_gn=LOC102899173;refGene_tx=XM_023244752.1;refGene_gn=LOC102899173;refGene_tx=XM_023244753.1;refGene_gn=LOC102899173;refGene_tx=XM_023244754.1
E1 46746542 46746644
refGene_gn=LOC102899173;refGene_tx=XM_019818045.2;refGene_gn=LOC102899173;refGene_tx=XM_023244736.1;refGene_gn=LOC102899173;refGene_tx=XM_023244737.1;refGene_gn=LOC102899173;refGene_tx=XM_023244738.1;refGene_gn=LOC102899173;refGene_tx=XM_023244739.1;refGene_gn=LOC102899173;refGene_tx=XM_023244740.1;refGene_gn=LOC102899173;refGene_tx=XM_023244741.1;refGene_gn=LOC102899173;refGene_tx=XM_023244742.1;refGene_gn=LOC102899173;refGene_tx=XM_023244743.1;refGene_gn=LOC102899173;refGene_tx=XM_023244744.1;refGene_gn=LOC102899173;refGene_tx=XM_023244745.1;refGene_gn=LOC102899173;refGene_tx=XM_023244746.1;refGene_gn=LOC102899173;refGene_tx=XM_023244747.1;refGene_gn=LOC102899173;refGene_tx=XM_023244748.1;refGene_gn=LOC102899173;refGene_tx=XM_023244749.1;refGene_gn=LOC102899173;refGene_tx=XM_023244750.1;refGene_gn=LOC102899173;refGene_tx=XM_023244751.1;refGene_gn=LOC102899173;refGene_tx=XM_023244752.1;refGene_gn=LOC102899173;refGene_tx=XM_023244753.1;refGene_gn=LOC102899173;refGene_tx=XM_023244754.1
E1 46747152 46747206
refGene_gn=LOC102899173;refGene_tx=XM_019818045.2;refGene_gn=LOC102899173;refGene_tx=XM_023244736.1;refGene_gn=LOC102899173;refGene_tx=XM_023244737.1;refGene_gn=LOC102899173;refGene_tx=XM_023244738.1;refGene_gn=LOC102899173;refGene_tx=XM_023244739.1;refGene_gn=LOC102899173;refGene_tx=XM_023244740.1;refGene_gn=LOC102899173;refGene_tx=XM_023244741.1;refGene_gn=LOC102899173;refGene_tx=XM_023244742.1;refGene_gn=LOC102899173;refGene_tx=XM_023244743.1;refGene_gn=LOC102899173;refGene_tx=XM_023244744.1;refGene_gn=LOC102899173;refGene_tx=XM_023244745.1;refGene_gn=LOC102899173;refGene_tx=XM_023244746.1;refGene_gn=LOC102899173;refGene_tx=XM_023244747.1;refGene_gn=LOC102899173;refGene_tx=XM_023244748.1;

refGene_gn=LOC102899173;refGene_tx=XM_023244749.1;refGene_gn=LOC102899173;refGene_tx=XM_023244750.1;refGene_gn=LOC102899173;refGene_tx=XM_023244751.1;refGene_gn=LOC102899173;refGene_tx=XM_023244752.1;refGene_gn=LOC102899173;refGene_tx=XM_023244753.1;refGene_gn=LOC102899173;refGene_tx=XM_023244754.1
E1 46747895 46748000
refGene_gn=LOC102899173;refGene_tx=XM_019818045.2;refGene_gn=LOC102899173;refGene_tx=XM_023244736.1;refGene_gn=LOC102899173;refGene_tx=XM_023244737.1;refGene_gn=LOC102899173;refGene_tx=XM_023244738.1;refGene_gn=LOC102899173;refGene_tx=XM_023244739.1;refGene_gn=LOC102899173;refGene_tx=XM_023244740.1;refGene_gn=LOC102899173;refGene_tx=XM_023244741.1;refGene_gn=LOC102899173;refGene_tx=XM_023244742.1;refGene_gn=LOC102899173;refGene_tx=XM_023244743.1;refGene_gn=LOC102899173;refGene_tx=XM_023244744.1;refGene_gn=LOC102899173;refGene_tx=XM_023244745.1;refGene_gn=LOC102899173;refGene_tx=XM_023244746.1;refGene_gn=LOC102899173;refGene_tx=XM_023244747.1;refGene_gn=LOC102899173;refGene_tx=XM_023244748.1;refGene_gn=LOC102899173;refGene_tx=XM_023244749.1;refGene_gn=LOC102899173;refGene_tx=XM_023244750.1;refGene_gn=LOC102899173;refGene_tx=XM_023244751.1;refGene_gn=LOC102899173;refGene_tx=XM_023244752.1;refGene_gn=LOC102899173;refGene_tx=XM_023244753.1;refGene_gn=LOC102899173;refGene_tx=XM_023244754.1
E1 46748788 46748842
refGene_gn=LOC102899173;refGene_tx=XM_019818045.2;refGene_gn=LOC102899173;refGene_tx=XM_023244736.1;refGene_gn=LOC102899173;refGene_tx=XM_023244737.1;refGene_gn=LOC102899173;refGene_tx=XM_023244738.1;refGene_gn=LOC102899173;refGene_tx=XM_023244739.1;refGene_gn=LOC102899173;refGene_tx=XM_023244740.1;refGene_gn=LOC102899173;refGene_tx=XM_023244741.1;refGene_gn=LOC102899173;refGene_tx=XM_023244742.1;refGene_gn=LOC102899173;refGene_tx=XM_023244743.1;refGene_gn=LOC102899173;refGene_tx=XM_023244744.1;refGene_gn=LOC102899173;refGene_tx=XM_023244745.1;refGene_gn=LOC102899173;refGene_tx=XM_023244746.1;refGene_gn=LOC102899173;refGene_tx=XM_023244747.1;refGene_gn=LOC102899173;refGene_tx=XM_023244748.1;refGene_gn=LOC102899173;refGene_tx=XM_023244749.1;refGene_gn=LOC102899173;refGene_tx=XM_023244750.1;refGene_gn=LOC102899173;refGene_tx=XM_023244751.1;refGene_gn=LOC102899173;refGene_tx=XM_023244752.1;refGene_gn=LOC102899173;refGene_tx=XM_023244753.1;refGene_gn=LOC102899173;refGene_tx=XM_023244754.1
E1 46750760 46750859
refGene_gn=LOC102899173;refGene_tx=XM_019818045.2;refGene_gn=LOC102899173;refGene_tx=XM_023244736.1;refGene_gn=LOC102899173;refGene_tx=XM_023244737.1;refGene_gn=LOC102899173;refGene_tx=XM_023244738.1;refGene_gn=LOC102899173;refGene_tx=XM_023244739.1;refGene_gn=LOC102899173;refGene_tx=XM_023244740.1;refGene_gn=LOC102899173;refGene_tx=XM_023244741.1;refGene_gn=LOC102899173;refGene_tx=XM_023244742.1;refGene_gn=LOC102899173;refGene_tx=XM_023244743.1;refGene_gn=LOC102899173;refGene_tx=XM_023244744.1;refGene_gn=LOC102899173;refGene_tx=XM_023244745.1;refGene_gn=LOC102899173;refGene_tx=XM_023244746.1;refGene_gn=LOC102899173;refGene_tx=XM_023244747.1;refGene_gn=LOC102899173;refGene_tx=XM_023244748.1;refGene_gn=LOC102899173;refGene_tx=XM_023244749.1;refGene_gn=LOC102899173;refGene_tx=XM_023244750.1;refGene_gn=LOC102899173;refGene_tx=XM_02

3244751.1;refGene_gn=LOC102899173;refGene_tx=XM_023244752.1;refGene_gn
=LOC102899173;refGene_tx=XM_023244753.1;refGene_gn=LOC102899173;refGen
e_tx=XM_023244754.1
E1 46753401 46753455
refGene_gn=LOC102899173;refGene_tx=XM_019818045.2;refGene_gn=LOC102899
173;refGene_tx=XM_023244736.1;refGene_gn=LOC102899173;refGene_tx=XM_02
3244737.1;refGene_gn=LOC102899173;refGene_tx=XM_023244738.1;refGene_gn
=LOC102899173;refGene_tx=XM_023244739.1;refGene_gn=LOC102899173;refGen
e_tx=XM_023244740.1;refGene_gn=LOC102899173;refGene_tx=XM_023244741.1;
refGene_gn=LOC102899173;refGene_tx=XM_023244742.1;refGene_gn=LOC102899
173;refGene_tx=XM_023244743.1;refGene_gn=LOC102899173;refGene_tx=XM_02
3244744.1;refGene_gn=LOC102899173;refGene_tx=XM_023244745.1;refGene_gn
=LOC102899173;refGene_tx=XM_023244746.1;refGene_gn=LOC102899173;refGen
e_tx=XM_023244747.1;refGene_gn=LOC102899173;refGene_tx=XM_023244748.1;
refGene_gn=LOC102899173;refGene_tx=XM_023244749.1;refGene_gn=LOC102899
173;refGene_tx=XM_023244750.1;refGene_gn=LOC102899173;refGene_tx=XM_02
3244751.1;refGene_gn=LOC102899173;refGene_tx=XM_023244752.1;refGene_gn
=LOC102899173;refGene_tx=XM_023244753.1;refGene_gn=LOC102899173;refGen
e_tx=XM_023244754.1
E1 46754508 46754613
refGene_gn=LOC102899173;refGene_tx=XM_019818045.2;refGene_gn=LOC102899
173;refGene_tx=XM_023244736.1;refGene_gn=LOC102899173;refGene_tx=XM_02
3244737.1;refGene_gn=LOC102899173;refGene_tx=XM_023244738.1;refGene_gn
=LOC102899173;refGene_tx=XM_023244739.1;refGene_gn=LOC102899173;refGen
e_tx=XM_023244740.1;refGene_gn=LOC102899173;refGene_tx=XM_023244741.1;
refGene_gn=LOC102899173;refGene_tx=XM_023244742.1;refGene_gn=LOC102899
173;refGene_tx=XM_023244743.1;refGene_gn=LOC102899173;refGene_tx=XM_02
3244744.1;refGene_gn=LOC102899173;refGene_tx=XM_023244745.1;refGene_gn
=LOC102899173;refGene_tx=XM_023244746.1;refGene_gn=LOC102899173;refGen
e_tx=XM_023244747.1;refGene_gn=LOC102899173;refGene_tx=XM_023244748.1;
refGene_gn=LOC102899173;refGene_tx=XM_023244749.1;refGene_gn=LOC102899
173;refGene_tx=XM_023244750.1;refGene_gn=LOC102899173;refGene_tx=XM_02
3244751.1;refGene_gn=LOC102899173;refGene_tx=XM_023244752.1;refGene_gn
=LOC102899173;refGene_tx=XM_023244753.1;refGene_gn=LOC102899173;refGen
e_tx=XM_023244754.1
E1 46755142 46755193
refGene_gn=LOC102899173;refGene_tx=XM_019818045.2;refGene_gn=LOC102899
173;refGene_tx=XM_023244736.1;refGene_gn=LOC102899173;refGene_tx=XM_02
3244737.1;refGene_gn=LOC102899173;refGene_tx=XM_023244738.1;refGene_gn
=LOC102899173;refGene_tx=XM_023244739.1;refGene_gn=LOC102899173;refGen
e_tx=XM_023244740.1;refGene_gn=LOC102899173;refGene_tx=XM_023244741.1;
refGene_gn=LOC102899173;refGene_tx=XM_023244742.1;refGene_gn=LOC102899
173;refGene_tx=XM_023244743.1;refGene_gn=LOC102899173;refGene_tx=XM_02
3244744.1;refGene_gn=LOC102899173;refGene_tx=XM_023244745.1;refGene_gn
=LOC102899173;refGene_tx=XM_023244746.1;refGene_gn=LOC102899173;refGen
e_tx=XM_023244747.1;refGene_gn=LOC102899173;refGene_tx=XM_023244748.1;
refGene_gn=LOC102899173;refGene_tx=XM_023244749.1;refGene_gn=LOC102899
173;refGene_tx=XM_023244750.1;refGene_gn=LOC102899173;refGene_tx=XM_02
3244751.1;refGene_gn=LOC102899173;refGene_tx=XM_023244752.1;refGene_gn
=LOC102899173;refGene_tx=XM_023244753.1;refGene_gn=LOC102899173;refGen

e_tx=XM_023244754.1
E1 46756326 46756431
refGene_gn=LOC102899173;refGene_tx=XM_019818045.2;refGene_gn=LOC102899173;refGene_tx=XM_023244736.1;refGene_gn=LOC102899173;refGene_tx=XM_023244737.1;refGene_gn=LOC102899173;refGene_tx=XM_023244738.1;refGene_gn=LOC102899173;refGene_tx=XM_023244739.1;refGene_gn=LOC102899173;refGene_tx=XM_023244740.1;refGene_gn=LOC102899173;refGene_tx=XM_023244741.1;refGene_gn=LOC102899173;refGene_tx=XM_023244742.1;refGene_gn=LOC102899173;refGene_tx=XM_023244743.1;refGene_gn=LOC102899173;refGene_tx=XM_023244744.1;refGene_gn=LOC102899173;refGene_tx=XM_023244745.1;refGene_gn=LOC102899173;refGene_tx=XM_023244746.1;refGene_gn=LOC102899173;refGene_tx=XM_023244747.1;refGene_gn=LOC102899173;refGene_tx=XM_023244748.1;refGene_gn=LOC102899173;refGene_tx=XM_023244749.1;refGene_gn=LOC102899173;refGene_tx=XM_023244750.1;refGene_gn=LOC102899173;refGene_tx=XM_023244751.1;refGene_gn=LOC102899173;refGene_tx=XM_023244752.1;refGene_gn=LOC102899173;refGene_tx=XM_023244753.1;refGene_gn=LOC102899173;refGene_tx=XM_023244754.1
E1 46768046 46768100
refGene_gn=LOC102899173;refGene_tx=XM_019818045.2;refGene_gn=LOC102899173;refGene_tx=XM_023244736.1;refGene_gn=LOC102899173;refGene_tx=XM_023244737.1;refGene_gn=LOC102899173;refGene_tx=XM_023244738.1;refGene_gn=LOC102899173;refGene_tx=XM_023244739.1;refGene_gn=LOC102899173;refGene_tx=XM_023244740.1;refGene_gn=LOC102899173;refGene_tx=XM_023244741.1;refGene_gn=LOC102899173;refGene_tx=XM_023244742.1;refGene_gn=LOC102899173;refGene_tx=XM_023244743.1;refGene_gn=LOC102899173;refGene_tx=XM_023244744.1;refGene_gn=LOC102899173;refGene_tx=XM_023244745.1;refGene_gn=LOC102899173;refGene_tx=XM_023244746.1;refGene_gn=LOC102899173;refGene_tx=XM_023244747.1;refGene_gn=LOC102899173;refGene_tx=XM_023244748.1;refGene_gn=LOC102899173;refGene_tx=XM_023244749.1;refGene_gn=LOC102899173;refGene_tx=XM_023244750.1;refGene_gn=LOC102899173;refGene_tx=XM_023244751.1;refGene_gn=LOC102899173;refGene_tx=XM_023244752.1;refGene_gn=LOC102899173;refGene_tx=XM_023244753.1;refGene_gn=LOC102899173;refGene_tx=XM_023244754.1
E1 46768917 46769022
refGene_gn=LOC102899173;refGene_tx=XM_019818045.2;refGene_gn=LOC102899173;refGene_tx=XM_023244736.1;refGene_gn=LOC102899173;refGene_tx=XM_023244737.1;refGene_gn=LOC102899173;refGene_tx=XM_023244738.1;refGene_gn=LOC102899173;refGene_tx=XM_023244739.1;refGene_gn=LOC102899173;refGene_tx=XM_023244740.1;refGene_gn=LOC102899173;refGene_tx=XM_023244741.1;refGene_gn=LOC102899173;refGene_tx=XM_023244742.1;refGene_gn=LOC102899173;refGene_tx=XM_023244743.1;refGene_gn=LOC102899173;refGene_tx=XM_023244744.1;refGene_gn=LOC102899173;refGene_tx=XM_023244745.1;refGene_gn=LOC102899173;refGene_tx=XM_023244746.1;refGene_gn=LOC102899173;refGene_tx=XM_023244747.1;refGene_gn=LOC102899173;refGene_tx=XM_023244748.1;refGene_gn=LOC102899173;refGene_tx=XM_023244749.1;refGene_gn=LOC102899173;refGene_tx=XM_023244750.1;refGene_gn=LOC102899173;refGene_tx=XM_023244751.1;refGene_gn=LOC102899173;refGene_tx=XM_023244752.1;refGene_gn=LOC102899173;refGene_tx=XM_023244753.1;refGene_gn=LOC102899173;refGene_tx=XM_023244754.1
E1 46770172 46770226

refGene_gn=LOC102899173;refGene_tx=XM_019818045.2;refGene_gn=LOC102899
173;refGene_tx=XM_023244736.1;refGene_gn=LOC102899173;refGene_tx=XM_02
3244737.1;refGene_gn=LOC102899173;refGene_tx=XM_023244738.1;refGene_gn
=LOC102899173;refGene_tx=XM_023244739.1;refGene_gn=LOC102899173;refGen
e_tx=XM_023244740.1;refGene_gn=LOC102899173;refGene_tx=XM_023244741.1;
refGene_gn=LOC102899173;refGene_tx=XM_023244742.1;refGene_gn=LOC102899
173;refGene_tx=XM_023244743.1;refGene_gn=LOC102899173;refGene_tx=XM_02
3244744.1;refGene_gn=LOC102899173;refGene_tx=XM_023244745.1;refGene_gn
=LOC102899173;refGene_tx=XM_023244746.1;refGene_gn=LOC102899173;refGen
e_tx=XM_023244747.1;refGene_gn=LOC102899173;refGene_tx=XM_023244748.1;
refGene_gn=LOC102899173;refGene_tx=XM_023244749.1;refGene_gn=LOC102899
173;refGene_tx=XM_023244750.1;refGene_gn=LOC102899173;refGene_tx=XM_02
3244751.1;refGene_gn=LOC102899173;refGene_tx=XM_023244752.1;refGene_gn
=LOC102899173;refGene_tx=XM_023244753.1;refGene_gn=LOC102899173;refGen
e_tx=XM_023244754.1
E1 46770820 46770925
refGene_gn=LOC102899173;refGene_tx=XM_019818045.2;refGene_gn=LOC102899
173;refGene_tx=XM_023244736.1;refGene_gn=LOC102899173;refGene_tx=XM_02
3244737.1;refGene_gn=LOC102899173;refGene_tx=XM_023244738.1;refGene_gn
=LOC102899173;refGene_tx=XM_023244739.1;refGene_gn=LOC102899173;refGen
e_tx=XM_023244740.1;refGene_gn=LOC102899173;refGene_tx=XM_023244741.1;
refGene_gn=LOC102899173;refGene_tx=XM_023244742.1;refGene_gn=LOC102899
173;refGene_tx=XM_023244743.1;refGene_gn=LOC102899173;refGene_tx=XM_02
3244744.1;refGene_gn=LOC102899173;refGene_tx=XM_023244745.1;refGene_gn
=LOC102899173;refGene_tx=XM_023244746.1;refGene_gn=LOC102899173;refGen
e_tx=XM_023244747.1;refGene_gn=LOC102899173;refGene_tx=XM_023244748.1;
refGene_gn=LOC102899173;refGene_tx=XM_023244749.1;refGene_gn=LOC102899
173;refGene_tx=XM_023244750.1;refGene_gn=LOC102899173;refGene_tx=XM_02
3244751.1;refGene_gn=LOC102899173;refGene_tx=XM_023244752.1;refGene_gn
=LOC102899173;refGene_tx=XM_023244753.1;refGene_gn=LOC102899173;refGen
e_tx=XM_023244754.1
E1 46776263 46776317
refGene_gn=LOC102899173;refGene_tx=XM_019818045.2;refGene_gn=LOC102899
173;refGene_tx=XM_023244736.1;refGene_gn=LOC102899173;refGene_tx=XM_02
3244737.1;refGene_gn=LOC102899173;refGene_tx=XM_023244738.1;refGene_gn
=LOC102899173;refGene_tx=XM_023244739.1;refGene_gn=LOC102899173;refGen
e_tx=XM_023244740.1;refGene_gn=LOC102899173;refGene_tx=XM_023244741.1;
refGene_gn=LOC102899173;refGene_tx=XM_023244742.1;refGene_gn=LOC102899
173;refGene_tx=XM_023244743.1;refGene_gn=LOC102899173;refGene_tx=XM_02
3244744.1;refGene_gn=LOC102899173;refGene_tx=XM_023244745.1;refGene_gn
=LOC102899173;refGene_tx=XM_023244746.1;refGene_gn=LOC102899173;refGen
e_tx=XM_023244747.1;refGene_gn=LOC102899173;refGene_tx=XM_023244748.1;
refGene_gn=LOC102899173;refGene_tx=XM_023244749.1;refGene_gn=LOC102899
173;refGene_tx=XM_023244750.1;refGene_gn=LOC102899173;refGene_tx=XM_02
3244751.1;refGene_gn=LOC102899173;refGene_tx=XM_023244752.1;refGene_gn
=LOC102899173;refGene_tx=XM_023244753.1;refGene_gn=LOC102899173;refGen
e_tx=XM_023244754.1
E1 46784044 46784149
refGene_gn=LOC102899173;refGene_tx=XM_019818045.2;refGene_gn=LOC102899
173;refGene_tx=XM_023244736.1;refGene_gn=LOC102899173;refGene_tx=XM_02

3244737.1;refGene_gn=LOC102899173;refGene_tx=XM_023244738.1;refGene_gn
=LOC102899173;refGene_tx=XM_023244739.1;refGene_gn=LOC102899173;refGen
e_tx=XM_023244740.1;refGene_gn=LOC102899173;refGene_tx=XM_023244741.1;
refGene_gn=LOC102899173;refGene_tx=XM_023244742.1;refGene_gn=LOC102899
173;refGene_tx=XM_023244743.1;refGene_gn=LOC102899173;refGene_tx=XM_02
3244744.1;refGene_gn=LOC102899173;refGene_tx=XM_023244745.1;refGene_gn
=LOC102899173;refGene_tx=XM_023244746.1;refGene_gn=LOC102899173;refGen
e_tx=XM_023244747.1;refGene_gn=LOC102899173;refGene_tx=XM_023244748.1;
refGene_gn=LOC102899173;refGene_tx=XM_023244749.1;refGene_gn=LOC102899
173;refGene_tx=XM_023244750.1;refGene_gn=LOC102899173;refGene_tx=XM_02
3244751.1;refGene_gn=LOC102899173;refGene_tx=XM_023244752.1;refGene_gn
=LOC102899173;refGene_tx=XM_023244753.1;refGene_gn=LOC102899173;refGen
e_tx=XM_023244754.1
E1 46784801 46784855
refGene_gn=LOC102899173;refGene_tx=XM_019818045.2;refGene_gn=LOC102899
173;refGene_tx=XM_023244736.1;refGene_gn=LOC102899173;refGene_tx=XM_02
3244737.1;refGene_gn=LOC102899173;refGene_tx=XM_023244738.1;refGene_gn
=LOC102899173;refGene_tx=XM_023244739.1;refGene_gn=LOC102899173;refGen
e_tx=XM_023244740.1;refGene_gn=LOC102899173;refGene_tx=XM_023244741.1;
refGene_gn=LOC102899173;refGene_tx=XM_023244742.1;refGene_gn=LOC102899
173;refGene_tx=XM_023244743.1;refGene_gn=LOC102899173;refGene_tx=XM_02
3244744.1;refGene_gn=LOC102899173;refGene_tx=XM_023244745.1;refGene_gn
=LOC102899173;refGene_tx=XM_023244746.1;refGene_gn=LOC102899173;refGen
e_tx=XM_023244747.1;refGene_gn=LOC102899173;refGene_tx=XM_023244748.1;
refGene_gn=LOC102899173;refGene_tx=XM_023244749.1;refGene_gn=LOC102899
173;refGene_tx=XM_023244750.1;refGene_gn=LOC102899173;refGene_tx=XM_02
3244751.1;refGene_gn=LOC102899173;refGene_tx=XM_023244752.1;refGene_gn
=LOC102899173;refGene_tx=XM_023244753.1;refGene_gn=LOC102899173;refGen
e_tx=XM_023244754.1
E1 46785229 46785334
refGene_gn=LOC102899173;refGene_tx=XM_019818045.2;refGene_gn=LOC102899
173;refGene_tx=XM_023244736.1;refGene_gn=LOC102899173;refGene_tx=XM_02
3244737.1;refGene_gn=LOC102899173;refGene_tx=XM_023244738.1;refGene_gn
=LOC102899173;refGene_tx=XM_023244739.1;refGene_gn=LOC102899173;refGen
e_tx=XM_023244740.1;refGene_gn=LOC102899173;refGene_tx=XM_023244741.1;
refGene_gn=LOC102899173;refGene_tx=XM_023244742.1;refGene_gn=LOC102899
173;refGene_tx=XM_023244743.1;refGene_gn=LOC102899173;refGene_tx=XM_02
3244744.1;refGene_gn=LOC102899173;refGene_tx=XM_023244745.1;refGene_gn
=LOC102899173;refGene_tx=XM_023244746.1;refGene_gn=LOC102899173;refGen
e_tx=XM_023244747.1;refGene_gn=LOC102899173;refGene_tx=XM_023244748.1;
refGene_gn=LOC102899173;refGene_tx=XM_023244749.1;refGene_gn=LOC102899
173;refGene_tx=XM_023244750.1;refGene_gn=LOC102899173;refGene_tx=XM_02
3244751.1;refGene_gn=LOC102899173;refGene_tx=XM_023244752.1;refGene_gn
=LOC102899173;refGene_tx=XM_023244753.1;refGene_gn=LOC102899173;refGen
e_tx=XM_023244754.1
E1 46793931 46793985
refGene_gn=LOC102899173;refGene_tx=XM_019818045.2;refGene_gn=LOC102899
173;refGene_tx=XM_023244736.1;refGene_gn=LOC102899173;refGene_tx=XM_02
3244737.1;refGene_gn=LOC102899173;refGene_tx=XM_023244738.1;refGene_gn
=LOC102899173;refGene_tx=XM_023244739.1;refGene_gn=LOC102899173;refGen

e_tx=XM_023244740.1;refGene_gn=LOC102899173;refGene_tx=XM_023244741.1;
refGene_gn=LOC102899173;refGene_tx=XM_023244742.1;refGene_gn=LOC102899
173;refGene_tx=XM_023244743.1;refGene_gn=LOC102899173;refGene_tx=XM_02
3244744.1;refGene_gn=LOC102899173;refGene_tx=XM_023244745.1;refGene_gn
=LOC102899173;refGene_tx=XM_023244746.1;refGene_gn=LOC102899173;refGen
e_tx=XM_023244747.1;refGene_gn=LOC102899173;refGene_tx=XM_023244748.1;
refGene_gn=LOC102899173;refGene_tx=XM_023244749.1;refGene_gn=LOC102899
173;refGene_tx=XM_023244750.1;refGene_gn=LOC102899173;refGene_tx=XM_02
3244751.1;refGene_gn=LOC102899173;refGene_tx=XM_023244752.1;refGene_gn
=LOC102899173;refGene_tx=XM_023244753.1;refGene_gn=LOC102899173;refGen
e_tx=XM_023244754.1
E1 46794691 46794796
refGene_gn=LOC102899173;refGene_tx=XM_019818045.2;refGene_gn=LOC102899
173;refGene_tx=XM_023244736.1;refGene_gn=LOC102899173;refGene_tx=XM_02
3244737.1;refGene_gn=LOC102899173;refGene_tx=XM_023244738.1;refGene_gn
=LOC102899173;refGene_tx=XM_023244739.1;refGene_gn=LOC102899173;refGen
e_tx=XM_023244740.1;refGene_gn=LOC102899173;refGene_tx=XM_023244741.1;
refGene_gn=LOC102899173;refGene_tx=XM_023244742.1;refGene_gn=LOC102899
173;refGene_tx=XM_023244743.1;refGene_gn=LOC102899173;refGene_tx=XM_02
3244744.1;refGene_gn=LOC102899173;refGene_tx=XM_023244745.1;refGene_gn
=LOC102899173;refGene_tx=XM_023244746.1;refGene_gn=LOC102899173;refGen
e_tx=XM_023244747.1;refGene_gn=LOC102899173;refGene_tx=XM_023244748.1;
refGene_gn=LOC102899173;refGene_tx=XM_023244749.1;refGene_gn=LOC102899
173;refGene_tx=XM_023244750.1;refGene_gn=LOC102899173;refGene_tx=XM_02
3244751.1;refGene_gn=LOC102899173;refGene_tx=XM_023244752.1;refGene_gn
=LOC102899173;refGene_tx=XM_023244753.1;refGene_gn=LOC102899173;refGen
e_tx=XM_023244754.1
E1 46803732 46803786
refGene_gn=LOC102899173;refGene_tx=XM_019818045.2;refGene_gn=LOC102899
173;refGene_tx=XM_023244736.1;refGene_gn=LOC102899173;refGene_tx=XM_02
3244737.1;refGene_gn=LOC102899173;refGene_tx=XM_023244738.1;refGene_gn
=LOC102899173;refGene_tx=XM_023244739.1;refGene_gn=LOC102899173;refGen
e_tx=XM_023244740.1;refGene_gn=LOC102899173;refGene_tx=XM_023244741.1;
refGene_gn=LOC102899173;refGene_tx=XM_023244742.1;refGene_gn=LOC102899
173;refGene_tx=XM_023244743.1;refGene_gn=LOC102899173;refGene_tx=XM_02
3244744.1;refGene_gn=LOC102899173;refGene_tx=XM_023244745.1;refGene_gn
=LOC102899173;refGene_tx=XM_023244746.1;refGene_gn=LOC102899173;refGen
e_tx=XM_023244747.1;refGene_gn=LOC102899173;refGene_tx=XM_023244748.1;
refGene_gn=LOC102899173;refGene_tx=XM_023244749.1;refGene_gn=LOC102899
173;refGene_tx=XM_023244750.1;refGene_gn=LOC102899173;refGene_tx=XM_02
3244751.1;refGene_gn=LOC102899173;refGene_tx=XM_023244752.1;refGene_gn
=LOC102899173;refGene_tx=XM_023244753.1;refGene_gn=LOC102899173;refGen
e_tx=XM_023244754.1
E1 46805341 46805446
refGene_gn=LOC102899173;refGene_tx=XM_019818045.2;refGene_gn=LOC102899
173;refGene_tx=XM_023244736.1;refGene_gn=LOC102899173;refGene_tx=XM_02
3244737.1;refGene_gn=LOC102899173;refGene_tx=XM_023244738.1;refGene_gn
=LOC102899173;refGene_tx=XM_023244739.1;refGene_gn=LOC102899173;refGen
e_tx=XM_023244740.1;refGene_gn=LOC102899173;refGene_tx=XM_023244741.1;
refGene_gn=LOC102899173;refGene_tx=XM_023244742.1;refGene_gn=LOC102899

173;refGene_tx=XM_023244743.1;refGene_gn=LOC102899173;refGene_tx=XM_023244744.1;refGene_gn=LOC102899173;refGene_tx=XM_023244745.1;refGene_gn=LOC102899173;refGene_tx=XM_023244746.1;refGene_gn=LOC102899173;refGene_tx=XM_023244747.1;refGene_gn=LOC102899173;refGene_tx=XM_023244748.1;refGene_gn=LOC102899173;refGene_tx=XM_023244749.1;refGene_gn=LOC102899173;refGene_tx=XM_023244750.1;refGene_gn=LOC102899173;refGene_tx=XM_023244751.1;refGene_gn=LOC102899173;refGene_tx=XM_023244752.1;refGene_gn=LOC102899173;refGene_tx=XM_023244753.1;refGene_gn=LOC102899173;refGene_tx=XM_023244754.1
E1 46806475 46806529
refGene_gn=LOC102899173;refGene_tx=XM_019818045.2;refGene_gn=LOC102899173;refGene_tx=XM_023244736.1;refGene_gn=LOC102899173;refGene_tx=XM_023244737.1;refGene_gn=LOC102899173;refGene_tx=XM_023244738.1;refGene_gn=LOC102899173;refGene_tx=XM_023244739.1;refGene_gn=LOC102899173;refGene_tx=XM_023244740.1;refGene_gn=LOC102899173;refGene_tx=XM_023244741.1;refGene_gn=LOC102899173;refGene_tx=XM_023244742.1;refGene_gn=LOC102899173;refGene_tx=XM_023244743.1;refGene_gn=LOC102899173;refGene_tx=XM_023244744.1;refGene_gn=LOC102899173;refGene_tx=XM_023244745.1;refGene_gn=LOC102899173;refGene_tx=XM_023244746.1;refGene_gn=LOC102899173;refGene_tx=XM_023244747.1;refGene_gn=LOC102899173;refGene_tx=XM_023244748.1;refGene_gn=LOC102899173;refGene_tx=XM_023244749.1;refGene_gn=LOC102899173;refGene_tx=XM_023244750.1;refGene_gn=LOC102899173;refGene_tx=XM_023244751.1;refGene_gn=LOC102899173;refGene_tx=XM_023244752.1;refGene_gn=LOC102899173;refGene_tx=XM_023244753.1;refGene_gn=LOC102899173;refGene_tx=XM_023244754.1
E1 46807833 46807938
refGene_gn=LOC102899173;refGene_tx=XM_019818045.2;refGene_gn=LOC102899173;refGene_tx=XM_023244736.1;refGene_gn=LOC102899173;refGene_tx=XM_023244737.1;refGene_gn=LOC102899173;refGene_tx=XM_023244738.1;refGene_gn=LOC102899173;refGene_tx=XM_023244739.1;refGene_gn=LOC102899173;refGene_tx=XM_023244740.1;refGene_gn=LOC102899173;refGene_tx=XM_023244741.1;refGene_gn=LOC102899173;refGene_tx=XM_023244742.1;refGene_gn=LOC102899173;refGene_tx=XM_023244743.1;refGene_gn=LOC102899173;refGene_tx=XM_023244744.1;refGene_gn=LOC102899173;refGene_tx=XM_023244745.1;refGene_gn=LOC102899173;refGene_tx=XM_023244746.1;refGene_gn=LOC102899173;refGene_tx=XM_023244747.1;refGene_gn=LOC102899173;refGene_tx=XM_023244748.1;refGene_gn=LOC102899173;refGene_tx=XM_023244749.1;refGene_gn=LOC102899173;refGene_tx=XM_023244750.1;refGene_gn=LOC102899173;refGene_tx=XM_023244751.1;refGene_gn=LOC102899173;refGene_tx=XM_023244752.1;refGene_gn=LOC102899173;refGene_tx=XM_023244753.1;refGene_gn=LOC102899173;refGene_tx=XM_023244754.1
E1 46809607 46809661
refGene_gn=LOC102899173;refGene_tx=XM_019818045.2;refGene_gn=LOC102899173;refGene_tx=XM_023244736.1;refGene_gn=LOC102899173;refGene_tx=XM_023244737.1;refGene_gn=LOC102899173;refGene_tx=XM_023244738.1;refGene_gn=LOC102899173;refGene_tx=XM_023244739.1;refGene_gn=LOC102899173;refGene_tx=XM_023244740.1;refGene_gn=LOC102899173;refGene_tx=XM_023244741.1;refGene_gn=LOC102899173;refGene_tx=XM_023244742.1;refGene_gn=LOC102899173;refGene_tx=XM_023244743.1;refGene_gn=LOC102899173;refGene_tx=XM_023244744.1;refGene_gn=LOC102899173;refGene_tx=XM_023244745.1;refGene_gn

=LOC102899173;refGene_tx=XM_023244746.1;refGene_gn=LOC102899173;refGene_tx=XM_023244747.1;refGene_gn=LOC102899173;refGene_tx=XM_023244748.1;refGene_gn=LOC102899173;refGene_tx=XM_023244749.1;refGene_gn=LOC102899173;refGene_tx=XM_023244750.1;refGene_gn=LOC102899173;refGene_tx=XM_023244751.1;refGene_gn=LOC102899173;refGene_tx=XM_023244752.1;refGene_gn=LOC102899173;refGene_tx=XM_023244753.1;refGene_gn=LOC102899173;refGene_tx=XM_023244754.1

E1 46810490 46810595

refGene_gn=LOC102899173;refGene_tx=XM_019818045.2;refGene_gn=LOC102899173;refGene_tx=XM_023244736.1;refGene_gn=LOC102899173;refGene_tx=XM_023244737.1;refGene_gn=LOC102899173;refGene_tx=XM_023244738.1;refGene_gn=LOC102899173;refGene_tx=XM_023244739.1;refGene_gn=LOC102899173;refGene_tx=XM_023244740.1;refGene_gn=LOC102899173;refGene_tx=XM_023244741.1;refGene_gn=LOC102899173;refGene_tx=XM_023244742.1;refGene_gn=LOC102899173;refGene_tx=XM_023244743.1;refGene_gn=LOC102899173;refGene_tx=XM_023244744.1;refGene_gn=LOC102899173;refGene_tx=XM_023244745.1;refGene_gn=LOC102899173;refGene_tx=XM_023244746.1;refGene_gn=LOC102899173;refGene_tx=XM_023244747.1;refGene_gn=LOC102899173;refGene_tx=XM_023244748.1;refGene_gn=LOC102899173;refGene_tx=XM_023244749.1;refGene_gn=LOC102899173;refGene_tx=XM_023244750.1;refGene_gn=LOC102899173;refGene_tx=XM_023244751.1;refGene_gn=LOC102899173;refGene_tx=XM_023244752.1;refGene_gn=LOC102899173;refGene_tx=XM_023244753.1;refGene_gn=LOC102899173;refGene_tx=XM_023244754.1

E1 46815065 46815119

refGene_gn=LOC102899173;refGene_tx=XM_019818045.2;refGene_gn=LOC102899173;refGene_tx=XM_023244736.1;refGene_gn=LOC102899173;refGene_tx=XM_023244737.1;refGene_gn=LOC102899173;refGene_tx=XM_023244738.1;refGene_gn=LOC102899173;refGene_tx=XM_023244739.1;refGene_gn=LOC102899173;refGene_tx=XM_023244740.1;refGene_gn=LOC102899173;refGene_tx=XM_023244741.1;refGene_gn=LOC102899173;refGene_tx=XM_023244742.1;refGene_gn=LOC102899173;refGene_tx=XM_023244743.1;refGene_gn=LOC102899173;refGene_tx=XM_023244744.1;refGene_gn=LOC102899173;refGene_tx=XM_023244745.1;refGene_gn=LOC102899173;refGene_tx=XM_023244746.1;refGene_gn=LOC102899173;refGene_tx=XM_023244747.1;refGene_gn=LOC102899173;refGene_tx=XM_023244748.1;refGene_gn=LOC102899173;refGene_tx=XM_023244749.1;refGene_gn=LOC102899173;refGene_tx=XM_023244750.1;refGene_gn=LOC102899173;refGene_tx=XM_023244751.1;refGene_gn=LOC102899173;refGene_tx=XM_023244752.1;refGene_gn=LOC102899173;refGene_tx=XM_023244753.1;refGene_gn=LOC102899173;refGene_tx=XM_023244754.1

E1 46815775 46815880

refGene_gn=LOC102899173;refGene_tx=XM_019818045.2;refGene_gn=LOC102899173;refGene_tx=XM_023244736.1;refGene_gn=LOC102899173;refGene_tx=XM_023244737.1;refGene_gn=LOC102899173;refGene_tx=XM_023244738.1;refGene_gn=LOC102899173;refGene_tx=XM_023244739.1;refGene_gn=LOC102899173;refGene_tx=XM_023244740.1;refGene_gn=LOC102899173;refGene_tx=XM_023244741.1;refGene_gn=LOC102899173;refGene_tx=XM_023244742.1;refGene_gn=LOC102899173;refGene_tx=XM_023244743.1;refGene_gn=LOC102899173;refGene_tx=XM_023244744.1;refGene_gn=LOC102899173;refGene_tx=XM_023244745.1;refGene_gn=LOC102899173;refGene_tx=XM_023244746.1;refGene_gn=LOC102899173;refGene_tx=XM_023244747.1;refGene_gn=LOC102899173;refGene_tx=XM_023244748.1;

refGene_gn=LOC102899173;refGene_tx=XM_023244749.1;refGene_gn=LOC102899
173;refGene_tx=XM_023244750.1;refGene_gn=LOC102899173;refGene_tx=XM_02
3244751.1;refGene_gn=LOC102899173;refGene_tx=XM_023244752.1;refGene_gn
=LOC102899173;refGene_tx=XM_023244753.1;refGene_gn=LOC102899173;refGen
e_tx=XM_023244754.1
E1 46816427 46816481
refGene_gn=LOC102899173;refGene_tx=XM_019818045.2;refGene_gn=LOC102899
173;refGene_tx=XM_023244736.1;refGene_gn=LOC102899173;refGene_tx=XM_02
3244737.1;refGene_gn=LOC102899173;refGene_tx=XM_023244738.1;refGene_gn
=LOC102899173;refGene_tx=XM_023244739.1;refGene_gn=LOC102899173;refGen
e_tx=XM_023244740.1;refGene_gn=LOC102899173;refGene_tx=XM_023244741.1;
refGene_gn=LOC102899173;refGene_tx=XM_023244742.1;refGene_gn=LOC102899
173;refGene_tx=XM_023244743.1;refGene_gn=LOC102899173;refGene_tx=XM_02
3244744.1;refGene_gn=LOC102899173;refGene_tx=XM_023244745.1;refGene_gn
=LOC102899173;refGene_tx=XM_023244746.1;refGene_gn=LOC102899173;refGen
e_tx=XM_023244747.1;refGene_gn=LOC102899173;refGene_tx=XM_023244748.1;
refGene_gn=LOC102899173;refGene_tx=XM_023244749.1;refGene_gn=LOC102899
173;refGene_tx=XM_023244750.1;refGene_gn=LOC102899173;refGene_tx=XM_02
3244751.1;refGene_gn=LOC102899173;refGene_tx=XM_023244752.1;refGene_gn
=LOC102899173;refGene_tx=XM_023244753.1;refGene_gn=LOC102899173;refGen
e_tx=XM_023244754.1
E1 46817315 46817420
refGene_gn=LOC102899173;refGene_tx=XM_019818045.2;refGene_gn=LOC102899
173;refGene_tx=XM_023244736.1;refGene_gn=LOC102899173;refGene_tx=XM_02
3244737.1;refGene_gn=LOC102899173;refGene_tx=XM_023244738.1;refGene_gn
=LOC102899173;refGene_tx=XM_023244739.1;refGene_gn=LOC102899173;refGen
e_tx=XM_023244740.1;refGene_gn=LOC102899173;refGene_tx=XM_023244741.1;
refGene_gn=LOC102899173;refGene_tx=XM_023244742.1;refGene_gn=LOC102899
173;refGene_tx=XM_023244743.1;refGene_gn=LOC102899173;refGene_tx=XM_02
3244744.1;refGene_gn=LOC102899173;refGene_tx=XM_023244745.1;refGene_gn
=LOC102899173;refGene_tx=XM_023244746.1;refGene_gn=LOC102899173;refGen
e_tx=XM_023244747.1;refGene_gn=LOC102899173;refGene_tx=XM_023244748.1;
refGene_gn=LOC102899173;refGene_tx=XM_023244749.1;refGene_gn=LOC102899
173;refGene_tx=XM_023244750.1;refGene_gn=LOC102899173;refGene_tx=XM_02
3244751.1;refGene_gn=LOC102899173;refGene_tx=XM_023244752.1;refGene_gn
=LOC102899173;refGene_tx=XM_023244753.1;refGene_gn=LOC102899173;refGen
e_tx=XM_023244754.1
E1 46820508 46820559
refGene_gn=LOC102899173;refGene_tx=XM_019818045.2;refGene_gn=LOC102899
173;refGene_tx=XM_023244736.1;refGene_gn=LOC102899173;refGene_tx=XM_02
3244737.1;refGene_gn=LOC102899173;refGene_tx=XM_023244738.1;refGene_gn
=LOC102899173;refGene_tx=XM_023244739.1;refGene_gn=LOC102899173;refGen
e_tx=XM_023244740.1;refGene_gn=LOC102899173;refGene_tx=XM_023244741.1;
refGene_gn=LOC102899173;refGene_tx=XM_023244742.1;refGene_gn=LOC102899
173;refGene_tx=XM_023244743.1;refGene_gn=LOC102899173;refGene_tx=XM_02
3244744.1;refGene_gn=LOC102899173;refGene_tx=XM_023244745.1;refGene_gn
=LOC102899173;refGene_tx=XM_023244746.1;refGene_gn=LOC102899173;refGen
e_tx=XM_023244747.1;refGene_gn=LOC102899173;refGene_tx=XM_023244748.1;
refGene_gn=LOC102899173;refGene_tx=XM_023244749.1;refGene_gn=LOC102899
173;refGene_tx=XM_023244750.1;refGene_gn=LOC102899173;refGene_tx=XM_02

3244751.1;refGene_gn=LOC102899173;refGene_tx=XM_023244752.1;refGene_gn=LOC102899173;refGene_tx=XM_023244753.1;refGene_gn=LOC102899173;refGene_tx=XM_023244754.1
E1 46821201 46821306
refGene_gn=LOC102899173;refGene_tx=XM_019818045.2;refGene_gn=LOC102899173;refGene_tx=XM_023244736.1;refGene_gn=LOC102899173;refGene_tx=XM_023244737.1;refGene_gn=LOC102899173;refGene_tx=XM_023244738.1;refGene_gn=LOC102899173;refGene_tx=XM_023244739.1;refGene_gn=LOC102899173;refGene_tx=XM_023244740.1;refGene_gn=LOC102899173;refGene_tx=XM_023244741.1;refGene_gn=LOC102899173;refGene_tx=XM_023244742.1;refGene_gn=LOC102899173;refGene_tx=XM_023244743.1;refGene_gn=LOC102899173;refGene_tx=XM_023244744.1;refGene_gn=LOC102899173;refGene_tx=XM_023244745.1;refGene_gn=LOC102899173;refGene_tx=XM_023244746.1;refGene_gn=LOC102899173;refGene_tx=XM_023244747.1;refGene_gn=LOC102899173;refGene_tx=XM_023244748.1;refGene_gn=LOC102899173;refGene_tx=XM_023244749.1;refGene_gn=LOC102899173;refGene_tx=XM_023244750.1;refGene_gn=LOC102899173;refGene_tx=XM_023244751.1;refGene_gn=LOC102899173;refGene_tx=XM_023244752.1;refGene_gn=LOC102899173;refGene_tx=XM_023244753.1;refGene_gn=LOC102899173;refGene_tx=XM_023244754.1
E1 46822124 46822178
refGene_gn=LOC102899173;refGene_tx=XM_019818045.2;refGene_gn=LOC102899173;refGene_tx=XM_023244736.1;refGene_gn=LOC102899173;refGene_tx=XM_023244737.1;refGene_gn=LOC102899173;refGene_tx=XM_023244738.1;refGene_gn=LOC102899173;refGene_tx=XM_023244739.1;refGene_gn=LOC102899173;refGene_tx=XM_023244740.1;refGene_gn=LOC102899173;refGene_tx=XM_023244741.1;refGene_gn=LOC102899173;refGene_tx=XM_023244742.1;refGene_gn=LOC102899173;refGene_tx=XM_023244743.1;refGene_gn=LOC102899173;refGene_tx=XM_023244744.1;refGene_gn=LOC102899173;refGene_tx=XM_023244745.1;refGene_gn=LOC102899173;refGene_tx=XM_023244746.1;refGene_gn=LOC102899173;refGene_tx=XM_023244747.1;refGene_gn=LOC102899173;refGene_tx=XM_023244748.1;refGene_gn=LOC102899173;refGene_tx=XM_023244749.1;refGene_gn=LOC102899173;refGene_tx=XM_023244750.1;refGene_gn=LOC102899173;refGene_tx=XM_023244751.1;refGene_gn=LOC102899173;refGene_tx=XM_023244752.1;refGene_gn=LOC102899173;refGene_tx=XM_023244753.1;refGene_gn=LOC102899173;refGene_tx=XM_023244754.1
E1 46822859 46822964
refGene_gn=LOC102899173;refGene_tx=XM_019818045.2;refGene_gn=LOC102899173;refGene_tx=XM_023244736.1;refGene_gn=LOC102899173;refGene_tx=XM_023244737.1;refGene_gn=LOC102899173;refGene_tx=XM_023244738.1;refGene_gn=LOC102899173;refGene_tx=XM_023244739.1;refGene_gn=LOC102899173;refGene_tx=XM_023244740.1;refGene_gn=LOC102899173;refGene_tx=XM_023244741.1;refGene_gn=LOC102899173;refGene_tx=XM_023244742.1;refGene_gn=LOC102899173;refGene_tx=XM_023244743.1;refGene_gn=LOC102899173;refGene_tx=XM_023244744.1;refGene_gn=LOC102899173;refGene_tx=XM_023244745.1;refGene_gn=LOC102899173;refGene_tx=XM_023244746.1;refGene_gn=LOC102899173;refGene_tx=XM_023244747.1;refGene_gn=LOC102899173;refGene_tx=XM_023244748.1;refGene_gn=LOC102899173;refGene_tx=XM_023244749.1;refGene_gn=LOC102899173;refGene_tx=XM_023244750.1;refGene_gn=LOC102899173;refGene_tx=XM_023244751.1;refGene_gn=LOC102899173;refGene_tx=XM_023244752.1;refGene_gn=LOC102899173;refGene_tx=XM_023244753.1;refGene_gn=LOC102899173;refGen

e_tx=XM_023244754.1
E1 46823643 46823697
refGene_gn=LOC102899173;refGene_tx=XM_019818045.2;refGene_gn=LOC102899173;refGene_tx=XM_023244736.1;refGene_gn=LOC102899173;refGene_tx=XM_023244737.1;refGene_gn=LOC102899173;refGene_tx=XM_023244738.1;refGene_gn=LOC102899173;refGene_tx=XM_023244739.1;refGene_gn=LOC102899173;refGene_tx=XM_023244740.1;refGene_gn=LOC102899173;refGene_tx=XM_023244741.1;refGene_gn=LOC102899173;refGene_tx=XM_023244742.1;refGene_gn=LOC102899173;refGene_tx=XM_023244743.1;refGene_gn=LOC102899173;refGene_tx=XM_023244744.1;refGene_gn=LOC102899173;refGene_tx=XM_023244745.1;refGene_gn=LOC102899173;refGene_tx=XM_023244746.1;refGene_gn=LOC102899173;refGene_tx=XM_023244747.1;refGene_gn=LOC102899173;refGene_tx=XM_023244748.1;refGene_gn=LOC102899173;refGene_tx=XM_023244749.1;refGene_gn=LOC102899173;refGene_tx=XM_023244750.1;refGene_gn=LOC102899173;refGene_tx=XM_023244751.1;refGene_gn=LOC102899173;refGene_tx=XM_023244752.1;refGene_gn=LOC102899173;refGene_tx=XM_023244753.1;refGene_gn=LOC102899173;refGene_tx=XM_023244754.1
E1 46824383 46824488
refGene_gn=LOC102899173;refGene_tx=XM_019818045.2;refGene_gn=LOC102899173;refGene_tx=XM_023244736.1;refGene_gn=LOC102899173;refGene_tx=XM_023244737.1;refGene_gn=LOC102899173;refGene_tx=XM_023244738.1;refGene_gn=LOC102899173;refGene_tx=XM_023244739.1;refGene_gn=LOC102899173;refGene_tx=XM_023244740.1;refGene_gn=LOC102899173;refGene_tx=XM_023244741.1;refGene_gn=LOC102899173;refGene_tx=XM_023244742.1;refGene_gn=LOC102899173;refGene_tx=XM_023244743.1;refGene_gn=LOC102899173;refGene_tx=XM_023244744.1;refGene_gn=LOC102899173;refGene_tx=XM_023244745.1;refGene_gn=LOC102899173;refGene_tx=XM_023244746.1;refGene_gn=LOC102899173;refGene_tx=XM_023244747.1;refGene_gn=LOC102899173;refGene_tx=XM_023244748.1;refGene_gn=LOC102899173;refGene_tx=XM_023244749.1;refGene_gn=LOC102899173;refGene_tx=XM_023244750.1;refGene_gn=LOC102899173;refGene_tx=XM_023244751.1;refGene_gn=LOC102899173;refGene_tx=XM_023244752.1;refGene_gn=LOC102899173;refGene_tx=XM_023244753.1;refGene_gn=LOC102899173;refGene_tx=XM_023244754.1
E1 46836595 46836649
refGene_gn=LOC102899173;refGene_tx=XM_019818045.2;refGene_gn=LOC102899173;refGene_tx=XM_023244736.1;refGene_gn=LOC102899173;refGene_tx=XM_023244737.1;refGene_gn=LOC102899173;refGene_tx=XM_023244738.1;refGene_gn=LOC102899173;refGene_tx=XM_023244739.1;refGene_gn=LOC102899173;refGene_tx=XM_023244740.1;refGene_gn=LOC102899173;refGene_tx=XM_023244741.1;refGene_gn=LOC102899173;refGene_tx=XM_023244742.1;refGene_gn=LOC102899173;refGene_tx=XM_023244743.1;refGene_gn=LOC102899173;refGene_tx=XM_023244744.1;refGene_gn=LOC102899173;refGene_tx=XM_023244745.1;refGene_gn=LOC102899173;refGene_tx=XM_023244746.1;refGene_gn=LOC102899173;refGene_tx=XM_023244747.1;refGene_gn=LOC102899173;refGene_tx=XM_023244748.1;refGene_gn=LOC102899173;refGene_tx=XM_023244749.1;refGene_gn=LOC102899173;refGene_tx=XM_023244750.1;refGene_gn=LOC102899173;refGene_tx=XM_023244751.1;refGene_gn=LOC102899173;refGene_tx=XM_023244752.1;refGene_gn=LOC102899173;refGene_tx=XM_023244753.1;refGene_gn=LOC102899173;refGene_tx=XM_023244754.1
E1 46837146 46837251

refGene_gn=LOC102899173;refGene_tx=XM_019818045.2;refGene_gn=LOC102899173;refGene_tx=XM_023244736.1;refGene_gn=LOC102899173;refGene_tx=XM_023244737.1;refGene_gn=LOC102899173;refGene_tx=XM_023244738.1;refGene_gn=LOC102899173;refGene_tx=XM_023244739.1;refGene_gn=LOC102899173;refGene_tx=XM_023244740.1;refGene_gn=LOC102899173;refGene_tx=XM_023244741.1;refGene_gn=LOC102899173;refGene_tx=XM_023244742.1;refGene_gn=LOC102899173;refGene_tx=XM_023244743.1;refGene_gn=LOC102899173;refGene_tx=XM_023244744.1;refGene_gn=LOC102899173;refGene_tx=XM_023244745.1;refGene_gn=LOC102899173;refGene_tx=XM_023244746.1;refGene_gn=LOC102899173;refGene_tx=XM_023244747.1;refGene_gn=LOC102899173;refGene_tx=XM_023244748.1;refGene_gn=LOC102899173;refGene_tx=XM_023244749.1;refGene_gn=LOC102899173;refGene_tx=XM_023244750.1;refGene_gn=LOC102899173;refGene_tx=XM_023244751.1;refGene_gn=LOC102899173;refGene_tx=XM_023244752.1;refGene_gn=LOC102899173;refGene_tx=XM_023244753.1;refGene_gn=LOC102899173;refGene_tx=XM_023244754.1
E1 46848033 46848087
refGene_gn=LOC102899173;refGene_tx=XM_019818045.2;refGene_gn=LOC102899173;refGene_tx=XM_023244736.1;refGene_gn=LOC102899173;refGene_tx=XM_023244737.1;refGene_gn=LOC102899173;refGene_tx=XM_023244738.1;refGene_gn=LOC102899173;refGene_tx=XM_023244739.1;refGene_gn=LOC102899173;refGene_tx=XM_023244740.1;refGene_gn=LOC102899173;refGene_tx=XM_023244741.1;refGene_gn=LOC102899173;refGene_tx=XM_023244742.1;refGene_gn=LOC102899173;refGene_tx=XM_023244743.1;refGene_gn=LOC102899173;refGene_tx=XM_023244744.1;refGene_gn=LOC102899173;refGene_tx=XM_023244745.1;refGene_gn=LOC102899173;refGene_tx=XM_023244746.1;refGene_gn=LOC102899173;refGene_tx=XM_023244747.1;refGene_gn=LOC102899173;refGene_tx=XM_023244748.1;refGene_gn=LOC102899173;refGene_tx=XM_023244749.1;refGene_gn=LOC102899173;refGene_tx=XM_023244750.1;refGene_gn=LOC102899173;refGene_tx=XM_023244751.1;refGene_gn=LOC102899173;refGene_tx=XM_023244752.1;refGene_gn=LOC102899173;refGene_tx=XM_023244753.1;refGene_gn=LOC102899173;refGene_tx=XM_023244754.1
E1 46849346 46849451
refGene_gn=LOC102899173;refGene_tx=XM_019818045.2;refGene_gn=LOC102899173;refGene_tx=XM_023244736.1;refGene_gn=LOC102899173;refGene_tx=XM_023244737.1;refGene_gn=LOC102899173;refGene_tx=XM_023244738.1;refGene_gn=LOC102899173;refGene_tx=XM_023244739.1;refGene_gn=LOC102899173;refGene_tx=XM_023244740.1;refGene_gn=LOC102899173;refGene_tx=XM_023244741.1;refGene_gn=LOC102899173;refGene_tx=XM_023244742.1;refGene_gn=LOC102899173;refGene_tx=XM_023244743.1;refGene_gn=LOC102899173;refGene_tx=XM_023244744.1;refGene_gn=LOC102899173;refGene_tx=XM_023244745.1;refGene_gn=LOC102899173;refGene_tx=XM_023244746.1;refGene_gn=LOC102899173;refGene_tx=XM_023244747.1;refGene_gn=LOC102899173;refGene_tx=XM_023244748.1;refGene_gn=LOC102899173;refGene_tx=XM_023244749.1;refGene_gn=LOC102899173;refGene_tx=XM_023244750.1;refGene_gn=LOC102899173;refGene_tx=XM_023244751.1;refGene_gn=LOC102899173;refGene_tx=XM_023244752.1;refGene_gn=LOC102899173;refGene_tx=XM_023244753.1;refGene_gn=LOC102899173;refGene_tx=XM_023244754.1
E1 46859233 46859287
refGene_gn=LOC102899173;refGene_tx=XM_019818045.2;refGene_gn=LOC102899173;refGene_tx=XM_023244736.1;refGene_gn=LOC102899173;refGene_tx=XM_02

3244737.1;refGene_gn=LOC102899173;refGene_tx=XM_023244738.1;refGene_gn
=LOC102899173;refGene_tx=XM_023244739.1;refGene_gn=LOC102899173;refGen
e_tx=XM_023244740.1;refGene_gn=LOC102899173;refGene_tx=XM_023244741.1;
refGene_gn=LOC102899173;refGene_tx=XM_023244742.1;refGene_gn=LOC102899
173;refGene_tx=XM_023244743.1;refGene_gn=LOC102899173;refGene_tx=XM_02
3244744.1;refGene_gn=LOC102899173;refGene_tx=XM_023244745.1;refGene_gn
=LOC102899173;refGene_tx=XM_023244746.1;refGene_gn=LOC102899173;refGen
e_tx=XM_023244747.1;refGene_gn=LOC102899173;refGene_tx=XM_023244748.1;
refGene_gn=LOC102899173;refGene_tx=XM_023244749.1;refGene_gn=LOC102899
173;refGene_tx=XM_023244750.1;refGene_gn=LOC102899173;refGene_tx=XM_02
3244751.1;refGene_gn=LOC102899173;refGene_tx=XM_023244752.1;refGene_gn
=LOC102899173;refGene_tx=XM_023244753.1;refGene_gn=LOC102899173;refGen
e_tx=XM_023244754.1
E1 46859739 46859844
refGene_gn=LOC102899173;refGene_tx=XM_019818045.2;refGene_gn=LOC102899
173;refGene_tx=XM_023244736.1;refGene_gn=LOC102899173;refGene_tx=XM_02
3244737.1;refGene_gn=LOC102899173;refGene_tx=XM_023244738.1;refGene_gn
=LOC102899173;refGene_tx=XM_023244739.1;refGene_gn=LOC102899173;refGen
e_tx=XM_023244740.1;refGene_gn=LOC102899173;refGene_tx=XM_023244741.1;
refGene_gn=LOC102899173;refGene_tx=XM_023244742.1;refGene_gn=LOC102899
173;refGene_tx=XM_023244743.1;refGene_gn=LOC102899173;refGene_tx=XM_02
3244744.1;refGene_gn=LOC102899173;refGene_tx=XM_023244745.1;refGene_gn
=LOC102899173;refGene_tx=XM_023244746.1;refGene_gn=LOC102899173;refGen
e_tx=XM_023244747.1;refGene_gn=LOC102899173;refGene_tx=XM_023244748.1;
refGene_gn=LOC102899173;refGene_tx=XM_023244749.1;refGene_gn=LOC102899
173;refGene_tx=XM_023244750.1;refGene_gn=LOC102899173;refGene_tx=XM_02
3244751.1;refGene_gn=LOC102899173;refGene_tx=XM_023244752.1;refGene_gn
=LOC102899173;refGene_tx=XM_023244753.1;refGene_gn=LOC102899173;refGen
e_tx=XM_023244754.1
E1 46863753 46863807
refGene_gn=LOC102899173;refGene_tx=XM_019818045.2;refGene_gn=LOC102899
173;refGene_tx=XM_023244736.1;refGene_gn=LOC102899173;refGene_tx=XM_02
3244737.1;refGene_gn=LOC102899173;refGene_tx=XM_023244738.1;refGene_gn
=LOC102899173;refGene_tx=XM_023244739.1;refGene_gn=LOC102899173;refGen
e_tx=XM_023244740.1;refGene_gn=LOC102899173;refGene_tx=XM_023244741.1;
refGene_gn=LOC102899173;refGene_tx=XM_023244742.1;refGene_gn=LOC102899
173;refGene_tx=XM_023244743.1;refGene_gn=LOC102899173;refGene_tx=XM_02
3244744.1;refGene_gn=LOC102899173;refGene_tx=XM_023244745.1;refGene_gn
=LOC102899173;refGene_tx=XM_023244746.1;refGene_gn=LOC102899173;refGen
e_tx=XM_023244747.1;refGene_gn=LOC102899173;refGene_tx=XM_023244748.1;
refGene_gn=LOC102899173;refGene_tx=XM_023244749.1;refGene_gn=LOC102899
173;refGene_tx=XM_023244750.1;refGene_gn=LOC102899173;refGene_tx=XM_02
3244751.1;refGene_gn=LOC102899173;refGene_tx=XM_023244752.1;refGene_gn
=LOC102899173;refGene_tx=XM_023244753.1;refGene_gn=LOC102899173;refGen
e_tx=XM_023244754.1
E1 46869026 46869131
refGene_gn=LOC102899173;refGene_tx=XM_019818045.2;refGene_gn=LOC102899
173;refGene_tx=XM_023244736.1;refGene_gn=LOC102899173;refGene_tx=XM_02
3244737.1;refGene_gn=LOC102899173;refGene_tx=XM_023244738.1;refGene_gn
=LOC102899173;refGene_tx=XM_023244739.1;refGene_gn=LOC102899173;refGen

e_tx=XM_023244740.1;refGene_gn=LOC102899173;refGene_tx=XM_023244741.1;
refGene_gn=LOC102899173;refGene_tx=XM_023244742.1;refGene_gn=LOC102899
173;refGene_tx=XM_023244743.1;refGene_gn=LOC102899173;refGene_tx=XM_02
3244744.1;refGene_gn=LOC102899173;refGene_tx=XM_023244745.1;refGene_gn
=LOC102899173;refGene_tx=XM_023244746.1;refGene_gn=LOC102899173;refGen
e_tx=XM_023244747.1;refGene_gn=LOC102899173;refGene_tx=XM_023244748.1;
refGene_gn=LOC102899173;refGene_tx=XM_023244749.1;refGene_gn=LOC102899
173;refGene_tx=XM_023244750.1;refGene_gn=LOC102899173;refGene_tx=XM_02
3244751.1;refGene_gn=LOC102899173;refGene_tx=XM_023244752.1;refGene_gn
=LOC102899173;refGene_tx=XM_023244753.1;refGene_gn=LOC102899173;refGen
e_tx=XM_023244754.1
E1 46869893 46869947
refGene_gn=LOC102899173;refGene_tx=XM_019818045.2;refGene_gn=LOC102899
173;refGene_tx=XM_023244736.1;refGene_gn=LOC102899173;refGene_tx=XM_02
3244737.1;refGene_gn=LOC102899173;refGene_tx=XM_023244738.1;refGene_gn
=LOC102899173;refGene_tx=XM_023244739.1;refGene_gn=LOC102899173;refGen
e_tx=XM_023244740.1;refGene_gn=LOC102899173;refGene_tx=XM_023244741.1;
refGene_gn=LOC102899173;refGene_tx=XM_023244742.1;refGene_gn=LOC102899
173;refGene_tx=XM_023244743.1;refGene_gn=LOC102899173;refGene_tx=XM_02
3244744.1;refGene_gn=LOC102899173;refGene_tx=XM_023244745.1;refGene_gn
=LOC102899173;refGene_tx=XM_023244746.1;refGene_gn=LOC102899173;refGen
e_tx=XM_023244747.1;refGene_gn=LOC102899173;refGene_tx=XM_023244748.1;
refGene_gn=LOC102899173;refGene_tx=XM_023244749.1;refGene_gn=LOC102899
173;refGene_tx=XM_023244750.1;refGene_gn=LOC102899173;refGene_tx=XM_02
3244751.1;refGene_gn=LOC102899173;refGene_tx=XM_023244752.1;refGene_gn
=LOC102899173;refGene_tx=XM_023244753.1;refGene_gn=LOC102899173;refGen
e_tx=XM_023244754.1
E1 46871786 46871891
refGene_gn=LOC102899173;refGene_tx=XM_019818045.2;refGene_gn=LOC102899
173;refGene_tx=XM_023244736.1;refGene_gn=LOC102899173;refGene_tx=XM_02
3244737.1;refGene_gn=LOC102899173;refGene_tx=XM_023244738.1;refGene_gn
=LOC102899173;refGene_tx=XM_023244739.1;refGene_gn=LOC102899173;refGen
e_tx=XM_023244740.1;refGene_gn=LOC102899173;refGene_tx=XM_023244741.1;
refGene_gn=LOC102899173;refGene_tx=XM_023244742.1;refGene_gn=LOC102899
173;refGene_tx=XM_023244743.1;refGene_gn=LOC102899173;refGene_tx=XM_02
3244744.1;refGene_gn=LOC102899173;refGene_tx=XM_023244745.1;refGene_gn
=LOC102899173;refGene_tx=XM_023244746.1;refGene_gn=LOC102899173;refGen
e_tx=XM_023244747.1;refGene_gn=LOC102899173;refGene_tx=XM_023244748.1;
refGene_gn=LOC102899173;refGene_tx=XM_023244749.1;refGene_gn=LOC102899
173;refGene_tx=XM_023244750.1;refGene_gn=LOC102899173;refGene_tx=XM_02
3244751.1;refGene_gn=LOC102899173;refGene_tx=XM_023244752.1;refGene_gn
=LOC102899173;refGene_tx=XM_023244753.1;refGene_gn=LOC102899173;refGen
e_tx=XM_023244754.1
E1 46874224 46874278
refGene_gn=LOC102899173;refGene_tx=XM_019818045.2;refGene_gn=LOC102899
173;refGene_tx=XM_023244736.1;refGene_gn=LOC102899173;refGene_tx=XM_02
3244737.1;refGene_gn=LOC102899173;refGene_tx=XM_023244738.1;refGene_gn
=LOC102899173;refGene_tx=XM_023244739.1;refGene_gn=LOC102899173;refGen
e_tx=XM_023244740.1;refGene_gn=LOC102899173;refGene_tx=XM_023244741.1;
refGene_gn=LOC102899173;refGene_tx=XM_023244742.1;refGene_gn=LOC102899

173;refGene_tx=XM_023244743.1;refGene_gn=LOC102899173;refGene_tx=XM_02
3244744.1;refGene_gn=LOC102899173;refGene_tx=XM_023244745.1;refGene_gn
=LOC102899173;refGene_tx=XM_023244746.1;refGene_gn=LOC102899173;refGen
e_tx=XM_023244747.1;refGene_gn=LOC102899173;refGene_tx=XM_023244748.1;
refGene_gn=LOC102899173;refGene_tx=XM_023244749.1;refGene_gn=LOC102899
173;refGene_tx=XM_023244750.1;refGene_gn=LOC102899173;refGene_tx=XM_02
3244751.1;refGene_gn=LOC102899173;refGene_tx=XM_023244752.1;refGene_gn
=LOC102899173;refGene_tx=XM_023244753.1;refGene_gn=LOC102899173;refGen
e_tx=XM_023244754.1
E1 46880248 46880353
refGene_gn=LOC102899173;refGene_tx=XM_019818045.2;refGene_gn=LOC102899
173;refGene_tx=XM_023244736.1;refGene_gn=LOC102899173;refGene_tx=XM_02
3244737.1;refGene_gn=LOC102899173;refGene_tx=XM_023244738.1;refGene_gn
=LOC102899173;refGene_tx=XM_023244739.1;refGene_gn=LOC102899173;refGen
e_tx=XM_023244740.1;refGene_gn=LOC102899173;refGene_tx=XM_023244741.1;
refGene_gn=LOC102899173;refGene_tx=XM_023244742.1;refGene_gn=LOC102899
173;refGene_tx=XM_023244743.1;refGene_gn=LOC102899173;refGene_tx=XM_02
3244744.1;refGene_gn=LOC102899173;refGene_tx=XM_023244745.1;refGene_gn
=LOC102899173;refGene_tx=XM_023244746.1;refGene_gn=LOC102899173;refGen
e_tx=XM_023244747.1;refGene_gn=LOC102899173;refGene_tx=XM_023244748.1;
refGene_gn=LOC102899173;refGene_tx=XM_023244749.1;refGene_gn=LOC102899
173;refGene_tx=XM_023244750.1;refGene_gn=LOC102899173;refGene_tx=XM_02
3244751.1;refGene_gn=LOC102899173;refGene_tx=XM_023244752.1;refGene_gn
=LOC102899173;refGene_tx=XM_023244753.1;refGene_gn=LOC102899173;refGen
e_tx=XM_023244754.1
E1 46881014 46881068
refGene_gn=LOC102899173;refGene_tx=XM_019818045.2;refGene_gn=LOC102899
173;refGene_tx=XM_023244736.1;refGene_gn=LOC102899173;refGene_tx=XM_02
3244737.1;refGene_gn=LOC102899173;refGene_tx=XM_023244738.1;refGene_gn
=LOC102899173;refGene_tx=XM_023244739.1;refGene_gn=LOC102899173;refGen
e_tx=XM_023244740.1;refGene_gn=LOC102899173;refGene_tx=XM_023244741.1;
refGene_gn=LOC102899173;refGene_tx=XM_023244742.1;refGene_gn=LOC102899
173;refGene_tx=XM_023244743.1;refGene_gn=LOC102899173;refGene_tx=XM_02
3244744.1;refGene_gn=LOC102899173;refGene_tx=XM_023244745.1;refGene_gn
=LOC102899173;refGene_tx=XM_023244746.1;refGene_gn=LOC102899173;refGen
e_tx=XM_023244747.1;refGene_gn=LOC102899173;refGene_tx=XM_023244748.1;
refGene_gn=LOC102899173;refGene_tx=XM_023244749.1;refGene_gn=LOC102899
173;refGene_tx=XM_023244750.1;refGene_gn=LOC102899173;refGene_tx=XM_02
3244751.1;refGene_gn=LOC102899173;refGene_tx=XM_023244752.1;refGene_gn
=LOC102899173;refGene_tx=XM_023244753.1;refGene_gn=LOC102899173;refGen
e_tx=XM_023244754.1
E1 46882966 46883071
refGene_gn=LOC102899173;refGene_tx=XM_019818045.2;refGene_gn=LOC102899
173;refGene_tx=XM_023244736.1;refGene_gn=LOC102899173;refGene_tx=XM_02
3244737.1;refGene_gn=LOC102899173;refGene_tx=XM_023244738.1;refGene_gn
=LOC102899173;refGene_tx=XM_023244739.1;refGene_gn=LOC102899173;refGen
e_tx=XM_023244740.1;refGene_gn=LOC102899173;refGene_tx=XM_023244741.1;
refGene_gn=LOC102899173;refGene_tx=XM_023244742.1;refGene_gn=LOC102899
173;refGene_tx=XM_023244743.1;refGene_gn=LOC102899173;refGene_tx=XM_02
3244744.1;refGene_gn=LOC102899173;refGene_tx=XM_023244745.1;refGene_gn

=LOC102899173;refGene_tx=XM_023244746.1;refGene_gn=LOC102899173;refGene_tx=XM_023244747.1;refGene_gn=LOC102899173;refGene_tx=XM_023244748.1;refGene_gn=LOC102899173;refGene_tx=XM_023244749.1;refGene_gn=LOC102899173;refGene_tx=XM_023244750.1;refGene_gn=LOC102899173;refGene_tx=XM_023244751.1;refGene_gn=LOC102899173;refGene_tx=XM_023244752.1;refGene_gn=LOC102899173;refGene_tx=XM_023244753.1;refGene_gn=LOC102899173;refGene_tx=XM_023244754.1
E1 46890533 46890587
refGene_gn=LOC102899173;refGene_tx=XM_019818045.2;refGene_gn=LOC102899173;refGene_tx=XM_023244736.1;refGene_gn=LOC102899173;refGene_tx=XM_023244737.1;refGene_gn=LOC102899173;refGene_tx=XM_023244738.1;refGene_gn=LOC102899173;refGene_tx=XM_023244739.1;refGene_gn=LOC102899173;refGene_tx=XM_023244740.1;refGene_gn=LOC102899173;refGene_tx=XM_023244741.1;refGene_gn=LOC102899173;refGene_tx=XM_023244742.1;refGene_gn=LOC102899173;refGene_tx=XM_023244743.1;refGene_gn=LOC102899173;refGene_tx=XM_023244744.1;refGene_gn=LOC102899173;refGene_tx=XM_023244745.1;refGene_gn=LOC102899173;refGene_tx=XM_023244746.1;refGene_gn=LOC102899173;refGene_tx=XM_023244747.1;refGene_gn=LOC102899173;refGene_tx=XM_023244748.1;refGene_gn=LOC102899173;refGene_tx=XM_023244749.1;refGene_gn=LOC102899173;refGene_tx=XM_023244750.1;refGene_gn=LOC102899173;refGene_tx=XM_023244751.1;refGene_gn=LOC102899173;refGene_tx=XM_023244752.1;refGene_gn=LOC102899173;refGene_tx=XM_023244753.1;refGene_gn=LOC102899173;refGene_tx=XM_023244754.1
E1 46891228 46891333
refGene_gn=LOC102899173;refGene_tx=XM_019818045.2;refGene_gn=LOC102899173;refGene_tx=XM_023244736.1;refGene_gn=LOC102899173;refGene_tx=XM_023244737.1;refGene_gn=LOC102899173;refGene_tx=XM_023244738.1;refGene_gn=LOC102899173;refGene_tx=XM_023244739.1;refGene_gn=LOC102899173;refGene_tx=XM_023244740.1;refGene_gn=LOC102899173;refGene_tx=XM_023244741.1;refGene_gn=LOC102899173;refGene_tx=XM_023244742.1;refGene_gn=LOC102899173;refGene_tx=XM_023244743.1;refGene_gn=LOC102899173;refGene_tx=XM_023244744.1;refGene_gn=LOC102899173;refGene_tx=XM_023244745.1;refGene_gn=LOC102899173;refGene_tx=XM_023244746.1;refGene_gn=LOC102899173;refGene_tx=XM_023244747.1;refGene_gn=LOC102899173;refGene_tx=XM_023244748.1;refGene_gn=LOC102899173;refGene_tx=XM_023244749.1;refGene_gn=LOC102899173;refGene_tx=XM_023244750.1;refGene_gn=LOC102899173;refGene_tx=XM_023244751.1;refGene_gn=LOC102899173;refGene_tx=XM_023244752.1;refGene_gn=LOC102899173;refGene_tx=XM_023244753.1;refGene_gn=LOC102899173;refGene_tx=XM_023244754.1
E1 46892902 46892956
refGene_gn=LOC102899173;refGene_tx=XM_019818045.2;refGene_gn=LOC102899173;refGene_tx=XM_023244736.1;refGene_gn=LOC102899173;refGene_tx=XM_023244737.1;refGene_gn=LOC102899173;refGene_tx=XM_023244738.1;refGene_gn=LOC102899173;refGene_tx=XM_023244739.1;refGene_gn=LOC102899173;refGene_tx=XM_023244740.1;refGene_gn=LOC102899173;refGene_tx=XM_023244741.1;refGene_gn=LOC102899173;refGene_tx=XM_023244742.1;refGene_gn=LOC102899173;refGene_tx=XM_023244743.1;refGene_gn=LOC102899173;refGene_tx=XM_023244744.1;refGene_gn=LOC102899173;refGene_tx=XM_023244745.1;refGene_gn=LOC102899173;refGene_tx=XM_023244746.1;refGene_gn=LOC102899173;refGene_tx=XM_023244747.1;refGene_gn=LOC102899173;refGene_tx=XM_023244748.1;

refGene_gn=LOC102899173;refGene_tx=XM_023244749.1;refGene_gn=LOC102899
173;refGene_tx=XM_023244750.1;refGene_gn=LOC102899173;refGene_tx=XM_02
3244751.1;refGene_gn=LOC102899173;refGene_tx=XM_023244752.1;refGene_gn
=LOC102899173;refGene_tx=XM_023244753.1;refGene_gn=LOC102899173;refGen
e_tx=XM_023244754.1
E1 46893851 46893956
refGene_gn=LOC102899173;refGene_tx=XM_019818045.2;refGene_gn=LOC102899
173;refGene_tx=XM_023244736.1;refGene_gn=LOC102899173;refGene_tx=XM_02
3244737.1;refGene_gn=LOC102899173;refGene_tx=XM_023244738.1;refGene_gn
=LOC102899173;refGene_tx=XM_023244739.1;refGene_gn=LOC102899173;refGen
e_tx=XM_023244740.1;refGene_gn=LOC102899173;refGene_tx=XM_023244741.1;
refGene_gn=LOC102899173;refGene_tx=XM_023244742.1;refGene_gn=LOC102899
173;refGene_tx=XM_023244743.1;refGene_gn=LOC102899173;refGene_tx=XM_02
3244744.1;refGene_gn=LOC102899173;refGene_tx=XM_023244746.1;refGene_gn
=LOC102899173;refGene_tx=XM_023244747.1;refGene_gn=LOC102899173;refGen
e_tx=XM_023244748.1;refGene_gn=LOC102899173;refGene_tx=XM_023244749.1;
refGene_gn=LOC102899173;refGene_tx=XM_023244750.1;refGene_gn=LOC102899
173;refGene_tx=XM_023244751.1;refGene_gn=LOC102899173;refGene_tx=XM_02
3244752.1;refGene_gn=LOC102899173;refGene_tx=XM_023244753.1;refGene_gn
=LOC102899173;refGene_tx=XM_023244754.1
E1 46896344 46896398
refGene_gn=LOC102899173;refGene_tx=XM_019818045.2;refGene_gn=LOC102899
173;refGene_tx=XM_023244736.1;refGene_gn=LOC102899173;refGene_tx=XM_02
3244737.1;refGene_gn=LOC102899173;refGene_tx=XM_023244738.1;refGene_gn
=LOC102899173;refGene_tx=XM_023244739.1;refGene_gn=LOC102899173;refGen
e_tx=XM_023244740.1;refGene_gn=LOC102899173;refGene_tx=XM_023244741.1;
refGene_gn=LOC102899173;refGene_tx=XM_023244742.1;refGene_gn=LOC102899
173;refGene_tx=XM_023244743.1;refGene_gn=LOC102899173;refGene_tx=XM_02
3244744.1;refGene_gn=LOC102899173;refGene_tx=XM_023244745.1;refGene_gn
=LOC102899173;refGene_tx=XM_023244746.1;refGene_gn=LOC102899173;refGen
e_tx=XM_023244747.1;refGene_gn=LOC102899173;refGene_tx=XM_023244748.1;
refGene_gn=LOC102899173;refGene_tx=XM_023244749.1;refGene_gn=LOC102899
173;refGene_tx=XM_023244750.1;refGene_gn=LOC102899173;refGene_tx=XM_02
3244751.1;refGene_gn=LOC102899173;refGene_tx=XM_023244752.1;refGene_gn
=LOC102899173;refGene_tx=XM_023244753.1;refGene_gn=LOC102899173;refGen
e_tx=XM_023244754.1
E1 46897247 46897352
refGene_gn=LOC102899173;refGene_tx=XM_019818045.2;refGene_gn=LOC102899
173;refGene_tx=XM_023244736.1;refGene_gn=LOC102899173;refGene_tx=XM_02
3244737.1;refGene_gn=LOC102899173;refGene_tx=XM_023244738.1;refGene_gn
=LOC102899173;refGene_tx=XM_023244739.1;refGene_gn=LOC102899173;refGen
e_tx=XM_023244740.1;refGene_gn=LOC102899173;refGene_tx=XM_023244741.1;
refGene_gn=LOC102899173;refGene_tx=XM_023244742.1;refGene_gn=LOC102899
173;refGene_tx=XM_023244743.1;refGene_gn=LOC102899173;refGene_tx=XM_02
3244744.1;refGene_gn=LOC102899173;refGene_tx=XM_023244745.1;refGene_gn
=LOC102899173;refGene_tx=XM_023244746.1;refGene_gn=LOC102899173;refGen
e_tx=XM_023244747.1;refGene_gn=LOC102899173;refGene_tx=XM_023244748.1;
refGene_gn=LOC102899173;refGene_tx=XM_023244749.1;refGene_gn=LOC102899
173;refGene_tx=XM_023244750.1;refGene_gn=LOC102899173;refGene_tx=XM_02
3244751.1;refGene_gn=LOC102899173;refGene_tx=XM_023244752.1;refGene_gn

=LOC102899173;refGene_tx=XM_023244753.1;refGene_gn=LOC102899173;refGene_tx=XM_023244754.1
E1 46898128 46898182
refGene_gn=LOC102899173;refGene_tx=XM_019818045.2;refGene_gn=LOC102899173;refGene_tx=XM_023244736.1;refGene_gn=LOC102899173;refGene_tx=XM_023244737.1;refGene_gn=LOC102899173;refGene_tx=XM_023244738.1;refGene_gn=LOC102899173;refGene_tx=XM_023244739.1;refGene_gn=LOC102899173;refGene_tx=XM_023244740.1;refGene_gn=LOC102899173;refGene_tx=XM_023244741.1;refGene_gn=LOC102899173;refGene_tx=XM_023244742.1;refGene_gn=LOC102899173;refGene_tx=XM_023244743.1;refGene_gn=LOC102899173;refGene_tx=XM_023244744.1;refGene_gn=LOC102899173;refGene_tx=XM_023244745.1;refGene_gn=LOC102899173;refGene_tx=XM_023244746.1;refGene_gn=LOC102899173;refGene_tx=XM_023244747.1;refGene_gn=LOC102899173;refGene_tx=XM_023244748.1;refGene_gn=LOC102899173;refGene_tx=XM_023244749.1;refGene_gn=LOC102899173;refGene_tx=XM_023244750.1;refGene_gn=LOC102899173;refGene_tx=XM_023244751.1;refGene_gn=LOC102899173;refGene_tx=XM_023244752.1;refGene_gn=LOC102899173;refGene_tx=XM_023244753.1;refGene_gn=LOC102899173;refGene_tx=XM_023244754.1
E1 46899794 46899899
refGene_gn=LOC102899173;refGene_tx=XM_019818045.2;refGene_gn=LOC102899173;refGene_tx=XM_023244736.1;refGene_gn=LOC102899173;refGene_tx=XM_023244737.1;refGene_gn=LOC102899173;refGene_tx=XM_023244738.1;refGene_gn=LOC102899173;refGene_tx=XM_023244739.1;refGene_gn=LOC102899173;refGene_tx=XM_023244740.1;refGene_gn=LOC102899173;refGene_tx=XM_023244741.1;refGene_gn=LOC102899173;refGene_tx=XM_023244742.1;refGene_gn=LOC102899173;refGene_tx=XM_023244743.1;refGene_gn=LOC102899173;refGene_tx=XM_023244744.1;refGene_gn=LOC102899173;refGene_tx=XM_023244745.1;refGene_gn=LOC102899173;refGene_tx=XM_023244746.1;refGene_gn=LOC102899173;refGene_tx=XM_023244747.1;refGene_gn=LOC102899173;refGene_tx=XM_023244748.1;refGene_gn=LOC102899173;refGene_tx=XM_023244749.1;refGene_gn=LOC102899173;refGene_tx=XM_023244750.1;refGene_gn=LOC102899173;refGene_tx=XM_023244751.1;refGene_gn=LOC102899173;refGene_tx=XM_023244752.1;refGene_gn=LOC102899173;refGene_tx=XM_023244753.1;refGene_gn=LOC102899173;refGene_tx=XM_023244754.1
E1 46907183 46907237
refGene_gn=LOC102899173;refGene_tx=XM_019818045.2;refGene_gn=LOC102899173;refGene_tx=XM_023244736.1;refGene_gn=LOC102899173;refGene_tx=XM_023244737.1;refGene_gn=LOC102899173;refGene_tx=XM_023244738.1;refGene_gn=LOC102899173;refGene_tx=XM_023244739.1;refGene_gn=LOC102899173;refGene_tx=XM_023244740.1;refGene_gn=LOC102899173;refGene_tx=XM_023244741.1;refGene_gn=LOC102899173;refGene_tx=XM_023244742.1;refGene_gn=LOC102899173;refGene_tx=XM_023244743.1;refGene_gn=LOC102899173;refGene_tx=XM_023244744.1;refGene_gn=LOC102899173;refGene_tx=XM_023244745.1;refGene_gn=LOC102899173;refGene_tx=XM_023244746.1;refGene_gn=LOC102899173;refGene_tx=XM_023244747.1;refGene_gn=LOC102899173;refGene_tx=XM_023244748.1;refGene_gn=LOC102899173;refGene_tx=XM_023244749.1;refGene_gn=LOC102899173;refGene_tx=XM_023244750.1;refGene_gn=LOC102899173;refGene_tx=XM_023244751.1;refGene_gn=LOC102899173;refGene_tx=XM_023244752.1;refGene_gn=LOC102899173;refGene_tx=XM_023244753.1;refGene_gn=LOC102899173;refGene_tx=XM_023244754.1

E1 46908362 46908467
refGene_gn=LOC102899173;refGene_tx=XM_019818045.2;refGene_gn=LOC102899173;refGene_tx=XM_023244736.1;refGene_gn=LOC102899173;refGene_tx=XM_023244737.1;refGene_gn=LOC102899173;refGene_tx=XM_023244738.1;refGene_gn=LOC102899173;refGene_tx=XM_023244739.1;refGene_gn=LOC102899173;refGene_tx=XM_023244740.1;refGene_gn=LOC102899173;refGene_tx=XM_023244741.1;refGene_gn=LOC102899173;refGene_tx=XM_023244742.1;refGene_gn=LOC102899173;refGene_tx=XM_023244743.1;refGene_gn=LOC102899173;refGene_tx=XM_023244744.1;refGene_gn=LOC102899173;refGene_tx=XM_023244745.1;refGene_gn=LOC102899173;refGene_tx=XM_023244746.1;refGene_gn=LOC102899173;refGene_tx=XM_023244747.1;refGene_gn=LOC102899173;refGene_tx=XM_023244748.1;refGene_gn=LOC102899173;refGene_tx=XM_023244749.1;refGene_gn=LOC102899173;refGene_tx=XM_023244750.1;refGene_gn=LOC102899173;refGene_tx=XM_023244751.1;refGene_gn=LOC102899173;refGene_tx=XM_023244752.1;refGene_gn=LOC102899173;refGene_tx=XM_023244753.1;refGene_gn=LOC102899173;refGene_tx=XM_023244754.1
E1 46912709 46912763
refGene_gn=LOC102899173;refGene_tx=XM_019818045.2;refGene_gn=LOC102899173;refGene_tx=XM_023244736.1;refGene_gn=LOC102899173;refGene_tx=XM_023244737.1;refGene_gn=LOC102899173;refGene_tx=XM_023244738.1;refGene_gn=LOC102899173;refGene_tx=XM_023244739.1;refGene_gn=LOC102899173;refGene_tx=XM_023244740.1;refGene_gn=LOC102899173;refGene_tx=XM_023244741.1;refGene_gn=LOC102899173;refGene_tx=XM_023244742.1;refGene_gn=LOC102899173;refGene_tx=XM_023244743.1;refGene_gn=LOC102899173;refGene_tx=XM_023244744.1;refGene_gn=LOC102899173;refGene_tx=XM_023244745.1;refGene_gn=LOC102899173;refGene_tx=XM_023244746.1;refGene_gn=LOC102899173;refGene_tx=XM_023244747.1;refGene_gn=LOC102899173;refGene_tx=XM_023244748.1;refGene_gn=LOC102899173;refGene_tx=XM_023244749.1;refGene_gn=LOC102899173;refGene_tx=XM_023244750.1;refGene_gn=LOC102899173;refGene_tx=XM_023244751.1;refGene_gn=LOC102899173;refGene_tx=XM_023244752.1;refGene_gn=LOC102899173;refGene_tx=XM_023244753.1;refGene_gn=LOC102899173;refGene_tx=XM_023244754.1
E1 46917605 46917659
refGene_gn=LOC102899173;refGene_tx=XM_019818045.2;refGene_gn=LOC102899173;refGene_tx=XM_023244736.1;refGene_gn=LOC102899173;refGene_tx=XM_023244737.1;refGene_gn=LOC102899173;refGene_tx=XM_023244738.1;refGene_gn=LOC102899173;refGene_tx=XM_023244739.1;refGene_gn=LOC102899173;refGene_tx=XM_023244740.1;refGene_gn=LOC102899173;refGene_tx=XM_023244741.1;refGene_gn=LOC102899173;refGene_tx=XM_023244742.1;refGene_gn=LOC102899173;refGene_tx=XM_023244744.1;refGene_gn=LOC102899173;refGene_tx=XM_023244745.1;refGene_gn=LOC102899173;refGene_tx=XM_023244746.1;refGene_gn=LOC102899173;refGene_tx=XM_023244747.1;refGene_gn=LOC102899173;refGene_tx=XM_023244748.1;refGene_gn=LOC102899173;refGene_tx=XM_023244749.1;refGene_gn=LOC102899173;refGene_tx=XM_023244750.1;refGene_gn=LOC102899173;refGene_tx=XM_023244751.1;refGene_gn=LOC102899173;refGene_tx=XM_023244752.1;refGene_gn=LOC102899173;refGene_tx=XM_023244753.1;refGene_gn=LOC102899173;refGene_tx=XM_023244754.1
E1 46929593 46929684
refGene_gn=LOC102899173;refGene_tx=XM_006940473.4;refGene_gn=LOC102899173;refGene_tx=XM_019818045.2;refGene_gn=LOC102899173;refGene_tx=XM_02

3244736.1;refGene_gn=LOC102899173;refGene_tx=XM_023244737.1;refGene_gn
=LOC102899173;refGene_tx=XM_023244738.1;refGene_gn=LOC102899173;refGen
e_tx=XM_023244739.1;refGene_gn=LOC102899173;refGene_tx=XM_023244740.1;
refGene_gn=LOC102899173;refGene_tx=XM_023244741.1;refGene_gn=LOC102899
173;refGene_tx=XM_023244742.1;refGene_gn=LOC102899173;refGene_tx=XM_02
3244743.1;refGene_gn=LOC102899173;refGene_tx=XM_023244744.1;refGene_gn
=LOC102899173;refGene_tx=XM_023244745.1;refGene_gn=LOC102899173;refGen
e_tx=XM_023244746.1;refGene_gn=LOC102899173;refGene_tx=XM_023244747.1;
refGene_gn=LOC102899173;refGene_tx=XM_023244748.1;refGene_gn=LOC102899
173;refGene_tx=XM_023244749.1;refGene_gn=LOC102899173;refGene_tx=XM_02
3244750.1;refGene_gn=LOC102899173;refGene_tx=XM_023244751.1;refGene_gn
=LOC102899173;refGene_tx=XM_023244752.1;refGene_gn=LOC102899173;refGen
e_tx=XM_023244753.1;refGene_gn=LOC102899173;refGene_tx=XM_023244754.1
E1 46933603 46933680
refGene_gn=LOC102899173;refGene_tx=XM_006940473.4;refGene_gn=LOC102899
173;refGene_tx=XM_019818045.2;refGene_gn=LOC102899173;refGene_tx=XM_02
3244736.1;refGene_gn=LOC102899173;refGene_tx=XM_023244737.1;refGene_gn
=LOC102899173;refGene_tx=XM_023244738.1;refGene_gn=LOC102899173;refGen
e_tx=XM_023244739.1;refGene_gn=LOC102899173;refGene_tx=XM_023244740.1;
refGene_gn=LOC102899173;refGene_tx=XM_023244741.1;refGene_gn=LOC102899
173;refGene_tx=XM_023244742.1;refGene_gn=LOC102899173;refGene_tx=XM_02
3244743.1;refGene_gn=LOC102899173;refGene_tx=XM_023244744.1;refGene_gn
=LOC102899173;refGene_tx=XM_023244745.1;refGene_gn=LOC102899173;refGen
e_tx=XM_023244746.1;refGene_gn=LOC102899173;refGene_tx=XM_023244747.1;
refGene_gn=LOC102899173;refGene_tx=XM_023244748.1;refGene_gn=LOC102899
173;refGene_tx=XM_023244749.1;refGene_gn=LOC102899173;refGene_tx=XM_02
3244750.1;refGene_gn=LOC102899173;refGene_tx=XM_023244751.1;refGene_gn
=LOC102899173;refGene_tx=XM_023244752.1;refGene_gn=LOC102899173;refGen
e_tx=XM_023244753.1;refGene_gn=LOC102899173;refGene_tx=XM_023244754.1
E1 46940694 46940838
refGene_gn=LOC102899173;refGene_tx=XM_006940473.4;refGene_gn=LOC102899
173;refGene_tx=XM_019818045.2;refGene_gn=LOC102899173;refGene_tx=XM_02
3244736.1;refGene_gn=LOC102899173;refGene_tx=XM_023244737.1;refGene_gn
=LOC102899173;refGene_tx=XM_023244738.1;refGene_gn=LOC102899173;refGen
e_tx=XM_023244739.1;refGene_gn=LOC102899173;refGene_tx=XM_023244740.1;
refGene_gn=LOC102899173;refGene_tx=XM_023244741.1;refGene_gn=LOC102899
173;refGene_tx=XM_023244742.1;refGene_gn=LOC102899173;refGene_tx=XM_02
3244743.1;refGene_gn=LOC102899173;refGene_tx=XM_023244744.1;refGene_gn
=LOC102899173;refGene_tx=XM_023244745.1;refGene_gn=LOC102899173;refGen
e_tx=XM_023244746.1;refGene_gn=LOC102899173;refGene_tx=XM_023244747.1;
refGene_gn=LOC102899173;refGene_tx=XM_023244748.1;refGene_gn=LOC102899
173;refGene_tx=XM_023244749.1;refGene_gn=LOC102899173;refGene_tx=XM_02
3244750.1;refGene_gn=LOC102899173;refGene_tx=XM_023244751.1;refGene_gn
=LOC102899173;refGene_tx=XM_023244752.1;refGene_gn=LOC102899173;refGen
e_tx=XM_023244753.1;refGene_gn=LOC102899173;refGene_tx=XM_023244754.1
E1 46942081 46942153
refGene_gn=LOC102899173;refGene_tx=XM_006940473.4;refGene_gn=LOC102899
173;refGene_tx=XM_019818045.2;refGene_gn=LOC102899173;refGene_tx=XM_02
3244736.1;refGene_gn=LOC102899173;refGene_tx=XM_023244737.1;refGene_gn
=LOC102899173;refGene_tx=XM_023244738.1;refGene_gn=LOC102899173;refGen

e_tx=XM_023244739.1;refGene_gn=LOC102899173;refGene_tx=XM_023244740.1;
refGene_gn=LOC102899173;refGene_tx=XM_023244741.1;refGene_gn=LOC102899
173;refGene_tx=XM_023244742.1;refGene_gn=LOC102899173;refGene_tx=XM_02
3244743.1;refGene_gn=LOC102899173;refGene_tx=XM_023244744.1;refGene_gn
=LOC102899173;refGene_tx=XM_023244745.1;refGene_gn=LOC102899173;refGen
e_tx=XM_023244746.1;refGene_gn=LOC102899173;refGene_tx=XM_023244747.1;
refGene_gn=LOC102899173;refGene_tx=XM_023244748.1;refGene_gn=LOC102899
173;refGene_tx=XM_023244749.1;refGene_gn=LOC102899173;refGene_tx=XM_02
3244750.1;refGene_gn=LOC102899173;refGene_tx=XM_023244751.1;refGene_gn
=LOC102899173;refGene_tx=XM_023244752.1;refGene_gn=LOC102899173;refGen
e_tx=XM_023244753.1;refGene_gn=LOC102899173;refGene_tx=XM_023244754.1
E1 46943690 46943811
refGene_gn=LOC102899173;refGene_tx=XM_006940473.4;refGene_gn=LOC102899
173;refGene_tx=XM_019818045.2;refGene_gn=LOC102899173;refGene_tx=XM_02
3244736.1;refGene_gn=LOC102899173;refGene_tx=XM_023244737.1;refGene_gn
=LOC102899173;refGene_tx=XM_023244738.1;refGene_gn=LOC102899173;refGen
e_tx=XM_023244739.1;refGene_gn=LOC102899173;refGene_tx=XM_023244740.1;
refGene_gn=LOC102899173;refGene_tx=XM_023244741.1;refGene_gn=LOC102899
173;refGene_tx=XM_023244742.1;refGene_gn=LOC102899173;refGene_tx=XM_02
3244743.1;refGene_gn=LOC102899173;refGene_tx=XM_023244744.1;refGene_gn
=LOC102899173;refGene_tx=XM_023244745.1;refGene_gn=LOC102899173;refGen
e_tx=XM_023244746.1;refGene_gn=LOC102899173;refGene_tx=XM_023244747.1;
refGene_gn=LOC102899173;refGene_tx=XM_023244748.1;refGene_gn=LOC102899
173;refGene_tx=XM_023244749.1;refGene_gn=LOC102899173;refGene_tx=XM_02
3244750.1;refGene_gn=LOC102899173;refGene_tx=XM_023244751.1;refGene_gn
=LOC102899173;refGene_tx=XM_023244752.1;refGene_gn=LOC102899173;refGen
e_tx=XM_023244753.1;refGene_gn=LOC102899173;refGene_tx=XM_023244754.1
E1 46947403 46947591
refGene_gn=LOC102899173;refGene_tx=XM_006940473.4;refGene_gn=LOC102899
173;refGene_tx=XM_019818045.2;refGene_gn=LOC102899173;refGene_tx=XM_02
3244736.1;refGene_gn=LOC102899173;refGene_tx=XM_023244737.1;refGene_gn
=LOC102899173;refGene_tx=XM_023244738.1;refGene_gn=LOC102899173;refGen
e_tx=XM_023244739.1;refGene_gn=LOC102899173;refGene_tx=XM_023244740.1;
refGene_gn=LOC102899173;refGene_tx=XM_023244741.1;refGene_gn=LOC102899
173;refGene_tx=XM_023244742.1;refGene_gn=LOC102899173;refGene_tx=XM_02
3244743.1;refGene_gn=LOC102899173;refGene_tx=XM_023244744.1;refGene_gn
=LOC102899173;refGene_tx=XM_023244745.1;refGene_gn=LOC102899173;refGen
e_tx=XM_023244746.1;refGene_gn=LOC102899173;refGene_tx=XM_023244747.1;
refGene_gn=LOC102899173;refGene_tx=XM_023244748.1;refGene_gn=LOC102899
173;refGene_tx=XM_023244749.1;refGene_gn=LOC102899173;refGene_tx=XM_02
3244750.1;refGene_gn=LOC102899173;refGene_tx=XM_023244751.1;refGene_gn
=LOC102899173;refGene_tx=XM_023244752.1;refGene_gn=LOC102899173;refGen
e_tx=XM_023244753.1;refGene_gn=LOC102899173;refGene_tx=XM_023244754.1
E1 46947938 46948123
refGene_gn=LOC102899173;refGene_tx=XM_006940473.4;refGene_gn=LOC102899
173;refGene_tx=XM_019818045.2;refGene_gn=LOC102899173;refGene_tx=XM_02
3244736.1;refGene_gn=LOC102899173;refGene_tx=XM_023244737.1;refGene_gn
=LOC102899173;refGene_tx=XM_023244738.1;refGene_gn=LOC102899173;refGen
e_tx=XM_023244739.1;refGene_gn=LOC102899173;refGene_tx=XM_023244740.1;
refGene_gn=LOC102899173;refGene_tx=XM_023244741.1;refGene_gn=LOC102899

173;refGene_tx=XM_023244742.1;refGene_gn=LOC102899173;refGene_tx=XM_023244743.1;refGene_gn=LOC102899173;refGene_tx=XM_023244744.1;refGene_gn=LOC102899173;refGene_tx=XM_023244745.1;refGene_gn=LOC102899173;refGene_tx=XM_023244746.1;refGene_gn=LOC102899173;refGene_tx=XM_023244747.1;refGene_gn=LOC102899173;refGene_tx=XM_023244748.1;refGene_gn=LOC102899173;refGene_tx=XM_023244749.1;refGene_gn=LOC102899173;refGene_tx=XM_023244750.1;refGene_gn=LOC102899173;refGene_tx=XM_023244751.1;refGene_gn=LOC102899173;refGene_tx=XM_023244752.1;refGene_gn=LOC102899173;refGene_tx=XM_023244753.1;refGene_gn=LOC102899173;refGene_tx=XM_023244754.1
E1 46953862 46953985
refGene_gn=LOC102899173;refGene_tx=XM_006940473.4;refGene_gn=LOC102899173;refGene_tx=XM_019818045.2;refGene_gn=LOC102899173;refGene_tx=XM_023244736.1;refGene_gn=LOC102899173;refGene_tx=XM_023244737.1;refGene_gn=LOC102899173;refGene_tx=XM_023244738.1;refGene_gn=LOC102899173;refGene_tx=XM_023244739.1;refGene_gn=LOC102899173;refGene_tx=XM_023244740.1;refGene_gn=LOC102899173;refGene_tx=XM_023244741.1;refGene_gn=LOC102899173;refGene_tx=XM_023244742.1;refGene_gn=LOC102899173;refGene_tx=XM_023244743.1;refGene_gn=LOC102899173;refGene_tx=XM_023244744.1;refGene_gn=LOC102899173;refGene_tx=XM_023244745.1;refGene_gn=LOC102899173;refGene_tx=XM_023244746.1;refGene_gn=LOC102899173;refGene_tx=XM_023244747.1;refGene_gn=LOC102899173;refGene_tx=XM_023244748.1;refGene_gn=LOC102899173;refGene_tx=XM_023244749.1;refGene_gn=LOC102899173;refGene_tx=XM_023244750.1;refGene_gn=LOC102899173;refGene_tx=XM_023244751.1;refGene_gn=LOC102899173;refGene_tx=XM_023244752.1;refGene_gn=LOC102899173;refGene_tx=XM_023244753.1;refGene_gn=LOC102899173;refGene_tx=XM_023244754.1
E1 46956776 46957103
refGene_gn=LOC102899173;refGene_tx=XM_006940473.4;refGene_gn=LOC102899173;refGene_tx=XM_019818045.2;refGene_gn=LOC102899173;refGene_tx=XM_023244736.1;refGene_gn=LOC102899173;refGene_tx=XM_023244737.1;refGene_gn=LOC102899173;refGene_tx=XM_023244738.1;refGene_gn=LOC102899173;refGene_tx=XM_023244739.1;refGene_gn=LOC102899173;refGene_tx=XM_023244740.1;refGene_gn=LOC102899173;refGene_tx=XM_023244741.1;refGene_gn=LOC102899173;refGene_tx=XM_023244742.1;refGene_gn=LOC102899173;refGene_tx=XM_023244743.1;refGene_gn=LOC102899173;refGene_tx=XM_023244744.1;refGene_gn=LOC102899173;refGene_tx=XM_023244745.1;refGene_gn=LOC102899173;refGene_tx=XM_023244746.1;refGene_gn=LOC102899173;refGene_tx=XM_023244747.1;refGene_gn=LOC102899173;refGene_tx=XM_023244748.1;refGene_gn=LOC102899173;refGene_tx=XM_023244749.1;refGene_gn=LOC102899173;refGene_tx=XM_023244750.1;refGene_gn=LOC102899173;refGene_tx=XM_023244751.1;refGene_gn=LOC102899173;refGene_tx=XM_023244752.1;refGene_gn=LOC102899173;refGene_tx=XM_023244753.1;refGene_gn=LOC102899173;refGene_tx=XM_023244754.1
E1 46961586 46961696
refGene_gn=METTL2A;refGene_tx=XM_004001403.5;refGene_gn=METTL2A;refGene_tx=XM_019818053.2;refGene_gn=METTL2A;refGene_tx=XM_019818054.1
E1 46961903 46962005
refGene_gn=METTL2A;refGene_tx=XM_004001403.5;refGene_gn=METTL2A;refGene_tx=XM_019818053.2;refGene_gn=METTL2A;refGene_tx=XM_019818054.1
E1 46963710 46964066
refGene_gn=METTL2A;refGene_tx=XM_004001403.5;refGene_gn=METTL2A;refGene_tx=XM_019818053.2;refGene_gn=METTL2A;refGene_tx=XM_019818054.1

E1 46965121 46965171
refGene_gn=METTL2A;refGene_tx=XM_004001403.5;refGene_gn=METTL2A;refGene_tx=XM_019818053.2;refGene_gn=METTL2A;refGene_tx=XM_019818054.1
E1 46965505 46965566
refGene_gn=METTL2A;refGene_tx=XM_004001403.5;refGene_gn=METTL2A;refGene_tx=XM_019818053.2;refGene_gn=METTL2A;refGene_tx=XM_019818054.1
E1 46967512 46967652
refGene_gn=METTL2A;refGene_tx=XM_004001403.5;refGene_gn=METTL2A;refGene_tx=XM_019818053.2;refGene_gn=METTL2A;refGene_tx=XM_019818054.1
E1 46969450 46969557
refGene_gn=METTL2A;refGene_tx=XM_004001403.5;refGene_gn=METTL2A;refGene_tx=XM_019818053.2;refGene_gn=METTL2A;refGene_tx=XM_019818054.1
E1 46971426 46971631
refGene_gn=METTL2A;refGene_tx=XM_004001403.5;refGene_gn=METTL2A;refGene_tx=XM_019818053.2;refGene_gn=METTL2A;refGene_tx=XM_019818054.1
E1 46972446 46972601
refGene_gn=METTL2A;refGene_tx=XM_004001403.5;refGene_gn=METTL2A;refGene_tx=XM_019818054.1
E1 47004119 47004200
refGene_gn=TLK2;refGene_tx=XM_019818056.2;refGene_gn=TLK2;refGene_tx=XM_019818057.2;refGene_gn=TLK2;refGene_tx=XM_019818059.2;refGene_gn=TLK2;refGene_tx=XM_023244579.1;refGene_gn=TLK2;refGene_tx=XM_023244580.1;refGene_gn=TLK2;refGene_tx=XM_023244581.1;refGene_gn=TLK2;refGene_tx=XM_023244582.1
E1 47026693 47026699 refGene_gn=TLK2;refGene_tx=XM_019818058.2
E1 47040965 47041037
refGene_gn=TLK2;refGene_tx=XM_019818056.2;refGene_gn=TLK2;refGene_tx=XM_019818057.2;refGene_gn=TLK2;refGene_tx=XM_019818058.2;refGene_gn=TLK2;refGene_tx=XM_019818059.2;refGene_gn=TLK2;refGene_tx=XM_023244579.1;refGene_gn=TLK2;refGene_tx=XM_023244580.1
E1 47041232 47041302
refGene_gn=TLK2;refGene_tx=XM_019818056.2;refGene_gn=TLK2;refGene_tx=XM_019818057.2;refGene_gn=TLK2;refGene_tx=XM_019818058.2;refGene_gn=TLK2;refGene_tx=XM_019818059.2;refGene_gn=TLK2;refGene_tx=XM_023244579.1;refGene_gn=TLK2;refGene_tx=XM_023244580.1
E1 47042327 47042371
refGene_gn=TLK2;refGene_tx=XM_019818056.2;refGene_gn=TLK2;refGene_tx=XM_019818057.2;refGene_gn=TLK2;refGene_tx=XM_019818058.2;refGene_gn=TLK2;refGene_tx=XM_019818059.2;refGene_gn=TLK2;refGene_tx=XM_023244579.1;refGene_gn=TLK2;refGene_tx=XM_023244580.1
E1 47043104 47043200
refGene_gn=TLK2;refGene_tx=XM_019818056.2;refGene_gn=TLK2;refGene_tx=XM_019818058.2;refGene_gn=TLK2;refGene_tx=XM_023244579.1;refGene_gn=TLK2;refGene_tx=XM_023244580.1
E1 47052663 47052831
refGene_gn=TLK2;refGene_tx=XM_019818056.2;refGene_gn=TLK2;refGene_tx=XM_019818057.2;refGene_gn=TLK2;refGene_tx=XM_019818058.2;refGene_gn=TLK2;refGene_tx=XM_019818059.2;refGene_gn=TLK2;refGene_tx=XM_023244579.1;refGene_gn=TLK2;refGene_tx=XM_023244580.1;refGene_gn=TLK2;refGene_tx=XM_023244581.1;refGene_gn=TLK2;refGene_tx=XM_023244582.1

E1 47064679 47064775
refGene_gn=TLK2;refGene_tx=XM_019818056.2;refGene_gn=TLK2;refGene_tx=XM_019818057.2;refGene_gn=TLK2;refGene_tx=XM_019818058.2;refGene_gn=TLK2;refGene_tx=XM_019818059.2;refGene_gn=TLK2;refGene_tx=XM_023244579.1;refGene_gn=TLK2;refGene_tx=XM_023244580.1;refGene_gn=TLK2;refGene_tx=XM_023244581.1;refGene_gn=TLK2;refGene_tx=XM_023244582.1
E1 47066231 47066324
refGene_gn=TLK2;refGene_tx=XM_019818056.2;refGene_gn=TLK2;refGene_tx=XM_019818057.2;refGene_gn=TLK2;refGene_tx=XM_019818058.2;refGene_gn=TLK2;refGene_tx=XM_019818059.2;refGene_gn=TLK2;refGene_tx=XM_023244579.1;refGene_gn=TLK2;refGene_tx=XM_023244580.1;refGene_gn=TLK2;refGene_tx=XM_023244581.1;refGene_gn=TLK2;refGene_tx=XM_023244582.1
E1 47071868 47071979
refGene_gn=TLK2;refGene_tx=XM_019818056.2;refGene_gn=TLK2;refGene_tx=XM_019818057.2;refGene_gn=TLK2;refGene_tx=XM_019818058.2;refGene_gn=TLK2;refGene_tx=XM_019818059.2;refGene_gn=TLK2;refGene_tx=XM_023244579.1;refGene_gn=TLK2;refGene_tx=XM_023244580.1;refGene_gn=TLK2;refGene_tx=XM_023244581.1;refGene_gn=TLK2;refGene_tx=XM_023244582.1
E1 47073140 47073277
refGene_gn=TLK2;refGene_tx=XM_019818056.2;refGene_gn=TLK2;refGene_tx=XM_019818057.2;refGene_gn=TLK2;refGene_tx=XM_019818058.2;refGene_gn=TLK2;refGene_tx=XM_019818059.2;refGene_gn=TLK2;refGene_tx=XM_023244579.1;refGene_gn=TLK2;refGene_tx=XM_023244580.1;refGene_gn=TLK2;refGene_tx=XM_023244581.1;refGene_gn=TLK2;refGene_tx=XM_023244582.1
E1 47080641 47080794
refGene_gn=TLK2;refGene_tx=XM_019818056.2;refGene_gn=TLK2;refGene_tx=XM_019818057.2;refGene_gn=TLK2;refGene_tx=XM_019818058.2;refGene_gn=TLK2;refGene_tx=XM_019818059.2;refGene_gn=TLK2;refGene_tx=XM_023244579.1;refGene_gn=TLK2;refGene_tx=XM_023244580.1;refGene_gn=TLK2;refGene_tx=XM_023244581.1;refGene_gn=TLK2;refGene_tx=XM_023244582.1
E1 47085296 47085363
refGene_gn=TLK2;refGene_tx=XM_019818056.2;refGene_gn=TLK2;refGene_tx=XM_019818057.2;refGene_gn=TLK2;refGene_tx=XM_019818058.2;refGene_gn=TLK2;refGene_tx=XM_019818059.2;refGene_gn=TLK2;refGene_tx=XM_023244579.1;refGene_gn=TLK2;refGene_tx=XM_023244580.1;refGene_gn=TLK2;refGene_tx=XM_023244581.1;refGene_gn=TLK2;refGene_tx=XM_023244582.1
E1 47086337 47086435
refGene_gn=TLK2;refGene_tx=XM_019818056.2;refGene_gn=TLK2;refGene_tx=XM_019818057.2;refGene_gn=TLK2;refGene_tx=XM_019818058.2;refGene_gn=TLK2;refGene_tx=XM_019818059.2;refGene_gn=TLK2;refGene_tx=XM_023244579.1;refGene_gn=TLK2;refGene_tx=XM_023244580.1;refGene_gn=TLK2;refGene_tx=XM_023244581.1;refGene_gn=TLK2;refGene_tx=XM_023244582.1
E1 47088441 47088523
refGene_gn=TLK2;refGene_tx=XM_019818056.2;refGene_gn=TLK2;refGene_tx=XM_019818057.2;refGene_gn=TLK2;refGene_tx=XM_019818058.2;refGene_gn=TLK2;refGene_tx=XM_019818059.2;refGene_gn=TLK2;refGene_tx=XM_023244579.1;refGene_gn=TLK2;refGene_tx=XM_023244580.1;refGene_gn=TLK2;refGene_tx=XM_023244581.1;refGene_gn=TLK2;refGene_tx=XM_023244582.1
E1 47093719 47093811
refGene_gn=TLK2;refGene_tx=XM_019818056.2;refGene_gn=TLK2;refGene_tx=X

M_019818057.2;refGene_gn=TLK2;refGene_tx=XM_019818058.2;refGene_gn=TLK2;refGene_tx=XM_019818059.2;refGene_gn=TLK2;refGene_tx=XM_023244579.1;refGene_gn=TLK2;refGene_tx=XM_023244580.1;refGene_gn=TLK2;refGene_tx=XM_023244581.1;refGene_gn=TLK2;refGene_tx=XM_023244582.1
E1 47098590 47098680
refGene_gn=TLK2;refGene_tx=XM_019818056.2;refGene_gn=TLK2;refGene_tx=XM_019818057.2;refGene_gn=TLK2;refGene_tx=XM_019818058.2;refGene_gn=TLK2;refGene_tx=XM_019818059.2;refGene_gn=TLK2;refGene_tx=XM_023244579.1;refGene_gn=TLK2;refGene_tx=XM_023244580.1;refGene_gn=TLK2;refGene_tx=XM_023244581.1;refGene_gn=TLK2;refGene_tx=XM_023244582.1
E1 47101926 47102096
refGene_gn=TLK2;refGene_tx=XM_019818056.2;refGene_gn=TLK2;refGene_tx=XM_019818057.2;refGene_gn=TLK2;refGene_tx=XM_019818058.2;refGene_gn=TLK2;refGene_tx=XM_019818059.2;refGene_gn=TLK2;refGene_tx=XM_023244579.1;refGene_gn=TLK2;refGene_tx=XM_023244580.1;refGene_gn=TLK2;refGene_tx=XM_023244581.1;refGene_gn=TLK2;refGene_tx=XM_023244582.1
E1 47103545 47103684
refGene_gn=TLK2;refGene_tx=XM_019818056.2;refGene_gn=TLK2;refGene_tx=XM_019818057.2;refGene_gn=TLK2;refGene_tx=XM_019818058.2;refGene_gn=TLK2;refGene_tx=XM_019818059.2;refGene_gn=TLK2;refGene_tx=XM_023244579.1;refGene_gn=TLK2;refGene_tx=XM_023244580.1;refGene_gn=TLK2;refGene_tx=XM_023244581.1;refGene_gn=TLK2;refGene_tx=XM_023244582.1
E1 47107664 47107776
refGene_gn=TLK2;refGene_tx=XM_019818056.2;refGene_gn=TLK2;refGene_tx=XM_019818057.2;refGene_gn=TLK2;refGene_tx=XM_019818058.2;refGene_gn=TLK2;refGene_tx=XM_019818059.2;refGene_gn=TLK2;refGene_tx=XM_023244579.1;refGene_gn=TLK2;refGene_tx=XM_023244580.1;refGene_gn=TLK2;refGene_tx=XM_023244581.1;refGene_gn=TLK2;refGene_tx=XM_023244582.1
E1 47109737 47110040
refGene_gn=TLK2;refGene_tx=XM_019818056.2;refGene_gn=TLK2;refGene_tx=XM_019818057.2;refGene_gn=TLK2;refGene_tx=XM_019818058.2;refGene_gn=TLK2;refGene_tx=XM_019818059.2;refGene_gn=TLK2;refGene_tx=XM_023244579.1;refGene_gn=TLK2;refGene_tx=XM_023244580.1;refGene_gn=TLK2;refGene_tx=XM_023244581.1;refGene_gn=TLK2;refGene_tx=XM_023244582.1
E1 47112663 47112837
refGene_gn=TLK2;refGene_tx=XM_019818056.2;refGene_gn=TLK2;refGene_tx=XM_019818059.2
E1 47125389 47125516
refGene_gn=MRC2;refGene_tx=XM_023244718.1;refGene_gn=MRC2;refGene_tx=XM_023244719.1
E1 47153988 47154390
refGene_gn=MRC2;refGene_tx=XM_023244718.1;refGene_gn=MRC2;refGene_tx=XM_023244719.1
E1 47155083 47155257
refGene_gn=MRC2;refGene_tx=XM_023244718.1;refGene_gn=MRC2;refGene_tx=XM_023244719.1
E1 47155455 47155620
refGene_gn=MRC2;refGene_tx=XM_023244718.1;refGene_gn=MRC2;refGene_tx=XM_023244719.1
E1 47155828 47155942

refGene_gn=MRC2;refGene_tx=XM_023244718.1;refGene_gn=MRC2;refGene_tx=XM_023244719.1
E1 47156441 47156585
refGene_gn=MRC2;refGene_tx=XM_023244718.1;refGene_gn=MRC2;refGene_tx=XM_023244719.1
E1 47159177 47159366
refGene_gn=MRC2;refGene_tx=XM_023244718.1;refGene_gn=MRC2;refGene_tx=XM_023244719.1
E1 47159526 47159681
refGene_gn=MRC2;refGene_tx=XM_023244718.1;refGene_gn=MRC2;refGene_tx=XM_023244719.1
E1 47160618 47160726
refGene_gn=MRC2;refGene_tx=XM_023244718.1;refGene_gn=MRC2;refGene_tx=XM_023244719.1
E1 47162663 47162779
refGene_gn=MRC2;refGene_tx=XM_023244718.1;refGene_gn=MRC2;refGene_tx=XM_023244719.1
E1 47163250 47163399
refGene_gn=MRC2;refGene_tx=XM_023244718.1;refGene_gn=MRC2;refGene_tx=XM_023244719.1
E1 47163972 47164190
refGene_gn=MRC2;refGene_tx=XM_023244718.1;refGene_gn=MRC2;refGene_tx=XM_023244719.1
E1 47164874 47165017
refGene_gn=MRC2;refGene_tx=XM_023244718.1;refGene_gn=MRC2;refGene_tx=XM_023244719.1
E1 47166048 47166151
refGene_gn=MRC2;refGene_tx=XM_023244718.1;refGene_gn=MRC2;refGene_tx=XM_023244719.1
E1 47166418 47166557
refGene_gn=MRC2;refGene_tx=XM_023244718.1;refGene_gn=MRC2;refGene_tx=XM_023244719.1
E1 47166671 47166707
refGene_gn=MRC2;refGene_tx=XM_023244718.1;refGene_gn=MRC2;refGene_tx=XM_023244719.1
E1 47167020 47167181
refGene_gn=MRC2;refGene_tx=XM_023244718.1;refGene_gn=MRC2;refGene_tx=XM_023244719.1
E1 47167267 47167335
refGene_gn=MRC2;refGene_tx=XM_023244718.1;refGene_gn=MRC2;refGene_tx=XM_023244719.1
E1 47168323 47168424
refGene_gn=MRC2;refGene_tx=XM_023244718.1;refGene_gn=MRC2;refGene_tx=XM_023244719.1
E1 47168722 47168865
refGene_gn=MRC2;refGene_tx=XM_023244718.1;refGene_gn=MRC2;refGene_tx=XM_023244719.1
E1 47172514 47172629
refGene_gn=MRC2;refGene_tx=XM_023244718.1;refGene_gn=MRC2;refGene_tx=XM_023244719.1

E1 47172760 47172924
refGene_gn=MRC2;refGene_tx=XM_023244718.1;refGene_gn=MRC2;refGene_tx=XM_023244719.1
E1 47173088 47173197
refGene_gn=MRC2;refGene_tx=XM_023244718.1;refGene_gn=MRC2;refGene_tx=XM_023244719.1
E1 47173731 47173970
refGene_gn=MRC2;refGene_tx=XM_023244718.1;refGene_gn=MRC2;refGene_tx=XM_023244719.1
E1 47174136 47174305
refGene_gn=MRC2;refGene_tx=XM_023244718.1;refGene_gn=MRC2;refGene_tx=XM_023244719.1
E1 47174399 47174549
refGene_gn=MRC2;refGene_tx=XM_023244718.1;refGene_gn=MRC2;refGene_tx=XM_023244719.1
E1 47174886 47175006
refGene_gn=MRC2;refGene_tx=XM_023244718.1;refGene_gn=MRC2;refGene_tx=XM_023244719.1
E1 47175200 47175380
refGene_gn=MRC2;refGene_tx=XM_023244718.1;refGene_gn=MRC2;refGene_tx=XM_023244719.1
E1 47175868 47175895
refGene_gn=MRC2;refGene_tx=XM_023244718.1;refGene_gn=MRC2;refGene_tx=XM_023244719.1
E1 47176017 47176241
refGene_gn=MRC2;refGene_tx=XM_023244718.1;refGene_gn=MRC2;refGene_tx=XM_023244719.1
E1 47186941 47187088
refGene_gn=MARCH10;refGene_tx=XM_006940477.3;refGene_gn=MARCH10;refGene_tx=XM_006940478.4;refGene_gn=MARCH10;refGene_tx=XM_011289289.3;refGene_gn=MARCH10;refGene_tx=XM_019818061.2;refGene_gn=MARCH10;refGene_tx=XM_019818062.2;refGene_gn=MARCH10;refGene_tx=XM_019818063.2;refGene_gn=MARCH10;refGene_tx=XM_019818064.2;refGene_gn=MARCH10;refGene_tx=XM_019818065.2;refGene_gn=MARCH10;refGene_tx=XM_019818066.2;refGene_gn=MARCH10;refGene_tx=XM_019818067.2;refGene_gn=MARCH10;refGene_tx=XM_019818068.2;refGene_gn=MARCH10;refGene_tx=XM_019818069.2;refGene_gn=MARCH10;refGene_tx=XM_019818070.2;refGene_gn=MARCH10;refGene_tx=XM_019818071.2
E1 47190456 47190499
refGene_gn=MARCH10;refGene_tx=XM_006940478.4;refGene_gn=MARCH10;refGene_tx=XM_011289289.3;refGene_gn=MARCH10;refGene_tx=XM_019818070.2
E1 47195747 47195861
refGene_gn=MARCH10;refGene_tx=XM_006940477.3;refGene_gn=MARCH10;refGene_tx=XM_006940478.4;refGene_gn=MARCH10;refGene_tx=XM_011289289.3;refGene_gn=MARCH10;refGene_tx=XM_019818061.2;refGene_gn=MARCH10;refGene_tx=XM_019818062.2;refGene_gn=MARCH10;refGene_tx=XM_019818063.2;refGene_gn=MARCH10;refGene_tx=XM_019818064.2;refGene_gn=MARCH10;refGene_tx=XM_019818065.2;refGene_gn=MARCH10;refGene_tx=XM_019818066.2;refGene_gn=MARCH10;refGene_tx=XM_019818067.2;refGene_gn=MARCH10;refGene_tx=XM_019818069.2;refGene_gn=MARCH10;refGene_tx=XM_019818071.2
E1 47205403 47205513

refGene_gn=MARCH10;refGene_tx=XM_006940477.3;refGene_gn=MARCH10;refGene_tx=XM_006940478.4;refGene_gn=MARCH10;refGene_tx=XM_011289289.3;refGene_gn=MARCH10;refGene_tx=XM_019818061.2;refGene_gn=MARCH10;refGene_tx=XM_019818062.2;refGene_gn=MARCH10;refGene_tx=XM_019818063.2;refGene_gn=MARCH10;refGene_tx=XM_019818064.2;refGene_gn=MARCH10;refGene_tx=XM_019818065.2;refGene_gn=MARCH10;refGene_tx=XM_019818066.2;refGene_gn=MARCH10;refGene_tx=XM_019818067.2;refGene_gn=MARCH10;refGene_tx=XM_019818068.2;refGene_gn=MARCH10;refGene_tx=XM_019818069.2;refGene_gn=MARCH10;refGene_tx=XM_019818070.2;refGene_gn=MARCH10;refGene_tx=XM_019818071.2
E1 47207878 47208045
refGene_gn=MARCH10;refGene_tx=XM_006940477.3;refGene_gn=MARCH10;refGene_tx=XM_006940478.4;refGene_gn=MARCH10;refGene_tx=XM_011289289.3;refGene_gn=MARCH10;refGene_tx=XM_019818061.2;refGene_gn=MARCH10;refGene_tx=XM_019818062.2;refGene_gn=MARCH10;refGene_tx=XM_019818063.2;refGene_gn=MARCH10;refGene_tx=XM_019818064.2;refGene_gn=MARCH10;refGene_tx=XM_019818065.2;refGene_gn=MARCH10;refGene_tx=XM_019818066.2;refGene_gn=MARCH10;refGene_tx=XM_019818067.2;refGene_gn=MARCH10;refGene_tx=XM_019818068.2;refGene_gn=MARCH10;refGene_tx=XM_019818069.2;refGene_gn=MARCH10;refGene_tx=XM_019818070.2
E1 47211724 47213144
refGene_gn=MARCH10;refGene_tx=XM_006940477.3;refGene_gn=MARCH10;refGene_tx=XM_006940478.4;refGene_gn=MARCH10;refGene_tx=XM_011289289.3;refGene_gn=MARCH10;refGene_tx=XM_019818061.2;refGene_gn=MARCH10;refGene_tx=XM_019818062.2;refGene_gn=MARCH10;refGene_tx=XM_019818063.2;refGene_gn=MARCH10;refGene_tx=XM_019818064.2;refGene_gn=MARCH10;refGene_tx=XM_019818065.2;refGene_gn=MARCH10;refGene_tx=XM_019818066.2;refGene_gn=MARCH10;refGene_tx=XM_019818067.2;refGene_gn=MARCH10;refGene_tx=XM_019818068.2;refGene_gn=MARCH10;refGene_tx=XM_019818069.2;refGene_gn=MARCH10;refGene_tx=XM_019818070.2
E1 47219386 47219539
refGene_gn=MARCH10;refGene_tx=XM_006940477.3;refGene_gn=MARCH10;refGene_tx=XM_006940478.4;refGene_gn=MARCH10;refGene_tx=XM_011289289.3;refGene_gn=MARCH10;refGene_tx=XM_019818061.2;refGene_gn=MARCH10;refGene_tx=XM_019818062.2;refGene_gn=MARCH10;refGene_tx=XM_019818063.2;refGene_gn=MARCH10;refGene_tx=XM_019818064.2;refGene_gn=MARCH10;refGene_tx=XM_019818065.2;refGene_gn=MARCH10;refGene_tx=XM_019818066.2;refGene_gn=MARCH10;refGene_tx=XM_019818067.2;refGene_gn=MARCH10;refGene_tx=XM_019818068.2;refGene_gn=MARCH10;refGene_tx=XM_019818069.2;refGene_gn=MARCH10;refGene_tx=XM_019818070.2;refGene_gn=MARCH10;refGene_tx=XM_019818071.2
E1 47230923 47231095
refGene_gn=MARCH10;refGene_tx=XM_006940477.3;refGene_gn=MARCH10;refGene_tx=XM_006940478.4;refGene_gn=MARCH10;refGene_tx=XM_011289289.3;refGene_gn=MARCH10;refGene_tx=XM_019818061.2;refGene_gn=MARCH10;refGene_tx=XM_019818062.2;refGene_gn=MARCH10;refGene_tx=XM_019818063.2;refGene_gn=MARCH10;refGene_tx=XM_019818064.2;refGene_gn=MARCH10;refGene_tx=XM_019818065.2;refGene_gn=MARCH10;refGene_tx=XM_019818066.2;refGene_gn=MARCH10;refGene_tx=XM_019818067.2;refGene_gn=MARCH10;refGene_tx=XM_019818068.2;refGene_gn=MARCH10;refGene_tx=XM_019818069.2;refGene_gn=MARCH10;refGene_tx=XM_019818070.2;refGene_gn=MARCH10;refGene_tx=XM_019818071.2
E1 47252503 47252623

refGene_gn=MARCH10;refGene_tx=XM_006940477.3;refGene_gn=MARCH10;refGene_tx=XM_006940478.4;refGene_gn=MARCH10;refGene_tx=XM_011289289.3;refGene_gn=MARCH10;refGene_tx=XM_019818061.2;refGene_gn=MARCH10;refGene_tx=XM_019818062.2;refGene_gn=MARCH10;refGene_tx=XM_019818063.2;refGene_gn=MARCH10;refGene_tx=XM_019818064.2;refGene_gn=MARCH10;refGene_tx=XM_019818065.2;refGene_gn=MARCH10;refGene_tx=XM_019818066.2;refGene_gn=MARCH10;refGene_tx=XM_019818067.2;refGene_gn=MARCH10;refGene_tx=XM_019818068.2;refGene_gn=MARCH10;refGene_tx=XM_019818070.2;refGene_gn=MARCH10;refGene_tx=XM_019818071.2
E1 47262981 47263071
refGene_gn=MARCH10;refGene_tx=XM_006940477.3;refGene_gn=MARCH10;refGene_tx=XM_006940478.4;refGene_gn=MARCH10;refGene_tx=XM_011289289.3;refGene_gn=MARCH10;refGene_tx=XM_019818061.2;refGene_gn=MARCH10;refGene_tx=XM_019818062.2;refGene_gn=MARCH10;refGene_tx=XM_019818063.2;refGene_gn=MARCH10;refGene_tx=XM_019818064.2;refGene_gn=MARCH10;refGene_tx=XM_019818065.2;refGene_gn=MARCH10;refGene_tx=XM_019818066.2;refGene_gn=MARCH10;refGene_tx=XM_019818067.2;refGene_gn=MARCH10;refGene_tx=XM_019818068.2;refGene_gn=MARCH10;refGene_tx=XM_019818069.2;refGene_gn=MARCH10;refGene_tx=XM_019818070.2;refGene_gn=MARCH10;refGene_tx=XM_019818071.2
E1 47465138 47465205
refGene_gn=TANC2;refGene_tx=XM_019818072.2;refGene_gn=TANC2;refGene_tx=XM_019818073.2;refGene_gn=TANC2;refGene_tx=XM_019818074.2;refGene_gn=TANC2;refGene_tx=XM_019818075.2;refGene_gn=TANC2;refGene_tx=XM_019818076.2;refGene_gn=TANC2;refGene_tx=XM_019818079.2;refGene_gn=TANC2;refGene_tx=XM_019818080.2
E1 47516884 47516956
refGene_gn=TANC2;refGene_tx=XM_019818072.2;refGene_gn=TANC2;refGene_tx=XM_019818073.2;refGene_gn=TANC2;refGene_tx=XM_019818074.2;refGene_gn=TANC2;refGene_tx=XM_019818075.2;refGene_gn=TANC2;refGene_tx=XM_019818076.2;refGene_gn=TANC2;refGene_tx=XM_019818079.2;refGene_gn=TANC2;refGene_tx=XM_019818080.2
E1 47546043 47546226
refGene_gn=TANC2;refGene_tx=XM_019818072.2;refGene_gn=TANC2;refGene_tx=XM_019818073.2;refGene_gn=TANC2;refGene_tx=XM_019818074.2;refGene_gn=TANC2;refGene_tx=XM_019818075.2;refGene_gn=TANC2;refGene_tx=XM_019818076.2;refGene_gn=TANC2;refGene_tx=XM_019818077.2;refGene_gn=TANC2;refGene_tx=XM_019818079.2;refGene_gn=TANC2;refGene_tx=XM_019818080.2
E1 47597604 47597715
refGene_gn=TANC2;refGene_tx=XM_019818072.2;refGene_gn=TANC2;refGene_tx=XM_019818073.2;refGene_gn=TANC2;refGene_tx=XM_019818074.2;refGene_gn=TANC2;refGene_tx=XM_019818076.2;refGene_gn=TANC2;refGene_tx=XM_019818077.2
E1 47628092 47628241
refGene_gn=TANC2;refGene_tx=XM_019818072.2;refGene_gn=TANC2;refGene_tx=XM_019818073.2;refGene_gn=TANC2;refGene_tx=XM_019818074.2;refGene_gn=TANC2;refGene_tx=XM_019818075.2;refGene_gn=TANC2;refGene_tx=XM_019818076.2;refGene_gn=TANC2;refGene_tx=XM_019818077.2;refGene_gn=TANC2;refGene_tx=XM_019818079.2;refGene_gn=TANC2;refGene_tx=XM_019818080.2;refGene_gn=TANC2;refGene_tx=XM_019818081.2
E1 47631932 47632119

refGene_gn=TANC2;refGene_tx=XM_019818072.2;refGene_gn=TANC2;refGene_tx=XM_019818073.2;refGene_gn=TANC2;refGene_tx=XM_019818074.2;refGene_gn=TANC2;refGene_tx=XM_019818075.2;refGene_gn=TANC2;refGene_tx=XM_019818076.2;refGene_gn=TANC2;refGene_tx=XM_019818077.2;refGene_gn=TANC2;refGene_tx=XM_019818079.2;refGene_gn=TANC2;refGene_tx=XM_019818080.2;refGene_gn=TANC2;refGene_tx=XM_019818081.2
E1 47646848 47647112
refGene_gn=TANC2;refGene_tx=XM_019818072.2;refGene_gn=TANC2;refGene_tx=XM_019818073.2;refGene_gn=TANC2;refGene_tx=XM_019818074.2;refGene_gn=TANC2;refGene_tx=XM_019818075.2;refGene_gn=TANC2;refGene_tx=XM_019818076.2;refGene_gn=TANC2;refGene_tx=XM_019818077.2;refGene_gn=TANC2;refGene_tx=XM_019818079.2;refGene_gn=TANC2;refGene_tx=XM_019818080.2;refGene_gn=TANC2;refGene_tx=XM_019818081.2
E1 47656462 47656588
refGene_gn=TANC2;refGene_tx=XM_019818072.2;refGene_gn=TANC2;refGene_tx=XM_019818073.2;refGene_gn=TANC2;refGene_tx=XM_019818074.2;refGene_gn=TANC2;refGene_tx=XM_019818075.2;refGene_gn=TANC2;refGene_tx=XM_019818076.2;refGene_gn=TANC2;refGene_tx=XM_019818077.2;refGene_gn=TANC2;refGene_tx=XM_019818079.2;refGene_gn=TANC2;refGene_tx=XM_019818080.2;refGene_gn=TANC2;refGene_tx=XM_019818081.2
E1 47679405 47679687
refGene_gn=TANC2;refGene_tx=XM_019818072.2;refGene_gn=TANC2;refGene_tx=XM_019818073.2;refGene_gn=TANC2;refGene_tx=XM_019818074.2;refGene_gn=TANC2;refGene_tx=XM_019818075.2;refGene_gn=TANC2;refGene_tx=XM_019818076.2;refGene_gn=TANC2;refGene_tx=XM_019818077.2;refGene_gn=TANC2;refGene_tx=XM_019818079.2;refGene_gn=TANC2;refGene_tx=XM_019818080.2;refGene_gn=TANC2;refGene_tx=XM_019818081.2
E1 47684096 47684230
refGene_gn=TANC2;refGene_tx=XM_019818072.2;refGene_gn=TANC2;refGene_tx=XM_019818073.2;refGene_gn=TANC2;refGene_tx=XM_019818074.2;refGene_gn=TANC2;refGene_tx=XM_019818075.2;refGene_gn=TANC2;refGene_tx=XM_019818076.2;refGene_gn=TANC2;refGene_tx=XM_019818077.2;refGene_gn=TANC2;refGene_tx=XM_019818079.2;refGene_gn=TANC2;refGene_tx=XM_019818080.2;refGene_gn=TANC2;refGene_tx=XM_019818081.2
E1 47692858 47693090
refGene_gn=TANC2;refGene_tx=XM_019818072.2;refGene_gn=TANC2;refGene_tx=XM_019818073.2;refGene_gn=TANC2;refGene_tx=XM_019818074.2;refGene_gn=TANC2;refGene_tx=XM_019818075.2;refGene_gn=TANC2;refGene_tx=XM_019818076.2;refGene_gn=TANC2;refGene_tx=XM_019818077.2;refGene_gn=TANC2;refGene_tx=XM_019818079.2;refGene_gn=TANC2;refGene_tx=XM_019818080.2;refGene_gn=TANC2;refGene_tx=XM_019818081.2
E1 47705687 47705854
refGene_gn=TANC2;refGene_tx=XM_019818072.2;refGene_gn=TANC2;refGene_tx=XM_019818073.2;refGene_gn=TANC2;refGene_tx=XM_019818074.2;refGene_gn=TANC2;refGene_tx=XM_019818075.2;refGene_gn=TANC2;refGene_tx=XM_019818076.2;refGene_gn=TANC2;refGene_tx=XM_019818077.2;refGene_gn=TANC2;refGene_tx=XM_019818079.2;refGene_gn=TANC2;refGene_tx=XM_019818080.2;refGene_gn=TANC2;refGene_tx=XM_019818081.2
E1 47711278 47711886
refGene_gn=TANC2;refGene_tx=XM_019818072.2;refGene_gn=TANC2;refGene_tx

=XM_019818073.2;refGene_gn=TANC2;refGene_tx=XM_019818074.2;refGene_gn=TANC2;refGene_tx=XM_019818075.2;refGene_gn=TANC2;refGene_tx=XM_019818076.2;refGene_gn=TANC2;refGene_tx=XM_019818077.2;refGene_gn=TANC2;refGene_tx=XM_019818079.2;refGene_gn=TANC2;refGene_tx=XM_019818080.2;refGene_gn=TANC2;refGene_tx=XM_019818081.2
E1 47727728 47727837
refGene_gn=TANC2;refGene_tx=XM_019818072.2;refGene_gn=TANC2;refGene_tx=XM_019818073.2;refGene_gn=TANC2;refGene_tx=XM_019818074.2;refGene_gn=TANC2;refGene_tx=XM_019818075.2;refGene_gn=TANC2;refGene_tx=XM_019818076.2;refGene_gn=TANC2;refGene_tx=XM_019818077.2;refGene_gn=TANC2;refGene_tx=XM_019818079.2;refGene_gn=TANC2;refGene_tx=XM_019818080.2;refGene_gn=TANC2;refGene_tx=XM_019818081.2
E1 47745491 47745614
refGene_gn=TANC2;refGene_tx=XM_019818072.2;refGene_gn=TANC2;refGene_tx=XM_019818073.2;refGene_gn=TANC2;refGene_tx=XM_019818074.2;refGene_gn=TANC2;refGene_tx=XM_019818075.2;refGene_gn=TANC2;refGene_tx=XM_019818076.2;refGene_gn=TANC2;refGene_tx=XM_019818077.2;refGene_gn=TANC2;refGene_tx=XM_019818079.2;refGene_gn=TANC2;refGene_tx=XM_019818080.2;refGene_gn=TANC2;refGene_tx=XM_019818081.2
E1 47746228 47746465
refGene_gn=TANC2;refGene_tx=XM_019818072.2;refGene_gn=TANC2;refGene_tx=XM_019818073.2;refGene_gn=TANC2;refGene_tx=XM_019818074.2;refGene_gn=TANC2;refGene_tx=XM_019818075.2;refGene_gn=TANC2;refGene_tx=XM_019818076.2;refGene_gn=TANC2;refGene_tx=XM_019818077.2;refGene_gn=TANC2;refGene_tx=XM_019818079.2;refGene_gn=TANC2;refGene_tx=XM_019818080.2;refGene_gn=TANC2;refGene_tx=XM_019818081.2
E1 47751123 47751309
refGene_gn=TANC2;refGene_tx=XM_019818072.2;refGene_gn=TANC2;refGene_tx=XM_019818073.2;refGene_gn=TANC2;refGene_tx=XM_019818074.2;refGene_gn=TANC2;refGene_tx=XM_019818075.2;refGene_gn=TANC2;refGene_tx=XM_019818076.2;refGene_gn=TANC2;refGene_tx=XM_019818077.2;refGene_gn=TANC2;refGene_tx=XM_019818079.2;refGene_gn=TANC2;refGene_tx=XM_019818080.2;refGene_gn=TANC2;refGene_tx=XM_019818081.2
E1 47753540 47753634
refGene_gn=TANC2;refGene_tx=XM_019818072.2;refGene_gn=TANC2;refGene_tx=XM_019818073.2;refGene_gn=TANC2;refGene_tx=XM_019818074.2;refGene_gn=TANC2;refGene_tx=XM_019818075.2;refGene_gn=TANC2;refGene_tx=XM_019818076.2;refGene_gn=TANC2;refGene_tx=XM_019818077.2;refGene_gn=TANC2;refGene_tx=XM_019818079.2;refGene_gn=TANC2;refGene_tx=XM_019818080.2;refGene_gn=TANC2;refGene_tx=XM_019818081.2
E1 47760056 47760190
refGene_gn=TANC2;refGene_tx=XM_019818072.2;refGene_gn=TANC2;refGene_tx=XM_019818073.2;refGene_gn=TANC2;refGene_tx=XM_019818074.2;refGene_gn=TANC2;refGene_tx=XM_019818075.2;refGene_gn=TANC2;refGene_tx=XM_019818076.2;refGene_gn=TANC2;refGene_tx=XM_019818077.2;refGene_gn=TANC2;refGene_tx=XM_019818079.2;refGene_gn=TANC2;refGene_tx=XM_019818080.2;refGene_gn=TANC2;refGene_tx=XM_019818081.2
E1 47761380 47761504
refGene_gn=TANC2;refGene_tx=XM_019818072.2;refGene_gn=TANC2;refGene_tx=XM_019818073.2;refGene_gn=TANC2;refGene_tx=XM_019818074.2;refGene_gn=

TANC2;refGene_tx=XM_019818075.2;refGene_gn=TANC2;refGene_tx=XM_0198180
76.2;refGene_gn=TANC2;refGene_tx=XM_019818077.2;refGene_gn=TANC2;refGe
ne_tx=XM_019818079.2;refGene_gn=TANC2;refGene_tx=XM_019818080.2;refGen
e_gn=TANC2;refGene_tx=XM_019818081.2
E1 47765689 47765865
refGene_gn=TANC2;refGene_tx=XM_019818072.2;refGene_gn=TANC2;refGene_tx
=XM_019818073.2;refGene_gn=TANC2;refGene_tx=XM_019818074.2;refGene_gn=
TANC2;refGene_tx=XM_019818075.2;refGene_gn=TANC2;refGene_tx=XM_0198180
76.2;refGene_gn=TANC2;refGene_tx=XM_019818077.2;refGene_gn=TANC2;refGe
ne_tx=XM_019818079.2;refGene_gn=TANC2;refGene_tx=XM_019818080.2;refGen
e_gn=TANC2;refGene_tx=XM_019818081.2
E1 47766180 47766313
refGene_gn=TANC2;refGene_tx=XM_019818072.2;refGene_gn=TANC2;refGene_tx
=XM_019818073.2;refGene_gn=TANC2;refGene_tx=XM_019818074.2;refGene_gn=
TANC2;refGene_tx=XM_019818075.2;refGene_gn=TANC2;refGene_tx=XM_0198180
76.2;refGene_gn=TANC2;refGene_tx=XM_019818077.2;refGene_gn=TANC2;refGe
ne_tx=XM_019818079.2;refGene_gn=TANC2;refGene_tx=XM_019818080.2;refGen
e_gn=TANC2;refGene_tx=XM_019818081.2
E1 47766870 47766900
refGene_gn=TANC2;refGene_tx=XM_019818072.2;refGene_gn=TANC2;refGene_tx
=XM_019818073.2;refGene_gn=TANC2;refGene_tx=XM_019818075.2;refGene_gn=
TANC2;refGene_tx=XM_019818076.2;refGene_gn=TANC2;refGene_tx=XM_0198180
77.2;refGene_gn=TANC2;refGene_tx=XM_019818079.2;refGene_gn=TANC2;refGe
ne_tx=XM_019818081.2
E1 47767734 47767826
refGene_gn=TANC2;refGene_tx=XM_019818072.2;refGene_gn=TANC2;refGene_tx
=XM_019818073.2;refGene_gn=TANC2;refGene_tx=XM_019818074.2;refGene_gn=
TANC2;refGene_tx=XM_019818075.2;refGene_gn=TANC2;refGene_tx=XM_0198180
76.2;refGene_gn=TANC2;refGene_tx=XM_019818077.2;refGene_gn=TANC2;refGe
ne_tx=XM_019818079.2;refGene_gn=TANC2;refGene_tx=XM_019818080.2;refGen
e_gn=TANC2;refGene_tx=XM_019818081.2
E1 47769837 47769984
refGene_gn=TANC2;refGene_tx=XM_019818072.2;refGene_gn=TANC2;refGene_tx
=XM_019818073.2;refGene_gn=TANC2;refGene_tx=XM_019818074.2;refGene_gn=
TANC2;refGene_tx=XM_019818075.2;refGene_gn=TANC2;refGene_tx=XM_0198180
76.2;refGene_gn=TANC2;refGene_tx=XM_019818077.2;refGene_gn=TANC2;refGe
ne_tx=XM_019818079.2;refGene_gn=TANC2;refGene_tx=XM_019818080.2;refGen
e_gn=TANC2;refGene_tx=XM_019818081.2
E1 47772467 47772568
refGene_gn=TANC2;refGene_tx=XM_019818072.2;refGene_gn=TANC2;refGene_tx
=XM_019818073.2;refGene_gn=TANC2;refGene_tx=XM_019818074.2;refGene_gn=
TANC2;refGene_tx=XM_019818075.2;refGene_gn=TANC2;refGene_tx=XM_0198180
76.2;refGene_gn=TANC2;refGene_tx=XM_019818077.2;refGene_gn=TANC2;refGe
ne_tx=XM_019818079.2;refGene_gn=TANC2;refGene_tx=XM_019818080.2;refGen
e_gn=TANC2;refGene_tx=XM_019818081.2
E1 47774163 47776117
refGene_gn=TANC2;refGene_tx=XM_019818072.2;refGene_gn=TANC2;refGene_tx
=XM_019818073.2;refGene_gn=TANC2;refGene_tx=XM_019818074.2;refGene_gn=
TANC2;refGene_tx=XM_019818075.2;refGene_gn=TANC2;refGene_tx=XM_0198180
76.2;refGene_gn=TANC2;refGene_tx=XM_019818077.2;refGene_gn=TANC2;refGe

ne_tx=XM_019818079.2;refGene_gn=TANC2;refGene_tx=XM_019818080.2;refGene_gn=TANC2;refGene_tx=XM_019818081.2
E1 47785507 47785700
refGene_gn=LOC101093813;refGene_tx=XM_023244577.1;refGene_gn=LOC101093813;refGene_tx=XM_023244578.1
E1 47785944 47786102
refGene_gn=LOC101093813;refGene_tx=XM_023244577.1;refGene_gn=LOC101093813;refGene_tx=XM_023244578.1
E1 47786583 47786687
refGene_gn=LOC101093813;refGene_tx=XM_023244577.1;refGene_gn=LOC101093813;refGene_tx=XM_023244578.1
E1 47786973 47787072
refGene_gn=LOC101093813;refGene_tx=XM_023244577.1;refGene_gn=LOC101093813;refGene_tx=XM_023244578.1
E1 47788019 47788224
refGene_gn=LOC101093813;refGene_tx=XM_023244577.1;refGene_gn=LOC101093813;refGene_tx=XM_023244578.1
E1 47818054 47818330 refGene_gn=ACE;refGene_tx=XM_023244575.1
E1 47818923 47819091 refGene_gn=ACE;refGene_tx=XM_023244575.1
E1 47819687 47819784 refGene_gn=ACE;refGene_tx=XM_023244575.1
E1 47820522 47820666 refGene_gn=ACE;refGene_tx=XM_023244575.1
E1 47821127 47821319 refGene_gn=ACE;refGene_tx=XM_023244575.1
E1 47821902 47822000 refGene_gn=ACE;refGene_tx=XM_023244575.1
E1 47822419 47822592 refGene_gn=ACE;refGene_tx=XM_023244575.1
E1 47822963 47823187 refGene_gn=ACE;refGene_tx=XM_023244575.1
E1 47823541 47823686 refGene_gn=ACE;refGene_tx=XM_023244575.1
E1 47823976 47824075 refGene_gn=ACE;refGene_tx=XM_023244575.1
E1 47824291 47824414 refGene_gn=ACE;refGene_tx=XM_023244575.1
E1 47824757 47824969 refGene_gn=ACE;refGene_tx=XM_023244575.1
E1 47825312 47825511 refGene_gn=ACE;refGene_tx=XM_023244576.1
E1 47825648 47825785
refGene_gn=ACE;refGene_tx=XM_023244575.1;refGene_gn=ACE;refGene_tx=XM_023244576.1
E1 47827205 47827364
refGene_gn=ACE;refGene_tx=XM_023244575.1;refGene_gn=ACE;refGene_tx=XM_023244576.1
E1 47827588 47827676
refGene_gn=ACE;refGene_tx=XM_023244575.1;refGene_gn=ACE;refGene_tx=XM_023244576.1
E1 47829879 47830023
refGene_gn=ACE;refGene_tx=XM_023244575.1;refGene_gn=ACE;refGene_tx=XM_023244576.1
E1 47830177 47830369
refGene_gn=ACE;refGene_tx=XM_023244575.1;refGene_gn=ACE;refGene_tx=XM_023244576.1
E1 47831727 47831825
refGene_gn=ACE;refGene_tx=XM_023244575.1;refGene_gn=ACE;refGene_tx=XM_023244576.1
E1 47832027 47832200
refGene_gn=ACE;refGene_tx=XM_023244575.1;refGene_gn=ACE;refGene_tx=XM_

023244576.1
E1	47834091	47834315
refGene_gn=ACE;refGene_tx=XM_023244575.1;refGene_gn=ACE;refGene_tx=XM_
023244576.1
E1	47834628	47834773
refGene_gn=ACE;refGene_tx=XM_023244575.1;refGene_gn=ACE;refGene_tx=XM_
023244576.1
E1	47835023	47835122
refGene_gn=ACE;refGene_tx=XM_023244575.1;refGene_gn=ACE;refGene_tx=XM_
023244576.1
E1	47837308	47837431
refGene_gn=ACE;refGene_tx=XM_023244575.1;refGene_gn=ACE;refGene_tx=XM_
023244576.1
E1	47837777	47837965
refGene_gn=ACE;refGene_tx=XM_023244575.1;refGene_gn=ACE;refGene_tx=XM_
023244576.1
E1	47838148	47838378
refGene_gn=ACE;refGene_tx=XM_023244575.1;refGene_gn=ACE;refGene_tx=XM_
023244576.1
E1	47846363	47846463
refGene_gn=LOC101083124;refGene_tx=XM_023244023.1
E1	47846560	47846697
refGene_gn=LOC101083124;refGene_tx=XM_023244023.1
E1	47847076	47847225
refGene_gn=LOC101083124;refGene_tx=XM_023244023.1
E1	47847477	47847575
refGene_gn=LOC101083124;refGene_tx=XM_023244023.1
E1	47847781	47847925
refGene_gn=LOC101083124;refGene_tx=XM_023244023.1
E1	47848709	47848739
refGene_gn=LOC101083124;refGene_tx=XM_023244023.1
E1	47848741	47848897
refGene_gn=LOC101083124;refGene_tx=XM_023244023.1
E1	47849009	47849107
refGene_gn=LOC101083124;refGene_tx=XM_023244023.1
E1	47849185	47849358
refGene_gn=LOC101083124;refGene_tx=XM_023244023.1
E1	47849661	47849885
refGene_gn=LOC101083124;refGene_tx=XM_023244023.1
E1	47850008	47850153
refGene_gn=LOC101083124;refGene_tx=XM_023244023.1
E1	47851994	47852114
refGene_gn=LOC101083124;refGene_tx=XM_023244023.1
E1	47852585	47852702
refGene_gn=LOC101083124;refGene_tx=XM_023244023.1
E1	47856343	47856528
refGene_gn=LOC101083124;refGene_tx=XM_023244023.1
E1	47857057	47857162
refGene_gn=LOC101083124;refGene_tx=XM_023244023.1
E1	47857164	47857451

refGene_gn=LOC101083124;refGene_tx=XM_023244023.1
E1 47860539 47860615
refGene_gn=KCNH6;refGene_tx=XM_019818085.2;refGene_gn=KCNH6;refGene_tx=XM_019818086.2;refGene_gn=KCNH6;refGene_tx=XM_019818088.2
E1 47861426 47861657
refGene_gn=KCNH6;refGene_tx=XM_019818085.2;refGene_gn=KCNH6;refGene_tx=XM_019818086.2;refGene_gn=KCNH6;refGene_tx=XM_019818088.2
E1 47866763 47866925
refGene_gn=KCNH6;refGene_tx=XM_019818085.2;refGene_gn=KCNH6;refGene_tx=XM_019818086.2;refGene_gn=KCNH6;refGene_tx=XM_019818088.2
E1 47867027 47867233
refGene_gn=KCNH6;refGene_tx=XM_019818085.2;refGene_gn=KCNH6;refGene_tx=XM_019818086.2;refGene_gn=KCNH6;refGene_tx=XM_019818088.2
E1 47870664 47871090
refGene_gn=KCNH6;refGene_tx=XM_019818085.2;refGene_gn=KCNH6;refGene_tx=XM_019818086.2;refGene_gn=KCNH6;refGene_tx=XM_019818088.2
E1 47872270 47872670
refGene_gn=KCNH6;refGene_tx=XM_019818085.2;refGene_gn=KCNH6;refGene_tx=XM_019818086.2;refGene_gn=KCNH6;refGene_tx=XM_019818088.2
E1 47873709 47873909
refGene_gn=KCNH6;refGene_tx=XM_019818085.2;refGene_gn=KCNH6;refGene_tx=XM_019818086.2;refGene_gn=KCNH6;refGene_tx=XM_019818088.2
E1 47874093 47874346
refGene_gn=KCNH6;refGene_tx=XM_019818085.2;refGene_gn=KCNH6;refGene_tx=XM_019818086.2;refGene_gn=KCNH6;refGene_tx=XM_019818088.2
E1 47877457 47877651
refGene_gn=KCNH6;refGene_tx=XM_019818085.2;refGene_gn=KCNH6;refGene_tx=XM_019818086.2;refGene_gn=KCNH6;refGene_tx=XM_019818088.2
E1 47879178 47879263
refGene_gn=KCNH6;refGene_tx=XM_019818085.2;refGene_gn=KCNH6;refGene_tx=XM_019818086.2;refGene_gn=KCNH6;refGene_tx=XM_019818088.2
E1 47879826 47879995
refGene_gn=KCNH6;refGene_tx=XM_019818085.2;refGene_gn=KCNH6;refGene_tx=XM_019818086.2;refGene_gn=KCNH6;refGene_tx=XM_019818088.2
E1 47880732 47880919
refGene_gn=KCNH6;refGene_tx=XM_019818085.2;refGene_gn=KCNH6;refGene_tx=XM_019818086.2
E1 47881196 47881481
refGene_gn=KCNH6;refGene_tx=XM_019818085.2;refGene_gn=KCNH6;refGene_tx=XM_019818086.2
E1 47881808 47882006 refGene_gn=KCNH6;refGene_tx=XM_019818085.2
E1 47885415 47885553 refGene_gn=DCAF7;refGene_tx=XM_019818090.2
E1 47903160 47903319 refGene_gn=DCAF7;refGene_tx=XM_019818090.2
E1 47904057 47904169 refGene_gn=DCAF7;refGene_tx=XM_019818090.2
E1 47904530 47904649 refGene_gn=DCAF7;refGene_tx=XM_019818090.2
E1 47907463 47907673 refGene_gn=DCAF7;refGene_tx=XM_019818090.2
E1 47908622 47908740 refGene_gn=DCAF7;refGene_tx=XM_019818090.2
E1 47910919 47911092 refGene_gn=DCAF7;refGene_tx=XM_019818090.2
E1 47920375 47920655
refGene_gn=LOC101095057;refGene_tx=XM_003997047.5;refGene_gn=LOC101095

057;refGene_tx=XM_006940481.3
E1 47922359 47922466
refGene_gn=LOC101095057;refGene_tx=XM_003997047.5;refGene_gn=LOC101095
057;refGene_tx=XM_006940481.3
E1 47923780 47923908
refGene_gn=LOC101095057;refGene_tx=XM_003997047.5;refGene_gn=LOC101095
057;refGene_tx=XM_006940481.3
E1 47924417 47924595
refGene_gn=LOC101095057;refGene_tx=XM_003997047.5;refGene_gn=LOC101095
057;refGene_tx=XM_006940481.3
E1 47924923 47925124
refGene_gn=LOC101095057;refGene_tx=XM_003997047.5;refGene_gn=LOC101095
057;refGene_tx=XM_006940481.3
E1 47934195 47934199
refGene_gn=MAP3K3;refGene_tx=XM_023244333.1;refGene_gn=MAP3K3;refGene_
tx=XM_023244334.1
E1 47943428 47943550
refGene_gn=MAP3K3;refGene_tx=XM_023244333.1;refGene_gn=MAP3K3;refGene_
tx=XM_023244334.1
E1 47955776 47955817
refGene_gn=MAP3K3;refGene_tx=XM_023244333.1;refGene_gn=MAP3K3;refGene_
tx=XM_023244334.1
E1 47961601 47961701
refGene_gn=MAP3K3;refGene_tx=XM_023244333.1;refGene_gn=MAP3K3;refGene_
tx=XM_023244334.1
E1 47963943 47963970 refGene_gn=MAP3K3;refGene_tx=XM_023244333.1
E1 47966245 47966359
refGene_gn=MAP3K3;refGene_tx=XM_023244333.1;refGene_gn=MAP3K3;refGene_
tx=XM_023244334.1
E1 47973822 47973943
refGene_gn=MAP3K3;refGene_tx=XM_023244333.1;refGene_gn=MAP3K3;refGene_
tx=XM_023244334.1
E1 47981948 47982082
refGene_gn=MAP3K3;refGene_tx=XM_023244333.1;refGene_gn=MAP3K3;refGene_
tx=XM_023244334.1
E1 47984887 47984961
refGene_gn=MAP3K3;refGene_tx=XM_023244333.1;refGene_gn=MAP3K3;refGene_
tx=XM_023244334.1
E1 47986904 47986972
refGene_gn=MAP3K3;refGene_tx=XM_023244333.1;refGene_gn=MAP3K3;refGene_
tx=XM_023244334.1
E1 47987178 47987271
refGene_gn=MAP3K3;refGene_tx=XM_023244333.1;refGene_gn=MAP3K3;refGene_
tx=XM_023244334.1
E1 47987881 47988073
refGene_gn=MAP3K3;refGene_tx=XM_023244333.1;refGene_gn=MAP3K3;refGene_
tx=XM_023244334.1
E1 47988791 47988940
refGene_gn=MAP3K3;refGene_tx=XM_023244333.1;refGene_gn=MAP3K3;refGene_
tx=XM_023244334.1

E1 47989589 47989721
refGene_gn=MAP3K3;refGene_tx=XM_023244333.1;refGene_gn=MAP3K3;refGene_tx=XM_023244334.1
E1 47990184 47990314
refGene_gn=MAP3K3;refGene_tx=XM_023244333.1;refGene_gn=MAP3K3;refGene_tx=XM_023244334.1
E1 47990644 47990822
refGene_gn=MAP3K3;refGene_tx=XM_023244333.1;refGene_gn=MAP3K3;refGene_tx=XM_023244334.1
E1 47991934 47992163
refGene_gn=MAP3K3;refGene_tx=XM_023244333.1;refGene_gn=MAP3K3;refGene_tx=XM_023244334.1
E1 47997034 47997194
refGene_gn=LIMD2;refGene_tx=XM_003997048.5;refGene_gn=LIMD2;refGene_tx=XM_023244343.1;refGene_gn=LIMD2;refGene_tx=XM_023244344.1
E1 47997269 47997409
refGene_gn=LIMD2;refGene_tx=XM_003997048.5;refGene_gn=LIMD2;refGene_tx=XM_023244343.1;refGene_gn=LIMD2;refGene_tx=XM_023244344.1
E1 47997488 47997533
refGene_gn=LIMD2;refGene_tx=XM_003997048.5;refGene_gn=LIMD2;refGene_tx=XM_023244343.1;refGene_gn=LIMD2;refGene_tx=XM_023244344.1
E1 47997724 47997766
refGene_gn=LIMD2;refGene_tx=XM_003997048.5;refGene_gn=LIMD2;refGene_tx=XM_023244343.1;refGene_gn=LIMD2;refGene_tx=XM_023244344.1
E1 48002035 48002188
refGene_gn=STRADA;refGene_tx=XM_003997049.5;refGene_gn=STRADA;refGene_tx=XM_011289304.3;refGene_gn=STRADA;refGene_tx=XM_011289306.3;refGene_gn=STRADA;refGene_tx=XM_019818097.2;refGene_gn=STRADA;refGene_tx=XM_019818103.2;refGene_gn=STRADA;refGene_tx=XM_019818104.2;refGene_gn=STRADA;refGene_tx=XM_023244335.1;refGene_gn=STRADA;refGene_tx=XM_023244336.1;refGene_gn=STRADA;refGene_tx=XM_023244337.1;refGene_gn=STRADA;refGene_tx=XM_023244338.1;refGene_gn=STRADA;refGene_tx=XM_023244339.1;refGene_gn=STRADA;refGene_tx=XM_023244340.1;refGene_gn=STRADA;refGene_tx=XM_023244341.1;refGene_gn=STRADA;refGene_tx=XM_023244342.1
E1 48002471 48002514
refGene_gn=STRADA;refGene_tx=XM_003997049.5;refGene_gn=STRADA;refGene_tx=XM_011289304.3;refGene_gn=STRADA;refGene_tx=XM_011289306.3;refGene_gn=STRADA;refGene_tx=XM_019818097.2;refGene_gn=STRADA;refGene_tx=XM_019818103.2;refGene_gn=STRADA;refGene_tx=XM_019818104.2;refGene_gn=STRADA;refGene_tx=XM_023244335.1;refGene_gn=STRADA;refGene_tx=XM_023244336.1;refGene_gn=STRADA;refGene_tx=XM_023244337.1;refGene_gn=STRADA;refGene_tx=XM_023244338.1;refGene_gn=STRADA;refGene_tx=XM_023244339.1;refGene_gn=STRADA;refGene_tx=XM_023244340.1;refGene_gn=STRADA;refGene_tx=XM_023244341.1;refGene_gn=STRADA;refGene_tx=XM_023244342.1
E1 48002819 48003061
refGene_gn=STRADA;refGene_tx=XM_003997049.5;refGene_gn=STRADA;refGene_tx=XM_011289304.3;refGene_gn=STRADA;refGene_tx=XM_011289306.3;refGene_gn=STRADA;refGene_tx=XM_019818097.2;refGene_gn=STRADA;refGene_tx=XM_019818103.2;refGene_gn=STRADA;refGene_tx=XM_019818104.2;refGene_gn=STRADA;refGene_tx=XM_023244335.1;refGene_gn=STRADA;refGene_tx=XM_023244336.

1;refGene_gn=STRADA;refGene_tx=XM_023244337.1;refGene_gn=STRADA;refGene_tx=XM_023244338.1;refGene_gn=STRADA;refGene_tx=XM_023244339.1;refGene_gn=STRADA;refGene_tx=XM_023244340.1;refGene_gn=STRADA;refGene_tx=XM_023244341.1;refGene_gn=STRADA;refGene_tx=XM_023244342.1
E1 48004974 48005079
refGene_gn=STRADA;refGene_tx=XM_003997049.5;refGene_gn=STRADA;refGene_tx=XM_011289304.3;refGene_gn=STRADA;refGene_tx=XM_011289306.3;refGene_gn=STRADA;refGene_tx=XM_019818097.2;refGene_gn=STRADA;refGene_tx=XM_019818103.2;refGene_gn=STRADA;refGene_tx=XM_019818104.2;refGene_gn=STRADA;refGene_tx=XM_023244335.1;refGene_gn=STRADA;refGene_tx=XM_023244336.1;refGene_gn=STRADA;refGene_tx=XM_023244337.1;refGene_gn=STRADA;refGene_tx=XM_023244338.1;refGene_gn=STRADA;refGene_tx=XM_023244339.1;refGene_gn=STRADA;refGene_tx=XM_023244340.1;refGene_gn=STRADA;refGene_tx=XM_023244341.1;refGene_gn=STRADA;refGene_tx=XM_023244342.1
E1 48005601 48005773
refGene_gn=STRADA;refGene_tx=XM_003997049.5;refGene_gn=STRADA;refGene_tx=XM_011289304.3;refGene_gn=STRADA;refGene_tx=XM_011289306.3;refGene_gn=STRADA;refGene_tx=XM_019818097.2;refGene_gn=STRADA;refGene_tx=XM_019818103.2;refGene_gn=STRADA;refGene_tx=XM_019818104.2;refGene_gn=STRADA;refGene_tx=XM_023244335.1;refGene_gn=STRADA;refGene_tx=XM_023244336.1;refGene_gn=STRADA;refGene_tx=XM_023244337.1;refGene_gn=STRADA;refGene_tx=XM_023244338.1;refGene_gn=STRADA;refGene_tx=XM_023244339.1;refGene_gn=STRADA;refGene_tx=XM_023244340.1;refGene_gn=STRADA;refGene_tx=XM_023244341.1;refGene_gn=STRADA;refGene_tx=XM_023244342.1
E1 48007284 48007408
refGene_gn=STRADA;refGene_tx=XM_003997049.5;refGene_gn=STRADA;refGene_tx=XM_011289304.3;refGene_gn=STRADA;refGene_tx=XM_011289306.3;refGene_gn=STRADA;refGene_tx=XM_019818097.2;refGene_gn=STRADA;refGene_tx=XM_019818103.2;refGene_gn=STRADA;refGene_tx=XM_019818104.2;refGene_gn=STRADA;refGene_tx=XM_023244335.1;refGene_gn=STRADA;refGene_tx=XM_023244336.1;refGene_gn=STRADA;refGene_tx=XM_023244337.1;refGene_gn=STRADA;refGene_tx=XM_023244338.1;refGene_gn=STRADA;refGene_tx=XM_023244339.1;refGene_gn=STRADA;refGene_tx=XM_023244340.1;refGene_gn=STRADA;refGene_tx=XM_023244341.1;refGene_gn=STRADA;refGene_tx=XM_023244342.1
E1 48007520 48007629
refGene_gn=STRADA;refGene_tx=XM_003997049.5;refGene_gn=STRADA;refGene_tx=XM_011289304.3;refGene_gn=STRADA;refGene_tx=XM_011289306.3;refGene_gn=STRADA;refGene_tx=XM_019818097.2;refGene_gn=STRADA;refGene_tx=XM_019818103.2;refGene_gn=STRADA;refGene_tx=XM_019818104.2;refGene_gn=STRADA;refGene_tx=XM_023244335.1;refGene_gn=STRADA;refGene_tx=XM_023244336.1;refGene_gn=STRADA;refGene_tx=XM_023244337.1;refGene_gn=STRADA;refGene_tx=XM_023244338.1;refGene_gn=STRADA;refGene_tx=XM_023244339.1;refGene_gn=STRADA;refGene_tx=XM_023244340.1;refGene_gn=STRADA;refGene_tx=XM_023244341.1;refGene_gn=STRADA;refGene_tx=XM_023244342.1
E1 48009543 48009665
refGene_gn=STRADA;refGene_tx=XM_003997049.5;refGene_gn=STRADA;refGene_tx=XM_011289304.3;refGene_gn=STRADA;refGene_tx=XM_011289306.3;refGene_gn=STRADA;refGene_tx=XM_019818097.2;refGene_gn=STRADA;refGene_tx=XM_019818103.2;refGene_gn=STRADA;refGene_tx=XM_019818104.2;refGene_gn=STRADA;refGene_tx=XM_023244335.1;refGene_gn=STRADA;refGene_tx=XM_023244336.

1;refGene_gn=STRADA;refGene_tx=XM_023244337.1;refGene_gn=STRADA;refGene_tx=XM_023244338.1;refGene_gn=STRADA;refGene_tx=XM_023244339.1;refGene_gn=STRADA;refGene_tx=XM_023244340.1;refGene_gn=STRADA;refGene_tx=XM_023244341.1;refGene_gn=STRADA;refGene_tx=XM_023244342.1
E1 48010570 48010673
refGene_gn=STRADA;refGene_tx=XM_003997049.5;refGene_gn=STRADA;refGene_tx=XM_011289304.3;refGene_gn=STRADA;refGene_tx=XM_011289306.3;refGene_gn=STRADA;refGene_tx=XM_019818097.2;refGene_gn=STRADA;refGene_tx=XM_019818103.2;refGene_gn=STRADA;refGene_tx=XM_019818104.2;refGene_gn=STRADA;refGene_tx=XM_023244335.1;refGene_gn=STRADA;refGene_tx=XM_023244336.1;refGene_gn=STRADA;refGene_tx=XM_023244337.1;refGene_gn=STRADA;refGene_tx=XM_023244338.1;refGene_gn=STRADA;refGene_tx=XM_023244339.1;refGene_gn=STRADA;refGene_tx=XM_023244340.1;refGene_gn=STRADA;refGene_tx=XM_023244341.1;refGene_gn=STRADA;refGene_tx=XM_023244342.1
E1 48016388 48016417
refGene_gn=STRADA;refGene_tx=XM_003997049.5;refGene_gn=STRADA;refGene_tx=XM_011289304.3;refGene_gn=STRADA;refGene_tx=XM_019818097.2;refGene_gn=STRADA;refGene_tx=XM_023244335.1;refGene_gn=STRADA;refGene_tx=XM_023244336.1;refGene_gn=STRADA;refGene_tx=XM_023244337.1;refGene_gn=STRADA;refGene_tx=XM_023244338.1
E1 48018703 48018761
refGene_gn=STRADA;refGene_tx=XM_003997049.5;refGene_gn=STRADA;refGene_tx=XM_011289304.3;refGene_gn=STRADA;refGene_tx=XM_019818097.2;refGene_gn=STRADA;refGene_tx=XM_023244335.1;refGene_gn=STRADA;refGene_tx=XM_023244336.1;refGene_gn=STRADA;refGene_tx=XM_023244337.1;refGene_gn=STRADA;refGene_tx=XM_023244338.1
E1 48020440 48020476
refGene_gn=STRADA;refGene_tx=XM_003997049.5;refGene_gn=STRADA;refGene_tx=XM_011289304.3;refGene_gn=STRADA;refGene_tx=XM_019818097.2;refGene_gn=STRADA;refGene_tx=XM_023244335.1;refGene_gn=STRADA;refGene_tx=XM_023244336.1;refGene_gn=STRADA;refGene_tx=XM_023244337.1;refGene_gn=STRADA;refGene_tx=XM_023244338.1;refGene_gn=STRADA;refGene_tx=XM_023244339.1;refGene_gn=STRADA;refGene_tx=XM_023244340.1;refGene_gn=STRADA;refGene_tx=XM_023244341.1
E1 48034344 48034425 refGene_gn=CCDC47;refGene_tx=XM_003997055.5
E1 48036415 48036565 refGene_gn=CCDC47;refGene_tx=XM_003997055.5
E1 48036566 48036584 refGene_gn=CCDC47;refGene_tx=XM_003997055.5
E1 48036974 48037084 refGene_gn=CCDC47;refGene_tx=XM_003997055.5
E1 48037401 48037460 refGene_gn=CCDC47;refGene_tx=XM_003997055.5
E1 48038776 48038862 refGene_gn=CCDC47;refGene_tx=XM_003997055.5
E1 48039522 48039633 refGene_gn=CCDC47;refGene_tx=XM_003997055.5
E1 48039832 48039934 refGene_gn=CCDC47;refGene_tx=XM_003997055.5
E1 48042475 48042541 refGene_gn=CCDC47;refGene_tx=XM_003997055.5
E1 48042770 48042892 refGene_gn=CCDC47;refGene_tx=XM_003997055.5
E1 48044528 48044703 refGene_gn=CCDC47;refGene_tx=XM_003997055.5
E1 48045212 48045320 refGene_gn=CCDC47;refGene_tx=XM_003997055.5
E1 48046000 48046264 refGene_gn=CCDC47;refGene_tx=XM_003997055.5
E1 48065766 48065987 refGene_gn=DDX42;refGene_tx=XM_006940486.4
E1 48068037 48068188 refGene_gn=DDX42;refGene_tx=XM_006940486.4
E1 48072864 48072926 refGene_gn=DDX42;refGene_tx=XM_006940486.4

E1 48074292 48074329 refGene_gn=DDX42;refGene_tx=XM_006940486.4
E1 48075132 48075282
refGene_gn=DDX42;refGene_tx=XM_006940486.4;refGene_gn=DDX42;refGene_tx
=XM_019818093.1
E1 48078847 48078952
refGene_gn=DDX42;refGene_tx=XM_006940486.4;refGene_gn=DDX42;refGene_tx
=XM_019818093.1
E1 48081013 48081133
refGene_gn=DDX42;refGene_tx=XM_006940486.4;refGene_gn=DDX42;refGene_tx
=XM_019818093.1
E1 48082000 48082177
refGene_gn=DDX42;refGene_tx=XM_006940486.4;refGene_gn=DDX42;refGene_tx
=XM_019818093.1
E1 48082879 48083008
refGene_gn=DDX42;refGene_tx=XM_006940486.4;refGene_gn=DDX42;refGene_tx
=XM_019818093.1
E1 48083603 48083703
refGene_gn=DDX42;refGene_tx=XM_006940486.4;refGene_gn=DDX42;refGene_tx
=XM_019818093.1
E1 48084274 48084322
refGene_gn=DDX42;refGene_tx=XM_006940486.4;refGene_gn=DDX42;refGene_tx
=XM_019818093.1
E1 48084844 48084942
refGene_gn=DDX42;refGene_tx=XM_006940486.4;refGene_gn=DDX42;refGene_tx
=XM_019818093.1
E1 48086318 48086595
refGene_gn=DDX42;refGene_tx=XM_006940486.4;refGene_gn=DDX42;refGene_tx
=XM_019818093.1
E1 48087523 48087750
refGene_gn=DDX42;refGene_tx=XM_006940486.4;refGene_gn=DDX42;refGene_tx
=XM_019818093.1
E1 48089405 48089516
refGene_gn=DDX42;refGene_tx=XM_006940486.4;refGene_gn=DDX42;refGene_tx
=XM_019818093.1
E1 48090646 48090748
refGene_gn=DDX42;refGene_tx=XM_006940486.4;refGene_gn=DDX42;refGene_tx
=XM_019818093.1
E1 48091404 48092094
refGene_gn=DDX42;refGene_tx=XM_006940486.4;refGene_gn=DDX42;refGene_tx
=XM_019818093.1
E1 48093432 48093625 refGene_gn=FTSJ3;refGene_tx=XM_003997057.5
E1 48093706 48093801 refGene_gn=FTSJ3;refGene_tx=XM_003997057.5
E1 48093891 48094075 refGene_gn=FTSJ3;refGene_tx=XM_003997057.5
E1 48094190 48094290 refGene_gn=FTSJ3;refGene_tx=XM_003997057.5
E1 48094379 48094465 refGene_gn=FTSJ3;refGene_tx=XM_003997057.5
E1 48094722 48094988 refGene_gn=FTSJ3;refGene_tx=XM_003997057.5
E1 48095075 48095196 refGene_gn=FTSJ3;refGene_tx=XM_003997057.5
E1 48095344 48095523 refGene_gn=FTSJ3;refGene_tx=XM_003997057.5
E1 48096877 48097013 refGene_gn=FTSJ3;refGene_tx=XM_003997057.5
E1 48097150 48097297 refGene_gn=FTSJ3;refGene_tx=XM_003997057.5

E1 48097420 48097501 refGene_gn=FTSJ3;refGene_tx=XM_003997057.5
E1 48097723 48097824 refGene_gn=FTSJ3;refGene_tx=XM_003997057.5
E1 48097909 48098014 refGene_gn=FTSJ3;refGene_tx=XM_003997057.5
E1 48098131 48098247 refGene_gn=FTSJ3;refGene_tx=XM_003997057.5
E1 48098326 48098521 refGene_gn=FTSJ3;refGene_tx=XM_003997057.5
E1 48098613 48098713 refGene_gn=FTSJ3;refGene_tx=XM_003997057.5
E1 48099191 48099271 refGene_gn=FTSJ3;refGene_tx=XM_003997057.5
E1 48099397 48099444 refGene_gn=FTSJ3;refGene_tx=XM_003997057.5
E1 48099962 48100068 refGene_gn=FTSJ3;refGene_tx=XM_003997057.5
E1 48100242 48100309 refGene_gn=FTSJ3;refGene_tx=XM_003997057.5
E1 48100799 48100946
refGene_gn=PSMC5;refGene_tx=XM_003997058.4;refGene_gn=PSMC5;refGene_tx
=XM_006940485.3
E1 48101414 48101486
refGene_gn=PSMC5;refGene_tx=XM_003997058.4;refGene_gn=PSMC5;refGene_tx
=XM_006940485.3;refGene_gn=PSMC5;refGene_tx=XM_019818096.2
E1 48102541 48102611
refGene_gn=PSMC5;refGene_tx=XM_003997058.4;refGene_gn=PSMC5;refGene_tx
=XM_006940485.3;refGene_gn=PSMC5;refGene_tx=XM_019818096.2
E1 48102934 48103032
refGene_gn=PSMC5;refGene_tx=XM_003997058.4;refGene_gn=PSMC5;refGene_tx
=XM_006940485.3;refGene_gn=PSMC5;refGene_tx=XM_019818096.2
E1 48103206 48103263
refGene_gn=PSMC5;refGene_tx=XM_003997058.4;refGene_gn=PSMC5;refGene_tx
=XM_006940485.3;refGene_gn=PSMC5;refGene_tx=XM_019818096.2
E1 48103334 48103565
refGene_gn=PSMC5;refGene_tx=XM_003997058.4;refGene_gn=PSMC5;refGene_tx
=XM_006940485.3;refGene_gn=PSMC5;refGene_tx=XM_019818096.2
E1 48103878 48104005
refGene_gn=PSMC5;refGene_tx=XM_003997058.4;refGene_gn=PSMC5;refGene_tx
=XM_006940485.3;refGene_gn=PSMC5;refGene_tx=XM_019818096.2
E1 48104101 48104292
refGene_gn=PSMC5;refGene_tx=XM_003997058.4;refGene_gn=PSMC5;refGene_tx
=XM_006940485.3;refGene_gn=PSMC5;refGene_tx=XM_019818096.2
E1 48104390 48104489
refGene_gn=PSMC5;refGene_tx=XM_003997058.4;refGene_gn=PSMC5;refGene_tx
=XM_006940485.3;refGene_gn=PSMC5;refGene_tx=XM_019818096.2
E1 48104575 48104686
refGene_gn=PSMC5;refGene_tx=XM_003997058.4;refGene_gn=PSMC5;refGene_tx
=XM_006940485.3;refGene_gn=PSMC5;refGene_tx=XM_019818096.2
E1 48104780 48104867
refGene_gn=PSMC5;refGene_tx=XM_003997058.4;refGene_gn=PSMC5;refGene_tx
=XM_006940485.3;refGene_gn=PSMC5;refGene_tx=XM_019818096.2
E1 48104969 48105023
refGene_gn=PSMC5;refGene_tx=XM_003997058.4;refGene_gn=PSMC5;refGene_tx
=XM_006940485.3;refGene_gn=PSMC5;refGene_tx=XM_019818096.2
E1 48105998 48106052
refGene_gn=SMARCD2;refGene_tx=XM_011289308.2;refGene_gn=SMARCD2;refGen
e_tx=XM_023243554.1
E1 48106128 48106230

refGene_gn=SMARCD2;refGene_tx=XM_011289308.2;refGene_gn=SMARCD2;refGene_tx=XM_023243554.1
E1	48106359	48106482	refGene_gn=SMARCD2;refGene_tx=XM_011289308.2;refGene_gn=SMARCD2;refGene_tx=XM_023243554.1
E1	48106667	48106803	refGene_gn=SMARCD2;refGene_tx=XM_011289308.2;refGene_gn=SMARCD2;refGene_tx=XM_023243554.1
E1	48107203	48107301	refGene_gn=SMARCD2;refGene_tx=XM_011289308.2;refGene_gn=SMARCD2;refGene_tx=XM_023243554.1
E1	48107471	48107633	refGene_gn=SMARCD2;refGene_tx=XM_011289308.2;refGene_gn=SMARCD2;refGene_tx=XM_023243554.1
E1	48107774	48107876	refGene_gn=SMARCD2;refGene_tx=XM_011289308.2;refGene_gn=SMARCD2;refGene_tx=XM_023243554.1
E1	48108019	48108115	refGene_gn=SMARCD2;refGene_tx=XM_011289308.2;refGene_gn=SMARCD2;refGene_tx=XM_023243554.1
E1	48108514	48108670	refGene_gn=SMARCD2;refGene_tx=XM_011289308.2;refGene_gn=SMARCD2;refGene_tx=XM_023243554.1
E1	48109685	48109808	refGene_gn=SMARCD2;refGene_tx=XM_011289308.2;refGene_gn=SMARCD2;refGene_tx=XM_023243554.1
E1	48109954	48109997	refGene_gn=SMARCD2;refGene_tx=XM_011289308.2;refGene_gn=SMARCD2;refGene_tx=XM_023243554.1
E1	48110193	48110378	refGene_gn=SMARCD2;refGene_tx=XM_011289308.2;refGene_gn=SMARCD2;refGene_tx=XM_023243554.1
E1	48114774	48114990	refGene_gn=SMARCD2;refGene_tx=XM_011289308.2
E1	48128273	48128327	refGene_gn=LOC101097853;refGene_tx=XM_019818094.2
E1	48132718	48132794	refGene_gn=LOC101097853;refGene_tx=XM_003997060.4;refGene_gn=LOC101097853;refGene_tx=XM_006940488.3;refGene_gn=LOC101097853;refGene_tx=XM_019818094.2
E1	48134094	48134361	refGene_gn=LOC101097853;refGene_tx=XM_003997060.4;refGene_gn=LOC101097853;refGene_tx=XM_006940488.3;refGene_gn=LOC101097853;refGene_tx=XM_019818094.2
E1	48135225	48135546	refGene_gn=LOC101097853;refGene_tx=XM_003997060.4;refGene_gn=LOC101097853;refGene_tx=XM_006940488.3;refGene_gn=LOC101097853;refGene_tx=XM_019818094.2
E1	48135803	48136082	refGene_gn=LOC101097853;refGene_tx=XM_003997060.4;refGene_gn=LOC101097853;refGene_tx=XM_006940488.3;refGene_gn=LOC101097853;refGene_tx=XM_01

9818094.2
E1 48136611 48136881
refGene_gn=LOC101097853;refGene_tx=XM_003997060.4;refGene_gn=LOC101097
853;refGene_tx=XM_006940488.3;refGene_gn=LOC101097853;refGene_tx=XM_01
9818094.2
E1 48137552 48137795
refGene_gn=LOC101097853;refGene_tx=XM_003997060.4;refGene_gn=LOC101097
853;refGene_tx=XM_006940488.3;refGene_gn=LOC101097853;refGene_tx=XM_01
9818094.2
E1 48137994 48138203
refGene_gn=LOC101097853;refGene_tx=XM_003997060.4;refGene_gn=LOC101097
853;refGene_tx=XM_006940488.3;refGene_gn=LOC101097853;refGene_tx=XM_01
9818094.2
E1 48150241 48150442 refGene_gn=GH1;refGene_tx=NM_001009337.1
E1 48150717 48150879 refGene_gn=GH1;refGene_tx=NM_001009337.1
E1 48151038 48151155 refGene_gn=GH1;refGene_tx=NM_001009337.1
E1 48151370 48151531 refGene_gn=GH1;refGene_tx=NM_001009337.1
E1 48151779 48151789 refGene_gn=GH1;refGene_tx=NM_001009337.1
E1 48160527 48160626
refGene_gn=CD79B;refGene_tx=XM_003997062.3;refGene_gn=CD79B;refGene_tx
=XM_006940490.2
E1 48160737 48160779
refGene_gn=CD79B;refGene_tx=XM_003997062.3;refGene_gn=CD79B;refGene_tx
=XM_006940490.2
E1 48161053 48161172
refGene_gn=CD79B;refGene_tx=XM_003997062.3;refGene_gn=CD79B;refGene_tx
=XM_006940490.2
E1 48161406 48161709
refGene_gn=CD79B;refGene_tx=XM_003997062.3;refGene_gn=CD79B;refGene_tx
=XM_006940490.2
E1 48162549 48162597
refGene_gn=CD79B;refGene_tx=XM_003997062.3;refGene_gn=CD79B;refGene_tx
=XM_006940490.2
E1 48163274 48163341
refGene_gn=CD79B;refGene_tx=XM_003997062.3;refGene_gn=CD79B;refGene_tx
=XM_006940490.2
E1 48169957 48171183
refGene_gn=SCN4A;refGene_tx=XM_003997064.5;refGene_gn=SCN4A;refGene_tx
=XM_006940491.3;refGene_gn=SCN4A;refGene_tx=XM_006940492.3
E1 48171949 48172220
refGene_gn=SCN4A;refGene_tx=XM_003997064.5;refGene_gn=SCN4A;refGene_tx
=XM_006940491.3;refGene_gn=SCN4A;refGene_tx=XM_006940492.3
E1 48172856 48172961
refGene_gn=SCN4A;refGene_tx=XM_003997064.5;refGene_gn=SCN4A;refGene_tx
=XM_006940491.3;refGene_gn=SCN4A;refGene_tx=XM_006940492.3
E1 48173467 48173605
refGene_gn=SCN4A;refGene_tx=XM_003997064.5;refGene_gn=SCN4A;refGene_tx
=XM_006940491.3;refGene_gn=SCN4A;refGene_tx=XM_006940492.3
E1 48173877 48173931
refGene_gn=SCN4A;refGene_tx=XM_003997064.5;refGene_gn=SCN4A;refGene_tx

=XM_006940491.3;refGene_gn=SCN4A;refGene_tx=XM_006940492.3
E1	48174201	48174480
refGene_gn=SCN4A;refGene_tx=XM_003997064.5;refGene_gn=SCN4A;refGene_tx=XM_006940491.3;refGene_gn=SCN4A;refGene_tx=XM_006940492.3
E1	48175777	48175900
refGene_gn=SCN4A;refGene_tx=XM_003997064.5;refGene_gn=SCN4A;refGene_tx=XM_006940491.3;refGene_gn=SCN4A;refGene_tx=XM_006940492.3
E1	48176024	48176198
refGene_gn=SCN4A;refGene_tx=XM_003997064.5;refGene_gn=SCN4A;refGene_tx=XM_006940491.3;refGene_gn=SCN4A;refGene_tx=XM_006940492.3
E1	48176731	48176886
refGene_gn=SCN4A;refGene_tx=XM_003997064.5;refGene_gn=SCN4A;refGene_tx=XM_006940491.3;refGene_gn=SCN4A;refGene_tx=XM_006940492.3
E1	48177519	48177652
refGene_gn=SCN4A;refGene_tx=XM_003997064.5;refGene_gn=SCN4A;refGene_tx=XM_006940491.3;refGene_gn=SCN4A;refGene_tx=XM_006940492.3
E1	48179546	48180035
refGene_gn=SCN4A;refGene_tx=XM_003997064.5;refGene_gn=SCN4A;refGene_tx=XM_006940491.3;refGene_gn=SCN4A;refGene_tx=XM_006940492.3
E1	48184556	48184913
refGene_gn=SCN4A;refGene_tx=XM_003997064.5;refGene_gn=SCN4A;refGene_tx=XM_006940491.3;refGene_gn=SCN4A;refGene_tx=XM_006940492.3
E1	48185645	48185819
refGene_gn=SCN4A;refGene_tx=XM_003997064.5;refGene_gn=SCN4A;refGene_tx=XM_006940491.3;refGene_gn=SCN4A;refGene_tx=XM_006940492.3
E1	48187565	48187804
refGene_gn=SCN4A;refGene_tx=XM_003997064.5;refGene_gn=SCN4A;refGene_tx=XM_006940491.3;refGene_gn=SCN4A;refGene_tx=XM_006940492.3
E1	48189745	48189899
refGene_gn=SCN4A;refGene_tx=XM_003997064.5;refGene_gn=SCN4A;refGene_tx=XM_006940491.3;refGene_gn=SCN4A;refGene_tx=XM_006940492.3
E1	48190514	48190724
refGene_gn=SCN4A;refGene_tx=XM_003997064.5;refGene_gn=SCN4A;refGene_tx=XM_006940491.3;refGene_gn=SCN4A;refGene_tx=XM_006940492.3
E1	48191505	48191647
refGene_gn=SCN4A;refGene_tx=XM_003997064.5;refGene_gn=SCN4A;refGene_tx=XM_006940491.3;refGene_gn=SCN4A;refGene_tx=XM_006940492.3
E1	48191886	48191950
refGene_gn=SCN4A;refGene_tx=XM_003997064.5;refGene_gn=SCN4A;refGene_tx=XM_006940491.3;refGene_gn=SCN4A;refGene_tx=XM_006940492.3
E1	48192798	48193113
refGene_gn=SCN4A;refGene_tx=XM_003997064.5;refGene_gn=SCN4A;refGene_tx=XM_006940491.3;refGene_gn=SCN4A;refGene_tx=XM_006940492.3
E1	48194705	48194797
refGene_gn=SCN4A;refGene_tx=XM_003997064.5;refGene_gn=SCN4A;refGene_tx=XM_006940491.3;refGene_gn=SCN4A;refGene_tx=XM_006940492.3
E1	48195270	48195399
refGene_gn=SCN4A;refGene_tx=XM_003997064.5;refGene_gn=SCN4A;refGene_tx=XM_006940491.3;refGene_gn=SCN4A;refGene_tx=XM_006940492.3
E1	48195683	48195773

refGene_gn=SCN4A;refGene_tx=XM_003997064.5;refGene_gn=SCN4A;refGene_tx=XM_006940491.3;refGene_gn=SCN4A;refGene_tx=XM_006940492.3
E1 48195911 48196030
refGene_gn=SCN4A;refGene_tx=XM_003997064.5;refGene_gn=SCN4A;refGene_tx=XM_006940491.3;refGene_gn=SCN4A;refGene_tx=XM_006940492.3
E1 48196120 48196393
refGene_gn=SCN4A;refGene_tx=XM_003997064.5;refGene_gn=SCN4A;refGene_tx=XM_006940491.3;refGene_gn=SCN4A;refGene_tx=XM_006940492.3
E1 48219249 48219333
refGene_gn=PRR29;refGene_tx=XM_011289426.3;refGene_gn=PRR29;refGene_tx=XM_023243626.1;refGene_gn=PRR29;refGene_tx=XM_023243627.1;refGene_gn=PRR29;refGene_tx=XM_023243628.1;refGene_gn=PRR29;refGene_tx=XM_023243629.1;refGene_gn=PRR29;refGene_tx=XM_023243631.1
E1 48219555 48219631
refGene_gn=PRR29;refGene_tx=XM_011289426.3;refGene_gn=PRR29;refGene_tx=XM_023243626.1;refGene_gn=PRR29;refGene_tx=XM_023243627.1;refGene_gn=PRR29;refGene_tx=XM_023243628.1;refGene_gn=PRR29;refGene_tx=XM_023243629.1;refGene_gn=PRR29;refGene_tx=XM_023243631.1
E1 48219823 48219960
refGene_gn=PRR29;refGene_tx=XM_011289426.3;refGene_gn=PRR29;refGene_tx=XM_023243626.1;refGene_gn=PRR29;refGene_tx=XM_023243627.1;refGene_gn=PRR29;refGene_tx=XM_023243628.1;refGene_gn=PRR29;refGene_tx=XM_023243629.1;refGene_gn=PRR29;refGene_tx=XM_023243630.1;refGene_gn=PRR29;refGene_tx=XM_023243631.1
E1 48220884 48221111
refGene_gn=PRR29;refGene_tx=XM_011289426.3;refGene_gn=PRR29;refGene_tx=XM_023243626.1;refGene_gn=PRR29;refGene_tx=XM_023243627.1;refGene_gn=PRR29;refGene_tx=XM_023243628.1;refGene_gn=PRR29;refGene_tx=XM_023243629.1;refGene_gn=PRR29;refGene_tx=XM_023243630.1;refGene_gn=PRR29;refGene_tx=XM_023243631.1
E1 48221255 48221326
refGene_gn=PRR29;refGene_tx=XM_011289426.3;refGene_gn=PRR29;refGene_tx=XM_023243626.1;refGene_gn=PRR29;refGene_tx=XM_023243627.1;refGene_gn=PRR29;refGene_tx=XM_023243628.1;refGene_gn=PRR29;refGene_tx=XM_023243629.1;refGene_gn=PRR29;refGene_tx=XM_023243630.1;refGene_gn=PRR29;refGene_tx=XM_023243631.1
E1 48221475 48221549
refGene_gn=PRR29;refGene_tx=XM_011289426.3;refGene_gn=PRR29;refGene_tx=XM_023243626.1;refGene_gn=PRR29;refGene_tx=XM_023243627.1;refGene_gn=PRR29;refGene_tx=XM_023243628.1;refGene_gn=PRR29;refGene_tx=XM_023243629.1;refGene_gn=PRR29;refGene_tx=XM_023243630.1;refGene_gn=PRR29;refGene_tx=XM_023243631.1
E1 48222550 48222699 refGene_gn=ICAM2;refGene_tx=XM_019818106.2
E1 48223362 48223683 refGene_gn=ICAM2;refGene_tx=XM_019818106.2
E1 48224872 48225136 refGene_gn=ICAM2;refGene_tx=XM_019818106.2
E1 48226918 48227111 refGene_gn=ICAM2;refGene_tx=XM_019818106.2
E1 48245299 48245302 refGene_gn=ERN1;refGene_tx=XM_019818108.2
E1 48252644 48252857
refGene_gn=ERN1;refGene_tx=XM_003997087.5;refGene_gn=ERN1;refGene_tx=XM_023244605.1

E1 48253400 48253468
refGene_gn=ERN1;refGene_tx=XM_003997087.5;refGene_gn=ERN1;refGene_tx=XM_019818108.2;refGene_gn=ERN1;refGene_tx=XM_023244605.1
E1 48253922 48254046
refGene_gn=ERN1;refGene_tx=XM_003997087.5;refGene_gn=ERN1;refGene_tx=XM_019818108.2;refGene_gn=ERN1;refGene_tx=XM_023244605.1
E1 48256253 48256381
refGene_gn=ERN1;refGene_tx=XM_003997087.5;refGene_gn=ERN1;refGene_tx=XM_019818108.2;refGene_gn=ERN1;refGene_tx=XM_023244605.1
E1 48258125 48258273
refGene_gn=ERN1;refGene_tx=XM_003997087.5;refGene_gn=ERN1;refGene_tx=XM_019818108.2;refGene_gn=ERN1;refGene_tx=XM_023244605.1
E1 48258684 48258884
refGene_gn=ERN1;refGene_tx=XM_003997087.5;refGene_gn=ERN1;refGene_tx=XM_019818108.2;refGene_gn=ERN1;refGene_tx=XM_023244605.1
E1 48259175 48259275
refGene_gn=ERN1;refGene_tx=XM_003997087.5;refGene_gn=ERN1;refGene_tx=XM_019818108.2;refGene_gn=ERN1;refGene_tx=XM_023244605.1
E1 48260045 48260235
refGene_gn=ERN1;refGene_tx=XM_003997087.5;refGene_gn=ERN1;refGene_tx=XM_019818108.2;refGene_gn=ERN1;refGene_tx=XM_023244605.1
E1 48260525 48260616
refGene_gn=ERN1;refGene_tx=XM_003997087.5;refGene_gn=ERN1;refGene_tx=XM_019818108.2;refGene_gn=ERN1;refGene_tx=XM_023244605.1
E1 48261353 48261621
refGene_gn=ERN1;refGene_tx=XM_003997087.5;refGene_gn=ERN1;refGene_tx=XM_019818108.2;refGene_gn=ERN1;refGene_tx=XM_023244605.1
E1 48263376 48263568
refGene_gn=ERN1;refGene_tx=XM_003997087.5;refGene_gn=ERN1;refGene_tx=XM_019818108.2;refGene_gn=ERN1;refGene_tx=XM_023244605.1
E1 48266292 48266411
refGene_gn=ERN1;refGene_tx=XM_003997087.5;refGene_gn=ERN1;refGene_tx=XM_019818108.2;refGene_gn=ERN1;refGene_tx=XM_023244605.1
E1 48270252 48270418
refGene_gn=ERN1;refGene_tx=XM_003997087.5;refGene_gn=ERN1;refGene_tx=XM_019818108.2;refGene_gn=ERN1;refGene_tx=XM_023244605.1
E1 48271595 48271674
refGene_gn=ERN1;refGene_tx=XM_003997087.5;refGene_gn=ERN1;refGene_tx=XM_019818108.2;refGene_gn=ERN1;refGene_tx=XM_023244605.1
E1 48272906 48273168
refGene_gn=ERN1;refGene_tx=XM_003997087.5;refGene_gn=ERN1;refGene_tx=XM_019818108.2;refGene_gn=ERN1;refGene_tx=XM_023244605.1
E1 48273995 48274097
refGene_gn=ERN1;refGene_tx=XM_003997087.5;refGene_gn=ERN1;refGene_tx=XM_019818108.2;refGene_gn=ERN1;refGene_tx=XM_023244605.1
E1 48275468 48275591
refGene_gn=ERN1;refGene_tx=XM_003997087.5;refGene_gn=ERN1;refGene_tx=XM_019818108.2;refGene_gn=ERN1;refGene_tx=XM_023244605.1
E1 48277925 48277998
refGene_gn=ERN1;refGene_tx=XM_003997087.5;refGene_gn=ERN1;refGene_tx=X

M_019818108.2;refGene_gn=ERN1;refGene_tx=XM_023244605.1
E1 48281170 48281243
refGene_gn=ERN1;refGene_tx=XM_003997087.5;refGene_gn=ERN1;refGene_tx=XM_019818108.2;refGene_gn=ERN1;refGene_tx=XM_023244605.1
E1 48282242 48282276
refGene_gn=ERN1;refGene_tx=XM_003997087.5;refGene_gn=ERN1;refGene_tx=XM_019818108.2;refGene_gn=ERN1;refGene_tx=XM_023244605.1
E1 48300449 48300570
refGene_gn=ERN1;refGene_tx=XM_003997087.5;refGene_gn=ERN1;refGene_tx=XM_019818108.2;refGene_gn=ERN1;refGene_tx=XM_023244605.1
E1 48300693 48300741
refGene_gn=ERN1;refGene_tx=XM_003997087.5;refGene_gn=ERN1;refGene_tx=XM_019818108.2
E1 48327592 48327637 refGene_gn=ERN1;refGene_tx=XM_023244605.1
E1 48343169 48343292
refGene_gn=TEX2;refGene_tx=XM_006940494.4;refGene_gn=TEX2;refGene_tx=XM_006940495.4;refGene_gn=TEX2;refGene_tx=XM_019818109.2;refGene_gn=TEX2;refGene_tx=XM_023244502.1;refGene_gn=TEX2;refGene_tx=XM_023244503.1;refGene_gn=TEX2;refGene_tx=XM_023244504.1;refGene_gn=TEX2;refGene_tx=XM_023244505.1
E1 48345100 48345221
refGene_gn=TEX2;refGene_tx=XM_006940494.4;refGene_gn=TEX2;refGene_tx=XM_006940495.4;refGene_gn=TEX2;refGene_tx=XM_019818109.2;refGene_gn=TEX2;refGene_tx=XM_023244502.1;refGene_gn=TEX2;refGene_tx=XM_023244503.1;refGene_gn=TEX2;refGene_tx=XM_023244504.1;refGene_gn=TEX2;refGene_tx=XM_023244505.1
E1 48347438 48347648
refGene_gn=TEX2;refGene_tx=XM_006940494.4;refGene_gn=TEX2;refGene_tx=XM_006940495.4;refGene_gn=TEX2;refGene_tx=XM_019818109.2;refGene_gn=TEX2;refGene_tx=XM_023244502.1;refGene_gn=TEX2;refGene_tx=XM_023244503.1;refGene_gn=TEX2;refGene_tx=XM_023244504.1;refGene_gn=TEX2;refGene_tx=XM_023244505.1
E1 48349105 48349231
refGene_gn=TEX2;refGene_tx=XM_006940494.4;refGene_gn=TEX2;refGene_tx=XM_006940495.4;refGene_gn=TEX2;refGene_tx=XM_019818109.2;refGene_gn=TEX2;refGene_tx=XM_023244502.1;refGene_gn=TEX2;refGene_tx=XM_023244503.1;refGene_gn=TEX2;refGene_tx=XM_023244504.1;refGene_gn=TEX2;refGene_tx=XM_023244505.1
E1 48349266 48349273 refGene_gn=TEX2;refGene_tx=XM_023244506.1
E1 48355068 48355201
refGene_gn=TEX2;refGene_tx=XM_006940494.4;refGene_gn=TEX2;refGene_tx=XM_006940495.4;refGene_gn=TEX2;refGene_tx=XM_019818109.2;refGene_gn=TEX2;refGene_tx=XM_023244502.1;refGene_gn=TEX2;refGene_tx=XM_023244503.1;refGene_gn=TEX2;refGene_tx=XM_023244504.1;refGene_gn=TEX2;refGene_tx=XM_023244505.1;refGene_gn=TEX2;refGene_tx=XM_023244506.1
E1 48364495 48364595
refGene_gn=TEX2;refGene_tx=XM_006940494.4;refGene_gn=TEX2;refGene_tx=XM_006940495.4;refGene_gn=TEX2;refGene_tx=XM_019818109.2;refGene_gn=TEX2;refGene_tx=XM_023244502.1;refGene_gn=TEX2;refGene_tx=XM_023244503.1;refGene_gn=TEX2;refGene_tx=XM_023244504.1;refGene_gn=TEX2;refGene_tx=X

M_023244505.1;refGene_gn=TEX2;refGene_tx=XM_023244506.1
E1 48368822 48368969
refGene_gn=TEX2;refGene_tx=XM_006940494.4;refGene_gn=TEX2;refGene_tx=XM_006940495.4;refGene_gn=TEX2;refGene_tx=XM_019818109.2;refGene_gn=TEX2;refGene_tx=XM_023244502.1;refGene_gn=TEX2;refGene_tx=XM_023244503.1;refGene_gn=TEX2;refGene_tx=XM_023244504.1;refGene_gn=TEX2;refGene_tx=XM_023244505.1;refGene_gn=TEX2;refGene_tx=XM_023244506.1
E1 48380297 48380545
refGene_gn=TEX2;refGene_tx=XM_006940494.4;refGene_gn=TEX2;refGene_tx=XM_006940495.4;refGene_gn=TEX2;refGene_tx=XM_019818109.2;refGene_gn=TEX2;refGene_tx=XM_023244502.1;refGene_gn=TEX2;refGene_tx=XM_023244503.1;refGene_gn=TEX2;refGene_tx=XM_023244504.1;refGene_gn=TEX2;refGene_tx=XM_023244505.1;refGene_gn=TEX2;refGene_tx=XM_023244506.1
E1 48381804 48382144
refGene_gn=TEX2;refGene_tx=XM_006940494.4;refGene_gn=TEX2;refGene_tx=XM_006940495.4;refGene_gn=TEX2;refGene_tx=XM_019818109.2;refGene_gn=TEX2;refGene_tx=XM_023244502.1;refGene_gn=TEX2;refGene_tx=XM_023244503.1;refGene_gn=TEX2;refGene_tx=XM_023244504.1;refGene_gn=TEX2;refGene_tx=XM_023244505.1;refGene_gn=TEX2;refGene_tx=XM_023244506.1
E1 48383282 48383483
refGene_gn=TEX2;refGene_tx=XM_006940494.4;refGene_gn=TEX2;refGene_tx=XM_006940495.4;refGene_gn=TEX2;refGene_tx=XM_019818109.2;refGene_gn=TEX2;refGene_tx=XM_023244502.1;refGene_gn=TEX2;refGene_tx=XM_023244503.1;refGene_gn=TEX2;refGene_tx=XM_023244504.1;refGene_gn=TEX2;refGene_tx=XM_023244505.1;refGene_gn=TEX2;refGene_tx=XM_023244506.1
E1 48400960 48402601
refGene_gn=TEX2;refGene_tx=XM_006940494.4;refGene_gn=TEX2;refGene_tx=XM_006940495.4;refGene_gn=TEX2;refGene_tx=XM_019818109.2;refGene_gn=TEX2;refGene_tx=XM_023244502.1;refGene_gn=TEX2;refGene_tx=XM_023244503.1;refGene_gn=TEX2;refGene_tx=XM_023244504.1;refGene_gn=TEX2;refGene_tx=XM_023244505.1;refGene_gn=TEX2;refGene_tx=XM_023244506.1
E1 48488879 48488909
refGene_gn=PECAM1;refGene_tx=XM_006940496.4;refGene_gn=PECAM1;refGene_tx=XM_006940497.4;refGene_gn=PECAM1;refGene_tx=XM_006940498.4;refGene_gn=PECAM1;refGene_tx=XM_019818114.2;refGene_gn=PECAM1;refGene_tx=XM_023244610.1
E1 48494696 48494719
refGene_gn=PECAM1;refGene_tx=XM_006940496.4;refGene_gn=PECAM1;refGene_tx=XM_006940497.4;refGene_gn=PECAM1;refGene_tx=XM_006940498.4;refGene_gn=PECAM1;refGene_tx=XM_019818114.2;refGene_gn=PECAM1;refGene_tx=XM_023244610.1
E1 48505187 48505244
refGene_gn=PECAM1;refGene_tx=XM_006940496.4;refGene_gn=PECAM1;refGene_tx=XM_006940497.4;refGene_gn=PECAM1;refGene_tx=XM_019818114.2;refGene_gn=PECAM1;refGene_tx=XM_023244610.1
E1 48507780 48507855
refGene_gn=PECAM1;refGene_tx=XM_006940496.4;refGene_gn=PECAM1;refGene_tx=XM_023244610.1
E1 48510560 48510623
refGene_gn=PECAM1;refGene_tx=XM_006940496.4;refGene_gn=PECAM1;refGene_

tx=XM_006940497.4;refGene_gn=PECAM1;refGene_tx=XM_006940498.4;refGene_gn=PECAM1;refGene_tx=XM_023244610.1
E1 48512075 48512129
refGene_gn=PECAM1;refGene_tx=XM_006940496.4;refGene_gn=PECAM1;refGene_tx=XM_006940497.4;refGene_gn=PECAM1;refGene_tx=XM_006940498.4;refGene_gn=PECAM1;refGene_tx=XM_019818114.2;refGene_gn=PECAM1;refGene_tx=XM_023244610.1
E1 48514787 48514864
refGene_gn=PECAM1;refGene_tx=XM_006940496.4;refGene_gn=PECAM1;refGene_tx=XM_006940497.4;refGene_gn=PECAM1;refGene_tx=XM_006940498.4;refGene_gn=PECAM1;refGene_tx=XM_019818114.2;refGene_gn=PECAM1;refGene_tx=XM_023244610.1
E1 48515631 48515659
refGene_gn=PECAM1;refGene_tx=XM_006940496.4;refGene_gn=PECAM1;refGene_tx=XM_006940497.4;refGene_gn=PECAM1;refGene_tx=XM_006940498.4;refGene_gn=PECAM1;refGene_tx=XM_019818114.2;refGene_gn=PECAM1;refGene_tx=XM_023244610.1
E1 48516756 48516864
refGene_gn=PECAM1;refGene_tx=XM_006940496.4;refGene_gn=PECAM1;refGene_tx=XM_006940497.4;refGene_gn=PECAM1;refGene_tx=XM_006940498.4;refGene_gn=PECAM1;refGene_tx=XM_019818114.2;refGene_gn=PECAM1;refGene_tx=XM_023244610.1
E1 48517568 48517856
refGene_gn=PECAM1;refGene_tx=XM_006940496.4;refGene_gn=PECAM1;refGene_tx=XM_006940497.4;refGene_gn=PECAM1;refGene_tx=XM_006940498.4;refGene_gn=PECAM1;refGene_tx=XM_019818114.2;refGene_gn=PECAM1;refGene_tx=XM_023244610.1
E1 48519703 48519979
refGene_gn=PECAM1;refGene_tx=XM_006940496.4;refGene_gn=PECAM1;refGene_tx=XM_006940497.4;refGene_gn=PECAM1;refGene_tx=XM_006940498.4;refGene_gn=PECAM1;refGene_tx=XM_019818114.2;refGene_gn=PECAM1;refGene_tx=XM_023244610.1
E1 48521734 48521983
refGene_gn=PECAM1;refGene_tx=XM_006940496.4;refGene_gn=PECAM1;refGene_tx=XM_006940497.4;refGene_gn=PECAM1;refGene_tx=XM_006940498.4;refGene_gn=PECAM1;refGene_tx=XM_019818114.2;refGene_gn=PECAM1;refGene_tx=XM_023244610.1
E1 48526431 48526707
refGene_gn=PECAM1;refGene_tx=XM_006940496.4;refGene_gn=PECAM1;refGene_tx=XM_006940497.4;refGene_gn=PECAM1;refGene_tx=XM_006940498.4;refGene_gn=PECAM1;refGene_tx=XM_019818114.2;refGene_gn=PECAM1;refGene_tx=XM_023244610.1
E1 48531618 48531924
refGene_gn=PECAM1;refGene_tx=XM_006940496.4;refGene_gn=PECAM1;refGene_tx=XM_006940497.4;refGene_gn=PECAM1;refGene_tx=XM_006940498.4;refGene_gn=PECAM1;refGene_tx=XM_019818114.2;refGene_gn=PECAM1;refGene_tx=XM_023244610.1
E1 48535507 48535801
refGene_gn=PECAM1;refGene_tx=XM_006940496.4;refGene_gn=PECAM1;refGene_tx=XM_006940497.4;refGene_gn=PECAM1;refGene_tx=XM_006940498.4;refGene_

gn=PECAM1;refGene_tx=XM_019818114.2;refGene_gn=PECAM1;refGene_tx=XM_023244610.1
E1 48547456 48547483 refGene_gn=PECAM1;refGene_tx=XM_006940496.4;refGene_gn=PECAM1;refGene_tx=XM_006940497.4;refGene_gn=PECAM1;refGene_tx=XM_006940498.4;refGene_gn=PECAM1;refGene_tx=XM_019818114.2;refGene_gn=PECAM1;refGene_tx=XM_023244610.1
E1 48547569 48547633 refGene_gn=PECAM1;refGene_tx=XM_006940496.4;refGene_gn=PECAM1;refGene_tx=XM_006940497.4;refGene_gn=PECAM1;refGene_tx=XM_006940498.4;refGene_gn=PECAM1;refGene_tx=XM_019818114.2;refGene_gn=PECAM1;refGene_tx=XM_023244610.1
E1 48573261 48573391 refGene_gn=MILR1;refGene_tx=XM_006940499.3;refGene_gn=MILR1;refGene_tx=XM_006940501.4;refGene_gn=MILR1;refGene_tx=XM_019818120.2
E1 48580925 48581201 refGene_gn=MILR1;refGene_tx=XM_006940499.3;refGene_gn=MILR1;refGene_tx=XM_006940501.4;refGene_gn=MILR1;refGene_tx=XM_019818120.2
E1 48586360 48586648 refGene_gn=MILR1;refGene_tx=XM_006940499.3;refGene_gn=MILR1;refGene_tx=XM_006940501.4;refGene_gn=MILR1;refGene_tx=XM_019818120.2
E1 48588294 48588405 refGene_gn=MILR1;refGene_tx=XM_006940499.3;refGene_gn=MILR1;refGene_tx=XM_006940501.4;refGene_gn=MILR1;refGene_tx=XM_019818120.2
E1 48590253 48590343 refGene_gn=MILR1;refGene_tx=XM_006940499.3;refGene_gn=MILR1;refGene_tx=XM_006940501.4;refGene_gn=MILR1;refGene_tx=XM_019818120.2
E1 48590869 48590920 refGene_gn=MILR1;refGene_tx=XM_006940499.3;refGene_gn=MILR1;refGene_tx=XM_006940501.4;refGene_gn=MILR1;refGene_tx=XM_019818120.2
E1 48591052 48591121 refGene_gn=MILR1;refGene_tx=XM_006940499.3;refGene_gn=MILR1;refGene_tx=XM_006940501.4;refGene_gn=MILR1;refGene_tx=XM_019818120.2
E1 48591565 48591621 refGene_gn=MILR1;refGene_tx=XM_006940499.3;refGene_gn=MILR1;refGene_tx=XM_006940501.4;refGene_gn=MILR1;refGene_tx=XM_019818120.2
E1 48594422 48594582 refGene_gn=POLG2;refGene_tx=XM_019818116.2
E1 48602191 48602357 refGene_gn=POLG2;refGene_tx=XM_011289309.3;refGene_gn=POLG2;refGene_tx=XM_011289310.3;refGene_gn=POLG2;refGene_tx=XM_011289311.3;refGene_gn=POLG2;refGene_tx=XM_011289312.3;refGene_gn=POLG2;refGene_tx=XM_011289313.3;refGene_gn=POLG2;refGene_tx=XM_019818117.2;refGene_gn=POLG2;refGene_tx=XM_019818118.2;refGene_gn=POLG2;refGene_tx=XM_019818119.2
E1 48604533 48604634 refGene_gn=POLG2;refGene_tx=XM_011289309.3;refGene_gn=POLG2;refGene_tx=XM_011289310.3;refGene_gn=POLG2;refGene_tx=XM_011289311.3;refGene_gn=POLG2;refGene_tx=XM_011289312.3;refGene_gn=POLG2;refGene_tx=XM_011289313.3;refGene_gn=POLG2;refGene_tx=XM_019818116.2;refGene_gn=POLG2;refGene_tx=XM_019818117.2;refGene_gn=POLG2;refGene_tx=XM_019818118.2;refGene_gn=POLG2;refGene_tx=XM_019818119.2

E1	48607251	48607332
refGene_gn=POLG2;refGene_tx=XM_011289309.3;refGene_gn=POLG2;refGene_tx=XM_011289311.3;refGene_gn=POLG2;refGene_tx=XM_011289312.3;refGene_gn=POLG2;refGene_tx=XM_019818116.2;refGene_gn=POLG2;refGene_tx=XM_019818117.2;refGene_gn=POLG2;refGene_tx=XM_019818118.2
E1	48609503	48609644
refGene_gn=POLG2;refGene_tx=XM_011289309.3;refGene_gn=POLG2;refGene_tx=XM_011289310.3;refGene_gn=POLG2;refGene_tx=XM_011289312.3;refGene_gn=POLG2;refGene_tx=XM_019818116.2;refGene_gn=POLG2;refGene_tx=XM_019818117.2;refGene_gn=POLG2;refGene_tx=XM_019818118.2;refGene_gn=POLG2;refGene_tx=XM_019818119.2
E1	48615408	48615582
refGene_gn=POLG2;refGene_tx=XM_011289309.3;refGene_gn=POLG2;refGene_tx=XM_011289310.3;refGene_gn=POLG2;refGene_tx=XM_011289311.3;refGene_gn=POLG2;refGene_tx=XM_011289312.3;refGene_gn=POLG2;refGene_tx=XM_011289313.3;refGene_gn=POLG2;refGene_tx=XM_019818116.2;refGene_gn=POLG2;refGene_tx=XM_019818117.2;refGene_gn=POLG2;refGene_tx=XM_019818119.2
E1	48616815	48616921
refGene_gn=POLG2;refGene_tx=XM_011289309.3;refGene_gn=POLG2;refGene_tx=XM_011289310.3;refGene_gn=POLG2;refGene_tx=XM_011289311.3;refGene_gn=POLG2;refGene_tx=XM_011289312.3;refGene_gn=POLG2;refGene_tx=XM_011289313.3;refGene_gn=POLG2;refGene_tx=XM_019818116.2;refGene_gn=POLG2;refGene_tx=XM_019818117.2;refGene_gn=POLG2;refGene_tx=XM_019818118.2;refGene_gn=POLG2;refGene_tx=XM_019818119.2
E1	48617046	48617173
refGene_gn=POLG2;refGene_tx=XM_011289309.3;refGene_gn=POLG2;refGene_tx=XM_011289310.3;refGene_gn=POLG2;refGene_tx=XM_011289311.3;refGene_gn=POLG2;refGene_tx=XM_011289312.3;refGene_gn=POLG2;refGene_tx=XM_011289313.3;refGene_gn=POLG2;refGene_tx=XM_019818116.2;refGene_gn=POLG2;refGene_tx=XM_019818117.2;refGene_gn=POLG2;refGene_tx=XM_019818118.2;refGene_gn=POLG2;refGene_tx=XM_019818119.2
E1	48620606	48621168
refGene_gn=POLG2;refGene_tx=XM_011289309.3;refGene_gn=POLG2;refGene_tx=XM_011289310.3;refGene_gn=POLG2;refGene_tx=XM_011289311.3;refGene_gn=POLG2;refGene_tx=XM_011289312.3;refGene_gn=POLG2;refGene_tx=XM_011289313.3;refGene_gn=POLG2;refGene_tx=XM_019818116.2;refGene_gn=POLG2;refGene_tx=XM_019818117.2;refGene_gn=POLG2;refGene_tx=XM_019818118.2;refGene_gn=POLG2;refGene_tx=XM_019818119.2
E1	48624053	48624457
refGene_gn=DDX5;refGene_tx=XM_003997067.3;refGene_gn=DDX5;refGene_tx=XM_006940502.2;refGene_gn=DDX5;refGene_tx=XM_006940503.3;refGene_gn=DDX5;refGene_tx=XM_019818115.2
E1	48624679	48624904
refGene_gn=DDX5;refGene_tx=XM_003997067.3;refGene_gn=DDX5;refGene_tx=XM_006940502.2;refGene_gn=DDX5;refGene_tx=XM_006940503.3;refGene_gn=DDX5;refGene_tx=XM_019818115.2
E1	48626147	48626207
refGene_gn=DDX5;refGene_tx=XM_003997067.3;refGene_gn=DDX5;refGene_tx=XM_006940502.2;refGene_gn=DDX5;refGene_tx=XM_006940503.3;refGene_gn=DDX5;refGene_tx=XM_019818115.2

E1 48626285 48626347
refGene_gn=DDX5;refGene_tx=XM_003997067.3;refGene_gn=DDX5;refGene_tx=X
M_006940502.2;refGene_gn=DDX5;refGene_tx=XM_006940503.3;refGene_gn=DDX
5;refGene_tx=XM_019818115.2
E1 48626573 48626684
refGene_gn=DDX5;refGene_tx=XM_003997067.3;refGene_gn=DDX5;refGene_tx=X
M_006940502.2;refGene_gn=DDX5;refGene_tx=XM_006940503.3;refGene_gn=DDX
5;refGene_tx=XM_019818115.2
E1 48627053 48627226
refGene_gn=DDX5;refGene_tx=XM_003997067.3;refGene_gn=DDX5;refGene_tx=X
M_006940502.2;refGene_gn=DDX5;refGene_tx=XM_006940503.3;refGene_gn=DDX
5;refGene_tx=XM_019818115.2
E1 48627318 48627479
refGene_gn=DDX5;refGene_tx=XM_003997067.3;refGene_gn=DDX5;refGene_tx=X
M_006940502.2;refGene_gn=DDX5;refGene_tx=XM_006940503.3;refGene_gn=DDX
5;refGene_tx=XM_019818115.2
E1 48627560 48627702
refGene_gn=DDX5;refGene_tx=XM_003997067.3;refGene_gn=DDX5;refGene_tx=X
M_006940502.2;refGene_gn=DDX5;refGene_tx=XM_006940503.3;refGene_gn=DDX
5;refGene_tx=XM_019818115.2
E1 48627946 48628012
refGene_gn=DDX5;refGene_tx=XM_003997067.3;refGene_gn=DDX5;refGene_tx=X
M_006940502.2;refGene_gn=DDX5;refGene_tx=XM_006940503.3;refGene_gn=DDX
5;refGene_tx=XM_019818115.2
E1 48628134 48628268
refGene_gn=DDX5;refGene_tx=XM_003997067.3;refGene_gn=DDX5;refGene_tx=X
M_006940502.2;refGene_gn=DDX5;refGene_tx=XM_006940503.3;refGene_gn=DDX
5;refGene_tx=XM_019818115.2
E1 48628361 48628458
refGene_gn=DDX5;refGene_tx=XM_003997067.3;refGene_gn=DDX5;refGene_tx=X
M_006940502.2;refGene_gn=DDX5;refGene_tx=XM_006940503.3;refGene_gn=DDX
5;refGene_tx=XM_019818115.2
E1 48628805 48628971
refGene_gn=DDX5;refGene_tx=XM_003997067.3;refGene_gn=DDX5;refGene_tx=X
M_006940502.2;refGene_gn=DDX5;refGene_tx=XM_006940503.3;refGene_gn=DDX
5;refGene_tx=XM_019818115.2
E1 48629456 48629638 refGene_gn=DDX5;refGene_tx=XM_019818115.2
E1 48630183 48630227
refGene_gn=DDX5;refGene_tx=XM_003997067.3;refGene_gn=DDX5;refGene_tx=X
M_006940502.2;refGene_gn=DDX5;refGene_tx=XM_006940503.3
E1 48631251 48631270 refGene_gn=CEP95;refGene_tx=XM_003997069.4
E1 48632744 48632873 refGene_gn=CEP95;refGene_tx=XM_003997069.4
E1 48636420 48636528 refGene_gn=CEP95;refGene_tx=XM_003997069.4
E1 48640146 48640257 refGene_gn=CEP95;refGene_tx=XM_003997069.4
E1 48643240 48643346 refGene_gn=CEP95;refGene_tx=XM_003997069.4
E1 48646768 48646884 refGene_gn=CEP95;refGene_tx=XM_003997069.4
E1 48654760 48654880 refGene_gn=CEP95;refGene_tx=XM_003997069.4
E1 48655429 48655623 refGene_gn=CEP95;refGene_tx=XM_003997069.4
E1 48657843 48657956 refGene_gn=CEP95;refGene_tx=XM_003997069.4
E1 48658131 48658261 refGene_gn=CEP95;refGene_tx=XM_003997069.4

E1 48659257 48659411 refGene_gn=CEP95;refGene_tx=XM_003997069.4
E1 48669451 48669591 refGene_gn=CEP95;refGene_tx=XM_003997069.4
E1 48670342 48670432 refGene_gn=CEP95;refGene_tx=XM_003997069.4
E1 48671241 48671374 refGene_gn=CEP95;refGene_tx=XM_003997069.4
E1 48672221 48672391 refGene_gn=CEP95;refGene_tx=XM_003997069.4
E1 48672505 48672580 refGene_gn=CEP95;refGene_tx=XM_003997069.4
E1 48673445 48673598 refGene_gn=CEP95;refGene_tx=XM_003997069.4
E1 48675124 48675271 refGene_gn=CEP95;refGene_tx=XM_003997069.4
E1 48675767 48675839 refGene_gn=CEP95;refGene_tx=XM_003997069.4
E1 48676124 48676301 refGene_gn=CEP95;refGene_tx=XM_003997069.4
E1 48685394 48685494 refGene_gn=SMURF2;refGene_tx=XM_023243700.1;refGene_gn=SMURF2;refGene_tx=XM_023243701.1
E1 48685804 48685880 refGene_gn=SMURF2;refGene_tx=XM_023243700.1;refGene_gn=SMURF2;refGene_tx=XM_023243701.1
E1 48687145 48687347 refGene_gn=SMURF2;refGene_tx=XM_023243700.1
E1 48689417 48689538 refGene_gn=SMURF2;refGene_tx=XM_023243700.1;refGene_gn=SMURF2;refGene_tx=XM_023243701.1
E1 48692902 48693040 refGene_gn=SMURF2;refGene_tx=XM_023243700.1;refGene_gn=SMURF2;refGene_tx=XM_023243701.1
E1 48694089 48694268 refGene_gn=SMURF2;refGene_tx=XM_023243700.1;refGene_gn=SMURF2;refGene_tx=XM_023243701.1
E1 48695903 48696018 refGene_gn=SMURF2;refGene_tx=XM_023243700.1;refGene_gn=SMURF2;refGene_tx=XM_023243701.1
E1 48698975 48699079 refGene_gn=SMURF2;refGene_tx=XM_023243700.1;refGene_gn=SMURF2;refGene_tx=XM_023243701.1
E1 48701425 48701621 refGene_gn=SMURF2;refGene_tx=XM_023243700.1;refGene_gn=SMURF2;refGene_tx=XM_023243701.1
E1 48706549 48706708 refGene_gn=SMURF2;refGene_tx=XM_023243700.1;refGene_gn=SMURF2;refGene_tx=XM_023243701.1
E1 48711523 48711608 refGene_gn=SMURF2;refGene_tx=XM_023243700.1;refGene_gn=SMURF2;refGene_tx=XM_023243701.1
E1 48713932 48714135 refGene_gn=SMURF2;refGene_tx=XM_023243700.1;refGene_gn=SMURF2;refGene_tx=XM_023243701.1
E1 48716675 48716759 refGene_gn=SMURF2;refGene_tx=XM_023243700.1;refGene_gn=SMURF2;refGene_tx=XM_023243701.1
E1 48718785 48718870 refGene_gn=SMURF2;refGene_tx=XM_023243700.1;refGene_gn=SMURF2;refGene_tx=XM_023243701.1

E1 48723402 48723468
refGene_gn=SMURF2;refGene_tx=XM_023243700.1;refGene_gn=SMURF2;refGene_tx=XM_023243701.1
E1 48726918 48727052
refGene_gn=SMURF2;refGene_tx=XM_023243700.1;refGene_gn=SMURF2;refGene_tx=XM_023243701.1
E1 48736651 48736760
refGene_gn=SMURF2;refGene_tx=XM_023243700.1;refGene_gn=SMURF2;refGene_tx=XM_023243701.1
E1 48742474 48742513
refGene_gn=SMURF2;refGene_tx=XM_023243700.1;refGene_gn=SMURF2;refGene_tx=XM_023243701.1
E1 48803830 48803882
refGene_gn=SMURF2;refGene_tx=XM_023243700.1;refGene_gn=SMURF2;refGene_tx=XM_023243701.1
E1 48880324 48880417 refGene_gn=KPNA2;refGene_tx=XM_023243470.1
E1 48881526 48881676 refGene_gn=KPNA2;refGene_tx=XM_023243470.1
E1 48883233 48883416 refGene_gn=KPNA2;refGene_tx=XM_023243470.1
E1 48883675 48883909 refGene_gn=KPNA2;refGene_tx=XM_023243470.1
E1 48884625 48884889 refGene_gn=KPNA2;refGene_tx=XM_023243470.1
E1 48884991 48885091 refGene_gn=KPNA2;refGene_tx=XM_023243470.1
E1 48886594 48886683 refGene_gn=KPNA2;refGene_tx=XM_023243470.1
E1 48888645 48888783 refGene_gn=KPNA2;refGene_tx=XM_023243470.1
E1 48888939 48889014 refGene_gn=KPNA2;refGene_tx=XM_023243470.1
E1 48915992 48916095
refGene_gn=CE1H17orf58;refGene_tx=XM_003997070.5
E1 48917081 48917272
refGene_gn=CE1H17orf58;refGene_tx=XM_003997070.5
E1 48923129 48923166
refGene_gn=BPTF;refGene_tx=XM_023243975.1;refGene_gn=BPTF;refGene_tx=XM_023243976.1;refGene_gn=BPTF;refGene_tx=XM_023243979.1;refGene_gn=BPTF;refGene_tx=XM_023243980.1;refGene_gn=BPTF;refGene_tx=XM_023243981.1;refGene_gn=BPTF;refGene_tx=XM_023243982.1;refGene_gn=BPTF;refGene_tx=XM_023243983.1;refGene_gn=BPTF;refGene_tx=XM_023243984.1;refGene_gn=BPTF;refGene_tx=XM_023243985.1;refGene_gn=BPTF;refGene_tx=XM_023243986.1;refGene_gn=BPTF;refGene_tx=XM_023243987.1;refGene_gn=BPTF;refGene_tx=XM_023243988.1;refGene_gn=BPTF;refGene_tx=XM_023243989.1
E1 48924005 48924012 refGene_gn=BPTF;refGene_tx=XM_023243978.1
E1 48928393 48928580
refGene_gn=BPTF;refGene_tx=XM_023243975.1;refGene_gn=BPTF;refGene_tx=XM_023243976.1;refGene_gn=BPTF;refGene_tx=XM_023243978.1;refGene_gn=BPTF;refGene_tx=XM_023243979.1;refGene_gn=BPTF;refGene_tx=XM_023243980.1;refGene_gn=BPTF;refGene_tx=XM_023243981.1;refGene_gn=BPTF;refGene_tx=XM_023243982.1;refGene_gn=BPTF;refGene_tx=XM_023243983.1;refGene_gn=BPTF;refGene_tx=XM_023243984.1;refGene_gn=BPTF;refGene_tx=XM_023243985.1;refGene_gn=BPTF;refGene_tx=XM_023243986.1;refGene_gn=BPTF;refGene_tx=XM_023243987.1;refGene_gn=BPTF;refGene_tx=XM_023243988.1;refGene_gn=BPTF;refGene_tx=XM_023243989.1
E1 48934977 48935062
refGene_gn=BPTF;refGene_tx=XM_023243975.1;refGene_gn=BPTF;refGene_tx=X

M_023243976.1;refGene_gn=BPTF;refGene_tx=XM_023243978.1;refGene_gn=BPTF;refGene_tx=XM_023243979.1;refGene_gn=BPTF;refGene_tx=XM_023243980.1;refGene_gn=BPTF;refGene_tx=XM_023243981.1;refGene_gn=BPTF;refGene_tx=XM_023243982.1;refGene_gn=BPTF;refGene_tx=XM_023243983.1;refGene_gn=BPTF;refGene_tx=XM_023243984.1;refGene_gn=BPTF;refGene_tx=XM_023243985.1;refGene_gn=BPTF;refGene_tx=XM_023243986.1;refGene_gn=BPTF;refGene_tx=XM_023243987.1;refGene_gn=BPTF;refGene_tx=XM_023243988.1;refGene_gn=BPTF;refGene_tx=XM_023243989.1

E1 48935896 48936089 refGene_gn=BPTF;refGene_tx=XM_023243975.1;refGene_gn=BPTF;refGene_tx=XM_023243976.1;refGene_gn=BPTF;refGene_tx=XM_023243978.1;refGene_gn=BPTF;refGene_tx=XM_023243979.1;refGene_gn=BPTF;refGene_tx=XM_023243980.1;refGene_gn=BPTF;refGene_tx=XM_023243981.1;refGene_gn=BPTF;refGene_tx=XM_023243982.1;refGene_gn=BPTF;refGene_tx=XM_023243983.1;refGene_gn=BPTF;refGene_tx=XM_023243984.1;refGene_gn=BPTF;refGene_tx=XM_023243985.1;refGene_gn=BPTF;refGene_tx=XM_023243986.1;refGene_gn=BPTF;refGene_tx=XM_023243987.1;refGene_gn=BPTF;refGene_tx=XM_023243988.1;refGene_gn=BPTF;refGene_tx=XM_023243989.1

E1 48936835 48937009 refGene_gn=BPTF;refGene_tx=XM_023243975.1;refGene_gn=BPTF;refGene_tx=XM_023243976.1;refGene_gn=BPTF;refGene_tx=XM_023243978.1;refGene_gn=BPTF;refGene_tx=XM_023243979.1;refGene_gn=BPTF;refGene_tx=XM_023243980.1;refGene_gn=BPTF;refGene_tx=XM_023243982.1;refGene_gn=BPTF;refGene_tx=XM_023243983.1;refGene_gn=BPTF;refGene_tx=XM_023243984.1;refGene_gn=BPTF;refGene_tx=XM_023243985.1;refGene_gn=BPTF;refGene_tx=XM_023243987.1

E1 48939861 48940202 refGene_gn=BPTF;refGene_tx=XM_023243975.1;refGene_gn=BPTF;refGene_tx=XM_023243976.1;refGene_gn=BPTF;refGene_tx=XM_023243978.1;refGene_gn=BPTF;refGene_tx=XM_023243979.1;refGene_gn=BPTF;refGene_tx=XM_023243980.1;refGene_gn=BPTF;refGene_tx=XM_023243981.1;refGene_gn=BPTF;refGene_tx=XM_023243982.1;refGene_gn=BPTF;refGene_tx=XM_023243983.1;refGene_gn=BPTF;refGene_tx=XM_023243984.1;refGene_gn=BPTF;refGene_tx=XM_023243985.1;refGene_gn=BPTF;refGene_tx=XM_023243986.1;refGene_gn=BPTF;refGene_tx=XM_023243987.1;refGene_gn=BPTF;refGene_tx=XM_023243988.1;refGene_gn=BPTF;refGene_tx=XM_023243989.1

E1 48946042 48946268 refGene_gn=BPTF;refGene_tx=XM_023243975.1;refGene_gn=BPTF;refGene_tx=XM_023243976.1;refGene_gn=BPTF;refGene_tx=XM_023243978.1;refGene_gn=BPTF;refGene_tx=XM_023243979.1;refGene_gn=BPTF;refGene_tx=XM_023243980.1;refGene_gn=BPTF;refGene_tx=XM_023243981.1;refGene_gn=BPTF;refGene_tx=XM_023243982.1;refGene_gn=BPTF;refGene_tx=XM_023243983.1;refGene_gn=BPTF;refGene_tx=XM_023243984.1;refGene_gn=BPTF;refGene_tx=XM_023243985.1;refGene_gn=BPTF;refGene_tx=XM_023243986.1;refGene_gn=BPTF;refGene_tx=XM_023243987.1;refGene_gn=BPTF;refGene_tx=XM_023243988.1;refGene_gn=BPTF;refGene_tx=XM_023243989.1

E1 48946581 48946664 refGene_gn=BPTF;refGene_tx=XM_023243975.1;refGene_gn=BPTF;refGene_tx=XM_023243976.1;refGene_gn=BPTF;refGene_tx=XM_023243978.1;refGene_gn=BPTF;refGene_tx=XM_023243979.1;refGene_gn=BPTF;refGene_tx=XM_023243980.1;refGene_gn=BPTF;refGene_tx=XM_023243981.1;refGene_gn=BPTF;refGene_tx=X

M_023243982.1;refGene_gn=BPTF;refGene_tx=XM_023243983.1;refGene_gn=BPTF;refGene_tx=XM_023243984.1;refGene_gn=BPTF;refGene_tx=XM_023243985.1;refGene_gn=BPTF;refGene_tx=XM_023243986.1;refGene_gn=BPTF;refGene_tx=XM_023243987.1;refGene_gn=BPTF;refGene_tx=XM_023243988.1;refGene_gn=BPTF;refGene_tx=XM_023243989.1
E1 48948259 48949176
refGene_gn=BPTF;refGene_tx=XM_023243975.1;refGene_gn=BPTF;refGene_tx=XM_023243976.1;refGene_gn=BPTF;refGene_tx=XM_023243978.1;refGene_gn=BPTF;refGene_tx=XM_023243979.1;refGene_gn=BPTF;refGene_tx=XM_023243980.1;refGene_gn=BPTF;refGene_tx=XM_023243981.1;refGene_gn=BPTF;refGene_tx=XM_023243982.1;refGene_gn=BPTF;refGene_tx=XM_023243983.1;refGene_gn=BPTF;refGene_tx=XM_023243984.1;refGene_gn=BPTF;refGene_tx=XM_023243985.1;refGene_gn=BPTF;refGene_tx=XM_023243986.1;refGene_gn=BPTF;refGene_tx=XM_023243987.1;refGene_gn=BPTF;refGene_tx=XM_023243988.1;refGene_gn=BPTF;refGene_tx=XM_023243989.1
E1 48949853 48950076
refGene_gn=BPTF;refGene_tx=XM_023243975.1;refGene_gn=BPTF;refGene_tx=XM_023243976.1;refGene_gn=BPTF;refGene_tx=XM_023243978.1;refGene_gn=BPTF;refGene_tx=XM_023243979.1;refGene_gn=BPTF;refGene_tx=XM_023243980.1;refGene_gn=BPTF;refGene_tx=XM_023243981.1;refGene_gn=BPTF;refGene_tx=XM_023243982.1;refGene_gn=BPTF;refGene_tx=XM_023243983.1;refGene_gn=BPTF;refGene_tx=XM_023243984.1;refGene_gn=BPTF;refGene_tx=XM_023243985.1;refGene_gn=BPTF;refGene_tx=XM_023243986.1;refGene_gn=BPTF;refGene_tx=XM_023243987.1;refGene_gn=BPTF;refGene_tx=XM_023243988.1;refGene_gn=BPTF;refGene_tx=XM_023243989.1
E1 48952939 48953157
refGene_gn=BPTF;refGene_tx=XM_023243975.1;refGene_gn=BPTF;refGene_tx=XM_023243976.1;refGene_gn=BPTF;refGene_tx=XM_023243978.1;refGene_gn=BPTF;refGene_tx=XM_023243979.1;refGene_gn=BPTF;refGene_tx=XM_023243980.1;refGene_gn=BPTF;refGene_tx=XM_023243981.1;refGene_gn=BPTF;refGene_tx=XM_023243982.1;refGene_gn=BPTF;refGene_tx=XM_023243983.1;refGene_gn=BPTF;refGene_tx=XM_023243984.1;refGene_gn=BPTF;refGene_tx=XM_023243985.1;refGene_gn=BPTF;refGene_tx=XM_023243986.1;refGene_gn=BPTF;refGene_tx=XM_023243987.1;refGene_gn=BPTF;refGene_tx=XM_023243988.1;refGene_gn=BPTF;refGene_tx=XM_023243989.1
E1 48959160 48959269
refGene_gn=BPTF;refGene_tx=XM_023243975.1;refGene_gn=BPTF;refGene_tx=XM_023243976.1;refGene_gn=BPTF;refGene_tx=XM_023243978.1;refGene_gn=BPTF;refGene_tx=XM_023243979.1;refGene_gn=BPTF;refGene_tx=XM_023243980.1;refGene_gn=BPTF;refGene_tx=XM_023243981.1;refGene_gn=BPTF;refGene_tx=XM_023243982.1;refGene_gn=BPTF;refGene_tx=XM_023243983.1;refGene_gn=BPTF;refGene_tx=XM_023243984.1;refGene_gn=BPTF;refGene_tx=XM_023243985.1;refGene_gn=BPTF;refGene_tx=XM_023243986.1;refGene_gn=BPTF;refGene_tx=XM_023243987.1;refGene_gn=BPTF;refGene_tx=XM_023243988.1;refGene_gn=BPTF;refGene_tx=XM_023243989.1
E1 48960183 48960335
refGene_gn=BPTF;refGene_tx=XM_023243975.1;refGene_gn=BPTF;refGene_tx=XM_023243976.1;refGene_gn=BPTF;refGene_tx=XM_023243978.1;refGene_gn=BPTF;refGene_tx=XM_023243979.1;refGene_gn=BPTF;refGene_tx=XM_023243980.1;refGene_gn=BPTF;refGene_tx=XM_023243981.1;refGene_gn=BPTF;refGene_tx=X

M_023243982.1;refGene_gn=BPTF;refGene_tx=XM_023243983.1;refGene_gn=BPT
F;refGene_tx=XM_023243984.1;refGene_gn=BPTF;refGene_tx=XM_023243985.1;
refGene_gn=BPTF;refGene_tx=XM_023243986.1;refGene_gn=BPTF;refGene_tx=X
M_023243987.1;refGene_gn=BPTF;refGene_tx=XM_023243988.1;refGene_gn=BPT
F;refGene_tx=XM_023243989.1
E1 48960552 48960714
refGene_gn=BPTF;refGene_tx=XM_023243975.1;refGene_gn=BPTF;refGene_tx=X
M_023243976.1;refGene_gn=BPTF;refGene_tx=XM_023243978.1;refGene_gn=BPT
F;refGene_tx=XM_023243979.1;refGene_gn=BPTF;refGene_tx=XM_023243981.1;
refGene_gn=BPTF;refGene_tx=XM_023243982.1;refGene_gn=BPTF;refGene_tx=X
M_023243983.1;refGene_gn=BPTF;refGene_tx=XM_023243984.1;refGene_gn=BPT
F;refGene_tx=XM_023243985.1;refGene_gn=BPTF;refGene_tx=XM_023243987.1
E1 48961076 48961323
refGene_gn=BPTF;refGene_tx=XM_023243975.1;refGene_gn=BPTF;refGene_tx=X
M_023243976.1;refGene_gn=BPTF;refGene_tx=XM_023243978.1;refGene_gn=BPT
F;refGene_tx=XM_023243979.1;refGene_gn=BPTF;refGene_tx=XM_023243980.1;
refGene_gn=BPTF;refGene_tx=XM_023243981.1;refGene_gn=BPTF;refGene_tx=X
M_023243982.1;refGene_gn=BPTF;refGene_tx=XM_023243983.1;refGene_gn=BPT
F;refGene_tx=XM_023243984.1;refGene_gn=BPTF;refGene_tx=XM_023243985.1;
refGene_gn=BPTF;refGene_tx=XM_023243986.1;refGene_gn=BPTF;refGene_tx=X
M_023243987.1;refGene_gn=BPTF;refGene_tx=XM_023243988.1;refGene_gn=BPT
F;refGene_tx=XM_023243989.1
E1 48962826 48962950
refGene_gn=BPTF;refGene_tx=XM_023243975.1;refGene_gn=BPTF;refGene_tx=X
M_023243976.1;refGene_gn=BPTF;refGene_tx=XM_023243978.1;refGene_gn=BPT
F;refGene_tx=XM_023243979.1;refGene_gn=BPTF;refGene_tx=XM_023243980.1;
refGene_gn=BPTF;refGene_tx=XM_023243981.1;refGene_gn=BPTF;refGene_tx=X
M_023243982.1;refGene_gn=BPTF;refGene_tx=XM_023243983.1;refGene_gn=BPT
F;refGene_tx=XM_023243984.1;refGene_gn=BPTF;refGene_tx=XM_023243985.1;
refGene_gn=BPTF;refGene_tx=XM_023243986.1;refGene_gn=BPTF;refGene_tx=X
M_023243987.1;refGene_gn=BPTF;refGene_tx=XM_023243988.1;refGene_gn=BPT
F;refGene_tx=XM_023243989.1
E1 48963289 48963440
refGene_gn=BPTF;refGene_tx=XM_023243975.1;refGene_gn=BPTF;refGene_tx=X
M_023243976.1;refGene_gn=BPTF;refGene_tx=XM_023243978.1;refGene_gn=BPT
F;refGene_tx=XM_023243979.1;refGene_gn=BPTF;refGene_tx=XM_023243980.1;
refGene_gn=BPTF;refGene_tx=XM_023243981.1;refGene_gn=BPTF;refGene_tx=X
M_023243982.1;refGene_gn=BPTF;refGene_tx=XM_023243983.1;refGene_gn=BPT
F;refGene_tx=XM_023243984.1;refGene_gn=BPTF;refGene_tx=XM_023243985.1;
refGene_gn=BPTF;refGene_tx=XM_023243986.1;refGene_gn=BPTF;refGene_tx=X
M_023243987.1;refGene_gn=BPTF;refGene_tx=XM_023243988.1;refGene_gn=BPT
F;refGene_tx=XM_023243989.1
E1 48965197 48965326
refGene_gn=BPTF;refGene_tx=XM_023243975.1;refGene_gn=BPTF;refGene_tx=X
M_023243976.1;refGene_gn=BPTF;refGene_tx=XM_023243978.1;refGene_gn=BPT
F;refGene_tx=XM_023243979.1;refGene_gn=BPTF;refGene_tx=XM_023243980.1;
refGene_gn=BPTF;refGene_tx=XM_023243981.1;refGene_gn=BPTF;refGene_tx=X
M_023243982.1;refGene_gn=BPTF;refGene_tx=XM_023243983.1;refGene_gn=BPT
F;refGene_tx=XM_023243984.1;refGene_gn=BPTF;refGene_tx=XM_023243985.1;
refGene_gn=BPTF;refGene_tx=XM_023243986.1;refGene_gn=BPTF;refGene_tx=X

M_023243987.1;refGene_gn=BPTF;refGene_tx=XM_023243988.1;refGene_gn=BPT
F;refGene_tx=XM_023243989.1
E1 48966637 48966762
refGene_gn=BPTF;refGene_tx=XM_023243975.1;refGene_gn=BPTF;refGene_tx=X
M_023243976.1;refGene_gn=BPTF;refGene_tx=XM_023243978.1;refGene_gn=BPT
F;refGene_tx=XM_023243979.1;refGene_gn=BPTF;refGene_tx=XM_023243980.1;
refGene_gn=BPTF;refGene_tx=XM_023243981.1;refGene_gn=BPTF;refGene_tx=X
M_023243982.1;refGene_gn=BPTF;refGene_tx=XM_023243983.1;refGene_gn=BPT
F;refGene_tx=XM_023243984.1;refGene_gn=BPTF;refGene_tx=XM_023243985.1;
refGene_gn=BPTF;refGene_tx=XM_023243986.1;refGene_gn=BPTF;refGene_tx=X
M_023243987.1;refGene_gn=BPTF;refGene_tx=XM_023243988.1;refGene_gn=BPT
F;refGene_tx=XM_023243989.1
E1 48971074 48973385
refGene_gn=BPTF;refGene_tx=XM_023243975.1;refGene_gn=BPTF;refGene_tx=X
M_023243976.1;refGene_gn=BPTF;refGene_tx=XM_023243978.1;refGene_gn=BPT
F;refGene_tx=XM_023243979.1;refGene_gn=BPTF;refGene_tx=XM_023243980.1;
refGene_gn=BPTF;refGene_tx=XM_023243981.1;refGene_gn=BPTF;refGene_tx=X
M_023243982.1;refGene_gn=BPTF;refGene_tx=XM_023243983.1;refGene_gn=BPT
F;refGene_tx=XM_023243984.1;refGene_gn=BPTF;refGene_tx=XM_023243985.1;
refGene_gn=BPTF;refGene_tx=XM_023243986.1;refGene_gn=BPTF;refGene_tx=X
M_023243987.1;refGene_gn=BPTF;refGene_tx=XM_023243988.1;refGene_gn=BPT
F;refGene_tx=XM_023243989.1
E1 48974439 48974619
refGene_gn=BPTF;refGene_tx=XM_023243975.1;refGene_gn=BPTF;refGene_tx=X
M_023243976.1;refGene_gn=BPTF;refGene_tx=XM_023243978.1;refGene_gn=BPT
F;refGene_tx=XM_023243979.1;refGene_gn=BPTF;refGene_tx=XM_023243980.1;
refGene_gn=BPTF;refGene_tx=XM_023243981.1;refGene_gn=BPTF;refGene_tx=X
M_023243982.1;refGene_gn=BPTF;refGene_tx=XM_023243983.1;refGene_gn=BPT
F;refGene_tx=XM_023243984.1;refGene_gn=BPTF;refGene_tx=XM_023243985.1;
refGene_gn=BPTF;refGene_tx=XM_023243986.1;refGene_gn=BPTF;refGene_tx=X
M_023243987.1;refGene_gn=BPTF;refGene_tx=XM_023243988.1;refGene_gn=BPT
F;refGene_tx=XM_023243989.1
E1 48977932 48978071
refGene_gn=BPTF;refGene_tx=XM_023243975.1;refGene_gn=BPTF;refGene_tx=X
M_023243976.1;refGene_gn=BPTF;refGene_tx=XM_023243978.1;refGene_gn=BPT
F;refGene_tx=XM_023243979.1;refGene_gn=BPTF;refGene_tx=XM_023243980.1;
refGene_gn=BPTF;refGene_tx=XM_023243981.1;refGene_gn=BPTF;refGene_tx=X
M_023243982.1;refGene_gn=BPTF;refGene_tx=XM_023243983.1;refGene_gn=BPT
F;refGene_tx=XM_023243984.1;refGene_gn=BPTF;refGene_tx=XM_023243985.1;
refGene_gn=BPTF;refGene_tx=XM_023243986.1;refGene_gn=BPTF;refGene_tx=X
M_023243987.1;refGene_gn=BPTF;refGene_tx=XM_023243988.1;refGene_gn=BPT
F;refGene_tx=XM_023243989.1
E1 48978821 48978951
refGene_gn=BPTF;refGene_tx=XM_023243975.1;refGene_gn=BPTF;refGene_tx=X
M_023243976.1;refGene_gn=BPTF;refGene_tx=XM_023243978.1;refGene_gn=BPT
F;refGene_tx=XM_023243979.1;refGene_gn=BPTF;refGene_tx=XM_023243980.1;
refGene_gn=BPTF;refGene_tx=XM_023243981.1;refGene_gn=BPTF;refGene_tx=X
M_023243982.1;refGene_gn=BPTF;refGene_tx=XM_023243983.1;refGene_gn=BPT
F;refGene_tx=XM_023243984.1;refGene_gn=BPTF;refGene_tx=XM_023243985.1;
refGene_gn=BPTF;refGene_tx=XM_023243986.1;refGene_gn=BPTF;refGene_tx=X

M_023243987.1;refGene_gn=BPTF;refGene_tx=XM_023243988.1;refGene_gn=BPTF;refGene_tx=XM_023243989.1
E1 48991889 48992021
refGene_gn=BPTF;refGene_tx=XM_023243975.1;refGene_gn=BPTF;refGene_tx=XM_023243976.1;refGene_gn=BPTF;refGene_tx=XM_023243978.1;refGene_gn=BPTF;refGene_tx=XM_023243979.1;refGene_gn=BPTF;refGene_tx=XM_023243980.1;refGene_gn=BPTF;refGene_tx=XM_023243981.1;refGene_gn=BPTF;refGene_tx=XM_023243982.1;refGene_gn=BPTF;refGene_tx=XM_023243983.1;refGene_gn=BPTF;refGene_tx=XM_023243984.1;refGene_gn=BPTF;refGene_tx=XM_023243985.1;refGene_gn=BPTF;refGene_tx=XM_023243986.1;refGene_gn=BPTF;refGene_tx=XM_023243987.1;refGene_gn=BPTF;refGene_tx=XM_023243988.1;refGene_gn=BPTF;refGene_tx=XM_023243989.1
E1 48992494 48992850
refGene_gn=BPTF;refGene_tx=XM_023243975.1;refGene_gn=BPTF;refGene_tx=XM_023243976.1;refGene_gn=BPTF;refGene_tx=XM_023243978.1;refGene_gn=BPTF;refGene_tx=XM_023243979.1;refGene_gn=BPTF;refGene_tx=XM_023243980.1;refGene_gn=BPTF;refGene_tx=XM_023243981.1;refGene_gn=BPTF;refGene_tx=XM_023243982.1;refGene_gn=BPTF;refGene_tx=XM_023243983.1;refGene_gn=BPTF;refGene_tx=XM_023243984.1;refGene_gn=BPTF;refGene_tx=XM_023243985.1;refGene_gn=BPTF;refGene_tx=XM_023243986.1;refGene_gn=BPTF;refGene_tx=XM_023243987.1;refGene_gn=BPTF;refGene_tx=XM_023243988.1;refGene_gn=BPTF;refGene_tx=XM_023243989.1
E1 48995007 48995195
refGene_gn=BPTF;refGene_tx=XM_023243975.1;refGene_gn=BPTF;refGene_tx=XM_023243976.1;refGene_gn=BPTF;refGene_tx=XM_023243978.1;refGene_gn=BPTF;refGene_tx=XM_023243979.1;refGene_gn=BPTF;refGene_tx=XM_023243980.1;refGene_gn=BPTF;refGene_tx=XM_023243981.1;refGene_gn=BPTF;refGene_tx=XM_023243982.1;refGene_gn=BPTF;refGene_tx=XM_023243983.1;refGene_gn=BPTF;refGene_tx=XM_023243984.1;refGene_gn=BPTF;refGene_tx=XM_023243985.1;refGene_gn=BPTF;refGene_tx=XM_023243986.1;refGene_gn=BPTF;refGene_tx=XM_023243987.1;refGene_gn=BPTF;refGene_tx=XM_023243988.1;refGene_gn=BPTF;refGene_tx=XM_023243989.1
E1 49000365 49000554
refGene_gn=BPTF;refGene_tx=XM_023243975.1;refGene_gn=BPTF;refGene_tx=XM_023243976.1;refGene_gn=BPTF;refGene_tx=XM_023243978.1;refGene_gn=BPTF;refGene_tx=XM_023243979.1;refGene_gn=BPTF;refGene_tx=XM_023243980.1;refGene_gn=BPTF;refGene_tx=XM_023243981.1;refGene_gn=BPTF;refGene_tx=XM_023243983.1;refGene_gn=BPTF;refGene_tx=XM_023243984.1;refGene_gn=BPTF;refGene_tx=XM_023243986.1;refGene_gn=BPTF;refGene_tx=XM_023243987.1;refGene_gn=BPTF;refGene_tx=XM_023243989.1
E1 49015970 49016159
refGene_gn=BPTF;refGene_tx=XM_023243975.1;refGene_gn=BPTF;refGene_tx=XM_023243976.1;refGene_gn=BPTF;refGene_tx=XM_023243978.1;refGene_gn=BPTF;refGene_tx=XM_023243979.1;refGene_gn=BPTF;refGene_tx=XM_023243980.1;refGene_gn=BPTF;refGene_tx=XM_023243981.1;refGene_gn=BPTF;refGene_tx=XM_023243982.1;refGene_gn=BPTF;refGene_tx=XM_023243986.1;refGene_gn=BPTF;refGene_tx=XM_023243987.1;refGene_gn=BPTF;refGene_tx=XM_023243989.1
E1 49016701 49016908
refGene_gn=BPTF;refGene_tx=XM_023243975.1;refGene_gn=BPTF;refGene_tx=XM_023243976.1;refGene_gn=BPTF;refGene_tx=XM_023243978.1;refGene_gn=BPT

F;refGene_tx=XM_023243979.1;refGene_gn=BPTF;refGene_tx=XM_023243980.1;refGene_gn=BPTF;refGene_tx=XM_023243981.1;refGene_gn=BPTF;refGene_tx=XM_023243982.1;refGene_gn=BPTF;refGene_tx=XM_023243983.1;refGene_gn=BPTF;refGene_tx=XM_023243984.1;refGene_gn=BPTF;refGene_tx=XM_023243985.1;refGene_gn=BPTF;refGene_tx=XM_023243986.1;refGene_gn=BPTF;refGene_tx=XM_023243987.1;refGene_gn=BPTF;refGene_tx=XM_023243988.1;refGene_gn=BPTF;refGene_tx=XM_023243989.1
E1 49023163 49023387
refGene_gn=BPTF;refGene_tx=XM_023243975.1;refGene_gn=BPTF;refGene_tx=XM_023243976.1;refGene_gn=BPTF;refGene_tx=XM_023243978.1;refGene_gn=BPTF;refGene_tx=XM_023243979.1;refGene_gn=BPTF;refGene_tx=XM_023243980.1;refGene_gn=BPTF;refGene_tx=XM_023243981.1;refGene_gn=BPTF;refGene_tx=XM_023243982.1;refGene_gn=BPTF;refGene_tx=XM_023243983.1;refGene_gn=BPTF;refGene_tx=XM_023243984.1;refGene_gn=BPTF;refGene_tx=XM_023243985.1;refGene_gn=BPTF;refGene_tx=XM_023243986.1;refGene_gn=BPTF;refGene_tx=XM_023243987.1;refGene_gn=BPTF;refGene_tx=XM_023243988.1;refGene_gn=BPTF;refGene_tx=XM_023243989.1
E1 49031921 49032744
refGene_gn=BPTF;refGene_tx=XM_023243975.1;refGene_gn=BPTF;refGene_tx=XM_023243976.1;refGene_gn=BPTF;refGene_tx=XM_023243978.1;refGene_gn=BPTF;refGene_tx=XM_023243979.1;refGene_gn=BPTF;refGene_tx=XM_023243980.1;refGene_gn=BPTF;refGene_tx=XM_023243981.1;refGene_gn=BPTF;refGene_tx=XM_023243982.1;refGene_gn=BPTF;refGene_tx=XM_023243983.1;refGene_gn=BPTF;refGene_tx=XM_023243984.1;refGene_gn=BPTF;refGene_tx=XM_023243985.1;refGene_gn=BPTF;refGene_tx=XM_023243986.1;refGene_gn=BPTF;refGene_tx=XM_023243987.1;refGene_gn=BPTF;refGene_tx=XM_023243988.1;refGene_gn=BPTF;refGene_tx=XM_023243989.1
E1 49071357 49071988
refGene_gn=BPTF;refGene_tx=XM_023243975.1;refGene_gn=BPTF;refGene_tx=XM_023243976.1;refGene_gn=BPTF;refGene_tx=XM_023243978.1;refGene_gn=BPTF;refGene_tx=XM_023243979.1;refGene_gn=BPTF;refGene_tx=XM_023243980.1;refGene_gn=BPTF;refGene_tx=XM_023243981.1;refGene_gn=BPTF;refGene_tx=XM_023243982.1;refGene_gn=BPTF;refGene_tx=XM_023243983.1;refGene_gn=BPTF;refGene_tx=XM_023243984.1;refGene_gn=BPTF;refGene_tx=XM_023243985.1;refGene_gn=BPTF;refGene_tx=XM_023243986.1;refGene_gn=BPTF;refGene_tx=XM_023243987.1;refGene_gn=BPTF;refGene_tx=XM_023243988.1;refGene_gn=BPTF;refGene_tx=XM_023243989.1
E1 49124831 49124948
refGene_gn=NOL11;refGene_tx=XM_011289317.3;refGene_gn=NOL11;refGene_tx=XM_019818123.2;refGene_gn=NOL11;refGene_tx=XM_019818124.1
E1 49125113 49125221
refGene_gn=NOL11;refGene_tx=XM_011289317.3;refGene_gn=NOL11;refGene_tx=XM_019818123.2;refGene_gn=NOL11;refGene_tx=XM_019818124.1
E1 49129946 49130039
refGene_gn=NOL11;refGene_tx=XM_011289317.3;refGene_gn=NOL11;refGene_tx=XM_019818123.2;refGene_gn=NOL11;refGene_tx=XM_019818124.1
E1 49130682 49130761
refGene_gn=NOL11;refGene_tx=XM_011289317.3;refGene_gn=NOL11;refGene_tx=XM_019818123.2;refGene_gn=NOL11;refGene_tx=XM_019818124.1
E1 49131196 49131424

refGene_gn=NOL11;refGene_tx=XM_011289317.3;refGene_gn=NOL11;refGene_tx=XM_019818123.2;refGene_gn=NOL11;refGene_tx=XM_019818124.1
E1 49131577 49131703
refGene_gn=NOL11;refGene_tx=XM_011289317.3;refGene_gn=NOL11;refGene_tx=XM_019818123.2;refGene_gn=NOL11;refGene_tx=XM_019818124.1
E1 49131855 49132040
refGene_gn=NOL11;refGene_tx=XM_011289317.3;refGene_gn=NOL11;refGene_tx=XM_019818123.2;refGene_gn=NOL11;refGene_tx=XM_019818124.1
E1 49132760 49132829
refGene_gn=NOL11;refGene_tx=XM_011289317.3;refGene_gn=NOL11;refGene_tx=XM_019818123.2;refGene_gn=NOL11;refGene_tx=XM_019818124.1
E1 49133411 49133500
refGene_gn=NOL11;refGene_tx=XM_011289317.3;refGene_gn=NOL11;refGene_tx=XM_019818123.2;refGene_gn=NOL11;refGene_tx=XM_019818124.1
E1 49134091 49134215
refGene_gn=NOL11;refGene_tx=XM_011289317.3;refGene_gn=NOL11;refGene_tx=XM_019818123.2;refGene_gn=NOL11;refGene_tx=XM_019818124.1
E1 49136706 49136783
refGene_gn=NOL11;refGene_tx=XM_011289317.3;refGene_gn=NOL11;refGene_tx=XM_019818123.2;refGene_gn=NOL11;refGene_tx=XM_019818124.1
E1 49138477 49138666
refGene_gn=NOL11;refGene_tx=XM_011289317.3;refGene_gn=NOL11;refGene_tx=XM_019818123.2;refGene_gn=NOL11;refGene_tx=XM_019818124.1
E1 49140878 49141023
refGene_gn=NOL11;refGene_tx=XM_011289317.3;refGene_gn=NOL11;refGene_tx=XM_019818123.2;refGene_gn=NOL11;refGene_tx=XM_019818124.1
E1 49141342 49141400
refGene_gn=NOL11;refGene_tx=XM_011289317.3;refGene_gn=NOL11;refGene_tx=XM_019818123.2;refGene_gn=NOL11;refGene_tx=XM_019818124.1
E1 49142771 49142920
refGene_gn=NOL11;refGene_tx=XM_011289317.3;refGene_gn=NOL11;refGene_tx=XM_019818123.2
E1 49145988 49146045
refGene_gn=NOL11;refGene_tx=XM_011289317.3;refGene_gn=NOL11;refGene_tx=XM_019818123.2
E1 49146170 49146284
refGene_gn=NOL11;refGene_tx=XM_011289317.3;refGene_gn=NOL11;refGene_tx=XM_019818123.2
E1 49148254 49148395
refGene_gn=NOL11;refGene_tx=XM_011289317.3;refGene_gn=NOL11;refGene_tx=XM_019818123.2
E1 49166588 49166905
refGene_gn=PITPNC1;refGene_tx=XM_019818126.2;refGene_gn=PITPNC1;refGene_tx=XM_019818127.2;refGene_gn=PITPNC1;refGene_tx=XM_019818128.2;refGene_gn=PITPNC1;refGene_tx=XM_019818129.2;refGene_gn=PITPNC1;refGene_tx=XM_019818130.2;refGene_gn=PITPNC1;refGene_tx=XM_023244106.1
E1 49170953 49171072 refGene_gn=PITPNC1;refGene_tx=XM_023244106.1
E1 49182365 49182429
refGene_gn=PITPNC1;refGene_tx=XM_019818126.2;refGene_gn=PITPNC1;refGene_tx=XM_019818127.2;refGene_gn=PITPNC1;refGene_tx=XM_019818128.2;refGe

ne_gn=PITPNC1;refGene_tx=XM_019818129.2;refGene_gn=PITPNC1;refGene_tx=XM_019818130.2;refGene_gn=PITPNC1;refGene_tx=XM_023244106.1
E1 49185636 49185792 refGene_gn=PITPNC1;refGene_tx=XM_019818126.2;refGene_gn=PITPNC1;refGene_tx=XM_019818127.2;refGene_gn=PITPNC1;refGene_tx=XM_019818128.2;refGene_gn=PITPNC1;refGene_tx=XM_019818129.2;refGene_gn=PITPNC1;refGene_tx=XM_019818130.2;refGene_gn=PITPNC1;refGene_tx=XM_023244106.1
E1 49213439 49213535 refGene_gn=PITPNC1;refGene_tx=XM_019818126.2;refGene_gn=PITPNC1;refGene_tx=XM_019818127.2;refGene_gn=PITPNC1;refGene_tx=XM_019818128.2;refGene_gn=PITPNC1;refGene_tx=XM_019818129.2;refGene_gn=PITPNC1;refGene_tx=XM_019818130.2;refGene_gn=PITPNC1;refGene_tx=XM_023244106.1
E1 49256276 49256348 refGene_gn=PITPNC1;refGene_tx=XM_019818126.2;refGene_gn=PITPNC1;refGene_tx=XM_019818127.2;refGene_gn=PITPNC1;refGene_tx=XM_019818128.2;refGene_gn=PITPNC1;refGene_tx=XM_019818129.2;refGene_gn=PITPNC1;refGene_tx=XM_019818130.2;refGene_gn=PITPNC1;refGene_tx=XM_023244106.1
E1 49279231 49279239 refGene_gn=PITPNC1;refGene_tx=XM_019818126.2;refGene_gn=PITPNC1;refGene_tx=XM_019818127.2;refGene_gn=PITPNC1;refGene_tx=XM_019818128.2;refGene_gn=PITPNC1;refGene_tx=XM_019818129.2;refGene_gn=PITPNC1;refGene_tx=XM_019818130.2;refGene_gn=PITPNC1;refGene_tx=XM_023244106.1
E1 49280326 49280415 refGene_gn=PITPNC1;refGene_tx=XM_019818126.2;refGene_gn=PITPNC1;refGene_tx=XM_019818127.2;refGene_gn=PITPNC1;refGene_tx=XM_019818128.2;refGene_gn=PITPNC1;refGene_tx=XM_019818129.2;refGene_gn=PITPNC1;refGene_tx=XM_019818130.2;refGene_gn=PITPNC1;refGene_tx=XM_023244106.1
E1 49297906 49298055 refGene_gn=PITPNC1;refGene_tx=XM_019818126.2;refGene_gn=PITPNC1;refGene_tx=XM_019818127.2;refGene_gn=PITPNC1;refGene_tx=XM_019818128.2;refGene_gn=PITPNC1;refGene_tx=XM_019818129.2;refGene_gn=PITPNC1;refGene_tx=XM_019818130.2;refGene_gn=PITPNC1;refGene_tx=XM_023244106.1
E1 49352543 49352576 refGene_gn=PITPNC1;refGene_tx=XM_019818127.2
E1 49427557 49427605 refGene_gn=PITPNC1;refGene_tx=XM_019818126.2;refGene_gn=PITPNC1;refGene_tx=XM_023244106.1
E1 49438973 49439081 refGene_gn=PSMD12;refGene_tx=XM_003997073.4
E1 49448144 49448204 refGene_gn=PSMD12;refGene_tx=XM_003997073.4
E1 49448291 49448420 refGene_gn=PSMD12;refGene_tx=XM_003997073.4
E1 49453245 49453353 refGene_gn=PSMD12;refGene_tx=XM_003997073.4
E1 49455385 49455490 refGene_gn=PSMD12;refGene_tx=XM_003997073.4
E1 49458581 49458731 refGene_gn=PSMD12;refGene_tx=XM_003997073.4
E1 49458827 49458962 refGene_gn=PSMD12;refGene_tx=XM_003997073.4
E1 49459922 49460035 refGene_gn=PSMD12;refGene_tx=XM_003997073.4
E1 49460721 49460896 refGene_gn=PSMD12;refGene_tx=XM_003997073.4
E1 49463766 49463844 refGene_gn=PSMD12;refGene_tx=XM_003997073.4
E1 49464806 49465016 refGene_gn=PSMD12;refGene_tx=XM_003997073.4
E1 49563794 49564004 refGene_gn=HELZ;refGene_tx=XM_023244319.1;refGene_gn=HELZ;refGene_tx=XM_023244320.1;refGene_gn=HELZ;refGene_tx=XM_023244321.1;refGene_gn=HEL

Z;refGene_tx=XM_023244322.1;refGene_gn=HELZ;refGene_tx=XM_023244323.1
E1 49566343 49566380
refGene_gn=HELZ;refGene_tx=XM_023244319.1;refGene_gn=HELZ;refGene_tx=X
M_023244320.1;refGene_gn=HELZ;refGene_tx=XM_023244321.1;refGene_gn=HEL
Z;refGene_tx=XM_023244322.1;refGene_gn=HELZ;refGene_tx=XM_023244323.1
E1 49575648 49575773
refGene_gn=HELZ;refGene_tx=XM_023244319.1;refGene_gn=HELZ;refGene_tx=X
M_023244320.1;refGene_gn=HELZ;refGene_tx=XM_023244321.1;refGene_gn=HEL
Z;refGene_tx=XM_023244322.1;refGene_gn=HELZ;refGene_tx=XM_023244323.1
E1 49578489 49578546
refGene_gn=HELZ;refGene_tx=XM_023244319.1;refGene_gn=HELZ;refGene_tx=X
M_023244320.1;refGene_gn=HELZ;refGene_tx=XM_023244321.1;refGene_gn=HEL
Z;refGene_tx=XM_023244322.1;refGene_gn=HELZ;refGene_tx=XM_023244323.1
E1 49583560 49583612
refGene_gn=HELZ;refGene_tx=XM_023244319.1;refGene_gn=HELZ;refGene_tx=X
M_023244320.1;refGene_gn=HELZ;refGene_tx=XM_023244321.1;refGene_gn=HEL
Z;refGene_tx=XM_023244322.1;refGene_gn=HELZ;refGene_tx=XM_023244323.1
E1 49584972 49585048
refGene_gn=HELZ;refGene_tx=XM_023244319.1;refGene_gn=HELZ;refGene_tx=X
M_023244320.1;refGene_gn=HELZ;refGene_tx=XM_023244321.1;refGene_gn=HEL
Z;refGene_tx=XM_023244322.1;refGene_gn=HELZ;refGene_tx=XM_023244323.1
E1 49587102 49587301
refGene_gn=HELZ;refGene_tx=XM_023244319.1;refGene_gn=HELZ;refGene_tx=X
M_023244320.1;refGene_gn=HELZ;refGene_tx=XM_023244321.1;refGene_gn=HEL
Z;refGene_tx=XM_023244322.1;refGene_gn=HELZ;refGene_tx=XM_023244323.1
E1 49587742 49587850
refGene_gn=HELZ;refGene_tx=XM_023244319.1;refGene_gn=HELZ;refGene_tx=X
M_023244320.1;refGene_gn=HELZ;refGene_tx=XM_023244321.1;refGene_gn=HEL
Z;refGene_tx=XM_023244322.1;refGene_gn=HELZ;refGene_tx=XM_023244323.1
E1 49588905 49589203
refGene_gn=HELZ;refGene_tx=XM_023244319.1;refGene_gn=HELZ;refGene_tx=X
M_023244320.1;refGene_gn=HELZ;refGene_tx=XM_023244321.1;refGene_gn=HEL
Z;refGene_tx=XM_023244322.1;refGene_gn=HELZ;refGene_tx=XM_023244323.1
E1 49601267 49601535
refGene_gn=HELZ;refGene_tx=XM_023244319.1;refGene_gn=HELZ;refGene_tx=X
M_023244320.1;refGene_gn=HELZ;refGene_tx=XM_023244321.1;refGene_gn=HEL
Z;refGene_tx=XM_023244322.1;refGene_gn=HELZ;refGene_tx=XM_023244323.1
E1 49611295 49611629
refGene_gn=HELZ;refGene_tx=XM_023244319.1;refGene_gn=HELZ;refGene_tx=X
M_023244320.1;refGene_gn=HELZ;refGene_tx=XM_023244321.1;refGene_gn=HEL
Z;refGene_tx=XM_023244322.1;refGene_gn=HELZ;refGene_tx=XM_023244323.1
E1 49613246 49613377
refGene_gn=HELZ;refGene_tx=XM_023244319.1;refGene_gn=HELZ;refGene_tx=X
M_023244320.1;refGene_gn=HELZ;refGene_tx=XM_023244321.1;refGene_gn=HEL
Z;refGene_tx=XM_023244322.1;refGene_gn=HELZ;refGene_tx=XM_023244323.1
E1 49618307 49618487
refGene_gn=HELZ;refGene_tx=XM_023244319.1;refGene_gn=HELZ;refGene_tx=X
M_023244320.1;refGene_gn=HELZ;refGene_tx=XM_023244321.1;refGene_gn=HEL
Z;refGene_tx=XM_023244322.1;refGene_gn=HELZ;refGene_tx=XM_023244323.1
E1 49618988 49619093

refGene_gn=HELZ;refGene_tx=XM_023244319.1;refGene_gn=HELZ;refGene_tx=XM_023244320.1;refGene_gn=HELZ;refGene_tx=XM_023244321.1;refGene_gn=HELZ;refGene_tx=XM_023244322.1;refGene_gn=HELZ;refGene_tx=XM_023244323.1
E1 49625785 49625964
refGene_gn=HELZ;refGene_tx=XM_023244319.1;refGene_gn=HELZ;refGene_tx=XM_023244320.1;refGene_gn=HELZ;refGene_tx=XM_023244321.1;refGene_gn=HELZ;refGene_tx=XM_023244322.1;refGene_gn=HELZ;refGene_tx=XM_023244323.1
E1 49628802 49628921
refGene_gn=HELZ;refGene_tx=XM_023244319.1;refGene_gn=HELZ;refGene_tx=XM_023244320.1;refGene_gn=HELZ;refGene_tx=XM_023244321.1;refGene_gn=HELZ;refGene_tx=XM_023244322.1;refGene_gn=HELZ;refGene_tx=XM_023244323.1
E1 49630249 49630395
refGene_gn=HELZ;refGene_tx=XM_023244319.1;refGene_gn=HELZ;refGene_tx=XM_023244320.1;refGene_gn=HELZ;refGene_tx=XM_023244321.1;refGene_gn=HELZ;refGene_tx=XM_023244322.1;refGene_gn=HELZ;refGene_tx=XM_023244323.1
E1 49632114 49632262
refGene_gn=HELZ;refGene_tx=XM_023244319.1;refGene_gn=HELZ;refGene_tx=XM_023244320.1;refGene_gn=HELZ;refGene_tx=XM_023244321.1;refGene_gn=HELZ;refGene_tx=XM_023244322.1;refGene_gn=HELZ;refGene_tx=XM_023244323.1
E1 49636109 49636293
refGene_gn=HELZ;refGene_tx=XM_023244319.1;refGene_gn=HELZ;refGene_tx=XM_023244320.1;refGene_gn=HELZ;refGene_tx=XM_023244321.1;refGene_gn=HELZ;refGene_tx=XM_023244322.1;refGene_gn=HELZ;refGene_tx=XM_023244323.1
E1 49638376 49638605
refGene_gn=HELZ;refGene_tx=XM_023244319.1;refGene_gn=HELZ;refGene_tx=XM_023244320.1;refGene_gn=HELZ;refGene_tx=XM_023244321.1;refGene_gn=HELZ;refGene_tx=XM_023244322.1;refGene_gn=HELZ;refGene_tx=XM_023244323.1
E1 49645549 49645754
refGene_gn=HELZ;refGene_tx=XM_023244319.1;refGene_gn=HELZ;refGene_tx=XM_023244320.1;refGene_gn=HELZ;refGene_tx=XM_023244321.1;refGene_gn=HELZ;refGene_tx=XM_023244322.1;refGene_gn=HELZ;refGene_tx=XM_023244323.1
E1 49652577 49652629
refGene_gn=HELZ;refGene_tx=XM_023244319.1;refGene_gn=HELZ;refGene_tx=XM_023244320.1;refGene_gn=HELZ;refGene_tx=XM_023244321.1;refGene_gn=HELZ;refGene_tx=XM_023244322.1;refGene_gn=HELZ;refGene_tx=XM_023244323.1
E1 49653368 49653559
refGene_gn=HELZ;refGene_tx=XM_023244319.1;refGene_gn=HELZ;refGene_tx=XM_023244320.1;refGene_gn=HELZ;refGene_tx=XM_023244321.1;refGene_gn=HELZ;refGene_tx=XM_023244322.1;refGene_gn=HELZ;refGene_tx=XM_023244323.1
E1 49655850 49656058
refGene_gn=HELZ;refGene_tx=XM_023244319.1;refGene_gn=HELZ;refGene_tx=XM_023244320.1;refGene_gn=HELZ;refGene_tx=XM_023244321.1;refGene_gn=HELZ;refGene_tx=XM_023244322.1;refGene_gn=HELZ;refGene_tx=XM_023244323.1
E1 49663134 49663214
refGene_gn=HELZ;refGene_tx=XM_023244319.1;refGene_gn=HELZ;refGene_tx=XM_023244320.1;refGene_gn=HELZ;refGene_tx=XM_023244321.1;refGene_gn=HELZ;refGene_tx=XM_023244322.1;refGene_gn=HELZ;refGene_tx=XM_023244323.1
E1 49667494 49668065
refGene_gn=HELZ;refGene_tx=XM_023244319.1;refGene_gn=HELZ;refGene_tx=XM_023244320.1;refGene_gn=HELZ;refGene_tx=XM_023244321.1;refGene_gn=HEL

Z;refGene_tx=XM_023244322.1;refGene_gn=HELZ;refGene_tx=XM_023244323.1
E1 49668467 49668684
refGene_gn=HELZ;refGene_tx=XM_023244319.1;refGene_gn=HELZ;refGene_tx=XM_023244320.1;refGene_gn=HELZ;refGene_tx=XM_023244321.1;refGene_gn=HELZ;refGene_tx=XM_023244322.1;refGene_gn=HELZ;refGene_tx=XM_023244323.1
E1 49669846 49670363
refGene_gn=HELZ;refGene_tx=XM_023244319.1;refGene_gn=HELZ;refGene_tx=XM_023244320.1;refGene_gn=HELZ;refGene_tx=XM_023244321.1;refGene_gn=HELZ;refGene_tx=XM_023244322.1;refGene_gn=HELZ;refGene_tx=XM_023244323.1
E1 49692854 49693107
refGene_gn=HELZ;refGene_tx=XM_023244319.1;refGene_gn=HELZ;refGene_tx=XM_023244320.1;refGene_gn=HELZ;refGene_tx=XM_023244321.1;refGene_gn=HELZ;refGene_tx=XM_023244322.1;refGene_gn=HELZ;refGene_tx=XM_023244323.1
E1 49698690 49699028
refGene_gn=HELZ;refGene_tx=XM_023244319.1;refGene_gn=HELZ;refGene_tx=XM_023244320.1;refGene_gn=HELZ;refGene_tx=XM_023244321.1;refGene_gn=HELZ;refGene_tx=XM_023244322.1;refGene_gn=HELZ;refGene_tx=XM_023244323.1
E1 49718959 49719186 refGene_gn=CACNG1;refGene_tx=XM_003997076.5
E1 49719369 49719507 refGene_gn=CACNG1;refGene_tx=XM_003997076.5
E1 49720506 49720581 refGene_gn=CACNG1;refGene_tx=XM_003997076.5
E1 49729769 49730004 refGene_gn=CACNG1;refGene_tx=XM_003997076.5
E1 49743212 49743751
refGene_gn=CACNG4;refGene_tx=XM_003997094.5;refGene_gn=CACNG4;refGene_tx=XM_019818133.2
E1 49746542 49746683
refGene_gn=CACNG4;refGene_tx=XM_003997094.5;refGene_gn=CACNG4;refGene_tx=XM_019818133.2
E1 49751560 49751644
refGene_gn=CACNG4;refGene_tx=XM_003997094.5;refGene_gn=CACNG4;refGene_tx=XM_019818133.2
E1 49763475 49763638 refGene_gn=CACNG4;refGene_tx=XM_019818133.2
E1 49800580 49800800 refGene_gn=CACNG4;refGene_tx=XM_003997094.5
E1 49873877 49874135 refGene_gn=CACNG5;refGene_tx=XM_023243849.1
E1 49874444 49874590 refGene_gn=CACNG5;refGene_tx=XM_023243849.1
E1 49877381 49877522 refGene_gn=CACNG5;refGene_tx=XM_023243849.1
E1 49879224 49879311 refGene_gn=CACNG5;refGene_tx=XM_023243849.1
E1 49880750 49880946 refGene_gn=CACNG5;refGene_tx=XM_023243849.1
E1 49941158 49941323
refGene_gn=PRKCA;refGene_tx=XM_023244468.1;refGene_gn=PRKCA;refGene_tx=XM_023244469.1
E1 49953671 49953812
refGene_gn=PRKCA;refGene_tx=XM_023244468.1;refGene_gn=PRKCA;refGene_tx=XM_023244469.1
E1 49956319 49956427
refGene_gn=PRKCA;refGene_tx=XM_023244468.1;refGene_gn=PRKCA;refGene_tx=XM_023244469.1
E1 49964996 49965077
refGene_gn=PRKCA;refGene_tx=XM_023244468.1;refGene_gn=PRKCA;refGene_tx=XM_023244469.1
E1 49989203 49989342

refGene_gn=PRKCA;refGene_tx=XM_023244468.1;refGene_gn=PRKCA;refGene_tx=XM_023244469.1
E1 49990228 49990291
refGene_gn=PRKCA;refGene_tx=XM_023244468.1;refGene_gn=PRKCA;refGene_tx=XM_023244469.1
E1 49992794 49992886
refGene_gn=PRKCA;refGene_tx=XM_023244468.1;refGene_gn=PRKCA;refGene_tx=XM_023244469.1
E1 49994880 49995054
refGene_gn=PRKCA;refGene_tx=XM_023244468.1;refGene_gn=PRKCA;refGene_tx=XM_023244469.1
E1 49996104 49996242
refGene_gn=PRKCA;refGene_tx=XM_023244468.1;refGene_gn=PRKCA;refGene_tx=XM_023244469.1
E1 50012083 50012107 refGene_gn=PRKCA;refGene_tx=XM_023244468.1
E1 50025603 50025700
refGene_gn=PRKCA;refGene_tx=XM_023244468.1;refGene_gn=PRKCA;refGene_tx=XM_023244469.1
E1 50025953 50026088
refGene_gn=PRKCA;refGene_tx=XM_023244468.1;refGene_gn=PRKCA;refGene_tx=XM_023244469.1
E1 50027161 50027318
refGene_gn=PRKCA;refGene_tx=XM_023244468.1;refGene_gn=PRKCA;refGene_tx=XM_023244469.1
E1 50062653 50062782
refGene_gn=PRKCA;refGene_tx=XM_023244468.1;refGene_gn=PRKCA;refGene_tx=XM_023244469.1
E1 50066210 50066322
refGene_gn=PRKCA;refGene_tx=XM_023244468.1;refGene_gn=PRKCA;refGene_tx=XM_023244469.1
E1 50182625 50182708
refGene_gn=PRKCA;refGene_tx=XM_023244468.1;refGene_gn=PRKCA;refGene_tx=XM_023244469.1
E1 50336748 50336780
refGene_gn=PRKCA;refGene_tx=XM_023244468.1;refGene_gn=PRKCA;refGene_tx=XM_023244469.1
E1 50339690 50339863
refGene_gn=PRKCA;refGene_tx=XM_023244468.1;refGene_gn=PRKCA;refGene_tx=XM_023244469.1
E1 50396379 50396443 refGene_gn=APOH;refGene_tx=XM_019818134.1
E1 50396838 50397015 refGene_gn=APOH;refGene_tx=XM_019818134.1
E1 50399029 50399126 refGene_gn=APOH;refGene_tx=XM_019818134.1
E1 50400237 50400314 refGene_gn=APOH;refGene_tx=XM_019818134.1
E1 50402406 50402595 refGene_gn=APOH;refGene_tx=XM_019818134.1
E1 50404585 50404765 refGene_gn=APOH;refGene_tx=XM_019818134.1
E1 50406407 50406605 refGene_gn=APOH;refGene_tx=XM_019818134.1
E1 50408246 50408302 refGene_gn=APOH;refGene_tx=XM_019818134.1
E1 50429476 50429582
refGene_gn=CEP112;refGene_tx=XM_023244533.1;refGene_gn=CEP112;refGene_tx=XM_023244534.1;refGene_gn=CEP112;refGene_tx=XM_023244536.1;refGene_

gn=CEP112;refGene_tx=XM_023244537.1
E1 50432895 50433086
refGene_gn=CEP112;refGene_tx=XM_023244533.1;refGene_gn=CEP112;refGene_tx=XM_023244534.1;refGene_gn=CEP112;refGene_tx=XM_023244536.1;refGene_gn=CEP112;refGene_tx=XM_023244537.1;refGene_gn=CEP112;refGene_tx=XM_023244538.1
E1 50435672 50435845
refGene_gn=CEP112;refGene_tx=XM_023244533.1;refGene_gn=CEP112;refGene_tx=XM_023244534.1;refGene_gn=CEP112;refGene_tx=XM_023244536.1;refGene_gn=CEP112;refGene_tx=XM_023244537.1;refGene_gn=CEP112;refGene_tx=XM_023244538.1;refGene_gn=CEP112;refGene_tx=XM_023244539.1
E1 50464991 50465085
refGene_gn=CEP112;refGene_tx=XM_023244533.1;refGene_gn=CEP112;refGene_tx=XM_023244534.1;refGene_gn=CEP112;refGene_tx=XM_023244536.1;refGene_gn=CEP112;refGene_tx=XM_023244537.1;refGene_gn=CEP112;refGene_tx=XM_023244538.1;refGene_gn=CEP112;refGene_tx=XM_023244539.1
E1 50467083 50467161
refGene_gn=CEP112;refGene_tx=XM_023244533.1;refGene_gn=CEP112;refGene_tx=XM_023244534.1;refGene_gn=CEP112;refGene_tx=XM_023244536.1;refGene_gn=CEP112;refGene_tx=XM_023244538.1;refGene_gn=CEP112;refGene_tx=XM_023244539.1
E1 50478124 50478172
refGene_gn=CEP112;refGene_tx=XM_023244533.1;refGene_gn=CEP112;refGene_tx=XM_023244534.1;refGene_gn=CEP112;refGene_tx=XM_023244536.1;refGene_gn=CEP112;refGene_tx=XM_023244537.1;refGene_gn=CEP112;refGene_tx=XM_023244538.1;refGene_gn=CEP112;refGene_tx=XM_023244539.1
E1 50478679 50478757
refGene_gn=CEP112;refGene_tx=XM_023244533.1;refGene_gn=CEP112;refGene_tx=XM_023244534.1;refGene_gn=CEP112;refGene_tx=XM_023244536.1;refGene_gn=CEP112;refGene_tx=XM_023244537.1;refGene_gn=CEP112;refGene_tx=XM_023244538.1;refGene_gn=CEP112;refGene_tx=XM_023244539.1;refGene_gn=CEP112;refGene_tx=XM_023244540.1;refGene_gn=CEP112;refGene_tx=XM_023244541.1;refGene_gn=CEP112;refGene_tx=XM_023244542.1;refGene_gn=CEP112;refGene_tx=XM_023244543.1
E1 50505628 50505715
refGene_gn=CEP112;refGene_tx=XM_023244533.1;refGene_gn=CEP112;refGene_tx=XM_023244534.1;refGene_gn=CEP112;refGene_tx=XM_023244536.1;refGene_gn=CEP112;refGene_tx=XM_023244537.1;refGene_gn=CEP112;refGene_tx=XM_023244538.1;refGene_gn=CEP112;refGene_tx=XM_023244539.1;refGene_gn=CEP112;refGene_tx=XM_023244540.1;refGene_gn=CEP112;refGene_tx=XM_023244541.1;refGene_gn=CEP112;refGene_tx=XM_023244542.1;refGene_gn=CEP112;refGene_tx=XM_023244543.1
E1 50508140 50508240
refGene_gn=CEP112;refGene_tx=XM_023244533.1;refGene_gn=CEP112;refGene_tx=XM_023244534.1;refGene_gn=CEP112;refGene_tx=XM_023244536.1;refGene_gn=CEP112;refGene_tx=XM_023244537.1;refGene_gn=CEP112;refGene_tx=XM_023244538.1;refGene_gn=CEP112;refGene_tx=XM_023244539.1;refGene_gn=CEP112;refGene_tx=XM_023244540.1;refGene_gn=CEP112;refGene_tx=XM_023244541.1;refGene_gn=CEP112;refGene_tx=XM_023244542.1;refGene_gn=CEP112;refGene_tx=XM_023244543.1

E1 50511631 50511750
refGene_gn=CEP112;refGene_tx=XM_023244533.1;refGene_gn=CEP112;refGene_tx=XM_023244534.1;refGene_gn=CEP112;refGene_tx=XM_023244536.1;refGene_gn=CEP112;refGene_tx=XM_023244537.1;refGene_gn=CEP112;refGene_tx=XM_023244538.1;refGene_gn=CEP112;refGene_tx=XM_023244539.1;refGene_gn=CEP112;refGene_tx=XM_023244540.1;refGene_gn=CEP112;refGene_tx=XM_023244541.1;refGene_gn=CEP112;refGene_tx=XM_023244542.1;refGene_gn=CEP112;refGene_tx=XM_023244543.1
E1 50517328 50517472
refGene_gn=CEP112;refGene_tx=XM_023244533.1;refGene_gn=CEP112;refGene_tx=XM_023244534.1;refGene_gn=CEP112;refGene_tx=XM_023244536.1;refGene_gn=CEP112;refGene_tx=XM_023244537.1;refGene_gn=CEP112;refGene_tx=XM_023244538.1;refGene_gn=CEP112;refGene_tx=XM_023244539.1;refGene_gn=CEP112;refGene_tx=XM_023244540.1;refGene_gn=CEP112;refGene_tx=XM_023244541.1;refGene_gn=CEP112;refGene_tx=XM_023244542.1;refGene_gn=CEP112;refGene_tx=XM_023244543.1
E1 50529610 50529765
refGene_gn=CEP112;refGene_tx=XM_023244533.1;refGene_gn=CEP112;refGene_tx=XM_023244534.1;refGene_gn=CEP112;refGene_tx=XM_023244536.1;refGene_gn=CEP112;refGene_tx=XM_023244537.1;refGene_gn=CEP112;refGene_tx=XM_023244538.1;refGene_gn=CEP112;refGene_tx=XM_023244539.1;refGene_gn=CEP112;refGene_tx=XM_023244540.1;refGene_gn=CEP112;refGene_tx=XM_023244541.1;refGene_gn=CEP112;refGene_tx=XM_023244542.1;refGene_gn=CEP112;refGene_tx=XM_023244543.1
E1 50530310 50530440
refGene_gn=CEP112;refGene_tx=XM_023244533.1;refGene_gn=CEP112;refGene_tx=XM_023244534.1;refGene_gn=CEP112;refGene_tx=XM_023244536.1;refGene_gn=CEP112;refGene_tx=XM_023244537.1;refGene_gn=CEP112;refGene_tx=XM_023244538.1;refGene_gn=CEP112;refGene_tx=XM_023244539.1;refGene_gn=CEP112;refGene_tx=XM_023244540.1;refGene_gn=CEP112;refGene_tx=XM_023244541.1;refGene_gn=CEP112;refGene_tx=XM_023244542.1;refGene_gn=CEP112;refGene_tx=XM_023244543.1
E1 50531479 50531572
refGene_gn=CEP112;refGene_tx=XM_023244533.1;refGene_gn=CEP112;refGene_tx=XM_023244534.1;refGene_gn=CEP112;refGene_tx=XM_023244536.1;refGene_gn=CEP112;refGene_tx=XM_023244537.1;refGene_gn=CEP112;refGene_tx=XM_023244538.1;refGene_gn=CEP112;refGene_tx=XM_023244539.1;refGene_gn=CEP112;refGene_tx=XM_023244540.1;refGene_gn=CEP112;refGene_tx=XM_023244541.1;refGene_gn=CEP112;refGene_tx=XM_023244542.1;refGene_gn=CEP112;refGene_tx=XM_023244543.1
E1 50532443 50532503
refGene_gn=CEP112;refGene_tx=XM_023244533.1;refGene_gn=CEP112;refGene_tx=XM_023244534.1;refGene_gn=CEP112;refGene_tx=XM_023244536.1;refGene_gn=CEP112;refGene_tx=XM_023244537.1;refGene_gn=CEP112;refGene_tx=XM_023244538.1;refGene_gn=CEP112;refGene_tx=XM_023244539.1;refGene_gn=CEP112;refGene_tx=XM_023244540.1;refGene_gn=CEP112;refGene_tx=XM_023244541.1;refGene_gn=CEP112;refGene_tx=XM_023244542.1;refGene_gn=CEP112;refGene_tx=XM_023244543.1
E1 50541423 50541503
refGene_gn=CEP112;refGene_tx=XM_023244533.1;refGene_gn=CEP112;refGene_

tx=XM_023244534.1;refGene_gn=CEP112;refGene_tx=XM_023244536.1;refGene_gn=CEP112;refGene_tx=XM_023244537.1;refGene_gn=CEP112;refGene_tx=XM_023244538.1;refGene_gn=CEP112;refGene_tx=XM_023244539.1;refGene_gn=CEP112;refGene_tx=XM_023244540.1;refGene_gn=CEP112;refGene_tx=XM_023244541.1;refGene_gn=CEP112;refGene_tx=XM_023244542.1;refGene_gn=CEP112;refGene_tx=XM_023244543.1
E1 50588013 50588149
refGene_gn=CEP112;refGene_tx=XM_023244533.1;refGene_gn=CEP112;refGene_tx=XM_023244534.1;refGene_gn=CEP112;refGene_tx=XM_023244536.1;refGene_gn=CEP112;refGene_tx=XM_023244537.1;refGene_gn=CEP112;refGene_tx=XM_023244538.1;refGene_gn=CEP112;refGene_tx=XM_023244539.1;refGene_gn=CEP112;refGene_tx=XM_023244540.1;refGene_gn=CEP112;refGene_tx=XM_023244541.1;refGene_gn=CEP112;refGene_tx=XM_023244542.1;refGene_gn=CEP112;refGene_tx=XM_023244543.1
E1 50605892 50606000
refGene_gn=CEP112;refGene_tx=XM_023244533.1;refGene_gn=CEP112;refGene_tx=XM_023244534.1;refGene_gn=CEP112;refGene_tx=XM_023244536.1;refGene_gn=CEP112;refGene_tx=XM_023244537.1;refGene_gn=CEP112;refGene_tx=XM_023244538.1;refGene_gn=CEP112;refGene_tx=XM_023244539.1;refGene_gn=CEP112;refGene_tx=XM_023244540.1;refGene_gn=CEP112;refGene_tx=XM_023244541.1;refGene_gn=CEP112;refGene_tx=XM_023244542.1;refGene_gn=CEP112;refGene_tx=XM_023244543.1
E1 50625044 50625227
refGene_gn=CEP112;refGene_tx=XM_023244533.1;refGene_gn=CEP112;refGene_tx=XM_023244534.1;refGene_gn=CEP112;refGene_tx=XM_023244536.1;refGene_gn=CEP112;refGene_tx=XM_023244537.1;refGene_gn=CEP112;refGene_tx=XM_023244538.1;refGene_gn=CEP112;refGene_tx=XM_023244539.1;refGene_gn=CEP112;refGene_tx=XM_023244540.1;refGene_gn=CEP112;refGene_tx=XM_023244541.1;refGene_gn=CEP112;refGene_tx=XM_023244542.1;refGene_gn=CEP112;refGene_tx=XM_023244543.1
E1 50676781 50677012
refGene_gn=CEP112;refGene_tx=XM_023244533.1;refGene_gn=CEP112;refGene_tx=XM_023244534.1;refGene_gn=CEP112;refGene_tx=XM_023244536.1;refGene_gn=CEP112;refGene_tx=XM_023244537.1;refGene_gn=CEP112;refGene_tx=XM_023244538.1;refGene_gn=CEP112;refGene_tx=XM_023244539.1;refGene_gn=CEP112;refGene_tx=XM_023244540.1;refGene_gn=CEP112;refGene_tx=XM_023244541.1;refGene_gn=CEP112;refGene_tx=XM_023244542.1;refGene_gn=CEP112;refGene_tx=XM_023244543.1
E1 50742869 50742932
refGene_gn=CEP112;refGene_tx=XM_023244533.1;refGene_gn=CEP112;refGene_tx=XM_023244534.1;refGene_gn=CEP112;refGene_tx=XM_023244536.1;refGene_gn=CEP112;refGene_tx=XM_023244537.1;refGene_gn=CEP112;refGene_tx=XM_023244538.1;refGene_gn=CEP112;refGene_tx=XM_023244539.1;refGene_gn=CEP112;refGene_tx=XM_023244540.1;refGene_gn=CEP112;refGene_tx=XM_023244541.1;refGene_gn=CEP112;refGene_tx=XM_023244542.1;refGene_gn=CEP112;refGene_tx=XM_023244543.1
E1 50749404 50749554
refGene_gn=CEP112;refGene_tx=XM_023244533.1;refGene_gn=CEP112;refGene_tx=XM_023244534.1;refGene_gn=CEP112;refGene_tx=XM_023244536.1;refGene_gn=CEP112;refGene_tx=XM_023244537.1;refGene_gn=CEP112;refGene_tx=XM_02

3244538.1;refGene_gn=CEP112;refGene_tx=XM_023244539.1;refGene_gn=CEP11
2;refGene_tx=XM_023244540.1;refGene_gn=CEP112;refGene_tx=XM_023244541.
1;refGene_gn=CEP112;refGene_tx=XM_023244542.1;refGene_gn=CEP112;refGen
e_tx=XM_023244543.1
E1 50794625 50794715
refGene_gn=CEP112;refGene_tx=XM_023244533.1;refGene_gn=CEP112;refGene_
tx=XM_023244534.1;refGene_gn=CEP112;refGene_tx=XM_023244536.1;refGene_
gn=CEP112;refGene_tx=XM_023244537.1;refGene_gn=CEP112;refGene_tx=XM_02
3244538.1;refGene_gn=CEP112;refGene_tx=XM_023244539.1;refGene_gn=CEP11
2;refGene_tx=XM_023244540.1;refGene_gn=CEP112;refGene_tx=XM_023244541.
1;refGene_gn=CEP112;refGene_tx=XM_023244542.1;refGene_gn=CEP112;refGen
e_tx=XM_023244543.1
E1 50832930 50833032
refGene_gn=CEP112;refGene_tx=XM_023244533.1;refGene_gn=CEP112;refGene_
tx=XM_023244534.1;refGene_gn=CEP112;refGene_tx=XM_023244536.1;refGene_
gn=CEP112;refGene_tx=XM_023244537.1;refGene_gn=CEP112;refGene_tx=XM_02
3244538.1;refGene_gn=CEP112;refGene_tx=XM_023244539.1;refGene_gn=CEP11
2;refGene_tx=XM_023244540.1;refGene_gn=CEP112;refGene_tx=XM_023244541.
1;refGene_gn=CEP112;refGene_tx=XM_023244542.1;refGene_gn=CEP112;refGen
e_tx=XM_023244543.1
E1 50837014 50837079
refGene_gn=CEP112;refGene_tx=XM_023244533.1;refGene_gn=CEP112;refGene_
tx=XM_023244534.1;refGene_gn=CEP112;refGene_tx=XM_023244536.1;refGene_
gn=CEP112;refGene_tx=XM_023244537.1;refGene_gn=CEP112;refGene_tx=XM_02
3244538.1;refGene_gn=CEP112;refGene_tx=XM_023244539.1;refGene_gn=CEP11
2;refGene_tx=XM_023244540.1;refGene_gn=CEP112;refGene_tx=XM_023244541.
1;refGene_gn=CEP112;refGene_tx=XM_023244542.1;refGene_gn=CEP112;refGen
e_tx=XM_023244543.1
E1 50838341 50838345
refGene_gn=CEP112;refGene_tx=XM_023244533.1;refGene_gn=CEP112;refGene_
tx=XM_023244534.1;refGene_gn=CEP112;refGene_tx=XM_023244536.1;refGene_
gn=CEP112;refGene_tx=XM_023244537.1;refGene_gn=CEP112;refGene_tx=XM_02
3244538.1;refGene_gn=CEP112;refGene_tx=XM_023244539.1;refGene_gn=CEP11
2;refGene_tx=XM_023244540.1;refGene_gn=CEP112;refGene_tx=XM_023244541.
1;refGene_gn=CEP112;refGene_tx=XM_023244542.1;refGene_gn=CEP112;refGen
e_tx=XM_023244543.1
E1 50889396 50890211
refGene_gn=AXIN2;refGene_tx=XM_011289320.3;refGene_gn=AXIN2;refGene_tx
=XM_019818136.2
E1 50898357 50898498
refGene_gn=AXIN2;refGene_tx=XM_011289320.3;refGene_gn=AXIN2;refGene_tx
=XM_019818136.2
E1 50907548 50907651
refGene_gn=AXIN2;refGene_tx=XM_011289320.3;refGene_gn=AXIN2;refGene_tx
=XM_019818136.2
E1 50910103 50910244
refGene_gn=AXIN2;refGene_tx=XM_011289320.3;refGene_gn=AXIN2;refGene_tx
=XM_019818136.2
E1 50910482 50910991
refGene_gn=AXIN2;refGene_tx=XM_011289320.3;refGene_gn=AXIN2;refGene_tx

=XM_019818136.2
E1 50911197 50911389
refGene_gn=AXIN2;refGene_tx=XM_011289320.3;refGene_gn=AXIN2;refGene_tx
=XM_019818136.2
E1 50911696 50911930
refGene_gn=AXIN2;refGene_tx=XM_011289320.3;refGene_gn=AXIN2;refGene_tx
=XM_019818136.2
E1 50912466 50912562
refGene_gn=AXIN2;refGene_tx=XM_011289320.3;refGene_gn=AXIN2;refGene_tx
=XM_019818136.2
E1 50914007 50914175
refGene_gn=AXIN2;refGene_tx=XM_011289320.3;refGene_gn=AXIN2;refGene_tx
=XM_019818136.2
E1 50917677 50917804
refGene_gn=AXIN2;refGene_tx=XM_011289320.3;refGene_gn=AXIN2;refGene_tx
=XM_019818136.2
E1 51151820 51151971
refGene_gn=RGS9;refGene_tx=XM_003997097.3;refGene_gn=RGS9;refGene_tx=X
M_019818137.2
E1 51153155 51153613
refGene_gn=RGS9;refGene_tx=XM_003997097.3;refGene_gn=RGS9;refGene_tx=X
M_019818137.2
E1 51161878 51161996
refGene_gn=RGS9;refGene_tx=XM_003997097.3;refGene_gn=RGS9;refGene_tx=X
M_019818137.2
E1 51164474 51164560
refGene_gn=RGS9;refGene_tx=XM_003997097.3;refGene_gn=RGS9;refGene_tx=X
M_019818137.2
E1 51167040 51167179
refGene_gn=RGS9;refGene_tx=XM_003997097.3;refGene_gn=RGS9;refGene_tx=X
M_019818137.2
E1 51169406 51169494
refGene_gn=RGS9;refGene_tx=XM_003997097.3;refGene_gn=RGS9;refGene_tx=X
M_019818137.2
E1 51173035 51173151
refGene_gn=RGS9;refGene_tx=XM_003997097.3;refGene_gn=RGS9;refGene_tx=X
M_019818137.2
E1 51175459 51175573
refGene_gn=RGS9;refGene_tx=XM_003997097.3;refGene_gn=RGS9;refGene_tx=X
M_019818137.2
E1 51177523 51177585
refGene_gn=RGS9;refGene_tx=XM_003997097.3;refGene_gn=RGS9;refGene_tx=X
M_019818137.2
E1 51178317 51178347
refGene_gn=RGS9;refGene_tx=XM_003997097.3;refGene_gn=RGS9;refGene_tx=X
M_019818137.2
E1 51187225 51187297
refGene_gn=RGS9;refGene_tx=XM_003997097.3;refGene_gn=RGS9;refGene_tx=X
M_019818137.2
E1 51194197 51194279

refGene_gn=RGS9;refGene_tx=XM_003997097.3;refGene_gn=RGS9;refGene_tx=XM_019818137.2
E1 51200667 51200744
refGene_gn=RGS9;refGene_tx=XM_003997097.3;refGene_gn=RGS9;refGene_tx=XM_019818137.2
E1 51202786 51202845
refGene_gn=RGS9;refGene_tx=XM_003997097.3;refGene_gn=RGS9;refGene_tx=XM_019818137.2
E1 51203106 51203158
refGene_gn=RGS9;refGene_tx=XM_003997097.3;refGene_gn=RGS9;refGene_tx=XM_019818137.2
E1 51203375 51203482
refGene_gn=RGS9;refGene_tx=XM_003997097.3;refGene_gn=RGS9;refGene_tx=XM_019818137.2
E1 51204908 51204959
refGene_gn=RGS9;refGene_tx=XM_003997097.3;refGene_gn=RGS9;refGene_tx=XM_019818137.2
E1 51210261 51210358
refGene_gn=RGS9;refGene_tx=XM_003997097.3;refGene_gn=RGS9;refGene_tx=XM_019818137.2
E1 51221941 51221998
refGene_gn=RGS9;refGene_tx=XM_003997097.3;refGene_gn=RGS9;refGene_tx=XM_019818137.2
E1 51278082 51278365 refGene_gn=GNA13;refGene_tx=XM_003997099.4
E1 51280778 51281005 refGene_gn=GNA13;refGene_tx=XM_003997099.4
E1 51310468 51310519 refGene_gn=GNA13;refGene_tx=XM_003997099.4
E1 51316465 51317038 refGene_gn=GNA13;refGene_tx=XM_003997099.4
E1 51351475 51351758 refGene_gn=AMZ2;refGene_tx=XM_011289324.3
E1 51352540 51352608 refGene_gn=AMZ2;refGene_tx=XM_011289324.3
E1 51352609 51352715 refGene_gn=AMZ2;refGene_tx=XM_011289324.3
E1 51352828 51352957 refGene_gn=AMZ2;refGene_tx=XM_011289324.3
E1 51356338 51356502 refGene_gn=AMZ2;refGene_tx=XM_011289324.3
E1 51357848 51358025 refGene_gn=AMZ2;refGene_tx=XM_011289324.3
E1 51358971 51359124 refGene_gn=AMZ2;refGene_tx=XM_011289324.3
E1 51369679 51369930
refGene_gn=SLC16A6;refGene_tx=XM_019818145.1;refGene_gn=SLC16A6;refGene_tx=XM_023244625.1;refGene_gn=SLC16A6;refGene_tx=XM_023244626.1
E1 51370808 51371618
refGene_gn=SLC16A6;refGene_tx=XM_019818145.1;refGene_gn=SLC16A6;refGene_tx=XM_023244625.1;refGene_gn=SLC16A6;refGene_tx=XM_023244626.1
E1 51374097 51374226
refGene_gn=SLC16A6;refGene_tx=XM_019818145.1;refGene_gn=SLC16A6;refGene_tx=XM_023244625.1
E1 51376279 51376423
refGene_gn=SLC16A6;refGene_tx=XM_019818145.1;refGene_gn=SLC16A6;refGene_tx=XM_023244625.1;refGene_gn=SLC16A6;refGene_tx=XM_023244626.1
E1 51379261 51379493
refGene_gn=SLC16A6;refGene_tx=XM_023244625.1;refGene_gn=SLC16A6;refGene_tx=XM_023244626.1
E1 51403131 51403349

refGene_gn=ARSG;refGene_tx=XM_023244620.1;refGene_gn=ARSG;refGene_tx=XM_023244621.1;refGene_gn=ARSG;refGene_tx=XM_023244622.1;refGene_gn=ARSG;refGene_tx=XM_023244623.1;refGene_gn=ARSG;refGene_tx=XM_023244624.1
E1 51425162 51425350
refGene_gn=ARSG;refGene_tx=XM_023244620.1;refGene_gn=ARSG;refGene_tx=XM_023244621.1;refGene_gn=ARSG;refGene_tx=XM_023244622.1;refGene_gn=ARSG;refGene_tx=XM_023244623.1;refGene_gn=ARSG;refGene_tx=XM_023244624.1
E1 51428703 51428751
refGene_gn=ARSG;refGene_tx=XM_023244620.1;refGene_gn=ARSG;refGene_tx=XM_023244621.1;refGene_gn=ARSG;refGene_tx=XM_023244622.1;refGene_gn=ARSG;refGene_tx=XM_023244623.1;refGene_gn=ARSG;refGene_tx=XM_023244624.1
E1 51430609 51430727
refGene_gn=ARSG;refGene_tx=XM_023244620.1;refGene_gn=ARSG;refGene_tx=XM_023244621.1;refGene_gn=ARSG;refGene_tx=XM_023244622.1;refGene_gn=ARSG;refGene_tx=XM_023244623.1;refGene_gn=ARSG;refGene_tx=XM_023244624.1
E1 51432985 51433159
refGene_gn=ARSG;refGene_tx=XM_023244620.1;refGene_gn=ARSG;refGene_tx=XM_023244621.1;refGene_gn=ARSG;refGene_tx=XM_023244623.1;refGene_gn=ARSG;refGene_tx=XM_023244624.1
E1 51441549 51441746
refGene_gn=ARSG;refGene_tx=XM_023244620.1;refGene_gn=ARSG;refGene_tx=XM_023244621.1;refGene_gn=ARSG;refGene_tx=XM_023244622.1;refGene_gn=ARSG;refGene_tx=XM_023244623.1;refGene_gn=ARSG;refGene_tx=XM_023244624.1
E1 51442807 51442888
refGene_gn=ARSG;refGene_tx=XM_023244620.1;refGene_gn=ARSG;refGene_tx=XM_023244621.1;refGene_gn=ARSG;refGene_tx=XM_023244622.1;refGene_gn=ARSG;refGene_tx=XM_023244623.1;refGene_gn=ARSG;refGene_tx=XM_023244624.1
E1 51454963 51455072
refGene_gn=ARSG;refGene_tx=XM_023244620.1;refGene_gn=ARSG;refGene_tx=XM_023244621.1;refGene_gn=ARSG;refGene_tx=XM_023244622.1;refGene_gn=ARSG;refGene_tx=XM_023244623.1;refGene_gn=ARSG;refGene_tx=XM_023244624.1
E1 51462631 51462752
refGene_gn=ARSG;refGene_tx=XM_023244620.1;refGene_gn=ARSG;refGene_tx=XM_023244621.1;refGene_gn=ARSG;refGene_tx=XM_023244622.1;refGene_gn=ARSG;refGene_tx=XM_023244623.1
E1 51467418 51467509
refGene_gn=ARSG;refGene_tx=XM_023244620.1;refGene_gn=ARSG;refGene_tx=XM_023244621.1;refGene_gn=ARSG;refGene_tx=XM_023244622.1;refGene_gn=ARSG;refGene_tx=XM_023244623.1;refGene_gn=ARSG;refGene_tx=XM_023244624.1
E1 51470304 51470312 refGene_gn=ARSG;refGene_tx=XM_023244623.1
E1 51470829 51471104
refGene_gn=ARSG;refGene_tx=XM_023244620.1;refGene_gn=ARSG;refGene_tx=XM_023244621.1;refGene_gn=ARSG;refGene_tx=XM_023244622.1;refGene_gn=ARSG;refGene_tx=XM_023244624.1
E1 51472202 51472250
refGene_gn=WIPI1;refGene_tx=XM_023244627.1;refGene_gn=WIPI1;refGene_tx=XM_023244628.1;refGene_gn=WIPI1;refGene_tx=XM_023244629.1
E1 51474652 51474768
refGene_gn=WIPI1;refGene_tx=XM_023244627.1;refGene_gn=WIPI1;refGene_tx=XM_023244629.1

E1 51475131 51475232
refGene_gn=WIPI1;refGene_tx=XM_023244627.1;refGene_gn=WIPI1;refGene_tx=XM_023244628.1;refGene_gn=WIPI1;refGene_tx=XM_023244629.1
E1 51475688 51475807
refGene_gn=WIPI1;refGene_tx=XM_023244627.1;refGene_gn=WIPI1;refGene_tx=XM_023244628.1;refGene_gn=WIPI1;refGene_tx=XM_023244629.1
E1 51476724 51476832
refGene_gn=WIPI1;refGene_tx=XM_023244627.1;refGene_gn=WIPI1;refGene_tx=XM_023244628.1;refGene_gn=WIPI1;refGene_tx=XM_023244629.1
E1 51477583 51477748
refGene_gn=WIPI1;refGene_tx=XM_023244627.1;refGene_gn=WIPI1;refGene_tx=XM_023244628.1;refGene_gn=WIPI1;refGene_tx=XM_023244629.1
E1 51478995 51479103
refGene_gn=WIPI1;refGene_tx=XM_023244627.1;refGene_gn=WIPI1;refGene_tx=XM_023244628.1;refGene_gn=WIPI1;refGene_tx=XM_023244629.1
E1 51479930 51480001
refGene_gn=WIPI1;refGene_tx=XM_023244627.1;refGene_gn=WIPI1;refGene_tx=XM_023244628.1;refGene_gn=WIPI1;refGene_tx=XM_023244629.1
E1 51480699 51480792
refGene_gn=WIPI1;refGene_tx=XM_023244627.1;refGene_gn=WIPI1;refGene_tx=XM_023244628.1;refGene_gn=WIPI1;refGene_tx=XM_023244629.1
E1 51481401 51481499
refGene_gn=WIPI1;refGene_tx=XM_023244627.1;refGene_gn=WIPI1;refGene_tx=XM_023244628.1;refGene_gn=WIPI1;refGene_tx=XM_023244629.1
E1 51490814 51490911
refGene_gn=WIPI1;refGene_tx=XM_023244627.1;refGene_gn=WIPI1;refGene_tx=XM_023244628.1;refGene_gn=WIPI1;refGene_tx=XM_023244629.1
E1 51495592 51495762
refGene_gn=WIPI1;refGene_tx=XM_023244627.1;refGene_gn=WIPI1;refGene_tx=XM_023244628.1;refGene_gn=WIPI1;refGene_tx=XM_023244629.1
E1 51497145 51497228
refGene_gn=WIPI1;refGene_tx=XM_023244627.1;refGene_gn=WIPI1;refGene_tx=XM_023244628.1
E1 51502916 51502996
refGene_gn=WIPI1;refGene_tx=XM_023244627.1;refGene_gn=WIPI1;refGene_tx=XM_023244628.1;refGene_gn=WIPI1;refGene_tx=XM_023244629.1
E1 51544002 51544176
refGene_gn=PRKAR1A;refGene_tx=XM_006940522.4;refGene_gn=PRKAR1A;refGene_tx=XM_019818149.2
E1 51549376 51549547
refGene_gn=PRKAR1A;refGene_tx=XM_006940522.4;refGene_gn=PRKAR1A;refGene_tx=XM_019818149.2
E1 51550344 51550436
refGene_gn=PRKAR1A;refGene_tx=XM_006940522.4;refGene_gn=PRKAR1A;refGene_tx=XM_019818149.2
E1 51550637 51550699
refGene_gn=PRKAR1A;refGene_tx=XM_006940522.4;refGene_gn=PRKAR1A;refGene_tx=XM_019818149.2
E1 51554751 51554798
refGene_gn=PRKAR1A;refGene_tx=XM_006940522.4;refGene_gn=PRKAR1A;refGen

e_tx=XM_019818149.2
E1 51555455 51555614
refGene_gn=PRKAR1A;refGene_tx=XM_006940522.4;refGene_gn=PRKAR1A;refGen
e_tx=XM_019818149.2
E1 51556658 51556719
refGene_gn=PRKAR1A;refGene_tx=XM_006940522.4;refGene_gn=PRKAR1A;refGen
e_tx=XM_019818149.2
E1 51557824 51557946
refGene_gn=PRKAR1A;refGene_tx=XM_006940522.4;refGene_gn=PRKAR1A;refGen
e_tx=XM_019818149.2
E1 51558914 51558996
refGene_gn=PRKAR1A;refGene_tx=XM_006940522.4;refGene_gn=PRKAR1A;refGen
e_tx=XM_019818149.2
E1 51559327 51559500
refGene_gn=PRKAR1A;refGene_tx=XM_006940522.4;refGene_gn=PRKAR1A;refGen
e_tx=XM_019818149.2
E1 51566133 51566398 refGene_gn=FAM20A;refGene_tx=XM_023243542.1
E1 51566820 51566839 refGene_gn=FAM20A;refGene_tx=XM_023243543.1
E1 51567607 51567667
refGene_gn=FAM20A;refGene_tx=XM_023243542.1;refGene_gn=FAM20A;refGene_
tx=XM_023243543.1
E1 51568130 51568212
refGene_gn=FAM20A;refGene_tx=XM_023243542.1;refGene_gn=FAM20A;refGene_
tx=XM_023243543.1
E1 51568591 51568701
refGene_gn=FAM20A;refGene_tx=XM_023243542.1;refGene_gn=FAM20A;refGene_
tx=XM_023243543.1
E1 51569688 51569869
refGene_gn=FAM20A;refGene_tx=XM_023243542.1;refGene_gn=FAM20A;refGene_
tx=XM_023243543.1
E1 51570549 51570665
refGene_gn=FAM20A;refGene_tx=XM_023243542.1;refGene_gn=FAM20A;refGene_
tx=XM_023243543.1
E1 51571438 51571531
refGene_gn=FAM20A;refGene_tx=XM_023243542.1;refGene_gn=FAM20A;refGene_
tx=XM_023243543.1
E1 51576644 51576723
refGene_gn=FAM20A;refGene_tx=XM_023243542.1;refGene_gn=FAM20A;refGene_
tx=XM_023243543.1
E1 51578264 51578315
refGene_gn=FAM20A;refGene_tx=XM_023243542.1;refGene_gn=FAM20A;refGene_
tx=XM_023243543.1
E1 51579045 51579230
refGene_gn=FAM20A;refGene_tx=XM_023243542.1;refGene_gn=FAM20A;refGene_
tx=XM_023243543.1
E1 51617773 51618165
refGene_gn=FAM20A;refGene_tx=XM_023243542.1;refGene_gn=FAM20A;refGene_
tx=XM_023243543.1
E1 51633029 51633200
refGene_gn=LOC109494090;refGene_tx=XM_019816962.1

E1 51686697 51686928
refGene_gn=LOC109494430;refGene_tx=XM_019818150.2;refGene_gn=LOC109494
430;refGene_tx=XM_019818151.2;refGene_gn=LOC109494430;refGene_tx=XM_01
9818152.2
E1 51689076 51689250
refGene_gn=LOC109494430;refGene_tx=XM_019818150.2;refGene_gn=LOC109494
430;refGene_tx=XM_019818151.2;refGene_gn=LOC109494430;refGene_tx=XM_01
9818152.2
E1 51691883 51691999
refGene_gn=LOC109494430;refGene_tx=XM_019818150.2;refGene_gn=LOC109494
430;refGene_tx=XM_019818151.2;refGene_gn=LOC109494430;refGene_tx=XM_01
9818152.2
E1 51701871 51702041
refGene_gn=LOC109494430;refGene_tx=XM_019818150.2;refGene_gn=LOC109494
430;refGene_tx=XM_019818151.2;refGene_gn=LOC109494430;refGene_tx=XM_01
9818152.2
E1 51711774 51711842
refGene_gn=LOC109494430;refGene_tx=XM_019818150.2;refGene_gn=LOC109494
430;refGene_tx=XM_019818151.2
E1 51790883 51790982
refGene_gn=ABCA8;refGene_tx=XM_011289325.3;refGene_gn=ABCA8;refGene_tx
=XM_011289327.3;refGene_gn=ABCA8;refGene_tx=XM_019818154.1;refGene_gn=
ABCA8;refGene_tx=XM_019818155.2;refGene_gn=ABCA8;refGene_tx=XM_0198181
57.1;refGene_gn=ABCA8;refGene_tx=XM_019818158.1;refGene_gn=ABCA8;refGe
ne_tx=XM_019818159.1
E1 51791113 51791169
refGene_gn=ABCA8;refGene_tx=XM_011289325.3;refGene_gn=ABCA8;refGene_tx
=XM_011289327.3;refGene_gn=ABCA8;refGene_tx=XM_019818154.1;refGene_gn=
ABCA8;refGene_tx=XM_019818155.2;refGene_gn=ABCA8;refGene_tx=XM_0198181
57.1;refGene_gn=ABCA8;refGene_tx=XM_019818158.1;refGene_gn=ABCA8;refGe
ne_tx=XM_019818159.1
E1 51792952 51793032
refGene_gn=ABCA8;refGene_tx=XM_011289325.3;refGene_gn=ABCA8;refGene_tx
=XM_011289327.3;refGene_gn=ABCA8;refGene_tx=XM_019818154.1;refGene_gn=
ABCA8;refGene_tx=XM_019818155.2;refGene_gn=ABCA8;refGene_tx=XM_0198181
57.1;refGene_gn=ABCA8;refGene_tx=XM_019818158.1;refGene_gn=ABCA8;refGe
ne_tx=XM_019818159.1
E1 51794759 51794900
refGene_gn=ABCA8;refGene_tx=XM_011289325.3;refGene_gn=ABCA8;refGene_tx
=XM_011289327.3;refGene_gn=ABCA8;refGene_tx=XM_019818154.1;refGene_gn=
ABCA8;refGene_tx=XM_019818155.2;refGene_gn=ABCA8;refGene_tx=XM_0198181
57.1;refGene_gn=ABCA8;refGene_tx=XM_019818158.1;refGene_gn=ABCA8;refGe
ne_tx=XM_019818159.1
E1 51795102 51795222
refGene_gn=ABCA8;refGene_tx=XM_011289325.3;refGene_gn=ABCA8;refGene_tx
=XM_011289327.3;refGene_gn=ABCA8;refGene_tx=XM_019818154.1;refGene_gn=
ABCA8;refGene_tx=XM_019818155.2;refGene_gn=ABCA8;refGene_tx=XM_0198181
57.1;refGene_gn=ABCA8;refGene_tx=XM_019818158.1;refGene_gn=ABCA8;refGe
ne_tx=XM_019818159.1
E1 51795866 51795961

refGene_gn=ABCA8;refGene_tx=XM_011289325.3;refGene_gn=ABCA8;refGene_tx=XM_011289327.3;refGene_gn=ABCA8;refGene_tx=XM_019818154.1;refGene_gn=ABCA8;refGene_tx=XM_019818155.2;refGene_gn=ABCA8;refGene_tx=XM_019818157.1;refGene_gn=ABCA8;refGene_tx=XM_019818158.1;refGene_gn=ABCA8;refGene_tx=XM_019818159.1
E1 51796032 51796108
refGene_gn=ABCA8;refGene_tx=XM_011289325.3;refGene_gn=ABCA8;refGene_tx=XM_011289327.3;refGene_gn=ABCA8;refGene_tx=XM_019818154.1;refGene_gn=ABCA8;refGene_tx=XM_019818155.2;refGene_gn=ABCA8;refGene_tx=XM_019818157.1;refGene_gn=ABCA8;refGene_tx=XM_019818158.1;refGene_gn=ABCA8;refGene_tx=XM_019818159.1
E1 51797277 51797438
refGene_gn=ABCA8;refGene_tx=XM_011289325.3;refGene_gn=ABCA8;refGene_tx=XM_011289327.3;refGene_gn=ABCA8;refGene_tx=XM_019818154.1;refGene_gn=ABCA8;refGene_tx=XM_019818155.2;refGene_gn=ABCA8;refGene_tx=XM_019818157.1;refGene_gn=ABCA8;refGene_tx=XM_019818158.1;refGene_gn=ABCA8;refGene_tx=XM_019818159.1
E1 51800447 51800539
refGene_gn=ABCA8;refGene_tx=XM_011289325.3;refGene_gn=ABCA8;refGene_tx=XM_011289327.3;refGene_gn=ABCA8;refGene_tx=XM_019818154.1;refGene_gn=ABCA8;refGene_tx=XM_019818155.2;refGene_gn=ABCA8;refGene_tx=XM_019818157.1;refGene_gn=ABCA8;refGene_tx=XM_019818158.1;refGene_gn=ABCA8;refGene_tx=XM_019818159.1
E1 51801223 51801341
refGene_gn=ABCA8;refGene_tx=XM_011289325.3;refGene_gn=ABCA8;refGene_tx=XM_011289327.3;refGene_gn=ABCA8;refGene_tx=XM_019818154.1;refGene_gn=ABCA8;refGene_tx=XM_019818155.2;refGene_gn=ABCA8;refGene_tx=XM_019818157.1;refGene_gn=ABCA8;refGene_tx=XM_019818158.1;refGene_gn=ABCA8;refGene_tx=XM_019818159.1
E1 51802208 51802329
refGene_gn=ABCA8;refGene_tx=XM_011289325.3;refGene_gn=ABCA8;refGene_tx=XM_011289327.3;refGene_gn=ABCA8;refGene_tx=XM_019818154.1;refGene_gn=ABCA8;refGene_tx=XM_019818155.2;refGene_gn=ABCA8;refGene_tx=XM_019818157.1;refGene_gn=ABCA8;refGene_tx=XM_019818158.1;refGene_gn=ABCA8;refGene_tx=XM_019818159.1
E1 51803972 51804064
refGene_gn=ABCA8;refGene_tx=XM_011289325.3;refGene_gn=ABCA8;refGene_tx=XM_011289327.3;refGene_gn=ABCA8;refGene_tx=XM_019818154.1;refGene_gn=ABCA8;refGene_tx=XM_019818155.2;refGene_gn=ABCA8;refGene_tx=XM_019818157.1;refGene_gn=ABCA8;refGene_tx=XM_019818158.1;refGene_gn=ABCA8;refGene_tx=XM_019818159.1
E1 51804514 51804580
refGene_gn=ABCA8;refGene_tx=XM_011289325.3;refGene_gn=ABCA8;refGene_tx=XM_011289327.3;refGene_gn=ABCA8;refGene_tx=XM_019818154.1;refGene_gn=ABCA8;refGene_tx=XM_019818155.2;refGene_gn=ABCA8;refGene_tx=XM_019818157.1;refGene_gn=ABCA8;refGene_tx=XM_019818158.1;refGene_gn=ABCA8;refGene_tx=XM_019818159.1
E1 51805444 51805567
refGene_gn=ABCA8;refGene_tx=XM_011289325.3;refGene_gn=ABCA8;refGene_tx=XM_011289327.3;refGene_gn=ABCA8;refGene_tx=XM_019818154.1;refGene_gn=

ABCA8;refGene_tx=XM_019818155.2;refGene_gn=ABCA8;refGene_tx=XM_0198181
58.1;refGene_gn=ABCA8;refGene_tx=XM_019818159.1
E1 51808740 51808854
refGene_gn=ABCA8;refGene_tx=XM_011289325.3;refGene_gn=ABCA8;refGene_tx
=XM_011289327.3;refGene_gn=ABCA8;refGene_tx=XM_019818154.1;refGene_gn=
ABCA8;refGene_tx=XM_019818155.2;refGene_gn=ABCA8;refGene_tx=XM_0198181
57.1;refGene_gn=ABCA8;refGene_tx=XM_019818158.1;refGene_gn=ABCA8;refGe
ne_tx=XM_019818159.1
E1 51809045 51809219
refGene_gn=ABCA8;refGene_tx=XM_011289325.3;refGene_gn=ABCA8;refGene_tx
=XM_011289327.3;refGene_gn=ABCA8;refGene_tx=XM_019818154.1;refGene_gn=
ABCA8;refGene_tx=XM_019818155.2;refGene_gn=ABCA8;refGene_tx=XM_0198181
57.1;refGene_gn=ABCA8;refGene_tx=XM_019818158.1;refGene_gn=ABCA8;refGe
ne_tx=XM_019818159.1
E1 51813960 51814068
refGene_gn=ABCA8;refGene_tx=XM_011289325.3;refGene_gn=ABCA8;refGene_tx
=XM_011289327.3;refGene_gn=ABCA8;refGene_tx=XM_019818154.1;refGene_gn=
ABCA8;refGene_tx=XM_019818155.2;refGene_gn=ABCA8;refGene_tx=XM_0198181
57.1;refGene_gn=ABCA8;refGene_tx=XM_019818158.1;refGene_gn=ABCA8;refGe
ne_tx=XM_019818159.1
E1 51816771 51816909
refGene_gn=ABCA8;refGene_tx=XM_011289325.3;refGene_gn=ABCA8;refGene_tx
=XM_011289327.3;refGene_gn=ABCA8;refGene_tx=XM_019818154.1;refGene_gn=
ABCA8;refGene_tx=XM_019818155.2;refGene_gn=ABCA8;refGene_tx=XM_0198181
57.1;refGene_gn=ABCA8;refGene_tx=XM_019818158.1;refGene_gn=ABCA8;refGe
ne_tx=XM_019818159.1
E1 51817686 51817820
refGene_gn=ABCA8;refGene_tx=XM_011289325.3;refGene_gn=ABCA8;refGene_tx
=XM_011289327.3;refGene_gn=ABCA8;refGene_tx=XM_019818154.1;refGene_gn=
ABCA8;refGene_tx=XM_019818155.2;refGene_gn=ABCA8;refGene_tx=XM_0198181
57.1;refGene_gn=ABCA8;refGene_tx=XM_019818158.1;refGene_gn=ABCA8;refGe
ne_tx=XM_019818159.1
E1 51821726 51821893
refGene_gn=ABCA8;refGene_tx=XM_011289325.3;refGene_gn=ABCA8;refGene_tx
=XM_011289327.3;refGene_gn=ABCA8;refGene_tx=XM_019818154.1;refGene_gn=
ABCA8;refGene_tx=XM_019818155.2;refGene_gn=ABCA8;refGene_tx=XM_0198181
57.1;refGene_gn=ABCA8;refGene_tx=XM_019818158.1;refGene_gn=ABCA8;refGe
ne_tx=XM_019818159.1
E1 51822618 51822817
refGene_gn=ABCA8;refGene_tx=XM_011289325.3;refGene_gn=ABCA8;refGene_tx
=XM_011289327.3;refGene_gn=ABCA8;refGene_tx=XM_019818154.1;refGene_gn=
ABCA8;refGene_tx=XM_019818155.2;refGene_gn=ABCA8;refGene_tx=XM_0198181
57.1;refGene_gn=ABCA8;refGene_tx=XM_019818158.1;refGene_gn=ABCA8;refGe
ne_tx=XM_019818159.1
E1 51826086 51826206
refGene_gn=ABCA8;refGene_tx=XM_011289325.3;refGene_gn=ABCA8;refGene_tx
=XM_011289327.3;refGene_gn=ABCA8;refGene_tx=XM_019818154.1;refGene_gn=
ABCA8;refGene_tx=XM_019818155.2;refGene_gn=ABCA8;refGene_tx=XM_0198181
57.1;refGene_gn=ABCA8;refGene_tx=XM_019818158.1;refGene_gn=ABCA8;refGe
ne_tx=XM_019818159.1

E1 51829702 51829842
refGene_gn=ABCA8;refGene_tx=XM_011289325.3;refGene_gn=ABCA8;refGene_tx=XM_011289327.3;refGene_gn=ABCA8;refGene_tx=XM_019818154.1;refGene_gn=ABCA8;refGene_tx=XM_019818155.2;refGene_gn=ABCA8;refGene_tx=XM_019818157.1;refGene_gn=ABCA8;refGene_tx=XM_019818158.1;refGene_gn=ABCA8;refGene_tx=XM_019818159.1
E1 51832995 51833086
refGene_gn=ABCA8;refGene_tx=XM_011289325.3;refGene_gn=ABCA8;refGene_tx=XM_011289327.3;refGene_gn=ABCA8;refGene_tx=XM_019818154.1;refGene_gn=ABCA8;refGene_tx=XM_019818155.2;refGene_gn=ABCA8;refGene_tx=XM_019818157.1;refGene_gn=ABCA8;refGene_tx=XM_019818158.1;refGene_gn=ABCA8;refGene_tx=XM_019818159.1
E1 51833881 51834020
refGene_gn=ABCA8;refGene_tx=XM_011289325.3;refGene_gn=ABCA8;refGene_tx=XM_011289327.3;refGene_gn=ABCA8;refGene_tx=XM_019818154.1;refGene_gn=ABCA8;refGene_tx=XM_019818155.2;refGene_gn=ABCA8;refGene_tx=XM_019818157.1;refGene_gn=ABCA8;refGene_tx=XM_019818158.1;refGene_gn=ABCA8;refGene_tx=XM_019818159.1
E1 51834265 51834385
refGene_gn=ABCA8;refGene_tx=XM_011289325.3;refGene_gn=ABCA8;refGene_tx=XM_011289327.3;refGene_gn=ABCA8;refGene_tx=XM_019818154.1;refGene_gn=ABCA8;refGene_tx=XM_019818155.2;refGene_gn=ABCA8;refGene_tx=XM_019818157.1;refGene_gn=ABCA8;refGene_tx=XM_019818158.1;refGene_gn=ABCA8;refGene_tx=XM_019818159.1
E1 51834830 51835006
refGene_gn=ABCA8;refGene_tx=XM_011289325.3;refGene_gn=ABCA8;refGene_tx=XM_011289327.3;refGene_gn=ABCA8;refGene_tx=XM_019818154.1;refGene_gn=ABCA8;refGene_tx=XM_019818155.2;refGene_gn=ABCA8;refGene_tx=XM_019818157.1;refGene_gn=ABCA8;refGene_tx=XM_019818158.1;refGene_gn=ABCA8;refGene_tx=XM_019818159.1
E1 51836051 51836162
refGene_gn=ABCA8;refGene_tx=XM_011289325.3;refGene_gn=ABCA8;refGene_tx=XM_011289327.3;refGene_gn=ABCA8;refGene_tx=XM_019818154.1;refGene_gn=ABCA8;refGene_tx=XM_019818155.2;refGene_gn=ABCA8;refGene_tx=XM_019818157.1;refGene_gn=ABCA8;refGene_tx=XM_019818158.1;refGene_gn=ABCA8;refGene_tx=XM_019818159.1
E1 51837358 51837417
refGene_gn=ABCA8;refGene_tx=XM_011289325.3;refGene_gn=ABCA8;refGene_tx=XM_011289327.3;refGene_gn=ABCA8;refGene_tx=XM_019818154.1;refGene_gn=ABCA8;refGene_tx=XM_019818155.2;refGene_gn=ABCA8;refGene_tx=XM_019818157.1;refGene_gn=ABCA8;refGene_tx=XM_019818158.1;refGene_gn=ABCA8;refGene_tx=XM_019818159.1
E1 51839742 51839911
refGene_gn=ABCA8;refGene_tx=XM_011289325.3;refGene_gn=ABCA8;refGene_tx=XM_011289327.3;refGene_gn=ABCA8;refGene_tx=XM_019818154.1;refGene_gn=ABCA8;refGene_tx=XM_019818155.2;refGene_gn=ABCA8;refGene_tx=XM_019818157.1;refGene_gn=ABCA8;refGene_tx=XM_019818158.1;refGene_gn=ABCA8;refGene_tx=XM_019818159.1
E1 51843090 51843238
refGene_gn=ABCA8;refGene_tx=XM_011289325.3;refGene_gn=ABCA8;refGene_tx

=XM_011289327.3;refGene_gn=ABCA8;refGene_tx=XM_019818154.1;refGene_gn=ABCA8;refGene_tx=XM_019818155.2;refGene_gn=ABCA8;refGene_tx=XM_019818157.1;refGene_gn=ABCA8;refGene_tx=XM_019818158.1;refGene_gn=ABCA8;refGene_tx=XM_019818159.1
E1 51844497 51844683
refGene_gn=ABCA8;refGene_tx=XM_011289325.3;refGene_gn=ABCA8;refGene_tx=XM_011289327.3;refGene_gn=ABCA8;refGene_tx=XM_019818154.1;refGene_gn=ABCA8;refGene_tx=XM_019818155.2;refGene_gn=ABCA8;refGene_tx=XM_019818157.1;refGene_gn=ABCA8;refGene_tx=XM_019818158.1;refGene_gn=ABCA8;refGene_tx=XM_019818159.1
E1 51844987 51845129
refGene_gn=ABCA8;refGene_tx=XM_011289325.3;refGene_gn=ABCA8;refGene_tx=XM_011289327.3;refGene_gn=ABCA8;refGene_tx=XM_019818154.1;refGene_gn=ABCA8;refGene_tx=XM_019818155.2;refGene_gn=ABCA8;refGene_tx=XM_019818157.1;refGene_gn=ABCA8;refGene_tx=XM_019818158.1
E1 51847181 51847408
refGene_gn=ABCA8;refGene_tx=XM_011289325.3;refGene_gn=ABCA8;refGene_tx=XM_011289327.3;refGene_gn=ABCA8;refGene_tx=XM_019818154.1;refGene_gn=ABCA8;refGene_tx=XM_019818155.2;refGene_gn=ABCA8;refGene_tx=XM_019818157.1;refGene_gn=ABCA8;refGene_tx=XM_019818158.1
E1 51848315 51848422
refGene_gn=ABCA8;refGene_tx=XM_011289325.3;refGene_gn=ABCA8;refGene_tx=XM_011289327.3;refGene_gn=ABCA8;refGene_tx=XM_019818154.1;refGene_gn=ABCA8;refGene_tx=XM_019818155.2;refGene_gn=ABCA8;refGene_tx=XM_019818157.1
E1 51852539 51852704
refGene_gn=ABCA8;refGene_tx=XM_011289325.3;refGene_gn=ABCA8;refGene_tx=XM_011289327.3;refGene_gn=ABCA8;refGene_tx=XM_019818154.1;refGene_gn=ABCA8;refGene_tx=XM_019818155.2;refGene_gn=ABCA8;refGene_tx=XM_019818157.1
E1 51856230 51856435
refGene_gn=ABCA8;refGene_tx=XM_011289325.3;refGene_gn=ABCA8;refGene_tx=XM_011289327.3;refGene_gn=ABCA8;refGene_tx=XM_019818154.1;refGene_gn=ABCA8;refGene_tx=XM_019818155.2;refGene_gn=ABCA8;refGene_tx=XM_019818157.1
E1 51857594 51857690
refGene_gn=ABCA8;refGene_tx=XM_011289325.3;refGene_gn=ABCA8;refGene_tx=XM_019818154.1;refGene_gn=ABCA8;refGene_tx=XM_019818155.2;refGene_gn=ABCA8;refGene_tx=XM_019818157.1
E1 51895144 51895243
refGene_gn=ABCA9;refGene_tx=XM_003997105.4;refGene_gn=ABCA9;refGene_tx=XM_019818161.2;refGene_gn=ABCA9;refGene_tx=XM_019818162.1;refGene_gn=ABCA9;refGene_tx=XM_019818163.2;refGene_gn=ABCA9;refGene_tx=XM_019818164.1
E1 51895374 51895430
refGene_gn=ABCA9;refGene_tx=XM_003997105.4;refGene_gn=ABCA9;refGene_tx=XM_019818161.2;refGene_gn=ABCA9;refGene_tx=XM_019818162.1;refGene_gn=ABCA9;refGene_tx=XM_019818163.2;refGene_gn=ABCA9;refGene_tx=XM_019818164.1
E1 51898223 51898303

refGene_gn=ABCA9;refGene_tx=XM_003997105.4;refGene_gn=ABCA9;refGene_tx=XM_019818161.2;refGene_gn=ABCA9;refGene_tx=XM_019818162.1;refGene_gn=ABCA9;refGene_tx=XM_019818163.2;refGene_gn=ABCA9;refGene_tx=XM_019818164.1
E1 51899137 51899278
refGene_gn=ABCA9;refGene_tx=XM_003997105.4;refGene_gn=ABCA9;refGene_tx=XM_019818161.2;refGene_gn=ABCA9;refGene_tx=XM_019818162.1;refGene_gn=ABCA9;refGene_tx=XM_019818163.2;refGene_gn=ABCA9;refGene_tx=XM_019818164.1
E1 51899479 51899599
refGene_gn=ABCA9;refGene_tx=XM_003997105.4;refGene_gn=ABCA9;refGene_tx=XM_019818161.2;refGene_gn=ABCA9;refGene_tx=XM_019818162.1;refGene_gn=ABCA9;refGene_tx=XM_019818163.2;refGene_gn=ABCA9;refGene_tx=XM_019818164.1
E1 51900115 51900210
refGene_gn=ABCA9;refGene_tx=XM_003997105.4;refGene_gn=ABCA9;refGene_tx=XM_019818161.2;refGene_gn=ABCA9;refGene_tx=XM_019818162.1;refGene_gn=ABCA9;refGene_tx=XM_019818163.2;refGene_gn=ABCA9;refGene_tx=XM_019818164.1
E1 51900281 51900357
refGene_gn=ABCA9;refGene_tx=XM_003997105.4;refGene_gn=ABCA9;refGene_tx=XM_019818161.2;refGene_gn=ABCA9;refGene_tx=XM_019818162.1;refGene_gn=ABCA9;refGene_tx=XM_019818163.2;refGene_gn=ABCA9;refGene_tx=XM_019818164.1
E1 51901190 51901351
refGene_gn=ABCA9;refGene_tx=XM_003997105.4;refGene_gn=ABCA9;refGene_tx=XM_019818161.2;refGene_gn=ABCA9;refGene_tx=XM_019818162.1;refGene_gn=ABCA9;refGene_tx=XM_019818163.2;refGene_gn=ABCA9;refGene_tx=XM_019818164.1
E1 51903514 51903606
refGene_gn=ABCA9;refGene_tx=XM_003997105.4;refGene_gn=ABCA9;refGene_tx=XM_019818161.2;refGene_gn=ABCA9;refGene_tx=XM_019818162.1;refGene_gn=ABCA9;refGene_tx=XM_019818163.2;refGene_gn=ABCA9;refGene_tx=XM_019818164.1
E1 51904352 51904470
refGene_gn=ABCA9;refGene_tx=XM_003997105.4;refGene_gn=ABCA9;refGene_tx=XM_019818161.2;refGene_gn=ABCA9;refGene_tx=XM_019818162.1;refGene_gn=ABCA9;refGene_tx=XM_019818163.2;refGene_gn=ABCA9;refGene_tx=XM_019818164.1
E1 51905070 51905191
refGene_gn=ABCA9;refGene_tx=XM_003997105.4;refGene_gn=ABCA9;refGene_tx=XM_019818161.2;refGene_gn=ABCA9;refGene_tx=XM_019818162.1;refGene_gn=ABCA9;refGene_tx=XM_019818163.2;refGene_gn=ABCA9;refGene_tx=XM_019818164.1
E1 51906416 51906508
refGene_gn=ABCA9;refGene_tx=XM_003997105.4;refGene_gn=ABCA9;refGene_tx=XM_019818161.2;refGene_gn=ABCA9;refGene_tx=XM_019818162.1;refGene_gn=ABCA9;refGene_tx=XM_019818163.2;refGene_gn=ABCA9;refGene_tx=XM_019818164.1
E1 51906994 51907063

refGene_gn=ABCA9;refGene_tx=XM_003997105.4;refGene_gn=ABCA9;refGene_tx=XM_019818161.2;refGene_gn=ABCA9;refGene_tx=XM_019818162.1;refGene_gn=ABCA9;refGene_tx=XM_019818163.2;refGene_gn=ABCA9;refGene_tx=XM_019818164.1
E1 51909460 51909580
refGene_gn=ABCA9;refGene_tx=XM_003997105.4;refGene_gn=ABCA9;refGene_tx=XM_019818161.2;refGene_gn=ABCA9;refGene_tx=XM_019818162.1;refGene_gn=ABCA9;refGene_tx=XM_019818163.2;refGene_gn=ABCA9;refGene_tx=XM_019818164.1
E1 51912353 51912467
refGene_gn=ABCA9;refGene_tx=XM_003997105.4;refGene_gn=ABCA9;refGene_tx=XM_019818161.2;refGene_gn=ABCA9;refGene_tx=XM_019818162.1;refGene_gn=ABCA9;refGene_tx=XM_019818163.2;refGene_gn=ABCA9;refGene_tx=XM_019818164.1
E1 51912654 51912828
refGene_gn=ABCA9;refGene_tx=XM_003997105.4;refGene_gn=ABCA9;refGene_tx=XM_019818161.2;refGene_gn=ABCA9;refGene_tx=XM_019818162.1;refGene_gn=ABCA9;refGene_tx=XM_019818163.2;refGene_gn=ABCA9;refGene_tx=XM_019818164.1
E1 51915688 51915796
refGene_gn=ABCA9;refGene_tx=XM_003997105.4;refGene_gn=ABCA9;refGene_tx=XM_019818161.2;refGene_gn=ABCA9;refGene_tx=XM_019818162.1;refGene_gn=ABCA9;refGene_tx=XM_019818163.2;refGene_gn=ABCA9;refGene_tx=XM_019818164.1
E1 51920934 51921072
refGene_gn=ABCA9;refGene_tx=XM_003997105.4;refGene_gn=ABCA9;refGene_tx=XM_019818161.2;refGene_gn=ABCA9;refGene_tx=XM_019818162.1;refGene_gn=ABCA9;refGene_tx=XM_019818163.2;refGene_gn=ABCA9;refGene_tx=XM_019818164.1
E1 51923701 51923835
refGene_gn=ABCA9;refGene_tx=XM_003997105.4;refGene_gn=ABCA9;refGene_tx=XM_019818161.2;refGene_gn=ABCA9;refGene_tx=XM_019818162.1;refGene_gn=ABCA9;refGene_tx=XM_019818163.2;refGene_gn=ABCA9;refGene_tx=XM_019818164.1
E1 51924478 51924645
refGene_gn=ABCA9;refGene_tx=XM_003997105.4;refGene_gn=ABCA9;refGene_tx=XM_019818161.2;refGene_gn=ABCA9;refGene_tx=XM_019818162.1;refGene_gn=ABCA9;refGene_tx=XM_019818163.2;refGene_gn=ABCA9;refGene_tx=XM_019818164.1
E1 51926420 51926619
refGene_gn=ABCA9;refGene_tx=XM_003997105.4;refGene_gn=ABCA9;refGene_tx=XM_019818161.2;refGene_gn=ABCA9;refGene_tx=XM_019818162.1;refGene_gn=ABCA9;refGene_tx=XM_019818163.2;refGene_gn=ABCA9;refGene_tx=XM_019818164.1
E1 51928664 51928784
refGene_gn=ABCA9;refGene_tx=XM_003997105.4;refGene_gn=ABCA9;refGene_tx=XM_019818161.2;refGene_gn=ABCA9;refGene_tx=XM_019818162.1;refGene_gn=ABCA9;refGene_tx=XM_019818163.2;refGene_gn=ABCA9;refGene_tx=XM_019818164.1
E1 51931488 51931628

refGene_gn=ABCA9;refGene_tx=XM_003997105.4;refGene_gn=ABCA9;refGene_tx=XM_019818161.2;refGene_gn=ABCA9;refGene_tx=XM_019818162.1;refGene_gn=ABCA9;refGene_tx=XM_019818163.2;refGene_gn=ABCA9;refGene_tx=XM_019818164.1
E1 51934376 51934467
refGene_gn=ABCA9;refGene_tx=XM_003997105.4;refGene_gn=ABCA9;refGene_tx=XM_019818161.2;refGene_gn=ABCA9;refGene_tx=XM_019818162.1;refGene_gn=ABCA9;refGene_tx=XM_019818163.2;refGene_gn=ABCA9;refGene_tx=XM_019818164.1
E1 51935097 51935236
refGene_gn=ABCA9;refGene_tx=XM_003997105.4;refGene_gn=ABCA9;refGene_tx=XM_019818161.2;refGene_gn=ABCA9;refGene_tx=XM_019818162.1;refGene_gn=ABCA9;refGene_tx=XM_019818163.2;refGene_gn=ABCA9;refGene_tx=XM_019818164.1
E1 51935449 51935569
refGene_gn=ABCA9;refGene_tx=XM_003997105.4;refGene_gn=ABCA9;refGene_tx=XM_019818161.2;refGene_gn=ABCA9;refGene_tx=XM_019818162.1;refGene_gn=ABCA9;refGene_tx=XM_019818163.2;refGene_gn=ABCA9;refGene_tx=XM_019818164.1
E1 51936371 51936547
refGene_gn=ABCA9;refGene_tx=XM_003997105.4;refGene_gn=ABCA9;refGene_tx=XM_019818161.2;refGene_gn=ABCA9;refGene_tx=XM_019818162.1;refGene_gn=ABCA9;refGene_tx=XM_019818163.2;refGene_gn=ABCA9;refGene_tx=XM_019818164.1
E1 51937174 51937285
refGene_gn=ABCA9;refGene_tx=XM_003997105.4;refGene_gn=ABCA9;refGene_tx=XM_019818161.2;refGene_gn=ABCA9;refGene_tx=XM_019818162.1;refGene_gn=ABCA9;refGene_tx=XM_019818163.2;refGene_gn=ABCA9;refGene_tx=XM_019818164.1
E1 51938499 51938561
refGene_gn=ABCA9;refGene_tx=XM_003997105.4;refGene_gn=ABCA9;refGene_tx=XM_019818161.2;refGene_gn=ABCA9;refGene_tx=XM_019818162.1;refGene_gn=ABCA9;refGene_tx=XM_019818163.2;refGene_gn=ABCA9;refGene_tx=XM_019818164.1
E1 51939320 51939492
refGene_gn=ABCA9;refGene_tx=XM_003997105.4;refGene_gn=ABCA9;refGene_tx=XM_019818161.2;refGene_gn=ABCA9;refGene_tx=XM_019818162.1;refGene_gn=ABCA9;refGene_tx=XM_019818163.2;refGene_gn=ABCA9;refGene_tx=XM_019818164.1
E1 51940108 51940256
refGene_gn=ABCA9;refGene_tx=XM_003997105.4;refGene_gn=ABCA9;refGene_tx=XM_019818161.2;refGene_gn=ABCA9;refGene_tx=XM_019818162.1;refGene_gn=ABCA9;refGene_tx=XM_019818163.2;refGene_gn=ABCA9;refGene_tx=XM_019818164.1
E1 51941363 51941549
refGene_gn=ABCA9;refGene_tx=XM_003997105.4;refGene_gn=ABCA9;refGene_tx=XM_019818161.2;refGene_gn=ABCA9;refGene_tx=XM_019818162.1;refGene_gn=ABCA9;refGene_tx=XM_019818163.2;refGene_gn=ABCA9;refGene_tx=XM_019818164.1
E1 51941840 51941982

refGene_gn=ABCA9;refGene_tx=XM_003997105.4;refGene_gn=ABCA9;refGene_tx=XM_019818161.2;refGene_gn=ABCA9;refGene_tx=XM_019818162.1;refGene_gn=ABCA9;refGene_tx=XM_019818163.2;refGene_gn=ABCA9;refGene_tx=XM_019818164.1
E1 51946677 51946904
refGene_gn=ABCA9;refGene_tx=XM_003997105.4;refGene_gn=ABCA9;refGene_tx=XM_019818161.2;refGene_gn=ABCA9;refGene_tx=XM_019818162.1;refGene_gn=ABCA9;refGene_tx=XM_019818163.2;refGene_gn=ABCA9;refGene_tx=XM_019818164.1
E1 51947506 51947610
refGene_gn=ABCA9;refGene_tx=XM_003997105.4;refGene_gn=ABCA9;refGene_tx=XM_019818161.2;refGene_gn=ABCA9;refGene_tx=XM_019818162.1;refGene_gn=ABCA9;refGene_tx=XM_019818163.2;refGene_gn=ABCA9;refGene_tx=XM_019818164.1
E1 51948458 51948623
refGene_gn=ABCA9;refGene_tx=XM_003997105.4;refGene_gn=ABCA9;refGene_tx=XM_019818161.2;refGene_gn=ABCA9;refGene_tx=XM_019818162.1;refGene_gn=ABCA9;refGene_tx=XM_019818163.2;refGene_gn=ABCA9;refGene_tx=XM_019818164.1
E1 51950073 51950281
refGene_gn=ABCA9;refGene_tx=XM_003997105.4;refGene_gn=ABCA9;refGene_tx=XM_019818161.2;refGene_gn=ABCA9;refGene_tx=XM_019818162.1;refGene_gn=ABCA9;refGene_tx=XM_019818163.2;refGene_gn=ABCA9;refGene_tx=XM_019818164.1
E1 51952078 51952174
refGene_gn=ABCA9;refGene_tx=XM_003997105.4;refGene_gn=ABCA9;refGene_tx=XM_019818161.2;refGene_gn=ABCA9;refGene_tx=XM_019818162.1;refGene_gn=ABCA9;refGene_tx=XM_019818163.2
E1 51980945 51981047 refGene_gn=ABCA6;refGene_tx=XM_006940527.4
E1 51981178 51981234 refGene_gn=ABCA6;refGene_tx=XM_006940527.4
E1 51983470 51983550 refGene_gn=ABCA6;refGene_tx=XM_006940527.4
E1 51985403 51985544 refGene_gn=ABCA6;refGene_tx=XM_006940527.4
E1 51986006 51986126 refGene_gn=ABCA6;refGene_tx=XM_006940527.4
E1 51986783 51986878 refGene_gn=ABCA6;refGene_tx=XM_006940527.4
E1 51986955 51987031 refGene_gn=ABCA6;refGene_tx=XM_006940527.4
E1 51987497 51987664 refGene_gn=ABCA6;refGene_tx=XM_006940527.4
E1 51988144 51988236 refGene_gn=ABCA6;refGene_tx=XM_006940527.4
E1 51989588 51989706 refGene_gn=ABCA6;refGene_tx=XM_006940527.4
E1 51990784 51990905 refGene_gn=ABCA6;refGene_tx=XM_006940527.4
E1 51991608 51991700 refGene_gn=ABCA6;refGene_tx=XM_006940527.4
E1 51993225 51993303 refGene_gn=ABCA6;refGene_tx=XM_006940527.4
E1 51995183 51995303 refGene_gn=ABCA6;refGene_tx=XM_006940527.4
E1 51998838 51998952 refGene_gn=ABCA6;refGene_tx=XM_006940527.4
E1 51999169 51999343 refGene_gn=ABCA6;refGene_tx=XM_006940527.4
E1 52000251 52000359 refGene_gn=ABCA6;refGene_tx=XM_006940527.4
E1 52002966 52003104 refGene_gn=ABCA6;refGene_tx=XM_006940527.4
E1 52003870 52004004 refGene_gn=ABCA6;refGene_tx=XM_006940527.4
E1 52006341 52006508 refGene_gn=ABCA6;refGene_tx=XM_006940527.4
E1 52006896 52006955 refGene_gn=ABCA6;refGene_tx=XM_006940527.4
E1 52006956 52007081 refGene_gn=ABCA6;refGene_tx=XM_006940527.4

| | | | |
|---|---|---|---|
| E1 | 52008039 | 52008156 | refGene_gn=ABCA6;refGene_tx=XM_006940527.4 |
| E1 | 52011582 | 52011722 | refGene_gn=ABCA6;refGene_tx=XM_006940527.4 |
| E1 | 52012639 | 52012730 | refGene_gn=ABCA6;refGene_tx=XM_006940527.4 |
| E1 | 52013258 | 52013397 | refGene_gn=ABCA6;refGene_tx=XM_006940527.4 |
| E1 | 52013653 | 52013773 | refGene_gn=ABCA6;refGene_tx=XM_006940527.4 |
| E1 | 52013984 | 52014160 | refGene_gn=ABCA6;refGene_tx=XM_006940527.4 |
| E1 | 52014560 | 52014671 | refGene_gn=ABCA6;refGene_tx=XM_006940527.4 |
| E1 | 52018038 | 52018097 | refGene_gn=ABCA6;refGene_tx=XM_006940527.4 |
| E1 | 52022195 | 52022361 | refGene_gn=ABCA6;refGene_tx=XM_006940527.4 |
| E1 | 52023930 | 52024078 | refGene_gn=ABCA6;refGene_tx=XM_006940527.4 |
| E1 | 52028280 | 52028466 | refGene_gn=ABCA6;refGene_tx=XM_006940527.4 |
| E1 | 52029258 | 52029400 | refGene_gn=ABCA6;refGene_tx=XM_006940527.4 |
| E1 | 52032657 | 52032884 | refGene_gn=ABCA6;refGene_tx=XM_006940527.4 |
| E1 | 52033288 | 52033392 | refGene_gn=ABCA6;refGene_tx=XM_006940527.4 |
| E1 | 52035076 | 52035238 | refGene_gn=ABCA6;refGene_tx=XM_006940527.4 |
| E1 | 52036252 | 52036454 | refGene_gn=ABCA6;refGene_tx=XM_006940527.4 |
| E1 | 52038230 | 52038326 | refGene_gn=ABCA6;refGene_tx=XM_006940527.4 |
| E1 | 52051832 | 52051931 | refGene_gn=ABCA10;refGene_tx=XM_023243471.1 |
| E1 | 52052050 | 52052106 | refGene_gn=ABCA10;refGene_tx=XM_023243471.1 |
| E1 | 52052998 | 52053078 | refGene_gn=ABCA10;refGene_tx=XM_023243471.1 |
| E1 | 52055551 | 52055692 | refGene_gn=ABCA10;refGene_tx=XM_023243471.1 |
| E1 | 52055910 | 52056030 | refGene_gn=ABCA10;refGene_tx=XM_023243471.1 |
| E1 | 52056733 | 52056828 | refGene_gn=ABCA10;refGene_tx=XM_023243471.1 |
| E1 | 52056907 | 52056983 | refGene_gn=ABCA10;refGene_tx=XM_023243471.1 |
| E1 | 52057262 | 52057624 | refGene_gn=ABCA10;refGene_tx=XM_023243471.1 |
| E1 | 52059962 | 52060121 | refGene_gn=ABCA10;refGene_tx=XM_023243471.1 |
| E1 | 52060982 | 52061103 | refGene_gn=ABCA10;refGene_tx=XM_023243471.1 |
| E1 | 52061804 | 52061896 | refGene_gn=ABCA10;refGene_tx=XM_023243471.1 |
| E1 | 52064658 | 52064745 | refGene_gn=ABCA10;refGene_tx=XM_023243471.1 |
| E1 | 52066901 | 52067021 | refGene_gn=ABCA10;refGene_tx=XM_023243471.1 |
| E1 | 52073427 | 52073541 | refGene_gn=ABCA10;refGene_tx=XM_023243471.1 |
| E1 | 52073735 | 52073906 | refGene_gn=ABCA10;refGene_tx=XM_023243471.1 |
| E1 | 52074687 | 52074795 | refGene_gn=ABCA10;refGene_tx=XM_023243471.1 |
| E1 | 52078031 | 52078169 | refGene_gn=ABCA10;refGene_tx=XM_023243471.1 |
| E1 | 52079797 | 52079931 | refGene_gn=ABCA10;refGene_tx=XM_023243471.1 |
| E1 | 52080652 | 52080819 | refGene_gn=ABCA10;refGene_tx=XM_023243471.1 |
| E1 | 52082665 | 52082864 | refGene_gn=ABCA10;refGene_tx=XM_023243471.1 |
| E1 | 52084150 | 52084270 | refGene_gn=ABCA10;refGene_tx=XM_023243471.1 |
| E1 | 52085091 | 52085233 | refGene_gn=ABCA10;refGene_tx=XM_023243471.1 |
| E1 | 52087236 | 52087325 | refGene_gn=ABCA10;refGene_tx=XM_023243471.1 |
| E1 | 52088650 | 52088789 | refGene_gn=ABCA10;refGene_tx=XM_023243471.1 |
| E1 | 52089025 | 52089145 | refGene_gn=ABCA10;refGene_tx=XM_023243471.1 |
| E1 | 52089563 | 52089739 | refGene_gn=ABCA10;refGene_tx=XM_023243471.1 |
| E1 | 52090304 | 52090415 | refGene_gn=ABCA10;refGene_tx=XM_023243471.1 |
| E1 | 52092718 | 52092777 | refGene_gn=ABCA10;refGene_tx=XM_023243471.1 |
| E1 | 52097515 | 52097684 | refGene_gn=ABCA10;refGene_tx=XM_023243471.1 |
| E1 | 52104584 | 52104732 | refGene_gn=ABCA10;refGene_tx=XM_023243471.1 |
| E1 | 52105893 | 52106079 | refGene_gn=ABCA10;refGene_tx=XM_023243471.1 |
| E1 | 52106296 | 52106438 | refGene_gn=ABCA10;refGene_tx=XM_023243471.1 |
| E1 | 52114149 | 52114376 | refGene_gn=ABCA10;refGene_tx=XM_023243471.1 |

E1 52116681 52116785 refGene_gn=ABCA10;refGene_tx=XM_023243471.1
E1 52117133 52117298 refGene_gn=ABCA10;refGene_tx=XM_023243471.1
E1 52119122 52119327 refGene_gn=ABCA10;refGene_tx=XM_023243471.1
E1 52121118 52121214 refGene_gn=ABCA10;refGene_tx=XM_023243471.1
E1 52144802 52144910 refGene_gn=ABCA5;refGene_tx=XM_023244816.1
E1 52145420 52145476 refGene_gn=ABCA5;refGene_tx=XM_023244816.1
E1 52146556 52146636 refGene_gn=ABCA5;refGene_tx=XM_023244816.1
E1 52147017 52147167 refGene_gn=ABCA5;refGene_tx=XM_023244816.1
E1 52148299 52148419 refGene_gn=ABCA5;refGene_tx=XM_023244816.1
E1 52149395 52149490 refGene_gn=ABCA5;refGene_tx=XM_023244816.1
E1 52149615 52149691 refGene_gn=ABCA5;refGene_tx=XM_023244816.1
E1 52150507 52150683 refGene_gn=ABCA5;refGene_tx=XM_023244816.1
E1 52151684 52151776 refGene_gn=ABCA5;refGene_tx=XM_023244816.1
E1 52151872 52151990 refGene_gn=ABCA5;refGene_tx=XM_023244816.1
E1 52152300 52152427 refGene_gn=ABCA5;refGene_tx=XM_023244816.1
E1 52154485 52154577 refGene_gn=ABCA5;refGene_tx=XM_023244816.1
E1 52155609 52155684 refGene_gn=ABCA5;refGene_tx=XM_023244816.1
E1 52157166 52157301 refGene_gn=ABCA5;refGene_tx=XM_023244816.1
E1 52157698 52157812 refGene_gn=ABCA5;refGene_tx=XM_023244816.1
E1 52159267 52159438 refGene_gn=ABCA5;refGene_tx=XM_023244816.1
E1 52162368 52162482 refGene_gn=ABCA5;refGene_tx=XM_023244816.1
E1 52165518 52165656 refGene_gn=ABCA5;refGene_tx=XM_023244816.1
E1 52166272 52166400 refGene_gn=ABCA5;refGene_tx=XM_023244816.1
E1 52168649 52168819 refGene_gn=ABCA5;refGene_tx=XM_023244816.1
E1 52171721 52171923 refGene_gn=ABCA5;refGene_tx=XM_023244816.1
E1 52176128 52176248 refGene_gn=ABCA5;refGene_tx=XM_023244816.1
E1 52177198 52177338 refGene_gn=ABCA5;refGene_tx=XM_023244816.1
E1 52177528 52177619 refGene_gn=ABCA5;refGene_tx=XM_023244816.1
E1 52178585 52178724 refGene_gn=ABCA5;refGene_tx=XM_023244816.1
E1 52179830 52179950 refGene_gn=ABCA5;refGene_tx=XM_023244816.1
E1 52180469 52180645 refGene_gn=ABCA5;refGene_tx=XM_023244816.1
E1 52181835 52181946 refGene_gn=ABCA5;refGene_tx=XM_023244816.1
E1 52188318 52188377 refGene_gn=ABCA5;refGene_tx=XM_023244816.1
E1 52189733 52189902 refGene_gn=ABCA5;refGene_tx=XM_023244816.1
E1 52191817 52191965 refGene_gn=ABCA5;refGene_tx=XM_023244816.1
E1 52193424 52193613 refGene_gn=ABCA5;refGene_tx=XM_023244816.1
E1 52194688 52194830 refGene_gn=ABCA5;refGene_tx=XM_023244816.1
E1 52196803 52197033 refGene_gn=ABCA5;refGene_tx=XM_023244816.1
E1 52199044 52199133 refGene_gn=ABCA5;refGene_tx=XM_023244816.1
E1 52200356 52200518 refGene_gn=ABCA5;refGene_tx=XM_023244816.1
E1 52202405 52202610 refGene_gn=ABCA5;refGene_tx=XM_023244816.1
E1 52203853 52203955 refGene_gn=ABCA5;refGene_tx=XM_023244816.1
E1 52281235 52281251 refGene_gn=MAP2K6;refGene_tx=XM_003997108.5
E1 52358882 52358949 refGene_gn=MAP2K6;refGene_tx=XM_003997108.5
E1 52368070 52368119 refGene_gn=MAP2K6;refGene_tx=XM_003997108.5
E1 52368932 52369046
refGene_gn=MAP2K6;refGene_tx=XM_003997108.5;refGene_gn=MAP2K6;refGene_tx=XM_019818165.2;refGene_gn=MAP2K6;refGene_tx=XM_019818166.2
E1 52370895 52371015
refGene_gn=MAP2K6;refGene_tx=XM_003997108.5;refGene_gn=MAP2K6;refGene_

tx=XM_019818165.2;refGene_gn=MAP2K6;refGene_tx=XM_019818166.2
E1 52372322 52372439
refGene_gn=MAP2K6;refGene_tx=XM_003997108.5;refGene_gn=MAP2K6;refGene_tx=XM_019818165.2;refGene_gn=MAP2K6;refGene_tx=XM_019818166.2
E1 52373098 52373150
refGene_gn=MAP2K6;refGene_tx=XM_003997108.5;refGene_gn=MAP2K6;refGene_tx=XM_019818165.2;refGene_gn=MAP2K6;refGene_tx=XM_019818166.2
E1 52375430 52375558
refGene_gn=MAP2K6;refGene_tx=XM_003997108.5;refGene_gn=MAP2K6;refGene_tx=XM_019818165.2;refGene_gn=MAP2K6;refGene_tx=XM_019818166.2
E1 52376470 52376548
refGene_gn=MAP2K6;refGene_tx=XM_003997108.5;refGene_gn=MAP2K6;refGene_tx=XM_019818165.2;refGene_gn=MAP2K6;refGene_tx=XM_019818166.2
E1 52378156 52378296
refGene_gn=MAP2K6;refGene_tx=XM_003997108.5;refGene_gn=MAP2K6;refGene_tx=XM_019818165.2;refGene_gn=MAP2K6;refGene_tx=XM_019818166.2
E1 52386108 52386154
refGene_gn=MAP2K6;refGene_tx=XM_003997108.5;refGene_gn=MAP2K6;refGene_tx=XM_019818165.2;refGene_gn=MAP2K6;refGene_tx=XM_019818166.2
E1 52392517 52392595
refGene_gn=MAP2K6;refGene_tx=XM_003997108.5;refGene_gn=MAP2K6;refGene_tx=XM_019818165.2;refGene_gn=MAP2K6;refGene_tx=XM_019818166.2
E1 52888258 52889518
refGene_gn=KCNJ16;refGene_tx=XM_003997109.5;refGene_gn=KCNJ16;refGene_tx=XM_019818167.2;refGene_gn=KCNJ16;refGene_tx=XM_019818168.2;refGene_gn=KCNJ16;refGene_tx=XM_019818169.2;refGene_gn=KCNJ16;refGene_tx=XM_019818170.2;refGene_gn=KCNJ16;refGene_tx=XM_019818171.2;refGene_gn=KCNJ16;refGene_tx=XM_019818173.2;refGene_gn=KCNJ16;refGene_tx=XM_019818175.2
E1 52926700 52927984
refGene_gn=KCNJ2;refGene_tx=XM_019818177.2;refGene_gn=KCNJ2;refGene_tx=XM_023243719.1;refGene_gn=KCNJ2;refGene_tx=XM_023243720.1
E1 54366784 54366820
refGene_gn=LOC109494092;refGene_tx=XM_023243472.1
E1 54368839 54369164
refGene_gn=LOC109494092;refGene_tx=XM_023243472.1
E1 54369233 54369924
refGene_gn=LOC109494092;refGene_tx=XM_023243472.1
E1 54370237 54370394
refGene_gn=LOC109494092;refGene_tx=XM_023243472.1
E1 54646418 54646849 refGene_gn=SOX9;refGene_tx=XM_023243815.1
E1 54647797 54648054 refGene_gn=SOX9;refGene_tx=XM_023243815.1
E1 54648656 54649504 refGene_gn=SOX9;refGene_tx=XM_023243815.1
E1 55078140 55078219
refGene_gn=SLC39A11;refGene_tx=XM_019818185.2;refGene_gn=SLC39A11;refGene_tx=XM_019818186.2;refGene_gn=SLC39A11;refGene_tx=XM_023243819.1;refGene_gn=SLC39A11;refGene_tx=XM_023243820.1;refGene_gn=SLC39A11;refGene_tx=XM_023243821.1;refGene_gn=SLC39A11;refGene_tx=XM_023243822.1
E1 55079254 55079413
refGene_gn=SLC39A11;refGene_tx=XM_019818185.2;refGene_gn=SLC39A11;refG

ene_tx=XM_019818186.2;refGene_gn=SLC39A11;refGene_tx=XM_023243819.1;refGene_gn=SLC39A11;refGene_tx=XM_023243820.1;refGene_gn=SLC39A11;refGene_tx=XM_023243821.1;refGene_gn=SLC39A11;refGene_tx=XM_023243822.1
E1 55082364 55082463
refGene_gn=SLC39A11;refGene_tx=XM_019818185.2;refGene_gn=SLC39A11;refGene_tx=XM_019818186.2;refGene_gn=SLC39A11;refGene_tx=XM_023243819.1;refGene_gn=SLC39A11;refGene_tx=XM_023243820.1;refGene_gn=SLC39A11;refGene_tx=XM_023243821.1;refGene_gn=SLC39A11;refGene_tx=XM_023243822.1
E1 55144577 55144647
refGene_gn=SLC39A11;refGene_tx=XM_019818185.2;refGene_gn=SLC39A11;refGene_tx=XM_019818186.2;refGene_gn=SLC39A11;refGene_tx=XM_023243819.1;refGene_gn=SLC39A11;refGene_tx=XM_023243820.1;refGene_gn=SLC39A11;refGene_tx=XM_023243821.1;refGene_gn=SLC39A11;refGene_tx=XM_023243822.1
E1 55225010 55225181
refGene_gn=SLC39A11;refGene_tx=XM_019818185.2;refGene_gn=SLC39A11;refGene_tx=XM_019818186.2;refGene_gn=SLC39A11;refGene_tx=XM_023243819.1;refGene_gn=SLC39A11;refGene_tx=XM_023243820.1;refGene_gn=SLC39A11;refGene_tx=XM_023243821.1;refGene_gn=SLC39A11;refGene_tx=XM_023243822.1
E1 55324698 55324840
refGene_gn=SLC39A11;refGene_tx=XM_019818185.2;refGene_gn=SLC39A11;refGene_tx=XM_019818186.2;refGene_gn=SLC39A11;refGene_tx=XM_023243819.1;refGene_gn=SLC39A11;refGene_tx=XM_023243820.1;refGene_gn=SLC39A11;refGene_tx=XM_023243821.1;refGene_gn=SLC39A11;refGene_tx=XM_023243822.1
E1 55378293 55378392
refGene_gn=SLC39A11;refGene_tx=XM_023243819.1;refGene_gn=SLC39A11;refGene_tx=XM_023243820.1;refGene_gn=SLC39A11;refGene_tx=XM_023243821.1
E1 55382454 55382613
refGene_gn=SLC39A11;refGene_tx=XM_019818185.2;refGene_gn=SLC39A11;refGene_tx=XM_019818186.2;refGene_gn=SLC39A11;refGene_tx=XM_023243819.1;refGene_gn=SLC39A11;refGene_tx=XM_023243820.1;refGene_gn=SLC39A11;refGene_tx=XM_023243821.1
E1 55419454 55419493
refGene_gn=SLC39A11;refGene_tx=XM_019818185.2;refGene_gn=SLC39A11;refGene_tx=XM_019818186.2;refGene_gn=SLC39A11;refGene_tx=XM_023243819.1;refGene_gn=SLC39A11;refGene_tx=XM_023243820.1;refGene_gn=SLC39A11;refGene_tx=XM_023243821.1;refGene_gn=SLC39A11;refGene_tx=XM_023243822.1
E1 55424128 55424247
refGene_gn=SLC39A11;refGene_tx=XM_019818185.2;refGene_gn=SLC39A11;refGene_tx=XM_019818186.2;refGene_gn=SLC39A11;refGene_tx=XM_023243819.1;refGene_gn=SLC39A11;refGene_tx=XM_023243820.1;refGene_gn=SLC39A11;refGene_tx=XM_023243821.1;refGene_gn=SLC39A11;refGene_tx=XM_023243822.1
E1 55485433 55486543
refGene_gn=SSTR2;refGene_tx=NM_001309046.1;refGene_gn=SSTR2;refGene_tx=XM_011289337.3;refGene_gn=SSTR2;refGene_tx=XM_019816878.2;refGene_gn=SSTR2;refGene_tx=XM_019816879.2;refGene_gn=SSTR2;refGene_tx=XM_019816880.2
E1 55507812 55508127
refGene_gn=COG1;refGene_tx=XM_003997114.5;refGene_gn=COG1;refGene_tx=XM_019818180.2;refGene_gn=COG1;refGene_tx=XM_019818181.2
E1 55508221 55508315

refGene_gn=SLC39A11;refGene_tx=XM_019818185.2;refGene_gn=SLC39A11;refGene_tx=XM_019818186.2;refGene_gn=SLC39A11;refGene_tx=XM_023243819.1;refGene_gn=SLC39A11;refGene_tx=XM_023243822.1
E1 55510696 55510789
refGene_gn=COG1;refGene_tx=XM_019818180.2;refGene_gn=COG1;refGene_tx=XM_019818181.2
E1 55511001 55511246
refGene_gn=COG1;refGene_tx=XM_003997114.5;refGene_gn=COG1;refGene_tx=XM_019818180.2;refGene_gn=COG1;refGene_tx=XM_019818181.2
E1 55511396 55511578
refGene_gn=COG1;refGene_tx=XM_003997114.5;refGene_gn=COG1;refGene_tx=XM_019818180.2;refGene_gn=COG1;refGene_tx=XM_019818181.2
E1 55511723 55511894
refGene_gn=COG1;refGene_tx=XM_003997114.5;refGene_gn=COG1;refGene_tx=XM_019818180.2;refGene_gn=COG1;refGene_tx=XM_019818181.2
E1 55513717 55513874
refGene_gn=COG1;refGene_tx=XM_003997114.5;refGene_gn=COG1;refGene_tx=XM_019818180.2;refGene_gn=COG1;refGene_tx=XM_019818181.2
E1 55514419 55514630
refGene_gn=COG1;refGene_tx=XM_003997114.5;refGene_gn=COG1;refGene_tx=XM_019818180.2;refGene_gn=COG1;refGene_tx=XM_019818181.2
E1 55514997 55515789
refGene_gn=COG1;refGene_tx=XM_003997114.5;refGene_gn=COG1;refGene_tx=XM_019818180.2;refGene_gn=COG1;refGene_tx=XM_019818181.2
E1 55516339 55516486
refGene_gn=COG1;refGene_tx=XM_003997114.5;refGene_gn=COG1;refGene_tx=XM_019818180.2;refGene_gn=COG1;refGene_tx=XM_019818181.2
E1 55516935 55517097
refGene_gn=COG1;refGene_tx=XM_003997114.5;refGene_gn=COG1;refGene_tx=XM_019818180.2;refGene_gn=COG1;refGene_tx=XM_019818181.2
E1 55518127 55518255
refGene_gn=COG1;refGene_tx=XM_003997114.5;refGene_gn=COG1;refGene_tx=XM_019818180.2;refGene_gn=COG1;refGene_tx=XM_019818181.2
E1 55518707 55518816
refGene_gn=COG1;refGene_tx=XM_003997114.5;refGene_gn=COG1;refGene_tx=XM_019818180.2;refGene_gn=COG1;refGene_tx=XM_019818181.2
E1 55519268 55519378
refGene_gn=COG1;refGene_tx=XM_003997114.5;refGene_gn=COG1;refGene_tx=XM_019818180.2;refGene_gn=COG1;refGene_tx=XM_019818181.2
E1 55519458 55519537
refGene_gn=COG1;refGene_tx=XM_003997114.5;refGene_gn=COG1;refGene_tx=XM_019818180.2;refGene_gn=COG1;refGene_tx=XM_019818181.2
E1 55520616 55520754
refGene_gn=COG1;refGene_tx=XM_003997114.5;refGene_gn=COG1;refGene_tx=XM_019818180.2;refGene_gn=COG1;refGene_tx=XM_019818181.2
E1 55521827 55522058
refGene_gn=FAM104A;refGene_tx=XM_003997115.5;refGene_gn=FAM104A;refGene_tx=XM_003997116.5;refGene_gn=FAM104A;refGene_tx=XM_019818191.2;refGene_gn=FAM104A;refGene_tx=XM_023243830.1
E1 55523696 55523759

refGene_gn=FAM104A;refGene_tx=XM_003997116.5;refGene_gn=FAM104A;refGene_tx=XM_019818191.2;refGene_gn=FAM104A;refGene_tx=XM_023243830.1
E1 55532384 55532484
refGene_gn=FAM104A;refGene_tx=XM_003997115.5;refGene_gn=FAM104A;refGene_tx=XM_003997116.5;refGene_gn=FAM104A;refGene_tx=XM_019818191.2;refGene_gn=FAM104A;refGene_tx=XM_023243830.1
E1 55532839 55532868 refGene_gn=FAM104A;refGene_tx=XM_023243830.1
E1 55536764 55536796 refGene_gn=FAM104A;refGene_tx=XM_019818191.2
E1 55536938 55537159
refGene_gn=FAM104A;refGene_tx=XM_003997115.5;refGene_gn=FAM104A;refGene_tx=XM_003997116.5
E1 55539751 55541219
refGene_gn=CE1H17orf80;refGene_tx=XM_003997118.5;refGene_gn=CE1H17orf80;refGene_tx=XM_006940530.3;refGene_gn=CE1H17orf80;refGene_tx=XM_006940531.4;refGene_gn=CE1H17orf80;refGene_tx=XM_019818182.2;refGene_gn=CE1H17orf80;refGene_tx=XM_019818183.2;refGene_gn=CE1H17orf80;refGene_tx=XM_023243817.1
E1 55545703 55545811
refGene_gn=CE1H17orf80;refGene_tx=XM_003997118.5;refGene_gn=CE1H17orf80;refGene_tx=XM_006940530.3;refGene_gn=CE1H17orf80;refGene_tx=XM_019818182.2;refGene_gn=CE1H17orf80;refGene_tx=XM_019818183.2;refGene_gn=CE1H17orf80;refGene_tx=XM_023243817.1
E1 55546372 55546477
refGene_gn=CE1H17orf80;refGene_tx=XM_003997118.5;refGene_gn=CE1H17orf80;refGene_tx=XM_006940530.3;refGene_gn=CE1H17orf80;refGene_tx=XM_006940531.4;refGene_gn=CE1H17orf80;refGene_tx=XM_019818182.2;refGene_gn=CE1H17orf80;refGene_tx=XM_019818183.2;refGene_gn=CE1H17orf80;refGene_tx=XM_023243817.1
E1 55547658 55547681
refGene_gn=CE1H17orf80;refGene_tx=XM_003997118.5;refGene_gn=CE1H17orf80;refGene_tx=XM_006940530.3;refGene_gn=CE1H17orf80;refGene_tx=XM_006940531.4;refGene_gn=CE1H17orf80;refGene_tx=XM_019818182.2;refGene_gn=CE1H17orf80;refGene_tx=XM_019818183.2;refGene_gn=CE1H17orf80;refGene_tx=XM_023243817.1
E1 55552603 55552693
refGene_gn=CPSF4L;refGene_tx=XM_023243823.1;refGene_gn=CPSF4L;refGene_tx=XM_023243824.1;refGene_gn=CPSF4L;refGene_tx=XM_023243825.1;refGene_gn=CPSF4L;refGene_tx=XM_023243827.1;refGene_gn=CPSF4L;refGene_tx=XM_023243828.1
E1 55553667 55553835
refGene_gn=CPSF4L;refGene_tx=XM_023243823.1;refGene_gn=CPSF4L;refGene_tx=XM_023243824.1;refGene_gn=CPSF4L;refGene_tx=XM_023243825.1;refGene_gn=CPSF4L;refGene_tx=XM_023243827.1;refGene_gn=CPSF4L;refGene_tx=XM_023243828.1
E1 55555013 55555053
refGene_gn=CPSF4L;refGene_tx=XM_023243823.1;refGene_gn=CPSF4L;refGene_tx=XM_023243824.1;refGene_gn=CPSF4L;refGene_tx=XM_023243825.1;refGene_gn=CPSF4L;refGene_tx=XM_023243827.1;refGene_gn=CPSF4L;refGene_tx=XM_023243828.1
E1 55555529 55555630

refGene_gn=CPSF4L;refGene_tx=XM_023243823.1;refGene_gn=CPSF4L;refGene_tx=XM_023243824.1;refGene_gn=CPSF4L;refGene_tx=XM_023243825.1;refGene_gn=CPSF4L;refGene_tx=XM_023243827.1;refGene_gn=CPSF4L;refGene_tx=XM_023243828.1;refGene_gn=CPSF4L;refGene_tx=XM_023243829.1
E1 55556562 55556658
refGene_gn=CPSF4L;refGene_tx=XM_023243823.1;refGene_gn=CPSF4L;refGene_tx=XM_023243824.1;refGene_gn=CPSF4L;refGene_tx=XM_023243825.1;refGene_gn=CPSF4L;refGene_tx=XM_023243827.1;refGene_gn=CPSF4L;refGene_tx=XM_023243828.1;refGene_gn=CPSF4L;refGene_tx=XM_023243829.1
E1 55559606 55559841
refGene_gn=CPSF4L;refGene_tx=XM_023243823.1;refGene_gn=CPSF4L;refGene_tx=XM_023243824.1;refGene_gn=CPSF4L;refGene_tx=XM_023243825.1;refGene_gn=CPSF4L;refGene_tx=XM_023243829.1
E1 55560509 55560560
refGene_gn=CPSF4L;refGene_tx=XM_023243827.1;refGene_gn=CPSF4L;refGene_tx=XM_023243828.1;refGene_gn=CPSF4L;refGene_tx=XM_023243829.1
E1 55561087 55561190
refGene_gn=CPSF4L;refGene_tx=XM_023243827.1;refGene_gn=CPSF4L;refGene_tx=XM_023243828.1;refGene_gn=CPSF4L;refGene_tx=XM_023243829.1
E1 55574432 55575482
refGene_gn=CDC42EP4;refGene_tx=XM_006940534.4;refGene_gn=CDC42EP4;refGene_tx=XM_023243835.1;refGene_gn=CDC42EP4;refGene_tx=XM_023243836.1;refGene_gn=CDC42EP4;refGene_tx=XM_023243837.1;refGene_gn=CDC42EP4;refGene_tx=XM_023243838.1;refGene_gn=CDC42EP4;refGene_tx=XM_023243839.1;refGene_gn=CDC42EP4;refGene_tx=XM_023243841.1;refGene_gn=CDC42EP4;refGene_tx=XM_023243842.1
E1 55614433 55614787 refGene_gn=SDK2;refGene_tx=XM_011289338.3
E1 55624025 55624152 refGene_gn=SDK2;refGene_tx=XM_011289338.3
E1 55626590 55626729 refGene_gn=SDK2;refGene_tx=XM_011289338.3
E1 55627000 55627141 refGene_gn=SDK2;refGene_tx=XM_011289338.3
E1 55628958 55629123 refGene_gn=SDK2;refGene_tx=XM_011289338.3
E1 55632799 55632925 refGene_gn=SDK2;refGene_tx=XM_011289338.3
E1 55635850 55636012 refGene_gn=SDK2;refGene_tx=XM_011289338.3
E1 55637541 55637679 refGene_gn=SDK2;refGene_tx=XM_011289338.3
E1 55640408 55640601 refGene_gn=SDK2;refGene_tx=XM_011289338.3
E1 55646279 55646395 refGene_gn=SDK2;refGene_tx=XM_011289338.3
E1 55646538 55646640 refGene_gn=SDK2;refGene_tx=XM_011289338.3
E1 55648198 55648255 refGene_gn=SDK2;refGene_tx=XM_011289338.3
E1 55649647 55649783 refGene_gn=SDK2;refGene_tx=XM_011289338.3
E1 55651261 55651332 refGene_gn=SDK2;refGene_tx=XM_011289338.3
E1 55651775 55651879 refGene_gn=SDK2;refGene_tx=XM_011289338.3
E1 55652781 55652983 refGene_gn=SDK2;refGene_tx=XM_011289338.3
E1 55653995 55654190 refGene_gn=SDK2;refGene_tx=XM_011289338.3
E1 55655605 55655704 refGene_gn=SDK2;refGene_tx=XM_011289338.3
E1 55656948 55657138 refGene_gn=SDK2;refGene_tx=XM_011289338.3
E1 55657794 55657910 refGene_gn=SDK2;refGene_tx=XM_011289338.3
E1 55658631 55658869 refGene_gn=SDK2;refGene_tx=XM_011289338.3
E1 55661405 55661556 refGene_gn=SDK2;refGene_tx=XM_011289338.3
E1 55661680 55661790 refGene_gn=SDK2;refGene_tx=XM_011289338.3
E1 55662428 55662476 refGene_gn=SDK2;refGene_tx=XM_011289338.3

E1 55662477 55662551 refGene_gn=SDK2;refGene_tx=XM_011289338.3
E1 55664102 55664294 refGene_gn=SDK2;refGene_tx=XM_011289338.3
E1 55664666 55664765 refGene_gn=SDK2;refGene_tx=XM_011289338.3
E1 55664969 55665165 refGene_gn=SDK2;refGene_tx=XM_011289338.3
E1 55673315 55673431 refGene_gn=SDK2;refGene_tx=XM_011289338.3
E1 55673972 55674154 refGene_gn=SDK2;refGene_tx=XM_011289338.3
E1 55675635 55675776 refGene_gn=SDK2;refGene_tx=XM_011289338.3
E1 55680559 55680707 refGene_gn=SDK2;refGene_tx=XM_011289338.3
E1 55681289 55681426 refGene_gn=SDK2;refGene_tx=XM_011289338.3
E1 55681674 55681851 refGene_gn=SDK2;refGene_tx=XM_011289338.3
E1 55688438 55688541 refGene_gn=SDK2;refGene_tx=XM_011289338.3
E1 55689493 55689661 refGene_gn=SDK2;refGene_tx=XM_011289338.3
E1 55691249 55691366 refGene_gn=SDK2;refGene_tx=XM_011289338.3
E1 55692582 55692777 refGene_gn=SDK2;refGene_tx=XM_011289338.3
E1 55694882 55694966 refGene_gn=SDK2;refGene_tx=XM_011289338.3
E1 55695133 55695324 refGene_gn=SDK2;refGene_tx=XM_011289338.3
E1 55696963 55697075 refGene_gn=SDK2;refGene_tx=XM_011289338.3
E1 55704207 55704341 refGene_gn=SDK2;refGene_tx=XM_011289338.3
E1 55710006 55710154 refGene_gn=SDK2;refGene_tx=XM_011289338.3
E1 55721327 55721434 refGene_gn=SDK2;refGene_tx=XM_011289338.3
E1 55749780 55749940 refGene_gn=SDK2;refGene_tx=XM_011289338.3
E1 55876165 55876229 refGene_gn=SDK2;refGene_tx=XM_011289338.3
E1 55876996 55877639
refGene_gn=LOC111557722;refGene_tx=XM_023243473.1
E1 55877837 55877995
refGene_gn=LOC111557722;refGene_tx=XM_023243473.1
E1 56333258 56333387
refGene_gn=TTYH2;refGene_tx=XM_003997129.4;refGene_gn=TTYH2;refGene_tx
=XM_019818194.1
E1 56340003 56340009 refGene_gn=TTYH2;refGene_tx=XM_019818193.1
E1 56340274 56340447
refGene_gn=TTYH2;refGene_tx=XM_003997129.4;refGene_gn=TTYH2;refGene_tx
=XM_019818193.1
E1 56344994 56345106
refGene_gn=TTYH2;refGene_tx=XM_003997129.4;refGene_gn=TTYH2;refGene_tx
=XM_019818193.1;refGene_gn=TTYH2;refGene_tx=XM_019818194.1
E1 56349933 56350154
refGene_gn=TTYH2;refGene_tx=XM_003997129.4;refGene_gn=TTYH2;refGene_tx
=XM_019818193.1;refGene_gn=TTYH2;refGene_tx=XM_019818194.1
E1 56354665 56354761
refGene_gn=TTYH2;refGene_tx=XM_003997129.4;refGene_gn=TTYH2;refGene_tx
=XM_019818193.1;refGene_gn=TTYH2;refGene_tx=XM_019818194.1
E1 56355329 56355402
refGene_gn=TTYH2;refGene_tx=XM_003997129.4;refGene_gn=TTYH2;refGene_tx
=XM_019818193.1;refGene_gn=TTYH2;refGene_tx=XM_019818194.1
E1 56359844 56359914
refGene_gn=TTYH2;refGene_tx=XM_003997129.4;refGene_gn=TTYH2;refGene_tx
=XM_019818193.1;refGene_gn=TTYH2;refGene_tx=XM_019818194.1
E1 56360172 56360228
refGene_gn=TTYH2;refGene_tx=XM_003997129.4;refGene_gn=TTYH2;refGene_tx

=XM_019818193.1;refGene_gn=TTYH2;refGene_tx=XM_019818194.1
E1 56360522 56360615
refGene_gn=TTYH2;refGene_tx=XM_003997129.4;refGene_gn=TTYH2;refGene_tx
=XM_019818193.1;refGene_gn=TTYH2;refGene_tx=XM_019818194.1
E1 56360894 56360987
refGene_gn=TTYH2;refGene_tx=XM_003997129.4;refGene_gn=TTYH2;refGene_tx
=XM_019818193.1;refGene_gn=TTYH2;refGene_tx=XM_019818194.1
E1 56362769 56362912
refGene_gn=TTYH2;refGene_tx=XM_003997129.4;refGene_gn=TTYH2;refGene_tx
=XM_019818193.1;refGene_gn=TTYH2;refGene_tx=XM_019818194.1
E1 56363596 56363782
refGene_gn=TTYH2;refGene_tx=XM_003997129.4;refGene_gn=TTYH2;refGene_tx
=XM_019818193.1;refGene_gn=TTYH2;refGene_tx=XM_019818194.1
E1 56364289 56364368
refGene_gn=TTYH2;refGene_tx=XM_003997129.4;refGene_gn=TTYH2;refGene_tx
=XM_019818193.1;refGene_gn=TTYH2;refGene_tx=XM_019818194.1
E1 56369209 56369290
refGene_gn=TTYH2;refGene_tx=XM_003997129.4;refGene_gn=TTYH2;refGene_tx
=XM_019818193.1;refGene_gn=TTYH2;refGene_tx=XM_019818194.1
E1 56384551 56384734
refGene_gn=DNAI2;refGene_tx=XM_003997188.4;refGene_gn=DNAI2;refGene_tx
=XM_019818199.2;refGene_gn=DNAI2;refGene_tx=XM_019818201.2
E1 56386716 56386878
refGene_gn=DNAI2;refGene_tx=XM_003997188.4;refGene_gn=DNAI2;refGene_tx
=XM_019818199.2;refGene_gn=DNAI2;refGene_tx=XM_019818201.2
E1 56387844 56387966
refGene_gn=DNAI2;refGene_tx=XM_003997188.4;refGene_gn=DNAI2;refGene_tx
=XM_019818199.2;refGene_gn=DNAI2;refGene_tx=XM_019818201.2
E1 56391199 56391342
refGene_gn=DNAI2;refGene_tx=XM_003997188.4;refGene_gn=DNAI2;refGene_tx
=XM_019818199.2;refGene_gn=DNAI2;refGene_tx=XM_019818201.2
E1 56392305 56392419
refGene_gn=DNAI2;refGene_tx=XM_003997188.4;refGene_gn=DNAI2;refGene_tx
=XM_019818199.2;refGene_gn=DNAI2;refGene_tx=XM_019818201.2
E1 56395872 56396012
refGene_gn=DNAI2;refGene_tx=XM_003997188.4;refGene_gn=DNAI2;refGene_tx
=XM_019818199.2;refGene_gn=DNAI2;refGene_tx=XM_019818201.2
E1 56397112 56397235
refGene_gn=DNAI2;refGene_tx=XM_003997188.4;refGene_gn=DNAI2;refGene_tx
=XM_019818199.2;refGene_gn=DNAI2;refGene_tx=XM_019818201.2
E1 56399255 56399479
refGene_gn=DNAI2;refGene_tx=XM_003997188.4;refGene_gn=DNAI2;refGene_tx
=XM_019818199.2;refGene_gn=DNAI2;refGene_tx=XM_019818201.2
E1 56403035 56403171
refGene_gn=DNAI2;refGene_tx=XM_003997188.4;refGene_gn=DNAI2;refGene_tx
=XM_019818199.2;refGene_gn=DNAI2;refGene_tx=XM_019818201.2
E1 56403693 56403840
refGene_gn=DNAI2;refGene_tx=XM_003997188.4;refGene_gn=DNAI2;refGene_tx
=XM_019818199.2;refGene_gn=DNAI2;refGene_tx=XM_019818201.2
E1 56404598 56404823

refGene_gn=DNAI2;refGene_tx=XM_003997188.4;refGene_gn=DNAI2;refGene_tx=XM_019818199.2
E1 56406479 56406566
refGene_gn=DNAI2;refGene_tx=XM_003997188.4;refGene_gn=DNAI2;refGene_tx=XM_019818199.2;refGene_gn=DNAI2;refGene_tx=XM_019818201.2
E1 56412872 56412911
refGene_gn=KIF19;refGene_tx=XM_019818195.2;refGene_gn=KIF19;refGene_tx=XM_023243900.1;refGene_gn=KIF19;refGene_tx=XM_023243901.1
E1 56414572 56414653
refGene_gn=KIF19;refGene_tx=XM_019818195.2;refGene_gn=KIF19;refGene_tx=XM_023243900.1;refGene_gn=KIF19;refGene_tx=XM_023243901.1
E1 56425719 56425830
refGene_gn=KIF19;refGene_tx=XM_019818195.2;refGene_gn=KIF19;refGene_tx=XM_023243900.1;refGene_gn=KIF19;refGene_tx=XM_023243901.1
E1 56426453 56426541
refGene_gn=KIF19;refGene_tx=XM_019818195.2;refGene_gn=KIF19;refGene_tx=XM_023243900.1;refGene_gn=KIF19;refGene_tx=XM_023243901.1
E1 56426815 56426952
refGene_gn=KIF19;refGene_tx=XM_019818195.2;refGene_gn=KIF19;refGene_tx=XM_023243900.1;refGene_gn=KIF19;refGene_tx=XM_023243901.1
E1 56428215 56428341
refGene_gn=KIF19;refGene_tx=XM_019818195.2;refGene_gn=KIF19;refGene_tx=XM_023243900.1
E1 56428688 56428883
refGene_gn=KIF19;refGene_tx=XM_019818195.2;refGene_gn=KIF19;refGene_tx=XM_023243900.1;refGene_gn=KIF19;refGene_tx=XM_023243901.1
E1 56430091 56430238
refGene_gn=KIF19;refGene_tx=XM_019818195.2;refGene_gn=KIF19;refGene_tx=XM_023243900.1;refGene_gn=KIF19;refGene_tx=XM_023243901.1
E1 56431444 56431567
refGene_gn=KIF19;refGene_tx=XM_019818195.2;refGene_gn=KIF19;refGene_tx=XM_023243900.1;refGene_gn=KIF19;refGene_tx=XM_023243901.1
E1 56432995 56433161
refGene_gn=KIF19;refGene_tx=XM_019818195.2;refGene_gn=KIF19;refGene_tx=XM_023243900.1;refGene_gn=KIF19;refGene_tx=XM_023243901.1
E1 56433878 56434053
refGene_gn=KIF19;refGene_tx=XM_019818195.2;refGene_gn=KIF19;refGene_tx=XM_023243900.1;refGene_gn=KIF19;refGene_tx=XM_023243901.1
E1 56434187 56434386
refGene_gn=KIF19;refGene_tx=XM_019818195.2;refGene_gn=KIF19;refGene_tx=XM_023243900.1;refGene_gn=KIF19;refGene_tx=XM_023243901.1
E1 56434997 56435268
refGene_gn=KIF19;refGene_tx=XM_019818195.2;refGene_gn=KIF19;refGene_tx=XM_023243900.1;refGene_gn=KIF19;refGene_tx=XM_023243901.1
E1 56435362 56435484
refGene_gn=KIF19;refGene_tx=XM_019818195.2;refGene_gn=KIF19;refGene_tx=XM_023243900.1;refGene_gn=KIF19;refGene_tx=XM_023243901.1
E1 56435961 56436095
refGene_gn=KIF19;refGene_tx=XM_019818195.2;refGene_gn=KIF19;refGene_tx=XM_023243900.1;refGene_gn=KIF19;refGene_tx=XM_023243901.1

E1 56436336 56436442
refGene_gn=KIF19;refGene_tx=XM_019818195.2;refGene_gn=KIF19;refGene_tx=XM_023243900.1;refGene_gn=KIF19;refGene_tx=XM_023243901.1
E1 56436633 56436730
refGene_gn=KIF19;refGene_tx=XM_019818195.2;refGene_gn=KIF19;refGene_tx=XM_023243900.1;refGene_gn=KIF19;refGene_tx=XM_023243901.1
E1 56437224 56437619
refGene_gn=KIF19;refGene_tx=XM_019818195.2;refGene_gn=KIF19;refGene_tx=XM_023243900.1;refGene_gn=KIF19;refGene_tx=XM_023243901.1
E1 56437803 56437963
refGene_gn=KIF19;refGene_tx=XM_019818195.2;refGene_gn=KIF19;refGene_tx=XM_023243900.1;refGene_gn=KIF19;refGene_tx=XM_023243901.1
E1 56438199 56438330
refGene_gn=KIF19;refGene_tx=XM_019818195.2;refGene_gn=KIF19;refGene_tx=XM_023243900.1;refGene_gn=KIF19;refGene_tx=XM_023243901.1
E1 56439631 56440706
refGene_gn=BTBD17;refGene_tx=XM_023243902.1;refGene_gn=BTBD17;refGene_tx=XM_023243903.1
E1 56442062 56442339
refGene_gn=BTBD17;refGene_tx=XM_023243902.1;refGene_gn=BTBD17;refGene_tx=XM_023243903.1
E1 56443771 56443856 refGene_gn=BTBD17;refGene_tx=XM_023243903.1
E1 56445030 56445514 refGene_gn=BTBD17;refGene_tx=XM_023243902.1
E1 56448961 56448986 refGene_gn=GPR142;refGene_tx=XM_023243904.1
E1 56449835 56449994 refGene_gn=GPR142;refGene_tx=XM_023243904.1
E1 56450898 56451770 refGene_gn=GPR142;refGene_tx=XM_023243904.1
E1 56493428 56493696
refGene_gn=GPRC5C;refGene_tx=XM_019818202.2;refGene_gn=GPRC5C;refGene_tx=XM_023243918.1
E1 56499303 56500386
refGene_gn=GPRC5C;refGene_tx=XM_006940544.4;refGene_gn=GPRC5C;refGene_tx=XM_006940545.3;refGene_gn=GPRC5C;refGene_tx=XM_006940546.4;refGene_gn=GPRC5C;refGene_tx=XM_019818202.2;refGene_gn=GPRC5C;refGene_tx=XM_023243918.1
E1 56503438 56503533
refGene_gn=GPRC5C;refGene_tx=XM_006940544.4;refGene_gn=GPRC5C;refGene_tx=XM_006940545.3;refGene_gn=GPRC5C;refGene_tx=XM_006940546.4;refGene_gn=GPRC5C;refGene_tx=XM_019818202.2;refGene_gn=GPRC5C;refGene_tx=XM_023243918.1
E1 56506019 56506199
refGene_gn=GPRC5C;refGene_tx=XM_006940544.4;refGene_gn=GPRC5C;refGene_tx=XM_006940545.3;refGene_gn=GPRC5C;refGene_tx=XM_006940546.4;refGene_gn=GPRC5C;refGene_tx=XM_019818202.2;refGene_gn=GPRC5C;refGene_tx=XM_023243918.1
E1 56508094 56508127
refGene_gn=GPRC5C;refGene_tx=XM_006940544.4;refGene_gn=GPRC5C;refGene_tx=XM_006940546.4;refGene_gn=GPRC5C;refGene_tx=XM_019818202.2
E1 56527506 56527567
refGene_gn=LOC101088701;refGene_tx=XM_006940550.3;refGene_gn=LOC101088701;refGene_tx=XM_006940551.2;refGene_gn=LOC101088701;refGene_tx=XM_01

1289340.2;refGene_gn=LOC101088701;refGene_tx=XM_019818205.1;refGene_gn=LOC101088701;refGene_tx=XM_019818206.1
E1 56529553 56529889
refGene_gn=LOC101088701;refGene_tx=XM_006940550.3;refGene_gn=LOC101088701;refGene_tx=XM_006940551.2;refGene_gn=LOC101088701;refGene_tx=XM_011289340.2;refGene_gn=LOC101088701;refGene_tx=XM_019818205.1;refGene_gn=LOC101088701;refGene_tx=XM_019818206.1;refGene_gn=LOC101088701;refGene_tx=XM_019818207.1
E1 56530504 56530735
refGene_gn=LOC101088701;refGene_tx=XM_006940550.3;refGene_gn=LOC101088701;refGene_tx=XM_006940551.2;refGene_gn=LOC101088701;refGene_tx=XM_011289340.2;refGene_gn=LOC101088701;refGene_tx=XM_019818205.1;refGene_gn=LOC101088701;refGene_tx=XM_019818206.1;refGene_gn=LOC101088701;refGene_tx=XM_019818207.1
E1 56533547 56533642
refGene_gn=LOC101088701;refGene_tx=XM_006940550.3;refGene_gn=LOC101088701;refGene_tx=XM_006940551.2;refGene_gn=LOC101088701;refGene_tx=XM_011289340.2;refGene_gn=LOC101088701;refGene_tx=XM_019818205.1;refGene_gn=LOC101088701;refGene_tx=XM_019818206.1;refGene_gn=LOC101088701;refGene_tx=XM_019818207.1
E1 56537489 56537527
refGene_gn=LOC101088701;refGene_tx=XM_006940550.3;refGene_gn=LOC101088701;refGene_tx=XM_006940551.2;refGene_gn=LOC101088701;refGene_tx=XM_011289340.2;refGene_gn=LOC101088701;refGene_tx=XM_019818205.1;refGene_gn=LOC101088701;refGene_tx=XM_019818206.1;refGene_gn=LOC101088701;refGene_tx=XM_019818207.1
E1 56537986 56538094
refGene_gn=LOC101088701;refGene_tx=XM_006940550.3;refGene_gn=LOC101088701;refGene_tx=XM_006940551.2;refGene_gn=LOC101088701;refGene_tx=XM_011289340.2;refGene_gn=LOC101088701;refGene_tx=XM_019818205.1;refGene_gn=LOC101088701;refGene_tx=XM_019818206.1;refGene_gn=LOC101088701;refGene_tx=XM_019818207.1
E1 56538927 56539065
refGene_gn=LOC101088701;refGene_tx=XM_006940550.3;refGene_gn=LOC101088701;refGene_tx=XM_006940551.2
E1 56539307 56539616
refGene_gn=LOC101088701;refGene_tx=XM_019818205.1;refGene_gn=LOC101088701;refGene_tx=XM_019818206.1;refGene_gn=LOC101088701;refGene_tx=XM_019818207.1
E1 56551282 56551456
refGene_gn=LOC101088701;refGene_tx=XM_011289340.2
E1 56564411 56564517
refGene_gn=LOC101088964;refGene_tx=XM_006940554.2
E1 56565327 56565415
refGene_gn=LOC101088964;refGene_tx=XM_006940554.2
E1 56566618 56566948
refGene_gn=LOC101088964;refGene_tx=XM_006940554.2
E1 56573740 56573780
refGene_gn=LOC101088964;refGene_tx=XM_006940554.2
E1 56577438 56577571

refGene_gn=LOC101089223;refGene_tx=XM_006940555.3
E1 56578673 56578797
refGene_gn=LOC101089223;refGene_tx=XM_006940555.3
E1 56580321 56580657
refGene_gn=LOC101089223;refGene_tx=XM_006940555.3
E1 56581189 56581250
refGene_gn=LOC101089223;refGene_tx=XM_006940555.3
E1 56595082 56595344
refGene_gn=LOC101089477;refGene_tx=XM_006940556.4
E1 56598019 56598377
refGene_gn=LOC101089477;refGene_tx=XM_006940556.4
E1 56603748 56603788
refGene_gn=LOC101089477;refGene_tx=XM_006940556.4
E1 56619855 56619895
refGene_gn=LOC109494122;refGene_tx=XM_019816998.1
E1 56621310 56621640
refGene_gn=LOC109494122;refGene_tx=XM_019816998.1
E1 56623073 56623200
refGene_gn=LOC109494122;refGene_tx=XM_019816998.1
E1 56625627 56625781
refGene_gn=LOC109494122;refGene_tx=XM_019816998.1
E1 56652803 56652843
refGene_gn=LOC109494123;refGene_tx=XM_019816999.2
E1 56654581 56654911
refGene_gn=LOC109494123;refGene_tx=XM_019816999.2
E1 56656371 56656495
refGene_gn=LOC109494123;refGene_tx=XM_019816999.2
E1 56658861 56658964
refGene_gn=LOC109494123;refGene_tx=XM_019816999.2
E1 56663311 56663490
refGene_gn=LOC102899494;refGene_tx=XM_011289341.2
E1 56664971 56665304
refGene_gn=LOC102899494;refGene_tx=XM_011289341.2
E1 56669587 56669627
refGene_gn=LOC102899494;refGene_tx=XM_011289341.2
E1 56693868 56694047
refGene_gn=LOC101089971;refGene_tx=XM_006940557.3
E1 56695831 56696164
refGene_gn=LOC101089971;refGene_tx=XM_006940557.3
E1 56700434 56700474
refGene_gn=LOC101089971;refGene_tx=XM_006940557.3
E1 56711648 56711688
refGene_gn=LOC109494125;refGene_tx=XM_019817001.2;refGene_gn=LOC109494125;refGene_tx=XM_019817002.2
E1 56713044 56713374
refGene_gn=LOC109494125;refGene_tx=XM_019817001.2;refGene_gn=LOC109494125;refGene_tx=XM_019817002.2
E1 56713708 56713832
refGene_gn=LOC109494125;refGene_tx=XM_019817001.2;refGene_gn=LOC109494125;refGene_tx=XM_019817002.2

E1 56716210 56716313
refGene_gn=LOC109494125;refGene_tx=XM_019817002.2
E1 56716366 56716511
refGene_gn=LOC109494125;refGene_tx=XM_019817001.2
E1 56722477 56722745 refGene_gn=CD300E;refGene_tx=XM_023244065.1
E1 56723450 56723562 refGene_gn=CD300E;refGene_tx=XM_023244065.1
E1 56725589 56725934 refGene_gn=CD300E;refGene_tx=XM_023244065.1
E1 56728271 56728311 refGene_gn=CD300E;refGene_tx=XM_023244065.1
E1 56741411 56741523
refGene_gn=LOC111557803;refGene_tx=XM_023244129.1
E1 56742805 56742902
refGene_gn=LOC111557803;refGene_tx=XM_023244129.1
E1 56744404 56744737
refGene_gn=LOC111557803;refGene_tx=XM_023244129.1
E1 56745992 56746032
refGene_gn=LOC111557803;refGene_tx=XM_023244129.1
E1 56794869 56794935 refGene_gn=RAB37;refGene_tx=XM_019818214.2
E1 56814981 56815137
refGene_gn=CD300LF;refGene_tx=XM_006940563.4;refGene_gn=CD300LF;refGene_tx=XM_019818212.2
E1 56815627 56815777
refGene_gn=CD300LF;refGene_tx=XM_006940563.4;refGene_gn=CD300LF;refGene_tx=XM_019818212.2
E1 56816057 56816080
refGene_gn=CD300LF;refGene_tx=XM_006940563.4;refGene_gn=CD300LF;refGene_tx=XM_019818212.2
E1 56817152 56817265
refGene_gn=CD300LF;refGene_tx=XM_006940563.4;refGene_gn=CD300LF;refGene_tx=XM_019818212.2
E1 56818744 56818847
refGene_gn=CD300LF;refGene_tx=XM_006940563.4;refGene_gn=CD300LF;refGene_tx=XM_019818212.2
E1 56819749 56820079
refGene_gn=CD300LF;refGene_tx=XM_006940563.4;refGene_gn=CD300LF;refGene_tx=XM_019818212.2
E1 56829760 56830040
refGene_gn=CD300LF;refGene_tx=XM_006940563.4;refGene_gn=CD300LF;refGene_tx=XM_019818212.2
E1 56838863 56839016
refGene_gn=RAB37;refGene_tx=XM_019818213.2;refGene_gn=RAB37;refGene_tx=XM_019818214.2
E1 56842377 56842461
refGene_gn=RAB37;refGene_tx=XM_011289343.3;refGene_gn=RAB37;refGene_tx=XM_023243947.1
E1 56845245 56845356
refGene_gn=RAB37;refGene_tx=XM_011289343.3;refGene_gn=RAB37;refGene_tx=XM_019818214.2;refGene_gn=RAB37;refGene_tx=XM_019818216.2;refGene_gn=RAB37;refGene_tx=XM_023243947.1
E1 56846408 56846450
refGene_gn=RAB37;refGene_tx=XM_011289343.3;refGene_gn=RAB37;refGene_tx

=XM_019818213.2;refGene_gn=RAB37;refGene_tx=XM_019818214.2;refGene_gn=RAB37;refGene_tx=XM_019818216.2;refGene_gn=RAB37;refGene_tx=XM_023243947.1
E1 56846959 56847026
refGene_gn=RAB37;refGene_tx=XM_011289343.3;refGene_gn=RAB37;refGene_tx=XM_019818213.2;refGene_gn=RAB37;refGene_tx=XM_019818214.2;refGene_gn=RAB37;refGene_tx=XM_019818216.2;refGene_gn=RAB37;refGene_tx=XM_023243947.1
E1 56847105 56847158
refGene_gn=RAB37;refGene_tx=XM_011289343.3;refGene_gn=RAB37;refGene_tx=XM_019818213.2;refGene_gn=RAB37;refGene_tx=XM_019818214.2;refGene_gn=RAB37;refGene_tx=XM_019818216.2;refGene_gn=RAB37;refGene_tx=XM_023243947.1
E1 56848110 56848176
refGene_gn=RAB37;refGene_tx=XM_011289343.3;refGene_gn=RAB37;refGene_tx=XM_019818213.2;refGene_gn=RAB37;refGene_tx=XM_019818214.2;refGene_gn=RAB37;refGene_tx=XM_019818216.2;refGene_gn=RAB37;refGene_tx=XM_023243947.1
E1 56848781 56848838
refGene_gn=RAB37;refGene_tx=XM_011289343.3;refGene_gn=RAB37;refGene_tx=XM_019818213.2;refGene_gn=RAB37;refGene_tx=XM_019818214.2;refGene_gn=RAB37;refGene_tx=XM_019818216.2;refGene_gn=RAB37;refGene_tx=XM_023243947.1
E1 56848915 56848992
refGene_gn=RAB37;refGene_tx=XM_011289343.3;refGene_gn=RAB37;refGene_tx=XM_019818213.2;refGene_gn=RAB37;refGene_tx=XM_019818214.2;refGene_gn=RAB37;refGene_tx=XM_019818216.2;refGene_gn=RAB37;refGene_tx=XM_023243947.1
E1 56849185 56849291
refGene_gn=RAB37;refGene_tx=XM_011289343.3;refGene_gn=RAB37;refGene_tx=XM_019818213.2;refGene_gn=RAB37;refGene_tx=XM_019818214.2;refGene_gn=RAB37;refGene_tx=XM_019818216.2;refGene_gn=RAB37;refGene_tx=XM_023243947.1
E1 56852078 56852540 refGene_gn=SLC9A3R1;refGene_tx=XM_006940564.4
E1 56863977 56864139 refGene_gn=SLC9A3R1;refGene_tx=XM_006940564.4
E1 56864998 56865152 refGene_gn=SLC9A3R1;refGene_tx=XM_006940564.4
E1 56867436 56867477 refGene_gn=SLC9A3R1;refGene_tx=XM_006940564.4
E1 56868633 56868723 refGene_gn=SLC9A3R1;refGene_tx=XM_006940564.4
E1 56868969 56869161 refGene_gn=SLC9A3R1;refGene_tx=XM_006940564.4
E1 56872110 56872245
refGene_gn=NAT9;refGene_tx=XM_011289345.3;refGene_gn=NAT9;refGene_tx=XM_019818222.1;refGene_gn=NAT9;refGene_tx=XM_023243948.1
E1 56872350 56872445
refGene_gn=NAT9;refGene_tx=XM_011289345.3;refGene_gn=NAT9;refGene_tx=XM_019818222.1;refGene_gn=NAT9;refGene_tx=XM_023243948.1;refGene_gn=NAT9;refGene_tx=XM_023243949.1
E1 56872665 56872725
refGene_gn=NAT9;refGene_tx=XM_011289345.3;refGene_gn=NAT9;refGene_tx=XM_019818222.1;refGene_gn=NAT9;refGene_tx=XM_023243948.1;refGene_gn=NAT9;refGene_tx=XM_023243949.1

E1 56873311 56873455
refGene_gn=NAT9;refGene_tx=XM_011289345.3;refGene_gn=NAT9;refGene_tx=XM_019818222.1;refGene_gn=NAT9;refGene_tx=XM_023243948.1;refGene_gn=NAT9;refGene_tx=XM_023243949.1
E1 56873979 56874089
refGene_gn=NAT9;refGene_tx=XM_011289345.3;refGene_gn=NAT9;refGene_tx=XM_019818222.1;refGene_gn=NAT9;refGene_tx=XM_023243948.1;refGene_gn=NAT9;refGene_tx=XM_023243949.1
E1 56875014 56875091
refGene_gn=NAT9;refGene_tx=XM_011289345.3;refGene_gn=NAT9;refGene_tx=XM_019818222.1;refGene_gn=NAT9;refGene_tx=XM_023243948.1;refGene_gn=NAT9;refGene_tx=XM_023243949.1
E1 56875948 56876057
refGene_gn=TMEM104;refGene_tx=XM_023243944.1;refGene_gn=TMEM104;refGene_tx=XM_023243945.1
E1 56876664 56876786
refGene_gn=TMEM104;refGene_tx=XM_023243944.1;refGene_gn=TMEM104;refGene_tx=XM_023243945.1;refGene_gn=TMEM104;refGene_tx=XM_023243946.1
E1 56885165 56885287
refGene_gn=TMEM104;refGene_tx=XM_023243944.1;refGene_gn=TMEM104;refGene_tx=XM_023243945.1;refGene_gn=TMEM104;refGene_tx=XM_023243946.1
E1 56887471 56887553
refGene_gn=TMEM104;refGene_tx=XM_023243944.1;refGene_gn=TMEM104;refGene_tx=XM_023243945.1;refGene_gn=TMEM104;refGene_tx=XM_023243946.1
E1 56888034 56888131
refGene_gn=TMEM104;refGene_tx=XM_023243944.1;refGene_gn=TMEM104;refGene_tx=XM_023243945.1;refGene_gn=TMEM104;refGene_tx=XM_023243946.1
E1 56888865 56888946
refGene_gn=TMEM104;refGene_tx=XM_023243944.1;refGene_gn=TMEM104;refGene_tx=XM_023243945.1;refGene_gn=TMEM104;refGene_tx=XM_023243946.1
E1 56892176 56892276
refGene_gn=TMEM104;refGene_tx=XM_023243944.1;refGene_gn=TMEM104;refGene_tx=XM_023243945.1;refGene_gn=TMEM104;refGene_tx=XM_023243946.1
E1 56892697 56892806
refGene_gn=TMEM104;refGene_tx=XM_023243944.1;refGene_gn=TMEM104;refGene_tx=XM_023243945.1;refGene_gn=TMEM104;refGene_tx=XM_023243946.1
E1 56916520 56916614
refGene_gn=TMEM104;refGene_tx=XM_023243944.1;refGene_gn=TMEM104;refGene_tx=XM_023243945.1;refGene_gn=TMEM104;refGene_tx=XM_023243946.1
E1 56932414 56933175
refGene_gn=TMEM104;refGene_tx=XM_023243944.1;refGene_gn=TMEM104;refGene_tx=XM_023243945.1;refGene_gn=TMEM104;refGene_tx=XM_023243946.1
E1 56939249 56940344
refGene_gn=GRIN2C;refGene_tx=XM_023243950.1;refGene_gn=GRIN2C;refGene_tx=XM_023243951.1;refGene_gn=GRIN2C;refGene_tx=XM_023243952.1;refGene_gn=GRIN2C;refGene_tx=XM_023243953.1;refGene_gn=GRIN2C;refGene_tx=XM_023243954.1
E1 56940766 56940999
refGene_gn=GRIN2C;refGene_tx=XM_023243950.1;refGene_gn=GRIN2C;refGene_tx=XM_023243951.1;refGene_gn=GRIN2C;refGene_tx=XM_023243952.1;refGene_

gn=GRIN2C;refGene_tx=XM_023243953.1;refGene_gn=GRIN2C;refGene_tx=XM_023243954.1
E1 56941934 56942122
refGene_gn=GRIN2C;refGene_tx=XM_023243950.1;refGene_gn=GRIN2C;refGene_tx=XM_023243951.1;refGene_gn=GRIN2C;refGene_tx=XM_023243952.1;refGene_gn=GRIN2C;refGene_tx=XM_023243953.1;refGene_gn=GRIN2C;refGene_tx=XM_023243954.1
E1 56942655 56942816
refGene_gn=GRIN2C;refGene_tx=XM_023243950.1;refGene_gn=GRIN2C;refGene_tx=XM_023243951.1;refGene_gn=GRIN2C;refGene_tx=XM_023243952.1;refGene_gn=GRIN2C;refGene_tx=XM_023243953.1;refGene_gn=GRIN2C;refGene_tx=XM_023243954.1
E1 56943199 56943429
refGene_gn=GRIN2C;refGene_tx=XM_023243950.1;refGene_gn=GRIN2C;refGene_tx=XM_023243951.1;refGene_gn=GRIN2C;refGene_tx=XM_023243952.1;refGene_gn=GRIN2C;refGene_tx=XM_023243953.1;refGene_gn=GRIN2C;refGene_tx=XM_023243954.1
E1 56943776 56943902
refGene_gn=GRIN2C;refGene_tx=XM_023243950.1;refGene_gn=GRIN2C;refGene_tx=XM_023243951.1;refGene_gn=GRIN2C;refGene_tx=XM_023243952.1;refGene_gn=GRIN2C;refGene_tx=XM_023243953.1;refGene_gn=GRIN2C;refGene_tx=XM_023243954.1
E1 56945245 56945399
refGene_gn=GRIN2C;refGene_tx=XM_023243950.1;refGene_gn=GRIN2C;refGene_tx=XM_023243951.1;refGene_gn=GRIN2C;refGene_tx=XM_023243952.1;refGene_gn=GRIN2C;refGene_tx=XM_023243953.1;refGene_gn=GRIN2C;refGene_tx=XM_023243954.1
E1 56945736 56945902
refGene_gn=GRIN2C;refGene_tx=XM_023243950.1;refGene_gn=GRIN2C;refGene_tx=XM_023243951.1;refGene_gn=GRIN2C;refGene_tx=XM_023243952.1;refGene_gn=GRIN2C;refGene_tx=XM_023243953.1;refGene_gn=GRIN2C;refGene_tx=XM_023243954.1
E1 56946055 56946267
refGene_gn=GRIN2C;refGene_tx=XM_023243950.1;refGene_gn=GRIN2C;refGene_tx=XM_023243951.1;refGene_gn=GRIN2C;refGene_tx=XM_023243952.1;refGene_gn=GRIN2C;refGene_tx=XM_023243953.1;refGene_gn=GRIN2C;refGene_tx=XM_023243954.1
E1 56947202 56947317
refGene_gn=GRIN2C;refGene_tx=XM_023243950.1;refGene_gn=GRIN2C;refGene_tx=XM_023243951.1;refGene_gn=GRIN2C;refGene_tx=XM_023243952.1;refGene_gn=GRIN2C;refGene_tx=XM_023243953.1;refGene_gn=GRIN2C;refGene_tx=XM_023243954.1
E1 56947647 56948246
refGene_gn=GRIN2C;refGene_tx=XM_023243950.1;refGene_gn=GRIN2C;refGene_tx=XM_023243951.1;refGene_gn=GRIN2C;refGene_tx=XM_023243952.1;refGene_gn=GRIN2C;refGene_tx=XM_023243953.1;refGene_gn=GRIN2C;refGene_tx=XM_023243954.1
E1 56950505 56951207
refGene_gn=GRIN2C;refGene_tx=XM_023243950.1;refGene_gn=GRIN2C;refGene_tx=XM_023243951.1;refGene_gn=GRIN2C;refGene_tx=XM_023243952.1;refGene_

gn=GRIN2C;refGene_tx=XM_023243953.1;refGene_gn=GRIN2C;refGene_tx=XM_023243954.1
E1 56955874 56955937 refGene_gn=GRIN2C;refGene_tx=XM_023243951.1
E1 56957070 56957386
refGene_gn=FDXR;refGene_tx=XM_003997131.4;refGene_gn=FDXR;refGene_tx=XM_011289349.2;refGene_gn=FDXR;refGene_tx=XM_011289350.2;refGene_gn=FDXR;refGene_tx=XM_011289352.2;refGene_gn=FDXR;refGene_tx=XM_019818224.1
E1 56957510 56957694
refGene_gn=FDXR;refGene_tx=XM_003997131.4;refGene_gn=FDXR;refGene_tx=XM_011289349.2;refGene_gn=FDXR;refGene_tx=XM_011289350.2;refGene_gn=FDXR;refGene_tx=XM_011289352.2;refGene_gn=FDXR;refGene_tx=XM_019818224.1
E1 56958539 56958711
refGene_gn=FDXR;refGene_tx=XM_003997131.4;refGene_gn=FDXR;refGene_tx=XM_011289349.2;refGene_gn=FDXR;refGene_tx=XM_011289350.2;refGene_gn=FDXR;refGene_tx=XM_011289352.2;refGene_gn=FDXR;refGene_tx=XM_019818224.1
E1 56958793 56958993
refGene_gn=FDXR;refGene_tx=XM_003997131.4;refGene_gn=FDXR;refGene_tx=XM_011289349.2;refGene_gn=FDXR;refGene_tx=XM_011289350.2;refGene_gn=FDXR;refGene_tx=XM_011289352.2;refGene_gn=FDXR;refGene_tx=XM_019818224.1
E1 56959125 56959210
refGene_gn=FDXR;refGene_tx=XM_003997131.4;refGene_gn=FDXR;refGene_tx=XM_011289349.2;refGene_gn=FDXR;refGene_tx=XM_011289350.2;refGene_gn=FDXR;refGene_tx=XM_011289352.2;refGene_gn=FDXR;refGene_tx=XM_019818224.1
E1 56959459 56959567
refGene_gn=FDXR;refGene_tx=XM_003997131.4;refGene_gn=FDXR;refGene_tx=XM_011289349.2;refGene_gn=FDXR;refGene_tx=XM_011289350.2;refGene_gn=FDXR;refGene_tx=XM_011289352.2;refGene_gn=FDXR;refGene_tx=XM_019818224.1
E1 56960408 56960510
refGene_gn=FDXR;refGene_tx=XM_003997131.4;refGene_gn=FDXR;refGene_tx=XM_011289349.2;refGene_gn=FDXR;refGene_tx=XM_011289350.2;refGene_gn=FDXR;refGene_tx=XM_011289352.2
E1 56960864 56960978
refGene_gn=FDXR;refGene_tx=XM_003997131.4;refGene_gn=FDXR;refGene_tx=XM_011289349.2;refGene_gn=FDXR;refGene_tx=XM_011289350.2;refGene_gn=FDXR;refGene_tx=XM_011289352.2;refGene_gn=FDXR;refGene_tx=XM_019818224.1
E1 56961157 56961280
refGene_gn=FDXR;refGene_tx=XM_003997131.4;refGene_gn=FDXR;refGene_tx=XM_011289349.2;refGene_gn=FDXR;refGene_tx=XM_011289350.2;refGene_gn=FDXR;refGene_tx=XM_011289352.2;refGene_gn=FDXR;refGene_tx=XM_019818224.1
E1 56961489 56961585
refGene_gn=FDXR;refGene_tx=XM_003997131.4;refGene_gn=FDXR;refGene_tx=XM_011289349.2;refGene_gn=FDXR;refGene_tx=XM_011289350.2;refGene_gn=FDXR;refGene_tx=XM_011289352.2;refGene_gn=FDXR;refGene_tx=XM_019818224.1
E1 56967845 56967943
refGene_gn=FDXR;refGene_tx=XM_003997131.4;refGene_gn=FDXR;refGene_tx=XM_011289349.2;refGene_gn=FDXR;refGene_tx=XM_011289350.2;refGene_gn=FDXR;refGene_tx=XM_011289352.2;refGene_gn=FDXR;refGene_tx=XM_019818224.1
E1 56968635 56968780
refGene_gn=FDXR;refGene_tx=XM_003997131.4;refGene_gn=FDXR;refGene_tx=XM_011289349.2;refGene_gn=FDXR;refGene_tx=XM_011289350.2;refGene_gn=FDX

R;refGene_tx=XM_011289352.2;refGene_gn=FDXR;refGene_tx=XM_019818224.1
E1 56973517 56973667 refGene_gn=FADS6;refGene_tx=XM_011289353.3
E1 56974278 56974305 refGene_gn=FADS6;refGene_tx=XM_023243956.1
E1 56974411 56974591
refGene_gn=FADS6;refGene_tx=XM_011289353.3;refGene_gn=FADS6;refGene_tx
=XM_023243956.1
E1 56975819 56976007
refGene_gn=FADS6;refGene_tx=XM_011289353.3;refGene_gn=FADS6;refGene_tx
=XM_023243956.1
E1 56976604 56976785
refGene_gn=FADS6;refGene_tx=XM_011289353.3;refGene_gn=FADS6;refGene_tx
=XM_023243956.1
E1 56986308 56986475
refGene_gn=FADS6;refGene_tx=XM_011289353.3;refGene_gn=FADS6;refGene_tx
=XM_023243956.1
E1 56986988 56987151
refGene_gn=FADS6;refGene_tx=XM_011289353.3;refGene_gn=FADS6;refGene_tx
=XM_023243956.1
E1 57002599 57002603 refGene_gn=USH1G;refGene_tx=XM_006940565.4
E1 57003492 57004714
refGene_gn=USH1G;refGene_tx=XM_003997136.5;refGene_gn=USH1G;refGene_tx
=XM_006940565.4
E1 57006943 57007107
refGene_gn=USH1G;refGene_tx=XM_003997136.5;refGene_gn=USH1G;refGene_tx
=XM_006940565.4
E1 57008573 57008886
refGene_gn=OTOP2;refGene_tx=XM_006940566.3;refGene_gn=OTOP2;refGene_tx
=XM_019818225.2;refGene_gn=OTOP2;refGene_tx=XM_023243957.1
E1 57009579 57009716
refGene_gn=OTOP2;refGene_tx=XM_006940566.3;refGene_gn=OTOP2;refGene_tx
=XM_019818225.2;refGene_gn=OTOP2;refGene_tx=XM_023243957.1
E1 57010764 57010823
refGene_gn=OTOP2;refGene_tx=XM_006940566.3;refGene_gn=OTOP2;refGene_tx
=XM_019818225.2;refGene_gn=OTOP2;refGene_tx=XM_023243957.1
E1 57011299 57011433
refGene_gn=OTOP2;refGene_tx=XM_006940566.3;refGene_gn=OTOP2;refGene_tx
=XM_019818225.2;refGene_gn=OTOP2;refGene_tx=XM_023243957.1
E1 57013045 57013923
refGene_gn=OTOP2;refGene_tx=XM_006940566.3;refGene_gn=OTOP2;refGene_tx
=XM_019818225.2;refGene_gn=OTOP2;refGene_tx=XM_023243957.1
E1 57015794 57015965
refGene_gn=OTOP2;refGene_tx=XM_006940566.3;refGene_gn=OTOP2;refGene_tx
=XM_019818225.2;refGene_gn=OTOP2;refGene_tx=XM_023243957.1
E1 57017778 57017797 refGene_gn=OTOP3;refGene_tx=XM_011289436.3
E1 57023927 57024341 refGene_gn=OTOP3;refGene_tx=XM_011289436.3
E1 57024430 57024567 refGene_gn=OTOP3;refGene_tx=XM_011289436.3
E1 57025152 57025211 refGene_gn=OTOP3;refGene_tx=XM_011289436.3
E1 57025486 57025605 refGene_gn=OTOP3;refGene_tx=XM_011289436.3
E1 57027335 57028135 refGene_gn=OTOP3;refGene_tx=XM_011289436.3
E1 57029307 57029478 refGene_gn=OTOP3;refGene_tx=XM_011289436.3

E1 57031682 57031746
refGene_gn=HID1;refGene_tx=XM_023243959.1;refGene_gn=HID1;refGene_tx=XM_023243960.1;refGene_gn=HID1;refGene_tx=XM_023243961.1
E1 57032008 57032167
refGene_gn=HID1;refGene_tx=XM_023243959.1;refGene_gn=HID1;refGene_tx=XM_023243960.1;refGene_gn=HID1;refGene_tx=XM_023243961.1
E1 57032310 57032402
refGene_gn=HID1;refGene_tx=XM_023243959.1;refGene_gn=HID1;refGene_tx=XM_023243960.1;refGene_gn=HID1;refGene_tx=XM_023243961.1
E1 57033002 57033083
refGene_gn=HID1;refGene_tx=XM_023243959.1;refGene_gn=HID1;refGene_tx=XM_023243960.1;refGene_gn=HID1;refGene_tx=XM_023243961.1
E1 57033587 57033694
refGene_gn=HID1;refGene_tx=XM_023243959.1;refGene_gn=HID1;refGene_tx=XM_023243960.1;refGene_gn=HID1;refGene_tx=XM_023243961.1
E1 57034036 57034264
refGene_gn=HID1;refGene_tx=XM_023243959.1;refGene_gn=HID1;refGene_tx=XM_023243960.1;refGene_gn=HID1;refGene_tx=XM_023243961.1
E1 57035431 57035596
refGene_gn=HID1;refGene_tx=XM_023243959.1;refGene_gn=HID1;refGene_tx=XM_023243960.1;refGene_gn=HID1;refGene_tx=XM_023243961.1
E1 57037433 57037512
refGene_gn=HID1;refGene_tx=XM_023243959.1;refGene_gn=HID1;refGene_tx=XM_023243960.1;refGene_gn=HID1;refGene_tx=XM_023243961.1
E1 57037618 57037770
refGene_gn=HID1;refGene_tx=XM_023243959.1;refGene_gn=HID1;refGene_tx=XM_023243960.1;refGene_gn=HID1;refGene_tx=XM_023243961.1
E1 57037994 57038085
refGene_gn=HID1;refGene_tx=XM_023243959.1;refGene_gn=HID1;refGene_tx=XM_023243960.1;refGene_gn=HID1;refGene_tx=XM_023243961.1
E1 57038266 57038407
refGene_gn=HID1;refGene_tx=XM_023243959.1;refGene_gn=HID1;refGene_tx=XM_023243960.1;refGene_gn=HID1;refGene_tx=XM_023243961.1
E1 57039224 57039281
refGene_gn=HID1;refGene_tx=XM_023243959.1;refGene_gn=HID1;refGene_tx=XM_023243960.1;refGene_gn=HID1;refGene_tx=XM_023243961.1
E1 57039450 57039673
refGene_gn=HID1;refGene_tx=XM_023243959.1;refGene_gn=HID1;refGene_tx=XM_023243960.1;refGene_gn=HID1;refGene_tx=XM_023243961.1
E1 57041051 57041168
refGene_gn=HID1;refGene_tx=XM_023243959.1;refGene_gn=HID1;refGene_tx=XM_023243960.1;refGene_gn=HID1;refGene_tx=XM_023243961.1
E1 57041432 57041539
refGene_gn=HID1;refGene_tx=XM_023243959.1;refGene_gn=HID1;refGene_tx=XM_023243960.1;refGene_gn=HID1;refGene_tx=XM_023243961.1
E1 57042133 57042250
refGene_gn=HID1;refGene_tx=XM_023243959.1;refGene_gn=HID1;refGene_tx=XM_023243960.1;refGene_gn=HID1;refGene_tx=XM_023243961.1
E1 57042780 57042951
refGene_gn=HID1;refGene_tx=XM_023243959.1;refGene_gn=HID1;refGene_tx=X

M_023243960.1;refGene_gn=HID1;refGene_tx=XM_023243961.1
E1 57043482 57043632
refGene_gn=HID1;refGene_tx=XM_023243959.1;refGene_gn=HID1;refGene_tx=XM_023243960.1;refGene_gn=HID1;refGene_tx=XM_023243961.1
E1 57048875 57048941
refGene_gn=HID1;refGene_tx=XM_023243959.1;refGene_gn=HID1;refGene_tx=XM_023243960.1;refGene_gn=HID1;refGene_tx=XM_023243961.1
E1 57062598 57062674 refGene_gn=CDR2L;refGene_tx=XM_023243962.1
E1 57071210 57071323 refGene_gn=CDR2L;refGene_tx=XM_023243962.1
E1 57071616 57071765 refGene_gn=CDR2L;refGene_tx=XM_023243962.1
E1 57072340 57072505 refGene_gn=CDR2L;refGene_tx=XM_023243962.1
E1 57073436 57074328 refGene_gn=CDR2L;refGene_tx=XM_023243962.1
E1 57079782 57079968
refGene_gn=MRPL58;refGene_tx=XM_003997137.4;refGene_gn=MRPL58;refGene_tx=XM_019818237.1
E1 57082971 57083008
refGene_gn=MRPL58;refGene_tx=XM_003997137.4;refGene_gn=MRPL58;refGene_tx=XM_019818237.1
E1 57085594 57085654 refGene_gn=MRPL58;refGene_tx=XM_003997137.4
E1 57086208 57086291
refGene_gn=MRPL58;refGene_tx=XM_003997137.4;refGene_gn=MRPL58;refGene_tx=XM_019818237.1
E1 57086384 57086554
refGene_gn=MRPL58;refGene_tx=XM_003997137.4;refGene_gn=MRPL58;refGene_tx=XM_019818237.1
E1 57086812 57086897
refGene_gn=MRPL58;refGene_tx=XM_003997137.4;refGene_gn=MRPL58;refGene_tx=XM_019818237.1
E1 57100313 57100445 refGene_gn=ATP5PD;refGene_tx=XM_003997138.5
E1 57100589 57100652 refGene_gn=ATP5PD;refGene_tx=XM_003997138.5
E1 57101387 57101459 refGene_gn=ATP5PD;refGene_tx=XM_003997138.5
E1 57102657 57102754 refGene_gn=ATP5PD;refGene_tx=XM_003997138.5
E1 57103029 57103151 refGene_gn=ATP5PD;refGene_tx=XM_003997138.5
E1 57105880 57106219 refGene_gn=KCTD2;refGene_tx=XM_003997210.4
E1 57108032 57108141 refGene_gn=KCTD2;refGene_tx=XM_003997210.4
E1 57111355 57111447 refGene_gn=KCTD2;refGene_tx=XM_003997210.4
E1 57116509 57116605 refGene_gn=KCTD2;refGene_tx=XM_003997210.4
E1 57118656 57118782 refGene_gn=KCTD2;refGene_tx=XM_003997210.4
E1 57119530 57119560 refGene_gn=KCTD2;refGene_tx=XM_003997210.4
E1 57125955 57125971
refGene_gn=LOC101093474;refGene_tx=XM_006940568.3;refGene_gn=LOC101093474;refGene_tx=XM_023243967.1;refGene_gn=LOC101093474;refGene_tx=XM_023243968.1;refGene_gn=LOC101093474;refGene_tx=XM_023243971.1;refGene_gn=LOC101093474;refGene_tx=XM_023243972.1
E1 57126186 57126944
refGene_gn=LOC101093474;refGene_tx=XM_006940568.3;refGene_gn=LOC101093474;refGene_tx=XM_023243967.1;refGene_gn=LOC101093474;refGene_tx=XM_023243968.1;refGene_gn=LOC101093474;refGene_tx=XM_023243969.1;refGene_gn=LOC101093474;refGene_tx=XM_023243970.1;refGene_gn=LOC101093474;refGene_tx=XM_023243971.1;refGene_gn=LOC101093474;refGene_tx=XM_023243972.1;

refGene_gn=LOC101093474;refGene_tx=XM_023243973.1;refGene_gn=LOC101093474;refGene_tx=XM_023243974.1
E1 57129346 57129442
refGene_gn=LOC101093474;refGene_tx=XM_006940568.3;refGene_gn=LOC101093474;refGene_tx=XM_023243967.1;refGene_gn=LOC101093474;refGene_tx=XM_023243968.1;refGene_gn=LOC101093474;refGene_tx=XM_023243969.1;refGene_gn=LOC101093474;refGene_tx=XM_023243970.1;refGene_gn=LOC101093474;refGene_tx=XM_023243971.1;refGene_gn=LOC101093474;refGene_tx=XM_023243972.1;refGene_gn=LOC101093474;refGene_tx=XM_023243973.1;refGene_gn=LOC101093474;refGene_tx=XM_023243974.1
E1 57129654 57129888
refGene_gn=LOC101093474;refGene_tx=XM_006940568.3;refGene_gn=LOC101093474;refGene_tx=XM_023243967.1;refGene_gn=LOC101093474;refGene_tx=XM_023243968.1;refGene_gn=LOC101093474;refGene_tx=XM_023243969.1;refGene_gn=LOC101093474;refGene_tx=XM_023243970.1;refGene_gn=LOC101093474;refGene_tx=XM_023243971.1;refGene_gn=LOC101093474;refGene_tx=XM_023243972.1;refGene_gn=LOC101093474;refGene_tx=XM_023243973.1;refGene_gn=LOC101093474;refGene_tx=XM_023243974.1
E1 57130781 57130944
refGene_gn=LOC101093474;refGene_tx=XM_006940568.3;refGene_gn=LOC101093474;refGene_tx=XM_023243967.1;refGene_gn=LOC101093474;refGene_tx=XM_023243968.1;refGene_gn=LOC101093474;refGene_tx=XM_023243969.1;refGene_gn=LOC101093474;refGene_tx=XM_023243970.1;refGene_gn=LOC101093474;refGene_tx=XM_023243971.1;refGene_gn=LOC101093474;refGene_tx=XM_023243972.1;refGene_gn=LOC101093474;refGene_tx=XM_023243973.1;refGene_gn=LOC101093474;refGene_tx=XM_023243974.1
E1 57131074 57131227
refGene_gn=LOC101093474;refGene_tx=XM_006940568.3;refGene_gn=LOC101093474;refGene_tx=XM_023243967.1;refGene_gn=LOC101093474;refGene_tx=XM_023243968.1;refGene_gn=LOC101093474;refGene_tx=XM_023243969.1;refGene_gn=LOC101093474;refGene_tx=XM_023243970.1;refGene_gn=LOC101093474;refGene_tx=XM_023243971.1;refGene_gn=LOC101093474;refGene_tx=XM_023243972.1;refGene_gn=LOC101093474;refGene_tx=XM_023243973.1;refGene_gn=LOC101093474;refGene_tx=XM_023243974.1
E1 57132122 57132685
refGene_gn=LOC101093474;refGene_tx=XM_006940568.3;refGene_gn=LOC101093474;refGene_tx=XM_023243967.1;refGene_gn=LOC101093474;refGene_tx=XM_023243968.1;refGene_gn=LOC101093474;refGene_tx=XM_023243969.1;refGene_gn=LOC101093474;refGene_tx=XM_023243970.1;refGene_gn=LOC101093474;refGene_tx=XM_023243971.1;refGene_gn=LOC101093474;refGene_tx=XM_023243972.1;refGene_gn=LOC101093474;refGene_tx=XM_023243973.1;refGene_gn=LOC101093474;refGene_tx=XM_023243974.1
E1 57149125 57149294
refGene_gn=SLC16A5;refGene_tx=XM_006940574.3;refGene_gn=SLC16A5;refGene_tx=XM_006940575.3
E1 57152893 57153092
refGene_gn=SLC16A5;refGene_tx=XM_003997212.3;refGene_gn=SLC16A5;refGene_tx=XM_006940572.2;refGene_gn=SLC16A5;refGene_tx=XM_019818238.1
E1 57155484 57155628
refGene_gn=SLC16A5;refGene_tx=XM_003997212.3;refGene_gn=SLC16A5;refGen

e_tx=XM_006940572.2;refGene_gn=SLC16A5;refGene_tx=XM_006940574.3;refGene_gn=SLC16A5;refGene_tx=XM_019818238.1
E1 57160419 57161229
refGene_gn=SLC16A5;refGene_tx=XM_003997212.3;refGene_gn=SLC16A5;refGene_tx=XM_006940572.2;refGene_gn=SLC16A5;refGene_tx=XM_006940574.3;refGene_gn=SLC16A5;refGene_tx=XM_006940575.3;refGene_gn=SLC16A5;refGene_tx=XM_019818238.1
E1 57162438 57162686
refGene_gn=SLC16A5;refGene_tx=XM_003997212.3;refGene_gn=SLC16A5;refGene_tx=XM_006940572.2;refGene_gn=SLC16A5;refGene_tx=XM_006940574.3;refGene_gn=SLC16A5;refGene_tx=XM_006940575.3;refGene_gn=SLC16A5;refGene_tx=XM_019818238.1
E1 57165997 57166088
refGene_gn=ARMC7;refGene_tx=XM_003997140.3;refGene_gn=ARMC7;refGene_tx=XM_011289357.2;refGene_gn=ARMC7;refGene_tx=XM_019818243.1;refGene_gn=ARMC7;refGene_tx=XM_019818245.2;refGene_gn=ARMC7;refGene_tx=XM_023243964.1
E1 57166182 57166326
refGene_gn=ARMC7;refGene_tx=XM_003997140.3;refGene_gn=ARMC7;refGene_tx=XM_011289357.2;refGene_gn=ARMC7;refGene_tx=XM_019818243.1;refGene_gn=ARMC7;refGene_tx=XM_019818245.2;refGene_gn=ARMC7;refGene_tx=XM_023243964.1
E1 57170967 57171123 refGene_gn=ARMC7;refGene_tx=XM_023243964.1
E1 57172746 57172873 refGene_gn=ARMC7;refGene_tx=XM_019818244.2
E1 57181417 57181704 refGene_gn=ARMC7;refGene_tx=XM_019818243.1
E1 57183733 57184095
refGene_gn=ARMC7;refGene_tx=XM_003997140.3;refGene_gn=ARMC7;refGene_tx=XM_011289357.2;refGene_gn=ARMC7;refGene_tx=XM_019818244.2
E1 57185587 57185742 refGene_gn=NT5C;refGene_tx=XM_011289437.3
E1 57185859 57185974 refGene_gn=NT5C;refGene_tx=XM_011289437.3
E1 57186319 57186380 refGene_gn=NT5C;refGene_tx=XM_011289437.3
E1 57186463 57186564 refGene_gn=NT5C;refGene_tx=XM_011289437.3
E1 57186975 57187161 refGene_gn=NT5C;refGene_tx=XM_011289437.3
E1 57190209 57190358 refGene_gn=JPT1;refGene_tx=XM_003997141.5
E1 57195773 57195792 refGene_gn=JPT1;refGene_tx=XM_003997141.5
E1 57196702 57196800 refGene_gn=JPT1;refGene_tx=XM_003997141.5
E1 57197129 57197272 refGene_gn=JPT1;refGene_tx=XM_003997141.5
E1 57204873 57204893 refGene_gn=ARMC7;refGene_tx=XM_019818245.2
E1 57206854 57206910 refGene_gn=JPT1;refGene_tx=XM_003997141.5
E1 57216648 57216711
refGene_gn=SUMO2;refGene_tx=XM_003997214.5;refGene_gn=SUMO2;refGene_tx=XM_023243965.1
E1 57219225 57219297
refGene_gn=SUMO2;refGene_tx=XM_003997214.5;refGene_gn=SUMO2;refGene_tx=XM_023243965.1
E1 57224918 57225050
refGene_gn=SUMO2;refGene_tx=XM_003997214.5;refGene_gn=SUMO2;refGene_tx=XM_023243965.1
E1 57225943 57225991 refGene_gn=SUMO2;refGene_tx=XM_003997214.5
E1 57226664 57226685 refGene_gn=SUMO2;refGene_tx=XM_023243965.1

E1 57240084 57240117
refGene_gn=NUP85;refGene_tx=XM_003997142.4;refGene_gn=NUP85;refGene_tx=XM_011289359.2
E1 57244199 57244293
refGene_gn=NUP85;refGene_tx=XM_003997142.4;refGene_gn=NUP85;refGene_tx=XM_011289359.2
E1 57247276 57247439
refGene_gn=NUP85;refGene_tx=XM_003997142.4;refGene_gn=NUP85;refGene_tx=XM_011289359.2
E1 57253304 57253441
refGene_gn=NUP85;refGene_tx=XM_003997142.4;refGene_gn=NUP85;refGene_tx=XM_011289359.2
E1 57253973 57254017
refGene_gn=NUP85;refGene_tx=XM_003997142.4;refGene_gn=NUP85;refGene_tx=XM_011289359.2
E1 57255026 57255096
refGene_gn=NUP85;refGene_tx=XM_003997142.4;refGene_gn=NUP85;refGene_tx=XM_011289359.2
E1 57256271 57256393
refGene_gn=NUP85;refGene_tx=XM_003997142.4;refGene_gn=NUP85;refGene_tx=XM_011289359.2
E1 57261071 57261206
refGene_gn=NUP85;refGene_tx=XM_003997142.4;refGene_gn=NUP85;refGene_tx=XM_011289359.2
E1 57261319 57261442
refGene_gn=NUP85;refGene_tx=XM_003997142.4;refGene_gn=NUP85;refGene_tx=XM_011289359.2
E1 57261642 57261774
refGene_gn=NUP85;refGene_tx=XM_003997142.4;refGene_gn=NUP85;refGene_tx=XM_011289359.2
E1 57262425 57262532
refGene_gn=NUP85;refGene_tx=XM_003997142.4;refGene_gn=NUP85;refGene_tx=XM_011289359.2
E1 57274235 57274319
refGene_gn=NUP85;refGene_tx=XM_003997142.4;refGene_gn=NUP85;refGene_tx=XM_011289359.2
E1 57274459 57274525
refGene_gn=NUP85;refGene_tx=XM_003997142.4;refGene_gn=NUP85;refGene_tx=XM_011289359.2
E1 57274694 57274846
refGene_gn=NUP85;refGene_tx=XM_003997142.4;refGene_gn=NUP85;refGene_tx=XM_011289359.2
E1 57275593 57275711
refGene_gn=NUP85;refGene_tx=XM_003997142.4;refGene_gn=NUP85;refGene_tx=XM_011289359.2
E1 57275820 57275921
refGene_gn=NUP85;refGene_tx=XM_003997142.4;refGene_gn=NUP85;refGene_tx=XM_011289359.2
E1 57276823 57276975
refGene_gn=NUP85;refGene_tx=XM_003997142.4;refGene_gn=NUP85;refGene_tx

=XM_011289359.2
E1 57277380 57277482
refGene_gn=NUP85;refGene_tx=XM_003997142.4;refGene_gn=NUP85;refGene_tx=XM_011289359.2
E1 57277855 57277954
refGene_gn=NUP85;refGene_tx=XM_003997142.4;refGene_gn=NUP85;refGene_tx=XM_011289359.2
E1 57280110 57280221
refGene_gn=GGA3;refGene_tx=XM_003997143.3;refGene_gn=GGA3;refGene_tx=XM_019818239.2;refGene_gn=GGA3;refGene_tx=XM_019818240.1
E1 57280437 57280548
refGene_gn=GGA3;refGene_tx=XM_003997143.3;refGene_gn=GGA3;refGene_tx=XM_019818239.2;refGene_gn=GGA3;refGene_tx=XM_019818240.1
E1 57280721 57280891
refGene_gn=GGA3;refGene_tx=XM_003997143.3;refGene_gn=GGA3;refGene_tx=XM_019818239.2;refGene_gn=GGA3;refGene_tx=XM_019818240.1
E1 57281194 57281403
refGene_gn=GGA3;refGene_tx=XM_003997143.3;refGene_gn=GGA3;refGene_tx=XM_019818239.2;refGene_gn=GGA3;refGene_tx=XM_019818240.1
E1 57281648 57281962
refGene_gn=GGA3;refGene_tx=XM_003997143.3;refGene_gn=GGA3;refGene_tx=XM_019818239.2;refGene_gn=GGA3;refGene_tx=XM_019818240.1
E1 57282265 57282336
refGene_gn=GGA3;refGene_tx=XM_003997143.3;refGene_gn=GGA3;refGene_tx=XM_019818239.2;refGene_gn=GGA3;refGene_tx=XM_019818240.1
E1 57282725 57282968
refGene_gn=GGA3;refGene_tx=XM_003997143.3;refGene_gn=GGA3;refGene_tx=XM_019818239.2
E1 57283298 57283415
refGene_gn=GGA3;refGene_tx=XM_003997143.3;refGene_gn=GGA3;refGene_tx=XM_019818239.2;refGene_gn=GGA3;refGene_tx=XM_019818240.1
E1 57283527 57283609
refGene_gn=GGA3;refGene_tx=XM_003997143.3;refGene_gn=GGA3;refGene_tx=XM_019818239.2;refGene_gn=GGA3;refGene_tx=XM_019818240.1
E1 57284152 57284290
refGene_gn=GGA3;refGene_tx=XM_003997143.3;refGene_gn=GGA3;refGene_tx=XM_019818239.2;refGene_gn=GGA3;refGene_tx=XM_019818240.1
E1 57284679 57284760
refGene_gn=GGA3;refGene_tx=XM_003997143.3;refGene_gn=GGA3;refGene_tx=XM_019818239.2;refGene_gn=GGA3;refGene_tx=XM_019818240.1
E1 57284894 57284998
refGene_gn=GGA3;refGene_tx=XM_003997143.3;refGene_gn=GGA3;refGene_tx=XM_019818239.2;refGene_gn=GGA3;refGene_tx=XM_019818240.1
E1 57285253 57285377
refGene_gn=GGA3;refGene_tx=XM_003997143.3;refGene_gn=GGA3;refGene_tx=XM_019818239.2;refGene_gn=GGA3;refGene_tx=XM_019818240.1
E1 57286073 57286172
refGene_gn=GGA3;refGene_tx=XM_003997143.3;refGene_gn=GGA3;refGene_tx=XM_019818239.2;refGene_gn=GGA3;refGene_tx=XM_019818240.1
E1 57287563 57287639

refGene_gn=GGA3;refGene_tx=XM_003997143.3;refGene_gn=GGA3;refGene_tx=XM_019818240.1
E1 57287765 57287850
refGene_gn=GGA3;refGene_tx=XM_003997143.3;refGene_gn=GGA3;refGene_tx=XM_019818240.1
E1 57294445 57294485
refGene_gn=GGA3;refGene_tx=XM_003997143.3;refGene_gn=GGA3;refGene_tx=XM_019818240.1
E1 57294617 57294664 refGene_gn=MRPS7;refGene_tx=XM_006940577.3
E1 57294798 57294881 refGene_gn=MRPS7;refGene_tx=XM_003997146.5
E1 57295420 57295612
refGene_gn=MRPS7;refGene_tx=XM_003997146.5;refGene_gn=MRPS7;refGene_tx=XM_006940577.3
E1 57295733 57295797
refGene_gn=MRPS7;refGene_tx=XM_003997146.5;refGene_gn=MRPS7;refGene_tx=XM_006940577.3
E1 57296168 57296336
refGene_gn=MRPS7;refGene_tx=XM_003997146.5;refGene_gn=MRPS7;refGene_tx=XM_006940577.3
E1 57297434 57297656
refGene_gn=MRPS7;refGene_tx=XM_003997146.5;refGene_gn=MRPS7;refGene_tx=XM_006940577.3
E1 57298481 57298709
refGene_gn=MIF4GD;refGene_tx=XM_003997148.5;refGene_gn=MIF4GD;refGene_tx=XM_019818242.2
E1 57299249 57299342
refGene_gn=MIF4GD;refGene_tx=XM_003997148.5;refGene_gn=MIF4GD;refGene_tx=XM_019818242.2
E1 57299431 57299587
refGene_gn=MIF4GD;refGene_tx=XM_003997148.5;refGene_gn=MIF4GD;refGene_tx=XM_019818242.2
E1 57299785 57299895
refGene_gn=MIF4GD;refGene_tx=XM_003997148.5;refGene_gn=MIF4GD;refGene_tx=XM_019818242.2
E1 57300763 57300851
refGene_gn=MIF4GD;refGene_tx=XM_003997148.5;refGene_gn=MIF4GD;refGene_tx=XM_019818242.2
E1 57304299 57304482
refGene_gn=SLC25A19;refGene_tx=XM_003997147.5;refGene_gn=SLC25A19;refGene_tx=XM_006940578.4;refGene_gn=SLC25A19;refGene_tx=XM_019818241.2
E1 57306041 57306172
refGene_gn=SLC25A19;refGene_tx=XM_003997147.5;refGene_gn=SLC25A19;refGene_tx=XM_006940578.4;refGene_gn=SLC25A19;refGene_tx=XM_019818241.2
E1 57306751 57306935
refGene_gn=SLC25A19;refGene_tx=XM_003997147.5;refGene_gn=SLC25A19;refGene_tx=XM_006940578.4;refGene_gn=SLC25A19;refGene_tx=XM_019818241.2
E1 57311595 57311766
refGene_gn=SLC25A19;refGene_tx=XM_003997147.5;refGene_gn=SLC25A19;refGene_tx=XM_006940578.4;refGene_gn=SLC25A19;refGene_tx=XM_019818241.2
E1 57313121 57313277

refGene_gn=SLC25A19;refGene_tx=XM_003997147.5;refGene_gn=SLC25A19;refGene_tx=XM_006940578.4;refGene_gn=SLC25A19;refGene_tx=XM_019818241.2
E1 57313445 57313577
refGene_gn=SLC25A19;refGene_tx=XM_003997147.5;refGene_gn=SLC25A19;refGene_tx=XM_006940578.4;refGene_gn=SLC25A19;refGene_tx=XM_019818241.2
E1 57339746 57339932
refGene_gn=GRB2;refGene_tx=XM_023243990.1;refGene_gn=GRB2;refGene_tx=XM_023243991.1
E1 57341117 57341286
refGene_gn=GRB2;refGene_tx=XM_023243990.1;refGene_gn=GRB2;refGene_tx=XM_023243991.1
E1 57344610 57344733
refGene_gn=GRB2;refGene_tx=XM_023243990.1;refGene_gn=GRB2;refGene_tx=XM_023243991.1
E1 57350814 57350912
refGene_gn=GRB2;refGene_tx=XM_023243990.1;refGene_gn=GRB2;refGene_tx=XM_023243991.1
E1 57355426 57355429 refGene_gn=GRB2;refGene_tx=XM_023243991.1
E1 57401585 57401663 refGene_gn=GRB2;refGene_tx=XM_023243990.1
E1 57461735 57461759 refGene_gn=TMEM94;refGene_tx=XM_011289373.3
E1 57470945 57471047 refGene_gn=TMEM94;refGene_tx=XM_011289373.3
E1 57471048 57471066 refGene_gn=TMEM94;refGene_tx=XM_011289373.3
E1 57471398 57471458 refGene_gn=TMEM94;refGene_tx=XM_011289373.3
E1 57471459 57471527 refGene_gn=TMEM94;refGene_tx=XM_011289373.3
E1 57471745 57471882 refGene_gn=TMEM94;refGene_tx=XM_011289373.3
E1 57473548 57473751 refGene_gn=TMEM94;refGene_tx=XM_011289373.3
E1 57474124 57474276 refGene_gn=TMEM94;refGene_tx=XM_011289373.3
E1 57474652 57474755 refGene_gn=TMEM94;refGene_tx=XM_011289373.3
E1 57474977 57475064 refGene_gn=TMEM94;refGene_tx=XM_011289373.3
E1 57475338 57475455 refGene_gn=TMEM94;refGene_tx=XM_011289373.3
E1 57475766 57475823 refGene_gn=TMEM94;refGene_tx=XM_011289373.3
E1 57476078 57476183 refGene_gn=TMEM94;refGene_tx=XM_011289373.3
E1 57476325 57476490 refGene_gn=TMEM94;refGene_tx=XM_011289373.3
E1 57476711 57476921 refGene_gn=TMEM94;refGene_tx=XM_011289373.3
E1 57477373 57477689 refGene_gn=TMEM94;refGene_tx=XM_011289373.3
E1 57478177 57478351 refGene_gn=TMEM94;refGene_tx=XM_011289373.3
E1 57478746 57478849 refGene_gn=TMEM94;refGene_tx=XM_011289373.3
E1 57478973 57479191 refGene_gn=TMEM94;refGene_tx=XM_011289373.3
E1 57479892 57480074 refGene_gn=TMEM94;refGene_tx=XM_011289373.3
E1 57480163 57480302 refGene_gn=TMEM94;refGene_tx=XM_011289373.3
E1 57480561 57480677 refGene_gn=TMEM94;refGene_tx=XM_011289373.3
E1 57480805 57480905 refGene_gn=TMEM94;refGene_tx=XM_011289373.3
E1 57481157 57481266 refGene_gn=TMEM94;refGene_tx=XM_011289373.3
E1 57481461 57481651 refGene_gn=TMEM94;refGene_tx=XM_011289373.3
E1 57481876 57481954 refGene_gn=TMEM94;refGene_tx=XM_011289373.3
E1 57482241 57482327 refGene_gn=TMEM94;refGene_tx=XM_011289373.3
E1 57482588 57482670 refGene_gn=TMEM94;refGene_tx=XM_011289373.3
E1 57482919 57483068 refGene_gn=TMEM94;refGene_tx=XM_011289373.3
E1 57483155 57483250 refGene_gn=TMEM94;refGene_tx=XM_011289373.3
E1 57483358 57483452 refGene_gn=TMEM94;refGene_tx=XM_011289373.3

E1 57483634 57483805 refGene_gn=TMEM94;refGene_tx=XM_011289373.3
E1 57483971 57484044 refGene_gn=TMEM94;refGene_tx=XM_011289373.3
E1 57485745 57485836
refGene_gn=CASKIN2;refGene_tx=XM_006940582.2;refGene_gn=CASKIN2;refGene_tx=XM_006940583.2;refGene_gn=CASKIN2;refGene_tx=XM_006940584.3
E1 57486261 57486484
refGene_gn=CASKIN2;refGene_tx=XM_006940582.2;refGene_gn=CASKIN2;refGene_tx=XM_006940583.2;refGene_gn=CASKIN2;refGene_tx=XM_006940584.3
E1 57486573 57488049
refGene_gn=CASKIN2;refGene_tx=XM_006940582.2;refGene_gn=CASKIN2;refGene_tx=XM_006940583.2;refGene_gn=CASKIN2;refGene_tx=XM_006940584.3
E1 57488206 57488345
refGene_gn=CASKIN2;refGene_tx=XM_006940582.2;refGene_gn=CASKIN2;refGene_tx=XM_006940583.2;refGene_gn=CASKIN2;refGene_tx=XM_006940584.3
E1 57488457 57488559
refGene_gn=CASKIN2;refGene_tx=XM_006940582.2;refGene_gn=CASKIN2;refGene_tx=XM_006940583.2;refGene_gn=CASKIN2;refGene_tx=XM_006940584.3
E1 57488637 57488748
refGene_gn=CASKIN2;refGene_tx=XM_006940582.2;refGene_gn=CASKIN2;refGene_tx=XM_006940583.2;refGene_gn=CASKIN2;refGene_tx=XM_006940584.3
E1 57488949 57489041
refGene_gn=CASKIN2;refGene_tx=XM_006940582.2;refGene_gn=CASKIN2;refGene_tx=XM_006940583.2;refGene_gn=CASKIN2;refGene_tx=XM_006940584.3
E1 57489118 57489179
refGene_gn=CASKIN2;refGene_tx=XM_006940582.2;refGene_gn=CASKIN2;refGene_tx=XM_006940583.2;refGene_gn=CASKIN2;refGene_tx=XM_006940584.3
E1 57489270 57489392
refGene_gn=CASKIN2;refGene_tx=XM_006940582.2;refGene_gn=CASKIN2;refGene_tx=XM_006940583.2;refGene_gn=CASKIN2;refGene_tx=XM_006940584.3
E1 57489511 57489773
refGene_gn=CASKIN2;refGene_tx=XM_006940582.2;refGene_gn=CASKIN2;refGene_tx=XM_006940583.2;refGene_gn=CASKIN2;refGene_tx=XM_006940584.3
E1 57489960 57490055
refGene_gn=CASKIN2;refGene_tx=XM_006940582.2;refGene_gn=CASKIN2;refGene_tx=XM_006940583.2;refGene_gn=CASKIN2;refGene_tx=XM_006940584.3
E1 57490299 57490408
refGene_gn=CASKIN2;refGene_tx=XM_006940582.2;refGene_gn=CASKIN2;refGene_tx=XM_006940583.2;refGene_gn=CASKIN2;refGene_tx=XM_006940584.3
E1 57490701 57490810
refGene_gn=CASKIN2;refGene_tx=XM_006940582.2;refGene_gn=CASKIN2;refGene_tx=XM_006940583.2;refGene_gn=CASKIN2;refGene_tx=XM_006940584.3
E1 57490992 57491123
refGene_gn=CASKIN2;refGene_tx=XM_006940582.2;refGene_gn=CASKIN2;refGene_tx=XM_006940583.2;refGene_gn=CASKIN2;refGene_tx=XM_006940584.3
E1 57491209 57491305
refGene_gn=CASKIN2;refGene_tx=XM_006940582.2;refGene_gn=CASKIN2;refGene_tx=XM_006940583.2;refGene_gn=CASKIN2;refGene_tx=XM_006940584.3
E1 57491404 57491550
refGene_gn=CASKIN2;refGene_tx=XM_006940582.2;refGene_gn=CASKIN2;refGene_tx=XM_006940583.2;refGene_gn=CASKIN2;refGene_tx=XM_006940584.3

E1	57491948 57492046
refGene_gn=CASKIN2;refGene_tx=XM_006940582.2;refGene_gn=CASKIN2;refGen
e_tx=XM_006940583.2;refGene_gn=CASKIN2;refGene_tx=XM_006940584.3
E1	57492577 57492629
refGene_gn=CASKIN2;refGene_tx=XM_006940582.2;refGene_gn=CASKIN2;refGen
e_tx=XM_006940583.2;refGene_gn=CASKIN2;refGene_tx=XM_006940584.3
E1	57492733 57492764 refGene_gn=CASKIN2;refGene_tx=XM_006940584.3
E1	57496759 57496853
refGene_gn=CASKIN2;refGene_tx=XM_006940582.2;refGene_gn=CASKIN2;refGen
e_tx=XM_006940583.2
E1	57499686 57499736
refGene_gn=TSEN54;refGene_tx=XM_003997216.5;refGene_gn=TSEN54;refGene_
tx=XM_023243994.1
E1	57499861 57500026
refGene_gn=TSEN54;refGene_tx=XM_003997216.5;refGene_gn=TSEN54;refGene_
tx=XM_023243994.1
E1	57500127 57500191
refGene_gn=TSEN54;refGene_tx=XM_003997216.5;refGene_gn=TSEN54;refGene_
tx=XM_023243994.1
E1	57500274 57500358
refGene_gn=TSEN54;refGene_tx=XM_003997216.5;refGene_gn=TSEN54;refGene_
tx=XM_023243994.1
E1	57500682 57500781
refGene_gn=TSEN54;refGene_tx=XM_003997216.5;refGene_gn=TSEN54;refGene_
tx=XM_023243994.1
E1	57501793 57501846
refGene_gn=TSEN54;refGene_tx=XM_003997216.5;refGene_gn=TSEN54;refGene_
tx=XM_023243994.1
E1	57503313 57503415
refGene_gn=TSEN54;refGene_tx=XM_003997216.5;refGene_gn=TSEN54;refGene_
tx=XM_023243994.1
E1	57503551 57504282
refGene_gn=TSEN54;refGene_tx=XM_003997216.5;refGene_gn=TSEN54;refGene_
tx=XM_023243994.1
E1	57505349 57505410
refGene_gn=TSEN54;refGene_tx=XM_003997216.5;refGene_gn=TSEN54;refGene_
tx=XM_023243994.1
E1	57505762 57505879
refGene_gn=TSEN54;refGene_tx=XM_003997216.5;refGene_gn=TSEN54;refGene_
tx=XM_023243994.1
E1	57506394 57506545
refGene_gn=TSEN54;refGene_tx=XM_003997216.5;refGene_gn=TSEN54;refGene_
tx=XM_023243994.1
E1	57519671 57519746
refGene_gn=LLGL2;refGene_tx=XM_023243996.1;refGene_gn=LLGL2;refGene_tx
=XM_023243997.1;refGene_gn=LLGL2;refGene_tx=XM_023243998.1;refGene_gn=
LLGL2;refGene_tx=XM_023243999.1;refGene_gn=LLGL2;refGene_tx=XM_0232440
00.1;refGene_gn=LLGL2;refGene_tx=XM_023244001.1
E1	57529001 57529099
refGene_gn=LLGL2;refGene_tx=XM_023243996.1;refGene_gn=LLGL2;refGene_tx

=XM_023243997.1;refGene_gn=LLGL2;refGene_tx=XM_023243998.1;refGene_gn=LLGL2;refGene_tx=XM_023243999.1;refGene_gn=LLGL2;refGene_tx=XM_023244000.1;refGene_gn=LLGL2;refGene_tx=XM_023244001.1
E1 57530338 57530420
refGene_gn=LLGL2;refGene_tx=XM_023243996.1;refGene_gn=LLGL2;refGene_tx=XM_023243997.1;refGene_gn=LLGL2;refGene_tx=XM_023243998.1;refGene_gn=LLGL2;refGene_tx=XM_023243999.1;refGene_gn=LLGL2;refGene_tx=XM_023244000.1;refGene_gn=LLGL2;refGene_tx=XM_023244001.1
E1 57530711 57530827
refGene_gn=LLGL2;refGene_tx=XM_023243996.1;refGene_gn=LLGL2;refGene_tx=XM_023243997.1;refGene_gn=LLGL2;refGene_tx=XM_023243998.1;refGene_gn=LLGL2;refGene_tx=XM_023243999.1;refGene_gn=LLGL2;refGene_tx=XM_023244000.1;refGene_gn=LLGL2;refGene_tx=XM_023244001.1
E1 57531265 57531424
refGene_gn=LLGL2;refGene_tx=XM_023243996.1;refGene_gn=LLGL2;refGene_tx=XM_023243997.1;refGene_gn=LLGL2;refGene_tx=XM_023243998.1;refGene_gn=LLGL2;refGene_tx=XM_023243999.1;refGene_gn=LLGL2;refGene_tx=XM_023244000.1;refGene_gn=LLGL2;refGene_tx=XM_023244001.1
E1 57533168 57533331
refGene_gn=LLGL2;refGene_tx=XM_023243996.1;refGene_gn=LLGL2;refGene_tx=XM_023243997.1;refGene_gn=LLGL2;refGene_tx=XM_023243998.1;refGene_gn=LLGL2;refGene_tx=XM_023243999.1;refGene_gn=LLGL2;refGene_tx=XM_023244000.1;refGene_gn=LLGL2;refGene_tx=XM_023244001.1
E1 57533473 57533606
refGene_gn=LLGL2;refGene_tx=XM_023243996.1;refGene_gn=LLGL2;refGene_tx=XM_023243997.1;refGene_gn=LLGL2;refGene_tx=XM_023243998.1;refGene_gn=LLGL2;refGene_tx=XM_023243999.1;refGene_gn=LLGL2;refGene_tx=XM_023244000.1;refGene_gn=LLGL2;refGene_tx=XM_023244001.1
E1 57533917 57533972
refGene_gn=LLGL2;refGene_tx=XM_023243996.1;refGene_gn=LLGL2;refGene_tx=XM_023243997.1;refGene_gn=LLGL2;refGene_tx=XM_023243998.1;refGene_gn=LLGL2;refGene_tx=XM_023243999.1;refGene_gn=LLGL2;refGene_tx=XM_023244000.1;refGene_gn=LLGL2;refGene_tx=XM_023244001.1
E1 57534569 57534724
refGene_gn=LLGL2;refGene_tx=XM_023243996.1;refGene_gn=LLGL2;refGene_tx=XM_023243997.1;refGene_gn=LLGL2;refGene_tx=XM_023243998.1;refGene_gn=LLGL2;refGene_tx=XM_023243999.1;refGene_gn=LLGL2;refGene_tx=XM_023244000.1;refGene_gn=LLGL2;refGene_tx=XM_023244001.1
E1 57536784 57537028
refGene_gn=LLGL2;refGene_tx=XM_023243996.1;refGene_gn=LLGL2;refGene_tx=XM_023243997.1;refGene_gn=LLGL2;refGene_tx=XM_023243998.1;refGene_gn=LLGL2;refGene_tx=XM_023243999.1;refGene_gn=LLGL2;refGene_tx=XM_023244000.1;refGene_gn=LLGL2;refGene_tx=XM_023244001.1
E1 57537100 57537168
refGene_gn=LLGL2;refGene_tx=XM_023243996.1;refGene_gn=LLGL2;refGene_tx=XM_023243997.1;refGene_gn=LLGL2;refGene_tx=XM_023243998.1;refGene_gn=LLGL2;refGene_tx=XM_023243999.1;refGene_gn=LLGL2;refGene_tx=XM_023244000.1
E1 57537314 57537468
refGene_gn=LLGL2;refGene_tx=XM_023243996.1;refGene_gn=LLGL2;refGene_tx

=XM_023243997.1;refGene_gn=LLGL2;refGene_tx=XM_023243998.1;refGene_gn=
LLGL2;refGene_tx=XM_023243999.1;refGene_gn=LLGL2;refGene_tx=XM_0232440
00.1
E1 57537557 57537662
refGene_gn=LLGL2;refGene_tx=XM_023243996.1;refGene_gn=LLGL2;refGene_tx
=XM_023243997.1;refGene_gn=LLGL2;refGene_tx=XM_023243998.1;refGene_gn=
LLGL2;refGene_tx=XM_023243999.1;refGene_gn=LLGL2;refGene_tx=XM_0232440
00.1
E1 57538080 57538373
refGene_gn=LLGL2;refGene_tx=XM_023243996.1;refGene_gn=LLGL2;refGene_tx
=XM_023243997.1;refGene_gn=LLGL2;refGene_tx=XM_023243998.1;refGene_gn=
LLGL2;refGene_tx=XM_023243999.1;refGene_gn=LLGL2;refGene_tx=XM_0232440
00.1
E1 57538473 57538624
refGene_gn=LLGL2;refGene_tx=XM_023243996.1;refGene_gn=LLGL2;refGene_tx
=XM_023243997.1;refGene_gn=LLGL2;refGene_tx=XM_023243998.1;refGene_gn=
LLGL2;refGene_tx=XM_023243999.1;refGene_gn=LLGL2;refGene_tx=XM_0232440
00.1
E1 57538938 57539089
refGene_gn=LLGL2;refGene_tx=XM_023243996.1;refGene_gn=LLGL2;refGene_tx
=XM_023243997.1;refGene_gn=LLGL2;refGene_tx=XM_023243998.1;refGene_gn=
LLGL2;refGene_tx=XM_023243999.1;refGene_gn=LLGL2;refGene_tx=XM_0232440
00.1
E1 57539721 57539838
refGene_gn=LLGL2;refGene_tx=XM_023243996.1;refGene_gn=LLGL2;refGene_tx
=XM_023243997.1;refGene_gn=LLGL2;refGene_tx=XM_023243998.1;refGene_gn=
LLGL2;refGene_tx=XM_023243999.1;refGene_gn=LLGL2;refGene_tx=XM_0232440
00.1
E1 57539961 57540128
refGene_gn=LLGL2;refGene_tx=XM_023243996.1;refGene_gn=LLGL2;refGene_tx
=XM_023243997.1;refGene_gn=LLGL2;refGene_tx=XM_023243998.1;refGene_gn=
LLGL2;refGene_tx=XM_023243999.1;refGene_gn=LLGL2;refGene_tx=XM_0232440
00.1
E1 57540524 57540789
refGene_gn=LLGL2;refGene_tx=XM_023243996.1;refGene_gn=LLGL2;refGene_tx
=XM_023243997.1;refGene_gn=LLGL2;refGene_tx=XM_023243998.1;refGene_gn=
LLGL2;refGene_tx=XM_023243999.1;refGene_gn=LLGL2;refGene_tx=XM_0232440
00.1
E1 57541057 57541211
refGene_gn=LLGL2;refGene_tx=XM_023243996.1;refGene_gn=LLGL2;refGene_tx
=XM_023243997.1;refGene_gn=LLGL2;refGene_tx=XM_023243998.1;refGene_gn=
LLGL2;refGene_tx=XM_023243999.1;refGene_gn=LLGL2;refGene_tx=XM_0232440
00.1
E1 57541467 57541496
refGene_gn=LLGL2;refGene_tx=XM_023243996.1;refGene_gn=LLGL2;refGene_tx
=XM_023243997.1;refGene_gn=LLGL2;refGene_tx=XM_023243998.1;refGene_gn=
LLGL2;refGene_tx=XM_023243999.1;refGene_gn=LLGL2;refGene_tx=XM_0232440
00.1
E1 57541740 57541788
refGene_gn=LLGL2;refGene_tx=XM_023243996.1;refGene_gn=LLGL2;refGene_tx

=XM_023243997.1;refGene_gn=LLGL2;refGene_tx=XM_023243998.1;refGene_gn=LLGL2;refGene_tx=XM_023243999.1
E1 57541981 57542024
refGene_gn=LLGL2;refGene_tx=XM_023243996.1;refGene_gn=LLGL2;refGene_tx=XM_023243998.1;refGene_gn=LLGL2;refGene_tx=XM_023244000.1
E1 57542150 57542212
refGene_gn=LLGL2;refGene_tx=XM_023243996.1;refGene_gn=LLGL2;refGene_tx=XM_023243997.1;refGene_gn=LLGL2;refGene_tx=XM_023243998.1;refGene_gn=LLGL2;refGene_tx=XM_023243999.1;refGene_gn=LLGL2;refGene_tx=XM_023244000.1
E1 57542381 57542432
refGene_gn=LLGL2;refGene_tx=XM_023243996.1;refGene_gn=LLGL2;refGene_tx=XM_023243997.1;refGene_gn=LLGL2;refGene_tx=XM_023243998.1;refGene_gn=LLGL2;refGene_tx=XM_023243999.1;refGene_gn=LLGL2;refGene_tx=XM_023244000.1
E1 57546107 57548164 refGene_gn=MYO15B;refGene_tx=XM_023243474.1
E1 57548568 57548632 refGene_gn=MYO15B;refGene_tx=XM_023243474.1
E1 57548882 57548992 refGene_gn=MYO15B;refGene_tx=XM_023243474.1
E1 57549162 57549237 refGene_gn=MYO15B;refGene_tx=XM_023243474.1
E1 57549570 57549667 refGene_gn=MYO15B;refGene_tx=XM_023243474.1
E1 57549929 57550033 refGene_gn=MYO15B;refGene_tx=XM_023243474.1
E1 57550211 57550275 refGene_gn=MYO15B;refGene_tx=XM_023243474.1
E1 57550467 57550581 refGene_gn=MYO15B;refGene_tx=XM_023243474.1
E1 57550726 57550888 refGene_gn=MYO15B;refGene_tx=XM_023243474.1
E1 57551708 57551822 refGene_gn=MYO15B;refGene_tx=XM_023243474.1
E1 57551940 57552207 refGene_gn=MYO15B;refGene_tx=XM_023243474.1
E1 57553175 57553271 refGene_gn=MYO15B;refGene_tx=XM_023243474.1
E1 57553523 57553655 refGene_gn=MYO15B;refGene_tx=XM_023243474.1
E1 57554075 57554201 refGene_gn=MYO15B;refGene_tx=XM_023243474.1
E1 57556887 57556965 refGene_gn=MYO15B;refGene_tx=XM_023243474.1
E1 57557262 57557378 refGene_gn=MYO15B;refGene_tx=XM_023243474.1
E1 57557467 57557513 refGene_gn=MYO15B;refGene_tx=XM_023243474.1
E1 57557633 57557758 refGene_gn=MYO15B;refGene_tx=XM_023243474.1
E1 57558167 57558285 refGene_gn=MYO15B;refGene_tx=XM_023243474.1
E1 57558491 57558661 refGene_gn=MYO15B;refGene_tx=XM_023243474.1
E1 57562106 57562200 refGene_gn=MYO15B;refGene_tx=XM_023243474.1
E1 57562754 57562811 refGene_gn=MYO15B;refGene_tx=XM_023243474.1
E1 57563115 57563173 refGene_gn=MYO15B;refGene_tx=XM_023243474.1
E1 57563390 57563521 refGene_gn=MYO15B;refGene_tx=XM_023243474.1
E1 57564236 57564332 refGene_gn=MYO15B;refGene_tx=XM_023243474.1
E1 57564445 57564681 refGene_gn=MYO15B;refGene_tx=XM_023243474.1
E1 57564764 57564943 refGene_gn=MYO15B;refGene_tx=XM_023243474.1
E1 57565095 57565168 refGene_gn=MYO15B;refGene_tx=XM_023243474.1
E1 57565517 57565679 refGene_gn=MYO15B;refGene_tx=XM_023243474.1
E1 57565926 57566027 refGene_gn=MYO15B;refGene_tx=XM_023243474.1
E1 57566128 57566205 refGene_gn=MYO15B;refGene_tx=XM_023243474.1
E1 57566322 57566407 refGene_gn=MYO15B;refGene_tx=XM_023243474.1
E1 57566570 57566669 refGene_gn=MYO15B;refGene_tx=XM_023243474.1
E1 57566793 57566903 refGene_gn=MYO15B;refGene_tx=XM_023243474.1
E1 57567002 57567182 refGene_gn=MYO15B;refGene_tx=XM_023243474.1

E1 57567408 57567487 refGene_gn=MYO15B;refGene_tx=XM_023243474.1
E1 57567648 57567753 refGene_gn=MYO15B;refGene_tx=XM_023243474.1
E1 57567835 57568094 refGene_gn=MYO15B;refGene_tx=XM_023243474.1
E1 57568172 57568263 refGene_gn=MYO15B;refGene_tx=XM_023243474.1
E1 57568379 57568434 refGene_gn=MYO15B;refGene_tx=XM_023243474.1
E1 57568436 57568516 refGene_gn=MYO15B;refGene_tx=XM_023243474.1
E1 57568517 57568581 refGene_gn=MYO15B;refGene_tx=XM_023243474.1
E1 57569035 57569148 refGene_gn=MYO15B;refGene_tx=XM_023243474.1
E1 57569356 57569416 refGene_gn=MYO15B;refGene_tx=XM_023243474.1
E1 57569983 57570059 refGene_gn=MYO15B;refGene_tx=XM_023243474.1
E1 57570174 57570293 refGene_gn=MYO15B;refGene_tx=XM_023243474.1
E1 57570476 57570595 refGene_gn=MYO15B;refGene_tx=XM_023243474.1
E1 57570675 57570817 refGene_gn=MYO15B;refGene_tx=XM_023243474.1
E1 57571023 57571135 refGene_gn=MYO15B;refGene_tx=XM_023243474.1
E1 57571241 57571408 refGene_gn=MYO15B;refGene_tx=XM_023243474.1
E1 57571805 57571957 refGene_gn=MYO15B;refGene_tx=XM_023243474.1
E1 57572125 57572186 refGene_gn=MYO15B;refGene_tx=XM_023243474.1
E1 57572272 57572345 refGene_gn=MYO15B;refGene_tx=XM_023243474.1
E1 57572758 57572835 refGene_gn=MYO15B;refGene_tx=XM_023243474.1
E1 57574077 57574151 refGene_gn=MYO15B;refGene_tx=XM_023243474.1
E1 57574244 57574360 refGene_gn=MYO15B;refGene_tx=XM_023243474.1
E1 57574452 57574547 refGene_gn=MYO15B;refGene_tx=XM_023243474.1
E1 57574661 57574739 refGene_gn=MYO15B;refGene_tx=XM_023243474.1
E1 57574852 57574949 refGene_gn=MYO15B;refGene_tx=XM_023243474.1
E1 57575130 57575273 refGene_gn=MYO15B;refGene_tx=XM_023243474.1
E1 57575470 57575586 refGene_gn=MYO15B;refGene_tx=XM_023243474.1
E1 57575921 57576055 refGene_gn=MYO15B;refGene_tx=XM_023243474.1
E1 57576231 57576365 refGene_gn=MYO15B;refGene_tx=XM_023243474.1
E1 57576479 57576620 refGene_gn=MYO15B;refGene_tx=XM_023243474.1
E1 57576739 57576799 refGene_gn=MYO15B;refGene_tx=XM_023243474.1
E1 57577820 57577921 refGene_gn=RECQL5;refGene_tx=XM_003997154.4
E1 57578013 57578083 refGene_gn=RECQL5;refGene_tx=XM_003997154.4
E1 57578306 57578531 refGene_gn=RECQL5;refGene_tx=XM_003997154.4
E1 57578756 57578847 refGene_gn=RECQL5;refGene_tx=XM_003997154.4
E1 57578999 57579535 refGene_gn=RECQL5;refGene_tx=XM_003997154.4
E1 57579853 57579988 refGene_gn=RECQL5;refGene_tx=XM_003997154.4
E1 57580273 57580367 refGene_gn=RECQL5;refGene_tx=XM_003997154.4
E1 57580735 57580809 refGene_gn=RECQL5;refGene_tx=XM_003997154.4
E1 57580889 57580948 refGene_gn=RECQL5;refGene_tx=XM_003997154.4
E1 57581070 57581107 refGene_gn=RECQL5;refGene_tx=XM_003997154.4
E1 57581243 57581343 refGene_gn=RECQL5;refGene_tx=XM_003997154.4
E1 57581596 57581815 refGene_gn=RECQL5;refGene_tx=XM_003997154.4
E1 57590172 57590299 refGene_gn=SMIM5;refGene_tx=XM_006940590.4
E1 57590760 57590870 refGene_gn=SMIM5;refGene_tx=XM_006940590.4
E1 57596004 57596187
refGene_gn=SMIM6;refGene_tx=XM_006940589.3;refGene_gn=SMIM6;refGene_tx=XM_019818254.1;refGene_gn=SMIM6;refGene_tx=XM_019818255.1;refGene_gn=SMIM6;refGene_tx=XM_019818256.2;refGene_gn=SMIM6;refGene_tx=XM_019818257.1
E1 57598551 57598631 refGene_gn=RECQL5;refGene_tx=XM_003997154.4

E1	57606478	57606641	refGene_gn=RECQL5;refGene_tx=XM_003997154.4
E1	57607839	57607951	refGene_gn=RECQL5;refGene_tx=XM_003997154.4
E1	57608651	57608757	refGene_gn=RECQL5;refGene_tx=XM_003997154.4
E1	57609466	57609985	refGene_gn=RECQL5;refGene_tx=XM_003997154.4
E1	57610933	57611055	refGene_gn=RECQL5;refGene_tx=XM_003997154.4
E1	57612097	57612218	refGene_gn=RECQL5;refGene_tx=XM_003997154.4
E1	57613056	57613162
refGene_gn=SAP30BP;refGene_tx=XM_003997156.3;refGene_gn=SAP30BP;refGene_tx=XM_019818252.1;refGene_gn=SAP30BP;refGene_tx=XM_019818253.1
E1	57615033	57615143
refGene_gn=SAP30BP;refGene_tx=XM_003997156.3;refGene_gn=SAP30BP;refGene_tx=XM_019818252.1;refGene_gn=SAP30BP;refGene_tx=XM_019818253.1
E1	57617385	57617433
refGene_gn=SAP30BP;refGene_tx=XM_003997156.3;refGene_gn=SAP30BP;refGene_tx=XM_019818252.1
E1	57635050	57635173	refGene_gn=SAP30BP;refGene_tx=XM_019818252.1
E1	57635779	57635822
refGene_gn=SAP30BP;refGene_tx=XM_003997156.3;refGene_gn=SAP30BP;refGene_tx=XM_019818252.1;refGene_gn=SAP30BP;refGene_tx=XM_019818253.1
E1	57641891	57641980
refGene_gn=SAP30BP;refGene_tx=XM_003997156.3;refGene_gn=SAP30BP;refGene_tx=XM_019818252.1;refGene_gn=SAP30BP;refGene_tx=XM_019818253.1
E1	57644227	57644319
refGene_gn=SAP30BP;refGene_tx=XM_003997156.3;refGene_gn=SAP30BP;refGene_tx=XM_019818252.1;refGene_gn=SAP30BP;refGene_tx=XM_019818253.1
E1	57645113	57645174
refGene_gn=SAP30BP;refGene_tx=XM_003997156.3;refGene_gn=SAP30BP;refGene_tx=XM_019818252.1;refGene_gn=SAP30BP;refGene_tx=XM_019818253.1
E1	57645648	57645700
refGene_gn=SAP30BP;refGene_tx=XM_003997156.3;refGene_gn=SAP30BP;refGene_tx=XM_019818252.1;refGene_gn=SAP30BP;refGene_tx=XM_019818253.1
E1	57646569	57646628
refGene_gn=SAP30BP;refGene_tx=XM_003997156.3;refGene_gn=SAP30BP;refGene_tx=XM_019818252.1;refGene_gn=SAP30BP;refGene_tx=XM_019818253.1
E1	57647620	57647705
refGene_gn=SAP30BP;refGene_tx=XM_003997156.3;refGene_gn=SAP30BP;refGene_tx=XM_019818252.1;refGene_gn=SAP30BP;refGene_tx=XM_019818253.1
E1	57647956	57648138
refGene_gn=SAP30BP;refGene_tx=XM_003997156.3;refGene_gn=SAP30BP;refGene_tx=XM_019818252.1;refGene_gn=SAP30BP;refGene_tx=XM_019818253.1
E1	57662234	57662313
refGene_gn=ITGB4;refGene_tx=XM_019818258.1;refGene_gn=ITGB4;refGene_tx=XM_019818259.1;refGene_gn=ITGB4;refGene_tx=XM_023244004.1;refGene_gn=ITGB4;refGene_tx=XM_023244005.1
E1	57663187	57663270
refGene_gn=ITGB4;refGene_tx=XM_019818258.1;refGene_gn=ITGB4;refGene_tx=XM_019818259.1;refGene_gn=ITGB4;refGene_tx=XM_023244004.1;refGene_gn=ITGB4;refGene_tx=XM_023244005.1
E1	57663514	57663616
refGene_gn=ITGB4;refGene_tx=XM_019818258.1;refGene_gn=ITGB4;refGene_tx

=XM_019818259.1;refGene_gn=ITGB4;refGene_tx=XM_023244004.1;refGene_gn=ITGB4;refGene_tx=XM_023244005.1
E1 57663769 57663974
refGene_gn=ITGB4;refGene_tx=XM_019818258.1;refGene_gn=ITGB4;refGene_tx=XM_019818259.1;refGene_gn=ITGB4;refGene_tx=XM_023244004.1;refGene_gn=ITGB4;refGene_tx=XM_023244005.1
E1 57664548 57664645
refGene_gn=ITGB4;refGene_tx=XM_019818258.1;refGene_gn=ITGB4;refGene_tx=XM_019818259.1;refGene_gn=ITGB4;refGene_tx=XM_023244004.1;refGene_gn=ITGB4;refGene_tx=XM_023244005.1
E1 57664823 57664995
refGene_gn=ITGB4;refGene_tx=XM_019818258.1;refGene_gn=ITGB4;refGene_tx=XM_019818259.1;refGene_gn=ITGB4;refGene_tx=XM_023244004.1;refGene_gn=ITGB4;refGene_tx=XM_023244005.1
E1 57665996 57666260
refGene_gn=ITGB4;refGene_tx=XM_019818258.1;refGene_gn=ITGB4;refGene_tx=XM_019818259.1;refGene_gn=ITGB4;refGene_tx=XM_023244004.1;refGene_gn=ITGB4;refGene_tx=XM_023244005.1
E1 57666599 57666689
refGene_gn=ITGB4;refGene_tx=XM_019818258.1;refGene_gn=ITGB4;refGene_tx=XM_019818259.1;refGene_gn=ITGB4;refGene_tx=XM_023244004.1;refGene_gn=ITGB4;refGene_tx=XM_023244005.1
E1 57667002 57667125
refGene_gn=ITGB4;refGene_tx=XM_019818258.1;refGene_gn=ITGB4;refGene_tx=XM_019818259.1;refGene_gn=ITGB4;refGene_tx=XM_023244004.1;refGene_gn=ITGB4;refGene_tx=XM_023244005.1
E1 57667700 57667865
refGene_gn=ITGB4;refGene_tx=XM_019818258.1;refGene_gn=ITGB4;refGene_tx=XM_019818259.1;refGene_gn=ITGB4;refGene_tx=XM_023244004.1;refGene_gn=ITGB4;refGene_tx=XM_023244005.1
E1 57668040 57668117
refGene_gn=ITGB4;refGene_tx=XM_019818258.1;refGene_gn=ITGB4;refGene_tx=XM_019818259.1;refGene_gn=ITGB4;refGene_tx=XM_023244004.1;refGene_gn=ITGB4;refGene_tx=XM_023244005.1
E1 57669207 57669410
refGene_gn=ITGB4;refGene_tx=XM_019818258.1;refGene_gn=ITGB4;refGene_tx=XM_019818259.1;refGene_gn=ITGB4;refGene_tx=XM_023244004.1;refGene_gn=ITGB4;refGene_tx=XM_023244005.1
E1 57671437 57671541
refGene_gn=ITGB4;refGene_tx=XM_019818258.1;refGene_gn=ITGB4;refGene_tx=XM_019818259.1;refGene_gn=ITGB4;refGene_tx=XM_023244004.1;refGene_gn=ITGB4;refGene_tx=XM_023244005.1
E1 57671661 57671760
refGene_gn=ITGB4;refGene_tx=XM_019818258.1;refGene_gn=ITGB4;refGene_tx=XM_019818259.1;refGene_gn=ITGB4;refGene_tx=XM_023244004.1;refGene_gn=ITGB4;refGene_tx=XM_023244005.1
E1 57671862 57671992
refGene_gn=ITGB4;refGene_tx=XM_019818258.1;refGene_gn=ITGB4;refGene_tx=XM_019818259.1;refGene_gn=ITGB4;refGene_tx=XM_023244004.1;refGene_gn=ITGB4;refGene_tx=XM_023244005.1

E1 57672596 57672719
refGene_gn=ITGB4;refGene_tx=XM_019818258.1;refGene_gn=ITGB4;refGene_tx=XM_019818259.1;refGene_gn=ITGB4;refGene_tx=XM_023244004.1;refGene_gn=ITGB4;refGene_tx=XM_023244005.1
E1 57672813 57672920
refGene_gn=ITGB4;refGene_tx=XM_019818258.1;refGene_gn=ITGB4;refGene_tx=XM_019818259.1;refGene_gn=ITGB4;refGene_tx=XM_023244004.1;refGene_gn=ITGB4;refGene_tx=XM_023244005.1
E1 57674062 57674096
refGene_gn=ITGB4;refGene_tx=XM_019818258.1;refGene_gn=ITGB4;refGene_tx=XM_019818259.1;refGene_gn=ITGB4;refGene_tx=XM_023244004.1;refGene_gn=ITGB4;refGene_tx=XM_023244005.1
E1 57674273 57674465
refGene_gn=ITGB4;refGene_tx=XM_019818258.1;refGene_gn=ITGB4;refGene_tx=XM_019818259.1;refGene_gn=ITGB4;refGene_tx=XM_023244004.1;refGene_gn=ITGB4;refGene_tx=XM_023244005.1
E1 57674749 57674853
refGene_gn=ITGB4;refGene_tx=XM_019818258.1;refGene_gn=ITGB4;refGene_tx=XM_019818259.1;refGene_gn=ITGB4;refGene_tx=XM_023244004.1;refGene_gn=ITGB4;refGene_tx=XM_023244005.1
E1 57675155 57675214
refGene_gn=ITGB4;refGene_tx=XM_019818258.1;refGene_gn=ITGB4;refGene_tx=XM_019818259.1;refGene_gn=ITGB4;refGene_tx=XM_023244004.1;refGene_gn=ITGB4;refGene_tx=XM_023244005.1
E1 57675350 57675374
refGene_gn=ITGB4;refGene_tx=XM_019818258.1;refGene_gn=ITGB4;refGene_tx=XM_019818259.1;refGene_gn=ITGB4;refGene_tx=XM_023244004.1;refGene_gn=ITGB4;refGene_tx=XM_023244005.1
E1 57676042 57676191
refGene_gn=ITGB4;refGene_tx=XM_019818258.1;refGene_gn=ITGB4;refGene_tx=XM_019818259.1;refGene_gn=ITGB4;refGene_tx=XM_023244004.1;refGene_gn=ITGB4;refGene_tx=XM_023244005.1
E1 57676286 57676466
refGene_gn=ITGB4;refGene_tx=XM_019818258.1;refGene_gn=ITGB4;refGene_tx=XM_019818259.1;refGene_gn=ITGB4;refGene_tx=XM_023244004.1;refGene_gn=ITGB4;refGene_tx=XM_023244005.1
E1 57677260 57677409
refGene_gn=ITGB4;refGene_tx=XM_019818258.1;refGene_gn=ITGB4;refGene_tx=XM_019818259.1;refGene_gn=ITGB4;refGene_tx=XM_023244004.1;refGene_gn=ITGB4;refGene_tx=XM_023244005.1
E1 57680139 57680344
refGene_gn=ITGB4;refGene_tx=XM_019818258.1;refGene_gn=ITGB4;refGene_tx=XM_019818259.1;refGene_gn=ITGB4;refGene_tx=XM_023244004.1;refGene_gn=ITGB4;refGene_tx=XM_023244005.1
E1 57681176 57681334
refGene_gn=ITGB4;refGene_tx=XM_019818258.1;refGene_gn=ITGB4;refGene_tx=XM_019818259.1;refGene_gn=ITGB4;refGene_tx=XM_023244004.1;refGene_gn=ITGB4;refGene_tx=XM_023244005.1
E1 57681718 57681899
refGene_gn=ITGB4;refGene_tx=XM_019818258.1;refGene_gn=ITGB4;refGene_tx

=XM_019818259.1;refGene_gn=ITGB4;refGene_tx=XM_023244004.1;refGene_gn=ITGB4;refGene_tx=XM_023244005.1
E1 57682040 57682178
refGene_gn=ITGB4;refGene_tx=XM_019818258.1;refGene_gn=ITGB4;refGene_tx=XM_019818259.1;refGene_gn=ITGB4;refGene_tx=XM_023244004.1;refGene_gn=ITGB4;refGene_tx=XM_023244005.1
E1 57684297 57684480
refGene_gn=ITGB4;refGene_tx=XM_019818258.1;refGene_gn=ITGB4;refGene_tx=XM_019818259.1;refGene_gn=ITGB4;refGene_tx=XM_023244004.1;refGene_gn=ITGB4;refGene_tx=XM_023244005.1
E1 57684572 57684704
refGene_gn=ITGB4;refGene_tx=XM_019818258.1;refGene_gn=ITGB4;refGene_tx=XM_019818259.1;refGene_gn=ITGB4;refGene_tx=XM_023244004.1;refGene_gn=ITGB4;refGene_tx=XM_023244005.1
E1 57685282 57685324 refGene_gn=GALK1;refGene_tx=XM_023244007.1
E1 57685923 57686142
refGene_gn=ITGB4;refGene_tx=XM_023244004.1;refGene_gn=ITGB4;refGene_tx=XM_023244005.1
E1 57686765 57687011
refGene_gn=ITGB4;refGene_tx=XM_019818258.1;refGene_gn=ITGB4;refGene_tx=XM_019818259.1;refGene_gn=ITGB4;refGene_tx=XM_023244004.1;refGene_gn=ITGB4;refGene_tx=XM_023244005.1
E1 57687150 57687309
refGene_gn=ITGB4;refGene_tx=XM_019818258.1;refGene_gn=ITGB4;refGene_tx=XM_023244004.1
E1 57687733 57687883
refGene_gn=ITGB4;refGene_tx=XM_019818258.1;refGene_gn=ITGB4;refGene_tx=XM_019818259.1;refGene_gn=ITGB4;refGene_tx=XM_023244004.1;refGene_gn=ITGB4;refGene_tx=XM_023244005.1
E1 57688436 57688625
refGene_gn=ITGB4;refGene_tx=XM_019818258.1;refGene_gn=ITGB4;refGene_tx=XM_019818259.1;refGene_gn=ITGB4;refGene_tx=XM_023244004.1;refGene_gn=ITGB4;refGene_tx=XM_023244005.1
E1 57688709 57688865
refGene_gn=ITGB4;refGene_tx=XM_019818258.1;refGene_gn=ITGB4;refGene_tx=XM_019818259.1;refGene_gn=ITGB4;refGene_tx=XM_023244004.1;refGene_gn=ITGB4;refGene_tx=XM_023244005.1
E1 57688948 57689113
refGene_gn=ITGB4;refGene_tx=XM_019818258.1;refGene_gn=ITGB4;refGene_tx=XM_019818259.1;refGene_gn=ITGB4;refGene_tx=XM_023244004.1;refGene_gn=ITGB4;refGene_tx=XM_023244005.1
E1 57689207 57689318
refGene_gn=ITGB4;refGene_tx=XM_019818258.1;refGene_gn=ITGB4;refGene_tx=XM_019818259.1;refGene_gn=ITGB4;refGene_tx=XM_023244004.1;refGene_gn=ITGB4;refGene_tx=XM_023244005.1
E1 57689425 57689565
refGene_gn=ITGB4;refGene_tx=XM_019818258.1;refGene_gn=ITGB4;refGene_tx=XM_019818259.1;refGene_gn=ITGB4;refGene_tx=XM_023244004.1;refGene_gn=ITGB4;refGene_tx=XM_023244005.1
E1 57690130 57690202

refGene_gn=GALK1;refGene_tx=XM_011289439.3;refGene_gn=GALK1;refGene_tx=XM_023244008.1
E1 57690294 57690457 refGene_gn=GALK1;refGene_tx=XM_011289439.3;refGene_gn=GALK1;refGene_tx=XM_023244007.1;refGene_gn=GALK1;refGene_tx=XM_023244008.1
E1 57690556 57690707 refGene_gn=GALK1;refGene_tx=XM_011289439.3;refGene_gn=GALK1;refGene_tx=XM_023244007.1;refGene_gn=GALK1;refGene_tx=XM_023244008.1
E1 57693156 57693338 refGene_gn=GALK1;refGene_tx=XM_011289439.3;refGene_gn=GALK1;refGene_tx=XM_023244007.1;refGene_gn=GALK1;refGene_tx=XM_023244008.1
E1 57693462 57693598 refGene_gn=GALK1;refGene_tx=XM_011289439.3;refGene_gn=GALK1;refGene_tx=XM_023244007.1;refGene_gn=GALK1;refGene_tx=XM_023244008.1
E1 57693785 57693905 refGene_gn=GALK1;refGene_tx=XM_011289439.3;refGene_gn=GALK1;refGene_tx=XM_023244007.1;refGene_gn=GALK1;refGene_tx=XM_023244008.1
E1 57694312 57694502 refGene_gn=GALK1;refGene_tx=XM_011289439.3;refGene_gn=GALK1;refGene_tx=XM_023244007.1;refGene_gn=GALK1;refGene_tx=XM_023244008.1
E1 57695410 57695575 refGene_gn=GALK1;refGene_tx=XM_011289439.3;refGene_gn=GALK1;refGene_tx=XM_023244007.1;refGene_gn=GALK1;refGene_tx=XM_023244008.1
E1 57707183 57707312 refGene_gn=LOC101080845;refGene_tx=XM_003997160.4
E1 57707406 57707560 refGene_gn=LOC101080845;refGene_tx=XM_003997160.4
E1 57707663 57707791 refGene_gn=LOC101080845;refGene_tx=XM_003997160.4
E1 57712432 57712536 refGene_gn=UNK;refGene_tx=XM_003997161.5;refGene_gn=UNK;refGene_tx=XM_006940594.3
E1 57716983 57716992 refGene_gn=UNK;refGene_tx=XM_006940593.3;refGene_gn=UNK;refGene_tx=XM_019818262.2;refGene_gn=UNK;refGene_tx=XM_019818263.2
E1 57717100 57717177 refGene_gn=UNK;refGene_tx=XM_006940593.3;refGene_gn=UNK;refGene_tx=XM_019818262.2
E1 57730346 57730390 refGene_gn=UNC13D;refGene_tx=XM_019818260.2
E1 57731395 57731605 refGene_gn=UNK;refGene_tx=XM_003997161.5;refGene_gn=UNK;refGene_tx=XM_006940593.3;refGene_gn=UNK;refGene_tx=XM_006940594.3;refGene_gn=UNK;refGene_tx=XM_019818262.2;refGene_gn=UNK;refGene_tx=XM_019818263.2
E1 57732438 57732615 refGene_gn=UNK;refGene_tx=XM_003997161.5;refGene_gn=UNK;refGene_tx=XM_006940593.3;refGene_gn=UNK;refGene_tx=XM_006940594.3;refGene_gn=UNK;refGene_tx=XM_019818262.2;refGene_gn=UNK;refGene_tx=XM_019818263.2
E1 57732753 57732884 refGene_gn=UNK;refGene_tx=XM_003997161.5;refGene_gn=UNK;refGene_tx=XM_006940593.3;refGene_gn=UNK;refGene_tx=XM_006940594.3;refGene_gn=UNK;re

fGene_tx=XM_019818262.2;refGene_gn=UNK;refGene_tx=XM_019818263.2
E1 57733342 57733478
refGene_gn=UNK;refGene_tx=XM_003997161.5;refGene_gn=UNK;refGene_tx=XM_006940593.3;refGene_gn=UNK;refGene_tx=XM_006940594.3;refGene_gn=UNK;refGene_tx=XM_019818262.2;refGene_gn=UNK;refGene_tx=XM_019818263.2
E1 57734068 57734186
refGene_gn=UNK;refGene_tx=XM_003997161.5;refGene_gn=UNK;refGene_tx=XM_006940593.3;refGene_gn=UNK;refGene_tx=XM_006940594.3;refGene_gn=UNK;refGene_tx=XM_019818262.2;refGene_gn=UNK;refGene_tx=XM_019818263.2
E1 57735150 57735235
refGene_gn=UNK;refGene_tx=XM_003997161.5;refGene_gn=UNK;refGene_tx=XM_006940593.3;refGene_gn=UNK;refGene_tx=XM_006940594.3;refGene_gn=UNK;refGene_tx=XM_019818262.2;refGene_gn=UNK;refGene_tx=XM_019818263.2
E1 57736484 57736627
refGene_gn=UNK;refGene_tx=XM_003997161.5;refGene_gn=UNK;refGene_tx=XM_006940593.3;refGene_gn=UNK;refGene_tx=XM_006940594.3;refGene_gn=UNK;refGene_tx=XM_019818262.2;refGene_gn=UNK;refGene_tx=XM_019818263.2
E1 57737052 57737253
refGene_gn=UNK;refGene_tx=XM_003997161.5;refGene_gn=UNK;refGene_tx=XM_006940593.3;refGene_gn=UNK;refGene_tx=XM_019818262.2;refGene_gn=UNK;refGene_tx=XM_019818263.2
E1 57737835 57737901
refGene_gn=UNK;refGene_tx=XM_003997161.5;refGene_gn=UNK;refGene_tx=XM_006940593.3;refGene_gn=UNK;refGene_tx=XM_006940594.3;refGene_gn=UNK;refGene_tx=XM_019818262.2;refGene_gn=UNK;refGene_tx=XM_019818263.2
E1 57738385 57738560
refGene_gn=UNK;refGene_tx=XM_003997161.5;refGene_gn=UNK;refGene_tx=XM_006940593.3;refGene_gn=UNK;refGene_tx=XM_006940594.3;refGene_gn=UNK;refGene_tx=XM_019818262.2;refGene_gn=UNK;refGene_tx=XM_019818263.2
E1 57739397 57739499
refGene_gn=UNK;refGene_tx=XM_003997161.5;refGene_gn=UNK;refGene_tx=XM_006940593.3;refGene_gn=UNK;refGene_tx=XM_006940594.3;refGene_gn=UNK;refGene_tx=XM_019818262.2;refGene_gn=UNK;refGene_tx=XM_019818263.2
E1 57739645 57739834
refGene_gn=UNK;refGene_tx=XM_003997161.5;refGene_gn=UNK;refGene_tx=XM_006940593.3;refGene_gn=UNK;refGene_tx=XM_006940594.3;refGene_gn=UNK;refGene_tx=XM_019818262.2;refGene_gn=UNK;refGene_tx=XM_019818263.2
E1 57741945 57742127
refGene_gn=UNK;refGene_tx=XM_003997161.5;refGene_gn=UNK;refGene_tx=XM_006940593.3;refGene_gn=UNK;refGene_tx=XM_006940594.3;refGene_gn=UNK;refGene_tx=XM_019818262.2;refGene_gn=UNK;refGene_tx=XM_019818263.2
E1 57742742 57743000
refGene_gn=UNK;refGene_tx=XM_003997161.5;refGene_gn=UNK;refGene_tx=XM_006940593.3;refGene_gn=UNK;refGene_tx=XM_006940594.3;refGene_gn=UNK;refGene_tx=XM_019818262.2;refGene_gn=UNK;refGene_tx=XM_019818263.2
E1 57743807 57743963
refGene_gn=UNK;refGene_tx=XM_003997161.5;refGene_gn=UNK;refGene_tx=XM_006940593.3;refGene_gn=UNK;refGene_tx=XM_006940594.3;refGene_gn=UNK;refGene_tx=XM_019818262.2;refGene_gn=UNK;refGene_tx=XM_019818263.2
E1 57746874 57746996

refGene_gn=UNC13D;refGene_tx=XM_003997162.4;refGene_gn=UNC13D;refGene_tx=XM_006940595.3;refGene_gn=UNC13D;refGene_tx=XM_019818261.1
E1 57747764 57747961
refGene_gn=UNC13D;refGene_tx=XM_003997162.4;refGene_gn=UNC13D;refGene_tx=XM_006940595.3;refGene_gn=UNC13D;refGene_tx=XM_019818261.1
E1 57748397 57748521
refGene_gn=UNC13D;refGene_tx=XM_003997162.4;refGene_gn=UNC13D;refGene_tx=XM_006940595.3;refGene_gn=UNC13D;refGene_tx=XM_019818261.1
E1 57748747 57748868
refGene_gn=UNC13D;refGene_tx=XM_003997162.4;refGene_gn=UNC13D;refGene_tx=XM_006940595.3;refGene_gn=UNC13D;refGene_tx=XM_019818260.2;refGene_gn=UNC13D;refGene_tx=XM_019818261.1
E1 57748964 57749048
refGene_gn=UNC13D;refGene_tx=XM_003997162.4;refGene_gn=UNC13D;refGene_tx=XM_006940595.3;refGene_gn=UNC13D;refGene_tx=XM_019818260.2;refGene_gn=UNC13D;refGene_tx=XM_019818261.1
E1 57749455 57749527
refGene_gn=UNC13D;refGene_tx=XM_003997162.4;refGene_gn=UNC13D;refGene_tx=XM_006940595.3;refGene_gn=UNC13D;refGene_tx=XM_019818260.2;refGene_gn=UNC13D;refGene_tx=XM_019818261.1
E1 57749601 57749707
refGene_gn=UNC13D;refGene_tx=XM_003997162.4;refGene_gn=UNC13D;refGene_tx=XM_006940595.3;refGene_gn=UNC13D;refGene_tx=XM_019818260.2;refGene_gn=UNC13D;refGene_tx=XM_019818261.1
E1 57751277 57751357
refGene_gn=UNC13D;refGene_tx=XM_003997162.4;refGene_gn=UNC13D;refGene_tx=XM_006940595.3;refGene_gn=UNC13D;refGene_tx=XM_019818260.2;refGene_gn=UNC13D;refGene_tx=XM_019818261.1
E1 57752228 57752297
refGene_gn=UNC13D;refGene_tx=XM_003997162.4;refGene_gn=UNC13D;refGene_tx=XM_006940595.3;refGene_gn=UNC13D;refGene_tx=XM_019818260.2;refGene_gn=UNC13D;refGene_tx=XM_019818261.1
E1 57752459 57752666
refGene_gn=UNC13D;refGene_tx=XM_003997162.4;refGene_gn=UNC13D;refGene_tx=XM_006940595.3;refGene_gn=UNC13D;refGene_tx=XM_019818260.2;refGene_gn=UNC13D;refGene_tx=XM_019818261.1
E1 57752769 57752868
refGene_gn=UNC13D;refGene_tx=XM_003997162.4;refGene_gn=UNC13D;refGene_tx=XM_006940595.3;refGene_gn=UNC13D;refGene_tx=XM_019818260.2;refGene_gn=UNC13D;refGene_tx=XM_019818261.1
E1 57753067 57753211
refGene_gn=UNC13D;refGene_tx=XM_003997162.4;refGene_gn=UNC13D;refGene_tx=XM_006940595.3;refGene_gn=UNC13D;refGene_tx=XM_019818260.2;refGene_gn=UNC13D;refGene_tx=XM_019818261.1
E1 57753557 57753678
refGene_gn=UNC13D;refGene_tx=XM_003997162.4;refGene_gn=UNC13D;refGene_tx=XM_006940595.3;refGene_gn=UNC13D;refGene_tx=XM_019818260.2;refGene_gn=UNC13D;refGene_tx=XM_019818261.1
E1 57753756 57753890
refGene_gn=UNC13D;refGene_tx=XM_003997162.4;refGene_gn=UNC13D;refGene_

tx=XM_006940595.3;refGene_gn=UNC13D;refGene_tx=XM_019818260.2;refGene_gn=UNC13D;refGene_tx=XM_019818261.1
E1 57753968 57754020
refGene_gn=UNC13D;refGene_tx=XM_003997162.4;refGene_gn=UNC13D;refGene_tx=XM_006940595.3;refGene_gn=UNC13D;refGene_tx=XM_019818260.2;refGene_gn=UNC13D;refGene_tx=XM_019818261.1
E1 57754125 57754223
refGene_gn=UNC13D;refGene_tx=XM_003997162.4;refGene_gn=UNC13D;refGene_tx=XM_006940595.3;refGene_gn=UNC13D;refGene_tx=XM_019818260.2;refGene_gn=UNC13D;refGene_tx=XM_019818261.1
E1 57754302 57754359
refGene_gn=UNC13D;refGene_tx=XM_003997162.4;refGene_gn=UNC13D;refGene_tx=XM_006940595.3;refGene_gn=UNC13D;refGene_tx=XM_019818260.2;refGene_gn=UNC13D;refGene_tx=XM_019818261.1
E1 57754441 57754532
refGene_gn=UNC13D;refGene_tx=XM_003997162.4;refGene_gn=UNC13D;refGene_tx=XM_006940595.3;refGene_gn=UNC13D;refGene_tx=XM_019818260.2;refGene_gn=UNC13D;refGene_tx=XM_019818261.1
E1 57754668 57754793
refGene_gn=UNC13D;refGene_tx=XM_003997162.4;refGene_gn=UNC13D;refGene_tx=XM_006940595.3;refGene_gn=UNC13D;refGene_tx=XM_019818260.2;refGene_gn=UNC13D;refGene_tx=XM_019818261.1
E1 57754897 57755015
refGene_gn=UNC13D;refGene_tx=XM_003997162.4;refGene_gn=UNC13D;refGene_tx=XM_006940595.3;refGene_gn=UNC13D;refGene_tx=XM_019818260.2;refGene_gn=UNC13D;refGene_tx=XM_019818261.1
E1 57756594 57756698
refGene_gn=UNC13D;refGene_tx=XM_003997162.4;refGene_gn=UNC13D;refGene_tx=XM_006940595.3;refGene_gn=UNC13D;refGene_tx=XM_019818260.2;refGene_gn=UNC13D;refGene_tx=XM_019818261.1
E1 57756772 57756865
refGene_gn=UNC13D;refGene_tx=XM_003997162.4;refGene_gn=UNC13D;refGene_tx=XM_006940595.3;refGene_gn=UNC13D;refGene_tx=XM_019818260.2;refGene_gn=UNC13D;refGene_tx=XM_019818261.1
E1 57756975 57757080
refGene_gn=UNC13D;refGene_tx=XM_003997162.4;refGene_gn=UNC13D;refGene_tx=XM_006940595.3;refGene_gn=UNC13D;refGene_tx=XM_019818260.2;refGene_gn=UNC13D;refGene_tx=XM_019818261.1
E1 57757269 57757339
refGene_gn=UNC13D;refGene_tx=XM_003997162.4;refGene_gn=UNC13D;refGene_tx=XM_006940595.3;refGene_gn=UNC13D;refGene_tx=XM_019818260.2;refGene_gn=UNC13D;refGene_tx=XM_019818261.1
E1 57757514 57757583
refGene_gn=UNC13D;refGene_tx=XM_003997162.4;refGene_gn=UNC13D;refGene_tx=XM_006940595.3;refGene_gn=UNC13D;refGene_tx=XM_019818260.2;refGene_gn=UNC13D;refGene_tx=XM_019818261.1
E1 57757720 57757765
refGene_gn=UNC13D;refGene_tx=XM_003997162.4;refGene_gn=UNC13D;refGene_tx=XM_006940595.3;refGene_gn=UNC13D;refGene_tx=XM_019818260.2;refGene_gn=UNC13D;refGene_tx=XM_019818261.1

E1 57758112 57758293
refGene_gn=UNC13D;refGene_tx=XM_003997162.4;refGene_gn=UNC13D;refGene_tx=XM_006940595.3;refGene_gn=UNC13D;refGene_tx=XM_019818260.2
E1 57758538 57758605
refGene_gn=UNC13D;refGene_tx=XM_003997162.4;refGene_gn=UNC13D;refGene_tx=XM_006940595.3;refGene_gn=UNC13D;refGene_tx=XM_019818260.2
E1 57758696 57758756
refGene_gn=UNC13D;refGene_tx=XM_003997162.4;refGene_gn=UNC13D;refGene_tx=XM_006940595.3;refGene_gn=UNC13D;refGene_tx=XM_019818260.2
E1 57758873 57758981
refGene_gn=UNC13D;refGene_tx=XM_003997162.4;refGene_gn=UNC13D;refGene_tx=XM_006940595.3;refGene_gn=UNC13D;refGene_tx=XM_019818260.2
E1 57759256 57759292
refGene_gn=UNC13D;refGene_tx=XM_003997162.4;refGene_gn=UNC13D;refGene_tx=XM_006940595.3;refGene_gn=UNC13D;refGene_tx=XM_019818260.2
E1 57759427 57759508 refGene_gn=UNC13D;refGene_tx=XM_006940595.3
E1 57759994 57760111
refGene_gn=UNC13D;refGene_tx=XM_003997162.4;refGene_gn=UNC13D;refGene_tx=XM_019818260.2
E1 57762410 57762464
refGene_gn=WBP2;refGene_tx=XM_003997163.5;refGene_gn=WBP2;refGene_tx=XM_006940596.4;refGene_gn=WBP2;refGene_tx=XM_019818264.1
E1 57762570 57762647
refGene_gn=WBP2;refGene_tx=XM_003997163.5;refGene_gn=WBP2;refGene_tx=XM_006940596.4;refGene_gn=WBP2;refGene_tx=XM_019818264.1
E1 57763111 57763234
refGene_gn=WBP2;refGene_tx=XM_003997163.5;refGene_gn=WBP2;refGene_tx=XM_006940596.4;refGene_gn=WBP2;refGene_tx=XM_019818264.1
E1 57763426 57763561
refGene_gn=WBP2;refGene_tx=XM_003997163.5;refGene_gn=WBP2;refGene_tx=XM_006940596.4
E1 57764210 57764303
refGene_gn=WBP2;refGene_tx=XM_003997163.5;refGene_gn=WBP2;refGene_tx=XM_006940596.4;refGene_gn=WBP2;refGene_tx=XM_019818264.1
E1 57765173 57765309
refGene_gn=WBP2;refGene_tx=XM_003997163.5;refGene_gn=WBP2;refGene_tx=XM_006940596.4;refGene_gn=WBP2;refGene_tx=XM_019818264.1
E1 57766440 57766549
refGene_gn=WBP2;refGene_tx=XM_003997163.5;refGene_gn=WBP2;refGene_tx=XM_006940596.4;refGene_gn=WBP2;refGene_tx=XM_019818264.1
E1 57768640 57768699
refGene_gn=WBP2;refGene_tx=XM_003997163.5;refGene_gn=WBP2;refGene_tx=XM_019818264.1
E1 57782722 57783363 refGene_gn=TRIM47;refGene_tx=XM_023244012.1
E1 57783617 57783692 refGene_gn=TRIM47;refGene_tx=XM_023244012.1
E1 57784170 57784339 refGene_gn=TRIM47;refGene_tx=XM_023244012.1
E1 57784520 57784751 refGene_gn=TRIM47;refGene_tx=XM_023244012.1
E1 57785036 57785132 refGene_gn=TRIM47;refGene_tx=XM_023244012.1
E1 57786208 57786895 refGene_gn=TRIM47;refGene_tx=XM_023244012.1
E1 57794183 57794758

refGene_gn=TRIM65;refGene_tx=XM_003997220.5;refGene_gn=TRIM65;refGene_tx=XM_006940597.4;refGene_gn=TRIM65;refGene_tx=XM_006940598.4;refGene_gn=TRIM65;refGene_tx=XM_006940599.4;refGene_gn=TRIM65;refGene_tx=XM_006940600.4
E1 57795292 57795665
refGene_gn=TRIM65;refGene_tx=XM_003997220.5;refGene_gn=TRIM65;refGene_tx=XM_006940597.4;refGene_gn=TRIM65;refGene_tx=XM_006940598.4;refGene_gn=TRIM65;refGene_tx=XM_006940599.4;refGene_gn=TRIM65;refGene_tx=XM_006940600.4
E1 57795785 57796019
refGene_gn=TRIM65;refGene_tx=XM_003997220.5;refGene_gn=TRIM65;refGene_tx=XM_006940597.4;refGene_gn=TRIM65;refGene_tx=XM_006940598.4;refGene_gn=TRIM65;refGene_tx=XM_006940599.4;refGene_gn=TRIM65;refGene_tx=XM_006940600.4
E1 57796271 57796367
refGene_gn=TRIM65;refGene_tx=XM_003997220.5;refGene_gn=TRIM65;refGene_tx=XM_006940597.4;refGene_gn=TRIM65;refGene_tx=XM_006940598.4;refGene_gn=TRIM65;refGene_tx=XM_006940599.4;refGene_gn=TRIM65;refGene_tx=XM_006940600.4
E1 57797279 57797303 refGene_gn=TRIM65;refGene_tx=XM_006940600.4
E1 57797622 57798066
refGene_gn=TRIM65;refGene_tx=XM_003997220.5;refGene_gn=TRIM65;refGene_tx=XM_006940597.4;refGene_gn=TRIM65;refGene_tx=XM_006940598.4;refGene_gn=TRIM65;refGene_tx=XM_006940599.4
E1 57799806 57799947
refGene_gn=MRPL38;refGene_tx=XM_003997165.3;refGene_gn=MRPL38;refGene_tx=XM_006940601.2
E1 57800161 57800298
refGene_gn=MRPL38;refGene_tx=XM_003997165.3;refGene_gn=MRPL38;refGene_tx=XM_006940601.2
E1 57800537 57800696
refGene_gn=MRPL38;refGene_tx=XM_003997165.3;refGene_gn=MRPL38;refGene_tx=XM_006940601.2
E1 57801675 57801721
refGene_gn=MRPL38;refGene_tx=XM_003997165.3;refGene_gn=MRPL38;refGene_tx=XM_006940601.2
E1 57801855 57801928
refGene_gn=MRPL38;refGene_tx=XM_003997165.3;refGene_gn=MRPL38;refGene_tx=XM_006940601.2
E1 57802463 57802672
refGene_gn=MRPL38;refGene_tx=XM_003997165.3;refGene_gn=MRPL38;refGene_tx=XM_006940601.2
E1 57802757 57802892
refGene_gn=MRPL38;refGene_tx=XM_003997165.3;refGene_gn=MRPL38;refGene_tx=XM_006940601.2
E1 57804264 57804444
refGene_gn=MRPL38;refGene_tx=XM_003997165.3;refGene_gn=MRPL38;refGene_tx=XM_006940601.2
E1 57804550 57804617
refGene_gn=MRPL38;refGene_tx=XM_003997165.3;refGene_gn=MRPL38;refGene_

tx=XM_006940601.2
E1	57807446	57807530	refGene_gn=FBF1;refGene_tx=XM_019818272.2
E1	57808122	57808232	refGene_gn=FBF1;refGene_tx=XM_019818272.2
E1	57809132	57809250	refGene_gn=FBF1;refGene_tx=XM_019818272.2
E1	57809340	57809478	refGene_gn=FBF1;refGene_tx=XM_019818272.2
E1	57809570	57809747	refGene_gn=FBF1;refGene_tx=XM_019818272.2
E1	57810391	57810577	refGene_gn=FBF1;refGene_tx=XM_019818272.2
E1	57810662	57810785	refGene_gn=FBF1;refGene_tx=XM_019818272.2
E1	57811793	57811833	refGene_gn=FBF1;refGene_tx=XM_019818272.2
E1	57811834	57811880	refGene_gn=FBF1;refGene_tx=XM_019818272.2
E1	57811967	57812107	refGene_gn=FBF1;refGene_tx=XM_019818272.2
E1	57812186	57812294	refGene_gn=FBF1;refGene_tx=XM_019818272.2
E1	57813064	57813271	refGene_gn=FBF1;refGene_tx=XM_019818272.2
E1	57813396	57813497	refGene_gn=FBF1;refGene_tx=XM_019818272.2
E1	57813620	57813776	refGene_gn=FBF1;refGene_tx=XM_019818272.2
E1	57814586	57814645	refGene_gn=FBF1;refGene_tx=XM_019818272.2
E1	57814808	57814897	refGene_gn=FBF1;refGene_tx=XM_019818272.2
E1	57815255	57815354	refGene_gn=FBF1;refGene_tx=XM_019818272.2
E1	57816665	57817124	refGene_gn=FBF1;refGene_tx=XM_019818272.2
E1	57818177	57818277	refGene_gn=FBF1;refGene_tx=XM_019818272.2
E1	57819344	57819478	refGene_gn=FBF1;refGene_tx=XM_019818272.2
E1	57819614	57819753	refGene_gn=FBF1;refGene_tx=XM_019818272.2
E1	57820095	57820215	refGene_gn=FBF1;refGene_tx=XM_019818272.2
E1	57820776	57820854	refGene_gn=FBF1;refGene_tx=XM_019818272.2
E1	57821332	57821474	refGene_gn=FBF1;refGene_tx=XM_019818272.2
E1	57822999	57823050	refGene_gn=FBF1;refGene_tx=XM_019818272.2
E1	57823753	57823814	refGene_gn=FBF1;refGene_tx=XM_019818272.2
E1	57824583	57824677	refGene_gn=FBF1;refGene_tx=XM_019818272.2
E1	57825951	57825993	refGene_gn=FBF1;refGene_tx=XM_019818272.2
E1	57827221	57827249	refGene_gn=FBF1;refGene_tx=XM_019818272.2
E1	57827793	57827796	refGene_gn=FBF1;refGene_tx=XM_019818272.2
E1	57833384	57833435
refGene_gn=ACOX1;refGene_tx=XM_003997166.5;refGene_gn=ACOX1;refGene_tx
=XM_003997167.5;refGene_gn=ACOX1;refGene_tx=XM_019818277.1
E1	57834291	57834498
refGene_gn=ACOX1;refGene_tx=XM_003997166.5;refGene_gn=ACOX1;refGene_tx
=XM_003997167.5;refGene_gn=ACOX1;refGene_tx=XM_019818277.1
E1	57834643	57834787
refGene_gn=ACOX1;refGene_tx=XM_003997166.5;refGene_gn=ACOX1;refGene_tx
=XM_003997167.5;refGene_gn=ACOX1;refGene_tx=XM_019818277.1
E1	57834960	57835066
refGene_gn=ACOX1;refGene_tx=XM_003997166.5;refGene_gn=ACOX1;refGene_tx
=XM_003997167.5;refGene_gn=ACOX1;refGene_tx=XM_019818277.1
E1	57835202	57835382
refGene_gn=ACOX1;refGene_tx=XM_003997166.5;refGene_gn=ACOX1;refGene_tx
=XM_003997167.5;refGene_gn=ACOX1;refGene_tx=XM_019818277.1
E1	57835642	57835833
refGene_gn=ACOX1;refGene_tx=XM_003997166.5;refGene_gn=ACOX1;refGene_tx
=XM_003997167.5;refGene_gn=ACOX1;refGene_tx=XM_019818277.1
E1	57836272	57836435

refGene_gn=ACOX1;refGene_tx=XM_003997166.5;refGene_gn=ACOX1;refGene_tx=XM_003997167.5;refGene_gn=ACOX1;refGene_tx=XM_019818277.1
E1 57837548 57837718
refGene_gn=ACOX1;refGene_tx=XM_003997166.5;refGene_gn=ACOX1;refGene_tx=XM_003997167.5;refGene_gn=ACOX1;refGene_tx=XM_019818277.1
E1 57838108 57838224
refGene_gn=ACOX1;refGene_tx=XM_003997166.5;refGene_gn=ACOX1;refGene_tx=XM_003997167.5;refGene_gn=ACOX1;refGene_tx=XM_019818277.1
E1 57838370 57838490
refGene_gn=ACOX1;refGene_tx=XM_003997166.5;refGene_gn=ACOX1;refGene_tx=XM_003997167.5;refGene_gn=ACOX1;refGene_tx=XM_019818277.1
E1 57839523 57839631
refGene_gn=ACOX1;refGene_tx=XM_003997166.5;refGene_gn=ACOX1;refGene_tx=XM_003997167.5;refGene_gn=ACOX1;refGene_tx=XM_019818277.1
E1 57841799 57841960
refGene_gn=ACOX1;refGene_tx=XM_003997166.5;refGene_gn=ACOX1;refGene_tx=XM_019818277.1
E1 57848963 57849124
refGene_gn=ACOX1;refGene_tx=XM_003997167.5;refGene_gn=ACOX1;refGene_tx=XM_019818277.1
E1 57853316 57853476
refGene_gn=ACOX1;refGene_tx=XM_003997166.5;refGene_gn=ACOX1;refGene_tx=XM_003997167.5
E1 57853768 57853877
refGene_gn=ACOX1;refGene_tx=XM_003997166.5;refGene_gn=ACOX1;refGene_tx=XM_003997167.5
E1 57854432 57854437 refGene_gn=CDK3;refGene_tx=XM_019818281.2
E1 57863453 57863542
refGene_gn=CDK3;refGene_tx=XM_019818278.2;refGene_gn=CDK3;refGene_tx=XM_019818279.2
E1 57867893 57868048
refGene_gn=CDK3;refGene_tx=XM_019818278.2;refGene_gn=CDK3;refGene_tx=XM_019818279.2;refGene_gn=CDK3;refGene_tx=XM_019818281.2;refGene_gn=CDK3;refGene_tx=XM_019818282.2
E1 57877187 57877278
refGene_gn=CDK3;refGene_tx=XM_003997169.5;refGene_gn=CDK3;refGene_tx=XM_019818278.2;refGene_gn=CDK3;refGene_tx=XM_019818279.2;refGene_gn=CDK3;refGene_tx=XM_019818281.2;refGene_gn=CDK3;refGene_tx=XM_019818282.2
E1 57878234 57878360
refGene_gn=CDK3;refGene_tx=XM_003997169.5;refGene_gn=CDK3;refGene_tx=XM_019818278.2;refGene_gn=CDK3;refGene_tx=XM_019818279.2;refGene_gn=CDK3;refGene_tx=XM_019818281.2;refGene_gn=CDK3;refGene_tx=XM_019818282.2
E1 57878677 57878755
refGene_gn=CDK3;refGene_tx=XM_003997169.5;refGene_gn=CDK3;refGene_tx=XM_019818278.2;refGene_gn=CDK3;refGene_tx=XM_019818279.2;refGene_gn=CDK3;refGene_tx=XM_019818281.2;refGene_gn=CDK3;refGene_tx=XM_019818282.2
E1 57878837 57878958
refGene_gn=CDK3;refGene_tx=XM_003997169.5;refGene_gn=CDK3;refGene_tx=XM_019818278.2;refGene_gn=CDK3;refGene_tx=XM_019818279.2;refGene_gn=CDK3;refGene_tx=XM_019818281.2;refGene_gn=CDK3;refGene_tx=XM_019818282.2

E1 57879050 57879221
refGene_gn=CDK3;refGene_tx=XM_003997169.5;refGene_gn=CDK3;refGene_tx=XM_019818278.2;refGene_gn=CDK3;refGene_tx=XM_019818279.2;refGene_gn=CDK3;refGene_tx=XM_019818281.2;refGene_gn=CDK3;refGene_tx=XM_019818282.2
E1 57879311 57879413
refGene_gn=CDK3;refGene_tx=XM_003997169.5;refGene_gn=CDK3;refGene_tx=XM_019818278.2;refGene_gn=CDK3;refGene_tx=XM_019818279.2;refGene_gn=CDK3;refGene_tx=XM_019818281.2;refGene_gn=CDK3;refGene_tx=XM_019818282.2
E1 57879689 57879893
refGene_gn=CDK3;refGene_tx=XM_003997169.5;refGene_gn=CDK3;refGene_tx=XM_019818278.2;refGene_gn=CDK3;refGene_tx=XM_019818279.2;refGene_gn=CDK3;refGene_tx=XM_019818281.2;refGene_gn=CDK3;refGene_tx=XM_019818282.2
E1 57880941 57881067
refGene_gn=CDK3;refGene_tx=XM_003997169.5;refGene_gn=CDK3;refGene_tx=XM_019818278.2;refGene_gn=CDK3;refGene_tx=XM_019818279.2;refGene_gn=CDK3;refGene_tx=XM_019818281.2;refGene_gn=CDK3;refGene_tx=XM_019818282.2
E1 57882247 57882318 refGene_gn=EVPL;refGene_tx=XM_023244014.1
E1 57882320 57882489 refGene_gn=EVPL;refGene_tx=XM_023244014.1
E1 57882490 57885694 refGene_gn=EVPL;refGene_tx=XM_023244014.1
E1 57886382 57886475 refGene_gn=EVPL;refGene_tx=XM_023244014.1
E1 57886582 57886693 refGene_gn=EVPL;refGene_tx=XM_023244014.1
E1 57886796 57886880 refGene_gn=EVPL;refGene_tx=XM_023244014.1
E1 57888176 57888327 refGene_gn=EVPL;refGene_tx=XM_023244014.1
E1 57888767 57888961 refGene_gn=EVPL;refGene_tx=XM_023244014.1
E1 57889067 57889206 refGene_gn=EVPL;refGene_tx=XM_023244014.1
E1 57889303 57889482 refGene_gn=EVPL;refGene_tx=XM_023244014.1
E1 57890810 57890983 refGene_gn=EVPL;refGene_tx=XM_023244014.1
E1 57891290 57891388 refGene_gn=EVPL;refGene_tx=XM_023244014.1
E1 57891558 57891713 refGene_gn=EVPL;refGene_tx=XM_023244014.1
E1 57891939 57892086 refGene_gn=EVPL;refGene_tx=XM_023244014.1
E1 57892562 57892688 refGene_gn=EVPL;refGene_tx=XM_023244014.1
E1 57893669 57893765 refGene_gn=EVPL;refGene_tx=XM_023244014.1
E1 57893944 57894052 refGene_gn=EVPL;refGene_tx=XM_023244014.1
E1 57894145 57894307 refGene_gn=EVPL;refGene_tx=XM_023244014.1
E1 57894444 57894483 refGene_gn=EVPL;refGene_tx=XM_023244014.1
E1 57894687 57894813 refGene_gn=EVPL;refGene_tx=XM_023244014.1
E1 57895646 57895773 refGene_gn=EVPL;refGene_tx=XM_023244014.1
E1 57895880 57896035 refGene_gn=EVPL;refGene_tx=XM_023244014.1
E1 57896356 57896456 refGene_gn=EVPL;refGene_tx=XM_023244014.1
E1 57899207 57899305 refGene_gn=EVPL;refGene_tx=XM_023244014.1
E1 57908293 57908521 refGene_gn=SRP68;refGene_tx=XM_003997170.4
E1 57909005 57909058 refGene_gn=SRP68;refGene_tx=XM_003997170.4
E1 57909477 57909553 refGene_gn=SRP68;refGene_tx=XM_003997170.4
E1 57911114 57911244 refGene_gn=SRP68;refGene_tx=XM_003997170.4
E1 57912579 57912674 refGene_gn=SRP68;refGene_tx=XM_003997170.4
E1 57913305 57913462 refGene_gn=SRP68;refGene_tx=XM_003997170.4
E1 57917487 57917552 refGene_gn=SRP68;refGene_tx=XM_003997170.4
E1 57919851 57919950 refGene_gn=SRP68;refGene_tx=XM_003997170.4
E1 57933560 57933701 refGene_gn=SRP68;refGene_tx=XM_003997170.4
E1 57936259 57936342 refGene_gn=SRP68;refGene_tx=XM_003997170.4

E1 57936697 57936807 refGene_gn=SRP68;refGene_tx=XM_003997170.4
E1 57937110 57937193 refGene_gn=SRP68;refGene_tx=XM_003997170.4
E1 57938057 57938253 refGene_gn=SRP68;refGene_tx=XM_003997170.4
E1 57939985 57940099 refGene_gn=SRP68;refGene_tx=XM_003997170.4
E1 57942933 57943000 refGene_gn=SRP68;refGene_tx=XM_003997170.4
E1 57944547 57944710 refGene_gn=SRP68;refGene_tx=XM_003997170.4
E1 57949348 57949716 refGene_gn=GALR2;refGene_tx=XM_011289441.3
E1 57951054 57951430 refGene_gn=GALR2;refGene_tx=XM_011289441.3
E1 57951431 57951854 refGene_gn=GALR2;refGene_tx=XM_011289441.3
E1 57953209 57953306 refGene_gn=ZACN;refGene_tx=XM_011289384.3
E1 57953391 57953516 refGene_gn=ZACN;refGene_tx=XM_011289384.3
E1 57953644 57953689 refGene_gn=ZACN;refGene_tx=XM_011289384.3
E1 57953828 57953935 refGene_gn=ZACN;refGene_tx=XM_011289384.3
E1 57954160 57954330 refGene_gn=ZACN;refGene_tx=XM_011289384.3
E1 57954871 57954996 refGene_gn=ZACN;refGene_tx=XM_011289384.3
E1 57955128 57955339 refGene_gn=ZACN;refGene_tx=XM_011289384.3
E1 57955477 57955642 refGene_gn=ZACN;refGene_tx=XM_011289384.3
E1 57956102 57956290 refGene_gn=ZACN;refGene_tx=XM_011289384.3
E1 57957333 57957436
refGene_gn=EXOC7;refGene_tx=XM_003997173.5;refGene_gn=EXOC7;refGene_tx=XM_003997174.5;refGene_gn=EXOC7;refGene_tx=XM_006940606.4;refGene_gn=EXOC7;refGene_tx=XM_006940607.4;refGene_gn=EXOC7;refGene_tx=XM_011289385.3;refGene_gn=EXOC7;refGene_tx=XM_019818283.2;refGene_gn=EXOC7;refGene_tx=XM_019818284.2;refGene_gn=EXOC7;refGene_tx=XM_019818286.2;refGene_gn=EXOC7;refGene_tx=XM_019818287.1
E1 57957675 57957809
refGene_gn=EXOC7;refGene_tx=XM_003997173.5;refGene_gn=EXOC7;refGene_tx=XM_003997174.5;refGene_gn=EXOC7;refGene_tx=XM_006940606.4;refGene_gn=EXOC7;refGene_tx=XM_006940607.4;refGene_gn=EXOC7;refGene_tx=XM_011289385.3;refGene_gn=EXOC7;refGene_tx=XM_019818283.2;refGene_gn=EXOC7;refGene_tx=XM_019818284.2;refGene_gn=EXOC7;refGene_tx=XM_019818286.2;refGene_gn=EXOC7;refGene_tx=XM_019818287.1
E1 57957917 57957959
refGene_gn=EXOC7;refGene_tx=XM_003997173.5;refGene_gn=EXOC7;refGene_tx=XM_003997174.5;refGene_gn=EXOC7;refGene_tx=XM_006940606.4;refGene_gn=EXOC7;refGene_tx=XM_006940607.4;refGene_gn=EXOC7;refGene_tx=XM_011289385.3;refGene_gn=EXOC7;refGene_tx=XM_019818283.2;refGene_gn=EXOC7;refGene_tx=XM_019818284.2;refGene_gn=EXOC7;refGene_tx=XM_019818286.2;refGene_gn=EXOC7;refGene_tx=XM_019818287.1
E1 57958167 57958231
refGene_gn=EXOC7;refGene_tx=XM_003997173.5;refGene_gn=EXOC7;refGene_tx=XM_003997174.5;refGene_gn=EXOC7;refGene_tx=XM_006940606.4;refGene_gn=EXOC7;refGene_tx=XM_006940607.4;refGene_gn=EXOC7;refGene_tx=XM_011289385.3;refGene_gn=EXOC7;refGene_tx=XM_019818283.2;refGene_gn=EXOC7;refGene_tx=XM_019818284.2;refGene_gn=EXOC7;refGene_tx=XM_019818286.2;refGene_gn=EXOC7;refGene_tx=XM_019818287.1
E1 57959137 57959233
refGene_gn=EXOC7;refGene_tx=XM_003997173.5;refGene_gn=EXOC7;refGene_tx=XM_003997174.5;refGene_gn=EXOC7;refGene_tx=XM_006940606.4;refGene_gn=EXOC7;refGene_tx=XM_006940607.4;refGene_gn=EXOC7;refGene_tx=XM_0112893

85.3;refGene_gn=EXOC7;refGene_tx=XM_019818283.2;refGene_gn=EXOC7;refGene_tx=XM_019818284.2;refGene_gn=EXOC7;refGene_tx=XM_019818286.2;refGene_gn=EXOC7;refGene_tx=XM_019818287.1
E1 57959464 57959585
refGene_gn=EXOC7;refGene_tx=XM_003997173.5;refGene_gn=EXOC7;refGene_tx=XM_003997174.5;refGene_gn=EXOC7;refGene_tx=XM_006940606.4;refGene_gn=EXOC7;refGene_tx=XM_006940607.4;refGene_gn=EXOC7;refGene_tx=XM_011289385.3;refGene_gn=EXOC7;refGene_tx=XM_019818283.2;refGene_gn=EXOC7;refGene_tx=XM_019818284.2;refGene_gn=EXOC7;refGene_tx=XM_019818286.2;refGene_gn=EXOC7;refGene_tx=XM_019818287.1
E1 57959879 57959945
refGene_gn=EXOC7;refGene_tx=XM_003997173.5;refGene_gn=EXOC7;refGene_tx=XM_003997174.5;refGene_gn=EXOC7;refGene_tx=XM_006940606.4;refGene_gn=EXOC7;refGene_tx=XM_006940607.4;refGene_gn=EXOC7;refGene_tx=XM_011289385.3;refGene_gn=EXOC7;refGene_tx=XM_019818283.2;refGene_gn=EXOC7;refGene_tx=XM_019818284.2;refGene_gn=EXOC7;refGene_tx=XM_019818286.2;refGene_gn=EXOC7;refGene_tx=XM_019818287.1
E1 57960647 57960686
refGene_gn=EXOC7;refGene_tx=XM_006940606.4;refGene_gn=EXOC7;refGene_tx=XM_006940607.4;refGene_gn=EXOC7;refGene_tx=XM_019818283.2;refGene_gn=EXOC7;refGene_tx=XM_019818284.2;refGene_gn=EXOC7;refGene_tx=XM_019818287.1
E1 57961353 57961420
refGene_gn=EXOC7;refGene_tx=XM_003997173.5;refGene_gn=EXOC7;refGene_tx=XM_003997174.5;refGene_gn=EXOC7;refGene_tx=XM_006940606.4;refGene_gn=EXOC7;refGene_tx=XM_006940607.4;refGene_gn=EXOC7;refGene_tx=XM_011289385.3;refGene_gn=EXOC7;refGene_tx=XM_019818283.2;refGene_gn=EXOC7;refGene_tx=XM_019818284.2;refGene_gn=EXOC7;refGene_tx=XM_019818286.2;refGene_gn=EXOC7;refGene_tx=XM_019818287.1
E1 57961662 57961725
refGene_gn=EXOC7;refGene_tx=XM_003997173.5;refGene_gn=EXOC7;refGene_tx=XM_003997174.5;refGene_gn=EXOC7;refGene_tx=XM_006940606.4;refGene_gn=EXOC7;refGene_tx=XM_006940607.4;refGene_gn=EXOC7;refGene_tx=XM_011289385.3;refGene_gn=EXOC7;refGene_tx=XM_019818283.2;refGene_gn=EXOC7;refGene_tx=XM_019818284.2;refGene_gn=EXOC7;refGene_tx=XM_019818286.2;refGene_gn=EXOC7;refGene_tx=XM_019818287.1
E1 57962036 57962135
refGene_gn=EXOC7;refGene_tx=XM_003997173.5;refGene_gn=EXOC7;refGene_tx=XM_003997174.5;refGene_gn=EXOC7;refGene_tx=XM_006940606.4;refGene_gn=EXOC7;refGene_tx=XM_006940607.4;refGene_gn=EXOC7;refGene_tx=XM_011289385.3;refGene_gn=EXOC7;refGene_tx=XM_019818283.2;refGene_gn=EXOC7;refGene_tx=XM_019818284.2;refGene_gn=EXOC7;refGene_tx=XM_019818286.2;refGene_gn=EXOC7;refGene_tx=XM_019818287.1
E1 57962385 57962538
refGene_gn=EXOC7;refGene_tx=XM_003997173.5;refGene_gn=EXOC7;refGene_tx=XM_003997174.5;refGene_gn=EXOC7;refGene_tx=XM_006940606.4;refGene_gn=EXOC7;refGene_tx=XM_006940607.4;refGene_gn=EXOC7;refGene_tx=XM_011289385.3;refGene_gn=EXOC7;refGene_tx=XM_019818283.2;refGene_gn=EXOC7;refGene_tx=XM_019818284.2;refGene_gn=EXOC7;refGene_tx=XM_019818286.2;refGene_gn=EXOC7;refGene_tx=XM_019818287.1

E1 57962804 57962950
refGene_gn=EXOC7;refGene_tx=XM_003997173.5;refGene_gn=EXOC7;refGene_tx=XM_003997174.5;refGene_gn=EXOC7;refGene_tx=XM_006940606.4;refGene_gn=EXOC7;refGene_tx=XM_006940607.4;refGene_gn=EXOC7;refGene_tx=XM_011289385.3;refGene_gn=EXOC7;refGene_tx=XM_019818283.2;refGene_gn=EXOC7;refGene_tx=XM_019818284.2;refGene_gn=EXOC7;refGene_tx=XM_019818286.2;refGene_gn=EXOC7;refGene_tx=XM_019818287.1
E1 57963911 57963980
refGene_gn=EXOC7;refGene_tx=XM_011289385.3;refGene_gn=EXOC7;refGene_tx=XM_019818283.2;refGene_gn=EXOC7;refGene_tx=XM_019818284.2;refGene_gn=EXOC7;refGene_tx=XM_019818286.2;refGene_gn=EXOC7;refGene_tx=XM_019818287.1
E1 57964900 57964993
refGene_gn=EXOC7;refGene_tx=XM_003997173.5;refGene_gn=EXOC7;refGene_tx=XM_006940606.4;refGene_gn=EXOC7;refGene_tx=XM_011289385.3;refGene_gn=EXOC7;refGene_tx=XM_019818283.2;refGene_gn=EXOC7;refGene_tx=XM_019818287.1
E1 57967431 57967599
refGene_gn=EXOC7;refGene_tx=XM_003997173.5;refGene_gn=EXOC7;refGene_tx=XM_003997174.5;refGene_gn=EXOC7;refGene_tx=XM_006940606.4;refGene_gn=EXOC7;refGene_tx=XM_006940607.4;refGene_gn=EXOC7;refGene_tx=XM_011289385.3;refGene_gn=EXOC7;refGene_tx=XM_019818283.2;refGene_gn=EXOC7;refGene_tx=XM_019818284.2;refGene_gn=EXOC7;refGene_tx=XM_019818286.2;refGene_gn=EXOC7;refGene_tx=XM_019818287.1
E1 57969667 57969890
refGene_gn=EXOC7;refGene_tx=XM_003997173.5;refGene_gn=EXOC7;refGene_tx=XM_003997174.5;refGene_gn=EXOC7;refGene_tx=XM_006940606.4;refGene_gn=EXOC7;refGene_tx=XM_006940607.4;refGene_gn=EXOC7;refGene_tx=XM_011289385.3;refGene_gn=EXOC7;refGene_tx=XM_019818283.2;refGene_gn=EXOC7;refGene_tx=XM_019818284.2;refGene_gn=EXOC7;refGene_tx=XM_019818286.2;refGene_gn=EXOC7;refGene_tx=XM_019818287.1
E1 57972030 57972136
refGene_gn=EXOC7;refGene_tx=XM_003997173.5;refGene_gn=EXOC7;refGene_tx=XM_003997174.5;refGene_gn=EXOC7;refGene_tx=XM_006940606.4;refGene_gn=EXOC7;refGene_tx=XM_006940607.4;refGene_gn=EXOC7;refGene_tx=XM_011289385.3;refGene_gn=EXOC7;refGene_tx=XM_019818283.2;refGene_gn=EXOC7;refGene_tx=XM_019818284.2;refGene_gn=EXOC7;refGene_tx=XM_019818286.2;refGene_gn=EXOC7;refGene_tx=XM_019818287.1
E1 57972328 57972513
refGene_gn=EXOC7;refGene_tx=XM_003997173.5;refGene_gn=EXOC7;refGene_tx=XM_003997174.5;refGene_gn=EXOC7;refGene_tx=XM_006940606.4;refGene_gn=EXOC7;refGene_tx=XM_006940607.4;refGene_gn=EXOC7;refGene_tx=XM_011289385.3;refGene_gn=EXOC7;refGene_tx=XM_019818283.2;refGene_gn=EXOC7;refGene_tx=XM_019818284.2;refGene_gn=EXOC7;refGene_tx=XM_019818286.2
E1 57973625 57973691
refGene_gn=EXOC7;refGene_tx=XM_003997173.5;refGene_gn=EXOC7;refGene_tx=XM_003997174.5;refGene_gn=EXOC7;refGene_tx=XM_006940606.4;refGene_gn=EXOC7;refGene_tx=XM_006940607.4;refGene_gn=EXOC7;refGene_tx=XM_011289385.3;refGene_gn=EXOC7;refGene_tx=XM_019818283.2;refGene_gn=EXOC7;refGene_tx=XM_019818284.2;refGene_gn=EXOC7;refGene_tx=XM_019818286.2

E1 57973885 57973945
refGene_gn=EXOC7;refGene_tx=XM_003997173.5;refGene_gn=EXOC7;refGene_tx=XM_003997174.5;refGene_gn=EXOC7;refGene_tx=XM_006940606.4;refGene_gn=EXOC7;refGene_tx=XM_006940607.4;refGene_gn=EXOC7;refGene_tx=XM_011289385.3;refGene_gn=EXOC7;refGene_tx=XM_019818283.2;refGene_gn=EXOC7;refGene_tx=XM_019818284.2;refGene_gn=EXOC7;refGene_tx=XM_019818286.2
E1 57997525 57998281 refGene_gn=FOXJ1;refGene_tx=XM_003997176.5
E1 58000087 58000585 refGene_gn=FOXJ1;refGene_tx=XM_003997176.5
E1 58005213 58005335
refGene_gn=RNF157;refGene_tx=XM_011289386.3;refGene_gn=RNF157;refGene_tx=XM_023244024.1;refGene_gn=RNF157;refGene_tx=XM_023244027.1
E1 58006999 58007040
refGene_gn=RNF157;refGene_tx=XM_023244025.1;refGene_gn=RNF157;refGene_tx=XM_023244028.1
E1 58010038 58010061 refGene_gn=RNF157;refGene_tx=XM_023244026.1
E1 58010777 58010888
refGene_gn=RNF157;refGene_tx=XM_011289386.3;refGene_gn=RNF157;refGene_tx=XM_023244024.1;refGene_gn=RNF157;refGene_tx=XM_023244025.1;refGene_gn=RNF157;refGene_tx=XM_023244026.1
E1 58012688 58012737
refGene_gn=RNF157;refGene_tx=XM_011289386.3;refGene_gn=RNF157;refGene_tx=XM_023244024.1;refGene_gn=RNF157;refGene_tx=XM_023244025.1;refGene_gn=RNF157;refGene_tx=XM_023244026.1;refGene_gn=RNF157;refGene_tx=XM_023244027.1;refGene_gn=RNF157;refGene_tx=XM_023244028.1
E1 58013754 58013820
refGene_gn=RNF157;refGene_tx=XM_011289386.3;refGene_gn=RNF157;refGene_tx=XM_023244024.1;refGene_gn=RNF157;refGene_tx=XM_023244025.1;refGene_gn=RNF157;refGene_tx=XM_023244026.1;refGene_gn=RNF157;refGene_tx=XM_023244027.1;refGene_gn=RNF157;refGene_tx=XM_023244028.1
E1 58014094 58014282
refGene_gn=RNF157;refGene_tx=XM_011289386.3;refGene_gn=RNF157;refGene_tx=XM_023244024.1;refGene_gn=RNF157;refGene_tx=XM_023244025.1;refGene_gn=RNF157;refGene_tx=XM_023244026.1;refGene_gn=RNF157;refGene_tx=XM_023244027.1;refGene_gn=RNF157;refGene_tx=XM_023244028.1
E1 58014721 58014833
refGene_gn=RNF157;refGene_tx=XM_011289386.3;refGene_gn=RNF157;refGene_tx=XM_023244024.1;refGene_gn=RNF157;refGene_tx=XM_023244025.1;refGene_gn=RNF157;refGene_tx=XM_023244026.1;refGene_gn=RNF157;refGene_tx=XM_023244027.1;refGene_gn=RNF157;refGene_tx=XM_023244028.1
E1 58016727 58016836
refGene_gn=RNF157;refGene_tx=XM_011289386.3;refGene_gn=RNF157;refGene_tx=XM_023244024.1;refGene_gn=RNF157;refGene_tx=XM_023244025.1;refGene_gn=RNF157;refGene_tx=XM_023244026.1;refGene_gn=RNF157;refGene_tx=XM_023244027.1;refGene_gn=RNF157;refGene_tx=XM_023244028.1
E1 58017557 58017796
refGene_gn=RNF157;refGene_tx=XM_011289386.3;refGene_gn=RNF157;refGene_tx=XM_023244024.1;refGene_gn=RNF157;refGene_tx=XM_023244025.1;refGene_gn=RNF157;refGene_tx=XM_023244026.1;refGene_gn=RNF157;refGene_tx=XM_023244027.1;refGene_gn=RNF157;refGene_tx=XM_023244028.1
E1 58020376 58020489

refGene_gn=RNF157;refGene_tx=XM_011289386.3;refGene_gn=RNF157;refGene_tx=XM_023244024.1;refGene_gn=RNF157;refGene_tx=XM_023244025.1;refGene_gn=RNF157;refGene_tx=XM_023244026.1;refGene_gn=RNF157;refGene_tx=XM_023244027.1;refGene_gn=RNF157;refGene_tx=XM_023244028.1
E1 58020643 58020803
refGene_gn=RNF157;refGene_tx=XM_011289386.3;refGene_gn=RNF157;refGene_tx=XM_023244024.1;refGene_gn=RNF157;refGene_tx=XM_023244025.1;refGene_gn=RNF157;refGene_tx=XM_023244026.1;refGene_gn=RNF157;refGene_tx=XM_023244027.1;refGene_gn=RNF157;refGene_tx=XM_023244028.1
E1 58021521 58021593
refGene_gn=RNF157;refGene_tx=XM_011289386.3;refGene_gn=RNF157;refGene_tx=XM_023244024.1;refGene_gn=RNF157;refGene_tx=XM_023244025.1;refGene_gn=RNF157;refGene_tx=XM_023244026.1;refGene_gn=RNF157;refGene_tx=XM_023244027.1;refGene_gn=RNF157;refGene_tx=XM_023244028.1
E1 58023539 58023587
refGene_gn=RNF157;refGene_tx=XM_011289386.3;refGene_gn=RNF157;refGene_tx=XM_023244024.1;refGene_gn=RNF157;refGene_tx=XM_023244025.1;refGene_gn=RNF157;refGene_tx=XM_023244026.1;refGene_gn=RNF157;refGene_tx=XM_023244027.1;refGene_gn=RNF157;refGene_tx=XM_023244028.1
E1 58023964 58024008
refGene_gn=RNF157;refGene_tx=XM_011289386.3;refGene_gn=RNF157;refGene_tx=XM_023244024.1;refGene_gn=RNF157;refGene_tx=XM_023244025.1;refGene_gn=RNF157;refGene_tx=XM_023244026.1;refGene_gn=RNF157;refGene_tx=XM_023244027.1;refGene_gn=RNF157;refGene_tx=XM_023244028.1
E1 58024311 58024378
refGene_gn=RNF157;refGene_tx=XM_011289386.3;refGene_gn=RNF157;refGene_tx=XM_023244024.1;refGene_gn=RNF157;refGene_tx=XM_023244025.1;refGene_gn=RNF157;refGene_tx=XM_023244026.1;refGene_gn=RNF157;refGene_tx=XM_023244027.1;refGene_gn=RNF157;refGene_tx=XM_023244028.1
E1 58024808 58024926
refGene_gn=RNF157;refGene_tx=XM_011289386.3;refGene_gn=RNF157;refGene_tx=XM_023244024.1;refGene_gn=RNF157;refGene_tx=XM_023244025.1;refGene_gn=RNF157;refGene_tx=XM_023244026.1;refGene_gn=RNF157;refGene_tx=XM_023244027.1;refGene_gn=RNF157;refGene_tx=XM_023244028.1
E1 58025363 58025510
refGene_gn=RNF157;refGene_tx=XM_011289386.3;refGene_gn=RNF157;refGene_tx=XM_023244024.1;refGene_gn=RNF157;refGene_tx=XM_023244025.1;refGene_gn=RNF157;refGene_tx=XM_023244026.1;refGene_gn=RNF157;refGene_tx=XM_023244027.1;refGene_gn=RNF157;refGene_tx=XM_023244028.1
E1 58031587 58031676
refGene_gn=RNF157;refGene_tx=XM_011289386.3;refGene_gn=RNF157;refGene_tx=XM_023244024.1;refGene_gn=RNF157;refGene_tx=XM_023244025.1;refGene_gn=RNF157;refGene_tx=XM_023244026.1;refGene_gn=RNF157;refGene_tx=XM_023244027.1;refGene_gn=RNF157;refGene_tx=XM_023244028.1
E1 58061628 58061747
refGene_gn=RNF157;refGene_tx=XM_011289386.3;refGene_gn=RNF157;refGene_tx=XM_023244024.1;refGene_gn=RNF157;refGene_tx=XM_023244025.1;refGene_gn=RNF157;refGene_tx=XM_023244026.1;refGene_gn=RNF157;refGene_tx=XM_023244027.1;refGene_gn=RNF157;refGene_tx=XM_023244028.1
E1 58084556 58084644

refGene_gn=RNF157;refGene_tx=XM_023244024.1;refGene_gn=RNF157;refGene_tx=XM_023244025.1;refGene_gn=RNF157;refGene_tx=XM_023244026.1;refGene_gn=RNF157;refGene_tx=XM_023244027.1;refGene_gn=RNF157;refGene_tx=XM_023244028.1
E1 58101163 58101283 refGene_gn=UBALD2;refGene_tx=XM_023244029.1;refGene_gn=UBALD2;refGene_tx=XM_023244030.1
E1 58101523 58101586 refGene_gn=UBALD2;refGene_tx=XM_023244029.1;refGene_gn=UBALD2;refGene_tx=XM_023244030.1
E1 58105452 58105773 refGene_gn=UBALD2;refGene_tx=XM_023244029.1;refGene_gn=UBALD2;refGene_tx=XM_023244030.1
E1 58109219 58109409 refGene_gn=QRICH2;refGene_tx=XM_023244032.1
E1 58110671 58110783 refGene_gn=QRICH2;refGene_tx=XM_023244032.1
E1 58110885 58111026 refGene_gn=QRICH2;refGene_tx=XM_023244032.1
E1 58111746 58111834 refGene_gn=QRICH2;refGene_tx=XM_023244032.1
E1 58112222 58112370 refGene_gn=QRICH2;refGene_tx=XM_023244032.1
E1 58112726 58112930 refGene_gn=QRICH2;refGene_tx=XM_023244032.1
E1 58113722 58113824 refGene_gn=QRICH2;refGene_tx=XM_023244032.1
E1 58114107 58114173 refGene_gn=QRICH2;refGene_tx=XM_023244032.1
E1 58114808 58114930 refGene_gn=QRICH2;refGene_tx=XM_023244032.1
E1 58115035 58115200 refGene_gn=QRICH2;refGene_tx=XM_023244032.1
E1 58115408 58115483 refGene_gn=QRICH2;refGene_tx=XM_023244032.1
E1 58115573 58115696 refGene_gn=QRICH2;refGene_tx=XM_023244032.1
E1 58116452 58116707 refGene_gn=QRICH2;refGene_tx=XM_023244032.1
E1 58118655 58118770 refGene_gn=QRICH2;refGene_tx=XM_023244032.1
E1 58119264 58119362 refGene_gn=QRICH2;refGene_tx=XM_023244032.1
E1 58119919 58120005 refGene_gn=QRICH2;refGene_tx=XM_023244032.1
E1 58120611 58121457 refGene_gn=QRICH2;refGene_tx=XM_023244032.1
E1 58121862 58124809 refGene_gn=QRICH2;refGene_tx=XM_023244032.1
E1 58140326 58140437 refGene_gn=QRICH2;refGene_tx=XM_023244032.1
E1 58140789 58140849 refGene_gn=QRICH2;refGene_tx=XM_023244032.1
E1 58142026 58142107 refGene_gn=QRICH2;refGene_tx=XM_023244032.1
E1 58142217 58142727 refGene_gn=QRICH2;refGene_tx=XM_023244032.1
E1 58146713 58146872 refGene_gn=PRPSAP1;refGene_tx=XM_019818291.2;refGene_gn=PRPSAP1;refGene_tx=XM_023244033.1;refGene_gn=PRPSAP1;refGene_tx=XM_023244034.1;refGene_gn=PRPSAP1;refGene_tx=XM_023244035.1
E1 58147418 58147565 refGene_gn=PRPSAP1;refGene_tx=XM_019818291.2;refGene_gn=PRPSAP1;refGene_tx=XM_023244033.1;refGene_gn=PRPSAP1;refGene_tx=XM_023244034.1;refGene_gn=PRPSAP1;refGene_tx=XM_023244035.1
E1 58148301 58148372 refGene_gn=PRPSAP1;refGene_tx=XM_019818291.2;refGene_gn=PRPSAP1;refGene_tx=XM_023244033.1;refGene_gn=PRPSAP1;refGene_tx=XM_023244034.1;refGene_gn=PRPSAP1;refGene_tx=XM_023244035.1
E1 58156173 58156319 refGene_gn=PRPSAP1;refGene_tx=XM_019818291.2;refGene_gn=PRPSAP1;refGene_tx=XM_023244033.1;refGene_gn=PRPSAP1;refGene_tx=XM_023244034.1;refGe

ne_gn=PRPSAP1;refGene_tx=XM_023244035.1
E1 58157103 58157159
refGene_gn=PRPSAP1;refGene_tx=XM_019818291.2;refGene_gn=PRPSAP1;refGene_tx=XM_023244033.1;refGene_gn=PRPSAP1;refGene_tx=XM_023244034.1;refGene_gn=PRPSAP1;refGene_tx=XM_023244035.1
E1 58157613 58157729
refGene_gn=PRPSAP1;refGene_tx=XM_019818291.2;refGene_gn=PRPSAP1;refGene_tx=XM_023244033.1;refGene_gn=PRPSAP1;refGene_tx=XM_023244034.1;refGene_gn=PRPSAP1;refGene_tx=XM_023244035.1
E1 58159070 58159243
refGene_gn=PRPSAP1;refGene_tx=XM_019818291.2;refGene_gn=PRPSAP1;refGene_tx=XM_023244033.1;refGene_gn=PRPSAP1;refGene_tx=XM_023244034.1;refGene_gn=PRPSAP1;refGene_tx=XM_023244035.1
E1 58166207 58166274 refGene_gn=PRPSAP1;refGene_tx=XM_023244033.1
E1 58169355 58169408 refGene_gn=PRPSAP1;refGene_tx=XM_023244033.1
E1 58173734 58173904 refGene_gn=PRPSAP1;refGene_tx=XM_023244033.1
E1 58197054 58197364
refGene_gn=SPHK1;refGene_tx=XM_011289395.3;refGene_gn=SPHK1;refGene_tx=XM_019818292.2;refGene_gn=SPHK1;refGene_tx=XM_023244037.1
E1 58197732 58197885
refGene_gn=SPHK1;refGene_tx=XM_011289395.3;refGene_gn=SPHK1;refGene_tx=XM_019818292.2;refGene_gn=SPHK1;refGene_tx=XM_023244037.1
E1 58197971 58198066
refGene_gn=SPHK1;refGene_tx=XM_011289395.3;refGene_gn=SPHK1;refGene_tx=XM_019818292.2;refGene_gn=SPHK1;refGene_tx=XM_023244037.1
E1 58198146 58198262
refGene_gn=SPHK1;refGene_tx=XM_011289395.3;refGene_gn=SPHK1;refGene_tx=XM_019818292.2;refGene_gn=SPHK1;refGene_tx=XM_023244037.1
E1 58198565 58199355
refGene_gn=SPHK1;refGene_tx=XM_011289395.3;refGene_gn=SPHK1;refGene_tx=XM_019818292.2;refGene_gn=SPHK1;refGene_tx=XM_023244037.1
E1 58202268 58202936 refGene_gn=UBE2O;refGene_tx=XM_023243476.1
E1 58203100 58203158 refGene_gn=UBE2O;refGene_tx=XM_023243476.1
E1 58203274 58203479 refGene_gn=UBE2O;refGene_tx=XM_023243476.1
E1 58206167 58206303 refGene_gn=UBE2O;refGene_tx=XM_023243476.1
E1 58206573 58207270 refGene_gn=UBE2O;refGene_tx=XM_023243476.1
E1 58208330 58208420 refGene_gn=UBE2O;refGene_tx=XM_023243476.1
E1 58209019 58209148 refGene_gn=UBE2O;refGene_tx=XM_023243476.1
E1 58209240 58209336 refGene_gn=UBE2O;refGene_tx=XM_023243476.1
E1 58209337 58209354 refGene_gn=UBE2O;refGene_tx=XM_023243476.1
E1 58209625 58209780 refGene_gn=UBE2O;refGene_tx=XM_023243476.1
E1 58210160 58210633 refGene_gn=UBE2O;refGene_tx=XM_023243476.1
E1 58210876 58211027 refGene_gn=UBE2O;refGene_tx=XM_023243476.1
E1 58211156 58211266 refGene_gn=UBE2O;refGene_tx=XM_023243476.1
E1 58211696 58211840 refGene_gn=UBE2O;refGene_tx=XM_023243476.1
E1 58212514 58212578 refGene_gn=UBE2O;refGene_tx=XM_023243476.1
E1 58212891 58212989 refGene_gn=UBE2O;refGene_tx=XM_023243476.1
E1 58215039 58215150 refGene_gn=UBE2O;refGene_tx=XM_023243476.1
E1 58215253 58215313 refGene_gn=UBE2O;refGene_tx=XM_023243476.1
E1 58257751 58258174 refGene_gn=UBE2O;refGene_tx=XM_023243476.1

E1 58274611 58274774 refGene_gn=AANAT;refGene_tx=XM_011289442.3
E1 58275049 58275204 refGene_gn=AANAT;refGene_tx=XM_011289442.3
E1 58275429 58275735 refGene_gn=AANAT;refGene_tx=XM_011289442.3
E1 58277458 58277878
refGene_gn=RHBDF2;refGene_tx=XM_023244038.1;refGene_gn=RHBDF2;refGene_tx=XM_023244039.1;refGene_gn=RHBDF2;refGene_tx=XM_023244040.1;refGene_gn=RHBDF2;refGene_tx=XM_023244041.1;refGene_gn=RHBDF2;refGene_tx=XM_023244042.1;refGene_gn=RHBDF2;refGene_tx=XM_023244043.1;refGene_gn=RHBDF2;refGene_tx=XM_023244044.1
E1 58278591 58278745
refGene_gn=RHBDF2;refGene_tx=XM_023244038.1;refGene_gn=RHBDF2;refGene_tx=XM_023244039.1;refGene_gn=RHBDF2;refGene_tx=XM_023244040.1;refGene_gn=RHBDF2;refGene_tx=XM_023244041.1;refGene_gn=RHBDF2;refGene_tx=XM_023244042.1;refGene_gn=RHBDF2;refGene_tx=XM_023244043.1;refGene_gn=RHBDF2;refGene_tx=XM_023244044.1
E1 58278965 58279066
refGene_gn=RHBDF2;refGene_tx=XM_023244038.1;refGene_gn=RHBDF2;refGene_tx=XM_023244039.1;refGene_gn=RHBDF2;refGene_tx=XM_023244040.1;refGene_gn=RHBDF2;refGene_tx=XM_023244041.1;refGene_gn=RHBDF2;refGene_tx=XM_023244042.1;refGene_gn=RHBDF2;refGene_tx=XM_023244043.1;refGene_gn=RHBDF2;refGene_tx=XM_023244044.1
E1 58279235 58279311
refGene_gn=RHBDF2;refGene_tx=XM_023244038.1;refGene_gn=RHBDF2;refGene_tx=XM_023244039.1;refGene_gn=RHBDF2;refGene_tx=XM_023244040.1;refGene_gn=RHBDF2;refGene_tx=XM_023244041.1;refGene_gn=RHBDF2;refGene_tx=XM_023244042.1;refGene_gn=RHBDF2;refGene_tx=XM_023244043.1;refGene_gn=RHBDF2;refGene_tx=XM_023244044.1
E1 58279588 58279683
refGene_gn=RHBDF2;refGene_tx=XM_023244038.1;refGene_gn=RHBDF2;refGene_tx=XM_023244039.1;refGene_gn=RHBDF2;refGene_tx=XM_023244040.1;refGene_gn=RHBDF2;refGene_tx=XM_023244041.1;refGene_gn=RHBDF2;refGene_tx=XM_023244042.1;refGene_gn=RHBDF2;refGene_tx=XM_023244043.1;refGene_gn=RHBDF2;refGene_tx=XM_023244044.1
E1 58279772 58279836
refGene_gn=RHBDF2;refGene_tx=XM_023244038.1;refGene_gn=RHBDF2;refGene_tx=XM_023244039.1;refGene_gn=RHBDF2;refGene_tx=XM_023244040.1;refGene_gn=RHBDF2;refGene_tx=XM_023244041.1;refGene_gn=RHBDF2;refGene_tx=XM_023244042.1;refGene_gn=RHBDF2;refGene_tx=XM_023244043.1;refGene_gn=RHBDF2;refGene_tx=XM_023244044.1
E1 58279991 58280101
refGene_gn=RHBDF2;refGene_tx=XM_023244038.1;refGene_gn=RHBDF2;refGene_tx=XM_023244039.1;refGene_gn=RHBDF2;refGene_tx=XM_023244040.1;refGene_gn=RHBDF2;refGene_tx=XM_023244041.1;refGene_gn=RHBDF2;refGene_tx=XM_023244042.1;refGene_gn=RHBDF2;refGene_tx=XM_023244043.1;refGene_gn=RHBDF2;refGene_tx=XM_023244044.1
E1 58280388 58280550
refGene_gn=RHBDF2;refGene_tx=XM_023244038.1;refGene_gn=RHBDF2;refGene_tx=XM_023244039.1;refGene_gn=RHBDF2;refGene_tx=XM_023244040.1;refGene_gn=RHBDF2;refGene_tx=XM_023244041.1;refGene_gn=RHBDF2;refGene_tx=XM_023244042.1;refGene_gn=RHBDF2;refGene_tx=XM_023244043.1;refGene_gn=RHBDF

2;refGene_tx=XM_023244044.1
E1 58280727 58280802
refGene_gn=RHBDF2;refGene_tx=XM_023244038.1;refGene_gn=RHBDF2;refGene_tx=XM_023244039.1;refGene_gn=RHBDF2;refGene_tx=XM_023244040.1;refGene_gn=RHBDF2;refGene_tx=XM_023244041.1;refGene_gn=RHBDF2;refGene_tx=XM_023244042.1;refGene_gn=RHBDF2;refGene_tx=XM_023244043.1;refGene_gn=RHBDF2;refGene_tx=XM_023244044.1
E1 58281004 58281116
refGene_gn=RHBDF2;refGene_tx=XM_023244038.1;refGene_gn=RHBDF2;refGene_tx=XM_023244039.1;refGene_gn=RHBDF2;refGene_tx=XM_023244040.1;refGene_gn=RHBDF2;refGene_tx=XM_023244041.1;refGene_gn=RHBDF2;refGene_tx=XM_023244042.1;refGene_gn=RHBDF2;refGene_tx=XM_023244043.1;refGene_gn=RHBDF2;refGene_tx=XM_023244044.1
E1 58282341 58282539
refGene_gn=RHBDF2;refGene_tx=XM_023244038.1;refGene_gn=RHBDF2;refGene_tx=XM_023244039.1;refGene_gn=RHBDF2;refGene_tx=XM_023244040.1;refGene_gn=RHBDF2;refGene_tx=XM_023244041.1;refGene_gn=RHBDF2;refGene_tx=XM_023244042.1;refGene_gn=RHBDF2;refGene_tx=XM_023244043.1;refGene_gn=RHBDF2;refGene_tx=XM_023244044.1
E1 58282681 58282800
refGene_gn=RHBDF2;refGene_tx=XM_023244038.1;refGene_gn=RHBDF2;refGene_tx=XM_023244039.1;refGene_gn=RHBDF2;refGene_tx=XM_023244040.1;refGene_gn=RHBDF2;refGene_tx=XM_023244041.1;refGene_gn=RHBDF2;refGene_tx=XM_023244042.1;refGene_gn=RHBDF2;refGene_tx=XM_023244043.1;refGene_gn=RHBDF2;refGene_tx=XM_023244044.1
E1 58283154 58283283
refGene_gn=RHBDF2;refGene_tx=XM_023244038.1;refGene_gn=RHBDF2;refGene_tx=XM_023244039.1;refGene_gn=RHBDF2;refGene_tx=XM_023244040.1;refGene_gn=RHBDF2;refGene_tx=XM_023244041.1;refGene_gn=RHBDF2;refGene_tx=XM_023244042.1;refGene_gn=RHBDF2;refGene_tx=XM_023244043.1;refGene_gn=RHBDF2;refGene_tx=XM_023244044.1
E1 58284088 58284292
refGene_gn=RHBDF2;refGene_tx=XM_023244038.1;refGene_gn=RHBDF2;refGene_tx=XM_023244039.1;refGene_gn=RHBDF2;refGene_tx=XM_023244040.1;refGene_gn=RHBDF2;refGene_tx=XM_023244041.1;refGene_gn=RHBDF2;refGene_tx=XM_023244042.1;refGene_gn=RHBDF2;refGene_tx=XM_023244043.1;refGene_gn=RHBDF2;refGene_tx=XM_023244044.1
E1 58284389 58284585
refGene_gn=RHBDF2;refGene_tx=XM_023244038.1;refGene_gn=RHBDF2;refGene_tx=XM_023244039.1;refGene_gn=RHBDF2;refGene_tx=XM_023244040.1;refGene_gn=RHBDF2;refGene_tx=XM_023244041.1;refGene_gn=RHBDF2;refGene_tx=XM_023244042.1;refGene_gn=RHBDF2;refGene_tx=XM_023244043.1;refGene_gn=RHBDF2;refGene_tx=XM_023244044.1
E1 58285125 58285253
refGene_gn=RHBDF2;refGene_tx=XM_023244038.1;refGene_gn=RHBDF2;refGene_tx=XM_023244039.1;refGene_gn=RHBDF2;refGene_tx=XM_023244040.1;refGene_gn=RHBDF2;refGene_tx=XM_023244041.1;refGene_gn=RHBDF2;refGene_tx=XM_023244042.1;refGene_gn=RHBDF2;refGene_tx=XM_023244043.1;refGene_gn=RHBDF2;refGene_tx=XM_023244044.1
E1 58285993 58286137

refGene_gn=RHBDF2;refGene_tx=XM_023244038.1;refGene_gn=RHBDF2;refGene_tx=XM_023244039.1;refGene_gn=RHBDF2;refGene_tx=XM_023244040.1;refGene_gn=RHBDF2;refGene_tx=XM_023244041.1;refGene_gn=RHBDF2;refGene_tx=XM_023244042.1;refGene_gn=RHBDF2;refGene_tx=XM_023244043.1;refGene_gn=RHBDF2;refGene_tx=XM_023244044.1
E1 58313787 58314156
refGene_gn=CYGB;refGene_tx=XM_011289397.3;refGene_gn=CYGB;refGene_tx=XM_011289398.3
E1 58316563 58316731
refGene_gn=CYGB;refGene_tx=XM_011289397.3;refGene_gn=CYGB;refGene_tx=XM_011289398.3;refGene_gn=CYGB;refGene_tx=XM_011289399.3
E1 58317050 58317282
refGene_gn=CYGB;refGene_tx=XM_011289397.3;refGene_gn=CYGB;refGene_tx=XM_011289398.3;refGene_gn=CYGB;refGene_tx=XM_011289399.3
E1 58322063 58322206
refGene_gn=CYGB;refGene_tx=XM_011289397.3;refGene_gn=CYGB;refGene_tx=XM_011289398.3;refGene_gn=CYGB;refGene_tx=XM_011289399.3
E1 58325387 58325461
refGene_gn=PRCD;refGene_tx=XM_023244047.1;refGene_gn=PRCD;refGene_tx=XM_023244048.1;refGene_gn=PRCD;refGene_tx=XM_023244049.1;refGene_gn=PRCD;refGene_tx=XM_023244051.1
E1 58325756 58325825
refGene_gn=PRCD;refGene_tx=XM_023244047.1;refGene_gn=PRCD;refGene_tx=XM_023244048.1;refGene_gn=PRCD;refGene_tx=XM_023244049.1;refGene_gn=PRCD;refGene_tx=XM_023244051.1
E1 58328174 58328196
refGene_gn=PRCD;refGene_tx=XM_023244047.1;refGene_gn=PRCD;refGene_tx=XM_023244048.1;refGene_gn=PRCD;refGene_tx=XM_023244049.1;refGene_gn=PRCD;refGene_tx=XM_023244051.1
E1 58333814 58333904
refGene_gn=LOC111557723;refGene_tx=XM_023243477.1
E1 58340307 58340453
refGene_gn=LOC111557723;refGene_tx=XM_023243477.1
E1 58340904 58340987
refGene_gn=LOC111557723;refGene_tx=XM_023243477.1
E1 58343843 58343929
refGene_gn=LOC111557723;refGene_tx=XM_023243477.1
E1 58344055 58344094
refGene_gn=LOC111557723;refGene_tx=XM_023243477.1
E1 58345628 58345817
refGene_gn=LOC111557723;refGene_tx=XM_023243477.1
E1 58347853 58348021
refGene_gn=ST6GALNAC2;refGene_tx=XM_023244046.1
E1 58349108 58349208
refGene_gn=ST6GALNAC2;refGene_tx=XM_023244046.1
E1 58350346 58350430
refGene_gn=ST6GALNAC2;refGene_tx=XM_023244046.1
E1 58352143 58352247
refGene_gn=ST6GALNAC2;refGene_tx=XM_023244046.1
E1 58353557 58353696

refGene_gn=ST6GALNAC2;refGene_tx=XM_023244046.1
E1 58354065 58354234
refGene_gn=ST6GALNAC2;refGene_tx=XM_023244046.1
E1 58354872 58355047
refGene_gn=ST6GALNAC2;refGene_tx=XM_023244046.1
E1 58359291 58359352
refGene_gn=ST6GALNAC2;refGene_tx=XM_023244046.1
E1 58362777 58362902
refGene_gn=ST6GALNAC2;refGene_tx=XM_023244046.1
E1 58383084 58383276
refGene_gn=ST6GALNAC1;refGene_tx=XM_003997179.4
E1 58383514 58383614
refGene_gn=ST6GALNAC1;refGene_tx=XM_003997179.4
E1 58383701 58383791
refGene_gn=ST6GALNAC1;refGene_tx=XM_003997179.4
E1 58383987 58384091
refGene_gn=ST6GALNAC1;refGene_tx=XM_003997179.4
E1 58384337 58384476
refGene_gn=ST6GALNAC1;refGene_tx=XM_003997179.4
E1 58384705 58384877
refGene_gn=ST6GALNAC1;refGene_tx=XM_003997179.4
E1 58385047 58385216
refGene_gn=ST6GALNAC1;refGene_tx=XM_003997179.4
E1 58385629 58386281
refGene_gn=ST6GALNAC1;refGene_tx=XM_003997179.4
E1 58401766 58401897
refGene_gn=ST6GALNAC1;refGene_tx=XM_003997179.4
E1 58431294 58431409 refGene_gn=MXRA7;refGene_tx=XM_023244060.1
E1 58434282 58434295 refGene_gn=MXRA7;refGene_tx=XM_023244062.1
E1 58437265 58437299 refGene_gn=MXRA7;refGene_tx=XM_023244061.1
E1 58438379 58438476
refGene_gn=MXRA7;refGene_tx=XM_023244060.1;refGene_gn=MXRA7;refGene_tx
=XM_023244061.1;refGene_gn=MXRA7;refGene_tx=XM_023244062.1
E1 58441811 58441875
refGene_gn=MXRA7;refGene_tx=XM_023244060.1;refGene_gn=MXRA7;refGene_tx
=XM_023244061.1;refGene_gn=MXRA7;refGene_tx=XM_023244062.1
E1 58459329 58459608
refGene_gn=MXRA7;refGene_tx=XM_023244060.1;refGene_gn=MXRA7;refGene_tx
=XM_023244061.1;refGene_gn=MXRA7;refGene_tx=XM_023244062.1
E1 58464688 58464725 refGene_gn=JMJD6;refGene_tx=XM_003997181.5
E1 58466007 58466135 refGene_gn=JMJD6;refGene_tx=XM_003997181.5
E1 58466971 58467110 refGene_gn=JMJD6;refGene_tx=XM_003997181.5
E1 58467901 58467968 refGene_gn=JMJD6;refGene_tx=XM_019818305.2
E1 58468456 58468592
refGene_gn=JMJD6;refGene_tx=XM_003997181.5;refGene_gn=JMJD6;refGene_tx
=XM_019818305.2
E1 58469731 58470018
refGene_gn=JMJD6;refGene_tx=XM_003997181.5;refGene_gn=JMJD6;refGene_tx
=XM_019818305.2
E1 58470729 58471118

refGene_gn=JMJD6;refGene_tx=XM_003997181.5;refGene_gn=JMJD6;refGene_tx=XM_019818305.2
E1 58471653 58471782 refGene_gn=JMJD6;refGene_tx=XM_003997181.5;refGene_gn=JMJD6;refGene_tx=XM_019818305.2
E1 58472473 58472559 refGene_gn=METTL23;refGene_tx=XM_023244057.1
E1 58474792 58474897 refGene_gn=METTL23;refGene_tx=XM_003997182.4;refGene_gn=METTL23;refGene_tx=XM_023244057.1
E1 58476154 58476392 refGene_gn=METTL23;refGene_tx=XM_003997182.4;refGene_gn=METTL23;refGene_tx=XM_023244057.1
E1 58476472 58476557 refGene_gn=METTL23;refGene_tx=XM_003997182.4;refGene_gn=METTL23;refGene_tx=XM_023244057.1
E1 58476814 58476983 refGene_gn=METTL23;refGene_tx=XM_003997182.4;refGene_gn=METTL23;refGene_tx=XM_023244057.1
E1 58479624 58479928 refGene_gn=SRSF2;refGene_tx=XM_023244058.1;refGene_gn=SRSF2;refGene_tx=XM_023244059.1
E1 58480146 58480508 refGene_gn=SRSF2;refGene_tx=XM_023244058.1;refGene_gn=SRSF2;refGene_tx=XM_023244059.1
E1 58481738 58481834 refGene_gn=MFSD11;refGene_tx=XM_003997183.5;refGene_gn=MFSD11;refGene_tx=XM_011289401.3;refGene_gn=MFSD11;refGene_tx=XM_019818304.2;refGene_gn=MFSD11;refGene_tx=XM_023244053.1;refGene_gn=MFSD11;refGene_tx=XM_023244054.1;refGene_gn=MFSD11;refGene_tx=XM_023244055.1;refGene_gn=MFSD11;refGene_tx=XM_023244056.1
E1 58482390 58482446 refGene_gn=MFSD11;refGene_tx=XM_003997183.5;refGene_gn=MFSD11;refGene_tx=XM_011289401.3;refGene_gn=MFSD11;refGene_tx=XM_023244053.1
E1 58483961 58484069 refGene_gn=MFSD11;refGene_tx=XM_003997183.5;refGene_gn=MFSD11;refGene_tx=XM_011289401.3;refGene_gn=MFSD11;refGene_tx=XM_023244053.1;refGene_gn=MFSD11;refGene_tx=XM_023244054.1;refGene_gn=MFSD11;refGene_tx=XM_023244055.1
E1 58484983 58485063 refGene_gn=MFSD11;refGene_tx=XM_003997183.5;refGene_gn=MFSD11;refGene_tx=XM_011289401.3;refGene_gn=MFSD11;refGene_tx=XM_019818304.2;refGene_gn=MFSD11;refGene_tx=XM_023244053.1;refGene_gn=MFSD11;refGene_tx=XM_023244054.1;refGene_gn=MFSD11;refGene_tx=XM_023244055.1;refGene_gn=MFSD11;refGene_tx=XM_023244056.1
E1 58485181 58485278 refGene_gn=MFSD11;refGene_tx=XM_003997183.5;refGene_gn=MFSD11;refGene_tx=XM_011289401.3;refGene_gn=MFSD11;refGene_tx=XM_019818304.2;refGene_gn=MFSD11;refGene_tx=XM_023244053.1;refGene_gn=MFSD11;refGene_tx=XM_023244054.1;refGene_gn=MFSD11;refGene_tx=XM_023244055.1;refGene_gn=MFSD11;refGene_tx=XM_023244056.1

E1 58486521 58486580
refGene_gn=MFSD11;refGene_tx=XM_003997183.5;refGene_gn=MFSD11;refGene_tx=XM_011289401.3;refGene_gn=MFSD11;refGene_tx=XM_019818304.2;refGene_gn=MFSD11;refGene_tx=XM_023244053.1;refGene_gn=MFSD11;refGene_tx=XM_023244054.1;refGene_gn=MFSD11;refGene_tx=XM_023244055.1;refGene_gn=MFSD11;refGene_tx=XM_023244056.1
E1 58487072 58487217
refGene_gn=MFSD11;refGene_tx=XM_003997183.5;refGene_gn=MFSD11;refGene_tx=XM_011289401.3;refGene_gn=MFSD11;refGene_tx=XM_019818304.2;refGene_gn=MFSD11;refGene_tx=XM_023244053.1;refGene_gn=MFSD11;refGene_tx=XM_023244054.1;refGene_gn=MFSD11;refGene_tx=XM_023244055.1;refGene_gn=MFSD11;refGene_tx=XM_023244056.1
E1 58492264 58492305
refGene_gn=MFSD11;refGene_tx=XM_003997183.5;refGene_gn=MFSD11;refGene_tx=XM_011289401.3;refGene_gn=MFSD11;refGene_tx=XM_019818304.2;refGene_gn=MFSD11;refGene_tx=XM_023244053.1;refGene_gn=MFSD11;refGene_tx=XM_023244054.1;refGene_gn=MFSD11;refGene_tx=XM_023244055.1;refGene_gn=MFSD11;refGene_tx=XM_023244056.1
E1 58499401 58499467
refGene_gn=MFSD11;refGene_tx=XM_003997183.5;refGene_gn=MFSD11;refGene_tx=XM_011289401.3;refGene_gn=MFSD11;refGene_tx=XM_019818304.2;refGene_gn=MFSD11;refGene_tx=XM_023244053.1;refGene_gn=MFSD11;refGene_tx=XM_023244054.1;refGene_gn=MFSD11;refGene_tx=XM_023244055.1;refGene_gn=MFSD11;refGene_tx=XM_023244056.1
E1 58501101 58501227
refGene_gn=MFSD11;refGene_tx=XM_003997183.5;refGene_gn=MFSD11;refGene_tx=XM_011289401.3;refGene_gn=MFSD11;refGene_tx=XM_019818304.2;refGene_gn=MFSD11;refGene_tx=XM_023244053.1;refGene_gn=MFSD11;refGene_tx=XM_023244054.1;refGene_gn=MFSD11;refGene_tx=XM_023244055.1;refGene_gn=MFSD11;refGene_tx=XM_023244056.1
E1 58504323 58504498
refGene_gn=MFSD11;refGene_tx=XM_003997183.5;refGene_gn=MFSD11;refGene_tx=XM_011289401.3;refGene_gn=MFSD11;refGene_tx=XM_019818304.2;refGene_gn=MFSD11;refGene_tx=XM_023244053.1;refGene_gn=MFSD11;refGene_tx=XM_023244054.1;refGene_gn=MFSD11;refGene_tx=XM_023244055.1;refGene_gn=MFSD11;refGene_tx=XM_023244056.1
E1 58505411 58505547
refGene_gn=MFSD11;refGene_tx=XM_003997183.5;refGene_gn=MFSD11;refGene_tx=XM_011289401.3;refGene_gn=MFSD11;refGene_tx=XM_019818304.2;refGene_gn=MFSD11;refGene_tx=XM_023244053.1;refGene_gn=MFSD11;refGene_tx=XM_023244054.1;refGene_gn=MFSD11;refGene_tx=XM_023244055.1;refGene_gn=MFSD11;refGene_tx=XM_023244056.1
E1 58506505 58506670
refGene_gn=MFSD11;refGene_tx=XM_003997183.5;refGene_gn=MFSD11;refGene_tx=XM_011289401.3;refGene_gn=MFSD11;refGene_tx=XM_019818304.2;refGene_gn=MFSD11;refGene_tx=XM_023244053.1;refGene_gn=MFSD11;refGene_tx=XM_023244054.1;refGene_gn=MFSD11;refGene_tx=XM_023244055.1;refGene_gn=MFSD11;refGene_tx=XM_023244056.1
E1 58558579 58558647
refGene_gn=MGAT5B;refGene_tx=XM_011289402.3;refGene_gn=MGAT5B;refGene_

tx=XM_011289403.3
E1 58562396 58562509
refGene_gn=MGAT5B;refGene_tx=XM_011289402.3;refGene_gn=MGAT5B;refGene_
tx=XM_011289403.3
E1 58570268 58570416
refGene_gn=MGAT5B;refGene_tx=XM_011289402.3;refGene_gn=MGAT5B;refGene_
tx=XM_011289403.3
E1 58584507 58584623
refGene_gn=MGAT5B;refGene_tx=XM_011289402.3;refGene_gn=MGAT5B;refGene_
tx=XM_011289403.3
E1 58585148 58585222
refGene_gn=MGAT5B;refGene_tx=XM_011289402.3;refGene_gn=MGAT5B;refGene_
tx=XM_011289403.3
E1 58586078 58586249
refGene_gn=MGAT5B;refGene_tx=XM_011289402.3;refGene_gn=MGAT5B;refGene_
tx=XM_011289403.3
E1 58586969 58587134
refGene_gn=MGAT5B;refGene_tx=XM_011289402.3;refGene_gn=MGAT5B;refGene_
tx=XM_011289403.3
E1 58588410 58588580
refGene_gn=MGAT5B;refGene_tx=XM_011289402.3;refGene_gn=MGAT5B;refGene_
tx=XM_011289403.3
E1 58604336 58604468
refGene_gn=MGAT5B;refGene_tx=XM_011289402.3;refGene_gn=MGAT5B;refGene_
tx=XM_011289403.3
E1 58605860 58605994
refGene_gn=MGAT5B;refGene_tx=XM_011289402.3;refGene_gn=MGAT5B;refGene_
tx=XM_011289403.3
E1 58610097 58610228
refGene_gn=MGAT5B;refGene_tx=XM_011289402.3;refGene_gn=MGAT5B;refGene_
tx=XM_011289403.3
E1 58610668 58610674 refGene_gn=MGAT5B;refGene_tx=XM_011289402.3
E1 58621587 58621743
refGene_gn=MGAT5B;refGene_tx=XM_011289402.3;refGene_gn=MGAT5B;refGene_
tx=XM_011289403.3
E1 58622528 58622675
refGene_gn=MGAT5B;refGene_tx=XM_011289402.3;refGene_gn=MGAT5B;refGene_
tx=XM_011289403.3
E1 58622838 58622955
refGene_gn=MGAT5B;refGene_tx=XM_011289402.3;refGene_gn=MGAT5B;refGene_
tx=XM_011289403.3
E1 58627722 58627797
refGene_gn=MGAT5B;refGene_tx=XM_011289402.3;refGene_gn=MGAT5B;refGene_
tx=XM_011289403.3
E1 58628945 58629202
refGene_gn=MGAT5B;refGene_tx=XM_011289402.3;refGene_gn=MGAT5B;refGene_
tx=XM_011289403.3
E1 58629558 58629757
refGene_gn=MGAT5B;refGene_tx=XM_011289402.3;refGene_gn=MGAT5B;refGene_
tx=XM_011289403.3

E1 58769881 58769944 refGene_gn=SEC14L1;refGene_tx=XM_023244091.1
E1 58800718 58800868 refGene_gn=SEC14L1;refGene_tx=XM_023244091.1
E1 58800978 58801110 refGene_gn=SEC14L1;refGene_tx=XM_023244091.1
E1 58803263 58803392 refGene_gn=SEC14L1;refGene_tx=XM_023244091.1
E1 58804177 58804412 refGene_gn=SEC14L1;refGene_tx=XM_023244091.1
E1 58805010 58805120 refGene_gn=SEC14L1;refGene_tx=XM_023244091.1
E1 58807539 58807729 refGene_gn=SEC14L1;refGene_tx=XM_023244091.1
E1 58809158 58809247 refGene_gn=SEC14L1;refGene_tx=XM_023244091.1
E1 58810337 58810408 refGene_gn=SEC14L1;refGene_tx=XM_023244091.1
E1 58811314 58811486 refGene_gn=SEC14L1;refGene_tx=XM_023244091.1
E1 58811819 58811954 refGene_gn=SEC14L1;refGene_tx=XM_023244091.1
E1 58813806 58813833 refGene_gn=SEC14L1;refGene_tx=XM_023244091.1
E1 58813834 58813942 refGene_gn=SEC14L1;refGene_tx=XM_023244091.1
E1 58816041 58816293 refGene_gn=SEC14L1;refGene_tx=XM_023244091.1
E1 58817547 58817726 refGene_gn=SEC14L1;refGene_tx=XM_023244091.1
E1 58817912 58818018 refGene_gn=SEC14L1;refGene_tx=XM_023244091.1
E1 58844239 58844360
refGene_gn=LOC111557724;refGene_tx=XM_023243478.1
E1 58844593 58845227
refGene_gn=LOC111557724;refGene_tx=XM_023243478.1
E1 58845267 58845403
refGene_gn=LOC111557724;refGene_tx=XM_023243478.1
E1 58846294 58846597
refGene_gn=LOC111557724;refGene_tx=XM_023243478.1
E1 58861584 58861603
refGene_gn=SEPT9;refGene_tx=XM_023244093.1;refGene_gn=SEPT9;refGene_tx=XM_023244095.1
E1 58878222 58878279 refGene_gn=SEPT9;refGene_tx=XM_023244093.1
E1 58887570 58887592 refGene_gn=SEPT9;refGene_tx=XM_011289408.3
E1 58921745 58921800 refGene_gn=SEPT9;refGene_tx=XM_023244094.1
E1 58944949 58945591
refGene_gn=SEPT9;refGene_tx=XM_011289408.3;refGene_gn=SEPT9;refGene_tx=XM_023244093.1;refGene_gn=SEPT9;refGene_tx=XM_023244094.1;refGene_gn=SEPT9;refGene_tx=XM_023244095.1
E1 59002129 59002321
refGene_gn=SEPT9;refGene_tx=XM_003997238.5;refGene_gn=SEPT9;refGene_tx=XM_011289408.3;refGene_gn=SEPT9;refGene_tx=XM_019818314.2;refGene_gn=SEPT9;refGene_tx=XM_019818315.2;refGene_gn=SEPT9;refGene_tx=XM_019818317.2;refGene_gn=SEPT9;refGene_tx=XM_023244093.1;refGene_gn=SEPT9;refGene_tx=XM_023244094.1;refGene_gn=SEPT9;refGene_tx=XM_023244095.1
E1 59004277 59004406
refGene_gn=SEPT9;refGene_tx=XM_003997238.5;refGene_gn=SEPT9;refGene_tx=XM_011289408.3;refGene_gn=SEPT9;refGene_tx=XM_019818314.2;refGene_gn=SEPT9;refGene_tx=XM_019818315.2;refGene_gn=SEPT9;refGene_tx=XM_019818317.2;refGene_gn=SEPT9;refGene_tx=XM_023244093.1;refGene_gn=SEPT9;refGene_tx=XM_023244094.1;refGene_gn=SEPT9;refGene_tx=XM_023244095.1
E1 59004790 59004872
refGene_gn=SEPT9;refGene_tx=XM_003997238.5;refGene_gn=SEPT9;refGene_tx=XM_011289408.3;refGene_gn=SEPT9;refGene_tx=XM_019818314.2;refGene_gn=SEPT9;refGene_tx=XM_019818315.2;refGene_gn=SEPT9;refGene_tx=XM_0198183

17.2;refGene_gn=SEPT9;refGene_tx=XM_023244093.1;refGene_gn=SEPT9;refGene_tx=XM_023244094.1;refGene_gn=SEPT9;refGene_tx=XM_023244095.1
E1 59005190 59005328
refGene_gn=SEPT9;refGene_tx=XM_003997238.5;refGene_gn=SEPT9;refGene_tx=XM_011289408.3;refGene_gn=SEPT9;refGene_tx=XM_019818314.2;refGene_gn=SEPT9;refGene_tx=XM_019818315.2;refGene_gn=SEPT9;refGene_tx=XM_019818317.2;refGene_gn=SEPT9;refGene_tx=XM_023244093.1;refGene_gn=SEPT9;refGene_tx=XM_023244094.1;refGene_gn=SEPT9;refGene_tx=XM_023244095.1
E1 59005959 59006077
refGene_gn=SEPT9;refGene_tx=XM_003997238.5;refGene_gn=SEPT9;refGene_tx=XM_011289408.3;refGene_gn=SEPT9;refGene_tx=XM_019818314.2;refGene_gn=SEPT9;refGene_tx=XM_019818315.2;refGene_gn=SEPT9;refGene_tx=XM_019818317.2;refGene_gn=SEPT9;refGene_tx=XM_023244093.1;refGene_gn=SEPT9;refGene_tx=XM_023244094.1;refGene_gn=SEPT9;refGene_tx=XM_023244095.1
E1 59006930 59007026
refGene_gn=SEPT9;refGene_tx=XM_003997238.5;refGene_gn=SEPT9;refGene_tx=XM_011289408.3;refGene_gn=SEPT9;refGene_tx=XM_019818314.2;refGene_gn=SEPT9;refGene_tx=XM_019818315.2;refGene_gn=SEPT9;refGene_tx=XM_019818317.2;refGene_gn=SEPT9;refGene_tx=XM_023244093.1;refGene_gn=SEPT9;refGene_tx=XM_023244094.1;refGene_gn=SEPT9;refGene_tx=XM_023244095.1
E1 59007308 59007405
refGene_gn=SEPT9;refGene_tx=XM_003997238.5;refGene_gn=SEPT9;refGene_tx=XM_011289408.3;refGene_gn=SEPT9;refGene_tx=XM_019818314.2;refGene_gn=SEPT9;refGene_tx=XM_019818315.2;refGene_gn=SEPT9;refGene_tx=XM_019818317.2;refGene_gn=SEPT9;refGene_tx=XM_023244093.1;refGene_gn=SEPT9;refGene_tx=XM_023244094.1;refGene_gn=SEPT9;refGene_tx=XM_023244095.1
E1 59010390 59010442
refGene_gn=SEPT9;refGene_tx=XM_003997238.5;refGene_gn=SEPT9;refGene_tx=XM_011289408.3;refGene_gn=SEPT9;refGene_tx=XM_019818314.2;refGene_gn=SEPT9;refGene_tx=XM_019818315.2;refGene_gn=SEPT9;refGene_tx=XM_019818317.2;refGene_gn=SEPT9;refGene_tx=XM_023244093.1;refGene_gn=SEPT9;refGene_tx=XM_023244094.1;refGene_gn=SEPT9;refGene_tx=XM_023244095.1
E1 59011478 59011608
refGene_gn=SEPT9;refGene_tx=XM_003997238.5;refGene_gn=SEPT9;refGene_tx=XM_011289408.3;refGene_gn=SEPT9;refGene_tx=XM_019818314.2;refGene_gn=SEPT9;refGene_tx=XM_019818315.2;refGene_gn=SEPT9;refGene_tx=XM_019818317.2;refGene_gn=SEPT9;refGene_tx=XM_023244093.1;refGene_gn=SEPT9;refGene_tx=XM_023244094.1;refGene_gn=SEPT9;refGene_tx=XM_023244095.1
E1 59029403 59029761
refGene_gn=LOC109494474;refGene_tx=XM_023244101.1
E1 59030461 59030577
refGene_gn=LOC109494474;refGene_tx=XM_023244101.1
E1 59030911 59031074
refGene_gn=LOC109494474;refGene_tx=XM_023244101.1
E1 59031655 59031759
refGene_gn=LOC109494474;refGene_tx=XM_023244101.1
E1 59067792 59068095
refGene_gn=LOC109494475;refGene_tx=XM_019818319.1;refGene_gn=LOC109494475;refGene_tx=XM_019818324.2
E1 59069523 59069586

refGene_gn=LOC109494475;refGene_tx=XM_019818319.1;refGene_gn=LOC109494475;refGene_tx=XM_019818324.2
E1 59070420 59070435
refGene_gn=LOC109494475;refGene_tx=XM_019818324.2
E1 59074730 59075009
refGene_gn=LOC109494475;refGene_tx=XM_019818319.1
E1 59394297 59394373 refGene_gn=TNRC6C;refGene_tx=XM_023244125.1
E1 59405591 59405918 refGene_gn=TNRC6C;refGene_tx=XM_023244125.1
E1 59426977 59429587
refGene_gn=TNRC6C;refGene_tx=XM_019818326.2;refGene_gn=TNRC6C;refGene_tx=XM_023244123.1;refGene_gn=TNRC6C;refGene_tx=XM_023244124.1;refGene_gn=TNRC6C;refGene_tx=XM_023244125.1
E1 59441099 59441315
refGene_gn=TNRC6C;refGene_tx=XM_019818326.2;refGene_gn=TNRC6C;refGene_tx=XM_023244123.1;refGene_gn=TNRC6C;refGene_tx=XM_023244124.1;refGene_gn=TNRC6C;refGene_tx=XM_023244125.1
E1 59443031 59443198
refGene_gn=TNRC6C;refGene_tx=XM_019818326.2;refGene_gn=TNRC6C;refGene_tx=XM_023244123.1;refGene_gn=TNRC6C;refGene_tx=XM_023244124.1;refGene_gn=TNRC6C;refGene_tx=XM_023244125.1
E1 59446713 59446794
refGene_gn=TNRC6C;refGene_tx=XM_019818326.2;refGene_gn=TNRC6C;refGene_tx=XM_023244123.1;refGene_gn=TNRC6C;refGene_tx=XM_023244124.1;refGene_gn=TNRC6C;refGene_tx=XM_023244125.1
E1 59448100 59448158
refGene_gn=TNRC6C;refGene_tx=XM_019818326.2;refGene_gn=TNRC6C;refGene_tx=XM_023244123.1;refGene_gn=TNRC6C;refGene_tx=XM_023244124.1;refGene_gn=TNRC6C;refGene_tx=XM_023244125.1
E1 59455792 59455935
refGene_gn=TNRC6C;refGene_tx=XM_019818326.2;refGene_gn=TNRC6C;refGene_tx=XM_023244123.1;refGene_gn=TNRC6C;refGene_tx=XM_023244124.1;refGene_gn=TNRC6C;refGene_tx=XM_023244125.1
E1 59457354 59457504
refGene_gn=TNRC6C;refGene_tx=XM_019818326.2;refGene_gn=TNRC6C;refGene_tx=XM_023244123.1;refGene_gn=TNRC6C;refGene_tx=XM_023244124.1;refGene_gn=TNRC6C;refGene_tx=XM_023244125.1
E1 59458922 59459069
refGene_gn=TNRC6C;refGene_tx=XM_019818326.2;refGene_gn=TNRC6C;refGene_tx=XM_023244123.1;refGene_gn=TNRC6C;refGene_tx=XM_023244124.1;refGene_gn=TNRC6C;refGene_tx=XM_023244125.1
E1 59460632 59460752
refGene_gn=TNRC6C;refGene_tx=XM_019818326.2;refGene_gn=TNRC6C;refGene_tx=XM_023244123.1;refGene_gn=TNRC6C;refGene_tx=XM_023244124.1;refGene_gn=TNRC6C;refGene_tx=XM_023244125.1
E1 59463908 59463992
refGene_gn=TNRC6C;refGene_tx=XM_019818326.2;refGene_gn=TNRC6C;refGene_tx=XM_023244123.1;refGene_gn=TNRC6C;refGene_tx=XM_023244124.1;refGene_gn=TNRC6C;refGene_tx=XM_023244125.1
E1 59464224 59464456
refGene_gn=TNRC6C;refGene_tx=XM_019818326.2;refGene_gn=TNRC6C;refGene_

tx=XM_023244123.1;refGene_gn=TNRC6C;refGene_tx=XM_023244124.1;refGene_gn=TNRC6C;refGene_tx=XM_023244125.1
E1 59467625 59467793 refGene_gn=TNRC6C;refGene_tx=XM_019818326.2;refGene_gn=TNRC6C;refGene_tx=XM_023244123.1;refGene_gn=TNRC6C;refGene_tx=XM_023244124.1;refGene_gn=TNRC6C;refGene_tx=XM_023244125.1
E1 59469373 59469565 refGene_gn=TNRC6C;refGene_tx=XM_019818326.2;refGene_gn=TNRC6C;refGene_tx=XM_023244123.1;refGene_gn=TNRC6C;refGene_tx=XM_023244124.1;refGene_gn=TNRC6C;refGene_tx=XM_023244125.1
E1 59470021 59470165 refGene_gn=TNRC6C;refGene_tx=XM_019818326.2;refGene_gn=TNRC6C;refGene_tx=XM_023244123.1;refGene_gn=TNRC6C;refGene_tx=XM_023244124.1;refGene_gn=TNRC6C;refGene_tx=XM_023244125.1
E1 59472362 59472479 refGene_gn=TNRC6C;refGene_tx=XM_019818326.2;refGene_gn=TNRC6C;refGene_tx=XM_023244123.1;refGene_gn=TNRC6C;refGene_tx=XM_023244124.1;refGene_gn=TNRC6C;refGene_tx=XM_023244125.1
E1 59474416 59474611 refGene_gn=TNRC6C;refGene_tx=XM_019818326.2;refGene_gn=TNRC6C;refGene_tx=XM_023244123.1;refGene_gn=TNRC6C;refGene_tx=XM_023244124.1;refGene_gn=TNRC6C;refGene_tx=XM_023244125.1
E1 59475383 59475454 refGene_gn=TNRC6C;refGene_tx=XM_019818326.2;refGene_gn=TNRC6C;refGene_tx=XM_023244123.1;refGene_gn=TNRC6C;refGene_tx=XM_023244124.1;refGene_gn=TNRC6C;refGene_tx=XM_023244125.1
E1 59475930 59476070 refGene_gn=TNRC6C;refGene_tx=XM_019818326.2;refGene_gn=TNRC6C;refGene_tx=XM_023244123.1;refGene_gn=TNRC6C;refGene_tx=XM_023244124.1;refGene_gn=TNRC6C;refGene_tx=XM_023244125.1
E1 59480550 59480911 refGene_gn=TNRC6C;refGene_tx=XM_019818326.2;refGene_gn=TNRC6C;refGene_tx=XM_023244123.1;refGene_gn=TNRC6C;refGene_tx=XM_023244124.1;refGene_gn=TNRC6C;refGene_tx=XM_023244125.1
E1 59484529 59484713 refGene_gn=TMC6;refGene_tx=XM_023243479.1
E1 59488459 59488527 refGene_gn=TMC6;refGene_tx=XM_023243479.1
E1 59490205 59490284 refGene_gn=TMC6;refGene_tx=XM_023243479.1
E1 59490416 59490593 refGene_gn=TMC6;refGene_tx=XM_023243479.1
E1 59490695 59490829 refGene_gn=TMC6;refGene_tx=XM_023243479.1
E1 59491611 59491687 refGene_gn=TMC6;refGene_tx=XM_023243479.1
E1 59491952 59492048 refGene_gn=TMC6;refGene_tx=XM_023243479.1
E1 59492855 59492953 refGene_gn=TMC6;refGene_tx=XM_023243479.1
E1 59493177 59493259 refGene_gn=TMC6;refGene_tx=XM_023243479.1
E1 59493373 59493525 refGene_gn=TMC6;refGene_tx=XM_023243479.1
E1 59493836 59493992 refGene_gn=TMC6;refGene_tx=XM_023243479.1
E1 59494797 59495058 refGene_gn=TMC6;refGene_tx=XM_023243479.1
E1 59496675 59496811 refGene_gn=TMC6;refGene_tx=XM_023243479.1
E1 59496964 59497070 refGene_gn=TMC6;refGene_tx=XM_023243479.1
E1 59498283 59498442 refGene_gn=TMC6;refGene_tx=XM_023243479.1
E1 59498689 59498785 refGene_gn=TMC6;refGene_tx=XM_023243479.1

E1 59498951 59499091 refGene_gn=TMC6;refGene_tx=XM_023243479.1
E1 59499268 59499324 refGene_gn=TMC6;refGene_tx=XM_023243479.1
E1 59504060 59504221
refGene_gn=TMC8;refGene_tx=XM_006940636.4;refGene_gn=TMC8;refGene_tx=XM_006940637.4
E1 59504413 59504562
refGene_gn=TMC8;refGene_tx=XM_006940636.4;refGene_gn=TMC8;refGene_tx=XM_006940637.4
E1 59504922 59505072
refGene_gn=TMC8;refGene_tx=XM_006940636.4;refGene_gn=TMC8;refGene_tx=XM_006940637.4
E1 59505364 59505447
refGene_gn=TMC8;refGene_tx=XM_006940636.4;refGene_gn=TMC8;refGene_tx=XM_006940637.4
E1 59505952 59506089
refGene_gn=TMC8;refGene_tx=XM_006940636.4;refGene_gn=TMC8;refGene_tx=XM_006940637.4
E1 59506407 59506555
refGene_gn=TMC8;refGene_tx=XM_006940636.4;refGene_gn=TMC8;refGene_tx=XM_006940637.4
E1 59506911 59507082
refGene_gn=TMC8;refGene_tx=XM_006940636.4;refGene_gn=TMC8;refGene_tx=XM_006940637.4
E1 59507376 59507516
refGene_gn=TMC8;refGene_tx=XM_006940636.4;refGene_gn=TMC8;refGene_tx=XM_006940637.4
E1 59508902 59509026
refGene_gn=TMC8;refGene_tx=XM_006940636.4;refGene_gn=TMC8;refGene_tx=XM_006940637.4
E1 59509325 59509423
refGene_gn=TMC8;refGene_tx=XM_006940636.4;refGene_gn=TMC8;refGene_tx=XM_006940637.4
E1 59509607 59509791
refGene_gn=TMC8;refGene_tx=XM_006940636.4;refGene_gn=TMC8;refGene_tx=XM_006940637.4
E1 59509961 59510092
refGene_gn=TMC8;refGene_tx=XM_006940636.4;refGene_gn=TMC8;refGene_tx=XM_006940637.4
E1 59510186 59510345
refGene_gn=TMC8;refGene_tx=XM_006940636.4;refGene_gn=TMC8;refGene_tx=XM_006940637.4
E1 59510772 59510851
refGene_gn=TMC8;refGene_tx=XM_006940636.4;refGene_gn=TMC8;refGene_tx=XM_006940637.4
E1 59511234 59511492
refGene_gn=TMC8;refGene_tx=XM_006940636.4;refGene_gn=TMC8;refGene_tx=XM_006940637.4
E1 59524989 59525088
refGene_gn=SYNGR2;refGene_tx=XM_023244126.1;refGene_gn=SYNGR2;refGene_tx=XM_023244127.1

E1 59526896 59527134
refGene_gn=SYNGR2;refGene_tx=XM_023244126.1;refGene_gn=SYNGR2;refGene_tx=XM_023244127.1
E1 59527513 59527653 refGene_gn=SYNGR2;refGene_tx=XM_023244126.1
E1 59527741 59527939
refGene_gn=SYNGR2;refGene_tx=XM_023244126.1;refGene_gn=SYNGR2;refGene_tx=XM_023244127.1
E1 59530617 59530818 refGene_gn=TK1;refGene_tx=XM_003997237.5
E1 59530917 59531037 refGene_gn=TK1;refGene_tx=XM_003997237.5
E1 59531416 59531506 refGene_gn=TK1;refGene_tx=XM_003997237.5
E1 59536517 59536611 refGene_gn=TK1;refGene_tx=XM_003997237.5
E1 59539011 59539122 refGene_gn=TK1;refGene_tx=XM_003997237.5
E1 59540448 59540480 refGene_gn=TK1;refGene_tx=XM_003997237.5
E1 59540584 59540650 refGene_gn=TK1;refGene_tx=XM_003997237.5
E1 59540956 59541016
refGene_gn=AFMID;refGene_tx=XM_006940639.4;refGene_gn=AFMID;refGene_tx=XM_019818330.2;refGene_gn=AFMID;refGene_tx=XM_019818331.2
E1 59542635 59542726
refGene_gn=AFMID;refGene_tx=XM_006940639.4;refGene_gn=AFMID;refGene_tx=XM_019818330.2;refGene_gn=AFMID;refGene_tx=XM_019818333.2
E1 59553252 59553361
refGene_gn=AFMID;refGene_tx=XM_006940639.4;refGene_gn=AFMID;refGene_tx=XM_019818330.2;refGene_gn=AFMID;refGene_tx=XM_019818331.2;refGene_gn=AFMID;refGene_tx=XM_019818333.2
E1 59553442 59553491
refGene_gn=AFMID;refGene_tx=XM_006940639.4;refGene_gn=AFMID;refGene_tx=XM_019818330.2;refGene_gn=AFMID;refGene_tx=XM_019818331.2;refGene_gn=AFMID;refGene_tx=XM_019818333.2
E1 59553911 59553997
refGene_gn=AFMID;refGene_tx=XM_006940639.4;refGene_gn=AFMID;refGene_tx=XM_019818330.2;refGene_gn=AFMID;refGene_tx=XM_019818331.2;refGene_gn=AFMID;refGene_tx=XM_019818333.2
E1 59554092 59554165
refGene_gn=AFMID;refGene_tx=XM_006940639.4;refGene_gn=AFMID;refGene_tx=XM_019818330.2;refGene_gn=AFMID;refGene_tx=XM_019818331.2;refGene_gn=AFMID;refGene_tx=XM_019818333.2
E1 59554356 59554454
refGene_gn=AFMID;refGene_tx=XM_006940639.4;refGene_gn=AFMID;refGene_tx=XM_019818330.2;refGene_gn=AFMID;refGene_tx=XM_019818331.2;refGene_gn=AFMID;refGene_tx=XM_019818333.2
E1 59554738 59554817
refGene_gn=AFMID;refGene_tx=XM_006940639.4;refGene_gn=AFMID;refGene_tx=XM_019818330.2;refGene_gn=AFMID;refGene_tx=XM_019818331.2;refGene_gn=AFMID;refGene_tx=XM_019818333.2
E1 59554936 59555078
refGene_gn=AFMID;refGene_tx=XM_006940639.4;refGene_gn=AFMID;refGene_tx=XM_019818330.2;refGene_gn=AFMID;refGene_tx=XM_019818331.2;refGene_gn=AFMID;refGene_tx=XM_019818333.2
E1 59555267 59555417
refGene_gn=AFMID;refGene_tx=XM_006940639.4;refGene_gn=AFMID;refGene_tx

=XM_019818330.2;refGene_gn=AFMID;refGene_tx=XM_019818331.2;refGene_gn=AFMID;refGene_tx=XM_019818333.2
E1 59557684 59557717 refGene_gn=AFMID;refGene_tx=XM_006940639.4
E1 59562228 59562339
refGene_gn=BIRC5;refGene_tx=NM_001009280.1;refGene_gn=BIRC5;refGene_tx=XM_019816885.2
E1 59562826 59562936
refGene_gn=BIRC5;refGene_tx=NM_001009280.1;refGene_gn=BIRC5;refGene_tx=XM_019816885.2
E1 59565411 59565529
refGene_gn=BIRC5;refGene_tx=NM_001009280.1;refGene_gn=BIRC5;refGene_tx=XM_019816885.2
E1 59569936 59569966 refGene_gn=BIRC5;refGene_tx=XM_019816885.2
E1 59570406 59570496 refGene_gn=BIRC5;refGene_tx=NM_001009280.1
E1 59577143 59577270 refGene_gn=TMEM235;refGene_tx=XM_023244514.1
E1 59577272 59577296 refGene_gn=TMEM235;refGene_tx=XM_023244514.1
E1 59578629 59578710 refGene_gn=TMEM235;refGene_tx=XM_023244514.1
E1 59578877 59579015 refGene_gn=TMEM235;refGene_tx=XM_023244514.1
E1 59581897 59581981 refGene_gn=TMEM235;refGene_tx=XM_023244514.1
E1 59581983 59582129 refGene_gn=TMEM235;refGene_tx=XM_023244514.1
E1 59584912 59585208
refGene_gn=LOC101081589;refGene_tx=XM_023244128.1
E1 59585765 59585958
refGene_gn=LOC101081589;refGene_tx=XM_023244128.1
E1 59586205 59586343
refGene_gn=LOC101081589;refGene_tx=XM_023244128.1
E1 59587027 59587213
refGene_gn=LOC101081589;refGene_tx=XM_023244128.1
E1 59587631 59587720
refGene_gn=LOC101081589;refGene_tx=XM_023244128.1
E1 59588268 59588352
refGene_gn=LOC101081589;refGene_tx=XM_023244128.1
E1 59589860 59590062
refGene_gn=LOC101081589;refGene_tx=XM_023244128.1
E1 59662083 59662761 refGene_gn=SOCS3;refGene_tx=XM_023244150.1
E1 59676123 59676266
refGene_gn=PGS1;refGene_tx=XM_023244152.1;refGene_gn=PGS1;refGene_tx=XM_023244153.1
E1 59684920 59685110
refGene_gn=PGS1;refGene_tx=XM_023244152.1;refGene_gn=PGS1;refGene_tx=XM_023244153.1
E1 59687744 59687822
refGene_gn=PGS1;refGene_tx=XM_023244152.1;refGene_gn=PGS1;refGene_tx=XM_023244153.1
E1 59689326 59689426
refGene_gn=PGS1;refGene_tx=XM_023244152.1;refGene_gn=PGS1;refGene_tx=XM_023244153.1
E1 59690179 59690369
refGene_gn=PGS1;refGene_tx=XM_023244152.1;refGene_gn=PGS1;refGene_tx=XM_023244153.1

E1 59691011 59691190
refGene_gn=PGS1;refGene_tx=XM_023244152.1;refGene_gn=PGS1;refGene_tx=X
M_023244153.1
E1 59693081 59693603
refGene_gn=PGS1;refGene_tx=XM_023244152.1;refGene_gn=PGS1;refGene_tx=X
M_023244153.1
E1 59701731 59701880
refGene_gn=PGS1;refGene_tx=XM_023244152.1;refGene_gn=PGS1;refGene_tx=X
M_023244153.1
E1 59706501 59706621 refGene_gn=PGS1;refGene_tx=XM_023244152.1
E1 59710506 59710754 refGene_gn=DNAH17;refGene_tx=XM_023244151.1
E1 59711750 59711976 refGene_gn=DNAH17;refGene_tx=XM_023244151.1
E1 59712459 59712603 refGene_gn=DNAH17;refGene_tx=XM_023244151.1
E1 59712929 59713112 refGene_gn=DNAH17;refGene_tx=XM_023244151.1
E1 59713998 59714181 refGene_gn=DNAH17;refGene_tx=XM_023244151.1
E1 59714510 59714690 refGene_gn=DNAH17;refGene_tx=XM_023244151.1
E1 59717515 59717707 refGene_gn=DNAH17;refGene_tx=XM_023244151.1
E1 59718773 59719001 refGene_gn=DNAH17;refGene_tx=XM_023244151.1
E1 59719698 59719826 refGene_gn=DNAH17;refGene_tx=XM_023244151.1
E1 59720568 59720717 refGene_gn=DNAH17;refGene_tx=XM_023244151.1
E1 59723532 59723726 refGene_gn=DNAH17;refGene_tx=XM_023244151.1
E1 59724249 59724372 refGene_gn=DNAH17;refGene_tx=XM_023244151.1
E1 59725893 59726064 refGene_gn=DNAH17;refGene_tx=XM_023244151.1
E1 59726323 59726464 refGene_gn=DNAH17;refGene_tx=XM_023244151.1
E1 59726667 59726832 refGene_gn=DNAH17;refGene_tx=XM_023244151.1
E1 59727287 59727353 refGene_gn=DNAH17;refGene_tx=XM_023244151.1
E1 59727354 59727493 refGene_gn=DNAH17;refGene_tx=XM_023244151.1
E1 59729213 59729336 refGene_gn=DNAH17;refGene_tx=XM_023244151.1
E1 59729997 59730233 refGene_gn=DNAH17;refGene_tx=XM_023244151.1
E1 59730744 59730937 refGene_gn=DNAH17;refGene_tx=XM_023244151.1
E1 59732384 59732500 refGene_gn=DNAH17;refGene_tx=XM_023244151.1
E1 59732795 59733003 refGene_gn=DNAH17;refGene_tx=XM_023244151.1
E1 59734289 59734507 refGene_gn=DNAH17;refGene_tx=XM_023244151.1
E1 59734661 59734757 refGene_gn=DNAH17;refGene_tx=XM_023244151.1
E1 59736219 59736384 refGene_gn=DNAH17;refGene_tx=XM_023244151.1
E1 59736754 59736880 refGene_gn=DNAH17;refGene_tx=XM_023244151.1
E1 59736881 59736989 refGene_gn=DNAH17;refGene_tx=XM_023244151.1
E1 59738435 59738594 refGene_gn=DNAH17;refGene_tx=XM_023244151.1
E1 59739598 59739865 refGene_gn=DNAH17;refGene_tx=XM_023244151.1
E1 59744313 59744505 refGene_gn=DNAH17;refGene_tx=XM_023244151.1
E1 59744727 59744892 refGene_gn=DNAH17;refGene_tx=XM_023244151.1
E1 59745384 59745546 refGene_gn=DNAH17;refGene_tx=XM_023244151.1
E1 59746176 59746268 refGene_gn=DNAH17;refGene_tx=XM_023244151.1
E1 59746654 59746802 refGene_gn=DNAH17;refGene_tx=XM_023244151.1
E1 59747866 59747969 refGene_gn=DNAH17;refGene_tx=XM_023244151.1
E1 59748749 59748915 refGene_gn=DNAH17;refGene_tx=XM_023244151.1
E1 59751071 59751279 refGene_gn=DNAH17;refGene_tx=XM_023244151.1
E1 59751486 59751660 refGene_gn=DNAH17;refGene_tx=XM_023244151.1
E1 59751759 59752042 refGene_gn=DNAH17;refGene_tx=XM_023244151.1
E1 59754371 59754520 refGene_gn=DNAH17;refGene_tx=XM_023244151.1

E1 59755130 59755258 refGene_gn=DNAH17;refGene_tx=XM_023244151.1
E1 59756004 59756137 refGene_gn=DNAH17;refGene_tx=XM_023244151.1
E1 59757298 59757436 refGene_gn=DNAH17;refGene_tx=XM_023244151.1
E1 59757809 59757971 refGene_gn=DNAH17;refGene_tx=XM_023244151.1
E1 59757972 59758038 refGene_gn=DNAH17;refGene_tx=XM_023244151.1
E1 59758162 59758301 refGene_gn=DNAH17;refGene_tx=XM_023244151.1
E1 59758730 59758888 refGene_gn=DNAH17;refGene_tx=XM_023244151.1
E1 59761074 59761179 refGene_gn=DNAH17;refGene_tx=XM_023244151.1
E1 59762267 59762424 refGene_gn=DNAH17;refGene_tx=XM_023244151.1
E1 59763055 59763216 refGene_gn=DNAH17;refGene_tx=XM_023244151.1
E1 59763500 59763632 refGene_gn=DNAH17;refGene_tx=XM_023244151.1
E1 59764830 59764938 refGene_gn=DNAH17;refGene_tx=XM_023244151.1
E1 59765084 59765210 refGene_gn=DNAH17;refGene_tx=XM_023244151.1
E1 59766437 59766590 refGene_gn=DNAH17;refGene_tx=XM_023244151.1
E1 59767222 59767349 refGene_gn=DNAH17;refGene_tx=XM_023244151.1
E1 59767685 59767777 refGene_gn=DNAH17;refGene_tx=XM_023244151.1
E1 59767883 59768231 refGene_gn=DNAH17;refGene_tx=XM_023244151.1
E1 59769500 59769623 refGene_gn=DNAH17;refGene_tx=XM_023244151.1
E1 59771451 59771700 refGene_gn=DNAH17;refGene_tx=XM_023244151.1
E1 59773575 59773728 refGene_gn=DNAH17;refGene_tx=XM_023244151.1
E1 59774750 59774837 refGene_gn=DNAH17;refGene_tx=XM_023244151.1
E1 59774917 59775034 refGene_gn=DNAH17;refGene_tx=XM_023244151.1
E1 59776181 59776404 refGene_gn=DNAH17;refGene_tx=XM_023244151.1
E1 59776982 59777152 refGene_gn=DNAH17;refGene_tx=XM_023244151.1
E1 59778162 59778417 refGene_gn=DNAH17;refGene_tx=XM_023244151.1
E1 59781371 59781554 refGene_gn=DNAH17;refGene_tx=XM_023244151.1
E1 59784217 59784361 refGene_gn=DNAH17;refGene_tx=XM_023244151.1
E1 59786924 59787065 refGene_gn=DNAH17;refGene_tx=XM_023244151.1
E1 59790557 59790661 refGene_gn=DNAH17;refGene_tx=XM_023244151.1
E1 59791628 59791737 refGene_gn=DNAH17;refGene_tx=XM_023244151.1
E1 59796901 59797048 refGene_gn=DNAH17;refGene_tx=XM_023244151.1
E1 59798922 59799118 refGene_gn=DNAH17;refGene_tx=XM_023244151.1
E1 59799846 59800112 refGene_gn=DNAH17;refGene_tx=XM_023244151.1
E1 59804510 59804627 refGene_gn=DNAH17;refGene_tx=XM_023244151.1
E1 59804849 59805017 refGene_gn=DNAH17;refGene_tx=XM_023244151.1
E1 59806638 59806725 refGene_gn=DNAH17;refGene_tx=XM_023244151.1
E1 59806836 59806989 refGene_gn=DNAH17;refGene_tx=XM_023244151.1
E1 59807651 59807777 refGene_gn=DNAH17;refGene_tx=XM_023244151.1
E1 59807957 59808043 refGene_gn=DNAH17;refGene_tx=XM_023244151.1
E1 59808182 59808282 refGene_gn=DNAH17;refGene_tx=XM_023244151.1
E1 59808491 59808684 refGene_gn=DNAH17;refGene_tx=XM_023244151.1
E1 59809565 59809759 refGene_gn=DNAH17;refGene_tx=XM_023244151.1
E1 59810828 59811173 refGene_gn=DNAH17;refGene_tx=XM_023244151.1
E1 59846915 59846994
refGene_gn=CYTH1;refGene_tx=XM_019818339.2;refGene_gn=CYTH1;refGene_tx
=XM_023244156.1;refGene_gn=CYTH1;refGene_tx=XM_023244157.1
E1 59850468 59850623
refGene_gn=CYTH1;refGene_tx=XM_019818339.2;refGene_gn=CYTH1;refGene_tx
=XM_023244156.1;refGene_gn=CYTH1;refGene_tx=XM_023244157.1
E1 59851263 59851335

refGene_gn=CYTH1;refGene_tx=XM_019818339.2;refGene_gn=CYTH1;refGene_tx
=XM_023244156.1;refGene_gn=CYTH1;refGene_tx=XM_023244157.1
E1 59854728 59854807
refGene_gn=CYTH1;refGene_tx=XM_019818339.2;refGene_gn=CYTH1;refGene_tx
=XM_023244156.1;refGene_gn=CYTH1;refGene_tx=XM_023244157.1
E1 59860019 59860132
refGene_gn=CYTH1;refGene_tx=XM_019818339.2;refGene_gn=CYTH1;refGene_tx
=XM_023244156.1;refGene_gn=CYTH1;refGene_tx=XM_023244157.1
E1 59860510 59860659
refGene_gn=CYTH1;refGene_tx=XM_019818339.2;refGene_gn=CYTH1;refGene_tx
=XM_023244156.1;refGene_gn=CYTH1;refGene_tx=XM_023244157.1
E1 59861580 59861693
refGene_gn=CYTH1;refGene_tx=XM_019818339.2;refGene_gn=CYTH1;refGene_tx
=XM_023244156.1;refGene_gn=CYTH1;refGene_tx=XM_023244157.1
E1 59862100 59862181
refGene_gn=CYTH1;refGene_tx=XM_019818339.2;refGene_gn=CYTH1;refGene_tx
=XM_023244156.1;refGene_gn=CYTH1;refGene_tx=XM_023244157.1
E1 59862500 59862619
refGene_gn=CYTH1;refGene_tx=XM_019818339.2;refGene_gn=CYTH1;refGene_tx
=XM_023244156.1;refGene_gn=CYTH1;refGene_tx=XM_023244157.1
E1 59862911 59862978
refGene_gn=CYTH1;refGene_tx=XM_019818339.2;refGene_gn=CYTH1;refGene_tx
=XM_023244156.1;refGene_gn=CYTH1;refGene_tx=XM_023244157.1
E1 59864526 59864591
refGene_gn=CYTH1;refGene_tx=XM_023244156.1;refGene_gn=CYTH1;refGene_tx
=XM_023244157.1
E1 59866103 59866186
refGene_gn=CYTH1;refGene_tx=XM_023244156.1;refGene_gn=CYTH1;refGene_tx
=XM_023244157.1
E1 59885712 59885740 refGene_gn=CYTH1;refGene_tx=XM_023244156.1
E1 59925025 59925047 refGene_gn=CYTH1;refGene_tx=XM_023244157.1
E1 59938234 59938366
refGene_gn=USP36;refGene_tx=XM_019818340.2;refGene_gn=USP36;refGene_tx
=XM_019818341.2;refGene_gn=USP36;refGene_tx=XM_023244158.1
E1 59938772 59938888
refGene_gn=USP36;refGene_tx=XM_019818340.2;refGene_gn=USP36;refGene_tx
=XM_019818341.2;refGene_gn=USP36;refGene_tx=XM_023244158.1
E1 59939347 59939449
refGene_gn=USP36;refGene_tx=XM_019818340.2;refGene_gn=USP36;refGene_tx
=XM_019818341.2;refGene_gn=USP36;refGene_tx=XM_023244158.1
E1 59941032 59941244
refGene_gn=USP36;refGene_tx=XM_019818340.2;refGene_gn=USP36;refGene_tx
=XM_019818341.2;refGene_gn=USP36;refGene_tx=XM_023244158.1
E1 59941691 59942249
refGene_gn=USP36;refGene_tx=XM_019818340.2;refGene_gn=USP36;refGene_tx
=XM_019818341.2;refGene_gn=USP36;refGene_tx=XM_023244158.1
E1 59944095 59944223
refGene_gn=USP36;refGene_tx=XM_019818340.2;refGene_gn=USP36;refGene_tx
=XM_019818341.2;refGene_gn=USP36;refGene_tx=XM_023244158.1
E1 59944844 59945504

refGene_gn=USP36;refGene_tx=XM_019818340.2;refGene_gn=USP36;refGene_tx=XM_019818341.2;refGene_gn=USP36;refGene_tx=XM_023244158.1
E1 59948070 59948209 refGene_gn=USP36;refGene_tx=XM_019818340.2;refGene_gn=USP36;refGene_tx=XM_019818341.2;refGene_gn=USP36;refGene_tx=XM_023244158.1
E1 59948808 59948909 refGene_gn=USP36;refGene_tx=XM_019818340.2;refGene_gn=USP36;refGene_tx=XM_019818341.2;refGene_gn=USP36;refGene_tx=XM_023244158.1
E1 59949852 59949993 refGene_gn=USP36;refGene_tx=XM_019818340.2;refGene_gn=USP36;refGene_tx=XM_019818341.2;refGene_gn=USP36;refGene_tx=XM_023244158.1
E1 59953938 59954050 refGene_gn=USP36;refGene_tx=XM_019818340.2;refGene_gn=USP36;refGene_tx=XM_019818341.2;refGene_gn=USP36;refGene_tx=XM_023244158.1
E1 59955058 59955141 refGene_gn=USP36;refGene_tx=XM_019818340.2;refGene_gn=USP36;refGene_tx=XM_019818341.2;refGene_gn=USP36;refGene_tx=XM_023244158.1
E1 59955779 59955850 refGene_gn=USP36;refGene_tx=XM_019818340.2;refGene_gn=USP36;refGene_tx=XM_019818341.2;refGene_gn=USP36;refGene_tx=XM_023244158.1
E1 59956306 59956374 refGene_gn=USP36;refGene_tx=XM_019818340.2;refGene_gn=USP36;refGene_tx=XM_019818341.2;refGene_gn=USP36;refGene_tx=XM_023244158.1
E1 59960335 59960438 refGene_gn=USP36;refGene_tx=XM_019818340.2;refGene_gn=USP36;refGene_tx=XM_019818341.2;refGene_gn=USP36;refGene_tx=XM_023244158.1
E1 59961014 59961125 refGene_gn=USP36;refGene_tx=XM_019818340.2;refGene_gn=USP36;refGene_tx=XM_019818341.2;refGene_gn=USP36;refGene_tx=XM_023244158.1
E1 59967916 59968138 refGene_gn=USP36;refGene_tx=XM_019818340.2;refGene_gn=USP36;refGene_tx=XM_019818341.2;refGene_gn=USP36;refGene_tx=XM_023244158.1
E1 59968673 59968926 refGene_gn=USP36;refGene_tx=XM_019818340.2;refGene_gn=USP36;refGene_tx=XM_019818341.2;refGene_gn=USP36;refGene_tx=XM_023244158.1
E1 59984694 59984750 refGene_gn=TIMP2;refGene_tx=XM_003997248.4
E1 59984752 59984891 refGene_gn=TIMP2;refGene_tx=XM_003997248.4
E1 59986196 59986321 refGene_gn=TIMP2;refGene_tx=XM_003997248.4
E1 59992342 59992451 refGene_gn=TIMP2;refGene_tx=XM_003997248.4
E1 59994396 59994497 refGene_gn=TIMP2;refGene_tx=XM_003997248.4
E1 60006332 60007349 refGene_gn=CEP295NL;refGene_tx=XM_011289417.3
E1 60030198 60030331 refGene_gn=TIMP2;refGene_tx=XM_003997248.4
E1 60059407 60060461 refGene_gn=LOC105261197;refGene_tx=XM_003997251.5;refGene_gn=LOC105261197;refGene_tx=XM_006940645.3;refGene_gn=LOC105261197;refGene_tx=XM_019818343.2
E1 60063029 60063288 refGene_gn=LOC105261197;refGene_tx=XM_003997251.5;refGene_gn=LOC105261197;refGene_tx=XM_006940645.3;refGene_gn=LOC105261197;refGene_tx=XM_019818343.2

E1 60064725 60064857
refGene_gn=LOC105261197;refGene_tx=XM_003997251.5;refGene_gn=LOC105261197;refGene_tx=XM_006940645.3;refGene_gn=LOC105261197;refGene_tx=XM_019818343.2
E1 60065432 60065624
refGene_gn=LOC105261197;refGene_tx=XM_003997251.5;refGene_gn=LOC105261197;refGene_tx=XM_006940645.3;refGene_gn=LOC105261197;refGene_tx=XM_019818343.2
E1 60066587 60066664
refGene_gn=LOC105261197;refGene_tx=XM_003997251.5;refGene_gn=LOC105261197;refGene_tx=XM_006940645.3;refGene_gn=LOC105261197;refGene_tx=XM_019818343.2
E1 60075914 60076260
refGene_gn=LOC105261197;refGene_tx=XM_019818343.2
E1 60077106 60077310
refGene_gn=LOC105261197;refGene_tx=XM_019818343.2
E1 60079218 60079849
refGene_gn=LOC105261197;refGene_tx=XM_019818343.2
E1 60113069 60113218
refGene_gn=C1QTNF1;refGene_tx=XM_011289419.3;refGene_gn=C1QTNF1;refGene_tx=XM_023244194.1;refGene_gn=C1QTNF1;refGene_tx=XM_023244195.1
E1 60116084 60116224
refGene_gn=C1QTNF1;refGene_tx=XM_011289419.3;refGene_gn=C1QTNF1;refGene_tx=XM_023244194.1;refGene_gn=C1QTNF1;refGene_tx=XM_023244195.1
E1 60117026 60117577
refGene_gn=C1QTNF1;refGene_tx=XM_011289419.3;refGene_gn=C1QTNF1;refGene_tx=XM_023244194.1;refGene_gn=C1QTNF1;refGene_tx=XM_023244195.1
E1 60135952 60136059
refGene_gn=ENGASE;refGene_tx=XM_019816973.2;refGene_gn=ENGASE;refGene_tx=XM_023244197.1;refGene_gn=ENGASE;refGene_tx=XM_023244198.1;refGene_gn=ENGASE;refGene_tx=XM_023244199.1
E1 60137595 60137663
refGene_gn=ENGASE;refGene_tx=XM_019816973.2;refGene_gn=ENGASE;refGene_tx=XM_023244197.1;refGene_gn=ENGASE;refGene_tx=XM_023244198.1;refGene_gn=ENGASE;refGene_tx=XM_023244199.1
E1 60137815 60138017
refGene_gn=ENGASE;refGene_tx=XM_019816973.2;refGene_gn=ENGASE;refGene_tx=XM_023244197.1;refGene_gn=ENGASE;refGene_tx=XM_023244198.1;refGene_gn=ENGASE;refGene_tx=XM_023244199.1;refGene_gn=ENGASE;refGene_tx=XM_023244200.1;refGene_gn=ENGASE;refGene_tx=XM_023244201.1
E1 60138780 60138929
refGene_gn=ENGASE;refGene_tx=XM_019816973.2;refGene_gn=ENGASE;refGene_tx=XM_023244197.1;refGene_gn=ENGASE;refGene_tx=XM_023244198.1;refGene_gn=ENGASE;refGene_tx=XM_023244199.1;refGene_gn=ENGASE;refGene_tx=XM_023244200.1;refGene_gn=ENGASE;refGene_tx=XM_023244201.1
E1 60139659 60139817
refGene_gn=ENGASE;refGene_tx=XM_019816973.2;refGene_gn=ENGASE;refGene_tx=XM_023244197.1;refGene_gn=ENGASE;refGene_tx=XM_023244198.1;refGene_gn=ENGASE;refGene_tx=XM_023244199.1;refGene_gn=ENGASE;refGene_tx=XM_023244200.1;refGene_gn=ENGASE;refGene_tx=XM_023244201.1

E1 60140365 60140514
refGene_gn=ENGASE;refGene_tx=XM_019816973.2;refGene_gn=ENGASE;refGene_tx=XM_023244197.1;refGene_gn=ENGASE;refGene_tx=XM_023244198.1;refGene_gn=ENGASE;refGene_tx=XM_023244199.1;refGene_gn=ENGASE;refGene_tx=XM_023244200.1;refGene_gn=ENGASE;refGene_tx=XM_023244201.1
E1 60140801 60140967
refGene_gn=ENGASE;refGene_tx=XM_019816973.2;refGene_gn=ENGASE;refGene_tx=XM_023244197.1;refGene_gn=ENGASE;refGene_tx=XM_023244198.1;refGene_gn=ENGASE;refGene_tx=XM_023244199.1;refGene_gn=ENGASE;refGene_tx=XM_023244200.1;refGene_gn=ENGASE;refGene_tx=XM_023244201.1
E1 60141863 60141967
refGene_gn=ENGASE;refGene_tx=XM_019816973.2;refGene_gn=ENGASE;refGene_tx=XM_023244197.1;refGene_gn=ENGASE;refGene_tx=XM_023244198.1;refGene_gn=ENGASE;refGene_tx=XM_023244199.1;refGene_gn=ENGASE;refGene_tx=XM_023244200.1;refGene_gn=ENGASE;refGene_tx=XM_023244201.1
E1 60142283 60142392
refGene_gn=ENGASE;refGene_tx=XM_019816973.2;refGene_gn=ENGASE;refGene_tx=XM_023244197.1;refGene_gn=ENGASE;refGene_tx=XM_023244198.1;refGene_gn=ENGASE;refGene_tx=XM_023244199.1;refGene_gn=ENGASE;refGene_tx=XM_023244200.1;refGene_gn=ENGASE;refGene_tx=XM_023244201.1
E1 60142560 60142751
refGene_gn=ENGASE;refGene_tx=XM_019816973.2;refGene_gn=ENGASE;refGene_tx=XM_023244197.1;refGene_gn=ENGASE;refGene_tx=XM_023244198.1;refGene_gn=ENGASE;refGene_tx=XM_023244199.1;refGene_gn=ENGASE;refGene_tx=XM_023244200.1;refGene_gn=ENGASE;refGene_tx=XM_023244201.1
E1 60143292 60143441
refGene_gn=ENGASE;refGene_tx=XM_019816973.2;refGene_gn=ENGASE;refGene_tx=XM_023244197.1;refGene_gn=ENGASE;refGene_tx=XM_023244198.1;refGene_gn=ENGASE;refGene_tx=XM_023244200.1;refGene_gn=ENGASE;refGene_tx=XM_023244201.1
E1 60143904 60144013
refGene_gn=ENGASE;refGene_tx=XM_019816973.2;refGene_gn=ENGASE;refGene_tx=XM_023244197.1;refGene_gn=ENGASE;refGene_tx=XM_023244198.1;refGene_gn=ENGASE;refGene_tx=XM_023244199.1;refGene_gn=ENGASE;refGene_tx=XM_023244200.1;refGene_gn=ENGASE;refGene_tx=XM_023244201.1
E1 60144304 60144419
refGene_gn=ENGASE;refGene_tx=XM_019816973.2;refGene_gn=ENGASE;refGene_tx=XM_023244197.1;refGene_gn=ENGASE;refGene_tx=XM_023244198.1;refGene_gn=ENGASE;refGene_tx=XM_023244199.1;refGene_gn=ENGASE;refGene_tx=XM_023244200.1;refGene_gn=ENGASE;refGene_tx=XM_023244201.1
E1 60144541 60144993
refGene_gn=ENGASE;refGene_tx=XM_019816973.2;refGene_gn=ENGASE;refGene_tx=XM_023244197.1;refGene_gn=ENGASE;refGene_tx=XM_023244198.1;refGene_gn=ENGASE;refGene_tx=XM_023244199.1;refGene_gn=ENGASE;refGene_tx=XM_023244200.1;refGene_gn=ENGASE;refGene_tx=XM_023244201.1
E1 60149136 60149139 refGene_gn=RBFOX3;refGene_tx=XM_023244204.1
E1 60150957 60151002
refGene_gn=RBFOX3;refGene_tx=XM_019818344.2;refGene_gn=RBFOX3;refGene_tx=XM_023244202.1;refGene_gn=RBFOX3;refGene_tx=XM_023244203.1;refGene_gn=RBFOX3;refGene_tx=XM_023244205.1;refGene_gn=RBFOX3;refGene_tx=XM_02

3244206.1;refGene_gn=RBFOX3;refGene_tx=XM_023244207.1;refGene_gn=RBFOX3;refGene_tx=XM_023244208.1
E1 60152643 60152722
refGene_gn=RBFOX3;refGene_tx=XM_019818344.2;refGene_gn=RBFOX3;refGene_tx=XM_023244202.1;refGene_gn=RBFOX3;refGene_tx=XM_023244203.1;refGene_gn=RBFOX3;refGene_tx=XM_023244204.1;refGene_gn=RBFOX3;refGene_tx=XM_023244205.1;refGene_gn=RBFOX3;refGene_tx=XM_023244206.1;refGene_gn=RBFOX3;refGene_tx=XM_023244207.1;refGene_gn=RBFOX3;refGene_tx=XM_023244208.1
E1 60153824 60153886
refGene_gn=RBFOX3;refGene_tx=XM_019818344.2;refGene_gn=RBFOX3;refGene_tx=XM_023244202.1;refGene_gn=RBFOX3;refGene_tx=XM_023244203.1;refGene_gn=RBFOX3;refGene_tx=XM_023244204.1;refGene_gn=RBFOX3;refGene_tx=XM_023244205.1;refGene_gn=RBFOX3;refGene_tx=XM_023244206.1;refGene_gn=RBFOX3;refGene_tx=XM_023244207.1;refGene_gn=RBFOX3;refGene_tx=XM_023244208.1
E1 60154796 60154959
refGene_gn=RBFOX3;refGene_tx=XM_019818344.2;refGene_gn=RBFOX3;refGene_tx=XM_023244202.1;refGene_gn=RBFOX3;refGene_tx=XM_023244203.1;refGene_gn=RBFOX3;refGene_tx=XM_023244204.1;refGene_gn=RBFOX3;refGene_tx=XM_023244205.1;refGene_gn=RBFOX3;refGene_tx=XM_023244206.1;refGene_gn=RBFOX3;refGene_tx=XM_023244207.1;refGene_gn=RBFOX3;refGene_tx=XM_023244208.1
E1 60155454 60155587
refGene_gn=RBFOX3;refGene_tx=XM_019818344.2;refGene_gn=RBFOX3;refGene_tx=XM_023244202.1;refGene_gn=RBFOX3;refGene_tx=XM_023244203.1;refGene_gn=RBFOX3;refGene_tx=XM_023244204.1;refGene_gn=RBFOX3;refGene_tx=XM_023244205.1;refGene_gn=RBFOX3;refGene_tx=XM_023244206.1;refGene_gn=RBFOX3;refGene_tx=XM_023244207.1;refGene_gn=RBFOX3;refGene_tx=XM_023244208.1
E1 60155789 60155843
refGene_gn=RBFOX3;refGene_tx=XM_019818344.2;refGene_gn=RBFOX3;refGene_tx=XM_023244202.1;refGene_gn=RBFOX3;refGene_tx=XM_023244203.1;refGene_gn=RBFOX3;refGene_tx=XM_023244204.1;refGene_gn=RBFOX3;refGene_tx=XM_023244205.1;refGene_gn=RBFOX3;refGene_tx=XM_023244206.1;refGene_gn=RBFOX3;refGene_tx=XM_023244207.1;refGene_gn=RBFOX3;refGene_tx=XM_023244208.1
E1 60158986 60159047
refGene_gn=RBFOX3;refGene_tx=XM_019818344.2;refGene_gn=RBFOX3;refGene_tx=XM_023244202.1;refGene_gn=RBFOX3;refGene_tx=XM_023244203.1;refGene_gn=RBFOX3;refGene_tx=XM_023244204.1;refGene_gn=RBFOX3;refGene_tx=XM_023244205.1;refGene_gn=RBFOX3;refGene_tx=XM_023244206.1;refGene_gn=RBFOX3;refGene_tx=XM_023244207.1;refGene_gn=RBFOX3;refGene_tx=XM_023244208.1
E1 60160527 60160620
refGene_gn=RBFOX3;refGene_tx=XM_019818344.2;refGene_gn=RBFOX3;refGene_tx=XM_023244202.1;refGene_gn=RBFOX3;refGene_tx=XM_023244203.1;refGene_gn=RBFOX3;refGene_tx=XM_023244204.1;refGene_gn=RBFOX3;refGene_tx=XM_023244205.1;refGene_gn=RBFOX3;refGene_tx=XM_023244206.1;refGene_gn=RBFOX3;refGene_tx=XM_023244207.1;refGene_gn=RBFOX3;refGene_tx=XM_023244208.

1
E1 60161444 60161498
refGene_gn=RBFOX3;refGene_tx=XM_019818344.2;refGene_gn=RBFOX3;refGene_tx=XM_023244202.1;refGene_gn=RBFOX3;refGene_tx=XM_023244203.1;refGene_gn=RBFOX3;refGene_tx=XM_023244204.1;refGene_gn=RBFOX3;refGene_tx=XM_023244205.1;refGene_gn=RBFOX3;refGene_tx=XM_023244206.1;refGene_gn=RBFOX3;refGene_tx=XM_023244207.1;refGene_gn=RBFOX3;refGene_tx=XM_023244208.1
E1 60163463 60163601
refGene_gn=RBFOX3;refGene_tx=XM_019818344.2;refGene_gn=RBFOX3;refGene_tx=XM_023244202.1;refGene_gn=RBFOX3;refGene_tx=XM_023244203.1;refGene_gn=RBFOX3;refGene_tx=XM_023244204.1;refGene_gn=RBFOX3;refGene_tx=XM_023244205.1;refGene_gn=RBFOX3;refGene_tx=XM_023244206.1;refGene_gn=RBFOX3;refGene_tx=XM_023244207.1;refGene_gn=RBFOX3;refGene_tx=XM_023244208.1
E1 60169042 60169297
refGene_gn=RBFOX3;refGene_tx=XM_019818344.2;refGene_gn=RBFOX3;refGene_tx=XM_023244202.1;refGene_gn=RBFOX3;refGene_tx=XM_023244203.1;refGene_gn=RBFOX3;refGene_tx=XM_023244204.1;refGene_gn=RBFOX3;refGene_tx=XM_023244205.1;refGene_gn=RBFOX3;refGene_tx=XM_023244206.1;refGene_gn=RBFOX3;refGene_tx=XM_023244207.1;refGene_gn=RBFOX3;refGene_tx=XM_023244208.1
E1 60226615 60226720
refGene_gn=RBFOX3;refGene_tx=XM_023244202.1;refGene_gn=RBFOX3;refGene_tx=XM_023244203.1;refGene_gn=RBFOX3;refGene_tx=XM_023244204.1;refGene_gn=RBFOX3;refGene_tx=XM_023244205.1;refGene_gn=RBFOX3;refGene_tx=XM_023244206.1
E1 60651696 60651949 refGene_gn=ENPP7;refGene_tx=XM_023243480.1
E1 60653113 60653259 refGene_gn=ENPP7;refGene_tx=XM_023243480.1
E1 60654419 60655046 refGene_gn=ENPP7;refGene_tx=XM_023243480.1
E1 60655687 60655997 refGene_gn=ENPP7;refGene_tx=XM_023243480.1
E1 60656080 60656239 refGene_gn=ENPP7;refGene_tx=XM_023243480.1
E1 60659071 60659202 refGene_gn=ENPP7;refGene_tx=XM_023243480.1
E1 60691789 60691861 refGene_gn=CBX2;refGene_tx=XM_023244232.1
E1 60691933 60691977 refGene_gn=CBX2;refGene_tx=XM_023244232.1
E1 60692988 60693054 refGene_gn=CBX2;refGene_tx=XM_023244232.1
E1 60695417 60695520 refGene_gn=CBX2;refGene_tx=XM_023244232.1
E1 60696913 60698218 refGene_gn=CBX2;refGene_tx=XM_023244232.1
E1 60707212 60708112 refGene_gn=CBX8;refGene_tx=XM_003997256.5
E1 60708561 60708628 refGene_gn=CBX8;refGene_tx=XM_003997256.5
E1 60708742 60708808 refGene_gn=CBX8;refGene_tx=XM_003997256.5
E1 60708993 60709037 refGene_gn=CBX8;refGene_tx=XM_003997256.5
E1 60709428 60709497 refGene_gn=CBX8;refGene_tx=XM_003997256.5
E1 60743824 60745240 refGene_gn=CBX4;refGene_tx=XM_023244233.1
E1 60745506 60745573 refGene_gn=CBX4;refGene_tx=XM_023244233.1
E1 60747820 60747886 refGene_gn=CBX4;refGene_tx=XM_023244233.1
E1 60749043 60749087 refGene_gn=CBX4;refGene_tx=XM_023244233.1
E1 60749233 60749302 refGene_gn=CBX4;refGene_tx=XM_023244233.1
E1 60817528 60817777 refGene_gn=TBC1D16;refGene_tx=XM_023244259.1
E1 60818863 60819010 refGene_gn=TBC1D16;refGene_tx=XM_023244259.1

E1 60821580 60821760 refGene_gn=TBC1D16;refGene_tx=XM_023244259.1
E1 60823520 60823707 refGene_gn=TBC1D16;refGene_tx=XM_023244259.1
E1 60824467 60824501 refGene_gn=TBC1D16;refGene_tx=XM_023244259.1
E1 60824503 60824601 refGene_gn=TBC1D16;refGene_tx=XM_023244259.1
E1 60828241 60828390 refGene_gn=TBC1D16;refGene_tx=XM_023244259.1
E1 60828885 60829050 refGene_gn=TBC1D16;refGene_tx=XM_023244259.1
E1 60829993 60830141 refGene_gn=TBC1D16;refGene_tx=XM_023244259.1
E1 60831100 60831262 refGene_gn=TBC1D16;refGene_tx=XM_023244259.1
E1 60873275 60873864 refGene_gn=TBC1D16;refGene_tx=XM_023244259.1
E1 60876742 60876920 refGene_gn=TBC1D16;refGene_tx=XM_023244259.1
E1 60890677 60890706
refGene_gn=CCDC40;refGene_tx=XM_023244257.1;refGene_gn=CCDC40;refGene_tx=XM_023244258.1
E1 60891352 60891375 refGene_gn=CCDC40;refGene_tx=XM_019818349.2
E1 60892277 60892362
refGene_gn=CCDC40;refGene_tx=XM_019818349.2;refGene_gn=CCDC40;refGene_tx=XM_023244257.1;refGene_gn=CCDC40;refGene_tx=XM_023244258.1
E1 60892791 60893241
refGene_gn=CCDC40;refGene_tx=XM_019818349.2;refGene_gn=CCDC40;refGene_tx=XM_023244257.1;refGene_gn=CCDC40;refGene_tx=XM_023244258.1
E1 60899797 60899942
refGene_gn=CCDC40;refGene_tx=XM_019818349.2;refGene_gn=CCDC40;refGene_tx=XM_023244257.1;refGene_gn=CCDC40;refGene_tx=XM_023244258.1
E1 60901235 60901411
refGene_gn=CCDC40;refGene_tx=XM_019818349.2;refGene_gn=CCDC40;refGene_tx=XM_023244257.1;refGene_gn=CCDC40;refGene_tx=XM_023244258.1
E1 60902212 60902296
refGene_gn=CCDC40;refGene_tx=XM_019818349.2;refGene_gn=CCDC40;refGene_tx=XM_023244257.1;refGene_gn=CCDC40;refGene_tx=XM_023244258.1
E1 60902376 60902596
refGene_gn=CCDC40;refGene_tx=XM_019818349.2;refGene_gn=CCDC40;refGene_tx=XM_023244257.1;refGene_gn=CCDC40;refGene_tx=XM_023244258.1
E1 60910238 60910396
refGene_gn=CCDC40;refGene_tx=XM_019818349.2;refGene_gn=CCDC40;refGene_tx=XM_023244257.1;refGene_gn=CCDC40;refGene_tx=XM_023244258.1
E1 60910595 60910718
refGene_gn=CCDC40;refGene_tx=XM_019818349.2;refGene_gn=CCDC40;refGene_tx=XM_023244257.1;refGene_gn=CCDC40;refGene_tx=XM_023244258.1
E1 60913707 60913829
refGene_gn=CCDC40;refGene_tx=XM_019818349.2;refGene_gn=CCDC40;refGene_tx=XM_023244257.1;refGene_gn=CCDC40;refGene_tx=XM_023244258.1
E1 60918550 60918794
refGene_gn=CCDC40;refGene_tx=XM_019818349.2;refGene_gn=CCDC40;refGene_tx=XM_023244257.1;refGene_gn=CCDC40;refGene_tx=XM_023244258.1
E1 60918873 60919056
refGene_gn=CCDC40;refGene_tx=XM_019818349.2;refGene_gn=CCDC40;refGene_tx=XM_023244257.1;refGene_gn=CCDC40;refGene_tx=XM_023244258.1
E1 60922144 60922390
refGene_gn=CCDC40;refGene_tx=XM_019818349.2;refGene_gn=CCDC40;refGene_tx=XM_023244257.1;refGene_gn=CCDC40;refGene_tx=XM_023244258.1

E1 60923474 60923688
refGene_gn=CCDC40;refGene_tx=XM_019818349.2;refGene_gn=CCDC40;refGene_tx=XM_023244257.1;refGene_gn=CCDC40;refGene_tx=XM_023244258.1
E1 60924857 60925027
refGene_gn=CCDC40;refGene_tx=XM_019818349.2;refGene_gn=CCDC40;refGene_tx=XM_023244257.1;refGene_gn=CCDC40;refGene_tx=XM_023244258.1
E1 60925129 60925221
refGene_gn=CCDC40;refGene_tx=XM_019818349.2;refGene_gn=CCDC40;refGene_tx=XM_023244257.1;refGene_gn=CCDC40;refGene_tx=XM_023244258.1
E1 60927807 60927928
refGene_gn=CCDC40;refGene_tx=XM_019818349.2;refGene_gn=CCDC40;refGene_tx=XM_023244257.1
E1 60929223 60929344 refGene_gn=CCDC40;refGene_tx=XM_023244258.1
E1 60932778 60932967
refGene_gn=CCDC40;refGene_tx=XM_019818349.2;refGene_gn=CCDC40;refGene_tx=XM_023244257.1;refGene_gn=CCDC40;refGene_tx=XM_023244258.1
E1 60934000 60934159
refGene_gn=CCDC40;refGene_tx=XM_019818349.2;refGene_gn=CCDC40;refGene_tx=XM_023244257.1;refGene_gn=CCDC40;refGene_tx=XM_023244258.1
E1 60935199 60935448
refGene_gn=CCDC40;refGene_tx=XM_019818349.2;refGene_gn=CCDC40;refGene_tx=XM_023244257.1;refGene_gn=CCDC40;refGene_tx=XM_023244258.1
E1 60939173 60939719
refGene_gn=GAA;refGene_tx=XM_003997259.5;refGene_gn=GAA;refGene_tx=XM_006940651.4;refGene_gn=GAA;refGene_tx=XM_006940652.3
E1 60940535 60940611 refGene_gn=GAA;refGene_tx=XM_006940653.3
E1 60940688 60940834
refGene_gn=GAA;refGene_tx=XM_003997259.5;refGene_gn=GAA;refGene_tx=XM_006940651.4;refGene_gn=GAA;refGene_tx=XM_006940652.3;refGene_gn=GAA;refGene_tx=XM_006940653.3
E1 60942639 60942805
refGene_gn=GAA;refGene_tx=XM_003997259.5;refGene_gn=GAA;refGene_tx=XM_006940651.4;refGene_gn=GAA;refGene_tx=XM_006940652.3;refGene_gn=GAA;refGene_tx=XM_006940653.3
E1 60942883 60942980
refGene_gn=GAA;refGene_tx=XM_003997259.5;refGene_gn=GAA;refGene_tx=XM_006940651.4;refGene_gn=GAA;refGene_tx=XM_006940652.3;refGene_gn=GAA;refGene_tx=XM_006940653.3
E1 60943395 60943515
refGene_gn=GAA;refGene_tx=XM_003997259.5;refGene_gn=GAA;refGene_tx=XM_006940651.4;refGene_gn=GAA;refGene_tx=XM_006940652.3;refGene_gn=GAA;refGene_tx=XM_006940653.3
E1 60943642 60943761
refGene_gn=GAA;refGene_tx=XM_003997259.5;refGene_gn=GAA;refGene_tx=XM_006940651.4;refGene_gn=GAA;refGene_tx=XM_006940652.3;refGene_gn=GAA;refGene_tx=XM_006940653.3
E1 60943837 60943969
refGene_gn=GAA;refGene_tx=XM_003997259.5;refGene_gn=GAA;refGene_tx=XM_006940651.4;refGene_gn=GAA;refGene_tx=XM_006940652.3;refGene_gn=GAA;refGene_tx=XM_006940653.3

E1 60944893 60945004
refGene_gn=GAA;refGene_tx=XM_003997259.5;refGene_gn=GAA;refGene_tx=XM_006940651.4;refGene_gn=GAA;refGene_tx=XM_006940652.3;refGene_gn=GAA;refGene_tx=XM_006940653.3
E1 60945560 60945674
refGene_gn=GAA;refGene_tx=XM_003997259.5;refGene_gn=GAA;refGene_tx=XM_006940651.4;refGene_gn=GAA;refGene_tx=XM_006940652.3;refGene_gn=GAA;refGene_tx=XM_006940653.3
E1 60945752 60945837
refGene_gn=GAA;refGene_tx=XM_003997259.5;refGene_gn=GAA;refGene_tx=XM_006940651.4;refGene_gn=GAA;refGene_tx=XM_006940652.3;refGene_gn=GAA;refGene_tx=XM_006940653.3
E1 60946308 60946426
refGene_gn=GAA;refGene_tx=XM_003997259.5;refGene_gn=GAA;refGene_tx=XM_006940651.4;refGene_gn=GAA;refGene_tx=XM_006940652.3;refGene_gn=GAA;refGene_tx=XM_006940653.3
E1 60946692 60946826
refGene_gn=GAA;refGene_tx=XM_003997259.5;refGene_gn=GAA;refGene_tx=XM_006940651.4;refGene_gn=GAA;refGene_tx=XM_006940652.3;refGene_gn=GAA;refGene_tx=XM_006940653.3
E1 60948610 60948762
refGene_gn=GAA;refGene_tx=XM_003997259.5;refGene_gn=GAA;refGene_tx=XM_006940651.4;refGene_gn=GAA;refGene_tx=XM_006940652.3;refGene_gn=GAA;refGene_tx=XM_006940653.3
E1 60948908 60949057
refGene_gn=GAA;refGene_tx=XM_003997259.5;refGene_gn=GAA;refGene_tx=XM_006940651.4;refGene_gn=GAA;refGene_tx=XM_006940652.3;refGene_gn=GAA;refGene_tx=XM_006940653.3
E1 60949876 60950018
refGene_gn=GAA;refGene_tx=XM_003997259.5;refGene_gn=GAA;refGene_tx=XM_006940651.4;refGene_gn=GAA;refGene_tx=XM_006940652.3;refGene_gn=GAA;refGene_tx=XM_006940653.3
E1 60950333 60950483
refGene_gn=GAA;refGene_tx=XM_003997259.5;refGene_gn=GAA;refGene_tx=XM_006940651.4;refGene_gn=GAA;refGene_tx=XM_006940652.3;refGene_gn=GAA;refGene_tx=XM_006940653.3
E1 60950851 60951016
refGene_gn=GAA;refGene_tx=XM_003997259.5;refGene_gn=GAA;refGene_tx=XM_006940651.4;refGene_gn=GAA;refGene_tx=XM_006940652.3;refGene_gn=GAA;refGene_tx=XM_006940653.3
E1 60951218 60951371
refGene_gn=GAA;refGene_tx=XM_003997259.5;refGene_gn=GAA;refGene_tx=XM_006940651.4;refGene_gn=GAA;refGene_tx=XM_006940652.3;refGene_gn=GAA;refGene_tx=XM_006940653.3
E1 60951751 60951811
refGene_gn=GAA;refGene_tx=XM_003997259.5;refGene_gn=GAA;refGene_tx=XM_006940651.4;refGene_gn=GAA;refGene_tx=XM_006940652.3;refGene_gn=GAA;refGene_tx=XM_006940653.3
E1 60957455 60957472 refGene_gn=EIF4A3;refGene_tx=XM_023244263.1
E1 60957662 60957790 refGene_gn=EIF4A3;refGene_tx=XM_023244263.1

| | | | |
|---|---|---|---|
| E1 | 60957883 | 60957991 | refGene_gn=EIF4A3;refGene_tx=XM_023244263.1 |
| E1 | 60960415 | 60960531 | refGene_gn=EIF4A3;refGene_tx=XM_023244263.1 |
| E1 | 60961282 | 60961421 | refGene_gn=EIF4A3;refGene_tx=XM_023244263.1 |
| E1 | 60961903 | 60962045 | refGene_gn=EIF4A3;refGene_tx=XM_023244263.1 |
| E1 | 60964223 | 60964304 | refGene_gn=EIF4A3;refGene_tx=XM_023244263.1 |
| E1 | 60965002 | 60965135 | refGene_gn=EIF4A3;refGene_tx=XM_023244263.1 |
| E1 | 60965721 | 60965784 | refGene_gn=EIF4A3;refGene_tx=XM_023244263.1 |
| E1 | 60966192 | 60966259 | refGene_gn=EIF4A3;refGene_tx=XM_023244263.1 |
| E1 | 60966839 | 60966912 | refGene_gn=EIF4A3;refGene_tx=XM_023244263.1 |
| E1 | 60968752 | 60968921 | refGene_gn=EIF4A3;refGene_tx=XM_023244263.1 |
| E1 | 60981431 | 60981642 | refGene_gn=CARD14;refGene_tx=XM_023244261.1 |
| E1 | 60984898 | 60984934 | refGene_gn=CARD14;refGene_tx=XM_023244261.1 |
| E1 | 60984936 | 60985035 | refGene_gn=CARD14;refGene_tx=XM_023244261.1 |
| E1 | 60986821 | 60987126 | refGene_gn=CARD14;refGene_tx=XM_023244261.1 |
| E1 | 60991433 | 60991601 | refGene_gn=CARD14;refGene_tx=XM_023244261.1 |
| E1 | 60992439 | 60992559 | refGene_gn=CARD14;refGene_tx=XM_023244261.1 |
| E1 | 60993256 | 60993382 | refGene_gn=CARD14;refGene_tx=XM_023244261.1 |
| E1 | 60993509 | 60993659 | refGene_gn=CARD14;refGene_tx=XM_023244261.1 |
| E1 | 60994703 | 60994811 | refGene_gn=CARD14;refGene_tx=XM_023244261.1 |
| E1 | 60995289 | 60995432 | refGene_gn=CARD14;refGene_tx=XM_023244261.1 |
| E1 | 60995570 | 60995665 | refGene_gn=CARD14;refGene_tx=XM_023244261.1 |
| E1 | 60996935 | 60996993 | refGene_gn=CARD14;refGene_tx=XM_023244261.1 |
| E1 | 60997209 | 60997402 | refGene_gn=CARD14;refGene_tx=XM_023244261.1 |
| E1 | 60999861 | 60999988 | refGene_gn=CARD14;refGene_tx=XM_023244261.1 |
| E1 | 61000291 | 61000529 | refGene_gn=CARD14;refGene_tx=XM_023244261.1 |
| E1 | 61001348 | 61001400 | refGene_gn=CARD14;refGene_tx=XM_023244261.1 |
| E1 | 61002090 | 61002205 | refGene_gn=CARD14;refGene_tx=XM_023244261.1 |
| E1 | 61002772 | 61002946 | refGene_gn=CARD14;refGene_tx=XM_023244261.1 |
| E1 | 61004941 | 61005063 | refGene_gn=CARD14;refGene_tx=XM_023244261.1 |
| E1 | 61005750 | 61005866 | refGene_gn=CARD14;refGene_tx=XM_023244261.1 |
| E1 | 61006387 | 61006595 | refGene_gn=CARD14;refGene_tx=XM_023244261.1 |
| E1 | 61008423 | 61008983 | refGene_gn=SGSH;refGene_tx=XM_023244262.1 |
| E1 | 61009725 | 61009929 | refGene_gn=SGSH;refGene_tx=XM_023244262.1 |
| E1 | 61010682 | 61010764 | refGene_gn=SGSH;refGene_tx=XM_023244262.1 |
| E1 | 61011107 | 61011264 | refGene_gn=SGSH;refGene_tx=XM_023244262.1 |
| E1 | 61011558 | 61011709 | refGene_gn=SGSH;refGene_tx=XM_023244262.1 |
| E1 | 61011954 | 61012060 | refGene_gn=SGSH;refGene_tx=XM_023244262.1 |
| E1 | 61013883 | 61014044 | refGene_gn=SGSH;refGene_tx=XM_023244262.1 |
| E1 | 61015157 | 61015254 | refGene_gn=SGSH;refGene_tx=XM_023244262.1 |
| E1 | 61016452 | 61016686 | refGene_gn=SLC26A11;refGene_tx=XM_003997294.5 |
| E1 | 61017227 | 61017420 | refGene_gn=SLC26A11;refGene_tx=XM_003997294.5 |
| E1 | 61017784 | 61017870 | refGene_gn=SLC26A11;refGene_tx=XM_003997294.5 |
| E1 | 61020313 | 61020393 | refGene_gn=SLC26A11;refGene_tx=XM_003997294.5 |
| E1 | 61021509 | 61021652 | refGene_gn=SLC26A11;refGene_tx=XM_003997294.5 |
| E1 | 61022110 | 61022286 | refGene_gn=SLC26A11;refGene_tx=XM_003997294.5 |
| E1 | 61022707 | 61022780 | refGene_gn=SLC26A11;refGene_tx=XM_003997294.5 |
| E1 | 61025136 | 61025187 | refGene_gn=SLC26A11;refGene_tx=XM_003997294.5 |
| E1 | 61026968 | 61027029 | refGene_gn=SLC26A11;refGene_tx=XM_003997294.5 |
| E1 | 61028516 | 61028572 | refGene_gn=SLC26A11;refGene_tx=XM_003997294.5 |
| E1 | 61028655 | 61028796 | refGene_gn=SLC26A11;refGene_tx=XM_003997294.5 |

E1	61031441 61031569 refGene_gn=SLC26A11;refGene_tx=XM_003997294.5
E1	61031935 61032035 refGene_gn=SLC26A11;refGene_tx=XM_003997294.5
E1	61032434 61032568 refGene_gn=SLC26A11;refGene_tx=XM_003997294.5
E1	61033030 61033103 refGene_gn=SLC26A11;refGene_tx=XM_003997294.5
E1	61033892 61033984 refGene_gn=SLC26A11;refGene_tx=XM_003997294.5
E1	61043902 61043999
refGene_gn=RNF213;refGene_tx=XM_019818352.2;refGene_gn=RNF213;refGene_tx=XM_023244265.1;refGene_gn=RNF213;refGene_tx=XM_023244266.1;refGene_gn=RNF213;refGene_tx=XM_023244267.1;refGene_gn=RNF213;refGene_tx=XM_023244268.1
E1	61049632 61049817
refGene_gn=RNF213;refGene_tx=XM_019818352.2;refGene_gn=RNF213;refGene_tx=XM_023244265.1;refGene_gn=RNF213;refGene_tx=XM_023244266.1;refGene_gn=RNF213;refGene_tx=XM_023244267.1;refGene_gn=RNF213;refGene_tx=XM_023244268.1
E1	61053637 61053781
refGene_gn=RNF213;refGene_tx=XM_019818352.2;refGene_gn=RNF213;refGene_tx=XM_023244265.1;refGene_gn=RNF213;refGene_tx=XM_023244267.1;refGene_gn=RNF213;refGene_tx=XM_023244268.1
E1	61062664 61063171
refGene_gn=RNF213;refGene_tx=XM_019818352.2;refGene_gn=RNF213;refGene_tx=XM_023244265.1;refGene_gn=RNF213;refGene_tx=XM_023244266.1;refGene_gn=RNF213;refGene_tx=XM_023244267.1;refGene_gn=RNF213;refGene_tx=XM_023244268.1
E1	61063399 61063522
refGene_gn=RNF213;refGene_tx=XM_019818352.2;refGene_gn=RNF213;refGene_tx=XM_023244265.1;refGene_gn=RNF213;refGene_tx=XM_023244266.1;refGene_gn=RNF213;refGene_tx=XM_023244267.1;refGene_gn=RNF213;refGene_tx=XM_023244268.1
E1	61064165 61064341
refGene_gn=RNF213;refGene_tx=XM_019818352.2;refGene_gn=RNF213;refGene_tx=XM_023244265.1;refGene_gn=RNF213;refGene_tx=XM_023244266.1;refGene_gn=RNF213;refGene_tx=XM_023244267.1;refGene_gn=RNF213;refGene_tx=XM_023244268.1
E1	61064864 61065023
refGene_gn=RNF213;refGene_tx=XM_019818352.2;refGene_gn=RNF213;refGene_tx=XM_023244265.1;refGene_gn=RNF213;refGene_tx=XM_023244266.1;refGene_gn=RNF213;refGene_tx=XM_023244267.1;refGene_gn=RNF213;refGene_tx=XM_023244268.1
E1	61065357 61065560
refGene_gn=RNF213;refGene_tx=XM_019818352.2;refGene_gn=RNF213;refGene_tx=XM_023244265.1;refGene_gn=RNF213;refGene_tx=XM_023244266.1;refGene_gn=RNF213;refGene_tx=XM_023244267.1;refGene_gn=RNF213;refGene_tx=XM_023244268.1
E1	61067926 61068210
refGene_gn=RNF213;refGene_tx=XM_019818352.2;refGene_gn=RNF213;refGene_tx=XM_023244265.1;refGene_gn=RNF213;refGene_tx=XM_023244266.1;refGene_gn=RNF213;refGene_tx=XM_023244267.1;refGene_gn=RNF213;refGene_tx=XM_023244268.1
E1	61069881 61070138

refGene_gn=RNF213;refGene_tx=XM_019818352.2;refGene_gn=RNF213;refGene_tx=XM_023244265.1;refGene_gn=RNF213;refGene_tx=XM_023244266.1;refGene_gn=RNF213;refGene_tx=XM_023244267.1;refGene_gn=RNF213;refGene_tx=XM_023244268.1
E1 61073496 61073694 refGene_gn=RNF213;refGene_tx=XM_019818352.2;refGene_gn=RNF213;refGene_tx=XM_023244265.1;refGene_gn=RNF213;refGene_tx=XM_023244266.1;refGene_gn=RNF213;refGene_tx=XM_023244267.1;refGene_gn=RNF213;refGene_tx=XM_023244268.1
E1 61075785 61076008 refGene_gn=RNF213;refGene_tx=XM_019818352.2;refGene_gn=RNF213;refGene_tx=XM_023244265.1;refGene_gn=RNF213;refGene_tx=XM_023244266.1;refGene_gn=RNF213;refGene_tx=XM_023244267.1;refGene_gn=RNF213;refGene_tx=XM_023244268.1
E1 61076532 61076588 refGene_gn=RNF213;refGene_tx=XM_019818352.2;refGene_gn=RNF213;refGene_tx=XM_023244265.1;refGene_gn=RNF213;refGene_tx=XM_023244266.1;refGene_gn=RNF213;refGene_tx=XM_023244267.1;refGene_gn=RNF213;refGene_tx=XM_023244268.1
E1 61077330 61077484 refGene_gn=RNF213;refGene_tx=XM_019818352.2;refGene_gn=RNF213;refGene_tx=XM_023244265.1;refGene_gn=RNF213;refGene_tx=XM_023244266.1;refGene_gn=RNF213;refGene_tx=XM_023244267.1;refGene_gn=RNF213;refGene_tx=XM_023244268.1
E1 61080274 61080448 refGene_gn=RNF213;refGene_tx=XM_019818352.2;refGene_gn=RNF213;refGene_tx=XM_023244265.1;refGene_gn=RNF213;refGene_tx=XM_023244266.1;refGene_gn=RNF213;refGene_tx=XM_023244267.1;refGene_gn=RNF213;refGene_tx=XM_023244268.1
E1 61082844 61082934 refGene_gn=RNF213;refGene_tx=XM_019818352.2;refGene_gn=RNF213;refGene_tx=XM_023244265.1;refGene_gn=RNF213;refGene_tx=XM_023244266.1;refGene_gn=RNF213;refGene_tx=XM_023244267.1
E1 61084042 61084165 refGene_gn=RNF213;refGene_tx=XM_019818352.2;refGene_gn=RNF213;refGene_tx=XM_023244266.1;refGene_gn=RNF213;refGene_tx=XM_023244267.1
E1 61086588 61086769 refGene_gn=RNF213;refGene_tx=XM_019818352.2;refGene_gn=RNF213;refGene_tx=XM_023244265.1;refGene_gn=RNF213;refGene_tx=XM_023244266.1;refGene_gn=RNF213;refGene_tx=XM_023244267.1;refGene_gn=RNF213;refGene_tx=XM_023244268.1
E1 61088493 61088667 refGene_gn=RNF213;refGene_tx=XM_019818352.2;refGene_gn=RNF213;refGene_tx=XM_023244265.1;refGene_gn=RNF213;refGene_tx=XM_023244266.1;refGene_gn=RNF213;refGene_tx=XM_023244267.1;refGene_gn=RNF213;refGene_tx=XM_023244268.1
E1 61089506 61089656 refGene_gn=RNF213;refGene_tx=XM_019818352.2;refGene_gn=RNF213;refGene_tx=XM_023244265.1;refGene_gn=RNF213;refGene_tx=XM_023244266.1;refGene_gn=RNF213;refGene_tx=XM_023244268.1

E1 61091568 61092158
refGene_gn=RNF213;refGene_tx=XM_019818352.2;refGene_gn=RNF213;refGene_tx=XM_023244265.1;refGene_gn=RNF213;refGene_tx=XM_023244266.1;refGene_gn=RNF213;refGene_tx=XM_023244267.1;refGene_gn=RNF213;refGene_tx=XM_023244268.1
E1 61093225 61093391
refGene_gn=RNF213;refGene_tx=XM_019818352.2;refGene_gn=RNF213;refGene_tx=XM_023244265.1;refGene_gn=RNF213;refGene_tx=XM_023244266.1;refGene_gn=RNF213;refGene_tx=XM_023244267.1;refGene_gn=RNF213;refGene_tx=XM_023244268.1
E1 61094377 61094595
refGene_gn=RNF213;refGene_tx=XM_019818352.2;refGene_gn=RNF213;refGene_tx=XM_023244265.1;refGene_gn=RNF213;refGene_tx=XM_023244266.1;refGene_gn=RNF213;refGene_tx=XM_023244267.1;refGene_gn=RNF213;refGene_tx=XM_023244268.1
E1 61096457 61096598
refGene_gn=RNF213;refGene_tx=XM_019818352.2;refGene_gn=RNF213;refGene_tx=XM_023244265.1;refGene_gn=RNF213;refGene_tx=XM_023244266.1;refGene_gn=RNF213;refGene_tx=XM_023244267.1;refGene_gn=RNF213;refGene_tx=XM_023244268.1
E1 61096712 61096877
refGene_gn=RNF213;refGene_tx=XM_019818352.2;refGene_gn=RNF213;refGene_tx=XM_023244265.1;refGene_gn=RNF213;refGene_tx=XM_023244266.1;refGene_gn=RNF213;refGene_tx=XM_023244267.1;refGene_gn=RNF213;refGene_tx=XM_023244268.1
E1 61097515 61098671
refGene_gn=RNF213;refGene_tx=XM_019818352.2;refGene_gn=RNF213;refGene_tx=XM_023244265.1;refGene_gn=RNF213;refGene_tx=XM_023244266.1;refGene_gn=RNF213;refGene_tx=XM_023244267.1;refGene_gn=RNF213;refGene_tx=XM_023244268.1
E1 61098934 61099128
refGene_gn=RNF213;refGene_tx=XM_019818352.2;refGene_gn=RNF213;refGene_tx=XM_023244265.1;refGene_gn=RNF213;refGene_tx=XM_023244266.1;refGene_gn=RNF213;refGene_tx=XM_023244267.1;refGene_gn=RNF213;refGene_tx=XM_023244268.1
E1 61099301 61099457
refGene_gn=RNF213;refGene_tx=XM_019818352.2;refGene_gn=RNF213;refGene_tx=XM_023244265.1;refGene_gn=RNF213;refGene_tx=XM_023244266.1;refGene_gn=RNF213;refGene_tx=XM_023244267.1;refGene_gn=RNF213;refGene_tx=XM_023244268.1
E1 61100509 61104130
refGene_gn=RNF213;refGene_tx=XM_019818352.2;refGene_gn=RNF213;refGene_tx=XM_023244265.1;refGene_gn=RNF213;refGene_tx=XM_023244266.1;refGene_gn=RNF213;refGene_tx=XM_023244267.1;refGene_gn=RNF213;refGene_tx=XM_023244268.1
E1 61106013 61106150
refGene_gn=RNF213;refGene_tx=XM_019818352.2;refGene_gn=RNF213;refGene_tx=XM_023244265.1;refGene_gn=RNF213;refGene_tx=XM_023244266.1;refGene_gn=RNF213;refGene_tx=XM_023244267.1;refGene_gn=RNF213;refGene_tx=XM_023244268.1

E1 61106546 61106642
refGene_gn=RNF213;refGene_tx=XM_019818352.2;refGene_gn=RNF213;refGene_tx=XM_023244265.1;refGene_gn=RNF213;refGene_tx=XM_023244266.1;refGene_gn=RNF213;refGene_tx=XM_023244267.1;refGene_gn=RNF213;refGene_tx=XM_023244268.1
E1 61108602 61108721
refGene_gn=RNF213;refGene_tx=XM_019818352.2;refGene_gn=RNF213;refGene_tx=XM_023244265.1;refGene_gn=RNF213;refGene_tx=XM_023244266.1;refGene_gn=RNF213;refGene_tx=XM_023244267.1;refGene_gn=RNF213;refGene_tx=XM_023244268.1
E1 61109995 61110115
refGene_gn=RNF213;refGene_tx=XM_019818352.2;refGene_gn=RNF213;refGene_tx=XM_023244265.1;refGene_gn=RNF213;refGene_tx=XM_023244266.1;refGene_gn=RNF213;refGene_tx=XM_023244267.1;refGene_gn=RNF213;refGene_tx=XM_023244268.1
E1 61110544 61110684
refGene_gn=RNF213;refGene_tx=XM_019818352.2;refGene_gn=RNF213;refGene_tx=XM_023244265.1;refGene_gn=RNF213;refGene_tx=XM_023244266.1;refGene_gn=RNF213;refGene_tx=XM_023244267.1;refGene_gn=RNF213;refGene_tx=XM_023244268.1
E1 61110905 61111050
refGene_gn=RNF213;refGene_tx=XM_019818352.2;refGene_gn=RNF213;refGene_tx=XM_023244265.1;refGene_gn=RNF213;refGene_tx=XM_023244266.1;refGene_gn=RNF213;refGene_tx=XM_023244267.1;refGene_gn=RNF213;refGene_tx=XM_023244268.1
E1 61111247 61111386
refGene_gn=RNF213;refGene_tx=XM_019818352.2;refGene_gn=RNF213;refGene_tx=XM_023244265.1;refGene_gn=RNF213;refGene_tx=XM_023244266.1;refGene_gn=RNF213;refGene_tx=XM_023244267.1;refGene_gn=RNF213;refGene_tx=XM_023244268.1
E1 61114020 61114212
refGene_gn=RNF213;refGene_tx=XM_019818352.2;refGene_gn=RNF213;refGene_tx=XM_023244265.1;refGene_gn=RNF213;refGene_tx=XM_023244266.1;refGene_gn=RNF213;refGene_tx=XM_023244267.1;refGene_gn=RNF213;refGene_tx=XM_023244268.1
E1 61115280 61115426
refGene_gn=RNF213;refGene_tx=XM_019818352.2;refGene_gn=RNF213;refGene_tx=XM_023244265.1;refGene_gn=RNF213;refGene_tx=XM_023244266.1;refGene_gn=RNF213;refGene_tx=XM_023244267.1;refGene_gn=RNF213;refGene_tx=XM_023244268.1
E1 61117068 61117223
refGene_gn=RNF213;refGene_tx=XM_019818352.2;refGene_gn=RNF213;refGene_tx=XM_023244265.1;refGene_gn=RNF213;refGene_tx=XM_023244266.1;refGene_gn=RNF213;refGene_tx=XM_023244267.1;refGene_gn=RNF213;refGene_tx=XM_023244268.1
E1 61119207 61119426
refGene_gn=RNF213;refGene_tx=XM_019818352.2;refGene_gn=RNF213;refGene_tx=XM_023244265.1;refGene_gn=RNF213;refGene_tx=XM_023244266.1;refGene_gn=RNF213;refGene_tx=XM_023244267.1;refGene_gn=RNF213;refGene_tx=XM_023244268.1

E1 61119775 61119957
refGene_gn=RNF213;refGene_tx=XM_019818352.2;refGene_gn=RNF213;refGene_tx=XM_023244265.1;refGene_gn=RNF213;refGene_tx=XM_023244266.1;refGene_gn=RNF213;refGene_tx=XM_023244267.1;refGene_gn=RNF213;refGene_tx=XM_023244268.1
E1 61120521 61120642
refGene_gn=RNF213;refGene_tx=XM_019818352.2;refGene_gn=RNF213;refGene_tx=XM_023244265.1;refGene_gn=RNF213;refGene_tx=XM_023244266.1;refGene_gn=RNF213;refGene_tx=XM_023244267.1;refGene_gn=RNF213;refGene_tx=XM_023244268.1
E1 61124363 61124464
refGene_gn=RNF213;refGene_tx=XM_019818352.2;refGene_gn=RNF213;refGene_tx=XM_023244265.1;refGene_gn=RNF213;refGene_tx=XM_023244266.1;refGene_gn=RNF213;refGene_tx=XM_023244267.1;refGene_gn=RNF213;refGene_tx=XM_023244268.1
E1 61124555 61124738
refGene_gn=RNF213;refGene_tx=XM_019818352.2;refGene_gn=RNF213;refGene_tx=XM_023244265.1;refGene_gn=RNF213;refGene_tx=XM_023244266.1;refGene_gn=RNF213;refGene_tx=XM_023244267.1;refGene_gn=RNF213;refGene_tx=XM_023244268.1
E1 61125724 61125894
refGene_gn=RNF213;refGene_tx=XM_019818352.2;refGene_gn=RNF213;refGene_tx=XM_023244265.1;refGene_gn=RNF213;refGene_tx=XM_023244266.1;refGene_gn=RNF213;refGene_tx=XM_023244267.1;refGene_gn=RNF213;refGene_tx=XM_023244268.1
E1 61125999 61126099
refGene_gn=RNF213;refGene_tx=XM_019818352.2;refGene_gn=RNF213;refGene_tx=XM_023244265.1;refGene_gn=RNF213;refGene_tx=XM_023244266.1;refGene_gn=RNF213;refGene_tx=XM_023244267.1;refGene_gn=RNF213;refGene_tx=XM_023244268.1
E1 61127210 61127322
refGene_gn=RNF213;refGene_tx=XM_019818352.2;refGene_gn=RNF213;refGene_tx=XM_023244265.1;refGene_gn=RNF213;refGene_tx=XM_023244266.1;refGene_gn=RNF213;refGene_tx=XM_023244267.1;refGene_gn=RNF213;refGene_tx=XM_023244268.1
E1 61128646 61128857
refGene_gn=RNF213;refGene_tx=XM_019818352.2;refGene_gn=RNF213;refGene_tx=XM_023244265.1;refGene_gn=RNF213;refGene_tx=XM_023244266.1;refGene_gn=RNF213;refGene_tx=XM_023244267.1;refGene_gn=RNF213;refGene_tx=XM_023244268.1
E1 61129095 61129286
refGene_gn=RNF213;refGene_tx=XM_019818352.2;refGene_gn=RNF213;refGene_tx=XM_023244265.1;refGene_gn=RNF213;refGene_tx=XM_023244266.1;refGene_gn=RNF213;refGene_tx=XM_023244267.1;refGene_gn=RNF213;refGene_tx=XM_023244268.1
E1 61129968 61130091
refGene_gn=RNF213;refGene_tx=XM_019818352.2;refGene_gn=RNF213;refGene_tx=XM_023244265.1;refGene_gn=RNF213;refGene_tx=XM_023244266.1;refGene_gn=RNF213;refGene_tx=XM_023244267.1;refGene_gn=RNF213;refGene_tx=XM_023244268.1

E1 61130846 61130957
refGene_gn=RNF213;refGene_tx=XM_019818352.2;refGene_gn=RNF213;refGene_tx=XM_023244265.1;refGene_gn=RNF213;refGene_tx=XM_023244266.1;refGene_gn=RNF213;refGene_tx=XM_023244267.1;refGene_gn=RNF213;refGene_tx=XM_023244268.1
E1 61131499 61131742
refGene_gn=RNF213;refGene_tx=XM_019818352.2;refGene_gn=RNF213;refGene_tx=XM_023244265.1;refGene_gn=RNF213;refGene_tx=XM_023244266.1;refGene_gn=RNF213;refGene_tx=XM_023244267.1;refGene_gn=RNF213;refGene_tx=XM_023244268.1
E1 61132027 61132109
refGene_gn=RNF213;refGene_tx=XM_019818352.2;refGene_gn=RNF213;refGene_tx=XM_023244265.1;refGene_gn=RNF213;refGene_tx=XM_023244266.1;refGene_gn=RNF213;refGene_tx=XM_023244267.1;refGene_gn=RNF213;refGene_tx=XM_023244268.1
E1 61133367 61133402
refGene_gn=RNF213;refGene_tx=XM_019818352.2;refGene_gn=RNF213;refGene_tx=XM_023244265.1;refGene_gn=RNF213;refGene_tx=XM_023244266.1;refGene_gn=RNF213;refGene_tx=XM_023244267.1;refGene_gn=RNF213;refGene_tx=XM_023244268.1
E1 61134493 61134588
refGene_gn=RNF213;refGene_tx=XM_019818352.2;refGene_gn=RNF213;refGene_tx=XM_023244265.1;refGene_gn=RNF213;refGene_tx=XM_023244266.1;refGene_gn=RNF213;refGene_tx=XM_023244267.1;refGene_gn=RNF213;refGene_tx=XM_023244268.1
E1 61136141 61136301
refGene_gn=RNF213;refGene_tx=XM_019818352.2;refGene_gn=RNF213;refGene_tx=XM_023244265.1;refGene_gn=RNF213;refGene_tx=XM_023244266.1;refGene_gn=RNF213;refGene_tx=XM_023244267.1;refGene_gn=RNF213;refGene_tx=XM_023244268.1
E1 61137590 61137771
refGene_gn=RNF213;refGene_tx=XM_019818352.2;refGene_gn=RNF213;refGene_tx=XM_023244265.1;refGene_gn=RNF213;refGene_tx=XM_023244266.1;refGene_gn=RNF213;refGene_tx=XM_023244267.1;refGene_gn=RNF213;refGene_tx=XM_023244268.1
E1 61138479 61138571
refGene_gn=RNF213;refGene_tx=XM_019818352.2;refGene_gn=RNF213;refGene_tx=XM_023244265.1;refGene_gn=RNF213;refGene_tx=XM_023244266.1;refGene_gn=RNF213;refGene_tx=XM_023244267.1;refGene_gn=RNF213;refGene_tx=XM_023244268.1
E1 61139177 61139435
refGene_gn=RNF213;refGene_tx=XM_019818352.2;refGene_gn=RNF213;refGene_tx=XM_023244265.1;refGene_gn=RNF213;refGene_tx=XM_023244266.1;refGene_gn=RNF213;refGene_tx=XM_023244267.1;refGene_gn=RNF213;refGene_tx=XM_023244268.1
E1 61140760 61140893
refGene_gn=RNF213;refGene_tx=XM_019818352.2;refGene_gn=RNF213;refGene_tx=XM_023244265.1;refGene_gn=RNF213;refGene_tx=XM_023244266.1;refGene_gn=RNF213;refGene_tx=XM_023244267.1;refGene_gn=RNF213;refGene_tx=XM_023244268.1

E1 61141197 61141281
refGene_gn=RNF213;refGene_tx=XM_019818352.2;refGene_gn=RNF213;refGene_tx=XM_023244265.1;refGene_gn=RNF213;refGene_tx=XM_023244266.1;refGene_gn=RNF213;refGene_tx=XM_023244267.1;refGene_gn=RNF213;refGene_tx=XM_023244268.1
E1 61141610 61141791
refGene_gn=RNF213;refGene_tx=XM_019818352.2;refGene_gn=RNF213;refGene_tx=XM_023244265.1;refGene_gn=RNF213;refGene_tx=XM_023244266.1;refGene_gn=RNF213;refGene_tx=XM_023244267.1;refGene_gn=RNF213;refGene_tx=XM_023244268.1
E1 61141979 61142181
refGene_gn=RNF213;refGene_tx=XM_019818352.2;refGene_gn=RNF213;refGene_tx=XM_023244265.1;refGene_gn=RNF213;refGene_tx=XM_023244266.1;refGene_gn=RNF213;refGene_tx=XM_023244267.1;refGene_gn=RNF213;refGene_tx=XM_023244268.1
E1 61143181 61143259
refGene_gn=RNF213;refGene_tx=XM_019818352.2;refGene_gn=RNF213;refGene_tx=XM_023244265.1;refGene_gn=RNF213;refGene_tx=XM_023244266.1;refGene_gn=RNF213;refGene_tx=XM_023244267.1;refGene_gn=RNF213;refGene_tx=XM_023244268.1
E1 61143820 61144015
refGene_gn=RNF213;refGene_tx=XM_019818352.2;refGene_gn=RNF213;refGene_tx=XM_023244265.1;refGene_gn=RNF213;refGene_tx=XM_023244266.1;refGene_gn=RNF213;refGene_tx=XM_023244267.1;refGene_gn=RNF213;refGene_tx=XM_023244268.1
E1 61144417 61144507
refGene_gn=RNF213;refGene_tx=XM_019818352.2;refGene_gn=RNF213;refGene_tx=XM_023244265.1;refGene_gn=RNF213;refGene_tx=XM_023244266.1;refGene_gn=RNF213;refGene_tx=XM_023244267.1;refGene_gn=RNF213;refGene_tx=XM_023244268.1
E1 61144594 61144773
refGene_gn=RNF213;refGene_tx=XM_019818352.2;refGene_gn=RNF213;refGene_tx=XM_023244265.1;refGene_gn=RNF213;refGene_tx=XM_023244266.1;refGene_gn=RNF213;refGene_tx=XM_023244267.1;refGene_gn=RNF213;refGene_tx=XM_023244268.1
E1 61147745 61147899
refGene_gn=RNF213;refGene_tx=XM_019818352.2;refGene_gn=RNF213;refGene_tx=XM_023244265.1;refGene_gn=RNF213;refGene_tx=XM_023244266.1;refGene_gn=RNF213;refGene_tx=XM_023244267.1;refGene_gn=RNF213;refGene_tx=XM_023244268.1
E1 61152908 61152964
refGene_gn=ENDOV;refGene_tx=XM_023244269.1;refGene_gn=ENDOV;refGene_tx=XM_023244270.1
E1 61153269 61153441
refGene_gn=ENDOV;refGene_tx=XM_023244269.1;refGene_gn=ENDOV;refGene_tx=XM_023244270.1
E1 61157968 61158103 refGene_gn=ENDOV;refGene_tx=XM_023244269.1
E1 61158339 61158379
refGene_gn=ENDOV;refGene_tx=XM_023244269.1;refGene_gn=ENDOV;refGene_tx=XM_023244270.1

E1 61159271 61159384
refGene_gn=ENDOV;refGene_tx=XM_023244269.1;refGene_gn=ENDOV;refGene_tx=XM_023244270.1
E1 61160501 61160570
refGene_gn=ENDOV;refGene_tx=XM_023244269.1;refGene_gn=ENDOV;refGene_tx=XM_023244270.1
E1 61160845 61160974
refGene_gn=ENDOV;refGene_tx=XM_023244269.1;refGene_gn=ENDOV;refGene_tx=XM_023244270.1
E1 61163522 61163596
refGene_gn=ENDOV;refGene_tx=XM_023244269.1;refGene_gn=ENDOV;refGene_tx=XM_023244270.1
E1 61164230 61164384
refGene_gn=ENDOV;refGene_tx=XM_023244269.1;refGene_gn=ENDOV;refGene_tx=XM_023244270.1
E1 61195813 61196035 refGene_gn=NPTX1;refGene_tx=XM_023244271.1
E1 61196680 61196860 refGene_gn=NPTX1;refGene_tx=XM_023244271.1
E1 61198076 61198321 refGene_gn=NPTX1;refGene_tx=XM_023244271.1
E1 61200482 61200690 refGene_gn=NPTX1;refGene_tx=XM_023244271.1
E1 61201013 61201472 refGene_gn=NPTX1;refGene_tx=XM_023244271.1
E1 61253459 61253621
refGene_gn=RPTOR;refGene_tx=XM_011289455.3;refGene_gn=RPTOR;refGene_tx=XM_023244284.1
E1 61326279 61326382
refGene_gn=RPTOR;refGene_tx=XM_011289455.3;refGene_gn=RPTOR;refGene_tx=XM_023244284.1
E1 61335147 61335199 refGene_gn=RPTOR;refGene_tx=XM_019818372.2
E1 61337803 61337886
refGene_gn=RPTOR;refGene_tx=XM_011289455.3;refGene_gn=RPTOR;refGene_tx=XM_019818372.2;refGene_gn=RPTOR;refGene_tx=XM_019818373.2;refGene_gn=RPTOR;refGene_tx=XM_023244284.1
E1 61392252 61392411
refGene_gn=RPTOR;refGene_tx=XM_011289455.3;refGene_gn=RPTOR;refGene_tx=XM_019818372.2;refGene_gn=RPTOR;refGene_tx=XM_019818373.2;refGene_gn=RPTOR;refGene_tx=XM_023244284.1
E1 61404310 61404457
refGene_gn=RPTOR;refGene_tx=XM_011289455.3;refGene_gn=RPTOR;refGene_tx=XM_019818372.2;refGene_gn=RPTOR;refGene_tx=XM_019818373.2;refGene_gn=RPTOR;refGene_tx=XM_019818375.2;refGene_gn=RPTOR;refGene_tx=XM_023244284.1;refGene_gn=RPTOR;refGene_tx=XM_023244285.1
E1 61418737 61418913
refGene_gn=RPTOR;refGene_tx=XM_011289455.3;refGene_gn=RPTOR;refGene_tx=XM_019818372.2;refGene_gn=RPTOR;refGene_tx=XM_019818373.2;refGene_gn=RPTOR;refGene_tx=XM_019818375.2;refGene_gn=RPTOR;refGene_tx=XM_023244284.1;refGene_gn=RPTOR;refGene_tx=XM_023244285.1
E1 61452012 61452072
refGene_gn=RPTOR;refGene_tx=XM_011289455.3;refGene_gn=RPTOR;refGene_tx=XM_019818372.2;refGene_gn=RPTOR;refGene_tx=XM_019818373.2;refGene_gn=RPTOR;refGene_tx=XM_019818375.2;refGene_gn=RPTOR;refGene_tx=XM_023244284.1;refGene_gn=RPTOR;refGene_tx=XM_023244285.1

E1 61476716 61476817
refGene_gn=RPTOR;refGene_tx=XM_011289455.3;refGene_gn=RPTOR;refGene_tx=XM_019818372.2;refGene_gn=RPTOR;refGene_tx=XM_019818373.2;refGene_gn=RPTOR;refGene_tx=XM_019818375.2;refGene_gn=RPTOR;refGene_tx=XM_023244284.1;refGene_gn=RPTOR;refGene_tx=XM_023244285.1
E1 61477377 61477522
refGene_gn=RPTOR;refGene_tx=XM_011289455.3;refGene_gn=RPTOR;refGene_tx=XM_019818372.2;refGene_gn=RPTOR;refGene_tx=XM_019818373.2;refGene_gn=RPTOR;refGene_tx=XM_019818375.2;refGene_gn=RPTOR;refGene_tx=XM_023244284.1;refGene_gn=RPTOR;refGene_tx=XM_023244285.1
E1 61485487 61485563
refGene_gn=RPTOR;refGene_tx=XM_011289455.3;refGene_gn=RPTOR;refGene_tx=XM_019818372.2;refGene_gn=RPTOR;refGene_tx=XM_019818373.2;refGene_gn=RPTOR;refGene_tx=XM_019818375.2;refGene_gn=RPTOR;refGene_tx=XM_023244284.1;refGene_gn=RPTOR;refGene_tx=XM_023244285.1
E1 61492851 61492953
refGene_gn=RPTOR;refGene_tx=XM_011289455.3;refGene_gn=RPTOR;refGene_tx=XM_019818372.2;refGene_gn=RPTOR;refGene_tx=XM_019818373.2;refGene_gn=RPTOR;refGene_tx=XM_019818375.2;refGene_gn=RPTOR;refGene_tx=XM_023244284.1;refGene_gn=RPTOR;refGene_tx=XM_023244285.1
E1 61498957 61499041
refGene_gn=RPTOR;refGene_tx=XM_011289455.3;refGene_gn=RPTOR;refGene_tx=XM_019818372.2;refGene_gn=RPTOR;refGene_tx=XM_019818373.2;refGene_gn=RPTOR;refGene_tx=XM_019818375.2;refGene_gn=RPTOR;refGene_tx=XM_023244284.1;refGene_gn=RPTOR;refGene_tx=XM_023244285.1
E1 61500309 61500420
refGene_gn=RPTOR;refGene_tx=XM_011289455.3;refGene_gn=RPTOR;refGene_tx=XM_019818372.2;refGene_gn=RPTOR;refGene_tx=XM_019818373.2;refGene_gn=RPTOR;refGene_tx=XM_019818375.2;refGene_gn=RPTOR;refGene_tx=XM_023244284.1;refGene_gn=RPTOR;refGene_tx=XM_023244285.1
E1 61523369 61523444
refGene_gn=RPTOR;refGene_tx=XM_011289455.3;refGene_gn=RPTOR;refGene_tx=XM_019818372.2;refGene_gn=RPTOR;refGene_tx=XM_019818373.2;refGene_gn=RPTOR;refGene_tx=XM_019818375.2;refGene_gn=RPTOR;refGene_tx=XM_023244284.1;refGene_gn=RPTOR;refGene_tx=XM_023244285.1
E1 61526787 61526853
refGene_gn=RPTOR;refGene_tx=XM_011289455.3;refGene_gn=RPTOR;refGene_tx=XM_019818372.2;refGene_gn=RPTOR;refGene_tx=XM_019818373.2;refGene_gn=RPTOR;refGene_tx=XM_019818375.2;refGene_gn=RPTOR;refGene_tx=XM_023244284.1;refGene_gn=RPTOR;refGene_tx=XM_023244285.1
E1 61527136 61527328
refGene_gn=RPTOR;refGene_tx=XM_011289455.3;refGene_gn=RPTOR;refGene_tx=XM_019818372.2;refGene_gn=RPTOR;refGene_tx=XM_019818373.2;refGene_gn=RPTOR;refGene_tx=XM_019818375.2;refGene_gn=RPTOR;refGene_tx=XM_023244284.1;refGene_gn=RPTOR;refGene_tx=XM_023244285.1
E1 61527473 61527614
refGene_gn=RPTOR;refGene_tx=XM_011289455.3;refGene_gn=RPTOR;refGene_tx=XM_019818372.2;refGene_gn=RPTOR;refGene_tx=XM_019818373.2;refGene_gn=RPTOR;refGene_tx=XM_019818375.2;refGene_gn=RPTOR;refGene_tx=XM_023244285.1

E1 61532694 61532812
refGene_gn=RPTOR;refGene_tx=XM_011289455.3;refGene_gn=RPTOR;refGene_tx=XM_019818372.2;refGene_gn=RPTOR;refGene_tx=XM_019818373.2;refGene_gn=RPTOR;refGene_tx=XM_019818375.2;refGene_gn=RPTOR;refGene_tx=XM_023244284.1;refGene_gn=RPTOR;refGene_tx=XM_023244285.1
E1 61533671 61533812
refGene_gn=RPTOR;refGene_tx=XM_011289455.3;refGene_gn=RPTOR;refGene_tx=XM_019818372.2;refGene_gn=RPTOR;refGene_tx=XM_019818373.2;refGene_gn=RPTOR;refGene_tx=XM_019818375.2;refGene_gn=RPTOR;refGene_tx=XM_023244284.1;refGene_gn=RPTOR;refGene_tx=XM_023244285.1
E1 61534135 61534294
refGene_gn=RPTOR;refGene_tx=XM_011289455.3;refGene_gn=RPTOR;refGene_tx=XM_019818372.2;refGene_gn=RPTOR;refGene_tx=XM_019818373.2;refGene_gn=RPTOR;refGene_tx=XM_019818375.2;refGene_gn=RPTOR;refGene_tx=XM_023244284.1;refGene_gn=RPTOR;refGene_tx=XM_023244285.1
E1 61542992 61543111
refGene_gn=RPTOR;refGene_tx=XM_011289455.3;refGene_gn=RPTOR;refGene_tx=XM_019818372.2;refGene_gn=RPTOR;refGene_tx=XM_019818373.2;refGene_gn=RPTOR;refGene_tx=XM_019818375.2;refGene_gn=RPTOR;refGene_tx=XM_023244284.1;refGene_gn=RPTOR;refGene_tx=XM_023244285.1
E1 61549171 61549275
refGene_gn=RPTOR;refGene_tx=XM_011289455.3;refGene_gn=RPTOR;refGene_tx=XM_019818372.2;refGene_gn=RPTOR;refGene_tx=XM_019818373.2;refGene_gn=RPTOR;refGene_tx=XM_019818375.2;refGene_gn=RPTOR;refGene_tx=XM_023244284.1;refGene_gn=RPTOR;refGene_tx=XM_023244285.1
E1 61549714 61549898
refGene_gn=RPTOR;refGene_tx=XM_011289455.3;refGene_gn=RPTOR;refGene_tx=XM_019818372.2;refGene_gn=RPTOR;refGene_tx=XM_019818373.2;refGene_gn=RPTOR;refGene_tx=XM_019818375.2;refGene_gn=RPTOR;refGene_tx=XM_023244284.1;refGene_gn=RPTOR;refGene_tx=XM_023244285.1
E1 61551300 61551411
refGene_gn=RPTOR;refGene_tx=XM_011289455.3;refGene_gn=RPTOR;refGene_tx=XM_019818372.2;refGene_gn=RPTOR;refGene_tx=XM_019818373.2;refGene_gn=RPTOR;refGene_tx=XM_019818375.2;refGene_gn=RPTOR;refGene_tx=XM_023244284.1;refGene_gn=RPTOR;refGene_tx=XM_023244285.1
E1 61562482 61562588
refGene_gn=RPTOR;refGene_tx=XM_011289455.3;refGene_gn=RPTOR;refGene_tx=XM_019818372.2;refGene_gn=RPTOR;refGene_tx=XM_019818373.2;refGene_gn=RPTOR;refGene_tx=XM_019818375.2;refGene_gn=RPTOR;refGene_tx=XM_023244284.1;refGene_gn=RPTOR;refGene_tx=XM_023244285.1
E1 61564926 61565041
refGene_gn=RPTOR;refGene_tx=XM_011289455.3;refGene_gn=RPTOR;refGene_tx=XM_019818372.2;refGene_gn=RPTOR;refGene_tx=XM_019818373.2;refGene_gn=RPTOR;refGene_tx=XM_019818375.2;refGene_gn=RPTOR;refGene_tx=XM_023244284.1;refGene_gn=RPTOR;refGene_tx=XM_023244285.1
E1 61566544 61566669
refGene_gn=RPTOR;refGene_tx=XM_011289455.3;refGene_gn=RPTOR;refGene_tx=XM_019818372.2;refGene_gn=RPTOR;refGene_tx=XM_019818373.2;refGene_gn=RPTOR;refGene_tx=XM_019818375.2;refGene_gn=RPTOR;refGene_tx=XM_023244284.1;refGene_gn=RPTOR;refGene_tx=XM_023244285.1

E1 61568700 61568805
refGene_gn=RPTOR;refGene_tx=XM_011289455.3;refGene_gn=RPTOR;refGene_tx=XM_019818372.2;refGene_gn=RPTOR;refGene_tx=XM_019818373.2;refGene_gn=RPTOR;refGene_tx=XM_019818375.2;refGene_gn=RPTOR;refGene_tx=XM_023244284.1;refGene_gn=RPTOR;refGene_tx=XM_023244285.1
E1 61578183 61578290
refGene_gn=RPTOR;refGene_tx=XM_011289455.3;refGene_gn=RPTOR;refGene_tx=XM_019818372.2;refGene_gn=RPTOR;refGene_tx=XM_019818373.2;refGene_gn=RPTOR;refGene_tx=XM_019818375.2;refGene_gn=RPTOR;refGene_tx=XM_023244284.1;refGene_gn=RPTOR;refGene_tx=XM_023244285.1
E1 61579615 61579743
refGene_gn=RPTOR;refGene_tx=XM_011289455.3;refGene_gn=RPTOR;refGene_tx=XM_019818372.2;refGene_gn=RPTOR;refGene_tx=XM_019818373.2;refGene_gn=RPTOR;refGene_tx=XM_019818375.2;refGene_gn=RPTOR;refGene_tx=XM_023244284.1;refGene_gn=RPTOR;refGene_tx=XM_023244285.1
E1 61581100 61581187
refGene_gn=RPTOR;refGene_tx=XM_011289455.3;refGene_gn=RPTOR;refGene_tx=XM_019818372.2;refGene_gn=RPTOR;refGene_tx=XM_019818373.2;refGene_gn=RPTOR;refGene_tx=XM_019818375.2;refGene_gn=RPTOR;refGene_tx=XM_023244284.1;refGene_gn=RPTOR;refGene_tx=XM_023244285.1
E1 61581991 61582108
refGene_gn=RPTOR;refGene_tx=XM_011289455.3;refGene_gn=RPTOR;refGene_tx=XM_019818372.2;refGene_gn=RPTOR;refGene_tx=XM_019818373.2;refGene_gn=RPTOR;refGene_tx=XM_019818375.2;refGene_gn=RPTOR;refGene_tx=XM_023244284.1;refGene_gn=RPTOR;refGene_tx=XM_023244285.1
E1 61582453 61582583
refGene_gn=RPTOR;refGene_tx=XM_011289455.3;refGene_gn=RPTOR;refGene_tx=XM_019818372.2;refGene_gn=RPTOR;refGene_tx=XM_019818373.2;refGene_gn=RPTOR;refGene_tx=XM_019818375.2;refGene_gn=RPTOR;refGene_tx=XM_023244284.1;refGene_gn=RPTOR;refGene_tx=XM_023244285.1
E1 61583255 61583324
refGene_gn=RPTOR;refGene_tx=XM_011289455.3;refGene_gn=RPTOR;refGene_tx=XM_019818372.2;refGene_gn=RPTOR;refGene_tx=XM_019818373.2;refGene_gn=RPTOR;refGene_tx=XM_019818375.2;refGene_gn=RPTOR;refGene_tx=XM_023244284.1;refGene_gn=RPTOR;refGene_tx=XM_023244285.1
E1 61600161 61600224 refGene_gn=CHMP6;refGene_tx=XM_023244291.1
E1 61602348 61602458 refGene_gn=CHMP6;refGene_tx=XM_023244291.1
E1 61602588 61602676 refGene_gn=CHMP6;refGene_tx=XM_023244291.1
E1 61602764 61602851 refGene_gn=CHMP6;refGene_tx=XM_023244291.1
E1 61603618 61603684 refGene_gn=CHMP6;refGene_tx=XM_023244291.1
E1 61603906 61603987 refGene_gn=CHMP6;refGene_tx=XM_023244291.1
E1 61604993 61605048 refGene_gn=CHMP6;refGene_tx=XM_023244291.1
E1 61605574 61605630 refGene_gn=CHMP6;refGene_tx=XM_023244291.1
E1 61623693 61623747
refGene_gn=BAIAP2;refGene_tx=XM_003997263.5;refGene_gn=BAIAP2;refGene_tx=XM_003997264.5;refGene_gn=BAIAP2;refGene_tx=XM_006940657.4;refGene_gn=BAIAP2;refGene_tx=XM_019818377.2
E1 61637915 61637991
refGene_gn=BAIAP2;refGene_tx=XM_003997263.5;refGene_gn=BAIAP2;refGene_tx=XM_003997264.5;refGene_gn=BAIAP2;refGene_tx=XM_006940657.4;refGene_

gn=BAIAP2;refGene_tx=XM_019818377.2
E1 61640616 61640703
refGene_gn=BAIAP2;refGene_tx=XM_003997263.5;refGene_gn=BAIAP2;refGene_tx=XM_003997264.5;refGene_gn=BAIAP2;refGene_tx=XM_006940657.4;refGene_gn=BAIAP2;refGene_tx=XM_019818377.2
E1 61661891 61661953
refGene_gn=BAIAP2;refGene_tx=XM_003997263.5;refGene_gn=BAIAP2;refGene_tx=XM_003997264.5;refGene_gn=BAIAP2;refGene_tx=XM_006940657.4;refGene_gn=BAIAP2;refGene_tx=XM_019818377.2
E1 61662727 61662799
refGene_gn=BAIAP2;refGene_tx=XM_003997263.5;refGene_gn=BAIAP2;refGene_tx=XM_003997264.5;refGene_gn=BAIAP2;refGene_tx=XM_006940657.4;refGene_gn=BAIAP2;refGene_tx=XM_019818377.2
E1 61663367 61663505
refGene_gn=BAIAP2;refGene_tx=XM_003997263.5;refGene_gn=BAIAP2;refGene_tx=XM_003997264.5;refGene_gn=BAIAP2;refGene_tx=XM_006940657.4;refGene_gn=BAIAP2;refGene_tx=XM_019818377.2
E1 61672728 61672881
refGene_gn=BAIAP2;refGene_tx=XM_003997263.5;refGene_gn=BAIAP2;refGene_tx=XM_003997264.5;refGene_gn=BAIAP2;refGene_tx=XM_006940657.4;refGene_gn=BAIAP2;refGene_tx=XM_019818377.2
E1 61675213 61675429
refGene_gn=BAIAP2;refGene_tx=XM_003997263.5;refGene_gn=BAIAP2;refGene_tx=XM_003997264.5;refGene_gn=BAIAP2;refGene_tx=XM_006940657.4;refGene_gn=BAIAP2;refGene_tx=XM_019818377.2
E1 61675569 61675771
refGene_gn=BAIAP2;refGene_tx=XM_003997263.5;refGene_gn=BAIAP2;refGene_tx=XM_003997264.5;refGene_gn=BAIAP2;refGene_tx=XM_006940657.4;refGene_gn=BAIAP2;refGene_tx=XM_019818377.2
E1 61676344 61676546
refGene_gn=BAIAP2;refGene_tx=XM_003997263.5;refGene_gn=BAIAP2;refGene_tx=XM_003997264.5;refGene_gn=BAIAP2;refGene_tx=XM_006940657.4;refGene_gn=BAIAP2;refGene_tx=XM_019818377.2
E1 61677513 61677582
refGene_gn=BAIAP2;refGene_tx=XM_003997263.5;refGene_gn=BAIAP2;refGene_tx=XM_003997264.5;refGene_gn=BAIAP2;refGene_tx=XM_006940657.4;refGene_gn=BAIAP2;refGene_tx=XM_019818377.2
E1 61677983 61678146
refGene_gn=BAIAP2;refGene_tx=XM_003997263.5;refGene_gn=BAIAP2;refGene_tx=XM_003997264.5;refGene_gn=BAIAP2;refGene_tx=XM_006940657.4;refGene_gn=BAIAP2;refGene_tx=XM_019818377.2
E1 61679830 61679865
refGene_gn=BAIAP2;refGene_tx=XM_003997263.5;refGene_gn=BAIAP2;refGene_tx=XM_003997264.5;refGene_gn=BAIAP2;refGene_tx=XM_006940657.4;refGene_gn=BAIAP2;refGene_tx=XM_019818377.2
E1 61680395 61680426 refGene_gn=BAIAP2;refGene_tx=XM_003997263.5
E1 61682299 61682327
refGene_gn=BAIAP2;refGene_tx=XM_006940657.4;refGene_gn=BAIAP2;refGene_tx=XM_019818377.2
E1 61685985 61686055 refGene_gn=BAIAP2;refGene_tx=XM_003997264.5

E1 61688370 61688411
refGene_gn=AATK;refGene_tx=XM_019818379.2;refGene_gn=AATK;refGene_tx=XM_023244279.1;refGene_gn=AATK;refGene_tx=XM_023244280.1;refGene_gn=AATK;refGene_tx=XM_023244281.1;refGene_gn=AATK;refGene_tx=XM_023244282.1;refGene_gn=AATK;refGene_tx=XM_023244283.1
E1 61689413 61689599
refGene_gn=AATK;refGene_tx=XM_019818379.2;refGene_gn=AATK;refGene_tx=XM_023244279.1;refGene_gn=AATK;refGene_tx=XM_023244280.1;refGene_gn=AATK;refGene_tx=XM_023244281.1;refGene_gn=AATK;refGene_tx=XM_023244282.1;refGene_gn=AATK;refGene_tx=XM_023244283.1
E1 61689729 61689877
refGene_gn=AATK;refGene_tx=XM_019818379.2;refGene_gn=AATK;refGene_tx=XM_023244279.1;refGene_gn=AATK;refGene_tx=XM_023244280.1;refGene_gn=AATK;refGene_tx=XM_023244281.1;refGene_gn=AATK;refGene_tx=XM_023244282.1;refGene_gn=AATK;refGene_tx=XM_023244283.1
E1 61690009 61692617
refGene_gn=AATK;refGene_tx=XM_019818379.2;refGene_gn=AATK;refGene_tx=XM_023244279.1;refGene_gn=AATK;refGene_tx=XM_023244280.1;refGene_gn=AATK;refGene_tx=XM_023244281.1;refGene_gn=AATK;refGene_tx=XM_023244282.1;refGene_gn=AATK;refGene_tx=XM_023244283.1
E1 61693068 61693218
refGene_gn=AATK;refGene_tx=XM_019818379.2;refGene_gn=AATK;refGene_tx=XM_023244279.1;refGene_gn=AATK;refGene_tx=XM_023244280.1;refGene_gn=AATK;refGene_tx=XM_023244281.1;refGene_gn=AATK;refGene_tx=XM_023244282.1;refGene_gn=AATK;refGene_tx=XM_023244283.1
E1 61694492 61694614
refGene_gn=AATK;refGene_tx=XM_019818379.2;refGene_gn=AATK;refGene_tx=XM_023244279.1;refGene_gn=AATK;refGene_tx=XM_023244280.1;refGene_gn=AATK;refGene_tx=XM_023244281.1;refGene_gn=AATK;refGene_tx=XM_023244282.1;refGene_gn=AATK;refGene_tx=XM_023244283.1
E1 61694706 61694791
refGene_gn=AATK;refGene_tx=XM_019818379.2;refGene_gn=AATK;refGene_tx=XM_023244279.1;refGene_gn=AATK;refGene_tx=XM_023244280.1;refGene_gn=AATK;refGene_tx=XM_023244281.1;refGene_gn=AATK;refGene_tx=XM_023244282.1;refGene_gn=AATK;refGene_tx=XM_023244283.1
E1 61696580 61696714
refGene_gn=AATK;refGene_tx=XM_019818379.2;refGene_gn=AATK;refGene_tx=XM_023244279.1;refGene_gn=AATK;refGene_tx=XM_023244280.1;refGene_gn=AATK;refGene_tx=XM_023244281.1;refGene_gn=AATK;refGene_tx=XM_023244282.1;refGene_gn=AATK;refGene_tx=XM_023244283.1
E1 61697551 61697717
refGene_gn=AATK;refGene_tx=XM_019818379.2;refGene_gn=AATK;refGene_tx=XM_023244279.1;refGene_gn=AATK;refGene_tx=XM_023244280.1;refGene_gn=AATK;refGene_tx=XM_023244281.1;refGene_gn=AATK;refGene_tx=XM_023244282.1;refGene_gn=AATK;refGene_tx=XM_023244283.1
E1 61697833 61697952
refGene_gn=AATK;refGene_tx=XM_023244279.1;refGene_gn=AATK;refGene_tx=XM_023244280.1;refGene_gn=AATK;refGene_tx=XM_023244281.1;refGene_gn=AATK;refGene_tx=XM_023244282.1;refGene_gn=AATK;refGene_tx=XM_023244283.1
E1 61698428 61698508

refGene_gn=AATK;refGene_tx=XM_023244279.1;refGene_gn=AATK;refGene_tx=XM_023244280.1;refGene_gn=AATK;refGene_tx=XM_023244281.1;refGene_gn=AATK;refGene_tx=XM_023244282.1;refGene_gn=AATK;refGene_tx=XM_023244283.1
E1 61700809 61700957
refGene_gn=AATK;refGene_tx=XM_023244279.1;refGene_gn=AATK;refGene_tx=XM_023244280.1;refGene_gn=AATK;refGene_tx=XM_023244281.1;refGene_gn=AATK;refGene_tx=XM_023244282.1;refGene_gn=AATK;refGene_tx=XM_023244283.1
E1 61702338 61702545 refGene_gn=AATK;refGene_tx=XM_023244280.1
E1 61703850 61703984
refGene_gn=AATK;refGene_tx=XM_023244279.1;refGene_gn=AATK;refGene_tx=XM_023244281.1
E1 61719703 61719785
refGene_gn=AATK;refGene_tx=XM_023244279.1;refGene_gn=AATK;refGene_tx=XM_023244281.1
E1 61730100 61730206
refGene_gn=LOC105261202;refGene_tx=XM_023244298.1
E1 61730436 61730572
refGene_gn=LOC105261202;refGene_tx=XM_023244298.1
E1 61730870 61730986
refGene_gn=LOC105261202;refGene_tx=XM_023244298.1
E1 61731967 61732014
refGene_gn=LOC105261202;refGene_tx=XM_023244298.1
E1 61737182 61737257 refGene_gn=CEP131;refGene_tx=XM_023244299.1
E1 61737334 61737395 refGene_gn=CEP131;refGene_tx=XM_023244299.1
E1 61738317 61738481 refGene_gn=CEP131;refGene_tx=XM_023244299.1
E1 61738560 61738738 refGene_gn=CEP131;refGene_tx=XM_023244299.1
E1 61738815 61738958 refGene_gn=CEP131;refGene_tx=XM_023244299.1
E1 61739826 61739901 refGene_gn=CEP131;refGene_tx=XM_023244299.1
E1 61740281 61740399 refGene_gn=CEP131;refGene_tx=XM_023244299.1
E1 61740480 61740588 refGene_gn=CEP131;refGene_tx=XM_023244299.1
E1 61740729 61740931 refGene_gn=CEP131;refGene_tx=XM_023244299.1
E1 61741663 61741766 refGene_gn=CEP131;refGene_tx=XM_023244299.1
E1 61742581 61742698 refGene_gn=CEP131;refGene_tx=XM_023244299.1
E1 61743076 61743202 refGene_gn=CEP131;refGene_tx=XM_023244299.1
E1 61743374 61743494 refGene_gn=CEP131;refGene_tx=XM_023244299.1
E1 61743568 61743745 refGene_gn=CEP131;refGene_tx=XM_023244299.1
E1 61743999 61744182 refGene_gn=CEP131;refGene_tx=XM_023244299.1
E1 61744608 61744703 refGene_gn=CEP131;refGene_tx=XM_023244299.1
E1 61745179 61745345 refGene_gn=CEP131;refGene_tx=XM_023244299.1
E1 61745593 61745699 refGene_gn=CEP131;refGene_tx=XM_023244299.1
E1 61746030 61746159 refGene_gn=CEP131;refGene_tx=XM_023244299.1
E1 61747418 61747574 refGene_gn=CEP131;refGene_tx=XM_023244299.1
E1 61748432 61748546 refGene_gn=CEP131;refGene_tx=XM_023244299.1
E1 61752403 61752531 refGene_gn=CEP131;refGene_tx=XM_023244299.1
E1 61752741 61752856 refGene_gn=CEP131;refGene_tx=XM_023244299.1
E1 61753364 61753474 refGene_gn=CEP131;refGene_tx=XM_023244299.1
E1 61754087 61754273 refGene_gn=CEP131;refGene_tx=XM_023244299.1
E1 61765831 61766344 refGene_gn=TEPSIN;refGene_tx=XM_023244290.1
E1 61767310 61767445 refGene_gn=TEPSIN;refGene_tx=XM_023244290.1
E1 61767682 61767761 refGene_gn=TEPSIN;refGene_tx=XM_023244290.1

E1 61767841 61767955 refGene_gn=TEPSIN;refGene_tx=XM_023244290.1
E1 61768041 61768219 refGene_gn=TEPSIN;refGene_tx=XM_023244290.1
E1 61769003 61769213 refGene_gn=TEPSIN;refGene_tx=XM_023244290.1
E1 61769940 61770012 refGene_gn=TEPSIN;refGene_tx=XM_023244290.1
E1 61770153 61770232 refGene_gn=TEPSIN;refGene_tx=XM_023244290.1
E1 61770395 61770463 refGene_gn=TEPSIN;refGene_tx=XM_023244290.1
E1 61772241 61772335 refGene_gn=TEPSIN;refGene_tx=XM_023244290.1
E1 61772472 61772564 refGene_gn=TEPSIN;refGene_tx=XM_023244290.1
E1 61772851 61772924 refGene_gn=TEPSIN;refGene_tx=XM_023244290.1
E1 61774003 61774051 refGene_gn=TEPSIN;refGene_tx=XM_023244290.1
E1 61774410 61774494
refGene_gn=NDUFAF8;refGene_tx=XM_023244293.1;refGene_gn=NDUFAF8;refGene_tx=XM_023244294.1;refGene_gn=NDUFAF8;refGene_tx=XM_023244295.1;refGene_gn=NDUFAF8;refGene_tx=XM_023244296.1;refGene_gn=NDUFAF8;refGene_tx=XM_023244297.1
E1 61774604 61774715
refGene_gn=NDUFAF8;refGene_tx=XM_023244293.1;refGene_gn=NDUFAF8;refGene_tx=XM_023244294.1;refGene_gn=NDUFAF8;refGene_tx=XM_023244295.1;refGene_gn=NDUFAF8;refGene_tx=XM_023244296.1;refGene_gn=NDUFAF8;refGene_tx=XM_023244297.1
E1 61776206 61776263
refGene_gn=NDUFAF8;refGene_tx=XM_023244293.1;refGene_gn=NDUFAF8;refGene_tx=XM_023244294.1;refGene_gn=NDUFAF8;refGene_tx=XM_023244297.1
E1 61776386 61776498
refGene_gn=NDUFAF8;refGene_tx=XM_023244293.1;refGene_gn=NDUFAF8;refGene_tx=XM_023244294.1;refGene_gn=NDUFAF8;refGene_tx=XM_023244295.1;refGene_gn=NDUFAF8;refGene_tx=XM_023244296.1
E1 61776799 61776876
refGene_gn=NDUFAF8;refGene_tx=XM_023244293.1;refGene_gn=NDUFAF8;refGene_tx=XM_023244294.1;refGene_gn=NDUFAF8;refGene_tx=XM_023244295.1;refGene_gn=NDUFAF8;refGene_tx=XM_023244296.1;refGene_gn=NDUFAF8;refGene_tx=XM_023244297.1
E1 61779743 61780798
refGene_gn=SLC38A10;refGene_tx=XM_023244286.1;refGene_gn=SLC38A10;refGene_tx=XM_023244287.1;refGene_gn=SLC38A10;refGene_tx=XM_023244288.1;refGene_gn=SLC38A10;refGene_tx=XM_023244289.1
E1 61781019 61781181
refGene_gn=SLC38A10;refGene_tx=XM_023244286.1;refGene_gn=SLC38A10;refGene_tx=XM_023244287.1;refGene_gn=SLC38A10;refGene_tx=XM_023244288.1;refGene_gn=SLC38A10;refGene_tx=XM_023244289.1
E1 61783247 61783271
refGene_gn=SLC38A10;refGene_tx=XM_023244286.1;refGene_gn=SLC38A10;refGene_tx=XM_023244287.1;refGene_gn=SLC38A10;refGene_tx=XM_023244289.1
E1 61784434 61784551
refGene_gn=SLC38A10;refGene_tx=XM_023244286.1;refGene_gn=SLC38A10;refGene_tx=XM_023244287.1;refGene_gn=SLC38A10;refGene_tx=XM_023244288.1;refGene_gn=SLC38A10;refGene_tx=XM_023244289.1
E1 61785190 61785646
refGene_gn=SLC38A10;refGene_tx=XM_023244286.1;refGene_gn=SLC38A10;refGene_tx=XM_023244287.1;refGene_gn=SLC38A10;refGene_tx=XM_023244288.1;re

fGene_gn=SLC38A10;refGene_tx=XM_023244289.1
E1 61785948 61786116
refGene_gn=SLC38A10;refGene_tx=XM_023244286.1;refGene_gn=SLC38A10;refGene_tx=XM_023244287.1;refGene_gn=SLC38A10;refGene_tx=XM_023244288.1;refGene_gn=SLC38A10;refGene_tx=XM_023244289.1
E1 61791351 61791508
refGene_gn=SLC38A10;refGene_tx=XM_023244286.1;refGene_gn=SLC38A10;refGene_tx=XM_023244287.1;refGene_gn=SLC38A10;refGene_tx=XM_023244288.1;refGene_gn=SLC38A10;refGene_tx=XM_023244289.1
E1 61802537 61802644
refGene_gn=SLC38A10;refGene_tx=XM_023244286.1;refGene_gn=SLC38A10;refGene_tx=XM_023244287.1;refGene_gn=SLC38A10;refGene_tx=XM_023244288.1;refGene_gn=SLC38A10;refGene_tx=XM_023244289.1
E1 61803717 61803829
refGene_gn=SLC38A10;refGene_tx=XM_023244286.1;refGene_gn=SLC38A10;refGene_tx=XM_023244287.1;refGene_gn=SLC38A10;refGene_tx=XM_023244288.1;refGene_gn=SLC38A10;refGene_tx=XM_023244289.1
E1 61807645 61807828
refGene_gn=SLC38A10;refGene_tx=XM_023244286.1;refGene_gn=SLC38A10;refGene_tx=XM_023244287.1;refGene_gn=SLC38A10;refGene_tx=XM_023244288.1;refGene_gn=SLC38A10;refGene_tx=XM_023244289.1
E1 61808745 61808848
refGene_gn=SLC38A10;refGene_tx=XM_023244286.1;refGene_gn=SLC38A10;refGene_tx=XM_023244287.1;refGene_gn=SLC38A10;refGene_tx=XM_023244288.1;refGene_gn=SLC38A10;refGene_tx=XM_023244289.1
E1 61811644 61811769
refGene_gn=SLC38A10;refGene_tx=XM_023244286.1;refGene_gn=SLC38A10;refGene_tx=XM_023244287.1;refGene_gn=SLC38A10;refGene_tx=XM_023244288.1;refGene_gn=SLC38A10;refGene_tx=XM_023244289.1
E1 61812989 61813133
refGene_gn=SLC38A10;refGene_tx=XM_023244286.1;refGene_gn=SLC38A10;refGene_tx=XM_023244287.1;refGene_gn=SLC38A10;refGene_tx=XM_023244288.1;refGene_gn=SLC38A10;refGene_tx=XM_023244289.1
E1 61813338 61813432
refGene_gn=SLC38A10;refGene_tx=XM_023244286.1;refGene_gn=SLC38A10;refGene_tx=XM_023244287.1;refGene_gn=SLC38A10;refGene_tx=XM_023244288.1;refGene_gn=SLC38A10;refGene_tx=XM_023244289.1
E1 61814190 61814236
refGene_gn=SLC38A10;refGene_tx=XM_023244286.1;refGene_gn=SLC38A10;refGene_tx=XM_023244287.1;refGene_gn=SLC38A10;refGene_tx=XM_023244288.1;refGene_gn=SLC38A10;refGene_tx=XM_023244289.1
E1 61818874 61818992
refGene_gn=SLC38A10;refGene_tx=XM_023244286.1;refGene_gn=SLC38A10;refGene_tx=XM_023244287.1;refGene_gn=SLC38A10;refGene_tx=XM_023244288.1;refGene_gn=SLC38A10;refGene_tx=XM_023244289.1
E1 61823445 61823544
refGene_gn=SLC38A10;refGene_tx=XM_023244286.1;refGene_gn=SLC38A10;refGene_tx=XM_023244287.1;refGene_gn=SLC38A10;refGene_tx=XM_023244288.1;refGene_gn=SLC38A10;refGene_tx=XM_023244289.1
E1 61914003 61914190 refGene_gn=BAHCC1;refGene_tx=XM_023244301.1

E1 61936796 61936976
refGene_gn=BAHCC1;refGene_tx=XM_023244301.1;refGene_gn=BAHCC1;refGene_tx=XM_023244302.1
E1 61946853 61946976
refGene_gn=BAHCC1;refGene_tx=XM_023244301.1;refGene_gn=BAHCC1;refGene_tx=XM_023244302.1
E1 61950063 61951827
refGene_gn=BAHCC1;refGene_tx=XM_023244301.1;refGene_gn=BAHCC1;refGene_tx=XM_023244302.1
E1 61952139 61952248
refGene_gn=BAHCC1;refGene_tx=XM_023244301.1;refGene_gn=BAHCC1;refGene_tx=XM_023244302.1
E1 61952578 61952784
refGene_gn=BAHCC1;refGene_tx=XM_023244301.1;refGene_gn=BAHCC1;refGene_tx=XM_023244302.1
E1 61952960 61953119
refGene_gn=BAHCC1;refGene_tx=XM_023244301.1;refGene_gn=BAHCC1;refGene_tx=XM_023244302.1
E1 61953370 61953555
refGene_gn=BAHCC1;refGene_tx=XM_023244301.1;refGene_gn=BAHCC1;refGene_tx=XM_023244302.1
E1 61953659 61953996
refGene_gn=BAHCC1;refGene_tx=XM_023244301.1;refGene_gn=BAHCC1;refGene_tx=XM_023244302.1
E1 61955063 61955891
refGene_gn=BAHCC1;refGene_tx=XM_023244301.1;refGene_gn=BAHCC1;refGene_tx=XM_023244302.1
E1 61961590 61961790
refGene_gn=BAHCC1;refGene_tx=XM_023244301.1;refGene_gn=BAHCC1;refGene_tx=XM_023244302.1
E1 61961922 61962059
refGene_gn=BAHCC1;refGene_tx=XM_023244301.1;refGene_gn=BAHCC1;refGene_tx=XM_023244302.1
E1 61962401 61962530
refGene_gn=BAHCC1;refGene_tx=XM_023244301.1;refGene_gn=BAHCC1;refGene_tx=XM_023244302.1
E1 61964550 61964668
refGene_gn=BAHCC1;refGene_tx=XM_023244301.1;refGene_gn=BAHCC1;refGene_tx=XM_023244302.1
E1 61965150 61965442
refGene_gn=BAHCC1;refGene_tx=XM_023244301.1;refGene_gn=BAHCC1;refGene_tx=XM_023244302.1
E1 61966141 61966324
refGene_gn=BAHCC1;refGene_tx=XM_023244301.1;refGene_gn=BAHCC1;refGene_tx=XM_023244302.1
E1 61966416 61966718
refGene_gn=BAHCC1;refGene_tx=XM_023244301.1;refGene_gn=BAHCC1;refGene_tx=XM_023244302.1
E1 61966964 61967072
refGene_gn=BAHCC1;refGene_tx=XM_023244301.1;refGene_gn=BAHCC1;refGene_

tx=XM_023244302.1
E1 61967162 61967334
refGene_gn=BAHCC1;refGene_tx=XM_023244301.1;refGene_gn=BAHCC1;refGene_
tx=XM_023244302.1
E1 61967399 61967514
refGene_gn=BAHCC1;refGene_tx=XM_023244301.1;refGene_gn=BAHCC1;refGene_
tx=XM_023244302.1
E1 61967616 61967692
refGene_gn=BAHCC1;refGene_tx=XM_023244301.1;refGene_gn=BAHCC1;refGene_
tx=XM_023244302.1
E1 61968027 61968136
refGene_gn=BAHCC1;refGene_tx=XM_023244301.1;refGene_gn=BAHCC1;refGene_
tx=XM_023244302.1
E1 61968349 61968469
refGene_gn=BAHCC1;refGene_tx=XM_023244301.1;refGene_gn=BAHCC1;refGene_
tx=XM_023244302.1
E1 61968561 61968738
refGene_gn=BAHCC1;refGene_tx=XM_023244301.1;refGene_gn=BAHCC1;refGene_
tx=XM_023244302.1
E1 61968864 61970069
refGene_gn=BAHCC1;refGene_tx=XM_023244301.1;refGene_gn=BAHCC1;refGene_
tx=XM_023244302.1
E1 61971098 61971335
refGene_gn=BAHCC1;refGene_tx=XM_023244301.1;refGene_gn=BAHCC1;refGene_
tx=XM_023244302.1
E1 61971887 61972100
refGene_gn=BAHCC1;refGene_tx=XM_023244301.1;refGene_gn=BAHCC1;refGene_
tx=XM_023244302.1
E1 62017381 62017525 refGene_gn=ACTG1;refGene_tx=XM_023244304.1
E1 62017614 62017796 refGene_gn=ACTG1;refGene_tx=XM_023244304.1
E1 62017888 62018327 refGene_gn=ACTG1;refGene_tx=XM_023244304.1
E1 62018610 62018973 refGene_gn=ACTG1;refGene_tx=XM_023244304.1
E1 62034067 62034893 refGene_gn=FSCN2;refGene_tx=XM_003997267.5
E1 62037826 62037983 refGene_gn=FSCN2;refGene_tx=XM_003997267.5
E1 62038405 62038527 refGene_gn=FSCN2;refGene_tx=XM_003997267.5
E1 62038910 62039078 refGene_gn=FSCN2;refGene_tx=XM_003997267.5
E1 62039179 62039385 refGene_gn=FSCN2;refGene_tx=XM_003997267.5
E1 62042669 62042801 refGene_gn=FAAP100;refGene_tx=XM_023244305.1
E1 62043085 62043172 refGene_gn=FAAP100;refGene_tx=XM_023244305.1
E1 62045140 62045257 refGene_gn=FAAP100;refGene_tx=XM_023244305.1
E1 62046491 62046628 refGene_gn=FAAP100;refGene_tx=XM_023244305.1
E1 62046725 62047498 refGene_gn=FAAP100;refGene_tx=XM_023244305.1
E1 62048353 62048510 refGene_gn=FAAP100;refGene_tx=XM_023244305.1
E1 62049633 62050592 refGene_gn=FAAP100;refGene_tx=XM_023244305.1
E1 62051425 62051544 refGene_gn=FAAP100;refGene_tx=XM_023244305.1
E1 62051869 62052034 refGene_gn=FAAP100;refGene_tx=XM_023244305.1
E1 62056876 62057034
refGene_gn=NPLOC4;refGene_tx=XM_023244307.1;refGene_gn=NPLOC4;refGene_
tx=XM_023244308.1
E1 62059002 62059105

refGene_gn=NPLOC4;refGene_tx=XM_023244307.1;refGene_gn=NPLOC4;refGene_tx=XM_023244308.1
E1 62060654 62060771
refGene_gn=NPLOC4;refGene_tx=XM_023244307.1;refGene_gn=NPLOC4;refGene_tx=XM_023244308.1
E1 62061823 62061919
refGene_gn=NPLOC4;refGene_tx=XM_023244307.1;refGene_gn=NPLOC4;refGene_tx=XM_023244308.1
E1 62065133 62065205
refGene_gn=NPLOC4;refGene_tx=XM_023244307.1;refGene_gn=NPLOC4;refGene_tx=XM_023244308.1
E1 62085696 62085857
refGene_gn=NPLOC4;refGene_tx=XM_023244307.1;refGene_gn=NPLOC4;refGene_tx=XM_023244308.1
E1 62089929 62090056
refGene_gn=NPLOC4;refGene_tx=XM_023244307.1;refGene_gn=NPLOC4;refGene_tx=XM_023244308.1
E1 62090427 62090499
refGene_gn=NPLOC4;refGene_tx=XM_023244307.1;refGene_gn=NPLOC4;refGene_tx=XM_023244308.1
E1 62093072 62093159
refGene_gn=NPLOC4;refGene_tx=XM_023244307.1;refGene_gn=NPLOC4;refGene_tx=XM_023244308.1
E1 62096312 62096492
refGene_gn=NPLOC4;refGene_tx=XM_023244307.1;refGene_gn=NPLOC4;refGene_tx=XM_023244308.1
E1 62097569 62097693
refGene_gn=NPLOC4;refGene_tx=XM_023244307.1;refGene_gn=NPLOC4;refGene_tx=XM_023244308.1
E1 62099344 62099439
refGene_gn=NPLOC4;refGene_tx=XM_023244307.1;refGene_gn=NPLOC4;refGene_tx=XM_023244308.1
E1 62100204 62100253
refGene_gn=NPLOC4;refGene_tx=XM_023244307.1;refGene_gn=NPLOC4;refGene_tx=XM_023244308.1
E1 62102178 62102355
refGene_gn=NPLOC4;refGene_tx=XM_023244307.1;refGene_gn=NPLOC4;refGene_tx=XM_023244308.1
E1 62106731 62106844
refGene_gn=NPLOC4;refGene_tx=XM_023244307.1;refGene_gn=NPLOC4;refGene_tx=XM_023244308.1
E1 62116090 62116099 refGene_gn=NPLOC4;refGene_tx=XM_023244308.1
E1 62116458 62116539 refGene_gn=NPLOC4;refGene_tx=XM_023244307.1
E1 62119949 62119964 refGene_gn=NPLOC4;refGene_tx=XM_023244307.1
E1 62124994 62125030
refGene_gn=TSPAN10;refGene_tx=XM_006940660.4;refGene_gn=TSPAN10;refGene_tx=XM_006940661.4;refGene_gn=TSPAN10;refGene_tx=XM_023244309.1
E1 62126370 62126945
refGene_gn=TSPAN10;refGene_tx=XM_006940660.4;refGene_gn=TSPAN10;refGene_tx=XM_006940661.4;refGene_gn=TSPAN10;refGene_tx=XM_023244309.1

E1 62128013 62128509
refGene_gn=TSPAN10;refGene_tx=XM_006940660.4;refGene_gn=TSPAN10;refGene_tx=XM_006940661.4;refGene_gn=TSPAN10;refGene_tx=XM_023244309.1
E1 62130373 62130450 refGene_gn=PDE6G;refGene_tx=XM_019818382.1
E1 62130836 62130877 refGene_gn=PDE6G;refGene_tx=XM_019818382.1
E1 62131284 62131430 refGene_gn=PDE6G;refGene_tx=XM_019818382.1
E1 62135412 62135832 refGene_gn=OXLD1;refGene_tx=XM_006940664.4
E1 62136640 62136766 refGene_gn=OXLD1;refGene_tx=XM_006940664.4
E1 62136906 62137037 refGene_gn=CCDC137;refGene_tx=XM_023244310.1
E1 62137398 62137532 refGene_gn=CCDC137;refGene_tx=XM_023244310.1
E1 62138830 62139053 refGene_gn=CCDC137;refGene_tx=XM_023244310.1
E1 62140337 62140420 refGene_gn=CCDC137;refGene_tx=XM_023244310.1
E1 62140550 62140630 refGene_gn=CCDC137;refGene_tx=XM_023244310.1
E1 62140800 62141025 refGene_gn=CCDC137;refGene_tx=XM_023244310.1
E1 62143888 62144081
refGene_gn=ARL16;refGene_tx=XM_003997309.5;refGene_gn=ARL16;refGene_tx=XM_019818380.2
E1 62144246 62144362
refGene_gn=ARL16;refGene_tx=XM_003997309.5;refGene_gn=ARL16;refGene_tx=XM_019818380.2
E1 62144936 62145050
refGene_gn=ARL16;refGene_tx=XM_003997309.5;refGene_gn=ARL16;refGene_tx=XM_019818380.2
E1 62145527 62145586 refGene_gn=ARL16;refGene_tx=XM_003997309.5
E1 62145744 62145805 refGene_gn=ARL16;refGene_tx=XM_003997309.5
E1 62146102 62146139 refGene_gn=HGS;refGene_tx=XM_003997269.5
E1 62147598 62147683 refGene_gn=HGS;refGene_tx=XM_003997269.5
E1 62148004 62148080 refGene_gn=HGS;refGene_tx=XM_003997269.5
E1 62148469 62148562 refGene_gn=HGS;refGene_tx=XM_003997269.5
E1 62150225 62150349
refGene_gn=HGS;refGene_tx=XM_003997269.5;refGene_gn=HGS;refGene_tx=XM_023244306.1
E1 62151608 62151661
refGene_gn=HGS;refGene_tx=XM_003997269.5;refGene_gn=HGS;refGene_tx=XM_023244306.1
E1 62152071 62152140
refGene_gn=HGS;refGene_tx=XM_003997269.5;refGene_gn=HGS;refGene_tx=XM_023244306.1
E1 62152841 62152966
refGene_gn=HGS;refGene_tx=XM_003997269.5;refGene_gn=HGS;refGene_tx=XM_023244306.1
E1 62153844 62153923
refGene_gn=HGS;refGene_tx=XM_003997269.5;refGene_gn=HGS;refGene_tx=XM_023244306.1
E1 62154058 62154157
refGene_gn=HGS;refGene_tx=XM_003997269.5;refGene_gn=HGS;refGene_tx=XM_023244306.1
E1 62154281 62154383
refGene_gn=HGS;refGene_tx=XM_003997269.5;refGene_gn=HGS;refGene_tx=XM_023244306.1

E1 62154988 62155027
refGene_gn=HGS;refGene_tx=XM_003997269.5;refGene_gn=HGS;refGene_tx=XM_023244306.1
E1 62155111 62155255
refGene_gn=HGS;refGene_tx=XM_003997269.5;refGene_gn=HGS;refGene_tx=XM_023244306.1
E1 62155347 62155407
refGene_gn=HGS;refGene_tx=XM_003997269.5;refGene_gn=HGS;refGene_tx=XM_023244306.1
E1 62155932 62156143
refGene_gn=HGS;refGene_tx=XM_003997269.5;refGene_gn=HGS;refGene_tx=XM_023244306.1
E1 62156511 62156684
refGene_gn=HGS;refGene_tx=XM_003997269.5;refGene_gn=HGS;refGene_tx=XM_023244306.1
E1 62156772 62156913
refGene_gn=HGS;refGene_tx=XM_003997269.5;refGene_gn=HGS;refGene_tx=XM_023244306.1
E1 62157027 62157205
refGene_gn=HGS;refGene_tx=XM_003997269.5;refGene_gn=HGS;refGene_tx=XM_023244306.1
E1 62159018 62159152
refGene_gn=HGS;refGene_tx=XM_003997269.5;refGene_gn=HGS;refGene_tx=XM_023244306.1
E1 62159227 62159347
refGene_gn=HGS;refGene_tx=XM_003997269.5;refGene_gn=HGS;refGene_tx=XM_023244306.1
E1 62159539 62159629
refGene_gn=HGS;refGene_tx=XM_003997269.5;refGene_gn=HGS;refGene_tx=XM_023244306.1
E1 62160016 62160127
refGene_gn=HGS;refGene_tx=XM_003997269.5;refGene_gn=HGS;refGene_tx=XM_023244306.1
E1 62162171 62162245 refGene_gn=MRPL12;refGene_tx=XM_003997270.4
E1 62162868 62163055 refGene_gn=MRPL12;refGene_tx=XM_003997270.4
E1 62163213 62163297 refGene_gn=MRPL12;refGene_tx=XM_003997270.4
E1 62165202 62165337 refGene_gn=MRPL12;refGene_tx=XM_003997270.4
E1 62165465 62165582 refGene_gn=MRPL12;refGene_tx=XM_003997270.4
E1 62170236 62170329 refGene_gn=SLC25A10;refGene_tx=XM_023244311.1
E1 62172805 62172925 refGene_gn=SLC25A10;refGene_tx=XM_023244311.1
E1 62173519 62173634 refGene_gn=SLC25A10;refGene_tx=XM_023244311.1
E1 62173722 62173771 refGene_gn=SLC25A10;refGene_tx=XM_023244311.1
E1 62173989 62174031 refGene_gn=SLC25A10;refGene_tx=XM_023244311.1
E1 62174736 62174780 refGene_gn=SLC25A10;refGene_tx=XM_023244311.1
E1 62174939 62175010 refGene_gn=SLC25A10;refGene_tx=XM_023244311.1
E1 62175131 62175224 refGene_gn=SLC25A10;refGene_tx=XM_023244311.1
E1 62175563 62175641 refGene_gn=SLC25A10;refGene_tx=XM_023244311.1
E1 62176385 62176442 refGene_gn=SLC25A10;refGene_tx=XM_023244311.1
E1 62176534 62176636 refGene_gn=SLC25A10;refGene_tx=XM_023244311.1
E1 62204933 62204999

refGene_gn=GCGR;refGene_tx=XM_023244312.1;refGene_gn=GCGR;refGene_tx=XM_023244313.1
E1 62205108 62205211
refGene_gn=GCGR;refGene_tx=XM_023244312.1;refGene_gn=GCGR;refGene_tx=XM_023244313.1
E1 62205920 62206028
refGene_gn=GCGR;refGene_tx=XM_023244312.1;refGene_gn=GCGR;refGene_tx=XM_023244313.1
E1 62206101 62206223
refGene_gn=GCGR;refGene_tx=XM_023244312.1;refGene_gn=GCGR;refGene_tx=XM_023244313.1
E1 62206318 62206425
refGene_gn=GCGR;refGene_tx=XM_023244312.1;refGene_gn=GCGR;refGene_tx=XM_023244313.1
E1 62206512 62206669
refGene_gn=GCGR;refGene_tx=XM_023244312.1;refGene_gn=GCGR;refGene_tx=XM_023244313.1
E1 62206757 62206917
refGene_gn=GCGR;refGene_tx=XM_023244312.1;refGene_gn=GCGR;refGene_tx=XM_023244313.1
E1 62206993 62207054
refGene_gn=GCGR;refGene_tx=XM_023244312.1;refGene_gn=GCGR;refGene_tx=XM_023244313.1
E1 62207274 62207344
refGene_gn=GCGR;refGene_tx=XM_023244312.1;refGene_gn=GCGR;refGene_tx=XM_023244313.1
E1 62207565 62207654
refGene_gn=GCGR;refGene_tx=XM_023244312.1;refGene_gn=GCGR;refGene_tx=XM_023244313.1
E1 62207774 62207913
refGene_gn=GCGR;refGene_tx=XM_023244312.1;refGene_gn=GCGR;refGene_tx=XM_023244313.1
E1 62207988 62208030
refGene_gn=GCGR;refGene_tx=XM_023244312.1;refGene_gn=GCGR;refGene_tx=XM_023244313.1
E1 62208377 62208608
refGene_gn=GCGR;refGene_tx=XM_023244312.1;refGene_gn=GCGR;refGene_tx=XM_023244313.1
E1 62214454 62214519
refGene_gn=MCRIP1;refGene_tx=XM_023244314.1;refGene_gn=MCRIP1;refGene_tx=XM_023244315.1
E1 62214728 62214830
refGene_gn=MCRIP1;refGene_tx=XM_023244314.1;refGene_gn=MCRIP1;refGene_tx=XM_023244315.1
E1 62215349 62215468
refGene_gn=MCRIP1;refGene_tx=XM_023244314.1;refGene_gn=MCRIP1;refGene_tx=XM_023244315.1
E1 62215561 62215569
refGene_gn=MCRIP1;refGene_tx=XM_023244314.1;refGene_gn=MCRIP1;refGene_tx=XM_023244315.1

E1 62221160 62221284 refGene_gn=PPP1R27;refGene_tx=XM_003997272.4
E1 62221604 62221755 refGene_gn=PPP1R27;refGene_tx=XM_003997272.4
E1 62221840 62222030 refGene_gn=PPP1R27;refGene_tx=XM_003997272.4
E1 62226803 62226884
refGene_gn=P4HB;refGene_tx=XM_023244316.1;refGene_gn=P4HB;refGene_tx=XM_023244317.1
E1 62227520 62227607
refGene_gn=P4HB;refGene_tx=XM_023244316.1;refGene_gn=P4HB;refGene_tx=XM_023244317.1
E1 62227722 62227904
refGene_gn=P4HB;refGene_tx=XM_023244316.1;refGene_gn=P4HB;refGene_tx=XM_023244317.1
E1 62228017 62228138
refGene_gn=P4HB;refGene_tx=XM_023244316.1;refGene_gn=P4HB;refGene_tx=XM_023244317.1
E1 62228465 62228666
refGene_gn=P4HB;refGene_tx=XM_023244316.1;refGene_gn=P4HB;refGene_tx=XM_023244317.1
E1 62228847 62228973
refGene_gn=P4HB;refGene_tx=XM_023244316.1;refGene_gn=P4HB;refGene_tx=XM_023244317.1
E1 62229157 62229262
refGene_gn=P4HB;refGene_tx=XM_023244316.1;refGene_gn=P4HB;refGene_tx=XM_023244317.1
E1 62232403 62232541
refGene_gn=P4HB;refGene_tx=XM_023244316.1;refGene_gn=P4HB;refGene_tx=XM_023244317.1
E1 62232678 62232812
refGene_gn=P4HB;refGene_tx=XM_023244316.1;refGene_gn=P4HB;refGene_tx=XM_023244317.1
E1 62234229 62234436
refGene_gn=P4HB;refGene_tx=XM_023244316.1;refGene_gn=P4HB;refGene_tx=XM_023244317.1
E1 62235337 62235488 refGene_gn=P4HB;refGene_tx=XM_023244316.1
E1 62237244 62237246 refGene_gn=P4HB;refGene_tx=XM_023244317.1
E1 62240864 62240880 refGene_gn=P4HB;refGene_tx=XM_023244317.1
E1 62241351 62241551
refGene_gn=ARHGDIA;refGene_tx=XM_003997273.5;refGene_gn=ARHGDIA;refGene_tx=XM_023244318.1
E1 62241711 62241775
refGene_gn=ARHGDIA;refGene_tx=XM_003997273.5;refGene_gn=ARHGDIA;refGene_tx=XM_023244318.1
E1 62241863 62241940
refGene_gn=ARHGDIA;refGene_tx=XM_003997273.5;refGene_gn=ARHGDIA;refGene_tx=XM_023244318.1
E1 62242236 62242320
refGene_gn=ARHGDIA;refGene_tx=XM_003997273.5;refGene_gn=ARHGDIA;refGene_tx=XM_023244318.1
E1 62242437 62242627
refGene_gn=ARHGDIA;refGene_tx=XM_003997273.5;refGene_gn=ARHGDIA;refGen

e_tx=XM_023244318.1
E1 62263259 62263274 refGene_gn=ALYREF;refGene_tx=XM_023243481.1
E1 62263375 62263553 refGene_gn=ALYREF;refGene_tx=XM_023243481.1
E1 62263636 62263700 refGene_gn=ALYREF;refGene_tx=XM_023243481.1
E1 62264310 62264458 refGene_gn=ALYREF;refGene_tx=XM_023243481.1
E1 62265888 62266020 refGene_gn=ALYREF;refGene_tx=XM_023243481.1
E1 62266469 62267075 refGene_gn=ALYREF;refGene_tx=XM_023243481.1
E1 62267412 62267538 refGene_gn=ALYREF;refGene_tx=XM_023243481.1
E1 62267911 62268020
refGene_gn=ANAPC11;refGene_tx=XM_003997275.5;refGene_gn=ANAPC11;refGene_tx=XM_006940666.4;refGene_gn=ANAPC11;refGene_tx=XM_023244326.1
E1 62272778 62272924
refGene_gn=ANAPC11;refGene_tx=XM_003997275.5;refGene_gn=ANAPC11;refGene_tx=XM_006940666.4;refGene_gn=ANAPC11;refGene_tx=XM_023244326.1
E1 62275394 62275637 refGene_gn=NPB;refGene_tx=XM_023244325.1
E1 62275746 62275875 refGene_gn=NPB;refGene_tx=XM_023244325.1
E1 62277589 62277701
refGene_gn=PCYT2;refGene_tx=XM_006940667.4;refGene_gn=PCYT2;refGene_tx=XM_006940668.4;refGene_gn=PCYT2;refGene_tx=XM_019818384.2;refGene_gn=PCYT2;refGene_tx=XM_023244324.1
E1 62277790 62277879
refGene_gn=PCYT2;refGene_tx=XM_006940667.4;refGene_gn=PCYT2;refGene_tx=XM_006940668.4;refGene_gn=PCYT2;refGene_tx=XM_019818384.2;refGene_gn=PCYT2;refGene_tx=XM_023244324.1
E1 62278078 62278144
refGene_gn=PCYT2;refGene_tx=XM_006940667.4;refGene_gn=PCYT2;refGene_tx=XM_006940668.4;refGene_gn=PCYT2;refGene_tx=XM_019818384.2;refGene_gn=PCYT2;refGene_tx=XM_023244324.1
E1 62278287 62278353
refGene_gn=PCYT2;refGene_tx=XM_006940667.4;refGene_gn=PCYT2;refGene_tx=XM_006940668.4;refGene_gn=PCYT2;refGene_tx=XM_019818384.2;refGene_gn=PCYT2;refGene_tx=XM_023244324.1
E1 62278556 62278634
refGene_gn=PCYT2;refGene_tx=XM_006940667.4;refGene_gn=PCYT2;refGene_tx=XM_006940668.4;refGene_gn=PCYT2;refGene_tx=XM_019818384.2;refGene_gn=PCYT2;refGene_tx=XM_023244324.1
E1 62278866 62278949
refGene_gn=PCYT2;refGene_tx=XM_006940667.4;refGene_gn=PCYT2;refGene_tx=XM_006940668.4;refGene_gn=PCYT2;refGene_tx=XM_019818384.2;refGene_gn=PCYT2;refGene_tx=XM_023244324.1
E1 62279139 62279278
refGene_gn=PCYT2;refGene_tx=XM_006940667.4;refGene_gn=PCYT2;refGene_tx=XM_006940668.4;refGene_gn=PCYT2;refGene_tx=XM_019818384.2;refGene_gn=PCYT2;refGene_tx=XM_023244324.1
E1 62279916 62279961
refGene_gn=PCYT2;refGene_tx=XM_006940667.4;refGene_gn=PCYT2;refGene_tx=XM_006940668.4;refGene_gn=PCYT2;refGene_tx=XM_019818384.2;refGene_gn=PCYT2;refGene_tx=XM_023244324.1
E1 62280147 62280232
refGene_gn=PCYT2;refGene_tx=XM_006940667.4;refGene_gn=PCYT2;refGene_tx

=XM_006940668.4;refGene_gn=PCYT2;refGene_tx=XM_019818384.2;refGene_gn=PCYT2;refGene_tx=XM_023244324.1
E1 62280758 62280825
refGene_gn=PCYT2;refGene_tx=XM_006940667.4;refGene_gn=PCYT2;refGene_tx=XM_006940668.4;refGene_gn=PCYT2;refGene_tx=XM_019818384.2;refGene_gn=PCYT2;refGene_tx=XM_023244324.1
E1 62281031 62281193
refGene_gn=PCYT2;refGene_tx=XM_006940667.4;refGene_gn=PCYT2;refGene_tx=XM_006940668.4;refGene_gn=PCYT2;refGene_tx=XM_019818384.2;refGene_gn=PCYT2;refGene_tx=XM_023244324.1
E1 62281731 62281820
refGene_gn=PCYT2;refGene_tx=XM_006940667.4;refGene_gn=PCYT2;refGene_tx=XM_023244324.1
E1 62283548 62283637 refGene_gn=PCYT2;refGene_tx=XM_023244324.1
E1 62284528 62284727 refGene_gn=SIRT7;refGene_tx=XM_023244327.1
E1 62285843 62285950 refGene_gn=SIRT7;refGene_tx=XM_023244327.1
E1 62286093 62286174 refGene_gn=SIRT7;refGene_tx=XM_023244327.1
E1 62286317 62286554 refGene_gn=SIRT7;refGene_tx=XM_023244327.1
E1 62286628 62286727 refGene_gn=SIRT7;refGene_tx=XM_023244327.1
E1 62287405 62287478 refGene_gn=SIRT7;refGene_tx=XM_023244327.1
E1 62287576 62287647 refGene_gn=SIRT7;refGene_tx=XM_023244327.1
E1 62289265 62289370 refGene_gn=SIRT7;refGene_tx=XM_023244327.1
E1 62289483 62289621 refGene_gn=SIRT7;refGene_tx=XM_023244327.1
E1 62289691 62289784 refGene_gn=SIRT7;refGene_tx=XM_023244327.1
E1 62294288 62294741
refGene_gn=MAFG;refGene_tx=XM_006940671.4;refGene_gn=MAFG;refGene_tx=XM_019818385.2;refGene_gn=MAFG;refGene_tx=XM_019818386.2;refGene_gn=MAFG;refGene_tx=XM_019818387.2;refGene_gn=MAFG;refGene_tx=XM_023244328.1;refGene_gn=MAFG;refGene_tx=XM_023244329.1;refGene_gn=MAFG;refGene_tx=XM_023244330.1
E1 62294837 62294900
refGene_gn=MAFG;refGene_tx=XM_006940671.4;refGene_gn=MAFG;refGene_tx=XM_019818385.2;refGene_gn=MAFG;refGene_tx=XM_019818386.2;refGene_gn=MAFG;refGene_tx=XM_019818387.2;refGene_gn=MAFG;refGene_tx=XM_023244328.1;refGene_gn=MAFG;refGene_tx=XM_023244329.1;refGene_gn=MAFG;refGene_tx=XM_023244330.1
E1 62296120 62296141 refGene_gn=MAFG;refGene_tx=XM_019818385.2
E1 62304095 62304261
refGene_gn=PYCR1;refGene_tx=XM_003997277.5;refGene_gn=PYCR1;refGene_tx=XM_019818383.2;refGene_gn=PYCR1;refGene_tx=XM_023244331.1;refGene_gn=PYCR1;refGene_tx=XM_023244332.1
E1 62305120 62305284
refGene_gn=PYCR1;refGene_tx=XM_003997277.5;refGene_gn=PYCR1;refGene_tx=XM_019818383.2;refGene_gn=PYCR1;refGene_tx=XM_023244331.1;refGene_gn=PYCR1;refGene_tx=XM_023244332.1
E1 62305431 62305524
refGene_gn=PYCR1;refGene_tx=XM_003997277.5;refGene_gn=PYCR1;refGene_tx=XM_023244331.1;refGene_gn=PYCR1;refGene_tx=XM_023244332.1
E1 62305699 62305921
refGene_gn=PYCR1;refGene_tx=XM_003997277.5;refGene_gn=PYCR1;refGene_tx

=XM_019818383.2;refGene_gn=PYCR1;refGene_tx=XM_023244331.1;refGene_gn=PYCR1;refGene_tx=XM_023244332.1
E1 62306126 62306306
refGene_gn=PYCR1;refGene_tx=XM_003997277.5;refGene_gn=PYCR1;refGene_tx=XM_019818383.2;refGene_gn=PYCR1;refGene_tx=XM_023244331.1;refGene_gn=PYCR1;refGene_tx=XM_023244332.1
E1 62306998 62307069
refGene_gn=PYCR1;refGene_tx=XM_003997277.5;refGene_gn=PYCR1;refGene_tx=XM_019818383.2;refGene_gn=PYCR1;refGene_tx=XM_023244331.1
E1 62307277 62307512
refGene_gn=PYCR1;refGene_tx=XM_003997277.5;refGene_gn=PYCR1;refGene_tx=XM_019818383.2;refGene_gn=PYCR1;refGene_tx=XM_023244331.1
E1 62311866 62312790 refGene_gn=MYADML2;refGene_tx=XM_006940673.2
E1 62320711 62321018 refGene_gn=NOTUM;refGene_tx=XM_023244345.1
E1 62321248 62321296 refGene_gn=NOTUM;refGene_tx=XM_023244345.1
E1 62322446 62322594 refGene_gn=NOTUM;refGene_tx=XM_023244345.1
E1 62323136 62323237 refGene_gn=NOTUM;refGene_tx=XM_023244345.1
E1 62323366 62323558 refGene_gn=NOTUM;refGene_tx=XM_023244345.1
E1 62324273 62324376 refGene_gn=NOTUM;refGene_tx=XM_023244345.1
E1 62324744 62324803 refGene_gn=NOTUM;refGene_tx=XM_023244345.1
E1 62325192 62325253 refGene_gn=NOTUM;refGene_tx=XM_023244345.1
E1 62325718 62325814 refGene_gn=NOTUM;refGene_tx=XM_023244345.1
E1 62325894 62325947 refGene_gn=NOTUM;refGene_tx=XM_023244345.1
E1 62327047 62327382 refGene_gn=NOTUM;refGene_tx=XM_023244345.1
E1 62342208 62342310
refGene_gn=ASPSCR1;refGene_tx=XM_023244347.1;refGene_gn=ASPSCR1;refGene_tx=XM_023244348.1;refGene_gn=ASPSCR1;refGene_tx=XM_023244349.1;refGene_gn=ASPSCR1;refGene_tx=XM_023244350.1
E1 62342920 62342932 refGene_gn=ASPSCR1;refGene_tx=XM_019818389.2
E1 62343272 62343328
refGene_gn=ASPSCR1;refGene_tx=XM_019818389.2;refGene_gn=ASPSCR1;refGene_tx=XM_023244347.1;refGene_gn=ASPSCR1;refGene_tx=XM_023244348.1;refGene_gn=ASPSCR1;refGene_tx=XM_023244349.1;refGene_gn=ASPSCR1;refGene_tx=XM_023244350.1
E1 62344903 62345018
refGene_gn=ASPSCR1;refGene_tx=XM_019818389.2;refGene_gn=ASPSCR1;refGene_tx=XM_019818391.2;refGene_gn=ASPSCR1;refGene_tx=XM_019818392.2;refGene_gn=ASPSCR1;refGene_tx=XM_019818396.2;refGene_gn=ASPSCR1;refGene_tx=XM_023244347.1;refGene_gn=ASPSCR1;refGene_tx=XM_023244348.1;refGene_gn=ASPSCR1;refGene_tx=XM_023244349.1;refGene_gn=ASPSCR1;refGene_tx=XM_023244350.1;refGene_gn=ASPSCR1;refGene_tx=XM_023244351.1;refGene_gn=ASPSCR1;refGene_tx=XM_023244352.1
E1 62346317 62346418
refGene_gn=ASPSCR1;refGene_tx=XM_019818389.2;refGene_gn=ASPSCR1;refGene_tx=XM_019818391.2;refGene_gn=ASPSCR1;refGene_tx=XM_019818392.2;refGene_gn=ASPSCR1;refGene_tx=XM_019818396.2;refGene_gn=ASPSCR1;refGene_tx=XM_023244347.1;refGene_gn=ASPSCR1;refGene_tx=XM_023244348.1;refGene_gn=ASPSCR1;refGene_tx=XM_023244349.1;refGene_gn=ASPSCR1;refGene_tx=XM_023244350.1;refGene_gn=ASPSCR1;refGene_tx=XM_023244351.1;refGene_gn=ASPSCR1;refGene_tx=XM_023244352.1

E1 62353648 62353706
refGene_gn=ASPSCR1;refGene_tx=XM_019818389.2;refGene_gn=ASPSCR1;refGene_tx=XM_019818391.2;refGene_gn=ASPSCR1;refGene_tx=XM_019818392.2;refGene_gn=ASPSCR1;refGene_tx=XM_019818396.2;refGene_gn=ASPSCR1;refGene_tx=XM_023244347.1;refGene_gn=ASPSCR1;refGene_tx=XM_023244348.1;refGene_gn=ASPSCR1;refGene_tx=XM_023244349.1;refGene_gn=ASPSCR1;refGene_tx=XM_023244350.1;refGene_gn=ASPSCR1;refGene_tx=XM_023244351.1;refGene_gn=ASPSCR1;refGene_tx=XM_023244352.1
E1 62354619 62354693
refGene_gn=ASPSCR1;refGene_tx=XM_019818389.2;refGene_gn=ASPSCR1;refGene_tx=XM_019818391.2;refGene_gn=ASPSCR1;refGene_tx=XM_019818392.2;refGene_gn=ASPSCR1;refGene_tx=XM_019818396.2;refGene_gn=ASPSCR1;refGene_tx=XM_023244347.1;refGene_gn=ASPSCR1;refGene_tx=XM_023244348.1;refGene_gn=ASPSCR1;refGene_tx=XM_023244349.1;refGene_gn=ASPSCR1;refGene_tx=XM_023244350.1;refGene_gn=ASPSCR1;refGene_tx=XM_023244351.1;refGene_gn=ASPSCR1;refGene_tx=XM_023244352.1
E1 62355039 62355439
refGene_gn=ASPSCR1;refGene_tx=XM_019818389.2;refGene_gn=ASPSCR1;refGene_tx=XM_019818391.2;refGene_gn=ASPSCR1;refGene_tx=XM_019818392.2;refGene_gn=ASPSCR1;refGene_tx=XM_019818396.2;refGene_gn=ASPSCR1;refGene_tx=XM_023244347.1;refGene_gn=ASPSCR1;refGene_tx=XM_023244348.1;refGene_gn=ASPSCR1;refGene_tx=XM_023244349.1;refGene_gn=ASPSCR1;refGene_tx=XM_023244350.1;refGene_gn=ASPSCR1;refGene_tx=XM_023244351.1;refGene_gn=ASPSCR1;refGene_tx=XM_023244352.1
E1 62361904 62362044
refGene_gn=ASPSCR1;refGene_tx=XM_019818389.2;refGene_gn=ASPSCR1;refGene_tx=XM_019818391.2;refGene_gn=ASPSCR1;refGene_tx=XM_019818392.2;refGene_gn=ASPSCR1;refGene_tx=XM_019818396.2;refGene_gn=ASPSCR1;refGene_tx=XM_023244347.1;refGene_gn=ASPSCR1;refGene_tx=XM_023244348.1;refGene_gn=ASPSCR1;refGene_tx=XM_023244349.1;refGene_gn=ASPSCR1;refGene_tx=XM_023244350.1;refGene_gn=ASPSCR1;refGene_tx=XM_023244351.1;refGene_gn=ASPSCR1;refGene_tx=XM_023244352.1
E1 62362173 62362255
refGene_gn=ASPSCR1;refGene_tx=XM_019818389.2;refGene_gn=ASPSCR1;refGene_tx=XM_019818391.2;refGene_gn=ASPSCR1;refGene_tx=XM_019818392.2;refGene_gn=ASPSCR1;refGene_tx=XM_019818396.2;refGene_gn=ASPSCR1;refGene_tx=XM_023244347.1;refGene_gn=ASPSCR1;refGene_tx=XM_023244348.1;refGene_gn=ASPSCR1;refGene_tx=XM_023244349.1;refGene_gn=ASPSCR1;refGene_tx=XM_023244350.1;refGene_gn=ASPSCR1;refGene_tx=XM_023244351.1;refGene_gn=ASPSCR1;refGene_tx=XM_023244352.1
E1 62362627 62362694
refGene_gn=ASPSCR1;refGene_tx=XM_019818389.2;refGene_gn=ASPSCR1;refGene_tx=XM_019818391.2;refGene_gn=ASPSCR1;refGene_tx=XM_019818392.2;refGene_gn=ASPSCR1;refGene_tx=XM_019818396.2;refGene_gn=ASPSCR1;refGene_tx=XM_023244347.1;refGene_gn=ASPSCR1;refGene_tx=XM_023244348.1;refGene_gn=ASPSCR1;refGene_tx=XM_023244349.1;refGene_gn=ASPSCR1;refGene_tx=XM_023244350.1;refGene_gn=ASPSCR1;refGene_tx=XM_023244351.1;refGene_gn=ASPSCR1;refGene_tx=XM_023244352.1
E1 62363467 62363530
refGene_gn=ASPSCR1;refGene_tx=XM_019818389.2;refGene_gn=ASPSCR1;refGen

e_tx=XM_019818391.2;refGene_gn=ASPSCR1;refGene_tx=XM_019818392.2;refGene_gn=ASPSCR1;refGene_tx=XM_019818396.2;refGene_gn=ASPSCR1;refGene_tx=XM_023244347.1;refGene_gn=ASPSCR1;refGene_tx=XM_023244348.1;refGene_gn=ASPSCR1;refGene_tx=XM_023244349.1;refGene_gn=ASPSCR1;refGene_tx=XM_023244350.1;refGene_gn=ASPSCR1;refGene_tx=XM_023244351.1;refGene_gn=ASPSCR1;refGene_tx=XM_023244352.1
E1 62364087 62364140
refGene_gn=ASPSCR1;refGene_tx=XM_019818389.2;refGene_gn=ASPSCR1;refGene_tx=XM_019818391.2;refGene_gn=ASPSCR1;refGene_tx=XM_019818392.2;refGene_gn=ASPSCR1;refGene_tx=XM_019818396.2;refGene_gn=ASPSCR1;refGene_tx=XM_023244347.1;refGene_gn=ASPSCR1;refGene_tx=XM_023244348.1;refGene_gn=ASPSCR1;refGene_tx=XM_023244349.1;refGene_gn=ASPSCR1;refGene_tx=XM_023244350.1;refGene_gn=ASPSCR1;refGene_tx=XM_023244351.1;refGene_gn=ASPSCR1;refGene_tx=XM_023244352.1
E1 62367370 62367422
refGene_gn=ASPSCR1;refGene_tx=XM_019818389.2;refGene_gn=ASPSCR1;refGene_tx=XM_019818391.2;refGene_gn=ASPSCR1;refGene_tx=XM_019818392.2;refGene_gn=ASPSCR1;refGene_tx=XM_019818396.2;refGene_gn=ASPSCR1;refGene_tx=XM_023244347.1;refGene_gn=ASPSCR1;refGene_tx=XM_023244348.1;refGene_gn=ASPSCR1;refGene_tx=XM_023244351.1;refGene_gn=ASPSCR1;refGene_tx=XM_023244352.1
E1 62367556 62367626
refGene_gn=ASPSCR1;refGene_tx=XM_019818389.2;refGene_gn=ASPSCR1;refGene_tx=XM_019818391.2;refGene_gn=ASPSCR1;refGene_tx=XM_019818392.2;refGene_gn=ASPSCR1;refGene_tx=XM_019818396.2;refGene_gn=ASPSCR1;refGene_tx=XM_023244347.1;refGene_gn=ASPSCR1;refGene_tx=XM_023244348.1;refGene_gn=ASPSCR1;refGene_tx=XM_023244349.1;refGene_gn=ASPSCR1;refGene_tx=XM_023244350.1;refGene_gn=ASPSCR1;refGene_tx=XM_023244351.1;refGene_gn=ASPSCR1;refGene_tx=XM_023244352.1
E1 62367705 62367881
refGene_gn=ASPSCR1;refGene_tx=XM_019818389.2;refGene_gn=ASPSCR1;refGene_tx=XM_019818391.2;refGene_gn=ASPSCR1;refGene_tx=XM_019818392.2;refGene_gn=ASPSCR1;refGene_tx=XM_019818396.2;refGene_gn=ASPSCR1;refGene_tx=XM_023244347.1;refGene_gn=ASPSCR1;refGene_tx=XM_023244348.1;refGene_gn=ASPSCR1;refGene_tx=XM_023244349.1;refGene_gn=ASPSCR1;refGene_tx=XM_023244350.1;refGene_gn=ASPSCR1;refGene_tx=XM_023244351.1;refGene_gn=ASPSCR1;refGene_tx=XM_023244352.1
E1 62368014 62368028 refGene_gn=ASPSCR1;refGene_tx=XM_023244348.1
E1 62368139 62368285
refGene_gn=ASPSCR1;refGene_tx=XM_019818389.2;refGene_gn=ASPSCR1;refGene_tx=XM_019818391.2;refGene_gn=ASPSCR1;refGene_tx=XM_019818392.2;refGene_gn=ASPSCR1;refGene_tx=XM_019818396.2;refGene_gn=ASPSCR1;refGene_tx=XM_023244347.1;refGene_gn=ASPSCR1;refGene_tx=XM_023244351.1;refGene_gn=ASPSCR1;refGene_tx=XM_023244352.1
E1 62369615 62369630 refGene_gn=CENPX;refGene_tx=XM_003997318.5
E1 62369725 62369814 refGene_gn=CENPX;refGene_tx=XM_003997318.5
E1 62370057 62370111 refGene_gn=CENPX;refGene_tx=XM_003997318.5
E1 62370281 62370333 refGene_gn=CENPX;refGene_tx=XM_003997318.5
E1 62372709 62372739 refGene_gn=CENPX;refGene_tx=XM_003997318.5
E1 62373409 62373629

refGene_gn=LRRC45;refGene_tx=XM_003997319.5;refGene_gn=LRRC45;refGene_tx=XM_011289465.3;refGene_gn=LRRC45;refGene_tx=XM_019818388.2
E1 62374180 62374242
refGene_gn=LRRC45;refGene_tx=XM_003997319.5;refGene_gn=LRRC45;refGene_tx=XM_011289465.3;refGene_gn=LRRC45;refGene_tx=XM_019818388.2
E1 62374675 62374746
refGene_gn=LRRC45;refGene_tx=XM_003997319.5;refGene_gn=LRRC45;refGene_tx=XM_011289465.3;refGene_gn=LRRC45;refGene_tx=XM_019818388.2
E1 62374996 62375175
refGene_gn=LRRC45;refGene_tx=XM_003997319.5;refGene_gn=LRRC45;refGene_tx=XM_011289465.3;refGene_gn=LRRC45;refGene_tx=XM_019818388.2
E1 62375428 62375557
refGene_gn=LRRC45;refGene_tx=XM_003997319.5;refGene_gn=LRRC45;refGene_tx=XM_011289465.3
E1 62376624 62376737
refGene_gn=LRRC45;refGene_tx=XM_003997319.5;refGene_gn=LRRC45;refGene_tx=XM_011289465.3;refGene_gn=LRRC45;refGene_tx=XM_019818388.2
E1 62377066 62377125
refGene_gn=LRRC45;refGene_tx=XM_003997319.5;refGene_gn=LRRC45;refGene_tx=XM_011289465.3;refGene_gn=LRRC45;refGene_tx=XM_019818388.2
E1 62377376 62377454
refGene_gn=LRRC45;refGene_tx=XM_003997319.5;refGene_gn=LRRC45;refGene_tx=XM_011289465.3;refGene_gn=LRRC45;refGene_tx=XM_019818388.2
E1 62377729 62377865
refGene_gn=LRRC45;refGene_tx=XM_003997319.5;refGene_gn=LRRC45;refGene_tx=XM_011289465.3;refGene_gn=LRRC45;refGene_tx=XM_019818388.2
E1 62377949 62378027
refGene_gn=LRRC45;refGene_tx=XM_003997319.5;refGene_gn=LRRC45;refGene_tx=XM_011289465.3;refGene_gn=LRRC45;refGene_tx=XM_019818388.2
E1 62378117 62378229
refGene_gn=LRRC45;refGene_tx=XM_003997319.5;refGene_gn=LRRC45;refGene_tx=XM_011289465.3;refGene_gn=LRRC45;refGene_tx=XM_019818388.2
E1 62378342 62378413
refGene_gn=LRRC45;refGene_tx=XM_003997319.5;refGene_gn=LRRC45;refGene_tx=XM_011289465.3;refGene_gn=LRRC45;refGene_tx=XM_019818388.2
E1 62378794 62378887
refGene_gn=LRRC45;refGene_tx=XM_003997319.5;refGene_gn=LRRC45;refGene_tx=XM_011289465.3;refGene_gn=LRRC45;refGene_tx=XM_019818388.2
E1 62379191 62379284
refGene_gn=LRRC45;refGene_tx=XM_003997319.5;refGene_gn=LRRC45;refGene_tx=XM_011289465.3;refGene_gn=LRRC45;refGene_tx=XM_019818388.2
E1 62379830 62380004
refGene_gn=LRRC45;refGene_tx=XM_003997319.5;refGene_gn=LRRC45;refGene_tx=XM_011289465.3;refGene_gn=LRRC45;refGene_tx=XM_019818388.2
E1 62380073 62380221
refGene_gn=LRRC45;refGene_tx=XM_003997319.5;refGene_gn=LRRC45;refGene_tx=XM_011289465.3;refGene_gn=LRRC45;refGene_tx=XM_019818388.2
E1 62380430 62380627
refGene_gn=LRRC45;refGene_tx=XM_003997319.5;refGene_gn=LRRC45;refGene_tx=XM_011289465.3;refGene_gn=LRRC45;refGene_tx=XM_019818388.2

E1 62381674 62381709
refGene_gn=RAC3;refGene_tx=XM_023244353.1;refGene_gn=RAC3;refGene_tx=XM_023244354.1
E1 62382258 62382330
refGene_gn=RAC3;refGene_tx=XM_023244353.1;refGene_gn=RAC3;refGene_tx=XM_023244354.1
E1 62382535 62382653
refGene_gn=RAC3;refGene_tx=XM_023244353.1;refGene_gn=RAC3;refGene_tx=XM_023244354.1
E1 62382782 62382845
refGene_gn=RAC3;refGene_tx=XM_023244353.1;refGene_gn=RAC3;refGene_tx=XM_023244354.1
E1 62383257 62383421
refGene_gn=RAC3;refGene_tx=XM_023244353.1;refGene_gn=RAC3;refGene_tx=XM_023244354.1
E1 62383654 62383785
refGene_gn=RAC3;refGene_tx=XM_023244353.1;refGene_gn=RAC3;refGene_tx=XM_023244354.1
E1 62385724 62385828 refGene_gn=DCXR;refGene_tx=XM_003997279.5
E1 62385955 62386073 refGene_gn=DCXR;refGene_tx=XM_003997279.5
E1 62386163 62386228 refGene_gn=DCXR;refGene_tx=XM_003997279.5
E1 62386316 62386416 refGene_gn=DCXR;refGene_tx=XM_003997279.5
E1 62386490 62386533 refGene_gn=DCXR;refGene_tx=XM_003997279.5
E1 62386626 62386781 refGene_gn=DCXR;refGene_tx=XM_003997279.5
E1 62387237 62387335 refGene_gn=DCXR;refGene_tx=XM_003997279.5
E1 62387410 62387462 refGene_gn=DCXR;refGene_tx=XM_003997279.5
E1 62392798 62392902
refGene_gn=LOC101091418;refGene_tx=XM_003997280.5
E1 62392991 62393109
refGene_gn=LOC101091418;refGene_tx=XM_003997280.5
E1 62393344 62393409
refGene_gn=LOC101091418;refGene_tx=XM_003997280.5
E1 62393525 62393625
refGene_gn=LOC101091418;refGene_tx=XM_003997280.5
E1 62393712 62393755
refGene_gn=LOC101091418;refGene_tx=XM_003997280.5
E1 62393848 62394003
refGene_gn=LOC101091418;refGene_tx=XM_003997280.5
E1 62394106 62394204
refGene_gn=LOC101091418;refGene_tx=XM_003997280.5
E1 62394688 62394740
refGene_gn=LOC101091418;refGene_tx=XM_003997280.5
E1 62409950 62410032
refGene_gn=RFNG;refGene_tx=XM_006940701.4;refGene_gn=RFNG;refGene_tx=XM_023244363.1
E1 62410113 62410199
refGene_gn=RFNG;refGene_tx=XM_006940701.4;refGene_gn=RFNG;refGene_tx=XM_023244363.1
E1 62410356 62410503 refGene_gn=RFNG;refGene_tx=XM_023244363.1
E1 62410682 62410848

refGene_gn=RFNG;refGene_tx=XM_006940701.4;refGene_gn=RFNG;refGene_tx=XM_023244363.1
E1 62410940 62411029
refGene_gn=RFNG;refGene_tx=XM_006940701.4;refGene_gn=RFNG;refGene_tx=XM_023244363.1
E1 62411276 62411430
refGene_gn=RFNG;refGene_tx=XM_006940701.4;refGene_gn=RFNG;refGene_tx=XM_023244363.1
E1 62411565 62411668
refGene_gn=RFNG;refGene_tx=XM_006940701.4;refGene_gn=RFNG;refGene_tx=XM_023244363.1
E1 62412212 62412261
refGene_gn=RFNG;refGene_tx=XM_006940701.4;refGene_gn=RFNG;refGene_tx=XM_023244363.1
E1 62412694 62412976
refGene_gn=RFNG;refGene_tx=XM_006940701.4;refGene_gn=RFNG;refGene_tx=XM_023244363.1
E1 62413127 62413160
refGene_gn=GPS1;refGene_tx=XM_023244355.1;refGene_gn=GPS1;refGene_tx=XM_023244356.1;refGene_gn=GPS1;refGene_tx=XM_023244357.1;refGene_gn=GPS1;refGene_tx=XM_023244358.1
E1 62413344 62413548
refGene_gn=GPS1;refGene_tx=XM_023244355.1;refGene_gn=GPS1;refGene_tx=XM_023244356.1
E1 62414120 62414213
refGene_gn=GPS1;refGene_tx=XM_023244355.1;refGene_gn=GPS1;refGene_tx=XM_023244356.1;refGene_gn=GPS1;refGene_tx=XM_023244357.1;refGene_gn=GPS1;refGene_tx=XM_023244358.1;refGene_gn=GPS1;refGene_tx=XM_023244359.1
E1 62414738 62414920
refGene_gn=GPS1;refGene_tx=XM_023244355.1;refGene_gn=GPS1;refGene_tx=XM_023244356.1;refGene_gn=GPS1;refGene_tx=XM_023244357.1;refGene_gn=GPS1;refGene_tx=XM_023244358.1;refGene_gn=GPS1;refGene_tx=XM_023244359.1
E1 62415373 62415538
refGene_gn=GPS1;refGene_tx=XM_023244355.1;refGene_gn=GPS1;refGene_tx=XM_023244356.1;refGene_gn=GPS1;refGene_tx=XM_023244357.1;refGene_gn=GPS1;refGene_tx=XM_023244358.1;refGene_gn=GPS1;refGene_tx=XM_023244359.1
E1 62415619 62415755
refGene_gn=GPS1;refGene_tx=XM_023244355.1;refGene_gn=GPS1;refGene_tx=XM_023244356.1;refGene_gn=GPS1;refGene_tx=XM_023244357.1;refGene_gn=GPS1;refGene_tx=XM_023244358.1;refGene_gn=GPS1;refGene_tx=XM_023244359.1
E1 62415865 62415943
refGene_gn=GPS1;refGene_tx=XM_023244355.1;refGene_gn=GPS1;refGene_tx=XM_023244356.1;refGene_gn=GPS1;refGene_tx=XM_023244357.1;refGene_gn=GPS1;refGene_tx=XM_023244358.1;refGene_gn=GPS1;refGene_tx=XM_023244359.1
E1 62416123 62416184
refGene_gn=GPS1;refGene_tx=XM_023244355.1;refGene_gn=GPS1;refGene_tx=XM_023244356.1;refGene_gn=GPS1;refGene_tx=XM_023244357.1;refGene_gn=GPS1;refGene_tx=XM_023244358.1;refGene_gn=GPS1;refGene_tx=XM_023244359.1
E1 62416611 62416697
refGene_gn=GPS1;refGene_tx=XM_023244355.1;refGene_gn=GPS1;refGene_tx=X

M_023244356.1;refGene_gn=GPS1;refGene_tx=XM_023244357.1;refGene_gn=GPS1;refGene_tx=XM_023244358.1;refGene_gn=GPS1;refGene_tx=XM_023244359.1
E1 62416936 62417031
refGene_gn=GPS1;refGene_tx=XM_023244355.1;refGene_gn=GPS1;refGene_tx=XM_023244356.1;refGene_gn=GPS1;refGene_tx=XM_023244357.1;refGene_gn=GPS1;refGene_tx=XM_023244358.1;refGene_gn=GPS1;refGene_tx=XM_023244359.1
E1 62417267 62417373
refGene_gn=GPS1;refGene_tx=XM_023244355.1;refGene_gn=GPS1;refGene_tx=XM_023244356.1;refGene_gn=GPS1;refGene_tx=XM_023244357.1;refGene_gn=GPS1;refGene_tx=XM_023244358.1;refGene_gn=GPS1;refGene_tx=XM_023244359.1
E1 62417452 62417533
refGene_gn=GPS1;refGene_tx=XM_023244355.1;refGene_gn=GPS1;refGene_tx=XM_023244356.1;refGene_gn=GPS1;refGene_tx=XM_023244357.1;refGene_gn=GPS1;refGene_tx=XM_023244358.1;refGene_gn=GPS1;refGene_tx=XM_023244359.1
E1 62417622 62417760
refGene_gn=GPS1;refGene_tx=XM_023244355.1;refGene_gn=GPS1;refGene_tx=XM_023244356.1;refGene_gn=GPS1;refGene_tx=XM_023244357.1;refGene_gn=GPS1;refGene_tx=XM_023244358.1;refGene_gn=GPS1;refGene_tx=XM_023244359.1
E1 62417845 62417980
refGene_gn=GPS1;refGene_tx=XM_023244355.1;refGene_gn=GPS1;refGene_tx=XM_023244356.1;refGene_gn=GPS1;refGene_tx=XM_023244357.1;refGene_gn=GPS1;refGene_tx=XM_023244358.1;refGene_gn=GPS1;refGene_tx=XM_023244359.1
E1 62418067 62418142
refGene_gn=GPS1;refGene_tx=XM_023244355.1;refGene_gn=GPS1;refGene_tx=XM_023244356.1;refGene_gn=GPS1;refGene_tx=XM_023244357.1;refGene_gn=GPS1;refGene_tx=XM_023244358.1;refGene_gn=GPS1;refGene_tx=XM_023244359.1
E1 62419010 62419081 refGene_gn=DUS1L;refGene_tx=XM_023244361.1
E1 62419183 62419259 refGene_gn=DUS1L;refGene_tx=XM_023244361.1
E1 62419503 62419541
refGene_gn=DUS1L;refGene_tx=XM_023244361.1;refGene_gn=DUS1L;refGene_tx=XM_023244362.1
E1 62419878 62420024
refGene_gn=DUS1L;refGene_tx=XM_023244361.1;refGene_gn=DUS1L;refGene_tx=XM_023244362.1
E1 62420432 62420521
refGene_gn=DUS1L;refGene_tx=XM_023244361.1;refGene_gn=DUS1L;refGene_tx=XM_023244362.1
E1 62420606 62420707
refGene_gn=DUS1L;refGene_tx=XM_023244361.1;refGene_gn=DUS1L;refGene_tx=XM_023244362.1
E1 62420822 62420967
refGene_gn=DUS1L;refGene_tx=XM_023244361.1;refGene_gn=DUS1L;refGene_tx=XM_023244362.1
E1 62421061 62421165
refGene_gn=DUS1L;refGene_tx=XM_023244361.1;refGene_gn=DUS1L;refGene_tx=XM_023244362.1
E1 62421396 62421479
refGene_gn=DUS1L;refGene_tx=XM_023244361.1;refGene_gn=DUS1L;refGene_tx=XM_023244362.1
E1 62422009 62422122

refGene_gn=DUS1L;refGene_tx=XM_023244361.1;refGene_gn=DUS1L;refGene_tx=XM_023244362.1
E1 62422393 62422444
refGene_gn=DUS1L;refGene_tx=XM_023244361.1;refGene_gn=DUS1L;refGene_tx=XM_023244362.1
E1 62422926 62423035
refGene_gn=DUS1L;refGene_tx=XM_023244361.1;refGene_gn=DUS1L;refGene_tx=XM_023244362.1
E1 62423482 62423719
refGene_gn=DUS1L;refGene_tx=XM_023244361.1;refGene_gn=DUS1L;refGene_tx=XM_023244362.1
E1 62442717 62443107
refGene_gn=FASN;refGene_tx=XM_023244364.1;refGene_gn=FASN;refGene_tx=XM_023244365.1;refGene_gn=FASN;refGene_tx=XM_023244367.1
E1 62443272 62443371
refGene_gn=FASN;refGene_tx=XM_023244364.1;refGene_gn=FASN;refGene_tx=XM_023244365.1;refGene_gn=FASN;refGene_tx=XM_023244367.1
E1 62443462 62443686
refGene_gn=FASN;refGene_tx=XM_023244364.1;refGene_gn=FASN;refGene_tx=XM_023244365.1;refGene_gn=FASN;refGene_tx=XM_023244367.1
E1 62443809 62444040
refGene_gn=FASN;refGene_tx=XM_023244364.1;refGene_gn=FASN;refGene_tx=XM_023244365.1;refGene_gn=FASN;refGene_tx=XM_023244367.1
E1 62444351 62444540
refGene_gn=FASN;refGene_tx=XM_023244364.1;refGene_gn=FASN;refGene_tx=XM_023244365.1;refGene_gn=FASN;refGene_tx=XM_023244367.1
E1 62444780 62445026
refGene_gn=FASN;refGene_tx=XM_023244364.1;refGene_gn=FASN;refGene_tx=XM_023244365.1;refGene_gn=FASN;refGene_tx=XM_023244367.1
E1 62445117 62445269
refGene_gn=FASN;refGene_tx=XM_023244364.1;refGene_gn=FASN;refGene_tx=XM_023244365.1;refGene_gn=FASN;refGene_tx=XM_023244367.1
E1 62445512 62445604
refGene_gn=FASN;refGene_tx=XM_023244364.1;refGene_gn=FASN;refGene_tx=XM_023244365.1;refGene_gn=FASN;refGene_tx=XM_023244367.1
E1 62445714 62445866
refGene_gn=FASN;refGene_tx=XM_023244364.1;refGene_gn=FASN;refGene_tx=XM_023244365.1;refGene_gn=FASN;refGene_tx=XM_023244367.1
E1 62445972 62446174
refGene_gn=FASN;refGene_tx=XM_023244364.1;refGene_gn=FASN;refGene_tx=XM_023244365.1;refGene_gn=FASN;refGene_tx=XM_023244367.1
E1 62446267 62446491
refGene_gn=FASN;refGene_tx=XM_023244364.1;refGene_gn=FASN;refGene_tx=XM_023244365.1;refGene_gn=FASN;refGene_tx=XM_023244367.1
E1 62446623 62446746
refGene_gn=FASN;refGene_tx=XM_023244364.1;refGene_gn=FASN;refGene_tx=XM_023244365.1;refGene_gn=FASN;refGene_tx=XM_023244367.1
E1 62446919 62447039
refGene_gn=FASN;refGene_tx=XM_023244364.1;refGene_gn=FASN;refGene_tx=XM_023244365.1;refGene_gn=FASN;refGene_tx=XM_023244367.1

E1 62447141 62447320
refGene_gn=FASN;refGene_tx=XM_023244364.1;refGene_gn=FASN;refGene_tx=XM_023244365.1;refGene_gn=FASN;refGene_tx=XM_023244367.1
E1 62447403 62447554
refGene_gn=FASN;refGene_tx=XM_023244364.1;refGene_gn=FASN;refGene_tx=XM_023244365.1;refGene_gn=FASN;refGene_tx=XM_023244367.1
E1 62447735 62447939
refGene_gn=FASN;refGene_tx=XM_023244364.1;refGene_gn=FASN;refGene_tx=XM_023244365.1;refGene_gn=FASN;refGene_tx=XM_023244367.1
E1 62448064 62448219
refGene_gn=FASN;refGene_tx=XM_023244364.1;refGene_gn=FASN;refGene_tx=XM_023244365.1;refGene_gn=FASN;refGene_tx=XM_023244367.1
E1 62448326 62448445
refGene_gn=FASN;refGene_tx=XM_023244364.1;refGene_gn=FASN;refGene_tx=XM_023244365.1;refGene_gn=FASN;refGene_tx=XM_023244367.1
E1 62448527 62448692
refGene_gn=FASN;refGene_tx=XM_023244364.1;refGene_gn=FASN;refGene_tx=XM_023244365.1;refGene_gn=FASN;refGene_tx=XM_023244367.1
E1 62448794 62449187
refGene_gn=FASN;refGene_tx=XM_023244364.1;refGene_gn=FASN;refGene_tx=XM_023244365.1;refGene_gn=FASN;refGene_tx=XM_023244367.1
E1 62449486 62449791
refGene_gn=FASN;refGene_tx=XM_023244364.1;refGene_gn=FASN;refGene_tx=XM_023244365.1;refGene_gn=FASN;refGene_tx=XM_023244367.1
E1 62450266 62450470
refGene_gn=FASN;refGene_tx=XM_023244364.1;refGene_gn=FASN;refGene_tx=XM_023244365.1;refGene_gn=FASN;refGene_tx=XM_023244367.1
E1 62450553 62450733
refGene_gn=FASN;refGene_tx=XM_023244364.1;refGene_gn=FASN;refGene_tx=XM_023244365.1;refGene_gn=FASN;refGene_tx=XM_023244367.1
E1 62450963 62451143
refGene_gn=FASN;refGene_tx=XM_023244364.1;refGene_gn=FASN;refGene_tx=XM_023244365.1;refGene_gn=FASN;refGene_tx=XM_023244367.1
E1 62451261 62451342
refGene_gn=FASN;refGene_tx=XM_023244364.1;refGene_gn=FASN;refGene_tx=XM_023244365.1;refGene_gn=FASN;refGene_tx=XM_023244367.1
E1 62451425 62451617
refGene_gn=FASN;refGene_tx=XM_023244364.1;refGene_gn=FASN;refGene_tx=XM_023244365.1;refGene_gn=FASN;refGene_tx=XM_023244367.1
E1 62451699 62451875
refGene_gn=FASN;refGene_tx=XM_023244364.1;refGene_gn=FASN;refGene_tx=XM_023244365.1;refGene_gn=FASN;refGene_tx=XM_023244367.1
E1 62452138 62452254
refGene_gn=FASN;refGene_tx=XM_023244364.1;refGene_gn=FASN;refGene_tx=XM_023244365.1;refGene_gn=FASN;refGene_tx=XM_023244367.1
E1 62452438 62452642
refGene_gn=FASN;refGene_tx=XM_023244364.1;refGene_gn=FASN;refGene_tx=XM_023244365.1;refGene_gn=FASN;refGene_tx=XM_023244367.1
E1 62452726 62452861
refGene_gn=FASN;refGene_tx=XM_023244364.1;refGene_gn=FASN;refGene_tx=X

M_023244365.1;refGene_gn=FASN;refGene_tx=XM_023244367.1
E1 62452971 62453066
refGene_gn=FASN;refGene_tx=XM_023244364.1;refGene_gn=FASN;refGene_tx=X
M_023244365.1;refGene_gn=FASN;refGene_tx=XM_023244367.1
E1 62453801 62453991
refGene_gn=FASN;refGene_tx=XM_023244364.1;refGene_gn=FASN;refGene_tx=X
M_023244365.1;refGene_gn=FASN;refGene_tx=XM_023244367.1
E1 62454162 62454350
refGene_gn=FASN;refGene_tx=XM_023244364.1;refGene_gn=FASN;refGene_tx=X
M_023244365.1;refGene_gn=FASN;refGene_tx=XM_023244367.1
E1 62454441 62454904
refGene_gn=FASN;refGene_tx=XM_023244364.1;refGene_gn=FASN;refGene_tx=X
M_023244365.1;refGene_gn=FASN;refGene_tx=XM_023244367.1
E1 62455534 62455669
refGene_gn=FASN;refGene_tx=XM_023244364.1;refGene_gn=FASN;refGene_tx=X
M_023244365.1;refGene_gn=FASN;refGene_tx=XM_023244367.1
E1 62455782 62455898
refGene_gn=FASN;refGene_tx=XM_023244364.1;refGene_gn=FASN;refGene_tx=X
M_023244365.1;refGene_gn=FASN;refGene_tx=XM_023244367.1
E1 62455984 62456107
refGene_gn=FASN;refGene_tx=XM_023244364.1;refGene_gn=FASN;refGene_tx=X
M_023244365.1;refGene_gn=FASN;refGene_tx=XM_023244367.1
E1 62456231 62456432
refGene_gn=FASN;refGene_tx=XM_023244364.1;refGene_gn=FASN;refGene_tx=X
M_023244365.1;refGene_gn=FASN;refGene_tx=XM_023244367.1
E1 62456532 62456706
refGene_gn=FASN;refGene_tx=XM_023244364.1;refGene_gn=FASN;refGene_tx=X
M_023244365.1;refGene_gn=FASN;refGene_tx=XM_023244367.1
E1 62457853 62458006
refGene_gn=FASN;refGene_tx=XM_023244364.1;refGene_gn=FASN;refGene_tx=X
M_023244365.1;refGene_gn=FASN;refGene_tx=XM_023244367.1
E1 62458736 62458863
refGene_gn=FASN;refGene_tx=XM_023244364.1;refGene_gn=FASN;refGene_tx=X
M_023244365.1;refGene_gn=FASN;refGene_tx=XM_023244367.1
E1 62463962 62464165 refGene_gn=CCDC57;refGene_tx=XM_023244368.1
E1 62492736 62492953 refGene_gn=CCDC57;refGene_tx=XM_023244368.1
E1 62493389 62493521 refGene_gn=CCDC57;refGene_tx=XM_023244368.1
E1 62501059 62501151 refGene_gn=CCDC57;refGene_tx=XM_023244368.1
E1 62509989 62510176 refGene_gn=CCDC57;refGene_tx=XM_023244368.1
E1 62514151 62514322 refGene_gn=CCDC57;refGene_tx=XM_023244368.1
E1 62517074 62517178 refGene_gn=CCDC57;refGene_tx=XM_023244368.1
E1 62523563 62523716 refGene_gn=CCDC57;refGene_tx=XM_023244368.1
E1 62524380 62524603 refGene_gn=CCDC57;refGene_tx=XM_023244368.1
E1 62531672 62531801 refGene_gn=CCDC57;refGene_tx=XM_023244368.1
E1 62532407 62532570 refGene_gn=CCDC57;refGene_tx=XM_023244368.1
E1 62536626 62536785 refGene_gn=CCDC57;refGene_tx=XM_023244368.1
E1 62540439 62540640 refGene_gn=CCDC57;refGene_tx=XM_023244368.1
E1 62544350 62544425 refGene_gn=CCDC57;refGene_tx=XM_023244368.1
E1 62544526 62544684 refGene_gn=CCDC57;refGene_tx=XM_023244368.1
E1 62546041 62546143 refGene_gn=CCDC57;refGene_tx=XM_023244368.1

E1 62552914 62553023 refGene_gn=CCDC57;refGene_tx=XM_023244368.1
E1 62555531 62555898 refGene_gn=CCDC57;refGene_tx=XM_023244368.1
E1 62555900 62555940 refGene_gn=CCDC57;refGene_tx=XM_023244368.1
E1 62584786 62585009
refGene_gn=SLC16A3;refGene_tx=XM_003997281.4;refGene_gn=SLC16A3;refGene_tx=XM_023244375.1;refGene_gn=SLC16A3;refGene_tx=XM_023244376.1;refGene_gn=SLC16A3;refGene_tx=XM_023244377.1;refGene_gn=SLC16A3;refGene_tx=XM_023244378.1;refGene_gn=SLC16A3;refGene_tx=XM_023244379.1;refGene_gn=SLC16A3;refGene_tx=XM_023244380.1
E1 62585472 62585676
refGene_gn=SLC16A3;refGene_tx=XM_003997281.4;refGene_gn=SLC16A3;refGene_tx=XM_023244375.1;refGene_gn=SLC16A3;refGene_tx=XM_023244376.1;refGene_gn=SLC16A3;refGene_tx=XM_023244377.1;refGene_gn=SLC16A3;refGene_tx=XM_023244378.1;refGene_gn=SLC16A3;refGene_tx=XM_023244379.1;refGene_gn=SLC16A3;refGene_tx=XM_023244380.1
E1 62585948 62586707
refGene_gn=SLC16A3;refGene_tx=XM_003997281.4;refGene_gn=SLC16A3;refGene_tx=XM_023244375.1;refGene_gn=SLC16A3;refGene_tx=XM_023244376.1;refGene_gn=SLC16A3;refGene_tx=XM_023244377.1;refGene_gn=SLC16A3;refGene_tx=XM_023244378.1;refGene_gn=SLC16A3;refGene_tx=XM_023244379.1;refGene_gn=SLC16A3;refGene_tx=XM_023244380.1
E1 62587413 62587670
refGene_gn=SLC16A3;refGene_tx=XM_003997281.4;refGene_gn=SLC16A3;refGene_tx=XM_023244375.1;refGene_gn=SLC16A3;refGene_tx=XM_023244376.1;refGene_gn=SLC16A3;refGene_tx=XM_023244377.1;refGene_gn=SLC16A3;refGene_tx=XM_023244378.1;refGene_gn=SLC16A3;refGene_tx=XM_023244379.1;refGene_gn=SLC16A3;refGene_tx=XM_023244380.1
E1 62588634 62588724
refGene_gn=CSNK1D;refGene_tx=XM_023244370.1;refGene_gn=CSNK1D;refGene_tx=XM_023244371.1
E1 62593089 62593140 refGene_gn=CSNK1D;refGene_tx=XM_023244372.1
E1 62594314 62594347 refGene_gn=CSNK1D;refGene_tx=XM_023244373.1
E1 62595635 62596004
refGene_gn=CSNK1D;refGene_tx=XM_019818399.2;refGene_gn=CSNK1D;refGene_tx=XM_023244369.1;refGene_gn=CSNK1D;refGene_tx=XM_023244370.1
E1 62597262 62597402
refGene_gn=CSNK1D;refGene_tx=XM_019818399.2;refGene_gn=CSNK1D;refGene_tx=XM_023244369.1;refGene_gn=CSNK1D;refGene_tx=XM_023244370.1;refGene_gn=CSNK1D;refGene_tx=XM_023244371.1;refGene_gn=CSNK1D;refGene_tx=XM_023244372.1;refGene_gn=CSNK1D;refGene_tx=XM_023244373.1
E1 62597924 62598096
refGene_gn=CSNK1D;refGene_tx=XM_019818399.2;refGene_gn=CSNK1D;refGene_tx=XM_023244369.1;refGene_gn=CSNK1D;refGene_tx=XM_023244370.1;refGene_gn=CSNK1D;refGene_tx=XM_023244371.1;refGene_gn=CSNK1D;refGene_tx=XM_023244372.1;refGene_gn=CSNK1D;refGene_tx=XM_023244373.1
E1 62599832 62599981
refGene_gn=CSNK1D;refGene_tx=XM_019818399.2;refGene_gn=CSNK1D;refGene_tx=XM_023244369.1;refGene_gn=CSNK1D;refGene_tx=XM_023244370.1;refGene_gn=CSNK1D;refGene_tx=XM_023244371.1;refGene_gn=CSNK1D;refGene_tx=XM_023244372.1;refGene_gn=CSNK1D;refGene_tx=XM_023244373.1

E1 62600410 62600581
refGene_gn=CSNK1D;refGene_tx=XM_019818399.2;refGene_gn=CSNK1D;refGene_tx=XM_023244369.1;refGene_gn=CSNK1D;refGene_tx=XM_023244370.1;refGene_gn=CSNK1D;refGene_tx=XM_023244371.1;refGene_gn=CSNK1D;refGene_tx=XM_023244372.1;refGene_gn=CSNK1D;refGene_tx=XM_023244373.1
E1 62601018 62601247
refGene_gn=CSNK1D;refGene_tx=XM_019818399.2;refGene_gn=CSNK1D;refGene_tx=XM_023244369.1;refGene_gn=CSNK1D;refGene_tx=XM_023244370.1;refGene_gn=CSNK1D;refGene_tx=XM_023244371.1;refGene_gn=CSNK1D;refGene_tx=XM_023244372.1;refGene_gn=CSNK1D;refGene_tx=XM_023244373.1
E1 62601743 62601892
refGene_gn=CSNK1D;refGene_tx=XM_019818399.2;refGene_gn=CSNK1D;refGene_tx=XM_023244369.1;refGene_gn=CSNK1D;refGene_tx=XM_023244370.1;refGene_gn=CSNK1D;refGene_tx=XM_023244371.1;refGene_gn=CSNK1D;refGene_tx=XM_023244372.1;refGene_gn=CSNK1D;refGene_tx=XM_023244373.1
E1 62613381 62613492
refGene_gn=CSNK1D;refGene_tx=XM_019818399.2;refGene_gn=CSNK1D;refGene_tx=XM_023244369.1;refGene_gn=CSNK1D;refGene_tx=XM_023244370.1;refGene_gn=CSNK1D;refGene_tx=XM_023244371.1;refGene_gn=CSNK1D;refGene_tx=XM_023244372.1;refGene_gn=CSNK1D;refGene_tx=XM_023244373.1
E1 62619561 62619619 refGene_gn=CSNK1D;refGene_tx=XM_019818399.2
E1 62620337 62620413
refGene_gn=CSNK1D;refGene_tx=XM_023244369.1;refGene_gn=CSNK1D;refGene_tx=XM_023244370.1;refGene_gn=CSNK1D;refGene_tx=XM_023244371.1;refGene_gn=CSNK1D;refGene_tx=XM_023244372.1;refGene_gn=CSNK1D;refGene_tx=XM_023244373.1
E1 62653829 62653937
refGene_gn=CD7;refGene_tx=NM_001009296.1;refGene_gn=CD7;refGene_tx=XM_019816886.2
E1 62654639 62654797
refGene_gn=CD7;refGene_tx=NM_001009296.1;refGene_gn=CD7;refGene_tx=XM_019816886.2
E1 62655022 62655337
refGene_gn=CD7;refGene_tx=NM_001009296.1;refGene_gn=CD7;refGene_tx=XM_019816886.2
E1 62656250 62656524
refGene_gn=CD7;refGene_tx=NM_001009296.1;refGene_gn=CD7;refGene_tx=XM_019816886.2
E1 62657964 62658119 refGene_gn=SECTM1;refGene_tx=XM_019818404.2
E1 62658283 62658346 refGene_gn=SECTM1;refGene_tx=XM_019818404.2
E1 62658869 62659181 refGene_gn=SECTM1;refGene_tx=XM_019818404.2
E1 62659483 62659615 refGene_gn=SECTM1;refGene_tx=XM_019818404.2
E1 62660963 62661098 refGene_gn=SECTM1;refGene_tx=XM_019818404.2
E1 62677497 62677647 refGene_gn=SECTM1;refGene_tx=XM_019818404.2
E1 62682498 62682748 refGene_gn=SECTM1;refGene_tx=XM_019818404.2
E1 62687584 62687641 refGene_gn=SECTM1;refGene_tx=XM_019818404.2
E1 62697521 62698443
refGene_gn=LOC111557819;refGene_tx=XM_023244382.1
E1 62698444 62698746
refGene_gn=LOC111557819;refGene_tx=XM_023244382.1

E1 62889709 62890026 refGene_gn=METRNL;refGene_tx=XM_023244384.1
E1 62890855 62890915 refGene_gn=METRNL;refGene_tx=XM_023244384.1
E1 62898182 62898571 refGene_gn=METRNL;refGene_tx=XM_023244384.1
E1 62903749 62903916 refGene_gn=METRNL;refGene_tx=XM_023244384.1
E1 62927722 62927761
refGene_gn=B3GNTL1;refGene_tx=XM_023244387.1;refGene_gn=B3GNTL1;refGen
e_tx=XM_023244388.1
E1 62929463 62929580
refGene_gn=B3GNTL1;refGene_tx=XM_023244387.1;refGene_gn=B3GNTL1;refGen
e_tx=XM_023244388.1
E1 62929679 62929779
refGene_gn=B3GNTL1;refGene_tx=XM_023244387.1;refGene_gn=B3GNTL1;refGen
e_tx=XM_023244388.1
E1 62944386 62944451
refGene_gn=B3GNTL1;refGene_tx=XM_023244387.1;refGene_gn=B3GNTL1;refGen
e_tx=XM_023244388.1
E1 62972909 62972975
refGene_gn=B3GNTL1;refGene_tx=XM_023244387.1;refGene_gn=B3GNTL1;refGen
e_tx=XM_023244388.1
E1 62982729 62982825
refGene_gn=B3GNTL1;refGene_tx=XM_023244387.1;refGene_gn=B3GNTL1;refGen
e_tx=XM_023244388.1
E1 63004936 63005032
refGene_gn=B3GNTL1;refGene_tx=XM_023244387.1;refGene_gn=B3GNTL1;refGen
e_tx=XM_023244388.1
E1 63011380 63011505
refGene_gn=B3GNTL1;refGene_tx=XM_023244387.1;refGene_gn=B3GNTL1;refGen
e_tx=XM_023244388.1
E1 63016505 63016641
refGene_gn=B3GNTL1;refGene_tx=XM_023244387.1;refGene_gn=B3GNTL1;refGen
e_tx=XM_023244388.1
E1 63016729 63016798
refGene_gn=B3GNTL1;refGene_tx=XM_023244387.1;refGene_gn=B3GNTL1;refGen
e_tx=XM_023244388.1
E1 63017143 63017218
refGene_gn=B3GNTL1;refGene_tx=XM_023244387.1;refGene_gn=B3GNTL1;refGen
e_tx=XM_023244388.1
E1 63023854 63023935 refGene_gn=B3GNTL1;refGene_tx=XM_023244387.1
E1 63026161 63026170 refGene_gn=B3GNTL1;refGene_tx=XM_023244388.1
E1 63027014 63027029
refGene_gn=TBCD;refGene_tx=XM_023244385.1;refGene_gn=TBCD;refGene_tx=X
M_023244386.1
E1 63031069 63031154
refGene_gn=TBCD;refGene_tx=XM_023244385.1;refGene_gn=TBCD;refGene_tx=X
M_023244386.1
E1 63033524 63033634
refGene_gn=TBCD;refGene_tx=XM_023244385.1;refGene_gn=TBCD;refGene_tx=X
M_023244386.1
E1 63034562 63034650
refGene_gn=TBCD;refGene_tx=XM_023244385.1;refGene_gn=TBCD;refGene_tx=X

M_023244386.1
E1	63034966	63035056
refGene_gn=TBCD;refGene_tx=XM_023244385.1;refGene_gn=TBCD;refGene_tx=X
M_023244386.1
E1	63039831	63039909
refGene_gn=TBCD;refGene_tx=XM_023244385.1;refGene_gn=TBCD;refGene_tx=X
M_023244386.1
E1	63041986	63042108
refGene_gn=TBCD;refGene_tx=XM_023244385.1;refGene_gn=TBCD;refGene_tx=X
M_023244386.1
E1	63044009	63044148
refGene_gn=TBCD;refGene_tx=XM_023244385.1;refGene_gn=TBCD;refGene_tx=X
M_023244386.1
E1	63044221	63044380
refGene_gn=TBCD;refGene_tx=XM_023244385.1;refGene_gn=TBCD;refGene_tx=X
M_023244386.1
E1	63044659	63044743
refGene_gn=TBCD;refGene_tx=XM_023244385.1;refGene_gn=TBCD;refGene_tx=X
M_023244386.1
E1	63047645	63047786
refGene_gn=TBCD;refGene_tx=XM_023244385.1;refGene_gn=TBCD;refGene_tx=X
M_023244386.1
E1	63048327	63048419
refGene_gn=TBCD;refGene_tx=XM_023244385.1;refGene_gn=TBCD;refGene_tx=X
M_023244386.1
E1	63048906	63049025
refGene_gn=TBCD;refGene_tx=XM_023244385.1;refGene_gn=TBCD;refGene_tx=X
M_023244386.1
E1	63050843	63050925
refGene_gn=TBCD;refGene_tx=XM_023244385.1;refGene_gn=TBCD;refGene_tx=X
M_023244386.1
E1	63052512	63052589
refGene_gn=TBCD;refGene_tx=XM_023244385.1;refGene_gn=TBCD;refGene_tx=X
M_023244386.1
E1	63053433	63053496
refGene_gn=TBCD;refGene_tx=XM_023244385.1;refGene_gn=TBCD;refGene_tx=X
M_023244386.1
E1	63057883	63057915
refGene_gn=TBCD;refGene_tx=XM_023244385.1;refGene_gn=TBCD;refGene_tx=X
M_023244386.1
E1	63062197	63062220
refGene_gn=TBCD;refGene_tx=XM_023244385.1;refGene_gn=TBCD;refGene_tx=X
M_023244386.1
E1	63063425	63063486
refGene_gn=TBCD;refGene_tx=XM_023244385.1;refGene_gn=TBCD;refGene_tx=X
M_023244386.1
E1	63064900	63065018
refGene_gn=TBCD;refGene_tx=XM_023244385.1;refGene_gn=TBCD;refGene_tx=X
M_023244386.1
E1	63066474	63066548

refGene_gn=TBCD;refGene_tx=XM_023244385.1;refGene_gn=TBCD;refGene_tx=XM_023244386.1
E1 63068396 63068477
refGene_gn=TBCD;refGene_tx=XM_023244385.1;refGene_gn=TBCD;refGene_tx=XM_023244386.1
E1 63074562 63074648
refGene_gn=TBCD;refGene_tx=XM_023244385.1;refGene_gn=TBCD;refGene_tx=XM_023244386.1
E1 63076942 63076972
refGene_gn=TBCD;refGene_tx=XM_023244385.1;refGene_gn=TBCD;refGene_tx=XM_023244386.1
E1 63085087 63085145
refGene_gn=TBCD;refGene_tx=XM_023244385.1;refGene_gn=TBCD;refGene_tx=XM_023244386.1
E1 63095876 63096033
refGene_gn=TBCD;refGene_tx=XM_023244385.1;refGene_gn=TBCD;refGene_tx=XM_023244386.1
E1 63133528 63134967 refGene_gn=ZNF750;refGene_tx=XM_003997289.3
E1 63135109 63135818 refGene_gn=ZNF750;refGene_tx=XM_003997289.3
E1 63143986 63144081
refGene_gn=TBCD;refGene_tx=XM_023244385.1;refGene_gn=TBCD;refGene_tx=XM_023244386.1
E1 63148519 63148594
refGene_gn=TBCD;refGene_tx=XM_023244385.1;refGene_gn=TBCD;refGene_tx=XM_023244386.1
E1 63149474 63149535
refGene_gn=TBCD;refGene_tx=XM_023244385.1;refGene_gn=TBCD;refGene_tx=XM_023244386.1
E1 63150586 63150723
refGene_gn=TBCD;refGene_tx=XM_023244385.1;refGene_gn=TBCD;refGene_tx=XM_023244386.1
E1 63156836 63156969
refGene_gn=TBCD;refGene_tx=XM_023244385.1;refGene_gn=TBCD;refGene_tx=XM_023244386.1
E1 63158351 63158397
refGene_gn=TBCD;refGene_tx=XM_023244385.1;refGene_gn=TBCD;refGene_tx=XM_023244386.1
E1 63168624 63168757
refGene_gn=TBCD;refGene_tx=XM_023244385.1;refGene_gn=TBCD;refGene_tx=XM_023244386.1
E1 63171925 63171981
refGene_gn=TBCD;refGene_tx=XM_023244385.1;refGene_gn=TBCD;refGene_tx=XM_023244386.1
E1 63174141 63174288
refGene_gn=TBCD;refGene_tx=XM_023244385.1;refGene_gn=TBCD;refGene_tx=XM_023244386.1
E1 63176744 63176846 refGene_gn=TBCD;refGene_tx=XM_023244385.1
E1 63177902 63178000
refGene_gn=TBCD;refGene_tx=XM_023244385.1;refGene_gn=TBCD;refGene_tx=XM_023244386.1

E1 63186029 63186080
refGene_gn=TBCD;refGene_tx=XM_023244385.1;refGene_gn=TBCD;refGene_tx=XM_023244386.1
E1 63187256 63187440
refGene_gn=TBCD;refGene_tx=XM_023244385.1;refGene_gn=TBCD;refGene_tx=XM_023244386.1
E1 63189231 63189570 refGene_gn=FN3K;refGene_tx=XM_003997288.4
E1 63190423 63190546 refGene_gn=FN3K;refGene_tx=XM_003997288.4
E1 63193900 63193983 refGene_gn=FN3K;refGene_tx=XM_003997288.4
E1 63194293 63194385 refGene_gn=FN3K;refGene_tx=XM_003997288.4
E1 63196081 63196233 refGene_gn=FN3K;refGene_tx=XM_003997288.4
E1 63198882 63199023 refGene_gn=FN3K;refGene_tx=XM_003997288.4
E1 63206327 63206666 refGene_gn=FN3KRP;refGene_tx=XM_003997287.4
E1 63206865 63206988 refGene_gn=FN3KRP;refGene_tx=XM_003997287.4
E1 63208040 63208123 refGene_gn=FN3KRP;refGene_tx=XM_003997287.4
E1 63209062 63209154 refGene_gn=FN3KRP;refGene_tx=XM_003997287.4
E1 63209527 63209679 refGene_gn=FN3KRP;refGene_tx=XM_003997287.4
E1 63213816 63213897 refGene_gn=FN3KRP;refGene_tx=XM_003997287.4
E1 63213898 63213958 refGene_gn=FN3KRP;refGene_tx=XM_003997287.4
E1 63226795 63226967
refGene_gn=RAB40B;refGene_tx=XM_023244389.1;refGene_gn=RAB40B;refGene_tx=XM_023244390.1
E1 63244722 63244783
refGene_gn=RAB40B;refGene_tx=XM_023244389.1;refGene_gn=RAB40B;refGene_tx=XM_023244390.1
E1 63246555 63246616
refGene_gn=RAB40B;refGene_tx=XM_023244389.1;refGene_gn=RAB40B;refGene_tx=XM_023244390.1
E1 63247255 63247333
refGene_gn=RAB40B;refGene_tx=XM_023244389.1;refGene_gn=RAB40B;refGene_tx=XM_023244390.1
E1 63247996 63248219
refGene_gn=RAB40B;refGene_tx=XM_023244389.1;refGene_gn=RAB40B;refGene_tx=XM_023244390.1
E1 63249390 63249593 refGene_gn=RAB40B;refGene_tx=XM_023244390.1
E1 63249734 63250006 refGene_gn=RAB40B;refGene_tx=XM_023244389.1
E1 63253482 63253802
refGene_gn=LOC109494096;refGene_tx=XM_023243482.1
E1 63256131 63256488
refGene_gn=LOC109494096;refGene_tx=XM_023243482.1
E1 63257705 63257850
refGene_gn=LOC109494096;refGene_tx=XM_023243482.1
E1 63257888 63257955
refGene_gn=WDR45B;refGene_tx=XM_023244391.1;refGene_gn=WDR45B;refGene_tx=XM_023244392.1;refGene_gn=WDR45B;refGene_tx=XM_023244393.1
E1 63265657 63265732
refGene_gn=WDR45B;refGene_tx=XM_023244391.1;refGene_gn=WDR45B;refGene_tx=XM_023244393.1
E1 63271555 63271657
refGene_gn=WDR45B;refGene_tx=XM_023244391.1;refGene_gn=WDR45B;refGene_

tx=XM_023244392.1;refGene_gn=WDR45B;refGene_tx=XM_023244393.1
E1 63275000 63275088
refGene_gn=WDR45B;refGene_tx=XM_023244391.1;refGene_gn=WDR45B;refGene_tx=XM_023244392.1
E1 63276113 63276208
refGene_gn=WDR45B;refGene_tx=XM_023244391.1;refGene_gn=WDR45B;refGene_tx=XM_023244392.1
E1 63278769 63278960
refGene_gn=WDR45B;refGene_tx=XM_023244391.1;refGene_gn=WDR45B;refGene_tx=XM_023244392.1;refGene_gn=WDR45B;refGene_tx=XM_023244393.1
E1 63279509 63279595
refGene_gn=WDR45B;refGene_tx=XM_023244391.1;refGene_gn=WDR45B;refGene_tx=XM_023244392.1;refGene_gn=WDR45B;refGene_tx=XM_023244393.1
E1 63280240 63280342
refGene_gn=WDR45B;refGene_tx=XM_023244391.1;refGene_gn=WDR45B;refGene_tx=XM_023244392.1;refGene_gn=WDR45B;refGene_tx=XM_023244393.1
E1 63281053 63281175
refGene_gn=WDR45B;refGene_tx=XM_023244391.1;refGene_gn=WDR45B;refGene_tx=XM_023244392.1;refGene_gn=WDR45B;refGene_tx=XM_023244393.1
E1 63281533 63281640
refGene_gn=WDR45B;refGene_tx=XM_023244391.1;refGene_gn=WDR45B;refGene_tx=XM_023244392.1;refGene_gn=WDR45B;refGene_tx=XM_023244393.1
E1 63303637 63303834 refGene_gn=FOXK2;refGene_tx=XM_023244395.1
E1 63314033 63314243 refGene_gn=FOXK2;refGene_tx=XM_023244395.1
E1 63314755 63315052 refGene_gn=FOXK2;refGene_tx=XM_023244395.1
E1 63316314 63316490 refGene_gn=FOXK2;refGene_tx=XM_023244395.1
E1 63317299 63317493 refGene_gn=FOXK2;refGene_tx=XM_023244395.1
E1 63323040 63323187 refGene_gn=FOXK2;refGene_tx=XM_023244395.1
E1 63325758 63325906 refGene_gn=FOXK2;refGene_tx=XM_023244395.1
E1 63329269 63329464 refGene_gn=FOXK2;refGene_tx=XM_023244395.1
E1 63365885 63366298 refGene_gn=FOXK2;refGene_tx=XM_023244395.1
E1 63391586 63391828
refGene_gn=NARF;refGene_tx=XM_023244397.1;refGene_gn=NARF;refGene_tx=XM_023244398.1;refGene_gn=NARF;refGene_tx=XM_023244399.1
E1 63393073 63393231
refGene_gn=NARF;refGene_tx=XM_023244397.1;refGene_gn=NARF;refGene_tx=XM_023244398.1;refGene_gn=NARF;refGene_tx=XM_023244399.1
E1 63393525 63393663
refGene_gn=NARF;refGene_tx=XM_023244397.1;refGene_gn=NARF;refGene_tx=XM_023244398.1;refGene_gn=NARF;refGene_tx=XM_023244399.1
E1 63394352 63394416
refGene_gn=NARF;refGene_tx=XM_023244397.1;refGene_gn=NARF;refGene_tx=XM_023244398.1;refGene_gn=NARF;refGene_tx=XM_023244399.1
E1 63395383 63395513
refGene_gn=NARF;refGene_tx=XM_023244397.1;refGene_gn=NARF;refGene_tx=XM_023244398.1;refGene_gn=NARF;refGene_tx=XM_023244399.1
E1 63396759 63396878
refGene_gn=NARF;refGene_tx=XM_023244397.1;refGene_gn=NARF;refGene_tx=XM_023244398.1;refGene_gn=NARF;refGene_tx=XM_023244399.1
E1 63398945 63399080

refGene_gn=NARF;refGene_tx=XM_023244397.1;refGene_gn=NARF;refGene_tx=XM_023244398.1;refGene_gn=NARF;refGene_tx=XM_023244399.1
E1 63401074 63401207
refGene_gn=NARF;refGene_tx=XM_023244397.1;refGene_gn=NARF;refGene_tx=XM_023244398.1;refGene_gn=NARF;refGene_tx=XM_023244399.1
E1 63404267 63404411
refGene_gn=NARF;refGene_tx=XM_023244397.1;refGene_gn=NARF;refGene_tx=XM_023244398.1;refGene_gn=NARF;refGene_tx=XM_023244399.1
E1 63406340 63406421
refGene_gn=NARF;refGene_tx=XM_023244397.1;refGene_gn=NARF;refGene_tx=XM_023244398.1;refGene_gn=NARF;refGene_tx=XM_023244399.1
E1 63407684 63407711
refGene_gn=NARF;refGene_tx=XM_023244397.1;refGene_gn=NARF;refGene_tx=XM_023244398.1;refGene_gn=NARF;refGene_tx=XM_023244399.1
E1 63414933 63415018
refGene_gn=CE1H17orf62;refGene_tx=XM_011289469.3
E1 63415893 63415935
refGene_gn=CE1H17orf62;refGene_tx=XM_011289469.3
E1 63416477 63416551
refGene_gn=CE1H17orf62;refGene_tx=XM_011289469.3
E1 63417222 63417319
refGene_gn=CE1H17orf62;refGene_tx=XM_011289469.3
E1 63417828 63417973
refGene_gn=CE1H17orf62;refGene_tx=XM_011289469.3
E1 63418304 63418425
refGene_gn=CE1H17orf62;refGene_tx=XM_011289469.3
E1 63419835 63420034
refGene_gn=HEXDC;refGene_tx=XM_006940683.4;refGene_gn=HEXDC;refGene_tx=XM_006940684.4
E1 63420707 63420797 refGene_gn=HEXDC;refGene_tx=XM_006940683.4
E1 63420993 63421095
refGene_gn=HEXDC;refGene_tx=XM_006940683.4;refGene_gn=HEXDC;refGene_tx=XM_006940684.4
E1 63421172 63421251
refGene_gn=HEXDC;refGene_tx=XM_006940683.4;refGene_gn=HEXDC;refGene_tx=XM_006940684.4
E1 63422264 63422347
refGene_gn=HEXDC;refGene_tx=XM_006940683.4;refGene_gn=HEXDC;refGene_tx=XM_006940684.4
E1 63422815 63423011
refGene_gn=HEXDC;refGene_tx=XM_006940683.4;refGene_gn=HEXDC;refGene_tx=XM_006940684.4
E1 63423327 63423399
refGene_gn=HEXDC;refGene_tx=XM_006940683.4;refGene_gn=HEXDC;refGene_tx=XM_006940684.4
E1 63424475 63424659
refGene_gn=HEXDC;refGene_tx=XM_006940683.4;refGene_gn=HEXDC;refGene_tx=XM_006940684.4
E1 63425168 63425333
refGene_gn=HEXDC;refGene_tx=XM_006940683.4;refGene_gn=HEXDC;refGene_tx

=XM_006940684.4
E1	63429760	63429848
refGene_gn=HEXDC;refGene_tx=XM_006940683.4;refGene_gn=HEXDC;refGene_tx
=XM_006940684.4
E1	63431785	63431895
refGene_gn=HEXDC;refGene_tx=XM_006940683.4;refGene_gn=HEXDC;refGene_tx
=XM_006940684.4
E1	63437263	63437362
refGene_gn=HEXDC;refGene_tx=XM_006940683.4;refGene_gn=HEXDC;refGene_tx
=XM_006940684.4
E1	63439401	63439475
refGene_gn=OGFOD3;refGene_tx=XM_023244400.1;refGene_gn=OGFOD3;refGene_
tx=XM_023244401.1;refGene_gn=OGFOD3;refGene_tx=XM_023244402.1
E1	63440413	63440631
refGene_gn=OGFOD3;refGene_tx=XM_023244400.1;refGene_gn=OGFOD3;refGene_
tx=XM_023244401.1;refGene_gn=OGFOD3;refGene_tx=XM_023244402.1
E1	63444506	63444582
refGene_gn=OGFOD3;refGene_tx=XM_023244400.1;refGene_gn=OGFOD3;refGene_
tx=XM_023244401.1;refGene_gn=OGFOD3;refGene_tx=XM_023244402.1
E1	63446027	63446070
refGene_gn=OGFOD3;refGene_tx=XM_023244400.1;refGene_gn=OGFOD3;refGene_
tx=XM_023244401.1;refGene_gn=OGFOD3;refGene_tx=XM_023244402.1
E1	63448459	63448524
refGene_gn=OGFOD3;refGene_tx=XM_023244400.1;refGene_gn=OGFOD3;refGene_
tx=XM_023244401.1;refGene_gn=OGFOD3;refGene_tx=XM_023244402.1
E1	63449069	63449126
refGene_gn=OGFOD3;refGene_tx=XM_023244400.1;refGene_gn=OGFOD3;refGene_
tx=XM_023244401.1;refGene_gn=OGFOD3;refGene_tx=XM_023244402.1
E1	63453978	63454132
refGene_gn=OGFOD3;refGene_tx=XM_023244400.1;refGene_gn=OGFOD3;refGene_
tx=XM_023244401.1;refGene_gn=OGFOD3;refGene_tx=XM_023244402.1
E1	63454420	63454561	refGene_gn=OGFOD3;refGene_tx=XM_023244402.1
E1	63457406	63457530
refGene_gn=OGFOD3;refGene_tx=XM_023244400.1;refGene_gn=OGFOD3;refGene_
tx=XM_023244401.1
E1	63459926	63460108	refGene_gn=OGFOD3;refGene_tx=XM_023244400.1
E1	63461551	63461688	refGene_gn=OGFOD3;refGene_tx=XM_023244401.1
E1	63474661	63475786
refGene_gn=UTS2R;refGene_tx=NM_001048011.1;refGene_gn=UTS2R;refGene_tx
=XM_023243407.1
E1	63486827	63486890
refGene_gn=LOC111557725;refGene_tx=XM_023243483.1
E1	63487468	63487788
refGene_gn=LOC111557725;refGene_tx=XM_023243483.1
E1	63487790	63488013
refGene_gn=LOC111557725;refGene_tx=XM_023243483.1
E1	63488014	63488161
refGene_gn=LOC111557725;refGene_tx=XM_023243483.1
E2	234365	234411
refGene_gn=LOC101088444;refGene_tx=XM_023245753.1;refGene_gn=LOC101088

444;refGene_tx=XM_023245754.1;refGene_gn=LOC101088444;refGene_tx=XM_02
3245755.1
E2 240834 240892
refGene_gn=LOC101088444;refGene_tx=XM_023245744.1;refGene_gn=LOC101088
444;refGene_tx=XM_023245745.1;refGene_gn=LOC101088444;refGene_tx=XM_02
3245746.1;refGene_gn=LOC101088444;refGene_tx=XM_023245747.1;refGene_gn
=LOC101088444;refGene_tx=XM_023245748.1;refGene_gn=LOC101088444;refGen
e_tx=XM_023245749.1;refGene_gn=LOC101088444;refGene_tx=XM_023245750.1;
refGene_gn=LOC101088444;refGene_tx=XM_023245751.1;refGene_gn=LOC101088
444;refGene_tx=XM_023245752.1;refGene_gn=LOC101088444;refGene_tx=XM_02
3245756.1;refGene_gn=LOC101088444;refGene_tx=XM_023245757.1
E2 241896 242011
refGene_gn=LOC101088444;refGene_tx=XM_023245744.1;refGene_gn=LOC101088
444;refGene_tx=XM_023245745.1;refGene_gn=LOC101088444;refGene_tx=XM_02
3245746.1;refGene_gn=LOC101088444;refGene_tx=XM_023245756.1
E2 243281 243361
refGene_gn=LOC101088444;refGene_tx=XM_023245744.1;refGene_gn=LOC101088
444;refGene_tx=XM_023245745.1;refGene_gn=LOC101088444;refGene_tx=XM_02
3245746.1;refGene_gn=LOC101088444;refGene_tx=XM_023245747.1;refGene_gn
=LOC101088444;refGene_tx=XM_023245748.1;refGene_gn=LOC101088444;refGen
e_tx=XM_023245749.1;refGene_gn=LOC101088444;refGene_tx=XM_023245750.1;
refGene_gn=LOC101088444;refGene_tx=XM_023245751.1;refGene_gn=LOC101088
444;refGene_tx=XM_023245752.1;refGene_gn=LOC101088444;refGene_tx=XM_02
3245753.1;refGene_gn=LOC101088444;refGene_tx=XM_023245754.1;refGene_gn
=LOC101088444;refGene_tx=XM_023245755.1;refGene_gn=LOC101088444;refGen
e_tx=XM_023245756.1;refGene_gn=LOC101088444;refGene_tx=XM_023245757.1
E2 251543 253331
refGene_gn=LOC101088444;refGene_tx=XM_023245744.1;refGene_gn=LOC101088
444;refGene_tx=XM_023245745.1;refGene_gn=LOC101088444;refGene_tx=XM_02
3245746.1;refGene_gn=LOC101088444;refGene_tx=XM_023245747.1;refGene_gn
=LOC101088444;refGene_tx=XM_023245748.1;refGene_gn=LOC101088444;refGen
e_tx=XM_023245749.1;refGene_gn=LOC101088444;refGene_tx=XM_023245750.1;
refGene_gn=LOC101088444;refGene_tx=XM_023245751.1;refGene_gn=LOC101088
444;refGene_tx=XM_023245752.1;refGene_gn=LOC101088444;refGene_tx=XM_02
3245753.1;refGene_gn=LOC101088444;refGene_tx=XM_023245754.1;refGene_gn
=LOC101088444;refGene_tx=XM_023245755.1;refGene_gn=LOC101088444;refGen
e_tx=XM_023245756.1;refGene_gn=LOC101088444;refGene_tx=XM_023245757.1
E2 258012 258049
refGene_gn=MZF1;refGene_tx=XM_003997334.5;refGene_gn=MZF1;refGene_tx=X
M_019818447.2
E2 258209 258333 refGene_gn=MZF1;refGene_tx=XM_006940713.4
E2 258398 258581 refGene_gn=MZF1;refGene_tx=XM_019818447.2
E2 259349 259786
refGene_gn=MZF1;refGene_tx=XM_003997334.5;refGene_gn=MZF1;refGene_tx=X
M_006940713.4;refGene_gn=MZF1;refGene_tx=XM_006940715.4;refGene_gn=MZF
1;refGene_tx=XM_006940716.4;refGene_gn=MZF1;refGene_tx=XM_019818447.2
E2 260230 260414
refGene_gn=MZF1;refGene_tx=XM_003997334.5;refGene_gn=MZF1;refGene_tx=X
M_006940713.4;refGene_gn=MZF1;refGene_tx=XM_006940715.4;refGene_gn=MZF
1;refGene_tx=XM_006940716.4;refGene_gn=MZF1;refGene_tx=XM_019818447.2

E2	261122	261193	refGene_gn=MZF1;refGene_tx=XM_003997334.5;refGene_gn=MZF1;refGene_tx=XM_006940713.4;refGene_gn=MZF1;refGene_tx=XM_006940715.4;refGene_gn=MZF1;refGene_tx=XM_006940716.4;refGene_gn=MZF1;refGene_tx=XM_019818447.2
E2	261313	261434	refGene_gn=MZF1;refGene_tx=XM_003997334.5;refGene_gn=MZF1;refGene_tx=XM_006940713.4;refGene_gn=MZF1;refGene_tx=XM_006940715.4;refGene_gn=MZF1;refGene_tx=XM_006940716.4;refGene_gn=MZF1;refGene_tx=XM_019818447.2
E2	265781	267214	refGene_gn=MZF1;refGene_tx=XM_003997334.5;refGene_gn=MZF1;refGene_tx=XM_006940713.4;refGene_gn=MZF1;refGene_tx=XM_006940715.4;refGene_gn=MZF1;refGene_tx=XM_006940716.4;refGene_gn=MZF1;refGene_tx=XM_019818447.2
E2	272191	272300	refGene_gn=UBE2M;refGene_tx=XM_023245780.1
E2	273363	273458	refGene_gn=UBE2M;refGene_tx=XM_023245780.1
E2	273598	273637	refGene_gn=UBE2M;refGene_tx=XM_023245780.1
E2	273747	273851	refGene_gn=UBE2M;refGene_tx=XM_023245780.1
E2	274045	274109	refGene_gn=UBE2M;refGene_tx=XM_023245780.1
E2	274202	274343	refGene_gn=UBE2M;refGene_tx=XM_023245780.1
E2	276348	276516	refGene_gn=CHMP2A;refGene_tx=XM_011289525.3;refGene_gn=CHMP2A;refGene_tx=XM_019818480.1;refGene_gn=CHMP2A;refGene_tx=XM_019818481.2;refGene_gn=CHMP2A;refGene_tx=XM_019818482.2;refGene_gn=CHMP2A;refGene_tx=XM_023245778.1
E2	277589	277769	refGene_gn=CHMP2A;refGene_tx=XM_011289525.3;refGene_gn=CHMP2A;refGene_tx=XM_019818480.1;refGene_gn=CHMP2A;refGene_tx=XM_019818481.2;refGene_gn=CHMP2A;refGene_tx=XM_019818482.2;refGene_gn=CHMP2A;refGene_tx=XM_023245778.1
E2	277856	277987	refGene_gn=CHMP2A;refGene_tx=XM_011289525.3;refGene_gn=CHMP2A;refGene_tx=XM_019818480.1;refGene_gn=CHMP2A;refGene_tx=XM_019818481.2;refGene_gn=CHMP2A;refGene_tx=XM_019818482.2;refGene_gn=CHMP2A;refGene_tx=XM_023245778.1
E2	278077	278139	refGene_gn=CHMP2A;refGene_tx=XM_011289525.3;refGene_gn=CHMP2A;refGene_tx=XM_019818480.1;refGene_gn=CHMP2A;refGene_tx=XM_019818481.2;refGene_gn=CHMP2A;refGene_tx=XM_019818482.2;refGene_gn=CHMP2A;refGene_tx=XM_023245778.1
E2	278255	278383	refGene_gn=CHMP2A;refGene_tx=XM_011289525.3;refGene_gn=CHMP2A;refGene_tx=XM_019818480.1;refGene_gn=CHMP2A;refGene_tx=XM_019818481.2;refGene_gn=CHMP2A;refGene_tx=XM_019818482.2;refGene_gn=CHMP2A;refGene_tx=XM_023245778.1
E2	279343	279520	refGene_gn=TRIM28;refGene_tx=XM_019818467.2
E2	279612	279750	refGene_gn=TRIM28;refGene_tx=XM_019818467.2
E2	279828	279867	refGene_gn=TRIM28;refGene_tx=XM_019818467.2
E2	279868	279916	refGene_gn=TRIM28;refGene_tx=XM_019818467.2
E2	279999	280123	refGene_gn=TRIM28;refGene_tx=XM_019818467.2
E2	280208	280528	refGene_gn=TRIM28;refGene_tx=XM_019818467.2
E2	280607	280860	refGene_gn=TRIM28;refGene_tx=XM_019818467.2

E2	280934	280980	refGene_gn=TRIM28;refGene_tx=XM_019818467.2
E2	281056	281093	refGene_gn=TRIM28;refGene_tx=XM_019818467.2
E2	281186	281293	refGene_gn=TRIM28;refGene_tx=XM_019818467.2
E2	281374	281489	refGene_gn=TRIM28;refGene_tx=XM_019818467.2
E2	281593	281740	refGene_gn=TRIM28;refGene_tx=XM_019818467.2
E2	281893	282008	refGene_gn=TRIM28;refGene_tx=XM_019818467.2
E2	282091	282208	refGene_gn=TRIM28;refGene_tx=XM_019818467.2
E2	282285	282421	refGene_gn=TRIM28;refGene_tx=XM_019818467.2
E2	284141	284274	refGene_gn=TRIM28;refGene_tx=XM_019818467.2
E2	284470	284583	refGene_gn=TRIM28;refGene_tx=XM_019818467.2
E2	284919	285253	refGene_gn=TRIM28;refGene_tx=XM_019818467.2
E2	301445	301481	refGene_gn=ZBTB45;refGene_tx=XM_006940722.3
E2	301596	301725	refGene_gn=ZBTB45;refGene_tx=XM_003997337.5
E2	303458	304740
refGene_gn=ZBTB45;refGene_tx=XM_003997337.5;refGene_gn=ZBTB45;refGene_tx=XM_006940722.3;refGene_gn=ZBTB45;refGene_tx=XM_006940723.4;refGene_gn=ZBTB45;refGene_tx=XM_006940724.4
E2	306592	306849
refGene_gn=ZBTB45;refGene_tx=XM_003997337.5;refGene_gn=ZBTB45;refGene_tx=XM_006940722.3;refGene_gn=ZBTB45;refGene_tx=XM_006940723.4;refGene_gn=ZBTB45;refGene_tx=XM_006940724.4
E2	308685	309364	refGene_gn=SLC27A5;refGene_tx=XM_003997338.5
E2	309658	309868	refGene_gn=SLC27A5;refGene_tx=XM_003997338.5
E2	310231	310390	refGene_gn=SLC27A5;refGene_tx=XM_003997338.5
E2	312251	312376	refGene_gn=SLC27A5;refGene_tx=XM_003997338.5
E2	313190	313385	refGene_gn=SLC27A5;refGene_tx=XM_003997338.5
E2	313464	313555	refGene_gn=SLC27A5;refGene_tx=XM_003997338.5
E2	314274	314473	refGene_gn=SLC27A5;refGene_tx=XM_003997338.5
E2	314715	314813	refGene_gn=SLC27A5;refGene_tx=XM_003997338.5
E2	315010	315141	refGene_gn=SLC27A5;refGene_tx=XM_003997338.5
E2	315225	315402	refGene_gn=SLC27A5;refGene_tx=XM_003997338.5
E2	337047	338015
refGene_gn=ZNF446;refGene_tx=XM_011289518.3;refGene_gn=ZNF446;refGene_tx=XM_011289519.3;refGene_gn=ZNF446;refGene_tx=XM_011289520.3;refGene_gn=ZNF446;refGene_tx=XM_011289521.3;refGene_gn=ZNF446;refGene_tx=XM_011289522.3;refGene_gn=ZNF446;refGene_tx=XM_011289523.3;refGene_gn=ZNF446;refGene_tx=XM_019818471.2;refGene_gn=ZNF446;refGene_tx=XM_019818472.2
E2	338241	338326
refGene_gn=ZNF446;refGene_tx=XM_011289518.3;refGene_gn=ZNF446;refGene_tx=XM_011289519.3;refGene_gn=ZNF446;refGene_tx=XM_011289520.3;refGene_gn=ZNF446;refGene_tx=XM_011289521.3;refGene_gn=ZNF446;refGene_tx=XM_011289522.3;refGene_gn=ZNF446;refGene_tx=XM_011289523.3;refGene_gn=ZNF446;refGene_tx=XM_019818471.2;refGene_gn=ZNF446;refGene_tx=XM_019818472.2
E2	339912	340007
refGene_gn=ZNF446;refGene_tx=XM_011289518.3;refGene_gn=ZNF446;refGene_tx=XM_011289519.3;refGene_gn=ZNF446;refGene_tx=XM_011289520.3;refGene_gn=ZNF446;refGene_tx=XM_011289521.3;refGene_gn=ZNF446;refGene_tx=XM_011289522.3;refGene_gn=ZNF446;refGene_tx=XM_011289523.3;refGene_gn=ZNF44

6;refGene_tx=XM_019818471.2;refGene_gn=ZNF446;refGene_tx=XM_019818472.2
E2 340320 340522 refGene_gn=ZNF446;refGene_tx=XM_011289518.3;refGene_gn=ZNF446;refGene_tx=XM_011289519.3;refGene_gn=ZNF446;refGene_tx=XM_011289520.3;refGene_gn=ZNF446;refGene_tx=XM_011289521.3;refGene_gn=ZNF446;refGene_tx=XM_011289522.3;refGene_gn=ZNF446;refGene_tx=XM_011289523.3;refGene_gn=ZNF446;refGene_tx=XM_019818471.2;refGene_gn=ZNF446;refGene_tx=XM_019818472.2
E2 340598 340979 refGene_gn=ZNF446;refGene_tx=XM_011289518.3;refGene_gn=ZNF446;refGene_tx=XM_011289519.3;refGene_gn=ZNF446;refGene_tx=XM_011289520.3;refGene_gn=ZNF446;refGene_tx=XM_011289521.3;refGene_gn=ZNF446;refGene_tx=XM_011289522.3;refGene_gn=ZNF446;refGene_tx=XM_011289523.3;refGene_gn=ZNF446;refGene_tx=XM_019818471.2
E2 342283 342388 refGene_gn=ZNF446;refGene_tx=XM_011289518.3;refGene_gn=ZNF446;refGene_tx=XM_011289519.3;refGene_gn=ZNF446;refGene_tx=XM_011289522.3;refGene_gn=ZNF446;refGene_tx=XM_011289523.3;refGene_gn=ZNF446;refGene_tx=XM_019818471.2;refGene_gn=ZNF446;refGene_tx=XM_019818472.2
E2 346099 347538 refGene_gn=ZNF324;refGene_tx=XM_023245758.1;refGene_gn=ZNF324;refGene_tx=XM_023245759.1;refGene_gn=ZNF324;refGene_tx=XM_023245760.1
E2 347945 348062 refGene_gn=ZNF324;refGene_tx=XM_023245758.1;refGene_gn=ZNF324;refGene_tx=XM_023245759.1;refGene_gn=ZNF324;refGene_tx=XM_023245760.1
E2 348390 348517 refGene_gn=ZNF324;refGene_tx=XM_023245758.1;refGene_gn=ZNF324;refGene_tx=XM_023245759.1;refGene_gn=ZNF324;refGene_tx=XM_023245760.1
E2 349473 349482 refGene_gn=ZNF324;refGene_tx=XM_023245759.1
E2 350008 350041 refGene_gn=ZNF324;refGene_tx=XM_023245758.1
E2 370065 370752 refGene_gn=LOC101089225;refGene_tx=XM_019818484.2
E2 370753 370903 refGene_gn=LOC101089225;refGene_tx=XM_019818484.2
E2 385506 385605 refGene_gn=LOC101089479;refGene_tx=XM_006940727.4
E2 385768 385783 refGene_gn=LOC101089479;refGene_tx=XM_006940728.3
E2 387916 388085 refGene_gn=LOC101089479;refGene_tx=XM_006940727.4;refGene_gn=LOC101089479;refGene_tx=XM_006940728.3
E2 389447 391378 refGene_gn=LOC101089479;refGene_tx=XM_006940727.4;refGene_gn=LOC101089479;refGene_tx=XM_006940728.3
E2 410499 411506 refGene_gn=ZNF584;refGene_tx=XM_006940729.4
E2 412116 412239 refGene_gn=ZNF584;refGene_tx=XM_006940729.4
E2 417304 417431 refGene_gn=ZNF584;refGene_tx=XM_006940729.4
E2 418329 418347 refGene_gn=ZNF584;refGene_tx=XM_006940729.4
E2 439815 440823 refGene_gn=RNF225;refGene_tx=XM_023245771.1

E2	442426	442495	refGene_gn=RPS5;refGene_tx=XM_003997339.4
E2	442576	442675	refGene_gn=RPS5;refGene_tx=XM_003997339.4
E2	445700	445829	refGene_gn=RPS5;refGene_tx=XM_003997339.4
E2	445990	446200	refGene_gn=RPS5;refGene_tx=XM_003997339.4
E2	448415	448523	refGene_gn=RPS5;refGene_tx=XM_003997339.4
E2	467064	467133
refGene_gn=LOC111558093;refGene_tx=XM_023245853.1
E2	467250	467426
refGene_gn=LOC111558093;refGene_tx=XM_023245853.1
E2	467550	467662
refGene_gn=LOC111558093;refGene_tx=XM_023245853.1
E2	467715	467778
refGene_gn=LOC111558093;refGene_tx=XM_023245853.1
E2	468270	468361
refGene_gn=LOC111558093;refGene_tx=XM_023245853.1
E2	468453	468553
refGene_gn=LOC111558093;refGene_tx=XM_023245853.1
E2	469009	469175
refGene_gn=LOC111558093;refGene_tx=XM_023245853.1
E2	469291	469311
refGene_gn=LOC111558093;refGene_tx=XM_023245853.1
E2	469407	469485
refGene_gn=LOC111558093;refGene_tx=XM_023245853.1
E2	469605	469807
refGene_gn=LOC111558093;refGene_tx=XM_023245853.1
E2	476566	476600	refGene_gn=A1BG;refGene_tx=XM_023245762.1
E2	476685	476724	refGene_gn=A1BG;refGene_tx=XM_023245762.1
E2	476815	477094
refGene_gn=A1BG;refGene_tx=XM_023245762.1;refGene_gn=A1BG;refGene_tx=XM_023245763.1;refGene_gn=A1BG;refGene_tx=XM_023245764.1;refGene_gn=A1BG;refGene_tx=XM_023245765.1;refGene_gn=A1BG;refGene_tx=XM_023245766.1;refGene_gn=A1BG;refGene_tx=XM_023245767.1;refGene_gn=A1BG;refGene_tx=XM_023245768.1;refGene_gn=A1BG;refGene_tx=XM_023245769.1
E2	477517	477790
refGene_gn=A1BG;refGene_tx=XM_023245762.1;refGene_gn=A1BG;refGene_tx=XM_023245763.1;refGene_gn=A1BG;refGene_tx=XM_023245764.1;refGene_gn=A1BG;refGene_tx=XM_023245765.1;refGene_gn=A1BG;refGene_tx=XM_023245766.1;refGene_gn=A1BG;refGene_tx=XM_023245767.1;refGene_gn=A1BG;refGene_tx=XM_023245768.1;refGene_gn=A1BG;refGene_tx=XM_023245769.1
E2	478076	478373
refGene_gn=A1BG;refGene_tx=XM_023245762.1;refGene_gn=A1BG;refGene_tx=XM_023245763.1;refGene_gn=A1BG;refGene_tx=XM_023245764.1;refGene_gn=A1BG;refGene_tx=XM_023245765.1;refGene_gn=A1BG;refGene_tx=XM_023245766.1;refGene_gn=A1BG;refGene_tx=XM_023245767.1;refGene_gn=A1BG;refGene_tx=XM_023245768.1;refGene_gn=A1BG;refGene_tx=XM_023245769.1
E2	479024	479321
refGene_gn=A1BG;refGene_tx=XM_023245762.1;refGene_gn=A1BG;refGene_tx=XM_023245763.1;refGene_gn=A1BG;refGene_tx=XM_023245764.1;refGene_gn=A1BG;refGene_tx=XM_023245765.1;refGene_gn=A1BG;refGene_tx=XM_023245766.1;refGene_gn=A1BG;refGene_tx=XM_023245767.1;refGene_gn=A1BG;refGene_tx=X

M_023245768.1;refGene_gn=A1BG;refGene_tx=XM_023245769.1
E2 479786 480074
refGene_gn=A1BG;refGene_tx=XM_023245762.1;refGene_gn=A1BG;refGene_tx=XM_023245763.1;refGene_gn=A1BG;refGene_tx=XM_023245764.1;refGene_gn=A1BG;refGene_tx=XM_023245765.1;refGene_gn=A1BG;refGene_tx=XM_023245766.1;refGene_gn=A1BG;refGene_tx=XM_023245767.1;refGene_gn=A1BG;refGene_tx=XM_023245768.1;refGene_gn=A1BG;refGene_tx=XM_023245769.1
E2 480449 480457
refGene_gn=A1BG;refGene_tx=XM_023245762.1;refGene_gn=A1BG;refGene_tx=XM_023245763.1;refGene_gn=A1BG;refGene_tx=XM_023245764.1;refGene_gn=A1BG;refGene_tx=XM_023245765.1;refGene_gn=A1BG;refGene_tx=XM_023245766.1;refGene_gn=A1BG;refGene_tx=XM_023245767.1;refGene_gn=A1BG;refGene_tx=XM_023245768.1;refGene_gn=A1BG;refGene_tx=XM_023245769.1
E2 488001 489077
refGene_gn=ZSCAN22;refGene_tx=XM_011289515.3;refGene_gn=ZSCAN22;refGene_tx=XM_019818477.2;refGene_gn=ZSCAN22;refGene_tx=XM_019818478.2
E2 491861 492261
refGene_gn=ZSCAN22;refGene_tx=XM_011289515.3;refGene_gn=ZSCAN22;refGene_tx=XM_019818477.2;refGene_gn=ZSCAN22;refGene_tx=XM_019818478.2
E2 508277 509396
refGene_gn=LOC102901988;refGene_tx=XM_023245781.1
E2 523192 524631 refGene_gn=ZNF8;refGene_tx=XM_006940741.4
E2 528036 528132 refGene_gn=ZNF8;refGene_tx=XM_006940741.4
E2 528430 528557 refGene_gn=ZNF8;refGene_tx=XM_006940741.4
E2 533823 533889 refGene_gn=ZNF8;refGene_tx=XM_006940741.4
E2 548261 550282
refGene_gn=ZNF544;refGene_tx=XM_019818448.2;refGene_gn=ZNF544;refGene_tx=XM_019818450.2;refGene_gn=ZNF544;refGene_tx=XM_019818451.2;refGene_gn=ZNF544;refGene_tx=XM_019818452.2;refGene_gn=ZNF544;refGene_tx=XM_019818453.2;refGene_gn=ZNF544;refGene_tx=XM_019818454.2;refGene_gn=ZNF544;refGene_tx=XM_019818456.2;refGene_gn=ZNF544;refGene_tx=XM_019818457.2;refGene_gn=ZNF544;refGene_tx=XM_019818458.2;refGene_gn=ZNF544;refGene_tx=XM_019818459.2
E2 553710 553794
refGene_gn=ZNF544;refGene_tx=XM_019818448.2;refGene_gn=ZNF544;refGene_tx=XM_019818450.2;refGene_gn=ZNF544;refGene_tx=XM_019818451.2;refGene_gn=ZNF544;refGene_tx=XM_019818452.2;refGene_gn=ZNF544;refGene_tx=XM_019818453.2;refGene_gn=ZNF544;refGene_tx=XM_019818454.2;refGene_gn=ZNF544;refGene_tx=XM_019818456.2;refGene_gn=ZNF544;refGene_tx=XM_019818457.2;refGene_gn=ZNF544;refGene_tx=XM_019818458.2;refGene_gn=ZNF544;refGene_tx=XM_019818459.2
E2 554099 554238
refGene_gn=ZNF544;refGene_tx=XM_019818448.2;refGene_gn=ZNF544;refGene_tx=XM_019818450.2;refGene_gn=ZNF544;refGene_tx=XM_019818451.2;refGene_gn=ZNF544;refGene_tx=XM_019818452.2;refGene_gn=ZNF544;refGene_tx=XM_019818453.2;refGene_gn=ZNF544;refGene_tx=XM_019818454.2;refGene_gn=ZNF544;refGene_tx=XM_019818456.2;refGene_gn=ZNF544;refGene_tx=XM_019818457.2;refGene_gn=ZNF544;refGene_tx=XM_019818458.2;refGene_gn=ZNF544;refGene_tx=XM_019818459.2
E2 554345 554405 refGene_gn=ZNF544;refGene_tx=XM_019818456.2

E2	555786	555878
refGene_gn=ZNF544;refGene_tx=XM_019818448.2;refGene_gn=ZNF544;refGene_tx=XM_019818450.2;refGene_gn=ZNF544;refGene_tx=XM_019818451.2;refGene_gn=ZNF544;refGene_tx=XM_019818452.2;refGene_gn=ZNF544;refGene_tx=XM_019818453.2;refGene_gn=ZNF544;refGene_tx=XM_019818454.2;refGene_gn=ZNF544;refGene_tx=XM_019818457.2;refGene_gn=ZNF544;refGene_tx=XM_019818458.2
E2	557181	557224
refGene_gn=ZNF544;refGene_tx=XM_019818448.2;refGene_gn=ZNF544;refGene_tx=XM_019818450.2;refGene_gn=ZNF544;refGene_tx=XM_019818451.2;refGene_gn=ZNF544;refGene_tx=XM_019818452.2;refGene_gn=ZNF544;refGene_tx=XM_019818453.2;refGene_gn=ZNF544;refGene_tx=XM_019818454.2
E2	575124	576026
refGene_gn=ZNF274;refGene_tx=XM_006940743.4;refGene_gn=ZNF274;refGene_tx=XM_006940744.4;refGene_gn=ZNF274;refGene_tx=XM_006940745.4;refGene_gn=ZNF274;refGene_tx=XM_006940746.4;refGene_gn=ZNF274;refGene_tx=XM_019818468.2;refGene_gn=ZNF274;refGene_tx=XM_019818469.2;refGene_gn=ZNF274;refGene_tx=XM_019818470.2
E2	576498	576625
refGene_gn=ZNF274;refGene_tx=XM_006940743.4;refGene_gn=ZNF274;refGene_tx=XM_006940744.4;refGene_gn=ZNF274;refGene_tx=XM_006940745.4;refGene_gn=ZNF274;refGene_tx=XM_006940746.4;refGene_gn=ZNF274;refGene_tx=XM_019818468.2;refGene_gn=ZNF274;refGene_tx=XM_019818469.2;refGene_gn=ZNF274;refGene_tx=XM_019818470.2
E2	577784	577885
refGene_gn=ZNF274;refGene_tx=XM_006940743.4;refGene_gn=ZNF274;refGene_tx=XM_006940744.4;refGene_gn=ZNF274;refGene_tx=XM_006940745.4;refGene_gn=ZNF274;refGene_tx=XM_006940746.4;refGene_gn=ZNF274;refGene_tx=XM_019818468.2;refGene_gn=ZNF274;refGene_tx=XM_019818469.2;refGene_gn=ZNF274;refGene_tx=XM_019818470.2
E2	586992	587475
refGene_gn=ZNF274;refGene_tx=XM_006940743.4;refGene_gn=ZNF274;refGene_tx=XM_006940744.4;refGene_gn=ZNF274;refGene_tx=XM_006940745.4;refGene_gn=ZNF274;refGene_tx=XM_006940746.4;refGene_gn=ZNF274;refGene_tx=XM_019818468.2;refGene_gn=ZNF274;refGene_tx=XM_019818469.2;refGene_gn=ZNF274;refGene_tx=XM_019818470.2
E2	602410	602506
refGene_gn=ZNF274;refGene_tx=XM_006940743.4;refGene_gn=ZNF274;refGene_tx=XM_006940745.4;refGene_gn=ZNF274;refGene_tx=XM_019818468.2;refGene_gn=ZNF274;refGene_tx=XM_019818469.2;refGene_gn=ZNF274;refGene_tx=XM_019818470.2
E2	603269	603396
refGene_gn=ZNF274;refGene_tx=XM_006940743.4;refGene_gn=ZNF274;refGene_tx=XM_006940744.4;refGene_gn=ZNF274;refGene_tx=XM_006940745.4;refGene_gn=ZNF274;refGene_tx=XM_019818468.2;refGene_gn=ZNF274;refGene_tx=XM_019818469.2;refGene_gn=ZNF274;refGene_tx=XM_019818470.2
E2	604161	604235
refGene_gn=ZNF274;refGene_tx=XM_006940743.4;refGene_gn=ZNF274;refGene_tx=XM_006940744.4;refGene_gn=ZNF274;refGene_tx=XM_006940745.4;refGene_gn=ZNF274;refGene_tx=XM_019818468.2

E2	604721	604845
refGene_gn=ZNF274;refGene_tx=XM_006940743.4;refGene_gn=ZNF274;refGene_tx=XM_006940744.4
E2	650115	651741	refGene_gn=ZNF329;refGene_tx=XM_003997342.4
E2	681945	682348
refGene_gn=ZSCAN18;refGene_tx=XM_006940747.4;refGene_gn=ZSCAN18;refGene_tx=XM_006940749.4;refGene_gn=ZSCAN18;refGene_tx=XM_019818474.2;refGene_gn=ZSCAN18;refGene_tx=XM_019818475.2
E2	682936	682999
refGene_gn=ZSCAN18;refGene_tx=XM_006940747.4;refGene_gn=ZSCAN18;refGene_tx=XM_006940749.4;refGene_gn=ZSCAN18;refGene_tx=XM_019818474.2;refGene_gn=ZSCAN18;refGene_tx=XM_019818475.2
E2	684038	684130
refGene_gn=ZSCAN18;refGene_tx=XM_006940747.4;refGene_gn=ZSCAN18;refGene_tx=XM_006940749.4;refGene_gn=ZSCAN18;refGene_tx=XM_019818474.2;refGene_gn=ZSCAN18;refGene_tx=XM_019818475.2
E2	685168	685268
refGene_gn=ZSCAN18;refGene_tx=XM_006940747.4;refGene_gn=ZSCAN18;refGene_tx=XM_006940749.4;refGene_gn=ZSCAN18;refGene_tx=XM_019818474.2;refGene_gn=ZSCAN18;refGene_tx=XM_019818475.2
E2	686687	686780
refGene_gn=ZSCAN18;refGene_tx=XM_006940747.4;refGene_gn=ZSCAN18;refGene_tx=XM_006940749.4;refGene_gn=ZSCAN18;refGene_tx=XM_019818474.2;refGene_gn=ZSCAN18;refGene_tx=XM_019818475.2
E2	688023	688766
refGene_gn=ZSCAN18;refGene_tx=XM_006940747.4;refGene_gn=ZSCAN18;refGene_tx=XM_006940749.4;refGene_gn=ZSCAN18;refGene_tx=XM_019818474.2;refGene_gn=ZSCAN18;refGene_tx=XM_019818475.2
E2	696309	696798	refGene_gn=ZNF135;refGene_tx=XM_023245770.1
E2	696799	697584	refGene_gn=ZNF135;refGene_tx=XM_023245770.1
E2	700936	701063	refGene_gn=ZNF135;refGene_tx=XM_023245770.1
E2	703710	703743	refGene_gn=ZNF135;refGene_tx=XM_023245770.1
E2	713106	713610
refGene_gn=LOC111558085;refGene_tx=XM_023245782.1
E2	714240	714380	refGene_gn=ZSCAN1;refGene_tx=XM_023245777.1
E2	714579	714691	refGene_gn=ZSCAN1;refGene_tx=XM_023245777.1
E2	721263	721343	refGene_gn=ZSCAN1;refGene_tx=XM_023245777.1
E2	727653	728652
refGene_gn=ZSCAN1;refGene_tx=XM_023245772.1;refGene_gn=ZSCAN1;refGene_tx=XM_023245773.1;refGene_gn=ZSCAN1;refGene_tx=XM_023245774.1;refGene_gn=ZSCAN1;refGene_tx=XM_023245775.1;refGene_gn=ZSCAN1;refGene_tx=XM_023245776.1;refGene_gn=ZSCAN1;refGene_tx=XM_023245777.1
E2	731124	731273
refGene_gn=LOC102901032;refGene_tx=XM_023245855.1
E2	731358	731832
refGene_gn=LOC102901032;refGene_tx=XM_023245855.1
E2	736265	736348
refGene_gn=LOC102901032;refGene_tx=XM_023245855.1
E2	738579	738723
refGene_gn=LOC102901032;refGene_tx=XM_023245855.1

E2	739168	739173
refGene_gn=LOC102901032;refGene_tx=XM_023245855.1
E2	769316	769347
refGene_gn=ZNF606;refGene_tx=XM_006940752.4;refGene_gn=ZNF606;refGene_tx=XM_006940754.4;refGene_gn=ZNF606;refGene_tx=XM_006940757.4;refGene_gn=ZNF606;refGene_tx=XM_019818460.2;refGene_gn=ZNF606;refGene_tx=XM_019818461.2;refGene_gn=ZNF606;refGene_tx=XM_019818462.2;refGene_gn=ZNF606;refGene_tx=XM_019818465.2;refGene_gn=ZNF606;refGene_tx=XM_023245739.1
E2	770628	770685
refGene_gn=ZNF606;refGene_tx=XM_006940752.4;refGene_gn=ZNF606;refGene_tx=XM_006940754.4;refGene_gn=ZNF606;refGene_tx=XM_006940757.4;refGene_gn=ZNF606;refGene_tx=XM_019818460.2;refGene_gn=ZNF606;refGene_tx=XM_019818461.2;refGene_gn=ZNF606;refGene_tx=XM_019818465.2;refGene_gn=ZNF606;refGene_tx=XM_023245739.1
E2	771220	771309
refGene_gn=ZNF606;refGene_tx=XM_006940752.4;refGene_gn=ZNF606;refGene_tx=XM_006940754.4;refGene_gn=ZNF606;refGene_tx=XM_006940757.4;refGene_gn=ZNF606;refGene_tx=XM_019818460.2;refGene_gn=ZNF606;refGene_tx=XM_019818461.2;refGene_gn=ZNF606;refGene_tx=XM_019818462.2;refGene_gn=ZNF606;refGene_tx=XM_019818465.2;refGene_gn=ZNF606;refGene_tx=XM_023245739.1
E2	780621	780748
refGene_gn=ZNF606;refGene_tx=XM_006940752.4;refGene_gn=ZNF606;refGene_tx=XM_006940754.4;refGene_gn=ZNF606;refGene_tx=XM_006940757.4;refGene_gn=ZNF606;refGene_tx=XM_019818460.2;refGene_gn=ZNF606;refGene_tx=XM_019818461.2;refGene_gn=ZNF606;refGene_tx=XM_019818462.2;refGene_gn=ZNF606;refGene_tx=XM_019818465.2;refGene_gn=ZNF606;refGene_tx=XM_023245739.1;refGene_gn=ZNF606;refGene_tx=XM_023245740.1;refGene_gn=ZNF606;refGene_tx=XM_023245741.1;refGene_gn=ZNF606;refGene_tx=XM_023245742.1;refGene_gn=ZNF606;refGene_tx=XM_023245743.1
E2	781225	781321
refGene_gn=ZNF606;refGene_tx=XM_006940752.4;refGene_gn=ZNF606;refGene_tx=XM_006940754.4;refGene_gn=ZNF606;refGene_tx=XM_006940757.4;refGene_gn=ZNF606;refGene_tx=XM_019818460.2;refGene_gn=ZNF606;refGene_tx=XM_019818461.2;refGene_gn=ZNF606;refGene_tx=XM_019818462.2;refGene_gn=ZNF606;refGene_tx=XM_019818465.2;refGene_gn=ZNF606;refGene_tx=XM_023245739.1;refGene_gn=ZNF606;refGene_tx=XM_023245740.1;refGene_gn=ZNF606;refGene_tx=XM_023245741.1;refGene_gn=ZNF606;refGene_tx=XM_023245742.1;refGene_gn=ZNF606;refGene_tx=XM_023245743.1
E2	783195	783262
refGene_gn=ZNF606;refGene_tx=XM_023245740.1;refGene_gn=ZNF606;refGene_tx=XM_023245743.1
E2	783950	784051	refGene_gn=ZNF606;refGene_tx=XM_023245743.1
E2	786899	788875
refGene_gn=ZNF606;refGene_tx=XM_006940752.4;refGene_gn=ZNF606;refGene_tx=XM_019818460.2;refGene_gn=ZNF606;refGene_tx=XM_019818461.2;refGene_gn=ZNF606;refGene_tx=XM_019818462.2;refGene_gn=ZNF606;refGene_tx=XM_023245739.1;refGene_gn=ZNF606;refGene_tx=XM_023245740.1;refGene_gn=ZNF606;refGene_tx=XM_023245741.1;refGene_gn=ZNF606;refGene_tx=XM_023245742.

1
E2 795468 795725
refGene_gn=ZNF606;refGene_tx=XM_006940754.4;refGene_gn=ZNF606;refGene_tx=XM_006940757.4;refGene_gn=ZNF606;refGene_tx=XM_019818465.2
E2 801851 801930
refGene_gn=CE2H19orf18;refGene_tx=XM_006940751.4;refGene_gn=CE2H19orf18;refGene_tx=XM_023245779.1
E2 802053 802119
refGene_gn=CE2H19orf18;refGene_tx=XM_006940751.4;refGene_gn=CE2H19orf18;refGene_tx=XM_023245779.1
E2 803063 803105
refGene_gn=CE2H19orf18;refGene_tx=XM_006940751.4
E2 805893 805986
refGene_gn=CE2H19orf18;refGene_tx=XM_006940751.4;refGene_gn=CE2H19orf18;refGene_tx=XM_023245779.1
E2 806853 806956
refGene_gn=CE2H19orf18;refGene_tx=XM_006940751.4;refGene_gn=CE2H19orf18;refGene_tx=XM_023245779.1
E2 810959 811126
refGene_gn=CE2H19orf18;refGene_tx=XM_006940751.4;refGene_gn=CE2H19orf18;refGene_tx=XM_023245779.1
E2 812500 812616
refGene_gn=CE2H19orf18;refGene_tx=XM_006940751.4;refGene_gn=CE2H19orf18;refGene_tx=XM_023245779.1
E2 828468 828507
refGene_gn=LOC109494649;refGene_tx=XM_023245856.1
E2 828935 829580
refGene_gn=LOC109494649;refGene_tx=XM_023245856.1
E2 853346 853552
refGene_gn=LOC102902115;refGene_tx=XM_023245783.1
E2 853554 853610
refGene_gn=LOC102902115;refGene_tx=XM_023245783.1
E2 853611 854473
refGene_gn=LOC102902115;refGene_tx=XM_023245783.1
E2 854475 854733
refGene_gn=LOC102902115;refGene_tx=XM_023245783.1
E2 856001 856128
refGene_gn=LOC102902115;refGene_tx=XM_023245783.1
E2 856597 856717
refGene_gn=LOC102902115;refGene_tx=XM_023245783.1
E2 860328 860653
refGene_gn=ZNF671;refGene_tx=XM_019818989.2;refGene_gn=ZNF671;refGene_tx=XM_023245247.1;refGene_gn=ZNF671;refGene_tx=XM_023245248.1;refGene_gn=ZNF671;refGene_tx=XM_023245249.1
E2 871933 872060
refGene_gn=ZNF671;refGene_tx=XM_019818989.2;refGene_gn=ZNF671;refGene_tx=XM_023245247.1;refGene_gn=ZNF671;refGene_tx=XM_023245249.1
E2 872543 872666
refGene_gn=ZNF671;refGene_tx=XM_019818989.2;refGene_gn=ZNF671;refGene_tx=XM_023245247.1;refGene_gn=ZNF671;refGene_tx=XM_023245248.1;refGene_

gn=ZNF671;refGene_tx=XM_023245249.1
E2 873273 874481
refGene_gn=ZNF671;refGene_tx=XM_019818989.2;refGene_gn=ZNF671;refGene_tx=XM_023245247.1;refGene_gn=ZNF671;refGene_tx=XM_023245248.1;refGene_gn=ZNF671;refGene_tx=XM_023245249.1
E2 901896 901929
refGene_gn=LOC101080879;refGene_tx=XM_023245667.1
E2 904225 904255
refGene_gn=LOC101080879;refGene_tx=XM_023245668.1
E2 904360 904487
refGene_gn=LOC101080879;refGene_tx=XM_023245667.1;refGene_gn=LOC101080879;refGene_tx=XM_023245668.1
E2 908475 910892
refGene_gn=LOC101080879;refGene_tx=XM_023245667.1;refGene_gn=LOC101080879;refGene_tx=XM_023245668.1
E2 934826 934859
refGene_gn=LOC101091237;refGene_tx=XM_023245669.1
E2 937314 937441
refGene_gn=LOC101091237;refGene_tx=XM_023245669.1
E2 940458 942080
refGene_gn=LOC101091237;refGene_tx=XM_023245669.1
E2 942081 942696
refGene_gn=LOC101091237;refGene_tx=XM_023245669.1
E2 961253 961286
refGene_gn=LOC105261205;refGene_tx=XM_011289510.3
E2 963724 963851
refGene_gn=LOC105261205;refGene_tx=XM_011289510.3
E2 967398 970559
refGene_gn=LOC105261205;refGene_tx=XM_011289510.3
E2 1010898 1010931
refGene_gn=LOC101091741;refGene_tx=XM_023246037.1
E2 1012803 1012930
refGene_gn=LOC101091741;refGene_tx=XM_023246037.1
E2 1018917 1020497
refGene_gn=LOC101091741;refGene_tx=XM_023246037.1
E2 1020498 1021167
refGene_gn=LOC101091741;refGene_tx=XM_023246037.1
E2 1021168 1021230
refGene_gn=LOC101091741;refGene_tx=XM_023246037.1
E2 1052927 1053033
refGene_gn=LOC101091741;refGene_tx=XM_023246037.1
E2 1053034 1053958
refGene_gn=LOC101091741;refGene_tx=XM_023246037.1
E2 1053959 1054394
refGene_gn=LOC101091741;refGene_tx=XM_023246037.1
E2 1054395 1054632
refGene_gn=LOC101091741;refGene_tx=XM_023246037.1
E2 1054633 1054870
refGene_gn=LOC101091741;refGene_tx=XM_023246037.1
E2 1129366 1129399

refGene_gn=LOC101092471;refGene_tx=XM_019818993.2
E2 1130958 1131085
refGene_gn=LOC101092471;refGene_tx=XM_019818993.2
E2 1131947 1133233
refGene_gn=LOC101092471;refGene_tx=XM_019818993.2
E2 1168390 1169874
refGene_gn=LOC101092972;refGene_tx=XM_003997358.5
E2 1171170 1171297
refGene_gn=LOC101092972;refGene_tx=XM_003997358.5
E2 1175400 1175433
refGene_gn=LOC101092972;refGene_tx=XM_003997358.5
E2 1185415 1185448
refGene_gn=LOC105261204;refGene_tx=XM_011289507.3
E2 1188377 1188504
refGene_gn=LOC105261204;refGene_tx=XM_011289507.3
E2 1189826 1189892
refGene_gn=LOC105261204;refGene_tx=XM_011289507.3
E2 1189894 1191213
refGene_gn=LOC105261204;refGene_tx=XM_011289507.3
E2 1211571 1213356
refGene_gn=LOC111556192;refGene_tx=XM_019818994.2;refGene_gn=LOC111556
192;refGene_tx=XM_019818995.2;refGene_gn=LOC111556192;refGene_tx=XM_01
9818997.2
E2 1214767 1214891
refGene_gn=LOC111556192;refGene_tx=XM_019818994.2
E2 1217647 1217680
refGene_gn=LOC111556192;refGene_tx=XM_019818994.2
E2 1235621 1237009
refGene_gn=LOC111557975;refGene_tx=XM_023244875.1;refGene_gn=LOC111557
975;refGene_tx=XM_023244876.1;refGene_gn=LOC111557975;refGene_tx=XM_02
3244877.1;refGene_gn=LOC111557975;refGene_tx=XM_023244878.1;refGene_gn
=LOC111557975;refGene_tx=XM_023244879.1
E2 1238331 1238557
refGene_gn=LOC111557975;refGene_tx=XM_023244875.1;refGene_gn=LOC111557
975;refGene_tx=XM_023244876.1;refGene_gn=LOC111557975;refGene_tx=XM_02
3244877.1;refGene_gn=LOC111557975;refGene_tx=XM_023244878.1;refGene_gn
=LOC111557975;refGene_tx=XM_023244879.1
E2 1241366 1241399
refGene_gn=LOC111557975;refGene_tx=XM_023244876.1;refGene_gn=LOC111557
975;refGene_tx=XM_023244877.1
E2 1251852 1251978
refGene_gn=LOC102901659;refGene_tx=XM_019819002.2
E2 1251979 1252596
refGene_gn=LOC102901659;refGene_tx=XM_019819002.2
E2 1252687 1252853
refGene_gn=LOC102901659;refGene_tx=XM_019819002.2
E2 1253373 1253766
refGene_gn=LOC102901659;refGene_tx=XM_019819002.2
E2 1253841 1253873 refGene_gn=ZNF211;refGene_tx=XM_023244874.1
E2 1272073 1273542 refGene_gn=ZNF211;refGene_tx=XM_023244873.1

E2 1281087 1281214
refGene_gn=ZNF211;refGene_tx=XM_023244873.1;refGene_gn=ZNF211;refGene_tx=XM_023244874.1
E2 1281814 1281853
refGene_gn=ZNF211;refGene_tx=XM_023244873.1;refGene_gn=ZNF211;refGene_tx=XM_023244874.1
E2 1282393 1282426
refGene_gn=ZNF211;refGene_tx=XM_023244873.1;refGene_gn=ZNF211;refGene_tx=XM_023244874.1
E2 1314421 1315656
refGene_gn=ZNF134;refGene_tx=XM_019819004.2;refGene_gn=ZNF134;refGene_tx=XM_019819005.2;refGene_gn=ZNF134;refGene_tx=XM_019819006.1;refGene_gn=ZNF134;refGene_tx=XM_019819007.2;refGene_gn=ZNF134;refGene_tx=XM_019819008.2;refGene_gn=ZNF134;refGene_tx=XM_023246011.1
E2 1316291 1316331
refGene_gn=ZNF134;refGene_tx=XM_019819005.2;refGene_gn=ZNF134;refGene_tx=XM_019819006.1;refGene_gn=ZNF134;refGene_tx=XM_019819007.2;refGene_gn=ZNF134;refGene_tx=XM_019819008.2;refGene_gn=ZNF134;refGene_tx=XM_023246011.1
E2 1320427 1320476 refGene_gn=ZNF134;refGene_tx=XM_019819004.2
E2 1328099 1328900
refGene_gn=FELCATV1R10;refGene_tx=NM_001167482.1;refGene_gn=FELCATV1R10;refGene_tx=XM_023244830.1
E2 1335233 1335243
refGene_gn=FELCATV1R10;refGene_tx=XM_023244830.1
E2 1342320 1342574
refGene_gn=LOC101096860;refGene_tx=XM_023246012.1
E2 1342576 1342970
refGene_gn=LOC101096860;refGene_tx=XM_023246012.1
E2 1347381 1347392
refGene_gn=LOC101096860;refGene_tx=XM_023246012.1
E2 1373309 1374659
refGene_gn=LOC101096860;refGene_tx=XM_023246012.1
E2 1376099 1376226
refGene_gn=LOC101096860;refGene_tx=XM_023246012.1
E2 1378830 1378863
refGene_gn=LOC101096860;refGene_tx=XM_023246012.1
E2 1386924 1387529
refGene_gn=LOC101097113;refGene_tx=XM_023245895.1
E2 1387531 1387847
refGene_gn=LOC101097113;refGene_tx=XM_023245895.1
E2 1402954 1403697
refGene_gn=LOC101097361;refGene_tx=XM_023245336.1
E2 1403788 1403954
refGene_gn=LOC101097361;refGene_tx=XM_023245336.1
E2 1404385 1404781
refGene_gn=LOC101097361;refGene_tx=XM_023245336.1
E2 1441799 1442717
refGene_gn=LOC111558028;refGene_tx=XM_023245337.1
E2 1443093 1444011

refGene_gn=FELCATV1R11;refGene_tx=NM_001167483.1
E2	1447680	1447713
refGene_gn=LOC111556196;refGene_tx=XM_019819014.2
E2	1450422	1450549
refGene_gn=LOC111556196;refGene_tx=XM_019819014.2
E2	1454892	1456472
refGene_gn=LOC111556196;refGene_tx=XM_019819014.2
E2	1467446	1467473	refGene_gn=ZNF550;refGene_tx=XM_006940780.4
E2	1469165	1469189
refGene_gn=ZNF550;refGene_tx=XM_019819017.2;refGene_gn=ZNF550;refGene_tx=XM_019819018.2;refGene_gn=ZNF550;refGene_tx=XM_019819019.2;refGene_gn=ZNF550;refGene_tx=XM_019819020.2;refGene_gn=ZNF550;refGene_tx=XM_019819021.2
E2	1469632	1469759
refGene_gn=ZNF550;refGene_tx=XM_006940780.4;refGene_gn=ZNF550;refGene_tx=XM_006940783.4;refGene_gn=ZNF550;refGene_tx=XM_019819017.2;refGene_gn=ZNF550;refGene_tx=XM_019819018.2;refGene_gn=ZNF550;refGene_tx=XM_019819019.2;refGene_gn=ZNF550;refGene_tx=XM_019819020.2;refGene_gn=ZNF550;refGene_tx=XM_019819021.2;refGene_gn=ZNF550;refGene_tx=XM_023245153.1;refGene_gn=ZNF550;refGene_tx=XM_023245154.1;refGene_gn=ZNF550;refGene_tx=XM_023245155.1
E2	1471374	1471470
refGene_gn=ZNF550;refGene_tx=XM_006940780.4;refGene_gn=ZNF550;refGene_tx=XM_006940783.4;refGene_gn=ZNF550;refGene_tx=XM_019819017.2;refGene_gn=ZNF550;refGene_tx=XM_019819018.2;refGene_gn=ZNF550;refGene_tx=XM_019819019.2;refGene_gn=ZNF550;refGene_tx=XM_019819020.2;refGene_gn=ZNF550;refGene_tx=XM_019819021.2;refGene_gn=ZNF550;refGene_tx=XM_023245153.1;refGene_gn=ZNF550;refGene_tx=XM_023245154.1;refGene_gn=ZNF550;refGene_tx=XM_023245155.1
E2	1475045	1476058
refGene_gn=ZNF550;refGene_tx=XM_006940780.4;refGene_gn=ZNF550;refGene_tx=XM_006940783.4;refGene_gn=ZNF550;refGene_tx=XM_019819017.2;refGene_gn=ZNF550;refGene_tx=XM_019819018.2;refGene_gn=ZNF550;refGene_tx=XM_019819019.2;refGene_gn=ZNF550;refGene_tx=XM_019819020.2;refGene_gn=ZNF550;refGene_tx=XM_019819021.2;refGene_gn=ZNF550;refGene_tx=XM_023245153.1;refGene_gn=ZNF550;refGene_tx=XM_023245154.1;refGene_gn=ZNF550;refGene_tx=XM_023245155.1
E2	1491882	1492812
refGene_gn=FELCATV1R4;refGene_tx=NM_001167476.1
E2	1503971	1505299
refGene_gn=LOC101097855;refGene_tx=XM_019819030.2;refGene_gn=LOC101097855;refGene_tx=XM_019819032.2;refGene_gn=LOC101097855;refGene_tx=XM_019819033.2
E2	1506489	1506616
refGene_gn=LOC101097855;refGene_tx=XM_019819030.2;refGene_gn=LOC101097855;refGene_tx=XM_019819032.2;refGene_gn=LOC101097855;refGene_tx=XM_019819033.2
E2	1509088	1509127
refGene_gn=LOC101097855;refGene_tx=XM_019819030.2;refGene_gn=LOC101097855;refGene_tx=XM_019819033.2

E2 1509941 1510064
refGene_gn=LOC101097855;refGene_tx=XM_019819030.2;refGene_gn=LOC101097
855;refGene_tx=XM_019819032.2
E2 1563719 1563838
refGene_gn=LOC101098592;refGene_tx=XM_019819037.2
E2 1576470 1577567
refGene_gn=LOC101098592;refGene_tx=XM_006940784.4
E2 1583096 1583195
refGene_gn=LOC101098592;refGene_tx=XM_006940784.4;refGene_gn=LOC101098
592;refGene_tx=XM_019819037.2
E2 1583699 1583826
refGene_gn=LOC101098592;refGene_tx=XM_006940784.4;refGene_gn=LOC101098
592;refGene_tx=XM_019819037.2
E2 1587869 1587902
refGene_gn=LOC101098592;refGene_tx=XM_006940784.4;refGene_gn=LOC101098
592;refGene_tx=XM_019819037.2
E2 1593462 1593540
refGene_gn=ZNF772;refGene_tx=XM_006940785.4;refGene_gn=ZNF772;refGene_
tx=XM_006940786.4;refGene_gn=ZNF772;refGene_tx=XM_006940787.4
E2 1594149 1594188
refGene_gn=ZNF772;refGene_tx=XM_006940785.4;refGene_gn=ZNF772;refGene_
tx=XM_006940786.4;refGene_gn=ZNF772;refGene_tx=XM_006940787.4
E2 1595532 1595659
refGene_gn=ZNF772;refGene_tx=XM_006940785.4;refGene_gn=ZNF772;refGene_
tx=XM_006940786.4;refGene_gn=ZNF772;refGene_tx=XM_006940787.4
E2 1596016 1596124
refGene_gn=ZNF772;refGene_tx=XM_006940785.4;refGene_gn=ZNF772;refGene_
tx=XM_006940786.4;refGene_gn=ZNF772;refGene_tx=XM_006940789.3;refGene_
gn=ZNF772;refGene_tx=XM_019819034.1
E2 1596716 1597861
refGene_gn=ZNF772;refGene_tx=XM_006940785.4;refGene_gn=ZNF772;refGene_
tx=XM_006940786.4;refGene_gn=ZNF772;refGene_tx=XM_006940787.4;refGene_
gn=ZNF772;refGene_tx=XM_006940789.3;refGene_gn=ZNF772;refGene_tx=XM_01
9819034.1
E2 1611220 1612533
refGene_gn=LOC101099091;refGene_tx=XM_019819035.2
E2 1613323 1613450
refGene_gn=LOC101099091;refGene_tx=XM_019819035.2
E2 1615210 1615225
refGene_gn=LOC101099091;refGene_tx=XM_019819035.2
E2 1635078 1636604
refGene_gn=LOC102899789;refGene_tx=XM_023245858.1
E2 1638410 1638537
refGene_gn=LOC102899789;refGene_tx=XM_023245858.1
E2 1641068 1641110
refGene_gn=LOC102899789;refGene_tx=XM_023245858.1
E2 1643080 1643249
refGene_gn=LOC102899789;refGene_tx=XM_023245858.1
E2 1653165 1653248
refGene_gn=LOC102899789;refGene_tx=XM_023245858.1

E2 1653362 1653427
refGene_gn=LOC102899789;refGene_tx=XM_023245858.1
E2 1654586 1654628
refGene_gn=LOC102899789;refGene_tx=XM_023245858.1
E2 1656057 1656284
refGene_gn=LOC102899789;refGene_tx=XM_023245858.1
E2 1661019 1662463
refGene_gn=LOC102899789;refGene_tx=XM_023245858.1
E2 1663614 1663741
refGene_gn=LOC102899789;refGene_tx=XM_023245858.1
E2 1665581 1665614
refGene_gn=LOC102899789;refGene_tx=XM_023245858.1
E2 1681151 1682971 refGene_gn=ZNF304;refGene_tx=XM_006940799.2
E2 1685149 1685276 refGene_gn=ZNF304;refGene_tx=XM_006940799.2
E2 1686933 1686966 refGene_gn=ZNF304;refGene_tx=XM_006940799.2
E2 1721565 1721928
refGene_gn=LOC101095625;refGene_tx=XM_019818843.2
E2 1721929 1721980
refGene_gn=LOC101095625;refGene_tx=XM_019818843.2
E2 1721981 1723621
refGene_gn=LOC101095625;refGene_tx=XM_019818843.2
E2 1729105 1729201
refGene_gn=LOC101095625;refGene_tx=XM_019818843.2
E2 1735263 1735390
refGene_gn=LOC101095625;refGene_tx=XM_019818843.2
E2 1737877 1737907
refGene_gn=LOC101095625;refGene_tx=XM_019818843.2
E2 1756588 1758207
refGene_gn=LOC102900013;refGene_tx=XM_011289494.3
E2 1766329 1766353
refGene_gn=LOC102900013;refGene_tx=XM_011289494.3
E2 1766355 1766424
refGene_gn=LOC102900013;refGene_tx=XM_011289494.3
E2 1771206 1771333
refGene_gn=LOC102900013;refGene_tx=XM_011289494.3
E2 1773207 1773240
refGene_gn=LOC102900013;refGene_tx=XM_011289494.3
E2 1777108 1777279
refGene_gn=AURKC;refGene_tx=XM_006940800.4;refGene_gn=AURKC;refGene_tx
=XM_019819079.2
E2 1777452 1777627
refGene_gn=AURKC;refGene_tx=XM_006940800.4;refGene_gn=AURKC;refGene_tx
=XM_019819079.2
E2 1778193 1778342
refGene_gn=AURKC;refGene_tx=XM_006940800.4;refGene_gn=AURKC;refGene_tx
=XM_019819079.2
E2 1779265 1779404
refGene_gn=AURKC;refGene_tx=XM_006940800.4;refGene_gn=AURKC;refGene_tx
=XM_019819079.2
E2 1779753 1779945

refGene_gn=AURKC;refGene_tx=XM_006940800.4;refGene_gn=AURKC;refGene_tx=XM_019819079.2
E2 1780145 1780185
refGene_gn=AURKC;refGene_tx=XM_006940800.4;refGene_gn=AURKC;refGene_tx=XM_019819079.2
E2 1780266 1780330
refGene_gn=AURKC;refGene_tx=XM_006940800.4;refGene_gn=AURKC;refGene_tx=XM_019819079.2
E2 1794072 1795706
refGene_gn=LOC101099850;refGene_tx=XM_011289492.3;refGene_gn=LOC101099850;refGene_tx=XM_019819067.2
E2 1801764 1801860
refGene_gn=LOC101099850;refGene_tx=XM_011289492.3;refGene_gn=LOC101099850;refGene_tx=XM_019819067.2
E2 1810947 1811074
refGene_gn=LOC101099850;refGene_tx=XM_011289492.3;refGene_gn=LOC101099850;refGene_tx=XM_019819067.2
E2 1811397 1811433
refGene_gn=LOC101099850;refGene_tx=XM_019819067.2
E2 1812668 1812698
refGene_gn=LOC101099850;refGene_tx=XM_011289492.3
E2 1826868 1826893 refGene_gn=DUXA;refGene_tx=XM_019818844.1
E2 1836129 1836284 refGene_gn=DUXA;refGene_tx=XM_019818844.1
E2 1836774 1836877 refGene_gn=DUXA;refGene_tx=XM_019818844.1
E2 1837299 1837445 refGene_gn=DUXA;refGene_tx=XM_019818844.1
E2 1842541 1842643 refGene_gn=DUXA;refGene_tx=XM_019818844.1
E2 2086714 2087108
refGene_gn=PEG3;refGene_tx=XM_011289488.3;refGene_gn=PEG3;refGene_tx=XM_011289490.3;refGene_gn=PEG3;refGene_tx=XM_019819039.2;refGene_gn=PEG3;refGene_tx=XM_019819040.2
E2 2087712 2087802
refGene_gn=PEG3;refGene_tx=XM_011289488.3;refGene_gn=PEG3;refGene_tx=XM_011289490.3;refGene_gn=PEG3;refGene_tx=XM_019819039.2;refGene_gn=PEG3;refGene_tx=XM_019819040.2
E2 2088503 2088653
refGene_gn=PEG3;refGene_tx=XM_011289488.3;refGene_gn=PEG3;refGene_tx=XM_011289490.3;refGene_gn=PEG3;refGene_tx=XM_019819039.2;refGene_gn=PEG3;refGene_tx=XM_019819040.2
E2 2088737 2088847
refGene_gn=PEG3;refGene_tx=XM_011289488.3;refGene_gn=PEG3;refGene_tx=XM_011289490.3;refGene_gn=PEG3;refGene_tx=XM_019819039.2;refGene_gn=PEG3;refGene_tx=XM_019819040.2
E2 2092764 2092867
refGene_gn=PEG3;refGene_tx=XM_011289488.3;refGene_gn=PEG3;refGene_tx=XM_011289490.3;refGene_gn=PEG3;refGene_tx=XM_019819039.2;refGene_gn=PEG3;refGene_tx=XM_019819040.2
E2 2093931 2094024
refGene_gn=PEG3;refGene_tx=XM_011289488.3;refGene_gn=PEG3;refGene_tx=XM_011289490.3;refGene_gn=PEG3;refGene_tx=XM_019819039.2;refGene_gn=PEG3;refGene_tx=XM_019819040.2

E2	2094192	2098217
refGene_gn=PEG3;refGene_tx=XM_011289488.3;refGene_gn=PEG3;refGene_tx=XM_011289490.3;refGene_gn=PEG3;refGene_tx=XM_019819039.2;refGene_gn=PEG3;refGene_tx=XM_019819040.2
E2	2119574	2119627	refGene_gn=ZIM2;refGene_tx=XM_023245859.1
E2	2128209	2128712	refGene_gn=ZIM2;refGene_tx=XM_023245859.1
E2	2130857	2130957	refGene_gn=ZIM2;refGene_tx=XM_023245859.1
E2	2131662	2131717	refGene_gn=ZIM2;refGene_tx=XM_023245859.1
E2	2132601	2132721	refGene_gn=ZIM2;refGene_tx=XM_023245859.1
E2	2132821	2132948	refGene_gn=ZIM2;refGene_tx=XM_023245859.1
E2	2135818	2135911	refGene_gn=ZIM2;refGene_tx=XM_023245859.1
E2	2138282	2139265	refGene_gn=ZIM2;refGene_tx=XM_023245859.1
E2	2160467	2161394
refGene_gn=LOC111558094;refGene_tx=XM_023245860.1
E2	2206213	2207152
refGene_gn=LOC111558120;refGene_tx=XM_023246084.1
E2	2225478	2226453
refGene_gn=LOC111556214;refGene_tx=XM_023246083.1
E2	2235747	2236686
refGene_gn=LOC101101097;refGene_tx=XM_003997378.2
E2	2276723	2277707
refGene_gn=LOC102900225;refGene_tx=XM_023246082.1
E2	2284840	2284951	refGene_gn=SMIM17;refGene_tx=XM_006940803.4
E2	2291625	2291702	refGene_gn=SMIM17;refGene_tx=XM_006940803.4
E2	2293230	2293393	refGene_gn=SMIM17;refGene_tx=XM_006940803.4
E2	2332855	2334312
refGene_gn=ZNF71;refGene_tx=XM_006940804.4;refGene_gn=ZNF71;refGene_tx=XM_006940806.4;refGene_gn=ZNF71;refGene_tx=XM_019819071.2;refGene_gn=ZNF71;refGene_tx=XM_023246073.1;refGene_gn=ZNF71;refGene_tx=XM_023246075.1;refGene_gn=ZNF71;refGene_tx=XM_023246076.1;refGene_gn=ZNF71;refGene_tx=XM_023246077.1
E2	2340527	2340654
refGene_gn=ZNF71;refGene_tx=XM_006940804.4;refGene_gn=ZNF71;refGene_tx=XM_006940806.4;refGene_gn=ZNF71;refGene_tx=XM_023246073.1;refGene_gn=ZNF71;refGene_tx=XM_023246075.1
E2	2349507	2349574
refGene_gn=ZNF71;refGene_tx=XM_006940804.4;refGene_gn=ZNF71;refGene_tx=XM_006940806.4;refGene_gn=ZNF71;refGene_tx=XM_023246073.1;refGene_gn=ZNF71;refGene_tx=XM_023246075.1
E2	2349760	2349765	refGene_gn=ZNF71;refGene_tx=XM_006940804.4
E2	2353771	2353815	refGene_gn=ZNF71;refGene_tx=XM_023246073.1
E2	2363947	2365821
refGene_gn=LOC101095388;refGene_tx=XM_023246063.1;refGene_gn=LOC101095388;refGene_tx=XM_023246064.1
E2	2368402	2368498
refGene_gn=LOC101095388;refGene_tx=XM_023246063.1;refGene_gn=LOC101095388;refGene_tx=XM_023246064.1
E2	2368679	2368806
refGene_gn=LOC101095388;refGene_tx=XM_023246063.1;refGene_gn=LOC101095388;refGene_tx=XM_023246064.1

E2 2370700 2370703
refGene_gn=LOC101095388;refGene_tx=XM_023246064.1
E2 2375192 2375225
refGene_gn=LOC101095388;refGene_tx=XM_023246063.1
E2 2387949 2389652
refGene_gn=LOC101095388;refGene_tx=XM_023246060.1;refGene_gn=LOC101095
388;refGene_tx=XM_023246061.1
E2 2396529 2396625
refGene_gn=LOC101095388;refGene_tx=XM_023246060.1;refGene_gn=LOC101095
388;refGene_tx=XM_023246061.1
E2 2397432 2397547
refGene_gn=LOC101095388;refGene_tx=XM_023246060.1;refGene_gn=LOC101095
388;refGene_tx=XM_023246061.1
E2 2397969 2398059
refGene_gn=LOC101095388;refGene_tx=XM_023246065.1
E2 2398172 2398336
refGene_gn=LOC101095388;refGene_tx=XM_023246060.1;refGene_gn=LOC101095
388;refGene_tx=XM_023246061.1;refGene_gn=LOC101095388;refGene_tx=XM_02
3246065.1
E2 2400133 2400229
refGene_gn=LOC101095388;refGene_tx=XM_023246060.1;refGene_gn=LOC101095
388;refGene_tx=XM_023246061.1;refGene_gn=LOC101095388;refGene_tx=XM_02
3246065.1
E2 2400410 2400537
refGene_gn=LOC101095388;refGene_tx=XM_023246060.1;refGene_gn=LOC101095
388;refGene_tx=XM_023246061.1;refGene_gn=LOC101095388;refGene_tx=XM_02
3246065.1
E2 2409410 2409502
refGene_gn=LOC101095388;refGene_tx=XM_023246060.1;refGene_gn=LOC101095
388;refGene_tx=XM_023246065.1
E2 2410009 2410229
refGene_gn=LOC101095388;refGene_tx=XM_023246060.1;refGene_gn=LOC101095
388;refGene_tx=XM_023246065.1
E2 2418517 2420532
refGene_gn=LOC101101602;refGene_tx=XM_006940809.4;refGene_gn=LOC101101
602;refGene_tx=XM_019819041.2;refGene_gn=LOC101101602;refGene_tx=XM_01
9819045.2;refGene_gn=LOC101101602;refGene_tx=XM_019819046.1;refGene_gn
=LOC101101602;refGene_tx=XM_019819048.1;refGene_gn=LOC101101602;refGen
e_tx=XM_019819049.2
E2 2424184 2424274
refGene_gn=LOC101101602;refGene_tx=XM_006940809.4;refGene_gn=LOC101101
602;refGene_tx=XM_019819041.2;refGene_gn=LOC101101602;refGene_tx=XM_01
9819049.2
E2 2426011 2426138
refGene_gn=LOC101101602;refGene_tx=XM_006940809.4;refGene_gn=LOC101101
602;refGene_tx=XM_019819041.2;refGene_gn=LOC101101602;refGene_tx=XM_01
9819045.2;refGene_gn=LOC101101602;refGene_tx=XM_019819049.2
E2 2432178 2432211
refGene_gn=LOC101101602;refGene_tx=XM_006940809.4;refGene_gn=LOC101101
602;refGene_tx=XM_019819041.2;refGene_gn=LOC101101602;refGene_tx=XM_01

9819045.2;refGene_gn=LOC101101602;refGene_tx=XM_019819049.2
E2	2474208	2475518	refGene_gn=ZNF582;refGene_tx=XM_019819072.2
E2	2480842	2480862	refGene_gn=ZNF582;refGene_tx=XM_019819075.2
E2	2482187	2482283
refGene_gn=ZNF582;refGene_tx=XM_019819072.2;refGene_gn=ZNF582;refGene_tx=XM_019819075.2
E2	2482513	2482640
refGene_gn=ZNF582;refGene_tx=XM_019819072.2;refGene_gn=ZNF582;refGene_tx=XM_019819075.2
E2	2483874	2483883
refGene_gn=ZNF582;refGene_tx=XM_019819072.2;refGene_gn=ZNF582;refGene_tx=XM_019819075.2
E2	2499014	2499023	refGene_gn=ZNF583;refGene_tx=XM_023246072.1
E2	2505199	2505326	refGene_gn=ZNF583;refGene_tx=XM_023246072.1
E2	2505618	2505717	refGene_gn=ZNF583;refGene_tx=XM_023246072.1
E2	2510791	2511306	refGene_gn=ZNF583;refGene_tx=XM_023246072.1
E2	2511307	2512270	refGene_gn=ZNF583;refGene_tx=XM_023246072.1
E2	2525521	2527101
refGene_gn=ZNF667;refGene_tx=XM_023246067.1;refGene_gn=ZNF667;refGene_tx=XM_023246068.1;refGene_gn=ZNF667;refGene_tx=XM_023246069.1;refGene_gn=ZNF667;refGene_tx=XM_023246070.1;refGene_gn=ZNF667;refGene_tx=XM_023246071.1
E2	2537892	2537985
refGene_gn=ZNF667;refGene_tx=XM_023246067.1;refGene_gn=ZNF667;refGene_tx=XM_023246068.1;refGene_gn=ZNF667;refGene_tx=XM_023246069.1;refGene_gn=ZNF667;refGene_tx=XM_023246070.1;refGene_gn=ZNF667;refGene_tx=XM_023246071.1
E2	2539536	2539663
refGene_gn=ZNF667;refGene_tx=XM_023246067.1;refGene_gn=ZNF667;refGene_tx=XM_023246068.1;refGene_gn=ZNF667;refGene_tx=XM_023246069.1;refGene_gn=ZNF667;refGene_tx=XM_023246070.1;refGene_gn=ZNF667;refGene_tx=XM_023246071.1
E2	2542074	2542134
refGene_gn=ZNF667;refGene_tx=XM_023246067.1;refGene_gn=ZNF667;refGene_tx=XM_023246068.1;refGene_gn=ZNF667;refGene_tx=XM_023246069.1;refGene_gn=ZNF667;refGene_tx=XM_023246070.1;refGene_gn=ZNF667;refGene_tx=XM_023246071.1
E2	2592827	2592852
refGene_gn=LOC111558121;refGene_tx=XM_023246088.1
E2	2594090	2594174
refGene_gn=LOC111558121;refGene_tx=XM_023246087.1;refGene_gn=LOC111558121;refGene_tx=XM_023246088.1
E2	2595188	2595368
refGene_gn=LOC111558121;refGene_tx=XM_023246085.1;refGene_gn=LOC111558121;refGene_tx=XM_023246086.1;refGene_gn=LOC111558121;refGene_tx=XM_023246089.1
E2	2596060	2596188
refGene_gn=LOC111558121;refGene_tx=XM_023246085.1;refGene_gn=LOC111558121;refGene_tx=XM_023246086.1;refGene_gn=LOC111558121;refGene_tx=XM_023246087.1;refGene_gn=LOC111558121;refGene_tx=XM_023246088.1;refGene_gn

=LOC111558121;refGene_tx=XM_023246089.1
E2 2598295 2598322
refGene_gn=LOC111558121;refGene_tx=XM_023246085.1;refGene_gn=LOC111558121;refGene_tx=XM_023246086.1;refGene_gn=LOC111558121;refGene_tx=XM_023246087.1;refGene_gn=LOC111558121;refGene_tx=XM_023246088.1;refGene_gn=LOC111558121;refGene_tx=XM_023246089.1
E2 2599231 2599258
refGene_gn=LOC111558121;refGene_tx=XM_023246085.1;refGene_gn=LOC111558121;refGene_tx=XM_023246086.1;refGene_gn=LOC111558121;refGene_tx=XM_023246087.1;refGene_gn=LOC111558121;refGene_tx=XM_023246088.1;refGene_gn=LOC111558121;refGene_tx=XM_023246089.1
E2 2599600 2599609
refGene_gn=LOC111558121;refGene_tx=XM_023246085.1;refGene_gn=LOC111558121;refGene_tx=XM_023246086.1;refGene_gn=LOC111558121;refGene_tx=XM_023246087.1;refGene_gn=LOC111558121;refGene_tx=XM_023246088.1;refGene_gn=LOC111558121;refGene_tx=XM_023246089.1
E2 2600198 2600207
refGene_gn=LOC111558121;refGene_tx=XM_023246085.1;refGene_gn=LOC111558121;refGene_tx=XM_023246086.1;refGene_gn=LOC111558121;refGene_tx=XM_023246087.1;refGene_gn=LOC111558121;refGene_tx=XM_023246088.1;refGene_gn=LOC111558121;refGene_tx=XM_023246089.1
E2 2601001 2601007
refGene_gn=LOC111558121;refGene_tx=XM_023246085.1;refGene_gn=LOC111558121;refGene_tx=XM_023246086.1;refGene_gn=LOC111558121;refGene_tx=XM_023246087.1;refGene_gn=LOC111558121;refGene_tx=XM_023246088.1;refGene_gn=LOC111558121;refGene_tx=XM_023246089.1
E2 2602654 2602672
refGene_gn=LOC111558121;refGene_tx=XM_023246085.1;refGene_gn=LOC111558121;refGene_tx=XM_023246087.1;refGene_gn=LOC111558121;refGene_tx=XM_023246088.1;refGene_gn=LOC111558121;refGene_tx=XM_023246089.1
E2 2605681 2605708
refGene_gn=LOC111558121;refGene_tx=XM_023246085.1;refGene_gn=LOC111558121;refGene_tx=XM_023246087.1;refGene_gn=LOC111558121;refGene_tx=XM_023246088.1
E2 2606102 2606111
refGene_gn=LOC111558121;refGene_tx=XM_023246085.1;refGene_gn=LOC111558121;refGene_tx=XM_023246086.1;refGene_gn=LOC111558121;refGene_tx=XM_023246087.1;refGene_gn=LOC111558121;refGene_tx=XM_023246088.1
E2 2606293 2606366
refGene_gn=LOC111558121;refGene_tx=XM_023246085.1;refGene_gn=LOC111558121;refGene_tx=XM_023246086.1;refGene_gn=LOC111558121;refGene_tx=XM_023246087.1;refGene_gn=LOC111558121;refGene_tx=XM_023246088.1
E2 2626724 2627398 refGene_gn=ZSCAN5B;refGene_tx=XM_023245862.1
E2 2627661 2627812 refGene_gn=ZSCAN5B;refGene_tx=XM_023245862.1
E2 2628733 2628937 refGene_gn=ZSCAN5B;refGene_tx=XM_023245862.1
E2 2630392 2630728 refGene_gn=ZSCAN5B;refGene_tx=XM_023245862.1
E2 2658752 2659033 refGene_gn=ZNF444;refGene_tx=XM_023246081.1
E2 2659035 2659338 refGene_gn=ZNF444;refGene_tx=XM_023246081.1
E2 2661000 2661085 refGene_gn=ZNF444;refGene_tx=XM_023246081.1
E2 2670083 2670425 refGene_gn=ZNF444;refGene_tx=XM_023246081.1

E2	2674654	2674666	refGene_gn=ZNF444;refGene_tx=XM_023246081.1
E2	2701240	2701319	refGene_gn=ZNF787;refGene_tx=XM_023246078.1;refGene_gn=ZNF787;refGene_tx=XM_023246080.1
E2	2707479	2707480	refGene_gn=ZNF787;refGene_tx=XM_023246079.1
E2	2711409	2712485	refGene_gn=ZNF787;refGene_tx=XM_023246078.1;refGene_gn=ZNF787;refGene_tx=XM_023246079.1;refGene_gn=ZNF787;refGene_tx=XM_023246080.1
E2	2726527	2726660	refGene_gn=NLRP5;refGene_tx=XM_023245863.1
E2	2731483	2731654	refGene_gn=NLRP5;refGene_tx=XM_023245863.1
E2	2733189	2733360	refGene_gn=NLRP5;refGene_tx=XM_023245863.1
E2	2735453	2735624	refGene_gn=NLRP5;refGene_tx=XM_023245863.1
E2	2738464	2738635	refGene_gn=NLRP5;refGene_tx=XM_023245863.1
E2	2740593	2740764	refGene_gn=NLRP5;refGene_tx=XM_023245863.1
E2	2742572	2742743	refGene_gn=NLRP5;refGene_tx=XM_023245863.1
E2	2744686	2744857	refGene_gn=NLRP5;refGene_tx=XM_023245863.1
E2	2745661	2745864	refGene_gn=NLRP5;refGene_tx=XM_023245863.1
E2	2745865	2747264	refGene_gn=NLRP5;refGene_tx=XM_023245863.1
E2	2751236	2751293	refGene_gn=NLRP5;refGene_tx=XM_023245863.1
E2	2755040	2755115	refGene_gn=NLRP5;refGene_tx=XM_023245863.1
E2	2757986	2758391	refGene_gn=NLRP5;refGene_tx=XM_023245863.1
E2	2767627	2767652	refGene_gn=NLRP5;refGene_tx=XM_023245863.1
E2	2768283	2768453	refGene_gn=NLRP8;refGene_tx=XM_023245865.1
E2	2770888	2770893	refGene_gn=NLRP8;refGene_tx=XM_023245865.1
E2	2771645	2771816	refGene_gn=NLRP8;refGene_tx=XM_023245865.1
E2	2773932	2774043	refGene_gn=NLRP8;refGene_tx=XM_023245865.1
E2	2775366	2775516	refGene_gn=NLRP8;refGene_tx=XM_023245865.1
E2	2778909	2779077	refGene_gn=NLRP8;refGene_tx=XM_023245865.1
E2	2781239	2781410	refGene_gn=NLRP8;refGene_tx=XM_023245865.1
E2	2783750	2785314	refGene_gn=NLRP8;refGene_tx=XM_023245865.1
E2	2787185	2787260	refGene_gn=NLRP8;refGene_tx=XM_023245865.1
E2	2788451	2788743	refGene_gn=NLRP8;refGene_tx=XM_023245865.1
E2	2803314	2803615	refGene_gn=NLRP13;refGene_tx=XM_023245269.1;refGene_gn=NLRP13;refGene_tx=XM_023245270.1;refGene_gn=NLRP13;refGene_tx=XM_023245271.1;refGene_gn=NLRP13;refGene_tx=XM_023245272.1;refGene_gn=NLRP13;refGene_tx=XM_023245273.1;refGene_gn=NLRP13;refGene_tx=XM_023245274.1
E2	2809150	2809219	refGene_gn=NLRP13;refGene_tx=XM_023245269.1;refGene_gn=NLRP13;refGene_tx=XM_023245270.1;refGene_gn=NLRP13;refGene_tx=XM_023245271.1;refGene_gn=NLRP13;refGene_tx=XM_023245273.1;refGene_gn=NLRP13;refGene_tx=XM_023245274.1
E2	2814310	2815898	refGene_gn=NLRP13;refGene_tx=XM_023245269.1;refGene_gn=NLRP13;refGene_tx=XM_023245270.1;refGene_gn=NLRP13;refGene_tx=XM_023245271.1;refGene_gn=NLRP13;refGene_tx=XM_023245272.1;refGene_gn=NLRP13;refGene_tx=XM_023245273.1;refGene_gn=NLRP13;refGene_tx=XM_023245274.1
E2	2817298	2817463	refGene_gn=NLRP13;refGene_tx=XM_023245269.1;refGene_gn=NLRP13;refGene_tx=XM_023245270.1;refGene_gn=NLRP13;refGene_tx=XM_023245271.1;refGene_

gn=NLRP13;refGene_tx=XM_023245272.1;refGene_gn=NLRP13;refGene_tx=XM_02
3245273.1;refGene_gn=NLRP13;refGene_tx=XM_023245274.1
E2 2818944 2819112
refGene_gn=NLRP13;refGene_tx=XM_023245269.1;refGene_gn=NLRP13;refGene_
tx=XM_023245270.1;refGene_gn=NLRP13;refGene_tx=XM_023245271.1;refGene_
gn=NLRP13;refGene_tx=XM_023245272.1;refGene_gn=NLRP13;refGene_tx=XM_02
3245273.1;refGene_gn=NLRP13;refGene_tx=XM_023245274.1
E2 2819560 2819731
refGene_gn=NLRP13;refGene_tx=XM_023245269.1;refGene_gn=NLRP13;refGene_
tx=XM_023245270.1;refGene_gn=NLRP13;refGene_tx=XM_023245271.1;refGene_
gn=NLRP13;refGene_tx=XM_023245272.1;refGene_gn=NLRP13;refGene_tx=XM_02
3245273.1
E2 2820310 2820481
refGene_gn=NLRP13;refGene_tx=XM_023245269.1;refGene_gn=NLRP13;refGene_
tx=XM_023245270.1;refGene_gn=NLRP13;refGene_tx=XM_023245271.1;refGene_
gn=NLRP13;refGene_tx=XM_023245272.1
E2 2823095 2823266
refGene_gn=NLRP13;refGene_tx=XM_023245269.1;refGene_gn=NLRP13;refGene_
tx=XM_023245270.1;refGene_gn=NLRP13;refGene_tx=XM_023245271.1;refGene_
gn=NLRP13;refGene_tx=XM_023245272.1;refGene_gn=NLRP13;refGene_tx=XM_02
3245273.1;refGene_gn=NLRP13;refGene_tx=XM_023245274.1
E2 2823879 2824050
refGene_gn=NLRP13;refGene_tx=XM_023245269.1;refGene_gn=NLRP13;refGene_
tx=XM_023245270.1;refGene_gn=NLRP13;refGene_tx=XM_023245271.1;refGene_
gn=NLRP13;refGene_tx=XM_023245272.1;refGene_gn=NLRP13;refGene_tx=XM_02
3245273.1;refGene_gn=NLRP13;refGene_tx=XM_023245274.1
E2 2825972 2826143
refGene_gn=NLRP13;refGene_tx=XM_023245269.1;refGene_gn=NLRP13;refGene_
tx=XM_023245270.1;refGene_gn=NLRP13;refGene_tx=XM_023245271.1;refGene_
gn=NLRP13;refGene_tx=XM_023245272.1;refGene_gn=NLRP13;refGene_tx=XM_02
3245273.1;refGene_gn=NLRP13;refGene_tx=XM_023245274.1
E2 2827359 2827483
refGene_gn=NLRP13;refGene_tx=XM_023245269.1;refGene_gn=NLRP13;refGene_
tx=XM_023245270.1;refGene_gn=NLRP13;refGene_tx=XM_023245271.1;refGene_
gn=NLRP13;refGene_tx=XM_023245272.1;refGene_gn=NLRP13;refGene_tx=XM_02
3245273.1;refGene_gn=NLRP13;refGene_tx=XM_023245274.1
E2 2829601 2829603 refGene_gn=NLRP13;refGene_tx=XM_023245270.1
E2 2832806 2832900
refGene_gn=LOC101087284;refGene_tx=XM_011289547.2
E2 2836377 2836548
refGene_gn=LOC101087284;refGene_tx=XM_011289547.2
E2 2837436 2837607
refGene_gn=LOC101087284;refGene_tx=XM_011289547.2
E2 2843302 2843473
refGene_gn=LOC101087284;refGene_tx=XM_011289547.2
E2 2846237 2846405
refGene_gn=LOC101087284;refGene_tx=XM_011289547.2
E2 2851074 2851242
refGene_gn=LOC101087284;refGene_tx=XM_011289547.2
E2 2852840 2853002

refGene_gn=LOC101087284;refGene_tx=XM_011289547.2
E2 2854057 2855630
refGene_gn=LOC101087284;refGene_tx=XM_011289547.2
E2 2862194 2862474
refGene_gn=LOC101087284;refGene_tx=XM_011289547.2
E2 2989243 2990152
refGene_gn=FELCATV1R5;refGene_tx=NM_001167477.1
E2 2994533 2994666 refGene_gn=NLRP9;refGene_tx=XM_019819093.2
E2 2995609 2995780 refGene_gn=NLRP9;refGene_tx=XM_019819093.2
E2 2996241 2996412 refGene_gn=NLRP9;refGene_tx=XM_019819093.2
E2 2997822 2997993 refGene_gn=NLRP9;refGene_tx=XM_019819093.2
E2 3003125 3003296 refGene_gn=NLRP9;refGene_tx=XM_019819093.2
E2 3006376 3006544 refGene_gn=NLRP9;refGene_tx=XM_019819093.2
E2 3007501 3007663 refGene_gn=NLRP9;refGene_tx=XM_019819093.2
E2 3009684 3011245 refGene_gn=NLRP9;refGene_tx=XM_019819093.2
E2 3016651 3016931 refGene_gn=NLRP9;refGene_tx=XM_019819093.2
E2 3027883 3028211
refGene_gn=RFPL4A;refGene_tx=XM_006940824.3;refGene_gn=RFPL4A;refGene_tx=XM_019819095.2;refGene_gn=RFPL4A;refGene_tx=XM_023245011.1
E2 3029799 3030377
refGene_gn=RFPL4A;refGene_tx=XM_006940824.3;refGene_gn=RFPL4A;refGene_tx=XM_019819095.2;refGene_gn=RFPL4A;refGene_tx=XM_023245011.1
E2 3033939 3034149
refGene_gn=EPN1;refGene_tx=XM_023245008.1;refGene_gn=EPN1;refGene_tx=XM_023245009.1;refGene_gn=EPN1;refGene_tx=XM_023245010.1
E2 3034241 3034502
refGene_gn=EPN1;refGene_tx=XM_023245008.1;refGene_gn=EPN1;refGene_tx=XM_023245009.1;refGene_gn=EPN1;refGene_tx=XM_023245010.1
E2 3035458 3035542
refGene_gn=EPN1;refGene_tx=XM_023245008.1;refGene_gn=EPN1;refGene_tx=XM_023245009.1;refGene_gn=EPN1;refGene_tx=XM_023245010.1
E2 3035697 3035808
refGene_gn=EPN1;refGene_tx=XM_023245008.1;refGene_gn=EPN1;refGene_tx=XM_023245009.1;refGene_gn=EPN1;refGene_tx=XM_023245010.1
E2 3036413 3036705
refGene_gn=EPN1;refGene_tx=XM_023245008.1;refGene_gn=EPN1;refGene_tx=XM_023245009.1;refGene_gn=EPN1;refGene_tx=XM_023245010.1
E2 3038096 3038180
refGene_gn=EPN1;refGene_tx=XM_023245008.1;refGene_gn=EPN1;refGene_tx=XM_023245009.1;refGene_gn=EPN1;refGene_tx=XM_023245010.1
E2 3038632 3038707
refGene_gn=EPN1;refGene_tx=XM_023245008.1;refGene_gn=EPN1;refGene_tx=XM_023245010.1
E2 3039722 3039847
refGene_gn=EPN1;refGene_tx=XM_023245008.1;refGene_gn=EPN1;refGene_tx=XM_023245009.1;refGene_gn=EPN1;refGene_tx=XM_023245010.1
E2 3040806 3041056
refGene_gn=EPN1;refGene_tx=XM_023245008.1;refGene_gn=EPN1;refGene_tx=XM_023245009.1;refGene_gn=EPN1;refGene_tx=XM_023245010.1
E2 3045314 3045542

refGene_gn=EPN1;refGene_tx=XM_023245008.1;refGene_gn=EPN1;refGene_tx=XM_023245009.1;refGene_gn=EPN1;refGene_tx=XM_023245010.1
E2 3049860 3049995
refGene_gn=U2AF2;refGene_tx=XM_011289549.3;refGene_gn=U2AF2;refGene_tx=XM_011289550.2
E2 3052850 3053099
refGene_gn=U2AF2;refGene_tx=XM_011289549.3;refGene_gn=U2AF2;refGene_tx=XM_011289550.2
E2 3053311 3053410
refGene_gn=U2AF2;refGene_tx=XM_011289549.3;refGene_gn=U2AF2;refGene_tx=XM_011289550.2
E2 3053720 3053843
refGene_gn=U2AF2;refGene_tx=XM_011289549.3;refGene_gn=U2AF2;refGene_tx=XM_011289550.2
E2 3053925 3054005
refGene_gn=U2AF2;refGene_tx=XM_011289549.3;refGene_gn=U2AF2;refGene_tx=XM_011289550.2
E2 3058729 3058868
refGene_gn=U2AF2;refGene_tx=XM_011289549.3;refGene_gn=U2AF2;refGene_tx=XM_011289550.2
E2 3059012 3059129
refGene_gn=U2AF2;refGene_tx=XM_011289549.3;refGene_gn=U2AF2;refGene_tx=XM_011289550.2
E2 3060408 3060560
refGene_gn=U2AF2;refGene_tx=XM_011289549.3;refGene_gn=U2AF2;refGene_tx=XM_011289550.2
E2 3060848 3060952
refGene_gn=U2AF2;refGene_tx=XM_011289549.3;refGene_gn=U2AF2;refGene_tx=XM_011289550.2
E2 3061223 3061268
refGene_gn=U2AF2;refGene_tx=XM_011289549.3;refGene_gn=U2AF2;refGene_tx=XM_011289550.2
E2 3062012 3062148
refGene_gn=U2AF2;refGene_tx=XM_011289549.3;refGene_gn=U2AF2;refGene_tx=XM_011289550.2
E2 3065752 3065801
refGene_gn=U2AF2;refGene_tx=XM_011289549.3;refGene_gn=U2AF2;refGene_tx=XM_011289550.2
E2 3068016 3068333
refGene_gn=CCDC106;refGene_tx=XM_003997389.5;refGene_gn=CCDC106;refGene_tx=XM_019819096.2;refGene_gn=CCDC106;refGene_tx=XM_023245012.1;refGene_gn=CCDC106;refGene_tx=XM_023245013.1;refGene_gn=CCDC106;refGene_tx=XM_023245014.1;refGene_gn=CCDC106;refGene_tx=XM_023245015.1;refGene_gn=CCDC106;refGene_tx=XM_023245016.1;refGene_gn=CCDC106;refGene_tx=XM_023245017.1
E2 3068693 3068906
refGene_gn=CCDC106;refGene_tx=XM_003997389.5;refGene_gn=CCDC106;refGene_tx=XM_019819096.2;refGene_gn=CCDC106;refGene_tx=XM_023245012.1;refGene_gn=CCDC106;refGene_tx=XM_023245013.1;refGene_gn=CCDC106;refGene_tx=XM_023245014.1;refGene_gn=CCDC106;refGene_tx=XM_023245015.1;refGene_gn

=CCDC106;refGene_tx=XM_023245016.1;refGene_gn=CCDC106;refGene_tx=XM_023245017.1
E2 3069971 3070148
refGene_gn=CCDC106;refGene_tx=XM_003997389.5;refGene_gn=CCDC106;refGene_tx=XM_019819096.2;refGene_gn=CCDC106;refGene_tx=XM_023245012.1;refGene_gn=CCDC106;refGene_tx=XM_023245013.1;refGene_gn=CCDC106;refGene_tx=XM_023245014.1;refGene_gn=CCDC106;refGene_tx=XM_023245015.1;refGene_gn=CCDC106;refGene_tx=XM_023245016.1;refGene_gn=CCDC106;refGene_tx=XM_023245017.1
E2 3070242 3070350
refGene_gn=CCDC106;refGene_tx=XM_003997389.5;refGene_gn=CCDC106;refGene_tx=XM_019819096.2;refGene_gn=CCDC106;refGene_tx=XM_023245012.1;refGene_gn=CCDC106;refGene_tx=XM_023245013.1;refGene_gn=CCDC106;refGene_tx=XM_023245014.1;refGene_gn=CCDC106;refGene_tx=XM_023245015.1;refGene_gn=CCDC106;refGene_tx=XM_023245016.1
E2 3070497 3070528
refGene_gn=CCDC106;refGene_tx=XM_003997389.5;refGene_gn=CCDC106;refGene_tx=XM_019819096.2;refGene_gn=CCDC106;refGene_tx=XM_023245012.1;refGene_gn=CCDC106;refGene_tx=XM_023245013.1;refGene_gn=CCDC106;refGene_tx=XM_023245014.1;refGene_gn=CCDC106;refGene_tx=XM_023245015.1;refGene_gn=CCDC106;refGene_tx=XM_023245016.1;refGene_gn=CCDC106;refGene_tx=XM_023245017.1
E2 3073464 3074061
refGene_gn=ZNF581;refGene_tx=XM_003997391.5;refGene_gn=ZNF581;refGene_tx=XM_006940827.3;refGene_gn=ZNF581;refGene_tx=XM_019819097.2
E2 3075602 3076133
refGene_gn=ZNF580;refGene_tx=XM_011289552.3;refGene_gn=ZNF580;refGene_tx=XM_019819098.1;refGene_gn=ZNF580;refGene_tx=XM_023245018.1;refGene_gn=ZNF580;refGene_tx=XM_023245019.1
E2 3076231 3076279 refGene_gn=ZNF580;refGene_tx=XM_011289552.3
E2 3097755 3097833 refGene_gn=ZNF784;refGene_tx=XM_003997410.4
E2 3099048 3099939 refGene_gn=ZNF784;refGene_tx=XM_003997410.4
E2 3103501 3106734 refGene_gn=ZNF865;refGene_tx=XM_023245867.1
E2 3112000 3112076 refGene_gn=ZNF865;refGene_tx=XM_023245867.1
E2 3112786 3112880 refGene_gn=ZNF865;refGene_tx=XM_023245867.1
E2 3113128 3113139 refGene_gn=ZNF865;refGene_tx=XM_023245867.1
E2 3114413 3115205 refGene_gn=ZNF524;refGene_tx=XM_003997392.5
E2 3118511 3118663 refGene_gn=FIZ1;refGene_tx=XM_023245332.1
E2 3119273 3119628
refGene_gn=FIZ1;refGene_tx=XM_023245332.1;refGene_gn=FIZ1;refGene_tx=XM_023245333.1;refGene_gn=FIZ1;refGene_tx=XM_023245334.1
E2 3120595 3121783
refGene_gn=FIZ1;refGene_tx=XM_023245332.1;refGene_gn=FIZ1;refGene_tx=XM_023245333.1;refGene_gn=FIZ1;refGene_tx=XM_023245334.1
E2 3130937 3132626 refGene_gn=ZNF579;refGene_tx=XM_023245344.1
E2 3148237 3148339 refGene_gn=SBK3;refGene_tx=XM_019818850.2
E2 3155044 3155195 refGene_gn=SBK3;refGene_tx=XM_019818850.2
E2 3155707 3155910 refGene_gn=SBK3;refGene_tx=XM_019818850.2
E2 3157844 3158552 refGene_gn=SBK3;refGene_tx=XM_019818850.2
E2 3161839 3162092

refGene_gn=SBK2;refGene_tx=XM_023245131.1;refGene_gn=SBK2;refGene_tx=XM_023245132.1;refGene_gn=SBK2;refGene_tx=XM_023245133.1;refGene_gn=SBK2;refGene_tx=XM_023245134.1;refGene_gn=SBK2;refGene_tx=XM_023245135.1
E2 3163919 3164122
refGene_gn=SBK2;refGene_tx=XM_023245131.1;refGene_gn=SBK2;refGene_tx=XM_023245132.1;refGene_gn=SBK2;refGene_tx=XM_023245133.1;refGene_gn=SBK2;refGene_tx=XM_023245134.1;refGene_gn=SBK2;refGene_tx=XM_023245135.1
E2 3164535 3165129
refGene_gn=SBK2;refGene_tx=XM_023245131.1;refGene_gn=SBK2;refGene_tx=XM_023245132.1;refGene_gn=SBK2;refGene_tx=XM_023245133.1;refGene_gn=SBK2;refGene_tx=XM_023245134.1;refGene_gn=SBK2;refGene_tx=XM_023245135.1
E2 3171973 3173646 refGene_gn=SSC5D;refGene_tx=XM_023245004.1
E2 3176491 3176638 refGene_gn=SSC5D;refGene_tx=XM_023245004.1
E2 3183510 3183678 refGene_gn=SSC5D;refGene_tx=XM_023245004.1
E2 3183800 3184115 refGene_gn=SSC5D;refGene_tx=XM_023245004.1
E2 3184188 3184583 refGene_gn=SSC5D;refGene_tx=XM_023245004.1
E2 3184584 3184791 refGene_gn=SSC5D;refGene_tx=XM_023245004.1
E2 3186176 3186519 refGene_gn=SSC5D;refGene_tx=XM_023245004.1
E2 3189139 3189319 refGene_gn=SSC5D;refGene_tx=XM_023245004.1
E2 3189556 3189874 refGene_gn=SSC5D;refGene_tx=XM_023245004.1
E2 3192244 3192553 refGene_gn=SSC5D;refGene_tx=XM_023245004.1
E2 3192903 3193017 refGene_gn=SSC5D;refGene_tx=XM_023245004.1
E2 3193254 3193374 refGene_gn=SSC5D;refGene_tx=XM_023245004.1
E2 3193561 3193870 refGene_gn=SSC5D;refGene_tx=XM_023245004.1
E2 3194166 3194193 refGene_gn=SSC5D;refGene_tx=XM_023245004.1
E2 3194731 3194756 refGene_gn=SSC5D;refGene_tx=XM_023245004.1
E2 3196307 3196856
refGene_gn=NAT14;refGene_tx=XM_003997393.5;refGene_gn=NAT14;refGene_tx=XM_006940828.4;refGene_gn=NAT14;refGene_tx=XM_006940829.4;refGene_gn=NAT14;refGene_tx=XM_011289555.3
E2 3197500 3197572
refGene_gn=NAT14;refGene_tx=XM_003997393.5;refGene_gn=NAT14;refGene_tx=XM_006940828.4;refGene_gn=NAT14;refGene_tx=XM_006940829.4;refGene_gn=NAT14;refGene_tx=XM_011289555.3
E2 3199134 3199443 refGene_gn=ZNF628;refGene_tx=XM_023245868.1
E2 3199444 3201243 refGene_gn=ZNF628;refGene_tx=XM_023245868.1
E2 3201245 3202442 refGene_gn=ZNF628;refGene_tx=XM_023245868.1
E2 3206501 3206714 refGene_gn=ZNF628;refGene_tx=XM_023245868.1
E2 3206759 3206802 refGene_gn=ZNF628;refGene_tx=XM_023245868.1
E2 3224778 3224913
refGene_gn=ISOC2;refGene_tx=XM_006940830.4;refGene_gn=ISOC2;refGene_tx=XM_019819100.2
E2 3226855 3227065
refGene_gn=ISOC2;refGene_tx=XM_006940830.4;refGene_gn=ISOC2;refGene_tx=XM_019819100.2
E2 3227341 3227412
refGene_gn=ISOC2;refGene_tx=XM_006940830.4;refGene_gn=ISOC2;refGene_tx=XM_019819100.2
E2 3227546 3227664
refGene_gn=ISOC2;refGene_tx=XM_006940830.4;refGene_gn=ISOC2;refGene_tx

=XM_019819100.2
E2 3229149 3229230
refGene_gn=ISOC2;refGene_tx=XM_006940830.4;refGene_gn=ISOC2;refGene_tx
=XM_019819100.2
E2 3229537 3229885 refGene_gn=SHISA7;refGene_tx=XM_023246188.1
E2 3236801 3237532 refGene_gn=SHISA7;refGene_tx=XM_023246188.1
E2 3238765 3238920 refGene_gn=SHISA7;refGene_tx=XM_023246188.1
E2 3240479 3240545 refGene_gn=SHISA7;refGene_tx=XM_023246188.1
E2 3241024 3241174 refGene_gn=SHISA7;refGene_tx=XM_023246188.1
E2 3243463 3244107 refGene_gn=SHISA7;refGene_tx=XM_023246188.1
E2 3258887 3258890 refGene_gn=UBE2S;refGene_tx=XM_019819105.2
E2 3259504 3259652 refGene_gn=UBE2S;refGene_tx=XM_019819105.2
E2 3260983 3261174 refGene_gn=UBE2S;refGene_tx=XM_019819105.2
E2 3262854 3263175 refGene_gn=UBE2S;refGene_tx=XM_019819105.2
E2 3269947 3270037 refGene_gn=RPL28;refGene_tx=XM_003997396.5
E2 3270128 3270247 refGene_gn=RPL28;refGene_tx=XM_003997396.5
E2 3271789 3271913 refGene_gn=RPL28;refGene_tx=XM_003997396.5
E2 3272008 3272097 refGene_gn=RPL28;refGene_tx=XM_003997396.5
E2 3272462 3272538 refGene_gn=RPL28;refGene_tx=XM_003997396.5
E2 3274899 3275433 refGene_gn=TMEM238;refGene_tx=XM_023245233.1
E2 3279096 3279337 refGene_gn=TMEM190;refGene_tx=XM_003997397.4
E2 3279430 3279515 refGene_gn=TMEM190;refGene_tx=XM_003997397.4
E2 3279602 3279728 refGene_gn=TMEM190;refGene_tx=XM_003997397.4
E2 3280793 3280829 refGene_gn=TMEM190;refGene_tx=XM_003997397.4
E2 3280921 3280979 refGene_gn=TMEM190;refGene_tx=XM_003997397.4
E2 3281227 3281310 refGene_gn=IL11;refGene_tx=XM_023245413.1
E2 3283568 3283579 refGene_gn=IL11;refGene_tx=XM_023245413.1
E2 3286783 3286790 refGene_gn=IL11;refGene_tx=XM_003997398.4
E2 3287933 3288106
refGene_gn=IL11;refGene_tx=XM_003997398.4;refGene_gn=IL11;refGene_tx=X
M_023245413.1
E2 3288271 3288358
refGene_gn=IL11;refGene_tx=XM_003997398.4;refGene_gn=IL11;refGene_tx=X
M_023245413.1
E2 3288488 3288650
refGene_gn=IL11;refGene_tx=XM_003997398.4;refGene_gn=IL11;refGene_tx=X
M_023245413.1
E2 3289416 3289587
refGene_gn=IL11;refGene_tx=XM_003997398.4;refGene_gn=IL11;refGene_tx=X
M_023245413.1
E2 3291672 3291822
refGene_gn=FAM71E2;refGene_tx=XM_019819114.2;refGene_gn=FAM71E2;refGen
e_tx=XM_023245991.1;refGene_gn=FAM71E2;refGene_tx=XM_023245993.1
E2 3292040 3292195
refGene_gn=FAM71E2;refGene_tx=XM_019819114.2;refGene_gn=FAM71E2;refGen
e_tx=XM_023245991.1;refGene_gn=FAM71E2;refGene_tx=XM_023245993.1
E2 3292332 3292591
refGene_gn=FAM71E2;refGene_tx=XM_019819114.2;refGene_gn=FAM71E2;refGen
e_tx=XM_023245991.1;refGene_gn=FAM71E2;refGene_tx=XM_023245993.1
E2 3293038 3293063

refGene_gn=FAM71E2;refGene_tx=XM_019819114.2;refGene_gn=FAM71E2;refGene_tx=XM_023245991.1;refGene_gn=FAM71E2;refGene_tx=XM_023245993.1
E2 3293224 3293289
refGene_gn=FAM71E2;refGene_tx=XM_019819114.2;refGene_gn=FAM71E2;refGene_tx=XM_023245991.1;refGene_gn=FAM71E2;refGene_tx=XM_023245993.1
E2 3293381 3293485
refGene_gn=FAM71E2;refGene_tx=XM_019819114.2;refGene_gn=FAM71E2;refGene_tx=XM_023245991.1;refGene_gn=FAM71E2;refGene_tx=XM_023245993.1
E2 3293576 3293629
refGene_gn=FAM71E2;refGene_tx=XM_019819114.2;refGene_gn=FAM71E2;refGene_tx=XM_023245991.1;refGene_gn=FAM71E2;refGene_tx=XM_023245993.1
E2 3293713 3293790
refGene_gn=FAM71E2;refGene_tx=XM_019819114.2;refGene_gn=FAM71E2;refGene_tx=XM_023245991.1;refGene_gn=FAM71E2;refGene_tx=XM_023245993.1
E2 3295377 3297191
refGene_gn=FAM71E2;refGene_tx=XM_019819114.2;refGene_gn=FAM71E2;refGene_tx=XM_023245991.1;refGene_gn=FAM71E2;refGene_tx=XM_023245993.1
E2 3297702 3297736
refGene_gn=FAM71E2;refGene_tx=XM_023245991.1;refGene_gn=FAM71E2;refGene_tx=XM_023245993.1
E2 3299525 3299553 refGene_gn=FAM71E2;refGene_tx=XM_019819114.2
E2 3300320 3300432
refGene_gn=LOC105261211;refGene_tx=XM_023246002.1
E2 3301197 3301298
refGene_gn=LOC105261211;refGene_tx=XM_023246002.1
E2 3301420 3301474
refGene_gn=LOC105261211;refGene_tx=XM_023246002.1
E2 3306485 3306985
refGene_gn=KMT5C;refGene_tx=XM_023245996.1;refGene_gn=KMT5C;refGene_tx=XM_023245997.1;refGene_gn=KMT5C;refGene_tx=XM_023245998.1
E2 3307162 3307338
refGene_gn=KMT5C;refGene_tx=XM_023245996.1;refGene_gn=KMT5C;refGene_tx=XM_023245997.1;refGene_gn=KMT5C;refGene_tx=XM_023245998.1
E2 3307696 3307833
refGene_gn=KMT5C;refGene_tx=XM_023245996.1;refGene_gn=KMT5C;refGene_tx=XM_023245997.1;refGene_gn=KMT5C;refGene_tx=XM_023245998.1
E2 3309674 3309694
refGene_gn=KMT5C;refGene_tx=XM_023245996.1;refGene_gn=KMT5C;refGene_tx=XM_023245997.1;refGene_gn=KMT5C;refGene_tx=XM_023245998.1
E2 3309818 3309982
refGene_gn=KMT5C;refGene_tx=XM_023245996.1;refGene_gn=KMT5C;refGene_tx=XM_023245997.1;refGene_gn=KMT5C;refGene_tx=XM_023245998.1
E2 3310810 3310920
refGene_gn=KMT5C;refGene_tx=XM_023245996.1;refGene_gn=KMT5C;refGene_tx=XM_023245997.1;refGene_gn=KMT5C;refGene_tx=XM_023245998.1
E2 3311310 3311476
refGene_gn=KMT5C;refGene_tx=XM_023245996.1;refGene_gn=KMT5C;refGene_tx=XM_023245997.1;refGene_gn=KMT5C;refGene_tx=XM_023245998.1
E2 3311630 3311869
refGene_gn=KMT5C;refGene_tx=XM_023245996.1;refGene_gn=KMT5C;refGene_tx

=XM_023245997.1;refGene_gn=KMT5C;refGene_tx=XM_023245998.1
E2 3328953 3329031 refGene_gn=KMT5C;refGene_tx=XM_023245997.1
E2 3330912 3331169
refGene_gn=TMEM150B;refGene_tx=XM_019819122.2;refGene_gn=TMEM150B;refGene_tx=XM_023245999.1;refGene_gn=TMEM150B;refGene_tx=XM_023246001.1
E2 3331805 3331854
refGene_gn=TMEM150B;refGene_tx=XM_019819122.2;refGene_gn=TMEM150B;refGene_tx=XM_023245999.1;refGene_gn=TMEM150B;refGene_tx=XM_023246001.1
E2 3332114 3332275
refGene_gn=TMEM150B;refGene_tx=XM_011289560.3;refGene_gn=TMEM150B;refGene_tx=XM_019819122.2;refGene_gn=TMEM150B;refGene_tx=XM_019819126.2;refGene_gn=TMEM150B;refGene_tx=XM_019819127.2;refGene_gn=TMEM150B;refGene_tx=XM_023245999.1;refGene_gn=TMEM150B;refGene_tx=XM_023246000.1;refGene_gn=TMEM150B;refGene_tx=XM_023246001.1
E2 3333140 3333200
refGene_gn=TMEM150B;refGene_tx=XM_011289560.3;refGene_gn=TMEM150B;refGene_tx=XM_019819122.2;refGene_gn=TMEM150B;refGene_tx=XM_019819126.2;refGene_gn=TMEM150B;refGene_tx=XM_019819127.2;refGene_gn=TMEM150B;refGene_tx=XM_023245999.1;refGene_gn=TMEM150B;refGene_tx=XM_023246000.1;refGene_gn=TMEM150B;refGene_tx=XM_023246001.1
E2 3333300 3333508
refGene_gn=TMEM150B;refGene_tx=XM_011289560.3;refGene_gn=TMEM150B;refGene_tx=XM_019819122.2;refGene_gn=TMEM150B;refGene_tx=XM_019819126.2;refGene_gn=TMEM150B;refGene_tx=XM_019819127.2;refGene_gn=TMEM150B;refGene_tx=XM_023245999.1;refGene_gn=TMEM150B;refGene_tx=XM_023246000.1;refGene_gn=TMEM150B;refGene_tx=XM_023246001.1
E2 3333539 3333710
refGene_gn=TMEM150B;refGene_tx=XM_011289560.3;refGene_gn=TMEM150B;refGene_tx=XM_019819122.2;refGene_gn=TMEM150B;refGene_tx=XM_019819126.2;refGene_gn=TMEM150B;refGene_tx=XM_019819127.2;refGene_gn=TMEM150B;refGene_tx=XM_023246000.1;refGene_gn=TMEM150B;refGene_tx=XM_023246001.1
E2 3335896 3336080
refGene_gn=TMEM150B;refGene_tx=XM_011289560.3;refGene_gn=TMEM150B;refGene_tx=XM_019819122.2;refGene_gn=TMEM150B;refGene_tx=XM_019819126.2;refGene_gn=TMEM150B;refGene_tx=XM_019819127.2;refGene_gn=TMEM150B;refGene_tx=XM_023246000.1
E2 3338275 3338472
refGene_gn=TMEM150B;refGene_tx=XM_011289560.3;refGene_gn=TMEM150B;refGene_tx=XM_019819122.2;refGene_gn=TMEM150B;refGene_tx=XM_019819126.2;refGene_gn=TMEM150B;refGene_tx=XM_019819127.2;refGene_gn=TMEM150B;refGene_tx=XM_023246000.1
E2 3339095 3339253
refGene_gn=BRSK1;refGene_tx=XM_023245994.1;refGene_gn=BRSK1;refGene_tx=XM_023245995.1
E2 3339987 3340077
refGene_gn=BRSK1;refGene_tx=XM_023245994.1;refGene_gn=BRSK1;refGene_tx=XM_023245995.1
E2 3341199 3341398
refGene_gn=BRSK1;refGene_tx=XM_023245994.1;refGene_gn=BRSK1;refGene_tx=XM_023245995.1

E2 3341956 3342080
refGene_gn=BRSK1;refGene_tx=XM_023245994.1;refGene_gn=BRSK1;refGene_tx=XM_023245995.1
E2 3342171 3342220
refGene_gn=BRSK1;refGene_tx=XM_023245994.1;refGene_gn=BRSK1;refGene_tx=XM_023245995.1
E2 3342665 3343035
refGene_gn=BRSK1;refGene_tx=XM_023245994.1;refGene_gn=BRSK1;refGene_tx=XM_023245995.1
E2 3343487 3343548
refGene_gn=BRSK1;refGene_tx=XM_023245994.1;refGene_gn=BRSK1;refGene_tx=XM_023245995.1
E2 3343781 3343941
refGene_gn=BRSK1;refGene_tx=XM_023245994.1;refGene_gn=BRSK1;refGene_tx=XM_023245995.1
E2 3344152 3344250
refGene_gn=BRSK1;refGene_tx=XM_023245994.1;refGene_gn=BRSK1;refGene_tx=XM_023245995.1
E2 3344771 3344942
refGene_gn=BRSK1;refGene_tx=XM_023245994.1;refGene_gn=BRSK1;refGene_tx=XM_023245995.1
E2 3345140 3345172
refGene_gn=BRSK1;refGene_tx=XM_023245994.1;refGene_gn=BRSK1;refGene_tx=XM_023245995.1
E2 3345538 3345685
refGene_gn=BRSK1;refGene_tx=XM_023245994.1;refGene_gn=BRSK1;refGene_tx=XM_023245995.1
E2 3348889 3348958
refGene_gn=BRSK1;refGene_tx=XM_023245994.1;refGene_gn=BRSK1;refGene_tx=XM_023245995.1
E2 3349038 3349072
refGene_gn=BRSK1;refGene_tx=XM_023245994.1;refGene_gn=BRSK1;refGene_tx=XM_023245995.1
E2 3349152 3349269
refGene_gn=BRSK1;refGene_tx=XM_023245994.1;refGene_gn=BRSK1;refGene_tx=XM_023245995.1
E2 3352590 3352731
refGene_gn=BRSK1;refGene_tx=XM_023245994.1;refGene_gn=BRSK1;refGene_tx=XM_023245995.1
E2 3354601 3354691
refGene_gn=BRSK1;refGene_tx=XM_023245994.1;refGene_gn=BRSK1;refGene_tx=XM_023245995.1
E2 3354786 3354881 refGene_gn=BRSK1;refGene_tx=XM_023245994.1
E2 3357464 3357600 refGene_gn=BRSK1;refGene_tx=XM_023245994.1
E2 3362198 3362537
refGene_gn=HSPBP1;refGene_tx=XM_006940834.4;refGene_gn=HSPBP1;refGene_tx=XM_011289561.3;refGene_gn=HSPBP1;refGene_tx=XM_011289562.3
E2 3364252 3364457
refGene_gn=HSPBP1;refGene_tx=XM_006940834.4;refGene_gn=HSPBP1;refGene_tx=XM_011289561.3;refGene_gn=HSPBP1;refGene_tx=XM_011289562.3

E2	3366831	3367056
refGene_gn=HSPBP1;refGene_tx=XM_006940834.4;refGene_gn=HSPBP1;refGene_tx=XM_011289561.3;refGene_gn=HSPBP1;refGene_tx=XM_011289562.3
E2	3369846	3370002
refGene_gn=HSPBP1;refGene_tx=XM_006940834.4;refGene_gn=HSPBP1;refGene_tx=XM_011289561.3;refGene_gn=HSPBP1;refGene_tx=XM_011289562.3
E2	3370112	3370209
refGene_gn=HSPBP1;refGene_tx=XM_006940834.4;refGene_gn=HSPBP1;refGene_tx=XM_011289561.3;refGene_gn=HSPBP1;refGene_tx=XM_011289562.3
E2	3370749	3370861
refGene_gn=HSPBP1;refGene_tx=XM_006940834.4;refGene_gn=HSPBP1;refGene_tx=XM_011289561.3;refGene_gn=HSPBP1;refGene_tx=XM_011289562.3
E2	3373952	3374027
refGene_gn=HSPBP1;refGene_tx=XM_006940834.4;refGene_gn=HSPBP1;refGene_tx=XM_011289561.3;refGene_gn=HSPBP1;refGene_tx=XM_011289562.3
E2	3386310	3386537	refGene_gn=PPP6R1;refGene_tx=XM_023245869.1
E2	3386810	3386999	refGene_gn=PPP6R1;refGene_tx=XM_023245869.1
E2	3387116	3387123	refGene_gn=PPP6R1;refGene_tx=XM_023245869.1
E2	3387265	3387915	refGene_gn=PPP6R1;refGene_tx=XM_023245869.1
E2	3388000	3388114	refGene_gn=PPP6R1;refGene_tx=XM_023245869.1
E2	3388491	3388654	refGene_gn=PPP6R1;refGene_tx=XM_023245869.1
E2	3388738	3388891	refGene_gn=PPP6R1;refGene_tx=XM_023245869.1
E2	3389012	3389147	refGene_gn=PPP6R1;refGene_tx=XM_023245869.1
E2	3389475	3389540	refGene_gn=PPP6R1;refGene_tx=XM_023245869.1
E2	3389631	3389747	refGene_gn=PPP6R1;refGene_tx=XM_023245869.1
E2	3389833	3389919	refGene_gn=PPP6R1;refGene_tx=XM_023245869.1
E2	3390075	3390165	refGene_gn=PPP6R1;refGene_tx=XM_023245869.1
E2	3390250	3390348	refGene_gn=PPP6R1;refGene_tx=XM_023245869.1
E2	3392353	3392411	refGene_gn=PPP6R1;refGene_tx=XM_023245869.1
E2	3392505	3392687	refGene_gn=PPP6R1;refGene_tx=XM_023245869.1
E2	3395329	3395454	refGene_gn=PPP6R1;refGene_tx=XM_023245869.1
E2	3395550	3395698	refGene_gn=PPP6R1;refGene_tx=XM_023245869.1
E2	3395796	3395867	refGene_gn=PPP6R1;refGene_tx=XM_023245869.1
E2	3395969	3396051	refGene_gn=PPP6R1;refGene_tx=XM_023245869.1
E2	3396425	3396536	refGene_gn=PPP6R1;refGene_tx=XM_023245869.1
E2	3396617	3396689	refGene_gn=PPP6R1;refGene_tx=XM_023245869.1
E2	3396799	3396803	refGene_gn=PPP6R1;refGene_tx=XM_023245869.1
E2	3398822	3398873
refGene_gn=TMEM86B;refGene_tx=XM_003997424.4;refGene_gn=TMEM86B;refGene_tx=XM_019819128.2
E2	3399117	3399364
refGene_gn=TMEM86B;refGene_tx=XM_003997424.4;refGene_gn=TMEM86B;refGene_tx=XM_019819128.2
E2	3399911	3400294
refGene_gn=TMEM86B;refGene_tx=XM_003997424.4;refGene_gn=TMEM86B;refGene_tx=XM_019819128.2
E2	3409533	3409584	refGene_gn=PTPRH;refGene_tx=XM_023245253.1
E2	3411150	3411184	refGene_gn=PTPRH;refGene_tx=XM_023245253.1
E2	3411401	3411668	refGene_gn=PTPRH;refGene_tx=XM_023245253.1
E2	3414495	3414750	refGene_gn=PTPRH;refGene_tx=XM_023245253.1

E2 3415892 3416162 refGene_gn=PTPRH;refGene_tx=XM_023245253.1
E2 3417105 3417405 refGene_gn=PTPRH;refGene_tx=XM_023245253.1
E2 3417751 3418018 refGene_gn=PTPRH;refGene_tx=XM_023245253.1
E2 3419828 3419907 refGene_gn=PTPRH;refGene_tx=XM_023245253.1
E2 3420028 3420076 refGene_gn=PTPRH;refGene_tx=XM_023245253.1
E2 3423209 3423300 refGene_gn=PTPRH;refGene_tx=XM_023245253.1
E2 3423674 3423765 refGene_gn=PTPRH;refGene_tx=XM_023245253.1
E2 3424037 3424114 refGene_gn=PTPRH;refGene_tx=XM_023245253.1
E2 3424207 3424342 refGene_gn=PTPRH;refGene_tx=XM_023245253.1
E2 3424589 3424712 refGene_gn=PTPRH;refGene_tx=XM_023245253.1
E2 3424927 3425088 refGene_gn=PTPRH;refGene_tx=XM_023245253.1
E2 3426431 3426567 refGene_gn=PTPRH;refGene_tx=XM_023245253.1
E2 3426674 3426785 refGene_gn=PTPRH;refGene_tx=XM_023245253.1
E2 3429078 3429157
refGene_gn=SYT5;refGene_tx=XM_003997401.5;refGene_gn=SYT5;refGene_tx=X
M_006940841.4
E2 3429713 3429886
refGene_gn=SYT5;refGene_tx=XM_003997401.5;refGene_gn=SYT5;refGene_tx=X
M_006940841.4
E2 3431483 3431603
refGene_gn=SYT5;refGene_tx=XM_003997401.5;refGene_gn=SYT5;refGene_tx=X
M_006940841.4
E2 3431716 3431884
refGene_gn=SYT5;refGene_tx=XM_003997401.5;refGene_gn=SYT5;refGene_tx=X
M_006940841.4
E2 3432245 3432413
refGene_gn=SYT5;refGene_tx=XM_003997401.5;refGene_gn=SYT5;refGene_tx=X
M_006940841.4
E2 3432686 3432804
refGene_gn=SYT5;refGene_tx=XM_003997401.5;refGene_gn=SYT5;refGene_tx=X
M_006940841.4
E2 3433073 3433207
refGene_gn=SYT5;refGene_tx=XM_003997401.5;refGene_gn=SYT5;refGene_tx=X
M_006940841.4
E2 3433964 3434165
refGene_gn=SYT5;refGene_tx=XM_003997401.5;refGene_gn=SYT5;refGene_tx=X
M_006940841.4
E2 3440053 3440138
refGene_gn=TNNI3;refGene_tx=XM_006940842.4;refGene_gn=TNNI3;refGene_tx
=XM_019819119.2;refGene_gn=TNNI3;refGene_tx=XM_023244836.1
E2 3440429 3440614
refGene_gn=TNNI3;refGene_tx=XM_006940842.4;refGene_gn=TNNI3;refGene_tx
=XM_019819119.2;refGene_gn=TNNI3;refGene_tx=XM_023244836.1
E2 3440747 3440841
refGene_gn=TNNI3;refGene_tx=XM_006940842.4;refGene_gn=TNNI3;refGene_tx
=XM_019819119.2;refGene_gn=TNNI3;refGene_tx=XM_023244836.1;refGene_gn=
TNNI3;refGene_tx=XM_023244837.1
E2 3442580 3442738
refGene_gn=TNNI3;refGene_tx=XM_006940842.4;refGene_gn=TNNI3;refGene_tx
=XM_019819119.2;refGene_gn=TNNI3;refGene_tx=XM_023244836.1;refGene_gn=

TNNI3;refGene_tx=XM_023244837.1
E2 3443010 3443193
refGene_gn=TNNI3;refGene_tx=XM_006940842.4;refGene_gn=TNNI3;refGene_tx=XM_019819119.2;refGene_gn=TNNI3;refGene_tx=XM_023244836.1;refGene_gn=TNNI3;refGene_tx=XM_023244837.1
E2 3443475 3443601
refGene_gn=TNNI3;refGene_tx=XM_006940842.4;refGene_gn=TNNI3;refGene_tx=XM_019819119.2;refGene_gn=TNNI3;refGene_tx=XM_023244836.1;refGene_gn=TNNI3;refGene_tx=XM_023244837.1
E2 3443707 3443830
refGene_gn=TNNI3;refGene_tx=XM_006940842.4;refGene_gn=TNNI3;refGene_tx=XM_019819119.2;refGene_gn=TNNI3;refGene_tx=XM_023244836.1;refGene_gn=TNNI3;refGene_tx=XM_023244837.1
E2 3444027 3444157
refGene_gn=TNNI3;refGene_tx=XM_006940842.4;refGene_gn=TNNI3;refGene_tx=XM_019819119.2;refGene_gn=TNNI3;refGene_tx=XM_023244836.1;refGene_gn=TNNI3;refGene_tx=XM_023244837.1
E2 3444954 3445066
refGene_gn=TNNI3;refGene_tx=XM_006940842.4;refGene_gn=TNNI3;refGene_tx=XM_019819119.2;refGene_gn=TNNI3;refGene_tx=XM_023244836.1;refGene_gn=TNNI3;refGene_tx=XM_023244837.1
E2 3445229 3445304
refGene_gn=TNNI3;refGene_tx=XM_006940842.4;refGene_gn=TNNI3;refGene_tx=XM_019819119.2;refGene_gn=TNNI3;refGene_tx=XM_023244836.1;refGene_gn=TNNI3;refGene_tx=XM_023244837.1
E2 3445388 3445842
refGene_gn=TNNI3;refGene_tx=XM_006940842.4;refGene_gn=TNNI3;refGene_tx=XM_019819119.2;refGene_gn=TNNI3;refGene_tx=XM_023244836.1;refGene_gn=TNNI3;refGene_tx=XM_023244837.1
E2 3446714 3446866
refGene_gn=TNNI3;refGene_tx=NM_001009237.1;refGene_gn=TNNI3;refGene_tx=XM_019819119.2;refGene_gn=TNNI3;refGene_tx=XM_023244836.1;refGene_gn=TNNI3;refGene_tx=XM_023244837.1
E2 3447130 3447140
refGene_gn=TNNI3;refGene_tx=NM_001009237.1;refGene_gn=TNNI3;refGene_tx=XM_019819119.2;refGene_gn=TNNI3;refGene_tx=XM_023244836.1;refGene_gn=TNNI3;refGene_tx=XM_023244837.1
E2 3447285 3447372
refGene_gn=TNNI3;refGene_tx=NM_001009237.1;refGene_gn=TNNI3;refGene_tx=XM_019819119.2;refGene_gn=TNNI3;refGene_tx=XM_023244836.1;refGene_gn=TNNI3;refGene_tx=XM_023244837.1
E2 3447778 3447820
refGene_gn=TNNI3;refGene_tx=NM_001009237.1;refGene_gn=TNNI3;refGene_tx=XM_019819119.2;refGene_gn=TNNI3;refGene_tx=XM_023244836.1;refGene_gn=TNNI3;refGene_tx=XM_023244837.1
E2 3448096 3448228
refGene_gn=TNNI3;refGene_tx=NM_001009237.1;refGene_gn=TNNI3;refGene_tx=XM_019819119.2;refGene_gn=TNNI3;refGene_tx=XM_023244836.1;refGene_gn=TNNI3;refGene_tx=XM_023244837.1
E2 3448618 3448708

refGene_gn=TNNI3;refGene_tx=NM_001009237.1;refGene_gn=TNNI3;refGene_tx=XM_019819119.2;refGene_gn=TNNI3;refGene_tx=XM_023244836.1;refGene_gn=TNNI3;refGene_tx=XM_023244837.1
E2 3449258 3449435 refGene_gn=TNNI3;refGene_tx=NM_001009237.1;refGene_gn=TNNI3;refGene_tx=XM_019819119.2;refGene_gn=TNNI3;refGene_tx=XM_023244836.1;refGene_gn=TNNI3;refGene_tx=XM_023244837.1
E2 3449901 3449985 refGene_gn=TNNI3;refGene_tx=NM_001009237.1;refGene_gn=TNNI3;refGene_tx=XM_019819119.2;refGene_gn=TNNI3;refGene_tx=XM_023244836.1;refGene_gn=TNNI3;refGene_tx=XM_023244837.1
E2 3454409 3454441 refGene_gn=TNNT1;refGene_tx=XM_006940843.3;refGene_gn=TNNT1;refGene_tx=XM_006940844.3
E2 3454548 3454562 refGene_gn=TNNT1;refGene_tx=XM_006940843.3;refGene_gn=TNNT1;refGene_tx=XM_006940844.3
E2 3454738 3454765 refGene_gn=TNNT1;refGene_tx=XM_006940843.3;refGene_gn=TNNT1;refGene_tx=XM_006940844.3
E2 3455006 3455039 refGene_gn=TNNT1;refGene_tx=XM_006940843.3
E2 3455774 3455796 refGene_gn=TNNT1;refGene_tx=XM_006940843.3;refGene_gn=TNNT1;refGene_tx=XM_006940844.3
E2 3457915 3457976 refGene_gn=TNNT1;refGene_tx=XM_006940843.3;refGene_gn=TNNT1;refGene_tx=XM_006940844.3
E2 3458254 3458371 refGene_gn=TNNT1;refGene_tx=XM_006940843.3;refGene_gn=TNNT1;refGene_tx=XM_006940844.3
E2 3458577 3458655 refGene_gn=TNNT1;refGene_tx=XM_006940843.3;refGene_gn=TNNT1;refGene_tx=XM_006940844.3
E2 3460397 3460511 refGene_gn=TNNT1;refGene_tx=XM_006940843.3;refGene_gn=TNNT1;refGene_tx=XM_006940844.3
E2 3461124 3461234 refGene_gn=TNNT1;refGene_tx=XM_006940843.3;refGene_gn=TNNT1;refGene_tx=XM_006940844.3
E2 3463840 3463931 refGene_gn=TNNT1;refGene_tx=XM_006940843.3;refGene_gn=TNNT1;refGene_tx=XM_006940844.3
E2 3464426 3464467 refGene_gn=TNNT1;refGene_tx=XM_006940843.3;refGene_gn=TNNT1;refGene_tx=XM_006940844.3
E2 3465178 3465224 refGene_gn=TNNT1;refGene_tx=XM_006940843.3;refGene_gn=TNNT1;refGene_tx=XM_006940844.3
E2 3472614 3472935 refGene_gn=PPP1R12C;refGene_tx=XM_011289595.2
E2 3476705 3476836 refGene_gn=PPP1R12C;refGene_tx=XM_011289595.2

E2	3476927	3477046	refGene_gn=PPP1R12C;refGene_tx=XM_011289595.2
E2	3482281	3482441	refGene_gn=PPP1R12C;refGene_tx=XM_011289595.2
E2	3485455	3485487	refGene_gn=PPP1R12C;refGene_tx=XM_011289595.2
E2	3485489	3485599	refGene_gn=PPP1R12C;refGene_tx=XM_011289595.2
E2	3485694	3485769	refGene_gn=PPP1R12C;refGene_tx=XM_011289595.2
E2	3487000	3487074	refGene_gn=PPP1R12C;refGene_tx=XM_011289595.2
E2	3487149	3487277	refGene_gn=PPP1R12C;refGene_tx=XM_011289595.2
E2	3488286	3488360	refGene_gn=PPP1R12C;refGene_tx=XM_011289595.2
E2	3488616	3488775	refGene_gn=PPP1R12C;refGene_tx=XM_011289595.2
E2	3488854	3488922	refGene_gn=PPP1R12C;refGene_tx=XM_011289595.2
E2	3489535	3489670	refGene_gn=PPP1R12C;refGene_tx=XM_011289595.2
E2	3489903	3489994	refGene_gn=PPP1R12C;refGene_tx=XM_011289595.2
E2	3490785	3490860	refGene_gn=PPP1R12C;refGene_tx=XM_011289595.2
E2	3490938	3490999	refGene_gn=PPP1R12C;refGene_tx=XM_011289595.2
E2	3491158	3491244	refGene_gn=PPP1R12C;refGene_tx=XM_011289595.2
E2	3491345	3491386	refGene_gn=PPP1R12C;refGene_tx=XM_011289595.2
E2	3491469	3491557	refGene_gn=PPP1R12C;refGene_tx=XM_011289595.2
E2	3491717	3491822	refGene_gn=PPP1R12C;refGene_tx=XM_011289595.2
E2	3491990	3492041	refGene_gn=PPP1R12C;refGene_tx=XM_011289595.2
E2	3492193	3492244	refGene_gn=PPP1R12C;refGene_tx=XM_011289595.2
E2	3492333	3492420	refGene_gn=PPP1R12C;refGene_tx=XM_011289595.2
E2	3495390	3495477
refGene_gn=EPS8L1;refGene_tx=XM_011289566.3;refGene_gn=EPS8L1;refGene_tx=XM_019819107.2;refGene_gn=EPS8L1;refGene_tx=XM_019819108.2;refGene_gn=EPS8L1;refGene_tx=XM_019819109.2;refGene_gn=EPS8L1;refGene_tx=XM_019819110.2;refGene_gn=EPS8L1;refGene_tx=XM_023246015.1
E2	3495558	3495691
refGene_gn=EPS8L1;refGene_tx=XM_011289566.3;refGene_gn=EPS8L1;refGene_tx=XM_019819107.2;refGene_gn=EPS8L1;refGene_tx=XM_019819108.2;refGene_gn=EPS8L1;refGene_tx=XM_019819109.2;refGene_gn=EPS8L1;refGene_tx=XM_019819110.2;refGene_gn=EPS8L1;refGene_tx=XM_023246015.1
E2	3496046	3496221
refGene_gn=EPS8L1;refGene_tx=XM_011289566.3;refGene_gn=EPS8L1;refGene_tx=XM_019819107.2;refGene_gn=EPS8L1;refGene_tx=XM_019819108.2;refGene_gn=EPS8L1;refGene_tx=XM_019819109.2;refGene_gn=EPS8L1;refGene_tx=XM_019819110.2;refGene_gn=EPS8L1;refGene_tx=XM_023246015.1
E2	3496400	3496500
refGene_gn=EPS8L1;refGene_tx=XM_011289566.3;refGene_gn=EPS8L1;refGene_tx=XM_019819107.2;refGene_gn=EPS8L1;refGene_tx=XM_019819108.2;refGene_gn=EPS8L1;refGene_tx=XM_019819109.2;refGene_gn=EPS8L1;refGene_tx=XM_019819110.2;refGene_gn=EPS8L1;refGene_tx=XM_023246015.1
E2	3496658	3496772
refGene_gn=EPS8L1;refGene_tx=XM_011289566.3;refGene_gn=EPS8L1;refGene_tx=XM_019819107.2;refGene_gn=EPS8L1;refGene_tx=XM_019819108.2;refGene_gn=EPS8L1;refGene_tx=XM_019819109.2;refGene_gn=EPS8L1;refGene_tx=XM_019819110.2;refGene_gn=EPS8L1;refGene_tx=XM_023246015.1
E2	3496859	3496986
refGene_gn=EPS8L1;refGene_tx=XM_011289566.3;refGene_gn=EPS8L1;refGene_tx=XM_019819107.2;refGene_gn=EPS8L1;refGene_tx=XM_019819108.2;refGene_gn=EPS8L1;refGene_tx=XM_019819109.2;refGene_gn=EPS8L1;refGene_tx=XM_01

9819110.2;refGene_gn=EPS8L1;refGene_tx=XM_023246015.1
E2	3498814	3498843
refGene_gn=EPS8L1;refGene_tx=XM_011289566.3;refGene_gn=EPS8L1;refGene_tx=XM_019819107.2;refGene_gn=EPS8L1;refGene_tx=XM_019819108.2;refGene_gn=EPS8L1;refGene_tx=XM_019819109.2;refGene_gn=EPS8L1;refGene_tx=XM_019819110.2;refGene_gn=EPS8L1;refGene_tx=XM_023246015.1
E2	3498949	3499091
refGene_gn=EPS8L1;refGene_tx=XM_011289566.3;refGene_gn=EPS8L1;refGene_tx=XM_019819107.2;refGene_gn=EPS8L1;refGene_tx=XM_019819108.2;refGene_gn=EPS8L1;refGene_tx=XM_019819109.2;refGene_gn=EPS8L1;refGene_tx=XM_019819110.2;refGene_gn=EPS8L1;refGene_tx=XM_023246015.1
E2	3499654	3499803
refGene_gn=EPS8L1;refGene_tx=XM_011289566.3;refGene_gn=EPS8L1;refGene_tx=XM_019819107.2;refGene_gn=EPS8L1;refGene_tx=XM_019819108.2;refGene_gn=EPS8L1;refGene_tx=XM_019819109.2;refGene_gn=EPS8L1;refGene_tx=XM_019819110.2;refGene_gn=EPS8L1;refGene_tx=XM_023246015.1
E2	3499906	3499981
refGene_gn=EPS8L1;refGene_tx=XM_011289566.3;refGene_gn=EPS8L1;refGene_tx=XM_019819107.2;refGene_gn=EPS8L1;refGene_tx=XM_019819108.2;refGene_gn=EPS8L1;refGene_tx=XM_019819109.2;refGene_gn=EPS8L1;refGene_tx=XM_019819110.2;refGene_gn=EPS8L1;refGene_tx=XM_023246015.1
E2	3500076	3500165
refGene_gn=EPS8L1;refGene_tx=XM_011289566.3;refGene_gn=EPS8L1;refGene_tx=XM_019819107.2;refGene_gn=EPS8L1;refGene_tx=XM_019819108.2;refGene_gn=EPS8L1;refGene_tx=XM_019819109.2;refGene_gn=EPS8L1;refGene_tx=XM_019819110.2;refGene_gn=EPS8L1;refGene_tx=XM_023246015.1
E2	3500382	3500509
refGene_gn=EPS8L1;refGene_tx=XM_011289566.3;refGene_gn=EPS8L1;refGene_tx=XM_019819107.2;refGene_gn=EPS8L1;refGene_tx=XM_019819108.2;refGene_gn=EPS8L1;refGene_tx=XM_019819109.2;refGene_gn=EPS8L1;refGene_tx=XM_019819110.2;refGene_gn=EPS8L1;refGene_tx=XM_023246015.1
E2	3500766	3501028
refGene_gn=EPS8L1;refGene_tx=XM_011289566.3;refGene_gn=EPS8L1;refGene_tx=XM_019819107.2;refGene_gn=EPS8L1;refGene_tx=XM_019819108.2;refGene_gn=EPS8L1;refGene_tx=XM_019819109.2;refGene_gn=EPS8L1;refGene_tx=XM_019819110.2;refGene_gn=EPS8L1;refGene_tx=XM_023246015.1
E2	3501408	3501491
refGene_gn=EPS8L1;refGene_tx=XM_011289566.3;refGene_gn=EPS8L1;refGene_tx=XM_019819107.2;refGene_gn=EPS8L1;refGene_tx=XM_019819108.2;refGene_gn=EPS8L1;refGene_tx=XM_019819109.2;refGene_gn=EPS8L1;refGene_tx=XM_019819110.2;refGene_gn=EPS8L1;refGene_tx=XM_023246015.1
E2	3502107	3502257
refGene_gn=EPS8L1;refGene_tx=XM_011289566.3;refGene_gn=EPS8L1;refGene_tx=XM_019819107.2;refGene_gn=EPS8L1;refGene_tx=XM_019819108.2;refGene_gn=EPS8L1;refGene_tx=XM_019819109.2;refGene_gn=EPS8L1;refGene_tx=XM_019819110.2;refGene_gn=EPS8L1;refGene_tx=XM_023246015.1
E2	3502545	3502707
refGene_gn=EPS8L1;refGene_tx=XM_011289566.3;refGene_gn=EPS8L1;refGene_tx=XM_019819107.2;refGene_gn=EPS8L1;refGene_tx=XM_019819108.2;refGene_gn=EPS8L1;refGene_tx=XM_019819109.2;refGene_gn=EPS8L1;refGene_tx=XM_01

9819110.2;refGene_gn=EPS8L1;refGene_tx=XM_023246015.1
E2 3503274 3503333
refGene_gn=EPS8L1;refGene_tx=XM_011289566.3;refGene_gn=EPS8L1;refGene_tx=XM_019819107.2;refGene_gn=EPS8L1;refGene_tx=XM_019819108.2;refGene_gn=EPS8L1;refGene_tx=XM_019819109.2;refGene_gn=EPS8L1;refGene_tx=XM_019819110.2;refGene_gn=EPS8L1;refGene_tx=XM_023246015.1
E2 3503695 3503736
refGene_gn=EPS8L1;refGene_tx=XM_011289566.3;refGene_gn=EPS8L1;refGene_tx=XM_019819107.2;refGene_gn=EPS8L1;refGene_tx=XM_019819108.2;refGene_gn=EPS8L1;refGene_tx=XM_019819109.2;refGene_gn=EPS8L1;refGene_tx=XM_019819110.2;refGene_gn=EPS8L1;refGene_tx=XM_023246015.1
E2 3504905 3505007
refGene_gn=EPS8L1;refGene_tx=XM_011289566.3;refGene_gn=EPS8L1;refGene_tx=XM_019819107.2;refGene_gn=EPS8L1;refGene_tx=XM_019819108.2;refGene_gn=EPS8L1;refGene_tx=XM_019819109.2;refGene_gn=EPS8L1;refGene_tx=XM_019819110.2;refGene_gn=EPS8L1;refGene_tx=XM_023246015.1
E2 3506015 3506143
refGene_gn=EPS8L1;refGene_tx=XM_019819107.2;refGene_gn=EPS8L1;refGene_tx=XM_019819109.2
E2 3506157 3506261 refGene_gn=EPS8L1;refGene_tx=XM_019819108.2
E2 3517855 3517920 refGene_gn=RDH13;refGene_tx=XM_006940848.3
E2 3520903 3521022 refGene_gn=RDH13;refGene_tx=XM_006940848.3
E2 3522586 3522742 refGene_gn=RDH13;refGene_tx=XM_006940848.3
E2 3525284 3525389 refGene_gn=RDH13;refGene_tx=XM_006940848.3
E2 3526317 3526530 refGene_gn=RDH13;refGene_tx=XM_006940848.3
E2 3527360 3527462 refGene_gn=RDH13;refGene_tx=XM_006940848.3
E2 3528724 3528951 refGene_gn=RDH13;refGene_tx=XM_006940848.3
E2 3550106 3550140
refGene_gn=LOC101093312;refGene_tx=XM_006940863.3
E2 3550372 3550408
refGene_gn=LOC101093312;refGene_tx=XM_006940863.3
E2 3552851 3553136
refGene_gn=LOC101093312;refGene_tx=XM_006940863.3
E2 3553558 3553846
refGene_gn=LOC101093312;refGene_tx=XM_006940863.3
E2 3555772 3555892
refGene_gn=LOC101093312;refGene_tx=XM_006940863.3
E2 3555973 3556016
refGene_gn=LOC101093312;refGene_tx=XM_006940863.3
E2 3556101 3556139
refGene_gn=LOC101093312;refGene_tx=XM_006940863.3
E2 3556409 3556462
refGene_gn=LOC101093312;refGene_tx=XM_006940863.3
E2 3556548 3556695
refGene_gn=LOC101093312;refGene_tx=XM_006940863.3
E2 3560073 3560107
refGene_gn=LOC101093562;refGene_tx=XM_003997430.5
E2 3564362 3564398
refGene_gn=LOC101093562;refGene_tx=XM_003997430.5
E2 3564526 3564784

refGene_gn=LOC101093562;refGene_tx=XM_003997430.5
E2 3567007 3567292
refGene_gn=LOC101093562;refGene_tx=XM_003997430.5
E2 3574354 3574408
refGene_gn=LOC101093562;refGene_tx=XM_003997430.5
E2 3577466 3577526
refGene_gn=LOC101093562;refGene_tx=XM_003997430.5
E2 3581129 3581180
refGene_gn=LOC101093562;refGene_tx=XM_003997430.5
E2 3581699 3581935
refGene_gn=LOC101093562;refGene_tx=XM_003997430.5
E2 3584403 3584542 refGene_gn=NLRP2;refGene_tx=XM_023245870.1
E2 3587567 3587738 refGene_gn=NLRP2;refGene_tx=XM_023245870.1
E2 3594510 3594681 refGene_gn=NLRP2;refGene_tx=XM_023245870.1
E2 3597038 3597209 refGene_gn=NLRP2;refGene_tx=XM_023245870.1
E2 3597551 3597722 refGene_gn=NLRP2;refGene_tx=XM_023245870.1
E2 3599420 3599585 refGene_gn=NLRP2;refGene_tx=XM_023245870.1
E2 3601593 3601764 refGene_gn=NLRP2;refGene_tx=XM_023245870.1
E2 3609054 3610705 refGene_gn=NLRP2;refGene_tx=XM_023245870.1
E2 3616204 3616484 refGene_gn=NLRP2;refGene_tx=XM_023245870.1
E2 3639456 3640284 refGene_gn=NLRP7;refGene_tx=XM_023245871.1
E2 3640285 3640754 refGene_gn=NLRP7;refGene_tx=XM_023245871.1
E2 3640756 3641015 refGene_gn=NLRP7;refGene_tx=XM_023245871.1
E2 3643174 3643345 refGene_gn=NLRP7;refGene_tx=XM_023245871.1
E2 3646154 3646325 refGene_gn=NLRP7;refGene_tx=XM_023245871.1
E2 3647801 3647985 refGene_gn=NLRP7;refGene_tx=XM_023245871.1
E2 3648293 3648457 refGene_gn=NLRP7;refGene_tx=XM_023245871.1
E2 3651081 3651249 refGene_gn=NLRP7;refGene_tx=XM_023245871.1
E2 3654455 3654765 refGene_gn=NLRP7;refGene_tx=XM_023245871.1
E2 3668653 3668715
refGene_gn=LOC101094307;refGene_tx=XM_023246047.1
E2 3668717 3668834
refGene_gn=LOC101094307;refGene_tx=XM_023246047.1
E2 3668951 3668999
refGene_gn=LOC101094307;refGene_tx=XM_023246047.1
E2 3669964 3670015
refGene_gn=LOC101094307;refGene_tx=XM_023246047.1
E2 3670276 3670555
refGene_gn=LOC101094307;refGene_tx=XM_023246047.1
E2 3672329 3672614
refGene_gn=LOC101094307;refGene_tx=XM_023246047.1
E2 3672797 3672833
refGene_gn=LOC101094307;refGene_tx=XM_023246047.1
E2 3672916 3672950
refGene_gn=LOC101094307;refGene_tx=XM_023246047.1
E2 3676486 3676544 refGene_gn=FCAR;refGene_tx=XM_023246048.1
E2 3677436 3677644 refGene_gn=FCAR;refGene_tx=XM_023246048.1
E2 3680037 3680325 refGene_gn=FCAR;refGene_tx=XM_023246048.1
E2 3682160 3682445 refGene_gn=FCAR;refGene_tx=XM_023246048.1
E2 3685374 3685410 refGene_gn=FCAR;refGene_tx=XM_023246048.1

E2	3685823	3685857	refGene_gn=FCAR;refGene_tx=XM_023246048.1
E2	3690476	3690632
refGene_gn=LOC101084434;refGene_tx=XM_023245873.1
E2	3690729	3690782
refGene_gn=LOC101084434;refGene_tx=XM_023245873.1
E2	3691772	3691848
refGene_gn=LOC101084434;refGene_tx=XM_023245873.1
E2	3691849	3691878
refGene_gn=LOC101084434;refGene_tx=XM_023245873.1
E2	3695511	3695562
refGene_gn=LOC101084434;refGene_tx=XM_023245873.1
E2	3696158	3696393
refGene_gn=LOC101084434;refGene_tx=XM_023245873.1
E2	3696395	3696460
refGene_gn=LOC101084434;refGene_tx=XM_023245873.1
E2	3697933	3698233
refGene_gn=LOC101084434;refGene_tx=XM_023245873.1
E2	3699478	3699961
refGene_gn=LOC101084434;refGene_tx=XM_023245873.1
E2	3700138	3700229
refGene_gn=LOC101084434;refGene_tx=XM_023245873.1
E2	3704941	3705087
refGene_gn=LOC101094553;refGene_tx=XM_023246054.1
E2	3705231	3705282
refGene_gn=LOC101094553;refGene_tx=XM_023246054.1
E2	3710027	3710330
refGene_gn=LOC101094553;refGene_tx=XM_023246054.1
E2	3710547	3710565
refGene_gn=LOC101094553;refGene_tx=XM_023246054.1
E2	3710567	3710843
refGene_gn=LOC101094553;refGene_tx=XM_023246054.1
E2	3711067	3711370
refGene_gn=LOC101094553;refGene_tx=XM_023246054.1
E2	3712859	3713156
refGene_gn=LOC101094553;refGene_tx=XM_023246054.1
E2	3713380	3713683
refGene_gn=LOC101094553;refGene_tx=XM_023246054.1
E2	3713834	3714119
refGene_gn=LOC101094553;refGene_tx=XM_023246054.1
E2	3714307	3714343
refGene_gn=LOC101094553;refGene_tx=XM_023246054.1
E2	3714501	3714643
refGene_gn=LOC101094553;refGene_tx=XM_023246054.1
E2	3728036	3728278
refGene_gn=LOC101094804;refGene_tx=XM_023245874.1
E2	3728280	3728409
refGene_gn=LOC101094804;refGene_tx=XM_023245874.1
E2	3729577	3729667
refGene_gn=LOC101094804;refGene_tx=XM_023245874.1
E2	3729669	3729738

refGene_gn=LOC101094804;refGene_tx=XM_023245874.1
E2 3729740 3729818
refGene_gn=LOC101094804;refGene_tx=XM_023245874.1
E2 3729820 3729886
refGene_gn=LOC101094804;refGene_tx=XM_023245874.1
E2 3730103 3730235
refGene_gn=LOC101094804;refGene_tx=XM_023245874.1
E2 3730237 3730389
refGene_gn=LOC101094804;refGene_tx=XM_023245874.1
E2 3730643 3730938
refGene_gn=LOC101094804;refGene_tx=XM_023245874.1
E2 3731084 3731369
refGene_gn=LOC101094804;refGene_tx=XM_023245874.1
E2 3731551 3731587
refGene_gn=LOC101094804;refGene_tx=XM_023245874.1
E2 3731744 3731778
refGene_gn=LOC101094804;refGene_tx=XM_023245874.1
E2 3767605 3768067
refGene_gn=CDC42EP5;refGene_tx=XM_023246049.1;refGene_gn=CDC42EP5;refGene_tx=XM_023246050.1;refGene_gn=CDC42EP5;refGene_tx=XM_023246051.1;refGene_gn=CDC42EP5;refGene_tx=XM_023246052.1
E2 3769342 3770552 refGene_gn=LENG9;refGene_tx=XM_011289599.3
E2 3770554 3770985 refGene_gn=LENG9;refGene_tx=XM_011289599.3
E2 3772192 3772355 refGene_gn=LENG8;refGene_tx=XM_023246053.1
E2 3774340 3774548 refGene_gn=LENG8;refGene_tx=XM_023246053.1
E2 3774630 3774760 refGene_gn=LENG8;refGene_tx=XM_023246053.1
E2 3774850 3774919 refGene_gn=LENG8;refGene_tx=XM_023246053.1
E2 3774998 3775100 refGene_gn=LENG8;refGene_tx=XM_023246053.1
E2 3775818 3776104 refGene_gn=LENG8;refGene_tx=XM_023246053.1
E2 3776247 3776388 refGene_gn=LENG8;refGene_tx=XM_023246053.1
E2 3776476 3776756 refGene_gn=LENG8;refGene_tx=XM_023246053.1
E2 3777132 3777338 refGene_gn=LENG8;refGene_tx=XM_023246053.1
E2 3777595 3777737 refGene_gn=LENG8;refGene_tx=XM_023246053.1
E2 3777945 3778198 refGene_gn=LENG8;refGene_tx=XM_023246053.1
E2 3779002 3779113 refGene_gn=LENG8;refGene_tx=XM_023246053.1
E2 3779882 3779984 refGene_gn=LENG8;refGene_tx=XM_023246053.1
E2 3780366 3780541 refGene_gn=LENG8;refGene_tx=XM_023246053.1
E2 3781232 3781270 refGene_gn=LENG8;refGene_tx=XM_023246053.1
E2 3787233 3787471
refGene_gn=TTYH1;refGene_tx=XM_023246042.1;refGene_gn=TTYH1;refGene_tx=XM_023246043.1
E2 3787560 3787606 refGene_gn=TTYH1;refGene_tx=XM_023246042.1
E2 3787725 3787868
refGene_gn=TTYH1;refGene_tx=XM_023246042.1;refGene_gn=TTYH1;refGene_tx=XM_023246043.1
E2 3790599 3790692
refGene_gn=TTYH1;refGene_tx=XM_023246042.1;refGene_gn=TTYH1;refGene_tx=XM_023246043.1
E2 3790834 3790927
refGene_gn=TTYH1;refGene_tx=XM_023246042.1;refGene_gn=TTYH1;refGene_tx

=XM_023246043.1
E2 3791069 3791125
refGene_gn=TTYH1;refGene_tx=XM_023246042.1;refGene_gn=TTYH1;refGene_tx
=XM_023246043.1
E2 3791593 3791669
refGene_gn=TTYH1;refGene_tx=XM_023246042.1;refGene_gn=TTYH1;refGene_tx
=XM_023246043.1
E2 3792031 3792104
refGene_gn=TTYH1;refGene_tx=XM_023246042.1;refGene_gn=TTYH1;refGene_tx
=XM_023246043.1
E2 3793368 3793464
refGene_gn=TTYH1;refGene_tx=XM_023246042.1;refGene_gn=TTYH1;refGene_tx
=XM_023246043.1
E2 3795947 3796168
refGene_gn=TTYH1;refGene_tx=XM_023246042.1;refGene_gn=TTYH1;refGene_tx
=XM_023246043.1
E2 3796631 3796743
refGene_gn=TTYH1;refGene_tx=XM_023246042.1;refGene_gn=TTYH1;refGene_tx
=XM_023246043.1
E2 3798642 3798821
refGene_gn=TTYH1;refGene_tx=XM_023246042.1;refGene_gn=TTYH1;refGene_tx
=XM_023246043.1
E2 3801418 3801544
refGene_gn=TTYH1;refGene_tx=XM_023246042.1;refGene_gn=TTYH1;refGene_tx
=XM_023246043.1
E2 3839008 3839042
refGene_gn=LAIR1;refGene_tx=XM_006940880.3;refGene_gn=LAIR1;refGene_tx
=XM_019819146.2
E2 3839419 3839455
refGene_gn=LAIR1;refGene_tx=XM_006940880.3;refGene_gn=LAIR1;refGene_tx
=XM_019819146.2
E2 3841962 3842244
refGene_gn=LAIR1;refGene_tx=XM_006940880.3;refGene_gn=LAIR1;refGene_tx
=XM_019819146.2
E2 3842402 3842453 refGene_gn=LAIR1;refGene_tx=XM_006940880.3
E2 3844633 3844672
refGene_gn=LAIR1;refGene_tx=XM_006940880.3;refGene_gn=LAIR1;refGene_tx
=XM_019819146.2
E2 3844961 3845072
refGene_gn=LAIR1;refGene_tx=XM_006940880.3;refGene_gn=LAIR1;refGene_tx
=XM_019819146.2
E2 3845169 3845212
refGene_gn=LAIR1;refGene_tx=XM_006940880.3;refGene_gn=LAIR1;refGene_tx
=XM_019819146.2
E2 3845299 3845337
refGene_gn=LAIR1;refGene_tx=XM_006940880.3;refGene_gn=LAIR1;refGene_tx
=XM_019819146.2
E2 3845503 3845557
refGene_gn=LAIR1;refGene_tx=XM_006940880.3;refGene_gn=LAIR1;refGene_tx
=XM_019819146.2

E2 3846087 3846200
refGene_gn=LAIR1;refGene_tx=XM_006940880.3;refGene_gn=LAIR1;refGene_tx=XM_019819146.2
E2 3864549 3864583
refGene_gn=LOC101082108;refGene_tx=XM_006940890.4;refGene_gn=LOC101082108;refGene_tx=XM_006940891.4;refGene_gn=LOC101082108;refGene_tx=XM_019819153.2;refGene_gn=LOC101082108;refGene_tx=XM_023246045.1
E2 3864740 3864776
refGene_gn=LOC101082108;refGene_tx=XM_006940890.4;refGene_gn=LOC101082108;refGene_tx=XM_006940891.4;refGene_gn=LOC101082108;refGene_tx=XM_019819153.2;refGene_gn=LOC101082108;refGene_tx=XM_023246045.1
E2 3864957 3865242
refGene_gn=LOC101082108;refGene_tx=XM_006940890.4;refGene_gn=LOC101082108;refGene_tx=XM_006940891.4;refGene_gn=LOC101082108;refGene_tx=XM_019819153.2;refGene_gn=LOC101082108;refGene_tx=XM_023246045.1
E2 3865392 3865695
refGene_gn=LOC101082108;refGene_tx=XM_006940890.4;refGene_gn=LOC101082108;refGene_tx=XM_006940891.4;refGene_gn=LOC101082108;refGene_tx=XM_019819153.2;refGene_gn=LOC101082108;refGene_tx=XM_023246045.1
E2 3865997 3866048
refGene_gn=LOC101082108;refGene_tx=XM_006940890.4;refGene_gn=LOC101082108;refGene_tx=XM_019819153.2;refGene_gn=LOC101082108;refGene_tx=XM_023246045.1
E2 3867430 3867550
refGene_gn=LOC101082108;refGene_tx=XM_006940890.4;refGene_gn=LOC101082108;refGene_tx=XM_006940891.4;refGene_gn=LOC101082108;refGene_tx=XM_019819153.2;refGene_gn=LOC101082108;refGene_tx=XM_023246045.1
E2 3869683 3869726
refGene_gn=LOC101082108;refGene_tx=XM_006940890.4;refGene_gn=LOC101082108;refGene_tx=XM_006940891.4;refGene_gn=LOC101082108;refGene_tx=XM_019819153.2;refGene_gn=LOC101082108;refGene_tx=XM_023246045.1
E2 3869817 3869855
refGene_gn=LOC101082108;refGene_tx=XM_006940890.4;refGene_gn=LOC101082108;refGene_tx=XM_006940891.4;refGene_gn=LOC101082108;refGene_tx=XM_019819153.2;refGene_gn=LOC101082108;refGene_tx=XM_023246045.1
E2 3870060 3870113
refGene_gn=LOC101082108;refGene_tx=XM_006940890.4;refGene_gn=LOC101082108;refGene_tx=XM_006940891.4;refGene_gn=LOC101082108;refGene_tx=XM_019819153.2
E2 3870998 3871196
refGene_gn=LOC101082108;refGene_tx=XM_006940890.4;refGene_gn=LOC101082108;refGene_tx=XM_006940891.4;refGene_gn=LOC101082108;refGene_tx=XM_019819153.2;refGene_gn=LOC101082108;refGene_tx=XM_023246045.1
E2 3875140 3875174
refGene_gn=LOC102899310;refGene_tx=XM_006940887.3;refGene_gn=LOC102899310;refGene_tx=XM_006940889.3;refGene_gn=LOC102899310;refGene_tx=XM_019819155.2;refGene_gn=LOC102899310;refGene_tx=XM_019819156.2;refGene_gn=LOC102899310;refGene_tx=XM_019819157.2;refGene_gn=LOC102899310;refGene_tx=XM_019819158.2;refGene_gn=LOC102899310;refGene_tx=XM_019819159.2;refGene_gn=LOC102899310;refGene_tx=XM_023246046.1

E2 3875333 3875369
refGene_gn=LOC102899310;refGene_tx=XM_006940887.3;refGene_gn=LOC102899310;refGene_tx=XM_006940889.3;refGene_gn=LOC102899310;refGene_tx=XM_019819155.2;refGene_gn=LOC102899310;refGene_tx=XM_019819156.2;refGene_gn=LOC102899310;refGene_tx=XM_019819157.2;refGene_gn=LOC102899310;refGene_tx=XM_019819158.2;refGene_gn=LOC102899310;refGene_tx=XM_019819159.2;refGene_gn=LOC102899310;refGene_tx=XM_023246046.1
E2 3875550 3875835
refGene_gn=LOC102899310;refGene_tx=XM_006940887.3;refGene_gn=LOC102899310;refGene_tx=XM_006940889.3;refGene_gn=LOC102899310;refGene_tx=XM_019819155.2;refGene_gn=LOC102899310;refGene_tx=XM_019819156.2;refGene_gn=LOC102899310;refGene_tx=XM_019819157.2;refGene_gn=LOC102899310;refGene_tx=XM_023246046.1
E2 3875981 3876284
refGene_gn=LOC102899310;refGene_tx=XM_006940887.3;refGene_gn=LOC102899310;refGene_tx=XM_006940889.3;refGene_gn=LOC102899310;refGene_tx=XM_019819155.2;refGene_gn=LOC102899310;refGene_tx=XM_019819156.2;refGene_gn=LOC102899310;refGene_tx=XM_019819157.2;refGene_gn=LOC102899310;refGene_tx=XM_019819158.2;refGene_gn=LOC102899310;refGene_tx=XM_019819159.2;refGene_gn=LOC102899310;refGene_tx=XM_023246046.1
E2 3876878 3876998
refGene_gn=LOC102899310;refGene_tx=XM_019819155.2;refGene_gn=LOC102899310;refGene_tx=XM_019819156.2;refGene_gn=LOC102899310;refGene_tx=XM_019819157.2;refGene_gn=LOC102899310;refGene_tx=XM_019819158.2;refGene_gn=LOC102899310;refGene_tx=XM_019819159.2;refGene_gn=LOC102899310;refGene_tx=XM_023246046.1
E2 3880508 3880559
refGene_gn=LOC102899310;refGene_tx=XM_006940887.3;refGene_gn=LOC102899310;refGene_tx=XM_006940889.3;refGene_gn=LOC102899310;refGene_tx=XM_019819155.2;refGene_gn=LOC102899310;refGene_tx=XM_019819156.2;refGene_gn=LOC102899310;refGene_tx=XM_019819157.2;refGene_gn=LOC102899310;refGene_tx=XM_019819158.2;refGene_gn=LOC102899310;refGene_tx=XM_019819159.2;refGene_gn=LOC102899310;refGene_tx=XM_023246046.1
E2 3880705 3880991
refGene_gn=LOC102899310;refGene_tx=XM_006940887.3;refGene_gn=LOC102899310;refGene_tx=XM_006940889.3;refGene_gn=LOC102899310;refGene_tx=XM_019819155.2;refGene_gn=LOC102899310;refGene_tx=XM_019819156.2;refGene_gn=LOC102899310;refGene_tx=XM_019819157.2;refGene_gn=LOC102899310;refGene_tx=XM_019819158.2;refGene_gn=LOC102899310;refGene_tx=XM_019819159.2;refGene_gn=LOC102899310;refGene_tx=XM_023246046.1
E2 3883943 3883992
refGene_gn=LOC111558116;refGene_tx=XM_023246044.1
E2 3884197 3884233
refGene_gn=LOC111558116;refGene_tx=XM_023246044.1
E2 3884399 3884684
refGene_gn=LOC111558116;refGene_tx=XM_023246044.1
E2 3884829 3885041
refGene_gn=LOC111558116;refGene_tx=XM_023246044.1
E2 3885042 3885130
refGene_gn=LOC111558116;refGene_tx=XM_023246044.1

E2 3885367 3885649
refGene_gn=LOC111558116;refGene_tx=XM_023246044.1
E2 3885953 3886253
refGene_gn=LOC111558116;refGene_tx=XM_023246044.1
E2 3891356 3891493
refGene_gn=LOC111558116;refGene_tx=XM_023246044.1
E2 3906947 3906981
refGene_gn=LOC102899271;refGene_tx=XM_023245728.1;refGene_gn=LOC102899271;refGene_tx=XM_023245729.1
E2 3907138 3907174
refGene_gn=LOC102899271;refGene_tx=XM_023245728.1;refGene_gn=LOC102899271;refGene_tx=XM_023245729.1
E2 3907356 3907641
refGene_gn=LOC102899271;refGene_tx=XM_023245728.1;refGene_gn=LOC102899271;refGene_tx=XM_023245729.1
E2 3907790 3908102
refGene_gn=LOC102899271;refGene_tx=XM_023245728.1;refGene_gn=LOC102899271;refGene_tx=XM_023245729.1
E2 3908327 3908615
refGene_gn=LOC102899271;refGene_tx=XM_023245729.1
E2 3908833 3909136
refGene_gn=LOC102899271;refGene_tx=XM_023245729.1
E2 3909448 3909499
refGene_gn=LOC102899271;refGene_tx=XM_023245728.1;refGene_gn=LOC102899271;refGene_tx=XM_023245729.1
E2 3910410 3910527
refGene_gn=LOC102899271;refGene_tx=XM_023245728.1;refGene_gn=LOC102899271;refGene_tx=XM_023245729.1
E2 3910855 3910898
refGene_gn=LOC102899271;refGene_tx=XM_023245728.1;refGene_gn=LOC102899271;refGene_tx=XM_023245729.1
E2 3910975 3911013
refGene_gn=LOC102899271;refGene_tx=XM_023245728.1;refGene_gn=LOC102899271;refGene_tx=XM_023245729.1
E2 3911194 3911250
refGene_gn=LOC102899271;refGene_tx=XM_023245728.1;refGene_gn=LOC102899271;refGene_tx=XM_023245729.1
E2 3912017 3912173
refGene_gn=LOC102899271;refGene_tx=XM_023245728.1;refGene_gn=LOC102899271;refGene_tx=XM_023245729.1
E2 3912251 3912398
refGene_gn=LOC102899271;refGene_tx=XM_023245728.1;refGene_gn=LOC102899271;refGene_tx=XM_023245729.1
E2 3916375 3916472 refGene_gn=RPS9;refGene_tx=XM_006940885.3
E2 3916700 3916823 refGene_gn=RPS9;refGene_tx=XM_006940885.3
E2 3921306 3921493 refGene_gn=RPS9;refGene_tx=XM_006940885.3
E2 3922348 3922526 refGene_gn=RPS9;refGene_tx=XM_006940885.3
E2 3923418 3923607
refGene_gn=LOC101082866;refGene_tx=XM_023245879.1
E2 3924316 3924369

refGene_gn=LOC101082866;refGene_tx=XM_023245879.1
E2 3924587 3924628
refGene_gn=LOC101082866;refGene_tx=XM_023245879.1
E2 3924706 3924755
refGene_gn=LOC101082866;refGene_tx=XM_023245879.1
E2 3925140 3925257
refGene_gn=LOC101082866;refGene_tx=XM_023245879.1
E2 3926207 3926243
refGene_gn=LOC101082866;refGene_tx=XM_023245879.1
E2 3926734 3927034
refGene_gn=LOC101082866;refGene_tx=XM_023245879.1
E2 3927333 3927615
refGene_gn=LOC101082866;refGene_tx=XM_023245879.1
E2 3927852 3928063
refGene_gn=LOC101082866;refGene_tx=XM_023245879.1
E2 3928065 3928153
refGene_gn=LOC101082866;refGene_tx=XM_023245879.1
E2 3928303 3928580
refGene_gn=LOC101082866;refGene_tx=XM_023245879.1
E2 3928987 3929036
refGene_gn=LOC101082866;refGene_tx=XM_023245879.1
E2 3950997 3951185
refGene_gn=TSEN34;refGene_tx=XM_003997444.5;refGene_gn=TSEN34;refGene_tx=XM_006940883.3;refGene_gn=TSEN34;refGene_tx=XM_023245730.1
E2 3954193 3954451
refGene_gn=TSEN34;refGene_tx=XM_003997444.5;refGene_gn=TSEN34;refGene_tx=XM_006940883.3;refGene_gn=TSEN34;refGene_tx=XM_023245730.1
E2 3954601 3954863
refGene_gn=TSEN34;refGene_tx=XM_003997444.5;refGene_gn=TSEN34;refGene_tx=XM_006940883.3;refGene_gn=TSEN34;refGene_tx=XM_023245730.1
E2 3955113 3955356
refGene_gn=TSEN34;refGene_tx=XM_003997444.5;refGene_gn=TSEN34;refGene_tx=XM_006940883.3;refGene_gn=TSEN34;refGene_tx=XM_023245730.1
E2 3957730 3957806 refGene_gn=MBOAT7;refGene_tx=XM_023245727.1
E2 3957908 3958038
refGene_gn=MBOAT7;refGene_tx=XM_011289620.3;refGene_gn=MBOAT7;refGene_tx=XM_023245727.1
E2 3958599 3958726 refGene_gn=MBOAT7;refGene_tx=XM_023245727.1
E2 3960585 3960745
refGene_gn=MBOAT7;refGene_tx=XM_011289620.3;refGene_gn=MBOAT7;refGene_tx=XM_023245727.1
E2 3962764 3963125
refGene_gn=MBOAT7;refGene_tx=XM_011289620.3;refGene_gn=MBOAT7;refGene_tx=XM_023245727.1
E2 3964798 3964975
refGene_gn=MBOAT7;refGene_tx=XM_011289620.3;refGene_gn=MBOAT7;refGene_tx=XM_023245727.1
E2 3968460 3968848
refGene_gn=MBOAT7;refGene_tx=XM_011289620.3;refGene_gn=MBOAT7;refGene_tx=XM_023245727.1

E2	3969717	3969796	refGene_gn=TMC4;refGene_tx=XM_011289615.3
E2	3970361	3970575	refGene_gn=TMC4;refGene_tx=XM_011289615.3
E2	3971791	3971937	refGene_gn=TMC4;refGene_tx=XM_011289615.3
E2	3972639	3972872	refGene_gn=TMC4;refGene_tx=XM_011289615.3
E2	3972973	3973095	refGene_gn=TMC4;refGene_tx=XM_011289615.3
E2	3976924	3977072	refGene_gn=TMC4;refGene_tx=XM_011289615.3
E2	3977569	3977737	refGene_gn=TMC4;refGene_tx=XM_011289615.3
E2	3977953	3978117	refGene_gn=TMC4;refGene_tx=XM_011289615.3
E2	3978384	3978511	refGene_gn=TMC4;refGene_tx=XM_011289615.3
E2	3978767	3978865	refGene_gn=TMC4;refGene_tx=XM_011289615.3
E2	3979113	3979297	refGene_gn=TMC4;refGene_tx=XM_011289615.3
E2	3979584	3979715	refGene_gn=TMC4;refGene_tx=XM_011289615.3
E2	3979817	3979973	refGene_gn=TMC4;refGene_tx=XM_011289615.3
E2	3980256	3980335	refGene_gn=TMC4;refGene_tx=XM_011289615.3
E2	3980431	3980500	refGene_gn=TMC4;refGene_tx=XM_011289615.3
E2	3981079	3981211
refGene_gn=LENG1;refGene_tx=XM_003997538.5;refGene_gn=LENG1;refGene_tx
=XM_011289616.3;refGene_gn=LENG1;refGene_tx=XM_011289618.3
E2	3982033	3982213
refGene_gn=LENG1;refGene_tx=XM_003997538.5;refGene_gn=LENG1;refGene_tx
=XM_011289616.3;refGene_gn=LENG1;refGene_tx=XM_011289618.3
E2	3982883	3983143
refGene_gn=LENG1;refGene_tx=XM_003997538.5;refGene_gn=LENG1;refGene_tx
=XM_011289616.3;refGene_gn=LENG1;refGene_tx=XM_011289618.3
E2	3983732	3984018
refGene_gn=LENG1;refGene_tx=XM_003997538.5;refGene_gn=LENG1;refGene_tx
=XM_011289616.3;refGene_gn=LENG1;refGene_tx=XM_011289618.3
E2	3984226	3984325
refGene_gn=CNOT3;refGene_tx=XM_003997441.5;refGene_gn=CNOT3;refGene_tx
=XM_006940882.4;refGene_gn=CNOT3;refGene_tx=XM_019819147.2
E2	3985691	3985817
refGene_gn=CNOT3;refGene_tx=XM_003997441.5;refGene_gn=CNOT3;refGene_tx
=XM_006940882.4;refGene_gn=CNOT3;refGene_tx=XM_019819147.2
E2	3986394	3986527
refGene_gn=CNOT3;refGene_tx=XM_003997441.5;refGene_gn=CNOT3;refGene_tx
=XM_006940882.4;refGene_gn=CNOT3;refGene_tx=XM_019819147.2
E2	3986790	3986989
refGene_gn=CNOT3;refGene_tx=XM_003997441.5;refGene_gn=CNOT3;refGene_tx
=XM_006940882.4;refGene_gn=CNOT3;refGene_tx=XM_019819147.2
E2	3987078	3987178
refGene_gn=CNOT3;refGene_tx=XM_003997441.5;refGene_gn=CNOT3;refGene_tx
=XM_006940882.4;refGene_gn=CNOT3;refGene_tx=XM_019819147.2
E2	3988682	3988881
refGene_gn=CNOT3;refGene_tx=XM_003997441.5;refGene_gn=CNOT3;refGene_tx
=XM_006940882.4;refGene_gn=CNOT3;refGene_tx=XM_019819147.2
E2	3989540	3989664
refGene_gn=CNOT3;refGene_tx=XM_003997441.5;refGene_gn=CNOT3;refGene_tx
=XM_006940882.4;refGene_gn=CNOT3;refGene_tx=XM_019819147.2
E2	3989740	3990107
refGene_gn=CNOT3;refGene_tx=XM_003997441.5;refGene_gn=CNOT3;refGene_tx

=XM_006940882.4;refGene_gn=CNOT3;refGene_tx=XM_019819147.2
E2 3991173 3991230
refGene_gn=CNOT3;refGene_tx=XM_003997441.5;refGene_gn=CNOT3;refGene_tx=XM_006940882.4;refGene_gn=CNOT3;refGene_tx=XM_019819147.2
E2 3991539 3991673
refGene_gn=CNOT3;refGene_tx=XM_003997441.5;refGene_gn=CNOT3;refGene_tx=XM_006940882.4;refGene_gn=CNOT3;refGene_tx=XM_019819147.2
E2 3991767 3991990
refGene_gn=CNOT3;refGene_tx=XM_003997441.5;refGene_gn=CNOT3;refGene_tx=XM_006940882.4;refGene_gn=CNOT3;refGene_tx=XM_019819147.2
E2 3992957 3993053
refGene_gn=CNOT3;refGene_tx=XM_003997441.5;refGene_gn=CNOT3;refGene_tx=XM_006940882.4;refGene_gn=CNOT3;refGene_tx=XM_019819147.2
E2 3993152 3993281
refGene_gn=CNOT3;refGene_tx=XM_003997441.5;refGene_gn=CNOT3;refGene_tx=XM_006940882.4;refGene_gn=CNOT3;refGene_tx=XM_019819147.2
E2 3993403 3993493
refGene_gn=CNOT3;refGene_tx=XM_003997441.5;refGene_gn=CNOT3;refGene_tx=XM_006940882.4;refGene_gn=CNOT3;refGene_tx=XM_019819147.2
E2 3993616 3993691
refGene_gn=CNOT3;refGene_tx=XM_003997441.5;refGene_gn=CNOT3;refGene_tx=XM_006940882.4;refGene_gn=CNOT3;refGene_tx=XM_019819147.2
E2 3993923 3993991
refGene_gn=CNOT3;refGene_tx=XM_003997441.5;refGene_gn=CNOT3;refGene_tx=XM_006940882.4;refGene_gn=CNOT3;refGene_tx=XM_019819147.2
E2 3994104 3994129
refGene_gn=CNOT3;refGene_tx=XM_003997441.5;refGene_gn=CNOT3;refGene_tx=XM_006940882.4;refGene_gn=CNOT3;refGene_tx=XM_019819147.2
E2 4001815 4001941
refGene_gn=PRPF31;refGene_tx=XM_003997437.5;refGene_gn=PRPF31;refGene_tx=XM_019819150.2
E2 4003023 4003122
refGene_gn=PRPF31;refGene_tx=XM_003997437.5;refGene_gn=PRPF31;refGene_tx=XM_019819150.2
E2 4003217 4003346
refGene_gn=PRPF31;refGene_tx=XM_003997437.5;refGene_gn=PRPF31;refGene_tx=XM_019819150.2
E2 4003815 4003888
refGene_gn=PRPF31;refGene_tx=XM_003997437.5;refGene_gn=PRPF31;refGene_tx=XM_019819150.2
E2 4003968 4004096
refGene_gn=PRPF31;refGene_tx=XM_003997437.5;refGene_gn=PRPF31;refGene_tx=XM_019819150.2
E2 4005431 4005521
refGene_gn=PRPF31;refGene_tx=XM_003997437.5;refGene_gn=PRPF31;refGene_tx=XM_019819150.2
E2 4007006 4007164
refGene_gn=PRPF31;refGene_tx=XM_003997437.5;refGene_gn=PRPF31;refGene_tx=XM_019819150.2
E2 4009000 4009170

refGene_gn=PRPF31;refGene_tx=XM_003997437.5;refGene_gn=PRPF31;refGene_tx=XM_019819150.2
E2 4010967 4011074
refGene_gn=PRPF31;refGene_tx=XM_003997437.5;refGene_gn=PRPF31;refGene_tx=XM_019819150.2
E2 4011983 4012081
refGene_gn=PRPF31;refGene_tx=XM_003997437.5;refGene_gn=PRPF31;refGene_tx=XM_019819150.2
E2 4012676 4012760
refGene_gn=PRPF31;refGene_tx=XM_003997437.5;refGene_gn=PRPF31;refGene_tx=XM_019819150.2
E2 4015617 4015678
refGene_gn=PRPF31;refGene_tx=XM_003997437.5;refGene_gn=PRPF31;refGene_tx=XM_019819150.2
E2 4015780 4015957
refGene_gn=PRPF31;refGene_tx=XM_003997437.5;refGene_gn=PRPF31;refGene_tx=XM_019819150.2
E2 4017712 4017735 refGene_gn=TFPT;refGene_tx=XM_011289614.2
E2 4018141 4018400
refGene_gn=TFPT;refGene_tx=XM_011289614.2;refGene_gn=TFPT;refGene_tx=XM_019819162.2
E2 4020930 4021001
refGene_gn=TFPT;refGene_tx=XM_011289614.2;refGene_gn=TFPT;refGene_tx=XM_019819162.2
E2 4023844 4023914
refGene_gn=TFPT;refGene_tx=XM_011289614.2;refGene_gn=TFPT;refGene_tx=XM_019819162.2
E2 4023993 4024212
refGene_gn=TFPT;refGene_tx=XM_011289614.2;refGene_gn=TFPT;refGene_tx=XM_019819162.2
E2 4024657 4024777
refGene_gn=TFPT;refGene_tx=XM_011289614.2;refGene_gn=TFPT;refGene_tx=XM_019819162.2
E2 4024929 4025021
refGene_gn=NDUFA3;refGene_tx=XM_003997445.3;refGene_gn=NDUFA3;refGene_tx=XM_011289613.2
E2 4025465 4025543
refGene_gn=NDUFA3;refGene_tx=XM_003997445.3;refGene_gn=NDUFA3;refGene_tx=XM_011289613.2
E2 4026828 4026930
refGene_gn=NDUFA3;refGene_tx=XM_003997445.3;refGene_gn=NDUFA3;refGene_tx=XM_011289613.2
E2 4027125 4027228
refGene_gn=NDUFA3;refGene_tx=XM_003997445.3;refGene_gn=NDUFA3;refGene_tx=XM_011289613.2
E2 4028213 4028250 refGene_gn=OSCAR;refGene_tx=XM_023245734.1
E2 4028819 4028835 refGene_gn=OSCAR;refGene_tx=XM_023245735.1
E2 4028954 4028987
refGene_gn=OSCAR;refGene_tx=XM_023245731.1;refGene_gn=OSCAR;refGene_tx=XM_023245732.1;refGene_gn=OSCAR;refGene_tx=XM_023245733.1;refGene_gn=

OSCAR;refGene_tx=XM_023245734.1;refGene_gn=OSCAR;refGene_tx=XM_0232457
35.1
E2 4032825 4033234
refGene_gn=OSCAR;refGene_tx=XM_023245731.1;refGene_gn=OSCAR;refGene_tx
=XM_023245732.1;refGene_gn=OSCAR;refGene_tx=XM_023245733.1;refGene_gn=
OSCAR;refGene_tx=XM_023245734.1;refGene_gn=OSCAR;refGene_tx=XM_0232457
35.1
E2 4034232 4034514
refGene_gn=OSCAR;refGene_tx=XM_023245731.1;refGene_gn=OSCAR;refGene_tx
=XM_023245732.1;refGene_gn=OSCAR;refGene_tx=XM_023245733.1;refGene_gn=
OSCAR;refGene_tx=XM_023245734.1;refGene_gn=OSCAR;refGene_tx=XM_0232457
35.1
E2 4035420 4035557
refGene_gn=OSCAR;refGene_tx=XM_023245731.1;refGene_gn=OSCAR;refGene_tx
=XM_023245732.1;refGene_gn=OSCAR;refGene_tx=XM_023245733.1;refGene_gn=
OSCAR;refGene_tx=XM_023245734.1;refGene_gn=OSCAR;refGene_tx=XM_0232457
35.1
E2 4047936 4048233
refGene_gn=LOC101083635;refGene_tx=XM_019818860.2
E2 4048234 4048430
refGene_gn=LOC101083635;refGene_tx=XM_019818860.2
E2 4048683 4049120
refGene_gn=LOC101083635;refGene_tx=XM_019818860.2
E2 4065333 4065367
refGene_gn=LOC101083890;refGene_tx=XM_006941038.3
E2 4070985 4071021
refGene_gn=LOC101083890;refGene_tx=XM_006941038.3
E2 4071152 4071443
refGene_gn=LOC101083890;refGene_tx=XM_006941038.3
E2 4071696 4071993
refGene_gn=LOC101083890;refGene_tx=XM_006941038.3
E2 4073327 4073470
refGene_gn=LOC101083890;refGene_tx=XM_006941038.3
E2 4098585 4098728
refGene_gn=LOC101084150;refGene_tx=XM_023245880.1
E2 4100063 4100349
refGene_gn=LOC101084150;refGene_tx=XM_023245880.1
E2 4109645 4109679
refGene_gn=LOC101084405;refGene_tx=XM_006940893.4;refGene_gn=LOC101084
405;refGene_tx=XM_019819166.2;refGene_gn=LOC101084405;refGene_tx=XM_01
9819167.2
E2 4114992 4115439
refGene_gn=LOC101084405;refGene_tx=XM_006940893.4;refGene_gn=LOC101084
405;refGene_tx=XM_019819166.2;refGene_gn=LOC101084405;refGene_tx=XM_01
9819167.2
E2 4118625 4118664
refGene_gn=LOC101084405;refGene_tx=XM_006940893.4;refGene_gn=LOC101084
405;refGene_tx=XM_019819166.2;refGene_gn=LOC101084405;refGene_tx=XM_01
9819167.2
E2 4123275 4123368

refGene_gn=LOC101084405;refGene_tx=XM_006940893.4;refGene_gn=LOC101084405;refGene_tx=XM_019819166.2;refGene_gn=LOC101084405;refGene_tx=XM_019819167.2
E2 4123504 4123532
refGene_gn=LOC101084405;refGene_tx=XM_006940893.4;refGene_gn=LOC101084405;refGene_tx=XM_019819166.2;refGene_gn=LOC101084405;refGene_tx=XM_019819167.2
E2 4123704 4123750
refGene_gn=LOC101084405;refGene_tx=XM_006940893.4;refGene_gn=LOC101084405;refGene_tx=XM_019819166.2;refGene_gn=LOC101084405;refGene_tx=XM_019819167.2
E2 4126247 4126252
refGene_gn=LOC101084405;refGene_tx=XM_019819167.2
E2 4144820 4144960 refGene_gn=CACNG6;refGene_tx=XM_011289609.3
E2 4149306 4149545 refGene_gn=CACNG6;refGene_tx=XM_011289608.2
E2 4160006 4160144
refGene_gn=CACNG6;refGene_tx=XM_011289608.2;refGene_gn=CACNG6;refGene_tx=XM_011289609.3
E2 4161921 4161996
refGene_gn=CACNG6;refGene_tx=XM_011289608.2;refGene_gn=CACNG6;refGene_tx=XM_011289609.3
E2 4166343 4166680
refGene_gn=CACNG6;refGene_tx=XM_011289608.2;refGene_gn=CACNG6;refGene_tx=XM_011289609.3
E2 4174219 4174983 refGene_gn=CACNG8;refGene_tx=XM_023245883.1
E2 4177230 4177371 refGene_gn=CACNG8;refGene_tx=XM_023245883.1
E2 4178405 4178489 refGene_gn=CACNG8;refGene_tx=XM_023245883.1
E2 4187616 4188089 refGene_gn=CACNG8;refGene_tx=XM_023245883.1
E2 4188178 4188213 refGene_gn=CACNG8;refGene_tx=XM_023245883.1
E2 4202872 4203130 refGene_gn=CACNG7;refGene_tx=XM_003997446.5
E2 4203433 4203579 refGene_gn=CACNG7;refGene_tx=XM_003997446.5
E2 4218721 4218862 refGene_gn=CACNG7;refGene_tx=XM_003997446.5
E2 4219249 4219336 refGene_gn=CACNG7;refGene_tx=XM_003997446.5
E2 4219655 4219851 refGene_gn=CACNG7;refGene_tx=XM_003997446.5
E2 4225318 4225507 refGene_gn=PRKCG;refGene_tx=XM_006940894.4
E2 4225737 4225878 refGene_gn=PRKCG;refGene_tx=XM_006940894.4
E2 4226829 4226937 refGene_gn=PRKCG;refGene_tx=XM_006940894.4
E2 4228513 4228594 refGene_gn=PRKCG;refGene_tx=XM_006940894.4
E2 4229850 4229989 refGene_gn=PRKCG;refGene_tx=XM_006940894.4
E2 4230445 4230508 refGene_gn=PRKCG;refGene_tx=XM_006940894.4
E2 4230588 4230680 refGene_gn=PRKCG;refGene_tx=XM_006940894.4
E2 4231810 4231999 refGene_gn=PRKCG;refGene_tx=XM_006940894.4
E2 4232317 4232470 refGene_gn=PRKCG;refGene_tx=XM_006940894.4
E2 4233885 4233915 refGene_gn=PRKCG;refGene_tx=XM_006940894.4
E2 4234215 4234303 refGene_gn=PRKCG;refGene_tx=XM_006940894.4
E2 4234602 4234737 refGene_gn=PRKCG;refGene_tx=XM_006940894.4
E2 4235416 4235573 refGene_gn=PRKCG;refGene_tx=XM_006940894.4
E2 4236557 4236689 refGene_gn=PRKCG;refGene_tx=XM_006940894.4
E2 4236856 4236968 refGene_gn=PRKCG;refGene_tx=XM_006940894.4
E2 4241162 4241245 refGene_gn=PRKCG;refGene_tx=XM_006940894.4

E2 4242205 4242237 refGene_gn=PRKCG;refGene_tx=XM_006940894.4
E2 4242741 4242911 refGene_gn=PRKCG;refGene_tx=XM_006940894.4
E2 4247473 4248442
refGene_gn=MYADM;refGene_tx=XM_003997448.5;refGene_gn=MYADM;refGene_tx=XM_006940895.4
E2 4295833 4296137 refGene_gn=NLRP12;refGene_tx=XM_023245577.1
E2 4304793 4304811 refGene_gn=NLRP12;refGene_tx=XM_023245577.1
E2 4305455 4306876 refGene_gn=NLRP12;refGene_tx=XM_023245577.1
E2 4306877 4307206 refGene_gn=NLRP12;refGene_tx=XM_023245577.1
E2 4312940 4313111 refGene_gn=NLRP12;refGene_tx=XM_023245577.1
E2 4314615 4314786 refGene_gn=NLRP12;refGene_tx=XM_023245577.1
E2 4317151 4317322 refGene_gn=NLRP12;refGene_tx=XM_023245577.1
E2 4320002 4320173 refGene_gn=NLRP12;refGene_tx=XM_023245577.1
E2 4324743 4324911 refGene_gn=NLRP12;refGene_tx=XM_023245577.1
E2 4326599 4326770 refGene_gn=NLRP12;refGene_tx=XM_023245577.1
E2 4327214 4327302 refGene_gn=NLRP12;refGene_tx=XM_023245577.1
E2 4371442 4371461
refGene_gn=LOC105261224;refGene_tx=XM_011289644.2
E2 4372780 4372932
refGene_gn=LOC105261224;refGene_tx=XM_011289644.2
E2 4374167 4374323
refGene_gn=LOC105261224;refGene_tx=XM_011289644.2
E2 4393693 4394083 refGene_gn=DPRX;refGene_tx=XM_023245884.1
E2 4395174 4395329 refGene_gn=DPRX;refGene_tx=XM_023245884.1
E2 4397079 4397119 refGene_gn=DPRX;refGene_tx=XM_023245884.1
E2 4420290 4421627
refGene_gn=ZNF331;refGene_tx=XM_006940899.4;refGene_gn=ZNF331;refGene_tx=XM_011289604.3;refGene_gn=ZNF331;refGene_tx=XM_011289605.3;refGene_gn=ZNF331;refGene_tx=XM_011289606.3
E2 4424230 4424357
refGene_gn=ZNF331;refGene_tx=XM_006940899.4;refGene_gn=ZNF331;refGene_tx=XM_011289604.3;refGene_gn=ZNF331;refGene_tx=XM_011289605.3;refGene_gn=ZNF331;refGene_tx=XM_011289606.3
E2 4425554 4425563
refGene_gn=ZNF331;refGene_tx=XM_006940899.4;refGene_gn=ZNF331;refGene_tx=XM_011289604.3;refGene_gn=ZNF331;refGene_tx=XM_011289605.3;refGene_gn=ZNF331;refGene_tx=XM_011289606.3
E2 4470047 4470992
refGene_gn=FELCATV1R17;refGene_tx=NM_001167489.1
E2 4594606 4594621
refGene_gn=LOC101085936;refGene_tx=XM_019819176.2
E2 4597913 4598040
refGene_gn=LOC101085936;refGene_tx=XM_019819176.2
E2 4598522 4598618
refGene_gn=LOC101085936;refGene_tx=XM_019819176.2
E2 4613371 4615344
refGene_gn=LOC101085936;refGene_tx=XM_019819176.2
E2 4769064 4769413
refGene_gn=LOC111557977;refGene_tx=XM_023244887.1
E2 4769415 4769786

refGene_gn=LOC111557977;refGene_tx=XM_023244887.1
E2 4778627 4778737
refGene_gn=LOC105261214;refGene_tx=XM_023245885.1
E2 4779239 4779384
refGene_gn=LOC105261214;refGene_tx=XM_023245885.1
E2 4790475 4790502
refGene_gn=LOC105261213;refGene_tx=XM_023244882.1
E2 4803180 4805114
refGene_gn=LOC105261213;refGene_tx=XM_023244882.1
E2 4853992 4854102
refGene_gn=LOC105261223;refGene_tx=XM_023245886.1
E2 4854599 4854674
refGene_gn=LOC105261223;refGene_tx=XM_023245886.1
E2 4854676 4854725
refGene_gn=LOC105261223;refGene_tx=XM_023245886.1
E2 4857764 4857779
refGene_gn=LOC105261223;refGene_tx=XM_023245886.1
E2 4865927 4865942
refGene_gn=LOC102901312;refGene_tx=XM_023245887.1
E2 4870315 4870442
refGene_gn=LOC102901312;refGene_tx=XM_023245887.1
E2 4870638 4870642
refGene_gn=LOC102901312;refGene_tx=XM_023245887.1
E2 4882411 4882477
refGene_gn=LOC102901312;refGene_tx=XM_023245887.1
E2 4882478 4882631
refGene_gn=LOC102901312;refGene_tx=XM_023245887.1
E2 4882632 4882798
refGene_gn=LOC102901312;refGene_tx=XM_023245887.1
E2 4882799 4883615
refGene_gn=LOC102901312;refGene_tx=XM_023245887.1
E2 4883695 4883785
refGene_gn=LOC102901312;refGene_tx=XM_023245887.1
E2 4883786 4883884
refGene_gn=LOC102901312;refGene_tx=XM_023245887.1
E2 4898545 4898612
refGene_gn=LOC102901312;refGene_tx=XM_023245887.1
E2 4899837 4899952
refGene_gn=LOC102901312;refGene_tx=XM_023245887.1
E2 4903519 4903644
refGene_gn=LOC102901312;refGene_tx=XM_023245887.1
E2 4904147 4904245
refGene_gn=LOC102901312;refGene_tx=XM_023245887.1
E2 4904444 4904575
refGene_gn=LOC102901312;refGene_tx=XM_023245887.1
E2 4914535 4916358
refGene_gn=LOC102901312;refGene_tx=XM_023245887.1
E2 4916359 4916878
refGene_gn=LOC102901312;refGene_tx=XM_023245887.1
E2 4943827 4944013

refGene_gn=LOC101086687;refGene_tx=XM_023244880.1
E2	4944014	4946320
refGene_gn=LOC101086687;refGene_tx=XM_023244880.1
E2	4965145	4965238
refGene_gn=LOC101086687;refGene_tx=XM_023244880.1
E2	4965740	4965867
refGene_gn=LOC101086687;refGene_tx=XM_023244880.1
E2	4969168	4969183
refGene_gn=LOC101086687;refGene_tx=XM_023244880.1
E2	5027770	5029500
refGene_gn=LOC101087185;refGene_tx=XM_023244884.1
E2	5041044	5041140
refGene_gn=LOC101087185;refGene_tx=XM_023244884.1
E2	5041636	5041763
refGene_gn=LOC101087185;refGene_tx=XM_023244884.1
E2	5044909	5044924
refGene_gn=LOC101087185;refGene_tx=XM_023244884.1
E2	5114318	5114490
refGene_gn=LOC111558092;refGene_tx=XM_023245851.1
E2	5116015	5116411
refGene_gn=LOC111558092;refGene_tx=XM_023245851.1
E2	5116413	5117401
refGene_gn=LOC111558092;refGene_tx=XM_023245851.1
E2	5117427	5117565
refGene_gn=LOC111558092;refGene_tx=XM_023245851.1
E2	5144328	5144426
refGene_gn=LOC111558092;refGene_tx=XM_023245851.1
E2	5144567	5144927
refGene_gn=LOC111558092;refGene_tx=XM_023245851.1
E2	5146192	5146216
refGene_gn=LOC111558092;refGene_tx=XM_023245851.1
E2	5266416	5266588
refGene_gn=LOC105261231;refGene_tx=XM_023244883.1
E2	5266589	5268312
refGene_gn=LOC105261231;refGene_tx=XM_023244883.1
E2	5272814	5272941
refGene_gn=LOC105261231;refGene_tx=XM_023244883.1
E2	5278829	5278844
refGene_gn=LOC105261231;refGene_tx=XM_023244883.1
E2	5305489	5305762
refGene_gn=LOC102900382;refGene_tx=XM_023244881.1
E2	5305763	5307775
refGene_gn=LOC102900382;refGene_tx=XM_023244881.1
E2	5308638	5308707
refGene_gn=LOC102900382;refGene_tx=XM_023244881.1
E2	5312744	5312871
refGene_gn=LOC102900382;refGene_tx=XM_023244881.1
E2	5322616	5322631
refGene_gn=LOC102900382;refGene_tx=XM_023244881.1
E2	5339991	5340048

refGene_gn=LOC109494583;refGene_tx=XM_023245888.1
E2 5342932 5343237
refGene_gn=LOC109494583;refGene_tx=XM_023245888.1
E2 5343239 5344589
refGene_gn=LOC109494583;refGene_tx=XM_023245888.1
E2 5344591 5345245
refGene_gn=LOC109494583;refGene_tx=XM_023245888.1
E2 5350483 5350636
refGene_gn=LOC109494583;refGene_tx=XM_023245888.1
E2 5359145 5359206
refGene_gn=LOC109494583;refGene_tx=XM_023245888.1
E2 5423843 5424004
refGene_gn=LOC109494585;refGene_tx=XM_023245889.1
E2 5424869 5425047
refGene_gn=LOC109494585;refGene_tx=XM_023245889.1
E2 5425496 5425599
refGene_gn=LOC109494585;refGene_tx=XM_023245889.1
E2 5425991 5426260
refGene_gn=LOC109494585;refGene_tx=XM_023245889.1
E2 5427152 5427644
refGene_gn=LOC109494585;refGene_tx=XM_023245889.1
E2 5444070 5445003
refGene_gn=FELCATV1R6;refGene_tx=NM_001167478.1
E2 5463622 5464816
refGene_gn=LOC109494586;refGene_tx=XM_023244885.1
E2 5500538 5500553
refGene_gn=LOC101083469;refGene_tx=XM_019818494.2
E2 5504487 5504653
refGene_gn=LOC101083469;refGene_tx=XM_019818494.2
E2 5512002 5513813
refGene_gn=LOC101083469;refGene_tx=XM_019818494.2
E2 5513814 5514250
refGene_gn=LOC101083469;refGene_tx=XM_019818494.2
E2 5514252 5514480
refGene_gn=LOC101083469;refGene_tx=XM_019818494.2
E2 5514482 5515396
refGene_gn=LOC101083469;refGene_tx=XM_019818494.2
E2 5532271 5532349
refGene_gn=LOC111557978;refGene_tx=XM_023244888.1
E2 5533692 5533911
refGene_gn=LOC111557978;refGene_tx=XM_023244888.1
E2 5580628 5581549
refGene_gn=LOC111557976;refGene_tx=XM_023244886.1
E2 5617544 5617683
refGene_gn=LOC105261237;refGene_tx=XM_023245890.1
E2 5618184 5618277
refGene_gn=LOC105261237;refGene_tx=XM_023245890.1
E2 5671240 5671293
refGene_gn=LOC105261237;refGene_tx=XM_023245890.1
E2 5679196 5679546

refGene_gn=LOC105259655;refGene_tx=XM_023245891.1
E2 5679631 5679751
refGene_gn=LOC105259655;refGene_tx=XM_023245891.1
E2 5679825 5684628
refGene_gn=LOC105259655;refGene_tx=XM_023245891.1
E2 5698104 5698240
refGene_gn=LOC105259655;refGene_tx=XM_023245891.1
E2 5704641 5704775
refGene_gn=LOC105259655;refGene_tx=XM_023245891.1
E2 5728152 5728270
refGene_gn=LOC105259655;refGene_tx=XM_023245891.1
E2 5729149 5730425
refGene_gn=LOC111558098;refGene_tx=XM_023245892.1
E2 5730655 5730874
refGene_gn=LOC111558098;refGene_tx=XM_023245892.1
E2 5730906 5731404
refGene_gn=LOC111558098;refGene_tx=XM_023245892.1
E2 5731538 5732170
refGene_gn=LOC111558098;refGene_tx=XM_023245892.1
E2 5790494 5790509
refGene_gn=LOC101088445;refGene_tx=XM_019819183.2
E2 5795211 5795338
refGene_gn=LOC101088445;refGene_tx=XM_019819183.2
E2 5795835 5795931
refGene_gn=LOC101088445;refGene_tx=XM_019819183.2
E2 5808018 5810003
refGene_gn=LOC101088445;refGene_tx=XM_019819183.2;refGene_gn=LOC101088445;refGene_tx=XM_023244964.1
E2 5831570 5832422
refGene_gn=LOC111557987;refGene_tx=XM_023244983.1
E2 5873458 5873473
refGene_gn=LOC101087952;refGene_tx=XM_023244965.1
E2 5878517 5878644
refGene_gn=LOC101087952;refGene_tx=XM_023244965.1
E2 5879135 5879231
refGene_gn=LOC101087952;refGene_tx=XM_023244965.1
E2 5899543 5901336
refGene_gn=LOC101087952;refGene_tx=XM_023244965.1
E2 5910546 5910573
refGene_gn=LOC105259643;refGene_tx=XM_019818864.2
E2 5916174 5916301
refGene_gn=LOC105259643;refGene_tx=XM_019818864.2
E2 5916783 5916890
refGene_gn=LOC105259643;refGene_tx=XM_019818864.2
E2 5963024 5963118
refGene_gn=LOC102901357;refGene_tx=XM_023244975.1
E2 5966814 5968334
refGene_gn=LOC102901357;refGene_tx=XM_023244975.1
E2 5971193 5971223
refGene_gn=LOC111558099;refGene_tx=XM_023245893.1

E2 5972585 5972712
refGene_gn=LOC111558099;refGene_tx=XM_023245893.1
E2 5973630 5973737
refGene_gn=LOC111558099;refGene_tx=XM_023245893.1
E2 6032888 6032995
refGene_gn=LOC105261241;refGene_tx=XM_023245894.1
E2 6033500 6033627
refGene_gn=LOC105261241;refGene_tx=XM_023245894.1
E2 6033747 6033768
refGene_gn=LOC105261241;refGene_tx=XM_023245894.1
E2 6099102 6099753
refGene_gn=CRX;refGene_tx=NM_001173443.1;refGene_gn=CRX;refGene_tx=XM_023244833.1
E2 6100917 6101068
refGene_gn=CRX;refGene_tx=NM_001173443.1;refGene_gn=CRX;refGene_tx=XM_023244833.1
E2 6101955 6102059
refGene_gn=CRX;refGene_tx=NM_001173443.1;refGene_gn=CRX;refGene_tx=XM_023244833.1
E2 6117906 6118848
refGene_gn=LOC105261236;refGene_tx=XM_023244982.1
E2 6131920 6132049
refGene_gn=LOC111558100;refGene_tx=XM_023245896.1
E2 6132498 6132695
refGene_gn=LOC111558100;refGene_tx=XM_023245896.1
E2 6132796 6132870
refGene_gn=LOC111558100;refGene_tx=XM_023245896.1
E2 6132993 6133262
refGene_gn=LOC111558100;refGene_tx=XM_023245896.1
E2 6134153 6134645
refGene_gn=LOC111558100;refGene_tx=XM_023245896.1
E2 6199180 6199470
refGene_gn=LOC102902178;refGene_tx=XM_023244968.1
E2 6199472 6199924
refGene_gn=LOC102902178;refGene_tx=XM_023244968.1
E2 6199925 6200048
refGene_gn=LOC102902178;refGene_tx=XM_023244968.1
E2 6200049 6200983
refGene_gn=LOC102902178;refGene_tx=XM_023244968.1
E2 6221134 6221261
refGene_gn=LOC102902178;refGene_tx=XM_023244968.1
E2 6227134 6227167
refGene_gn=LOC102902178;refGene_tx=XM_023244968.1
E2 6251333 6252745
refGene_gn=LOC101090728;refGene_tx=XM_019818583.2
E2 6257605 6257732
refGene_gn=LOC101090728;refGene_tx=XM_019818583.2
E2 6259288 6259303
refGene_gn=LOC101090728;refGene_tx=XM_019818583.2
E2 6277939 6278068

refGene_gn=LOC101090977;refGene_tx=XM_019818524.2
E2 6285512 6285678
refGene_gn=LOC101090977;refGene_tx=XM_019818524.2
E2 6291816 6292415
refGene_gn=LOC101090977;refGene_tx=XM_019818524.2
E2 6292416 6292857
refGene_gn=LOC101090977;refGene_tx=XM_019818524.2
E2 6292858 6293875
refGene_gn=LOC101090977;refGene_tx=XM_019818524.2
E2 6300747 6301692
refGene_gn=FELCATV1R9;refGene_tx=NM_001167481.1
E2 6330769 6331720
refGene_gn=LOC101091239;refGene_tx=XM_023244981.1
E2 6361818 6362778
refGene_gn=FELCATV1R12;refGene_tx=NM_001167484.1
E2 6375188 6376181
refGene_gn=FELCATV1R7;refGene_tx=NM_001167479.1;refGene_gn=FELCATV1R7;refGene_tx=XM_023244828.1;refGene_gn=FELCATV1R7;refGene_tx=XM_023244829.1
E2 6402103 6403054
refGene_gn=FELCATV1R13;refGene_tx=NM_001167485.1
E2 6428084 6428101 refGene_gn=PPP2R1A;refGene_tx=XM_003997451.5
E2 6428254 6428346 refGene_gn=PPP2R1A;refGene_tx=XM_003997451.5
E2 6430833 6430976 refGene_gn=PPP2R1A;refGene_tx=XM_003997451.5
E2 6431959 6432114 refGene_gn=PPP2R1A;refGene_tx=XM_003997451.5
E2 6432711 6432772 refGene_gn=PPP2R1A;refGene_tx=XM_003997451.5
E2 6433047 6433221 refGene_gn=PPP2R1A;refGene_tx=XM_003997451.5
E2 6439562 6439697 refGene_gn=PPP2R1A;refGene_tx=XM_003997451.5
E2 6440134 6440205 refGene_gn=PPP2R1A;refGene_tx=XM_003997451.5
E2 6440309 6440424 refGene_gn=PPP2R1A;refGene_tx=XM_003997451.5
E2 6442390 6442546 refGene_gn=PPP2R1A;refGene_tx=XM_003997451.5
E2 6442648 6442796 refGene_gn=PPP2R1A;refGene_tx=XM_003997451.5
E2 6443699 6443932 refGene_gn=PPP2R1A;refGene_tx=XM_003997451.5
E2 6447965 6448066 refGene_gn=PPP2R1A;refGene_tx=XM_003997451.5
E2 6450892 6450983 refGene_gn=PPP2R1A;refGene_tx=XM_003997451.5
E2 6464323 6464401 refGene_gn=PPP2R1A;refGene_tx=XM_003997451.5
E2 6496026 6497052
refGene_gn=LOC101091742;refGene_tx=XM_003997553.4
E2 6569578 6570939
refGene_gn=ZNF350;refGene_tx=XM_019818578.2;refGene_gn=ZNF350;refGene_tx=XM_023244976.1;refGene_gn=ZNF350;refGene_tx=XM_023244977.1;refGene_gn=ZNF350;refGene_tx=XM_023244978.1;refGene_gn=ZNF350;refGene_tx=XM_023244979.1
E2 6574520 6574616
refGene_gn=ZNF350;refGene_tx=XM_019818578.2;refGene_gn=ZNF350;refGene_tx=XM_023244976.1;refGene_gn=ZNF350;refGene_tx=XM_023244977.1;refGene_gn=ZNF350;refGene_tx=XM_023244978.1
E2 6576026 6576030 refGene_gn=ZNF350;refGene_tx=XM_023244978.1
E2 6576161 6576288
refGene_gn=ZNF350;refGene_tx=XM_019818578.2;refGene_gn=ZNF350;refGene_

tx=XM_023244976.1;refGene_gn=ZNF350;refGene_tx=XM_023244977.1
E2 6576668 6576683
refGene_gn=ZNF350;refGene_tx=XM_019818578.2;refGene_gn=ZNF350;refGene_tx=XM_023244976.1;refGene_gn=ZNF350;refGene_tx=XM_023244977.1
E2 6589740 6590127 refGene_gn=ZNF615;refGene_tx=XM_019818509.2
E2 6590128 6591702 refGene_gn=ZNF615;refGene_tx=XM_019818509.2
E2 6595387 6595480 refGene_gn=ZNF615;refGene_tx=XM_019818509.2
E2 6595979 6596106 refGene_gn=ZNF615;refGene_tx=XM_019818509.2
E2 6598033 6598048 refGene_gn=ZNF615;refGene_tx=XM_019818509.2
E2 6611845 6612019
refGene_gn=LOC101083984;refGene_tx=XM_023244984.1
E2 6612020 6612075
refGene_gn=LOC101083984;refGene_tx=XM_023244984.1
E2 6612076 6612911
refGene_gn=LOC101083984;refGene_tx=XM_023244984.1
E2 6630542 6632276
refGene_gn=LOC101083984;refGene_tx=XM_023244984.1
E2 6637353 6637449
refGene_gn=LOC101083984;refGene_tx=XM_023244984.1
E2 6638788 6638915
refGene_gn=LOC101083984;refGene_tx=XM_023244984.1
E2 6641963 6641978
refGene_gn=LOC101083984;refGene_tx=XM_023244984.1
E2 6660200 6661918
refGene_gn=ZNF613;refGene_tx=XM_019818539.2;refGene_gn=ZNF613;refGene_tx=XM_019818540.2;refGene_gn=ZNF613;refGene_tx=XM_019818542.2;refGene_gn=ZNF613;refGene_tx=XM_019818544.2;refGene_gn=ZNF613;refGene_tx=XM_019818546.2;refGene_gn=ZNF613;refGene_tx=XM_019818547.2;refGene_gn=ZNF613;refGene_tx=XM_023244966.1
E2 6666720 6666864
refGene_gn=ZNF613;refGene_tx=XM_019818539.2;refGene_gn=ZNF613;refGene_tx=XM_019818540.2;refGene_gn=ZNF613;refGene_tx=XM_019818542.2;refGene_gn=ZNF613;refGene_tx=XM_019818544.2;refGene_gn=ZNF613;refGene_tx=XM_019818546.2;refGene_gn=ZNF613;refGene_tx=XM_019818547.2;refGene_gn=ZNF613;refGene_tx=XM_023244966.1
E2 6667105 6667232
refGene_gn=ZNF613;refGene_tx=XM_019818539.2;refGene_gn=ZNF613;refGene_tx=XM_019818540.2;refGene_gn=ZNF613;refGene_tx=XM_019818542.2;refGene_gn=ZNF613;refGene_tx=XM_019818544.2;refGene_gn=ZNF613;refGene_tx=XM_019818546.2;refGene_gn=ZNF613;refGene_tx=XM_023244966.1
E2 6669990 6670008
refGene_gn=ZNF613;refGene_tx=XM_019818539.2;refGene_gn=ZNF613;refGene_tx=XM_019818540.2;refGene_gn=ZNF613;refGene_tx=XM_023244966.1
E2 6670079 6670094
refGene_gn=ZNF613;refGene_tx=XM_019818542.2;refGene_gn=ZNF613;refGene_tx=XM_019818544.2
E2 6687629 6687644
refGene_gn=LOC109494588;refGene_tx=XM_023244969.1;refGene_gn=LOC109494588;refGene_tx=XM_023244970.1;refGene_gn=LOC109494588;refGene_tx=XM_023244971.1;refGene_gn=LOC109494588;refGene_tx=XM_023244972.1

E2	6692567	6692694
refGene_gn=LOC109494588;refGene_tx=XM_023244969.1;refGene_gn=LOC109494588;refGene_tx=XM_023244970.1;refGene_gn=LOC109494588;refGene_tx=XM_023244971.1;refGene_gn=LOC109494588;refGene_tx=XM_023244972.1
E2	6692995	6693088
refGene_gn=LOC109494588;refGene_tx=XM_023244969.1;refGene_gn=LOC109494588;refGene_tx=XM_023244970.1;refGene_gn=LOC109494588;refGene_tx=XM_023244971.1
E2	6695260	6696825
refGene_gn=LOC109494588;refGene_tx=XM_023244969.1;refGene_gn=LOC109494588;refGene_tx=XM_023244970.1;refGene_gn=LOC109494588;refGene_tx=XM_023244971.1;refGene_gn=LOC109494588;refGene_tx=XM_023244972.1;refGene_gn=LOC109494588;refGene_tx=XM_023244973.1
E2	6704893	6705289
refGene_gn=LOC102901153;refGene_tx=XM_019818497.2
E2	6712426	6712553
refGene_gn=LOC102901153;refGene_tx=XM_019818497.2
E2	6712814	6712910
refGene_gn=LOC102901153;refGene_tx=XM_019818497.2
E2	6720355	6721953
refGene_gn=LOC102901153;refGene_tx=XM_019818497.2
E2	6729733	6729867
refGene_gn=LOC102901153;refGene_tx=XM_019818497.2
E2	6731161	6731216
refGene_gn=LOC102901153;refGene_tx=XM_019818497.2
E2	6739209	6739230
refGene_gn=LOC102901153;refGene_tx=XM_019818497.2
E2	6741844	6741971
refGene_gn=LOC102901153;refGene_tx=XM_019818497.2
E2	6742241	6742337
refGene_gn=LOC102901153;refGene_tx=XM_019818497.2
E2	6748731	6750065
refGene_gn=LOC102901153;refGene_tx=XM_019818497.2
E2	6767223	6768135
refGene_gn=LOC111558102;refGene_tx=XM_023245901.1
E2	6797373	6797397
refGene_gn=LOC101092973;refGene_tx=XM_006940918.4;refGene_gn=LOC101092973;refGene_tx=XM_019818526.2;refGene_gn=LOC101092973;refGene_tx=XM_019818527.2;refGene_gn=LOC101092973;refGene_tx=XM_019818528.2;refGene_gn=LOC101092973;refGene_tx=XM_019818529.2;refGene_gn=LOC101092973;refGene_tx=XM_019818530.2;refGene_gn=LOC101092973;refGene_tx=XM_019818531.2;refGene_gn=LOC101092973;refGene_tx=XM_019818532.2;refGene_gn=LOC101092973;refGene_tx=XM_019818533.2
E2	6800128	6800267
refGene_gn=LOC101092973;refGene_tx=XM_006940918.4;refGene_gn=LOC101092973;refGene_tx=XM_019818526.2;refGene_gn=LOC101092973;refGene_tx=XM_019818527.2;refGene_gn=LOC101092973;refGene_tx=XM_019818528.2;refGene_gn=LOC101092973;refGene_tx=XM_019818529.2;refGene_gn=LOC101092973;refGene_tx=XM_019818530.2;refGene_gn=LOC101092973;refGene_tx=XM_019818531.2;refGene_gn=LOC101092973;refGene_tx=XM_019818532.2;refGene_gn=LOC101092

973;refGene_tx=XM_019818533.2;refGene_gn=LOC101092973;refGene_tx=XM_01
9818536.2;refGene_gn=LOC101092973;refGene_tx=XM_023245071.1;refGene_gn
=LOC101092973;refGene_tx=XM_023245072.1
E2 6800546 6800639
refGene_gn=LOC101092973;refGene_tx=XM_006940918.4;refGene_gn=LOC101092
973;refGene_tx=XM_019818526.2;refGene_gn=LOC101092973;refGene_tx=XM_01
9818527.2;refGene_gn=LOC101092973;refGene_tx=XM_019818528.2;refGene_gn
=LOC101092973;refGene_tx=XM_019818529.2;refGene_gn=LOC101092973;refGen
e_tx=XM_019818530.2;refGene_gn=LOC101092973;refGene_tx=XM_019818531.2;
refGene_gn=LOC101092973;refGene_tx=XM_019818532.2;refGene_gn=LOC101092
973;refGene_tx=XM_019818533.2;refGene_gn=LOC101092973;refGene_tx=XM_01
9818536.2;refGene_gn=LOC101092973;refGene_tx=XM_023245071.1;refGene_gn
=LOC101092973;refGene_tx=XM_023245072.1
E2 6801555 6803414
refGene_gn=LOC101092973;refGene_tx=XM_006940918.4;refGene_gn=LOC101092
973;refGene_tx=XM_019818526.2;refGene_gn=LOC101092973;refGene_tx=XM_01
9818527.2;refGene_gn=LOC101092973;refGene_tx=XM_019818528.2;refGene_gn
=LOC101092973;refGene_tx=XM_019818529.2;refGene_gn=LOC101092973;refGen
e_tx=XM_019818530.2;refGene_gn=LOC101092973;refGene_tx=XM_019818531.2;
refGene_gn=LOC101092973;refGene_tx=XM_019818532.2;refGene_gn=LOC101092
973;refGene_tx=XM_019818533.2;refGene_gn=LOC101092973;refGene_tx=XM_01
9818536.2;refGene_gn=LOC101092973;refGene_tx=XM_023245071.1;refGene_gn
=LOC101092973;refGene_tx=XM_023245072.1
E2 6835016 6836072
refGene_gn=FPR2;refGene_tx=XM_019818608.2;refGene_gn=FPR2;refGene_tx=X
M_023245239.1
E2 6865661 6865670 refGene_gn=HAS1;refGene_tx=XM_011289756.3
E2 6869204 6869912 refGene_gn=HAS1;refGene_tx=XM_011289756.3
E2 6871639 6871865 refGene_gn=HAS1;refGene_tx=XM_011289756.3
E2 6872458 6872591 refGene_gn=HAS1;refGene_tx=XM_011289756.3
E2 6874281 6874957 refGene_gn=HAS1;refGene_tx=XM_011289756.3
E2 6881382 6881383 refGene_gn=SPACA6;refGene_tx=XM_023245026.1
E2 6881455 6881489
refGene_gn=SPACA6;refGene_tx=XM_006940919.4;refGene_gn=SPACA6;refGene_
tx=XM_019818615.1;refGene_gn=SPACA6;refGene_tx=XM_023245023.1;refGene_
gn=SPACA6;refGene_tx=XM_023245024.1;refGene_gn=SPACA6;refGene_tx=XM_02
3245025.1;refGene_gn=SPACA6;refGene_tx=XM_023245027.1;refGene_gn=SPACA
6;refGene_tx=XM_023245028.1;refGene_gn=SPACA6;refGene_tx=XM_023245029.
1
E2 6881817 6882028
refGene_gn=SPACA6;refGene_tx=XM_006940919.4;refGene_gn=SPACA6;refGene_
tx=XM_019818615.1;refGene_gn=SPACA6;refGene_tx=XM_023245023.1;refGene_
gn=SPACA6;refGene_tx=XM_023245024.1;refGene_gn=SPACA6;refGene_tx=XM_02
3245025.1;refGene_gn=SPACA6;refGene_tx=XM_023245026.1;refGene_gn=SPACA
6;refGene_tx=XM_023245027.1;refGene_gn=SPACA6;refGene_tx=XM_023245028.
1;refGene_gn=SPACA6;refGene_tx=XM_023245029.1
E2 6882107 6882264
refGene_gn=SPACA6;refGene_tx=XM_006940919.4;refGene_gn=SPACA6;refGene_
tx=XM_019818615.1;refGene_gn=SPACA6;refGene_tx=XM_023245023.1;refGene_
gn=SPACA6;refGene_tx=XM_023245024.1;refGene_gn=SPACA6;refGene_tx=XM_02

3245025.1;refGene_gn=SPACA6;refGene_tx=XM_023245026.1;refGene_gn=SPACA6;refGene_tx=XM_023245027.1;refGene_gn=SPACA6;refGene_tx=XM_023245028.1;refGene_gn=SPACA6;refGene_tx=XM_023245029.1
E2 6882697 6882807
refGene_gn=SPACA6;refGene_tx=XM_006940919.4;refGene_gn=SPACA6;refGene_tx=XM_019818615.1;refGene_gn=SPACA6;refGene_tx=XM_023245023.1;refGene_gn=SPACA6;refGene_tx=XM_023245024.1;refGene_gn=SPACA6;refGene_tx=XM_023245025.1;refGene_gn=SPACA6;refGene_tx=XM_023245026.1;refGene_gn=SPACA6;refGene_tx=XM_023245027.1;refGene_gn=SPACA6;refGene_tx=XM_023245028.1;refGene_gn=SPACA6;refGene_tx=XM_023245029.1
E2 6882894 6883004
refGene_gn=SPACA6;refGene_tx=XM_006940919.4;refGene_gn=SPACA6;refGene_tx=XM_019818615.1;refGene_gn=SPACA6;refGene_tx=XM_023245023.1;refGene_gn=SPACA6;refGene_tx=XM_023245024.1;refGene_gn=SPACA6;refGene_tx=XM_023245025.1;refGene_gn=SPACA6;refGene_tx=XM_023245026.1;refGene_gn=SPACA6;refGene_tx=XM_023245027.1;refGene_gn=SPACA6;refGene_tx=XM_023245028.1;refGene_gn=SPACA6;refGene_tx=XM_023245029.1
E2 6883350 6883374
refGene_gn=SPACA6;refGene_tx=XM_006940919.4;refGene_gn=SPACA6;refGene_tx=XM_019818615.1;refGene_gn=SPACA6;refGene_tx=XM_023245023.1;refGene_gn=SPACA6;refGene_tx=XM_023245025.1;refGene_gn=SPACA6;refGene_tx=XM_023245026.1
E2 6884050 6884119
refGene_gn=SPACA6;refGene_tx=XM_006940919.4;refGene_gn=SPACA6;refGene_tx=XM_019818615.1;refGene_gn=SPACA6;refGene_tx=XM_023245023.1;refGene_gn=SPACA6;refGene_tx=XM_023245024.1;refGene_gn=SPACA6;refGene_tx=XM_023245025.1;refGene_gn=SPACA6;refGene_tx=XM_023245026.1;refGene_gn=SPACA6;refGene_tx=XM_023245027.1
E2 6890190 6890294
refGene_gn=SPACA6;refGene_tx=XM_006940919.4;refGene_gn=SPACA6;refGene_tx=XM_019818615.1;refGene_gn=SPACA6;refGene_tx=XM_023245023.1;refGene_gn=SPACA6;refGene_tx=XM_023245024.1;refGene_gn=SPACA6;refGene_tx=XM_023245025.1;refGene_gn=SPACA6;refGene_tx=XM_023245026.1;refGene_gn=SPACA6;refGene_tx=XM_023245027.1;refGene_gn=SPACA6;refGene_tx=XM_023245028.1;refGene_gn=SPACA6;refGene_tx=XM_023245029.1
E2 6890752 6890966
refGene_gn=SPACA6;refGene_tx=XM_006940919.4;refGene_gn=SPACA6;refGene_tx=XM_023245023.1;refGene_gn=SPACA6;refGene_tx=XM_023245024.1;refGene_gn=SPACA6;refGene_tx=XM_023245025.1;refGene_gn=SPACA6;refGene_tx=XM_023245026.1;refGene_gn=SPACA6;refGene_tx=XM_023245027.1;refGene_gn=SPACA6;refGene_tx=XM_023245028.1;refGene_gn=SPACA6;refGene_tx=XM_023245029.1
E2 6892536 6892580 refGene_gn=SPACA6;refGene_tx=XM_019818615.1
E2 6935310 6935347
refGene_gn=LOC101093728;refGene_tx=XM_023245222.1
E2 6935424 6935564
refGene_gn=LOC101093728;refGene_tx=XM_023245222.1
E2 6935565 6935800
refGene_gn=LOC101093728;refGene_tx=XM_023245222.1
E2 6936023 6936302

refGene_gn=LOC101093728;refGene_tx=XM_023245222.1
E2 6936694 6936742
refGene_gn=LOC101093728;refGene_tx=XM_023245222.1
E2 6939157 6939415
refGene_gn=LOC101093728;refGene_tx=XM_023245222.1
E2 6939525 6939661
refGene_gn=LOC101093728;refGene_tx=XM_023245222.1
E2 6939803 6939873
refGene_gn=LOC101093728;refGene_tx=XM_023245222.1
E2 6949586 6949629
refGene_gn=SIGLEC5;refGene_tx=XM_023245219.1;refGene_gn=SIGLEC5;refGene_tx=XM_023245221.1
E2 6951322 6951436
refGene_gn=SIGLEC5;refGene_tx=XM_023245219.1;refGene_gn=SIGLEC5;refGene_tx=XM_023245221.1
E2 6952096 6952133 refGene_gn=SIGLEC5;refGene_tx=XM_023245220.1
E2 6952210 6952585
refGene_gn=SIGLEC5;refGene_tx=XM_023245219.1;refGene_gn=SIGLEC5;refGene_tx=XM_023245220.1;refGene_gn=SIGLEC5;refGene_tx=XM_023245221.1
E2 6952808 6953087
refGene_gn=SIGLEC5;refGene_tx=XM_023245219.1;refGene_gn=SIGLEC5;refGene_tx=XM_023245220.1;refGene_gn=SIGLEC5;refGene_tx=XM_023245221.1
E2 6953733 6953778
refGene_gn=SIGLEC5;refGene_tx=XM_023245219.1;refGene_gn=SIGLEC5;refGene_tx=XM_023245220.1;refGene_gn=SIGLEC5;refGene_tx=XM_023245221.1
E2 6954313 6954571
refGene_gn=SIGLEC5;refGene_tx=XM_023245219.1;refGene_gn=SIGLEC5;refGene_tx=XM_023245220.1;refGene_gn=SIGLEC5;refGene_tx=XM_023245221.1
E2 6954662 6954953
refGene_gn=SIGLEC5;refGene_tx=XM_023245219.1;refGene_gn=SIGLEC5;refGene_tx=XM_023245220.1;refGene_gn=SIGLEC5;refGene_tx=XM_023245221.1
E2 6955189 6955289
refGene_gn=SIGLEC5;refGene_tx=XM_023245219.1;refGene_gn=SIGLEC5;refGene_tx=XM_023245220.1;refGene_gn=SIGLEC5;refGene_tx=XM_023245221.1
E2 6955877 6955959
refGene_gn=SIGLEC5;refGene_tx=XM_023245219.1;refGene_gn=SIGLEC5;refGene_tx=XM_023245220.1;refGene_gn=SIGLEC5;refGene_tx=XM_023245221.1
E2 6956215 6956704
refGene_gn=SIGLEC5;refGene_tx=XM_023245219.1;refGene_gn=SIGLEC5;refGene_tx=XM_023245220.1
E2 6962636 6962828 refGene_gn=SIGLEC5;refGene_tx=XM_023245221.1
E2 6970190 6970765
refGene_gn=LOC111558053;refGene_tx=XM_023245506.1;refGene_gn=LOC111558053;refGene_tx=XM_023245507.1
E2 6972346 6972854
refGene_gn=LOC111558053;refGene_tx=XM_023245506.1;refGene_gn=LOC111558053;refGene_tx=XM_023245507.1
E2 6987562 6989412
refGene_gn=ZNF175;refGene_tx=XM_006940922.4;refGene_gn=ZNF175;refGene_tx=XM_019818508.2

E2 6995573 6995669
refGene_gn=ZNF175;refGene_tx=XM_006940922.4;refGene_gn=ZNF175;refGene_tx=XM_019818508.2
E2 6995951 6996078
refGene_gn=ZNF175;refGene_tx=XM_006940922.4;refGene_gn=ZNF175;refGene_tx=XM_019818508.2
E2 7019687 7019837 refGene_gn=ZNF175;refGene_tx=XM_006940922.4
E2 7034196 7034227
refGene_gn=LOC101094717;refGene_tx=XM_011289725.3
E2 7034307 7034667
refGene_gn=LOC101094717;refGene_tx=XM_011289725.3
E2 7034835 7035114
refGene_gn=LOC101094717;refGene_tx=XM_011289725.3
E2 7035311 7035359
refGene_gn=LOC101094717;refGene_tx=XM_011289725.3
E2 7035771 7036029
refGene_gn=LOC101094717;refGene_tx=XM_011289725.3
E2 7036118 7036406
refGene_gn=LOC101094717;refGene_tx=XM_011289725.3
E2 7037164 7037261
refGene_gn=LOC101094717;refGene_tx=XM_011289725.3
E2 7037599 7037678
refGene_gn=LOC101094717;refGene_tx=XM_011289725.3
E2 7041744 7041924
refGene_gn=LOC101094717;refGene_tx=XM_011289725.3
E2 7044663 7045105
refGene_gn=LOC105261234;refGene_tx=XM_019818618.2
E2 7045286 7045565
refGene_gn=LOC105261234;refGene_tx=XM_019818618.2
E2 7045790 7045838
refGene_gn=LOC105261234;refGene_tx=XM_019818618.2
E2 7047760 7047849
refGene_gn=LOC105261234;refGene_tx=XM_019818618.2
E2 7051943 7051980
refGene_gn=LOC101094969;refGene_tx=XM_011289724.3;refGene_gn=LOC101094969;refGene_tx=XM_019818520.2
E2 7052045 7052429
refGene_gn=LOC101094969;refGene_tx=XM_011289724.3;refGene_gn=LOC101094969;refGene_tx=XM_019818520.2
E2 7052588 7052867
refGene_gn=LOC101094969;refGene_tx=XM_019818520.2
E2 7053068 7053116
refGene_gn=LOC101094969;refGene_tx=XM_019818520.2
E2 7054258 7054516
refGene_gn=LOC101094969;refGene_tx=XM_019818520.2
E2 7054611 7054747
refGene_gn=LOC101094969;refGene_tx=XM_019818520.2
E2 7054875 7054918
refGene_gn=LOC101094969;refGene_tx=XM_019818520.2
E2 7072928 7072965

refGene_gn=LOC101094969;refGene_tx=XM_019818517.2;refGene_gn=LOC101094969;refGene_tx=XM_019818518.2;refGene_gn=LOC101094969;refGene_tx=XM_019818519.2;refGene_gn=LOC101094969;refGene_tx=XM_019818522.2;refGene_gn=LOC101094969;refGene_tx=XM_019818523.2;refGene_gn=LOC101094969;refGene_tx=XM_023245314.1;refGene_gn=LOC101094969;refGene_tx=XM_023245315.1;refGene_gn=LOC101094969;refGene_tx=XM_023245316.1;refGene_gn=LOC101094969;refGene_tx=XM_023245317.1
E2 7073045 7073426
refGene_gn=LOC101094969;refGene_tx=XM_011289724.3;refGene_gn=LOC101094969;refGene_tx=XM_019818517.2;refGene_gn=LOC101094969;refGene_tx=XM_019818518.2;refGene_gn=LOC101094969;refGene_tx=XM_019818519.2;refGene_gn=LOC101094969;refGene_tx=XM_019818522.2;refGene_gn=LOC101094969;refGene_tx=XM_019818523.2;refGene_gn=LOC101094969;refGene_tx=XM_023245314.1;refGene_gn=LOC101094969;refGene_tx=XM_023245315.1;refGene_gn=LOC101094969;refGene_tx=XM_023245316.1;refGene_gn=LOC101094969;refGene_tx=XM_023245317.1
E2 7073586 7073871
refGene_gn=LOC101094969;refGene_tx=XM_011289724.3;refGene_gn=LOC101094969;refGene_tx=XM_019818517.2;refGene_gn=LOC101094969;refGene_tx=XM_019818518.2;refGene_gn=LOC101094969;refGene_tx=XM_019818519.2;refGene_gn=LOC101094969;refGene_tx=XM_019818522.2;refGene_gn=LOC101094969;refGene_tx=XM_019818523.2;refGene_gn=LOC101094969;refGene_tx=XM_023245314.1;refGene_gn=LOC101094969;refGene_tx=XM_023245315.1;refGene_gn=LOC101094969;refGene_tx=XM_023245316.1;refGene_gn=LOC101094969;refGene_tx=XM_023245317.1
E2 7074148 7074196
refGene_gn=LOC101094969;refGene_tx=XM_011289724.3;refGene_gn=LOC101094969;refGene_tx=XM_019818517.2;refGene_gn=LOC101094969;refGene_tx=XM_019818518.2;refGene_gn=LOC101094969;refGene_tx=XM_019818519.2;refGene_gn=LOC101094969;refGene_tx=XM_019818522.2;refGene_gn=LOC101094969;refGene_tx=XM_019818523.2;refGene_gn=LOC101094969;refGene_tx=XM_023245314.1;refGene_gn=LOC101094969;refGene_tx=XM_023245315.1;refGene_gn=LOC101094969;refGene_tx=XM_023245316.1;refGene_gn=LOC101094969;refGene_tx=XM_023245317.1
E2 7074310 7074574
refGene_gn=LOC101094969;refGene_tx=XM_011289724.3;refGene_gn=LOC101094969;refGene_tx=XM_019818517.2;refGene_gn=LOC101094969;refGene_tx=XM_019818518.2;refGene_gn=LOC101094969;refGene_tx=XM_019818519.2;refGene_gn=LOC101094969;refGene_tx=XM_019818522.2;refGene_gn=LOC101094969;refGene_tx=XM_019818523.2;refGene_gn=LOC101094969;refGene_tx=XM_023245314.1;refGene_gn=LOC101094969;refGene_tx=XM_023245315.1;refGene_gn=LOC101094969;refGene_tx=XM_023245316.1;refGene_gn=LOC101094969;refGene_tx=XM_023245317.1
E2 7075871 7075919
refGene_gn=LOC101094969;refGene_tx=XM_011289724.3;refGene_gn=LOC101094969;refGene_tx=XM_019818517.2;refGene_gn=LOC101094969;refGene_tx=XM_019818518.2;refGene_gn=LOC101094969;refGene_tx=XM_019818519.2;refGene_gn=LOC101094969;refGene_tx=XM_019818522.2;refGene_gn=LOC101094969;refGene_tx=XM_019818523.2;refGene_gn=LOC101094969;refGene_tx=XM_023245314.1;refGene_gn=LOC101094969;refGene_tx=XM_023245315.1;refGene_gn=LOC101094

969;refGene_tx=XM_023245316.1;refGene_gn=LOC101094969;refGene_tx=XM_023245317.1
E2 7076049 7076307
refGene_gn=LOC101094969;refGene_tx=XM_011289724.3;refGene_gn=LOC101094969;refGene_tx=XM_019818517.2;refGene_gn=LOC101094969;refGene_tx=XM_019818518.2;refGene_gn=LOC101094969;refGene_tx=XM_019818519.2;refGene_gn=LOC101094969;refGene_tx=XM_019818522.2;refGene_gn=LOC101094969;refGene_tx=XM_019818523.2;refGene_gn=LOC101094969;refGene_tx=XM_023245314.1;refGene_gn=LOC101094969;refGene_tx=XM_023245315.1;refGene_gn=LOC101094969;refGene_tx=XM_023245316.1;refGene_gn=LOC101094969;refGene_tx=XM_023245317.1
E2 7076390 7076675
refGene_gn=LOC101094969;refGene_tx=XM_011289724.3;refGene_gn=LOC101094969;refGene_tx=XM_019818517.2;refGene_gn=LOC101094969;refGene_tx=XM_019818518.2;refGene_gn=LOC101094969;refGene_tx=XM_019818519.2;refGene_gn=LOC101094969;refGene_tx=XM_023245314.1;refGene_gn=LOC101094969;refGene_tx=XM_023245315.1
E2 7076994 7077088
refGene_gn=LOC101094969;refGene_tx=XM_011289724.3;refGene_gn=LOC101094969;refGene_tx=XM_019818517.2;refGene_gn=LOC101094969;refGene_tx=XM_019818518.2;refGene_gn=LOC101094969;refGene_tx=XM_019818519.2;refGene_gn=LOC101094969;refGene_tx=XM_019818522.2;refGene_gn=LOC101094969;refGene_tx=XM_019818523.2;refGene_gn=LOC101094969;refGene_tx=XM_023245314.1;refGene_gn=LOC101094969;refGene_tx=XM_023245315.1;refGene_gn=LOC101094969;refGene_tx=XM_023245316.1;refGene_gn=LOC101094969;refGene_tx=XM_023245317.1
E2 7077183 7077349
refGene_gn=LOC101094969;refGene_tx=XM_011289724.3;refGene_gn=LOC101094969;refGene_tx=XM_019818517.2;refGene_gn=LOC101094969;refGene_tx=XM_019818518.2;refGene_gn=LOC101094969;refGene_tx=XM_019818519.2;refGene_gn=LOC101094969;refGene_tx=XM_019818522.2;refGene_gn=LOC101094969;refGene_tx=XM_019818523.2;refGene_gn=LOC101094969;refGene_tx=XM_023245314.1;refGene_gn=LOC101094969;refGene_tx=XM_023245315.1;refGene_gn=LOC101094969;refGene_tx=XM_023245316.1;refGene_gn=LOC101094969;refGene_tx=XM_023245317.1
E2 7079941 7080226
refGene_gn=LOC101094969;refGene_tx=XM_011289724.3;refGene_gn=LOC101094969;refGene_tx=XM_019818517.2;refGene_gn=LOC101094969;refGene_tx=XM_019818518.2;refGene_gn=LOC101094969;refGene_tx=XM_019818522.2;refGene_gn=LOC101094969;refGene_tx=XM_019818523.2
E2 7081126 7081690
refGene_gn=LOC101094969;refGene_tx=XM_019818519.2;refGene_gn=LOC101094969;refGene_tx=XM_023245314.1;refGene_gn=LOC101094969;refGene_tx=XM_023245315.1;refGene_gn=LOC101094969;refGene_tx=XM_023245316.1;refGene_gn=LOC101094969;refGene_tx=XM_023245317.1
E2 7091792 7091827
refGene_gn=CE2H19orf84;refGene_tx=XM_023245553.1
E2 7092683 7093212
refGene_gn=CE2H19orf84;refGene_tx=XM_023245553.1
E2 7094545 7094720 refGene_gn=LIM2;refGene_tx=XM_019818667.2

E2	7098946	7099096	refGene_gn=LIM2;refGene_tx=XM_019818667.2
E2	7100166	7100301	refGene_gn=LIM2;refGene_tx=XM_019818667.2
E2	7100531	7100593	refGene_gn=LIM2;refGene_tx=XM_019818667.2
E2	7107467	7107624	refGene_gn=NKG7;refGene_tx=XM_003997485.5
E2	7107704	7107851	refGene_gn=NKG7;refGene_tx=XM_003997485.5
E2	7107935	7108070	refGene_gn=NKG7;refGene_tx=XM_003997485.5
E2	7108161	7108220	refGene_gn=NKG7;refGene_tx=XM_003997485.5
E2	7110273	7110442
refGene_gn=CLDND2;refGene_tx=XM_006940927.3;refGene_gn=CLDND2;refGene_tx=XM_011289722.3;refGene_gn=CLDND2;refGene_tx=XM_019818660.2;refGene_gn=CLDND2;refGene_tx=XM_019818661.2;refGene_gn=CLDND2;refGene_tx=XM_023245528.1
E2	7110774	7110912
refGene_gn=CLDND2;refGene_tx=XM_006940927.3;refGene_gn=CLDND2;refGene_tx=XM_011289722.3;refGene_gn=CLDND2;refGene_tx=XM_019818660.2;refGene_gn=CLDND2;refGene_tx=XM_019818661.2;refGene_gn=CLDND2;refGene_tx=XM_023245528.1
E2	7111141	7111398
refGene_gn=CLDND2;refGene_tx=XM_006940927.3;refGene_gn=CLDND2;refGene_tx=XM_011289722.3;refGene_gn=CLDND2;refGene_tx=XM_019818660.2;refGene_gn=CLDND2;refGene_tx=XM_019818661.2;refGene_gn=CLDND2;refGene_tx=XM_023245528.1
E2	7111486	7111560	refGene_gn=CLDND2;refGene_tx=XM_023245528.1
E2	7112417	7112474	refGene_gn=ETFB;refGene_tx=XM_003997457.4
E2	7120003	7120162	refGene_gn=ETFB;refGene_tx=XM_003997457.4
E2	7120274	7120433	refGene_gn=ETFB;refGene_tx=XM_003997457.4
E2	7122535	7122598	refGene_gn=ETFB;refGene_tx=XM_003997457.4
E2	7124892	7125051	refGene_gn=ETFB;refGene_tx=XM_003997457.4
E2	7125659	7125830	refGene_gn=ETFB;refGene_tx=XM_003997457.4
E2	7129047	7129087	refGene_gn=VSIG10L;refGene_tx=XM_023245113.1
E2	7129167	7129998	refGene_gn=VSIG10L;refGene_tx=XM_023245113.1
E2	7130441	7130735	refGene_gn=VSIG10L;refGene_tx=XM_023245113.1
E2	7130894	7131179	refGene_gn=VSIG10L;refGene_tx=XM_023245113.1
E2	7132086	7132341	refGene_gn=VSIG10L;refGene_tx=XM_023245113.1
E2	7132774	7133053	refGene_gn=VSIG10L;refGene_tx=XM_023245113.1
E2	7133284	7133581	refGene_gn=VSIG10L;refGene_tx=XM_023245113.1
E2	7134435	7134549	refGene_gn=VSIG10L;refGene_tx=XM_023245113.1
E2	7134763	7134918	refGene_gn=VSIG10L;refGene_tx=XM_023245113.1
E2	7135881	7135953	refGene_gn=VSIG10L;refGene_tx=XM_023245113.1
E2	7141016	7141105	refGene_gn=IGLON5;refGene_tx=XM_023245114.1
E2	7142390	7142545	refGene_gn=IGLON5;refGene_tx=XM_023245114.1
E2	7143074	7143195	refGene_gn=IGLON5;refGene_tx=XM_023245114.1
E2	7143360	7143495	refGene_gn=IGLON5;refGene_tx=XM_023245114.1
E2	7144911	7145031	refGene_gn=IGLON5;refGene_tx=XM_023245114.1
E2	7146433	7146666	refGene_gn=IGLON5;refGene_tx=XM_023245114.1
E2	7147683	7147762	refGene_gn=IGLON5;refGene_tx=XM_023245114.1
E2	7155444	7155523	refGene_gn=IGLON5;refGene_tx=XM_023245114.1
E2	7164884	7165154
refGene_gn=LOC101095701;refGene_tx=XM_019818622.2;refGene_gn=LOC101095701;refGene_tx=XM_019818623.2

E2	7166019	7166158
refGene_gn=LOC101095701;refGene_tx=XM_019818622.2;refGene_gn=LOC101095
701;refGene_tx=XM_019818623.2
E2	7169368	7169450
refGene_gn=LOC101095701;refGene_tx=XM_006940928.4
E2	7171172	7171275
refGene_gn=LOC101095701;refGene_tx=XM_006940928.4;refGene_gn=LOC101095
701;refGene_tx=XM_019818622.2;refGene_gn=LOC101095701;refGene_tx=XM_01
9818623.2
E2	7171421	7171703
refGene_gn=LOC101095701;refGene_tx=XM_006940928.4;refGene_gn=LOC101095
701;refGene_tx=XM_019818622.2;refGene_gn=LOC101095701;refGene_tx=XM_01
9818623.2
E2	7172051	7172073
refGene_gn=LOC101095701;refGene_tx=XM_006940928.4;refGene_gn=LOC101095
701;refGene_tx=XM_019818622.2;refGene_gn=LOC101095701;refGene_tx=XM_01
9818623.2
E2	7180802	7180817
refGene_gn=SIGLECL1;refGene_tx=XM_006940930.4;refGene_gn=SIGLECL1;refG
ene_tx=XM_006940933.4;refGene_gn=SIGLECL1;refGene_tx=XM_023245499.1;re
fGene_gn=SIGLECL1;refGene_tx=XM_023245500.1;refGene_gn=SIGLECL1;refGen
e_tx=XM_023245501.1;refGene_gn=SIGLECL1;refGene_tx=XM_023245502.1;refG
ene_gn=SIGLECL1;refGene_tx=XM_023245503.1
E2	7181840	7181997
refGene_gn=SIGLECL1;refGene_tx=XM_006940930.4;refGene_gn=SIGLECL1;refG
ene_tx=XM_006940933.4;refGene_gn=SIGLECL1;refGene_tx=XM_023245499.1;re
fGene_gn=SIGLECL1;refGene_tx=XM_023245500.1;refGene_gn=SIGLECL1;refGen
e_tx=XM_023245501.1;refGene_gn=SIGLECL1;refGene_tx=XM_023245502.1;refG
ene_gn=SIGLECL1;refGene_tx=XM_023245503.1
E2	7183139	7183245
refGene_gn=SIGLECL1;refGene_tx=XM_006940930.4;refGene_gn=SIGLECL1;refG
ene_tx=XM_006940933.4;refGene_gn=SIGLECL1;refGene_tx=XM_023245499.1;re
fGene_gn=SIGLECL1;refGene_tx=XM_023245500.1;refGene_gn=SIGLECL1;refGen
e_tx=XM_023245501.1;refGene_gn=SIGLECL1;refGene_tx=XM_023245502.1;refG
ene_gn=SIGLECL1;refGene_tx=XM_023245503.1
E2	7183364	7183646
refGene_gn=SIGLECL1;refGene_tx=XM_006940930.4;refGene_gn=SIGLECL1;refG
ene_tx=XM_006940933.4;refGene_gn=SIGLECL1;refGene_tx=XM_023245499.1;re
fGene_gn=SIGLECL1;refGene_tx=XM_023245500.1;refGene_gn=SIGLECL1;refGen
e_tx=XM_023245501.1;refGene_gn=SIGLECL1;refGene_tx=XM_023245502.1;refG
ene_gn=SIGLECL1;refGene_tx=XM_023245503.1
E2	7185097	7185101
refGene_gn=SIGLECL1;refGene_tx=XM_006940930.4;refGene_gn=SIGLECL1;refG
ene_tx=XM_006940933.4;refGene_gn=SIGLECL1;refGene_tx=XM_023245499.1;re
fGene_gn=SIGLECL1;refGene_tx=XM_023245500.1;refGene_gn=SIGLECL1;refGen
e_tx=XM_023245501.1;refGene_gn=SIGLECL1;refGene_tx=XM_023245502.1;refG
ene_gn=SIGLECL1;refGene_tx=XM_023245503.1
E2	7205280	7205454
refGene_gn=LOC101096188;refGene_tx=XM_006940936.4
E2	7207138	7207220

refGene_gn=LOC101096188;refGene_tx=XM_006940936.4
E2	7207511	7207611
refGene_gn=LOC101096188;refGene_tx=XM_006940936.4
E2	7207927	7208212
refGene_gn=LOC101096188;refGene_tx=XM_006940936.4
E2	7208292	7208550
refGene_gn=LOC101096188;refGene_tx=XM_006940936.4
E2	7211747	7211795
refGene_gn=LOC101096188;refGene_tx=XM_006940936.4
E2	7211989	7212268
refGene_gn=LOC101096188;refGene_tx=XM_006940936.4
E2	7212441	7212846
refGene_gn=LOC101096188;refGene_tx=XM_006940936.4
E2	7212931	7212983
refGene_gn=LOC101096188;refGene_tx=XM_006940936.4
E2	7223908	7224073
refGene_gn=LOC102902160;refGene_tx=XM_006940934.3;refGene_gn=LOC102902
160;refGene_tx=XM_023245384.1
E2	7229240	7229322
refGene_gn=LOC102902160;refGene_tx=XM_006940934.3;refGene_gn=LOC102902
160;refGene_tx=XM_023245384.1
E2	7229610	7229707
refGene_gn=LOC102902160;refGene_tx=XM_006940934.3;refGene_gn=LOC102902
160;refGene_tx=XM_023245384.1
E2	7230941	7230989
refGene_gn=LOC102902160;refGene_tx=XM_006940934.3;refGene_gn=LOC102902
160;refGene_tx=XM_023245384.1
E2	7231218	7231497
refGene_gn=LOC102902160;refGene_tx=XM_006940934.3;refGene_gn=LOC102902
160;refGene_tx=XM_023245384.1
E2	7231637	7232115
refGene_gn=LOC102902160;refGene_tx=XM_006940934.3;refGene_gn=LOC102902
160;refGene_tx=XM_023245384.1
E2	7239414	7239476
refGene_gn=LOC101096432;refGene_tx=XM_023245902.1
E2	7239578	7239626
refGene_gn=LOC101096432;refGene_tx=XM_023245902.1
E2	7244132	7244358
refGene_gn=LOC101096432;refGene_tx=XM_023245902.1
E2	7244360	7244413
refGene_gn=LOC101096432;refGene_tx=XM_023245902.1
E2	7244558	7244868
refGene_gn=LOC101096432;refGene_tx=XM_023245902.1
E2	7244869	7244943
refGene_gn=LOC101096432;refGene_tx=XM_023245902.1
E2	7245008	7245045
refGene_gn=LOC101096432;refGene_tx=XM_023245902.1
E2	7258694	7258746	refGene_gn=CEACAM18;refGene_tx=XM_019818607.2
E2	7260558	7260915	refGene_gn=CEACAM18;refGene_tx=XM_019818607.2
E2	7262068	7262341	refGene_gn=CEACAM18;refGene_tx=XM_019818607.2

E2	7264800	7265082	refGene_gn=CEACAM18;refGene_tx=XM_019818607.2
E2	7266359	7266495	refGene_gn=CEACAM18;refGene_tx=XM_019818607.2
E2	7270752	7270929
refGene_gn=LOC109494587;refGene_tx=XM_023245545.1;refGene_gn=LOC109494
587;refGene_tx=XM_023245546.1
E2	7271745	7271764	refGene_gn=CEACAM18;refGene_tx=XM_019818607.2
E2	7276028	7276125
refGene_gn=LOC109494587;refGene_tx=XM_023245545.1;refGene_gn=LOC109494
587;refGene_tx=XM_023245546.1
E2	7276686	7276783
refGene_gn=LOC109494587;refGene_tx=XM_023245545.1;refGene_gn=LOC109494
587;refGene_tx=XM_023245546.1
E2	7277579	7277828
refGene_gn=LOC109494587;refGene_tx=XM_023245545.1;refGene_gn=LOC109494
587;refGene_tx=XM_023245546.1
E2	7278676	7278724
refGene_gn=LOC109494587;refGene_tx=XM_023245545.1;refGene_gn=LOC109494
587;refGene_tx=XM_023245546.1
E2	7278947	7279226
refGene_gn=LOC109494587;refGene_tx=XM_023245545.1;refGene_gn=LOC109494
587;refGene_tx=XM_023245546.1
E2	7279409	7279833
refGene_gn=LOC109494587;refGene_tx=XM_023245545.1;refGene_gn=LOC109494
587;refGene_tx=XM_023245546.1
E2	7308089	7308129
refGene_gn=KLK14;refGene_tx=XM_019818634.2;refGene_gn=KLK14;refGene_tx
=XM_019818635.2
E2	7308918	7309090
refGene_gn=KLK14;refGene_tx=XM_019818634.2;refGene_gn=KLK14;refGene_tx
=XM_019818635.2
E2	7310830	7311084
refGene_gn=KLK14;refGene_tx=XM_019818634.2;refGene_gn=KLK14;refGene_tx
=XM_019818635.2
E2	7311187	7311324
refGene_gn=KLK14;refGene_tx=XM_019818634.2;refGene_gn=KLK14;refGene_tx
=XM_019818635.2
E2	7311899	7312052
refGene_gn=KLK14;refGene_tx=XM_019818634.2;refGene_gn=KLK14;refGene_tx
=XM_019818635.2
E2	7327381	7327433
refGene_gn=KLK13;refGene_tx=XM_011289718.3;refGene_gn=KLK13;refGene_tx
=XM_011289719.3
E2	7330168	7330470
refGene_gn=KLK13;refGene_tx=XM_006940938.4;refGene_gn=KLK13;refGene_tx
=XM_011289718.3;refGene_gn=KLK13;refGene_tx=XM_011289719.3
E2	7330713	7330979
refGene_gn=KLK13;refGene_tx=XM_006940938.4;refGene_gn=KLK13;refGene_tx
=XM_011289718.3;refGene_gn=KLK13;refGene_tx=XM_011289719.3
E2	7332132	7332269
refGene_gn=KLK13;refGene_tx=XM_006940938.4;refGene_gn=KLK13;refGene_tx

=XM_011289718.3;refGene_gn=KLK13;refGene_tx=XM_011289719.3
E2 7333832 7334027
refGene_gn=KLK13;refGene_tx=XM_006940938.4;refGene_gn=KLK13;refGene_tx=XM_011289718.3;refGene_gn=KLK13;refGene_tx=XM_011289719.3
E2 7344331 7344360 refGene_gn=KLK12;refGene_tx=XM_019818620.1
E2 7344542 7344598 refGene_gn=KLK12;refGene_tx=XM_019818620.1
E2 7345322 7345482 refGene_gn=KLK12;refGene_tx=XM_019818620.1
E2 7346265 7346525 refGene_gn=KLK12;refGene_tx=XM_019818620.1
E2 7347426 7347560 refGene_gn=KLK12;refGene_tx=XM_019818620.1
E2 7348457 7348613 refGene_gn=KLK12;refGene_tx=XM_019818620.1
E2 7351929 7351969
refGene_gn=KLK11;refGene_tx=XM_006940939.2;refGene_gn=KLK11;refGene_tx=XM_019818638.1
E2 7352407 7352564
refGene_gn=KLK11;refGene_tx=XM_006940939.2;refGene_gn=KLK11;refGene_tx=XM_019818638.1
E2 7352938 7353204
refGene_gn=KLK11;refGene_tx=XM_006940939.2;refGene_gn=KLK11;refGene_tx=XM_019818638.1
E2 7353730 7354023
refGene_gn=KLK11;refGene_tx=XM_006940939.2;refGene_gn=KLK11;refGene_tx=XM_019818638.1
E2 7354283 7354436 refGene_gn=KLK11;refGene_tx=XM_006940939.2
E2 7357346 7357437 refGene_gn=KLK10;refGene_tx=XM_003997472.5
E2 7358131 7358312 refGene_gn=KLK10;refGene_tx=XM_003997472.5
E2 7358907 7359041 refGene_gn=KLK10;refGene_tx=XM_003997472.5
E2 7359043 7359181 refGene_gn=KLK10;refGene_tx=XM_003997472.5
E2 7359406 7359435 refGene_gn=KLK10;refGene_tx=XM_003997472.5
E2 7359436 7359541 refGene_gn=KLK10;refGene_tx=XM_003997472.5
E2 7360285 7360438 refGene_gn=KLK10;refGene_tx=XM_003997472.5
E2 7365371 7365414
refGene_gn=KLK9;refGene_tx=XM_023245237.1;refGene_gn=KLK9;refGene_tx=XM_023245238.1
E2 7365601 7365758
refGene_gn=KLK9;refGene_tx=XM_023245237.1;refGene_gn=KLK9;refGene_tx=XM_023245238.1
E2 7367964 7368230
refGene_gn=KLK9;refGene_tx=XM_023245237.1;refGene_gn=KLK9;refGene_tx=XM_023245238.1
E2 7369326 7369463
refGene_gn=KLK9;refGene_tx=XM_023245237.1;refGene_gn=KLK9;refGene_tx=XM_023245238.1
E2 7369899 7370052
refGene_gn=KLK9;refGene_tx=XM_023245237.1;refGene_gn=KLK9;refGene_tx=XM_023245238.1
E2 7371983 7372053
refGene_gn=KLK8;refGene_tx=XM_019818629.2;refGene_gn=KLK8;refGene_tx=XM_019818630.1;refGene_gn=KLK8;refGene_tx=XM_019818631.2
E2 7372530 7372690
refGene_gn=KLK8;refGene_tx=XM_019818629.2;refGene_gn=KLK8;refGene_tx=X

M_019818630.1;refGene_gn=KLK8;refGene_tx=XM_019818631.2
E2 7372846 7373106
refGene_gn=KLK8;refGene_tx=XM_019818629.2;refGene_gn=KLK8;refGene_tx=XM_019818630.1;refGene_gn=KLK8;refGene_tx=XM_019818631.2
E2 7375823 7375957
refGene_gn=KLK8;refGene_tx=XM_019818629.2;refGene_gn=KLK8;refGene_tx=XM_019818630.1;refGene_gn=KLK8;refGene_tx=XM_019818631.2
E2 7376185 7376323 refGene_gn=KLK8;refGene_tx=XM_019818631.2
E2 7377141 7377297
refGene_gn=KLK8;refGene_tx=XM_019818629.2;refGene_gn=KLK8;refGene_tx=XM_019818630.1
E2 7386711 7386784 refGene_gn=KLK7;refGene_tx=XM_003997473.3
E2 7387248 7387408 refGene_gn=KLK7;refGene_tx=XM_003997473.3
E2 7387589 7387837 refGene_gn=KLK7;refGene_tx=XM_003997473.3
E2 7388025 7388162 refGene_gn=KLK7;refGene_tx=XM_003997473.3
E2 7390156 7390312 refGene_gn=KLK7;refGene_tx=XM_003997473.3
E2 7396715 7396758
refGene_gn=KLK6;refGene_tx=XM_011289717.3;refGene_gn=KLK6;refGene_tx=XM_019818645.2
E2 7397060 7397217
refGene_gn=KLK6;refGene_tx=XM_011289717.3;refGene_gn=KLK6;refGene_tx=XM_019818645.2
E2 7400367 7400615
refGene_gn=KLK6;refGene_tx=XM_011289717.3;refGene_gn=KLK6;refGene_tx=XM_019818645.2
E2 7401748 7401885
refGene_gn=KLK6;refGene_tx=XM_011289717.3;refGene_gn=KLK6;refGene_tx=XM_019818645.2
E2 7403089 7403242
refGene_gn=KLK6;refGene_tx=XM_011289717.3;refGene_gn=KLK6;refGene_tx=XM_019818645.2
E2 7408381 7408454 refGene_gn=KLK5;refGene_tx=XM_003997480.3
E2 7410320 7410591 refGene_gn=KLK5;refGene_tx=XM_003997480.3
E2 7411115 7411372 refGene_gn=KLK5;refGene_tx=XM_003997480.3
E2 7411496 7411630 refGene_gn=KLK5;refGene_tx=XM_003997480.3
E2 7414949 7415105 refGene_gn=KLK5;refGene_tx=XM_003997480.3
E2 7441102 7441138 refGene_gn=KLK4;refGene_tx=XM_023245412.1
E2 7441139 7441164 refGene_gn=KLK4;refGene_tx=XM_023245412.1
E2 7442438 7442601 refGene_gn=KLK4;refGene_tx=XM_023245412.1
E2 7443018 7443269 refGene_gn=KLK4;refGene_tx=XM_023245412.1
E2 7443355 7443492 refGene_gn=KLK4;refGene_tx=XM_023245412.1
E2 7444082 7444235 refGene_gn=KLK4;refGene_tx=XM_023245412.1
E2 7451521 7451722
refGene_gn=LOC101097936;refGene_tx=XM_019818671.2
E2 7453459 7453596
refGene_gn=LOC101097936;refGene_tx=XM_019818671.2
E2 7453727 7454032
refGene_gn=LOC101097936;refGene_tx=XM_019818671.2
E2 7455264 7455424
refGene_gn=LOC101097936;refGene_tx=XM_019818671.2

E2 7456394 7456440
refGene_gn=LOC101097936;refGene_tx=XM_019818671.2
E2 7474617 7474713 refGene_gn=KLK15;refGene_tx=XM_023245903.1
E2 7475810 7475919 refGene_gn=KLK15;refGene_tx=XM_023245903.1
E2 7476137 7476235 refGene_gn=KLK15;refGene_tx=XM_023245903.1
E2 7484528 7484789 refGene_gn=KLK15;refGene_tx=XM_023245903.1
E2 7484888 7484953 refGene_gn=KLK15;refGene_tx=XM_023245903.1
E2 7485038 7485105 refGene_gn=KLK15;refGene_tx=XM_023245903.1
E2 7485107 7485339 refGene_gn=KLK15;refGene_tx=XM_023245903.1
E2 7485459 7485549 refGene_gn=KLK15;refGene_tx=XM_023245903.1
E2 7485551 7485595 refGene_gn=KLK15;refGene_tx=XM_023245903.1
E2 7486237 7486402 refGene_gn=KLK15;refGene_tx=XM_023245903.1
E2 7488198 7488244 refGene_gn=KLK1;refGene_tx=XM_019818628.1
E2 7489652 7489812 refGene_gn=KLK1;refGene_tx=XM_019818628.1
E2 7491059 7491346 refGene_gn=KLK1;refGene_tx=XM_019818628.1
E2 7491483 7491620 refGene_gn=KLK1;refGene_tx=XM_019818628.1
E2 7492228 7492384 refGene_gn=KLK1;refGene_tx=XM_019818628.1
E2 7493216 7493457
refGene_gn=LOC102899222;refGene_tx=XM_006940948.4;refGene_gn=LOC102899222;refGene_tx=XM_023244921.1;refGene_gn=LOC102899222;refGene_tx=XM_023244922.1;refGene_gn=LOC102899222;refGene_tx=XM_023244923.1;refGene_gn=LOC102899222;refGene_tx=XM_023244924.1
E2 7493642 7493776
refGene_gn=LOC102899222;refGene_tx=XM_006940948.4;refGene_gn=LOC102899222;refGene_tx=XM_023244921.1;refGene_gn=LOC102899222;refGene_tx=XM_023244922.1;refGene_gn=LOC102899222;refGene_tx=XM_023244923.1;refGene_gn=LOC102899222;refGene_tx=XM_023244924.1;refGene_gn=LOC102899222;refGene_tx=XM_023244925.1
E2 7494182 7494329
refGene_gn=LOC102899222;refGene_tx=XM_006940948.4;refGene_gn=LOC102899222;refGene_tx=XM_023244921.1;refGene_gn=LOC102899222;refGene_tx=XM_023244922.1;refGene_gn=LOC102899222;refGene_tx=XM_023244923.1;refGene_gn=LOC102899222;refGene_tx=XM_023244924.1;refGene_gn=LOC102899222;refGene_tx=XM_023244925.1
E2 7499981 7500182
refGene_gn=LOC102899222;refGene_tx=XM_006940948.4;refGene_gn=LOC102899222;refGene_tx=XM_023244921.1;refGene_gn=LOC102899222;refGene_tx=XM_023244922.1;refGene_gn=LOC102899222;refGene_tx=XM_023244923.1;refGene_gn=LOC102899222;refGene_tx=XM_023244924.1;refGene_gn=LOC102899222;refGene_tx=XM_023244925.1
E2 7500652 7500701
refGene_gn=LOC102899222;refGene_tx=XM_023244921.1;refGene_gn=LOC102899222;refGene_tx=XM_023244922.1;refGene_gn=LOC102899222;refGene_tx=XM_023244923.1;refGene_gn=LOC102899222;refGene_tx=XM_023244924.1;refGene_gn=LOC102899222;refGene_tx=XM_023244925.1
E2 7500964 7501043
refGene_gn=LOC102899222;refGene_tx=XM_023244921.1;refGene_gn=LOC102899222;refGene_tx=XM_023244922.1;refGene_gn=LOC102899222;refGene_tx=XM_023244923.1;refGene_gn=LOC102899222;refGene_tx=XM_023244924.1;refGene_gn=LOC102899222;refGene_tx=XM_023244925.1

E2 7502637 7502699
refGene_gn=LOC102899222;refGene_tx=XM_023244921.1;refGene_gn=LOC102899222;refGene_tx=XM_023244922.1;refGene_gn=LOC102899222;refGene_tx=XM_023244923.1;refGene_gn=LOC102899222;refGene_tx=XM_023244924.1;refGene_gn=LOC102899222;refGene_tx=XM_023244925.1
E2 7502946 7503824
refGene_gn=LOC102899129;refGene_tx=XM_006940945.4;refGene_gn=LOC102899129;refGene_tx=XM_019818586.2
E2 7504238 7504516
refGene_gn=LOC102899129;refGene_tx=XM_006940945.4;refGene_gn=LOC102899129;refGene_tx=XM_019818586.2
E2 7505076 7505223
refGene_gn=LOC102899129;refGene_tx=XM_006940945.4;refGene_gn=LOC102899129;refGene_tx=XM_019818586.2
E2 7507126 7507258
refGene_gn=LOC102899129;refGene_tx=XM_006940945.4
E2 7507554 7507622
refGene_gn=LOC102899129;refGene_tx=XM_006940945.4
E2 7511345 7511469
refGene_gn=LOC102899286;refGene_tx=XM_023244919.1
E2 7513378 7513421
refGene_gn=LOC102899286;refGene_tx=XM_023244919.1
E2 7514226 7515163
refGene_gn=LOC102899286;refGene_tx=XM_023244919.1;refGene_gn=LOC102899286;refGene_tx=XM_023244920.1
E2 7517733 7517849 refGene_gn=ACP4;refGene_tx=XM_003997468.4
E2 7517927 7518106 refGene_gn=ACP4;refGene_tx=XM_003997468.4
E2 7518349 7518474 refGene_gn=ACP4;refGene_tx=XM_003997468.4
E2 7518905 7518988 refGene_gn=ACP4;refGene_tx=XM_003997468.4
E2 7519072 7519205 refGene_gn=ACP4;refGene_tx=XM_003997468.4
E2 7519860 7519956 refGene_gn=ACP4;refGene_tx=XM_003997468.4
E2 7520059 7520158 refGene_gn=ACP4;refGene_tx=XM_003997468.4
E2 7520495 7520642 refGene_gn=ACP4;refGene_tx=XM_003997468.4
E2 7521688 7521775 refGene_gn=ACP4;refGene_tx=XM_003997468.4
E2 7521854 7521959 refGene_gn=ACP4;refGene_tx=XM_003997468.4
E2 7522030 7522047 refGene_gn=ACP4;refGene_tx=XM_003997468.4
E2 7522049 7522128 refGene_gn=ACP4;refGene_tx=XM_003997468.4
E2 7550816 7551262 refGene_gn=CLEC11A;refGene_tx=XM_003997469.5
E2 7551699 7551891 refGene_gn=CLEC11A;refGene_tx=XM_003997469.5
E2 7552202 7552392 refGene_gn=CLEC11A;refGene_tx=XM_003997469.5
E2 7552621 7552768 refGene_gn=CLEC11A;refGene_tx=XM_003997469.5
E2 7559503 7559758 refGene_gn=SHANK1;refGene_tx=XM_023244937.1
E2 7559961 7560165 refGene_gn=SHANK1;refGene_tx=XM_023244937.1
E2 7560673 7560745 refGene_gn=SHANK1;refGene_tx=XM_023244937.1
E2 7561837 7561946 refGene_gn=SHANK1;refGene_tx=XM_023244937.1
E2 7562037 7562189 refGene_gn=SHANK1;refGene_tx=XM_023244937.1
E2 7564049 7564217 refGene_gn=SHANK1;refGene_tx=XM_023244937.1
E2 7564645 7564762 refGene_gn=SHANK1;refGene_tx=XM_023244937.1
E2 7568677 7568755 refGene_gn=SHANK1;refGene_tx=XM_023244937.1
E2 7568992 7569059 refGene_gn=SHANK1;refGene_tx=XM_023244937.1

E2 7569295 7569626 refGene_gn=SHANK1;refGene_tx=XM_023244937.1
E2 7570526 7570720 refGene_gn=SHANK1;refGene_tx=XM_023244937.1
E2 7573970 7574084 refGene_gn=SHANK1;refGene_tx=XM_023244937.1
E2 7574263 7574339 refGene_gn=SHANK1;refGene_tx=XM_023244937.1
E2 7574776 7574806 refGene_gn=SHANK1;refGene_tx=XM_023244937.1
E2 7582538 7582621 refGene_gn=SHANK1;refGene_tx=XM_023244937.1
E2 7582831 7582956 refGene_gn=SHANK1;refGene_tx=XM_023244937.1
E2 7583525 7583661 refGene_gn=SHANK1;refGene_tx=XM_023244937.1
E2 7583938 7584019 refGene_gn=SHANK1;refGene_tx=XM_023244937.1
E2 7584797 7584866 refGene_gn=SHANK1;refGene_tx=XM_023244937.1
E2 7585312 7585431 refGene_gn=SHANK1;refGene_tx=XM_023244937.1
E2 7592573 7592670 refGene_gn=SHANK1;refGene_tx=XM_023244937.1
E2 7594572 7595588 refGene_gn=SHANK1;refGene_tx=XM_023244937.1
E2 7595589 7597721 refGene_gn=SHANK1;refGene_tx=XM_023244937.1
E2 7599643 7600358 refGene_gn=SHANK1;refGene_tx=XM_023244937.1
E2 7603269 7603465
refGene_gn=CE2H19orf81;refGene_tx=XM_011289752.3
E2 7604306 7604446
refGene_gn=CE2H19orf81;refGene_tx=XM_011289752.3
E2 7605983 7606104
refGene_gn=CE2H19orf81;refGene_tx=XM_011289752.3
E2 7606210 7606283
refGene_gn=CE2H19orf81;refGene_tx=XM_011289752.3
E2 7611718 7611914
refGene_gn=CE2H19orf81;refGene_tx=XM_011289752.3
E2 7621047 7621198
refGene_gn=SYT3;refGene_tx=XM_003997466.4;refGene_gn=SYT3;refGene_tx=XM_006940950.2;refGene_gn=SYT3;refGene_tx=XM_019818564.2;refGene_gn=SYT3;refGene_tx=XM_019818565.2;refGene_gn=SYT3;refGene_tx=XM_019818566.2;refGene_gn=SYT3;refGene_tx=XM_019818567.2;refGene_gn=SYT3;refGene_tx=XM_023245330.1;refGene_gn=SYT3;refGene_tx=XM_023245331.1
E2 7624505 7625031
refGene_gn=SYT3;refGene_tx=XM_003997466.4;refGene_gn=SYT3;refGene_tx=XM_006940950.2;refGene_gn=SYT3;refGene_tx=XM_019818564.2;refGene_gn=SYT3;refGene_tx=XM_019818565.2;refGene_gn=SYT3;refGene_tx=XM_019818566.2;refGene_gn=SYT3;refGene_tx=XM_019818567.2;refGene_gn=SYT3;refGene_tx=XM_023245330.1;refGene_gn=SYT3;refGene_tx=XM_023245331.1
E2 7627500 7627888
refGene_gn=SYT3;refGene_tx=XM_003997466.4;refGene_gn=SYT3;refGene_tx=XM_006940950.2;refGene_gn=SYT3;refGene_tx=XM_019818564.2;refGene_gn=SYT3;refGene_tx=XM_019818565.2;refGene_gn=SYT3;refGene_tx=XM_019818566.2;refGene_gn=SYT3;refGene_tx=XM_019818567.2;refGene_gn=SYT3;refGene_tx=XM_023245330.1;refGene_gn=SYT3;refGene_tx=XM_023245331.1
E2 7628161 7628380
refGene_gn=SYT3;refGene_tx=XM_003997466.4;refGene_gn=SYT3;refGene_tx=XM_006940950.2;refGene_gn=SYT3;refGene_tx=XM_019818564.2;refGene_gn=SYT3;refGene_tx=XM_019818565.2;refGene_gn=SYT3;refGene_tx=XM_019818566.2;refGene_gn=SYT3;refGene_tx=XM_019818567.2;refGene_gn=SYT3;refGene_tx=XM_023245330.1;refGene_gn=SYT3;refGene_tx=XM_023245331.1
E2 7630479 7630600

refGene_gn=SYT3;refGene_tx=XM_003997466.4;refGene_gn=SYT3;refGene_tx=XM_006940950.2;refGene_gn=SYT3;refGene_tx=XM_019818564.2;refGene_gn=SYT3;refGene_tx=XM_019818565.2;refGene_gn=SYT3;refGene_tx=XM_019818566.2;refGene_gn=SYT3;refGene_tx=XM_019818567.2;refGene_gn=SYT3;refGene_tx=XM_023245330.1;refGene_gn=SYT3;refGene_tx=XM_023245331.1
E2 7631203 7631375
refGene_gn=SYT3;refGene_tx=XM_003997466.4;refGene_gn=SYT3;refGene_tx=XM_006940950.2;refGene_gn=SYT3;refGene_tx=XM_019818564.2;refGene_gn=SYT3;refGene_tx=XM_019818565.2;refGene_gn=SYT3;refGene_tx=XM_019818566.2;refGene_gn=SYT3;refGene_tx=XM_019818567.2;refGene_gn=SYT3;refGene_tx=XM_023245330.1;refGene_gn=SYT3;refGene_tx=XM_023245331.1
E2 7631480 7631613
refGene_gn=SYT3;refGene_tx=XM_003997466.4;refGene_gn=SYT3;refGene_tx=XM_006940950.2;refGene_gn=SYT3;refGene_tx=XM_019818564.2;refGene_gn=SYT3;refGene_tx=XM_019818565.2;refGene_gn=SYT3;refGene_tx=XM_019818566.2;refGene_gn=SYT3;refGene_tx=XM_019818567.2;refGene_gn=SYT3;refGene_tx=XM_023245330.1;refGene_gn=SYT3;refGene_tx=XM_023245331.1
E2 7634257 7634323
refGene_gn=SYT3;refGene_tx=XM_003997466.4;refGene_gn=SYT3;refGene_tx=XM_006940950.2;refGene_gn=SYT3;refGene_tx=XM_019818564.2;refGene_gn=SYT3;refGene_tx=XM_019818565.2;refGene_gn=SYT3;refGene_tx=XM_019818566.2;refGene_gn=SYT3;refGene_tx=XM_019818567.2;refGene_gn=SYT3;refGene_tx=XM_023245330.1;refGene_gn=SYT3;refGene_tx=XM_023245331.1
E2 7680664 7680961
refGene_gn=LRRC4B;refGene_tx=XM_023245952.1;refGene_gn=LRRC4B;refGene_tx=XM_023245953.1;refGene_gn=LRRC4B;refGene_tx=XM_023245954.1
E2 7702262 7704113
refGene_gn=LRRC4B;refGene_tx=XM_023245952.1;refGene_gn=LRRC4B;refGene_tx=XM_023245953.1;refGene_gn=LRRC4B;refGene_tx=XM_023245954.1
E2 7706179 7706235
refGene_gn=ASPDH;refGene_tx=XM_023245957.1;refGene_gn=ASPDH;refGene_tx=XM_023245959.1;refGene_gn=ASPDH;refGene_tx=XM_023245961.1
E2 7706387 7706814
refGene_gn=ASPDH;refGene_tx=XM_019818594.2;refGene_gn=ASPDH;refGene_tx=XM_023245955.1;refGene_gn=ASPDH;refGene_tx=XM_023245956.1;refGene_gn=ASPDH;refGene_tx=XM_023245957.1;refGene_gn=ASPDH;refGene_tx=XM_023245960.1
E2 7707048 7707103
refGene_gn=ASPDH;refGene_tx=XM_019818593.2;refGene_gn=ASPDH;refGene_tx=XM_019818594.2;refGene_gn=ASPDH;refGene_tx=XM_023245956.1;refGene_gn=ASPDH;refGene_tx=XM_023245957.1;refGene_gn=ASPDH;refGene_tx=XM_023245959.1;refGene_gn=ASPDH;refGene_tx=XM_023245960.1;refGene_gn=ASPDH;refGene_tx=XM_023245961.1
E2 7707516 7707661
refGene_gn=ASPDH;refGene_tx=XM_019818593.2;refGene_gn=ASPDH;refGene_tx=XM_019818594.2;refGene_gn=ASPDH;refGene_tx=XM_023245955.1;refGene_gn=ASPDH;refGene_tx=XM_023245956.1;refGene_gn=ASPDH;refGene_tx=XM_023245957.1;refGene_gn=ASPDH;refGene_tx=XM_023245959.1;refGene_gn=ASPDH;refGene_tx=XM_023245960.1;refGene_gn=ASPDH;refGene_tx=XM_023245961.1
E2 7707903 7707988

refGene_gn=ASPDH;refGene_tx=XM_019818593.2;refGene_gn=ASPDH;refGene_tx=XM_019818594.2;refGene_gn=ASPDH;refGene_tx=XM_023245955.1;refGene_gn=ASPDH;refGene_tx=XM_023245956.1;refGene_gn=ASPDH;refGene_tx=XM_023245957.1;refGene_gn=ASPDH;refGene_tx=XM_023245959.1;refGene_gn=ASPDH;refGene_tx=XM_023245960.1;refGene_gn=ASPDH;refGene_tx=XM_023245961.1
E2 7708084 7708234
refGene_gn=ASPDH;refGene_tx=XM_019818593.2;refGene_gn=ASPDH;refGene_tx=XM_019818594.2;refGene_gn=ASPDH;refGene_tx=XM_023245955.1;refGene_gn=ASPDH;refGene_tx=XM_023245956.1;refGene_gn=ASPDH;refGene_tx=XM_023245957.1;refGene_gn=ASPDH;refGene_tx=XM_023245959.1;refGene_gn=ASPDH;refGene_tx=XM_023245960.1;refGene_gn=ASPDH;refGene_tx=XM_023245961.1
E2 7708317 7708538
refGene_gn=ASPDH;refGene_tx=XM_019818593.2;refGene_gn=ASPDH;refGene_tx=XM_019818594.2;refGene_gn=ASPDH;refGene_tx=XM_023245955.1;refGene_gn=ASPDH;refGene_tx=XM_023245956.1;refGene_gn=ASPDH;refGene_tx=XM_023245957.1;refGene_gn=ASPDH;refGene_tx=XM_023245959.1;refGene_gn=ASPDH;refGene_tx=XM_023245960.1;refGene_gn=ASPDH;refGene_tx=XM_023245961.1
E2 7708611 7708766
refGene_gn=ASPDH;refGene_tx=XM_019818593.2;refGene_gn=ASPDH;refGene_tx=XM_019818594.2;refGene_gn=ASPDH;refGene_tx=XM_023245955.1;refGene_gn=ASPDH;refGene_tx=XM_023245956.1;refGene_gn=ASPDH;refGene_tx=XM_023245957.1;refGene_gn=ASPDH;refGene_tx=XM_023245959.1;refGene_gn=ASPDH;refGene_tx=XM_023245960.1;refGene_gn=ASPDH;refGene_tx=XM_023245961.1
E2 7709152 7709196
refGene_gn=ASPDH;refGene_tx=XM_019818593.2;refGene_gn=ASPDH;refGene_tx=XM_019818594.2;refGene_gn=ASPDH;refGene_tx=XM_023245955.1;refGene_gn=ASPDH;refGene_tx=XM_023245956.1;refGene_gn=ASPDH;refGene_tx=XM_023245957.1;refGene_gn=ASPDH;refGene_tx=XM_023245959.1;refGene_gn=ASPDH;refGene_tx=XM_023245960.1;refGene_gn=ASPDH;refGene_tx=XM_023245961.1
E2 7710443 7710589
refGene_gn=JOSD2;refGene_tx=XM_006940951.4;refGene_gn=JOSD2;refGene_tx=XM_023245962.1;refGene_gn=JOSD2;refGene_tx=XM_023245963.1;refGene_gn=JOSD2;refGene_tx=XM_023245964.1;refGene_gn=JOSD2;refGene_tx=XM_023245965.1
E2 7711391 7711448
refGene_gn=JOSD2;refGene_tx=XM_023245962.1;refGene_gn=JOSD2;refGene_tx=XM_023245963.1;refGene_gn=JOSD2;refGene_tx=XM_023245964.1;refGene_gn=JOSD2;refGene_tx=XM_023245965.1
E2 7712090 7712216
refGene_gn=JOSD2;refGene_tx=XM_006940951.4;refGene_gn=JOSD2;refGene_tx=XM_023245962.1;refGene_gn=JOSD2;refGene_tx=XM_023245963.1;refGene_gn=JOSD2;refGene_tx=XM_023245964.1;refGene_gn=JOSD2;refGene_tx=XM_023245965.1
E2 7712865 7713060
refGene_gn=JOSD2;refGene_tx=XM_006940951.4;refGene_gn=JOSD2;refGene_tx=XM_023245962.1;refGene_gn=JOSD2;refGene_tx=XM_023245963.1;refGene_gn=JOSD2;refGene_tx=XM_023245964.1;refGene_gn=JOSD2;refGene_tx=XM_023245965.1
E2 7713163 7713263
refGene_gn=JOSD2;refGene_tx=XM_006940951.4;refGene_gn=JOSD2;refGene_tx

=XM_023245962.1;refGene_gn=JOSD2;refGene_tx=XM_023245963.1;refGene_gn=JOSD2;refGene_tx=XM_023245964.1;refGene_gn=JOSD2;refGene_tx=XM_023245965.1
E2 7729796 7729907 refGene_gn=EMC10;refGene_tx=XM_003997458.5
E2 7730131 7730218 refGene_gn=EMC10;refGene_tx=XM_003997459.4
E2 7731125 7731219 refGene_gn=EMC10;refGene_tx=XM_003997458.5;refGene_gn=EMC10;refGene_tx=XM_003997459.4
E2 7732336 7732518 refGene_gn=EMC10;refGene_tx=XM_003997458.5;refGene_gn=EMC10;refGene_tx=XM_003997459.4
E2 7732856 7732961 refGene_gn=EMC10;refGene_tx=XM_003997458.5;refGene_gn=EMC10;refGene_tx=XM_003997459.4
E2 7733886 7733996 refGene_gn=EMC10;refGene_tx=XM_003997458.5;refGene_gn=EMC10;refGene_tx=XM_003997459.4
E2 7734914 7734987 refGene_gn=EMC10;refGene_tx=XM_003997458.5;refGene_gn=EMC10;refGene_tx=XM_003997459.4
E2 7736142 7736256 refGene_gn=EMC10;refGene_tx=XM_003997458.5;refGene_gn=EMC10;refGene_tx=XM_003997459.4
E2 7736564 7736637 refGene_gn=FAM71E1;refGene_tx=XM_006940953.4;refGene_gn=FAM71E1;refGene_tx=XM_019818644.2
E2 7736744 7736944 refGene_gn=FAM71E1;refGene_tx=XM_006940953.4;refGene_gn=FAM71E1;refGene_tx=XM_019818644.2
E2 7737293 7737424 refGene_gn=FAM71E1;refGene_tx=XM_006940953.4;refGene_gn=FAM71E1;refGene_tx=XM_019818644.2
E2 7740575 7740858 refGene_gn=FAM71E1;refGene_tx=XM_006940953.4;refGene_gn=FAM71E1;refGene_tx=XM_019818644.2
E2 7741454 7741511 refGene_gn=FAM71E1;refGene_tx=XM_006940953.4;refGene_gn=FAM71E1;refGene_tx=XM_019818644.2
E2 7742232 7742243 refGene_gn=MYBPC2;refGene_tx=XM_006940954.4
E2 7743588 7743775 refGene_gn=MYBPC2;refGene_tx=XM_006940954.4
E2 7746046 7746183 refGene_gn=MYBPC2;refGene_tx=XM_006940954.4
E2 7746368 7746528 refGene_gn=MYBPC2;refGene_tx=XM_006940954.4
E2 7747705 7747845 refGene_gn=MYBPC2;refGene_tx=XM_006940954.4
E2 7748079 7748275 refGene_gn=MYBPC2;refGene_tx=XM_006940954.4
E2 7748367 7748456 refGene_gn=MYBPC2;refGene_tx=XM_006940954.4
E2 7748632 7748800 refGene_gn=MYBPC2;refGene_tx=XM_006940954.4
E2 7750375 7750510 refGene_gn=MYBPC2;refGene_tx=XM_006940954.4
E2 7750893 7751082 refGene_gn=MYBPC2;refGene_tx=XM_006940954.4
E2 7751533 7751638 refGene_gn=MYBPC2;refGene_tx=XM_006940954.4
E2 7751722 7751882 refGene_gn=MYBPC2;refGene_tx=XM_006940954.4

E2 7752709 7752849 refGene_gn=MYBPC2;refGene_tx=XM_006940954.4
E2 7753683 7753713 refGene_gn=MYBPC2;refGene_tx=XM_006940954.4
E2 7754034 7754141 refGene_gn=MYBPC2;refGene_tx=XM_006940954.4
E2 7755582 7755748 refGene_gn=MYBPC2;refGene_tx=XM_006940954.4
E2 7757967 7758140 refGene_gn=MYBPC2;refGene_tx=XM_006940954.4
E2 7759743 7759849 refGene_gn=MYBPC2;refGene_tx=XM_006940954.4
E2 7759939 7760064 refGene_gn=MYBPC2;refGene_tx=XM_006940954.4
E2 7762618 7762751 refGene_gn=MYBPC2;refGene_tx=XM_006940954.4
E2 7763021 7763215 refGene_gn=MYBPC2;refGene_tx=XM_006940954.4
E2 7764612 7764672 refGene_gn=MYBPC2;refGene_tx=XM_006940954.4
E2 7764839 7764888 refGene_gn=MYBPC2;refGene_tx=XM_006940954.4
E2 7765537 7765655 refGene_gn=MYBPC2;refGene_tx=XM_006940954.4
E2 7766048 7766197 refGene_gn=MYBPC2;refGene_tx=XM_006940954.4
E2 7766408 7766495 refGene_gn=MYBPC2;refGene_tx=XM_006940954.4
E2 7766993 7767083 refGene_gn=MYBPC2;refGene_tx=XM_006940954.4
E2 7768143 7768162 refGene_gn=MYBPC2;refGene_tx=XM_006940954.4
E2 7771897 7772196
refGene_gn=SPIB;refGene_tx=XM_011289709.3;refGene_gn=SPIB;refGene_tx=X
M_019818626.2
E2 7773377 7773528
refGene_gn=SPIB;refGene_tx=XM_011289709.3;refGene_gn=SPIB;refGene_tx=X
M_019818626.2
E2 7773915 7774130
refGene_gn=SPIB;refGene_tx=XM_011289709.3;refGene_gn=SPIB;refGene_tx=X
M_019818626.2
E2 7774408 7774481
refGene_gn=SPIB;refGene_tx=XM_011289709.3;refGene_gn=SPIB;refGene_tx=X
M_019818626.2
E2 7776108 7776162
refGene_gn=SPIB;refGene_tx=XM_011289709.3;refGene_gn=SPIB;refGene_tx=X
M_019818626.2
E2 7777090 7777113 refGene_gn=SPIB;refGene_tx=XM_011289709.3
E2 7778095 7778201 refGene_gn=POLD1;refGene_tx=XM_023245137.1
E2 7778819 7778917 refGene_gn=POLD1;refGene_tx=XM_023245137.1
E2 7778997 7779050 refGene_gn=POLD1;refGene_tx=XM_023245137.1
E2 7779405 7779519 refGene_gn=POLD1;refGene_tx=XM_023245137.1
E2 7779604 7779737 refGene_gn=POLD1;refGene_tx=XM_023245137.1
E2 7780501 7780604 refGene_gn=POLD1;refGene_tx=XM_023245137.1
E2 7780740 7780893 refGene_gn=POLD1;refGene_tx=XM_023245137.1
E2 7781164 7781340 refGene_gn=POLD1;refGene_tx=XM_023245137.1
E2 7782223 7782361 refGene_gn=POLD1;refGene_tx=XM_023245137.1
E2 7782468 7782564 refGene_gn=POLD1;refGene_tx=XM_023245137.1
E2 7785092 7785240 refGene_gn=POLD1;refGene_tx=XM_023245137.1
E2 7785562 7785676 refGene_gn=POLD1;refGene_tx=XM_023245137.1
E2 7785830 7785947 refGene_gn=POLD1;refGene_tx=XM_023245137.1
E2 7786030 7786119 refGene_gn=POLD1;refGene_tx=XM_023245137.1
E2 7786285 7786477 refGene_gn=POLD1;refGene_tx=XM_023245137.1
E2 7786904 7787015 refGene_gn=POLD1;refGene_tx=XM_023245137.1
E2 7787110 7787251 refGene_gn=POLD1;refGene_tx=XM_023245137.1
E2 7790508 7790613 refGene_gn=POLD1;refGene_tx=XM_023245137.1

E2 7790941 7791108 refGene_gn=POLD1;refGene_tx=XM_023245137.1
E2 7791616 7791746 refGene_gn=POLD1;refGene_tx=XM_023245137.1
E2 7791817 7791899 refGene_gn=POLD1;refGene_tx=XM_023245137.1
E2 7792000 7792169 refGene_gn=POLD1;refGene_tx=XM_023245137.1
E2 7792240 7792366 refGene_gn=POLD1;refGene_tx=XM_023245137.1
E2 7792450 7792597 refGene_gn=POLD1;refGene_tx=XM_023245137.1
E2 7794782 7794896 refGene_gn=POLD1;refGene_tx=XM_023245137.1
E2 7795258 7795460 refGene_gn=POLD1;refGene_tx=XM_023245137.1
E2 7810578 7810725 refGene_gn=NR1H2;refGene_tx=XM_006940956.4
E2 7811061 7811270 refGene_gn=NR1H2;refGene_tx=XM_006940956.4
E2 7813562 7813662 refGene_gn=NR1H2;refGene_tx=XM_006940956.4
E2 7814319 7814499 refGene_gn=NR1H2;refGene_tx=XM_006940956.4
E2 7814698 7814970 refGene_gn=NR1H2;refGene_tx=XM_006940956.4
E2 7815052 7815343 refGene_gn=NR1H2;refGene_tx=XM_006940956.4
E2 7815715 7815847 refGene_gn=NR1H2;refGene_tx=XM_006940956.4
E2 7815927 7815970 refGene_gn=NR1H2;refGene_tx=XM_006940956.4
E2 7823023 7823090
refGene_gn=LOC102899949;refGene_tx=XM_019818641.2
E2 7825412 7825489 refGene_gn=NAPSA;refGene_tx=XM_003997470.4
E2 7827042 7827058
refGene_gn=LOC102899949;refGene_tx=XM_023245070.1
E2 7828846 7828988 refGene_gn=NAPSA;refGene_tx=XM_003997470.4
E2 7829075 7829199 refGene_gn=NAPSA;refGene_tx=XM_003997470.4
E2 7829392 7829511 refGene_gn=NAPSA;refGene_tx=XM_003997470.4
E2 7829806 7830006 refGene_gn=NAPSA;refGene_tx=XM_003997470.4
E2 7830117 7830121
refGene_gn=LOC102899949;refGene_tx=XM_019818642.2
E2 7830503 7830592
refGene_gn=LOC102899949;refGene_tx=XM_019818641.2;refGene_gn=LOC102899
949;refGene_tx=XM_019818642.2;refGene_gn=LOC102899949;refGene_tx=XM_02
3245070.1
E2 7831594 7831717 refGene_gn=NAPSA;refGene_tx=XM_003997470.4
E2 7831958 7832103 refGene_gn=NAPSA;refGene_tx=XM_003997470.4
E2 7834470 7834553
refGene_gn=LOC102899949;refGene_tx=XM_019818641.2;refGene_gn=LOC102899
949;refGene_tx=XM_019818642.2;refGene_gn=LOC102899949;refGene_tx=XM_02
3245070.1
E2 7834702 7834801 refGene_gn=NAPSA;refGene_tx=XM_003997470.4
E2 7835003 7835243 refGene_gn=NAPSA;refGene_tx=XM_003997470.4
E2 7836761 7836806
refGene_gn=LOC102899949;refGene_tx=XM_019818641.2;refGene_gn=LOC102899
949;refGene_tx=XM_019818642.2;refGene_gn=LOC102899949;refGene_tx=XM_02
3245070.1
E2 7837150 7837212
refGene_gn=LOC102899949;refGene_tx=XM_019818641.2;refGene_gn=LOC102899
949;refGene_tx=XM_019818642.2;refGene_gn=LOC102899949;refGene_tx=XM_02
3245070.1
E2 7838001 7838276
refGene_gn=LOC102899949;refGene_tx=XM_019818641.2;refGene_gn=LOC102899
949;refGene_tx=XM_019818642.2;refGene_gn=LOC102899949;refGene_tx=XM_02

3245070.1
E2	7839600	7840485	refGene_gn=KCNC3;refGene_tx=XM_023245069.1
E2	7844660	7845771	refGene_gn=KCNC3;refGene_tx=XM_023245069.1
E2	7847225	7847417	refGene_gn=KCNC3;refGene_tx=XM_023245069.1
E2	7847669	7847689	refGene_gn=KCNC3;refGene_tx=XM_023245069.1
E2	7847690	7847796	refGene_gn=KCNC3;refGene_tx=XM_023245069.1
E2	7848419	7848573	refGene_gn=KCNC3;refGene_tx=XM_023245069.1
E2	7850583	7850643	refGene_gn=KCNC3;refGene_tx=XM_023245069.1
E2	7851740	7851912	refGene_gn=KCNC3;refGene_tx=XM_023245069.1
E2	7856431	7856591
refGene_gn=MYH14;refGene_tx=XM_023245063.1;refGene_gn=MYH14;refGene_tx=XM_023245064.1;refGene_gn=MYH14;refGene_tx=XM_023245065.1;refGene_gn=MYH14;refGene_tx=XM_023245066.1;refGene_gn=MYH14;refGene_tx=XM_023245067.1;refGene_gn=MYH14;refGene_tx=XM_023245068.1
E2	7856752	7856925
refGene_gn=MYH14;refGene_tx=XM_023245063.1;refGene_gn=MYH14;refGene_tx=XM_023245064.1;refGene_gn=MYH14;refGene_tx=XM_023245065.1;refGene_gn=MYH14;refGene_tx=XM_023245066.1;refGene_gn=MYH14;refGene_tx=XM_023245067.1;refGene_gn=MYH14;refGene_tx=XM_023245068.1
E2	7860758	7860867
refGene_gn=MYH14;refGene_tx=XM_023245063.1;refGene_gn=MYH14;refGene_tx=XM_023245064.1;refGene_gn=MYH14;refGene_tx=XM_023245065.1;refGene_gn=MYH14;refGene_tx=XM_023245066.1;refGene_gn=MYH14;refGene_tx=XM_023245067.1;refGene_gn=MYH14;refGene_tx=XM_023245068.1
E2	7863549	7863758
refGene_gn=MYH14;refGene_tx=XM_023245063.1;refGene_gn=MYH14;refGene_tx=XM_023245064.1;refGene_gn=MYH14;refGene_tx=XM_023245065.1;refGene_gn=MYH14;refGene_tx=XM_023245066.1;refGene_gn=MYH14;refGene_tx=XM_023245067.1;refGene_gn=MYH14;refGene_tx=XM_023245068.1
E2	7869363	7869487
refGene_gn=MYH14;refGene_tx=XM_023245063.1;refGene_gn=MYH14;refGene_tx=XM_023245064.1;refGene_gn=MYH14;refGene_tx=XM_023245065.1;refGene_gn=MYH14;refGene_tx=XM_023245066.1;refGene_gn=MYH14;refGene_tx=XM_023245067.1;refGene_gn=MYH14;refGene_tx=XM_023245068.1
E2	7869579	7869668
refGene_gn=MYH14;refGene_tx=XM_023245063.1;refGene_gn=MYH14;refGene_tx=XM_023245064.1;refGene_gn=MYH14;refGene_tx=XM_023245065.1;refGene_gn=MYH14;refGene_tx=XM_023245066.1;refGene_gn=MYH14;refGene_tx=XM_023245067.1;refGene_gn=MYH14;refGene_tx=XM_023245068.1
E2	7870277	7870406
refGene_gn=MYH14;refGene_tx=XM_023245063.1;refGene_gn=MYH14;refGene_tx=XM_023245064.1;refGene_gn=MYH14;refGene_tx=XM_023245065.1;refGene_gn=MYH14;refGene_tx=XM_023245066.1;refGene_gn=MYH14;refGene_tx=XM_023245067.1;refGene_gn=MYH14;refGene_tx=XM_023245068.1
E2	7872973	7873135
refGene_gn=MYH14;refGene_tx=XM_023245063.1;refGene_gn=MYH14;refGene_tx=XM_023245064.1;refGene_gn=MYH14;refGene_tx=XM_023245065.1;refGene_gn=MYH14;refGene_tx=XM_023245066.1;refGene_gn=MYH14;refGene_tx=XM_023245067.1;refGene_gn=MYH14;refGene_tx=XM_023245068.1
E2	7874011	7874224

refGene_gn=MYH14;refGene_tx=XM_023245063.1;refGene_gn=MYH14;refGene_tx=XM_023245064.1;refGene_gn=MYH14;refGene_tx=XM_023245065.1;refGene_gn=MYH14;refGene_tx=XM_023245066.1;refGene_gn=MYH14;refGene_tx=XM_023245067.1;refGene_gn=MYH14;refGene_tx=XM_023245068.1
E2 7876434 7876647
refGene_gn=MYH14;refGene_tx=XM_023245063.1;refGene_gn=MYH14;refGene_tx=XM_023245064.1;refGene_gn=MYH14;refGene_tx=XM_023245065.1;refGene_gn=MYH14;refGene_tx=XM_023245066.1;refGene_gn=MYH14;refGene_tx=XM_023245067.1;refGene_gn=MYH14;refGene_tx=XM_023245068.1
E2 7881839 7882088
refGene_gn=MYH14;refGene_tx=XM_023245063.1;refGene_gn=MYH14;refGene_tx=XM_023245064.1;refGene_gn=MYH14;refGene_tx=XM_023245065.1;refGene_gn=MYH14;refGene_tx=XM_023245066.1;refGene_gn=MYH14;refGene_tx=XM_023245067.1;refGene_gn=MYH14;refGene_tx=XM_023245068.1
E2 7883200 7883353
refGene_gn=MYH14;refGene_tx=XM_023245063.1;refGene_gn=MYH14;refGene_tx=XM_023245064.1;refGene_gn=MYH14;refGene_tx=XM_023245065.1;refGene_gn=MYH14;refGene_tx=XM_023245066.1;refGene_gn=MYH14;refGene_tx=XM_023245067.1;refGene_gn=MYH14;refGene_tx=XM_023245068.1
E2 7883448 7883553
refGene_gn=MYH14;refGene_tx=XM_023245063.1;refGene_gn=MYH14;refGene_tx=XM_023245064.1;refGene_gn=MYH14;refGene_tx=XM_023245065.1;refGene_gn=MYH14;refGene_tx=XM_023245066.1;refGene_gn=MYH14;refGene_tx=XM_023245067.1;refGene_gn=MYH14;refGene_tx=XM_023245068.1
E2 7885663 7885870
refGene_gn=MYH14;refGene_tx=XM_023245063.1;refGene_gn=MYH14;refGene_tx=XM_023245064.1;refGene_gn=MYH14;refGene_tx=XM_023245065.1;refGene_gn=MYH14;refGene_tx=XM_023245066.1;refGene_gn=MYH14;refGene_tx=XM_023245067.1;refGene_gn=MYH14;refGene_tx=XM_023245068.1
E2 7886391 7886536
refGene_gn=MYH14;refGene_tx=XM_023245063.1;refGene_gn=MYH14;refGene_tx=XM_023245064.1;refGene_gn=MYH14;refGene_tx=XM_023245065.1;refGene_gn=MYH14;refGene_tx=XM_023245066.1;refGene_gn=MYH14;refGene_tx=XM_023245067.1;refGene_gn=MYH14;refGene_tx=XM_023245068.1
E2 7887109 7887322
refGene_gn=MYH14;refGene_tx=XM_023245063.1;refGene_gn=MYH14;refGene_tx=XM_023245064.1;refGene_gn=MYH14;refGene_tx=XM_023245065.1;refGene_gn=MYH14;refGene_tx=XM_023245066.1;refGene_gn=MYH14;refGene_tx=XM_023245067.1;refGene_gn=MYH14;refGene_tx=XM_023245068.1
E2 7889680 7889852
refGene_gn=MYH14;refGene_tx=XM_023245063.1;refGene_gn=MYH14;refGene_tx=XM_023245064.1;refGene_gn=MYH14;refGene_tx=XM_023245065.1;refGene_gn=MYH14;refGene_tx=XM_023245066.1;refGene_gn=MYH14;refGene_tx=XM_023245067.1;refGene_gn=MYH14;refGene_tx=XM_023245068.1
E2 7890517 7890641
refGene_gn=MYH14;refGene_tx=XM_023245063.1;refGene_gn=MYH14;refGene_tx=XM_023245064.1;refGene_gn=MYH14;refGene_tx=XM_023245065.1;refGene_gn=MYH14;refGene_tx=XM_023245066.1;refGene_gn=MYH14;refGene_tx=XM_023245067.1;refGene_gn=MYH14;refGene_tx=XM_023245068.1
E2 7893024 7893162

refGene_gn=MYH14;refGene_tx=XM_023245063.1;refGene_gn=MYH14;refGene_tx=XM_023245064.1;refGene_gn=MYH14;refGene_tx=XM_023245065.1;refGene_gn=MYH14;refGene_tx=XM_023245066.1;refGene_gn=MYH14;refGene_tx=XM_023245067.1;refGene_gn=MYH14;refGene_tx=XM_023245068.1
E2 7895620 7895827
refGene_gn=MYH14;refGene_tx=XM_023245063.1;refGene_gn=MYH14;refGene_tx=XM_023245064.1;refGene_gn=MYH14;refGene_tx=XM_023245065.1;refGene_gn=MYH14;refGene_tx=XM_023245066.1;refGene_gn=MYH14;refGene_tx=XM_023245067.1;refGene_gn=MYH14;refGene_tx=XM_023245068.1
E2 7896926 7897058
refGene_gn=MYH14;refGene_tx=XM_023245063.1;refGene_gn=MYH14;refGene_tx=XM_023245064.1;refGene_gn=MYH14;refGene_tx=XM_023245065.1;refGene_gn=MYH14;refGene_tx=XM_023245066.1;refGene_gn=MYH14;refGene_tx=XM_023245067.1;refGene_gn=MYH14;refGene_tx=XM_023245068.1
E2 7899530 7899639
refGene_gn=MYH14;refGene_tx=XM_023245063.1;refGene_gn=MYH14;refGene_tx=XM_023245064.1;refGene_gn=MYH14;refGene_tx=XM_023245065.1;refGene_gn=MYH14;refGene_tx=XM_023245066.1;refGene_gn=MYH14;refGene_tx=XM_023245067.1;refGene_gn=MYH14;refGene_tx=XM_023245068.1
E2 7900940 7901101
refGene_gn=MYH14;refGene_tx=XM_023245063.1;refGene_gn=MYH14;refGene_tx=XM_023245064.1;refGene_gn=MYH14;refGene_tx=XM_023245065.1;refGene_gn=MYH14;refGene_tx=XM_023245066.1;refGene_gn=MYH14;refGene_tx=XM_023245067.1;refGene_gn=MYH14;refGene_tx=XM_023245068.1
E2 7901223 7901293
refGene_gn=MYH14;refGene_tx=XM_023245063.1;refGene_gn=MYH14;refGene_tx=XM_023245064.1;refGene_gn=MYH14;refGene_tx=XM_023245065.1;refGene_gn=MYH14;refGene_tx=XM_023245066.1;refGene_gn=MYH14;refGene_tx=XM_023245067.1;refGene_gn=MYH14;refGene_tx=XM_023245068.1
E2 7903476 7903598
refGene_gn=MYH14;refGene_tx=XM_023245063.1;refGene_gn=MYH14;refGene_tx=XM_023245064.1;refGene_gn=MYH14;refGene_tx=XM_023245065.1;refGene_gn=MYH14;refGene_tx=XM_023245066.1;refGene_gn=MYH14;refGene_tx=XM_023245067.1;refGene_gn=MYH14;refGene_tx=XM_023245068.1
E2 7908371 7908559
refGene_gn=MYH14;refGene_tx=XM_023245063.1;refGene_gn=MYH14;refGene_tx=XM_023245064.1;refGene_gn=MYH14;refGene_tx=XM_023245065.1;refGene_gn=MYH14;refGene_tx=XM_023245066.1;refGene_gn=MYH14;refGene_tx=XM_023245067.1;refGene_gn=MYH14;refGene_tx=XM_023245068.1
E2 7910661 7910760
refGene_gn=MYH14;refGene_tx=XM_023245063.1;refGene_gn=MYH14;refGene_tx=XM_023245064.1;refGene_gn=MYH14;refGene_tx=XM_023245065.1
E2 7912564 7912679
refGene_gn=MYH14;refGene_tx=XM_023245063.1;refGene_gn=MYH14;refGene_tx=XM_023245064.1;refGene_gn=MYH14;refGene_tx=XM_023245065.1;refGene_gn=MYH14;refGene_tx=XM_023245066.1;refGene_gn=MYH14;refGene_tx=XM_023245067.1;refGene_gn=MYH14;refGene_tx=XM_023245068.1
E2 7916970 7917144
refGene_gn=MYH14;refGene_tx=XM_023245063.1;refGene_gn=MYH14;refGene_tx=XM_023245064.1;refGene_gn=MYH14;refGene_tx=XM_023245065.1;refGene_gn=

MYH14;refGene_tx=XM_023245066.1;refGene_gn=MYH14;refGene_tx=XM_0232450
67.1;refGene_gn=MYH14;refGene_tx=XM_023245068.1
E2 7917422 7917596
refGene_gn=MYH14;refGene_tx=XM_023245063.1;refGene_gn=MYH14;refGene_tx
=XM_023245064.1;refGene_gn=MYH14;refGene_tx=XM_023245065.1;refGene_gn=
MYH14;refGene_tx=XM_023245066.1;refGene_gn=MYH14;refGene_tx=XM_0232450
67.1;refGene_gn=MYH14;refGene_tx=XM_023245068.1
E2 7919099 7919252
refGene_gn=MYH14;refGene_tx=XM_023245063.1;refGene_gn=MYH14;refGene_tx
=XM_023245064.1;refGene_gn=MYH14;refGene_tx=XM_023245065.1;refGene_gn=
MYH14;refGene_tx=XM_023245066.1;refGene_gn=MYH14;refGene_tx=XM_0232450
67.1;refGene_gn=MYH14;refGene_tx=XM_023245068.1
E2 7920509 7920628
refGene_gn=MYH14;refGene_tx=XM_023245063.1;refGene_gn=MYH14;refGene_tx
=XM_023245064.1;refGene_gn=MYH14;refGene_tx=XM_023245065.1;refGene_gn=
MYH14;refGene_tx=XM_023245066.1;refGene_gn=MYH14;refGene_tx=XM_0232450
67.1;refGene_gn=MYH14;refGene_tx=XM_023245068.1
E2 7921845 7921941
refGene_gn=MYH14;refGene_tx=XM_023245063.1;refGene_gn=MYH14;refGene_tx
=XM_023245064.1;refGene_gn=MYH14;refGene_tx=XM_023245065.1;refGene_gn=
MYH14;refGene_tx=XM_023245066.1;refGene_gn=MYH14;refGene_tx=XM_0232450
67.1;refGene_gn=MYH14;refGene_tx=XM_023245068.1
E2 7931786 7931927
refGene_gn=MYH14;refGene_tx=XM_023245063.1;refGene_gn=MYH14;refGene_tx
=XM_023245064.1;refGene_gn=MYH14;refGene_tx=XM_023245065.1;refGene_gn=
MYH14;refGene_tx=XM_023245066.1;refGene_gn=MYH14;refGene_tx=XM_0232450
67.1;refGene_gn=MYH14;refGene_tx=XM_023245068.1
E2 7932579 7932678
refGene_gn=MYH14;refGene_tx=XM_023245063.1;refGene_gn=MYH14;refGene_tx
=XM_023245064.1;refGene_gn=MYH14;refGene_tx=XM_023245065.1;refGene_gn=
MYH14;refGene_tx=XM_023245066.1;refGene_gn=MYH14;refGene_tx=XM_0232450
67.1;refGene_gn=MYH14;refGene_tx=XM_023245068.1
E2 7934877 7934941
refGene_gn=MYH14;refGene_tx=XM_023245063.1;refGene_gn=MYH14;refGene_tx
=XM_023245064.1;refGene_gn=MYH14;refGene_tx=XM_023245065.1;refGene_gn=
MYH14;refGene_tx=XM_023245066.1;refGene_gn=MYH14;refGene_tx=XM_0232450
67.1;refGene_gn=MYH14;refGene_tx=XM_023245068.1
E2 7936144 7936237
refGene_gn=MYH14;refGene_tx=XM_023245063.1;refGene_gn=MYH14;refGene_tx
=XM_023245064.1;refGene_gn=MYH14;refGene_tx=XM_023245065.1;refGene_gn=
MYH14;refGene_tx=XM_023245066.1;refGene_gn=MYH14;refGene_tx=XM_0232450
67.1;refGene_gn=MYH14;refGene_tx=XM_023245068.1
E2 7937523 7937544
refGene_gn=MYH14;refGene_tx=XM_023245063.1;refGene_gn=MYH14;refGene_tx
=XM_023245064.1;refGene_gn=MYH14;refGene_tx=XM_023245066.1
E2 7938175 7938278
refGene_gn=MYH14;refGene_tx=XM_023245063.1;refGene_gn=MYH14;refGene_tx
=XM_023245064.1;refGene_gn=MYH14;refGene_tx=XM_023245065.1;refGene_gn=
MYH14;refGene_tx=XM_023245066.1;refGene_gn=MYH14;refGene_tx=XM_0232450
67.1;refGene_gn=MYH14;refGene_tx=XM_023245068.1

E2	7938407	7938435
refGene_gn=MYH14;refGene_tx=XM_023245063.1;refGene_gn=MYH14;refGene_tx
=XM_023245064.1;refGene_gn=MYH14;refGene_tx=XM_023245065.1;refGene_gn=
MYH14;refGene_tx=XM_023245066.1;refGene_gn=MYH14;refGene_tx=XM_0232450
67.1;refGene_gn=MYH14;refGene_tx=XM_023245068.1
E2	7943726	7943883
refGene_gn=MYH14;refGene_tx=XM_023245063.1;refGene_gn=MYH14;refGene_tx
=XM_023245064.1;refGene_gn=MYH14;refGene_tx=XM_023245065.1;refGene_gn=
MYH14;refGene_tx=XM_023245066.1;refGene_gn=MYH14;refGene_tx=XM_0232450
67.1;refGene_gn=MYH14;refGene_tx=XM_023245068.1
E2	7949298	7949706
refGene_gn=MYH14;refGene_tx=XM_023245063.1;refGene_gn=MYH14;refGene_tx
=XM_023245065.1;refGene_gn=MYH14;refGene_tx=XM_023245066.1;refGene_gn=
MYH14;refGene_tx=XM_023245067.1
E2	7950631	7951039
refGene_gn=MYH14;refGene_tx=XM_023245064.1;refGene_gn=MYH14;refGene_tx
=XM_023245068.1
E2	7951481	7951499
refGene_gn=LOC111558104;refGene_tx=XM_023245904.1
E2	7951835	7952075
refGene_gn=LOC111558104;refGene_tx=XM_023245904.1
E2	7992071	7992303
refGene_gn=LOC101089310;refGene_tx=XM_003997475.4;refGene_gn=LOC101089
310;refGene_tx=XM_023245579.1
E2	7992515	7992590
refGene_gn=LOC101089310;refGene_tx=XM_003997475.4;refGene_gn=LOC101089
310;refGene_tx=XM_023245579.1
E2	7996174	7996261
refGene_gn=LOC101089310;refGene_tx=XM_003997475.4;refGene_gn=LOC101089
310;refGene_tx=XM_023245579.1
E2	7996484	7996505
refGene_gn=LOC101089310;refGene_tx=XM_003997475.4;refGene_gn=LOC101089
310;refGene_tx=XM_023245579.1
E2	7998012	7998093
refGene_gn=LOC101089310;refGene_tx=XM_003997475.4;refGene_gn=LOC101089
310;refGene_tx=XM_023245579.1
E2	8001263	8001390
refGene_gn=LOC101089310;refGene_tx=XM_003997475.4
E2	8002906	8002949
refGene_gn=LOC101089310;refGene_tx=XM_003997475.4
E2	8009135	8009201
refGene_gn=LOC101089310;refGene_tx=XM_006940960.4
E2	8013708	8013810
refGene_gn=LOC101089310;refGene_tx=XM_006940960.4;refGene_gn=LOC101089
310;refGene_tx=XM_019818507.2;refGene_gn=LOC101089310;refGene_tx=XM_02
3245579.1
E2	8015147	8015274
refGene_gn=LOC101089310;refGene_tx=XM_006940960.4;refGene_gn=LOC101089
310;refGene_tx=XM_019818507.2;refGene_gn=LOC101089310;refGene_tx=XM_02
3245579.1

E2	8017751	8017853
refGene_gn=LOC101089310;refGene_tx=XM_006940960.4;refGene_gn=LOC101089310;refGene_tx=XM_019818507.2;refGene_gn=LOC101089310;refGene_tx=XM_023245579.1
E2	8020899	8022794
refGene_gn=LOC101089310;refGene_tx=XM_006940960.4;refGene_gn=LOC101089310;refGene_tx=XM_019818507.2;refGene_gn=LOC101089310;refGene_tx=XM_023245579.1
E2	8033953	8036409
refGene_gn=ZNF473;refGene_tx=XM_011289703.3;refGene_gn=ZNF473;refGene_tx=XM_011289705.3;refGene_gn=ZNF473;refGene_tx=XM_019818492.2
E2	8037673	8037763
refGene_gn=ZNF473;refGene_tx=XM_011289703.3;refGene_gn=ZNF473;refGene_tx=XM_011289705.3;refGene_gn=ZNF473;refGene_tx=XM_019818492.2
E2	8039820	8039920
refGene_gn=ZNF473;refGene_tx=XM_011289703.3;refGene_gn=ZNF473;refGene_tx=XM_011289705.3;refGene_gn=ZNF473;refGene_tx=XM_019818492.2
E2	8053334	8053473
refGene_gn=VRK3;refGene_tx=XM_011289700.3;refGene_gn=VRK3;refGene_tx=XM_011289701.3
E2	8054046	8054196
refGene_gn=VRK3;refGene_tx=XM_011289700.3;refGene_gn=VRK3;refGene_tx=XM_011289701.3
E2	8055380	8055575
refGene_gn=VRK3;refGene_tx=XM_011289700.3;refGene_gn=VRK3;refGene_tx=XM_011289701.3
E2	8061316	8061381
refGene_gn=VRK3;refGene_tx=XM_011289700.3;refGene_gn=VRK3;refGene_tx=XM_011289701.3
E2	8062961	8063028
refGene_gn=VRK3;refGene_tx=XM_011289700.3;refGene_gn=VRK3;refGene_tx=XM_011289701.3
E2	8063906	8063991
refGene_gn=VRK3;refGene_tx=XM_011289700.3;refGene_gn=VRK3;refGene_tx=XM_011289701.3
E2	8064267	8064373
refGene_gn=VRK3;refGene_tx=XM_011289700.3;refGene_gn=VRK3;refGene_tx=XM_011289701.3
E2	8065814	8065907
refGene_gn=VRK3;refGene_tx=XM_011289700.3;refGene_gn=VRK3;refGene_tx=XM_011289701.3
E2	8067386	8067519
refGene_gn=VRK3;refGene_tx=XM_011289700.3;refGene_gn=VRK3;refGene_tx=XM_011289701.3
E2	8068617	8068738
refGene_gn=VRK3;refGene_tx=XM_011289700.3;refGene_gn=VRK3;refGene_tx=XM_011289701.3
E2	8076491	8076550
refGene_gn=VRK3;refGene_tx=XM_011289700.3;refGene_gn=VRK3;refGene_tx=XM_011289701.3

E2 8080874 8081023
refGene_gn=VRK3;refGene_tx=XM_011289700.3;refGene_gn=VRK3;refGene_tx=XM_011289701.3
E2 8089473 8089665
refGene_gn=SIGLEC11;refGene_tx=XM_011289698.3;refGene_gn=SIGLEC11;refGene_tx=XM_011289699.3;refGene_gn=SIGLEC11;refGene_tx=XM_019819184.2
E2 8091425 8091526
refGene_gn=SIGLEC11;refGene_tx=XM_011289698.3;refGene_gn=SIGLEC11;refGene_tx=XM_011289699.3;refGene_gn=SIGLEC11;refGene_tx=XM_019819184.2;refGene_gn=SIGLEC11;refGene_tx=XM_019819185.2
E2 8091793 8091890
refGene_gn=SIGLEC11;refGene_tx=XM_011289698.3;refGene_gn=SIGLEC11;refGene_tx=XM_011289699.3;refGene_gn=SIGLEC11;refGene_tx=XM_019819184.2;refGene_gn=SIGLEC11;refGene_tx=XM_019819185.2
E2 8092145 8092436
refGene_gn=SIGLEC11;refGene_tx=XM_011289698.3;refGene_gn=SIGLEC11;refGene_tx=XM_011289699.3;refGene_gn=SIGLEC11;refGene_tx=XM_019819184.2;refGene_gn=SIGLEC11;refGene_tx=XM_019819185.2
E2 8092530 8092788
refGene_gn=SIGLEC11;refGene_tx=XM_011289698.3;refGene_gn=SIGLEC11;refGene_tx=XM_011289699.3;refGene_gn=SIGLEC11;refGene_tx=XM_019819184.2;refGene_gn=SIGLEC11;refGene_tx=XM_019819185.2
E2 8096456 8096504
refGene_gn=SIGLEC11;refGene_tx=XM_011289698.3;refGene_gn=SIGLEC11;refGene_tx=XM_011289699.3;refGene_gn=SIGLEC11;refGene_tx=XM_019819184.2;refGene_gn=SIGLEC11;refGene_tx=XM_019819185.2
E2 8096703 8096988
refGene_gn=SIGLEC11;refGene_tx=XM_011289698.3;refGene_gn=SIGLEC11;refGene_tx=XM_011289699.3;refGene_gn=SIGLEC11;refGene_tx=XM_019819184.2;refGene_gn=SIGLEC11;refGene_tx=XM_019819185.2
E2 8097369 8097750
refGene_gn=SIGLEC11;refGene_tx=XM_011289698.3;refGene_gn=SIGLEC11;refGene_tx=XM_011289699.3;refGene_gn=SIGLEC11;refGene_tx=XM_019819184.2;refGene_gn=SIGLEC11;refGene_tx=XM_019819185.2
E2 8097839 8097876
refGene_gn=SIGLEC11;refGene_tx=XM_011289698.3;refGene_gn=SIGLEC11;refGene_tx=XM_011289699.3;refGene_gn=SIGLEC11;refGene_tx=XM_019819184.2;refGene_gn=SIGLEC11;refGene_tx=XM_019819185.2
E2 8115917 8116582 refGene_gn=ATF5;refGene_tx=XM_006940973.3
E2 8117526 8117704 refGene_gn=ATF5;refGene_tx=XM_006940973.3
E2 8139889 8141470
refGene_gn=NUP62;refGene_tx=XM_006940971.4;refGene_gn=NUP62;refGene_tx=XM_006940972.4
E2 8149648 8149661 refGene_gn=IL4I1;refGene_tx=XM_006940968.4
E2 8149764 8150003 refGene_gn=IL4I1;refGene_tx=XM_006940968.4
E2 8150658 8150771 refGene_gn=IL4I1;refGene_tx=XM_006940968.4
E2 8151203 8151405 refGene_gn=IL4I1;refGene_tx=XM_006940968.4
E2 8153077 8153146 refGene_gn=IL4I1;refGene_tx=XM_006940968.4
E2 8153470 8153607 refGene_gn=IL4I1;refGene_tx=XM_006940968.4
E2 8154097 8154164 refGene_gn=IL4I1;refGene_tx=XM_006940968.4

E2	8154165	8155020	refGene_gn=IL4I1;refGene_tx=XM_006940968.4
E2	8155795	8156014	refGene_gn=TBC1D17;refGene_tx=XM_003997491.5
E2	8156103	8156190	refGene_gn=TBC1D17;refGene_tx=XM_003997491.5
E2	8156716	8156833	refGene_gn=TBC1D17;refGene_tx=XM_003997491.5
E2	8156993	8157091	refGene_gn=TBC1D17;refGene_tx=XM_003997491.5
E2	8158275	8158375	refGene_gn=TBC1D17;refGene_tx=XM_003997491.5
E2	8158455	8158556	refGene_gn=TBC1D17;refGene_tx=XM_003997491.5
E2	8158659	8158776	refGene_gn=TBC1D17;refGene_tx=XM_003997491.5
E2	8159209	8159304	refGene_gn=TBC1D17;refGene_tx=XM_003997491.5
E2	8160131	8160235	refGene_gn=TBC1D17;refGene_tx=XM_003997491.5
E2	8160330	8160459	refGene_gn=TBC1D17;refGene_tx=XM_003997491.5
E2	8160789	8160948	refGene_gn=TBC1D17;refGene_tx=XM_003997491.5
E2	8161058	8161170	refGene_gn=TBC1D17;refGene_tx=XM_003997491.5
E2	8161593	8161801	refGene_gn=TBC1D17;refGene_tx=XM_003997491.5
E2	8162513	8162637	refGene_gn=TBC1D17;refGene_tx=XM_003997491.5
E2	8166709	8166784	refGene_gn=TBC1D17;refGene_tx=XM_003997491.5
E2	8167013	8167112	refGene_gn=TBC1D17;refGene_tx=XM_003997491.5
E2	8167392	8167413	refGene_gn=TBC1D17;refGene_tx=XM_003997491.5
E2	8168504	8168557	refGene_gn=AKT1S1;refGene_tx=XM_019819195.2
E2	8171869	8172255
refGene_gn=AKT1S1;refGene_tx=XM_019819195.2;refGene_gn=AKT1S1;refGene_tx=XM_023245725.1
E2	8173290	8173368
refGene_gn=AKT1S1;refGene_tx=XM_019819195.2;refGene_gn=AKT1S1;refGene_tx=XM_023245725.1
E2	8173450	8173620
refGene_gn=AKT1S1;refGene_tx=XM_019819195.2;refGene_gn=AKT1S1;refGene_tx=XM_023245725.1
E2	8175009	8175153
refGene_gn=AKT1S1;refGene_tx=XM_019819195.2;refGene_gn=AKT1S1;refGene_tx=XM_023245725.1
E2	8176781	8176972
refGene_gn=PNKP;refGene_tx=XM_023245718.1;refGene_gn=PNKP;refGene_tx=XM_023245719.1;refGene_gn=PNKP;refGene_tx=XM_023245720.1;refGene_gn=PNKP;refGene_tx=XM_023245722.1
E2	8177253	8177498
refGene_gn=PNKP;refGene_tx=XM_023245718.1;refGene_gn=PNKP;refGene_tx=XM_023245719.1;refGene_gn=PNKP;refGene_tx=XM_023245720.1;refGene_gn=PNKP;refGene_tx=XM_023245722.1
E2	8177962	8178118
refGene_gn=PNKP;refGene_tx=XM_023245718.1;refGene_gn=PNKP;refGene_tx=XM_023245719.1;refGene_gn=PNKP;refGene_tx=XM_023245720.1;refGene_gn=PNKP;refGene_tx=XM_023245721.1;refGene_gn=PNKP;refGene_tx=XM_023245722.1;refGene_gn=PNKP;refGene_tx=XM_023245723.1
E2	8178454	8178757
refGene_gn=PNKP;refGene_tx=XM_023245718.1;refGene_gn=PNKP;refGene_tx=XM_023245719.1;refGene_gn=PNKP;refGene_tx=XM_023245720.1;refGene_gn=PNKP;refGene_tx=XM_023245721.1;refGene_gn=PNKP;refGene_tx=XM_023245722.1;refGene_gn=PNKP;refGene_tx=XM_023245723.1
E2	8179656	8179736

refGene_gn=PNKP;refGene_tx=XM_023245718.1;refGene_gn=PNKP;refGene_tx=XM_023245719.1;refGene_gn=PNKP;refGene_tx=XM_023245720.1;refGene_gn=PNKP;refGene_tx=XM_023245721.1;refGene_gn=PNKP;refGene_tx=XM_023245722.1;refGene_gn=PNKP;refGene_tx=XM_023245723.1
E2 8179818 8179876
refGene_gn=PNKP;refGene_tx=XM_023245718.1;refGene_gn=PNKP;refGene_tx=XM_023245719.1;refGene_gn=PNKP;refGene_tx=XM_023245720.1;refGene_gn=PNKP;refGene_tx=XM_023245721.1;refGene_gn=PNKP;refGene_tx=XM_023245722.1;refGene_gn=PNKP;refGene_tx=XM_023245723.1
E2 8179966 8180074
refGene_gn=PNKP;refGene_tx=XM_023245718.1;refGene_gn=PNKP;refGene_tx=XM_023245719.1;refGene_gn=PNKP;refGene_tx=XM_023245720.1;refGene_gn=PNKP;refGene_tx=XM_023245721.1;refGene_gn=PNKP;refGene_tx=XM_023245722.1;refGene_gn=PNKP;refGene_tx=XM_023245723.1
E2 8180378 8180711
refGene_gn=PNKP;refGene_tx=XM_023245718.1;refGene_gn=PNKP;refGene_tx=XM_023245719.1;refGene_gn=PNKP;refGene_tx=XM_023245720.1;refGene_gn=PNKP;refGene_tx=XM_023245721.1;refGene_gn=PNKP;refGene_tx=XM_023245722.1;refGene_gn=PNKP;refGene_tx=XM_023245723.1
E2 8181391 8181440
refGene_gn=PNKP;refGene_tx=XM_023245718.1;refGene_gn=PNKP;refGene_tx=XM_023245719.1;refGene_gn=PNKP;refGene_tx=XM_023245720.1;refGene_gn=PNKP;refGene_tx=XM_023245721.1;refGene_gn=PNKP;refGene_tx=XM_023245722.1;refGene_gn=PNKP;refGene_tx=XM_023245723.1
E2 8181547 8181618
refGene_gn=PNKP;refGene_tx=XM_023245718.1;refGene_gn=PNKP;refGene_tx=XM_023245719.1;refGene_gn=PNKP;refGene_tx=XM_023245720.1;refGene_gn=PNKP;refGene_tx=XM_023245721.1;refGene_gn=PNKP;refGene_tx=XM_023245722.1;refGene_gn=PNKP;refGene_tx=XM_023245723.1
E2 8181704 8181797
refGene_gn=PNKP;refGene_tx=XM_023245718.1;refGene_gn=PNKP;refGene_tx=XM_023245719.1;refGene_gn=PNKP;refGene_tx=XM_023245720.1;refGene_gn=PNKP;refGene_tx=XM_023245721.1;refGene_gn=PNKP;refGene_tx=XM_023245722.1;refGene_gn=PNKP;refGene_tx=XM_023245723.1
E2 8181884 8181981
refGene_gn=PNKP;refGene_tx=XM_023245718.1;refGene_gn=PNKP;refGene_tx=XM_023245719.1;refGene_gn=PNKP;refGene_tx=XM_023245720.1;refGene_gn=PNKP;refGene_tx=XM_023245721.1;refGene_gn=PNKP;refGene_tx=XM_023245722.1;refGene_gn=PNKP;refGene_tx=XM_023245723.1
E2 8182068 8182130
refGene_gn=PNKP;refGene_tx=XM_023245718.1;refGene_gn=PNKP;refGene_tx=XM_023245719.1;refGene_gn=PNKP;refGene_tx=XM_023245720.1;refGene_gn=PNKP;refGene_tx=XM_023245721.1;refGene_gn=PNKP;refGene_tx=XM_023245722.1;refGene_gn=PNKP;refGene_tx=XM_023245723.1
E2 8182368 8182478
refGene_gn=PNKP;refGene_tx=XM_023245718.1;refGene_gn=PNKP;refGene_tx=XM_023245719.1;refGene_gn=PNKP;refGene_tx=XM_023245720.1;refGene_gn=PNKP;refGene_tx=XM_023245721.1;refGene_gn=PNKP;refGene_tx=XM_023245722.1;refGene_gn=PNKP;refGene_tx=XM_023245723.1
E2 8182556 8182644

refGene_gn=PNKP;refGene_tx=XM_023245718.1;refGene_gn=PNKP;refGene_tx=XM_023245719.1;refGene_gn=PNKP;refGene_tx=XM_023245720.1;refGene_gn=PNKP;refGene_tx=XM_023245721.1;refGene_gn=PNKP;refGene_tx=XM_023245722.1;refGene_gn=PNKP;refGene_tx=XM_023245723.1
E2	8182723	8182785
refGene_gn=PNKP;refGene_tx=XM_023245718.1;refGene_gn=PNKP;refGene_tx=XM_023245719.1;refGene_gn=PNKP;refGene_tx=XM_023245720.1;refGene_gn=PNKP;refGene_tx=XM_023245721.1;refGene_gn=PNKP;refGene_tx=XM_023245722.1;refGene_gn=PNKP;refGene_tx=XM_023245723.1
E2	8182872	8182990
refGene_gn=PNKP;refGene_tx=XM_023245718.1;refGene_gn=PNKP;refGene_tx=XM_023245719.1;refGene_gn=PNKP;refGene_tx=XM_023245720.1;refGene_gn=PNKP;refGene_tx=XM_023245721.1;refGene_gn=PNKP;refGene_tx=XM_023245722.1;refGene_gn=PNKP;refGene_tx=XM_023245723.1
E2	8184300	8184312	refGene_gn=PTOV1;refGene_tx=XM_023245724.1
E2	8184399	8184597	refGene_gn=PTOV1;refGene_tx=XM_023245724.1
E2	8186052	8186157	refGene_gn=PTOV1;refGene_tx=XM_023245724.1
E2	8186432	8186490	refGene_gn=PTOV1;refGene_tx=XM_023245724.1
E2	8186566	8186640	refGene_gn=PTOV1;refGene_tx=XM_023245724.1
E2	8186741	8186828	refGene_gn=PTOV1;refGene_tx=XM_023245724.1
E2	8187311	8187467	refGene_gn=PTOV1;refGene_tx=XM_023245724.1
E2	8189010	8189106	refGene_gn=PTOV1;refGene_tx=XM_023245724.1
E2	8189107	8189119	refGene_gn=PTOV1;refGene_tx=XM_023245724.1
E2	8189206	8189264	refGene_gn=PTOV1;refGene_tx=XM_023245724.1
E2	8189345	8189428	refGene_gn=PTOV1;refGene_tx=XM_023245724.1
E2	8189549	8189687	refGene_gn=PTOV1;refGene_tx=XM_023245724.1
E2	8192242	8192413	refGene_gn=PTOV1;refGene_tx=XM_023245724.1
E2	8204792	8205037
refGene_gn=MED25;refGene_tx=XM_006940974.4;refGene_gn=MED25;refGene_tx=XM_006940975.4;refGene_gn=MED25;refGene_tx=XM_019819186.2;refGene_gn=MED25;refGene_tx=XM_019819188.2;refGene_gn=MED25;refGene_tx=XM_019819189.2;refGene_gn=MED25;refGene_tx=XM_019819191.2
E2	8206773	8206871
refGene_gn=MED25;refGene_tx=XM_003997494.5;refGene_gn=MED25;refGene_tx=XM_019819190.2
E2	8207345	8207526
refGene_gn=MED25;refGene_tx=XM_003997494.5;refGene_gn=MED25;refGene_tx=XM_006940974.4;refGene_gn=MED25;refGene_tx=XM_006940975.4;refGene_gn=MED25;refGene_tx=XM_019819186.2;refGene_gn=MED25;refGene_tx=XM_019819188.2;refGene_gn=MED25;refGene_tx=XM_019819189.2;refGene_gn=MED25;refGene_tx=XM_019819190.2;refGene_gn=MED25;refGene_tx=XM_019819191.2
E2	8207833	8208052
refGene_gn=MED25;refGene_tx=XM_003997494.5;refGene_gn=MED25;refGene_tx=XM_006940974.4;refGene_gn=MED25;refGene_tx=XM_006940975.4;refGene_gn=MED25;refGene_tx=XM_019819186.2;refGene_gn=MED25;refGene_tx=XM_019819188.2;refGene_gn=MED25;refGene_tx=XM_019819189.2;refGene_gn=MED25;refGene_tx=XM_019819190.2;refGene_gn=MED25;refGene_tx=XM_019819191.2
E2	8208184	8208256
refGene_gn=MED25;refGene_tx=XM_003997494.5;refGene_gn=MED25;refGene_tx=XM_006940974.4;refGene_gn=MED25;refGene_tx=XM_006940975.4;refGene_gn=

MED25;refGene_tx=XM_019819186.2;refGene_gn=MED25;refGene_tx=XM_0198191
89.2;refGene_gn=MED25;refGene_tx=XM_019819190.2;refGene_gn=MED25;refGe
ne_tx=XM_019819191.2
E2 8208635 8208824
refGene_gn=MED25;refGene_tx=XM_003997494.5;refGene_gn=MED25;refGene_tx
=XM_006940974.4;refGene_gn=MED25;refGene_tx=XM_006940975.4;refGene_gn=
MED25;refGene_tx=XM_019819186.2;refGene_gn=MED25;refGene_tx=XM_0198191
88.2;refGene_gn=MED25;refGene_tx=XM_019819189.2;refGene_gn=MED25;refGe
ne_tx=XM_019819190.2;refGene_gn=MED25;refGene_tx=XM_019819191.2
E2 8208945 8209053
refGene_gn=MED25;refGene_tx=XM_003997494.5;refGene_gn=MED25;refGene_tx
=XM_006940974.4;refGene_gn=MED25;refGene_tx=XM_006940975.4;refGene_gn=
MED25;refGene_tx=XM_019819186.2;refGene_gn=MED25;refGene_tx=XM_0198191
88.2;refGene_gn=MED25;refGene_tx=XM_019819189.2;refGene_gn=MED25;refGe
ne_tx=XM_019819190.2;refGene_gn=MED25;refGene_tx=XM_019819191.2
E2 8209157 8209215
refGene_gn=MED25;refGene_tx=XM_003997494.5;refGene_gn=MED25;refGene_tx
=XM_006940974.4;refGene_gn=MED25;refGene_tx=XM_006940975.4;refGene_gn=
MED25;refGene_tx=XM_019819186.2;refGene_gn=MED25;refGene_tx=XM_0198191
88.2;refGene_gn=MED25;refGene_tx=XM_019819189.2;refGene_gn=MED25;refGe
ne_tx=XM_019819190.2;refGene_gn=MED25;refGene_tx=XM_019819191.2
E2 8209293 8209379
refGene_gn=MED25;refGene_tx=XM_003997494.5;refGene_gn=MED25;refGene_tx
=XM_006940974.4;refGene_gn=MED25;refGene_tx=XM_006940975.4;refGene_gn=
MED25;refGene_tx=XM_019819186.2;refGene_gn=MED25;refGene_tx=XM_0198191
88.2;refGene_gn=MED25;refGene_tx=XM_019819189.2;refGene_gn=MED25;refGe
ne_tx=XM_019819190.2;refGene_gn=MED25;refGene_tx=XM_019819191.2
E2 8209758 8209887
refGene_gn=MED25;refGene_tx=XM_003997494.5;refGene_gn=MED25;refGene_tx
=XM_006940974.4;refGene_gn=MED25;refGene_tx=XM_006940975.4;refGene_gn=
MED25;refGene_tx=XM_019819186.2;refGene_gn=MED25;refGene_tx=XM_0198191
88.2;refGene_gn=MED25;refGene_tx=XM_019819189.2;refGene_gn=MED25;refGe
ne_tx=XM_019819190.2;refGene_gn=MED25;refGene_tx=XM_019819191.2
E2 8210306 8210500
refGene_gn=MED25;refGene_tx=XM_003997494.5;refGene_gn=MED25;refGene_tx
=XM_006940974.4;refGene_gn=MED25;refGene_tx=XM_006940975.4;refGene_gn=
MED25;refGene_tx=XM_019819186.2;refGene_gn=MED25;refGene_tx=XM_0198191
88.2;refGene_gn=MED25;refGene_tx=XM_019819189.2;refGene_gn=MED25;refGe
ne_tx=XM_019819190.2;refGene_gn=MED25;refGene_tx=XM_019819191.2
E2 8210592 8210680
refGene_gn=MED25;refGene_tx=XM_003997494.5;refGene_gn=MED25;refGene_tx
=XM_006940974.4;refGene_gn=MED25;refGene_tx=XM_006940975.4;refGene_gn=
MED25;refGene_tx=XM_019819186.2;refGene_gn=MED25;refGene_tx=XM_0198191
88.2;refGene_gn=MED25;refGene_tx=XM_019819189.2;refGene_gn=MED25;refGe
ne_tx=XM_019819190.2;refGene_gn=MED25;refGene_tx=XM_019819191.2
E2 8210972 8211103
refGene_gn=MED25;refGene_tx=XM_003997494.5;refGene_gn=MED25;refGene_tx
=XM_006940974.4;refGene_gn=MED25;refGene_tx=XM_006940975.4;refGene_gn=
MED25;refGene_tx=XM_019819186.2;refGene_gn=MED25;refGene_tx=XM_0198191
88.2;refGene_gn=MED25;refGene_tx=XM_019819189.2;refGene_gn=MED25;refGe

ne_tx=XM_019819190.2;refGene_gn=MED25;refGene_tx=XM_019819191.2
E2	8211228	8211391
refGene_gn=MED25;refGene_tx=XM_003997494.5;refGene_gn=MED25;refGene_tx=XM_006940974.4;refGene_gn=MED25;refGene_tx=XM_006940975.4;refGene_gn=MED25;refGene_tx=XM_019819186.2;refGene_gn=MED25;refGene_tx=XM_019819188.2;refGene_gn=MED25;refGene_tx=XM_019819189.2;refGene_gn=MED25;refGene_tx=XM_019819190.2;refGene_gn=MED25;refGene_tx=XM_019819191.2
E2	8211666	8211832
refGene_gn=MED25;refGene_tx=XM_003997494.5;refGene_gn=MED25;refGene_tx=XM_006940974.4;refGene_gn=MED25;refGene_tx=XM_006940975.4;refGene_gn=MED25;refGene_tx=XM_019819186.2;refGene_gn=MED25;refGene_tx=XM_019819188.2;refGene_gn=MED25;refGene_tx=XM_019819189.2;refGene_gn=MED25;refGene_tx=XM_019819190.2;refGene_gn=MED25;refGene_tx=XM_019819191.2
E2	8212188	8212287
refGene_gn=MED25;refGene_tx=XM_003997494.5;refGene_gn=MED25;refGene_tx=XM_006940974.4;refGene_gn=MED25;refGene_tx=XM_006940975.4;refGene_gn=MED25;refGene_tx=XM_019819186.2;refGene_gn=MED25;refGene_tx=XM_019819188.2;refGene_gn=MED25;refGene_tx=XM_019819189.2;refGene_gn=MED25;refGene_tx=XM_019819190.2;refGene_gn=MED25;refGene_tx=XM_019819191.2
E2	8219643	8219768
refGene_gn=MED25;refGene_tx=XM_003997494.5;refGene_gn=MED25;refGene_tx=XM_006940974.4;refGene_gn=MED25;refGene_tx=XM_019819186.2;refGene_gn=MED25;refGene_tx=XM_019819188.2;refGene_gn=MED25;refGene_tx=XM_019819189.2;refGene_gn=MED25;refGene_tx=XM_019819190.2
E2	8219959	8220005
refGene_gn=MED25;refGene_tx=XM_003997494.5;refGene_gn=MED25;refGene_tx=XM_006940974.4;refGene_gn=MED25;refGene_tx=XM_019819186.2;refGene_gn=MED25;refGene_tx=XM_019819188.2;refGene_gn=MED25;refGene_tx=XM_019819189.2;refGene_gn=MED25;refGene_tx=XM_019819190.2
E2	8220101	8220235
refGene_gn=MED25;refGene_tx=XM_003997494.5;refGene_gn=MED25;refGene_tx=XM_006940974.4;refGene_gn=MED25;refGene_tx=XM_019819186.2;refGene_gn=MED25;refGene_tx=XM_019819188.2;refGene_gn=MED25;refGene_tx=XM_019819189.2;refGene_gn=MED25;refGene_tx=XM_019819190.2
E2	8227925	8228036	refGene_gn=FUZ;refGene_tx=XM_003997495.5
E2	8228302	8228424	refGene_gn=FUZ;refGene_tx=XM_003997495.5
E2	8228676	8228761	refGene_gn=FUZ;refGene_tx=XM_003997495.5
E2	8228880	8228949	refGene_gn=FUZ;refGene_tx=XM_003997495.5
E2	8229147	8229252	refGene_gn=FUZ;refGene_tx=XM_003997495.5
E2	8230051	8230249	refGene_gn=FUZ;refGene_tx=XM_003997495.5
E2	8230377	8230473	refGene_gn=FUZ;refGene_tx=XM_003997495.5
E2	8230784	8230891	refGene_gn=FUZ;refGene_tx=XM_003997495.5
E2	8230969	8231034	refGene_gn=FUZ;refGene_tx=XM_003997495.5
E2	8231114	8231189	refGene_gn=FUZ;refGene_tx=XM_003997495.5
E2	8232129	8232161	refGene_gn=FUZ;refGene_tx=XM_003997495.5
E2	8232162	8232357	refGene_gn=FUZ;refGene_tx=XM_003997495.5
E2	8232755	8232833
refGene_gn=AP2A1;refGene_tx=XM_011289690.3;refGene_gn=AP2A1;refGene_tx=XM_011289691.3
E2	8233145	8233280

refGene_gn=AP2A1;refGene_tx=XM_011289690.3;refGene_gn=AP2A1;refGene_tx=XM_011289691.3
E2 8233478 8233548
refGene_gn=AP2A1;refGene_tx=XM_011289690.3;refGene_gn=AP2A1;refGene_tx=XM_011289691.3
E2 8233657 8233774
refGene_gn=AP2A1;refGene_tx=XM_011289690.3;refGene_gn=AP2A1;refGene_tx=XM_011289691.3
E2 8233873 8233997
refGene_gn=AP2A1;refGene_tx=XM_011289690.3;refGene_gn=AP2A1;refGene_tx=XM_011289691.3
E2 8235135 8235225
refGene_gn=AP2A1;refGene_tx=XM_011289690.3;refGene_gn=AP2A1;refGene_tx=XM_011289691.3
E2 8235357 8235440
refGene_gn=AP2A1;refGene_tx=XM_011289690.3;refGene_gn=AP2A1;refGene_tx=XM_011289691.3
E2 8235550 8235607
refGene_gn=AP2A1;refGene_tx=XM_011289690.3;refGene_gn=AP2A1;refGene_tx=XM_011289691.3
E2 8235953 8236019 refGene_gn=AP2A1;refGene_tx=XM_011289690.3
E2 8236332 8236496
refGene_gn=AP2A1;refGene_tx=XM_011289690.3;refGene_gn=AP2A1;refGene_tx=XM_011289691.3
E2 8236575 8236743
refGene_gn=AP2A1;refGene_tx=XM_011289690.3;refGene_gn=AP2A1;refGene_tx=XM_011289691.3
E2 8236850 8237082
refGene_gn=AP2A1;refGene_tx=XM_011289690.3;refGene_gn=AP2A1;refGene_tx=XM_011289691.3
E2 8237287 8237385
refGene_gn=AP2A1;refGene_tx=XM_011289690.3;refGene_gn=AP2A1;refGene_tx=XM_011289691.3
E2 8237877 8238060
refGene_gn=AP2A1;refGene_tx=XM_011289690.3;refGene_gn=AP2A1;refGene_tx=XM_011289691.3
E2 8238237 8238375
refGene_gn=AP2A1;refGene_tx=XM_011289690.3;refGene_gn=AP2A1;refGene_tx=XM_011289691.3
E2 8238481 8238650
refGene_gn=AP2A1;refGene_tx=XM_011289690.3;refGene_gn=AP2A1;refGene_tx=XM_011289691.3
E2 8238968 8239119
refGene_gn=AP2A1;refGene_tx=XM_011289690.3;refGene_gn=AP2A1;refGene_tx=XM_011289691.3
E2 8242565 8242674
refGene_gn=AP2A1;refGene_tx=XM_011289690.3;refGene_gn=AP2A1;refGene_tx=XM_011289691.3
E2 8244044 8244146
refGene_gn=AP2A1;refGene_tx=XM_011289690.3;refGene_gn=AP2A1;refGene_tx

=XM_011289691.3
E2 8244526 8244656
refGene_gn=AP2A1;refGene_tx=XM_011289690.3;refGene_gn=AP2A1;refGene_tx
=XM_011289691.3
E2 8250735 8250929
refGene_gn=AP2A1;refGene_tx=XM_011289690.3;refGene_gn=AP2A1;refGene_tx
=XM_011289691.3
E2 8251118 8251261
refGene_gn=AP2A1;refGene_tx=XM_011289690.3;refGene_gn=AP2A1;refGene_tx
=XM_011289691.3
E2 8251410 8251479
refGene_gn=AP2A1;refGene_tx=XM_011289690.3;refGene_gn=AP2A1;refGene_tx
=XM_011289691.3
E2 8261875 8261942
refGene_gn=AP2A1;refGene_tx=XM_011289690.3;refGene_gn=AP2A1;refGene_tx
=XM_011289691.3
E2 8266458 8266628
refGene_gn=TSKS;refGene_tx=XM_003997497.5;refGene_gn=TSKS;refGene_tx=X
M_003997498.5;refGene_gn=TSKS;refGene_tx=XM_006940980.4;refGene_gn=TSK
S;refGene_tx=XM_006940981.4;refGene_gn=TSKS;refGene_tx=XM_006940982.4;
refGene_gn=TSKS;refGene_tx=XM_006940983.4;refGene_gn=TSKS;refGene_tx=X
M_006940984.4;refGene_gn=TSKS;refGene_tx=XM_006940985.4;refGene_gn=TSK
S;refGene_tx=XM_006940986.4
E2 8266985 8267211
refGene_gn=TSKS;refGene_tx=XM_003997497.5;refGene_gn=TSKS;refGene_tx=X
M_003997498.5;refGene_gn=TSKS;refGene_tx=XM_006940980.4;refGene_gn=TSK
S;refGene_tx=XM_006940981.4;refGene_gn=TSKS;refGene_tx=XM_006940982.4;
refGene_gn=TSKS;refGene_tx=XM_006940983.4;refGene_gn=TSKS;refGene_tx=X
M_006940984.4;refGene_gn=TSKS;refGene_tx=XM_006940985.4;refGene_gn=TSK
S;refGene_tx=XM_006940986.4
E2 8276053 8276149
refGene_gn=TSKS;refGene_tx=XM_003997497.5;refGene_gn=TSKS;refGene_tx=X
M_003997498.5;refGene_gn=TSKS;refGene_tx=XM_006940980.4;refGene_gn=TSK
S;refGene_tx=XM_006940981.4;refGene_gn=TSKS;refGene_tx=XM_006940982.4;
refGene_gn=TSKS;refGene_tx=XM_006940983.4;refGene_gn=TSKS;refGene_tx=X
M_006940984.4;refGene_gn=TSKS;refGene_tx=XM_006940985.4;refGene_gn=TSK
S;refGene_tx=XM_006940986.4
E2 8276353 8276437
refGene_gn=TSKS;refGene_tx=XM_003997497.5;refGene_gn=TSKS;refGene_tx=X
M_003997498.5;refGene_gn=TSKS;refGene_tx=XM_006940980.4;refGene_gn=TSK
S;refGene_tx=XM_006940981.4;refGene_gn=TSKS;refGene_tx=XM_006940982.4;
refGene_gn=TSKS;refGene_tx=XM_006940983.4;refGene_gn=TSKS;refGene_tx=X
M_006940984.4;refGene_gn=TSKS;refGene_tx=XM_006940985.4;refGene_gn=TSK
S;refGene_tx=XM_006940986.4
E2 8276702 8276813
refGene_gn=TSKS;refGene_tx=XM_003997497.5;refGene_gn=TSKS;refGene_tx=X
M_003997498.5;refGene_gn=TSKS;refGene_tx=XM_006940980.4;refGene_gn=TSK
S;refGene_tx=XM_006940981.4;refGene_gn=TSKS;refGene_tx=XM_006940982.4;
refGene_gn=TSKS;refGene_tx=XM_006940983.4;refGene_gn=TSKS;refGene_tx=X
M_006940984.4;refGene_gn=TSKS;refGene_tx=XM_006940985.4;refGene_gn=TSK

S;refGene_tx=XM_006940986.4
E2 8277377 8277664
refGene_gn=TSKS;refGene_tx=XM_003997497.5;refGene_gn=TSKS;refGene_tx=XM_003997498.5;refGene_gn=TSKS;refGene_tx=XM_006940980.4;refGene_gn=TSKS;refGene_tx=XM_006940981.4;refGene_gn=TSKS;refGene_tx=XM_006940982.4;refGene_gn=TSKS;refGene_tx=XM_006940983.4;refGene_gn=TSKS;refGene_tx=XM_006940984.4;refGene_gn=TSKS;refGene_tx=XM_006940985.4;refGene_gn=TSKS;refGene_tx=XM_006940986.4
E2 8278126 8278321
refGene_gn=TSKS;refGene_tx=XM_003997497.5;refGene_gn=TSKS;refGene_tx=XM_003997498.5;refGene_gn=TSKS;refGene_tx=XM_006940980.4;refGene_gn=TSKS;refGene_tx=XM_006940981.4;refGene_gn=TSKS;refGene_tx=XM_006940982.4;refGene_gn=TSKS;refGene_tx=XM_006940983.4;refGene_gn=TSKS;refGene_tx=XM_006940984.4;refGene_gn=TSKS;refGene_tx=XM_006940985.4;refGene_gn=TSKS;refGene_tx=XM_006940986.4
E2 8278754 8278928
refGene_gn=TSKS;refGene_tx=XM_003997497.5;refGene_gn=TSKS;refGene_tx=XM_003997498.5;refGene_gn=TSKS;refGene_tx=XM_006940980.4;refGene_gn=TSKS;refGene_tx=XM_006940981.4;refGene_gn=TSKS;refGene_tx=XM_006940982.4;refGene_gn=TSKS;refGene_tx=XM_006940983.4;refGene_gn=TSKS;refGene_tx=XM_006940984.4;refGene_gn=TSKS;refGene_tx=XM_006940985.4;refGene_gn=TSKS;refGene_tx=XM_006940986.4
E2 8279637 8279773
refGene_gn=TSKS;refGene_tx=XM_003997497.5;refGene_gn=TSKS;refGene_tx=XM_003997498.5;refGene_gn=TSKS;refGene_tx=XM_006940980.4;refGene_gn=TSKS;refGene_tx=XM_006940981.4;refGene_gn=TSKS;refGene_tx=XM_006940982.4;refGene_gn=TSKS;refGene_tx=XM_006940983.4;refGene_gn=TSKS;refGene_tx=XM_006940984.4;refGene_gn=TSKS;refGene_tx=XM_006940985.4;refGene_gn=TSKS;refGene_tx=XM_006940986.4
E2 8280808 8281008
refGene_gn=TSKS;refGene_tx=XM_003997497.5;refGene_gn=TSKS;refGene_tx=XM_003997498.5;refGene_gn=TSKS;refGene_tx=XM_006940980.4;refGene_gn=TSKS;refGene_tx=XM_006940981.4;refGene_gn=TSKS;refGene_tx=XM_006940982.4;refGene_gn=TSKS;refGene_tx=XM_006940983.4;refGene_gn=TSKS;refGene_tx=XM_006940984.4;refGene_gn=TSKS;refGene_tx=XM_006940985.4;refGene_gn=TSKS;refGene_tx=XM_006940986.4
E2 8281108 8281265
refGene_gn=TSKS;refGene_tx=XM_003997497.5;refGene_gn=TSKS;refGene_tx=XM_003997498.5;refGene_gn=TSKS;refGene_tx=XM_006940980.4;refGene_gn=TSKS;refGene_tx=XM_006940981.4;refGene_gn=TSKS;refGene_tx=XM_006940982.4;refGene_gn=TSKS;refGene_tx=XM_006940983.4;refGene_gn=TSKS;refGene_tx=XM_006940984.4
E2 8306490 8306676
refGene_gn=CPT1C;refGene_tx=XM_011289687.3;refGene_gn=CPT1C;refGene_tx=XM_019819199.2;refGene_gn=CPT1C;refGene_tx=XM_019819200.2;refGene_gn=CPT1C;refGene_tx=XM_019819201.2;refGene_gn=CPT1C;refGene_tx=XM_019819202.2
E2 8307170 8307263
refGene_gn=CPT1C;refGene_tx=XM_011289687.3;refGene_gn=CPT1C;refGene_tx=XM_019819199.2;refGene_gn=CPT1C;refGene_tx=XM_019819200.2;refGene_gn=

CPT1C;refGene_tx=XM_019819201.2;refGene_gn=CPT1C;refGene_tx=XM_0198192
02.2
E2	8307393	8307507
refGene_gn=CPT1C;refGene_tx=XM_011289687.3;refGene_gn=CPT1C;refGene_tx
=XM_019819199.2;refGene_gn=CPT1C;refGene_tx=XM_019819200.2;refGene_gn=
CPT1C;refGene_tx=XM_019819201.2;refGene_gn=CPT1C;refGene_tx=XM_0198192
02.2
E2	8307982	8308135
refGene_gn=CPT1C;refGene_tx=XM_011289687.3;refGene_gn=CPT1C;refGene_tx
=XM_019819199.2;refGene_gn=CPT1C;refGene_tx=XM_019819200.2;refGene_gn=
CPT1C;refGene_tx=XM_019819201.2;refGene_gn=CPT1C;refGene_tx=XM_0198192
02.2
E2	8308782	8308917
refGene_gn=CPT1C;refGene_tx=XM_011289687.3;refGene_gn=CPT1C;refGene_tx
=XM_019819199.2;refGene_gn=CPT1C;refGene_tx=XM_019819200.2;refGene_gn=
CPT1C;refGene_tx=XM_019819201.2
E2	8309175	8309340
refGene_gn=CPT1C;refGene_tx=XM_011289687.3;refGene_gn=CPT1C;refGene_tx
=XM_019819199.2;refGene_gn=CPT1C;refGene_tx=XM_019819200.2;refGene_gn=
CPT1C;refGene_tx=XM_019819201.2;refGene_gn=CPT1C;refGene_tx=XM_0198192
02.2
E2	8309699	8309816
refGene_gn=CPT1C;refGene_tx=XM_011289687.3;refGene_gn=CPT1C;refGene_tx
=XM_019819199.2;refGene_gn=CPT1C;refGene_tx=XM_019819200.2;refGene_gn=
CPT1C;refGene_tx=XM_019819201.2;refGene_gn=CPT1C;refGene_tx=XM_0198192
02.2
E2	8311441	8311547
refGene_gn=CPT1C;refGene_tx=XM_011289687.3;refGene_gn=CPT1C;refGene_tx
=XM_019819199.2;refGene_gn=CPT1C;refGene_tx=XM_019819200.2;refGene_gn=
CPT1C;refGene_tx=XM_019819201.2;refGene_gn=CPT1C;refGene_tx=XM_0198192
02.2
E2	8312006	8312189
refGene_gn=CPT1C;refGene_tx=XM_011289687.3;refGene_gn=CPT1C;refGene_tx
=XM_019819199.2;refGene_gn=CPT1C;refGene_tx=XM_019819200.2;refGene_gn=
CPT1C;refGene_tx=XM_019819201.2;refGene_gn=CPT1C;refGene_tx=XM_0198192
02.2
E2	8312367	8312563
refGene_gn=CPT1C;refGene_tx=XM_011289687.3;refGene_gn=CPT1C;refGene_tx
=XM_019819199.2;refGene_gn=CPT1C;refGene_tx=XM_019819200.2;refGene_gn=
CPT1C;refGene_tx=XM_019819201.2;refGene_gn=CPT1C;refGene_tx=XM_0198192
02.2
E2	8312879	8312964
refGene_gn=CPT1C;refGene_tx=XM_011289687.3;refGene_gn=CPT1C;refGene_tx
=XM_019819199.2;refGene_gn=CPT1C;refGene_tx=XM_019819200.2;refGene_gn=
CPT1C;refGene_tx=XM_019819201.2;refGene_gn=CPT1C;refGene_tx=XM_0198192
02.2
E2	8313067	8313175
refGene_gn=CPT1C;refGene_tx=XM_011289687.3;refGene_gn=CPT1C;refGene_tx
=XM_019819199.2;refGene_gn=CPT1C;refGene_tx=XM_019819200.2;refGene_gn=
CPT1C;refGene_tx=XM_019819201.2;refGene_gn=CPT1C;refGene_tx=XM_0198192

02.2
E2	8313402	8313480
refGene_gn=CPT1C;refGene_tx=XM_011289687.3;refGene_gn=CPT1C;refGene_tx=XM_019819199.2;refGene_gn=CPT1C;refGene_tx=XM_019819200.2;refGene_gn=CPT1C;refGene_tx=XM_019819201.2;refGene_gn=CPT1C;refGene_tx=XM_019819202.2
E2	8317491	8317629
refGene_gn=CPT1C;refGene_tx=XM_011289687.3;refGene_gn=CPT1C;refGene_tx=XM_019819199.2;refGene_gn=CPT1C;refGene_tx=XM_019819200.2;refGene_gn=CPT1C;refGene_tx=XM_019819201.2;refGene_gn=CPT1C;refGene_tx=XM_019819202.2
E2	8317716	8317818
refGene_gn=CPT1C;refGene_tx=XM_011289687.3;refGene_gn=CPT1C;refGene_tx=XM_019819199.2;refGene_gn=CPT1C;refGene_tx=XM_019819200.2;refGene_gn=CPT1C;refGene_tx=XM_019819201.2;refGene_gn=CPT1C;refGene_tx=XM_019819202.2
E2	8318051	8318223
refGene_gn=CPT1C;refGene_tx=XM_011289687.3;refGene_gn=CPT1C;refGene_tx=XM_019819199.2;refGene_gn=CPT1C;refGene_tx=XM_019819200.2;refGene_gn=CPT1C;refGene_tx=XM_019819201.2;refGene_gn=CPT1C;refGene_tx=XM_019819202.2
E2	8319664	8319804
refGene_gn=CPT1C;refGene_tx=XM_011289687.3;refGene_gn=CPT1C;refGene_tx=XM_019819199.2;refGene_gn=CPT1C;refGene_tx=XM_019819200.2;refGene_gn=CPT1C;refGene_tx=XM_019819201.2;refGene_gn=CPT1C;refGene_tx=XM_019819202.2
E2	8324213	8324354
refGene_gn=CPT1C;refGene_tx=XM_011289687.3;refGene_gn=CPT1C;refGene_tx=XM_019819199.2;refGene_gn=CPT1C;refGene_tx=XM_019819200.2;refGene_gn=CPT1C;refGene_tx=XM_019819201.2;refGene_gn=CPT1C;refGene_tx=XM_019819202.2
E2	8326076	8326332	refGene_gn=ADM5;refGene_tx=XM_004001414.3
E2	8326541	8326609	refGene_gn=ADM5;refGene_tx=XM_004001414.3
E2	8328115	8328199
refGene_gn=PRMT1;refGene_tx=XM_003997500.5;refGene_gn=PRMT1;refGene_tx=XM_003997501.5;refGene_gn=PRMT1;refGene_tx=XM_019819203.2;refGene_gn=PRMT1;refGene_tx=XM_019819204.2;refGene_gn=PRMT1;refGene_tx=XM_019819205.2
E2	8329225	8329347
refGene_gn=PRMT1;refGene_tx=XM_003997500.5;refGene_gn=PRMT1;refGene_tx=XM_003997501.5;refGene_gn=PRMT1;refGene_tx=XM_019819203.2;refGene_gn=PRMT1;refGene_tx=XM_019819204.2;refGene_gn=PRMT1;refGene_tx=XM_019819205.2
E2	8329641	8329792
refGene_gn=PRMT1;refGene_tx=XM_003997500.5;refGene_gn=PRMT1;refGene_tx=XM_003997501.5;refGene_gn=PRMT1;refGene_tx=XM_019819203.2;refGene_gn=PRMT1;refGene_tx=XM_019819204.2;refGene_gn=PRMT1;refGene_tx=XM_019819205.2
E2	8330415	8330531
refGene_gn=PRMT1;refGene_tx=XM_003997500.5;refGene_gn=PRMT1;refGene_tx

=XM_003997501.5;refGene_gn=PRMT1;refGene_tx=XM_019819203.2;refGene_gn=PRMT1;refGene_tx=XM_019819204.2;refGene_gn=PRMT1;refGene_tx=XM_019819205.2
E2 8330625 8330713 refGene_gn=PRMT1;refGene_tx=XM_003997500.5;refGene_gn=PRMT1;refGene_tx=XM_003997501.5;refGene_gn=PRMT1;refGene_tx=XM_019819203.2;refGene_gn=PRMT1;refGene_tx=XM_019819204.2;refGene_gn=PRMT1;refGene_tx=XM_019819205.2
E2 8331478 8331621 refGene_gn=PRMT1;refGene_tx=XM_003997500.5;refGene_gn=PRMT1;refGene_tx=XM_003997501.5;refGene_gn=PRMT1;refGene_tx=XM_019819203.2;refGene_gn=PRMT1;refGene_tx=XM_019819204.2;refGene_gn=PRMT1;refGene_tx=XM_019819205.2
E2 8333285 8333349 refGene_gn=PRMT1;refGene_tx=XM_003997500.5;refGene_gn=PRMT1;refGene_tx=XM_003997501.5;refGene_gn=PRMT1;refGene_tx=XM_019819203.2;refGene_gn=PRMT1;refGene_tx=XM_019819204.2;refGene_gn=PRMT1;refGene_tx=XM_019819205.2
E2 8333490 8333646 refGene_gn=PRMT1;refGene_tx=XM_003997500.5;refGene_gn=PRMT1;refGene_tx=XM_003997501.5;refGene_gn=PRMT1;refGene_tx=XM_019819203.2;refGene_gn=PRMT1;refGene_tx=XM_019819204.2;refGene_gn=PRMT1;refGene_tx=XM_019819205.2
E2 8334452 8334554 refGene_gn=PRMT1;refGene_tx=XM_003997500.5;refGene_gn=PRMT1;refGene_tx=XM_003997501.5;refGene_gn=PRMT1;refGene_tx=XM_019819203.2;refGene_gn=PRMT1;refGene_tx=XM_019819204.2;refGene_gn=PRMT1;refGene_tx=XM_019819205.2
E2 8334768 8334786 refGene_gn=PRMT1;refGene_tx=XM_019819204.2;refGene_gn=PRMT1;refGene_tx=XM_019819205.2
E2 8335087 8335141 refGene_gn=PRMT1;refGene_tx=XM_003997501.5;refGene_gn=PRMT1;refGene_tx=XM_019819203.2
E2 8336399 8336525 refGene_gn=PRMT1;refGene_tx=XM_019819203.2
E2 8337335 8337371 refGene_gn=PRMT1;refGene_tx=XM_003997500.5;refGene_gn=PRMT1;refGene_tx=XM_003997501.5
E2 8341536 8341587 refGene_gn=BCL2L12;refGene_tx=XM_006940994.4;refGene_gn=BCL2L12;refGene_tx=XM_019819211.2
E2 8343302 8343590 refGene_gn=BCL2L12;refGene_tx=XM_006940994.4;refGene_gn=BCL2L12;refGene_tx=XM_019819211.2
E2 8345458 8345550 refGene_gn=BCL2L12;refGene_tx=XM_006940994.4;refGene_gn=BCL2L12;refGene_tx=XM_019819211.2
E2 8345638 8345725 refGene_gn=BCL2L12;refGene_tx=XM_006940994.4;refGene_gn=BCL2L12;refGene_tx=XM_019819211.2

E2 8346589 8346732
refGene_gn=BCL2L12;refGene_tx=XM_006940994.4;refGene_gn=BCL2L12;refGen
e_tx=XM_019819211.2
E2 8346946 8347107
refGene_gn=BCL2L12;refGene_tx=XM_006940994.4;refGene_gn=BCL2L12;refGen
e_tx=XM_019819211.2
E2 8348988 8349198
refGene_gn=IRF3;refGene_tx=XM_003997503.5;refGene_gn=IRF3;refGene_tx=X
M_019819206.2;refGene_gn=IRF3;refGene_tx=XM_019819207.2;refGene_gn=IRF
3;refGene_tx=XM_019819208.2;refGene_gn=IRF3;refGene_tx=XM_019819209.1
E2 8350076 8350248
refGene_gn=IRF3;refGene_tx=XM_003997503.5;refGene_gn=IRF3;refGene_tx=X
M_019819206.2;refGene_gn=IRF3;refGene_tx=XM_019819207.2;refGene_gn=IRF
3;refGene_tx=XM_019819208.2;refGene_gn=IRF3;refGene_tx=XM_019819209.1
E2 8350333 8350404
refGene_gn=IRF3;refGene_tx=XM_003997503.5;refGene_gn=IRF3;refGene_tx=X
M_019819206.2;refGene_gn=IRF3;refGene_tx=XM_019819207.2;refGene_gn=IRF
3;refGene_tx=XM_019819208.2;refGene_gn=IRF3;refGene_tx=XM_019819209.1
E2 8351086 8351279
refGene_gn=IRF3;refGene_tx=XM_003997503.5;refGene_gn=IRF3;refGene_tx=X
M_019819206.2;refGene_gn=IRF3;refGene_tx=XM_019819207.2;refGene_gn=IRF
3;refGene_tx=XM_019819208.2;refGene_gn=IRF3;refGene_tx=XM_019819209.1;
refGene_gn=IRF3;refGene_tx=XM_019819210.2
E2 8351377 8351755
refGene_gn=IRF3;refGene_tx=XM_003997503.5;refGene_gn=IRF3;refGene_tx=X
M_019819206.2;refGene_gn=IRF3;refGene_tx=XM_019819207.2;refGene_gn=IRF
3;refGene_tx=XM_019819208.2;refGene_gn=IRF3;refGene_tx=XM_019819209.1;
refGene_gn=IRF3;refGene_tx=XM_019819210.2
E2 8352585 8352701
refGene_gn=IRF3;refGene_tx=XM_003997503.5;refGene_gn=IRF3;refGene_tx=X
M_019819206.2;refGene_gn=IRF3;refGene_tx=XM_019819207.2;refGene_gn=IRF
3;refGene_tx=XM_019819208.2;refGene_gn=IRF3;refGene_tx=XM_019819209.1;
refGene_gn=IRF3;refGene_tx=XM_019819210.2
E2 8353209 8353377
refGene_gn=IRF3;refGene_tx=XM_003997503.5;refGene_gn=IRF3;refGene_tx=X
M_019819206.2;refGene_gn=IRF3;refGene_tx=XM_019819207.2;refGene_gn=IRF
3;refGene_tx=XM_019819208.2;refGene_gn=IRF3;refGene_tx=XM_019819209.1;
refGene_gn=IRF3;refGene_tx=XM_019819210.2
E2 8354810 8355002 refGene_gn=SCAF1;refGene_tx=XM_023245677.1
E2 8355300 8355429 refGene_gn=SCAF1;refGene_tx=XM_023245677.1
E2 8356462 8356681 refGene_gn=SCAF1;refGene_tx=XM_023245677.1
E2 8356910 8356993 refGene_gn=SCAF1;refGene_tx=XM_023245677.1
E2 8357643 8360514 refGene_gn=SCAF1;refGene_tx=XM_023245677.1
E2 8363243 8363362 refGene_gn=SCAF1;refGene_tx=XM_023245677.1
E2 8363452 8363553 refGene_gn=SCAF1;refGene_tx=XM_023245677.1
E2 8363802 8363897 refGene_gn=SCAF1;refGene_tx=XM_023245677.1
E2 8364637 8364695 refGene_gn=SCAF1;refGene_tx=XM_023245677.1
E2 8364942 8365056 refGene_gn=SCAF1;refGene_tx=XM_023245677.1
E2 8365516 8365528 refGene_gn=SCAF1;refGene_tx=XM_023245677.1
E2 8369470 8369623

refGene_gn=RRAS;refGene_tx=XM_011289683.2;refGene_gn=RRAS;refGene_tx=XM_011289684.2
E2 8371559 8371662
refGene_gn=RRAS;refGene_tx=XM_011289683.2;refGene_gn=RRAS;refGene_tx=XM_011289684.2
E2 8371780 8371883
refGene_gn=RRAS;refGene_tx=XM_011289683.2;refGene_gn=RRAS;refGene_tx=XM_011289684.2
E2 8371974 8372083
refGene_gn=RRAS;refGene_tx=XM_011289683.2;refGene_gn=RRAS;refGene_tx=XM_011289684.2
E2 8372870 8372989
refGene_gn=RRAS;refGene_tx=XM_011289683.2;refGene_gn=RRAS;refGene_tx=XM_011289684.2
E2 8373066 8373151
refGene_gn=RRAS;refGene_tx=XM_011289683.2;refGene_gn=RRAS;refGene_tx=XM_011289684.2
E2 8378297 8378444 refGene_gn=PRR12;refGene_tx=XM_023246139.1
E2 8378739 8378835 refGene_gn=PRR12;refGene_tx=XM_023246139.1
E2 8378932 8379079 refGene_gn=PRR12;refGene_tx=XM_023246139.1
E2 8380170 8380268 refGene_gn=PRR12;refGene_tx=XM_023246139.1
E2 8380374 8380500 refGene_gn=PRR12;refGene_tx=XM_023246139.1
E2 8385841 8386338 refGene_gn=PRR12;refGene_tx=XM_023246139.1
E2 8386643 8386777 refGene_gn=PRR12;refGene_tx=XM_023246139.1
E2 8387032 8387149 refGene_gn=PRR12;refGene_tx=XM_023246139.1
E2 8394845 8395237 refGene_gn=PRR12;refGene_tx=XM_023246139.1
E2 8397838 8398499 refGene_gn=PRR12;refGene_tx=XM_023246139.1
E2 8399044 8402361 refGene_gn=PRR12;refGene_tx=XM_023246139.1
E2 8402445 8402607 refGene_gn=PRR12;refGene_tx=XM_023246139.1
E2 8403640 8403753 refGene_gn=PRR12;refGene_tx=XM_023246139.1
E2 8405292 8405378 refGene_gn=PRR12;refGene_tx=XM_023246139.1
E2 8406644 8406663
refGene_gn=PRRG2;refGene_tx=XM_003997505.4;refGene_gn=PRRG2;refGene_tx=XM_019819214.2
E2 8406981 8407128
refGene_gn=PRRG2;refGene_tx=XM_003997505.4;refGene_gn=PRRG2;refGene_tx=XM_019819214.2
E2 8407971 8408110
refGene_gn=PRRG2;refGene_tx=XM_003997505.4;refGene_gn=PRRG2;refGene_tx=XM_019819214.2
E2 8410655 8410695
refGene_gn=PRRG2;refGene_tx=XM_003997505.4;refGene_gn=PRRG2;refGene_tx=XM_019819214.2
E2 8410846 8411022
refGene_gn=PRRG2;refGene_tx=XM_003997505.4;refGene_gn=PRRG2;refGene_tx=XM_019819214.2
E2 8411210 8411295
refGene_gn=PRRG2;refGene_tx=XM_003997505.4;refGene_gn=PRRG2;refGene_tx=XM_019819214.2
E2 8422785 8422855

refGene_gn=NOSIP;refGene_tx=XM_003997506.5;refGene_gn=NOSIP;refGene_tx=XM_019819212.2;refGene_gn=NOSIP;refGene_tx=XM_019819213.2
E2 8423404 8423510
refGene_gn=NOSIP;refGene_tx=XM_003997506.5;refGene_gn=NOSIP;refGene_tx=XM_019819212.2;refGene_gn=NOSIP;refGene_tx=XM_019819213.2
E2 8424547 8424629
refGene_gn=NOSIP;refGene_tx=XM_003997506.5;refGene_gn=NOSIP;refGene_tx=XM_019819212.2;refGene_gn=NOSIP;refGene_tx=XM_019819213.2
E2 8427028 8427188
refGene_gn=NOSIP;refGene_tx=XM_003997506.5;refGene_gn=NOSIP;refGene_tx=XM_019819212.2;refGene_gn=NOSIP;refGene_tx=XM_019819213.2
E2 8427279 8427398
refGene_gn=NOSIP;refGene_tx=XM_003997506.5;refGene_gn=NOSIP;refGene_tx=XM_019819212.2;refGene_gn=NOSIP;refGene_tx=XM_019819213.2
E2 8427553 8427741
refGene_gn=NOSIP;refGene_tx=XM_003997506.5;refGene_gn=NOSIP;refGene_tx=XM_019819212.2;refGene_gn=NOSIP;refGene_tx=XM_019819213.2
E2 8427864 8427973
refGene_gn=NOSIP;refGene_tx=XM_003997506.5;refGene_gn=NOSIP;refGene_tx=XM_019819212.2;refGene_gn=NOSIP;refGene_tx=XM_019819213.2
E2 8428167 8428239
refGene_gn=NOSIP;refGene_tx=XM_003997506.5;refGene_gn=NOSIP;refGene_tx=XM_019819212.2;refGene_gn=NOSIP;refGene_tx=XM_019819213.2
E2 8438804 8438912 refGene_gn=RCN3;refGene_tx=XM_011289682.3
E2 8439138 8439338 refGene_gn=RCN3;refGene_tx=XM_011289682.3
E2 8442697 8442758 refGene_gn=RCN3;refGene_tx=XM_011289682.3
E2 8447642 8447815 refGene_gn=RCN3;refGene_tx=XM_011289682.3
E2 8448953 8449156 refGene_gn=RCN3;refGene_tx=XM_011289682.3
E2 8450984 8451226 refGene_gn=RCN3;refGene_tx=XM_011289682.3
E2 8452905 8452985
refGene_gn=FCGRT;refGene_tx=XM_023245230.1;refGene_gn=FCGRT;refGene_tx=XM_023245231.1;refGene_gn=FCGRT;refGene_tx=XM_023245232.1
E2 8453285 8453402
refGene_gn=FCGRT;refGene_tx=XM_023245230.1;refGene_gn=FCGRT;refGene_tx=XM_023245231.1;refGene_gn=FCGRT;refGene_tx=XM_023245232.1
E2 8454103 8454373
refGene_gn=FCGRT;refGene_tx=XM_023245230.1;refGene_gn=FCGRT;refGene_tx=XM_023245231.1;refGene_gn=FCGRT;refGene_tx=XM_023245232.1
E2 8459157 8459433
refGene_gn=FCGRT;refGene_tx=XM_023245230.1;refGene_gn=FCGRT;refGene_tx=XM_023245231.1;refGene_gn=FCGRT;refGene_tx=XM_023245232.1
E2 8459706 8459961
refGene_gn=FCGRT;refGene_tx=XM_023245230.1;refGene_gn=FCGRT;refGene_tx=XM_023245231.1;refGene_gn=FCGRT;refGene_tx=XM_023245232.1
E2 8460480 8460550
refGene_gn=FCGRT;refGene_tx=XM_023245230.1;refGene_gn=FCGRT;refGene_tx=XM_023245231.1;refGene_gn=FCGRT;refGene_tx=XM_023245232.1
E2 8472402 8472526 refGene_gn=RPS11;refGene_tx=XM_003997507.5
E2 8472962 8473092 refGene_gn=RPS11;refGene_tx=XM_003997507.5
E2 8473354 8473430 refGene_gn=RPS11;refGene_tx=XM_003997507.5

E2	8473607	8473739	refGene_gn=RPS11;refGene_tx=XM_003997507.5
E2	8474388	8474403	refGene_gn=RPS11;refGene_tx=XM_003997507.5
E2	8481688	8481775	refGene_gn=RPL13A;refGene_tx=XM_011289680.3
E2	8481959	8482082	refGene_gn=RPL13A;refGene_tx=XM_011289680.3
E2	8482334	8482394	refGene_gn=RPL13A;refGene_tx=XM_011289680.3
E2	8482571	8482657	refGene_gn=RPL13A;refGene_tx=XM_011289680.3
E2	8482847	8482949	refGene_gn=RPL13A;refGene_tx=XM_011289680.3
E2	8483129	8483195	refGene_gn=RPL13A;refGene_tx=XM_011289680.3
E2	8483614	8483687	refGene_gn=RPL13A;refGene_tx=XM_011289680.3
E2	8485319	8485334	refGene_gn=RPL13A;refGene_tx=XM_011289680.3
E2	8486354	8486567
refGene_gn=FLT3LG;refGene_tx=NM_001009842.1;refGene_gn=FLT3LG;refGene_tx=XM_011289647.3;refGene_gn=FLT3LG;refGene_tx=XM_011289648.3;refGene_gn=FLT3LG;refGene_tx=XM_019818430.2;refGene_gn=FLT3LG;refGene_tx=XM_019818431.2;refGene_gn=FLT3LG;refGene_tx=XM_019818432.2
E2	8489781	8489966
refGene_gn=FLT3LG;refGene_tx=NM_001009842.1;refGene_gn=FLT3LG;refGene_tx=XM_011289647.3;refGene_gn=FLT3LG;refGene_tx=XM_011289648.3;refGene_gn=FLT3LG;refGene_tx=XM_019818430.2;refGene_gn=FLT3LG;refGene_tx=XM_019818431.2;refGene_gn=FLT3LG;refGene_tx=XM_019818432.2
E2	8492095	8492234
refGene_gn=FLT3LG;refGene_tx=NM_001009842.1;refGene_gn=FLT3LG;refGene_tx=XM_011289647.3;refGene_gn=FLT3LG;refGene_tx=XM_011289648.3;refGene_gn=FLT3LG;refGene_tx=XM_019818430.2;refGene_gn=FLT3LG;refGene_tx=XM_019818431.2;refGene_gn=FLT3LG;refGene_tx=XM_019818432.2
E2	8492711	8492855
refGene_gn=FLT3LG;refGene_tx=NM_001009842.1;refGene_gn=FLT3LG;refGene_tx=XM_011289647.3;refGene_gn=FLT3LG;refGene_tx=XM_011289648.3;refGene_gn=FLT3LG;refGene_tx=XM_019818430.2;refGene_gn=FLT3LG;refGene_tx=XM_019818431.2;refGene_gn=FLT3LG;refGene_tx=XM_019818432.2
E2	8493068	8493122
refGene_gn=FLT3LG;refGene_tx=NM_001009842.1;refGene_gn=FLT3LG;refGene_tx=XM_011289647.3;refGene_gn=FLT3LG;refGene_tx=XM_011289648.3;refGene_gn=FLT3LG;refGene_tx=XM_019818430.2;refGene_gn=FLT3LG;refGene_tx=XM_019818431.2;refGene_gn=FLT3LG;refGene_tx=XM_019818432.2
E2	8493329	8493440
refGene_gn=FLT3LG;refGene_tx=NM_001009842.1;refGene_gn=FLT3LG;refGene_tx=XM_011289647.3;refGene_gn=FLT3LG;refGene_tx=XM_011289648.3;refGene_gn=FLT3LG;refGene_tx=XM_019818430.2;refGene_gn=FLT3LG;refGene_tx=XM_019818431.2
E2	8494215	8494248
refGene_gn=FLT3LG;refGene_tx=NM_001009842.1;refGene_gn=FLT3LG;refGene_tx=XM_011289647.3;refGene_gn=FLT3LG;refGene_tx=XM_011289648.3;refGene_gn=FLT3LG;refGene_tx=XM_019818430.2;refGene_gn=FLT3LG;refGene_tx=XM_019818431.2;refGene_gn=FLT3LG;refGene_tx=XM_019818432.2
E2	8500738	8500900
refGene_gn=ALDH16A1;refGene_tx=XM_011289678.3;refGene_gn=ALDH16A1;refGene_tx=XM_019819217.2
E2	8501569	8501692
refGene_gn=ALDH16A1;refGene_tx=XM_011289678.3;refGene_gn=ALDH16A1;refG

ene_tx=XM_019819217.2
E2 8501971 8502157
refGene_gn=ALDH16A1;refGene_tx=XM_011289678.3;refGene_gn=ALDH16A1;refGene_tx=XM_019819217.2
E2 8504969 8505171
refGene_gn=ALDH16A1;refGene_tx=XM_011289678.3;refGene_gn=ALDH16A1;refGene_tx=XM_019819217.2
E2 8505437 8505605
refGene_gn=ALDH16A1;refGene_tx=XM_011289678.3;refGene_gn=ALDH16A1;refGene_tx=XM_019819217.2
E2 8505898 8506029
refGene_gn=ALDH16A1;refGene_tx=XM_011289678.3;refGene_gn=ALDH16A1;refGene_tx=XM_019819217.2
E2 8506119 8506225
refGene_gn=ALDH16A1;refGene_tx=XM_011289678.3;refGene_gn=ALDH16A1;refGene_tx=XM_019819217.2
E2 8506424 8506561
refGene_gn=ALDH16A1;refGene_tx=XM_011289678.3;refGene_gn=ALDH16A1;refGene_tx=XM_019819217.2
E2 8506751 8506847
refGene_gn=ALDH16A1;refGene_tx=XM_011289678.3;refGene_gn=ALDH16A1;refGene_tx=XM_019819217.2
E2 8507401 8507587
refGene_gn=ALDH16A1;refGene_tx=XM_011289678.3;refGene_gn=ALDH16A1;refGene_tx=XM_019819217.2
E2 8508064 8508217
refGene_gn=ALDH16A1;refGene_tx=XM_011289678.3;refGene_gn=ALDH16A1;refGene_tx=XM_019819217.2
E2 8508299 8508481
refGene_gn=ALDH16A1;refGene_tx=XM_011289678.3;refGene_gn=ALDH16A1;refGene_tx=XM_019819217.2
E2 8509106 8509184
refGene_gn=ALDH16A1;refGene_tx=XM_011289678.3;refGene_gn=ALDH16A1;refGene_tx=XM_019819217.2
E2 8509270 8509449
refGene_gn=ALDH16A1;refGene_tx=XM_011289678.3;refGene_gn=ALDH16A1;refGene_tx=XM_019819217.2
E2 8510164 8510291
refGene_gn=ALDH16A1;refGene_tx=XM_011289678.3;refGene_gn=ALDH16A1;refGene_tx=XM_019819217.2
E2 8510812 8510915
refGene_gn=ALDH16A1;refGene_tx=XM_011289678.3;refGene_gn=ALDH16A1;refGene_tx=XM_019819217.2
E2 8511286 8511298 refGene_gn=ALDH16A1;refGene_tx=XM_019819217.2
E2 8516076 8516166 refGene_gn=ALDH16A1;refGene_tx=XM_011289678.3
E2 8517105 8517195
refGene_gn=PIH1D1;refGene_tx=XM_003997508.5;refGene_gn=PIH1D1;refGene_tx=XM_006940997.4
E2 8517555 8517622
refGene_gn=PIH1D1;refGene_tx=XM_003997508.5;refGene_gn=PIH1D1;refGene_

tx=XM_006940997.4
E2	8517967	8518147
refGene_gn=PIH1D1;refGene_tx=XM_003997508.5;refGene_gn=PIH1D1;refGene_tx=XM_006940997.4
E2	8519260	8519322
refGene_gn=PIH1D1;refGene_tx=XM_003997508.5;refGene_gn=PIH1D1;refGene_tx=XM_006940997.4
E2	8519410	8519492
refGene_gn=PIH1D1;refGene_tx=XM_003997508.5;refGene_gn=PIH1D1;refGene_tx=XM_006940997.4
E2	8519952	8520082
refGene_gn=PIH1D1;refGene_tx=XM_003997508.5;refGene_gn=PIH1D1;refGene_tx=XM_006940997.4
E2	8520327	8520403
refGene_gn=PIH1D1;refGene_tx=XM_003997508.5;refGene_gn=PIH1D1;refGene_tx=XM_006940997.4
E2	8520692	8520836
refGene_gn=PIH1D1;refGene_tx=XM_003997508.5;refGene_gn=PIH1D1;refGene_tx=XM_006940997.4
E2	8520938	8520980
refGene_gn=PIH1D1;refGene_tx=XM_003997508.5;refGene_gn=PIH1D1;refGene_tx=XM_006940997.4
E2	8524321	8524383	refGene_gn=SLC17A7;refGene_tx=XM_023245257.1
E2	8528403	8528656	refGene_gn=SLC17A7;refGene_tx=XM_023245257.1
E2	8529833	8529952	refGene_gn=SLC17A7;refGene_tx=XM_023245257.1
E2	8530238	8530353	refGene_gn=SLC17A7;refGene_tx=XM_023245257.1
E2	8530431	8530519	refGene_gn=SLC17A7;refGene_tx=XM_023245257.1
E2	8530674	8530761	refGene_gn=SLC17A7;refGene_tx=XM_023245257.1
E2	8530851	8530994	refGene_gn=SLC17A7;refGene_tx=XM_023245257.1
E2	8531657	8531807	refGene_gn=SLC17A7;refGene_tx=XM_023245257.1
E2	8531978	8532111	refGene_gn=SLC17A7;refGene_tx=XM_023245257.1
E2	8532579	8532690	refGene_gn=SLC17A7;refGene_tx=XM_023245257.1
E2	8532925	8533053	refGene_gn=SLC17A7;refGene_tx=XM_023245257.1
E2	8533287	8533581	refGene_gn=SLC17A7;refGene_tx=XM_023245257.1
E2	8535007	8535207	refGene_gn=GFY;refGene_tx=XM_023245258.1
E2	8535710	8535884	refGene_gn=GFY;refGene_tx=XM_023245258.1
E2	8536101	8537284	refGene_gn=GFY;refGene_tx=XM_023245258.1
E2	8539718	8539843	refGene_gn=PTH2;refGene_tx=XM_003997510.4
E2	8540365	8540540	refGene_gn=PTH2;refGene_tx=XM_003997510.4
E2	8544347	8544489
refGene_gn=CCDC155;refGene_tx=XM_023245188.1;refGene_gn=CCDC155;refGene_tx=XM_023245189.1;refGene_gn=CCDC155;refGene_tx=XM_023245190.1;refGene_gn=CCDC155;refGene_tx=XM_023245191.1;refGene_gn=CCDC155;refGene_tx=XM_023245192.1;refGene_gn=CCDC155;refGene_tx=XM_023245193.1;refGene_gn=CCDC155;refGene_tx=XM_023245194.1;refGene_gn=CCDC155;refGene_tx=XM_023245195.1;refGene_gn=CCDC155;refGene_tx=XM_023245196.1
E2	8544607	8544727
refGene_gn=CCDC155;refGene_tx=XM_023245188.1;refGene_gn=CCDC155;refGene_tx=XM_023245189.1;refGene_gn=CCDC155;refGene_tx=XM_023245190.1;refGene_gn=CCDC155;refGene_tx=XM_023245191.1;refGene_gn=CCDC155;refGene_tx=

XM_023245192.1;refGene_gn=CCDC155;refGene_tx=XM_023245193.1;refGene_gn=CCDC155;refGene_tx=XM_023245194.1;refGene_gn=CCDC155;refGene_tx=XM_023245195.1;refGene_gn=CCDC155;refGene_tx=XM_023245196.1
E2 8544809 8544874
refGene_gn=CCDC155;refGene_tx=XM_023245188.1;refGene_gn=CCDC155;refGene_tx=XM_023245189.1;refGene_gn=CCDC155;refGene_tx=XM_023245190.1;refGene_gn=CCDC155;refGene_tx=XM_023245191.1;refGene_gn=CCDC155;refGene_tx=XM_023245192.1;refGene_gn=CCDC155;refGene_tx=XM_023245193.1;refGene_gn=CCDC155;refGene_tx=XM_023245194.1;refGene_gn=CCDC155;refGene_tx=XM_023245195.1;refGene_gn=CCDC155;refGene_tx=XM_023245196.1
E2 8546588 8546634
refGene_gn=CCDC155;refGene_tx=XM_023245188.1;refGene_gn=CCDC155;refGene_tx=XM_023245189.1;refGene_gn=CCDC155;refGene_tx=XM_023245190.1;refGene_gn=CCDC155;refGene_tx=XM_023245191.1;refGene_gn=CCDC155;refGene_tx=XM_023245192.1;refGene_gn=CCDC155;refGene_tx=XM_023245193.1;refGene_gn=CCDC155;refGene_tx=XM_023245194.1;refGene_gn=CCDC155;refGene_tx=XM_023245195.1;refGene_gn=CCDC155;refGene_tx=XM_023245196.1
E2 8548399 8548458
refGene_gn=CCDC155;refGene_tx=XM_023245188.1;refGene_gn=CCDC155;refGene_tx=XM_023245189.1;refGene_gn=CCDC155;refGene_tx=XM_023245190.1;refGene_gn=CCDC155;refGene_tx=XM_023245191.1;refGene_gn=CCDC155;refGene_tx=XM_023245192.1;refGene_gn=CCDC155;refGene_tx=XM_023245193.1;refGene_gn=CCDC155;refGene_tx=XM_023245194.1;refGene_gn=CCDC155;refGene_tx=XM_023245195.1;refGene_gn=CCDC155;refGene_tx=XM_023245196.1
E2 8551289 8551418
refGene_gn=CCDC155;refGene_tx=XM_023245188.1;refGene_gn=CCDC155;refGene_tx=XM_023245189.1;refGene_gn=CCDC155;refGene_tx=XM_023245190.1;refGene_gn=CCDC155;refGene_tx=XM_023245191.1;refGene_gn=CCDC155;refGene_tx=XM_023245192.1;refGene_gn=CCDC155;refGene_tx=XM_023245193.1;refGene_gn=CCDC155;refGene_tx=XM_023245194.1;refGene_gn=CCDC155;refGene_tx=XM_023245195.1;refGene_gn=CCDC155;refGene_tx=XM_023245196.1
E2 8551659 8551747
refGene_gn=CCDC155;refGene_tx=XM_023245188.1;refGene_gn=CCDC155;refGene_tx=XM_023245189.1;refGene_gn=CCDC155;refGene_tx=XM_023245190.1;refGene_gn=CCDC155;refGene_tx=XM_023245191.1;refGene_gn=CCDC155;refGene_tx=XM_023245192.1;refGene_gn=CCDC155;refGene_tx=XM_023245193.1;refGene_gn=CCDC155;refGene_tx=XM_023245194.1;refGene_gn=CCDC155;refGene_tx=XM_023245195.1;refGene_gn=CCDC155;refGene_tx=XM_023245196.1
E2 8551911 8551976
refGene_gn=CCDC155;refGene_tx=XM_023245188.1;refGene_gn=CCDC155;refGene_tx=XM_023245189.1;refGene_gn=CCDC155;refGene_tx=XM_023245190.1;refGene_gn=CCDC155;refGene_tx=XM_023245191.1;refGene_gn=CCDC155;refGene_tx=XM_023245192.1;refGene_gn=CCDC155;refGene_tx=XM_023245193.1;refGene_gn=CCDC155;refGene_tx=XM_023245194.1;refGene_gn=CCDC155;refGene_tx=XM_023245195.1;refGene_gn=CCDC155;refGene_tx=XM_023245196.1
E2 8552644 8552704
refGene_gn=CCDC155;refGene_tx=XM_023245188.1;refGene_gn=CCDC155;refGene_tx=XM_023245189.1;refGene_gn=CCDC155;refGene_tx=XM_023245190.1;refGene_gn=CCDC155;refGene_tx=XM_023245191.1;refGene_gn=CCDC155;refGene_tx=XM_023245192.1;refGene_gn=CCDC155;refGene_tx=XM_023245193.1;refGene_gn

=CCDC155;refGene_tx=XM_023245194.1;refGene_gn=CCDC155;refGene_tx=XM_023245195.1;refGene_gn=CCDC155;refGene_tx=XM_023245196.1
E2 8553010 8553067
refGene_gn=CCDC155;refGene_tx=XM_023245188.1;refGene_gn=CCDC155;refGene_tx=XM_023245189.1;refGene_gn=CCDC155;refGene_tx=XM_023245190.1;refGene_gn=CCDC155;refGene_tx=XM_023245191.1;refGene_gn=CCDC155;refGene_tx=XM_023245192.1;refGene_gn=CCDC155;refGene_tx=XM_023245193.1;refGene_gn=CCDC155;refGene_tx=XM_023245194.1;refGene_gn=CCDC155;refGene_tx=XM_023245196.1
E2 8553274 8553403
refGene_gn=CCDC155;refGene_tx=XM_023245188.1;refGene_gn=CCDC155;refGene_tx=XM_023245189.1;refGene_gn=CCDC155;refGene_tx=XM_023245190.1;refGene_gn=CCDC155;refGene_tx=XM_023245191.1;refGene_gn=CCDC155;refGene_tx=XM_023245192.1;refGene_gn=CCDC155;refGene_tx=XM_023245193.1;refGene_gn=CCDC155;refGene_tx=XM_023245194.1;refGene_gn=CCDC155;refGene_tx=XM_023245195.1;refGene_gn=CCDC155;refGene_tx=XM_023245196.1
E2 8558370 8558421
refGene_gn=CCDC155;refGene_tx=XM_023245188.1;refGene_gn=CCDC155;refGene_tx=XM_023245189.1;refGene_gn=CCDC155;refGene_tx=XM_023245190.1;refGene_gn=CCDC155;refGene_tx=XM_023245191.1;refGene_gn=CCDC155;refGene_tx=XM_023245192.1;refGene_gn=CCDC155;refGene_tx=XM_023245193.1;refGene_gn=CCDC155;refGene_tx=XM_023245194.1;refGene_gn=CCDC155;refGene_tx=XM_023245195.1;refGene_gn=CCDC155;refGene_tx=XM_023245196.1
E2 8558753 8558871
refGene_gn=CCDC155;refGene_tx=XM_023245188.1;refGene_gn=CCDC155;refGene_tx=XM_023245189.1;refGene_gn=CCDC155;refGene_tx=XM_023245190.1;refGene_gn=CCDC155;refGene_tx=XM_023245191.1;refGene_gn=CCDC155;refGene_tx=XM_023245192.1;refGene_gn=CCDC155;refGene_tx=XM_023245193.1;refGene_gn=CCDC155;refGene_tx=XM_023245194.1;refGene_gn=CCDC155;refGene_tx=XM_023245195.1;refGene_gn=CCDC155;refGene_tx=XM_023245196.1
E2 8559313 8559475
refGene_gn=CCDC155;refGene_tx=XM_023245188.1;refGene_gn=CCDC155;refGene_tx=XM_023245189.1;refGene_gn=CCDC155;refGene_tx=XM_023245190.1;refGene_gn=CCDC155;refGene_tx=XM_023245191.1;refGene_gn=CCDC155;refGene_tx=XM_023245192.1;refGene_gn=CCDC155;refGene_tx=XM_023245193.1;refGene_gn=CCDC155;refGene_tx=XM_023245194.1;refGene_gn=CCDC155;refGene_tx=XM_023245195.1;refGene_gn=CCDC155;refGene_tx=XM_023245196.1
E2 8559708 8559775
refGene_gn=CCDC155;refGene_tx=XM_023245188.1;refGene_gn=CCDC155;refGene_tx=XM_023245189.1;refGene_gn=CCDC155;refGene_tx=XM_023245190.1;refGene_gn=CCDC155;refGene_tx=XM_023245191.1;refGene_gn=CCDC155;refGene_tx=XM_023245192.1;refGene_gn=CCDC155;refGene_tx=XM_023245193.1;refGene_gn=CCDC155;refGene_tx=XM_023245194.1;refGene_gn=CCDC155;refGene_tx=XM_023245195.1;refGene_gn=CCDC155;refGene_tx=XM_023245196.1
E2 8560988 8561053
refGene_gn=CCDC155;refGene_tx=XM_023245188.1;refGene_gn=CCDC155;refGene_tx=XM_023245189.1;refGene_gn=CCDC155;refGene_tx=XM_023245190.1;refGene_gn=CCDC155;refGene_tx=XM_023245191.1;refGene_gn=CCDC155;refGene_tx=XM_023245192.1;refGene_gn=CCDC155;refGene_tx=XM_023245193.1;refGene_gn=CCDC155;refGene_tx=XM_023245194.1;refGene_gn=CCDC155;refGene_tx=XM_02

3245195.1;refGene_gn=CCDC155;refGene_tx=XM_023245196.1
E2 8561541 8561728
refGene_gn=CCDC155;refGene_tx=XM_023245188.1;refGene_gn=CCDC155;refGene_tx=XM_023245189.1;refGene_gn=CCDC155;refGene_tx=XM_023245190.1;refGene_gn=CCDC155;refGene_tx=XM_023245191.1;refGene_gn=CCDC155;refGene_tx=XM_023245192.1;refGene_gn=CCDC155;refGene_tx=XM_023245193.1;refGene_gn=CCDC155;refGene_tx=XM_023245194.1;refGene_gn=CCDC155;refGene_tx=XM_023245195.1;refGene_gn=CCDC155;refGene_tx=XM_023245196.1
E2 8562233 8562338
refGene_gn=CCDC155;refGene_tx=XM_023245188.1;refGene_gn=CCDC155;refGene_tx=XM_023245189.1;refGene_gn=CCDC155;refGene_tx=XM_023245190.1;refGene_gn=CCDC155;refGene_tx=XM_023245191.1;refGene_gn=CCDC155;refGene_tx=XM_023245192.1;refGene_gn=CCDC155;refGene_tx=XM_023245193.1;refGene_gn=CCDC155;refGene_tx=XM_023245194.1;refGene_gn=CCDC155;refGene_tx=XM_023245195.1;refGene_gn=CCDC155;refGene_tx=XM_023245196.1
E2 8565753 8565891
refGene_gn=CCDC155;refGene_tx=XM_023245188.1;refGene_gn=CCDC155;refGene_tx=XM_023245189.1;refGene_gn=CCDC155;refGene_tx=XM_023245190.1;refGene_gn=CCDC155;refGene_tx=XM_023245191.1;refGene_gn=CCDC155;refGene_tx=XM_023245192.1;refGene_gn=CCDC155;refGene_tx=XM_023245193.1;refGene_gn=CCDC155;refGene_tx=XM_023245194.1;refGene_gn=CCDC155;refGene_tx=XM_023245195.1;refGene_gn=CCDC155;refGene_tx=XM_023245196.1
E2 8568981 8569261
refGene_gn=CCDC155;refGene_tx=XM_023245188.1;refGene_gn=CCDC155;refGene_tx=XM_023245189.1;refGene_gn=CCDC155;refGene_tx=XM_023245190.1;refGene_gn=CCDC155;refGene_tx=XM_023245191.1;refGene_gn=CCDC155;refGene_tx=XM_023245192.1;refGene_gn=CCDC155;refGene_tx=XM_023245193.1;refGene_gn=CCDC155;refGene_tx=XM_023245194.1;refGene_gn=CCDC155;refGene_tx=XM_023245195.1
E2 8574764 8575076
refGene_gn=DKKL1;refGene_tx=XM_006941000.4;refGene_gn=DKKL1;refGene_tx=XM_006941001.2;refGene_gn=DKKL1;refGene_tx=XM_006941002.4;refGene_gn=DKKL1;refGene_tx=XM_019819220.1
E2 8576956 8577049
refGene_gn=DKKL1;refGene_tx=XM_006941000.4;refGene_gn=DKKL1;refGene_tx=XM_006941001.2;refGene_gn=DKKL1;refGene_tx=XM_006941002.4;refGene_gn=DKKL1;refGene_tx=XM_019819220.1
E2 8577171 8577312
refGene_gn=DKKL1;refGene_tx=XM_006941000.4;refGene_gn=DKKL1;refGene_tx=XM_006941001.2;refGene_gn=DKKL1;refGene_tx=XM_006941002.4;refGene_gn=DKKL1;refGene_tx=XM_019819220.1
E2 8577862 8578037
refGene_gn=DKKL1;refGene_tx=XM_006941000.4;refGene_gn=DKKL1;refGene_tx=XM_006941001.2;refGene_gn=DKKL1;refGene_tx=XM_006941002.4
E2 8578442 8578489 refGene_gn=DKKL1;refGene_tx=XM_006941000.4
E2 8581460 8581692
refGene_gn=TEAD2;refGene_tx=XM_003997513.4;refGene_gn=TEAD2;refGene_tx=XM_006941008.3;refGene_gn=TEAD2;refGene_tx=XM_023246190.1
E2 8582073 8582138
refGene_gn=TEAD2;refGene_tx=XM_003997513.4;refGene_gn=TEAD2;refGene_tx

=XM_006941008.3;refGene_gn=TEAD2;refGene_tx=XM_023246190.1
E2 8582560 8582623
refGene_gn=TEAD2;refGene_tx=XM_003997513.4;refGene_gn=TEAD2;refGene_tx=XM_006941008.3;refGene_gn=TEAD2;refGene_tx=XM_023246190.1
E2 8583847 8583859
refGene_gn=TEAD2;refGene_tx=XM_003997513.4;refGene_gn=TEAD2;refGene_tx=XM_006941008.3
E2 8584087 8584195
refGene_gn=TEAD2;refGene_tx=XM_003997513.4;refGene_gn=TEAD2;refGene_tx=XM_006941008.3;refGene_gn=TEAD2;refGene_tx=XM_023246190.1
E2 8584281 8584340
refGene_gn=TEAD2;refGene_tx=XM_003997513.4;refGene_gn=TEAD2;refGene_tx=XM_006941008.3;refGene_gn=TEAD2;refGene_tx=XM_023246190.1
E2 8587723 8587788
refGene_gn=TEAD2;refGene_tx=XM_003997513.4;refGene_gn=TEAD2;refGene_tx=XM_006941008.3;refGene_gn=TEAD2;refGene_tx=XM_023246190.1
E2 8588821 8588970
refGene_gn=TEAD2;refGene_tx=XM_003997513.4;refGene_gn=TEAD2;refGene_tx=XM_006941008.3;refGene_gn=TEAD2;refGene_tx=XM_023246190.1
E2 8592022 8592196
refGene_gn=TEAD2;refGene_tx=XM_003997513.4;refGene_gn=TEAD2;refGene_tx=XM_006941008.3;refGene_gn=TEAD2;refGene_tx=XM_023246190.1
E2 8593794 8593956
refGene_gn=TEAD2;refGene_tx=XM_003997513.4;refGene_gn=TEAD2;refGene_tx=XM_006941008.3;refGene_gn=TEAD2;refGene_tx=XM_023246190.1
E2 8594638 8594791
refGene_gn=TEAD2;refGene_tx=XM_003997513.4;refGene_gn=TEAD2;refGene_tx=XM_006941008.3;refGene_gn=TEAD2;refGene_tx=XM_023246190.1
E2 8595231 8595345
refGene_gn=TEAD2;refGene_tx=XM_003997513.4;refGene_gn=TEAD2;refGene_tx=XM_006941008.3;refGene_gn=TEAD2;refGene_tx=XM_023246190.1
E2 8596211 8596218 refGene_gn=CD37;refGene_tx=XM_006941003.4
E2 8596313 8596391
refGene_gn=CD37;refGene_tx=XM_003997517.5;refGene_gn=CD37;refGene_tx=XM_006941003.4;refGene_gn=CD37;refGene_tx=XM_006941004.4;refGene_gn=CD37;refGene_tx=XM_006941005.4;refGene_gn=CD37;refGene_tx=XM_006941006.4;refGene_gn=CD37;refGene_tx=XM_006941007.4
E2 8597397 8597481
refGene_gn=CD37;refGene_tx=XM_003997517.5;refGene_gn=CD37;refGene_tx=XM_006941003.4;refGene_gn=CD37;refGene_tx=XM_006941004.4;refGene_gn=CD37;refGene_tx=XM_006941005.4;refGene_gn=CD37;refGene_tx=XM_006941006.4;refGene_gn=CD37;refGene_tx=XM_006941007.4
E2 8597861 8598098
refGene_gn=CD37;refGene_tx=XM_003997517.5;refGene_gn=CD37;refGene_tx=XM_006941003.4;refGene_gn=CD37;refGene_tx=XM_006941004.4;refGene_gn=CD37;refGene_tx=XM_006941005.4;refGene_gn=CD37;refGene_tx=XM_006941006.4;refGene_gn=CD37;refGene_tx=XM_006941007.4
E2 8598770 8598875
refGene_gn=CD37;refGene_tx=XM_003997517.5;refGene_gn=CD37;refGene_tx=XM_006941003.4;refGene_gn=CD37;refGene_tx=XM_006941004.4;refGene_gn=CD3

7;refGene_tx=XM_006941005.4;refGene_gn=CD37;refGene_tx=XM_006941006.4;
refGene_gn=CD37;refGene_tx=XM_006941007.4
E2 8599424 8599499
refGene_gn=CD37;refGene_tx=XM_003997517.5;refGene_gn=CD37;refGene_tx=X
M_006941003.4;refGene_gn=CD37;refGene_tx=XM_006941004.4;refGene_gn=CD3
7;refGene_tx=XM_006941005.4;refGene_gn=CD37;refGene_tx=XM_006941006.4;
refGene_gn=CD37;refGene_tx=XM_006941007.4
E2 8599594 8599719
refGene_gn=CD37;refGene_tx=XM_003997517.5;refGene_gn=CD37;refGene_tx=X
M_006941003.4;refGene_gn=CD37;refGene_tx=XM_006941004.4;refGene_gn=CD3
7;refGene_tx=XM_006941005.4;refGene_gn=CD37;refGene_tx=XM_006941006.4;
refGene_gn=CD37;refGene_tx=XM_006941007.4
E2 8600606 8600679
refGene_gn=CD37;refGene_tx=XM_003997517.5;refGene_gn=CD37;refGene_tx=X
M_006941003.4;refGene_gn=CD37;refGene_tx=XM_006941004.4;refGene_gn=CD3
7;refGene_tx=XM_006941005.4
E2 8600790 8600865
refGene_gn=CD37;refGene_tx=XM_003997517.5;refGene_gn=CD37;refGene_tx=X
M_006941003.4;refGene_gn=CD37;refGene_tx=XM_006941004.4;refGene_gn=CD3
7;refGene_tx=XM_006941005.4
E2 8601096 8601159
refGene_gn=CD37;refGene_tx=XM_003997517.5;refGene_gn=CD37;refGene_tx=X
M_006941003.4;refGene_gn=CD37;refGene_tx=XM_006941005.4
E2 8627883 8628079 refGene_gn=SLC6A16;refGene_tx=XM_023245905.1
E2 8628468 8628626 refGene_gn=SLC6A16;refGene_tx=XM_023245905.1
E2 8628815 8628942 refGene_gn=SLC6A16;refGene_tx=XM_023245905.1
E2 8629143 8629322 refGene_gn=SLC6A16;refGene_tx=XM_023245905.1
E2 8629545 8629601 refGene_gn=SLC6A16;refGene_tx=XM_023245905.1
E2 8629602 8629652 refGene_gn=SLC6A16;refGene_tx=XM_023245905.1
E2 8629818 8630059 refGene_gn=SLC6A16;refGene_tx=XM_023245905.1
E2 8637008 8637095 refGene_gn=SLC6A16;refGene_tx=XM_023245905.1
E2 8637096 8637193 refGene_gn=SLC6A16;refGene_tx=XM_023245905.1
E2 8637525 8637724 refGene_gn=SLC6A16;refGene_tx=XM_023245905.1
E2 8637863 8638065 refGene_gn=SLC6A16;refGene_tx=XM_023245905.1
E2 8639300 8639456 refGene_gn=SLC6A16;refGene_tx=XM_023245905.1
E2 8639556 8639926 refGene_gn=SLC6A16;refGene_tx=XM_023245905.1
E2 8668587 8668927
refGene_gn=LOC101082794;refGene_tx=XM_011289743.3;refGene_gn=LOC101082
794;refGene_tx=XM_019819226.2;refGene_gn=LOC101082794;refGene_tx=XM_01
9819227.2;refGene_gn=LOC101082794;refGene_tx=XM_019819229.2;refGene_gn
=LOC101082794;refGene_tx=XM_019819230.2;refGene_gn=LOC101082794;refGen
e_tx=XM_019819233.1;refGene_gn=LOC101082794;refGene_tx=XM_023245245.1
E2 8669361 8669519
refGene_gn=LOC101082794;refGene_tx=XM_011289743.3;refGene_gn=LOC101082
794;refGene_tx=XM_019819226.2;refGene_gn=LOC101082794;refGene_tx=XM_01
9819227.2;refGene_gn=LOC101082794;refGene_tx=XM_019819229.2;refGene_gn
=LOC101082794;refGene_tx=XM_019819230.2;refGene_gn=LOC101082794;refGen
e_tx=XM_019819233.1;refGene_gn=LOC101082794;refGene_tx=XM_023245245.1
E2 8669709 8669836
refGene_gn=LOC101082794;refGene_tx=XM_011289743.3;refGene_gn=LOC101082

794;refGene_tx=XM_019819226.2;refGene_gn=LOC101082794;refGene_tx=XM_01
9819227.2;refGene_gn=LOC101082794;refGene_tx=XM_019819229.2;refGene_gn
=LOC101082794;refGene_tx=XM_019819230.2;refGene_gn=LOC101082794;refGen
e_tx=XM_019819233.1;refGene_gn=LOC101082794;refGene_tx=XM_023245245.1
E2 8670030 8670206
refGene_gn=LOC101082794;refGene_tx=XM_011289743.3;refGene_gn=LOC101082
794;refGene_tx=XM_019819226.2;refGene_gn=LOC101082794;refGene_tx=XM_01
9819227.2;refGene_gn=LOC101082794;refGene_tx=XM_019819229.2;refGene_gn
=LOC101082794;refGene_tx=XM_019819230.2;refGene_gn=LOC101082794;refGen
e_tx=XM_019819233.1;refGene_gn=LOC101082794;refGene_tx=XM_023245245.1
E2 8670402 8670513
refGene_gn=LOC101082794;refGene_tx=XM_011289743.3;refGene_gn=LOC101082
794;refGene_tx=XM_019819226.2;refGene_gn=LOC101082794;refGene_tx=XM_01
9819227.2;refGene_gn=LOC101082794;refGene_tx=XM_019819229.2;refGene_gn
=LOC101082794;refGene_tx=XM_019819230.2;refGene_gn=LOC101082794;refGen
e_tx=XM_019819233.1;refGene_gn=LOC101082794;refGene_tx=XM_023245245.1
E2 8670689 8670931
refGene_gn=LOC101082794;refGene_tx=XM_011289743.3;refGene_gn=LOC101082
794;refGene_tx=XM_019819226.2;refGene_gn=LOC101082794;refGene_tx=XM_01
9819227.2;refGene_gn=LOC101082794;refGene_tx=XM_019819229.2;refGene_gn
=LOC101082794;refGene_tx=XM_019819230.2;refGene_gn=LOC101082794;refGen
e_tx=XM_019819233.1;refGene_gn=LOC101082794;refGene_tx=XM_023245245.1
E2 8672302 8672489
refGene_gn=LOC101082794;refGene_tx=XM_011289743.3;refGene_gn=LOC101082
794;refGene_tx=XM_019819226.2;refGene_gn=LOC101082794;refGene_tx=XM_01
9819227.2;refGene_gn=LOC101082794;refGene_tx=XM_019819229.2;refGene_gn
=LOC101082794;refGene_tx=XM_019819230.2;refGene_gn=LOC101082794;refGen
e_tx=XM_019819233.1;refGene_gn=LOC101082794;refGene_tx=XM_023245245.1
E2 8672889 8673091
refGene_gn=LOC101082794;refGene_tx=XM_011289743.3;refGene_gn=LOC101082
794;refGene_tx=XM_019819226.2;refGene_gn=LOC101082794;refGene_tx=XM_01
9819227.2;refGene_gn=LOC101082794;refGene_tx=XM_019819229.2;refGene_gn
=LOC101082794;refGene_tx=XM_019819230.2;refGene_gn=LOC101082794;refGen
e_tx=XM_019819233.1;refGene_gn=LOC101082794;refGene_tx=XM_023245245.1
E2 8673210 8673370
refGene_gn=LOC101082794;refGene_tx=XM_011289743.3;refGene_gn=LOC101082
794;refGene_tx=XM_019819226.2;refGene_gn=LOC101082794;refGene_tx=XM_01
9819227.2;refGene_gn=LOC101082794;refGene_tx=XM_019819229.2;refGene_gn
=LOC101082794;refGene_tx=XM_019819230.2;refGene_gn=LOC101082794;refGen
e_tx=XM_019819233.1;refGene_gn=LOC101082794;refGene_tx=XM_023245245.1
E2 8674498 8674661
refGene_gn=LOC101082794;refGene_tx=XM_011289743.3;refGene_gn=LOC101082
794;refGene_tx=XM_019819226.2;refGene_gn=LOC101082794;refGene_tx=XM_01
9819227.2;refGene_gn=LOC101082794;refGene_tx=XM_019819229.2;refGene_gn
=LOC101082794;refGene_tx=XM_019819230.2;refGene_gn=LOC101082794;refGen
e_tx=XM_023245245.1
E2 8674872 8675118
refGene_gn=LOC101082794;refGene_tx=XM_011289743.3;refGene_gn=LOC101082
794;refGene_tx=XM_019819226.2;refGene_gn=LOC101082794;refGene_tx=XM_01
9819227.2;refGene_gn=LOC101082794;refGene_tx=XM_019819229.2;refGene_gn

=LOC101082794;refGene_tx=XM_019819230.2;refGene_gn=LOC101082794;refGene_tx=XM_019819233.1;refGene_gn=LOC101082794;refGene_tx=XM_023245245.1
E2 8690945 8691168
refGene_gn=LOC102899166;refGene_tx=XM_006941009.4;refGene_gn=LOC102899166;refGene_tx=XM_011289670.3;refGene_gn=LOC102899166;refGene_tx=XM_011289671.3;refGene_gn=LOC102899166;refGene_tx=XM_011289672.3;refGene_gn=LOC102899166;refGene_tx=XM_011289673.3;refGene_gn=LOC102899166;refGene_tx=XM_011289674.3;refGene_gn=LOC102899166;refGene_tx=XM_019819222.2;refGene_gn=LOC102899166;refGene_tx=XM_019819223.2
E2 8691344 8691502
refGene_gn=LOC102899166;refGene_tx=XM_006941009.4;refGene_gn=LOC102899166;refGene_tx=XM_011289670.3;refGene_gn=LOC102899166;refGene_tx=XM_011289671.3;refGene_gn=LOC102899166;refGene_tx=XM_011289672.3;refGene_gn=LOC102899166;refGene_tx=XM_011289673.3;refGene_gn=LOC102899166;refGene_tx=XM_011289674.3;refGene_gn=LOC102899166;refGene_tx=XM_019819222.2;refGene_gn=LOC102899166;refGene_tx=XM_019819223.2
E2 8691609 8691736
refGene_gn=LOC102899166;refGene_tx=XM_006941009.4;refGene_gn=LOC102899166;refGene_tx=XM_011289670.3;refGene_gn=LOC102899166;refGene_tx=XM_011289671.3;refGene_gn=LOC102899166;refGene_tx=XM_011289672.3;refGene_gn=LOC102899166;refGene_tx=XM_011289673.3;refGene_gn=LOC102899166;refGene_tx=XM_011289674.3;refGene_gn=LOC102899166;refGene_tx=XM_019819222.2;refGene_gn=LOC102899166;refGene_tx=XM_019819223.2;refGene_gn=LOC102899166;refGene_tx=XM_019819225.2
E2 8691888 8692067
refGene_gn=LOC102899166;refGene_tx=XM_006941009.4;refGene_gn=LOC102899166;refGene_tx=XM_011289670.3;refGene_gn=LOC102899166;refGene_tx=XM_011289671.3;refGene_gn=LOC102899166;refGene_tx=XM_011289672.3;refGene_gn=LOC102899166;refGene_tx=XM_011289673.3;refGene_gn=LOC102899166;refGene_tx=XM_011289674.3;refGene_gn=LOC102899166;refGene_tx=XM_019819222.2;refGene_gn=LOC102899166;refGene_tx=XM_019819223.2;refGene_gn=LOC102899166;refGene_tx=XM_019819225.2
E2 8693063 8693270
refGene_gn=LOC102899166;refGene_tx=XM_006941009.4;refGene_gn=LOC102899166;refGene_tx=XM_011289670.3;refGene_gn=LOC102899166;refGene_tx=XM_011289671.3;refGene_gn=LOC102899166;refGene_tx=XM_011289672.3;refGene_gn=LOC102899166;refGene_tx=XM_011289673.3;refGene_gn=LOC102899166;refGene_tx=XM_011289674.3;refGene_gn=LOC102899166;refGene_tx=XM_019819222.2;refGene_gn=LOC102899166;refGene_tx=XM_019819223.2;refGene_gn=LOC102899166;refGene_tx=XM_019819225.2
E2 8693471 8693710
refGene_gn=LOC102899166;refGene_tx=XM_006941009.4;refGene_gn=LOC102899166;refGene_tx=XM_011289670.3;refGene_gn=LOC102899166;refGene_tx=XM_011289671.3;refGene_gn=LOC102899166;refGene_tx=XM_011289672.3;refGene_gn=LOC102899166;refGene_tx=XM_011289673.3;refGene_gn=LOC102899166;refGene_tx=XM_011289674.3;refGene_gn=LOC102899166;refGene_tx=XM_019819222.2;refGene_gn=LOC102899166;refGene_tx=XM_019819223.2;refGene_gn=LOC102899166;refGene_tx=XM_019819225.2
E2 8697286 8697473
refGene_gn=LOC102899166;refGene_tx=XM_006941009.4;refGene_gn=LOC102899

166;refGene_tx=XM_011289670.3;refGene_gn=LOC102899166;refGene_tx=XM_01
1289671.3;refGene_gn=LOC102899166;refGene_tx=XM_011289672.3;refGene_gn
=LOC102899166;refGene_tx=XM_011289673.3;refGene_gn=LOC102899166;refGen
e_tx=XM_011289674.3;refGene_gn=LOC102899166;refGene_tx=XM_019819222.2;
refGene_gn=LOC102899166;refGene_tx=XM_019819223.2;refGene_gn=LOC102899
166;refGene_tx=XM_019819225.2
E2 8697892 8698094
refGene_gn=LOC102899166;refGene_tx=XM_006941009.4;refGene_gn=LOC102899
166;refGene_tx=XM_011289670.3;refGene_gn=LOC102899166;refGene_tx=XM_01
1289671.3;refGene_gn=LOC102899166;refGene_tx=XM_011289672.3;refGene_gn
=LOC102899166;refGene_tx=XM_011289673.3;refGene_gn=LOC102899166;refGen
e_tx=XM_011289674.3;refGene_gn=LOC102899166;refGene_tx=XM_019819222.2;
refGene_gn=LOC102899166;refGene_tx=XM_019819223.2;refGene_gn=LOC102899
166;refGene_tx=XM_019819225.2
E2 8698235 8698395
refGene_gn=LOC102899166;refGene_tx=XM_006941009.4;refGene_gn=LOC102899
166;refGene_tx=XM_011289670.3;refGene_gn=LOC102899166;refGene_tx=XM_01
1289671.3;refGene_gn=LOC102899166;refGene_tx=XM_011289672.3;refGene_gn
=LOC102899166;refGene_tx=XM_011289673.3;refGene_gn=LOC102899166;refGen
e_tx=XM_011289674.3;refGene_gn=LOC102899166;refGene_tx=XM_019819222.2;
refGene_gn=LOC102899166;refGene_tx=XM_019819223.2;refGene_gn=LOC102899
166;refGene_tx=XM_019819225.2
E2 8698708 8698874
refGene_gn=LOC102899166;refGene_tx=XM_006941009.4;refGene_gn=LOC102899
166;refGene_tx=XM_011289670.3;refGene_gn=LOC102899166;refGene_tx=XM_01
1289671.3;refGene_gn=LOC102899166;refGene_tx=XM_011289672.3;refGene_gn
=LOC102899166;refGene_tx=XM_011289673.3;refGene_gn=LOC102899166;refGen
e_tx=XM_011289674.3;refGene_gn=LOC102899166;refGene_tx=XM_019819222.2;
refGene_gn=LOC102899166;refGene_tx=XM_019819223.2;refGene_gn=LOC102899
166;refGene_tx=XM_019819225.2
E2 8699087 8699570
refGene_gn=LOC102899166;refGene_tx=XM_006941009.4;refGene_gn=LOC102899
166;refGene_tx=XM_011289670.3;refGene_gn=LOC102899166;refGene_tx=XM_01
1289671.3;refGene_gn=LOC102899166;refGene_tx=XM_011289672.3;refGene_gn
=LOC102899166;refGene_tx=XM_011289674.3;refGene_gn=LOC102899166;refGen
e_tx=XM_019819222.2;refGene_gn=LOC102899166;refGene_tx=XM_019819223.2;
refGene_gn=LOC102899166;refGene_tx=XM_019819225.2
E2 8700966 8701110
refGene_gn=LOC102899166;refGene_tx=XM_011289673.3
E2 8705725 8705730
refGene_gn=TRPM4;refGene_tx=XM_023245982.1;refGene_gn=TRPM4;refGene_tx
=XM_023245983.1
E2 8705979 8706085
refGene_gn=TRPM4;refGene_tx=XM_023245982.1;refGene_gn=TRPM4;refGene_tx
=XM_023245983.1
E2 8706170 8706243
refGene_gn=TRPM4;refGene_tx=XM_023245982.1;refGene_gn=TRPM4;refGene_tx
=XM_023245983.1
E2 8706478 8706611
refGene_gn=TRPM4;refGene_tx=XM_023245982.1;refGene_gn=TRPM4;refGene_tx

=XM_023245983.1
E2 8706830 8707024
refGene_gn=TRPM4;refGene_tx=XM_023245982.1;refGene_gn=TRPM4;refGene_tx
=XM_023245983.1
E2 8710620 8710798
refGene_gn=TRPM4;refGene_tx=XM_023245982.1;refGene_gn=TRPM4;refGene_tx
=XM_023245983.1
E2 8711057 8711232
refGene_gn=TRPM4;refGene_tx=XM_023245982.1;refGene_gn=TRPM4;refGene_tx
=XM_023245983.1
E2 8711387 8711520
refGene_gn=TRPM4;refGene_tx=XM_023245982.1;refGene_gn=TRPM4;refGene_tx
=XM_023245983.1
E2 8717279 8717711
refGene_gn=TRPM4;refGene_tx=XM_023245982.1;refGene_gn=TRPM4;refGene_tx
=XM_023245983.1
E2 8722308 8722386
refGene_gn=TRPM4;refGene_tx=XM_023245982.1;refGene_gn=TRPM4;refGene_tx
=XM_023245983.1
E2 8722779 8722892
refGene_gn=TRPM4;refGene_tx=XM_023245982.1;refGene_gn=TRPM4;refGene_tx
=XM_023245983.1
E2 8724186 8724332
refGene_gn=TRPM4;refGene_tx=XM_023245982.1;refGene_gn=TRPM4;refGene_tx
=XM_023245983.1
E2 8724506 8724636
refGene_gn=TRPM4;refGene_tx=XM_023245982.1;refGene_gn=TRPM4;refGene_tx
=XM_023245983.1
E2 8729241 8729376
refGene_gn=TRPM4;refGene_tx=XM_023245982.1;refGene_gn=TRPM4;refGene_tx
=XM_023245983.1
E2 8729531 8729873
refGene_gn=TRPM4;refGene_tx=XM_023245982.1;refGene_gn=TRPM4;refGene_tx
=XM_023245983.1
E2 8730190 8730303
refGene_gn=TRPM4;refGene_tx=XM_023245982.1;refGene_gn=TRPM4;refGene_tx
=XM_023245983.1
E2 8734562 8734662
refGene_gn=TRPM4;refGene_tx=XM_023245982.1;refGene_gn=TRPM4;refGene_tx
=XM_023245983.1
E2 8734952 8735144
refGene_gn=TRPM4;refGene_tx=XM_023245982.1;refGene_gn=TRPM4;refGene_tx
=XM_023245983.1
E2 8735496 8735558
refGene_gn=TRPM4;refGene_tx=XM_023245982.1;refGene_gn=TRPM4;refGene_tx
=XM_023245983.1
E2 8738717 8738901
refGene_gn=TRPM4;refGene_tx=XM_023245982.1;refGene_gn=TRPM4;refGene_tx
=XM_023245983.1
E2 8739066 8739230

refGene_gn=TRPM4;refGene_tx=XM_023245982.1;refGene_gn=TRPM4;refGene_tx=XM_023245983.1
E2 8739383 8739564 refGene_gn=TRPM4;refGene_tx=XM_023245982.1
E2 8739911 8740086 refGene_gn=TRPM4;refGene_tx=XM_023245982.1
E2 8744453 8744521
refGene_gn=TRPM4;refGene_tx=XM_023245982.1;refGene_gn=TRPM4;refGene_tx=XM_023245983.1
E2 8744817 8744841 refGene_gn=TRPM4;refGene_tx=XM_023245982.1
E2 8745959 8746018 refGene_gn=TRPM4;refGene_tx=XM_023245983.1
E2 8746406 8748765 refGene_gn=HRC;refGene_tx=XM_023245990.1
E2 8748840 8748911 refGene_gn=HRC;refGene_tx=XM_023245990.1
E2 8749477 8749546 refGene_gn=HRC;refGene_tx=XM_023245990.1
E2 8749799 8749854 refGene_gn=HRC;refGene_tx=XM_023245990.1
E2 8750327 8750364 refGene_gn=HRC;refGene_tx=XM_023245990.1
E2 8750553 8750590 refGene_gn=HRC;refGene_tx=XM_023245990.1
E2 8751756 8751815
refGene_gn=PPFIA3;refGene_tx=XM_023245985.1;refGene_gn=PPFIA3;refGene_tx=XM_023245986.1;refGene_gn=PPFIA3;refGene_tx=XM_023245987.1;refGene_gn=PPFIA3;refGene_tx=XM_023245988.1;refGene_gn=PPFIA3;refGene_tx=XM_023245989.1
E2 8752292 8752464
refGene_gn=PPFIA3;refGene_tx=XM_023245985.1;refGene_gn=PPFIA3;refGene_tx=XM_023245986.1;refGene_gn=PPFIA3;refGene_tx=XM_023245987.1;refGene_gn=PPFIA3;refGene_tx=XM_023245988.1;refGene_gn=PPFIA3;refGene_tx=XM_023245989.1
E2 8752684 8752753
refGene_gn=PPFIA3;refGene_tx=XM_023245985.1;refGene_gn=PPFIA3;refGene_tx=XM_023245986.1;refGene_gn=PPFIA3;refGene_tx=XM_023245987.1;refGene_gn=PPFIA3;refGene_tx=XM_023245988.1;refGene_gn=PPFIA3;refGene_tx=XM_023245989.1
E2 8752845 8753021
refGene_gn=PPFIA3;refGene_tx=XM_023245985.1;refGene_gn=PPFIA3;refGene_tx=XM_023245986.1;refGene_gn=PPFIA3;refGene_tx=XM_023245987.1;refGene_gn=PPFIA3;refGene_tx=XM_023245988.1;refGene_gn=PPFIA3;refGene_tx=XM_023245989.1
E2 8753229 8753327
refGene_gn=PPFIA3;refGene_tx=XM_023245985.1;refGene_gn=PPFIA3;refGene_tx=XM_023245986.1;refGene_gn=PPFIA3;refGene_tx=XM_023245987.1;refGene_gn=PPFIA3;refGene_tx=XM_023245988.1;refGene_gn=PPFIA3;refGene_tx=XM_023245989.1
E2 8753740 8753916
refGene_gn=PPFIA3;refGene_tx=XM_023245985.1;refGene_gn=PPFIA3;refGene_tx=XM_023245986.1;refGene_gn=PPFIA3;refGene_tx=XM_023245987.1;refGene_gn=PPFIA3;refGene_tx=XM_023245988.1;refGene_gn=PPFIA3;refGene_tx=XM_023245989.1
E2 8754956 8754983
refGene_gn=PPFIA3;refGene_tx=XM_023245985.1;refGene_gn=PPFIA3;refGene_tx=XM_023245986.1;refGene_gn=PPFIA3;refGene_tx=XM_023245989.1
E2 8755150 8755213
refGene_gn=PPFIA3;refGene_tx=XM_023245985.1;refGene_gn=PPFIA3;refGene_

tx=XM_023245986.1;refGene_gn=PPFIA3;refGene_tx=XM_023245987.1;refGene_gn=PPFIA3;refGene_tx=XM_023245988.1;refGene_gn=PPFIA3;refGene_tx=XM_023245989.1
E2 8756884 8757085
refGene_gn=PPFIA3;refGene_tx=XM_023245985.1;refGene_gn=PPFIA3;refGene_tx=XM_023245986.1;refGene_gn=PPFIA3;refGene_tx=XM_023245987.1;refGene_gn=PPFIA3;refGene_tx=XM_023245988.1;refGene_gn=PPFIA3;refGene_tx=XM_023245989.1
E2 8757806 8757888
refGene_gn=PPFIA3;refGene_tx=XM_023245985.1;refGene_gn=PPFIA3;refGene_tx=XM_023245986.1;refGene_gn=PPFIA3;refGene_tx=XM_023245987.1;refGene_gn=PPFIA3;refGene_tx=XM_023245988.1;refGene_gn=PPFIA3;refGene_tx=XM_023245989.1
E2 8758342 8758436
refGene_gn=PPFIA3;refGene_tx=XM_023245985.1;refGene_gn=PPFIA3;refGene_tx=XM_023245986.1;refGene_gn=PPFIA3;refGene_tx=XM_023245987.1;refGene_gn=PPFIA3;refGene_tx=XM_023245988.1;refGene_gn=PPFIA3;refGene_tx=XM_023245989.1
E2 8759478 8759606
refGene_gn=PPFIA3;refGene_tx=XM_023245985.1;refGene_gn=PPFIA3;refGene_tx=XM_023245986.1;refGene_gn=PPFIA3;refGene_tx=XM_023245987.1;refGene_gn=PPFIA3;refGene_tx=XM_023245988.1;refGene_gn=PPFIA3;refGene_tx=XM_023245989.1
E2 8759736 8759900
refGene_gn=PPFIA3;refGene_tx=XM_023245985.1;refGene_gn=PPFIA3;refGene_tx=XM_023245986.1;refGene_gn=PPFIA3;refGene_tx=XM_023245987.1;refGene_gn=PPFIA3;refGene_tx=XM_023245988.1;refGene_gn=PPFIA3;refGene_tx=XM_023245989.1
E2 8760530 8760753
refGene_gn=PPFIA3;refGene_tx=XM_023245985.1;refGene_gn=PPFIA3;refGene_tx=XM_023245986.1;refGene_gn=PPFIA3;refGene_tx=XM_023245987.1;refGene_gn=PPFIA3;refGene_tx=XM_023245988.1;refGene_gn=PPFIA3;refGene_tx=XM_023245989.1
E2 8761875 8762063
refGene_gn=PPFIA3;refGene_tx=XM_023245985.1;refGene_gn=PPFIA3;refGene_tx=XM_023245986.1;refGene_gn=PPFIA3;refGene_tx=XM_023245987.1;refGene_gn=PPFIA3;refGene_tx=XM_023245988.1;refGene_gn=PPFIA3;refGene_tx=XM_023245989.1
E2 8762573 8762718
refGene_gn=PPFIA3;refGene_tx=XM_023245985.1;refGene_gn=PPFIA3;refGene_tx=XM_023245986.1;refGene_gn=PPFIA3;refGene_tx=XM_023245987.1;refGene_gn=PPFIA3;refGene_tx=XM_023245988.1;refGene_gn=PPFIA3;refGene_tx=XM_023245989.1
E2 8762991 8763071
refGene_gn=PPFIA3;refGene_tx=XM_023245985.1;refGene_gn=PPFIA3;refGene_tx=XM_023245986.1;refGene_gn=PPFIA3;refGene_tx=XM_023245987.1;refGene_gn=PPFIA3;refGene_tx=XM_023245988.1;refGene_gn=PPFIA3;refGene_tx=XM_023245989.1
E2 8763218 8763281
refGene_gn=PPFIA3;refGene_tx=XM_023245985.1;refGene_gn=PPFIA3;refGene_

tx=XM_023245986.1;refGene_gn=PPFIA3;refGene_tx=XM_023245987.1;refGene_
gn=PPFIA3;refGene_tx=XM_023245988.1;refGene_gn=PPFIA3;refGene_tx=XM_02
3245989.1
E2 8763746 8763878
refGene_gn=PPFIA3;refGene_tx=XM_023245985.1;refGene_gn=PPFIA3;refGene_
tx=XM_023245986.1;refGene_gn=PPFIA3;refGene_tx=XM_023245987.1;refGene_
gn=PPFIA3;refGene_tx=XM_023245988.1;refGene_gn=PPFIA3;refGene_tx=XM_02
3245989.1
E2 8764051 8764135
refGene_gn=PPFIA3;refGene_tx=XM_023245985.1;refGene_gn=PPFIA3;refGene_
tx=XM_023245986.1;refGene_gn=PPFIA3;refGene_tx=XM_023245987.1;refGene_
gn=PPFIA3;refGene_tx=XM_023245988.1;refGene_gn=PPFIA3;refGene_tx=XM_02
3245989.1
E2 8764527 8764662
refGene_gn=PPFIA3;refGene_tx=XM_023245985.1;refGene_gn=PPFIA3;refGene_
tx=XM_023245986.1;refGene_gn=PPFIA3;refGene_tx=XM_023245987.1;refGene_
gn=PPFIA3;refGene_tx=XM_023245988.1;refGene_gn=PPFIA3;refGene_tx=XM_02
3245989.1
E2 8764761 8764908
refGene_gn=PPFIA3;refGene_tx=XM_023245985.1;refGene_gn=PPFIA3;refGene_
tx=XM_023245986.1;refGene_gn=PPFIA3;refGene_tx=XM_023245987.1;refGene_
gn=PPFIA3;refGene_tx=XM_023245988.1;refGene_gn=PPFIA3;refGene_tx=XM_02
3245989.1
E2 8766301 8766523
refGene_gn=PPFIA3;refGene_tx=XM_023245985.1;refGene_gn=PPFIA3;refGene_
tx=XM_023245986.1;refGene_gn=PPFIA3;refGene_tx=XM_023245987.1;refGene_
gn=PPFIA3;refGene_tx=XM_023245988.1;refGene_gn=PPFIA3;refGene_tx=XM_02
3245989.1
E2 8766838 8766913
refGene_gn=PPFIA3;refGene_tx=XM_023245985.1;refGene_gn=PPFIA3;refGene_
tx=XM_023245986.1;refGene_gn=PPFIA3;refGene_tx=XM_023245987.1;refGene_
gn=PPFIA3;refGene_tx=XM_023245988.1;refGene_gn=PPFIA3;refGene_tx=XM_02
3245989.1
E2 8767225 8767300
refGene_gn=PPFIA3;refGene_tx=XM_023245985.1;refGene_gn=PPFIA3;refGene_
tx=XM_023245986.1;refGene_gn=PPFIA3;refGene_tx=XM_023245987.1;refGene_
gn=PPFIA3;refGene_tx=XM_023245988.1;refGene_gn=PPFIA3;refGene_tx=XM_02
3245989.1
E2 8767630 8767795
refGene_gn=PPFIA3;refGene_tx=XM_023245985.1;refGene_gn=PPFIA3;refGene_
tx=XM_023245986.1;refGene_gn=PPFIA3;refGene_tx=XM_023245987.1;refGene_
gn=PPFIA3;refGene_tx=XM_023245988.1;refGene_gn=PPFIA3;refGene_tx=XM_02
3245989.1
E2 8768168 8768270
refGene_gn=PPFIA3;refGene_tx=XM_023245985.1;refGene_gn=PPFIA3;refGene_
tx=XM_023245986.1;refGene_gn=PPFIA3;refGene_tx=XM_023245987.1;refGene_
gn=PPFIA3;refGene_tx=XM_023245988.1;refGene_gn=PPFIA3;refGene_tx=XM_02
3245989.1
E2 8768535 8768775
refGene_gn=PPFIA3;refGene_tx=XM_023245985.1;refGene_gn=PPFIA3;refGene_

tx=XM_023245986.1;refGene_gn=PPFIA3;refGene_tx=XM_023245987.1;refGene_gn=PPFIA3;refGene_tx=XM_023245988.1;refGene_gn=PPFIA3;refGene_tx=XM_023245989.1
E2 8774225 8774247 refGene_gn=LIN7B;refGene_tx=XM_003997519.4
E2 8774589 8774753 refGene_gn=LIN7B;refGene_tx=XM_003997519.4
E2 8775747 8775957 refGene_gn=LIN7B;refGene_tx=XM_003997519.4
E2 8776584 8776656 refGene_gn=LIN7B;refGene_tx=XM_003997519.4
E2 8776931 8777050 refGene_gn=LIN7B;refGene_tx=XM_003997519.4
E2 8777461 8777498 refGene_gn=LIN7B;refGene_tx=XM_003997519.4
E2 8783105 8783313 refGene_gn=SNRNP70;refGene_tx=XM_019819238.2
E2 8783314 8783402 refGene_gn=SNRNP70;refGene_tx=XM_019819238.2
E2 8783404 8783757 refGene_gn=SNRNP70;refGene_tx=XM_019819238.2
E2 8783839 8783927 refGene_gn=SNRNP70;refGene_tx=XM_019819238.2
E2 8787686 8787788 refGene_gn=SNRNP70;refGene_tx=XM_019819238.2
E2 8791036 8791118 refGene_gn=SNRNP70;refGene_tx=XM_019819238.2
E2 8791779 8791842 refGene_gn=SNRNP70;refGene_tx=XM_019819238.2
E2 8792010 8792075 refGene_gn=SNRNP70;refGene_tx=XM_019819238.2
E2 8796593 8796648 refGene_gn=SNRNP70;refGene_tx=XM_019819238.2
E2 8796761 8796824 refGene_gn=SNRNP70;refGene_tx=XM_019819238.2
E2 8798601 8798748 refGene_gn=SNRNP70;refGene_tx=XM_019819238.2
E2 8808418 8809240 refGene_gn=KCNA7;refGene_tx=XM_023245393.1
E2 8810193 8811009 refGene_gn=KCNA7;refGene_tx=XM_023245393.1
E2 8817210 8817843
refGene_gn=NTF4;refGene_tx=XM_006941016.4;refGene_gn=NTF4;refGene_tx=XM_023245586.1
E2 8819815 8819971
refGene_gn=LOC109494735;refGene_tx=XM_019819243.2
E2 8820234 8820435
refGene_gn=LOC109494735;refGene_tx=XM_019819243.2
E2 8820552 8820714
refGene_gn=LOC109494735;refGene_tx=XM_019819243.2
E2 8821444 8821572
refGene_gn=LOC109494735;refGene_tx=XM_019819243.2
E2 8822091 8822431
refGene_gn=LOC109494735;refGene_tx=XM_019819243.2
E2 8822748 8822856 refGene_gn=LHB;refGene_tx=XM_019818433.2
E2 8823384 8823399 refGene_gn=LHB;refGene_tx=NM_001009277.1
E2 8823757 8823931
refGene_gn=LHB;refGene_tx=NM_001009277.1;refGene_gn=LHB;refGene_tx=XM_019818433.2
E2 8824225 8824468
refGene_gn=LHB;refGene_tx=NM_001009277.1;refGene_gn=LHB;refGene_tx=XM_019818433.2
E2 8824743 8824769 refGene_gn=RUVBL2;refGene_tx=XM_023245201.1
E2 8824874 8824989 refGene_gn=RUVBL2;refGene_tx=XM_023245201.1
E2 8825397 8825527 refGene_gn=RUVBL2;refGene_tx=XM_023245201.1
E2 8825901 8826021 refGene_gn=RUVBL2;refGene_tx=XM_023245201.1
E2 8827798 8827917 refGene_gn=RUVBL2;refGene_tx=XM_023245201.1
E2 8828008 8828103 refGene_gn=RUVBL2;refGene_tx=XM_023245201.1
E2 8828330 8828454 refGene_gn=RUVBL2;refGene_tx=XM_023245201.1

E2 8828818 8828912 refGene_gn=RUVBL2;refGene_tx=XM_023245201.1
E2 8829000 8829107 refGene_gn=RUVBL2;refGene_tx=XM_023245201.1
E2 8832466 8832533 refGene_gn=RUVBL2;refGene_tx=XM_023245201.1
E2 8832693 8832823 refGene_gn=RUVBL2;refGene_tx=XM_023245201.1
E2 8834051 8834193 refGene_gn=RUVBL2;refGene_tx=XM_023245201.1
E2 8834716 8834772 refGene_gn=RUVBL2;refGene_tx=XM_023245201.1
E2 8836765 8836820 refGene_gn=RUVBL2;refGene_tx=XM_023245201.1
E2 8840551 8840563 refGene_gn=RUVBL2;refGene_tx=XM_023245201.1
E2 8841355 8841473 refGene_gn=GYS1;refGene_tx=XM_003997522.5
E2 8842965 8843147 refGene_gn=GYS1;refGene_tx=XM_003997522.5
E2 8845183 8845375 refGene_gn=GYS1;refGene_tx=XM_003997522.5
E2 8846280 8846466 refGene_gn=GYS1;refGene_tx=XM_003997522.5
E2 8846719 8846864 refGene_gn=GYS1;refGene_tx=XM_003997522.5
E2 8847917 8848035 refGene_gn=GYS1;refGene_tx=XM_003997522.5
E2 8848353 8848474 refGene_gn=GYS1;refGene_tx=XM_003997522.5
E2 8848759 8848866 refGene_gn=GYS1;refGene_tx=XM_003997522.5
E2 8850172 8850232 refGene_gn=GYS1;refGene_tx=XM_003997522.5
E2 8850320 8850399 refGene_gn=GYS1;refGene_tx=XM_003997522.5
E2 8852682 8852796 refGene_gn=GYS1;refGene_tx=XM_003997522.5
E2 8853059 8853186 refGene_gn=GYS1;refGene_tx=XM_003997522.5
E2 8855002 8855098 refGene_gn=GYS1;refGene_tx=XM_003997522.5
E2 8855264 8855428 refGene_gn=GYS1;refGene_tx=XM_003997522.5
E2 8855844 8855925 refGene_gn=GYS1;refGene_tx=XM_003997522.5
E2 8856080 8856404 refGene_gn=GYS1;refGene_tx=XM_003997522.5
E2 8858743 8858896 refGene_gn=FTL;refGene_tx=NM_001048150.1
E2 8859012 8859138 refGene_gn=FTL;refGene_tx=NM_001048150.1
E2 8859409 8859556 refGene_gn=FTL;refGene_tx=NM_001048150.1
E2 8859726 8859828 refGene_gn=FTL;refGene_tx=NM_001048150.1
E2 8862820 8862925
refGene_gn=BAX;refGene_tx=NM_001009282.2;refGene_gn=BAX;refGene_tx=XM_
011289649.3
E2 8863372 8863477
refGene_gn=BAX;refGene_tx=NM_001009282.2;refGene_gn=BAX;refGene_tx=XM_
011289649.3
E2 8864949 8865085
refGene_gn=BAX;refGene_tx=NM_001009282.2;refGene_gn=BAX;refGene_tx=XM_
011289649.3
E2 8865484 8865631
refGene_gn=BAX;refGene_tx=NM_001009282.2;refGene_gn=BAX;refGene_tx=XM_
011289649.3
E2 8865721 8865773 refGene_gn=BAX;refGene_tx=NM_001009282.2
E2 8866350 8866384 refGene_gn=BAX;refGene_tx=NM_001009282.2
E2 8870432 8870542 refGene_gn=DHDH;refGene_tx=XM_003997525.4
E2 8870791 8870945 refGene_gn=DHDH;refGene_tx=XM_003997525.4
E2 8871770 8871895 refGene_gn=DHDH;refGene_tx=XM_003997525.4
E2 8874970 8875223 refGene_gn=DHDH;refGene_tx=XM_003997525.4
E2 8877295 8877459 refGene_gn=DHDH;refGene_tx=XM_003997525.4
E2 8877975 8878087 refGene_gn=DHDH;refGene_tx=XM_003997525.4
E2 8878481 8878571 refGene_gn=DHDH;refGene_tx=XM_003997525.4
E2 8887571 8887684

refGene_gn=NUCB1;refGene_tx=XM_003997526.5;refGene_gn=NUCB1;refGene_tx=XM_006941017.4;refGene_gn=NUCB1;refGene_tx=XM_011289663.3
E2 8887788 8887885
refGene_gn=NUCB1;refGene_tx=XM_003997526.5;refGene_gn=NUCB1;refGene_tx=XM_006941017.4;refGene_gn=NUCB1;refGene_tx=XM_011289663.3
E2 8888353 8888524
refGene_gn=NUCB1;refGene_tx=XM_003997526.5;refGene_gn=NUCB1;refGene_tx=XM_006941017.4;refGene_gn=NUCB1;refGene_tx=XM_011289663.3
E2 8888863 8888956
refGene_gn=NUCB1;refGene_tx=XM_003997526.5;refGene_gn=NUCB1;refGene_tx=XM_006941017.4;refGene_gn=NUCB1;refGene_tx=XM_011289663.3
E2 8889041 8889134
refGene_gn=NUCB1;refGene_tx=XM_003997526.5;refGene_gn=NUCB1;refGene_tx=XM_006941017.4;refGene_gn=NUCB1;refGene_tx=XM_011289663.3
E2 8889394 8889453
refGene_gn=NUCB1;refGene_tx=XM_003997526.5;refGene_gn=NUCB1;refGene_tx=XM_006941017.4;refGene_gn=NUCB1;refGene_tx=XM_011289663.3
E2 8892064 8892155
refGene_gn=NUCB1;refGene_tx=XM_003997526.5;refGene_gn=NUCB1;refGene_tx=XM_006941017.4;refGene_gn=NUCB1;refGene_tx=XM_011289663.3
E2 8892437 8892623
refGene_gn=NUCB1;refGene_tx=XM_003997526.5;refGene_gn=NUCB1;refGene_tx=XM_006941017.4;refGene_gn=NUCB1;refGene_tx=XM_011289663.3
E2 8896173 8896277
refGene_gn=NUCB1;refGene_tx=XM_003997526.5;refGene_gn=NUCB1;refGene_tx=XM_006941017.4;refGene_gn=NUCB1;refGene_tx=XM_011289663.3
E2 8899767 8899900
refGene_gn=NUCB1;refGene_tx=XM_003997526.5;refGene_gn=NUCB1;refGene_tx=XM_006941017.4;refGene_gn=NUCB1;refGene_tx=XM_011289663.3
E2 8902327 8902435
refGene_gn=NUCB1;refGene_tx=XM_003997526.5;refGene_gn=NUCB1;refGene_tx=XM_006941017.4;refGene_gn=NUCB1;refGene_tx=XM_011289663.3
E2 8904170 8904317
refGene_gn=NUCB1;refGene_tx=XM_003997526.5;refGene_gn=NUCB1;refGene_tx=XM_006941017.4;refGene_gn=NUCB1;refGene_tx=XM_011289663.3
E2 8905894 8905926
refGene_gn=TULP2;refGene_tx=XM_006941019.4;refGene_gn=TULP2;refGene_tx=XM_006941020.4;refGene_gn=TULP2;refGene_tx=XM_006941021.4;refGene_gn=TULP2;refGene_tx=XM_006941022.4;refGene_gn=TULP2;refGene_tx=XM_019819253.2
E2 8906320 8906372
refGene_gn=TULP2;refGene_tx=XM_006941019.4;refGene_gn=TULP2;refGene_tx=XM_006941020.4;refGene_gn=TULP2;refGene_tx=XM_006941021.4;refGene_gn=TULP2;refGene_tx=XM_006941022.4;refGene_gn=TULP2;refGene_tx=XM_019819253.2
E2 8906779 8906906
refGene_gn=TULP2;refGene_tx=XM_006941019.4;refGene_gn=TULP2;refGene_tx=XM_006941020.4;refGene_gn=TULP2;refGene_tx=XM_006941021.4;refGene_gn=TULP2;refGene_tx=XM_006941022.4;refGene_gn=TULP2;refGene_tx=XM_019819253.2

E2 8907497 8907635
refGene_gn=TULP2;refGene_tx=XM_006941019.4;refGene_gn=TULP2;refGene_tx=XM_006941020.4;refGene_gn=TULP2;refGene_tx=XM_006941021.4;refGene_gn=TULP2;refGene_tx=XM_006941022.4;refGene_gn=TULP2;refGene_tx=XM_019819253.2
E2 8907885 8908002
refGene_gn=TULP2;refGene_tx=XM_006941019.4;refGene_gn=TULP2;refGene_tx=XM_006941020.4;refGene_gn=TULP2;refGene_tx=XM_019819253.2
E2 8908963 8909088
refGene_gn=TULP2;refGene_tx=XM_006941019.4;refGene_gn=TULP2;refGene_tx=XM_006941020.4;refGene_gn=TULP2;refGene_tx=XM_006941021.4;refGene_gn=TULP2;refGene_tx=XM_006941022.4;refGene_gn=TULP2;refGene_tx=XM_019819253.2
E2 8909774 8910083
refGene_gn=TULP2;refGene_tx=XM_006941019.4;refGene_gn=TULP2;refGene_tx=XM_006941020.4;refGene_gn=TULP2;refGene_tx=XM_006941021.4;refGene_gn=TULP2;refGene_tx=XM_006941022.4;refGene_gn=TULP2;refGene_tx=XM_019819253.2
E2 8911043 8911156
refGene_gn=TULP2;refGene_tx=XM_006941019.4;refGene_gn=TULP2;refGene_tx=XM_006941020.4;refGene_gn=TULP2;refGene_tx=XM_006941021.4;refGene_gn=TULP2;refGene_tx=XM_006941022.4;refGene_gn=TULP2;refGene_tx=XM_019819253.2
E2 8912127 8912242
refGene_gn=TULP2;refGene_tx=XM_006941019.4;refGene_gn=TULP2;refGene_tx=XM_006941020.4;refGene_gn=TULP2;refGene_tx=XM_006941021.4;refGene_gn=TULP2;refGene_tx=XM_006941022.4;refGene_gn=TULP2;refGene_tx=XM_019819253.2
E2 8912326 8912425
refGene_gn=TULP2;refGene_tx=XM_006941019.4;refGene_gn=TULP2;refGene_tx=XM_006941020.4;refGene_gn=TULP2;refGene_tx=XM_006941021.4;refGene_gn=TULP2;refGene_tx=XM_006941022.4;refGene_gn=TULP2;refGene_tx=XM_019819253.2
E2 8912665 8912837
refGene_gn=TULP2;refGene_tx=XM_006941019.4;refGene_gn=TULP2;refGene_tx=XM_006941020.4;refGene_gn=TULP2;refGene_tx=XM_006941021.4;refGene_gn=TULP2;refGene_tx=XM_006941022.4;refGene_gn=TULP2;refGene_tx=XM_019819253.2
E2 8913129 8913263
refGene_gn=TULP2;refGene_tx=XM_006941019.4;refGene_gn=TULP2;refGene_tx=XM_006941020.4;refGene_gn=TULP2;refGene_tx=XM_006941021.4;refGene_gn=TULP2;refGene_tx=XM_006941022.4;refGene_gn=TULP2;refGene_tx=XM_019819253.2
E2 8916032 8916380
refGene_gn=PPP1R15A;refGene_tx=XM_003997602.5;refGene_gn=PPP1R15A;refGene_tx=XM_011289661.3
E2 8916813 8918481
refGene_gn=PPP1R15A;refGene_tx=XM_003997602.5;refGene_gn=PPP1R15A;refGene_tx=XM_011289661.3
E2 8921990 8922074

refGene_gn=PLEKHA4;refGene_tx=XM_011289655.3;refGene_gn=PLEKHA4;refGene_tx=XM_011289656.3;refGene_gn=PLEKHA4;refGene_tx=XM_011289657.3;refGene_gn=PLEKHA4;refGene_tx=XM_011289658.3;refGene_gn=PLEKHA4;refGene_tx=XM_019819245.2;refGene_gn=PLEKHA4;refGene_tx=XM_019819249.2;refGene_gn=PLEKHA4;refGene_tx=XM_019819250.2;refGene_gn=PLEKHA4;refGene_tx=XM_019819251.2
E2 8923163 8923271
refGene_gn=PLEKHA4;refGene_tx=XM_011289655.3;refGene_gn=PLEKHA4;refGene_tx=XM_011289656.3;refGene_gn=PLEKHA4;refGene_tx=XM_011289657.3;refGene_gn=PLEKHA4;refGene_tx=XM_011289658.3;refGene_gn=PLEKHA4;refGene_tx=XM_019819245.2;refGene_gn=PLEKHA4;refGene_tx=XM_019819249.2;refGene_gn=PLEKHA4;refGene_tx=XM_019819250.2;refGene_gn=PLEKHA4;refGene_tx=XM_019819251.2
E2 8925028 8925101
refGene_gn=PLEKHA4;refGene_tx=XM_011289655.3;refGene_gn=PLEKHA4;refGene_tx=XM_011289656.3;refGene_gn=PLEKHA4;refGene_tx=XM_011289657.3;refGene_gn=PLEKHA4;refGene_tx=XM_011289658.3;refGene_gn=PLEKHA4;refGene_tx=XM_019819245.2;refGene_gn=PLEKHA4;refGene_tx=XM_019819249.2;refGene_gn=PLEKHA4;refGene_tx=XM_019819250.2;refGene_gn=PLEKHA4;refGene_tx=XM_019819251.2
E2 8925247 8925348
refGene_gn=PLEKHA4;refGene_tx=XM_011289655.3;refGene_gn=PLEKHA4;refGene_tx=XM_011289656.3;refGene_gn=PLEKHA4;refGene_tx=XM_011289657.3;refGene_gn=PLEKHA4;refGene_tx=XM_011289658.3;refGene_gn=PLEKHA4;refGene_tx=XM_019819245.2;refGene_gn=PLEKHA4;refGene_tx=XM_019819249.2;refGene_gn=PLEKHA4;refGene_tx=XM_019819250.2;refGene_gn=PLEKHA4;refGene_tx=XM_019819251.2
E2 8925501 8925611
refGene_gn=PLEKHA4;refGene_tx=XM_011289655.3;refGene_gn=PLEKHA4;refGene_tx=XM_011289656.3;refGene_gn=PLEKHA4;refGene_tx=XM_011289657.3;refGene_gn=PLEKHA4;refGene_tx=XM_011289658.3;refGene_gn=PLEKHA4;refGene_tx=XM_019819245.2;refGene_gn=PLEKHA4;refGene_tx=XM_019819249.2;refGene_gn=PLEKHA4;refGene_tx=XM_019819250.2;refGene_gn=PLEKHA4;refGene_tx=XM_019819251.2
E2 8926202 8926418
refGene_gn=PLEKHA4;refGene_tx=XM_011289655.3;refGene_gn=PLEKHA4;refGene_tx=XM_011289656.3;refGene_gn=PLEKHA4;refGene_tx=XM_011289657.3;refGene_gn=PLEKHA4;refGene_tx=XM_011289658.3;refGene_gn=PLEKHA4;refGene_tx=XM_019819245.2;refGene_gn=PLEKHA4;refGene_tx=XM_019819249.2;refGene_gn=PLEKHA4;refGene_tx=XM_019819250.2;refGene_gn=PLEKHA4;refGene_tx=XM_019819251.2
E2 8926838 8927130
refGene_gn=PLEKHA4;refGene_tx=XM_011289655.3;refGene_gn=PLEKHA4;refGene_tx=XM_011289656.3;refGene_gn=PLEKHA4;refGene_tx=XM_011289657.3;refGene_gn=PLEKHA4;refGene_tx=XM_011289658.3;refGene_gn=PLEKHA4;refGene_tx=XM_019819245.2;refGene_gn=PLEKHA4;refGene_tx=XM_019819249.2;refGene_gn=PLEKHA4;refGene_tx=XM_019819250.2;refGene_gn=PLEKHA4;refGene_tx=XM_019819251.2
E2 8927753 8927825
refGene_gn=PLEKHA4;refGene_tx=XM_011289655.3;refGene_gn=PLEKHA4;refGen

e_tx=XM_011289656.3;refGene_gn=PLEKHA4;refGene_tx=XM_011289657.3;refGene_gn=PLEKHA4;refGene_tx=XM_011289658.3;refGene_gn=PLEKHA4;refGene_tx=XM_019819245.2;refGene_gn=PLEKHA4;refGene_tx=XM_019819249.2;refGene_gn=PLEKHA4;refGene_tx=XM_019819250.2;refGene_gn=PLEKHA4;refGene_tx=XM_019819251.2
E2 8929322 8929370
refGene_gn=PLEKHA4;refGene_tx=XM_011289655.3;refGene_gn=PLEKHA4;refGene_tx=XM_011289656.3;refGene_gn=PLEKHA4;refGene_tx=XM_019819245.2;refGene_gn=PLEKHA4;refGene_tx=XM_019819249.2
E2 8929520 8929601
refGene_gn=PLEKHA4;refGene_tx=XM_011289655.3;refGene_gn=PLEKHA4;refGene_tx=XM_011289656.3;refGene_gn=PLEKHA4;refGene_tx=XM_019819245.2;refGene_gn=PLEKHA4;refGene_tx=XM_019819249.2
E2 8929757 8929907
refGene_gn=PLEKHA4;refGene_tx=XM_011289655.3;refGene_gn=PLEKHA4;refGene_tx=XM_011289656.3;refGene_gn=PLEKHA4;refGene_tx=XM_019819245.2;refGene_gn=PLEKHA4;refGene_tx=XM_019819249.2
E2 8931397 8931496
refGene_gn=PLEKHA4;refGene_tx=XM_011289655.3;refGene_gn=PLEKHA4;refGene_tx=XM_011289656.3;refGene_gn=PLEKHA4;refGene_tx=XM_011289657.3;refGene_gn=PLEKHA4;refGene_tx=XM_011289658.3;refGene_gn=PLEKHA4;refGene_tx=XM_019819245.2
E2 8933961 8934102
refGene_gn=PLEKHA4;refGene_tx=XM_011289655.3;refGene_gn=PLEKHA4;refGene_tx=XM_011289656.3;refGene_gn=PLEKHA4;refGene_tx=XM_011289657.3;refGene_gn=PLEKHA4;refGene_tx=XM_011289658.3;refGene_gn=PLEKHA4;refGene_tx=XM_019819245.2;refGene_gn=PLEKHA4;refGene_tx=XM_019819249.2;refGene_gn=PLEKHA4;refGene_tx=XM_019819250.2;refGene_gn=PLEKHA4;refGene_tx=XM_019819251.2
E2 8934753 8934853
refGene_gn=PLEKHA4;refGene_tx=XM_011289655.3;refGene_gn=PLEKHA4;refGene_tx=XM_011289656.3;refGene_gn=PLEKHA4;refGene_tx=XM_011289657.3;refGene_gn=PLEKHA4;refGene_tx=XM_011289658.3;refGene_gn=PLEKHA4;refGene_tx=XM_019819245.2;refGene_gn=PLEKHA4;refGene_tx=XM_019819249.2;refGene_gn=PLEKHA4;refGene_tx=XM_019819250.2;refGene_gn=PLEKHA4;refGene_tx=XM_019819251.2
E2 8934925 8935002
refGene_gn=PLEKHA4;refGene_tx=XM_011289655.3;refGene_gn=PLEKHA4;refGene_tx=XM_011289656.3;refGene_gn=PLEKHA4;refGene_tx=XM_011289657.3;refGene_gn=PLEKHA4;refGene_tx=XM_011289658.3;refGene_gn=PLEKHA4;refGene_tx=XM_019819245.2;refGene_gn=PLEKHA4;refGene_tx=XM_019819249.2;refGene_gn=PLEKHA4;refGene_tx=XM_019819250.2;refGene_gn=PLEKHA4;refGene_tx=XM_019819251.2
E2 8942611 8942773
refGene_gn=PLEKHA4;refGene_tx=XM_011289655.3;refGene_gn=PLEKHA4;refGene_tx=XM_011289656.3;refGene_gn=PLEKHA4;refGene_tx=XM_011289657.3;refGene_gn=PLEKHA4;refGene_tx=XM_011289658.3;refGene_gn=PLEKHA4;refGene_tx=XM_019819245.2;refGene_gn=PLEKHA4;refGene_tx=XM_019819249.2;refGene_gn=PLEKHA4;refGene_tx=XM_019819250.2;refGene_gn=PLEKHA4;refGene_tx=XM_019819251.2

E2	8943668	8943727
refGene_gn=PLEKHA4;refGene_tx=XM_011289655.3;refGene_gn=PLEKHA4;refGene_tx=XM_011289656.3;refGene_gn=PLEKHA4;refGene_tx=XM_011289657.3;refGene_gn=PLEKHA4;refGene_tx=XM_011289658.3;refGene_gn=PLEKHA4;refGene_tx=XM_019819245.2;refGene_gn=PLEKHA4;refGene_tx=XM_019819249.2;refGene_gn=PLEKHA4;refGene_tx=XM_019819250.2;refGene_gn=PLEKHA4;refGene_tx=XM_019819251.2
E2	8944098	8944211
refGene_gn=PLEKHA4;refGene_tx=XM_011289655.3;refGene_gn=PLEKHA4;refGene_tx=XM_011289656.3;refGene_gn=PLEKHA4;refGene_tx=XM_011289657.3;refGene_gn=PLEKHA4;refGene_tx=XM_011289658.3;refGene_gn=PLEKHA4;refGene_tx=XM_019819245.2;refGene_gn=PLEKHA4;refGene_tx=XM_019819249.2;refGene_gn=PLEKHA4;refGene_tx=XM_019819250.2;refGene_gn=PLEKHA4;refGene_tx=XM_019819251.2
E2	8944485	8944754
refGene_gn=PLEKHA4;refGene_tx=XM_011289655.3;refGene_gn=PLEKHA4;refGene_tx=XM_011289656.3;refGene_gn=PLEKHA4;refGene_tx=XM_011289657.3;refGene_gn=PLEKHA4;refGene_tx=XM_011289658.3;refGene_gn=PLEKHA4;refGene_tx=XM_019819245.2;refGene_gn=PLEKHA4;refGene_tx=XM_019819249.2;refGene_gn=PLEKHA4;refGene_tx=XM_019819250.2;refGene_gn=PLEKHA4;refGene_tx=XM_019819251.2
E2	8945288	8945376	refGene_gn=HSD17B14;refGene_tx=XM_023246028.1
E2	8945877	8945916	refGene_gn=HSD17B14;refGene_tx=XM_023246028.1
E2	8950547	8950703	refGene_gn=HSD17B14;refGene_tx=XM_023246028.1
E2	8953298	8953390	refGene_gn=HSD17B14;refGene_tx=XM_023246028.1
E2	8962131	8962236	refGene_gn=HSD17B14;refGene_tx=XM_023246028.1
E2	8962474	8962542	refGene_gn=HSD17B14;refGene_tx=XM_023246028.1
E2	8962637	8962734	refGene_gn=HSD17B14;refGene_tx=XM_023246028.1
E2	8962835	8963009	refGene_gn=HSD17B14;refGene_tx=XM_023246028.1
E2	8965267	8965291
refGene_gn=BCAT2;refGene_tx=XM_023246024.1;refGene_gn=BCAT2;refGene_tx=XM_023246025.1
E2	8968894	8968969
refGene_gn=BCAT2;refGene_tx=XM_023246024.1;refGene_gn=BCAT2;refGene_tx=XM_023246025.1
E2	8969335	8969539
refGene_gn=BCAT2;refGene_tx=XM_023246024.1;refGene_gn=BCAT2;refGene_tx=XM_023246025.1;refGene_gn=BCAT2;refGene_tx=XM_023246027.1
E2	8970712	8970826
refGene_gn=BCAT2;refGene_tx=XM_023246024.1;refGene_gn=BCAT2;refGene_tx=XM_023246025.1;refGene_gn=BCAT2;refGene_tx=XM_023246027.1
E2	8970909	8971029
refGene_gn=BCAT2;refGene_tx=XM_023246024.1;refGene_gn=BCAT2;refGene_tx=XM_023246025.1;refGene_gn=BCAT2;refGene_tx=XM_023246027.1
E2	8971175	8971339
refGene_gn=BCAT2;refGene_tx=XM_023246024.1;refGene_gn=BCAT2;refGene_tx=XM_023246025.1;refGene_gn=BCAT2;refGene_tx=XM_023246027.1
E2	8972381	8972524
refGene_gn=BCAT2;refGene_tx=XM_023246024.1;refGene_gn=BCAT2;refGene_tx=XM_023246025.1;refGene_gn=BCAT2;refGene_tx=XM_023246027.1

E2 8972687 8972773
refGene_gn=BCAT2;refGene_tx=XM_023246024.1;refGene_gn=BCAT2;refGene_tx=XM_023246025.1;refGene_gn=BCAT2;refGene_tx=XM_023246027.1
E2 8972990 8973131
refGene_gn=BCAT2;refGene_tx=XM_023246024.1;refGene_gn=BCAT2;refGene_tx=XM_023246025.1;refGene_gn=BCAT2;refGene_tx=XM_023246027.1
E2 8973209 8973284
refGene_gn=BCAT2;refGene_tx=XM_023246024.1;refGene_gn=BCAT2;refGene_tx=XM_023246025.1;refGene_gn=BCAT2;refGene_tx=XM_023246027.1
E2 8974154 8974193
refGene_gn=BCAT2;refGene_tx=XM_023246024.1;refGene_gn=BCAT2;refGene_tx=XM_023246025.1;refGene_gn=BCAT2;refGene_tx=XM_023246027.1
E2 8989414 8989705 refGene_gn=FGF21;refGene_tx=XM_003997528.3
E2 8990307 8990411 refGene_gn=FGF21;refGene_tx=XM_003997528.3
E2 8990532 8990770 refGene_gn=FGF21;refGene_tx=XM_003997528.3
E2 8992738 8992874 refGene_gn=FUT1;refGene_tx=XM_003997529.3
E2 8993437 8994519 refGene_gn=FUT1;refGene_tx=XM_003997529.3
E2 8997893 8998128
refGene_gn=LOC101085336;refGene_tx=XM_003997605.5;refGene_gn=LOC101085336;refGene_tx=XM_006941027.4;refGene_gn=LOC101085336;refGene_tx=XM_006941028.4;refGene_gn=LOC101085336;refGene_tx=XM_006941029.4;refGene_gn=LOC101085336;refGene_tx=XM_019819265.2
E2 8998512 8998587
refGene_gn=LOC101085336;refGene_tx=XM_006941027.4;refGene_gn=LOC101085336;refGene_tx=XM_006941028.4;refGene_gn=LOC101085336;refGene_tx=XM_006941029.4
E2 8998923 8999010
refGene_gn=LOC101085336;refGene_tx=XM_003997605.5;refGene_gn=LOC101085336;refGene_tx=XM_006941027.4;refGene_gn=LOC101085336;refGene_tx=XM_006941028.4;refGene_gn=LOC101085336;refGene_tx=XM_006941029.4;refGene_gn=LOC101085336;refGene_tx=XM_019819265.2
E2 8999235 8999256
refGene_gn=LOC101085336;refGene_tx=XM_003997605.5;refGene_gn=LOC101085336;refGene_tx=XM_006941027.4;refGene_gn=LOC101085336;refGene_tx=XM_006941029.4
E2 8999609 8999690
refGene_gn=LOC101085336;refGene_tx=XM_003997605.5;refGene_gn=LOC101085336;refGene_tx=XM_006941027.4;refGene_gn=LOC101085336;refGene_tx=XM_006941028.4;refGene_gn=LOC101085336;refGene_tx=XM_006941029.4;refGene_gn=LOC101085336;refGene_tx=XM_019819265.2
E2 8999876 8999977
refGene_gn=LOC101085336;refGene_tx=XM_003997605.5;refGene_gn=LOC101085336;refGene_tx=XM_006941027.4;refGene_gn=LOC101085336;refGene_tx=XM_006941028.4;refGene_gn=LOC101085336;refGene_tx=XM_006941029.4;refGene_gn=LOC101085336;refGene_tx=XM_019819265.2
E2 9000237 9000391
refGene_gn=LOC101085336;refGene_tx=XM_003997605.5;refGene_gn=LOC101085336;refGene_tx=XM_006941027.4;refGene_gn=LOC101085336;refGene_tx=XM_006941028.4;refGene_gn=LOC101085336;refGene_tx=XM_006941029.4;refGene_gn=LOC101085336;refGene_tx=XM_019819265.2

E2	9000674	9000849
refGene_gn=LOC101085336;refGene_tx=XM_003997605.5;refGene_gn=LOC101085336;refGene_tx=XM_006941027.4;refGene_gn=LOC101085336;refGene_tx=XM_006941028.4;refGene_gn=LOC101085336;refGene_tx=XM_006941029.4;refGene_gn=LOC101085336;refGene_tx=XM_019819265.2
E2	9001063	9001216
refGene_gn=LOC101085336;refGene_tx=XM_003997605.5;refGene_gn=LOC101085336;refGene_tx=XM_006941027.4;refGene_gn=LOC101085336;refGene_tx=XM_006941028.4;refGene_gn=LOC101085336;refGene_tx=XM_006941029.4;refGene_gn=LOC101085336;refGene_tx=XM_019819265.2
E2	9002340	9002575
refGene_gn=LOC101085591;refGene_tx=XM_019819268.2;refGene_gn=LOC101085591;refGene_tx=XM_023245684.1
E2	9002967	9003042
refGene_gn=LOC101085591;refGene_tx=XM_019819268.2;refGene_gn=LOC101085591;refGene_tx=XM_023245684.1
E2	9003421	9003508
refGene_gn=LOC101085591;refGene_tx=XM_019819268.2;refGene_gn=LOC101085591;refGene_tx=XM_023245684.1
E2	9003779	9003800
refGene_gn=LOC101085591;refGene_tx=XM_019819268.2;refGene_gn=LOC101085591;refGene_tx=XM_023245684.1
E2	9004152	9004233
refGene_gn=LOC101085591;refGene_tx=XM_019819268.2;refGene_gn=LOC101085591;refGene_tx=XM_023245684.1
E2	9004423	9004524
refGene_gn=LOC101085591;refGene_tx=XM_019819268.2;refGene_gn=LOC101085591;refGene_tx=XM_023245684.1
E2	9004775	9004929
refGene_gn=LOC101085591;refGene_tx=XM_019819268.2;refGene_gn=LOC101085591;refGene_tx=XM_023245684.1
E2	9005222	9005397
refGene_gn=LOC101085591;refGene_tx=XM_023245684.1
E2	9005621	9005817
refGene_gn=LOC101085591;refGene_tx=XM_019819268.2;refGene_gn=LOC101085591;refGene_tx=XM_023245684.1
E2	9006999	9007234
refGene_gn=LOC102900416;refGene_tx=XM_006941030.4;refGene_gn=LOC102900416;refGene_tx=XM_023245690.1;refGene_gn=LOC102900416;refGene_tx=XM_023245691.1
E2	9007613	9007688
refGene_gn=LOC102900416;refGene_tx=XM_006941030.4;refGene_gn=LOC102900416;refGene_tx=XM_023245690.1;refGene_gn=LOC102900416;refGene_tx=XM_023245691.1
E2	9008027	9008114
refGene_gn=LOC102900416;refGene_tx=XM_006941030.4;refGene_gn=LOC102900416;refGene_tx=XM_023245690.1
E2	9008374	9008395
refGene_gn=LOC102900416;refGene_tx=XM_006941030.4
E2	9008745	9008826

```
refGene_gn=LOC102900416;refGene_tx=XM_006941030.4;refGene_gn=LOC102900
416;refGene_tx=XM_023245690.1;refGene_gn=LOC102900416;refGene_tx=XM_02
3245691.1
E2	9009011	9009112
refGene_gn=LOC102900416;refGene_tx=XM_006941030.4;refGene_gn=LOC102900
416;refGene_tx=XM_023245690.1;refGene_gn=LOC102900416;refGene_tx=XM_02
3245691.1
E2	9009363	9009517
refGene_gn=LOC102900416;refGene_tx=XM_006941030.4;refGene_gn=LOC102900
416;refGene_tx=XM_023245690.1;refGene_gn=LOC102900416;refGene_tx=XM_02
3245691.1
E2	9009756	9009991
refGene_gn=LOC102900416;refGene_tx=XM_006941030.4;refGene_gn=LOC102900
416;refGene_tx=XM_023245690.1;refGene_gn=LOC102900416;refGene_tx=XM_02
3245691.1
E2	9010209	9010288
refGene_gn=LOC102900416;refGene_tx=XM_006941030.4;refGene_gn=LOC102900
416;refGene_tx=XM_023245690.1;refGene_gn=LOC102900416;refGene_tx=XM_02
3245691.1
E2	9021935	9022170
refGene_gn=LOC102900001;refGene_tx=XM_023245685.1;refGene_gn=LOC102900
001;refGene_tx=XM_023245686.1;refGene_gn=LOC102900001;refGene_tx=XM_02
3245688.1;refGene_gn=LOC102900001;refGene_tx=XM_023245689.1
E2	9022628	9022703
refGene_gn=LOC102900001;refGene_tx=XM_023245685.1;refGene_gn=LOC102900
001;refGene_tx=XM_023245686.1;refGene_gn=LOC102900001;refGene_tx=XM_02
3245688.1;refGene_gn=LOC102900001;refGene_tx=XM_023245689.1
E2	9023071	9023158
refGene_gn=LOC102900001;refGene_tx=XM_023245685.1;refGene_gn=LOC102900
001;refGene_tx=XM_023245686.1
E2	9023446	9023467
refGene_gn=LOC102900001;refGene_tx=XM_023245685.1;refGene_gn=LOC102900
001;refGene_tx=XM_023245686.1;refGene_gn=LOC102900001;refGene_tx=XM_02
3245688.1
E2	9023816	9023897
refGene_gn=LOC102900001;refGene_tx=XM_023245685.1;refGene_gn=LOC102900
001;refGene_tx=XM_023245686.1;refGene_gn=LOC102900001;refGene_tx=XM_02
3245688.1;refGene_gn=LOC102900001;refGene_tx=XM_023245689.1
E2	9024072	9024173
refGene_gn=LOC102900001;refGene_tx=XM_023245685.1;refGene_gn=LOC102900
001;refGene_tx=XM_023245686.1;refGene_gn=LOC102900001;refGene_tx=XM_02
3245688.1;refGene_gn=LOC102900001;refGene_tx=XM_023245689.1
E2	9024433	9024587
refGene_gn=LOC102900001;refGene_tx=XM_023245685.1;refGene_gn=LOC102900
001;refGene_tx=XM_023245686.1;refGene_gn=LOC102900001;refGene_tx=XM_02
3245688.1;refGene_gn=LOC102900001;refGene_tx=XM_023245689.1
E2	9024893	9025152
refGene_gn=LOC102900001;refGene_tx=XM_023245686.1
E2	9025382	9025476
refGene_gn=LOC102900001;refGene_tx=XM_023245686.1
```

E2 9026754 9026989
refGene_gn=LOC105260116;refGene_tx=XM_023245682.1
E2 9027368 9027443
refGene_gn=LOC105260116;refGene_tx=XM_023245682.1
E2 9027780 9027864
refGene_gn=LOC105260116;refGene_tx=XM_023245682.1
E2 9028119 9028146
refGene_gn=LOC105260116;refGene_tx=XM_023245682.1
E2 9028501 9028582
refGene_gn=LOC105260116;refGene_tx=XM_023245682.1
E2 9028766 9028867
refGene_gn=LOC105260116;refGene_tx=XM_023245682.1
E2 9029118 9029272
refGene_gn=LOC105260116;refGene_tx=XM_023245682.1
E2 9029556 9029824
refGene_gn=LOC105260116;refGene_tx=XM_023245682.1
E2 9030046 9030239
refGene_gn=LOC105260116;refGene_tx=XM_023245682.1
E2 9034330 9034505
refGene_gn=LOC102900001;refGene_tx=XM_023245685.1;refGene_gn=LOC102900001;refGene_tx=XM_023245688.1;refGene_gn=LOC102900001;refGene_tx=XM_023245689.1
E2 9034730 9034923
refGene_gn=LOC102900001;refGene_tx=XM_023245685.1;refGene_gn=LOC102900001;refGene_tx=XM_023245688.1;refGene_gn=LOC102900001;refGene_tx=XM_023245689.1
E2 9035535 9035672 refGene_gn=RASIP1;refGene_tx=XM_023245906.1
E2 9036340 9036971 refGene_gn=RASIP1;refGene_tx=XM_023245906.1
E2 9038848 9039322 refGene_gn=RASIP1;refGene_tx=XM_023245906.1
E2 9042162 9042816 refGene_gn=RASIP1;refGene_tx=XM_023245906.1
E2 9044023 9044061 refGene_gn=RASIP1;refGene_tx=XM_023245906.1
E2 9044288 9044440 refGene_gn=RASIP1;refGene_tx=XM_023245906.1
E2 9044556 9044660 refGene_gn=RASIP1;refGene_tx=XM_023245906.1
E2 9046427 9046671 refGene_gn=RASIP1;refGene_tx=XM_023245906.1
E2 9046877 9047050 refGene_gn=RASIP1;refGene_tx=XM_023245906.1
E2 9048062 9048210 refGene_gn=RASIP1;refGene_tx=XM_023245906.1
E2 9048771 9048971 refGene_gn=RASIP1;refGene_tx=XM_023245906.1
E2 9050681 9050739
refGene_gn=MAMSTR;refGene_tx=XM_006944257.4;refGene_gn=MAMSTR;refGene_tx=XM_023245679.1;refGene_gn=MAMSTR;refGene_tx=XM_023245680.1;refGene_gn=MAMSTR;refGene_tx=XM_023245681.1
E2 9052117 9052156
refGene_gn=MAMSTR;refGene_tx=XM_006944257.4;refGene_gn=MAMSTR;refGene_tx=XM_023245679.1;refGene_gn=MAMSTR;refGene_tx=XM_023245680.1;refGene_gn=MAMSTR;refGene_tx=XM_023245681.1
E2 9052745 9052912
refGene_gn=MAMSTR;refGene_tx=XM_006944257.4;refGene_gn=MAMSTR;refGene_tx=XM_023245679.1;refGene_gn=MAMSTR;refGene_tx=XM_023245680.1;refGene_gn=MAMSTR;refGene_tx=XM_023245681.1
E2 9053098 9053283

refGene_gn=MAMSTR;refGene_tx=XM_006944257.4;refGene_gn=MAMSTR;refGene_tx=XM_023245679.1;refGene_gn=MAMSTR;refGene_tx=XM_023245680.1;refGene_gn=MAMSTR;refGene_tx=XM_023245681.1
E2 9053637 9053740 refGene_gn=MAMSTR;refGene_tx=XM_006944257.4;refGene_gn=MAMSTR;refGene_tx=XM_023245679.1;refGene_gn=MAMSTR;refGene_tx=XM_023245680.1;refGene_gn=MAMSTR;refGene_tx=XM_023245681.1
E2 9053947 9054142 refGene_gn=MAMSTR;refGene_tx=XM_006944257.4;refGene_gn=MAMSTR;refGene_tx=XM_023245679.1;refGene_gn=MAMSTR;refGene_tx=XM_023245680.1;refGene_gn=MAMSTR;refGene_tx=XM_023245681.1
E2 9054419 9054587 refGene_gn=MAMSTR;refGene_tx=XM_006944257.4;refGene_gn=MAMSTR;refGene_tx=XM_023245679.1;refGene_gn=MAMSTR;refGene_tx=XM_023245680.1;refGene_gn=MAMSTR;refGene_tx=XM_023245681.1
E2 9054665 9054720 refGene_gn=MAMSTR;refGene_tx=XM_006944257.4;refGene_gn=MAMSTR;refGene_tx=XM_023245679.1;refGene_gn=MAMSTR;refGene_tx=XM_023245680.1;refGene_gn=MAMSTR;refGene_tx=XM_023245681.1
E2 9055152 9055436 refGene_gn=MAMSTR;refGene_tx=XM_006944257.4
E2 9062796 9062840 refGene_gn=MAMSTR;refGene_tx=XM_023245681.1
E2 9062916 9063062 refGene_gn=MAMSTR;refGene_tx=XM_023245679.1;refGene_gn=MAMSTR;refGene_tx=XM_023245680.1
E2 9086255 9087280 refGene_gn=LOC101085847;refGene_tx=XM_023245916.1
E2 9088965 9088970 refGene_gn=LOC101085847;refGene_tx=XM_023245916.1
E2 9093638 9094269 refGene_gn=NTN5;refGene_tx=XM_023245164.1;refGene_gn=NTN5;refGene_tx=XM_023245165.1
E2 9096879 9097068 refGene_gn=NTN5;refGene_tx=XM_023245164.1;refGene_gn=NTN5;refGene_tx=XM_023245165.1
E2 9097210 9097360 refGene_gn=NTN5;refGene_tx=XM_023245164.1;refGene_gn=NTN5;refGene_tx=XM_023245165.1
E2 9097698 9097752 refGene_gn=NTN5;refGene_tx=XM_023245164.1;refGene_gn=NTN5;refGene_tx=XM_023245165.1
E2 9097973 9098054 refGene_gn=NTN5;refGene_tx=XM_023245164.1;refGene_gn=NTN5;refGene_tx=XM_023245165.1
E2 9099300 9099665 refGene_gn=NTN5;refGene_tx=XM_023245164.1;refGene_gn=NTN5;refGene_tx=XM_023245165.1
E2 9109034 9109101 refGene_gn=CA11;refGene_tx=XM_003997530.5
E2 9109186 9109261 refGene_gn=CA11;refGene_tx=XM_003997530.5
E2 9109989 9110132 refGene_gn=CA11;refGene_tx=XM_003997530.5
E2 9111497 9111683 refGene_gn=CA11;refGene_tx=XM_003997530.5

E2 9111901 9111997 refGene_gn=CA11;refGene_tx=XM_003997530.5
E2 9112141 9112214 refGene_gn=CA11;refGene_tx=XM_003997530.5
E2 9112317 9112472 refGene_gn=CA11;refGene_tx=XM_003997530.5
E2 9112722 9112888 refGene_gn=CA11;refGene_tx=XM_003997530.5
E2 9113702 9113728 refGene_gn=CA11;refGene_tx=XM_003997530.5
E2 9114845 9114984 refGene_gn=DBP;refGene_tx=XM_023245972.1
E2 9115888 9116299 refGene_gn=DBP;refGene_tx=XM_023245972.1
E2 9117918 9118130 refGene_gn=DBP;refGene_tx=XM_023245972.1
E2 9119649 9119865 refGene_gn=DBP;refGene_tx=XM_023245972.1
E2 9120947 9122052
refGene_gn=SPHK2;refGene_tx=XM_006941031.3;refGene_gn=SPHK2;refGene_tx=XM_006941032.4;refGene_gn=SPHK2;refGene_tx=XM_006941033.3;refGene_gn=SPHK2;refGene_tx=XM_006941035.3;refGene_gn=SPHK2;refGene_tx=XM_019819270.2;refGene_gn=SPHK2;refGene_tx=XM_019819271.2;refGene_gn=SPHK2;refGene_tx=XM_019819272.2
E2 9122499 9122615
refGene_gn=SPHK2;refGene_tx=XM_006941031.3;refGene_gn=SPHK2;refGene_tx=XM_006941032.4;refGene_gn=SPHK2;refGene_tx=XM_006941033.3;refGene_gn=SPHK2;refGene_tx=XM_006941035.3;refGene_gn=SPHK2;refGene_tx=XM_019819270.2;refGene_gn=SPHK2;refGene_tx=XM_019819271.2;refGene_gn=SPHK2;refGene_tx=XM_019819272.2
E2 9122700 9122795
refGene_gn=SPHK2;refGene_tx=XM_006941031.3;refGene_gn=SPHK2;refGene_tx=XM_006941032.4;refGene_gn=SPHK2;refGene_tx=XM_006941033.3;refGene_gn=SPHK2;refGene_tx=XM_006941035.3;refGene_gn=SPHK2;refGene_tx=XM_019819270.2;refGene_gn=SPHK2;refGene_tx=XM_019819271.2;refGene_gn=SPHK2;refGene_tx=XM_019819272.2
E2 9122933 9123083
refGene_gn=SPHK2;refGene_tx=XM_006941031.3;refGene_gn=SPHK2;refGene_tx=XM_006941032.4;refGene_gn=SPHK2;refGene_tx=XM_006941033.3;refGene_gn=SPHK2;refGene_tx=XM_006941035.3;refGene_gn=SPHK2;refGene_tx=XM_019819270.2;refGene_gn=SPHK2;refGene_tx=XM_019819271.2;refGene_gn=SPHK2;refGene_tx=XM_019819272.2
E2 9123992 9124470
refGene_gn=SPHK2;refGene_tx=XM_006941031.3;refGene_gn=SPHK2;refGene_tx=XM_006941032.4;refGene_gn=SPHK2;refGene_tx=XM_006941033.3;refGene_gn=SPHK2;refGene_tx=XM_019819270.2;refGene_gn=SPHK2;refGene_tx=XM_019819271.2;refGene_gn=SPHK2;refGene_tx=XM_019819272.2
E2 9126064 9126163 refGene_gn=SPHK2;refGene_tx=XM_019819270.2
E2 9127641 9127680
refGene_gn=SPHK2;refGene_tx=XM_006941031.3;refGene_gn=SPHK2;refGene_tx=XM_006941032.4;refGene_gn=SPHK2;refGene_tx=XM_006941033.3
E2 9129206 9129209 refGene_gn=RPL18;refGene_tx=XM_003997532.5
E2 9130543 9130630 refGene_gn=RPL18;refGene_tx=XM_003997532.5
E2 9130972 9131080 refGene_gn=RPL18;refGene_tx=XM_003997532.5
E2 9131534 9131633 refGene_gn=RPL18;refGene_tx=XM_003997532.5
E2 9132380 9132504 refGene_gn=RPL18;refGene_tx=XM_003997532.5
E2 9132587 9132657 refGene_gn=RPL18;refGene_tx=XM_003997532.5
E2 9133125 9133201 refGene_gn=RPL18;refGene_tx=XM_003997532.5
E2 9135095 9135569 refGene_gn=FAM83E;refGene_tx=XM_023245235.1

```
E2	9136757	9136925	refGene_gn=FAM83E;refGene_tx=XM_023245235.1
E2	9137311	9137436	refGene_gn=FAM83E;refGene_tx=XM_023245235.1
E2	9139097	9139478	refGene_gn=SPACA4;refGene_tx=XM_006941036.4
E2	9140717	9141162	refGene_gn=FAM83E;refGene_tx=XM_023245235.1
E2	9142559	9142835	refGene_gn=FAM83E;refGene_tx=XM_023245235.1
E2	9143629	9143811	refGene_gn=SULT2B1;refGene_tx=XM_023245236.1
E2	9145528	9145709	refGene_gn=SULT2B1;refGene_tx=XM_023245236.1
E2	9149462	9149557	refGene_gn=SULT2B1;refGene_tx=XM_023245236.1
E2	9150075	9150202	refGene_gn=SULT2B1;refGene_tx=XM_023245236.1
E2	9151830	9152039	refGene_gn=SULT2B1;refGene_tx=XM_023245236.1
E2	9156264	9156404	refGene_gn=SULT2B1;refGene_tx=XM_023245236.1
E2	9175509	9175577	refGene_gn=SULT2B1;refGene_tx=XM_023245236.1
E2	9205897	9206133	refGene_gn=LMTK3;refGene_tx=XM_023245254.1
E2	9206255	9206368	refGene_gn=LMTK3;refGene_tx=XM_023245254.1
E2	9207301	9207435	refGene_gn=LMTK3;refGene_tx=XM_023245254.1
E2	9207690	9207841	refGene_gn=LMTK3;refGene_tx=XM_023245254.1
E2	9208358	9208435	refGene_gn=LMTK3;refGene_tx=XM_023245254.1
E2	9208915	9209034	refGene_gn=LMTK3;refGene_tx=XM_023245254.1
E2	9211332	9211420	refGene_gn=LMTK3;refGene_tx=XM_023245254.1
E2	9211764	9211913	refGene_gn=LMTK3;refGene_tx=XM_023245254.1
E2	9212311	9212396	refGene_gn=LMTK3;refGene_tx=XM_023245254.1
E2	9212487	9212609	refGene_gn=LMTK3;refGene_tx=XM_023245254.1
E2	9212762	9212912	refGene_gn=LMTK3;refGene_tx=XM_023245254.1
E2	9213659	9216169	refGene_gn=LMTK3;refGene_tx=XM_023245254.1
E2	9218806	9219009	refGene_gn=LMTK3;refGene_tx=XM_023245254.1
E2	9219011	9219236	refGene_gn=LMTK3;refGene_tx=XM_023245254.1
E2	9220465	9220601	refGene_gn=LMTK3;refGene_tx=XM_023245254.1
E2	9221624	9221762	refGene_gn=LMTK3;refGene_tx=XM_023245254.1
E2	9224439	9224456	refGene_gn=LMTK3;refGene_tx=XM_023245254.1
E2	9230201	9230289
refGene_gn=CYTH2;refGene_tx=XM_023245250.1;refGene_gn=CYTH2;refGene_tx
=XM_023245252.1
E2	9230384	9230539
refGene_gn=CYTH2;refGene_tx=XM_023245250.1;refGene_gn=CYTH2;refGene_tx
=XM_023245252.1
E2	9230664	9230736
refGene_gn=CYTH2;refGene_tx=XM_023245250.1;refGene_gn=CYTH2;refGene_tx
=XM_023245252.1
E2	9231026	9231105
refGene_gn=CYTH2;refGene_tx=XM_023245250.1;refGene_gn=CYTH2;refGene_tx
=XM_023245252.1
E2	9233269	9233382
refGene_gn=CYTH2;refGene_tx=XM_023245250.1;refGene_gn=CYTH2;refGene_tx
=XM_023245252.1
E2	9233869	9234018
refGene_gn=CYTH2;refGene_tx=XM_023245250.1;refGene_gn=CYTH2;refGene_tx
=XM_023245252.1
E2	9234113	9234226
refGene_gn=CYTH2;refGene_tx=XM_023245250.1;refGene_gn=CYTH2;refGene_tx
=XM_023245252.1
```

E2	9234666	9234747
refGene_gn=CYTH2;refGene_tx=XM_023245250.1;refGene_gn=CYTH2;refGene_tx
=XM_023245252.1
E2	9235736	9235855
refGene_gn=CYTH2;refGene_tx=XM_023245250.1;refGene_gn=CYTH2;refGene_tx
=XM_023245252.1
E2	9236301	9236368
refGene_gn=CYTH2;refGene_tx=XM_023245250.1;refGene_gn=CYTH2;refGene_tx
=XM_023245252.1
E2	9236457	9236605
refGene_gn=CYTH2;refGene_tx=XM_023245250.1;refGene_gn=CYTH2;refGene_tx
=XM_023245252.1
E2	9236883	9236968	refGene_gn=CYTH2;refGene_tx=XM_023245252.1
E2	9237421	9237440	refGene_gn=CYTH2;refGene_tx=XM_023245250.1
E2	9241169	9241766	refGene_gn=KCNJ14;refGene_tx=XM_003997721.5
E2	9243217	9243919	refGene_gn=KCNJ14;refGene_tx=XM_003997721.5
E2	9249841	9250159	refGene_gn=GRWD1;refGene_tx=XM_003997698.5
E2	9251265	9251406	refGene_gn=GRWD1;refGene_tx=XM_003997698.5
E2	9251407	9251464	refGene_gn=GRWD1;refGene_tx=XM_003997698.5
E2	9251927	9252070	refGene_gn=GRWD1;refGene_tx=XM_003997698.5
E2	9252213	9252427	refGene_gn=GRWD1;refGene_tx=XM_003997698.5
E2	9254392	9254555	refGene_gn=GRWD1;refGene_tx=XM_003997698.5
E2	9254709	9254827	refGene_gn=GRWD1;refGene_tx=XM_003997698.5
E2	9255036	9255223	refGene_gn=GRWD1;refGene_tx=XM_003997698.5
E2	9256838	9258719
refGene_gn=GRIN2D;refGene_tx=XM_023245297.1;refGene_gn=GRIN2D;refGene_
tx=XM_023245298.1
E2	9258902	9259135
refGene_gn=GRIN2D;refGene_tx=XM_023245297.1;refGene_gn=GRIN2D;refGene_
tx=XM_023245298.1
E2	9259359	9259547
refGene_gn=GRIN2D;refGene_tx=XM_023245297.1;refGene_gn=GRIN2D;refGene_
tx=XM_023245298.1
E2	9275570	9275731
refGene_gn=GRIN2D;refGene_tx=XM_023245297.1;refGene_gn=GRIN2D;refGene_
tx=XM_023245298.1
E2	9277077	9277307
refGene_gn=GRIN2D;refGene_tx=XM_023245297.1;refGene_gn=GRIN2D;refGene_
tx=XM_023245298.1
E2	9277502	9277628
refGene_gn=GRIN2D;refGene_tx=XM_023245297.1;refGene_gn=GRIN2D;refGene_
tx=XM_023245298.1
E2	9279978	9280132
refGene_gn=GRIN2D;refGene_tx=XM_023245297.1;refGene_gn=GRIN2D;refGene_
tx=XM_023245298.1
E2	9280621	9280790
refGene_gn=GRIN2D;refGene_tx=XM_023245297.1;refGene_gn=GRIN2D;refGene_
tx=XM_023245298.1
E2	9281046	9281258
refGene_gn=GRIN2D;refGene_tx=XM_023245297.1;refGene_gn=GRIN2D;refGene_

```
tx=XM_023245298.1
E2	9281623	9281738	
refGene_gn=GRIN2D;refGene_tx=XM_023245297.1;refGene_gn=GRIN2D;refGene_
tx=XM_023245298.1
E2	9285841	9286461	
refGene_gn=GRIN2D;refGene_tx=XM_023245297.1;refGene_gn=GRIN2D;refGene_
tx=XM_023245298.1
E2	9290330	9290795	refGene_gn=GRIN2D;refGene_tx=XM_023245297.1
E2	9297237	9297328	refGene_gn=KDELR1;refGene_tx=XM_003997697.5
E2	9298033	9298134	refGene_gn=KDELR1;refGene_tx=XM_003997697.5
E2	9298421	9298580	refGene_gn=KDELR1;refGene_tx=XM_003997697.5
E2	9302114	9302367	refGene_gn=KDELR1;refGene_tx=XM_003997697.5
E2	9303436	9303471	refGene_gn=KDELR1;refGene_tx=XM_003997697.5
E2	9307031	9307265	
refGene_gn=SYNGR4;refGene_tx=XM_003997696.5;refGene_gn=SYNGR4;refGene_
tx=XM_023245302.1
E2	9307586	9307726	
refGene_gn=SYNGR4;refGene_tx=XM_003997696.5;refGene_gn=SYNGR4;refGene_
tx=XM_023245302.1
E2	9307871	9308077	refGene_gn=SYNGR4;refGene_tx=XM_019819281.2
E2	9308951	9309189	
refGene_gn=SYNGR4;refGene_tx=XM_003997696.5;refGene_gn=SYNGR4;refGene_
tx=XM_019819281.2;refGene_gn=SYNGR4;refGene_tx=XM_023245302.1
E2	9314994	9315194	
refGene_gn=SYNGR4;refGene_tx=XM_003997696.5;refGene_gn=SYNGR4;refGene_
tx=XM_019819281.2;refGene_gn=SYNGR4;refGene_tx=XM_023245302.1
E2	9316161	9316594	refGene_gn=SYNGR4;refGene_tx=XM_019819281.2
E2	9316649	9316688	
refGene_gn=TMEM143;refGene_tx=XM_023245300.1;refGene_gn=TMEM143;refGen
e_tx=XM_023245301.1
E2	9317036	9317277	refGene_gn=TMEM143;refGene_tx=XM_023245300.1
E2	9318586	9318691	
refGene_gn=TMEM143;refGene_tx=XM_023245300.1;refGene_gn=TMEM143;refGen
e_tx=XM_023245301.1
E2	9327350	9327545	
refGene_gn=TMEM143;refGene_tx=XM_023245300.1;refGene_gn=TMEM143;refGen
e_tx=XM_023245301.1
E2	9328369	9328497	
refGene_gn=TMEM143;refGene_tx=XM_023245300.1;refGene_gn=TMEM143;refGen
e_tx=XM_023245301.1
E2	9329479	9329759	
refGene_gn=TMEM143;refGene_tx=XM_023245300.1;refGene_gn=TMEM143;refGen
e_tx=XM_023245301.1
E2	9334318	9334508	
refGene_gn=TMEM143;refGene_tx=XM_023245300.1;refGene_gn=TMEM143;refGen
e_tx=XM_023245301.1
E2	9335159	9335374	
refGene_gn=TMEM143;refGene_tx=XM_023245300.1;refGene_gn=TMEM143;refGen
e_tx=XM_023245301.1
E2	9337255	9337425
```

refGene_gn=EMP3;refGene_tx=XM_003997695.4;refGene_gn=EMP3;refGene_tx=XM_019819284.2;refGene_gn=EMP3;refGene_tx=XM_019819285.1
E2 9338216 9338357
refGene_gn=EMP3;refGene_tx=XM_003997695.4;refGene_gn=EMP3;refGene_tx=XM_019819284.2;refGene_gn=EMP3;refGene_tx=XM_019819285.1
E2 9339477 9339580
refGene_gn=EMP3;refGene_tx=XM_003997695.4;refGene_gn=EMP3;refGene_tx=XM_019819284.2;refGene_gn=EMP3;refGene_tx=XM_019819285.1
E2 9339881 9339959
refGene_gn=EMP3;refGene_tx=XM_003997695.4;refGene_gn=EMP3;refGene_tx=XM_019819284.2;refGene_gn=EMP3;refGene_tx=XM_019819285.1
E2 9345258 9345328
refGene_gn=CCDC114;refGene_tx=XM_023245183.1;refGene_gn=CCDC114;refGene_tx=XM_023245184.1;refGene_gn=CCDC114;refGene_tx=XM_023245185.1
E2 9346735 9346835
refGene_gn=CCDC114;refGene_tx=XM_023245183.1;refGene_gn=CCDC114;refGene_tx=XM_023245184.1;refGene_gn=CCDC114;refGene_tx=XM_023245185.1
E2 9347005 9347195
refGene_gn=CCDC114;refGene_tx=XM_023245183.1;refGene_gn=CCDC114;refGene_tx=XM_023245184.1;refGene_gn=CCDC114;refGene_tx=XM_023245185.1
E2 9358239 9358362
refGene_gn=CCDC114;refGene_tx=XM_023245183.1;refGene_gn=CCDC114;refGene_tx=XM_023245184.1;refGene_gn=CCDC114;refGene_tx=XM_023245185.1
E2 9358701 9358815
refGene_gn=CCDC114;refGene_tx=XM_023245183.1;refGene_gn=CCDC114;refGene_tx=XM_023245184.1;refGene_gn=CCDC114;refGene_tx=XM_023245185.1
E2 9362527 9362595
refGene_gn=CCDC114;refGene_tx=XM_023245183.1;refGene_gn=CCDC114;refGene_tx=XM_023245184.1;refGene_gn=CCDC114;refGene_tx=XM_023245185.1
E2 9365059 9365247
refGene_gn=CCDC114;refGene_tx=XM_023245183.1;refGene_gn=CCDC114;refGene_tx=XM_023245184.1;refGene_gn=CCDC114;refGene_tx=XM_023245185.1
E2 9365355 9365490
refGene_gn=CCDC114;refGene_tx=XM_023245183.1;refGene_gn=CCDC114;refGene_tx=XM_023245184.1;refGene_gn=CCDC114;refGene_tx=XM_023245185.1
E2 9366179 9366262
refGene_gn=CCDC114;refGene_tx=XM_023245183.1;refGene_gn=CCDC114;refGene_tx=XM_023245184.1;refGene_gn=CCDC114;refGene_tx=XM_023245185.1
E2 9366396 9366565
refGene_gn=CCDC114;refGene_tx=XM_023245183.1;refGene_gn=CCDC114;refGene_tx=XM_023245184.1;refGene_gn=CCDC114;refGene_tx=XM_023245185.1
E2 9368397 9368567
refGene_gn=CCDC114;refGene_tx=XM_023245183.1;refGene_gn=CCDC114;refGene_tx=XM_023245184.1;refGene_gn=CCDC114;refGene_tx=XM_023245185.1
E2 9368675 9368779
refGene_gn=CCDC114;refGene_tx=XM_023245183.1;refGene_gn=CCDC114;refGene_tx=XM_023245184.1;refGene_gn=CCDC114;refGene_tx=XM_023245185.1
E2 9369100 9369179
refGene_gn=CCDC114;refGene_tx=XM_023245183.1;refGene_gn=CCDC114;refGene_tx=XM_023245184.1;refGene_gn=CCDC114;refGene_tx=XM_023245185.1

E2	9369269	9369749
refGene_gn=CCDC114;refGene_tx=XM_023245183.1;refGene_gn=CCDC114;refGene_tx=XM_023245184.1;refGene_gn=CCDC114;refGene_tx=XM_023245185.1
E2	9380262	9381365
refGene_gn=ZNF114;refGene_tx=XM_006941152.4;refGene_gn=ZNF114;refGene_tx=XM_019819286.2;refGene_gn=ZNF114;refGene_tx=XM_023245379.1
E2	9382970	9383097
refGene_gn=ZNF114;refGene_tx=XM_006941152.4;refGene_gn=ZNF114;refGene_tx=XM_019819286.2;refGene_gn=ZNF114;refGene_tx=XM_023245379.1
E2	9391049	9391106
refGene_gn=ZNF114;refGene_tx=XM_006941152.4;refGene_gn=ZNF114;refGene_tx=XM_019819286.2
E2	9428569	9428628
refGene_gn=CARD8;refGene_tx=XM_006941145.3;refGene_gn=CARD8;refGene_tx=XM_006941146.3;refGene_gn=CARD8;refGene_tx=XM_006941147.3;refGene_gn=CARD8;refGene_tx=XM_006941148.3;refGene_gn=CARD8;refGene_tx=XM_011289783.3
E2	9430118	9430277
refGene_gn=CARD8;refGene_tx=XM_006941145.3;refGene_gn=CARD8;refGene_tx=XM_006941146.3;refGene_gn=CARD8;refGene_tx=XM_006941147.3;refGene_gn=CARD8;refGene_tx=XM_006941148.3;refGene_gn=CARD8;refGene_tx=XM_006941150.4;refGene_gn=CARD8;refGene_tx=XM_006941151.4;refGene_gn=CARD8;refGene_tx=XM_011289783.3
E2	9432152	9432317
refGene_gn=CARD8;refGene_tx=XM_006941145.3;refGene_gn=CARD8;refGene_tx=XM_006941146.3;refGene_gn=CARD8;refGene_tx=XM_006941147.3;refGene_gn=CARD8;refGene_tx=XM_006941148.3;refGene_gn=CARD8;refGene_tx=XM_006941150.4;refGene_gn=CARD8;refGene_tx=XM_006941151.4;refGene_gn=CARD8;refGene_tx=XM_011289783.3
E2	9436821	9436877
refGene_gn=CARD8;refGene_tx=XM_006941145.3;refGene_gn=CARD8;refGene_tx=XM_006941146.3;refGene_gn=CARD8;refGene_tx=XM_006941147.3;refGene_gn=CARD8;refGene_tx=XM_006941148.3;refGene_gn=CARD8;refGene_tx=XM_006941150.4;refGene_gn=CARD8;refGene_tx=XM_006941151.4;refGene_gn=CARD8;refGene_tx=XM_011289783.3
E2	9438921	9439072
refGene_gn=CARD8;refGene_tx=XM_006941145.3;refGene_gn=CARD8;refGene_tx=XM_006941146.3;refGene_gn=CARD8;refGene_tx=XM_006941147.3;refGene_gn=CARD8;refGene_tx=XM_006941148.3;refGene_gn=CARD8;refGene_tx=XM_006941150.4;refGene_gn=CARD8;refGene_tx=XM_006941151.4;refGene_gn=CARD8;refGene_tx=XM_011289783.3
E2	9439882	9440112
refGene_gn=CARD8;refGene_tx=XM_006941145.3;refGene_gn=CARD8;refGene_tx=XM_006941146.3;refGene_gn=CARD8;refGene_tx=XM_006941147.3;refGene_gn=CARD8;refGene_tx=XM_006941148.3;refGene_gn=CARD8;refGene_tx=XM_006941150.4;refGene_gn=CARD8;refGene_tx=XM_006941151.4;refGene_gn=CARD8;refGene_tx=XM_011289783.3
E2	9440186	9440449
refGene_gn=CARD8;refGene_tx=XM_006941145.3;refGene_gn=CARD8;refGene_tx=XM_006941146.3;refGene_gn=CARD8;refGene_tx=XM_006941147.3;refGene_gn=

CARD8;refGene_tx=XM_006941148.3;refGene_gn=CARD8;refGene_tx=XM_0069411
50.4;refGene_gn=CARD8;refGene_tx=XM_006941151.4;refGene_gn=CARD8;refGe
ne_tx=XM_011289783.3
E2 9441711 9441837
refGene_gn=CARD8;refGene_tx=XM_006941145.3;refGene_gn=CARD8;refGene_tx
=XM_006941146.3;refGene_gn=CARD8;refGene_tx=XM_006941147.3;refGene_gn=
CARD8;refGene_tx=XM_006941148.3;refGene_gn=CARD8;refGene_tx=XM_0069411
50.4;refGene_gn=CARD8;refGene_tx=XM_006941151.4;refGene_gn=CARD8;refGe
ne_tx=XM_011289783.3
E2 9442167 9442309
refGene_gn=CARD8;refGene_tx=XM_006941145.3;refGene_gn=CARD8;refGene_tx
=XM_006941146.3;refGene_gn=CARD8;refGene_tx=XM_006941147.3;refGene_gn=
CARD8;refGene_tx=XM_006941148.3;refGene_gn=CARD8;refGene_tx=XM_0069411
50.4;refGene_gn=CARD8;refGene_tx=XM_006941151.4;refGene_gn=CARD8;refGe
ne_tx=XM_011289783.3
E2 9443946 9443991
refGene_gn=CARD8;refGene_tx=XM_006941145.3;refGene_gn=CARD8;refGene_tx
=XM_006941146.3;refGene_gn=CARD8;refGene_tx=XM_006941147.3;refGene_gn=
CARD8;refGene_tx=XM_006941150.4;refGene_gn=CARD8;refGene_tx=XM_0069411
51.4;refGene_gn=CARD8;refGene_tx=XM_011289783.3
E2 9447800 9448072
refGene_gn=CARD8;refGene_tx=XM_006941145.3;refGene_gn=CARD8;refGene_tx
=XM_006941146.3;refGene_gn=CARD8;refGene_tx=XM_006941147.3;refGene_gn=
CARD8;refGene_tx=XM_006941148.3;refGene_gn=CARD8;refGene_tx=XM_0069411
50.4;refGene_gn=CARD8;refGene_tx=XM_006941151.4;refGene_gn=CARD8;refGe
ne_tx=XM_011289783.3
E2 9459234 9461364
refGene_gn=ZSWIM9;refGene_tx=XM_023246152.1;refGene_gn=ZSWIM9;refGene_
tx=XM_023246153.1;refGene_gn=ZSWIM9;refGene_tx=XM_023246154.1;refGene_
gn=ZSWIM9;refGene_tx=XM_023246155.1;refGene_gn=ZSWIM9;refGene_tx=XM_02
3246156.1;refGene_gn=ZSWIM9;refGene_tx=XM_023246157.1
E2 9470613 9471057
refGene_gn=ZSWIM9;refGene_tx=XM_023246152.1;refGene_gn=ZSWIM9;refGene_
tx=XM_023246153.1;refGene_gn=ZSWIM9;refGene_tx=XM_023246154.1;refGene_
gn=ZSWIM9;refGene_tx=XM_023246155.1;refGene_gn=ZSWIM9;refGene_tx=XM_02
3246157.1
E2 9477807 9478080
refGene_gn=ZSWIM9;refGene_tx=XM_023246152.1;refGene_gn=ZSWIM9;refGene_
tx=XM_023246153.1;refGene_gn=ZSWIM9;refGene_tx=XM_023246154.1;refGene_
gn=ZSWIM9;refGene_tx=XM_023246155.1;refGene_gn=ZSWIM9;refGene_tx=XM_02
3246156.1;refGene_gn=ZSWIM9;refGene_tx=XM_023246157.1
E2 9478540 9478803
refGene_gn=ZSWIM9;refGene_tx=XM_023246152.1;refGene_gn=ZSWIM9;refGene_
tx=XM_023246153.1;refGene_gn=ZSWIM9;refGene_tx=XM_023246156.1;refGene_
gn=ZSWIM9;refGene_tx=XM_023246157.1
E2 9485285 9485302
refGene_gn=LIG1;refGene_tx=XM_023246158.1;refGene_gn=LIG1;refGene_tx=X
M_023246159.1;refGene_gn=LIG1;refGene_tx=XM_023246160.1
E2 9487254 9487338
refGene_gn=LIG1;refGene_tx=XM_023246158.1;refGene_gn=LIG1;refGene_tx=X

M_023246159.1;refGene_gn=LIG1;refGene_tx=XM_023246160.1
E2 9487988 9488124
refGene_gn=LIG1;refGene_tx=XM_023246158.1;refGene_gn=LIG1;refGene_tx=XM_023246159.1;refGene_gn=LIG1;refGene_tx=XM_023246160.1
E2 9490424 9490551
refGene_gn=LIG1;refGene_tx=XM_023246158.1;refGene_gn=LIG1;refGene_tx=XM_023246159.1;refGene_gn=LIG1;refGene_tx=XM_023246160.1
E2 9492985 9493078
refGene_gn=LIG1;refGene_tx=XM_023246158.1;refGene_gn=LIG1;refGene_tx=XM_023246159.1;refGene_gn=LIG1;refGene_tx=XM_023246160.1
E2 9495081 9495183
refGene_gn=LIG1;refGene_tx=XM_023246158.1;refGene_gn=LIG1;refGene_tx=XM_023246159.1;refGene_gn=LIG1;refGene_tx=XM_023246160.1
E2 9496084 9496204
refGene_gn=LIG1;refGene_tx=XM_023246158.1;refGene_gn=LIG1;refGene_tx=XM_023246159.1;refGene_gn=LIG1;refGene_tx=XM_023246160.1
E2 9496397 9496473
refGene_gn=LIG1;refGene_tx=XM_023246158.1;refGene_gn=LIG1;refGene_tx=XM_023246159.1;refGene_gn=LIG1;refGene_tx=XM_023246160.1
E2 9500414 9500495
refGene_gn=LIG1;refGene_tx=XM_023246158.1;refGene_gn=LIG1;refGene_tx=XM_023246159.1;refGene_gn=LIG1;refGene_tx=XM_023246160.1
E2 9500705 9500762
refGene_gn=LIG1;refGene_tx=XM_023246158.1;refGene_gn=LIG1;refGene_tx=XM_023246159.1;refGene_gn=LIG1;refGene_tx=XM_023246160.1
E2 9503391 9503564
refGene_gn=LIG1;refGene_tx=XM_023246158.1;refGene_gn=LIG1;refGene_tx=XM_023246159.1;refGene_gn=LIG1;refGene_tx=XM_023246160.1
E2 9503913 9504080
refGene_gn=LIG1;refGene_tx=XM_023246158.1;refGene_gn=LIG1;refGene_tx=XM_023246159.1;refGene_gn=LIG1;refGene_tx=XM_023246160.1
E2 9504457 9504534
refGene_gn=LIG1;refGene_tx=XM_023246158.1;refGene_gn=LIG1;refGene_tx=XM_023246159.1;refGene_gn=LIG1;refGene_tx=XM_023246160.1
E2 9505281 9505373
refGene_gn=LIG1;refGene_tx=XM_023246158.1;refGene_gn=LIG1;refGene_tx=XM_023246159.1;refGene_gn=LIG1;refGene_tx=XM_023246160.1
E2 9505563 9505663
refGene_gn=LIG1;refGene_tx=XM_023246158.1;refGene_gn=LIG1;refGene_tx=XM_023246159.1;refGene_gn=LIG1;refGene_tx=XM_023246160.1
E2 9506411 9506497
refGene_gn=LIG1;refGene_tx=XM_023246158.1;refGene_gn=LIG1;refGene_tx=XM_023246159.1;refGene_gn=LIG1;refGene_tx=XM_023246160.1
E2 9507227 9507343
refGene_gn=LIG1;refGene_tx=XM_023246158.1;refGene_gn=LIG1;refGene_tx=XM_023246159.1;refGene_gn=LIG1;refGene_tx=XM_023246160.1
E2 9509884 9509980
refGene_gn=LIG1;refGene_tx=XM_023246158.1;refGene_gn=LIG1;refGene_tx=XM_023246159.1;refGene_gn=LIG1;refGene_tx=XM_023246160.1
E2 9512272 9512383

refGene_gn=LIG1;refGene_tx=XM_023246158.1;refGene_gn=LIG1;refGene_tx=XM_023246159.1;refGene_gn=LIG1;refGene_tx=XM_023246160.1
E2 9512939 9513011
refGene_gn=LIG1;refGene_tx=XM_023246158.1;refGene_gn=LIG1;refGene_tx=XM_023246159.1;refGene_gn=LIG1;refGene_tx=XM_023246160.1
E2 9515375 9515520
refGene_gn=LIG1;refGene_tx=XM_023246158.1;refGene_gn=LIG1;refGene_tx=XM_023246159.1;refGene_gn=LIG1;refGene_tx=XM_023246160.1
E2 9515643 9515726
refGene_gn=LIG1;refGene_tx=XM_023246158.1;refGene_gn=LIG1;refGene_tx=XM_023246159.1;refGene_gn=LIG1;refGene_tx=XM_023246160.1
E2 9516947 9517100
refGene_gn=LIG1;refGene_tx=XM_023246158.1;refGene_gn=LIG1;refGene_tx=XM_023246159.1;refGene_gn=LIG1;refGene_tx=XM_023246160.1
E2 9518892 9518946
refGene_gn=LIG1;refGene_tx=XM_023246158.1;refGene_gn=LIG1;refGene_tx=XM_023246159.1;refGene_gn=LIG1;refGene_tx=XM_023246160.1
E2 9519790 9519934
refGene_gn=LIG1;refGene_tx=XM_023246158.1;refGene_gn=LIG1;refGene_tx=XM_023246159.1;refGene_gn=LIG1;refGene_tx=XM_023246160.1
E2 9523987 9524080
refGene_gn=LIG1;refGene_tx=XM_023246158.1;refGene_gn=LIG1;refGene_tx=XM_023246159.1;refGene_gn=LIG1;refGene_tx=XM_023246160.1
E2 9524203 9524287
refGene_gn=LIG1;refGene_tx=XM_023246158.1;refGene_gn=LIG1;refGene_tx=XM_023246159.1;refGene_gn=LIG1;refGene_tx=XM_023246160.1
E2 9526588 9526651 refGene_gn=CABP5;refGene_tx=XM_003997694.4
E2 9527568 9527599 refGene_gn=CABP5;refGene_tx=XM_003997694.4
E2 9528551 9528695 refGene_gn=CABP5;refGene_tx=XM_003997694.4
E2 9529631 9529741 refGene_gn=CABP5;refGene_tx=XM_003997694.4
E2 9533375 9533523 refGene_gn=CABP5;refGene_tx=XM_003997694.4
E2 9535146 9535172 refGene_gn=CABP5;refGene_tx=XM_003997694.4
E2 9540333 9540491
refGene_gn=ELSPBP1;refGene_tx=XM_023245560.1;refGene_gn=ELSPBP1;refGene_tx=XM_023245561.1;refGene_gn=ELSPBP1;refGene_tx=XM_023245562.1;refGene_gn=ELSPBP1;refGene_tx=XM_023245563.1
E2 9542487 9542646
refGene_gn=ELSPBP1;refGene_tx=XM_023245560.1;refGene_gn=ELSPBP1;refGene_tx=XM_023245561.1;refGene_gn=ELSPBP1;refGene_tx=XM_023245562.1;refGene_gn=ELSPBP1;refGene_tx=XM_023245563.1
E2 9544625 9544772
refGene_gn=ELSPBP1;refGene_tx=XM_023245560.1;refGene_gn=ELSPBP1;refGene_tx=XM_023245561.1;refGene_gn=ELSPBP1;refGene_tx=XM_023245562.1;refGene_gn=ELSPBP1;refGene_tx=XM_023245563.1
E2 9550123 9550261
refGene_gn=ELSPBP1;refGene_tx=XM_023245560.1;refGene_gn=ELSPBP1;refGene_tx=XM_023245561.1;refGene_gn=ELSPBP1;refGene_tx=XM_023245562.1;refGene_gn=ELSPBP1;refGene_tx=XM_023245563.1
E2 9556512 9556579
refGene_gn=ELSPBP1;refGene_tx=XM_023245560.1;refGene_gn=ELSPBP1;refGen

e_tx=XM_023245561.1;refGene_gn=ELSPBP1;refGene_tx=XM_023245562.1;refGene_gn=ELSPBP1;refGene_tx=XM_023245563.1
E2 9611947 9612553 refGene_gn=TPRX1;refGene_tx=XM_023245917.1
E2 9613072 9613240 refGene_gn=TPRX1;refGene_tx=XM_023245917.1
E2 9614223 9614367 refGene_gn=TPRX1;refGene_tx=XM_023245917.1
E2 9614758 9614797 refGene_gn=TPRX1;refGene_tx=XM_023245917.1
E2 9624594 9625242 refGene_gn=CRX;refGene_tx=XM_023245410.1
E2 9626407 9626559 refGene_gn=CRX;refGene_tx=XM_023245410.1
E2 9627453 9627553 refGene_gn=CRX;refGene_tx=XM_023245410.1
E2 9634781 9634787 refGene_gn=LOC101092810;refGene_tx=XM_019818980.2
E2 9636526 9636609 refGene_gn=LOC101092810;refGene_tx=XM_019818980.2
E2 9646231 9646397 refGene_gn=LOC101092810;refGene_tx=XM_019818980.2
E2 9646921 9647377 refGene_gn=LOC101092810;refGene_tx=XM_019818980.2
E2 9660894 9660975 refGene_gn=SELENOW;refGene_tx=XM_004001320.5
E2 9661074 9661149 refGene_gn=SELENOW;refGene_tx=XM_004001320.5
E2 9661328 9661382 refGene_gn=SELENOW;refGene_tx=XM_004001320.5
E2 9661488 9661513 refGene_gn=SELENOW;refGene_tx=XM_004001320.5
E2 9662951 9662998 refGene_gn=SELENOW;refGene_tx=XM_004001320.5
E2 9678786 9678793 refGene_gn=NOP53;refGene_tx=XM_023246013.1
E2 9679137 9679198 refGene_gn=NOP53;refGene_tx=XM_023246013.1;refGene_gn=NOP53;refGene_tx=XM_023246014.1
E2 9679294 9679371 refGene_gn=NOP53;refGene_tx=XM_023246013.1;refGene_gn=NOP53;refGene_tx=XM_023246014.1
E2 9679982 9680049 refGene_gn=NOP53;refGene_tx=XM_023246013.1;refGene_gn=NOP53;refGene_tx=XM_023246014.1
E2 9680212 9680388 refGene_gn=NOP53;refGene_tx=XM_023246013.1;refGene_gn=NOP53;refGene_tx=XM_023246014.1
E2 9680770 9680974 refGene_gn=NOP53;refGene_tx=XM_023246013.1;refGene_gn=NOP53;refGene_tx=XM_023246014.1
E2 9681084 9681189 refGene_gn=NOP53;refGene_tx=XM_023246013.1;refGene_gn=NOP53;refGene_tx=XM_023246014.1
E2 9682432 9682528 refGene_gn=NOP53;refGene_tx=XM_023246013.1;refGene_gn=NOP53;refGene_tx=XM_023246014.1
E2 9683800 9683871 refGene_gn=NOP53;refGene_tx=XM_023246013.1;refGene_gn=NOP53;refGene_tx=XM_023246014.1
E2 9684250 9684450 refGene_gn=NOP53;refGene_tx=XM_023246013.1;refGene_gn=NOP53;refGene_tx=XM_023246014.1

E2 9685111 9685220
refGene_gn=NOP53;refGene_tx=XM_023246013.1;refGene_gn=NOP53;refGene_tx
=XM_023246014.1
E2 9687488 9687553
refGene_gn=NOP53;refGene_tx=XM_023246013.1;refGene_gn=NOP53;refGene_tx
=XM_023246014.1
E2 9688035 9688259
refGene_gn=NOP53;refGene_tx=XM_023246013.1;refGene_gn=NOP53;refGene_tx
=XM_023246014.1
E2 9691668 9692220 refGene_gn=EHD2;refGene_tx=XM_003997613.5
E2 9695001 9695166 refGene_gn=EHD2;refGene_tx=XM_003997613.5
E2 9699357 9699770 refGene_gn=EHD2;refGene_tx=XM_003997613.5
E2 9703775 9703873 refGene_gn=EHD2;refGene_tx=XM_003997613.5
E2 9704621 9705025 refGene_gn=EHD2;refGene_tx=XM_003997613.5
E2 9712646 9713306 refGene_gn=BICRA;refGene_tx=XM_023245881.1
E2 9713307 9713780 refGene_gn=BICRA;refGene_tx=XM_023245881.1
E2 9714342 9714445 refGene_gn=BICRA;refGene_tx=XM_023245881.1
E2 9714659 9714754 refGene_gn=BICRA;refGene_tx=XM_023245881.1
E2 9714893 9715042 refGene_gn=BICRA;refGene_tx=XM_023245881.1
E2 9716086 9716139 refGene_gn=BICRA;refGene_tx=XM_023245881.1
E2 9716867 9716977 refGene_gn=BICRA;refGene_tx=XM_023245881.1
E2 9717256 9717437 refGene_gn=BICRA;refGene_tx=XM_023245881.1
E2 9717590 9718166 refGene_gn=BICRA;refGene_tx=XM_023245881.1
E2 9728514 9728691 refGene_gn=BICRA;refGene_tx=XM_023245881.1
E2 9729261 9731208 refGene_gn=BICRA;refGene_tx=XM_023245881.1
E2 9733215 9733281 refGene_gn=BICRA;refGene_tx=XM_023245881.1
E2 9735145 9735188 refGene_gn=BICRA;refGene_tx=XM_023245881.1
E2 9735278 9735319 refGene_gn=BICRA;refGene_tx=XM_023245881.1
E2 9833463 9833770
refGene_gn=ZNF541;refGene_tx=XM_023245225.1;refGene_gn=ZNF541;refGene_
tx=XM_023245226.1;refGene_gn=ZNF541;refGene_tx=XM_023245227.1;refGene_
gn=ZNF541;refGene_tx=XM_023245228.1;refGene_gn=ZNF541;refGene_tx=XM_02
3245229.1
E2 9838014 9838258
refGene_gn=ZNF541;refGene_tx=XM_023245225.1;refGene_gn=ZNF541;refGene_
tx=XM_023245226.1;refGene_gn=ZNF541;refGene_tx=XM_023245227.1;refGene_
gn=ZNF541;refGene_tx=XM_023245228.1;refGene_gn=ZNF541;refGene_tx=XM_02
3245229.1
E2 9840110 9841944
refGene_gn=ZNF541;refGene_tx=XM_023245225.1;refGene_gn=ZNF541;refGene_
tx=XM_023245226.1;refGene_gn=ZNF541;refGene_tx=XM_023245227.1;refGene_
gn=ZNF541;refGene_tx=XM_023245228.1;refGene_gn=ZNF541;refGene_tx=XM_02
3245229.1
E2 9843810 9843869
refGene_gn=ZNF541;refGene_tx=XM_023245225.1;refGene_gn=ZNF541;refGene_
tx=XM_023245226.1;refGene_gn=ZNF541;refGene_tx=XM_023245227.1;refGene_
gn=ZNF541;refGene_tx=XM_023245228.1;refGene_gn=ZNF541;refGene_tx=XM_02
3245229.1
E2 9844163 9844323
refGene_gn=ZNF541;refGene_tx=XM_023245225.1;refGene_gn=ZNF541;refGene_

tx=XM_023245226.1;refGene_gn=ZNF541;refGene_tx=XM_023245227.1;refGene_gn=ZNF541;refGene_tx=XM_023245228.1;refGene_gn=ZNF541;refGene_tx=XM_023245229.1
E2 9844518 9844692
refGene_gn=ZNF541;refGene_tx=XM_023245225.1;refGene_gn=ZNF541;refGene_tx=XM_023245226.1;refGene_gn=ZNF541;refGene_tx=XM_023245227.1;refGene_gn=ZNF541;refGene_tx=XM_023245228.1;refGene_gn=ZNF541;refGene_tx=XM_023245229.1
E2 9845583 9845920
refGene_gn=ZNF541;refGene_tx=XM_023245225.1;refGene_gn=ZNF541;refGene_tx=XM_023245226.1;refGene_gn=ZNF541;refGene_tx=XM_023245227.1;refGene_gn=ZNF541;refGene_tx=XM_023245228.1;refGene_gn=ZNF541;refGene_tx=XM_023245229.1
E2 9852746 9852810
refGene_gn=ZNF541;refGene_tx=XM_023245225.1;refGene_gn=ZNF541;refGene_tx=XM_023245226.1;refGene_gn=ZNF541;refGene_tx=XM_023245227.1;refGene_gn=ZNF541;refGene_tx=XM_023245228.1;refGene_gn=ZNF541;refGene_tx=XM_023245229.1
E2 9853444 9853587
refGene_gn=ZNF541;refGene_tx=XM_023245225.1;refGene_gn=ZNF541;refGene_tx=XM_023245226.1;refGene_gn=ZNF541;refGene_tx=XM_023245227.1;refGene_gn=ZNF541;refGene_tx=XM_023245228.1
E2 9853921 9854025
refGene_gn=ZNF541;refGene_tx=XM_023245225.1;refGene_gn=ZNF541;refGene_tx=XM_023245226.1;refGene_gn=ZNF541;refGene_tx=XM_023245227.1;refGene_gn=ZNF541;refGene_tx=XM_023245228.1;refGene_gn=ZNF541;refGene_tx=XM_023245229.1
E2 9855879 9855955
refGene_gn=ZNF541;refGene_tx=XM_023245225.1;refGene_gn=ZNF541;refGene_tx=XM_023245226.1;refGene_gn=ZNF541;refGene_tx=XM_023245227.1;refGene_gn=ZNF541;refGene_tx=XM_023245228.1;refGene_gn=ZNF541;refGene_tx=XM_023245229.1
E2 9856470 9856559
refGene_gn=ZNF541;refGene_tx=XM_023245225.1;refGene_gn=ZNF541;refGene_tx=XM_023245226.1;refGene_gn=ZNF541;refGene_tx=XM_023245227.1;refGene_gn=ZNF541;refGene_tx=XM_023245228.1;refGene_gn=ZNF541;refGene_tx=XM_023245229.1
E2 9862498 9862639
refGene_gn=ZNF541;refGene_tx=XM_023245225.1;refGene_gn=ZNF541;refGene_tx=XM_023245226.1;refGene_gn=ZNF541;refGene_tx=XM_023245227.1;refGene_gn=ZNF541;refGene_tx=XM_023245229.1
E2 9862804 9862986
refGene_gn=ZNF541;refGene_tx=XM_023245225.1;refGene_gn=ZNF541;refGene_tx=XM_023245226.1;refGene_gn=ZNF541;refGene_tx=XM_023245227.1;refGene_gn=ZNF541;refGene_tx=XM_023245228.1
E2 9863087 9863250
refGene_gn=ZNF541;refGene_tx=XM_023245225.1;refGene_gn=ZNF541;refGene_tx=XM_023245226.1;refGene_gn=ZNF541;refGene_tx=XM_023245227.1;refGene_gn=ZNF541;refGene_tx=XM_023245228.1
E2 9868612 9868710 refGene_gn=NAPA;refGene_tx=XM_003997614.4

E2 9874519 9874545 refGene_gn=NAPA;refGene_tx=XM_006941073.4
E2 9877266 9877346
refGene_gn=NAPA;refGene_tx=XM_003997614.4;refGene_gn=NAPA;refGene_tx=XM_006941073.4
E2 9879934 9880051
refGene_gn=NAPA;refGene_tx=XM_003997614.4;refGene_gn=NAPA;refGene_tx=XM_006941073.4
E2 9884676 9884723
refGene_gn=NAPA;refGene_tx=XM_003997614.4;refGene_gn=NAPA;refGene_tx=XM_006941073.4
E2 9886080 9886158
refGene_gn=NAPA;refGene_tx=XM_003997614.4;refGene_gn=NAPA;refGene_tx=XM_006941073.4
E2 9886383 9886439
refGene_gn=NAPA;refGene_tx=XM_003997614.4;refGene_gn=NAPA;refGene_tx=XM_006941073.4
E2 9886556 9886641
refGene_gn=NAPA;refGene_tx=XM_003997614.4;refGene_gn=NAPA;refGene_tx=XM_006941073.4
E2 9887377 9887482
refGene_gn=NAPA;refGene_tx=XM_003997614.4;refGene_gn=NAPA;refGene_tx=XM_006941073.4
E2 9888560 9888629
refGene_gn=NAPA;refGene_tx=XM_003997614.4;refGene_gn=NAPA;refGene_tx=XM_006941073.4
E2 9889032 9889083
refGene_gn=NAPA;refGene_tx=XM_003997614.4;refGene_gn=NAPA;refGene_tx=XM_006941073.4
E2 9889983 9890085
refGene_gn=NAPA;refGene_tx=XM_003997614.4;refGene_gn=NAPA;refGene_tx=XM_006941073.4
E2 9893972 9894195
refGene_gn=KPTN;refGene_tx=XM_006941074.4;refGene_gn=KPTN;refGene_tx=XM_006941075.4;refGene_gn=KPTN;refGene_tx=XM_006941077.4
E2 9894532 9894615
refGene_gn=KPTN;refGene_tx=XM_006941074.4;refGene_gn=KPTN;refGene_tx=XM_006941075.4;refGene_gn=KPTN;refGene_tx=XM_006941077.4
E2 9894737 9894822
refGene_gn=KPTN;refGene_tx=XM_006941074.4;refGene_gn=KPTN;refGene_tx=XM_006941075.4;refGene_gn=KPTN;refGene_tx=XM_006941077.4
E2 9894916 9894971
refGene_gn=KPTN;refGene_tx=XM_006941074.4;refGene_gn=KPTN;refGene_tx=XM_006941075.4;refGene_gn=KPTN;refGene_tx=XM_006941077.4
E2 9896170 9896306
refGene_gn=KPTN;refGene_tx=XM_006941074.4;refGene_gn=KPTN;refGene_tx=XM_006941075.4;refGene_gn=KPTN;refGene_tx=XM_006941077.4
E2 9896436 9896510
refGene_gn=KPTN;refGene_tx=XM_006941074.4;refGene_gn=KPTN;refGene_tx=XM_006941075.4;refGene_gn=KPTN;refGene_tx=XM_006941077.4
E2 9896831 9896941

refGene_gn=KPTN;refGene_tx=XM_006941074.4;refGene_gn=KPTN;refGene_tx=XM_006941075.4;refGene_gn=KPTN;refGene_tx=XM_006941077.4
E2 9897021 9897099
refGene_gn=KPTN;refGene_tx=XM_006941074.4;refGene_gn=KPTN;refGene_tx=XM_006941075.4;refGene_gn=KPTN;refGene_tx=XM_006941077.4
E2 9897712 9897788
refGene_gn=KPTN;refGene_tx=XM_006941074.4;refGene_gn=KPTN;refGene_tx=XM_006941075.4;refGene_gn=KPTN;refGene_tx=XM_006941077.4
E2 9898474 9898610
refGene_gn=KPTN;refGene_tx=XM_006941074.4;refGene_gn=KPTN;refGene_tx=XM_006941075.4;refGene_gn=KPTN;refGene_tx=XM_006941077.4
E2 9898700 9898883
refGene_gn=KPTN;refGene_tx=XM_006941074.4;refGene_gn=KPTN;refGene_tx=XM_006941075.4;refGene_gn=KPTN;refGene_tx=XM_006941077.4
E2 9900154 9900200 refGene_gn=KPTN;refGene_tx=XM_006941074.4
E2 9900632 9900764
refGene_gn=KPTN;refGene_tx=XM_006941074.4;refGene_gn=KPTN;refGene_tx=XM_006941075.4;refGene_gn=KPTN;refGene_tx=XM_006941077.4
E2 9907518 9908193
refGene_gn=SLC8A2;refGene_tx=XM_023245973.1;refGene_gn=SLC8A2;refGene_tx=XM_023245974.1
E2 9911738 9912403
refGene_gn=SLC8A2;refGene_tx=XM_023245973.1;refGene_gn=SLC8A2;refGene_tx=XM_023245974.1
E2 9919562 9919985
refGene_gn=SLC8A2;refGene_tx=XM_023245973.1;refGene_gn=SLC8A2;refGene_tx=XM_023245974.1
E2 9924026 9924130
refGene_gn=SLC8A2;refGene_tx=XM_023245973.1;refGene_gn=SLC8A2;refGene_tx=XM_023245974.1
E2 9924291 9924309
refGene_gn=SLC8A2;refGene_tx=XM_023245973.1;refGene_gn=SLC8A2;refGene_tx=XM_023245974.1
E2 9925560 9925685
refGene_gn=SLC8A2;refGene_tx=XM_023245973.1;refGene_gn=SLC8A2;refGene_tx=XM_023245974.1
E2 9925957 9926057
refGene_gn=SLC8A2;refGene_tx=XM_023245973.1;refGene_gn=SLC8A2;refGene_tx=XM_023245974.1
E2 9930022 9930301
refGene_gn=SLC8A2;refGene_tx=XM_023245973.1;refGene_gn=SLC8A2;refGene_tx=XM_023245974.1
E2 9931451 9931828
refGene_gn=SLC8A2;refGene_tx=XM_023245973.1;refGene_gn=SLC8A2;refGene_tx=XM_023245974.1
E2 9948590 9948602
refGene_gn=MEIS3;refGene_tx=XM_023245975.1;refGene_gn=MEIS3;refGene_tx=XM_023245976.1
E2 9950275 9950457
refGene_gn=MEIS3;refGene_tx=XM_023245975.1;refGene_gn=MEIS3;refGene_tx

=XM_023245976.1
E2	9950629	9950789
refGene_gn=MEIS3;refGene_tx=XM_023245975.1;refGene_gn=MEIS3;refGene_tx=XM_023245976.1
E2	9950884	9950935
refGene_gn=MEIS3;refGene_tx=XM_023245975.1;refGene_gn=MEIS3;refGene_tx=XM_023245976.1
E2	9951890	9951941
refGene_gn=MEIS3;refGene_tx=XM_023245975.1;refGene_gn=MEIS3;refGene_tx=XM_023245976.1
E2	9952126	9952276
refGene_gn=MEIS3;refGene_tx=XM_023245975.1;refGene_gn=MEIS3;refGene_tx=XM_023245976.1
E2	9954563	9954675
refGene_gn=MEIS3;refGene_tx=XM_023245975.1;refGene_gn=MEIS3;refGene_tx=XM_023245976.1
E2	9954860	9955006
refGene_gn=MEIS3;refGene_tx=XM_023245975.1;refGene_gn=MEIS3;refGene_tx=XM_023245976.1
E2	9955908	9955985
refGene_gn=MEIS3;refGene_tx=XM_023245975.1;refGene_gn=MEIS3;refGene_tx=XM_023245976.1
E2	9956202	9956261
refGene_gn=MEIS3;refGene_tx=XM_023245975.1;refGene_gn=MEIS3;refGene_tx=XM_023245976.1
E2	9956368	9956452
refGene_gn=MEIS3;refGene_tx=XM_023245975.1;refGene_gn=MEIS3;refGene_tx=XM_023245976.1
E2	9956814	9956864
refGene_gn=MEIS3;refGene_tx=XM_023245975.1;refGene_gn=MEIS3;refGene_tx=XM_023245976.1
E2	9968922	9969056
refGene_gn=DHX34;refGene_tx=XM_019819292.2;refGene_gn=DHX34;refGene_tx=XM_019819293.2
E2	9969719	9969858
refGene_gn=DHX34;refGene_tx=XM_019819292.2;refGene_gn=DHX34;refGene_tx=XM_019819293.2
E2	9970088	9970265
refGene_gn=DHX34;refGene_tx=XM_019819292.2;refGene_gn=DHX34;refGene_tx=XM_019819293.2
E2	9971069	9971360
refGene_gn=DHX34;refGene_tx=XM_019819292.2;refGene_gn=DHX34;refGene_tx=XM_019819293.2
E2	9973430	9973537
refGene_gn=DHX34;refGene_tx=XM_019819292.2;refGene_gn=DHX34;refGene_tx=XM_019819293.2
E2	9975419	9975537
refGene_gn=DHX34;refGene_tx=XM_019819292.2;refGene_gn=DHX34;refGene_tx=XM_019819293.2
E2	9975910	9976084

refGene_gn=DHX34;refGene_tx=XM_019819292.2;refGene_gn=DHX34;refGene_tx=XM_019819293.2
E2 9976298 9976523
refGene_gn=DHX34;refGene_tx=XM_019819292.2;refGene_gn=DHX34;refGene_tx=XM_019819293.2
E2 9977548 9977650
refGene_gn=DHX34;refGene_tx=XM_019819292.2;refGene_gn=DHX34;refGene_tx=XM_019819293.2
E2 9978325 9978519
refGene_gn=DHX34;refGene_tx=XM_019819292.2;refGene_gn=DHX34;refGene_tx=XM_019819293.2
E2 9980213 9980388
refGene_gn=DHX34;refGene_tx=XM_019819292.2;refGene_gn=DHX34;refGene_tx=XM_019819293.2
E2 9982359 9982577
refGene_gn=DHX34;refGene_tx=XM_019819292.2;refGene_gn=DHX34;refGene_tx=XM_019819293.2
E2 9983783 9983886
refGene_gn=DHX34;refGene_tx=XM_019819292.2;refGene_gn=DHX34;refGene_tx=XM_019819293.2
E2 9984957 9985212
refGene_gn=DHX34;refGene_tx=XM_019819292.2;refGene_gn=DHX34;refGene_tx=XM_019819293.2
E2 9986649 9986961
refGene_gn=DHX34;refGene_tx=XM_019819292.2;refGene_gn=DHX34;refGene_tx=XM_019819293.2
E2 9988201 9988903
refGene_gn=DHX34;refGene_tx=XM_019819292.2;refGene_gn=DHX34;refGene_tx=XM_019819293.2
E2 9998274 9998310 refGene_gn=C5AR2;refGene_tx=XM_006941078.4
E2 9998311 9999304 refGene_gn=C5AR2;refGene_tx=XM_006941078.4
E2 10004385 10004484 refGene_gn=C5AR2;refGene_tx=XM_006941078.4
E2 10012028 10013081
refGene_gn=C5AR1;refGene_tx=XM_006941080.3;refGene_gn=C5AR1;refGene_tx=XM_023245544.1
E2 10020315 10020318 refGene_gn=C5AR1;refGene_tx=XM_006941080.3
E2 10042556 10042724 refGene_gn=INAFM1;refGene_tx=XM_023244947.1
E2 10042725 10042992 refGene_gn=INAFM1;refGene_tx=XM_023244947.1
E2 10046183 10046615
refGene_gn=CCDC9;refGene_tx=XM_023244945.1;refGene_gn=CCDC9;refGene_tx=XM_023244946.1
E2 10046702 10046793
refGene_gn=CCDC9;refGene_tx=XM_023244945.1;refGene_gn=CCDC9;refGene_tx=XM_023244946.1
E2 10047246 10047379
refGene_gn=CCDC9;refGene_tx=XM_023244945.1;refGene_gn=CCDC9;refGene_tx=XM_023244946.1
E2 10047487 10047534
refGene_gn=CCDC9;refGene_tx=XM_023244945.1;refGene_gn=CCDC9;refGene_tx=XM_023244946.1

E2 10049287 10049469
refGene_gn=CCDC9;refGene_tx=XM_023244945.1;refGene_gn=CCDC9;refGene_tx=XM_023244946.1
E2 10050276 10050450
refGene_gn=CCDC9;refGene_tx=XM_023244945.1;refGene_gn=CCDC9;refGene_tx=XM_023244946.1
E2 10050544 10050628
refGene_gn=CCDC9;refGene_tx=XM_023244945.1;refGene_gn=CCDC9;refGene_tx=XM_023244946.1
E2 10052276 10052525
refGene_gn=CCDC9;refGene_tx=XM_023244945.1;refGene_gn=CCDC9;refGene_tx=XM_023244946.1
E2 10052738 10052840
refGene_gn=CCDC9;refGene_tx=XM_023244945.1;refGene_gn=CCDC9;refGene_tx=XM_023244946.1
E2 10054646 10054798
refGene_gn=CCDC9;refGene_tx=XM_023244945.1;refGene_gn=CCDC9;refGene_tx=XM_023244946.1
E2 10054945 10055019
refGene_gn=CCDC9;refGene_tx=XM_023244945.1;refGene_gn=CCDC9;refGene_tx=XM_023244946.1
E2 10055454 10055468 refGene_gn=CCDC9;refGene_tx=XM_023244945.1
E2 10077932 10078206
refGene_gn=BBC3;refGene_tx=XM_023245381.1;refGene_gn=BBC3;refGene_tx=XM_023245383.1
E2 10079678 10079869
refGene_gn=BBC3;refGene_tx=XM_023245381.1;refGene_gn=BBC3;refGene_tx=XM_023245383.1
E2 10082197 10082314
refGene_gn=BBC3;refGene_tx=XM_023245381.1;refGene_gn=BBC3;refGene_tx=XM_023245383.1
E2 10091573 10091666
refGene_gn=SAE1;refGene_tx=XM_003997616.5;refGene_gn=SAE1;refGene_tx=XM_019819295.2;refGene_gn=SAE1;refGene_tx=XM_019819296.2
E2 10095913 10095983
refGene_gn=SAE1;refGene_tx=XM_003997616.5;refGene_gn=SAE1;refGene_tx=XM_019819296.2
E2 10107143 10107288
refGene_gn=SAE1;refGene_tx=XM_003997616.5;refGene_gn=SAE1;refGene_tx=XM_019819295.2;refGene_gn=SAE1;refGene_tx=XM_019819296.2
E2 10130827 10130933
refGene_gn=SAE1;refGene_tx=XM_003997616.5;refGene_gn=SAE1;refGene_tx=XM_019819295.2;refGene_gn=SAE1;refGene_tx=XM_019819296.2
E2 10142644 10142860 refGene_gn=SAE1;refGene_tx=XM_019819297.2
E2 10143117 10143217
refGene_gn=SAE1;refGene_tx=XM_003997616.5;refGene_gn=SAE1;refGene_tx=XM_019819295.2;refGene_gn=SAE1;refGene_tx=XM_019819296.2;refGene_gn=SAE1;refGene_tx=XM_019819297.2
E2 10145484 10145627
refGene_gn=SAE1;refGene_tx=XM_003997616.5;refGene_gn=SAE1;refGene_tx=X

M_019819295.2;refGene_gn=SAE1;refGene_tx=XM_019819296.2;refGene_gn=SAE1;refGene_tx=XM_019819297.2
E2 10147728 10147902
refGene_gn=SAE1;refGene_tx=XM_003997616.5;refGene_gn=SAE1;refGene_tx=XM_019819295.2;refGene_gn=SAE1;refGene_tx=XM_019819297.2
E2 10153839 10153951
refGene_gn=SAE1;refGene_tx=XM_003997616.5;refGene_gn=SAE1;refGene_tx=XM_019819295.2;refGene_gn=SAE1;refGene_tx=XM_019819296.2;refGene_gn=SAE1;refGene_tx=XM_019819297.2
E2 10166806 10166904
refGene_gn=SAE1;refGene_tx=XM_003997616.5;refGene_gn=SAE1;refGene_tx=XM_019819295.2;refGene_gn=SAE1;refGene_tx=XM_019819296.2;refGene_gn=SAE1;refGene_tx=XM_019819297.2
E2 10187095 10187256
refGene_gn=ZC3H4;refGene_tx=XM_023245812.1;refGene_gn=ZC3H4;refGene_tx=XM_023245813.1
E2 10187330 10187516
refGene_gn=ZC3H4;refGene_tx=XM_023245811.1;refGene_gn=ZC3H4;refGene_tx=XM_023245814.1;refGene_gn=ZC3H4;refGene_tx=XM_023245818.1
E2 10188695 10188757
refGene_gn=ZC3H4;refGene_tx=XM_023245811.1;refGene_gn=ZC3H4;refGene_tx=XM_023245816.1;refGene_gn=ZC3H4;refGene_tx=XM_023245818.1
E2 10189023 10189046 refGene_gn=ZC3H4;refGene_tx=XM_023245817.1
E2 10204368 10204588
refGene_gn=ZC3H4;refGene_tx=XM_023245811.1;refGene_gn=ZC3H4;refGene_tx=XM_023245812.1;refGene_gn=ZC3H4;refGene_tx=XM_023245813.1;refGene_gn=ZC3H4;refGene_tx=XM_023245814.1;refGene_gn=ZC3H4;refGene_tx=XM_023245816.1;refGene_gn=ZC3H4;refGene_tx=XM_023245817.1;refGene_gn=ZC3H4;refGene_tx=XM_023245818.1
E2 10204897 10205008
refGene_gn=ZC3H4;refGene_tx=XM_023245811.1;refGene_gn=ZC3H4;refGene_tx=XM_023245812.1;refGene_gn=ZC3H4;refGene_tx=XM_023245813.1;refGene_gn=ZC3H4;refGene_tx=XM_023245814.1;refGene_gn=ZC3H4;refGene_tx=XM_023245816.1;refGene_gn=ZC3H4;refGene_tx=XM_023245817.1;refGene_gn=ZC3H4;refGene_tx=XM_023245818.1
E2 10208140 10208363
refGene_gn=ZC3H4;refGene_tx=XM_023245811.1;refGene_gn=ZC3H4;refGene_tx=XM_023245812.1;refGene_gn=ZC3H4;refGene_tx=XM_023245813.1;refGene_gn=ZC3H4;refGene_tx=XM_023245814.1;refGene_gn=ZC3H4;refGene_tx=XM_023245816.1;refGene_gn=ZC3H4;refGene_tx=XM_023245817.1;refGene_gn=ZC3H4;refGene_tx=XM_023245818.1
E2 10210961 10211113
refGene_gn=ZC3H4;refGene_tx=XM_023245811.1;refGene_gn=ZC3H4;refGene_tx=XM_023245812.1;refGene_gn=ZC3H4;refGene_tx=XM_023245813.1;refGene_gn=ZC3H4;refGene_tx=XM_023245814.1;refGene_gn=ZC3H4;refGene_tx=XM_023245816.1;refGene_gn=ZC3H4;refGene_tx=XM_023245817.1;refGene_gn=ZC3H4;refGene_tx=XM_023245818.1
E2 10211779 10211876
refGene_gn=ZC3H4;refGene_tx=XM_023245811.1;refGene_gn=ZC3H4;refGene_tx=XM_023245812.1;refGene_gn=ZC3H4;refGene_tx=XM_023245813.1;refGene_gn=

ZC3H4;refGene_tx=XM_023245814.1;refGene_gn=ZC3H4;refGene_tx=XM_0232458
16.1;refGene_gn=ZC3H4;refGene_tx=XM_023245817.1;refGene_gn=ZC3H4;refGe
ne_tx=XM_023245818.1
E2 10211988 10212128
refGene_gn=ZC3H4;refGene_tx=XM_023245811.1;refGene_gn=ZC3H4;refGene_tx
=XM_023245812.1;refGene_gn=ZC3H4;refGene_tx=XM_023245813.1;refGene_gn=
ZC3H4;refGene_tx=XM_023245814.1;refGene_gn=ZC3H4;refGene_tx=XM_0232458
16.1;refGene_gn=ZC3H4;refGene_tx=XM_023245817.1;refGene_gn=ZC3H4;refGe
ne_tx=XM_023245818.1
E2 10212688 10212796
refGene_gn=ZC3H4;refGene_tx=XM_023245811.1;refGene_gn=ZC3H4;refGene_tx
=XM_023245812.1;refGene_gn=ZC3H4;refGene_tx=XM_023245813.1;refGene_gn=
ZC3H4;refGene_tx=XM_023245814.1;refGene_gn=ZC3H4;refGene_tx=XM_0232458
16.1;refGene_gn=ZC3H4;refGene_tx=XM_023245817.1;refGene_gn=ZC3H4;refGe
ne_tx=XM_023245818.1
E2 10214157 10214269
refGene_gn=ZC3H4;refGene_tx=XM_023245811.1;refGene_gn=ZC3H4;refGene_tx
=XM_023245812.1;refGene_gn=ZC3H4;refGene_tx=XM_023245813.1;refGene_gn=
ZC3H4;refGene_tx=XM_023245814.1;refGene_gn=ZC3H4;refGene_tx=XM_0232458
16.1;refGene_gn=ZC3H4;refGene_tx=XM_023245817.1;refGene_gn=ZC3H4;refGe
ne_tx=XM_023245818.1
E2 10214786 10214896
refGene_gn=ZC3H4;refGene_tx=XM_023245811.1;refGene_gn=ZC3H4;refGene_tx
=XM_023245812.1;refGene_gn=ZC3H4;refGene_tx=XM_023245813.1;refGene_gn=
ZC3H4;refGene_tx=XM_023245814.1;refGene_gn=ZC3H4;refGene_tx=XM_0232458
16.1;refGene_gn=ZC3H4;refGene_tx=XM_023245817.1;refGene_gn=ZC3H4;refGe
ne_tx=XM_023245818.1
E2 10221381 10221746
refGene_gn=ZC3H4;refGene_tx=XM_023245811.1;refGene_gn=ZC3H4;refGene_tx
=XM_023245812.1;refGene_gn=ZC3H4;refGene_tx=XM_023245813.1;refGene_gn=
ZC3H4;refGene_tx=XM_023245814.1;refGene_gn=ZC3H4;refGene_tx=XM_0232458
16.1;refGene_gn=ZC3H4;refGene_tx=XM_023245817.1;refGene_gn=ZC3H4;refGe
ne_tx=XM_023245818.1
E2 10221943 10222275
refGene_gn=ZC3H4;refGene_tx=XM_023245811.1;refGene_gn=ZC3H4;refGene_tx
=XM_023245812.1;refGene_gn=ZC3H4;refGene_tx=XM_023245813.1;refGene_gn=
ZC3H4;refGene_tx=XM_023245814.1;refGene_gn=ZC3H4;refGene_tx=XM_0232458
16.1;refGene_gn=ZC3H4;refGene_tx=XM_023245817.1;refGene_gn=ZC3H4;refGe
ne_tx=XM_023245818.1
E2 10225169 10225421
refGene_gn=ZC3H4;refGene_tx=XM_023245811.1;refGene_gn=ZC3H4;refGene_tx
=XM_023245812.1;refGene_gn=ZC3H4;refGene_tx=XM_023245813.1;refGene_gn=
ZC3H4;refGene_tx=XM_023245814.1;refGene_gn=ZC3H4;refGene_tx=XM_0232458
16.1;refGene_gn=ZC3H4;refGene_tx=XM_023245817.1
E2 10226609 10228129
refGene_gn=ZC3H4;refGene_tx=XM_023245811.1;refGene_gn=ZC3H4;refGene_tx
=XM_023245812.1;refGene_gn=ZC3H4;refGene_tx=XM_023245813.1;refGene_gn=
ZC3H4;refGene_tx=XM_023245814.1;refGene_gn=ZC3H4;refGene_tx=XM_0232458
16.1;refGene_gn=ZC3H4;refGene_tx=XM_023245817.1;refGene_gn=ZC3H4;refGe
ne_tx=XM_023245818.1

E2 10245347 10245555 refGene_gn=TMEM160;refGene_tx=XM_003997619.3
E2 10246841 10246934 refGene_gn=TMEM160;refGene_tx=XM_003997619.3
E2 10247602 10247868 refGene_gn=TMEM160;refGene_tx=XM_003997619.3
E2 10248184 10248648
refGene_gn=NPAS1;refGene_tx=XM_011289788.3;refGene_gn=NPAS1;refGene_tx
=XM_011289789.3;refGene_gn=NPAS1;refGene_tx=XM_011289791.3
E2 10248978 10249073
refGene_gn=NPAS1;refGene_tx=XM_011289788.3;refGene_gn=NPAS1;refGene_tx
=XM_011289789.3;refGene_gn=NPAS1;refGene_tx=XM_011289791.3
E2 10249733 10249881
refGene_gn=NPAS1;refGene_tx=XM_011289788.3;refGene_gn=NPAS1;refGene_tx
=XM_011289789.3;refGene_gn=NPAS1;refGene_tx=XM_011289791.3
E2 10250202 10250309
refGene_gn=NPAS1;refGene_tx=XM_011289788.3;refGene_gn=NPAS1;refGene_tx
=XM_011289789.3;refGene_gn=NPAS1;refGene_tx=XM_011289791.3
E2 10250758 10250916
refGene_gn=NPAS1;refGene_tx=XM_011289788.3;refGene_gn=NPAS1;refGene_tx
=XM_011289789.3;refGene_gn=NPAS1;refGene_tx=XM_011289791.3
E2 10251086 10251202
refGene_gn=NPAS1;refGene_tx=XM_011289788.3;refGene_gn=NPAS1;refGene_tx
=XM_011289789.3;refGene_gn=NPAS1;refGene_tx=XM_011289791.3
E2 10252573 10252745
refGene_gn=NPAS1;refGene_tx=XM_011289788.3;refGene_gn=NPAS1;refGene_tx
=XM_011289789.3;refGene_gn=NPAS1;refGene_tx=XM_011289791.3
E2 10254636 10254726
refGene_gn=NPAS1;refGene_tx=XM_011289788.3;refGene_gn=NPAS1;refGene_tx
=XM_011289789.3;refGene_gn=NPAS1;refGene_tx=XM_011289791.3
E2 10254985 10255059
refGene_gn=NPAS1;refGene_tx=XM_011289788.3;refGene_gn=NPAS1;refGene_tx
=XM_011289789.3;refGene_gn=NPAS1;refGene_tx=XM_011289791.3
E2 10261164 10261400
refGene_gn=NPAS1;refGene_tx=XM_011289788.3;refGene_gn=NPAS1;refGene_tx
=XM_011289789.3;refGene_gn=NPAS1;refGene_tx=XM_011289791.3
E2 10261828 10261950
refGene_gn=NPAS1;refGene_tx=XM_011289788.3;refGene_gn=NPAS1;refGene_tx
=XM_011289789.3;refGene_gn=NPAS1;refGene_tx=XM_011289791.3
E2 10277747 10278105 refGene_gn=ARHGAP35;refGene_tx=XM_023245327.1
E2 10278887 10278993 refGene_gn=ARHGAP35;refGene_tx=XM_023245327.1
E2 10285976 10286108 refGene_gn=ARHGAP35;refGene_tx=XM_023245327.1
E2 10287718 10287796 refGene_gn=ARHGAP35;refGene_tx=XM_023245327.1
E2 10337930 10338075 refGene_gn=ARHGAP35;refGene_tx=XM_023245327.1
E2 10349657 10353341 refGene_gn=ARHGAP35;refGene_tx=XM_023245327.1
E2 10411312 10411315 refGene_gn=AP2S1;refGene_tx=XM_023245338.1
E2 10415435 10415585 refGene_gn=AP2S1;refGene_tx=XM_023245338.1
E2 10421361 10421475 refGene_gn=AP2S1;refGene_tx=XM_023245338.1
E2 10421827 10421887 refGene_gn=AP2S1;refGene_tx=XM_023245338.1
E2 10422072 10422174 refGene_gn=AP2S1;refGene_tx=XM_023245338.1
E2 10453498 10454067 refGene_gn=SLC1A5;refGene_tx=NM_001290234.1
E2 10456950 10456993 refGene_gn=SLC1A5;refGene_tx=NM_001290234.1
E2 10457426 10457474 refGene_gn=SLC1A5;refGene_tx=NM_001290234.1

E2 10458360 10458527 refGene_gn=SLC1A5;refGene_tx=NM_001290234.1
E2 10461099 10461333 refGene_gn=SLC1A5;refGene_tx=NM_001290234.1
E2 10462112 10462307 refGene_gn=SLC1A5;refGene_tx=NM_001290234.1
E2 10462450 10462585 refGene_gn=SLC1A5;refGene_tx=NM_001290234.1
E2 10463781 10464016 refGene_gn=SLC1A5;refGene_tx=NM_001290234.1
E2 10470847 10472335 refGene_gn=FKRP;refGene_tx=XM_023245662.1
E2 10478778 10479060
refGene_gn=STRN4;refGene_tx=XM_023245655.1;refGene_gn=STRN4;refGene_tx
=XM_023245656.1;refGene_gn=STRN4;refGene_tx=XM_023245657.1;refGene_gn=
STRN4;refGene_tx=XM_023245658.1
E2 10485604 10485613 refGene_gn=STRN4;refGene_tx=XM_023245659.1
E2 10486458 10486562
refGene_gn=STRN4;refGene_tx=XM_023245655.1;refGene_gn=STRN4;refGene_tx
=XM_023245656.1;refGene_gn=STRN4;refGene_tx=XM_023245657.1;refGene_gn=
STRN4;refGene_tx=XM_023245658.1;refGene_gn=STRN4;refGene_tx=XM_0232456
59.1;refGene_gn=STRN4;refGene_tx=XM_023245660.1
E2 10487089 10487163
refGene_gn=STRN4;refGene_tx=XM_023245655.1;refGene_gn=STRN4;refGene_tx
=XM_023245656.1;refGene_gn=STRN4;refGene_tx=XM_023245657.1;refGene_gn=
STRN4;refGene_tx=XM_023245658.1;refGene_gn=STRN4;refGene_tx=XM_0232456
59.1;refGene_gn=STRN4;refGene_tx=XM_023245660.1
E2 10488201 10488280
refGene_gn=STRN4;refGene_tx=XM_023245655.1;refGene_gn=STRN4;refGene_tx
=XM_023245656.1;refGene_gn=STRN4;refGene_tx=XM_023245657.1;refGene_gn=
STRN4;refGene_tx=XM_023245658.1;refGene_gn=STRN4;refGene_tx=XM_0232456
59.1;refGene_gn=STRN4;refGene_tx=XM_023245660.1
E2 10491080 10491278
refGene_gn=STRN4;refGene_tx=XM_023245655.1;refGene_gn=STRN4;refGene_tx
=XM_023245656.1;refGene_gn=STRN4;refGene_tx=XM_023245657.1;refGene_gn=
STRN4;refGene_tx=XM_023245658.1;refGene_gn=STRN4;refGene_tx=XM_0232456
59.1;refGene_gn=STRN4;refGene_tx=XM_023245660.1
E2 10493179 10493321
refGene_gn=STRN4;refGene_tx=XM_023245655.1;refGene_gn=STRN4;refGene_tx
=XM_023245656.1;refGene_gn=STRN4;refGene_tx=XM_023245657.1;refGene_gn=
STRN4;refGene_tx=XM_023245658.1;refGene_gn=STRN4;refGene_tx=XM_0232456
59.1;refGene_gn=STRN4;refGene_tx=XM_023245660.1
E2 10496513 10496673
refGene_gn=STRN4;refGene_tx=XM_023245655.1;refGene_gn=STRN4;refGene_tx
=XM_023245656.1;refGene_gn=STRN4;refGene_tx=XM_023245657.1;refGene_gn=
STRN4;refGene_tx=XM_023245658.1;refGene_gn=STRN4;refGene_tx=XM_0232456
59.1;refGene_gn=STRN4;refGene_tx=XM_023245660.1
E2 10497226 10497361
refGene_gn=STRN4;refGene_tx=XM_023245655.1;refGene_gn=STRN4;refGene_tx
=XM_023245656.1;refGene_gn=STRN4;refGene_tx=XM_023245657.1;refGene_gn=
STRN4;refGene_tx=XM_023245658.1;refGene_gn=STRN4;refGene_tx=XM_0232456
59.1;refGene_gn=STRN4;refGene_tx=XM_023245660.1
E2 10497695 10497811
refGene_gn=STRN4;refGene_tx=XM_023245655.1;refGene_gn=STRN4;refGene_tx
=XM_023245656.1;refGene_gn=STRN4;refGene_tx=XM_023245657.1;refGene_gn=
STRN4;refGene_tx=XM_023245658.1;refGene_gn=STRN4;refGene_tx=XM_0232456

59.1;refGene_gn=STRN4;refGene_tx=XM_023245660.1
E2 10499360 10499536
refGene_gn=STRN4;refGene_tx=XM_023245655.1;refGene_gn=STRN4;refGene_tx=XM_023245656.1;refGene_gn=STRN4;refGene_tx=XM_023245657.1;refGene_gn=STRN4;refGene_tx=XM_023245658.1;refGene_gn=STRN4;refGene_tx=XM_023245659.1;refGene_gn=STRN4;refGene_tx=XM_023245660.1
E2 10499613 10499661
refGene_gn=STRN4;refGene_tx=XM_023245655.1;refGene_gn=STRN4;refGene_tx=XM_023245656.1;refGene_gn=STRN4;refGene_tx=XM_023245657.1;refGene_gn=STRN4;refGene_tx=XM_023245658.1;refGene_gn=STRN4;refGene_tx=XM_023245659.1;refGene_gn=STRN4;refGene_tx=XM_023245660.1
E2 10499971 10500093
refGene_gn=STRN4;refGene_tx=XM_023245655.1;refGene_gn=STRN4;refGene_tx=XM_023245656.1;refGene_gn=STRN4;refGene_tx=XM_023245657.1;refGene_gn=STRN4;refGene_tx=XM_023245658.1;refGene_gn=STRN4;refGene_tx=XM_023245659.1;refGene_gn=STRN4;refGene_tx=XM_023245660.1
E2 10501279 10501450
refGene_gn=STRN4;refGene_tx=XM_023245655.1;refGene_gn=STRN4;refGene_tx=XM_023245656.1;refGene_gn=STRN4;refGene_tx=XM_023245657.1;refGene_gn=STRN4;refGene_tx=XM_023245658.1;refGene_gn=STRN4;refGene_tx=XM_023245659.1;refGene_gn=STRN4;refGene_tx=XM_023245660.1
E2 10501608 10501749
refGene_gn=STRN4;refGene_tx=XM_023245655.1;refGene_gn=STRN4;refGene_tx=XM_023245656.1;refGene_gn=STRN4;refGene_tx=XM_023245657.1;refGene_gn=STRN4;refGene_tx=XM_023245658.1;refGene_gn=STRN4;refGene_tx=XM_023245659.1;refGene_gn=STRN4;refGene_tx=XM_023245660.1
E2 10502190 10502289
refGene_gn=STRN4;refGene_tx=XM_023245655.1;refGene_gn=STRN4;refGene_tx=XM_023245656.1;refGene_gn=STRN4;refGene_tx=XM_023245657.1;refGene_gn=STRN4;refGene_tx=XM_023245658.1;refGene_gn=STRN4;refGene_tx=XM_023245659.1;refGene_gn=STRN4;refGene_tx=XM_023245660.1
E2 10502401 10502488
refGene_gn=STRN4;refGene_tx=XM_023245655.1;refGene_gn=STRN4;refGene_tx=XM_023245656.1;refGene_gn=STRN4;refGene_tx=XM_023245657.1;refGene_gn=STRN4;refGene_tx=XM_023245658.1;refGene_gn=STRN4;refGene_tx=XM_023245659.1;refGene_gn=STRN4;refGene_tx=XM_023245660.1
E2 10503789 10503959
refGene_gn=STRN4;refGene_tx=XM_023245655.1;refGene_gn=STRN4;refGene_tx=XM_023245656.1;refGene_gn=STRN4;refGene_tx=XM_023245657.1;refGene_gn=STRN4;refGene_tx=XM_023245658.1;refGene_gn=STRN4;refGene_tx=XM_023245659.1;refGene_gn=STRN4;refGene_tx=XM_023245660.1
E2 10508515 10508755
refGene_gn=PRKD2;refGene_tx=XM_011289800.3;refGene_gn=PRKD2;refGene_tx=XM_019819304.2
E2 10510858 10510997
refGene_gn=PRKD2;refGene_tx=XM_011289800.3;refGene_gn=PRKD2;refGene_tx=XM_019819304.2
E2 10515773 10515905
refGene_gn=PRKD2;refGene_tx=XM_011289800.3;refGene_gn=PRKD2;refGene_tx=XM_019819304.2;refGene_gn=PRKD2;refGene_tx=XM_019819305.2

E2 10518971 10519126
refGene_gn=PRKD2;refGene_tx=XM_011289800.3;refGene_gn=PRKD2;refGene_tx=XM_019819304.2;refGene_gn=PRKD2;refGene_tx=XM_019819305.2
E2 10519222 10519445
refGene_gn=PRKD2;refGene_tx=XM_011289800.3;refGene_gn=PRKD2;refGene_tx=XM_019819304.2;refGene_gn=PRKD2;refGene_tx=XM_019819305.2
E2 10521090 10521168
refGene_gn=PRKD2;refGene_tx=XM_011289800.3;refGene_gn=PRKD2;refGene_tx=XM_019819304.2;refGene_gn=PRKD2;refGene_tx=XM_019819305.2
E2 10521246 10521400
refGene_gn=PRKD2;refGene_tx=XM_011289800.3;refGene_gn=PRKD2;refGene_tx=XM_019819304.2;refGene_gn=PRKD2;refGene_tx=XM_019819305.2
E2 10525307 10525425
refGene_gn=PRKD2;refGene_tx=XM_011289800.3;refGene_gn=PRKD2;refGene_tx=XM_019819304.2;refGene_gn=PRKD2;refGene_tx=XM_019819305.2
E2 10526168 10526246
refGene_gn=PRKD2;refGene_tx=XM_011289800.3;refGene_gn=PRKD2;refGene_tx=XM_019819304.2;refGene_gn=PRKD2;refGene_tx=XM_019819305.2
E2 10527050 10527309
refGene_gn=PRKD2;refGene_tx=XM_011289800.3;refGene_gn=PRKD2;refGene_tx=XM_019819304.2;refGene_gn=PRKD2;refGene_tx=XM_019819305.2
E2 10528891 10528944
refGene_gn=PRKD2;refGene_tx=XM_011289800.3;refGene_gn=PRKD2;refGene_tx=XM_019819304.2;refGene_gn=PRKD2;refGene_tx=XM_019819305.2
E2 10529083 10529156
refGene_gn=PRKD2;refGene_tx=XM_011289800.3;refGene_gn=PRKD2;refGene_tx=XM_019819304.2;refGene_gn=PRKD2;refGene_tx=XM_019819305.2
E2 10529618 10529725
refGene_gn=PRKD2;refGene_tx=XM_011289800.3;refGene_gn=PRKD2;refGene_tx=XM_019819304.2;refGene_gn=PRKD2;refGene_tx=XM_019819305.2
E2 10530930 10531092
refGene_gn=PRKD2;refGene_tx=XM_011289800.3;refGene_gn=PRKD2;refGene_tx=XM_019819304.2;refGene_gn=PRKD2;refGene_tx=XM_019819305.2
E2 10536759 10536858
refGene_gn=PRKD2;refGene_tx=XM_011289800.3;refGene_gn=PRKD2;refGene_tx=XM_019819304.2;refGene_gn=PRKD2;refGene_tx=XM_019819305.2
E2 10540535 10540827
refGene_gn=PRKD2;refGene_tx=XM_011289800.3;refGene_gn=PRKD2;refGene_tx=XM_019819304.2;refGene_gn=PRKD2;refGene_tx=XM_019819305.2
E2 10544312 10544398
refGene_gn=PRKD2;refGene_tx=XM_011289800.3;refGene_gn=PRKD2;refGene_tx=XM_019819304.2;refGene_gn=PRKD2;refGene_tx=XM_019819305.2
E2 10544715 10544928
refGene_gn=PRKD2;refGene_tx=XM_011289800.3;refGene_gn=PRKD2;refGene_tx=XM_019819304.2;refGene_gn=PRKD2;refGene_tx=XM_019819305.2
E2 10551907 10552156 refGene_gn=DACT3;refGene_tx=XM_023245286.1
E2 10557628 10557725 refGene_gn=DACT3;refGene_tx=XM_023245286.1
E2 10557865 10558018 refGene_gn=DACT3;refGene_tx=XM_023245286.1
E2 10558998 10560395 refGene_gn=DACT3;refGene_tx=XM_023245286.1
E2 10567457 10567626 refGene_gn=GNG8;refGene_tx=XM_019819306.2

E2 10571156 10571242
refGene_gn=GNG8;refGene_tx=XM_003997624.5;refGene_gn=GNG8;refGene_tx=XM_006941086.4;refGene_gn=GNG8;refGene_tx=XM_006941087.4;refGene_gn=GNG8;refGene_tx=XM_019819306.2;refGene_gn=GNG8;refGene_tx=XM_019819309.2;refGene_gn=GNG8;refGene_tx=XM_019819310.2
E2 10571472 10571601
refGene_gn=GNG8;refGene_tx=XM_003997624.5;refGene_gn=GNG8;refGene_tx=XM_006941086.4;refGene_gn=GNG8;refGene_tx=XM_006941087.4;refGene_gn=GNG8;refGene_tx=XM_019819306.2;refGene_gn=GNG8;refGene_tx=XM_019819309.2;refGene_gn=GNG8;refGene_tx=XM_019819310.2
E2 10578003 10578771 refGene_gn=PTGIR;refGene_tx=XM_006941088.4
E2 10579817 10580204 refGene_gn=PTGIR;refGene_tx=XM_006941088.4
E2 10588346 10588375
refGene_gn=CALM3;refGene_tx=XM_011289830.3;refGene_gn=CALM3;refGene_tx=XM_023245109.1
E2 10588516 10588652
refGene_gn=CALM3;refGene_tx=XM_011289830.3;refGene_gn=CALM3;refGene_tx=XM_023245109.1
E2 10588889 10588996
refGene_gn=CALM3;refGene_tx=XM_011289830.3;refGene_gn=CALM3;refGene_tx=XM_023245109.1
E2 10589145 10589289
refGene_gn=CALM3;refGene_tx=XM_011289830.3;refGene_gn=CALM3;refGene_tx=XM_023245109.1
E2 10591572 10591603 refGene_gn=CALM3;refGene_tx=XM_023245109.1
E2 10596047 10596050 refGene_gn=CALM3;refGene_tx=XM_023245109.1
E2 10633005 10633025 refGene_gn=PNMA8B;refGene_tx=XM_023245918.1
E2 10633646 10634023 refGene_gn=PNMA8B;refGene_tx=XM_023245918.1
E2 10634025 10634396 refGene_gn=PNMA8B;refGene_tx=XM_023245918.1
E2 10634398 10634540 refGene_gn=PNMA8B;refGene_tx=XM_023245918.1
E2 10634542 10634581 refGene_gn=PNMA8B;refGene_tx=XM_023245918.1
E2 10634583 10634796 refGene_gn=PNMA8B;refGene_tx=XM_023245918.1
E2 10634798 10635076 refGene_gn=PNMA8B;refGene_tx=XM_023245918.1
E2 10642536 10643979 refGene_gn=CCDC8;refGene_tx=XM_003997664.4
E2 10660118 10661270
refGene_gn=LOC105261245;refGene_tx=XM_023245106.1;refGene_gn=LOC105261245;refGene_tx=XM_023245107.1;refGene_gn=LOC105261245;refGene_tx=XM_023245108.1
E2 10687441 10688896
refGene_gn=LOC101091323;refGene_tx=XM_006941089.4
E2 10729587 10729650 refGene_gn=PPP5C;refGene_tx=XM_011289833.3
E2 10729838 10729920 refGene_gn=PPP5C;refGene_tx=XM_011289833.3
E2 10730883 10731062 refGene_gn=PPP5C;refGene_tx=XM_011289833.3
E2 10731150 10731191 refGene_gn=PPP5C;refGene_tx=XM_011289833.3
E2 10731948 10732036 refGene_gn=PPP5C;refGene_tx=XM_011289833.3
E2 10732175 10732318 refGene_gn=PPP5C;refGene_tx=XM_011289833.3
E2 10733086 10733192 refGene_gn=PPP5C;refGene_tx=XM_011289833.3
E2 10733990 10734089 refGene_gn=PPP5C;refGene_tx=XM_011289833.3
E2 10734406 10734472 refGene_gn=PPP5C;refGene_tx=XM_011289833.3
E2 10737294 10737416 refGene_gn=PPP5C;refGene_tx=XM_011289833.3

E2 10738085 10738233 refGene_gn=PPP5C;refGene_tx=XM_011289833.3
E2 10745891 10746133 refGene_gn=PPP5C;refGene_tx=XM_011289833.3
E2 10751909 10752030 refGene_gn=PPP5C;refGene_tx=XM_011289833.3
E2 10756720 10756818
refGene_gn=HIF3A;refGene_tx=XM_019819313.2;refGene_gn=HIF3A;refGene_tx=XM_023244996.1;refGene_gn=HIF3A;refGene_tx=XM_023244998.1
E2 10759599 10759681
refGene_gn=HIF3A;refGene_tx=XM_019819313.2;refGene_gn=HIF3A;refGene_tx=XM_023244996.1;refGene_gn=HIF3A;refGene_tx=XM_023244998.1;refGene_gn=HIF3A;refGene_tx=XM_023244999.1
E2 10759823 10759940 refGene_gn=HIF3A;refGene_tx=XM_023244997.1
E2 10761161 10761279
refGene_gn=HIF3A;refGene_tx=XM_019819313.2;refGene_gn=HIF3A;refGene_tx=XM_023244996.1;refGene_gn=HIF3A;refGene_tx=XM_023244997.1;refGene_gn=HIF3A;refGene_tx=XM_023244998.1;refGene_gn=HIF3A;refGene_tx=XM_023244999.1
E2 10762872 10763144
refGene_gn=HIF3A;refGene_tx=XM_019819313.2;refGene_gn=HIF3A;refGene_tx=XM_023244996.1;refGene_gn=HIF3A;refGene_tx=XM_023244997.1;refGene_gn=HIF3A;refGene_tx=XM_023244998.1
E2 10768019 10768124
refGene_gn=HIF3A;refGene_tx=XM_019819313.2;refGene_gn=HIF3A;refGene_tx=XM_023244996.1;refGene_gn=HIF3A;refGene_tx=XM_023244997.1;refGene_gn=HIF3A;refGene_tx=XM_023244998.1;refGene_gn=HIF3A;refGene_tx=XM_023244999.1
E2 10768662 10768853
refGene_gn=HIF3A;refGene_tx=XM_019819313.2;refGene_gn=HIF3A;refGene_tx=XM_023244996.1;refGene_gn=HIF3A;refGene_tx=XM_023244997.1;refGene_gn=HIF3A;refGene_tx=XM_023244998.1;refGene_gn=HIF3A;refGene_tx=XM_023244999.1
E2 10769875 10769994
refGene_gn=HIF3A;refGene_tx=XM_019819313.2;refGene_gn=HIF3A;refGene_tx=XM_023244996.1;refGene_gn=HIF3A;refGene_tx=XM_023244997.1;refGene_gn=HIF3A;refGene_tx=XM_023244998.1;refGene_gn=HIF3A;refGene_tx=XM_023244999.1
E2 10773897 10774045
refGene_gn=HIF3A;refGene_tx=XM_019819313.2;refGene_gn=HIF3A;refGene_tx=XM_023244996.1;refGene_gn=HIF3A;refGene_tx=XM_023244997.1;refGene_gn=HIF3A;refGene_tx=XM_023244998.1;refGene_gn=HIF3A;refGene_tx=XM_023244999.1
E2 10774259 10774366
refGene_gn=HIF3A;refGene_tx=XM_019819313.2;refGene_gn=HIF3A;refGene_tx=XM_023244996.1;refGene_gn=HIF3A;refGene_tx=XM_023244997.1;refGene_gn=HIF3A;refGene_tx=XM_023244998.1;refGene_gn=HIF3A;refGene_tx=XM_023244999.1
E2 10775960 10776169
refGene_gn=HIF3A;refGene_tx=XM_019819313.2;refGene_gn=HIF3A;refGene_tx=XM_023244996.1;refGene_gn=HIF3A;refGene_tx=XM_023244997.1;refGene_gn=HIF3A;refGene_tx=XM_023244998.1;refGene_gn=HIF3A;refGene_tx=XM_023244999.1

E2 10776589 10776702
refGene_gn=HIF3A;refGene_tx=XM_019819313.2;refGene_gn=HIF3A;refGene_tx=XM_023244996.1;refGene_gn=HIF3A;refGene_tx=XM_023244997.1;refGene_gn=HIF3A;refGene_tx=XM_023244998.1;refGene_gn=HIF3A;refGene_tx=XM_023244999.1
E2 10776954 10777039
refGene_gn=HIF3A;refGene_tx=XM_019819313.2;refGene_gn=HIF3A;refGene_tx=XM_023244996.1;refGene_gn=HIF3A;refGene_tx=XM_023244997.1;refGene_gn=HIF3A;refGene_tx=XM_023244998.1;refGene_gn=HIF3A;refGene_tx=XM_023244999.1
E2 10778722 10778868
refGene_gn=HIF3A;refGene_tx=XM_023244996.1;refGene_gn=HIF3A;refGene_tx=XM_023244997.1;refGene_gn=HIF3A;refGene_tx=XM_023244999.1
E2 10779891 10780082
refGene_gn=HIF3A;refGene_tx=XM_019819313.2;refGene_gn=HIF3A;refGene_tx=XM_023244996.1;refGene_gn=HIF3A;refGene_tx=XM_023244997.1;refGene_gn=HIF3A;refGene_tx=XM_023244998.1;refGene_gn=HIF3A;refGene_tx=XM_023244999.1
E2 10785067 10785093
refGene_gn=HIF3A;refGene_tx=XM_023244996.1;refGene_gn=HIF3A;refGene_tx=XM_023244997.1;refGene_gn=HIF3A;refGene_tx=XM_023244999.1
E2 10810227 10810246
refGene_gn=LOC101101352;refGene_tx=XM_003997666.4
E2 10817785 10817839
refGene_gn=LOC101101352;refGene_tx=XM_003997666.4
E2 10817975 10818243
refGene_gn=LOC101101352;refGene_tx=XM_003997666.4
E2 10818429 10818463
refGene_gn=LOC101101352;refGene_tx=XM_003997666.4
E2 10836381 10836406
refGene_gn=LOC101101603;refGene_tx=XM_006941094.4;refGene_gn=LOC101101603;refGene_tx=XM_019819315.2
E2 10837754 10837808
refGene_gn=LOC101101603;refGene_tx=XM_006941094.4;refGene_gn=LOC101101603;refGene_tx=XM_019819315.2
E2 10837884 10838164
refGene_gn=LOC101101603;refGene_tx=XM_006941094.4;refGene_gn=LOC101101603;refGene_tx=XM_019819315.2
E2 10838332 10838336
refGene_gn=LOC101101603;refGene_tx=XM_006941094.4;refGene_gn=LOC101101603;refGene_tx=XM_019819315.2
E2 10857852 10857957 refGene_gn=PGLYRP1;refGene_tx=XM_003997668.4
E2 10857958 10858006 refGene_gn=PGLYRP1;refGene_tx=XM_003997668.4
E2 10858007 10858144 refGene_gn=PGLYRP1;refGene_tx=XM_003997668.4
E2 10860786 10860908 refGene_gn=PGLYRP1;refGene_tx=XM_003997668.4
E2 10861460 10861639 refGene_gn=PGLYRP1;refGene_tx=XM_003997668.4
E2 10862451 10862549 refGene_gn=CCDC61;refGene_tx=XM_003997628.5
E2 10862656 10862732 refGene_gn=CCDC61;refGene_tx=XM_003997628.5
E2 10863231 10863283 refGene_gn=CCDC61;refGene_tx=XM_003997628.5
E2 10863460 10863539 refGene_gn=CCDC61;refGene_tx=XM_003997628.5

E2 10863626 10863769 refGene_gn=CCDC61;refGene_tx=XM_003997628.5
E2 10864120 10864196 refGene_gn=CCDC61;refGene_tx=XM_003997628.5
E2 10864318 10864488 refGene_gn=CCDC61;refGene_tx=XM_003997628.5
E2 10865212 10865295 refGene_gn=CCDC61;refGene_tx=XM_003997628.5
E2 10865391 10865602 refGene_gn=CCDC61;refGene_tx=XM_003997628.5
E2 10873429 10873575 refGene_gn=CCDC61;refGene_tx=XM_003997628.5
E2 10873576 10873592 refGene_gn=CCDC61;refGene_tx=XM_003997628.5
E2 10874713 10874871 refGene_gn=CCDC61;refGene_tx=XM_003997628.5
E2 10879385 10879468 refGene_gn=CCDC61;refGene_tx=XM_003997628.5
E2 10879719 10879878 refGene_gn=CCDC61;refGene_tx=XM_003997628.5
E2 10884111 10884124 refGene_gn=CCDC61;refGene_tx=XM_003997628.5
E2 10900981 10901066 refGene_gn=NOVA2;refGene_tx=XM_023244858.1
E2 10910133 10910277 refGene_gn=NOVA2;refGene_tx=XM_023244858.1
E2 10915770 10915937 refGene_gn=NOVA2;refGene_tx=XM_023244858.1
E2 10925149 10925197 refGene_gn=NOVA2;refGene_tx=XM_023244858.1
E2 10925198 10926221 refGene_gn=NOVA2;refGene_tx=XM_023244858.1
E2 10955591 10956008 refGene_gn=NANOS2;refGene_tx=XM_003997629.2
E2 10967153 10967652 refGene_gn=MYPOP;refGene_tx=XM_023245198.1
E2 10972830 10973540 refGene_gn=MYPOP;refGene_tx=XM_023245198.1
E2 10978464 10980219 refGene_gn=IRF2BP1;refGene_tx=XM_003997630.4
E2 10989485 10990469 refGene_gn=FOXA3;refGene_tx=XM_023245573.1
E2 10996383 10996452 refGene_gn=FOXA3;refGene_tx=XM_023245573.1
E2 11001575 11001680
refGene_gn=SYMPK;refGene_tx=XM_003997631.5;refGene_gn=SYMPK;refGene_tx
=XM_006941097.4;refGene_gn=SYMPK;refGene_tx=XM_019819318.2
E2 11001824 11001890
refGene_gn=SYMPK;refGene_tx=XM_003997631.5;refGene_gn=SYMPK;refGene_tx
=XM_006941097.4;refGene_gn=SYMPK;refGene_tx=XM_019819318.2
E2 11003362 11003416
refGene_gn=SYMPK;refGene_tx=XM_003997631.5;refGene_gn=SYMPK;refGene_tx
=XM_006941097.4;refGene_gn=SYMPK;refGene_tx=XM_019819318.2
E2 11003507 11003581
refGene_gn=SYMPK;refGene_tx=XM_003997631.5;refGene_gn=SYMPK;refGene_tx
=XM_006941097.4;refGene_gn=SYMPK;refGene_tx=XM_019819318.2
E2 11006394 11006521
refGene_gn=SYMPK;refGene_tx=XM_003997631.5;refGene_gn=SYMPK;refGene_tx
=XM_006941097.4;refGene_gn=SYMPK;refGene_tx=XM_019819318.2
E2 11007211 11007461
refGene_gn=SYMPK;refGene_tx=XM_003997631.5;refGene_gn=SYMPK;refGene_tx
=XM_006941097.4;refGene_gn=SYMPK;refGene_tx=XM_019819318.2
E2 11008023 11008194
refGene_gn=SYMPK;refGene_tx=XM_003997631.5;refGene_gn=SYMPK;refGene_tx
=XM_006941097.4;refGene_gn=SYMPK;refGene_tx=XM_019819318.2
E2 11008881 11009011 refGene_gn=SYMPK;refGene_tx=XM_006941098.3
E2 11009420 11009660
refGene_gn=SYMPK;refGene_tx=XM_003997631.5;refGene_gn=SYMPK;refGene_tx
=XM_006941097.4;refGene_gn=SYMPK;refGene_tx=XM_006941098.3;refGene_gn=
SYMPK;refGene_tx=XM_019819318.2
E2 11012957 11013112
refGene_gn=SYMPK;refGene_tx=XM_003997631.5;refGene_gn=SYMPK;refGene_tx

=XM_006941097.4;refGene_gn=SYMPK;refGene_tx=XM_006941098.3;refGene_gn=SYMPK;refGene_tx=XM_019819318.2
E2 11015503 11015654
refGene_gn=SYMPK;refGene_tx=XM_003997631.5;refGene_gn=SYMPK;refGene_tx=XM_006941097.4;refGene_gn=SYMPK;refGene_tx=XM_006941098.3;refGene_gn=SYMPK;refGene_tx=XM_019819318.2
E2 11017032 11017237
refGene_gn=SYMPK;refGene_tx=XM_003997631.5;refGene_gn=SYMPK;refGene_tx=XM_006941097.4;refGene_gn=SYMPK;refGene_tx=XM_006941098.3;refGene_gn=SYMPK;refGene_tx=XM_019819318.2
E2 11018108 11018259
refGene_gn=SYMPK;refGene_tx=XM_003997631.5;refGene_gn=SYMPK;refGene_tx=XM_006941097.4;refGene_gn=SYMPK;refGene_tx=XM_006941098.3;refGene_gn=SYMPK;refGene_tx=XM_019819318.2
E2 11019099 11019335
refGene_gn=SYMPK;refGene_tx=XM_003997631.5;refGene_gn=SYMPK;refGene_tx=XM_006941097.4;refGene_gn=SYMPK;refGene_tx=XM_006941098.3;refGene_gn=SYMPK;refGene_tx=XM_019819318.2
E2 11020348 11020430
refGene_gn=SYMPK;refGene_tx=XM_003997631.5;refGene_gn=SYMPK;refGene_tx=XM_006941097.4;refGene_gn=SYMPK;refGene_tx=XM_006941098.3;refGene_gn=SYMPK;refGene_tx=XM_019819318.2
E2 11020647 11020761
refGene_gn=SYMPK;refGene_tx=XM_003997631.5;refGene_gn=SYMPK;refGene_tx=XM_006941097.4;refGene_gn=SYMPK;refGene_tx=XM_006941098.3;refGene_gn=SYMPK;refGene_tx=XM_019819318.2
E2 11021785 11021933
refGene_gn=SYMPK;refGene_tx=XM_003997631.5;refGene_gn=SYMPK;refGene_tx=XM_006941097.4;refGene_gn=SYMPK;refGene_tx=XM_006941098.3;refGene_gn=SYMPK;refGene_tx=XM_019819318.2
E2 11023237 11023398
refGene_gn=SYMPK;refGene_tx=XM_003997631.5;refGene_gn=SYMPK;refGene_tx=XM_006941097.4;refGene_gn=SYMPK;refGene_tx=XM_006941098.3;refGene_gn=SYMPK;refGene_tx=XM_019819318.2
E2 11024891 11025000
refGene_gn=SYMPK;refGene_tx=XM_003997631.5;refGene_gn=SYMPK;refGene_tx=XM_006941097.4;refGene_gn=SYMPK;refGene_tx=XM_006941098.3;refGene_gn=SYMPK;refGene_tx=XM_019819318.2
E2 11025250 11025351
refGene_gn=SYMPK;refGene_tx=XM_003997631.5;refGene_gn=SYMPK;refGene_tx=XM_006941097.4;refGene_gn=SYMPK;refGene_tx=XM_006941098.3;refGene_gn=SYMPK;refGene_tx=XM_019819318.2
E2 11026013 11026104
refGene_gn=SYMPK;refGene_tx=XM_003997631.5;refGene_gn=SYMPK;refGene_tx=XM_006941097.4;refGene_gn=SYMPK;refGene_tx=XM_006941098.3;refGene_gn=SYMPK;refGene_tx=XM_019819318.2
E2 11026735 11026837
refGene_gn=SYMPK;refGene_tx=XM_003997631.5;refGene_gn=SYMPK;refGene_tx=XM_006941097.4;refGene_gn=SYMPK;refGene_tx=XM_006941098.3;refGene_gn=SYMPK;refGene_tx=XM_019819318.2

E2 11030335 11030523
refGene_gn=SYMPK;refGene_tx=XM_003997631.5;refGene_gn=SYMPK;refGene_tx=XM_006941097.4;refGene_gn=SYMPK;refGene_tx=XM_006941098.3;refGene_gn=SYMPK;refGene_tx=XM_019819318.2
E2 11031393 11031570
refGene_gn=SYMPK;refGene_tx=XM_003997631.5;refGene_gn=SYMPK;refGene_tx=XM_006941097.4;refGene_gn=SYMPK;refGene_tx=XM_006941098.3;refGene_gn=SYMPK;refGene_tx=XM_019819318.2
E2 11031746 11031842
refGene_gn=SYMPK;refGene_tx=XM_003997631.5;refGene_gn=SYMPK;refGene_tx=XM_006941097.4;refGene_gn=SYMPK;refGene_tx=XM_006941098.3;refGene_gn=SYMPK;refGene_tx=XM_019819318.2
E2 11032094 11032427
refGene_gn=SYMPK;refGene_tx=XM_003997631.5;refGene_gn=SYMPK;refGene_tx=XM_006941097.4;refGene_gn=SYMPK;refGene_tx=XM_006941098.3;refGene_gn=SYMPK;refGene_tx=XM_019819318.2
E2 11032596 11032731
refGene_gn=SYMPK;refGene_tx=XM_003997631.5;refGene_gn=SYMPK;refGene_tx=XM_006941097.4;refGene_gn=SYMPK;refGene_tx=XM_006941098.3;refGene_gn=SYMPK;refGene_tx=XM_019819318.2
E2 11033145 11033813
refGene_gn=RSPH6A;refGene_tx=XM_003997672.4;refGene_gn=RSPH6A;refGene_tx=XM_019819320.2;refGene_gn=RSPH6A;refGene_tx=XM_023245940.1
E2 11037127 11037362
refGene_gn=RSPH6A;refGene_tx=XM_003997672.4;refGene_gn=RSPH6A;refGene_tx=XM_019819320.2;refGene_gn=RSPH6A;refGene_tx=XM_023245940.1
E2 11042020 11042785
refGene_gn=RSPH6A;refGene_tx=XM_003997672.4;refGene_gn=RSPH6A;refGene_tx=XM_019819320.2;refGene_gn=RSPH6A;refGene_tx=XM_023245940.1
E2 11044037 11044182
refGene_gn=RSPH6A;refGene_tx=XM_003997672.4;refGene_gn=RSPH6A;refGene_tx=XM_019819320.2;refGene_gn=RSPH6A;refGene_tx=XM_023245940.1
E2 11044912 11045030
refGene_gn=RSPH6A;refGene_tx=XM_003997672.4;refGene_gn=RSPH6A;refGene_tx=XM_023245940.1
E2 11048227 11048330 refGene_gn=RSPH6A;refGene_tx=XM_023245940.1
E2 11049485 11049723
refGene_gn=RSPH6A;refGene_tx=XM_003997672.4;refGene_gn=RSPH6A;refGene_tx=XM_019819320.2
E2 11051894 11052335
refGene_gn=DMWD;refGene_tx=XM_023245914.1;refGene_gn=DMWD;refGene_tx=XM_023245915.1
E2 11053609 11053792
refGene_gn=DMWD;refGene_tx=XM_023245914.1;refGene_gn=DMWD;refGene_tx=XM_023245915.1
E2 11055653 11056928
refGene_gn=DMWD;refGene_tx=XM_023245914.1;refGene_gn=DMWD;refGene_tx=XM_023245915.1
E2 11057792 11057867 refGene_gn=DMWD;refGene_tx=XM_023245914.1
E2 11058210 11058258

refGene_gn=DMWD;refGene_tx=XM_023245914.1;refGene_gn=DMWD;refGene_tx=XM_023245915.1
E2 11060107 11060267
refGene_gn=DMPK;refGene_tx=XM_023245908.1;refGene_gn=DMPK;refGene_tx=XM_023245909.1;refGene_gn=DMPK;refGene_tx=XM_023245910.1;refGene_gn=DMPK;refGene_tx=XM_023245911.1;refGene_gn=DMPK;refGene_tx=XM_023245912.1;refGene_gn=DMPK;refGene_tx=XM_023245913.1
E2 11062523 11062615
refGene_gn=DMPK;refGene_tx=XM_023245908.1;refGene_gn=DMPK;refGene_tx=XM_023245909.1;refGene_gn=DMPK;refGene_tx=XM_023245910.1;refGene_gn=DMPK;refGene_tx=XM_023245911.1;refGene_gn=DMPK;refGene_tx=XM_023245912.1;refGene_gn=DMPK;refGene_tx=XM_023245913.1
E2 11062917 11063001
refGene_gn=DMPK;refGene_tx=XM_023245908.1;refGene_gn=DMPK;refGene_tx=XM_023245909.1;refGene_gn=DMPK;refGene_tx=XM_023245910.1;refGene_gn=DMPK;refGene_tx=XM_023245911.1;refGene_gn=DMPK;refGene_tx=XM_023245912.1;refGene_gn=DMPK;refGene_tx=XM_023245913.1
E2 11063090 11063186
refGene_gn=DMPK;refGene_tx=XM_023245908.1;refGene_gn=DMPK;refGene_tx=XM_023245909.1;refGene_gn=DMPK;refGene_tx=XM_023245910.1;refGene_gn=DMPK;refGene_tx=XM_023245911.1;refGene_gn=DMPK;refGene_tx=XM_023245912.1;refGene_gn=DMPK;refGene_tx=XM_023245913.1
E2 11063742 11063891
refGene_gn=DMPK;refGene_tx=XM_023245908.1;refGene_gn=DMPK;refGene_tx=XM_023245909.1;refGene_gn=DMPK;refGene_tx=XM_023245910.1;refGene_gn=DMPK;refGene_tx=XM_023245911.1;refGene_gn=DMPK;refGene_tx=XM_023245912.1;refGene_gn=DMPK;refGene_tx=XM_023245913.1
E2 11064255 11064349
refGene_gn=DMPK;refGene_tx=XM_023245908.1;refGene_gn=DMPK;refGene_tx=XM_023245909.1;refGene_gn=DMPK;refGene_tx=XM_023245910.1;refGene_gn=DMPK;refGene_tx=XM_023245911.1;refGene_gn=DMPK;refGene_tx=XM_023245912.1;refGene_gn=DMPK;refGene_tx=XM_023245913.1
E2 11064638 11064845
refGene_gn=DMPK;refGene_tx=XM_023245908.1;refGene_gn=DMPK;refGene_tx=XM_023245909.1;refGene_gn=DMPK;refGene_tx=XM_023245910.1;refGene_gn=DMPK;refGene_tx=XM_023245911.1;refGene_gn=DMPK;refGene_tx=XM_023245912.1;refGene_gn=DMPK;refGene_tx=XM_023245913.1
E2 11064929 11065193
refGene_gn=DMPK;refGene_tx=XM_023245908.1;refGene_gn=DMPK;refGene_tx=XM_023245909.1;refGene_gn=DMPK;refGene_tx=XM_023245910.1;refGene_gn=DMPK;refGene_tx=XM_023245911.1;refGene_gn=DMPK;refGene_tx=XM_023245912.1;refGene_gn=DMPK;refGene_tx=XM_023245913.1
E2 11065913 11065999
refGene_gn=DMPK;refGene_tx=XM_023245908.1;refGene_gn=DMPK;refGene_tx=XM_023245909.1;refGene_gn=DMPK;refGene_tx=XM_023245910.1;refGene_gn=DMPK;refGene_tx=XM_023245911.1;refGene_gn=DMPK;refGene_tx=XM_023245912.1;refGene_gn=DMPK;refGene_tx=XM_023245913.1
E2 11067556 11067668
refGene_gn=DMPK;refGene_tx=XM_023245908.1;refGene_gn=DMPK;refGene_tx=XM_023245909.1;refGene_gn=DMPK;refGene_tx=XM_023245910.1;refGene_gn=DMP

K;refGene_tx=XM_023245911.1;refGene_gn=DMPK;refGene_tx=XM_023245912.1;
refGene_gn=DMPK;refGene_tx=XM_023245913.1
E2 11068321 11068497
refGene_gn=DMPK;refGene_tx=XM_023245908.1;refGene_gn=DMPK;refGene_tx=X
M_023245909.1;refGene_gn=DMPK;refGene_tx=XM_023245910.1;refGene_gn=DMP
K;refGene_tx=XM_023245911.1;refGene_gn=DMPK;refGene_tx=XM_023245912.1;
refGene_gn=DMPK;refGene_tx=XM_023245913.1
E2 11068592 11068690
refGene_gn=DMPK;refGene_tx=XM_023245908.1;refGene_gn=DMPK;refGene_tx=X
M_023245909.1;refGene_gn=DMPK;refGene_tx=XM_023245910.1;refGene_gn=DMP
K;refGene_tx=XM_023245911.1;refGene_gn=DMPK;refGene_tx=XM_023245912.1;
refGene_gn=DMPK;refGene_tx=XM_023245913.1
E2 11068853 11068900
refGene_gn=DMPK;refGene_tx=XM_023245908.1;refGene_gn=DMPK;refGene_tx=X
M_023245909.1;refGene_gn=DMPK;refGene_tx=XM_023245910.1;refGene_gn=DMP
K;refGene_tx=XM_023245911.1
E2 11069192 11069282
refGene_gn=DMPK;refGene_tx=XM_023245908.1;refGene_gn=DMPK;refGene_tx=X
M_023245909.1;refGene_gn=DMPK;refGene_tx=XM_023245910.1;refGene_gn=DMP
K;refGene_tx=XM_023245911.1;refGene_gn=DMPK;refGene_tx=XM_023245912.1
E2 11069625 11069785
refGene_gn=DMPK;refGene_tx=XM_023245908.1;refGene_gn=DMPK;refGene_tx=X
M_023245909.1;refGene_gn=DMPK;refGene_tx=XM_023245910.1;refGene_gn=DMP
K;refGene_tx=XM_023245911.1;refGene_gn=DMPK;refGene_tx=XM_023245912.1;
refGene_gn=DMPK;refGene_tx=XM_023245913.1
E2 11071394 11072191 refGene_gn=SIX5;refGene_tx=XM_023245907.1
E2 11073152 11073967 refGene_gn=SIX5;refGene_tx=XM_023245907.1
E2 11074110 11074730 refGene_gn=SIX5;refGene_tx=XM_023245907.1
E2 11076219 11076385 refGene_gn=BHMG1;refGene_tx=XM_019819329.2
E2 11076975 11077065 refGene_gn=BHMG1;refGene_tx=XM_019819329.2
E2 11077870 11078130 refGene_gn=BHMG1;refGene_tx=XM_019819329.2
E2 11078205 11078370 refGene_gn=BHMG1;refGene_tx=XM_019819329.2
E2 11079213 11079287 refGene_gn=BHMG1;refGene_tx=XM_019819329.2
E2 11079827 11079983 refGene_gn=BHMG1;refGene_tx=XM_019819329.2
E2 11080440 11080541 refGene_gn=BHMG1;refGene_tx=XM_019819329.2
E2 11081414 11081523 refGene_gn=BHMG1;refGene_tx=XM_019819329.2
E2 11082255 11082501 refGene_gn=BHMG1;refGene_tx=XM_019819329.2
E2 11083796 11083931 refGene_gn=BHMG1;refGene_tx=XM_019819329.2
E2 11085438 11085550 refGene_gn=BHMG1;refGene_tx=XM_019819329.2
E2 11088524 11088615 refGene_gn=BHMG1;refGene_tx=XM_019819329.2
E2 11094312 11094383 refGene_gn=BHMG1;refGene_tx=XM_019819329.2
E2 11110311 11112123 refGene_gn=FBXO46;refGene_tx=XM_006941100.4
E2 11118020 11118166 refGene_gn=QPCTL;refGene_tx=XM_003997633.5
E2 11121750 11121867 refGene_gn=QPCTL;refGene_tx=XM_003997633.5
E2 11122368 11122468 refGene_gn=QPCTL;refGene_tx=XM_003997633.5
E2 11122554 11122707 refGene_gn=QPCTL;refGene_tx=XM_003997633.5
E2 11124287 11124569 refGene_gn=QPCTL;refGene_tx=XM_003997633.5
E2 11125334 11125478 refGene_gn=QPCTL;refGene_tx=XM_003997633.5
E2 11125971 11126475 refGene_gn=QPCTL;refGene_tx=XM_003997633.5
E2 11126991 11126993 refGene_gn=SNRPD2;refGene_tx=XM_003997635.5

E2 11128956 11129136 refGene_gn=SNRPD2;refGene_tx=XM_003997635.5
E2 11133317 11133492 refGene_gn=SNRPD2;refGene_tx=XM_003997635.5
E2 11134487 11134547 refGene_gn=GIPR;refGene_tx=XM_023245806.1
E2 11137010 11137217
refGene_gn=GIPR;refGene_tx=XM_023245804.1;refGene_gn=GIPR;refGene_tx=XM_023245805.1
E2 11137302 11137344 refGene_gn=GIPR;refGene_tx=XM_023245804.1
E2 11139378 11139517
refGene_gn=GIPR;refGene_tx=XM_023245804.1;refGene_gn=GIPR;refGene_tx=XM_023245805.1;refGene_gn=GIPR;refGene_tx=XM_023245806.1
E2 11139631 11139720
refGene_gn=GIPR;refGene_tx=XM_023245804.1;refGene_gn=GIPR;refGene_tx=XM_023245805.1;refGene_gn=GIPR;refGene_tx=XM_023245806.1
E2 11139838 11139908
refGene_gn=GIPR;refGene_tx=XM_023245804.1;refGene_gn=GIPR;refGene_tx=XM_023245805.1;refGene_gn=GIPR;refGene_tx=XM_023245806.1
E2 11140240 11140301
refGene_gn=GIPR;refGene_tx=XM_023245804.1;refGene_gn=GIPR;refGene_tx=XM_023245805.1;refGene_gn=GIPR;refGene_tx=XM_023245806.1
E2 11140510 11140670
refGene_gn=GIPR;refGene_tx=XM_023245804.1;refGene_gn=GIPR;refGene_tx=XM_023245805.1;refGene_gn=GIPR;refGene_tx=XM_023245806.1
E2 11141472 11141617
refGene_gn=GIPR;refGene_tx=XM_023245804.1;refGene_gn=GIPR;refGene_tx=XM_023245805.1;refGene_gn=GIPR;refGene_tx=XM_023245806.1
E2 11141813 11141917
refGene_gn=GIPR;refGene_tx=XM_023245804.1;refGene_gn=GIPR;refGene_tx=XM_023245805.1;refGene_gn=GIPR;refGene_tx=XM_023245806.1
E2 11142609 11142713
refGene_gn=GIPR;refGene_tx=XM_023245804.1;refGene_gn=GIPR;refGene_tx=XM_023245805.1;refGene_gn=GIPR;refGene_tx=XM_023245806.1
E2 11143102 11143210
refGene_gn=GIPR;refGene_tx=XM_023245804.1;refGene_gn=GIPR;refGene_tx=XM_023245805.1;refGene_gn=GIPR;refGene_tx=XM_023245806.1
E2 11143650 11143750
refGene_gn=GIPR;refGene_tx=XM_023245804.1;refGene_gn=GIPR;refGene_tx=XM_023245805.1;refGene_gn=GIPR;refGene_tx=XM_023245806.1
E2 11144139 11144214
refGene_gn=GIPR;refGene_tx=XM_023245804.1;refGene_gn=GIPR;refGene_tx=XM_023245805.1;refGene_gn=GIPR;refGene_tx=XM_023245806.1
E2 11152751 11152923 refGene_gn=EML2;refGene_tx=XM_023245803.1
E2 11153144 11153202 refGene_gn=EML2;refGene_tx=XM_019819333.2
E2 11154560 11154570 refGene_gn=EML2;refGene_tx=XM_019819332.2
E2 11155482 11155701
refGene_gn=EML2;refGene_tx=XM_006941102.4;refGene_gn=EML2;refGene_tx=XM_006941103.4;refGene_gn=EML2;refGene_tx=XM_019819332.2;refGene_gn=EML2;refGene_tx=XM_019819333.2;refGene_gn=EML2;refGene_tx=XM_023245803.1
E2 11155792 11155916
refGene_gn=EML2;refGene_tx=XM_006941102.4;refGene_gn=EML2;refGene_tx=XM_006941103.4;refGene_gn=EML2;refGene_tx=XM_019819332.2;refGene_gn=EML

2;refGene_tx=XM_019819333.2;refGene_gn=EML2;refGene_tx=XM_023245803.1
E2 11156112 11156217
refGene_gn=EML2;refGene_tx=XM_006941102.4;refGene_gn=EML2;refGene_tx=XM_006941103.4;refGene_gn=EML2;refGene_tx=XM_019819332.2;refGene_gn=EML2;refGene_tx=XM_019819333.2;refGene_gn=EML2;refGene_tx=XM_023245803.1
E2 11157565 11157585 refGene_gn=EML2;refGene_tx=XM_011289803.3
E2 11158094 11158123
refGene_gn=EML2;refGene_tx=XM_006941102.4;refGene_gn=EML2;refGene_tx=XM_006941103.4;refGene_gn=EML2;refGene_tx=XM_011289803.3;refGene_gn=EML2;refGene_tx=XM_011289804.3;refGene_gn=EML2;refGene_tx=XM_019819332.2;refGene_gn=EML2;refGene_tx=XM_019819333.2;refGene_gn=EML2;refGene_tx=XM_023245803.1
E2 11158535 11158665
refGene_gn=EML2;refGene_tx=XM_006941102.4;refGene_gn=EML2;refGene_tx=XM_006941103.4;refGene_gn=EML2;refGene_tx=XM_011289803.3;refGene_gn=EML2;refGene_tx=XM_011289804.3;refGene_gn=EML2;refGene_tx=XM_019819332.2;refGene_gn=EML2;refGene_tx=XM_019819333.2;refGene_gn=EML2;refGene_tx=XM_023245803.1
E2 11161916 11162066
refGene_gn=EML2;refGene_tx=XM_006941102.4;refGene_gn=EML2;refGene_tx=XM_006941103.4;refGene_gn=EML2;refGene_tx=XM_011289803.3;refGene_gn=EML2;refGene_tx=XM_011289804.3;refGene_gn=EML2;refGene_tx=XM_019819332.2;refGene_gn=EML2;refGene_tx=XM_019819333.2;refGene_gn=EML2;refGene_tx=XM_023245803.1
E2 11162612 11162682
refGene_gn=EML2;refGene_tx=XM_006941102.4;refGene_gn=EML2;refGene_tx=XM_006941103.4;refGene_gn=EML2;refGene_tx=XM_011289803.3;refGene_gn=EML2;refGene_tx=XM_011289804.3;refGene_gn=EML2;refGene_tx=XM_019819332.2;refGene_gn=EML2;refGene_tx=XM_019819333.2;refGene_gn=EML2;refGene_tx=XM_023245803.1
E2 11162781 11162892
refGene_gn=EML2;refGene_tx=XM_006941102.4;refGene_gn=EML2;refGene_tx=XM_006941103.4;refGene_gn=EML2;refGene_tx=XM_011289803.3;refGene_gn=EML2;refGene_tx=XM_011289804.3;refGene_gn=EML2;refGene_tx=XM_019819332.2;refGene_gn=EML2;refGene_tx=XM_019819333.2;refGene_gn=EML2;refGene_tx=XM_023245803.1
E2 11164099 11164195
refGene_gn=EML2;refGene_tx=XM_006941102.4;refGene_gn=EML2;refGene_tx=XM_006941103.4;refGene_gn=EML2;refGene_tx=XM_011289803.3;refGene_gn=EML2;refGene_tx=XM_011289804.3;refGene_gn=EML2;refGene_tx=XM_019819332.2;refGene_gn=EML2;refGene_tx=XM_019819333.2;refGene_gn=EML2;refGene_tx=XM_023245803.1
E2 11165077 11165212
refGene_gn=EML2;refGene_tx=XM_006941102.4;refGene_gn=EML2;refGene_tx=XM_006941103.4;refGene_gn=EML2;refGene_tx=XM_011289803.3;refGene_gn=EML2;refGene_tx=XM_011289804.3;refGene_gn=EML2;refGene_tx=XM_019819332.2;refGene_gn=EML2;refGene_tx=XM_019819333.2;refGene_gn=EML2;refGene_tx=XM_023245803.1
E2 11166206 11166306
refGene_gn=EML2;refGene_tx=XM_006941102.4;refGene_gn=EML2;refGene_tx=X

M_006941103.4;refGene_gn=EML2;refGene_tx=XM_011289803.3;refGene_gn=EML2;refGene_tx=XM_011289804.3;refGene_gn=EML2;refGene_tx=XM_019819332.2;refGene_gn=EML2;refGene_tx=XM_019819333.2;refGene_gn=EML2;refGene_tx=XM_023245803.1
E2 11169015 11169170
refGene_gn=EML2;refGene_tx=XM_006941102.4;refGene_gn=EML2;refGene_tx=XM_006941103.4;refGene_gn=EML2;refGene_tx=XM_011289803.3;refGene_gn=EML2;refGene_tx=XM_011289804.3;refGene_gn=EML2;refGene_tx=XM_019819332.2;refGene_gn=EML2;refGene_tx=XM_019819333.2;refGene_gn=EML2;refGene_tx=XM_023245803.1
E2 11169324 11169450
refGene_gn=EML2;refGene_tx=XM_006941102.4;refGene_gn=EML2;refGene_tx=XM_006941103.4;refGene_gn=EML2;refGene_tx=XM_011289803.3;refGene_gn=EML2;refGene_tx=XM_011289804.3;refGene_gn=EML2;refGene_tx=XM_019819332.2;refGene_gn=EML2;refGene_tx=XM_019819333.2;refGene_gn=EML2;refGene_tx=XM_023245803.1
E2 11171027 11171159
refGene_gn=EML2;refGene_tx=XM_006941102.4;refGene_gn=EML2;refGene_tx=XM_006941103.4;refGene_gn=EML2;refGene_tx=XM_011289803.3;refGene_gn=EML2;refGene_tx=XM_011289804.3;refGene_gn=EML2;refGene_tx=XM_019819332.2;refGene_gn=EML2;refGene_tx=XM_019819333.2;refGene_gn=EML2;refGene_tx=XM_023245803.1
E2 11174329 11174397
refGene_gn=EML2;refGene_tx=XM_006941102.4;refGene_gn=EML2;refGene_tx=XM_006941103.4;refGene_gn=EML2;refGene_tx=XM_011289803.3;refGene_gn=EML2;refGene_tx=XM_011289804.3;refGene_gn=EML2;refGene_tx=XM_019819332.2;refGene_gn=EML2;refGene_tx=XM_019819333.2;refGene_gn=EML2;refGene_tx=XM_023245803.1
E2 11174692 11174781
refGene_gn=EML2;refGene_tx=XM_006941102.4;refGene_gn=EML2;refGene_tx=XM_006941103.4;refGene_gn=EML2;refGene_tx=XM_011289803.3;refGene_gn=EML2;refGene_tx=XM_011289804.3;refGene_gn=EML2;refGene_tx=XM_019819332.2;refGene_gn=EML2;refGene_tx=XM_019819333.2;refGene_gn=EML2;refGene_tx=XM_023245803.1
E2 11175020 11175118
refGene_gn=EML2;refGene_tx=XM_006941102.4;refGene_gn=EML2;refGene_tx=XM_006941103.4;refGene_gn=EML2;refGene_tx=XM_011289803.3;refGene_gn=EML2;refGene_tx=XM_011289804.3;refGene_gn=EML2;refGene_tx=XM_019819332.2;refGene_gn=EML2;refGene_tx=XM_019819333.2;refGene_gn=EML2;refGene_tx=XM_023245803.1
E2 11175554 11175642
refGene_gn=EML2;refGene_tx=XM_006941102.4;refGene_gn=EML2;refGene_tx=XM_006941103.4;refGene_gn=EML2;refGene_tx=XM_011289803.3;refGene_gn=EML2;refGene_tx=XM_011289804.3;refGene_gn=EML2;refGene_tx=XM_019819332.2;refGene_gn=EML2;refGene_tx=XM_019819333.2;refGene_gn=EML2;refGene_tx=XM_023245803.1
E2 11176299 11176395
refGene_gn=EML2;refGene_tx=XM_006941102.4;refGene_gn=EML2;refGene_tx=XM_006941103.4;refGene_gn=EML2;refGene_tx=XM_011289803.3;refGene_gn=EML2;refGene_tx=XM_011289804.3;refGene_gn=EML2;refGene_tx=XM_019819332.2;

refGene_gn=EML2;refGene_tx=XM_019819333.2;refGene_gn=EML2;refGene_tx=XM_023245803.1
E2 11177034 11177165
refGene_gn=EML2;refGene_tx=XM_006941102.4;refGene_gn=EML2;refGene_tx=XM_006941103.4;refGene_gn=EML2;refGene_tx=XM_011289803.3;refGene_gn=EML2;refGene_tx=XM_011289804.3;refGene_gn=EML2;refGene_tx=XM_019819332.2;refGene_gn=EML2;refGene_tx=XM_019819333.2;refGene_gn=EML2;refGene_tx=XM_023245803.1
E2 11177494 11177620
refGene_gn=EML2;refGene_tx=XM_006941102.4;refGene_gn=EML2;refGene_tx=XM_006941103.4;refGene_gn=EML2;refGene_tx=XM_011289803.3;refGene_gn=EML2;refGene_tx=XM_011289804.3;refGene_gn=EML2;refGene_tx=XM_019819332.2;refGene_gn=EML2;refGene_tx=XM_019819333.2;refGene_gn=EML2;refGene_tx=XM_023245803.1
E2 11185747 11185979
refGene_gn=LOC102899082;refGene_tx=XM_023245654.1
E2 11189920 11190579
refGene_gn=LOC102899082;refGene_tx=XM_023245654.1
E2 11200519 11201608 refGene_gn=GPR4;refGene_tx=XM_023245540.1
E2 11205583 11205725 refGene_gn=OPA3;refGene_tx=XM_003997677.4
E2 11253657 11254115 refGene_gn=OPA3;refGene_tx=XM_003997677.4
E2 11281298 11281336
refGene_gn=VASP;refGene_tx=XM_006941107.4;refGene_gn=VASP;refGene_tx=XM_006941108.4;refGene_gn=VASP;refGene_tx=XM_019819344.2;refGene_gn=VASP;refGene_tx=XM_023245541.1
E2 11281457 11281515
refGene_gn=VASP;refGene_tx=XM_006941107.4;refGene_gn=VASP;refGene_tx=XM_006941108.4;refGene_gn=VASP;refGene_tx=XM_019819344.2;refGene_gn=VASP;refGene_tx=XM_023245541.1
E2 11282101 11282192
refGene_gn=VASP;refGene_tx=XM_006941107.4;refGene_gn=VASP;refGene_tx=XM_006941108.4;refGene_gn=VASP;refGene_tx=XM_019819344.2;refGene_gn=VASP;refGene_tx=XM_023245541.1
E2 11282346 11282392
refGene_gn=VASP;refGene_tx=XM_006941107.4;refGene_gn=VASP;refGene_tx=XM_006941108.4;refGene_gn=VASP;refGene_tx=XM_019819344.2;refGene_gn=VASP;refGene_tx=XM_023245541.1
E2 11282619 11282656
refGene_gn=VASP;refGene_tx=XM_006941107.4;refGene_gn=VASP;refGene_tx=XM_006941108.4;refGene_gn=VASP;refGene_tx=XM_019819344.2;refGene_gn=VASP;refGene_tx=XM_023245541.1
E2 11282813 11282865
refGene_gn=VASP;refGene_tx=XM_006941107.4;refGene_gn=VASP;refGene_tx=XM_006941108.4;refGene_gn=VASP;refGene_tx=XM_019819344.2;refGene_gn=VASP;refGene_tx=XM_023245541.1
E2 11283441 11283542
refGene_gn=VASP;refGene_tx=XM_006941107.4;refGene_gn=VASP;refGene_tx=XM_006941108.4;refGene_gn=VASP;refGene_tx=XM_019819344.2;refGene_gn=VASP;refGene_tx=XM_023245541.1
E2 11283713 11283955

refGene_gn=VASP;refGene_tx=XM_006941107.4;refGene_gn=VASP;refGene_tx=XM_006941108.4;refGene_gn=VASP;refGene_tx=XM_019819344.2;refGene_gn=VASP;refGene_tx=XM_023245541.1
E2 11284080 11284130
refGene_gn=VASP;refGene_tx=XM_006941107.4;refGene_gn=VASP;refGene_tx=XM_006941108.4;refGene_gn=VASP;refGene_tx=XM_019819344.2;refGene_gn=VASP;refGene_tx=XM_023245541.1
E2 11284614 11284708
refGene_gn=VASP;refGene_tx=XM_006941107.4;refGene_gn=VASP;refGene_tx=XM_006941108.4;refGene_gn=VASP;refGene_tx=XM_019819344.2;refGene_gn=VASP;refGene_tx=XM_023245541.1
E2 11285972 11286138
refGene_gn=VASP;refGene_tx=XM_006941107.4;refGene_gn=VASP;refGene_tx=XM_006941108.4;refGene_gn=VASP;refGene_tx=XM_019819344.2;refGene_gn=VASP;refGene_tx=XM_023245541.1
E2 11286232 11286404
refGene_gn=VASP;refGene_tx=XM_006941107.4;refGene_gn=VASP;refGene_tx=XM_006941108.4;refGene_gn=VASP;refGene_tx=XM_019819344.2;refGene_gn=VASP;refGene_tx=XM_023245541.1
E2 11293174 11293179
refGene_gn=VASP;refGene_tx=XM_006941107.4;refGene_gn=VASP;refGene_tx=XM_006941108.4;refGene_gn=VASP;refGene_tx=XM_019819344.2;refGene_gn=VASP;refGene_tx=XM_023245541.1
E2 11295481 11295619 refGene_gn=PPM1N;refGene_tx=XM_019819339.2
E2 11296476 11296545 refGene_gn=PPM1N;refGene_tx=XM_006941106.3
E2 11297672 11297733
refGene_gn=PPM1N;refGene_tx=XM_006941106.3;refGene_gn=PPM1N;refGene_tx=XM_019819339.2
E2 11297818 11297924
refGene_gn=PPM1N;refGene_tx=XM_006941106.3;refGene_gn=PPM1N;refGene_tx=XM_019819339.2
E2 11298002 11298120
refGene_gn=PPM1N;refGene_tx=XM_006941106.3;refGene_gn=PPM1N;refGene_tx=XM_019819339.2
E2 11298707 11299625
refGene_gn=PPM1N;refGene_tx=XM_006941106.3;refGene_gn=PPM1N;refGene_tx=XM_019819339.2
E2 11301061 11301095 refGene_gn=RTN2;refGene_tx=XM_019819343.2
E2 11302972 11303017 refGene_gn=RTN2;refGene_tx=XM_019819343.2
E2 11303140 11303611 refGene_gn=RTN2;refGene_tx=XM_019819343.2
E2 11303717 11303975 refGene_gn=RTN2;refGene_tx=XM_019819343.2
E2 11304673 11304686 refGene_gn=RTN2;refGene_tx=XM_006941109.2
E2 11306888 11307096
refGene_gn=RTN2;refGene_tx=XM_006941109.2;refGene_gn=RTN2;refGene_tx=XM_019819343.2
E2 11307435 11307574
refGene_gn=RTN2;refGene_tx=XM_006941109.2;refGene_gn=RTN2;refGene_tx=XM_019819343.2
E2 11307727 11307797
refGene_gn=RTN2;refGene_tx=XM_006941109.2;refGene_gn=RTN2;refGene_tx=X

M_019819343.2
E2	11307923	11307970
refGene_gn=RTN2;refGene_tx=XM_006941109.2;refGene_gn=RTN2;refGene_tx=XM_019819343.2
E2	11308750	11308809
refGene_gn=RTN2;refGene_tx=XM_006941109.2;refGene_gn=RTN2;refGene_tx=XM_019819343.2
E2	11309074	11309156
refGene_gn=RTN2;refGene_tx=XM_006941109.2;refGene_gn=RTN2;refGene_tx=XM_019819343.2
E2	11317775	11318237
refGene_gn=FOSB;refGene_tx=XM_003997640.5;refGene_gn=FOSB;refGene_tx=XM_019819354.2
E2	11319224	11319332
refGene_gn=FOSB;refGene_tx=XM_003997640.5;refGene_gn=FOSB;refGene_tx=XM_019819354.2
E2	11319547	11319868
refGene_gn=FOSB;refGene_tx=XM_003997640.5;refGene_gn=FOSB;refGene_tx=XM_019819354.2
E2	11321808	11321934
refGene_gn=FOSB;refGene_tx=XM_003997640.5;refGene_gn=FOSB;refGene_tx=XM_019819354.2
E2	11356967	11357060	refGene_gn=ERCC1;refGene_tx=XM_006941110.2
E2	11358032	11358248	refGene_gn=ERCC1;refGene_tx=XM_006941110.2
E2	11360088	11360192	refGene_gn=ERCC1;refGene_tx=XM_006941110.2
E2	11361047	11361147	refGene_gn=ERCC1;refGene_tx=XM_006941110.2
E2	11362126	11362203	refGene_gn=ERCC1;refGene_tx=XM_006941110.2
E2	11370689	11370789	refGene_gn=ERCC1;refGene_tx=XM_006941110.2
E2	11371797	11371869	refGene_gn=ERCC1;refGene_tx=XM_006941110.2
E2	11372060	11372129	refGene_gn=ERCC1;refGene_tx=XM_006941110.2
E2	11372448	11372502	refGene_gn=ERCC1;refGene_tx=XM_006941110.2
E2	11372900	11374173	refGene_gn=CD3EAP;refGene_tx=XM_003997680.4
E2	11374345	11374487	refGene_gn=CD3EAP;refGene_tx=XM_003997680.4
E2	11374853	11374875	refGene_gn=CD3EAP;refGene_tx=XM_003997680.4
E2	11380874	11380929
refGene_gn=PPP1R13L;refGene_tx=XM_023246172.1;refGene_gn=PPP1R13L;refGene_tx=XM_023246173.1;refGene_gn=PPP1R13L;refGene_tx=XM_023246174.1;refGene_gn=PPP1R13L;refGene_tx=XM_023246175.1
E2	11381035	11381178
refGene_gn=PPP1R13L;refGene_tx=XM_023246172.1;refGene_gn=PPP1R13L;refGene_tx=XM_023246173.1;refGene_gn=PPP1R13L;refGene_tx=XM_023246174.1;refGene_gn=PPP1R13L;refGene_tx=XM_023246175.1;refGene_gn=PPP1R13L;refGene_tx=XM_023246177.1
E2	11381555	11382069
refGene_gn=PPP1R13L;refGene_tx=XM_023246172.1;refGene_gn=PPP1R13L;refGene_tx=XM_023246173.1;refGene_gn=PPP1R13L;refGene_tx=XM_023246174.1;refGene_gn=PPP1R13L;refGene_tx=XM_023246175.1;refGene_gn=PPP1R13L;refGene_tx=XM_023246177.1
E2	11382158	11382257
refGene_gn=PPP1R13L;refGene_tx=XM_023246172.1;refGene_gn=PPP1R13L;refG

ene_tx=XM_023246173.1;refGene_gn=PPP1R13L;refGene_tx=XM_023246174.1;refGene_gn=PPP1R13L;refGene_tx=XM_023246175.1;refGene_gn=PPP1R13L;refGene_tx=XM_023246177.1
E2 11382340 11382432
refGene_gn=PPP1R13L;refGene_tx=XM_023246172.1;refGene_gn=PPP1R13L;refGene_tx=XM_023246173.1;refGene_gn=PPP1R13L;refGene_tx=XM_023246174.1;refGene_gn=PPP1R13L;refGene_tx=XM_023246175.1;refGene_gn=PPP1R13L;refGene_tx=XM_023246177.1
E2 11382733 11383196
refGene_gn=PPP1R13L;refGene_tx=XM_023246172.1;refGene_gn=PPP1R13L;refGene_tx=XM_023246173.1;refGene_gn=PPP1R13L;refGene_tx=XM_023246174.1;refGene_gn=PPP1R13L;refGene_tx=XM_023246175.1;refGene_gn=PPP1R13L;refGene_tx=XM_023246177.1
E2 11384345 11384809
refGene_gn=PPP1R13L;refGene_tx=XM_023246172.1;refGene_gn=PPP1R13L;refGene_tx=XM_023246173.1;refGene_gn=PPP1R13L;refGene_tx=XM_023246174.1;refGene_gn=PPP1R13L;refGene_tx=XM_023246175.1;refGene_gn=PPP1R13L;refGene_tx=XM_023246177.1
E2 11388399 11388531
refGene_gn=PPP1R13L;refGene_tx=XM_023246172.1;refGene_gn=PPP1R13L;refGene_tx=XM_023246173.1;refGene_gn=PPP1R13L;refGene_tx=XM_023246174.1;refGene_gn=PPP1R13L;refGene_tx=XM_023246175.1;refGene_gn=PPP1R13L;refGene_tx=XM_023246177.1
E2 11388627 11388761
refGene_gn=PPP1R13L;refGene_tx=XM_023246172.1;refGene_gn=PPP1R13L;refGene_tx=XM_023246173.1;refGene_gn=PPP1R13L;refGene_tx=XM_023246174.1;refGene_gn=PPP1R13L;refGene_tx=XM_023246175.1;refGene_gn=PPP1R13L;refGene_tx=XM_023246177.1
E2 11388882 11389049
refGene_gn=PPP1R13L;refGene_tx=XM_023246172.1;refGene_gn=PPP1R13L;refGene_tx=XM_023246173.1;refGene_gn=PPP1R13L;refGene_tx=XM_023246174.1;refGene_gn=PPP1R13L;refGene_tx=XM_023246175.1;refGene_gn=PPP1R13L;refGene_tx=XM_023246177.1
E2 11390608 11390808
refGene_gn=PPP1R13L;refGene_tx=XM_023246172.1;refGene_gn=PPP1R13L;refGene_tx=XM_023246173.1;refGene_gn=PPP1R13L;refGene_tx=XM_023246174.1;refGene_gn=PPP1R13L;refGene_tx=XM_023246175.1;refGene_gn=PPP1R13L;refGene_tx=XM_023246177.1
E2 11391818 11391857
refGene_gn=PPP1R13L;refGene_tx=XM_023246172.1;refGene_gn=PPP1R13L;refGene_tx=XM_023246173.1;refGene_gn=PPP1R13L;refGene_tx=XM_023246174.1;refGene_gn=PPP1R13L;refGene_tx=XM_023246175.1;refGene_gn=PPP1R13L;refGene_tx=XM_023246177.1
E2 11402140 11402145 refGene_gn=ERCC2;refGene_tx=XM_023244940.1
E2 11402464 11402564
refGene_gn=ERCC2;refGene_tx=XM_023244940.1;refGene_gn=ERCC2;refGene_tx=XM_023244941.1
E2 11403897 11403975
refGene_gn=ERCC2;refGene_tx=XM_023244940.1;refGene_gn=ERCC2;refGene_tx=XM_023244941.1

E2 11404059 11404122
refGene_gn=ERCC2;refGene_tx=XM_023244940.1;refGene_gn=ERCC2;refGene_tx=XM_023244941.1
E2 11404293 11404407
refGene_gn=ERCC2;refGene_tx=XM_023244940.1;refGene_gn=ERCC2;refGene_tx=XM_023244941.1
E2 11407506 11407623
refGene_gn=ERCC2;refGene_tx=XM_023244940.1;refGene_gn=ERCC2;refGene_tx=XM_023244941.1;refGene_gn=ERCC2;refGene_tx=XM_023244942.1
E2 11407716 11407833
refGene_gn=ERCC2;refGene_tx=XM_023244940.1;refGene_gn=ERCC2;refGene_tx=XM_023244941.1;refGene_gn=ERCC2;refGene_tx=XM_023244942.1
E2 11408053 11408177
refGene_gn=ERCC2;refGene_tx=XM_023244940.1;refGene_gn=ERCC2;refGene_tx=XM_023244941.1;refGene_gn=ERCC2;refGene_tx=XM_023244942.1
E2 11408259 11408356
refGene_gn=ERCC2;refGene_tx=XM_023244940.1;refGene_gn=ERCC2;refGene_tx=XM_023244941.1;refGene_gn=ERCC2;refGene_tx=XM_023244942.1
E2 11408544 11408678
refGene_gn=ERCC2;refGene_tx=XM_023244940.1;refGene_gn=ERCC2;refGene_tx=XM_023244941.1;refGene_gn=ERCC2;refGene_tx=XM_023244942.1
E2 11408758 11408927
refGene_gn=ERCC2;refGene_tx=XM_023244940.1;refGene_gn=ERCC2;refGene_tx=XM_023244941.1;refGene_gn=ERCC2;refGene_tx=XM_023244942.1
E2 11410588 11410707
refGene_gn=ERCC2;refGene_tx=XM_023244940.1;refGene_gn=ERCC2;refGene_tx=XM_023244941.1;refGene_gn=ERCC2;refGene_tx=XM_023244942.1
E2 11412979 11413049
refGene_gn=ERCC2;refGene_tx=XM_023244940.1;refGene_gn=ERCC2;refGene_tx=XM_023244941.1;refGene_gn=ERCC2;refGene_tx=XM_023244942.1
E2 11413132 11413202
refGene_gn=ERCC2;refGene_tx=XM_023244940.1;refGene_gn=ERCC2;refGene_tx=XM_023244941.1;refGene_gn=ERCC2;refGene_tx=XM_023244942.1
E2 11413295 11413397
refGene_gn=ERCC2;refGene_tx=XM_023244940.1;refGene_gn=ERCC2;refGene_tx=XM_023244941.1;refGene_gn=ERCC2;refGene_tx=XM_023244942.1
E2 11414939 11415003
refGene_gn=ERCC2;refGene_tx=XM_023244940.1;refGene_gn=ERCC2;refGene_tx=XM_023244941.1;refGene_gn=ERCC2;refGene_tx=XM_023244942.1
E2 11415629 11415751
refGene_gn=ERCC2;refGene_tx=XM_023244940.1;refGene_gn=ERCC2;refGene_tx=XM_023244941.1;refGene_gn=ERCC2;refGene_tx=XM_023244942.1
E2 11416869 11416962
refGene_gn=ERCC2;refGene_tx=XM_023244940.1;refGene_gn=ERCC2;refGene_tx=XM_023244941.1;refGene_gn=ERCC2;refGene_tx=XM_023244942.1
E2 11417062 11417135
refGene_gn=ERCC2;refGene_tx=XM_023244940.1;refGene_gn=ERCC2;refGene_tx=XM_023244941.1;refGene_gn=ERCC2;refGene_tx=XM_023244942.1
E2 11417373 11417444
refGene_gn=ERCC2;refGene_tx=XM_023244940.1;refGene_gn=ERCC2;refGene_tx

=XM_023244941.1;refGene_gn=ERCC2;refGene_tx=XM_023244942.1
E2 11417557 11417701
refGene_gn=ERCC2;refGene_tx=XM_023244940.1;refGene_gn=ERCC2;refGene_tx
=XM_023244941.1;refGene_gn=ERCC2;refGene_tx=XM_023244942.1
E2 11417831 11417975
refGene_gn=ERCC2;refGene_tx=XM_023244940.1;refGene_gn=ERCC2;refGene_tx
=XM_023244941.1;refGene_gn=ERCC2;refGene_tx=XM_023244942.1
E2 11418316 11418409
refGene_gn=ERCC2;refGene_tx=XM_023244940.1;refGene_gn=ERCC2;refGene_tx
=XM_023244941.1;refGene_gn=ERCC2;refGene_tx=XM_023244942.1
E2 11418670 11418742 refGene_gn=KLC3;refGene_tx=XM_023244943.1
E2 11418928 11418992 refGene_gn=KLC3;refGene_tx=XM_023244943.1
E2 11419181 11419288 refGene_gn=KLC3;refGene_tx=XM_023244943.1
E2 11419371 11419412 refGene_gn=KLC3;refGene_tx=XM_023244943.1
E2 11419489 11419580 refGene_gn=KLC3;refGene_tx=XM_023244943.1
E2 11420103 11420277 refGene_gn=KLC3;refGene_tx=XM_023244943.1
E2 11420715 11420817 refGene_gn=KLC3;refGene_tx=XM_023244943.1
E2 11420890 11420978 refGene_gn=KLC3;refGene_tx=XM_023244943.1
E2 11421321 11421541 refGene_gn=KLC3;refGene_tx=XM_023244943.1
E2 11421920 11421990 refGene_gn=KLC3;refGene_tx=XM_023244943.1
E2 11422296 11422527 refGene_gn=KLC3;refGene_tx=XM_023244943.1
E2 11423019 11423277 refGene_gn=KLC3;refGene_tx=XM_023244943.1
E2 11439087 11439280 refGene_gn=CKM;refGene_tx=XM_003997644.5
E2 11440047 11440202 refGene_gn=CKM;refGene_tx=XM_003997644.5
E2 11440585 11440718 refGene_gn=CKM;refGene_tx=XM_003997644.5
E2 11443384 11443556 refGene_gn=CKM;refGene_tx=XM_003997644.5
E2 11444568 11444692 refGene_gn=CKM;refGene_tx=XM_003997644.5
E2 11445596 11445786 refGene_gn=CKM;refGene_tx=XM_003997644.5
E2 11446302 11446481 refGene_gn=CKM;refGene_tx=XM_003997644.5
E2 11450572 11450909
refGene_gn=MARK4;refGene_tx=XM_023246016.1;refGene_gn=MARK4;refGene_tx
=XM_023246017.1
E2 11452605 11452650
refGene_gn=MARK4;refGene_tx=XM_023246016.1;refGene_gn=MARK4;refGene_tx
=XM_023246017.1
E2 11453503 11453583 refGene_gn=MARK4;refGene_tx=XM_023246017.1
E2 11453780 11454059
refGene_gn=MARK4;refGene_tx=XM_023246016.1;refGene_gn=MARK4;refGene_tx
=XM_023246017.1
E2 11456156 11456260
refGene_gn=MARK4;refGene_tx=XM_023246016.1;refGene_gn=MARK4;refGene_tx
=XM_023246017.1
E2 11459685 11459903
refGene_gn=MARK4;refGene_tx=XM_023246016.1;refGene_gn=MARK4;refGene_tx
=XM_023246017.1
E2 11462525 11462685
refGene_gn=MARK4;refGene_tx=XM_023246016.1;refGene_gn=MARK4;refGene_tx
=XM_023246017.1
E2 11462793 11462903
refGene_gn=MARK4;refGene_tx=XM_023246016.1;refGene_gn=MARK4;refGene_tx

=XM_023246017.1
E2 11464249 11464349
refGene_gn=MARK4;refGene_tx=XM_023246016.1;refGene_gn=MARK4;refGene_tx
=XM_023246017.1
E2 11464827 11464947
refGene_gn=MARK4;refGene_tx=XM_023246016.1;refGene_gn=MARK4;refGene_tx
=XM_023246017.1
E2 11468181 11468418
refGene_gn=MARK4;refGene_tx=XM_023246016.1;refGene_gn=MARK4;refGene_tx
=XM_023246017.1
E2 11470452 11470509
refGene_gn=MARK4;refGene_tx=XM_023246016.1;refGene_gn=MARK4;refGene_tx
=XM_023246017.1
E2 11471139 11471210
refGene_gn=MARK4;refGene_tx=XM_023246016.1;refGene_gn=MARK4;refGene_tx
=XM_023246017.1
E2 11471287 11471353
refGene_gn=MARK4;refGene_tx=XM_023246016.1;refGene_gn=MARK4;refGene_tx
=XM_023246017.1
E2 11472063 11472112
refGene_gn=MARK4;refGene_tx=XM_023246016.1;refGene_gn=MARK4;refGene_tx
=XM_023246017.1
E2 11472215 11472269
refGene_gn=MARK4;refGene_tx=XM_023246016.1;refGene_gn=MARK4;refGene_tx
=XM_023246017.1
E2 11474749 11474950
refGene_gn=MARK4;refGene_tx=XM_023246016.1;refGene_gn=MARK4;refGene_tx
=XM_023246017.1
E2 11479559 11479610
refGene_gn=MARK4;refGene_tx=XM_023246016.1;refGene_gn=MARK4;refGene_tx
=XM_023246017.1
E2 11488230 11488753 refGene_gn=EXOC3L2;refGene_tx=XM_023245234.1
E2 11491623 11492257 refGene_gn=EXOC3L2;refGene_tx=XM_023245234.1
E2 11494558 11494670 refGene_gn=EXOC3L2;refGene_tx=XM_023245234.1
E2 11495853 11495955 refGene_gn=EXOC3L2;refGene_tx=XM_023245234.1
E2 11496071 11496172 refGene_gn=EXOC3L2;refGene_tx=XM_023245234.1
E2 11496400 11496511 refGene_gn=EXOC3L2;refGene_tx=XM_023245234.1
E2 11498099 11498235 refGene_gn=EXOC3L2;refGene_tx=XM_023245234.1
E2 11501563 11501686 refGene_gn=EXOC3L2;refGene_tx=XM_023245234.1
E2 11502125 11502281 refGene_gn=EXOC3L2;refGene_tx=XM_023245234.1
E2 11503019 11503141 refGene_gn=EXOC3L2;refGene_tx=XM_023245234.1
E2 11504800 11505089 refGene_gn=EXOC3L2;refGene_tx=XM_023245234.1
E2 11531559 11532156 refGene_gn=BLOC1S3;refGene_tx=XM_023244956.1
E2 11533225 11533309
refGene_gn=TRAPPC6A;refGene_tx=XM_003997645.5;refGene_gn=TRAPPC6A;refG
ene_tx=XM_006941115.4;refGene_gn=TRAPPC6A;refGene_tx=XM_006941116.4
E2 11541080 11541148
refGene_gn=TRAPPC6A;refGene_tx=XM_003997645.5;refGene_gn=TRAPPC6A;refG
ene_tx=XM_006941115.4;refGene_gn=TRAPPC6A;refGene_tx=XM_019819369.2
E2 11541327 11541464

refGene_gn=TRAPPC6A;refGene_tx=XM_003997645.5;refGene_gn=TRAPPC6A;refGene_tx=XM_006941115.4;refGene_gn=TRAPPC6A;refGene_tx=XM_006941116.4;refGene_gn=TRAPPC6A;refGene_tx=XM_019819369.2
E2 11542088 11542172
refGene_gn=TRAPPC6A;refGene_tx=XM_003997645.5;refGene_gn=TRAPPC6A;refGene_tx=XM_006941115.4;refGene_gn=TRAPPC6A;refGene_tx=XM_006941116.4;refGene_gn=TRAPPC6A;refGene_tx=XM_019819369.2
E2 11542377 11542471
refGene_gn=TRAPPC6A;refGene_tx=XM_003997645.5;refGene_gn=TRAPPC6A;refGene_tx=XM_006941115.4;refGene_gn=TRAPPC6A;refGene_tx=XM_006941116.4;refGene_gn=TRAPPC6A;refGene_tx=XM_019819369.2
E2 11543289 11543321
refGene_gn=TRAPPC6A;refGene_tx=XM_003997645.5;refGene_gn=TRAPPC6A;refGene_tx=XM_006941115.4;refGene_gn=TRAPPC6A;refGene_tx=XM_006941116.4;refGene_gn=TRAPPC6A;refGene_tx=XM_019819369.2
E2 11546597 11546697
refGene_gn=NKPD1;refGene_tx=XM_023244949.1;refGene_gn=NKPD1;refGene_tx=XM_023244950.1;refGene_gn=NKPD1;refGene_tx=XM_023244951.1;refGene_gn=NKPD1;refGene_tx=XM_023244952.1;refGene_gn=NKPD1;refGene_tx=XM_023244955.1
E2 11547468 11547861
refGene_gn=NKPD1;refGene_tx=XM_023244949.1;refGene_gn=NKPD1;refGene_tx=XM_023244950.1;refGene_gn=NKPD1;refGene_tx=XM_023244951.1;refGene_gn=NKPD1;refGene_tx=XM_023244952.1;refGene_gn=NKPD1;refGene_tx=XM_023244953.1;refGene_gn=NKPD1;refGene_tx=XM_023244954.1
E2 11550053 11550185
refGene_gn=NKPD1;refGene_tx=XM_023244949.1;refGene_gn=NKPD1;refGene_tx=XM_023244950.1;refGene_gn=NKPD1;refGene_tx=XM_023244951.1;refGene_gn=NKPD1;refGene_tx=XM_023244952.1;refGene_gn=NKPD1;refGene_tx=XM_023244953.1;refGene_gn=NKPD1;refGene_tx=XM_023244954.1;refGene_gn=NKPD1;refGene_tx=XM_023244955.1
E2 11551554 11553404
refGene_gn=NKPD1;refGene_tx=XM_023244949.1;refGene_gn=NKPD1;refGene_tx=XM_023244950.1;refGene_gn=NKPD1;refGene_tx=XM_023244951.1;refGene_gn=NKPD1;refGene_tx=XM_023244952.1;refGene_gn=NKPD1;refGene_tx=XM_023244953.1;refGene_gn=NKPD1;refGene_tx=XM_023244954.1;refGene_gn=NKPD1;refGene_tx=XM_023244955.1
E2 11556593 11556676
refGene_gn=PPP1R37;refGene_tx=XM_011289843.3;refGene_gn=PPP1R37;refGene_tx=XM_023244856.1;refGene_gn=PPP1R37;refGene_tx=XM_023244857.1
E2 11557008 11557735
refGene_gn=PPP1R37;refGene_tx=XM_011289843.3;refGene_gn=PPP1R37;refGene_tx=XM_023244856.1;refGene_gn=PPP1R37;refGene_tx=XM_023244857.1
E2 11557877 11558044
refGene_gn=PPP1R37;refGene_tx=XM_011289843.3;refGene_gn=PPP1R37;refGene_tx=XM_023244856.1;refGene_gn=PPP1R37;refGene_tx=XM_023244857.1
E2 11558132 11558274
refGene_gn=PPP1R37;refGene_tx=XM_011289843.3;refGene_gn=PPP1R37;refGene_tx=XM_023244856.1;refGene_gn=PPP1R37;refGene_tx=XM_023244857.1
E2 11559088 11559201

refGene_gn=PPP1R37;refGene_tx=XM_011289843.3;refGene_gn=PPP1R37;refGene_tx=XM_023244856.1;refGene_gn=PPP1R37;refGene_tx=XM_023244857.1
E2 11560226 11560382
refGene_gn=PPP1R37;refGene_tx=XM_011289843.3;refGene_gn=PPP1R37;refGene_tx=XM_023244856.1;refGene_gn=PPP1R37;refGene_tx=XM_023244857.1
E2 11560476 11560627
refGene_gn=PPP1R37;refGene_tx=XM_011289843.3;refGene_gn=PPP1R37;refGene_tx=XM_023244856.1;refGene_gn=PPP1R37;refGene_tx=XM_023244857.1
E2 11561091 11561211
refGene_gn=PPP1R37;refGene_tx=XM_011289843.3;refGene_gn=PPP1R37;refGene_tx=XM_023244856.1;refGene_gn=PPP1R37;refGene_tx=XM_023244857.1
E2 11561903 11562004
refGene_gn=PPP1R37;refGene_tx=XM_011289843.3;refGene_gn=PPP1R37;refGene_tx=XM_023244856.1;refGene_gn=PPP1R37;refGene_tx=XM_023244857.1
E2 11562240 11562286
refGene_gn=PPP1R37;refGene_tx=XM_011289843.3;refGene_gn=PPP1R37;refGene_tx=XM_023244856.1;refGene_gn=PPP1R37;refGene_tx=XM_023244857.1
E2 11564016 11564114
refGene_gn=PPP1R37;refGene_tx=XM_011289843.3;refGene_gn=PPP1R37;refGene_tx=XM_023244856.1;refGene_gn=PPP1R37;refGene_tx=XM_023244857.1
E2 11578378 11578409 refGene_gn=PPP1R37;refGene_tx=XM_023244857.1
E2 11596405 11596607
refGene_gn=PPP1R37;refGene_tx=XM_011289843.3;refGene_gn=PPP1R37;refGene_tx=XM_023244856.1
E2 11598967 11599371
refGene_gn=GEMIN7;refGene_tx=XM_019819371.2;refGene_gn=GEMIN7;refGene_tx=XM_019819372.2;refGene_gn=GEMIN7;refGene_tx=XM_019819373.2;refGene_gn=GEMIN7;refGene_tx=XM_019819374.2;refGene_gn=GEMIN7;refGene_tx=XM_019819375.2;refGene_gn=GEMIN7;refGene_tx=XM_023246008.1;refGene_gn=GEMIN7;refGene_tx=XM_023246010.1
E2 11606110 11606150
refGene_gn=GEMIN7;refGene_tx=XM_019819371.2;refGene_gn=GEMIN7;refGene_tx=XM_019819372.2;refGene_gn=GEMIN7;refGene_tx=XM_019819373.2;refGene_gn=GEMIN7;refGene_tx=XM_019819374.2;refGene_gn=GEMIN7;refGene_tx=XM_019819375.2;refGene_gn=GEMIN7;refGene_tx=XM_023246008.1
E2 11611573 11611871 refGene_gn=ZNF296;refGene_tx=XM_011289809.3
E2 11612091 11612241 refGene_gn=ZNF296;refGene_tx=XM_011289809.3
E2 11614107 11615081 refGene_gn=ZNF296;refGene_tx=XM_011289809.3
E2 11615708 11616053
refGene_gn=CLASRP;refGene_tx=XM_023246003.1;refGene_gn=CLASRP;refGene_tx=XM_023246004.1;refGene_gn=CLASRP;refGene_tx=XM_023246005.1;refGene_gn=CLASRP;refGene_tx=XM_023246007.1
E2 11616151 11616386
refGene_gn=CLASRP;refGene_tx=XM_023246003.1;refGene_gn=CLASRP;refGene_tx=XM_023246004.1;refGene_gn=CLASRP;refGene_tx=XM_023246005.1;refGene_gn=CLASRP;refGene_tx=XM_023246007.1
E2 11617705 11617709 refGene_gn=CLASRP;refGene_tx=XM_023246006.1
E2 11618099 11618146
refGene_gn=CLASRP;refGene_tx=XM_023246003.1;refGene_gn=CLASRP;refGene_tx=XM_023246004.1;refGene_gn=CLASRP;refGene_tx=XM_023246005.1;refGene_

gn=CLASRP;refGene_tx=XM_023246006.1;refGene_gn=CLASRP;refGene_tx=XM_023246007.1
E2 11618223 11618282
refGene_gn=CLASRP;refGene_tx=XM_023246003.1;refGene_gn=CLASRP;refGene_tx=XM_023246004.1;refGene_gn=CLASRP;refGene_tx=XM_023246005.1;refGene_gn=CLASRP;refGene_tx=XM_023246006.1;refGene_gn=CLASRP;refGene_tx=XM_023246007.1
E2 11618505 11618566
refGene_gn=CLASRP;refGene_tx=XM_023246003.1;refGene_gn=CLASRP;refGene_tx=XM_023246004.1;refGene_gn=CLASRP;refGene_tx=XM_023246005.1;refGene_gn=CLASRP;refGene_tx=XM_023246006.1;refGene_gn=CLASRP;refGene_tx=XM_023246007.1
E2 11618910 11618950
refGene_gn=CLASRP;refGene_tx=XM_023246003.1;refGene_gn=CLASRP;refGene_tx=XM_023246004.1;refGene_gn=CLASRP;refGene_tx=XM_023246005.1;refGene_gn=CLASRP;refGene_tx=XM_023246006.1;refGene_gn=CLASRP;refGene_tx=XM_023246007.1
E2 11619272 11619539
refGene_gn=CLASRP;refGene_tx=XM_023246003.1;refGene_gn=CLASRP;refGene_tx=XM_023246004.1;refGene_gn=CLASRP;refGene_tx=XM_023246005.1;refGene_gn=CLASRP;refGene_tx=XM_023246006.1;refGene_gn=CLASRP;refGene_tx=XM_023246007.1
E2 11621173 11621464
refGene_gn=CLASRP;refGene_tx=XM_023246003.1;refGene_gn=CLASRP;refGene_tx=XM_023246004.1;refGene_gn=CLASRP;refGene_tx=XM_023246005.1;refGene_gn=CLASRP;refGene_tx=XM_023246006.1;refGene_gn=CLASRP;refGene_tx=XM_023246007.1
E2 11621580 11621796
refGene_gn=CLASRP;refGene_tx=XM_023246003.1;refGene_gn=CLASRP;refGene_tx=XM_023246004.1;refGene_gn=CLASRP;refGene_tx=XM_023246005.1;refGene_gn=CLASRP;refGene_tx=XM_023246006.1;refGene_gn=CLASRP;refGene_tx=XM_023246007.1
E2 11622643 11622685
refGene_gn=CLASRP;refGene_tx=XM_023246003.1;refGene_gn=CLASRP;refGene_tx=XM_023246004.1;refGene_gn=CLASRP;refGene_tx=XM_023246005.1;refGene_gn=CLASRP;refGene_tx=XM_023246006.1;refGene_gn=CLASRP;refGene_tx=XM_023246007.1
E2 11623964 11624038
refGene_gn=CLASRP;refGene_tx=XM_023246003.1;refGene_gn=CLASRP;refGene_tx=XM_023246004.1;refGene_gn=CLASRP;refGene_tx=XM_023246005.1;refGene_gn=CLASRP;refGene_tx=XM_023246006.1;refGene_gn=CLASRP;refGene_tx=XM_023246007.1
E2 11624120 11624199
refGene_gn=CLASRP;refGene_tx=XM_023246003.1;refGene_gn=CLASRP;refGene_tx=XM_023246004.1;refGene_gn=CLASRP;refGene_tx=XM_023246005.1;refGene_gn=CLASRP;refGene_tx=XM_023246006.1;refGene_gn=CLASRP;refGene_tx=XM_023246007.1
E2 11625194 11625291
refGene_gn=CLASRP;refGene_tx=XM_023246003.1;refGene_gn=CLASRP;refGene_tx=XM_023246004.1;refGene_gn=CLASRP;refGene_tx=XM_023246005.1;refGene_

gn=CLASRP;refGene_tx=XM_023246006.1;refGene_gn=CLASRP;refGene_tx=XM_023246007.1
E2 11626124 11626273 refGene_gn=CLASRP;refGene_tx=XM_023246003.1;refGene_gn=CLASRP;refGene_tx=XM_023246004.1;refGene_gn=CLASRP;refGene_tx=XM_023246005.1;refGene_gn=CLASRP;refGene_tx=XM_023246006.1
E2 11627448 11627533 refGene_gn=CLASRP;refGene_tx=XM_023246003.1;refGene_gn=CLASRP;refGene_tx=XM_023246004.1;refGene_gn=CLASRP;refGene_tx=XM_023246005.1;refGene_gn=CLASRP;refGene_tx=XM_023246006.1
E2 11629825 11629905 refGene_gn=CLASRP;refGene_tx=XM_023246003.1;refGene_gn=CLASRP;refGene_tx=XM_023246004.1;refGene_gn=CLASRP;refGene_tx=XM_023246005.1;refGene_gn=CLASRP;refGene_tx=XM_023246006.1
E2 11630119 11630221 refGene_gn=CLASRP;refGene_tx=XM_023246003.1;refGene_gn=CLASRP;refGene_tx=XM_023246004.1;refGene_gn=CLASRP;refGene_tx=XM_023246005.1;refGene_gn=CLASRP;refGene_tx=XM_023246006.1
E2 11630810 11630908 refGene_gn=CLASRP;refGene_tx=XM_023246003.1;refGene_gn=CLASRP;refGene_tx=XM_023246004.1;refGene_gn=CLASRP;refGene_tx=XM_023246005.1;refGene_gn=CLASRP;refGene_tx=XM_023246006.1
E2 11638250 11638349 refGene_gn=CLASRP;refGene_tx=XM_023246003.1;refGene_gn=CLASRP;refGene_tx=XM_023246004.1;refGene_gn=CLASRP;refGene_tx=XM_023246005.1;refGene_gn=CLASRP;refGene_tx=XM_023246006.1
E2 11640083 11640472 refGene_gn=RELB;refGene_tx=XM_023245919.1
E2 11642482 11642560 refGene_gn=RELB;refGene_tx=XM_023245919.1
E2 11642681 11642750 refGene_gn=RELB;refGene_tx=XM_023245919.1
E2 11643482 11643698 refGene_gn=RELB;refGene_tx=XM_023245919.1
E2 11647660 11647765 refGene_gn=RELB;refGene_tx=XM_023245919.1
E2 11649527 11649659 refGene_gn=RELB;refGene_tx=XM_023245919.1
E2 11649846 11649938 refGene_gn=RELB;refGene_tx=XM_023245919.1
E2 11651719 11651877 refGene_gn=RELB;refGene_tx=XM_023245919.1
E2 11662160 11662373 refGene_gn=RELB;refGene_tx=XM_023245919.1
E2 11662374 11662508 refGene_gn=RELB;refGene_tx=XM_023245919.1
E2 11664214 11664223 refGene_gn=RELB;refGene_tx=XM_023245919.1
E2 11667659 11667707 refGene_gn=RELB;refGene_tx=XM_023245919.1
E2 11668615 11668924 refGene_gn=RELB;refGene_tx=XM_023245919.1
E2 11669576 11669619 refGene_gn=RELB;refGene_tx=XM_023245919.1
E2 11673962 11674261 refGene_gn=CLPTM1;refGene_tx=XM_023244957.1
E2 11674397 11674565 refGene_gn=CLPTM1;refGene_tx=XM_023244957.1
E2 11675172 11675308 refGene_gn=CLPTM1;refGene_tx=XM_023244957.1
E2 11675602 11675698 refGene_gn=CLPTM1;refGene_tx=XM_023244957.1
E2 11675946 11676137 refGene_gn=CLPTM1;refGene_tx=XM_023244957.1
E2 11678109 11678203 refGene_gn=CLPTM1;refGene_tx=XM_023244957.1
E2 11678923 11679168 refGene_gn=CLPTM1;refGene_tx=XM_023244957.1
E2 11679739 11679860 refGene_gn=CLPTM1;refGene_tx=XM_023244957.1
E2 11680705 11680791 refGene_gn=CLPTM1;refGene_tx=XM_023244957.1
E2 11685024 11685142 refGene_gn=CLPTM1;refGene_tx=XM_023244957.1

E2 11687092 11687251 refGene_gn=CLPTM1;refGene_tx=XM_023244957.1
E2 11688407 11688531 refGene_gn=CLPTM1;refGene_tx=XM_023244957.1
E2 11699079 11699192 refGene_gn=CLPTM1;refGene_tx=XM_023244957.1
E2 11705164 11705236 refGene_gn=CLPTM1;refGene_tx=XM_023244957.1
E2 11709895 11709986 refGene_gn=APOC2;refGene_tx=XM_003997648.5
E2 11710225 11710385 refGene_gn=APOC2;refGene_tx=XM_003997648.5
E2 11710555 11710610 refGene_gn=APOC2;refGene_tx=XM_003997648.5
E2 11712298 11712464
refGene_gn=APOC4;refGene_tx=XM_003997689.4;refGene_gn=APOC4;refGene_tx=XM_006941120.4;refGene_gn=APOC4;refGene_tx=XM_019819378.2;refGene_gn=APOC4;refGene_tx=XM_019819379.1;refGene_gn=APOC4;refGene_tx=XM_019819380.2
E2 11712726 11712856
refGene_gn=APOC4;refGene_tx=XM_003997689.4;refGene_gn=APOC4;refGene_tx=XM_006941120.4;refGene_gn=APOC4;refGene_tx=XM_019819378.2;refGene_gn=APOC4;refGene_tx=XM_019819379.1;refGene_gn=APOC4;refGene_tx=XM_019819380.2
E2 11712991 11713031 refGene_gn=APOC4;refGene_tx=XM_019819379.1
E2 11713967 11714043 refGene_gn=APOC4;refGene_tx=XM_003997689.4
E2 11714459 11714587
refGene_gn=APOC4;refGene_tx=XM_006941120.4;refGene_gn=APOC4;refGene_tx=XM_019819378.2;refGene_gn=APOC4;refGene_tx=XM_019819380.2
E2 11717027 11717081 refGene_gn=APOC4;refGene_tx=XM_006941120.4
E2 11723694 11723752 refGene_gn=APOC1;refGene_tx=XM_019819381.2
E2 11725258 11725379 refGene_gn=APOC1;refGene_tx=XM_019819381.2
E2 11725910 11725968 refGene_gn=APOC1;refGene_tx=XM_019819381.2
E2 11729751 11730436 refGene_gn=APOE;refGene_tx=XM_023245793.1
E2 11730768 11730955 refGene_gn=APOE;refGene_tx=XM_023245793.1
E2 11731442 11731485 refGene_gn=APOE;refGene_tx=XM_023245793.1
E2 11734512 11734652 refGene_gn=TOMM40;refGene_tx=XM_023245792.1
E2 11735628 11735731 refGene_gn=TOMM40;refGene_tx=XM_023245792.1
E2 11735845 11735922 refGene_gn=TOMM40;refGene_tx=XM_023245792.1
E2 11736063 11736186 refGene_gn=TOMM40;refGene_tx=XM_023245792.1
E2 11741175 11741281 refGene_gn=TOMM40;refGene_tx=XM_023245792.1
E2 11741361 11741463 refGene_gn=TOMM40;refGene_tx=XM_023245792.1
E2 11742531 11742624 refGene_gn=TOMM40;refGene_tx=XM_023245792.1
E2 11743102 11743170 refGene_gn=TOMM40;refGene_tx=XM_023245792.1
E2 11743925 11744199 refGene_gn=TOMM40;refGene_tx=XM_023245792.1
E2 11746625 11746889 refGene_gn=NECTIN2;refGene_tx=XM_003997692.4
E2 11747442 11747529 refGene_gn=NECTIN2;refGene_tx=XM_003997692.4
E2 11747672 11747736 refGene_gn=NECTIN2;refGene_tx=XM_003997692.4
E2 11749927 11750081 refGene_gn=NECTIN2;refGene_tx=XM_003997692.4
E2 11757112 11757261 refGene_gn=NECTIN2;refGene_tx=XM_003997692.4
E2 11758779 11758897 refGene_gn=NECTIN2;refGene_tx=XM_003997692.4
E2 11760005 11760302 refGene_gn=NECTIN2;refGene_tx=XM_003997692.4
E2 11765622 11766018 refGene_gn=NECTIN2;refGene_tx=XM_003997692.4
E2 11777219 11777310 refGene_gn=NECTIN2;refGene_tx=XM_003997692.4
E2 11784777 11784783 refGene_gn=BCAM;refGene_tx=XM_023245130.1
E2 11784899 11785017 refGene_gn=BCAM;refGene_tx=XM_023245130.1
E2 11785857 11786002 refGene_gn=BCAM;refGene_tx=XM_023245130.1

E2 11786107 11786252 refGene_gn=BCAM;refGene_tx=XM_023245130.1
E2 11786435 11786572 refGene_gn=BCAM;refGene_tx=XM_023245130.1
E2 11786764 11786906 refGene_gn=BCAM;refGene_tx=XM_023245130.1
E2 11787014 11787130 refGene_gn=BCAM;refGene_tx=XM_023245130.1
E2 11789158 11789315 refGene_gn=BCAM;refGene_tx=XM_023245130.1
E2 11789682 11789819 refGene_gn=BCAM;refGene_tx=XM_023245130.1
E2 11790285 11790468 refGene_gn=BCAM;refGene_tx=XM_023245130.1
E2 11790563 11790660 refGene_gn=BCAM;refGene_tx=XM_023245130.1
E2 11790878 11790949 refGene_gn=BCAM;refGene_tx=XM_023245130.1
E2 11791040 11791269 refGene_gn=BCAM;refGene_tx=XM_023245130.1
E2 11791865 11791987 refGene_gn=BCAM;refGene_tx=XM_023245130.1
E2 11793944 11794026 refGene_gn=BCAM;refGene_tx=XM_023245130.1
E2 11799847 11799898
refGene_gn=CBLC;refGene_tx=XM_023245372.1;refGene_gn=CBLC;refGene_tx=X
M_023245373.1;refGene_gn=CBLC;refGene_tx=XM_023245374.1
E2 11800022 11800138
refGene_gn=CBLC;refGene_tx=XM_023245372.1;refGene_gn=CBLC;refGene_tx=X
M_023245373.1
E2 11802701 11802761
refGene_gn=CBLC;refGene_tx=XM_023245372.1;refGene_gn=CBLC;refGene_tx=X
M_023245374.1
E2 11803085 11803247
refGene_gn=CBLC;refGene_tx=XM_023245372.1;refGene_gn=CBLC;refGene_tx=X
M_023245373.1;refGene_gn=CBLC;refGene_tx=XM_023245374.1
E2 11804987 11805119
refGene_gn=CBLC;refGene_tx=XM_023245372.1;refGene_gn=CBLC;refGene_tx=X
M_023245373.1;refGene_gn=CBLC;refGene_tx=XM_023245374.1
E2 11805806 11805894
refGene_gn=CBLC;refGene_tx=XM_023245372.1;refGene_gn=CBLC;refGene_tx=X
M_023245373.1;refGene_gn=CBLC;refGene_tx=XM_023245374.1
E2 11807765 11807903
refGene_gn=CBLC;refGene_tx=XM_023245372.1;refGene_gn=CBLC;refGene_tx=X
M_023245373.1;refGene_gn=CBLC;refGene_tx=XM_023245374.1
E2 11808039 11808161
refGene_gn=CBLC;refGene_tx=XM_023245372.1;refGene_gn=CBLC;refGene_tx=X
M_023245373.1;refGene_gn=CBLC;refGene_tx=XM_023245374.1
E2 11808396 11808553
refGene_gn=CBLC;refGene_tx=XM_023245372.1;refGene_gn=CBLC;refGene_tx=X
M_023245373.1;refGene_gn=CBLC;refGene_tx=XM_023245374.1
E2 11811084 11811231
refGene_gn=CBLC;refGene_tx=XM_023245372.1;refGene_gn=CBLC;refGene_tx=X
M_023245373.1;refGene_gn=CBLC;refGene_tx=XM_023245374.1
E2 11813098 11813451
refGene_gn=CBLC;refGene_tx=XM_023245372.1;refGene_gn=CBLC;refGene_tx=X
M_023245373.1;refGene_gn=CBLC;refGene_tx=XM_023245374.1
E2 11824388 11824576 refGene_gn=BCL3;refGene_tx=XM_023245186.1
E2 11826593 11826711 refGene_gn=BCL3;refGene_tx=XM_023245186.1
E2 11826949 11827117 refGene_gn=BCL3;refGene_tx=XM_023245186.1
E2 11827769 11827847 refGene_gn=BCL3;refGene_tx=XM_023245186.1
E2 11828063 11828152 refGene_gn=BCL3;refGene_tx=XM_023245186.1

E2 11828250 11828455 refGene_gn=BCL3;refGene_tx=XM_023245186.1
E2 11828684 11828793 refGene_gn=BCL3;refGene_tx=XM_023245186.1
E2 11833622 11833776 refGene_gn=BCL3;refGene_tx=XM_023245186.1
E2 11835469 11835725 refGene_gn=BCL3;refGene_tx=XM_023245186.1
E2 11859199 11859264 refGene_gn=CEACAM16;refGene_tx=XM_023245275.1
E2 11861505 11861832 refGene_gn=CEACAM16;refGene_tx=XM_023245275.1
E2 11863986 11864218 refGene_gn=CEACAM16;refGene_tx=XM_023245275.1
E2 11864219 11864266 refGene_gn=CEACAM16;refGene_tx=XM_023245275.1
E2 11865272 11865423 refGene_gn=CEACAM16;refGene_tx=XM_023245275.1
E2 11865424 11865549 refGene_gn=CEACAM16;refGene_tx=XM_023245275.1
E2 11865871 11866216 refGene_gn=CEACAM16;refGene_tx=XM_023245275.1
E2 11867429 11867466 refGene_gn=CEACAM16;refGene_tx=XM_023245275.1
E2 11877592 11877646
refGene_gn=CEACAM19;refGene_tx=XM_019819394.2;refGene_gn=CEACAM19;refGene_tx=XM_019819395.2
E2 11878071 11878104
refGene_gn=CEACAM19;refGene_tx=XM_019819394.2;refGene_gn=CEACAM19;refGene_tx=XM_019819395.2
E2 11878605 11878691
refGene_gn=CEACAM19;refGene_tx=XM_019819394.2;refGene_gn=CEACAM19;refGene_tx=XM_019819395.2
E2 11879283 11879330
refGene_gn=CEACAM19;refGene_tx=XM_019819394.2;refGene_gn=CEACAM19;refGene_tx=XM_019819395.2
E2 11879907 11879991 refGene_gn=CEACAM19;refGene_tx=XM_019819394.2
E2 11882395 11882546
refGene_gn=CEACAM19;refGene_tx=XM_019819394.2;refGene_gn=CEACAM19;refGene_tx=XM_019819395.2
E2 11894889 11895258
refGene_gn=CEACAM19;refGene_tx=XM_019819394.2;refGene_gn=CEACAM19;refGene_tx=XM_019819395.2
E2 11896146 11896201
refGene_gn=CEACAM19;refGene_tx=XM_019819394.2;refGene_gn=CEACAM19;refGene_tx=XM_019819395.2
E2 11906863 11906935 refGene_gn=PVR;refGene_tx=XM_003997727.5
E2 11907426 11907458 refGene_gn=PVR;refGene_tx=XM_003997727.5
E2 11910585 11910714 refGene_gn=PVR;refGene_tx=XM_003997727.5
E2 11911580 11911729 refGene_gn=PVR;refGene_tx=XM_003997727.5
E2 11915217 11915335 refGene_gn=PVR;refGene_tx=XM_003997727.5
E2 11918105 11918300 refGene_gn=PVR;refGene_tx=XM_003997727.5
E2 11918302 11918404 refGene_gn=PVR;refGene_tx=XM_003997727.5
E2 11920525 11920885 refGene_gn=PVR;refGene_tx=XM_003997727.5
E2 11924383 11924444 refGene_gn=PVR;refGene_tx=XM_003997727.5
E2 11935018 11935052 refGene_gn=IGSF23;refGene_tx=XM_023245296.1
E2 11936680 11936834 refGene_gn=IGSF23;refGene_tx=XM_023245296.1
E2 11938785 11939058 refGene_gn=IGSF23;refGene_tx=XM_023245296.1
E2 11940597 11940927 refGene_gn=IGSF23;refGene_tx=XM_023245296.1
E2 11944753 11944817 refGene_gn=IGSF23;refGene_tx=XM_023245296.1
E2 11992398 11992518 refGene_gn=CEACAM20;refGene_tx=XM_023245920.1
E2 11993457 11993740 refGene_gn=CEACAM20;refGene_tx=XM_023245920.1

E2 11999811 11999907 refGene_gn=CEACAM20;refGene_tx=XM_023245920.1
E2 11999909 11999960 refGene_gn=CEACAM20;refGene_tx=XM_023245920.1
E2 12000491 12000767 refGene_gn=CEACAM20;refGene_tx=XM_023245920.1
E2 12001222 12001504 refGene_gn=CEACAM20;refGene_tx=XM_023245920.1
E2 12002360 12002485 refGene_gn=CEACAM20;refGene_tx=XM_023245920.1
E2 12003990 12004144 refGene_gn=CEACAM20;refGene_tx=XM_023245920.1
E2 12007001 12007280 refGene_gn=CEACAM20;refGene_tx=XM_023245920.1
E2 12008419 12008534 refGene_gn=CEACAM20;refGene_tx=XM_023245920.1
E2 12008743 12008832 refGene_gn=CEACAM20;refGene_tx=XM_023245920.1
E2 12009566 12009628 refGene_gn=CEACAM20;refGene_tx=XM_023245920.1
E2 12009991 12010027 refGene_gn=CEACAM20;refGene_tx=XM_023245920.1
E2 12010527 12010650 refGene_gn=CEACAM20;refGene_tx=XM_023245920.1
E2 12022584 12022638 refGene_gn=CEACAM20;refGene_tx=XM_023245920.1
E2 12029359 12029392
refGene_gn=ZNF180;refGene_tx=XM_019819413.2;refGene_gn=ZNF180;refGene_tx=XM_019819415.2
E2 12029756 12029795
refGene_gn=ZNF180;refGene_tx=XM_019819412.1;refGene_gn=ZNF180;refGene_tx=XM_019819414.1
E2 12032305 12032395
refGene_gn=ZNF180;refGene_tx=XM_006941156.3;refGene_gn=ZNF180;refGene_tx=XM_019819412.1;refGene_gn=ZNF180;refGene_tx=XM_019819413.2;refGene_gn=ZNF180;refGene_tx=XM_019819414.1;refGene_gn=ZNF180;refGene_tx=XM_019819415.2;refGene_gn=ZNF180;refGene_tx=XM_019819416.1
E2 12039852 12039927
refGene_gn=ZNF180;refGene_tx=XM_006941156.3;refGene_gn=ZNF180;refGene_tx=XM_019819412.1;refGene_gn=ZNF180;refGene_tx=XM_019819413.2
E2 12044240 12044367
refGene_gn=ZNF180;refGene_tx=XM_006941156.3;refGene_gn=ZNF180;refGene_tx=XM_019819412.1;refGene_gn=ZNF180;refGene_tx=XM_019819413.2;refGene_gn=ZNF180;refGene_tx=XM_019819414.1;refGene_gn=ZNF180;refGene_tx=XM_019819415.2;refGene_gn=ZNF180;refGene_tx=XM_019819416.1
E2 12045702 12047438
refGene_gn=ZNF180;refGene_tx=XM_006941156.3;refGene_gn=ZNF180;refGene_tx=XM_019819412.1;refGene_gn=ZNF180;refGene_tx=XM_019819413.2;refGene_gn=ZNF180;refGene_tx=XM_019819414.1;refGene_gn=ZNF180;refGene_tx=XM_019819415.2;refGene_gn=ZNF180;refGene_tx=XM_019819416.1
E2 12087285 12087450 refGene_gn=ZNF229;refGene_tx=XM_019819405.2
E2 12093439 12093566 refGene_gn=ZNF229;refGene_tx=XM_019819405.2
E2 12095008 12096390 refGene_gn=ZNF229;refGene_tx=XM_019819405.2
E2 12096391 12096892 refGene_gn=ZNF229;refGene_tx=XM_019819405.2
E2 12096893 12097280 refGene_gn=ZNF229;refGene_tx=XM_019819405.2
E2 12102484 12102537 refGene_gn=ZNF285;refGene_tx=XM_023245161.1
E2 12113515 12113573 refGene_gn=ZNF285;refGene_tx=XM_023245161.1
E2 12120888 12120939 refGene_gn=ZNF285;refGene_tx=XM_023245161.1
E2 12121154 12121233 refGene_gn=ZNF285;refGene_tx=XM_023245161.1
E2 12123374 12125026
refGene_gn=ZNF285;refGene_tx=XM_019819419.2;refGene_gn=ZNF285;refGene_tx=XM_019819420.2;refGene_gn=ZNF285;refGene_tx=XM_023245161.1
E2 12171113 12171128 refGene_gn=ZNF112;refGene_tx=XM_003997732.5

E2 12171232 12171250 refGene_gn=ZNF112;refGene_tx=XM_019819424.2
E2 12174003 12174130
refGene_gn=ZNF112;refGene_tx=XM_003997732.5;refGene_gn=ZNF112;refGene_tx=XM_019819424.2
E2 12179070 12179166
refGene_gn=ZNF112;refGene_tx=XM_003997732.5;refGene_gn=ZNF112;refGene_tx=XM_019819424.2;refGene_gn=ZNF112;refGene_tx=XM_019819425.2;refGene_gn=ZNF112;refGene_tx=XM_019819426.2
E2 12185096 12187597
refGene_gn=ZNF112;refGene_tx=XM_003997732.5;refGene_gn=ZNF112;refGene_tx=XM_019819424.2;refGene_gn=ZNF112;refGene_tx=XM_019819425.2;refGene_gn=ZNF112;refGene_tx=XM_019819426.2
E2 12224644 12224659
refGene_gn=LOC101090223;refGene_tx=XM_019819427.2;refGene_gn=LOC101090223;refGene_tx=XM_019819428.2
E2 12227470 12227597
refGene_gn=LOC101090223;refGene_tx=XM_019819427.2;refGene_gn=LOC101090223;refGene_tx=XM_019819428.2
E2 12228566 12228662
refGene_gn=LOC101090223;refGene_tx=XM_019819427.2;refGene_gn=LOC101090223;refGene_tx=XM_019819428.2
E2 12255308 12257245
refGene_gn=LOC101090223;refGene_tx=XM_019819427.2;refGene_gn=LOC101090223;refGene_tx=XM_019819428.2
E2 12310318 12310620
refGene_gn=LOC101098504;refGene_tx=XM_023245283.1;refGene_gn=LOC101098504;refGene_tx=XM_023245284.1
E2 12310666 12311243
refGene_gn=LOC101098504;refGene_tx=XM_023245283.1;refGene_gn=LOC101098504;refGene_tx=XM_023245284.1
E2 12341702 12343819
refGene_gn=LOC101099005;refGene_tx=XM_006941164.4;refGene_gn=LOC101099005;refGene_tx=XM_011289935.3;refGene_gn=LOC101099005;refGene_tx=XM_019819429.2;refGene_gn=LOC101099005;refGene_tx=XM_019819432.2;refGene_gn=LOC101099005;refGene_tx=XM_019819436.2;refGene_gn=LOC101099005;refGene_tx=XM_023245277.1
E2 12347296 12347419
refGene_gn=LOC101099005;refGene_tx=XM_006941164.4;refGene_gn=LOC101099005;refGene_tx=XM_011289935.3;refGene_gn=LOC101099005;refGene_tx=XM_019819429.2;refGene_gn=LOC101099005;refGene_tx=XM_019819432.2;refGene_gn=LOC101099005;refGene_tx=XM_019819436.2;refGene_gn=LOC101099005;refGene_tx=XM_023245277.1
E2 12348468 12348595
refGene_gn=LOC101099005;refGene_tx=XM_006941164.4;refGene_gn=LOC101099005;refGene_tx=XM_011289935.3;refGene_gn=LOC101099005;refGene_tx=XM_019819429.2;refGene_gn=LOC101099005;refGene_tx=XM_019819432.2;refGene_gn=LOC101099005;refGene_tx=XM_019819436.2;refGene_gn=LOC101099005;refGene_tx=XM_023245277.1
E2 12350705 12350765
refGene_gn=LOC101099005;refGene_tx=XM_006941164.4;refGene_gn=LOC101099

005;refGene_tx=XM_019819429.2;refGene_gn=LOC101099005;refGene_tx=XM_019819432.2
E2 12372673 12374859
refGene_gn=LOC111556191;refGene_tx=XM_006941166.3;refGene_gn=LOC111556191;refGene_tx=XM_011289932.3;refGene_gn=LOC111556191;refGene_tx=XM_023245279.1;refGene_gn=LOC111556191;refGene_tx=XM_023245280.1;refGene_gn=LOC111556191;refGene_tx=XM_023245281.1;refGene_gn=LOC111556191;refGene_tx=XM_023245282.1
E2 12378042 12378135
refGene_gn=LOC111556191;refGene_tx=XM_006941166.3;refGene_gn=LOC111556191;refGene_tx=XM_011289932.3;refGene_gn=LOC111556191;refGene_tx=XM_023245279.1;refGene_gn=LOC111556191;refGene_tx=XM_023245280.1;refGene_gn=LOC111556191;refGene_tx=XM_023245281.1
E2 12378976 12379103
refGene_gn=LOC111556191;refGene_tx=XM_006941166.3;refGene_gn=LOC111556191;refGene_tx=XM_023245279.1;refGene_gn=LOC111556191;refGene_tx=XM_023245280.1
E2 12380200 12380215
refGene_gn=LOC111556191;refGene_tx=XM_023245279.1
E2 12398266 12399912
refGene_gn=LOC111556191;refGene_tx=XM_023245282.1
E2 12407803 12407930
refGene_gn=LOC111556191;refGene_tx=XM_023245282.1
E2 12410585 12410594
refGene_gn=LOC101099005;refGene_tx=XM_023245277.1;refGene_gn=LOC111556191;refGene_tx=XM_023245282.1
E2 12424100 12425494
refGene_gn=LOC105259641;refGene_tx=XM_023245921.1
E2 12433168 12433261
refGene_gn=LOC105259641;refGene_tx=XM_023245921.1
E2 12433777 12433904
refGene_gn=LOC105259641;refGene_tx=XM_023245921.1
E2 12436278 12436302
refGene_gn=LOC105259641;refGene_tx=XM_023245921.1
E2 12459894 12461726
refGene_gn=LOC105259641;refGene_tx=XM_023245921.1
E2 12466439 12466535
refGene_gn=LOC105259641;refGene_tx=XM_023245921.1
E2 12467854 12467981
refGene_gn=LOC105259641;refGene_tx=XM_023245921.1
E2 12472626 12472641
refGene_gn=LOC105259641;refGene_tx=XM_023245921.1
E2 12514104 12514113
refGene_gn=LOC101089974;refGene_tx=XM_003997700.4
E2 12516911 12517038
refGene_gn=LOC101089974;refGene_tx=XM_003997700.4
E2 12523842 12523967
refGene_gn=LOC101089974;refGene_tx=XM_003997700.4
E2 12523968 12524481
refGene_gn=LOC101089974;refGene_tx=XM_003997700.4

E2 12524482 12525367
refGene_gn=LOC101089974;refGene_tx=XM_003997700.4
E2 12545073 12545108
refGene_gn=LOC101100265;refGene_tx=XM_019819441.2
E2 12545110 12548116
refGene_gn=LOC101100265;refGene_tx=XM_019819441.2
E2 12554171 12554298
refGene_gn=LOC101100265;refGene_tx=XM_019819441.2
E2 12556129 12556138
refGene_gn=LOC101100265;refGene_tx=XM_019819441.2
E2 12577984 12578893
refGene_gn=FELCATV1R1;refGene_tx=NM_001167473.1
E2 12613648 12613670 refGene_gn=LYPD5;refGene_tx=XM_011289963.3
E2 12613958 12614186 refGene_gn=LYPD5;refGene_tx=XM_011289963.3
E2 12615923 12616052 refGene_gn=LYPD5;refGene_tx=XM_011289963.3
E2 12616535 12616712 refGene_gn=LYPD5;refGene_tx=XM_011289963.3
E2 12616919 12617066 refGene_gn=LYPD5;refGene_tx=XM_011289963.3
E2 12617408 12617671 refGene_gn=LYPD5;refGene_tx=XM_011289963.3
E2 12632004 12632163 refGene_gn=KCNN4;refGene_tx=XM_006941172.4
E2 12635031 12635127 refGene_gn=KCNN4;refGene_tx=XM_006941172.4
E2 12636361 12636789 refGene_gn=KCNN4;refGene_tx=XM_006941172.4
E2 12638623 12638759 refGene_gn=KCNN4;refGene_tx=XM_006941172.4
E2 12640940 12641051 refGene_gn=KCNN4;refGene_tx=XM_006941172.4
E2 12641226 12641345 refGene_gn=KCNN4;refGene_tx=XM_006941172.4
E2 12641683 12641753 refGene_gn=KCNN4;refGene_tx=XM_006941172.4
E2 12642867 12643032 refGene_gn=KCNN4;refGene_tx=XM_006941172.4
E2 12652858 12652930 refGene_gn=SMG9;refGene_tx=XM_023246038.1
E2 12654783 12654939
refGene_gn=SMG9;refGene_tx=XM_006941173.4;refGene_gn=SMG9;refGene_tx=XM_023246038.1;refGene_gn=SMG9;refGene_tx=XM_023246039.1;refGene_gn=SMG9;refGene_tx=XM_023246040.1;refGene_gn=SMG9;refGene_tx=XM_023246041.1
E2 12657227 12657302
refGene_gn=SMG9;refGene_tx=XM_006941173.4;refGene_gn=SMG9;refGene_tx=XM_006941176.4;refGene_gn=SMG9;refGene_tx=XM_023246038.1;refGene_gn=SMG9;refGene_tx=XM_023246039.1;refGene_gn=SMG9;refGene_tx=XM_023246040.1;refGene_gn=SMG9;refGene_tx=XM_023246041.1
E2 12657391 12657656
refGene_gn=SMG9;refGene_tx=XM_006941173.4;refGene_gn=SMG9;refGene_tx=XM_006941176.4;refGene_gn=SMG9;refGene_tx=XM_023246038.1;refGene_gn=SMG9;refGene_tx=XM_023246039.1;refGene_gn=SMG9;refGene_tx=XM_023246040.1;refGene_gn=SMG9;refGene_tx=XM_023246041.1
E2 12657760 12657858
refGene_gn=SMG9;refGene_tx=XM_006941173.4;refGene_gn=SMG9;refGene_tx=XM_006941176.4;refGene_gn=SMG9;refGene_tx=XM_023246038.1;refGene_gn=SMG9;refGene_tx=XM_023246039.1;refGene_gn=SMG9;refGene_tx=XM_023246040.1;refGene_gn=SMG9;refGene_tx=XM_023246041.1
E2 12659722 12659835
refGene_gn=SMG9;refGene_tx=XM_006941173.4;refGene_gn=SMG9;refGene_tx=XM_006941176.4;refGene_gn=SMG9;refGene_tx=XM_023246038.1;refGene_gn=SMG9;refGene_tx=XM_023246039.1;refGene_gn=SMG9;refGene_tx=XM_023246040.1;

refGene_gn=SMG9;refGene_tx=XM_023246041.1
E2 12663136 12663248
refGene_gn=SMG9;refGene_tx=XM_006941173.4;refGene_gn=SMG9;refGene_tx=XM_006941176.4;refGene_gn=SMG9;refGene_tx=XM_023246038.1;refGene_gn=SMG9;refGene_tx=XM_023246039.1;refGene_gn=SMG9;refGene_tx=XM_023246040.1;refGene_gn=SMG9;refGene_tx=XM_023246041.1
E2 12665179 12665275
refGene_gn=SMG9;refGene_tx=XM_006941173.4;refGene_gn=SMG9;refGene_tx=XM_006941176.4;refGene_gn=SMG9;refGene_tx=XM_023246038.1;refGene_gn=SMG9;refGene_tx=XM_023246039.1;refGene_gn=SMG9;refGene_tx=XM_023246040.1;refGene_gn=SMG9;refGene_tx=XM_023246041.1
E2 12665711 12665797
refGene_gn=SMG9;refGene_tx=XM_006941173.4;refGene_gn=SMG9;refGene_tx=XM_006941176.4;refGene_gn=SMG9;refGene_tx=XM_023246038.1;refGene_gn=SMG9;refGene_tx=XM_023246039.1;refGene_gn=SMG9;refGene_tx=XM_023246040.1;refGene_gn=SMG9;refGene_tx=XM_023246041.1
E2 12668622 12668729
refGene_gn=SMG9;refGene_tx=XM_006941173.4;refGene_gn=SMG9;refGene_tx=XM_006941176.4;refGene_gn=SMG9;refGene_tx=XM_023246038.1;refGene_gn=SMG9;refGene_tx=XM_023246039.1;refGene_gn=SMG9;refGene_tx=XM_023246040.1;refGene_gn=SMG9;refGene_tx=XM_023246041.1
E2 12669465 12669573
refGene_gn=SMG9;refGene_tx=XM_006941173.4;refGene_gn=SMG9;refGene_tx=XM_006941176.4;refGene_gn=SMG9;refGene_tx=XM_023246038.1;refGene_gn=SMG9;refGene_tx=XM_023246039.1;refGene_gn=SMG9;refGene_tx=XM_023246040.1;refGene_gn=SMG9;refGene_tx=XM_023246041.1
E2 12669746 12669875
refGene_gn=SMG9;refGene_tx=XM_006941173.4;refGene_gn=SMG9;refGene_tx=XM_006941176.4;refGene_gn=SMG9;refGene_tx=XM_023246038.1;refGene_gn=SMG9;refGene_tx=XM_023246039.1;refGene_gn=SMG9;refGene_tx=XM_023246040.1;refGene_gn=SMG9;refGene_tx=XM_023246041.1
E2 12670188 12670333
refGene_gn=SMG9;refGene_tx=XM_006941173.4;refGene_gn=SMG9;refGene_tx=XM_006941176.4;refGene_gn=SMG9;refGene_tx=XM_023246038.1;refGene_gn=SMG9;refGene_tx=XM_023246039.1;refGene_gn=SMG9;refGene_tx=XM_023246040.1;refGene_gn=SMG9;refGene_tx=XM_023246041.1
E2 12671404 12671483
refGene_gn=SMG9;refGene_tx=XM_006941173.4;refGene_gn=SMG9;refGene_tx=XM_006941176.4;refGene_gn=SMG9;refGene_tx=XM_023246038.1;refGene_gn=SMG9;refGene_tx=XM_023246039.1;refGene_gn=SMG9;refGene_tx=XM_023246040.1;refGene_gn=SMG9;refGene_tx=XM_023246041.1
E2 12673364 12674008
refGene_gn=LOC101101273;refGene_tx=XM_003997740.4
E2 12674612 12675216
refGene_gn=LOC101101273;refGene_tx=XM_003997740.4
E2 12677830 12679225 refGene_gn=IFGGE;refGene_tx=XM_006941177.3
E2 12720709 12720767
refGene_gn=PLAUR;refGene_tx=XM_006941178.4;refGene_gn=PLAUR;refGene_tx=XM_006941179.3
E2 12722798 12722909

refGene_gn=PLAUR;refGene_tx=XM_006941178.4;refGene_gn=PLAUR;refGene_tx=XM_006941179.3;refGene_gn=PLAUR;refGene_tx=XM_019819445.2
E2 12724436 12724580
refGene_gn=PLAUR;refGene_tx=XM_006941178.4;refGene_gn=PLAUR;refGene_tx=XM_006941179.3;refGene_gn=PLAUR;refGene_tx=XM_019819445.2
E2 12730184 12730343
refGene_gn=PLAUR;refGene_tx=XM_006941178.4;refGene_gn=PLAUR;refGene_tx=XM_006941179.3;refGene_gn=PLAUR;refGene_tx=XM_019819445.2
E2 12730909 12731044
refGene_gn=PLAUR;refGene_tx=XM_006941178.4;refGene_gn=PLAUR;refGene_tx=XM_006941179.3;refGene_gn=PLAUR;refGene_tx=XM_019819445.2
E2 12732166 12732313
refGene_gn=PLAUR;refGene_tx=XM_006941178.4;refGene_gn=PLAUR;refGene_tx=XM_006941179.3;refGene_gn=PLAUR;refGene_tx=XM_019819445.2
E2 12734542 12734796
refGene_gn=PLAUR;refGene_tx=XM_006941178.4;refGene_gn=PLAUR;refGene_tx=XM_006941179.3;refGene_gn=PLAUR;refGene_tx=XM_019819445.2
E2 12741911 12741975
refGene_gn=CADM4;refGene_tx=XM_023245350.1;refGene_gn=CADM4;refGene_tx=XM_023245351.1
E2 12751380 12751527
refGene_gn=CADM4;refGene_tx=XM_023245350.1;refGene_gn=CADM4;refGene_tx=XM_023245351.1
E2 12751853 12752006
refGene_gn=CADM4;refGene_tx=XM_023245350.1;refGene_gn=CADM4;refGene_tx=XM_023245351.1
E2 12752256 12752391
refGene_gn=CADM4;refGene_tx=XM_023245350.1;refGene_gn=CADM4;refGene_tx=XM_023245351.1
E2 12752774 12752939
refGene_gn=CADM4;refGene_tx=XM_023245350.1;refGene_gn=CADM4;refGene_tx=XM_023245351.1
E2 12753067 12753158
refGene_gn=CADM4;refGene_tx=XM_023245350.1;refGene_gn=CADM4;refGene_tx=XM_023245351.1
E2 12753532 12753705
refGene_gn=CADM4;refGene_tx=XM_023245350.1;refGene_gn=CADM4;refGene_tx=XM_023245351.1
E2 12753889 12753897 refGene_gn=CADM4;refGene_tx=XM_023245351.1
E2 12754543 12754672 refGene_gn=CADM4;refGene_tx=XM_023245350.1
E2 12755215 12755325 refGene_gn=CADM4;refGene_tx=XM_023245350.1
E2 12761225 12762940
refGene_gn=SRRM5;refGene_tx=XM_023245287.1;refGene_gn=ZNF428;refGene_tx=XM_023245288.1
E2 12763047 12763446 refGene_gn=SRRM5;refGene_tx=XM_023245287.1
E2 12763569 12763633 refGene_gn=ZNF428;refGene_tx=XM_023245289.1
E2 12765723 12766226
refGene_gn=ZNF428;refGene_tx=XM_023245288.1;refGene_gn=ZNF428;refGene_tx=XM_023245289.1
E2 12773644 12774072

refGene_gn=ZNF576;refGene_tx=XM_006941181.4;refGene_gn=ZNF576;refGene_tx=XM_006941182.4;refGene_gn=ZNF576;refGene_tx=XM_019819448.2
E2 12775502 12775587
refGene_gn=ZNF576;refGene_tx=XM_006941181.4;refGene_gn=ZNF576;refGene_tx=XM_006941182.4;refGene_gn=ZNF576;refGene_tx=XM_019819448.2
E2 12777038 12777568
refGene_gn=IRGQ;refGene_tx=XM_023246178.1;refGene_gn=IRGQ;refGene_tx=XM_023246179.1;refGene_gn=IRGQ;refGene_tx=XM_023246180.1
E2 12778780 12780140
refGene_gn=IRGQ;refGene_tx=XM_023246178.1;refGene_gn=IRGQ;refGene_tx=XM_023246179.1;refGene_gn=IRGQ;refGene_tx=XM_023246180.1
E2 12786710 12786865
refGene_gn=PINLYP;refGene_tx=XM_019819450.2;refGene_gn=PINLYP;refGene_tx=XM_019819451.2
E2 12787046 12787190
refGene_gn=PINLYP;refGene_tx=XM_019819450.2;refGene_gn=PINLYP;refGene_tx=XM_019819451.2
E2 12787415 12787568
refGene_gn=PINLYP;refGene_tx=XM_019819450.2;refGene_gn=PINLYP;refGene_tx=XM_019819451.2
E2 12789361 12789478
refGene_gn=PINLYP;refGene_tx=XM_019819450.2;refGene_gn=PINLYP;refGene_tx=XM_019819451.2
E2 12790300 12790370
refGene_gn=PINLYP;refGene_tx=XM_019819450.2;refGene_gn=PINLYP;refGene_tx=XM_019819451.2
E2 12791734 12791785
refGene_gn=XRCC1;refGene_tx=XM_003997708.5;refGene_gn=XRCC1;refGene_tx=XM_006941184.4
E2 12792135 12792228
refGene_gn=XRCC1;refGene_tx=XM_003997708.5;refGene_gn=XRCC1;refGene_tx=XM_006941184.4
E2 12802037 12802148 refGene_gn=XRCC1;refGene_tx=XM_003997708.5
E2 12812353 12812512
refGene_gn=XRCC1;refGene_tx=XM_003997708.5;refGene_gn=XRCC1;refGene_tx=XM_006941184.4
E2 12813384 12813459
refGene_gn=XRCC1;refGene_tx=XM_003997708.5;refGene_gn=XRCC1;refGene_tx=XM_006941184.4
E2 12813554 12813660
refGene_gn=XRCC1;refGene_tx=XM_003997708.5;refGene_gn=XRCC1;refGene_tx=XM_006941184.4
E2 12813878 12813988
refGene_gn=XRCC1;refGene_tx=XM_003997708.5;refGene_gn=XRCC1;refGene_tx=XM_006941184.4
E2 12814063 12814175
refGene_gn=XRCC1;refGene_tx=XM_003997708.5;refGene_gn=XRCC1;refGene_tx=XM_006941184.4
E2 12814401 12814660
refGene_gn=XRCC1;refGene_tx=XM_003997708.5;refGene_gn=XRCC1;refGene_tx

=XM_006941184.4
E2 12814958 12815075
refGene_gn=XRCC1;refGene_tx=XM_003997708.5;refGene_gn=XRCC1;refGene_tx
=XM_006941184.4
E2 12815636 12815730
refGene_gn=XRCC1;refGene_tx=XM_003997708.5;refGene_gn=XRCC1;refGene_tx
=XM_006941184.4
E2 12815864 12815997
refGene_gn=XRCC1;refGene_tx=XM_003997708.5;refGene_gn=XRCC1;refGene_tx
=XM_006941184.4
E2 12816298 12816353
refGene_gn=XRCC1;refGene_tx=XM_003997708.5;refGene_gn=XRCC1;refGene_tx
=XM_006941184.4
E2 12816445 12816585
refGene_gn=XRCC1;refGene_tx=XM_003997708.5;refGene_gn=XRCC1;refGene_tx
=XM_006941184.4
E2 12817636 12817727
refGene_gn=XRCC1;refGene_tx=XM_003997708.5;refGene_gn=XRCC1;refGene_tx
=XM_006941184.4
E2 12818114 12818190
refGene_gn=XRCC1;refGene_tx=XM_003997708.5;refGene_gn=XRCC1;refGene_tx
=XM_006941184.4
E2 12818294 12818408
refGene_gn=XRCC1;refGene_tx=XM_003997708.5;refGene_gn=XRCC1;refGene_tx
=XM_006941184.4
E2 12823472 12824131
refGene_gn=ZNF575;refGene_tx=XM_011289920.3;refGene_gn=ZNF575;refGene_
tx=XM_011289921.3;refGene_gn=ZNF575;refGene_tx=XM_019819452.2;refGene_
gn=ZNF575;refGene_tx=XM_019819453.1;refGene_gn=ZNF575;refGene_tx=XM_01
9819455.1;refGene_gn=ZNF575;refGene_tx=XM_019819456.2;refGene_gn=ZNF57
5;refGene_tx=XM_019819457.1
E2 12824603 12824685
refGene_gn=ZNF575;refGene_tx=XM_011289920.3;refGene_gn=ZNF575;refGene_
tx=XM_011289921.3;refGene_gn=ZNF575;refGene_tx=XM_019819452.2;refGene_
gn=ZNF575;refGene_tx=XM_019819453.1;refGene_gn=ZNF575;refGene_tx=XM_01
9819455.1;refGene_gn=ZNF575;refGene_tx=XM_019819456.2;refGene_gn=ZNF57
5;refGene_tx=XM_019819457.1
E2 12830664 12830745 refGene_gn=ETHE1;refGene_tx=XM_023245255.1
E2 12831127 12831272
refGene_gn=ETHE1;refGene_tx=XM_023245255.1;refGene_gn=ETHE1;refGene_tx
=XM_023245256.1
E2 12831381 12831530 refGene_gn=ETHE1;refGene_tx=XM_023245255.1
E2 12844992 12845122
refGene_gn=ETHE1;refGene_tx=XM_023245255.1;refGene_gn=ETHE1;refGene_tx
=XM_023245256.1
E2 12847048 12847138
refGene_gn=ETHE1;refGene_tx=XM_023245255.1;refGene_gn=ETHE1;refGene_tx
=XM_023245256.1
E2 12847539 12847656
refGene_gn=ETHE1;refGene_tx=XM_023245255.1;refGene_gn=ETHE1;refGene_tx

=XM_023245256.1
E2 12847821 12847874
refGene_gn=ETHE1;refGene_tx=XM_023245255.1;refGene_gn=ETHE1;refGene_tx
=XM_023245256.1
E2 12849927 12850188 refGene_gn=PHLDB3;refGene_tx=XM_006941187.3
E2 12850860 12851043 refGene_gn=PHLDB3;refGene_tx=XM_006941187.3
E2 12851264 12851402 refGene_gn=PHLDB3;refGene_tx=XM_006941187.3
E2 12852751 12852880 refGene_gn=PHLDB3;refGene_tx=XM_006941187.3
E2 12853108 12853270 refGene_gn=PHLDB3;refGene_tx=XM_006941187.3
E2 12854252 12854363 refGene_gn=PHLDB3;refGene_tx=XM_006941187.3
E2 12854524 12854608 refGene_gn=PHLDB3;refGene_tx=XM_006941187.3
E2 12854964 12855075 refGene_gn=PHLDB3;refGene_tx=XM_006941187.3
E2 12864719 12864819 refGene_gn=PHLDB3;refGene_tx=XM_006941187.3
E2 12864984 12865075 refGene_gn=PHLDB3;refGene_tx=XM_006941187.3
E2 12865157 12865245 refGene_gn=PHLDB3;refGene_tx=XM_006941187.3
E2 12865972 12866029 refGene_gn=PHLDB3;refGene_tx=XM_006941187.3
E2 12870313 12870533 refGene_gn=PHLDB3;refGene_tx=XM_006941187.3
E2 12871152 12871238 refGene_gn=PHLDB3;refGene_tx=XM_006941187.3
E2 12872505 12872640 refGene_gn=PHLDB3;refGene_tx=XM_006941187.3
E2 12878734 12878819 refGene_gn=LYPD3;refGene_tx=XM_006941191.4
E2 12879385 12879517 refGene_gn=LYPD3;refGene_tx=XM_006941191.4
E2 12880311 12880482 refGene_gn=LYPD3;refGene_tx=XM_006941191.4
E2 12880771 12880933 refGene_gn=LYPD3;refGene_tx=XM_006941191.4
E2 12881758 12882240 refGene_gn=LYPD3;refGene_tx=XM_006941191.4
E2 12918689 12918964
refGene_gn=TEX101;refGene_tx=XM_006941189.4;refGene_gn=TEX101;refGene_
tx=XM_006941190.4;refGene_gn=TEX101;refGene_tx=XM_023244872.1
E2 12919122 12919251
refGene_gn=TEX101;refGene_tx=XM_006941189.4;refGene_gn=TEX101;refGene_
tx=XM_006941190.4;refGene_gn=TEX101;refGene_tx=XM_023244872.1
E2 12920811 12921006
refGene_gn=TEX101;refGene_tx=XM_006941189.4;refGene_gn=TEX101;refGene_
tx=XM_006941190.4;refGene_gn=TEX101;refGene_tx=XM_023244872.1
E2 12921122 12921266
refGene_gn=TEX101;refGene_tx=XM_006941189.4;refGene_gn=TEX101;refGene_
tx=XM_006941190.4;refGene_gn=TEX101;refGene_tx=XM_023244872.1
E2 12921388 12921584
refGene_gn=TEX101;refGene_tx=XM_006941189.4;refGene_gn=TEX101;refGene_
tx=XM_006941190.4;refGene_gn=TEX101;refGene_tx=XM_023244872.1
E2 12921702 12921888
refGene_gn=TEX101;refGene_tx=XM_006941189.4;refGene_gn=TEX101;refGene_
tx=XM_006941190.4
E2 12935139 12935176
refGene_gn=CD177;refGene_tx=XM_023244862.1;refGene_gn=CD177;refGene_tx
=XM_023244863.1;refGene_gn=CD177;refGene_tx=XM_023244864.1;refGene_gn=
CD177;refGene_tx=XM_023244865.1;refGene_gn=CD177;refGene_tx=XM_0232448
66.1;refGene_gn=CD177;refGene_tx=XM_023244867.1;refGene_gn=CD177;refGe
ne_tx=XM_023244868.1;refGene_gn=CD177;refGene_tx=XM_023244869.1
E2 12935653 12935705
refGene_gn=CD177;refGene_tx=XM_023244861.1;refGene_gn=CD177;refGene_tx

=XM_023244871.1
E2 12935779 12935920
refGene_gn=CD177;refGene_tx=XM_023244861.1;refGene_gn=CD177;refGene_tx=XM_023244862.1;refGene_gn=CD177;refGene_tx=XM_023244863.1;refGene_gn=CD177;refGene_tx=XM_023244864.1;refGene_gn=CD177;refGene_tx=XM_023244865.1;refGene_gn=CD177;refGene_tx=XM_023244866.1;refGene_gn=CD177;refGene_tx=XM_023244867.1;refGene_gn=CD177;refGene_tx=XM_023244868.1;refGene_gn=CD177;refGene_tx=XM_023244869.1;refGene_gn=CD177;refGene_tx=XM_023244870.1;refGene_gn=CD177;refGene_tx=XM_023244871.1
E2 12936385 12936571
refGene_gn=CD177;refGene_tx=XM_023244861.1;refGene_gn=CD177;refGene_tx=XM_023244862.1;refGene_gn=CD177;refGene_tx=XM_023244863.1;refGene_gn=CD177;refGene_tx=XM_023244864.1;refGene_gn=CD177;refGene_tx=XM_023244865.1;refGene_gn=CD177;refGene_tx=XM_023244866.1;refGene_gn=CD177;refGene_tx=XM_023244867.1;refGene_gn=CD177;refGene_tx=XM_023244868.1;refGene_gn=CD177;refGene_tx=XM_023244869.1;refGene_gn=CD177;refGene_tx=XM_023244870.1;refGene_gn=CD177;refGene_tx=XM_023244871.1
E2 12937563 12937683
refGene_gn=CD177;refGene_tx=XM_023244861.1;refGene_gn=CD177;refGene_tx=XM_023244862.1;refGene_gn=CD177;refGene_tx=XM_023244863.1;refGene_gn=CD177;refGene_tx=XM_023244864.1;refGene_gn=CD177;refGene_tx=XM_023244865.1;refGene_gn=CD177;refGene_tx=XM_023244866.1;refGene_gn=CD177;refGene_tx=XM_023244867.1;refGene_gn=CD177;refGene_tx=XM_023244868.1;refGene_gn=CD177;refGene_tx=XM_023244869.1;refGene_gn=CD177;refGene_tx=XM_023244870.1;refGene_gn=CD177;refGene_tx=XM_023244871.1
E2 12937879 12937996
refGene_gn=CD177;refGene_tx=XM_023244861.1;refGene_gn=CD177;refGene_tx=XM_023244862.1;refGene_gn=CD177;refGene_tx=XM_023244863.1;refGene_gn=CD177;refGene_tx=XM_023244864.1;refGene_gn=CD177;refGene_tx=XM_023244865.1;refGene_gn=CD177;refGene_tx=XM_023244866.1;refGene_gn=CD177;refGene_tx=XM_023244867.1;refGene_gn=CD177;refGene_tx=XM_023244868.1;refGene_gn=CD177;refGene_tx=XM_023244869.1;refGene_gn=CD177;refGene_tx=XM_023244870.1;refGene_gn=CD177;refGene_tx=XM_023244871.1
E2 12939705 12939843
refGene_gn=CD177;refGene_tx=XM_023244861.1;refGene_gn=CD177;refGene_tx=XM_023244862.1;refGene_gn=CD177;refGene_tx=XM_023244863.1;refGene_gn=CD177;refGene_tx=XM_023244864.1;refGene_gn=CD177;refGene_tx=XM_023244865.1;refGene_gn=CD177;refGene_tx=XM_023244866.1;refGene_gn=CD177;refGene_tx=XM_023244867.1;refGene_gn=CD177;refGene_tx=XM_023244868.1;refGene_gn=CD177;refGene_tx=XM_023244869.1;refGene_gn=CD177;refGene_tx=XM_023244870.1;refGene_gn=CD177;refGene_tx=XM_023244871.1
E2 12940510 12940696
refGene_gn=CD177;refGene_tx=XM_023244861.1;refGene_gn=CD177;refGene_tx=XM_023244862.1;refGene_gn=CD177;refGene_tx=XM_023244863.1;refGene_gn=CD177;refGene_tx=XM_023244864.1;refGene_gn=CD177;refGene_tx=XM_023244865.1;refGene_gn=CD177;refGene_tx=XM_023244866.1;refGene_gn=CD177;refGene_tx=XM_023244867.1;refGene_gn=CD177;refGene_tx=XM_023244868.1;refGene_gn=CD177;refGene_tx=XM_023244869.1;refGene_gn=CD177;refGene_tx=XM_023244870.1;refGene_gn=CD177;refGene_tx=XM_023244871.1
E2 12940829 12940964

refGene_gn=CD177;refGene_tx=XM_023244861.1;refGene_gn=CD177;refGene_tx=XM_023244862.1;refGene_gn=CD177;refGene_tx=XM_023244863.1;refGene_gn=CD177;refGene_tx=XM_023244864.1;refGene_gn=CD177;refGene_tx=XM_023244865.1;refGene_gn=CD177;refGene_tx=XM_023244866.1;refGene_gn=CD177;refGene_tx=XM_023244867.1;refGene_gn=CD177;refGene_tx=XM_023244868.1;refGene_gn=CD177;refGene_tx=XM_023244869.1;refGene_gn=CD177;refGene_tx=XM_023244870.1
E2 12941217 12941438
refGene_gn=CD177;refGene_tx=XM_023244861.1;refGene_gn=CD177;refGene_tx=XM_023244862.1;refGene_gn=CD177;refGene_tx=XM_023244863.1;refGene_gn=CD177;refGene_tx=XM_023244864.1;refGene_gn=CD177;refGene_tx=XM_023244865.1;refGene_gn=CD177;refGene_tx=XM_023244866.1;refGene_gn=CD177;refGene_tx=XM_023244867.1;refGene_gn=CD177;refGene_tx=XM_023244868.1;refGene_gn=CD177;refGene_tx=XM_023244869.1;refGene_gn=CD177;refGene_tx=XM_023244870.1
E2 12958521 12958585
refGene_gn=LOC101082109;refGene_tx=XM_023245941.1
E2 12959287 12959647
refGene_gn=LOC101082109;refGene_tx=XM_023245941.1
E2 12963893 12964172
refGene_gn=LOC101082109;refGene_tx=XM_023245941.1
E2 12965653 12965774
refGene_gn=LOC101082109;refGene_tx=XM_023245941.1
E2 12966752 12966805
refGene_gn=LOC101082109;refGene_tx=XM_023245941.1
E2 12967100 12967132
refGene_gn=LOC101082109;refGene_tx=XM_023245941.1
E2 12967557 12967599
refGene_gn=LOC101082109;refGene_tx=XM_023245941.1
E2 12967857 12967911
refGene_gn=LOC101082109;refGene_tx=XM_023245941.1
E2 12975517 12975696
refGene_gn=LOC101082109;refGene_tx=XM_023245941.1
E2 12975782 12975892
refGene_gn=LOC101082109;refGene_tx=XM_023245941.1
E2 12977778 12978138
refGene_gn=LOC101082109;refGene_tx=XM_023245941.1
E2 12980149 12980428
refGene_gn=LOC101082109;refGene_tx=XM_023245941.1
E2 12980664 12980919
refGene_gn=LOC101082109;refGene_tx=XM_023245941.1
E2 12983617 12983896
refGene_gn=LOC101082109;refGene_tx=XM_023245941.1
E2 12985366 12985487
refGene_gn=LOC101082109;refGene_tx=XM_023245941.1
E2 12986465 12986518
refGene_gn=LOC101082109;refGene_tx=XM_023245941.1
E2 12986813 12986845
refGene_gn=LOC101082109;refGene_tx=XM_023245941.1
E2 12987272 12987314

refGene_gn=LOC101082109;refGene_tx=XM_023245941.1
E2 12987572 12987638
refGene_gn=LOC101082109;refGene_tx=XM_023245941.1
E2 12995455 12995658
refGene_gn=LYPD4;refGene_tx=XM_019819465.2;refGene_gn=LYPD4;refGene_tx=XM_019819466.1;refGene_gn=LYPD4;refGene_tx=XM_019819467.2;refGene_gn=LYPD4;refGene_tx=XM_019819468.1;refGene_gn=LYPD4;refGene_tx=XM_019819469.2;refGene_gn=LYPD4;refGene_tx=XM_023245362.1;refGene_gn=LYPD4;refGene_tx=XM_023245364.1
E2 12995896 12996223
refGene_gn=LYPD4;refGene_tx=XM_019819465.2;refGene_gn=LYPD4;refGene_tx=XM_019819466.1;refGene_gn=LYPD4;refGene_tx=XM_019819467.2;refGene_gn=LYPD4;refGene_tx=XM_019819468.1;refGene_gn=LYPD4;refGene_tx=XM_019819469.2;refGene_gn=LYPD4;refGene_tx=XM_023245362.1;refGene_gn=LYPD4;refGene_tx=XM_023245364.1
E2 12996549 12996699
refGene_gn=LYPD4;refGene_tx=XM_019819465.2;refGene_gn=LYPD4;refGene_tx=XM_019819466.1;refGene_gn=LYPD4;refGene_tx=XM_019819467.2;refGene_gn=LYPD4;refGene_tx=XM_019819468.1;refGene_gn=LYPD4;refGene_tx=XM_019819469.2;refGene_gn=LYPD4;refGene_tx=XM_023245362.1
E2 12996896 12996963
refGene_gn=LYPD4;refGene_tx=XM_019819465.2;refGene_gn=LYPD4;refGene_tx=XM_019819466.1;refGene_gn=LYPD4;refGene_tx=XM_019819467.2;refGene_gn=LYPD4;refGene_tx=XM_019819468.1;refGene_gn=LYPD4;refGene_tx=XM_019819469.2;refGene_gn=LYPD4;refGene_tx=XM_023245362.1;refGene_gn=LYPD4;refGene_tx=XM_023245364.1
E2 13002016 13002132
refGene_gn=DMRTC2;refGene_tx=XM_019819461.2;refGene_gn=DMRTC2;refGene_tx=XM_023245359.1
E2 13002329 13002557
refGene_gn=DMRTC2;refGene_tx=XM_006941206.4;refGene_gn=DMRTC2;refGene_tx=XM_019819461.2;refGene_gn=DMRTC2;refGene_tx=XM_019819463.2;refGene_gn=DMRTC2;refGene_tx=XM_023245359.1;refGene_gn=DMRTC2;refGene_tx=XM_023245360.1;refGene_gn=DMRTC2;refGene_tx=XM_023245361.1
E2 13002685 13002843
refGene_gn=DMRTC2;refGene_tx=XM_006941206.4;refGene_gn=DMRTC2;refGene_tx=XM_019819461.2;refGene_gn=DMRTC2;refGene_tx=XM_019819463.2;refGene_gn=DMRTC2;refGene_tx=XM_023245359.1;refGene_gn=DMRTC2;refGene_tx=XM_023245360.1;refGene_gn=DMRTC2;refGene_tx=XM_023245361.1
E2 13003117 13003194
refGene_gn=DMRTC2;refGene_tx=XM_006941206.4;refGene_gn=DMRTC2;refGene_tx=XM_019819461.2;refGene_gn=DMRTC2;refGene_tx=XM_019819463.2;refGene_gn=DMRTC2;refGene_tx=XM_023245359.1;refGene_gn=DMRTC2;refGene_tx=XM_023245360.1;refGene_gn=DMRTC2;refGene_tx=XM_023245361.1
E2 13003467 13003651
refGene_gn=DMRTC2;refGene_tx=XM_006941206.4;refGene_gn=DMRTC2;refGene_tx=XM_019819461.2;refGene_gn=DMRTC2;refGene_tx=XM_019819463.2;refGene_gn=DMRTC2;refGene_tx=XM_023245359.1;refGene_gn=DMRTC2;refGene_tx=XM_023245360.1;refGene_gn=DMRTC2;refGene_tx=XM_023245361.1
E2 13003792 13003919

refGene_gn=DMRTC2;refGene_tx=XM_006941206.4;refGene_gn=DMRTC2;refGene_tx=XM_019819461.2;refGene_gn=DMRTC2;refGene_tx=XM_019819463.2;refGene_gn=DMRTC2;refGene_tx=XM_023245359.1;refGene_gn=DMRTC2;refGene_tx=XM_023245360.1;refGene_gn=DMRTC2;refGene_tx=XM_023245361.1
E2 13004691 13004752
refGene_gn=DMRTC2;refGene_tx=XM_006941206.4;refGene_gn=DMRTC2;refGene_tx=XM_019819461.2;refGene_gn=DMRTC2;refGene_tx=XM_019819463.2;refGene_gn=DMRTC2;refGene_tx=XM_023245359.1;refGene_gn=DMRTC2;refGene_tx=XM_023245360.1;refGene_gn=DMRTC2;refGene_tx=XM_023245361.1
E2 13004907 13005082
refGene_gn=DMRTC2;refGene_tx=XM_006941206.4;refGene_gn=DMRTC2;refGene_tx=XM_019819461.2;refGene_gn=DMRTC2;refGene_tx=XM_019819463.2;refGene_gn=DMRTC2;refGene_tx=XM_023245359.1;refGene_gn=DMRTC2;refGene_tx=XM_023245360.1;refGene_gn=DMRTC2;refGene_tx=XM_023245361.1
E2 13005743 13005790 refGene_gn=DMRTC2;refGene_tx=XM_023245359.1
E2 13006112 13006225
refGene_gn=DMRTC2;refGene_tx=XM_006941206.4;refGene_gn=DMRTC2;refGene_tx=XM_019819461.2;refGene_gn=DMRTC2;refGene_tx=XM_019819463.2;refGene_gn=DMRTC2;refGene_tx=XM_023245360.1;refGene_gn=DMRTC2;refGene_tx=XM_023245361.1
E2 13013977 13014048
refGene_gn=RPS19;refGene_tx=XM_003997752.5;refGene_gn=RPS19;refGene_tx=XM_006941215.4;refGene_gn=RPS19;refGene_tx=XM_019819470.2;refGene_gn=RPS19;refGene_tx=XM_019819471.2;refGene_gn=RPS19;refGene_tx=XM_019819472.2
E2 13014339 13014440
refGene_gn=RPS19;refGene_tx=XM_003997752.5;refGene_gn=RPS19;refGene_tx=XM_006941215.4;refGene_gn=RPS19;refGene_tx=XM_019819470.2;refGene_gn=RPS19;refGene_tx=XM_019819471.2;refGene_gn=RPS19;refGene_tx=XM_019819472.2
E2 13019939 13020195
refGene_gn=RPS19;refGene_tx=XM_003997752.5;refGene_gn=RPS19;refGene_tx=XM_006941215.4;refGene_gn=RPS19;refGene_tx=XM_019819470.2;refGene_gn=RPS19;refGene_tx=XM_019819471.2;refGene_gn=RPS19;refGene_tx=XM_019819472.2
E2 13020300 13020403
refGene_gn=RPS19;refGene_tx=XM_006941215.4;refGene_gn=RPS19;refGene_tx=XM_019819470.2
E2 13020725 13020780
refGene_gn=RPS19;refGene_tx=XM_003997752.5;refGene_gn=RPS19;refGene_tx=XM_019819471.2;refGene_gn=RPS19;refGene_tx=XM_019819472.2
E2 13021550 13021571 refGene_gn=RPS19;refGene_tx=XM_019819472.2
E2 13021733 13021760
refGene_gn=RPS19;refGene_tx=XM_003997752.5;refGene_gn=RPS19;refGene_tx=XM_019819471.2
E2 13025767 13025846 refGene_gn=CD79A;refGene_tx=XM_023245568.1
E2 13027423 13027747 refGene_gn=CD79A;refGene_tx=XM_023245568.1
E2 13027997 13028116 refGene_gn=CD79A;refGene_tx=XM_023245568.1
E2 13028712 13028781 refGene_gn=CD79A;refGene_tx=XM_023245568.1
E2 13028909 13029029 refGene_gn=CD79A;refGene_tx=XM_023245568.1

E2 13033266 13033280
refGene_gn=ARHGEF1;refGene_tx=XM_023244917.1;refGene_gn=ARHGEF1;refGene_tx=XM_023244918.1
E2 13034816 13034859
refGene_gn=ARHGEF1;refGene_tx=XM_023244917.1;refGene_gn=ARHGEF1;refGene_tx=XM_023244918.1
E2 13034941 13035025
refGene_gn=ARHGEF1;refGene_tx=XM_023244917.1;refGene_gn=ARHGEF1;refGene_tx=XM_023244918.1
E2 13035484 13035598
refGene_gn=ARHGEF1;refGene_tx=XM_023244917.1;refGene_gn=ARHGEF1;refGene_tx=XM_023244918.1
E2 13037793 13037892
refGene_gn=ARHGEF1;refGene_tx=XM_023244917.1;refGene_gn=ARHGEF1;refGene_tx=XM_023244918.1
E2 13038092 13038135
refGene_gn=ARHGEF1;refGene_tx=XM_023244917.1;refGene_gn=ARHGEF1;refGene_tx=XM_023244918.1
E2 13038332 13038579
refGene_gn=ARHGEF1;refGene_tx=XM_023244917.1;refGene_gn=ARHGEF1;refGene_tx=XM_023244918.1
E2 13039014 13039044
refGene_gn=ARHGEF1;refGene_tx=XM_023244917.1;refGene_gn=ARHGEF1;refGene_tx=XM_023244918.1
E2 13039320 13039420
refGene_gn=ARHGEF1;refGene_tx=XM_023244917.1;refGene_gn=ARHGEF1;refGene_tx=XM_023244918.1
E2 13039565 13039662
refGene_gn=ARHGEF1;refGene_tx=XM_023244917.1;refGene_gn=ARHGEF1;refGene_tx=XM_023244918.1
E2 13039738 13039774
refGene_gn=ARHGEF1;refGene_tx=XM_023244917.1;refGene_gn=ARHGEF1;refGene_tx=XM_023244918.1
E2 13039907 13040045
refGene_gn=ARHGEF1;refGene_tx=XM_023244917.1;refGene_gn=ARHGEF1;refGene_tx=XM_023244918.1
E2 13040739 13040845
refGene_gn=ARHGEF1;refGene_tx=XM_023244917.1;refGene_gn=ARHGEF1;refGene_tx=XM_023244918.1
E2 13041329 13041341 refGene_gn=ARHGEF1;refGene_tx=XM_023244917.1
E2 13041887 13042055 refGene_gn=ARHGEF1;refGene_tx=XM_023244917.1
E2 13042337 13042483
refGene_gn=ARHGEF1;refGene_tx=XM_023244917.1;refGene_gn=ARHGEF1;refGene_tx=XM_023244918.1
E2 13043927 13044074
refGene_gn=ARHGEF1;refGene_tx=XM_023244917.1;refGene_gn=ARHGEF1;refGene_tx=XM_023244918.1
E2 13044459 13044542
refGene_gn=ARHGEF1;refGene_tx=XM_023244917.1;refGene_gn=ARHGEF1;refGene_tx=XM_023244918.1

E2 13044768 13044894
refGene_gn=ARHGEF1;refGene_tx=XM_023244917.1;refGene_gn=ARHGEF1;refGene_tx=XM_023244918.1
E2 13044983 13045098
refGene_gn=ARHGEF1;refGene_tx=XM_023244917.1;refGene_gn=ARHGEF1;refGene_tx=XM_023244918.1
E2 13045193 13045294
refGene_gn=ARHGEF1;refGene_tx=XM_023244917.1;refGene_gn=ARHGEF1;refGene_tx=XM_023244918.1
E2 13045488 13045566
refGene_gn=ARHGEF1;refGene_tx=XM_023244917.1;refGene_gn=ARHGEF1;refGene_tx=XM_023244918.1
E2 13045831 13045907
refGene_gn=ARHGEF1;refGene_tx=XM_023244917.1;refGene_gn=ARHGEF1;refGene_tx=XM_023244918.1
E2 13045987 13046155
refGene_gn=ARHGEF1;refGene_tx=XM_023244917.1;refGene_gn=ARHGEF1;refGene_tx=XM_023244918.1
E2 13046671 13046759
refGene_gn=ARHGEF1;refGene_tx=XM_023244917.1;refGene_gn=ARHGEF1;refGene_tx=XM_023244918.1
E2 13046894 13046981
refGene_gn=ARHGEF1;refGene_tx=XM_023244917.1;refGene_gn=ARHGEF1;refGene_tx=XM_023244918.1
E2 13047258 13047329
refGene_gn=ARHGEF1;refGene_tx=XM_023244917.1;refGene_gn=ARHGEF1;refGene_tx=XM_023244918.1
E2 13047417 13047504
refGene_gn=ARHGEF1;refGene_tx=XM_023244917.1;refGene_gn=ARHGEF1;refGene_tx=XM_023244918.1
E2 13047781 13047951
refGene_gn=ARHGEF1;refGene_tx=XM_023244917.1;refGene_gn=ARHGEF1;refGene_tx=XM_023244918.1
E2 13048051 13048135
refGene_gn=ARHGEF1;refGene_tx=XM_023244917.1;refGene_gn=ARHGEF1;refGene_tx=XM_023244918.1
E2 13049510 13049959
refGene_gn=LOC101084152;refGene_tx=XM_019819473.2
E2 13050337 13050455
refGene_gn=LOC101084152;refGene_tx=XM_019819473.2
E2 13050573 13050769
refGene_gn=LOC101084152;refGene_tx=XM_019819473.2
E2 13051109 13051344
refGene_gn=LOC101084152;refGene_tx=XM_019819473.2
E2 13054150 13054230
refGene_gn=LOC101084152;refGene_tx=XM_019819473.2
E2 13068903 13069071
refGene_gn=LOC101084152;refGene_tx=XM_019819473.2
E2 13069191 13069391
refGene_gn=LOC101084152;refGene_tx=XM_019819473.2

E2 13070037 13070073
refGene_gn=LOC101084152;refGene_tx=XM_019819473.2
E2 13077024 13077112
refGene_gn=LOC101084152;refGene_tx=XM_019819473.2
E2 13077396 13077404
refGene_gn=LOC101084152;refGene_tx=XM_019819473.2
E2 13090816 13090905 refGene_gn=RABAC1;refGene_tx=XM_003997754.5
E2 13090987 13091089 refGene_gn=RABAC1;refGene_tx=XM_003997754.5
E2 13092593 13092691 refGene_gn=RABAC1;refGene_tx=XM_003997754.5
E2 13092985 13093198 refGene_gn=RABAC1;refGene_tx=XM_003997754.5
E2 13093514 13093570 refGene_gn=RABAC1;refGene_tx=XM_003997754.5
E2 13098565 13098594
refGene_gn=ATP1A3;refGene_tx=XM_023245179.1;refGene_gn=ATP1A3;refGene_tx=XM_023245180.1;refGene_gn=ATP1A3;refGene_tx=XM_023245181.1;refGene_gn=ATP1A3;refGene_tx=XM_023245182.1
E2 13098833 13098925
refGene_gn=ATP1A3;refGene_tx=XM_023245179.1;refGene_gn=ATP1A3;refGene_tx=XM_023245180.1;refGene_gn=ATP1A3;refGene_tx=XM_023245181.1;refGene_gn=ATP1A3;refGene_tx=XM_023245182.1
E2 13099404 13099506
refGene_gn=ATP1A3;refGene_tx=XM_023245179.1;refGene_gn=ATP1A3;refGene_tx=XM_023245180.1;refGene_gn=ATP1A3;refGene_tx=XM_023245181.1;refGene_gn=ATP1A3;refGene_tx=XM_023245182.1
E2 13100151 13100282
refGene_gn=ATP1A3;refGene_tx=XM_023245179.1;refGene_gn=ATP1A3;refGene_tx=XM_023245180.1;refGene_gn=ATP1A3;refGene_tx=XM_023245181.1;refGene_gn=ATP1A3;refGene_tx=XM_023245182.1
E2 13100788 13100934
refGene_gn=ATP1A3;refGene_tx=XM_023245179.1;refGene_gn=ATP1A3;refGene_tx=XM_023245180.1;refGene_gn=ATP1A3;refGene_tx=XM_023245181.1;refGene_gn=ATP1A3;refGene_tx=XM_023245182.1
E2 13101727 13101851
refGene_gn=ATP1A3;refGene_tx=XM_023245179.1;refGene_gn=ATP1A3;refGene_tx=XM_023245180.1;refGene_gn=ATP1A3;refGene_tx=XM_023245181.1;refGene_gn=ATP1A3;refGene_tx=XM_023245182.1
E2 13101927 13102082
refGene_gn=ATP1A3;refGene_tx=XM_023245179.1;refGene_gn=ATP1A3;refGene_tx=XM_023245180.1;refGene_gn=ATP1A3;refGene_tx=XM_023245181.1;refGene_gn=ATP1A3;refGene_tx=XM_023245182.1
E2 13104470 13104639
refGene_gn=ATP1A3;refGene_tx=XM_023245179.1;refGene_gn=ATP1A3;refGene_tx=XM_023245180.1;refGene_gn=ATP1A3;refGene_tx=XM_023245181.1;refGene_gn=ATP1A3;refGene_tx=XM_023245182.1
E2 13104744 13104895
refGene_gn=ATP1A3;refGene_tx=XM_023245179.1;refGene_gn=ATP1A3;refGene_tx=XM_023245180.1;refGene_gn=ATP1A3;refGene_tx=XM_023245181.1;refGene_gn=ATP1A3;refGene_tx=XM_023245182.1
E2 13105226 13105363
refGene_gn=ATP1A3;refGene_tx=XM_023245179.1;refGene_gn=ATP1A3;refGene_tx=XM_023245180.1;refGene_gn=ATP1A3;refGene_tx=XM_023245181.1;refGene_

gn=ATP1A3;refGene_tx=XM_023245182.1
E2 13105448 13105623
refGene_gn=ATP1A3;refGene_tx=XM_023245179.1;refGene_gn=ATP1A3;refGene_tx=XM_023245180.1;refGene_gn=ATP1A3;refGene_tx=XM_023245181.1;refGene_gn=ATP1A3;refGene_tx=XM_023245182.1
E2 13105944 13106138
refGene_gn=ATP1A3;refGene_tx=XM_023245179.1;refGene_gn=ATP1A3;refGene_tx=XM_023245180.1;refGene_gn=ATP1A3;refGene_tx=XM_023245181.1;refGene_gn=ATP1A3;refGene_tx=XM_023245182.1
E2 13108516 13108651
refGene_gn=ATP1A3;refGene_tx=XM_023245179.1;refGene_gn=ATP1A3;refGene_tx=XM_023245180.1;refGene_gn=ATP1A3;refGene_tx=XM_023245181.1;refGene_gn=ATP1A3;refGene_tx=XM_023245182.1
E2 13108734 13108844
refGene_gn=ATP1A3;refGene_tx=XM_023245179.1;refGene_gn=ATP1A3;refGene_tx=XM_023245180.1;refGene_gn=ATP1A3;refGene_tx=XM_023245181.1;refGene_gn=ATP1A3;refGene_tx=XM_023245182.1
E2 13108929 13109128
refGene_gn=ATP1A3;refGene_tx=XM_023245179.1;refGene_gn=ATP1A3;refGene_tx=XM_023245180.1;refGene_gn=ATP1A3;refGene_tx=XM_023245181.1;refGene_gn=ATP1A3;refGene_tx=XM_023245182.1
E2 13110346 13110615
refGene_gn=ATP1A3;refGene_tx=XM_023245179.1;refGene_gn=ATP1A3;refGene_tx=XM_023245180.1;refGene_gn=ATP1A3;refGene_tx=XM_023245181.1;refGene_gn=ATP1A3;refGene_tx=XM_023245182.1
E2 13110755 13110873
refGene_gn=ATP1A3;refGene_tx=XM_023245179.1;refGene_gn=ATP1A3;refGene_tx=XM_023245180.1;refGene_gn=ATP1A3;refGene_tx=XM_023245181.1;refGene_gn=ATP1A3;refGene_tx=XM_023245182.1
E2 13112212 13112347
refGene_gn=ATP1A3;refGene_tx=XM_023245179.1;refGene_gn=ATP1A3;refGene_tx=XM_023245180.1;refGene_gn=ATP1A3;refGene_tx=XM_023245181.1;refGene_gn=ATP1A3;refGene_tx=XM_023245182.1
E2 13112459 13112573
refGene_gn=ATP1A3;refGene_tx=XM_023245179.1;refGene_gn=ATP1A3;refGene_tx=XM_023245180.1;refGene_gn=ATP1A3;refGene_tx=XM_023245181.1;refGene_gn=ATP1A3;refGene_tx=XM_023245182.1
E2 13113931 13114135
refGene_gn=ATP1A3;refGene_tx=XM_023245179.1;refGene_gn=ATP1A3;refGene_tx=XM_023245180.1;refGene_gn=ATP1A3;refGene_tx=XM_023245181.1;refGene_gn=ATP1A3;refGene_tx=XM_023245182.1
E2 13114259 13114319
refGene_gn=ATP1A3;refGene_tx=XM_023245179.1;refGene_gn=ATP1A3;refGene_tx=XM_023245180.1;refGene_gn=ATP1A3;refGene_tx=XM_023245181.1;refGene_gn=ATP1A3;refGene_tx=XM_023245182.1
E2 13114451 13114580
refGene_gn=ATP1A3;refGene_tx=XM_023245179.1;refGene_gn=ATP1A3;refGene_tx=XM_023245180.1;refGene_gn=ATP1A3;refGene_tx=XM_023245181.1;refGene_gn=ATP1A3;refGene_tx=XM_023245182.1
E2 13117631 13117649

refGene_gn=ATP1A3;refGene_tx=XM_023245179.1;refGene_gn=ATP1A3;refGene_tx=XM_023245180.1
E2 13118652 13118658
refGene_gn=ATP1A3;refGene_tx=XM_023245181.1;refGene_gn=ATP1A3;refGene_tx=XM_023245182.1
E2 13123497 13123926
refGene_gn=GRIK5;refGene_tx=XM_023245922.1;refGene_gn=GRIK5;refGene_tx=XM_023245923.1;refGene_gn=GRIK5;refGene_tx=XM_023245924.1;refGene_gn=GRIK5;refGene_tx=XM_023245925.1;refGene_gn=GRIK5;refGene_tx=XM_023245926.1;refGene_gn=GRIK5;refGene_tx=XM_023245927.1
E2 13124693 13124843 refGene_gn=GRIK5;refGene_tx=XM_023245928.1
E2 13127195 13127317
refGene_gn=GRIK5;refGene_tx=XM_023245922.1;refGene_gn=GRIK5;refGene_tx=XM_023245923.1;refGene_gn=GRIK5;refGene_tx=XM_023245924.1;refGene_gn=GRIK5;refGene_tx=XM_023245925.1;refGene_gn=GRIK5;refGene_tx=XM_023245926.1;refGene_gn=GRIK5;refGene_tx=XM_023245927.1;refGene_gn=GRIK5;refGene_tx=XM_023245928.1
E2 13127411 13127540
refGene_gn=GRIK5;refGene_tx=XM_023245922.1;refGene_gn=GRIK5;refGene_tx=XM_023245923.1;refGene_gn=GRIK5;refGene_tx=XM_023245924.1;refGene_gn=GRIK5;refGene_tx=XM_023245925.1;refGene_gn=GRIK5;refGene_tx=XM_023245926.1;refGene_gn=GRIK5;refGene_tx=XM_023245927.1;refGene_gn=GRIK5;refGene_tx=XM_023245928.1
E2 13129331 13129557
refGene_gn=GRIK5;refGene_tx=XM_023245922.1;refGene_gn=GRIK5;refGene_tx=XM_023245923.1;refGene_gn=GRIK5;refGene_tx=XM_023245924.1;refGene_gn=GRIK5;refGene_tx=XM_023245925.1;refGene_gn=GRIK5;refGene_tx=XM_023245926.1;refGene_gn=GRIK5;refGene_tx=XM_023245927.1;refGene_gn=GRIK5;refGene_tx=XM_023245928.1
E2 13130850 13131016
refGene_gn=GRIK5;refGene_tx=XM_023245922.1;refGene_gn=GRIK5;refGene_tx=XM_023245923.1;refGene_gn=GRIK5;refGene_tx=XM_023245924.1;refGene_gn=GRIK5;refGene_tx=XM_023245925.1;refGene_gn=GRIK5;refGene_tx=XM_023245926.1;refGene_gn=GRIK5;refGene_tx=XM_023245927.1;refGene_gn=GRIK5;refGene_tx=XM_023245928.1
E2 13141095 13141269
refGene_gn=GRIK5;refGene_tx=XM_023245922.1;refGene_gn=GRIK5;refGene_tx=XM_023245923.1;refGene_gn=GRIK5;refGene_tx=XM_023245924.1;refGene_gn=GRIK5;refGene_tx=XM_023245925.1;refGene_gn=GRIK5;refGene_tx=XM_023245926.1;refGene_gn=GRIK5;refGene_tx=XM_023245927.1;refGene_gn=GRIK5;refGene_tx=XM_023245928.1
E2 13141651 13141761
refGene_gn=GRIK5;refGene_tx=XM_023245922.1;refGene_gn=GRIK5;refGene_tx=XM_023245923.1;refGene_gn=GRIK5;refGene_tx=XM_023245924.1;refGene_gn=GRIK5;refGene_tx=XM_023245925.1;refGene_gn=GRIK5;refGene_tx=XM_023245926.1;refGene_gn=GRIK5;refGene_tx=XM_023245927.1;refGene_gn=GRIK5;refGene_tx=XM_023245928.1
E2 13141957 13142074
refGene_gn=GRIK5;refGene_tx=XM_023245922.1;refGene_gn=GRIK5;refGene_tx=XM_023245923.1;refGene_gn=GRIK5;refGene_tx=XM_023245924.1;refGene_gn=

GRIK5;refGene_tx=XM_023245925.1;refGene_gn=GRIK5;refGene_tx=XM_023245926.1;refGene_gn=GRIK5;refGene_tx=XM_023245927.1;refGene_gn=GRIK5;refGene_tx=XM_023245928.1
E2 13143332 13143461
refGene_gn=GRIK5;refGene_tx=XM_023245922.1;refGene_gn=GRIK5;refGene_tx=XM_023245923.1;refGene_gn=GRIK5;refGene_tx=XM_023245924.1;refGene_gn=GRIK5;refGene_tx=XM_023245925.1;refGene_gn=GRIK5;refGene_tx=XM_023245928.1
E2 13156134 13156338
refGene_gn=GRIK5;refGene_tx=XM_023245922.1;refGene_gn=GRIK5;refGene_tx=XM_023245923.1;refGene_gn=GRIK5;refGene_tx=XM_023245924.1;refGene_gn=GRIK5;refGene_tx=XM_023245925.1;refGene_gn=GRIK5;refGene_tx=XM_023245926.1;refGene_gn=GRIK5;refGene_tx=XM_023245927.1;refGene_gn=GRIK5;refGene_tx=XM_023245928.1
E2 13163619 13163727
refGene_gn=GRIK5;refGene_tx=XM_023245922.1;refGene_gn=GRIK5;refGene_tx=XM_023245923.1;refGene_gn=GRIK5;refGene_tx=XM_023245924.1;refGene_gn=GRIK5;refGene_tx=XM_023245925.1;refGene_gn=GRIK5;refGene_tx=XM_023245926.1;refGene_gn=GRIK5;refGene_tx=XM_023245927.1;refGene_gn=GRIK5;refGene_tx=XM_023245928.1
E2 13163810 13163915
refGene_gn=GRIK5;refGene_tx=XM_023245922.1;refGene_gn=GRIK5;refGene_tx=XM_023245923.1;refGene_gn=GRIK5;refGene_tx=XM_023245924.1;refGene_gn=GRIK5;refGene_tx=XM_023245925.1;refGene_gn=GRIK5;refGene_tx=XM_023245926.1;refGene_gn=GRIK5;refGene_tx=XM_023245927.1;refGene_gn=GRIK5;refGene_tx=XM_023245928.1
E2 13164304 13164457
refGene_gn=GRIK5;refGene_tx=XM_023245922.1;refGene_gn=GRIK5;refGene_tx=XM_023245923.1;refGene_gn=GRIK5;refGene_tx=XM_023245924.1;refGene_gn=GRIK5;refGene_tx=XM_023245925.1;refGene_gn=GRIK5;refGene_tx=XM_023245926.1;refGene_gn=GRIK5;refGene_tx=XM_023245927.1;refGene_gn=GRIK5;refGene_tx=XM_023245928.1
E2 13166040 13166202
refGene_gn=GRIK5;refGene_tx=XM_023245922.1;refGene_gn=GRIK5;refGene_tx=XM_023245923.1;refGene_gn=GRIK5;refGene_tx=XM_023245924.1;refGene_gn=GRIK5;refGene_tx=XM_023245925.1;refGene_gn=GRIK5;refGene_tx=XM_023245926.1;refGene_gn=GRIK5;refGene_tx=XM_023245927.1;refGene_gn=GRIK5;refGene_tx=XM_023245928.1
E2 13166304 13166358
refGene_gn=GRIK5;refGene_tx=XM_023245922.1;refGene_gn=GRIK5;refGene_tx=XM_023245924.1;refGene_gn=GRIK5;refGene_tx=XM_023245925.1;refGene_gn=GRIK5;refGene_tx=XM_023245926.1;refGene_gn=GRIK5;refGene_tx=XM_023245927.1;refGene_gn=GRIK5;refGene_tx=XM_023245928.1
E2 13167827 13168006
refGene_gn=GRIK5;refGene_tx=XM_023245922.1;refGene_gn=GRIK5;refGene_tx=XM_023245923.1;refGene_gn=GRIK5;refGene_tx=XM_023245924.1;refGene_gn=GRIK5;refGene_tx=XM_023245925.1;refGene_gn=GRIK5;refGene_tx=XM_023245926.1;refGene_gn=GRIK5;refGene_tx=XM_023245927.1;refGene_gn=GRIK5;refGene_tx=XM_023245928.1
E2 13170003 13170169

refGene_gn=GRIK5;refGene_tx=XM_023245922.1;refGene_gn=GRIK5;refGene_tx=XM_023245923.1;refGene_gn=GRIK5;refGene_tx=XM_023245924.1;refGene_gn=GRIK5;refGene_tx=XM_023245925.1;refGene_gn=GRIK5;refGene_tx=XM_023245926.1;refGene_gn=GRIK5;refGene_tx=XM_023245927.1;refGene_gn=GRIK5;refGene_tx=XM_023245928.1
E2 13170265 13170363
refGene_gn=GRIK5;refGene_tx=XM_023245922.1;refGene_gn=GRIK5;refGene_tx=XM_023245923.1;refGene_gn=GRIK5;refGene_tx=XM_023245924.1;refGene_gn=GRIK5;refGene_tx=XM_023245925.1;refGene_gn=GRIK5;refGene_tx=XM_023245926.1;refGene_gn=GRIK5;refGene_tx=XM_023245927.1;refGene_gn=GRIK5;refGene_tx=XM_023245928.1
E2 13172624 13172789
refGene_gn=GRIK5;refGene_tx=XM_023245922.1;refGene_gn=GRIK5;refGene_tx=XM_023245923.1;refGene_gn=GRIK5;refGene_tx=XM_023245924.1;refGene_gn=GRIK5;refGene_tx=XM_023245925.1;refGene_gn=GRIK5;refGene_tx=XM_023245926.1;refGene_gn=GRIK5;refGene_tx=XM_023245927.1;refGene_gn=GRIK5;refGene_tx=XM_023245928.1
E2 13173094 13173223
refGene_gn=GRIK5;refGene_tx=XM_023245922.1;refGene_gn=GRIK5;refGene_tx=XM_023245923.1;refGene_gn=GRIK5;refGene_tx=XM_023245924.1;refGene_gn=GRIK5;refGene_tx=XM_023245925.1;refGene_gn=GRIK5;refGene_tx=XM_023245926.1;refGene_gn=GRIK5;refGene_tx=XM_023245927.1;refGene_gn=GRIK5;refGene_tx=XM_023245928.1
E2 13176337 13176479 refGene_gn=ZNF574;refGene_tx=XM_023245929.1
E2 13179657 13179709
refGene_gn=GRIK5;refGene_tx=XM_023245922.1;refGene_gn=GRIK5;refGene_tx=XM_023245923.1;refGene_gn=GRIK5;refGene_tx=XM_023245926.1;refGene_gn=GRIK5;refGene_tx=XM_023245927.1;refGene_gn=GRIK5;refGene_tx=XM_023245928.1
E2 13183832 13186543
refGene_gn=ZNF574;refGene_tx=XM_023245929.1;refGene_gn=ZNF574;refGene_tx=XM_023245930.1;refGene_gn=ZNF574;refGene_tx=XM_023245931.1;refGene_gn=ZNF574;refGene_tx=XM_023245933.1;refGene_gn=ZNF574;refGene_tx=XM_023245934.1;refGene_gn=ZNF574;refGene_tx=XM_023245935.1;refGene_gn=ZNF574;refGene_tx=XM_023245936.1;refGene_gn=ZNF574;refGene_tx=XM_023245937.1;refGene_gn=ZNF574;refGene_tx=XM_023245938.1
E2 13196146 13196592
refGene_gn=POU2F2;refGene_tx=XM_003997848.5;refGene_gn=POU2F2;refGene_tx=XM_023245172.1;refGene_gn=POU2F2;refGene_tx=XM_023245173.1;refGene_gn=POU2F2;refGene_tx=XM_023245174.1;refGene_gn=POU2F2;refGene_tx=XM_023245175.1;refGene_gn=POU2F2;refGene_tx=XM_023245176.1;refGene_gn=POU2F2;refGene_tx=XM_023245177.1;refGene_gn=POU2F2;refGene_tx=XM_023245178.1
E2 13196850 13196924
refGene_gn=POU2F2;refGene_tx=XM_023245172.1;refGene_gn=POU2F2;refGene_tx=XM_023245173.1;refGene_gn=POU2F2;refGene_tx=XM_023245174.1;refGene_gn=POU2F2;refGene_tx=XM_023245175.1;refGene_gn=POU2F2;refGene_tx=XM_023245176.1
E2 13197064 13197266
refGene_gn=POU2F2;refGene_tx=XM_003997848.5;refGene_gn=POU2F2;refGene_

tx=XM_023245172.1;refGene_gn=POU2F2;refGene_tx=XM_023245173.1;refGene_
gn=POU2F2;refGene_tx=XM_023245174.1;refGene_gn=POU2F2;refGene_tx=XM_02
3245175.1;refGene_gn=POU2F2;refGene_tx=XM_023245176.1;refGene_gn=POU2F
2;refGene_tx=XM_023245177.1;refGene_gn=POU2F2;refGene_tx=XM_023245178.
1
E2 13198491 13198558
refGene_gn=POU2F2;refGene_tx=XM_003997848.5;refGene_gn=POU2F2;refGene_
tx=XM_023245172.1;refGene_gn=POU2F2;refGene_tx=XM_023245173.1;refGene_
gn=POU2F2;refGene_tx=XM_023245174.1;refGene_gn=POU2F2;refGene_tx=XM_02
3245175.1;refGene_gn=POU2F2;refGene_tx=XM_023245176.1;refGene_gn=POU2F
2;refGene_tx=XM_023245177.1;refGene_gn=POU2F2;refGene_tx=XM_023245178.
1
E2 13199830 13200007
refGene_gn=POU2F2;refGene_tx=XM_003997848.5;refGene_gn=POU2F2;refGene_
tx=XM_023245172.1;refGene_gn=POU2F2;refGene_tx=XM_023245173.1;refGene_
gn=POU2F2;refGene_tx=XM_023245174.1;refGene_gn=POU2F2;refGene_tx=XM_02
3245175.1;refGene_gn=POU2F2;refGene_tx=XM_023245176.1;refGene_gn=POU2F
2;refGene_tx=XM_023245177.1;refGene_gn=POU2F2;refGene_tx=XM_023245178.
1
E2 13200108 13200257
refGene_gn=POU2F2;refGene_tx=XM_003997848.5;refGene_gn=POU2F2;refGene_
tx=XM_023245172.1;refGene_gn=POU2F2;refGene_tx=XM_023245173.1;refGene_
gn=POU2F2;refGene_tx=XM_023245174.1;refGene_gn=POU2F2;refGene_tx=XM_02
3245175.1;refGene_gn=POU2F2;refGene_tx=XM_023245176.1;refGene_gn=POU2F
2;refGene_tx=XM_023245177.1;refGene_gn=POU2F2;refGene_tx=XM_023245178.
1
E2 13200349 13200491
refGene_gn=POU2F2;refGene_tx=XM_003997848.5;refGene_gn=POU2F2;refGene_
tx=XM_023245172.1;refGene_gn=POU2F2;refGene_tx=XM_023245173.1;refGene_
gn=POU2F2;refGene_tx=XM_023245174.1;refGene_gn=POU2F2;refGene_tx=XM_02
3245175.1;refGene_gn=POU2F2;refGene_tx=XM_023245176.1;refGene_gn=POU2F
2;refGene_tx=XM_023245177.1;refGene_gn=POU2F2;refGene_tx=XM_023245178.
1
E2 13200725 13200887
refGene_gn=POU2F2;refGene_tx=XM_003997848.5;refGene_gn=POU2F2;refGene_
tx=XM_023245172.1;refGene_gn=POU2F2;refGene_tx=XM_023245173.1;refGene_
gn=POU2F2;refGene_tx=XM_023245174.1;refGene_gn=POU2F2;refGene_tx=XM_02
3245175.1;refGene_gn=POU2F2;refGene_tx=XM_023245176.1;refGene_gn=POU2F
2;refGene_tx=XM_023245177.1;refGene_gn=POU2F2;refGene_tx=XM_023245178.
1
E2 13202490 13202582
refGene_gn=POU2F2;refGene_tx=XM_003997848.5;refGene_gn=POU2F2;refGene_
tx=XM_023245172.1;refGene_gn=POU2F2;refGene_tx=XM_023245173.1;refGene_
gn=POU2F2;refGene_tx=XM_023245174.1;refGene_gn=POU2F2;refGene_tx=XM_02
3245175.1;refGene_gn=POU2F2;refGene_tx=XM_023245176.1;refGene_gn=POU2F
2;refGene_tx=XM_023245177.1;refGene_gn=POU2F2;refGene_tx=XM_023245178.
1
E2 13202670 13202776
refGene_gn=POU2F2;refGene_tx=XM_003997848.5;refGene_gn=POU2F2;refGene_
tx=XM_023245172.1;refGene_gn=POU2F2;refGene_tx=XM_023245173.1;refGene_

gn=POU2F2;refGene_tx=XM_023245174.1;refGene_gn=POU2F2;refGene_tx=XM_02
3245175.1;refGene_gn=POU2F2;refGene_tx=XM_023245176.1;refGene_gn=POU2F
2;refGene_tx=XM_023245177.1;refGene_gn=POU2F2;refGene_tx=XM_023245178.
1
E2 13215608 13215887
refGene_gn=POU2F2;refGene_tx=XM_023245172.1;refGene_gn=POU2F2;refGene_
tx=XM_023245173.1;refGene_gn=POU2F2;refGene_tx=XM_023245174.1;refGene_
gn=POU2F2;refGene_tx=XM_023245177.1
E2 13216151 13216268
refGene_gn=POU2F2;refGene_tx=XM_003997848.5;refGene_gn=POU2F2;refGene_
tx=XM_023245172.1;refGene_gn=POU2F2;refGene_tx=XM_023245173.1;refGene_
gn=POU2F2;refGene_tx=XM_023245174.1;refGene_gn=POU2F2;refGene_tx=XM_02
3245175.1;refGene_gn=POU2F2;refGene_tx=XM_023245176.1;refGene_gn=POU2F
2;refGene_tx=XM_023245177.1;refGene_gn=POU2F2;refGene_tx=XM_023245178.
1
E2 13219984 13220041
refGene_gn=POU2F2;refGene_tx=XM_003997848.5;refGene_gn=POU2F2;refGene_
tx=XM_023245172.1;refGene_gn=POU2F2;refGene_tx=XM_023245173.1;refGene_
gn=POU2F2;refGene_tx=XM_023245174.1;refGene_gn=POU2F2;refGene_tx=XM_02
3245175.1;refGene_gn=POU2F2;refGene_tx=XM_023245176.1;refGene_gn=POU2F
2;refGene_tx=XM_023245177.1;refGene_gn=POU2F2;refGene_tx=XM_023245178.
1
E2 13220180 13220215
refGene_gn=POU2F2;refGene_tx=XM_003997848.5;refGene_gn=POU2F2;refGene_
tx=XM_023245172.1;refGene_gn=POU2F2;refGene_tx=XM_023245173.1;refGene_
gn=POU2F2;refGene_tx=XM_023245174.1;refGene_gn=POU2F2;refGene_tx=XM_02
3245175.1;refGene_gn=POU2F2;refGene_tx=XM_023245176.1;refGene_gn=POU2F
2;refGene_tx=XM_023245177.1;refGene_gn=POU2F2;refGene_tx=XM_023245178.
1
E2 13220319 13220385
refGene_gn=POU2F2;refGene_tx=XM_003997848.5;refGene_gn=POU2F2;refGene_
tx=XM_023245172.1;refGene_gn=POU2F2;refGene_tx=XM_023245173.1;refGene_
gn=POU2F2;refGene_tx=XM_023245174.1;refGene_gn=POU2F2;refGene_tx=XM_02
3245175.1;refGene_gn=POU2F2;refGene_tx=XM_023245176.1;refGene_gn=POU2F
2;refGene_tx=XM_023245177.1;refGene_gn=POU2F2;refGene_tx=XM_023245178.
1
E2 13227500 13227567
refGene_gn=POU2F2;refGene_tx=XM_003997848.5;refGene_gn=POU2F2;refGene_
tx=XM_023245172.1;refGene_gn=POU2F2;refGene_tx=XM_023245173.1;refGene_
gn=POU2F2;refGene_tx=XM_023245175.1;refGene_gn=POU2F2;refGene_tx=XM_02
3245176.1;refGene_gn=POU2F2;refGene_tx=XM_023245177.1;refGene_gn=POU2F
2;refGene_tx=XM_023245178.1
E2 13289206 13289598
refGene_gn=DEDD2;refGene_tx=XM_003997756.4;refGene_gn=DEDD2;refGene_tx
=XM_019819481.2;refGene_gn=DEDD2;refGene_tx=XM_019819482.2
E2 13297613 13297754
refGene_gn=DEDD2;refGene_tx=XM_003997756.4;refGene_gn=DEDD2;refGene_tx
=XM_019819481.2;refGene_gn=DEDD2;refGene_tx=XM_019819482.2
E2 13301880 13302003
refGene_gn=DEDD2;refGene_tx=XM_003997756.4;refGene_gn=DEDD2;refGene_tx

=XM_019819481.2;refGene_gn=DEDD2;refGene_tx=XM_019819482.2
E2 13303495 13303823
refGene_gn=DEDD2;refGene_tx=XM_003997756.4;refGene_gn=DEDD2;refGene_tx=XM_019819481.2;refGene_gn=DEDD2;refGene_tx=XM_019819482.2
E2 13306213 13306306 refGene_gn=ZNF526;refGene_tx=XM_019819480.2
E2 13308381 13310427
refGene_gn=ZNF526;refGene_tx=XM_019819480.2;refGene_gn=ZNF526;refGene_tx=XM_023245819.1;refGene_gn=ZNF526;refGene_tx=XM_023245820.1;refGene_gn=ZNF526;refGene_tx=XM_023245821.1;refGene_gn=ZNF526;refGene_tx=XM_023245822.1;refGene_gn=ZNF526;refGene_tx=XM_023245823.1;refGene_gn=ZNF526;refGene_tx=XM_023245824.1
E2 13313857 13313931 refGene_gn=GSK3A;refGene_tx=XM_003997757.5
E2 13314497 13314590 refGene_gn=GSK3A;refGene_tx=XM_003997757.5
E2 13314919 13315106 refGene_gn=GSK3A;refGene_tx=XM_003997757.5
E2 13315522 13315618 refGene_gn=GSK3A;refGene_tx=XM_003997757.5
E2 13315700 13315798 refGene_gn=GSK3A;refGene_tx=XM_003997757.5
E2 13316746 13316853 refGene_gn=GSK3A;refGene_tx=XM_003997757.5
E2 13316947 13317078 refGene_gn=GSK3A;refGene_tx=XM_003997757.5
E2 13318958 13319069 refGene_gn=GSK3A;refGene_tx=XM_003997757.5
E2 13319233 13319317 refGene_gn=GSK3A;refGene_tx=XM_003997757.5
E2 13320656 13320844 refGene_gn=GSK3A;refGene_tx=XM_003997757.5
E2 13322735 13323018 refGene_gn=GSK3A;refGene_tx=XM_003997757.5
E2 13327999 13329288 refGene_gn=ERF;refGene_tx=XM_011289905.3
E2 13329394 13329510 refGene_gn=ERF;refGene_tx=XM_011289905.3
E2 13329887 13330122 refGene_gn=ERF;refGene_tx=XM_011289905.3
E2 13334530 13334552 refGene_gn=ERF;refGene_tx=XM_011289905.3
E2 13348188 13348437 refGene_gn=CIC;refGene_tx=XM_019819484.2
E2 13348438 13350992 refGene_gn=CIC;refGene_tx=XM_019819484.2
E2 13362851 13363001 refGene_gn=CIC;refGene_tx=XM_019819484.2
E2 13363088 13363323 refGene_gn=CIC;refGene_tx=XM_019819484.2
E2 13363403 13363533 refGene_gn=CIC;refGene_tx=XM_019819484.2
E2 13363627 13363810 refGene_gn=CIC;refGene_tx=XM_019819484.2
E2 13363892 13364058 refGene_gn=CIC;refGene_tx=XM_019819484.2
E2 13364901 13365104 refGene_gn=CIC;refGene_tx=XM_019819484.2
E2 13365186 13365412 refGene_gn=CIC;refGene_tx=XM_019819484.2
E2 13365829 13365933 refGene_gn=CIC;refGene_tx=XM_019819484.2
E2 13366183 13367417 refGene_gn=CIC;refGene_tx=XM_019819484.2
E2 13367496 13367684 refGene_gn=CIC;refGene_tx=XM_019819484.2
E2 13368058 13368180 refGene_gn=CIC;refGene_tx=XM_019819484.2
E2 13368265 13368432 refGene_gn=CIC;refGene_tx=XM_019819484.2
E2 13368531 13368822 refGene_gn=CIC;refGene_tx=XM_019819484.2
E2 13368918 13369244 refGene_gn=CIC;refGene_tx=XM_019819484.2
E2 13369540 13369794 refGene_gn=CIC;refGene_tx=XM_019819484.2
E2 13369894 13370049 refGene_gn=CIC;refGene_tx=XM_019819484.2
E2 13370127 13370259 refGene_gn=CIC;refGene_tx=XM_019819484.2
E2 13370542 13370674 refGene_gn=CIC;refGene_tx=XM_019819484.2
E2 13370770 13371138 refGene_gn=CIC;refGene_tx=XM_019819484.2
E2 13372652 13372940
refGene_gn=PAFAH1B3;refGene_tx=XM_006941221.4;refGene_gn=PAFAH1B3;refGene_tx=XM_006941223.3;refGene_gn=PAFAH1B3;refGene_tx=XM_006941224.4;re

fGene_gn=PAFAH1B3;refGene_tx=XM_023245826.1
E2 13373485 13373608
refGene_gn=PAFAH1B3;refGene_tx=XM_006941221.4;refGene_gn=PAFAH1B3;refGene_tx=XM_006941223.3;refGene_gn=PAFAH1B3;refGene_tx=XM_006941224.4;refGene_gn=PAFAH1B3;refGene_tx=XM_023245826.1
E2 13373686 13373803
refGene_gn=PAFAH1B3;refGene_tx=XM_006941221.4;refGene_gn=PAFAH1B3;refGene_tx=XM_006941223.3;refGene_gn=PAFAH1B3;refGene_tx=XM_006941224.4;refGene_gn=PAFAH1B3;refGene_tx=XM_023245826.1
E2 13374459 13374549
refGene_gn=PAFAH1B3;refGene_tx=XM_006941221.4;refGene_gn=PAFAH1B3;refGene_tx=XM_006941223.3;refGene_gn=PAFAH1B3;refGene_tx=XM_006941224.4;refGene_gn=PAFAH1B3;refGene_tx=XM_023245826.1
E2 13374736 13374814
refGene_gn=PAFAH1B3;refGene_tx=XM_006941221.4;refGene_gn=PAFAH1B3;refGene_tx=XM_006941223.3;refGene_gn=PAFAH1B3;refGene_tx=XM_006941224.4;refGene_gn=PAFAH1B3;refGene_tx=XM_023245826.1
E2 13375732 13375822 refGene_gn=PRR19;refGene_tx=XM_003997762.4
E2 13377965 13378572 refGene_gn=PRR19;refGene_tx=XM_003997762.4
E2 13378658 13379131 refGene_gn=PRR19;refGene_tx=XM_003997762.4
E2 13381168 13381288
refGene_gn=TMEM145;refGene_tx=XM_019818908.2;refGene_gn=TMEM145;refGene_tx=XM_023245825.1
E2 13382036 13382111
refGene_gn=TMEM145;refGene_tx=XM_019818908.2;refGene_gn=TMEM145;refGene_tx=XM_023245825.1
E2 13382227 13382305
refGene_gn=TMEM145;refGene_tx=XM_019818908.2;refGene_gn=TMEM145;refGene_tx=XM_023245825.1
E2 13382387 13382474
refGene_gn=TMEM145;refGene_tx=XM_019818908.2;refGene_gn=TMEM145;refGene_tx=XM_023245825.1
E2 13382558 13382660
refGene_gn=TMEM145;refGene_tx=XM_019818908.2;refGene_gn=TMEM145;refGene_tx=XM_023245825.1
E2 13382765 13382850
refGene_gn=TMEM145;refGene_tx=XM_019818908.2;refGene_gn=TMEM145;refGene_tx=XM_023245825.1
E2 13382965 13383046
refGene_gn=TMEM145;refGene_tx=XM_019818908.2;refGene_gn=TMEM145;refGene_tx=XM_023245825.1
E2 13383146 13383215
refGene_gn=TMEM145;refGene_tx=XM_019818908.2;refGene_gn=TMEM145;refGene_tx=XM_023245825.1
E2 13383578 13383659
refGene_gn=TMEM145;refGene_tx=XM_019818908.2;refGene_gn=TMEM145;refGene_tx=XM_023245825.1
E2 13383742 13383821
refGene_gn=TMEM145;refGene_tx=XM_019818908.2;refGene_gn=TMEM145;refGene_tx=XM_023245825.1

E2 13383968 13384062
refGene_gn=TMEM145;refGene_tx=XM_019818908.2;refGene_gn=TMEM145;refGene_tx=XM_023245825.1
E2 13384826 13384999 refGene_gn=TMEM145;refGene_tx=XM_019818908.2
E2 13385843 13385964
refGene_gn=TMEM145;refGene_tx=XM_019818908.2;refGene_gn=TMEM145;refGene_tx=XM_023245825.1
E2 13388031 13388609
refGene_gn=TMEM145;refGene_tx=XM_019818908.2;refGene_gn=TMEM145;refGene_tx=XM_023245825.1
E2 13389208 13389392 refGene_gn=TMEM145;refGene_tx=XM_019818908.2
E2 13390709 13390896
refGene_gn=MEGF8;refGene_tx=XM_006941225.4;refGene_gn=MEGF8;refGene_tx=XM_019819489.2;refGene_gn=MEGF8;refGene_tx=XM_019819490.2
E2 13395619 13395783
refGene_gn=MEGF8;refGene_tx=XM_006941225.4;refGene_gn=MEGF8;refGene_tx=XM_019819489.2;refGene_gn=MEGF8;refGene_tx=XM_019819490.2
E2 13396038 13396245
refGene_gn=MEGF8;refGene_tx=XM_006941225.4;refGene_gn=MEGF8;refGene_tx=XM_019819489.2;refGene_gn=MEGF8;refGene_tx=XM_019819490.2
E2 13397191 13397372
refGene_gn=MEGF8;refGene_tx=XM_006941225.4;refGene_gn=MEGF8;refGene_tx=XM_019819489.2;refGene_gn=MEGF8;refGene_tx=XM_019819490.2
E2 13397451 13397540
refGene_gn=MEGF8;refGene_tx=XM_006941225.4;refGene_gn=MEGF8;refGene_tx=XM_019819489.2;refGene_gn=MEGF8;refGene_tx=XM_019819490.2
E2 13398058 13398474
refGene_gn=MEGF8;refGene_tx=XM_006941225.4;refGene_gn=MEGF8;refGene_tx=XM_019819489.2;refGene_gn=MEGF8;refGene_tx=XM_019819490.2
E2 13398904 13399050
refGene_gn=MEGF8;refGene_tx=XM_006941225.4;refGene_gn=MEGF8;refGene_tx=XM_019819489.2;refGene_gn=MEGF8;refGene_tx=XM_019819490.2
E2 13399175 13399298
refGene_gn=MEGF8;refGene_tx=XM_006941225.4;refGene_gn=MEGF8;refGene_tx=XM_019819489.2;refGene_gn=MEGF8;refGene_tx=XM_019819490.2
E2 13401394 13401549
refGene_gn=MEGF8;refGene_tx=XM_006941225.4;refGene_gn=MEGF8;refGene_tx=XM_019819489.2;refGene_gn=MEGF8;refGene_tx=XM_019819490.2
E2 13401857 13401977
refGene_gn=MEGF8;refGene_tx=XM_006941225.4;refGene_gn=MEGF8;refGene_tx=XM_019819489.2;refGene_gn=MEGF8;refGene_tx=XM_019819490.2
E2 13402329 13402474
refGene_gn=MEGF8;refGene_tx=XM_006941225.4;refGene_gn=MEGF8;refGene_tx=XM_019819489.2;refGene_gn=MEGF8;refGene_tx=XM_019819490.2
E2 13402557 13402721
refGene_gn=MEGF8;refGene_tx=XM_006941225.4;refGene_gn=MEGF8;refGene_tx=XM_019819489.2;refGene_gn=MEGF8;refGene_tx=XM_019819490.2
E2 13405964 13406165
refGene_gn=MEGF8;refGene_tx=XM_006941225.4;refGene_gn=MEGF8;refGene_tx=XM_019819489.2;refGene_gn=MEGF8;refGene_tx=XM_019819490.2

E2	13406549	13406750
refGene_gn=MEGF8;refGene_tx=XM_006941225.4;refGene_gn=MEGF8;refGene_tx=XM_019819489.2;refGene_gn=MEGF8;refGene_tx=XM_019819490.2
E2	13407202	13407439
refGene_gn=MEGF8;refGene_tx=XM_006941225.4;refGene_gn=MEGF8;refGene_tx=XM_019819489.2;refGene_gn=MEGF8;refGene_tx=XM_019819490.2
E2	13408335	13408454
refGene_gn=MEGF8;refGene_tx=XM_006941225.4;refGene_gn=MEGF8;refGene_tx=XM_019819489.2;refGene_gn=MEGF8;refGene_tx=XM_019819490.2
E2	13408536	13408668
refGene_gn=MEGF8;refGene_tx=XM_006941225.4;refGene_gn=MEGF8;refGene_tx=XM_019819489.2;refGene_gn=MEGF8;refGene_tx=XM_019819490.2
E2	13408762	13408876
refGene_gn=MEGF8;refGene_tx=XM_006941225.4;refGene_gn=MEGF8;refGene_tx=XM_019819489.2;refGene_gn=MEGF8;refGene_tx=XM_019819490.2
E2	13409182	13409431
refGene_gn=MEGF8;refGene_tx=XM_006941225.4;refGene_gn=MEGF8;refGene_tx=XM_019819489.2;refGene_gn=MEGF8;refGene_tx=XM_019819490.2
E2	13409934	13410134
refGene_gn=MEGF8;refGene_tx=XM_006941225.4;refGene_gn=MEGF8;refGene_tx=XM_019819489.2;refGene_gn=MEGF8;refGene_tx=XM_019819490.2
E2	13410416	13410627
refGene_gn=MEGF8;refGene_tx=XM_006941225.4;refGene_gn=MEGF8;refGene_tx=XM_019819489.2;refGene_gn=MEGF8;refGene_tx=XM_019819490.2
E2	13410725	13410975
refGene_gn=MEGF8;refGene_tx=XM_006941225.4;refGene_gn=MEGF8;refGene_tx=XM_019819489.2;refGene_gn=MEGF8;refGene_tx=XM_019819490.2
E2	13411513	13411646
refGene_gn=MEGF8;refGene_tx=XM_006941225.4;refGene_gn=MEGF8;refGene_tx=XM_019819489.2;refGene_gn=MEGF8;refGene_tx=XM_019819490.2
E2	13413555	13413803
refGene_gn=MEGF8;refGene_tx=XM_006941225.4;refGene_gn=MEGF8;refGene_tx=XM_019819489.2;refGene_gn=MEGF8;refGene_tx=XM_019819490.2
E2	13413895	13414006
refGene_gn=MEGF8;refGene_tx=XM_006941225.4;refGene_gn=MEGF8;refGene_tx=XM_019819489.2;refGene_gn=MEGF8;refGene_tx=XM_019819490.2
E2	13414168	13414287
refGene_gn=MEGF8;refGene_tx=XM_006941225.4;refGene_gn=MEGF8;refGene_tx=XM_019819489.2;refGene_gn=MEGF8;refGene_tx=XM_019819490.2
E2	13414652	13414860
refGene_gn=MEGF8;refGene_tx=XM_006941225.4;refGene_gn=MEGF8;refGene_tx=XM_019819489.2;refGene_gn=MEGF8;refGene_tx=XM_019819490.2
E2	13415402	13415583
refGene_gn=MEGF8;refGene_tx=XM_006941225.4;refGene_gn=MEGF8;refGene_tx=XM_019819489.2;refGene_gn=MEGF8;refGene_tx=XM_019819490.2
E2	13415896	13415905
refGene_gn=MEGF8;refGene_tx=XM_019819489.2;refGene_gn=MEGF8;refGene_tx=XM_019819490.2
E2	13416116	13416280
refGene_gn=MEGF8;refGene_tx=XM_006941225.4;refGene_gn=MEGF8;refGene_tx

=XM_019819489.2;refGene_gn=MEGF8;refGene_tx=XM_019819490.2
E2 13417033 13417178
refGene_gn=MEGF8;refGene_tx=XM_006941225.4;refGene_gn=MEGF8;refGene_tx=XM_019819489.2;refGene_gn=MEGF8;refGene_tx=XM_019819490.2
E2 13418791 13419023
refGene_gn=MEGF8;refGene_tx=XM_006941225.4;refGene_gn=MEGF8;refGene_tx=XM_019819489.2;refGene_gn=MEGF8;refGene_tx=XM_019819490.2
E2 13420211 13420335
refGene_gn=MEGF8;refGene_tx=XM_006941225.4;refGene_gn=MEGF8;refGene_tx=XM_019819489.2;refGene_gn=MEGF8;refGene_tx=XM_019819490.2
E2 13420462 13420676
refGene_gn=MEGF8;refGene_tx=XM_006941225.4;refGene_gn=MEGF8;refGene_tx=XM_019819489.2;refGene_gn=MEGF8;refGene_tx=XM_019819490.2
E2 13420801 13421016
refGene_gn=MEGF8;refGene_tx=XM_006941225.4;refGene_gn=MEGF8;refGene_tx=XM_019819489.2;refGene_gn=MEGF8;refGene_tx=XM_019819490.2
E2 13424861 13425069
refGene_gn=MEGF8;refGene_tx=XM_006941225.4;refGene_gn=MEGF8;refGene_tx=XM_019819489.2;refGene_gn=MEGF8;refGene_tx=XM_019819490.2
E2 13425232 13425392
refGene_gn=MEGF8;refGene_tx=XM_006941225.4;refGene_gn=MEGF8;refGene_tx=XM_019819489.2;refGene_gn=MEGF8;refGene_tx=XM_019819490.2
E2 13425705 13425898
refGene_gn=MEGF8;refGene_tx=XM_006941225.4;refGene_gn=MEGF8;refGene_tx=XM_019819489.2;refGene_gn=MEGF8;refGene_tx=XM_019819490.2
E2 13426401 13426572
refGene_gn=MEGF8;refGene_tx=XM_006941225.4;refGene_gn=MEGF8;refGene_tx=XM_019819489.2;refGene_gn=MEGF8;refGene_tx=XM_019819490.2
E2 13426669 13426800
refGene_gn=MEGF8;refGene_tx=XM_006941225.4;refGene_gn=MEGF8;refGene_tx=XM_019819489.2;refGene_gn=MEGF8;refGene_tx=XM_019819490.2
E2 13427319 13427452
refGene_gn=MEGF8;refGene_tx=XM_006941225.4;refGene_gn=MEGF8;refGene_tx=XM_019819489.2;refGene_gn=MEGF8;refGene_tx=XM_019819490.2
E2 13428386 13429655
refGene_gn=MEGF8;refGene_tx=XM_006941225.4;refGene_gn=MEGF8;refGene_tx=XM_019819489.2;refGene_gn=MEGF8;refGene_tx=XM_019819490.2
E2 13435278 13435368
refGene_gn=CNFN;refGene_tx=XM_006941226.2;refGene_gn=CNFN;refGene_tx=XM_019819493.1;refGene_gn=CNFN;refGene_tx=XM_019819494.1;refGene_gn=CNFN;refGene_tx=XM_019819495.1
E2 13435471 13435608
refGene_gn=CNFN;refGene_tx=XM_006941226.2;refGene_gn=CNFN;refGene_tx=XM_019819493.1;refGene_gn=CNFN;refGene_tx=XM_019819494.1;refGene_gn=CNFN;refGene_tx=XM_019819495.1
E2 13436018 13436129
refGene_gn=CNFN;refGene_tx=XM_006941226.2;refGene_gn=CNFN;refGene_tx=XM_019819493.1;refGene_gn=CNFN;refGene_tx=XM_019819494.1;refGene_gn=CNFN;refGene_tx=XM_019819495.1
E2 13436580 13436617

refGene_gn=CNFN;refGene_tx=XM_019819493.1;refGene_gn=CNFN;refGene_tx=XM_019819494.1;refGene_gn=CNFN;refGene_tx=XM_019819495.1
E2 13446061 13446322
refGene_gn=LIPE;refGene_tx=XM_019818909.2;refGene_gn=LIPE;refGene_tx=XM_023245841.1;refGene_gn=LIPE;refGene_tx=XM_023245842.1;refGene_gn=LIPE;refGene_tx=XM_023245844.1;refGene_gn=LIPE;refGene_tx=XM_023245845.1;refGene_gn=LIPE;refGene_tx=XM_023245846.1
E2 13446746 13447165
refGene_gn=LIPE;refGene_tx=XM_019818909.2;refGene_gn=LIPE;refGene_tx=XM_023245841.1;refGene_gn=LIPE;refGene_tx=XM_023245842.1;refGene_gn=LIPE;refGene_tx=XM_023245844.1;refGene_gn=LIPE;refGene_tx=XM_023245845.1;refGene_gn=LIPE;refGene_tx=XM_023245846.1
E2 13449441 13449618
refGene_gn=LIPE;refGene_tx=XM_019818909.2;refGene_gn=LIPE;refGene_tx=XM_023245841.1;refGene_gn=LIPE;refGene_tx=XM_023245842.1;refGene_gn=LIPE;refGene_tx=XM_023245844.1;refGene_gn=LIPE;refGene_tx=XM_023245845.1;refGene_gn=LIPE;refGene_tx=XM_023245846.1
E2 13449832 13450060
refGene_gn=LIPE;refGene_tx=XM_019818909.2;refGene_gn=LIPE;refGene_tx=XM_023245841.1;refGene_gn=LIPE;refGene_tx=XM_023245842.1;refGene_gn=LIPE;refGene_tx=XM_023245844.1;refGene_gn=LIPE;refGene_tx=XM_023245845.1;refGene_gn=LIPE;refGene_tx=XM_023245846.1
E2 13450860 13451155
refGene_gn=LIPE;refGene_tx=XM_019818909.2;refGene_gn=LIPE;refGene_tx=XM_023245841.1;refGene_gn=LIPE;refGene_tx=XM_023245842.1;refGene_gn=LIPE;refGene_tx=XM_023245844.1;refGene_gn=LIPE;refGene_tx=XM_023245845.1;refGene_gn=LIPE;refGene_tx=XM_023245846.1
E2 13451325 13451511
refGene_gn=LIPE;refGene_tx=XM_019818909.2;refGene_gn=LIPE;refGene_tx=XM_023245841.1;refGene_gn=LIPE;refGene_tx=XM_023245842.1;refGene_gn=LIPE;refGene_tx=XM_023245844.1;refGene_gn=LIPE;refGene_tx=XM_023245845.1;refGene_gn=LIPE;refGene_tx=XM_023245846.1
E2 13451714 13451860
refGene_gn=LIPE;refGene_tx=XM_019818909.2;refGene_gn=LIPE;refGene_tx=XM_023245841.1;refGene_gn=LIPE;refGene_tx=XM_023245842.1;refGene_gn=LIPE;refGene_tx=XM_023245844.1;refGene_gn=LIPE;refGene_tx=XM_023245845.1;refGene_gn=LIPE;refGene_tx=XM_023245846.1
E2 13451982 13452073
refGene_gn=LIPE;refGene_tx=XM_019818909.2;refGene_gn=LIPE;refGene_tx=XM_023245841.1;refGene_gn=LIPE;refGene_tx=XM_023245842.1;refGene_gn=LIPE;refGene_tx=XM_023245844.1;refGene_gn=LIPE;refGene_tx=XM_023245845.1;refGene_gn=LIPE;refGene_tx=XM_023245846.1
E2 13454196 13454732
refGene_gn=LIPE;refGene_tx=XM_019818909.2;refGene_gn=LIPE;refGene_tx=XM_023245841.1;refGene_gn=LIPE;refGene_tx=XM_023245842.1;refGene_gn=LIPE;refGene_tx=XM_023245844.1;refGene_gn=LIPE;refGene_tx=XM_023245845.1;refGene_gn=LIPE;refGene_tx=XM_023245846.1
E2 13455002 13455150 refGene_gn=LIPE;refGene_tx=XM_019818909.2
E2 13455888 13455949 refGene_gn=LIPE;refGene_tx=XM_023245845.1
E2 13460935 13461053 refGene_gn=LIPE;refGene_tx=XM_023245844.1

E2 13463048 13463973 refGene_gn=LIPE;refGene_tx=XM_023245841.1
E2 13467210 13467308 refGene_gn=CXCL17;refGene_tx=XM_003997765.5
E2 13470987 13471089 refGene_gn=CXCL17;refGene_tx=XM_003997765.5
E2 13471704 13471785 refGene_gn=CXCL17;refGene_tx=XM_003997765.5
E2 13480496 13480575 refGene_gn=CXCL17;refGene_tx=XM_003997765.5
E2 13522113 13522233
refGene_gn=LOC101086271;refGene_tx=XM_011289901.3
E2 13523525 13523557
refGene_gn=LOC101086271;refGene_tx=XM_011289901.3;refGene_gn=LOC101086271;refGene_tx=XM_011289902.3
E2 13524275 13524328
refGene_gn=LOC101086271;refGene_tx=XM_011289901.3
E2 13525045 13525172
refGene_gn=LOC101086271;refGene_tx=XM_011289901.3;refGene_gn=LOC101086271;refGene_tx=XM_011289902.3
E2 13529479 13529758
refGene_gn=LOC101086271;refGene_tx=XM_011289901.3;refGene_gn=LOC101086271;refGene_tx=XM_011289902.3
E2 13530971 13531226
refGene_gn=LOC101086271;refGene_tx=XM_011289901.3;refGene_gn=LOC101086271;refGene_tx=XM_011289902.3
E2 13531864 13532143
refGene_gn=LOC101086271;refGene_tx=XM_011289901.3;refGene_gn=LOC101086271;refGene_tx=XM_011289902.3
E2 13535822 13536182
refGene_gn=LOC101086271;refGene_tx=XM_011289901.3;refGene_gn=LOC101086271;refGene_tx=XM_011289902.3
E2 13537040 13537104
refGene_gn=LOC101086271;refGene_tx=XM_011289901.3;refGene_gn=LOC101086271;refGene_tx=XM_011289902.3
E2 13619585 13619813 refGene_gn=ERICH4;refGene_tx=XM_003997854.3
E2 13621166 13621340 refGene_gn=ERICH4;refGene_tx=XM_003997854.3
E2 13625296 13625314 refGene_gn=DMAC2;refGene_tx=XM_003997767.5
E2 13625444 13625489 refGene_gn=DMAC2;refGene_tx=XM_006941227.4
E2 13626393 13626812
refGene_gn=DMAC2;refGene_tx=XM_003997767.5;refGene_gn=DMAC2;refGene_tx=XM_006941227.4;refGene_gn=DMAC2;refGene_tx=XM_023245848.1
E2 13628193 13628274
refGene_gn=DMAC2;refGene_tx=XM_003997767.5;refGene_gn=DMAC2;refGene_tx=XM_006941227.4;refGene_gn=DMAC2;refGene_tx=XM_023245848.1
E2 13628798 13628935
refGene_gn=DMAC2;refGene_tx=XM_003997767.5;refGene_gn=DMAC2;refGene_tx=XM_006941227.4;refGene_gn=DMAC2;refGene_tx=XM_023245848.1
E2 13629029 13629192
refGene_gn=DMAC2;refGene_tx=XM_003997767.5;refGene_gn=DMAC2;refGene_tx=XM_006941227.4;refGene_gn=DMAC2;refGene_tx=XM_023245848.1
E2 13629598 13629776
refGene_gn=DMAC2;refGene_tx=XM_003997767.5;refGene_gn=DMAC2;refGene_tx=XM_006941227.4;refGene_gn=DMAC2;refGene_tx=XM_023245848.1
E2 13633972 13635166

refGene_gn=B3GNT8;refGene_tx=XM_011289900.2;refGene_gn=B3GNT8;refGene_tx=XM_019819491.1
E2	13636127	13636298	refGene_gn=BCKDHA;refGene_tx=XM_003997770.5
E2	13637028	13637200	refGene_gn=BCKDHA;refGene_tx=XM_003997770.5
E2	13637405	13637547	refGene_gn=BCKDHA;refGene_tx=XM_003997770.5
E2	13637743	13637950	refGene_gn=BCKDHA;refGene_tx=XM_003997770.5
E2	13639563	13639725	refGene_gn=BCKDHA;refGene_tx=XM_003997770.5
E2	13642269	13642378	refGene_gn=BCKDHA;refGene_tx=XM_003997770.5
E2	13646104	13646191	refGene_gn=BCKDHA;refGene_tx=XM_003997770.5
E2	13646297	13646480	refGene_gn=BCKDHA;refGene_tx=XM_003997770.5
E2	13654402	13654510	refGene_gn=BCKDHA;refGene_tx=XM_003997770.5
E2	13655007	13655155	refGene_gn=EXOSC5;refGene_tx=XM_003997771.5
E2	13657939	13658053	refGene_gn=EXOSC5;refGene_tx=XM_003997771.5
E2	13658474	13658596	refGene_gn=EXOSC5;refGene_tx=XM_003997771.5
E2	13660060	13660201	refGene_gn=EXOSC5;refGene_tx=XM_003997771.5
E2	13661293	13661383	refGene_gn=EXOSC5;refGene_tx=XM_003997771.5
E2	13662018	13662111	refGene_gn=EXOSC5;refGene_tx=XM_003997771.5
E2	13664153	13664312
refGene_gn=TMEM91;refGene_tx=XM_003997772.5;refGene_gn=TMEM91;refGene_tx=XM_006941229.4;refGene_gn=TMEM91;refGene_tx=XM_006941230.4;refGene_gn=TMEM91;refGene_tx=XM_019819492.2
E2	13664823	13664973
refGene_gn=TMEM91;refGene_tx=XM_003997772.5;refGene_gn=TMEM91;refGene_tx=XM_006941229.4;refGene_gn=TMEM91;refGene_tx=XM_006941230.4;refGene_gn=TMEM91;refGene_tx=XM_019819492.2
E2	13667665	13667904
refGene_gn=TMEM91;refGene_tx=XM_003997772.5;refGene_gn=TMEM91;refGene_tx=XM_006941229.4;refGene_gn=TMEM91;refGene_tx=XM_006941230.4;refGene_gn=TMEM91;refGene_tx=XM_019819492.2
E2	13668881	13668894	refGene_gn=TMEM91;refGene_tx=XM_003997772.5
E2	13671024	13671109
refGene_gn=TMEM91;refGene_tx=XM_006941229.4;refGene_gn=TMEM91;refGene_tx=XM_019819492.2
E2	13675725	13675813
refGene_gn=B9D2;refGene_tx=XM_003997773.4;refGene_gn=B9D2;refGene_tx=XM_006941231.3
E2	13678768	13678894
refGene_gn=B9D2;refGene_tx=XM_003997773.4;refGene_gn=B9D2;refGene_tx=XM_006941231.3
E2	13681153	13681467
refGene_gn=B9D2;refGene_tx=XM_003997773.4;refGene_gn=B9D2;refGene_tx=XM_006941231.3
E2	13683299	13683654
refGene_gn=TGFB1;refGene_tx=XM_003997774.5;refGene_gn=TGFB1;refGene_tx=XM_006941232.4
E2	13686698	13686859
refGene_gn=TGFB1;refGene_tx=XM_003997774.5;refGene_gn=TGFB1;refGene_tx=XM_006941232.4
E2	13688061	13688179
refGene_gn=TGFB1;refGene_tx=XM_003997774.5;refGene_gn=TGFB1;refGene_tx

=XM_006941232.4
E2	13689769 13689847
refGene_gn=TGFB1;refGene_tx=XM_003997774.5;refGene_gn=TGFB1;refGene_tx
=XM_006941232.4
E2	13689995 13690146
refGene_gn=TGFB1;refGene_tx=XM_003997774.5;refGene_gn=TGFB1;refGene_tx
=XM_006941232.4
E2	13695743 13695897
refGene_gn=TGFB1;refGene_tx=XM_003997774.5;refGene_gn=TGFB1;refGene_tx
=XM_006941232.4
E2	13696385 13696544
refGene_gn=TGFB1;refGene_tx=XM_003997774.5;refGene_gn=TGFB1;refGene_tx
=XM_006941232.4
E2	13703683 13703804 refGene_gn=CCDC97;refGene_tx=XM_003997775.5
E2	13705638 13705768 refGene_gn=CCDC97;refGene_tx=XM_003997775.5
E2	13706207 13706483 refGene_gn=CCDC97;refGene_tx=XM_003997775.5
E2	13708070 13708526 refGene_gn=CCDC97;refGene_tx=XM_003997775.5
E2	13711923 13711969 refGene_gn=CCDC97;refGene_tx=XM_003997775.5
E2	13715243 13715360
refGene_gn=HNRNPUL1;refGene_tx=XM_003997776.5;refGene_gn=HNRNPUL1;refG
ene_tx=XM_006941234.4;refGene_gn=HNRNPUL1;refGene_tx=XM_019819497.2;re
fGene_gn=HNRNPUL1;refGene_tx=XM_023245211.1
E2	13715903 13716095
refGene_gn=HNRNPUL1;refGene_tx=XM_003997776.5;refGene_gn=HNRNPUL1;refG
ene_tx=XM_006941234.4;refGene_gn=HNRNPUL1;refGene_tx=XM_019819497.2;re
fGene_gn=HNRNPUL1;refGene_tx=XM_023245211.1
E2	13717389 13717679
refGene_gn=HNRNPUL1;refGene_tx=XM_003997776.5;refGene_gn=HNRNPUL1;refG
ene_tx=XM_006941234.4;refGene_gn=HNRNPUL1;refGene_tx=XM_019819497.2;re
fGene_gn=HNRNPUL1;refGene_tx=XM_023245211.1
E2	13718767 13719052
refGene_gn=HNRNPUL1;refGene_tx=XM_003997776.5;refGene_gn=HNRNPUL1;refG
ene_tx=XM_006941234.4;refGene_gn=HNRNPUL1;refGene_tx=XM_019819497.2;re
fGene_gn=HNRNPUL1;refGene_tx=XM_023245211.1
E2	13719656 13719825
refGene_gn=HNRNPUL1;refGene_tx=XM_003997776.5;refGene_gn=HNRNPUL1;refG
ene_tx=XM_006941234.4;refGene_gn=HNRNPUL1;refGene_tx=XM_019819497.2;re
fGene_gn=HNRNPUL1;refGene_tx=XM_023245211.1
E2	13728141 13728270
refGene_gn=HNRNPUL1;refGene_tx=XM_003997776.5;refGene_gn=HNRNPUL1;refG
ene_tx=XM_006941234.4;refGene_gn=HNRNPUL1;refGene_tx=XM_019819497.2;re
fGene_gn=HNRNPUL1;refGene_tx=XM_023245211.1
E2	13728368 13728491
refGene_gn=HNRNPUL1;refGene_tx=XM_003997776.5;refGene_gn=HNRNPUL1;refG
ene_tx=XM_006941234.4;refGene_gn=HNRNPUL1;refGene_tx=XM_019819497.2;re
fGene_gn=HNRNPUL1;refGene_tx=XM_023245211.1
E2	13729731 13729998
refGene_gn=HNRNPUL1;refGene_tx=XM_003997776.5;refGene_gn=HNRNPUL1;refG
ene_tx=XM_006941234.4;refGene_gn=HNRNPUL1;refGene_tx=XM_019819497.2;re
fGene_gn=HNRNPUL1;refGene_tx=XM_023245211.1

E2 13735072 13735185
refGene_gn=HNRNPUL1;refGene_tx=XM_003997776.5;refGene_gn=HNRNPUL1;refGene_tx=XM_006941234.4;refGene_gn=HNRNPUL1;refGene_tx=XM_019819497.2;refGene_gn=HNRNPUL1;refGene_tx=XM_023245211.1
E2 13737084 13737184
refGene_gn=HNRNPUL1;refGene_tx=XM_003997776.5;refGene_gn=HNRNPUL1;refGene_tx=XM_006941234.4;refGene_gn=HNRNPUL1;refGene_tx=XM_019819497.2;refGene_gn=HNRNPUL1;refGene_tx=XM_023245211.1
E2 13739067 13739207
refGene_gn=HNRNPUL1;refGene_tx=XM_003997776.5;refGene_gn=HNRNPUL1;refGene_tx=XM_006941234.4;refGene_gn=HNRNPUL1;refGene_tx=XM_019819497.2;refGene_gn=HNRNPUL1;refGene_tx=XM_023245211.1
E2 13740689 13740763
refGene_gn=HNRNPUL1;refGene_tx=XM_003997776.5;refGene_gn=HNRNPUL1;refGene_tx=XM_006941234.4;refGene_gn=HNRNPUL1;refGene_tx=XM_019819497.2;refGene_gn=HNRNPUL1;refGene_tx=XM_023245211.1
E2 13742533 13742690
refGene_gn=HNRNPUL1;refGene_tx=XM_003997776.5;refGene_gn=HNRNPUL1;refGene_tx=XM_006941234.4;refGene_gn=HNRNPUL1;refGene_tx=XM_019819497.2;refGene_gn=HNRNPUL1;refGene_tx=XM_023245211.1
E2 13745303 13745426
refGene_gn=HNRNPUL1;refGene_tx=XM_003997776.5;refGene_gn=HNRNPUL1;refGene_tx=XM_006941234.4;refGene_gn=HNRNPUL1;refGene_tx=XM_019819497.2;refGene_gn=HNRNPUL1;refGene_tx=XM_023245211.1
E2 13748938 13749233
refGene_gn=HNRNPUL1;refGene_tx=XM_003997776.5;refGene_gn=HNRNPUL1;refGene_tx=XM_006941234.4
E2 13753216 13753568 refGene_gn=AXL;refGene_tx=XM_023245942.1
E2 13754365 13754502 refGene_gn=AXL;refGene_tx=XM_023245942.1
E2 13755017 13755177 refGene_gn=AXL;refGene_tx=XM_023245942.1
E2 13758089 13758199 refGene_gn=AXL;refGene_tx=XM_023245942.1
E2 13758840 13758962 refGene_gn=AXL;refGene_tx=XM_023245942.1
E2 13759373 13759466 refGene_gn=AXL;refGene_tx=XM_023245942.1
E2 13760907 13760985 refGene_gn=AXL;refGene_tx=XM_023245942.1
E2 13761127 13761279 refGene_gn=AXL;refGene_tx=XM_023245942.1
E2 13766006 13766139 refGene_gn=AXL;refGene_tx=XM_023245942.1
E2 13767490 13767517 refGene_gn=AXL;refGene_tx=XM_023245942.1
E2 13767718 13767869 refGene_gn=AXL;refGene_tx=XM_023245942.1
E2 13768527 13768667 refGene_gn=AXL;refGene_tx=XM_023245942.1
E2 13768978 13769189 refGene_gn=AXL;refGene_tx=XM_023245942.1
E2 13772486 13772602 refGene_gn=AXL;refGene_tx=XM_023245942.1
E2 13772742 13772823 refGene_gn=AXL;refGene_tx=XM_023245942.1
E2 13777727 13777904 refGene_gn=AXL;refGene_tx=XM_023245942.1
E2 13778490 13778591 refGene_gn=AXL;refGene_tx=XM_023245942.1
E2 13778933 13779156 refGene_gn=AXL;refGene_tx=XM_023245942.1
E2 13780293 13780357 refGene_gn=AXL;refGene_tx=XM_023245942.1
E2 13789952 13790258
refGene_gn=LOC101086936;refGene_tx=XM_011289895.3
E2 13790259 13790432
refGene_gn=LOC101086936;refGene_tx=XM_011289895.3

E2	13790526	13790668
refGene_gn=LOC101086936;refGene_tx=XM_011289895.3
E2	13792554	13792742
refGene_gn=LOC101086936;refGene_tx=XM_011289895.3
E2	13793859	13794001
refGene_gn=LOC101086936;refGene_tx=XM_011289895.3
E2	13796146	13796326
refGene_gn=LOC101086936;refGene_tx=XM_011289895.3
E2	13796461	13796622
refGene_gn=LOC101086936;refGene_tx=XM_011289895.3
E2	13797077	13797227
refGene_gn=LOC101086936;refGene_tx=XM_011289895.3
E2	13799102	13799268
refGene_gn=LOC101086936;refGene_tx=XM_011289895.3
E2	13800395	13800566
refGene_gn=LOC101086936;refGene_tx=XM_011289895.3
E2	13845108	13845396
refGene_gn=LOC105259649;refGene_tx=XM_023245329.1
E2	13848702	13848865
refGene_gn=LOC105259649;refGene_tx=XM_023245329.1
E2	13849011	13849161
refGene_gn=LOC105259649;refGene_tx=XM_023245329.1
E2	13850820	13850981
refGene_gn=LOC105259649;refGene_tx=XM_023245329.1
E2	13852522	13852699
refGene_gn=LOC105259649;refGene_tx=XM_023245329.1
E2	13852993	13853135
refGene_gn=LOC105259649;refGene_tx=XM_023245329.1
E2	13854877	13855065
refGene_gn=LOC105259649;refGene_tx=XM_023245329.1
E2	13855269	13855411
refGene_gn=LOC105259649;refGene_tx=XM_023245329.1
E2	13858640	13858700
refGene_gn=LOC105259649;refGene_tx=XM_023245329.1
E2	13858702	13858739
refGene_gn=LOC105259649;refGene_tx=XM_023245329.1
E2	13858740	13858828
refGene_gn=LOC105259649;refGene_tx=XM_023245329.1
E2	13904023	13904203
refGene_gn=LOC101089311;refGene_tx=XM_003997779.3;refGene_gn=LOC101089
311;refGene_tx=XM_019819500.2
E2	13904801	13904964
refGene_gn=LOC101089311;refGene_tx=XM_003997779.3
E2	13906492	13906652
refGene_gn=LOC101089311;refGene_tx=XM_003997779.3;refGene_gn=LOC101089
311;refGene_tx=XM_019819500.2
E2	13908126	13908287
refGene_gn=LOC101089311;refGene_tx=XM_003997779.3;refGene_gn=LOC101089
311;refGene_tx=XM_019819500.2
E2	13908537	13908714

refGene_gn=LOC101089311;refGene_tx=XM_003997779.3;refGene_gn=LOC101089311;refGene_tx=XM_019819500.2
E2 13909430 13909572
refGene_gn=LOC101089311;refGene_tx=XM_003997779.3;refGene_gn=LOC101089311;refGene_tx=XM_019819500.2
E2 13911037 13911225
refGene_gn=LOC101089311;refGene_tx=XM_003997779.3;refGene_gn=LOC101089311;refGene_tx=XM_019819500.2
E2 13911339 13911481
refGene_gn=LOC101089311;refGene_tx=XM_003997779.3;refGene_gn=LOC101089311;refGene_tx=XM_019819500.2
E2 13913588 13913770
refGene_gn=LOC101089311;refGene_tx=XM_003997779.3;refGene_gn=LOC101089311;refGene_tx=XM_019819500.2
E2 13926977 13927157
refGene_gn=LOC101089561;refGene_tx=XM_011289955.3
E2 13927435 13927598
refGene_gn=LOC101089561;refGene_tx=XM_011289955.3
E2 13928095 13928245
refGene_gn=LOC101089561;refGene_tx=XM_011289955.3
E2 13928483 13928644
refGene_gn=LOC101089561;refGene_tx=XM_011289955.3
E2 13929815 13929992
refGene_gn=LOC101089561;refGene_tx=XM_011289955.3
E2 13930473 13930615
refGene_gn=LOC101089561;refGene_tx=XM_011289955.3
E2 13931067 13931255
refGene_gn=LOC101089561;refGene_tx=XM_011289955.3
E2 13931886 13932028
refGene_gn=LOC101089561;refGene_tx=XM_011289955.3
E2 13932567 13932749
refGene_gn=LOC101089561;refGene_tx=XM_011289955.3
E2 13943986 13944168
refGene_gn=LOC101089816;refGene_tx=XM_003997781.5
E2 13944708 13944850
refGene_gn=LOC101089816;refGene_tx=XM_003997781.5
E2 13945471 13945659
refGene_gn=LOC101089816;refGene_tx=XM_003997781.5
E2 13946110 13946252
refGene_gn=LOC101089816;refGene_tx=XM_003997781.5
E2 13946733 13946910
refGene_gn=LOC101089816;refGene_tx=XM_003997781.5
E2 13948044 13948205
refGene_gn=LOC101089816;refGene_tx=XM_003997781.5
E2 13948448 13948598
refGene_gn=LOC101089816;refGene_tx=XM_003997781.5
E2 13949354 13949517
refGene_gn=LOC101089816;refGene_tx=XM_003997781.5
E2 13950036 13950216
refGene_gn=LOC101089816;refGene_tx=XM_003997781.5

E2 13979050 13979106
refGene_gn=LOC101087438;refGene_tx=XM_023245335.1
E2 13979211 13979276
refGene_gn=LOC101087438;refGene_tx=XM_023245335.1
E2 13980492 13980684
refGene_gn=LOC101087438;refGene_tx=XM_006941235.4
E2 13980783 13980946
refGene_gn=LOC101087438;refGene_tx=XM_006941235.4
E2 13983697 13983847
refGene_gn=LOC101087438;refGene_tx=XM_006941235.4;refGene_gn=LOC101087438;refGene_tx=XM_023245335.1
E2 13986171 13986332
refGene_gn=LOC101087438;refGene_tx=XM_006941235.4;refGene_gn=LOC101087438;refGene_tx=XM_023245335.1
E2 13986673 13986850
refGene_gn=LOC101087438;refGene_tx=XM_006941235.4;refGene_gn=LOC101087438;refGene_tx=XM_023245335.1
E2 13987259 13987401
refGene_gn=LOC101087438;refGene_tx=XM_006941235.4;refGene_gn=LOC101087438;refGene_tx=XM_023245335.1
E2 13988242 13988430
refGene_gn=LOC101087438;refGene_tx=XM_006941235.4;refGene_gn=LOC101087438;refGene_tx=XM_023245335.1
E2 13989021 13989163
refGene_gn=LOC101087438;refGene_tx=XM_006941235.4;refGene_gn=LOC101087438;refGene_tx=XM_023245335.1
E2 13990628 13990810
refGene_gn=LOC101087438;refGene_tx=XM_006941235.4;refGene_gn=LOC101087438;refGene_tx=XM_023245335.1
E2 13994194 13994395
refGene_gn=LOC101087695;refGene_tx=XM_023245944.1
E2 13995031 13995194
refGene_gn=LOC101087695;refGene_tx=XM_023245944.1
E2 13995406 13995766
refGene_gn=LOC101087695;refGene_tx=XM_023245944.1
E2 13996003 13996164
refGene_gn=LOC101087695;refGene_tx=XM_023245944.1
E2 13996528 13996705
refGene_gn=LOC101087695;refGene_tx=XM_023245944.1
E2 13996789 13996931
refGene_gn=LOC101087695;refGene_tx=XM_023245944.1
E2 13997024 13997099
refGene_gn=LOC101087695;refGene_tx=XM_023245944.1
E2 13997101 13997199
refGene_gn=LOC101087695;refGene_tx=XM_023245944.1
E2 13997405 13997547
refGene_gn=LOC101087695;refGene_tx=XM_023245944.1
E2 13997642 13997824
refGene_gn=LOC101087695;refGene_tx=XM_023245944.1
E2 13998516 13998613

refGene_gn=EGLN2;refGene_tx=XM_003997782.4;refGene_gn=EGLN2;refGene_tx=XM_011289893.2;refGene_gn=EGLN2;refGene_tx=XM_019819502.2;refGene_gn=EGLN2;refGene_tx=XM_019819503.2
E2 13998894 13998962
refGene_gn=EGLN2;refGene_tx=XM_003997782.4;refGene_gn=EGLN2;refGene_tx=XM_011289893.2;refGene_gn=EGLN2;refGene_tx=XM_019819502.2
E2 13999183 13999320
refGene_gn=EGLN2;refGene_tx=XM_003997782.4;refGene_gn=EGLN2;refGene_tx=XM_011289893.2;refGene_gn=EGLN2;refGene_tx=XM_019819502.2;refGene_gn=EGLN2;refGene_tx=XM_019819503.2
E2 13999808 13999928
refGene_gn=EGLN2;refGene_tx=XM_003997782.4;refGene_gn=EGLN2;refGene_tx=XM_011289893.2;refGene_gn=EGLN2;refGene_tx=XM_019819502.2;refGene_gn=EGLN2;refGene_tx=XM_019819503.2
E2 14004815 14005652
refGene_gn=EGLN2;refGene_tx=XM_003997782.4;refGene_gn=EGLN2;refGene_tx=XM_011289893.2;refGene_gn=EGLN2;refGene_tx=XM_019819502.2;refGene_gn=EGLN2;refGene_tx=XM_019819503.2
E2 14012222 14012338
refGene_gn=RAB4B;refGene_tx=XM_003997783.5;refGene_gn=RAB4B;refGene_tx=XM_019819504.2
E2 14012433 14012529
refGene_gn=RAB4B;refGene_tx=XM_003997783.5;refGene_gn=RAB4B;refGene_tx=XM_019819504.2
E2 14013579 14013734
refGene_gn=RAB4B;refGene_tx=XM_003997783.5;refGene_gn=RAB4B;refGene_tx=XM_019819504.2
E2 14013822 14013885
refGene_gn=RAB4B;refGene_tx=XM_003997783.5;refGene_gn=RAB4B;refGene_tx=XM_019819504.2
E2 14015225 14015340
refGene_gn=RAB4B;refGene_tx=XM_003997783.5;refGene_gn=RAB4B;refGene_tx=XM_019819504.2
E2 14015595 14015676
refGene_gn=RAB4B;refGene_tx=XM_003997783.5;refGene_gn=RAB4B;refGene_tx=XM_019819504.2
E2 14017084 14017100
refGene_gn=RAB4B;refGene_tx=XM_003997783.5;refGene_gn=RAB4B;refGene_tx=XM_019819504.2
E2 14018115 14018139 refGene_gn=MIA;refGene_tx=XM_006941240.4
E2 14018450 14018561 refGene_gn=MIA;refGene_tx=XM_006941240.4
E2 14019035 14019169 refGene_gn=MIA;refGene_tx=XM_006941240.4
E2 14019262 14019389 refGene_gn=MIA;refGene_tx=XM_006941240.4
E2 14023566 14023726
refGene_gn=SNRPA;refGene_tx=XM_003997785.5;refGene_gn=SNRPA;refGene_tx=XM_006941241.4
E2 14024867 14024956
refGene_gn=SNRPA;refGene_tx=XM_003997785.5;refGene_gn=SNRPA;refGene_tx=XM_006941241.4
E2 14025404 14025578

refGene_gn=SNRPA;refGene_tx=XM_003997785.5;refGene_gn=SNRPA;refGene_tx=XM_006941241.4
E2 14027517 14027697
refGene_gn=SNRPA;refGene_tx=XM_003997785.5;refGene_gn=SNRPA;refGene_tx=XM_006941241.4
E2 14030386 14030559
refGene_gn=SNRPA;refGene_tx=XM_003997785.5;refGene_gn=SNRPA;refGene_tx=XM_006941241.4
E2 14033996 14034069
refGene_gn=SNRPA;refGene_tx=XM_003997785.5;refGene_gn=SNRPA;refGene_tx=XM_006941241.4
E2 14035646 14035990
refGene_gn=CE2H19orf54;refGene_tx=XM_003997858.5;refGene_gn=CE2H19orf54;refGene_tx=XM_023245040.1
E2 14037922 14037930
refGene_gn=CE2H19orf54;refGene_tx=XM_023245033.1;refGene_gn=CE2H19orf54;refGene_tx=XM_023245034.1
E2 14038464 14038533
refGene_gn=CE2H19orf54;refGene_tx=XM_003997858.5;refGene_gn=CE2H19orf54;refGene_tx=XM_023245033.1;refGene_gn=CE2H19orf54;refGene_tx=XM_023245034.1;refGene_gn=CE2H19orf54;refGene_tx=XM_023245040.1
E2 14039019 14039153
refGene_gn=CE2H19orf54;refGene_tx=XM_003997858.5;refGene_gn=CE2H19orf54;refGene_tx=XM_023245033.1;refGene_gn=CE2H19orf54;refGene_tx=XM_023245034.1;refGene_gn=CE2H19orf54;refGene_tx=XM_023245035.1;refGene_gn=CE2H19orf54;refGene_tx=XM_023245036.1;refGene_gn=CE2H19orf54;refGene_tx=XM_023245037.1;refGene_gn=CE2H19orf54;refGene_tx=XM_023245038.1;refGene_gn=CE2H19orf54;refGene_tx=XM_023245039.1;refGene_gn=CE2H19orf54;refGene_tx=XM_023245040.1
E2 14039454 14039584
refGene_gn=CE2H19orf54;refGene_tx=XM_003997858.5;refGene_gn=CE2H19orf54;refGene_tx=XM_023245033.1;refGene_gn=CE2H19orf54;refGene_tx=XM_023245034.1;refGene_gn=CE2H19orf54;refGene_tx=XM_023245035.1;refGene_gn=CE2H19orf54;refGene_tx=XM_023245036.1;refGene_gn=CE2H19orf54;refGene_tx=XM_023245037.1;refGene_gn=CE2H19orf54;refGene_tx=XM_023245038.1;refGene_gn=CE2H19orf54;refGene_tx=XM_023245039.1
E2 14039675 14039746
refGene_gn=CE2H19orf54;refGene_tx=XM_003997858.5;refGene_gn=CE2H19orf54;refGene_tx=XM_023245033.1;refGene_gn=CE2H19orf54;refGene_tx=XM_023245034.1;refGene_gn=CE2H19orf54;refGene_tx=XM_023245035.1;refGene_gn=CE2H19orf54;refGene_tx=XM_023245036.1;refGene_gn=CE2H19orf54;refGene_tx=XM_023245037.1;refGene_gn=CE2H19orf54;refGene_tx=XM_023245038.1;refGene_gn=CE2H19orf54;refGene_tx=XM_023245039.1;refGene_gn=CE2H19orf54;refGene_tx=XM_023245040.1
E2 14040507 14040845
refGene_gn=CE2H19orf54;refGene_tx=XM_003997858.5;refGene_gn=CE2H19orf54;refGene_tx=XM_023245033.1;refGene_gn=CE2H19orf54;refGene_tx=XM_023245034.1;refGene_gn=CE2H19orf54;refGene_tx=XM_023245035.1;refGene_gn=CE2H19orf54;refGene_tx=XM_023245036.1;refGene_gn=CE2H19orf54;refGene_tx=XM_023245037.1;refGene_gn=CE2H19orf54;refGene_tx=XM_023245038.1;refGene

_gn=CE2H19orf54;refGene_tx=XM_023245039.1
E2	14043987	14044191	refGene_gn=ITPKC;refGene_tx=XM_006941342.4
E2	14046038	14046110	refGene_gn=ITPKC;refGene_tx=XM_006941342.4
E2	14046752	14046854	refGene_gn=ITPKC;refGene_tx=XM_006941342.4
E2	14049037	14049242	refGene_gn=ITPKC;refGene_tx=XM_006941342.4
E2	14050460	14050674	refGene_gn=ITPKC;refGene_tx=XM_006941342.4
E2	14053049	14053149	refGene_gn=ITPKC;refGene_tx=XM_006941342.4
E2	14057223	14058330	refGene_gn=ITPKC;refGene_tx=XM_006941342.4
E2	14061955	14062057	refGene_gn=COQ8B;refGene_tx=XM_011289889.3
E2	14062205	14062325	refGene_gn=COQ8B;refGene_tx=XM_011289889.3
E2	14062461	14062528	refGene_gn=COQ8B;refGene_tx=XM_011289889.3
E2	14064308	14064386	refGene_gn=COQ8B;refGene_tx=XM_011289889.3
E2	14067903	14068026	refGene_gn=COQ8B;refGene_tx=XM_011289889.3
E2	14068172	14068258	refGene_gn=COQ8B;refGene_tx=XM_011289889.3
E2	14070207	14070348	refGene_gn=COQ8B;refGene_tx=XM_011289889.3
E2	14070436	14070518	refGene_gn=COQ8B;refGene_tx=XM_011289889.3
E2	14071221	14071315	refGene_gn=COQ8B;refGene_tx=XM_011289889.3
E2	14073193	14073335	refGene_gn=COQ8B;refGene_tx=XM_011289889.3
E2	14073482	14073590	refGene_gn=COQ8B;refGene_tx=XM_011289889.3
E2	14076048	14076114	refGene_gn=COQ8B;refGene_tx=XM_011289889.3
E2	14078761	14078806	refGene_gn=COQ8B;refGene_tx=XM_011289889.3
E2	14078807	14078849	refGene_gn=COQ8B;refGene_tx=XM_011289889.3
E2	14079260	14079542	refGene_gn=COQ8B;refGene_tx=XM_011289889.3
E2	14081245	14081269
refGene_gn=NUMBL;refGene_tx=XM_003997787.5;refGene_gn=NUMBL;refGene_tx
=XM_006941242.4;refGene_gn=NUMBL;refGene_tx=XM_006941244.4;refGene_gn=
NUMBL;refGene_tx=XM_019819506.2
E2	14084481	14084566
refGene_gn=NUMBL;refGene_tx=XM_003997787.5;refGene_gn=NUMBL;refGene_tx
=XM_006941242.4;refGene_gn=NUMBL;refGene_tx=XM_006941244.4;refGene_gn=
NUMBL;refGene_tx=XM_019819506.2
E2	14086022	14086162
refGene_gn=NUMBL;refGene_tx=XM_003997787.5;refGene_gn=NUMBL;refGene_tx
=XM_006941242.4;refGene_gn=NUMBL;refGene_tx=XM_006941244.4;refGene_gn=
NUMBL;refGene_tx=XM_019819505.2;refGene_gn=NUMBL;refGene_tx=XM_0198195
06.2;refGene_gn=NUMBL;refGene_tx=XM_019819507.2;refGene_gn=NUMBL;refGe
ne_tx=XM_019819508.2
E2	14087145	14087220
refGene_gn=NUMBL;refGene_tx=XM_003997787.5;refGene_gn=NUMBL;refGene_tx
=XM_006941242.4;refGene_gn=NUMBL;refGene_tx=XM_006941244.4;refGene_gn=
NUMBL;refGene_tx=XM_019819505.2;refGene_gn=NUMBL;refGene_tx=XM_0198195
06.2;refGene_gn=NUMBL;refGene_tx=XM_019819507.2;refGene_gn=NUMBL;refGe
ne_tx=XM_019819508.2
E2	14087308	14087383
refGene_gn=NUMBL;refGene_tx=XM_003997787.5;refGene_gn=NUMBL;refGene_tx
=XM_006941242.4;refGene_gn=NUMBL;refGene_tx=XM_006941244.4;refGene_gn=
NUMBL;refGene_tx=XM_019819505.2;refGene_gn=NUMBL;refGene_tx=XM_0198195
06.2;refGene_gn=NUMBL;refGene_tx=XM_019819507.2;refGene_gn=NUMBL;refGe
ne_tx=XM_019819508.2
E2	14088581	14088722

refGene_gn=NUMBL;refGene_tx=XM_003997787.5;refGene_gn=NUMBL;refGene_tx=XM_006941242.4;refGene_gn=NUMBL;refGene_tx=XM_006941244.4;refGene_gn=NUMBL;refGene_tx=XM_019819505.2;refGene_gn=NUMBL;refGene_tx=XM_019819506.2;refGene_gn=NUMBL;refGene_tx=XM_019819507.2;refGene_gn=NUMBL;refGene_tx=XM_019819508.2
E2 14097068 14097258
refGene_gn=NUMBL;refGene_tx=XM_003997787.5;refGene_gn=NUMBL;refGene_tx=XM_006941242.4;refGene_gn=NUMBL;refGene_tx=XM_006941244.4;refGene_gn=NUMBL;refGene_tx=XM_019819505.2;refGene_gn=NUMBL;refGene_tx=XM_019819506.2;refGene_gn=NUMBL;refGene_tx=XM_019819507.2;refGene_gn=NUMBL;refGene_tx=XM_019819508.2
E2 14101526 14101832
refGene_gn=NUMBL;refGene_tx=XM_003997787.5;refGene_gn=NUMBL;refGene_tx=XM_006941242.4;refGene_gn=NUMBL;refGene_tx=XM_006941244.4;refGene_gn=NUMBL;refGene_tx=XM_019819505.2;refGene_gn=NUMBL;refGene_tx=XM_019819506.2;refGene_gn=NUMBL;refGene_tx=XM_019819507.2;refGene_gn=NUMBL;refGene_tx=XM_019819508.2
E2 14103071 14103155
refGene_gn=NUMBL;refGene_tx=XM_006941242.4;refGene_gn=NUMBL;refGene_tx=XM_019819505.2;refGene_gn=NUMBL;refGene_tx=XM_019819506.2;refGene_gn=NUMBL;refGene_tx=XM_019819507.2;refGene_gn=NUMBL;refGene_tx=XM_019819508.2
E2 14104749 14104872
refGene_gn=NUMBL;refGene_tx=XM_003997787.5;refGene_gn=NUMBL;refGene_tx=XM_006941242.4;refGene_gn=NUMBL;refGene_tx=XM_006941244.4;refGene_gn=NUMBL;refGene_tx=XM_019819505.2;refGene_gn=NUMBL;refGene_tx=XM_019819506.2;refGene_gn=NUMBL;refGene_tx=XM_019819507.2;refGene_gn=NUMBL;refGene_tx=XM_019819508.2
E2 14105751 14106386
refGene_gn=NUMBL;refGene_tx=XM_003997787.5;refGene_gn=NUMBL;refGene_tx=XM_006941242.4;refGene_gn=NUMBL;refGene_tx=XM_006941244.4;refGene_gn=NUMBL;refGene_tx=XM_019819505.2;refGene_gn=NUMBL;refGene_tx=XM_019819506.2;refGene_gn=NUMBL;refGene_tx=XM_019819507.2;refGene_gn=NUMBL;refGene_tx=XM_019819508.2
E2 14123124 14123279
refGene_gn=LTBP4;refGene_tx=XM_023245977.1;refGene_gn=LTBP4;refGene_tx=XM_023245978.1;refGene_gn=LTBP4;refGene_tx=XM_023245979.1
E2 14124318 14124471
refGene_gn=LTBP4;refGene_tx=XM_023245977.1;refGene_gn=LTBP4;refGene_tx=XM_023245978.1;refGene_gn=LTBP4;refGene_tx=XM_023245979.1
E2 14124879 14125260
refGene_gn=LTBP4;refGene_tx=XM_023245977.1;refGene_gn=LTBP4;refGene_tx=XM_023245978.1;refGene_gn=LTBP4;refGene_tx=XM_023245979.1
E2 14125992 14126145
refGene_gn=LTBP4;refGene_tx=XM_023245977.1;refGene_gn=LTBP4;refGene_tx=XM_023245978.1;refGene_gn=LTBP4;refGene_tx=XM_023245979.1
E2 14126995 14127142
refGene_gn=LTBP4;refGene_tx=XM_023245977.1;refGene_gn=LTBP4;refGene_tx=XM_023245978.1;refGene_gn=LTBP4;refGene_tx=XM_023245979.1
E2 14127348 14127477

refGene_gn=LTBP4;refGene_tx=XM_023245977.1;refGene_gn=LTBP4;refGene_tx=XM_023245978.1;refGene_gn=LTBP4;refGene_tx=XM_023245979.1
E2 14127766 14127838
refGene_gn=LTBP4;refGene_tx=XM_023245977.1;refGene_gn=LTBP4;refGene_tx=XM_023245978.1;refGene_gn=LTBP4;refGene_tx=XM_023245979.1
E2 14128140 14128407
refGene_gn=LTBP4;refGene_tx=XM_023245977.1;refGene_gn=LTBP4;refGene_tx=XM_023245978.1;refGene_gn=LTBP4;refGene_tx=XM_023245979.1
E2 14129737 14129884
refGene_gn=LTBP4;refGene_tx=XM_023245977.1;refGene_gn=LTBP4;refGene_tx=XM_023245978.1;refGene_gn=LTBP4;refGene_tx=XM_023245979.1
E2 14132278 14132404
refGene_gn=LTBP4;refGene_tx=XM_023245977.1;refGene_gn=LTBP4;refGene_tx=XM_023245978.1;refGene_gn=LTBP4;refGene_tx=XM_023245979.1
E2 14132483 14132615
refGene_gn=LTBP4;refGene_tx=XM_023245977.1;refGene_gn=LTBP4;refGene_tx=XM_023245978.1;refGene_gn=LTBP4;refGene_tx=XM_023245979.1
E2 14134587 14134719
refGene_gn=LTBP4;refGene_tx=XM_023245977.1;refGene_gn=LTBP4;refGene_tx=XM_023245978.1;refGene_gn=LTBP4;refGene_tx=XM_023245979.1
E2 14134909 14135032
refGene_gn=LTBP4;refGene_tx=XM_023245977.1;refGene_gn=LTBP4;refGene_tx=XM_023245978.1;refGene_gn=LTBP4;refGene_tx=XM_023245979.1
E2 14135433 14135559
refGene_gn=LTBP4;refGene_tx=XM_023245977.1;refGene_gn=LTBP4;refGene_tx=XM_023245978.1;refGene_gn=LTBP4;refGene_tx=XM_023245979.1
E2 14135766 14135895
refGene_gn=LTBP4;refGene_tx=XM_023245977.1;refGene_gn=LTBP4;refGene_tx=XM_023245978.1;refGene_gn=LTBP4;refGene_tx=XM_023245979.1
E2 14137028 14137148
refGene_gn=LTBP4;refGene_tx=XM_023245977.1;refGene_gn=LTBP4;refGene_tx=XM_023245978.1;refGene_gn=LTBP4;refGene_tx=XM_023245979.1
E2 14137236 14137362
refGene_gn=LTBP4;refGene_tx=XM_023245977.1;refGene_gn=LTBP4;refGene_tx=XM_023245978.1;refGene_gn=LTBP4;refGene_tx=XM_023245979.1
E2 14137842 14138085
refGene_gn=LTBP4;refGene_tx=XM_023245977.1;refGene_gn=LTBP4;refGene_tx=XM_023245978.1;refGene_gn=LTBP4;refGene_tx=XM_023245979.1
E2 14138607 14138733
refGene_gn=LTBP4;refGene_tx=XM_023245977.1;refGene_gn=LTBP4;refGene_tx=XM_023245978.1;refGene_gn=LTBP4;refGene_tx=XM_023245979.1
E2 14139492 14139618
refGene_gn=LTBP4;refGene_tx=XM_023245977.1;refGene_gn=LTBP4;refGene_tx=XM_023245978.1;refGene_gn=LTBP4;refGene_tx=XM_023245979.1
E2 14139702 14139834
refGene_gn=LTBP4;refGene_tx=XM_023245977.1;refGene_gn=LTBP4;refGene_tx=XM_023245978.1;refGene_gn=LTBP4;refGene_tx=XM_023245979.1
E2 14140446 14140554
refGene_gn=LTBP4;refGene_tx=XM_023245977.1;refGene_gn=LTBP4;refGene_tx=XM_023245978.1;refGene_gn=LTBP4;refGene_tx=XM_023245979.1

E2 14140689 14140836
refGene_gn=LTBP4;refGene_tx=XM_023245977.1;refGene_gn=LTBP4;refGene_tx=XM_023245978.1;refGene_gn=LTBP4;refGene_tx=XM_023245979.1
E2 14141448 14141613
refGene_gn=LTBP4;refGene_tx=XM_023245977.1;refGene_gn=LTBP4;refGene_tx=XM_023245978.1;refGene_gn=LTBP4;refGene_tx=XM_023245979.1
E2 14142168 14142291
refGene_gn=LTBP4;refGene_tx=XM_023245977.1;refGene_gn=LTBP4;refGene_tx=XM_023245978.1;refGene_gn=LTBP4;refGene_tx=XM_023245979.1
E2 14142389 14142464
refGene_gn=LTBP4;refGene_tx=XM_023245977.1;refGene_gn=LTBP4;refGene_tx=XM_023245978.1;refGene_gn=LTBP4;refGene_tx=XM_023245979.1
E2 14142860 14142963
refGene_gn=LTBP4;refGene_tx=XM_023245977.1;refGene_gn=LTBP4;refGene_tx=XM_023245978.1;refGene_gn=LTBP4;refGene_tx=XM_023245979.1
E2 14143040 14143288
refGene_gn=LTBP4;refGene_tx=XM_023245977.1;refGene_gn=LTBP4;refGene_tx=XM_023245978.1;refGene_gn=LTBP4;refGene_tx=XM_023245979.1
E2 14143473 14143665
refGene_gn=LTBP4;refGene_tx=XM_023245977.1;refGene_gn=LTBP4;refGene_tx=XM_023245978.1;refGene_gn=LTBP4;refGene_tx=XM_023245979.1
E2 14146978 14147219 refGene_gn=LTBP4;refGene_tx=XM_023245979.1
E2 14148409 14148519
refGene_gn=LTBP4;refGene_tx=XM_023245977.1;refGene_gn=LTBP4;refGene_tx=XM_023245978.1
E2 14149077 14149216
refGene_gn=LTBP4;refGene_tx=XM_023245977.1;refGene_gn=LTBP4;refGene_tx=XM_023245978.1
E2 14149379 14149438
refGene_gn=LTBP4;refGene_tx=XM_023245977.1;refGene_gn=LTBP4;refGene_tx=XM_023245978.1
E2 14151303 14151540
refGene_gn=LTBP4;refGene_tx=XM_023245977.1;refGene_gn=LTBP4;refGene_tx=XM_023245978.1
E2 14155319 14155539
refGene_gn=SHKBP1;refGene_tx=XM_003997791.4;refGene_gn=SHKBP1;refGene_tx=XM_006941245.3
E2 14155651 14155766
refGene_gn=SHKBP1;refGene_tx=XM_003997791.4;refGene_gn=SHKBP1;refGene_tx=XM_006941245.3
E2 14156012 14156191
refGene_gn=SHKBP1;refGene_tx=XM_003997791.4;refGene_gn=SHKBP1;refGene_tx=XM_006941245.3
E2 14157218 14157315
refGene_gn=SHKBP1;refGene_tx=XM_003997791.4;refGene_gn=SHKBP1;refGene_tx=XM_006941245.3
E2 14157687 14157843
refGene_gn=SHKBP1;refGene_tx=XM_003997791.4;refGene_gn=SHKBP1;refGene_tx=XM_006941245.3
E2 14158809 14158980

refGene_gn=SHKBP1;refGene_tx=XM_003997791.4;refGene_gn=SHKBP1;refGene_tx=XM_006941245.3
E2 14159902 14160019
refGene_gn=SHKBP1;refGene_tx=XM_003997791.4;refGene_gn=SHKBP1;refGene_tx=XM_006941245.3
E2 14160125 14160213
refGene_gn=SHKBP1;refGene_tx=XM_003997791.4;refGene_gn=SHKBP1;refGene_tx=XM_006941245.3
E2 14161014 14161130
refGene_gn=SHKBP1;refGene_tx=XM_003997791.4;refGene_gn=SHKBP1;refGene_tx=XM_006941245.3
E2 14161898 14162089
refGene_gn=SHKBP1;refGene_tx=XM_003997791.4;refGene_gn=SHKBP1;refGene_tx=XM_006941245.3
E2 14162178 14162269
refGene_gn=SHKBP1;refGene_tx=XM_003997791.4;refGene_gn=SHKBP1;refGene_tx=XM_006941245.3
E2 14162352 14162514
refGene_gn=SHKBP1;refGene_tx=XM_003997791.4;refGene_gn=SHKBP1;refGene_tx=XM_006941245.3
E2 14164330 14164411
refGene_gn=SHKBP1;refGene_tx=XM_003997791.4;refGene_gn=SHKBP1;refGene_tx=XM_006941245.3
E2 14164629 14164688
refGene_gn=SHKBP1;refGene_tx=XM_003997791.4;refGene_gn=SHKBP1;refGene_tx=XM_006941245.3
E2 14164971 14165045
refGene_gn=SHKBP1;refGene_tx=XM_003997791.4;refGene_gn=SHKBP1;refGene_tx=XM_006941245.3
E2 14165150 14165196
refGene_gn=SHKBP1;refGene_tx=XM_003997791.4;refGene_gn=SHKBP1;refGene_tx=XM_006941245.3
E2 14165346 14165400
refGene_gn=SHKBP1;refGene_tx=XM_003997791.4;refGene_gn=SHKBP1;refGene_tx=XM_006941245.3
E2 14165623 14165709
refGene_gn=SHKBP1;refGene_tx=XM_003997791.4;refGene_gn=SHKBP1;refGene_tx=XM_006941245.3
E2 14167035 14167194
refGene_gn=SPTBN4;refGene_tx=XM_023246182.1;refGene_gn=SPTBN4;refGene_tx=XM_023246183.1
E2 14168409 14168452
refGene_gn=SPTBN4;refGene_tx=XM_023246182.1;refGene_gn=SPTBN4;refGene_tx=XM_023246183.1
E2 14168603 14168777
refGene_gn=SPTBN4;refGene_tx=XM_023246182.1;refGene_gn=SPTBN4;refGene_tx=XM_023246183.1
E2 14169939 14170232
refGene_gn=SPTBN4;refGene_tx=XM_023246182.1;refGene_gn=SPTBN4;refGene_tx=XM_023246183.1

E2 14170846 14170916
refGene_gn=SPTBN4;refGene_tx=XM_023246182.1;refGene_gn=SPTBN4;refGene_tx=XM_023246183.1
E2 14171931 14172551
refGene_gn=SPTBN4;refGene_tx=XM_023246182.1;refGene_gn=SPTBN4;refGene_tx=XM_023246183.1
E2 14173129 14173326
refGene_gn=SPTBN4;refGene_tx=XM_023246182.1;refGene_gn=SPTBN4;refGene_tx=XM_023246183.1
E2 14173674 14173759
refGene_gn=SPTBN4;refGene_tx=XM_023246182.1;refGene_gn=SPTBN4;refGene_tx=XM_023246183.1
E2 14173852 14173991
refGene_gn=SPTBN4;refGene_tx=XM_023246182.1;refGene_gn=SPTBN4;refGene_tx=XM_023246183.1
E2 14176879 14177124
refGene_gn=SPTBN4;refGene_tx=XM_023246182.1;refGene_gn=SPTBN4;refGene_tx=XM_023246183.1
E2 14178258 14178639
refGene_gn=SPTBN4;refGene_tx=XM_023246182.1;refGene_gn=SPTBN4;refGene_tx=XM_023246183.1
E2 14179027 14179232
refGene_gn=SPTBN4;refGene_tx=XM_023246182.1;refGene_gn=SPTBN4;refGene_tx=XM_023246183.1
E2 14180122 14180253
refGene_gn=SPTBN4;refGene_tx=XM_023246182.1;refGene_gn=SPTBN4;refGene_tx=XM_023246183.1
E2 14180345 14180624
refGene_gn=SPTBN4;refGene_tx=XM_023246182.1;refGene_gn=SPTBN4;refGene_tx=XM_023246183.1
E2 14183230 14183320
refGene_gn=SPTBN4;refGene_tx=XM_023246182.1;refGene_gn=SPTBN4;refGene_tx=XM_023246183.1
E2 14184348 14184573
refGene_gn=SPTBN4;refGene_tx=XM_023246182.1;refGene_gn=SPTBN4;refGene_tx=XM_023246183.1
E2 14194590 14194854
refGene_gn=SPTBN4;refGene_tx=XM_023246182.1;refGene_gn=SPTBN4;refGene_tx=XM_023246183.1
E2 14196079 14196226
refGene_gn=SPTBN4;refGene_tx=XM_023246182.1;refGene_gn=SPTBN4;refGene_tx=XM_023246183.1
E2 14199676 14199767
refGene_gn=SPTBN4;refGene_tx=XM_023246182.1;refGene_gn=SPTBN4;refGene_tx=XM_023246183.1
E2 14203771 14203974
refGene_gn=SPTBN4;refGene_tx=XM_023246182.1;refGene_gn=SPTBN4;refGene_tx=XM_023246183.1
E2 14205425 14206176
refGene_gn=SPTBN4;refGene_tx=XM_023246182.1;refGene_gn=SPTBN4;refGene_

tx=XM_023246183.1
E2 14209457 14209595
refGene_gn=SPTBN4;refGene_tx=XM_023246182.1;refGene_gn=SPTBN4;refGene_tx=XM_023246183.1
E2 14210544 14211502
refGene_gn=SPTBN4;refGene_tx=XM_023246182.1;refGene_gn=SPTBN4;refGene_tx=XM_023246183.1
E2 14215265 14215416
refGene_gn=SPTBN4;refGene_tx=XM_023246182.1;refGene_gn=SPTBN4;refGene_tx=XM_023246183.1
E2 14216403 14216706
refGene_gn=SPTBN4;refGene_tx=XM_023246182.1;refGene_gn=SPTBN4;refGene_tx=XM_023246183.1
E2 14217338 14217497
refGene_gn=SPTBN4;refGene_tx=XM_023246182.1;refGene_gn=SPTBN4;refGene_tx=XM_023246183.1
E2 14217768 14217886
refGene_gn=SPTBN4;refGene_tx=XM_023246182.1;refGene_gn=SPTBN4;refGene_tx=XM_023246183.1
E2 14217960 14218148
refGene_gn=SPTBN4;refGene_tx=XM_023246182.1;refGene_gn=SPTBN4;refGene_tx=XM_023246183.1
E2 14218244 14218357
refGene_gn=SPTBN4;refGene_tx=XM_023246182.1;refGene_gn=SPTBN4;refGene_tx=XM_023246183.1
E2 14220299 14220415
refGene_gn=SPTBN4;refGene_tx=XM_023246182.1;refGene_gn=SPTBN4;refGene_tx=XM_023246183.1
E2 14221560 14221641
refGene_gn=SPTBN4;refGene_tx=XM_023246182.1;refGene_gn=SPTBN4;refGene_tx=XM_023246183.1
E2 14222824 14222916
refGene_gn=SPTBN4;refGene_tx=XM_023246182.1;refGene_gn=SPTBN4;refGene_tx=XM_023246183.1
E2 14226323 14226497
refGene_gn=SPTBN4;refGene_tx=XM_023246182.1;refGene_gn=SPTBN4;refGene_tx=XM_023246183.1
E2 14227795 14227947
refGene_gn=SPTBN4;refGene_tx=XM_023246182.1;refGene_gn=SPTBN4;refGene_tx=XM_023246183.1
E2 14242794 14242966
refGene_gn=SPTBN4;refGene_tx=XM_023246182.1;refGene_gn=SPTBN4;refGene_tx=XM_023246183.1
E2 14247232 14247311 refGene_gn=BLVRB;refGene_tx=XM_003997792.5
E2 14257270 14257435 refGene_gn=BLVRB;refGene_tx=XM_003997792.5
E2 14257571 14257661 refGene_gn=BLVRB;refGene_tx=XM_003997792.5
E2 14259027 14259156 refGene_gn=BLVRB;refGene_tx=XM_003997792.5
E2 14262699 14262857 refGene_gn=BLVRB;refGene_tx=XM_003997792.5
E2 14267382 14267979
refGene_gn=SERTAD3;refGene_tx=XM_006941247.4;refGene_gn=SERTAD3;refGen

e_tx=XM_019819513.2;refGene_gn=SERTAD3;refGene_tx=XM_019819514.2;refGene_gn=SERTAD3;refGene_tx=XM_019819515.2;refGene_gn=SERTAD3;refGene_tx=XM_019819516.2
E2 14278833 14279562 refGene_gn=SERTAD1;refGene_tx=XM_003997794.3
E2 14285637 14285680 refGene_gn=PRX;refGene_tx=XM_019819517.2
E2 14289451 14289521 refGene_gn=PRX;refGene_tx=XM_011289887.3
E2 14289922 14289966
refGene_gn=PRX;refGene_tx=XM_011289887.3;refGene_gn=PRX;refGene_tx=XM_019819517.2
E2 14290049 14290148
refGene_gn=PRX;refGene_tx=XM_011289887.3;refGene_gn=PRX;refGene_tx=XM_019819517.2
E2 14290271 14290397
refGene_gn=PRX;refGene_tx=XM_003997796.4;refGene_gn=PRX;refGene_tx=XM_011289887.3;refGene_gn=PRX;refGene_tx=XM_019819517.2;refGene_gn=PRX;refGene_tx=XM_019819518.2
E2 14292397 14292554
refGene_gn=PRX;refGene_tx=XM_003997796.4;refGene_gn=PRX;refGene_tx=XM_011289887.3;refGene_gn=PRX;refGene_tx=XM_019819517.2;refGene_gn=PRX;refGene_tx=XM_019819518.2
E2 14295273 14295470
refGene_gn=PRX;refGene_tx=XM_003997796.4;refGene_gn=PRX;refGene_tx=XM_011289887.3;refGene_gn=PRX;refGene_tx=XM_019819517.2;refGene_gn=PRX;refGene_tx=XM_019819518.2
E2 14296110 14299812
refGene_gn=PRX;refGene_tx=XM_003997796.4;refGene_gn=PRX;refGene_tx=XM_011289887.3;refGene_gn=PRX;refGene_tx=XM_019819517.2;refGene_gn=PRX;refGene_tx=XM_019819518.2
E2 14302324 14302789 refGene_gn=HIPK4;refGene_tx=XM_003997797.5
E2 14305481 14305838 refGene_gn=HIPK4;refGene_tx=XM_003997797.5
E2 14306586 14307435 refGene_gn=HIPK4;refGene_tx=XM_003997797.5
E2 14308172 14308352 refGene_gn=HIPK4;refGene_tx=XM_003997797.5
E2 14309710 14309898
refGene_gn=PLD3;refGene_tx=XM_006941250.3;refGene_gn=PLD3;refGene_tx=XM_006941251.4;refGene_gn=PLD3;refGene_tx=XM_019819521.1;refGene_gn=PLD3;refGene_tx=XM_019819522.1;refGene_gn=PLD3;refGene_tx=XM_019819523.1
E2 14309982 14310082
refGene_gn=PLD3;refGene_tx=XM_006941250.3;refGene_gn=PLD3;refGene_tx=XM_006941251.4;refGene_gn=PLD3;refGene_tx=XM_019819521.1;refGene_gn=PLD3;refGene_tx=XM_019819522.1;refGene_gn=PLD3;refGene_tx=XM_019819523.1
E2 14310595 14310761
refGene_gn=PLD3;refGene_tx=XM_006941250.3;refGene_gn=PLD3;refGene_tx=XM_006941251.4;refGene_gn=PLD3;refGene_tx=XM_019819521.1;refGene_gn=PLD3;refGene_tx=XM_019819522.1;refGene_gn=PLD3;refGene_tx=XM_019819523.1
E2 14311246 14311386
refGene_gn=PLD3;refGene_tx=XM_006941250.3;refGene_gn=PLD3;refGene_tx=XM_006941251.4;refGene_gn=PLD3;refGene_tx=XM_019819521.1;refGene_gn=PLD3;refGene_tx=XM_019819522.1;refGene_gn=PLD3;refGene_tx=XM_019819523.1
E2 14313118 14313319
refGene_gn=PLD3;refGene_tx=XM_006941250.3;refGene_gn=PLD3;refGene_tx=X

M_006941251.4;refGene_gn=PLD3;refGene_tx=XM_019819521.1;refGene_gn=PLD3;refGene_tx=XM_019819522.1;refGene_gn=PLD3;refGene_tx=XM_019819523.1
E2 14314431 14314559
refGene_gn=PLD3;refGene_tx=XM_006941250.3;refGene_gn=PLD3;refGene_tx=XM_006941251.4;refGene_gn=PLD3;refGene_tx=XM_019819521.1;refGene_gn=PLD3;refGene_tx=XM_019819522.1;refGene_gn=PLD3;refGene_tx=XM_019819523.1
E2 14314636 14314757
refGene_gn=PLD3;refGene_tx=XM_006941250.3;refGene_gn=PLD3;refGene_tx=XM_006941251.4;refGene_gn=PLD3;refGene_tx=XM_019819521.1;refGene_gn=PLD3;refGene_tx=XM_019819522.1;refGene_gn=PLD3;refGene_tx=XM_019819523.1
E2 14316119 14316303
refGene_gn=PLD3;refGene_tx=XM_006941250.3;refGene_gn=PLD3;refGene_tx=XM_006941251.4;refGene_gn=PLD3;refGene_tx=XM_019819521.1;refGene_gn=PLD3;refGene_tx=XM_019819522.1;refGene_gn=PLD3;refGene_tx=XM_019819523.1
E2 14316787 14316930
refGene_gn=PLD3;refGene_tx=XM_006941250.3;refGene_gn=PLD3;refGene_tx=XM_006941251.4;refGene_gn=PLD3;refGene_tx=XM_019819521.1;refGene_gn=PLD3;refGene_tx=XM_019819522.1;refGene_gn=PLD3;refGene_tx=XM_019819523.1
E2 14317018 14317093
refGene_gn=PLD3;refGene_tx=XM_006941250.3;refGene_gn=PLD3;refGene_tx=XM_006941251.4;refGene_gn=PLD3;refGene_tx=XM_019819521.1;refGene_gn=PLD3;refGene_tx=XM_019819522.1;refGene_gn=PLD3;refGene_tx=XM_019819523.1
E2 14317193 14317220
refGene_gn=PLD3;refGene_tx=XM_006941250.3;refGene_gn=PLD3;refGene_tx=XM_006941251.4;refGene_gn=PLD3;refGene_tx=XM_019819521.1;refGene_gn=PLD3;refGene_tx=XM_019819522.1;refGene_gn=PLD3;refGene_tx=XM_019819523.1
E2 14332412 14332431
refGene_gn=CE2H19orf47;refGene_tx=XM_023246059.1
E2 14336411 14336499
refGene_gn=CE2H19orf47;refGene_tx=XM_023246059.1
E2 14336662 14336777
refGene_gn=CE2H19orf47;refGene_tx=XM_023246059.1
E2 14338649 14338731
refGene_gn=CE2H19orf47;refGene_tx=XM_023246059.1
E2 14343187 14343325
refGene_gn=CE2H19orf47;refGene_tx=XM_023246059.1
E2 14346208 14346318
refGene_gn=CE2H19orf47;refGene_tx=XM_023246059.1
E2 14346319 14346360
refGene_gn=CE2H19orf47;refGene_tx=XM_023246059.1
E2 14347733 14347806
refGene_gn=CE2H19orf47;refGene_tx=XM_023246059.1
E2 14349450 14349945
refGene_gn=CE2H19orf47;refGene_tx=XM_023246059.1
E2 14385882 14385928
refGene_gn=AKT2;refGene_tx=XM_003997800.5;refGene_gn=AKT2;refGene_tx=XM_006941257.4;refGene_gn=AKT2;refGene_tx=XM_006941258.4;refGene_gn=AKT2;refGene_tx=XM_023246057.1
E2 14394214 14394343
refGene_gn=AKT2;refGene_tx=XM_003997800.5;refGene_gn=AKT2;refGene_tx=X

M_006941257.4;refGene_gn=AKT2;refGene_tx=XM_006941258.4;refGene_gn=AKT2;refGene_tx=XM_023246057.1
E2 14395968 14396080
refGene_gn=AKT2;refGene_tx=XM_003997800.5;refGene_gn=AKT2;refGene_tx=XM_006941257.4;refGene_gn=AKT2;refGene_tx=XM_006941258.4;refGene_gn=AKT2;refGene_tx=XM_023246057.1;refGene_gn=AKT2;refGene_tx=XM_023246058.1
E2 14406346 14406500
refGene_gn=AKT2;refGene_tx=XM_003997800.5;refGene_gn=AKT2;refGene_tx=XM_006941257.4;refGene_gn=AKT2;refGene_tx=XM_006941258.4;refGene_gn=AKT2;refGene_tx=XM_023246057.1;refGene_gn=AKT2;refGene_tx=XM_023246058.1
E2 14406912 14407044
refGene_gn=AKT2;refGene_tx=XM_003997800.5;refGene_gn=AKT2;refGene_tx=XM_006941257.4;refGene_gn=AKT2;refGene_tx=XM_006941258.4;refGene_gn=AKT2;refGene_tx=XM_023246057.1;refGene_gn=AKT2;refGene_tx=XM_023246058.1
E2 14408499 14408565
refGene_gn=AKT2;refGene_tx=XM_003997800.5;refGene_gn=AKT2;refGene_tx=XM_006941257.4;refGene_gn=AKT2;refGene_tx=XM_006941258.4;refGene_gn=AKT2;refGene_tx=XM_023246057.1;refGene_gn=AKT2;refGene_tx=XM_023246058.1
E2 14409572 14409641
refGene_gn=AKT2;refGene_tx=XM_003997800.5;refGene_gn=AKT2;refGene_tx=XM_006941257.4;refGene_gn=AKT2;refGene_tx=XM_006941258.4;refGene_gn=AKT2;refGene_tx=XM_023246057.1;refGene_gn=AKT2;refGene_tx=XM_023246058.1
E2 14410463 14410586
refGene_gn=AKT2;refGene_tx=XM_003997800.5;refGene_gn=AKT2;refGene_tx=XM_006941257.4;refGene_gn=AKT2;refGene_tx=XM_006941258.4;refGene_gn=AKT2;refGene_tx=XM_023246057.1;refGene_gn=AKT2;refGene_tx=XM_023246058.1
E2 14412043 14412172
refGene_gn=AKT2;refGene_tx=XM_003997800.5;refGene_gn=AKT2;refGene_tx=XM_006941257.4;refGene_gn=AKT2;refGene_tx=XM_006941258.4;refGene_gn=AKT2;refGene_tx=XM_023246057.1;refGene_gn=AKT2;refGene_tx=XM_023246058.1
E2 14412398 14412613
refGene_gn=AKT2;refGene_tx=XM_003997800.5;refGene_gn=AKT2;refGene_tx=XM_006941257.4;refGene_gn=AKT2;refGene_tx=XM_006941258.4;refGene_gn=AKT2;refGene_tx=XM_023246057.1;refGene_gn=AKT2;refGene_tx=XM_023246058.1
E2 14413090 14413178
refGene_gn=AKT2;refGene_tx=XM_003997800.5;refGene_gn=AKT2;refGene_tx=XM_006941257.4;refGene_gn=AKT2;refGene_tx=XM_006941258.4;refGene_gn=AKT2;refGene_tx=XM_023246057.1;refGene_gn=AKT2;refGene_tx=XM_023246058.1
E2 14413280 14413383
refGene_gn=AKT2;refGene_tx=XM_003997800.5;refGene_gn=AKT2;refGene_tx=XM_006941257.4;refGene_gn=AKT2;refGene_tx=XM_006941258.4;refGene_gn=AKT2;refGene_tx=XM_023246057.1;refGene_gn=AKT2;refGene_tx=XM_023246058.1
E2 14414357 14414437
refGene_gn=AKT2;refGene_tx=XM_003997800.5;refGene_gn=AKT2;refGene_tx=XM_006941257.4;refGene_gn=AKT2;refGene_tx=XM_006941258.4;refGene_gn=AKT2;refGene_tx=XM_023246057.1;refGene_gn=AKT2;refGene_tx=XM_023246058.1
E2 14420887 14420944 refGene_gn=CNTD2;refGene_tx=XM_023245945.1
E2 14421080 14421281 refGene_gn=CNTD2;refGene_tx=XM_023245945.1
E2 14422517 14422607 refGene_gn=CNTD2;refGene_tx=XM_023245945.1
E2 14422684 14422840 refGene_gn=CNTD2;refGene_tx=XM_023245945.1

E2 14423371 14423530 refGene_gn=CNTD2;refGene_tx=XM_023245945.1
E2 14423606 14423831 refGene_gn=CNTD2;refGene_tx=XM_023245945.1
E2 14427189 14427604 refGene_gn=TTC9B;refGene_tx=XM_003997802.3
E2 14428152 14428335 refGene_gn=TTC9B;refGene_tx=XM_003997802.3
E2 14429206 14429334 refGene_gn=TTC9B;refGene_tx=XM_003997802.3
E2 14429885 14430208 refGene_gn=MAP3K10;refGene_tx=XM_023245041.1
E2 14431192 14431846 refGene_gn=MAP3K10;refGene_tx=XM_023245041.1
E2 14432214 14432327 refGene_gn=MAP3K10;refGene_tx=XM_023245041.1
E2 14432431 14432603 refGene_gn=MAP3K10;refGene_tx=XM_023245041.1
E2 14434436 14434553 refGene_gn=MAP3K10;refGene_tx=XM_023245041.1
E2 14436144 14436391 refGene_gn=MAP3K10;refGene_tx=XM_023245041.1
E2 14437448 14437624 refGene_gn=MAP3K10;refGene_tx=XM_023245041.1
E2 14438077 14438226 refGene_gn=MAP3K10;refGene_tx=XM_023245041.1
E2 14441235 14441416 refGene_gn=MAP3K10;refGene_tx=XM_023245041.1
E2 14445185 14445867 refGene_gn=MAP3K10;refGene_tx=XM_023245041.1
E2 14460766 14460903
refGene_gn=LOC101089975;refGene_tx=XM_019819525.2;refGene_gn=LOC101089975;refGene_tx=XM_019819526.2
E2 14488157 14488230
refGene_gn=LOC101089975;refGene_tx=XM_019819525.2;refGene_gn=LOC101089975;refGene_tx=XM_019819526.2
E2 14491387 14491423
refGene_gn=LOC101089975;refGene_tx=XM_019819530.2;refGene_gn=LOC101089975;refGene_tx=XM_019819531.2;refGene_gn=LOC101089975;refGene_tx=XM_019819533.2;refGene_gn=LOC101089975;refGene_tx=XM_023245157.1;refGene_gn=LOC101089975;refGene_tx=XM_023245158.1;refGene_gn=LOC101089975;refGene_tx=XM_023245160.1
E2 14491674 14491807
refGene_gn=LOC101089975;refGene_tx=XM_019819525.2;refGene_gn=LOC101089975;refGene_tx=XM_019819526.2;refGene_gn=LOC101089975;refGene_tx=XM_019819530.2;refGene_gn=LOC101089975;refGene_tx=XM_019819531.2;refGene_gn=LOC101089975;refGene_tx=XM_019819533.2;refGene_gn=LOC101089975;refGene_tx=XM_019819538.2;refGene_gn=LOC101089975;refGene_tx=XM_023245157.1;refGene_gn=LOC101089975;refGene_tx=XM_023245158.1;refGene_gn=LOC101089975;refGene_tx=XM_023245160.1
E2 14493671 14493770
refGene_gn=LOC101089975;refGene_tx=XM_019819525.2;refGene_gn=LOC101089975;refGene_tx=XM_019819526.2;refGene_gn=LOC101089975;refGene_tx=XM_019819530.2;refGene_gn=LOC101089975;refGene_tx=XM_019819531.2;refGene_gn=LOC101089975;refGene_tx=XM_019819533.2;refGene_gn=LOC101089975;refGene_tx=XM_019819538.2;refGene_gn=LOC101089975;refGene_tx=XM_023245157.1;refGene_gn=LOC101089975;refGene_tx=XM_023245158.1;refGene_gn=LOC101089975;refGene_tx=XM_023245160.1
E2 14503012 14505876
refGene_gn=LOC101089975;refGene_tx=XM_019819525.2;refGene_gn=LOC101089975;refGene_tx=XM_019819526.2;refGene_gn=LOC101089975;refGene_tx=XM_019819530.2;refGene_gn=LOC101089975;refGene_tx=XM_019819531.2;refGene_gn=LOC101089975;refGene_tx=XM_019819533.2;refGene_gn=LOC101089975;refGene_tx=XM_019819538.2;refGene_gn=LOC101089975;refGene_tx=XM_023245157.1;refGene_gn=LOC101089975;refGene_tx=XM_023245158.1;refGene_gn=LOC101089

975;refGene_tx=XM_023245160.1
E2 14517615 14517665 refGene_gn=PSMC4;refGene_tx=XM_023245171.1
E2 14521071 14521185 refGene_gn=PSMC4;refGene_tx=XM_011289885.2
E2 14521362 14521418 refGene_gn=PSMC4;refGene_tx=XM_011289885.2
E2 14521614 14521783
refGene_gn=PSMC4;refGene_tx=XM_011289885.2;refGene_gn=PSMC4;refGene_tx
=XM_023245171.1
E2 14521926 14522003
refGene_gn=PSMC4;refGene_tx=XM_011289885.2;refGene_gn=PSMC4;refGene_tx
=XM_023245171.1
E2 14522086 14522254
refGene_gn=PSMC4;refGene_tx=XM_011289885.2;refGene_gn=PSMC4;refGene_tx
=XM_023245171.1
E2 14526086 14526180
refGene_gn=PSMC4;refGene_tx=XM_011289885.2;refGene_gn=PSMC4;refGene_tx
=XM_023245171.1
E2 14526276 14526386
refGene_gn=PSMC4;refGene_tx=XM_011289885.2;refGene_gn=PSMC4;refGene_tx
=XM_023245171.1
E2 14526474 14526621
refGene_gn=PSMC4;refGene_tx=XM_011289885.2;refGene_gn=PSMC4;refGene_tx
=XM_023245171.1
E2 14528470 14528657
refGene_gn=PSMC4;refGene_tx=XM_011289885.2;refGene_gn=PSMC4;refGene_tx
=XM_023245171.1
E2 14528793 14528892
refGene_gn=PSMC4;refGene_tx=XM_011289885.2;refGene_gn=PSMC4;refGene_tx
=XM_023245171.1
E2 14529701 14529737
refGene_gn=PSMC4;refGene_tx=XM_011289885.2;refGene_gn=PSMC4;refGene_tx
=XM_023245171.1
E2 14560611 14560666 refGene_gn=FCGBP;refGene_tx=XM_023245830.1
E2 14561943 14563191 refGene_gn=FCGBP;refGene_tx=XM_023245830.1
E2 14565288 14565635 refGene_gn=FCGBP;refGene_tx=XM_023245830.1
E2 14569418 14569989 refGene_gn=FCGBP;refGene_tx=XM_023245830.1
E2 14572080 14572685 refGene_gn=FCGBP;refGene_tx=XM_023245830.1
E2 14574853 14575421 refGene_gn=FCGBP;refGene_tx=XM_023245830.1
E2 14579270 14579863 refGene_gn=FCGBP;refGene_tx=XM_023245830.1
E2 14582114 14582706 refGene_gn=FCGBP;refGene_tx=XM_023245830.1
E2 14582846 14583046 refGene_gn=FCGBP;refGene_tx=XM_023245830.1
E2 14584436 14584578 refGene_gn=FCGBP;refGene_tx=XM_023245830.1
E2 14585848 14586186 refGene_gn=FCGBP;refGene_tx=XM_023245830.1
E2 14587185 14587759 refGene_gn=FCGBP;refGene_tx=XM_023245830.1
E2 14588114 14588731 refGene_gn=FCGBP;refGene_tx=XM_023245830.1
E2 14589160 14589704 refGene_gn=FCGBP;refGene_tx=XM_023245830.1
E2 14590724 14591296 refGene_gn=FCGBP;refGene_tx=XM_023245830.1
E2 14591616 14592190 refGene_gn=FCGBP;refGene_tx=XM_023245830.1
E2 14593122 14593322 refGene_gn=FCGBP;refGene_tx=XM_023245830.1
E2 14596302 14596686 refGene_gn=FCGBP;refGene_tx=XM_023245830.1
E2 14599672 14599941 refGene_gn=FCGBP;refGene_tx=XM_023245830.1

E2 14600065 14600075 refGene_gn=FCGBP;refGene_tx=XM_023245830.1
E2 14604679 14604689 refGene_gn=FBL;refGene_tx=XM_003997803.5
E2 14610348 14610501 refGene_gn=FBL;refGene_tx=XM_003997803.5
E2 14610607 14610709 refGene_gn=FBL;refGene_tx=XM_003997803.5
E2 14610797 14610892 refGene_gn=FBL;refGene_tx=XM_003997803.5
E2 14612034 14612205 refGene_gn=FBL;refGene_tx=XM_003997803.5
E2 14612923 14613056 refGene_gn=FBL;refGene_tx=XM_003997803.5
E2 14614637 14614750 refGene_gn=FBL;refGene_tx=XM_003997803.5
E2 14615521 14615667 refGene_gn=FBL;refGene_tx=XM_003997803.5
E2 14615778 14615803 refGene_gn=FBL;refGene_tx=XM_003997803.5
E2 14617037 14617116
refGene_gn=DYRK1B;refGene_tx=XM_023245831.1;refGene_gn=DYRK1B;refGene_tx=XM_023245832.1;refGene_gn=DYRK1B;refGene_tx=XM_023245833.1
E2 14617260 14617300 refGene_gn=DYRK1B;refGene_tx=XM_023245834.1
E2 14618372 14618536
refGene_gn=DYRK1B;refGene_tx=XM_003997804.5;refGene_gn=DYRK1B;refGene_tx=XM_023245831.1;refGene_gn=DYRK1B;refGene_tx=XM_023245832.1;refGene_gn=DYRK1B;refGene_tx=XM_023245833.1;refGene_gn=DYRK1B;refGene_tx=XM_023245834.1
E2 14619573 14619693
refGene_gn=DYRK1B;refGene_tx=XM_003997804.5;refGene_gn=DYRK1B;refGene_tx=XM_023245831.1;refGene_gn=DYRK1B;refGene_tx=XM_023245832.1;refGene_gn=DYRK1B;refGene_tx=XM_023245833.1;refGene_gn=DYRK1B;refGene_tx=XM_023245834.1
E2 14619780 14619969
refGene_gn=DYRK1B;refGene_tx=XM_003997804.5;refGene_gn=DYRK1B;refGene_tx=XM_023245831.1;refGene_gn=DYRK1B;refGene_tx=XM_023245832.1;refGene_gn=DYRK1B;refGene_tx=XM_023245833.1;refGene_gn=DYRK1B;refGene_tx=XM_023245834.1
E2 14620133 14620281
refGene_gn=DYRK1B;refGene_tx=XM_003997804.5;refGene_gn=DYRK1B;refGene_tx=XM_023245831.1;refGene_gn=DYRK1B;refGene_tx=XM_023245832.1;refGene_gn=DYRK1B;refGene_tx=XM_023245833.1;refGene_gn=DYRK1B;refGene_tx=XM_023245834.1
E2 14621145 14621432
refGene_gn=DYRK1B;refGene_tx=XM_003997804.5;refGene_gn=DYRK1B;refGene_tx=XM_023245831.1;refGene_gn=DYRK1B;refGene_tx=XM_023245832.1;refGene_gn=DYRK1B;refGene_tx=XM_023245833.1;refGene_gn=DYRK1B;refGene_tx=XM_023245834.1
E2 14622153 14622300
refGene_gn=DYRK1B;refGene_tx=XM_003997804.5;refGene_gn=DYRK1B;refGene_tx=XM_023245831.1;refGene_gn=DYRK1B;refGene_tx=XM_023245832.1;refGene_gn=DYRK1B;refGene_tx=XM_023245833.1;refGene_gn=DYRK1B;refGene_tx=XM_023245834.1
E2 14622386 14622527
refGene_gn=DYRK1B;refGene_tx=XM_003997804.5;refGene_gn=DYRK1B;refGene_tx=XM_023245831.1;refGene_gn=DYRK1B;refGene_tx=XM_023245832.1;refGene_gn=DYRK1B;refGene_tx=XM_023245833.1;refGene_gn=DYRK1B;refGene_tx=XM_023245834.1
E2 14622838 14623154

refGene_gn=DYRK1B;refGene_tx=XM_003997804.5;refGene_gn=DYRK1B;refGene_tx=XM_023245831.1;refGene_gn=DYRK1B;refGene_tx=XM_023245832.1;refGene_gn=DYRK1B;refGene_tx=XM_023245833.1;refGene_gn=DYRK1B;refGene_tx=XM_023245834.1
E2 14623548 14623655
refGene_gn=DYRK1B;refGene_tx=XM_003997804.5;refGene_gn=DYRK1B;refGene_tx=XM_023245831.1;refGene_gn=DYRK1B;refGene_tx=XM_023245832.1;refGene_gn=DYRK1B;refGene_tx=XM_023245833.1;refGene_gn=DYRK1B;refGene_tx=XM_023245834.1
E2 14623749 14624121
refGene_gn=DYRK1B;refGene_tx=XM_003997804.5;refGene_gn=DYRK1B;refGene_tx=XM_023245831.1;refGene_gn=DYRK1B;refGene_tx=XM_023245832.1;refGene_gn=DYRK1B;refGene_tx=XM_023245833.1;refGene_gn=DYRK1B;refGene_tx=XM_023245834.1
E2 14660146 14660605 refGene_gn=LEUTX;refGene_tx=XM_023245946.1
E2 14661320 14661389 refGene_gn=LEUTX;refGene_tx=XM_023245946.1
E2 14685915 14686479
refGene_gn=LOC102902632;refGene_tx=XM_023245348.1
E2 14701395 14701473 refGene_gn=SELENOV;refGene_tx=XM_011289881.3
E2 14701588 14701663 refGene_gn=SELENOV;refGene_tx=XM_011289881.3
E2 14701778 14701832 refGene_gn=SELENOV;refGene_tx=XM_011289881.3
E2 14701961 14701986 refGene_gn=SELENOV;refGene_tx=XM_011289881.3
E2 14703894 14704889 refGene_gn=SELENOV;refGene_tx=XM_011289881.3
E2 14711205 14711211 refGene_gn=DLL3;refGene_tx=XM_003997806.5
E2 14711559 14711644 refGene_gn=DLL3;refGene_tx=XM_003997806.5
E2 14711885 14712477 refGene_gn=DLL3;refGene_tx=XM_003997806.5
E2 14713932 14714155 refGene_gn=DLL3;refGene_tx=XM_003997806.5
E2 14715077 14715292 refGene_gn=DLL3;refGene_tx=XM_003997806.5
E2 14716331 14716574 refGene_gn=DLL3;refGene_tx=XM_003997806.5
E2 14717717 14717772 refGene_gn=DLL3;refGene_tx=XM_003997806.5
E2 14718645 14718927 refGene_gn=DLL3;refGene_tx=XM_003997806.5
E2 14719056 14719125 refGene_gn=DLL3;refGene_tx=XM_003997806.5
E2 14724681 14724783
refGene_gn=TIMM50;refGene_tx=XM_011289880.3;refGene_gn=TIMM50;refGene_tx=XM_019819544.2
E2 14725666 14725773
refGene_gn=TIMM50;refGene_tx=XM_011289880.3;refGene_gn=TIMM50;refGene_tx=XM_019819544.2
E2 14726084 14726241
refGene_gn=TIMM50;refGene_tx=XM_011289880.3;refGene_gn=TIMM50;refGene_tx=XM_019819544.2
E2 14726596 14726695
refGene_gn=TIMM50;refGene_tx=XM_011289880.3;refGene_gn=TIMM50;refGene_tx=XM_019819544.2
E2 14726793 14726898
refGene_gn=TIMM50;refGene_tx=XM_011289880.3;refGene_gn=TIMM50;refGene_tx=XM_019819544.2
E2 14727309 14727429
refGene_gn=TIMM50;refGene_tx=XM_011289880.3;refGene_gn=TIMM50;refGene_tx=XM_019819544.2

E2 14727520 14727579 refGene_gn=TIMM50;refGene_tx=XM_011289880.3
E2 14729616 14729638 refGene_gn=TIMM50;refGene_tx=XM_011289880.3
E2 14729852 14729884 refGene_gn=TIMM50;refGene_tx=XM_011289880.3
E2 14730424 14730581 refGene_gn=TIMM50;refGene_tx=XM_011289880.3
E2 14731412 14731535 refGene_gn=TIMM50;refGene_tx=XM_011289880.3
E2 14733589 14733733
refGene_gn=SUPT5H;refGene_tx=XM_006941276.4;refGene_gn=SUPT5H;refGene_tx=XM_006941277.4;refGene_gn=SUPT5H;refGene_tx=XM_019819546.2
E2 14733805 14733901
refGene_gn=SUPT5H;refGene_tx=XM_006941276.4;refGene_gn=SUPT5H;refGene_tx=XM_006941277.4;refGene_gn=SUPT5H;refGene_tx=XM_019819546.2
E2 14735655 14735859
refGene_gn=SUPT5H;refGene_tx=XM_006941276.4;refGene_gn=SUPT5H;refGene_tx=XM_006941277.4;refGene_gn=SUPT5H;refGene_tx=XM_019819546.2
E2 14735971 14736140
refGene_gn=SUPT5H;refGene_tx=XM_006941276.4;refGene_gn=SUPT5H;refGene_tx=XM_006941277.4;refGene_gn=SUPT5H;refGene_tx=XM_019819546.2
E2 14736257 14736416
refGene_gn=SUPT5H;refGene_tx=XM_006941276.4;refGene_gn=SUPT5H;refGene_tx=XM_006941277.4;refGene_gn=SUPT5H;refGene_tx=XM_019819546.2
E2 14737073 14737179
refGene_gn=SUPT5H;refGene_tx=XM_006941276.4;refGene_gn=SUPT5H;refGene_tx=XM_006941277.4;refGene_gn=SUPT5H;refGene_tx=XM_019819546.2
E2 14737279 14737407
refGene_gn=SUPT5H;refGene_tx=XM_006941276.4;refGene_gn=SUPT5H;refGene_tx=XM_006941277.4;refGene_gn=SUPT5H;refGene_tx=XM_019819546.2
E2 14737489 14737592
refGene_gn=SUPT5H;refGene_tx=XM_006941276.4;refGene_gn=SUPT5H;refGene_tx=XM_006941277.4;refGene_gn=SUPT5H;refGene_tx=XM_019819546.2
E2 14737674 14737794
refGene_gn=SUPT5H;refGene_tx=XM_006941276.4;refGene_gn=SUPT5H;refGene_tx=XM_006941277.4;refGene_gn=SUPT5H;refGene_tx=XM_019819546.2
E2 14738134 14738219
refGene_gn=SUPT5H;refGene_tx=XM_006941276.4;refGene_gn=SUPT5H;refGene_tx=XM_006941277.4;refGene_gn=SUPT5H;refGene_tx=XM_019819546.2
E2 14739067 14739193
refGene_gn=SUPT5H;refGene_tx=XM_006941276.4;refGene_gn=SUPT5H;refGene_tx=XM_006941277.4;refGene_gn=SUPT5H;refGene_tx=XM_019819546.2
E2 14739283 14739430
refGene_gn=SUPT5H;refGene_tx=XM_006941276.4;refGene_gn=SUPT5H;refGene_tx=XM_006941277.4;refGene_gn=SUPT5H;refGene_tx=XM_019819546.2
E2 14740262 14740409
refGene_gn=SUPT5H;refGene_tx=XM_006941276.4;refGene_gn=SUPT5H;refGene_tx=XM_006941277.4;refGene_gn=SUPT5H;refGene_tx=XM_019819546.2
E2 14740513 14740669
refGene_gn=SUPT5H;refGene_tx=XM_006941276.4;refGene_gn=SUPT5H;refGene_tx=XM_006941277.4;refGene_gn=SUPT5H;refGene_tx=XM_019819546.2
E2 14741078 14741215
refGene_gn=SUPT5H;refGene_tx=XM_006941276.4;refGene_gn=SUPT5H;refGene_tx=XM_006941277.4;refGene_gn=SUPT5H;refGene_tx=XM_019819546.2

E2	14741306	14741400
refGene_gn=SUPT5H;refGene_tx=XM_006941276.4;refGene_gn=SUPT5H;refGene_tx=XM_006941277.4;refGene_gn=SUPT5H;refGene_tx=XM_019819546.2
E2	14741615	14741721
refGene_gn=SUPT5H;refGene_tx=XM_006941276.4;refGene_gn=SUPT5H;refGene_tx=XM_006941277.4;refGene_gn=SUPT5H;refGene_tx=XM_019819546.2
E2	14743234	14743305
refGene_gn=SUPT5H;refGene_tx=XM_006941276.4;refGene_gn=SUPT5H;refGene_tx=XM_006941277.4;refGene_gn=SUPT5H;refGene_tx=XM_019819546.2
E2	14743412	14743502
refGene_gn=SUPT5H;refGene_tx=XM_006941276.4;refGene_gn=SUPT5H;refGene_tx=XM_006941277.4;refGene_gn=SUPT5H;refGene_tx=XM_019819546.2
E2	14744638	14744890
refGene_gn=SUPT5H;refGene_tx=XM_006941276.4;refGene_gn=SUPT5H;refGene_tx=XM_006941277.4;refGene_gn=SUPT5H;refGene_tx=XM_019819546.2
E2	14748010	14748079
refGene_gn=SUPT5H;refGene_tx=XM_006941276.4;refGene_gn=SUPT5H;refGene_tx=XM_006941277.4;refGene_gn=SUPT5H;refGene_tx=XM_019819546.2
E2	14748430	14748461
refGene_gn=SUPT5H;refGene_tx=XM_006941276.4;refGene_gn=SUPT5H;refGene_tx=XM_006941277.4;refGene_gn=SUPT5H;refGene_tx=XM_019819546.2
E2	14748707	14748773
refGene_gn=SUPT5H;refGene_tx=XM_006941276.4;refGene_gn=SUPT5H;refGene_tx=XM_006941277.4;refGene_gn=SUPT5H;refGene_tx=XM_019819546.2
E2	14748871	14748940
refGene_gn=SUPT5H;refGene_tx=XM_006941276.4;refGene_gn=SUPT5H;refGene_tx=XM_006941277.4;refGene_gn=SUPT5H;refGene_tx=XM_019819546.2
E2	14749067	14749137
refGene_gn=SUPT5H;refGene_tx=XM_006941276.4;refGene_gn=SUPT5H;refGene_tx=XM_006941277.4;refGene_gn=SUPT5H;refGene_tx=XM_019819546.2
E2	14749628	14749640
refGene_gn=SUPT5H;refGene_tx=XM_006941276.4;refGene_gn=SUPT5H;refGene_tx=XM_006941277.4
E2	14750180	14750246
refGene_gn=SUPT5H;refGene_tx=XM_006941276.4;refGene_gn=SUPT5H;refGene_tx=XM_006941277.4;refGene_gn=SUPT5H;refGene_tx=XM_019819546.2
E2	14753737	14753903
refGene_gn=SUPT5H;refGene_tx=XM_006941276.4;refGene_gn=SUPT5H;refGene_tx=XM_006941277.4;refGene_gn=SUPT5H;refGene_tx=XM_019819546.2
E2	14758928	14759003
refGene_gn=SUPT5H;refGene_tx=XM_006941276.4;refGene_gn=SUPT5H;refGene_tx=XM_006941277.4;refGene_gn=SUPT5H;refGene_tx=XM_019819546.2
E2	14765604	14765652	refGene_gn=RPS16;refGene_tx=XM_003997810.3
E2	14765788	14765890	refGene_gn=RPS16;refGene_tx=XM_003997810.3
E2	14767828	14767925	refGene_gn=RPS16;refGene_tx=XM_003997810.3
E2	14768031	14768079	refGene_gn=RPS16;refGene_tx=XM_003997810.3
E2	14768208	14768354	refGene_gn=RPS16;refGene_tx=XM_003997810.3
E2	14772457	14774082
refGene_gn=PLEKHG2;refGene_tx=XM_006941278.4;refGene_gn=PLEKHG2;refGene_tx=XM_006941279.4;refGene_gn=PLEKHG2;refGene_tx=XM_019819553.2;refGe

ne_gn=PLEKHG2;refGene_tx=XM_019819554.2;refGene_gn=PLEKHG2;refGene_tx=XM_019819555.2;refGene_gn=PLEKHG2;refGene_tx=XM_023245074.1
E2 14774161 14775086
refGene_gn=PLEKHG2;refGene_tx=XM_006941278.4;refGene_gn=PLEKHG2;refGene_tx=XM_006941279.4;refGene_gn=PLEKHG2;refGene_tx=XM_019819553.2;refGene_gn=PLEKHG2;refGene_tx=XM_019819554.2;refGene_gn=PLEKHG2;refGene_tx=XM_019819555.2;refGene_gn=PLEKHG2;refGene_tx=XM_023245074.1
E2 14775192 14775369
refGene_gn=PLEKHG2;refGene_tx=XM_006941278.4;refGene_gn=PLEKHG2;refGene_tx=XM_006941279.4;refGene_gn=PLEKHG2;refGene_tx=XM_019819553.2;refGene_gn=PLEKHG2;refGene_tx=XM_019819554.2;refGene_gn=PLEKHG2;refGene_tx=XM_019819555.2;refGene_gn=PLEKHG2;refGene_tx=XM_023245074.1
E2 14775476 14775493
refGene_gn=PLEKHG2;refGene_tx=XM_006941278.4;refGene_gn=PLEKHG2;refGene_tx=XM_006941279.4;refGene_gn=PLEKHG2;refGene_tx=XM_019819553.2;refGene_gn=PLEKHG2;refGene_tx=XM_019819554.2;refGene_gn=PLEKHG2;refGene_tx=XM_019819555.2;refGene_gn=PLEKHG2;refGene_tx=XM_023245074.1
E2 14775657 14775696
refGene_gn=PLEKHG2;refGene_tx=XM_006941278.4;refGene_gn=PLEKHG2;refGene_tx=XM_006941279.4;refGene_gn=PLEKHG2;refGene_tx=XM_019819553.2;refGene_gn=PLEKHG2;refGene_tx=XM_019819554.2;refGene_gn=PLEKHG2;refGene_tx=XM_019819555.2;refGene_gn=PLEKHG2;refGene_tx=XM_023245074.1
E2 14775775 14775823
refGene_gn=PLEKHG2;refGene_tx=XM_006941278.4;refGene_gn=PLEKHG2;refGene_tx=XM_006941279.4;refGene_gn=PLEKHG2;refGene_tx=XM_019819553.2;refGene_gn=PLEKHG2;refGene_tx=XM_019819554.2;refGene_gn=PLEKHG2;refGene_tx=XM_019819555.2;refGene_gn=PLEKHG2;refGene_tx=XM_023245074.1
E2 14775941 14776043
refGene_gn=PLEKHG2;refGene_tx=XM_006941278.4;refGene_gn=PLEKHG2;refGene_tx=XM_006941279.4;refGene_gn=PLEKHG2;refGene_tx=XM_019819553.2;refGene_gn=PLEKHG2;refGene_tx=XM_019819554.2;refGene_gn=PLEKHG2;refGene_tx=XM_019819555.2;refGene_gn=PLEKHG2;refGene_tx=XM_023245074.1
E2 14776135 14776169
refGene_gn=PLEKHG2;refGene_tx=XM_006941278.4;refGene_gn=PLEKHG2;refGene_tx=XM_006941279.4;refGene_gn=PLEKHG2;refGene_tx=XM_019819553.2;refGene_gn=PLEKHG2;refGene_tx=XM_019819554.2;refGene_gn=PLEKHG2;refGene_tx=XM_019819555.2;refGene_gn=PLEKHG2;refGene_tx=XM_023245074.1
E2 14777001 14777268
refGene_gn=PLEKHG2;refGene_tx=XM_006941278.4;refGene_gn=PLEKHG2;refGene_tx=XM_006941279.4;refGene_gn=PLEKHG2;refGene_tx=XM_019819553.2;refGene_gn=PLEKHG2;refGene_tx=XM_019819554.2;refGene_gn=PLEKHG2;refGene_tx=XM_019819555.2;refGene_gn=PLEKHG2;refGene_tx=XM_023245074.1
E2 14777725 14777926
refGene_gn=PLEKHG2;refGene_tx=XM_006941278.4;refGene_gn=PLEKHG2;refGene_tx=XM_006941279.4;refGene_gn=PLEKHG2;refGene_tx=XM_019819553.2;refGene_gn=PLEKHG2;refGene_tx=XM_019819554.2;refGene_gn=PLEKHG2;refGene_tx=XM_019819555.2;refGene_gn=PLEKHG2;refGene_tx=XM_023245074.1
E2 14778093 14778231
refGene_gn=PLEKHG2;refGene_tx=XM_006941278.4;refGene_gn=PLEKHG2;refGene_tx=XM_006941279.4;refGene_gn=PLEKHG2;refGene_tx=XM_019819553.2;refGe

ne_gn=PLEKHG2;refGene_tx=XM_019819554.2;refGene_gn=PLEKHG2;refGene_tx=XM_019819555.2;refGene_gn=PLEKHG2;refGene_tx=XM_023245074.1
E2 14778530 14778681 refGene_gn=PLEKHG2;refGene_tx=XM_006941278.4;refGene_gn=PLEKHG2;refGene_tx=XM_006941279.4;refGene_gn=PLEKHG2;refGene_tx=XM_019819553.2;refGene_gn=PLEKHG2;refGene_tx=XM_019819554.2;refGene_gn=PLEKHG2;refGene_tx=XM_019819555.2;refGene_gn=PLEKHG2;refGene_tx=XM_023245074.1
E2 14778921 14778968 refGene_gn=PLEKHG2;refGene_tx=XM_006941278.4;refGene_gn=PLEKHG2;refGene_tx=XM_006941279.4;refGene_gn=PLEKHG2;refGene_tx=XM_019819553.2;refGene_gn=PLEKHG2;refGene_tx=XM_019819554.2;refGene_gn=PLEKHG2;refGene_tx=XM_019819555.2;refGene_gn=PLEKHG2;refGene_tx=XM_023245074.1
E2 14779073 14779140 refGene_gn=PLEKHG2;refGene_tx=XM_006941278.4;refGene_gn=PLEKHG2;refGene_tx=XM_006941279.4;refGene_gn=PLEKHG2;refGene_tx=XM_019819553.2;refGene_gn=PLEKHG2;refGene_tx=XM_019819554.2;refGene_gn=PLEKHG2;refGene_tx=XM_019819555.2;refGene_gn=PLEKHG2;refGene_tx=XM_023245074.1
E2 14779925 14780026 refGene_gn=PLEKHG2;refGene_tx=XM_006941278.4;refGene_gn=PLEKHG2;refGene_tx=XM_006941279.4;refGene_gn=PLEKHG2;refGene_tx=XM_019819553.2;refGene_gn=PLEKHG2;refGene_tx=XM_019819554.2;refGene_gn=PLEKHG2;refGene_tx=XM_019819555.2;refGene_gn=PLEKHG2;refGene_tx=XM_023245074.1
E2 14780107 14780376 refGene_gn=PLEKHG2;refGene_tx=XM_006941278.4;refGene_gn=PLEKHG2;refGene_tx=XM_006941279.4;refGene_gn=PLEKHG2;refGene_tx=XM_019819553.2;refGene_gn=PLEKHG2;refGene_tx=XM_019819554.2;refGene_gn=PLEKHG2;refGene_tx=XM_019819555.2;refGene_gn=PLEKHG2;refGene_tx=XM_023245074.1
E2 14781198 14781307 refGene_gn=PLEKHG2;refGene_tx=XM_006941278.4;refGene_gn=PLEKHG2;refGene_tx=XM_006941279.4;refGene_gn=PLEKHG2;refGene_tx=XM_019819553.2;refGene_gn=PLEKHG2;refGene_tx=XM_019819554.2;refGene_gn=PLEKHG2;refGene_tx=XM_019819555.2;refGene_gn=PLEKHG2;refGene_tx=XM_023245074.1
E2 14786880 14787834 refGene_gn=ZFP36;refGene_tx=XM_003997812.4
E2 14788665 14788689 refGene_gn=ZFP36;refGene_tx=XM_003997812.4
E2 14789303 14789570 refGene_gn=MED29;refGene_tx=XM_019819557.2
E2 14792245 14792247 refGene_gn=MED29;refGene_tx=XM_019819557.2
E2 14794360 14794603 refGene_gn=MED29;refGene_tx=XM_003997813.5
E2 14796254 14796339 refGene_gn=MED29;refGene_tx=XM_003997813.5;refGene_gn=MED29;refGene_tx=XM_019819557.2
E2 14796600 14796659 refGene_gn=MED29;refGene_tx=XM_003997813.5;refGene_gn=MED29;refGene_tx=XM_019819557.2
E2 14797199 14797415 refGene_gn=MED29;refGene_tx=XM_003997813.5;refGene_gn=MED29;refGene_tx=XM_019819557.2
E2 14797972 14798019 refGene_gn=PAF1;refGene_tx=XM_003997814.5
E2 14798612 14798642 refGene_gn=PAF1;refGene_tx=XM_003997814.5
E2 14798739 14798832 refGene_gn=PAF1;refGene_tx=XM_003997814.5
E2 14799165 14799287 refGene_gn=PAF1;refGene_tx=XM_003997814.5

E2 14799375 14799442 refGene_gn=PAF1;refGene_tx=XM_003997814.5
E2 14799534 14799636 refGene_gn=PAF1;refGene_tx=XM_003997814.5
E2 14799715 14799821 refGene_gn=PAF1;refGene_tx=XM_003997814.5
E2 14799921 14799990 refGene_gn=PAF1;refGene_tx=XM_003997814.5
E2 14800074 14800178 refGene_gn=PAF1;refGene_tx=XM_003997814.5
E2 14800258 14800375 refGene_gn=PAF1;refGene_tx=XM_003997814.5
E2 14800467 14800596 refGene_gn=PAF1;refGene_tx=XM_003997814.5
E2 14801553 14801659 refGene_gn=PAF1;refGene_tx=XM_003997814.5
E2 14801767 14801858 refGene_gn=PAF1;refGene_tx=XM_003997814.5
E2 14801944 14802348 refGene_gn=PAF1;refGene_tx=XM_003997814.5
E2 14804823 14804852
refGene_gn=SAMD4B;refGene_tx=XM_006941285.4;refGene_gn=SAMD4B;refGene_tx=XM_023245402.1
E2 14805061 14805145
refGene_gn=SAMD4B;refGene_tx=XM_006941285.4;refGene_gn=SAMD4B;refGene_tx=XM_023245402.1
E2 14807340 14807464
refGene_gn=SAMD4B;refGene_tx=XM_006941285.4;refGene_gn=SAMD4B;refGene_tx=XM_023245402.1
E2 14807655 14807854
refGene_gn=SAMD4B;refGene_tx=XM_006941285.4;refGene_gn=SAMD4B;refGene_tx=XM_023245402.1
E2 14808421 14808540
refGene_gn=SAMD4B;refGene_tx=XM_006941285.4;refGene_gn=SAMD4B;refGene_tx=XM_023245402.1
E2 14809028 14809114
refGene_gn=SAMD4B;refGene_tx=XM_006941285.4;refGene_gn=SAMD4B;refGene_tx=XM_023245402.1
E2 14809753 14810096
refGene_gn=SAMD4B;refGene_tx=XM_006941285.4;refGene_gn=SAMD4B;refGene_tx=XM_023245402.1
E2 14810801 14810888
refGene_gn=SAMD4B;refGene_tx=XM_006941285.4;refGene_gn=SAMD4B;refGene_tx=XM_023245402.1
E2 14811027 14811137
refGene_gn=SAMD4B;refGene_tx=XM_006941285.4;refGene_gn=SAMD4B;refGene_tx=XM_023245402.1
E2 14811699 14811939
refGene_gn=SAMD4B;refGene_tx=XM_006941285.4;refGene_gn=SAMD4B;refGene_tx=XM_023245402.1
E2 14816282 14816753
refGene_gn=SAMD4B;refGene_tx=XM_006941285.4;refGene_gn=SAMD4B;refGene_tx=XM_023245402.1
E2 14825890 14826086
refGene_gn=SAMD4B;refGene_tx=XM_006941285.4;refGene_gn=SAMD4B;refGene_tx=XM_023245402.1
E2 14841033 14841036 refGene_gn=GMFG;refGene_tx=XM_006941286.4
E2 14845259 14845262 refGene_gn=GMFG;refGene_tx=XM_003997816.5
E2 14845514 14845517 refGene_gn=GMFG;refGene_tx=XM_006941287.4
E2 14845686 14845783

refGene_gn=GMFG;refGene_tx=XM_003997816.5;refGene_gn=GMFG;refGene_tx=XM_006941286.4;refGene_gn=GMFG;refGene_tx=XM_006941287.4;refGene_gn=GMFG;refGene_tx=XM_011289879.3;refGene_gn=GMFG;refGene_tx=XM_019819565.2
E2 14845910 14845960
refGene_gn=GMFG;refGene_tx=XM_003997816.5;refGene_gn=GMFG;refGene_tx=XM_006941286.4;refGene_gn=GMFG;refGene_tx=XM_006941287.4;refGene_gn=GMFG;refGene_tx=XM_011289879.3;refGene_gn=GMFG;refGene_tx=XM_019819565.2
E2 14846780 14846830
refGene_gn=GMFG;refGene_tx=XM_003997816.5;refGene_gn=GMFG;refGene_tx=XM_006941286.4;refGene_gn=GMFG;refGene_tx=XM_006941287.4;refGene_gn=GMFG;refGene_tx=XM_011289879.3;refGene_gn=GMFG;refGene_tx=XM_019819565.2
E2 14848735 14848818
refGene_gn=GMFG;refGene_tx=XM_003997816.5;refGene_gn=GMFG;refGene_tx=XM_006941286.4;refGene_gn=GMFG;refGene_tx=XM_006941287.4;refGene_gn=GMFG;refGene_tx=XM_011289879.3;refGene_gn=GMFG;refGene_tx=XM_019819565.2
E2 14849330 14849404
refGene_gn=GMFG;refGene_tx=XM_003997816.5;refGene_gn=GMFG;refGene_tx=XM_006941286.4;refGene_gn=GMFG;refGene_tx=XM_006941287.4;refGene_gn=GMFG;refGene_tx=XM_011289879.3;refGene_gn=GMFG;refGene_tx=XM_019819565.2
E2 14849586 14849658
refGene_gn=GMFG;refGene_tx=XM_003997816.5;refGene_gn=GMFG;refGene_tx=XM_006941286.4;refGene_gn=GMFG;refGene_tx=XM_006941287.4;refGene_gn=GMFG;refGene_tx=XM_011289879.3;refGene_gn=GMFG;refGene_tx=XM_019819565.2
E2 14860511 14861917
refGene_gn=LRFN1;refGene_tx=XM_019819558.2;refGene_gn=LRFN1;refGene_tx=XM_023245394.1;refGene_gn=LRFN1;refGene_tx=XM_023245395.1;refGene_gn=LRFN1;refGene_tx=XM_023245396.1;refGene_gn=LRFN1;refGene_tx=XM_023245397.1;refGene_gn=LRFN1;refGene_tx=XM_023245399.1;refGene_gn=LRFN1;refGene_tx=XM_023245400.1;refGene_gn=LRFN1;refGene_tx=XM_023245401.1
E2 14865222 14866129
refGene_gn=LRFN1;refGene_tx=XM_019819558.2;refGene_gn=LRFN1;refGene_tx=XM_023245394.1;refGene_gn=LRFN1;refGene_tx=XM_023245395.1;refGene_gn=LRFN1;refGene_tx=XM_023245396.1;refGene_gn=LRFN1;refGene_tx=XM_023245397.1;refGene_gn=LRFN1;refGene_tx=XM_023245399.1;refGene_gn=LRFN1;refGene_tx=XM_023245400.1;refGene_gn=LRFN1;refGene_tx=XM_023245401.1
E2 14874707 14874806
refGene_gn=IFNL1;refGene_tx=XM_006941288.4;refGene_gn=IFNL1;refGene_tx=XM_019818982.2
E2 14874899 14874983
refGene_gn=IFNL1;refGene_tx=XM_006941288.4;refGene_gn=IFNL1;refGene_tx=XM_019818982.2
E2 14875088 14875232
refGene_gn=IFNL1;refGene_tx=XM_006941288.4;refGene_gn=IFNL1;refGene_tx=XM_019818982.2
E2 14875797 14875869
refGene_gn=IFNL1;refGene_tx=XM_006941288.4;refGene_gn=IFNL1;refGene_tx=XM_019818982.2
E2 14876066 14876237
refGene_gn=IFNL1;refGene_tx=XM_006941288.4;refGene_gn=IFNL1;refGene_tx=XM_019818982.2

E2 14885422 14885450 refGene_gn=IFNL1;refGene_tx=XM_019818982.2
E2 14905247 14905285
refGene_gn=LOC102899096;refGene_tx=XM_023245947.1
E2 14905287 14905374
refGene_gn=LOC102899096;refGene_tx=XM_023245947.1
E2 14905483 14905582
refGene_gn=LOC102899096;refGene_tx=XM_023245947.1
E2 14905809 14905984
refGene_gn=LOC102899096;refGene_tx=XM_023245947.1
E2 14906373 14906541
refGene_gn=LOC102899096;refGene_tx=XM_023245947.1
E2 14922085 14922092
refGene_gn=LOC101091743;refGene_tx=XM_023245948.1
E2 14922497 14922674
refGene_gn=LOC101091743;refGene_tx=XM_023245948.1
E2 14922894 14923065
refGene_gn=LOC101091743;refGene_tx=XM_023245948.1
E2 14923299 14923361
refGene_gn=LOC101091743;refGene_tx=XM_023245948.1
E2 14923490 14923631
refGene_gn=LOC101091743;refGene_tx=XM_023245948.1
E2 14923712 14923800
refGene_gn=LOC101091743;refGene_tx=XM_023245948.1
E2 14923873 14923924
refGene_gn=LOC101091743;refGene_tx=XM_023245948.1
E2 14923926 14923970
refGene_gn=LOC101091743;refGene_tx=XM_023245948.1
E2 14931060 14931094
refGene_gn=LOC101091983;refGene_tx=XM_006941349.1
E2 14938918 14939046
refGene_gn=LOC101091983;refGene_tx=XM_006941349.1
E2 14939170 14939325
refGene_gn=LOC101091983;refGene_tx=XM_006941349.1
E2 14939407 14939480
refGene_gn=LOC101091983;refGene_tx=XM_006941349.1
E2 14939566 14939662
refGene_gn=LOC101091983;refGene_tx=XM_006941349.1
E2 14953313 14953708 refGene_gn=SYCN;refGene_tx=XM_003997817.4
E2 14954529 14954539 refGene_gn=SYCN;refGene_tx=XM_003997817.4
E2 14955937 14956078 refGene_gn=NCCRP1;refGene_tx=XM_019819571.2
E2 14956210 14956349 refGene_gn=NCCRP1;refGene_tx=XM_019819571.2
E2 14956940 14957036 refGene_gn=NCCRP1;refGene_tx=XM_019819571.2
E2 14957611 14957663 refGene_gn=NCCRP1;refGene_tx=XM_019819571.2
E2 14957747 14957810 refGene_gn=NCCRP1;refGene_tx=XM_019819571.2
E2 14958413 14958672 refGene_gn=NCCRP1;refGene_tx=XM_019819571.2
E2 14974241 14974397
refGene_gn=PAK4;refGene_tx=XM_011289876.3;refGene_gn=PAK4;refGene_tx=XM_011289877.3;refGene_gn=PAK4;refGene_tx=XM_011289878.3;refGene_gn=PAK4;refGene_tx=XM_019819566.2;refGene_gn=PAK4;refGene_tx=XM_019819567.2;refGene_gn=PAK4;refGene_tx=XM_019819568.2;refGene_gn=PAK4;refGene_tx=X

M_019819570.2
E2 14974828 14974963
refGene_gn=PAK4;refGene_tx=XM_011289876.3;refGene_gn=PAK4;refGene_tx=XM_011289877.3;refGene_gn=PAK4;refGene_tx=XM_011289878.3;refGene_gn=PAK4;refGene_tx=XM_019819566.2;refGene_gn=PAK4;refGene_tx=XM_019819567.2;refGene_gn=PAK4;refGene_tx=XM_019819568.2;refGene_gn=PAK4;refGene_tx=XM_019819570.2
E2 14975795 14975921
refGene_gn=PAK4;refGene_tx=XM_011289876.3;refGene_gn=PAK4;refGene_tx=XM_011289877.3;refGene_gn=PAK4;refGene_tx=XM_011289878.3;refGene_gn=PAK4;refGene_tx=XM_019819566.2;refGene_gn=PAK4;refGene_tx=XM_019819567.2;refGene_gn=PAK4;refGene_tx=XM_019819568.2;refGene_gn=PAK4;refGene_tx=XM_019819570.2
E2 14977069 14977196
refGene_gn=PAK4;refGene_tx=XM_011289876.3;refGene_gn=PAK4;refGene_tx=XM_011289877.3;refGene_gn=PAK4;refGene_tx=XM_011289878.3;refGene_gn=PAK4;refGene_tx=XM_019819566.2;refGene_gn=PAK4;refGene_tx=XM_019819567.2;refGene_gn=PAK4;refGene_tx=XM_019819568.2;refGene_gn=PAK4;refGene_tx=XM_019819570.2
E2 14977414 14977548
refGene_gn=PAK4;refGene_tx=XM_011289876.3;refGene_gn=PAK4;refGene_tx=XM_011289877.3;refGene_gn=PAK4;refGene_tx=XM_011289878.3;refGene_gn=PAK4;refGene_tx=XM_019819566.2;refGene_gn=PAK4;refGene_tx=XM_019819567.2;refGene_gn=PAK4;refGene_tx=XM_019819568.2;refGene_gn=PAK4;refGene_tx=XM_019819570.2
E2 14978879 14979317
refGene_gn=PAK4;refGene_tx=XM_011289876.3;refGene_gn=PAK4;refGene_tx=XM_011289877.3;refGene_gn=PAK4;refGene_tx=XM_011289878.3;refGene_gn=PAK4;refGene_tx=XM_019819566.2;refGene_gn=PAK4;refGene_tx=XM_019819567.2;refGene_gn=PAK4;refGene_tx=XM_019819568.2;refGene_gn=PAK4;refGene_tx=XM_019819570.2
E2 14979531 14979990
refGene_gn=PAK4;refGene_tx=XM_011289876.3;refGene_gn=PAK4;refGene_tx=XM_011289877.3;refGene_gn=PAK4;refGene_tx=XM_011289878.3;refGene_gn=PAK4;refGene_tx=XM_019819566.2;refGene_gn=PAK4;refGene_tx=XM_019819567.2;refGene_gn=PAK4;refGene_tx=XM_019819568.2
E2 14982741 14982945
refGene_gn=PAK4;refGene_tx=XM_011289876.3;refGene_gn=PAK4;refGene_tx=XM_011289877.3;refGene_gn=PAK4;refGene_tx=XM_011289878.3;refGene_gn=PAK4;refGene_tx=XM_019819566.2;refGene_gn=PAK4;refGene_tx=XM_019819567.2;refGene_gn=PAK4;refGene_tx=XM_019819568.2;refGene_gn=PAK4;refGene_tx=XM_019819570.2
E2 15026691 15026757 refGene_gn=ACP7;refGene_tx=XM_023245575.1
E2 15028797 15028935 refGene_gn=ACP7;refGene_tx=XM_023245575.1
E2 15031676 15031748 refGene_gn=ACP7;refGene_tx=XM_023245575.1
E2 15031858 15031926 refGene_gn=ACP7;refGene_tx=XM_023245575.1
E2 15032004 15032062 refGene_gn=ACP7;refGene_tx=XM_023245575.1
E2 15032152 15032260 refGene_gn=ACP7;refGene_tx=XM_023245575.1
E2 15032350 15032465 refGene_gn=ACP7;refGene_tx=XM_023245575.1
E2 15032562 15032625 refGene_gn=ACP7;refGene_tx=XM_023245575.1

E2 15032838 15032962 refGene_gn=ACP7;refGene_tx=XM_023245575.1
E2 15033833 15034016 refGene_gn=ACP7;refGene_tx=XM_023245575.1
E2 15034323 15034524 refGene_gn=ACP7;refGene_tx=XM_023245575.1
E2 15040370 15040491 refGene_gn=ACP7;refGene_tx=XM_023245575.1
E2 15085732 15086084
refGene_gn=FBXO27;refGene_tx=XM_023245387.1;refGene_gn=FBXO27;refGene_tx=XM_023245388.1
E2 15086239 15086351
refGene_gn=FBXO27;refGene_tx=XM_023245387.1;refGene_gn=FBXO27;refGene_tx=XM_023245388.1
E2 15086433 15086529
refGene_gn=FBXO27;refGene_tx=XM_023245387.1;refGene_gn=FBXO27;refGene_tx=XM_023245388.1
E2 15090516 15090652
refGene_gn=FBXO27;refGene_tx=XM_023245387.1;refGene_gn=FBXO27;refGene_tx=XM_023245388.1
E2 15091430 15091574 refGene_gn=FBXO27;refGene_tx=XM_023245388.1
E2 15091617 15091785 refGene_gn=FBXO27;refGene_tx=XM_023245387.1
E2 15146368 15146714
refGene_gn=FBXO17;refGene_tx=XM_023245292.1;refGene_gn=FBXO17;refGene_tx=XM_023245293.1;refGene_gn=FBXO17;refGene_tx=XM_023245294.1;refGene_gn=FBXO17;refGene_tx=XM_023245295.1
E2 15147516 15147628
refGene_gn=FBXO17;refGene_tx=XM_023245292.1;refGene_gn=FBXO17;refGene_tx=XM_023245293.1;refGene_gn=FBXO17;refGene_tx=XM_023245294.1;refGene_gn=FBXO17;refGene_tx=XM_023245295.1
E2 15149329 15149425
refGene_gn=FBXO17;refGene_tx=XM_023245292.1;refGene_gn=FBXO17;refGene_tx=XM_023245293.1;refGene_gn=FBXO17;refGene_tx=XM_023245294.1;refGene_gn=FBXO17;refGene_tx=XM_023245295.1
E2 15150205 15150341
refGene_gn=FBXO17;refGene_tx=XM_023245292.1;refGene_gn=FBXO17;refGene_tx=XM_023245293.1;refGene_gn=FBXO17;refGene_tx=XM_023245294.1;refGene_gn=FBXO17;refGene_tx=XM_023245295.1
E2 15151137 15151281
refGene_gn=FBXO17;refGene_tx=XM_023245292.1;refGene_gn=FBXO17;refGene_tx=XM_023245293.1;refGene_gn=FBXO17;refGene_tx=XM_023245294.1;refGene_gn=FBXO17;refGene_tx=XM_023245295.1
E2 15153333 15153704
refGene_gn=MRPS12;refGene_tx=XM_003997820.5;refGene_gn=MRPS12;refGene_tx=XM_023245197.1
E2 15157105 15157154
refGene_gn=MRPS12;refGene_tx=XM_003997820.5;refGene_gn=MRPS12;refGene_tx=XM_023245197.1
E2 15157745 15158012 refGene_gn=SARS2;refGene_tx=XM_023245949.1
E2 15160664 15160760 refGene_gn=SARS2;refGene_tx=XM_023245949.1
E2 15161884 15161914 refGene_gn=SARS2;refGene_tx=XM_023245949.1
E2 15162445 15162625 refGene_gn=SARS2;refGene_tx=XM_023245949.1
E2 15162668 15162710 refGene_gn=SARS2;refGene_tx=XM_023245949.1
E2 15163582 15163896 refGene_gn=SARS2;refGene_tx=XM_023245949.1

E2	15164110	15164216	refGene_gn=SARS2;refGene_tx=XM_023245949.1
E2	15164882	15164930	refGene_gn=SARS2;refGene_tx=XM_023245949.1
E2	15165141	15165250	refGene_gn=SARS2;refGene_tx=XM_023245949.1
E2	15165806	15165852	refGene_gn=SARS2;refGene_tx=XM_023245949.1
E2	15166198	15166286	refGene_gn=SARS2;refGene_tx=XM_023245949.1
E2	15166370	15166480	refGene_gn=SARS2;refGene_tx=XM_023245949.1
E2	15167185	15167279	refGene_gn=SARS2;refGene_tx=XM_023245949.1
E2	15167369	15167462	refGene_gn=SARS2;refGene_tx=XM_023245949.1
E2	15167582	15167648	refGene_gn=SARS2;refGene_tx=XM_023245949.1
E2	15167734	15167878	refGene_gn=SARS2;refGene_tx=XM_023245949.1
E2	15170178	15170254	refGene_gn=CCER2;refGene_tx=XM_023245204.1
E2	15170490	15170531	refGene_gn=CCER2;refGene_tx=XM_023245204.1
E2	15170685	15170773	refGene_gn=CCER2;refGene_tx=XM_023245204.1
E2	15171101	15171631	refGene_gn=CCER2;refGene_tx=XM_023245204.1
E2	15172274	15172364	refGene_gn=CCER2;refGene_tx=XM_023245204.1
E2	15172719	15172815
refGene_gn=NFKBIB;refGene_tx=XM_004001636.5;refGene_gn=NFKBIB;refGene_tx=XM_023245203.1
E2	15173533	15173794
refGene_gn=NFKBIB;refGene_tx=XM_004001636.5;refGene_gn=NFKBIB;refGene_tx=XM_023245203.1
E2	15173883	15173972
refGene_gn=NFKBIB;refGene_tx=XM_004001636.5;refGene_gn=NFKBIB;refGene_tx=XM_023245203.1
E2	15174853	15175187
refGene_gn=NFKBIB;refGene_tx=XM_004001636.5;refGene_gn=NFKBIB;refGene_tx=XM_023245203.1
E2	15175266	15175372
refGene_gn=NFKBIB;refGene_tx=XM_004001636.5;refGene_gn=NFKBIB;refGene_tx=XM_023245203.1
E2	15181444	15181623
refGene_gn=NFKBIB;refGene_tx=XM_004001636.5;refGene_gn=NFKBIB;refGene_tx=XM_023245203.1
E2	15181965	15182091
refGene_gn=SIRT2;refGene_tx=XM_003997822.5;refGene_gn=SIRT2;refGene_tx=XM_006941289.4;refGene_gn=SIRT2;refGene_tx=XM_006941290.3;refGene_gn=SIRT2;refGene_tx=XM_006941291.4;refGene_gn=SIRT2;refGene_tx=XM_011289873.2;refGene_gn=SIRT2;refGene_tx=XM_011289874.2
E2	15182968	15183015
refGene_gn=SIRT2;refGene_tx=XM_003997822.5;refGene_gn=SIRT2;refGene_tx=XM_006941289.4;refGene_gn=SIRT2;refGene_tx=XM_006941291.4
E2	15185588	15185657
refGene_gn=SIRT2;refGene_tx=XM_006941289.4;refGene_gn=SIRT2;refGene_tx=XM_006941290.3;refGene_gn=SIRT2;refGene_tx=XM_011289873.2
E2	15185971	15186020
refGene_gn=SIRT2;refGene_tx=XM_003997822.5;refGene_gn=SIRT2;refGene_tx=XM_006941289.4;refGene_gn=SIRT2;refGene_tx=XM_006941290.3;refGene_gn=SIRT2;refGene_tx=XM_006941291.4;refGene_gn=SIRT2;refGene_tx=XM_006941292.3;refGene_gn=SIRT2;refGene_tx=XM_011289873.2;refGene_gn=SIRT2;refGene_tx=XM_011289874.2

E2 15186248 15186362
refGene_gn=SIRT2;refGene_tx=XM_003997822.5;refGene_gn=SIRT2;refGene_tx=XM_006941289.4;refGene_gn=SIRT2;refGene_tx=XM_006941290.3;refGene_gn=SIRT2;refGene_tx=XM_006941291.4;refGene_gn=SIRT2;refGene_tx=XM_006941292.3;refGene_gn=SIRT2;refGene_tx=XM_011289873.2;refGene_gn=SIRT2;refGene_tx=XM_011289874.2
E2 15189083 15189125
refGene_gn=SIRT2;refGene_tx=XM_003997822.5;refGene_gn=SIRT2;refGene_tx=XM_006941289.4;refGene_gn=SIRT2;refGene_tx=XM_006941290.3;refGene_gn=SIRT2;refGene_tx=XM_006941291.4;refGene_gn=SIRT2;refGene_tx=XM_006941292.3;refGene_gn=SIRT2;refGene_tx=XM_011289873.2;refGene_gn=SIRT2;refGene_tx=XM_011289874.2
E2 15189278 15189385
refGene_gn=SIRT2;refGene_tx=XM_003997822.5;refGene_gn=SIRT2;refGene_tx=XM_006941289.4;refGene_gn=SIRT2;refGene_tx=XM_006941290.3;refGene_gn=SIRT2;refGene_tx=XM_006941291.4;refGene_gn=SIRT2;refGene_tx=XM_006941292.3;refGene_gn=SIRT2;refGene_tx=XM_011289873.2;refGene_gn=SIRT2;refGene_tx=XM_011289874.2
E2 15189487 15189544
refGene_gn=SIRT2;refGene_tx=XM_003997822.5;refGene_gn=SIRT2;refGene_tx=XM_006941289.4;refGene_gn=SIRT2;refGene_tx=XM_006941290.3;refGene_gn=SIRT2;refGene_tx=XM_006941291.4;refGene_gn=SIRT2;refGene_tx=XM_006941292.3;refGene_gn=SIRT2;refGene_tx=XM_011289873.2;refGene_gn=SIRT2;refGene_tx=XM_011289874.2
E2 15190042 15190111
refGene_gn=SIRT2;refGene_tx=XM_003997822.5;refGene_gn=SIRT2;refGene_tx=XM_006941289.4;refGene_gn=SIRT2;refGene_tx=XM_006941290.3;refGene_gn=SIRT2;refGene_tx=XM_006941291.4;refGene_gn=SIRT2;refGene_tx=XM_006941292.3;refGene_gn=SIRT2;refGene_tx=XM_011289873.2;refGene_gn=SIRT2;refGene_tx=XM_011289874.2
E2 15194799 15194929
refGene_gn=SIRT2;refGene_tx=XM_003997822.5;refGene_gn=SIRT2;refGene_tx=XM_006941289.4;refGene_gn=SIRT2;refGene_tx=XM_006941290.3;refGene_gn=SIRT2;refGene_tx=XM_006941291.4;refGene_gn=SIRT2;refGene_tx=XM_006941292.3;refGene_gn=SIRT2;refGene_tx=XM_011289873.2;refGene_gn=SIRT2;refGene_tx=XM_011289874.2
E2 15195436 15195511
refGene_gn=SIRT2;refGene_tx=XM_006941289.4;refGene_gn=SIRT2;refGene_tx=XM_006941291.4;refGene_gn=SIRT2;refGene_tx=XM_006941292.3;refGene_gn=SIRT2;refGene_tx=XM_011289873.2;refGene_gn=SIRT2;refGene_tx=XM_011289874.2
E2 15196637 15196697
refGene_gn=SIRT2;refGene_tx=XM_003997822.5;refGene_gn=SIRT2;refGene_tx=XM_006941289.4;refGene_gn=SIRT2;refGene_tx=XM_006941290.3;refGene_gn=SIRT2;refGene_tx=XM_006941291.4;refGene_gn=SIRT2;refGene_tx=XM_006941292.3;refGene_gn=SIRT2;refGene_tx=XM_011289873.2;refGene_gn=SIRT2;refGene_tx=XM_011289874.2
E2 15196963 15197019
refGene_gn=SIRT2;refGene_tx=XM_003997822.5;refGene_gn=SIRT2;refGene_tx=XM_006941289.4;refGene_gn=SIRT2;refGene_tx=XM_006941290.3;refGene_gn=

SIRT2;refGene_tx=XM_006941291.4;refGene_gn=SIRT2;refGene_tx=XM_0069412
92.3;refGene_gn=SIRT2;refGene_tx=XM_011289873.2;refGene_gn=SIRT2;refGe
ne_tx=XM_011289874.2
E2 15197199 15197276
refGene_gn=SIRT2;refGene_tx=XM_003997822.5;refGene_gn=SIRT2;refGene_tx
=XM_006941289.4;refGene_gn=SIRT2;refGene_tx=XM_006941290.3;refGene_gn=
SIRT2;refGene_tx=XM_006941291.4;refGene_gn=SIRT2;refGene_tx=XM_0069412
92.3;refGene_gn=SIRT2;refGene_tx=XM_011289873.2;refGene_gn=SIRT2;refGe
ne_tx=XM_011289874.2
E2 15197362 15197414
refGene_gn=SIRT2;refGene_tx=XM_003997822.5;refGene_gn=SIRT2;refGene_tx
=XM_006941289.4;refGene_gn=SIRT2;refGene_tx=XM_006941290.3;refGene_gn=
SIRT2;refGene_tx=XM_006941291.4;refGene_gn=SIRT2;refGene_tx=XM_0069412
92.3;refGene_gn=SIRT2;refGene_tx=XM_011289873.2;refGene_gn=SIRT2;refGe
ne_tx=XM_011289874.2
E2 15198611 15198682
refGene_gn=SIRT2;refGene_tx=XM_003997822.5;refGene_gn=SIRT2;refGene_tx
=XM_006941289.4;refGene_gn=SIRT2;refGene_tx=XM_006941290.3;refGene_gn=
SIRT2;refGene_tx=XM_006941291.4;refGene_gn=SIRT2;refGene_tx=XM_0069412
92.3;refGene_gn=SIRT2;refGene_tx=XM_011289873.2;refGene_gn=SIRT2;refGe
ne_tx=XM_011289874.2
E2 15198804 15198871
refGene_gn=SIRT2;refGene_tx=XM_003997822.5;refGene_gn=SIRT2;refGene_tx
=XM_006941289.4;refGene_gn=SIRT2;refGene_tx=XM_006941290.3;refGene_gn=
SIRT2;refGene_tx=XM_006941291.4;refGene_gn=SIRT2;refGene_tx=XM_0069412
92.3;refGene_gn=SIRT2;refGene_tx=XM_011289873.2;refGene_gn=SIRT2;refGe
ne_tx=XM_011289874.2
E2 15198990 15199149
refGene_gn=SIRT2;refGene_tx=XM_003997822.5;refGene_gn=SIRT2;refGene_tx
=XM_006941289.4;refGene_gn=SIRT2;refGene_tx=XM_006941290.3;refGene_gn=
SIRT2;refGene_tx=XM_006941291.4;refGene_gn=SIRT2;refGene_tx=XM_0069412
92.3;refGene_gn=SIRT2;refGene_tx=XM_011289873.2;refGene_gn=SIRT2;refGe
ne_tx=XM_011289874.2
E2 15201854 15201904
refGene_gn=RINL;refGene_tx=XM_019819575.2;refGene_gn=RINL;refGene_tx=X
M_019819576.2
E2 15202107 15202267
refGene_gn=RINL;refGene_tx=XM_019819575.2;refGene_gn=RINL;refGene_tx=X
M_019819576.2
E2 15203593 15203696
refGene_gn=RINL;refGene_tx=XM_019819575.2;refGene_gn=RINL;refGene_tx=X
M_019819576.2
E2 15203961 15204034
refGene_gn=RINL;refGene_tx=XM_019819575.2;refGene_gn=RINL;refGene_tx=X
M_019819576.2
E2 15204118 15204183
refGene_gn=RINL;refGene_tx=XM_019819575.2;refGene_gn=RINL;refGene_tx=X
M_019819576.2
E2 15204440 15204590
refGene_gn=RINL;refGene_tx=XM_019819575.2;refGene_gn=RINL;refGene_tx=X

M_019819576.2
E2	15204681	15205107
refGene_gn=RINL;refGene_tx=XM_019819575.2;refGene_gn=RINL;refGene_tx=XM_019819576.2
E2	15205480	15205774
refGene_gn=RINL;refGene_tx=XM_019819575.2;refGene_gn=RINL;refGene_tx=XM_019819576.2
E2	15206029	15206161
refGene_gn=RINL;refGene_tx=XM_019819575.2;refGene_gn=RINL;refGene_tx=XM_019819576.2
E2	15206309	15206479
refGene_gn=RINL;refGene_tx=XM_019819575.2;refGene_gn=RINL;refGene_tx=XM_019819576.2
E2	15206561	15206621
refGene_gn=RINL;refGene_tx=XM_019819575.2;refGene_gn=RINL;refGene_tx=XM_019819576.2
E2	15237541	15237805
refGene_gn=HNRNPL;refGene_tx=XM_003997824.3;refGene_gn=HNRNPL;refGene_tx=XM_006941293.2;refGene_gn=HNRNPL;refGene_tx=XM_019819573.1
E2	15239785	15239904
refGene_gn=HNRNPL;refGene_tx=XM_003997824.3;refGene_gn=HNRNPL;refGene_tx=XM_006941293.2;refGene_gn=HNRNPL;refGene_tx=XM_019819573.1
E2	15240516	15240754
refGene_gn=HNRNPL;refGene_tx=XM_003997824.3;refGene_gn=HNRNPL;refGene_tx=XM_006941293.2;refGene_gn=HNRNPL;refGene_tx=XM_019819573.1;refGene_gn=HNRNPL;refGene_tx=XM_019819574.2;refGene_gn=HNRNPL;refGene_tx=XM_023245202.1
E2	15240878	15240964
refGene_gn=HNRNPL;refGene_tx=XM_003997824.3;refGene_gn=HNRNPL;refGene_tx=XM_006941293.2;refGene_gn=HNRNPL;refGene_tx=XM_019819573.1;refGene_gn=HNRNPL;refGene_tx=XM_019819574.2;refGene_gn=HNRNPL;refGene_tx=XM_023245202.1
E2	15241788	15241885
refGene_gn=HNRNPL;refGene_tx=XM_003997824.3;refGene_gn=HNRNPL;refGene_tx=XM_006941293.2;refGene_gn=HNRNPL;refGene_tx=XM_019819573.1;refGene_gn=HNRNPL;refGene_tx=XM_019819574.2;refGene_gn=HNRNPL;refGene_tx=XM_023245202.1
E2	15241968	15242041
refGene_gn=HNRNPL;refGene_tx=XM_003997824.3;refGene_gn=HNRNPL;refGene_tx=XM_006941293.2;refGene_gn=HNRNPL;refGene_tx=XM_019819573.1;refGene_gn=HNRNPL;refGene_tx=XM_019819574.2;refGene_gn=HNRNPL;refGene_tx=XM_023245202.1
E2	15245439	15245903
refGene_gn=HNRNPL;refGene_tx=XM_003997824.3;refGene_gn=HNRNPL;refGene_tx=XM_006941293.2;refGene_gn=HNRNPL;refGene_tx=XM_019819573.1;refGene_gn=HNRNPL;refGene_tx=XM_019819574.2;refGene_gn=HNRNPL;refGene_tx=XM_023245202.1
E2	15246973	15247095
refGene_gn=HNRNPL;refGene_tx=XM_003997824.3;refGene_gn=HNRNPL;refGene_tx=XM_006941293.2;refGene_gn=HNRNPL;refGene_tx=XM_019819573.1;refGene_

gn=HNRNPL;refGene_tx=XM_019819574.2;refGene_gn=HNRNPL;refGene_tx=XM_02
3245202.1
E2	15247399	15247601
refGene_gn=HNRNPL;refGene_tx=XM_003997824.3;refGene_gn=HNRNPL;refGene_
tx=XM_006941293.2;refGene_gn=HNRNPL;refGene_tx=XM_019819573.1;refGene_
gn=HNRNPL;refGene_tx=XM_019819574.2;refGene_gn=HNRNPL;refGene_tx=XM_02
3245202.1
E2	15248307	15248365
refGene_gn=HNRNPL;refGene_tx=XM_003997824.3;refGene_gn=HNRNPL;refGene_
tx=XM_006941293.2;refGene_gn=HNRNPL;refGene_tx=XM_019819573.1;refGene_
gn=HNRNPL;refGene_tx=XM_019819574.2;refGene_gn=HNRNPL;refGene_tx=XM_02
3245202.1
E2	15248474	15248570
refGene_gn=HNRNPL;refGene_tx=XM_003997824.3;refGene_gn=HNRNPL;refGene_
tx=XM_006941293.2;refGene_gn=HNRNPL;refGene_tx=XM_019819573.1;refGene_
gn=HNRNPL;refGene_tx=XM_019819574.2;refGene_gn=HNRNPL;refGene_tx=XM_02
3245202.1
E2	15249146	15249205
refGene_gn=HNRNPL;refGene_tx=XM_003997824.3;refGene_gn=HNRNPL;refGene_
tx=XM_006941293.2;refGene_gn=HNRNPL;refGene_tx=XM_019819573.1;refGene_
gn=HNRNPL;refGene_tx=XM_019819574.2;refGene_gn=HNRNPL;refGene_tx=XM_02
3245202.1
E2	15252755	15252795	refGene_gn=ECH1;refGene_tx=XM_003997877.3
E2	15253452	15253648	refGene_gn=ECH1;refGene_tx=XM_003997877.3
E2	15253786	15253875	refGene_gn=ECH1;refGene_tx=XM_003997877.3
E2	15258571	15258696	refGene_gn=ECH1;refGene_tx=XM_003997877.3
E2	15258783	15258832	refGene_gn=ECH1;refGene_tx=XM_003997877.3
E2	15259032	15259097	refGene_gn=ECH1;refGene_tx=XM_003997877.3
E2	15259731	15259802	refGene_gn=ECH1;refGene_tx=XM_003997877.3
E2	15259894	15259966	refGene_gn=ECH1;refGene_tx=XM_003997877.3
E2	15260214	15260365	refGene_gn=ECH1;refGene_tx=XM_003997877.3
E2	15260545	15260650	refGene_gn=ECH1;refGene_tx=XM_003997877.3
E2	15262295	15262340	refGene_gn=LGALS4;refGene_tx=XM_003997825.3
E2	15262640	15262729	refGene_gn=LGALS4;refGene_tx=XM_003997825.3
E2	15264052	15264257	refGene_gn=LGALS4;refGene_tx=XM_003997825.3
E2	15266331	15266466	refGene_gn=LGALS4;refGene_tx=XM_003997825.3
E2	15267624	15267663	refGene_gn=LGALS4;refGene_tx=XM_003997825.3
E2	15267854	15267884	refGene_gn=LGALS4;refGene_tx=XM_003997825.3
E2	15268162	15268251	refGene_gn=LGALS4;refGene_tx=XM_003997825.3
E2	15268387	15268553	refGene_gn=LGALS4;refGene_tx=XM_003997825.3
E2	15268634	15268781	refGene_gn=LGALS4;refGene_tx=XM_003997825.3
E2	15278026	15278140
refGene_gn=LOC101100430;refGene_tx=XM_023245548.1
E2	15278695	15279157
refGene_gn=LOC101100430;refGene_tx=XM_023245548.1
E2	15302006	15302243	refGene_gn=CAPN12;refGene_tx=XM_023245391.1
E2	15302566	15302636	refGene_gn=CAPN12;refGene_tx=XM_023245391.1
E2	15302767	15302886	refGene_gn=CAPN12;refGene_tx=XM_023245391.1
E2	15303050	15303184	refGene_gn=CAPN12;refGene_tx=XM_023245391.1
E2	15303929	15304098	refGene_gn=CAPN12;refGene_tx=XM_023245391.1

E2 15305127 15305202 refGene_gn=CAPN12;refGene_tx=XM_023245391.1
E2 15305272 15305358 refGene_gn=CAPN12;refGene_tx=XM_023245391.1
E2 15305428 15305503 refGene_gn=CAPN12;refGene_tx=XM_023245391.1
E2 15306128 15306232 refGene_gn=CAPN12;refGene_tx=XM_023245391.1
E2 15306234 15306291 refGene_gn=CAPN12;refGene_tx=XM_023245391.1
E2 15306387 15306620 refGene_gn=CAPN12;refGene_tx=XM_023245391.1
E2 15306907 15306919 refGene_gn=CAPN12;refGene_tx=XM_023245391.1
E2 15307049 15307258 refGene_gn=CAPN12;refGene_tx=XM_023245391.1
E2 15307393 15307436 refGene_gn=CAPN12;refGene_tx=XM_023245391.1
E2 15307532 15307592 refGene_gn=CAPN12;refGene_tx=XM_023245391.1
E2 15308102 15308160 refGene_gn=CAPN12;refGene_tx=XM_023245391.1
E2 15308537 15308608 refGene_gn=CAPN12;refGene_tx=XM_023245391.1
E2 15308708 15308771 refGene_gn=CAPN12;refGene_tx=XM_023245391.1
E2 15309380 15309459 refGene_gn=CAPN12;refGene_tx=XM_023245391.1
E2 15312414 15312531 refGene_gn=CAPN12;refGene_tx=XM_023245391.1
E2 15312617 15312676 refGene_gn=CAPN12;refGene_tx=XM_023245391.1
E2 15312769 15312796 refGene_gn=CAPN12;refGene_tx=XM_023245391.1
E2 15314499 15314658
refGene_gn=ACTN4;refGene_tx=XM_011289869.2;refGene_gn=ACTN4;refGene_tx=XM_019819580.2;refGene_gn=ACTN4;refGene_tx=XM_019819581.2;refGene_gn=ACTN4;refGene_tx=XM_019819582.2;refGene_gn=ACTN4;refGene_tx=XM_019819583.2
E2 15314795 15314954
refGene_gn=ACTN4;refGene_tx=XM_011289869.2;refGene_gn=ACTN4;refGene_tx=XM_019819580.2;refGene_gn=ACTN4;refGene_tx=XM_019819581.2;refGene_gn=ACTN4;refGene_tx=XM_019819582.2;refGene_gn=ACTN4;refGene_tx=XM_019819583.2
E2 15315536 15315602
refGene_gn=ACTN4;refGene_tx=XM_019819581.2;refGene_gn=ACTN4;refGene_tx=XM_019819582.2;refGene_gn=ACTN4;refGene_tx=XM_019819583.2
E2 15315909 15315990
refGene_gn=ACTN4;refGene_tx=XM_011289869.2;refGene_gn=ACTN4;refGene_tx=XM_019819580.2;refGene_gn=ACTN4;refGene_tx=XM_019819583.2
E2 15316827 15316974
refGene_gn=ACTN4;refGene_tx=XM_011289869.2;refGene_gn=ACTN4;refGene_tx=XM_019819580.2;refGene_gn=ACTN4;refGene_tx=XM_019819581.2;refGene_gn=ACTN4;refGene_tx=XM_019819582.2;refGene_gn=ACTN4;refGene_tx=XM_019819583.2
E2 15317682 15317862
refGene_gn=ACTN4;refGene_tx=XM_011289869.2;refGene_gn=ACTN4;refGene_tx=XM_019819580.2;refGene_gn=ACTN4;refGene_tx=XM_019819581.2;refGene_gn=ACTN4;refGene_tx=XM_019819582.2;refGene_gn=ACTN4;refGene_tx=XM_019819583.2
E2 15318621 15318756
refGene_gn=ACTN4;refGene_tx=XM_011289869.2;refGene_gn=ACTN4;refGene_tx=XM_019819580.2;refGene_gn=ACTN4;refGene_tx=XM_019819581.2;refGene_gn=ACTN4;refGene_tx=XM_019819582.2;refGene_gn=ACTN4;refGene_tx=XM_019819583.2
E2 15318866 15319049
refGene_gn=ACTN4;refGene_tx=XM_011289869.2;refGene_gn=ACTN4;refGene_tx

=XM_019819580.2;refGene_gn=ACTN4;refGene_tx=XM_019819581.2;refGene_gn=ACTN4;refGene_tx=XM_019819582.2;refGene_gn=ACTN4;refGene_tx=XM_019819583.2
E2 15319123 15319264
refGene_gn=ACTN4;refGene_tx=XM_011289869.2;refGene_gn=ACTN4;refGene_tx=XM_019819580.2;refGene_gn=ACTN4;refGene_tx=XM_019819581.2;refGene_gn=ACTN4;refGene_tx=XM_019819582.2;refGene_gn=ACTN4;refGene_tx=XM_019819583.2
E2 15319452 15319561
refGene_gn=ACTN4;refGene_tx=XM_011289869.2;refGene_gn=ACTN4;refGene_tx=XM_019819580.2;refGene_gn=ACTN4;refGene_tx=XM_019819581.2;refGene_gn=ACTN4;refGene_tx=XM_019819582.2;refGene_gn=ACTN4;refGene_tx=XM_019819583.2
E2 15322075 15322226
refGene_gn=ACTN4;refGene_tx=XM_011289869.2;refGene_gn=ACTN4;refGene_tx=XM_019819580.2;refGene_gn=ACTN4;refGene_tx=XM_019819581.2;refGene_gn=ACTN4;refGene_tx=XM_019819582.2;refGene_gn=ACTN4;refGene_tx=XM_019819583.2
E2 15325545 15325693
refGene_gn=ACTN4;refGene_tx=XM_011289869.2;refGene_gn=ACTN4;refGene_tx=XM_019819580.2;refGene_gn=ACTN4;refGene_tx=XM_019819581.2;refGene_gn=ACTN4;refGene_tx=XM_019819582.2;refGene_gn=ACTN4;refGene_tx=XM_019819583.2
E2 15326269 15326500
refGene_gn=ACTN4;refGene_tx=XM_011289869.2;refGene_gn=ACTN4;refGene_tx=XM_019819580.2;refGene_gn=ACTN4;refGene_tx=XM_019819581.2;refGene_gn=ACTN4;refGene_tx=XM_019819582.2;refGene_gn=ACTN4;refGene_tx=XM_019819583.2
E2 15328122 15328215
refGene_gn=ACTN4;refGene_tx=XM_011289869.2;refGene_gn=ACTN4;refGene_tx=XM_019819580.2;refGene_gn=ACTN4;refGene_tx=XM_019819581.2;refGene_gn=ACTN4;refGene_tx=XM_019819582.2;refGene_gn=ACTN4;refGene_tx=XM_019819583.2
E2 15331458 15331544
refGene_gn=ACTN4;refGene_tx=XM_019819580.2;refGene_gn=ACTN4;refGene_tx=XM_019819582.2
E2 15332776 15332862
refGene_gn=ACTN4;refGene_tx=XM_011289869.2;refGene_gn=ACTN4;refGene_tx=XM_019819581.2;refGene_gn=ACTN4;refGene_tx=XM_019819583.2
E2 15333636 15333718
refGene_gn=ACTN4;refGene_tx=XM_011289869.2;refGene_gn=ACTN4;refGene_tx=XM_019819580.2;refGene_gn=ACTN4;refGene_tx=XM_019819581.2;refGene_gn=ACTN4;refGene_tx=XM_019819582.2;refGene_gn=ACTN4;refGene_tx=XM_019819583.2
E2 15334907 15334986
refGene_gn=ACTN4;refGene_tx=XM_011289869.2;refGene_gn=ACTN4;refGene_tx=XM_019819580.2;refGene_gn=ACTN4;refGene_tx=XM_019819581.2;refGene_gn=ACTN4;refGene_tx=XM_019819582.2;refGene_gn=ACTN4;refGene_tx=XM_019819583.2
E2 15335603 15335691

refGene_gn=ACTN4;refGene_tx=XM_011289869.2;refGene_gn=ACTN4;refGene_tx=XM_019819580.2;refGene_gn=ACTN4;refGene_tx=XM_019819581.2;refGene_gn=ACTN4;refGene_tx=XM_019819582.2;refGene_gn=ACTN4;refGene_tx=XM_019819583.2
E2 15336726 15336813
refGene_gn=ACTN4;refGene_tx=XM_011289869.2;refGene_gn=ACTN4;refGene_tx=XM_019819580.2;refGene_gn=ACTN4;refGene_tx=XM_019819581.2;refGene_gn=ACTN4;refGene_tx=XM_019819582.2;refGene_gn=ACTN4;refGene_tx=XM_019819583.2
E2 15340253 15340373
refGene_gn=ACTN4;refGene_tx=XM_011289869.2;refGene_gn=ACTN4;refGene_tx=XM_019819580.2;refGene_gn=ACTN4;refGene_tx=XM_019819581.2;refGene_gn=ACTN4;refGene_tx=XM_019819582.2;refGene_gn=ACTN4;refGene_tx=XM_019819583.2
E2 15340630 15340745
refGene_gn=ACTN4;refGene_tx=XM_011289869.2;refGene_gn=ACTN4;refGene_tx=XM_019819580.2;refGene_gn=ACTN4;refGene_tx=XM_019819581.2;refGene_gn=ACTN4;refGene_tx=XM_019819582.2;refGene_gn=ACTN4;refGene_tx=XM_019819583.2
E2 15384052 15384214
refGene_gn=ACTN4;refGene_tx=XM_011289869.2;refGene_gn=ACTN4;refGene_tx=XM_019819580.2;refGene_gn=ACTN4;refGene_tx=XM_019819581.2;refGene_gn=ACTN4;refGene_tx=XM_019819582.2;refGene_gn=ACTN4;refGene_tx=XM_019819583.2
E2 15389736 15389768
refGene_gn=EIF3K;refGene_tx=XM_003997829.5;refGene_gn=EIF3K;refGene_tx=XM_006941299.3
E2 15390968 15391402
refGene_gn=EIF3K;refGene_tx=XM_003997829.5;refGene_gn=EIF3K;refGene_tx=XM_006941299.3;refGene_gn=EIF3K;refGene_tx=XM_019819584.2;refGene_gn=EIF3K;refGene_tx=XM_019819585.2
E2 15393410 15393488
refGene_gn=EIF3K;refGene_tx=XM_003997829.5;refGene_gn=EIF3K;refGene_tx=XM_006941299.3;refGene_gn=EIF3K;refGene_tx=XM_019819584.2;refGene_gn=EIF3K;refGene_tx=XM_019819585.2
E2 15393591 15393658
refGene_gn=EIF3K;refGene_tx=XM_003997829.5;refGene_gn=EIF3K;refGene_tx=XM_006941299.3;refGene_gn=EIF3K;refGene_tx=XM_019819584.2;refGene_gn=EIF3K;refGene_tx=XM_019819585.2
E2 15396110 15396191
refGene_gn=EIF3K;refGene_tx=XM_003997829.5;refGene_gn=EIF3K;refGene_tx=XM_006941299.3;refGene_gn=EIF3K;refGene_tx=XM_019819584.2;refGene_gn=EIF3K;refGene_tx=XM_019819585.2
E2 15397859 15397980
refGene_gn=EIF3K;refGene_tx=XM_003997829.5;refGene_gn=EIF3K;refGene_tx=XM_006941299.3;refGene_gn=EIF3K;refGene_tx=XM_019819584.2;refGene_gn=EIF3K;refGene_tx=XM_019819585.2
E2 15399815 15399914
refGene_gn=EIF3K;refGene_tx=XM_003997829.5;refGene_gn=EIF3K;refGene_tx=XM_006941299.3;refGene_gn=EIF3K;refGene_tx=XM_019819584.2;refGene_gn=

EIF3K;refGene_tx=XM_019819585.2
E2	15400763	15400822
refGene_gn=EIF3K;refGene_tx=XM_003997829.5;refGene_gn=EIF3K;refGene_tx
=XM_006941299.3;refGene_gn=EIF3K;refGene_tx=XM_019819584.2;refGene_gn=
EIF3K;refGene_tx=XM_019819585.2
E2	15402153	15402252	refGene_gn=MAP4K1;refGene_tx=XM_011289867.3
E2	15402421	15402479	refGene_gn=MAP4K1;refGene_tx=XM_011289867.3
E2	15402598	15402689	refGene_gn=MAP4K1;refGene_tx=XM_011289867.3
E2	15403409	15403474	refGene_gn=MAP4K1;refGene_tx=XM_011289867.3
E2	15405021	15405077	refGene_gn=MAP4K1;refGene_tx=XM_011289867.3
E2	15405166	15405214	refGene_gn=MAP4K1;refGene_tx=XM_011289867.3
E2	15405397	15405440	refGene_gn=MAP4K1;refGene_tx=XM_011289867.3
E2	15405530	15405603	refGene_gn=MAP4K1;refGene_tx=XM_011289867.3
E2	15406219	15406351	refGene_gn=MAP4K1;refGene_tx=XM_011289867.3
E2	15406773	15406836	refGene_gn=MAP4K1;refGene_tx=XM_011289867.3
E2	15406943	15407025	refGene_gn=MAP4K1;refGene_tx=XM_011289867.3
E2	15407443	15407458	refGene_gn=MAP4K1;refGene_tx=XM_011289867.3
E2	15407459	15407561	refGene_gn=MAP4K1;refGene_tx=XM_011289867.3
E2	15407807	15407886	refGene_gn=MAP4K1;refGene_tx=XM_011289867.3
E2	15408414	15408473	refGene_gn=MAP4K1;refGene_tx=XM_011289867.3
E2	15408549	15408593	refGene_gn=MAP4K1;refGene_tx=XM_011289867.3
E2	15408673	15408718	refGene_gn=MAP4K1;refGene_tx=XM_011289867.3
E2	15409360	15409403	refGene_gn=MAP4K1;refGene_tx=XM_011289867.3
E2	15410081	15410244	refGene_gn=MAP4K1;refGene_tx=XM_011289867.3
E2	15410331	15410414	refGene_gn=MAP4K1;refGene_tx=XM_011289867.3
E2	15410807	15410892	refGene_gn=MAP4K1;refGene_tx=XM_011289867.3
E2	15412087	15412164	refGene_gn=MAP4K1;refGene_tx=XM_011289867.3
E2	15412253	15412314	refGene_gn=MAP4K1;refGene_tx=XM_011289867.3
E2	15413105	15413214	refGene_gn=MAP4K1;refGene_tx=XM_011289867.3
E2	15413298	15413357	refGene_gn=MAP4K1;refGene_tx=XM_011289867.3
E2	15413566	15413670	refGene_gn=MAP4K1;refGene_tx=XM_011289867.3
E2	15414246	15414421	refGene_gn=MAP4K1;refGene_tx=XM_011289867.3
E2	15414683	15414746	refGene_gn=MAP4K1;refGene_tx=XM_011289867.3
E2	15414938	15415028	refGene_gn=MAP4K1;refGene_tx=XM_011289867.3
E2	15415111	15415182	refGene_gn=MAP4K1;refGene_tx=XM_011289867.3
E2	15415855	15415911	refGene_gn=MAP4K1;refGene_tx=XM_011289867.3
E2	15417676	15417746	refGene_gn=MAP4K1;refGene_tx=XM_011289867.3
E2	15418093	15418189	refGene_gn=RYR1;refGene_tx=XM_023245508.1
E2	15418331	15418383	refGene_gn=RYR1;refGene_tx=XM_023245508.1
E2	15418545	15418646	refGene_gn=RYR1;refGene_tx=XM_023245508.1
E2	15418734	15418799	refGene_gn=RYR1;refGene_tx=XM_023245508.1
E2	15419226	15419383	refGene_gn=RYR1;refGene_tx=XM_023245508.1
E2	15421372	15421507	refGene_gn=RYR1;refGene_tx=XM_023245508.1
E2	15421747	15421894	refGene_gn=RYR1;refGene_tx=XM_023245508.1
E2	15422358	15422419	refGene_gn=RYR1;refGene_tx=XM_023245508.1
E2	15422524	15422655	refGene_gn=RYR1;refGene_tx=XM_023245508.1
E2	15423887	15423930	refGene_gn=RYR1;refGene_tx=XM_023245508.1
E2	15425988	15426119	refGene_gn=RYR1;refGene_tx=XM_023245508.1
E2	15426691	15426943	refGene_gn=RYR1;refGene_tx=XM_023245508.1
E2	15427761	15427848	refGene_gn=RYR1;refGene_tx=XM_023245508.1

| | | | |
|---|---|---|---|
| E2 | 15428849 | 15428957 | refGene_gn=RYR1;refGene_tx=XM_023245508.1 |
| E2 | 15428958 | 15429004 | refGene_gn=RYR1;refGene_tx=XM_023245508.1 |
| E2 | 15429523 | 15429600 | refGene_gn=RYR1;refGene_tx=XM_023245508.1 |
| E2 | 15430289 | 15431108 | refGene_gn=RYR1;refGene_tx=XM_023245508.1 |
| E2 | 15433490 | 15433832 | refGene_gn=RYR1;refGene_tx=XM_023245508.1 |
| E2 | 15445405 | 15445593 | refGene_gn=RYR1;refGene_tx=XM_023245508.1 |
| E2 | 15448775 | 15448857 | refGene_gn=RYR1;refGene_tx=XM_023245508.1 |
| E2 | 15451015 | 15451120 | refGene_gn=RYR1;refGene_tx=XM_023245508.1 |
| E2 | 15451219 | 15451348 | refGene_gn=RYR1;refGene_tx=XM_023245508.1 |
| E2 | 15451443 | 15451532 | refGene_gn=RYR1;refGene_tx=XM_023245508.1 |
| E2 | 15456278 | 15456359 | refGene_gn=RYR1;refGene_tx=XM_023245508.1 |
| E2 | 15457474 | 15457492 | refGene_gn=RYR1;refGene_tx=XM_023245508.1 |
| E2 | 15458150 | 15458224 | refGene_gn=RYR1;refGene_tx=XM_023245508.1 |
| E2 | 15459043 | 15459120 | refGene_gn=RYR1;refGene_tx=XM_023245508.1 |
| E2 | 15459213 | 15459293 | refGene_gn=RYR1;refGene_tx=XM_023245508.1 |
| E2 | 15459574 | 15459674 | refGene_gn=RYR1;refGene_tx=XM_023245508.1 |
| E2 | 15461322 | 15461391 | refGene_gn=RYR1;refGene_tx=XM_023245508.1 |
| E2 | 15461525 | 15461577 | refGene_gn=RYR1;refGene_tx=XM_023245508.1 |
| E2 | 15463108 | 15463215 | refGene_gn=RYR1;refGene_tx=XM_023245508.1 |
| E2 | 15463463 | 15463560 | refGene_gn=RYR1;refGene_tx=XM_023245508.1 |
| E2 | 15463762 | 15463875 | refGene_gn=RYR1;refGene_tx=XM_023245508.1 |
| E2 | 15464428 | 15464566 | refGene_gn=RYR1;refGene_tx=XM_023245508.1 |
| E2 | 15464850 | 15464910 | refGene_gn=RYR1;refGene_tx=XM_023245508.1 |
| E2 | 15466458 | 15466629 | refGene_gn=RYR1;refGene_tx=XM_023245508.1 |
| E2 | 15467825 | 15467840 | refGene_gn=RYR1;refGene_tx=XM_023245508.1 |
| E2 | 15468829 | 15468922 | refGene_gn=RYR1;refGene_tx=XM_023245508.1 |
| E2 | 15469001 | 15469089 | refGene_gn=RYR1;refGene_tx=XM_023245508.1 |
| E2 | 15471661 | 15471902 | refGene_gn=RYR1;refGene_tx=XM_023245508.1 |
| E2 | 15472678 | 15472937 | refGene_gn=RYR1;refGene_tx=XM_023245508.1 |
| E2 | 15472938 | 15473012 | refGene_gn=RYR1;refGene_tx=XM_023245508.1 |
| E2 | 15473233 | 15473364 | refGene_gn=RYR1;refGene_tx=XM_023245508.1 |
| E2 | 15473908 | 15473990 | refGene_gn=RYR1;refGene_tx=XM_023245508.1 |
| E2 | 15476457 | 15476696 | refGene_gn=RYR1;refGene_tx=XM_023245508.1 |
| E2 | 15476802 | 15476863 | refGene_gn=RYR1;refGene_tx=XM_023245508.1 |
| E2 | 15477277 | 15477327 | refGene_gn=RYR1;refGene_tx=XM_023245508.1 |
| E2 | 15477808 | 15477930 | refGene_gn=RYR1;refGene_tx=XM_023245508.1 |
| E2 | 15478035 | 15478103 | refGene_gn=RYR1;refGene_tx=XM_023245508.1 |
| E2 | 15479672 | 15479788 | refGene_gn=RYR1;refGene_tx=XM_023245508.1 |
| E2 | 15480329 | 15480453 | refGene_gn=RYR1;refGene_tx=XM_023245508.1 |
| E2 | 15480758 | 15480834 | refGene_gn=RYR1;refGene_tx=XM_023245508.1 |
| E2 | 15480933 | 15481008 | refGene_gn=RYR1;refGene_tx=XM_023245508.1 |
| E2 | 15481408 | 15481549 | refGene_gn=RYR1;refGene_tx=XM_023245508.1 |
| E2 | 15482024 | 15482114 | refGene_gn=RYR1;refGene_tx=XM_023245508.1 |
| E2 | 15482386 | 15482465 | refGene_gn=RYR1;refGene_tx=XM_023245508.1 |
| E2 | 15482582 | 15482746 | refGene_gn=RYR1;refGene_tx=XM_023245508.1 |
| E2 | 15483147 | 15483288 | refGene_gn=RYR1;refGene_tx=XM_023245508.1 |
| E2 | 15484163 | 15484254 | refGene_gn=RYR1;refGene_tx=XM_023245508.1 |
| E2 | 15484346 | 15484567 | refGene_gn=RYR1;refGene_tx=XM_023245508.1 |
| E2 | 15485640 | 15485810 | refGene_gn=RYR1;refGene_tx=XM_023245508.1 |
| E2 | 15485906 | 15486027 | refGene_gn=RYR1;refGene_tx=XM_023245508.1 |

E2 15486427 15486536 refGene_gn=RYR1;refGene_tx=XM_023245508.1
E2 15486626 15486813 refGene_gn=RYR1;refGene_tx=XM_023245508.1
E2 15487052 15487188 refGene_gn=RYR1;refGene_tx=XM_023245508.1
E2 15488437 15488532 refGene_gn=RYR1;refGene_tx=XM_023245508.1
E2 15488939 15489072 refGene_gn=RYR1;refGene_tx=XM_023245508.1
E2 15489157 15489272 refGene_gn=RYR1;refGene_tx=XM_023245508.1
E2 15489991 15490265 refGene_gn=RYR1;refGene_tx=XM_023245508.1
E2 15492502 15492649 refGene_gn=RYR1;refGene_tx=XM_023245508.1
E2 15494109 15494221 refGene_gn=RYR1;refGene_tx=XM_023245508.1
E2 15494561 15494762 refGene_gn=RYR1;refGene_tx=XM_023245508.1
E2 15495105 15495360 refGene_gn=RYR1;refGene_tx=XM_023245508.1
E2 15496856 15497469 refGene_gn=RYR1;refGene_tx=XM_023245508.1
E2 15498787 15499014 refGene_gn=RYR1;refGene_tx=XM_023245508.1
E2 15499174 15499261 refGene_gn=RYR1;refGene_tx=XM_023245508.1
E2 15501352 15501518 refGene_gn=RYR1;refGene_tx=XM_023245508.1
E2 15501947 15502108 refGene_gn=RYR1;refGene_tx=XM_023245508.1
E2 15502960 15503093 refGene_gn=RYR1;refGene_tx=XM_023245508.1
E2 15504039 15504434 refGene_gn=RYR1;refGene_tx=XM_023245508.1
E2 15507309 15507518 refGene_gn=RYR1;refGene_tx=XM_023245508.1
E2 15507682 15507857 refGene_gn=RYR1;refGene_tx=XM_023245508.1
E2 15508821 15509024 refGene_gn=RYR1;refGene_tx=XM_023245508.1
E2 15509922 15510230 refGene_gn=RYR1;refGene_tx=XM_023245508.1
E2 15511031 15511115 refGene_gn=RYR1;refGene_tx=XM_023245508.1
E2 15511508 15511612 refGene_gn=RYR1;refGene_tx=XM_023245508.1
E2 15511826 15511931 refGene_gn=RYR1;refGene_tx=XM_023245508.1
E2 15513489 15513706 refGene_gn=RYR1;refGene_tx=XM_023245508.1
E2 15514930 15515123 refGene_gn=RYR1;refGene_tx=XM_023245508.1
E2 15515680 15515922 refGene_gn=RYR1;refGene_tx=XM_023245508.1
E2 15516430 15516564 refGene_gn=RYR1;refGene_tx=XM_023245508.1
E2 15517705 15517824 refGene_gn=RYR1;refGene_tx=XM_023245508.1
E2 15517907 15518003 refGene_gn=RYR1;refGene_tx=XM_023245508.1
E2 15518085 15518221 refGene_gn=RYR1;refGene_tx=XM_023245508.1
E2 15522247 15522443 refGene_gn=RYR1;refGene_tx=XM_023245508.1
E2 15522926 15523048 refGene_gn=RYR1;refGene_tx=XM_023245508.1
E2 15524952 15525117 refGene_gn=RYR1;refGene_tx=XM_023245508.1
E2 15525243 15525400 refGene_gn=RYR1;refGene_tx=XM_023245508.1
E2 15525884 15525959 refGene_gn=RYR1;refGene_tx=XM_023245508.1
E2 15526114 15526208 refGene_gn=RYR1;refGene_tx=XM_023245508.1
E2 15527244 15527338 refGene_gn=RYR1;refGene_tx=XM_023245508.1
E2 15527648 15527761 refGene_gn=RYR1;refGene_tx=XM_023245508.1
E2 15528023 15528102 refGene_gn=RYR1;refGene_tx=XM_023245508.1
E2 15528180 15528255 refGene_gn=RYR1;refGene_tx=XM_023245508.1
E2 15529325 15529430 refGene_gn=RYR1;refGene_tx=XM_023245508.1
E2 15530548 15530668 refGene_gn=RYR1;refGene_tx=XM_023245508.1
E2 15535471 15535519 refGene_gn=RYR1;refGene_tx=XM_023245508.1
E2 15540646 15540669
refGene_gn=RASGRP4;refGene_tx=XM_006941300.3;refGene_gn=RASGRP4;refGen
e_tx=XM_006941301.3;refGene_gn=RASGRP4;refGene_tx=XM_006941302.3
E2 15543175 15543360
refGene_gn=RASGRP4;refGene_tx=XM_006941300.3;refGene_gn=RASGRP4;refGen

e_tx=XM_006941301.3;refGene_gn=RASGRP4;refGene_tx=XM_006941302.3
E2 15543785 15543891
refGene_gn=RASGRP4;refGene_tx=XM_006941300.3;refGene_gn=RASGRP4;refGen
e_tx=XM_006941301.3;refGene_gn=RASGRP4;refGene_tx=XM_006941302.3
E2 15544005 15544068
refGene_gn=RASGRP4;refGene_tx=XM_006941300.3;refGene_gn=RASGRP4;refGen
e_tx=XM_006941301.3;refGene_gn=RASGRP4;refGene_tx=XM_006941302.3
E2 15544654 15544786
refGene_gn=RASGRP4;refGene_tx=XM_006941300.3;refGene_gn=RASGRP4;refGen
e_tx=XM_006941301.3;refGene_gn=RASGRP4;refGene_tx=XM_006941302.3
E2 15544885 15545039
refGene_gn=RASGRP4;refGene_tx=XM_006941300.3;refGene_gn=RASGRP4;refGen
e_tx=XM_006941301.3;refGene_gn=RASGRP4;refGene_tx=XM_006941302.3
E2 15546202 15546376
refGene_gn=RASGRP4;refGene_tx=XM_006941300.3;refGene_gn=RASGRP4;refGen
e_tx=XM_006941301.3;refGene_gn=RASGRP4;refGene_tx=XM_006941302.3
E2 15547640 15547757
refGene_gn=RASGRP4;refGene_tx=XM_006941300.3;refGene_gn=RASGRP4;refGen
e_tx=XM_006941301.3;refGene_gn=RASGRP4;refGene_tx=XM_006941302.3
E2 15548560 15548836
refGene_gn=RASGRP4;refGene_tx=XM_006941300.3;refGene_gn=RASGRP4;refGen
e_tx=XM_006941301.3;refGene_gn=RASGRP4;refGene_tx=XM_006941302.3
E2 15549949 15550030
refGene_gn=RASGRP4;refGene_tx=XM_006941300.3;refGene_gn=RASGRP4;refGen
e_tx=XM_006941301.3;refGene_gn=RASGRP4;refGene_tx=XM_006941302.3
E2 15550108 15550213
refGene_gn=RASGRP4;refGene_tx=XM_006941300.3;refGene_gn=RASGRP4;refGen
e_tx=XM_006941301.3;refGene_gn=RASGRP4;refGene_tx=XM_006941302.3
E2 15550325 15550444
refGene_gn=RASGRP4;refGene_tx=XM_006941300.3;refGene_gn=RASGRP4;refGen
e_tx=XM_006941301.3;refGene_gn=RASGRP4;refGene_tx=XM_006941302.3
E2 15550555 15550700
refGene_gn=RASGRP4;refGene_tx=XM_006941300.3;refGene_gn=RASGRP4;refGen
e_tx=XM_006941301.3;refGene_gn=RASGRP4;refGene_tx=XM_006941302.3
E2 15551494 15551531
refGene_gn=RASGRP4;refGene_tx=XM_006941300.3;refGene_gn=RASGRP4;refGen
e_tx=XM_006941301.3;refGene_gn=RASGRP4;refGene_tx=XM_006941302.3
E2 15551635 15551770
refGene_gn=RASGRP4;refGene_tx=XM_006941300.3;refGene_gn=RASGRP4;refGen
e_tx=XM_006941301.3;refGene_gn=RASGRP4;refGene_tx=XM_006941302.3
E2 15551888 15552109
refGene_gn=RASGRP4;refGene_tx=XM_006941300.3;refGene_gn=RASGRP4;refGen
e_tx=XM_006941301.3;refGene_gn=RASGRP4;refGene_tx=XM_006941302.3
E2 15552723 15552768 refGene_gn=RASGRP4;refGene_tx=XM_006941302.3
E2 15554225 15554369 refGene_gn=FAM98C;refGene_tx=XM_023244927.1
E2 15554480 15554648 refGene_gn=FAM98C;refGene_tx=XM_023244927.1
E2 15554993 15555076 refGene_gn=FAM98C;refGene_tx=XM_023244927.1
E2 15555078 15555109 refGene_gn=FAM98C;refGene_tx=XM_023244927.1
E2 15555199 15555277 refGene_gn=FAM98C;refGene_tx=XM_023244927.1
E2 15555516 15555722 refGene_gn=FAM98C;refGene_tx=XM_023244927.1

E2 15556132 15556267 refGene_gn=FAM98C;refGene_tx=XM_023244927.1
E2 15556346 15556495 refGene_gn=FAM98C;refGene_tx=XM_023244927.1
E2 15556608 15556673 refGene_gn=FAM98C;refGene_tx=XM_023244927.1
E2 15561438 15562104 refGene_gn=SPRED3;refGene_tx=XM_023244926.1
E2 15562869 15563013 refGene_gn=SPRED3;refGene_tx=XM_023244926.1
E2 15565272 15565349 refGene_gn=SPRED3;refGene_tx=XM_023244926.1
E2 15565450 15565619 refGene_gn=SPRED3;refGene_tx=XM_023244926.1
E2 15566709 15566718 refGene_gn=SPRED3;refGene_tx=XM_023244926.1
E2 15566719 15566887 refGene_gn=SPRED3;refGene_tx=XM_023244926.1
E2 15570310 15571148
refGene_gn=GGN;refGene_tx=XM_023245143.1;refGene_gn=GGN;refGene_tx=XM_023245144.1;refGene_gn=GGN;refGene_tx=XM_023245145.1
E2 15571184 15572238
refGene_gn=GGN;refGene_tx=XM_023245143.1;refGene_gn=GGN;refGene_tx=XM_023245144.1;refGene_gn=GGN;refGene_tx=XM_023245145.1
E2 15573173 15573291
refGene_gn=GGN;refGene_tx=XM_023245143.1;refGene_gn=GGN;refGene_tx=XM_023245144.1;refGene_gn=GGN;refGene_tx=XM_023245145.1
E2 15574292 15574430 refGene_gn=PSMD8;refGene_tx=XM_011289865.3
E2 15575158 15575270 refGene_gn=PSMD8;refGene_tx=XM_011289865.3
E2 15576469 15576570 refGene_gn=PSMD8;refGene_tx=XM_011289865.3
E2 15577591 15577757 refGene_gn=PSMD8;refGene_tx=XM_011289865.3
E2 15579604 15579707 refGene_gn=PSMD8;refGene_tx=XM_011289865.3
E2 15579815 15579888 refGene_gn=PSMD8;refGene_tx=XM_011289865.3
E2 15580892 15581255 refGene_gn=PSMD8;refGene_tx=XM_011289865.3
E2 15589168 15589453
refGene_gn=CATSPERG;refGene_tx=XM_023245086.1;refGene_gn=CATSPERG;refGene_tx=XM_023245088.1;refGene_gn=CATSPERG;refGene_tx=XM_023245089.1;refGene_gn=CATSPERG;refGene_tx=XM_023245090.1;refGene_gn=CATSPERG;refGene_tx=XM_023245091.1;refGene_gn=CATSPERG;refGene_tx=XM_023245092.1;refGene_gn=CATSPERG;refGene_tx=XM_023245093.1;refGene_gn=CATSPERG;refGene_tx=XM_023245094.1;refGene_gn=CATSPERG;refGene_tx=XM_023245095.1;refGene_gn=CATSPERG;refGene_tx=XM_023245100.1
E2 15589535 15589658
refGene_gn=CATSPERG;refGene_tx=XM_023245086.1;refGene_gn=CATSPERG;refGene_tx=XM_023245088.1;refGene_gn=CATSPERG;refGene_tx=XM_023245089.1;refGene_gn=CATSPERG;refGene_tx=XM_023245090.1;refGene_gn=CATSPERG;refGene_tx=XM_023245091.1;refGene_gn=CATSPERG;refGene_tx=XM_023245092.1;refGene_gn=CATSPERG;refGene_tx=XM_023245093.1;refGene_gn=CATSPERG;refGene_tx=XM_023245094.1;refGene_gn=CATSPERG;refGene_tx=XM_023245095.1;refGene_gn=CATSPERG;refGene_tx=XM_023245100.1
E2 15589724 15589817
refGene_gn=CATSPERG;refGene_tx=XM_023245086.1;refGene_gn=CATSPERG;refGene_tx=XM_023245088.1;refGene_gn=CATSPERG;refGene_tx=XM_023245089.1;refGene_gn=CATSPERG;refGene_tx=XM_023245090.1;refGene_gn=CATSPERG;refGene_tx=XM_023245091.1;refGene_gn=CATSPERG;refGene_tx=XM_023245092.1;refGene_gn=CATSPERG;refGene_tx=XM_023245093.1;refGene_gn=CATSPERG;refGene_tx=XM_023245094.1;refGene_gn=CATSPERG;refGene_tx=XM_023245095.1;refGene_gn=CATSPERG;refGene_tx=XM_023245098.1;refGene_gn=CATSPERG;refGene_tx=XM_023245099.1;refGene_gn=CATSPERG;refGene_tx=XM_023245100.1

E2 15590362 15590475
refGene_gn=CATSPERG;refGene_tx=XM_023245096.1;refGene_gn=CATSPERG;refGene_tx=XM_023245097.1
E2 15590839 15590929
refGene_gn=CATSPERG;refGene_tx=XM_023245086.1;refGene_gn=CATSPERG;refGene_tx=XM_023245088.1;refGene_gn=CATSPERG;refGene_tx=XM_023245089.1;refGene_gn=CATSPERG;refGene_tx=XM_023245090.1;refGene_gn=CATSPERG;refGene_tx=XM_023245091.1;refGene_gn=CATSPERG;refGene_tx=XM_023245092.1;refGene_gn=CATSPERG;refGene_tx=XM_023245093.1;refGene_gn=CATSPERG;refGene_tx=XM_023245094.1;refGene_gn=CATSPERG;refGene_tx=XM_023245095.1;refGene_gn=CATSPERG;refGene_tx=XM_023245096.1;refGene_gn=CATSPERG;refGene_tx=XM_023245097.1;refGene_gn=CATSPERG;refGene_tx=XM_023245098.1;refGene_gn=CATSPERG;refGene_tx=XM_023245099.1;refGene_gn=CATSPERG;refGene_tx=XM_023245100.1
E2 15591228 15591324
refGene_gn=CATSPERG;refGene_tx=XM_023245086.1;refGene_gn=CATSPERG;refGene_tx=XM_023245088.1;refGene_gn=CATSPERG;refGene_tx=XM_023245089.1;refGene_gn=CATSPERG;refGene_tx=XM_023245090.1;refGene_gn=CATSPERG;refGene_tx=XM_023245091.1;refGene_gn=CATSPERG;refGene_tx=XM_023245092.1;refGene_gn=CATSPERG;refGene_tx=XM_023245093.1;refGene_gn=CATSPERG;refGene_tx=XM_023245094.1;refGene_gn=CATSPERG;refGene_tx=XM_023245095.1;refGene_gn=CATSPERG;refGene_tx=XM_023245096.1;refGene_gn=CATSPERG;refGene_tx=XM_023245100.1
E2 15591473 15591537
refGene_gn=CATSPERG;refGene_tx=XM_023245086.1;refGene_gn=CATSPERG;refGene_tx=XM_023245088.1;refGene_gn=CATSPERG;refGene_tx=XM_023245089.1;refGene_gn=CATSPERG;refGene_tx=XM_023245090.1;refGene_gn=CATSPERG;refGene_tx=XM_023245091.1;refGene_gn=CATSPERG;refGene_tx=XM_023245092.1;refGene_gn=CATSPERG;refGene_tx=XM_023245093.1;refGene_gn=CATSPERG;refGene_tx=XM_023245094.1;refGene_gn=CATSPERG;refGene_tx=XM_023245095.1;refGene_gn=CATSPERG;refGene_tx=XM_023245096.1;refGene_gn=CATSPERG;refGene_tx=XM_023245100.1
E2 15591740 15591918
refGene_gn=CATSPERG;refGene_tx=XM_023245086.1;refGene_gn=CATSPERG;refGene_tx=XM_023245088.1;refGene_gn=CATSPERG;refGene_tx=XM_023245089.1;refGene_gn=CATSPERG;refGene_tx=XM_023245090.1;refGene_gn=CATSPERG;refGene_tx=XM_023245091.1;refGene_gn=CATSPERG;refGene_tx=XM_023245092.1;refGene_gn=CATSPERG;refGene_tx=XM_023245093.1;refGene_gn=CATSPERG;refGene_tx=XM_023245094.1;refGene_gn=CATSPERG;refGene_tx=XM_023245095.1;refGene_gn=CATSPERG;refGene_tx=XM_023245096.1;refGene_gn=CATSPERG;refGene_tx=XM_023245097.1;refGene_gn=CATSPERG;refGene_tx=XM_023245098.1;refGene_gn=CATSPERG;refGene_tx=XM_023245099.1;refGene_gn=CATSPERG;refGene_tx=XM_023245100.1;refGene_gn=CATSPERG;refGene_tx=XM_023245102.1;refGene_gn=CATSPERG;refGene_tx=XM_023245103.1
E2 15596550 15596613 refGene_gn=CATSPERG;refGene_tx=XM_023245101.1
E2 15596862 15596919
refGene_gn=CATSPERG;refGene_tx=XM_023245086.1;refGene_gn=CATSPERG;refGene_tx=XM_023245088.1;refGene_gn=CATSPERG;refGene_tx=XM_023245089.1;refGene_gn=CATSPERG;refGene_tx=XM_023245090.1;refGene_gn=CATSPERG;refGene_tx=XM_023245091.1;refGene_gn=CATSPERG;refGene_tx=XM_023245092.1;refG

ene_gn=CATSPERG;refGene_tx=XM_023245093.1;refGene_gn=CATSPERG;refGene_tx=XM_023245094.1;refGene_gn=CATSPERG;refGene_tx=XM_023245095.1;refGene_gn=CATSPERG;refGene_tx=XM_023245096.1;refGene_gn=CATSPERG;refGene_tx=XM_023245097.1;refGene_gn=CATSPERG;refGene_tx=XM_023245098.1;refGene_gn=CATSPERG;refGene_tx=XM_023245099.1;refGene_gn=CATSPERG;refGene_tx=XM_023245100.1;refGene_gn=CATSPERG;refGene_tx=XM_023245101.1;refGene_gn=CATSPERG;refGene_tx=XM_023245102.1;refGene_gn=CATSPERG;refGene_tx=XM_023245103.1
E2 15597009 15597090
refGene_gn=CATSPERG;refGene_tx=XM_023245086.1;refGene_gn=CATSPERG;refGene_tx=XM_023245088.1;refGene_gn=CATSPERG;refGene_tx=XM_023245089.1;refGene_gn=CATSPERG;refGene_tx=XM_023245090.1;refGene_gn=CATSPERG;refGene_tx=XM_023245091.1;refGene_gn=CATSPERG;refGene_tx=XM_023245092.1;refGene_gn=CATSPERG;refGene_tx=XM_023245093.1;refGene_gn=CATSPERG;refGene_tx=XM_023245094.1;refGene_gn=CATSPERG;refGene_tx=XM_023245095.1;refGene_gn=CATSPERG;refGene_tx=XM_023245096.1;refGene_gn=CATSPERG;refGene_tx=XM_023245097.1;refGene_gn=CATSPERG;refGene_tx=XM_023245098.1;refGene_gn=CATSPERG;refGene_tx=XM_023245099.1;refGene_gn=CATSPERG;refGene_tx=XM_023245100.1;refGene_gn=CATSPERG;refGene_tx=XM_023245101.1;refGene_gn=CATSPERG;refGene_tx=XM_023245102.1;refGene_gn=CATSPERG;refGene_tx=XM_023245103.1
E2 15597214 15597336
refGene_gn=CATSPERG;refGene_tx=XM_023245086.1;refGene_gn=CATSPERG;refGene_tx=XM_023245088.1;refGene_gn=CATSPERG;refGene_tx=XM_023245089.1;refGene_gn=CATSPERG;refGene_tx=XM_023245090.1;refGene_gn=CATSPERG;refGene_tx=XM_023245091.1;refGene_gn=CATSPERG;refGene_tx=XM_023245092.1;refGene_gn=CATSPERG;refGene_tx=XM_023245093.1;refGene_gn=CATSPERG;refGene_tx=XM_023245094.1;refGene_gn=CATSPERG;refGene_tx=XM_023245095.1;refGene_gn=CATSPERG;refGene_tx=XM_023245096.1;refGene_gn=CATSPERG;refGene_tx=XM_023245097.1;refGene_gn=CATSPERG;refGene_tx=XM_023245098.1;refGene_gn=CATSPERG;refGene_tx=XM_023245099.1;refGene_gn=CATSPERG;refGene_tx=XM_023245100.1;refGene_gn=CATSPERG;refGene_tx=XM_023245101.1;refGene_gn=CATSPERG;refGene_tx=XM_023245102.1;refGene_gn=CATSPERG;refGene_tx=XM_023245103.1
E2 15597452 15597651
refGene_gn=CATSPERG;refGene_tx=XM_023245086.1;refGene_gn=CATSPERG;refGene_tx=XM_023245088.1;refGene_gn=CATSPERG;refGene_tx=XM_023245089.1;refGene_gn=CATSPERG;refGene_tx=XM_023245090.1;refGene_gn=CATSPERG;refGene_tx=XM_023245091.1;refGene_gn=CATSPERG;refGene_tx=XM_023245092.1;refGene_gn=CATSPERG;refGene_tx=XM_023245093.1;refGene_gn=CATSPERG;refGene_tx=XM_023245094.1;refGene_gn=CATSPERG;refGene_tx=XM_023245095.1;refGene_gn=CATSPERG;refGene_tx=XM_023245096.1;refGene_gn=CATSPERG;refGene_tx=XM_023245097.1;refGene_gn=CATSPERG;refGene_tx=XM_023245098.1;refGene_gn=CATSPERG;refGene_tx=XM_023245099.1;refGene_gn=CATSPERG;refGene_tx=XM_023245100.1;refGene_gn=CATSPERG;refGene_tx=XM_023245101.1;refGene_gn=CATSPERG;refGene_tx=XM_023245102.1;refGene_gn=CATSPERG;refGene_tx=XM_023245103.1
E2 15597733 15597793
refGene_gn=CATSPERG;refGene_tx=XM_023245086.1;refGene_gn=CATSPERG;refGene_tx=XM_023245088.1;refGene_gn=CATSPERG;refGene_tx=XM_023245089.1;re

fGene_gn=CATSPERG;refGene_tx=XM_023245090.1;refGene_gn=CATSPERG;refGene_tx=XM_023245091.1;refGene_gn=CATSPERG;refGene_tx=XM_023245092.1;refGene_gn=CATSPERG;refGene_tx=XM_023245093.1;refGene_gn=CATSPERG;refGene_tx=XM_023245094.1;refGene_gn=CATSPERG;refGene_tx=XM_023245095.1;refGene_gn=CATSPERG;refGene_tx=XM_023245096.1;refGene_gn=CATSPERG;refGene_tx=XM_023245097.1;refGene_gn=CATSPERG;refGene_tx=XM_023245098.1;refGene_gn=CATSPERG;refGene_tx=XM_023245099.1;refGene_gn=CATSPERG;refGene_tx=XM_023245100.1;refGene_gn=CATSPERG;refGene_tx=XM_023245101.1;refGene_gn=CATSPERG;refGene_tx=XM_023245102.1;refGene_gn=CATSPERG;refGene_tx=XM_023245103.1
E2 15598054 15598307
refGene_gn=CATSPERG;refGene_tx=XM_023245086.1;refGene_gn=CATSPERG;refGene_tx=XM_023245088.1;refGene_gn=CATSPERG;refGene_tx=XM_023245089.1;refGene_gn=CATSPERG;refGene_tx=XM_023245090.1;refGene_gn=CATSPERG;refGene_tx=XM_023245091.1;refGene_gn=CATSPERG;refGene_tx=XM_023245092.1;refGene_gn=CATSPERG;refGene_tx=XM_023245093.1;refGene_gn=CATSPERG;refGene_tx=XM_023245094.1;refGene_gn=CATSPERG;refGene_tx=XM_023245095.1;refGene_gn=CATSPERG;refGene_tx=XM_023245096.1;refGene_gn=CATSPERG;refGene_tx=XM_023245097.1;refGene_gn=CATSPERG;refGene_tx=XM_023245098.1;refGene_gn=CATSPERG;refGene_tx=XM_023245099.1;refGene_gn=CATSPERG;refGene_tx=XM_023245100.1;refGene_gn=CATSPERG;refGene_tx=XM_023245101.1;refGene_gn=CATSPERG;refGene_tx=XM_023245102.1;refGene_gn=CATSPERG;refGene_tx=XM_023245103.1
E2 15599055 15599168
refGene_gn=CATSPERG;refGene_tx=XM_023245086.1;refGene_gn=CATSPERG;refGene_tx=XM_023245088.1;refGene_gn=CATSPERG;refGene_tx=XM_023245089.1;refGene_gn=CATSPERG;refGene_tx=XM_023245090.1;refGene_gn=CATSPERG;refGene_tx=XM_023245091.1;refGene_gn=CATSPERG;refGene_tx=XM_023245092.1;refGene_gn=CATSPERG;refGene_tx=XM_023245093.1;refGene_gn=CATSPERG;refGene_tx=XM_023245094.1;refGene_gn=CATSPERG;refGene_tx=XM_023245095.1;refGene_gn=CATSPERG;refGene_tx=XM_023245096.1;refGene_gn=CATSPERG;refGene_tx=XM_023245097.1;refGene_gn=CATSPERG;refGene_tx=XM_023245098.1;refGene_gn=CATSPERG;refGene_tx=XM_023245099.1;refGene_gn=CATSPERG;refGene_tx=XM_023245100.1;refGene_gn=CATSPERG;refGene_tx=XM_023245101.1;refGene_gn=CATSPERG;refGene_tx=XM_023245102.1;refGene_gn=CATSPERG;refGene_tx=XM_023245103.1
E2 15599262 15599421
refGene_gn=CATSPERG;refGene_tx=XM_023245086.1;refGene_gn=CATSPERG;refGene_tx=XM_023245088.1;refGene_gn=CATSPERG;refGene_tx=XM_023245089.1;refGene_gn=CATSPERG;refGene_tx=XM_023245090.1;refGene_gn=CATSPERG;refGene_tx=XM_023245091.1;refGene_gn=CATSPERG;refGene_tx=XM_023245092.1;refGene_gn=CATSPERG;refGene_tx=XM_023245093.1;refGene_gn=CATSPERG;refGene_tx=XM_023245094.1;refGene_gn=CATSPERG;refGene_tx=XM_023245095.1;refGene_gn=CATSPERG;refGene_tx=XM_023245096.1;refGene_gn=CATSPERG;refGene_tx=XM_023245097.1;refGene_gn=CATSPERG;refGene_tx=XM_023245098.1;refGene_gn=CATSPERG;refGene_tx=XM_023245099.1;refGene_gn=CATSPERG;refGene_tx=XM_023245100.1;refGene_gn=CATSPERG;refGene_tx=XM_023245101.1;refGene_gn=CATSPERG;refGene_tx=XM_023245102.1;refGene_gn=CATSPERG;refGene_tx=XM_023245103.1
E2 15600385 15600497

refGene_gn=CATSPERG;refGene_tx=XM_023245086.1;refGene_gn=CATSPERG;refG
ene_tx=XM_023245088.1;refGene_gn=CATSPERG;refGene_tx=XM_023245089.1;re
fGene_gn=CATSPERG;refGene_tx=XM_023245090.1;refGene_gn=CATSPERG;refGen
e_tx=XM_023245091.1;refGene_gn=CATSPERG;refGene_tx=XM_023245092.1;refG
ene_gn=CATSPERG;refGene_tx=XM_023245093.1;refGene_gn=CATSPERG;refGene_
tx=XM_023245094.1;refGene_gn=CATSPERG;refGene_tx=XM_023245095.1;refGen
e_gn=CATSPERG;refGene_tx=XM_023245096.1;refGene_gn=CATSPERG;refGene_tx
=XM_023245097.1;refGene_gn=CATSPERG;refGene_tx=XM_023245098.1;refGene_
gn=CATSPERG;refGene_tx=XM_023245099.1;refGene_gn=CATSPERG;refGene_tx=X
M_023245100.1;refGene_gn=CATSPERG;refGene_tx=XM_023245101.1;refGene_gn
=CATSPERG;refGene_tx=XM_023245102.1;refGene_gn=CATSPERG;refGene_tx=XM_
023245103.1
E2 15601006 15601136
refGene_gn=CATSPERG;refGene_tx=XM_023245086.1;refGene_gn=CATSPERG;refG
ene_tx=XM_023245088.1;refGene_gn=CATSPERG;refGene_tx=XM_023245089.1;re
fGene_gn=CATSPERG;refGene_tx=XM_023245090.1;refGene_gn=CATSPERG;refGen
e_tx=XM_023245091.1;refGene_gn=CATSPERG;refGene_tx=XM_023245092.1;refG
ene_gn=CATSPERG;refGene_tx=XM_023245093.1;refGene_gn=CATSPERG;refGene_
tx=XM_023245094.1;refGene_gn=CATSPERG;refGene_tx=XM_023245095.1;refGen
e_gn=CATSPERG;refGene_tx=XM_023245096.1;refGene_gn=CATSPERG;refGene_tx
=XM_023245097.1;refGene_gn=CATSPERG;refGene_tx=XM_023245098.1;refGene_
gn=CATSPERG;refGene_tx=XM_023245099.1;refGene_gn=CATSPERG;refGene_tx=X
M_023245100.1;refGene_gn=CATSPERG;refGene_tx=XM_023245101.1;refGene_gn
=CATSPERG;refGene_tx=XM_023245102.1;refGene_gn=CATSPERG;refGene_tx=XM_
023245103.1
E2 15601240 15601345
refGene_gn=CATSPERG;refGene_tx=XM_023245086.1;refGene_gn=CATSPERG;refG
ene_tx=XM_023245088.1;refGene_gn=CATSPERG;refGene_tx=XM_023245089.1;re
fGene_gn=CATSPERG;refGene_tx=XM_023245090.1;refGene_gn=CATSPERG;refGen
e_tx=XM_023245091.1;refGene_gn=CATSPERG;refGene_tx=XM_023245092.1;refG
ene_gn=CATSPERG;refGene_tx=XM_023245093.1;refGene_gn=CATSPERG;refGene_
tx=XM_023245094.1;refGene_gn=CATSPERG;refGene_tx=XM_023245095.1;refGen
e_gn=CATSPERG;refGene_tx=XM_023245096.1;refGene_gn=CATSPERG;refGene_tx
=XM_023245097.1;refGene_gn=CATSPERG;refGene_tx=XM_023245098.1;refGene_
gn=CATSPERG;refGene_tx=XM_023245099.1;refGene_gn=CATSPERG;refGene_tx=X
M_023245100.1;refGene_gn=CATSPERG;refGene_tx=XM_023245101.1;refGene_gn
=CATSPERG;refGene_tx=XM_023245102.1;refGene_gn=CATSPERG;refGene_tx=XM_
023245103.1
E2 15601814 15601934
refGene_gn=CATSPERG;refGene_tx=XM_023245086.1;refGene_gn=CATSPERG;refG
ene_tx=XM_023245088.1;refGene_gn=CATSPERG;refGene_tx=XM_023245089.1;re
fGene_gn=CATSPERG;refGene_tx=XM_023245090.1;refGene_gn=CATSPERG;refGen
e_tx=XM_023245091.1;refGene_gn=CATSPERG;refGene_tx=XM_023245092.1;refG
ene_gn=CATSPERG;refGene_tx=XM_023245094.1;refGene_gn=CATSPERG;refGene_
tx=XM_023245096.1;refGene_gn=CATSPERG;refGene_tx=XM_023245097.1;refGen
e_gn=CATSPERG;refGene_tx=XM_023245098.1;refGene_gn=CATSPERG;refGene_tx
=XM_023245099.1;refGene_gn=CATSPERG;refGene_tx=XM_023245100.1;refGene_
gn=CATSPERG;refGene_tx=XM_023245101.1;refGene_gn=CATSPERG;refGene_tx=X
M_023245102.1;refGene_gn=CATSPERG;refGene_tx=XM_023245103.1
E2 15602132 15602192

refGene_gn=CATSPERG;refGene_tx=XM_023245086.1;refGene_gn=CATSPERG;refGene_tx=XM_023245088.1;refGene_gn=CATSPERG;refGene_tx=XM_023245089.1;refGene_gn=CATSPERG;refGene_tx=XM_023245090.1;refGene_gn=CATSPERG;refGene_tx=XM_023245091.1;refGene_gn=CATSPERG;refGene_tx=XM_023245092.1;refGene_gn=CATSPERG;refGene_tx=XM_023245094.1;refGene_gn=CATSPERG;refGene_tx=XM_023245096.1;refGene_gn=CATSPERG;refGene_tx=XM_023245097.1;refGene_gn=CATSPERG;refGene_tx=XM_023245098.1;refGene_gn=CATSPERG;refGene_tx=XM_023245099.1;refGene_gn=CATSPERG;refGene_tx=XM_023245100.1;refGene_gn=CATSPERG;refGene_tx=XM_023245101.1;refGene_gn=CATSPERG;refGene_tx=XM_023245102.1;refGene_gn=CATSPERG;refGene_tx=XM_023245103.1
E2 15603207 15603345
refGene_gn=CATSPERG;refGene_tx=XM_023245086.1;refGene_gn=CATSPERG;refGene_tx=XM_023245088.1;refGene_gn=CATSPERG;refGene_tx=XM_023245089.1;refGene_gn=CATSPERG;refGene_tx=XM_023245090.1;refGene_gn=CATSPERG;refGene_tx=XM_023245091.1;refGene_gn=CATSPERG;refGene_tx=XM_023245092.1;refGene_gn=CATSPERG;refGene_tx=XM_023245093.1;refGene_gn=CATSPERG;refGene_tx=XM_023245094.1;refGene_gn=CATSPERG;refGene_tx=XM_023245095.1;refGene_gn=CATSPERG;refGene_tx=XM_023245096.1;refGene_gn=CATSPERG;refGene_tx=XM_023245097.1;refGene_gn=CATSPERG;refGene_tx=XM_023245098.1;refGene_gn=CATSPERG;refGene_tx=XM_023245099.1;refGene_gn=CATSPERG;refGene_tx=XM_023245100.1;refGene_gn=CATSPERG;refGene_tx=XM_023245101.1;refGene_gn=CATSPERG;refGene_tx=XM_023245102.1;refGene_gn=CATSPERG;refGene_tx=XM_023245103.1
E2 15605311 15605483
refGene_gn=CATSPERG;refGene_tx=XM_023245086.1;refGene_gn=CATSPERG;refGene_tx=XM_023245088.1;refGene_gn=CATSPERG;refGene_tx=XM_023245089.1;refGene_gn=CATSPERG;refGene_tx=XM_023245090.1;refGene_gn=CATSPERG;refGene_tx=XM_023245091.1;refGene_gn=CATSPERG;refGene_tx=XM_023245092.1;refGene_gn=CATSPERG;refGene_tx=XM_023245093.1;refGene_gn=CATSPERG;refGene_tx=XM_023245094.1;refGene_gn=CATSPERG;refGene_tx=XM_023245095.1;refGene_gn=CATSPERG;refGene_tx=XM_023245096.1;refGene_gn=CATSPERG;refGene_tx=XM_023245097.1;refGene_gn=CATSPERG;refGene_tx=XM_023245098.1;refGene_gn=CATSPERG;refGene_tx=XM_023245099.1;refGene_gn=CATSPERG;refGene_tx=XM_023245100.1;refGene_gn=CATSPERG;refGene_tx=XM_023245101.1;refGene_gn=CATSPERG;refGene_tx=XM_023245102.1;refGene_gn=CATSPERG;refGene_tx=XM_023245103.1
E2 15607219 15607375
refGene_gn=CATSPERG;refGene_tx=XM_023245086.1;refGene_gn=CATSPERG;refGene_tx=XM_023245088.1;refGene_gn=CATSPERG;refGene_tx=XM_023245089.1;refGene_gn=CATSPERG;refGene_tx=XM_023245090.1;refGene_gn=CATSPERG;refGene_tx=XM_023245091.1;refGene_gn=CATSPERG;refGene_tx=XM_023245092.1;refGene_gn=CATSPERG;refGene_tx=XM_023245093.1;refGene_gn=CATSPERG;refGene_tx=XM_023245094.1;refGene_gn=CATSPERG;refGene_tx=XM_023245095.1;refGene_gn=CATSPERG;refGene_tx=XM_023245096.1;refGene_gn=CATSPERG;refGene_tx=XM_023245097.1;refGene_gn=CATSPERG;refGene_tx=XM_023245098.1;refGene_gn=CATSPERG;refGene_tx=XM_023245099.1;refGene_gn=CATSPERG;refGene_tx=XM_023245100.1;refGene_gn=CATSPERG;refGene_tx=XM_023245101.1;refGene_gn=CATSPERG;refGene_tx=XM_023245102.1;refGene_gn=CATSPERG;refGene_tx=XM_023245103.1
E2 15608516 15608589

refGene_gn=CATSPERG;refGene_tx=XM_023245086.1;refGene_gn=CATSPERG;refG
ene_tx=XM_023245088.1;refGene_gn=CATSPERG;refGene_tx=XM_023245089.1;re
fGene_gn=CATSPERG;refGene_tx=XM_023245090.1;refGene_gn=CATSPERG;refGen
e_tx=XM_023245091.1;refGene_gn=CATSPERG;refGene_tx=XM_023245092.1;refG
ene_gn=CATSPERG;refGene_tx=XM_023245093.1;refGene_gn=CATSPERG;refGene_
tx=XM_023245094.1;refGene_gn=CATSPERG;refGene_tx=XM_023245095.1;refGen
e_gn=CATSPERG;refGene_tx=XM_023245096.1;refGene_gn=CATSPERG;refGene_tx
=XM_023245097.1;refGene_gn=CATSPERG;refGene_tx=XM_023245098.1;refGene_
gn=CATSPERG;refGene_tx=XM_023245099.1;refGene_gn=CATSPERG;refGene_tx=X
M_023245100.1;refGene_gn=CATSPERG;refGene_tx=XM_023245101.1;refGene_gn
=CATSPERG;refGene_tx=XM_023245102.1;refGene_gn=CATSPERG;refGene_tx=XM_
023245103.1
E2 15608759 15608886
refGene_gn=CATSPERG;refGene_tx=XM_023245086.1;refGene_gn=CATSPERG;refG
ene_tx=XM_023245088.1;refGene_gn=CATSPERG;refGene_tx=XM_023245089.1;re
fGene_gn=CATSPERG;refGene_tx=XM_023245090.1;refGene_gn=CATSPERG;refGen
e_tx=XM_023245091.1;refGene_gn=CATSPERG;refGene_tx=XM_023245092.1;refG
ene_gn=CATSPERG;refGene_tx=XM_023245093.1;refGene_gn=CATSPERG;refGene_
tx=XM_023245094.1;refGene_gn=CATSPERG;refGene_tx=XM_023245095.1;refGen
e_gn=CATSPERG;refGene_tx=XM_023245096.1;refGene_gn=CATSPERG;refGene_tx
=XM_023245097.1;refGene_gn=CATSPERG;refGene_tx=XM_023245098.1;refGene_
gn=CATSPERG;refGene_tx=XM_023245099.1;refGene_gn=CATSPERG;refGene_tx=X
M_023245100.1;refGene_gn=CATSPERG;refGene_tx=XM_023245101.1;refGene_gn
=CATSPERG;refGene_tx=XM_023245102.1;refGene_gn=CATSPERG;refGene_tx=XM_
023245103.1
E2 15609141 15609286
refGene_gn=CATSPERG;refGene_tx=XM_023245086.1;refGene_gn=CATSPERG;refG
ene_tx=XM_023245088.1;refGene_gn=CATSPERG;refGene_tx=XM_023245089.1;re
fGene_gn=CATSPERG;refGene_tx=XM_023245090.1;refGene_gn=CATSPERG;refGen
e_tx=XM_023245091.1;refGene_gn=CATSPERG;refGene_tx=XM_023245092.1;refG
ene_gn=CATSPERG;refGene_tx=XM_023245093.1;refGene_gn=CATSPERG;refGene_
tx=XM_023245094.1;refGene_gn=CATSPERG;refGene_tx=XM_023245095.1;refGen
e_gn=CATSPERG;refGene_tx=XM_023245096.1;refGene_gn=CATSPERG;refGene_tx
=XM_023245097.1;refGene_gn=CATSPERG;refGene_tx=XM_023245098.1;refGene_
gn=CATSPERG;refGene_tx=XM_023245099.1;refGene_gn=CATSPERG;refGene_tx=X
M_023245100.1;refGene_gn=CATSPERG;refGene_tx=XM_023245101.1;refGene_gn
=CATSPERG;refGene_tx=XM_023245102.1;refGene_gn=CATSPERG;refGene_tx=XM_
023245103.1
E2 15612798 15612852
refGene_gn=CATSPERG;refGene_tx=XM_023245086.1;refGene_gn=CATSPERG;refG
ene_tx=XM_023245088.1;refGene_gn=CATSPERG;refGene_tx=XM_023245089.1;re
fGene_gn=CATSPERG;refGene_tx=XM_023245090.1;refGene_gn=CATSPERG;refGen
e_tx=XM_023245091.1;refGene_gn=CATSPERG;refGene_tx=XM_023245092.1;refG
ene_gn=CATSPERG;refGene_tx=XM_023245093.1;refGene_gn=CATSPERG;refGene_
tx=XM_023245094.1;refGene_gn=CATSPERG;refGene_tx=XM_023245095.1;refGen
e_gn=CATSPERG;refGene_tx=XM_023245096.1;refGene_gn=CATSPERG;refGene_tx
=XM_023245097.1;refGene_gn=CATSPERG;refGene_tx=XM_023245098.1;refGene_
gn=CATSPERG;refGene_tx=XM_023245099.1;refGene_gn=CATSPERG;refGene_tx=X
M_023245100.1;refGene_gn=CATSPERG;refGene_tx=XM_023245101.1;refGene_gn
=CATSPERG;refGene_tx=XM_023245102.1;refGene_gn=CATSPERG;refGene_tx=XM_

023245103.1
E2 15612939 15613223
refGene_gn=CATSPERG;refGene_tx=XM_023245086.1;refGene_gn=CATSPERG;refGene_tx=XM_023245088.1;refGene_gn=CATSPERG;refGene_tx=XM_023245089.1;refGene_gn=CATSPERG;refGene_tx=XM_023245090.1;refGene_gn=CATSPERG;refGene_tx=XM_023245091.1;refGene_gn=CATSPERG;refGene_tx=XM_023245092.1;refGene_gn=CATSPERG;refGene_tx=XM_023245093.1;refGene_gn=CATSPERG;refGene_tx=XM_023245094.1;refGene_gn=CATSPERG;refGene_tx=XM_023245095.1;refGene_gn=CATSPERG;refGene_tx=XM_023245096.1;refGene_gn=CATSPERG;refGene_tx=XM_023245097.1;refGene_gn=CATSPERG;refGene_tx=XM_023245098.1;refGene_gn=CATSPERG;refGene_tx=XM_023245099.1;refGene_gn=CATSPERG;refGene_tx=XM_023245100.1;refGene_gn=CATSPERG;refGene_tx=XM_023245101.1;refGene_gn=CATSPERG;refGene_tx=XM_023245102.1;refGene_gn=CATSPERG;refGene_tx=XM_023245103.1
E2 15613779 15614380
refGene_gn=CATSPERG;refGene_tx=XM_023245086.1;refGene_gn=CATSPERG;refGene_tx=XM_023245088.1;refGene_gn=CATSPERG;refGene_tx=XM_023245089.1;refGene_gn=CATSPERG;refGene_tx=XM_023245090.1;refGene_gn=CATSPERG;refGene_tx=XM_023245091.1;refGene_gn=CATSPERG;refGene_tx=XM_023245092.1;refGene_gn=CATSPERG;refGene_tx=XM_023245093.1;refGene_gn=CATSPERG;refGene_tx=XM_023245094.1;refGene_gn=CATSPERG;refGene_tx=XM_023245095.1;refGene_gn=CATSPERG;refGene_tx=XM_023245096.1;refGene_gn=CATSPERG;refGene_tx=XM_023245097.1;refGene_gn=CATSPERG;refGene_tx=XM_023245098.1;refGene_gn=CATSPERG;refGene_tx=XM_023245099.1;refGene_gn=CATSPERG;refGene_tx=XM_023245101.1;refGene_gn=CATSPERG;refGene_tx=XM_023245102.1;refGene_gn=CATSPERG;refGene_tx=XM_023245103.1
E2 15620968 15621192 refGene_gn=KCNK6;refGene_tx=XM_019818921.2
E2 15621314 15621710 refGene_gn=KCNK6;refGene_tx=XM_019818921.2
E2 15629203 15629525 refGene_gn=KCNK6;refGene_tx=XM_019818921.2
E2 15632243 15632295
refGene_gn=YIF1B;refGene_tx=XM_011289940.3;refGene_gn=YIF1B;refGene_tx=XM_023245223.1
E2 15635541 15635575 refGene_gn=YIF1B;refGene_tx=XM_023245224.1
E2 15636378 15636617
refGene_gn=YIF1B;refGene_tx=XM_011289940.3;refGene_gn=YIF1B;refGene_tx=XM_023245223.1;refGene_gn=YIF1B;refGene_tx=XM_023245224.1
E2 15636699 15636804
refGene_gn=YIF1B;refGene_tx=XM_011289940.3;refGene_gn=YIF1B;refGene_tx=XM_023245223.1;refGene_gn=YIF1B;refGene_tx=XM_023245224.1
E2 15637005 15637084
refGene_gn=YIF1B;refGene_tx=XM_011289940.3;refGene_gn=YIF1B;refGene_tx=XM_023245223.1;refGene_gn=YIF1B;refGene_tx=XM_023245224.1
E2 15637218 15637276
refGene_gn=YIF1B;refGene_tx=XM_011289940.3;refGene_gn=YIF1B;refGene_tx=XM_023245223.1;refGene_gn=YIF1B;refGene_tx=XM_023245224.1
E2 15638332 15638488
refGene_gn=YIF1B;refGene_tx=XM_011289940.3;refGene_gn=YIF1B;refGene_tx=XM_023245223.1;refGene_gn=YIF1B;refGene_tx=XM_023245224.1
E2 15638569 15638663
refGene_gn=YIF1B;refGene_tx=XM_011289940.3;refGene_gn=YIF1B;refGene_tx

=XM_023245223.1;refGene_gn=YIF1B;refGene_tx=XM_023245224.1
E2 15639542 15639698
refGene_gn=YIF1B;refGene_tx=XM_011289940.3;refGene_gn=YIF1B;refGene_tx
=XM_023245223.1;refGene_gn=YIF1B;refGene_tx=XM_023245224.1
E2 15640082 15640193
refGene_gn=CE2H19orf33;refGene_tx=XM_011289859.3
E2 15640322 15640400
refGene_gn=CE2H19orf33;refGene_tx=XM_011289859.3
E2 15640542 15640655
refGene_gn=CE2H19orf33;refGene_tx=XM_011289859.3
E2 15640773 15640876
refGene_gn=CE2H19orf33;refGene_tx=XM_011289859.3
E2 15650209 15650376 refGene_gn=SPINT2;refGene_tx=XM_003997831.5
E2 15651760 15651799 refGene_gn=SPINT2;refGene_tx=XM_003997831.5
E2 15652113 15652275 refGene_gn=SPINT2;refGene_tx=XM_003997831.5
E2 15652822 15652876 refGene_gn=SPINT2;refGene_tx=XM_003997831.5
E2 15654177 15654237 refGene_gn=SPINT2;refGene_tx=XM_003997831.5
E2 15657109 15657280 refGene_gn=SPINT2;refGene_tx=XM_003997831.5
E2 15678215 15678321 refGene_gn=SPINT2;refGene_tx=XM_003997831.5
E2 15686586 15686787 refGene_gn=PPP1R14A;refGene_tx=XM_023245504.1
E2 15689014 15689095 refGene_gn=PPP1R14A;refGene_tx=XM_023245504.1
E2 15689707 15689740 refGene_gn=PPP1R14A;refGene_tx=XM_023245504.1
E2 15690701 15690830 refGene_gn=PPP1R14A;refGene_tx=XM_023245504.1
E2 15704327 15704359
refGene_gn=DPF1;refGene_tx=XM_019819594.2;refGene_gn=DPF1;refGene_tx=X
M_019819595.2;refGene_gn=DPF1;refGene_tx=XM_019819597.2
E2 15708634 15708768
refGene_gn=DPF1;refGene_tx=XM_003997832.4;refGene_gn=DPF1;refGene_tx=X
M_011289857.3;refGene_gn=DPF1;refGene_tx=XM_019819590.2;refGene_gn=DPF
1;refGene_tx=XM_019819592.2;refGene_gn=DPF1;refGene_tx=XM_019819593.2;
refGene_gn=DPF1;refGene_tx=XM_023245529.1;refGene_gn=DPF1;refGene_tx=X
M_023245530.1;refGene_gn=DPF1;refGene_tx=XM_023245531.1;refGene_gn=DPF
1;refGene_tx=XM_023245532.1;refGene_gn=DPF1;refGene_tx=XM_023245533.1
E2 15710150 15710311
refGene_gn=DPF1;refGene_tx=XM_003997832.4;refGene_gn=DPF1;refGene_tx=X
M_011289857.3;refGene_gn=DPF1;refGene_tx=XM_019819590.2;refGene_gn=DPF
1;refGene_tx=XM_019819592.2;refGene_gn=DPF1;refGene_tx=XM_019819593.2;
refGene_gn=DPF1;refGene_tx=XM_019819594.2;refGene_gn=DPF1;refGene_tx=X
M_019819595.2;refGene_gn=DPF1;refGene_tx=XM_019819597.2;refGene_gn=DPF
1;refGene_tx=XM_023245529.1;refGene_gn=DPF1;refGene_tx=XM_023245530.1;
refGene_gn=DPF1;refGene_tx=XM_023245531.1;refGene_gn=DPF1;refGene_tx=X
M_023245532.1;refGene_gn=DPF1;refGene_tx=XM_023245533.1
E2 15710404 15710512
refGene_gn=DPF1;refGene_tx=XM_003997832.4;refGene_gn=DPF1;refGene_tx=X
M_011289857.3;refGene_gn=DPF1;refGene_tx=XM_019819590.2;refGene_gn=DPF
1;refGene_tx=XM_019819592.2;refGene_gn=DPF1;refGene_tx=XM_019819593.2;
refGene_gn=DPF1;refGene_tx=XM_019819594.2;refGene_gn=DPF1;refGene_tx=X
M_019819595.2;refGene_gn=DPF1;refGene_tx=XM_019819597.2;refGene_gn=DPF
1;refGene_tx=XM_023245529.1;refGene_gn=DPF1;refGene_tx=XM_023245530.1;
refGene_gn=DPF1;refGene_tx=XM_023245531.1;refGene_gn=DPF1;refGene_tx=X

M_023245532.1;refGene_gn=DPF1;refGene_tx=XM_023245533.1
E2 15711975 15712103
refGene_gn=DPF1;refGene_tx=XM_003997832.4;refGene_gn=DPF1;refGene_tx=XM_011289857.3;refGene_gn=DPF1;refGene_tx=XM_019819590.2;refGene_gn=DPF1;refGene_tx=XM_019819592.2;refGene_gn=DPF1;refGene_tx=XM_019819593.2;refGene_gn=DPF1;refGene_tx=XM_019819594.2;refGene_gn=DPF1;refGene_tx=XM_019819595.2;refGene_gn=DPF1;refGene_tx=XM_019819597.2;refGene_gn=DPF1;refGene_tx=XM_023245529.1;refGene_gn=DPF1;refGene_tx=XM_023245530.1;refGene_gn=DPF1;refGene_tx=XM_023245531.1;refGene_gn=DPF1;refGene_tx=XM_023245532.1;refGene_gn=DPF1;refGene_tx=XM_023245533.1
E2 15712361 15712451
refGene_gn=DPF1;refGene_tx=XM_003997832.4;refGene_gn=DPF1;refGene_tx=XM_011289857.3;refGene_gn=DPF1;refGene_tx=XM_019819590.2;refGene_gn=DPF1;refGene_tx=XM_019819592.2;refGene_gn=DPF1;refGene_tx=XM_019819593.2;refGene_gn=DPF1;refGene_tx=XM_019819594.2;refGene_gn=DPF1;refGene_tx=XM_019819595.2;refGene_gn=DPF1;refGene_tx=XM_019819597.2;refGene_gn=DPF1;refGene_tx=XM_023245529.1;refGene_gn=DPF1;refGene_tx=XM_023245530.1;refGene_gn=DPF1;refGene_tx=XM_023245531.1;refGene_gn=DPF1;refGene_tx=XM_023245532.1;refGene_gn=DPF1;refGene_tx=XM_023245533.1
E2 15712829 15712908
refGene_gn=DPF1;refGene_tx=XM_003997832.4;refGene_gn=DPF1;refGene_tx=XM_011289857.3;refGene_gn=DPF1;refGene_tx=XM_019819590.2;refGene_gn=DPF1;refGene_tx=XM_019819592.2;refGene_gn=DPF1;refGene_tx=XM_019819593.2;refGene_gn=DPF1;refGene_tx=XM_019819594.2;refGene_gn=DPF1;refGene_tx=XM_019819595.2;refGene_gn=DPF1;refGene_tx=XM_019819597.2;refGene_gn=DPF1;refGene_tx=XM_023245529.1;refGene_gn=DPF1;refGene_tx=XM_023245530.1;refGene_gn=DPF1;refGene_tx=XM_023245531.1;refGene_gn=DPF1;refGene_tx=XM_023245532.1;refGene_gn=DPF1;refGene_tx=XM_023245533.1
E2 15713106 15713238
refGene_gn=DPF1;refGene_tx=XM_003997832.4;refGene_gn=DPF1;refGene_tx=XM_011289857.3;refGene_gn=DPF1;refGene_tx=XM_019819590.2;refGene_gn=DPF1;refGene_tx=XM_019819592.2;refGene_gn=DPF1;refGene_tx=XM_019819593.2;refGene_gn=DPF1;refGene_tx=XM_019819594.2;refGene_gn=DPF1;refGene_tx=XM_019819595.2;refGene_gn=DPF1;refGene_tx=XM_019819597.2;refGene_gn=DPF1;refGene_tx=XM_023245529.1;refGene_gn=DPF1;refGene_tx=XM_023245530.1;refGene_gn=DPF1;refGene_tx=XM_023245531.1;refGene_gn=DPF1;refGene_tx=XM_023245532.1;refGene_gn=DPF1;refGene_tx=XM_023245533.1
E2 15714120 15714171
refGene_gn=DPF1;refGene_tx=XM_003997832.4;refGene_gn=DPF1;refGene_tx=XM_011289857.3;refGene_gn=DPF1;refGene_tx=XM_019819590.2;refGene_gn=DPF1;refGene_tx=XM_019819592.2;refGene_gn=DPF1;refGene_tx=XM_019819593.2;refGene_gn=DPF1;refGene_tx=XM_019819594.2;refGene_gn=DPF1;refGene_tx=XM_019819595.2;refGene_gn=DPF1;refGene_tx=XM_023245529.1;refGene_gn=DPF1;refGene_tx=XM_023245530.1;refGene_gn=DPF1;refGene_tx=XM_023245533.1
E2 15714268 15714388
refGene_gn=DPF1;refGene_tx=XM_003997832.4;refGene_gn=DPF1;refGene_tx=XM_011289857.3;refGene_gn=DPF1;refGene_tx=XM_019819590.2;refGene_gn=DPF1;refGene_tx=XM_019819592.2;refGene_gn=DPF1;refGene_tx=XM_019819593.2;refGene_gn=DPF1;refGene_tx=XM_019819594.2;refGene_gn=DPF1;refGene_tx=XM_019819595.2;refGene_gn=DPF1;refGene_tx=XM_019819597.2;refGene_gn=DPF

1;refGene_tx=XM_023245529.1;refGene_gn=DPF1;refGene_tx=XM_023245530.1;
refGene_gn=DPF1;refGene_tx=XM_023245531.1;refGene_gn=DPF1;refGene_tx=X
M_023245532.1;refGene_gn=DPF1;refGene_tx=XM_023245533.1
E2 15715509 15715622
refGene_gn=DPF1;refGene_tx=XM_003997832.4;refGene_gn=DPF1;refGene_tx=X
M_011289857.3;refGene_gn=DPF1;refGene_tx=XM_019819590.2;refGene_gn=DPF
1;refGene_tx=XM_019819592.2;refGene_gn=DPF1;refGene_tx=XM_019819593.2;
refGene_gn=DPF1;refGene_tx=XM_019819594.2;refGene_gn=DPF1;refGene_tx=X
M_019819595.2;refGene_gn=DPF1;refGene_tx=XM_019819597.2;refGene_gn=DPF
1;refGene_tx=XM_023245529.1;refGene_gn=DPF1;refGene_tx=XM_023245530.1;
refGene_gn=DPF1;refGene_tx=XM_023245531.1;refGene_gn=DPF1;refGene_tx=X
M_023245532.1;refGene_gn=DPF1;refGene_tx=XM_023245533.1
E2 15715766 15715793 refGene_gn=DPF1;refGene_tx=XM_023245533.1
E2 15716201 15716325
refGene_gn=DPF1;refGene_tx=XM_003997832.4;refGene_gn=DPF1;refGene_tx=X
M_011289857.3;refGene_gn=DPF1;refGene_tx=XM_019819590.2;refGene_gn=DPF
1;refGene_tx=XM_019819592.2;refGene_gn=DPF1;refGene_tx=XM_019819593.2;
refGene_gn=DPF1;refGene_tx=XM_019819594.2;refGene_gn=DPF1;refGene_tx=X
M_019819595.2;refGene_gn=DPF1;refGene_tx=XM_019819597.2;refGene_gn=DPF
1;refGene_tx=XM_023245529.1;refGene_gn=DPF1;refGene_tx=XM_023245530.1;
refGene_gn=DPF1;refGene_tx=XM_023245531.1;refGene_gn=DPF1;refGene_tx=X
M_023245532.1
E2 15716467 15716538
refGene_gn=DPF1;refGene_tx=XM_003997832.4;refGene_gn=DPF1;refGene_tx=X
M_011289857.3;refGene_gn=DPF1;refGene_tx=XM_019819590.2;refGene_gn=DPF
1;refGene_tx=XM_019819592.2;refGene_gn=DPF1;refGene_tx=XM_019819593.2;
refGene_gn=DPF1;refGene_tx=XM_019819594.2;refGene_gn=DPF1;refGene_tx=X
M_019819595.2;refGene_gn=DPF1;refGene_tx=XM_019819597.2;refGene_gn=DPF
1;refGene_tx=XM_023245529.1;refGene_gn=DPF1;refGene_tx=XM_023245530.1;
refGene_gn=DPF1;refGene_tx=XM_023245531.1;refGene_gn=DPF1;refGene_tx=X
M_023245532.1
E2 15722124 15722268
refGene_gn=SIPA1L3;refGene_tx=XM_023245055.1;refGene_gn=SIPA1L3;refGen
e_tx=XM_023245056.1;refGene_gn=SIPA1L3;refGene_tx=XM_023245057.1;refGe
ne_gn=SIPA1L3;refGene_tx=XM_023245058.1;refGene_gn=SIPA1L3;refGene_tx=
XM_023245059.1
E2 15723504 15723586
refGene_gn=SIPA1L3;refGene_tx=XM_023245055.1;refGene_gn=SIPA1L3;refGen
e_tx=XM_023245056.1;refGene_gn=SIPA1L3;refGene_tx=XM_023245057.1;refGe
ne_gn=SIPA1L3;refGene_tx=XM_023245058.1;refGene_gn=SIPA1L3;refGene_tx=
XM_023245059.1
E2 15725282 15725418
refGene_gn=SIPA1L3;refGene_tx=XM_023245055.1;refGene_gn=SIPA1L3;refGen
e_tx=XM_023245056.1;refGene_gn=SIPA1L3;refGene_tx=XM_023245057.1;refGe
ne_gn=SIPA1L3;refGene_tx=XM_023245058.1;refGene_gn=SIPA1L3;refGene_tx=
XM_023245059.1
E2 15729400 15729544
refGene_gn=SIPA1L3;refGene_tx=XM_023245055.1;refGene_gn=SIPA1L3;refGen
e_tx=XM_023245056.1;refGene_gn=SIPA1L3;refGene_tx=XM_023245057.1;refGe
ne_gn=SIPA1L3;refGene_tx=XM_023245058.1;refGene_gn=SIPA1L3;refGene_tx=

XM_023245059.1
E2 15731836 15732086
refGene_gn=SIPA1L3;refGene_tx=XM_023245055.1;refGene_gn=SIPA1L3;refGene_tx=XM_023245056.1;refGene_gn=SIPA1L3;refGene_tx=XM_023245057.1;refGene_gn=SIPA1L3;refGene_tx=XM_023245058.1;refGene_gn=SIPA1L3;refGene_tx=XM_023245059.1
E2 15732826 15732992
refGene_gn=SIPA1L3;refGene_tx=XM_023245055.1;refGene_gn=SIPA1L3;refGene_tx=XM_023245056.1;refGene_gn=SIPA1L3;refGene_tx=XM_023245057.1;refGene_gn=SIPA1L3;refGene_tx=XM_023245058.1;refGene_gn=SIPA1L3;refGene_tx=XM_023245059.1
E2 15743818 15744040
refGene_gn=SIPA1L3;refGene_tx=XM_023245055.1;refGene_gn=SIPA1L3;refGene_tx=XM_023245056.1;refGene_gn=SIPA1L3;refGene_tx=XM_023245057.1;refGene_gn=SIPA1L3;refGene_tx=XM_023245058.1;refGene_gn=SIPA1L3;refGene_tx=XM_023245059.1
E2 15758441 15758869
refGene_gn=SIPA1L3;refGene_tx=XM_023245055.1;refGene_gn=SIPA1L3;refGene_tx=XM_023245056.1;refGene_gn=SIPA1L3;refGene_tx=XM_023245057.1;refGene_gn=SIPA1L3;refGene_tx=XM_023245058.1;refGene_gn=SIPA1L3;refGene_tx=XM_023245059.1
E2 15761103 15761222
refGene_gn=SIPA1L3;refGene_tx=XM_023245055.1;refGene_gn=SIPA1L3;refGene_tx=XM_023245056.1;refGene_gn=SIPA1L3;refGene_tx=XM_023245057.1;refGene_gn=SIPA1L3;refGene_tx=XM_023245058.1;refGene_gn=SIPA1L3;refGene_tx=XM_023245059.1
E2 15767548 15767676
refGene_gn=SIPA1L3;refGene_tx=XM_023245055.1;refGene_gn=SIPA1L3;refGene_tx=XM_023245056.1;refGene_gn=SIPA1L3;refGene_tx=XM_023245057.1;refGene_gn=SIPA1L3;refGene_tx=XM_023245058.1;refGene_gn=SIPA1L3;refGene_tx=XM_023245059.1
E2 15777801 15777939
refGene_gn=SIPA1L3;refGene_tx=XM_023245055.1;refGene_gn=SIPA1L3;refGene_tx=XM_023245056.1;refGene_gn=SIPA1L3;refGene_tx=XM_023245057.1;refGene_gn=SIPA1L3;refGene_tx=XM_023245058.1;refGene_gn=SIPA1L3;refGene_tx=XM_023245059.1
E2 15778701 15778953
refGene_gn=SIPA1L3;refGene_tx=XM_023245055.1;refGene_gn=SIPA1L3;refGene_tx=XM_023245056.1;refGene_gn=SIPA1L3;refGene_tx=XM_023245057.1;refGene_gn=SIPA1L3;refGene_tx=XM_023245058.1;refGene_gn=SIPA1L3;refGene_tx=XM_023245059.1
E2 15786262 15786537
refGene_gn=SIPA1L3;refGene_tx=XM_023245055.1;refGene_gn=SIPA1L3;refGene_tx=XM_023245056.1;refGene_gn=SIPA1L3;refGene_tx=XM_023245057.1;refGene_gn=SIPA1L3;refGene_tx=XM_023245058.1;refGene_gn=SIPA1L3;refGene_tx=XM_023245059.1
E2 15793274 15793851
refGene_gn=SIPA1L3;refGene_tx=XM_023245055.1;refGene_gn=SIPA1L3;refGene_tx=XM_023245056.1;refGene_gn=SIPA1L3;refGene_tx=XM_023245057.1;refGene_gn=SIPA1L3;refGene_tx=XM_023245058.1;refGene_gn=SIPA1L3;refGene_tx=

XM_023245059.1
E2	15800058	15800216
refGene_gn=SIPA1L3;refGene_tx=XM_023245055.1;refGene_gn=SIPA1L3;refGene_tx=XM_023245056.1;refGene_gn=SIPA1L3;refGene_tx=XM_023245057.1;refGene_gn=SIPA1L3;refGene_tx=XM_023245058.1;refGene_gn=SIPA1L3;refGene_tx=XM_023245059.1
E2	15801980	15802084
refGene_gn=SIPA1L3;refGene_tx=XM_023245055.1;refGene_gn=SIPA1L3;refGene_tx=XM_023245056.1;refGene_gn=SIPA1L3;refGene_tx=XM_023245057.1;refGene_gn=SIPA1L3;refGene_tx=XM_023245058.1;refGene_gn=SIPA1L3;refGene_tx=XM_023245059.1
E2	15805059	15805234
refGene_gn=SIPA1L3;refGene_tx=XM_023245055.1;refGene_gn=SIPA1L3;refGene_tx=XM_023245056.1;refGene_gn=SIPA1L3;refGene_tx=XM_023245057.1;refGene_gn=SIPA1L3;refGene_tx=XM_023245058.1;refGene_gn=SIPA1L3;refGene_tx=XM_023245059.1
E2	15806074	15806263
refGene_gn=SIPA1L3;refGene_tx=XM_023245055.1;refGene_gn=SIPA1L3;refGene_tx=XM_023245056.1;refGene_gn=SIPA1L3;refGene_tx=XM_023245057.1;refGene_gn=SIPA1L3;refGene_tx=XM_023245058.1;refGene_gn=SIPA1L3;refGene_tx=XM_023245059.1
E2	15814193	15814324
refGene_gn=SIPA1L3;refGene_tx=XM_023245055.1;refGene_gn=SIPA1L3;refGene_tx=XM_023245056.1;refGene_gn=SIPA1L3;refGene_tx=XM_023245057.1;refGene_gn=SIPA1L3;refGene_tx=XM_023245058.1;refGene_gn=SIPA1L3;refGene_tx=XM_023245059.1
E2	15818256	15819769
refGene_gn=SIPA1L3;refGene_tx=XM_023245055.1;refGene_gn=SIPA1L3;refGene_tx=XM_023245056.1;refGene_gn=SIPA1L3;refGene_tx=XM_023245057.1;refGene_gn=SIPA1L3;refGene_tx=XM_023245058.1;refGene_gn=SIPA1L3;refGene_tx=XM_023245059.1
E2	15964812	15964887	refGene_gn=WDR87;refGene_tx=XM_023245060.1
E2	15965903	15966071	refGene_gn=WDR87;refGene_tx=XM_023245060.1
E2	15966453	15969320	refGene_gn=WDR87;refGene_tx=XM_023245060.1
E2	15969771	15970040	refGene_gn=WDR87;refGene_tx=XM_023245060.1
E2	15970421	15971995	refGene_gn=WDR87;refGene_tx=XM_023245060.1
E2	15971996	15973919	refGene_gn=WDR87;refGene_tx=XM_023245060.1
E2	15974075	15974114	refGene_gn=WDR87;refGene_tx=XM_023245060.1
E2	15974627	15976451	refGene_gn=WDR87;refGene_tx=XM_023245060.1
E2	16006609	16006720
refGene_gn=LOC101097194;refGene_tx=XM_023245951.1
E2	16008707	16008875
refGene_gn=LOC101097194;refGene_tx=XM_023245951.1
E2	16009287	16012163
refGene_gn=LOC101097194;refGene_tx=XM_023245951.1
E2	16012625	16012885
refGene_gn=LOC101097194;refGene_tx=XM_023245951.1
E2	16013408	16015243
refGene_gn=LOC101097194;refGene_tx=XM_023245951.1
E2	16015411	16015550

refGene_gn=LOC101097194;refGene_tx=XM_023245951.1
E2 16015642 16016717
refGene_gn=LOC101097194;refGene_tx=XM_023245951.1
E2 16017536 16019337
refGene_gn=LOC101097194;refGene_tx=XM_023245951.1
E2 16038563 16040410
refGene_gn=LOC111556200;refGene_tx=XM_006941310.3;refGene_gn=LOC111556200;refGene_tx=XM_011289854.3;refGene_gn=LOC111556200;refGene_tx=XM_019818744.2
E2 16048188 16048287
refGene_gn=LOC111556200;refGene_tx=XM_006941310.3;refGene_gn=LOC111556200;refGene_tx=XM_011289854.3;refGene_gn=LOC111556200;refGene_tx=XM_019818744.2
E2 16050851 16050978
refGene_gn=LOC111556200;refGene_tx=XM_006941310.3;refGene_gn=LOC111556200;refGene_tx=XM_011289854.3;refGene_gn=LOC111556200;refGene_tx=XM_019818744.2
E2 16052710 16052719
refGene_gn=LOC111556200;refGene_tx=XM_006941310.3;refGene_gn=LOC111556200;refGene_tx=XM_011289854.3;refGene_gn=LOC111556200;refGene_tx=XM_019818744.2
E2 16067423 16069135
refGene_gn=LOC102900592;refGene_tx=XM_023245446.1;refGene_gn=LOC102900592;refGene_tx=XM_023245447.1
E2 16091870 16091966
refGene_gn=LOC102900592;refGene_tx=XM_023245446.1;refGene_gn=LOC102900592;refGene_tx=XM_023245447.1
E2 16092842 16092873
refGene_gn=LOC102900592;refGene_tx=XM_023245446.1
E2 16126298 16128634 refGene_gn=ZNF585A;refGene_tx=XM_006941311.4
E2 16130365 16130458 refGene_gn=ZNF585A;refGene_tx=XM_006941311.4
E2 16130717 16130844 refGene_gn=ZNF585A;refGene_tx=XM_006941311.4
E2 16140542 16140620 refGene_gn=ZNF585A;refGene_tx=XM_006941311.4
E2 16181666 16181675
refGene_gn=LOC101097690;refGene_tx=XM_023245458.1
E2 16184059 16184186
refGene_gn=LOC101097690;refGene_tx=XM_023245458.1
E2 16184494 16184590
refGene_gn=LOC101097690;refGene_tx=XM_023245458.1
E2 16189596 16189721
refGene_gn=LOC101097690;refGene_tx=XM_023245458.1
E2 16189722 16190541
refGene_gn=LOC101097690;refGene_tx=XM_023245458.1
E2 16190542 16190794
refGene_gn=LOC101097690;refGene_tx=XM_023245458.1
E2 16225593 16225626
refGene_gn=LOC109491353;refGene_tx=XM_006941318.3
E2 16238045 16238172
refGene_gn=LOC109491353;refGene_tx=XM_006941318.3
E2 16238855 16238951

refGene_gn=LOC109491353;refGene_tx=XM_006941318.3
E2 16248021 16249056
refGene_gn=LOC109491353;refGene_tx=XM_006941318.3
E2 16249058 16249144
refGene_gn=LOC109491353;refGene_tx=XM_006941318.3
E2 16249145 16249784
refGene_gn=LOC109491353;refGene_tx=XM_006941318.3
E2 16268549 16268582 refGene_gn=ZNF527;refGene_tx=XM_003997838.4
E2 16273305 16273432
refGene_gn=ZNF527;refGene_tx=XM_003997838.4;refGene_gn=ZNF527;refGene_tx=XM_019818747.1
E2 16274527 16274623
refGene_gn=ZNF527;refGene_tx=XM_003997838.4;refGene_gn=ZNF527;refGene_tx=XM_019818747.1
E2 16291595 16291617 refGene_gn=ZNF527;refGene_tx=XM_006941320.3
E2 16295016 16296590
refGene_gn=ZNF527;refGene_tx=XM_003997838.4;refGene_gn=ZNF527;refGene_tx=XM_006941320.3;refGene_gn=ZNF527;refGene_tx=XM_019818747.1
E2 16358700 16359374
refGene_gn=LOC102901034;refGene_tx=XM_006941321.4
E2 16359376 16360525
refGene_gn=LOC102901034;refGene_tx=XM_006941321.4
E2 16365869 16365965
refGene_gn=LOC102901034;refGene_tx=XM_006941321.4
E2 16366277 16366404
refGene_gn=LOC102901034;refGene_tx=XM_006941321.4
E2 16378713 16378728
refGene_gn=LOC102901034;refGene_tx=XM_006941321.4
E2 16394468 16394471
refGene_gn=ZNF570;refGene_tx=XM_019818769.2;refGene_gn=ZNF570;refGene_tx=XM_023245449.1
E2 16401784 16401868
refGene_gn=ZNF570;refGene_tx=XM_019818769.2;refGene_gn=ZNF570;refGene_tx=XM_023245449.1;refGene_gn=ZNF570;refGene_tx=XM_023245450.1;refGene_gn=ZNF570;refGene_tx=XM_023245451.1;refGene_gn=ZNF570;refGene_tx=XM_023245452.1
E2 16407895 16408022
refGene_gn=ZNF570;refGene_tx=XM_019818769.2;refGene_gn=ZNF570;refGene_tx=XM_023245449.1;refGene_gn=ZNF570;refGene_tx=XM_023245450.1;refGene_gn=ZNF570;refGene_tx=XM_023245451.1;refGene_gn=ZNF570;refGene_tx=XM_023245452.1
E2 16408346 16408442
refGene_gn=ZNF570;refGene_tx=XM_019818769.2;refGene_gn=ZNF570;refGene_tx=XM_023245449.1;refGene_gn=ZNF570;refGene_tx=XM_023245450.1;refGene_gn=ZNF570;refGene_tx=XM_023245451.1;refGene_gn=ZNF570;refGene_tx=XM_023245452.1
E2 16423686 16423705 refGene_gn=ZNF570;refGene_tx=XM_023245453.1
E2 16428168 16429526
refGene_gn=ZNF570;refGene_tx=XM_019818769.2;refGene_gn=ZNF570;refGene_tx=XM_023245449.1;refGene_gn=ZNF570;refGene_tx=XM_023245450.1;refGene_

gn=ZNF570;refGene_tx=XM_023245451.1;refGene_gn=ZNF570;refGene_tx=XM_02
3245452.1;refGene_gn=ZNF570;refGene_tx=XM_023245453.1
E2 16455986 16457917
refGene_gn=LOC102901416;refGene_tx=XM_011289849.2;refGene_gn=LOC102901
416;refGene_tx=XM_011289850.2;refGene_gn=LOC102901416;refGene_tx=XM_01
9818717.1;refGene_gn=LOC102901416;refGene_tx=XM_019818718.1;refGene_gn
=LOC102901416;refGene_tx=XM_019818719.1;refGene_gn=LOC102901416;refGen
e_tx=XM_019818720.2
E2 16473185 16473315
refGene_gn=LOC102901416;refGene_tx=XM_011289849.2
E2 16474600 16474609
refGene_gn=LOC102901416;refGene_tx=XM_011289849.2
E2 16515817 16517514
refGene_gn=LOC101082283;refGene_tx=XM_011289938.3;refGene_gn=LOC101082
283;refGene_tx=XM_019818728.2
E2 16536079 16536178
refGene_gn=LOC101082283;refGene_tx=XM_019818728.2
E2 16536456 16536583
refGene_gn=LOC101082283;refGene_tx=XM_011289938.3;refGene_gn=LOC101082
283;refGene_tx=XM_019818728.2
E2 16537958 16537967
refGene_gn=LOC101082283;refGene_tx=XM_011289938.3;refGene_gn=LOC101082
283;refGene_tx=XM_019818728.2
E2 16611536 16612400
refGene_gn=LOC102901341;refGene_tx=XM_019818788.2
E2 16647265 16647288
refGene_gn=LOC102901341;refGene_tx=XM_019818788.2
E2 16655986 16658138
refGene_gn=LOC102901341;refGene_tx=XM_019818788.2
E2 16702192 16702201
refGene_gn=ZNF790;refGene_tx=XM_003997835.5;refGene_gn=ZNF790;refGene_
tx=XM_011290061.3;refGene_gn=ZNF790;refGene_tx=XM_019818737.2;refGene_
gn=ZNF790;refGene_tx=XM_019818742.2;refGene_gn=ZNF790;refGene_tx=XM_02
3245442.1;refGene_gn=ZNF790;refGene_tx=XM_023245443.1
E2 16703221 16703348
refGene_gn=ZNF790;refGene_tx=XM_003997835.5;refGene_gn=ZNF790;refGene_
tx=XM_011290061.3;refGene_gn=ZNF790;refGene_tx=XM_019818737.2;refGene_
gn=ZNF790;refGene_tx=XM_019818738.2;refGene_gn=ZNF790;refGene_tx=XM_01
9818739.2;refGene_gn=ZNF790;refGene_tx=XM_019818742.2;refGene_gn=ZNF79
0;refGene_tx=XM_023245442.1;refGene_gn=ZNF790;refGene_tx=XM_023245443.
1
E2 16703644 16703740
refGene_gn=ZNF790;refGene_tx=XM_003997835.5;refGene_gn=ZNF790;refGene_
tx=XM_011290061.3;refGene_gn=ZNF790;refGene_tx=XM_019818737.2;refGene_
gn=ZNF790;refGene_tx=XM_019818738.2;refGene_gn=ZNF790;refGene_tx=XM_01
9818739.2;refGene_gn=ZNF790;refGene_tx=XM_019818742.2;refGene_gn=ZNF79
0;refGene_tx=XM_023245442.1;refGene_gn=ZNF790;refGene_tx=XM_023245443.
1
E2 16711370 16713028
refGene_gn=ZNF790;refGene_tx=XM_003997835.5;refGene_gn=ZNF790;refGene_

tx=XM_011290061.3;refGene_gn=ZNF790;refGene_tx=XM_019818737.2;refGene_gn=ZNF790;refGene_tx=XM_019818738.2;refGene_gn=ZNF790;refGene_tx=XM_019818739.2;refGene_gn=ZNF790;refGene_tx=XM_019818742.2;refGene_gn=ZNF790;refGene_tx=XM_023245442.1;refGene_gn=ZNF790;refGene_tx=XM_023245443.1
E2 16715885 16716046
refGene_gn=ZNF790;refGene_tx=XM_019818742.2;refGene_gn=ZNF790;refGene_tx=XM_023245442.1;refGene_gn=ZNF790;refGene_tx=XM_023245443.1
E2 16716866 16716873 refGene_gn=ZNF790;refGene_tx=XM_019818742.2
E2 16737190 16738884
refGene_gn=LOC102900191;refGene_tx=XM_011290066.3;refGene_gn=LOC102900191;refGene_tx=XM_019818702.2;refGene_gn=LOC102900191;refGene_tx=XM_019818703.2;refGene_gn=LOC102900191;refGene_tx=XM_019818705.2;refGene_gn=LOC102900191;refGene_tx=XM_019818706.2;refGene_gn=LOC102900191;refGene_tx=XM_019818709.2;refGene_gn=LOC102900191;refGene_tx=XM_019818710.2;refGene_gn=LOC102900191;refGene_tx=XM_023245423.1;refGene_gn=LOC102900191;refGene_tx=XM_023245424.1;refGene_gn=LOC102900191;refGene_tx=XM_023245425.1
E2 16745185 16745284
refGene_gn=LOC102900191;refGene_tx=XM_011290066.3;refGene_gn=LOC102900191;refGene_tx=XM_019818702.2;refGene_gn=LOC102900191;refGene_tx=XM_019818703.2;refGene_gn=LOC102900191;refGene_tx=XM_019818705.2;refGene_gn=LOC102900191;refGene_tx=XM_019818706.2;refGene_gn=LOC102900191;refGene_tx=XM_019818709.2;refGene_gn=LOC102900191;refGene_tx=XM_019818710.2;refGene_gn=LOC102900191;refGene_tx=XM_023245423.1;refGene_gn=LOC102900191;refGene_tx=XM_023245424.1;refGene_gn=LOC102900191;refGene_tx=XM_023245425.1
E2 16746575 16746702
refGene_gn=LOC102900191;refGene_tx=XM_011290066.3;refGene_gn=LOC102900191;refGene_tx=XM_019818702.2;refGene_gn=LOC102900191;refGene_tx=XM_019818703.2;refGene_gn=LOC102900191;refGene_tx=XM_019818705.2;refGene_gn=LOC102900191;refGene_tx=XM_019818706.2;refGene_gn=LOC102900191;refGene_tx=XM_019818709.2;refGene_gn=LOC102900191;refGene_tx=XM_019818710.2;refGene_gn=LOC102900191;refGene_tx=XM_023245423.1;refGene_gn=LOC102900191;refGene_tx=XM_023245424.1;refGene_gn=LOC102900191;refGene_tx=XM_023245425.1
E2 16747547 16747596 refGene_gn=ZNF790;refGene_tx=XM_023245443.1
E2 16748982 16749068
refGene_gn=LOC102900191;refGene_tx=XM_011290066.3;refGene_gn=LOC102900191;refGene_tx=XM_019818702.2;refGene_gn=LOC102900191;refGene_tx=XM_019818703.2;refGene_gn=LOC102900191;refGene_tx=XM_019818705.2;refGene_gn=LOC102900191;refGene_tx=XM_019818706.2;refGene_gn=LOC102900191;refGene_tx=XM_019818709.2;refGene_gn=LOC102900191;refGene_tx=XM_019818710.2;refGene_gn=LOC102900191;refGene_tx=XM_023245423.1;refGene_gn=LOC102900191;refGene_tx=XM_023245424.1;refGene_gn=LOC102900191;refGene_tx=XM_023245425.1
E2 16755962 16755990
refGene_gn=LOC102900191;refGene_tx=XM_011290066.3
E2 16756262 16756443
refGene_gn=LOC102900191;refGene_tx=XM_023245425.1

E2 16775309 16776679
refGene_gn=LOC102900191;refGene_tx=XM_023245426.1;refGene_gn=LOC102900191;refGene_tx=XM_023245427.1
E2 16783070 16783169
refGene_gn=LOC102900191;refGene_tx=XM_019818702.2;refGene_gn=LOC102900191;refGene_tx=XM_019818703.2;refGene_gn=LOC102900191;refGene_tx=XM_019818705.2;refGene_gn=LOC102900191;refGene_tx=XM_019818706.2;refGene_gn=LOC102900191;refGene_tx=XM_023245423.1;refGene_gn=LOC102900191;refGene_tx=XM_023245424.1;refGene_gn=LOC102900191;refGene_tx=XM_023245426.1;refGene_gn=LOC102900191;refGene_tx=XM_023245427.1
E2 16784148 16784293
refGene_gn=LOC102900191;refGene_tx=XM_019818702.2;refGene_gn=LOC102900191;refGene_tx=XM_019818703.2;refGene_gn=LOC102900191;refGene_tx=XM_019818705.2;refGene_gn=LOC102900191;refGene_tx=XM_019818706.2;refGene_gn=LOC102900191;refGene_tx=XM_023245423.1;refGene_gn=LOC102900191;refGene_tx=XM_023245424.1;refGene_gn=LOC102900191;refGene_tx=XM_023245426.1;refGene_gn=LOC102900191;refGene_tx=XM_023245427.1
E2 16786407 16786416
refGene_gn=LOC102900191;refGene_tx=XM_019818702.2;refGene_gn=LOC102900191;refGene_tx=XM_019818703.2;refGene_gn=LOC102900191;refGene_tx=XM_019818705.2;refGene_gn=LOC102900191;refGene_tx=XM_019818706.2
E2 16836185 16836226
refGene_gn=ZNF793;refGene_tx=XM_019818804.1;refGene_gn=ZNF793;refGene_tx=XM_019818807.1
E2 16850937 16850980
refGene_gn=ZNF793;refGene_tx=XM_011290056.2;refGene_gn=ZNF793;refGene_tx=XM_019818804.1;refGene_gn=ZNF793;refGene_tx=XM_019818807.1;refGene_gn=ZNF793;refGene_tx=XM_019818808.1
E2 16856204 16856334
refGene_gn=ZNF793;refGene_tx=XM_011290056.2;refGene_gn=ZNF793;refGene_tx=XM_019818804.1;refGene_gn=ZNF793;refGene_tx=XM_019818807.1;refGene_gn=ZNF793;refGene_tx=XM_019818808.1
E2 16856862 16856958
refGene_gn=ZNF793;refGene_tx=XM_011290056.2;refGene_gn=ZNF793;refGene_tx=XM_019818804.1;refGene_gn=ZNF793;refGene_tx=XM_019818807.1;refGene_gn=ZNF793;refGene_tx=XM_019818808.1
E2 16858699 16859652
refGene_gn=ZNF793;refGene_tx=XM_011290056.2;refGene_gn=ZNF793;refGene_tx=XM_019818804.1;refGene_gn=ZNF793;refGene_tx=XM_019818807.1;refGene_gn=ZNF793;refGene_tx=XM_019818808.1
E2 16880498 16881478
refGene_gn=ZNF829;refGene_tx=XM_006941332.4;refGene_gn=ZNF829;refGene_tx=XM_006941333.4;refGene_gn=ZNF829;refGene_tx=XM_011290048.3;refGene_gn=ZNF829;refGene_tx=XM_019818795.2;refGene_gn=ZNF829;refGene_tx=XM_019818796.2;refGene_gn=ZNF829;refGene_tx=XM_019818797.2;refGene_gn=ZNF829;refGene_tx=XM_019818798.2;refGene_gn=ZNF829;refGene_tx=XM_019818799.2;refGene_gn=ZNF829;refGene_tx=XM_019818800.2;refGene_gn=ZNF829;refGene_tx=XM_023245459.1;refGene_gn=ZNF829;refGene_tx=XM_023245460.1;refGene_gn=ZNF829;refGene_tx=XM_023245461.1;refGene_gn=ZNF829;refGene_tx=XM_023245462.1;refGene_gn=ZNF829;refGene_tx=XM_023245463.1;refGene_gn=ZNF

829;refGene_tx=XM_023245464.1;refGene_gn=ZNF829;refGene_tx=XM_023245465.1;refGene_gn=ZNF829;refGene_tx=XM_023245466.1;refGene_gn=ZNF829;refGene_tx=XM_023245467.1;refGene_gn=ZNF829;refGene_tx=XM_023245468.1;refGene_gn=ZNF829;refGene_tx=XM_023245469.1
E2 16884962 16885026
refGene_gn=ZNF829;refGene_tx=XM_019818795.2;refGene_gn=ZNF829;refGene_tx=XM_019818797.2;refGene_gn=ZNF829;refGene_tx=XM_019818798.2;refGene_gn=ZNF829;refGene_tx=XM_019818799.2;refGene_gn=ZNF829;refGene_tx=XM_019818800.2;refGene_gn=ZNF829;refGene_tx=XM_023245462.1
E2 16885185 16885249
refGene_gn=ZNF829;refGene_tx=XM_019818795.2;refGene_gn=ZNF829;refGene_tx=XM_019818797.2;refGene_gn=ZNF829;refGene_tx=XM_019818798.2;refGene_gn=ZNF829;refGene_tx=XM_019818799.2;refGene_gn=ZNF829;refGene_tx=XM_019818800.2
E2 16894057 16894153
refGene_gn=ZNF829;refGene_tx=XM_006941332.4;refGene_gn=ZNF829;refGene_tx=XM_006941333.4;refGene_gn=ZNF829;refGene_tx=XM_011290048.3;refGene_gn=ZNF829;refGene_tx=XM_019818796.2;refGene_gn=ZNF829;refGene_tx=XM_023245459.1;refGene_gn=ZNF829;refGene_tx=XM_023245460.1;refGene_gn=ZNF829;refGene_tx=XM_023245461.1
E2 16894451 16894578
refGene_gn=ZNF829;refGene_tx=XM_006941332.4;refGene_gn=ZNF829;refGene_tx=XM_006941333.4;refGene_gn=ZNF829;refGene_tx=XM_011290048.3;refGene_gn=ZNF829;refGene_tx=XM_019818796.2
E2 16904877 16904946
refGene_gn=ZNF829;refGene_tx=XM_006941332.4;refGene_gn=ZNF829;refGene_tx=XM_006941333.4;refGene_gn=ZNF829;refGene_tx=XM_011290048.3;refGene_gn=ZNF829;refGene_tx=XM_019818795.2;refGene_gn=ZNF829;refGene_tx=XM_019818796.2;refGene_gn=ZNF829;refGene_tx=XM_019818797.2;refGene_gn=ZNF829;refGene_tx=XM_019818798.2;refGene_gn=ZNF829;refGene_tx=XM_019818799.2;refGene_gn=ZNF829;refGene_tx=XM_019818800.2
E2 16905191 16905376
refGene_gn=ZNF829;refGene_tx=XM_019818795.2;refGene_gn=ZNF829;refGene_tx=XM_019818796.2
E2 16905624 16905629 refGene_gn=ZNF829;refGene_tx=XM_019818797.2
E2 16905993 16906098
refGene_gn=LOC101098182;refGene_tx=XM_019818831.2
E2 16910267 16910448
refGene_gn=LOC101098182;refGene_tx=XM_019818831.2
E2 16912871 16912930
refGene_gn=LOC101098182;refGene_tx=XM_019818831.2
E2 16920927 16920942
refGene_gn=LOC101098182;refGene_tx=XM_019818831.2
E2 16934035 16934162
refGene_gn=LOC101098182;refGene_tx=XM_019818831.2
E2 16934411 16934507
refGene_gn=LOC101098182;refGene_tx=XM_019818831.2
E2 16938275 16939852
refGene_gn=LOC101098182;refGene_tx=XM_019818831.2
E2 16942562 16942574

refGene_gn=LOC101098182;refGene_tx=XM_019818831.2
E2	16983023	16984160
refGene_gn=LOC101098182;refGene_tx=XM_019818831.2
E2	17007627	17008994
refGene_gn=ZFP14;refGene_tx=XM_019818749.2;refGene_gn=ZFP14;refGene_tx=XM_019818750.2;refGene_gn=ZFP14;refGene_tx=XM_019818751.2;refGene_gn=ZFP14;refGene_tx=XM_019818752.2;refGene_gn=ZFP14;refGene_tx=XM_019818753.2
E2	17021942	17022041
refGene_gn=ZFP14;refGene_tx=XM_019818749.2;refGene_gn=ZFP14;refGene_tx=XM_019818750.2;refGene_gn=ZFP14;refGene_tx=XM_019818751.2;refGene_gn=ZFP14;refGene_tx=XM_019818753.2
E2	17023087	17023214
refGene_gn=ZFP14;refGene_tx=XM_019818749.2;refGene_gn=ZFP14;refGene_tx=XM_019818750.2;refGene_gn=ZFP14;refGene_tx=XM_019818751.2;refGene_gn=ZFP14;refGene_tx=XM_019818752.2;refGene_gn=ZFP14;refGene_tx=XM_019818753.2
E2	17026648	17026734
refGene_gn=ZFP14;refGene_tx=XM_019818749.2;refGene_gn=ZFP14;refGene_tx=XM_019818750.2;refGene_gn=ZFP14;refGene_tx=XM_019818751.2;refGene_gn=ZFP14;refGene_tx=XM_019818752.2;refGene_gn=ZFP14;refGene_tx=XM_019818753.2
E2	17029732	17029817
refGene_gn=ZFP14;refGene_tx=XM_019818749.2;refGene_gn=ZFP14;refGene_tx=XM_019818750.2;refGene_gn=ZFP14;refGene_tx=XM_019818751.2;refGene_gn=ZFP14;refGene_tx=XM_019818752.2
E2	17049816	17051171
refGene_gn=LOC101085512;refGene_tx=XM_003997926.5;refGene_gn=LOC101085512;refGene_tx=XM_011290046.3;refGene_gn=LOC101085512;refGene_tx=XM_011290047.2;refGene_gn=LOC101085512;refGene_tx=XM_023245456.1;refGene_gn=LOC101085512;refGene_tx=XM_023245457.1
E2	17055585	17055670
refGene_gn=LOC101085512;refGene_tx=XM_011290047.2
E2	17060826	17060919
refGene_gn=LOC101085512;refGene_tx=XM_003997926.5;refGene_gn=LOC101085512;refGene_tx=XM_011290046.3
E2	17062458	17062585
refGene_gn=LOC101085512;refGene_tx=XM_003997926.5;refGene_gn=LOC101085512;refGene_tx=XM_011290046.3
E2	17063898	17063907
refGene_gn=LOC101085512;refGene_tx=XM_003997926.5
E2	17068371	17068376
refGene_gn=LOC101085512;refGene_tx=XM_023245457.1
E2	17081672	17081695
refGene_gn=LOC111558045;refGene_tx=XM_023245496.1;refGene_gn=LOC111558045;refGene_tx=XM_023245497.1
E2	17090447	17090546
refGene_gn=LOC111558045;refGene_tx=XM_023245496.1;refGene_gn=LOC111558045;refGene_tx=XM_023245497.1
E2	17090832	17090950

refGene_gn=LOC111558045;refGene_tx=XM_023245496.1;refGene_gn=LOC111558045;refGene_tx=XM_023245497.1
E2 17093500 17094525
refGene_gn=LOC101085512;refGene_tx=XM_019818756.1;refGene_gn=LOC101085512;refGene_tx=XM_023245455.1;refGene_gn=LOC101085512;refGene_tx=XM_023245456.1;refGene_gn=LOC101085512;refGene_tx=XM_023245457.1
E2 17104840 17104939
refGene_gn=LOC101085512;refGene_tx=XM_023245455.1;refGene_gn=LOC101085512;refGene_tx=XM_023245456.1;refGene_gn=LOC101085512;refGene_tx=XM_023245457.1
E2 17105225 17105343
refGene_gn=LOC101085512;refGene_tx=XM_023245455.1;refGene_gn=LOC101085512;refGene_tx=XM_023245456.1;refGene_gn=LOC101085512;refGene_tx=XM_023245457.1
E2 17166568 17167798 refGene_gn=ZNF260;refGene_tx=XM_006941369.4
E2 17199922 17201337 refGene_gn=ZNF529;refGene_tx=XM_023245454.1
E2 17204837 17204964 refGene_gn=ZNF529;refGene_tx=XM_023245454.1
E2 17205885 17205939 refGene_gn=ZNF529;refGene_tx=XM_023245454.1
E2 17280777 17280786
refGene_gn=ZNF382;refGene_tx=XM_006941373.2;refGene_gn=ZNF382;refGene_tx=XM_006941374.2;refGene_gn=ZNF382;refGene_tx=XM_019818777.2;refGene_gn=ZNF382;refGene_tx=XM_019818778.1;refGene_gn=ZNF382;refGene_tx=XM_019818779.1
E2 17284247 17284374
refGene_gn=ZNF382;refGene_tx=XM_006941373.2;refGene_gn=ZNF382;refGene_tx=XM_006941374.2;refGene_gn=ZNF382;refGene_tx=XM_006941376.2;refGene_gn=ZNF382;refGene_tx=XM_019818777.2;refGene_gn=ZNF382;refGene_tx=XM_019818778.1;refGene_gn=ZNF382;refGene_tx=XM_019818779.1;refGene_gn=ZNF382;refGene_tx=XM_019818780.1
E2 17284646 17284742
refGene_gn=ZNF382;refGene_tx=XM_006941373.2;refGene_gn=ZNF382;refGene_tx=XM_006941374.2;refGene_gn=ZNF382;refGene_tx=XM_006941376.2;refGene_gn=ZNF382;refGene_tx=XM_019818777.2;refGene_gn=ZNF382;refGene_tx=XM_019818778.1;refGene_gn=ZNF382;refGene_tx=XM_019818779.1;refGene_gn=ZNF382;refGene_tx=XM_019818780.1
E2 17302325 17303746
refGene_gn=ZNF382;refGene_tx=XM_006941373.2;refGene_gn=ZNF382;refGene_tx=XM_006941374.2;refGene_gn=ZNF382;refGene_tx=XM_006941376.2;refGene_gn=ZNF382;refGene_tx=XM_019818777.2;refGene_gn=ZNF382;refGene_tx=XM_019818778.1;refGene_gn=ZNF382;refGene_tx=XM_019818779.1;refGene_gn=ZNF382;refGene_tx=XM_019818780.1
E2 17325687 17327075
refGene_gn=ZNF461;refGene_tx=XM_006941444.4;refGene_gn=ZNF461;refGene_tx=XM_006941445.2;refGene_gn=ZNF461;refGene_tx=XM_011290040.3;refGene_gn=ZNF461;refGene_tx=XM_011290041.3;refGene_gn=ZNF461;refGene_tx=XM_011290042.2;refGene_gn=ZNF461;refGene_tx=XM_019818782.2
E2 17341859 17341955
refGene_gn=ZNF461;refGene_tx=XM_006941444.4;refGene_gn=ZNF461;refGene_tx=XM_006941445.2;refGene_gn=ZNF461;refGene_tx=XM_011290040.3;refGene_gn=ZNF461;refGene_tx=XM_011290041.3;refGene_gn=ZNF461;refGene_tx=XM_01

1290042.2
E2 17342796 17342923
refGene_gn=ZNF461;refGene_tx=XM_006941444.4;refGene_gn=ZNF461;refGene_tx=XM_006941445.2;refGene_gn=ZNF461;refGene_tx=XM_011290040.3;refGene_gn=ZNF461;refGene_tx=XM_011290041.3;refGene_gn=ZNF461;refGene_tx=XM_011290042.2
E2 17349059 17349062 refGene_gn=ZNF461;refGene_tx=XM_006941445.2
E2 17356675 17356684 refGene_gn=ZNF461;refGene_tx=XM_006941444.4
E2 17364283 17364292 refGene_gn=ZNF461;refGene_tx=XM_011290040.3
E2 17384106 17384128 refGene_gn=ZNF567;refGene_tx=XM_019818721.2
E2 17387293 17387367 refGene_gn=ZNF567;refGene_tx=XM_019818721.2
E2 17397440 17397567 refGene_gn=ZNF567;refGene_tx=XM_019818721.2
E2 17397831 17397918 refGene_gn=ZNF567;refGene_tx=XM_019818721.2
E2 17401815 17403536 refGene_gn=ZNF567;refGene_tx=XM_019818721.2
E2 17449845 17450724
refGene_gn=ZNF146;refGene_tx=XM_003997927.5;refGene_gn=ZNF146;refGene_tx=XM_011290035.3;refGene_gn=ZNF146;refGene_tx=XM_011290036.3;refGene_gn=ZNF146;refGene_tx=XM_011290038.3;refGene_gn=ZNF146;refGene_tx=XM_019818815.2
E2 17490255 17490264
refGene_gn=ZNF565;refGene_tx=XM_006941440.4;refGene_gn=ZNF565;refGene_tx=XM_006941443.4;refGene_gn=ZNF565;refGene_tx=XM_019818783.2;refGene_gn=ZNF565;refGene_tx=XM_019818785.2;refGene_gn=ZNF565;refGene_tx=XM_019818787.2
E2 17494497 17494624
refGene_gn=ZNF565;refGene_tx=XM_006941440.4;refGene_gn=ZNF565;refGene_tx=XM_006941443.4;refGene_gn=ZNF565;refGene_tx=XM_019818783.2;refGene_gn=ZNF565;refGene_tx=XM_019818785.2;refGene_gn=ZNF565;refGene_tx=XM_019818787.2
E2 17495318 17495414
refGene_gn=ZNF565;refGene_tx=XM_006941440.4;refGene_gn=ZNF565;refGene_tx=XM_006941443.4;refGene_gn=ZNF565;refGene_tx=XM_019818783.2;refGene_gn=ZNF565;refGene_tx=XM_019818785.2;refGene_gn=ZNF565;refGene_tx=XM_019818787.2
E2 17504182 17505453
refGene_gn=ZNF565;refGene_tx=XM_006941440.4;refGene_gn=ZNF565;refGene_tx=XM_006941443.4;refGene_gn=ZNF565;refGene_tx=XM_019818783.2;refGene_gn=ZNF565;refGene_tx=XM_019818785.2;refGene_gn=ZNF565;refGene_tx=XM_019818787.2
E2 17529652 17529765
refGene_gn=LOC101083819;refGene_tx=XM_003997919.3;refGene_gn=LOC101083819;refGene_tx=XM_019818828.1
E2 17530395 17530482
refGene_gn=LOC101083819;refGene_tx=XM_003997919.3;refGene_gn=LOC101083819;refGene_tx=XM_019818828.1
E2 17530618 17530703
refGene_gn=LOC101083819;refGene_tx=XM_003997919.3;refGene_gn=LOC101083819;refGene_tx=XM_019818828.1
E2 17531177 17531233
refGene_gn=LOC101083819;refGene_tx=XM_003997919.3;refGene_gn=LOC101083

819;refGene_tx=XM_019818828.1
E2	17532553 17532580 refGene_gn=CAPNS1;refGene_tx=XM_003997895.5
E2	17532763 17532822 refGene_gn=CAPNS1;refGene_tx=XM_003997895.5
E2	17535082 17535199 refGene_gn=CAPNS1;refGene_tx=XM_003997895.5
E2	17535341 17535420 refGene_gn=CAPNS1;refGene_tx=XM_003997895.5
E2	17535520 17535589 refGene_gn=CAPNS1;refGene_tx=XM_003997895.5
E2	17537570 17537635 refGene_gn=CAPNS1;refGene_tx=XM_003997895.5
E2	17537736 17537794 refGene_gn=CAPNS1;refGene_tx=XM_003997895.5
E2	17537947 17538037 refGene_gn=CAPNS1;refGene_tx=XM_003997895.5
E2	17538319 17538353 refGene_gn=CAPNS1;refGene_tx=XM_003997895.5
E2	17539339 17539539 refGene_gn=CAPNS1;refGene_tx=XM_003997895.5
E2	17558617 17558732
refGene_gn=TBCB;refGene_tx=XM_003997911.5;refGene_gn=TBCB;refGene_tx=XM_019818818.1
E2	17558904 17558977
refGene_gn=TBCB;refGene_tx=XM_003997911.5;refGene_gn=TBCB;refGene_tx=XM_019818818.1
E2	17561978 17562170
refGene_gn=TBCB;refGene_tx=XM_003997911.5;refGene_gn=TBCB;refGene_tx=XM_019818818.1
E2	17562842 17562939
refGene_gn=TBCB;refGene_tx=XM_003997911.5;refGene_gn=TBCB;refGene_tx=XM_019818818.1
E2	17566293 17566437
refGene_gn=TBCB;refGene_tx=XM_003997911.5;refGene_gn=TBCB;refGene_tx=XM_019818818.1
E2	17566707 17566725 refGene_gn=TBCB;refGene_tx=XM_019818818.1
E2	17566916 17567030 refGene_gn=TBCB;refGene_tx=XM_003997911.5
E2	17567690 17567749 refGene_gn=POLR2I;refGene_tx=XM_003997912.5
E2	17567827 17567882 refGene_gn=POLR2I;refGene_tx=XM_003997912.5
E2	17568227 17568301 refGene_gn=POLR2I;refGene_tx=XM_003997912.5
E2	17568389 17568464 refGene_gn=POLR2I;refGene_tx=XM_003997912.5
E2	17568566 17568618 refGene_gn=POLR2I;refGene_tx=XM_003997912.5
E2	17568831 17568894 refGene_gn=POLR2I;refGene_tx=XM_003997912.5
E2	17568960 17569190 refGene_gn=OVOL3;refGene_tx=XM_019818820.2
E2	17569643 17569845 refGene_gn=OVOL3;refGene_tx=XM_019818820.2
E2	17570589 17570654 refGene_gn=OVOL3;refGene_tx=XM_019818820.2
E2	17570730 17570824 refGene_gn=OVOL3;refGene_tx=XM_019818820.2
E2	17575432 17575708
refGene_gn=WDR62;refGene_tx=XM_006941438.3;refGene_gn=WDR62;refGene_tx=XM_011290033.3;refGene_gn=WDR62;refGene_tx=XM_019818674.2
E2	17575788 17575946
refGene_gn=WDR62;refGene_tx=XM_006941438.3;refGene_gn=WDR62;refGene_tx=XM_011290033.3;refGene_gn=WDR62;refGene_tx=XM_019818674.2
E2	17576504 17577158
refGene_gn=WDR62;refGene_tx=XM_006941438.3;refGene_gn=WDR62;refGene_tx=XM_011290033.3;refGene_gn=WDR62;refGene_tx=XM_019818674.2
E2	17577321 17577382
refGene_gn=WDR62;refGene_tx=XM_006941438.3;refGene_gn=WDR62;refGene_tx=XM_011290033.3;refGene_gn=WDR62;refGene_tx=XM_019818674.2

E2	17577462	17577604
refGene_gn=WDR62;refGene_tx=XM_006941438.3;refGene_gn=WDR62;refGene_tx=XM_011290033.3;refGene_gn=WDR62;refGene_tx=XM_019818674.2
E2	17577715	17577827
refGene_gn=WDR62;refGene_tx=XM_006941438.3;refGene_gn=WDR62;refGene_tx=XM_011290033.3;refGene_gn=WDR62;refGene_tx=XM_019818674.2
E2	17578100	17578235
refGene_gn=WDR62;refGene_tx=XM_006941438.3;refGene_gn=WDR62;refGene_tx=XM_011290033.3;refGene_gn=WDR62;refGene_tx=XM_019818674.2
E2	17578519	17578630
refGene_gn=WDR62;refGene_tx=XM_006941438.3;refGene_gn=WDR62;refGene_tx=XM_011290033.3;refGene_gn=WDR62;refGene_tx=XM_019818674.2
E2	17578982	17579086
refGene_gn=WDR62;refGene_tx=XM_006941438.3;refGene_gn=WDR62;refGene_tx=XM_011290033.3;refGene_gn=WDR62;refGene_tx=XM_019818674.2
E2	17579410	17579538
refGene_gn=WDR62;refGene_tx=XM_006941438.3;refGene_gn=WDR62;refGene_tx=XM_011290033.3;refGene_gn=WDR62;refGene_tx=XM_019818674.2
E2	17580712	17580931
refGene_gn=WDR62;refGene_tx=XM_006941438.3;refGene_gn=WDR62;refGene_tx=XM_011290033.3;refGene_gn=WDR62;refGene_tx=XM_019818674.2
E2	17583271	17583324
refGene_gn=WDR62;refGene_tx=XM_006941438.3;refGene_gn=WDR62;refGene_tx=XM_011290033.3;refGene_gn=WDR62;refGene_tx=XM_019818674.2
E2	17585067	17585201
refGene_gn=WDR62;refGene_tx=XM_006941438.3;refGene_gn=WDR62;refGene_tx=XM_011290033.3;refGene_gn=WDR62;refGene_tx=XM_019818674.2
E2	17586260	17586386
refGene_gn=WDR62;refGene_tx=XM_006941438.3;refGene_gn=WDR62;refGene_tx=XM_011290033.3;refGene_gn=WDR62;refGene_tx=XM_019818674.2
E2	17587272	17587336
refGene_gn=WDR62;refGene_tx=XM_006941438.3;refGene_gn=WDR62;refGene_tx=XM_011290033.3;refGene_gn=WDR62;refGene_tx=XM_019818674.2
E2	17587431	17587543
refGene_gn=WDR62;refGene_tx=XM_006941438.3;refGene_gn=WDR62;refGene_tx=XM_011290033.3;refGene_gn=WDR62;refGene_tx=XM_019818674.2
E2	17588290	17588366
refGene_gn=WDR62;refGene_tx=XM_006941438.3;refGene_gn=WDR62;refGene_tx=XM_011290033.3;refGene_gn=WDR62;refGene_tx=XM_019818674.2
E2	17589266	17589388
refGene_gn=WDR62;refGene_tx=XM_006941438.3;refGene_gn=WDR62;refGene_tx=XM_011290033.3;refGene_gn=WDR62;refGene_tx=XM_019818674.2
E2	17589469	17589537
refGene_gn=WDR62;refGene_tx=XM_006941438.3;refGene_gn=WDR62;refGene_tx=XM_011290033.3;refGene_gn=WDR62;refGene_tx=XM_019818674.2
E2	17592132	17592258
refGene_gn=WDR62;refGene_tx=XM_006941438.3;refGene_gn=WDR62;refGene_tx=XM_011290033.3;refGene_gn=WDR62;refGene_tx=XM_019818674.2
E2	17594388	17594480
refGene_gn=WDR62;refGene_tx=XM_006941438.3;refGene_gn=WDR62;refGene_tx

=XM_011290033.3;refGene_gn=WDR62;refGene_tx=XM_019818674.2
E2 17595880 17596059
refGene_gn=WDR62;refGene_tx=XM_006941438.3;refGene_gn=WDR62;refGene_tx=XM_011290033.3;refGene_gn=WDR62;refGene_tx=XM_019818674.2
E2 17597543 17597681
refGene_gn=WDR62;refGene_tx=XM_006941438.3;refGene_gn=WDR62;refGene_tx=XM_011290033.3;refGene_gn=WDR62;refGene_tx=XM_019818674.2
E2 17606077 17606267
refGene_gn=WDR62;refGene_tx=XM_006941438.3;refGene_gn=WDR62;refGene_tx=XM_011290033.3;refGene_gn=WDR62;refGene_tx=XM_019818674.2
E2 17607459 17607620
refGene_gn=WDR62;refGene_tx=XM_006941438.3;refGene_gn=WDR62;refGene_tx=XM_011290033.3;refGene_gn=WDR62;refGene_tx=XM_019818674.2
E2 17610062 17610245
refGene_gn=WDR62;refGene_tx=XM_006941438.3;refGene_gn=WDR62;refGene_tx=XM_011290033.3;refGene_gn=WDR62;refGene_tx=XM_019818674.2
E2 17610669 17610807
refGene_gn=WDR62;refGene_tx=XM_006941438.3;refGene_gn=WDR62;refGene_tx=XM_011290033.3;refGene_gn=WDR62;refGene_tx=XM_019818674.2
E2 17611565 17611736
refGene_gn=WDR62;refGene_tx=XM_006941438.3;refGene_gn=WDR62;refGene_tx=XM_011290033.3;refGene_gn=WDR62;refGene_tx=XM_019818674.2
E2 17611965 17612023
refGene_gn=WDR62;refGene_tx=XM_006941438.3;refGene_gn=WDR62;refGene_tx=XM_011290033.3;refGene_gn=WDR62;refGene_tx=XM_019818674.2
E2 17615034 17615097
refGene_gn=WDR62;refGene_tx=XM_006941438.3;refGene_gn=WDR62;refGene_tx=XM_011290033.3;refGene_gn=WDR62;refGene_tx=XM_019818674.2
E2 17615973 17616065
refGene_gn=WDR62;refGene_tx=XM_006941438.3;refGene_gn=WDR62;refGene_tx=XM_011290033.3;refGene_gn=WDR62;refGene_tx=XM_019818674.2
E2 17619241 17619418
refGene_gn=WDR62;refGene_tx=XM_006941438.3;refGene_gn=WDR62;refGene_tx=XM_011290033.3;refGene_gn=WDR62;refGene_tx=XM_019818674.2
E2 17620154 17620237 refGene_gn=THAP8;refGene_tx=XM_006941437.4
E2 17630072 17630265 refGene_gn=THAP8;refGene_tx=XM_006941437.4
E2 17630497 17630893 refGene_gn=THAP8;refGene_tx=XM_006941437.4
E2 17634921 17635087 refGene_gn=CLIP3;refGene_tx=XM_003997905.4
E2 17638045 17638185 refGene_gn=CLIP3;refGene_tx=XM_003997905.4
E2 17638264 17638358 refGene_gn=CLIP3;refGene_tx=XM_003997905.4
E2 17638569 17638731 refGene_gn=CLIP3;refGene_tx=XM_003997905.4
E2 17639184 17639303 refGene_gn=CLIP3;refGene_tx=XM_003997905.4
E2 17640201 17640438 refGene_gn=CLIP3;refGene_tx=XM_003997905.4
E2 17642746 17642882 refGene_gn=CLIP3;refGene_tx=XM_003997905.4
E2 17643030 17643159 refGene_gn=CLIP3;refGene_tx=XM_003997905.4
E2 17644070 17644214 refGene_gn=CLIP3;refGene_tx=XM_003997905.4
E2 17644292 17644416 refGene_gn=CLIP3;refGene_tx=XM_003997905.4
E2 17644608 17644673 refGene_gn=CLIP3;refGene_tx=XM_003997905.4
E2 17645056 17645129 refGene_gn=CLIP3;refGene_tx=XM_003997905.4
E2 17645467 17645522 refGene_gn=CLIP3;refGene_tx=XM_003997905.4

E2 17647898 17648173 refGene_gn=ALKBH6;refGene_tx=XM_011290031.3
E2 17648289 17648368 refGene_gn=ALKBH6;refGene_tx=XM_011290031.3
E2 17648678 17648747 refGene_gn=ALKBH6;refGene_tx=XM_011290031.3
E2 17650508 17650569 refGene_gn=ALKBH6;refGene_tx=XM_011290031.3
E2 17650933 17651085 refGene_gn=ALKBH6;refGene_tx=XM_011290031.3
E2 17651330 17651396 refGene_gn=ALKBH6;refGene_tx=XM_011290031.3
E2 17652672 17652743
refGene_gn=SYNE4;refGene_tx=XM_006941432.2;refGene_gn=SYNE4;refGene_tx=XM_019818802.1
E2 17653435 17653573
refGene_gn=SYNE4;refGene_tx=XM_006941432.2;refGene_gn=SYNE4;refGene_tx=XM_019818802.1;refGene_gn=SYNE4;refGene_tx=XM_023245477.1;refGene_gn=SYNE4;refGene_tx=XM_023245478.1
E2 17653764 17653915
refGene_gn=SYNE4;refGene_tx=XM_006941432.2;refGene_gn=SYNE4;refGene_tx=XM_019818802.1;refGene_gn=SYNE4;refGene_tx=XM_023245477.1
E2 17654404 17654548
refGene_gn=SYNE4;refGene_tx=XM_006941432.2;refGene_gn=SYNE4;refGene_tx=XM_019818802.1;refGene_gn=SYNE4;refGene_tx=XM_023245477.1;refGene_gn=SYNE4;refGene_tx=XM_023245478.1
E2 17654753 17655279
refGene_gn=SYNE4;refGene_tx=XM_006941432.2;refGene_gn=SYNE4;refGene_tx=XM_019818802.1;refGene_gn=SYNE4;refGene_tx=XM_023245477.1;refGene_gn=SYNE4;refGene_tx=XM_023245478.1
E2 17655987 17656065
refGene_gn=SYNE4;refGene_tx=XM_006941432.2;refGene_gn=SYNE4;refGene_tx=XM_019818802.1;refGene_gn=SYNE4;refGene_tx=XM_023245477.1;refGene_gn=SYNE4;refGene_tx=XM_023245478.1
E2 17656774 17656830
refGene_gn=SYNE4;refGene_tx=XM_006941432.2;refGene_gn=SYNE4;refGene_tx=XM_019818802.1;refGene_gn=SYNE4;refGene_tx=XM_023245477.1;refGene_gn=SYNE4;refGene_tx=XM_023245478.1
E2 17656923 17657124
refGene_gn=SYNE4;refGene_tx=XM_006941432.2;refGene_gn=SYNE4;refGene_tx=XM_019818802.1;refGene_gn=SYNE4;refGene_tx=XM_023245477.1;refGene_gn=SYNE4;refGene_tx=XM_023245478.1
E2 17664185 17664566 refGene_gn=SDHAF1;refGene_tx=XM_006941427.4
E2 17665146 17665195
refGene_gn=LOC109494605;refGene_tx=XM_019818794.2
E2 17665329 17665619
refGene_gn=LOC109494605;refGene_tx=XM_019818794.2
E2 17668128 17668524
refGene_gn=LOC109494605;refGene_tx=XM_019818794.2
E2 17670695 17670895
refGene_gn=LOC109494605;refGene_tx=XM_019818794.2
E2 17671402 17671949
refGene_gn=LOC109494605;refGene_tx=XM_019818794.2
E2 17673543 17674145
refGene_gn=LOC109494605;refGene_tx=XM_019818794.2
E2 17674513 17674547

refGene_gn=LOC109494605;refGene_tx=XM_019818794.2
E2 17674599 17675163
refGene_gn=LOC111556290;refGene_tx=XM_023245966.1
E2 17676325 17677008
refGene_gn=LOC111556290;refGene_tx=XM_023245966.1
E2 17677132 17677274
refGene_gn=LOC111556290;refGene_tx=XM_023245966.1
E2 17677393 17677593
refGene_gn=LOC111556290;refGene_tx=XM_023245966.1
E2 17679640 17680118
refGene_gn=LOC111556290;refGene_tx=XM_023245966.1
E2 17681061 17681439
refGene_gn=LOC111556290;refGene_tx=XM_023245966.1
E2 17681748 17681781
refGene_gn=LOC111556290;refGene_tx=XM_023245966.1
E2 17682105 17682771
refGene_gn=LOC109494656;refGene_tx=XM_023245418.1
E2 17685156 17685755
refGene_gn=LOC109494656;refGene_tx=XM_023245418.1
E2 17686402 17686958
refGene_gn=LOC109494656;refGene_tx=XM_023245418.1
E2 17689917 17690799
refGene_gn=LOC109494656;refGene_tx=XM_023245418.1
E2 17702975 17703441 refGene_gn=LRFN3;refGene_tx=XM_003997932.3
E2 17705473 17706900 refGene_gn=LRFN3;refGene_tx=XM_003997932.3
E2 17736036 17736097
refGene_gn=TYROBP;refGene_tx=XM_003997915.3;refGene_gn=TYROBP;refGene_
tx=XM_006941426.2
E2 17736484 17736511
refGene_gn=TYROBP;refGene_tx=XM_003997915.3;refGene_gn=TYROBP;refGene_
tx=XM_006941426.2
E2 17736660 17736795
refGene_gn=TYROBP;refGene_tx=XM_003997915.3;refGene_gn=TYROBP;refGene_
tx=XM_006941426.2
E2 17736964 17737011
refGene_gn=TYROBP;refGene_tx=XM_003997915.3;refGene_gn=TYROBP;refGene_
tx=XM_006941426.2
E2 17738436 17738502
refGene_gn=TYROBP;refGene_tx=XM_003997915.3;refGene_gn=TYROBP;refGene_
tx=XM_006941426.2
E2 17743959 17743974
refGene_gn=NFKBID;refGene_tx=XM_006941420.3;refGene_gn=NFKBID;refGene_
tx=XM_006941421.4
E2 17744640 17744646
refGene_gn=NFKBID;refGene_tx=XM_006941422.3;refGene_gn=NFKBID;refGene_
tx=XM_006941423.3;refGene_gn=NFKBID;refGene_tx=XM_023245488.1
E2 17744881 17745072
refGene_gn=NFKBID;refGene_tx=XM_006941420.3;refGene_gn=NFKBID;refGene_
tx=XM_006941421.4;refGene_gn=NFKBID;refGene_tx=XM_006941422.3;refGene_
gn=NFKBID;refGene_tx=XM_006941423.3;refGene_gn=NFKBID;refGene_tx=XM_02

3245488.1
E2 17745154 17745260
refGene_gn=NFKBID;refGene_tx=XM_006941420.3;refGene_gn=NFKBID;refGene_tx=XM_006941421.4;refGene_gn=NFKBID;refGene_tx=XM_006941422.3;refGene_gn=NFKBID;refGene_tx=XM_006941423.3;refGene_gn=NFKBID;refGene_tx=XM_023245488.1
E2 17745357 17745504
refGene_gn=NFKBID;refGene_tx=XM_006941420.3;refGene_gn=NFKBID;refGene_tx=XM_006941421.4;refGene_gn=NFKBID;refGene_tx=XM_006941422.3;refGene_gn=NFKBID;refGene_tx=XM_006941423.3;refGene_gn=NFKBID;refGene_tx=XM_023245488.1
E2 17745619 17745671
refGene_gn=NFKBID;refGene_tx=XM_006941420.3;refGene_gn=NFKBID;refGene_tx=XM_006941421.4;refGene_gn=NFKBID;refGene_tx=XM_006941422.3;refGene_gn=NFKBID;refGene_tx=XM_006941423.3;refGene_gn=NFKBID;refGene_tx=XM_023245488.1
E2 17745754 17745903
refGene_gn=NFKBID;refGene_tx=XM_006941420.3;refGene_gn=NFKBID;refGene_tx=XM_006941421.4;refGene_gn=NFKBID;refGene_tx=XM_006941422.3;refGene_gn=NFKBID;refGene_tx=XM_006941423.3;refGene_gn=NFKBID;refGene_tx=XM_023245488.1
E2 17747786 17747910
refGene_gn=NFKBID;refGene_tx=XM_006941420.3;refGene_gn=NFKBID;refGene_tx=XM_006941421.4;refGene_gn=NFKBID;refGene_tx=XM_006941422.3;refGene_gn=NFKBID;refGene_tx=XM_006941423.3;refGene_gn=NFKBID;refGene_tx=XM_023245488.1
E2 17748251 17748415
refGene_gn=NFKBID;refGene_tx=XM_006941420.3;refGene_gn=NFKBID;refGene_tx=XM_006941421.4;refGene_gn=NFKBID;refGene_tx=XM_006941422.3;refGene_gn=NFKBID;refGene_tx=XM_006941423.3;refGene_gn=NFKBID;refGene_tx=XM_023245488.1
E2 17749048 17749102
refGene_gn=NFKBID;refGene_tx=XM_006941420.3;refGene_gn=NFKBID;refGene_tx=XM_006941421.4;refGene_gn=NFKBID;refGene_tx=XM_006941422.3;refGene_gn=NFKBID;refGene_tx=XM_006941423.3;refGene_gn=NFKBID;refGene_tx=XM_023245488.1
E2 17753993 17754092
refGene_gn=APLP1;refGene_tx=XM_019818928.2;refGene_gn=APLP1;refGene_tx=XM_023245430.1;refGene_gn=APLP1;refGene_tx=XM_023245431.1;refGene_gn=APLP1;refGene_tx=XM_023245432.1;refGene_gn=APLP1;refGene_tx=XM_023245433.1;refGene_gn=APLP1;refGene_tx=XM_023245434.1;refGene_gn=APLP1;refGene_tx=XM_023245435.1;refGene_gn=APLP1;refGene_tx=XM_023245436.1
E2 17754210 17754354
refGene_gn=APLP1;refGene_tx=XM_019818928.2;refGene_gn=APLP1;refGene_tx=XM_023245430.1;refGene_gn=APLP1;refGene_tx=XM_023245431.1;refGene_gn=APLP1;refGene_tx=XM_023245432.1;refGene_gn=APLP1;refGene_tx=XM_023245433.1;refGene_gn=APLP1;refGene_tx=XM_023245434.1;refGene_gn=APLP1;refGene_tx=XM_023245435.1;refGene_gn=APLP1;refGene_tx=XM_023245436.1
E2 17754485 17754548
refGene_gn=APLP1;refGene_tx=XM_019818928.2;refGene_gn=APLP1;refGene_tx

=XM_023245430.1;refGene_gn=APLP1;refGene_tx=XM_023245432.1;refGene_gn=APLP1;refGene_tx=XM_023245435.1;refGene_gn=APLP1;refGene_tx=XM_023245436.1
E2 17754780 17754851
refGene_gn=APLP1;refGene_tx=XM_019818928.2;refGene_gn=APLP1;refGene_tx=XM_023245430.1;refGene_gn=APLP1;refGene_tx=XM_023245431.1;refGene_gn=APLP1;refGene_tx=XM_023245432.1;refGene_gn=APLP1;refGene_tx=XM_023245433.1;refGene_gn=APLP1;refGene_tx=XM_023245434.1;refGene_gn=APLP1;refGene_tx=XM_023245435.1;refGene_gn=APLP1;refGene_tx=XM_023245436.1
E2 17755040 17755148
refGene_gn=APLP1;refGene_tx=XM_019818928.2;refGene_gn=APLP1;refGene_tx=XM_023245430.1;refGene_gn=APLP1;refGene_tx=XM_023245431.1;refGene_gn=APLP1;refGene_tx=XM_023245432.1;refGene_gn=APLP1;refGene_tx=XM_023245433.1;refGene_gn=APLP1;refGene_tx=XM_023245434.1;refGene_gn=APLP1;refGene_tx=XM_023245435.1;refGene_gn=APLP1;refGene_tx=XM_023245436.1
E2 17755323 17755431
refGene_gn=APLP1;refGene_tx=XM_019818928.2;refGene_gn=APLP1;refGene_tx=XM_023245430.1;refGene_gn=APLP1;refGene_tx=XM_023245431.1;refGene_gn=APLP1;refGene_tx=XM_023245432.1;refGene_gn=APLP1;refGene_tx=XM_023245433.1;refGene_gn=APLP1;refGene_tx=XM_023245434.1
E2 17756403 17756503
refGene_gn=APLP1;refGene_tx=XM_019818928.2;refGene_gn=APLP1;refGene_tx=XM_023245430.1;refGene_gn=APLP1;refGene_tx=XM_023245431.1;refGene_gn=APLP1;refGene_tx=XM_023245432.1;refGene_gn=APLP1;refGene_tx=XM_023245433.1;refGene_gn=APLP1;refGene_tx=XM_023245434.1;refGene_gn=APLP1;refGene_tx=XM_023245435.1;refGene_gn=APLP1;refGene_tx=XM_023245436.1
E2 17757889 17758018
refGene_gn=APLP1;refGene_tx=XM_019818928.2;refGene_gn=APLP1;refGene_tx=XM_023245430.1;refGene_gn=APLP1;refGene_tx=XM_023245431.1;refGene_gn=APLP1;refGene_tx=XM_023245432.1;refGene_gn=APLP1;refGene_tx=XM_023245433.1;refGene_gn=APLP1;refGene_tx=XM_023245434.1;refGene_gn=APLP1;refGene_tx=XM_023245435.1;refGene_gn=APLP1;refGene_tx=XM_023245436.1
E2 17758095 17758254
refGene_gn=APLP1;refGene_tx=XM_019818928.2;refGene_gn=APLP1;refGene_tx=XM_023245430.1;refGene_gn=APLP1;refGene_tx=XM_023245431.1;refGene_gn=APLP1;refGene_tx=XM_023245432.1;refGene_gn=APLP1;refGene_tx=XM_023245433.1;refGene_gn=APLP1;refGene_tx=XM_023245434.1;refGene_gn=APLP1;refGene_tx=XM_023245435.1;refGene_gn=APLP1;refGene_tx=XM_023245436.1
E2 17759119 17759194
refGene_gn=APLP1;refGene_tx=XM_019818928.2;refGene_gn=APLP1;refGene_tx=XM_023245430.1;refGene_gn=APLP1;refGene_tx=XM_023245431.1;refGene_gn=APLP1;refGene_tx=XM_023245432.1;refGene_gn=APLP1;refGene_tx=XM_023245433.1;refGene_gn=APLP1;refGene_tx=XM_023245434.1;refGene_gn=APLP1;refGene_tx=XM_023245435.1;refGene_gn=APLP1;refGene_tx=XM_023245436.1
E2 17759569 17759700
refGene_gn=APLP1;refGene_tx=XM_019818928.2;refGene_gn=APLP1;refGene_tx=XM_023245430.1;refGene_gn=APLP1;refGene_tx=XM_023245431.1;refGene_gn=APLP1;refGene_tx=XM_023245432.1;refGene_gn=APLP1;refGene_tx=XM_023245433.1;refGene_gn=APLP1;refGene_tx=XM_023245434.1;refGene_gn=APLP1;refGene_tx=XM_023245435.1;refGene_gn=APLP1;refGene_tx=XM_023245436.1

E2 17760156 17760341
refGene_gn=APLP1;refGene_tx=XM_019818928.2;refGene_gn=APLP1;refGene_tx
=XM_023245430.1;refGene_gn=APLP1;refGene_tx=XM_023245431.1;refGene_gn=
APLP1;refGene_tx=XM_023245432.1;refGene_gn=APLP1;refGene_tx=XM_0232454
33.1;refGene_gn=APLP1;refGene_tx=XM_023245434.1;refGene_gn=APLP1;refGe
ne_tx=XM_023245435.1;refGene_gn=APLP1;refGene_tx=XM_023245436.1
E2 17760450 17760584
refGene_gn=APLP1;refGene_tx=XM_019818928.2;refGene_gn=APLP1;refGene_tx
=XM_023245430.1;refGene_gn=APLP1;refGene_tx=XM_023245431.1;refGene_gn=
APLP1;refGene_tx=XM_023245432.1;refGene_gn=APLP1;refGene_tx=XM_0232454
33.1;refGene_gn=APLP1;refGene_tx=XM_023245434.1;refGene_gn=APLP1;refGe
ne_tx=XM_023245435.1;refGene_gn=APLP1;refGene_tx=XM_023245436.1
E2 17760815 17760928
refGene_gn=APLP1;refGene_tx=XM_019818928.2;refGene_gn=APLP1;refGene_tx
=XM_023245430.1;refGene_gn=APLP1;refGene_tx=XM_023245431.1;refGene_gn=
APLP1;refGene_tx=XM_023245432.1;refGene_gn=APLP1;refGene_tx=XM_0232454
33.1;refGene_gn=APLP1;refGene_tx=XM_023245434.1;refGene_gn=APLP1;refGe
ne_tx=XM_023245435.1;refGene_gn=APLP1;refGene_tx=XM_023245436.1
E2 17761138 17761271
refGene_gn=APLP1;refGene_tx=XM_019818928.2;refGene_gn=APLP1;refGene_tx
=XM_023245430.1;refGene_gn=APLP1;refGene_tx=XM_023245431.1;refGene_gn=
APLP1;refGene_tx=XM_023245432.1;refGene_gn=APLP1;refGene_tx=XM_0232454
33.1;refGene_gn=APLP1;refGene_tx=XM_023245434.1;refGene_gn=APLP1;refGe
ne_tx=XM_023245435.1;refGene_gn=APLP1;refGene_tx=XM_023245436.1
E2 17762020 17762164
refGene_gn=APLP1;refGene_tx=XM_019818928.2;refGene_gn=APLP1;refGene_tx
=XM_023245430.1;refGene_gn=APLP1;refGene_tx=XM_023245431.1;refGene_gn=
APLP1;refGene_tx=XM_023245432.1;refGene_gn=APLP1;refGene_tx=XM_0232454
33.1;refGene_gn=APLP1;refGene_tx=XM_023245434.1;refGene_gn=APLP1;refGe
ne_tx=XM_023245435.1;refGene_gn=APLP1;refGene_tx=XM_023245436.1
E2 17763055 17763202
refGene_gn=APLP1;refGene_tx=XM_019818928.2;refGene_gn=APLP1;refGene_tx
=XM_023245430.1;refGene_gn=APLP1;refGene_tx=XM_023245431.1;refGene_gn=
APLP1;refGene_tx=XM_023245432.1;refGene_gn=APLP1;refGene_tx=XM_0232454
33.1;refGene_gn=APLP1;refGene_tx=XM_023245434.1;refGene_gn=APLP1;refGe
ne_tx=XM_023245435.1;refGene_gn=APLP1;refGene_tx=XM_023245436.1
E2 17764986 17765322
refGene_gn=KIRREL2;refGene_tx=XM_019818695.2;refGene_gn=KIRREL2;refGen
e_tx=XM_019818696.2;refGene_gn=KIRREL2;refGene_tx=XM_019818697.2
E2 17765397 17765409 refGene_gn=KIRREL2;refGene_tx=XM_019818698.2
E2 17766484 17766550
refGene_gn=KIRREL2;refGene_tx=XM_019818695.2;refGene_gn=KIRREL2;refGen
e_tx=XM_019818696.2;refGene_gn=KIRREL2;refGene_tx=XM_019818697.2;refGe
ne_gn=KIRREL2;refGene_tx=XM_019818698.2
E2 17766897 17767001
refGene_gn=KIRREL2;refGene_tx=XM_019818695.2;refGene_gn=KIRREL2;refGen
e_tx=XM_019818696.2;refGene_gn=KIRREL2;refGene_tx=XM_019818697.2;refGe
ne_gn=KIRREL2;refGene_tx=XM_019818698.2
E2 17767292 17767397
refGene_gn=KIRREL2;refGene_tx=XM_019818695.2;refGene_gn=KIRREL2;refGen

e_tx=XM_019818697.2;refGene_gn=KIRREL2;refGene_tx=XM_019818698.2
E2 17767899 17768119
refGene_gn=KIRREL2;refGene_tx=XM_019818695.2;refGene_gn=KIRREL2;refGene_tx=XM_019818696.2;refGene_gn=KIRREL2;refGene_tx=XM_019818697.2;refGene_gn=KIRREL2;refGene_tx=XM_019818698.2
E2 17768283 17768384
refGene_gn=KIRREL2;refGene_tx=XM_019818695.2;refGene_gn=KIRREL2;refGene_tx=XM_019818696.2;refGene_gn=KIRREL2;refGene_tx=XM_019818697.2;refGene_gn=KIRREL2;refGene_tx=XM_019818698.2
E2 17768690 17768823
refGene_gn=KIRREL2;refGene_tx=XM_019818695.2;refGene_gn=KIRREL2;refGene_tx=XM_019818696.2;refGene_gn=KIRREL2;refGene_tx=XM_019818697.2;refGene_gn=KIRREL2;refGene_tx=XM_019818698.2
E2 17768914 17769042
refGene_gn=KIRREL2;refGene_tx=XM_019818695.2;refGene_gn=KIRREL2;refGene_tx=XM_019818696.2;refGene_gn=KIRREL2;refGene_tx=XM_019818697.2;refGene_gn=KIRREL2;refGene_tx=XM_019818698.2
E2 17769276 17769425
refGene_gn=KIRREL2;refGene_tx=XM_019818695.2;refGene_gn=KIRREL2;refGene_tx=XM_019818696.2;refGene_gn=KIRREL2;refGene_tx=XM_019818697.2;refGene_gn=KIRREL2;refGene_tx=XM_019818698.2
E2 17769538 17769644
refGene_gn=KIRREL2;refGene_tx=XM_019818695.2;refGene_gn=KIRREL2;refGene_tx=XM_019818696.2;refGene_gn=KIRREL2;refGene_tx=XM_019818697.2;refGene_gn=KIRREL2;refGene_tx=XM_019818698.2
E2 17770175 17770326
refGene_gn=KIRREL2;refGene_tx=XM_019818695.2;refGene_gn=KIRREL2;refGene_tx=XM_019818696.2;refGene_gn=KIRREL2;refGene_tx=XM_019818697.2;refGene_gn=KIRREL2;refGene_tx=XM_019818698.2
E2 17770719 17770880
refGene_gn=KIRREL2;refGene_tx=XM_019818695.2;refGene_gn=KIRREL2;refGene_tx=XM_019818696.2;refGene_gn=KIRREL2;refGene_tx=XM_019818697.2;refGene_gn=KIRREL2;refGene_tx=XM_019818698.2
E2 17771039 17771189
refGene_gn=KIRREL2;refGene_tx=XM_019818695.2;refGene_gn=KIRREL2;refGene_tx=XM_019818696.2;refGene_gn=KIRREL2;refGene_tx=XM_019818698.2
E2 17771731 17771881
refGene_gn=KIRREL2;refGene_tx=XM_019818695.2;refGene_gn=KIRREL2;refGene_tx=XM_019818696.2;refGene_gn=KIRREL2;refGene_tx=XM_019818697.2;refGene_gn=KIRREL2;refGene_tx=XM_019818698.2
E2 17772081 17772592
refGene_gn=KIRREL2;refGene_tx=XM_019818695.2;refGene_gn=KIRREL2;refGene_tx=XM_019818696.2;refGene_gn=KIRREL2;refGene_tx=XM_019818697.2;refGene_gn=KIRREL2;refGene_tx=XM_019818698.2
E2 17775832 17775890
refGene_gn=NPHS1;refGene_tx=XM_003997902.4;refGene_gn=NPHS1;refGene_tx=XM_019818675.2
E2 17775923 17776215
refGene_gn=NPHS1;refGene_tx=XM_003997902.4;refGene_gn=NPHS1;refGene_tx=XM_019818675.2;refGene_gn=NPHS1;refGene_tx=XM_023245415.1;refGene_gn=

NPHS1;refGene_tx=XM_023245416.1
E2 17776298 17776421
refGene_gn=NPHS1;refGene_tx=XM_003997902.4;refGene_gn=NPHS1;refGene_tx=XM_019818675.2;refGene_gn=NPHS1;refGene_tx=XM_023245415.1;refGene_gn=NPHS1;refGene_tx=XM_023245416.1
E2 17776618 17776747
refGene_gn=NPHS1;refGene_tx=XM_003997902.4;refGene_gn=NPHS1;refGene_tx=XM_019818675.2;refGene_gn=NPHS1;refGene_tx=XM_023245415.1;refGene_gn=NPHS1;refGene_tx=XM_023245416.1
E2 17777184 17777354
refGene_gn=NPHS1;refGene_tx=XM_003997902.4;refGene_gn=NPHS1;refGene_tx=XM_019818675.2;refGene_gn=NPHS1;refGene_tx=XM_019818679.2;refGene_gn=NPHS1;refGene_tx=XM_019818680.2;refGene_gn=NPHS1;refGene_tx=XM_023245415.1;refGene_gn=NPHS1;refGene_tx=XM_023245416.1;refGene_gn=NPHS1;refGene_tx=XM_023245417.1
E2 17777694 17777835
refGene_gn=NPHS1;refGene_tx=XM_003997902.4;refGene_gn=NPHS1;refGene_tx=XM_019818675.2;refGene_gn=NPHS1;refGene_tx=XM_019818679.2;refGene_gn=NPHS1;refGene_tx=XM_019818680.2;refGene_gn=NPHS1;refGene_tx=XM_023245415.1;refGene_gn=NPHS1;refGene_tx=XM_023245416.1;refGene_gn=NPHS1;refGene_tx=XM_023245417.1
E2 17778019 17778147
refGene_gn=NPHS1;refGene_tx=XM_003997902.4;refGene_gn=NPHS1;refGene_tx=XM_019818675.2;refGene_gn=NPHS1;refGene_tx=XM_019818679.2;refGene_gn=NPHS1;refGene_tx=XM_019818680.2;refGene_gn=NPHS1;refGene_tx=XM_023245415.1;refGene_gn=NPHS1;refGene_tx=XM_023245416.1;refGene_gn=NPHS1;refGene_tx=XM_023245417.1
E2 17778230 17778402
refGene_gn=NPHS1;refGene_tx=XM_003997902.4;refGene_gn=NPHS1;refGene_tx=XM_019818675.2;refGene_gn=NPHS1;refGene_tx=XM_019818679.2;refGene_gn=NPHS1;refGene_tx=XM_019818680.2;refGene_gn=NPHS1;refGene_tx=XM_023245415.1;refGene_gn=NPHS1;refGene_tx=XM_023245416.1;refGene_gn=NPHS1;refGene_tx=XM_023245417.1
E2 17778585 17778743
refGene_gn=NPHS1;refGene_tx=XM_003997902.4;refGene_gn=NPHS1;refGene_tx=XM_019818675.2;refGene_gn=NPHS1;refGene_tx=XM_019818679.2;refGene_gn=NPHS1;refGene_tx=XM_019818680.2;refGene_gn=NPHS1;refGene_tx=XM_023245415.1;refGene_gn=NPHS1;refGene_tx=XM_023245416.1;refGene_gn=NPHS1;refGene_tx=XM_023245417.1
E2 17779001 17779146
refGene_gn=NPHS1;refGene_tx=XM_003997902.4;refGene_gn=NPHS1;refGene_tx=XM_019818675.2;refGene_gn=NPHS1;refGene_tx=XM_019818679.2;refGene_gn=NPHS1;refGene_tx=XM_019818680.2;refGene_gn=NPHS1;refGene_tx=XM_023245415.1;refGene_gn=NPHS1;refGene_tx=XM_023245416.1;refGene_gn=NPHS1;refGene_tx=XM_023245417.1
E2 17779266 17779391
refGene_gn=NPHS1;refGene_tx=XM_003997902.4;refGene_gn=NPHS1;refGene_tx=XM_019818675.2;refGene_gn=NPHS1;refGene_tx=XM_019818679.2;refGene_gn=NPHS1;refGene_tx=XM_019818680.2;refGene_gn=NPHS1;refGene_tx=XM_023245415.1;refGene_gn=NPHS1;refGene_tx=XM_023245416.1;refGene_gn=NPHS1;refGe

ne_tx=XM_023245417.1
E2 17780597 17780784
refGene_gn=NPHS1;refGene_tx=XM_003997902.4;refGene_gn=NPHS1;refGene_tx=XM_019818675.2;refGene_gn=NPHS1;refGene_tx=XM_019818679.2;refGene_gn=NPHS1;refGene_tx=XM_019818680.2;refGene_gn=NPHS1;refGene_tx=XM_023245415.1;refGene_gn=NPHS1;refGene_tx=XM_023245416.1;refGene_gn=NPHS1;refGene_tx=XM_023245417.1
E2 17780958 17781088
refGene_gn=NPHS1;refGene_tx=XM_003997902.4;refGene_gn=NPHS1;refGene_tx=XM_019818675.2;refGene_gn=NPHS1;refGene_tx=XM_019818679.2;refGene_gn=NPHS1;refGene_tx=XM_019818680.2;refGene_gn=NPHS1;refGene_tx=XM_023245415.1;refGene_gn=NPHS1;refGene_tx=XM_023245416.1;refGene_gn=NPHS1;refGene_tx=XM_023245417.1
E2 17781257 17781430
refGene_gn=NPHS1;refGene_tx=XM_003997902.4;refGene_gn=NPHS1;refGene_tx=XM_019818675.2;refGene_gn=NPHS1;refGene_tx=XM_019818679.2;refGene_gn=NPHS1;refGene_tx=XM_019818680.2;refGene_gn=NPHS1;refGene_tx=XM_023245415.1;refGene_gn=NPHS1;refGene_tx=XM_023245416.1;refGene_gn=NPHS1;refGene_tx=XM_023245417.1
E2 17781863 17782004
refGene_gn=NPHS1;refGene_tx=XM_003997902.4;refGene_gn=NPHS1;refGene_tx=XM_019818675.2;refGene_gn=NPHS1;refGene_tx=XM_019818679.2;refGene_gn=NPHS1;refGene_tx=XM_019818680.2;refGene_gn=NPHS1;refGene_tx=XM_023245415.1;refGene_gn=NPHS1;refGene_tx=XM_023245416.1;refGene_gn=NPHS1;refGene_tx=XM_023245417.1
E2 17782086 17782227
refGene_gn=NPHS1;refGene_tx=XM_003997902.4;refGene_gn=NPHS1;refGene_tx=XM_019818675.2;refGene_gn=NPHS1;refGene_tx=XM_019818679.2;refGene_gn=NPHS1;refGene_tx=XM_019818680.2;refGene_gn=NPHS1;refGene_tx=XM_023245415.1;refGene_gn=NPHS1;refGene_tx=XM_023245416.1;refGene_gn=NPHS1;refGene_tx=XM_023245417.1
E2 17782770 17782892
refGene_gn=NPHS1;refGene_tx=XM_003997902.4;refGene_gn=NPHS1;refGene_tx=XM_019818675.2;refGene_gn=NPHS1;refGene_tx=XM_019818679.2;refGene_gn=NPHS1;refGene_tx=XM_019818680.2;refGene_gn=NPHS1;refGene_tx=XM_023245415.1;refGene_gn=NPHS1;refGene_tx=XM_023245416.1;refGene_gn=NPHS1;refGene_tx=XM_023245417.1
E2 17783878 17784050
refGene_gn=NPHS1;refGene_tx=XM_003997902.4;refGene_gn=NPHS1;refGene_tx=XM_019818675.2;refGene_gn=NPHS1;refGene_tx=XM_019818679.2;refGene_gn=NPHS1;refGene_tx=XM_019818680.2;refGene_gn=NPHS1;refGene_tx=XM_023245415.1;refGene_gn=NPHS1;refGene_tx=XM_023245416.1;refGene_gn=NPHS1;refGene_tx=XM_023245417.1
E2 17784148 17784305
refGene_gn=NPHS1;refGene_tx=XM_003997902.4;refGene_gn=NPHS1;refGene_tx=XM_019818675.2;refGene_gn=NPHS1;refGene_tx=XM_019818679.2;refGene_gn=NPHS1;refGene_tx=XM_019818680.2;refGene_gn=NPHS1;refGene_tx=XM_023245415.1;refGene_gn=NPHS1;refGene_tx=XM_023245416.1;refGene_gn=NPHS1;refGene_tx=XM_023245417.1
E2 17784400 17784552

refGene_gn=NPHS1;refGene_tx=XM_003997902.4;refGene_gn=NPHS1;refGene_tx
=XM_019818675.2;refGene_gn=NPHS1;refGene_tx=XM_019818679.2;refGene_gn=
NPHS1;refGene_tx=XM_019818680.2;refGene_gn=NPHS1;refGene_tx=XM_0232454
15.1;refGene_gn=NPHS1;refGene_tx=XM_023245416.1;refGene_gn=NPHS1;refGe
ne_tx=XM_023245417.1
E2 17785956 17786068
refGene_gn=NPHS1;refGene_tx=XM_003997902.4;refGene_gn=NPHS1;refGene_tx
=XM_019818675.2;refGene_gn=NPHS1;refGene_tx=XM_019818679.2;refGene_gn=
NPHS1;refGene_tx=XM_019818680.2;refGene_gn=NPHS1;refGene_tx=XM_0232454
15.1;refGene_gn=NPHS1;refGene_tx=XM_023245416.1;refGene_gn=NPHS1;refGe
ne_tx=XM_023245417.1
E2 17786151 17786333
refGene_gn=NPHS1;refGene_tx=XM_003997902.4;refGene_gn=NPHS1;refGene_tx
=XM_019818675.2;refGene_gn=NPHS1;refGene_tx=XM_019818679.2;refGene_gn=
NPHS1;refGene_tx=XM_019818680.2;refGene_gn=NPHS1;refGene_tx=XM_0232454
15.1;refGene_gn=NPHS1;refGene_tx=XM_023245416.1;refGene_gn=NPHS1;refGe
ne_tx=XM_023245417.1
E2 17788381 17788438
refGene_gn=NPHS1;refGene_tx=XM_003997902.4;refGene_gn=NPHS1;refGene_tx
=XM_019818675.2;refGene_gn=NPHS1;refGene_tx=XM_019818679.2;refGene_gn=
NPHS1;refGene_tx=XM_019818680.2;refGene_gn=NPHS1;refGene_tx=XM_0232454
15.1;refGene_gn=NPHS1;refGene_tx=XM_023245416.1;refGene_gn=NPHS1;refGe
ne_tx=XM_023245417.1
E2 17790140 17790260
refGene_gn=NPHS1;refGene_tx=XM_003997902.4;refGene_gn=NPHS1;refGene_tx
=XM_019818675.2;refGene_gn=NPHS1;refGene_tx=XM_019818679.2;refGene_gn=
NPHS1;refGene_tx=XM_019818680.2;refGene_gn=NPHS1;refGene_tx=XM_0232454
15.1;refGene_gn=NPHS1;refGene_tx=XM_023245416.1;refGene_gn=NPHS1;refGe
ne_tx=XM_023245417.1
E2 17790399 17790424
refGene_gn=NPHS1;refGene_tx=XM_003997902.4;refGene_gn=NPHS1;refGene_tx
=XM_019818675.2;refGene_gn=NPHS1;refGene_tx=XM_019818679.2;refGene_gn=
NPHS1;refGene_tx=XM_019818680.2;refGene_gn=NPHS1;refGene_tx=XM_0232454
15.1;refGene_gn=NPHS1;refGene_tx=XM_023245416.1;refGene_gn=NPHS1;refGe
ne_tx=XM_023245417.1
E2 17790530 17790606
refGene_gn=NPHS1;refGene_tx=XM_003997902.4;refGene_gn=NPHS1;refGene_tx
=XM_019818675.2;refGene_gn=NPHS1;refGene_tx=XM_019818679.2;refGene_gn=
NPHS1;refGene_tx=XM_019818680.2;refGene_gn=NPHS1;refGene_tx=XM_0232454
15.1;refGene_gn=NPHS1;refGene_tx=XM_023245416.1;refGene_gn=NPHS1;refGe
ne_tx=XM_023245417.1
E2 17790777 17790871
refGene_gn=NPHS1;refGene_tx=XM_003997902.4;refGene_gn=NPHS1;refGene_tx
=XM_019818675.2;refGene_gn=NPHS1;refGene_tx=XM_019818679.2;refGene_gn=
NPHS1;refGene_tx=XM_019818680.2;refGene_gn=NPHS1;refGene_tx=XM_0232454
15.1;refGene_gn=NPHS1;refGene_tx=XM_023245416.1;refGene_gn=NPHS1;refGe
ne_tx=XM_023245417.1
E2 17790967 17791080
refGene_gn=NPHS1;refGene_tx=XM_003997902.4;refGene_gn=NPHS1;refGene_tx
=XM_019818675.2;refGene_gn=NPHS1;refGene_tx=XM_019818679.2;refGene_gn=

NPHS1;refGene_tx=XM_019818680.2;refGene_gn=NPHS1;refGene_tx=XM_0232454
15.1;refGene_gn=NPHS1;refGene_tx=XM_023245416.1;refGene_gn=NPHS1;refGe
ne_tx=XM_023245417.1
E2 17791233 17791242
refGene_gn=NPHS1;refGene_tx=XM_019818675.2;refGene_gn=NPHS1;refGene_tx
=XM_019818679.2;refGene_gn=NPHS1;refGene_tx=XM_019818680.2;refGene_gn=
NPHS1;refGene_tx=XM_023245415.1;refGene_gn=NPHS1;refGene_tx=XM_0232454
17.1
E2 17793334 17793466
refGene_gn=NPHS1;refGene_tx=XM_003997902.4;refGene_gn=NPHS1;refGene_tx
=XM_019818675.2;refGene_gn=NPHS1;refGene_tx=XM_019818679.2;refGene_gn=
NPHS1;refGene_tx=XM_019818680.2;refGene_gn=NPHS1;refGene_tx=XM_0232454
15.1;refGene_gn=NPHS1;refGene_tx=XM_023245416.1;refGene_gn=NPHS1;refGe
ne_tx=XM_023245417.1
E2 17808833 17809010
refGene_gn=PRODH2;refGene_tx=XM_003997934.4;refGene_gn=PRODH2;refGene_
tx=XM_019818793.2
E2 17809086 17809283
refGene_gn=PRODH2;refGene_tx=XM_003997934.4;refGene_gn=PRODH2;refGene_
tx=XM_019818793.2
E2 17809363 17809502
refGene_gn=PRODH2;refGene_tx=XM_003997934.4;refGene_gn=PRODH2;refGene_
tx=XM_019818793.2
E2 17809584 17809671
refGene_gn=PRODH2;refGene_tx=XM_003997934.4;refGene_gn=PRODH2;refGene_
tx=XM_019818793.2
E2 17813454 17813535
refGene_gn=PRODH2;refGene_tx=XM_003997934.4;refGene_gn=PRODH2;refGene_
tx=XM_019818793.2
E2 17813612 17813768
refGene_gn=PRODH2;refGene_tx=XM_003997934.4;refGene_gn=PRODH2;refGene_
tx=XM_019818793.2
E2 17813844 17814011
refGene_gn=PRODH2;refGene_tx=XM_003997934.4;refGene_gn=PRODH2;refGene_
tx=XM_019818793.2
E2 17817579 17817690
refGene_gn=PRODH2;refGene_tx=XM_003997934.4;refGene_gn=PRODH2;refGene_
tx=XM_019818793.2
E2 17818088 17818174
refGene_gn=PRODH2;refGene_tx=XM_003997934.4;refGene_gn=PRODH2;refGene_
tx=XM_019818793.2
E2 17819096 17819281
refGene_gn=PRODH2;refGene_tx=XM_003997934.4;refGene_gn=PRODH2;refGene_
tx=XM_019818793.2
E2 17821522 17821655
refGene_gn=LOC102899712;refGene_tx=XM_006941415.2;refGene_gn=LOC102899
712;refGene_tx=XM_011290026.3;refGene_gn=LOC102899712;refGene_tx=XM_02
3245437.1;refGene_gn=LOC102899712;refGene_tx=XM_023245438.1;refGene_gn
=LOC102899712;refGene_tx=XM_023245439.1;refGene_gn=LOC102899712;refGen
e_tx=XM_023245440.1

E2 17823877 17823992
refGene_gn=LOC102899712;refGene_tx=XM_006941415.2;refGene_gn=LOC102899712;refGene_tx=XM_011290026.3;refGene_gn=LOC102899712;refGene_tx=XM_023245437.1;refGene_gn=LOC102899712;refGene_tx=XM_023245438.1;refGene_gn=LOC102899712;refGene_tx=XM_023245439.1;refGene_gn=LOC102899712;refGene_tx=XM_023245440.1
E2 17824367 17824533
refGene_gn=LOC102899712;refGene_tx=XM_006941415.2;refGene_gn=LOC102899712;refGene_tx=XM_011290026.3;refGene_gn=LOC102899712;refGene_tx=XM_023245437.1;refGene_gn=LOC102899712;refGene_tx=XM_023245438.1;refGene_gn=LOC102899712;refGene_tx=XM_023245440.1
E2 17824612 17824767
refGene_gn=LOC102899712;refGene_tx=XM_006941415.2;refGene_gn=LOC102899712;refGene_tx=XM_011290026.3;refGene_gn=LOC102899712;refGene_tx=XM_023245437.1;refGene_gn=LOC102899712;refGene_tx=XM_023245438.1;refGene_gn=LOC102899712;refGene_tx=XM_023245440.1
E2 17824847 17824908
refGene_gn=LOC102899712;refGene_tx=XM_006941415.2;refGene_gn=LOC102899712;refGene_tx=XM_011290026.3;refGene_gn=LOC102899712;refGene_tx=XM_023245437.1;refGene_gn=LOC102899712;refGene_tx=XM_023245438.1;refGene_gn=LOC102899712;refGene_tx=XM_023245439.1;refGene_gn=LOC102899712;refGene_tx=XM_023245440.1
E2 17824984 17825109
refGene_gn=LOC102899712;refGene_tx=XM_006941415.2;refGene_gn=LOC102899712;refGene_tx=XM_011290026.3;refGene_gn=LOC102899712;refGene_tx=XM_023245437.1;refGene_gn=LOC102899712;refGene_tx=XM_023245438.1;refGene_gn=LOC102899712;refGene_tx=XM_023245439.1;refGene_gn=LOC102899712;refGene_tx=XM_023245440.1
E2 17825246 17825339
refGene_gn=LOC102899712;refGene_tx=XM_006941415.2;refGene_gn=LOC102899712;refGene_tx=XM_011290026.3;refGene_gn=LOC102899712;refGene_tx=XM_023245437.1;refGene_gn=LOC102899712;refGene_tx=XM_023245438.1;refGene_gn=LOC102899712;refGene_tx=XM_023245439.1;refGene_gn=LOC102899712;refGene_tx=XM_023245440.1
E2 17825769 17826105
refGene_gn=LOC102899712;refGene_tx=XM_006941415.2;refGene_gn=LOC102899712;refGene_tx=XM_011290026.3;refGene_gn=LOC102899712;refGene_tx=XM_023245437.1;refGene_gn=LOC102899712;refGene_tx=XM_023245438.1;refGene_gn=LOC102899712;refGene_tx=XM_023245439.1;refGene_gn=LOC102899712;refGene_tx=XM_023245440.1
E2 17826184 17826342
refGene_gn=LOC102899712;refGene_tx=XM_006941415.2;refGene_gn=LOC102899712;refGene_tx=XM_011290026.3;refGene_gn=LOC102899712;refGene_tx=XM_023245437.1;refGene_gn=LOC102899712;refGene_tx=XM_023245438.1;refGene_gn=LOC102899712;refGene_tx=XM_023245439.1;refGene_gn=LOC102899712;refGene_tx=XM_023245440.1
E2 17826418 17826612
refGene_gn=LOC102899712;refGene_tx=XM_006941415.2;refGene_gn=LOC102899712;refGene_tx=XM_011290026.3;refGene_gn=LOC102899712;refGene_tx=XM_023245437.1;refGene_gn=LOC102899712;refGene_tx=XM_023245438.1;refGene_gn

=LOC102899712;refGene_tx=XM_023245439.1;refGene_gn=LOC102899712;refGene_tx=XM_023245440.1
E2 17826694 17826767 refGene_gn=LOC102899712;refGene_tx=XM_006941415.2;refGene_gn=LOC102899712;refGene_tx=XM_011290026.3;refGene_gn=LOC102899712;refGene_tx=XM_023245437.1;refGene_gn=LOC102899712;refGene_tx=XM_023245438.1;refGene_gn=LOC102899712;refGene_tx=XM_023245439.1;refGene_gn=LOC102899712;refGene_tx=XM_023245440.1
E2 17826850 17826939 refGene_gn=LOC102899712;refGene_tx=XM_006941415.2;refGene_gn=LOC102899712;refGene_tx=XM_011290026.3;refGene_gn=LOC102899712;refGene_tx=XM_023245437.1;refGene_gn=LOC102899712;refGene_tx=XM_023245439.1
E2 17827028 17827199 refGene_gn=LOC102899712;refGene_tx=XM_006941415.2;refGene_gn=LOC102899712;refGene_tx=XM_011290026.3;refGene_gn=LOC102899712;refGene_tx=XM_023245437.1;refGene_gn=LOC102899712;refGene_tx=XM_023245439.1;refGene_gn=LOC102899712;refGene_tx=XM_023245440.1
E2 17827715 17827765 refGene_gn=LOC102899712;refGene_tx=XM_006941415.2;refGene_gn=LOC102899712;refGene_tx=XM_011290026.3;refGene_gn=LOC102899712;refGene_tx=XM_023245438.1;refGene_gn=LOC102899712;refGene_tx=XM_023245439.1
E2 17827852 17827949 refGene_gn=LOC102899712;refGene_tx=XM_006941415.2;refGene_gn=LOC102899712;refGene_tx=XM_011290026.3;refGene_gn=LOC102899712;refGene_tx=XM_023245437.1;refGene_gn=LOC102899712;refGene_tx=XM_023245438.1;refGene_gn=LOC102899712;refGene_tx=XM_023245439.1
E2 17828489 17829275 refGene_gn=ARHGAP33;refGene_tx=XM_023245414.1
E2 17829368 17829760 refGene_gn=ARHGAP33;refGene_tx=XM_023245414.1
E2 17829852 17830527 refGene_gn=ARHGAP33;refGene_tx=XM_023245414.1
E2 17831585 17831660 refGene_gn=ARHGAP33;refGene_tx=XM_023245414.1
E2 17831739 17831885 refGene_gn=ARHGAP33;refGene_tx=XM_023245414.1
E2 17831969 17832123 refGene_gn=ARHGAP33;refGene_tx=XM_023245414.1
E2 17832761 17832907 refGene_gn=ARHGAP33;refGene_tx=XM_023245414.1
E2 17834248 17834356 refGene_gn=ARHGAP33;refGene_tx=XM_023245414.1
E2 17834439 17834522 refGene_gn=ARHGAP33;refGene_tx=XM_023245414.1
E2 17834602 17834747 refGene_gn=ARHGAP33;refGene_tx=XM_023245414.1
E2 17835908 17836011 refGene_gn=ARHGAP33;refGene_tx=XM_023245414.1
E2 17836100 17836253 refGene_gn=ARHGAP33;refGene_tx=XM_023245414.1
E2 17836342 17836394 refGene_gn=ARHGAP33;refGene_tx=XM_023245414.1
E2 17836395 17836403 refGene_gn=ARHGAP33;refGene_tx=XM_023245414.1
E2 17836485 17836552 refGene_gn=ARHGAP33;refGene_tx=XM_023245414.1
E2 17836636 17836714 refGene_gn=ARHGAP33;refGene_tx=XM_023245414.1
E2 17836802 17836925 refGene_gn=ARHGAP33;refGene_tx=XM_023245414.1
E2 17837495 17837588 refGene_gn=ARHGAP33;refGene_tx=XM_023245414.1
E2 17838012 17838150 refGene_gn=ARHGAP33;refGene_tx=XM_023245414.1
E2 17838257 17838338 refGene_gn=ARHGAP33;refGene_tx=XM_023245414.1
E2 17838692 17838777 refGene_gn=ARHGAP33;refGene_tx=XM_023245414.1
E2 17838856 17838954 refGene_gn=ARHGAP33;refGene_tx=XM_023245414.1
E2 17840863 17840869 refGene_gn=ARHGAP33;refGene_tx=XM_023245414.1
E2 17843794 17843936 refGene_gn=PROSER3;refGene_tx=XM_023245476.1

E2 17844047 17844129 refGene_gn=PROSER3;refGene_tx=XM_023245476.1
E2 17844217 17844503 refGene_gn=PROSER3;refGene_tx=XM_023245476.1
E2 17844774 17845170
refGene_gn=PROSER3;refGene_tx=XM_023245470.1;refGene_gn=PROSER3;refGen
e_tx=XM_023245471.1;refGene_gn=PROSER3;refGene_tx=XM_023245472.1;refGe
ne_gn=PROSER3;refGene_tx=XM_023245473.1;refGene_gn=PROSER3;refGene_tx=
XM_023245474.1;refGene_gn=PROSER3;refGene_tx=XM_023245475.1
E2 17845287 17845463
refGene_gn=PROSER3;refGene_tx=XM_023245470.1;refGene_gn=PROSER3;refGen
e_tx=XM_023245471.1;refGene_gn=PROSER3;refGene_tx=XM_023245472.1;refGe
ne_gn=PROSER3;refGene_tx=XM_023245473.1;refGene_gn=PROSER3;refGene_tx=
XM_023245474.1;refGene_gn=PROSER3;refGene_tx=XM_023245475.1;refGene_gn
=PROSER3;refGene_tx=XM_023245476.1
E2 17848531 17848671
refGene_gn=PROSER3;refGene_tx=XM_023245470.1;refGene_gn=PROSER3;refGen
e_tx=XM_023245471.1;refGene_gn=PROSER3;refGene_tx=XM_023245472.1;refGe
ne_gn=PROSER3;refGene_tx=XM_023245473.1;refGene_gn=PROSER3;refGene_tx=
XM_023245474.1;refGene_gn=PROSER3;refGene_tx=XM_023245475.1;refGene_gn
=PROSER3;refGene_tx=XM_023245476.1
E2 17848769 17848852
refGene_gn=PROSER3;refGene_tx=XM_023245470.1;refGene_gn=PROSER3;refGen
e_tx=XM_023245471.1;refGene_gn=PROSER3;refGene_tx=XM_023245472.1;refGe
ne_gn=PROSER3;refGene_tx=XM_023245473.1;refGene_gn=PROSER3;refGene_tx=
XM_023245474.1;refGene_gn=PROSER3;refGene_tx=XM_023245475.1;refGene_gn
=PROSER3;refGene_tx=XM_023245476.1
E2 17849537 17849665
refGene_gn=PROSER3;refGene_tx=XM_023245470.1;refGene_gn=PROSER3;refGen
e_tx=XM_023245471.1;refGene_gn=PROSER3;refGene_tx=XM_023245472.1;refGe
ne_gn=PROSER3;refGene_tx=XM_023245473.1;refGene_gn=PROSER3;refGene_tx=
XM_023245474.1;refGene_gn=PROSER3;refGene_tx=XM_023245475.1;refGene_gn
=PROSER3;refGene_tx=XM_023245476.1
E2 17849754 17849876
refGene_gn=PROSER3;refGene_tx=XM_023245470.1;refGene_gn=PROSER3;refGen
e_tx=XM_023245471.1;refGene_gn=PROSER3;refGene_tx=XM_023245472.1;refGe
ne_gn=PROSER3;refGene_tx=XM_023245473.1;refGene_gn=PROSER3;refGene_tx=
XM_023245474.1;refGene_gn=PROSER3;refGene_tx=XM_023245475.1;refGene_gn
=PROSER3;refGene_tx=XM_023245476.1
E2 17850703 17850906
refGene_gn=PROSER3;refGene_tx=XM_023245470.1;refGene_gn=PROSER3;refGen
e_tx=XM_023245471.1;refGene_gn=PROSER3;refGene_tx=XM_023245472.1;refGe
ne_gn=PROSER3;refGene_tx=XM_023245473.1;refGene_gn=PROSER3;refGene_tx=
XM_023245474.1;refGene_gn=PROSER3;refGene_tx=XM_023245475.1;refGene_gn
=PROSER3;refGene_tx=XM_023245476.1
E2 17851216 17851313
refGene_gn=PROSER3;refGene_tx=XM_023245470.1;refGene_gn=PROSER3;refGen
e_tx=XM_023245471.1;refGene_gn=PROSER3;refGene_tx=XM_023245472.1;refGe
ne_gn=PROSER3;refGene_tx=XM_023245473.1;refGene_gn=PROSER3;refGene_tx=
XM_023245474.1;refGene_gn=PROSER3;refGene_tx=XM_023245475.1;refGene_gn
=PROSER3;refGene_tx=XM_023245476.1
E2 17852411 17852422

refGene_gn=PROSER3;refGene_tx=XM_023245470.1;refGene_gn=PROSER3;refGene_tx=XM_023245471.1;refGene_gn=PROSER3;refGene_tx=XM_023245472.1;refGene_gn=PROSER3;refGene_tx=XM_023245473.1;refGene_gn=PROSER3;refGene_tx=XM_023245474.1;refGene_gn=PROSER3;refGene_tx=XM_023245475.1;refGene_gn=PROSER3;refGene_tx=XM_023245476.1
E2 17853623 17853806 refGene_gn=HSPB6;refGene_tx=XM_011290021.3
E2 17854828 17854951 refGene_gn=HSPB6;refGene_tx=XM_011290021.3
E2 17855060 17855240 refGene_gn=HSPB6;refGene_tx=XM_011290021.3
E2 17856285 17856367 refGene_gn=LIN37;refGene_tx=XM_003997910.5;refGene_gn=LIN37;refGene_tx=XM_011290020.3
E2 17856451 17856525 refGene_gn=LIN37;refGene_tx=XM_003997910.5;refGene_gn=LIN37;refGene_tx=XM_011290020.3
E2 17856612 17856753 refGene_gn=LIN37;refGene_tx=XM_003997910.5;refGene_gn=LIN37;refGene_tx=XM_011290020.3
E2 17857298 17857465 refGene_gn=LIN37;refGene_tx=XM_003997910.5;refGene_gn=LIN37;refGene_tx=XM_011290020.3
E2 17857550 17857636 refGene_gn=LIN37;refGene_tx=XM_003997910.5;refGene_gn=LIN37;refGene_tx=XM_011290020.3
E2 17857717 17857747 refGene_gn=LIN37;refGene_tx=XM_003997910.5;refGene_gn=LIN37;refGene_tx=XM_011290020.3
E2 17858065 17858119 refGene_gn=LIN37;refGene_tx=XM_003997910.5;refGene_gn=LIN37;refGene_tx=XM_011290020.3
E2 17858276 17858352 refGene_gn=LIN37;refGene_tx=XM_003997910.5;refGene_gn=LIN37;refGene_tx=XM_011290020.3
E2 17861300 17861334 refGene_gn=LIN37;refGene_tx=XM_003997910.5;refGene_gn=LIN37;refGene_tx=XM_011290020.3
E2 17862991 17863131 refGene_gn=PSENEN;refGene_tx=XM_003997920.4;refGene_gn=PSENEN;refGene_tx=XM_006941413.4
E2 17863290 17863395 refGene_gn=PSENEN;refGene_tx=XM_003997920.4;refGene_gn=PSENEN;refGene_tx=XM_006941413.4
E2 17863815 17863973 refGene_gn=PSENEN;refGene_tx=XM_003997920.4;refGene_gn=PSENEN;refGene_tx=XM_006941413.4
E2 17864107 17864148 refGene_gn=PSENEN;refGene_tx=XM_003997920.4
E2 17864415 17864459 refGene_gn=U2AF1L4;refGene_tx=XM_006941411.4
E2 17864605 17864693 refGene_gn=U2AF1L4;refGene_tx=XM_006941411.4
E2 17865064 17865131 refGene_gn=U2AF1L4;refGene_tx=XM_006941411.4
E2 17865213 17865263 refGene_gn=U2AF1L4;refGene_tx=XM_006941411.4
E2 17865338 17865437 refGene_gn=U2AF1L4;refGene_tx=XM_006941411.4

E2 17865602 17865736 refGene_gn=U2AF1L4;refGene_tx=XM_006941411.4
E2 17865841 17865937 refGene_gn=U2AF1L4;refGene_tx=XM_006941411.4
E2 17866301 17866386 refGene_gn=U2AF1L4;refGene_tx=XM_006941411.4
E2 17867378 17867535
refGene_gn=IGFLR1;refGene_tx=XM_003997976.5;refGene_gn=IGFLR1;refGene_tx=XM_023245484.1
E2 17868005 17868190
refGene_gn=IGFLR1;refGene_tx=XM_003997976.5;refGene_gn=IGFLR1;refGene_tx=XM_023245484.1
E2 17868494 17868873
refGene_gn=IGFLR1;refGene_tx=XM_003997976.5;refGene_gn=IGFLR1;refGene_tx=XM_023245484.1
E2 17868956 17869282
refGene_gn=IGFLR1;refGene_tx=XM_003997976.5;refGene_gn=IGFLR1;refGene_tx=XM_023245484.1
E2 17870003 17870279 refGene_gn=KMT2B;refGene_tx=XM_023245969.1
E2 17870369 17870499 refGene_gn=KMT2B;refGene_tx=XM_023245969.1
E2 17870618 17870702 refGene_gn=KMT2B;refGene_tx=XM_023245969.1
E2 17870817 17870925 refGene_gn=KMT2B;refGene_tx=XM_023245969.1
E2 17871202 17871380 refGene_gn=KMT2B;refGene_tx=XM_023245969.1
E2 17871478 17871553 refGene_gn=KMT2B;refGene_tx=XM_023245969.1
E2 17871634 17871772 refGene_gn=KMT2B;refGene_tx=XM_023245969.1
E2 17874551 17874661 refGene_gn=KMT2B;refGene_tx=XM_023245969.1
E2 17874738 17874828 refGene_gn=KMT2B;refGene_tx=XM_023245969.1
E2 17874910 17876207 refGene_gn=KMT2B;refGene_tx=XM_023245969.1
E2 17876290 17876518 refGene_gn=KMT2B;refGene_tx=XM_023245969.1
E2 17877581 17877742 refGene_gn=KMT2B;refGene_tx=XM_023245969.1
E2 17877843 17877922 refGene_gn=KMT2B;refGene_tx=XM_023245969.1
E2 17877999 17878120 refGene_gn=KMT2B;refGene_tx=XM_023245969.1
E2 17878348 17878507 refGene_gn=KMT2B;refGene_tx=XM_023245969.1
E2 17879352 17879490 refGene_gn=KMT2B;refGene_tx=XM_023245969.1
E2 17879572 17879664 refGene_gn=KMT2B;refGene_tx=XM_023245969.1
E2 17879760 17879876 refGene_gn=KMT2B;refGene_tx=XM_023245969.1
E2 17880474 17880548 refGene_gn=KMT2B;refGene_tx=XM_023245969.1
E2 17880681 17880786 refGene_gn=KMT2B;refGene_tx=XM_023245969.1
E2 17880876 17880966 refGene_gn=KMT2B;refGene_tx=XM_023245969.1
E2 17881039 17881224 refGene_gn=KMT2B;refGene_tx=XM_023245969.1
E2 17881403 17881517 refGene_gn=KMT2B;refGene_tx=XM_023245969.1
E2 17882044 17882162 refGene_gn=KMT2B;refGene_tx=XM_023245969.1
E2 17882633 17882729 refGene_gn=KMT2B;refGene_tx=XM_023245969.1
E2 17882832 17882979 refGene_gn=KMT2B;refGene_tx=XM_023245969.1
E2 17883119 17883233 refGene_gn=KMT2B;refGene_tx=XM_023245969.1
E2 17883369 17883468 refGene_gn=KMT2B;refGene_tx=XM_023245969.1
E2 17883748 17883843 refGene_gn=KMT2B;refGene_tx=XM_023245969.1
E2 17884181 17884460 refGene_gn=KMT2B;refGene_tx=XM_023245969.1
E2 17884708 17884764 refGene_gn=KMT2B;refGene_tx=XM_023245969.1
E2 17884925 17885205 refGene_gn=KMT2B;refGene_tx=XM_023245969.1
E2 17885471 17885622 refGene_gn=KMT2B;refGene_tx=XM_023245969.1
E2 17885714 17885828 refGene_gn=KMT2B;refGene_tx=XM_023245969.1
E2 17886069 17887287 refGene_gn=KMT2B;refGene_tx=XM_023245969.1

E2 17888119 17888295 refGene_gn=KMT2B;refGene_tx=XM_023245969.1
E2 17888539 17888612 refGene_gn=KMT2B;refGene_tx=XM_023245969.1
E2 17889738 17890098 refGene_gn=KMT2B;refGene_tx=XM_023245969.1
E2 17891387 17891665
refGene_gn=ZBTB32;refGene_tx=XM_003997977.5;refGene_gn=ZBTB32;refGene_tx=XM_006941410.4;refGene_gn=ZBTB32;refGene_tx=XM_019818760.2;refGene_gn=ZBTB32;refGene_tx=XM_019818761.2;refGene_gn=ZBTB32;refGene_tx=XM_019818765.2;refGene_gn=ZBTB32;refGene_tx=XM_019818766.2
E2 17891843 17892005
refGene_gn=ZBTB32;refGene_tx=XM_003997977.5;refGene_gn=ZBTB32;refGene_tx=XM_006941410.4;refGene_gn=ZBTB32;refGene_tx=XM_019818760.2;refGene_gn=ZBTB32;refGene_tx=XM_019818761.2;refGene_gn=ZBTB32;refGene_tx=XM_019818765.2;refGene_gn=ZBTB32;refGene_tx=XM_019818766.2
E2 17892121 17892190
refGene_gn=ZBTB32;refGene_tx=XM_003997977.5;refGene_gn=ZBTB32;refGene_tx=XM_006941410.4;refGene_gn=ZBTB32;refGene_tx=XM_019818760.2;refGene_gn=ZBTB32;refGene_tx=XM_019818761.2;refGene_gn=ZBTB32;refGene_tx=XM_019818765.2;refGene_gn=ZBTB32;refGene_tx=XM_019818766.2
E2 17892300 17892374
refGene_gn=ZBTB32;refGene_tx=XM_003997977.5;refGene_gn=ZBTB32;refGene_tx=XM_006941410.4;refGene_gn=ZBTB32;refGene_tx=XM_019818760.2;refGene_gn=ZBTB32;refGene_tx=XM_019818761.2;refGene_gn=ZBTB32;refGene_tx=XM_019818765.2
E2 17892627 17893508
refGene_gn=ZBTB32;refGene_tx=XM_003997977.5;refGene_gn=ZBTB32;refGene_tx=XM_006941410.4;refGene_gn=ZBTB32;refGene_tx=XM_019818760.2;refGene_gn=ZBTB32;refGene_tx=XM_019818761.2;refGene_gn=ZBTB32;refGene_tx=XM_019818765.2
E2 17894720 17894821 refGene_gn=ZBTB32;refGene_tx=XM_019818765.2
E2 17911722 17911767 refGene_gn=UPK1A;refGene_tx=XM_023245490.1
E2 17911849 17911933 refGene_gn=UPK1A;refGene_tx=XM_023245490.1
E2 17913236 17913416 refGene_gn=UPK1A;refGene_tx=XM_023245490.1
E2 17914876 17914984 refGene_gn=UPK1A;refGene_tx=XM_023245490.1
E2 17915123 17915170 refGene_gn=UPK1A;refGene_tx=XM_023245490.1
E2 17915172 17915197 refGene_gn=UPK1A;refGene_tx=XM_023245490.1
E2 17918832 17919033 refGene_gn=UPK1A;refGene_tx=XM_023245490.1
E2 17920521 17920602 refGene_gn=UPK1A;refGene_tx=XM_023245490.1
E2 17940367 17940421
refGene_gn=LOC101100266;refGene_tx=XM_003997899.5;refGene_gn=LOC101100266;refGene_tx=XM_006941409.4
E2 17945159 17945260
refGene_gn=LOC101100266;refGene_tx=XM_003997899.5;refGene_gn=LOC101100266;refGene_tx=XM_006941409.4
E2 17947005 17947111
refGene_gn=LOC101100266;refGene_tx=XM_003997899.5;refGene_gn=LOC101100266;refGene_tx=XM_006941409.4
E2 17951083 17951284
refGene_gn=ETV2;refGene_tx=XM_006941408.3;refGene_gn=ETV2;refGene_tx=XM_023245483.1
E2 17951542 17951655

refGene_gn=ETV2;refGene_tx=XM_006941408.3;refGene_gn=ETV2;refGene_tx=XM_023245483.1
E2 17952144 17952624
refGene_gn=ETV2;refGene_tx=XM_006941408.3;refGene_gn=ETV2;refGene_tx=XM_023245483.1
E2 17952868 17952928
refGene_gn=ETV2;refGene_tx=XM_006941408.3;refGene_gn=ETV2;refGene_tx=XM_023245483.1
E2 17953188 17953326
refGene_gn=ETV2;refGene_tx=XM_006941408.3;refGene_gn=ETV2;refGene_tx=XM_023245483.1
E2 17953396 17953466
refGene_gn=ETV2;refGene_tx=XM_006941408.3;refGene_gn=ETV2;refGene_tx=XM_023245483.1
E2 17957202 17957315
refGene_gn=RBM42;refGene_tx=XM_003997930.3;refGene_gn=RBM42;refGene_tx=XM_003997931.3;refGene_gn=RBM42;refGene_tx=XM_006941406.3;refGene_gn=RBM42;refGene_tx=XM_006941407.3
E2 17957407 17957602
refGene_gn=RBM42;refGene_tx=XM_003997930.3;refGene_gn=RBM42;refGene_tx=XM_003997931.3;refGene_gn=RBM42;refGene_tx=XM_006941406.3;refGene_gn=RBM42;refGene_tx=XM_006941407.3
E2 17959801 17959919
refGene_gn=RBM42;refGene_tx=XM_003997930.3;refGene_gn=RBM42;refGene_tx=XM_003997931.3;refGene_gn=RBM42;refGene_tx=XM_006941406.3;refGene_gn=RBM42;refGene_tx=XM_006941407.3
E2 17960250 17960583
refGene_gn=RBM42;refGene_tx=XM_003997930.3;refGene_gn=RBM42;refGene_tx=XM_003997931.3;refGene_gn=RBM42;refGene_tx=XM_006941406.3;refGene_gn=RBM42;refGene_tx=XM_006941407.3
E2 17961019 17961195
refGene_gn=RBM42;refGene_tx=XM_003997930.3;refGene_gn=RBM42;refGene_tx=XM_003997931.3;refGene_gn=RBM42;refGene_tx=XM_006941406.3;refGene_gn=RBM42;refGene_tx=XM_006941407.3
E2 17961274 17961340
refGene_gn=RBM42;refGene_tx=XM_003997930.3;refGene_gn=RBM42;refGene_tx=XM_006941406.3
E2 17963138 17963213
refGene_gn=RBM42;refGene_tx=XM_003997930.3;refGene_gn=RBM42;refGene_tx=XM_003997931.3;refGene_gn=RBM42;refGene_tx=XM_006941406.3;refGene_gn=RBM42;refGene_tx=XM_006941407.3
E2 17963318 17963403
refGene_gn=RBM42;refGene_tx=XM_003997930.3;refGene_gn=RBM42;refGene_tx=XM_003997931.3;refGene_gn=RBM42;refGene_tx=XM_006941406.3;refGene_gn=RBM42;refGene_tx=XM_006941407.3
E2 17964613 17964758
refGene_gn=RBM42;refGene_tx=XM_003997930.3;refGene_gn=RBM42;refGene_tx=XM_003997931.3;refGene_gn=RBM42;refGene_tx=XM_006941406.3;refGene_gn=RBM42;refGene_tx=XM_006941407.3
E2 17964991 17965119

refGene_gn=RBM42;refGene_tx=XM_003997930.3;refGene_gn=RBM42;refGene_tx=XM_003997931.3;refGene_gn=RBM42;refGene_tx=XM_006941406.3;refGene_gn=RBM42;refGene_tx=XM_006941407.3
E2 17968960 17969078
refGene_gn=HAUS5;refGene_tx=XM_003997978.5;refGene_gn=HAUS5;refGene_tx=XM_019818730.2;refGene_gn=HAUS5;refGene_tx=XM_019818731.1;refGene_gn=HAUS5;refGene_tx=XM_019818732.1;refGene_gn=HAUS5;refGene_tx=XM_019818733.2;refGene_gn=HAUS5;refGene_tx=XM_019818734.1
E2 17969241 17969374
refGene_gn=HAUS5;refGene_tx=XM_003997978.5;refGene_gn=HAUS5;refGene_tx=XM_019818730.2;refGene_gn=HAUS5;refGene_tx=XM_019818731.1;refGene_gn=HAUS5;refGene_tx=XM_019818732.1;refGene_gn=HAUS5;refGene_tx=XM_019818733.2;refGene_gn=HAUS5;refGene_tx=XM_019818734.1
E2 17971369 17971502
refGene_gn=HAUS5;refGene_tx=XM_003997978.5;refGene_gn=HAUS5;refGene_tx=XM_019818730.2;refGene_gn=HAUS5;refGene_tx=XM_019818731.1;refGene_gn=HAUS5;refGene_tx=XM_019818732.1;refGene_gn=HAUS5;refGene_tx=XM_019818733.2;refGene_gn=HAUS5;refGene_tx=XM_019818734.1
E2 17971579 17971691
refGene_gn=HAUS5;refGene_tx=XM_003997978.5;refGene_gn=HAUS5;refGene_tx=XM_019818730.2;refGene_gn=HAUS5;refGene_tx=XM_019818731.1;refGene_gn=HAUS5;refGene_tx=XM_019818732.1;refGene_gn=HAUS5;refGene_tx=XM_019818733.2;refGene_gn=HAUS5;refGene_tx=XM_019818734.1
E2 17971951 17972097
refGene_gn=HAUS5;refGene_tx=XM_003997978.5;refGene_gn=HAUS5;refGene_tx=XM_019818730.2;refGene_gn=HAUS5;refGene_tx=XM_019818731.1;refGene_gn=HAUS5;refGene_tx=XM_019818732.1;refGene_gn=HAUS5;refGene_tx=XM_019818733.2;refGene_gn=HAUS5;refGene_tx=XM_019818734.1
E2 17972198 17972279
refGene_gn=HAUS5;refGene_tx=XM_003997978.5;refGene_gn=HAUS5;refGene_tx=XM_019818730.2;refGene_gn=HAUS5;refGene_tx=XM_019818731.1;refGene_gn=HAUS5;refGene_tx=XM_019818732.1;refGene_gn=HAUS5;refGene_tx=XM_019818733.2;refGene_gn=HAUS5;refGene_tx=XM_019818734.1
E2 17972495 17972658
refGene_gn=HAUS5;refGene_tx=XM_003997978.5;refGene_gn=HAUS5;refGene_tx=XM_019818730.2;refGene_gn=HAUS5;refGene_tx=XM_019818731.1;refGene_gn=HAUS5;refGene_tx=XM_019818732.1;refGene_gn=HAUS5;refGene_tx=XM_019818733.2;refGene_gn=HAUS5;refGene_tx=XM_019818734.1
E2 17972880 17973011
refGene_gn=HAUS5;refGene_tx=XM_003997978.5;refGene_gn=HAUS5;refGene_tx=XM_019818730.2;refGene_gn=HAUS5;refGene_tx=XM_019818731.1;refGene_gn=HAUS5;refGene_tx=XM_019818732.1;refGene_gn=HAUS5;refGene_tx=XM_019818733.2;refGene_gn=HAUS5;refGene_tx=XM_019818734.1
E2 17973122 17973186
refGene_gn=HAUS5;refGene_tx=XM_003997978.5;refGene_gn=HAUS5;refGene_tx=XM_019818730.2;refGene_gn=HAUS5;refGene_tx=XM_019818731.1;refGene_gn=HAUS5;refGene_tx=XM_019818732.1;refGene_gn=HAUS5;refGene_tx=XM_019818733.2;refGene_gn=HAUS5;refGene_tx=XM_019818734.1
E2 17973403 17973528
refGene_gn=HAUS5;refGene_tx=XM_003997978.5;refGene_gn=HAUS5;refGene_tx

=XM_019818730.2;refGene_gn=HAUS5;refGene_tx=XM_019818731.1;refGene_gn=HAUS5;refGene_tx=XM_019818732.1;refGene_gn=HAUS5;refGene_tx=XM_019818733.2;refGene_gn=HAUS5;refGene_tx=XM_019818734.1
E2 17973680 17973738
refGene_gn=HAUS5;refGene_tx=XM_003997978.5;refGene_gn=HAUS5;refGene_tx=XM_019818730.2;refGene_gn=HAUS5;refGene_tx=XM_019818731.1;refGene_gn=HAUS5;refGene_tx=XM_019818732.1;refGene_gn=HAUS5;refGene_tx=XM_019818733.2;refGene_gn=HAUS5;refGene_tx=XM_019818734.1
E2 17974263 17974346
refGene_gn=HAUS5;refGene_tx=XM_003997978.5;refGene_gn=HAUS5;refGene_tx=XM_019818730.2;refGene_gn=HAUS5;refGene_tx=XM_019818731.1;refGene_gn=HAUS5;refGene_tx=XM_019818732.1;refGene_gn=HAUS5;refGene_tx=XM_019818733.2;refGene_gn=HAUS5;refGene_tx=XM_019818734.1
E2 17974447 17974517
refGene_gn=HAUS5;refGene_tx=XM_003997978.5;refGene_gn=HAUS5;refGene_tx=XM_019818730.2;refGene_gn=HAUS5;refGene_tx=XM_019818731.1;refGene_gn=HAUS5;refGene_tx=XM_019818732.1;refGene_gn=HAUS5;refGene_tx=XM_019818733.2;refGene_gn=HAUS5;refGene_tx=XM_019818734.1
E2 17975604 17975764
refGene_gn=HAUS5;refGene_tx=XM_003997978.5;refGene_gn=HAUS5;refGene_tx=XM_019818730.2;refGene_gn=HAUS5;refGene_tx=XM_019818731.1;refGene_gn=HAUS5;refGene_tx=XM_019818732.1;refGene_gn=HAUS5;refGene_tx=XM_019818733.2;refGene_gn=HAUS5;refGene_tx=XM_019818734.1
E2 17975889 17975995
refGene_gn=HAUS5;refGene_tx=XM_003997978.5;refGene_gn=HAUS5;refGene_tx=XM_019818733.2;refGene_gn=HAUS5;refGene_tx=XM_019818734.1
E2 17976604 17976629 refGene_gn=HAUS5;refGene_tx=XM_003997978.5
E2 17976761 17976794
refGene_gn=HAUS5;refGene_tx=XM_003997978.5;refGene_gn=HAUS5;refGene_tx=XM_019818730.2;refGene_gn=HAUS5;refGene_tx=XM_019818731.1;refGene_gn=HAUS5;refGene_tx=XM_019818732.1
E2 17976886 17976949
refGene_gn=HAUS5;refGene_tx=XM_003997978.5;refGene_gn=HAUS5;refGene_tx=XM_019818730.2;refGene_gn=HAUS5;refGene_tx=XM_019818731.1;refGene_gn=HAUS5;refGene_tx=XM_019818732.1
E2 17977709 17977807 refGene_gn=HAUS5;refGene_tx=XM_003997978.5
E2 18004260 18004389
refGene_gn=LOC109494610;refGene_tx=XM_019818824.2
E2 18004564 18004861
refGene_gn=LOC109494610;refGene_tx=XM_019818824.2
E2 18048997 18049126
refGene_gn=ATP4A;refGene_tx=XM_023245419.1;refGene_gn=ATP4A;refGene_tx=XM_023245420.1;refGene_gn=ATP4A;refGene_tx=XM_023245421.1
E2 18049212 18049272
refGene_gn=ATP4A;refGene_tx=XM_023245419.1;refGene_gn=ATP4A;refGene_tx=XM_023245420.1;refGene_gn=ATP4A;refGene_tx=XM_023245421.1
E2 18049688 18049892
refGene_gn=ATP4A;refGene_tx=XM_023245419.1;refGene_gn=ATP4A;refGene_tx=XM_023245420.1;refGene_gn=ATP4A;refGene_tx=XM_023245421.1
E2 18051044 18051158

refGene_gn=ATP4A;refGene_tx=XM_023245419.1;refGene_gn=ATP4A;refGene_tx=XM_023245420.1;refGene_gn=ATP4A;refGene_tx=XM_023245421.1
E2 18051386 18051639
refGene_gn=ATP4A;refGene_tx=XM_023245419.1;refGene_gn=ATP4A;refGene_tx=XM_023245420.1;refGene_gn=ATP4A;refGene_tx=XM_023245421.1
E2 18051883 18052152
refGene_gn=ATP4A;refGene_tx=XM_023245419.1;refGene_gn=ATP4A;refGene_tx=XM_023245420.1;refGene_gn=ATP4A;refGene_tx=XM_023245421.1
E2 18052787 18052986
refGene_gn=ATP4A;refGene_tx=XM_023245419.1;refGene_gn=ATP4A;refGene_tx=XM_023245420.1;refGene_gn=ATP4A;refGene_tx=XM_023245421.1
E2 18053372 18053482
refGene_gn=ATP4A;refGene_tx=XM_023245419.1;refGene_gn=ATP4A;refGene_tx=XM_023245420.1;refGene_gn=ATP4A;refGene_tx=XM_023245421.1
E2 18053562 18053697
refGene_gn=ATP4A;refGene_tx=XM_023245419.1;refGene_gn=ATP4A;refGene_tx=XM_023245420.1;refGene_gn=ATP4A;refGene_tx=XM_023245421.1
E2 18054179 18054372
refGene_gn=ATP4A;refGene_tx=XM_023245419.1;refGene_gn=ATP4A;refGene_tx=XM_023245420.1;refGene_gn=ATP4A;refGene_tx=XM_023245421.1
E2 18055144 18055320
refGene_gn=ATP4A;refGene_tx=XM_023245419.1;refGene_gn=ATP4A;refGene_tx=XM_023245420.1;refGene_gn=ATP4A;refGene_tx=XM_023245421.1
E2 18056467 18056604
refGene_gn=ATP4A;refGene_tx=XM_023245419.1;refGene_gn=ATP4A;refGene_tx=XM_023245420.1;refGene_gn=ATP4A;refGene_tx=XM_023245421.1
E2 18056691 18056842
refGene_gn=ATP4A;refGene_tx=XM_023245419.1;refGene_gn=ATP4A;refGene_tx=XM_023245420.1;refGene_gn=ATP4A;refGene_tx=XM_023245421.1
E2 18056935 18057104
refGene_gn=ATP4A;refGene_tx=XM_023245419.1;refGene_gn=ATP4A;refGene_tx=XM_023245420.1;refGene_gn=ATP4A;refGene_tx=XM_023245421.1
E2 18057195 18057350
refGene_gn=ATP4A;refGene_tx=XM_023245419.1;refGene_gn=ATP4A;refGene_tx=XM_023245420.1;refGene_gn=ATP4A;refGene_tx=XM_023245421.1
E2 18058475 18058599
refGene_gn=ATP4A;refGene_tx=XM_023245419.1;refGene_gn=ATP4A;refGene_tx=XM_023245420.1;refGene_gn=ATP4A;refGene_tx=XM_023245421.1
E2 18059023 18059169
refGene_gn=ATP4A;refGene_tx=XM_023245419.1;refGene_gn=ATP4A;refGene_tx=XM_023245420.1;refGene_gn=ATP4A;refGene_tx=XM_023245421.1
E2 18060469 18060603
refGene_gn=ATP4A;refGene_tx=XM_023245419.1;refGene_gn=ATP4A;refGene_tx=XM_023245420.1;refGene_gn=ATP4A;refGene_tx=XM_023245421.1
E2 18060964 18061066
refGene_gn=ATP4A;refGene_tx=XM_023245419.1;refGene_gn=ATP4A;refGene_tx=XM_023245420.1;refGene_gn=ATP4A;refGene_tx=XM_023245421.1
E2 18061157 18061249
refGene_gn=ATP4A;refGene_tx=XM_023245419.1;refGene_gn=ATP4A;refGene_tx=XM_023245420.1;refGene_gn=ATP4A;refGene_tx=XM_023245421.1

E2 18061485 18061514
refGene_gn=ATP4A;refGene_tx=XM_023245419.1;refGene_gn=ATP4A;refGene_tx
=XM_023245420.1;refGene_gn=ATP4A;refGene_tx=XM_023245421.1
E2 18064701 18064825 refGene_gn=TMEM147;refGene_tx=XM_006941405.2
E2 18064908 18065030 refGene_gn=TMEM147;refGene_tx=XM_006941405.2
E2 18065116 18065201 refGene_gn=TMEM147;refGene_tx=XM_006941405.2
E2 18065330 18065467 refGene_gn=TMEM147;refGene_tx=XM_006941405.2
E2 18065552 18065612 refGene_gn=TMEM147;refGene_tx=XM_006941405.2
E2 18066056 18066126 refGene_gn=TMEM147;refGene_tx=XM_006941405.2
E2 18066194 18066271 refGene_gn=TMEM147;refGene_tx=XM_006941405.2
E2 18066699 18066772 refGene_gn=GAPDHS;refGene_tx=XM_023245481.1
E2 18066861 18066959 refGene_gn=GAPDHS;refGene_tx=XM_023245481.1
E2 18068024 18068187 refGene_gn=GAPDHS;refGene_tx=XM_023245481.1
E2 18068350 18068502 refGene_gn=GAPDHS;refGene_tx=XM_023245481.1
E2 18068810 18068892 refGene_gn=GAPDHS;refGene_tx=XM_023245481.1
E2 18069261 18069380 refGene_gn=GAPDHS;refGene_tx=XM_023245481.1
E2 18069467 18069558 refGene_gn=GAPDHS;refGene_tx=XM_023245481.1
E2 18072626 18072733 refGene_gn=GAPDHS;refGene_tx=XM_023245481.1
E2 18072915 18073012 refGene_gn=GAPDHS;refGene_tx=XM_023245481.1
E2 18073744 18073913 refGene_gn=GAPDHS;refGene_tx=XM_023245481.1
E2 18076673 18076740 refGene_gn=GAPDHS;refGene_tx=XM_023245481.1
E2 18081490 18082101
refGene_gn=SBSN;refGene_tx=XM_006941404.2;refGene_gn=SBSN;refGene_tx=X
M_023245444.1;refGene_gn=SBSN;refGene_tx=XM_023245445.1
E2 18082666 18083344
refGene_gn=SBSN;refGene_tx=XM_006941404.2;refGene_gn=SBSN;refGene_tx=X
M_023245444.1;refGene_gn=SBSN;refGene_tx=XM_023245445.1
E2 18084601 18084667
refGene_gn=SBSN;refGene_tx=XM_006941404.2;refGene_gn=SBSN;refGene_tx=X
M_023245444.1;refGene_gn=SBSN;refGene_tx=XM_023245445.1
E2 18084762 18084807
refGene_gn=SBSN;refGene_tx=XM_006941404.2;refGene_gn=SBSN;refGene_tx=X
M_023245444.1;refGene_gn=SBSN;refGene_tx=XM_023245445.1
E2 18085655 18085679
refGene_gn=SBSN;refGene_tx=XM_006941404.2;refGene_gn=SBSN;refGene_tx=X
M_023245444.1;refGene_gn=SBSN;refGene_tx=XM_023245445.1
E2 18093604 18093982 refGene_gn=DMKN;refGene_tx=XM_011290009.3
E2 18094239 18094455 refGene_gn=DMKN;refGene_tx=XM_011290009.3
E2 18094579 18094639 refGene_gn=DMKN;refGene_tx=XM_011290009.3
E2 18094933 18094984 refGene_gn=DMKN;refGene_tx=XM_011290009.3
E2 18095165 18095360 refGene_gn=DMKN;refGene_tx=XM_011290009.3
E2 18096045 18096102 refGene_gn=DMKN;refGene_tx=XM_011290009.3
E2 18096471 18096498 refGene_gn=DMKN;refGene_tx=XM_011290009.3
E2 18096499 18096538 refGene_gn=DMKN;refGene_tx=XM_011290009.3
E2 18096802 18096853 refGene_gn=DMKN;refGene_tx=XM_011290009.3
E2 18098335 18098383 refGene_gn=DMKN;refGene_tx=XM_011290009.3
E2 18099037 18099091 refGene_gn=DMKN;refGene_tx=XM_011290009.3
E2 18100055 18100100 refGene_gn=DMKN;refGene_tx=XM_011290009.3
E2 18100246 18100294 refGene_gn=DMKN;refGene_tx=XM_011290009.3
E2 18100493 18100541 refGene_gn=DMKN;refGene_tx=XM_011290009.3

E2 18102963 18103020 refGene_gn=DMKN;refGene_tx=XM_011290009.3
E2 18103314 18103362 refGene_gn=DMKN;refGene_tx=XM_011290009.3
E2 18105238 18105286 refGene_gn=DMKN;refGene_tx=XM_011290009.3
E2 18105402 18105444 refGene_gn=DMKN;refGene_tx=XM_011290009.3
E2 18105772 18105841 refGene_gn=DMKN;refGene_tx=XM_011290009.3
E2 18107264 18107288 refGene_gn=DMKN;refGene_tx=XM_011290009.3
E2 18107396 18107444 refGene_gn=DMKN;refGene_tx=XM_011290009.3
E2 18118620 18118707 refGene_gn=KRTDAP;refGene_tx=XM_003997917.4
E2 18120097 18120136 refGene_gn=KRTDAP;refGene_tx=XM_003997917.4
E2 18120236 18120278 refGene_gn=KRTDAP;refGene_tx=XM_003997917.4
E2 18120447 18120492 refGene_gn=KRTDAP;refGene_tx=XM_003997917.4
E2 18121177 18121225 refGene_gn=KRTDAP;refGene_tx=XM_003997917.4
E2 18121503 18121542 refGene_gn=KRTDAP;refGene_tx=XM_003997917.4
E2 18151241 18152231 refGene_gn=FFAR2;refGene_tx=XM_003997914.4
E2 18188042 18189029
refGene_gn=LOC101100526;refGene_tx=XM_003997981.3
E2 18213669 18213771 refGene_gn=FFAR3;refGene_tx=XM_023245482.1
E2 18213773 18214769 refGene_gn=FFAR3;refGene_tx=XM_023245482.1
E2 18218595 18219558 refGene_gn=FFAR1;refGene_tx=NM_001319275.1
E2 18224026 18224158
refGene_gn=CD22;refGene_tx=XM_003997907.5;refGene_gn=CD22;refGene_tx=X
M_019818690.1;refGene_gn=CD22;refGene_tx=XM_019818692.1;refGene_gn=CD2
2;refGene_tx=XM_019818693.1;refGene_gn=CD22;refGene_tx=XM_019818694.2
E2 18224455 18224540
refGene_gn=CD22;refGene_tx=XM_003997907.5;refGene_gn=CD22;refGene_tx=X
M_019818690.1;refGene_gn=CD22;refGene_tx=XM_019818692.1;refGene_gn=CD2
2;refGene_tx=XM_019818693.1;refGene_gn=CD22;refGene_tx=XM_019818694.2
E2 18224718 18224843
refGene_gn=CD22;refGene_tx=XM_003997907.5;refGene_gn=CD22;refGene_tx=X
M_019818690.1;refGene_gn=CD22;refGene_tx=XM_019818692.1;refGene_gn=CD2
2;refGene_tx=XM_019818693.1;refGene_gn=CD22;refGene_tx=XM_019818694.2
E2 18225012 18225088
refGene_gn=CD22;refGene_tx=XM_003997907.5;refGene_gn=CD22;refGene_tx=X
M_019818690.1;refGene_gn=CD22;refGene_tx=XM_019818692.1;refGene_gn=CD2
2;refGene_tx=XM_019818693.1;refGene_gn=CD22;refGene_tx=XM_019818694.2
E2 18225232 18225329
refGene_gn=CD22;refGene_tx=XM_003997907.5;refGene_gn=CD22;refGene_tx=X
M_019818690.1;refGene_gn=CD22;refGene_tx=XM_019818692.1;refGene_gn=CD2
2;refGene_tx=XM_019818693.1;refGene_gn=CD22;refGene_tx=XM_019818694.2
E2 18226817 18227081
refGene_gn=CD22;refGene_tx=XM_003997907.5;refGene_gn=CD22;refGene_tx=X
M_019818690.1;refGene_gn=CD22;refGene_tx=XM_019818692.1;refGene_gn=CD2
2;refGene_tx=XM_019818693.1;refGene_gn=CD22;refGene_tx=XM_019818694.2
E2 18227158 18227422
refGene_gn=CD22;refGene_tx=XM_003997907.5;refGene_gn=CD22;refGene_tx=X
M_019818690.1;refGene_gn=CD22;refGene_tx=XM_019818692.1;refGene_gn=CD2
2;refGene_tx=XM_019818693.1;refGene_gn=CD22;refGene_tx=XM_019818694.2
E2 18227632 18227890
refGene_gn=CD22;refGene_tx=XM_003997907.5;refGene_gn=CD22;refGene_tx=X
M_019818690.1;refGene_gn=CD22;refGene_tx=XM_019818692.1;refGene_gn=CD2

2;refGene_tx=XM_019818693.1;refGene_gn=CD22;refGene_tx=XM_019818694.2
E2 18228916 18229180
refGene_gn=CD22;refGene_tx=XM_003997907.5;refGene_gn=CD22;refGene_tx=XM_019818690.1;refGene_gn=CD22;refGene_tx=XM_019818692.1
E2 18229308 18229575
refGene_gn=CD22;refGene_tx=XM_003997907.5;refGene_gn=CD22;refGene_tx=XM_019818690.1
E2 18230683 18230977
refGene_gn=CD22;refGene_tx=XM_003997907.5;refGene_gn=CD22;refGene_tx=XM_019818690.1;refGene_gn=CD22;refGene_tx=XM_019818692.1;refGene_gn=CD22;refGene_tx=XM_019818693.1;refGene_gn=CD22;refGene_tx=XM_019818694.2
E2 18231916 18232294
refGene_gn=CD22;refGene_tx=XM_003997907.5;refGene_gn=CD22;refGene_tx=XM_019818690.1;refGene_gn=CD22;refGene_tx=XM_019818692.1;refGene_gn=CD22;refGene_tx=XM_019818693.1;refGene_gn=CD22;refGene_tx=XM_019818694.2
E2 18232438 18232547
refGene_gn=CD22;refGene_tx=XM_003997907.5;refGene_gn=CD22;refGene_tx=XM_019818690.1;refGene_gn=CD22;refGene_tx=XM_019818692.1;refGene_gn=CD22;refGene_tx=XM_019818693.1;refGene_gn=CD22;refGene_tx=XM_019818694.2
E2 18250492 18250657
refGene_gn=MAG;refGene_tx=XM_006941397.3;refGene_gn=MAG;refGene_tx=XM_019818743.1
E2 18251314 18251341 refGene_gn=MAG;refGene_tx=XM_006941398.3
E2 18251597 18251697
refGene_gn=MAG;refGene_tx=XM_006941397.3;refGene_gn=MAG;refGene_tx=XM_006941398.3;refGene_gn=MAG;refGene_tx=XM_019818743.1
E2 18252491 18252588
refGene_gn=MAG;refGene_tx=XM_006941397.3;refGene_gn=MAG;refGene_tx=XM_006941398.3;refGene_gn=MAG;refGene_tx=XM_019818743.1
E2 18252931 18253219
refGene_gn=MAG;refGene_tx=XM_006941397.3;refGene_gn=MAG;refGene_tx=XM_006941398.3;refGene_gn=MAG;refGene_tx=XM_019818743.1
E2 18257471 18257732
refGene_gn=MAG;refGene_tx=XM_006941397.3;refGene_gn=MAG;refGene_tx=XM_006941398.3;refGene_gn=MAG;refGene_tx=XM_019818743.1
E2 18258842 18259100
refGene_gn=MAG;refGene_tx=XM_006941397.3;refGene_gn=MAG;refGene_tx=XM_006941398.3;refGene_gn=MAG;refGene_tx=XM_019818743.1
E2 18259208 18259505
refGene_gn=MAG;refGene_tx=XM_006941397.3;refGene_gn=MAG;refGene_tx=XM_006941398.3;refGene_gn=MAG;refGene_tx=XM_019818743.1
E2 18260970 18261339
refGene_gn=MAG;refGene_tx=XM_006941397.3;refGene_gn=MAG;refGene_tx=XM_006941398.3;refGene_gn=MAG;refGene_tx=XM_019818743.1
E2 18261509 18261555
refGene_gn=MAG;refGene_tx=XM_006941397.3;refGene_gn=MAG;refGene_tx=XM_006941398.3;refGene_gn=MAG;refGene_tx=XM_019818743.1
E2 18267925 18268030 refGene_gn=HAMP;refGene_tx=XM_003997921.2
E2 18268114 18268174 refGene_gn=HAMP;refGene_tx=XM_003997921.2
E2 18269232 18269319 refGene_gn=HAMP;refGene_tx=XM_003997921.2

E2 18272140 18272230
refGene_gn=USF2;refGene_tx=XM_023245485.1;refGene_gn=USF2;refGene_tx=XM_023245486.1
E2 18272336 18272465
refGene_gn=USF2;refGene_tx=XM_023245485.1;refGene_gn=USF2;refGene_tx=XM_023245486.1
E2 18272629 18272724
refGene_gn=USF2;refGene_tx=XM_023245485.1;refGene_gn=USF2;refGene_tx=XM_023245486.1
E2 18278981 18279040
refGene_gn=USF2;refGene_tx=XM_023245485.1;refGene_gn=USF2;refGene_tx=XM_023245486.1
E2 18279331 18279419
refGene_gn=USF2;refGene_tx=XM_023245485.1;refGene_gn=USF2;refGene_tx=XM_023245486.1
E2 18279541 18279692
refGene_gn=USF2;refGene_tx=XM_023245485.1;refGene_gn=USF2;refGene_tx=XM_023245486.1
E2 18280115 18280316
refGene_gn=USF2;refGene_tx=XM_023245485.1;refGene_gn=USF2;refGene_tx=XM_023245486.1
E2 18280451 18280570
refGene_gn=USF2;refGene_tx=XM_023245485.1;refGene_gn=USF2;refGene_tx=XM_023245486.1
E2 18280681 18280728
refGene_gn=USF2;refGene_tx=XM_023245485.1;refGene_gn=USF2;refGene_tx=XM_023245486.1
E2 18280992 18281054
refGene_gn=USF2;refGene_tx=XM_023245485.1;refGene_gn=USF2;refGene_tx=XM_023245486.1
E2 18282349 18282385 refGene_gn=LSR;refGene_tx=XM_019818801.2
E2 18282475 18283101 refGene_gn=LSR;refGene_tx=XM_019818801.2
E2 18283394 18283526 refGene_gn=LSR;refGene_tx=XM_019818801.2
E2 18283625 18283685 refGene_gn=LSR;refGene_tx=XM_019818801.2
E2 18283837 18283999 refGene_gn=LSR;refGene_tx=XM_019818801.2
E2 18286279 18286426 refGene_gn=LSR;refGene_tx=XM_019818801.2
E2 18287000 18287057 refGene_gn=LSR;refGene_tx=XM_019818801.2
E2 18288768 18288888 refGene_gn=LSR;refGene_tx=XM_019818801.2
E2 18294114 18294459 refGene_gn=LSR;refGene_tx=XM_019818801.2
E2 18295649 18295915 refGene_gn=LSR;refGene_tx=XM_019818801.2
E2 18297511 18297519 refGene_gn=LSR;refGene_tx=XM_019818801.2
E2 18305914 18306631
refGene_gn=LOC111558044;refGene_tx=XM_023245491.1
E2 18323939 18324709
refGene_gn=LOC101088130;refGene_tx=XM_003997937.5
E2 18327459 18327694
refGene_gn=LOC101088130;refGene_tx=XM_003997937.5
E2 18327696 18327846
refGene_gn=LOC101088130;refGene_tx=XM_003997937.5
E2 18365987 18366037

refGene_gn=FXYD5;refGene_tx=XM_011290003.2;refGene_gn=FXYD5;refGene_tx=XM_011290004.3;refGene_gn=FXYD5;refGene_tx=XM_011290005.2;refGene_gn=FXYD5;refGene_tx=XM_011290006.2;refGene_gn=FXYD5;refGene_tx=XM_019818816.1;refGene_gn=FXYD5;refGene_tx=XM_019818817.1
E2 18367924 18367999 refGene_gn=FXYD5;refGene_tx=XM_011290003.2;refGene_gn=FXYD5;refGene_tx=XM_011290004.3;refGene_gn=FXYD5;refGene_tx=XM_011290005.2;refGene_gn=FXYD5;refGene_tx=XM_011290006.2;refGene_gn=FXYD5;refGene_tx=XM_019818816.1;refGene_gn=FXYD5;refGene_tx=XM_019818817.1
E2 18368077 18368107 refGene_gn=FXYD5;refGene_tx=XM_011290003.2;refGene_gn=FXYD5;refGene_tx=XM_011290004.3;refGene_gn=FXYD5;refGene_tx=XM_011290005.2;refGene_gn=FXYD5;refGene_tx=XM_011290006.2;refGene_gn=FXYD5;refGene_tx=XM_019818816.1;refGene_gn=FXYD5;refGene_tx=XM_019818817.1
E2 18369153 18369249 refGene_gn=FXYD5;refGene_tx=XM_011290003.2;refGene_gn=FXYD5;refGene_tx=XM_011290004.3;refGene_gn=FXYD5;refGene_tx=XM_011290005.2;refGene_gn=FXYD5;refGene_tx=XM_011290006.2;refGene_gn=FXYD5;refGene_tx=XM_019818816.1;refGene_gn=FXYD5;refGene_tx=XM_019818817.1
E2 18372036 18372138 refGene_gn=FXYD5;refGene_tx=XM_011290003.2;refGene_gn=FXYD5;refGene_tx=XM_011290004.3;refGene_gn=FXYD5;refGene_tx=XM_011290005.2;refGene_gn=FXYD5;refGene_tx=XM_011290006.2;refGene_gn=FXYD5;refGene_tx=XM_019818816.1;refGene_gn=FXYD5;refGene_tx=XM_019818817.1
E2 18373546 18373600 refGene_gn=FXYD5;refGene_tx=XM_011290003.2;refGene_gn=FXYD5;refGene_tx=XM_011290004.3;refGene_gn=FXYD5;refGene_tx=XM_011290005.2;refGene_gn=FXYD5;refGene_tx=XM_011290006.2;refGene_gn=FXYD5;refGene_tx=XM_019818816.1;refGene_gn=FXYD5;refGene_tx=XM_019818817.1
E2 18374341 18374422 refGene_gn=FXYD5;refGene_tx=XM_011290003.2;refGene_gn=FXYD5;refGene_tx=XM_011290004.3;refGene_gn=FXYD5;refGene_tx=XM_011290005.2;refGene_gn=FXYD5;refGene_tx=XM_011290006.2;refGene_gn=FXYD5;refGene_tx=XM_019818816.1;refGene_gn=FXYD5;refGene_tx=XM_019818817.1
E2 18374752 18374821 refGene_gn=FXYD5;refGene_tx=XM_011290003.2;refGene_gn=FXYD5;refGene_tx=XM_011290005.2;refGene_gn=FXYD5;refGene_tx=XM_011290006.2;refGene_gn=FXYD5;refGene_tx=XM_019818816.1
E2 18382814 18382953 refGene_gn=FXYD5;refGene_tx=XM_019818816.1
E2 18383008 18383069 refGene_gn=FXYD5;refGene_tx=XM_011290003.2;refGene_gn=FXYD5;refGene_tx=XM_011290004.3;refGene_gn=FXYD5;refGene_tx=XM_011290005.2;refGene_gn=FXYD5;refGene_tx=XM_011290006.2;refGene_gn=FXYD5;refGene_tx=XM_019818817.1
E2 18383814 18383937 refGene_gn=FXYD5;refGene_tx=XM_011290003.2;refGene_gn=FXYD5;refGene_tx=XM_011290004.3
E2 18384760 18384783 refGene_gn=FXYD7;refGene_tx=XM_019818821.1;refGene_gn=FXYD7;refGene_tx=XM_019818822.2

E2 18386753 18386794
refGene_gn=FXYD7;refGene_tx=XM_019818821.1;refGene_gn=FXYD7;refGene_tx=XM_019818822.2
E2 18386953 18386996
refGene_gn=FXYD7;refGene_tx=XM_019818821.1;refGene_gn=FXYD7;refGene_tx=XM_019818822.2
E2 18387144 18387219
refGene_gn=FXYD7;refGene_tx=XM_019818821.1;refGene_gn=FXYD7;refGene_tx=XM_019818822.2
E2 18389391 18389421
refGene_gn=FXYD7;refGene_tx=XM_019818821.1;refGene_gn=FXYD7;refGene_tx=XM_019818822.2
E2 18394143 18394174
refGene_gn=FXYD7;refGene_tx=XM_019818821.1;refGene_gn=FXYD7;refGene_tx=XM_019818822.2
E2 18394735 18394758 refGene_gn=FXYD1;refGene_tx=XM_011290002.3
E2 18394951 18395001 refGene_gn=FXYD1;refGene_tx=XM_011290002.3
E2 18395915 18395952 refGene_gn=FXYD1;refGene_tx=XM_011290002.3
E2 18396216 18396291 refGene_gn=FXYD1;refGene_tx=XM_011290002.3
E2 18396866 18396899 refGene_gn=FXYD1;refGene_tx=XM_011290002.3
E2 18397354 18397419 refGene_gn=FXYD1;refGene_tx=XM_011290002.3
E2 18398261 18398618 refGene_gn=FXYD1;refGene_tx=XM_011290002.3
E2 18400400 18400426 refGene_gn=FXYD1;refGene_tx=XM_011290002.3
E2 18402406 18402582
refGene_gn=LGI4;refGene_tx=XM_003997904.5;refGene_gn=LGI4;refGene_tx=XM_006941392.4
E2 18402990 18403062
refGene_gn=LGI4;refGene_tx=XM_003997904.5;refGene_gn=LGI4;refGene_tx=XM_006941392.4
E2 18403315 18403387
refGene_gn=LGI4;refGene_tx=XM_003997904.5;refGene_gn=LGI4;refGene_tx=XM_006941392.4
E2 18403957 18404270
refGene_gn=LOC109494611;refGene_tx=XM_019818825.2
E2 18404408 18404480
refGene_gn=LGI4;refGene_tx=XM_003997904.5;refGene_gn=LGI4;refGene_tx=XM_006941392.4
E2 18404614 18404686
refGene_gn=LGI4;refGene_tx=XM_003997904.5;refGene_gn=LGI4;refGene_tx=XM_006941392.4
E2 18404923 18405093 refGene_gn=LGI4;refGene_tx=XM_003997904.5
E2 18405186 18405266
refGene_gn=LOC109494611;refGene_tx=XM_019818825.2
E2 18406940 18407233
refGene_gn=LGI4;refGene_tx=XM_003997904.5;refGene_gn=LGI4;refGene_tx=XM_006941392.4
E2 18407352 18407858
refGene_gn=LGI4;refGene_tx=XM_003997904.5;refGene_gn=LGI4;refGene_tx=XM_006941392.4
E2 18408694 18409009

refGene_gn=LGI4;refGene_tx=XM_003997904.5;refGene_gn=LGI4;refGene_tx=XM_006941392.4
E2 18410704 18410724 refGene_gn=FXYD3;refGene_tx=XM_019818823.1
E2 18410853 18410891 refGene_gn=FXYD3;refGene_tx=XM_019818823.1
E2 18411200 18411314 refGene_gn=FXYD3;refGene_tx=XM_019818823.1
E2 18411315 18411397 refGene_gn=FXYD3;refGene_tx=XM_019818823.1
E2 18412121 18412145 refGene_gn=FXYD3;refGene_tx=XM_019818823.1
E2 18412255 18412288 refGene_gn=FXYD3;refGene_tx=XM_019818823.1
E2 18413871 18413928 refGene_gn=FXYD3;refGene_tx=XM_019818823.1
E2 18414154 18414230 refGene_gn=FXYD3;refGene_tx=XM_019818823.1
E2 18445695 18445734 refGene_gn=HPN;refGene_tx=XM_019818811.2
E2 18445952 18446117 refGene_gn=HPN;refGene_tx=XM_019818811.2
E2 18446299 18446442 refGene_gn=HPN;refGene_tx=XM_019818811.2
E2 18446635 18446731 refGene_gn=HPN;refGene_tx=XM_019818811.2
E2 18449744 18449935 refGene_gn=HPN;refGene_tx=XM_019818811.2
E2 18450024 18450190 refGene_gn=HPN;refGene_tx=XM_019818811.2
E2 18450446 18450487 refGene_gn=HPN;refGene_tx=XM_019818811.2
E2 18450581 18450704 refGene_gn=HPN;refGene_tx=XM_019818811.2
E2 18450777 18450907 refGene_gn=HPN;refGene_tx=XM_019818811.2
E2 18456006 18456048 refGene_gn=HPN;refGene_tx=XM_019818811.2
E2 18456136 18456238 refGene_gn=HPN;refGene_tx=XM_019818811.2
E2 18461928 18461944 refGene_gn=HPN;refGene_tx=XM_019818811.2
E2 18464774 18464841
refGene_gn=SCN1B;refGene_tx=XM_023245492.1;refGene_gn=SCN1B;refGene_tx=XM_023245493.1
E2 18465184 18465326
refGene_gn=SCN1B;refGene_tx=XM_023245492.1;refGene_gn=SCN1B;refGene_tx=XM_023245493.1
E2 18469718 18469959
refGene_gn=SCN1B;refGene_tx=XM_023245492.1;refGene_gn=SCN1B;refGene_tx=XM_023245493.1
E2 18470748 18470915
refGene_gn=SCN1B;refGene_tx=XM_023245492.1;refGene_gn=SCN1B;refGene_tx=XM_023245493.1
E2 18472651 18472691
refGene_gn=SCN1B;refGene_tx=XM_023245492.1;refGene_gn=SCN1B;refGene_tx=XM_023245493.1
E2 18476290 18476383 refGene_gn=GRAMD1A;refGene_tx=XM_023245428.1
E2 18478088 18478209 refGene_gn=GRAMD1A;refGene_tx=XM_023245428.1
E2 18478292 18478400 refGene_gn=GRAMD1A;refGene_tx=XM_023245428.1
E2 18478729 18478741 refGene_gn=GRAMD1A;refGene_tx=XM_023245428.1
E2 18479575 18479640 refGene_gn=GRAMD1A;refGene_tx=XM_023245428.1
E2 18479641 18479664 refGene_gn=GRAMD1A;refGene_tx=XM_023245428.1
E2 18479761 18479935 refGene_gn=GRAMD1A;refGene_tx=XM_023245428.1
E2 18480020 18480124 refGene_gn=GRAMD1A;refGene_tx=XM_023245428.1
E2 18480826 18480969 refGene_gn=GRAMD1A;refGene_tx=XM_023245428.1
E2 18481051 18481170 refGene_gn=GRAMD1A;refGene_tx=XM_023245428.1
E2 18482743 18482887 refGene_gn=GRAMD1A;refGene_tx=XM_023245428.1
E2 18483771 18483970 refGene_gn=GRAMD1A;refGene_tx=XM_023245428.1
E2 18484519 18484670 refGene_gn=GRAMD1A;refGene_tx=XM_023245428.1

E2	18484840 18484953 refGene_gn=GRAMD1A;refGene_tx=XM_023245428.1
E2	18488147 18488228 refGene_gn=GRAMD1A;refGene_tx=XM_023245428.1
E2	18488962 18489058 refGene_gn=GRAMD1A;refGene_tx=XM_023245428.1
E2	18489135 18489239 refGene_gn=GRAMD1A;refGene_tx=XM_023245428.1
E2	18489377 18489462 refGene_gn=GRAMD1A;refGene_tx=XM_023245428.1
E2	18489927 18489948 refGene_gn=GRAMD1A;refGene_tx=XM_023245428.1
E2	18490036 18490247 refGene_gn=GRAMD1A;refGene_tx=XM_023245428.1
E2	18496726 18496734 refGene_gn=GRAMD1A;refGene_tx=XM_023245428.1
E2	18543274 18543310
refGene_gn=ZNF792;refGene_tx=XM_003997987.5;refGene_gn=ZNF792;refGene_tx=XM_019818735.2
E2	18546482 18546609
refGene_gn=ZNF792;refGene_tx=XM_003997987.5;refGene_gn=ZNF792;refGene_tx=XM_019818735.2
E2	18547091 18547214
refGene_gn=ZNF792;refGene_tx=XM_003997987.5;refGene_gn=ZNF792;refGene_tx=XM_019818735.2
E2	18548092 18549708
refGene_gn=ZNF792;refGene_tx=XM_003997987.5;refGene_gn=ZNF792;refGene_tx=XM_019818735.2
E2	18572182 18572961 refGene_gn=ZNF599;refGene_tx=XM_023245480.1
E2	18622796 18624322
refGene_gn=ZNF599;refGene_tx=XM_023245479.1;refGene_gn=ZNF599;refGene_tx=XM_023245480.1
E2	18632405 18632501
refGene_gn=ZNF599;refGene_tx=XM_023245479.1;refGene_gn=ZNF599;refGene_tx=XM_023245480.1
E2	18632851 18632978
refGene_gn=ZNF599;refGene_tx=XM_023245479.1;refGene_gn=ZNF599;refGene_tx=XM_023245480.1
E2	18635670 18635688 refGene_gn=ZNF599;refGene_tx=XM_023245480.1
E2	18792980 18793121 refGene_gn=WTIP;refGene_tx=XM_019819599.2
E2	18794567 18794636 refGene_gn=WTIP;refGene_tx=XM_019819599.2
E2	18795297 18795349 refGene_gn=WTIP;refGene_tx=XM_019819599.2
E2	18796161 18796292 refGene_gn=WTIP;refGene_tx=XM_019819599.2
E2	18796456 18796519 refGene_gn=WTIP;refGene_tx=XM_019819599.2
E2	18796674 18796742 refGene_gn=WTIP;refGene_tx=XM_019819599.2
E2	18806860 18806962 refGene_gn=WTIP;refGene_tx=XM_019819599.2
E2	18812174 18812850 refGene_gn=WTIP;refGene_tx=XM_019819599.2
E2	18820310 18820492
refGene_gn=UBA2;refGene_tx=XM_011289986.2;refGene_gn=UBA2;refGene_tx=XM_019819600.2;refGene_gn=UBA2;refGene_tx=XM_019819601.2
E2	18821941 18822078
refGene_gn=UBA2;refGene_tx=XM_011289986.2;refGene_gn=UBA2;refGene_tx=XM_019819600.2;refGene_gn=UBA2;refGene_tx=XM_019819601.2
E2	18822591 18822628 refGene_gn=UBA2;refGene_tx=XM_019819602.2
E2	18823652 18823758
refGene_gn=UBA2;refGene_tx=XM_011289986.2;refGene_gn=UBA2;refGene_tx=XM_019819600.2;refGene_gn=UBA2;refGene_tx=XM_019819601.2;refGene_gn=UBA2;refGene_tx=XM_019819602.2

E2 18828008 18828105
refGene_gn=UBA2;refGene_tx=XM_011289986.2;refGene_gn=UBA2;refGene_tx=XM_019819600.2;refGene_gn=UBA2;refGene_tx=XM_019819601.2;refGene_gn=UBA2;refGene_tx=XM_019819602.2
E2 18829914 18830070
refGene_gn=UBA2;refGene_tx=XM_011289986.2;refGene_gn=UBA2;refGene_tx=XM_019819600.2;refGene_gn=UBA2;refGene_tx=XM_019819601.2;refGene_gn=UBA2;refGene_tx=XM_019819602.2
E2 18835657 18835770
refGene_gn=UBA2;refGene_tx=XM_011289986.2;refGene_gn=UBA2;refGene_tx=XM_019819600.2;refGene_gn=UBA2;refGene_tx=XM_019819602.2
E2 18835857 18835951
refGene_gn=UBA2;refGene_tx=XM_011289986.2;refGene_gn=UBA2;refGene_tx=XM_019819600.2;refGene_gn=UBA2;refGene_tx=XM_019819602.2
E2 18838569 18838736
refGene_gn=UBA2;refGene_tx=XM_011289986.2;refGene_gn=UBA2;refGene_tx=XM_019819600.2;refGene_gn=UBA2;refGene_tx=XM_019819601.2;refGene_gn=UBA2;refGene_tx=XM_019819602.2
E2 18839977 18840077
refGene_gn=UBA2;refGene_tx=XM_011289986.2;refGene_gn=UBA2;refGene_tx=XM_019819600.2;refGene_gn=UBA2;refGene_tx=XM_019819601.2;refGene_gn=UBA2;refGene_tx=XM_019819602.2
E2 18843579 18843701
refGene_gn=UBA2;refGene_tx=XM_011289986.2;refGene_gn=UBA2;refGene_tx=XM_019819600.2;refGene_gn=UBA2;refGene_tx=XM_019819601.2;refGene_gn=UBA2;refGene_tx=XM_019819602.2
E2 18844298 18844366
refGene_gn=UBA2;refGene_tx=XM_011289986.2;refGene_gn=UBA2;refGene_tx=XM_019819600.2;refGene_gn=UBA2;refGene_tx=XM_019819601.2;refGene_gn=UBA2;refGene_tx=XM_019819602.2
E2 18850321 18850443
refGene_gn=UBA2;refGene_tx=XM_011289986.2;refGene_gn=UBA2;refGene_tx=XM_019819600.2;refGene_gn=UBA2;refGene_tx=XM_019819601.2;refGene_gn=UBA2;refGene_tx=XM_019819602.2
E2 18852368 18852469
refGene_gn=UBA2;refGene_tx=XM_011289986.2;refGene_gn=UBA2;refGene_tx=XM_019819600.2;refGene_gn=UBA2;refGene_tx=XM_019819601.2;refGene_gn=UBA2;refGene_tx=XM_019819602.2
E2 18853636 18853701
refGene_gn=UBA2;refGene_tx=XM_011289986.2;refGene_gn=UBA2;refGene_tx=XM_019819600.2;refGene_gn=UBA2;refGene_tx=XM_019819601.2;refGene_gn=UBA2;refGene_tx=XM_019819602.2
E2 18855141 18855212
refGene_gn=UBA2;refGene_tx=XM_011289986.2;refGene_gn=UBA2;refGene_tx=XM_019819600.2;refGene_gn=UBA2;refGene_tx=XM_019819601.2;refGene_gn=UBA2;refGene_tx=XM_019819602.2
E2 18856336 18856420
refGene_gn=UBA2;refGene_tx=XM_011289986.2;refGene_gn=UBA2;refGene_tx=XM_019819600.2;refGene_gn=UBA2;refGene_tx=XM_019819601.2;refGene_gn=UBA2;refGene_tx=XM_019819602.2

E2 18859028 18859166
refGene_gn=UBA2;refGene_tx=XM_011289986.2;refGene_gn=UBA2;refGene_tx=XM_019819600.2;refGene_gn=UBA2;refGene_tx=XM_019819601.2;refGene_gn=UBA2;refGene_tx=XM_019819602.2
E2 18864453 18864584
refGene_gn=PDCD2L;refGene_tx=XM_006941452.3;refGene_gn=PDCD2L;refGene_tx=XM_006941453.3
E2 18874513 18874662
refGene_gn=PDCD2L;refGene_tx=XM_006941452.3;refGene_gn=PDCD2L;refGene_tx=XM_006941453.3
E2 18884540 18884651
refGene_gn=PDCD2L;refGene_tx=XM_006941452.3;refGene_gn=PDCD2L;refGene_tx=XM_006941453.3
E2 18885850 18886218
refGene_gn=PDCD2L;refGene_tx=XM_006941452.3;refGene_gn=PDCD2L;refGene_tx=XM_006941453.3
E2 18887415 18887476
refGene_gn=PDCD2L;refGene_tx=XM_006941452.3;refGene_gn=PDCD2L;refGene_tx=XM_006941453.3
E2 18887599 18887766
refGene_gn=PDCD2L;refGene_tx=XM_006941452.3;refGene_gn=PDCD2L;refGene_tx=XM_006941453.3
E2 18887867 18887975
refGene_gn=PDCD2L;refGene_tx=XM_006941452.3;refGene_gn=PDCD2L;refGene_tx=XM_006941453.3
E2 18895315 18895451 refGene_gn=GPI;refGene_tx=XM_023245645.1
E2 18895556 18895623 refGene_gn=GPI;refGene_tx=XM_023245645.1
E2 18895708 18895784 refGene_gn=GPI;refGene_tx=XM_023245645.1
E2 18896001 18896130 refGene_gn=GPI;refGene_tx=XM_023245645.1
E2 18897181 18897258 refGene_gn=GPI;refGene_tx=XM_023245645.1
E2 18897413 18897543 refGene_gn=GPI;refGene_tx=XM_023245645.1
E2 18898738 18898891 refGene_gn=GPI;refGene_tx=XM_023245645.1
E2 18898995 18899039 refGene_gn=GPI;refGene_tx=XM_023245645.1
E2 18899455 18899516 refGene_gn=GPI;refGene_tx=XM_023245645.1
E2 18910058 18910112 refGene_gn=GPI;refGene_tx=XM_023245645.1
E2 18911425 18911470 refGene_gn=GPI;refGene_tx=XM_023245645.1
E2 18911972 18912044 refGene_gn=GPI;refGene_tx=XM_023245645.1
E2 18912920 18913067 refGene_gn=GPI;refGene_tx=XM_023245645.1
E2 18913196 18913280 refGene_gn=GPI;refGene_tx=XM_023245645.1
E2 18917851 18917971 refGene_gn=GPI;refGene_tx=XM_023245645.1
E2 18919400 18919469 refGene_gn=GPI;refGene_tx=XM_023245645.1
E2 18919826 18919917 refGene_gn=GPI;refGene_tx=XM_023245645.1
E2 18920962 18921084 refGene_gn=GPI;refGene_tx=XM_023245645.1
E2 18932458 18932767
refGene_gn=KIAA0355;refGene_tx=XM_003997945.5;refGene_gn=KIAA0355;refGene_tx=XM_023245827.1;refGene_gn=KIAA0355;refGene_tx=XM_023245828.1
E2 18933216 18933295
refGene_gn=KIAA0355;refGene_tx=XM_003997945.5;refGene_gn=KIAA0355;refGene_tx=XM_023245827.1;refGene_gn=KIAA0355;refGene_tx=XM_023245828.1
E2 18934814 18934952

refGene_gn=KIAA0355;refGene_tx=XM_003997945.5;refGene_gn=KIAA0355;refGene_tx=XM_023245827.1;refGene_gn=KIAA0355;refGene_tx=XM_023245828.1
E2 18935709 18935875
refGene_gn=KIAA0355;refGene_tx=XM_003997945.5;refGene_gn=KIAA0355;refGene_tx=XM_023245827.1;refGene_gn=KIAA0355;refGene_tx=XM_023245828.1
E2 18939143 18940177
refGene_gn=KIAA0355;refGene_tx=XM_003997945.5;refGene_gn=KIAA0355;refGene_tx=XM_023245827.1;refGene_gn=KIAA0355;refGene_tx=XM_023245828.1
E2 18941258 18941384
refGene_gn=KIAA0355;refGene_tx=XM_003997945.5;refGene_gn=KIAA0355;refGene_tx=XM_023245827.1;refGene_gn=KIAA0355;refGene_tx=XM_023245828.1
E2 18946885 18946983
refGene_gn=KIAA0355;refGene_tx=XM_003997945.5;refGene_gn=KIAA0355;refGene_tx=XM_023245827.1;refGene_gn=KIAA0355;refGene_tx=XM_023245828.1
E2 18949265 18949424
refGene_gn=KIAA0355;refGene_tx=XM_003997945.5;refGene_gn=KIAA0355;refGene_tx=XM_023245827.1;refGene_gn=KIAA0355;refGene_tx=XM_023245828.1
E2 18950311 18950473
refGene_gn=KIAA0355;refGene_tx=XM_003997945.5;refGene_gn=KIAA0355;refGene_tx=XM_023245827.1;refGene_gn=KIAA0355;refGene_tx=XM_023245828.1
E2 18950598 18950694
refGene_gn=KIAA0355;refGene_tx=XM_003997945.5;refGene_gn=KIAA0355;refGene_tx=XM_023245827.1;refGene_gn=KIAA0355;refGene_tx=XM_023245828.1
E2 18950897 18951038
refGene_gn=KIAA0355;refGene_tx=XM_003997945.5;refGene_gn=KIAA0355;refGene_tx=XM_023245827.1;refGene_gn=KIAA0355;refGene_tx=XM_023245828.1
E2 18957640 18957847
refGene_gn=KIAA0355;refGene_tx=XM_003997945.5;refGene_gn=KIAA0355;refGene_tx=XM_023245827.1;refGene_gn=KIAA0355;refGene_tx=XM_023245828.1
E2 18983297 18983792
refGene_gn=KIAA0355;refGene_tx=XM_003997945.5;refGene_gn=KIAA0355;refGene_tx=XM_023245827.1;refGene_gn=KIAA0355;refGene_tx=XM_023245828.1
E2 19042553 19042577
refGene_gn=LSM14A;refGene_tx=XM_023246136.1;refGene_gn=LSM14A;refGene_tx=XM_023246138.1
E2 19043547 19043571 refGene_gn=LSM14A;refGene_tx=XM_023246137.1
E2 19048326 19048558
refGene_gn=LSM14A;refGene_tx=XM_023246136.1;refGene_gn=LSM14A;refGene_tx=XM_023246137.1;refGene_gn=LSM14A;refGene_tx=XM_023246138.1
E2 19049905 19050077
refGene_gn=LSM14A;refGene_tx=XM_023246136.1;refGene_gn=LSM14A;refGene_tx=XM_023246137.1;refGene_gn=LSM14A;refGene_tx=XM_023246138.1
E2 19050181 19050364
refGene_gn=LSM14A;refGene_tx=XM_023246136.1;refGene_gn=LSM14A;refGene_tx=XM_023246137.1;refGene_gn=LSM14A;refGene_tx=XM_023246138.1
E2 19052826 19052892
refGene_gn=LSM14A;refGene_tx=XM_023246136.1;refGene_gn=LSM14A;refGene_tx=XM_023246137.1;refGene_gn=LSM14A;refGene_tx=XM_023246138.1
E2 19053176 19053353
refGene_gn=LSM14A;refGene_tx=XM_023246136.1;refGene_gn=LSM14A;refGene_

tx=XM_023246137.1
E2 19058696 19058819
refGene_gn=LSM14A;refGene_tx=XM_023246136.1;refGene_gn=LSM14A;refGene_tx=XM_023246137.1;refGene_gn=LSM14A;refGene_tx=XM_023246138.1
E2 19064467 19064597
refGene_gn=LSM14A;refGene_tx=XM_023246136.1;refGene_gn=LSM14A;refGene_tx=XM_023246137.1;refGene_gn=LSM14A;refGene_tx=XM_023246138.1
E2 19067470 19067634
refGene_gn=LSM14A;refGene_tx=XM_023246136.1;refGene_gn=LSM14A;refGene_tx=XM_023246137.1;refGene_gn=LSM14A;refGene_tx=XM_023246138.1
E2 19090931 19091052
refGene_gn=LSM14A;refGene_tx=XM_023246136.1;refGene_gn=LSM14A;refGene_tx=XM_023246137.1;refGene_gn=LSM14A;refGene_tx=XM_023246138.1
E2 19411183 19411342
refGene_gn=KCTD15;refGene_tx=XM_023245084.1;refGene_gn=KCTD15;refGene_tx=XM_023245085.1
E2 19412554 19412860
refGene_gn=KCTD15;refGene_tx=XM_023245084.1;refGene_gn=KCTD15;refGene_tx=XM_023245085.1
E2 19415243 19415388
refGene_gn=KCTD15;refGene_tx=XM_023245084.1;refGene_gn=KCTD15;refGene_tx=XM_023245085.1
E2 19420303 19420479
refGene_gn=KCTD15;refGene_tx=XM_023245084.1;refGene_gn=KCTD15;refGene_tx=XM_023245085.1
E2 19421113 19421179
refGene_gn=KCTD15;refGene_tx=XM_023245084.1;refGene_gn=KCTD15;refGene_tx=XM_023245085.1
E2 19453234 19454344
refGene_gn=CHST8;refGene_tx=XM_003997947.5;refGene_gn=CHST8;refGene_tx=XM_019819604.2;refGene_gn=CHST8;refGene_tx=XM_023245166.1
E2 19454897 19454935
refGene_gn=CHST8;refGene_tx=XM_003997947.5;refGene_gn=CHST8;refGene_tx=XM_019819604.2;refGene_gn=CHST8;refGene_tx=XM_023245166.1
E2 19517368 19517498
refGene_gn=CHST8;refGene_tx=XM_003997947.5;refGene_gn=CHST8;refGene_tx=XM_019819604.2;refGene_gn=CHST8;refGene_tx=XM_023245166.1
E2 19656596 19656613 refGene_gn=PEPD;refGene_tx=XM_023245695.1
E2 19664166 19664264 refGene_gn=PEPD;refGene_tx=XM_023245694.1
E2 19665091 19665275
refGene_gn=PEPD;refGene_tx=XM_023245694.1;refGene_gn=PEPD;refGene_tx=XM_023245695.1
E2 19667256 19667384
refGene_gn=PEPD;refGene_tx=XM_023245694.1;refGene_gn=PEPD;refGene_tx=XM_023245695.1
E2 19675302 19675366
refGene_gn=PEPD;refGene_tx=XM_023245694.1;refGene_gn=PEPD;refGene_tx=XM_023245695.1
E2 19681839 19681887
refGene_gn=PEPD;refGene_tx=XM_023245694.1;refGene_gn=PEPD;refGene_tx=X

M_023245695.1
E2 19683490 19683552
refGene_gn=PEPD;refGene_tx=XM_023245694.1;refGene_gn=PEPD;refGene_tx=XM_023245695.1
E2 19691688 19691733
refGene_gn=PEPD;refGene_tx=XM_023245694.1;refGene_gn=PEPD;refGene_tx=XM_023245695.1
E2 19700747 19700823
refGene_gn=PEPD;refGene_tx=XM_023245694.1;refGene_gn=PEPD;refGene_tx=XM_023245695.1
E2 19701752 19701799
refGene_gn=PEPD;refGene_tx=XM_023245694.1;refGene_gn=PEPD;refGene_tx=XM_023245695.1
E2 19746057 19746126
refGene_gn=PEPD;refGene_tx=XM_023245694.1;refGene_gn=PEPD;refGene_tx=XM_023245695.1
E2 19747796 19747874
refGene_gn=PEPD;refGene_tx=XM_023245694.1;refGene_gn=PEPD;refGene_tx=XM_023245695.1
E2 19757707 19757856
refGene_gn=PEPD;refGene_tx=XM_023245694.1;refGene_gn=PEPD;refGene_tx=XM_023245695.1
E2 19765404 19765589
refGene_gn=PEPD;refGene_tx=XM_023245694.1;refGene_gn=PEPD;refGene_tx=XM_023245695.1
E2 19768560 19768752
refGene_gn=PEPD;refGene_tx=XM_023245694.1;refGene_gn=PEPD;refGene_tx=XM_023245695.1
E2 19769126 19769264
refGene_gn=PEPD;refGene_tx=XM_023245694.1;refGene_gn=PEPD;refGene_tx=XM_023245695.1
E2 19775907 19776351
refGene_gn=CEBPG;refGene_tx=XM_003997948.5;refGene_gn=CEBPG;refGene_tx=XM_023245696.1;refGene_gn=CEBPG;refGene_tx=XM_023245697.1
E2 19846594 19847674 refGene_gn=CEBPA;refGene_tx=XM_023245285.1
E2 19908344 19908495
refGene_gn=SLC7A10;refGene_tx=XM_003997949.5;refGene_gn=SLC7A10;refGene_tx=XM_019819607.2
E2 19918001 19918206
refGene_gn=SLC7A10;refGene_tx=XM_003997949.5;refGene_gn=SLC7A10;refGene_tx=XM_019819607.2
E2 19920163 19920315
refGene_gn=SLC7A10;refGene_tx=XM_003997949.5;refGene_gn=SLC7A10;refGene_tx=XM_019819607.2
E2 19920531 19920657
refGene_gn=SLC7A10;refGene_tx=XM_003997949.5;refGene_gn=SLC7A10;refGene_tx=XM_019819607.2
E2 19920727 19920881
refGene_gn=SLC7A10;refGene_tx=XM_003997949.5;refGene_gn=SLC7A10;refGene_tx=XM_019819607.2

E2 19921626 19921750
refGene_gn=SLC7A10;refGene_tx=XM_003997949.5;refGene_gn=SLC7A10;refGene_tx=XM_019819607.2
E2 19921830 19921934
refGene_gn=SLC7A10;refGene_tx=XM_003997949.5;refGene_gn=SLC7A10;refGene_tx=XM_019819607.2
E2 19922278 19922375
refGene_gn=SLC7A10;refGene_tx=XM_003997949.5;refGene_gn=SLC7A10;refGene_tx=XM_019819607.2
E2 19922591 19922741 refGene_gn=SLC7A10;refGene_tx=XM_003997949.5
E2 19923453 19923631
refGene_gn=SLC7A10;refGene_tx=XM_003997949.5;refGene_gn=SLC7A10;refGene_tx=XM_019819607.2
E2 19923872 19924003
refGene_gn=SLC7A10;refGene_tx=XM_003997949.5;refGene_gn=SLC7A10;refGene_tx=XM_019819607.2
E2 19925365 19925969
refGene_gn=LRP3;refGene_tx=XM_023245205.1;refGene_gn=LRP3;refGene_tx=XM_023245206.1
E2 19926191 19926324
refGene_gn=LRP3;refGene_tx=XM_023245205.1;refGene_gn=LRP3;refGene_tx=XM_023245206.1
E2 19926540 19927657
refGene_gn=LRP3;refGene_tx=XM_023245205.1;refGene_gn=LRP3;refGene_tx=XM_023245206.1
E2 19928043 19928258
refGene_gn=LRP3;refGene_tx=XM_023245205.1;refGene_gn=LRP3;refGene_tx=XM_023245206.1
E2 19929595 19929734
refGene_gn=LRP3;refGene_tx=XM_023245205.1;refGene_gn=LRP3;refGene_tx=XM_023245206.1
E2 19934919 19934967
refGene_gn=LRP3;refGene_tx=XM_023245205.1;refGene_gn=LRP3;refGene_tx=XM_023245206.1
E2 19936869 19936942
refGene_gn=LRP3;refGene_tx=XM_023245205.1;refGene_gn=LRP3;refGene_tx=XM_023245206.1
E2 19968165 19968357
refGene_gn=WDR88;refGene_tx=XM_003997994.5;refGene_gn=WDR88;refGene_tx=XM_011289983.3
E2 19970250 19970343
refGene_gn=WDR88;refGene_tx=XM_003997994.5;refGene_gn=WDR88;refGene_tx=XM_011289983.3
E2 19976471 19976540
refGene_gn=WDR88;refGene_tx=XM_003997994.5;refGene_gn=WDR88;refGene_tx=XM_011289983.3
E2 19978092 19978175
refGene_gn=WDR88;refGene_tx=XM_003997994.5;refGene_gn=WDR88;refGene_tx=XM_011289983.3
E2 19980147 19980335 refGene_gn=WDR88;refGene_tx=XM_003997994.5

E2 19986928 19987058 refGene_gn=WDR88;refGene_tx=XM_003997994.5
E2 19988502 19988641
refGene_gn=WDR88;refGene_tx=XM_003997994.5;refGene_gn=WDR88;refGene_tx=XM_011289983.3
E2 19991174 19991238
refGene_gn=WDR88;refGene_tx=XM_003997994.5;refGene_gn=WDR88;refGene_tx=XM_011289983.3
E2 19992450 19992539
refGene_gn=WDR88;refGene_tx=XM_003997994.5;refGene_gn=WDR88;refGene_tx=XM_011289983.3
E2 19997939 19998050
refGene_gn=WDR88;refGene_tx=XM_003997994.5;refGene_gn=WDR88;refGene_tx=XM_011289983.3
E2 20013476 20013788
refGene_gn=WDR88;refGene_tx=XM_003997994.5;refGene_gn=WDR88;refGene_tx=XM_011289983.3
E2 20015077 20015108
refGene_gn=GPATCH1;refGene_tx=XM_003997950.5;refGene_gn=GPATCH1;refGene_tx=XM_006941457.4;refGene_gn=GPATCH1;refGene_tx=XM_023245081.1
E2 20017678 20017827
refGene_gn=GPATCH1;refGene_tx=XM_003997950.5;refGene_gn=GPATCH1;refGene_tx=XM_006941457.4;refGene_gn=GPATCH1;refGene_tx=XM_023245081.1
E2 20019360 20019458
refGene_gn=GPATCH1;refGene_tx=XM_003997950.5;refGene_gn=GPATCH1;refGene_tx=XM_006941457.4;refGene_gn=GPATCH1;refGene_tx=XM_023245081.1
E2 20021782 20021890
refGene_gn=GPATCH1;refGene_tx=XM_003997950.5;refGene_gn=GPATCH1;refGene_tx=XM_006941457.4;refGene_gn=GPATCH1;refGene_tx=XM_023245081.1
E2 20023146 20023363
refGene_gn=GPATCH1;refGene_tx=XM_003997950.5;refGene_gn=GPATCH1;refGene_tx=XM_006941457.4;refGene_gn=GPATCH1;refGene_tx=XM_023245081.1
E2 20027312 20027479
refGene_gn=GPATCH1;refGene_tx=XM_003997950.5;refGene_gn=GPATCH1;refGene_tx=XM_006941457.4;refGene_gn=GPATCH1;refGene_tx=XM_023245081.1
E2 20027806 20027943
refGene_gn=GPATCH1;refGene_tx=XM_003997950.5;refGene_gn=GPATCH1;refGene_tx=XM_006941457.4;refGene_gn=GPATCH1;refGene_tx=XM_023245081.1
E2 20028602 20028730
refGene_gn=GPATCH1;refGene_tx=XM_003997950.5;refGene_gn=GPATCH1;refGene_tx=XM_006941457.4;refGene_gn=GPATCH1;refGene_tx=XM_023245081.1
E2 20028975 20029154
refGene_gn=GPATCH1;refGene_tx=XM_003997950.5;refGene_gn=GPATCH1;refGene_tx=XM_006941457.4;refGene_gn=GPATCH1;refGene_tx=XM_023245081.1
E2 20030377 20030674
refGene_gn=GPATCH1;refGene_tx=XM_003997950.5;refGene_gn=GPATCH1;refGene_tx=XM_006941457.4;refGene_gn=GPATCH1;refGene_tx=XM_023245081.1
E2 20032407 20032612
refGene_gn=GPATCH1;refGene_tx=XM_003997950.5;refGene_gn=GPATCH1;refGene_tx=XM_006941457.4;refGene_gn=GPATCH1;refGene_tx=XM_023245081.1
E2 20035675 20035755

refGene_gn=GPATCH1;refGene_tx=XM_003997950.5;refGene_gn=GPATCH1;refGene_tx=XM_006941457.4;refGene_gn=GPATCH1;refGene_tx=XM_023245081.1
E2 20039497 20039645 refGene_gn=GPATCH1;refGene_tx=XM_003997950.5;refGene_gn=GPATCH1;refGene_tx=XM_006941457.4;refGene_gn=GPATCH1;refGene_tx=XM_023245081.1
E2 20040562 20040805 refGene_gn=GPATCH1;refGene_tx=XM_003997950.5;refGene_gn=GPATCH1;refGene_tx=XM_006941457.4;refGene_gn=GPATCH1;refGene_tx=XM_023245081.1
E2 20041167 20041226 refGene_gn=GPATCH1;refGene_tx=XM_003997950.5;refGene_gn=GPATCH1;refGene_tx=XM_006941457.4;refGene_gn=GPATCH1;refGene_tx=XM_023245081.1
E2 20043093 20043191 refGene_gn=GPATCH1;refGene_tx=XM_003997950.5;refGene_gn=GPATCH1;refGene_tx=XM_006941457.4;refGene_gn=GPATCH1;refGene_tx=XM_023245081.1
E2 20043837 20043998 refGene_gn=GPATCH1;refGene_tx=XM_003997950.5;refGene_gn=GPATCH1;refGene_tx=XM_006941457.4;refGene_gn=GPATCH1;refGene_tx=XM_023245081.1
E2 20045940 20046026 refGene_gn=GPATCH1;refGene_tx=XM_003997950.5;refGene_gn=GPATCH1;refGene_tx=XM_006941457.4;refGene_gn=GPATCH1;refGene_tx=XM_023245081.1
E2 20048431 20048566 refGene_gn=GPATCH1;refGene_tx=XM_003997950.5;refGene_gn=GPATCH1;refGene_tx=XM_006941457.4
E2 20053810 20053883 refGene_gn=GPATCH1;refGene_tx=XM_003997950.5;refGene_gn=GPATCH1;refGene_tx=XM_006941457.4
E2 20061766 20061835 refGene_gn=RHPN2;refGene_tx=XM_003997951.5
E2 20074940 20075056 refGene_gn=RHPN2;refGene_tx=XM_003997951.5
E2 20089270 20089399 refGene_gn=RHPN2;refGene_tx=XM_003997951.5
E2 20091890 20091966 refGene_gn=RHPN2;refGene_tx=XM_003997951.5
E2 20100029 20100107 refGene_gn=RHPN2;refGene_tx=XM_003997951.5
E2 20100483 20100608 refGene_gn=RHPN2;refGene_tx=XM_003997951.5
E2 20102334 20102501 refGene_gn=RHPN2;refGene_tx=XM_003997951.5
E2 20104809 20104997 refGene_gn=RHPN2;refGene_tx=XM_003997951.5
E2 20105402 20105559 refGene_gn=RHPN2;refGene_tx=XM_003997951.5
E2 20106415 20106957 refGene_gn=LOC102899819;refGene_tx=XM_019819617.1
E2 20106959 20107062 refGene_gn=LOC102899819;refGene_tx=XM_019819617.1
E2 20108701 20108840 refGene_gn=LOC102899819;refGene_tx=XM_019819617.1;refGene_gn=RHPN2;refGene_tx=XM_003997951.5
E2 20110494 20110689 refGene_gn=RHPN2;refGene_tx=XM_003997951.5
E2 20113070 20113147 refGene_gn=RHPN2;refGene_tx=XM_003997951.5
E2 20114383 20114530 refGene_gn=RHPN2;refGene_tx=XM_003997951.5
E2 20114845 20115001 refGene_gn=RHPN2;refGene_tx=XM_003997951.5
E2 20120038 20120299 refGene_gn=RHPN2;refGene_tx=XM_003997951.5
E2 20122509 20122758 refGene_gn=FAAP24;refGene_tx=XM_003997952.5;refGene_gn=FAAP24;refGene_tx=XM_006941458.4;refGene_gn=FAAP24;refGene_tx=XM_019819615.2;refGene_

gn=FAAP24;refGene_tx=XM_019819616.2
E2 20128384 20128537
refGene_gn=FAAP24;refGene_tx=XM_003997952.5;refGene_gn=FAAP24;refGene_tx=XM_006941458.4;refGene_gn=FAAP24;refGene_tx=XM_019819615.2;refGene_gn=FAAP24;refGene_tx=XM_019819616.2
E2 20128722 20128862
refGene_gn=FAAP24;refGene_tx=XM_003997952.5;refGene_gn=FAAP24;refGene_tx=XM_006941458.4;refGene_gn=FAAP24;refGene_tx=XM_019819615.2
E2 20128976 20129095
refGene_gn=FAAP24;refGene_tx=XM_003997952.5;refGene_gn=FAAP24;refGene_tx=XM_006941458.4;refGene_gn=FAAP24;refGene_tx=XM_019819615.2
E2 20129330 20129500 refGene_gn=FAAP24;refGene_tx=XM_003997952.5
E2 20129845 20129884 refGene_gn=CEP89;refGene_tx=XM_006941460.3
E2 20132413 20132520 refGene_gn=CEP89;refGene_tx=XM_006941460.3
E2 20138918 20139077 refGene_gn=CEP89;refGene_tx=XM_006941460.3
E2 20143669 20143850 refGene_gn=CEP89;refGene_tx=XM_006941460.3
E2 20152977 20153011 refGene_gn=CEP89;refGene_tx=XM_006941460.3
E2 20153013 20153079 refGene_gn=CEP89;refGene_tx=XM_006941460.3
E2 20154213 20154242 refGene_gn=CEP89;refGene_tx=XM_006941460.3
E2 20155545 20155588 refGene_gn=CEP89;refGene_tx=XM_006941460.3
E2 20157064 20157283 refGene_gn=CEP89;refGene_tx=XM_006941460.3
E2 20158722 20158865 refGene_gn=CEP89;refGene_tx=XM_006941460.3
E2 20164761 20164812 refGene_gn=CEP89;refGene_tx=XM_006941460.3
E2 20164976 20165060 refGene_gn=CEP89;refGene_tx=XM_006941460.3
E2 20167468 20167572 refGene_gn=CEP89;refGene_tx=XM_006941460.3
E2 20171103 20171219 refGene_gn=CEP89;refGene_tx=XM_006941460.3
E2 20173259 20173440 refGene_gn=CEP89;refGene_tx=XM_006941460.3
E2 20185891 20186059 refGene_gn=CEP89;refGene_tx=XM_006941460.3
E2 20186655 20186797 refGene_gn=CEP89;refGene_tx=XM_006941460.3
E2 20190455 20190545 refGene_gn=CEP89;refGene_tx=XM_006941460.3
E2 20197684 20197854 refGene_gn=CEP89;refGene_tx=XM_006941460.3
E2 20203384 20203541 refGene_gn=CEP89;refGene_tx=XM_006941460.3
E2 20214823 20214919
refGene_gn=SLC7A9;refGene_tx=XM_003997953.4;refGene_gn=SLC7A9;refGene_tx=XM_019819626.2;refGene_gn=SLC7A9;refGene_tx=XM_019819627.2;refGene_gn=SLC7A9;refGene_tx=XM_019819629.2;refGene_gn=SLC7A9;refGene_tx=XM_019819632.2;refGene_gn=SLC7A9;refGene_tx=XM_019819633.2
E2 20216701 20216849
refGene_gn=SLC7A9;refGene_tx=XM_003997953.4;refGene_gn=SLC7A9;refGene_tx=XM_019819626.2;refGene_gn=SLC7A9;refGene_tx=XM_019819627.2;refGene_gn=SLC7A9;refGene_tx=XM_019819629.2;refGene_gn=SLC7A9;refGene_tx=XM_019819632.2;refGene_gn=SLC7A9;refGene_tx=XM_019819633.2
E2 20217129 20217372
refGene_gn=SLC7A9;refGene_tx=XM_003997953.4;refGene_gn=SLC7A9;refGene_tx=XM_019819626.2;refGene_gn=SLC7A9;refGene_tx=XM_019819627.2;refGene_gn=SLC7A9;refGene_tx=XM_019819629.2;refGene_gn=SLC7A9;refGene_tx=XM_019819632.2;refGene_gn=SLC7A9;refGene_tx=XM_019819633.2;refGene_gn=SLC7A9;refGene_tx=XM_019819634.2
E2 20217746 20217872
refGene_gn=SLC7A9;refGene_tx=XM_003997953.4;refGene_gn=SLC7A9;refGene_

tx=XM_019819626.2;refGene_gn=SLC7A9;refGene_tx=XM_019819627.2;refGene_gn=SLC7A9;refGene_tx=XM_019819629.2;refGene_gn=SLC7A9;refGene_tx=XM_019819632.2;refGene_gn=SLC7A9;refGene_tx=XM_019819633.2;refGene_gn=SLC7A9;refGene_tx=XM_019819634.2
E2 20218103 20218203
refGene_gn=SLC7A9;refGene_tx=XM_003997953.4;refGene_gn=SLC7A9;refGene_tx=XM_019819626.2;refGene_gn=SLC7A9;refGene_tx=XM_019819627.2;refGene_gn=SLC7A9;refGene_tx=XM_019819629.2;refGene_gn=SLC7A9;refGene_tx=XM_019819632.2;refGene_gn=SLC7A9;refGene_tx=XM_019819633.2;refGene_gn=SLC7A9;refGene_tx=XM_019819634.2
E2 20218925 20218970
refGene_gn=SLC7A9;refGene_tx=XM_003997953.4;refGene_gn=SLC7A9;refGene_tx=XM_019819626.2;refGene_gn=SLC7A9;refGene_tx=XM_019819627.2;refGene_gn=SLC7A9;refGene_tx=XM_019819629.2;refGene_gn=SLC7A9;refGene_tx=XM_019819632.2;refGene_gn=SLC7A9;refGene_tx=XM_019819633.2;refGene_gn=SLC7A9;refGene_tx=XM_019819634.2
E2 20219432 20219556
refGene_gn=SLC7A9;refGene_tx=XM_003997953.4;refGene_gn=SLC7A9;refGene_tx=XM_019819626.2;refGene_gn=SLC7A9;refGene_tx=XM_019819627.2;refGene_gn=SLC7A9;refGene_tx=XM_019819629.2;refGene_gn=SLC7A9;refGene_tx=XM_019819632.2;refGene_gn=SLC7A9;refGene_tx=XM_019819633.2;refGene_gn=SLC7A9;refGene_tx=XM_019819634.2
E2 20221112 20221216
refGene_gn=SLC7A9;refGene_tx=XM_003997953.4;refGene_gn=SLC7A9;refGene_tx=XM_019819626.2;refGene_gn=SLC7A9;refGene_tx=XM_019819627.2;refGene_gn=SLC7A9;refGene_tx=XM_019819629.2;refGene_gn=SLC7A9;refGene_tx=XM_019819632.2;refGene_gn=SLC7A9;refGene_tx=XM_019819633.2;refGene_gn=SLC7A9;refGene_tx=XM_019819634.2
E2 20227455 20227552
refGene_gn=SLC7A9;refGene_tx=XM_003997953.4;refGene_gn=SLC7A9;refGene_tx=XM_019819626.2;refGene_gn=SLC7A9;refGene_tx=XM_019819627.2;refGene_gn=SLC7A9;refGene_tx=XM_019819629.2;refGene_gn=SLC7A9;refGene_tx=XM_019819632.2;refGene_gn=SLC7A9;refGene_tx=XM_019819633.2;refGene_gn=SLC7A9;refGene_tx=XM_019819634.2
E2 20228630 20228780
refGene_gn=SLC7A9;refGene_tx=XM_003997953.4;refGene_gn=SLC7A9;refGene_tx=XM_019819626.2;refGene_gn=SLC7A9;refGene_tx=XM_019819627.2;refGene_gn=SLC7A9;refGene_tx=XM_019819629.2;refGene_gn=SLC7A9;refGene_tx=XM_019819632.2;refGene_gn=SLC7A9;refGene_tx=XM_019819633.2;refGene_gn=SLC7A9;refGene_tx=XM_019819634.2
E2 20237510 20237685
refGene_gn=SLC7A9;refGene_tx=XM_003997953.4;refGene_gn=SLC7A9;refGene_tx=XM_019819626.2;refGene_gn=SLC7A9;refGene_tx=XM_019819627.2;refGene_gn=SLC7A9;refGene_tx=XM_019819629.2;refGene_gn=SLC7A9;refGene_tx=XM_019819632.2;refGene_gn=SLC7A9;refGene_tx=XM_019819633.2;refGene_gn=SLC7A9;refGene_tx=XM_019819634.2
E2 20239431 20239496
refGene_gn=SLC7A9;refGene_tx=XM_003997953.4;refGene_gn=SLC7A9;refGene_tx=XM_019819632.2;refGene_gn=SLC7A9;refGene_tx=XM_019819633.2
E2 20241013 20241318

refGene_gn=SLC7A9;refGene_tx=XM_019819626.2;refGene_gn=SLC7A9;refGene_tx=XM_019819627.2;refGene_gn=SLC7A9;refGene_tx=XM_019819629.2;refGene_gn=SLC7A9;refGene_tx=XM_019819634.2
E2 20242102 20242194 refGene_gn=TDRD12;refGene_tx=XM_006941462.4
E2 20243474 20243544 refGene_gn=TDRD12;refGene_tx=XM_006941462.4
E2 20243628 20243782 refGene_gn=TDRD12;refGene_tx=XM_006941462.4
E2 20243865 20243979 refGene_gn=TDRD12;refGene_tx=XM_006941462.4
E2 20244067 20244128
refGene_gn=TDRD12;refGene_tx=XM_006941462.4;refGene_gn=TDRD12;refGene_tx=XM_019819620.2
E2 20245600 20245710 refGene_gn=TDRD12;refGene_tx=XM_006941462.4
E2 20252857 20253030
refGene_gn=TDRD12;refGene_tx=XM_006941462.4;refGene_gn=TDRD12;refGene_tx=XM_019819620.2
E2 20254892 20254985
refGene_gn=TDRD12;refGene_tx=XM_006941462.4;refGene_gn=TDRD12;refGene_tx=XM_019819620.2
E2 20255981 20256192
refGene_gn=TDRD12;refGene_tx=XM_006941462.4;refGene_gn=TDRD12;refGene_tx=XM_019819620.2
E2 20256953 20257141
refGene_gn=TDRD12;refGene_tx=XM_006941462.4;refGene_gn=TDRD12;refGene_tx=XM_019819620.2
E2 20262916 20262928 refGene_gn=TDRD12;refGene_tx=XM_019819621.2
E2 20264257 20264357
refGene_gn=TDRD12;refGene_tx=XM_006941462.4;refGene_gn=TDRD12;refGene_tx=XM_019819620.2;refGene_gn=TDRD12;refGene_tx=XM_019819621.2
E2 20266793 20267014
refGene_gn=TDRD12;refGene_tx=XM_006941462.4;refGene_gn=TDRD12;refGene_tx=XM_019819620.2;refGene_gn=TDRD12;refGene_tx=XM_019819621.2;refGene_gn=TDRD12;refGene_tx=XM_019819622.2
E2 20267147 20267287
refGene_gn=TDRD12;refGene_tx=XM_006941462.4;refGene_gn=TDRD12;refGene_tx=XM_019819620.2;refGene_gn=TDRD12;refGene_tx=XM_019819621.2;refGene_gn=TDRD12;refGene_tx=XM_019819622.2
E2 20267943 20268061
refGene_gn=TDRD12;refGene_tx=XM_006941462.4;refGene_gn=TDRD12;refGene_tx=XM_019819620.2;refGene_gn=TDRD12;refGene_tx=XM_019819621.2;refGene_gn=TDRD12;refGene_tx=XM_019819622.2
E2 20268604 20268652 refGene_gn=TDRD12;refGene_tx=XM_019819623.2
E2 20268947 20269076
refGene_gn=TDRD12;refGene_tx=XM_006941462.4;refGene_gn=TDRD12;refGene_tx=XM_019819620.2;refGene_gn=TDRD12;refGene_tx=XM_019819621.2;refGene_gn=TDRD12;refGene_tx=XM_019819622.2;refGene_gn=TDRD12;refGene_tx=XM_019819623.2
E2 20269325 20269517
refGene_gn=TDRD12;refGene_tx=XM_006941462.4;refGene_gn=TDRD12;refGene_tx=XM_019819620.2;refGene_gn=TDRD12;refGene_tx=XM_019819621.2;refGene_gn=TDRD12;refGene_tx=XM_019819622.2;refGene_gn=TDRD12;refGene_tx=XM_019819623.2

E2 20270904 20271032
refGene_gn=TDRD12;refGene_tx=XM_006941462.4;refGene_gn=TDRD12;refGene_tx=XM_019819620.2;refGene_gn=TDRD12;refGene_tx=XM_019819621.2;refGene_gn=TDRD12;refGene_tx=XM_019819622.2;refGene_gn=TDRD12;refGene_tx=XM_019819623.2
E2 20271494 20271651
refGene_gn=TDRD12;refGene_tx=XM_006941462.4;refGene_gn=TDRD12;refGene_tx=XM_019819620.2;refGene_gn=TDRD12;refGene_tx=XM_019819621.2;refGene_gn=TDRD12;refGene_tx=XM_019819622.2;refGene_gn=TDRD12;refGene_tx=XM_019819623.2
E2 20273218 20273404
refGene_gn=TDRD12;refGene_tx=XM_006941462.4;refGene_gn=TDRD12;refGene_tx=XM_019819620.2;refGene_gn=TDRD12;refGene_tx=XM_019819621.2;refGene_gn=TDRD12;refGene_tx=XM_019819622.2;refGene_gn=TDRD12;refGene_tx=XM_019819623.2
E2 20275074 20275182
refGene_gn=TDRD12;refGene_tx=XM_006941462.4;refGene_gn=TDRD12;refGene_tx=XM_019819620.2;refGene_gn=TDRD12;refGene_tx=XM_019819621.2;refGene_gn=TDRD12;refGene_tx=XM_019819622.2;refGene_gn=TDRD12;refGene_tx=XM_019819623.2
E2 20275512 20275573
refGene_gn=TDRD12;refGene_tx=XM_006941462.4;refGene_gn=TDRD12;refGene_tx=XM_019819620.2;refGene_gn=TDRD12;refGene_tx=XM_019819621.2;refGene_gn=TDRD12;refGene_tx=XM_019819622.2;refGene_gn=TDRD12;refGene_tx=XM_019819623.2
E2 20277345 20277426
refGene_gn=TDRD12;refGene_tx=XM_006941462.4;refGene_gn=TDRD12;refGene_tx=XM_019819620.2;refGene_gn=TDRD12;refGene_tx=XM_019819621.2;refGene_gn=TDRD12;refGene_tx=XM_019819622.2;refGene_gn=TDRD12;refGene_tx=XM_019819623.2
E2 20279867 20279944
refGene_gn=TDRD12;refGene_tx=XM_006941462.4;refGene_gn=TDRD12;refGene_tx=XM_019819620.2;refGene_gn=TDRD12;refGene_tx=XM_019819621.2;refGene_gn=TDRD12;refGene_tx=XM_019819622.2;refGene_gn=TDRD12;refGene_tx=XM_019819623.2
E2 20280753 20280851
refGene_gn=TDRD12;refGene_tx=XM_006941462.4;refGene_gn=TDRD12;refGene_tx=XM_019819620.2;refGene_gn=TDRD12;refGene_tx=XM_019819621.2;refGene_gn=TDRD12;refGene_tx=XM_019819622.2;refGene_gn=TDRD12;refGene_tx=XM_019819623.2
E2 20289343 20289436
refGene_gn=TDRD12;refGene_tx=XM_006941462.4;refGene_gn=TDRD12;refGene_tx=XM_019819620.2;refGene_gn=TDRD12;refGene_tx=XM_019819621.2;refGene_gn=TDRD12;refGene_tx=XM_019819622.2;refGene_gn=TDRD12;refGene_tx=XM_019819623.2
E2 20290221 20290411
refGene_gn=TDRD12;refGene_tx=XM_006941462.4;refGene_gn=TDRD12;refGene_tx=XM_019819620.2;refGene_gn=TDRD12;refGene_tx=XM_019819621.2;refGene_gn=TDRD12;refGene_tx=XM_019819622.2;refGene_gn=TDRD12;refGene_tx=XM_019819623.2

E2 20295619 20295705
refGene_gn=TDRD12;refGene_tx=XM_006941462.4;refGene_gn=TDRD12;refGene_tx=XM_019819620.2;refGene_gn=TDRD12;refGene_tx=XM_019819621.2;refGene_gn=TDRD12;refGene_tx=XM_019819622.2;refGene_gn=TDRD12;refGene_tx=XM_019819623.2
E2 20296912 20296968
refGene_gn=TDRD12;refGene_tx=XM_006941462.4;refGene_gn=TDRD12;refGene_tx=XM_019819620.2;refGene_gn=TDRD12;refGene_tx=XM_019819621.2;refGene_gn=TDRD12;refGene_tx=XM_019819622.2;refGene_gn=TDRD12;refGene_tx=XM_019819623.2
E2 20299645 20299765
refGene_gn=TDRD12;refGene_tx=XM_006941462.4;refGene_gn=TDRD12;refGene_tx=XM_019819620.2;refGene_gn=TDRD12;refGene_tx=XM_019819621.2;refGene_gn=TDRD12;refGene_tx=XM_019819622.2;refGene_gn=TDRD12;refGene_tx=XM_019819623.2
E2 20301405 20301542
refGene_gn=TDRD12;refGene_tx=XM_006941462.4;refGene_gn=TDRD12;refGene_tx=XM_019819620.2;refGene_gn=TDRD12;refGene_tx=XM_019819621.2;refGene_gn=TDRD12;refGene_tx=XM_019819622.2;refGene_gn=TDRD12;refGene_tx=XM_019819623.2
E2 20303709 20303868
refGene_gn=TDRD12;refGene_tx=XM_006941462.4;refGene_gn=TDRD12;refGene_tx=XM_019819620.2;refGene_gn=TDRD12;refGene_tx=XM_019819621.2;refGene_gn=TDRD12;refGene_tx=XM_019819622.2;refGene_gn=TDRD12;refGene_tx=XM_019819623.2
E2 20311779 20311803
refGene_gn=TDRD12;refGene_tx=XM_006941462.4;refGene_gn=TDRD12;refGene_tx=XM_019819620.2;refGene_gn=TDRD12;refGene_tx=XM_019819621.2;refGene_gn=TDRD12;refGene_tx=XM_019819622.2;refGene_gn=TDRD12;refGene_tx=XM_019819623.2
E2 20324762 20324968
refGene_gn=NUDT19;refGene_tx=XM_023245168.1;refGene_gn=NUDT19;refGene_tx=XM_023245169.1
E2 20326381 20326589
refGene_gn=NUDT19;refGene_tx=XM_023245168.1;refGene_gn=NUDT19;refGene_tx=XM_023245170.1
E2 20331823 20331878
refGene_gn=NUDT19;refGene_tx=XM_023245169.1;refGene_gn=NUDT19;refGene_tx=XM_023245170.1
E2 20332250 20332952
refGene_gn=NUDT19;refGene_tx=XM_023245168.1;refGene_gn=NUDT19;refGene_tx=XM_023245169.1;refGene_gn=NUDT19;refGene_tx=XM_023245170.1
E2 20345505 20346213 refGene_gn=RGS9BP;refGene_tx=XM_023245676.1
E2 20347680 20347713
refGene_gn=ANKRD27;refGene_tx=XM_019819635.2;refGene_gn=ANKRD27;refGene_tx=XM_019819636.2;refGene_gn=ANKRD27;refGene_tx=XM_019819638.2;refGene_gn=ANKRD27;refGene_tx=XM_019819639.2;refGene_gn=ANKRD27;refGene_tx=XM_023245675.1
E2 20358198 20358330
refGene_gn=ANKRD27;refGene_tx=XM_003997998.4;refGene_gn=ANKRD27;refGen

e_tx=XM_019819635.2;refGene_gn=ANKRD27;refGene_tx=XM_019819636.2;refGene_gn=ANKRD27;refGene_tx=XM_019819638.2;refGene_gn=ANKRD27;refGene_tx=XM_019819639.2;refGene_gn=ANKRD27;refGene_tx=XM_023245675.1
E2 20364370 20364481
refGene_gn=ANKRD27;refGene_tx=XM_003997998.4;refGene_gn=ANKRD27;refGene_tx=XM_019819635.2;refGene_gn=ANKRD27;refGene_tx=XM_019819636.2;refGene_gn=ANKRD27;refGene_tx=XM_019819638.2;refGene_gn=ANKRD27;refGene_tx=XM_019819639.2;refGene_gn=ANKRD27;refGene_tx=XM_023245675.1
E2 20366322 20366479
refGene_gn=ANKRD27;refGene_tx=XM_003997998.4;refGene_gn=ANKRD27;refGene_tx=XM_019819635.2;refGene_gn=ANKRD27;refGene_tx=XM_019819636.2;refGene_gn=ANKRD27;refGene_tx=XM_019819638.2;refGene_gn=ANKRD27;refGene_tx=XM_019819639.2;refGene_gn=ANKRD27;refGene_tx=XM_023245675.1
E2 20367443 20367598
refGene_gn=ANKRD27;refGene_tx=XM_003997998.4;refGene_gn=ANKRD27;refGene_tx=XM_019819635.2;refGene_gn=ANKRD27;refGene_tx=XM_019819636.2;refGene_gn=ANKRD27;refGene_tx=XM_019819638.2;refGene_gn=ANKRD27;refGene_tx=XM_019819639.2;refGene_gn=ANKRD27;refGene_tx=XM_023245675.1
E2 20368030 20368090
refGene_gn=ANKRD27;refGene_tx=XM_003997998.4;refGene_gn=ANKRD27;refGene_tx=XM_019819635.2;refGene_gn=ANKRD27;refGene_tx=XM_019819636.2;refGene_gn=ANKRD27;refGene_tx=XM_019819638.2;refGene_gn=ANKRD27;refGene_tx=XM_019819639.2;refGene_gn=ANKRD27;refGene_tx=XM_023245675.1
E2 20368179 20368233
refGene_gn=ANKRD27;refGene_tx=XM_003997998.4;refGene_gn=ANKRD27;refGene_tx=XM_019819635.2;refGene_gn=ANKRD27;refGene_tx=XM_019819636.2;refGene_gn=ANKRD27;refGene_tx=XM_019819638.2;refGene_gn=ANKRD27;refGene_tx=XM_019819639.2;refGene_gn=ANKRD27;refGene_tx=XM_023245675.1
E2 20368336 20368402
refGene_gn=ANKRD27;refGene_tx=XM_003997998.4;refGene_gn=ANKRD27;refGene_tx=XM_019819635.2;refGene_gn=ANKRD27;refGene_tx=XM_019819636.2;refGene_gn=ANKRD27;refGene_tx=XM_019819638.2;refGene_gn=ANKRD27;refGene_tx=XM_019819639.2;refGene_gn=ANKRD27;refGene_tx=XM_023245675.1
E2 20368490 20368567
refGene_gn=ANKRD27;refGene_tx=XM_003997998.4;refGene_gn=ANKRD27;refGene_tx=XM_019819635.2;refGene_gn=ANKRD27;refGene_tx=XM_019819636.2;refGene_gn=ANKRD27;refGene_tx=XM_019819638.2;refGene_gn=ANKRD27;refGene_tx=XM_019819639.2;refGene_gn=ANKRD27;refGene_tx=XM_023245675.1
E2 20369237 20369359
refGene_gn=ANKRD27;refGene_tx=XM_003997998.4;refGene_gn=ANKRD27;refGene_tx=XM_019819635.2;refGene_gn=ANKRD27;refGene_tx=XM_019819636.2;refGene_gn=ANKRD27;refGene_tx=XM_019819638.2;refGene_gn=ANKRD27;refGene_tx=XM_019819639.2;refGene_gn=ANKRD27;refGene_tx=XM_023245675.1
E2 20372494 20372573
refGene_gn=ANKRD27;refGene_tx=XM_003997998.4;refGene_gn=ANKRD27;refGene_tx=XM_019819635.2;refGene_gn=ANKRD27;refGene_tx=XM_019819636.2;refGene_gn=ANKRD27;refGene_tx=XM_019819638.2;refGene_gn=ANKRD27;refGene_tx=XM_019819639.2;refGene_gn=ANKRD27;refGene_tx=XM_023245675.1
E2 20373069 20373202
refGene_gn=ANKRD27;refGene_tx=XM_003997998.4;refGene_gn=ANKRD27;refGen

e_tx=XM_019819635.2;refGene_gn=ANKRD27;refGene_tx=XM_019819636.2;refGene_gn=ANKRD27;refGene_tx=XM_019819638.2;refGene_gn=ANKRD27;refGene_tx=XM_019819639.2;refGene_gn=ANKRD27;refGene_tx=XM_023245675.1
E2 20377496 20377589
refGene_gn=ANKRD27;refGene_tx=XM_003997998.4;refGene_gn=ANKRD27;refGene_tx=XM_019819635.2;refGene_gn=ANKRD27;refGene_tx=XM_019819636.2;refGene_gn=ANKRD27;refGene_tx=XM_019819638.2;refGene_gn=ANKRD27;refGene_tx=XM_019819639.2;refGene_gn=ANKRD27;refGene_tx=XM_023245675.1
E2 20378688 20378816
refGene_gn=ANKRD27;refGene_tx=XM_003997998.4;refGene_gn=ANKRD27;refGene_tx=XM_019819635.2;refGene_gn=ANKRD27;refGene_tx=XM_019819636.2;refGene_gn=ANKRD27;refGene_tx=XM_019819638.2;refGene_gn=ANKRD27;refGene_tx=XM_019819639.2;refGene_gn=ANKRD27;refGene_tx=XM_023245675.1
E2 20379375 20379458
refGene_gn=ANKRD27;refGene_tx=XM_003997998.4;refGene_gn=ANKRD27;refGene_tx=XM_019819635.2;refGene_gn=ANKRD27;refGene_tx=XM_019819636.2;refGene_gn=ANKRD27;refGene_tx=XM_019819638.2;refGene_gn=ANKRD27;refGene_tx=XM_019819639.2;refGene_gn=ANKRD27;refGene_tx=XM_023245675.1
E2 20379809 20379925
refGene_gn=ANKRD27;refGene_tx=XM_003997998.4;refGene_gn=ANKRD27;refGene_tx=XM_019819635.2;refGene_gn=ANKRD27;refGene_tx=XM_019819636.2;refGene_gn=ANKRD27;refGene_tx=XM_019819638.2;refGene_gn=ANKRD27;refGene_tx=XM_019819639.2;refGene_gn=ANKRD27;refGene_tx=XM_023245675.1
E2 20380325 20380418
refGene_gn=ANKRD27;refGene_tx=XM_003997998.4;refGene_gn=ANKRD27;refGene_tx=XM_019819635.2;refGene_gn=ANKRD27;refGene_tx=XM_019819636.2;refGene_gn=ANKRD27;refGene_tx=XM_019819638.2;refGene_gn=ANKRD27;refGene_tx=XM_019819639.2;refGene_gn=ANKRD27;refGene_tx=XM_023245675.1
E2 20381997 20382195
refGene_gn=ANKRD27;refGene_tx=XM_003997998.4;refGene_gn=ANKRD27;refGene_tx=XM_019819635.2;refGene_gn=ANKRD27;refGene_tx=XM_019819636.2;refGene_gn=ANKRD27;refGene_tx=XM_019819638.2;refGene_gn=ANKRD27;refGene_tx=XM_019819639.2;refGene_gn=ANKRD27;refGene_tx=XM_023245675.1
E2 20383845 20383905
refGene_gn=ANKRD27;refGene_tx=XM_003997998.4;refGene_gn=ANKRD27;refGene_tx=XM_019819635.2;refGene_gn=ANKRD27;refGene_tx=XM_019819636.2;refGene_gn=ANKRD27;refGene_tx=XM_019819638.2;refGene_gn=ANKRD27;refGene_tx=XM_019819639.2;refGene_gn=ANKRD27;refGene_tx=XM_023245675.1
E2 20384001 20384118
refGene_gn=ANKRD27;refGene_tx=XM_003997998.4;refGene_gn=ANKRD27;refGene_tx=XM_019819635.2;refGene_gn=ANKRD27;refGene_tx=XM_019819636.2;refGene_gn=ANKRD27;refGene_tx=XM_019819638.2;refGene_gn=ANKRD27;refGene_tx=XM_019819639.2;refGene_gn=ANKRD27;refGene_tx=XM_023245675.1
E2 20385666 20385711
refGene_gn=ANKRD27;refGene_tx=XM_003997998.4;refGene_gn=ANKRD27;refGene_tx=XM_019819635.2;refGene_gn=ANKRD27;refGene_tx=XM_019819636.2;refGene_gn=ANKRD27;refGene_tx=XM_019819638.2;refGene_gn=ANKRD27;refGene_tx=XM_019819639.2;refGene_gn=ANKRD27;refGene_tx=XM_023245675.1
E2 20387641 20387764
refGene_gn=ANKRD27;refGene_tx=XM_003997998.4;refGene_gn=ANKRD27;refGen

e_tx=XM_019819635.2;refGene_gn=ANKRD27;refGene_tx=XM_019819636.2;refGene_gn=ANKRD27;refGene_tx=XM_019819638.2;refGene_gn=ANKRD27;refGene_tx=XM_019819639.2;refGene_gn=ANKRD27;refGene_tx=XM_023245675.1
E2 20393033 20393231
refGene_gn=ANKRD27;refGene_tx=XM_003997998.4;refGene_gn=ANKRD27;refGene_tx=XM_019819635.2;refGene_gn=ANKRD27;refGene_tx=XM_019819636.2;refGene_gn=ANKRD27;refGene_tx=XM_019819638.2;refGene_gn=ANKRD27;refGene_tx=XM_019819639.2;refGene_gn=ANKRD27;refGene_tx=XM_023245675.1
E2 20395231 20395351
refGene_gn=ANKRD27;refGene_tx=XM_003997998.4;refGene_gn=ANKRD27;refGene_tx=XM_019819635.2;refGene_gn=ANKRD27;refGene_tx=XM_019819636.2;refGene_gn=ANKRD27;refGene_tx=XM_019819638.2;refGene_gn=ANKRD27;refGene_tx=XM_019819639.2;refGene_gn=ANKRD27;refGene_tx=XM_023245675.1
E2 20395450 20395612
refGene_gn=ANKRD27;refGene_tx=XM_003997998.4;refGene_gn=ANKRD27;refGene_tx=XM_019819635.2;refGene_gn=ANKRD27;refGene_tx=XM_019819636.2;refGene_gn=ANKRD27;refGene_tx=XM_019819638.2;refGene_gn=ANKRD27;refGene_tx=XM_019819639.2;refGene_gn=ANKRD27;refGene_tx=XM_023245675.1
E2 20396861 20396973
refGene_gn=ANKRD27;refGene_tx=XM_003997998.4;refGene_gn=ANKRD27;refGene_tx=XM_019819635.2;refGene_gn=ANKRD27;refGene_tx=XM_019819636.2;refGene_gn=ANKRD27;refGene_tx=XM_019819638.2;refGene_gn=ANKRD27;refGene_tx=XM_019819639.2;refGene_gn=ANKRD27;refGene_tx=XM_023245675.1
E2 20398889 20398968
refGene_gn=ANKRD27;refGene_tx=XM_003997998.4;refGene_gn=ANKRD27;refGene_tx=XM_019819635.2;refGene_gn=ANKRD27;refGene_tx=XM_019819636.2;refGene_gn=ANKRD27;refGene_tx=XM_019819638.2;refGene_gn=ANKRD27;refGene_tx=XM_019819639.2;refGene_gn=ANKRD27;refGene_tx=XM_023245675.1
E2 20399155 20399228
refGene_gn=ANKRD27;refGene_tx=XM_003997998.4;refGene_gn=ANKRD27;refGene_tx=XM_019819635.2;refGene_gn=ANKRD27;refGene_tx=XM_019819636.2;refGene_gn=ANKRD27;refGene_tx=XM_019819638.2;refGene_gn=ANKRD27;refGene_tx=XM_019819639.2;refGene_gn=ANKRD27;refGene_tx=XM_023245675.1
E2 20399861 20400046
refGene_gn=ANKRD27;refGene_tx=XM_019819636.2;refGene_gn=ANKRD27;refGene_tx=XM_019819638.2
E2 20400456 20400474 refGene_gn=ANKRD27;refGene_tx=XM_019819639.2
E2 20400492 20400729
refGene_gn=ANKRD27;refGene_tx=XM_003997998.4;refGene_gn=ANKRD27;refGene_tx=XM_019819635.2;refGene_gn=ANKRD27;refGene_tx=XM_023245675.1
E2 20404123 20404171 refGene_gn=PDCD5;refGene_tx=XM_003997954.5
E2 20404489 20404561 refGene_gn=PDCD5;refGene_tx=XM_003997954.5
E2 20405491 20405583 refGene_gn=PDCD5;refGene_tx=XM_003997954.5
E2 20406399 20406461 refGene_gn=PDCD5;refGene_tx=XM_003997954.5
E2 20409789 20409827 refGene_gn=PDCD5;refGene_tx=XM_003997954.5
E2 20410685 20410751 refGene_gn=PDCD5;refGene_tx=XM_003997954.5
E2 20494305 20494467
refGene_gn=DPY19L3;refGene_tx=XM_019819640.2;refGene_gn=DPY19L3;refGene_tx=XM_019819641.2;refGene_gn=DPY19L3;refGene_tx=XM_023245665.1;refGene_gn=DPY19L3;refGene_tx=XM_023245666.1

E2 20496313 20496472
refGene_gn=DPY19L3;refGene_tx=XM_019819640.2;refGene_gn=DPY19L3;refGene_tx=XM_019819641.2;refGene_gn=DPY19L3;refGene_tx=XM_023245665.1;refGene_gn=DPY19L3;refGene_tx=XM_023245666.1
E2 20498932 20499065
refGene_gn=DPY19L3;refGene_tx=XM_019819640.2;refGene_gn=DPY19L3;refGene_tx=XM_019819641.2;refGene_gn=DPY19L3;refGene_tx=XM_023245665.1;refGene_gn=DPY19L3;refGene_tx=XM_023245666.1
E2 20507443 20507526
refGene_gn=DPY19L3;refGene_tx=XM_019819640.2;refGene_gn=DPY19L3;refGene_tx=XM_019819641.2;refGene_gn=DPY19L3;refGene_tx=XM_023245665.1;refGene_gn=DPY19L3;refGene_tx=XM_023245666.1
E2 20511388 20511445
refGene_gn=DPY19L3;refGene_tx=XM_019819640.2;refGene_gn=DPY19L3;refGene_tx=XM_019819641.2;refGene_gn=DPY19L3;refGene_tx=XM_023245665.1;refGene_gn=DPY19L3;refGene_tx=XM_023245666.1
E2 20512203 20512315
refGene_gn=DPY19L3;refGene_tx=XM_019819640.2;refGene_gn=DPY19L3;refGene_tx=XM_019819641.2;refGene_gn=DPY19L3;refGene_tx=XM_023245665.1;refGene_gn=DPY19L3;refGene_tx=XM_023245666.1
E2 20512694 20512817
refGene_gn=DPY19L3;refGene_tx=XM_019819640.2;refGene_gn=DPY19L3;refGene_tx=XM_019819641.2;refGene_gn=DPY19L3;refGene_tx=XM_023245665.1;refGene_gn=DPY19L3;refGene_tx=XM_023245666.1
E2 20515669 20515828
refGene_gn=DPY19L3;refGene_tx=XM_019819640.2;refGene_gn=DPY19L3;refGene_tx=XM_019819641.2;refGene_gn=DPY19L3;refGene_tx=XM_023245665.1;refGene_gn=DPY19L3;refGene_tx=XM_023245666.1
E2 20515997 20516071
refGene_gn=DPY19L3;refGene_tx=XM_019819640.2;refGene_gn=DPY19L3;refGene_tx=XM_019819641.2;refGene_gn=DPY19L3;refGene_tx=XM_023245665.1;refGene_gn=DPY19L3;refGene_tx=XM_023245666.1
E2 20517805 20517907
refGene_gn=DPY19L3;refGene_tx=XM_019819640.2;refGene_gn=DPY19L3;refGene_tx=XM_019819641.2;refGene_gn=DPY19L3;refGene_tx=XM_023245665.1;refGene_gn=DPY19L3;refGene_tx=XM_023245666.1
E2 20519279 20519411
refGene_gn=DPY19L3;refGene_tx=XM_019819640.2;refGene_gn=DPY19L3;refGene_tx=XM_019819641.2;refGene_gn=DPY19L3;refGene_tx=XM_023245665.1;refGene_gn=DPY19L3;refGene_tx=XM_023245666.1
E2 20531402 20531537
refGene_gn=DPY19L3;refGene_tx=XM_019819640.2;refGene_gn=DPY19L3;refGene_tx=XM_019819641.2;refGene_gn=DPY19L3;refGene_tx=XM_023245665.1;refGene_gn=DPY19L3;refGene_tx=XM_023245666.1
E2 20532908 20533032
refGene_gn=DPY19L3;refGene_tx=XM_019819640.2;refGene_gn=DPY19L3;refGene_tx=XM_019819641.2;refGene_gn=DPY19L3;refGene_tx=XM_023245665.1;refGene_gn=DPY19L3;refGene_tx=XM_023245666.1
E2 20534503 20534649
refGene_gn=DPY19L3;refGene_tx=XM_019819640.2;refGene_gn=DPY19L3;refGen

e_tx=XM_019819641.2;refGene_gn=DPY19L3;refGene_tx=XM_023245665.1;refGene_gn=DPY19L3;refGene_tx=XM_023245666.1
E2 20535628 20535750
refGene_gn=DPY19L3;refGene_tx=XM_019819640.2;refGene_gn=DPY19L3;refGene_tx=XM_019819641.2;refGene_gn=DPY19L3;refGene_tx=XM_023245665.1;refGene_gn=DPY19L3;refGene_tx=XM_023245666.1
E2 20541443 20541532 refGene_gn=DPY19L3;refGene_tx=XM_019819640.2
E2 20541738 20541829
refGene_gn=DPY19L3;refGene_tx=XM_019819640.2;refGene_gn=DPY19L3;refGene_tx=XM_023245665.1;refGene_gn=DPY19L3;refGene_tx=XM_023245666.1
E2 20564693 20564827
refGene_gn=DPY19L3;refGene_tx=XM_023245665.1;refGene_gn=DPY19L3;refGene_tx=XM_023245666.1
E2 20567195 20567298 refGene_gn=DPY19L3;refGene_tx=XM_023245665.1
E2 20586145 20586521
refGene_gn=ZNF507;refGene_tx=XM_011289976.3;refGene_gn=ZNF507;refGene_tx=XM_011289977.3;refGene_gn=ZNF507;refGene_tx=XM_019819643.2
E2 20586636 20586771
refGene_gn=ZNF507;refGene_tx=XM_011289976.3;refGene_gn=ZNF507;refGene_tx=XM_011289977.3;refGene_gn=ZNF507;refGene_tx=XM_019819643.2
E2 20612348 20612463
refGene_gn=ZNF507;refGene_tx=XM_011289976.3;refGene_gn=ZNF507;refGene_tx=XM_011289977.3;refGene_gn=ZNF507;refGene_tx=XM_019819643.2
E2 20616318 20616436
refGene_gn=ZNF507;refGene_tx=XM_011289976.3;refGene_gn=ZNF507;refGene_tx=XM_011289977.3;refGene_gn=ZNF507;refGene_tx=XM_019819643.2
E2 20617793 20619923
refGene_gn=ZNF507;refGene_tx=XM_011289976.3;refGene_gn=ZNF507;refGene_tx=XM_011289977.3;refGene_gn=ZNF507;refGene_tx=XM_019819643.2
E2 21469023 21469063 refGene_gn=TSHZ3;refGene_tx=XM_023245049.1
E2 21533050 21533208 refGene_gn=TSHZ3;refGene_tx=XM_023245049.1
E2 21533209 21536266 refGene_gn=TSHZ3;refGene_tx=XM_023245049.1
E2 22075853 22076098
refGene_gn=ZNF536;refGene_tx=XM_019819656.2;refGene_gn=ZNF536;refGene_tx=XM_023245646.1;refGene_gn=ZNF536;refGene_tx=XM_023245648.1;refGene_gn=ZNF536;refGene_tx=XM_023245649.1;refGene_gn=ZNF536;refGene_tx=XM_023245650.1;refGene_gn=ZNF536;refGene_tx=XM_023245651.1;refGene_gn=ZNF536;refGene_tx=XM_023245652.1
E2 22213448 22213456 refGene_gn=ZNF536;refGene_tx=XM_023245653.1
E2 22220840 22222412
refGene_gn=ZNF536;refGene_tx=XM_019819656.2;refGene_gn=ZNF536;refGene_tx=XM_023245646.1;refGene_gn=ZNF536;refGene_tx=XM_023245648.1;refGene_gn=ZNF536;refGene_tx=XM_023245649.1;refGene_gn=ZNF536;refGene_tx=XM_023245650.1;refGene_gn=ZNF536;refGene_tx=XM_023245651.1;refGene_gn=ZNF536;refGene_tx=XM_023245652.1;refGene_gn=ZNF536;refGene_tx=XM_023245653.1
E2 22229240 22229294
refGene_gn=ZNF536;refGene_tx=XM_019819656.2;refGene_gn=ZNF536;refGene_tx=XM_023245646.1;refGene_gn=ZNF536;refGene_tx=XM_023245648.1;refGene_gn=ZNF536;refGene_tx=XM_023245649.1;refGene_gn=ZNF536;refGene_tx=XM_02

3245650.1;refGene_gn=ZNF536;refGene_tx=XM_023245651.1;refGene_gn=ZNF53
6;refGene_tx=XM_023245653.1
E2 22234620 22234773
refGene_gn=ZNF536;refGene_tx=XM_019819656.2;refGene_gn=ZNF536;refGene_
tx=XM_023245646.1;refGene_gn=ZNF536;refGene_tx=XM_023245648.1;refGene_
gn=ZNF536;refGene_tx=XM_023245649.1;refGene_gn=ZNF536;refGene_tx=XM_02
3245650.1;refGene_gn=ZNF536;refGene_tx=XM_023245651.1;refGene_gn=ZNF53
6;refGene_tx=XM_023245652.1;refGene_gn=ZNF536;refGene_tx=XM_023245653.
1
E2 22309783 22311953
refGene_gn=ZNF536;refGene_tx=XM_019819656.2;refGene_gn=ZNF536;refGene_
tx=XM_023245646.1;refGene_gn=ZNF536;refGene_tx=XM_023245648.1;refGene_
gn=ZNF536;refGene_tx=XM_023245649.1;refGene_gn=ZNF536;refGene_tx=XM_02
3245650.1;refGene_gn=ZNF536;refGene_tx=XM_023245651.1;refGene_gn=ZNF53
6;refGene_tx=XM_023245652.1;refGene_gn=ZNF536;refGene_tx=XM_023245653.
1
E2 22685649 22685832
refGene_gn=URI1;refGene_tx=XM_011289974.3;refGene_gn=URI1;refGene_tx=X
M_019819663.2;refGene_gn=URI1;refGene_tx=XM_019819664.2;refGene_gn=URI
1;refGene_tx=XM_023245900.1
E2 22689228 22689478
refGene_gn=URI1;refGene_tx=XM_011289974.3;refGene_gn=URI1;refGene_tx=X
M_019819663.2;refGene_gn=URI1;refGene_tx=XM_019819664.2;refGene_gn=URI
1;refGene_tx=XM_023245900.1
E2 22690265 22690408
refGene_gn=URI1;refGene_tx=XM_011289974.3;refGene_gn=URI1;refGene_tx=X
M_019819663.2;refGene_gn=URI1;refGene_tx=XM_019819664.2;refGene_gn=URI
1;refGene_tx=XM_023245900.1
E2 22692351 22692697
refGene_gn=URI1;refGene_tx=XM_011289974.3;refGene_gn=URI1;refGene_tx=X
M_019819663.2;refGene_gn=URI1;refGene_tx=XM_019819664.2;refGene_gn=URI
1;refGene_tx=XM_023245900.1
E2 22694128 22694297
refGene_gn=URI1;refGene_tx=XM_011289974.3;refGene_gn=URI1;refGene_tx=X
M_019819663.2;refGene_gn=URI1;refGene_tx=XM_019819664.2;refGene_gn=URI
1;refGene_tx=XM_023245900.1
E2 22696163 22696221
refGene_gn=URI1;refGene_tx=XM_011289974.3;refGene_gn=URI1;refGene_tx=X
M_019819663.2;refGene_gn=URI1;refGene_tx=XM_019819664.2;refGene_gn=URI
1;refGene_tx=XM_023245900.1
E2 22696409 22696501
refGene_gn=URI1;refGene_tx=XM_011289974.3;refGene_gn=URI1;refGene_tx=X
M_019819663.2;refGene_gn=URI1;refGene_tx=XM_019819664.2;refGene_gn=URI
1;refGene_tx=XM_023245900.1
E2 22722873 22723009
refGene_gn=URI1;refGene_tx=XM_011289974.3;refGene_gn=URI1;refGene_tx=X
M_019819663.2;refGene_gn=URI1;refGene_tx=XM_019819664.2;refGene_gn=URI
1;refGene_tx=XM_023245900.1
E2 22724205 22724284
refGene_gn=URI1;refGene_tx=XM_011289974.3;refGene_gn=URI1;refGene_tx=X

M_019819663.2;refGene_gn=URI1;refGene_tx=XM_019819664.2;refGene_gn=URI1;refGene_tx=XM_023245900.1
E2 22738290 22738325
refGene_gn=URI1;refGene_tx=XM_011289974.3;refGene_gn=URI1;refGene_tx=XM_019819663.2;refGene_gn=URI1;refGene_tx=XM_019819664.2;refGene_gn=URI1;refGene_tx=XM_023245900.1
E2 22768497 22768608 refGene_gn=URI1;refGene_tx=XM_011289974.3
E2 22769065 22769086
refGene_gn=URI1;refGene_tx=XM_019819664.2;refGene_gn=URI1;refGene_tx=XM_023245900.1
E2 22786104 22786167 refGene_gn=URI1;refGene_tx=XM_019819663.2
E2 22871318 22871441 refGene_gn=CCNE1;refGene_tx=XM_023245346.1
E2 22871733 22871891 refGene_gn=CCNE1;refGene_tx=XM_023245346.1
E2 22871970 22872082 refGene_gn=CCNE1;refGene_tx=XM_023245346.1
E2 22872189 22872324 refGene_gn=CCNE1;refGene_tx=XM_023245346.1
E2 22872538 22872634 refGene_gn=CCNE1;refGene_tx=XM_023245346.1
E2 22873183 22873330 refGene_gn=CCNE1;refGene_tx=XM_023245346.1
E2 22875817 22875953 refGene_gn=CCNE1;refGene_tx=XM_023245346.1
E2 22876072 22876218 refGene_gn=CCNE1;refGene_tx=XM_023245346.1
E2 22879854 22879923 refGene_gn=CCNE1;refGene_tx=XM_023245346.1
E2 22880145 22880233 refGene_gn=CCNE1;refGene_tx=XM_023245346.1
E2 22880345 22880368 refGene_gn=CCNE1;refGene_tx=XM_023245346.1
E2 22957542 22957715
refGene_gn=CE2H19orf12;refGene_tx=XM_023245110.1
E2 22957999 22958041
refGene_gn=CE2H19orf12;refGene_tx=XM_006941471.3;refGene_gn=CE2H19orf12;refGene_tx=XM_019819667.2
E2 22959342 22959401
refGene_gn=CE2H19orf12;refGene_tx=XM_006941471.3;refGene_gn=CE2H19orf12;refGene_tx=XM_019819667.2
E2 22962788 22962958
refGene_gn=CE2H19orf12;refGene_tx=XM_003997958.5;refGene_gn=CE2H19orf12;refGene_tx=XM_006941471.3;refGene_gn=CE2H19orf12;refGene_tx=XM_006941472.2;refGene_gn=CE2H19orf12;refGene_tx=XM_006941473.2;refGene_gn=CE2H19orf12;refGene_tx=XM_019819667.2;refGene_gn=CE2H19orf12;refGene_tx=XM_023245110.1;refGene_gn=CE2H19orf12;refGene_tx=XM_023245111.1;refGene_gn=CE2H19orf12;refGene_tx=XM_023245112.1
E2 22966343 22966609
refGene_gn=CE2H19orf12;refGene_tx=XM_003997958.5;refGene_gn=CE2H19orf12;refGene_tx=XM_006941471.3;refGene_gn=CE2H19orf12;refGene_tx=XM_006941472.2;refGene_gn=CE2H19orf12;refGene_tx=XM_006941473.2;refGene_gn=CE2H19orf12;refGene_tx=XM_019819667.2;refGene_gn=CE2H19orf12;refGene_tx=XM_023245110.1;refGene_gn=CE2H19orf12;refGene_tx=XM_023245111.1;refGene_gn=CE2H19orf12;refGene_tx=XM_023245112.1
E2 22988346 22989189
refGene_gn=PLEKHF1;refGene_tx=XM_006941474.4;refGene_gn=PLEKHF1;refGene_tx=XM_006941475.4;refGene_gn=PLEKHF1;refGene_tx=XM_006941476.4;refGene_gn=PLEKHF1;refGene_tx=XM_006941477.4
E2 23030788 23030925
refGene_gn=POP4;refGene_tx=XM_003997960.3;refGene_gn=POP4;refGene_tx=X

M_019819669.2
E2 23032081 23032183
refGene_gn=POP4;refGene_tx=XM_003997960.3;refGene_gn=POP4;refGene_tx=X
M_019819669.2
E2 23033789 23033851
refGene_gn=POP4;refGene_tx=XM_003997960.3;refGene_gn=POP4;refGene_tx=X
M_019819669.2
E2 23034017 23034095
refGene_gn=POP4;refGene_tx=XM_003997960.3;refGene_gn=POP4;refGene_tx=X
M_019819669.2
E2 23035519 23035743
refGene_gn=POP4;refGene_tx=XM_003997960.3;refGene_gn=POP4;refGene_tx=X
M_019819669.2
E2 23037585 23037638 refGene_gn=POP4;refGene_tx=XM_003997960.3
E2 23039631 23039638 refGene_gn=POP4;refGene_tx=XM_003997960.3
E2 23079602 23079691 refGene_gn=VSTM2B;refGene_tx=XM_023245380.1
E2 23101677 23102158 refGene_gn=VSTM2B;refGene_tx=XM_023245380.1
E2 23103512 23103542 refGene_gn=VSTM2B;refGene_tx=XM_023245380.1
E2 23104458 23104643 refGene_gn=VSTM2B;refGene_tx=XM_023245380.1
E2 23105197 23105279 refGene_gn=VSTM2B;refGene_tx=XM_023245380.1
E2 23366578 23366792
refGene_gn=LOC101094311;refGene_tx=XM_003997961.4
E2 23371250 23371861
refGene_gn=LOC101094311;refGene_tx=XM_003997961.4
E2 26893316 26893642 refGene_gn=SHCBP1;refGene_tx=XM_003998003.4
E2 26895725 26895867 refGene_gn=SHCBP1;refGene_tx=XM_003998003.4
E2 26896246 26896333 refGene_gn=SHCBP1;refGene_tx=XM_003998003.4
E2 26901924 26902043 refGene_gn=SHCBP1;refGene_tx=XM_003998003.4
E2 26902685 26902817 refGene_gn=SHCBP1;refGene_tx=XM_003998003.4
E2 26905603 26905724 refGene_gn=SHCBP1;refGene_tx=XM_003998003.4
E2 26906033 26906202 refGene_gn=SHCBP1;refGene_tx=XM_003998003.4
E2 26906292 26906526 refGene_gn=SHCBP1;refGene_tx=XM_003998003.4
E2 26909729 26909822 refGene_gn=SHCBP1;refGene_tx=XM_003998003.4
E2 26914502 26914711 refGene_gn=SHCBP1;refGene_tx=XM_003998003.4
E2 26916127 26916243 refGene_gn=SHCBP1;refGene_tx=XM_003998003.4
E2 26916422 26916590 refGene_gn=SHCBP1;refGene_tx=XM_003998003.4
E2 26919199 26919308 refGene_gn=SHCBP1;refGene_tx=XM_003998003.4
E2 26964231 26964244
refGene_gn=LOC101091491;refGene_tx=XM_019818941.1
E2 26964491 26965082
refGene_gn=LOC101091491;refGene_tx=XM_019818941.1
E2 26968104 26968679
refGene_gn=LOC101091491;refGene_tx=XM_019818941.1
E2 26973754 26973934
refGene_gn=VPS35;refGene_tx=XM_003998004.5;refGene_gn=VPS35;refGene_tx
=XM_019819672.2
E2 26974589 26974733
refGene_gn=VPS35;refGene_tx=XM_003998004.5;refGene_gn=VPS35;refGene_tx
=XM_019819672.2
E2 26975862 26976102

refGene_gn=VPS35;refGene_tx=XM_003998004.5;refGene_gn=VPS35;refGene_tx=XM_019819672.2
E2	26976597	26976777
refGene_gn=VPS35;refGene_tx=XM_003998004.5;refGene_gn=VPS35;refGene_tx=XM_019819672.2
E2	26978626	26978749
refGene_gn=VPS35;refGene_tx=XM_003998004.5;refGene_gn=VPS35;refGene_tx=XM_019819672.2
E2	26981299	26981455
refGene_gn=VPS35;refGene_tx=XM_003998004.5;refGene_gn=VPS35;refGene_tx=XM_019819672.2
E2	26982204	26982412
refGene_gn=VPS35;refGene_tx=XM_003998004.5;refGene_gn=VPS35;refGene_tx=XM_019819672.2
E2	26984386	26984535
refGene_gn=VPS35;refGene_tx=XM_003998004.5;refGene_gn=VPS35;refGene_tx=XM_019819672.2
E2	26984633	26984730
refGene_gn=VPS35;refGene_tx=XM_003998004.5;refGene_gn=VPS35;refGene_tx=XM_019819672.2
E2	26988479	26988589
refGene_gn=VPS35;refGene_tx=XM_003998004.5;refGene_gn=VPS35;refGene_tx=XM_019819672.2
E2	26988991	26989075
refGene_gn=VPS35;refGene_tx=XM_003998004.5;refGene_gn=VPS35;refGene_tx=XM_019819672.2
E2	26990250	26990464
refGene_gn=VPS35;refGene_tx=XM_003998004.5;refGene_gn=VPS35;refGene_tx=XM_019819672.2
E2	26992101	26992284
refGene_gn=VPS35;refGene_tx=XM_003998004.5;refGene_gn=VPS35;refGene_tx=XM_019819672.2
E2	26993524	26993648
refGene_gn=VPS35;refGene_tx=XM_003998004.5;refGene_gn=VPS35;refGene_tx=XM_019819672.2
E2	26994729	26994826
refGene_gn=VPS35;refGene_tx=XM_003998004.5;refGene_gn=VPS35;refGene_tx=XM_019819672.2
E2	26996128	26996227
refGene_gn=VPS35;refGene_tx=XM_003998004.5;refGene_gn=VPS35;refGene_tx=XM_019819672.2
E2	27002338	27002341	refGene_gn=VPS35;refGene_tx=XM_003998004.5
E2	27002543	27002747	refGene_gn=VPS35;refGene_tx=XM_019819672.2
E2	27002909	27002974
refGene_gn=ORC6;refGene_tx=XM_003998005.3;refGene_gn=ORC6;refGene_tx=XM_019819674.1
E2	27004177	27004307
refGene_gn=ORC6;refGene_tx=XM_003998005.3;refGene_gn=ORC6;refGene_tx=XM_019819674.1;refGene_gn=ORC6;refGene_tx=XM_019819675.2
E2	27005511	27005675

refGene_gn=ORC6;refGene_tx=XM_003998005.3;refGene_gn=ORC6;refGene_tx=XM_019819674.1;refGene_gn=ORC6;refGene_tx=XM_019819675.2
E2 27005894 27005984
refGene_gn=ORC6;refGene_tx=XM_003998005.3;refGene_gn=ORC6;refGene_tx=XM_019819675.2
E2 27007523 27007636
refGene_gn=ORC6;refGene_tx=XM_003998005.3;refGene_gn=ORC6;refGene_tx=XM_019819674.1;refGene_gn=ORC6;refGene_tx=XM_019819675.2
E2 27008101 27008170
refGene_gn=ORC6;refGene_tx=XM_003998005.3;refGene_gn=ORC6;refGene_tx=XM_019819674.1;refGene_gn=ORC6;refGene_tx=XM_019819675.2
E2 27008781 27008909
refGene_gn=ORC6;refGene_tx=XM_003998005.3;refGene_gn=ORC6;refGene_tx=XM_019819674.1;refGene_gn=ORC6;refGene_tx=XM_019819675.2
E2 27036381 27036441
refGene_gn=MYLK3;refGene_tx=XM_011290185.2;refGene_gn=MYLK3;refGene_tx=XM_011290186.3;refGene_gn=MYLK3;refGene_tx=XM_019819678.1;refGene_gn=MYLK3;refGene_tx=XM_019819679.1
E2 27037719 27037852
refGene_gn=MYLK3;refGene_tx=XM_011290185.2;refGene_gn=MYLK3;refGene_tx=XM_011290186.3;refGene_gn=MYLK3;refGene_tx=XM_019819678.1;refGene_gn=MYLK3;refGene_tx=XM_019819679.1
E2 27038800 27038953
refGene_gn=MYLK3;refGene_tx=XM_011290185.2;refGene_gn=MYLK3;refGene_tx=XM_011290186.3;refGene_gn=MYLK3;refGene_tx=XM_019819678.1;refGene_gn=MYLK3;refGene_tx=XM_019819679.1
E2 27040149 27040278
refGene_gn=MYLK3;refGene_tx=XM_011290185.2;refGene_gn=MYLK3;refGene_tx=XM_011290186.3;refGene_gn=MYLK3;refGene_tx=XM_019819678.1;refGene_gn=MYLK3;refGene_tx=XM_019819679.1
E2 27044717 27044788
refGene_gn=MYLK3;refGene_tx=XM_011290185.2;refGene_gn=MYLK3;refGene_tx=XM_011290186.3;refGene_gn=MYLK3;refGene_tx=XM_019819678.1;refGene_gn=MYLK3;refGene_tx=XM_019819679.1
E2 27052628 27052770
refGene_gn=MYLK3;refGene_tx=XM_011290185.2;refGene_gn=MYLK3;refGene_tx=XM_011290186.3;refGene_gn=MYLK3;refGene_tx=XM_019819678.1;refGene_gn=MYLK3;refGene_tx=XM_019819679.1
E2 27053633 27053743
refGene_gn=MYLK3;refGene_tx=XM_011290185.2;refGene_gn=MYLK3;refGene_tx=XM_011290186.3;refGene_gn=MYLK3;refGene_tx=XM_019819678.1;refGene_gn=MYLK3;refGene_tx=XM_019819679.1
E2 27054164 27054258
refGene_gn=MYLK3;refGene_tx=XM_011290185.2;refGene_gn=MYLK3;refGene_tx=XM_011290186.3;refGene_gn=MYLK3;refGene_tx=XM_019819678.1;refGene_gn=MYLK3;refGene_tx=XM_019819679.1
E2 27054812 27054918
refGene_gn=MYLK3;refGene_tx=XM_011290185.2;refGene_gn=MYLK3;refGene_tx=XM_011290186.3;refGene_gn=MYLK3;refGene_tx=XM_019819678.1;refGene_gn=MYLK3;refGene_tx=XM_019819679.1

E2 27056957 27057409
refGene_gn=MYLK3;refGene_tx=XM_011290185.2;refGene_gn=MYLK3;refGene_tx=XM_011290186.3;refGene_gn=MYLK3;refGene_tx=XM_019819678.1;refGene_gn=MYLK3;refGene_tx=XM_019819679.1
E2 27061468 27061853
refGene_gn=MYLK3;refGene_tx=XM_011290185.2;refGene_gn=MYLK3;refGene_tx=XM_011290186.3;refGene_gn=MYLK3;refGene_tx=XM_019819678.1;refGene_gn=MYLK3;refGene_tx=XM_019819679.1
E2 27063114 27063205
refGene_gn=MYLK3;refGene_tx=XM_011290185.2;refGene_gn=MYLK3;refGene_tx=XM_011290186.3
E2 27068979 27069456 refGene_gn=MYLK3;refGene_tx=XM_011290185.2
E2 27080250 27080268 refGene_gn=MYLK3;refGene_tx=XM_011290186.3
E2 27113722 27113841
refGene_gn=CE2H16orf87;refGene_tx=XM_023246021.1;refGene_gn=CE2H16orf87;refGene_tx=XM_023246022.1
E2 27115087 27115128
refGene_gn=CE2H16orf87;refGene_tx=XM_023246020.1
E2 27117017 27117109
refGene_gn=CE2H16orf87;refGene_tx=XM_023246019.1
E2 27121022 27121205
refGene_gn=CE2H16orf87;refGene_tx=XM_023246019.1;refGene_gn=CE2H16orf87;refGene_tx=XM_023246020.1;refGene_gn=CE2H16orf87;refGene_tx=XM_023246021.1
E2 27137609 27137706
refGene_gn=CE2H16orf87;refGene_tx=XM_023246019.1;refGene_gn=CE2H16orf87;refGene_tx=XM_023246020.1;refGene_gn=CE2H16orf87;refGene_tx=XM_023246021.1;refGene_gn=CE2H16orf87;refGene_tx=XM_023246022.1
E2 27146263 27146329
refGene_gn=CE2H16orf87;refGene_tx=XM_023246019.1;refGene_gn=CE2H16orf87;refGene_tx=XM_023246020.1;refGene_gn=CE2H16orf87;refGene_tx=XM_023246021.1;refGene_gn=CE2H16orf87;refGene_tx=XM_023246022.1
E2 27146648 27146795
refGene_gn=LOC111556304;refGene_tx=XM_023245116.1
E2 27154425 27154781
refGene_gn=LOC111556304;refGene_tx=XM_023245116.1
E2 27200738 27201263
refGene_gn=LOC111556304;refGene_tx=XM_023245116.1
E2 27213969 27214087
refGene_gn=LOC111556304;refGene_tx=XM_023245116.1
E2 27249930 27249938
refGene_gn=LOC109494660;refGene_tx=XM_019818943.1
E2 27254861 27255213
refGene_gn=LOC109494660;refGene_tx=XM_019818943.1
E2 27260382 27260625 refGene_gn=GPT2;refGene_tx=XM_003998006.4
E2 27268182 27268272 refGene_gn=GPT2;refGene_tx=XM_003998006.4
E2 27270364 27270473 refGene_gn=GPT2;refGene_tx=XM_003998006.4
E2 27274852 27274986 refGene_gn=GPT2;refGene_tx=XM_003998006.4
E2 27277092 27277336 refGene_gn=GPT2;refGene_tx=XM_003998006.4
E2 27281168 27281248 refGene_gn=GPT2;refGene_tx=XM_003998006.4

E2 27283355 27283492 refGene_gn=GPT2;refGene_tx=XM_003998006.4
E2 27286699 27286874 refGene_gn=GPT2;refGene_tx=XM_003998006.4
E2 27288349 27288505 refGene_gn=GPT2;refGene_tx=XM_003998006.4
E2 27289593 27289706 refGene_gn=GPT2;refGene_tx=XM_003998006.4
E2 27291206 27291297 refGene_gn=GPT2;refGene_tx=XM_003998006.4
E2 27316740 27316932 refGene_gn=DNAJA2;refGene_tx=XM_023245082.1
E2 27317527 27317655 refGene_gn=DNAJA2;refGene_tx=XM_023245082.1
E2 27317785 27317930 refGene_gn=DNAJA2;refGene_tx=XM_023245082.1
E2 27326298 27326495 refGene_gn=DNAJA2;refGene_tx=XM_023245082.1
E2 27328209 27328343 refGene_gn=DNAJA2;refGene_tx=XM_023245082.1
E2 27328594 27328675 refGene_gn=DNAJA2;refGene_tx=XM_023245082.1
E2 27331647 27331871 refGene_gn=DNAJA2;refGene_tx=XM_023245082.1
E2 27332155 27332215 refGene_gn=DNAJA2;refGene_tx=XM_023245082.1
E2 27333935 27334013 refGene_gn=DNAJA2;refGene_tx=XM_023245082.1
E2 27405160 27405741
refGene_gn=NETO2;refGene_tx=XM_023245020.1;refGene_gn=NETO2;refGene_tx=XM_023245021.1;refGene_gn=NETO2;refGene_tx=XM_023245022.1
E2 27407531 27407645
refGene_gn=NETO2;refGene_tx=XM_023245020.1;refGene_gn=NETO2;refGene_tx=XM_023245021.1;refGene_gn=NETO2;refGene_tx=XM_023245022.1
E2 27422569 27422798
refGene_gn=NETO2;refGene_tx=XM_023245020.1;refGene_gn=NETO2;refGene_tx=XM_023245021.1;refGene_gn=NETO2;refGene_tx=XM_023245022.1
E2 27439296 27439424
refGene_gn=NETO2;refGene_tx=XM_023245020.1;refGene_gn=NETO2;refGene_tx=XM_023245021.1;refGene_gn=NETO2;refGene_tx=XM_023245022.1
E2 27439506 27439551
refGene_gn=NETO2;refGene_tx=XM_023245020.1;refGene_gn=NETO2;refGene_tx=XM_023245021.1;refGene_gn=NETO2;refGene_tx=XM_023245022.1
E2 27443385 27443634
refGene_gn=NETO2;refGene_tx=XM_023245020.1;refGene_gn=NETO2;refGene_tx=XM_023245021.1;refGene_gn=NETO2;refGene_tx=XM_023245022.1
E2 27444524 27444665
refGene_gn=NETO2;refGene_tx=XM_023245020.1;refGene_gn=NETO2;refGene_tx=XM_023245021.1;refGene_gn=NETO2;refGene_tx=XM_023245022.1
E2 27447415 27447472
refGene_gn=NETO2;refGene_tx=XM_023245020.1;refGene_gn=NETO2;refGene_tx=XM_023245021.1
E2 27459736 27459770
refGene_gn=NETO2;refGene_tx=XM_023245020.1;refGene_gn=NETO2;refGene_tx=XM_023245021.1;refGene_gn=NETO2;refGene_tx=XM_023245022.1
E2 27471621 27471681 refGene_gn=ITFG1;refGene_tx=XM_023245671.1
E2 27474939 27475057 refGene_gn=ITFG1;refGene_tx=XM_023245671.1
E2 27478320 27478403 refGene_gn=ITFG1;refGene_tx=XM_023245671.1
E2 27484905 27485030
refGene_gn=ITFG1;refGene_tx=XM_023245671.1;refGene_gn=ITFG1;refGene_tx=XM_023245672.1
E2 27535756 27535835
refGene_gn=ITFG1;refGene_tx=XM_023245671.1;refGene_gn=ITFG1;refGene_tx=XM_023245672.1

E2 27565285 27565329 refGene_gn=ITFG1;refGene_tx=XM_023245671.1
E2 27604819 27604928
refGene_gn=ITFG1;refGene_tx=XM_023245671.1;refGene_gn=ITFG1;refGene_tx
=XM_023245672.1
E2 27607685 27607836
refGene_gn=ITFG1;refGene_tx=XM_023245671.1;refGene_gn=ITFG1;refGene_tx
=XM_023245672.1
E2 27664794 27664967
refGene_gn=ITFG1;refGene_tx=XM_023245671.1;refGene_gn=ITFG1;refGene_tx
=XM_023245672.1
E2 27667437 27667532
refGene_gn=ITFG1;refGene_tx=XM_023245671.1;refGene_gn=ITFG1;refGene_tx
=XM_023245672.1
E2 27720522 27720604
refGene_gn=ITFG1;refGene_tx=XM_023245671.1;refGene_gn=ITFG1;refGene_tx
=XM_023245672.1
E2 27736080 27736142
refGene_gn=ITFG1;refGene_tx=XM_023245671.1;refGene_gn=ITFG1;refGene_tx
=XM_023245672.1
E2 27784526 27784621
refGene_gn=ITFG1;refGene_tx=XM_023245671.1;refGene_gn=ITFG1;refGene_tx
=XM_023245672.1
E2 27808995 27809070
refGene_gn=ITFG1;refGene_tx=XM_023245671.1;refGene_gn=ITFG1;refGene_tx
=XM_023245672.1
E2 27810487 27810545
refGene_gn=ITFG1;refGene_tx=XM_023245671.1;refGene_gn=ITFG1;refGene_tx
=XM_023245672.1
E2 27812612 27812758
refGene_gn=ITFG1;refGene_tx=XM_023245671.1;refGene_gn=ITFG1;refGene_tx
=XM_023245672.1
E2 27818936 27819009
refGene_gn=ITFG1;refGene_tx=XM_023245671.1;refGene_gn=ITFG1;refGene_tx
=XM_023245672.1
E2 27820712 27820920
refGene_gn=ITFG1;refGene_tx=XM_023245671.1;refGene_gn=ITFG1;refGene_tx
=XM_023245672.1
E2 27821230 27821306
refGene_gn=PHKB;refGene_tx=XM_011290177.2;refGene_gn=PHKB;refGene_tx=X
M_011290178.2
E2 27823536 27823591
refGene_gn=PHKB;refGene_tx=XM_019819688.1;refGene_gn=PHKB;refGene_tx=X
M_019819689.1;refGene_gn=PHKB;refGene_tx=XM_019819690.2;refGene_gn=PHK
B;refGene_tx=XM_019819691.2;refGene_gn=PHKB;refGene_tx=XM_023245670.1
E2 27866255 27866345
refGene_gn=PHKB;refGene_tx=XM_011290177.2;refGene_gn=PHKB;refGene_tx=X
M_011290178.2;refGene_gn=PHKB;refGene_tx=XM_019819688.1;refGene_gn=PHK
B;refGene_tx=XM_019819689.1;refGene_gn=PHKB;refGene_tx=XM_019819690.2;
refGene_gn=PHKB;refGene_tx=XM_019819691.2;refGene_gn=PHKB;refGene_tx=X
M_023245670.1

E2 27868114 27868253
refGene_gn=PHKB;refGene_tx=XM_011290177.2;refGene_gn=PHKB;refGene_tx=XM_011290178.2;refGene_gn=PHKB;refGene_tx=XM_019819688.1;refGene_gn=PHKB;refGene_tx=XM_019819689.1;refGene_gn=PHKB;refGene_tx=XM_019819690.2;refGene_gn=PHKB;refGene_tx=XM_019819691.2;refGene_gn=PHKB;refGene_tx=XM_023245670.1
E2 27884291 27884391
refGene_gn=PHKB;refGene_tx=XM_011290177.2;refGene_gn=PHKB;refGene_tx=XM_011290178.2;refGene_gn=PHKB;refGene_tx=XM_019819688.1;refGene_gn=PHKB;refGene_tx=XM_019819689.1;refGene_gn=PHKB;refGene_tx=XM_019819690.2;refGene_gn=PHKB;refGene_tx=XM_019819691.2;refGene_gn=PHKB;refGene_tx=XM_023245670.1
E2 27889070 27889178
refGene_gn=PHKB;refGene_tx=XM_011290177.2;refGene_gn=PHKB;refGene_tx=XM_011290178.2;refGene_gn=PHKB;refGene_tx=XM_019819688.1;refGene_gn=PHKB;refGene_tx=XM_019819689.1;refGene_gn=PHKB;refGene_tx=XM_019819690.2;refGene_gn=PHKB;refGene_tx=XM_019819691.2;refGene_gn=PHKB;refGene_tx=XM_023245670.1
E2 27892805 27892886
refGene_gn=PHKB;refGene_tx=XM_011290177.2;refGene_gn=PHKB;refGene_tx=XM_011290178.2;refGene_gn=PHKB;refGene_tx=XM_019819688.1;refGene_gn=PHKB;refGene_tx=XM_019819689.1;refGene_gn=PHKB;refGene_tx=XM_019819690.2;refGene_gn=PHKB;refGene_tx=XM_019819691.2;refGene_gn=PHKB;refGene_tx=XM_023245670.1
E2 27935716 27935832
refGene_gn=PHKB;refGene_tx=XM_011290177.2;refGene_gn=PHKB;refGene_tx=XM_011290178.2;refGene_gn=PHKB;refGene_tx=XM_019819688.1;refGene_gn=PHKB;refGene_tx=XM_019819689.1;refGene_gn=PHKB;refGene_tx=XM_019819690.2;refGene_gn=PHKB;refGene_tx=XM_019819691.2;refGene_gn=PHKB;refGene_tx=XM_023245670.1
E2 27973711 27973775
refGene_gn=PHKB;refGene_tx=XM_011290177.2;refGene_gn=PHKB;refGene_tx=XM_011290178.2;refGene_gn=PHKB;refGene_tx=XM_019819688.1;refGene_gn=PHKB;refGene_tx=XM_019819689.1;refGene_gn=PHKB;refGene_tx=XM_019819690.2;refGene_gn=PHKB;refGene_tx=XM_019819691.2;refGene_gn=PHKB;refGene_tx=XM_023245670.1
E2 27982565 27982661
refGene_gn=PHKB;refGene_tx=XM_011290177.2;refGene_gn=PHKB;refGene_tx=XM_011290178.2;refGene_gn=PHKB;refGene_tx=XM_019819688.1;refGene_gn=PHKB;refGene_tx=XM_019819689.1;refGene_gn=PHKB;refGene_tx=XM_019819690.2;refGene_gn=PHKB;refGene_tx=XM_019819691.2;refGene_gn=PHKB;refGene_tx=XM_023245670.1
E2 27983932 27984130
refGene_gn=PHKB;refGene_tx=XM_011290177.2;refGene_gn=PHKB;refGene_tx=XM_011290178.2;refGene_gn=PHKB;refGene_tx=XM_019819688.1;refGene_gn=PHKB;refGene_tx=XM_019819689.1;refGene_gn=PHKB;refGene_tx=XM_019819690.2;refGene_gn=PHKB;refGene_tx=XM_019819691.2;refGene_gn=PHKB;refGene_tx=XM_023245670.1
E2 27987778 27987836
refGene_gn=PHKB;refGene_tx=XM_011290177.2;refGene_gn=PHKB;refGene_tx=X

M_011290178.2;refGene_gn=PHKB;refGene_tx=XM_019819688.1;refGene_gn=PHK
B;refGene_tx=XM_019819689.1;refGene_gn=PHKB;refGene_tx=XM_019819690.2;
refGene_gn=PHKB;refGene_tx=XM_019819691.2;refGene_gn=PHKB;refGene_tx=X
M_023245670.1
E2 27988381 27988459
refGene_gn=PHKB;refGene_tx=XM_011290177.2;refGene_gn=PHKB;refGene_tx=X
M_011290178.2;refGene_gn=PHKB;refGene_tx=XM_019819688.1;refGene_gn=PHK
B;refGene_tx=XM_019819689.1;refGene_gn=PHKB;refGene_tx=XM_019819690.2;
refGene_gn=PHKB;refGene_tx=XM_019819691.2;refGene_gn=PHKB;refGene_tx=X
M_023245670.1
E2 27990463 27990622
refGene_gn=PHKB;refGene_tx=XM_011290177.2;refGene_gn=PHKB;refGene_tx=X
M_011290178.2;refGene_gn=PHKB;refGene_tx=XM_019819688.1;refGene_gn=PHK
B;refGene_tx=XM_019819689.1;refGene_gn=PHKB;refGene_tx=XM_019819690.2;
refGene_gn=PHKB;refGene_tx=XM_019819691.2;refGene_gn=PHKB;refGene_tx=X
M_023245670.1
E2 28001009 28001104
refGene_gn=PHKB;refGene_tx=XM_011290177.2;refGene_gn=PHKB;refGene_tx=X
M_011290178.2;refGene_gn=PHKB;refGene_tx=XM_019819688.1;refGene_gn=PHK
B;refGene_tx=XM_019819689.1;refGene_gn=PHKB;refGene_tx=XM_019819690.2;
refGene_gn=PHKB;refGene_tx=XM_019819691.2;refGene_gn=PHKB;refGene_tx=X
M_023245670.1
E2 28041325 28041381
refGene_gn=PHKB;refGene_tx=XM_011290177.2;refGene_gn=PHKB;refGene_tx=X
M_011290178.2;refGene_gn=PHKB;refGene_tx=XM_019819688.1;refGene_gn=PHK
B;refGene_tx=XM_019819689.1;refGene_gn=PHKB;refGene_tx=XM_019819690.2;
refGene_gn=PHKB;refGene_tx=XM_019819691.2;refGene_gn=PHKB;refGene_tx=X
M_023245670.1
E2 28041899 28041993
refGene_gn=PHKB;refGene_tx=XM_011290177.2;refGene_gn=PHKB;refGene_tx=X
M_011290178.2;refGene_gn=PHKB;refGene_tx=XM_019819688.1;refGene_gn=PHK
B;refGene_tx=XM_019819689.1;refGene_gn=PHKB;refGene_tx=XM_019819690.2;
refGene_gn=PHKB;refGene_tx=XM_019819691.2;refGene_gn=PHKB;refGene_tx=X
M_023245670.1
E2 28048856 28048940
refGene_gn=PHKB;refGene_tx=XM_011290177.2;refGene_gn=PHKB;refGene_tx=X
M_011290178.2;refGene_gn=PHKB;refGene_tx=XM_019819688.1;refGene_gn=PHK
B;refGene_tx=XM_019819689.1;refGene_gn=PHKB;refGene_tx=XM_019819690.2;
refGene_gn=PHKB;refGene_tx=XM_019819691.2;refGene_gn=PHKB;refGene_tx=X
M_023245670.1
E2 28049459 28049564
refGene_gn=PHKB;refGene_tx=XM_011290177.2;refGene_gn=PHKB;refGene_tx=X
M_011290178.2;refGene_gn=PHKB;refGene_tx=XM_019819688.1;refGene_gn=PHK
B;refGene_tx=XM_019819689.1;refGene_gn=PHKB;refGene_tx=XM_019819690.2;
refGene_gn=PHKB;refGene_tx=XM_019819691.2;refGene_gn=PHKB;refGene_tx=X
M_023245670.1
E2 28051004 28051087
refGene_gn=PHKB;refGene_tx=XM_011290177.2;refGene_gn=PHKB;refGene_tx=X
M_011290178.2;refGene_gn=PHKB;refGene_tx=XM_019819688.1;refGene_gn=PHK
B;refGene_tx=XM_019819689.1;refGene_gn=PHKB;refGene_tx=XM_019819690.2;

refGene_gn=PHKB;refGene_tx=XM_019819691.2;refGene_gn=PHKB;refGene_tx=X
M_023245670.1
E2 28051290 28051381
refGene_gn=PHKB;refGene_tx=XM_011290177.2;refGene_gn=PHKB;refGene_tx=X
M_011290178.2;refGene_gn=PHKB;refGene_tx=XM_019819688.1;refGene_gn=PHK
B;refGene_tx=XM_019819689.1;refGene_gn=PHKB;refGene_tx=XM_019819690.2;
refGene_gn=PHKB;refGene_tx=XM_019819691.2;refGene_gn=PHKB;refGene_tx=X
M_023245670.1
E2 28056899 28056961
refGene_gn=PHKB;refGene_tx=XM_011290177.2;refGene_gn=PHKB;refGene_tx=X
M_011290178.2;refGene_gn=PHKB;refGene_tx=XM_019819688.1;refGene_gn=PHK
B;refGene_tx=XM_019819689.1;refGene_gn=PHKB;refGene_tx=XM_019819690.2;
refGene_gn=PHKB;refGene_tx=XM_019819691.2;refGene_gn=PHKB;refGene_tx=X
M_023245670.1
E2 28057047 28057210
refGene_gn=PHKB;refGene_tx=XM_011290177.2;refGene_gn=PHKB;refGene_tx=X
M_011290178.2;refGene_gn=PHKB;refGene_tx=XM_019819688.1;refGene_gn=PHK
B;refGene_tx=XM_019819689.1;refGene_gn=PHKB;refGene_tx=XM_019819690.2;
refGene_gn=PHKB;refGene_tx=XM_019819691.2;refGene_gn=PHKB;refGene_tx=X
M_023245670.1
E2 28058387 28058469
refGene_gn=PHKB;refGene_tx=XM_011290177.2;refGene_gn=PHKB;refGene_tx=X
M_011290178.2;refGene_gn=PHKB;refGene_tx=XM_019819688.1;refGene_gn=PHK
B;refGene_tx=XM_019819689.1;refGene_gn=PHKB;refGene_tx=XM_019819690.2;
refGene_gn=PHKB;refGene_tx=XM_019819691.2;refGene_gn=PHKB;refGene_tx=X
M_023245670.1
E2 28060366 28060424
refGene_gn=PHKB;refGene_tx=XM_011290177.2;refGene_gn=PHKB;refGene_tx=X
M_011290178.2;refGene_gn=PHKB;refGene_tx=XM_019819688.1;refGene_gn=PHK
B;refGene_tx=XM_019819689.1;refGene_gn=PHKB;refGene_tx=XM_019819690.2;
refGene_gn=PHKB;refGene_tx=XM_019819691.2;refGene_gn=PHKB;refGene_tx=X
M_023245670.1
E2 28061748 28061839 refGene_gn=PHKB;refGene_tx=XM_011290177.2
E2 28062772 28062863
refGene_gn=PHKB;refGene_tx=XM_011290178.2;refGene_gn=PHKB;refGene_tx=X
M_019819688.1;refGene_gn=PHKB;refGene_tx=XM_019819689.1;refGene_gn=PHK
B;refGene_tx=XM_019819690.2;refGene_gn=PHKB;refGene_tx=XM_019819691.2;
refGene_gn=PHKB;refGene_tx=XM_023245670.1
E2 28065686 28065889
refGene_gn=PHKB;refGene_tx=XM_011290177.2;refGene_gn=PHKB;refGene_tx=X
M_011290178.2;refGene_gn=PHKB;refGene_tx=XM_019819688.1;refGene_gn=PHK
B;refGene_tx=XM_019819689.1;refGene_gn=PHKB;refGene_tx=XM_019819690.2;
refGene_gn=PHKB;refGene_tx=XM_019819691.2;refGene_gn=PHKB;refGene_tx=X
M_023245670.1
E2 28078053 28078188
refGene_gn=PHKB;refGene_tx=XM_011290177.2;refGene_gn=PHKB;refGene_tx=X
M_011290178.2;refGene_gn=PHKB;refGene_tx=XM_019819688.1;refGene_gn=PHK
B;refGene_tx=XM_019819689.1;refGene_gn=PHKB;refGene_tx=XM_019819690.2;
refGene_gn=PHKB;refGene_tx=XM_019819691.2;refGene_gn=PHKB;refGene_tx=X
M_023245670.1

E2 28083548 28083678
refGene_gn=PHKB;refGene_tx=XM_011290177.2;refGene_gn=PHKB;refGene_tx=XM_011290178.2;refGene_gn=PHKB;refGene_tx=XM_019819688.1;refGene_gn=PHKB;refGene_tx=XM_019819689.1;refGene_gn=PHKB;refGene_tx=XM_019819690.2;refGene_gn=PHKB;refGene_tx=XM_019819691.2;refGene_gn=PHKB;refGene_tx=XM_023245670.1
E2 28086939 28087047
refGene_gn=PHKB;refGene_tx=XM_011290177.2;refGene_gn=PHKB;refGene_tx=XM_011290178.2;refGene_gn=PHKB;refGene_tx=XM_019819688.1;refGene_gn=PHKB;refGene_tx=XM_019819689.1;refGene_gn=PHKB;refGene_tx=XM_019819690.2;refGene_gn=PHKB;refGene_tx=XM_019819691.2;refGene_gn=PHKB;refGene_tx=XM_023245670.1
E2 28089614 28089755
refGene_gn=PHKB;refGene_tx=XM_011290177.2;refGene_gn=PHKB;refGene_tx=XM_011290178.2;refGene_gn=PHKB;refGene_tx=XM_019819688.1;refGene_gn=PHKB;refGene_tx=XM_019819689.1;refGene_gn=PHKB;refGene_tx=XM_019819690.2;refGene_gn=PHKB;refGene_tx=XM_019819691.2;refGene_gn=PHKB;refGene_tx=XM_023245670.1
E2 28090400 28090538
refGene_gn=PHKB;refGene_tx=XM_011290177.2;refGene_gn=PHKB;refGene_tx=XM_011290178.2;refGene_gn=PHKB;refGene_tx=XM_019819688.1;refGene_gn=PHKB;refGene_tx=XM_019819689.1;refGene_gn=PHKB;refGene_tx=XM_019819690.2;refGene_gn=PHKB;refGene_tx=XM_019819691.2;refGene_gn=PHKB;refGene_tx=XM_023245670.1
E2 28415177 28415270
refGene_gn=ABCC12;refGene_tx=XM_006941488.3;refGene_gn=ABCC12;refGene_tx=XM_019819692.2;refGene_gn=ABCC12;refGene_tx=XM_019819695.2;refGene_gn=ABCC12;refGene_tx=XM_023245345.1
E2 28415374 28415539
refGene_gn=ABCC12;refGene_tx=XM_006941488.3;refGene_gn=ABCC12;refGene_tx=XM_019819692.2;refGene_gn=ABCC12;refGene_tx=XM_019819695.2;refGene_gn=ABCC12;refGene_tx=XM_023245345.1
E2 28417056 28417242
refGene_gn=ABCC12;refGene_tx=XM_006941488.3;refGene_gn=ABCC12;refGene_tx=XM_019819692.2;refGene_gn=ABCC12;refGene_tx=XM_019819695.2;refGene_gn=ABCC12;refGene_tx=XM_023245345.1
E2 28418228 28418307
refGene_gn=ABCC12;refGene_tx=XM_006941488.3;refGene_gn=ABCC12;refGene_tx=XM_019819692.2;refGene_gn=ABCC12;refGene_tx=XM_019819695.2;refGene_gn=ABCC12;refGene_tx=XM_019819698.2;refGene_gn=ABCC12;refGene_tx=XM_023245345.1
E2 28419311 28419471
refGene_gn=ABCC12;refGene_tx=XM_006941488.3;refGene_gn=ABCC12;refGene_tx=XM_019819692.2;refGene_gn=ABCC12;refGene_tx=XM_019819695.2;refGene_gn=ABCC12;refGene_tx=XM_019819698.2;refGene_gn=ABCC12;refGene_tx=XM_023245345.1
E2 28419905 28420095
refGene_gn=ABCC12;refGene_tx=XM_006941488.3;refGene_gn=ABCC12;refGene_tx=XM_019819692.2;refGene_gn=ABCC12;refGene_tx=XM_019819695.2;refGene_gn=ABCC12;refGene_tx=XM_019819698.2;refGene_gn=ABCC12;refGene_tx=XM_02

3245345.1
E2 28421963 28422053
refGene_gn=ABCC12;refGene_tx=XM_006941488.3;refGene_gn=ABCC12;refGene_tx=XM_019819692.2;refGene_gn=ABCC12;refGene_tx=XM_019819695.2;refGene_gn=ABCC12;refGene_tx=XM_019819698.2;refGene_gn=ABCC12;refGene_tx=XM_023245345.1
E2 28426108 28426295
refGene_gn=ABCC12;refGene_tx=XM_006941488.3;refGene_gn=ABCC12;refGene_tx=XM_019819692.2;refGene_gn=ABCC12;refGene_tx=XM_019819695.2;refGene_gn=ABCC12;refGene_tx=XM_019819698.2;refGene_gn=ABCC12;refGene_tx=XM_023245345.1
E2 28430452 28430590
refGene_gn=ABCC12;refGene_tx=XM_006941488.3;refGene_gn=ABCC12;refGene_tx=XM_019819692.2;refGene_gn=ABCC12;refGene_tx=XM_019819695.2;refGene_gn=ABCC12;refGene_tx=XM_019819698.2;refGene_gn=ABCC12;refGene_tx=XM_023245345.1
E2 28434356 28434583
refGene_gn=ABCC12;refGene_tx=XM_006941488.3;refGene_gn=ABCC12;refGene_tx=XM_019819692.2;refGene_gn=ABCC12;refGene_tx=XM_019819695.2;refGene_gn=ABCC12;refGene_tx=XM_019819698.2;refGene_gn=ABCC12;refGene_tx=XM_023245345.1
E2 28435358 28435556
refGene_gn=ABCC12;refGene_tx=XM_006941488.3;refGene_gn=ABCC12;refGene_tx=XM_019819692.2;refGene_gn=ABCC12;refGene_tx=XM_019819695.2;refGene_gn=ABCC12;refGene_tx=XM_019819698.2;refGene_gn=ABCC12;refGene_tx=XM_023245345.1
E2 28437514 28437618
refGene_gn=ABCC12;refGene_tx=XM_006941488.3;refGene_gn=ABCC12;refGene_tx=XM_019819692.2;refGene_gn=ABCC12;refGene_tx=XM_019819695.2;refGene_gn=ABCC12;refGene_tx=XM_019819698.2;refGene_gn=ABCC12;refGene_tx=XM_023245345.1
E2 28438328 28438418
refGene_gn=ABCC12;refGene_tx=XM_006941488.3;refGene_gn=ABCC12;refGene_tx=XM_019819692.2;refGene_gn=ABCC12;refGene_tx=XM_019819695.2;refGene_gn=ABCC12;refGene_tx=XM_019819698.2;refGene_gn=ABCC12;refGene_tx=XM_023245345.1
E2 28441224 28441296
refGene_gn=ABCC12;refGene_tx=XM_006941488.3;refGene_gn=ABCC12;refGene_tx=XM_019819692.2;refGene_gn=ABCC12;refGene_tx=XM_019819695.2;refGene_gn=ABCC12;refGene_tx=XM_019819698.2;refGene_gn=ABCC12;refGene_tx=XM_023245345.1
E2 28441379 28441455
refGene_gn=ABCC12;refGene_tx=XM_006941488.3;refGene_gn=ABCC12;refGene_tx=XM_019819692.2;refGene_gn=ABCC12;refGene_tx=XM_019819695.2;refGene_gn=ABCC12;refGene_tx=XM_019819698.2;refGene_gn=ABCC12;refGene_tx=XM_023245345.1
E2 28441565 28441700
refGene_gn=ABCC12;refGene_tx=XM_006941488.3;refGene_gn=ABCC12;refGene_tx=XM_019819692.2;refGene_gn=ABCC12;refGene_tx=XM_019819695.2;refGene_gn=ABCC12;refGene_tx=XM_019819698.2;refGene_gn=ABCC12;refGene_tx=XM_02

3245345.1
E2 28445130 28445334
refGene_gn=ABCC12;refGene_tx=XM_006941488.3;refGene_gn=ABCC12;refGene_tx=XM_019819692.2;refGene_gn=ABCC12;refGene_tx=XM_019819695.2;refGene_gn=ABCC12;refGene_tx=XM_019819698.2
E2 28447046 28447119
refGene_gn=ABCC12;refGene_tx=XM_006941488.3;refGene_gn=ABCC12;refGene_tx=XM_019819692.2;refGene_gn=ABCC12;refGene_tx=XM_019819695.2;refGene_gn=ABCC12;refGene_tx=XM_019819698.2;refGene_gn=ABCC12;refGene_tx=XM_023245345.1
E2 28459051 28459176
refGene_gn=ABCC12;refGene_tx=XM_006941488.3;refGene_gn=ABCC12;refGene_tx=XM_019819692.2;refGene_gn=ABCC12;refGene_tx=XM_019819695.2;refGene_gn=ABCC12;refGene_tx=XM_019819698.2;refGene_gn=ABCC12;refGene_tx=XM_023245345.1
E2 28461577 28461649
refGene_gn=ABCC12;refGene_tx=XM_006941488.3;refGene_gn=ABCC12;refGene_tx=XM_019819692.2;refGene_gn=ABCC12;refGene_tx=XM_019819695.2;refGene_gn=ABCC12;refGene_tx=XM_019819698.2;refGene_gn=ABCC12;refGene_tx=XM_023245345.1
E2 28465803 28466079
refGene_gn=ABCC12;refGene_tx=XM_006941488.3;refGene_gn=ABCC12;refGene_tx=XM_019819692.2;refGene_gn=ABCC12;refGene_tx=XM_019819695.2;refGene_gn=ABCC12;refGene_tx=XM_019819698.2;refGene_gn=ABCC12;refGene_tx=XM_023245345.1
E2 28468117 28468225
refGene_gn=ABCC12;refGene_tx=XM_006941488.3;refGene_gn=ABCC12;refGene_tx=XM_019819692.2;refGene_gn=ABCC12;refGene_tx=XM_019819695.2;refGene_gn=ABCC12;refGene_tx=XM_019819698.2;refGene_gn=ABCC12;refGene_tx=XM_023245345.1
E2 28470641 28470790
refGene_gn=ABCC12;refGene_tx=XM_006941488.3;refGene_gn=ABCC12;refGene_tx=XM_019819692.2;refGene_gn=ABCC12;refGene_tx=XM_019819695.2;refGene_gn=ABCC12;refGene_tx=XM_019819698.2;refGene_gn=ABCC12;refGene_tx=XM_023245345.1
E2 28473995 28474143
refGene_gn=ABCC12;refGene_tx=XM_006941488.3;refGene_gn=ABCC12;refGene_tx=XM_019819692.2;refGene_gn=ABCC12;refGene_tx=XM_019819695.2;refGene_gn=ABCC12;refGene_tx=XM_019819698.2;refGene_gn=ABCC12;refGene_tx=XM_023245345.1
E2 28474597 28474771
refGene_gn=ABCC12;refGene_tx=XM_006941488.3;refGene_gn=ABCC12;refGene_tx=XM_019819692.2;refGene_gn=ABCC12;refGene_tx=XM_019819695.2;refGene_gn=ABCC12;refGene_tx=XM_019819698.2;refGene_gn=ABCC12;refGene_tx=XM_023245345.1
E2 28476028 28476262
refGene_gn=ABCC12;refGene_tx=XM_006941488.3;refGene_gn=ABCC12;refGene_tx=XM_019819692.2;refGene_gn=ABCC12;refGene_tx=XM_019819695.2;refGene_gn=ABCC12;refGene_tx=XM_019819698.2;refGene_gn=ABCC12;refGene_tx=XM_023245345.1

E2 28476551 28476699
refGene_gn=ABCC12;refGene_tx=XM_006941488.3;refGene_gn=ABCC12;refGene_tx=XM_019819692.2;refGene_gn=ABCC12;refGene_tx=XM_019819695.2;refGene_gn=ABCC12;refGene_tx=XM_019819698.2;refGene_gn=ABCC12;refGene_tx=XM_023245345.1
E2 28480681 28480837
refGene_gn=ABCC12;refGene_tx=XM_006941488.3;refGene_gn=ABCC12;refGene_tx=XM_019819692.2;refGene_gn=ABCC12;refGene_tx=XM_019819698.2;refGene_gn=ABCC12;refGene_tx=XM_023245345.1
E2 28490657 28490776
refGene_gn=ABCC12;refGene_tx=XM_006941488.3;refGene_gn=ABCC12;refGene_tx=XM_019819692.2;refGene_gn=ABCC12;refGene_tx=XM_019819695.2;refGene_gn=ABCC12;refGene_tx=XM_019819698.2;refGene_gn=ABCC12;refGene_tx=XM_023245345.1
E2 28518544 28518637
refGene_gn=ABCC11;refGene_tx=XM_003998010.5;refGene_gn=ABCC11;refGene_tx=XM_019819699.2;refGene_gn=ABCC11;refGene_tx=XM_019819701.2
E2 28518764 28518929
refGene_gn=ABCC11;refGene_tx=XM_003998010.5;refGene_gn=ABCC11;refGene_tx=XM_019819699.2;refGene_gn=ABCC11;refGene_tx=XM_019819701.2
E2 28521870 28521984
refGene_gn=ABCC11;refGene_tx=XM_003998010.5;refGene_gn=ABCC11;refGene_tx=XM_019819699.2;refGene_gn=ABCC11;refGene_tx=XM_019819701.2
E2 28522656 28522735
refGene_gn=ABCC11;refGene_tx=XM_003998010.5;refGene_gn=ABCC11;refGene_tx=XM_019819699.2;refGene_gn=ABCC11;refGene_tx=XM_019819701.2
E2 28525601 28525761
refGene_gn=ABCC11;refGene_tx=XM_003998010.5;refGene_gn=ABCC11;refGene_tx=XM_019819699.2;refGene_gn=ABCC11;refGene_tx=XM_019819701.2
E2 28527127 28527317
refGene_gn=ABCC11;refGene_tx=XM_003998010.5;refGene_gn=ABCC11;refGene_tx=XM_019819699.2;refGene_gn=ABCC11;refGene_tx=XM_019819701.2
E2 28528696 28528786
refGene_gn=ABCC11;refGene_tx=XM_003998010.5;refGene_gn=ABCC11;refGene_tx=XM_019819699.2;refGene_gn=ABCC11;refGene_tx=XM_019819701.2
E2 28533264 28533451
refGene_gn=ABCC11;refGene_tx=XM_003998010.5;refGene_gn=ABCC11;refGene_tx=XM_019819699.2;refGene_gn=ABCC11;refGene_tx=XM_019819701.2
E2 28534895 28535033
refGene_gn=ABCC11;refGene_tx=XM_003998010.5;refGene_gn=ABCC11;refGene_tx=XM_019819699.2;refGene_gn=ABCC11;refGene_tx=XM_019819701.2
E2 28535153 28535380
refGene_gn=ABCC11;refGene_tx=XM_003998010.5;refGene_gn=ABCC11;refGene_tx=XM_019819699.2;refGene_gn=ABCC11;refGene_tx=XM_019819701.2
E2 28537657 28537855
refGene_gn=ABCC11;refGene_tx=XM_003998010.5;refGene_gn=ABCC11;refGene_tx=XM_019819699.2;refGene_gn=ABCC11;refGene_tx=XM_019819701.2
E2 28540338 28540442
refGene_gn=ABCC11;refGene_tx=XM_003998010.5;refGene_gn=ABCC11;refGene_tx=XM_019819699.2;refGene_gn=ABCC11;refGene_tx=XM_019819701.2

E2 28542973 28543063
refGene_gn=ABCC11;refGene_tx=XM_003998010.5;refGene_gn=ABCC11;refGene_tx=XM_019819699.2;refGene_gn=ABCC11;refGene_tx=XM_019819701.2
E2 28544808 28544905
refGene_gn=ABCC11;refGene_tx=XM_003998010.5;refGene_gn=ABCC11;refGene_tx=XM_019819699.2;refGene_gn=ABCC11;refGene_tx=XM_019819701.2
E2 28544991 28545126
refGene_gn=ABCC11;refGene_tx=XM_003998010.5;refGene_gn=ABCC11;refGene_tx=XM_019819699.2;refGene_gn=ABCC11;refGene_tx=XM_019819701.2
E2 28546596 28546800
refGene_gn=ABCC11;refGene_tx=XM_003998010.5;refGene_gn=ABCC11;refGene_tx=XM_019819699.2;refGene_gn=ABCC11;refGene_tx=XM_019819701.2
E2 28548483 28548556
refGene_gn=ABCC11;refGene_tx=XM_003998010.5;refGene_gn=ABCC11;refGene_tx=XM_019819699.2;refGene_gn=ABCC11;refGene_tx=XM_019819701.2
E2 28550522 28550647
refGene_gn=ABCC11;refGene_tx=XM_003998010.5;refGene_gn=ABCC11;refGene_tx=XM_019819699.2;refGene_gn=ABCC11;refGene_tx=XM_019819701.2
E2 28553043 28553115
refGene_gn=ABCC11;refGene_tx=XM_003998010.5;refGene_gn=ABCC11;refGene_tx=XM_019819699.2;refGene_gn=ABCC11;refGene_tx=XM_019819701.2
E2 28554334 28554595
refGene_gn=ABCC11;refGene_tx=XM_003998010.5;refGene_gn=ABCC11;refGene_tx=XM_019819699.2;refGene_gn=ABCC11;refGene_tx=XM_019819701.2
E2 28558115 28558223
refGene_gn=ABCC11;refGene_tx=XM_003998010.5;refGene_gn=ABCC11;refGene_tx=XM_019819699.2
E2 28559548 28559697
refGene_gn=ABCC11;refGene_tx=XM_003998010.5;refGene_gn=ABCC11;refGene_tx=XM_019819699.2;refGene_gn=ABCC11;refGene_tx=XM_019819701.2
E2 28559854 28560002
refGene_gn=ABCC11;refGene_tx=XM_003998010.5;refGene_gn=ABCC11;refGene_tx=XM_019819699.2;refGene_gn=ABCC11;refGene_tx=XM_019819701.2
E2 28560830 28561004
refGene_gn=ABCC11;refGene_tx=XM_003998010.5;refGene_gn=ABCC11;refGene_tx=XM_019819699.2;refGene_gn=ABCC11;refGene_tx=XM_019819701.2
E2 28565411 28565645
refGene_gn=ABCC11;refGene_tx=XM_003998010.5;refGene_gn=ABCC11;refGene_tx=XM_019819699.2;refGene_gn=ABCC11;refGene_tx=XM_019819701.2
E2 28567798 28567946
refGene_gn=ABCC11;refGene_tx=XM_003998010.5;refGene_gn=ABCC11;refGene_tx=XM_019819699.2;refGene_gn=ABCC11;refGene_tx=XM_019819701.2
E2 28572308 28572467
refGene_gn=ABCC11;refGene_tx=XM_003998010.5;refGene_gn=ABCC11;refGene_tx=XM_019819699.2;refGene_gn=ABCC11;refGene_tx=XM_019819701.2
E2 28575471 28575608
refGene_gn=ABCC11;refGene_tx=XM_003998010.5;refGene_gn=ABCC11;refGene_tx=XM_019819699.2;refGene_gn=ABCC11;refGene_tx=XM_019819701.2
E2 28577368 28577467
refGene_gn=ABCC11;refGene_tx=XM_003998010.5;refGene_gn=ABCC11;refGene_

tx=XM_019819699.2;refGene_gn=ABCC11;refGene_tx=XM_019819701.2
E2	28581940	28582173
refGene_gn=LONP2;refGene_tx=XM_003998012.4;refGene_gn=LONP2;refGene_tx
=XM_023245149.1
E2	28588627	28588862
refGene_gn=LONP2;refGene_tx=XM_003998012.4;refGene_gn=LONP2;refGene_tx
=XM_023245149.1
E2	28592704	28592836
refGene_gn=LONP2;refGene_tx=XM_003998012.4;refGene_gn=LONP2;refGene_tx
=XM_023245149.1
E2	28594189	28594312
refGene_gn=LONP2;refGene_tx=XM_003998012.4;refGene_gn=LONP2;refGene_tx
=XM_023245149.1
E2	28599737	28599901
refGene_gn=LONP2;refGene_tx=XM_003998012.4;refGene_gn=LONP2;refGene_tx
=XM_023245149.1
E2	28601215	28601310
refGene_gn=LONP2;refGene_tx=XM_003998012.4;refGene_gn=LONP2;refGene_tx
=XM_023245149.1
E2	28607217	28607476
refGene_gn=LONP2;refGene_tx=XM_003998012.4;refGene_gn=LONP2;refGene_tx
=XM_023245149.1
E2	28615891	28616033
refGene_gn=LONP2;refGene_tx=XM_003998012.4;refGene_gn=LONP2;refGene_tx
=XM_023245149.1
E2	28637687	28637838
refGene_gn=LONP2;refGene_tx=XM_003998012.4;refGene_gn=LONP2;refGene_tx
=XM_023245149.1
E2	28640740	28640867
refGene_gn=LONP2;refGene_tx=XM_003998012.4;refGene_gn=LONP2;refGene_tx
=XM_023245149.1
E2	28650114	28650248
refGene_gn=LONP2;refGene_tx=XM_003998012.4;refGene_gn=LONP2;refGene_tx
=XM_023245149.1
E2	28672061	28672204
refGene_gn=LONP2;refGene_tx=XM_003998012.4;refGene_gn=LONP2;refGene_tx
=XM_023245149.1
E2	28682066	28682274
refGene_gn=LONP2;refGene_tx=XM_003998012.4;refGene_gn=LONP2;refGene_tx
=XM_023245149.1
E2	28682687	28682878
refGene_gn=LONP2;refGene_tx=XM_003998012.4;refGene_gn=LONP2;refGene_tx
=XM_023245149.1
E2	28683077	28683092	refGene_gn=LONP2;refGene_tx=XM_023245149.1
E2	28685729	28685951	refGene_gn=LONP2;refGene_tx=XM_003998012.4
E2	28692238	28693089
refGene_gn=SIAH1;refGene_tx=XM_003998013.5;refGene_gn=SIAH1;refGene_tx
=XM_023245150.1;refGene_gn=SIAH1;refGene_tx=XM_023245151.1
E2	28693629	28693642	refGene_gn=SIAH1;refGene_tx=XM_023245150.1
E2	28695615	28695706	refGene_gn=SIAH1;refGene_tx=XM_003998013.5

E2 28853457 28853815
refGene_gn=N4BP1;refGene_tx=XM_023245980.1;refGene_gn=N4BP1;refGene_tx
=XM_023245981.1
E2 28855448 28855556
refGene_gn=N4BP1;refGene_tx=XM_023245980.1;refGene_gn=N4BP1;refGene_tx
=XM_023245981.1
E2 28857570 28857678 refGene_gn=N4BP1;refGene_tx=XM_023245980.1
E2 28860609 28860706
refGene_gn=N4BP1;refGene_tx=XM_023245980.1;refGene_gn=N4BP1;refGene_tx
=XM_023245981.1
E2 28862244 28862375
refGene_gn=N4BP1;refGene_tx=XM_023245980.1;refGene_gn=N4BP1;refGene_tx
=XM_023245981.1
E2 28870724 28872415
refGene_gn=N4BP1;refGene_tx=XM_023245980.1;refGene_gn=N4BP1;refGene_tx
=XM_023245981.1
E2 28899949 28900147
refGene_gn=N4BP1;refGene_tx=XM_023245980.1;refGene_gn=N4BP1;refGene_tx
=XM_023245981.1
E2 29441081 29441279 refGene_gn=CBLN1;refGene_tx=XM_023245547.1
E2 29442559 29442679 refGene_gn=CBLN1;refGene_tx=XM_023245547.1
E2 29442820 29443084 refGene_gn=CBLN1;refGene_tx=XM_023245547.1
E2 29539347 29539497
refGene_gn=CE2H16orf78;refGene_tx=XM_003998026.4;refGene_gn=CE2H16orf7
8;refGene_tx=XM_023244928.1;refGene_gn=CE2H16orf78;refGene_tx=XM_02324
4929.1;refGene_gn=CE2H16orf78;refGene_tx=XM_023244930.1;refGene_gn=CE2
H16orf78;refGene_tx=XM_023244931.1;refGene_gn=CE2H16orf78;refGene_tx=X
M_023244932.1;refGene_gn=CE2H16orf78;refGene_tx=XM_023244933.1
E2 29542012 29542147
refGene_gn=CE2H16orf78;refGene_tx=XM_019819709.2
E2 29542462 29542570
refGene_gn=CE2H16orf78;refGene_tx=XM_003998026.4;refGene_gn=CE2H16orf7
8;refGene_tx=XM_019819709.2;refGene_gn=CE2H16orf78;refGene_tx=XM_02324
4928.1;refGene_gn=CE2H16orf78;refGene_tx=XM_023244929.1;refGene_gn=CE2
H16orf78;refGene_tx=XM_023244930.1;refGene_gn=CE2H16orf78;refGene_tx=X
M_023244931.1;refGene_gn=CE2H16orf78;refGene_tx=XM_023244932.1;refGene
_gn=CE2H16orf78;refGene_tx=XM_023244933.1
E2 29543116 29543231
refGene_gn=CE2H16orf78;refGene_tx=XM_003998026.4;refGene_gn=CE2H16orf7
8;refGene_tx=XM_019819709.2;refGene_gn=CE2H16orf78;refGene_tx=XM_02324
4928.1;refGene_gn=CE2H16orf78;refGene_tx=XM_023244929.1;refGene_gn=CE2
H16orf78;refGene_tx=XM_023244930.1;refGene_gn=CE2H16orf78;refGene_tx=X
M_023244931.1;refGene_gn=CE2H16orf78;refGene_tx=XM_023244932.1;refGene
_gn=CE2H16orf78;refGene_tx=XM_023244933.1
E2 29550901 29551157
refGene_gn=CE2H16orf78;refGene_tx=XM_003998026.4;refGene_gn=CE2H16orf7
8;refGene_tx=XM_019819709.2;refGene_gn=CE2H16orf78;refGene_tx=XM_02324
4928.1;refGene_gn=CE2H16orf78;refGene_tx=XM_023244929.1;refGene_gn=CE2
H16orf78;refGene_tx=XM_023244930.1;refGene_gn=CE2H16orf78;refGene_tx=X
M_023244931.1;refGene_gn=CE2H16orf78;refGene_tx=XM_023244932.1;refGene

_gn=CE2H16orf78;refGene_tx=XM_023244933.1
E2 29551264 29551322
refGene_gn=CE2H16orf78;refGene_tx=XM_023244930.1
E2 29551467 29551480
refGene_gn=CE2H16orf78;refGene_tx=XM_023244933.1
E2 29551665 29551729
refGene_gn=CE2H16orf78;refGene_tx=XM_023244929.1
E2 29552693 29552709
refGene_gn=CE2H16orf78;refGene_tx=XM_023244932.1
E2 29553281 29553429
refGene_gn=CE2H16orf78;refGene_tx=XM_003998026.4;refGene_gn=CE2H16orf78;refGene_tx=XM_019819709.2;refGene_gn=CE2H16orf78;refGene_tx=XM_023244928.1
E2 29554859 29554919
refGene_gn=CE2H16orf78;refGene_tx=XM_003998026.4;refGene_gn=CE2H16orf78;refGene_tx=XM_019819709.2;refGene_gn=CE2H16orf78;refGene_tx=XM_023244931.1
E2 29642300 29642330
refGene_gn=ZNF423;refGene_tx=XM_003998016.4;refGene_gn=ZNF423;refGene_tx=XM_023245564.1
E2 29670212 29670328
refGene_gn=ZNF423;refGene_tx=XM_003998016.4;refGene_gn=ZNF423;refGene_tx=XM_023245564.1
E2 29671974 29672106
refGene_gn=ZNF423;refGene_tx=XM_003998016.4;refGene_gn=ZNF423;refGene_tx=XM_023245564.1
E2 29745972 29746028 refGene_gn=ZNF423;refGene_tx=XM_023245565.1
E2 29772000 29772085
refGene_gn=ZNF423;refGene_tx=XM_003998016.4;refGene_gn=ZNF423;refGene_tx=XM_023245564.1;refGene_gn=ZNF423;refGene_tx=XM_023245565.1
E2 29779071 29782212
refGene_gn=ZNF423;refGene_tx=XM_003998016.4;refGene_gn=ZNF423;refGene_tx=XM_023245564.1;refGene_gn=ZNF423;refGene_tx=XM_023245565.1
E2 30059994 30060050
refGene_gn=LOC102902033;refGene_tx=XM_023245306.1
E2 30063395 30063542
refGene_gn=LOC102902033;refGene_tx=XM_019819721.2;refGene_gn=LOC102902033;refGene_tx=XM_019819722.2;refGene_gn=LOC102902033;refGene_tx=XM_019819723.2;refGene_gn=LOC102902033;refGene_tx=XM_023245304.1;refGene_gn=LOC102902033;refGene_tx=XM_023245306.1
E2 30065948 30066143
refGene_gn=LOC102902033;refGene_tx=XM_019819721.2;refGene_gn=LOC102902033;refGene_tx=XM_019819722.2;refGene_gn=LOC102902033;refGene_tx=XM_019819723.2;refGene_gn=LOC102902033;refGene_tx=XM_023245304.1;refGene_gn=LOC102902033;refGene_tx=XM_023245306.1
E2 30067301 30067577
refGene_gn=LOC102902033;refGene_tx=XM_019819721.2;refGene_gn=LOC102902033;refGene_tx=XM_019819722.2;refGene_gn=LOC102902033;refGene_tx=XM_019819723.2;refGene_gn=LOC102902033;refGene_tx=XM_023245304.1;refGene_gn=LOC102902033;refGene_tx=XM_023245306.1

E2 30070491 30070691
refGene_gn=LOC102902033;refGene_tx=XM_019819721.2;refGene_gn=LOC102902033;refGene_tx=XM_019819722.2;refGene_gn=LOC102902033;refGene_tx=XM_019819723.2;refGene_gn=LOC102902033;refGene_tx=XM_023245304.1;refGene_gn=LOC102902033;refGene_tx=XM_023245306.1
E2 30074593 30074720
refGene_gn=LOC102902033;refGene_tx=XM_019819721.2;refGene_gn=LOC102902033;refGene_tx=XM_019819722.2;refGene_gn=LOC102902033;refGene_tx=XM_019819723.2;refGene_gn=LOC102902033;refGene_tx=XM_023245304.1;refGene_gn=LOC102902033;refGene_tx=XM_023245306.1
E2 30074870 30074953
refGene_gn=LOC102902033;refGene_tx=XM_019819721.2;refGene_gn=LOC102902033;refGene_tx=XM_019819722.2;refGene_gn=LOC102902033;refGene_tx=XM_019819723.2;refGene_gn=LOC102902033;refGene_tx=XM_023245304.1;refGene_gn=LOC102902033;refGene_tx=XM_023245306.1
E2 30077484 30077565
refGene_gn=LOC102902033;refGene_tx=XM_019819721.2;refGene_gn=LOC102902033;refGene_tx=XM_019819722.2;refGene_gn=LOC102902033;refGene_tx=XM_019819723.2;refGene_gn=LOC102902033;refGene_tx=XM_023245304.1;refGene_gn=LOC102902033;refGene_tx=XM_023245306.1
E2 30097392 30097451
refGene_gn=LOC102902033;refGene_tx=XM_019819721.2;refGene_gn=LOC102902033;refGene_tx=XM_019819722.2;refGene_gn=LOC102902033;refGene_tx=XM_019819723.2;refGene_gn=LOC102902033;refGene_tx=XM_023245304.1;refGene_gn=LOC102902033;refGene_tx=XM_023245306.1
E2 30150247 30150272 refGene_gn=CNEP1R1;refGene_tx=XM_006941496.3
E2 30151367 30151439 refGene_gn=CNEP1R1;refGene_tx=XM_006941496.3
E2 30154393 30154467 refGene_gn=CNEP1R1;refGene_tx=XM_006941496.3
E2 30157498 30157608 refGene_gn=CNEP1R1;refGene_tx=XM_006941496.3
E2 30157724 30157779 refGene_gn=CNEP1R1;refGene_tx=XM_006941496.3
E2 30164070 30164112 refGene_gn=CNEP1R1;refGene_tx=XM_006941496.3
E2 30184568 30184700
refGene_gn=HEATR3;refGene_tx=XM_023244839.1;refGene_gn=HEATR3;refGene_tx=XM_023244840.1;refGene_gn=HEATR3;refGene_tx=XM_023244841.1;refGene_gn=HEATR3;refGene_tx=XM_023244842.1;refGene_gn=HEATR3;refGene_tx=XM_023244843.1;refGene_gn=HEATR3;refGene_tx=XM_023244844.1
E2 30184831 30185004
refGene_gn=HEATR3;refGene_tx=XM_023244839.1;refGene_gn=HEATR3;refGene_tx=XM_023244840.1;refGene_gn=HEATR3;refGene_tx=XM_023244841.1;refGene_gn=HEATR3;refGene_tx=XM_023244842.1;refGene_gn=HEATR3;refGene_tx=XM_023244843.1;refGene_gn=HEATR3;refGene_tx=XM_023244844.1
E2 30186973 30187061
refGene_gn=HEATR3;refGene_tx=XM_023244839.1;refGene_gn=HEATR3;refGene_tx=XM_023244840.1;refGene_gn=HEATR3;refGene_tx=XM_023244841.1;refGene_gn=HEATR3;refGene_tx=XM_023244842.1;refGene_gn=HEATR3;refGene_tx=XM_023244843.1;refGene_gn=HEATR3;refGene_tx=XM_023244844.1
E2 30188429 30188542
refGene_gn=HEATR3;refGene_tx=XM_023244839.1;refGene_gn=HEATR3;refGene_tx=XM_023244840.1;refGene_gn=HEATR3;refGene_tx=XM_023244841.1;refGene_gn=HEATR3;refGene_tx=XM_023244842.1;refGene_gn=HEATR3;refGene_tx=XM_02

3244843.1;refGene_gn=HEATR3;refGene_tx=XM_023244844.1;refGene_gn=HEATR3;refGene_tx=XM_023244845.1
E2 30190554 30190664
refGene_gn=HEATR3;refGene_tx=XM_023244839.1;refGene_gn=HEATR3;refGene_tx=XM_023244840.1;refGene_gn=HEATR3;refGene_tx=XM_023244841.1;refGene_gn=HEATR3;refGene_tx=XM_023244842.1;refGene_gn=HEATR3;refGene_tx=XM_023244843.1;refGene_gn=HEATR3;refGene_tx=XM_023244844.1
E2 30193365 30193506
refGene_gn=HEATR3;refGene_tx=XM_023244839.1;refGene_gn=HEATR3;refGene_tx=XM_023244840.1;refGene_gn=HEATR3;refGene_tx=XM_023244841.1;refGene_gn=HEATR3;refGene_tx=XM_023244842.1;refGene_gn=HEATR3;refGene_tx=XM_023244843.1;refGene_gn=HEATR3;refGene_tx=XM_023244844.1;refGene_gn=HEATR3;refGene_tx=XM_023244845.1
E2 30195234 30195512
refGene_gn=HEATR3;refGene_tx=XM_023244839.1;refGene_gn=HEATR3;refGene_tx=XM_023244840.1;refGene_gn=HEATR3;refGene_tx=XM_023244841.1;refGene_gn=HEATR3;refGene_tx=XM_023244842.1;refGene_gn=HEATR3;refGene_tx=XM_023244843.1;refGene_gn=HEATR3;refGene_tx=XM_023244844.1;refGene_gn=HEATR3;refGene_tx=XM_023244845.1
E2 30200058 30200149
refGene_gn=HEATR3;refGene_tx=XM_023244839.1;refGene_gn=HEATR3;refGene_tx=XM_023244840.1;refGene_gn=HEATR3;refGene_tx=XM_023244841.1;refGene_gn=HEATR3;refGene_tx=XM_023244842.1;refGene_gn=HEATR3;refGene_tx=XM_023244843.1;refGene_gn=HEATR3;refGene_tx=XM_023244844.1;refGene_gn=HEATR3;refGene_tx=XM_023244845.1
E2 30200256 30200414
refGene_gn=HEATR3;refGene_tx=XM_023244839.1;refGene_gn=HEATR3;refGene_tx=XM_023244840.1;refGene_gn=HEATR3;refGene_tx=XM_023244841.1;refGene_gn=HEATR3;refGene_tx=XM_023244842.1;refGene_gn=HEATR3;refGene_tx=XM_023244843.1;refGene_gn=HEATR3;refGene_tx=XM_023244844.1;refGene_gn=HEATR3;refGene_tx=XM_023244845.1
E2 30200681 30200764
refGene_gn=HEATR3;refGene_tx=XM_023244839.1;refGene_gn=HEATR3;refGene_tx=XM_023244840.1;refGene_gn=HEATR3;refGene_tx=XM_023244841.1;refGene_gn=HEATR3;refGene_tx=XM_023244842.1;refGene_gn=HEATR3;refGene_tx=XM_023244843.1;refGene_gn=HEATR3;refGene_tx=XM_023244844.1;refGene_gn=HEATR3;refGene_tx=XM_023244845.1
E2 30201688 30201825
refGene_gn=HEATR3;refGene_tx=XM_023244839.1;refGene_gn=HEATR3;refGene_tx=XM_023244840.1;refGene_gn=HEATR3;refGene_tx=XM_023244841.1;refGene_gn=HEATR3;refGene_tx=XM_023244842.1;refGene_gn=HEATR3;refGene_tx=XM_023244843.1;refGene_gn=HEATR3;refGene_tx=XM_023244844.1;refGene_gn=HEATR3;refGene_tx=XM_023244845.1
E2 30208340 30208429
refGene_gn=HEATR3;refGene_tx=XM_023244839.1;refGene_gn=HEATR3;refGene_tx=XM_023244840.1;refGene_gn=HEATR3;refGene_tx=XM_023244841.1;refGene_gn=HEATR3;refGene_tx=XM_023244842.1;refGene_gn=HEATR3;refGene_tx=XM_023244843.1;refGene_gn=HEATR3;refGene_tx=XM_023244844.1;refGene_gn=HEATR3;refGene_tx=XM_023244845.1
E2 30214949 30215093

refGene_gn=HEATR3;refGene_tx=XM_023244839.1;refGene_gn=HEATR3;refGene_tx=XM_023244841.1;refGene_gn=HEATR3;refGene_tx=XM_023244842.1;refGene_gn=HEATR3;refGene_tx=XM_023244843.1;refGene_gn=HEATR3;refGene_tx=XM_023244844.1;refGene_gn=HEATR3;refGene_tx=XM_023244845.1
E2 30218353 30218530
refGene_gn=HEATR3;refGene_tx=XM_023244839.1;refGene_gn=HEATR3;refGene_tx=XM_023244840.1;refGene_gn=HEATR3;refGene_tx=XM_023244841.1;refGene_gn=HEATR3;refGene_tx=XM_023244842.1;refGene_gn=HEATR3;refGene_tx=XM_023244843.1;refGene_gn=HEATR3;refGene_tx=XM_023244844.1;refGene_gn=HEATR3;refGene_tx=XM_023244845.1
E2 30223241 30223364 refGene_gn=HEATR3;refGene_tx=XM_023244842.1
E2 30227778 30227906
refGene_gn=HEATR3;refGene_tx=XM_023244839.1;refGene_gn=HEATR3;refGene_tx=XM_023244840.1;refGene_gn=HEATR3;refGene_tx=XM_023244841.1;refGene_gn=HEATR3;refGene_tx=XM_023244845.1
E2 30238795 30238886
refGene_gn=HEATR3;refGene_tx=XM_023244839.1;refGene_gn=HEATR3;refGene_tx=XM_023244840.1;refGene_gn=HEATR3;refGene_tx=XM_023244841.1;refGene_gn=HEATR3;refGene_tx=XM_023244843.1;refGene_gn=HEATR3;refGene_tx=XM_023244845.1
E2 30239186 30239384
refGene_gn=HEATR3;refGene_tx=XM_023244839.1;refGene_gn=HEATR3;refGene_tx=XM_023244840.1;refGene_gn=HEATR3;refGene_tx=XM_023244845.1
E2 30252253 30252259
refGene_gn=HEATR3;refGene_tx=XM_023244841.1;refGene_gn=HEATR3;refGene_tx=XM_023244844.1
E2 30255788 30256423
refGene_gn=PAPD5;refGene_tx=XM_023244906.1;refGene_gn=PAPD5;refGene_tx=XM_023244907.1;refGene_gn=PAPD5;refGene_tx=XM_023244908.1
E2 30310763 30310887
refGene_gn=PAPD5;refGene_tx=XM_023244906.1;refGene_gn=PAPD5;refGene_tx=XM_023244907.1;refGene_gn=PAPD5;refGene_tx=XM_023244908.1
E2 30312651 30312698
refGene_gn=PAPD5;refGene_tx=XM_023244906.1;refGene_gn=PAPD5;refGene_tx=XM_023244907.1;refGene_gn=PAPD5;refGene_tx=XM_023244908.1
E2 30314177 30314298
refGene_gn=PAPD5;refGene_tx=XM_023244906.1;refGene_gn=PAPD5;refGene_tx=XM_023244907.1;refGene_gn=PAPD5;refGene_tx=XM_023244908.1
E2 30316192 30316300
refGene_gn=PAPD5;refGene_tx=XM_023244906.1;refGene_gn=PAPD5;refGene_tx=XM_023244907.1;refGene_gn=PAPD5;refGene_tx=XM_023244908.1
E2 30318637 30318766
refGene_gn=PAPD5;refGene_tx=XM_023244906.1;refGene_gn=PAPD5;refGene_tx=XM_023244907.1;refGene_gn=PAPD5;refGene_tx=XM_023244908.1
E2 30319519 30319733
refGene_gn=PAPD5;refGene_tx=XM_023244906.1;refGene_gn=PAPD5;refGene_tx=XM_023244907.1;refGene_gn=PAPD5;refGene_tx=XM_023244908.1
E2 30321387 30321514
refGene_gn=PAPD5;refGene_tx=XM_023244906.1;refGene_gn=PAPD5;refGene_tx=XM_023244907.1;refGene_gn=PAPD5;refGene_tx=XM_023244908.1

E2 30321613 30321717
refGene_gn=PAPD5;refGene_tx=XM_023244906.1;refGene_gn=PAPD5;refGene_tx=XM_023244907.1;refGene_gn=PAPD5;refGene_tx=XM_023244908.1
E2 30321820 30322008
refGene_gn=PAPD5;refGene_tx=XM_023244906.1;refGene_gn=PAPD5;refGene_tx=XM_023244907.1;refGene_gn=PAPD5;refGene_tx=XM_023244908.1
E2 30324030 30324195
refGene_gn=PAPD5;refGene_tx=XM_023244906.1;refGene_gn=PAPD5;refGene_tx=XM_023244907.1;refGene_gn=PAPD5;refGene_tx=XM_023244908.1
E2 30325640 30325817
refGene_gn=PAPD5;refGene_tx=XM_023244906.1;refGene_gn=PAPD5;refGene_tx=XM_023244907.1;refGene_gn=PAPD5;refGene_tx=XM_023244908.1
E2 30331356 30331409 refGene_gn=PAPD5;refGene_tx=XM_023244907.1
E2 30374365 30374536
refGene_gn=ADCY7;refGene_tx=XM_019819728.2;refGene_gn=ADCY7;refGene_tx=XM_019819729.2
E2 30376318 30376522
refGene_gn=ADCY7;refGene_tx=XM_019819728.2;refGene_gn=ADCY7;refGene_tx=XM_019819729.2;refGene_gn=ADCY7;refGene_tx=XM_019819730.2
E2 30377393 30377560
refGene_gn=ADCY7;refGene_tx=XM_019819728.2;refGene_gn=ADCY7;refGene_tx=XM_019819729.2;refGene_gn=ADCY7;refGene_tx=XM_019819730.2
E2 30378292 30378442
refGene_gn=ADCY7;refGene_tx=XM_019819728.2;refGene_gn=ADCY7;refGene_tx=XM_019819729.2;refGene_gn=ADCY7;refGene_tx=XM_019819730.2
E2 30378763 30378912
refGene_gn=ADCY7;refGene_tx=XM_019819728.2;refGene_gn=ADCY7;refGene_tx=XM_019819729.2;refGene_gn=ADCY7;refGene_tx=XM_019819730.2
E2 30379711 30379823
refGene_gn=ADCY7;refGene_tx=XM_019819728.2;refGene_gn=ADCY7;refGene_tx=XM_019819729.2;refGene_gn=ADCY7;refGene_tx=XM_019819730.2
E2 30382767 30382895
refGene_gn=ADCY7;refGene_tx=XM_019819728.2;refGene_gn=ADCY7;refGene_tx=XM_019819729.2;refGene_gn=ADCY7;refGene_tx=XM_019819730.2
E2 30383065 30383224
refGene_gn=ADCY7;refGene_tx=XM_019819728.2;refGene_gn=ADCY7;refGene_tx=XM_019819729.2;refGene_gn=ADCY7;refGene_tx=XM_019819730.2
E2 30383439 30383572
refGene_gn=ADCY7;refGene_tx=XM_019819728.2;refGene_gn=ADCY7;refGene_tx=XM_019819729.2;refGene_gn=ADCY7;refGene_tx=XM_019819730.2
E2 30387036 30387228
refGene_gn=ADCY7;refGene_tx=XM_019819728.2;refGene_gn=ADCY7;refGene_tx=XM_019819729.2;refGene_gn=ADCY7;refGene_tx=XM_019819730.2
E2 30387362 30387397
refGene_gn=ADCY7;refGene_tx=XM_019819728.2;refGene_gn=ADCY7;refGene_tx=XM_019819729.2;refGene_gn=ADCY7;refGene_tx=XM_019819730.2
E2 30387898 30387982
refGene_gn=ADCY7;refGene_tx=XM_019819728.2;refGene_gn=ADCY7;refGene_tx=XM_019819729.2;refGene_gn=ADCY7;refGene_tx=XM_019819730.2
E2 30388208 30388281

refGene_gn=ADCY7;refGene_tx=XM_019819728.2;refGene_gn=ADCY7;refGene_tx=XM_019819729.2;refGene_gn=ADCY7;refGene_tx=XM_019819730.2
E2 30389639 30389737
refGene_gn=ADCY7;refGene_tx=XM_019819728.2;refGene_gn=ADCY7;refGene_tx=XM_019819729.2;refGene_gn=ADCY7;refGene_tx=XM_019819730.2
E2 30390451 30390536
refGene_gn=ADCY7;refGene_tx=XM_019819728.2;refGene_gn=ADCY7;refGene_tx=XM_019819729.2;refGene_gn=ADCY7;refGene_tx=XM_019819730.2
E2 30390921 30391047
refGene_gn=ADCY7;refGene_tx=XM_019819728.2;refGene_gn=ADCY7;refGene_tx=XM_019819729.2;refGene_gn=ADCY7;refGene_tx=XM_019819730.2
E2 30392002 30392104
refGene_gn=ADCY7;refGene_tx=XM_019819728.2;refGene_gn=ADCY7;refGene_tx=XM_019819729.2;refGene_gn=ADCY7;refGene_tx=XM_019819730.2
E2 30392727 30392921
refGene_gn=ADCY7;refGene_tx=XM_019819728.2;refGene_gn=ADCY7;refGene_tx=XM_019819729.2;refGene_gn=ADCY7;refGene_tx=XM_019819730.2
E2 30393766 30393860
refGene_gn=ADCY7;refGene_tx=XM_019819728.2;refGene_gn=ADCY7;refGene_tx=XM_019819729.2;refGene_gn=ADCY7;refGene_tx=XM_019819730.2
E2 30394269 30394425
refGene_gn=ADCY7;refGene_tx=XM_019819728.2;refGene_gn=ADCY7;refGene_tx=XM_019819729.2;refGene_gn=ADCY7;refGene_tx=XM_019819730.2
E2 30394982 30395129
refGene_gn=ADCY7;refGene_tx=XM_019819728.2;refGene_gn=ADCY7;refGene_tx=XM_019819729.2;refGene_gn=ADCY7;refGene_tx=XM_019819730.2
E2 30396072 30396177
refGene_gn=ADCY7;refGene_tx=XM_019819728.2;refGene_gn=ADCY7;refGene_tx=XM_019819729.2;refGene_gn=ADCY7;refGene_tx=XM_019819730.2
E2 30396423 30396538
refGene_gn=ADCY7;refGene_tx=XM_019819728.2;refGene_gn=ADCY7;refGene_tx=XM_019819729.2;refGene_gn=ADCY7;refGene_tx=XM_019819730.2
E2 30397153 30397278
refGene_gn=ADCY7;refGene_tx=XM_019819728.2;refGene_gn=ADCY7;refGene_tx=XM_019819729.2;refGene_gn=ADCY7;refGene_tx=XM_019819730.2
E2 30397484 30397631
refGene_gn=ADCY7;refGene_tx=XM_019819728.2;refGene_gn=ADCY7;refGene_tx=XM_019819729.2;refGene_gn=ADCY7;refGene_tx=XM_019819730.2
E2 30401451 30401507
refGene_gn=BRD7;refGene_tx=XM_023245808.1;refGene_gn=BRD7;refGene_tx=XM_023245809.1;refGene_gn=BRD7;refGene_tx=XM_023245810.1
E2 30401823 30401967
refGene_gn=BRD7;refGene_tx=XM_023245808.1;refGene_gn=BRD7;refGene_tx=XM_023245809.1;refGene_gn=BRD7;refGene_tx=XM_023245810.1
E2 30402501 30402645
refGene_gn=BRD7;refGene_tx=XM_023245808.1;refGene_gn=BRD7;refGene_tx=XM_023245809.1;refGene_gn=BRD7;refGene_tx=XM_023245810.1
E2 30402925 30403037
refGene_gn=BRD7;refGene_tx=XM_023245808.1;refGene_gn=BRD7;refGene_tx=XM_023245809.1;refGene_gn=BRD7;refGene_tx=XM_023245810.1

E2 30405863 30405920
refGene_gn=BRD7;refGene_tx=XM_023245808.1;refGene_gn=BRD7;refGene_tx=XM_023245809.1;refGene_gn=BRD7;refGene_tx=XM_023245810.1
E2 30408073 30408209
refGene_gn=BRD7;refGene_tx=XM_023245808.1;refGene_gn=BRD7;refGene_tx=XM_023245809.1;refGene_gn=BRD7;refGene_tx=XM_023245810.1
E2 30408673 30408809
refGene_gn=BRD7;refGene_tx=XM_023245808.1;refGene_gn=BRD7;refGene_tx=XM_023245809.1;refGene_gn=BRD7;refGene_tx=XM_023245810.1
E2 30409166 30409274
refGene_gn=BRD7;refGene_tx=XM_023245808.1;refGene_gn=BRD7;refGene_tx=XM_023245809.1;refGene_gn=BRD7;refGene_tx=XM_023245810.1
E2 30411892 30411968
refGene_gn=BRD7;refGene_tx=XM_023245808.1;refGene_gn=BRD7;refGene_tx=XM_023245809.1;refGene_gn=BRD7;refGene_tx=XM_023245810.1
E2 30412588 30412712
refGene_gn=BRD7;refGene_tx=XM_023245808.1;refGene_gn=BRD7;refGene_tx=XM_023245809.1;refGene_gn=BRD7;refGene_tx=XM_023245810.1
E2 30413688 30413873
refGene_gn=BRD7;refGene_tx=XM_023245808.1;refGene_gn=BRD7;refGene_tx=XM_023245809.1;refGene_gn=BRD7;refGene_tx=XM_023245810.1
E2 30418682 30418793
refGene_gn=BRD7;refGene_tx=XM_023245808.1;refGene_gn=BRD7;refGene_tx=XM_023245809.1;refGene_gn=BRD7;refGene_tx=XM_023245810.1
E2 30422549 30422694
refGene_gn=BRD7;refGene_tx=XM_023245808.1;refGene_gn=BRD7;refGene_tx=XM_023245809.1;refGene_gn=BRD7;refGene_tx=XM_023245810.1
E2 30425310 30425368
refGene_gn=BRD7;refGene_tx=XM_023245808.1;refGene_gn=BRD7;refGene_tx=XM_023245809.1;refGene_gn=BRD7;refGene_tx=XM_023245810.1
E2 30425715 30425845
refGene_gn=BRD7;refGene_tx=XM_023245808.1;refGene_gn=BRD7;refGene_tx=XM_023245809.1;refGene_gn=BRD7;refGene_tx=XM_023245810.1
E2 30441939 30442148
refGene_gn=BRD7;refGene_tx=XM_023245808.1;refGene_gn=BRD7;refGene_tx=XM_023245809.1;refGene_gn=BRD7;refGene_tx=XM_023245810.1
E2 30442634 30442683
refGene_gn=BRD7;refGene_tx=XM_023245808.1;refGene_gn=BRD7;refGene_tx=XM_023245809.1;refGene_gn=BRD7;refGene_tx=XM_023245810.1
E2 30605283 30605308 refGene_gn=NKD1;refGene_tx=XM_023244859.1
E2 30605439 30605472 refGene_gn=NKD1;refGene_tx=XM_023244859.1
E2 30606191 30606325 refGene_gn=NKD1;refGene_tx=XM_023244859.1
E2 30608751 30608754 refGene_gn=NKD1;refGene_tx=XM_023244860.1
E2 30659706 30659773
refGene_gn=NKD1;refGene_tx=XM_023244859.1;refGene_gn=NKD1;refGene_tx=XM_023244860.1
E2 30673557 30673664
refGene_gn=NKD1;refGene_tx=XM_023244859.1;refGene_gn=NKD1;refGene_tx=XM_023244860.1
E2 30676738 30676834

refGene_gn=NKD1;refGene_tx=XM_023244859.1;refGene_gn=NKD1;refGene_tx=XM_023244860.1
E2	30681989	30682137
refGene_gn=NKD1;refGene_tx=XM_023244859.1;refGene_gn=NKD1;refGene_tx=XM_023244860.1
E2	30682671	30682756
refGene_gn=NKD1;refGene_tx=XM_023244859.1;refGene_gn=NKD1;refGene_tx=XM_023244860.1
E2	30684133	30684261
refGene_gn=NKD1;refGene_tx=XM_023244859.1;refGene_gn=NKD1;refGene_tx=XM_023244860.1
E2	30684996	30685586
refGene_gn=NKD1;refGene_tx=XM_023244859.1;refGene_gn=NKD1;refGene_tx=XM_023244860.1
E2	30699810	30700002
refGene_gn=LOC109494662;refGene_tx=XM_023245117.1
E2	30701845	30702297
refGene_gn=LOC109494662;refGene_tx=XM_023245117.1
E2	30702821	30702934
refGene_gn=LOC109494662;refGene_tx=XM_023245117.1
E2	30712597	30712644
refGene_gn=LOC109494662;refGene_tx=XM_023245117.1
E2	30718813	30719482
refGene_gn=SNX20;refGene_tx=XM_006941504.4;refGene_gn=SNX20;refGene_tx=XM_006941505.4;refGene_gn=SNX20;refGene_tx=XM_019819735.2;refGene_gn=SNX20;refGene_tx=XM_019819736.2;refGene_gn=SNX20;refGene_tx=XM_023245371.1
E2	30721796	30721948
refGene_gn=SNX20;refGene_tx=XM_006941504.4;refGene_gn=SNX20;refGene_tx=XM_006941505.4;refGene_gn=SNX20;refGene_tx=XM_019819735.2;refGene_gn=SNX20;refGene_tx=XM_019819736.2;refGene_gn=SNX20;refGene_tx=XM_023245371.1
E2	30722512	30722593
refGene_gn=SNX20;refGene_tx=XM_006941504.4;refGene_gn=SNX20;refGene_tx=XM_006941505.4
E2	30723385	30723515
refGene_gn=SNX20;refGene_tx=XM_006941504.4;refGene_gn=SNX20;refGene_tx=XM_006941505.4;refGene_gn=SNX20;refGene_tx=XM_023245371.1
E2	30741896	30741906	refGene_gn=NOD2;refGene_tx=XM_019819738.2
E2	30743229	30743696
refGene_gn=NOD2;refGene_tx=XM_019819738.2;refGene_gn=NOD2;refGene_tx=XM_023245187.1
E2	30749482	30749588
refGene_gn=NOD2;refGene_tx=XM_019819738.2;refGene_gn=NOD2;refGene_tx=XM_023245187.1
E2	30752666	30754485
refGene_gn=NOD2;refGene_tx=XM_019819738.2;refGene_gn=NOD2;refGene_tx=XM_023245187.1
E2	30758652	30758736
refGene_gn=NOD2;refGene_tx=XM_019819738.2;refGene_gn=NOD2;refGene_tx=X

M_023245187.1
E2 30758946 30759030
refGene_gn=NOD2;refGene_tx=XM_019819738.2;refGene_gn=NOD2;refGene_tx=XM_023245187.1
E2 30761524 30761608
refGene_gn=NOD2;refGene_tx=XM_019819738.2;refGene_gn=NOD2;refGene_tx=XM_023245187.1
E2 30763468 30763552
refGene_gn=NOD2;refGene_tx=XM_019819738.2;refGene_gn=NOD2;refGene_tx=XM_023245187.1
E2 30764298 30764382
refGene_gn=NOD2;refGene_tx=XM_019819738.2;refGene_gn=NOD2;refGene_tx=XM_023245187.1
E2 30766344 30766428
refGene_gn=NOD2;refGene_tx=XM_019819738.2;refGene_gn=NOD2;refGene_tx=XM_023245187.1
E2 30769359 30769443
refGene_gn=NOD2;refGene_tx=XM_019819738.2;refGene_gn=NOD2;refGene_tx=XM_023245187.1
E2 30771378 30771451
refGene_gn=NOD2;refGene_tx=XM_019819738.2;refGene_gn=NOD2;refGene_tx=XM_023245187.1
E2 30786961 30787465
refGene_gn=CYLD;refGene_tx=XM_023244890.1;refGene_gn=CYLD;refGene_tx=XM_023244891.1;refGene_gn=CYLD;refGene_tx=XM_023244892.1;refGene_gn=CYLD;refGene_tx=XM_023244893.1;refGene_gn=CYLD;refGene_tx=XM_023244894.1;refGene_gn=CYLD;refGene_tx=XM_023244895.1
E2 30788691 30788994
refGene_gn=CYLD;refGene_tx=XM_023244890.1;refGene_gn=CYLD;refGene_tx=XM_023244891.1;refGene_gn=CYLD;refGene_tx=XM_023244892.1;refGene_gn=CYLD;refGene_tx=XM_023244893.1;refGene_gn=CYLD;refGene_tx=XM_023244894.1;refGene_gn=CYLD;refGene_tx=XM_023244895.1
E2 30790182 30790288
refGene_gn=CYLD;refGene_tx=XM_023244890.1;refGene_gn=CYLD;refGene_tx=XM_023244891.1;refGene_gn=CYLD;refGene_tx=XM_023244892.1;refGene_gn=CYLD;refGene_tx=XM_023244893.1;refGene_gn=CYLD;refGene_tx=XM_023244894.1;refGene_gn=CYLD;refGene_tx=XM_023244895.1
E2 30816341 30816350
refGene_gn=CYLD;refGene_tx=XM_023244890.1;refGene_gn=CYLD;refGene_tx=XM_023244891.1;refGene_gn=CYLD;refGene_tx=XM_023244892.1;refGene_gn=CYLD;refGene_tx=XM_023244893.1
E2 30817322 30817421
refGene_gn=CYLD;refGene_tx=XM_023244890.1;refGene_gn=CYLD;refGene_tx=XM_023244891.1;refGene_gn=CYLD;refGene_tx=XM_023244892.1;refGene_gn=CYLD;refGene_tx=XM_023244893.1;refGene_gn=CYLD;refGene_tx=XM_023244894.1
E2 30819002 30819119
refGene_gn=CYLD;refGene_tx=XM_023244890.1;refGene_gn=CYLD;refGene_tx=XM_023244891.1;refGene_gn=CYLD;refGene_tx=XM_023244892.1;refGene_gn=CYLD;refGene_tx=XM_023244893.1;refGene_gn=CYLD;refGene_tx=XM_023244894.1
E2 30826475 30826855

refGene_gn=CYLD;refGene_tx=XM_023244890.1;refGene_gn=CYLD;refGene_tx=XM_023244891.1;refGene_gn=CYLD;refGene_tx=XM_023244892.1;refGene_gn=CYLD;refGene_tx=XM_023244893.1;refGene_gn=CYLD;refGene_tx=XM_023244894.1;refGene_gn=CYLD;refGene_tx=XM_023244895.1
E2 30827745 30827911
refGene_gn=CYLD;refGene_tx=XM_023244890.1;refGene_gn=CYLD;refGene_tx=XM_023244891.1;refGene_gn=CYLD;refGene_tx=XM_023244892.1;refGene_gn=CYLD;refGene_tx=XM_023244893.1;refGene_gn=CYLD;refGene_tx=XM_023244894.1;refGene_gn=CYLD;refGene_tx=XM_023244895.1
E2 30829278 30829420
refGene_gn=CYLD;refGene_tx=XM_023244890.1;refGene_gn=CYLD;refGene_tx=XM_023244891.1;refGene_gn=CYLD;refGene_tx=XM_023244892.1;refGene_gn=CYLD;refGene_tx=XM_023244893.1;refGene_gn=CYLD;refGene_tx=XM_023244894.1;refGene_gn=CYLD;refGene_tx=XM_023244895.1
E2 30830684 30830807
refGene_gn=CYLD;refGene_tx=XM_023244890.1;refGene_gn=CYLD;refGene_tx=XM_023244891.1;refGene_gn=CYLD;refGene_tx=XM_023244892.1;refGene_gn=CYLD;refGene_tx=XM_023244893.1;refGene_gn=CYLD;refGene_tx=XM_023244894.1;refGene_gn=CYLD;refGene_tx=XM_023244895.1
E2 30833539 30833631
refGene_gn=CYLD;refGene_tx=XM_023244890.1;refGene_gn=CYLD;refGene_tx=XM_023244891.1;refGene_gn=CYLD;refGene_tx=XM_023244892.1;refGene_gn=CYLD;refGene_tx=XM_023244893.1;refGene_gn=CYLD;refGene_tx=XM_023244894.1;refGene_gn=CYLD;refGene_tx=XM_023244895.1
E2 30834510 30834577
refGene_gn=CYLD;refGene_tx=XM_023244890.1;refGene_gn=CYLD;refGene_tx=XM_023244891.1;refGene_gn=CYLD;refGene_tx=XM_023244892.1;refGene_gn=CYLD;refGene_tx=XM_023244893.1;refGene_gn=CYLD;refGene_tx=XM_023244894.1;refGene_gn=CYLD;refGene_tx=XM_023244895.1
E2 30839561 30839694
refGene_gn=CYLD;refGene_tx=XM_023244890.1;refGene_gn=CYLD;refGene_tx=XM_023244891.1;refGene_gn=CYLD;refGene_tx=XM_023244892.1;refGene_gn=CYLD;refGene_tx=XM_023244893.1;refGene_gn=CYLD;refGene_tx=XM_023244894.1;refGene_gn=CYLD;refGene_tx=XM_023244895.1
E2 30840964 30841073
refGene_gn=CYLD;refGene_tx=XM_023244890.1;refGene_gn=CYLD;refGene_tx=XM_023244891.1;refGene_gn=CYLD;refGene_tx=XM_023244892.1;refGene_gn=CYLD;refGene_tx=XM_023244893.1;refGene_gn=CYLD;refGene_tx=XM_023244894.1;refGene_gn=CYLD;refGene_tx=XM_023244895.1
E2 30843234 30843353
refGene_gn=CYLD;refGene_tx=XM_023244890.1;refGene_gn=CYLD;refGene_tx=XM_023244891.1;refGene_gn=CYLD;refGene_tx=XM_023244892.1;refGene_gn=CYLD;refGene_tx=XM_023244893.1;refGene_gn=CYLD;refGene_tx=XM_023244894.1;refGene_gn=CYLD;refGene_tx=XM_023244895.1
E2 30843645 30843862
refGene_gn=CYLD;refGene_tx=XM_023244890.1;refGene_gn=CYLD;refGene_tx=XM_023244891.1;refGene_gn=CYLD;refGene_tx=XM_023244892.1;refGene_gn=CYLD;refGene_tx=XM_023244893.1;refGene_gn=CYLD;refGene_tx=XM_023244894.1;refGene_gn=CYLD;refGene_tx=XM_023244895.1
E2 30845376 30845561

refGene_gn=CYLD;refGene_tx=XM_023244890.1;refGene_gn=CYLD;refGene_tx=XM_023244891.1;refGene_gn=CYLD;refGene_tx=XM_023244892.1;refGene_gn=CYLD;refGene_tx=XM_023244893.1;refGene_gn=CYLD;refGene_tx=XM_023244894.1;refGene_gn=CYLD;refGene_tx=XM_023244895.1
E2 31158228 31158669 refGene_gn=SALL1;refGene_tx=XM_019819743.2;refGene_gn=SALL1;refGene_tx=XM_019819744.2;refGene_gn=SALL1;refGene_tx=XM_019819745.2;refGene_gn=SALL1;refGene_tx=XM_023245148.1
E2 31159805 31163239 refGene_gn=SALL1;refGene_tx=XM_019819743.2;refGene_gn=SALL1;refGene_tx=XM_019819744.2;refGene_gn=SALL1;refGene_tx=XM_019819745.2;refGene_gn=SALL1;refGene_tx=XM_023245148.1
E2 31171332 31171417 refGene_gn=SALL1;refGene_tx=XM_019819743.2
E2 31172251 31172327 refGene_gn=SALL1;refGene_tx=XM_019819744.2;refGene_gn=SALL1;refGene_tx=XM_023245148.1
E2 32375858 32375993 refGene_gn=TOX3;refGene_tx=XM_023244962.1
E2 32375994 32376564 refGene_gn=TOX3;refGene_tx=XM_023244962.1
E2 32380471 32380552 refGene_gn=TOX3;refGene_tx=XM_023244962.1
E2 32382424 32382652 refGene_gn=TOX3;refGene_tx=XM_023244962.1
E2 32386726 32386996 refGene_gn=TOX3;refGene_tx=XM_023244962.1
E2 32400508 32400763 refGene_gn=TOX3;refGene_tx=XM_023244962.1
E2 32404726 32404792 refGene_gn=TOX3;refGene_tx=XM_023244962.1
E2 32481730 32481817 refGene_gn=TOX3;refGene_tx=XM_023244962.1
E2 33010241 33011699 refGene_gn=CHD9;refGene_tx=XM_023245104.1
E2 33057448 33057780 refGene_gn=CHD9;refGene_tx=XM_023245104.1
E2 33071428 33071540 refGene_gn=CHD9;refGene_tx=XM_023245104.1
E2 33075203 33075350 refGene_gn=CHD9;refGene_tx=XM_023245104.1
E2 33076316 33076385 refGene_gn=CHD9;refGene_tx=XM_023245104.1
E2 33076463 33076519 refGene_gn=CHD9;refGene_tx=XM_023245104.1
E2 33077406 33077524 refGene_gn=CHD9;refGene_tx=XM_023245104.1
E2 33080804 33080891 refGene_gn=CHD9;refGene_tx=XM_023245104.1
E2 33081081 33081219 refGene_gn=CHD9;refGene_tx=XM_023245104.1
E2 33089664 33089786 refGene_gn=CHD9;refGene_tx=XM_023245104.1
E2 33093469 33093713 refGene_gn=CHD9;refGene_tx=XM_023245104.1
E2 33097816 33097993 refGene_gn=CHD9;refGene_tx=XM_023245104.1
E2 33099131 33099275 refGene_gn=CHD9;refGene_tx=XM_023245104.1
E2 33099394 33099650 refGene_gn=CHD9;refGene_tx=XM_023245104.1
E2 33101190 33101401 refGene_gn=CHD9;refGene_tx=XM_023245104.1
E2 33103453 33103649 refGene_gn=CHD9;refGene_tx=XM_023245104.1
E2 33108139 33108307 refGene_gn=CHD9;refGene_tx=XM_023245104.1
E2 33109014 33109194 refGene_gn=CHD9;refGene_tx=XM_023245104.1
E2 33115212 33115323 refGene_gn=CHD9;refGene_tx=XM_023245104.1
E2 33120190 33120387 refGene_gn=CHD9;refGene_tx=XM_023245104.1
E2 33120855 33121055 refGene_gn=CHD9;refGene_tx=XM_023245104.1
E2 33125675 33125806 refGene_gn=CHD9;refGene_tx=XM_023245104.1
E2 33125807 33125836 refGene_gn=CHD9;refGene_tx=XM_023245104.1
E2 33126271 33126361 refGene_gn=CHD9;refGene_tx=XM_023245104.1
E2 33131240 33131344 refGene_gn=CHD9;refGene_tx=XM_023245104.1
E2 33131821 33131939 refGene_gn=CHD9;refGene_tx=XM_023245104.1

E2	33132805	33132863	refGene_gn=CHD9;refGene_tx=XM_023245104.1
E2	33136366	33136409	refGene_gn=CHD9;refGene_tx=XM_023245104.1
E2	33137591	33137811	refGene_gn=CHD9;refGene_tx=XM_023245104.1
E2	33140339	33140542	refGene_gn=CHD9;refGene_tx=XM_023245104.1
E2	33143812	33144721	refGene_gn=CHD9;refGene_tx=XM_023245104.1
E2	33145721	33145882	refGene_gn=CHD9;refGene_tx=XM_023245104.1
E2	33146986	33147259	refGene_gn=CHD9;refGene_tx=XM_023245104.1
E2	33148000	33148169	refGene_gn=CHD9;refGene_tx=XM_023245104.1
E2	33154293	33154433	refGene_gn=CHD9;refGene_tx=XM_023245104.1
E2	33154713	33154935	refGene_gn=CHD9;refGene_tx=XM_023245104.1
E2	33158899	33159028	refGene_gn=CHD9;refGene_tx=XM_023245104.1
E2	33162025	33162130	refGene_gn=CHD9;refGene_tx=XM_023245104.1
E2	33164837	33165716	refGene_gn=CHD9;refGene_tx=XM_023245104.1
E2	33185766	33185904	refGene_gn=RBL2;refGene_tx=XM_023245701.1
E2	33185905	33186007	refGene_gn=RBL2;refGene_tx=XM_023245701.1
E2	33188206	33188337	refGene_gn=RBL2;refGene_tx=XM_023245701.1
E2	33190968	33191169	refGene_gn=RBL2;refGene_tx=XM_023245701.1
E2	33194342	33194407	refGene_gn=RBL2;refGene_tx=XM_023245701.1
E2	33198205	33198334	refGene_gn=RBL2;refGene_tx=XM_023245701.1
E2	33202699	33202860	refGene_gn=RBL2;refGene_tx=XM_023245701.1
E2	33202952	33203017	refGene_gn=RBL2;refGene_tx=XM_023245701.1
E2	33203652	33203839	refGene_gn=RBL2;refGene_tx=XM_023245701.1
E2	33207821	33207988	refGene_gn=RBL2;refGene_tx=XM_023245701.1
E2	33211509	33211619	refGene_gn=RBL2;refGene_tx=XM_023245701.1
E2	33212287	33212391	refGene_gn=RBL2;refGene_tx=XM_023245701.1
E2	33214862	33215000	refGene_gn=RBL2;refGene_tx=XM_023245701.1
E2	33216225	33216390	refGene_gn=RBL2;refGene_tx=XM_023245701.1
E2	33216908	33217020	refGene_gn=RBL2;refGene_tx=XM_023245701.1
E2	33219632	33219902	refGene_gn=RBL2;refGene_tx=XM_023245701.1
E2	33220087	33220368	refGene_gn=RBL2;refGene_tx=XM_023245701.1
E2	33220458	33220635	refGene_gn=RBL2;refGene_tx=XM_023245701.1
E2	33228509	33228581	refGene_gn=RBL2;refGene_tx=XM_023245701.1
E2	33229239	33229345	refGene_gn=RBL2;refGene_tx=XM_023245701.1
E2	33229945	33230151	refGene_gn=RBL2;refGene_tx=XM_023245701.1
E2	33231254	33231419	refGene_gn=RBL2;refGene_tx=XM_023245701.1
E2	33242071	33242242	refGene_gn=RBL2;refGene_tx=XM_023245701.1
E2	33244299	33244410
refGene_gn=AKTIP;refGene_tx=XM_006941509.4;refGene_gn=AKTIP;refGene_tx
=XM_006941510.4;refGene_gn=AKTIP;refGene_tx=XM_019819752.2;refGene_gn=
AKTIP;refGene_tx=XM_023245703.1
E2	33244573	33244634
refGene_gn=AKTIP;refGene_tx=XM_006941509.4;refGene_gn=AKTIP;refGene_tx
=XM_006941510.4;refGene_gn=AKTIP;refGene_tx=XM_019819752.2;refGene_gn=
AKTIP;refGene_tx=XM_023245703.1
E2	33245457	33245565
refGene_gn=AKTIP;refGene_tx=XM_006941509.4;refGene_gn=AKTIP;refGene_tx
=XM_006941510.4;refGene_gn=AKTIP;refGene_tx=XM_019819752.2;refGene_gn=
AKTIP;refGene_tx=XM_023245703.1
E2	33245657	33245756
refGene_gn=AKTIP;refGene_tx=XM_006941509.4;refGene_gn=AKTIP;refGene_tx

=XM_006941510.4;refGene_gn=AKTIP;refGene_tx=XM_019819752.2;refGene_gn=AKTIP;refGene_tx=XM_023245703.1
E2 33245833 33245922
refGene_gn=AKTIP;refGene_tx=XM_006941509.4;refGene_gn=AKTIP;refGene_tx=XM_006941510.4;refGene_gn=AKTIP;refGene_tx=XM_019819752.2;refGene_gn=AKTIP;refGene_tx=XM_023245703.1
E2 33246426 33246527
refGene_gn=AKTIP;refGene_tx=XM_006941509.4;refGene_gn=AKTIP;refGene_tx=XM_006941510.4;refGene_gn=AKTIP;refGene_tx=XM_019819752.2;refGene_gn=AKTIP;refGene_tx=XM_023245703.1
E2 33246614 33246679
refGene_gn=AKTIP;refGene_tx=XM_006941509.4;refGene_gn=AKTIP;refGene_tx=XM_006941510.4;refGene_gn=AKTIP;refGene_tx=XM_019819752.2;refGene_gn=AKTIP;refGene_tx=XM_023245703.1
E2 33248870 33249076
refGene_gn=AKTIP;refGene_tx=XM_006941509.4;refGene_gn=AKTIP;refGene_tx=XM_006941510.4;refGene_gn=AKTIP;refGene_tx=XM_019819752.2;refGene_gn=AKTIP;refGene_tx=XM_023245703.1
E2 33250615 33250657
refGene_gn=AKTIP;refGene_tx=XM_006941509.4;refGene_gn=AKTIP;refGene_tx=XM_006941510.4;refGene_gn=AKTIP;refGene_tx=XM_019819752.2;refGene_gn=AKTIP;refGene_tx=XM_023245703.1
E2 33334638 33334814
refGene_gn=RPGRIP1L;refGene_tx=XM_011290158.3;refGene_gn=RPGRIP1L;refGene_tx=XM_019819754.2;refGene_gn=RPGRIP1L;refGene_tx=XM_019819755.2
E2 33337096 33337230
refGene_gn=RPGRIP1L;refGene_tx=XM_011290158.3;refGene_gn=RPGRIP1L;refGene_tx=XM_019819754.2;refGene_gn=RPGRIP1L;refGene_tx=XM_019819755.2
E2 33343568 33343653
refGene_gn=RPGRIP1L;refGene_tx=XM_011290158.3;refGene_gn=RPGRIP1L;refGene_tx=XM_019819754.2;refGene_gn=RPGRIP1L;refGene_tx=XM_019819755.2
E2 33350874 33351058
refGene_gn=RPGRIP1L;refGene_tx=XM_011290158.3;refGene_gn=RPGRIP1L;refGene_tx=XM_019819754.2;refGene_gn=RPGRIP1L;refGene_tx=XM_019819755.2
E2 33363826 33363900
refGene_gn=RPGRIP1L;refGene_tx=XM_011290158.3;refGene_gn=RPGRIP1L;refGene_tx=XM_019819754.2;refGene_gn=RPGRIP1L;refGene_tx=XM_019819755.2
E2 33365596 33365756
refGene_gn=RPGRIP1L;refGene_tx=XM_011290158.3;refGene_gn=RPGRIP1L;refGene_tx=XM_019819754.2;refGene_gn=RPGRIP1L;refGene_tx=XM_019819755.2
E2 33366202 33366304
refGene_gn=RPGRIP1L;refGene_tx=XM_011290158.3;refGene_gn=RPGRIP1L;refGene_tx=XM_019819754.2;refGene_gn=RPGRIP1L;refGene_tx=XM_019819755.2
E2 33369436 33369520
refGene_gn=RPGRIP1L;refGene_tx=XM_011290158.3;refGene_gn=RPGRIP1L;refGene_tx=XM_019819754.2;refGene_gn=RPGRIP1L;refGene_tx=XM_019819755.2
E2 33369641 33369826
refGene_gn=RPGRIP1L;refGene_tx=XM_011290158.3;refGene_gn=RPGRIP1L;refGene_tx=XM_019819754.2;refGene_gn=RPGRIP1L;refGene_tx=XM_019819755.2
E2 33372889 33373268

refGene_gn=RPGRIP1L;refGene_tx=XM_011290158.3;refGene_gn=RPGRIP1L;refGene_tx=XM_019819754.2;refGene_gn=RPGRIP1L;refGene_tx=XM_019819755.2
E2 33377687 33377839
refGene_gn=RPGRIP1L;refGene_tx=XM_011290158.3;refGene_gn=RPGRIP1L;refGene_tx=XM_019819754.2;refGene_gn=RPGRIP1L;refGene_tx=XM_019819755.2
E2 33381468 33381921
refGene_gn=RPGRIP1L;refGene_tx=XM_011290158.3;refGene_gn=RPGRIP1L;refGene_tx=XM_019819754.2;refGene_gn=RPGRIP1L;refGene_tx=XM_019819755.2
E2 33386500 33386618
refGene_gn=RPGRIP1L;refGene_tx=XM_011290158.3;refGene_gn=RPGRIP1L;refGene_tx=XM_019819754.2;refGene_gn=RPGRIP1L;refGene_tx=XM_019819755.2
E2 33387160 33387340
refGene_gn=RPGRIP1L;refGene_tx=XM_011290158.3;refGene_gn=RPGRIP1L;refGene_tx=XM_019819754.2;refGene_gn=RPGRIP1L;refGene_tx=XM_019819755.2
E2 33388022 33388073
refGene_gn=RPGRIP1L;refGene_tx=XM_011290158.3;refGene_gn=RPGRIP1L;refGene_tx=XM_019819754.2;refGene_gn=RPGRIP1L;refGene_tx=XM_019819755.2
E2 33388380 33388487
refGene_gn=RPGRIP1L;refGene_tx=XM_011290158.3;refGene_gn=RPGRIP1L;refGene_tx=XM_019819754.2;refGene_gn=RPGRIP1L;refGene_tx=XM_019819755.2
E2 33388820 33388832
refGene_gn=RPGRIP1L;refGene_tx=XM_019819754.2;refGene_gn=RPGRIP1L;refGene_tx=XM_019819755.2
E2 33393356 33393496
refGene_gn=RPGRIP1L;refGene_tx=XM_011290158.3;refGene_gn=RPGRIP1L;refGene_tx=XM_019819754.2;refGene_gn=RPGRIP1L;refGene_tx=XM_019819755.2
E2 33399708 33399782
refGene_gn=RPGRIP1L;refGene_tx=XM_011290158.3;refGene_gn=RPGRIP1L;refGene_tx=XM_019819754.2;refGene_gn=RPGRIP1L;refGene_tx=XM_019819755.2
E2 33401103 33401250
refGene_gn=RPGRIP1L;refGene_tx=XM_011290158.3;refGene_gn=RPGRIP1L;refGene_tx=XM_019819754.2;refGene_gn=RPGRIP1L;refGene_tx=XM_019819755.2
E2 33403549 33403655
refGene_gn=RPGRIP1L;refGene_tx=XM_011290158.3;refGene_gn=RPGRIP1L;refGene_tx=XM_019819754.2;refGene_gn=RPGRIP1L;refGene_tx=XM_019819755.2
E2 33412575 33412719
refGene_gn=RPGRIP1L;refGene_tx=XM_011290158.3;refGene_gn=RPGRIP1L;refGene_tx=XM_019819754.2;refGene_gn=RPGRIP1L;refGene_tx=XM_019819755.2
E2 33414082 33414185
refGene_gn=RPGRIP1L;refGene_tx=XM_011290158.3;refGene_gn=RPGRIP1L;refGene_tx=XM_019819754.2;refGene_gn=RPGRIP1L;refGene_tx=XM_019819755.2
E2 33417759 33418058
refGene_gn=RPGRIP1L;refGene_tx=XM_011290158.3;refGene_gn=RPGRIP1L;refGene_tx=XM_019819754.2;refGene_gn=RPGRIP1L;refGene_tx=XM_019819755.2
E2 33421348 33421493
refGene_gn=RPGRIP1L;refGene_tx=XM_011290158.3;refGene_gn=RPGRIP1L;refGene_tx=XM_019819754.2;refGene_gn=RPGRIP1L;refGene_tx=XM_019819755.2
E2 33423095 33423189
refGene_gn=RPGRIP1L;refGene_tx=XM_011290158.3;refGene_gn=RPGRIP1L;refGene_tx=XM_019819754.2;refGene_gn=RPGRIP1L;refGene_tx=XM_019819755.2

E2 33426009 33426267
refGene_gn=RPGRIP1L;refGene_tx=XM_011290158.3;refGene_gn=RPGRIP1L;refGene_tx=XM_019819755.2
E2 33426372 33426417 refGene_gn=FTO;refGene_tx=XM_003998041.4
E2 33536439 33536517 refGene_gn=FTO;refGene_tx=XM_003998041.4
E2 33553276 33553759 refGene_gn=FTO;refGene_tx=XM_003998041.4
E2 33553760 33553902 refGene_gn=FTO;refGene_tx=XM_003998041.4
E2 33572052 33572196 refGene_gn=FTO;refGene_tx=XM_003998041.4
E2 33604559 33604639 refGene_gn=FTO;refGene_tx=XM_003998041.4
E2 33611461 33611605 refGene_gn=FTO;refGene_tx=XM_003998041.4
E2 33619616 33619736 refGene_gn=FTO;refGene_tx=XM_003998041.4
E2 33661834 33661959 refGene_gn=FTO;refGene_tx=XM_003998041.4
E2 33821053 33821207 refGene_gn=FTO;refGene_tx=XM_003998041.4
E2 33961954 33962009 refGene_gn=IRX3;refGene_tx=XM_023245118.1
E2 33962490 33962557 refGene_gn=IRX3;refGene_tx=XM_023245118.1
E2 33962739 33963859 refGene_gn=IRX3;refGene_tx=XM_023245118.1
E2 33964045 33964521 refGene_gn=IRX3;refGene_tx=XM_023245118.1
E2 33965377 33965573 refGene_gn=IRX3;refGene_tx=XM_023245118.1
E2 33965779 33965797
refGene_gn=LOC111558002;refGene_tx=XM_023245119.1
E2 33967698 33967936
refGene_gn=LOC111558002;refGene_tx=XM_023245119.1
E2 33968410 33968895
refGene_gn=LOC111558002;refGene_tx=XM_023245119.1
E2 34545749 34545998
refGene_gn=IRX5;refGene_tx=XM_023245199.1;refGene_gn=IRX5;refGene_tx=XM_023245200.1
E2 34547064 34547470
refGene_gn=IRX5;refGene_tx=XM_023245199.1;refGene_gn=IRX5;refGene_tx=XM_023245200.1
E2 34547637 34548434
refGene_gn=IRX5;refGene_tx=XM_023245199.1;refGene_gn=IRX5;refGene_tx=XM_023245200.1
E2 34565253 34565318 refGene_gn=IRX5;refGene_tx=XM_023245199.1
E2 34878938 34878983
refGene_gn=IRX6;refGene_tx=XM_023245566.1;refGene_gn=IRX6;refGene_tx=XM_023245567.1
E2 34879942 34880200
refGene_gn=IRX6;refGene_tx=XM_023245566.1;refGene_gn=IRX6;refGene_tx=XM_023245567.1
E2 34880915 34881025
refGene_gn=IRX6;refGene_tx=XM_023245566.1;refGene_gn=IRX6;refGene_tx=XM_023245567.1
E2 34881226 34881534
refGene_gn=IRX6;refGene_tx=XM_023245566.1;refGene_gn=IRX6;refGene_tx=XM_023245567.1
E2 34882415 34883042
refGene_gn=IRX6;refGene_tx=XM_023245566.1;refGene_gn=IRX6;refGene_tx=XM_023245567.1
E2 34883981 34883989

refGene_gn=IRX6;refGene_tx=XM_023245566.1;refGene_gn=IRX6;refGene_tx=XM_023245567.1
E2 35010214 35010370 refGene_gn=MMP2;refGene_tx=XM_003998042.3
E2 35012953 35013180 refGene_gn=MMP2;refGene_tx=XM_003998042.3
E2 35014127 35014276 refGene_gn=MMP2;refGene_tx=XM_003998042.3
E2 35015316 35015445 refGene_gn=MMP2;refGene_tx=XM_003998042.3
E2 35015680 35015854 refGene_gn=MMP2;refGene_tx=XM_003998042.3
E2 35018269 35018443 refGene_gn=MMP2;refGene_tx=XM_003998042.3
E2 35019265 35019439 refGene_gn=MMP2;refGene_tx=XM_003998042.3
E2 35021113 35021269 refGene_gn=MMP2;refGene_tx=XM_003998042.3
E2 35022453 35022589 refGene_gn=MMP2;refGene_tx=XM_003998042.3
E2 35024784 35024921 refGene_gn=MMP2;refGene_tx=XM_003998042.3
E2 35026019 35026179 refGene_gn=MMP2;refGene_tx=XM_003998042.3
E2 35030776 35030886 refGene_gn=MMP2;refGene_tx=XM_003998042.3
E2 35032846 35032950 refGene_gn=MMP2;refGene_tx=XM_003998042.3
E2 35038105 35038276 refGene_gn=LPCAT2;refGene_tx=XM_023244935.1;refGene_gn=LPCAT2;refGene_tx=XM_023244936.1
E2 35052577 35052717 refGene_gn=LPCAT2;refGene_tx=XM_023244935.1;refGene_gn=LPCAT2;refGene_tx=XM_023244936.1
E2 35055778 35055996 refGene_gn=LPCAT2;refGene_tx=XM_023244935.1;refGene_gn=LPCAT2;refGene_tx=XM_023244936.1
E2 35057569 35057682 refGene_gn=LPCAT2;refGene_tx=XM_023244935.1;refGene_gn=LPCAT2;refGene_tx=XM_023244936.1
E2 35059578 35059639 refGene_gn=LPCAT2;refGene_tx=XM_023244935.1
E2 35060553 35060612 refGene_gn=LPCAT2;refGene_tx=XM_023244935.1
E2 35061870 35061905 refGene_gn=LPCAT2;refGene_tx=XM_023244935.1;refGene_gn=LPCAT2;refGene_tx=XM_023244936.1
E2 35065186 35065241 refGene_gn=LPCAT2;refGene_tx=XM_023244935.1;refGene_gn=LPCAT2;refGene_tx=XM_023244936.1
E2 35075626 35075709 refGene_gn=LPCAT2;refGene_tx=XM_023244935.1;refGene_gn=LPCAT2;refGene_tx=XM_023244936.1
E2 35078360 35078486 refGene_gn=LPCAT2;refGene_tx=XM_023244935.1;refGene_gn=LPCAT2;refGene_tx=XM_023244936.1
E2 35080270 35080424 refGene_gn=LPCAT2;refGene_tx=XM_023244935.1;refGene_gn=LPCAT2;refGene_tx=XM_023244936.1
E2 35091401 35092157 refGene_gn=CAPNS2;refGene_tx=XM_019819760.1
E2 35100773 35100872 refGene_gn=LPCAT2;refGene_tx=XM_023244935.1;refGene_gn=LPCAT2;refGene_tx=XM_023244936.1
E2 35104464 35104600 refGene_gn=LPCAT2;refGene_tx=XM_023244935.1;refGene_gn=LPCAT2;refGene_

tx=XM_023244936.1
E2 35108851 35109036
refGene_gn=LPCAT2;refGene_tx=XM_023244935.1;refGene_gn=LPCAT2;refGene_tx=XM_023244936.1
E2 35146836 35147110
refGene_gn=SLC6A2;refGene_tx=XM_023245162.1;refGene_gn=SLC6A2;refGene_tx=XM_023245163.1
E2 35159499 35159631
refGene_gn=SLC6A2;refGene_tx=XM_023245162.1;refGene_gn=SLC6A2;refGene_tx=XM_023245163.1
E2 35161556 35161794
refGene_gn=SLC6A2;refGene_tx=XM_023245162.1;refGene_gn=SLC6A2;refGene_tx=XM_023245163.1
E2 35173500 35173639
refGene_gn=SLC6A2;refGene_tx=XM_023245162.1;refGene_gn=SLC6A2;refGene_tx=XM_023245163.1
E2 35181343 35181478
refGene_gn=SLC6A2;refGene_tx=XM_023245162.1;refGene_gn=SLC6A2;refGene_tx=XM_023245163.1
E2 35183165 35183269
refGene_gn=SLC6A2;refGene_tx=XM_023245162.1;refGene_gn=SLC6A2;refGene_tx=XM_023245163.1
E2 35184992 35185117
refGene_gn=SLC6A2;refGene_tx=XM_023245162.1;refGene_gn=SLC6A2;refGene_tx=XM_023245163.1
E2 35185869 35185982
refGene_gn=SLC6A2;refGene_tx=XM_023245162.1;refGene_gn=SLC6A2;refGene_tx=XM_023245163.1
E2 35187103 35187232
refGene_gn=SLC6A2;refGene_tx=XM_023245162.1;refGene_gn=SLC6A2;refGene_tx=XM_023245163.1
E2 35187731 35187831
refGene_gn=SLC6A2;refGene_tx=XM_023245162.1;refGene_gn=SLC6A2;refGene_tx=XM_023245163.1
E2 35189408 35189509
refGene_gn=SLC6A2;refGene_tx=XM_023245162.1;refGene_gn=SLC6A2;refGene_tx=XM_023245163.1
E2 35189954 35190602
refGene_gn=SLC6A2;refGene_tx=XM_023245162.1;refGene_gn=SLC6A2;refGene_tx=XM_023245163.1
E2 35191533 35191605 refGene_gn=SLC6A2;refGene_tx=XM_023245163.1
E2 35192202 35192226 refGene_gn=SLC6A2;refGene_tx=XM_023245163.1
E2 35205829 35206013
refGene_gn=CES1;refGene_tx=NM_001009249.1;refGene_gn=CES1;refGene_tx=XM_011290080.2
E2 35206276 35206349
refGene_gn=CES1;refGene_tx=NM_001009249.1;refGene_gn=CES1;refGene_tx=XM_011290080.2
E2 35207548 35207680
refGene_gn=CES1;refGene_tx=NM_001009249.1;refGene_gn=CES1;refGene_tx=X

M_011290080.2
E2	35211795	35211943
refGene_gn=CES1;refGene_tx=NM_001009249.1;refGene_gn=CES1;refGene_tx=X
M_011290080.2
E2	35212188	35212269
refGene_gn=CES1;refGene_tx=NM_001009249.1;refGene_gn=CES1;refGene_tx=X
M_011290080.2
E2	35213860	35214001
refGene_gn=CES1;refGene_tx=NM_001009249.1;refGene_gn=CES1;refGene_tx=X
M_011290080.2
E2	35218173	35218212
refGene_gn=CES1;refGene_tx=NM_001009249.1;refGene_gn=CES1;refGene_tx=X
M_011290080.2
E2	35220806	35220911
refGene_gn=CES1;refGene_tx=NM_001009249.1;refGene_gn=CES1;refGene_tx=X
M_011290080.2
E2	35221881	35221986
refGene_gn=CES1;refGene_tx=NM_001009249.1;refGene_gn=CES1;refGene_tx=X
M_011290080.2
E2	35223523	35223677
refGene_gn=CES1;refGene_tx=NM_001009249.1;refGene_gn=CES1;refGene_tx=X
M_011290080.2
E2	35226071	35226205
refGene_gn=CES1;refGene_tx=NM_001009249.1;refGene_gn=CES1;refGene_tx=X
M_011290080.2
E2	35229008	35229153
refGene_gn=CES1;refGene_tx=NM_001009249.1;refGene_gn=CES1;refGene_tx=X
M_011290080.2
E2	35232835	35233043
refGene_gn=CES1;refGene_tx=NM_001009249.1;refGene_gn=CES1;refGene_tx=X
M_011290080.2
E2	35236129	35236184
refGene_gn=CES1;refGene_tx=NM_001009249.1;refGene_gn=CES1;refGene_tx=X
M_011290080.2
E2	35249757	35249899	refGene_gn=CES5A;refGene_tx=NM_001009188.1
E2	35249988	35250061	refGene_gn=CES5A;refGene_tx=NM_001009188.1
E2	35252366	35252516	refGene_gn=CES5A;refGene_tx=NM_001009188.1
E2	35259840	35259988	refGene_gn=CES5A;refGene_tx=NM_001009188.1
E2	35262582	35262648	refGene_gn=CES5A;refGene_tx=NM_001009188.1
E2	35265657	35265798	refGene_gn=CES5A;refGene_tx=NM_001009188.1
E2	35266864	35266969	refGene_gn=CES5A;refGene_tx=NM_001009188.1
E2	35268745	35268853	refGene_gn=CES5A;refGene_tx=NM_001009188.1
E2	35270608	35270762	refGene_gn=CES5A;refGene_tx=NM_001009188.1
E2	35271232	35271366	refGene_gn=CES5A;refGene_tx=NM_001009188.1
E2	35272718	35272857	refGene_gn=CES5A;refGene_tx=NM_001009188.1
E2	35274905	35275110	refGene_gn=CES5A;refGene_tx=NM_001009188.1
E2	35276234	35276307	refGene_gn=CES5A;refGene_tx=NM_001009188.1
E2	35537810	35537928
refGene_gn=GNAO1;refGene_tx=XM_011290154.3;refGene_gn=GNAO1;refGene_tx
=XM_011290155.3;refGene_gn=GNAO1;refGene_tx=XM_011290156.3

E2 35538149 35538192
refGene_gn=GNAO1;refGene_tx=XM_011290154.3;refGene_gn=GNAO1;refGene_tx=XM_011290155.3;refGene_gn=GNAO1;refGene_tx=XM_011290156.3
E2 35622310 35622452
refGene_gn=GNAO1;refGene_tx=XM_011290154.3;refGene_gn=GNAO1;refGene_tx=XM_011290155.3;refGene_gn=GNAO1;refGene_tx=XM_011290156.3
E2 35681736 35681897
refGene_gn=GNAO1;refGene_tx=XM_011290154.3;refGene_gn=GNAO1;refGene_tx=XM_011290155.3;refGene_gn=GNAO1;refGene_tx=XM_011290156.3
E2 35687202 35687331
refGene_gn=GNAO1;refGene_tx=XM_011290154.3;refGene_gn=GNAO1;refGene_tx=XM_011290155.3;refGene_gn=GNAO1;refGene_tx=XM_011290156.3
E2 35688542 35688672
refGene_gn=GNAO1;refGene_tx=XM_011290154.3;refGene_gn=GNAO1;refGene_tx=XM_011290155.3;refGene_gn=GNAO1;refGene_tx=XM_011290156.3
E2 35692668 35692822 refGene_gn=GNAO1;refGene_tx=XM_011290154.3
E2 35694880 35695068 refGene_gn=GNAO1;refGene_tx=XM_011290154.3
E2 35701680 35701834
refGene_gn=GNAO1;refGene_tx=XM_011290155.3;refGene_gn=GNAO1;refGene_tx=XM_011290156.3
E2 35705001 35705189
refGene_gn=GNAO1;refGene_tx=XM_011290155.3;refGene_gn=GNAO1;refGene_tx=XM_011290156.3
E2 35711545 35711693 refGene_gn=AMFR;refGene_tx=XM_023246140.1
E2 35712352 35712537 refGene_gn=AMFR;refGene_tx=XM_023246140.1
E2 35715552 35715636 refGene_gn=AMFR;refGene_tx=XM_023246140.1
E2 35717337 35717472 refGene_gn=AMFR;refGene_tx=XM_023246140.1
E2 35722435 35722539 refGene_gn=AMFR;refGene_tx=XM_023246140.1
E2 35724797 35724988 refGene_gn=AMFR;refGene_tx=XM_023246140.1
E2 35732589 35732700 refGene_gn=AMFR;refGene_tx=XM_023246140.1
E2 35733567 35733701 refGene_gn=AMFR;refGene_tx=XM_023246140.1
E2 35736157 35736290 refGene_gn=AMFR;refGene_tx=XM_023246140.1
E2 35736454 35736495 refGene_gn=AMFR;refGene_tx=XM_023246140.1
E2 35739265 35739418 refGene_gn=AMFR;refGene_tx=XM_023246140.1
E2 35740686 35740851 refGene_gn=AMFR;refGene_tx=XM_023246140.1
E2 35744408 35744501 refGene_gn=AMFR;refGene_tx=XM_023246140.1
E2 35753548 35753803 refGene_gn=AMFR;refGene_tx=XM_023246140.1
E2 35761542 35761564 refGene_gn=NUDT21;refGene_tx=XM_003998050.5
E2 35762850 35762965 refGene_gn=NUDT21;refGene_tx=XM_003998050.5
E2 35763586 35763662 refGene_gn=NUDT21;refGene_tx=XM_003998050.5
E2 35767703 35767793 refGene_gn=NUDT21;refGene_tx=XM_003998050.5
E2 35772502 35772566 refGene_gn=NUDT21;refGene_tx=XM_003998050.5
E2 35773469 35773670 refGene_gn=NUDT21;refGene_tx=XM_003998050.5
E2 35776118 35776234 refGene_gn=NUDT21;refGene_tx=XM_003998050.5
E2 35776646 35776800 refGene_gn=OGFOD1;refGene_tx=XM_003998051.5
E2 35778429 35778575 refGene_gn=OGFOD1;refGene_tx=XM_003998051.5
E2 35782392 35782439 refGene_gn=OGFOD1;refGene_tx=XM_003998051.5
E2 35785316 35785389 refGene_gn=OGFOD1;refGene_tx=XM_003998051.5
E2 35785390 35785418 refGene_gn=OGFOD1;refGene_tx=XM_003998051.5
E2 35788912 35789029 refGene_gn=OGFOD1;refGene_tx=XM_003998051.5

E2	35789607	35789699	refGene_gn=OGFOD1;refGene_tx=XM_003998051.5
E2	35789906	35790035	refGene_gn=OGFOD1;refGene_tx=XM_003998051.5
E2	35790315	35790429	refGene_gn=OGFOD1;refGene_tx=XM_003998051.5
E2	35791867	35791947	refGene_gn=OGFOD1;refGene_tx=XM_003998051.5
E2	35792213	35792527	refGene_gn=OGFOD1;refGene_tx=XM_003998051.5
E2	35796842	35796965	refGene_gn=OGFOD1;refGene_tx=XM_003998051.5
E2	35797461	35797520	refGene_gn=OGFOD1;refGene_tx=XM_003998051.5
E2	35798018	35798180	refGene_gn=OGFOD1;refGene_tx=XM_003998051.5
E2	35806214	35806321
refGene_gn=BBS2;refGene_tx=XM_003998052.5;refGene_gn=BBS2;refGene_tx=XM_006941525.4;refGene_gn=BBS2;refGene_tx=XM_019819765.2;refGene_gn=BBS2;refGene_tx=XM_019819766.1
E2	35807028	35807177
refGene_gn=BBS2;refGene_tx=XM_003998052.5;refGene_gn=BBS2;refGene_tx=XM_006941525.4;refGene_gn=BBS2;refGene_tx=XM_019819765.2;refGene_gn=BBS2;refGene_tx=XM_019819766.1
E2	35814205	35814318
refGene_gn=BBS2;refGene_tx=XM_003998052.5;refGene_gn=BBS2;refGene_tx=XM_006941525.4;refGene_gn=BBS2;refGene_tx=XM_019819765.2;refGene_gn=BBS2;refGene_tx=XM_019819766.1
E2	35815196	35815334
refGene_gn=BBS2;refGene_tx=XM_003998052.5;refGene_gn=BBS2;refGene_tx=XM_006941525.4;refGene_gn=BBS2;refGene_tx=XM_019819765.2;refGene_gn=BBS2;refGene_tx=XM_019819766.1
E2	35815819	35815951
refGene_gn=BBS2;refGene_tx=XM_003998052.5;refGene_gn=BBS2;refGene_tx=XM_006941525.4;refGene_gn=BBS2;refGene_tx=XM_019819765.2;refGene_gn=BBS2;refGene_tx=XM_019819766.1
E2	35817296	35817426
refGene_gn=BBS2;refGene_tx=XM_003998052.5;refGene_gn=BBS2;refGene_tx=XM_006941525.4;refGene_gn=BBS2;refGene_tx=XM_019819765.2;refGene_gn=BBS2;refGene_tx=XM_019819766.1
E2	35818062	35818234
refGene_gn=BBS2;refGene_tx=XM_003998052.5;refGene_gn=BBS2;refGene_tx=XM_006941525.4;refGene_gn=BBS2;refGene_tx=XM_019819765.2;refGene_gn=BBS2;refGene_tx=XM_019819766.1
E2	35818321	35818463
refGene_gn=BBS2;refGene_tx=XM_003998052.5;refGene_gn=BBS2;refGene_tx=XM_006941525.4;refGene_gn=BBS2;refGene_tx=XM_019819765.2;refGene_gn=BBS2;refGene_tx=XM_019819766.1
E2	35819591	35819731
refGene_gn=BBS2;refGene_tx=XM_003998052.5;refGene_gn=BBS2;refGene_tx=XM_006941525.4;refGene_gn=BBS2;refGene_tx=XM_019819765.2;refGene_gn=BBS2;refGene_tx=XM_019819766.1
E2	35819975	35820111
refGene_gn=BBS2;refGene_tx=XM_003998052.5;refGene_gn=BBS2;refGene_tx=XM_006941525.4;refGene_gn=BBS2;refGene_tx=XM_019819765.2;refGene_gn=BBS2;refGene_tx=XM_019819766.1
E2	35823130	35823217
refGene_gn=BBS2;refGene_tx=XM_003998052.5;refGene_gn=BBS2;refGene_tx=X

M_006941525.4;refGene_gn=BBS2;refGene_tx=XM_019819765.2;refGene_gn=BBS2;refGene_tx=XM_019819766.1
E2 35823300 35823405
refGene_gn=BBS2;refGene_tx=XM_003998052.5;refGene_gn=BBS2;refGene_tx=XM_006941525.4;refGene_gn=BBS2;refGene_tx=XM_019819765.2;refGene_gn=BBS2;refGene_tx=XM_019819766.1
E2 35826799 35826877
refGene_gn=BBS2;refGene_tx=XM_003998052.5;refGene_gn=BBS2;refGene_tx=XM_006941525.4;refGene_gn=BBS2;refGene_tx=XM_019819765.2;refGene_gn=BBS2;refGene_tx=XM_019819766.1
E2 35827678 35827741
refGene_gn=BBS2;refGene_tx=XM_003998052.5;refGene_gn=BBS2;refGene_tx=XM_006941525.4;refGene_gn=BBS2;refGene_tx=XM_019819765.2;refGene_gn=BBS2;refGene_tx=XM_019819766.1
E2 35828005 35828131
refGene_gn=BBS2;refGene_tx=XM_003998052.5;refGene_gn=BBS2;refGene_tx=XM_006941525.4;refGene_gn=BBS2;refGene_tx=XM_019819765.2;refGene_gn=BBS2;refGene_tx=XM_019819766.1
E2 35832505 35832733
refGene_gn=BBS2;refGene_tx=XM_003998052.5;refGene_gn=BBS2;refGene_tx=XM_006941525.4;refGene_gn=BBS2;refGene_tx=XM_019819765.2;refGene_gn=BBS2;refGene_tx=XM_019819766.1
E2 35837409 35837526
refGene_gn=BBS2;refGene_tx=XM_003998052.5;refGene_gn=BBS2;refGene_tx=XM_006941525.4;refGene_gn=BBS2;refGene_tx=XM_019819765.2
E2 35839154 35839181 refGene_gn=BBS2;refGene_tx=XM_019819766.1
E2 35869802 35869833
refGene_gn=LOC109494884;refGene_tx=XM_019819767.1
E2 35871237 35871303
refGene_gn=LOC109494884;refGene_tx=XM_019819767.1
E2 35871873 35871965
refGene_gn=LOC109494884;refGene_tx=XM_019819767.1
E2 35885355 35885853
refGene_gn=LOC109494886;refGene_tx=XM_023246150.1
E2 35886099 35886165
refGene_gn=LOC109494886;refGene_tx=XM_023246150.1
E2 35886391 35886456
refGene_gn=LOC109494886;refGene_tx=XM_023246150.1
E2 35886990 35887021
refGene_gn=LOC109494886;refGene_tx=XM_023246150.1
E2 35904246 35904274
refGene_gn=LOC101081428;refGene_tx=XM_003998055.5
E2 35904579 35904645
refGene_gn=LOC101081428;refGene_tx=XM_003998055.5
E2 35904843 35904935
refGene_gn=LOC101081428;refGene_tx=XM_003998055.5
E2 35923295 35923323
refGene_gn=LOC111558024;refGene_tx=XM_023245328.1
E2 35923897 35923963
refGene_gn=LOC111558024;refGene_tx=XM_023245328.1

E2 35924634 35924726
refGene_gn=LOC111558024;refGene_tx=XM_023245328.1
E2 35970647 35970826 refGene_gn=NUP93;refGene_tx=XM_006941526.4
E2 35980548 35980666 refGene_gn=NUP93;refGene_tx=XM_006941526.4
E2 36019370 36019433 refGene_gn=NUP93;refGene_tx=XM_006941526.4
E2 36024477 36024606
refGene_gn=NUP93;refGene_tx=XM_006941526.4;refGene_gn=NUP93;refGene_tx=XM_006941527.4;refGene_gn=NUP93;refGene_tx=XM_019819769.1
E2 36033128 36033203
refGene_gn=NUP93;refGene_tx=XM_006941526.4;refGene_gn=NUP93;refGene_tx=XM_006941527.4;refGene_gn=NUP93;refGene_tx=XM_019819769.1
E2 36035230 36035320
refGene_gn=NUP93;refGene_tx=XM_006941526.4;refGene_gn=NUP93;refGene_tx=XM_006941527.4;refGene_gn=NUP93;refGene_tx=XM_019819769.1
E2 36037172 36037312
refGene_gn=NUP93;refGene_tx=XM_006941526.4;refGene_gn=NUP93;refGene_tx=XM_006941527.4;refGene_gn=NUP93;refGene_tx=XM_019819769.1
E2 36041060 36041193
refGene_gn=NUP93;refGene_tx=XM_006941526.4;refGene_gn=NUP93;refGene_tx=XM_006941527.4;refGene_gn=NUP93;refGene_tx=XM_019819769.1
E2 36042635 36042793
refGene_gn=NUP93;refGene_tx=XM_006941526.4;refGene_gn=NUP93;refGene_tx=XM_006941527.4;refGene_gn=NUP93;refGene_tx=XM_019819769.1
E2 36043952 36044118
refGene_gn=NUP93;refGene_tx=XM_006941526.4;refGene_gn=NUP93;refGene_tx=XM_006941527.4;refGene_gn=NUP93;refGene_tx=XM_019819769.1
E2 36044387 36044481
refGene_gn=NUP93;refGene_tx=XM_006941526.4;refGene_gn=NUP93;refGene_tx=XM_006941527.4;refGene_gn=NUP93;refGene_tx=XM_019819769.1
E2 36045249 36045441
refGene_gn=NUP93;refGene_tx=XM_006941526.4;refGene_gn=NUP93;refGene_tx=XM_006941527.4;refGene_gn=NUP93;refGene_tx=XM_019819769.1
E2 36046038 36046165
refGene_gn=NUP93;refGene_tx=XM_006941526.4;refGene_gn=NUP93;refGene_tx=XM_006941527.4;refGene_gn=NUP93;refGene_tx=XM_019819769.1
E2 36046277 36046350
refGene_gn=NUP93;refGene_tx=XM_006941526.4;refGene_gn=NUP93;refGene_tx=XM_006941527.4;refGene_gn=NUP93;refGene_tx=XM_019819769.1
E2 36046656 36046701
refGene_gn=NUP93;refGene_tx=XM_006941526.4;refGene_gn=NUP93;refGene_tx=XM_006941527.4;refGene_gn=NUP93;refGene_tx=XM_019819769.1
E2 36048374 36048491
refGene_gn=NUP93;refGene_tx=XM_006941526.4;refGene_gn=NUP93;refGene_tx=XM_006941527.4;refGene_gn=NUP93;refGene_tx=XM_019819769.1
E2 36049262 36049381
refGene_gn=NUP93;refGene_tx=XM_006941526.4;refGene_gn=NUP93;refGene_tx=XM_006941527.4;refGene_gn=NUP93;refGene_tx=XM_019819769.1
E2 36050260 36050378
refGene_gn=NUP93;refGene_tx=XM_006941526.4;refGene_gn=NUP93;refGene_tx=XM_006941527.4;refGene_gn=NUP93;refGene_tx=XM_019819769.1

E2 36050822 36050906
refGene_gn=NUP93;refGene_tx=XM_006941526.4;refGene_gn=NUP93;refGene_tx=XM_006941527.4;refGene_gn=NUP93;refGene_tx=XM_019819769.1
E2 36052365 36052494
refGene_gn=NUP93;refGene_tx=XM_006941526.4;refGene_gn=NUP93;refGene_tx=XM_006941527.4;refGene_gn=NUP93;refGene_tx=XM_019819769.1
E2 36054956 36055067
refGene_gn=NUP93;refGene_tx=XM_006941526.4;refGene_gn=NUP93;refGene_tx=XM_006941527.4;refGene_gn=NUP93;refGene_tx=XM_019819769.1
E2 36070625 36070907
refGene_gn=SLC12A3;refGene_tx=XM_023245212.1;refGene_gn=SLC12A3;refGene_tx=XM_023245213.1;refGene_gn=SLC12A3;refGene_tx=XM_023245214.1;refGene_gn=SLC12A3;refGene_tx=XM_023245215.1;refGene_gn=SLC12A3;refGene_tx=XM_023245216.1;refGene_gn=SLC12A3;refGene_tx=XM_023245217.1;refGene_gn=SLC12A3;refGene_tx=XM_023245218.1
E2 36071343 36071490
refGene_gn=SLC12A3;refGene_tx=XM_023245212.1;refGene_gn=SLC12A3;refGene_tx=XM_023245213.1;refGene_gn=SLC12A3;refGene_tx=XM_023245214.1;refGene_gn=SLC12A3;refGene_tx=XM_023245215.1;refGene_gn=SLC12A3;refGene_tx=XM_023245216.1;refGene_gn=SLC12A3;refGene_tx=XM_023245217.1;refGene_gn=SLC12A3;refGene_tx=XM_023245218.1
E2 36072796 36072872
refGene_gn=SLC12A3;refGene_tx=XM_023245212.1;refGene_gn=SLC12A3;refGene_tx=XM_023245213.1;refGene_gn=SLC12A3;refGene_tx=XM_023245214.1;refGene_gn=SLC12A3;refGene_tx=XM_023245215.1;refGene_gn=SLC12A3;refGene_tx=XM_023245216.1;refGene_gn=SLC12A3;refGene_tx=XM_023245217.1;refGene_gn=SLC12A3;refGene_tx=XM_023245218.1
E2 36073825 36073921
refGene_gn=SLC12A3;refGene_tx=XM_023245212.1;refGene_gn=SLC12A3;refGene_tx=XM_023245213.1;refGene_gn=SLC12A3;refGene_tx=XM_023245214.1;refGene_gn=SLC12A3;refGene_tx=XM_023245215.1;refGene_gn=SLC12A3;refGene_tx=XM_023245216.1;refGene_gn=SLC12A3;refGene_tx=XM_023245217.1;refGene_gn=SLC12A3;refGene_tx=XM_023245218.1
E2 36074237 36074377
refGene_gn=SLC12A3;refGene_tx=XM_023245212.1;refGene_gn=SLC12A3;refGene_tx=XM_023245213.1;refGene_gn=SLC12A3;refGene_tx=XM_023245214.1;refGene_gn=SLC12A3;refGene_tx=XM_023245215.1;refGene_gn=SLC12A3;refGene_tx=XM_023245216.1;refGene_gn=SLC12A3;refGene_tx=XM_023245217.1;refGene_gn=SLC12A3;refGene_tx=XM_023245218.1
E2 36074718 36074829
refGene_gn=SLC12A3;refGene_tx=XM_023245212.1;refGene_gn=SLC12A3;refGene_tx=XM_023245213.1;refGene_gn=SLC12A3;refGene_tx=XM_023245214.1;refGene_gn=SLC12A3;refGene_tx=XM_023245215.1;refGene_gn=SLC12A3;refGene_tx=XM_023245216.1;refGene_gn=SLC12A3;refGene_tx=XM_023245217.1;refGene_gn=SLC12A3;refGene_tx=XM_023245218.1
E2 36075820 36075932
refGene_gn=SLC12A3;refGene_tx=XM_023245212.1;refGene_gn=SLC12A3;refGene_tx=XM_023245213.1;refGene_gn=SLC12A3;refGene_tx=XM_023245214.1;refGene_gn=SLC12A3;refGene_tx=XM_023245215.1;refGene_gn=SLC12A3;refGene_tx=XM_023245216.1;refGene_gn=SLC12A3;refGene_tx=XM_023245217.1;refGene_gn

=SLC12A3;refGene_tx=XM_023245218.1
E2 36076132 36076263
refGene_gn=SLC12A3;refGene_tx=XM_023245212.1;refGene_gn=SLC12A3;refGene_tx=XM_023245213.1;refGene_gn=SLC12A3;refGene_tx=XM_023245214.1;refGene_gn=SLC12A3;refGene_tx=XM_023245215.1;refGene_gn=SLC12A3;refGene_tx=XM_023245216.1;refGene_gn=SLC12A3;refGene_tx=XM_023245217.1;refGene_gn=SLC12A3;refGene_tx=XM_023245218.1
E2 36081099 36081184
refGene_gn=SLC12A3;refGene_tx=XM_023245212.1;refGene_gn=SLC12A3;refGene_tx=XM_023245213.1;refGene_gn=SLC12A3;refGene_tx=XM_023245214.1;refGene_gn=SLC12A3;refGene_tx=XM_023245215.1;refGene_gn=SLC12A3;refGene_tx=XM_023245216.1;refGene_gn=SLC12A3;refGene_tx=XM_023245217.1;refGene_gn=SLC12A3;refGene_tx=XM_023245218.1
E2 36082148 36082303
refGene_gn=SLC12A3;refGene_tx=XM_023245212.1;refGene_gn=SLC12A3;refGene_tx=XM_023245213.1;refGene_gn=SLC12A3;refGene_tx=XM_023245214.1;refGene_gn=SLC12A3;refGene_tx=XM_023245215.1;refGene_gn=SLC12A3;refGene_tx=XM_023245216.1;refGene_gn=SLC12A3;refGene_tx=XM_023245217.1;refGene_gn=SLC12A3;refGene_tx=XM_023245218.1
E2 36082595 36082703
refGene_gn=SLC12A3;refGene_tx=XM_023245212.1;refGene_gn=SLC12A3;refGene_tx=XM_023245213.1;refGene_gn=SLC12A3;refGene_tx=XM_023245214.1;refGene_gn=SLC12A3;refGene_tx=XM_023245215.1;refGene_gn=SLC12A3;refGene_tx=XM_023245216.1;refGene_gn=SLC12A3;refGene_tx=XM_023245217.1;refGene_gn=SLC12A3;refGene_tx=XM_023245218.1
E2 36083371 36083495
refGene_gn=SLC12A3;refGene_tx=XM_023245212.1;refGene_gn=SLC12A3;refGene_tx=XM_023245213.1;refGene_gn=SLC12A3;refGene_tx=XM_023245214.1;refGene_gn=SLC12A3;refGene_tx=XM_023245215.1;refGene_gn=SLC12A3;refGene_tx=XM_023245216.1;refGene_gn=SLC12A3;refGene_tx=XM_023245217.1;refGene_gn=SLC12A3;refGene_tx=XM_023245218.1
E2 36085474 36085576
refGene_gn=SLC12A3;refGene_tx=XM_023245212.1;refGene_gn=SLC12A3;refGene_tx=XM_023245213.1;refGene_gn=SLC12A3;refGene_tx=XM_023245214.1;refGene_gn=SLC12A3;refGene_tx=XM_023245215.1;refGene_gn=SLC12A3;refGene_tx=XM_023245216.1;refGene_gn=SLC12A3;refGene_tx=XM_023245217.1;refGene_gn=SLC12A3;refGene_tx=XM_023245218.1
E2 36086469 36086625
refGene_gn=SLC12A3;refGene_tx=XM_023245212.1;refGene_gn=SLC12A3;refGene_tx=XM_023245213.1;refGene_gn=SLC12A3;refGene_tx=XM_023245214.1;refGene_gn=SLC12A3;refGene_tx=XM_023245215.1;refGene_gn=SLC12A3;refGene_tx=XM_023245216.1;refGene_gn=SLC12A3;refGene_tx=XM_023245217.1;refGene_gn=SLC12A3;refGene_tx=XM_023245218.1
E2 36087322 36087422
refGene_gn=SLC12A3;refGene_tx=XM_023245212.1;refGene_gn=SLC12A3;refGene_tx=XM_023245213.1;refGene_gn=SLC12A3;refGene_tx=XM_023245214.1;refGene_gn=SLC12A3;refGene_tx=XM_023245215.1;refGene_gn=SLC12A3;refGene_tx=XM_023245216.1;refGene_gn=SLC12A3;refGene_tx=XM_023245217.1;refGene_gn=SLC12A3;refGene_tx=XM_023245218.1
E2 36088138 36088250

refGene_gn=SLC12A3;refGene_tx=XM_023245212.1;refGene_gn=SLC12A3;refGene_tx=XM_023245213.1;refGene_gn=SLC12A3;refGene_tx=XM_023245214.1;refGene_gn=SLC12A3;refGene_tx=XM_023245215.1;refGene_gn=SLC12A3;refGene_tx=XM_023245216.1;refGene_gn=SLC12A3;refGene_tx=XM_023245217.1;refGene_gn=SLC12A3;refGene_tx=XM_023245218.1
E2 36088674 36088815
refGene_gn=SLC12A3;refGene_tx=XM_023245212.1;refGene_gn=SLC12A3;refGene_tx=XM_023245213.1;refGene_gn=SLC12A3;refGene_tx=XM_023245214.1;refGene_gn=SLC12A3;refGene_tx=XM_023245215.1;refGene_gn=SLC12A3;refGene_tx=XM_023245216.1;refGene_gn=SLC12A3;refGene_tx=XM_023245217.1;refGene_gn=SLC12A3;refGene_tx=XM_023245218.1
E2 36089940 36090047
refGene_gn=SLC12A3;refGene_tx=XM_023245212.1;refGene_gn=SLC12A3;refGene_tx=XM_023245213.1;refGene_gn=SLC12A3;refGene_tx=XM_023245214.1;refGene_gn=SLC12A3;refGene_tx=XM_023245215.1;refGene_gn=SLC12A3;refGene_tx=XM_023245216.1;refGene_gn=SLC12A3;refGene_tx=XM_023245217.1;refGene_gn=SLC12A3;refGene_tx=XM_023245218.1
E2 36091420 36091503
refGene_gn=SLC12A3;refGene_tx=XM_023245212.1;refGene_gn=SLC12A3;refGene_tx=XM_023245213.1;refGene_gn=SLC12A3;refGene_tx=XM_023245214.1;refGene_gn=SLC12A3;refGene_tx=XM_023245215.1;refGene_gn=SLC12A3;refGene_tx=XM_023245216.1;refGene_gn=SLC12A3;refGene_tx=XM_023245217.1;refGene_gn=SLC12A3;refGene_tx=XM_023245218.1
E2 36092788 36092962
refGene_gn=SLC12A3;refGene_tx=XM_023245212.1;refGene_gn=SLC12A3;refGene_tx=XM_023245213.1;refGene_gn=SLC12A3;refGene_tx=XM_023245214.1;refGene_gn=SLC12A3;refGene_tx=XM_023245215.1;refGene_gn=SLC12A3;refGene_tx=XM_023245216.1
E2 36093536 36093698
refGene_gn=SLC12A3;refGene_tx=XM_023245212.1;refGene_gn=SLC12A3;refGene_tx=XM_023245213.1;refGene_gn=SLC12A3;refGene_tx=XM_023245214.1;refGene_gn=SLC12A3;refGene_tx=XM_023245215.1;refGene_gn=SLC12A3;refGene_tx=XM_023245216.1;refGene_gn=SLC12A3;refGene_tx=XM_023245217.1;refGene_gn=SLC12A3;refGene_tx=XM_023245218.1
E2 36095551 36095663
refGene_gn=SLC12A3;refGene_tx=XM_023245212.1;refGene_gn=SLC12A3;refGene_tx=XM_023245213.1;refGene_gn=SLC12A3;refGene_tx=XM_023245214.1;refGene_gn=SLC12A3;refGene_tx=XM_023245215.1;refGene_gn=SLC12A3;refGene_tx=XM_023245216.1;refGene_gn=SLC12A3;refGene_tx=XM_023245217.1;refGene_gn=SLC12A3;refGene_tx=XM_023245218.1
E2 36098026 36098113
refGene_gn=SLC12A3;refGene_tx=XM_023245212.1;refGene_gn=SLC12A3;refGene_tx=XM_023245213.1;refGene_gn=SLC12A3;refGene_tx=XM_023245214.1;refGene_gn=SLC12A3;refGene_tx=XM_023245215.1;refGene_gn=SLC12A3;refGene_tx=XM_023245216.1;refGene_gn=SLC12A3;refGene_tx=XM_023245217.1;refGene_gn=SLC12A3;refGene_tx=XM_023245218.1
E2 36101188 36101324
refGene_gn=SLC12A3;refGene_tx=XM_023245212.1;refGene_gn=SLC12A3;refGene_tx=XM_023245213.1;refGene_gn=SLC12A3;refGene_tx=XM_023245214.1;refGene_gn=SLC12A3;refGene_tx=XM_023245215.1;refGene_gn=SLC12A3;refGene_tx=

XM_023245216.1;refGene_gn=SLC12A3;refGene_tx=XM_023245217.1;refGene_gn=SLC12A3;refGene_tx=XM_023245218.1
E2 36103011 36103079
refGene_gn=SLC12A3;refGene_tx=XM_023245212.1;refGene_gn=SLC12A3;refGene_tx=XM_023245213.1;refGene_gn=SLC12A3;refGene_tx=XM_023245214.1;refGene_gn=SLC12A3;refGene_tx=XM_023245215.1;refGene_gn=SLC12A3;refGene_tx=XM_023245216.1;refGene_gn=SLC12A3;refGene_tx=XM_023245217.1;refGene_gn=SLC12A3;refGene_tx=XM_023245218.1
E2 36109107 36109249
refGene_gn=SLC12A3;refGene_tx=XM_023245212.1;refGene_gn=SLC12A3;refGene_tx=XM_023245213.1;refGene_gn=SLC12A3;refGene_tx=XM_023245214.1;refGene_gn=SLC12A3;refGene_tx=XM_023245215.1;refGene_gn=SLC12A3;refGene_tx=XM_023245216.1;refGene_gn=SLC12A3;refGene_tx=XM_023245217.1;refGene_gn=SLC12A3;refGene_tx=XM_023245218.1
E2 36122719 36122872
refGene_gn=HERPUD1;refGene_tx=XM_003998061.5;refGene_gn=HERPUD1;refGene_tx=XM_019819770.2;refGene_gn=HERPUD1;refGene_tx=XM_023245136.1
E2 36125528 36125606
refGene_gn=HERPUD1;refGene_tx=XM_003998061.5;refGene_gn=HERPUD1;refGene_tx=XM_019819770.2;refGene_gn=HERPUD1;refGene_tx=XM_023245136.1
E2 36125692 36125767
refGene_gn=HERPUD1;refGene_tx=XM_003998061.5;refGene_gn=HERPUD1;refGene_tx=XM_019819770.2;refGene_gn=HERPUD1;refGene_tx=XM_023245136.1
E2 36127015 36127146
refGene_gn=HERPUD1;refGene_tx=XM_003998061.5;refGene_gn=HERPUD1;refGene_tx=XM_019819770.2;refGene_gn=HERPUD1;refGene_tx=XM_023245136.1
E2 36129208 36129331
refGene_gn=HERPUD1;refGene_tx=XM_003998061.5;refGene_gn=HERPUD1;refGene_tx=XM_019819770.2;refGene_gn=HERPUD1;refGene_tx=XM_023245136.1
E2 36129827 36130178
refGene_gn=HERPUD1;refGene_tx=XM_003998061.5;refGene_gn=HERPUD1;refGene_tx=XM_019819770.2;refGene_gn=HERPUD1;refGene_tx=XM_023245136.1
E2 36131551 36131657
refGene_gn=HERPUD1;refGene_tx=XM_003998061.5;refGene_gn=HERPUD1;refGene_tx=XM_019819770.2;refGene_gn=HERPUD1;refGene_tx=XM_023245136.1
E2 36132955 36133120
refGene_gn=HERPUD1;refGene_tx=XM_003998061.5;refGene_gn=HERPUD1;refGene_tx=XM_019819770.2
E2 36200766 36201061
refGene_gn=NLRC5;refGene_tx=XM_003998063.4;refGene_gn=NLRC5;refGene_tx=XM_011290148.2;refGene_gn=NLRC5;refGene_tx=XM_011290149.2;refGene_gn=NLRC5;refGene_tx=XM_011290150.2;refGene_gn=NLRC5;refGene_tx=XM_011290151.2;refGene_gn=NLRC5;refGene_tx=XM_011290152.2;refGene_gn=NLRC5;refGene_tx=XM_019819771.2;refGene_gn=NLRC5;refGene_tx=XM_019819773.1;refGene_gn=NLRC5;refGene_tx=XM_019819775.1;refGene_gn=NLRC5;refGene_tx=XM_019819776.1;refGene_gn=NLRC5;refGene_tx=XM_019819777.1
E2 36201875 36201968
refGene_gn=NLRC5;refGene_tx=XM_003998063.4;refGene_gn=NLRC5;refGene_tx=XM_011290148.2;refGene_gn=NLRC5;refGene_tx=XM_011290149.2;refGene_gn=NLRC5;refGene_tx=XM_011290150.2;refGene_gn=NLRC5;refGene_tx=XM_0112901

51.2;refGene_gn=NLRC5;refGene_tx=XM_011290152.2;refGene_gn=NLRC5;refGene_tx=XM_019819771.2;refGene_gn=NLRC5;refGene_tx=XM_019819773.1;refGene_gn=NLRC5;refGene_tx=XM_019819775.1;refGene_gn=NLRC5;refGene_tx=XM_019819776.1;refGene_gn=NLRC5;refGene_tx=XM_019819777.1
E2 36205925 36206006
refGene_gn=NLRC5;refGene_tx=XM_003998063.4;refGene_gn=NLRC5;refGene_tx=XM_011290148.2;refGene_gn=NLRC5;refGene_tx=XM_011290149.2;refGene_gn=NLRC5;refGene_tx=XM_011290150.2;refGene_gn=NLRC5;refGene_tx=XM_011290151.2;refGene_gn=NLRC5;refGene_tx=XM_011290152.2;refGene_gn=NLRC5;refGene_tx=XM_019819771.2;refGene_gn=NLRC5;refGene_tx=XM_019819773.1;refGene_gn=NLRC5;refGene_tx=XM_019819775.1;refGene_gn=NLRC5;refGene_tx=XM_019819776.1;refGene_gn=NLRC5;refGene_tx=XM_019819777.1
E2 36207078 36208753
refGene_gn=NLRC5;refGene_tx=XM_003998063.4;refGene_gn=NLRC5;refGene_tx=XM_011290148.2;refGene_gn=NLRC5;refGene_tx=XM_011290149.2;refGene_gn=NLRC5;refGene_tx=XM_011290150.2;refGene_gn=NLRC5;refGene_tx=XM_011290151.2;refGene_gn=NLRC5;refGene_tx=XM_011290152.2;refGene_gn=NLRC5;refGene_tx=XM_019819771.2;refGene_gn=NLRC5;refGene_tx=XM_019819773.1;refGene_gn=NLRC5;refGene_tx=XM_019819775.1;refGene_gn=NLRC5;refGene_tx=XM_019819776.1;refGene_gn=NLRC5;refGene_tx=XM_019819777.1
E2 36209768 36209852
refGene_gn=NLRC5;refGene_tx=XM_003998063.4;refGene_gn=NLRC5;refGene_tx=XM_011290148.2;refGene_gn=NLRC5;refGene_tx=XM_011290149.2;refGene_gn=NLRC5;refGene_tx=XM_011290150.2;refGene_gn=NLRC5;refGene_tx=XM_011290151.2;refGene_gn=NLRC5;refGene_tx=XM_011290152.2;refGene_gn=NLRC5;refGene_tx=XM_019819771.2;refGene_gn=NLRC5;refGene_tx=XM_019819773.1;refGene_gn=NLRC5;refGene_tx=XM_019819775.1;refGene_gn=NLRC5;refGene_tx=XM_019819776.1;refGene_gn=NLRC5;refGene_tx=XM_019819777.1
E2 36209998 36210082
refGene_gn=NLRC5;refGene_tx=XM_003998063.4;refGene_gn=NLRC5;refGene_tx=XM_011290148.2;refGene_gn=NLRC5;refGene_tx=XM_011290149.2;refGene_gn=NLRC5;refGene_tx=XM_011290150.2;refGene_gn=NLRC5;refGene_tx=XM_011290151.2;refGene_gn=NLRC5;refGene_tx=XM_011290152.2;refGene_gn=NLRC5;refGene_tx=XM_019819771.2;refGene_gn=NLRC5;refGene_tx=XM_019819773.1;refGene_gn=NLRC5;refGene_tx=XM_019819775.1;refGene_gn=NLRC5;refGene_tx=XM_019819776.1;refGene_gn=NLRC5;refGene_tx=XM_019819777.1
E2 36211451 36211535
refGene_gn=NLRC5;refGene_tx=XM_003998063.4;refGene_gn=NLRC5;refGene_tx=XM_011290148.2;refGene_gn=NLRC5;refGene_tx=XM_011290149.2;refGene_gn=NLRC5;refGene_tx=XM_011290150.2;refGene_gn=NLRC5;refGene_tx=XM_011290151.2;refGene_gn=NLRC5;refGene_tx=XM_019819771.2;refGene_gn=NLRC5;refGene_tx=XM_019819773.1;refGene_gn=NLRC5;refGene_tx=XM_019819775.1;refGene_gn=NLRC5;refGene_tx=XM_019819776.1;refGene_gn=NLRC5;refGene_tx=XM_019819777.1
E2 36211655 36211745
refGene_gn=NLRC5;refGene_tx=XM_003998063.4;refGene_gn=NLRC5;refGene_tx=XM_011290148.2;refGene_gn=NLRC5;refGene_tx=XM_011290149.2;refGene_gn=NLRC5;refGene_tx=XM_011290150.2;refGene_gn=NLRC5;refGene_tx=XM_011290151.2;refGene_gn=NLRC5;refGene_tx=XM_019819771.2;refGene_gn=NLRC5;refGene_tx=XM_019819773.1;refGene_gn=NLRC5;refGene_tx=XM_019819775.1;refGen

e_gn=NLRC5;refGene_tx=XM_019819776.1
E2 36212731 36212791
refGene_gn=NLRC5;refGene_tx=XM_003998063.4;refGene_gn=NLRC5;refGene_tx=XM_011290148.2;refGene_gn=NLRC5;refGene_tx=XM_011290149.2;refGene_gn=NLRC5;refGene_tx=XM_011290150.2;refGene_gn=NLRC5;refGene_tx=XM_011290151.2;refGene_gn=NLRC5;refGene_tx=XM_011290152.2;refGene_gn=NLRC5;refGene_tx=XM_019819771.2;refGene_gn=NLRC5;refGene_tx=XM_019819773.1;refGene_gn=NLRC5;refGene_tx=XM_019819775.1;refGene_gn=NLRC5;refGene_tx=XM_019819776.1;refGene_gn=NLRC5;refGene_tx=XM_019819777.1
E2 36214519 36214585
refGene_gn=NLRC5;refGene_tx=XM_003998063.4;refGene_gn=NLRC5;refGene_tx=XM_011290148.2;refGene_gn=NLRC5;refGene_tx=XM_011290149.2;refGene_gn=NLRC5;refGene_tx=XM_011290150.2;refGene_gn=NLRC5;refGene_tx=XM_011290151.2;refGene_gn=NLRC5;refGene_tx=XM_011290152.2;refGene_gn=NLRC5;refGene_tx=XM_019819771.2;refGene_gn=NLRC5;refGene_tx=XM_019819773.1;refGene_gn=NLRC5;refGene_tx=XM_019819775.1;refGene_gn=NLRC5;refGene_tx=XM_019819776.1;refGene_gn=NLRC5;refGene_tx=XM_019819777.1
E2 36215051 36215135
refGene_gn=NLRC5;refGene_tx=XM_003998063.4;refGene_gn=NLRC5;refGene_tx=XM_011290148.2;refGene_gn=NLRC5;refGene_tx=XM_011290149.2;refGene_gn=NLRC5;refGene_tx=XM_011290150.2;refGene_gn=NLRC5;refGene_tx=XM_011290151.2;refGene_gn=NLRC5;refGene_tx=XM_011290152.2;refGene_gn=NLRC5;refGene_tx=XM_019819771.2;refGene_gn=NLRC5;refGene_tx=XM_019819773.1;refGene_gn=NLRC5;refGene_tx=XM_019819775.1;refGene_gn=NLRC5;refGene_tx=XM_019819777.1
E2 36216904 36216988
refGene_gn=NLRC5;refGene_tx=XM_003998063.4;refGene_gn=NLRC5;refGene_tx=XM_011290148.2;refGene_gn=NLRC5;refGene_tx=XM_011290149.2;refGene_gn=NLRC5;refGene_tx=XM_011290150.2;refGene_gn=NLRC5;refGene_tx=XM_011290151.2;refGene_gn=NLRC5;refGene_tx=XM_011290152.2;refGene_gn=NLRC5;refGene_tx=XM_019819771.2;refGene_gn=NLRC5;refGene_tx=XM_019819773.1;refGene_gn=NLRC5;refGene_tx=XM_019819775.1;refGene_gn=NLRC5;refGene_tx=XM_019819776.1;refGene_gn=NLRC5;refGene_tx=XM_019819777.1
E2 36217960 36218050
refGene_gn=NLRC5;refGene_tx=XM_003998063.4;refGene_gn=NLRC5;refGene_tx=XM_011290148.2;refGene_gn=NLRC5;refGene_tx=XM_011290149.2;refGene_gn=NLRC5;refGene_tx=XM_011290150.2;refGene_gn=NLRC5;refGene_tx=XM_011290151.2;refGene_gn=NLRC5;refGene_tx=XM_011290152.2;refGene_gn=NLRC5;refGene_tx=XM_019819771.2;refGene_gn=NLRC5;refGene_tx=XM_019819773.1;refGene_gn=NLRC5;refGene_tx=XM_019819775.1;refGene_gn=NLRC5;refGene_tx=XM_019819776.1;refGene_gn=NLRC5;refGene_tx=XM_019819777.1
E2 36219835 36219904
refGene_gn=NLRC5;refGene_tx=XM_003998063.4;refGene_gn=NLRC5;refGene_tx=XM_011290148.2;refGene_gn=NLRC5;refGene_tx=XM_011290149.2;refGene_gn=NLRC5;refGene_tx=XM_011290150.2;refGene_gn=NLRC5;refGene_tx=XM_011290151.2;refGene_gn=NLRC5;refGene_tx=XM_011290152.2;refGene_gn=NLRC5;refGene_tx=XM_019819771.2;refGene_gn=NLRC5;refGene_tx=XM_019819773.1;refGene_gn=NLRC5;refGene_tx=XM_019819775.1;refGene_gn=NLRC5;refGene_tx=XM_019819776.1;refGene_gn=NLRC5;refGene_tx=XM_019819777.1
E2 36220676 36220745

refGene_gn=NLRC5;refGene_tx=XM_003998063.4;refGene_gn=NLRC5;refGene_tx=XM_011290148.2;refGene_gn=NLRC5;refGene_tx=XM_011290149.2;refGene_gn=NLRC5;refGene_tx=XM_011290150.2;refGene_gn=NLRC5;refGene_tx=XM_011290151.2;refGene_gn=NLRC5;refGene_tx=XM_011290152.2;refGene_gn=NLRC5;refGene_tx=XM_019819771.2;refGene_gn=NLRC5;refGene_tx=XM_019819773.1;refGene_gn=NLRC5;refGene_tx=XM_019819775.1;refGene_gn=NLRC5;refGene_tx=XM_019819776.1;refGene_gn=NLRC5;refGene_tx=XM_019819777.1
E2 36222112 36222196
refGene_gn=NLRC5;refGene_tx=XM_003998063.4;refGene_gn=NLRC5;refGene_tx=XM_011290148.2;refGene_gn=NLRC5;refGene_tx=XM_011290149.2;refGene_gn=NLRC5;refGene_tx=XM_011290150.2;refGene_gn=NLRC5;refGene_tx=XM_011290151.2;refGene_gn=NLRC5;refGene_tx=XM_011290152.2;refGene_gn=NLRC5;refGene_tx=XM_019819771.2;refGene_gn=NLRC5;refGene_tx=XM_019819773.1;refGene_gn=NLRC5;refGene_tx=XM_019819775.1;refGene_gn=NLRC5;refGene_tx=XM_019819776.1;refGene_gn=NLRC5;refGene_tx=XM_019819777.1
E2 36222635 36222719
refGene_gn=NLRC5;refGene_tx=XM_003998063.4;refGene_gn=NLRC5;refGene_tx=XM_011290148.2;refGene_gn=NLRC5;refGene_tx=XM_011290149.2;refGene_gn=NLRC5;refGene_tx=XM_011290150.2;refGene_gn=NLRC5;refGene_tx=XM_011290151.2;refGene_gn=NLRC5;refGene_tx=XM_011290152.2;refGene_gn=NLRC5;refGene_tx=XM_019819771.2;refGene_gn=NLRC5;refGene_tx=XM_019819773.1;refGene_gn=NLRC5;refGene_tx=XM_019819775.1;refGene_gn=NLRC5;refGene_tx=XM_019819776.1;refGene_gn=NLRC5;refGene_tx=XM_019819777.1
E2 36224245 36224335
refGene_gn=NLRC5;refGene_tx=XM_003998063.4;refGene_gn=NLRC5;refGene_tx=XM_011290148.2;refGene_gn=NLRC5;refGene_tx=XM_011290149.2;refGene_gn=NLRC5;refGene_tx=XM_011290150.2;refGene_gn=NLRC5;refGene_tx=XM_011290151.2;refGene_gn=NLRC5;refGene_tx=XM_011290152.2;refGene_gn=NLRC5;refGene_tx=XM_019819771.2;refGene_gn=NLRC5;refGene_tx=XM_019819773.1;refGene_gn=NLRC5;refGene_tx=XM_019819775.1;refGene_gn=NLRC5;refGene_tx=XM_019819776.1;refGene_gn=NLRC5;refGene_tx=XM_019819777.1
E2 36225087 36225132
refGene_gn=NLRC5;refGene_tx=XM_003998063.4;refGene_gn=NLRC5;refGene_tx=XM_011290148.2;refGene_gn=NLRC5;refGene_tx=XM_011290149.2;refGene_gn=NLRC5;refGene_tx=XM_011290150.2;refGene_gn=NLRC5;refGene_tx=XM_011290151.2;refGene_gn=NLRC5;refGene_tx=XM_011290152.2;refGene_gn=NLRC5;refGene_tx=XM_019819771.2;refGene_gn=NLRC5;refGene_tx=XM_019819773.1;refGene_gn=NLRC5;refGene_tx=XM_019819775.1;refGene_gn=NLRC5;refGene_tx=XM_019819776.1;refGene_gn=NLRC5;refGene_tx=XM_019819777.1
E2 36226088 36226178
refGene_gn=NLRC5;refGene_tx=XM_003998063.4;refGene_gn=NLRC5;refGene_tx=XM_011290148.2;refGene_gn=NLRC5;refGene_tx=XM_011290149.2;refGene_gn=NLRC5;refGene_tx=XM_011290150.2;refGene_gn=NLRC5;refGene_tx=XM_011290151.2;refGene_gn=NLRC5;refGene_tx=XM_011290152.2;refGene_gn=NLRC5;refGene_tx=XM_019819771.2;refGene_gn=NLRC5;refGene_tx=XM_019819773.1;refGene_gn=NLRC5;refGene_tx=XM_019819775.1;refGene_gn=NLRC5;refGene_tx=XM_019819776.1;refGene_gn=NLRC5;refGene_tx=XM_019819777.1
E2 36227103 36227187
refGene_gn=NLRC5;refGene_tx=XM_003998063.4;refGene_gn=NLRC5;refGene_tx=XM_011290148.2;refGene_gn=NLRC5;refGene_tx=XM_011290149.2;refGene_gn=

NLRC5;refGene_tx=XM_011290150.2;refGene_gn=NLRC5;refGene_tx=XM_0112901
51.2;refGene_gn=NLRC5;refGene_tx=XM_011290152.2;refGene_gn=NLRC5;refGe
ne_tx=XM_019819771.2;refGene_gn=NLRC5;refGene_tx=XM_019819773.1;refGen
e_gn=NLRC5;refGene_tx=XM_019819775.1;refGene_gn=NLRC5;refGene_tx=XM_01
9819776.1;refGene_gn=NLRC5;refGene_tx=XM_019819777.1
E2 36231055 36231139
refGene_gn=NLRC5;refGene_tx=XM_003998063.4;refGene_gn=NLRC5;refGene_tx
=XM_011290148.2;refGene_gn=NLRC5;refGene_tx=XM_011290149.2;refGene_gn=
NLRC5;refGene_tx=XM_011290150.2;refGene_gn=NLRC5;refGene_tx=XM_0112901
51.2;refGene_gn=NLRC5;refGene_tx=XM_011290152.2;refGene_gn=NLRC5;refGe
ne_tx=XM_019819771.2;refGene_gn=NLRC5;refGene_tx=XM_019819773.1;refGen
e_gn=NLRC5;refGene_tx=XM_019819775.1;refGene_gn=NLRC5;refGene_tx=XM_01
9819776.1;refGene_gn=NLRC5;refGene_tx=XM_019819777.1
E2 36233510 36233600
refGene_gn=NLRC5;refGene_tx=XM_003998063.4;refGene_gn=NLRC5;refGene_tx
=XM_011290148.2;refGene_gn=NLRC5;refGene_tx=XM_011290149.2;refGene_gn=
NLRC5;refGene_tx=XM_011290150.2;refGene_gn=NLRC5;refGene_tx=XM_0112901
51.2;refGene_gn=NLRC5;refGene_tx=XM_011290152.2;refGene_gn=NLRC5;refGe
ne_tx=XM_019819771.2;refGene_gn=NLRC5;refGene_tx=XM_019819773.1;refGen
e_gn=NLRC5;refGene_tx=XM_019819775.1;refGene_gn=NLRC5;refGene_tx=XM_01
9819776.1;refGene_gn=NLRC5;refGene_tx=XM_019819777.1
E2 36233787 36233850
refGene_gn=NLRC5;refGene_tx=XM_003998063.4;refGene_gn=NLRC5;refGene_tx
=XM_011290148.2;refGene_gn=NLRC5;refGene_tx=XM_011290149.2;refGene_gn=
NLRC5;refGene_tx=XM_011290150.2;refGene_gn=NLRC5;refGene_tx=XM_0112901
51.2;refGene_gn=NLRC5;refGene_tx=XM_011290152.2;refGene_gn=NLRC5;refGe
ne_tx=XM_019819771.2;refGene_gn=NLRC5;refGene_tx=XM_019819773.1;refGen
e_gn=NLRC5;refGene_tx=XM_019819775.1;refGene_gn=NLRC5;refGene_tx=XM_01
9819776.1;refGene_gn=NLRC5;refGene_tx=XM_019819777.1
E2 36236552 36236639
refGene_gn=NLRC5;refGene_tx=XM_003998063.4;refGene_gn=NLRC5;refGene_tx
=XM_011290148.2;refGene_gn=NLRC5;refGene_tx=XM_011290149.2;refGene_gn=
NLRC5;refGene_tx=XM_011290150.2;refGene_gn=NLRC5;refGene_tx=XM_0112901
51.2;refGene_gn=NLRC5;refGene_tx=XM_011290152.2;refGene_gn=NLRC5;refGe
ne_tx=XM_019819771.2;refGene_gn=NLRC5;refGene_tx=XM_019819773.1;refGen
e_gn=NLRC5;refGene_tx=XM_019819775.1;refGene_gn=NLRC5;refGene_tx=XM_01
9819776.1;refGene_gn=NLRC5;refGene_tx=XM_019819777.1
E2 36238296 36238380
refGene_gn=NLRC5;refGene_tx=XM_003998063.4;refGene_gn=NLRC5;refGene_tx
=XM_011290148.2;refGene_gn=NLRC5;refGene_tx=XM_011290149.2;refGene_gn=
NLRC5;refGene_tx=XM_011290150.2;refGene_gn=NLRC5;refGene_tx=XM_0112901
51.2;refGene_gn=NLRC5;refGene_tx=XM_011290152.2;refGene_gn=NLRC5;refGe
ne_tx=XM_019819771.2;refGene_gn=NLRC5;refGene_tx=XM_019819773.1;refGen
e_gn=NLRC5;refGene_tx=XM_019819775.1;refGene_gn=NLRC5;refGene_tx=XM_01
9819776.1;refGene_gn=NLRC5;refGene_tx=XM_019819777.1
E2 36239026 36239116
refGene_gn=NLRC5;refGene_tx=XM_003998063.4;refGene_gn=NLRC5;refGene_tx
=XM_011290148.2;refGene_gn=NLRC5;refGene_tx=XM_011290149.2;refGene_gn=
NLRC5;refGene_tx=XM_011290150.2;refGene_gn=NLRC5;refGene_tx=XM_0112901
51.2;refGene_gn=NLRC5;refGene_tx=XM_011290152.2;refGene_gn=NLRC5;refGe

ne_tx=XM_019819771.2;refGene_gn=NLRC5;refGene_tx=XM_019819773.1;refGen
e_gn=NLRC5;refGene_tx=XM_019819775.1;refGene_gn=NLRC5;refGene_tx=XM_01
9819776.1;refGene_gn=NLRC5;refGene_tx=XM_019819777.1
E2 36239437 36239503
refGene_gn=NLRC5;refGene_tx=XM_003998063.4;refGene_gn=NLRC5;refGene_tx
=XM_011290148.2;refGene_gn=NLRC5;refGene_tx=XM_011290149.2;refGene_gn=
NLRC5;refGene_tx=XM_011290150.2;refGene_gn=NLRC5;refGene_tx=XM_0112901
51.2;refGene_gn=NLRC5;refGene_tx=XM_011290152.2;refGene_gn=NLRC5;refGe
ne_tx=XM_019819771.2;refGene_gn=NLRC5;refGene_tx=XM_019819773.1;refGen
e_gn=NLRC5;refGene_tx=XM_019819775.1;refGene_gn=NLRC5;refGene_tx=XM_01
9819776.1;refGene_gn=NLRC5;refGene_tx=XM_019819777.1
E2 36241104 36241188
refGene_gn=NLRC5;refGene_tx=XM_003998063.4;refGene_gn=NLRC5;refGene_tx
=XM_011290148.2;refGene_gn=NLRC5;refGene_tx=XM_011290149.2;refGene_gn=
NLRC5;refGene_tx=XM_011290150.2;refGene_gn=NLRC5;refGene_tx=XM_0112901
51.2;refGene_gn=NLRC5;refGene_tx=XM_011290152.2;refGene_gn=NLRC5;refGe
ne_tx=XM_019819771.2;refGene_gn=NLRC5;refGene_tx=XM_019819773.1;refGen
e_gn=NLRC5;refGene_tx=XM_019819775.1;refGene_gn=NLRC5;refGene_tx=XM_01
9819776.1;refGene_gn=NLRC5;refGene_tx=XM_019819777.1
E2 36241268 36241352
refGene_gn=NLRC5;refGene_tx=XM_003998063.4;refGene_gn=NLRC5;refGene_tx
=XM_011290148.2;refGene_gn=NLRC5;refGene_tx=XM_011290149.2;refGene_gn=
NLRC5;refGene_tx=XM_011290150.2;refGene_gn=NLRC5;refGene_tx=XM_0112901
51.2;refGene_gn=NLRC5;refGene_tx=XM_011290152.2;refGene_gn=NLRC5;refGe
ne_tx=XM_019819771.2;refGene_gn=NLRC5;refGene_tx=XM_019819773.1;refGen
e_gn=NLRC5;refGene_tx=XM_019819775.1;refGene_gn=NLRC5;refGene_tx=XM_01
9819776.1;refGene_gn=NLRC5;refGene_tx=XM_019819777.1
E2 36244676 36244763
refGene_gn=NLRC5;refGene_tx=XM_003998063.4;refGene_gn=NLRC5;refGene_tx
=XM_011290148.2;refGene_gn=NLRC5;refGene_tx=XM_011290149.2;refGene_gn=
NLRC5;refGene_tx=XM_011290150.2;refGene_gn=NLRC5;refGene_tx=XM_0112901
51.2;refGene_gn=NLRC5;refGene_tx=XM_011290152.2;refGene_gn=NLRC5;refGe
ne_tx=XM_019819771.2;refGene_gn=NLRC5;refGene_tx=XM_019819773.1;refGen
e_gn=NLRC5;refGene_tx=XM_019819775.1;refGene_gn=NLRC5;refGene_tx=XM_01
9819776.1;refGene_gn=NLRC5;refGene_tx=XM_019819777.1
E2 36245257 36245335
refGene_gn=NLRC5;refGene_tx=XM_003998063.4;refGene_gn=NLRC5;refGene_tx
=XM_011290148.2;refGene_gn=NLRC5;refGene_tx=XM_011290149.2;refGene_gn=
NLRC5;refGene_tx=XM_011290150.2;refGene_gn=NLRC5;refGene_tx=XM_0112901
51.2;refGene_gn=NLRC5;refGene_tx=XM_011290152.2;refGene_gn=NLRC5;refGe
ne_tx=XM_019819771.2;refGene_gn=NLRC5;refGene_tx=XM_019819773.1;refGen
e_gn=NLRC5;refGene_tx=XM_019819775.1;refGene_gn=NLRC5;refGene_tx=XM_01
9819776.1;refGene_gn=NLRC5;refGene_tx=XM_019819777.1
E2 36246134 36246218
refGene_gn=NLRC5;refGene_tx=XM_003998063.4;refGene_gn=NLRC5;refGene_tx
=XM_011290148.2;refGene_gn=NLRC5;refGene_tx=XM_011290149.2;refGene_gn=
NLRC5;refGene_tx=XM_011290150.2;refGene_gn=NLRC5;refGene_tx=XM_0112901
51.2;refGene_gn=NLRC5;refGene_tx=XM_011290152.2;refGene_gn=NLRC5;refGe
ne_tx=XM_019819771.2;refGene_gn=NLRC5;refGene_tx=XM_019819773.1;refGen
e_gn=NLRC5;refGene_tx=XM_019819775.1;refGene_gn=NLRC5;refGene_tx=XM_01

9819776.1;refGene_gn=NLRC5;refGene_tx=XM_019819777.1
E2 36246534 36246618
refGene_gn=NLRC5;refGene_tx=XM_003998063.4;refGene_gn=NLRC5;refGene_tx=XM_011290148.2;refGene_gn=NLRC5;refGene_tx=XM_011290149.2;refGene_gn=NLRC5;refGene_tx=XM_011290150.2;refGene_gn=NLRC5;refGene_tx=XM_011290151.2;refGene_gn=NLRC5;refGene_tx=XM_011290152.2;refGene_gn=NLRC5;refGene_tx=XM_019819771.2;refGene_gn=NLRC5;refGene_tx=XM_019819773.1;refGene_gn=NLRC5;refGene_tx=XM_019819775.1;refGene_gn=NLRC5;refGene_tx=XM_019819776.1;refGene_gn=NLRC5;refGene_tx=XM_019819777.1
E2 36248108 36248192
refGene_gn=NLRC5;refGene_tx=XM_003998063.4;refGene_gn=NLRC5;refGene_tx=XM_011290148.2;refGene_gn=NLRC5;refGene_tx=XM_011290149.2;refGene_gn=NLRC5;refGene_tx=XM_011290150.2;refGene_gn=NLRC5;refGene_tx=XM_011290151.2;refGene_gn=NLRC5;refGene_tx=XM_011290152.2;refGene_gn=NLRC5;refGene_tx=XM_019819771.2;refGene_gn=NLRC5;refGene_tx=XM_019819773.1;refGene_gn=NLRC5;refGene_tx=XM_019819775.1;refGene_gn=NLRC5;refGene_tx=XM_019819776.1;refGene_gn=NLRC5;refGene_tx=XM_019819777.1
E2 36248806 36248890
refGene_gn=NLRC5;refGene_tx=XM_003998063.4;refGene_gn=NLRC5;refGene_tx=XM_011290148.2;refGene_gn=NLRC5;refGene_tx=XM_011290149.2;refGene_gn=NLRC5;refGene_tx=XM_011290150.2;refGene_gn=NLRC5;refGene_tx=XM_011290151.2;refGene_gn=NLRC5;refGene_tx=XM_011290152.2;refGene_gn=NLRC5;refGene_tx=XM_019819771.2;refGene_gn=NLRC5;refGene_tx=XM_019819773.1;refGene_gn=NLRC5;refGene_tx=XM_019819775.1;refGene_gn=NLRC5;refGene_tx=XM_019819776.1;refGene_gn=NLRC5;refGene_tx=XM_019819777.1
E2 36250609 36250693
refGene_gn=NLRC5;refGene_tx=XM_003998063.4;refGene_gn=NLRC5;refGene_tx=XM_011290148.2;refGene_gn=NLRC5;refGene_tx=XM_011290149.2;refGene_gn=NLRC5;refGene_tx=XM_011290150.2;refGene_gn=NLRC5;refGene_tx=XM_011290151.2;refGene_gn=NLRC5;refGene_tx=XM_011290152.2;refGene_gn=NLRC5;refGene_tx=XM_019819771.2;refGene_gn=NLRC5;refGene_tx=XM_019819773.1;refGene_gn=NLRC5;refGene_tx=XM_019819775.1;refGene_gn=NLRC5;refGene_tx=XM_019819776.1;refGene_gn=NLRC5;refGene_tx=XM_019819777.1
E2 36252141 36252225 refGene_gn=NLRC5;refGene_tx=XM_003998063.4
E2 36252627 36252711
refGene_gn=NLRC5;refGene_tx=XM_003998063.4;refGene_gn=NLRC5;refGene_tx=XM_011290148.2;refGene_gn=NLRC5;refGene_tx=XM_011290149.2;refGene_gn=NLRC5;refGene_tx=XM_011290150.2;refGene_gn=NLRC5;refGene_tx=XM_011290151.2;refGene_gn=NLRC5;refGene_tx=XM_011290152.2;refGene_gn=NLRC5;refGene_tx=XM_019819771.2;refGene_gn=NLRC5;refGene_tx=XM_019819773.1;refGene_gn=NLRC5;refGene_tx=XM_019819775.1;refGene_gn=NLRC5;refGene_tx=XM_019819776.1;refGene_gn=NLRC5;refGene_tx=XM_019819777.1
E2 36253013 36253097
refGene_gn=NLRC5;refGene_tx=XM_003998063.4;refGene_gn=NLRC5;refGene_tx=XM_011290148.2;refGene_gn=NLRC5;refGene_tx=XM_011290149.2;refGene_gn=NLRC5;refGene_tx=XM_011290150.2;refGene_gn=NLRC5;refGene_tx=XM_011290151.2;refGene_gn=NLRC5;refGene_tx=XM_011290152.2;refGene_gn=NLRC5;refGene_tx=XM_019819771.2;refGene_gn=NLRC5;refGene_tx=XM_019819773.1;refGene_gn=NLRC5;refGene_tx=XM_019819775.1;refGene_gn=NLRC5;refGene_tx=XM_019819776.1;refGene_gn=NLRC5;refGene_tx=XM_019819777.1

E2 36253242 36253320
refGene_gn=NLRC5;refGene_tx=XM_003998063.4;refGene_gn=NLRC5;refGene_tx=XM_011290148.2;refGene_gn=NLRC5;refGene_tx=XM_011290149.2;refGene_gn=NLRC5;refGene_tx=XM_011290150.2;refGene_gn=NLRC5;refGene_tx=XM_011290151.2;refGene_gn=NLRC5;refGene_tx=XM_011290152.2;refGene_gn=NLRC5;refGene_tx=XM_019819771.2;refGene_gn=NLRC5;refGene_tx=XM_019819773.1;refGene_gn=NLRC5;refGene_tx=XM_019819775.1;refGene_gn=NLRC5;refGene_tx=XM_019819776.1;refGene_gn=NLRC5;refGene_tx=XM_019819777.1
E2 36254042 36254126
refGene_gn=NLRC5;refGene_tx=XM_003998063.4;refGene_gn=NLRC5;refGene_tx=XM_011290148.2;refGene_gn=NLRC5;refGene_tx=XM_011290149.2;refGene_gn=NLRC5;refGene_tx=XM_011290150.2;refGene_gn=NLRC5;refGene_tx=XM_011290151.2;refGene_gn=NLRC5;refGene_tx=XM_011290152.2;refGene_gn=NLRC5;refGene_tx=XM_019819771.2;refGene_gn=NLRC5;refGene_tx=XM_019819773.1;refGene_gn=NLRC5;refGene_tx=XM_019819775.1;refGene_gn=NLRC5;refGene_tx=XM_019819776.1;refGene_gn=NLRC5;refGene_tx=XM_019819777.1
E2 36254211 36254283
refGene_gn=NLRC5;refGene_tx=XM_003998063.4;refGene_gn=NLRC5;refGene_tx=XM_011290148.2;refGene_gn=NLRC5;refGene_tx=XM_011290149.2;refGene_gn=NLRC5;refGene_tx=XM_011290150.2;refGene_gn=NLRC5;refGene_tx=XM_011290151.2;refGene_gn=NLRC5;refGene_tx=XM_011290152.2;refGene_gn=NLRC5;refGene_tx=XM_019819771.2;refGene_gn=NLRC5;refGene_tx=XM_019819773.1;refGene_gn=NLRC5;refGene_tx=XM_019819775.1;refGene_gn=NLRC5;refGene_tx=XM_019819776.1;refGene_gn=NLRC5;refGene_tx=XM_019819777.1
E2 36254564 36254648
refGene_gn=NLRC5;refGene_tx=XM_003998063.4;refGene_gn=NLRC5;refGene_tx=XM_011290148.2;refGene_gn=NLRC5;refGene_tx=XM_011290149.2;refGene_gn=NLRC5;refGene_tx=XM_011290150.2;refGene_gn=NLRC5;refGene_tx=XM_011290151.2;refGene_gn=NLRC5;refGene_tx=XM_011290152.2;refGene_gn=NLRC5;refGene_tx=XM_019819771.2;refGene_gn=NLRC5;refGene_tx=XM_019819773.1;refGene_gn=NLRC5;refGene_tx=XM_019819775.1;refGene_gn=NLRC5;refGene_tx=XM_019819776.1;refGene_gn=NLRC5;refGene_tx=XM_019819777.1
E2 36256316 36256400
refGene_gn=NLRC5;refGene_tx=XM_003998063.4;refGene_gn=NLRC5;refGene_tx=XM_011290148.2;refGene_gn=NLRC5;refGene_tx=XM_011290149.2;refGene_gn=NLRC5;refGene_tx=XM_011290150.2;refGene_gn=NLRC5;refGene_tx=XM_011290151.2;refGene_gn=NLRC5;refGene_tx=XM_011290152.2;refGene_gn=NLRC5;refGene_tx=XM_019819771.2;refGene_gn=NLRC5;refGene_tx=XM_019819773.1;refGene_gn=NLRC5;refGene_tx=XM_019819775.1;refGene_gn=NLRC5;refGene_tx=XM_019819776.1;refGene_gn=NLRC5;refGene_tx=XM_019819777.1
E2 36256705 36256789
refGene_gn=NLRC5;refGene_tx=XM_003998063.4;refGene_gn=NLRC5;refGene_tx=XM_011290148.2;refGene_gn=NLRC5;refGene_tx=XM_011290149.2;refGene_gn=NLRC5;refGene_tx=XM_011290150.2;refGene_gn=NLRC5;refGene_tx=XM_011290151.2;refGene_gn=NLRC5;refGene_tx=XM_011290152.2;refGene_gn=NLRC5;refGene_tx=XM_019819771.2;refGene_gn=NLRC5;refGene_tx=XM_019819773.1;refGene_gn=NLRC5;refGene_tx=XM_019819775.1;refGene_gn=NLRC5;refGene_tx=XM_019819776.1;refGene_gn=NLRC5;refGene_tx=XM_019819777.1
E2 36257615 36257727
refGene_gn=NLRC5;refGene_tx=XM_003998063.4;refGene_gn=NLRC5;refGene_tx

=XM_011290148.2;refGene_gn=NLRC5;refGene_tx=XM_011290149.2;refGene_gn=NLRC5;refGene_tx=XM_011290150.2;refGene_gn=NLRC5;refGene_tx=XM_011290151.2;refGene_gn=NLRC5;refGene_tx=XM_011290152.2;refGene_gn=NLRC5;refGene_tx=XM_019819771.2;refGene_gn=NLRC5;refGene_tx=XM_019819773.1;refGene_gn=NLRC5;refGene_tx=XM_019819775.1;refGene_gn=NLRC5;refGene_tx=XM_019819776.1;refGene_gn=NLRC5;refGene_tx=XM_019819777.1
E2 36282094 36282274 refGene_gn=CPNE2;refGene_tx=XM_023245147.1
E2 36285017 36285197 refGene_gn=CPNE2;refGene_tx=XM_023245147.1
E2 36286916 36286991 refGene_gn=CPNE2;refGene_tx=XM_023245147.1
E2 36288847 36288919 refGene_gn=CPNE2;refGene_tx=XM_023245147.1
E2 36289792 36289876 refGene_gn=CPNE2;refGene_tx=XM_023245147.1
E2 36290151 36290241 refGene_gn=CPNE2;refGene_tx=XM_023245147.1
E2 36291594 36291693 refGene_gn=CPNE2;refGene_tx=XM_023245147.1
E2 36292240 36292327 refGene_gn=CPNE2;refGene_tx=XM_023245147.1
E2 36293795 36293855 refGene_gn=CPNE2;refGene_tx=XM_023245147.1
E2 36295717 36295851 refGene_gn=CPNE2;refGene_tx=XM_023245147.1
E2 36297970 36298025 refGene_gn=CPNE2;refGene_tx=XM_023245147.1
E2 36303180 36303232 refGene_gn=CPNE2;refGene_tx=XM_023245147.1
E2 36304595 36304729 refGene_gn=CPNE2;refGene_tx=XM_023245147.1
E2 36311486 36311723 refGene_gn=CPNE2;refGene_tx=XM_023245147.1
E2 36313003 36313111 refGene_gn=CPNE2;refGene_tx=XM_023245147.1
E2 36319728 36319946
refGene_gn=FAM192A;refGene_tx=XM_003998065.4;refGene_gn=FAM192A;refGene_tx=XM_006941530.4;refGene_gn=FAM192A;refGene_tx=XM_006941532.4;refGene_gn=FAM192A;refGene_tx=XM_006941533.4;refGene_gn=FAM192A;refGene_tx=XM_006941535.3;refGene_gn=FAM192A;refGene_tx=XM_006941536.4
E2 36327810 36327875
refGene_gn=FAM192A;refGene_tx=XM_003998065.4;refGene_gn=FAM192A;refGene_tx=XM_006941530.4;refGene_gn=FAM192A;refGene_tx=XM_006941532.4;refGene_gn=FAM192A;refGene_tx=XM_006941533.4;refGene_gn=FAM192A;refGene_tx=XM_006941535.3;refGene_gn=FAM192A;refGene_tx=XM_006941536.4
E2 36334804 36334938
refGene_gn=FAM192A;refGene_tx=XM_003998065.4;refGene_gn=FAM192A;refGene_tx=XM_006941530.4;refGene_gn=FAM192A;refGene_tx=XM_006941532.4;refGene_gn=FAM192A;refGene_tx=XM_006941533.4;refGene_gn=FAM192A;refGene_tx=XM_006941535.3;refGene_gn=FAM192A;refGene_tx=XM_006941536.4
E2 36339628 36339750
refGene_gn=FAM192A;refGene_tx=XM_003998065.4;refGene_gn=FAM192A;refGene_tx=XM_006941530.4;refGene_gn=FAM192A;refGene_tx=XM_006941532.4;refGene_gn=FAM192A;refGene_tx=XM_006941533.4;refGene_gn=FAM192A;refGene_tx=XM_006941535.3;refGene_gn=FAM192A;refGene_tx=XM_006941536.4
E2 36340151 36340250
refGene_gn=FAM192A;refGene_tx=XM_003998065.4;refGene_gn=FAM192A;refGene_tx=XM_006941530.4;refGene_gn=FAM192A;refGene_tx=XM_006941532.4;refGene_gn=FAM192A;refGene_tx=XM_006941533.4;refGene_gn=FAM192A;refGene_tx=XM_006941535.3;refGene_gn=FAM192A;refGene_tx=XM_006941536.4
E2 36340785 36340912
refGene_gn=FAM192A;refGene_tx=XM_003998065.4;refGene_gn=FAM192A;refGene_tx=XM_006941530.4;refGene_gn=FAM192A;refGene_tx=XM_006941532.4;refGene_gn=FAM192A;refGene_tx=XM_006941533.4;refGene_gn=FAM192A;refGene_tx=

XM_006941535.3;refGene_gn=FAM192A;refGene_tx=XM_006941536.4
E2 36364095 36364445
refGene_gn=RSPRY1;refGene_tx=XM_006941537.4;refGene_gn=RSPRY1;refGene_tx=XM_006941538.4;refGene_gn=RSPRY1;refGene_tx=XM_023245146.1
E2 36370068 36370121
refGene_gn=RSPRY1;refGene_tx=XM_006941537.4;refGene_gn=RSPRY1;refGene_tx=XM_006941538.4;refGene_gn=RSPRY1;refGene_tx=XM_023245146.1
E2 36370805 36370918
refGene_gn=RSPRY1;refGene_tx=XM_006941537.4;refGene_gn=RSPRY1;refGene_tx=XM_006941538.4;refGene_gn=RSPRY1;refGene_tx=XM_023245146.1
E2 36372226 36372353
refGene_gn=RSPRY1;refGene_tx=XM_006941537.4;refGene_gn=RSPRY1;refGene_tx=XM_006941538.4;refGene_gn=RSPRY1;refGene_tx=XM_023245146.1
E2 36373399 36373458
refGene_gn=RSPRY1;refGene_tx=XM_006941537.4;refGene_gn=RSPRY1;refGene_tx=XM_006941538.4;refGene_gn=RSPRY1;refGene_tx=XM_023245146.1
E2 36375351 36375418
refGene_gn=RSPRY1;refGene_tx=XM_006941537.4;refGene_gn=RSPRY1;refGene_tx=XM_006941538.4;refGene_gn=RSPRY1;refGene_tx=XM_023245146.1
E2 36376048 36376180
refGene_gn=RSPRY1;refGene_tx=XM_006941537.4;refGene_gn=RSPRY1;refGene_tx=XM_006941538.4;refGene_gn=RSPRY1;refGene_tx=XM_023245146.1
E2 36377854 36377970
refGene_gn=RSPRY1;refGene_tx=XM_006941537.4;refGene_gn=RSPRY1;refGene_tx=XM_006941538.4;refGene_gn=RSPRY1;refGene_tx=XM_023245146.1
E2 36379125 36379269
refGene_gn=RSPRY1;refGene_tx=XM_006941537.4;refGene_gn=RSPRY1;refGene_tx=XM_006941538.4;refGene_gn=RSPRY1;refGene_tx=XM_023245146.1
E2 36383889 36384001
refGene_gn=RSPRY1;refGene_tx=XM_006941537.4;refGene_gn=RSPRY1;refGene_tx=XM_006941538.4;refGene_gn=RSPRY1;refGene_tx=XM_023245146.1
E2 36385330 36385433
refGene_gn=RSPRY1;refGene_tx=XM_006941537.4;refGene_gn=RSPRY1;refGene_tx=XM_006941538.4;refGene_gn=RSPRY1;refGene_tx=XM_023245146.1
E2 36385771 36385924
refGene_gn=RSPRY1;refGene_tx=XM_006941537.4;refGene_gn=RSPRY1;refGene_tx=XM_006941538.4;refGene_gn=RSPRY1;refGene_tx=XM_023245146.1
E2 36390935 36391040
refGene_gn=RSPRY1;refGene_tx=XM_006941537.4;refGene_gn=RSPRY1;refGene_tx=XM_006941538.4;refGene_gn=RSPRY1;refGene_tx=XM_023245146.1
E2 36394205 36394302
refGene_gn=RSPRY1;refGene_tx=XM_006941537.4;refGene_gn=RSPRY1;refGene_tx=XM_006941538.4;refGene_gn=RSPRY1;refGene_tx=XM_023245146.1
E2 36400903 36400941
refGene_gn=ARL2BP;refGene_tx=XM_003998067.4;refGene_gn=ARL2BP;refGene_tx=XM_006941540.4
E2 36401640 36401702
refGene_gn=ARL2BP;refGene_tx=XM_003998067.4;refGene_gn=ARL2BP;refGene_tx=XM_006941540.4
E2 36404403 36404510

refGene_gn=ARL2BP;refGene_tx=XM_003998067.4;refGene_gn=ARL2BP;refGene_tx=XM_006941540.4
E2 36406879 36406965
refGene_gn=ARL2BP;refGene_tx=XM_003998067.4;refGene_gn=ARL2BP;refGene_tx=XM_006941540.4
E2 36407177 36407274
refGene_gn=ARL2BP;refGene_tx=XM_003998067.4;refGene_gn=ARL2BP;refGene_tx=XM_006941540.4
E2 36407718 36407869
refGene_gn=ARL2BP;refGene_tx=XM_003998067.4;refGene_gn=ARL2BP;refGene_tx=XM_006941540.4
E2 36411925 36412042 refGene_gn=PLLP;refGene_tx=XM_003998068.4
E2 36413174 36413297 refGene_gn=PLLP;refGene_tx=XM_003998068.4
E2 36415431 36415605 refGene_gn=PLLP;refGene_tx=XM_003998068.4
E2 36431376 36431511 refGene_gn=PLLP;refGene_tx=XM_003998068.4
E2 36482972 36483045 refGene_gn=CCL22;refGene_tx=XM_003998096.4
E2 36484451 36484575 refGene_gn=CCL22;refGene_tx=XM_003998096.4
E2 36487454 36487539 refGene_gn=CCL22;refGene_tx=XM_003998096.4
E2 36496550 36496620 refGene_gn=CX3CL1;refGene_tx=XM_023245001.1
E2 36503081 36503202 refGene_gn=CX3CL1;refGene_tx=XM_023245001.1
E2 36505101 36506227
refGene_gn=CX3CL1;refGene_tx=XM_023245001.1;refGene_gn=CX3CL1;refGene_tx=XM_023245002.1
E2 36523718 36523788 refGene_gn=CCL17;refGene_tx=NM_001009849.1
E2 36525219 36525337 refGene_gn=CCL17;refGene_tx=NM_001009849.1
E2 36525973 36526085 refGene_gn=CCL17;refGene_tx=NM_001009849.1
E2 36534446 36534557 refGene_gn=CIAPIN1;refGene_tx=XM_003998072.4
E2 36535079 36535161 refGene_gn=CIAPIN1;refGene_tx=XM_003998072.4
E2 36538159 36538275 refGene_gn=CIAPIN1;refGene_tx=XM_003998072.4
E2 36539155 36539229 refGene_gn=CIAPIN1;refGene_tx=XM_003998072.4
E2 36540666 36540835 refGene_gn=CIAPIN1;refGene_tx=XM_003998072.4
E2 36542503 36542580 refGene_gn=CIAPIN1;refGene_tx=XM_003998072.4
E2 36544023 36544176 refGene_gn=CIAPIN1;refGene_tx=XM_003998072.4
E2 36544624 36544781 refGene_gn=CIAPIN1;refGene_tx=XM_003998072.4
E2 36550475 36550650 refGene_gn=COQ9;refGene_tx=XM_003998073.4
E2 36553437 36553606 refGene_gn=COQ9;refGene_tx=XM_003998073.4
E2 36554986 36555122 refGene_gn=COQ9;refGene_tx=XM_003998073.4
E2 36559519 36559662 refGene_gn=COQ9;refGene_tx=XM_003998073.4
E2 36559945 36560030 refGene_gn=COQ9;refGene_tx=XM_003998073.4
E2 36561514 36561619 refGene_gn=COQ9;refGene_tx=XM_003998073.4
E2 36563120 36563276 refGene_gn=COQ9;refGene_tx=XM_003998073.4
E2 36563535 36563589 refGene_gn=COQ9;refGene_tx=XM_003998073.4
E2 36563914 36563950 refGene_gn=COQ9;refGene_tx=XM_003998073.4
E2 36566421 36566507 refGene_gn=POLR2C;refGene_tx=XM_003998097.5
E2 36566691 36566741 refGene_gn=POLR2C;refGene_tx=XM_003998097.5
E2 36569077 36569146 refGene_gn=POLR2C;refGene_tx=XM_003998097.5
E2 36569292 36569345 refGene_gn=POLR2C;refGene_tx=XM_003998097.5
E2 36572277 36572406 refGene_gn=POLR2C;refGene_tx=XM_003998097.5
E2 36572828 36572880 refGene_gn=POLR2C;refGene_tx=XM_003998097.5
E2 36573098 36573267 refGene_gn=POLR2C;refGene_tx=XM_003998097.5

E2 36573412 36573487 refGene_gn=POLR2C;refGene_tx=XM_003998097.5
E2 36574013 36574158 refGene_gn=POLR2C;refGene_tx=XM_003998097.5
E2 36576293 36576653
refGene_gn=DOK4;refGene_tx=XM_003998074.4;refGene_gn=DOK4;refGene_tx=XM_019819784.2;refGene_gn=DOK4;refGene_tx=XM_023244938.1;refGene_gn=DOK4;refGene_tx=XM_023244939.1
E2 36576809 36576948
refGene_gn=DOK4;refGene_tx=XM_003998074.4;refGene_gn=DOK4;refGene_tx=XM_019819784.2;refGene_gn=DOK4;refGene_tx=XM_023244938.1;refGene_gn=DOK4;refGene_tx=XM_023244939.1
E2 36577687 36577877
refGene_gn=DOK4;refGene_tx=XM_003998074.4;refGene_gn=DOK4;refGene_tx=XM_019819784.2;refGene_gn=DOK4;refGene_tx=XM_023244938.1;refGene_gn=DOK4;refGene_tx=XM_023244939.1
E2 36577967 36578087
refGene_gn=DOK4;refGene_tx=XM_003998074.4;refGene_gn=DOK4;refGene_tx=XM_019819784.2;refGene_gn=DOK4;refGene_tx=XM_023244938.1;refGene_gn=DOK4;refGene_tx=XM_023244939.1
E2 36578380 36578495
refGene_gn=DOK4;refGene_tx=XM_003998074.4;refGene_gn=DOK4;refGene_tx=XM_019819784.2;refGene_gn=DOK4;refGene_tx=XM_023244938.1;refGene_gn=DOK4;refGene_tx=XM_023244939.1
E2 36578661 36578769
refGene_gn=DOK4;refGene_tx=XM_003998074.4;refGene_gn=DOK4;refGene_tx=XM_019819784.2;refGene_gn=DOK4;refGene_tx=XM_023244938.1;refGene_gn=DOK4;refGene_tx=XM_023244939.1
E2 36582282 36582348
refGene_gn=DOK4;refGene_tx=XM_003998074.4;refGene_gn=DOK4;refGene_tx=XM_019819784.2;refGene_gn=DOK4;refGene_tx=XM_023244938.1;refGene_gn=DOK4;refGene_tx=XM_023244939.1
E2 36600553 36600727
refGene_gn=LOC111556252;refGene_tx=XM_023245210.1
E2 36602150 36602269
refGene_gn=LOC111556252;refGene_tx=XM_023245210.1
E2 36603505 36603675
refGene_gn=LOC111556252;refGene_tx=XM_023245210.1
E2 36604000 36604049
refGene_gn=LOC111556252;refGene_tx=XM_023245210.1
E2 36605847 36605958
refGene_gn=LOC111556252;refGene_tx=XM_023245210.1
E2 36606181 36606260
refGene_gn=LOC111556252;refGene_tx=XM_023245210.1
E2 36608898 36609028
refGene_gn=CCDC102A;refGene_tx=XM_023245207.1;refGene_gn=CCDC102A;refGene_tx=XM_023245208.1;refGene_gn=CCDC102A;refGene_tx=XM_023245209.1
E2 36611240 36611344
refGene_gn=CCDC102A;refGene_tx=XM_023245207.1;refGene_gn=CCDC102A;refGene_tx=XM_023245208.1;refGene_gn=CCDC102A;refGene_tx=XM_023245209.1
E2 36612395 36612566
refGene_gn=CCDC102A;refGene_tx=XM_023245207.1;refGene_gn=CCDC102A;refG

ene_tx=XM_023245208.1;refGene_gn=CCDC102A;refGene_tx=XM_023245209.1
E2 36612946 36613156
refGene_gn=CCDC102A;refGene_tx=XM_023245207.1;refGene_gn=CCDC102A;refGene_tx=XM_023245208.1;refGene_gn=CCDC102A;refGene_tx=XM_023245209.1
E2 36613515 36613632
refGene_gn=CCDC102A;refGene_tx=XM_023245207.1;refGene_gn=CCDC102A;refGene_tx=XM_023245208.1;refGene_gn=CCDC102A;refGene_tx=XM_023245209.1
E2 36616103 36616212
refGene_gn=CCDC102A;refGene_tx=XM_023245207.1;refGene_gn=CCDC102A;refGene_tx=XM_023245208.1;refGene_gn=CCDC102A;refGene_tx=XM_023245209.1
E2 36619134 36619361
refGene_gn=CCDC102A;refGene_tx=XM_023245207.1;refGene_gn=CCDC102A;refGene_tx=XM_023245208.1;refGene_gn=CCDC102A;refGene_tx=XM_023245209.1
E2 36621499 36622084
refGene_gn=CCDC102A;refGene_tx=XM_023245207.1;refGene_gn=CCDC102A;refGene_tx=XM_023245208.1;refGene_gn=CCDC102A;refGene_tx=XM_023245209.1
E2 36642158 36642222
refGene_gn=ADGRG5;refGene_tx=XM_006941542.2;refGene_gn=ADGRG5;refGene_tx=XM_023245318.1;refGene_gn=ADGRG5;refGene_tx=XM_023245319.1
E2 36642421 36642500
refGene_gn=ADGRG5;refGene_tx=XM_006941542.2;refGene_gn=ADGRG5;refGene_tx=XM_023245318.1;refGene_gn=ADGRG5;refGene_tx=XM_023245319.1
E2 36643132 36643289
refGene_gn=ADGRG5;refGene_tx=XM_006941542.2;refGene_gn=ADGRG5;refGene_tx=XM_023245318.1;refGene_gn=ADGRG5;refGene_tx=XM_023245319.1
E2 36643852 36643969
refGene_gn=ADGRG5;refGene_tx=XM_006941542.2;refGene_gn=ADGRG5;refGene_tx=XM_023245318.1;refGene_gn=ADGRG5;refGene_tx=XM_023245319.1
E2 36644482 36644596
refGene_gn=ADGRG5;refGene_tx=XM_006941542.2;refGene_gn=ADGRG5;refGene_tx=XM_023245318.1;refGene_gn=ADGRG5;refGene_tx=XM_023245319.1
E2 36646062 36646215
refGene_gn=ADGRG5;refGene_tx=XM_006941542.2;refGene_gn=ADGRG5;refGene_tx=XM_023245318.1;refGene_gn=ADGRG5;refGene_tx=XM_023245319.1
E2 36646954 36647076
refGene_gn=ADGRG5;refGene_tx=XM_006941542.2;refGene_gn=ADGRG5;refGene_tx=XM_023245318.1;refGene_gn=ADGRG5;refGene_tx=XM_023245319.1
E2 36647330 36647590
refGene_gn=ADGRG5;refGene_tx=XM_006941542.2;refGene_gn=ADGRG5;refGene_tx=XM_023245318.1;refGene_gn=ADGRG5;refGene_tx=XM_023245319.1
E2 36651154 36651272
refGene_gn=ADGRG5;refGene_tx=XM_006941542.2;refGene_gn=ADGRG5;refGene_tx=XM_023245318.1;refGene_gn=ADGRG5;refGene_tx=XM_023245319.1
E2 36653335 36653513 refGene_gn=ADGRG5;refGene_tx=XM_023245319.1
E2 36653933 36654211
refGene_gn=ADGRG5;refGene_tx=XM_006941542.2;refGene_gn=ADGRG5;refGene_tx=XM_023245318.1
E2 36654593 36654694
refGene_gn=ADGRG5;refGene_tx=XM_006941542.2;refGene_gn=ADGRG5;refGene_tx=XM_023245318.1

E2 36716495 36716559
refGene_gn=ADGRG1;refGene_tx=XM_003998075.5;refGene_gn=ADGRG1;refGene_tx=XM_006941543.3;refGene_gn=ADGRG1;refGene_tx=XM_006941544.4;refGene_gn=ADGRG1;refGene_tx=XM_006941545.3;refGene_gn=ADGRG1;refGene_tx=XM_019819790.2;refGene_gn=ADGRG1;refGene_tx=XM_019819791.2;refGene_gn=ADGRG1;refGene_tx=XM_019819792.2;refGene_gn=ADGRG1;refGene_tx=XM_019819793.2
E2 36717308 36717746
refGene_gn=ADGRG1;refGene_tx=XM_003998075.5;refGene_gn=ADGRG1;refGene_tx=XM_006941543.3;refGene_gn=ADGRG1;refGene_tx=XM_006941544.4;refGene_gn=ADGRG1;refGene_tx=XM_006941545.3;refGene_gn=ADGRG1;refGene_tx=XM_019819790.2;refGene_gn=ADGRG1;refGene_tx=XM_019819791.2;refGene_gn=ADGRG1;refGene_tx=XM_019819792.2;refGene_gn=ADGRG1;refGene_tx=XM_019819793.2
E2 36718382 36718515
refGene_gn=ADGRG1;refGene_tx=XM_003998075.5;refGene_gn=ADGRG1;refGene_tx=XM_006941543.3;refGene_gn=ADGRG1;refGene_tx=XM_006941544.4;refGene_gn=ADGRG1;refGene_tx=XM_006941545.3;refGene_gn=ADGRG1;refGene_tx=XM_019819790.2;refGene_gn=ADGRG1;refGene_tx=XM_019819791.2;refGene_gn=ADGRG1;refGene_tx=XM_019819792.2;refGene_gn=ADGRG1;refGene_tx=XM_019819793.2
E2 36719080 36719228
refGene_gn=ADGRG1;refGene_tx=XM_003998075.5;refGene_gn=ADGRG1;refGene_tx=XM_006941543.3;refGene_gn=ADGRG1;refGene_tx=XM_006941544.4;refGene_gn=ADGRG1;refGene_tx=XM_006941545.3;refGene_gn=ADGRG1;refGene_tx=XM_019819790.2;refGene_gn=ADGRG1;refGene_tx=XM_019819791.2;refGene_gn=ADGRG1;refGene_tx=XM_019819792.2;refGene_gn=ADGRG1;refGene_tx=XM_019819793.2
E2 36719770 36719902
refGene_gn=ADGRG1;refGene_tx=XM_003998075.5;refGene_gn=ADGRG1;refGene_tx=XM_006941543.3;refGene_gn=ADGRG1;refGene_tx=XM_006941544.4;refGene_gn=ADGRG1;refGene_tx=XM_006941545.3;refGene_gn=ADGRG1;refGene_tx=XM_019819790.2;refGene_gn=ADGRG1;refGene_tx=XM_019819791.2;refGene_gn=ADGRG1;refGene_tx=XM_019819792.2;refGene_gn=ADGRG1;refGene_tx=XM_019819793.2
E2 36720260 36720377
refGene_gn=ADGRG1;refGene_tx=XM_003998075.5;refGene_gn=ADGRG1;refGene_tx=XM_006941543.3;refGene_gn=ADGRG1;refGene_tx=XM_006941544.4;refGene_gn=ADGRG1;refGene_tx=XM_006941545.3;refGene_gn=ADGRG1;refGene_tx=XM_019819790.2;refGene_gn=ADGRG1;refGene_tx=XM_019819791.2;refGene_gn=ADGRG1;refGene_tx=XM_019819792.2;refGene_gn=ADGRG1;refGene_tx=XM_019819793.2
E2 36720587 36720633
refGene_gn=ADGRG1;refGene_tx=XM_003998075.5;refGene_gn=ADGRG1;refGene_tx=XM_006941543.3;refGene_gn=ADGRG1;refGene_tx=XM_006941544.4;refGene_gn=ADGRG1;refGene_tx=XM_006941545.3;refGene_gn=ADGRG1;refGene_tx=XM_019819790.2;refGene_gn=ADGRG1;refGene_tx=XM_019819791.2;refGene_gn=ADGRG1;refGene_tx=XM_019819792.2;refGene_gn=ADGRG1;refGene_tx=XM_019819793.2
E2 36720876 36720980

refGene_gn=ADGRG1;refGene_tx=XM_003998075.5;refGene_gn=ADGRG1;refGene_tx=XM_006941543.3;refGene_gn=ADGRG1;refGene_tx=XM_006941544.4;refGene_gn=ADGRG1;refGene_tx=XM_006941545.3;refGene_gn=ADGRG1;refGene_tx=XM_019819790.2;refGene_gn=ADGRG1;refGene_tx=XM_019819791.2;refGene_gn=ADGRG1;refGene_tx=XM_019819792.2;refGene_gn=ADGRG1;refGene_tx=XM_019819793.2
E2 36721754 36721873
refGene_gn=ADGRG1;refGene_tx=XM_003998075.5;refGene_gn=ADGRG1;refGene_tx=XM_006941543.3;refGene_gn=ADGRG1;refGene_tx=XM_006941544.4;refGene_gn=ADGRG1;refGene_tx=XM_006941545.3;refGene_gn=ADGRG1;refGene_tx=XM_019819790.2;refGene_gn=ADGRG1;refGene_tx=XM_019819791.2;refGene_gn=ADGRG1;refGene_tx=XM_019819792.2;refGene_gn=ADGRG1;refGene_tx=XM_019819793.2
E2 36723551 36723820
refGene_gn=ADGRG1;refGene_tx=XM_003998075.5;refGene_gn=ADGRG1;refGene_tx=XM_006941543.3;refGene_gn=ADGRG1;refGene_tx=XM_006941544.4;refGene_gn=ADGRG1;refGene_tx=XM_006941545.3;refGene_gn=ADGRG1;refGene_tx=XM_019819790.2;refGene_gn=ADGRG1;refGene_tx=XM_019819791.2;refGene_gn=ADGRG1;refGene_tx=XM_019819792.2;refGene_gn=ADGRG1;refGene_tx=XM_019819793.2
E2 36724784 36724893
refGene_gn=ADGRG1;refGene_tx=XM_003998075.5;refGene_gn=ADGRG1;refGene_tx=XM_006941543.3;refGene_gn=ADGRG1;refGene_tx=XM_006941544.4;refGene_gn=ADGRG1;refGene_tx=XM_006941545.3;refGene_gn=ADGRG1;refGene_tx=XM_019819790.2;refGene_gn=ADGRG1;refGene_tx=XM_019819791.2;refGene_gn=ADGRG1;refGene_tx=XM_019819792.2;refGene_gn=ADGRG1;refGene_tx=XM_019819793.2
E2 36725291 36725560
refGene_gn=ADGRG1;refGene_tx=XM_003998075.5;refGene_gn=ADGRG1;refGene_tx=XM_006941543.3;refGene_gn=ADGRG1;refGene_tx=XM_006941544.4;refGene_gn=ADGRG1;refGene_tx=XM_006941545.3;refGene_gn=ADGRG1;refGene_tx=XM_019819790.2;refGene_gn=ADGRG1;refGene_tx=XM_019819791.2;refGene_gn=ADGRG1;refGene_tx=XM_019819792.2;refGene_gn=ADGRG1;refGene_tx=XM_019819793.2
E2 36726765 36726896
refGene_gn=ADGRG1;refGene_tx=XM_003998075.5;refGene_gn=ADGRG1;refGene_tx=XM_006941543.3;refGene_gn=ADGRG1;refGene_tx=XM_006941544.4;refGene_gn=ADGRG1;refGene_tx=XM_006941545.3;refGene_gn=ADGRG1;refGene_tx=XM_019819790.2;refGene_gn=ADGRG1;refGene_tx=XM_019819791.2;refGene_gn=ADGRG1;refGene_tx=XM_019819792.2;refGene_gn=ADGRG1;refGene_tx=XM_019819793.2
E2 36731626 36731687
refGene_gn=ADGRG3;refGene_tx=XM_019819797.2;refGene_gn=ADGRG3;refGene_tx=XM_023245526.1;refGene_gn=ADGRG3;refGene_tx=XM_023245527.1
E2 36736459 36736619
refGene_gn=ADGRG3;refGene_tx=XM_019819797.2;refGene_gn=ADGRG3;refGene_tx=XM_023245526.1;refGene_gn=ADGRG3;refGene_tx=XM_023245527.1
E2 36748392 36748528
refGene_gn=ADGRG3;refGene_tx=XM_019819797.2;refGene_gn=ADGRG3;refGene_tx=XM_023245526.1;refGene_gn=ADGRG3;refGene_tx=XM_023245527.1

E2 36750074 36750221
refGene_gn=ADGRG3;refGene_tx=XM_019819797.2;refGene_gn=ADGRG3;refGene_tx=XM_023245526.1;refGene_gn=ADGRG3;refGene_tx=XM_023245527.1
E2 36751072 36751207
refGene_gn=ADGRG3;refGene_tx=XM_019819797.2;refGene_gn=ADGRG3;refGene_tx=XM_023245526.1;refGene_gn=ADGRG3;refGene_tx=XM_023245527.1
E2 36751746 36751786
refGene_gn=ADGRG3;refGene_tx=XM_019819797.2;refGene_gn=ADGRG3;refGene_tx=XM_023245526.1;refGene_gn=ADGRG3;refGene_tx=XM_023245527.1
E2 36751955 36752056
refGene_gn=ADGRG3;refGene_tx=XM_019819797.2;refGene_gn=ADGRG3;refGene_tx=XM_023245526.1;refGene_gn=ADGRG3;refGene_tx=XM_023245527.1
E2 36752189 36752302
refGene_gn=ADGRG3;refGene_tx=XM_019819797.2;refGene_gn=ADGRG3;refGene_tx=XM_023245526.1;refGene_gn=ADGRG3;refGene_tx=XM_023245527.1
E2 36753426 36753642
refGene_gn=ADGRG3;refGene_tx=XM_019819797.2;refGene_gn=ADGRG3;refGene_tx=XM_023245526.1
E2 36754335 36754616
refGene_gn=ADGRG3;refGene_tx=XM_019819797.2;refGene_gn=ADGRG3;refGene_tx=XM_023245526.1;refGene_gn=ADGRG3;refGene_tx=XM_023245527.1
E2 36754811 36754905
refGene_gn=ADGRG3;refGene_tx=XM_019819797.2;refGene_gn=ADGRG3;refGene_tx=XM_023245526.1;refGene_gn=ADGRG3;refGene_tx=XM_023245527.1
E2 36756216 36756500
refGene_gn=ADGRG3;refGene_tx=XM_019819797.2;refGene_gn=ADGRG3;refGene_tx=XM_023245526.1;refGene_gn=ADGRG3;refGene_tx=XM_023245527.1
E2 36758364 36758477
refGene_gn=ADGRG3;refGene_tx=XM_019819797.2;refGene_gn=ADGRG3;refGene_tx=XM_023245526.1;refGene_gn=ADGRG3;refGene_tx=XM_023245527.1
E2 36768276 36768473 refGene_gn=DRC7;refGene_tx=XM_003998076.4
E2 36768871 36769046 refGene_gn=DRC7;refGene_tx=XM_003998076.4
E2 36770068 36770194 refGene_gn=DRC7;refGene_tx=XM_003998076.4
E2 36770954 36771149 refGene_gn=DRC7;refGene_tx=XM_003998076.4
E2 36772834 36772993 refGene_gn=DRC7;refGene_tx=XM_003998076.4
E2 36773838 36774057 refGene_gn=DRC7;refGene_tx=XM_003998076.4
E2 36776714 36776843 refGene_gn=DRC7;refGene_tx=XM_003998076.4
E2 36779140 36779213 refGene_gn=DRC7;refGene_tx=XM_003998076.4
E2 36780026 36780155 refGene_gn=DRC7;refGene_tx=XM_003998076.4
E2 36780340 36780469 refGene_gn=DRC7;refGene_tx=XM_003998076.4
E2 36781727 36781948 refGene_gn=DRC7;refGene_tx=XM_003998076.4
E2 36782736 36782952 refGene_gn=DRC7;refGene_tx=XM_003998076.4
E2 36783530 36783641 refGene_gn=DRC7;refGene_tx=XM_003998076.4
E2 36783730 36783841 refGene_gn=DRC7;refGene_tx=XM_003998076.4
E2 36784736 36784931 refGene_gn=DRC7;refGene_tx=XM_003998076.4
E2 36785515 36785655 refGene_gn=DRC7;refGene_tx=XM_003998076.4
E2 36785747 36785841 refGene_gn=DRC7;refGene_tx=XM_003998076.4
E2 36790926 36790966
refGene_gn=KATNB1;refGene_tx=XM_003998077.5;refGene_gn=KATNB1;refGene_tx=XM_006941546.4

E2	36795129	36795260
refGene_gn=KATNB1;refGene_tx=XM_003998077.5;refGene_gn=KATNB1;refGene_tx=XM_006941546.4
E2	36798312	36798430
refGene_gn=KATNB1;refGene_tx=XM_003998077.5;refGene_gn=KATNB1;refGene_tx=XM_006941546.4
E2	36803705	36803806
refGene_gn=KATNB1;refGene_tx=XM_003998077.5;refGene_gn=KATNB1;refGene_tx=XM_006941546.4
E2	36804416	36804458
refGene_gn=KATNB1;refGene_tx=XM_003998077.5;refGene_gn=KATNB1;refGene_tx=XM_006941546.4
E2	36805078	36805162
refGene_gn=KATNB1;refGene_tx=XM_003998077.5;refGene_gn=KATNB1;refGene_tx=XM_006941546.4
E2	36805383	36805499
refGene_gn=KATNB1;refGene_tx=XM_003998077.5;refGene_gn=KATNB1;refGene_tx=XM_006941546.4
E2	36805926	36805998
refGene_gn=KATNB1;refGene_tx=XM_003998077.5;refGene_gn=KATNB1;refGene_tx=XM_006941546.4
E2	36806159	36806310
refGene_gn=KATNB1;refGene_tx=XM_003998077.5;refGene_gn=KATNB1;refGene_tx=XM_006941546.4
E2	36806410	36806601
refGene_gn=KATNB1;refGene_tx=XM_003998077.5;refGene_gn=KATNB1;refGene_tx=XM_006941546.4
E2	36806715	36806846
refGene_gn=KATNB1;refGene_tx=XM_003998077.5;refGene_gn=KATNB1;refGene_tx=XM_006941546.4
E2	36807257	36807308
refGene_gn=KATNB1;refGene_tx=XM_003998077.5;refGene_gn=KATNB1;refGene_tx=XM_006941546.4
E2	36808154	36808222
refGene_gn=KATNB1;refGene_tx=XM_003998077.5;refGene_gn=KATNB1;refGene_tx=XM_006941546.4
E2	36808361	36808481
refGene_gn=KATNB1;refGene_tx=XM_003998077.5;refGene_gn=KATNB1;refGene_tx=XM_006941546.4
E2	36808571	36808721
refGene_gn=KATNB1;refGene_tx=XM_003998077.5;refGene_gn=KATNB1;refGene_tx=XM_006941546.4
E2	36808984	36809061
refGene_gn=KATNB1;refGene_tx=XM_003998077.5;refGene_gn=KATNB1;refGene_tx=XM_006941546.4
E2	36809141	36809216
refGene_gn=KATNB1;refGene_tx=XM_003998077.5;refGene_gn=KATNB1;refGene_tx=XM_006941546.4
E2	36809476	36809593
refGene_gn=KATNB1;refGene_tx=XM_003998077.5;refGene_gn=KATNB1;refGene_

tx=XM_006941546.4
E2 36809872 36810005
refGene_gn=KATNB1;refGene_tx=XM_003998077.5;refGene_gn=KATNB1;refGene_
tx=XM_006941546.4
E2 36811946 36811972
refGene_gn=KIFC3;refGene_tx=XM_006941548.4;refGene_gn=KIFC3;refGene_tx
=XM_006941549.4;refGene_gn=KIFC3;refGene_tx=XM_006941550.4;refGene_gn=
KIFC3;refGene_tx=XM_019819801.2;refGene_gn=KIFC3;refGene_tx=XM_0232458
36.1;refGene_gn=KIFC3;refGene_tx=XM_023245837.1;refGene_gn=KIFC3;refGe
ne_tx=XM_023245838.1;refGene_gn=KIFC3;refGene_tx=XM_023245840.1
E2 36812185 36812190 refGene_gn=KIFC3;refGene_tx=XM_011290143.3
E2 36812772 36812881
refGene_gn=KIFC3;refGene_tx=XM_006941548.4;refGene_gn=KIFC3;refGene_tx
=XM_006941549.4;refGene_gn=KIFC3;refGene_tx=XM_006941550.4;refGene_gn=
KIFC3;refGene_tx=XM_011290143.3;refGene_gn=KIFC3;refGene_tx=XM_0198198
01.2;refGene_gn=KIFC3;refGene_tx=XM_023245836.1;refGene_gn=KIFC3;refGe
ne_tx=XM_023245837.1;refGene_gn=KIFC3;refGene_tx=XM_023245838.1;refGen
e_gn=KIFC3;refGene_tx=XM_023245840.1
E2 36813237 36813372
refGene_gn=KIFC3;refGene_tx=XM_006941548.4;refGene_gn=KIFC3;refGene_tx
=XM_006941549.4;refGene_gn=KIFC3;refGene_tx=XM_006941550.4;refGene_gn=
KIFC3;refGene_tx=XM_011290143.3;refGene_gn=KIFC3;refGene_tx=XM_0198198
01.2;refGene_gn=KIFC3;refGene_tx=XM_023245836.1;refGene_gn=KIFC3;refGe
ne_tx=XM_023245837.1;refGene_gn=KIFC3;refGene_tx=XM_023245838.1;refGen
e_gn=KIFC3;refGene_tx=XM_023245840.1
E2 36813593 36813823
refGene_gn=KIFC3;refGene_tx=XM_006941548.4;refGene_gn=KIFC3;refGene_tx
=XM_006941549.4;refGene_gn=KIFC3;refGene_tx=XM_006941550.4;refGene_gn=
KIFC3;refGene_tx=XM_011290143.3;refGene_gn=KIFC3;refGene_tx=XM_0198198
01.2;refGene_gn=KIFC3;refGene_tx=XM_023245836.1;refGene_gn=KIFC3;refGe
ne_tx=XM_023245837.1;refGene_gn=KIFC3;refGene_tx=XM_023245838.1;refGen
e_gn=KIFC3;refGene_tx=XM_023245840.1
E2 36813913 36814043
refGene_gn=KIFC3;refGene_tx=XM_006941548.4;refGene_gn=KIFC3;refGene_tx
=XM_006941549.4;refGene_gn=KIFC3;refGene_tx=XM_006941550.4;refGene_gn=
KIFC3;refGene_tx=XM_011290143.3;refGene_gn=KIFC3;refGene_tx=XM_0198198
01.2;refGene_gn=KIFC3;refGene_tx=XM_023245836.1;refGene_gn=KIFC3;refGe
ne_tx=XM_023245837.1;refGene_gn=KIFC3;refGene_tx=XM_023245838.1;refGen
e_gn=KIFC3;refGene_tx=XM_023245840.1
E2 36814246 36814370
refGene_gn=KIFC3;refGene_tx=XM_006941548.4;refGene_gn=KIFC3;refGene_tx
=XM_006941549.4;refGene_gn=KIFC3;refGene_tx=XM_006941550.4;refGene_gn=
KIFC3;refGene_tx=XM_011290143.3;refGene_gn=KIFC3;refGene_tx=XM_0198198
01.2;refGene_gn=KIFC3;refGene_tx=XM_023245836.1;refGene_gn=KIFC3;refGe
ne_tx=XM_023245837.1;refGene_gn=KIFC3;refGene_tx=XM_023245838.1;refGen
e_gn=KIFC3;refGene_tx=XM_023245840.1
E2 36814897 36815028
refGene_gn=KIFC3;refGene_tx=XM_006941548.4;refGene_gn=KIFC3;refGene_tx
=XM_006941549.4;refGene_gn=KIFC3;refGene_tx=XM_006941550.4;refGene_gn=
KIFC3;refGene_tx=XM_011290143.3;refGene_gn=KIFC3;refGene_tx=XM_0198198

01.2;refGene_gn=KIFC3;refGene_tx=XM_023245836.1;refGene_gn=KIFC3;refGene_tx=XM_023245837.1;refGene_gn=KIFC3;refGene_tx=XM_023245838.1;refGene_gn=KIFC3;refGene_tx=XM_023245840.1
E2 36816724 36816829
refGene_gn=KIFC3;refGene_tx=XM_006941548.4;refGene_gn=KIFC3;refGene_tx=XM_006941549.4;refGene_gn=KIFC3;refGene_tx=XM_006941550.4;refGene_gn=KIFC3;refGene_tx=XM_011290143.3;refGene_gn=KIFC3;refGene_tx=XM_019819801.2;refGene_gn=KIFC3;refGene_tx=XM_023245836.1;refGene_gn=KIFC3;refGene_tx=XM_023245837.1;refGene_gn=KIFC3;refGene_tx=XM_023245838.1;refGene_gn=KIFC3;refGene_tx=XM_023245840.1
E2 36817642 36817824
refGene_gn=KIFC3;refGene_tx=XM_006941548.4;refGene_gn=KIFC3;refGene_tx=XM_006941549.4;refGene_gn=KIFC3;refGene_tx=XM_006941550.4;refGene_gn=KIFC3;refGene_tx=XM_011290143.3;refGene_gn=KIFC3;refGene_tx=XM_019819801.2;refGene_gn=KIFC3;refGene_tx=XM_023245836.1;refGene_gn=KIFC3;refGene_tx=XM_023245837.1;refGene_gn=KIFC3;refGene_tx=XM_023245838.1;refGene_gn=KIFC3;refGene_tx=XM_023245840.1
E2 36819061 36819173
refGene_gn=KIFC3;refGene_tx=XM_006941548.4;refGene_gn=KIFC3;refGene_tx=XM_006941549.4;refGene_gn=KIFC3;refGene_tx=XM_006941550.4;refGene_gn=KIFC3;refGene_tx=XM_011290143.3;refGene_gn=KIFC3;refGene_tx=XM_019819801.2;refGene_gn=KIFC3;refGene_tx=XM_023245836.1;refGene_gn=KIFC3;refGene_tx=XM_023245837.1;refGene_gn=KIFC3;refGene_tx=XM_023245838.1;refGene_gn=KIFC3;refGene_tx=XM_023245840.1
E2 36820329 36820460
refGene_gn=KIFC3;refGene_tx=XM_006941548.4;refGene_gn=KIFC3;refGene_tx=XM_006941549.4;refGene_gn=KIFC3;refGene_tx=XM_006941550.4;refGene_gn=KIFC3;refGene_tx=XM_011290143.3;refGene_gn=KIFC3;refGene_tx=XM_019819801.2;refGene_gn=KIFC3;refGene_tx=XM_023245836.1;refGene_gn=KIFC3;refGene_tx=XM_023245837.1;refGene_gn=KIFC3;refGene_tx=XM_023245838.1;refGene_gn=KIFC3;refGene_tx=XM_023245840.1
E2 36820541 36820689
refGene_gn=KIFC3;refGene_tx=XM_006941548.4;refGene_gn=KIFC3;refGene_tx=XM_006941549.4;refGene_gn=KIFC3;refGene_tx=XM_006941550.4;refGene_gn=KIFC3;refGene_tx=XM_011290143.3;refGene_gn=KIFC3;refGene_tx=XM_019819801.2;refGene_gn=KIFC3;refGene_tx=XM_023245836.1;refGene_gn=KIFC3;refGene_tx=XM_023245837.1;refGene_gn=KIFC3;refGene_tx=XM_023245838.1;refGene_gn=KIFC3;refGene_tx=XM_023245840.1
E2 36821338 36821512
refGene_gn=KIFC3;refGene_tx=XM_006941548.4;refGene_gn=KIFC3;refGene_tx=XM_006941549.4;refGene_gn=KIFC3;refGene_tx=XM_006941550.4;refGene_gn=KIFC3;refGene_tx=XM_011290143.3;refGene_gn=KIFC3;refGene_tx=XM_019819801.2;refGene_gn=KIFC3;refGene_tx=XM_023245836.1;refGene_gn=KIFC3;refGene_tx=XM_023245837.1;refGene_gn=KIFC3;refGene_tx=XM_023245838.1;refGene_gn=KIFC3;refGene_tx=XM_023245840.1
E2 36822036 36822276
refGene_gn=KIFC3;refGene_tx=XM_006941548.4;refGene_gn=KIFC3;refGene_tx=XM_006941549.4;refGene_gn=KIFC3;refGene_tx=XM_006941550.4;refGene_gn=KIFC3;refGene_tx=XM_011290143.3;refGene_gn=KIFC3;refGene_tx=XM_019819801.2;refGene_gn=KIFC3;refGene_tx=XM_023245836.1;refGene_gn=KIFC3;refGe

ne_tx=XM_023245837.1;refGene_gn=KIFC3;refGene_tx=XM_023245838.1;refGene_gn=KIFC3;refGene_tx=XM_023245840.1
E2 36822413 36822557
refGene_gn=KIFC3;refGene_tx=XM_006941548.4;refGene_gn=KIFC3;refGene_tx=XM_006941549.4;refGene_gn=KIFC3;refGene_tx=XM_006941550.4;refGene_gn=KIFC3;refGene_tx=XM_011290143.3;refGene_gn=KIFC3;refGene_tx=XM_019819801.2;refGene_gn=KIFC3;refGene_tx=XM_023245836.1;refGene_gn=KIFC3;refGene_tx=XM_023245837.1;refGene_gn=KIFC3;refGene_tx=XM_023245838.1;refGene_gn=KIFC3;refGene_tx=XM_023245840.1
E2 36823084 36823150
refGene_gn=KIFC3;refGene_tx=XM_006941548.4;refGene_gn=KIFC3;refGene_tx=XM_006941549.4;refGene_gn=KIFC3;refGene_tx=XM_006941550.4;refGene_gn=KIFC3;refGene_tx=XM_011290143.3;refGene_gn=KIFC3;refGene_tx=XM_019819801.2;refGene_gn=KIFC3;refGene_tx=XM_023245836.1;refGene_gn=KIFC3;refGene_tx=XM_023245837.1;refGene_gn=KIFC3;refGene_tx=XM_023245838.1;refGene_gn=KIFC3;refGene_tx=XM_023245840.1
E2 36823288 36823297 refGene_gn=KIFC3;refGene_tx=XM_006941550.4
E2 36840490 36840693
refGene_gn=KIFC3;refGene_tx=XM_006941548.4;refGene_gn=KIFC3;refGene_tx=XM_006941549.4;refGene_gn=KIFC3;refGene_tx=XM_011290143.3;refGene_gn=KIFC3;refGene_tx=XM_019819801.2;refGene_gn=KIFC3;refGene_tx=XM_023245836.1;refGene_gn=KIFC3;refGene_tx=XM_023245838.1
E2 36843731 36843870
refGene_gn=KIFC3;refGene_tx=XM_011290143.3;refGene_gn=KIFC3;refGene_tx=XM_019819801.2;refGene_gn=KIFC3;refGene_tx=XM_023245836.1;refGene_gn=KIFC3;refGene_tx=XM_023245838.1
E2 36847701 36847844
refGene_gn=KIFC3;refGene_tx=XM_011290143.3;refGene_gn=KIFC3;refGene_tx=XM_023245836.1;refGene_gn=KIFC3;refGene_tx=XM_023245837.1;refGene_gn=KIFC3;refGene_tx=XM_023245838.1;refGene_gn=KIFC3;refGene_tx=XM_023245840.1
E2 36850410 36850618
refGene_gn=KIFC3;refGene_tx=XM_011290143.3;refGene_gn=KIFC3;refGene_tx=XM_023245836.1;refGene_gn=KIFC3;refGene_tx=XM_023245837.1;refGene_gn=KIFC3;refGene_tx=XM_023245838.1;refGene_gn=KIFC3;refGene_tx=XM_023245840.1
E2 36886458 36886689
refGene_gn=KIFC3;refGene_tx=XM_011290143.3;refGene_gn=KIFC3;refGene_tx=XM_023245836.1;refGene_gn=KIFC3;refGene_tx=XM_023245837.1
E2 36920039 36920528 refGene_gn=CNGB1;refGene_tx=XM_023245061.1
E2 36923591 36923808 refGene_gn=CNGB1;refGene_tx=XM_023245061.1
E2 36930595 36930742 refGene_gn=CNGB1;refGene_tx=XM_023245061.1
E2 36930977 36931096 refGene_gn=CNGB1;refGene_tx=XM_023245061.1
E2 36934138 36934222 refGene_gn=CNGB1;refGene_tx=XM_023245061.1
E2 36934321 36934419 refGene_gn=CNGB1;refGene_tx=XM_023245061.1
E2 36936117 36936277 refGene_gn=CNGB1;refGene_tx=XM_023245061.1
E2 36937010 36937152 refGene_gn=CNGB1;refGene_tx=XM_023245061.1
E2 36942180 36942303 refGene_gn=CNGB1;refGene_tx=XM_023245061.1
E2 36943146 36943205 refGene_gn=CNGB1;refGene_tx=XM_023245061.1
E2 36943206 36943212 refGene_gn=CNGB1;refGene_tx=XM_023245061.1

E2	36945139 36945226 refGene_gn=CNGB1;refGene_tx=XM_023245061.1
E2	36945488 36945539 refGene_gn=CNGB1;refGene_tx=XM_023245061.1
E2	36946505 36946714 refGene_gn=CNGB1;refGene_tx=XM_023245061.1
E2	36947708 36947867 refGene_gn=CNGB1;refGene_tx=XM_023245061.1
E2	36948917 36949075 refGene_gn=CNGB1;refGene_tx=XM_023245061.1
E2	36951185 36951293 refGene_gn=CNGB1;refGene_tx=XM_023245061.1
E2	36954256 36954416 refGene_gn=CNGB1;refGene_tx=XM_023245061.1
E2	36960369 36960406 refGene_gn=CNGB1;refGene_tx=XM_023245061.1
E2	36961100 36961188 refGene_gn=CNGB1;refGene_tx=XM_023245061.1
E2	36966965 36967049 refGene_gn=CNGB1;refGene_tx=XM_023245061.1
E2	36967973 36968136 refGene_gn=CNGB1;refGene_tx=XM_023245061.1
E2	36973927 36973964 refGene_gn=CNGB1;refGene_tx=XM_023245061.1
E2	36975525 36975601 refGene_gn=CNGB1;refGene_tx=XM_023245061.1
E2	36977048 36977208 refGene_gn=CNGB1;refGene_tx=XM_023245061.1
E2	36977587 36977627 refGene_gn=CNGB1;refGene_tx=XM_023245061.1
E2	36977916 36977992 refGene_gn=CNGB1;refGene_tx=XM_023245061.1
E2	36979590 36979627 refGene_gn=CNGB1;refGene_tx=XM_023245061.1
E2	36979825 36979868 refGene_gn=CNGB1;refGene_tx=XM_023245061.1
E2	36979968 36980089 refGene_gn=CNGB1;refGene_tx=XM_023245061.1
E2	36980910 36980983 refGene_gn=CNGB1;refGene_tx=XM_023245061.1
E2	36981280 36981338 refGene_gn=CNGB1;refGene_tx=XM_023245061.1
E2	36983388 36983556 refGene_gn=CNGB1;refGene_tx=XM_023245061.1
E2	36991390 36991613
refGene_gn=TEPP;refGene_tx=XM_011290142.3;refGene_gn=TEPP;refGene_tx=XM_019819810.2;refGene_gn=TEPP;refGene_tx=XM_019819811.2;refGene_gn=TEPP;refGene_tx=XM_023245062.1
E2	36994837 36994899
refGene_gn=TEPP;refGene_tx=XM_011290142.3;refGene_gn=TEPP;refGene_tx=XM_019819810.2;refGene_gn=TEPP;refGene_tx=XM_019819811.2;refGene_gn=TEPP;refGene_tx=XM_023245062.1
E2	36995163 36995338
refGene_gn=TEPP;refGene_tx=XM_011290142.3;refGene_gn=TEPP;refGene_tx=XM_019819810.2;refGene_gn=TEPP;refGene_tx=XM_019819811.2;refGene_gn=TEPP;refGene_tx=XM_023245062.1
E2	36995775 36995837
refGene_gn=TEPP;refGene_tx=XM_011290142.3;refGene_gn=TEPP;refGene_tx=XM_019819811.2
E2	36996068 36996133
refGene_gn=TEPP;refGene_tx=XM_011290142.3;refGene_gn=TEPP;refGene_tx=XM_019819810.2;refGene_gn=TEPP;refGene_tx=XM_019819811.2;refGene_gn=TEPP;refGene_tx=XM_023245062.1
E2	36996510 36996543
refGene_gn=TEPP;refGene_tx=XM_011290142.3;refGene_gn=TEPP;refGene_tx=XM_019819810.2;refGene_gn=TEPP;refGene_tx=XM_019819811.2;refGene_gn=TEPP;refGene_tx=XM_023245062.1
E2	36997675 36997804
refGene_gn=TEPP;refGene_tx=XM_011290142.3;refGene_gn=TEPP;refGene_tx=XM_019819810.2;refGene_gn=TEPP;refGene_tx=XM_019819811.2;refGene_gn=TEPP;refGene_tx=XM_023245062.1
E2	37007068 37008826 refGene_gn=ZNF319;refGene_tx=XM_023245708.1

E2 37012172 37012276 refGene_gn=USB1;refGene_tx=XM_003998082.5
E2 37013127 37013294
refGene_gn=USB1;refGene_tx=XM_003998082.5;refGene_gn=USB1;refGene_tx=XM_019819812.2
E2 37015386 37015570
refGene_gn=USB1;refGene_tx=XM_003998082.5;refGene_gn=USB1;refGene_tx=XM_019819812.2
E2 37018244 37018298
refGene_gn=USB1;refGene_tx=XM_003998082.5;refGene_gn=USB1;refGene_tx=XM_019819812.2
E2 37019090 37019196
refGene_gn=USB1;refGene_tx=XM_003998082.5;refGene_gn=USB1;refGene_tx=XM_019819812.2
E2 37020414 37020498
refGene_gn=USB1;refGene_tx=XM_003998082.5;refGene_gn=USB1;refGene_tx=XM_019819812.2
E2 37021542 37021647
refGene_gn=USB1;refGene_tx=XM_003998082.5;refGene_gn=USB1;refGene_tx=XM_019819812.2
E2 37035236 37035398 refGene_gn=MMP15;refGene_tx=XM_003998084.4
E2 37046492 37046641 refGene_gn=MMP15;refGene_tx=XM_003998084.4
E2 37047312 37047441 refGene_gn=MMP15;refGene_tx=XM_003998084.4
E2 37048745 37049053 refGene_gn=MMP15;refGene_tx=XM_003998084.4
E2 37049418 37049580 refGene_gn=MMP15;refGene_tx=XM_003998084.4
E2 37050526 37050780 refGene_gn=MMP15;refGene_tx=XM_003998084.4
E2 37051159 37051298 refGene_gn=MMP15;refGene_tx=XM_003998084.4
E2 37052188 37052339 refGene_gn=MMP15;refGene_tx=XM_003998084.4
E2 37052422 37052538 refGene_gn=MMP15;refGene_tx=XM_003998084.4
E2 37053895 37054362 refGene_gn=MMP15;refGene_tx=XM_003998084.4
E2 37096562 37096568 refGene_gn=CFAP20;refGene_tx=XM_006941553.2
E2 37097049 37097160 refGene_gn=CFAP20;refGene_tx=XM_006941553.2
E2 37097508 37097697 refGene_gn=CFAP20;refGene_tx=XM_006941553.2
E2 37098727 37098839 refGene_gn=CFAP20;refGene_tx=XM_006941553.2
E2 37099547 37099627 refGene_gn=CFAP20;refGene_tx=XM_006941553.2
E2 37108496 37108580 refGene_gn=CFAP20;refGene_tx=XM_006941553.2
E2 37123542 37123619
refGene_gn=CSNK2A2;refGene_tx=XM_023245638.1;refGene_gn=CSNK2A2;refGene_tx=XM_023245639.1;refGene_gn=CSNK2A2;refGene_tx=XM_023245640.1;refGene_gn=CSNK2A2;refGene_tx=XM_023245641.1
E2 37125266 37125415
refGene_gn=CSNK2A2;refGene_tx=XM_023245638.1;refGene_gn=CSNK2A2;refGene_tx=XM_023245639.1;refGene_gn=CSNK2A2;refGene_tx=XM_023245640.1;refGene_gn=CSNK2A2;refGene_tx=XM_023245641.1
E2 37126219 37126320
refGene_gn=CSNK2A2;refGene_tx=XM_023245638.1;refGene_gn=CSNK2A2;refGene_tx=XM_023245639.1;refGene_gn=CSNK2A2;refGene_tx=XM_023245640.1;refGene_gn=CSNK2A2;refGene_tx=XM_023245641.1
E2 37126840 37126942
refGene_gn=CSNK2A2;refGene_tx=XM_023245638.1;refGene_gn=CSNK2A2;refGene_tx=XM_023245639.1;refGene_gn=CSNK2A2;refGene_tx=XM_023245640.1;refGe

ne_gn=CSNK2A2;refGene_tx=XM_023245641.1
E2 37127207 37127318
refGene_gn=CSNK2A2;refGene_tx=XM_023245638.1;refGene_gn=CSNK2A2;refGene_tx=XM_023245639.1;refGene_gn=CSNK2A2;refGene_tx=XM_023245640.1;refGene_gn=CSNK2A2;refGene_tx=XM_023245641.1
E2 37128114 37128198
refGene_gn=CSNK2A2;refGene_tx=XM_023245638.1;refGene_gn=CSNK2A2;refGene_tx=XM_023245639.1;refGene_gn=CSNK2A2;refGene_tx=XM_023245640.1;refGene_gn=CSNK2A2;refGene_tx=XM_023245641.1
E2 37133501 37133561
refGene_gn=CSNK2A2;refGene_tx=XM_023245638.1;refGene_gn=CSNK2A2;refGene_tx=XM_023245639.1;refGene_gn=CSNK2A2;refGene_tx=XM_023245640.1;refGene_gn=CSNK2A2;refGene_tx=XM_023245641.1
E2 37142971 37143022
refGene_gn=CSNK2A2;refGene_tx=XM_023245638.1;refGene_gn=CSNK2A2;refGene_tx=XM_023245639.1;refGene_gn=CSNK2A2;refGene_tx=XM_023245640.1;refGene_gn=CSNK2A2;refGene_tx=XM_023245641.1
E2 37145882 37145984
refGene_gn=CSNK2A2;refGene_tx=XM_023245638.1;refGene_gn=CSNK2A2;refGene_tx=XM_023245639.1;refGene_gn=CSNK2A2;refGene_tx=XM_023245641.1
E2 37154773 37154885
refGene_gn=CSNK2A2;refGene_tx=XM_023245638.1;refGene_gn=CSNK2A2;refGene_tx=XM_023245639.1;refGene_gn=CSNK2A2;refGene_tx=XM_023245640.1;refGene_gn=CSNK2A2;refGene_tx=XM_023245641.1
E2 37155691 37155795
refGene_gn=CSNK2A2;refGene_tx=XM_023245638.1;refGene_gn=CSNK2A2;refGene_tx=XM_023245639.1;refGene_gn=CSNK2A2;refGene_tx=XM_023245640.1;refGene_gn=CSNK2A2;refGene_tx=XM_023245641.1
E2 37193908 37194009
refGene_gn=CCDC113;refGene_tx=XM_006941554.4;refGene_gn=CCDC113;refGene_tx=XM_019819818.2;refGene_gn=CCDC113;refGene_tx=XM_019819819.2
E2 37195820 37195947
refGene_gn=CCDC113;refGene_tx=XM_006941554.4;refGene_gn=CCDC113;refGene_tx=XM_019819818.2;refGene_gn=CCDC113;refGene_tx=XM_019819819.2
E2 37196921 37197083
refGene_gn=CCDC113;refGene_tx=XM_006941554.4;refGene_gn=CCDC113;refGene_tx=XM_019819818.2;refGene_gn=CCDC113;refGene_tx=XM_019819819.2
E2 37202206 37202362
refGene_gn=CCDC113;refGene_tx=XM_006941554.4;refGene_gn=CCDC113;refGene_tx=XM_019819818.2;refGene_gn=CCDC113;refGene_tx=XM_019819819.2
E2 37203145 37203226
refGene_gn=CCDC113;refGene_tx=XM_006941554.4;refGene_gn=CCDC113;refGene_tx=XM_019819818.2;refGene_gn=CCDC113;refGene_tx=XM_019819819.2
E2 37205687 37205843
refGene_gn=CCDC113;refGene_tx=XM_006941554.4;refGene_gn=CCDC113;refGene_tx=XM_019819818.2
E2 37210545 37210653
refGene_gn=CCDC113;refGene_tx=XM_006941554.4;refGene_gn=CCDC113;refGene_tx=XM_019819818.2;refGene_gn=CCDC113;refGene_tx=XM_019819819.2
E2 37221766 37221922

refGene_gn=CCDC113;refGene_tx=XM_006941554.4;refGene_gn=CCDC113;refGene_tx=XM_019819818.2;refGene_gn=CCDC113;refGene_tx=XM_019819819.2
E2 37223138 37223240
refGene_gn=CCDC113;refGene_tx=XM_019819818.2;refGene_gn=CCDC113;refGene_tx=XM_019819819.2
E2 37223359 37223446 refGene_gn=CCDC113;refGene_tx=XM_006941554.4
E2 37223905 37224445 refGene_gn=PRSS54;refGene_tx=XM_023245637.1
E2 37227442 37227574 refGene_gn=PRSS54;refGene_tx=XM_023245637.1
E2 37228167 37228426 refGene_gn=PRSS54;refGene_tx=XM_023245637.1
E2 37233928 37234100 refGene_gn=PRSS54;refGene_tx=XM_023245637.1
E2 37237314 37237363 refGene_gn=PRSS54;refGene_tx=XM_023245637.1
E2 37237364 37237400 refGene_gn=PRSS54;refGene_tx=XM_023245637.1
E2 37322199 37322385 refGene_gn=GINS3;refGene_tx=XM_023245353.1
E2 37326608 37326842 refGene_gn=GINS3;refGene_tx=XM_023245353.1
E2 37328959 37329190 refGene_gn=GINS3;refGene_tx=XM_023245353.1
E2 37384811 37384848
refGene_gn=NDRG4;refGene_tx=XM_006941560.4;refGene_gn=NDRG4;refGene_tx=XM_011290141.2;refGene_gn=NDRG4;refGene_tx=XM_023246168.1
E2 37404310 37404405
refGene_gn=NDRG4;refGene_tx=XM_006941560.4;refGene_gn=NDRG4;refGene_tx=XM_011290141.2;refGene_gn=NDRG4;refGene_tx=XM_023246168.1;refGene_gn=NDRG4;refGene_tx=XM_023246169.1
E2 37409129 37409174
refGene_gn=NDRG4;refGene_tx=XM_006941560.4;refGene_gn=NDRG4;refGene_tx=XM_011290141.2;refGene_gn=NDRG4;refGene_tx=XM_023246168.1;refGene_gn=NDRG4;refGene_tx=XM_023246169.1
E2 37413280 37413301
refGene_gn=NDRG4;refGene_tx=XM_003998089.5;refGene_gn=NDRG4;refGene_tx=XM_003998090.5;refGene_gn=NDRG4;refGene_tx=XM_023246170.1
E2 37416640 37416746
refGene_gn=NDRG4;refGene_tx=XM_003998089.5;refGene_gn=NDRG4;refGene_tx=XM_003998090.5;refGene_gn=NDRG4;refGene_tx=XM_006941560.4;refGene_gn=NDRG4;refGene_tx=XM_011290141.2;refGene_gn=NDRG4;refGene_tx=XM_023246168.1;refGene_gn=NDRG4;refGene_tx=XM_023246169.1;refGene_gn=NDRG4;refGene_tx=XM_023246170.1
E2 37416967 37417088
refGene_gn=NDRG4;refGene_tx=XM_003998089.5;refGene_gn=NDRG4;refGene_tx=XM_003998090.5;refGene_gn=NDRG4;refGene_tx=XM_006941560.4;refGene_gn=NDRG4;refGene_tx=XM_011290141.2;refGene_gn=NDRG4;refGene_tx=XM_023246168.1;refGene_gn=NDRG4;refGene_tx=XM_023246169.1;refGene_gn=NDRG4;refGene_tx=XM_023246170.1
E2 37417173 37417236
refGene_gn=NDRG4;refGene_tx=XM_003998089.5;refGene_gn=NDRG4;refGene_tx=XM_003998090.5;refGene_gn=NDRG4;refGene_tx=XM_006941560.4;refGene_gn=NDRG4;refGene_tx=XM_011290141.2;refGene_gn=NDRG4;refGene_tx=XM_023246168.1;refGene_gn=NDRG4;refGene_tx=XM_023246169.1;refGene_gn=NDRG4;refGene_tx=XM_023246170.1
E2 37417351 37417412
refGene_gn=NDRG4;refGene_tx=XM_003998089.5;refGene_gn=NDRG4;refGene_tx=XM_003998090.5;refGene_gn=NDRG4;refGene_tx=XM_006941560.4;refGene_gn=

NDRG4;refGene_tx=XM_011290141.2;refGene_gn=NDRG4;refGene_tx=XM_023246168.1;refGene_gn=NDRG4;refGene_tx=XM_023246169.1;refGene_gn=NDRG4;refGene_tx=XM_023246170.1
E2 37421225 37421312
refGene_gn=NDRG4;refGene_tx=XM_003998089.5;refGene_gn=NDRG4;refGene_tx=XM_003998090.5;refGene_gn=NDRG4;refGene_tx=XM_006941560.4;refGene_gn=NDRG4;refGene_tx=XM_011290141.2;refGene_gn=NDRG4;refGene_tx=XM_023246168.1;refGene_gn=NDRG4;refGene_tx=XM_023246169.1;refGene_gn=NDRG4;refGene_tx=XM_023246170.1
E2 37421391 37421448
refGene_gn=NDRG4;refGene_tx=XM_003998089.5;refGene_gn=NDRG4;refGene_tx=XM_003998090.5;refGene_gn=NDRG4;refGene_tx=XM_006941560.4;refGene_gn=NDRG4;refGene_tx=XM_011290141.2;refGene_gn=NDRG4;refGene_tx=XM_023246168.1;refGene_gn=NDRG4;refGene_tx=XM_023246169.1;refGene_gn=NDRG4;refGene_tx=XM_023246170.1
E2 37421761 37421865
refGene_gn=NDRG4;refGene_tx=XM_003998089.5;refGene_gn=NDRG4;refGene_tx=XM_003998090.5;refGene_gn=NDRG4;refGene_tx=XM_006941560.4;refGene_gn=NDRG4;refGene_tx=XM_011290141.2;refGene_gn=NDRG4;refGene_tx=XM_023246168.1;refGene_gn=NDRG4;refGene_tx=XM_023246169.1;refGene_gn=NDRG4;refGene_tx=XM_023246170.1
E2 37422745 37422802
refGene_gn=NDRG4;refGene_tx=XM_003998089.5;refGene_gn=NDRG4;refGene_tx=XM_003998090.5;refGene_gn=NDRG4;refGene_tx=XM_006941560.4;refGene_gn=NDRG4;refGene_tx=XM_011290141.2;refGene_gn=NDRG4;refGene_tx=XM_023246168.1;refGene_gn=NDRG4;refGene_tx=XM_023246169.1;refGene_gn=NDRG4;refGene_tx=XM_023246170.1
E2 37422879 37422931
refGene_gn=NDRG4;refGene_tx=XM_003998089.5;refGene_gn=NDRG4;refGene_tx=XM_003998090.5;refGene_gn=NDRG4;refGene_tx=XM_006941560.4;refGene_gn=NDRG4;refGene_tx=XM_011290141.2;refGene_gn=NDRG4;refGene_tx=XM_023246168.1;refGene_gn=NDRG4;refGene_tx=XM_023246169.1;refGene_gn=NDRG4;refGene_tx=XM_023246170.1
E2 37423670 37423718
refGene_gn=NDRG4;refGene_tx=XM_003998089.5;refGene_gn=NDRG4;refGene_tx=XM_003998090.5;refGene_gn=NDRG4;refGene_tx=XM_006941560.4;refGene_gn=NDRG4;refGene_tx=XM_011290141.2;refGene_gn=NDRG4;refGene_tx=XM_023246168.1;refGene_gn=NDRG4;refGene_tx=XM_023246169.1;refGene_gn=NDRG4;refGene_tx=XM_023246170.1
E2 37423911 37423947
refGene_gn=NDRG4;refGene_tx=XM_003998089.5;refGene_gn=NDRG4;refGene_tx=XM_003998090.5;refGene_gn=NDRG4;refGene_tx=XM_006941560.4;refGene_gn=NDRG4;refGene_tx=XM_011290141.2;refGene_gn=NDRG4;refGene_tx=XM_023246168.1;refGene_gn=NDRG4;refGene_tx=XM_023246169.1;refGene_gn=NDRG4;refGene_tx=XM_023246170.1
E2 37424055 37424107
refGene_gn=NDRG4;refGene_tx=XM_003998089.5;refGene_gn=NDRG4;refGene_tx=XM_003998090.5;refGene_gn=NDRG4;refGene_tx=XM_006941560.4;refGene_gn=NDRG4;refGene_tx=XM_011290141.2;refGene_gn=NDRG4;refGene_tx=XM_023246168.1;refGene_gn=NDRG4;refGene_tx=XM_023246169.1;refGene_gn=NDRG4;refGe

ne_tx=XM_023246170.1
E2 37425459 37425582
refGene_gn=NDRG4;refGene_tx=XM_003998090.5;refGene_gn=NDRG4;refGene_tx=XM_011290141.2;refGene_gn=NDRG4;refGene_tx=XM_023246168.1;refGene_gn=NDRG4;refGene_tx=XM_023246169.1;refGene_gn=NDRG4;refGene_tx=XM_023246170.1
E2 37426429 37426584
refGene_gn=NDRG4;refGene_tx=XM_003998089.5;refGene_gn=NDRG4;refGene_tx=XM_003998090.5;refGene_gn=NDRG4;refGene_tx=XM_006941560.4;refGene_gn=NDRG4;refGene_tx=XM_011290141.2;refGene_gn=NDRG4;refGene_tx=XM_023246168.1;refGene_gn=NDRG4;refGene_tx=XM_023246169.1;refGene_gn=NDRG4;refGene_tx=XM_023246170.1
E2 37429757 37429784 refGene_gn=SETD6;refGene_tx=XM_023246167.1
E2 37430018 37430123 refGene_gn=SETD6;refGene_tx=XM_023246167.1
E2 37430196 37430341 refGene_gn=SETD6;refGene_tx=XM_023246167.1
E2 37430419 37430561 refGene_gn=SETD6;refGene_tx=XM_023246167.1
E2 37430696 37430891 refGene_gn=SETD6;refGene_tx=XM_023246167.1
E2 37431021 37431142 refGene_gn=SETD6;refGene_tx=XM_023246167.1
E2 37432571 37432752 refGene_gn=SETD6;refGene_tx=XM_023246167.1
E2 37432926 37433069 refGene_gn=SETD6;refGene_tx=XM_023246167.1
E2 37433224 37433530 refGene_gn=SETD6;refGene_tx=XM_023246167.1
E2 37435458 37435537
refGene_gn=CNOT1;refGene_tx=XM_011290135.2;refGene_gn=CNOT1;refGene_tx=XM_011290136.3;refGene_gn=CNOT1;refGene_tx=XM_011290137.2;refGene_gn=CNOT1;refGene_tx=XM_011290138.2;refGene_gn=CNOT1;refGene_tx=XM_023246163.1;refGene_gn=CNOT1;refGene_tx=XM_023246164.1;refGene_gn=CNOT1;refGene_tx=XM_023246165.1
E2 37435691 37435826
refGene_gn=CNOT1;refGene_tx=XM_011290135.2;refGene_gn=CNOT1;refGene_tx=XM_011290136.3;refGene_gn=CNOT1;refGene_tx=XM_011290137.2;refGene_gn=CNOT1;refGene_tx=XM_011290138.2;refGene_gn=CNOT1;refGene_tx=XM_023246163.1;refGene_gn=CNOT1;refGene_tx=XM_023246164.1;refGene_gn=CNOT1;refGene_tx=XM_023246165.1
E2 37436741 37436874
refGene_gn=CNOT1;refGene_tx=XM_011290135.2;refGene_gn=CNOT1;refGene_tx=XM_011290136.3;refGene_gn=CNOT1;refGene_tx=XM_011290137.2;refGene_gn=CNOT1;refGene_tx=XM_011290138.2;refGene_gn=CNOT1;refGene_tx=XM_023246163.1;refGene_gn=CNOT1;refGene_tx=XM_023246164.1;refGene_gn=CNOT1;refGene_tx=XM_023246165.1
E2 37437480 37437661
refGene_gn=CNOT1;refGene_tx=XM_011290135.2;refGene_gn=CNOT1;refGene_tx=XM_011290136.3;refGene_gn=CNOT1;refGene_tx=XM_011290137.2;refGene_gn=CNOT1;refGene_tx=XM_011290138.2;refGene_gn=CNOT1;refGene_tx=XM_023246163.1;refGene_gn=CNOT1;refGene_tx=XM_023246164.1;refGene_gn=CNOT1;refGene_tx=XM_023246165.1
E2 37438613 37438763
refGene_gn=CNOT1;refGene_tx=XM_011290135.2;refGene_gn=CNOT1;refGene_tx=XM_011290136.3;refGene_gn=CNOT1;refGene_tx=XM_011290137.2;refGene_gn=CNOT1;refGene_tx=XM_011290138.2;refGene_gn=CNOT1;refGene_tx=XM_023246163.1;refGene_gn=CNOT1;refGene_tx=XM_023246164.1;refGene_gn=CNOT1;refGe

ne_tx=XM_023246165.1
E2 37439560 37439734
refGene_gn=CNOT1;refGene_tx=XM_011290135.2;refGene_gn=CNOT1;refGene_tx=XM_011290136.3;refGene_gn=CNOT1;refGene_tx=XM_011290137.2;refGene_gn=CNOT1;refGene_tx=XM_011290138.2;refGene_gn=CNOT1;refGene_tx=XM_023246163.1;refGene_gn=CNOT1;refGene_tx=XM_023246164.1;refGene_gn=CNOT1;refGene_tx=XM_023246165.1
E2 37442526 37442628
refGene_gn=CNOT1;refGene_tx=XM_011290135.2;refGene_gn=CNOT1;refGene_tx=XM_011290136.3;refGene_gn=CNOT1;refGene_tx=XM_011290137.2;refGene_gn=CNOT1;refGene_tx=XM_011290138.2;refGene_gn=CNOT1;refGene_tx=XM_023246163.1;refGene_gn=CNOT1;refGene_tx=XM_023246164.1;refGene_gn=CNOT1;refGene_tx=XM_023246165.1
E2 37444008 37444156
refGene_gn=CNOT1;refGene_tx=XM_011290135.2;refGene_gn=CNOT1;refGene_tx=XM_011290136.3;refGene_gn=CNOT1;refGene_tx=XM_011290137.2;refGene_gn=CNOT1;refGene_tx=XM_011290138.2;refGene_gn=CNOT1;refGene_tx=XM_023246163.1;refGene_gn=CNOT1;refGene_tx=XM_023246164.1;refGene_gn=CNOT1;refGene_tx=XM_023246165.1;refGene_gn=CNOT1;refGene_tx=XM_023246166.1
E2 37444350 37444514
refGene_gn=CNOT1;refGene_tx=XM_011290135.2;refGene_gn=CNOT1;refGene_tx=XM_011290136.3;refGene_gn=CNOT1;refGene_tx=XM_011290137.2;refGene_gn=CNOT1;refGene_tx=XM_011290138.2;refGene_gn=CNOT1;refGene_tx=XM_023246163.1;refGene_gn=CNOT1;refGene_tx=XM_023246164.1;refGene_gn=CNOT1;refGene_tx=XM_023246165.1;refGene_gn=CNOT1;refGene_tx=XM_023246166.1
E2 37445686 37445935
refGene_gn=CNOT1;refGene_tx=XM_011290135.2;refGene_gn=CNOT1;refGene_tx=XM_011290136.3;refGene_gn=CNOT1;refGene_tx=XM_011290137.2;refGene_gn=CNOT1;refGene_tx=XM_011290138.2;refGene_gn=CNOT1;refGene_tx=XM_023246163.1;refGene_gn=CNOT1;refGene_tx=XM_023246164.1;refGene_gn=CNOT1;refGene_tx=XM_023246165.1;refGene_gn=CNOT1;refGene_tx=XM_023246166.1
E2 37448139 37448392
refGene_gn=CNOT1;refGene_tx=XM_011290135.2;refGene_gn=CNOT1;refGene_tx=XM_011290136.3;refGene_gn=CNOT1;refGene_tx=XM_011290137.2;refGene_gn=CNOT1;refGene_tx=XM_011290138.2;refGene_gn=CNOT1;refGene_tx=XM_023246163.1;refGene_gn=CNOT1;refGene_tx=XM_023246164.1;refGene_gn=CNOT1;refGene_tx=XM_023246165.1;refGene_gn=CNOT1;refGene_tx=XM_023246166.1
E2 37449025 37449195
refGene_gn=CNOT1;refGene_tx=XM_011290135.2;refGene_gn=CNOT1;refGene_tx=XM_011290136.3;refGene_gn=CNOT1;refGene_tx=XM_011290137.2;refGene_gn=CNOT1;refGene_tx=XM_011290138.2;refGene_gn=CNOT1;refGene_tx=XM_023246163.1;refGene_gn=CNOT1;refGene_tx=XM_023246164.1;refGene_gn=CNOT1;refGene_tx=XM_023246165.1;refGene_gn=CNOT1;refGene_tx=XM_023246166.1
E2 37449293 37449402
refGene_gn=CNOT1;refGene_tx=XM_011290135.2;refGene_gn=CNOT1;refGene_tx=XM_011290136.3;refGene_gn=CNOT1;refGene_tx=XM_011290137.2;refGene_gn=CNOT1;refGene_tx=XM_011290138.2;refGene_gn=CNOT1;refGene_tx=XM_023246163.1;refGene_gn=CNOT1;refGene_tx=XM_023246164.1;refGene_gn=CNOT1;refGene_tx=XM_023246165.1;refGene_gn=CNOT1;refGene_tx=XM_023246166.1
E2 37449850 37449993

refGene_gn=CNOT1;refGene_tx=XM_011290135.2;refGene_gn=CNOT1;refGene_tx=XM_011290136.3;refGene_gn=CNOT1;refGene_tx=XM_011290137.2;refGene_gn=CNOT1;refGene_tx=XM_011290138.2;refGene_gn=CNOT1;refGene_tx=XM_023246163.1;refGene_gn=CNOT1;refGene_tx=XM_023246164.1;refGene_gn=CNOT1;refGene_tx=XM_023246165.1;refGene_gn=CNOT1;refGene_tx=XM_023246166.1
E2 37450805 37450997
refGene_gn=CNOT1;refGene_tx=XM_011290135.2;refGene_gn=CNOT1;refGene_tx=XM_011290136.3;refGene_gn=CNOT1;refGene_tx=XM_011290137.2;refGene_gn=CNOT1;refGene_tx=XM_011290138.2;refGene_gn=CNOT1;refGene_tx=XM_023246163.1;refGene_gn=CNOT1;refGene_tx=XM_023246164.1;refGene_gn=CNOT1;refGene_tx=XM_023246165.1;refGene_gn=CNOT1;refGene_tx=XM_023246166.1
E2 37452152 37452272
refGene_gn=CNOT1;refGene_tx=XM_011290135.2;refGene_gn=CNOT1;refGene_tx=XM_011290136.3;refGene_gn=CNOT1;refGene_tx=XM_011290137.2;refGene_gn=CNOT1;refGene_tx=XM_011290138.2;refGene_gn=CNOT1;refGene_tx=XM_023246163.1;refGene_gn=CNOT1;refGene_tx=XM_023246164.1;refGene_gn=CNOT1;refGene_tx=XM_023246165.1;refGene_gn=CNOT1;refGene_tx=XM_023246166.1
E2 37453018 37453123
refGene_gn=CNOT1;refGene_tx=XM_011290135.2;refGene_gn=CNOT1;refGene_tx=XM_011290136.3;refGene_gn=CNOT1;refGene_tx=XM_011290137.2;refGene_gn=CNOT1;refGene_tx=XM_011290138.2;refGene_gn=CNOT1;refGene_tx=XM_023246163.1;refGene_gn=CNOT1;refGene_tx=XM_023246164.1;refGene_gn=CNOT1;refGene_tx=XM_023246165.1;refGene_gn=CNOT1;refGene_tx=XM_023246166.1
E2 37453217 37453358
refGene_gn=CNOT1;refGene_tx=XM_011290135.2;refGene_gn=CNOT1;refGene_tx=XM_011290136.3;refGene_gn=CNOT1;refGene_tx=XM_011290137.2;refGene_gn=CNOT1;refGene_tx=XM_011290138.2;refGene_gn=CNOT1;refGene_tx=XM_023246163.1;refGene_gn=CNOT1;refGene_tx=XM_023246164.1;refGene_gn=CNOT1;refGene_tx=XM_023246165.1;refGene_gn=CNOT1;refGene_tx=XM_023246166.1
E2 37454036 37454333
refGene_gn=CNOT1;refGene_tx=XM_011290135.2;refGene_gn=CNOT1;refGene_tx=XM_011290136.3;refGene_gn=CNOT1;refGene_tx=XM_011290137.2;refGene_gn=CNOT1;refGene_tx=XM_011290138.2;refGene_gn=CNOT1;refGene_tx=XM_023246163.1;refGene_gn=CNOT1;refGene_tx=XM_023246164.1;refGene_gn=CNOT1;refGene_tx=XM_023246165.1;refGene_gn=CNOT1;refGene_tx=XM_023246166.1
E2 37456194 37456325
refGene_gn=CNOT1;refGene_tx=XM_011290135.2;refGene_gn=CNOT1;refGene_tx=XM_011290136.3;refGene_gn=CNOT1;refGene_tx=XM_011290137.2;refGene_gn=CNOT1;refGene_tx=XM_011290138.2;refGene_gn=CNOT1;refGene_tx=XM_023246163.1;refGene_gn=CNOT1;refGene_tx=XM_023246164.1;refGene_gn=CNOT1;refGene_tx=XM_023246165.1;refGene_gn=CNOT1;refGene_tx=XM_023246166.1
E2 37457179 37457357
refGene_gn=CNOT1;refGene_tx=XM_011290135.2;refGene_gn=CNOT1;refGene_tx=XM_011290136.3;refGene_gn=CNOT1;refGene_tx=XM_011290137.2;refGene_gn=CNOT1;refGene_tx=XM_011290138.2;refGene_gn=CNOT1;refGene_tx=XM_023246163.1;refGene_gn=CNOT1;refGene_tx=XM_023246164.1;refGene_gn=CNOT1;refGene_tx=XM_023246165.1;refGene_gn=CNOT1;refGene_tx=XM_023246166.1
E2 37457543 37457621
refGene_gn=CNOT1;refGene_tx=XM_011290135.2;refGene_gn=CNOT1;refGene_tx=XM_011290136.3;refGene_gn=CNOT1;refGene_tx=XM_011290137.2;refGene_gn=

CNOT1;refGene_tx=XM_011290138.2;refGene_gn=CNOT1;refGene_tx=XM_0232461
63.1;refGene_gn=CNOT1;refGene_tx=XM_023246164.1;refGene_gn=CNOT1;refGe
ne_tx=XM_023246165.1;refGene_gn=CNOT1;refGene_tx=XM_023246166.1
E2 37457934 37458045
refGene_gn=CNOT1;refGene_tx=XM_011290135.2;refGene_gn=CNOT1;refGene_tx
=XM_011290136.3;refGene_gn=CNOT1;refGene_tx=XM_011290137.2;refGene_gn=
CNOT1;refGene_tx=XM_011290138.2;refGene_gn=CNOT1;refGene_tx=XM_0232461
63.1;refGene_gn=CNOT1;refGene_tx=XM_023246164.1;refGene_gn=CNOT1;refGe
ne_tx=XM_023246165.1;refGene_gn=CNOT1;refGene_tx=XM_023246166.1
E2 37458381 37458498
refGene_gn=CNOT1;refGene_tx=XM_011290135.2;refGene_gn=CNOT1;refGene_tx
=XM_011290136.3;refGene_gn=CNOT1;refGene_tx=XM_011290137.2;refGene_gn=
CNOT1;refGene_tx=XM_011290138.2;refGene_gn=CNOT1;refGene_tx=XM_0232461
63.1;refGene_gn=CNOT1;refGene_tx=XM_023246164.1;refGene_gn=CNOT1;refGe
ne_tx=XM_023246165.1;refGene_gn=CNOT1;refGene_tx=XM_023246166.1
E2 37460031 37460211
refGene_gn=CNOT1;refGene_tx=XM_011290135.2;refGene_gn=CNOT1;refGene_tx
=XM_011290136.3;refGene_gn=CNOT1;refGene_tx=XM_011290137.2;refGene_gn=
CNOT1;refGene_tx=XM_011290138.2;refGene_gn=CNOT1;refGene_tx=XM_0232461
63.1;refGene_gn=CNOT1;refGene_tx=XM_023246164.1;refGene_gn=CNOT1;refGe
ne_tx=XM_023246165.1;refGene_gn=CNOT1;refGene_tx=XM_023246166.1
E2 37461139 37461280
refGene_gn=CNOT1;refGene_tx=XM_011290135.2;refGene_gn=CNOT1;refGene_tx
=XM_011290136.3;refGene_gn=CNOT1;refGene_tx=XM_011290137.2;refGene_gn=
CNOT1;refGene_tx=XM_011290138.2;refGene_gn=CNOT1;refGene_tx=XM_0232461
63.1;refGene_gn=CNOT1;refGene_tx=XM_023246164.1;refGene_gn=CNOT1;refGe
ne_tx=XM_023246165.1;refGene_gn=CNOT1;refGene_tx=XM_023246166.1
E2 37461593 37461824
refGene_gn=CNOT1;refGene_tx=XM_011290135.2;refGene_gn=CNOT1;refGene_tx
=XM_011290136.3;refGene_gn=CNOT1;refGene_tx=XM_011290137.2;refGene_gn=
CNOT1;refGene_tx=XM_011290138.2;refGene_gn=CNOT1;refGene_tx=XM_0232461
63.1;refGene_gn=CNOT1;refGene_tx=XM_023246164.1;refGene_gn=CNOT1;refGe
ne_tx=XM_023246165.1;refGene_gn=CNOT1;refGene_tx=XM_023246166.1
E2 37463752 37463831
refGene_gn=CNOT1;refGene_tx=XM_011290135.2;refGene_gn=CNOT1;refGene_tx
=XM_011290136.3;refGene_gn=CNOT1;refGene_tx=XM_011290137.2;refGene_gn=
CNOT1;refGene_tx=XM_011290138.2;refGene_gn=CNOT1;refGene_tx=XM_0232461
63.1;refGene_gn=CNOT1;refGene_tx=XM_023246164.1;refGene_gn=CNOT1;refGe
ne_tx=XM_023246165.1;refGene_gn=CNOT1;refGene_tx=XM_023246166.1
E2 37464868 37465155
refGene_gn=CNOT1;refGene_tx=XM_011290135.2;refGene_gn=CNOT1;refGene_tx
=XM_011290136.3;refGene_gn=CNOT1;refGene_tx=XM_011290137.2;refGene_gn=
CNOT1;refGene_tx=XM_011290138.2;refGene_gn=CNOT1;refGene_tx=XM_0232461
63.1;refGene_gn=CNOT1;refGene_tx=XM_023246164.1;refGene_gn=CNOT1;refGe
ne_tx=XM_023246165.1;refGene_gn=CNOT1;refGene_tx=XM_023246166.1
E2 37465583 37465708
refGene_gn=CNOT1;refGene_tx=XM_011290135.2;refGene_gn=CNOT1;refGene_tx
=XM_011290136.3;refGene_gn=CNOT1;refGene_tx=XM_011290137.2;refGene_gn=
CNOT1;refGene_tx=XM_011290138.2;refGene_gn=CNOT1;refGene_tx=XM_0232461
63.1;refGene_gn=CNOT1;refGene_tx=XM_023246164.1;refGene_gn=CNOT1;refGe

ne_tx=XM_023246165.1;refGene_gn=CNOT1;refGene_tx=XM_023246166.1
E2 37467097 37467244
refGene_gn=CNOT1;refGene_tx=XM_011290135.2;refGene_gn=CNOT1;refGene_tx=XM_011290136.3;refGene_gn=CNOT1;refGene_tx=XM_011290137.2;refGene_gn=CNOT1;refGene_tx=XM_011290138.2;refGene_gn=CNOT1;refGene_tx=XM_023246163.1;refGene_gn=CNOT1;refGene_tx=XM_023246164.1;refGene_gn=CNOT1;refGene_tx=XM_023246165.1;refGene_gn=CNOT1;refGene_tx=XM_023246166.1
E2 37468231 37468433
refGene_gn=CNOT1;refGene_tx=XM_011290135.2;refGene_gn=CNOT1;refGene_tx=XM_011290136.3;refGene_gn=CNOT1;refGene_tx=XM_011290137.2;refGene_gn=CNOT1;refGene_tx=XM_011290138.2;refGene_gn=CNOT1;refGene_tx=XM_023246163.1;refGene_gn=CNOT1;refGene_tx=XM_023246164.1;refGene_gn=CNOT1;refGene_tx=XM_023246165.1;refGene_gn=CNOT1;refGene_tx=XM_023246166.1
E2 37469638 37469789
refGene_gn=CNOT1;refGene_tx=XM_011290135.2;refGene_gn=CNOT1;refGene_tx=XM_011290136.3;refGene_gn=CNOT1;refGene_tx=XM_011290137.2;refGene_gn=CNOT1;refGene_tx=XM_011290138.2;refGene_gn=CNOT1;refGene_tx=XM_023246163.1;refGene_gn=CNOT1;refGene_tx=XM_023246164.1;refGene_gn=CNOT1;refGene_tx=XM_023246165.1;refGene_gn=CNOT1;refGene_tx=XM_023246166.1
E2 37475823 37475975
refGene_gn=CNOT1;refGene_tx=XM_011290135.2;refGene_gn=CNOT1;refGene_tx=XM_011290136.3;refGene_gn=CNOT1;refGene_tx=XM_011290137.2;refGene_gn=CNOT1;refGene_tx=XM_011290138.2;refGene_gn=CNOT1;refGene_tx=XM_023246163.1;refGene_gn=CNOT1;refGene_tx=XM_023246164.1;refGene_gn=CNOT1;refGene_tx=XM_023246165.1;refGene_gn=CNOT1;refGene_tx=XM_023246166.1
E2 37476185 37476308
refGene_gn=CNOT1;refGene_tx=XM_011290135.2;refGene_gn=CNOT1;refGene_tx=XM_011290136.3;refGene_gn=CNOT1;refGene_tx=XM_011290137.2;refGene_gn=CNOT1;refGene_tx=XM_011290138.2;refGene_gn=CNOT1;refGene_tx=XM_023246163.1;refGene_gn=CNOT1;refGene_tx=XM_023246164.1;refGene_gn=CNOT1;refGene_tx=XM_023246165.1;refGene_gn=CNOT1;refGene_tx=XM_023246166.1
E2 37477076 37477196
refGene_gn=CNOT1;refGene_tx=XM_011290135.2;refGene_gn=CNOT1;refGene_tx=XM_011290136.3;refGene_gn=CNOT1;refGene_tx=XM_011290137.2;refGene_gn=CNOT1;refGene_tx=XM_011290138.2;refGene_gn=CNOT1;refGene_tx=XM_023246163.1;refGene_gn=CNOT1;refGene_tx=XM_023246164.1;refGene_gn=CNOT1;refGene_tx=XM_023246165.1;refGene_gn=CNOT1;refGene_tx=XM_023246166.1
E2 37478754 37478995
refGene_gn=CNOT1;refGene_tx=XM_011290135.2;refGene_gn=CNOT1;refGene_tx=XM_011290136.3;refGene_gn=CNOT1;refGene_tx=XM_011290137.2;refGene_gn=CNOT1;refGene_tx=XM_011290138.2;refGene_gn=CNOT1;refGene_tx=XM_023246163.1;refGene_gn=CNOT1;refGene_tx=XM_023246164.1;refGene_gn=CNOT1;refGene_tx=XM_023246165.1;refGene_gn=CNOT1;refGene_tx=XM_023246166.1
E2 37481305 37481433
refGene_gn=CNOT1;refGene_tx=XM_011290135.2;refGene_gn=CNOT1;refGene_tx=XM_011290136.3;refGene_gn=CNOT1;refGene_tx=XM_011290137.2;refGene_gn=CNOT1;refGene_tx=XM_011290138.2;refGene_gn=CNOT1;refGene_tx=XM_023246163.1;refGene_gn=CNOT1;refGene_tx=XM_023246164.1;refGene_gn=CNOT1;refGene_tx=XM_023246165.1;refGene_gn=CNOT1;refGene_tx=XM_023246166.1
E2 37481853 37482024

refGene_gn=CNOT1;refGene_tx=XM_011290135.2;refGene_gn=CNOT1;refGene_tx
=XM_011290136.3;refGene_gn=CNOT1;refGene_tx=XM_011290137.2;refGene_gn=
CNOT1;refGene_tx=XM_011290138.2;refGene_gn=CNOT1;refGene_tx=XM_0232461
63.1;refGene_gn=CNOT1;refGene_tx=XM_023246164.1;refGene_gn=CNOT1;refGe
ne_tx=XM_023246165.1;refGene_gn=CNOT1;refGene_tx=XM_023246166.1
E2 37485156 37485267
refGene_gn=CNOT1;refGene_tx=XM_011290135.2;refGene_gn=CNOT1;refGene_tx
=XM_011290136.3;refGene_gn=CNOT1;refGene_tx=XM_011290137.2;refGene_gn=
CNOT1;refGene_tx=XM_011290138.2;refGene_gn=CNOT1;refGene_tx=XM_0232461
63.1;refGene_gn=CNOT1;refGene_tx=XM_023246164.1;refGene_gn=CNOT1;refGe
ne_tx=XM_023246165.1;refGene_gn=CNOT1;refGene_tx=XM_023246166.1
E2 37485414 37485541
refGene_gn=CNOT1;refGene_tx=XM_011290135.2;refGene_gn=CNOT1;refGene_tx
=XM_011290136.3;refGene_gn=CNOT1;refGene_tx=XM_011290137.2;refGene_gn=
CNOT1;refGene_tx=XM_011290138.2;refGene_gn=CNOT1;refGene_tx=XM_0232461
63.1;refGene_gn=CNOT1;refGene_tx=XM_023246164.1;refGene_gn=CNOT1;refGe
ne_tx=XM_023246165.1;refGene_gn=CNOT1;refGene_tx=XM_023246166.1
E2 37490145 37490314
refGene_gn=CNOT1;refGene_tx=XM_011290135.2;refGene_gn=CNOT1;refGene_tx
=XM_011290136.3;refGene_gn=CNOT1;refGene_tx=XM_011290137.2;refGene_gn=
CNOT1;refGene_tx=XM_011290138.2;refGene_gn=CNOT1;refGene_tx=XM_0232461
63.1;refGene_gn=CNOT1;refGene_tx=XM_023246164.1;refGene_gn=CNOT1;refGe
ne_tx=XM_023246165.1;refGene_gn=CNOT1;refGene_tx=XM_023246166.1
E2 37491390 37491594
refGene_gn=CNOT1;refGene_tx=XM_011290135.2;refGene_gn=CNOT1;refGene_tx
=XM_011290136.3;refGene_gn=CNOT1;refGene_tx=XM_011290137.2;refGene_gn=
CNOT1;refGene_tx=XM_011290138.2;refGene_gn=CNOT1;refGene_tx=XM_0232461
63.1;refGene_gn=CNOT1;refGene_tx=XM_023246164.1;refGene_gn=CNOT1;refGe
ne_tx=XM_023246165.1;refGene_gn=CNOT1;refGene_tx=XM_023246166.1
E2 37492310 37492365
refGene_gn=CNOT1;refGene_tx=XM_011290135.2;refGene_gn=CNOT1;refGene_tx
=XM_011290136.3;refGene_gn=CNOT1;refGene_tx=XM_011290137.2;refGene_gn=
CNOT1;refGene_tx=XM_011290138.2;refGene_gn=CNOT1;refGene_tx=XM_0232461
63.1;refGene_gn=CNOT1;refGene_tx=XM_023246164.1;refGene_gn=CNOT1;refGe
ne_tx=XM_023246165.1;refGene_gn=CNOT1;refGene_tx=XM_023246166.1
E2 37492452 37492521
refGene_gn=CNOT1;refGene_tx=XM_011290135.2;refGene_gn=CNOT1;refGene_tx
=XM_011290136.3;refGene_gn=CNOT1;refGene_tx=XM_011290137.2;refGene_gn=
CNOT1;refGene_tx=XM_011290138.2;refGene_gn=CNOT1;refGene_tx=XM_0232461
63.1;refGene_gn=CNOT1;refGene_tx=XM_023246164.1;refGene_gn=CNOT1;refGe
ne_tx=XM_023246165.1;refGene_gn=CNOT1;refGene_tx=XM_023246166.1
E2 37492895 37492994
refGene_gn=CNOT1;refGene_tx=XM_011290135.2;refGene_gn=CNOT1;refGene_tx
=XM_011290136.3;refGene_gn=CNOT1;refGene_tx=XM_011290137.2;refGene_gn=
CNOT1;refGene_tx=XM_011290138.2;refGene_gn=CNOT1;refGene_tx=XM_0232461
63.1;refGene_gn=CNOT1;refGene_tx=XM_023246164.1;refGene_gn=CNOT1;refGe
ne_tx=XM_023246165.1;refGene_gn=CNOT1;refGene_tx=XM_023246166.1
E2 37493821 37493929
refGene_gn=CNOT1;refGene_tx=XM_011290135.2;refGene_gn=CNOT1;refGene_tx
=XM_011290136.3;refGene_gn=CNOT1;refGene_tx=XM_011290137.2;refGene_gn=

CNOT1;refGene_tx=XM_011290138.2;refGene_gn=CNOT1;refGene_tx=XM_0232461
63.1;refGene_gn=CNOT1;refGene_tx=XM_023246164.1;refGene_gn=CNOT1;refGe
ne_tx=XM_023246165.1;refGene_gn=CNOT1;refGene_tx=XM_023246166.1
E2 37516469 37516571
refGene_gn=CNOT1;refGene_tx=XM_011290135.2;refGene_gn=CNOT1;refGene_tx
=XM_011290136.3;refGene_gn=CNOT1;refGene_tx=XM_011290137.2;refGene_gn=
CNOT1;refGene_tx=XM_011290138.2;refGene_gn=CNOT1;refGene_tx=XM_0232461
63.1;refGene_gn=CNOT1;refGene_tx=XM_023246164.1;refGene_gn=CNOT1;refGe
ne_tx=XM_023246165.1;refGene_gn=CNOT1;refGene_tx=XM_023246166.1
E2 37581291 37581394 refGene_gn=SLC38A7;refGene_tx=XM_019819830.2
E2 37583470 37583525 refGene_gn=SLC38A7;refGene_tx=XM_019819830.2
E2 37584389 37584589 refGene_gn=SLC38A7;refGene_tx=XM_019819830.2
E2 37584997 37585145 refGene_gn=SLC38A7;refGene_tx=XM_019819830.2
E2 37586481 37586596 refGene_gn=SLC38A7;refGene_tx=XM_019819830.2
E2 37586813 37586871 refGene_gn=SLC38A7;refGene_tx=XM_019819830.2
E2 37587448 37587547 refGene_gn=SLC38A7;refGene_tx=XM_019819830.2
E2 37588553 37588698 refGene_gn=SLC38A7;refGene_tx=XM_019819830.2
E2 37588934 37589133 refGene_gn=SLC38A7;refGene_tx=XM_019819830.2
E2 37589776 37590046 refGene_gn=SLC38A7;refGene_tx=XM_019819830.2
E2 37598238 37598361 refGene_gn=GOT2;refGene_tx=XM_003998093.4
E2 37598984 37599135 refGene_gn=GOT2;refGene_tx=XM_003998093.4
E2 37602855 37603021 refGene_gn=GOT2;refGene_tx=XM_003998093.4
E2 37603457 37603608 refGene_gn=GOT2;refGene_tx=XM_003998093.4
E2 37604702 37604807 refGene_gn=GOT2;refGene_tx=XM_003998093.4
E2 37605026 37605188 refGene_gn=GOT2;refGene_tx=XM_003998093.4
E2 37605995 37606055 refGene_gn=GOT2;refGene_tx=XM_003998093.4
E2 37607046 37607175 refGene_gn=GOT2;refGene_tx=XM_003998093.4
E2 37608788 37608945 refGene_gn=GOT2;refGene_tx=XM_003998093.4
E2 37618653 37618742 refGene_gn=GOT2;refGene_tx=XM_003998093.4
E2 40129399 40129893
refGene_gn=LOC101094809;refGene_tx=XM_003998105.5;refGene_gn=LOC101094
809;refGene_tx=XM_019819837.2;refGene_gn=LOC101094809;refGene_tx=XM_02
3244991.1;refGene_gn=LOC101094809;refGene_tx=XM_023244992.1;refGene_gn
=LOC101094809;refGene_tx=XM_023244993.1;refGene_gn=LOC101094809;refGen
e_tx=XM_023244995.1
E2 40131292 40131544
refGene_gn=LOC101094809;refGene_tx=XM_003998105.5;refGene_gn=LOC101094
809;refGene_tx=XM_019819837.2;refGene_gn=LOC101094809;refGene_tx=XM_02
3244991.1;refGene_gn=LOC101094809;refGene_tx=XM_023244992.1;refGene_gn
=LOC101094809;refGene_tx=XM_023244993.1;refGene_gn=LOC101094809;refGen
e_tx=XM_023244995.1
E2 40156350 40156507
refGene_gn=LOC101094809;refGene_tx=XM_023244995.1
E2 40182168 40182286
refGene_gn=LOC101094809;refGene_tx=XM_003998105.5;refGene_gn=LOC101094
809;refGene_tx=XM_019819837.2;refGene_gn=LOC101094809;refGene_tx=XM_02
3244991.1;refGene_gn=LOC101094809;refGene_tx=XM_023244992.1;refGene_gn
=LOC101094809;refGene_tx=XM_023244993.1
E2 40192121 40192376
refGene_gn=LOC101094809;refGene_tx=XM_019819837.2;refGene_gn=LOC101094

809;refGene_tx=XM_023244991.1;refGene_gn=LOC101094809;refGene_tx=XM_023244992.1;refGene_gn=LOC101094809;refGene_tx=XM_023244993.1
E2 40195802 40195924
refGene_gn=LOC101094809;refGene_tx=XM_003998105.5;refGene_gn=LOC101094809;refGene_tx=XM_019819837.2;refGene_gn=LOC101094809;refGene_tx=XM_023244991.1;refGene_gn=LOC101094809;refGene_tx=XM_023244992.1;refGene_gn=LOC101094809;refGene_tx=XM_023244993.1
E2 40267698 40267835
refGene_gn=LOC101094809;refGene_tx=XM_003998105.5;refGene_gn=LOC101094809;refGene_tx=XM_019819837.2;refGene_gn=LOC101094809;refGene_tx=XM_023244991.1;refGene_gn=LOC101094809;refGene_tx=XM_023244992.1;refGene_gn=LOC101094809;refGene_tx=XM_023244993.1
E2 40289455 40289709
refGene_gn=LOC101094809;refGene_tx=XM_003998105.5;refGene_gn=LOC101094809;refGene_tx=XM_019819837.2;refGene_gn=LOC101094809;refGene_tx=XM_023244991.1;refGene_gn=LOC101094809;refGene_tx=XM_023244992.1;refGene_gn=LOC101094809;refGene_tx=XM_023244993.1
E2 40292882 40293070
refGene_gn=LOC101094809;refGene_tx=XM_003998105.5;refGene_gn=LOC101094809;refGene_tx=XM_019819837.2;refGene_gn=LOC101094809;refGene_tx=XM_023244991.1;refGene_gn=LOC101094809;refGene_tx=XM_023244992.1;refGene_gn=LOC101094809;refGene_tx=XM_023244993.1
E2 40297320 40297488
refGene_gn=LOC101094809;refGene_tx=XM_003998105.5;refGene_gn=LOC101094809;refGene_tx=XM_019819837.2;refGene_gn=LOC101094809;refGene_tx=XM_023244991.1;refGene_gn=LOC101094809;refGene_tx=XM_023244992.1;refGene_gn=LOC101094809;refGene_tx=XM_023244993.1
E2 40331225 40331345
refGene_gn=LOC101094809;refGene_tx=XM_003998105.5;refGene_gn=LOC101094809;refGene_tx=XM_019819837.2;refGene_gn=LOC101094809;refGene_tx=XM_023244991.1;refGene_gn=LOC101094809;refGene_tx=XM_023244992.1
E2 40375255 40375550
refGene_gn=LOC101094809;refGene_tx=XM_003998105.5;refGene_gn=LOC101094809;refGene_tx=XM_019819837.2;refGene_gn=LOC101094809;refGene_tx=XM_023244991.1;refGene_gn=LOC101094809;refGene_tx=XM_023244992.1;refGene_gn=LOC101094809;refGene_tx=XM_023244993.1
E2 40476067 40476316
refGene_gn=LOC101094809;refGene_tx=XM_003998105.5;refGene_gn=LOC101094809;refGene_tx=XM_019819837.2;refGene_gn=LOC101094809;refGene_tx=XM_023244991.1;refGene_gn=LOC101094809;refGene_tx=XM_023244992.1;refGene_gn=LOC101094809;refGene_tx=XM_023244993.1
E2 42954302 42954799
refGene_gn=LOC101095064;refGene_tx=XM_023245550.1;refGene_gn=LOC101095064;refGene_tx=XM_023245551.1
E2 42957306 42957558
refGene_gn=LOC101095064;refGene_tx=XM_023245550.1;refGene_gn=LOC101095064;refGene_tx=XM_023245551.1
E2 42972657 42972775
refGene_gn=LOC101095064;refGene_tx=XM_023245550.1;refGene_gn=LOC101095064;refGene_tx=XM_023245551.1

E2 42973008 42973142
refGene_gn=LOC101095064;refGene_tx=XM_023245550.1;refGene_gn=LOC101095
064;refGene_tx=XM_023245551.1
E2 42973556 42973693
refGene_gn=LOC101095064;refGene_tx=XM_023245550.1;refGene_gn=LOC101095
064;refGene_tx=XM_023245551.1
E2 42979890 42980144
refGene_gn=LOC101095064;refGene_tx=XM_023245550.1;refGene_gn=LOC101095
064;refGene_tx=XM_023245551.1
E2 42986114 42986302
refGene_gn=LOC101095064;refGene_tx=XM_023245550.1;refGene_gn=LOC101095
064;refGene_tx=XM_023245551.1
E2 42991355 42991523
refGene_gn=LOC101095064;refGene_tx=XM_023245550.1;refGene_gn=LOC101095
064;refGene_tx=XM_023245551.1
E2 42993726 42993846
refGene_gn=LOC101095064;refGene_tx=XM_023245550.1;refGene_gn=LOC101095
064;refGene_tx=XM_023245551.1
E2 42996727 42997022
refGene_gn=LOC101095064;refGene_tx=XM_023245550.1;refGene_gn=LOC101095
064;refGene_tx=XM_023245551.1
E2 43001679 43001907
refGene_gn=LOC101095064;refGene_tx=XM_023245550.1;refGene_gn=LOC101095
064;refGene_tx=XM_023245551.1
E2 44269747 44269873 refGene_gn=CDH5;refGene_tx=XM_006941565.4
E2 44269874 44269943 refGene_gn=CDH5;refGene_tx=XM_006941565.4
E2 44274882 44275171 refGene_gn=CDH5;refGene_tx=XM_006941565.4
E2 44276239 44276356 refGene_gn=CDH5;refGene_tx=XM_006941565.4
E2 44277316 44277481 refGene_gn=CDH5;refGene_tx=XM_006941565.4
E2 44278366 44278416 refGene_gn=CDH5;refGene_tx=XM_006941565.4
E2 44278417 44278555 refGene_gn=CDH5;refGene_tx=XM_006941565.4
E2 44280168 44280419 refGene_gn=CDH5;refGene_tx=XM_006941565.4
E2 44284360 44284503 refGene_gn=CDH5;refGene_tx=XM_006941565.4
E2 44286523 44286648 refGene_gn=CDH5;refGene_tx=XM_006941565.4
E2 44287003 44287112 refGene_gn=CDH5;refGene_tx=XM_006941565.4
E2 44289434 44289680 refGene_gn=CDH5;refGene_tx=XM_006941565.4
E2 44291724 44292239 refGene_gn=CDH5;refGene_tx=XM_006941565.4
E2 44326441 44326466
refGene_gn=BEAN1;refGene_tx=XM_023245290.1;refGene_gn=BEAN1;refGene_tx
=XM_023245291.1
E2 44343283 44343547
refGene_gn=BEAN1;refGene_tx=XM_023245290.1;refGene_gn=BEAN1;refGene_tx
=XM_023245291.1
E2 44349944 44350092
refGene_gn=BEAN1;refGene_tx=XM_023245290.1;refGene_gn=BEAN1;refGene_tx
=XM_023245291.1
E2 44351043 44351371
refGene_gn=BEAN1;refGene_tx=XM_023245290.1;refGene_gn=BEAN1;refGene_tx
=XM_023245291.1
E2 44359550 44359892

refGene_gn=LOC102900682;refGene_tx=XM_006941641.3
E2	44360499	44361600
refGene_gn=LOC102900682;refGene_tx=XM_006941641.3
E2	44372415	44372514
refGene_gn=TK2;refGene_tx=XM_006941571.4;refGene_gn=TK2;refGene_tx=XM_011290128.3;refGene_gn=TK2;refGene_tx=XM_019819841.2;refGene_gn=TK2;refGene_tx=XM_019819843.2;refGene_gn=TK2;refGene_tx=XM_023245603.1;refGene_gn=TK2;refGene_tx=XM_023245604.1;refGene_gn=TK2;refGene_tx=XM_023245605.1;refGene_gn=TK2;refGene_tx=XM_023245606.1;refGene_gn=TK2;refGene_tx=XM_023245607.1
E2	44373200	44373266	refGene_gn=TK2;refGene_tx=XM_019819842.2
E2	44373663	44373744
refGene_gn=TK2;refGene_tx=XM_006941571.4;refGene_gn=TK2;refGene_tx=XM_011290128.3;refGene_gn=TK2;refGene_tx=XM_019819841.2;refGene_gn=TK2;refGene_tx=XM_019819842.2;refGene_gn=TK2;refGene_tx=XM_019819843.2;refGene_gn=TK2;refGene_tx=XM_023245603.1;refGene_gn=TK2;refGene_tx=XM_023245604.1;refGene_gn=TK2;refGene_tx=XM_023245605.1;refGene_gn=TK2;refGene_tx=XM_023245606.1;refGene_gn=TK2;refGene_tx=XM_023245607.1
E2	44376193	44376273
refGene_gn=TK2;refGene_tx=XM_006941571.4;refGene_gn=TK2;refGene_tx=XM_011290128.3;refGene_gn=TK2;refGene_tx=XM_019819841.2;refGene_gn=TK2;refGene_tx=XM_019819842.2;refGene_gn=TK2;refGene_tx=XM_019819843.2;refGene_gn=TK2;refGene_tx=XM_023245603.1;refGene_gn=TK2;refGene_tx=XM_023245604.1;refGene_gn=TK2;refGene_tx=XM_023245605.1;refGene_gn=TK2;refGene_tx=XM_023245606.1;refGene_gn=TK2;refGene_tx=XM_023245607.1
E2	44376876	44376965
refGene_gn=TK2;refGene_tx=XM_006941571.4;refGene_gn=TK2;refGene_tx=XM_011290128.3;refGene_gn=TK2;refGene_tx=XM_019819841.2;refGene_gn=TK2;refGene_tx=XM_019819842.2;refGene_gn=TK2;refGene_tx=XM_019819843.2;refGene_gn=TK2;refGene_tx=XM_023245603.1;refGene_gn=TK2;refGene_tx=XM_023245604.1;refGene_gn=TK2;refGene_tx=XM_023245605.1;refGene_gn=TK2;refGene_tx=XM_023245606.1;refGene_gn=TK2;refGene_tx=XM_023245607.1
E2	44387560	44387634
refGene_gn=TK2;refGene_tx=XM_006941571.4;refGene_gn=TK2;refGene_tx=XM_011290128.3;refGene_gn=TK2;refGene_tx=XM_019819841.2;refGene_gn=TK2;refGene_tx=XM_019819842.2;refGene_gn=TK2;refGene_tx=XM_019819843.2;refGene_gn=TK2;refGene_tx=XM_023245603.1;refGene_gn=TK2;refGene_tx=XM_023245604.1;refGene_gn=TK2;refGene_tx=XM_023245605.1;refGene_gn=TK2;refGene_tx=XM_023245606.1;refGene_gn=TK2;refGene_tx=XM_023245607.1
E2	44390232	44390322
refGene_gn=TK2;refGene_tx=XM_006941571.4;refGene_gn=TK2;refGene_tx=XM_011290128.3;refGene_gn=TK2;refGene_tx=XM_019819841.2;refGene_gn=TK2;refGene_tx=XM_019819842.2;refGene_gn=TK2;refGene_tx=XM_019819843.2;refGene_gn=TK2;refGene_tx=XM_023245603.1;refGene_gn=TK2;refGene_tx=XM_023245604.1;refGene_gn=TK2;refGene_tx=XM_023245605.1;refGene_gn=TK2;refGene_tx=XM_023245606.1;refGene_gn=TK2;refGene_tx=XM_023245607.1
E2	44391190	44391238	refGene_gn=TK2;refGene_tx=XM_019819843.2
E2	44392751	44392805
refGene_gn=TK2;refGene_tx=XM_006941571.4;refGene_gn=TK2;refGene_tx=XM_011290128.3;refGene_gn=TK2;refGene_tx=XM_019819841.2;refGene_gn=TK2;re

fGene_tx=XM_019819842.2;refGene_gn=TK2;refGene_tx=XM_023245603.1;refGene_gn=TK2;refGene_tx=XM_023245604.1;refGene_gn=TK2;refGene_tx=XM_023245605.1;refGene_gn=TK2;refGene_tx=XM_023245606.1;refGene_gn=TK2;refGene_tx=XM_023245607.1
E2 44395748 44395823
refGene_gn=TK2;refGene_tx=XM_006941571.4;refGene_gn=TK2;refGene_tx=XM_011290128.3;refGene_gn=TK2;refGene_tx=XM_019819841.2;refGene_gn=TK2;refGene_tx=XM_019819842.2;refGene_gn=TK2;refGene_tx=XM_023245603.1
E2 44399182 44399220
refGene_gn=TK2;refGene_tx=XM_006941571.4;refGene_gn=TK2;refGene_tx=XM_011290128.3;refGene_gn=TK2;refGene_tx=XM_019819841.2;refGene_gn=TK2;refGene_tx=XM_019819842.2;refGene_gn=TK2;refGene_tx=XM_023245603.1;refGene_gn=TK2;refGene_tx=XM_023245604.1;refGene_gn=TK2;refGene_tx=XM_023245605.1;refGene_gn=TK2;refGene_tx=XM_023245606.1;refGene_gn=TK2;refGene_tx=XM_023245607.1
E2 44399953 44400131
refGene_gn=TK2;refGene_tx=XM_011290128.3;refGene_gn=TK2;refGene_tx=XM_019819841.2;refGene_gn=TK2;refGene_tx=XM_019819842.2;refGene_gn=TK2;refGene_tx=XM_023245604.1;refGene_gn=TK2;refGene_tx=XM_023245605.1
E2 44400151 44400266
refGene_gn=TK2;refGene_tx=XM_023245603.1;refGene_gn=TK2;refGene_tx=XM_023245606.1
E2 44401934 44401992
refGene_gn=TK2;refGene_tx=XM_006941571.4;refGene_gn=TK2;refGene_tx=XM_023245607.1
E2 44402371 44402443
refGene_gn=LOC109491355;refGene_tx=XM_023245596.1;refGene_gn=LOC109491355;refGene_tx=XM_023245598.1;refGene_gn=LOC109491355;refGene_tx=XM_023245599.1;refGene_gn=LOC109491355;refGene_tx=XM_023245600.1;refGene_gn=LOC109491355;refGene_tx=XM_023245601.1;refGene_gn=LOC109491355;refGene_tx=XM_023245602.1
E2 44407515 44407674
refGene_gn=LOC109491355;refGene_tx=XM_023245596.1;refGene_gn=LOC109491355;refGene_tx=XM_023245598.1;refGene_gn=LOC109491355;refGene_tx=XM_023245599.1;refGene_gn=LOC109491355;refGene_tx=XM_023245600.1;refGene_gn=LOC109491355;refGene_tx=XM_023245601.1;refGene_gn=LOC109491355;refGene_tx=XM_023245602.1
E2 44417613 44417709
refGene_gn=LOC109491355;refGene_tx=XM_023245596.1;refGene_gn=LOC109491355;refGene_tx=XM_023245598.1;refGene_gn=LOC109491355;refGene_tx=XM_023245599.1;refGene_gn=LOC109491355;refGene_tx=XM_023245600.1;refGene_gn=LOC109491355;refGene_tx=XM_023245601.1;refGene_gn=LOC109491355;refGene_tx=XM_023245602.1
E2 44417964 44417970
refGene_gn=LOC109491355;refGene_tx=XM_023245602.1
E2 44420373 44420490
refGene_gn=LOC109491355;refGene_tx=XM_023245599.1
E2 44420569 44420584
refGene_gn=LOC109491355;refGene_tx=XM_023245601.1
E2 44420999 44421431

refGene_gn=LOC109491355;refGene_tx=XM_006941569.4;refGene_gn=LOC109491355;refGene_tx=XM_023245594.1;refGene_gn=LOC109491355;refGene_tx=XM_023245595.1
E2 44432859 44432958
refGene_gn=LOC109491355;refGene_tx=XM_006941569.4;refGene_gn=LOC109491355;refGene_tx=XM_023245594.1;refGene_gn=LOC109491355;refGene_tx=XM_023245596.1;refGene_gn=LOC109491355;refGene_tx=XM_023245598.1;refGene_gn=LOC109491355;refGene_tx=XM_023245600.1
E2 44433671 44433677
refGene_gn=LOC109491355;refGene_tx=XM_023245600.1
E2 44435014 44435122
refGene_gn=LOC109491355;refGene_tx=XM_006941569.4;refGene_gn=LOC109491355;refGene_tx=XM_023245598.1
E2 44435974 44436259
refGene_gn=LOC109491355;refGene_tx=XM_006941568.4;refGene_gn=LOC109491355;refGene_tx=XM_023245597.1
E2 44436396 44436555
refGene_gn=LOC109491355;refGene_tx=XM_006941568.4
E2 44449359 44449461
refGene_gn=LOC109491355;refGene_tx=XM_006941568.4;refGene_gn=LOC109491355;refGene_tx=XM_023245594.1;refGene_gn=LOC109491355;refGene_tx=XM_023245595.1;refGene_gn=LOC109491355;refGene_tx=XM_023245596.1;refGene_gn=LOC109491355;refGene_tx=XM_023245597.1
E2 44450234 44450468
refGene_gn=LOC109491355;refGene_tx=XM_006941568.4;refGene_gn=LOC109491355;refGene_tx=XM_023245594.1;refGene_gn=LOC109491355;refGene_tx=XM_023245595.1;refGene_gn=LOC109491355;refGene_tx=XM_023245596.1;refGene_gn=LOC109491355;refGene_tx=XM_023245597.1
E2 44464554 44464713
refGene_gn=CMTM3;refGene_tx=XM_023246029.1;refGene_gn=CMTM3;refGene_tx=XM_023246030.1
E2 44467725 44467881
refGene_gn=CMTM3;refGene_tx=XM_023246029.1;refGene_gn=CMTM3;refGene_tx=XM_023246030.1
E2 44468891 44468987
refGene_gn=CMTM3;refGene_tx=XM_023246029.1;refGene_gn=CMTM3;refGene_tx=XM_023246030.1
E2 44469392 44469513 refGene_gn=CMTM3;refGene_tx=XM_023246029.1
E2 44471022 44471109
refGene_gn=CMTM3;refGene_tx=XM_023246029.1;refGene_gn=CMTM3;refGene_tx=XM_023246030.1
E2 44480495 44480660
refGene_gn=CMTM4;refGene_tx=XM_011290194.3;refGene_gn=CMTM4;refGene_tx=XM_023246031.1
E2 44482266 44482365 refGene_gn=CMTM4;refGene_tx=XM_011290194.3
E2 44492487 44492664
refGene_gn=CMTM4;refGene_tx=XM_011290194.3;refGene_gn=CMTM4;refGene_tx=XM_023246031.1
E2 44542268 44542454
refGene_gn=CMTM4;refGene_tx=XM_011290194.3;refGene_gn=CMTM4;refGene_tx

=XM_023246031.1
E2	44561401 44561502 refGene_gn=DYNC1LI2;refGene_tx=XM_003998107.5
E2	44563093 44563210 refGene_gn=DYNC1LI2;refGene_tx=XM_003998107.5
E2	44565232 44565350 refGene_gn=DYNC1LI2;refGene_tx=XM_003998107.5
E2	44565744 44565786 refGene_gn=DYNC1LI2;refGene_tx=XM_003998107.5
E2	44566561 44566621 refGene_gn=DYNC1LI2;refGene_tx=XM_003998107.5
E2	44567202 44567314 refGene_gn=DYNC1LI2;refGene_tx=XM_003998107.5
E2	44569422 44569558 refGene_gn=DYNC1LI2;refGene_tx=XM_003998107.5
E2	44570516 44570610 refGene_gn=DYNC1LI2;refGene_tx=XM_003998107.5
E2	44571649 44571819 refGene_gn=DYNC1LI2;refGene_tx=XM_003998107.5
E2	44576787 44577018 refGene_gn=DYNC1LI2;refGene_tx=XM_003998107.5
E2	44581237 44581354 refGene_gn=DYNC1LI2;refGene_tx=XM_003998107.5
E2	44583855 44583929 refGene_gn=DYNC1LI2;refGene_tx=XM_003998107.5
E2	44584095 44584202 refGene_gn=DYNC1LI2;refGene_tx=XM_003998107.5
E2	44589411 44589599
refGene_gn=TERB1;refGene_tx=XM_023245625.1;refGene_gn=TERB1;refGene_tx=XM_023245627.1;refGene_gn=TERB1;refGene_tx=XM_023245630.1
E2	44589937 44589975
refGene_gn=TERB1;refGene_tx=XM_023245628.1;refGene_gn=TERB1;refGene_tx=XM_023245629.1;refGene_gn=TERB1;refGene_tx=XM_023245631.1
E2	44594356 44594422
refGene_gn=TERB1;refGene_tx=XM_023245625.1;refGene_gn=TERB1;refGene_tx=XM_023245627.1;refGene_gn=TERB1;refGene_tx=XM_023245628.1;refGene_gn=TERB1;refGene_tx=XM_023245629.1;refGene_gn=TERB1;refGene_tx=XM_023245630.1;refGene_gn=TERB1;refGene_tx=XM_023245631.1
E2	44594718 44594868
refGene_gn=TERB1;refGene_tx=XM_023245625.1;refGene_gn=TERB1;refGene_tx=XM_023245627.1;refGene_gn=TERB1;refGene_tx=XM_023245628.1;refGene_gn=TERB1;refGene_tx=XM_023245629.1
E2	44600944 44601040
refGene_gn=TERB1;refGene_tx=XM_023245625.1;refGene_gn=TERB1;refGene_tx=XM_023245627.1;refGene_gn=TERB1;refGene_tx=XM_023245628.1;refGene_gn=TERB1;refGene_tx=XM_023245629.1;refGene_gn=TERB1;refGene_tx=XM_023245630.1;refGene_gn=TERB1;refGene_tx=XM_023245631.1
E2	44601343 44601408
refGene_gn=TERB1;refGene_tx=XM_023245625.1;refGene_gn=TERB1;refGene_tx=XM_023245627.1;refGene_gn=TERB1;refGene_tx=XM_023245628.1;refGene_gn=TERB1;refGene_tx=XM_023245629.1;refGene_gn=TERB1;refGene_tx=XM_023245630.1;refGene_gn=TERB1;refGene_tx=XM_023245631.1
E2	44602665 44603015
refGene_gn=TERB1;refGene_tx=XM_023245625.1;refGene_gn=TERB1;refGene_tx=XM_023245627.1;refGene_gn=TERB1;refGene_tx=XM_023245628.1;refGene_gn=TERB1;refGene_tx=XM_023245629.1;refGene_gn=TERB1;refGene_tx=XM_023245630.1;refGene_gn=TERB1;refGene_tx=XM_023245631.1
E2	44606662 44606820
refGene_gn=TERB1;refGene_tx=XM_023245625.1;refGene_gn=TERB1;refGene_tx=XM_023245627.1;refGene_gn=TERB1;refGene_tx=XM_023245628.1;refGene_gn=TERB1;refGene_tx=XM_023245629.1;refGene_gn=TERB1;refGene_tx=XM_023245630.1;refGene_gn=TERB1;refGene_tx=XM_023245631.1
E2	44607921 44608047

refGene_gn=TERB1;refGene_tx=XM_023245625.1;refGene_gn=TERB1;refGene_tx=XM_023245627.1;refGene_gn=TERB1;refGene_tx=XM_023245628.1;refGene_gn=TERB1;refGene_tx=XM_023245629.1;refGene_gn=TERB1;refGene_tx=XM_023245630.1;refGene_gn=TERB1;refGene_tx=XM_023245631.1
E2 44608644 44608776
refGene_gn=TERB1;refGene_tx=XM_023245625.1;refGene_gn=TERB1;refGene_tx=XM_023245627.1;refGene_gn=TERB1;refGene_tx=XM_023245628.1;refGene_gn=TERB1;refGene_tx=XM_023245629.1;refGene_gn=TERB1;refGene_tx=XM_023245630.1;refGene_gn=TERB1;refGene_tx=XM_023245631.1
E2 44610711 44610864
refGene_gn=TERB1;refGene_tx=XM_023245625.1;refGene_gn=TERB1;refGene_tx=XM_023245627.1;refGene_gn=TERB1;refGene_tx=XM_023245628.1;refGene_gn=TERB1;refGene_tx=XM_023245629.1;refGene_gn=TERB1;refGene_tx=XM_023245630.1;refGene_gn=TERB1;refGene_tx=XM_023245631.1
E2 44618038 44618161
refGene_gn=TERB1;refGene_tx=XM_023245625.1;refGene_gn=TERB1;refGene_tx=XM_023245627.1;refGene_gn=TERB1;refGene_tx=XM_023245628.1;refGene_gn=TERB1;refGene_tx=XM_023245629.1;refGene_gn=TERB1;refGene_tx=XM_023245630.1;refGene_gn=TERB1;refGene_tx=XM_023245631.1
E2 44618260 44618376
refGene_gn=TERB1;refGene_tx=XM_023245625.1;refGene_gn=TERB1;refGene_tx=XM_023245627.1;refGene_gn=TERB1;refGene_tx=XM_023245628.1;refGene_gn=TERB1;refGene_tx=XM_023245629.1;refGene_gn=TERB1;refGene_tx=XM_023245630.1;refGene_gn=TERB1;refGene_tx=XM_023245631.1
E2 44618469 44618530
refGene_gn=TERB1;refGene_tx=XM_023245625.1;refGene_gn=TERB1;refGene_tx=XM_023245627.1;refGene_gn=TERB1;refGene_tx=XM_023245628.1;refGene_gn=TERB1;refGene_tx=XM_023245629.1;refGene_gn=TERB1;refGene_tx=XM_023245630.1;refGene_gn=TERB1;refGene_tx=XM_023245631.1
E2 44619602 44619734
refGene_gn=TERB1;refGene_tx=XM_023245625.1;refGene_gn=TERB1;refGene_tx=XM_023245627.1;refGene_gn=TERB1;refGene_tx=XM_023245628.1;refGene_gn=TERB1;refGene_tx=XM_023245629.1;refGene_gn=TERB1;refGene_tx=XM_023245630.1;refGene_gn=TERB1;refGene_tx=XM_023245631.1
E2 44621884 44622013
refGene_gn=TERB1;refGene_tx=XM_023245625.1;refGene_gn=TERB1;refGene_tx=XM_023245627.1;refGene_gn=TERB1;refGene_tx=XM_023245628.1;refGene_gn=TERB1;refGene_tx=XM_023245629.1;refGene_gn=TERB1;refGene_tx=XM_023245630.1;refGene_gn=TERB1;refGene_tx=XM_023245631.1
E2 44622197 44622308
refGene_gn=TERB1;refGene_tx=XM_023245625.1;refGene_gn=TERB1;refGene_tx=XM_023245627.1;refGene_gn=TERB1;refGene_tx=XM_023245628.1;refGene_gn=TERB1;refGene_tx=XM_023245629.1;refGene_gn=TERB1;refGene_tx=XM_023245630.1;refGene_gn=TERB1;refGene_tx=XM_023245631.1
E2 44633247 44633278
refGene_gn=TERB1;refGene_tx=XM_023245625.1;refGene_gn=TERB1;refGene_tx=XM_023245627.1;refGene_gn=TERB1;refGene_tx=XM_023245628.1;refGene_gn=TERB1;refGene_tx=XM_023245629.1;refGene_gn=TERB1;refGene_tx=XM_023245630.1;refGene_gn=TERB1;refGene_tx=XM_023245631.1
E2 44638518 44638628

refGene_gn=NAE1;refGene_tx=XM_011290125.2;refGene_gn=NAE1;refGene_tx=XM_011290126.2;refGene_gn=NAE1;refGene_tx=XM_011290127.2;refGene_gn=NAE1;refGene_tx=XM_019819852.1
E2 44641100 44641150
refGene_gn=NAE1;refGene_tx=XM_011290125.2;refGene_gn=NAE1;refGene_tx=XM_011290126.2;refGene_gn=NAE1;refGene_tx=XM_011290127.2;refGene_gn=NAE1;refGene_tx=XM_019819852.1
E2 44641230 44641345
refGene_gn=NAE1;refGene_tx=XM_011290125.2;refGene_gn=NAE1;refGene_tx=XM_011290126.2;refGene_gn=NAE1;refGene_tx=XM_011290127.2;refGene_gn=NAE1;refGene_tx=XM_019819852.1
E2 44644229 44644322
refGene_gn=NAE1;refGene_tx=XM_011290125.2;refGene_gn=NAE1;refGene_tx=XM_011290126.2;refGene_gn=NAE1;refGene_tx=XM_011290127.2;refGene_gn=NAE1;refGene_tx=XM_019819852.1
E2 44644832 44644919
refGene_gn=NAE1;refGene_tx=XM_011290125.2;refGene_gn=NAE1;refGene_tx=XM_011290126.2;refGene_gn=NAE1;refGene_tx=XM_011290127.2;refGene_gn=NAE1;refGene_tx=XM_019819852.1
E2 44646395 44646435
refGene_gn=NAE1;refGene_tx=XM_011290125.2;refGene_gn=NAE1;refGene_tx=XM_011290126.2;refGene_gn=NAE1;refGene_tx=XM_011290127.2;refGene_gn=NAE1;refGene_tx=XM_019819852.1
E2 44646740 44646816
refGene_gn=NAE1;refGene_tx=XM_011290125.2;refGene_gn=NAE1;refGene_tx=XM_011290126.2;refGene_gn=NAE1;refGene_tx=XM_011290127.2;refGene_gn=NAE1;refGene_tx=XM_019819852.1
E2 44648554 44648688
refGene_gn=NAE1;refGene_tx=XM_011290125.2;refGene_gn=NAE1;refGene_tx=XM_011290126.2;refGene_gn=NAE1;refGene_tx=XM_011290127.2;refGene_gn=NAE1;refGene_tx=XM_019819852.1
E2 44648765 44648825
refGene_gn=NAE1;refGene_tx=XM_011290125.2;refGene_gn=NAE1;refGene_tx=XM_011290127.2;refGene_gn=NAE1;refGene_tx=XM_019819852.1
E2 44650714 44650806
refGene_gn=NAE1;refGene_tx=XM_011290125.2;refGene_gn=NAE1;refGene_tx=XM_011290126.2;refGene_gn=NAE1;refGene_tx=XM_011290127.2;refGene_gn=NAE1;refGene_tx=XM_019819852.1
E2 44652056 44652120
refGene_gn=NAE1;refGene_tx=XM_011290125.2;refGene_gn=NAE1;refGene_tx=XM_011290126.2;refGene_gn=NAE1;refGene_tx=XM_011290127.2;refGene_gn=NAE1;refGene_tx=XM_019819852.1
E2 44652516 44652579
refGene_gn=NAE1;refGene_tx=XM_011290125.2;refGene_gn=NAE1;refGene_tx=XM_011290126.2;refGene_gn=NAE1;refGene_tx=XM_019819852.1
E2 44653668 44653778
refGene_gn=NAE1;refGene_tx=XM_011290125.2;refGene_gn=NAE1;refGene_tx=XM_011290126.2;refGene_gn=NAE1;refGene_tx=XM_011290127.2;refGene_gn=NAE1;refGene_tx=XM_019819852.1
E2 44656151 44656261

refGene_gn=NAE1;refGene_tx=XM_011290125.2;refGene_gn=NAE1;refGene_tx=XM_011290126.2;refGene_gn=NAE1;refGene_tx=XM_011290127.2;refGene_gn=NAE1;refGene_tx=XM_019819852.1
E2 44656839 44656919
refGene_gn=NAE1;refGene_tx=XM_011290125.2;refGene_gn=NAE1;refGene_tx=XM_011290126.2;refGene_gn=NAE1;refGene_tx=XM_011290127.2;refGene_gn=NAE1;refGene_tx=XM_019819852.1
E2 44657269 44657341
refGene_gn=NAE1;refGene_tx=XM_011290125.2;refGene_gn=NAE1;refGene_tx=XM_011290126.2;refGene_gn=NAE1;refGene_tx=XM_011290127.2;refGene_gn=NAE1;refGene_tx=XM_019819852.1
E2 44658031 44658062
refGene_gn=NAE1;refGene_tx=XM_011290125.2;refGene_gn=NAE1;refGene_tx=XM_011290126.2;refGene_gn=NAE1;refGene_tx=XM_011290127.2
E2 44659142 44659203
refGene_gn=NAE1;refGene_tx=XM_011290125.2;refGene_gn=NAE1;refGene_tx=XM_011290126.2;refGene_gn=NAE1;refGene_tx=XM_011290127.2
E2 44659290 44659394
refGene_gn=NAE1;refGene_tx=XM_011290125.2;refGene_gn=NAE1;refGene_tx=XM_011290126.2;refGene_gn=NAE1;refGene_tx=XM_011290127.2
E2 44662933 44662986
refGene_gn=NAE1;refGene_tx=XM_011290125.2;refGene_gn=NAE1;refGene_tx=XM_011290126.2;refGene_gn=NAE1;refGene_tx=XM_011290127.2
E2 44670085 44670125 refGene_gn=CA7;refGene_tx=XM_011290124.3
E2 44670577 44670629
refGene_gn=CA7;refGene_tx=XM_019819855.2;refGene_gn=CA7;refGene_tx=XM_019819856.2
E2 44672656 44672854
refGene_gn=CA7;refGene_tx=XM_011290124.3;refGene_gn=CA7;refGene_tx=XM_019819855.2;refGene_gn=CA7;refGene_tx=XM_019819856.2
E2 44675630 44675749
refGene_gn=CA7;refGene_tx=XM_011290124.3;refGene_gn=CA7;refGene_tx=XM_019819855.2;refGene_gn=CA7;refGene_tx=XM_019819856.2
E2 44676445 44676541
refGene_gn=CA7;refGene_tx=XM_011290124.3;refGene_gn=CA7;refGene_tx=XM_019819855.2
E2 44676644 44676707
refGene_gn=CA7;refGene_tx=XM_011290124.3;refGene_gn=CA7;refGene_tx=XM_019819855.2;refGene_gn=CA7;refGene_tx=XM_019819856.2
E2 44677234 44677390
refGene_gn=CA7;refGene_tx=XM_011290124.3;refGene_gn=CA7;refGene_tx=XM_019819855.2;refGene_gn=CA7;refGene_tx=XM_019819856.2
E2 44677849 44677972
refGene_gn=CA7;refGene_tx=XM_011290124.3;refGene_gn=CA7;refGene_tx=XM_019819855.2;refGene_gn=CA7;refGene_tx=XM_019819856.2
E2 44697312 44698905
refGene_gn=PDP2;refGene_tx=XM_006941574.4;refGene_gn=PDP2;refGene_tx=XM_006941575.4;refGene_gn=PDP2;refGene_tx=XM_023245632.1
E2 44705293 44705313
refGene_gn=CDH16;refGene_tx=XM_019819850.2;refGene_gn=CDH16;refGene_tx

=XM_019819851.2
E2 44715453 44715551
refGene_gn=CDH16;refGene_tx=XM_003998109.3;refGene_gn=CDH16;refGene_tx=XM_019819849.1
E2 44716381 44716498
refGene_gn=CDH16;refGene_tx=XM_003998109.3;refGene_gn=CDH16;refGene_tx=XM_019819849.1;refGene_gn=CDH16;refGene_tx=XM_019819850.2
E2 44716810 44716918
refGene_gn=CDH16;refGene_tx=XM_003998109.3;refGene_gn=CDH16;refGene_tx=XM_019819849.1;refGene_gn=CDH16;refGene_tx=XM_019819850.2;refGene_gn=CDH16;refGene_tx=XM_019819851.2
E2 44717066 44717309
refGene_gn=CDH16;refGene_tx=XM_003998109.3;refGene_gn=CDH16;refGene_tx=XM_019819849.1;refGene_gn=CDH16;refGene_tx=XM_019819850.2;refGene_gn=CDH16;refGene_tx=XM_019819851.2
E2 44718038 44718172
refGene_gn=CDH16;refGene_tx=XM_003998109.3;refGene_gn=CDH16;refGene_tx=XM_019819849.1;refGene_gn=CDH16;refGene_tx=XM_019819850.2;refGene_gn=CDH16;refGene_tx=XM_019819851.2
E2 44718722 44718964
refGene_gn=CDH16;refGene_tx=XM_003998109.3;refGene_gn=CDH16;refGene_tx=XM_019819849.1;refGene_gn=CDH16;refGene_tx=XM_019819850.2;refGene_gn=CDH16;refGene_tx=XM_019819851.2
E2 44719065 44719254
refGene_gn=CDH16;refGene_tx=XM_003998109.3;refGene_gn=CDH16;refGene_tx=XM_019819849.1;refGene_gn=CDH16;refGene_tx=XM_019819850.2;refGene_gn=CDH16;refGene_tx=XM_019819851.2
E2 44719328 44719405
refGene_gn=CDH16;refGene_tx=XM_003998109.3;refGene_gn=CDH16;refGene_tx=XM_019819849.1;refGene_gn=CDH16;refGene_tx=XM_019819850.2;refGene_gn=CDH16;refGene_tx=XM_019819851.2
E2 44719484 44719712
refGene_gn=CDH16;refGene_tx=XM_003998109.3;refGene_gn=CDH16;refGene_tx=XM_019819849.1;refGene_gn=CDH16;refGene_tx=XM_019819850.2;refGene_gn=CDH16;refGene_tx=XM_019819851.2
E2 44719928 44720079
refGene_gn=CDH16;refGene_tx=XM_003998109.3;refGene_gn=CDH16;refGene_tx=XM_019819849.1;refGene_gn=CDH16;refGene_tx=XM_019819850.2;refGene_gn=CDH16;refGene_tx=XM_019819851.2
E2 44720292 44720415
refGene_gn=CDH16;refGene_tx=XM_003998109.3;refGene_gn=CDH16;refGene_tx=XM_019819849.1;refGene_gn=CDH16;refGene_tx=XM_019819850.2;refGene_gn=CDH16;refGene_tx=XM_019819851.2
E2 44721044 44721241
refGene_gn=CDH16;refGene_tx=XM_003998109.3;refGene_gn=CDH16;refGene_tx=XM_019819849.1;refGene_gn=CDH16;refGene_tx=XM_019819850.2;refGene_gn=CDH16;refGene_tx=XM_019819851.2
E2 44722002 44722161
refGene_gn=CDH16;refGene_tx=XM_003998109.3;refGene_gn=CDH16;refGene_tx=XM_019819849.1;refGene_gn=CDH16;refGene_tx=XM_019819850.2;refGene_gn=

CDH16;refGene_tx=XM_019819851.2
E2 44722555 44722694
refGene_gn=CDH16;refGene_tx=XM_003998109.3;refGene_gn=CDH16;refGene_tx=XM_019819849.1;refGene_gn=CDH16;refGene_tx=XM_019819850.2;refGene_gn=CDH16;refGene_tx=XM_019819851.2
E2 44722764 44722920
refGene_gn=CDH16;refGene_tx=XM_003998109.3;refGene_gn=CDH16;refGene_tx=XM_019819849.1;refGene_gn=CDH16;refGene_tx=XM_019819850.2;refGene_gn=CDH16;refGene_tx=XM_019819851.2
E2 44723556 44723640
refGene_gn=CDH16;refGene_tx=XM_003998109.3;refGene_gn=CDH16;refGene_tx=XM_019819849.1;refGene_gn=CDH16;refGene_tx=XM_019819850.2;refGene_gn=CDH16;refGene_tx=XM_019819851.2
E2 44723904 44723949
refGene_gn=CDH16;refGene_tx=XM_003998109.3;refGene_gn=CDH16;refGene_tx=XM_019819849.1;refGene_gn=CDH16;refGene_tx=XM_019819850.2;refGene_gn=CDH16;refGene_tx=XM_019819851.2
E2 44727948 44728226
refGene_gn=RRAD;refGene_tx=XM_023245633.1;refGene_gn=RRAD;refGene_tx=XM_023245635.1
E2 44729069 44729274
refGene_gn=RRAD;refGene_tx=XM_023245633.1;refGene_gn=RRAD;refGene_tx=XM_023245635.1
E2 44729406 44729480
refGene_gn=RRAD;refGene_tx=XM_023245633.1;refGene_gn=RRAD;refGene_tx=XM_023245635.1
E2 44730131 44730501
refGene_gn=RRAD;refGene_tx=XM_023245633.1;refGene_gn=RRAD;refGene_tx=XM_023245635.1
E2 44736951 44737049 refGene_gn=FAM96B;refGene_tx=XM_003998113.5
E2 44737501 44737547 refGene_gn=FAM96B;refGene_tx=XM_003998113.5
E2 44738007 44738133 refGene_gn=FAM96B;refGene_tx=XM_003998113.5
E2 44738376 44738456 refGene_gn=FAM96B;refGene_tx=XM_003998113.5
E2 44738612 44738754 refGene_gn=FAM96B;refGene_tx=XM_003998113.5
E2 44739985 44740058 refGene_gn=CES2;refGene_tx=XM_019819862.1
E2 44745227 44745432 refGene_gn=CES2;refGene_tx=XM_019819862.1
E2 44746331 44746473 refGene_gn=CES2;refGene_tx=XM_019819862.1
E2 44747282 44747416 refGene_gn=CES2;refGene_tx=XM_019819862.1
E2 44747496 44747755 refGene_gn=CES2;refGene_tx=XM_019819862.1
E2 44748074 44748173 refGene_gn=CES2;refGene_tx=XM_019819862.1
E2 44748435 44748576 refGene_gn=CES2;refGene_tx=XM_019819862.1
E2 44748693 44748768 refGene_gn=CES2;refGene_tx=XM_019819862.1
E2 44749017 44749162 refGene_gn=CES2;refGene_tx=XM_019819862.1
E2 44749590 44749728 refGene_gn=CES2;refGene_tx=XM_019819862.1
E2 44750255 44750328 refGene_gn=CES2;refGene_tx=XM_019819862.1
E2 44750735 44750922 refGene_gn=CES2;refGene_tx=XM_019819862.1
E2 44754366 44754448 refGene_gn=CES3;refGene_tx=XM_023246092.1
E2 44755804 44756009 refGene_gn=CES3;refGene_tx=XM_023246092.1
E2 44756120 44756331 refGene_gn=CES3;refGene_tx=XM_023246092.1
E2 44756425 44756559 refGene_gn=CES3;refGene_tx=XM_023246092.1

E2 44757089 44757243 refGene_gn=CES3;refGene_tx=XM_023246092.1
E2 44757346 44757451 refGene_gn=CES3;refGene_tx=XM_023246092.1
E2 44758094 44758196 refGene_gn=CES3;refGene_tx=XM_023246092.1
E2 44758567 44758705 refGene_gn=CES3;refGene_tx=XM_023246092.1
E2 44762942 44763023 refGene_gn=CES3;refGene_tx=XM_023246092.1
E2 44763824 44763972 refGene_gn=CES3;refGene_tx=XM_023246092.1
E2 44764885 44765035 refGene_gn=CES3;refGene_tx=XM_023246092.1
E2 44765184 44765267 refGene_gn=CES3;refGene_tx=XM_023246092.1
E2 44765334 44765535 refGene_gn=CES3;refGene_tx=XM_023246092.1
E2 44773062 44773246
refGene_gn=CES4A;refGene_tx=XM_006941576.4;refGene_gn=CES4A;refGene_tx=XM_019819857.2;refGene_gn=CES4A;refGene_tx=XM_019819858.2;refGene_gn=CES4A;refGene_tx=XM_019819859.2;refGene_gn=CES4A;refGene_tx=XM_019819860.2;refGene_gn=CES4A;refGene_tx=XM_019819861.2
E2 44776960 44777162
refGene_gn=CES4A;refGene_tx=XM_006941576.4;refGene_gn=CES4A;refGene_tx=XM_019819857.2;refGene_gn=CES4A;refGene_tx=XM_019819858.2;refGene_gn=CES4A;refGene_tx=XM_019819859.2;refGene_gn=CES4A;refGene_tx=XM_019819861.2
E2 44779703 44779845
refGene_gn=CES4A;refGene_tx=XM_006941576.4;refGene_gn=CES4A;refGene_tx=XM_019819857.2;refGene_gn=CES4A;refGene_tx=XM_019819858.2;refGene_gn=CES4A;refGene_tx=XM_019819859.2;refGene_gn=CES4A;refGene_tx=XM_019819860.2;refGene_gn=CES4A;refGene_tx=XM_019819861.2
E2 44779925 44780059
refGene_gn=CES4A;refGene_tx=XM_006941576.4;refGene_gn=CES4A;refGene_tx=XM_019819857.2;refGene_gn=CES4A;refGene_tx=XM_019819858.2;refGene_gn=CES4A;refGene_tx=XM_019819859.2;refGene_gn=CES4A;refGene_tx=XM_019819860.2;refGene_gn=CES4A;refGene_tx=XM_019819861.2
E2 44780379 44780533
refGene_gn=CES4A;refGene_tx=XM_006941576.4;refGene_gn=CES4A;refGene_tx=XM_019819857.2;refGene_gn=CES4A;refGene_tx=XM_019819858.2;refGene_gn=CES4A;refGene_tx=XM_019819859.2;refGene_gn=CES4A;refGene_tx=XM_019819860.2;refGene_gn=CES4A;refGene_tx=XM_019819861.2
E2 44782021 44782126
refGene_gn=CES4A;refGene_tx=XM_006941576.4;refGene_gn=CES4A;refGene_tx=XM_019819857.2;refGene_gn=CES4A;refGene_tx=XM_019819859.2;refGene_gn=CES4A;refGene_tx=XM_019819860.2;refGene_gn=CES4A;refGene_tx=XM_019819861.2
E2 44782148 44782301
refGene_gn=CES4A;refGene_tx=XM_006941576.4;refGene_gn=CES4A;refGene_tx=XM_019819857.2;refGene_gn=CES4A;refGene_tx=XM_019819858.2;refGene_gn=CES4A;refGene_tx=XM_019819860.2;refGene_gn=CES4A;refGene_tx=XM_019819861.2
E2 44782446 44782485
refGene_gn=CES4A;refGene_tx=XM_006941576.4;refGene_gn=CES4A;refGene_tx=XM_019819857.2;refGene_gn=CES4A;refGene_tx=XM_019819858.2;refGene_gn=CES4A;refGene_tx=XM_019819859.2;refGene_gn=CES4A;refGene_tx=XM_019819860.2;refGene_gn=CES4A;refGene_tx=XM_019819861.2
E2 44782961 44783102

refGene_gn=CES4A;refGene_tx=XM_006941576.4;refGene_gn=CES4A;refGene_tx=XM_019819857.2;refGene_gn=CES4A;refGene_tx=XM_019819858.2;refGene_gn=CES4A;refGene_tx=XM_019819859.2;refGene_gn=CES4A;refGene_tx=XM_019819860.2;refGene_gn=CES4A;refGene_tx=XM_019819861.2
E2 44783633 44783714
refGene_gn=CES4A;refGene_tx=XM_006941576.4;refGene_gn=CES4A;refGene_tx=XM_019819857.2;refGene_gn=CES4A;refGene_tx=XM_019819858.2;refGene_gn=CES4A;refGene_tx=XM_019819859.2;refGene_gn=CES4A;refGene_tx=XM_019819860.2;refGene_gn=CES4A;refGene_tx=XM_019819861.2
E2 44784066 44784220
refGene_gn=CES4A;refGene_tx=XM_006941576.4;refGene_gn=CES4A;refGene_tx=XM_019819857.2;refGene_gn=CES4A;refGene_tx=XM_019819858.2;refGene_gn=CES4A;refGene_tx=XM_019819859.2;refGene_gn=CES4A;refGene_tx=XM_019819860.2
E2 44784821 44784968
refGene_gn=CES4A;refGene_tx=XM_006941576.4;refGene_gn=CES4A;refGene_tx=XM_019819857.2;refGene_gn=CES4A;refGene_tx=XM_019819858.2;refGene_gn=CES4A;refGene_tx=XM_019819859.2;refGene_gn=CES4A;refGene_tx=XM_019819860.2;refGene_gn=CES4A;refGene_tx=XM_019819861.2
E2 44785191 44785264
refGene_gn=CES4A;refGene_tx=XM_006941576.4;refGene_gn=CES4A;refGene_tx=XM_019819857.2;refGene_gn=CES4A;refGene_tx=XM_019819858.2;refGene_gn=CES4A;refGene_tx=XM_019819859.2;refGene_gn=CES4A;refGene_tx=XM_019819860.2;refGene_gn=CES4A;refGene_tx=XM_019819861.2
E2 44786232 44786374
refGene_gn=CES4A;refGene_tx=XM_006941576.4;refGene_gn=CES4A;refGene_tx=XM_019819857.2;refGene_gn=CES4A;refGene_tx=XM_019819858.2;refGene_gn=CES4A;refGene_tx=XM_019819859.2;refGene_gn=CES4A;refGene_tx=XM_019819860.2
E2 44803945 44804023
refGene_gn=CBFB;refGene_tx=NM_001305028.1;refGene_gn=CBFB;refGene_tx=XM_023244834.1
E2 44804238 44804325
refGene_gn=CBFB;refGene_tx=NM_001305028.1;refGene_gn=CBFB;refGene_tx=XM_023244834.1
E2 44808872 44808989
refGene_gn=CBFB;refGene_tx=NM_001305028.1;refGene_gn=CBFB;refGene_tx=XM_023244834.1
E2 44835123 44835240
refGene_gn=CBFB;refGene_tx=NM_001305028.1;refGene_gn=CBFB;refGene_tx=XM_023244834.1
E2 44845589 44845716
refGene_gn=CBFB;refGene_tx=NM_001305028.1;refGene_gn=CBFB;refGene_tx=XM_023244834.1
E2 44858698 44858767
refGene_gn=CBFB;refGene_tx=NM_001305028.1;refGene_gn=CBFB;refGene_tx=XM_023244834.1
E2 44868237 44868301
refGene_gn=CE2H16orf70;refGene_tx=XM_003998116.5;refGene_gn=CE2H16orf70;refGene_tx=XM_023246122.1;refGene_gn=CE2H16orf70;refGene_tx=XM_02324

6123.1
E2 44876414 44876497
refGene_gn=CE2H16orf70;refGene_tx=XM_003998116.5;refGene_gn=CE2H16orf70;refGene_tx=XM_023246122.1;refGene_gn=CE2H16orf70;refGene_tx=XM_023246123.1
E2 44881419 44881503
refGene_gn=CE2H16orf70;refGene_tx=XM_003998116.5;refGene_gn=CE2H16orf70;refGene_tx=XM_023246122.1;refGene_gn=CE2H16orf70;refGene_tx=XM_023246123.1
E2 44885725 44885769
refGene_gn=CE2H16orf70;refGene_tx=XM_003998116.5;refGene_gn=CE2H16orf70;refGene_tx=XM_023246122.1;refGene_gn=CE2H16orf70;refGene_tx=XM_023246123.1
E2 44888305 44888385
refGene_gn=CE2H16orf70;refGene_tx=XM_003998116.5;refGene_gn=CE2H16orf70;refGene_tx=XM_023246122.1;refGene_gn=CE2H16orf70;refGene_tx=XM_023246123.1
E2 44888659 44888754
refGene_gn=CE2H16orf70;refGene_tx=XM_003998116.5;refGene_gn=CE2H16orf70;refGene_tx=XM_023246122.1;refGene_gn=CE2H16orf70;refGene_tx=XM_023246123.1
E2 44889391 44889489
refGene_gn=CE2H16orf70;refGene_tx=XM_003998116.5;refGene_gn=CE2H16orf70;refGene_tx=XM_023246122.1;refGene_gn=CE2H16orf70;refGene_tx=XM_023246123.1
E2 44889575 44889688
refGene_gn=CE2H16orf70;refGene_tx=XM_003998116.5;refGene_gn=CE2H16orf70;refGene_tx=XM_023246122.1;refGene_gn=CE2H16orf70;refGene_tx=XM_023246123.1
E2 44892444 44892578
refGene_gn=CE2H16orf70;refGene_tx=XM_003998116.5;refGene_gn=CE2H16orf70;refGene_tx=XM_023246122.1;refGene_gn=CE2H16orf70;refGene_tx=XM_023246123.1
E2 44893045 44893129
refGene_gn=CE2H16orf70;refGene_tx=XM_003998116.5;refGene_gn=CE2H16orf70;refGene_tx=XM_023246122.1;refGene_gn=CE2H16orf70;refGene_tx=XM_023246123.1
E2 44894584 44894667
refGene_gn=CE2H16orf70;refGene_tx=XM_003998116.5;refGene_gn=CE2H16orf70;refGene_tx=XM_023246122.1;refGene_gn=CE2H16orf70;refGene_tx=XM_023246123.1
E2 44895047 44895091
refGene_gn=CE2H16orf70;refGene_tx=XM_003998116.5;refGene_gn=CE2H16orf70;refGene_tx=XM_023246122.1;refGene_gn=CE2H16orf70;refGene_tx=XM_023246123.1
E2 44895568 44895612
refGene_gn=CE2H16orf70;refGene_tx=XM_003998116.5;refGene_gn=CE2H16orf70;refGene_tx=XM_023246122.1;refGene_gn=CE2H16orf70;refGene_tx=XM_023246123.1
E2 44895763 44895822

refGene_gn=CE2H16orf70;refGene_tx=XM_003998116.5;refGene_gn=CE2H16orf70;refGene_tx=XM_023246122.1;refGene_gn=CE2H16orf70;refGene_tx=XM_023246123.1
E2 44896435 44896508
refGene_gn=CE2H16orf70;refGene_tx=XM_003998116.5;refGene_gn=CE2H16orf70;refGene_tx=XM_023246122.1;refGene_gn=CE2H16orf70;refGene_tx=XM_023246123.1
E2 44897262 44897349
refGene_gn=CE2H16orf70;refGene_tx=XM_003998116.5;refGene_gn=CE2H16orf70;refGene_tx=XM_023246122.1;refGene_gn=CE2H16orf70;refGene_tx=XM_023246123.1
E2 44899533 44900780
refGene_gn=B3GNT9;refGene_tx=XM_006941581.3;refGene_gn=B3GNT9;refGene_tx=XM_006941582.3
E2 44901154 44901317 refGene_gn=B3GNT9;refGene_tx=XM_006941581.3
E2 44903312 44903623
refGene_gn=TRADD;refGene_tx=XM_003998118.3;refGene_gn=TRADD;refGene_tx=XM_019819893.2;refGene_gn=TRADD;refGene_tx=XM_019819894.2;refGene_gn=TRADD;refGene_tx=XM_019819895.1
E2 44903770 44903966
refGene_gn=TRADD;refGene_tx=XM_003998118.3;refGene_gn=TRADD;refGene_tx=XM_019819893.2;refGene_gn=TRADD;refGene_tx=XM_019819894.2;refGene_gn=TRADD;refGene_tx=XM_019819895.1
E2 44904058 44904336
refGene_gn=TRADD;refGene_tx=XM_003998118.3;refGene_gn=TRADD;refGene_tx=XM_019819893.2;refGene_gn=TRADD;refGene_tx=XM_019819894.2;refGene_gn=TRADD;refGene_tx=XM_019819895.1
E2 44905184 44905359
refGene_gn=TRADD;refGene_tx=XM_003998118.3;refGene_gn=TRADD;refGene_tx=XM_019819893.2;refGene_gn=TRADD;refGene_tx=XM_019819894.2;refGene_gn=TRADD;refGene_tx=XM_019819895.1
E2 44910201 44910353
refGene_gn=FBXL8;refGene_tx=XM_003998119.5;refGene_gn=FBXL8;refGene_tx=XM_011290117.3
E2 44910948 44911921
refGene_gn=FBXL8;refGene_tx=XM_003998119.5;refGene_gn=FBXL8;refGene_tx=XM_011290117.3
E2 44912794 44912917
refGene_gn=HSF4;refGene_tx=XM_011290112.3;refGene_gn=HSF4;refGene_tx=XM_023246114.1
E2 44913632 44913741
refGene_gn=HSF4;refGene_tx=XM_011290112.3;refGene_gn=HSF4;refGene_tx=XM_019819878.1;refGene_gn=HSF4;refGene_tx=XM_023246114.1
E2 44913830 44913958
refGene_gn=HSF4;refGene_tx=XM_011290112.3;refGene_gn=HSF4;refGene_tx=XM_019819878.1;refGene_gn=HSF4;refGene_tx=XM_023246114.1
E2 44914057 44914182
refGene_gn=HSF4;refGene_tx=XM_011290112.3;refGene_gn=HSF4;refGene_tx=XM_019819878.1;refGene_gn=HSF4;refGene_tx=XM_023246114.1
E2 44914454 44914530

refGene_gn=HSF4;refGene_tx=XM_011290112.3;refGene_gn=HSF4;refGene_tx=XM_019819878.1;refGene_gn=HSF4;refGene_tx=XM_023246114.1
E2 44914671 44914736
refGene_gn=HSF4;refGene_tx=XM_011290112.3;refGene_gn=HSF4;refGene_tx=XM_019819878.1;refGene_gn=HSF4;refGene_tx=XM_023246114.1
E2 44915261 44915364
refGene_gn=HSF4;refGene_tx=XM_011290112.3;refGene_gn=HSF4;refGene_tx=XM_019819878.1;refGene_gn=HSF4;refGene_tx=XM_023246114.1
E2 44915618 44915746
refGene_gn=HSF4;refGene_tx=XM_011290112.3;refGene_gn=HSF4;refGene_tx=XM_019819878.1;refGene_gn=HSF4;refGene_tx=XM_023246114.1
E2 44915867 44916095
refGene_gn=HSF4;refGene_tx=XM_011290112.3;refGene_gn=HSF4;refGene_tx=XM_019819878.1;refGene_gn=HSF4;refGene_tx=XM_023246114.1
E2 44916569 44916735
refGene_gn=HSF4;refGene_tx=XM_011290112.3;refGene_gn=HSF4;refGene_tx=XM_019819878.1;refGene_gn=HSF4;refGene_tx=XM_023246114.1
E2 44916823 44916889
refGene_gn=HSF4;refGene_tx=XM_011290112.3;refGene_gn=HSF4;refGene_tx=XM_019819878.1;refGene_gn=HSF4;refGene_tx=XM_023246114.1
E2 44917012 44917082
refGene_gn=HSF4;refGene_tx=XM_011290112.3;refGene_gn=HSF4;refGene_tx=XM_019819878.1;refGene_gn=HSF4;refGene_tx=XM_023246114.1
E2 44917360 44917512
refGene_gn=HSF4;refGene_tx=XM_011290112.3;refGene_gn=HSF4;refGene_tx=XM_019819878.1;refGene_gn=HSF4;refGene_tx=XM_023246114.1
E2 44918582 44918625 refGene_gn=NOL3;refGene_tx=XM_019819896.2
E2 44920947 44921250
refGene_gn=NOL3;refGene_tx=XM_006941587.4;refGene_gn=NOL3;refGene_tx=XM_019819896.2;refGene_gn=NOL3;refGene_tx=XM_019819897.1;refGene_gn=NOL3;refGene_tx=XM_019819899.2
E2 44921397 44921646
refGene_gn=NOL3;refGene_tx=XM_006941587.4;refGene_gn=NOL3;refGene_tx=XM_019819896.2;refGene_gn=NOL3;refGene_tx=XM_019819897.1;refGene_gn=NOL3;refGene_tx=XM_019819899.2
E2 44921746 44921754
refGene_gn=NOL3;refGene_tx=XM_006941587.4;refGene_gn=NOL3;refGene_tx=XM_019819896.2;refGene_gn=NOL3;refGene_tx=XM_019819897.1;refGene_gn=NOL3;refGene_tx=XM_019819899.2
E2 44923539 44923704
refGene_gn=KIAA0895L;refGene_tx=XM_006941596.3;refGene_gn=KIAA0895L;refGene_tx=XM_006941597.3;refGene_gn=KIAA0895L;refGene_tx=XM_019819883.1;refGene_gn=KIAA0895L;refGene_tx=XM_019819884.1;refGene_gn=KIAA0895L;refGene_tx=XM_019819885.1;refGene_gn=KIAA0895L;refGene_tx=XM_019819886.1;refGene_gn=KIAA0895L;refGene_tx=XM_023246119.1;refGene_gn=KIAA0895L;refGene_tx=XM_023246120.1
E2 44924670 44924768
refGene_gn=KIAA0895L;refGene_tx=XM_006941596.3;refGene_gn=KIAA0895L;refGene_tx=XM_006941597.3;refGene_gn=KIAA0895L;refGene_tx=XM_019819883.1;refGene_gn=KIAA0895L;refGene_tx=XM_019819884.1;refGene_gn=KIAA0895L;r

efGene_tx=XM_019819885.1;refGene_gn=KIAA0895L;refGene_tx=XM_019819886.1;refGene_gn=KIAA0895L;refGene_tx=XM_023246119.1;refGene_gn=KIAA0895L;refGene_tx=XM_023246120.1
E2 44924900 44925206
refGene_gn=KIAA0895L;refGene_tx=XM_006941596.3;refGene_gn=KIAA0895L;refGene_tx=XM_006941597.3;refGene_gn=KIAA0895L;refGene_tx=XM_019819883.1;refGene_gn=KIAA0895L;refGene_tx=XM_019819884.1;refGene_gn=KIAA0895L;refGene_tx=XM_019819885.1;refGene_gn=KIAA0895L;refGene_tx=XM_019819886.1;refGene_gn=KIAA0895L;refGene_tx=XM_023246119.1;refGene_gn=KIAA0895L;refGene_tx=XM_023246120.1
E2 44926157 44926290
refGene_gn=KIAA0895L;refGene_tx=XM_006941596.3;refGene_gn=KIAA0895L;refGene_tx=XM_006941597.3;refGene_gn=KIAA0895L;refGene_tx=XM_019819883.1;refGene_gn=KIAA0895L;refGene_tx=XM_019819884.1;refGene_gn=KIAA0895L;refGene_tx=XM_019819885.1;refGene_gn=KIAA0895L;refGene_tx=XM_019819886.1;refGene_gn=KIAA0895L;refGene_tx=XM_023246119.1;refGene_gn=KIAA0895L;refGene_tx=XM_023246120.1
E2 44926558 44926702
refGene_gn=KIAA0895L;refGene_tx=XM_006941596.3;refGene_gn=KIAA0895L;refGene_tx=XM_006941597.3;refGene_gn=KIAA0895L;refGene_tx=XM_019819883.1;refGene_gn=KIAA0895L;refGene_tx=XM_019819884.1;refGene_gn=KIAA0895L;refGene_tx=XM_019819885.1;refGene_gn=KIAA0895L;refGene_tx=XM_019819886.1;refGene_gn=KIAA0895L;refGene_tx=XM_023246119.1;refGene_gn=KIAA0895L;refGene_tx=XM_023246120.1
E2 44926783 44927353
refGene_gn=KIAA0895L;refGene_tx=XM_006941596.3;refGene_gn=KIAA0895L;refGene_tx=XM_006941597.3;refGene_gn=KIAA0895L;refGene_tx=XM_019819883.1;refGene_gn=KIAA0895L;refGene_tx=XM_019819884.1;refGene_gn=KIAA0895L;refGene_tx=XM_019819885.1;refGene_gn=KIAA0895L;refGene_tx=XM_019819886.1;refGene_gn=KIAA0895L;refGene_tx=XM_023246119.1;refGene_gn=KIAA0895L;refGene_tx=XM_023246120.1
E2 44930634 44930824
refGene_gn=EXOC3L1;refGene_tx=XM_006941592.3;refGene_gn=EXOC3L1;refGene_tx=XM_006941593.2;refGene_gn=EXOC3L1;refGene_tx=XM_023246105.1;refGene_gn=EXOC3L1;refGene_tx=XM_023246106.1;refGene_gn=EXOC3L1;refGene_tx=XM_023246107.1
E2 44930904 44931029
refGene_gn=EXOC3L1;refGene_tx=XM_006941592.3;refGene_gn=EXOC3L1;refGene_tx=XM_006941593.2;refGene_gn=EXOC3L1;refGene_tx=XM_023246105.1;refGene_gn=EXOC3L1;refGene_tx=XM_023246106.1;refGene_gn=EXOC3L1;refGene_tx=XM_023246107.1
E2 44931120 44931276
refGene_gn=EXOC3L1;refGene_tx=XM_006941592.3;refGene_gn=EXOC3L1;refGene_tx=XM_006941593.2;refGene_gn=EXOC3L1;refGene_tx=XM_023246105.1;refGene_gn=EXOC3L1;refGene_tx=XM_023246106.1;refGene_gn=EXOC3L1;refGene_tx=XM_023246107.1
E2 44931353 44931476
refGene_gn=EXOC3L1;refGene_tx=XM_006941592.3;refGene_gn=EXOC3L1;refGene_tx=XM_006941593.2;refGene_gn=EXOC3L1;refGene_tx=XM_023246105.1;refGene_gn=EXOC3L1;refGene_tx=XM_023246106.1;refGene_gn=EXOC3L1;refGene_tx=

XM_023246107.1
E2 44931603 44931733
refGene_gn=EXOC3L1;refGene_tx=XM_006941592.3;refGene_gn=EXOC3L1;refGene_tx=XM_006941593.2;refGene_gn=EXOC3L1;refGene_tx=XM_023246105.1;refGene_gn=EXOC3L1;refGene_tx=XM_023246106.1;refGene_gn=EXOC3L1;refGene_tx=XM_023246107.1
E2 44932408 44932519
refGene_gn=EXOC3L1;refGene_tx=XM_006941592.3;refGene_gn=EXOC3L1;refGene_tx=XM_006941593.2;refGene_gn=EXOC3L1;refGene_tx=XM_023246105.1;refGene_gn=EXOC3L1;refGene_tx=XM_023246106.1;refGene_gn=EXOC3L1;refGene_tx=XM_023246107.1
E2 44932726 44932827
refGene_gn=EXOC3L1;refGene_tx=XM_006941592.3;refGene_gn=EXOC3L1;refGene_tx=XM_006941593.2;refGene_gn=EXOC3L1;refGene_tx=XM_023246105.1;refGene_gn=EXOC3L1;refGene_tx=XM_023246106.1;refGene_gn=EXOC3L1;refGene_tx=XM_023246107.1
E2 44932926 44933052
refGene_gn=EXOC3L1;refGene_tx=XM_006941592.3;refGene_gn=EXOC3L1;refGene_tx=XM_006941593.2;refGene_gn=EXOC3L1;refGene_tx=XM_023246105.1;refGene_gn=EXOC3L1;refGene_tx=XM_023246106.1;refGene_gn=EXOC3L1;refGene_tx=XM_023246107.1
E2 44933193 44933311
refGene_gn=EXOC3L1;refGene_tx=XM_006941592.3;refGene_gn=EXOC3L1;refGene_tx=XM_006941593.2;refGene_gn=EXOC3L1;refGene_tx=XM_023246105.1;refGene_gn=EXOC3L1;refGene_tx=XM_023246106.1;refGene_gn=EXOC3L1;refGene_tx=XM_023246107.1
E2 44933389 44934002
refGene_gn=EXOC3L1;refGene_tx=XM_006941592.3;refGene_gn=EXOC3L1;refGene_tx=XM_006941593.2;refGene_gn=EXOC3L1;refGene_tx=XM_023246105.1;refGene_gn=EXOC3L1;refGene_tx=XM_023246106.1;refGene_gn=EXOC3L1;refGene_tx=XM_023246107.1
E2 44934543 44934763
refGene_gn=EXOC3L1;refGene_tx=XM_006941592.3;refGene_gn=EXOC3L1;refGene_tx=XM_006941593.2;refGene_gn=EXOC3L1;refGene_tx=XM_023246105.1;refGene_gn=EXOC3L1;refGene_tx=XM_023246106.1;refGene_gn=EXOC3L1;refGene_tx=XM_023246107.1
E2 44934852 44935028
refGene_gn=EXOC3L1;refGene_tx=XM_006941592.3;refGene_gn=EXOC3L1;refGene_tx=XM_006941593.2;refGene_gn=EXOC3L1;refGene_tx=XM_023246105.1;refGene_gn=EXOC3L1;refGene_tx=XM_023246106.1;refGene_gn=EXOC3L1;refGene_tx=XM_023246107.1
E2 44935237 44935420
refGene_gn=EXOC3L1;refGene_tx=XM_006941592.3;refGene_gn=EXOC3L1;refGene_tx=XM_006941593.2;refGene_gn=EXOC3L1;refGene_tx=XM_023246105.1;refGene_gn=EXOC3L1;refGene_tx=XM_023246106.1;refGene_gn=EXOC3L1;refGene_tx=XM_023246107.1
E2 44936875 44936921
refGene_gn=EXOC3L1;refGene_tx=XM_006941592.3;refGene_gn=EXOC3L1;refGene_tx=XM_023246105.1
E2 44937288 44937423 refGene_gn=E2F4;refGene_tx=XM_003998123.5

E2 44937860 44937970 refGene_gn=E2F4;refGene_tx=XM_003998123.5
E2 44938111 44938273 refGene_gn=E2F4;refGene_tx=XM_003998123.5
E2 44938589 44938633 refGene_gn=E2F4;refGene_tx=XM_003998123.5
E2 44939349 44939411 refGene_gn=E2F4;refGene_tx=XM_003998123.5
E2 44939629 44939927 refGene_gn=E2F4;refGene_tx=XM_003998123.5
E2 44940398 44940599 refGene_gn=E2F4;refGene_tx=XM_003998123.5
E2 44942666 44942714 refGene_gn=E2F4;refGene_tx=XM_003998123.5
E2 44942919 44942964 refGene_gn=E2F4;refGene_tx=XM_003998123.5
E2 44943062 44943178 refGene_gn=E2F4;refGene_tx=XM_003998123.5
E2 44944405 44944483
refGene_gn=ELMO3;refGene_tx=XM_006941589.4;refGene_gn=ELMO3;refGene_tx=XM_006941590.4;refGene_gn=ELMO3;refGene_tx=XM_019819873.2;refGene_gn=ELMO3;refGene_tx=XM_019819874.2
E2 44944614 44944655
refGene_gn=ELMO3;refGene_tx=XM_006941589.4;refGene_gn=ELMO3;refGene_tx=XM_006941590.4;refGene_gn=ELMO3;refGene_tx=XM_019819873.2;refGene_gn=ELMO3;refGene_tx=XM_019819874.2
E2 44944755 44944828
refGene_gn=ELMO3;refGene_tx=XM_006941589.4;refGene_gn=ELMO3;refGene_tx=XM_006941590.4;refGene_gn=ELMO3;refGene_tx=XM_019819873.2;refGene_gn=ELMO3;refGene_tx=XM_019819874.2
E2 44945037 44945088
refGene_gn=ELMO3;refGene_tx=XM_006941589.4;refGene_gn=ELMO3;refGene_tx=XM_006941590.4;refGene_gn=ELMO3;refGene_tx=XM_019819874.2
E2 44945272 44945442
refGene_gn=ELMO3;refGene_tx=XM_006941589.4;refGene_gn=ELMO3;refGene_tx=XM_006941590.4;refGene_gn=ELMO3;refGene_tx=XM_019819873.2;refGene_gn=ELMO3;refGene_tx=XM_019819874.2
E2 44945527 44945627
refGene_gn=ELMO3;refGene_tx=XM_006941589.4;refGene_gn=ELMO3;refGene_tx=XM_006941590.4;refGene_gn=ELMO3;refGene_tx=XM_019819873.2;refGene_gn=ELMO3;refGene_tx=XM_019819874.2
E2 44945719 44945871
refGene_gn=ELMO3;refGene_tx=XM_006941589.4;refGene_gn=ELMO3;refGene_tx=XM_006941590.4;refGene_gn=ELMO3;refGene_tx=XM_019819873.2;refGene_gn=ELMO3;refGene_tx=XM_019819874.2
E2 44945961 44946040
refGene_gn=ELMO3;refGene_tx=XM_006941589.4;refGene_gn=ELMO3;refGene_tx=XM_006941590.4;refGene_gn=ELMO3;refGene_tx=XM_019819873.2;refGene_gn=ELMO3;refGene_tx=XM_019819874.2
E2 44946168 44946219
refGene_gn=ELMO3;refGene_tx=XM_006941589.4;refGene_gn=ELMO3;refGene_tx=XM_006941590.4;refGene_gn=ELMO3;refGene_tx=XM_019819873.2;refGene_gn=ELMO3;refGene_tx=XM_019819874.2
E2 44946307 44946430
refGene_gn=ELMO3;refGene_tx=XM_006941589.4;refGene_gn=ELMO3;refGene_tx=XM_006941590.4;refGene_gn=ELMO3;refGene_tx=XM_019819873.2;refGene_gn=ELMO3;refGene_tx=XM_019819874.2
E2 44946526 44946862
refGene_gn=ELMO3;refGene_tx=XM_006941589.4;refGene_gn=ELMO3;refGene_tx

=XM_006941590.4;refGene_gn=ELMO3;refGene_tx=XM_019819873.2;refGene_gn=ELMO3;refGene_tx=XM_019819874.2
E2 44946964 44947073
refGene_gn=ELMO3;refGene_tx=XM_006941589.4;refGene_gn=ELMO3;refGene_tx=XM_006941590.4;refGene_gn=ELMO3;refGene_tx=XM_019819873.2;refGene_gn=ELMO3;refGene_tx=XM_019819874.2
E2 44947180 44947317
refGene_gn=ELMO3;refGene_tx=XM_006941589.4;refGene_gn=ELMO3;refGene_tx=XM_006941590.4;refGene_gn=ELMO3;refGene_tx=XM_019819873.2;refGene_gn=ELMO3;refGene_tx=XM_019819874.2;refGene_gn=ELMO3;refGene_tx=XM_019819876.1
E2 44947412 44947576
refGene_gn=ELMO3;refGene_tx=XM_006941589.4;refGene_gn=ELMO3;refGene_tx=XM_006941590.4;refGene_gn=ELMO3;refGene_tx=XM_019819873.2;refGene_gn=ELMO3;refGene_tx=XM_019819874.2;refGene_gn=ELMO3;refGene_tx=XM_019819876.1
E2 44947685 44947798
refGene_gn=ELMO3;refGene_tx=XM_006941589.4;refGene_gn=ELMO3;refGene_tx=XM_006941590.4;refGene_gn=ELMO3;refGene_tx=XM_019819873.2;refGene_gn=ELMO3;refGene_tx=XM_019819874.2;refGene_gn=ELMO3;refGene_tx=XM_019819876.1
E2 44947902 44948007
refGene_gn=ELMO3;refGene_tx=XM_006941589.4;refGene_gn=ELMO3;refGene_tx=XM_006941590.4;refGene_gn=ELMO3;refGene_tx=XM_019819873.2;refGene_gn=ELMO3;refGene_tx=XM_019819874.2;refGene_gn=ELMO3;refGene_tx=XM_019819876.1
E2 44948093 44948190
refGene_gn=ELMO3;refGene_tx=XM_006941589.4;refGene_gn=ELMO3;refGene_tx=XM_006941590.4;refGene_gn=ELMO3;refGene_tx=XM_019819873.2;refGene_gn=ELMO3;refGene_tx=XM_019819874.2;refGene_gn=ELMO3;refGene_tx=XM_019819876.1
E2 44948281 44948368
refGene_gn=ELMO3;refGene_tx=XM_006941589.4;refGene_gn=ELMO3;refGene_tx=XM_006941590.4;refGene_gn=ELMO3;refGene_tx=XM_019819873.2;refGene_gn=ELMO3;refGene_tx=XM_019819876.1
E2 44948456 44948669
refGene_gn=ELMO3;refGene_tx=XM_006941589.4;refGene_gn=ELMO3;refGene_tx=XM_006941590.4;refGene_gn=ELMO3;refGene_tx=XM_019819873.2;refGene_gn=ELMO3;refGene_tx=XM_019819876.1
E2 44951401 44951570
refGene_gn=LOC111558130;refGene_tx=XM_023246108.1;refGene_gn=LOC111558130;refGene_tx=XM_023246109.1;refGene_gn=LOC111558130;refGene_tx=XM_023246110.1;refGene_gn=LOC111558130;refGene_tx=XM_023246111.1;refGene_gn=LOC111558130;refGene_tx=XM_023246112.1;refGene_gn=LOC111558130;refGene_tx=XM_023246113.1
E2 44951659 44951877
refGene_gn=LOC111558130;refGene_tx=XM_023246108.1;refGene_gn=LOC111558130;refGene_tx=XM_023246109.1;refGene_gn=LOC111558130;refGene_tx=XM_023246110.1;refGene_gn=LOC111558130;refGene_tx=XM_023246111.1;refGene_gn=LOC111558130;refGene_tx=XM_023246112.1;refGene_gn=LOC111558130;refGen

e_tx=XM_023246113.1
E2	44952071 44952269
refGene_gn=LOC111558130;refGene_tx=XM_023246108.1;refGene_gn=LOC111558130;refGene_tx=XM_023246109.1;refGene_gn=LOC111558130;refGene_tx=XM_023246110.1;refGene_gn=LOC111558130;refGene_tx=XM_023246111.1;refGene_gn=LOC111558130;refGene_tx=XM_023246112.1;refGene_gn=LOC111558130;refGene_tx=XM_023246113.1
E2	44952350 44952566
refGene_gn=LOC111558130;refGene_tx=XM_023246108.1;refGene_gn=LOC111558130;refGene_tx=XM_023246109.1;refGene_gn=LOC111558130;refGene_tx=XM_023246110.1;refGene_gn=LOC111558130;refGene_tx=XM_023246111.1;refGene_gn=LOC111558130;refGene_tx=XM_023246112.1
E2	44953692 44954685
refGene_gn=LOC111558130;refGene_tx=XM_023246108.1;refGene_gn=LOC111558130;refGene_tx=XM_023246109.1;refGene_gn=LOC111558130;refGene_tx=XM_023246110.1;refGene_gn=LOC111558130;refGene_tx=XM_023246111.1;refGene_gn=LOC111558130;refGene_tx=XM_023246112.1;refGene_gn=LOC111558130;refGene_tx=XM_023246113.1
E2	44962499 44962601
refGene_gn=LOC111558130;refGene_tx=XM_023246108.1;refGene_gn=LOC111558130;refGene_tx=XM_023246109.1;refGene_gn=LOC111558130;refGene_tx=XM_023246110.1;refGene_gn=LOC111558130;refGene_tx=XM_023246111.1;refGene_gn=LOC111558130;refGene_tx=XM_023246112.1;refGene_gn=LOC111558130;refGene_tx=XM_023246113.1
E2	44965738 44965762
refGene_gn=LOC111558130;refGene_tx=XM_023246108.1;refGene_gn=LOC111558130;refGene_tx=XM_023246112.1;refGene_gn=LOC111558130;refGene_tx=XM_023246113.1
E2	44965984 44965990
refGene_gn=TMEM208;refGene_tx=XM_003998124.4;refGene_gn=TMEM208;refGene_tx=XM_019819900.2
E2	44966617 44966713
refGene_gn=TMEM208;refGene_tx=XM_003998124.4;refGene_gn=TMEM208;refGene_tx=XM_019819900.2
E2	44967188 44967248
refGene_gn=TMEM208;refGene_tx=XM_003998124.4;refGene_gn=TMEM208;refGene_tx=XM_019819900.2
E2	44967326 44967463
refGene_gn=TMEM208;refGene_tx=XM_003998124.4;refGene_gn=TMEM208;refGene_tx=XM_019819900.2
E2	44967610 44967713
refGene_gn=TMEM208;refGene_tx=XM_003998124.4;refGene_gn=TMEM208;refGene_tx=XM_019819900.2
E2	44967812 44967950
refGene_gn=TMEM208;refGene_tx=XM_003998124.4;refGene_gn=TMEM208;refGene_tx=XM_019819900.2
E2	44968421 44968504
refGene_gn=FHOD1;refGene_tx=XM_006941602.4;refGene_gn=FHOD1;refGene_tx=XM_006941603.4;refGene_gn=FHOD1;refGene_tx=XM_019819868.1
E2	44968577 44968778

refGene_gn=FHOD1;refGene_tx=XM_006941602.4;refGene_gn=FHOD1;refGene_tx=XM_006941603.4;refGene_gn=FHOD1;refGene_tx=XM_019819868.1
E2 44968853 44969016
refGene_gn=FHOD1;refGene_tx=XM_006941602.4;refGene_gn=FHOD1;refGene_tx=XM_006941603.4;refGene_gn=FHOD1;refGene_tx=XM_019819868.1
E2 44969097 44969290
refGene_gn=FHOD1;refGene_tx=XM_006941602.4;refGene_gn=FHOD1;refGene_tx=XM_006941603.4;refGene_gn=FHOD1;refGene_tx=XM_019819868.1
E2 44969380 44969571
refGene_gn=FHOD1;refGene_tx=XM_006941602.4;refGene_gn=FHOD1;refGene_tx=XM_006941603.4;refGene_gn=FHOD1;refGene_tx=XM_019819868.1
E2 44969981 44970143
refGene_gn=FHOD1;refGene_tx=XM_006941602.4;refGene_gn=FHOD1;refGene_tx=XM_006941603.4;refGene_gn=FHOD1;refGene_tx=XM_019819868.1
E2 44970220 44970340
refGene_gn=FHOD1;refGene_tx=XM_006941602.4;refGene_gn=FHOD1;refGene_tx=XM_006941603.4;refGene_gn=FHOD1;refGene_tx=XM_019819868.1
E2 44970426 44970609
refGene_gn=FHOD1;refGene_tx=XM_006941602.4;refGene_gn=FHOD1;refGene_tx=XM_006941603.4;refGene_gn=FHOD1;refGene_tx=XM_019819868.1
E2 44970841 44970997
refGene_gn=FHOD1;refGene_tx=XM_006941602.4;refGene_gn=FHOD1;refGene_tx=XM_006941603.4;refGene_gn=FHOD1;refGene_tx=XM_019819868.1
E2 44971337 44971942
refGene_gn=FHOD1;refGene_tx=XM_006941602.4;refGene_gn=FHOD1;refGene_tx=XM_006941603.4;refGene_gn=FHOD1;refGene_tx=XM_019819868.1
E2 44972026 44972142
refGene_gn=FHOD1;refGene_tx=XM_006941602.4;refGene_gn=FHOD1;refGene_tx=XM_006941603.4;refGene_gn=FHOD1;refGene_tx=XM_019819868.1
E2 44973041 44973095
refGene_gn=FHOD1;refGene_tx=XM_006941602.4;refGene_gn=FHOD1;refGene_tx=XM_019819868.1
E2 44973477 44973501
refGene_gn=FHOD1;refGene_tx=XM_006941602.4;refGene_gn=FHOD1;refGene_tx=XM_019819868.1
E2 44973635 44973872
refGene_gn=FHOD1;refGene_tx=XM_006941602.4;refGene_gn=FHOD1;refGene_tx=XM_006941603.4;refGene_gn=FHOD1;refGene_tx=XM_019819868.1
E2 44973954 44974100
refGene_gn=FHOD1;refGene_tx=XM_006941602.4;refGene_gn=FHOD1;refGene_tx=XM_006941603.4;refGene_gn=FHOD1;refGene_tx=XM_019819868.1
E2 44974185 44974329
refGene_gn=FHOD1;refGene_tx=XM_006941602.4;refGene_gn=FHOD1;refGene_tx=XM_006941603.4;refGene_gn=FHOD1;refGene_tx=XM_019819868.1
E2 44974438 44974533
refGene_gn=FHOD1;refGene_tx=XM_006941602.4;refGene_gn=FHOD1;refGene_tx=XM_006941603.4;refGene_gn=FHOD1;refGene_tx=XM_019819868.1
E2 44974623 44974735
refGene_gn=FHOD1;refGene_tx=XM_006941602.4;refGene_gn=FHOD1;refGene_tx=XM_006941603.4;refGene_gn=FHOD1;refGene_tx=XM_019819868.1

E2 44975003 44975098
refGene_gn=FHOD1;refGene_tx=XM_006941602.4;refGene_gn=FHOD1;refGene_tx=XM_006941603.4;refGene_gn=FHOD1;refGene_tx=XM_019819868.1
E2 44975174 44975280
refGene_gn=FHOD1;refGene_tx=XM_006941602.4;refGene_gn=FHOD1;refGene_tx=XM_006941603.4;refGene_gn=FHOD1;refGene_tx=XM_019819868.1
E2 44975353 44975421
refGene_gn=FHOD1;refGene_tx=XM_006941602.4;refGene_gn=FHOD1;refGene_tx=XM_006941603.4
E2 44975718 44975783
refGene_gn=FHOD1;refGene_tx=XM_006941602.4;refGene_gn=FHOD1;refGene_tx=XM_006941603.4;refGene_gn=FHOD1;refGene_tx=XM_019819868.1
E2 44976158 44976265
refGene_gn=FHOD1;refGene_tx=XM_006941602.4;refGene_gn=FHOD1;refGene_tx=XM_006941603.4
E2 44984260 44984461
refGene_gn=FHOD1;refGene_tx=XM_006941602.4;refGene_gn=FHOD1;refGene_tx=XM_006941603.4
E2 44986102 44986295
refGene_gn=SLC9A5;refGene_tx=XM_023246098.1;refGene_gn=SLC9A5;refGene_tx=XM_023246099.1;refGene_gn=SLC9A5;refGene_tx=XM_023246100.1;refGene_gn=SLC9A5;refGene_tx=XM_023246101.1;refGene_gn=SLC9A5;refGene_tx=XM_023246102.1;refGene_gn=SLC9A5;refGene_tx=XM_023246103.1;refGene_gn=SLC9A5;refGene_tx=XM_023246104.1
E2 44989524 44989827
refGene_gn=SLC9A5;refGene_tx=XM_023246098.1;refGene_gn=SLC9A5;refGene_tx=XM_023246099.1;refGene_gn=SLC9A5;refGene_tx=XM_023246100.1;refGene_gn=SLC9A5;refGene_tx=XM_023246101.1;refGene_gn=SLC9A5;refGene_tx=XM_023246102.1;refGene_gn=SLC9A5;refGene_tx=XM_023246103.1;refGene_gn=SLC9A5;refGene_tx=XM_023246104.1
E2 44991574 44991738
refGene_gn=SLC9A5;refGene_tx=XM_023246098.1;refGene_gn=SLC9A5;refGene_tx=XM_023246099.1;refGene_gn=SLC9A5;refGene_tx=XM_023246100.1;refGene_gn=SLC9A5;refGene_tx=XM_023246101.1;refGene_gn=SLC9A5;refGene_tx=XM_023246102.1;refGene_gn=SLC9A5;refGene_tx=XM_023246103.1;refGene_gn=SLC9A5;refGene_tx=XM_023246104.1
E2 44991928 44992007
refGene_gn=SLC9A5;refGene_tx=XM_023246098.1;refGene_gn=SLC9A5;refGene_tx=XM_023246099.1;refGene_gn=SLC9A5;refGene_tx=XM_023246100.1;refGene_gn=SLC9A5;refGene_tx=XM_023246101.1;refGene_gn=SLC9A5;refGene_tx=XM_023246102.1;refGene_gn=SLC9A5;refGene_tx=XM_023246103.1;refGene_gn=SLC9A5;refGene_tx=XM_023246104.1
E2 44992277 44992455
refGene_gn=SLC9A5;refGene_tx=XM_023246098.1;refGene_gn=SLC9A5;refGene_tx=XM_023246099.1;refGene_gn=SLC9A5;refGene_tx=XM_023246100.1;refGene_gn=SLC9A5;refGene_tx=XM_023246101.1;refGene_gn=SLC9A5;refGene_tx=XM_023246102.1;refGene_gn=SLC9A5;refGene_tx=XM_023246103.1;refGene_gn=SLC9A5;refGene_tx=XM_023246104.1
E2 44993028 44993249
refGene_gn=SLC9A5;refGene_tx=XM_023246098.1;refGene_gn=SLC9A5;refGene_

tx=XM_023246099.1;refGene_gn=SLC9A5;refGene_tx=XM_023246100.1;refGene_gn=SLC9A5;refGene_tx=XM_023246101.1;refGene_gn=SLC9A5;refGene_tx=XM_023246102.1;refGene_gn=SLC9A5;refGene_tx=XM_023246103.1;refGene_gn=SLC9A5;refGene_tx=XM_023246104.1
E2 44993464 44993667
refGene_gn=SLC9A5;refGene_tx=XM_023246098.1;refGene_gn=SLC9A5;refGene_tx=XM_023246099.1;refGene_gn=SLC9A5;refGene_tx=XM_023246100.1;refGene_gn=SLC9A5;refGene_tx=XM_023246101.1;refGene_gn=SLC9A5;refGene_tx=XM_023246102.1;refGene_gn=SLC9A5;refGene_tx=XM_023246103.1;refGene_gn=SLC9A5;refGene_tx=XM_023246104.1
E2 44993887 44993977
refGene_gn=SLC9A5;refGene_tx=XM_023246098.1;refGene_gn=SLC9A5;refGene_tx=XM_023246099.1;refGene_gn=SLC9A5;refGene_tx=XM_023246100.1;refGene_gn=SLC9A5;refGene_tx=XM_023246101.1;refGene_gn=SLC9A5;refGene_tx=XM_023246102.1;refGene_gn=SLC9A5;refGene_tx=XM_023246103.1;refGene_gn=SLC9A5;refGene_tx=XM_023246104.1
E2 44994071 44994142
refGene_gn=SLC9A5;refGene_tx=XM_023246098.1;refGene_gn=SLC9A5;refGene_tx=XM_023246099.1;refGene_gn=SLC9A5;refGene_tx=XM_023246100.1;refGene_gn=SLC9A5;refGene_tx=XM_023246101.1;refGene_gn=SLC9A5;refGene_tx=XM_023246102.1;refGene_gn=SLC9A5;refGene_tx=XM_023246103.1;refGene_gn=SLC9A5;refGene_tx=XM_023246104.1
E2 44994866 44994996
refGene_gn=SLC9A5;refGene_tx=XM_023246098.1;refGene_gn=SLC9A5;refGene_tx=XM_023246099.1;refGene_gn=SLC9A5;refGene_tx=XM_023246100.1;refGene_gn=SLC9A5;refGene_tx=XM_023246101.1;refGene_gn=SLC9A5;refGene_tx=XM_023246102.1;refGene_gn=SLC9A5;refGene_tx=XM_023246103.1;refGene_gn=SLC9A5;refGene_tx=XM_023246104.1
E2 44995462 44995554
refGene_gn=SLC9A5;refGene_tx=XM_023246098.1;refGene_gn=SLC9A5;refGene_tx=XM_023246099.1;refGene_gn=SLC9A5;refGene_tx=XM_023246100.1;refGene_gn=SLC9A5;refGene_tx=XM_023246101.1;refGene_gn=SLC9A5;refGene_tx=XM_023246102.1;refGene_gn=SLC9A5;refGene_tx=XM_023246103.1;refGene_gn=SLC9A5;refGene_tx=XM_023246104.1
E2 44995716 44995843
refGene_gn=SLC9A5;refGene_tx=XM_023246098.1;refGene_gn=SLC9A5;refGene_tx=XM_023246099.1;refGene_gn=SLC9A5;refGene_tx=XM_023246100.1;refGene_gn=SLC9A5;refGene_tx=XM_023246101.1;refGene_gn=SLC9A5;refGene_tx=XM_023246102.1;refGene_gn=SLC9A5;refGene_tx=XM_023246103.1;refGene_gn=SLC9A5;refGene_tx=XM_023246104.1
E2 44997255 44997303
refGene_gn=SLC9A5;refGene_tx=XM_023246098.1;refGene_gn=SLC9A5;refGene_tx=XM_023246099.1;refGene_gn=SLC9A5;refGene_tx=XM_023246100.1
E2 44999665 44999836
refGene_gn=SLC9A5;refGene_tx=XM_023246098.1;refGene_gn=SLC9A5;refGene_tx=XM_023246099.1;refGene_gn=SLC9A5;refGene_tx=XM_023246100.1;refGene_gn=SLC9A5;refGene_tx=XM_023246101.1;refGene_gn=SLC9A5;refGene_tx=XM_023246102.1;refGene_gn=SLC9A5;refGene_tx=XM_023246103.1
E2 45000358 45000428
refGene_gn=SLC9A5;refGene_tx=XM_023246098.1;refGene_gn=SLC9A5;refGene_

tx=XM_023246099.1;refGene_gn=SLC9A5;refGene_tx=XM_023246100.1;refGene_gn=SLC9A5;refGene_tx=XM_023246101.1;refGene_gn=SLC9A5;refGene_tx=XM_023246102.1;refGene_gn=SLC9A5;refGene_tx=XM_023246103.1
E2 45001107 45001242
refGene_gn=SLC9A5;refGene_tx=XM_023246098.1;refGene_gn=SLC9A5;refGene_tx=XM_023246099.1;refGene_gn=SLC9A5;refGene_tx=XM_023246100.1;refGene_gn=SLC9A5;refGene_tx=XM_023246101.1;refGene_gn=SLC9A5;refGene_tx=XM_023246102.1;refGene_gn=SLC9A5;refGene_tx=XM_023246103.1
E2 45004168 45004641
refGene_gn=SLC9A5;refGene_tx=XM_023246098.1;refGene_gn=SLC9A5;refGene_tx=XM_023246099.1;refGene_gn=SLC9A5;refGene_tx=XM_023246100.1;refGene_gn=SLC9A5;refGene_tx=XM_023246101.1;refGene_gn=SLC9A5;refGene_tx=XM_023246102.1;refGene_gn=SLC9A5;refGene_tx=XM_023246103.1;refGene_gn=SLC9A5;refGene_tx=XM_023246104.1
E2 45011391 45011902
refGene_gn=PLEKHG4;refGene_tx=XM_006941609.4;refGene_gn=PLEKHG4;refGene_tx=XM_023246094.1;refGene_gn=PLEKHG4;refGene_tx=XM_023246095.1;refGene_gn=PLEKHG4;refGene_tx=XM_023246096.1
E2 45012059 45012155
refGene_gn=PLEKHG4;refGene_tx=XM_006941609.4;refGene_gn=PLEKHG4;refGene_tx=XM_023246094.1;refGene_gn=PLEKHG4;refGene_tx=XM_023246095.1;refGene_gn=PLEKHG4;refGene_tx=XM_023246096.1
E2 45012230 45012354
refGene_gn=PLEKHG4;refGene_tx=XM_006941609.4;refGene_gn=PLEKHG4;refGene_tx=XM_023246094.1;refGene_gn=PLEKHG4;refGene_tx=XM_023246095.1;refGene_gn=PLEKHG4;refGene_tx=XM_023246096.1
E2 45012440 45012534
refGene_gn=PLEKHG4;refGene_tx=XM_006941609.4;refGene_gn=PLEKHG4;refGene_tx=XM_023246094.1;refGene_gn=PLEKHG4;refGene_tx=XM_023246095.1;refGene_gn=PLEKHG4;refGene_tx=XM_023246096.1
E2 45012847 45012922
refGene_gn=PLEKHG4;refGene_tx=XM_006941609.4;refGene_gn=PLEKHG4;refGene_tx=XM_023246094.1;refGene_gn=PLEKHG4;refGene_tx=XM_023246095.1;refGene_gn=PLEKHG4;refGene_tx=XM_023246096.1
E2 45013002 45013119
refGene_gn=PLEKHG4;refGene_tx=XM_006941609.4;refGene_gn=PLEKHG4;refGene_tx=XM_023246094.1;refGene_gn=PLEKHG4;refGene_tx=XM_023246095.1;refGene_gn=PLEKHG4;refGene_tx=XM_023246096.1
E2 45013286 45013385
refGene_gn=PLEKHG4;refGene_tx=XM_006941609.4;refGene_gn=PLEKHG4;refGene_tx=XM_023246094.1;refGene_gn=PLEKHG4;refGene_tx=XM_023246095.1;refGene_gn=PLEKHG4;refGene_tx=XM_023246096.1
E2 45013476 45013625
refGene_gn=PLEKHG4;refGene_tx=XM_006941609.4;refGene_gn=PLEKHG4;refGene_tx=XM_023246094.1;refGene_gn=PLEKHG4;refGene_tx=XM_023246095.1;refGene_gn=PLEKHG4;refGene_tx=XM_023246096.1
E2 45013786 45013925
refGene_gn=PLEKHG4;refGene_tx=XM_006941609.4;refGene_gn=PLEKHG4;refGene_tx=XM_023246094.1;refGene_gn=PLEKHG4;refGene_tx=XM_023246095.1;refGene_gn=PLEKHG4;refGene_tx=XM_023246096.1

E2 45014019 45014136
refGene_gn=PLEKHG4;refGene_tx=XM_006941609.4;refGene_gn=PLEKHG4;refGene_tx=XM_023246094.1;refGene_gn=PLEKHG4;refGene_tx=XM_023246095.1;refGene_gn=PLEKHG4;refGene_tx=XM_023246096.1
E2 45015970 45016153
refGene_gn=PLEKHG4;refGene_tx=XM_006941609.4;refGene_gn=PLEKHG4;refGene_tx=XM_023246094.1;refGene_gn=PLEKHG4;refGene_tx=XM_023246095.1;refGene_gn=PLEKHG4;refGene_tx=XM_023246096.1
E2 45016410 45016925
refGene_gn=PLEKHG4;refGene_tx=XM_006941609.4;refGene_gn=PLEKHG4;refGene_tx=XM_023246094.1;refGene_gn=PLEKHG4;refGene_tx=XM_023246095.1;refGene_gn=PLEKHG4;refGene_tx=XM_023246096.1
E2 45017000 45017247
refGene_gn=PLEKHG4;refGene_tx=XM_006941609.4;refGene_gn=PLEKHG4;refGene_tx=XM_023246094.1;refGene_gn=PLEKHG4;refGene_tx=XM_023246095.1;refGene_gn=PLEKHG4;refGene_tx=XM_023246096.1
E2 45017904 45017994
refGene_gn=PLEKHG4;refGene_tx=XM_006941609.4;refGene_gn=PLEKHG4;refGene_tx=XM_023246094.1;refGene_gn=PLEKHG4;refGene_tx=XM_023246095.1;refGene_gn=PLEKHG4;refGene_tx=XM_023246096.1
E2 45018079 45018301
refGene_gn=PLEKHG4;refGene_tx=XM_006941609.4;refGene_gn=PLEKHG4;refGene_tx=XM_023246094.1;refGene_gn=PLEKHG4;refGene_tx=XM_023246095.1;refGene_gn=PLEKHG4;refGene_tx=XM_023246096.1
E2 45018393 45018564
refGene_gn=PLEKHG4;refGene_tx=XM_006941609.4;refGene_gn=PLEKHG4;refGene_tx=XM_023246094.1;refGene_gn=PLEKHG4;refGene_tx=XM_023246095.1;refGene_gn=PLEKHG4;refGene_tx=XM_023246096.1
E2 45018647 45018825
refGene_gn=PLEKHG4;refGene_tx=XM_006941609.4;refGene_gn=PLEKHG4;refGene_tx=XM_023246094.1;refGene_gn=PLEKHG4;refGene_tx=XM_023246095.1;refGene_gn=PLEKHG4;refGene_tx=XM_023246096.1
E2 45019210 45019327
refGene_gn=PLEKHG4;refGene_tx=XM_006941609.4;refGene_gn=PLEKHG4;refGene_tx=XM_023246094.1;refGene_gn=PLEKHG4;refGene_tx=XM_023246095.1;refGene_gn=PLEKHG4;refGene_tx=XM_023246096.1
E2 45019468 45019705
refGene_gn=PLEKHG4;refGene_tx=XM_006941609.4;refGene_gn=PLEKHG4;refGene_tx=XM_023246094.1;refGene_gn=PLEKHG4;refGene_tx=XM_023246095.1;refGene_gn=PLEKHG4;refGene_tx=XM_023246096.1
E2 45019789 45019905
refGene_gn=PLEKHG4;refGene_tx=XM_006941609.4;refGene_gn=PLEKHG4;refGene_tx=XM_023246094.1;refGene_gn=PLEKHG4;refGene_tx=XM_023246095.1;refGene_gn=PLEKHG4;refGene_tx=XM_023246096.1
E2 45020068 45020074 refGene_gn=PLEKHG4;refGene_tx=XM_023246096.1
E2 45020831 45020945 refGene_gn=KCTD19;refGene_tx=XM_023246097.1
E2 45021867 45021969 refGene_gn=KCTD19;refGene_tx=XM_023246097.1
E2 45022276 45022431 refGene_gn=KCTD19;refGene_tx=XM_023246097.1
E2 45022604 45022796 refGene_gn=KCTD19;refGene_tx=XM_023246097.1
E2 45024204 45024832 refGene_gn=KCTD19;refGene_tx=XM_023246097.1

E2 45025232 45025347 refGene_gn=KCTD19;refGene_tx=XM_023246097.1
E2 45025609 45025693 refGene_gn=KCTD19;refGene_tx=XM_023246097.1
E2 45025881 45026024 refGene_gn=KCTD19;refGene_tx=XM_023246097.1
E2 45026497 45026601 refGene_gn=KCTD19;refGene_tx=XM_023246097.1
E2 45027835 45027996 refGene_gn=KCTD19;refGene_tx=XM_023246097.1
E2 45029154 45029365 refGene_gn=KCTD19;refGene_tx=XM_023246097.1
E2 45031764 45031896 refGene_gn=KCTD19;refGene_tx=XM_023246097.1
E2 45033105 45033256 refGene_gn=KCTD19;refGene_tx=XM_023246097.1
E2 45033257 45033298 refGene_gn=KCTD19;refGene_tx=XM_023246097.1
E2 45034101 45034252 refGene_gn=KCTD19;refGene_tx=XM_023246097.1
E2 45045598 45045616
refGene_gn=PLEKHG4;refGene_tx=XM_006941609.4;refGene_gn=PLEKHG4;refGene_tx=XM_023246094.1;refGene_gn=PLEKHG4;refGene_tx=XM_023246095.1
E2 45046961 45047258 refGene_gn=KCTD19;refGene_tx=XM_023246097.1
E2 45049772 45049784 refGene_gn=KCTD19;refGene_tx=XM_023246097.1
E2 45052263 45052333
refGene_gn=LRRC36;refGene_tx=XM_003998167.3;refGene_gn=LRRC36;refGene_tx=XM_006941611.4;refGene_gn=LRRC36;refGene_tx=XM_006941612.3;refGene_gn=LRRC36;refGene_tx=XM_006941613.4
E2 45062300 45062428
refGene_gn=LRRC36;refGene_tx=XM_003998167.3;refGene_gn=LRRC36;refGene_tx=XM_006941611.4;refGene_gn=LRRC36;refGene_tx=XM_006941612.3;refGene_gn=LRRC36;refGene_tx=XM_006941613.4
E2 45065185 45065378
refGene_gn=LRRC36;refGene_tx=XM_003998167.3;refGene_gn=LRRC36;refGene_tx=XM_006941611.4;refGene_gn=LRRC36;refGene_tx=XM_006941612.3;refGene_gn=LRRC36;refGene_tx=XM_006941613.4
E2 45066174 45066271
refGene_gn=LRRC36;refGene_tx=XM_003998167.3;refGene_gn=LRRC36;refGene_tx=XM_006941611.4;refGene_gn=LRRC36;refGene_tx=XM_006941612.3;refGene_gn=LRRC36;refGene_tx=XM_006941613.4
E2 45069612 45069701
refGene_gn=LRRC36;refGene_tx=XM_003998167.3;refGene_gn=LRRC36;refGene_tx=XM_006941611.4;refGene_gn=LRRC36;refGene_tx=XM_006941612.3;refGene_gn=LRRC36;refGene_tx=XM_006941613.4
E2 45079466 45079594
refGene_gn=LRRC36;refGene_tx=XM_003998167.3;refGene_gn=LRRC36;refGene_tx=XM_006941611.4;refGene_gn=LRRC36;refGene_tx=XM_006941612.3;refGene_gn=LRRC36;refGene_tx=XM_006941613.4
E2 45082025 45082071
refGene_gn=LRRC36;refGene_tx=XM_003998167.3;refGene_gn=LRRC36;refGene_tx=XM_006941611.4;refGene_gn=LRRC36;refGene_tx=XM_006941612.3;refGene_gn=LRRC36;refGene_tx=XM_006941613.4
E2 45086017 45086458
refGene_gn=LRRC36;refGene_tx=XM_003998167.3;refGene_gn=LRRC36;refGene_tx=XM_006941611.4;refGene_gn=LRRC36;refGene_tx=XM_006941612.3;refGene_gn=LRRC36;refGene_tx=XM_006941613.4
E2 45088246 45088536
refGene_gn=LRRC36;refGene_tx=XM_003998167.3;refGene_gn=LRRC36;refGene_tx=XM_006941611.4;refGene_gn=LRRC36;refGene_tx=XM_006941612.3;refGene_

gn=LRRC36;refGene_tx=XM_006941613.4
E2 45093295 45093461
refGene_gn=LRRC36;refGene_tx=XM_003998167.3;refGene_gn=LRRC36;refGene_tx=XM_006941611.4;refGene_gn=LRRC36;refGene_tx=XM_006941612.3;refGene_gn=LRRC36;refGene_tx=XM_006941613.4
E2 45094618 45094764
refGene_gn=LRRC36;refGene_tx=XM_003998167.3;refGene_gn=LRRC36;refGene_tx=XM_006941611.4;refGene_gn=LRRC36;refGene_tx=XM_006941612.3;refGene_gn=LRRC36;refGene_tx=XM_006941613.4
E2 45095540 45095664
refGene_gn=LRRC36;refGene_tx=XM_003998167.3;refGene_gn=LRRC36;refGene_tx=XM_006941611.4;refGene_gn=LRRC36;refGene_tx=XM_006941612.3;refGene_gn=LRRC36;refGene_tx=XM_006941613.4
E2 45099269 45099384
refGene_gn=LRRC36;refGene_tx=XM_003998167.3;refGene_gn=LRRC36;refGene_tx=XM_006941611.4;refGene_gn=LRRC36;refGene_tx=XM_006941612.3;refGene_gn=LRRC36;refGene_tx=XM_006941613.4
E2 45101906 45102132
refGene_gn=LRRC36;refGene_tx=XM_003998167.3;refGene_gn=LRRC36;refGene_tx=XM_006941611.4;refGene_gn=LRRC36;refGene_tx=XM_006941612.3;refGene_gn=LRRC36;refGene_tx=XM_006941613.4
E2 45104204 45104299 refGene_gn=LRRC36;refGene_tx=XM_006941612.3
E2 45105405 45105548 refGene_gn=LRRC36;refGene_tx=XM_006941611.4
E2 45106857 45107046 refGene_gn=TPPP3;refGene_tx=XM_006941610.3
E2 45107165 45107319 refGene_gn=TPPP3;refGene_tx=XM_006941610.3
E2 45107434 45107622 refGene_gn=TPPP3;refGene_tx=XM_006941610.3
E2 45110025 45110037
refGene_gn=LRRC36;refGene_tx=XM_006941611.4;refGene_gn=LRRC36;refGene_tx=XM_006941613.4
E2 45111348 45111632
refGene_gn=ZDHHC1;refGene_tx=XM_011290103.3;refGene_gn=ZDHHC1;refGene_tx=XM_011290105.3;refGene_gn=ZDHHC1;refGene_tx=XM_011290107.3;refGene_gn=ZDHHC1;refGene_tx=XM_023246115.1;refGene_gn=ZDHHC1;refGene_tx=XM_023246116.1;refGene_gn=ZDHHC1;refGene_tx=XM_023246117.1;refGene_gn=ZDHHC1;refGene_tx=XM_023246118.1
E2 45111735 45111814
refGene_gn=ZDHHC1;refGene_tx=XM_011290103.3;refGene_gn=ZDHHC1;refGene_tx=XM_011290105.3;refGene_gn=ZDHHC1;refGene_tx=XM_011290107.3;refGene_gn=ZDHHC1;refGene_tx=XM_023246115.1;refGene_gn=ZDHHC1;refGene_tx=XM_023246116.1;refGene_gn=ZDHHC1;refGene_tx=XM_023246117.1;refGene_gn=ZDHHC1;refGene_tx=XM_023246118.1
E2 45111944 45112032
refGene_gn=ZDHHC1;refGene_tx=XM_011290103.3;refGene_gn=ZDHHC1;refGene_tx=XM_011290105.3;refGene_gn=ZDHHC1;refGene_tx=XM_011290107.3;refGene_gn=ZDHHC1;refGene_tx=XM_023246116.1
E2 45112275 45112358
refGene_gn=ZDHHC1;refGene_tx=XM_011290103.3;refGene_gn=ZDHHC1;refGene_tx=XM_011290105.3;refGene_gn=ZDHHC1;refGene_tx=XM_011290107.3;refGene_gn=ZDHHC1;refGene_tx=XM_023246115.1;refGene_gn=ZDHHC1;refGene_tx=XM_023246116.1;refGene_gn=ZDHHC1;refGene_tx=XM_023246117.1;refGene_gn=ZDHHC

1;refGene_tx=XM_023246118.1
E2 45116008 45116121
refGene_gn=ZDHHC1;refGene_tx=XM_011290103.3;refGene_gn=ZDHHC1;refGene_tx=XM_011290105.3;refGene_gn=ZDHHC1;refGene_tx=XM_011290107.3;refGene_gn=ZDHHC1;refGene_tx=XM_023246115.1;refGene_gn=ZDHHC1;refGene_tx=XM_023246116.1;refGene_gn=ZDHHC1;refGene_tx=XM_023246118.1
E2 45116390 45116567
refGene_gn=ZDHHC1;refGene_tx=XM_011290103.3;refGene_gn=ZDHHC1;refGene_tx=XM_011290105.3;refGene_gn=ZDHHC1;refGene_tx=XM_011290107.3;refGene_gn=ZDHHC1;refGene_tx=XM_023246115.1;refGene_gn=ZDHHC1;refGene_tx=XM_023246117.1;refGene_gn=ZDHHC1;refGene_tx=XM_023246118.1
E2 45116646 45116771
refGene_gn=ZDHHC1;refGene_tx=XM_011290103.3;refGene_gn=ZDHHC1;refGene_tx=XM_011290105.3;refGene_gn=ZDHHC1;refGene_tx=XM_011290107.3;refGene_gn=ZDHHC1;refGene_tx=XM_023246115.1;refGene_gn=ZDHHC1;refGene_tx=XM_023246116.1;refGene_gn=ZDHHC1;refGene_tx=XM_023246117.1;refGene_gn=ZDHHC1;refGene_tx=XM_023246118.1
E2 45117185 45117287
refGene_gn=ZDHHC1;refGene_tx=XM_011290103.3;refGene_gn=ZDHHC1;refGene_tx=XM_011290105.3;refGene_gn=ZDHHC1;refGene_tx=XM_011290107.3;refGene_gn=ZDHHC1;refGene_tx=XM_023246115.1;refGene_gn=ZDHHC1;refGene_tx=XM_023246116.1;refGene_gn=ZDHHC1;refGene_tx=XM_023246117.1;refGene_gn=ZDHHC1;refGene_tx=XM_023246118.1
E2 45118878 45119058
refGene_gn=ZDHHC1;refGene_tx=XM_011290103.3;refGene_gn=ZDHHC1;refGene_tx=XM_011290105.3;refGene_gn=ZDHHC1;refGene_tx=XM_011290107.3;refGene_gn=ZDHHC1;refGene_tx=XM_023246115.1;refGene_gn=ZDHHC1;refGene_tx=XM_023246116.1;refGene_gn=ZDHHC1;refGene_tx=XM_023246117.1;refGene_gn=ZDHHC1;refGene_tx=XM_023246118.1
E2 45123797 45124040
refGene_gn=ZDHHC1;refGene_tx=XM_011290103.3;refGene_gn=ZDHHC1;refGene_tx=XM_011290105.3;refGene_gn=ZDHHC1;refGene_tx=XM_011290107.3;refGene_gn=ZDHHC1;refGene_tx=XM_023246115.1;refGene_gn=ZDHHC1;refGene_tx=XM_023246116.1;refGene_gn=ZDHHC1;refGene_tx=XM_023246117.1;refGene_gn=ZDHHC1;refGene_tx=XM_023246118.1
E2 45125431 45125440
refGene_gn=ZDHHC1;refGene_tx=XM_011290103.3;refGene_gn=ZDHHC1;refGene_tx=XM_011290105.3;refGene_gn=ZDHHC1;refGene_tx=XM_023246115.1;refGene_gn=ZDHHC1;refGene_tx=XM_023246116.1;refGene_gn=ZDHHC1;refGene_tx=XM_023246117.1;refGene_gn=ZDHHC1;refGene_tx=XM_023246118.1
E2 45144269 45144534 refGene_gn=HSD11B2;refGene_tx=XM_023246056.1
E2 45147933 45148146 refGene_gn=HSD11B2;refGene_tx=XM_023246056.1
E2 45148269 45148455 refGene_gn=HSD11B2;refGene_tx=XM_023246056.1
E2 45148562 45148700 refGene_gn=HSD11B2;refGene_tx=XM_023246056.1
E2 45148877 45149293 refGene_gn=HSD11B2;refGene_tx=XM_023246056.1
E2 45150720 45150882 refGene_gn=ATP6V0D1;refGene_tx=XM_003998130.5
E2 45150983 45151061 refGene_gn=ATP6V0D1;refGene_tx=XM_003998130.5
E2 45151170 45151347 refGene_gn=ATP6V0D1;refGene_tx=XM_003998130.5
E2 45151473 45151551 refGene_gn=ATP6V0D1;refGene_tx=XM_003998130.5
E2 45154922 45155002 refGene_gn=ATP6V0D1;refGene_tx=XM_003998130.5

E2 45156512 45156691 refGene_gn=ATP6V0D1;refGene_tx=XM_003998130.5
E2 45165106 45165278 refGene_gn=ATP6V0D1;refGene_tx=XM_003998130.5
E2 45188879 45189009 refGene_gn=ATP6V0D1;refGene_tx=XM_003998130.5
E2 45189876 45190059
refGene_gn=AGRP;refGene_tx=XM_003998131.2;refGene_gn=AGRP;refGene_tx=XM_019819905.1
E2 45190279 45190365
refGene_gn=AGRP;refGene_tx=XM_003998131.2;refGene_gn=AGRP;refGene_tx=XM_019819905.1
E2 45190550 45190680
refGene_gn=AGRP;refGene_tx=XM_003998131.2;refGene_gn=AGRP;refGene_tx=XM_019819905.1
E2 45228595 45228614
refGene_gn=RIPOR1;refGene_tx=XM_006941616.3;refGene_gn=RIPOR1;refGene_tx=XM_019819901.2;refGene_gn=RIPOR1;refGene_tx=XM_019819902.2;refGene_gn=RIPOR1;refGene_tx=XM_023245340.1
E2 45229431 45229558
refGene_gn=RIPOR1;refGene_tx=XM_006941616.3;refGene_gn=RIPOR1;refGene_tx=XM_019819901.2;refGene_gn=RIPOR1;refGene_tx=XM_019819902.2;refGene_gn=RIPOR1;refGene_tx=XM_023245339.1;refGene_gn=RIPOR1;refGene_tx=XM_023245340.1
E2 45229674 45229827
refGene_gn=RIPOR1;refGene_tx=XM_006941616.3;refGene_gn=RIPOR1;refGene_tx=XM_019819901.2;refGene_gn=RIPOR1;refGene_tx=XM_019819902.2;refGene_gn=RIPOR1;refGene_tx=XM_023245339.1;refGene_gn=RIPOR1;refGene_tx=XM_023245340.1
E2 45229999 45230078
refGene_gn=RIPOR1;refGene_tx=XM_006941616.3;refGene_gn=RIPOR1;refGene_tx=XM_019819901.2;refGene_gn=RIPOR1;refGene_tx=XM_019819902.2;refGene_gn=RIPOR1;refGene_tx=XM_023245339.1;refGene_gn=RIPOR1;refGene_tx=XM_023245340.1
E2 45230685 45230709
refGene_gn=RIPOR1;refGene_tx=XM_006941616.3;refGene_gn=RIPOR1;refGene_tx=XM_019819901.2;refGene_gn=RIPOR1;refGene_tx=XM_019819902.2;refGene_gn=RIPOR1;refGene_tx=XM_023245339.1;refGene_gn=RIPOR1;refGene_tx=XM_023245340.1
E2 45230804 45230858
refGene_gn=RIPOR1;refGene_tx=XM_006941616.3;refGene_gn=RIPOR1;refGene_tx=XM_019819901.2;refGene_gn=RIPOR1;refGene_tx=XM_019819902.2;refGene_gn=RIPOR1;refGene_tx=XM_023245339.1;refGene_gn=RIPOR1;refGene_tx=XM_023245340.1
E2 45230991 45231144
refGene_gn=RIPOR1;refGene_tx=XM_006941616.3;refGene_gn=RIPOR1;refGene_tx=XM_019819901.2;refGene_gn=RIPOR1;refGene_tx=XM_019819902.2;refGene_gn=RIPOR1;refGene_tx=XM_023245339.1;refGene_gn=RIPOR1;refGene_tx=XM_023245340.1
E2 45231245 45231309
refGene_gn=RIPOR1;refGene_tx=XM_006941616.3;refGene_gn=RIPOR1;refGene_tx=XM_019819901.2;refGene_gn=RIPOR1;refGene_tx=XM_019819902.2;refGene_gn=RIPOR1;refGene_tx=XM_023245339.1;refGene_gn=RIPOR1;refGene_tx=XM_02

3245340.1
E2 45231400 45231444
refGene_gn=RIPOR1;refGene_tx=XM_006941616.3;refGene_gn=RIPOR1;refGene_tx=XM_019819901.2;refGene_gn=RIPOR1;refGene_tx=XM_019819902.2;refGene_gn=RIPOR1;refGene_tx=XM_023245339.1;refGene_gn=RIPOR1;refGene_tx=XM_023245340.1
E2 45231521 45231647
refGene_gn=RIPOR1;refGene_tx=XM_006941616.3;refGene_gn=RIPOR1;refGene_tx=XM_019819901.2;refGene_gn=RIPOR1;refGene_tx=XM_019819902.2;refGene_gn=RIPOR1;refGene_tx=XM_023245339.1;refGene_gn=RIPOR1;refGene_tx=XM_023245340.1
E2 45232083 45232232
refGene_gn=RIPOR1;refGene_tx=XM_006941616.3;refGene_gn=RIPOR1;refGene_tx=XM_019819901.2;refGene_gn=RIPOR1;refGene_tx=XM_019819902.2;refGene_gn=RIPOR1;refGene_tx=XM_023245339.1;refGene_gn=RIPOR1;refGene_tx=XM_023245340.1
E2 45232307 45232437
refGene_gn=RIPOR1;refGene_tx=XM_006941616.3;refGene_gn=RIPOR1;refGene_tx=XM_019819901.2;refGene_gn=RIPOR1;refGene_tx=XM_019819902.2;refGene_gn=RIPOR1;refGene_tx=XM_023245339.1;refGene_gn=RIPOR1;refGene_tx=XM_023245340.1
E2 45232520 45234041
refGene_gn=RIPOR1;refGene_tx=XM_006941616.3;refGene_gn=RIPOR1;refGene_tx=XM_019819901.2;refGene_gn=RIPOR1;refGene_tx=XM_019819902.2;refGene_gn=RIPOR1;refGene_tx=XM_023245339.1;refGene_gn=RIPOR1;refGene_tx=XM_023245340.1
E2 45234120 45234242
refGene_gn=RIPOR1;refGene_tx=XM_006941616.3;refGene_gn=RIPOR1;refGene_tx=XM_019819901.2;refGene_gn=RIPOR1;refGene_tx=XM_019819902.2;refGene_gn=RIPOR1;refGene_tx=XM_023245339.1;refGene_gn=RIPOR1;refGene_tx=XM_023245340.1
E2 45235127 45235260
refGene_gn=RIPOR1;refGene_tx=XM_006941616.3;refGene_gn=RIPOR1;refGene_tx=XM_019819901.2;refGene_gn=RIPOR1;refGene_tx=XM_019819902.2;refGene_gn=RIPOR1;refGene_tx=XM_023245339.1;refGene_gn=RIPOR1;refGene_tx=XM_023245340.1
E2 45235514 45235650
refGene_gn=RIPOR1;refGene_tx=XM_006941616.3;refGene_gn=RIPOR1;refGene_tx=XM_019819901.2;refGene_gn=RIPOR1;refGene_tx=XM_019819902.2;refGene_gn=RIPOR1;refGene_tx=XM_023245339.1;refGene_gn=RIPOR1;refGene_tx=XM_023245340.1
E2 45235758 45235920
refGene_gn=RIPOR1;refGene_tx=XM_006941616.3;refGene_gn=RIPOR1;refGene_tx=XM_019819901.2;refGene_gn=RIPOR1;refGene_tx=XM_019819902.2;refGene_gn=RIPOR1;refGene_tx=XM_023245339.1;refGene_gn=RIPOR1;refGene_tx=XM_023245340.1
E2 45236186 45236345
refGene_gn=RIPOR1;refGene_tx=XM_006941616.3;refGene_gn=RIPOR1;refGene_tx=XM_019819901.2;refGene_gn=RIPOR1;refGene_tx=XM_019819902.2;refGene_gn=RIPOR1;refGene_tx=XM_023245339.1

E2 45236475 45236672
refGene_gn=RIPOR1;refGene_tx=XM_006941616.3;refGene_gn=RIPOR1;refGene_tx=XM_019819901.2;refGene_gn=RIPOR1;refGene_tx=XM_019819902.2;refGene_gn=RIPOR1;refGene_tx=XM_023245339.1;refGene_gn=RIPOR1;refGene_tx=XM_023245340.1
E2 45236762 45236846
refGene_gn=RIPOR1;refGene_tx=XM_006941616.3;refGene_gn=RIPOR1;refGene_tx=XM_019819901.2;refGene_gn=RIPOR1;refGene_tx=XM_019819902.2;refGene_gn=RIPOR1;refGene_tx=XM_023245339.1;refGene_gn=RIPOR1;refGene_tx=XM_023245340.1
E2 45236959 45237050 refGene_gn=RIPOR1;refGene_tx=XM_006941616.3
E2 45237191 45237407
refGene_gn=RIPOR1;refGene_tx=XM_006941616.3;refGene_gn=RIPOR1;refGene_tx=XM_019819901.2;refGene_gn=RIPOR1;refGene_tx=XM_019819902.2;refGene_gn=RIPOR1;refGene_tx=XM_023245339.1;refGene_gn=RIPOR1;refGene_tx=XM_023245340.1
E2 45277922 45278703
refGene_gn=CTCF;refGene_tx=XM_023245341.1;refGene_gn=CTCF;refGene_tx=XM_023245342.1;refGene_gn=CTCF;refGene_tx=XM_023245343.1
E2 45279062 45279233
refGene_gn=CTCF;refGene_tx=XM_023245341.1;refGene_gn=CTCF;refGene_tx=XM_023245342.1;refGene_gn=CTCF;refGene_tx=XM_023245343.1
E2 45282966 45283100
refGene_gn=CTCF;refGene_tx=XM_023245341.1;refGene_gn=CTCF;refGene_tx=XM_023245342.1;refGene_gn=CTCF;refGene_tx=XM_023245343.1
E2 45284713 45284834
refGene_gn=CTCF;refGene_tx=XM_023245341.1;refGene_gn=CTCF;refGene_tx=XM_023245342.1;refGene_gn=CTCF;refGene_tx=XM_023245343.1
E2 45285523 45285673
refGene_gn=CTCF;refGene_tx=XM_023245341.1;refGene_gn=CTCF;refGene_tx=XM_023245342.1;refGene_gn=CTCF;refGene_tx=XM_023245343.1
E2 45290556 45290717
refGene_gn=CTCF;refGene_tx=XM_023245341.1;refGene_gn=CTCF;refGene_tx=XM_023245342.1;refGene_gn=CTCF;refGene_tx=XM_023245343.1
E2 45291246 45291429
refGene_gn=CTCF;refGene_tx=XM_023245341.1;refGene_gn=CTCF;refGene_tx=XM_023245342.1;refGene_gn=CTCF;refGene_tx=XM_023245343.1
E2 45291980 45292116
refGene_gn=CTCF;refGene_tx=XM_023245341.1;refGene_gn=CTCF;refGene_tx=XM_023245342.1;refGene_gn=CTCF;refGene_tx=XM_023245343.1
E2 45296235 45296397
refGene_gn=CTCF;refGene_tx=XM_023245341.1;refGene_gn=CTCF;refGene_tx=XM_023245342.1;refGene_gn=CTCF;refGene_tx=XM_023245343.1
E2 45297576 45297761
refGene_gn=CTCF;refGene_tx=XM_023245341.1;refGene_gn=CTCF;refGene_tx=XM_023245342.1;refGene_gn=CTCF;refGene_tx=XM_023245343.1
E2 45303488 45303528 refGene_gn=CARMIL2;refGene_tx=XM_019819916.2
E2 45303764 45303859 refGene_gn=CARMIL2;refGene_tx=XM_019819916.2
E2 45303953 45304004 refGene_gn=CARMIL2;refGene_tx=XM_019819916.2
E2 45304230 45304293 refGene_gn=CARMIL2;refGene_tx=XM_019819916.2

E2 45304392 45304517 refGene_gn=CARMIL2;refGene_tx=XM_019819916.2
E2 45304651 45304743 refGene_gn=CARMIL2;refGene_tx=XM_019819916.2
E2 45304942 45305013 refGene_gn=CARMIL2;refGene_tx=XM_019819916.2
E2 45305128 45305202 refGene_gn=CARMIL2;refGene_tx=XM_019819916.2
E2 45305335 45305411 refGene_gn=CARMIL2;refGene_tx=XM_019819916.2
E2 45305490 45305579 refGene_gn=CARMIL2;refGene_tx=XM_019819916.2
E2 45305707 45305802 refGene_gn=CARMIL2;refGene_tx=XM_019819916.2
E2 45305879 45305966 refGene_gn=CARMIL2;refGene_tx=XM_019819916.2
E2 45306037 45306150 refGene_gn=CARMIL2;refGene_tx=XM_019819916.2
E2 45306265 45306343 refGene_gn=CARMIL2;refGene_tx=XM_019819916.2
E2 45306473 45306550 refGene_gn=CARMIL2;refGene_tx=XM_019819916.2
E2 45306643 45306748 refGene_gn=CARMIL2;refGene_tx=XM_019819916.2
E2 45306937 45307007 refGene_gn=CARMIL2;refGene_tx=XM_019819916.2
E2 45307148 45307230 refGene_gn=CARMIL2;refGene_tx=XM_019819916.2
E2 45307322 45307419 refGene_gn=CARMIL2;refGene_tx=XM_019819916.2
E2 45307518 45307576 refGene_gn=CARMIL2;refGene_tx=XM_019819916.2
E2 45307684 45307857 refGene_gn=CARMIL2;refGene_tx=XM_019819916.2
E2 45308026 45308189 refGene_gn=CARMIL2;refGene_tx=XM_019819916.2
E2 45308269 45308371 refGene_gn=CARMIL2;refGene_tx=XM_019819916.2
E2 45309618 45309747 refGene_gn=CARMIL2;refGene_tx=XM_019819916.2
E2 45309837 45309951 refGene_gn=CARMIL2;refGene_tx=XM_019819916.2
E2 45310101 45310262 refGene_gn=CARMIL2;refGene_tx=XM_019819916.2
E2 45310347 45310435 refGene_gn=CARMIL2;refGene_tx=XM_019819916.2
E2 45310634 45310790 refGene_gn=CARMIL2;refGene_tx=XM_019819916.2
E2 45310869 45310936 refGene_gn=CARMIL2;refGene_tx=XM_019819916.2
E2 45311408 45311638 refGene_gn=CARMIL2;refGene_tx=XM_019819916.2
E2 45312502 45312603 refGene_gn=CARMIL2;refGene_tx=XM_019819916.2
E2 45312685 45313046 refGene_gn=CARMIL2;refGene_tx=XM_019819916.2
E2 45313127 45313250 refGene_gn=CARMIL2;refGene_tx=XM_019819916.2
E2 45313786 45313823 refGene_gn=CARMIL2;refGene_tx=XM_019819916.2
E2 45313983 45314055 refGene_gn=CARMIL2;refGene_tx=XM_019819916.2
E2 45314144 45314321 refGene_gn=CARMIL2;refGene_tx=XM_019819916.2
E2 45314323 45314365 refGene_gn=CARMIL2;refGene_tx=XM_019819916.2
E2 45314922 45315000 refGene_gn=CARMIL2;refGene_tx=XM_019819916.2
E2 45315088 45315201 refGene_gn=CARMIL2;refGene_tx=XM_019819916.2
E2 45315328 45315407
refGene_gn=ACD;refGene_tx=XM_003998170.5;refGene_gn=ACD;refGene_tx=XM_006941623.4;refGene_gn=ACD;refGene_tx=XM_023245006.1
E2 45315487 45315621
refGene_gn=ACD;refGene_tx=XM_003998170.5;refGene_gn=ACD;refGene_tx=XM_006941623.4;refGene_gn=ACD;refGene_tx=XM_019819917.2;refGene_gn=ACD;refGene_tx=XM_023245006.1
E2 45315715 45316089
refGene_gn=ACD;refGene_tx=XM_003998170.5;refGene_gn=ACD;refGene_tx=XM_006941623.4;refGene_gn=ACD;refGene_tx=XM_019819917.2;refGene_gn=ACD;refGene_tx=XM_023245006.1
E2 45316222 45316303
refGene_gn=ACD;refGene_tx=XM_003998170.5;refGene_gn=ACD;refGene_tx=XM_006941623.4;refGene_gn=ACD;refGene_tx=XM_019819917.2;refGene_gn=ACD;refGene_tx=XM_023245006.1

E2 45316385 45316482
refGene_gn=ACD;refGene_tx=XM_003998170.5;refGene_gn=ACD;refGene_tx=XM_006941623.4;refGene_gn=ACD;refGene_tx=XM_019819917.2;refGene_gn=ACD;refGene_tx=XM_023245006.1
E2 45316556 45316708
refGene_gn=ACD;refGene_tx=XM_003998170.5;refGene_gn=ACD;refGene_tx=XM_006941623.4;refGene_gn=ACD;refGene_tx=XM_019819917.2;refGene_gn=ACD;refGene_tx=XM_023245006.1
E2 45317126 45317161
refGene_gn=ACD;refGene_tx=XM_003998170.5;refGene_gn=ACD;refGene_tx=XM_006941623.4;refGene_gn=ACD;refGene_tx=XM_019819917.2;refGene_gn=ACD;refGene_tx=XM_023245006.1
E2 45317268 45317313
refGene_gn=ACD;refGene_tx=XM_003998170.5;refGene_gn=ACD;refGene_tx=XM_006941623.4;refGene_gn=ACD;refGene_tx=XM_019819917.2;refGene_gn=ACD;refGene_tx=XM_023245006.1
E2 45317446 45317523
refGene_gn=ACD;refGene_tx=XM_003998170.5;refGene_gn=ACD;refGene_tx=XM_006941623.4;refGene_gn=ACD;refGene_tx=XM_019819917.2;refGene_gn=ACD;refGene_tx=XM_023245006.1
E2 45317606 45317700
refGene_gn=ACD;refGene_tx=XM_003998170.5;refGene_gn=ACD;refGene_tx=XM_006941623.4;refGene_gn=ACD;refGene_tx=XM_019819917.2;refGene_gn=ACD;refGene_tx=XM_023245006.1
E2 45317838 45317975
refGene_gn=ACD;refGene_tx=XM_003998170.5;refGene_gn=ACD;refGene_tx=XM_006941623.4;refGene_gn=ACD;refGene_tx=XM_019819917.2;refGene_gn=ACD;refGene_tx=XM_023245006.1
E2 45318051 45318163
refGene_gn=ACD;refGene_tx=XM_003998170.5;refGene_gn=ACD;refGene_tx=XM_006941623.4;refGene_gn=ACD;refGene_tx=XM_019819917.2;refGene_gn=ACD;refGene_tx=XM_023245006.1
E2 45318369 45318407
refGene_gn=ACD;refGene_tx=XM_003998170.5;refGene_gn=ACD;refGene_tx=XM_006941623.4;refGene_gn=ACD;refGene_tx=XM_019819917.2
E2 45318936 45318999
refGene_gn=PARD6A;refGene_tx=XM_003998134.5;refGene_gn=PARD6A;refGene_tx=XM_006941620.4;refGene_gn=PARD6A;refGene_tx=XM_019819919.2
E2 45319374 45319597
refGene_gn=PARD6A;refGene_tx=XM_003998134.5;refGene_gn=PARD6A;refGene_tx=XM_006941620.4;refGene_gn=PARD6A;refGene_tx=XM_019819919.2
E2 45319840 45320592
refGene_gn=PARD6A;refGene_tx=XM_003998134.5;refGene_gn=PARD6A;refGene_tx=XM_006941620.4;refGene_gn=PARD6A;refGene_tx=XM_019819919.2
E2 45321116 45321277 refGene_gn=ENKD1;refGene_tx=XM_003998171.5
E2 45321374 45321511 refGene_gn=ENKD1;refGene_tx=XM_003998171.5
E2 45321611 45321775 refGene_gn=ENKD1;refGene_tx=XM_003998171.5
E2 45321877 45321925 refGene_gn=ENKD1;refGene_tx=XM_003998171.5
E2 45321926 45322004 refGene_gn=ENKD1;refGene_tx=XM_003998171.5
E2 45323009 45323182 refGene_gn=ENKD1;refGene_tx=XM_003998171.5

E2 45323737 45323932 refGene_gn=ENKD1;refGene_tx=XM_003998171.5
E2 45324028 45324113 refGene_gn=ENKD1;refGene_tx=XM_003998171.5
E2 45324945 45325169
refGene_gn=CE2H16orf86;refGene_tx=XM_019819920.2
E2 45325532 45325750
refGene_gn=CE2H16orf86;refGene_tx=XM_019819920.2
E2 45325826 45326030
refGene_gn=CE2H16orf86;refGene_tx=XM_019819920.2
E2 45326254 45326382
refGene_gn=CE2H16orf86;refGene_tx=XM_019819920.2
E2 45332296 45333195
refGene_gn=GFOD2;refGene_tx=XM_003998135.4;refGene_gn=GFOD2;refGene_tx
=XM_019819921.2
E2 45338271 45338530 refGene_gn=GFOD2;refGene_tx=XM_003998135.4
E2 45365838 45366046 refGene_gn=GFOD2;refGene_tx=XM_019819921.2
E2 45371545 45371676 refGene_gn=RANBP10;refGene_tx=NM_001134988.1
E2 45372128 45372240 refGene_gn=RANBP10;refGene_tx=NM_001134988.1
E2 45372540 45372686 refGene_gn=RANBP10;refGene_tx=NM_001134988.1
E2 45373180 45373302 refGene_gn=RANBP10;refGene_tx=NM_001134988.1
E2 45374043 45374248 refGene_gn=RANBP10;refGene_tx=NM_001134988.1
E2 45374441 45374590 refGene_gn=RANBP10;refGene_tx=NM_001134988.1
E2 45374700 45374809 refGene_gn=RANBP10;refGene_tx=NM_001134988.1
E2 45376245 45376358 refGene_gn=RANBP10;refGene_tx=NM_001134988.1
E2 45378779 45378964 refGene_gn=RANBP10;refGene_tx=NM_001134988.1
E2 45380931 45380954 refGene_gn=RANBP10;refGene_tx=NM_001134988.1
E2 45388848 45389016 refGene_gn=RANBP10;refGene_tx=NM_001134988.1
E2 45409144 45409197 refGene_gn=RANBP10;refGene_tx=NM_001134988.1
E2 45431217 45431329 refGene_gn=RANBP10;refGene_tx=NM_001134988.1
E2 45432126 45432361 refGene_gn=RANBP10;refGene_tx=NM_001134988.1
E2 45432943 45433018
refGene_gn=TSNAXIP1;refGene_tx=XM_023245307.1;refGene_gn=TSNAXIP1;refG
ene_tx=XM_023245308.1;refGene_gn=TSNAXIP1;refGene_tx=XM_023245309.1;re
fGene_gn=TSNAXIP1;refGene_tx=XM_023245310.1;refGene_gn=TSNAXIP1;refGen
e_tx=XM_023245311.1
E2 45435777 45435877
refGene_gn=TSNAXIP1;refGene_tx=XM_023245307.1;refGene_gn=TSNAXIP1;refG
ene_tx=XM_023245309.1;refGene_gn=TSNAXIP1;refGene_tx=XM_023245310.1
E2 45439883 45439996
refGene_gn=TSNAXIP1;refGene_tx=XM_023245307.1;refGene_gn=TSNAXIP1;refG
ene_tx=XM_023245308.1;refGene_gn=TSNAXIP1;refGene_tx=XM_023245309.1;re
fGene_gn=TSNAXIP1;refGene_tx=XM_023245310.1
E2 45440143 45440270
refGene_gn=TSNAXIP1;refGene_tx=XM_023245307.1;refGene_gn=TSNAXIP1;refG
ene_tx=XM_023245308.1;refGene_gn=TSNAXIP1;refGene_tx=XM_023245309.1;re
fGene_gn=TSNAXIP1;refGene_tx=XM_023245310.1
E2 45441408 45441502
refGene_gn=TSNAXIP1;refGene_tx=XM_023245307.1;refGene_gn=TSNAXIP1;refG
ene_tx=XM_023245308.1;refGene_gn=TSNAXIP1;refGene_tx=XM_023245309.1;re
fGene_gn=TSNAXIP1;refGene_tx=XM_023245310.1
E2 45442043 45442240

refGene_gn=TSNAXIP1;refGene_tx=XM_023245307.1;refGene_gn=TSNAXIP1;refGene_tx=XM_023245308.1;refGene_gn=TSNAXIP1;refGene_tx=XM_023245309.1;refGene_gn=TSNAXIP1;refGene_tx=XM_023245310.1
E2 45442649 45442785
refGene_gn=TSNAXIP1;refGene_tx=XM_023245307.1;refGene_gn=TSNAXIP1;refGene_tx=XM_023245308.1;refGene_gn=TSNAXIP1;refGene_tx=XM_023245309.1;refGene_gn=TSNAXIP1;refGene_tx=XM_023245310.1;refGene_gn=TSNAXIP1;refGene_tx=XM_023245311.1
E2 45443240 45443410
refGene_gn=TSNAXIP1;refGene_tx=XM_023245307.1;refGene_gn=TSNAXIP1;refGene_tx=XM_023245308.1;refGene_gn=TSNAXIP1;refGene_tx=XM_023245309.1;refGene_gn=TSNAXIP1;refGene_tx=XM_023245310.1;refGene_gn=TSNAXIP1;refGene_tx=XM_023245311.1
E2 45443490 45443650
refGene_gn=TSNAXIP1;refGene_tx=XM_023245307.1;refGene_gn=TSNAXIP1;refGene_tx=XM_023245308.1;refGene_gn=TSNAXIP1;refGene_tx=XM_023245309.1;refGene_gn=TSNAXIP1;refGene_tx=XM_023245310.1;refGene_gn=TSNAXIP1;refGene_tx=XM_023245311.1
E2 45443726 45443857
refGene_gn=TSNAXIP1;refGene_tx=XM_023245307.1;refGene_gn=TSNAXIP1;refGene_tx=XM_023245308.1;refGene_gn=TSNAXIP1;refGene_tx=XM_023245309.1;refGene_gn=TSNAXIP1;refGene_tx=XM_023245310.1;refGene_gn=TSNAXIP1;refGene_tx=XM_023245311.1
E2 45443991 45444117
refGene_gn=TSNAXIP1;refGene_tx=XM_023245307.1;refGene_gn=TSNAXIP1;refGene_tx=XM_023245308.1;refGene_gn=TSNAXIP1;refGene_tx=XM_023245309.1;refGene_gn=TSNAXIP1;refGene_tx=XM_023245310.1;refGene_gn=TSNAXIP1;refGene_tx=XM_023245311.1
E2 45444248 45444401
refGene_gn=TSNAXIP1;refGene_tx=XM_023245307.1;refGene_gn=TSNAXIP1;refGene_tx=XM_023245308.1;refGene_gn=TSNAXIP1;refGene_tx=XM_023245310.1;refGene_gn=TSNAXIP1;refGene_tx=XM_023245311.1
E2 45444504 45444614
refGene_gn=TSNAXIP1;refGene_tx=XM_023245307.1;refGene_gn=TSNAXIP1;refGene_tx=XM_023245308.1;refGene_gn=TSNAXIP1;refGene_tx=XM_023245309.1;refGene_gn=TSNAXIP1;refGene_tx=XM_023245310.1;refGene_gn=TSNAXIP1;refGene_tx=XM_023245311.1
E2 45444826 45444953
refGene_gn=TSNAXIP1;refGene_tx=XM_023245307.1;refGene_gn=TSNAXIP1;refGene_tx=XM_023245308.1;refGene_gn=TSNAXIP1;refGene_tx=XM_023245309.1;refGene_gn=TSNAXIP1;refGene_tx=XM_023245310.1;refGene_gn=TSNAXIP1;refGene_tx=XM_023245311.1
E2 45445069 45445176
refGene_gn=TSNAXIP1;refGene_tx=XM_023245307.1;refGene_gn=TSNAXIP1;refGene_tx=XM_023245308.1;refGene_gn=TSNAXIP1;refGene_tx=XM_023245309.1;refGene_gn=TSNAXIP1;refGene_tx=XM_023245311.1
E2 45445381 45445622
refGene_gn=TSNAXIP1;refGene_tx=XM_023245307.1;refGene_gn=TSNAXIP1;refGene_tx=XM_023245308.1;refGene_gn=TSNAXIP1;refGene_tx=XM_023245309.1;refGene_gn=TSNAXIP1;refGene_tx=XM_023245310.1;refGene_gn=TSNAXIP1;refGen

e_tx=XM_023245311.1
E2 45445906 45446030 refGene_gn=CENPT;refGene_tx=XM_006941628.3
E2 45446104 45446209 refGene_gn=CENPT;refGene_tx=XM_006941628.3
E2 45446298 45446475 refGene_gn=CENPT;refGene_tx=XM_006941628.3
E2 45447266 45447357 refGene_gn=CENPT;refGene_tx=XM_006941628.3
E2 45447586 45447892 refGene_gn=CENPT;refGene_tx=XM_006941628.3
E2 45448123 45448324 refGene_gn=CENPT;refGene_tx=XM_006941628.3
E2 45448894 45449043 refGene_gn=CENPT;refGene_tx=XM_006941628.3
E2 45449255 45449292 refGene_gn=CENPT;refGene_tx=XM_006941628.3
E2 45449414 45449551 refGene_gn=CENPT;refGene_tx=XM_006941628.3
E2 45449671 45449768 refGene_gn=CENPT;refGene_tx=XM_006941628.3
E2 45449876 45449964 refGene_gn=CENPT;refGene_tx=XM_006941628.3
E2 45450104 45450195 refGene_gn=CENPT;refGene_tx=XM_006941628.3
E2 45451729 45451854 refGene_gn=CENPT;refGene_tx=XM_006941628.3
E2 45455511 45456414 refGene_gn=THAP11;refGene_tx=XM_003998138.4
E2 45471187 45471286
refGene_gn=NUTF2;refGene_tx=XM_006941629.4;refGene_gn=NUTF2;refGene_tx
=XM_023245312.1;refGene_gn=NUTF2;refGene_tx=XM_023245313.1
E2 45473358 45473430
refGene_gn=NUTF2;refGene_tx=XM_006941629.4;refGene_gn=NUTF2;refGene_tx
=XM_023245312.1;refGene_gn=NUTF2;refGene_tx=XM_023245313.1
E2 45473514 45473613
refGene_gn=NUTF2;refGene_tx=XM_006941629.4;refGene_gn=NUTF2;refGene_tx
=XM_023245312.1;refGene_gn=NUTF2;refGene_tx=XM_023245313.1
E2 45475455 45475569
refGene_gn=NUTF2;refGene_tx=XM_006941629.4;refGene_gn=NUTF2;refGene_tx
=XM_023245312.1;refGene_gn=NUTF2;refGene_tx=XM_023245313.1
E2 45477897 45477951 refGene_gn=EDC4;refGene_tx=XM_011290101.1
E2 45477952 45477983 refGene_gn=EDC4;refGene_tx=XM_011290101.1
E2 45480560 45480717 refGene_gn=EDC4;refGene_tx=XM_011290101.1
E2 45481050 45481162 refGene_gn=EDC4;refGene_tx=XM_011290101.1
E2 45481459 45481559 refGene_gn=EDC4;refGene_tx=XM_011290101.1
E2 45481820 45482010 refGene_gn=EDC4;refGene_tx=XM_011290101.1
E2 45482117 45482265 refGene_gn=EDC4;refGene_tx=XM_011290101.1
E2 45482351 45482456 refGene_gn=EDC4;refGene_tx=XM_011290101.1
E2 45482783 45482893 refGene_gn=EDC4;refGene_tx=XM_011290101.1
E2 45482984 45483068 refGene_gn=EDC4;refGene_tx=XM_011290101.1
E2 45483161 45483257 refGene_gn=EDC4;refGene_tx=XM_011290101.1
E2 45483347 45483450 refGene_gn=EDC4;refGene_tx=XM_011290101.1
E2 45483567 45483748 refGene_gn=EDC4;refGene_tx=XM_011290101.1
E2 45483835 45483908 refGene_gn=EDC4;refGene_tx=XM_011290101.1
E2 45484010 45484102 refGene_gn=EDC4;refGene_tx=XM_011290101.1
E2 45484180 45484368 refGene_gn=EDC4;refGene_tx=XM_011290101.1
E2 45484462 45484584 refGene_gn=EDC4;refGene_tx=XM_011290101.1
E2 45484675 45484829 refGene_gn=EDC4;refGene_tx=XM_011290101.1
E2 45485155 45485589 refGene_gn=EDC4;refGene_tx=XM_011290101.1
E2 45485673 45485778 refGene_gn=EDC4;refGene_tx=XM_011290101.1
E2 45485862 45486015 refGene_gn=EDC4;refGene_tx=XM_011290101.1
E2 45486104 45486141 refGene_gn=EDC4;refGene_tx=XM_011290101.1
E2 45486272 45486450 refGene_gn=EDC4;refGene_tx=XM_011290101.1

E2 45486561 45486717 refGene_gn=EDC4;refGene_tx=XM_011290101.1
E2 45486821 45486937 refGene_gn=EDC4;refGene_tx=XM_011290101.1
E2 45487033 45487199 refGene_gn=EDC4;refGene_tx=XM_011290101.1
E2 45487286 45487473 refGene_gn=EDC4;refGene_tx=XM_011290101.1
E2 45487565 45487785 refGene_gn=EDC4;refGene_tx=XM_011290101.1
E2 45488171 45488335 refGene_gn=EDC4;refGene_tx=XM_011290101.1
E2 45488574 45488767 refGene_gn=EDC4;refGene_tx=XM_011290101.1
E2 45489471 45489562
refGene_gn=NRN1L;refGene_tx=XM_003998173.4;refGene_gn=NRN1L;refGene_tx
=XM_019819927.2
E2 45490322 45490455
refGene_gn=NRN1L;refGene_tx=XM_003998173.4;refGene_gn=NRN1L;refGene_tx
=XM_019819927.2
E2 45490589 45490875
refGene_gn=NRN1L;refGene_tx=XM_003998173.4;refGene_gn=NRN1L;refGene_tx
=XM_019819927.2
E2 45506016 45506973
refGene_gn=PSKH1;refGene_tx=XM_003998141.3;refGene_gn=PSKH1;refGene_tx
=XM_006941637.4
E2 45515441 45515548 refGene_gn=PSKH1;refGene_tx=XM_006941637.4
E2 45520877 45521195
refGene_gn=PSKH1;refGene_tx=XM_003998141.3;refGene_gn=PSKH1;refGene_tx
=XM_006941637.4
E2 45523334 45523496 refGene_gn=CTRL;refGene_tx=XM_006941636.3
E2 45523583 45523717 refGene_gn=CTRL;refGene_tx=XM_006941636.3
E2 45523803 45523984 refGene_gn=CTRL;refGene_tx=XM_006941636.3
E2 45524151 45524233 refGene_gn=CTRL;refGene_tx=XM_006941636.3
E2 45524345 45524425 refGene_gn=CTRL;refGene_tx=XM_006941636.3
E2 45524516 45524620 refGene_gn=CTRL;refGene_tx=XM_006941636.3
E2 45525219 45525271 refGene_gn=CTRL;refGene_tx=XM_006941636.3
E2 45527218 45527330 refGene_gn=PSMB10;refGene_tx=XM_003998142.4
E2 45527456 45527608 refGene_gn=PSMB10;refGene_tx=XM_003998142.4
E2 45528270 45528329 refGene_gn=PSMB10;refGene_tx=XM_003998142.4
E2 45528438 45528554 refGene_gn=PSMB10;refGene_tx=XM_003998142.4
E2 45528828 45528969 refGene_gn=PSMB10;refGene_tx=XM_003998142.4
E2 45529063 45529161 refGene_gn=PSMB10;refGene_tx=XM_003998142.4
E2 45529249 45529337 refGene_gn=PSMB10;refGene_tx=XM_003998142.4
E2 45529544 45529600 refGene_gn=PSMB10;refGene_tx=XM_003998142.4
E2 45531132 45531707 refGene_gn=LCAT;refGene_tx=XM_003998147.4
E2 45532630 45532855 refGene_gn=LCAT;refGene_tx=XM_003998147.4
E2 45532936 45533032 refGene_gn=LCAT;refGene_tx=XM_003998147.4
E2 45533123 45533239 refGene_gn=LCAT;refGene_tx=XM_003998147.4
E2 45533318 45533475 refGene_gn=LCAT;refGene_tx=XM_003998147.4
E2 45534110 45534264 refGene_gn=LCAT;refGene_tx=XM_003998147.4
E2 45534951 45535043
refGene_gn=SLC12A4;refGene_tx=XM_003998143.3;refGene_gn=SLC12A4;refGen
e_tx=XM_019819928.1
E2 45535215 45535349
refGene_gn=SLC12A4;refGene_tx=XM_003998143.3;refGene_gn=SLC12A4;refGen
e_tx=XM_019819928.1

E2	45535463	45535651
refGene_gn=SLC12A4;refGene_tx=XM_003998143.3;refGene_gn=SLC12A4;refGen
e_tx=XM_019819928.1
E2	45535783	45535891
refGene_gn=SLC12A4;refGene_tx=XM_003998143.3;refGene_gn=SLC12A4;refGen
e_tx=XM_019819928.1
E2	45535968	45536100
refGene_gn=SLC12A4;refGene_tx=XM_003998143.3;refGene_gn=SLC12A4;refGen
e_tx=XM_019819928.1
E2	45536190	45536360
refGene_gn=SLC12A4;refGene_tx=XM_003998143.3;refGene_gn=SLC12A4;refGen
e_tx=XM_019819928.1
E2	45536478	45536674
refGene_gn=SLC12A4;refGene_tx=XM_003998143.3;refGene_gn=SLC12A4;refGen
e_tx=XM_019819928.1
E2	45536956	45537125
refGene_gn=SLC12A4;refGene_tx=XM_003998143.3;refGene_gn=SLC12A4;refGen
e_tx=XM_019819928.1
E2	45537374	45537479
refGene_gn=SLC12A4;refGene_tx=XM_003998143.3;refGene_gn=SLC12A4;refGen
e_tx=XM_019819928.1
E2	45537686	45537806
refGene_gn=SLC12A4;refGene_tx=XM_003998143.3;refGene_gn=SLC12A4;refGen
e_tx=XM_019819928.1
E2	45537998	45538097
refGene_gn=SLC12A4;refGene_tx=XM_003998143.3;refGene_gn=SLC12A4;refGen
e_tx=XM_019819928.1
E2	45539744	45539863
refGene_gn=SLC12A4;refGene_tx=XM_003998143.3;refGene_gn=SLC12A4;refGen
e_tx=XM_019819928.1
E2	45540299	45540474
refGene_gn=SLC12A4;refGene_tx=XM_003998143.3;refGene_gn=SLC12A4;refGen
e_tx=XM_019819928.1
E2	45540836	45540894
refGene_gn=SLC12A4;refGene_tx=XM_003998143.3;refGene_gn=SLC12A4;refGen
e_tx=XM_019819928.1
E2	45541171	45541270
refGene_gn=SLC12A4;refGene_tx=XM_003998143.3;refGene_gn=SLC12A4;refGen
e_tx=XM_019819928.1
E2	45541346	45541511
refGene_gn=SLC12A4;refGene_tx=XM_003998143.3;refGene_gn=SLC12A4;refGen
e_tx=XM_019819928.1
E2	45542030	45542245
refGene_gn=SLC12A4;refGene_tx=XM_003998143.3;refGene_gn=SLC12A4;refGen
e_tx=XM_019819928.1
E2	45542389	45542631
refGene_gn=SLC12A4;refGene_tx=XM_003998143.3;refGene_gn=SLC12A4;refGen
e_tx=XM_019819928.1
E2	45544188	45544319
refGene_gn=SLC12A4;refGene_tx=XM_003998143.3;refGene_gn=SLC12A4;refGen

e_tx=XM_019819928.1
E2	45546098	45546153
refGene_gn=SLC12A4;refGene_tx=XM_003998143.3;refGene_gn=SLC12A4;refGene_tx=XM_019819928.1
E2	45546239	45546386
refGene_gn=SLC12A4;refGene_tx=XM_003998143.3;refGene_gn=SLC12A4;refGene_tx=XM_019819928.1
E2	45550109	45550241
refGene_gn=SLC12A4;refGene_tx=XM_003998143.3;refGene_gn=SLC12A4;refGene_tx=XM_019819928.1
E2	45551678	45551773
refGene_gn=SLC12A4;refGene_tx=XM_003998143.3;refGene_gn=SLC12A4;refGene_tx=XM_019819928.1
E2	45552152	45552249	refGene_gn=SLC12A4;refGene_tx=XM_019819928.1
E2	45556455	45556570	refGene_gn=SLC12A4;refGene_tx=XM_003998143.3
E2	45563898	45564132	refGene_gn=DPEP3;refGene_tx=XM_019819939.1
E2	45564223	45564359	refGene_gn=DPEP3;refGene_tx=XM_019819939.1
E2	45564842	45564918	refGene_gn=DPEP3;refGene_tx=XM_019819939.1
E2	45565403	45565488	refGene_gn=DPEP3;refGene_tx=XM_019819939.1
E2	45565794	45565971	refGene_gn=DPEP3;refGene_tx=XM_019819939.1
E2	45566087	45566157	refGene_gn=DPEP3;refGene_tx=XM_019819939.1
E2	45566441	45566583	refGene_gn=DPEP3;refGene_tx=XM_019819939.1
E2	45566672	45566802	refGene_gn=DPEP3;refGene_tx=XM_019819939.1
E2	45567302	45567429	refGene_gn=DPEP3;refGene_tx=XM_019819939.1
E2	45567775	45568038	refGene_gn=DPEP3;refGene_tx=XM_019819939.1
E2	45568238	45568274	refGene_gn=DPEP2;refGene_tx=XM_023245408.1
E2	45574663	45574921
refGene_gn=DPEP2;refGene_tx=XM_011290097.3;refGene_gn=DPEP2;refGene_tx=XM_019819938.2;refGene_gn=DPEP2;refGene_tx=XM_023245406.1;refGene_gn=DPEP2;refGene_tx=XM_023245407.1
E2	45575014	45575150
refGene_gn=DPEP2;refGene_tx=XM_011290097.3;refGene_gn=DPEP2;refGene_tx=XM_019819938.2;refGene_gn=DPEP2;refGene_tx=XM_023245406.1;refGene_gn=DPEP2;refGene_tx=XM_023245407.1;refGene_gn=DPEP2;refGene_tx=XM_023245408.1
E2	45576125	45576201
refGene_gn=DPEP2;refGene_tx=XM_011290097.3;refGene_gn=DPEP2;refGene_tx=XM_019819938.2;refGene_gn=DPEP2;refGene_tx=XM_023245406.1;refGene_gn=DPEP2;refGene_tx=XM_023245407.1;refGene_gn=DPEP2;refGene_tx=XM_023245408.1
E2	45576830	45576915
refGene_gn=DPEP2;refGene_tx=XM_011290097.3;refGene_gn=DPEP2;refGene_tx=XM_019819938.2;refGene_gn=DPEP2;refGene_tx=XM_023245406.1;refGene_gn=DPEP2;refGene_tx=XM_023245407.1;refGene_gn=DPEP2;refGene_tx=XM_023245408.1
E2	45577303	45577480
refGene_gn=DPEP2;refGene_tx=XM_011290097.3;refGene_gn=DPEP2;refGene_tx=XM_019819938.2;refGene_gn=DPEP2;refGene_tx=XM_023245406.1;refGene_gn=DPEP2;refGene_tx=XM_023245407.1;refGene_gn=DPEP2;refGene_tx=XM_023245408.1

E2 45577597 45577667
refGene_gn=DPEP2;refGene_tx=XM_011290097.3;refGene_gn=DPEP2;refGene_tx=XM_019819938.2;refGene_gn=DPEP2;refGene_tx=XM_023245406.1;refGene_gn=DPEP2;refGene_tx=XM_023245407.1;refGene_gn=DPEP2;refGene_tx=XM_023245408.1
E2 45577959 45578101
refGene_gn=DPEP2;refGene_tx=XM_011290097.3;refGene_gn=DPEP2;refGene_tx=XM_019819938.2;refGene_gn=DPEP2;refGene_tx=XM_023245406.1;refGene_gn=DPEP2;refGene_tx=XM_023245407.1;refGene_gn=DPEP2;refGene_tx=XM_023245408.1
E2 45578185 45578315
refGene_gn=DPEP2;refGene_tx=XM_011290097.3;refGene_gn=DPEP2;refGene_tx=XM_019819938.2;refGene_gn=DPEP2;refGene_tx=XM_023245406.1;refGene_gn=DPEP2;refGene_tx=XM_023245407.1;refGene_gn=DPEP2;refGene_tx=XM_023245408.1
E2 45578579 45578706
refGene_gn=DPEP2;refGene_tx=XM_011290097.3;refGene_gn=DPEP2;refGene_tx=XM_023245406.1;refGene_gn=DPEP2;refGene_tx=XM_023245408.1
E2 45579037 45579262
refGene_gn=DPEP2;refGene_tx=XM_011290097.3;refGene_gn=DPEP2;refGene_tx=XM_023245406.1;refGene_gn=DPEP2;refGene_tx=XM_023245407.1;refGene_gn=DPEP2;refGene_tx=XM_023245408.1
E2 45589035 45589334 refGene_gn=DPEP2NB;refGene_tx=XM_011290095.3
E2 45590469 45590536 refGene_gn=DPEP2NB;refGene_tx=XM_011290095.3
E2 45593343 45594963 refGene_gn=DDX28;refGene_tx=XM_003998148.5
E2 45606844 45606970
refGene_gn=DUS2;refGene_tx=XM_023246130.1;refGene_gn=DUS2;refGene_tx=XM_023246131.1;refGene_gn=DUS2;refGene_tx=XM_023246132.1;refGene_gn=DUS2;refGene_tx=XM_023246133.1;refGene_gn=DUS2;refGene_tx=XM_023246134.1
E2 45618809 45618855
refGene_gn=DUS2;refGene_tx=XM_023246130.1;refGene_gn=DUS2;refGene_tx=XM_023246131.1;refGene_gn=DUS2;refGene_tx=XM_023246132.1;refGene_gn=DUS2;refGene_tx=XM_023246133.1;refGene_gn=DUS2;refGene_tx=XM_023246134.1
E2 45621415 45621507
refGene_gn=DUS2;refGene_tx=XM_023246130.1;refGene_gn=DUS2;refGene_tx=XM_023246131.1;refGene_gn=DUS2;refGene_tx=XM_023246132.1;refGene_gn=DUS2;refGene_tx=XM_023246133.1;refGene_gn=DUS2;refGene_tx=XM_023246134.1
E2 45622537 45622581
refGene_gn=DUS2;refGene_tx=XM_023246130.1;refGene_gn=DUS2;refGene_tx=XM_023246131.1;refGene_gn=DUS2;refGene_tx=XM_023246132.1;refGene_gn=DUS2;refGene_tx=XM_023246133.1;refGene_gn=DUS2;refGene_tx=XM_023246134.1
E2 45624220 45624281
refGene_gn=DUS2;refGene_tx=XM_023246130.1;refGene_gn=DUS2;refGene_tx=XM_023246131.1;refGene_gn=DUS2;refGene_tx=XM_023246132.1;refGene_gn=DUS2;refGene_tx=XM_023246133.1;refGene_gn=DUS2;refGene_tx=XM_023246134.1
E2 45627578 45627626
refGene_gn=DUS2;refGene_tx=XM_023246130.1;refGene_gn=DUS2;refGene_tx=XM_023246131.1;refGene_gn=DUS2;refGene_tx=XM_023246132.1;refGene_gn=DUS2;refGene_tx=XM_023246133.1;refGene_gn=DUS2;refGene_tx=XM_023246134.1
E2 45632617 45632683

refGene_gn=DUS2;refGene_tx=XM_023246130.1;refGene_gn=DUS2;refGene_tx=XM_023246131.1;refGene_gn=DUS2;refGene_tx=XM_023246132.1;refGene_gn=DUS2;refGene_tx=XM_023246133.1;refGene_gn=DUS2;refGene_tx=XM_023246134.1
E2 45632868 45632939
refGene_gn=DUS2;refGene_tx=XM_023246130.1;refGene_gn=DUS2;refGene_tx=XM_023246131.1;refGene_gn=DUS2;refGene_tx=XM_023246132.1;refGene_gn=DUS2;refGene_tx=XM_023246133.1;refGene_gn=DUS2;refGene_tx=XM_023246134.1
E2 45635173 45635260
refGene_gn=DUS2;refGene_tx=XM_023246130.1;refGene_gn=DUS2;refGene_tx=XM_023246131.1;refGene_gn=DUS2;refGene_tx=XM_023246132.1;refGene_gn=DUS2;refGene_tx=XM_023246133.1;refGene_gn=DUS2;refGene_tx=XM_023246134.1
E2 45636612 45636781
refGene_gn=DUS2;refGene_tx=XM_023246130.1;refGene_gn=DUS2;refGene_tx=XM_023246131.1;refGene_gn=DUS2;refGene_tx=XM_023246132.1;refGene_gn=DUS2;refGene_tx=XM_023246133.1;refGene_gn=DUS2;refGene_tx=XM_023246134.1
E2 45639326 45639448
refGene_gn=DUS2;refGene_tx=XM_023246130.1;refGene_gn=DUS2;refGene_tx=XM_023246131.1;refGene_gn=DUS2;refGene_tx=XM_023246132.1;refGene_gn=DUS2;refGene_tx=XM_023246133.1;refGene_gn=DUS2;refGene_tx=XM_023246134.1
E2 45640989 45641139
refGene_gn=DUS2;refGene_tx=XM_023246130.1;refGene_gn=DUS2;refGene_tx=XM_023246131.1;refGene_gn=DUS2;refGene_tx=XM_023246132.1;refGene_gn=DUS2;refGene_tx=XM_023246133.1;refGene_gn=DUS2;refGene_tx=XM_023246134.1
E2 45642226 45642314
refGene_gn=DUS2;refGene_tx=XM_023246130.1;refGene_gn=DUS2;refGene_tx=XM_023246131.1;refGene_gn=DUS2;refGene_tx=XM_023246132.1;refGene_gn=DUS2;refGene_tx=XM_023246133.1;refGene_gn=DUS2;refGene_tx=XM_023246134.1
E2 45643454 45643528
refGene_gn=DUS2;refGene_tx=XM_023246130.1;refGene_gn=DUS2;refGene_tx=XM_023246131.1;refGene_gn=DUS2;refGene_tx=XM_023246132.1;refGene_gn=DUS2;refGene_tx=XM_023246133.1;refGene_gn=DUS2;refGene_tx=XM_023246134.1
E2 45643766 45644004
refGene_gn=DUS2;refGene_tx=XM_023246130.1;refGene_gn=DUS2;refGene_tx=XM_023246131.1;refGene_gn=DUS2;refGene_tx=XM_023246132.1;refGene_gn=DUS2;refGene_tx=XM_023246133.1;refGene_gn=DUS2;refGene_tx=XM_023246134.1
E2 45650000 45650103
refGene_gn=NFATC3;refGene_tx=XM_011290091.3;refGene_gn=NFATC3;refGene_tx=XM_011290092.3;refGene_gn=NFATC3;refGene_tx=XM_019819930.2
E2 45689960 45691095
refGene_gn=NFATC3;refGene_tx=XM_011290091.3;refGene_gn=NFATC3;refGene_tx=XM_011290092.3;refGene_gn=NFATC3;refGene_tx=XM_019819930.2
E2 45693805 45693968
refGene_gn=NFATC3;refGene_tx=XM_011290091.3;refGene_gn=NFATC3;refGene_tx=XM_011290092.3;refGene_gn=NFATC3;refGene_tx=XM_019819930.2
E2 45723873 45724073
refGene_gn=NFATC3;refGene_tx=XM_011290091.3;refGene_gn=NFATC3;refGene_tx=XM_011290092.3;refGene_gn=NFATC3;refGene_tx=XM_019819930.2
E2 45730311 45730484
refGene_gn=NFATC3;refGene_tx=XM_011290091.3;refGene_gn=NFATC3;refGene_tx=XM_011290092.3;refGene_gn=NFATC3;refGene_tx=XM_019819930.2

E2 45735587 45735728
refGene_gn=NFATC3;refGene_tx=XM_011290091.3;refGene_gn=NFATC3;refGene_tx=XM_011290092.3;refGene_gn=NFATC3;refGene_tx=XM_019819930.2
E2 45742782 45742838
refGene_gn=NFATC3;refGene_tx=XM_011290091.3;refGene_gn=NFATC3;refGene_tx=XM_011290092.3
E2 45743855 45743982
refGene_gn=NFATC3;refGene_tx=XM_011290091.3;refGene_gn=NFATC3;refGene_tx=XM_011290092.3
E2 45754652 45755660
refGene_gn=NFATC3;refGene_tx=XM_011290091.3;refGene_gn=NFATC3;refGene_tx=XM_011290092.3;refGene_gn=NFATC3;refGene_tx=XM_019819930.2
E2 45774314 45774415 refGene_gn=NFATC3;refGene_tx=XM_011290092.3
E2 45785666 45785788
refGene_gn=NFATC3;refGene_tx=XM_011290091.3;refGene_gn=NFATC3;refGene_tx=XM_019819930.2
E2 45789522 45789612
refGene_gn=ESRP2;refGene_tx=XM_006941652.3;refGene_gn=ESRP2;refGene_tx=XM_011290093.2;refGene_gn=ESRP2;refGene_tx=XM_019819932.1;refGene_gn=ESRP2;refGene_tx=XM_019819933.1
E2 45789688 45789854
refGene_gn=ESRP2;refGene_tx=XM_006941652.3;refGene_gn=ESRP2;refGene_tx=XM_011290093.2;refGene_gn=ESRP2;refGene_tx=XM_019819932.1;refGene_gn=ESRP2;refGene_tx=XM_019819933.1
E2 45790163 45790397
refGene_gn=ESRP2;refGene_tx=XM_006941652.3;refGene_gn=ESRP2;refGene_tx=XM_011290093.2;refGene_gn=ESRP2;refGene_tx=XM_019819932.1;refGene_gn=ESRP2;refGene_tx=XM_019819933.1;refGene_gn=ESRP2;refGene_tx=XM_019819934.1
E2 45790497 45790696
refGene_gn=ESRP2;refGene_tx=XM_006941652.3;refGene_gn=ESRP2;refGene_tx=XM_011290093.2;refGene_gn=ESRP2;refGene_tx=XM_019819932.1;refGene_gn=ESRP2;refGene_tx=XM_019819933.1;refGene_gn=ESRP2;refGene_tx=XM_019819934.1
E2 45790805 45791018
refGene_gn=ESRP2;refGene_tx=XM_006941652.3;refGene_gn=ESRP2;refGene_tx=XM_011290093.2;refGene_gn=ESRP2;refGene_tx=XM_019819932.1;refGene_gn=ESRP2;refGene_tx=XM_019819933.1;refGene_gn=ESRP2;refGene_tx=XM_019819934.1
E2 45791131 45791433
refGene_gn=ESRP2;refGene_tx=XM_006941652.3;refGene_gn=ESRP2;refGene_tx=XM_011290093.2;refGene_gn=ESRP2;refGene_tx=XM_019819932.1;refGene_gn=ESRP2;refGene_tx=XM_019819933.1;refGene_gn=ESRP2;refGene_tx=XM_019819934.1
E2 45791592 45791635
refGene_gn=ESRP2;refGene_tx=XM_006941652.3;refGene_gn=ESRP2;refGene_tx=XM_011290093.2;refGene_gn=ESRP2;refGene_tx=XM_019819932.1;refGene_gn=ESRP2;refGene_tx=XM_019819933.1;refGene_gn=ESRP2;refGene_tx=XM_019819934.1
E2 45791721 45791854

refGene_gn=ESRP2;refGene_tx=XM_006941652.3;refGene_gn=ESRP2;refGene_tx=XM_011290093.2;refGene_gn=ESRP2;refGene_tx=XM_019819932.1;refGene_gn=ESRP2;refGene_tx=XM_019819933.1;refGene_gn=ESRP2;refGene_tx=XM_019819934.1
E2 45791932 45792043
refGene_gn=ESRP2;refGene_tx=XM_006941652.3;refGene_gn=ESRP2;refGene_tx=XM_011290093.2;refGene_gn=ESRP2;refGene_tx=XM_019819932.1;refGene_gn=ESRP2;refGene_tx=XM_019819933.1;refGene_gn=ESRP2;refGene_tx=XM_019819934.1
E2 45792112 45792167
refGene_gn=ESRP2;refGene_tx=XM_006941652.3;refGene_gn=ESRP2;refGene_tx=XM_011290093.2;refGene_gn=ESRP2;refGene_tx=XM_019819932.1;refGene_gn=ESRP2;refGene_tx=XM_019819933.1;refGene_gn=ESRP2;refGene_tx=XM_019819934.1
E2 45792386 45792485
refGene_gn=ESRP2;refGene_tx=XM_006941652.3;refGene_gn=ESRP2;refGene_tx=XM_011290093.2;refGene_gn=ESRP2;refGene_tx=XM_019819932.1;refGene_gn=ESRP2;refGene_tx=XM_019819933.1;refGene_gn=ESRP2;refGene_tx=XM_019819934.1
E2 45792819 45792934
refGene_gn=ESRP2;refGene_tx=XM_006941652.3;refGene_gn=ESRP2;refGene_tx=XM_011290093.2;refGene_gn=ESRP2;refGene_tx=XM_019819932.1;refGene_gn=ESRP2;refGene_tx=XM_019819933.1;refGene_gn=ESRP2;refGene_tx=XM_019819934.1
E2 45793043 45793157
refGene_gn=ESRP2;refGene_tx=XM_006941652.3;refGene_gn=ESRP2;refGene_tx=XM_011290093.2;refGene_gn=ESRP2;refGene_tx=XM_019819932.1;refGene_gn=ESRP2;refGene_tx=XM_019819934.1
E2 45794611 45794740
refGene_gn=ESRP2;refGene_tx=XM_006941652.3;refGene_gn=ESRP2;refGene_tx=XM_011290093.2;refGene_gn=ESRP2;refGene_tx=XM_019819932.1;refGene_gn=ESRP2;refGene_tx=XM_019819933.1;refGene_gn=ESRP2;refGene_tx=XM_019819934.1
E2 45794828 45795026
refGene_gn=ESRP2;refGene_tx=XM_006941652.3;refGene_gn=ESRP2;refGene_tx=XM_011290093.2;refGene_gn=ESRP2;refGene_tx=XM_019819932.1;refGene_gn=ESRP2;refGene_tx=XM_019819933.1;refGene_gn=ESRP2;refGene_tx=XM_019819934.1;refGene_gn=PLA2G15;refGene_tx=XM_019819944.2
E2 45802660 45802775
refGene_gn=PLA2G15;refGene_tx=XM_003998181.5;refGene_gn=PLA2G15;refGene_tx=XM_019819945.2
E2 45804539 45804540
refGene_gn=PLA2G15;refGene_tx=XM_019819942.2;refGene_gn=PLA2G15;refGene_tx=XM_019819943.2
E2 45805608 45805765
refGene_gn=PLA2G15;refGene_tx=XM_003998181.5;refGene_gn=PLA2G15;refGene_tx=XM_019819942.2;refGene_gn=PLA2G15;refGene_tx=XM_019819943.2;refGene_gn=PLA2G15;refGene_tx=XM_019819944.2;refGene_gn=PLA2G15;refGene_tx=XM_019819945.2
E2 45811146 45811265

refGene_gn=PLA2G15;refGene_tx=XM_003998181.5;refGene_gn=PLA2G15;refGene_tx=XM_019819942.2;refGene_gn=PLA2G15;refGene_tx=XM_019819943.2;refGene_gn=PLA2G15;refGene_tx=XM_019819944.2;refGene_gn=PLA2G15;refGene_tx=XM_019819945.2
E2 45811509 45811608
refGene_gn=PLA2G15;refGene_tx=XM_003998181.5;refGene_gn=PLA2G15;refGene_tx=XM_019819942.2;refGene_gn=PLA2G15;refGene_tx=XM_019819943.2;refGene_gn=PLA2G15;refGene_tx=XM_019819944.2;refGene_gn=PLA2G15;refGene_tx=XM_019819945.2
E2 45812008 45812233
refGene_gn=PLA2G15;refGene_tx=XM_003998181.5;refGene_gn=PLA2G15;refGene_tx=XM_019819942.2;refGene_gn=PLA2G15;refGene_tx=XM_019819943.2;refGene_gn=PLA2G15;refGene_tx=XM_019819944.2
E2 45814000 45814512
refGene_gn=PLA2G15;refGene_tx=XM_003998181.5;refGene_gn=PLA2G15;refGene_tx=XM_019819942.2;refGene_gn=PLA2G15;refGene_tx=XM_019819943.2;refGene_gn=PLA2G15;refGene_tx=XM_019819944.2;refGene_gn=PLA2G15;refGene_tx=XM_019819945.2
E2 45830295 45830815
refGene_gn=SLC7A6;refGene_tx=XM_003998182.4;refGene_gn=SLC7A6;refGene_tx=XM_019819955.2;refGene_gn=SLC7A6;refGene_tx=XM_019819956.2;refGene_gn=SLC7A6;refGene_tx=XM_023245051.1;refGene_gn=SLC7A6;refGene_tx=XM_023245053.1;refGene_gn=SLC7A6;refGene_tx=XM_023245054.1
E2 45840949 45841075
refGene_gn=SLC7A6;refGene_tx=XM_003998182.4;refGene_gn=SLC7A6;refGene_tx=XM_019819955.2;refGene_gn=SLC7A6;refGene_tx=XM_019819956.2;refGene_gn=SLC7A6;refGene_tx=XM_023245051.1;refGene_gn=SLC7A6;refGene_tx=XM_023245053.1;refGene_gn=SLC7A6;refGene_tx=XM_023245054.1
E2 45842960 45843105
refGene_gn=SLC7A6;refGene_tx=XM_003998182.4;refGene_gn=SLC7A6;refGene_tx=XM_019819955.2;refGene_gn=SLC7A6;refGene_tx=XM_019819956.2;refGene_gn=SLC7A6;refGene_tx=XM_023245051.1;refGene_gn=SLC7A6;refGene_tx=XM_023245053.1;refGene_gn=SLC7A6;refGene_tx=XM_023245054.1
E2 45843794 45843918
refGene_gn=SLC7A6;refGene_tx=XM_003998182.4;refGene_gn=SLC7A6;refGene_tx=XM_019819955.2;refGene_gn=SLC7A6;refGene_tx=XM_019819956.2;refGene_gn=SLC7A6;refGene_tx=XM_023245051.1;refGene_gn=SLC7A6;refGene_tx=XM_023245053.1;refGene_gn=SLC7A6;refGene_tx=XM_023245054.1
E2 45844127 45844231
refGene_gn=SLC7A6;refGene_tx=XM_003998182.4;refGene_gn=SLC7A6;refGene_tx=XM_019819955.2;refGene_gn=SLC7A6;refGene_tx=XM_019819956.2;refGene_gn=SLC7A6;refGene_tx=XM_023245051.1;refGene_gn=SLC7A6;refGene_tx=XM_023245053.1;refGene_gn=SLC7A6;refGene_tx=XM_023245054.1
E2 45847983 45848080
refGene_gn=SLC7A6;refGene_tx=XM_003998182.4;refGene_gn=SLC7A6;refGene_tx=XM_019819955.2;refGene_gn=SLC7A6;refGene_tx=XM_019819956.2;refGene_gn=SLC7A6;refGene_tx=XM_023245051.1;refGene_gn=SLC7A6;refGene_tx=XM_023245053.1;refGene_gn=SLC7A6;refGene_tx=XM_023245054.1
E2 45850350 45850500
refGene_gn=SLC7A6;refGene_tx=XM_003998182.4;refGene_gn=SLC7A6;refGene_

tx=XM_019819955.2;refGene_gn=SLC7A6;refGene_tx=XM_019819956.2;refGene_gn=SLC7A6;refGene_tx=XM_023245051.1;refGene_gn=SLC7A6;refGene_tx=XM_023245053.1;refGene_gn=SLC7A6;refGene_tx=XM_023245054.1
E2 45850611 45850795
refGene_gn=SLC7A6;refGene_tx=XM_003998182.4;refGene_gn=SLC7A6;refGene_tx=XM_019819955.2;refGene_gn=SLC7A6;refGene_tx=XM_019819956.2;refGene_gn=SLC7A6;refGene_tx=XM_023245051.1;refGene_gn=SLC7A6;refGene_tx=XM_023245053.1;refGene_gn=SLC7A6;refGene_tx=XM_023245054.1
E2 45851233 45851328
refGene_gn=SLC7A6;refGene_tx=XM_003998182.4;refGene_gn=SLC7A6;refGene_tx=XM_019819955.2;refGene_gn=SLC7A6;refGene_tx=XM_019819956.2;refGene_gn=SLC7A6;refGene_tx=XM_023245051.1;refGene_gn=SLC7A6;refGene_tx=XM_023245053.1
E2 45855686 45855817 refGene_gn=SLC7A6OS;refGene_tx=XM_003998183.4
E2 45857104 45857225 refGene_gn=SLC7A6OS;refGene_tx=XM_003998183.4
E2 45859590 45859794 refGene_gn=SLC7A6OS;refGene_tx=XM_003998183.4
E2 45867891 45868185 refGene_gn=SLC7A6OS;refGene_tx=XM_003998183.4
E2 45868309 45868504 refGene_gn=SLC7A6OS;refGene_tx=XM_003998183.4
E2 45873318 45873413
refGene_gn=PRMT7;refGene_tx=XM_003998184.3;refGene_gn=PRMT7;refGene_tx=XM_006941653.2;refGene_gn=PRMT7;refGene_tx=XM_006941654.2;refGene_gn=PRMT7;refGene_tx=XM_006941655.2;refGene_gn=PRMT7;refGene_tx=XM_006941656.3;refGene_gn=PRMT7;refGene_tx=XM_006941657.2;refGene_gn=PRMT7;refGene_tx=XM_006941658.2;refGene_gn=PRMT7;refGene_tx=XM_006941659.2;refGene_gn=PRMT7;refGene_tx=XM_006941660.2;refGene_gn=PRMT7;refGene_tx=XM_006941661.2;refGene_gn=PRMT7;refGene_tx=XM_006941663.2
E2 45877024 45877061
refGene_gn=PRMT7;refGene_tx=XM_003998184.3;refGene_gn=PRMT7;refGene_tx=XM_006941653.2;refGene_gn=PRMT7;refGene_tx=XM_006941654.2;refGene_gn=PRMT7;refGene_tx=XM_006941655.2;refGene_gn=PRMT7;refGene_tx=XM_006941656.3;refGene_gn=PRMT7;refGene_tx=XM_006941657.2;refGene_gn=PRMT7;refGene_tx=XM_006941658.2;refGene_gn=PRMT7;refGene_tx=XM_006941659.2;refGene_gn=PRMT7;refGene_tx=XM_006941660.2;refGene_gn=PRMT7;refGene_tx=XM_006941661.2;refGene_gn=PRMT7;refGene_tx=XM_006941663.2
E2 45884458 45884608
refGene_gn=PRMT7;refGene_tx=XM_003998184.3;refGene_gn=PRMT7;refGene_tx=XM_006941653.2;refGene_gn=PRMT7;refGene_tx=XM_006941654.2;refGene_gn=PRMT7;refGene_tx=XM_006941655.2;refGene_gn=PRMT7;refGene_tx=XM_006941656.3;refGene_gn=PRMT7;refGene_tx=XM_006941657.2;refGene_gn=PRMT7;refGene_tx=XM_006941658.2;refGene_gn=PRMT7;refGene_tx=XM_006941659.2;refGene_gn=PRMT7;refGene_tx=XM_006941660.2;refGene_gn=PRMT7;refGene_tx=XM_006941662.2
E2 45890673 45890782
refGene_gn=PRMT7;refGene_tx=XM_003998184.3;refGene_gn=PRMT7;refGene_tx=XM_006941653.2;refGene_gn=PRMT7;refGene_tx=XM_006941654.2;refGene_gn=PRMT7;refGene_tx=XM_006941655.2;refGene_gn=PRMT7;refGene_tx=XM_006941656.3;refGene_gn=PRMT7;refGene_tx=XM_006941657.2;refGene_gn=PRMT7;refGene_tx=XM_006941658.2;refGene_gn=PRMT7;refGene_tx=XM_006941659.2;refGene_gn=PRMT7;refGene_tx=XM_006941660.2;refGene_gn=PRMT7;refGene_tx=XM_006941661.2;refGene_gn=PRMT7;refGene_tx=XM_006941662.2;refGene_gn=PRMT7;

refGene_tx=XM_006941663.2
E2 45897202 45897315
refGene_gn=PRMT7;refGene_tx=XM_003998184.3;refGene_gn=PRMT7;refGene_tx=XM_006941653.2;refGene_gn=PRMT7;refGene_tx=XM_006941654.2;refGene_gn=PRMT7;refGene_tx=XM_006941655.2;refGene_gn=PRMT7;refGene_tx=XM_006941656.3;refGene_gn=PRMT7;refGene_tx=XM_006941657.2;refGene_gn=PRMT7;refGene_tx=XM_006941658.2;refGene_gn=PRMT7;refGene_tx=XM_006941659.2;refGene_gn=PRMT7;refGene_tx=XM_006941660.2;refGene_gn=PRMT7;refGene_tx=XM_006941661.2;refGene_gn=PRMT7;refGene_tx=XM_006941662.2;refGene_gn=PRMT7;refGene_tx=XM_006941663.2;refGene_gn=PRMT7;refGene_tx=XM_006941664.2
E2 45901064 45901306
refGene_gn=PRMT7;refGene_tx=XM_003998184.3;refGene_gn=PRMT7;refGene_tx=XM_006941653.2;refGene_gn=PRMT7;refGene_tx=XM_006941654.2;refGene_gn=PRMT7;refGene_tx=XM_006941655.2;refGene_gn=PRMT7;refGene_tx=XM_006941656.3;refGene_gn=PRMT7;refGene_tx=XM_006941657.2;refGene_gn=PRMT7;refGene_tx=XM_006941658.2;refGene_gn=PRMT7;refGene_tx=XM_006941659.2;refGene_gn=PRMT7;refGene_tx=XM_006941660.2;refGene_gn=PRMT7;refGene_tx=XM_006941661.2;refGene_gn=PRMT7;refGene_tx=XM_006941662.2;refGene_gn=PRMT7;refGene_tx=XM_006941663.2;refGene_gn=PRMT7;refGene_tx=XM_006941664.2
E2 45901530 45901711
refGene_gn=PRMT7;refGene_tx=XM_003998184.3;refGene_gn=PRMT7;refGene_tx=XM_006941653.2;refGene_gn=PRMT7;refGene_tx=XM_006941654.2;refGene_gn=PRMT7;refGene_tx=XM_006941655.2;refGene_gn=PRMT7;refGene_tx=XM_006941656.3;refGene_gn=PRMT7;refGene_tx=XM_006941657.2;refGene_gn=PRMT7;refGene_tx=XM_006941658.2;refGene_gn=PRMT7;refGene_tx=XM_006941659.2;refGene_gn=PRMT7;refGene_tx=XM_006941660.2;refGene_gn=PRMT7;refGene_tx=XM_006941661.2;refGene_gn=PRMT7;refGene_tx=XM_006941662.2;refGene_gn=PRMT7;refGene_tx=XM_006941663.2;refGene_gn=PRMT7;refGene_tx=XM_006941664.2
E2 45905191 45905319
refGene_gn=PRMT7;refGene_tx=XM_003998184.3;refGene_gn=PRMT7;refGene_tx=XM_006941653.2;refGene_gn=PRMT7;refGene_tx=XM_006941654.2;refGene_gn=PRMT7;refGene_tx=XM_006941655.2;refGene_gn=PRMT7;refGene_tx=XM_006941656.3;refGene_gn=PRMT7;refGene_tx=XM_006941657.2;refGene_gn=PRMT7;refGene_tx=XM_006941658.2;refGene_gn=PRMT7;refGene_tx=XM_006941659.2;refGene_gn=PRMT7;refGene_tx=XM_006941660.2;refGene_gn=PRMT7;refGene_tx=XM_006941661.2;refGene_gn=PRMT7;refGene_tx=XM_006941662.2;refGene_gn=PRMT7;refGene_tx=XM_006941663.2;refGene_gn=PRMT7;refGene_tx=XM_006941664.2
E2 45905749 45905885
refGene_gn=PRMT7;refGene_tx=XM_003998184.3;refGene_gn=PRMT7;refGene_tx=XM_006941653.2;refGene_gn=PRMT7;refGene_tx=XM_006941654.2;refGene_gn=PRMT7;refGene_tx=XM_006941655.2;refGene_gn=PRMT7;refGene_tx=XM_006941656.3;refGene_gn=PRMT7;refGene_tx=XM_006941657.2;refGene_gn=PRMT7;refGene_tx=XM_006941658.2;refGene_gn=PRMT7;refGene_tx=XM_006941659.2;refGene_gn=PRMT7;refGene_tx=XM_006941660.2;refGene_gn=PRMT7;refGene_tx=XM_006941661.2;refGene_gn=PRMT7;refGene_tx=XM_006941662.2;refGene_gn=PRMT7;refGene_tx=XM_006941663.2;refGene_gn=PRMT7;refGene_tx=XM_006941664.2
E2 45906856 45906940
refGene_gn=PRMT7;refGene_tx=XM_003998184.3;refGene_gn=PRMT7;refGene_tx=XM_006941653.2;refGene_gn=PRMT7;refGene_tx=XM_006941654.2;refGene_gn=PRMT7;refGene_tx=XM_006941655.2;refGene_gn=PRMT7;refGene_tx=XM_0069416

56.3;refGene_gn=PRMT7;refGene_tx=XM_006941657.2;refGene_gn=PRMT7;refGene_tx=XM_006941658.2;refGene_gn=PRMT7;refGene_tx=XM_006941659.2;refGene_gn=PRMT7;refGene_tx=XM_006941660.2;refGene_gn=PRMT7;refGene_tx=XM_006941661.2;refGene_gn=PRMT7;refGene_tx=XM_006941662.2;refGene_gn=PRMT7;refGene_tx=XM_006941663.2;refGene_gn=PRMT7;refGene_tx=XM_006941664.2
E2 45907741 45907789
refGene_gn=PRMT7;refGene_tx=XM_003998184.3;refGene_gn=PRMT7;refGene_tx=XM_006941653.2;refGene_gn=PRMT7;refGene_tx=XM_006941654.2;refGene_gn=PRMT7;refGene_tx=XM_006941655.2;refGene_gn=PRMT7;refGene_tx=XM_006941656.3;refGene_gn=PRMT7;refGene_tx=XM_006941657.2;refGene_gn=PRMT7;refGene_tx=XM_006941658.2;refGene_gn=PRMT7;refGene_tx=XM_006941659.2;refGene_gn=PRMT7;refGene_tx=XM_006941660.2;refGene_gn=PRMT7;refGene_tx=XM_006941661.2;refGene_gn=PRMT7;refGene_tx=XM_006941662.2;refGene_gn=PRMT7;refGene_tx=XM_006941663.2;refGene_gn=PRMT7;refGene_tx=XM_006941664.2
E2 45908408 45908498
refGene_gn=PRMT7;refGene_tx=XM_003998184.3;refGene_gn=PRMT7;refGene_tx=XM_006941653.2;refGene_gn=PRMT7;refGene_tx=XM_006941654.2;refGene_gn=PRMT7;refGene_tx=XM_006941655.2;refGene_gn=PRMT7;refGene_tx=XM_006941656.3;refGene_gn=PRMT7;refGene_tx=XM_006941657.2;refGene_gn=PRMT7;refGene_tx=XM_006941658.2;refGene_gn=PRMT7;refGene_tx=XM_006941659.2;refGene_gn=PRMT7;refGene_tx=XM_006941661.2;refGene_gn=PRMT7;refGene_tx=XM_006941662.2;refGene_gn=PRMT7;refGene_tx=XM_006941663.2;refGene_gn=PRMT7;refGene_tx=XM_006941664.2
E2 45912478 45912640
refGene_gn=PRMT7;refGene_tx=XM_003998184.3;refGene_gn=PRMT7;refGene_tx=XM_006941653.2;refGene_gn=PRMT7;refGene_tx=XM_006941654.2;refGene_gn=PRMT7;refGene_tx=XM_006941655.2;refGene_gn=PRMT7;refGene_tx=XM_006941656.3;refGene_gn=PRMT7;refGene_tx=XM_006941657.2;refGene_gn=PRMT7;refGene_tx=XM_006941658.2;refGene_gn=PRMT7;refGene_tx=XM_006941659.2;refGene_gn=PRMT7;refGene_tx=XM_006941660.2;refGene_gn=PRMT7;refGene_tx=XM_006941661.2;refGene_gn=PRMT7;refGene_tx=XM_006941662.2;refGene_gn=PRMT7;refGene_tx=XM_006941663.2;refGene_gn=PRMT7;refGene_tx=XM_006941664.2
E2 45913757 45913832
refGene_gn=PRMT7;refGene_tx=XM_003998184.3;refGene_gn=PRMT7;refGene_tx=XM_006941653.2;refGene_gn=PRMT7;refGene_tx=XM_006941654.2;refGene_gn=PRMT7;refGene_tx=XM_006941655.2;refGene_gn=PRMT7;refGene_tx=XM_006941656.3;refGene_gn=PRMT7;refGene_tx=XM_006941657.2;refGene_gn=PRMT7;refGene_tx=XM_006941658.2;refGene_gn=PRMT7;refGene_tx=XM_006941659.2;refGene_gn=PRMT7;refGene_tx=XM_006941660.2;refGene_gn=PRMT7;refGene_tx=XM_006941661.2;refGene_gn=PRMT7;refGene_tx=XM_006941662.2;refGene_gn=PRMT7;refGene_tx=XM_006941663.2;refGene_gn=PRMT7;refGene_tx=XM_006941664.2
E2 45914545 45914760
refGene_gn=PRMT7;refGene_tx=XM_003998184.3;refGene_gn=PRMT7;refGene_tx=XM_006941653.2;refGene_gn=PRMT7;refGene_tx=XM_006941654.2;refGene_gn=PRMT7;refGene_tx=XM_006941655.2;refGene_gn=PRMT7;refGene_tx=XM_006941656.3;refGene_gn=PRMT7;refGene_tx=XM_006941657.2;refGene_gn=PRMT7;refGene_tx=XM_006941658.2;refGene_gn=PRMT7;refGene_tx=XM_006941659.2;refGene_gn=PRMT7;refGene_tx=XM_006941660.2;refGene_gn=PRMT7;refGene_tx=XM_006941661.2;refGene_gn=PRMT7;refGene_tx=XM_006941662.2;refGene_gn=PRMT7;refGene_tx=XM_006941663.2;refGene_gn=PRMT7;refGene_tx=XM_006941664.2

E2 45915461 45915572
refGene_gn=PRMT7;refGene_tx=XM_003998184.3;refGene_gn=PRMT7;refGene_tx=XM_006941653.2;refGene_gn=PRMT7;refGene_tx=XM_006941654.2;refGene_gn=PRMT7;refGene_tx=XM_006941655.2;refGene_gn=PRMT7;refGene_tx=XM_006941656.3;refGene_gn=PRMT7;refGene_tx=XM_006941657.2;refGene_gn=PRMT7;refGene_tx=XM_006941658.2;refGene_gn=PRMT7;refGene_tx=XM_006941659.2;refGene_gn=PRMT7;refGene_tx=XM_006941660.2;refGene_gn=PRMT7;refGene_tx=XM_006941661.2;refGene_gn=PRMT7;refGene_tx=XM_006941662.2;refGene_gn=PRMT7;refGene_tx=XM_006941663.2;refGene_gn=PRMT7;refGene_tx=XM_006941664.2
E2 45915753 45915961
refGene_gn=PRMT7;refGene_tx=XM_003998184.3;refGene_gn=PRMT7;refGene_tx=XM_006941653.2;refGene_gn=PRMT7;refGene_tx=XM_006941654.2;refGene_gn=PRMT7;refGene_tx=XM_006941655.2;refGene_gn=PRMT7;refGene_tx=XM_006941656.3;refGene_gn=PRMT7;refGene_tx=XM_006941657.2;refGene_gn=PRMT7;refGene_tx=XM_006941658.2;refGene_gn=PRMT7;refGene_tx=XM_006941659.2;refGene_gn=PRMT7;refGene_tx=XM_006941660.2;refGene_gn=PRMT7;refGene_tx=XM_006941661.2;refGene_gn=PRMT7;refGene_tx=XM_006941662.2;refGene_gn=PRMT7;refGene_tx=XM_006941663.2;refGene_gn=PRMT7;refGene_tx=XM_006941664.2
E2 45919819 45919921
refGene_gn=SMPD3;refGene_tx=XM_003998186.5;refGene_gn=SMPD3;refGene_tx=XM_006941665.4
E2 45920183 45920346
refGene_gn=SMPD3;refGene_tx=XM_003998186.5;refGene_gn=SMPD3;refGene_tx=XM_006941665.4
E2 45922272 45922336
refGene_gn=SMPD3;refGene_tx=XM_003998186.5;refGene_gn=SMPD3;refGene_tx=XM_006941665.4
E2 45922609 45922699
refGene_gn=SMPD3;refGene_tx=XM_003998186.5;refGene_gn=SMPD3;refGene_tx=XM_006941665.4
E2 45924216 45924372
refGene_gn=SMPD3;refGene_tx=XM_003998186.5;refGene_gn=SMPD3;refGene_tx=XM_006941665.4
E2 45924510 45924586
refGene_gn=SMPD3;refGene_tx=XM_003998186.5;refGene_gn=SMPD3;refGene_tx=XM_006941665.4
E2 45929394 45930708
refGene_gn=SMPD3;refGene_tx=XM_003998186.5;refGene_gn=SMPD3;refGene_tx=XM_006941665.4
E2 46040245 46040257
refGene_gn=ZFP90;refGene_tx=XM_019819951.2;refGene_gn=ZFP90;refGene_tx=XM_019819952.2
E2 46078412 46078445
refGene_gn=ZFP90;refGene_tx=XM_003998187.4;refGene_gn=ZFP90;refGene_tx=XM_019819948.1;refGene_gn=ZFP90;refGene_tx=XM_019819950.2;refGene_gn=ZFP90;refGene_tx=XM_019819953.2;refGene_gn=ZFP90;refGene_tx=XM_019819954.2
E2 46107746 46107897
refGene_gn=ZFP90;refGene_tx=XM_003998187.4;refGene_gn=ZFP90;refGene_tx=XM_019819948.1;refGene_gn=ZFP90;refGene_tx=XM_019819950.2;refGene_gn=

ZFP90;refGene_tx=XM_019819951.2;refGene_gn=ZFP90;refGene_tx=XM_0198199
52.2;refGene_gn=ZFP90;refGene_tx=XM_019819953.2;refGene_gn=ZFP90;refGe
ne_tx=XM_019819954.2
E2 46108235 46108331
refGene_gn=ZFP90;refGene_tx=XM_003998187.4;refGene_gn=ZFP90;refGene_tx
=XM_019819948.1;refGene_gn=ZFP90;refGene_tx=XM_019819950.2;refGene_gn=
ZFP90;refGene_tx=XM_019819951.2;refGene_gn=ZFP90;refGene_tx=XM_0198199
52.2;refGene_gn=ZFP90;refGene_tx=XM_019819953.2;refGene_gn=ZFP90;refGe
ne_tx=XM_019819954.2
E2 46111611 46113260
refGene_gn=ZFP90;refGene_tx=XM_003998187.4;refGene_gn=ZFP90;refGene_tx
=XM_019819948.1;refGene_gn=ZFP90;refGene_tx=XM_019819950.2;refGene_gn=
ZFP90;refGene_tx=XM_019819951.2;refGene_gn=ZFP90;refGene_tx=XM_0198199
52.2
E2 46113751 46113966 refGene_gn=ZFP90;refGene_tx=XM_019819953.2
E2 46114844 46115011 refGene_gn=ZFP90;refGene_tx=XM_019819954.2
E2 46176653 46176662 refGene_gn=CDH3;refGene_tx=XM_019819961.2
E2 46189992 46190052 refGene_gn=CDH3;refGene_tx=XM_006941669.3
E2 46190189 46190373
refGene_gn=CDH3;refGene_tx=XM_006941669.3;refGene_gn=CDH3;refGene_tx=X
M_019819960.2;refGene_gn=CDH3;refGene_tx=XM_019819961.2
E2 46219321 46219407
refGene_gn=CDH3;refGene_tx=XM_006941669.3;refGene_gn=CDH3;refGene_tx=X
M_019819960.2;refGene_gn=CDH3;refGene_tx=XM_019819961.2
E2 46220396 46220522
refGene_gn=CDH3;refGene_tx=XM_006941669.3;refGene_gn=CDH3;refGene_tx=X
M_019819960.2;refGene_gn=CDH3;refGene_tx=XM_019819961.2
E2 46220718 46220874
refGene_gn=CDH3;refGene_tx=XM_006941669.3;refGene_gn=CDH3;refGene_tx=X
M_019819960.2;refGene_gn=CDH3;refGene_tx=XM_019819961.2
E2 46220964 46221109
refGene_gn=CDH3;refGene_tx=XM_006941669.3;refGene_gn=CDH3;refGene_tx=X
M_019819960.2;refGene_gn=CDH3;refGene_tx=XM_019819961.2
E2 46222455 46222631
refGene_gn=CDH3;refGene_tx=XM_006941669.3;refGene_gn=CDH3;refGene_tx=X
M_019819960.2;refGene_gn=CDH3;refGene_tx=XM_019819961.2
E2 46223298 46223427
refGene_gn=CDH3;refGene_tx=XM_006941669.3;refGene_gn=CDH3;refGene_tx=X
M_019819960.2;refGene_gn=CDH3;refGene_tx=XM_019819961.2
E2 46224718 46224904
refGene_gn=CDH3;refGene_tx=XM_006941669.3;refGene_gn=CDH3;refGene_tx=X
M_019819960.2;refGene_gn=CDH3;refGene_tx=XM_019819961.2
E2 46226253 46226495
refGene_gn=CDH3;refGene_tx=XM_006941669.3;refGene_gn=CDH3;refGene_tx=X
M_019819960.2;refGene_gn=CDH3;refGene_tx=XM_019819961.2
E2 46226888 46227034
refGene_gn=CDH3;refGene_tx=XM_006941669.3;refGene_gn=CDH3;refGene_tx=X
M_019819960.2;refGene_gn=CDH3;refGene_tx=XM_019819961.2
E2 46227785 46228010
refGene_gn=CDH3;refGene_tx=XM_006941669.3;refGene_gn=CDH3;refGene_tx=X

M_019819960.2;refGene_gn=CDH3;refGene_tx=XM_019819961.2
E2 46231952 46232159
refGene_gn=CDH3;refGene_tx=XM_006941669.3;refGene_gn=CDH3;refGene_tx=X
M_019819960.2;refGene_gn=CDH3;refGene_tx=XM_019819961.2
E2 46234507 46234638
refGene_gn=CDH3;refGene_tx=XM_006941669.3;refGene_gn=CDH3;refGene_tx=X
M_019819960.2;refGene_gn=CDH3;refGene_tx=XM_019819961.2
E2 46235031 46235178
refGene_gn=CDH3;refGene_tx=XM_006941669.3;refGene_gn=CDH3;refGene_tx=X
M_019819960.2;refGene_gn=CDH3;refGene_tx=XM_019819961.2
E2 46237169 46237379
refGene_gn=CDH3;refGene_tx=XM_006941669.3;refGene_gn=CDH3;refGene_tx=X
M_019819960.2;refGene_gn=CDH3;refGene_tx=XM_019819961.2
E2 46274486 46274534 refGene_gn=CDH1;refGene_tx=XM_003998189.4
E2 46275417 46275532 refGene_gn=CDH1;refGene_tx=XM_003998189.4
E2 46333796 46334014 refGene_gn=CDH1;refGene_tx=XM_003998189.4
E2 46335701 46335845 refGene_gn=CDH1;refGene_tx=XM_003998189.4
E2 46335944 46336100 refGene_gn=CDH1;refGene_tx=XM_003998189.4
E2 46336847 46336992 refGene_gn=CDH1;refGene_tx=XM_003998189.4
E2 46338335 46338511 refGene_gn=CDH1;refGene_tx=XM_003998189.4
E2 46338771 46338900 refGene_gn=CDH1;refGene_tx=XM_003998189.4
E2 46340904 46341087 refGene_gn=CDH1;refGene_tx=XM_003998189.4
E2 46342358 46342603 refGene_gn=CDH1;refGene_tx=XM_003998189.4
E2 46346335 46346481 refGene_gn=CDH1;refGene_tx=XM_003998189.4
E2 46349473 46349698 refGene_gn=CDH1;refGene_tx=XM_003998189.4
E2 46350011 46350242 refGene_gn=CDH1;refGene_tx=XM_003998189.4
E2 46353071 46353202 refGene_gn=CDH1;refGene_tx=XM_003998189.4
E2 46355055 46355199 refGene_gn=CDH1;refGene_tx=XM_003998189.4
E2 46356127 46356337 refGene_gn=CDH1;refGene_tx=XM_003998189.4
E2 46368035 46368132 refGene_gn=TANGO6;refGene_tx=XM_003998190.5
E2 46383466 46384107 refGene_gn=TANGO6;refGene_tx=XM_003998190.5
E2 46388182 46388299 refGene_gn=TANGO6;refGene_tx=XM_003998190.5
E2 46390950 46391092 refGene_gn=TANGO6;refGene_tx=XM_003998190.5
E2 46397506 46397643 refGene_gn=TANGO6;refGene_tx=XM_003998190.5
E2 46400437 46400600 refGene_gn=TANGO6;refGene_tx=XM_003998190.5
E2 46407151 46407234 refGene_gn=TANGO6;refGene_tx=XM_003998190.5
E2 46423349 46423462 refGene_gn=TANGO6;refGene_tx=XM_003998190.5
E2 46424905 46425082 refGene_gn=TANGO6;refGene_tx=XM_003998190.5
E2 46428623 46428756 refGene_gn=TANGO6;refGene_tx=XM_003998190.5
E2 46431181 46431370 refGene_gn=TANGO6;refGene_tx=XM_003998190.5
E2 46438532 46438667 refGene_gn=TANGO6;refGene_tx=XM_003998190.5
E2 46446947 46447457 refGene_gn=TANGO6;refGene_tx=XM_003998190.5
E2 46451545 46451603 refGene_gn=TANGO6;refGene_tx=XM_003998190.5
E2 46482431 46482572 refGene_gn=TANGO6;refGene_tx=XM_003998190.5
E2 46516702 46516854 refGene_gn=TANGO6;refGene_tx=XM_003998190.5
E2 46527276 46527390 refGene_gn=TANGO6;refGene_tx=XM_003998190.5
E2 46562415 46562592 refGene_gn=TANGO6;refGene_tx=XM_003998190.5
E2 46582896 46583532
refGene_gn=HAS3;refGene_tx=XM_003998193.5;refGene_gn=HAS3;refGene_tx=X
M_023245706.1

E2 46586411 46586513
refGene_gn=HAS3;refGene_tx=XM_003998193.5;refGene_gn=HAS3;refGene_tx=XM_023245706.1
E2 46587319 46588243 refGene_gn=HAS3;refGene_tx=XM_023245706.1
E2 46591414 46591522 refGene_gn=HAS3;refGene_tx=XM_003998193.5
E2 46593545 46593770
refGene_gn=CHTF8;refGene_tx=XM_011290085.3;refGene_gn=CHTF8;refGene_tx=XM_023245707.1
E2 46594172 46594290
refGene_gn=CHTF8;refGene_tx=XM_011290085.3;refGene_gn=CHTF8;refGene_tx=XM_023245707.1
E2 46594551 46594574
refGene_gn=CHTF8;refGene_tx=XM_011290085.3;refGene_gn=CHTF8;refGene_tx=XM_023245707.1
E2 46603576 46603735 refGene_gn=UTP4;refGene_tx=XM_003998194.4
E2 46607408 46607600 refGene_gn=UTP4;refGene_tx=XM_003998194.4
E2 46608739 46608824 refGene_gn=UTP4;refGene_tx=XM_003998194.4
E2 46610566 46610656 refGene_gn=UTP4;refGene_tx=XM_003998194.4
E2 46613274 46613486 refGene_gn=UTP4;refGene_tx=XM_003998194.4
E2 46620284 46620456 refGene_gn=UTP4;refGene_tx=XM_003998194.4
E2 46620559 46620651 refGene_gn=UTP4;refGene_tx=XM_003998194.4
E2 46621850 46621947 refGene_gn=UTP4;refGene_tx=XM_003998194.4
E2 46622704 46622769 refGene_gn=UTP4;refGene_tx=XM_003998194.4
E2 46624029 46624152 refGene_gn=UTP4;refGene_tx=XM_003998194.4
E2 46624534 46624691 refGene_gn=UTP4;refGene_tx=XM_003998194.4
E2 46626706 46626813 refGene_gn=UTP4;refGene_tx=XM_003998194.4
E2 46627834 46627930 refGene_gn=UTP4;refGene_tx=XM_003998194.4
E2 46631022 46631208 refGene_gn=UTP4;refGene_tx=XM_003998194.4
E2 46632685 46632796 refGene_gn=UTP4;refGene_tx=XM_003998194.4
E2 46633926 46634043 refGene_gn=UTP4;refGene_tx=XM_003998194.4
E2 46644862 46645442 refGene_gn=SNTB2;refGene_tx=XM_023246128.1
E2 46691978 46692192 refGene_gn=SNTB2;refGene_tx=XM_023246128.1
E2 46706719 46706930 refGene_gn=SNTB2;refGene_tx=XM_023246128.1
E2 46718386 46718529 refGene_gn=SNTB2;refGene_tx=XM_023246128.1
E2 46736602 46736799 refGene_gn=SNTB2;refGene_tx=XM_023246128.1
E2 46743256 46743441 refGene_gn=SNTB2;refGene_tx=XM_023246128.1
E2 46745991 46746084 refGene_gn=SNTB2;refGene_tx=XM_023246128.1
E2 46759213 46759234
refGene_gn=VPS4A;refGene_tx=XM_023245075.1;refGene_gn=VPS4A;refGene_tx=XM_023245076.1
E2 46762631 46762743
refGene_gn=VPS4A;refGene_tx=XM_023245075.1;refGene_gn=VPS4A;refGene_tx=XM_023245076.1
E2 46762826 46762974
refGene_gn=VPS4A;refGene_tx=XM_023245075.1;refGene_gn=VPS4A;refGene_tx=XM_023245076.1
E2 46764336 46764398
refGene_gn=VPS4A;refGene_tx=XM_023245075.1;refGene_gn=VPS4A;refGene_tx=XM_023245076.1
E2 46764513 46764633

refGene_gn=VPS4A;refGene_tx=XM_023245075.1;refGene_gn=VPS4A;refGene_tx=XM_023245076.1
E2 46765120 46765277 refGene_gn=VPS4A;refGene_tx=XM_023245075.1;refGene_gn=VPS4A;refGene_tx=XM_023245076.1
E2 46765819 46765968 refGene_gn=VPS4A;refGene_tx=XM_023245075.1;refGene_gn=VPS4A;refGene_tx=XM_023245076.1
E2 46766320 46766402 refGene_gn=VPS4A;refGene_tx=XM_023245075.1;refGene_gn=VPS4A;refGene_tx=XM_023245076.1
E2 46766617 46766837 refGene_gn=VPS4A;refGene_tx=XM_023245075.1;refGene_gn=VPS4A;refGene_tx=XM_023245076.1
E2 46768301 46768442 refGene_gn=VPS4A;refGene_tx=XM_023245075.1;refGene_gn=VPS4A;refGene_tx=XM_023245076.1
E2 46769155 46769257 refGene_gn=VPS4A;refGene_tx=XM_023245075.1;refGene_gn=VPS4A;refGene_tx=XM_023245076.1
E2 46773471 46773629 refGene_gn=PDF;refGene_tx=XM_006941714.4
E2 46774507 46775099 refGene_gn=PDF;refGene_tx=XM_006941714.4
E2 46775405 46775680 refGene_gn=COG8;refGene_tx=XM_003998224.5
E2 46776460 46776629 refGene_gn=COG8;refGene_tx=XM_003998224.5
E2 46778463 46779291 refGene_gn=COG8;refGene_tx=XM_003998224.5
E2 46780113 46780321 refGene_gn=COG8;refGene_tx=XM_003998224.5
E2 46781484 46781891 refGene_gn=COG8;refGene_tx=XM_003998224.5
E2 46782169 46782225 refGene_gn=NIP7;refGene_tx=XM_003998195.5;refGene_gn=NIP7;refGene_tx=XM_006941672.4;refGene_gn=NIP7;refGene_tx=XM_023245591.1;refGene_gn=NIP7;refGene_tx=XM_023245592.1;refGene_gn=NIP7;refGene_tx=XM_023245593.1
E2 46782348 46782435 refGene_gn=NIP7;refGene_tx=XM_003998195.5;refGene_gn=NIP7;refGene_tx=XM_006941672.4;refGene_gn=NIP7;refGene_tx=XM_023245591.1;refGene_gn=NIP7;refGene_tx=XM_023245592.1;refGene_gn=NIP7;refGene_tx=XM_023245593.1
E2 46782530 46782669 refGene_gn=NIP7;refGene_tx=XM_003998195.5;refGene_gn=NIP7;refGene_tx=XM_006941672.4;refGene_gn=NIP7;refGene_tx=XM_023245591.1;refGene_gn=NIP7;refGene_tx=XM_023245592.1;refGene_gn=NIP7;refGene_tx=XM_023245593.1
E2 46783631 46783772 refGene_gn=NIP7;refGene_tx=XM_003998195.5;refGene_gn=NIP7;refGene_tx=XM_006941672.4;refGene_gn=NIP7;refGene_tx=XM_023245591.1;refGene_gn=NIP7;refGene_tx=XM_023245592.1;refGene_gn=NIP7;refGene_tx=XM_023245593.1
E2 46783988 46784108 refGene_gn=NIP7;refGene_tx=XM_003998195.5
E2 46787274 46787508 refGene_gn=TMED6;refGene_tx=XM_023245589.1;refGene_gn=TMED6;refGene_tx=XM_023245590.1
E2 46788642 46788794 refGene_gn=TMED6;refGene_tx=XM_023245589.1;refGene_gn=TMED6;refGene_tx=XM_023245590.1

E2 46790470 46790597
refGene_gn=TMED6;refGene_tx=XM_023245589.1;refGene_gn=TMED6;refGene_tx
=XM_023245590.1
E2 46791840 46792053
refGene_gn=TMED6;refGene_tx=XM_023245589.1;refGene_gn=TMED6;refGene_tx
=XM_023245590.1
E2 46795335 46795494
refGene_gn=TERF2;refGene_tx=XM_011290188.3;refGene_gn=TERF2;refGene_tx
=XM_023245587.1;refGene_gn=TERF2;refGene_tx=XM_023245588.1
E2 46796012 46796056
refGene_gn=TERF2;refGene_tx=XM_011290188.3;refGene_gn=TERF2;refGene_tx
=XM_023245587.1;refGene_gn=TERF2;refGene_tx=XM_023245588.1
E2 46798321 46798407
refGene_gn=TERF2;refGene_tx=XM_011290188.3;refGene_gn=TERF2;refGene_tx
=XM_023245587.1;refGene_gn=TERF2;refGene_tx=XM_023245588.1
E2 46802973 46803366
refGene_gn=TERF2;refGene_tx=XM_011290188.3;refGene_gn=TERF2;refGene_tx
=XM_023245587.1;refGene_gn=TERF2;refGene_tx=XM_023245588.1
E2 46804702 46804809
refGene_gn=TERF2;refGene_tx=XM_011290188.3;refGene_gn=TERF2;refGene_tx
=XM_023245587.1;refGene_gn=TERF2;refGene_tx=XM_023245588.1
E2 46805005 46805029 refGene_gn=NIP7;refGene_tx=XM_023245592.1
E2 46806789 46806936
refGene_gn=TERF2;refGene_tx=XM_011290188.3;refGene_gn=TERF2;refGene_tx
=XM_023245587.1;refGene_gn=TERF2;refGene_tx=XM_023245588.1
E2 46807990 46808077
refGene_gn=TERF2;refGene_tx=XM_011290188.3;refGene_gn=TERF2;refGene_tx
=XM_023245587.1
E2 46818021 46818152
refGene_gn=TERF2;refGene_tx=XM_011290188.3;refGene_gn=TERF2;refGene_tx
=XM_023245587.1;refGene_gn=TERF2;refGene_tx=XM_023245588.1
E2 46818410 46818557 refGene_gn=NIP7;refGene_tx=XM_023245591.1
E2 46818811 46818907
refGene_gn=TERF2;refGene_tx=XM_011290188.3;refGene_gn=TERF2;refGene_tx
=XM_023245587.1;refGene_gn=TERF2;refGene_tx=XM_023245588.1
E2 46819010 46819389
refGene_gn=TERF2;refGene_tx=XM_011290188.3;refGene_gn=TERF2;refGene_tx
=XM_023245587.1;refGene_gn=TERF2;refGene_tx=XM_023245588.1
E2 46833482 46833500 refGene_gn=NIP7;refGene_tx=XM_023245593.1
E2 46844626 46844686 refGene_gn=NIP7;refGene_tx=XM_006941672.4
E2 46856367 46856529
refGene_gn=LOC101083640;refGene_tx=XM_006941673.4
E2 46876695 46876824
refGene_gn=LOC101083640;refGene_tx=XM_006941673.4
E2 46877328 46877358
refGene_gn=LOC101083640;refGene_tx=XM_006941673.4
E2 46886446 46886475
refGene_gn=LOC101083640;refGene_tx=XM_006941673.4
E2 46890688 46890779
refGene_gn=LOC101083640;refGene_tx=XM_006941673.4

E2 46983303 46983376 refGene_gn=NFAT5;refGene_tx=XM_023245636.1
E2 46985185 46985239 refGene_gn=NFAT5;refGene_tx=XM_023245636.1
E2 47027923 47028049 refGene_gn=NFAT5;refGene_tx=XM_023245636.1
E2 47043126 47043685 refGene_gn=NFAT5;refGene_tx=XM_023245636.1
E2 47048456 47048649 refGene_gn=NFAT5;refGene_tx=XM_023245636.1
E2 47050677 47050868 refGene_gn=NFAT5;refGene_tx=XM_023245636.1
E2 47052995 47053168 refGene_gn=NFAT5;refGene_tx=XM_023245636.1
E2 47063954 47064089 refGene_gn=NFAT5;refGene_tx=XM_023245636.1
E2 47064208 47064261 refGene_gn=NFAT5;refGene_tx=XM_023245636.1
E2 47073372 47073505 refGene_gn=NFAT5;refGene_tx=XM_023245636.1
E2 47079661 47079745 refGene_gn=NFAT5;refGene_tx=XM_023245636.1
E2 47088536 47088682 refGene_gn=NFAT5;refGene_tx=XM_023245636.1
E2 47089577 47091407 refGene_gn=NFAT5;refGene_tx=XM_023245636.1
E2 47091408 47092072 refGene_gn=NFAT5;refGene_tx=XM_023245636.1
E2 47092722 47092958 refGene_gn=NFAT5;refGene_tx=XM_023245636.1
E2 47106786 47107092 refGene_gn=NQO1;refGene_tx=XM_011290082.3
E2 47108004 47108106 refGene_gn=NQO1;refGene_tx=XM_011290082.3
E2 47108966 47109080 refGene_gn=NQO1;refGene_tx=XM_011290082.3
E2 47111895 47112026 refGene_gn=NQO1;refGene_tx=XM_011290082.3
E2 47112139 47112248 refGene_gn=NQO1;refGene_tx=XM_011290082.3
E2 47139231 47139501 refGene_gn=NOB1;refGene_tx=XM_003998201.5
E2 47140304 47140449 refGene_gn=NOB1;refGene_tx=XM_003998201.5
E2 47143165 47143263 refGene_gn=NOB1;refGene_tx=XM_003998201.5
E2 47143806 47144010 refGene_gn=NOB1;refGene_tx=XM_003998201.5
E2 47144102 47144249 refGene_gn=NOB1;refGene_tx=XM_003998201.5
E2 47144446 47144518 refGene_gn=NOB1;refGene_tx=XM_003998201.5
E2 47146194 47146325 refGene_gn=NOB1;refGene_tx=XM_003998201.5
E2 47147538 47147671 refGene_gn=NOB1;refGene_tx=XM_003998201.5
E2 47147792 47147855 refGene_gn=NOB1;refGene_tx=XM_003998201.5
E2 47178634 47178704 refGene_gn=WWP2;refGene_tx=XM_023245698.1
E2 47184759 47184907 refGene_gn=WWP2;refGene_tx=XM_023245698.1
E2 47185252 47185374 refGene_gn=WWP2;refGene_tx=XM_023245698.1
E2 47213767 47213846 refGene_gn=WWP2;refGene_tx=XM_023245699.1
E2 47217737 47217738 refGene_gn=WWP2;refGene_tx=XM_023245700.1
E2 47217827 47217965
refGene_gn=WWP2;refGene_tx=XM_023245698.1;refGene_gn=WWP2;refGene_tx=XM_023245699.1;refGene_gn=WWP2;refGene_tx=XM_023245700.1
E2 47219796 47219893
refGene_gn=WWP2;refGene_tx=XM_023245698.1;refGene_gn=WWP2;refGene_tx=XM_023245699.1;refGene_gn=WWP2;refGene_tx=XM_023245700.1
E2 47245937 47246065
refGene_gn=WWP2;refGene_tx=XM_023245698.1;refGene_gn=WWP2;refGene_tx=XM_023245699.1;refGene_gn=WWP2;refGene_tx=XM_023245700.1
E2 47258164 47258375
refGene_gn=WWP2;refGene_tx=XM_023245698.1;refGene_gn=WWP2;refGene_tx=XM_023245699.1;refGene_gn=WWP2;refGene_tx=XM_023245700.1
E2 47272255 47272345
refGene_gn=WWP2;refGene_tx=XM_023245698.1;refGene_gn=WWP2;refGene_tx=XM_023245699.1;refGene_gn=WWP2;refGene_tx=XM_023245700.1
E2 47283546 47283721

refGene_gn=WWP2;refGene_tx=XM_023245698.1;refGene_gn=WWP2;refGene_tx=XM_023245699.1;refGene_gn=WWP2;refGene_tx=XM_023245700.1
E2 47290414 47290469 refGene_gn=WWP2;refGene_tx=XM_023245698.1;refGene_gn=WWP2;refGene_tx=XM_023245699.1;refGene_gn=WWP2;refGene_tx=XM_023245700.1
E2 47293980 47294062 refGene_gn=WWP2;refGene_tx=XM_023245698.1;refGene_gn=WWP2;refGene_tx=XM_023245699.1;refGene_gn=WWP2;refGene_tx=XM_023245700.1
E2 47294608 47294737 refGene_gn=WWP2;refGene_tx=XM_023245698.1;refGene_gn=WWP2;refGene_tx=XM_023245699.1;refGene_gn=WWP2;refGene_tx=XM_023245700.1
E2 47295115 47295191 refGene_gn=WWP2;refGene_tx=XM_023245698.1;refGene_gn=WWP2;refGene_tx=XM_023245699.1;refGene_gn=WWP2;refGene_tx=XM_023245700.1
E2 47295502 47295574 refGene_gn=WWP2;refGene_tx=XM_023245698.1;refGene_gn=WWP2;refGene_tx=XM_023245699.1;refGene_gn=WWP2;refGene_tx=XM_023245700.1
E2 47295796 47295885 refGene_gn=WWP2;refGene_tx=XM_023245698.1;refGene_gn=WWP2;refGene_tx=XM_023245699.1;refGene_gn=WWP2;refGene_tx=XM_023245700.1
E2 47297727 47297887 refGene_gn=WWP2;refGene_tx=XM_023245698.1;refGene_gn=WWP2;refGene_tx=XM_023245699.1;refGene_gn=WWP2;refGene_tx=XM_023245700.1
E2 47299137 47299271 refGene_gn=WWP2;refGene_tx=XM_023245698.1;refGene_gn=WWP2;refGene_tx=XM_023245699.1;refGene_gn=WWP2;refGene_tx=XM_023245700.1
E2 47299599 47299740 refGene_gn=WWP2;refGene_tx=XM_023245698.1;refGene_gn=WWP2;refGene_tx=XM_023245699.1;refGene_gn=WWP2;refGene_tx=XM_023245700.1
E2 47300334 47300455 refGene_gn=WWP2;refGene_tx=XM_023245698.1;refGene_gn=WWP2;refGene_tx=XM_023245699.1;refGene_gn=WWP2;refGene_tx=XM_023245700.1
E2 47300801 47300906 refGene_gn=WWP2;refGene_tx=XM_023245698.1;refGene_gn=WWP2;refGene_tx=XM_023245699.1;refGene_gn=WWP2;refGene_tx=XM_023245700.1
E2 47301835 47301932 refGene_gn=WWP2;refGene_tx=XM_023245698.1;refGene_gn=WWP2;refGene_tx=XM_023245699.1;refGene_gn=WWP2;refGene_tx=XM_023245700.1
E2 47302123 47302196 refGene_gn=WWP2;refGene_tx=XM_023245698.1;refGene_gn=WWP2;refGene_tx=XM_023245699.1;refGene_gn=WWP2;refGene_tx=XM_023245700.1
E2 47302683 47302783 refGene_gn=WWP2;refGene_tx=XM_023245698.1;refGene_gn=WWP2;refGene_tx=XM_023245699.1;refGene_gn=WWP2;refGene_tx=XM_023245700.1
E2 47318370 47318815 refGene_gn=PSMD7;refGene_tx=XM_003998205.4
E2 47319505 47319597 refGene_gn=PSMD7;refGene_tx=XM_003998205.4
E2 47321421 47321502 refGene_gn=PSMD7;refGene_tx=XM_003998205.4
E2 47322248 47322346 refGene_gn=PSMD7;refGene_tx=XM_003998205.4
E2 47323078 47323171 refGene_gn=PSMD7;refGene_tx=XM_003998205.4
E2 47324234 47324326 refGene_gn=PSMD7;refGene_tx=XM_003998205.4

E2 47326620 47326694 refGene_gn=PSMD7;refGene_tx=XM_003998205.4
E2 48410137 48410185
refGene_gn=CE2H16orf47;refGene_tx=XM_023245120.1
E2 48410186 48410222
refGene_gn=CE2H16orf47;refGene_tx=XM_023245120.1
E2 48410223 48410394
refGene_gn=CE2H16orf47;refGene_tx=XM_023245120.1
E2 48553302 48556015
refGene_gn=ZFHX3;refGene_tx=XM_006941677.4;refGene_gn=ZFHX3;refGene_tx=XM_019819974.2
E2 48560928 48561425
refGene_gn=ZFHX3;refGene_tx=XM_006941677.4;refGene_gn=ZFHX3;refGene_tx=XM_019819974.2
E2 48615360 48615592
refGene_gn=ZFHX3;refGene_tx=XM_006941677.4;refGene_gn=ZFHX3;refGene_tx=XM_019819974.2
E2 48668332 48668440
refGene_gn=ZFHX3;refGene_tx=XM_006941677.4;refGene_gn=ZFHX3;refGene_tx=XM_019819974.2
E2 48686822 48686956
refGene_gn=ZFHX3;refGene_tx=XM_006941677.4;refGene_gn=ZFHX3;refGene_tx=XM_019819974.2
E2 48687108 48687309
refGene_gn=ZFHX3;refGene_tx=XM_006941677.4;refGene_gn=ZFHX3;refGene_tx=XM_019819974.2
E2 48698871 48698974
refGene_gn=ZFHX3;refGene_tx=XM_006941677.4;refGene_gn=ZFHX3;refGene_tx=XM_019819974.2
E2 48700205 48705638
refGene_gn=ZFHX3;refGene_tx=XM_006941677.4;refGene_gn=ZFHX3;refGene_tx=XM_019819974.2
E2 48709362 48711032
refGene_gn=ZFHX3;refGene_tx=XM_006941677.4;refGene_gn=ZFHX3;refGene_tx=XM_019819974.2
E2 49333613 49333664
refGene_gn=PMFBP1;refGene_tx=XM_006941678.4;refGene_gn=PMFBP1;refGene_tx=XM_011290241.3;refGene_gn=PMFBP1;refGene_tx=XM_019819975.2;refGene_gn=PMFBP1;refGene_tx=XM_019819976.2;refGene_gn=PMFBP1;refGene_tx=XM_019819977.2;refGene_gn=PMFBP1;refGene_tx=XM_019819978.2;refGene_gn=PMFBP1;refGene_tx=XM_019819979.2
E2 49338766 49338922
refGene_gn=PMFBP1;refGene_tx=XM_006941678.4;refGene_gn=PMFBP1;refGene_tx=XM_011290241.3;refGene_gn=PMFBP1;refGene_tx=XM_019819975.2;refGene_gn=PMFBP1;refGene_tx=XM_019819976.2;refGene_gn=PMFBP1;refGene_tx=XM_019819977.2;refGene_gn=PMFBP1;refGene_tx=XM_019819978.2;refGene_gn=PMFBP1;refGene_tx=XM_019819979.2
E2 49347561 49347804
refGene_gn=PMFBP1;refGene_tx=XM_006941678.4;refGene_gn=PMFBP1;refGene_tx=XM_011290241.3;refGene_gn=PMFBP1;refGene_tx=XM_019819975.2;refGene_gn=PMFBP1;refGene_tx=XM_019819976.2;refGene_gn=PMFBP1;refGene_tx=XM_01

9819977.2;refGene_gn=PMFBP1;refGene_tx=XM_019819978.2;refGene_gn=PMFBP1;refGene_tx=XM_019819979.2
E2 49350361 49350583
refGene_gn=PMFBP1;refGene_tx=XM_006941678.4;refGene_gn=PMFBP1;refGene_tx=XM_011290241.3;refGene_gn=PMFBP1;refGene_tx=XM_019819975.2;refGene_gn=PMFBP1;refGene_tx=XM_019819976.2;refGene_gn=PMFBP1;refGene_tx=XM_019819977.2;refGene_gn=PMFBP1;refGene_tx=XM_019819978.2;refGene_gn=PMFBP1;refGene_tx=XM_019819979.2
E2 49353150 49353321
refGene_gn=PMFBP1;refGene_tx=XM_006941678.4;refGene_gn=PMFBP1;refGene_tx=XM_011290241.3;refGene_gn=PMFBP1;refGene_tx=XM_019819975.2;refGene_gn=PMFBP1;refGene_tx=XM_019819976.2;refGene_gn=PMFBP1;refGene_tx=XM_019819977.2;refGene_gn=PMFBP1;refGene_tx=XM_019819978.2;refGene_gn=PMFBP1;refGene_tx=XM_019819979.2
E2 49354299 49354410
refGene_gn=PMFBP1;refGene_tx=XM_006941678.4;refGene_gn=PMFBP1;refGene_tx=XM_011290241.3;refGene_gn=PMFBP1;refGene_tx=XM_019819975.2;refGene_gn=PMFBP1;refGene_tx=XM_019819976.2;refGene_gn=PMFBP1;refGene_tx=XM_019819977.2;refGene_gn=PMFBP1;refGene_tx=XM_019819978.2;refGene_gn=PMFBP1;refGene_tx=XM_019819979.2
E2 49358319 49358446
refGene_gn=PMFBP1;refGene_tx=XM_006941678.4;refGene_gn=PMFBP1;refGene_tx=XM_011290241.3;refGene_gn=PMFBP1;refGene_tx=XM_019819975.2;refGene_gn=PMFBP1;refGene_tx=XM_019819976.2;refGene_gn=PMFBP1;refGene_tx=XM_019819977.2;refGene_gn=PMFBP1;refGene_tx=XM_019819978.2;refGene_gn=PMFBP1;refGene_tx=XM_019819979.2
E2 49358537 49358695
refGene_gn=PMFBP1;refGene_tx=XM_006941678.4;refGene_gn=PMFBP1;refGene_tx=XM_011290241.3;refGene_gn=PMFBP1;refGene_tx=XM_019819975.2;refGene_gn=PMFBP1;refGene_tx=XM_019819976.2;refGene_gn=PMFBP1;refGene_tx=XM_019819977.2;refGene_gn=PMFBP1;refGene_tx=XM_019819978.2;refGene_gn=PMFBP1;refGene_tx=XM_019819979.2
E2 49360701 49360942
refGene_gn=PMFBP1;refGene_tx=XM_006941678.4;refGene_gn=PMFBP1;refGene_tx=XM_011290241.3;refGene_gn=PMFBP1;refGene_tx=XM_019819975.2;refGene_gn=PMFBP1;refGene_tx=XM_019819976.2;refGene_gn=PMFBP1;refGene_tx=XM_019819977.2;refGene_gn=PMFBP1;refGene_tx=XM_019819978.2;refGene_gn=PMFBP1;refGene_tx=XM_019819979.2
E2 49362921 49363111
refGene_gn=PMFBP1;refGene_tx=XM_006941678.4;refGene_gn=PMFBP1;refGene_tx=XM_011290241.3;refGene_gn=PMFBP1;refGene_tx=XM_019819975.2;refGene_gn=PMFBP1;refGene_tx=XM_019819976.2;refGene_gn=PMFBP1;refGene_tx=XM_019819977.2;refGene_gn=PMFBP1;refGene_tx=XM_019819978.2;refGene_gn=PMFBP1;refGene_tx=XM_019819979.2
E2 49363273 49363427
refGene_gn=PMFBP1;refGene_tx=XM_006941678.4;refGene_gn=PMFBP1;refGene_tx=XM_011290241.3;refGene_gn=PMFBP1;refGene_tx=XM_019819975.2;refGene_gn=PMFBP1;refGene_tx=XM_019819976.2;refGene_gn=PMFBP1;refGene_tx=XM_019819977.2;refGene_gn=PMFBP1;refGene_tx=XM_019819978.2;refGene_gn=PMFBP1;refGene_tx=XM_019819979.2

E2 49364053 49364221
refGene_gn=PMFBP1;refGene_tx=XM_006941678.4;refGene_gn=PMFBP1;refGene_tx=XM_011290241.3;refGene_gn=PMFBP1;refGene_tx=XM_019819975.2;refGene_gn=PMFBP1;refGene_tx=XM_019819976.2;refGene_gn=PMFBP1;refGene_tx=XM_019819977.2;refGene_gn=PMFBP1;refGene_tx=XM_019819978.2;refGene_gn=PMFBP1;refGene_tx=XM_019819979.2
E2 49364489 49364627
refGene_gn=PMFBP1;refGene_tx=XM_006941678.4;refGene_gn=PMFBP1;refGene_tx=XM_011290241.3;refGene_gn=PMFBP1;refGene_tx=XM_019819975.2;refGene_gn=PMFBP1;refGene_tx=XM_019819976.2;refGene_gn=PMFBP1;refGene_tx=XM_019819977.2;refGene_gn=PMFBP1;refGene_tx=XM_019819978.2;refGene_gn=PMFBP1;refGene_tx=XM_019819979.2
E2 49369508 49369673
refGene_gn=PMFBP1;refGene_tx=XM_006941678.4;refGene_gn=PMFBP1;refGene_tx=XM_011290241.3;refGene_gn=PMFBP1;refGene_tx=XM_019819975.2;refGene_gn=PMFBP1;refGene_tx=XM_019819976.2;refGene_gn=PMFBP1;refGene_tx=XM_019819977.2;refGene_gn=PMFBP1;refGene_tx=XM_019819978.2;refGene_gn=PMFBP1;refGene_tx=XM_019819979.2
E2 49370221 49370389
refGene_gn=PMFBP1;refGene_tx=XM_006941678.4;refGene_gn=PMFBP1;refGene_tx=XM_011290241.3;refGene_gn=PMFBP1;refGene_tx=XM_019819975.2;refGene_gn=PMFBP1;refGene_tx=XM_019819976.2;refGene_gn=PMFBP1;refGene_tx=XM_019819977.2;refGene_gn=PMFBP1;refGene_tx=XM_019819979.2
E2 49370699 49370867
refGene_gn=PMFBP1;refGene_tx=XM_006941678.4;refGene_gn=PMFBP1;refGene_tx=XM_011290241.3;refGene_gn=PMFBP1;refGene_tx=XM_019819975.2;refGene_gn=PMFBP1;refGene_tx=XM_019819976.2;refGene_gn=PMFBP1;refGene_tx=XM_019819977.2;refGene_gn=PMFBP1;refGene_tx=XM_019819978.2;refGene_gn=PMFBP1;refGene_tx=XM_019819979.2
E2 49371700 49371807
refGene_gn=PMFBP1;refGene_tx=XM_006941678.4;refGene_gn=PMFBP1;refGene_tx=XM_011290241.3;refGene_gn=PMFBP1;refGene_tx=XM_019819975.2;refGene_gn=PMFBP1;refGene_tx=XM_019819976.2;refGene_gn=PMFBP1;refGene_tx=XM_019819977.2;refGene_gn=PMFBP1;refGene_tx=XM_019819978.2
E2 49372295 49372370
refGene_gn=PMFBP1;refGene_tx=XM_006941678.4;refGene_gn=PMFBP1;refGene_tx=XM_011290241.3;refGene_gn=PMFBP1;refGene_tx=XM_019819975.2;refGene_gn=PMFBP1;refGene_tx=XM_019819976.2;refGene_gn=PMFBP1;refGene_tx=XM_019819977.2;refGene_gn=PMFBP1;refGene_tx=XM_019819978.2
E2 49374409 49374648
refGene_gn=PMFBP1;refGene_tx=XM_006941678.4;refGene_gn=PMFBP1;refGene_tx=XM_011290241.3;refGene_gn=PMFBP1;refGene_tx=XM_019819975.2;refGene_gn=PMFBP1;refGene_tx=XM_019819976.2;refGene_gn=PMFBP1;refGene_tx=XM_019819977.2;refGene_gn=PMFBP1;refGene_tx=XM_019819978.2;refGene_gn=PMFBP1;refGene_tx=XM_019819979.2
E2 49375211 49375228 refGene_gn=PMFBP1;refGene_tx=XM_006941678.4
E2 49375678 49375695 refGene_gn=PMFBP1;refGene_tx=XM_019819977.2
E2 49377589 49377693
refGene_gn=PMFBP1;refGene_tx=XM_019819975.2;refGene_gn=PMFBP1;refGene_tx=XM_019819976.2;refGene_gn=PMFBP1;refGene_tx=XM_019819978.2

E2	49379896	49379981	refGene_gn=DHX38;refGene_tx=XM_003998209.5
E2	49380688	49380705	refGene_gn=PMFBP1;refGene_tx=XM_011290241.3
E2	49381832	49381954	refGene_gn=DHX38;refGene_tx=XM_003998209.5
E2	49383376	49383472	refGene_gn=DHX38;refGene_tx=XM_003998209.5
E2	49384213	49384339	refGene_gn=DHX38;refGene_tx=XM_003998209.5
E2	49384540	49384675	refGene_gn=DHX38;refGene_tx=XM_003998209.5
E2	49384771	49384927	refGene_gn=DHX38;refGene_tx=XM_003998209.5
E2	49385355	49385510	refGene_gn=DHX38;refGene_tx=XM_003998209.5
E2	49385634	49385843	refGene_gn=DHX38;refGene_tx=XM_003998209.5
E2	49386570	49386683	refGene_gn=DHX38;refGene_tx=XM_003998209.5
E2	49387231	49387339	refGene_gn=DHX38;refGene_tx=XM_003998209.5
E2	49387522	49387639	refGene_gn=DHX38;refGene_tx=XM_003998209.5
E2	49387734	49387845	refGene_gn=DHX38;refGene_tx=XM_003998209.5
E2	49388194	49388335	refGene_gn=DHX38;refGene_tx=XM_003998209.5
E2	49388720	49388906	refGene_gn=DHX38;refGene_tx=XM_003998209.5
E2	49389041	49389228	refGene_gn=DHX38;refGene_tx=XM_003998209.5
E2	49389839	49389977	refGene_gn=DHX38;refGene_tx=XM_003998209.5
E2	49391535	49391648	refGene_gn=DHX38;refGene_tx=XM_003998209.5
E2	49391995	49392103	refGene_gn=DHX38;refGene_tx=XM_003998209.5
E2	49392293	49392455	refGene_gn=DHX38;refGene_tx=XM_003998209.5
E2	49392971	49393127	refGene_gn=DHX38;refGene_tx=XM_003998209.5
E2	49393593	49393670	refGene_gn=DHX38;refGene_tx=XM_003998209.5
E2	49393820	49393939	refGene_gn=DHX38;refGene_tx=XM_003998209.5
E2	49394093	49394241	refGene_gn=DHX38;refGene_tx=XM_003998209.5
E2	49395053	49395155	refGene_gn=DHX38;refGene_tx=XM_003998209.5
E2	49395802	49395990	refGene_gn=DHX38;refGene_tx=XM_003998209.5
E2	49396338	49396661	refGene_gn=DHX38;refGene_tx=XM_003998209.5
E2	49402416	49402548
refGene_gn=TXNL4B;refGene_tx=XM_003998210.5;refGene_gn=TXNL4B;refGene_tx=XM_006941679.3;refGene_gn=TXNL4B;refGene_tx=XM_006941681.4;refGene_gn=TXNL4B;refGene_tx=XM_019819983.1;refGene_gn=TXNL4B;refGene_tx=XM_023245663.1;refGene_gn=TXNL4B;refGene_tx=XM_023245664.1
E2	49403798	49403950
refGene_gn=TXNL4B;refGene_tx=XM_003998210.5;refGene_gn=TXNL4B;refGene_tx=XM_006941679.3;refGene_gn=TXNL4B;refGene_tx=XM_006941681.4;refGene_gn=TXNL4B;refGene_tx=XM_019819983.1;refGene_gn=TXNL4B;refGene_tx=XM_023245663.1;refGene_gn=TXNL4B;refGene_tx=XM_023245664.1
E2	49405717	49405883
refGene_gn=TXNL4B;refGene_tx=XM_003998210.5;refGene_gn=TXNL4B;refGene_tx=XM_006941679.3;refGene_gn=TXNL4B;refGene_tx=XM_006941681.4;refGene_gn=TXNL4B;refGene_tx=XM_023245663.1;refGene_gn=TXNL4B;refGene_tx=XM_023245664.1
E2	49409634	49410413	refGene_gn=HP;refGene_tx=NM_001198928.2
E2	49411325	49411400	refGene_gn=HP;refGene_tx=NM_001198928.2
E2	49412144	49412246	refGene_gn=HP;refGene_tx=NM_001198928.2
E2	49412553	49412636	refGene_gn=HP;refGene_tx=NM_001198928.2
E2	49413769	49413774	refGene_gn=HP;refGene_tx=NM_001198928.2
E2	49441270	49441376	refGene_gn=TXNL4B;refGene_tx=XM_019819983.1
E2	49446933	49447031
refGene_gn=DHODH;refGene_tx=XM_003998211.3;refGene_gn=DHODH;refGene_tx

=XM_019819981.1
E2 49447512 49447672 refGene_gn=DHODH;refGene_tx=XM_003998211.3
E2 49447845 49447999
refGene_gn=DHODH;refGene_tx=XM_003998211.3;refGene_gn=DHODH;refGene_tx
=XM_019819981.1
E2 49448882 49448996
refGene_gn=DHODH;refGene_tx=XM_003998211.3;refGene_gn=DHODH;refGene_tx
=XM_019819981.1
E2 49449662 49449850
refGene_gn=DHODH;refGene_tx=XM_003998211.3;refGene_gn=DHODH;refGene_tx
=XM_019819981.1
E2 49452472 49452555
refGene_gn=DHODH;refGene_tx=XM_003998211.3;refGene_gn=DHODH;refGene_tx
=XM_019819981.1
E2 49454509 49454709
refGene_gn=DHODH;refGene_tx=XM_003998211.3;refGene_gn=DHODH;refGene_tx
=XM_019819981.1
E2 49454829 49455042
refGene_gn=DHODH;refGene_tx=XM_003998211.3;refGene_gn=DHODH;refGene_tx
=XM_019819981.1
E2 49457135 49457156
refGene_gn=DHODH;refGene_tx=XM_003998211.3;refGene_gn=DHODH;refGene_tx
=XM_019819981.1
E2 49459144 49459339 refGene_gn=PKD1L3;refGene_tx=XM_023245121.1
E2 49459340 49459441 refGene_gn=PKD1L3;refGene_tx=XM_023245121.1
E2 49474386 49474543 refGene_gn=PKD1L3;refGene_tx=XM_023245121.1
E2 49478309 49478441 refGene_gn=PKD1L3;refGene_tx=XM_023245121.1
E2 49482031 49482202 refGene_gn=PKD1L3;refGene_tx=XM_023245121.1
E2 49483863 49483991 refGene_gn=PKD1L3;refGene_tx=XM_023245121.1
E2 49485098 49485181 refGene_gn=PKD1L3;refGene_tx=XM_023245121.1
E2 49485183 49485218 refGene_gn=PKD1L3;refGene_tx=XM_023245121.1
E2 49485527 49485759 refGene_gn=PKD1L3;refGene_tx=XM_023245121.1
E2 49487489 49487683 refGene_gn=PKD1L3;refGene_tx=XM_023245121.1
E2 49490707 49490746 refGene_gn=PKD1L3;refGene_tx=XM_023245121.1
E2 49490748 49490949 refGene_gn=PKD1L3;refGene_tx=XM_023245121.1
E2 49491934 49491972 refGene_gn=PKD1L3;refGene_tx=XM_023245121.1
E2 49491973 49492049 refGene_gn=PKD1L3;refGene_tx=XM_023245121.1
E2 49492300 49492372 refGene_gn=PKD1L3;refGene_tx=XM_023245121.1
E2 49492373 49492483 refGene_gn=PKD1L3;refGene_tx=XM_023245121.1
E2 49496324 49496471 refGene_gn=PKD1L3;refGene_tx=XM_023245121.1
E2 49501210 49501267 refGene_gn=PKD1L3;refGene_tx=XM_023245121.1
E2 49501268 49501387 refGene_gn=PKD1L3;refGene_tx=XM_023245121.1
E2 49501904 49501915 refGene_gn=PKD1L3;refGene_tx=XM_023245121.1
E2 49503105 49503279 refGene_gn=PKD1L3;refGene_tx=XM_023245121.1
E2 49503899 49504117 refGene_gn=PKD1L3;refGene_tx=XM_023245121.1
E2 49504219 49504454 refGene_gn=PKD1L3;refGene_tx=XM_023245121.1
E2 49511033 49511133 refGene_gn=PKD1L3;refGene_tx=XM_023245121.1
E2 49513380 49513518 refGene_gn=PKD1L3;refGene_tx=XM_023245121.1
E2 49514475 49514622 refGene_gn=PKD1L3;refGene_tx=XM_023245121.1
E2 49514624 49514657 refGene_gn=PKD1L3;refGene_tx=XM_023245121.1

E2 49528791 49528912 refGene_gn=PKD1L3;refGene_tx=XM_023245121.1
E2 49531193 49531327 refGene_gn=PKD1L3;refGene_tx=XM_023245121.1
E2 49533093 49533101 refGene_gn=PKD1L3;refGene_tx=XM_023245121.1
E2 49534595 49534767 refGene_gn=PKD1L3;refGene_tx=XM_023245121.1
E2 49534769 49534796 refGene_gn=PKD1L3;refGene_tx=XM_023245121.1
E2 49534798 49534863 refGene_gn=PKD1L3;refGene_tx=XM_023245121.1
E2 49535472 49535552 refGene_gn=PKD1L3;refGene_tx=XM_023245121.1
E2 49535553 49535631 refGene_gn=PKD1L3;refGene_tx=XM_023245121.1
E2 49536835 49537037 refGene_gn=PKD1L3;refGene_tx=XM_023245121.1
E2 49537532 49537634 refGene_gn=PKD1L3;refGene_tx=XM_023245121.1
E2 49539746 49539832 refGene_gn=PKD1L3;refGene_tx=XM_023245121.1
E2 49539833 49539872 refGene_gn=PKD1L3;refGene_tx=XM_023245121.1
E2 49540019 49540116 refGene_gn=PKD1L3;refGene_tx=XM_023245121.1
E2 49540245 49540320 refGene_gn=PKD1L3;refGene_tx=XM_023245121.1
E2 49542527 49542727
refGene_gn=IST1;refGene_tx=XM_023245042.1;refGene_gn=IST1;refGene_tx=XM_023245043.1;refGene_gn=IST1;refGene_tx=XM_023245044.1;refGene_gn=IST1;refGene_tx=XM_023245045.1
E2 49544160 49544223
refGene_gn=IST1;refGene_tx=XM_023245042.1;refGene_gn=IST1;refGene_tx=XM_023245043.1;refGene_gn=IST1;refGene_tx=XM_023245044.1;refGene_gn=IST1;refGene_tx=XM_023245045.1
E2 49545311 49545404
refGene_gn=IST1;refGene_tx=XM_023245042.1;refGene_gn=IST1;refGene_tx=XM_023245043.1;refGene_gn=IST1;refGene_tx=XM_023245044.1;refGene_gn=IST1;refGene_tx=XM_023245045.1
E2 49546042 49546249
refGene_gn=IST1;refGene_tx=XM_023245042.1;refGene_gn=IST1;refGene_tx=XM_023245043.1;refGene_gn=IST1;refGene_tx=XM_023245044.1;refGene_gn=IST1;refGene_tx=XM_023245045.1
E2 49547140 49547251
refGene_gn=IST1;refGene_tx=XM_023245042.1;refGene_gn=IST1;refGene_tx=XM_023245043.1;refGene_gn=IST1;refGene_tx=XM_023245044.1;refGene_gn=IST1;refGene_tx=XM_023245045.1
E2 49547759 49547843
refGene_gn=IST1;refGene_tx=XM_023245042.1;refGene_gn=IST1;refGene_tx=XM_023245043.1;refGene_gn=IST1;refGene_tx=XM_023245044.1;refGene_gn=IST1;refGene_tx=XM_023245045.1
E2 49550399 49550487
refGene_gn=IST1;refGene_tx=XM_023245042.1;refGene_gn=IST1;refGene_tx=XM_023245043.1;refGene_gn=IST1;refGene_tx=XM_023245044.1;refGene_gn=IST1;refGene_tx=XM_023245045.1
E2 49551670 49551851
refGene_gn=IST1;refGene_tx=XM_023245042.1;refGene_gn=IST1;refGene_tx=XM_023245043.1;refGene_gn=IST1;refGene_tx=XM_023245044.1;refGene_gn=IST1;refGene_tx=XM_023245045.1
E2 49552261 49552349
refGene_gn=IST1;refGene_tx=XM_023245042.1;refGene_gn=IST1;refGene_tx=XM_023245043.1;refGene_gn=IST1;refGene_tx=XM_023245044.1;refGene_gn=IST1;refGene_tx=XM_023245045.1

E2 49589655 49589695
refGene_gn=ZNF821;refGene_tx=XM_011290228.3;refGene_gn=ZNF821;refGene_tx=XM_011290229.3;refGene_gn=ZNF821;refGene_tx=XM_011290230.3
E2 49597372 49597498
refGene_gn=ZNF821;refGene_tx=XM_011290228.3;refGene_gn=ZNF821;refGene_tx=XM_011290229.3;refGene_gn=ZNF821;refGene_tx=XM_011290230.3;refGene_gn=ZNF821;refGene_tx=XM_011290232.3
E2 49600156 49600302
refGene_gn=ZNF821;refGene_tx=XM_011290228.3;refGene_gn=ZNF821;refGene_tx=XM_011290229.3;refGene_gn=ZNF821;refGene_tx=XM_011290230.3;refGene_gn=ZNF821;refGene_tx=XM_011290232.3
E2 49600469 49600484 refGene_gn=ZNF821;refGene_tx=XM_019819989.1
E2 49600938 49601043
refGene_gn=ZNF821;refGene_tx=XM_011290228.3;refGene_gn=ZNF821;refGene_tx=XM_011290229.3;refGene_gn=ZNF821;refGene_tx=XM_011290230.3;refGene_gn=ZNF821;refGene_tx=XM_011290232.3;refGene_gn=ZNF821;refGene_tx=XM_019819989.1
E2 49602017 49602184
refGene_gn=ZNF821;refGene_tx=XM_011290228.3;refGene_gn=ZNF821;refGene_tx=XM_011290229.3;refGene_gn=ZNF821;refGene_tx=XM_011290230.3;refGene_gn=ZNF821;refGene_tx=XM_011290232.3;refGene_gn=ZNF821;refGene_tx=XM_019819989.1;refGene_gn=ZNF821;refGene_tx=XM_019819990.1;refGene_gn=ZNF821;refGene_tx=XM_019819991.1
E2 49602959 49603614
refGene_gn=ZNF821;refGene_tx=XM_011290228.3;refGene_gn=ZNF821;refGene_tx=XM_011290229.3;refGene_gn=ZNF821;refGene_tx=XM_011290230.3;refGene_gn=ZNF821;refGene_tx=XM_011290232.3;refGene_gn=ZNF821;refGene_tx=XM_019819989.1;refGene_gn=ZNF821;refGene_tx=XM_019819990.1;refGene_gn=ZNF821;refGene_tx=XM_019819991.1
E2 49612324 49614391
refGene_gn=ATXN1L;refGene_tx=XM_006941688.4;refGene_gn=ATXN1L;refGene_tx=XM_011290234.3
E2 49670845 49671046
refGene_gn=AP1G1;refGene_tx=XM_003998215.5;refGene_gn=AP1G1;refGene_tx=XM_011290227.3
E2 49683118 49683243
refGene_gn=AP1G1;refGene_tx=XM_003998215.5;refGene_gn=AP1G1;refGene_tx=XM_011290227.3
E2 49692076 49692218
refGene_gn=AP1G1;refGene_tx=XM_003998215.5;refGene_gn=AP1G1;refGene_tx=XM_011290227.3
E2 49693743 49693840
refGene_gn=AP1G1;refGene_tx=XM_003998215.5;refGene_gn=AP1G1;refGene_tx=XM_011290227.3
E2 49695041 49695118
refGene_gn=AP1G1;refGene_tx=XM_003998215.5;refGene_gn=AP1G1;refGene_tx=XM_011290227.3
E2 49696015 49696024 refGene_gn=AP1G1;refGene_tx=XM_003998215.5
E2 49698345 49698441
refGene_gn=AP1G1;refGene_tx=XM_003998215.5;refGene_gn=AP1G1;refGene_tx

=XM_011290227.3
E2	49698909	49698990
refGene_gn=AP1G1;refGene_tx=XM_003998215.5;refGene_gn=AP1G1;refGene_tx
=XM_011290227.3
E2	49699195	49699294
refGene_gn=AP1G1;refGene_tx=XM_003998215.5;refGene_gn=AP1G1;refGene_tx
=XM_011290227.3
E2	49701800	49701856
refGene_gn=AP1G1;refGene_tx=XM_003998215.5;refGene_gn=AP1G1;refGene_tx
=XM_011290227.3
E2	49703443	49703557
refGene_gn=AP1G1;refGene_tx=XM_003998215.5;refGene_gn=AP1G1;refGene_tx
=XM_011290227.3
E2	49706608	49706749
refGene_gn=AP1G1;refGene_tx=XM_003998215.5;refGene_gn=AP1G1;refGene_tx
=XM_011290227.3
E2	49707550	49707605
refGene_gn=AP1G1;refGene_tx=XM_003998215.5;refGene_gn=AP1G1;refGene_tx
=XM_011290227.3
E2	49711635	49711758
refGene_gn=AP1G1;refGene_tx=XM_003998215.5;refGene_gn=AP1G1;refGene_tx
=XM_011290227.3
E2	49711923	49712013
refGene_gn=AP1G1;refGene_tx=XM_003998215.5;refGene_gn=AP1G1;refGene_tx
=XM_011290227.3
E2	49712614	49712742
refGene_gn=AP1G1;refGene_tx=XM_003998215.5;refGene_gn=AP1G1;refGene_tx
=XM_011290227.3
E2	49713682	49713787
refGene_gn=AP1G1;refGene_tx=XM_003998215.5;refGene_gn=AP1G1;refGene_tx
=XM_011290227.3
E2	49714431	49714573
refGene_gn=AP1G1;refGene_tx=XM_003998215.5;refGene_gn=AP1G1;refGene_tx
=XM_011290227.3
E2	49714771	49714898
refGene_gn=AP1G1;refGene_tx=XM_003998215.5;refGene_gn=AP1G1;refGene_tx
=XM_011290227.3
E2	49718792	49718900
refGene_gn=AP1G1;refGene_tx=XM_003998215.5;refGene_gn=AP1G1;refGene_tx
=XM_011290227.3
E2	49719007	49719168
refGene_gn=AP1G1;refGene_tx=XM_003998215.5;refGene_gn=AP1G1;refGene_tx
=XM_011290227.3
E2	49723957	49724056
refGene_gn=AP1G1;refGene_tx=XM_003998215.5;refGene_gn=AP1G1;refGene_tx
=XM_011290227.3
E2	49725492	49725594
refGene_gn=AP1G1;refGene_tx=XM_003998215.5;refGene_gn=AP1G1;refGene_tx
=XM_011290227.3
E2	49733796	49733895	refGene_gn=PHLPP2;refGene_tx=XM_023246184.1

E2 49734671 49734758
refGene_gn=PHLPP2;refGene_tx=XM_003998216.5;refGene_gn=PHLPP2;refGene_tx=XM_019819992.2
E2 49741150 49741413
refGene_gn=PHLPP2;refGene_tx=XM_003998216.5;refGene_gn=PHLPP2;refGene_tx=XM_019819992.2;refGene_gn=PHLPP2;refGene_tx=XM_023246184.1
E2 49753021 49753155
refGene_gn=PHLPP2;refGene_tx=XM_003998216.5;refGene_gn=PHLPP2;refGene_tx=XM_019819992.2;refGene_gn=PHLPP2;refGene_tx=XM_023246184.1;refGene_gn=PHLPP2;refGene_tx=XM_023246186.1
E2 49768325 49768516
refGene_gn=PHLPP2;refGene_tx=XM_003998216.5;refGene_gn=PHLPP2;refGene_tx=XM_019819992.2;refGene_gn=PHLPP2;refGene_tx=XM_023246184.1;refGene_gn=PHLPP2;refGene_tx=XM_023246186.1
E2 49775643 49775727 refGene_gn=PHLPP2;refGene_tx=XM_019819993.2
E2 49778802 49778928
refGene_gn=PHLPP2;refGene_tx=XM_003998216.5;refGene_gn=PHLPP2;refGene_tx=XM_019819992.2;refGene_gn=PHLPP2;refGene_tx=XM_019819993.2;refGene_gn=PHLPP2;refGene_tx=XM_023246184.1;refGene_gn=PHLPP2;refGene_tx=XM_023246186.1
E2 49780239 49780394
refGene_gn=PHLPP2;refGene_tx=XM_003998216.5;refGene_gn=PHLPP2;refGene_tx=XM_019819992.2;refGene_gn=PHLPP2;refGene_tx=XM_019819993.2;refGene_gn=PHLPP2;refGene_tx=XM_023246184.1;refGene_gn=PHLPP2;refGene_tx=XM_023246186.1
E2 49782958 49783105
refGene_gn=PHLPP2;refGene_tx=XM_003998216.5;refGene_gn=PHLPP2;refGene_tx=XM_019819992.2;refGene_gn=PHLPP2;refGene_tx=XM_019819993.2;refGene_gn=PHLPP2;refGene_tx=XM_023246184.1;refGene_gn=PHLPP2;refGene_tx=XM_023246186.1
E2 49783483 49783714
refGene_gn=PHLPP2;refGene_tx=XM_003998216.5;refGene_gn=PHLPP2;refGene_tx=XM_019819992.2;refGene_gn=PHLPP2;refGene_tx=XM_019819993.2;refGene_gn=PHLPP2;refGene_tx=XM_023246184.1;refGene_gn=PHLPP2;refGene_tx=XM_023246186.1
E2 49786581 49786784
refGene_gn=PHLPP2;refGene_tx=XM_003998216.5;refGene_gn=PHLPP2;refGene_tx=XM_019819992.2;refGene_gn=PHLPP2;refGene_tx=XM_019819993.2;refGene_gn=PHLPP2;refGene_tx=XM_023246184.1;refGene_gn=PHLPP2;refGene_tx=XM_023246186.1
E2 49790196 49790257
refGene_gn=PHLPP2;refGene_tx=XM_003998216.5;refGene_gn=PHLPP2;refGene_tx=XM_019819992.2;refGene_gn=PHLPP2;refGene_tx=XM_019819993.2;refGene_gn=PHLPP2;refGene_tx=XM_023246184.1;refGene_gn=PHLPP2;refGene_tx=XM_023246186.1
E2 49793955 49794051
refGene_gn=PHLPP2;refGene_tx=XM_003998216.5;refGene_gn=PHLPP2;refGene_tx=XM_019819992.2;refGene_gn=PHLPP2;refGene_tx=XM_019819993.2;refGene_gn=PHLPP2;refGene_tx=XM_023246184.1;refGene_gn=PHLPP2;refGene_tx=XM_023246186.1

E2 49796040 49796196
refGene_gn=PHLPP2;refGene_tx=XM_003998216.5;refGene_gn=PHLPP2;refGene_tx=XM_019819992.2;refGene_gn=PHLPP2;refGene_tx=XM_019819993.2;refGene_gn=PHLPP2;refGene_tx=XM_023246184.1;refGene_gn=PHLPP2;refGene_tx=XM_023246186.1
E2 49799173 49799374
refGene_gn=PHLPP2;refGene_tx=XM_003998216.5;refGene_gn=PHLPP2;refGene_tx=XM_019819992.2;refGene_gn=PHLPP2;refGene_tx=XM_019819993.2;refGene_gn=PHLPP2;refGene_tx=XM_023246184.1;refGene_gn=PHLPP2;refGene_tx=XM_023246186.1
E2 49803238 49803401
refGene_gn=PHLPP2;refGene_tx=XM_003998216.5;refGene_gn=PHLPP2;refGene_tx=XM_019819992.2;refGene_gn=PHLPP2;refGene_tx=XM_019819993.2;refGene_gn=PHLPP2;refGene_tx=XM_023246184.1;refGene_gn=PHLPP2;refGene_tx=XM_023246186.1
E2 49803622 49803753
refGene_gn=PHLPP2;refGene_tx=XM_003998216.5;refGene_gn=PHLPP2;refGene_tx=XM_019819992.2;refGene_gn=PHLPP2;refGene_tx=XM_019819993.2;refGene_gn=PHLPP2;refGene_tx=XM_023246184.1;refGene_gn=PHLPP2;refGene_tx=XM_023246186.1
E2 49804381 49804492
refGene_gn=PHLPP2;refGene_tx=XM_003998216.5;refGene_gn=PHLPP2;refGene_tx=XM_019819992.2;refGene_gn=PHLPP2;refGene_tx=XM_019819993.2;refGene_gn=PHLPP2;refGene_tx=XM_023246184.1;refGene_gn=PHLPP2;refGene_tx=XM_023246186.1
E2 49805314 49805509
refGene_gn=PHLPP2;refGene_tx=XM_003998216.5;refGene_gn=PHLPP2;refGene_tx=XM_019819992.2;refGene_gn=PHLPP2;refGene_tx=XM_019819993.2;refGene_gn=PHLPP2;refGene_tx=XM_023246184.1;refGene_gn=PHLPP2;refGene_tx=XM_023246186.1
E2 49808423 49808655
refGene_gn=PHLPP2;refGene_tx=XM_003998216.5;refGene_gn=PHLPP2;refGene_tx=XM_019819992.2;refGene_gn=PHLPP2;refGene_tx=XM_019819993.2;refGene_gn=PHLPP2;refGene_tx=XM_023246184.1;refGene_gn=PHLPP2;refGene_tx=XM_023246186.1
E2 49811658 49812813
refGene_gn=PHLPP2;refGene_tx=XM_003998216.5;refGene_gn=PHLPP2;refGene_tx=XM_019819992.2;refGene_gn=PHLPP2;refGene_tx=XM_019819993.2;refGene_gn=PHLPP2;refGene_tx=XM_023246184.1;refGene_gn=PHLPP2;refGene_tx=XM_023246186.1
E2 49819643 49820281 refGene_gn=MARVELD3;refGene_tx=XM_019819996.2
E2 49825834 49826445 refGene_gn=MARVELD3;refGene_tx=XM_003998228.4
E2 49831302 49831430
refGene_gn=MARVELD3;refGene_tx=XM_003998228.4;refGene_gn=MARVELD3;refGene_tx=XM_019819996.2
E2 49834354 49834740
refGene_gn=MARVELD3;refGene_tx=XM_003998228.4;refGene_gn=MARVELD3;refGene_tx=XM_019819996.2
E2 49839332 49839482
refGene_gn=TAT;refGene_tx=XM_019819994.2;refGene_gn=TAT;refGene_tx=XM_

019819995.2
E2	49874491 49874738
refGene_gn=TAT;refGene_tx=XM_006941692.3;refGene_gn=TAT;refGene_tx=XM_019819994.2;refGene_gn=TAT;refGene_tx=XM_019819995.2
E2	49874891 49874996
refGene_gn=TAT;refGene_tx=XM_006941692.3;refGene_gn=TAT;refGene_tx=XM_019819994.2;refGene_gn=TAT;refGene_tx=XM_019819995.2
E2	49877247 49877315
refGene_gn=TAT;refGene_tx=XM_006941692.3;refGene_gn=TAT;refGene_tx=XM_019819994.2;refGene_gn=TAT;refGene_tx=XM_019819995.2
E2	49878432 49878591
refGene_gn=TAT;refGene_tx=XM_006941692.3;refGene_gn=TAT;refGene_tx=XM_019819994.2;refGene_gn=TAT;refGene_tx=XM_019819995.2
E2	49878791 49878930
refGene_gn=TAT;refGene_tx=XM_006941692.3;refGene_gn=TAT;refGene_tx=XM_019819994.2;refGene_gn=TAT;refGene_tx=XM_019819995.2
E2	49879496 49879549
refGene_gn=TAT;refGene_tx=XM_006941692.3;refGene_gn=TAT;refGene_tx=XM_019819994.2;refGene_gn=TAT;refGene_tx=XM_019819995.2
E2	49880340 49880493
refGene_gn=TAT;refGene_tx=XM_006941692.3;refGene_gn=TAT;refGene_tx=XM_019819994.2;refGene_gn=TAT;refGene_tx=XM_019819995.2
E2	49881111 49881240
refGene_gn=TAT;refGene_tx=XM_006941692.3;refGene_gn=TAT;refGene_tx=XM_019819994.2;refGene_gn=TAT;refGene_tx=XM_019819995.2
E2	49881579 49881663
refGene_gn=TAT;refGene_tx=XM_006941692.3;refGene_gn=TAT;refGene_tx=XM_019819994.2;refGene_gn=TAT;refGene_tx=XM_019819995.2
E2	49882239 49882338
refGene_gn=TAT;refGene_tx=XM_006941692.3;refGene_gn=TAT;refGene_tx=XM_019819994.2;refGene_gn=TAT;refGene_tx=XM_019819995.2
E2	49883193 49883334
refGene_gn=TAT;refGene_tx=XM_006941692.3;refGene_gn=TAT;refGene_tx=XM_019819994.2;refGene_gn=TAT;refGene_tx=XM_019819995.2
E2	49909594 49910731 refGene_gn=CHST4;refGene_tx=XM_011290226.3
E2	49979245 49979278
refGene_gn=LOC101081348;refGene_tx=XM_019820023.1
E2	49982007 49982026
refGene_gn=LOC101081348;refGene_tx=XM_019820023.1
E2	49982028 49982133
refGene_gn=LOC101081348;refGene_tx=XM_019820023.1
E2	49982690 49982804
refGene_gn=LOC101081348;refGene_tx=XM_019820023.1
E2	49985911 49986678
refGene_gn=LOC101081348;refGene_tx=XM_019820023.1
E2	49988102 49988197 refGene_gn=ZNF19;refGene_tx=XM_023245787.1
E2	49989729 49989794
refGene_gn=ZNF19;refGene_tx=XM_006941705.3;refGene_gn=ZNF19;refGene_tx=XM_023245784.1;refGene_gn=ZNF19;refGene_tx=XM_023245785.1;refGene_gn=ZNF19;refGene_tx=XM_023245786.1;refGene_gn=ZNF19;refGene_tx=XM_0232457

87.1
E2	49992965 49993092
refGene_gn=ZNF19;refGene_tx=XM_006941705.3;refGene_gn=ZNF19;refGene_tx=XM_023245784.1;refGene_gn=ZNF19;refGene_tx=XM_023245785.1;refGene_gn=ZNF19;refGene_tx=XM_023245786.1;refGene_gn=ZNF19;refGene_tx=XM_023245788.1;refGene_gn=ZNF19;refGene_tx=XM_023245789.1;refGene_gn=ZNF19;refGene_tx=XM_023245790.1
E2	49993235 49993251 refGene_gn=ZNF19;refGene_tx=XM_023245791.1
E2	49993638 49993752
refGene_gn=ZNF19;refGene_tx=XM_006941705.3;refGene_gn=ZNF19;refGene_tx=XM_023245784.1;refGene_gn=ZNF19;refGene_tx=XM_023245785.1;refGene_gn=ZNF19;refGene_tx=XM_023245786.1;refGene_gn=ZNF19;refGene_tx=XM_023245787.1;refGene_gn=ZNF19;refGene_tx=XM_023245788.1;refGene_gn=ZNF19;refGene_tx=XM_023245789.1;refGene_gn=ZNF19;refGene_tx=XM_023245790.1;refGene_gn=ZNF19;refGene_tx=XM_023245791.1
E2	49997578 49998681
refGene_gn=ZNF19;refGene_tx=XM_006941705.3;refGene_gn=ZNF19;refGene_tx=XM_023245784.1;refGene_gn=ZNF19;refGene_tx=XM_023245785.1;refGene_gn=ZNF19;refGene_tx=XM_023245786.1;refGene_gn=ZNF19;refGene_tx=XM_023245787.1;refGene_gn=ZNF19;refGene_tx=XM_023245788.1;refGene_gn=ZNF19;refGene_tx=XM_023245789.1;refGene_gn=ZNF19;refGene_tx=XM_023245790.1;refGene_gn=ZNF19;refGene_tx=XM_023245791.1
E2	50019114 50019207 refGene_gn=ZNF23;refGene_tx=XM_011290222.3
E2	50021361 50021488 refGene_gn=ZNF23;refGene_tx=XM_011290222.3
E2	50022082 50022196 refGene_gn=ZNF23;refGene_tx=XM_011290222.3
E2	50027319 50027678 refGene_gn=ZNF23;refGene_tx=XM_011290222.3
E2	50027679 50028204 refGene_gn=ZNF23;refGene_tx=XM_011290222.3
E2	50028205 50028457 refGene_gn=ZNF23;refGene_tx=XM_011290222.3
E2	50028458 50029115 refGene_gn=ZNF23;refGene_tx=XM_011290222.3
E2	50041045 50041356
refGene_gn=LOC105261272;refGene_tx=XM_019820019.2;refGene_gn=LOC105261272;refGene_tx=XM_019820020.2;refGene_gn=LOC105261272;refGene_tx=XM_019820021.2
E2	50041457 50041480
refGene_gn=LOC105261272;refGene_tx=XM_019820019.2;refGene_gn=LOC105261272;refGene_tx=XM_019820020.2;refGene_gn=LOC105261272;refGene_tx=XM_019820021.2
E2	50041631 50041690
refGene_gn=LOC105261272;refGene_tx=XM_019820019.2;refGene_gn=LOC105261272;refGene_tx=XM_019820020.2;refGene_gn=LOC105261272;refGene_tx=XM_019820021.2
E2	50042207 50042420
refGene_gn=LOC105261272;refGene_tx=XM_019820019.2;refGene_gn=LOC105261272;refGene_tx=XM_019820020.2;refGene_gn=LOC105261272;refGene_tx=XM_019820021.2
E2	50042528 50042770
refGene_gn=LOC105261272;refGene_tx=XM_019820019.2;refGene_gn=LOC105261272;refGene_tx=XM_019820020.2;refGene_gn=LOC105261272;refGene_tx=XM_019820021.2
E2	50042976 50043118

refGene_gn=LOC105261272;refGene_tx=XM_019820019.2;refGene_gn=LOC105261
272;refGene_tx=XM_019820020.2;refGene_gn=LOC105261272;refGene_tx=XM_01
9820021.2
E2 50043504 50043658
refGene_gn=LOC105261272;refGene_tx=XM_019820019.2;refGene_gn=LOC105261
272;refGene_tx=XM_019820020.2;refGene_gn=LOC105261272;refGene_tx=XM_01
9820021.2
E2 50044217 50044291
refGene_gn=LOC105261272;refGene_tx=XM_019820019.2;refGene_gn=LOC105261
272;refGene_tx=XM_019820020.2
E2 50044604 50044956
refGene_gn=LOC105261272;refGene_tx=XM_019820019.2;refGene_gn=LOC105261
272;refGene_tx=XM_019820020.2;refGene_gn=LOC105261272;refGene_tx=XM_01
9820021.2
E2 50075323 50075440 refGene_gn=CALB2;refGene_tx=XM_011290221.2
E2 50079091 50079163 refGene_gn=CALB2;refGene_tx=XM_011290221.2
E2 50079752 50079806 refGene_gn=CALB2;refGene_tx=XM_011290221.2
E2 50080550 50080590 refGene_gn=CALB2;refGene_tx=XM_011290221.2
E2 50080903 50080959 refGene_gn=CALB2;refGene_tx=XM_011290221.2
E2 50081443 50081521 refGene_gn=CALB2;refGene_tx=XM_011290221.2
E2 50082084 50082141 refGene_gn=CALB2;refGene_tx=XM_011290221.2
E2 50085420 50085501 refGene_gn=CALB2;refGene_tx=XM_011290221.2
E2 50088660 50088750 refGene_gn=CALB2;refGene_tx=XM_011290221.2
E2 50091312 50091389 refGene_gn=CALB2;refGene_tx=XM_011290221.2
E2 50103459 50103553 refGene_gn=CALB2;refGene_tx=XM_011290221.2
E2 50185797 50188098
refGene_gn=CMTR2;refGene_tx=XM_003998219.5;refGene_gn=CMTR2;refGene_tx
=XM_019820024.2;refGene_gn=CMTR2;refGene_tx=XM_019820025.2;refGene_gn=
CMTR2;refGene_tx=XM_019820026.2;refGene_gn=CMTR2;refGene_tx=XM_0198200
27.2
E2 50280346 50280481
refGene_gn=HYDIN;refGene_tx=XM_023246143.1;refGene_gn=HYDIN;refGene_tx
=XM_023246144.1;refGene_gn=HYDIN;refGene_tx=XM_023246147.1;refGene_gn=
HYDIN;refGene_tx=XM_023246148.1;refGene_gn=HYDIN;refGene_tx=XM_0232461
49.1
E2 50282283 50282409
refGene_gn=HYDIN;refGene_tx=XM_023246143.1;refGene_gn=HYDIN;refGene_tx
=XM_023246144.1;refGene_gn=HYDIN;refGene_tx=XM_023246145.1;refGene_gn=
HYDIN;refGene_tx=XM_023246147.1;refGene_gn=HYDIN;refGene_tx=XM_0232461
48.1;refGene_gn=HYDIN;refGene_tx=XM_023246149.1
E2 50291758 50291878
refGene_gn=HYDIN;refGene_tx=XM_023246143.1;refGene_gn=HYDIN;refGene_tx
=XM_023246144.1;refGene_gn=HYDIN;refGene_tx=XM_023246145.1;refGene_gn=
HYDIN;refGene_tx=XM_023246147.1;refGene_gn=HYDIN;refGene_tx=XM_0232461
48.1;refGene_gn=HYDIN;refGene_tx=XM_023246149.1
E2 50295207 50295342
refGene_gn=HYDIN;refGene_tx=XM_023246143.1;refGene_gn=HYDIN;refGene_tx
=XM_023246144.1;refGene_gn=HYDIN;refGene_tx=XM_023246145.1;refGene_gn=
HYDIN;refGene_tx=XM_023246146.1;refGene_gn=HYDIN;refGene_tx=XM_0232461
47.1;refGene_gn=HYDIN;refGene_tx=XM_023246148.1;refGene_gn=HYDIN;refGe

ne_tx=XM_023246149.1
E2 50308496 50308696
refGene_gn=HYDIN;refGene_tx=XM_023246143.1;refGene_gn=HYDIN;refGene_tx=XM_023246144.1;refGene_gn=HYDIN;refGene_tx=XM_023246145.1;refGene_gn=HYDIN;refGene_tx=XM_023246146.1;refGene_gn=HYDIN;refGene_tx=XM_023246147.1;refGene_gn=HYDIN;refGene_tx=XM_023246148.1;refGene_gn=HYDIN;refGene_tx=XM_023246149.1
E2 50325128 50325253
refGene_gn=HYDIN;refGene_tx=XM_023246143.1;refGene_gn=HYDIN;refGene_tx=XM_023246144.1;refGene_gn=HYDIN;refGene_tx=XM_023246145.1;refGene_gn=HYDIN;refGene_tx=XM_023246146.1;refGene_gn=HYDIN;refGene_tx=XM_023246147.1;refGene_gn=HYDIN;refGene_tx=XM_023246148.1;refGene_gn=HYDIN;refGene_tx=XM_023246149.1
E2 50344182 50344384
refGene_gn=HYDIN;refGene_tx=XM_023246143.1;refGene_gn=HYDIN;refGene_tx=XM_023246144.1;refGene_gn=HYDIN;refGene_tx=XM_023246145.1;refGene_gn=HYDIN;refGene_tx=XM_023246146.1;refGene_gn=HYDIN;refGene_tx=XM_023246147.1;refGene_gn=HYDIN;refGene_tx=XM_023246148.1;refGene_gn=HYDIN;refGene_tx=XM_023246149.1
E2 50350530 50350714
refGene_gn=HYDIN;refGene_tx=XM_023246143.1;refGene_gn=HYDIN;refGene_tx=XM_023246144.1;refGene_gn=HYDIN;refGene_tx=XM_023246145.1;refGene_gn=HYDIN;refGene_tx=XM_023246146.1;refGene_gn=HYDIN;refGene_tx=XM_023246147.1;refGene_gn=HYDIN;refGene_tx=XM_023246148.1;refGene_gn=HYDIN;refGene_tx=XM_023246149.1
E2 50364227 50364327
refGene_gn=HYDIN;refGene_tx=XM_023246143.1;refGene_gn=HYDIN;refGene_tx=XM_023246144.1;refGene_gn=HYDIN;refGene_tx=XM_023246145.1;refGene_gn=HYDIN;refGene_tx=XM_023246146.1;refGene_gn=HYDIN;refGene_tx=XM_023246147.1;refGene_gn=HYDIN;refGene_tx=XM_023246148.1;refGene_gn=HYDIN;refGene_tx=XM_023246149.1
E2 50383714 50383833
refGene_gn=HYDIN;refGene_tx=XM_023246143.1;refGene_gn=HYDIN;refGene_tx=XM_023246144.1;refGene_gn=HYDIN;refGene_tx=XM_023246145.1;refGene_gn=HYDIN;refGene_tx=XM_023246146.1;refGene_gn=HYDIN;refGene_tx=XM_023246147.1;refGene_gn=HYDIN;refGene_tx=XM_023246148.1;refGene_gn=HYDIN;refGene_tx=XM_023246149.1
E2 50388388 50388612
refGene_gn=HYDIN;refGene_tx=XM_023246143.1;refGene_gn=HYDIN;refGene_tx=XM_023246144.1;refGene_gn=HYDIN;refGene_tx=XM_023246145.1;refGene_gn=HYDIN;refGene_tx=XM_023246146.1;refGene_gn=HYDIN;refGene_tx=XM_023246147.1;refGene_gn=HYDIN;refGene_tx=XM_023246148.1;refGene_gn=HYDIN;refGene_tx=XM_023246149.1
E2 50395958 50396026
refGene_gn=HYDIN;refGene_tx=XM_023246143.1;refGene_gn=HYDIN;refGene_tx=XM_023246144.1;refGene_gn=HYDIN;refGene_tx=XM_023246145.1;refGene_gn=HYDIN;refGene_tx=XM_023246146.1;refGene_gn=HYDIN;refGene_tx=XM_023246147.1;refGene_gn=HYDIN;refGene_tx=XM_023246148.1;refGene_gn=HYDIN;refGene_tx=XM_023246149.1
E2 50401320 50401556

refGene_gn=HYDIN;refGene_tx=XM_023246143.1;refGene_gn=HYDIN;refGene_tx=XM_023246144.1;refGene_gn=HYDIN;refGene_tx=XM_023246145.1;refGene_gn=HYDIN;refGene_tx=XM_023246146.1;refGene_gn=HYDIN;refGene_tx=XM_023246147.1;refGene_gn=HYDIN;refGene_tx=XM_023246148.1;refGene_gn=HYDIN;refGene_tx=XM_023246149.1
E2 50404164 50404265
refGene_gn=HYDIN;refGene_tx=XM_023246143.1;refGene_gn=HYDIN;refGene_tx=XM_023246144.1;refGene_gn=HYDIN;refGene_tx=XM_023246145.1;refGene_gn=HYDIN;refGene_tx=XM_023246146.1;refGene_gn=HYDIN;refGene_tx=XM_023246147.1;refGene_gn=HYDIN;refGene_tx=XM_023246148.1;refGene_gn=HYDIN;refGene_tx=XM_023246149.1
E2 50406884 50407020
refGene_gn=HYDIN;refGene_tx=XM_023246143.1;refGene_gn=HYDIN;refGene_tx=XM_023246144.1;refGene_gn=HYDIN;refGene_tx=XM_023246145.1;refGene_gn=HYDIN;refGene_tx=XM_023246146.1;refGene_gn=HYDIN;refGene_tx=XM_023246147.1;refGene_gn=HYDIN;refGene_tx=XM_023246148.1;refGene_gn=HYDIN;refGene_tx=XM_023246149.1
E2 50409850 50410015
refGene_gn=HYDIN;refGene_tx=XM_023246143.1;refGene_gn=HYDIN;refGene_tx=XM_023246144.1;refGene_gn=HYDIN;refGene_tx=XM_023246145.1;refGene_gn=HYDIN;refGene_tx=XM_023246146.1;refGene_gn=HYDIN;refGene_tx=XM_023246147.1;refGene_gn=HYDIN;refGene_tx=XM_023246148.1;refGene_gn=HYDIN;refGene_tx=XM_023246149.1
E2 50412068 50412221
refGene_gn=HYDIN;refGene_tx=XM_023246143.1;refGene_gn=HYDIN;refGene_tx=XM_023246144.1;refGene_gn=HYDIN;refGene_tx=XM_023246145.1;refGene_gn=HYDIN;refGene_tx=XM_023246146.1;refGene_gn=HYDIN;refGene_tx=XM_023246147.1;refGene_gn=HYDIN;refGene_tx=XM_023246148.1;refGene_gn=HYDIN;refGene_tx=XM_023246149.1
E2 50442425 50442664
refGene_gn=HYDIN;refGene_tx=XM_023246143.1;refGene_gn=HYDIN;refGene_tx=XM_023246144.1;refGene_gn=HYDIN;refGene_tx=XM_023246145.1;refGene_gn=HYDIN;refGene_tx=XM_023246146.1;refGene_gn=HYDIN;refGene_tx=XM_023246147.1;refGene_gn=HYDIN;refGene_tx=XM_023246148.1;refGene_gn=HYDIN;refGene_tx=XM_023246149.1
E2 50445515 50445789
refGene_gn=HYDIN;refGene_tx=XM_023246143.1;refGene_gn=HYDIN;refGene_tx=XM_023246144.1;refGene_gn=HYDIN;refGene_tx=XM_023246145.1;refGene_gn=HYDIN;refGene_tx=XM_023246146.1;refGene_gn=HYDIN;refGene_tx=XM_023246147.1;refGene_gn=HYDIN;refGene_tx=XM_023246148.1;refGene_gn=HYDIN;refGene_tx=XM_023246149.1
E2 50450389 50450533
refGene_gn=HYDIN;refGene_tx=XM_023246143.1;refGene_gn=HYDIN;refGene_tx=XM_023246144.1;refGene_gn=HYDIN;refGene_tx=XM_023246145.1;refGene_gn=HYDIN;refGene_tx=XM_023246146.1;refGene_gn=HYDIN;refGene_tx=XM_023246147.1;refGene_gn=HYDIN;refGene_tx=XM_023246148.1;refGene_gn=HYDIN;refGene_tx=XM_023246149.1
E2 50455386 50455530
refGene_gn=HYDIN;refGene_tx=XM_023246143.1;refGene_gn=HYDIN;refGene_tx=XM_023246144.1;refGene_gn=HYDIN;refGene_tx=XM_023246145.1;refGene_gn=

HYDIN;refGene_tx=XM_023246146.1;refGene_gn=HYDIN;refGene_tx=XM_0232461
47.1;refGene_gn=HYDIN;refGene_tx=XM_023246148.1;refGene_gn=HYDIN;refGe
ne_tx=XM_023246149.1
E2 50461412 50461726
refGene_gn=HYDIN;refGene_tx=XM_023246143.1;refGene_gn=HYDIN;refGene_tx
=XM_023246144.1;refGene_gn=HYDIN;refGene_tx=XM_023246145.1;refGene_gn=
HYDIN;refGene_tx=XM_023246146.1;refGene_gn=HYDIN;refGene_tx=XM_0232461
47.1;refGene_gn=HYDIN;refGene_tx=XM_023246148.1;refGene_gn=HYDIN;refGe
ne_tx=XM_023246149.1
E2 50478703 50478841
refGene_gn=HYDIN;refGene_tx=XM_023246143.1;refGene_gn=HYDIN;refGene_tx
=XM_023246144.1;refGene_gn=HYDIN;refGene_tx=XM_023246145.1;refGene_gn=
HYDIN;refGene_tx=XM_023246146.1;refGene_gn=HYDIN;refGene_tx=XM_0232461
47.1;refGene_gn=HYDIN;refGene_tx=XM_023246148.1;refGene_gn=HYDIN;refGe
ne_tx=XM_023246149.1
E2 50479547 50479629
refGene_gn=HYDIN;refGene_tx=XM_023246143.1;refGene_gn=HYDIN;refGene_tx
=XM_023246144.1;refGene_gn=HYDIN;refGene_tx=XM_023246145.1;refGene_gn=
HYDIN;refGene_tx=XM_023246146.1;refGene_gn=HYDIN;refGene_tx=XM_0232461
47.1;refGene_gn=HYDIN;refGene_tx=XM_023246148.1;refGene_gn=HYDIN;refGe
ne_tx=XM_023246149.1
E2 50482968 50483088
refGene_gn=HYDIN;refGene_tx=XM_023246143.1;refGene_gn=HYDIN;refGene_tx
=XM_023246144.1;refGene_gn=HYDIN;refGene_tx=XM_023246145.1;refGene_gn=
HYDIN;refGene_tx=XM_023246146.1;refGene_gn=HYDIN;refGene_tx=XM_0232461
47.1;refGene_gn=HYDIN;refGene_tx=XM_023246148.1;refGene_gn=HYDIN;refGe
ne_tx=XM_023246149.1
E2 50483355 50483565
refGene_gn=HYDIN;refGene_tx=XM_023246143.1;refGene_gn=HYDIN;refGene_tx
=XM_023246144.1;refGene_gn=HYDIN;refGene_tx=XM_023246145.1;refGene_gn=
HYDIN;refGene_tx=XM_023246146.1;refGene_gn=HYDIN;refGene_tx=XM_0232461
47.1;refGene_gn=HYDIN;refGene_tx=XM_023246148.1;refGene_gn=HYDIN;refGe
ne_tx=XM_023246149.1
E2 50485759 50485897
refGene_gn=HYDIN;refGene_tx=XM_023246143.1;refGene_gn=HYDIN;refGene_tx
=XM_023246144.1;refGene_gn=HYDIN;refGene_tx=XM_023246145.1;refGene_gn=
HYDIN;refGene_tx=XM_023246146.1;refGene_gn=HYDIN;refGene_tx=XM_0232461
47.1;refGene_gn=HYDIN;refGene_tx=XM_023246148.1;refGene_gn=HYDIN;refGe
ne_tx=XM_023246149.1
E2 50487225 50487403
refGene_gn=HYDIN;refGene_tx=XM_023246143.1;refGene_gn=HYDIN;refGene_tx
=XM_023246144.1;refGene_gn=HYDIN;refGene_tx=XM_023246145.1;refGene_gn=
HYDIN;refGene_tx=XM_023246146.1;refGene_gn=HYDIN;refGene_tx=XM_0232461
47.1;refGene_gn=HYDIN;refGene_tx=XM_023246148.1;refGene_gn=HYDIN;refGe
ne_tx=XM_023246149.1
E2 50489113 50489238
refGene_gn=HYDIN;refGene_tx=XM_023246143.1;refGene_gn=HYDIN;refGene_tx
=XM_023246144.1;refGene_gn=HYDIN;refGene_tx=XM_023246145.1;refGene_gn=
HYDIN;refGene_tx=XM_023246146.1;refGene_gn=HYDIN;refGene_tx=XM_0232461
47.1;refGene_gn=HYDIN;refGene_tx=XM_023246148.1;refGene_gn=HYDIN;refGe

ne_tx=XM_023246149.1
E2 50497163 50497291
refGene_gn=HYDIN;refGene_tx=XM_023246143.1;refGene_gn=HYDIN;refGene_tx=XM_023246144.1;refGene_gn=HYDIN;refGene_tx=XM_023246145.1;refGene_gn=HYDIN;refGene_tx=XM_023246146.1;refGene_gn=HYDIN;refGene_tx=XM_023246147.1;refGene_gn=HYDIN;refGene_tx=XM_023246148.1;refGene_gn=HYDIN;refGene_tx=XM_023246149.1
E2 50497733 50497870
refGene_gn=HYDIN;refGene_tx=XM_023246143.1;refGene_gn=HYDIN;refGene_tx=XM_023246144.1;refGene_gn=HYDIN;refGene_tx=XM_023246145.1;refGene_gn=HYDIN;refGene_tx=XM_023246146.1;refGene_gn=HYDIN;refGene_tx=XM_023246147.1;refGene_gn=HYDIN;refGene_tx=XM_023246148.1;refGene_gn=HYDIN;refGene_tx=XM_023246149.1
E2 50498121 50498249
refGene_gn=HYDIN;refGene_tx=XM_023246143.1;refGene_gn=HYDIN;refGene_tx=XM_023246144.1;refGene_gn=HYDIN;refGene_tx=XM_023246145.1;refGene_gn=HYDIN;refGene_tx=XM_023246146.1;refGene_gn=HYDIN;refGene_tx=XM_023246147.1;refGene_gn=HYDIN;refGene_tx=XM_023246148.1;refGene_gn=HYDIN;refGene_tx=XM_023246149.1
E2 50498399 50498591
refGene_gn=HYDIN;refGene_tx=XM_023246143.1;refGene_gn=HYDIN;refGene_tx=XM_023246144.1;refGene_gn=HYDIN;refGene_tx=XM_023246145.1;refGene_gn=HYDIN;refGene_tx=XM_023246146.1;refGene_gn=HYDIN;refGene_tx=XM_023246147.1;refGene_gn=HYDIN;refGene_tx=XM_023246148.1;refGene_gn=HYDIN;refGene_tx=XM_023246149.1
E2 50498926 50499076
refGene_gn=HYDIN;refGene_tx=XM_023246143.1;refGene_gn=HYDIN;refGene_tx=XM_023246144.1;refGene_gn=HYDIN;refGene_tx=XM_023246145.1;refGene_gn=HYDIN;refGene_tx=XM_023246146.1;refGene_gn=HYDIN;refGene_tx=XM_023246147.1;refGene_gn=HYDIN;refGene_tx=XM_023246148.1;refGene_gn=HYDIN;refGene_tx=XM_023246149.1
E2 50502302 50502542
refGene_gn=HYDIN;refGene_tx=XM_023246143.1;refGene_gn=HYDIN;refGene_tx=XM_023246144.1;refGene_gn=HYDIN;refGene_tx=XM_023246145.1;refGene_gn=HYDIN;refGene_tx=XM_023246146.1;refGene_gn=HYDIN;refGene_tx=XM_023246147.1;refGene_gn=HYDIN;refGene_tx=XM_023246148.1;refGene_gn=HYDIN;refGene_tx=XM_023246149.1
E2 50506832 50507001
refGene_gn=HYDIN;refGene_tx=XM_023246143.1;refGene_gn=HYDIN;refGene_tx=XM_023246144.1;refGene_gn=HYDIN;refGene_tx=XM_023246145.1;refGene_gn=HYDIN;refGene_tx=XM_023246146.1;refGene_gn=HYDIN;refGene_tx=XM_023246147.1;refGene_gn=HYDIN;refGene_tx=XM_023246148.1;refGene_gn=HYDIN;refGene_tx=XM_023246149.1
E2 50509682 50509862
refGene_gn=HYDIN;refGene_tx=XM_023246143.1;refGene_gn=HYDIN;refGene_tx=XM_023246144.1;refGene_gn=HYDIN;refGene_tx=XM_023246145.1;refGene_gn=HYDIN;refGene_tx=XM_023246146.1;refGene_gn=HYDIN;refGene_tx=XM_023246147.1;refGene_gn=HYDIN;refGene_tx=XM_023246148.1;refGene_gn=HYDIN;refGene_tx=XM_023246149.1
E2 50510343 50510517

refGene_gn=HYDIN;refGene_tx=XM_023246143.1;refGene_gn=HYDIN;refGene_tx=XM_023246144.1;refGene_gn=HYDIN;refGene_tx=XM_023246145.1;refGene_gn=HYDIN;refGene_tx=XM_023246146.1;refGene_gn=HYDIN;refGene_tx=XM_023246147.1;refGene_gn=HYDIN;refGene_tx=XM_023246148.1;refGene_gn=HYDIN;refGene_tx=XM_023246149.1
E2 50512653 50512827
refGene_gn=HYDIN;refGene_tx=XM_023246143.1;refGene_gn=HYDIN;refGene_tx=XM_023246144.1;refGene_gn=HYDIN;refGene_tx=XM_023246145.1;refGene_gn=HYDIN;refGene_tx=XM_023246146.1;refGene_gn=HYDIN;refGene_tx=XM_023246147.1;refGene_gn=HYDIN;refGene_tx=XM_023246148.1;refGene_gn=HYDIN;refGene_tx=XM_023246149.1
E2 50516646 50516858
refGene_gn=HYDIN;refGene_tx=XM_023246143.1;refGene_gn=HYDIN;refGene_tx=XM_023246144.1;refGene_gn=HYDIN;refGene_tx=XM_023246145.1;refGene_gn=HYDIN;refGene_tx=XM_023246146.1;refGene_gn=HYDIN;refGene_tx=XM_023246147.1;refGene_gn=HYDIN;refGene_tx=XM_023246148.1;refGene_gn=HYDIN;refGene_tx=XM_023246149.1
E2 50521476 50521614
refGene_gn=HYDIN;refGene_tx=XM_023246143.1;refGene_gn=HYDIN;refGene_tx=XM_023246144.1;refGene_gn=HYDIN;refGene_tx=XM_023246145.1;refGene_gn=HYDIN;refGene_tx=XM_023246146.1;refGene_gn=HYDIN;refGene_tx=XM_023246147.1;refGene_gn=HYDIN;refGene_tx=XM_023246148.1;refGene_gn=HYDIN;refGene_tx=XM_023246149.1
E2 50523337 50523521
refGene_gn=HYDIN;refGene_tx=XM_023246143.1;refGene_gn=HYDIN;refGene_tx=XM_023246144.1;refGene_gn=HYDIN;refGene_tx=XM_023246145.1;refGene_gn=HYDIN;refGene_tx=XM_023246146.1;refGene_gn=HYDIN;refGene_tx=XM_023246147.1;refGene_gn=HYDIN;refGene_tx=XM_023246148.1;refGene_gn=HYDIN;refGene_tx=XM_023246149.1
E2 50525818 50525960
refGene_gn=HYDIN;refGene_tx=XM_023246143.1;refGene_gn=HYDIN;refGene_tx=XM_023246144.1;refGene_gn=HYDIN;refGene_tx=XM_023246145.1;refGene_gn=HYDIN;refGene_tx=XM_023246146.1;refGene_gn=HYDIN;refGene_tx=XM_023246147.1;refGene_gn=HYDIN;refGene_tx=XM_023246148.1;refGene_gn=HYDIN;refGene_tx=XM_023246149.1
E2 50528591 50528754
refGene_gn=HYDIN;refGene_tx=XM_023246143.1;refGene_gn=HYDIN;refGene_tx=XM_023246144.1;refGene_gn=HYDIN;refGene_tx=XM_023246145.1;refGene_gn=HYDIN;refGene_tx=XM_023246146.1;refGene_gn=HYDIN;refGene_tx=XM_023246147.1;refGene_gn=HYDIN;refGene_tx=XM_023246148.1;refGene_gn=HYDIN;refGene_tx=XM_023246149.1
E2 50530475 50531099
refGene_gn=HYDIN;refGene_tx=XM_023246143.1;refGene_gn=HYDIN;refGene_tx=XM_023246144.1;refGene_gn=HYDIN;refGene_tx=XM_023246145.1;refGene_gn=HYDIN;refGene_tx=XM_023246146.1;refGene_gn=HYDIN;refGene_tx=XM_023246147.1;refGene_gn=HYDIN;refGene_tx=XM_023246148.1;refGene_gn=HYDIN;refGene_tx=XM_023246149.1
E2 50533086 50533308
refGene_gn=HYDIN;refGene_tx=XM_023246143.1;refGene_gn=HYDIN;refGene_tx=XM_023246144.1;refGene_gn=HYDIN;refGene_tx=XM_023246145.1;refGene_gn=

HYDIN;refGene_tx=XM_023246146.1;refGene_gn=HYDIN;refGene_tx=XM_0232461
47.1;refGene_gn=HYDIN;refGene_tx=XM_023246148.1;refGene_gn=HYDIN;refGe
ne_tx=XM_023246149.1
E2 50536075 50536284
refGene_gn=HYDIN;refGene_tx=XM_023246143.1;refGene_gn=HYDIN;refGene_tx
=XM_023246144.1;refGene_gn=HYDIN;refGene_tx=XM_023246145.1;refGene_gn=
HYDIN;refGene_tx=XM_023246146.1;refGene_gn=HYDIN;refGene_tx=XM_0232461
47.1;refGene_gn=HYDIN;refGene_tx=XM_023246148.1;refGene_gn=HYDIN;refGe
ne_tx=XM_023246149.1
E2 50536504 50536687
refGene_gn=HYDIN;refGene_tx=XM_023246143.1;refGene_gn=HYDIN;refGene_tx
=XM_023246144.1;refGene_gn=HYDIN;refGene_tx=XM_023246145.1;refGene_gn=
HYDIN;refGene_tx=XM_023246146.1;refGene_gn=HYDIN;refGene_tx=XM_0232461
47.1;refGene_gn=HYDIN;refGene_tx=XM_023246148.1;refGene_gn=HYDIN;refGe
ne_tx=XM_023246149.1
E2 50537574 50537694
refGene_gn=HYDIN;refGene_tx=XM_023246143.1;refGene_gn=HYDIN;refGene_tx
=XM_023246144.1;refGene_gn=HYDIN;refGene_tx=XM_023246145.1;refGene_gn=
HYDIN;refGene_tx=XM_023246146.1;refGene_gn=HYDIN;refGene_tx=XM_0232461
47.1;refGene_gn=HYDIN;refGene_tx=XM_023246148.1;refGene_gn=HYDIN;refGe
ne_tx=XM_023246149.1
E2 50540318 50540478
refGene_gn=HYDIN;refGene_tx=XM_023246143.1;refGene_gn=HYDIN;refGene_tx
=XM_023246144.1;refGene_gn=HYDIN;refGene_tx=XM_023246145.1;refGene_gn=
HYDIN;refGene_tx=XM_023246146.1;refGene_gn=HYDIN;refGene_tx=XM_0232461
47.1;refGene_gn=HYDIN;refGene_tx=XM_023246148.1;refGene_gn=HYDIN;refGe
ne_tx=XM_023246149.1
E2 50540582 50540755
refGene_gn=HYDIN;refGene_tx=XM_023246143.1;refGene_gn=HYDIN;refGene_tx
=XM_023246144.1;refGene_gn=HYDIN;refGene_tx=XM_023246145.1;refGene_gn=
HYDIN;refGene_tx=XM_023246146.1;refGene_gn=HYDIN;refGene_tx=XM_0232461
47.1;refGene_gn=HYDIN;refGene_tx=XM_023246148.1;refGene_gn=HYDIN;refGe
ne_tx=XM_023246149.1
E2 50541379 50541578
refGene_gn=HYDIN;refGene_tx=XM_023246143.1;refGene_gn=HYDIN;refGene_tx
=XM_023246144.1;refGene_gn=HYDIN;refGene_tx=XM_023246145.1;refGene_gn=
HYDIN;refGene_tx=XM_023246146.1;refGene_gn=HYDIN;refGene_tx=XM_0232461
47.1;refGene_gn=HYDIN;refGene_tx=XM_023246148.1;refGene_gn=HYDIN;refGe
ne_tx=XM_023246149.1
E2 50544144 50544244
refGene_gn=HYDIN;refGene_tx=XM_023246143.1;refGene_gn=HYDIN;refGene_tx
=XM_023246144.1;refGene_gn=HYDIN;refGene_tx=XM_023246145.1;refGene_gn=
HYDIN;refGene_tx=XM_023246146.1;refGene_gn=HYDIN;refGene_tx=XM_0232461
47.1;refGene_gn=HYDIN;refGene_tx=XM_023246148.1;refGene_gn=HYDIN;refGe
ne_tx=XM_023246149.1
E2 50546454 50546554
refGene_gn=HYDIN;refGene_tx=XM_023246143.1;refGene_gn=HYDIN;refGene_tx
=XM_023246144.1;refGene_gn=HYDIN;refGene_tx=XM_023246145.1;refGene_gn=
HYDIN;refGene_tx=XM_023246146.1;refGene_gn=HYDIN;refGene_tx=XM_0232461
47.1;refGene_gn=HYDIN;refGene_tx=XM_023246148.1;refGene_gn=HYDIN;refGe

ne_tx=XM_023246149.1
E2 50548638 50548807
refGene_gn=HYDIN;refGene_tx=XM_023246143.1;refGene_gn=HYDIN;refGene_tx=XM_023246144.1;refGene_gn=HYDIN;refGene_tx=XM_023246145.1;refGene_gn=HYDIN;refGene_tx=XM_023246146.1;refGene_gn=HYDIN;refGene_tx=XM_023246147.1;refGene_gn=HYDIN;refGene_tx=XM_023246148.1;refGene_gn=HYDIN;refGene_tx=XM_023246149.1
E2 50549267 50549506
refGene_gn=HYDIN;refGene_tx=XM_023246143.1;refGene_gn=HYDIN;refGene_tx=XM_023246144.1;refGene_gn=HYDIN;refGene_tx=XM_023246145.1;refGene_gn=HYDIN;refGene_tx=XM_023246146.1;refGene_gn=HYDIN;refGene_tx=XM_023246147.1;refGene_gn=HYDIN;refGene_tx=XM_023246148.1;refGene_gn=HYDIN;refGene_tx=XM_023246149.1
E2 50551360 50551472
refGene_gn=HYDIN;refGene_tx=XM_023246143.1;refGene_gn=HYDIN;refGene_tx=XM_023246144.1;refGene_gn=HYDIN;refGene_tx=XM_023246145.1;refGene_gn=HYDIN;refGene_tx=XM_023246146.1;refGene_gn=HYDIN;refGene_tx=XM_023246147.1;refGene_gn=HYDIN;refGene_tx=XM_023246148.1;refGene_gn=HYDIN;refGene_tx=XM_023246149.1
E2 50556602 50556807
refGene_gn=HYDIN;refGene_tx=XM_023246143.1;refGene_gn=HYDIN;refGene_tx=XM_023246144.1;refGene_gn=HYDIN;refGene_tx=XM_023246145.1;refGene_gn=HYDIN;refGene_tx=XM_023246146.1;refGene_gn=HYDIN;refGene_tx=XM_023246148.1;refGene_gn=HYDIN;refGene_tx=XM_023246149.1
E2 50557533 50557769
refGene_gn=HYDIN;refGene_tx=XM_023246143.1;refGene_gn=HYDIN;refGene_tx=XM_023246144.1;refGene_gn=HYDIN;refGene_tx=XM_023246145.1;refGene_gn=HYDIN;refGene_tx=XM_023246146.1;refGene_gn=HYDIN;refGene_tx=XM_023246148.1;refGene_gn=HYDIN;refGene_tx=XM_023246149.1
E2 50559689 50559841
refGene_gn=HYDIN;refGene_tx=XM_023246143.1;refGene_gn=HYDIN;refGene_tx=XM_023246144.1;refGene_gn=HYDIN;refGene_tx=XM_023246145.1;refGene_gn=HYDIN;refGene_tx=XM_023246146.1;refGene_gn=HYDIN;refGene_tx=XM_023246148.1
E2 50559953 50560143
refGene_gn=HYDIN;refGene_tx=XM_023246143.1;refGene_gn=HYDIN;refGene_tx=XM_023246144.1;refGene_gn=HYDIN;refGene_tx=XM_023246145.1;refGene_gn=HYDIN;refGene_tx=XM_023246146.1;refGene_gn=HYDIN;refGene_tx=XM_023246148.1;refGene_gn=HYDIN;refGene_tx=XM_023246149.1
E2 50563391 50563494
refGene_gn=HYDIN;refGene_tx=XM_023246143.1;refGene_gn=HYDIN;refGene_tx=XM_023246144.1;refGene_gn=HYDIN;refGene_tx=XM_023246145.1;refGene_gn=HYDIN;refGene_tx=XM_023246146.1;refGene_gn=HYDIN;refGene_tx=XM_023246147.1;refGene_gn=HYDIN;refGene_tx=XM_023246148.1
E2 50563722 50564001
refGene_gn=HYDIN;refGene_tx=XM_023246143.1;refGene_gn=HYDIN;refGene_tx=XM_023246144.1;refGene_gn=HYDIN;refGene_tx=XM_023246145.1;refGene_gn=HYDIN;refGene_tx=XM_023246146.1;refGene_gn=HYDIN;refGene_tx=XM_023246147.1;refGene_gn=HYDIN;refGene_tx=XM_023246148.1
E2 50565755 50565898

refGene_gn=HYDIN;refGene_tx=XM_023246143.1;refGene_gn=HYDIN;refGene_tx=XM_023246144.1;refGene_gn=HYDIN;refGene_tx=XM_023246145.1;refGene_gn=HYDIN;refGene_tx=XM_023246146.1;refGene_gn=HYDIN;refGene_tx=XM_023246147.1;refGene_gn=HYDIN;refGene_tx=XM_023246148.1
E2 50568278 50568497
refGene_gn=HYDIN;refGene_tx=XM_023246143.1;refGene_gn=HYDIN;refGene_tx=XM_023246144.1;refGene_gn=HYDIN;refGene_tx=XM_023246145.1;refGene_gn=HYDIN;refGene_tx=XM_023246146.1;refGene_gn=HYDIN;refGene_tx=XM_023246147.1;refGene_gn=HYDIN;refGene_tx=XM_023246148.1
E2 50571107 50571268
refGene_gn=HYDIN;refGene_tx=XM_023246143.1;refGene_gn=HYDIN;refGene_tx=XM_023246144.1;refGene_gn=HYDIN;refGene_tx=XM_023246145.1;refGene_gn=HYDIN;refGene_tx=XM_023246146.1;refGene_gn=HYDIN;refGene_tx=XM_023246147.1;refGene_gn=HYDIN;refGene_tx=XM_023246148.1
E2 50573521 50573619
refGene_gn=HYDIN;refGene_tx=XM_023246143.1;refGene_gn=HYDIN;refGene_tx=XM_023246144.1;refGene_gn=HYDIN;refGene_tx=XM_023246145.1;refGene_gn=HYDIN;refGene_tx=XM_023246146.1;refGene_gn=HYDIN;refGene_tx=XM_023246147.1;refGene_gn=HYDIN;refGene_tx=XM_023246148.1
E2 50574382 50574590
refGene_gn=HYDIN;refGene_tx=XM_023246143.1;refGene_gn=HYDIN;refGene_tx=XM_023246144.1;refGene_gn=HYDIN;refGene_tx=XM_023246145.1;refGene_gn=HYDIN;refGene_tx=XM_023246146.1;refGene_gn=HYDIN;refGene_tx=XM_023246147.1;refGene_gn=HYDIN;refGene_tx=XM_023246148.1
E2 50575769 50575982
refGene_gn=HYDIN;refGene_tx=XM_023246143.1;refGene_gn=HYDIN;refGene_tx=XM_023246144.1;refGene_gn=HYDIN;refGene_tx=XM_023246145.1;refGene_gn=HYDIN;refGene_tx=XM_023246146.1;refGene_gn=HYDIN;refGene_tx=XM_023246147.1;refGene_gn=HYDIN;refGene_tx=XM_023246148.1
E2 50576522 50576661
refGene_gn=HYDIN;refGene_tx=XM_023246143.1;refGene_gn=HYDIN;refGene_tx=XM_023246144.1;refGene_gn=HYDIN;refGene_tx=XM_023246145.1;refGene_gn=HYDIN;refGene_tx=XM_023246146.1;refGene_gn=HYDIN;refGene_tx=XM_023246147.1;refGene_gn=HYDIN;refGene_tx=XM_023246148.1
E2 50577781 50577947
refGene_gn=HYDIN;refGene_tx=XM_023246143.1;refGene_gn=HYDIN;refGene_tx=XM_023246144.1;refGene_gn=HYDIN;refGene_tx=XM_023246145.1;refGene_gn=HYDIN;refGene_tx=XM_023246146.1;refGene_gn=HYDIN;refGene_tx=XM_023246147.1;refGene_gn=HYDIN;refGene_tx=XM_023246148.1
E2 50580724 50580872
refGene_gn=HYDIN;refGene_tx=XM_023246143.1;refGene_gn=HYDIN;refGene_tx=XM_023246144.1;refGene_gn=HYDIN;refGene_tx=XM_023246145.1;refGene_gn=HYDIN;refGene_tx=XM_023246146.1;refGene_gn=HYDIN;refGene_tx=XM_023246147.1;refGene_gn=HYDIN;refGene_tx=XM_023246148.1
E2 50581289 50581497
refGene_gn=HYDIN;refGene_tx=XM_023246143.1;refGene_gn=HYDIN;refGene_tx=XM_023246144.1;refGene_gn=HYDIN;refGene_tx=XM_023246145.1;refGene_gn=HYDIN;refGene_tx=XM_023246146.1;refGene_gn=HYDIN;refGene_tx=XM_023246147.1;refGene_gn=HYDIN;refGene_tx=XM_023246148.1
E2 50582073 50582295

refGene_gn=HYDIN;refGene_tx=XM_023246143.1;refGene_gn=HYDIN;refGene_tx=XM_023246144.1;refGene_gn=HYDIN;refGene_tx=XM_023246145.1;refGene_gn=HYDIN;refGene_tx=XM_023246146.1;refGene_gn=HYDIN;refGene_tx=XM_023246147.1;refGene_gn=HYDIN;refGene_tx=XM_023246148.1
E2 50586278 50586448
refGene_gn=HYDIN;refGene_tx=XM_023246143.1;refGene_gn=HYDIN;refGene_tx=XM_023246144.1;refGene_gn=HYDIN;refGene_tx=XM_023246145.1;refGene_gn=HYDIN;refGene_tx=XM_023246146.1;refGene_gn=HYDIN;refGene_tx=XM_023246147.1;refGene_gn=HYDIN;refGene_tx=XM_023246148.1
E2 50588233 50588432
refGene_gn=HYDIN;refGene_tx=XM_023246143.1;refGene_gn=HYDIN;refGene_tx=XM_023246144.1;refGene_gn=HYDIN;refGene_tx=XM_023246145.1;refGene_gn=HYDIN;refGene_tx=XM_023246146.1;refGene_gn=HYDIN;refGene_tx=XM_023246147.1;refGene_gn=HYDIN;refGene_tx=XM_023246148.1
E2 50590189 50590348
refGene_gn=HYDIN;refGene_tx=XM_023246143.1;refGene_gn=HYDIN;refGene_tx=XM_023246144.1;refGene_gn=HYDIN;refGene_tx=XM_023246145.1;refGene_gn=HYDIN;refGene_tx=XM_023246146.1;refGene_gn=HYDIN;refGene_tx=XM_023246147.1;refGene_gn=HYDIN;refGene_tx=XM_023246148.1
E2 50592191 50592469
refGene_gn=HYDIN;refGene_tx=XM_023246143.1;refGene_gn=HYDIN;refGene_tx=XM_023246144.1;refGene_gn=HYDIN;refGene_tx=XM_023246145.1;refGene_gn=HYDIN;refGene_tx=XM_023246146.1;refGene_gn=HYDIN;refGene_tx=XM_023246147.1;refGene_gn=HYDIN;refGene_tx=XM_023246148.1
E2 50593632 50593852
refGene_gn=HYDIN;refGene_tx=XM_023246143.1;refGene_gn=HYDIN;refGene_tx=XM_023246144.1;refGene_gn=HYDIN;refGene_tx=XM_023246145.1;refGene_gn=HYDIN;refGene_tx=XM_023246146.1;refGene_gn=HYDIN;refGene_tx=XM_023246147.1;refGene_gn=HYDIN;refGene_tx=XM_023246148.1
E2 50595452 50595665
refGene_gn=HYDIN;refGene_tx=XM_023246143.1;refGene_gn=HYDIN;refGene_tx=XM_023246144.1;refGene_gn=HYDIN;refGene_tx=XM_023246145.1;refGene_gn=HYDIN;refGene_tx=XM_023246146.1;refGene_gn=HYDIN;refGene_tx=XM_023246147.1;refGene_gn=HYDIN;refGene_tx=XM_023246148.1
E2 50596899 50597063
refGene_gn=HYDIN;refGene_tx=XM_023246143.1;refGene_gn=HYDIN;refGene_tx=XM_023246144.1;refGene_gn=HYDIN;refGene_tx=XM_023246145.1;refGene_gn=HYDIN;refGene_tx=XM_023246146.1;refGene_gn=HYDIN;refGene_tx=XM_023246147.1;refGene_gn=HYDIN;refGene_tx=XM_023246148.1
E2 50597919 50598070
refGene_gn=HYDIN;refGene_tx=XM_023246143.1;refGene_gn=HYDIN;refGene_tx=XM_023246144.1;refGene_gn=HYDIN;refGene_tx=XM_023246145.1;refGene_gn=HYDIN;refGene_tx=XM_023246146.1;refGene_gn=HYDIN;refGene_tx=XM_023246147.1;refGene_gn=HYDIN;refGene_tx=XM_023246148.1
E2 50604716 50604722 refGene_gn=HYDIN;refGene_tx=XM_023246148.1
E2 50605319 50605550
refGene_gn=HYDIN;refGene_tx=XM_023246143.1;refGene_gn=HYDIN;refGene_tx=XM_023246144.1;refGene_gn=HYDIN;refGene_tx=XM_023246145.1;refGene_gn=HYDIN;refGene_tx=XM_023246146.1;refGene_gn=HYDIN;refGene_tx=XM_023246147.1

E2 50614155 50614380
refGene_gn=HYDIN;refGene_tx=XM_023246143.1;refGene_gn=HYDIN;refGene_tx=XM_023246144.1;refGene_gn=HYDIN;refGene_tx=XM_023246145.1;refGene_gn=HYDIN;refGene_tx=XM_023246146.1;refGene_gn=HYDIN;refGene_tx=XM_023246147.1
E2 50615924 50616407
refGene_gn=HYDIN;refGene_tx=XM_023246143.1;refGene_gn=HYDIN;refGene_tx=XM_023246144.1;refGene_gn=HYDIN;refGene_tx=XM_023246145.1;refGene_gn=HYDIN;refGene_tx=XM_023246146.1;refGene_gn=HYDIN;refGene_tx=XM_023246147.1
E2 50622720 50622824
refGene_gn=VAC14;refGene_tx=XM_006941719.3;refGene_gn=VAC14;refGene_tx=XM_011290219.2
E2 50635521 50635672
refGene_gn=VAC14;refGene_tx=XM_006941719.3;refGene_gn=VAC14;refGene_tx=XM_011290219.2
E2 50635987 50636155
refGene_gn=VAC14;refGene_tx=XM_006941719.3;refGene_gn=VAC14;refGene_tx=XM_011290219.2
E2 50636942 50637005
refGene_gn=VAC14;refGene_tx=XM_006941719.3;refGene_gn=VAC14;refGene_tx=XM_011290219.2
E2 50637602 50637710
refGene_gn=VAC14;refGene_tx=XM_006941719.3;refGene_gn=VAC14;refGene_tx=XM_011290219.2
E2 50638236 50638346
refGene_gn=VAC14;refGene_tx=XM_006941719.3;refGene_gn=VAC14;refGene_tx=XM_011290219.2
E2 50638633 50638740
refGene_gn=VAC14;refGene_tx=XM_006941719.3;refGene_gn=VAC14;refGene_tx=XM_011290219.2
E2 50639696 50639831
refGene_gn=VAC14;refGene_tx=XM_006941719.3;refGene_gn=VAC14;refGene_tx=XM_011290219.2
E2 50640298 50640457
refGene_gn=VAC14;refGene_tx=XM_006941719.3;refGene_gn=VAC14;refGene_tx=XM_011290219.2
E2 50646886 50646950
refGene_gn=VAC14;refGene_tx=XM_006941719.3;refGene_gn=VAC14;refGene_tx=XM_011290219.2
E2 50653832 50653977
refGene_gn=VAC14;refGene_tx=XM_006941719.3;refGene_gn=VAC14;refGene_tx=XM_011290219.2
E2 50654252 50654318
refGene_gn=VAC14;refGene_tx=XM_006941719.3;refGene_gn=VAC14;refGene_tx=XM_011290219.2
E2 50671830 50671987
refGene_gn=VAC14;refGene_tx=XM_006941719.3;refGene_gn=VAC14;refGene_tx=XM_011290219.2
E2 50683593 50683726

refGene_gn=VAC14;refGene_tx=XM_006941719.3;refGene_gn=VAC14;refGene_tx=XM_011290219.2
E2 50713351 50713526
refGene_gn=VAC14;refGene_tx=XM_006941719.3;refGene_gn=VAC14;refGene_tx=XM_011290219.2
E2 50714991 50715110
refGene_gn=VAC14;refGene_tx=XM_006941719.3;refGene_gn=VAC14;refGene_tx=XM_011290219.2
E2 50716262 50716342
refGene_gn=VAC14;refGene_tx=XM_006941719.3;refGene_gn=VAC14;refGene_tx=XM_011290219.2
E2 50718134 50718285
refGene_gn=VAC14;refGene_tx=XM_006941719.3;refGene_gn=VAC14;refGene_tx=XM_011290219.2
E2 50722896 50723059
refGene_gn=VAC14;refGene_tx=XM_006941719.3;refGene_gn=VAC14;refGene_tx=XM_011290219.2
E2 50725163 50725232
refGene_gn=MTSS1L;refGene_tx=XM_023245580.1;refGene_gn=MTSS1L;refGene_tx=XM_023245581.1;refGene_gn=MTSS1L;refGene_tx=XM_023245582.1;refGene_gn=MTSS1L;refGene_tx=XM_023245583.1;refGene_gn=MTSS1L;refGene_tx=XM_023245584.1
E2 50730162 50730224
refGene_gn=MTSS1L;refGene_tx=XM_023245580.1;refGene_gn=MTSS1L;refGene_tx=XM_023245581.1;refGene_gn=MTSS1L;refGene_tx=XM_023245582.1;refGene_gn=MTSS1L;refGene_tx=XM_023245583.1;refGene_gn=MTSS1L;refGene_tx=XM_023245584.1
E2 50730310 50730384
refGene_gn=MTSS1L;refGene_tx=XM_023245580.1;refGene_gn=MTSS1L;refGene_tx=XM_023245581.1;refGene_gn=MTSS1L;refGene_tx=XM_023245582.1;refGene_gn=MTSS1L;refGene_tx=XM_023245583.1;refGene_gn=MTSS1L;refGene_tx=XM_023245584.1
E2 50731196 50731281
refGene_gn=MTSS1L;refGene_tx=XM_023245580.1;refGene_gn=MTSS1L;refGene_tx=XM_023245581.1;refGene_gn=MTSS1L;refGene_tx=XM_023245582.1;refGene_gn=MTSS1L;refGene_tx=XM_023245583.1;refGene_gn=MTSS1L;refGene_tx=XM_023245584.1
E2 50731365 50731457
refGene_gn=MTSS1L;refGene_tx=XM_023245580.1;refGene_gn=MTSS1L;refGene_tx=XM_023245581.1;refGene_gn=MTSS1L;refGene_tx=XM_023245582.1;refGene_gn=MTSS1L;refGene_tx=XM_023245583.1;refGene_gn=MTSS1L;refGene_tx=XM_023245584.1
E2 50731542 50731617
refGene_gn=MTSS1L;refGene_tx=XM_023245580.1;refGene_gn=MTSS1L;refGene_tx=XM_023245581.1;refGene_gn=MTSS1L;refGene_tx=XM_023245582.1;refGene_gn=MTSS1L;refGene_tx=XM_023245583.1;refGene_gn=MTSS1L;refGene_tx=XM_023245584.1
E2 50731949 50731958
refGene_gn=MTSS1L;refGene_tx=XM_023245580.1;refGene_gn=MTSS1L;refGene_tx=XM_023245582.1;refGene_gn=MTSS1L;refGene_tx=XM_023245583.1

E2	50732894 50733052
refGene_gn=MTSS1L;refGene_tx=XM_023245580.1;refGene_gn=MTSS1L;refGene_tx=XM_023245581.1;refGene_gn=MTSS1L;refGene_tx=XM_023245582.1;refGene_gn=MTSS1L;refGene_tx=XM_023245583.1;refGene_gn=MTSS1L;refGene_tx=XM_023245584.1
E2	50733482 50733590
refGene_gn=MTSS1L;refGene_tx=XM_023245580.1;refGene_gn=MTSS1L;refGene_tx=XM_023245581.1;refGene_gn=MTSS1L;refGene_tx=XM_023245582.1;refGene_gn=MTSS1L;refGene_tx=XM_023245583.1;refGene_gn=MTSS1L;refGene_tx=XM_023245584.1
E2	50734471 50734569
refGene_gn=MTSS1L;refGene_tx=XM_023245580.1;refGene_gn=MTSS1L;refGene_tx=XM_023245581.1;refGene_gn=MTSS1L;refGene_tx=XM_023245582.1;refGene_gn=MTSS1L;refGene_tx=XM_023245583.1;refGene_gn=MTSS1L;refGene_tx=XM_023245584.1
E2	50736047 50736270
refGene_gn=MTSS1L;refGene_tx=XM_023245580.1;refGene_gn=MTSS1L;refGene_tx=XM_023245581.1;refGene_gn=MTSS1L;refGene_tx=XM_023245582.1;refGene_gn=MTSS1L;refGene_tx=XM_023245583.1;refGene_gn=MTSS1L;refGene_tx=XM_023245584.1
E2	50744693 50744768
refGene_gn=MTSS1L;refGene_tx=XM_023245580.1;refGene_gn=MTSS1L;refGene_tx=XM_023245581.1;refGene_gn=MTSS1L;refGene_tx=XM_023245583.1;refGene_gn=MTSS1L;refGene_tx=XM_023245584.1
E2	50745102 50745279
refGene_gn=MTSS1L;refGene_tx=XM_023245580.1;refGene_gn=MTSS1L;refGene_tx=XM_023245581.1;refGene_gn=MTSS1L;refGene_tx=XM_023245582.1;refGene_gn=MTSS1L;refGene_tx=XM_023245583.1;refGene_gn=MTSS1L;refGene_tx=XM_023245584.1
E2	50745458 50745624
refGene_gn=MTSS1L;refGene_tx=XM_023245580.1;refGene_gn=MTSS1L;refGene_tx=XM_023245581.1;refGene_gn=MTSS1L;refGene_tx=XM_023245582.1;refGene_gn=MTSS1L;refGene_tx=XM_023245583.1;refGene_gn=MTSS1L;refGene_tx=XM_023245584.1
E2	50745702 50746448
refGene_gn=MTSS1L;refGene_tx=XM_023245580.1;refGene_gn=MTSS1L;refGene_tx=XM_023245581.1;refGene_gn=MTSS1L;refGene_tx=XM_023245582.1;refGene_gn=MTSS1L;refGene_tx=XM_023245583.1;refGene_gn=MTSS1L;refGene_tx=XM_023245584.1
E2	50749739 50749912
refGene_gn=IL34;refGene_tx=XM_006941720.4;refGene_gn=IL34;refGene_tx=XM_019820031.2;refGene_gn=IL34;refGene_tx=XM_019820032.2;refGene_gn=IL34;refGene_tx=XM_023245585.1
E2	50750143 50750279
refGene_gn=IL34;refGene_tx=XM_006941720.4;refGene_gn=IL34;refGene_tx=XM_019820031.2;refGene_gn=IL34;refGene_tx=XM_019820032.2;refGene_gn=IL34;refGene_tx=XM_023245585.1
E2	50753176 50753338
refGene_gn=IL34;refGene_tx=XM_006941720.4;refGene_gn=IL34;refGene_tx=XM_019820031.2;refGene_gn=IL34;refGene_tx=XM_019820032.2;refGene_gn=IL3

4;refGene_tx=XM_023245585.1
E2 50753623 50753701
refGene_gn=IL34;refGene_tx=XM_006941720.4;refGene_gn=IL34;refGene_tx=XM_019820031.2;refGene_gn=IL34;refGene_tx=XM_019820032.2;refGene_gn=IL34;refGene_tx=XM_023245585.1
E2 50754488 50754622
refGene_gn=IL34;refGene_tx=XM_006941720.4;refGene_gn=IL34;refGene_tx=XM_019820031.2;refGene_gn=IL34;refGene_tx=XM_019820032.2;refGene_gn=IL34;refGene_tx=XM_023245585.1
E2 50761260 50761288
refGene_gn=IL34;refGene_tx=XM_006941720.4;refGene_gn=IL34;refGene_tx=XM_019820031.2;refGene_gn=IL34;refGene_tx=XM_023245585.1
E2 50839870 50840011
refGene_gn=SF3B3;refGene_tx=XM_003998231.4;refGene_gn=SF3B3;refGene_tx=XM_006941723.2
E2 50840423 50840528
refGene_gn=SF3B3;refGene_tx=XM_003998231.4;refGene_gn=SF3B3;refGene_tx=XM_006941723.2
E2 50841224 50841368
refGene_gn=SF3B3;refGene_tx=XM_003998231.4;refGene_gn=SF3B3;refGene_tx=XM_006941723.2
E2 50842077 50842176
refGene_gn=SF3B3;refGene_tx=XM_003998231.4;refGene_gn=SF3B3;refGene_tx=XM_006941723.2
E2 50843574 50843787
refGene_gn=SF3B3;refGene_tx=XM_003998231.4;refGene_gn=SF3B3;refGene_tx=XM_006941723.2
E2 50844201 50844327
refGene_gn=SF3B3;refGene_tx=XM_003998231.4;refGene_gn=SF3B3;refGene_tx=XM_006941723.2
E2 50845736 50845893
refGene_gn=SF3B3;refGene_tx=XM_003998231.4;refGene_gn=SF3B3;refGene_tx=XM_006941723.2
E2 50845994 50846200
refGene_gn=SF3B3;refGene_tx=XM_003998231.4;refGene_gn=SF3B3;refGene_tx=XM_006941723.2
E2 50846953 50847128
refGene_gn=SF3B3;refGene_tx=XM_003998231.4;refGene_gn=SF3B3;refGene_tx=XM_006941723.2
E2 50848348 50848503
refGene_gn=SF3B3;refGene_tx=XM_003998231.4;refGene_gn=SF3B3;refGene_tx=XM_006941723.2
E2 50849238 50849361
refGene_gn=SF3B3;refGene_tx=XM_003998231.4;refGene_gn=SF3B3;refGene_tx=XM_006941723.2
E2 50852356 50852500
refGene_gn=SF3B3;refGene_tx=XM_003998231.4;refGene_gn=SF3B3;refGene_tx=XM_006941723.2
E2 50853269 50853425
refGene_gn=SF3B3;refGene_tx=XM_003998231.4;refGene_gn=SF3B3;refGene_tx

=XM_006941723.2
E2 50854391 50854547
refGene_gn=SF3B3;refGene_tx=XM_003998231.4;refGene_gn=SF3B3;refGene_tx
=XM_006941723.2
E2 50855076 50855228
refGene_gn=SF3B3;refGene_tx=XM_003998231.4;refGene_gn=SF3B3;refGene_tx
=XM_006941723.2
E2 50860706 50860779
refGene_gn=SF3B3;refGene_tx=XM_003998231.4;refGene_gn=SF3B3;refGene_tx
=XM_006941723.2
E2 50864509 50864605
refGene_gn=SF3B3;refGene_tx=XM_003998231.4;refGene_gn=SF3B3;refGene_tx
=XM_006941723.2
E2 50867445 50867611
refGene_gn=SF3B3;refGene_tx=XM_003998231.4;refGene_gn=SF3B3;refGene_tx
=XM_006941723.2
E2 50873901 50874005
refGene_gn=SF3B3;refGene_tx=XM_003998231.4;refGene_gn=SF3B3;refGene_tx
=XM_006941723.2
E2 50875455 50875593
refGene_gn=SF3B3;refGene_tx=XM_003998231.4;refGene_gn=SF3B3;refGene_tx
=XM_006941723.2
E2 50878381 50878494
refGene_gn=SF3B3;refGene_tx=XM_003998231.4;refGene_gn=SF3B3;refGene_tx
=XM_006941723.2
E2 50880957 50881099
refGene_gn=SF3B3;refGene_tx=XM_003998231.4;refGene_gn=SF3B3;refGene_tx
=XM_006941723.2
E2 50881619 50881792
refGene_gn=SF3B3;refGene_tx=XM_003998231.4;refGene_gn=SF3B3;refGene_tx
=XM_006941723.2
E2 50882789 50883116
refGene_gn=SF3B3;refGene_tx=XM_003998231.4;refGene_gn=SF3B3;refGene_tx
=XM_006941723.2
E2 50884741 50884811
refGene_gn=SF3B3;refGene_tx=XM_003998231.4;refGene_gn=SF3B3;refGene_tx
=XM_006941723.2
E2 50888996 50889164 refGene_gn=COG4;refGene_tx=XM_003998232.3
E2 50891552 50891635 refGene_gn=COG4;refGene_tx=XM_003998232.3
E2 50892830 50892945 refGene_gn=COG4;refGene_tx=XM_003998232.3
E2 50894905 50895080 refGene_gn=COG4;refGene_tx=XM_003998232.3
E2 50895821 50896015 refGene_gn=COG4;refGene_tx=XM_003998232.3
E2 50898199 50898305 refGene_gn=COG4;refGene_tx=XM_003998232.3
E2 50898824 50898982 refGene_gn=COG4;refGene_tx=XM_003998232.3
E2 50899713 50899772 refGene_gn=COG4;refGene_tx=XM_003998232.3
E2 50903682 50903816 refGene_gn=COG4;refGene_tx=XM_003998232.3
E2 50905347 50905466 refGene_gn=COG4;refGene_tx=XM_003998232.3
E2 50906027 50906194 refGene_gn=COG4;refGene_tx=XM_003998232.3
E2 50906985 50907151 refGene_gn=COG4;refGene_tx=XM_003998232.3
E2 50911002 50911065 refGene_gn=COG4;refGene_tx=XM_003998232.3

E2 50913713 50913830 refGene_gn=COG4;refGene_tx=XM_003998232.3
E2 50914408 50914501 refGene_gn=COG4;refGene_tx=XM_003998232.3
E2 50915009 50915093 refGene_gn=COG4;refGene_tx=XM_003998232.3
E2 50915329 50915431 refGene_gn=COG4;refGene_tx=XM_003998232.3
E2 50915707 50915836 refGene_gn=COG4;refGene_tx=XM_003998232.3
E2 50916054 50916189 refGene_gn=COG4;refGene_tx=XM_003998232.3
E2 50917592 50917679
refGene_gn=FUK;refGene_tx=XM_019820028.2;refGene_gn=FUK;refGene_tx=XM_019820029.2;refGene_gn=FUK;refGene_tx=XM_019820030.2
E2 50917843 50918067
refGene_gn=FUK;refGene_tx=XM_019820028.2;refGene_gn=FUK;refGene_tx=XM_019820029.2;refGene_gn=FUK;refGene_tx=XM_019820030.2
E2 50918590 50918690
refGene_gn=FUK;refGene_tx=XM_019820028.2;refGene_gn=FUK;refGene_tx=XM_019820029.2;refGene_gn=FUK;refGene_tx=XM_019820030.2
E2 50918777 50918965
refGene_gn=FUK;refGene_tx=XM_019820028.2;refGene_gn=FUK;refGene_tx=XM_019820029.2;refGene_gn=FUK;refGene_tx=XM_019820030.2
E2 50920584 50920704
refGene_gn=FUK;refGene_tx=XM_019820028.2;refGene_gn=FUK;refGene_tx=XM_019820029.2;refGene_gn=FUK;refGene_tx=XM_019820030.2
E2 50920850 50920994
refGene_gn=FUK;refGene_tx=XM_019820028.2;refGene_gn=FUK;refGene_tx=XM_019820029.2;refGene_gn=FUK;refGene_tx=XM_019820030.2
E2 50921350 50921572
refGene_gn=FUK;refGene_tx=XM_019820028.2;refGene_gn=FUK;refGene_tx=XM_019820029.2;refGene_gn=FUK;refGene_tx=XM_019820030.2
E2 50921665 50921832
refGene_gn=FUK;refGene_tx=XM_019820028.2;refGene_gn=FUK;refGene_tx=XM_019820029.2;refGene_gn=FUK;refGene_tx=XM_019820030.2
E2 50922003 50922214
refGene_gn=FUK;refGene_tx=XM_019820028.2;refGene_gn=FUK;refGene_tx=XM_019820029.2;refGene_gn=FUK;refGene_tx=XM_019820030.2
E2 50922842 50923213
refGene_gn=FUK;refGene_tx=XM_019820028.2;refGene_gn=FUK;refGene_tx=XM_019820029.2;refGene_gn=FUK;refGene_tx=XM_019820030.2
E2 50923620 50923685
refGene_gn=FUK;refGene_tx=XM_019820028.2;refGene_gn=FUK;refGene_tx=XM_019820029.2;refGene_gn=FUK;refGene_tx=XM_019820030.2
E2 50924078 50924249
refGene_gn=FUK;refGene_tx=XM_019820028.2;refGene_gn=FUK;refGene_tx=XM_019820029.2;refGene_gn=FUK;refGene_tx=XM_019820030.2
E2 50924370 50924472
refGene_gn=FUK;refGene_tx=XM_019820028.2;refGene_gn=FUK;refGene_tx=XM_019820029.2;refGene_gn=FUK;refGene_tx=XM_019820030.2
E2 50925146 50925259
refGene_gn=FUK;refGene_tx=XM_019820028.2;refGene_gn=FUK;refGene_tx=XM_019820029.2;refGene_gn=FUK;refGene_tx=XM_019820030.2
E2 50925651 50925823
refGene_gn=FUK;refGene_tx=XM_019820028.2;refGene_gn=FUK;refGene_tx=XM_

019820029.2;refGene_gn=FUK;refGene_tx=XM_019820030.2
E2 50925986 50926106
refGene_gn=FUK;refGene_tx=XM_019820028.2;refGene_gn=FUK;refGene_tx=XM_019820029.2;refGene_gn=FUK;refGene_tx=XM_019820030.2
E2 50926686 50926767
refGene_gn=FUK;refGene_tx=XM_019820028.2;refGene_gn=FUK;refGene_tx=XM_019820029.2;refGene_gn=FUK;refGene_tx=XM_019820030.2
E2 50927156 50927254
refGene_gn=FUK;refGene_tx=XM_019820028.2;refGene_gn=FUK;refGene_tx=XM_019820029.2;refGene_gn=FUK;refGene_tx=XM_019820030.2
E2 50927662 50927735
refGene_gn=FUK;refGene_tx=XM_019820028.2;refGene_gn=FUK;refGene_tx=XM_019820029.2;refGene_gn=FUK;refGene_tx=XM_019820030.2
E2 50928100 50928226
refGene_gn=FUK;refGene_tx=XM_019820028.2;refGene_gn=FUK;refGene_tx=XM_019820029.2;refGene_gn=FUK;refGene_tx=XM_019820030.2
E2 50930250 50930301 refGene_gn=FUK;refGene_tx=XM_019820028.2
E2 50931068 50931220 refGene_gn=FUK;refGene_tx=XM_019820028.2
E2 50931999 50932156
refGene_gn=FUK;refGene_tx=XM_019820028.2;refGene_gn=FUK;refGene_tx=XM_019820029.2;refGene_gn=FUK;refGene_tx=XM_019820030.2
E2 50932280 50932313 refGene_gn=FUK;refGene_tx=XM_019820028.2
E2 50981038 50981377
refGene_gn=ST3GAL2;refGene_tx=XM_003998234.5;refGene_gn=ST3GAL2;refGene_tx=XM_023244896.1
E2 50984297 50984491
refGene_gn=ST3GAL2;refGene_tx=XM_003998234.5;refGene_gn=ST3GAL2;refGene_tx=XM_023244896.1
E2 50990394 50990574
refGene_gn=ST3GAL2;refGene_tx=XM_003998234.5;refGene_gn=ST3GAL2;refGene_tx=XM_023244896.1
E2 50992799 50992845
refGene_gn=ST3GAL2;refGene_tx=XM_003998234.5;refGene_gn=ST3GAL2;refGene_tx=XM_023244896.1
E2 50993136 50993256
refGene_gn=ST3GAL2;refGene_tx=XM_003998234.5;refGene_gn=ST3GAL2;refGene_tx=XM_023244896.1
E2 50994123 50994297
refGene_gn=ST3GAL2;refGene_tx=XM_003998234.5;refGene_gn=ST3GAL2;refGene_tx=XM_023244896.1
E2 51008679 51008741
refGene_gn=DDX19A;refGene_tx=XM_003998235.4;refGene_gn=DDX19A;refGene_tx=XM_019820037.1;refGene_gn=DDX19A;refGene_tx=XM_023245543.1
E2 51009127 51009319
refGene_gn=DDX19A;refGene_tx=XM_003998235.4;refGene_gn=DDX19A;refGene_tx=XM_019820037.1;refGene_gn=DDX19A;refGene_tx=XM_023245543.1
E2 51009805 51009968
refGene_gn=DDX19A;refGene_tx=XM_003998235.4;refGene_gn=DDX19A;refGene_tx=XM_019820037.1;refGene_gn=DDX19A;refGene_tx=XM_023245543.1
E2 51010518 51010756

refGene_gn=DDX19A;refGene_tx=XM_003998235.4;refGene_gn=DDX19A;refGene_tx=XM_019820037.1;refGene_gn=DDX19A;refGene_tx=XM_023245543.1
E2 51011136 51011314
refGene_gn=DDX19A;refGene_tx=XM_003998235.4;refGene_gn=DDX19A;refGene_tx=XM_019820037.1;refGene_gn=DDX19A;refGene_tx=XM_023245543.1
E2 51015937 51016052
refGene_gn=DDX19A;refGene_tx=XM_003998235.4;refGene_gn=DDX19A;refGene_tx=XM_019820037.1;refGene_gn=DDX19A;refGene_tx=XM_023245543.1
E2 51016667 51016770
refGene_gn=DDX19A;refGene_tx=XM_003998235.4;refGene_gn=DDX19A;refGene_tx=XM_019820037.1;refGene_gn=DDX19A;refGene_tx=XM_023245543.1
E2 51018361 51018454
refGene_gn=DDX19A;refGene_tx=XM_003998235.4;refGene_gn=DDX19A;refGene_tx=XM_019820037.1;refGene_gn=DDX19A;refGene_tx=XM_023245543.1
E2 51020453 51020476 refGene_gn=DDX19A;refGene_tx=XM_019820037.1
E2 51020811 51020947
refGene_gn=DDX19A;refGene_tx=XM_003998235.4;refGene_gn=DDX19A;refGene_tx=XM_023245543.1
E2 51022184 51022235 refGene_gn=DDX19A;refGene_tx=XM_023245543.1
E2 51022898 51022949 refGene_gn=DDX19A;refGene_tx=XM_003998235.4
E2 51025778 51025827
refGene_gn=DDX19A;refGene_tx=XM_003998235.4;refGene_gn=DDX19A;refGene_tx=XM_023245543.1
E2 51031466 51031523
refGene_gn=DDX19A;refGene_tx=XM_003998235.4;refGene_gn=DDX19A;refGene_tx=XM_023245543.1
E2 51044150 51044212
refGene_gn=LOC101087696;refGene_tx=XM_006941728.4;refGene_gn=LOC101087696;refGene_tx=XM_019820034.2;refGene_gn=LOC101087696;refGene_tx=XM_019820035.2;refGene_gn=LOC101087696;refGene_tx=XM_019820036.2;refGene_gn=LOC101087696;refGene_tx=XM_023245542.1
E2 51044598 51044790
refGene_gn=LOC101087696;refGene_tx=XM_006941728.4;refGene_gn=LOC101087696;refGene_tx=XM_019820034.2;refGene_gn=LOC101087696;refGene_tx=XM_019820035.2;refGene_gn=LOC101087696;refGene_tx=XM_019820036.2;refGene_gn=LOC101087696;refGene_tx=XM_023245542.1
E2 51045276 51045439
refGene_gn=LOC101087696;refGene_tx=XM_006941728.4;refGene_gn=LOC101087696;refGene_tx=XM_019820034.2;refGene_gn=LOC101087696;refGene_tx=XM_019820035.2;refGene_gn=LOC101087696;refGene_tx=XM_019820036.2;refGene_gn=LOC101087696;refGene_tx=XM_023245542.1
E2 51045990 51046228
refGene_gn=LOC101087696;refGene_tx=XM_006941728.4;refGene_gn=LOC101087696;refGene_tx=XM_019820034.2;refGene_gn=LOC101087696;refGene_tx=XM_019820035.2;refGene_gn=LOC101087696;refGene_tx=XM_019820036.2;refGene_gn=LOC101087696;refGene_tx=XM_023245542.1
E2 51046587 51046765
refGene_gn=LOC101087696;refGene_tx=XM_006941728.4;refGene_gn=LOC101087696;refGene_tx=XM_019820034.2;refGene_gn=LOC101087696;refGene_tx=XM_019820035.2;refGene_gn=LOC101087696;refGene_tx=XM_019820036.2;refGene_gn

=LOC101087696;refGene_tx=XM_023245542.1
E2	51049017	51049132
refGene_gn=LOC101087696;refGene_tx=XM_006941728.4;refGene_gn=LOC101087696;refGene_tx=XM_019820034.2;refGene_gn=LOC101087696;refGene_tx=XM_019820035.2;refGene_gn=LOC101087696;refGene_tx=XM_019820036.2;refGene_gn=LOC101087696;refGene_tx=XM_023245542.1
E2	51049669	51049772
refGene_gn=LOC101087696;refGene_tx=XM_006941728.4;refGene_gn=LOC101087696;refGene_tx=XM_019820034.2;refGene_gn=LOC101087696;refGene_tx=XM_019820035.2;refGene_gn=LOC101087696;refGene_tx=XM_019820036.2;refGene_gn=LOC101087696;refGene_tx=XM_023245542.1
E2	51053232	51053325
refGene_gn=LOC101087696;refGene_tx=XM_006941728.4;refGene_gn=LOC101087696;refGene_tx=XM_019820034.2;refGene_gn=LOC101087696;refGene_tx=XM_019820035.2;refGene_gn=LOC101087696;refGene_tx=XM_019820036.2;refGene_gn=LOC101087696;refGene_tx=XM_023245542.1
E2	51053641	51053777
refGene_gn=LOC101087696;refGene_tx=XM_006941728.4;refGene_gn=LOC101087696;refGene_tx=XM_019820034.2;refGene_gn=LOC101087696;refGene_tx=XM_019820035.2;refGene_gn=LOC101087696;refGene_tx=XM_019820036.2;refGene_gn=LOC101087696;refGene_tx=XM_023245542.1
E2	51054498	51054552
refGene_gn=LOC101087696;refGene_tx=XM_006941728.4;refGene_gn=LOC101087696;refGene_tx=XM_019820034.2;refGene_gn=LOC101087696;refGene_tx=XM_019820035.2;refGene_gn=LOC101087696;refGene_tx=XM_019820036.2;refGene_gn=LOC101087696;refGene_tx=XM_023245542.1
E2	51055451	51055500
refGene_gn=LOC101087696;refGene_tx=XM_006941728.4;refGene_gn=LOC101087696;refGene_tx=XM_019820034.2;refGene_gn=LOC101087696;refGene_tx=XM_019820035.2;refGene_gn=LOC101087696;refGene_tx=XM_019820036.2;refGene_gn=LOC101087696;refGene_tx=XM_023245542.1
E2	51062987	51063161
refGene_gn=LOC101087696;refGene_tx=XM_019820034.2;refGene_gn=LOC101087696;refGene_tx=XM_023245542.1
E2	51064578	51064626
refGene_gn=LOC101087696;refGene_tx=XM_019820035.2;refGene_gn=LOC101087696;refGene_tx=XM_019820036.2
E2	51065230	51065302
refGene_gn=LOC101087696;refGene_tx=XM_006941728.4
E2	51071945	51072089
refGene_gn=AARS;refGene_tx=XM_003998242.5;refGene_gn=AARS;refGene_tx=XM_019820033.2
E2	51075750	51075939
refGene_gn=AARS;refGene_tx=XM_003998242.5;refGene_gn=AARS;refGene_tx=XM_019820033.2
E2	51076286	51076432
refGene_gn=AARS;refGene_tx=XM_003998242.5;refGene_gn=AARS;refGene_tx=XM_019820033.2
E2	51078628	51078820
refGene_gn=AARS;refGene_tx=XM_003998242.5;refGene_gn=AARS;refGene_tx=X

M_019820033.2
E2 51079362 51079507
refGene_gn=AARS;refGene_tx=XM_003998242.5;refGene_gn=AARS;refGene_tx=X
M_019820033.2
E2 51079873 51080019
refGene_gn=AARS;refGene_tx=XM_003998242.5;refGene_gn=AARS;refGene_tx=X
M_019820033.2
E2 51080645 51080754
refGene_gn=AARS;refGene_tx=XM_003998242.5;refGene_gn=AARS;refGene_tx=X
M_019820033.2
E2 51080913 51081064
refGene_gn=AARS;refGene_tx=XM_003998242.5;refGene_gn=AARS;refGene_tx=X
M_019820033.2
E2 51082020 51082145
refGene_gn=AARS;refGene_tx=XM_003998242.5;refGene_gn=AARS;refGene_tx=X
M_019820033.2
E2 51082576 51082721
refGene_gn=AARS;refGene_tx=XM_003998242.5;refGene_gn=AARS;refGene_tx=X
M_019820033.2
E2 51083500 51083679
refGene_gn=AARS;refGene_tx=XM_003998242.5;refGene_gn=AARS;refGene_tx=X
M_019820033.2
E2 51084791 51084905
refGene_gn=AARS;refGene_tx=XM_003998242.5;refGene_gn=AARS;refGene_tx=X
M_019820033.2
E2 51086361 51086568
refGene_gn=AARS;refGene_tx=XM_003998242.5;refGene_gn=AARS;refGene_tx=X
M_019820033.2
E2 51087094 51087279
refGene_gn=AARS;refGene_tx=XM_003998242.5;refGene_gn=AARS;refGene_tx=X
M_019820033.2
E2 51089111 51089220
refGene_gn=AARS;refGene_tx=XM_003998242.5;refGene_gn=AARS;refGene_tx=X
M_019820033.2
E2 51089914 51090028
refGene_gn=AARS;refGene_tx=XM_003998242.5;refGene_gn=AARS;refGene_tx=X
M_019820033.2
E2 51090627 51090747
refGene_gn=AARS;refGene_tx=XM_003998242.5;refGene_gn=AARS;refGene_tx=X
M_019820033.2
E2 51090862 51090949
refGene_gn=AARS;refGene_tx=XM_003998242.5;refGene_gn=AARS;refGene_tx=X
M_019820033.2
E2 51091352 51091466
refGene_gn=AARS;refGene_tx=XM_003998242.5;refGene_gn=AARS;refGene_tx=X
M_019820033.2
E2 51091854 51092040
refGene_gn=AARS;refGene_tx=XM_003998242.5;refGene_gn=AARS;refGene_tx=X
M_019820033.2
E2 51093663 51094482 refGene_gn=EXOSC6;refGene_tx=XM_023245549.1

E2 51096362 51096401
refGene_gn=LOC102902362;refGene_tx=XM_006941731.3;refGene_gn=LOC102902362;refGene_tx=XM_019820040.2;refGene_gn=LOC102902362;refGene_tx=XM_019820041.2;refGene_gn=LOC102902362;refGene_tx=XM_019820042.2;refGene_gn=LOC102902362;refGene_tx=XM_019820044.2;refGene_gn=LOC102902362;refGene_tx=XM_019820045.2;refGene_gn=LOC102902362;refGene_tx=XM_019820046.2;refGene_gn=LOC102902362;refGene_tx=XM_019820047.2;refGene_gn=LOC102902362;refGene_tx=XM_023245555.1;refGene_gn=LOC102902362;refGene_tx=XM_023245556.1;refGene_gn=LOC102902362;refGene_tx=XM_023245557.1;refGene_gn=LOC102902362;refGene_tx=XM_023245558.1
E2 51097321 51097464
refGene_gn=LOC102902362;refGene_tx=XM_006941731.3;refGene_gn=LOC102902362;refGene_tx=XM_019820040.2;refGene_gn=LOC102902362;refGene_tx=XM_019820041.2;refGene_gn=LOC102902362;refGene_tx=XM_019820042.2;refGene_gn=LOC102902362;refGene_tx=XM_019820044.2;refGene_gn=LOC102902362;refGene_tx=XM_019820045.2;refGene_gn=LOC102902362;refGene_tx=XM_019820046.2;refGene_gn=LOC102902362;refGene_tx=XM_019820047.2;refGene_gn=LOC102902362;refGene_tx=XM_023245555.1;refGene_gn=LOC102902362;refGene_tx=XM_023245556.1;refGene_gn=LOC102902362;refGene_tx=XM_023245557.1;refGene_gn=LOC102902362;refGene_tx=XM_023245558.1
E2 51097785 51097938
refGene_gn=LOC102902362;refGene_tx=XM_006941731.3;refGene_gn=LOC102902362;refGene_tx=XM_019820040.2;refGene_gn=LOC102902362;refGene_tx=XM_019820041.2;refGene_gn=LOC102902362;refGene_tx=XM_019820042.2;refGene_gn=LOC102902362;refGene_tx=XM_019820044.2;refGene_gn=LOC102902362;refGene_tx=XM_019820045.2;refGene_gn=LOC102902362;refGene_tx=XM_019820046.2;refGene_gn=LOC102902362;refGene_tx=XM_019820047.2;refGene_gn=LOC102902362;refGene_tx=XM_023245555.1;refGene_gn=LOC102902362;refGene_tx=XM_023245556.1;refGene_gn=LOC102902362;refGene_tx=XM_023245557.1;refGene_gn=LOC102902362;refGene_tx=XM_023245558.1
E2 51098083 51098232
refGene_gn=LOC102902362;refGene_tx=XM_006941731.3;refGene_gn=LOC102902362;refGene_tx=XM_019820040.2;refGene_gn=LOC102902362;refGene_tx=XM_019820041.2;refGene_gn=LOC102902362;refGene_tx=XM_019820042.2;refGene_gn=LOC102902362;refGene_tx=XM_019820044.2;refGene_gn=LOC102902362;refGene_tx=XM_019820045.2;refGene_gn=LOC102902362;refGene_tx=XM_019820046.2;refGene_gn=LOC102902362;refGene_tx=XM_019820047.2;refGene_gn=LOC102902362;refGene_tx=XM_023245555.1;refGene_gn=LOC102902362;refGene_tx=XM_023245556.1;refGene_gn=LOC102902362;refGene_tx=XM_023245557.1;refGene_gn=LOC102902362;refGene_tx=XM_023245558.1
E2 51098312 51098518
refGene_gn=LOC102902362;refGene_tx=XM_006941731.3;refGene_gn=LOC102902362;refGene_tx=XM_019820040.2;refGene_gn=LOC102902362;refGene_tx=XM_019820041.2;refGene_gn=LOC102902362;refGene_tx=XM_019820042.2;refGene_gn=LOC102902362;refGene_tx=XM_019820044.2;refGene_gn=LOC102902362;refGene_tx=XM_019820045.2;refGene_gn=LOC102902362;refGene_tx=XM_019820046.2;refGene_gn=LOC102902362;refGene_tx=XM_019820047.2;refGene_gn=LOC102902362;refGene_tx=XM_023245555.1;refGene_gn=LOC102902362;refGene_tx=XM_023245556.1;refGene_gn=LOC102902362;refGene_tx=XM_023245557.1;refGene_gn=LOC102902362;refGene_tx=XM_023245558.1

E2 51102158 51102229
refGene_gn=LOC102902362;refGene_tx=XM_006941731.3;refGene_gn=LOC102902362;refGene_tx=XM_019820040.2;refGene_gn=LOC102902362;refGene_tx=XM_019820041.2;refGene_gn=LOC102902362;refGene_tx=XM_019820042.2;refGene_gn=LOC102902362;refGene_tx=XM_019820044.2;refGene_gn=LOC102902362;refGene_tx=XM_019820045.2;refGene_gn=LOC102902362;refGene_tx=XM_019820046.2;refGene_gn=LOC102902362;refGene_tx=XM_019820047.2;refGene_gn=LOC102902362;refGene_tx=XM_023245557.1;refGene_gn=LOC102902362;refGene_tx=XM_023245558.1
E2 51111916 51111954
refGene_gn=LOC101095392;refGene_tx=XM_006941732.4;refGene_gn=LOC101095392;refGene_tx=XM_011290215.3;refGene_gn=LOC101095392;refGene_tx=XM_019820048.2
E2 51112466 51112558
refGene_gn=LOC101095392;refGene_tx=XM_006941732.4;refGene_gn=LOC101095392;refGene_tx=XM_011290215.3;refGene_gn=LOC101095392;refGene_tx=XM_019820048.2
E2 51113758 51113855
refGene_gn=LOC101095392;refGene_tx=XM_006941732.4;refGene_gn=LOC101095392;refGene_tx=XM_011290215.3;refGene_gn=LOC101095392;refGene_tx=XM_019820048.2
E2 51114147 51114277
refGene_gn=LOC101095392;refGene_tx=XM_006941732.4;refGene_gn=LOC101095392;refGene_tx=XM_011290215.3;refGene_gn=LOC101095392;refGene_tx=XM_019820048.2
E2 51114886 51114993
refGene_gn=LOC101095392;refGene_tx=XM_006941732.4;refGene_gn=LOC101095392;refGene_tx=XM_011290215.3;refGene_gn=LOC101095392;refGene_tx=XM_019820048.2
E2 51115394 51115487
refGene_gn=LOC101095392;refGene_tx=XM_006941732.4;refGene_gn=LOC101095392;refGene_tx=XM_011290215.3;refGene_gn=LOC101095392;refGene_tx=XM_019820048.2
E2 51116068 51116176
refGene_gn=LOC101095392;refGene_tx=XM_006941732.4;refGene_gn=LOC101095392;refGene_tx=XM_011290215.3;refGene_gn=LOC101095392;refGene_tx=XM_019820048.2
E2 51116365 51116487
refGene_gn=LOC101095392;refGene_tx=XM_006941732.4;refGene_gn=LOC101095392;refGene_tx=XM_011290215.3;refGene_gn=LOC101095392;refGene_tx=XM_019820048.2
E2 51116876 51116974
refGene_gn=LOC101095392;refGene_tx=XM_006941732.4;refGene_gn=LOC101095392;refGene_tx=XM_011290215.3;refGene_gn=LOC101095392;refGene_tx=XM_019820048.2
E2 51121670 51121910
refGene_gn=LOC101095392;refGene_tx=XM_006941732.4;refGene_gn=LOC101095392;refGene_tx=XM_011290215.3;refGene_gn=LOC101095392;refGene_tx=XM_019820048.2
E2 51124011 51124130

refGene_gn=LOC101095392;refGene_tx=XM_006941732.4;refGene_gn=LOC101095392;refGene_tx=XM_011290215.3;refGene_gn=LOC101095392;refGene_tx=XM_019820048.2
E2 51124613 51124752
refGene_gn=LOC101095392;refGene_tx=XM_006941732.4;refGene_gn=LOC101095392;refGene_tx=XM_011290215.3;refGene_gn=LOC101095392;refGene_tx=XM_019820048.2
E2 51152718 51152942
refGene_gn=LOC102902613;refGene_tx=XM_006941735.4;refGene_gn=LOC102902613;refGene_tx=XM_006941736.4;refGene_gn=LOC102902613;refGene_tx=XM_006941737.3;refGene_gn=LOC102902613;refGene_tx=XM_019820052.2
E2 51154098 51154232
refGene_gn=LOC102902613;refGene_tx=XM_006941735.4;refGene_gn=LOC102902613;refGene_tx=XM_006941736.4;refGene_gn=LOC102902613;refGene_tx=XM_006941737.3;refGene_gn=LOC102902613;refGene_tx=XM_019820052.2
E2 51155761 51155952
refGene_gn=LOC102902613;refGene_tx=XM_006941735.4;refGene_gn=LOC102902613;refGene_tx=XM_019820052.2
E2 51158114 51158195
refGene_gn=LOC102902613;refGene_tx=XM_006941736.4
E2 51158628 51158651
refGene_gn=LOC102902613;refGene_tx=XM_006941736.4
E2 51160821 51161226
refGene_gn=LOC101088897;refGene_tx=XM_023245673.1;refGene_gn=LOC101088897;refGene_tx=XM_023245674.1
E2 51162810 51162993
refGene_gn=LOC101088897;refGene_tx=XM_023245673.1;refGene_gn=LOC101088897;refGene_tx=XM_023245674.1
E2 51163456 51163506
refGene_gn=LOC102902613;refGene_tx=XM_006941737.3
E2 51168076 51168166
refGene_gn=LOC101088897;refGene_tx=XM_023245673.1;refGene_gn=LOC101088897;refGene_tx=XM_023245674.1
E2 51169719 51169814
refGene_gn=LOC101088897;refGene_tx=XM_023245673.1;refGene_gn=LOC101088897;refGene_tx=XM_023245674.1
E2 51170895 51171008
refGene_gn=LOC101088897;refGene_tx=XM_023245673.1;refGene_gn=LOC101088897;refGene_tx=XM_023245674.1
E2 51171500 51171649
refGene_gn=LOC101088897;refGene_tx=XM_023245673.1;refGene_gn=LOC101088897;refGene_tx=XM_023245674.1
E2 51172278 51172412
refGene_gn=LOC101088897;refGene_tx=XM_023245673.1;refGene_gn=LOC101088897;refGene_tx=XM_023245674.1
E2 51172607 51172763
refGene_gn=LOC101088897;refGene_tx=XM_023245673.1;refGene_gn=LOC101088897;refGene_tx=XM_023245674.1
E2 51174601 51174726
refGene_gn=LOC101088897;refGene_tx=XM_023245673.1;refGene_gn=LOC101088

897;refGene_tx=XM_023245674.1
E2 51176412 51176605
refGene_gn=LOC101088897;refGene_tx=XM_023245673.1;refGene_gn=LOC101088897;refGene_tx=XM_023245674.1
E2 51177679 51177829
refGene_gn=LOC101088897;refGene_tx=XM_023245673.1
E2 51178560 51178678
refGene_gn=LOC101088897;refGene_tx=XM_023245673.1
E2 51179447 51179569
refGene_gn=LOC101088897;refGene_tx=XM_023245673.1
E2 51180264 51180428
refGene_gn=LOC101088897;refGene_tx=XM_023245673.1
E2 51180524 51180606
refGene_gn=LOC101088897;refGene_tx=XM_023245673.1
E2 51185989 51186050
refGene_gn=LOC101088897;refGene_tx=XM_023245673.1;refGene_gn=LOC101088897;refGene_tx=XM_023245674.1
E2 51206614 51206702
refGene_gn=LOC102899111;refGene_tx=XM_006941738.3;refGene_gn=LOC102899111;refGene_tx=XM_019820053.2
E2 51213796 51214156
refGene_gn=LOC102899111;refGene_tx=XM_006941738.3;refGene_gn=LOC102899111;refGene_tx=XM_019820053.2
E2 51214705 51214999
refGene_gn=LOC102899111;refGene_tx=XM_006941738.3;refGene_gn=LOC102899111;refGene_tx=XM_019820053.2
E2 51215325 51215382
refGene_gn=LOC102899111;refGene_tx=XM_006941738.3;refGene_gn=LOC102899111;refGene_tx=XM_019820053.2
E2 51216067 51216372
refGene_gn=LOC102899111;refGene_tx=XM_006941738.3;refGene_gn=LOC102899111;refGene_tx=XM_019820053.2
E2 51221667 51221743 refGene_gn=GLG1;refGene_tx=XM_023245357.1
E2 51222802 51222970
refGene_gn=GLG1;refGene_tx=XM_023245355.1;refGene_gn=GLG1;refGene_tx=XM_023245356.1;refGene_gn=GLG1;refGene_tx=XM_023245357.1
E2 51224945 51225052
refGene_gn=GLG1;refGene_tx=XM_023245355.1;refGene_gn=GLG1;refGene_tx=XM_023245356.1;refGene_gn=GLG1;refGene_tx=XM_023245357.1
E2 51225670 51225791
refGene_gn=GLG1;refGene_tx=XM_023245355.1;refGene_gn=GLG1;refGene_tx=XM_023245356.1;refGene_gn=GLG1;refGene_tx=XM_023245357.1
E2 51227370 51227478
refGene_gn=GLG1;refGene_tx=XM_023245355.1;refGene_gn=GLG1;refGene_tx=XM_023245356.1;refGene_gn=GLG1;refGene_tx=XM_023245357.1
E2 51228658 51228760
refGene_gn=GLG1;refGene_tx=XM_023245355.1;refGene_gn=GLG1;refGene_tx=XM_023245356.1;refGene_gn=GLG1;refGene_tx=XM_023245357.1
E2 51228950 51229093
refGene_gn=GLG1;refGene_tx=XM_023245355.1;refGene_gn=GLG1;refGene_tx=X

M_023245356.1;refGene_gn=GLG1;refGene_tx=XM_023245357.1
E2 51230003 51230127
refGene_gn=GLG1;refGene_tx=XM_023245355.1;refGene_gn=GLG1;refGene_tx=XM_023245356.1;refGene_gn=GLG1;refGene_tx=XM_023245357.1
E2 51232433 51232571
refGene_gn=GLG1;refGene_tx=XM_023245355.1;refGene_gn=GLG1;refGene_tx=XM_023245356.1;refGene_gn=GLG1;refGene_tx=XM_023245357.1
E2 51236152 51236245
refGene_gn=GLG1;refGene_tx=XM_023245355.1;refGene_gn=GLG1;refGene_tx=XM_023245356.1;refGene_gn=GLG1;refGene_tx=XM_023245357.1
E2 51237454 51237572
refGene_gn=GLG1;refGene_tx=XM_023245355.1;refGene_gn=GLG1;refGene_tx=XM_023245356.1;refGene_gn=GLG1;refGene_tx=XM_023245357.1
E2 51238698 51238787
refGene_gn=GLG1;refGene_tx=XM_023245355.1;refGene_gn=GLG1;refGene_tx=XM_023245356.1;refGene_gn=GLG1;refGene_tx=XM_023245357.1
E2 51239811 51239925
refGene_gn=GLG1;refGene_tx=XM_023245355.1;refGene_gn=GLG1;refGene_tx=XM_023245356.1;refGene_gn=GLG1;refGene_tx=XM_023245357.1
E2 51240642 51240705
refGene_gn=GLG1;refGene_tx=XM_023245355.1;refGene_gn=GLG1;refGene_tx=XM_023245356.1;refGene_gn=GLG1;refGene_tx=XM_023245357.1
E2 51241989 51242076
refGene_gn=GLG1;refGene_tx=XM_023245355.1;refGene_gn=GLG1;refGene_tx=XM_023245357.1
E2 51244301 51244439
refGene_gn=GLG1;refGene_tx=XM_023245355.1;refGene_gn=GLG1;refGene_tx=XM_023245356.1;refGene_gn=GLG1;refGene_tx=XM_023245357.1
E2 51245680 51245834
refGene_gn=GLG1;refGene_tx=XM_023245355.1;refGene_gn=GLG1;refGene_tx=XM_023245356.1;refGene_gn=GLG1;refGene_tx=XM_023245357.1
E2 51247411 51247513
refGene_gn=GLG1;refGene_tx=XM_023245355.1;refGene_gn=GLG1;refGene_tx=XM_023245356.1;refGene_gn=GLG1;refGene_tx=XM_023245357.1
E2 51248957 51249079
refGene_gn=GLG1;refGene_tx=XM_023245355.1;refGene_gn=GLG1;refGene_tx=XM_023245356.1;refGene_gn=GLG1;refGene_tx=XM_023245357.1
E2 51252952 51253167
refGene_gn=GLG1;refGene_tx=XM_023245355.1;refGene_gn=GLG1;refGene_tx=XM_023245356.1;refGene_gn=GLG1;refGene_tx=XM_023245357.1
E2 51256604 51256788
refGene_gn=GLG1;refGene_tx=XM_023245355.1;refGene_gn=GLG1;refGene_tx=XM_023245356.1;refGene_gn=GLG1;refGene_tx=XM_023245357.1
E2 51257700 51257772
refGene_gn=GLG1;refGene_tx=XM_023245355.1;refGene_gn=GLG1;refGene_tx=XM_023245356.1;refGene_gn=GLG1;refGene_tx=XM_023245357.1
E2 51258488 51258692
refGene_gn=GLG1;refGene_tx=XM_023245355.1;refGene_gn=GLG1;refGene_tx=XM_023245356.1;refGene_gn=GLG1;refGene_tx=XM_023245357.1
E2 51264359 51264575

refGene_gn=GLG1;refGene_tx=XM_023245355.1;refGene_gn=GLG1;refGene_tx=XM_023245356.1;refGene_gn=GLG1;refGene_tx=XM_023245357.1
E2 51267944 51268031
refGene_gn=GLG1;refGene_tx=XM_023245355.1;refGene_gn=GLG1;refGene_tx=XM_023245356.1;refGene_gn=GLG1;refGene_tx=XM_023245357.1
E2 51278766 51278799
refGene_gn=GLG1;refGene_tx=XM_023245355.1;refGene_gn=GLG1;refGene_tx=XM_023245356.1;refGene_gn=GLG1;refGene_tx=XM_023245357.1
E2 51367521 51367962
refGene_gn=GLG1;refGene_tx=XM_023245355.1;refGene_gn=GLG1;refGene_tx=XM_023245356.1;refGene_gn=GLG1;refGene_tx=XM_023245357.1
E2 51388659 51388803 refGene_gn=RFWD3;refGene_tx=XM_003998245.5
E2 51390639 51390851 refGene_gn=RFWD3;refGene_tx=XM_003998245.5
E2 51391728 51391943 refGene_gn=RFWD3;refGene_tx=XM_003998245.5
E2 51393984 51394161 refGene_gn=RFWD3;refGene_tx=XM_003998245.5
E2 51395446 51395597 refGene_gn=RFWD3;refGene_tx=XM_003998245.5
E2 51399822 51400051 refGene_gn=RFWD3;refGene_tx=XM_003998245.5
E2 51400546 51400661 refGene_gn=RFWD3;refGene_tx=XM_003998245.5
E2 51409172 51409264 refGene_gn=RFWD3;refGene_tx=XM_003998245.5
E2 51409360 51409552 refGene_gn=RFWD3;refGene_tx=XM_003998245.5
E2 51411167 51411238 refGene_gn=RFWD3;refGene_tx=XM_003998245.5
E2 51414251 51414448 refGene_gn=RFWD3;refGene_tx=XM_003998245.5
E2 51423599 51424069 refGene_gn=RFWD3;refGene_tx=XM_003998245.5
E2 51444111 51444191 refGene_gn=FA2H;refGene_tx=XM_003998246.3
E2 51446988 51447241 refGene_gn=FA2H;refGene_tx=XM_003998246.3
E2 51448547 51448720 refGene_gn=FA2H;refGene_tx=XM_003998246.3
E2 51456789 51456896 refGene_gn=FA2H;refGene_tx=XM_003998246.3
E2 51457707 51457850 refGene_gn=FA2H;refGene_tx=XM_003998246.3
E2 51469708 51469801 refGene_gn=FA2H;refGene_tx=XM_003998246.3
E2 51495749 51496019 refGene_gn=FA2H;refGene_tx=XM_003998246.3
E2 51564116 51564352
refGene_gn=WDR59;refGene_tx=XM_003998247.4;refGene_gn=WDR59;refGene_tx=XM_006941739.3;refGene_gn=WDR59;refGene_tx=XM_011290211.2;refGene_gn=WDR59;refGene_tx=XM_011290212.2
E2 51564638 51564757 refGene_gn=WDR59;refGene_tx=XM_011290210.3
E2 51571172 51571315
refGene_gn=WDR59;refGene_tx=XM_003998247.4;refGene_gn=WDR59;refGene_tx=XM_006941739.3;refGene_gn=WDR59;refGene_tx=XM_011290210.3;refGene_gn=WDR59;refGene_tx=XM_011290211.2;refGene_gn=WDR59;refGene_tx=XM_011290212.2
E2 51571469 51571596
refGene_gn=WDR59;refGene_tx=XM_003998247.4;refGene_gn=WDR59;refGene_tx=XM_006941739.3;refGene_gn=WDR59;refGene_tx=XM_011290210.3;refGene_gn=WDR59;refGene_tx=XM_011290211.2;refGene_gn=WDR59;refGene_tx=XM_011290212.2
E2 51572754 51572827
refGene_gn=WDR59;refGene_tx=XM_003998247.4;refGene_gn=WDR59;refGene_tx=XM_006941739.3;refGene_gn=WDR59;refGene_tx=XM_011290210.3;refGene_gn=WDR59;refGene_tx=XM_011290211.2;refGene_gn=WDR59;refGene_tx=XM_011290212.2

E2 51572922 51573073
refGene_gn=WDR59;refGene_tx=XM_003998247.4;refGene_gn=WDR59;refGene_tx=XM_006941739.3;refGene_gn=WDR59;refGene_tx=XM_011290210.3;refGene_gn=WDR59;refGene_tx=XM_011290211.2;refGene_gn=WDR59;refGene_tx=XM_011290212.2
E2 51574429 51574542
refGene_gn=WDR59;refGene_tx=XM_003998247.4;refGene_gn=WDR59;refGene_tx=XM_006941739.3;refGene_gn=WDR59;refGene_tx=XM_011290210.3;refGene_gn=WDR59;refGene_tx=XM_011290211.2;refGene_gn=WDR59;refGene_tx=XM_011290212.2
E2 51577952 51578034
refGene_gn=WDR59;refGene_tx=XM_003998247.4;refGene_gn=WDR59;refGene_tx=XM_006941739.3;refGene_gn=WDR59;refGene_tx=XM_011290210.3;refGene_gn=WDR59;refGene_tx=XM_011290211.2;refGene_gn=WDR59;refGene_tx=XM_011290212.2
E2 51579160 51579294
refGene_gn=WDR59;refGene_tx=XM_003998247.4;refGene_gn=WDR59;refGene_tx=XM_006941739.3;refGene_gn=WDR59;refGene_tx=XM_011290210.3;refGene_gn=WDR59;refGene_tx=XM_011290211.2;refGene_gn=WDR59;refGene_tx=XM_011290212.2
E2 51579317 51579344 refGene_gn=WDR59;refGene_tx=XM_019820055.2
E2 51584754 51584811
refGene_gn=WDR59;refGene_tx=XM_006941739.3;refGene_gn=WDR59;refGene_tx=XM_011290210.3;refGene_gn=WDR59;refGene_tx=XM_011290211.2;refGene_gn=WDR59;refGene_tx=XM_011290212.2
E2 51588389 51588543
refGene_gn=WDR59;refGene_tx=XM_003998247.4;refGene_gn=WDR59;refGene_tx=XM_006941739.3;refGene_gn=WDR59;refGene_tx=XM_011290210.3;refGene_gn=WDR59;refGene_tx=XM_011290211.2;refGene_gn=WDR59;refGene_tx=XM_011290212.2;refGene_gn=WDR59;refGene_tx=XM_019820055.2
E2 51592707 51592723 refGene_gn=WDR59;refGene_tx=XM_019820056.2
E2 51593269 51593339
refGene_gn=WDR59;refGene_tx=XM_003998247.4;refGene_gn=WDR59;refGene_tx=XM_006941739.3;refGene_gn=WDR59;refGene_tx=XM_011290210.3;refGene_gn=WDR59;refGene_tx=XM_011290211.2;refGene_gn=WDR59;refGene_tx=XM_011290212.2;refGene_gn=WDR59;refGene_tx=XM_019820055.2;refGene_gn=WDR59;refGene_tx=XM_019820056.2
E2 51593787 51593944
refGene_gn=WDR59;refGene_tx=XM_003998247.4;refGene_gn=WDR59;refGene_tx=XM_006941739.3;refGene_gn=WDR59;refGene_tx=XM_011290210.3;refGene_gn=WDR59;refGene_tx=XM_011290211.2;refGene_gn=WDR59;refGene_tx=XM_011290212.2;refGene_gn=WDR59;refGene_tx=XM_019820055.2;refGene_gn=WDR59;refGene_tx=XM_019820056.2
E2 51594257 51594353
refGene_gn=WDR59;refGene_tx=XM_003998247.4;refGene_gn=WDR59;refGene_tx=XM_006941739.3;refGene_gn=WDR59;refGene_tx=XM_011290210.3;refGene_gn=WDR59;refGene_tx=XM_011290211.2;refGene_gn=WDR59;refGene_tx=XM_011290212.2;refGene_gn=WDR59;refGene_tx=XM_019820055.2;refGene_gn=WDR59;refGene_tx=XM_019820056.2
E2 51596362 51596527

refGene_gn=WDR59;refGene_tx=XM_003998247.4;refGene_gn=WDR59;refGene_tx=XM_006941739.3;refGene_gn=WDR59;refGene_tx=XM_011290210.3;refGene_gn=WDR59;refGene_tx=XM_011290211.2;refGene_gn=WDR59;refGene_tx=XM_011290212.2;refGene_gn=WDR59;refGene_tx=XM_019820055.2;refGene_gn=WDR59;refGene_tx=XM_019820056.2
E2 51598793 51598918
refGene_gn=WDR59;refGene_tx=XM_003998247.4;refGene_gn=WDR59;refGene_tx=XM_006941739.3;refGene_gn=WDR59;refGene_tx=XM_011290210.3;refGene_gn=WDR59;refGene_tx=XM_011290211.2;refGene_gn=WDR59;refGene_tx=XM_011290212.2;refGene_gn=WDR59;refGene_tx=XM_019820055.2;refGene_gn=WDR59;refGene_tx=XM_019820056.2
E2 51599053 51599186
refGene_gn=WDR59;refGene_tx=XM_003998247.4;refGene_gn=WDR59;refGene_tx=XM_006941739.3;refGene_gn=WDR59;refGene_tx=XM_011290210.3;refGene_gn=WDR59;refGene_tx=XM_011290211.2;refGene_gn=WDR59;refGene_tx=XM_011290212.2;refGene_gn=WDR59;refGene_tx=XM_019820055.2;refGene_gn=WDR59;refGene_tx=XM_019820056.2
E2 51600351 51600431
refGene_gn=WDR59;refGene_tx=XM_003998247.4;refGene_gn=WDR59;refGene_tx=XM_006941739.3;refGene_gn=WDR59;refGene_tx=XM_011290210.3;refGene_gn=WDR59;refGene_tx=XM_011290211.2;refGene_gn=WDR59;refGene_tx=XM_011290212.2;refGene_gn=WDR59;refGene_tx=XM_019820055.2;refGene_gn=WDR59;refGene_tx=XM_019820056.2
E2 51603685 51603842
refGene_gn=WDR59;refGene_tx=XM_003998247.4;refGene_gn=WDR59;refGene_tx=XM_006941739.3;refGene_gn=WDR59;refGene_tx=XM_011290210.3;refGene_gn=WDR59;refGene_tx=XM_011290211.2;refGene_gn=WDR59;refGene_tx=XM_011290212.2;refGene_gn=WDR59;refGene_tx=XM_019820055.2;refGene_gn=WDR59;refGene_tx=XM_019820056.2
E2 51605968 51606046
refGene_gn=WDR59;refGene_tx=XM_003998247.4;refGene_gn=WDR59;refGene_tx=XM_006941739.3;refGene_gn=WDR59;refGene_tx=XM_011290210.3;refGene_gn=WDR59;refGene_tx=XM_011290211.2;refGene_gn=WDR59;refGene_tx=XM_011290212.2;refGene_gn=WDR59;refGene_tx=XM_019820055.2;refGene_gn=WDR59;refGene_tx=XM_019820056.2
E2 51610120 51610237
refGene_gn=WDR59;refGene_tx=XM_003998247.4;refGene_gn=WDR59;refGene_tx=XM_006941739.3;refGene_gn=WDR59;refGene_tx=XM_011290210.3;refGene_gn=WDR59;refGene_tx=XM_011290211.2;refGene_gn=WDR59;refGene_tx=XM_011290212.2;refGene_gn=WDR59;refGene_tx=XM_019820055.2;refGene_gn=WDR59;refGene_tx=XM_019820056.2
E2 51614041 51614130
refGene_gn=WDR59;refGene_tx=XM_003998247.4;refGene_gn=WDR59;refGene_tx=XM_006941739.3;refGene_gn=WDR59;refGene_tx=XM_011290210.3;refGene_gn=WDR59;refGene_tx=XM_011290211.2;refGene_gn=WDR59;refGene_tx=XM_011290212.2;refGene_gn=WDR59;refGene_tx=XM_019820055.2;refGene_gn=WDR59;refGene_tx=XM_019820056.2
E2 51617988 51618026
refGene_gn=WDR59;refGene_tx=XM_003998247.4;refGene_gn=WDR59;refGene_tx=XM_006941739.3;refGene_gn=WDR59;refGene_tx=XM_011290210.3;refGene_gn=

WDR59;refGene_tx=XM_011290211.2;refGene_gn=WDR59;refGene_tx=XM_0112902
12.2;refGene_gn=WDR59;refGene_tx=XM_019820055.2;refGene_gn=WDR59;refGe
ne_tx=XM_019820056.2
E2 51620145 51620226
refGene_gn=WDR59;refGene_tx=XM_003998247.4;refGene_gn=WDR59;refGene_tx
=XM_006941739.3;refGene_gn=WDR59;refGene_tx=XM_011290210.3;refGene_gn=
WDR59;refGene_tx=XM_011290211.2;refGene_gn=WDR59;refGene_tx=XM_0112902
12.2;refGene_gn=WDR59;refGene_tx=XM_019820055.2;refGene_gn=WDR59;refGe
ne_tx=XM_019820056.2
E2 51621227 51621313
refGene_gn=WDR59;refGene_tx=XM_003998247.4;refGene_gn=WDR59;refGene_tx
=XM_006941739.3;refGene_gn=WDR59;refGene_tx=XM_011290210.3;refGene_gn=
WDR59;refGene_tx=XM_011290211.2;refGene_gn=WDR59;refGene_tx=XM_0112902
12.2;refGene_gn=WDR59;refGene_tx=XM_019820055.2;refGene_gn=WDR59;refGe
ne_tx=XM_019820056.2
E2 51626476 51626612
refGene_gn=WDR59;refGene_tx=XM_003998247.4;refGene_gn=WDR59;refGene_tx
=XM_006941739.3;refGene_gn=WDR59;refGene_tx=XM_011290210.3;refGene_gn=
WDR59;refGene_tx=XM_011290211.2;refGene_gn=WDR59;refGene_tx=XM_0112902
12.2;refGene_gn=WDR59;refGene_tx=XM_019820055.2;refGene_gn=WDR59;refGe
ne_tx=XM_019820056.2
E2 51631106 51631156
refGene_gn=WDR59;refGene_tx=XM_003998247.4;refGene_gn=WDR59;refGene_tx
=XM_006941739.3;refGene_gn=WDR59;refGene_tx=XM_011290210.3;refGene_gn=
WDR59;refGene_tx=XM_011290211.2;refGene_gn=WDR59;refGene_tx=XM_0112902
12.2;refGene_gn=WDR59;refGene_tx=XM_019820055.2;refGene_gn=WDR59;refGe
ne_tx=XM_019820056.2
E2 51647927 51647981
refGene_gn=WDR59;refGene_tx=XM_003998247.4;refGene_gn=WDR59;refGene_tx
=XM_006941739.3;refGene_gn=WDR59;refGene_tx=XM_011290210.3;refGene_gn=
WDR59;refGene_tx=XM_019820055.2;refGene_gn=WDR59;refGene_tx=XM_0198200
56.2
E2 51659280 51659704
refGene_gn=ZNRF1;refGene_tx=XM_003998248.5;refGene_gn=ZNRF1;refGene_tx
=XM_006941741.4;refGene_gn=ZNRF1;refGene_tx=XM_006941742.4
E2 51750549 51750645
refGene_gn=ZNRF1;refGene_tx=XM_003998248.5;refGene_gn=ZNRF1;refGene_tx
=XM_006941741.4;refGene_gn=ZNRF1;refGene_tx=XM_006941742.4
E2 51760955 51761061
refGene_gn=ZNRF1;refGene_tx=XM_003998248.5;refGene_gn=ZNRF1;refGene_tx
=XM_006941741.4;refGene_gn=ZNRF1;refGene_tx=XM_006941742.4
E2 51762713 51762771
refGene_gn=ZNRF1;refGene_tx=XM_003998248.5;refGene_gn=ZNRF1;refGene_tx
=XM_006941741.4;refGene_gn=ZNRF1;refGene_tx=XM_006941742.4
E2 51768336 51768502
refGene_gn=LDHD;refGene_tx=XM_003998251.5;refGene_gn=LDHD;refGene_tx=X
M_019820057.2
E2 51768591 51768703
refGene_gn=LDHD;refGene_tx=XM_003998251.5;refGene_gn=LDHD;refGene_tx=X
M_019820057.2

E2 51768789 51768880
refGene_gn=LDHD;refGene_tx=XM_003998251.5;refGene_gn=LDHD;refGene_tx=XM_019820057.2
E2 51769455 51769583
refGene_gn=LDHD;refGene_tx=XM_003998251.5;refGene_gn=LDHD;refGene_tx=XM_019820057.2
E2 51769661 51769790
refGene_gn=LDHD;refGene_tx=XM_003998251.5;refGene_gn=LDHD;refGene_tx=XM_019820057.2
E2 51769883 51770083
refGene_gn=LDHD;refGene_tx=XM_003998251.5;refGene_gn=LDHD;refGene_tx=XM_019820057.2
E2 51770378 51770538
refGene_gn=LDHD;refGene_tx=XM_003998251.5;refGene_gn=LDHD;refGene_tx=XM_019820057.2
E2 51770663 51770805
refGene_gn=LDHD;refGene_tx=XM_003998251.5;refGene_gn=LDHD;refGene_tx=XM_019820057.2
E2 51771011 51771153
refGene_gn=LDHD;refGene_tx=XM_003998251.5;refGene_gn=LDHD;refGene_tx=XM_019820057.2
E2 51771382 51771495
refGene_gn=LDHD;refGene_tx=XM_003998251.5;refGene_gn=LDHD;refGene_tx=XM_019820057.2
E2 51772229 51772304 refGene_gn=LDHD;refGene_tx=XM_003998251.5
E2 51795167 51795182
refGene_gn=ZFP1;refGene_tx=XM_003998252.5;refGene_gn=ZFP1;refGene_tx=XM_011290247.3
E2 51806687 51806814
refGene_gn=ZFP1;refGene_tx=XM_003998252.5;refGene_gn=ZFP1;refGene_tx=XM_011290247.3
E2 51808382 51809458
refGene_gn=ZFP1;refGene_tx=XM_003998252.5;refGene_gn=ZFP1;refGene_tx=XM_011290247.3;refGene_gn=ZFP1;refGene_tx=XM_019820058.2;refGene_gn=ZFP1;refGene_tx=XM_019820059.1
E2 51835631 51835793
refGene_gn=LOC101096125;refGene_tx=XM_023245390.1
E2 51836144 51836278
refGene_gn=LOC101096125;refGene_tx=XM_023245390.1
E2 51837421 51837602
refGene_gn=LOC101096125;refGene_tx=XM_023245390.1
E2 51837852 51837931
refGene_gn=LOC101096125;refGene_tx=XM_023245390.1
E2 51838032 51838112
refGene_gn=LOC101096125;refGene_tx=XM_023245390.1
E2 51838226 51838330
refGene_gn=LOC101096125;refGene_tx=XM_023245390.1
E2 51840173 51840228
refGene_gn=LOC101096125;refGene_tx=XM_023245390.1
E2 51854768 51854820

refGene_gn=LOC101091168;refGene_tx=XM_011290248.2
E2 51855477 51855581
refGene_gn=LOC101091168;refGene_tx=XM_011290248.2
E2 51855695 51855775
refGene_gn=LOC101091168;refGene_tx=XM_011290248.2
E2 51856116 51856195
refGene_gn=LOC101091168;refGene_tx=XM_011290248.2
E2 51856478 51856659
refGene_gn=LOC101091168;refGene_tx=XM_011290248.2
E2 51856745 51856879
refGene_gn=LOC101091168;refGene_tx=XM_011290248.2
E2 51857526 51857688
refGene_gn=LOC101091168;refGene_tx=XM_011290248.2
E2 51858201 51858253
refGene_gn=LOC101096371;refGene_tx=XM_023246162.1
E2 51859053 51859157
refGene_gn=LOC101096371;refGene_tx=XM_023246162.1
E2 51859243 51859323
refGene_gn=LOC101096371;refGene_tx=XM_023246162.1
E2 51859687 51859766
refGene_gn=LOC101096371;refGene_tx=XM_023246162.1
E2 51860194 51860375
refGene_gn=LOC101096371;refGene_tx=XM_023246162.1
E2 51861471 51861605
refGene_gn=LOC101096371;refGene_tx=XM_023246162.1
E2 51861955 51862117
refGene_gn=LOC101096371;refGene_tx=XM_023246162.1
E2 51863508 51864021
refGene_gn=BCAR1;refGene_tx=XM_011290249.3;refGene_gn=BCAR1;refGene_tx=XM_019820063.2;refGene_gn=BCAR1;refGene_tx=XM_023246161.1
E2 51867194 51867284
refGene_gn=BCAR1;refGene_tx=XM_011290249.3;refGene_gn=BCAR1;refGene_tx=XM_019820063.2;refGene_gn=BCAR1;refGene_tx=XM_023246161.1
E2 51868209 51869307
refGene_gn=BCAR1;refGene_tx=XM_011290249.3;refGene_gn=BCAR1;refGene_tx=XM_019820063.2;refGene_gn=BCAR1;refGene_tx=XM_023246161.1
E2 51870479 51870596
refGene_gn=BCAR1;refGene_tx=XM_011290249.3;refGene_gn=BCAR1;refGene_tx=XM_019820063.2;refGene_gn=BCAR1;refGene_tx=XM_023246161.1
E2 51870767 51870929
refGene_gn=BCAR1;refGene_tx=XM_011290249.3;refGene_gn=BCAR1;refGene_tx=XM_019820063.2;refGene_gn=BCAR1;refGene_tx=XM_023246161.1
E2 51876545 51876584
refGene_gn=BCAR1;refGene_tx=XM_019820063.2;refGene_gn=BCAR1;refGene_tx=XM_023246161.1
E2 51876668 51877280
refGene_gn=BCAR1;refGene_tx=XM_011290249.3;refGene_gn=BCAR1;refGene_tx=XM_019820063.2;refGene_gn=BCAR1;refGene_tx=XM_023246161.1
E2 51887576 51887588 refGene_gn=BCAR1;refGene_tx=XM_023246161.1
E2 51900601 51900613

refGene_gn=BCAR1;refGene_tx=XM_011290249.3;refGene_gn=BCAR1;refGene_tx=XM_019820063.2
E2 51923616 51923707 refGene_gn=CFDP1;refGene_tx=XM_003998254.5;refGene_gn=CFDP1;refGene_tx=XM_019820079.2;refGene_gn=CFDP1;refGene_tx=XM_019820080.2
E2 51934109 51934268 refGene_gn=CFDP1;refGene_tx=XM_003998254.5;refGene_gn=CFDP1;refGene_tx=XM_019820079.2
E2 51995263 51995342 refGene_gn=CFDP1;refGene_tx=XM_023245704.1
E2 52021246 52021366 refGene_gn=CFDP1;refGene_tx=XM_003998254.5;refGene_gn=CFDP1;refGene_tx=XM_019820079.2;refGene_gn=CFDP1;refGene_tx=XM_019820080.2;refGene_gn=CFDP1;refGene_tx=XM_023245704.1
E2 52032667 52032795 refGene_gn=CFDP1;refGene_tx=XM_003998254.5;refGene_gn=CFDP1;refGene_tx=XM_019820079.2;refGene_gn=CFDP1;refGene_tx=XM_019820080.2;refGene_gn=CFDP1;refGene_tx=XM_023245704.1
E2 52033331 52033551 refGene_gn=CFDP1;refGene_tx=XM_003998254.5;refGene_gn=CFDP1;refGene_tx=XM_019820079.2;refGene_gn=CFDP1;refGene_tx=XM_019820080.2;refGene_gn=CFDP1;refGene_tx=XM_023245704.1
E2 52034829 52034947 refGene_gn=CFDP1;refGene_tx=XM_003998254.5;refGene_gn=CFDP1;refGene_tx=XM_019820079.2;refGene_gn=CFDP1;refGene_tx=XM_019820080.2;refGene_gn=CFDP1;refGene_tx=XM_023245704.1
E2 52038115 52038167 refGene_gn=CFDP1;refGene_tx=XM_019820079.2
E2 52044272 52044336 refGene_gn=CFDP1;refGene_tx=XM_003998254.5;refGene_gn=CFDP1;refGene_tx=XM_019820080.2;refGene_gn=CFDP1;refGene_tx=XM_023245704.1
E2 52054771 52054902 refGene_gn=TMEM170A;refGene_tx=XM_003998255.5;refGene_gn=TMEM170A;refGene_tx=XM_011290251.3;refGene_gn=TMEM170A;refGene_tx=XM_019820083.2;refGene_gn=TMEM170A;refGene_tx=XM_023245031.1
E2 52058117 52058288 refGene_gn=TMEM170A;refGene_tx=XM_003998255.5;refGene_gn=TMEM170A;refGene_tx=XM_011290251.3;refGene_gn=TMEM170A;refGene_tx=XM_019820083.2;refGene_gn=TMEM170A;refGene_tx=XM_023245031.1
E2 52070702 52070835 refGene_gn=TMEM170A;refGene_tx=XM_003998255.5;refGene_gn=TMEM170A;refGene_tx=XM_011290251.3
E2 52078762 52079950 refGene_gn=LOC101097117;refGene_tx=XM_023245030.1
E2 52094185 52094366 refGene_gn=TMEM231;refGene_tx=XM_003998257.5
E2 52094722 52094828 refGene_gn=TMEM231;refGene_tx=XM_003998257.5
E2 52095971 52096053 refGene_gn=TMEM231;refGene_tx=XM_003998257.5
E2 52098133 52098277 refGene_gn=TMEM231;refGene_tx=XM_003998257.5
E2 52098616 52098745 refGene_gn=TMEM231;refGene_tx=XM_003998257.5
E2 52111205 52111375 refGene_gn=TMEM231;refGene_tx=XM_003998257.5
E2 52111470 52111609 refGene_gn=TMEM231;refGene_tx=XM_003998257.5
E2 52118883 52118917

refGene_gn=GABARAPL2;refGene_tx=XM_006941744.2;refGene_gn=GABARAPL2;refGene_tx=XM_019820085.1
E2 52119249 52119305
refGene_gn=GABARAPL2;refGene_tx=XM_006941744.2;refGene_gn=GABARAPL2;refGene_tx=XM_019820085.1
E2 52120477 52120650
refGene_gn=GABARAPL2;refGene_tx=XM_006941744.2;refGene_gn=GABARAPL2;refGene_tx=XM_019820085.1
E2 52123937 52124054
refGene_gn=GABARAPL2;refGene_tx=XM_019820085.1
E2 52128994 52129085
refGene_gn=GABARAPL2;refGene_tx=XM_006941744.2;refGene_gn=GABARAPL2;refGene_tx=XM_019820085.1
E2 52129647 52129723 refGene_gn=ADAT1;refGene_tx=XM_019820070.2
E2 52139041 52139150 refGene_gn=ADAT1;refGene_tx=XM_011290252.3
E2 52146072 52146205
refGene_gn=ADAT1;refGene_tx=XM_003998258.4;refGene_gn=ADAT1;refGene_tx=XM_006941745.3;refGene_gn=ADAT1;refGene_tx=XM_019820068.2;refGene_gn=ADAT1;refGene_tx=XM_019820069.2;refGene_gn=ADAT1;refGene_tx=XM_019820074.1;refGene_gn=ADAT1;refGene_tx=XM_019820078.1
E2 52148800 52148887
refGene_gn=ADAT1;refGene_tx=XM_003998258.4;refGene_gn=ADAT1;refGene_tx=XM_006941745.3;refGene_gn=ADAT1;refGene_tx=XM_011290252.3;refGene_gn=ADAT1;refGene_tx=XM_019820068.2;refGene_gn=ADAT1;refGene_tx=XM_019820069.2;refGene_gn=ADAT1;refGene_tx=XM_019820070.2;refGene_gn=ADAT1;refGene_tx=XM_019820074.1;refGene_gn=ADAT1;refGene_tx=XM_019820075.2;refGene_gn=ADAT1;refGene_tx=XM_019820076.2;refGene_gn=ADAT1;refGene_tx=XM_019820077.2;refGene_gn=ADAT1;refGene_tx=XM_019820078.1
E2 52154395 52154495
refGene_gn=ADAT1;refGene_tx=XM_003998258.4;refGene_gn=ADAT1;refGene_tx=XM_006941745.3;refGene_gn=ADAT1;refGene_tx=XM_011290252.3;refGene_gn=ADAT1;refGene_tx=XM_019820068.2;refGene_gn=ADAT1;refGene_tx=XM_019820069.2;refGene_gn=ADAT1;refGene_tx=XM_019820070.2;refGene_gn=ADAT1;refGene_tx=XM_019820074.1;refGene_gn=ADAT1;refGene_tx=XM_019820078.1
E2 52156259 52156405
refGene_gn=ADAT1;refGene_tx=XM_003998258.4;refGene_gn=ADAT1;refGene_tx=XM_006941745.3;refGene_gn=ADAT1;refGene_tx=XM_011290252.3;refGene_gn=ADAT1;refGene_tx=XM_019820068.2;refGene_gn=ADAT1;refGene_tx=XM_019820069.2;refGene_gn=ADAT1;refGene_tx=XM_019820070.2;refGene_gn=ADAT1;refGene_tx=XM_019820074.1;refGene_gn=ADAT1;refGene_tx=XM_019820075.2;refGene_gn=ADAT1;refGene_tx=XM_019820076.2;refGene_gn=ADAT1;refGene_tx=XM_019820077.2;refGene_gn=ADAT1;refGene_tx=XM_019820078.1
E2 52159091 52159695
refGene_gn=ADAT1;refGene_tx=XM_003998258.4;refGene_gn=ADAT1;refGene_tx=XM_006941745.3;refGene_gn=ADAT1;refGene_tx=XM_011290252.3;refGene_gn=ADAT1;refGene_tx=XM_019820068.2;refGene_gn=ADAT1;refGene_tx=XM_019820069.2;refGene_gn=ADAT1;refGene_tx=XM_019820070.2;refGene_gn=ADAT1;refGene_tx=XM_019820074.1;refGene_gn=ADAT1;refGene_tx=XM_019820075.2;refGene_gn=ADAT1;refGene_tx=XM_019820076.2;refGene_gn=ADAT1;refGene_tx=XM_019820077.2;refGene_gn=ADAT1;refGene_tx=XM_019820078.1

E2 52162900 52163031
refGene_gn=ADAT1;refGene_tx=XM_003998258.4;refGene_gn=ADAT1;refGene_tx=XM_006941745.3;refGene_gn=ADAT1;refGene_tx=XM_011290252.3;refGene_gn=ADAT1;refGene_tx=XM_019820068.2;refGene_gn=ADAT1;refGene_tx=XM_019820069.2;refGene_gn=ADAT1;refGene_tx=XM_019820070.2;refGene_gn=ADAT1;refGene_tx=XM_019820074.1;refGene_gn=ADAT1;refGene_tx=XM_019820075.2;refGene_gn=ADAT1;refGene_tx=XM_019820076.2;refGene_gn=ADAT1;refGene_tx=XM_019820077.2
E2 52163477 52163532
refGene_gn=ADAT1;refGene_tx=XM_003998258.4;refGene_gn=ADAT1;refGene_tx=XM_006941745.3;refGene_gn=ADAT1;refGene_tx=XM_011290252.3;refGene_gn=ADAT1;refGene_tx=XM_019820068.2;refGene_gn=ADAT1;refGene_tx=XM_019820069.2;refGene_gn=ADAT1;refGene_tx=XM_019820070.2;refGene_gn=ADAT1;refGene_tx=XM_019820075.2;refGene_gn=ADAT1;refGene_tx=XM_019820076.2;refGene_gn=ADAT1;refGene_tx=XM_019820077.2
E2 52164643 52164712
refGene_gn=ADAT1;refGene_tx=XM_003998258.4;refGene_gn=ADAT1;refGene_tx=XM_006941745.3;refGene_gn=ADAT1;refGene_tx=XM_011290252.3;refGene_gn=ADAT1;refGene_tx=XM_019820068.2;refGene_gn=ADAT1;refGene_tx=XM_019820069.2;refGene_gn=ADAT1;refGene_tx=XM_019820070.2;refGene_gn=ADAT1;refGene_tx=XM_019820074.1;refGene_gn=ADAT1;refGene_tx=XM_019820075.2;refGene_gn=ADAT1;refGene_tx=XM_019820076.2;refGene_gn=ADAT1;refGene_tx=XM_019820077.2
E2 52165003 52165172
refGene_gn=ADAT1;refGene_tx=XM_003998258.4;refGene_gn=ADAT1;refGene_tx=XM_006941745.3;refGene_gn=ADAT1;refGene_tx=XM_011290252.3;refGene_gn=ADAT1;refGene_tx=XM_019820068.2;refGene_gn=ADAT1;refGene_tx=XM_019820069.2;refGene_gn=ADAT1;refGene_tx=XM_019820070.2;refGene_gn=ADAT1;refGene_tx=XM_019820075.2;refGene_gn=ADAT1;refGene_tx=XM_019820076.2;refGene_gn=ADAT1;refGene_tx=XM_019820077.2
E2 52172798 52172897
refGene_gn=KARS;refGene_tx=XM_003998259.4;refGene_gn=KARS;refGene_tx=XM_003998260.3
E2 52173385 52173529
refGene_gn=KARS;refGene_tx=XM_003998259.4;refGene_gn=KARS;refGene_tx=XM_003998260.3
E2 52174234 52174361
refGene_gn=KARS;refGene_tx=XM_003998259.4;refGene_gn=KARS;refGene_tx=XM_003998260.3
E2 52175010 52175096
refGene_gn=KARS;refGene_tx=XM_003998259.4;refGene_gn=KARS;refGene_tx=XM_003998260.3
E2 52175434 52175520
refGene_gn=KARS;refGene_tx=XM_003998259.4;refGene_gn=KARS;refGene_tx=XM_003998260.3
E2 52175678 52175852
refGene_gn=KARS;refGene_tx=XM_003998259.4;refGene_gn=KARS;refGene_tx=XM_003998260.3
E2 52175961 52176124
refGene_gn=KARS;refGene_tx=XM_003998259.4;refGene_gn=KARS;refGene_tx=X

M_003998260.3
E2 52177385 52177505
refGene_gn=KARS;refGene_tx=XM_003998259.4;refGene_gn=KARS;refGene_tx=XM_003998260.3
E2 52178419 52178545
refGene_gn=KARS;refGene_tx=XM_003998259.4;refGene_gn=KARS;refGene_tx=XM_003998260.3
E2 52178630 52178817
refGene_gn=KARS;refGene_tx=XM_003998259.4;refGene_gn=KARS;refGene_tx=XM_003998260.3
E2 52179185 52179279
refGene_gn=KARS;refGene_tx=XM_003998259.4;refGene_gn=KARS;refGene_tx=XM_003998260.3
E2 52181842 52182008
refGene_gn=KARS;refGene_tx=XM_003998259.4;refGene_gn=KARS;refGene_tx=XM_003998260.3
E2 52182962 52183122
refGene_gn=KARS;refGene_tx=XM_003998259.4;refGene_gn=KARS;refGene_tx=XM_003998260.3
E2 52184758 52184904 refGene_gn=KARS;refGene_tx=XM_003998260.3
E2 52187444 52187506 refGene_gn=KARS;refGene_tx=XM_003998259.4
E2 52187740 52188431 refGene_gn=TERF2IP;refGene_tx=XM_019820087.2
E2 52192839 52192964 refGene_gn=TERF2IP;refGene_tx=XM_019820087.2
E2 52195428 52195833 refGene_gn=TERF2IP;refGene_tx=XM_019820087.2
E2 52661673 52661758
refGene_gn=CNTNAP4;refGene_tx=XM_011290253.3;refGene_gn=CNTNAP4;refGene_tx=XM_019820088.2;refGene_gn=CNTNAP4;refGene_tx=XM_019820089.2;refGene_gn=CNTNAP4;refGene_tx=XM_023245046.1;refGene_gn=CNTNAP4;refGene_tx=XM_023245047.1
E2 52699156 52699267
refGene_gn=CNTNAP4;refGene_tx=XM_011290253.3;refGene_gn=CNTNAP4;refGene_tx=XM_019820088.2;refGene_gn=CNTNAP4;refGene_tx=XM_019820089.2;refGene_gn=CNTNAP4;refGene_tx=XM_023245046.1;refGene_gn=CNTNAP4;refGene_tx=XM_023245047.1
E2 52737801 52737995
refGene_gn=CNTNAP4;refGene_tx=XM_011290253.3;refGene_gn=CNTNAP4;refGene_tx=XM_019820088.2;refGene_gn=CNTNAP4;refGene_tx=XM_019820089.2;refGene_gn=CNTNAP4;refGene_tx=XM_023245046.1;refGene_gn=CNTNAP4;refGene_tx=XM_023245047.1
E2 52792386 52792534
refGene_gn=CNTNAP4;refGene_tx=XM_011290253.3;refGene_gn=CNTNAP4;refGene_tx=XM_019820088.2;refGene_gn=CNTNAP4;refGene_tx=XM_019820089.2;refGene_gn=CNTNAP4;refGene_tx=XM_023245046.1;refGene_gn=CNTNAP4;refGene_tx=XM_023245047.1
E2 52808119 52808323
refGene_gn=CNTNAP4;refGene_tx=XM_011290253.3;refGene_gn=CNTNAP4;refGene_tx=XM_019820088.2;refGene_gn=CNTNAP4;refGene_tx=XM_019820089.2;refGene_gn=CNTNAP4;refGene_tx=XM_023245046.1;refGene_gn=CNTNAP4;refGene_tx=XM_023245047.1
E2 52809085 52809270

refGene_gn=CNTNAP4;refGene_tx=XM_011290253.3;refGene_gn=CNTNAP4;refGene_tx=XM_019820088.2;refGene_gn=CNTNAP4;refGene_tx=XM_019820089.2;refGene_gn=CNTNAP4;refGene_tx=XM_023245046.1;refGene_gn=CNTNAP4;refGene_tx=XM_023245047.1
E2 52810078 52810222
refGene_gn=CNTNAP4;refGene_tx=XM_011290253.3;refGene_gn=CNTNAP4;refGene_tx=XM_019820088.2;refGene_gn=CNTNAP4;refGene_tx=XM_019820089.2;refGene_gn=CNTNAP4;refGene_tx=XM_023245046.1;refGene_gn=CNTNAP4;refGene_tx=XM_023245047.1
E2 52813252 52813514
refGene_gn=CNTNAP4;refGene_tx=XM_011290253.3;refGene_gn=CNTNAP4;refGene_tx=XM_019820088.2;refGene_gn=CNTNAP4;refGene_tx=XM_019820089.2;refGene_gn=CNTNAP4;refGene_tx=XM_023245046.1;refGene_gn=CNTNAP4;refGene_tx=XM_023245047.1
E2 52821354 52821504
refGene_gn=CNTNAP4;refGene_tx=XM_011290253.3;refGene_gn=CNTNAP4;refGene_tx=XM_019820088.2;refGene_gn=CNTNAP4;refGene_tx=XM_019820089.2;refGene_gn=CNTNAP4;refGene_tx=XM_023245046.1;refGene_gn=CNTNAP4;refGene_tx=XM_023245047.1
E2 52827141 52827313
refGene_gn=CNTNAP4;refGene_tx=XM_011290253.3;refGene_gn=CNTNAP4;refGene_tx=XM_019820088.2;refGene_gn=CNTNAP4;refGene_tx=XM_019820089.2;refGene_gn=CNTNAP4;refGene_tx=XM_023245046.1;refGene_gn=CNTNAP4;refGene_tx=XM_023245047.1
E2 52833781 52833888
refGene_gn=CNTNAP4;refGene_tx=XM_011290253.3;refGene_gn=CNTNAP4;refGene_tx=XM_019820088.2;refGene_gn=CNTNAP4;refGene_tx=XM_019820089.2;refGene_gn=CNTNAP4;refGene_tx=XM_023245046.1;refGene_gn=CNTNAP4;refGene_tx=XM_023245047.1
E2 52837236 52837356
refGene_gn=CNTNAP4;refGene_tx=XM_011290253.3;refGene_gn=CNTNAP4;refGene_tx=XM_019820088.2;refGene_gn=CNTNAP4;refGene_tx=XM_019820089.2;refGene_gn=CNTNAP4;refGene_tx=XM_023245046.1;refGene_gn=CNTNAP4;refGene_tx=XM_023245047.1
E2 52847904 52848102
refGene_gn=CNTNAP4;refGene_tx=XM_011290253.3;refGene_gn=CNTNAP4;refGene_tx=XM_019820088.2;refGene_gn=CNTNAP4;refGene_tx=XM_019820089.2;refGene_gn=CNTNAP4;refGene_tx=XM_023245046.1;refGene_gn=CNTNAP4;refGene_tx=XM_023245047.1
E2 52852608 52852765
refGene_gn=CNTNAP4;refGene_tx=XM_011290253.3;refGene_gn=CNTNAP4;refGene_tx=XM_019820088.2;refGene_gn=CNTNAP4;refGene_tx=XM_019820089.2;refGene_gn=CNTNAP4;refGene_tx=XM_023245046.1;refGene_gn=CNTNAP4;refGene_tx=XM_023245047.1
E2 52857133 52857261
refGene_gn=CNTNAP4;refGene_tx=XM_011290253.3;refGene_gn=CNTNAP4;refGene_tx=XM_019820088.2;refGene_gn=CNTNAP4;refGene_tx=XM_019820089.2;refGene_gn=CNTNAP4;refGene_tx=XM_023245046.1;refGene_gn=CNTNAP4;refGene_tx=XM_023245047.1
E2 52876282 52876453

refGene_gn=CNTNAP4;refGene_tx=XM_011290253.3;refGene_gn=CNTNAP4;refGene_tx=XM_019820088.2;refGene_gn=CNTNAP4;refGene_tx=XM_019820089.2;refGene_gn=CNTNAP4;refGene_tx=XM_023245046.1;refGene_gn=CNTNAP4;refGene_tx=XM_023245047.1
E2 52877154 52877373
refGene_gn=CNTNAP4;refGene_tx=XM_011290253.3;refGene_gn=CNTNAP4;refGene_tx=XM_019820088.2;refGene_gn=CNTNAP4;refGene_tx=XM_019820089.2;refGene_gn=CNTNAP4;refGene_tx=XM_023245046.1;refGene_gn=CNTNAP4;refGene_tx=XM_023245047.1
E2 52893676 52893916
refGene_gn=CNTNAP4;refGene_tx=XM_011290253.3;refGene_gn=CNTNAP4;refGene_tx=XM_019820089.2;refGene_gn=CNTNAP4;refGene_tx=XM_023245046.1
E2 52896289 52896514
refGene_gn=CNTNAP4;refGene_tx=XM_011290253.3;refGene_gn=CNTNAP4;refGene_tx=XM_019820088.2;refGene_gn=CNTNAP4;refGene_tx=XM_019820089.2;refGene_gn=CNTNAP4;refGene_tx=XM_023245046.1;refGene_gn=CNTNAP4;refGene_tx=XM_023245047.1
E2 52897902 52898036
refGene_gn=CNTNAP4;refGene_tx=XM_011290253.3;refGene_gn=CNTNAP4;refGene_tx=XM_019820088.2;refGene_gn=CNTNAP4;refGene_tx=XM_019820089.2;refGene_gn=CNTNAP4;refGene_tx=XM_023245046.1;refGene_gn=CNTNAP4;refGene_tx=XM_023245047.1
E2 52898774 52898862
refGene_gn=CNTNAP4;refGene_tx=XM_011290253.3;refGene_gn=CNTNAP4;refGene_tx=XM_019820088.2;refGene_gn=CNTNAP4;refGene_tx=XM_023245046.1;refGene_gn=CNTNAP4;refGene_tx=XM_023245047.1
E2 52909326 52909545
refGene_gn=CNTNAP4;refGene_tx=XM_011290253.3;refGene_gn=CNTNAP4;refGene_tx=XM_019820088.2;refGene_gn=CNTNAP4;refGene_tx=XM_023245046.1;refGene_gn=CNTNAP4;refGene_tx=XM_023245047.1
E2 52909863 52909935
refGene_gn=CNTNAP4;refGene_tx=XM_011290253.3;refGene_gn=CNTNAP4;refGene_tx=XM_019820088.2;refGene_gn=CNTNAP4;refGene_tx=XM_023245046.1;refGene_gn=CNTNAP4;refGene_tx=XM_023245047.1
E2 52914685 52914879
refGene_gn=CNTNAP4;refGene_tx=XM_011290253.3;refGene_gn=CNTNAP4;refGene_tx=XM_019820088.2;refGene_gn=CNTNAP4;refGene_tx=XM_023245046.1;refGene_gn=CNTNAP4;refGene_tx=XM_023245047.1
E2 52914989 52915086
refGene_gn=CNTNAP4;refGene_tx=XM_023245046.1;refGene_gn=CNTNAP4;refGene_tx=XM_023245047.1
E2 53442945 53443060
refGene_gn=LOC109494688;refGene_tx=XM_023245642.1;refGene_gn=LOC109494688;refGene_tx=XM_023245643.1;refGene_gn=LOC109494688;refGene_tx=XM_023245644.1
E2 53443243 53443317
refGene_gn=LOC109494688;refGene_tx=XM_023245642.1;refGene_gn=LOC109494688;refGene_tx=XM_023245643.1;refGene_gn=LOC109494688;refGene_tx=XM_023245644.1
E2 53443464 53443579

refGene_gn=LOC109494688;refGene_tx=XM_023245642.1;refGene_gn=LOC109494688;refGene_tx=XM_023245643.1;refGene_gn=LOC109494688;refGene_tx=XM_023245644.1
E2 53443837 53443959
refGene_gn=LOC109494688;refGene_tx=XM_023245642.1;refGene_gn=LOC109494688;refGene_tx=XM_023245643.1
E2 53444233 53444446
refGene_gn=LOC109494688;refGene_tx=XM_023245642.1;refGene_gn=LOC109494688;refGene_tx=XM_023245643.1;refGene_gn=LOC109494688;refGene_tx=XM_023245644.1
E2 53477773 53477921 refGene_gn=MON1B;refGene_tx=XM_003998261.5
E2 53479895 53480225 refGene_gn=MON1B;refGene_tx=XM_003998261.5
E2 53480721 53481541 refGene_gn=MON1B;refGene_tx=XM_003998261.5
E2 53481888 53482036 refGene_gn=MON1B;refGene_tx=XM_003998261.5
E2 53484147 53484357 refGene_gn=MON1B;refGene_tx=XM_003998261.5
E2 53485461 53485531
refGene_gn=SYCE1L;refGene_tx=XM_019820091.1;refGene_gn=SYCE1L;refGene_tx=XM_019820092.1;refGene_gn=SYCE1L;refGene_tx=XM_019820093.1;refGene_gn=SYCE1L;refGene_tx=XM_019820094.1;refGene_gn=SYCE1L;refGene_tx=XM_019820095.1;refGene_gn=SYCE1L;refGene_tx=XM_019820097.1;refGene_gn=SYCE1L;refGene_tx=XM_019820098.1;refGene_gn=SYCE1L;refGene_tx=XM_019820101.1;refGene_gn=SYCE1L;refGene_tx=XM_019820102.2;refGene_gn=SYCE1L;refGene_tx=XM_023244889.1
E2 53491558 53491621
refGene_gn=SYCE1L;refGene_tx=XM_019820091.1;refGene_gn=SYCE1L;refGene_tx=XM_019820092.1;refGene_gn=SYCE1L;refGene_tx=XM_019820094.1;refGene_gn=SYCE1L;refGene_tx=XM_019820095.1;refGene_gn=SYCE1L;refGene_tx=XM_019820096.1;refGene_gn=SYCE1L;refGene_tx=XM_019820097.1;refGene_gn=SYCE1L;refGene_tx=XM_019820098.1;refGene_gn=SYCE1L;refGene_tx=XM_019820101.1;refGene_gn=SYCE1L;refGene_tx=XM_019820102.2
E2 53493315 53493375
refGene_gn=SYCE1L;refGene_tx=XM_019820091.1;refGene_gn=SYCE1L;refGene_tx=XM_019820092.1;refGene_gn=SYCE1L;refGene_tx=XM_019820093.1;refGene_gn=SYCE1L;refGene_tx=XM_019820094.1;refGene_gn=SYCE1L;refGene_tx=XM_019820095.1;refGene_gn=SYCE1L;refGene_tx=XM_019820096.1;refGene_gn=SYCE1L;refGene_tx=XM_019820097.1;refGene_gn=SYCE1L;refGene_tx=XM_019820098.1;refGene_gn=SYCE1L;refGene_tx=XM_019820099.1;refGene_gn=SYCE1L;refGene_tx=XM_019820100.1;refGene_gn=SYCE1L;refGene_tx=XM_019820101.1;refGene_gn=SYCE1L;refGene_tx=XM_019820102.2;refGene_gn=SYCE1L;refGene_tx=XM_023244889.1
E2 53493563 53493638
refGene_gn=SYCE1L;refGene_tx=XM_019820091.1;refGene_gn=SYCE1L;refGene_tx=XM_019820092.1;refGene_gn=SYCE1L;refGene_tx=XM_019820093.1;refGene_gn=SYCE1L;refGene_tx=XM_019820094.1;refGene_gn=SYCE1L;refGene_tx=XM_019820095.1;refGene_gn=SYCE1L;refGene_tx=XM_019820096.1;refGene_gn=SYCE1L;refGene_tx=XM_019820097.1;refGene_gn=SYCE1L;refGene_tx=XM_019820098.1;refGene_gn=SYCE1L;refGene_tx=XM_019820099.1;refGene_gn=SYCE1L;refGene_tx=XM_019820100.1;refGene_gn=SYCE1L;refGene_tx=XM_019820101.1;refGene_gn=SYCE1L;refGene_tx=XM_019820102.2;refGene_gn=SYCE1L;refGene_tx=XM_023244889.1

E2 53494203 53494251
refGene_gn=SYCE1L;refGene_tx=XM_019820091.1;refGene_gn=SYCE1L;refGene_tx=XM_019820093.1;refGene_gn=SYCE1L;refGene_tx=XM_019820094.1;refGene_gn=SYCE1L;refGene_tx=XM_019820095.1;refGene_gn=SYCE1L;refGene_tx=XM_019820096.1;refGene_gn=SYCE1L;refGene_tx=XM_019820098.1;refGene_gn=SYCE1L;refGene_tx=XM_019820099.1;refGene_gn=SYCE1L;refGene_tx=XM_019820100.1;refGene_gn=SYCE1L;refGene_tx=XM_019820102.2;refGene_gn=SYCE1L;refGene_tx=XM_023244889.1
E2 53494524 53494579
refGene_gn=SYCE1L;refGene_tx=XM_019820091.1;refGene_gn=SYCE1L;refGene_tx=XM_019820092.1;refGene_gn=SYCE1L;refGene_tx=XM_019820093.1;refGene_gn=SYCE1L;refGene_tx=XM_019820094.1;refGene_gn=SYCE1L;refGene_tx=XM_019820095.1;refGene_gn=SYCE1L;refGene_tx=XM_019820096.1;refGene_gn=SYCE1L;refGene_tx=XM_019820097.1;refGene_gn=SYCE1L;refGene_tx=XM_019820098.1;refGene_gn=SYCE1L;refGene_tx=XM_019820099.1;refGene_gn=SYCE1L;refGene_tx=XM_019820100.1;refGene_gn=SYCE1L;refGene_tx=XM_019820101.1;refGene_gn=SYCE1L;refGene_tx=XM_019820102.2;refGene_gn=SYCE1L;refGene_tx=XM_023244889.1
E2 53495755 53495839
refGene_gn=SYCE1L;refGene_tx=XM_019820091.1;refGene_gn=SYCE1L;refGene_tx=XM_019820092.1;refGene_gn=SYCE1L;refGene_tx=XM_019820093.1;refGene_gn=SYCE1L;refGene_tx=XM_019820095.1;refGene_gn=SYCE1L;refGene_tx=XM_019820096.1;refGene_gn=SYCE1L;refGene_tx=XM_019820099.1;refGene_gn=SYCE1L;refGene_tx=XM_019820100.1
E2 53496095 53496429
refGene_gn=SYCE1L;refGene_tx=XM_019820091.1;refGene_gn=SYCE1L;refGene_tx=XM_019820092.1;refGene_gn=SYCE1L;refGene_tx=XM_019820093.1;refGene_gn=SYCE1L;refGene_tx=XM_019820094.1;refGene_gn=SYCE1L;refGene_tx=XM_019820095.1;refGene_gn=SYCE1L;refGene_tx=XM_019820096.1;refGene_gn=SYCE1L;refGene_tx=XM_019820097.1;refGene_gn=SYCE1L;refGene_tx=XM_019820099.1;refGene_gn=SYCE1L;refGene_tx=XM_019820100.1;refGene_gn=SYCE1L;refGene_tx=XM_019820102.2
E2 53497033 53497103
refGene_gn=SYCE1L;refGene_tx=XM_019820091.1;refGene_gn=SYCE1L;refGene_tx=XM_019820092.1;refGene_gn=SYCE1L;refGene_tx=XM_019820093.1;refGene_gn=SYCE1L;refGene_tx=XM_019820094.1;refGene_gn=SYCE1L;refGene_tx=XM_019820095.1;refGene_gn=SYCE1L;refGene_tx=XM_019820096.1;refGene_gn=SYCE1L;refGene_tx=XM_019820097.1;refGene_gn=SYCE1L;refGene_tx=XM_019820098.1;refGene_gn=SYCE1L;refGene_tx=XM_019820099.1;refGene_gn=SYCE1L;refGene_tx=XM_019820100.1;refGene_gn=SYCE1L;refGene_tx=XM_019820101.1;refGene_gn=SYCE1L;refGene_tx=XM_023244889.1
E2 53497192 53497280
refGene_gn=SYCE1L;refGene_tx=XM_019820091.1;refGene_gn=SYCE1L;refGene_tx=XM_019820092.1;refGene_gn=SYCE1L;refGene_tx=XM_019820093.1;refGene_gn=SYCE1L;refGene_tx=XM_019820094.1;refGene_gn=SYCE1L;refGene_tx=XM_019820095.1;refGene_gn=SYCE1L;refGene_tx=XM_019820096.1;refGene_gn=SYCE1L;refGene_tx=XM_019820097.1;refGene_gn=SYCE1L;refGene_tx=XM_019820098.1;refGene_gn=SYCE1L;refGene_tx=XM_019820099.1;refGene_gn=SYCE1L;refGene_tx=XM_019820100.1;refGene_gn=SYCE1L;refGene_tx=XM_019820101.1;refGene_gn=SYCE1L;refGene_tx=XM_023244889.1

E2 53497540 53497625
refGene_gn=SYCE1L;refGene_tx=XM_019820091.1;refGene_gn=SYCE1L;refGene_tx=XM_019820092.1;refGene_gn=SYCE1L;refGene_tx=XM_019820093.1;refGene_gn=SYCE1L;refGene_tx=XM_019820094.1;refGene_gn=SYCE1L;refGene_tx=XM_019820095.1;refGene_gn=SYCE1L;refGene_tx=XM_019820096.1;refGene_gn=SYCE1L;refGene_tx=XM_019820097.1;refGene_gn=SYCE1L;refGene_tx=XM_019820098.1;refGene_gn=SYCE1L;refGene_tx=XM_019820099.1;refGene_gn=SYCE1L;refGene_tx=XM_019820100.1;refGene_gn=SYCE1L;refGene_tx=XM_019820101.1;refGene_gn=SYCE1L;refGene_tx=XM_023244889.1
E2 53497824 53497996
refGene_gn=SYCE1L;refGene_tx=XM_019820091.1;refGene_gn=SYCE1L;refGene_tx=XM_019820092.1;refGene_gn=SYCE1L;refGene_tx=XM_019820093.1;refGene_gn=SYCE1L;refGene_tx=XM_019820094.1;refGene_gn=SYCE1L;refGene_tx=XM_019820095.1;refGene_gn=SYCE1L;refGene_tx=XM_019820096.1;refGene_gn=SYCE1L;refGene_tx=XM_019820097.1;refGene_gn=SYCE1L;refGene_tx=XM_019820098.1;refGene_gn=SYCE1L;refGene_tx=XM_019820099.1;refGene_gn=SYCE1L;refGene_tx=XM_019820100.1;refGene_gn=SYCE1L;refGene_tx=XM_019820101.1;refGene_gn=SYCE1L;refGene_tx=XM_023244889.1
E2 53554788 53554904 refGene_gn=ADAMTS18;refGene_tx=XM_023245122.1
E2 53560309 53560457 refGene_gn=ADAMTS18;refGene_tx=XM_023245122.1
E2 53563210 53563423 refGene_gn=ADAMTS18;refGene_tx=XM_023245122.1
E2 53565377 53565563 refGene_gn=ADAMTS18;refGene_tx=XM_023245122.1
E2 53568186 53568391 refGene_gn=ADAMTS18;refGene_tx=XM_023245122.1
E2 53571725 53571852 refGene_gn=ADAMTS18;refGene_tx=XM_023245122.1
E2 53574068 53574210 refGene_gn=ADAMTS18;refGene_tx=XM_023245122.1
E2 53593004 53593249 refGene_gn=ADAMTS18;refGene_tx=XM_023245122.1
E2 53593957 53594081 refGene_gn=ADAMTS18;refGene_tx=XM_023245122.1
E2 53594898 53595029 refGene_gn=ADAMTS18;refGene_tx=XM_023245122.1
E2 53599540 53599713 refGene_gn=ADAMTS18;refGene_tx=XM_023245122.1
E2 53606570 53606719 refGene_gn=ADAMTS18;refGene_tx=XM_023245122.1
E2 53611107 53611203 refGene_gn=ADAMTS18;refGene_tx=XM_023245122.1
E2 53619475 53619629 refGene_gn=ADAMTS18;refGene_tx=XM_023245122.1
E2 53621965 53622103 refGene_gn=ADAMTS18;refGene_tx=XM_023245122.1
E2 53624759 53624865 refGene_gn=ADAMTS18;refGene_tx=XM_023245122.1
E2 53626235 53626395 refGene_gn=ADAMTS18;refGene_tx=XM_023245122.1
E2 53628305 53628389 refGene_gn=ADAMTS18;refGene_tx=XM_023245122.1
E2 53628648 53628842 refGene_gn=ADAMTS18;refGene_tx=XM_023245122.1
E2 53631215 53631498 refGene_gn=ADAMTS18;refGene_tx=XM_023245122.1
E2 53691639 53691956 refGene_gn=ADAMTS18;refGene_tx=XM_023245122.1
E2 53694904 53694989 refGene_gn=ADAMTS18;refGene_tx=XM_023245122.1
E2 53695101 53695191 refGene_gn=ADAMTS18;refGene_tx=XM_023245122.1
E2 53947952 53947987 refGene_gn=NUDT7;refGene_tx=XM_003998282.3
E2 53950161 53950315 refGene_gn=NUDT7;refGene_tx=XM_003998282.3
E2 53956113 53956272 refGene_gn=NUDT7;refGene_tx=XM_003998282.3
E2 53962005 53962374 refGene_gn=NUDT7;refGene_tx=XM_003998282.3
E2 54002791 54003024 refGene_gn=VAT1L;refGene_tx=XM_023246187.1
E2 54026873 54027003 refGene_gn=VAT1L;refGene_tx=XM_023246187.1
E2 54031224 54031440 refGene_gn=VAT1L;refGene_tx=XM_023246187.1
E2 54065524 54065667 refGene_gn=VAT1L;refGene_tx=XM_023246187.1
E2 54075813 54075917 refGene_gn=VAT1L;refGene_tx=XM_023246187.1

E2 54078135 54078191 refGene_gn=VAT1L;refGene_tx=XM_023246187.1
E2 54081861 54082056 refGene_gn=VAT1L;refGene_tx=XM_023246187.1
E2 54146127 54146211 refGene_gn=VAT1L;refGene_tx=XM_023246187.1
E2 54150007 54150106 refGene_gn=VAT1L;refGene_tx=XM_023246187.1
E2 54186992 54187107 refGene_gn=CLEC3A;refGene_tx=XM_006941747.3
E2 54191353 54191437 refGene_gn=CLEC3A;refGene_tx=XM_006941747.3
E2 54193650 54194042 refGene_gn=CLEC3A;refGene_tx=XM_006941747.3
E2 54248386 54248493
refGene_gn=WWOX;refGene_tx=XM_003998263.5;refGene_gn=WWOX;refGene_tx=XM_019818834.2
E2 54254741 54254806
refGene_gn=WWOX;refGene_tx=XM_003998263.5;refGene_gn=WWOX;refGene_tx=XM_019818834.2
E2 54255167 54255225
refGene_gn=WWOX;refGene_tx=XM_003998263.5;refGene_gn=WWOX;refGene_tx=XM_019818834.2
E2 54257125 54257304
refGene_gn=WWOX;refGene_tx=XM_003998263.5;refGene_gn=WWOX;refGene_tx=XM_019818834.2
E2 54296946 54297053
refGene_gn=WWOX;refGene_tx=XM_003998263.5;refGene_gn=WWOX;refGene_tx=XM_019818834.2
E2 54501301 54501390
refGene_gn=WWOX;refGene_tx=XM_003998263.5;refGene_gn=WWOX;refGene_tx=XM_019818834.2
E2 54542791 54542977
refGene_gn=WWOX;refGene_tx=XM_003998263.5;refGene_gn=WWOX;refGene_tx=XM_019818834.2
E2 54548937 54549202
refGene_gn=WWOX;refGene_tx=XM_003998263.5;refGene_gn=WWOX;refGene_tx=XM_019818834.2
E2 54551358 54551376 refGene_gn=WWOX;refGene_tx=XM_019818834.2
E2 55231461 55231650 refGene_gn=WWOX;refGene_tx=XM_003998263.5
E2 55565921 55566015 refGene_gn=MAF;refGene_tx=XM_023245367.1
E2 55570402 55571553 refGene_gn=MAF;refGene_tx=XM_023245367.1
E2 56391977 56391980 refGene_gn=DYNLRB2;refGene_tx=XM_006941755.4
E2 56394628 56394704
refGene_gn=DYNLRB2;refGene_tx=XM_006941755.4;refGene_gn=DYNLRB2;refGene_tx=XM_019818835.2
E2 56400437 56400605
refGene_gn=DYNLRB2;refGene_tx=XM_006941755.4;refGene_gn=DYNLRB2;refGene_tx=XM_019818835.2
E2 56401373 56401417
refGene_gn=DYNLRB2;refGene_tx=XM_006941755.4;refGene_gn=DYNLRB2;refGene_tx=XM_019818835.2
E2 56445540 56445699 refGene_gn=CDYL2;refGene_tx=XM_023244944.1
E2 56448332 56448476 refGene_gn=CDYL2;refGene_tx=XM_023244944.1
E2 56452950 56453161 refGene_gn=CDYL2;refGene_tx=XM_023244944.1
E2 56459426 56459599 refGene_gn=CDYL2;refGene_tx=XM_023244944.1
E2 56467049 56467267 refGene_gn=CDYL2;refGene_tx=XM_023244944.1

E2 56494141 56494730 refGene_gn=CDYL2;refGene_tx=XM_023244944.1
E2 56607582 56607606 refGene_gn=CDYL2;refGene_tx=XM_023244944.1
E2 56745467 56745554
refGene_gn=CMC2;refGene_tx=XM_003998285.5;refGene_gn=CMC2;refGene_tx=XM_023244959.1;refGene_gn=CMC2;refGene_tx=XM_023244960.1;refGene_gn=CMC2;refGene_tx=XM_023244961.1
E2 56749318 56749390
refGene_gn=CMC2;refGene_tx=XM_003998285.5;refGene_gn=CMC2;refGene_tx=XM_023244959.1;refGene_gn=CMC2;refGene_tx=XM_023244960.1;refGene_gn=CMC2;refGene_tx=XM_023244961.1
E2 56753945 56754060
refGene_gn=CMC2;refGene_tx=XM_003998285.5;refGene_gn=CMC2;refGene_tx=XM_023244959.1;refGene_gn=CMC2;refGene_tx=XM_023244960.1;refGene_gn=CMC2;refGene_tx=XM_023244961.1
E2 56761275 56761457 refGene_gn=CMC2;refGene_tx=XM_003998285.5
E2 56766118 56766289
refGene_gn=CENPN;refGene_tx=XM_003998286.5;refGene_gn=CENPN;refGene_tx=XM_006941761.4;refGene_gn=CENPN;refGene_tx=XM_019820107.2;refGene_gn=CENPN;refGene_tx=XM_019820109.2
E2 56768173 56768219
refGene_gn=CENPN;refGene_tx=XM_003998286.5;refGene_gn=CENPN;refGene_tx=XM_006941761.4;refGene_gn=CENPN;refGene_tx=XM_019820107.2;refGene_gn=CENPN;refGene_tx=XM_019820109.2
E2 56770811 56770871
refGene_gn=CENPN;refGene_tx=XM_003998286.5;refGene_gn=CENPN;refGene_tx=XM_006941761.4;refGene_gn=CENPN;refGene_tx=XM_019820107.2;refGene_gn=CENPN;refGene_tx=XM_019820109.2
E2 56771588 56771665
refGene_gn=CENPN;refGene_tx=XM_003998286.5;refGene_gn=CENPN;refGene_tx=XM_006941761.4;refGene_gn=CENPN;refGene_tx=XM_019820107.2;refGene_gn=CENPN;refGene_tx=XM_019820109.2
E2 56778699 56778876
refGene_gn=CENPN;refGene_tx=XM_003998286.5;refGene_gn=CENPN;refGene_tx=XM_006941761.4;refGene_gn=CENPN;refGene_tx=XM_019820107.2;refGene_gn=CENPN;refGene_tx=XM_019820109.2
E2 56781703 56781805
refGene_gn=CENPN;refGene_tx=XM_003998286.5;refGene_gn=CENPN;refGene_tx=XM_006941761.4;refGene_gn=CENPN;refGene_tx=XM_019820107.2;refGene_gn=CENPN;refGene_tx=XM_019820109.2
E2 56782863 56782924
refGene_gn=CENPN;refGene_tx=XM_003998286.5;refGene_gn=CENPN;refGene_tx=XM_006941761.4;refGene_gn=CENPN;refGene_tx=XM_019820107.2
E2 56784439 56784552
refGene_gn=CENPN;refGene_tx=XM_003998286.5;refGene_gn=CENPN;refGene_tx=XM_006941761.4;refGene_gn=CENPN;refGene_tx=XM_019820107.2
E2 56787163 56787290
refGene_gn=CENPN;refGene_tx=XM_003998286.5;refGene_gn=CENPN;refGene_tx=XM_006941761.4;refGene_gn=CENPN;refGene_tx=XM_019820107.2;refGene_gn=CENPN;refGene_tx=XM_019820109.2
E2 56787524 56787607

refGene_gn=CENPN;refGene_tx=XM_003998286.5;refGene_gn=CENPN;refGene_tx=XM_006941761.4;refGene_gn=CENPN;refGene_tx=XM_019820107.2;refGene_gn=CENPN;refGene_tx=XM_019820109.2
E2 56791059 56791392 refGene_gn=ATMIN;refGene_tx=XM_011290261.2
E2 56800597 56800723 refGene_gn=ATMIN;refGene_tx=XM_011290261.2
E2 56801503 56801703 refGene_gn=ATMIN;refGene_tx=XM_011290261.2
E2 56802364 56804177 refGene_gn=ATMIN;refGene_tx=XM_011290261.2
E2 56812720 56813692
refGene_gn=CE2H16orf46;refGene_tx=XM_003998289.5;refGene_gn=CE2H16orf46;refGene_tx=XM_006941762.4;refGene_gn=CE2H16orf46;refGene_tx=XM_006941763.4;refGene_gn=CE2H16orf46;refGene_tx=XM_006941764.4;refGene_gn=CE2H16orf46;refGene_tx=XM_006941765.4;refGene_gn=CE2H16orf46;refGene_tx=XM_006941766.4;refGene_gn=CE2H16orf46;refGene_tx=XM_011290263.3;refGene_gn=CE2H16orf46;refGene_tx=XM_019820116.2;refGene_gn=CE2H16orf46;refGene_tx=XM_019820118.2;refGene_gn=CE2H16orf46;refGene_tx=XM_019820119.2
E2 56816420 56816630
refGene_gn=CE2H16orf46;refGene_tx=XM_003998289.5;refGene_gn=CE2H16orf46;refGene_tx=XM_006941762.4;refGene_gn=CE2H16orf46;refGene_tx=XM_006941763.4;refGene_gn=CE2H16orf46;refGene_tx=XM_006941764.4;refGene_gn=CE2H16orf46;refGene_tx=XM_006941765.4;refGene_gn=CE2H16orf46;refGene_tx=XM_006941766.4;refGene_gn=CE2H16orf46;refGene_tx=XM_011290263.3;refGene_gn=CE2H16orf46;refGene_tx=XM_019820116.2;refGene_gn=CE2H16orf46;refGene_tx=XM_019820118.2;refGene_gn=CE2H16orf46;refGene_tx=XM_019820119.2
E2 56829250 56829348 refGene_gn=GCSH;refGene_tx=XM_023244989.1
E2 56830963 56831095 refGene_gn=GCSH;refGene_tx=XM_023244989.1
E2 56836689 56836753 refGene_gn=GCSH;refGene_tx=XM_023244989.1
E2 56837943 56838023 refGene_gn=GCSH;refGene_tx=XM_023244989.1
E2 56841379 56841527 refGene_gn=GCSH;refGene_tx=XM_023244989.1
E2 56845737 56845929 refGene_gn=PKD1L2;refGene_tx=XM_023245123.1
E2 56847088 56847220 refGene_gn=PKD1L2;refGene_tx=XM_023245123.1
E2 56847997 56848099 refGene_gn=PKD1L2;refGene_tx=XM_023245123.1
E2 56849657 56849868 refGene_gn=PKD1L2;refGene_tx=XM_023245123.1
E2 56854306 56854467 refGene_gn=PKD1L2;refGene_tx=XM_023245123.1
E2 56857379 56857507 refGene_gn=PKD1L2;refGene_tx=XM_023245123.1
E2 56859310 56859444 refGene_gn=PKD1L2;refGene_tx=XM_023245123.1
E2 56861082 56861410 refGene_gn=PKD1L2;refGene_tx=XM_023245123.1
E2 56862334 56862520 refGene_gn=PKD1L2;refGene_tx=XM_023245123.1
E2 56864017 56864155 refGene_gn=PKD1L2;refGene_tx=XM_023245123.1
E2 56867098 56867195 refGene_gn=PKD1L2;refGene_tx=XM_023245123.1
E2 56868434 56868639 refGene_gn=PKD1L2;refGene_tx=XM_023245123.1
E2 56869993 56870381 refGene_gn=PKD1L2;refGene_tx=XM_023245123.1
E2 56873587 56873804 refGene_gn=PKD1L2;refGene_tx=XM_023245123.1
E2 56874538 56874756 refGene_gn=PKD1L2;refGene_tx=XM_023245123.1
E2 56875407 56875607 refGene_gn=PKD1L2;refGene_tx=XM_023245123.1
E2 56878524 56878671 refGene_gn=PKD1L2;refGene_tx=XM_023245123.1
E2 56880313 56880492 refGene_gn=PKD1L2;refGene_tx=XM_023245123.1
E2 56882605 56882707 refGene_gn=PKD1L2;refGene_tx=XM_023245123.1
E2 56885497 56885728 refGene_gn=PKD1L2;refGene_tx=XM_023245123.1
E2 56889842 56890036 refGene_gn=PKD1L2;refGene_tx=XM_023245123.1
E2 56890597 56890820 refGene_gn=PKD1L2;refGene_tx=XM_023245123.1

E2 56894620 56894752 refGene_gn=PKD1L2;refGene_tx=XM_023245123.1
E2 56895417 56895544 refGene_gn=PKD1L2;refGene_tx=XM_023245123.1
E2 56896543 56896668 refGene_gn=PKD1L2;refGene_tx=XM_023245123.1
E2 56898113 56898296 refGene_gn=PKD1L2;refGene_tx=XM_023245123.1
E2 56899658 56899790 refGene_gn=PKD1L2;refGene_tx=XM_023245123.1
E2 56902621 56902816 refGene_gn=PKD1L2;refGene_tx=XM_023245123.1
E2 56903492 56903638 refGene_gn=PKD1L2;refGene_tx=XM_023245123.1
E2 56904958 56905135 refGene_gn=PKD1L2;refGene_tx=XM_023245123.1
E2 56906373 56906515 refGene_gn=PKD1L2;refGene_tx=XM_023245123.1
E2 56907194 56907315 refGene_gn=PKD1L2;refGene_tx=XM_023245123.1
E2 56909001 56909209 refGene_gn=PKD1L2;refGene_tx=XM_023245123.1
E2 56910779 56910897 refGene_gn=PKD1L2;refGene_tx=XM_023245123.1
E2 56912645 56912695 refGene_gn=PKD1L2;refGene_tx=XM_023245123.1
E2 56915526 56915604 refGene_gn=PKD1L2;refGene_tx=XM_023245123.1
E2 56917639 56918041 refGene_gn=PKD1L2;refGene_tx=XM_023245123.1
E2 56919814 56920044 refGene_gn=PKD1L2;refGene_tx=XM_023245123.1
E2 56928795 56928943 refGene_gn=PKD1L2;refGene_tx=XM_023245123.1
E2 56932615 56932726 refGene_gn=PKD1L2;refGene_tx=XM_023245123.1
E2 56936904 56937090 refGene_gn=PKD1L2;refGene_tx=XM_023245123.1
E2 56939159 56939324 refGene_gn=PKD1L2;refGene_tx=XM_023245123.1
E2 56941384 56941685 refGene_gn=PKD1L2;refGene_tx=XM_023245123.1
E2 56958380 56958444
refGene_gn=BCO1;refGene_tx=XM_003998290.4;refGene_gn=BCO1;refGene_tx=XM_023244985.1;refGene_gn=BCO1;refGene_tx=XM_023244986.1;refGene_gn=BCO1;refGene_tx=XM_023244987.1;refGene_gn=BCO1;refGene_tx=XM_023244988.1
E2 56962922 56963051
refGene_gn=BCO1;refGene_tx=XM_003998290.4;refGene_gn=BCO1;refGene_tx=XM_023244985.1;refGene_gn=BCO1;refGene_tx=XM_023244986.1;refGene_gn=BCO1;refGene_tx=XM_023244987.1
E2 56973368 56973498
refGene_gn=BCO1;refGene_tx=XM_003998290.4;refGene_gn=BCO1;refGene_tx=XM_023244985.1;refGene_gn=BCO1;refGene_tx=XM_023244986.1;refGene_gn=BCO1;refGene_tx=XM_023244987.1;refGene_gn=BCO1;refGene_tx=XM_023244988.1
E2 56973678 56973686 refGene_gn=BCO1;refGene_tx=XM_019820114.2
E2 56975058 56975206
refGene_gn=BCO1;refGene_tx=XM_003998290.4;refGene_gn=BCO1;refGene_tx=XM_019820114.2;refGene_gn=BCO1;refGene_tx=XM_023244985.1;refGene_gn=BCO1;refGene_tx=XM_023244986.1;refGene_gn=BCO1;refGene_tx=XM_023244987.1;refGene_gn=BCO1;refGene_tx=XM_023244988.1
E2 56976830 56976978
refGene_gn=BCO1;refGene_tx=XM_003998290.4;refGene_gn=BCO1;refGene_tx=XM_019820114.2;refGene_gn=BCO1;refGene_tx=XM_023244985.1;refGene_gn=BCO1;refGene_tx=XM_023244986.1;refGene_gn=BCO1;refGene_tx=XM_023244987.1;refGene_gn=BCO1;refGene_tx=XM_023244988.1
E2 56979325 56979546
refGene_gn=BCO1;refGene_tx=XM_003998290.4;refGene_gn=BCO1;refGene_tx=XM_019820114.2;refGene_gn=BCO1;refGene_tx=XM_023244985.1;refGene_gn=BCO1;refGene_tx=XM_023244986.1;refGene_gn=BCO1;refGene_tx=XM_023244987.1;refGene_gn=BCO1;refGene_tx=XM_023244988.1
E2 56982523 56982781

refGene_gn=BCO1;refGene_tx=XM_003998290.4;refGene_gn=BCO1;refGene_tx=XM_019820114.2;refGene_gn=BCO1;refGene_tx=XM_023244985.1;refGene_gn=BCO1;refGene_tx=XM_023244986.1;refGene_gn=BCO1;refGene_tx=XM_023244987.1;refGene_gn=BCO1;refGene_tx=XM_023244988.1
E2 56990645 56990751
refGene_gn=BCO1;refGene_tx=XM_003998290.4;refGene_gn=BCO1;refGene_tx=XM_019820114.2;refGene_gn=BCO1;refGene_tx=XM_023244985.1;refGene_gn=BCO1;refGene_tx=XM_023244986.1;refGene_gn=BCO1;refGene_tx=XM_023244987.1;refGene_gn=BCO1;refGene_tx=XM_023244988.1
E2 56993773 56993868
refGene_gn=BCO1;refGene_tx=XM_003998290.4;refGene_gn=BCO1;refGene_tx=XM_019820114.2;refGene_gn=BCO1;refGene_tx=XM_023244985.1;refGene_gn=BCO1;refGene_tx=XM_023244986.1;refGene_gn=BCO1;refGene_tx=XM_023244987.1;refGene_gn=BCO1;refGene_tx=XM_023244988.1
E2 56995720 56995832
refGene_gn=BCO1;refGene_tx=XM_003998290.4;refGene_gn=BCO1;refGene_tx=XM_019820114.2;refGene_gn=BCO1;refGene_tx=XM_023244985.1;refGene_gn=BCO1;refGene_tx=XM_023244986.1;refGene_gn=BCO1;refGene_tx=XM_023244987.1;refGene_gn=BCO1;refGene_tx=XM_023244988.1
E2 56997795 56998025
refGene_gn=BCO1;refGene_tx=XM_003998290.4;refGene_gn=BCO1;refGene_tx=XM_019820114.2;refGene_gn=BCO1;refGene_tx=XM_023244985.1;refGene_gn=BCO1;refGene_tx=XM_023244986.1;refGene_gn=BCO1;refGene_tx=XM_023244987.1;refGene_gn=BCO1;refGene_tx=XM_023244988.1
E2 57020320 57020487 refGene_gn=GAN;refGene_tx=XM_023246127.1
E2 57048122 57048237 refGene_gn=GAN;refGene_tx=XM_023246127.1
E2 57050619 57050970 refGene_gn=GAN;refGene_tx=XM_023246127.1
E2 57053423 57053641 refGene_gn=GAN;refGene_tx=XM_023246127.1
E2 57053859 57053981 refGene_gn=GAN;refGene_tx=XM_023246127.1
E2 57057167 57057280 refGene_gn=GAN;refGene_tx=XM_023246127.1
E2 57058202 57058352 refGene_gn=GAN;refGene_tx=XM_023246127.1
E2 57059703 57059840 refGene_gn=GAN;refGene_tx=XM_023246127.1
E2 57060530 57060659 refGene_gn=GAN;refGene_tx=XM_023246127.1
E2 57069640 57069750 refGene_gn=GAN;refGene_tx=XM_023246127.1
E2 57069839 57070021 refGene_gn=GAN;refGene_tx=XM_023246127.1
E2 57115473 57115493
refGene_gn=LOC111557974;refGene_tx=XM_023244851.1
E2 57116229 57116347
refGene_gn=LOC111557974;refGene_tx=XM_023244848.1
E2 57117181 57117416
refGene_gn=LOC111557974;refGene_tx=XM_023244848.1;refGene_gn=LOC111557974;refGene_tx=XM_023244849.1;refGene_gn=LOC111557974;refGene_tx=XM_023244850.1;refGene_gn=LOC111557974;refGene_tx=XM_023244851.1;refGene_gn=LOC111557974;refGene_tx=XM_023244852.1;refGene_gn=LOC111557974;refGene_tx=XM_023244853.1
E2 57117940 57118021
refGene_gn=LOC111557974;refGene_tx=XM_023244848.1;refGene_gn=LOC111557974;refGene_tx=XM_023244849.1;refGene_gn=LOC111557974;refGene_tx=XM_023244850.1;refGene_gn=LOC111557974;refGene_tx=XM_023244851.1;refGene_gn=LOC111557974;refGene_tx=XM_023244852.1;refGene_gn=LOC111557974;refGen

e_tx=XM_023244853.1
E2 57119241 57119302
refGene_gn=LOC111557974;refGene_tx=XM_023244854.1;refGene_gn=LOC111557
974;refGene_tx=XM_023244855.1
E2 57119417 57120058
refGene_gn=LOC111557974;refGene_tx=XM_023244848.1;refGene_gn=LOC111557
974;refGene_tx=XM_023244849.1;refGene_gn=LOC111557974;refGene_tx=XM_02
3244850.1;refGene_gn=LOC111557974;refGene_tx=XM_023244851.1;refGene_gn
=LOC111557974;refGene_tx=XM_023244852.1;refGene_gn=LOC111557974;refGen
e_tx=XM_023244853.1;refGene_gn=LOC111557974;refGene_tx=XM_023244854.1;
refGene_gn=LOC111557974;refGene_tx=XM_023244855.1
E2 57120301 57120601 refGene_gn=CMIP;refGene_tx=XM_023244846.1
E2 57260113 57260167 refGene_gn=CMIP;refGene_tx=XM_011290265.3
E2 57262456 57262582
refGene_gn=CMIP;refGene_tx=XM_011290265.3;refGene_gn=CMIP;refGene_tx=X
M_023244846.1
E2 57274397 57274448
refGene_gn=CMIP;refGene_tx=XM_011290265.3;refGene_gn=CMIP;refGene_tx=X
M_023244846.1
E2 57294651 57294669 refGene_gn=CMIP;refGene_tx=XM_023244847.1
E2 57300408 57300570
refGene_gn=CMIP;refGene_tx=XM_011290265.3;refGene_gn=CMIP;refGene_tx=X
M_023244846.1;refGene_gn=CMIP;refGene_tx=XM_023244847.1
E2 57305356 57305398
refGene_gn=CMIP;refGene_tx=XM_011290265.3;refGene_gn=CMIP;refGene_tx=X
M_023244846.1;refGene_gn=CMIP;refGene_tx=XM_023244847.1
E2 57312138 57312201
refGene_gn=CMIP;refGene_tx=XM_011290265.3;refGene_gn=CMIP;refGene_tx=X
M_023244846.1;refGene_gn=CMIP;refGene_tx=XM_023244847.1
E2 57314618 57314699
refGene_gn=CMIP;refGene_tx=XM_011290265.3;refGene_gn=CMIP;refGene_tx=X
M_023244846.1;refGene_gn=CMIP;refGene_tx=XM_023244847.1
E2 57319747 57319851
refGene_gn=CMIP;refGene_tx=XM_011290265.3;refGene_gn=CMIP;refGene_tx=X
M_023244846.1;refGene_gn=CMIP;refGene_tx=XM_023244847.1
E2 57321356 57321461
refGene_gn=CMIP;refGene_tx=XM_011290265.3;refGene_gn=CMIP;refGene_tx=X
M_023244846.1;refGene_gn=CMIP;refGene_tx=XM_023244847.1
E2 57327618 57327972
refGene_gn=CMIP;refGene_tx=XM_011290265.3;refGene_gn=CMIP;refGene_tx=X
M_023244846.1;refGene_gn=CMIP;refGene_tx=XM_023244847.1
E2 57336954 57337020
refGene_gn=CMIP;refGene_tx=XM_011290265.3;refGene_gn=CMIP;refGene_tx=X
M_023244846.1;refGene_gn=CMIP;refGene_tx=XM_023244847.1
E2 57338248 57338275
refGene_gn=CMIP;refGene_tx=XM_011290265.3;refGene_gn=CMIP;refGene_tx=X
M_023244846.1;refGene_gn=CMIP;refGene_tx=XM_023244847.1
E2 57338513 57338562
refGene_gn=CMIP;refGene_tx=XM_011290265.3;refGene_gn=CMIP;refGene_tx=X
M_023244846.1;refGene_gn=CMIP;refGene_tx=XM_023244847.1

E2 57341105 57341213
refGene_gn=CMIP;refGene_tx=XM_011290265.3;refGene_gn=CMIP;refGene_tx=XM_023244846.1;refGene_gn=CMIP;refGene_tx=XM_023244847.1
E2 57343301 57343418
refGene_gn=CMIP;refGene_tx=XM_011290265.3;refGene_gn=CMIP;refGene_tx=XM_023244846.1;refGene_gn=CMIP;refGene_tx=XM_023244847.1
E2 57344911 57345052
refGene_gn=CMIP;refGene_tx=XM_011290265.3;refGene_gn=CMIP;refGene_tx=XM_023244846.1;refGene_gn=CMIP;refGene_tx=XM_023244847.1
E2 57345806 57345854
refGene_gn=CMIP;refGene_tx=XM_011290265.3;refGene_gn=CMIP;refGene_tx=XM_023244846.1;refGene_gn=CMIP;refGene_tx=XM_023244847.1
E2 57346973 57347120
refGene_gn=CMIP;refGene_tx=XM_011290265.3;refGene_gn=CMIP;refGene_tx=XM_023244846.1;refGene_gn=CMIP;refGene_tx=XM_023244847.1
E2 57348007 57348113
refGene_gn=CMIP;refGene_tx=XM_011290265.3;refGene_gn=CMIP;refGene_tx=XM_023244846.1;refGene_gn=CMIP;refGene_tx=XM_023244847.1
E2 57349428 57349499
refGene_gn=CMIP;refGene_tx=XM_011290265.3;refGene_gn=CMIP;refGene_tx=XM_023244846.1;refGene_gn=CMIP;refGene_tx=XM_023244847.1
E2 57352067 57352121
refGene_gn=CMIP;refGene_tx=XM_011290265.3;refGene_gn=CMIP;refGene_tx=XM_023244846.1;refGene_gn=CMIP;refGene_tx=XM_023244847.1
E2 57402278 57402471
refGene_gn=PLCG2;refGene_tx=XM_019820120.2;refGene_gn=PLCG2;refGene_tx=XM_019820121.2;refGene_gn=PLCG2;refGene_tx=XM_023245864.1
E2 57456540 57456684
refGene_gn=PLCG2;refGene_tx=XM_019820120.2;refGene_gn=PLCG2;refGene_tx=XM_019820121.2;refGene_gn=PLCG2;refGene_tx=XM_023245864.1
E2 57458898 57458992
refGene_gn=PLCG2;refGene_tx=XM_019820120.2;refGene_gn=PLCG2;refGene_tx=XM_019820121.2;refGene_gn=PLCG2;refGene_tx=XM_023245864.1
E2 57459746 57459794
refGene_gn=PLCG2;refGene_tx=XM_019820120.2;refGene_gn=PLCG2;refGene_tx=XM_019820121.2;refGene_gn=PLCG2;refGene_tx=XM_023245864.1
E2 57469976 57470061
refGene_gn=PLCG2;refGene_tx=XM_019820120.2;refGene_gn=PLCG2;refGene_tx=XM_019820121.2;refGene_gn=PLCG2;refGene_tx=XM_023245864.1
E2 57471720 57471804
refGene_gn=PLCG2;refGene_tx=XM_019820120.2;refGene_gn=PLCG2;refGene_tx=XM_019820121.2;refGene_gn=PLCG2;refGene_tx=XM_023245864.1
E2 57481105 57481149
refGene_gn=PLCG2;refGene_tx=XM_019820120.2;refGene_gn=PLCG2;refGene_tx=XM_019820121.2;refGene_gn=PLCG2;refGene_tx=XM_023245864.1
E2 57483821 57483894
refGene_gn=PLCG2;refGene_tx=XM_019820120.2;refGene_gn=PLCG2;refGene_tx=XM_019820121.2;refGene_gn=PLCG2;refGene_tx=XM_023245864.1
E2 57488374 57488476
refGene_gn=PLCG2;refGene_tx=XM_019820120.2;refGene_gn=PLCG2;refGene_tx

=XM_019820121.2;refGene_gn=PLCG2;refGene_tx=XM_023245864.1
E2 57489853 57489972
refGene_gn=PLCG2;refGene_tx=XM_019820120.2;refGene_gn=PLCG2;refGene_tx=XM_019820121.2;refGene_gn=PLCG2;refGene_tx=XM_023245864.1
E2 57491628 57491714
refGene_gn=PLCG2;refGene_tx=XM_019820120.2;refGene_gn=PLCG2;refGene_tx=XM_019820121.2;refGene_gn=PLCG2;refGene_tx=XM_023245864.1
E2 57493416 57493537
refGene_gn=PLCG2;refGene_tx=XM_019820120.2;refGene_gn=PLCG2;refGene_tx=XM_019820121.2;refGene_gn=PLCG2;refGene_tx=XM_023245864.1
E2 57493858 57494027
refGene_gn=PLCG2;refGene_tx=XM_019820120.2;refGene_gn=PLCG2;refGene_tx=XM_019820121.2;refGene_gn=PLCG2;refGene_tx=XM_023245864.1
E2 57497115 57497220
refGene_gn=PLCG2;refGene_tx=XM_019820120.2;refGene_gn=PLCG2;refGene_tx=XM_019820121.2;refGene_gn=PLCG2;refGene_tx=XM_023245864.1
E2 57498174 57498264
refGene_gn=PLCG2;refGene_tx=XM_019820120.2;refGene_gn=PLCG2;refGene_tx=XM_019820121.2;refGene_gn=PLCG2;refGene_tx=XM_023245864.1
E2 57498843 57499019
refGene_gn=PLCG2;refGene_tx=XM_019820120.2;refGene_gn=PLCG2;refGene_tx=XM_019820121.2;refGene_gn=PLCG2;refGene_tx=XM_023245864.1
E2 57500377 57500578
refGene_gn=PLCG2;refGene_tx=XM_019820120.2;refGene_gn=PLCG2;refGene_tx=XM_019820121.2;refGene_gn=PLCG2;refGene_tx=XM_023245864.1
E2 57501553 57501673
refGene_gn=PLCG2;refGene_tx=XM_019820120.2;refGene_gn=PLCG2;refGene_tx=XM_019820121.2;refGene_gn=PLCG2;refGene_tx=XM_023245864.1
E2 57506365 57506546
refGene_gn=PLCG2;refGene_tx=XM_019820120.2;refGene_gn=PLCG2;refGene_tx=XM_019820121.2;refGene_gn=PLCG2;refGene_tx=XM_023245864.1
E2 57507915 57507987
refGene_gn=PLCG2;refGene_tx=XM_019820120.2;refGene_gn=PLCG2;refGene_tx=XM_019820121.2;refGene_gn=PLCG2;refGene_tx=XM_023245864.1
E2 57510262 57510372
refGene_gn=PLCG2;refGene_tx=XM_019820120.2;refGene_gn=PLCG2;refGene_tx=XM_019820121.2;refGene_gn=PLCG2;refGene_tx=XM_023245864.1
E2 57514229 57514326
refGene_gn=PLCG2;refGene_tx=XM_019820120.2;refGene_gn=PLCG2;refGene_tx=XM_019820121.2;refGene_gn=PLCG2;refGene_tx=XM_023245864.1
E2 57515691 57515758
refGene_gn=PLCG2;refGene_tx=XM_019820120.2;refGene_gn=PLCG2;refGene_tx=XM_019820121.2;refGene_gn=PLCG2;refGene_tx=XM_023245864.1
E2 57516855 57517013
refGene_gn=PLCG2;refGene_tx=XM_019820120.2;refGene_gn=PLCG2;refGene_tx=XM_019820121.2;refGene_gn=PLCG2;refGene_tx=XM_023245864.1
E2 57520359 57520462
refGene_gn=PLCG2;refGene_tx=XM_019820120.2;refGene_gn=PLCG2;refGene_tx=XM_019820121.2;refGene_gn=PLCG2;refGene_tx=XM_023245864.1
E2 57521780 57521990

refGene_gn=PLCG2;refGene_tx=XM_019820120.2;refGene_gn=PLCG2;refGene_tx=XM_019820121.2;refGene_gn=PLCG2;refGene_tx=XM_023245864.1
E2 57523540 57523686 refGene_gn=PLCG2;refGene_tx=XM_019820120.2;refGene_gn=PLCG2;refGene_tx=XM_019820121.2;refGene_gn=PLCG2;refGene_tx=XM_023245864.1
E2 57524264 57524379 refGene_gn=PLCG2;refGene_tx=XM_019820120.2;refGene_gn=PLCG2;refGene_tx=XM_019820121.2;refGene_gn=PLCG2;refGene_tx=XM_023245864.1
E2 57525220 57525388 refGene_gn=PLCG2;refGene_tx=XM_019820120.2;refGene_gn=PLCG2;refGene_tx=XM_019820121.2;refGene_gn=PLCG2;refGene_tx=XM_023245864.1
E2 57530789 57530878 refGene_gn=PLCG2;refGene_tx=XM_019820120.2;refGene_gn=PLCG2;refGene_tx=XM_019820121.2;refGene_gn=PLCG2;refGene_tx=XM_023245864.1
E2 57548066 57548251 refGene_gn=PLCG2;refGene_tx=XM_019820120.2;refGene_gn=PLCG2;refGene_tx=XM_019820121.2;refGene_gn=PLCG2;refGene_tx=XM_023245864.1
E2 57549735 57549778 refGene_gn=PLCG2;refGene_tx=XM_019820120.2;refGene_gn=PLCG2;refGene_tx=XM_019820121.2;refGene_gn=PLCG2;refGene_tx=XM_023245864.1
E2 57578127 57579241 refGene_gn=SDR42E1;refGene_tx=XM_003998293.5;refGene_gn=SDR42E1;refGene_tx=XM_019820125.2;refGene_gn=SDR42E1;refGene_tx=XM_019820126.2;refGene_gn=SDR42E1;refGene_tx=XM_019820127.2
E2 57580080 57580148 refGene_gn=SDR42E1;refGene_tx=XM_003998293.5;refGene_gn=SDR42E1;refGene_tx=XM_019820125.2;refGene_gn=SDR42E1;refGene_tx=XM_019820126.2;refGene_gn=SDR42E1;refGene_tx=XM_019820127.2
E2 57602971 57603236 refGene_gn=HSD17B2;refGene_tx=XM_003998308.3
E2 57652197 57652410 refGene_gn=HSD17B2;refGene_tx=XM_003998308.3
E2 57658610 57658796 refGene_gn=HSD17B2;refGene_tx=XM_003998308.3
E2 57681856 57681994 refGene_gn=HSD17B2;refGene_tx=XM_003998308.3
E2 57694408 57694752 refGene_gn=HSD17B2;refGene_tx=XM_003998308.3
E2 57718331 57718467 refGene_gn=MPHOSPH6;refGene_tx=XM_003998294.5;refGene_gn=MPHOSPH6;refGene_tx=XM_023244934.1
E2 57718922 57719017 refGene_gn=MPHOSPH6;refGene_tx=XM_003998294.5;refGene_gn=MPHOSPH6;refGene_tx=XM_023244934.1
E2 57721717 57721808 refGene_gn=MPHOSPH6;refGene_tx=XM_003998294.5;refGene_gn=MPHOSPH6;refGene_tx=XM_023244934.1
E2 57733816 57733929 refGene_gn=MPHOSPH6;refGene_tx=XM_003998294.5;refGene_gn=MPHOSPH6;refGene_tx=XM_023244934.1
E2 57738954 57738990 refGene_gn=MPHOSPH6;refGene_tx=XM_023244934.1
E2 57739719 57739770 refGene_gn=MPHOSPH6;refGene_tx=XM_003998294.5
E2 58138038 58138083 refGene_gn=CDH13;refGene_tx=XM_003998309.4;refGene_gn=CDH13;refGene_tx=XM_011290267.3

E2 58360169 58360281
refGene_gn=CDH13;refGene_tx=XM_003998309.4;refGene_gn=CDH13;refGene_tx=XM_011290267.3
E2 58517566 58517775
refGene_gn=CDH13;refGene_tx=XM_003998309.4;refGene_gn=CDH13;refGene_tx=XM_011290267.3
E2 58591187 58591304
refGene_gn=CDH13;refGene_tx=XM_003998309.4;refGene_gn=CDH13;refGene_tx=XM_011290267.3
E2 58676031 58676184
refGene_gn=CDH13;refGene_tx=XM_003998309.4;refGene_gn=CDH13;refGene_tx=XM_011290267.3
E2 58679900 58679954 refGene_gn=CDH13;refGene_tx=XM_011290267.3
E2 58783510 58783655 refGene_gn=CDH13;refGene_tx=XM_003998309.4
E2 58924228 58924407 refGene_gn=CDH13;refGene_tx=XM_003998309.4
E2 59024097 59024238 refGene_gn=CDH13;refGene_tx=XM_003998309.4
E2 59071269 59071452 refGene_gn=CDH13;refGene_tx=XM_003998309.4
E2 59078581 59078835 refGene_gn=CDH13;refGene_tx=XM_003998309.4
E2 59139993 59140136 refGene_gn=CDH13;refGene_tx=XM_003998309.4
E2 59159189 59159423 refGene_gn=CDH13;refGene_tx=XM_003998309.4
E2 59162874 59163093 refGene_gn=CDH13;refGene_tx=XM_003998309.4
E2 59173288 59173296 refGene_gn=CDH13;refGene_tx=XM_003998309.4
E2 59263386 59263923 refGene_gn=MLYCD;refGene_tx=XM_004001386.3
E2 59271371 59271484 refGene_gn=MLYCD;refGene_tx=XM_004001386.3
E2 59272337 59272494 refGene_gn=MLYCD;refGene_tx=XM_004001386.3
E2 59275136 59275286 refGene_gn=MLYCD;refGene_tx=XM_004001386.3
E2 59277028 59277562 refGene_gn=MLYCD;refGene_tx=XM_004001386.3
E2 59312578 59312645
refGene_gn=OSGIN1;refGene_tx=XM_006941770.3;refGene_gn=OSGIN1;refGene_tx=XM_011290268.3;refGene_gn=OSGIN1;refGene_tx=XM_019820132.1;refGene_gn=OSGIN1;refGene_tx=XM_019820133.2;refGene_gn=OSGIN1;refGene_tx=XM_019820134.1;refGene_gn=OSGIN1;refGene_tx=XM_023245726.1
E2 59313989 59314126
refGene_gn=OSGIN1;refGene_tx=XM_006941770.3;refGene_gn=OSGIN1;refGene_tx=XM_011290268.3;refGene_gn=OSGIN1;refGene_tx=XM_019820132.1;refGene_gn=OSGIN1;refGene_tx=XM_019820133.2;refGene_gn=OSGIN1;refGene_tx=XM_019820134.1;refGene_gn=OSGIN1;refGene_tx=XM_023245726.1
E2 59314921 59315113
refGene_gn=OSGIN1;refGene_tx=XM_006941770.3;refGene_gn=OSGIN1;refGene_tx=XM_011290268.3;refGene_gn=OSGIN1;refGene_tx=XM_019820132.1;refGene_gn=OSGIN1;refGene_tx=XM_019820133.2;refGene_gn=OSGIN1;refGene_tx=XM_019820134.1;refGene_gn=OSGIN1;refGene_tx=XM_023245726.1
E2 59315379 59315471
refGene_gn=OSGIN1;refGene_tx=XM_006941770.3;refGene_gn=OSGIN1;refGene_tx=XM_011290268.3;refGene_gn=OSGIN1;refGene_tx=XM_019820132.1;refGene_gn=OSGIN1;refGene_tx=XM_019820133.2;refGene_gn=OSGIN1;refGene_tx=XM_019820134.1;refGene_gn=OSGIN1;refGene_tx=XM_023245726.1
E2 59317907 59318856
refGene_gn=OSGIN1;refGene_tx=XM_006941770.3;refGene_gn=OSGIN1;refGene_tx=XM_011290268.3;refGene_gn=OSGIN1;refGene_tx=XM_019820132.1;refGene_

gn=OSGIN1;refGene_tx=XM_019820133.2;refGene_gn=OSGIN1;refGene_tx=XM_019820134.1;refGene_gn=OSGIN1;refGene_tx=XM_023245726.1
E2 59320674 59320899 refGene_gn=NECAB2;refGene_tx=XM_023245243.1;refGene_gn=NECAB2;refGene_tx=XM_023245244.1
E2 59323411 59323436 refGene_gn=NECAB2;refGene_tx=XM_023245243.1;refGene_gn=NECAB2;refGene_tx=XM_023245244.1
E2 59328147 59328256 refGene_gn=NECAB2;refGene_tx=XM_023245243.1;refGene_gn=NECAB2;refGene_tx=XM_023245244.1
E2 59330352 59330378 refGene_gn=NECAB2;refGene_tx=XM_023245243.1;refGene_gn=NECAB2;refGene_tx=XM_023245244.1
E2 59330527 59330625 refGene_gn=NECAB2;refGene_tx=XM_023245243.1;refGene_gn=NECAB2;refGene_tx=XM_023245244.1
E2 59341056 59341193 refGene_gn=NECAB2;refGene_tx=XM_023245243.1;refGene_gn=NECAB2;refGene_tx=XM_023245244.1
E2 59343253 59343372 refGene_gn=NECAB2;refGene_tx=XM_023245243.1;refGene_gn=NECAB2;refGene_tx=XM_023245244.1
E2 59343534 59343614 refGene_gn=NECAB2;refGene_tx=XM_023245243.1;refGene_gn=NECAB2;refGene_tx=XM_023245244.1
E2 59346239 59346293 refGene_gn=NECAB2;refGene_tx=XM_023245243.1;refGene_gn=NECAB2;refGene_tx=XM_023245244.1
E2 59346847 59346960 refGene_gn=NECAB2;refGene_tx=XM_023245243.1;refGene_gn=NECAB2;refGene_tx=XM_023245244.1
E2 59348435 59348513 refGene_gn=NECAB2;refGene_tx=XM_023245243.1;refGene_gn=NECAB2;refGene_tx=XM_023245244.1
E2 59348931 59349127 refGene_gn=NECAB2;refGene_tx=XM_023245243.1;refGene_gn=NECAB2;refGene_tx=XM_023245244.1
E2 59349364 59349393 refGene_gn=NECAB2;refGene_tx=XM_023245244.1
E2 59355634 59355728 refGene_gn=SLC38A8;refGene_tx=XM_019820135.2;refGene_gn=SLC38A8;refGene_tx=XM_023245240.1;refGene_gn=SLC38A8;refGene_tx=XM_023245241.1;refGene_gn=SLC38A8;refGene_tx=XM_023245242.1
E2 59357876 59357928 refGene_gn=SLC38A8;refGene_tx=XM_019820135.2;refGene_gn=SLC38A8;refGene_tx=XM_023245240.1;refGene_gn=SLC38A8;refGene_tx=XM_023245241.1;refGene_gn=SLC38A8;refGene_tx=XM_023245242.1
E2 59358613 59358822 refGene_gn=SLC38A8;refGene_tx=XM_019820135.2;refGene_gn=SLC38A8;refGene_tx=XM_023245240.1;refGene_gn=SLC38A8;refGene_tx=XM_023245241.1;refGe

ne_gn=SLC38A8;refGene_tx=XM_023245242.1
E2 59359001 59359149
refGene_gn=SLC38A8;refGene_tx=XM_019820135.2;refGene_gn=SLC38A8;refGene_tx=XM_023245240.1;refGene_gn=SLC38A8;refGene_tx=XM_023245241.1;refGene_gn=SLC38A8;refGene_tx=XM_023245242.1
E2 59359940 59360055
refGene_gn=SLC38A8;refGene_tx=XM_019820135.2;refGene_gn=SLC38A8;refGene_tx=XM_023245240.1;refGene_gn=SLC38A8;refGene_tx=XM_023245241.1;refGene_gn=SLC38A8;refGene_tx=XM_023245242.1
E2 59363868 59363926
refGene_gn=SLC38A8;refGene_tx=XM_019820135.2;refGene_gn=SLC38A8;refGene_tx=XM_023245240.1;refGene_gn=SLC38A8;refGene_tx=XM_023245241.1;refGene_gn=SLC38A8;refGene_tx=XM_023245242.1
E2 59365999 59366101
refGene_gn=SLC38A8;refGene_tx=XM_019820135.2;refGene_gn=SLC38A8;refGene_tx=XM_023245240.1;refGene_gn=SLC38A8;refGene_tx=XM_023245241.1;refGene_gn=SLC38A8;refGene_tx=XM_023245242.1
E2 59367205 59367347
refGene_gn=SLC38A8;refGene_tx=XM_019820135.2;refGene_gn=SLC38A8;refGene_tx=XM_023245240.1;refGene_gn=SLC38A8;refGene_tx=XM_023245241.1;refGene_gn=SLC38A8;refGene_tx=XM_023245242.1
E2 59370511 59370710
refGene_gn=SLC38A8;refGene_tx=XM_019820135.2;refGene_gn=SLC38A8;refGene_tx=XM_023245240.1;refGene_gn=SLC38A8;refGene_tx=XM_023245241.1;refGene_gn=SLC38A8;refGene_tx=XM_023245242.1
E2 59371988 59372179
refGene_gn=SLC38A8;refGene_tx=XM_019820135.2;refGene_gn=SLC38A8;refGene_tx=XM_023245240.1;refGene_gn=SLC38A8;refGene_tx=XM_023245241.1;refGene_gn=SLC38A8;refGene_tx=XM_023245242.1
E2 59372434 59372474
refGene_gn=SLC38A8;refGene_tx=XM_019820135.2;refGene_gn=SLC38A8;refGene_tx=XM_023245240.1;refGene_gn=SLC38A8;refGene_tx=XM_023245241.1;refGene_gn=SLC38A8;refGene_tx=XM_023245242.1
E2 59382343 59382540 refGene_gn=MBTPS1;refGene_tx=XM_006941772.3
E2 59383774 59383905 refGene_gn=MBTPS1;refGene_tx=XM_006941772.3
E2 59386826 59386953 refGene_gn=MBTPS1;refGene_tx=XM_006941772.3
E2 59387934 59388066 refGene_gn=MBTPS1;refGene_tx=XM_006941772.3
E2 59389795 59389936 refGene_gn=MBTPS1;refGene_tx=XM_006941772.3
E2 59392305 59392383 refGene_gn=MBTPS1;refGene_tx=XM_006941772.3
E2 59393162 59393287 refGene_gn=MBTPS1;refGene_tx=XM_006941772.3
E2 59394026 59394183 refGene_gn=MBTPS1;refGene_tx=XM_006941772.3
E2 59394702 59394818 refGene_gn=MBTPS1;refGene_tx=XM_006941772.3
E2 59396124 59396297 refGene_gn=MBTPS1;refGene_tx=XM_006941772.3
E2 59396797 59396986 refGene_gn=MBTPS1;refGene_tx=XM_006941772.3
E2 59401525 59401670 refGene_gn=MBTPS1;refGene_tx=XM_006941772.3
E2 59405595 59405757 refGene_gn=MBTPS1;refGene_tx=XM_006941772.3
E2 59408642 59408794 refGene_gn=MBTPS1;refGene_tx=XM_006941772.3
E2 59410343 59410446 refGene_gn=MBTPS1;refGene_tx=XM_006941772.3
E2 59412863 59412931 refGene_gn=MBTPS1;refGene_tx=XM_006941772.3
E2 59413402 59413519 refGene_gn=MBTPS1;refGene_tx=XM_006941772.3

E2 59414798 59414908 refGene_gn=MBTPS1;refGene_tx=XM_006941772.3
E2 59415307 59415418 refGene_gn=MBTPS1;refGene_tx=XM_006941772.3
E2 59416963 59417155 refGene_gn=MBTPS1;refGene_tx=XM_006941772.3
E2 59420398 59420656 refGene_gn=MBTPS1;refGene_tx=XM_006941772.3
E2 59421364 59421533 refGene_gn=MBTPS1;refGene_tx=XM_006941772.3
E2 59439166 59439265 refGene_gn=HSDL1;refGene_tx=XM_023245878.1
E2 59445354 59445582 refGene_gn=HSDL1;refGene_tx=XM_023245878.1
E2 59445788 59446234 refGene_gn=HSDL1;refGene_tx=XM_023245878.1
E2 59446778 59446998 refGene_gn=HSDL1;refGene_tx=XM_023245878.1
E2 59458870 59458994
refGene_gn=DNAAF1;refGene_tx=XM_011290301.3;refGene_gn=DNAAF1;refGene_tx=XM_019820139.2;refGene_gn=DNAAF1;refGene_tx=XM_019820140.2
E2 59461108 59461244
refGene_gn=DNAAF1;refGene_tx=XM_011290301.3;refGene_gn=DNAAF1;refGene_tx=XM_019820139.2;refGene_gn=DNAAF1;refGene_tx=XM_019820140.2
E2 59461740 59461832
refGene_gn=DNAAF1;refGene_tx=XM_011290301.3;refGene_gn=DNAAF1;refGene_tx=XM_019820139.2;refGene_gn=DNAAF1;refGene_tx=XM_019820140.2
E2 59463835 59464057
refGene_gn=DNAAF1;refGene_tx=XM_011290301.3;refGene_gn=DNAAF1;refGene_tx=XM_019820139.2;refGene_gn=DNAAF1;refGene_tx=XM_019820140.2
E2 59464185 59464352
refGene_gn=DNAAF1;refGene_tx=XM_011290301.3;refGene_gn=DNAAF1;refGene_tx=XM_019820139.2;refGene_gn=DNAAF1;refGene_tx=XM_019820140.2
E2 59467355 59467477
refGene_gn=DNAAF1;refGene_tx=XM_011290301.3;refGene_gn=DNAAF1;refGene_tx=XM_019820139.2;refGene_gn=DNAAF1;refGene_tx=XM_019820140.2
E2 59470723 59470890
refGene_gn=DNAAF1;refGene_tx=XM_011290301.3;refGene_gn=DNAAF1;refGene_tx=XM_019820139.2;refGene_gn=DNAAF1;refGene_tx=XM_019820140.2
E2 59474558 59474825
refGene_gn=DNAAF1;refGene_tx=XM_011290301.3;refGene_gn=DNAAF1;refGene_tx=XM_019820139.2
E2 59478530 59478640
refGene_gn=DNAAF1;refGene_tx=XM_011290301.3;refGene_gn=DNAAF1;refGene_tx=XM_019820139.2;refGene_gn=DNAAF1;refGene_tx=XM_019820140.2
E2 59479894 59479951
refGene_gn=DNAAF1;refGene_tx=XM_011290301.3;refGene_gn=DNAAF1;refGene_tx=XM_019820139.2;refGene_gn=DNAAF1;refGene_tx=XM_019820140.2
E2 59480806 59481173
refGene_gn=DNAAF1;refGene_tx=XM_011290301.3;refGene_gn=DNAAF1;refGene_tx=XM_019820139.2;refGene_gn=DNAAF1;refGene_tx=XM_019820140.2
E2 59482191 59482286
refGene_gn=DNAAF1;refGene_tx=XM_011290301.3;refGene_gn=DNAAF1;refGene_tx=XM_019820139.2;refGene_gn=DNAAF1;refGene_tx=XM_019820140.2
E2 59483345 59484319
refGene_gn=TAF1C;refGene_tx=XM_023245875.1;refGene_gn=TAF1C;refGene_tx=XM_023245876.1
E2 59484406 59484544
refGene_gn=TAF1C;refGene_tx=XM_023245875.1;refGene_gn=TAF1C;refGene_tx

=XM_023245876.1
E2 59484619 59484794
refGene_gn=TAF1C;refGene_tx=XM_023245875.1;refGene_gn=TAF1C;refGene_tx
=XM_023245876.1
E2 59485396 59485405 refGene_gn=TAF1C;refGene_tx=XM_023245876.1
E2 59485591 59485735 refGene_gn=TAF1C;refGene_tx=XM_023245875.1
E2 59485903 59486039 refGene_gn=TAF1C;refGene_tx=XM_023245875.1
E2 59486168 59486240 refGene_gn=TAF1C;refGene_tx=XM_023245875.1
E2 59486321 59486439 refGene_gn=TAF1C;refGene_tx=XM_023245875.1
E2 59486531 59486648 refGene_gn=TAF1C;refGene_tx=XM_023245875.1
E2 59486787 59487026 refGene_gn=TAF1C;refGene_tx=XM_023245875.1
E2 59487669 59487743 refGene_gn=TAF1C;refGene_tx=XM_023245875.1
E2 59487833 59487923 refGene_gn=TAF1C;refGene_tx=XM_023245875.1
E2 59487995 59488093 refGene_gn=TAF1C;refGene_tx=XM_023245875.1
E2 59488256 59488338 refGene_gn=TAF1C;refGene_tx=XM_023245875.1
E2 59489240 59489378 refGene_gn=TAF1C;refGene_tx=XM_023245875.1
E2 59493641 59493852 refGene_gn=ADAD2;refGene_tx=XM_011290271.3
E2 59493960 59494104 refGene_gn=ADAD2;refGene_tx=XM_011290271.3
E2 59496296 59496432 refGene_gn=ADAD2;refGene_tx=XM_019820143.2
E2 59496660 59496801
refGene_gn=ADAD2;refGene_tx=XM_011290271.3;refGene_gn=ADAD2;refGene_tx
=XM_019820143.2
E2 59497122 59497170
refGene_gn=ADAD2;refGene_tx=XM_011290271.3;refGene_gn=ADAD2;refGene_tx
=XM_019820143.2
E2 59497267 59497393
refGene_gn=ADAD2;refGene_tx=XM_011290271.3;refGene_gn=ADAD2;refGene_tx
=XM_019820143.2
E2 59497485 59497636
refGene_gn=ADAD2;refGene_tx=XM_011290271.3;refGene_gn=ADAD2;refGene_tx
=XM_019820143.2
E2 59497729 59497897
refGene_gn=ADAD2;refGene_tx=XM_011290271.3;refGene_gn=ADAD2;refGene_tx
=XM_019820143.2
E2 59498047 59498277
refGene_gn=ADAD2;refGene_tx=XM_011290271.3;refGene_gn=ADAD2;refGene_tx
=XM_019820143.2
E2 59498391 59498620
refGene_gn=ADAD2;refGene_tx=XM_011290271.3;refGene_gn=ADAD2;refGene_tx
=XM_019820143.2
E2 59498876 59498997
refGene_gn=ADAD2;refGene_tx=XM_011290271.3;refGene_gn=ADAD2;refGene_tx
=XM_019820143.2
E2 59499080 59499185
refGene_gn=ADAD2;refGene_tx=XM_011290271.3;refGene_gn=ADAD2;refGene_tx
=XM_019820143.2
E2 59509574 59510345
refGene_gn=KCNG4;refGene_tx=XM_019820145.2;refGene_gn=KCNG4;refGene_tx
=XM_023245349.1
E2 59517929 59518692

refGene_gn=KCNG4;refGene_tx=XM_019820145.2;refGene_gn=KCNG4;refGene_tx=XM_023245349.1
E2 59519751 59519753 refGene_gn=KCNG4;refGene_tx=XM_019820145.2
E2 59555676 59555946 refGene_gn=WFDC1;refGene_tx=XM_023245048.1
E2 59566745 59566929 refGene_gn=WFDC1;refGene_tx=XM_023245048.1
E2 59569696 59569780 refGene_gn=WFDC1;refGene_tx=XM_023245048.1
E2 59570714 59570855 refGene_gn=WFDC1;refGene_tx=XM_023245048.1
E2 59574084 59574126 refGene_gn=WFDC1;refGene_tx=XM_023245048.1
E2 59576337 59576396 refGene_gn=WFDC1;refGene_tx=XM_023245048.1
E2 59598630 59598729 refGene_gn=ATP2C2;refGene_tx=XM_023246034.1;refGene_gn=ATP2C2;refGene_tx=XM_023246035.1
E2 59620631 59620742 refGene_gn=ATP2C2;refGene_tx=XM_023246034.1;refGene_gn=ATP2C2;refGene_tx=XM_023246035.1
E2 59622000 59622117 refGene_gn=ATP2C2;refGene_tx=XM_023246034.1;refGene_gn=ATP2C2;refGene_tx=XM_023246035.1
E2 59627256 59627346 refGene_gn=ATP2C2;refGene_tx=XM_023246034.1;refGene_gn=ATP2C2;refGene_tx=XM_023246035.1
E2 59628183 59628219 refGene_gn=ATP2C2;refGene_tx=XM_023246034.1;refGene_gn=ATP2C2;refGene_tx=XM_023246035.1
E2 59628321 59628383 refGene_gn=ATP2C2;refGene_tx=XM_023246034.1;refGene_gn=ATP2C2;refGene_tx=XM_023246035.1
E2 59633418 59633527 refGene_gn=ATP2C2;refGene_tx=XM_023246034.1;refGene_gn=ATP2C2;refGene_tx=XM_023246035.1
E2 59641032 59641182 refGene_gn=ATP2C2;refGene_tx=XM_023246034.1;refGene_gn=ATP2C2;refGene_tx=XM_023246035.1
E2 59641260 59641329 refGene_gn=ATP2C2;refGene_tx=XM_023246034.1;refGene_gn=ATP2C2;refGene_tx=XM_023246035.1
E2 59641889 59641965 refGene_gn=ATP2C2;refGene_tx=XM_023246034.1;refGene_gn=ATP2C2;refGene_tx=XM_023246035.1
E2 59644299 59644366 refGene_gn=ATP2C2;refGene_tx=XM_023246034.1;refGene_gn=ATP2C2;refGene_tx=XM_023246035.1
E2 59647764 59647889 refGene_gn=ATP2C2;refGene_tx=XM_023246034.1;refGene_gn=ATP2C2;refGene_tx=XM_023246035.1
E2 59648021 59648119 refGene_gn=ATP2C2;refGene_tx=XM_023246034.1;refGene_gn=ATP2C2;refGene_tx=XM_023246035.1
E2 59648999 59649101 refGene_gn=ATP2C2;refGene_tx=XM_023246034.1;refGene_gn=ATP2C2;refGene_

tx=XM_023246035.1
E2 59650362 59650452 refGene_gn=ATP2C2;refGene_tx=XM_023246034.1
E2 59653201 59653303
refGene_gn=ATP2C2;refGene_tx=XM_023246034.1;refGene_gn=ATP2C2;refGene_
tx=XM_023246035.1
E2 59654622 59654779
refGene_gn=ATP2C2;refGene_tx=XM_023246034.1;refGene_gn=ATP2C2;refGene_
tx=XM_023246035.1
E2 59655455 59655626
refGene_gn=ATP2C2;refGene_tx=XM_023246034.1;refGene_gn=ATP2C2;refGene_
tx=XM_023246035.1
E2 59656318 59656416
refGene_gn=ATP2C2;refGene_tx=XM_023246034.1;refGene_gn=ATP2C2;refGene_
tx=XM_023246035.1
E2 59656510 59656561
refGene_gn=ATP2C2;refGene_tx=XM_023246034.1;refGene_gn=ATP2C2;refGene_
tx=XM_023246035.1
E2 59657499 59657666
refGene_gn=ATP2C2;refGene_tx=XM_023246034.1;refGene_gn=ATP2C2;refGene_
tx=XM_023246035.1
E2 59658958 59659027
refGene_gn=ATP2C2;refGene_tx=XM_023246034.1;refGene_gn=ATP2C2;refGene_
tx=XM_023246035.1
E2 59659116 59659233
refGene_gn=ATP2C2;refGene_tx=XM_023246034.1;refGene_gn=ATP2C2;refGene_
tx=XM_023246035.1
E2 59660509 59660657
refGene_gn=ATP2C2;refGene_tx=XM_023246034.1;refGene_gn=ATP2C2;refGene_
tx=XM_023246035.1
E2 59661786 59661885
refGene_gn=ATP2C2;refGene_tx=XM_023246034.1;refGene_gn=ATP2C2;refGene_
tx=XM_023246035.1
E2 59662041 59662183
refGene_gn=ATP2C2;refGene_tx=XM_023246034.1;refGene_gn=ATP2C2;refGene_
tx=XM_023246035.1
E2 59663186 59663305
refGene_gn=ATP2C2;refGene_tx=XM_023246034.1;refGene_gn=ATP2C2;refGene_
tx=XM_023246035.1
E2 59673752 59673866
refGene_gn=TLDC1;refGene_tx=XM_006941773.3;refGene_gn=TLDC1;refGene_tx
=XM_019820148.2;refGene_gn=TLDC1;refGene_tx=XM_023246036.1
E2 59674535 59674715
refGene_gn=TLDC1;refGene_tx=XM_006941773.3;refGene_gn=TLDC1;refGene_tx
=XM_019820148.2
E2 59676180 59676299
refGene_gn=TLDC1;refGene_tx=XM_006941773.3;refGene_gn=TLDC1;refGene_tx
=XM_019820148.2;refGene_gn=TLDC1;refGene_tx=XM_023246036.1
E2 59680501 59680930
refGene_gn=TLDC1;refGene_tx=XM_006941773.3;refGene_gn=TLDC1;refGene_tx
=XM_019820148.2;refGene_gn=TLDC1;refGene_tx=XM_023246036.1

E2 59682816 59682961
refGene_gn=TLDC1;refGene_tx=XM_006941773.3;refGene_gn=TLDC1;refGene_tx=XM_019820148.2;refGene_gn=TLDC1;refGene_tx=XM_023246036.1
E2 59683247 59683478
refGene_gn=TLDC1;refGene_tx=XM_006941773.3;refGene_gn=TLDC1;refGene_tx=XM_019820148.2;refGene_gn=TLDC1;refGene_tx=XM_023246036.1
E2 59685650 59685803
refGene_gn=TLDC1;refGene_tx=XM_006941773.3;refGene_gn=TLDC1;refGene_tx=XM_019820148.2;refGene_gn=TLDC1;refGene_tx=XM_023246036.1
E2 59733102 59733213 refGene_gn=COTL1;refGene_tx=XM_003998318.4
E2 59746457 59746615 refGene_gn=COTL1;refGene_tx=XM_003998318.4
E2 59767530 59767613 refGene_gn=COTL1;refGene_tx=XM_003998318.4
E2 59767875 59767952 refGene_gn=COTL1;refGene_tx=XM_003998318.4
E2 59795225 59795288
refGene_gn=LOC101089411;refGene_tx=XM_023245534.1;refGene_gn=LOC101089411;refGene_tx=XM_023245535.1;refGene_gn=LOC101089411;refGene_tx=XM_023245538.1;refGene_gn=LOC101089411;refGene_tx=XM_023245539.1
E2 59799642 59800716
refGene_gn=LOC101089411;refGene_tx=XM_023245538.1;refGene_gn=LOC101089411;refGene_tx=XM_023245539.1
E2 59802298 59802456
refGene_gn=LOC101089411;refGene_tx=XM_023245538.1;refGene_gn=LOC101089411;refGene_tx=XM_023245539.1
E2 59804480 59805126
refGene_gn=LOC101089411;refGene_tx=XM_023245538.1;refGene_gn=LOC101089411;refGene_tx=XM_023245539.1
E2 59827389 59827410
refGene_gn=LOC101089411;refGene_tx=XM_023245536.1;refGene_gn=LOC101089411;refGene_tx=XM_023245537.1
E2 59856515 59856584
refGene_gn=LOC101089411;refGene_tx=XM_023245534.1;refGene_gn=LOC101089411;refGene_tx=XM_023245535.1;refGene_gn=LOC101089411;refGene_tx=XM_023245536.1
E2 59863776 59863837
refGene_gn=LOC101089411;refGene_tx=XM_023245534.1;refGene_gn=LOC101089411;refGene_tx=XM_023245535.1;refGene_gn=LOC101089411;refGene_tx=XM_023245536.1;refGene_gn=LOC101089411;refGene_tx=XM_023245537.1
E2 59868176 59869217
refGene_gn=LOC101089411;refGene_tx=XM_023245534.1;refGene_gn=LOC101089411;refGene_tx=XM_023245535.1;refGene_gn=LOC101089411;refGene_tx=XM_023245536.1;refGene_gn=LOC101089411;refGene_tx=XM_023245537.1
E2 59878980 59879072
refGene_gn=LOC101089411;refGene_tx=XM_023245534.1;refGene_gn=LOC101089411;refGene_tx=XM_023245535.1;refGene_gn=LOC101089411;refGene_tx=XM_023245536.1;refGene_gn=LOC101089411;refGene_tx=XM_023245537.1
E2 59879598 59879708
refGene_gn=LOC101089411;refGene_tx=XM_023245534.1;refGene_gn=LOC101089411;refGene_tx=XM_023245535.1;refGene_gn=LOC101089411;refGene_tx=XM_023245536.1;refGene_gn=LOC101089411;refGene_tx=XM_023245537.1
E2 59880120 59880176

refGene_gn=LOC101089411;refGene_tx=XM_023245534.1;refGene_gn=LOC101089
411;refGene_tx=XM_023245535.1;refGene_gn=LOC101089411;refGene_tx=XM_02
3245536.1;refGene_gn=LOC101089411;refGene_tx=XM_023245537.1
E2 59880437 59880541
refGene_gn=LOC101089411;refGene_tx=XM_023245534.1;refGene_gn=LOC101089
411;refGene_tx=XM_023245535.1;refGene_gn=LOC101089411;refGene_tx=XM_02
3245536.1;refGene_gn=LOC101089411;refGene_tx=XM_023245537.1
E2 59883410 59883510
refGene_gn=LOC101089411;refGene_tx=XM_023245534.1;refGene_gn=LOC101089
411;refGene_tx=XM_023245535.1;refGene_gn=LOC101089411;refGene_tx=XM_02
3245536.1;refGene_gn=LOC101089411;refGene_tx=XM_023245537.1
E2 59884284 59884462
refGene_gn=LOC101089411;refGene_tx=XM_023245534.1;refGene_gn=LOC101089
411;refGene_tx=XM_023245535.1;refGene_gn=LOC101089411;refGene_tx=XM_02
3245536.1;refGene_gn=LOC101089411;refGene_tx=XM_023245537.1
E2 59887189 59887355
refGene_gn=LOC101089411;refGene_tx=XM_023245534.1;refGene_gn=LOC101089
411;refGene_tx=XM_023245535.1;refGene_gn=LOC101089411;refGene_tx=XM_02
3245536.1;refGene_gn=LOC101089411;refGene_tx=XM_023245537.1
E2 59890562 59890707
refGene_gn=LOC101089411;refGene_tx=XM_023245534.1;refGene_gn=LOC101089
411;refGene_tx=XM_023245535.1;refGene_gn=LOC101089411;refGene_tx=XM_02
3245536.1;refGene_gn=LOC101089411;refGene_tx=XM_023245537.1
E2 59891825 59891891
refGene_gn=LOC101089411;refGene_tx=XM_023245534.1;refGene_gn=LOC101089
411;refGene_tx=XM_023245535.1;refGene_gn=LOC101089411;refGene_tx=XM_02
3245536.1;refGene_gn=LOC101089411;refGene_tx=XM_023245537.1
E2 59895529 59895717
refGene_gn=LOC101089411;refGene_tx=XM_023245534.1;refGene_gn=LOC101089
411;refGene_tx=XM_023245535.1;refGene_gn=LOC101089411;refGene_tx=XM_02
3245536.1;refGene_gn=LOC101089411;refGene_tx=XM_023245537.1
E2 59935516 59935756 refGene_gn=CRISPLD2;refGene_tx=XM_003998300.4
E2 59940997 59941116 refGene_gn=CRISPLD2;refGene_tx=XM_003998300.4
E2 59943771 59943904 refGene_gn=CRISPLD2;refGene_tx=XM_003998300.4
E2 59944230 59944346 refGene_gn=CRISPLD2;refGene_tx=XM_003998300.4
E2 59947020 59947121 refGene_gn=CRISPLD2;refGene_tx=XM_003998300.4
E2 59951108 59951252 refGene_gn=CRISPLD2;refGene_tx=XM_003998300.4
E2 59952963 59953024 refGene_gn=CRISPLD2;refGene_tx=XM_003998300.4
E2 59955889 59955956 refGene_gn=CRISPLD2;refGene_tx=XM_003998300.4
E2 59956409 59956540 refGene_gn=CRISPLD2;refGene_tx=XM_003998300.4
E2 59958276 59958320 refGene_gn=CRISPLD2;refGene_tx=XM_003998300.4
E2 59960851 59960924 refGene_gn=CRISPLD2;refGene_tx=XM_003998300.4
E2 59962493 59962569 refGene_gn=CRISPLD2;refGene_tx=XM_003998300.4
E2 59968249 59968383 refGene_gn=CRISPLD2;refGene_tx=XM_003998300.4
E2 59979058 59979113 refGene_gn=CRISPLD2;refGene_tx=XM_003998300.4
E2 60008677 60008762
refGene_gn=LOC111558005;refGene_tx=XM_023245124.1
E2 60031023 60031132
refGene_gn=LOC111558005;refGene_tx=XM_023245124.1
E2 60036371 60036532

refGene_gn=LOC111558005;refGene_tx=XM_023245124.1
E2 60037836 60038076
refGene_gn=LOC111558005;refGene_tx=XM_023245124.1
E2 60038966 60039037
refGene_gn=LOC111558005;refGene_tx=XM_023245124.1
E2 60039939 60040116
refGene_gn=ZDHHC7;refGene_tx=XM_003998301.5;refGene_gn=ZDHHC7;refGene_tx=XM_019820157.2;refGene_gn=ZDHHC7;refGene_tx=XM_023245897.1;refGene_gn=ZDHHC7;refGene_tx=XM_023245898.1;refGene_gn=ZDHHC7;refGene_tx=XM_023245899.1
E2 60040786 60040917
refGene_gn=ZDHHC7;refGene_tx=XM_003998301.5;refGene_gn=ZDHHC7;refGene_tx=XM_019820157.2;refGene_gn=ZDHHC7;refGene_tx=XM_023245897.1;refGene_gn=ZDHHC7;refGene_tx=XM_023245898.1;refGene_gn=ZDHHC7;refGene_tx=XM_023245899.1
E2 60041629 60041711
refGene_gn=ZDHHC7;refGene_tx=XM_003998301.5;refGene_gn=ZDHHC7;refGene_tx=XM_019820157.2;refGene_gn=ZDHHC7;refGene_tx=XM_023245897.1;refGene_gn=ZDHHC7;refGene_tx=XM_023245898.1;refGene_gn=ZDHHC7;refGene_tx=XM_023245899.1
E2 60042120 60042217
refGene_gn=ZDHHC7;refGene_tx=XM_003998301.5;refGene_gn=ZDHHC7;refGene_tx=XM_019820157.2;refGene_gn=ZDHHC7;refGene_tx=XM_023245897.1;refGene_gn=ZDHHC7;refGene_tx=XM_023245898.1;refGene_gn=ZDHHC7;refGene_tx=XM_023245899.1
E2 60043581 60043706
refGene_gn=ZDHHC7;refGene_tx=XM_003998301.5;refGene_gn=ZDHHC7;refGene_tx=XM_019820157.2;refGene_gn=ZDHHC7;refGene_tx=XM_023245897.1;refGene_gn=ZDHHC7;refGene_tx=XM_023245898.1;refGene_gn=ZDHHC7;refGene_tx=XM_023245899.1
E2 60047841 60048156
refGene_gn=ZDHHC7;refGene_tx=XM_003998301.5;refGene_gn=ZDHHC7;refGene_tx=XM_019820157.2;refGene_gn=ZDHHC7;refGene_tx=XM_023245897.1;refGene_gn=ZDHHC7;refGene_tx=XM_023245898.1;refGene_gn=ZDHHC7;refGene_tx=XM_023245899.1
E2 60111903 60112232
refGene_gn=KIAA0513;refGene_tx=XM_011290276.3;refGene_gn=KIAA0513;refGene_tx=XM_023246124.1;refGene_gn=KIAA0513;refGene_tx=XM_023246125.1;refGene_gn=KIAA0513;refGene_tx=XM_023246126.1
E2 60113651 60113751
refGene_gn=KIAA0513;refGene_tx=XM_011290276.3;refGene_gn=KIAA0513;refGene_tx=XM_023246124.1;refGene_gn=KIAA0513;refGene_tx=XM_023246125.1;refGene_gn=KIAA0513;refGene_tx=XM_023246126.1
E2 60115032 60115106
refGene_gn=KIAA0513;refGene_tx=XM_011290276.3;refGene_gn=KIAA0513;refGene_tx=XM_023246124.1;refGene_gn=KIAA0513;refGene_tx=XM_023246125.1;refGene_gn=KIAA0513;refGene_tx=XM_023246126.1
E2 60117883 60117954
refGene_gn=KIAA0513;refGene_tx=XM_011290276.3;refGene_gn=KIAA0513;refGene_tx=XM_023246124.1;refGene_gn=KIAA0513;refGene_tx=XM_023246125.1;re

fGene_gn=KIAA0513;refGene_tx=XM_023246126.1
E2 60119388 60119596
refGene_gn=KIAA0513;refGene_tx=XM_011290276.3;refGene_gn=KIAA0513;refGene_tx=XM_023246124.1;refGene_gn=KIAA0513;refGene_tx=XM_023246125.1;refGene_gn=KIAA0513;refGene_tx=XM_023246126.1
E2 60120362 60120403
refGene_gn=KIAA0513;refGene_tx=XM_011290276.3;refGene_gn=KIAA0513;refGene_tx=XM_023246124.1;refGene_gn=KIAA0513;refGene_tx=XM_023246125.1;refGene_gn=KIAA0513;refGene_tx=XM_023246126.1
E2 60120737 60120816
refGene_gn=KIAA0513;refGene_tx=XM_011290276.3;refGene_gn=KIAA0513;refGene_tx=XM_023246124.1;refGene_gn=KIAA0513;refGene_tx=XM_023246125.1;refGene_gn=KIAA0513;refGene_tx=XM_023246126.1
E2 60122561 60122639
refGene_gn=KIAA0513;refGene_tx=XM_011290276.3;refGene_gn=KIAA0513;refGene_tx=XM_023246124.1;refGene_gn=KIAA0513;refGene_tx=XM_023246125.1;refGene_gn=KIAA0513;refGene_tx=XM_023246126.1
E2 60123821 60123851
refGene_gn=KIAA0513;refGene_tx=XM_011290276.3;refGene_gn=KIAA0513;refGene_tx=XM_023246124.1;refGene_gn=KIAA0513;refGene_tx=XM_023246125.1
E2 60126895 60126976
refGene_gn=KIAA0513;refGene_tx=XM_011290276.3;refGene_gn=KIAA0513;refGene_tx=XM_023246124.1;refGene_gn=KIAA0513;refGene_tx=XM_023246125.1;refGene_gn=KIAA0513;refGene_tx=XM_023246126.1
E2 60127420 60127515
refGene_gn=KIAA0513;refGene_tx=XM_011290276.3;refGene_gn=KIAA0513;refGene_tx=XM_023246124.1;refGene_gn=KIAA0513;refGene_tx=XM_023246125.1;refGene_gn=KIAA0513;refGene_tx=XM_023246126.1
E2 60128605 60128655
refGene_gn=KIAA0513;refGene_tx=XM_011290276.3;refGene_gn=KIAA0513;refGene_tx=XM_023246124.1;refGene_gn=KIAA0513;refGene_tx=XM_023246125.1;refGene_gn=KIAA0513;refGene_tx=XM_023246126.1
E2 60138708 60138813 refGene_gn=FAM92B;refGene_tx=XM_019820158.2
E2 60139595 60139697 refGene_gn=FAM92B;refGene_tx=XM_019820158.2
E2 60141685 60141799 refGene_gn=FAM92B;refGene_tx=XM_019820158.2
E2 60144718 60144823 refGene_gn=FAM92B;refGene_tx=XM_019820158.2
E2 60145504 60145510 refGene_gn=FAM92B;refGene_tx=XM_019820158.2
E2 60145626 60145728 refGene_gn=FAM92B;refGene_tx=XM_019820158.2
E2 60145818 60145887 refGene_gn=FAM92B;refGene_tx=XM_019820158.2
E2 60146786 60147021 refGene_gn=FAM92B;refGene_tx=XM_019820158.2
E2 60148053 60148073 refGene_gn=FAM92B;refGene_tx=XM_019820158.2
E2 60198685 60198757
refGene_gn=LOC111558006;refGene_tx=XM_023245125.1
E2 60198853 60199357
refGene_gn=LOC111558006;refGene_tx=XM_023245125.1
E2 60199421 60199718
refGene_gn=LOC111558006;refGene_tx=XM_023245125.1
E2 60200016 60201012
refGene_gn=LOC101093230;refGene_tx=XM_023245320.1;refGene_gn=LOC101093230;refGene_tx=XM_023245321.1;refGene_gn=LOC101093230;refGene_tx=XM_02

3245322.1;refGene_gn=LOC101093230;refGene_tx=XM_023245323.1;refGene_gn=LOC101093230;refGene_tx=XM_023245324.1;refGene_gn=LOC101093230;refGene_tx=XM_023245325.1
E2 60342399 60342580
refGene_gn=LOC101093230;refGene_tx=XM_023245320.1;refGene_gn=LOC101093230;refGene_tx=XM_023245321.1;refGene_gn=LOC101093230;refGene_tx=XM_023245322.1;refGene_gn=LOC101093230;refGene_tx=XM_023245323.1;refGene_gn=LOC101093230;refGene_tx=XM_023245324.1;refGene_gn=LOC101093230;refGene_tx=XM_023245325.1
E2 60557394 60557614
refGene_gn=LOC101093230;refGene_tx=XM_023245320.1;refGene_gn=LOC101093230;refGene_tx=XM_023245321.1;refGene_gn=LOC101093230;refGene_tx=XM_023245322.1;refGene_gn=LOC101093230;refGene_tx=XM_023245323.1;refGene_gn=LOC101093230;refGene_tx=XM_023245325.1
E2 60568748 60568948
refGene_gn=LOC101093230;refGene_tx=XM_023245320.1;refGene_gn=LOC101093230;refGene_tx=XM_023245321.1;refGene_gn=LOC101093230;refGene_tx=XM_023245322.1;refGene_gn=LOC101093230;refGene_tx=XM_023245323.1;refGene_gn=LOC101093230;refGene_tx=XM_023245324.1;refGene_gn=LOC101093230;refGene_tx=XM_023245325.1
E2 60573200 60573376
refGene_gn=LOC101093230;refGene_tx=XM_023245320.1;refGene_gn=LOC101093230;refGene_tx=XM_023245321.1;refGene_gn=LOC101093230;refGene_tx=XM_023245322.1;refGene_gn=LOC101093230;refGene_tx=XM_023245323.1;refGene_gn=LOC101093230;refGene_tx=XM_023245324.1;refGene_gn=LOC101093230;refGene_tx=XM_023245325.1
E2 60573609 60573810
refGene_gn=LOC101093230;refGene_tx=XM_023245320.1;refGene_gn=LOC101093230;refGene_tx=XM_023245321.1;refGene_gn=LOC101093230;refGene_tx=XM_023245322.1;refGene_gn=LOC101093230;refGene_tx=XM_023245323.1;refGene_gn=LOC101093230;refGene_tx=XM_023245324.1;refGene_gn=LOC101093230;refGene_tx=XM_023245325.1
E2 60574348 60574540
refGene_gn=LOC101093230;refGene_tx=XM_023245320.1;refGene_gn=LOC101093230;refGene_tx=XM_023245321.1;refGene_gn=LOC101093230;refGene_tx=XM_023245322.1;refGene_gn=LOC101093230;refGene_tx=XM_023245323.1;refGene_gn=LOC101093230;refGene_tx=XM_023245324.1;refGene_gn=LOC101093230;refGene_tx=XM_023245325.1
E2 60574886 60575185
refGene_gn=LOC101093230;refGene_tx=XM_023245320.1;refGene_gn=LOC101093230;refGene_tx=XM_023245321.1;refGene_gn=LOC101093230;refGene_tx=XM_023245322.1;refGene_gn=LOC101093230;refGene_tx=XM_023245323.1;refGene_gn=LOC101093230;refGene_tx=XM_023245324.1;refGene_gn=LOC101093230;refGene_tx=XM_023245325.1
E2 60575643 60575968
refGene_gn=LOC101093230;refGene_tx=XM_023245320.1;refGene_gn=LOC101093230;refGene_tx=XM_023245321.1;refGene_gn=LOC101093230;refGene_tx=XM_023245322.1;refGene_gn=LOC101093230;refGene_tx=XM_023245323.1;refGene_gn=LOC101093230;refGene_tx=XM_023245324.1;refGene_gn=LOC101093230;refGene_tx=XM_023245325.1

E2 60578061 60578675
refGene_gn=LOC101093230;refGene_tx=XM_023245320.1;refGene_gn=LOC101093230;refGene_tx=XM_023245321.1;refGene_gn=LOC101093230;refGene_tx=XM_023245322.1;refGene_gn=LOC101093230;refGene_tx=XM_023245323.1;refGene_gn=LOC101093230;refGene_tx=XM_023245324.1;refGene_gn=LOC101093230;refGene_tx=XM_023245325.1
E2 60579759 60579872
refGene_gn=LOC101093230;refGene_tx=XM_023245320.1;refGene_gn=LOC101093230;refGene_tx=XM_023245321.1;refGene_gn=LOC101093230;refGene_tx=XM_023245322.1;refGene_gn=LOC101093230;refGene_tx=XM_023245323.1;refGene_gn=LOC101093230;refGene_tx=XM_023245324.1;refGene_gn=LOC101093230;refGene_tx=XM_023245325.1
E2 60580150 60580421
refGene_gn=LOC101093230;refGene_tx=XM_023245320.1;refGene_gn=LOC101093230;refGene_tx=XM_023245321.1;refGene_gn=LOC101093230;refGene_tx=XM_023245322.1;refGene_gn=LOC101093230;refGene_tx=XM_023245323.1;refGene_gn=LOC101093230;refGene_tx=XM_023245324.1;refGene_gn=LOC101093230;refGene_tx=XM_023245325.1
E2 60581626 60581718
refGene_gn=LOC101093230;refGene_tx=XM_023245325.1
E2 60581922 60582036
refGene_gn=LOC101093230;refGene_tx=XM_023245320.1;refGene_gn=LOC101093230;refGene_tx=XM_023245321.1;refGene_gn=LOC101093230;refGene_tx=XM_023245322.1;refGene_gn=LOC101093230;refGene_tx=XM_023245324.1
E2 60582902 60583256
refGene_gn=LOC101093230;refGene_tx=XM_023245320.1;refGene_gn=LOC101093230;refGene_tx=XM_023245321.1;refGene_gn=LOC101093230;refGene_tx=XM_023245322.1;refGene_gn=LOC101093230;refGene_tx=XM_023245323.1;refGene_gn=LOC101093230;refGene_tx=XM_023245324.1
E2 60584704 60584980
refGene_gn=LOC101093230;refGene_tx=XM_023245320.1;refGene_gn=LOC101093230;refGene_tx=XM_023245321.1;refGene_gn=LOC101093230;refGene_tx=XM_023245322.1;refGene_gn=LOC101093230;refGene_tx=XM_023245323.1;refGene_gn=LOC101093230;refGene_tx=XM_023245324.1
E2 60586885 60586989
refGene_gn=LOC101093230;refGene_tx=XM_023245320.1;refGene_gn=LOC101093230;refGene_tx=XM_023245321.1;refGene_gn=LOC101093230;refGene_tx=XM_023245322.1;refGene_gn=LOC101093230;refGene_tx=XM_023245323.1;refGene_gn=LOC101093230;refGene_tx=XM_023245324.1
E2 60587734 60587869
refGene_gn=LOC101093230;refGene_tx=XM_023245320.1;refGene_gn=LOC101093230;refGene_tx=XM_023245321.1;refGene_gn=LOC101093230;refGene_tx=XM_023245322.1;refGene_gn=LOC101093230;refGene_tx=XM_023245323.1;refGene_gn=LOC101093230;refGene_tx=XM_023245324.1
E2 60593460 60593586 refGene_gn=GINS2;refGene_tx=XM_003998320.5
E2 60593700 60593827 refGene_gn=GINS2;refGene_tx=XM_003998320.5
E2 60597016 60597116 refGene_gn=GINS2;refGene_tx=XM_003998320.5
E2 60601156 60601271 refGene_gn=GINS2;refGene_tx=XM_003998320.5
E2 60602493 60602583 refGene_gn=GINS2;refGene_tx=XM_003998320.5
E2 60670179 60670339

refGene_gn=EMC8;refGene_tx=XM_023245608.1;refGene_gn=EMC8;refGene_tx=XM_023245609.1;refGene_gn=EMC8;refGene_tx=XM_023245610.1
E2 60670813 60670908
refGene_gn=EMC8;refGene_tx=XM_023245608.1;refGene_gn=EMC8;refGene_tx=XM_023245609.1;refGene_gn=EMC8;refGene_tx=XM_023245610.1
E2 60671645 60671715
refGene_gn=EMC8;refGene_tx=XM_023245608.1;refGene_gn=EMC8;refGene_tx=XM_023245609.1
E2 60678796 60678873
refGene_gn=EMC8;refGene_tx=XM_023245608.1;refGene_gn=EMC8;refGene_tx=XM_023245609.1;refGene_gn=EMC8;refGene_tx=XM_023245610.1
E2 60686596 60686827 refGene_gn=EMC8;refGene_tx=XM_023245608.1
E2 60687582 60687930
refGene_gn=EMC8;refGene_tx=XM_023245609.1;refGene_gn=EMC8;refGene_tx=XM_023245610.1
E2 60688449 60688522
refGene_gn=LOC101089914;refGene_tx=XM_023245612.1
E2 60692718 60692886
refGene_gn=LOC101089914;refGene_tx=XM_023245612.1
E2 60693434 60693566
refGene_gn=LOC101089914;refGene_tx=XM_023245612.1
E2 60694416 60694553
refGene_gn=LOC101089914;refGene_tx=XM_023245612.1
E2 60766194 60766368
refGene_gn=IRF8;refGene_tx=XM_023245141.1;refGene_gn=IRF8;refGene_tx=XM_023245142.1
E2 60771237 60771421
refGene_gn=IRF8;refGene_tx=XM_023245141.1;refGene_gn=IRF8;refGene_tx=XM_023245142.1
E2 60773785 60773874
refGene_gn=IRF8;refGene_tx=XM_023245141.1;refGene_gn=IRF8;refGene_tx=XM_023245142.1
E2 60775302 60775408
refGene_gn=IRF8;refGene_tx=XM_023245141.1;refGene_gn=IRF8;refGene_tx=XM_023245142.1
E2 60776594 60776642
refGene_gn=IRF8;refGene_tx=XM_023245141.1;refGene_gn=IRF8;refGene_tx=XM_023245142.1
E2 60780875 60781262
refGene_gn=IRF8;refGene_tx=XM_023245141.1;refGene_gn=IRF8;refGene_tx=XM_023245142.1
E2 60782253 60782369
refGene_gn=IRF8;refGene_tx=XM_023245141.1;refGene_gn=IRF8;refGene_tx=XM_023245142.1
E2 60782974 60783151
refGene_gn=IRF8;refGene_tx=XM_023245141.1;refGene_gn=IRF8;refGene_tx=XM_023245142.1
E2 60842513 60842638
refGene_gn=LOC111558007;refGene_tx=XM_023245126.1
E2 60843801 60844155

refGene_gn=LOC111558007;refGene_tx=XM_023245126.1
E2 60857490 60857553
refGene_gn=LOC111558007;refGene_tx=XM_023245126.1
E2 60867260 60867414
refGene_gn=LOC111558007;refGene_tx=XM_023245126.1
E2 60873109 60873201
refGene_gn=LOC111558007;refGene_tx=XM_023245126.1
E2 60874095 60874270
refGene_gn=LOC111558007;refGene_tx=XM_023245126.1
E2 61211054 61211243
refGene_gn=LOC111556293;refGene_tx=XM_023245079.1;refGene_gn=LOC111556293;refGene_tx=XM_023245080.1
E2 61211384 61212365
refGene_gn=LOC111556293;refGene_tx=XM_023245079.1;refGene_gn=LOC111556293;refGene_tx=XM_023245080.1
E2 61213588 61214732 refGene_gn=FOXF1;refGene_tx=XM_023245078.1
E2 61216170 61216331 refGene_gn=FOXF1;refGene_tx=XM_023245078.1
E2 61229542 61230052
refGene_gn=MTHFSD;refGene_tx=XM_003998340.4;refGene_gn=MTHFSD;refGene_tx=XM_006941792.3;refGene_gn=MTHFSD;refGene_tx=XM_019820160.2;refGene_gn=MTHFSD;refGene_tx=XM_019820161.2
E2 61239537 61239663
refGene_gn=MTHFSD;refGene_tx=XM_003998340.4;refGene_gn=MTHFSD;refGene_tx=XM_006941792.3;refGene_gn=MTHFSD;refGene_tx=XM_019820160.2;refGene_gn=MTHFSD;refGene_tx=XM_019820161.2
E2 61239927 61240040
refGene_gn=MTHFSD;refGene_tx=XM_003998340.4;refGene_gn=MTHFSD;refGene_tx=XM_006941792.3;refGene_gn=MTHFSD;refGene_tx=XM_019820160.2;refGene_gn=MTHFSD;refGene_tx=XM_019820161.2
E2 61242377 61242468
refGene_gn=MTHFSD;refGene_tx=XM_003998340.4;refGene_gn=MTHFSD;refGene_tx=XM_006941792.3;refGene_gn=MTHFSD;refGene_tx=XM_019820160.2;refGene_gn=MTHFSD;refGene_tx=XM_019820161.2
E2 61244647 61244761
refGene_gn=MTHFSD;refGene_tx=XM_003998340.4;refGene_gn=MTHFSD;refGene_tx=XM_006941792.3;refGene_gn=MTHFSD;refGene_tx=XM_019820160.2;refGene_gn=MTHFSD;refGene_tx=XM_019820161.2
E2 61247463 61247577
refGene_gn=MTHFSD;refGene_tx=XM_003998340.4;refGene_gn=MTHFSD;refGene_tx=XM_019820160.2
E2 61247609 61247723 refGene_gn=MTHFSD;refGene_tx=XM_006941792.3
E2 61249152 61249256
refGene_gn=MTHFSD;refGene_tx=XM_003998340.4;refGene_gn=MTHFSD;refGene_tx=XM_006941792.3;refGene_gn=MTHFSD;refGene_tx=XM_019820160.2;refGene_gn=MTHFSD;refGene_tx=XM_019820161.2
E2 61253138 61253154
refGene_gn=MTHFSD;refGene_tx=XM_003998340.4;refGene_gn=MTHFSD;refGene_tx=XM_006941792.3;refGene_gn=MTHFSD;refGene_tx=XM_019820161.2
E2 61265437 61266958 refGene_gn=FOXC2;refGene_tx=XM_023245554.1
E2 61276426 61277521 refGene_gn=FOXL1;refGene_tx=XM_023245552.1

E2 61797871 61797890
refGene_gn=CE2H16orf95;refGene_tx=XM_023245385.1
E2 61800039 61800250
refGene_gn=CE2H16orf95;refGene_tx=XM_023245385.1
E2 61803560 61803597
refGene_gn=CE2H16orf95;refGene_tx=XM_023245385.1
E2 61804367 61804514
refGene_gn=CE2H16orf95;refGene_tx=XM_023245385.1
E2 61810299 61810437
refGene_gn=CE2H16orf95;refGene_tx=XM_023245385.1
E2 61824050 61824273
refGene_gn=FBXO31;refGene_tx=XM_011290281.2;refGene_gn=FBXO31;refGene_tx=XM_011290282.2
E2 61826957 61827358
refGene_gn=FBXO31;refGene_tx=XM_011290281.2;refGene_gn=FBXO31;refGene_tx=XM_011290282.2
E2 61828648 61828802
refGene_gn=FBXO31;refGene_tx=XM_011290281.2;refGene_gn=FBXO31;refGene_tx=XM_011290282.2
E2 61829858 61829968
refGene_gn=FBXO31;refGene_tx=XM_011290281.2;refGene_gn=FBXO31;refGene_tx=XM_011290282.2
E2 61836685 61836760
refGene_gn=FBXO31;refGene_tx=XM_011290281.2;refGene_gn=FBXO31;refGene_tx=XM_011290282.2
E2 61837421 61837589
refGene_gn=FBXO31;refGene_tx=XM_011290281.2;refGene_gn=FBXO31;refGene_tx=XM_011290282.2
E2 61842711 61842788
refGene_gn=FBXO31;refGene_tx=XM_011290281.2;refGene_gn=FBXO31;refGene_tx=XM_011290282.2
E2 61849587 61849674 refGene_gn=FBXO31;refGene_tx=XM_011290281.2
E2 61852763 61852835
refGene_gn=FBXO31;refGene_tx=XM_011290281.2;refGene_gn=FBXO31;refGene_tx=XM_011290282.2
E2 61864914 61865263
refGene_gn=FBXO31;refGene_tx=XM_011290281.2;refGene_gn=FBXO31;refGene_tx=XM_011290282.2
E2 61871219 61871259 refGene_gn=MAP1LC3B;refGene_tx=XM_023245115.1
E2 61877782 61877838 refGene_gn=MAP1LC3B;refGene_tx=XM_023245115.1
E2 61881684 61881791 refGene_gn=MAP1LC3B;refGene_tx=XM_023245115.1
E2 61882432 61882607 refGene_gn=MAP1LC3B;refGene_tx=XM_023245115.1
E2 61889887 61889943
refGene_gn=ZCCHC14;refGene_tx=XM_023245794.1;refGene_gn=ZCCHC14;refGene_tx=XM_023245795.1;refGene_gn=ZCCHC14;refGene_tx=XM_023245796.1
E2 61890556 61892035
refGene_gn=ZCCHC14;refGene_tx=XM_023245794.1;refGene_gn=ZCCHC14;refGene_tx=XM_023245795.1;refGene_gn=ZCCHC14;refGene_tx=XM_023245796.1
E2 61892119 61892260
refGene_gn=ZCCHC14;refGene_tx=XM_023245794.1;refGene_gn=ZCCHC14;refGen

e_tx=XM_023245795.1
E2 61893089 61893217
refGene_gn=ZCCHC14;refGene_tx=XM_023245794.1;refGene_gn=ZCCHC14;refGene_tx=XM_023245795.1;refGene_gn=ZCCHC14;refGene_tx=XM_023245796.1
E2 61893919 61894011
refGene_gn=ZCCHC14;refGene_tx=XM_023245794.1;refGene_gn=ZCCHC14;refGene_tx=XM_023245795.1;refGene_gn=ZCCHC14;refGene_tx=XM_023245796.1
E2 61895692 61895972
refGene_gn=ZCCHC14;refGene_tx=XM_023245794.1;refGene_gn=ZCCHC14;refGene_tx=XM_023245795.1;refGene_gn=ZCCHC14;refGene_tx=XM_023245796.1
E2 61896985 61897040
refGene_gn=ZCCHC14;refGene_tx=XM_023245794.1;refGene_gn=ZCCHC14;refGene_tx=XM_023245795.1;refGene_gn=ZCCHC14;refGene_tx=XM_023245796.1
E2 61897119 61897214
refGene_gn=ZCCHC14;refGene_tx=XM_023245794.1;refGene_gn=ZCCHC14;refGene_tx=XM_023245795.1;refGene_gn=ZCCHC14;refGene_tx=XM_023245796.1
E2 61897862 61897972
refGene_gn=ZCCHC14;refGene_tx=XM_023245794.1;refGene_gn=ZCCHC14;refGene_tx=XM_023245795.1;refGene_gn=ZCCHC14;refGene_tx=XM_023245796.1
E2 61900645 61900717
refGene_gn=ZCCHC14;refGene_tx=XM_023245794.1;refGene_gn=ZCCHC14;refGene_tx=XM_023245796.1
E2 61909314 61909388
refGene_gn=ZCCHC14;refGene_tx=XM_023245794.1;refGene_gn=ZCCHC14;refGene_tx=XM_023245795.1;refGene_gn=ZCCHC14;refGene_tx=XM_023245796.1
E2 61920632 61920756
refGene_gn=ZCCHC14;refGene_tx=XM_023245794.1;refGene_gn=ZCCHC14;refGene_tx=XM_023245795.1;refGene_gn=ZCCHC14;refGene_tx=XM_023245796.1
E2 61943358 61943937
refGene_gn=ZCCHC14;refGene_tx=XM_023245794.1;refGene_gn=ZCCHC14;refGene_tx=XM_023245795.1;refGene_gn=ZCCHC14;refGene_tx=XM_023245796.1
E2 61949546 61949700
refGene_gn=JPH3;refGene_tx=XM_023245797.1;refGene_gn=JPH3;refGene_tx=XM_023245798.1;refGene_gn=JPH3;refGene_tx=XM_023245799.1;refGene_gn=JPH3;refGene_tx=XM_023245801.1;refGene_gn=JPH3;refGene_tx=XM_023245802.1
E2 61955534 61955597
refGene_gn=JPH3;refGene_tx=XM_023245797.1;refGene_gn=JPH3;refGene_tx=XM_023245798.1;refGene_gn=JPH3;refGene_tx=XM_023245799.1;refGene_gn=JPH3;refGene_tx=XM_023245801.1;refGene_gn=JPH3;refGene_tx=XM_023245802.1
E2 62033209 62033591
refGene_gn=JPH3;refGene_tx=XM_003998322.5;refGene_gn=JPH3;refGene_tx=XM_019820170.2
E2 62065840 62066618
refGene_gn=JPH3;refGene_tx=XM_003998322.5;refGene_gn=JPH3;refGene_tx=XM_019820170.2;refGene_gn=JPH3;refGene_tx=XM_023245797.1;refGene_gn=JPH3;refGene_tx=XM_023245798.1;refGene_gn=JPH3;refGene_tx=XM_023245799.1;refGene_gn=JPH3;refGene_tx=XM_023245801.1;refGene_gn=JPH3;refGene_tx=XM_023245802.1
E2 62105684 62105809
refGene_gn=JPH3;refGene_tx=XM_003998322.5;refGene_gn=JPH3;refGene_tx=X

M_019820170.2;refGene_gn=JPH3;refGene_tx=XM_023245797.1;refGene_gn=JPH3;refGene_tx=XM_023245798.1;refGene_gn=JPH3;refGene_tx=XM_023245799.1;refGene_gn=JPH3;refGene_tx=XM_023245801.1;refGene_gn=JPH3;refGene_tx=XM_023245802.1
E2 62107279 62108178
refGene_gn=JPH3;refGene_tx=XM_003998322.5;refGene_gn=JPH3;refGene_tx=XM_019820170.2;refGene_gn=JPH3;refGene_tx=XM_023245797.1;refGene_gn=JPH3;refGene_tx=XM_023245798.1;refGene_gn=JPH3;refGene_tx=XM_023245799.1;refGene_gn=JPH3;refGene_tx=XM_023245801.1;refGene_gn=JPH3;refGene_tx=XM_023245802.1
E2 62109784 62109865
refGene_gn=JPH3;refGene_tx=XM_003998322.5;refGene_gn=JPH3;refGene_tx=XM_023245797.1;refGene_gn=JPH3;refGene_tx=XM_023245798.1;refGene_gn=JPH3;refGene_tx=XM_023245799.1;refGene_gn=JPH3;refGene_tx=XM_023245801.1;refGene_gn=JPH3;refGene_tx=XM_023245802.1
E2 62119219 62119413 refGene_gn=KLHDC4;refGene_tx=XM_006941796.3
E2 62121347 62121750 refGene_gn=KLHDC4;refGene_tx=XM_006941796.3
E2 62122257 62122466 refGene_gn=KLHDC4;refGene_tx=XM_006941796.3
E2 62123992 62124068 refGene_gn=KLHDC4;refGene_tx=XM_006941796.3
E2 62127101 62127261 refGene_gn=KLHDC4;refGene_tx=XM_006941796.3
E2 62129992 62130085 refGene_gn=KLHDC4;refGene_tx=XM_006941796.3
E2 62148799 62148936 refGene_gn=KLHDC4;refGene_tx=XM_006941796.3
E2 62160437 62160536 refGene_gn=KLHDC4;refGene_tx=XM_006941796.3
E2 62161272 62161351 refGene_gn=KLHDC4;refGene_tx=XM_006941796.3
E2 62165052 62165144 refGene_gn=KLHDC4;refGene_tx=XM_006941796.3
E2 62169424 62169523 refGene_gn=KLHDC4;refGene_tx=XM_006941796.3
E2 62211433 62211489 refGene_gn=SLC7A5;refGene_tx=XM_011290283.3
E2 62212576 62212754 refGene_gn=SLC7A5;refGene_tx=XM_011290283.3
E2 62214699 62214849 refGene_gn=SLC7A5;refGene_tx=XM_011290283.3
E2 62216146 62216243 refGene_gn=SLC7A5;refGene_tx=XM_011290283.3
E2 62216666 62216770 refGene_gn=SLC7A5;refGene_tx=XM_011290283.3
E2 62217690 62217814 refGene_gn=SLC7A5;refGene_tx=XM_011290283.3
E2 62218274 62218319 refGene_gn=SLC7A5;refGene_tx=XM_011290283.3
E2 62219276 62219382 refGene_gn=SLC7A5;refGene_tx=XM_011290283.3
E2 62228386 62228512 refGene_gn=SLC7A5;refGene_tx=XM_011290283.3
E2 62241333 62241877 refGene_gn=SLC7A5;refGene_tx=XM_011290283.3
E2 62252655 62253032
refGene_gn=CA5A;refGene_tx=XM_023245265.1;refGene_gn=CA5A;refGene_tx=XM_023245267.1
E2 62259032 62259176
refGene_gn=CA5A;refGene_tx=XM_019820177.2;refGene_gn=CA5A;refGene_tx=XM_019820178.2
E2 62261418 62261466 refGene_gn=CA5A;refGene_tx=XM_023245266.1
E2 62261999 62262155
refGene_gn=CA5A;refGene_tx=XM_019820177.2;refGene_gn=CA5A;refGene_tx=XM_019820178.2;refGene_gn=CA5A;refGene_tx=XM_023245265.1;refGene_gn=CA5A;refGene_tx=XM_023245266.1;refGene_gn=CA5A;refGene_tx=XM_023245267.1
E2 62265295 62265394 refGene_gn=CA5A;refGene_tx=XM_023245268.1
E2 62266063 62266126
refGene_gn=CA5A;refGene_tx=XM_019820177.2;refGene_gn=CA5A;refGene_tx=X

M_019820178.2;refGene_gn=CA5A;refGene_tx=XM_023245265.1;refGene_gn=CA5A;refGene_tx=XM_023245266.1;refGene_gn=CA5A;refGene_tx=XM_023245268.1
E2 62266271 62266367
refGene_gn=CA5A;refGene_tx=XM_019820177.2;refGene_gn=CA5A;refGene_tx=XM_019820178.2;refGene_gn=CA5A;refGene_tx=XM_023245265.1;refGene_gn=CA5A;refGene_tx=XM_023245266.1;refGene_gn=CA5A;refGene_tx=XM_023245268.1
E2 62268425 62268968
refGene_gn=CA5A;refGene_tx=XM_019820177.2;refGene_gn=CA5A;refGene_tx=XM_019820178.2;refGene_gn=CA5A;refGene_tx=XM_023245265.1;refGene_gn=CA5A;refGene_tx=XM_023245266.1;refGene_gn=CA5A;refGene_tx=XM_023245268.1
E2 62273424 62274408 refGene_gn=CA5A;refGene_tx=XM_023245264.1
E2 62281963 62282161
refGene_gn=CA5A;refGene_tx=XM_019820178.2;refGene_gn=CA5A;refGene_tx=XM_023245267.1
E2 62285450 62285607
refGene_gn=CA5A;refGene_tx=XM_019820178.2;refGene_gn=CA5A;refGene_tx=XM_023245267.1
E2 62324613 62324683
refGene_gn=BANP;refGene_tx=XM_011290284.3;refGene_gn=BANP;refGene_tx=XM_019820172.2;refGene_gn=BANP;refGene_tx=XM_023245259.1;refGene_gn=BANP;refGene_tx=XM_023245260.1;refGene_gn=BANP;refGene_tx=XM_023245262.1;refGene_gn=BANP;refGene_tx=XM_023245263.1
E2 62329330 62329422
refGene_gn=BANP;refGene_tx=XM_011290284.3;refGene_gn=BANP;refGene_tx=XM_019820172.2;refGene_gn=BANP;refGene_tx=XM_023245259.1;refGene_gn=BANP;refGene_tx=XM_023245260.1;refGene_gn=BANP;refGene_tx=XM_023245262.1;refGene_gn=BANP;refGene_tx=XM_023245263.1
E2 62330818 62331018
refGene_gn=BANP;refGene_tx=XM_011290284.3;refGene_gn=BANP;refGene_tx=XM_019820172.2;refGene_gn=BANP;refGene_tx=XM_023245259.1;refGene_gn=BANP;refGene_tx=XM_023245260.1;refGene_gn=BANP;refGene_tx=XM_023245262.1;refGene_gn=BANP;refGene_tx=XM_023245263.1
E2 62349344 62349461
refGene_gn=BANP;refGene_tx=XM_011290284.3;refGene_gn=BANP;refGene_tx=XM_019820172.2;refGene_gn=BANP;refGene_tx=XM_023245259.1;refGene_gn=BANP;refGene_tx=XM_023245260.1
E2 62351119 62351295
refGene_gn=BANP;refGene_tx=XM_011290284.3;refGene_gn=BANP;refGene_tx=XM_019820172.2;refGene_gn=BANP;refGene_tx=XM_023245259.1;refGene_gn=BANP;refGene_tx=XM_023245260.1;refGene_gn=BANP;refGene_tx=XM_023245262.1;refGene_gn=BANP;refGene_tx=XM_023245263.1
E2 62364333 62364573
refGene_gn=BANP;refGene_tx=XM_011290284.3;refGene_gn=BANP;refGene_tx=XM_019820172.2;refGene_gn=BANP;refGene_tx=XM_023245259.1;refGene_gn=BANP;refGene_tx=XM_023245260.1;refGene_gn=BANP;refGene_tx=XM_023245262.1;refGene_gn=BANP;refGene_tx=XM_023245263.1
E2 62373330 62373498
refGene_gn=BANP;refGene_tx=XM_011290284.3;refGene_gn=BANP;refGene_tx=XM_019820172.2;refGene_gn=BANP;refGene_tx=XM_023245259.1;refGene_gn=BANP;refGene_tx=XM_023245260.1;refGene_gn=BANP;refGene_tx=XM_023245262.1;

refGene_gn=BANP;refGene_tx=XM_023245263.1
E2 62377778 62377909
refGene_gn=BANP;refGene_tx=XM_011290284.3;refGene_gn=BANP;refGene_tx=XM_019820172.2;refGene_gn=BANP;refGene_tx=XM_023245259.1;refGene_gn=BANP;refGene_tx=XM_023245260.1;refGene_gn=BANP;refGene_tx=XM_023245262.1;refGene_gn=BANP;refGene_tx=XM_023245263.1
E2 62379486 62379558
refGene_gn=BANP;refGene_tx=XM_011290284.3;refGene_gn=BANP;refGene_tx=XM_019820172.2;refGene_gn=BANP;refGene_tx=XM_023245259.1;refGene_gn=BANP;refGene_tx=XM_023245260.1;refGene_gn=BANP;refGene_tx=XM_023245262.1;refGene_gn=BANP;refGene_tx=XM_023245263.1
E2 62380952 62380991
refGene_gn=BANP;refGene_tx=XM_011290284.3;refGene_gn=BANP;refGene_tx=XM_019820172.2;refGene_gn=BANP;refGene_tx=XM_023245259.1;refGene_gn=BANP;refGene_tx=XM_023245260.1;refGene_gn=BANP;refGene_tx=XM_023245262.1;refGene_gn=BANP;refGene_tx=XM_023245263.1
E2 62407428 62407572
refGene_gn=BANP;refGene_tx=XM_011290284.3;refGene_gn=BANP;refGene_tx=XM_019820172.2;refGene_gn=BANP;refGene_tx=XM_023245259.1;refGene_gn=BANP;refGene_tx=XM_023245260.1;refGene_gn=BANP;refGene_tx=XM_023245262.1;refGene_gn=BANP;refGene_tx=XM_023245263.1
E2 62411483 62411555
refGene_gn=BANP;refGene_tx=XM_011290284.3;refGene_gn=BANP;refGene_tx=XM_019820172.2;refGene_gn=BANP;refGene_tx=XM_023245259.1;refGene_gn=BANP;refGene_tx=XM_023245260.1;refGene_gn=BANP;refGene_tx=XM_023245262.1;refGene_gn=BANP;refGene_tx=XM_023245263.1
E2 62654122 62665474 refGene_gn=ZNF469;refGene_tx=XM_023245050.1
E2 62675104 62675144 refGene_gn=ZFPM1;refGene_tx=XM_023244948.1
E2 62703242 62703359 refGene_gn=ZFPM1;refGene_tx=XM_023244948.1
E2 62705667 62705790 refGene_gn=ZFPM1;refGene_tx=XM_023244948.1
E2 62727717 62727851 refGene_gn=ZFPM1;refGene_tx=XM_023244948.1
E2 62734824 62734927 refGene_gn=ZFPM1;refGene_tx=XM_023244948.1
E2 62736525 62736738 refGene_gn=ZFPM1;refGene_tx=XM_023244948.1
E2 62739565 62739796 refGene_gn=ZFPM1;refGene_tx=XM_023244948.1
E2 62740085 62740181 refGene_gn=ZFPM1;refGene_tx=XM_023244948.1
E2 62740251 62740398 refGene_gn=ZFPM1;refGene_tx=XM_023244948.1
E2 62740594 62742369 refGene_gn=ZFPM1;refGene_tx=XM_023244948.1
E2 62789608 62790154
refGene_gn=ZC3H18;refGene_tx=XM_023245970.1;refGene_gn=ZC3H18;refGene_tx=XM_023245971.1
E2 62800452 62800537
refGene_gn=ZC3H18;refGene_tx=XM_023245970.1;refGene_gn=ZC3H18;refGene_tx=XM_023245971.1
E2 62801218 62801290 refGene_gn=ZC3H18;refGene_tx=XM_023245970.1
E2 62811238 62811387
refGene_gn=ZC3H18;refGene_tx=XM_023245970.1;refGene_gn=ZC3H18;refGene_tx=XM_023245971.1
E2 62811710 62811803
refGene_gn=ZC3H18;refGene_tx=XM_023245970.1;refGene_gn=ZC3H18;refGene_tx=XM_023245971.1

E2 62812521 62812679
refGene_gn=ZC3H18;refGene_tx=XM_023245970.1;refGene_gn=ZC3H18;refGene_tx=XM_023245971.1
E2 62817373 62817491
refGene_gn=ZC3H18;refGene_tx=XM_023245970.1;refGene_gn=ZC3H18;refGene_tx=XM_023245971.1
E2 62818379 62818648
refGene_gn=ZC3H18;refGene_tx=XM_023245970.1;refGene_gn=ZC3H18;refGene_tx=XM_023245971.1
E2 62831097 62831289
refGene_gn=ZC3H18;refGene_tx=XM_023245970.1;refGene_gn=ZC3H18;refGene_tx=XM_023245971.1
E2 62831879 62832005
refGene_gn=ZC3H18;refGene_tx=XM_023245970.1;refGene_gn=ZC3H18;refGene_tx=XM_023245971.1
E2 62834435 62834540
refGene_gn=ZC3H18;refGene_tx=XM_023245970.1;refGene_gn=ZC3H18;refGene_tx=XM_023245971.1
E2 62835509 62835653
refGene_gn=ZC3H18;refGene_tx=XM_023245970.1;refGene_gn=ZC3H18;refGene_tx=XM_023245971.1
E2 62836018 62836078
refGene_gn=ZC3H18;refGene_tx=XM_023245970.1;refGene_gn=ZC3H18;refGene_tx=XM_023245971.1
E2 62837030 62837191
refGene_gn=ZC3H18;refGene_tx=XM_023245970.1;refGene_gn=ZC3H18;refGene_tx=XM_023245971.1
E2 62837421 62837618
refGene_gn=ZC3H18;refGene_tx=XM_023245970.1;refGene_gn=ZC3H18;refGene_tx=XM_023245971.1
E2 62838168 62838265
refGene_gn=ZC3H18;refGene_tx=XM_023245970.1;refGene_gn=ZC3H18;refGene_tx=XM_023245971.1
E2 62839208 62839305
refGene_gn=ZC3H18;refGene_tx=XM_023245970.1;refGene_gn=ZC3H18;refGene_tx=XM_023245971.1
E2 62839930 62840144
refGene_gn=ZC3H18;refGene_tx=XM_023245970.1;refGene_gn=ZC3H18;refGene_tx=XM_023245971.1
E2 62845306 62845309 refGene_gn=IL17C;refGene_tx=XM_023244897.1
E2 62845627 62845956 refGene_gn=IL17C;refGene_tx=XM_023244897.1
E2 62846285 62846547 refGene_gn=IL17C;refGene_tx=XM_023244897.1
E2 62847581 62847904
refGene_gn=LOC111558008;refGene_tx=XM_023245127.1
E2 62848104 62848366
refGene_gn=LOC111558008;refGene_tx=XM_023245127.1
E2 62850043 62850259
refGene_gn=LOC101091745;refGene_tx=XM_006941797.2
E2 62852038 62852120
refGene_gn=LOC101091745;refGene_tx=XM_006941797.2

E2 62852605 62852689
refGene_gn=LOC101091745;refGene_tx=XM_006941797.2
E2 62852886 62852961
refGene_gn=LOC101091745;refGene_tx=XM_006941797.2
E2 62853521 62853591
refGene_gn=LOC101091745;refGene_tx=XM_006941797.2
E2 62858588 62858646
refGene_gn=LOC101091745;refGene_tx=XM_006941797.2
E2 62860672 62860753
refGene_gn=MVD;refGene_tx=XM_023244901.1;refGene_gn=MVD;refGene_tx=XM_023244902.1;refGene_gn=MVD;refGene_tx=XM_023244903.1
E2 62861315 62861424
refGene_gn=MVD;refGene_tx=XM_023244901.1;refGene_gn=MVD;refGene_tx=XM_023244902.1;refGene_gn=MVD;refGene_tx=XM_023244903.1
E2 62862313 62862429
refGene_gn=MVD;refGene_tx=XM_023244901.1;refGene_gn=MVD;refGene_tx=XM_023244902.1;refGene_gn=MVD;refGene_tx=XM_023244903.1
E2 62862726 62862945
refGene_gn=MVD;refGene_tx=XM_023244901.1;refGene_gn=MVD;refGene_tx=XM_023244902.1;refGene_gn=MVD;refGene_tx=XM_023244903.1
E2 62863050 62863125
refGene_gn=MVD;refGene_tx=XM_023244901.1;refGene_gn=MVD;refGene_tx=XM_023244902.1;refGene_gn=MVD;refGene_tx=XM_023244903.1
E2 62863395 62863595
refGene_gn=MVD;refGene_tx=XM_023244901.1;refGene_gn=MVD;refGene_tx=XM_023244902.1;refGene_gn=MVD;refGene_tx=XM_023244903.1
E2 62864310 62864457
refGene_gn=MVD;refGene_tx=XM_023244901.1;refGene_gn=MVD;refGene_tx=XM_023244902.1;refGene_gn=MVD;refGene_tx=XM_023244903.1
E2 62864794 62864909
refGene_gn=MVD;refGene_tx=XM_023244901.1;refGene_gn=MVD;refGene_tx=XM_023244902.1;refGene_gn=MVD;refGene_tx=XM_023244903.1
E2 62865271 62865342
refGene_gn=MVD;refGene_tx=XM_023244901.1;refGene_gn=MVD;refGene_tx=XM_023244902.1;refGene_gn=MVD;refGene_tx=XM_023244903.1
E2 62868017 62868090 refGene_gn=MVD;refGene_tx=XM_023244901.1
E2 62868963 62869009
refGene_gn=MVD;refGene_tx=XM_023244902.1;refGene_gn=MVD;refGene_tx=XM_023244903.1
E2 62889832 62890014 refGene_gn=SNAI3;refGene_tx=XM_023244904.1
E2 62892022 62892580 refGene_gn=SNAI3;refGene_tx=XM_023244904.1
E2 62895773 62895849 refGene_gn=SNAI3;refGene_tx=XM_023244904.1
E2 62902252 62902318 refGene_gn=RNF166;refGene_tx=XM_023244905.1
E2 62903309 62903417 refGene_gn=RNF166;refGene_tx=XM_023244905.1
E2 62903503 62903618 refGene_gn=RNF166;refGene_tx=XM_023244905.1
E2 62903947 62904060 refGene_gn=RNF166;refGene_tx=XM_023244905.1
E2 62905440 62905534 refGene_gn=RNF166;refGene_tx=XM_023244905.1
E2 62905535 62905589 refGene_gn=RNF166;refGene_tx=XM_023244905.1
E2 62910516 62910671 refGene_gn=RNF166;refGene_tx=XM_023244905.1
E2 62910942 62911010 refGene_gn=CTU2;refGene_tx=XM_023244899.1

E2	62911615 62911699 refGene_gn=CTU2;refGene_tx=XM_023244899.1
E2	62913985 62914064 refGene_gn=CTU2;refGene_tx=XM_023244899.1
E2	62914239 62914299 refGene_gn=CTU2;refGene_tx=XM_023244899.1
E2	62915362 62915423 refGene_gn=CTU2;refGene_tx=XM_023244899.1
E2	62915462 62915553 refGene_gn=CTU2;refGene_tx=XM_023244900.1
E2	62915928 62916038
refGene_gn=CTU2;refGene_tx=XM_023244899.1;refGene_gn=CTU2;refGene_tx=XM_023244900.1
E2	62916161 62916457
refGene_gn=CTU2;refGene_tx=XM_023244899.1;refGene_gn=CTU2;refGene_tx=XM_023244900.1
E2	62916525 62916661
refGene_gn=CTU2;refGene_tx=XM_023244899.1;refGene_gn=CTU2;refGene_tx=XM_023244900.1
E2	62916817 62916949
refGene_gn=CTU2;refGene_tx=XM_023244899.1;refGene_gn=CTU2;refGene_tx=XM_023244900.1
E2	62917417 62917509
refGene_gn=CTU2;refGene_tx=XM_023244899.1;refGene_gn=CTU2;refGene_tx=XM_023244900.1
E2	62917610 62917705
refGene_gn=CTU2;refGene_tx=XM_023244899.1;refGene_gn=CTU2;refGene_tx=XM_023244900.1
E2	62917788 62917924
refGene_gn=CTU2;refGene_tx=XM_023244899.1;refGene_gn=CTU2;refGene_tx=XM_023244900.1
E2	62918036 62918103
refGene_gn=CTU2;refGene_tx=XM_023244899.1;refGene_gn=CTU2;refGene_tx=XM_023244900.1
E2	62918200 62918259
refGene_gn=CTU2;refGene_tx=XM_023244899.1;refGene_gn=CTU2;refGene_tx=XM_023244900.1
E2	62918384 62918454
refGene_gn=CTU2;refGene_tx=XM_023244899.1;refGene_gn=CTU2;refGene_tx=XM_023244900.1
E2	62918719 62918969 refGene_gn=PIEZO1;refGene_tx=XM_023244898.1
E2	62919050 62919240 refGene_gn=PIEZO1;refGene_tx=XM_023244898.1
E2	62919331 62919411 refGene_gn=PIEZO1;refGene_tx=XM_023244898.1
E2	62919596 62919719 refGene_gn=PIEZO1;refGene_tx=XM_023244898.1
E2	62919806 62919979 refGene_gn=PIEZO1;refGene_tx=XM_023244898.1
E2	62920065 62920158 refGene_gn=PIEZO1;refGene_tx=XM_023244898.1
E2	62920295 62920484 refGene_gn=PIEZO1;refGene_tx=XM_023244898.1
E2	62921855 62922003 refGene_gn=PIEZO1;refGene_tx=XM_023244898.1
E2	62922081 62922240 refGene_gn=PIEZO1;refGene_tx=XM_023244898.1
E2	62922332 62922547 refGene_gn=PIEZO1;refGene_tx=XM_023244898.1
E2	62922651 62922799 refGene_gn=PIEZO1;refGene_tx=XM_023244898.1
E2	62922886 62922983 refGene_gn=PIEZO1;refGene_tx=XM_023244898.1
E2	62923259 62923554 refGene_gn=PIEZO1;refGene_tx=XM_023244898.1
E2	62923689 62923878 refGene_gn=PIEZO1;refGene_tx=XM_023244898.1
E2	62923949 62924208 refGene_gn=PIEZO1;refGene_tx=XM_023244898.1

| | | | |
|---|---|---|---|
| E2 | 62924291 | 62924483 | refGene_gn=PIEZO1;refGene_tx=XM_023244898.1 |
| E2 | 62924650 | 62924757 | refGene_gn=PIEZO1;refGene_tx=XM_023244898.1 |
| E2 | 62924841 | 62925014 | refGene_gn=PIEZO1;refGene_tx=XM_023244898.1 |
| E2 | 62925107 | 62925164 | refGene_gn=PIEZO1;refGene_tx=XM_023244898.1 |
| E2 | 62925243 | 62925340 | refGene_gn=PIEZO1;refGene_tx=XM_023244898.1 |
| E2 | 62925849 | 62925950 | refGene_gn=PIEZO1;refGene_tx=XM_023244898.1 |
| E2 | 62926421 | 62926493 | refGene_gn=PIEZO1;refGene_tx=XM_023244898.1 |
| E2 | 62926766 | 62926861 | refGene_gn=PIEZO1;refGene_tx=XM_023244898.1 |
| E2 | 62926935 | 62927025 | refGene_gn=PIEZO1;refGene_tx=XM_023244898.1 |
| E2 | 62927451 | 62927623 | refGene_gn=PIEZO1;refGene_tx=XM_023244898.1 |
| E2 | 62927718 | 62927815 | refGene_gn=PIEZO1;refGene_tx=XM_023244898.1 |
| E2 | 62927903 | 62928147 | refGene_gn=PIEZO1;refGene_tx=XM_023244898.1 |
| E2 | 62928369 | 62928523 | refGene_gn=PIEZO1;refGene_tx=XM_023244898.1 |
| E2 | 62929090 | 62929195 | refGene_gn=PIEZO1;refGene_tx=XM_023244898.1 |
| E2 | 62931483 | 62931688 | refGene_gn=PIEZO1;refGene_tx=XM_023244898.1 |
| E2 | 62932128 | 62932329 | refGene_gn=PIEZO1;refGene_tx=XM_023244898.1 |
| E2 | 62932428 | 62932554 | refGene_gn=PIEZO1;refGene_tx=XM_023244898.1 |
| E2 | 62932821 | 62932928 | refGene_gn=PIEZO1;refGene_tx=XM_023244898.1 |
| E2 | 62932930 | 62932997 | refGene_gn=PIEZO1;refGene_tx=XM_023244898.1 |
| E2 | 62933148 | 62933309 | refGene_gn=PIEZO1;refGene_tx=XM_023244898.1 |
| E2 | 62933410 | 62933517 | refGene_gn=PIEZO1;refGene_tx=XM_023244898.1 |
| E2 | 62933896 | 62934079 | refGene_gn=PIEZO1;refGene_tx=XM_023244898.1 |
| E2 | 62934163 | 62934312 | refGene_gn=PIEZO1;refGene_tx=XM_023244898.1 |
| E2 | 62934395 | 62934574 | refGene_gn=PIEZO1;refGene_tx=XM_023244898.1 |
| E2 | 62934716 | 62934828 | refGene_gn=PIEZO1;refGene_tx=XM_023244898.1 |
| E2 | 62935354 | 62935615 | refGene_gn=PIEZO1;refGene_tx=XM_023244898.1 |
| E2 | 62935934 | 62936035 | refGene_gn=PIEZO1;refGene_tx=XM_023244898.1 |
| E2 | 62936751 | 62936845 | refGene_gn=PIEZO1;refGene_tx=XM_023244898.1 |
| E2 | 62936928 | 62937042 | refGene_gn=PIEZO1;refGene_tx=XM_023244898.1 |
| E2 | 62937142 | 62937314 | refGene_gn=PIEZO1;refGene_tx=XM_023244898.1 |
| E2 | 62937427 | 62937641 | refGene_gn=PIEZO1;refGene_tx=XM_023244898.1 |
| E2 | 62937773 | 62937942 | refGene_gn=PIEZO1;refGene_tx=XM_023244898.1 |
| E2 | 62940575 | 62940714 | refGene_gn=PIEZO1;refGene_tx=XM_023244898.1 |
| E2 | 62940840 | 62940883 | refGene_gn=PIEZO1;refGene_tx=XM_023244898.1 |
| E2 | 62941769 | 62942022 | refGene_gn=PIEZO1;refGene_tx=XM_023244898.1 |
| E2 | 62988389 | 62988620 | refGene_gn=CDT1;refGene_tx=XM_023245559.1 |
| E2 | 62989152 | 62989248 | refGene_gn=CDT1;refGene_tx=XM_023245559.1 |
| E2 | 62989321 | 62989476 | refGene_gn=CDT1;refGene_tx=XM_023245559.1 |
| E2 | 62989999 | 62990197 | refGene_gn=CDT1;refGene_tx=XM_023245559.1 |
| E2 | 62990291 | 62990437 | refGene_gn=CDT1;refGene_tx=XM_023245559.1 |
| E2 | 62990602 | 62990703 | refGene_gn=CDT1;refGene_tx=XM_023245559.1 |
| E2 | 62991150 | 62991339 | refGene_gn=CDT1;refGene_tx=XM_023245559.1 |
| E2 | 62991569 | 62991710 | refGene_gn=CDT1;refGene_tx=XM_023245559.1 |
| E2 | 62991782 | 62991984 | refGene_gn=CDT1;refGene_tx=XM_023245559.1 |
| E2 | 62992397 | 62992561 | refGene_gn=CDT1;refGene_tx=XM_023245559.1 |
| E2 | 62993372 | 62993515 | refGene_gn=APRT;refGene_tx=XM_023245716.1 |
| E2 | 62993753 | 62993832 | refGene_gn=APRT;refGene_tx=XM_023245716.1 |
| E2 | 62994062 | 62994196 | refGene_gn=APRT;refGene_tx=XM_023245716.1 |
| E2 | 62995288 | 62995395 | refGene_gn=APRT;refGene_tx=XM_023245716.1 |
| E2 | 62995514 | 62995594 | refGene_gn=APRT;refGene_tx=XM_023245716.1 |

E2 62996854 62996941 refGene_gn=GALNS;refGene_tx=XM_023245710.1
E2 62999082 62999200 refGene_gn=GALNS;refGene_tx=XM_023245710.1
E2 63000436 63000485 refGene_gn=GALNS;refGene_tx=XM_023245712.1
E2 63000889 63001011
refGene_gn=GALNS;refGene_tx=XM_023245710.1;refGene_gn=GALNS;refGene_tx=XM_023245712.1
E2 63005318 63005421
refGene_gn=GALNS;refGene_tx=XM_023245710.1;refGene_gn=GALNS;refGene_tx=XM_023245712.1
E2 63006094 63006231
refGene_gn=GALNS;refGene_tx=XM_023245710.1;refGene_gn=GALNS;refGene_tx=XM_023245712.1
E2 63009242 63009346
refGene_gn=GALNS;refGene_tx=XM_023245710.1;refGene_gn=GALNS;refGene_tx=XM_023245712.1
E2 63011073 63011213
refGene_gn=GALNS;refGene_tx=XM_023245710.1;refGene_gn=GALNS;refGene_tx=XM_023245712.1
E2 63011540 63011668
refGene_gn=GALNS;refGene_tx=XM_023245710.1;refGene_gn=GALNS;refGene_tx=XM_023245712.1
E2 63011752 63011819
refGene_gn=GALNS;refGene_tx=XM_023245710.1;refGene_gn=GALNS;refGene_tx=XM_023245712.1
E2 63012722 63012866
refGene_gn=GALNS;refGene_tx=XM_023245710.1;refGene_gn=GALNS;refGene_tx=XM_023245712.1
E2 63015426 63015529
refGene_gn=GALNS;refGene_tx=XM_023245710.1;refGene_gn=GALNS;refGene_tx=XM_023245712.1
E2 63016395 63016470
refGene_gn=GALNS;refGene_tx=XM_023245710.1;refGene_gn=GALNS;refGene_tx=XM_023245712.1
E2 63016756 63016880
refGene_gn=GALNS;refGene_tx=XM_023245710.1;refGene_gn=GALNS;refGene_tx=XM_023245712.1
E2 63021239 63021389
refGene_gn=GALNS;refGene_tx=XM_023245710.1;refGene_gn=GALNS;refGene_tx=XM_023245712.1
E2 63021517 63021550
refGene_gn=TRAPPC2L;refGene_tx=XM_023245713.1;refGene_gn=TRAPPC2L;refGene_tx=XM_023245715.1
E2 63023002 63023175
refGene_gn=TRAPPC2L;refGene_tx=XM_019820188.2;refGene_gn=TRAPPC2L;refGene_tx=XM_023245713.1;refGene_gn=TRAPPC2L;refGene_tx=XM_023245714.1;refGene_gn=TRAPPC2L;refGene_tx=XM_023245715.1
E2 63024172 63024260
refGene_gn=TRAPPC2L;refGene_tx=XM_019820188.2;refGene_gn=TRAPPC2L;refGene_tx=XM_023245713.1;refGene_gn=TRAPPC2L;refGene_tx=XM_023245714.1;refGene_gn=TRAPPC2L;refGene_tx=XM_023245715.1

E2 63024396 63024476
refGene_gn=TRAPPC2L;refGene_tx=XM_019820188.2;refGene_gn=TRAPPC2L;refGene_tx=XM_023245713.1;refGene_gn=TRAPPC2L;refGene_tx=XM_023245714.1;refGene_gn=TRAPPC2L;refGene_tx=XM_023245715.1
E2 63024578 63024700
refGene_gn=TRAPPC2L;refGene_tx=XM_019820188.2;refGene_gn=TRAPPC2L;refGene_tx=XM_023245713.1;refGene_gn=TRAPPC2L;refGene_tx=XM_023245714.1
E2 63025270 63025316
refGene_gn=TRAPPC2L;refGene_tx=XM_019820188.2;refGene_gn=TRAPPC2L;refGene_tx=XM_023245713.1;refGene_gn=TRAPPC2L;refGene_tx=XM_023245714.1;refGene_gn=TRAPPC2L;refGene_tx=XM_023245715.1
E2 63029175 63029215 refGene_gn=PABPN1L;refGene_tx=XM_003998332.2
E2 63029418 63029561 refGene_gn=PABPN1L;refGene_tx=XM_003998332.2
E2 63030069 63030157 refGene_gn=PABPN1L;refGene_tx=XM_003998332.2
E2 63030292 63030399 refGene_gn=PABPN1L;refGene_tx=XM_003998332.2
E2 63030822 63030890 refGene_gn=PABPN1L;refGene_tx=XM_003998332.2
E2 63031158 63031297 refGene_gn=PABPN1L;refGene_tx=XM_003998332.2
E2 63031621 63031870 refGene_gn=PABPN1L;refGene_tx=XM_003998332.2
E2 63039519 63039819
refGene_gn=CBFA2T3;refGene_tx=XM_023244909.1;refGene_gn=CBFA2T3;refGene_tx=XM_023244910.1;refGene_gn=CBFA2T3;refGene_tx=XM_023244911.1;refGene_gn=CBFA2T3;refGene_tx=XM_023244912.1;refGene_gn=CBFA2T3;refGene_tx=XM_023244913.1;refGene_gn=CBFA2T3;refGene_tx=XM_023244914.1;refGene_gn=CBFA2T3;refGene_tx=XM_023244915.1;refGene_gn=CBFA2T3;refGene_tx=XM_023244916.1
E2 63042324 63042515
refGene_gn=CBFA2T3;refGene_tx=XM_023244909.1;refGene_gn=CBFA2T3;refGene_tx=XM_023244910.1;refGene_gn=CBFA2T3;refGene_tx=XM_023244911.1;refGene_gn=CBFA2T3;refGene_tx=XM_023244912.1;refGene_gn=CBFA2T3;refGene_tx=XM_023244913.1;refGene_gn=CBFA2T3;refGene_tx=XM_023244914.1;refGene_gn=CBFA2T3;refGene_tx=XM_023244915.1;refGene_gn=CBFA2T3;refGene_tx=XM_023244916.1
E2 63043781 63043850
refGene_gn=CBFA2T3;refGene_tx=XM_023244909.1;refGene_gn=CBFA2T3;refGene_tx=XM_023244910.1;refGene_gn=CBFA2T3;refGene_tx=XM_023244911.1;refGene_gn=CBFA2T3;refGene_tx=XM_023244912.1;refGene_gn=CBFA2T3;refGene_tx=XM_023244913.1;refGene_gn=CBFA2T3;refGene_tx=XM_023244914.1;refGene_gn=CBFA2T3;refGene_tx=XM_023244915.1;refGene_gn=CBFA2T3;refGene_tx=XM_023244916.1
E2 63044319 63044518
refGene_gn=CBFA2T3;refGene_tx=XM_023244909.1;refGene_gn=CBFA2T3;refGene_tx=XM_023244910.1;refGene_gn=CBFA2T3;refGene_tx=XM_023244911.1;refGene_gn=CBFA2T3;refGene_tx=XM_023244912.1;refGene_gn=CBFA2T3;refGene_tx=XM_023244913.1;refGene_gn=CBFA2T3;refGene_tx=XM_023244914.1;refGene_gn=CBFA2T3;refGene_tx=XM_023244915.1;refGene_gn=CBFA2T3;refGene_tx=XM_023244916.1
E2 63045109 63045195
refGene_gn=CBFA2T3;refGene_tx=XM_023244909.1;refGene_gn=CBFA2T3;refGene_tx=XM_023244910.1;refGene_gn=CBFA2T3;refGene_tx=XM_023244911.1;refGene_gn=CBFA2T3;refGene_tx=XM_023244912.1;refGene_gn=CBFA2T3;refGene_tx=

XM_023244913.1;refGene_gn=CBFA2T3;refGene_tx=XM_023244914.1;refGene_gn
=CBFA2T3;refGene_tx=XM_023244915.1;refGene_gn=CBFA2T3;refGene_tx=XM_02
3244916.1
E2 63048195 63048422
refGene_gn=CBFA2T3;refGene_tx=XM_023244909.1;refGene_gn=CBFA2T3;refGen
e_tx=XM_023244910.1;refGene_gn=CBFA2T3;refGene_tx=XM_023244911.1;refGe
ne_gn=CBFA2T3;refGene_tx=XM_023244912.1;refGene_gn=CBFA2T3;refGene_tx=
XM_023244913.1;refGene_gn=CBFA2T3;refGene_tx=XM_023244914.1;refGene_gn
=CBFA2T3;refGene_tx=XM_023244915.1;refGene_gn=CBFA2T3;refGene_tx=XM_02
3244916.1
E2 63049060 63049242
refGene_gn=CBFA2T3;refGene_tx=XM_023244909.1;refGene_gn=CBFA2T3;refGen
e_tx=XM_023244910.1;refGene_gn=CBFA2T3;refGene_tx=XM_023244911.1;refGe
ne_gn=CBFA2T3;refGene_tx=XM_023244912.1;refGene_gn=CBFA2T3;refGene_tx=
XM_023244913.1;refGene_gn=CBFA2T3;refGene_tx=XM_023244914.1;refGene_gn
=CBFA2T3;refGene_tx=XM_023244915.1;refGene_gn=CBFA2T3;refGene_tx=XM_02
3244916.1
E2 63053446 63053536
refGene_gn=CBFA2T3;refGene_tx=XM_023244909.1;refGene_gn=CBFA2T3;refGen
e_tx=XM_023244910.1;refGene_gn=CBFA2T3;refGene_tx=XM_023244911.1;refGe
ne_gn=CBFA2T3;refGene_tx=XM_023244912.1;refGene_gn=CBFA2T3;refGene_tx=
XM_023244913.1;refGene_gn=CBFA2T3;refGene_tx=XM_023244914.1;refGene_gn
=CBFA2T3;refGene_tx=XM_023244915.1;refGene_gn=CBFA2T3;refGene_tx=XM_02
3244916.1
E2 63053644 63053883
refGene_gn=CBFA2T3;refGene_tx=XM_023244909.1;refGene_gn=CBFA2T3;refGen
e_tx=XM_023244910.1;refGene_gn=CBFA2T3;refGene_tx=XM_023244911.1;refGe
ne_gn=CBFA2T3;refGene_tx=XM_023244912.1;refGene_gn=CBFA2T3;refGene_tx=
XM_023244913.1;refGene_gn=CBFA2T3;refGene_tx=XM_023244914.1;refGene_gn
=CBFA2T3;refGene_tx=XM_023244915.1;refGene_gn=CBFA2T3;refGene_tx=XM_02
3244916.1
E2 63058154 63058232
refGene_gn=CBFA2T3;refGene_tx=XM_023244909.1;refGene_gn=CBFA2T3;refGen
e_tx=XM_023244911.1;refGene_gn=CBFA2T3;refGene_tx=XM_023244912.1;refGe
ne_gn=CBFA2T3;refGene_tx=XM_023244913.1;refGene_gn=CBFA2T3;refGene_tx=
XM_023244914.1;refGene_gn=CBFA2T3;refGene_tx=XM_023244915.1;refGene_gn
=CBFA2T3;refGene_tx=XM_023244916.1
E2 63061152 63061299
refGene_gn=CBFA2T3;refGene_tx=XM_023244909.1;refGene_gn=CBFA2T3;refGen
e_tx=XM_023244910.1;refGene_gn=CBFA2T3;refGene_tx=XM_023244911.1;refGe
ne_gn=CBFA2T3;refGene_tx=XM_023244912.1;refGene_gn=CBFA2T3;refGene_tx=
XM_023244913.1;refGene_gn=CBFA2T3;refGene_tx=XM_023244914.1;refGene_gn
=CBFA2T3;refGene_tx=XM_023244915.1;refGene_gn=CBFA2T3;refGene_tx=XM_02
3244916.1
E2 63123135 63123604
refGene_gn=CBFA2T3;refGene_tx=XM_023244909.1;refGene_gn=CBFA2T3;refGen
e_tx=XM_023244910.1
E2 63201903 63201997
refGene_gn=LOC111558009;refGene_tx=XM_023245128.1
E2 63202236 63202435

refGene_gn=LOC111558009;refGene_tx=XM_023245128.1
E2 63208790 63209444
refGene_gn=LOC111558009;refGene_tx=XM_023245128.1
E2 63211785 63211935
refGene_gn=LOC111558009;refGene_tx=XM_023245128.1
E2 63212542 63212765
refGene_gn=LOC111558009;refGene_tx=XM_023245128.1
E2 63214465 63215113
refGene_gn=ACSF3;refGene_tx=XM_006941804.3;refGene_gn=ACSF3;refGene_tx=XM_019820192.2;refGene_gn=ACSF3;refGene_tx=XM_019820193.2;refGene_gn=ACSF3;refGene_tx=XM_023245524.1
E2 63216467 63216623
refGene_gn=ACSF3;refGene_tx=XM_006941804.3;refGene_gn=ACSF3;refGene_tx=XM_019820192.2;refGene_gn=ACSF3;refGene_tx=XM_019820193.2;refGene_gn=ACSF3;refGene_tx=XM_023245524.1
E2 63222829 63222984
refGene_gn=ACSF3;refGene_tx=XM_006941804.3;refGene_gn=ACSF3;refGene_tx=XM_019820192.2;refGene_gn=ACSF3;refGene_tx=XM_019820193.2;refGene_gn=ACSF3;refGene_tx=XM_023245524.1
E2 63224460 63224606
refGene_gn=ACSF3;refGene_tx=XM_006941804.3;refGene_gn=ACSF3;refGene_tx=XM_019820192.2;refGene_gn=ACSF3;refGene_tx=XM_019820193.2;refGene_gn=ACSF3;refGene_tx=XM_023245524.1
E2 63228761 63228874
refGene_gn=ACSF3;refGene_tx=XM_006941804.3;refGene_gn=ACSF3;refGene_tx=XM_019820192.2;refGene_gn=ACSF3;refGene_tx=XM_019820193.2;refGene_gn=ACSF3;refGene_tx=XM_023245524.1
E2 63239330 63239457
refGene_gn=ACSF3;refGene_tx=XM_006941804.3;refGene_gn=ACSF3;refGene_tx=XM_019820192.2;refGene_gn=ACSF3;refGene_tx=XM_019820193.2;refGene_gn=ACSF3;refGene_tx=XM_023245524.1
E2 63241060 63241179 refGene_gn=ACSF3;refGene_tx=XM_019820193.2
E2 63249219 63249354
refGene_gn=ACSF3;refGene_tx=XM_006941804.3;refGene_gn=ACSF3;refGene_tx=XM_019820192.2;refGene_gn=ACSF3;refGene_tx=XM_023245524.1
E2 63249935 63250047
refGene_gn=ACSF3;refGene_tx=XM_006941804.3;refGene_gn=ACSF3;refGene_tx=XM_019820192.2;refGene_gn=ACSF3;refGene_tx=XM_023245524.1
E2 63252200 63252210 refGene_gn=ACSF3;refGene_tx=XM_019820192.2
E2 63253931 63254100 refGene_gn=ACSF3;refGene_tx=XM_006941804.3
E2 63263753 63263859 refGene_gn=ACSF3;refGene_tx=XM_023245524.1
E2 63263885 63264050 refGene_gn=CDH15;refGene_tx=XM_023245523.1
E2 63269681 63269837 refGene_gn=CDH15;refGene_tx=XM_011290290.3
E2 63276019 63276178
refGene_gn=CDH15;refGene_tx=XM_011290290.3;refGene_gn=CDH15;refGene_tx=XM_023245523.1
E2 63276371 63276527
refGene_gn=CDH15;refGene_tx=XM_011290290.3;refGene_gn=CDH15;refGene_tx=XM_023245523.1
E2 63277622 63277767

refGene_gn=CDH15;refGene_tx=XM_011290290.3;refGene_gn=CDH15;refGene_tx=XM_023245523.1
E2 63281619 63281780
refGene_gn=CDH15;refGene_tx=XM_011290290.3;refGene_gn=CDH15;refGene_tx=XM_023245523.1
E2 63282673 63282802
refGene_gn=CDH15;refGene_tx=XM_011290290.3;refGene_gn=CDH15;refGene_tx=XM_023245523.1
E2 63283282 63283468
refGene_gn=CDH15;refGene_tx=XM_011290290.3;refGene_gn=CDH15;refGene_tx=XM_023245523.1
E2 63284295 63284549
refGene_gn=CDH15;refGene_tx=XM_011290290.3;refGene_gn=CDH15;refGene_tx=XM_023245523.1
E2 63285137 63285280
refGene_gn=CDH15;refGene_tx=XM_011290290.3;refGene_gn=CDH15;refGene_tx=XM_023245523.1
E2 63285442 63285679
refGene_gn=CDH15;refGene_tx=XM_011290290.3;refGene_gn=CDH15;refGene_tx=XM_023245523.1
E2 63285956 63286196
refGene_gn=CDH15;refGene_tx=XM_011290290.3;refGene_gn=CDH15;refGene_tx=XM_023245523.1
E2 63286433 63286570
refGene_gn=CDH15;refGene_tx=XM_011290290.3;refGene_gn=CDH15;refGene_tx=XM_023245523.1
E2 63286809 63286977
refGene_gn=CDH15;refGene_tx=XM_011290290.3;refGene_gn=CDH15;refGene_tx=XM_023245523.1
E2 63287907 63288108
refGene_gn=CDH15;refGene_tx=XM_011290290.3;refGene_gn=CDH15;refGene_tx=XM_023245523.1
E2 63288666 63288901 refGene_gn=SLC22A31;refGene_tx=XM_023245525.1
E2 63289215 63289327 refGene_gn=SLC22A31;refGene_tx=XM_023245525.1
E2 63289614 63289829 refGene_gn=SLC22A31;refGene_tx=XM_023245525.1
E2 63289955 63290064 refGene_gn=SLC22A31;refGene_tx=XM_023245525.1
E2 63290148 63290294 refGene_gn=SLC22A31;refGene_tx=XM_023245525.1
E2 63290497 63290666 refGene_gn=SLC22A31;refGene_tx=XM_023245525.1
E2 63290921 63291076 refGene_gn=SLC22A31;refGene_tx=XM_023245525.1
E2 63291196 63291300 refGene_gn=SLC22A31;refGene_tx=XM_023245525.1
E2 63292782 63293160 refGene_gn=SLC22A31;refGene_tx=XM_023245525.1
E2 63321515 63322120
refGene_gn=ANKRD11;refGene_tx=XM_023245509.1;refGene_gn=ANKRD11;refGene_tx=XM_023245510.1;refGene_gn=ANKRD11;refGene_tx=XM_023245511.1;refGene_gn=ANKRD11;refGene_tx=XM_023245512.1;refGene_gn=ANKRD11;refGene_tx=XM_023245513.1;refGene_gn=ANKRD11;refGene_tx=XM_023245514.1;refGene_gn=ANKRD11;refGene_tx=XM_023245515.1;refGene_gn=ANKRD11;refGene_tx=XM_023245516.1;refGene_gn=ANKRD11;refGene_tx=XM_023245517.1;refGene_gn=ANKRD11;refGene_tx=XM_023245518.1;refGene_gn=ANKRD11;refGene_tx=XM_023245519.1;refGene_gn=ANKRD11;refGene_tx=XM_023245520.1;refGene_gn=ANKRD11;r

efGene_tx=XM_023245521.1;refGene_gn=ANKRD11;refGene_tx=XM_023245522.1
E2 63323303 63323469
refGene_gn=ANKRD11;refGene_tx=XM_023245509.1;refGene_gn=ANKRD11;refGene_tx=XM_023245510.1;refGene_gn=ANKRD11;refGene_tx=XM_023245511.1;refGene_gn=ANKRD11;refGene_tx=XM_023245512.1;refGene_gn=ANKRD11;refGene_tx=XM_023245513.1;refGene_gn=ANKRD11;refGene_tx=XM_023245514.1;refGene_gn=ANKRD11;refGene_tx=XM_023245515.1;refGene_gn=ANKRD11;refGene_tx=XM_023245516.1;refGene_gn=ANKRD11;refGene_tx=XM_023245517.1;refGene_gn=ANKRD11;refGene_tx=XM_023245518.1
E2 63323652 63323745
refGene_gn=ANKRD11;refGene_tx=XM_023245509.1;refGene_gn=ANKRD11;refGene_tx=XM_023245510.1;refGene_gn=ANKRD11;refGene_tx=XM_023245511.1;refGene_gn=ANKRD11;refGene_tx=XM_023245512.1;refGene_gn=ANKRD11;refGene_tx=XM_023245513.1;refGene_gn=ANKRD11;refGene_tx=XM_023245514.1;refGene_gn=ANKRD11;refGene_tx=XM_023245515.1;refGene_gn=ANKRD11;refGene_tx=XM_023245516.1;refGene_gn=ANKRD11;refGene_tx=XM_023245517.1;refGene_gn=ANKRD11;refGene_tx=XM_023245518.1;refGene_gn=ANKRD11;refGene_tx=XM_023245519.1;refGene_gn=ANKRD11;refGene_tx=XM_023245520.1;refGene_gn=ANKRD11;refGene_tx=XM_023245521.1;refGene_gn=ANKRD11;refGene_tx=XM_023245522.1
E2 63325704 63325848
refGene_gn=ANKRD11;refGene_tx=XM_023245509.1;refGene_gn=ANKRD11;refGene_tx=XM_023245510.1;refGene_gn=ANKRD11;refGene_tx=XM_023245511.1;refGene_gn=ANKRD11;refGene_tx=XM_023245512.1;refGene_gn=ANKRD11;refGene_tx=XM_023245513.1;refGene_gn=ANKRD11;refGene_tx=XM_023245514.1;refGene_gn=ANKRD11;refGene_tx=XM_023245515.1;refGene_gn=ANKRD11;refGene_tx=XM_023245516.1;refGene_gn=ANKRD11;refGene_tx=XM_023245517.1;refGene_gn=ANKRD11;refGene_tx=XM_023245518.1;refGene_gn=ANKRD11;refGene_tx=XM_023245519.1;refGene_gn=ANKRD11;refGene_tx=XM_023245520.1;refGene_gn=ANKRD11;refGene_tx=XM_023245521.1;refGene_gn=ANKRD11;refGene_tx=XM_023245522.1
E2 63325934 63326033
refGene_gn=ANKRD11;refGene_tx=XM_023245509.1;refGene_gn=ANKRD11;refGene_tx=XM_023245510.1;refGene_gn=ANKRD11;refGene_tx=XM_023245511.1;refGene_gn=ANKRD11;refGene_tx=XM_023245512.1;refGene_gn=ANKRD11;refGene_tx=XM_023245513.1;refGene_gn=ANKRD11;refGene_tx=XM_023245514.1;refGene_gn=ANKRD11;refGene_tx=XM_023245515.1;refGene_gn=ANKRD11;refGene_tx=XM_023245516.1;refGene_gn=ANKRD11;refGene_tx=XM_023245517.1;refGene_gn=ANKRD11;refGene_tx=XM_023245518.1;refGene_gn=ANKRD11;refGene_tx=XM_023245519.1;refGene_gn=ANKRD11;refGene_tx=XM_023245520.1;refGene_gn=ANKRD11;refGene_tx=XM_023245521.1;refGene_gn=ANKRD11;refGene_tx=XM_023245522.1
E2 63327402 63333914
refGene_gn=ANKRD11;refGene_tx=XM_023245509.1;refGene_gn=ANKRD11;refGene_tx=XM_023245510.1;refGene_gn=ANKRD11;refGene_tx=XM_023245511.1;refGene_gn=ANKRD11;refGene_tx=XM_023245512.1;refGene_gn=ANKRD11;refGene_tx=XM_023245513.1;refGene_gn=ANKRD11;refGene_tx=XM_023245514.1;refGene_gn=ANKRD11;refGene_tx=XM_023245515.1;refGene_gn=ANKRD11;refGene_tx=XM_023245516.1;refGene_gn=ANKRD11;refGene_tx=XM_023245517.1;refGene_gn=ANKRD11;refGene_tx=XM_023245518.1;refGene_gn=ANKRD11;refGene_tx=XM_023245519.1;refGene_gn=ANKRD11;refGene_tx=XM_023245520.1;refGene_gn=ANKRD11;refGene_tx=XM_023245521.1;refGene_gn=ANKRD11;refGene_tx=XM_023245522.1
E2 63334199 63334347

refGene_gn=ANKRD11;refGene_tx=XM_023245509.1;refGene_gn=ANKRD11;refGene_tx=XM_023245510.1;refGene_gn=ANKRD11;refGene_tx=XM_023245511.1;refGene_gn=ANKRD11;refGene_tx=XM_023245512.1;refGene_gn=ANKRD11;refGene_tx=XM_023245513.1;refGene_gn=ANKRD11;refGene_tx=XM_023245514.1;refGene_gn=ANKRD11;refGene_tx=XM_023245515.1;refGene_gn=ANKRD11;refGene_tx=XM_023245516.1;refGene_gn=ANKRD11;refGene_tx=XM_023245517.1;refGene_gn=ANKRD11;refGene_tx=XM_023245518.1;refGene_gn=ANKRD11;refGene_tx=XM_023245519.1;refGene_gn=ANKRD11;refGene_tx=XM_023245520.1;refGene_gn=ANKRD11;refGene_tx=XM_023245521.1;refGene_gn=ANKRD11;refGene_tx=XM_023245522.1
E2 63336502 63336645
refGene_gn=ANKRD11;refGene_tx=XM_023245509.1;refGene_gn=ANKRD11;refGene_tx=XM_023245510.1;refGene_gn=ANKRD11;refGene_tx=XM_023245511.1;refGene_gn=ANKRD11;refGene_tx=XM_023245512.1;refGene_gn=ANKRD11;refGene_tx=XM_023245513.1;refGene_gn=ANKRD11;refGene_tx=XM_023245514.1;refGene_gn=ANKRD11;refGene_tx=XM_023245515.1;refGene_gn=ANKRD11;refGene_tx=XM_023245516.1;refGene_gn=ANKRD11;refGene_tx=XM_023245517.1;refGene_gn=ANKRD11;refGene_tx=XM_023245518.1;refGene_gn=ANKRD11;refGene_tx=XM_023245519.1;refGene_gn=ANKRD11;refGene_tx=XM_023245520.1;refGene_gn=ANKRD11;refGene_tx=XM_023245521.1;refGene_gn=ANKRD11;refGene_tx=XM_023245522.1
E2 63337814 63338018
refGene_gn=ANKRD11;refGene_tx=XM_023245509.1;refGene_gn=ANKRD11;refGene_tx=XM_023245510.1;refGene_gn=ANKRD11;refGene_tx=XM_023245511.1;refGene_gn=ANKRD11;refGene_tx=XM_023245512.1;refGene_gn=ANKRD11;refGene_tx=XM_023245513.1;refGene_gn=ANKRD11;refGene_tx=XM_023245514.1;refGene_gn=ANKRD11;refGene_tx=XM_023245515.1;refGene_gn=ANKRD11;refGene_tx=XM_023245516.1;refGene_gn=ANKRD11;refGene_tx=XM_023245517.1;refGene_gn=ANKRD11;refGene_tx=XM_023245518.1;refGene_gn=ANKRD11;refGene_tx=XM_023245519.1;refGene_gn=ANKRD11;refGene_tx=XM_023245520.1;refGene_gn=ANKRD11;refGene_tx=XM_023245521.1;refGene_gn=ANKRD11;refGene_tx=XM_023245522.1
E2 63338118 63338289
refGene_gn=ANKRD11;refGene_tx=XM_023245509.1;refGene_gn=ANKRD11;refGene_tx=XM_023245510.1;refGene_gn=ANKRD11;refGene_tx=XM_023245511.1;refGene_gn=ANKRD11;refGene_tx=XM_023245512.1;refGene_gn=ANKRD11;refGene_tx=XM_023245513.1;refGene_gn=ANKRD11;refGene_tx=XM_023245514.1;refGene_gn=ANKRD11;refGene_tx=XM_023245515.1;refGene_gn=ANKRD11;refGene_tx=XM_023245516.1;refGene_gn=ANKRD11;refGene_tx=XM_023245517.1;refGene_gn=ANKRD11;refGene_tx=XM_023245518.1;refGene_gn=ANKRD11;refGene_tx=XM_023245519.1;refGene_gn=ANKRD11;refGene_tx=XM_023245520.1;refGene_gn=ANKRD11;refGene_tx=XM_023245521.1;refGene_gn=ANKRD11;refGene_tx=XM_023245522.1
E2 63338729 63338826
refGene_gn=ANKRD11;refGene_tx=XM_023245518.1;refGene_gn=ANKRD11;refGene_tx=XM_023245521.1
E2 63347984 63348123
refGene_gn=ANKRD11;refGene_tx=XM_023245509.1;refGene_gn=ANKRD11;refGene_tx=XM_023245510.1;refGene_gn=ANKRD11;refGene_tx=XM_023245511.1;refGene_gn=ANKRD11;refGene_tx=XM_023245512.1;refGene_gn=ANKRD11;refGene_tx=XM_023245513.1;refGene_gn=ANKRD11;refGene_tx=XM_023245514.1;refGene_gn=ANKRD11;refGene_tx=XM_023245515.1;refGene_gn=ANKRD11;refGene_tx=XM_023245516.1;refGene_gn=ANKRD11;refGene_tx=XM_023245517.1;refGene_gn=ANKRD11;refGene_tx=XM_023245518.1;refGene_gn=ANKRD11;refGene_tx=XM_0232455

19.1;refGene_gn=ANKRD11;refGene_tx=XM_023245520.1;refGene_gn=ANKRD11;refGene_tx=XM_023245521.1;refGene_gn=ANKRD11;refGene_tx=XM_023245522.1
E2 63354019 63354106
refGene_gn=ANKRD11;refGene_tx=XM_023245509.1;refGene_gn=ANKRD11;refGene_tx=XM_023245510.1;refGene_gn=ANKRD11;refGene_tx=XM_023245511.1;refGene_gn=ANKRD11;refGene_tx=XM_023245512.1;refGene_gn=ANKRD11;refGene_tx=XM_023245513.1;refGene_gn=ANKRD11;refGene_tx=XM_023245514.1;refGene_gn=ANKRD11;refGene_tx=XM_023245515.1;refGene_gn=ANKRD11;refGene_tx=XM_023245516.1;refGene_gn=ANKRD11;refGene_tx=XM_023245517.1;refGene_gn=ANKRD11;refGene_tx=XM_023245519.1;refGene_gn=ANKRD11;refGene_tx=XM_023245520.1;refGene_gn=ANKRD11;refGene_tx=XM_023245522.1
E2 63496664 63496707
refGene_gn=LOC111558010;refGene_tx=XM_023245129.1
E2 63499043 63499957
refGene_gn=LOC111558010;refGene_tx=XM_023245129.1
E2 63511516 63511699 refGene_gn=SPG7;refGene_tx=XM_011290291.3
E2 63514143 63514246 refGene_gn=SPG7;refGene_tx=XM_011290291.3
E2 63515953 63516043 refGene_gn=SPG7;refGene_tx=XM_011290291.3
E2 63523080 63523322 refGene_gn=SPG7;refGene_tx=XM_011290291.3
E2 63525306 63525446 refGene_gn=SPG7;refGene_tx=XM_011290291.3
E2 63526317 63526420 refGene_gn=SPG7;refGene_tx=XM_011290291.3
E2 63527967 63528093 refGene_gn=SPG7;refGene_tx=XM_011290291.3
E2 63528414 63528471 refGene_gn=SPG7;refGene_tx=XM_011290291.3
E2 63528472 63528578 refGene_gn=SPG7;refGene_tx=XM_011290291.3
E2 63528919 63529093 refGene_gn=SPG7;refGene_tx=XM_011290291.3
E2 63534861 63534986 refGene_gn=SPG7;refGene_tx=XM_011290291.3
E2 63535715 63535818 refGene_gn=SPG7;refGene_tx=XM_011290291.3
E2 63536501 63536612 refGene_gn=SPG7;refGene_tx=XM_011290291.3
E2 63538407 63538523 refGene_gn=SPG7;refGene_tx=XM_011290291.3
E2 63539984 63540141 refGene_gn=SPG7;refGene_tx=XM_011290291.3
E2 63540549 63540716 refGene_gn=SPG7;refGene_tx=XM_011290291.3
E2 63541313 63541391 refGene_gn=SPG7;refGene_tx=XM_011290291.3
E2 63543018 63543219 refGene_gn=SPG7;refGene_tx=XM_011290291.3
E2 63548784 63548888 refGene_gn=RPL13;refGene_tx=XM_023245576.1
E2 63549053 63549195 refGene_gn=RPL13;refGene_tx=XM_023245576.1
E2 63549390 63549564 refGene_gn=RPL13;refGene_tx=XM_023245576.1
E2 63549947 63550004 refGene_gn=RPL13;refGene_tx=XM_023245576.1
E2 63550632 63550791 refGene_gn=RPL13;refGene_tx=XM_023245576.1
E2 63562056 63562230
refGene_gn=CPNE7;refGene_tx=XM_023245375.1;refGene_gn=CPNE7;refGene_tx=XM_023245376.1;refGene_gn=CPNE7;refGene_tx=XM_023245378.1
E2 63564116 63564299
refGene_gn=CPNE7;refGene_tx=XM_023245375.1;refGene_gn=CPNE7;refGene_tx=XM_023245376.1;refGene_gn=CPNE7;refGene_tx=XM_023245378.1
E2 63566755 63566830
refGene_gn=CPNE7;refGene_tx=XM_023245375.1;refGene_gn=CPNE7;refGene_tx=XM_023245376.1;refGene_gn=CPNE7;refGene_tx=XM_023245378.1
E2 63566982 63567057
refGene_gn=CPNE7;refGene_tx=XM_023245375.1;refGene_gn=CPNE7;refGene_tx=XM_023245376.1;refGene_gn=CPNE7;refGene_tx=XM_023245378.1

E2 63567684 63567768
refGene_gn=CPNE7;refGene_tx=XM_023245375.1;refGene_gn=CPNE7;refGene_tx
=XM_023245376.1;refGene_gn=CPNE7;refGene_tx=XM_023245378.1
E2 63568439 63568529
refGene_gn=CPNE7;refGene_tx=XM_023245375.1;refGene_gn=CPNE7;refGene_tx
=XM_023245376.1;refGene_gn=CPNE7;refGene_tx=XM_023245378.1
E2 63568653 63568752
refGene_gn=CPNE7;refGene_tx=XM_023245375.1;refGene_gn=CPNE7;refGene_tx
=XM_023245376.1;refGene_gn=CPNE7;refGene_tx=XM_023245378.1
E2 63569091 63569178
refGene_gn=CPNE7;refGene_tx=XM_023245375.1;refGene_gn=CPNE7;refGene_tx
=XM_023245376.1;refGene_gn=CPNE7;refGene_tx=XM_023245378.1
E2 63569512 63569572
refGene_gn=CPNE7;refGene_tx=XM_023245375.1;refGene_gn=CPNE7;refGene_tx
=XM_023245376.1
E2 63570499 63570633
refGene_gn=CPNE7;refGene_tx=XM_023245375.1;refGene_gn=CPNE7;refGene_tx
=XM_023245376.1;refGene_gn=CPNE7;refGene_tx=XM_023245378.1
E2 63572223 63572278
refGene_gn=CPNE7;refGene_tx=XM_023245375.1;refGene_gn=CPNE7;refGene_tx
=XM_023245376.1;refGene_gn=CPNE7;refGene_tx=XM_023245378.1
E2 63572789 63572841
refGene_gn=CPNE7;refGene_tx=XM_023245375.1;refGene_gn=CPNE7;refGene_tx
=XM_023245376.1;refGene_gn=CPNE7;refGene_tx=XM_023245378.1
E2 63572929 63573063
refGene_gn=CPNE7;refGene_tx=XM_023245375.1;refGene_gn=CPNE7;refGene_tx
=XM_023245376.1;refGene_gn=CPNE7;refGene_tx=XM_023245378.1
E2 63576397 63576634
refGene_gn=CPNE7;refGene_tx=XM_023245375.1;refGene_gn=CPNE7;refGene_tx
=XM_023245376.1;refGene_gn=CPNE7;refGene_tx=XM_023245378.1
E2 63577287 63577473
refGene_gn=CPNE7;refGene_tx=XM_023245375.1;refGene_gn=CPNE7;refGene_tx
=XM_023245376.1;refGene_gn=CPNE7;refGene_tx=XM_023245378.1
E2 63607989 63608093
refGene_gn=DPEP1;refGene_tx=NM_001043356.1;refGene_gn=DPEP1;refGene_tx
=XM_019818435.1;refGene_gn=DPEP1;refGene_tx=XM_019818436.1;refGene_gn=
DPEP1;refGene_tx=XM_019818437.2;refGene_gn=DPEP1;refGene_tx=XM_0198184
38.1
E2 63611822 63611955
refGene_gn=DPEP1;refGene_tx=NM_001043356.1;refGene_gn=DPEP1;refGene_tx
=XM_019818435.1;refGene_gn=DPEP1;refGene_tx=XM_019818436.1;refGene_gn=
DPEP1;refGene_tx=XM_019818437.2;refGene_gn=DPEP1;refGene_tx=XM_0198184
38.1;refGene_gn=DPEP1;refGene_tx=XM_019818439.1
E2 63612202 63612335
refGene_gn=DPEP1;refGene_tx=NM_001043356.1;refGene_gn=DPEP1;refGene_tx
=XM_019818435.1;refGene_gn=DPEP1;refGene_tx=XM_019818436.1;refGene_gn=
DPEP1;refGene_tx=XM_019818437.2;refGene_gn=DPEP1;refGene_tx=XM_0198184
38.1;refGene_gn=DPEP1;refGene_tx=XM_019818439.1
E2 63612450 63612634
refGene_gn=DPEP1;refGene_tx=NM_001043356.1;refGene_gn=DPEP1;refGene_tx

=XM_019818435.1;refGene_gn=DPEP1;refGene_tx=XM_019818436.1;refGene_gn=DPEP1;refGene_tx=XM_019818437.2;refGene_gn=DPEP1;refGene_tx=XM_019818438.1;refGene_gn=DPEP1;refGene_tx=XM_019818439.1
E2 63612807 63612877
refGene_gn=DPEP1;refGene_tx=NM_001043356.1;refGene_gn=DPEP1;refGene_tx=XM_019818435.1;refGene_gn=DPEP1;refGene_tx=XM_019818436.1;refGene_gn=DPEP1;refGene_tx=XM_019818437.2;refGene_gn=DPEP1;refGene_tx=XM_019818438.1;refGene_gn=DPEP1;refGene_tx=XM_019818439.1
E2 63613215 63613392
refGene_gn=DPEP1;refGene_tx=NM_001043356.1;refGene_gn=DPEP1;refGene_tx=XM_019818435.1;refGene_gn=DPEP1;refGene_tx=XM_019818436.1;refGene_gn=DPEP1;refGene_tx=XM_019818437.2;refGene_gn=DPEP1;refGene_tx=XM_019818438.1;refGene_gn=DPEP1;refGene_tx=XM_019818439.1
E2 63613495 63613580
refGene_gn=DPEP1;refGene_tx=NM_001043356.1;refGene_gn=DPEP1;refGene_tx=XM_019818435.1;refGene_gn=DPEP1;refGene_tx=XM_019818436.1;refGene_gn=DPEP1;refGene_tx=XM_019818437.2;refGene_gn=DPEP1;refGene_tx=XM_019818438.1;refGene_gn=DPEP1;refGene_tx=XM_019818439.1
E2 63613660 63613736
refGene_gn=DPEP1;refGene_tx=NM_001043356.1;refGene_gn=DPEP1;refGene_tx=XM_019818435.1;refGene_gn=DPEP1;refGene_tx=XM_019818436.1;refGene_gn=DPEP1;refGene_tx=XM_019818437.2;refGene_gn=DPEP1;refGene_tx=XM_019818438.1;refGene_gn=DPEP1;refGene_tx=XM_019818439.1
E2 63613990 63614126
refGene_gn=DPEP1;refGene_tx=NM_001043356.1;refGene_gn=DPEP1;refGene_tx=XM_019818435.1;refGene_gn=DPEP1;refGene_tx=XM_019818436.1;refGene_gn=DPEP1;refGene_tx=XM_019818437.2;refGene_gn=DPEP1;refGene_tx=XM_019818438.1;refGene_gn=DPEP1;refGene_tx=XM_019818439.1
E2 63614204 63614372
refGene_gn=DPEP1;refGene_tx=NM_001043356.1;refGene_gn=DPEP1;refGene_tx=XM_019818435.1;refGene_gn=DPEP1;refGene_tx=XM_019818436.1;refGene_gn=DPEP1;refGene_tx=XM_019818437.2;refGene_gn=DPEP1;refGene_tx=XM_019818438.1;refGene_gn=DPEP1;refGene_tx=XM_019818439.1
E2 63618360 63618382 refGene_gn=CHMP1A;refGene_tx=XM_006941832.4
E2 63618930 63619118 refGene_gn=CHMP1A;refGene_tx=XM_006941832.4
E2 63619554 63619683 refGene_gn=CHMP1A;refGene_tx=XM_006941832.4
E2 63620036 63620183 refGene_gn=CHMP1A;refGene_tx=XM_006941832.4
E2 63621080 63621158 refGene_gn=CHMP1A;refGene_tx=XM_006941832.4
E2 63623288 63623308 refGene_gn=CHMP1A;refGene_tx=XM_006941832.4
E2 63625171 63625178 refGene_gn=CHMP1A;refGene_tx=XM_006941832.4
E2 63625472 63625509 refGene_gn=SPATA33;refGene_tx=XM_019820197.2
E2 63625845 63626010 refGene_gn=SPATA33;refGene_tx=XM_019820197.2
E2 63635389 63635595 refGene_gn=SPATA33;refGene_tx=XM_019820197.2
E2 63637938 63638025
refGene_gn=CDK10;refGene_tx=XM_003998365.4;refGene_gn=CDK10;refGene_tx=XM_006941827.3
E2 63640468 63640541
refGene_gn=CDK10;refGene_tx=XM_003998365.4;refGene_gn=CDK10;refGene_tx=XM_006941827.3
E2 63641240 63641312

refGene_gn=CDK10;refGene_tx=XM_003998365.4;refGene_gn=CDK10;refGene_tx=XM_006941827.3;refGene_gn=CDK10;refGene_tx=XM_006941828.3;refGene_gn=CDK10;refGene_tx=XM_006941829.3;refGene_gn=CDK10;refGene_tx=XM_006941830.2
E2 63646121 63646224
refGene_gn=CDK10;refGene_tx=XM_003998365.4;refGene_gn=CDK10;refGene_tx=XM_006941827.3;refGene_gn=CDK10;refGene_tx=XM_006941828.3;refGene_gn=CDK10;refGene_tx=XM_006941829.3;refGene_gn=CDK10;refGene_tx=XM_006941830.2
E2 63646362 63646444
refGene_gn=CDK10;refGene_tx=XM_003998365.4;refGene_gn=CDK10;refGene_tx=XM_006941827.3;refGene_gn=CDK10;refGene_tx=XM_006941828.3;refGene_gn=CDK10;refGene_tx=XM_006941829.3;refGene_gn=CDK10;refGene_tx=XM_006941830.2
E2 63646533 63646601
refGene_gn=CDK10;refGene_tx=XM_003998365.4;refGene_gn=CDK10;refGene_tx=XM_006941827.3;refGene_gn=CDK10;refGene_tx=XM_006941828.3;refGene_gn=CDK10;refGene_tx=XM_006941829.3;refGene_gn=CDK10;refGene_tx=XM_006941830.2
E2 63648356 63648409
refGene_gn=CDK10;refGene_tx=XM_003998365.4;refGene_gn=CDK10;refGene_tx=XM_006941827.3;refGene_gn=CDK10;refGene_tx=XM_006941828.3;refGene_gn=CDK10;refGene_tx=XM_006941829.3;refGene_gn=CDK10;refGene_tx=XM_006941830.2
E2 63648571 63648641
refGene_gn=CDK10;refGene_tx=XM_003998365.4;refGene_gn=CDK10;refGene_tx=XM_006941827.3;refGene_gn=CDK10;refGene_tx=XM_006941828.3;refGene_gn=CDK10;refGene_tx=XM_006941829.3;refGene_gn=CDK10;refGene_tx=XM_006941830.2
E2 63649161 63649221
refGene_gn=CDK10;refGene_tx=XM_003998365.4;refGene_gn=CDK10;refGene_tx=XM_006941827.3;refGene_gn=CDK10;refGene_tx=XM_006941828.3;refGene_gn=CDK10;refGene_tx=XM_006941829.3;refGene_gn=CDK10;refGene_tx=XM_006941830.2
E2 63649569 63649693
refGene_gn=CDK10;refGene_tx=XM_003998365.4;refGene_gn=CDK10;refGene_tx=XM_006941827.3;refGene_gn=CDK10;refGene_tx=XM_006941828.3;refGene_gn=CDK10;refGene_tx=XM_006941829.3;refGene_gn=CDK10;refGene_tx=XM_006941830.2
E2 63649790 63649930
refGene_gn=CDK10;refGene_tx=XM_003998365.4;refGene_gn=CDK10;refGene_tx=XM_006941827.3;refGene_gn=CDK10;refGene_tx=XM_006941828.3;refGene_gn=CDK10;refGene_tx=XM_006941829.3;refGene_gn=CDK10;refGene_tx=XM_006941830.2
E2 63650258 63650311
refGene_gn=CDK10;refGene_tx=XM_003998365.4;refGene_gn=CDK10;refGene_tx=XM_006941827.3;refGene_gn=CDK10;refGene_tx=XM_006941828.3;refGene_gn=CDK10;refGene_tx=XM_006941829.3;refGene_gn=CDK10;refGene_tx=XM_006941830.2
E2 63650670 63650768

refGene_gn=CDK10;refGene_tx=XM_003998365.4;refGene_gn=CDK10;refGene_tx=XM_006941827.3;refGene_gn=CDK10;refGene_tx=XM_006941828.3;refGene_gn=CDK10;refGene_tx=XM_006941829.3;refGene_gn=CDK10;refGene_tx=XM_006941830.2
E2 63652423 63653368 refGene_gn=SPATA2L;refGene_tx=XM_006941824.4;refGene_gn=SPATA2L;refGene_tx=XM_006941825.3;refGene_gn=SPATA2L;refGene_tx=XM_019820196.1
E2 63654882 63655272 refGene_gn=SPATA2L;refGene_tx=XM_006941824.4;refGene_gn=SPATA2L;refGene_tx=XM_006941825.3;refGene_gn=SPATA2L;refGene_tx=XM_019820196.1
E2 63655416 63655472 refGene_gn=SPATA2L;refGene_tx=XM_006941824.4
E2 63658264 63658358 refGene_gn=VPS9D1;refGene_tx=XM_023245615.1;refGene_gn=VPS9D1;refGene_tx=XM_023245616.1;refGene_gn=VPS9D1;refGene_tx=XM_023245617.1;refGene_gn=VPS9D1;refGene_tx=XM_023245618.1;refGene_gn=VPS9D1;refGene_tx=XM_023245619.1;refGene_gn=VPS9D1;refGene_tx=XM_023245620.1;refGene_gn=VPS9D1;refGene_tx=XM_023245621.1
E2 63658592 63658697 refGene_gn=VPS9D1;refGene_tx=XM_023245615.1;refGene_gn=VPS9D1;refGene_tx=XM_023245616.1;refGene_gn=VPS9D1;refGene_tx=XM_023245617.1;refGene_gn=VPS9D1;refGene_tx=XM_023245618.1;refGene_gn=VPS9D1;refGene_tx=XM_023245619.1;refGene_gn=VPS9D1;refGene_tx=XM_023245620.1;refGene_gn=VPS9D1;refGene_tx=XM_023245621.1
E2 63658750 63658838 refGene_gn=VPS9D1;refGene_tx=XM_023245622.1
E2 63659078 63659201 refGene_gn=VPS9D1;refGene_tx=XM_023245615.1;refGene_gn=VPS9D1;refGene_tx=XM_023245616.1;refGene_gn=VPS9D1;refGene_tx=XM_023245617.1;refGene_gn=VPS9D1;refGene_tx=XM_023245620.1;refGene_gn=VPS9D1;refGene_tx=XM_023245621.1
E2 63660083 63660180 refGene_gn=VPS9D1;refGene_tx=XM_023245615.1;refGene_gn=VPS9D1;refGene_tx=XM_023245616.1;refGene_gn=VPS9D1;refGene_tx=XM_023245617.1;refGene_gn=VPS9D1;refGene_tx=XM_023245618.1;refGene_gn=VPS9D1;refGene_tx=XM_023245619.1;refGene_gn=VPS9D1;refGene_tx=XM_023245620.1;refGene_gn=VPS9D1;refGene_tx=XM_023245621.1;refGene_gn=VPS9D1;refGene_tx=XM_023245622.1
E2 63660464 63660673 refGene_gn=VPS9D1;refGene_tx=XM_023245615.1;refGene_gn=VPS9D1;refGene_tx=XM_023245616.1;refGene_gn=VPS9D1;refGene_tx=XM_023245617.1;refGene_gn=VPS9D1;refGene_tx=XM_023245618.1;refGene_gn=VPS9D1;refGene_tx=XM_023245619.1;refGene_gn=VPS9D1;refGene_tx=XM_023245620.1;refGene_gn=VPS9D1;refGene_tx=XM_023245621.1;refGene_gn=VPS9D1;refGene_tx=XM_023245622.1
E2 63660898 63661028 refGene_gn=VPS9D1;refGene_tx=XM_023245615.1;refGene_gn=VPS9D1;refGene_tx=XM_023245616.1;refGene_gn=VPS9D1;refGene_tx=XM_023245617.1;refGene_gn=VPS9D1;refGene_tx=XM_023245618.1;refGene_gn=VPS9D1;refGene_tx=XM_023245619.1;refGene_gn=VPS9D1;refGene_tx=XM_023245620.1;refGene_gn=VPS9D1;refGene_tx=XM_023245621.1;refGene_gn=VPS9D1;refGene_tx=XM_023245622.1

E2 63662048 63662566
refGene_gn=VPS9D1;refGene_tx=XM_023245615.1;refGene_gn=VPS9D1;refGene_
tx=XM_023245616.1;refGene_gn=VPS9D1;refGene_tx=XM_023245617.1;refGene_
gn=VPS9D1;refGene_tx=XM_023245618.1;refGene_gn=VPS9D1;refGene_tx=XM_02
3245619.1;refGene_gn=VPS9D1;refGene_tx=XM_023245620.1;refGene_gn=VPS9D
1;refGene_tx=XM_023245621.1;refGene_gn=VPS9D1;refGene_tx=XM_023245622.
1
E2 63662930 63663016
refGene_gn=VPS9D1;refGene_tx=XM_023245615.1;refGene_gn=VPS9D1;refGene_
tx=XM_023245616.1;refGene_gn=VPS9D1;refGene_tx=XM_023245617.1;refGene_
gn=VPS9D1;refGene_tx=XM_023245618.1;refGene_gn=VPS9D1;refGene_tx=XM_02
3245619.1;refGene_gn=VPS9D1;refGene_tx=XM_023245620.1;refGene_gn=VPS9D
1;refGene_tx=XM_023245621.1;refGene_gn=VPS9D1;refGene_tx=XM_023245622.
1
E2 63663220 63663308
refGene_gn=VPS9D1;refGene_tx=XM_023245615.1;refGene_gn=VPS9D1;refGene_
tx=XM_023245616.1;refGene_gn=VPS9D1;refGene_tx=XM_023245617.1;refGene_
gn=VPS9D1;refGene_tx=XM_023245618.1;refGene_gn=VPS9D1;refGene_tx=XM_02
3245619.1;refGene_gn=VPS9D1;refGene_tx=XM_023245620.1;refGene_gn=VPS9D
1;refGene_tx=XM_023245621.1;refGene_gn=VPS9D1;refGene_tx=XM_023245622.
1
E2 63663412 63663465
refGene_gn=VPS9D1;refGene_tx=XM_023245615.1;refGene_gn=VPS9D1;refGene_
tx=XM_023245616.1;refGene_gn=VPS9D1;refGene_tx=XM_023245617.1;refGene_
gn=VPS9D1;refGene_tx=XM_023245618.1;refGene_gn=VPS9D1;refGene_tx=XM_02
3245619.1;refGene_gn=VPS9D1;refGene_tx=XM_023245620.1;refGene_gn=VPS9D
1;refGene_tx=XM_023245621.1;refGene_gn=VPS9D1;refGene_tx=XM_023245622.
1
E2 63663798 63663861
refGene_gn=VPS9D1;refGene_tx=XM_023245615.1;refGene_gn=VPS9D1;refGene_
tx=XM_023245616.1;refGene_gn=VPS9D1;refGene_tx=XM_023245617.1;refGene_
gn=VPS9D1;refGene_tx=XM_023245618.1;refGene_gn=VPS9D1;refGene_tx=XM_02
3245619.1;refGene_gn=VPS9D1;refGene_tx=XM_023245620.1;refGene_gn=VPS9D
1;refGene_tx=XM_023245621.1;refGene_gn=VPS9D1;refGene_tx=XM_023245622.
1
E2 63663935 63664047
refGene_gn=VPS9D1;refGene_tx=XM_023245615.1;refGene_gn=VPS9D1;refGene_
tx=XM_023245616.1;refGene_gn=VPS9D1;refGene_tx=XM_023245617.1;refGene_
gn=VPS9D1;refGene_tx=XM_023245618.1;refGene_gn=VPS9D1;refGene_tx=XM_02
3245619.1;refGene_gn=VPS9D1;refGene_tx=XM_023245620.1;refGene_gn=VPS9D
1;refGene_tx=XM_023245621.1;refGene_gn=VPS9D1;refGene_tx=XM_023245622.
1
E2 63665381 63665544
refGene_gn=VPS9D1;refGene_tx=XM_023245615.1;refGene_gn=VPS9D1;refGene_
tx=XM_023245616.1;refGene_gn=VPS9D1;refGene_tx=XM_023245617.1;refGene_
gn=VPS9D1;refGene_tx=XM_023245618.1;refGene_gn=VPS9D1;refGene_tx=XM_02
3245619.1;refGene_gn=VPS9D1;refGene_tx=XM_023245620.1;refGene_gn=VPS9D
1;refGene_tx=XM_023245621.1;refGene_gn=VPS9D1;refGene_tx=XM_023245622.
1
E2 63665642 63665735

refGene_gn=VPS9D1;refGene_tx=XM_023245615.1;refGene_gn=VPS9D1;refGene_tx=XM_023245616.1;refGene_gn=VPS9D1;refGene_tx=XM_023245617.1;refGene_gn=VPS9D1;refGene_tx=XM_023245618.1;refGene_gn=VPS9D1;refGene_tx=XM_023245619.1;refGene_gn=VPS9D1;refGene_tx=XM_023245620.1;refGene_gn=VPS9D1;refGene_tx=XM_023245621.1;refGene_gn=VPS9D1;refGene_tx=XM_023245622.1
E2	63667306	63667382	refGene_gn=VPS9D1;refGene_tx=XM_023245615.1;refGene_gn=VPS9D1;refGene_tx=XM_023245616.1;refGene_gn=VPS9D1;refGene_tx=XM_023245617.1;refGene_gn=VPS9D1;refGene_tx=XM_023245618.1;refGene_gn=VPS9D1;refGene_tx=XM_023245619.1;refGene_gn=VPS9D1;refGene_tx=XM_023245622.1
E2	63668677	63668776	refGene_gn=VPS9D1;refGene_tx=XM_023245615.1;refGene_gn=VPS9D1;refGene_tx=XM_023245616.1;refGene_gn=VPS9D1;refGene_tx=XM_023245617.1;refGene_gn=VPS9D1;refGene_tx=XM_023245618.1;refGene_gn=VPS9D1;refGene_tx=XM_023245619.1;refGene_gn=VPS9D1;refGene_tx=XM_023245622.1
E2	63669016	63669026	refGene_gn=ZNF276;refGene_tx=XM_023245614.1
E2	63669194	63669704	refGene_gn=ZNF276;refGene_tx=XM_023245614.1
E2	63673835	63674139	refGene_gn=ZNF276;refGene_tx=XM_023245614.1
E2	63674482	63674529	refGene_gn=ZNF276;refGene_tx=XM_023245614.1
E2	63674624	63675071	refGene_gn=ZNF276;refGene_tx=XM_023245614.1
E2	63676298	63676377	refGene_gn=ZNF276;refGene_tx=XM_023245614.1
E2	63677596	63677680	refGene_gn=ZNF276;refGene_tx=XM_023245614.1
E2	63680931	63681042	refGene_gn=ZNF276;refGene_tx=XM_023245614.1
E2	63681133	63681209	refGene_gn=ZNF276;refGene_tx=XM_023245614.1
E2	63681475	63681593	refGene_gn=ZNF276;refGene_tx=XM_023245614.1
E2	63683701	63683801	refGene_gn=ZNF276;refGene_tx=XM_023245614.1
E2	63683881	63684128	refGene_gn=ZNF276;refGene_tx=XM_023245614.1
E2	63684502	63684910	refGene_gn=FANCA;refGene_tx=XM_003998369.5;refGene_gn=FANCA;refGene_tx=XM_006941834.3;refGene_gn=FANCA;refGene_tx=XM_011290295.3;refGene_gn=FANCA;refGene_tx=XM_019820198.2;refGene_gn=FANCA;refGene_tx=XM_019820200.2;refGene_gn=FANCA;refGene_tx=XM_019820201.2;refGene_gn=FANCA;refGene_tx=XM_019820203.2
E2	63685057	63685218	refGene_gn=FANCA;refGene_tx=XM_003998369.5;refGene_gn=FANCA;refGene_tx=XM_006941834.3;refGene_gn=FANCA;refGene_tx=XM_011290295.3;refGene_gn=FANCA;refGene_tx=XM_019820198.2;refGene_gn=FANCA;refGene_tx=XM_019820200.2;refGene_gn=FANCA;refGene_tx=XM_019820201.2;refGene_gn=FANCA;refGene_tx=XM_019820203.2
E2	63685353	63685539	refGene_gn=FANCA;refGene_tx=XM_003998369.5;refGene_gn=FANCA;refGene_tx=XM_006941834.3;refGene_gn=FANCA;refGene_tx=XM_006941835.3;refGene_gn=FANCA;refGene_tx=XM_011290295.3;refGene_gn=FANCA;refGene_tx=XM_019820198.2;refGene_gn=FANCA;refGene_tx=XM_019820200.2;refGene_gn=FANCA;refGene_tx=XM_019820201.2;refGene_gn=FANCA;refGene_tx=XM_019820203.2
E2	63686008	63686111	refGene_gn=FANCA;refGene_tx=XM_003998369.5;refGene_gn=FANCA;refGene_tx=XM_006941834.3;refGene_gn=FANCA;refGene_tx=XM_011290295.3;refGene_gn=FANCA;refGene_tx=XM_019820198.2;refGene_gn=FANCA;refGene_tx=XM_0198202

00.2;refGene_gn=FANCA;refGene_tx=XM_019820201.2;refGene_gn=FANCA;refGene_tx=XM_019820203.2
E2 63686138 63686213 refGene_gn=FANCA;refGene_tx=XM_006941836.4
E2 63686400 63686478
refGene_gn=FANCA;refGene_tx=XM_003998369.5;refGene_gn=FANCA;refGene_tx=XM_006941834.3;refGene_gn=FANCA;refGene_tx=XM_006941835.3;refGene_gn=FANCA;refGene_tx=XM_006941836.4;refGene_gn=FANCA;refGene_tx=XM_011290295.3;refGene_gn=FANCA;refGene_tx=XM_019820198.2;refGene_gn=FANCA;refGene_tx=XM_019820200.2;refGene_gn=FANCA;refGene_tx=XM_019820201.2;refGene_gn=FANCA;refGene_tx=XM_019820203.2
E2 63687703 63687842
refGene_gn=FANCA;refGene_tx=XM_003998369.5;refGene_gn=FANCA;refGene_tx=XM_006941834.3;refGene_gn=FANCA;refGene_tx=XM_006941835.3;refGene_gn=FANCA;refGene_tx=XM_006941836.4;refGene_gn=FANCA;refGene_tx=XM_011290295.3;refGene_gn=FANCA;refGene_tx=XM_019820198.2;refGene_gn=FANCA;refGene_tx=XM_019820200.2;refGene_gn=FANCA;refGene_tx=XM_019820201.2;refGene_gn=FANCA;refGene_tx=XM_019820203.2
E2 63688741 63688854
refGene_gn=FANCA;refGene_tx=XM_003998369.5;refGene_gn=FANCA;refGene_tx=XM_006941834.3;refGene_gn=FANCA;refGene_tx=XM_006941835.3;refGene_gn=FANCA;refGene_tx=XM_006941836.4;refGene_gn=FANCA;refGene_tx=XM_011290295.3;refGene_gn=FANCA;refGene_tx=XM_019820198.2;refGene_gn=FANCA;refGene_tx=XM_019820200.2;refGene_gn=FANCA;refGene_tx=XM_019820201.2;refGene_gn=FANCA;refGene_tx=XM_019820203.2
E2 63689752 63689857
refGene_gn=FANCA;refGene_tx=XM_003998369.5;refGene_gn=FANCA;refGene_tx=XM_006941834.3;refGene_gn=FANCA;refGene_tx=XM_006941835.3;refGene_gn=FANCA;refGene_tx=XM_006941836.4;refGene_gn=FANCA;refGene_tx=XM_011290295.3;refGene_gn=FANCA;refGene_tx=XM_019820198.2;refGene_gn=FANCA;refGene_tx=XM_019820200.2;refGene_gn=FANCA;refGene_tx=XM_019820201.2;refGene_gn=FANCA;refGene_tx=XM_019820203.2
E2 63690006 63690063
refGene_gn=FANCA;refGene_tx=XM_003998369.5;refGene_gn=FANCA;refGene_tx=XM_006941834.3;refGene_gn=FANCA;refGene_tx=XM_006941835.3;refGene_gn=FANCA;refGene_tx=XM_006941836.4;refGene_gn=FANCA;refGene_tx=XM_019820198.2;refGene_gn=FANCA;refGene_tx=XM_019820200.2;refGene_gn=FANCA;refGene_tx=XM_019820201.2;refGene_gn=FANCA;refGene_tx=XM_019820203.2
E2 63691025 63691131
refGene_gn=FANCA;refGene_tx=XM_003998369.5;refGene_gn=FANCA;refGene_tx=XM_006941834.3;refGene_gn=FANCA;refGene_tx=XM_006941835.3;refGene_gn=FANCA;refGene_tx=XM_006941836.4;refGene_gn=FANCA;refGene_tx=XM_011290295.3;refGene_gn=FANCA;refGene_tx=XM_019820198.2;refGene_gn=FANCA;refGene_tx=XM_019820200.2;refGene_gn=FANCA;refGene_tx=XM_019820201.2;refGene_gn=FANCA;refGene_tx=XM_019820203.2
E2 63691653 63691820
refGene_gn=FANCA;refGene_tx=XM_003998369.5;refGene_gn=FANCA;refGene_tx=XM_006941834.3;refGene_gn=FANCA;refGene_tx=XM_006941835.3;refGene_gn=FANCA;refGene_tx=XM_006941836.4;refGene_gn=FANCA;refGene_tx=XM_011290295.3;refGene_gn=FANCA;refGene_tx=XM_019820198.2;refGene_gn=FANCA;refGene_tx=XM_019820200.2;refGene_gn=FANCA;refGene_tx=XM_019820201.2;refGen

e_gn=FANCA;refGene_tx=XM_019820203.2
E2 63693584 63693669
refGene_gn=FANCA;refGene_tx=XM_003998369.5;refGene_gn=FANCA;refGene_tx=XM_006941834.3;refGene_gn=FANCA;refGene_tx=XM_006941835.3;refGene_gn=FANCA;refGene_tx=XM_006941836.4;refGene_gn=FANCA;refGene_tx=XM_011290295.3;refGene_gn=FANCA;refGene_tx=XM_019820198.2;refGene_gn=FANCA;refGene_tx=XM_019820200.2;refGene_gn=FANCA;refGene_tx=XM_019820201.2;refGene_gn=FANCA;refGene_tx=XM_019820203.2
E2 63697416 63697545
refGene_gn=FANCA;refGene_tx=XM_003998369.5;refGene_gn=FANCA;refGene_tx=XM_006941834.3;refGene_gn=FANCA;refGene_tx=XM_006941835.3;refGene_gn=FANCA;refGene_tx=XM_006941836.4;refGene_gn=FANCA;refGene_tx=XM_011290295.3;refGene_gn=FANCA;refGene_tx=XM_019820198.2;refGene_gn=FANCA;refGene_tx=XM_019820201.2;refGene_gn=FANCA;refGene_tx=XM_019820203.2
E2 63697848 63697978 refGene_gn=FANCA;refGene_tx=XM_019820204.2
E2 63699104 63699178
refGene_gn=FANCA;refGene_tx=XM_003998369.5;refGene_gn=FANCA;refGene_tx=XM_006941834.3;refGene_gn=FANCA;refGene_tx=XM_006941835.3;refGene_gn=FANCA;refGene_tx=XM_006941836.4;refGene_gn=FANCA;refGene_tx=XM_011290295.3;refGene_gn=FANCA;refGene_tx=XM_019820198.2;refGene_gn=FANCA;refGene_tx=XM_019820200.2;refGene_gn=FANCA;refGene_tx=XM_019820201.2;refGene_gn=FANCA;refGene_tx=XM_019820203.2;refGene_gn=FANCA;refGene_tx=XM_019820204.2
E2 63700595 63700772
refGene_gn=FANCA;refGene_tx=XM_003998369.5;refGene_gn=FANCA;refGene_tx=XM_006941834.3;refGene_gn=FANCA;refGene_tx=XM_006941835.3;refGene_gn=FANCA;refGene_tx=XM_006941836.4;refGene_gn=FANCA;refGene_tx=XM_011290295.3;refGene_gn=FANCA;refGene_tx=XM_019820198.2;refGene_gn=FANCA;refGene_tx=XM_019820200.2;refGene_gn=FANCA;refGene_tx=XM_019820201.2;refGene_gn=FANCA;refGene_tx=XM_019820203.2;refGene_gn=FANCA;refGene_tx=XM_019820204.2
E2 63702198 63702295
refGene_gn=FANCA;refGene_tx=XM_003998369.5;refGene_gn=FANCA;refGene_tx=XM_006941834.3;refGene_gn=FANCA;refGene_tx=XM_006941835.3;refGene_gn=FANCA;refGene_tx=XM_006941836.4;refGene_gn=FANCA;refGene_tx=XM_011290295.3;refGene_gn=FANCA;refGene_tx=XM_019820198.2;refGene_gn=FANCA;refGene_tx=XM_019820200.2;refGene_gn=FANCA;refGene_tx=XM_019820201.2;refGene_gn=FANCA;refGene_tx=XM_019820203.2;refGene_gn=FANCA;refGene_tx=XM_019820204.2
E2 63703764 63703952
refGene_gn=FANCA;refGene_tx=XM_003998369.5;refGene_gn=FANCA;refGene_tx=XM_006941834.3;refGene_gn=FANCA;refGene_tx=XM_006941835.3;refGene_gn=FANCA;refGene_tx=XM_006941836.4;refGene_gn=FANCA;refGene_tx=XM_011290295.3;refGene_gn=FANCA;refGene_tx=XM_019820198.2;refGene_gn=FANCA;refGene_tx=XM_019820200.2;refGene_gn=FANCA;refGene_tx=XM_019820201.2;refGene_gn=FANCA;refGene_tx=XM_019820203.2;refGene_gn=FANCA;refGene_tx=XM_019820204.2
E2 63704032 63704129
refGene_gn=FANCA;refGene_tx=XM_003998369.5;refGene_gn=FANCA;refGene_tx=XM_006941834.3;refGene_gn=FANCA;refGene_tx=XM_006941835.3;refGene_gn=

FANCA;refGene_tx=XM_006941836.4;refGene_gn=FANCA;refGene_tx=XM_0112902
95.3;refGene_gn=FANCA;refGene_tx=XM_019820198.2;refGene_gn=FANCA;refGe
ne_tx=XM_019820200.2;refGene_gn=FANCA;refGene_tx=XM_019820201.2;refGen
e_gn=FANCA;refGene_tx=XM_019820203.2;refGene_gn=FANCA;refGene_tx=XM_01
9820204.2
E2 63704505 63704576
refGene_gn=FANCA;refGene_tx=XM_003998369.5;refGene_gn=FANCA;refGene_tx
=XM_006941834.3;refGene_gn=FANCA;refGene_tx=XM_006941835.3;refGene_gn=
FANCA;refGene_tx=XM_006941836.4;refGene_gn=FANCA;refGene_tx=XM_0112902
95.3;refGene_gn=FANCA;refGene_tx=XM_019820198.2;refGene_gn=FANCA;refGe
ne_tx=XM_019820200.2;refGene_gn=FANCA;refGene_tx=XM_019820201.2;refGen
e_gn=FANCA;refGene_tx=XM_019820203.2;refGene_gn=FANCA;refGene_tx=XM_01
9820204.2
E2 63705030 63705167
refGene_gn=FANCA;refGene_tx=XM_003998369.5;refGene_gn=FANCA;refGene_tx
=XM_006941834.3;refGene_gn=FANCA;refGene_tx=XM_006941835.3;refGene_gn=
FANCA;refGene_tx=XM_006941836.4;refGene_gn=FANCA;refGene_tx=XM_0112902
95.3;refGene_gn=FANCA;refGene_tx=XM_019820198.2;refGene_gn=FANCA;refGe
ne_tx=XM_019820200.2;refGene_gn=FANCA;refGene_tx=XM_019820201.2;refGen
e_gn=FANCA;refGene_tx=XM_019820203.2;refGene_gn=FANCA;refGene_tx=XM_01
9820204.2
E2 63706249 63706363
refGene_gn=FANCA;refGene_tx=XM_003998369.5;refGene_gn=FANCA;refGene_tx
=XM_006941834.3;refGene_gn=FANCA;refGene_tx=XM_006941835.3;refGene_gn=
FANCA;refGene_tx=XM_006941836.4;refGene_gn=FANCA;refGene_tx=XM_0112902
95.3;refGene_gn=FANCA;refGene_tx=XM_019820198.2;refGene_gn=FANCA;refGe
ne_tx=XM_019820200.2;refGene_gn=FANCA;refGene_tx=XM_019820201.2;refGen
e_gn=FANCA;refGene_tx=XM_019820203.2;refGene_gn=FANCA;refGene_tx=XM_01
9820204.2
E2 63708192 63708269
refGene_gn=FANCA;refGene_tx=XM_003998369.5;refGene_gn=FANCA;refGene_tx
=XM_006941834.3;refGene_gn=FANCA;refGene_tx=XM_006941835.3;refGene_gn=
FANCA;refGene_tx=XM_006941836.4;refGene_gn=FANCA;refGene_tx=XM_0112902
95.3;refGene_gn=FANCA;refGene_tx=XM_019820198.2;refGene_gn=FANCA;refGe
ne_tx=XM_019820200.2;refGene_gn=FANCA;refGene_tx=XM_019820201.2;refGen
e_gn=FANCA;refGene_tx=XM_019820203.2;refGene_gn=FANCA;refGene_tx=XM_01
9820204.2
E2 63715690 63715740
refGene_gn=FANCA;refGene_tx=XM_003998369.5;refGene_gn=FANCA;refGene_tx
=XM_006941834.3;refGene_gn=FANCA;refGene_tx=XM_006941835.3;refGene_gn=
FANCA;refGene_tx=XM_006941836.4;refGene_gn=FANCA;refGene_tx=XM_0112902
95.3;refGene_gn=FANCA;refGene_tx=XM_019820198.2;refGene_gn=FANCA;refGe
ne_tx=XM_019820200.2;refGene_gn=FANCA;refGene_tx=XM_019820201.2;refGen
e_gn=FANCA;refGene_tx=XM_019820203.2;refGene_gn=FANCA;refGene_tx=XM_01
9820204.2
E2 63715819 63715880
refGene_gn=FANCA;refGene_tx=XM_003998369.5;refGene_gn=FANCA;refGene_tx
=XM_006941834.3;refGene_gn=FANCA;refGene_tx=XM_006941835.3;refGene_gn=
FANCA;refGene_tx=XM_006941836.4;refGene_gn=FANCA;refGene_tx=XM_0112902
95.3;refGene_gn=FANCA;refGene_tx=XM_019820198.2;refGene_gn=FANCA;refGe

ne_tx=XM_019820200.2;refGene_gn=FANCA;refGene_tx=XM_019820201.2;refGene_gn=FANCA;refGene_tx=XM_019820203.2;refGene_gn=FANCA;refGene_tx=XM_019820204.2
E2 63716577 63716666
refGene_gn=FANCA;refGene_tx=XM_003998369.5;refGene_gn=FANCA;refGene_tx=XM_006941834.3;refGene_gn=FANCA;refGene_tx=XM_006941835.3;refGene_gn=FANCA;refGene_tx=XM_006941836.4;refGene_gn=FANCA;refGene_tx=XM_011290295.3;refGene_gn=FANCA;refGene_tx=XM_019820198.2;refGene_gn=FANCA;refGene_tx=XM_019820200.2;refGene_gn=FANCA;refGene_tx=XM_019820201.2;refGene_gn=FANCA;refGene_tx=XM_019820203.2;refGene_gn=FANCA;refGene_tx=XM_019820204.2
E2 63717518 63717578
refGene_gn=FANCA;refGene_tx=XM_003998369.5;refGene_gn=FANCA;refGene_tx=XM_006941834.3;refGene_gn=FANCA;refGene_tx=XM_006941835.3;refGene_gn=FANCA;refGene_tx=XM_006941836.4;refGene_gn=FANCA;refGene_tx=XM_011290295.3;refGene_gn=FANCA;refGene_tx=XM_019820198.2;refGene_gn=FANCA;refGene_tx=XM_019820200.2;refGene_gn=FANCA;refGene_tx=XM_019820201.2;refGene_gn=FANCA;refGene_tx=XM_019820203.2;refGene_gn=FANCA;refGene_tx=XM_019820204.2
E2 63717684 63717780
refGene_gn=FANCA;refGene_tx=XM_003998369.5;refGene_gn=FANCA;refGene_tx=XM_006941834.3;refGene_gn=FANCA;refGene_tx=XM_006941835.3;refGene_gn=FANCA;refGene_tx=XM_006941836.4;refGene_gn=FANCA;refGene_tx=XM_011290295.3;refGene_gn=FANCA;refGene_tx=XM_019820198.2;refGene_gn=FANCA;refGene_tx=XM_019820200.2;refGene_gn=FANCA;refGene_tx=XM_019820201.2;refGene_gn=FANCA;refGene_tx=XM_019820203.2;refGene_gn=FANCA;refGene_tx=XM_019820204.2
E2 63719671 63719782
refGene_gn=FANCA;refGene_tx=XM_003998369.5;refGene_gn=FANCA;refGene_tx=XM_006941834.3;refGene_gn=FANCA;refGene_tx=XM_006941835.3;refGene_gn=FANCA;refGene_tx=XM_006941836.4;refGene_gn=FANCA;refGene_tx=XM_011290295.3;refGene_gn=FANCA;refGene_tx=XM_019820198.2;refGene_gn=FANCA;refGene_tx=XM_019820200.2;refGene_gn=FANCA;refGene_tx=XM_019820201.2;refGene_gn=FANCA;refGene_tx=XM_019820203.2;refGene_gn=FANCA;refGene_tx=XM_019820204.2
E2 63725736 63725870
refGene_gn=FANCA;refGene_tx=XM_003998369.5;refGene_gn=FANCA;refGene_tx=XM_006941834.3;refGene_gn=FANCA;refGene_tx=XM_006941835.3;refGene_gn=FANCA;refGene_tx=XM_006941836.4;refGene_gn=FANCA;refGene_tx=XM_011290295.3;refGene_gn=FANCA;refGene_tx=XM_019820198.2;refGene_gn=FANCA;refGene_tx=XM_019820200.2;refGene_gn=FANCA;refGene_tx=XM_019820201.2;refGene_gn=FANCA;refGene_tx=XM_019820203.2;refGene_gn=FANCA;refGene_tx=XM_019820204.2
E2 63726207 63726349
refGene_gn=FANCA;refGene_tx=XM_003998369.5;refGene_gn=FANCA;refGene_tx=XM_006941834.3;refGene_gn=FANCA;refGene_tx=XM_006941835.3;refGene_gn=FANCA;refGene_tx=XM_006941836.4;refGene_gn=FANCA;refGene_tx=XM_011290295.3;refGene_gn=FANCA;refGene_tx=XM_019820198.2;refGene_gn=FANCA;refGene_tx=XM_019820200.2;refGene_gn=FANCA;refGene_tx=XM_019820201.2;refGene_gn=FANCA;refGene_tx=XM_019820203.2;refGene_gn=FANCA;refGene_tx=XM_01

9820204.2
E2 63726626 63726703
refGene_gn=FANCA;refGene_tx=XM_003998369.5;refGene_gn=FANCA;refGene_tx=XM_006941834.3;refGene_gn=FANCA;refGene_tx=XM_006941835.3;refGene_gn=FANCA;refGene_tx=XM_006941836.4;refGene_gn=FANCA;refGene_tx=XM_011290295.3;refGene_gn=FANCA;refGene_tx=XM_019820198.2;refGene_gn=FANCA;refGene_tx=XM_019820200.2;refGene_gn=FANCA;refGene_tx=XM_019820201.2;refGene_gn=FANCA;refGene_tx=XM_019820203.2;refGene_gn=FANCA;refGene_tx=XM_019820204.2
E2 63728564 63728677
refGene_gn=FANCA;refGene_tx=XM_003998369.5;refGene_gn=FANCA;refGene_tx=XM_006941834.3;refGene_gn=FANCA;refGene_tx=XM_006941835.3;refGene_gn=FANCA;refGene_tx=XM_006941836.4;refGene_gn=FANCA;refGene_tx=XM_011290295.3;refGene_gn=FANCA;refGene_tx=XM_019820198.2;refGene_gn=FANCA;refGene_tx=XM_019820200.2;refGene_gn=FANCA;refGene_tx=XM_019820201.2;refGene_gn=FANCA;refGene_tx=XM_019820203.2;refGene_gn=FANCA;refGene_tx=XM_019820204.2
E2 63729512 63729513 refGene_gn=FANCA;refGene_tx=XM_019820205.2
E2 63730091 63730095 refGene_gn=FANCA;refGene_tx=XM_006941837.3
E2 63730230 63730297
refGene_gn=FANCA;refGene_tx=XM_003998369.5;refGene_gn=FANCA;refGene_tx=XM_006941834.3;refGene_gn=FANCA;refGene_tx=XM_006941835.3;refGene_gn=FANCA;refGene_tx=XM_006941836.4;refGene_gn=FANCA;refGene_tx=XM_006941837.3;refGene_gn=FANCA;refGene_tx=XM_011290295.3;refGene_gn=FANCA;refGene_tx=XM_019820198.2;refGene_gn=FANCA;refGene_tx=XM_019820200.2;refGene_gn=FANCA;refGene_tx=XM_019820201.2;refGene_gn=FANCA;refGene_tx=XM_019820204.2;refGene_gn=FANCA;refGene_tx=XM_019820205.2
E2 63730723 63730757
refGene_gn=FANCA;refGene_tx=XM_003998369.5;refGene_gn=FANCA;refGene_tx=XM_006941834.3;refGene_gn=FANCA;refGene_tx=XM_006941835.3;refGene_gn=FANCA;refGene_tx=XM_006941836.4;refGene_gn=FANCA;refGene_tx=XM_006941837.3;refGene_gn=FANCA;refGene_tx=XM_011290295.3;refGene_gn=FANCA;refGene_tx=XM_019820198.2;refGene_gn=FANCA;refGene_tx=XM_019820200.2;refGene_gn=FANCA;refGene_tx=XM_019820201.2;refGene_gn=FANCA;refGene_tx=XM_019820204.2;refGene_gn=FANCA;refGene_tx=XM_019820205.2
E2 63731240 63731323
refGene_gn=FANCA;refGene_tx=XM_003998369.5;refGene_gn=FANCA;refGene_tx=XM_006941834.3;refGene_gn=FANCA;refGene_tx=XM_006941835.3;refGene_gn=FANCA;refGene_tx=XM_006941836.4;refGene_gn=FANCA;refGene_tx=XM_006941837.3;refGene_gn=FANCA;refGene_tx=XM_011290295.3;refGene_gn=FANCA;refGene_tx=XM_019820198.2;refGene_gn=FANCA;refGene_tx=XM_019820200.2;refGene_gn=FANCA;refGene_tx=XM_019820201.2;refGene_gn=FANCA;refGene_tx=XM_019820204.2;refGene_gn=FANCA;refGene_tx=XM_019820205.2
E2 63733194 63733304
refGene_gn=FANCA;refGene_tx=XM_003998369.5;refGene_gn=FANCA;refGene_tx=XM_006941834.3;refGene_gn=FANCA;refGene_tx=XM_006941835.3;refGene_gn=FANCA;refGene_tx=XM_006941836.4;refGene_gn=FANCA;refGene_tx=XM_006941837.3;refGene_gn=FANCA;refGene_tx=XM_011290295.3;refGene_gn=FANCA;refGene_tx=XM_019820198.2;refGene_gn=FANCA;refGene_tx=XM_019820200.2;refGene_gn=FANCA;refGene_tx=XM_019820201.2;refGene_gn=FANCA;refGene_tx=XM_01

9820204.2;refGene_gn=FANCA;refGene_tx=XM_019820205.2
E2 63737490 63737564
refGene_gn=FANCA;refGene_tx=XM_003998369.5;refGene_gn=FANCA;refGene_tx=XM_006941834.3;refGene_gn=FANCA;refGene_tx=XM_006941835.3;refGene_gn=FANCA;refGene_tx=XM_006941836.4;refGene_gn=FANCA;refGene_tx=XM_006941837.3;refGene_gn=FANCA;refGene_tx=XM_011290295.3;refGene_gn=FANCA;refGene_tx=XM_019820198.2;refGene_gn=FANCA;refGene_tx=XM_019820200.2;refGene_gn=FANCA;refGene_tx=XM_019820201.2;refGene_gn=FANCA;refGene_tx=XM_019820204.2;refGene_gn=FANCA;refGene_tx=XM_019820205.2
E2 63739915 63740011
refGene_gn=FANCA;refGene_tx=XM_003998369.5;refGene_gn=FANCA;refGene_tx=XM_006941834.3;refGene_gn=FANCA;refGene_tx=XM_006941835.3;refGene_gn=FANCA;refGene_tx=XM_006941836.4;refGene_gn=FANCA;refGene_tx=XM_006941837.3;refGene_gn=FANCA;refGene_tx=XM_011290295.3;refGene_gn=FANCA;refGene_tx=XM_019820198.2;refGene_gn=FANCA;refGene_tx=XM_019820200.2;refGene_gn=FANCA;refGene_tx=XM_019820201.2;refGene_gn=FANCA;refGene_tx=XM_019820204.2;refGene_gn=FANCA;refGene_tx=XM_019820205.2
E2 63740147 63740290
refGene_gn=FANCA;refGene_tx=XM_003998369.5;refGene_gn=FANCA;refGene_tx=XM_006941834.3;refGene_gn=FANCA;refGene_tx=XM_006941835.3;refGene_gn=FANCA;refGene_tx=XM_006941836.4;refGene_gn=FANCA;refGene_tx=XM_006941837.3;refGene_gn=FANCA;refGene_tx=XM_011290295.3;refGene_gn=FANCA;refGene_tx=XM_019820198.2;refGene_gn=FANCA;refGene_tx=XM_019820200.2;refGene_gn=FANCA;refGene_tx=XM_019820201.2;refGene_gn=FANCA;refGene_tx=XM_019820204.2;refGene_gn=FANCA;refGene_tx=XM_019820205.2
E2 63744338 63744423
refGene_gn=FANCA;refGene_tx=XM_003998369.5;refGene_gn=FANCA;refGene_tx=XM_006941834.3;refGene_gn=FANCA;refGene_tx=XM_006941835.3;refGene_gn=FANCA;refGene_tx=XM_006941836.4;refGene_gn=FANCA;refGene_tx=XM_006941837.3;refGene_gn=FANCA;refGene_tx=XM_011290295.3;refGene_gn=FANCA;refGene_tx=XM_019820198.2;refGene_gn=FANCA;refGene_tx=XM_019820200.2;refGene_gn=FANCA;refGene_tx=XM_019820201.2;refGene_gn=FANCA;refGene_tx=XM_019820204.2;refGene_gn=FANCA;refGene_tx=XM_019820205.2
E2 63744778 63744888
refGene_gn=FANCA;refGene_tx=XM_003998369.5;refGene_gn=FANCA;refGene_tx=XM_006941834.3;refGene_gn=FANCA;refGene_tx=XM_006941835.3;refGene_gn=FANCA;refGene_tx=XM_006941836.4;refGene_gn=FANCA;refGene_tx=XM_006941837.3;refGene_gn=FANCA;refGene_tx=XM_011290295.3;refGene_gn=FANCA;refGene_tx=XM_019820198.2;refGene_gn=FANCA;refGene_tx=XM_019820200.2;refGene_gn=FANCA;refGene_tx=XM_019820201.2;refGene_gn=FANCA;refGene_tx=XM_019820204.2;refGene_gn=FANCA;refGene_tx=XM_019820205.2
E2 63745738 63745817
refGene_gn=FANCA;refGene_tx=XM_003998369.5;refGene_gn=FANCA;refGene_tx=XM_006941834.3;refGene_gn=FANCA;refGene_tx=XM_006941835.3;refGene_gn=FANCA;refGene_tx=XM_006941836.4;refGene_gn=FANCA;refGene_tx=XM_006941837.3;refGene_gn=FANCA;refGene_tx=XM_011290295.3;refGene_gn=FANCA;refGene_tx=XM_019820198.2;refGene_gn=FANCA;refGene_tx=XM_019820200.2;refGene_gn=FANCA;refGene_tx=XM_019820201.2;refGene_gn=FANCA;refGene_tx=XM_019820204.2;refGene_gn=FANCA;refGene_tx=XM_019820205.2
E2 63762263 63762576

refGene_gn=SPIRE2;refGene_tx=XM_023245138.1;refGene_gn=SPIRE2;refGene_tx=XM_023245139.1
E2 63775105 63775149
refGene_gn=SPIRE2;refGene_tx=XM_023245138.1;refGene_gn=SPIRE2;refGene_tx=XM_023245139.1;refGene_gn=SPIRE2;refGene_tx=XM_023245140.1
E2 63780633 63780990
refGene_gn=SPIRE2;refGene_tx=XM_023245138.1;refGene_gn=SPIRE2;refGene_tx=XM_023245139.1;refGene_gn=SPIRE2;refGene_tx=XM_023245140.1
E2 63784765 63784846
refGene_gn=SPIRE2;refGene_tx=XM_023245138.1;refGene_gn=SPIRE2;refGene_tx=XM_023245139.1;refGene_gn=SPIRE2;refGene_tx=XM_023245140.1
E2 63784973 63785138
refGene_gn=SPIRE2;refGene_tx=XM_023245138.1;refGene_gn=SPIRE2;refGene_tx=XM_023245139.1;refGene_gn=SPIRE2;refGene_tx=XM_023245140.1
E2 63785878 63785965
refGene_gn=SPIRE2;refGene_tx=XM_023245138.1;refGene_gn=SPIRE2;refGene_tx=XM_023245139.1;refGene_gn=SPIRE2;refGene_tx=XM_023245140.1
E2 63786206 63786330
refGene_gn=SPIRE2;refGene_tx=XM_023245138.1;refGene_gn=SPIRE2;refGene_tx=XM_023245139.1;refGene_gn=SPIRE2;refGene_tx=XM_023245140.1
E2 63786417 63786587
refGene_gn=SPIRE2;refGene_tx=XM_023245138.1;refGene_gn=SPIRE2;refGene_tx=XM_023245139.1;refGene_gn=SPIRE2;refGene_tx=XM_023245140.1
E2 63787200 63787390
refGene_gn=SPIRE2;refGene_tx=XM_023245138.1;refGene_gn=SPIRE2;refGene_tx=XM_023245139.1;refGene_gn=SPIRE2;refGene_tx=XM_023245140.1
E2 63788540 63788662
refGene_gn=SPIRE2;refGene_tx=XM_023245138.1;refGene_gn=SPIRE2;refGene_tx=XM_023245139.1;refGene_gn=SPIRE2;refGene_tx=XM_023245140.1
E2 63791899 63792034
refGene_gn=SPIRE2;refGene_tx=XM_023245138.1;refGene_gn=SPIRE2;refGene_tx=XM_023245139.1;refGene_gn=SPIRE2;refGene_tx=XM_023245140.1
E2 63792163 63792231
refGene_gn=SPIRE2;refGene_tx=XM_023245138.1;refGene_gn=SPIRE2;refGene_tx=XM_023245139.1;refGene_gn=SPIRE2;refGene_tx=XM_023245140.1
E2 63795342 63795370
refGene_gn=SPIRE2;refGene_tx=XM_023245138.1;refGene_gn=SPIRE2;refGene_tx=XM_023245139.1;refGene_gn=SPIRE2;refGene_tx=XM_023245140.1
E2 63796436 63796577
refGene_gn=SPIRE2;refGene_tx=XM_023245138.1;refGene_gn=SPIRE2;refGene_tx=XM_023245139.1;refGene_gn=SPIRE2;refGene_tx=XM_023245140.1
E2 63796881 63797104
refGene_gn=SPIRE2;refGene_tx=XM_023245138.1;refGene_gn=SPIRE2;refGene_tx=XM_023245139.1;refGene_gn=SPIRE2;refGene_tx=XM_023245140.1
E2 63801186 63801378
refGene_gn=TCF25;refGene_tx=XM_003998371.5;refGene_gn=TCF25;refGene_tx=XM_006941839.4
E2 63808017 63808167
refGene_gn=TCF25;refGene_tx=XM_003998371.5;refGene_gn=TCF25;refGene_tx=XM_006941839.4

E2 63809145 63809220
refGene_gn=TCF25;refGene_tx=XM_003998371.5;refGene_gn=TCF25;refGene_tx=XM_006941839.4
E2 63809900 63810019
refGene_gn=TCF25;refGene_tx=XM_003998371.5;refGene_gn=TCF25;refGene_tx=XM_006941839.4
E2 63810591 63810657
refGene_gn=TCF25;refGene_tx=XM_003998371.5;refGene_gn=TCF25;refGene_tx=XM_006941839.4
E2 63814098 63814181
refGene_gn=TCF25;refGene_tx=XM_003998371.5;refGene_gn=TCF25;refGene_tx=XM_006941839.4
E2 63815255 63815386
refGene_gn=TCF25;refGene_tx=XM_003998371.5;refGene_gn=TCF25;refGene_tx=XM_006941839.4
E2 63815826 63815926
refGene_gn=TCF25;refGene_tx=XM_003998371.5;refGene_gn=TCF25;refGene_tx=XM_006941839.4
E2 63816715 63816809
refGene_gn=TCF25;refGene_tx=XM_003998371.5;refGene_gn=TCF25;refGene_tx=XM_006941839.4
E2 63817964 63818057
refGene_gn=TCF25;refGene_tx=XM_003998371.5;refGene_gn=TCF25;refGene_tx=XM_006941839.4
E2 63818179 63818285
refGene_gn=TCF25;refGene_tx=XM_003998371.5;refGene_gn=TCF25;refGene_tx=XM_006941839.4
E2 63819405 63819565
refGene_gn=TCF25;refGene_tx=XM_003998371.5;refGene_gn=TCF25;refGene_tx=XM_006941839.4
E2 63821026 63821114
refGene_gn=TCF25;refGene_tx=XM_003998371.5;refGene_gn=TCF25;refGene_tx=XM_006941839.4
E2 63822075 63822234
refGene_gn=TCF25;refGene_tx=XM_003998371.5;refGene_gn=TCF25;refGene_tx=XM_006941839.4
E2 63822385 63822476
refGene_gn=TCF25;refGene_tx=XM_003998371.5;refGene_gn=TCF25;refGene_tx=XM_006941839.4
E2 63822926 63823006
refGene_gn=TCF25;refGene_tx=XM_003998371.5;refGene_gn=TCF25;refGene_tx=XM_006941839.4
E2 63824558 63824631
refGene_gn=TCF25;refGene_tx=XM_003998371.5;refGene_gn=TCF25;refGene_tx=XM_006941839.4
E2 63824940 63825099 refGene_gn=TCF25;refGene_tx=XM_003998371.5
E2 63826372 63826414 refGene_gn=TCF25;refGene_tx=XM_006941839.4
E2 63829684 63830638 refGene_gn=MC1R;refGene_tx=NM_001009324.1
E2 63854887 63854946
refGene_gn=DEF8;refGene_tx=XM_011290297.2;refGene_gn=DEF8;refGene_tx=X

M_023245032.1
E2	63855535 63855669
refGene_gn=DEF8;refGene_tx=XM_006941841.4;refGene_gn=DEF8;refGene_tx=XM_011290297.2;refGene_gn=DEF8;refGene_tx=XM_011290299.1;refGene_gn=DEF8;refGene_tx=XM_023245032.1
E2	63856866 63856964
refGene_gn=DEF8;refGene_tx=XM_006941841.4;refGene_gn=DEF8;refGene_tx=XM_011290297.2;refGene_gn=DEF8;refGene_tx=XM_011290299.1;refGene_gn=DEF8;refGene_tx=XM_023245032.1
E2	63859450 63859600
refGene_gn=DEF8;refGene_tx=XM_006941841.4;refGene_gn=DEF8;refGene_tx=XM_011290297.2;refGene_gn=DEF8;refGene_tx=XM_011290299.1;refGene_gn=DEF8;refGene_tx=XM_023245032.1
E2	63860497 63860639
refGene_gn=DEF8;refGene_tx=XM_006941841.4;refGene_gn=DEF8;refGene_tx=XM_011290297.2;refGene_gn=DEF8;refGene_tx=XM_011290299.1;refGene_gn=DEF8;refGene_tx=XM_023245032.1
E2	63861942 63862107
refGene_gn=DEF8;refGene_tx=XM_006941841.4;refGene_gn=DEF8;refGene_tx=XM_011290297.2;refGene_gn=DEF8;refGene_tx=XM_011290299.1;refGene_gn=DEF8;refGene_tx=XM_023245032.1
E2	63862643 63862771
refGene_gn=DEF8;refGene_tx=XM_006941841.4;refGene_gn=DEF8;refGene_tx=XM_011290297.2;refGene_gn=DEF8;refGene_tx=XM_011290299.1;refGene_gn=DEF8;refGene_tx=XM_023245032.1
E2	63862955 63863069
refGene_gn=DEF8;refGene_tx=XM_006941841.4;refGene_gn=DEF8;refGene_tx=XM_011290297.2;refGene_gn=DEF8;refGene_tx=XM_011290299.1;refGene_gn=DEF8;refGene_tx=XM_023245032.1
E2	63863614 63863695
refGene_gn=DEF8;refGene_tx=XM_006941841.4;refGene_gn=DEF8;refGene_tx=XM_011290297.2;refGene_gn=DEF8;refGene_tx=XM_011290299.1;refGene_gn=DEF8;refGene_tx=XM_023245032.1
E2	63864490 63864631
refGene_gn=DEF8;refGene_tx=XM_006941841.4;refGene_gn=DEF8;refGene_tx=XM_011290297.2;refGene_gn=DEF8;refGene_tx=XM_011290299.1;refGene_gn=DEF8;refGene_tx=XM_023245032.1
E2	63864772 63864882
refGene_gn=DEF8;refGene_tx=XM_006941841.4;refGene_gn=DEF8;refGene_tx=XM_011290297.2;refGene_gn=DEF8;refGene_tx=XM_011290299.1;refGene_gn=DEF8;refGene_tx=XM_023245032.1
E2	63865948 63866051
refGene_gn=DEF8;refGene_tx=XM_006941841.4;refGene_gn=DEF8;refGene_tx=XM_011290297.2;refGene_gn=DEF8;refGene_tx=XM_011290299.1;refGene_gn=DEF8;refGene_tx=XM_023245032.1
E2	63873873 63875499 refGene_gn=CENPBD1;refGene_tx=XM_003998379.4
E2	63888012 63888113
refGene_gn=LOC101083987;refGene_tx=XM_019820216.2
E2	63902134 63902225
refGene_gn=LOC101083987;refGene_tx=XM_019820216.2

E2 63902449 63902599
refGene_gn=LOC101083987;refGene_tx=XM_019820216.2
E2 63903675 63903783
refGene_gn=LOC101083987;refGene_tx=XM_019820216.2
E2 63904583 63904769
refGene_gn=LOC101083987;refGene_tx=XM_019820216.2
E2 63908346 63908504
refGene_gn=DBNDD1;refGene_tx=XM_023245570.1;refGene_gn=DBNDD1;refGene_tx=XM_023245571.1;refGene_gn=DBNDD1;refGene_tx=XM_023245572.1
E2 63909436 63909577
refGene_gn=DBNDD1;refGene_tx=XM_023245570.1;refGene_gn=DBNDD1;refGene_tx=XM_023245571.1;refGene_gn=DBNDD1;refGene_tx=XM_023245572.1
E2 63909936 63910104
refGene_gn=DBNDD1;refGene_tx=XM_023245570.1;refGene_gn=DBNDD1;refGene_tx=XM_023245571.1;refGene_gn=DBNDD1;refGene_tx=XM_023245572.1
E2 63916857 63916888
refGene_gn=DBNDD1;refGene_tx=XM_023245570.1;refGene_gn=DBNDD1;refGene_tx=XM_023245571.1
E2 63921633 63921906
refGene_gn=GAS8;refGene_tx=XM_003998382.5;refGene_gn=GAS8;refGene_tx=XM_019820213.2;refGene_gn=GAS8;refGene_tx=XM_023245569.1
E2 63924251 63924338
refGene_gn=GAS8;refGene_tx=XM_003998382.5;refGene_gn=GAS8;refGene_tx=XM_019820213.2;refGene_gn=GAS8;refGene_tx=XM_023245569.1
E2 63927595 63927793
refGene_gn=GAS8;refGene_tx=XM_003998382.5;refGene_gn=GAS8;refGene_tx=XM_019820213.2;refGene_gn=GAS8;refGene_tx=XM_023245569.1
E2 63928756 63928963
refGene_gn=GAS8;refGene_tx=XM_003998382.5;refGene_gn=GAS8;refGene_tx=XM_019820213.2;refGene_gn=GAS8;refGene_tx=XM_023245569.1
E2 63930826 63930881
refGene_gn=GAS8;refGene_tx=XM_003998382.5;refGene_gn=GAS8;refGene_tx=XM_019820213.2;refGene_gn=GAS8;refGene_tx=XM_023245569.1
E2 63931752 63931958
refGene_gn=GAS8;refGene_tx=XM_003998382.5;refGene_gn=GAS8;refGene_tx=XM_019820213.2;refGene_gn=GAS8;refGene_tx=XM_019820215.1;refGene_gn=GAS8;refGene_tx=XM_023245569.1
E2 63932496 63932664
refGene_gn=GAS8;refGene_tx=XM_003998382.5;refGene_gn=GAS8;refGene_tx=XM_019820213.2;refGene_gn=GAS8;refGene_tx=XM_019820215.1;refGene_gn=GAS8;refGene_tx=XM_023245569.1
E2 63933126 63933213
refGene_gn=GAS8;refGene_tx=XM_003998382.5;refGene_gn=GAS8;refGene_tx=XM_019820213.2;refGene_gn=GAS8;refGene_tx=XM_019820215.1;refGene_gn=GAS8;refGene_tx=XM_023245569.1
E2 63936140 63936350
refGene_gn=GAS8;refGene_tx=XM_003998382.5;refGene_gn=GAS8;refGene_tx=XM_019820213.2;refGene_gn=GAS8;refGene_tx=XM_019820215.1;refGene_gn=GAS8;refGene_tx=XM_023245569.1
E2 63939063 63939129

refGene_gn=GAS8;refGene_tx=XM_003998382.5;refGene_gn=GAS8;refGene_tx=XM_019820213.2;refGene_gn=GAS8;refGene_tx=XM_019820215.1
E2 63940757 63940907
refGene_gn=GAS8;refGene_tx=XM_003998382.5;refGene_gn=GAS8;refGene_tx=XM_019820213.2;refGene_gn=GAS8;refGene_tx=XM_019820215.1;refGene_gn=GAS8;refGene_tx=XM_023245569.1
E2 63941328 63941418
refGene_gn=LOC102899897;refGene_tx=XM_019820217.1
E2 63941873 63942009
refGene_gn=LOC102899897;refGene_tx=XM_019820217.1
E2 63943246 63943386
refGene_gn=LOC102899897;refGene_tx=XM_019820217.1
E2 63943832 63943892
refGene_gn=LOC102899897;refGene_tx=XM_019820217.1
E2 63971587 63973143
refGene_gn=PRDM7;refGene_tx=XM_019820207.2;refGene_gn=PRDM7;refGene_tx=XM_019820210.2;refGene_gn=PRDM7;refGene_tx=XM_023245365.1;refGene_gn=PRDM7;refGene_tx=XM_023245366.1
E2 63975044 63975238
refGene_gn=PRDM7;refGene_tx=XM_019820207.2;refGene_gn=PRDM7;refGene_tx=XM_019820210.2;refGene_gn=PRDM7;refGene_tx=XM_023245365.1;refGene_gn=PRDM7;refGene_tx=XM_023245366.1
E2 63976426 63976494
refGene_gn=PRDM7;refGene_tx=XM_019820207.2;refGene_gn=PRDM7;refGene_tx=XM_019820210.2;refGene_gn=PRDM7;refGene_tx=XM_023245365.1;refGene_gn=PRDM7;refGene_tx=XM_023245366.1
E2 63976913 63977185
refGene_gn=PRDM7;refGene_tx=XM_019820207.2;refGene_gn=PRDM7;refGene_tx=XM_019820210.2;refGene_gn=PRDM7;refGene_tx=XM_023245365.1;refGene_gn=PRDM7;refGene_tx=XM_023245366.1
E2 63977399 63977501
refGene_gn=PRDM7;refGene_tx=XM_019820207.2;refGene_gn=PRDM7;refGene_tx=XM_019820210.2;refGene_gn=PRDM7;refGene_tx=XM_023245365.1;refGene_gn=PRDM7;refGene_tx=XM_023245366.1
E2 63978136 63978293
refGene_gn=PRDM7;refGene_tx=XM_019820207.2;refGene_gn=PRDM7;refGene_tx=XM_019820210.2;refGene_gn=PRDM7;refGene_tx=XM_023245365.1;refGene_gn=PRDM7;refGene_tx=XM_023245366.1
E2 63980986 63981036
refGene_gn=PRDM7;refGene_tx=XM_019820207.2;refGene_gn=PRDM7;refGene_tx=XM_019820210.2;refGene_gn=PRDM7;refGene_tx=XM_023245365.1;refGene_gn=PRDM7;refGene_tx=XM_023245366.1
E2 63986165 63986273
refGene_gn=PRDM7;refGene_tx=XM_019820207.2;refGene_gn=PRDM7;refGene_tx=XM_019820210.2;refGene_gn=PRDM7;refGene_tx=XM_023245365.1;refGene_gn=PRDM7;refGene_tx=XM_023245366.1
E2 63986591 63986715
refGene_gn=PRDM7;refGene_tx=XM_019820207.2;refGene_gn=PRDM7;refGene_tx=XM_019820210.2;refGene_gn=PRDM7;refGene_tx=XM_023245365.1;refGene_gn=PRDM7;refGene_tx=XM_023245366.1

E2 63987121 63987247
refGene_gn=PRDM7;refGene_tx=XM_019820207.2;refGene_gn=PRDM7;refGene_tx=XM_019820210.2;refGene_gn=PRDM7;refGene_tx=XM_023245365.1;refGene_gn=PRDM7;refGene_tx=XM_023245366.1
E3 398190 398798 refGene_gn=FAM20C;refGene_tx=XM_023247122.1
E3 400657 400836 refGene_gn=FAM20C;refGene_tx=XM_023247122.1
E3 411535 411614 refGene_gn=FAM20C;refGene_tx=XM_023247122.1
E3 433528 433621 refGene_gn=FAM20C;refGene_tx=XM_023247122.1
E3 434974 435090 refGene_gn=FAM20C;refGene_tx=XM_023247122.1
E3 441616 441797 refGene_gn=FAM20C;refGene_tx=XM_023247122.1
E3 442397 442507 refGene_gn=FAM20C;refGene_tx=XM_023247122.1
E3 442663 442745 refGene_gn=FAM20C;refGene_tx=XM_023247122.1
E3 443550 443610 refGene_gn=FAM20C;refGene_tx=XM_023247122.1
E3 444183 444433 refGene_gn=FAM20C;refGene_tx=XM_023247122.1
E3 460683 460790
refGene_gn=LOC101087030;refGene_tx=XM_003998391.4
E3 461177 461346
refGene_gn=LOC101087030;refGene_tx=XM_003998391.4
E3 461629 461818
refGene_gn=LOC101087030;refGene_tx=XM_003998391.4
E3 601632 601688
refGene_gn=PDGFA;refGene_tx=XM_023246417.1;refGene_gn=PDGFA;refGene_tx=XM_023246418.1
E3 602092 602219
refGene_gn=PDGFA;refGene_tx=XM_023246417.1;refGene_gn=PDGFA;refGene_tx=XM_023246418.1
E3 608749 608937
refGene_gn=PDGFA;refGene_tx=XM_023246417.1;refGene_gn=PDGFA;refGene_tx=XM_023246418.1
E3 611133 611238
refGene_gn=PDGFA;refGene_tx=XM_023246417.1;refGene_gn=PDGFA;refGene_tx=XM_023246418.1
E3 615890 615987
refGene_gn=PDGFA;refGene_tx=XM_023246417.1;refGene_gn=PDGFA;refGene_tx=XM_023246418.1
E3 616458 616503 refGene_gn=PDGFA;refGene_tx=XM_023246417.1
E3 617439 617502 refGene_gn=PDGFA;refGene_tx=XM_023246418.1
E3 642131 642304
refGene_gn=PRKAR1B;refGene_tx=XM_019820308.2;refGene_gn=PRKAR1B;refGene_tx=XM_023246635.1
E3 642643 642725
refGene_gn=PRKAR1B;refGene_tx=XM_019820308.2;refGene_gn=PRKAR1B;refGene_tx=XM_023246635.1
E3 654302 654424
refGene_gn=PRKAR1B;refGene_tx=XM_019820308.2;refGene_gn=PRKAR1B;refGene_tx=XM_023246635.1
E3 659588 659649
refGene_gn=PRKAR1B;refGene_tx=XM_019820308.2;refGene_gn=PRKAR1B;refGene_tx=XM_023246635.1
E3 665377 665536

refGene_gn=PRKAR1B;refGene_tx=XM_019820308.2;refGene_gn=PRKAR1B;refGene_tx=XM_023246635.1
E3 674057 674104
refGene_gn=PRKAR1B;refGene_tx=XM_019820308.2;refGene_gn=PRKAR1B;refGene_tx=XM_023246635.1
E3 674501 674563
refGene_gn=PRKAR1B;refGene_tx=XM_019820308.2;refGene_gn=PRKAR1B;refGene_tx=XM_023246635.1
E3 729735 729827
refGene_gn=PRKAR1B;refGene_tx=XM_019820308.2;refGene_gn=PRKAR1B;refGene_tx=XM_023246635.1
E3 732532 732691
refGene_gn=PRKAR1B;refGene_tx=XM_019820308.2;refGene_gn=PRKAR1B;refGene_tx=XM_023246635.1
E3 749036 749216
refGene_gn=PRKAR1B;refGene_tx=XM_019820308.2;refGene_gn=PRKAR1B;refGene_tx=XM_023246635.1
E3 763340 763968
refGene_gn=DNAAF5;refGene_tx=XM_023246633.1;refGene_gn=DNAAF5;refGene_tx=XM_023246634.1
E3 766178 766363
refGene_gn=DNAAF5;refGene_tx=XM_023246633.1;refGene_gn=DNAAF5;refGene_tx=XM_023246634.1
E3 774215 774340
refGene_gn=DNAAF5;refGene_tx=XM_023246633.1;refGene_gn=DNAAF5;refGene_tx=XM_023246634.1
E3 774722 774841
refGene_gn=DNAAF5;refGene_tx=XM_023246633.1;refGene_gn=DNAAF5;refGene_tx=XM_023246634.1
E3 786802 787035
refGene_gn=DNAAF5;refGene_tx=XM_023246633.1;refGene_gn=DNAAF5;refGene_tx=XM_023246634.1
E3 788014 788227
refGene_gn=DNAAF5;refGene_tx=XM_023246633.1;refGene_gn=DNAAF5;refGene_tx=XM_023246634.1
E3 793379 793523 refGene_gn=DNAAF5;refGene_tx=XM_023246633.1
E3 794789 794958
refGene_gn=DNAAF5;refGene_tx=XM_023246633.1;refGene_gn=DNAAF5;refGene_tx=XM_023246634.1
E3 796927 797075
refGene_gn=DNAAF5;refGene_tx=XM_023246633.1;refGene_gn=DNAAF5;refGene_tx=XM_023246634.1
E3 799404 799555
refGene_gn=DNAAF5;refGene_tx=XM_023246633.1;refGene_gn=DNAAF5;refGene_tx=XM_023246634.1
E3 801225 801382
refGene_gn=DNAAF5;refGene_tx=XM_023246633.1;refGene_gn=DNAAF5;refGene_tx=XM_023246634.1
E3 803765 803960
refGene_gn=DNAAF5;refGene_tx=XM_023246633.1;refGene_gn=DNAAF5;refGene_

tx=XM_023246634.1
E3 811694 811831
refGene_gn=DNAAF5;refGene_tx=XM_023246633.1;refGene_gn=DNAAF5;refGene_tx=XM_023246634.1
E3 845628 845705
refGene_gn=SUN1;refGene_tx=XM_011290457.2;refGene_gn=SUN1;refGene_tx=XM_011290458.2;refGene_gn=SUN1;refGene_tx=XM_019820309.1;refGene_gn=SUN1;refGene_tx=XM_019820310.1;refGene_gn=SUN1;refGene_tx=XM_019820312.1;refGene_gn=SUN1;refGene_tx=XM_019820313.1;refGene_gn=SUN1;refGene_tx=XM_019820314.1;refGene_gn=SUN1;refGene_tx=XM_019820315.1;refGene_gn=SUN1;refGene_tx=XM_019820316.2;refGene_gn=SUN1;refGene_tx=XM_023246231.1
E3 852081 852270
refGene_gn=SUN1;refGene_tx=XM_011290457.2;refGene_gn=SUN1;refGene_tx=XM_011290458.2;refGene_gn=SUN1;refGene_tx=XM_019820309.1;refGene_gn=SUN1;refGene_tx=XM_019820310.1;refGene_gn=SUN1;refGene_tx=XM_019820312.1;refGene_gn=SUN1;refGene_tx=XM_019820313.1;refGene_gn=SUN1;refGene_tx=XM_019820314.1;refGene_gn=SUN1;refGene_tx=XM_019820315.1;refGene_gn=SUN1;refGene_tx=XM_019820316.2;refGene_gn=SUN1;refGene_tx=XM_023246230.1;refGene_gn=SUN1;refGene_tx=XM_023246231.1
E3 853691 853876
refGene_gn=SUN1;refGene_tx=XM_011290457.2;refGene_gn=SUN1;refGene_tx=XM_011290458.2;refGene_gn=SUN1;refGene_tx=XM_019820309.1;refGene_gn=SUN1;refGene_tx=XM_019820310.1;refGene_gn=SUN1;refGene_tx=XM_019820312.1;refGene_gn=SUN1;refGene_tx=XM_019820313.1;refGene_gn=SUN1;refGene_tx=XM_019820314.1;refGene_gn=SUN1;refGene_tx=XM_019820315.1;refGene_gn=SUN1;refGene_tx=XM_019820316.2;refGene_gn=SUN1;refGene_tx=XM_023246230.1;refGene_gn=SUN1;refGene_tx=XM_023246231.1
E3 854863 854890
refGene_gn=SUN1;refGene_tx=XM_011290457.2;refGene_gn=SUN1;refGene_tx=XM_011290458.2;refGene_gn=SUN1;refGene_tx=XM_019820309.1;refGene_gn=SUN1;refGene_tx=XM_019820310.1;refGene_gn=SUN1;refGene_tx=XM_019820312.1;refGene_gn=SUN1;refGene_tx=XM_019820313.1;refGene_gn=SUN1;refGene_tx=XM_019820314.1;refGene_gn=SUN1;refGene_tx=XM_019820315.1;refGene_gn=SUN1;refGene_tx=XM_019820316.2;refGene_gn=SUN1;refGene_tx=XM_023246230.1;refGene_gn=SUN1;refGene_tx=XM_023246231.1
E3 854996 855194
refGene_gn=SUN1;refGene_tx=XM_011290457.2;refGene_gn=SUN1;refGene_tx=XM_011290458.2;refGene_gn=SUN1;refGene_tx=XM_019820309.1;refGene_gn=SUN1;refGene_tx=XM_019820310.1;refGene_gn=SUN1;refGene_tx=XM_019820312.1;refGene_gn=SUN1;refGene_tx=XM_019820313.1;refGene_gn=SUN1;refGene_tx=XM_019820314.1;refGene_gn=SUN1;refGene_tx=XM_019820315.1;refGene_gn=SUN1;refGene_tx=XM_019820316.2;refGene_gn=SUN1;refGene_tx=XM_023246230.1;refGene_gn=SUN1;refGene_tx=XM_023246231.1
E3 860727 860925
refGene_gn=SUN1;refGene_tx=XM_019820309.1;refGene_gn=SUN1;refGene_tx=XM_019820310.1;refGene_gn=SUN1;refGene_tx=XM_019820312.1;refGene_gn=SUN1;refGene_tx=XM_019820314.1;refGene_gn=SUN1;refGene_tx=XM_019820316.2;refGene_gn=SUN1;refGene_tx=XM_023246230.1;refGene_gn=SUN1;refGene_tx=XM_023246231.1
E3 861481 861565

refGene_gn=SUN1;refGene_tx=XM_019820309.1;refGene_gn=SUN1;refGene_tx=X
M_019820310.1;refGene_gn=SUN1;refGene_tx=XM_019820312.1;refGene_gn=SUN
1;refGene_tx=XM_019820313.1;refGene_gn=SUN1;refGene_tx=XM_019820315.1;
refGene_gn=SUN1;refGene_tx=XM_019820316.2;refGene_gn=SUN1;refGene_tx=X
M_023246230.1;refGene_gn=SUN1;refGene_tx=XM_023246231.1
E3 861873 861984
refGene_gn=SUN1;refGene_tx=XM_019820309.1;refGene_gn=SUN1;refGene_tx=X
M_019820313.1;refGene_gn=SUN1;refGene_tx=XM_019820316.2;refGene_gn=SUN
1;refGene_tx=XM_023246230.1;refGene_gn=SUN1;refGene_tx=XM_023246231.1
E3 863049 863148
refGene_gn=SUN1;refGene_tx=XM_011290457.2;refGene_gn=SUN1;refGene_tx=X
M_019820309.1;refGene_gn=SUN1;refGene_tx=XM_019820310.1;refGene_gn=SUN
1;refGene_tx=XM_019820313.1;refGene_gn=SUN1;refGene_tx=XM_019820316.2;
refGene_gn=SUN1;refGene_tx=XM_023246230.1;refGene_gn=SUN1;refGene_tx=X
M_023246231.1
E3 863697 863791
refGene_gn=SUN1;refGene_tx=XM_011290457.2;refGene_gn=SUN1;refGene_tx=X
M_011290458.2;refGene_gn=SUN1;refGene_tx=XM_019820309.1;refGene_gn=SUN
1;refGene_tx=XM_019820310.1;refGene_gn=SUN1;refGene_tx=XM_019820312.1;
refGene_gn=SUN1;refGene_tx=XM_019820313.1;refGene_gn=SUN1;refGene_tx=X
M_019820314.1;refGene_gn=SUN1;refGene_tx=XM_019820315.1;refGene_gn=SUN
1;refGene_tx=XM_019820316.2;refGene_gn=SUN1;refGene_tx=XM_023246230.1;
refGene_gn=SUN1;refGene_tx=XM_023246231.1
E3 865496 865555
refGene_gn=SUN1;refGene_tx=XM_011290457.2;refGene_gn=SUN1;refGene_tx=X
M_011290458.2;refGene_gn=SUN1;refGene_tx=XM_019820309.1;refGene_gn=SUN
1;refGene_tx=XM_019820310.1;refGene_gn=SUN1;refGene_tx=XM_019820312.1;
refGene_gn=SUN1;refGene_tx=XM_019820313.1;refGene_gn=SUN1;refGene_tx=X
M_019820314.1;refGene_gn=SUN1;refGene_tx=XM_019820315.1;refGene_gn=SUN
1;refGene_tx=XM_019820316.2;refGene_gn=SUN1;refGene_tx=XM_023246230.1;
refGene_gn=SUN1;refGene_tx=XM_023246231.1
E3 865699 865845
refGene_gn=SUN1;refGene_tx=XM_011290457.2;refGene_gn=SUN1;refGene_tx=X
M_011290458.2;refGene_gn=SUN1;refGene_tx=XM_019820309.1;refGene_gn=SUN
1;refGene_tx=XM_019820310.1;refGene_gn=SUN1;refGene_tx=XM_019820312.1;
refGene_gn=SUN1;refGene_tx=XM_019820313.1;refGene_gn=SUN1;refGene_tx=X
M_019820314.1;refGene_gn=SUN1;refGene_tx=XM_019820315.1;refGene_gn=SUN
1;refGene_tx=XM_019820316.2;refGene_gn=SUN1;refGene_tx=XM_023246230.1;
refGene_gn=SUN1;refGene_tx=XM_023246231.1
E3 866314 866491
refGene_gn=SUN1;refGene_tx=XM_011290457.2;refGene_gn=SUN1;refGene_tx=X
M_011290458.2;refGene_gn=SUN1;refGene_tx=XM_019820309.1;refGene_gn=SUN
1;refGene_tx=XM_019820310.1;refGene_gn=SUN1;refGene_tx=XM_019820312.1;
refGene_gn=SUN1;refGene_tx=XM_019820313.1;refGene_gn=SUN1;refGene_tx=X
M_019820314.1;refGene_gn=SUN1;refGene_tx=XM_019820315.1;refGene_gn=SUN
1;refGene_tx=XM_019820316.2;refGene_gn=SUN1;refGene_tx=XM_023246230.1;
refGene_gn=SUN1;refGene_tx=XM_023246231.1
E3 869271 869358
refGene_gn=SUN1;refGene_tx=XM_011290457.2;refGene_gn=SUN1;refGene_tx=X
M_011290458.2;refGene_gn=SUN1;refGene_tx=XM_019820309.1;refGene_gn=SUN

1;refGene_tx=XM_019820310.1;refGene_gn=SUN1;refGene_tx=XM_019820312.1;
refGene_gn=SUN1;refGene_tx=XM_019820313.1;refGene_gn=SUN1;refGene_tx=X
M_019820314.1;refGene_gn=SUN1;refGene_tx=XM_019820315.1;refGene_gn=SUN
1;refGene_tx=XM_019820316.2;refGene_gn=SUN1;refGene_tx=XM_023246230.1;
refGene_gn=SUN1;refGene_tx=XM_023246231.1
E3 870495 870539
refGene_gn=SUN1;refGene_tx=XM_011290457.2;refGene_gn=SUN1;refGene_tx=X
M_011290458.2;refGene_gn=SUN1;refGene_tx=XM_019820309.1;refGene_gn=SUN
1;refGene_tx=XM_019820310.1;refGene_gn=SUN1;refGene_tx=XM_019820312.1;
refGene_gn=SUN1;refGene_tx=XM_019820313.1;refGene_gn=SUN1;refGene_tx=X
M_019820314.1;refGene_gn=SUN1;refGene_tx=XM_019820315.1;refGene_gn=SUN
1;refGene_tx=XM_019820316.2;refGene_gn=SUN1;refGene_tx=XM_023246230.1;
refGene_gn=SUN1;refGene_tx=XM_023246231.1
E3 872179 872315
refGene_gn=SUN1;refGene_tx=XM_011290457.2;refGene_gn=SUN1;refGene_tx=X
M_011290458.2;refGene_gn=SUN1;refGene_tx=XM_019820309.1;refGene_gn=SUN
1;refGene_tx=XM_019820310.1;refGene_gn=SUN1;refGene_tx=XM_019820312.1;
refGene_gn=SUN1;refGene_tx=XM_019820313.1;refGene_gn=SUN1;refGene_tx=X
M_019820314.1;refGene_gn=SUN1;refGene_tx=XM_019820315.1;refGene_gn=SUN
1;refGene_tx=XM_019820316.2;refGene_gn=SUN1;refGene_tx=XM_023246230.1;
refGene_gn=SUN1;refGene_tx=XM_023246231.1
E3 873986 874238
refGene_gn=SUN1;refGene_tx=XM_011290457.2;refGene_gn=SUN1;refGene_tx=X
M_011290458.2;refGene_gn=SUN1;refGene_tx=XM_019820309.1;refGene_gn=SUN
1;refGene_tx=XM_019820310.1;refGene_gn=SUN1;refGene_tx=XM_019820312.1;
refGene_gn=SUN1;refGene_tx=XM_019820313.1;refGene_gn=SUN1;refGene_tx=X
M_019820314.1;refGene_gn=SUN1;refGene_tx=XM_019820315.1;refGene_gn=SUN
1;refGene_tx=XM_019820316.2;refGene_gn=SUN1;refGene_tx=XM_023246230.1;
refGene_gn=SUN1;refGene_tx=XM_023246231.1
E3 874686 874771
refGene_gn=SUN1;refGene_tx=XM_011290457.2;refGene_gn=SUN1;refGene_tx=X
M_011290458.2;refGene_gn=SUN1;refGene_tx=XM_019820309.1;refGene_gn=SUN
1;refGene_tx=XM_019820310.1;refGene_gn=SUN1;refGene_tx=XM_019820312.1;
refGene_gn=SUN1;refGene_tx=XM_019820313.1;refGene_gn=SUN1;refGene_tx=X
M_019820314.1;refGene_gn=SUN1;refGene_tx=XM_019820315.1;refGene_gn=SUN
1;refGene_tx=XM_019820316.2;refGene_gn=SUN1;refGene_tx=XM_023246230.1;
refGene_gn=SUN1;refGene_tx=XM_023246231.1
E3 877536 877652
refGene_gn=SUN1;refGene_tx=XM_011290457.2;refGene_gn=SUN1;refGene_tx=X
M_011290458.2;refGene_gn=SUN1;refGene_tx=XM_019820309.1;refGene_gn=SUN
1;refGene_tx=XM_019820310.1;refGene_gn=SUN1;refGene_tx=XM_019820312.1;
refGene_gn=SUN1;refGene_tx=XM_019820313.1;refGene_gn=SUN1;refGene_tx=X
M_019820314.1;refGene_gn=SUN1;refGene_tx=XM_019820315.1;refGene_gn=SUN
1;refGene_tx=XM_019820316.2;refGene_gn=SUN1;refGene_tx=XM_023246230.1;
refGene_gn=SUN1;refGene_tx=XM_023246231.1
E3 879986 880154
refGene_gn=SUN1;refGene_tx=XM_011290457.2;refGene_gn=SUN1;refGene_tx=X
M_011290458.2;refGene_gn=SUN1;refGene_tx=XM_019820309.1;refGene_gn=SUN
1;refGene_tx=XM_019820310.1;refGene_gn=SUN1;refGene_tx=XM_019820312.1;
refGene_gn=SUN1;refGene_tx=XM_019820313.1;refGene_gn=SUN1;refGene_tx=X

M_019820314.1;refGene_gn=SUN1;refGene_tx=XM_019820315.1;refGene_gn=SUN1;refGene_tx=XM_019820316.2;refGene_gn=SUN1;refGene_tx=XM_023246230.1;refGene_gn=SUN1;refGene_tx=XM_023246231.1
E3 881774 881867
refGene_gn=SUN1;refGene_tx=XM_011290457.2;refGene_gn=SUN1;refGene_tx=XM_011290458.2;refGene_gn=SUN1;refGene_tx=XM_019820309.1;refGene_gn=SUN1;refGene_tx=XM_019820310.1;refGene_gn=SUN1;refGene_tx=XM_019820312.1;refGene_gn=SUN1;refGene_tx=XM_019820313.1;refGene_gn=SUN1;refGene_tx=XM_019820314.1;refGene_gn=SUN1;refGene_tx=XM_019820315.1;refGene_gn=SUN1;refGene_tx=XM_019820316.2;refGene_gn=SUN1;refGene_tx=XM_023246230.1;refGene_gn=SUN1;refGene_tx=XM_023246231.1
E3 882286 882400
refGene_gn=SUN1;refGene_tx=XM_011290457.2;refGene_gn=SUN1;refGene_tx=XM_011290458.2;refGene_gn=SUN1;refGene_tx=XM_019820309.1;refGene_gn=SUN1;refGene_tx=XM_019820310.1;refGene_gn=SUN1;refGene_tx=XM_019820312.1;refGene_gn=SUN1;refGene_tx=XM_019820313.1;refGene_gn=SUN1;refGene_tx=XM_019820314.1;refGene_gn=SUN1;refGene_tx=XM_019820315.1;refGene_gn=SUN1;refGene_tx=XM_019820316.2;refGene_gn=SUN1;refGene_tx=XM_023246230.1;refGene_gn=SUN1;refGene_tx=XM_023246231.1
E3 885935 886090 refGene_gn=GET4;refGene_tx=XM_023246232.1
E3 890699 890778
refGene_gn=GET4;refGene_tx=XM_023246232.1;refGene_gn=GET4;refGene_tx=XM_023246233.1;refGene_gn=GET4;refGene_tx=XM_023246234.1
E3 891110 891192
refGene_gn=GET4;refGene_tx=XM_023246232.1;refGene_gn=GET4;refGene_tx=XM_023246233.1;refGene_gn=GET4;refGene_tx=XM_023246234.1
E3 891566 891716
refGene_gn=GET4;refGene_tx=XM_023246232.1;refGene_gn=GET4;refGene_tx=XM_023246233.1;refGene_gn=GET4;refGene_tx=XM_023246234.1
E3 897992 898131
refGene_gn=GET4;refGene_tx=XM_023246232.1;refGene_gn=GET4;refGene_tx=XM_023246233.1;refGene_gn=GET4;refGene_tx=XM_023246234.1
E3 899174 899315
refGene_gn=GET4;refGene_tx=XM_023246232.1;refGene_gn=GET4;refGene_tx=XM_023246233.1;refGene_gn=GET4;refGene_tx=XM_023246234.1
E3 899469 899545
refGene_gn=GET4;refGene_tx=XM_023246232.1;refGene_gn=GET4;refGene_tx=XM_023246233.1;refGene_gn=GET4;refGene_tx=XM_023246234.1
E3 899677 899750
refGene_gn=GET4;refGene_tx=XM_023246232.1;refGene_gn=GET4;refGene_tx=XM_023246233.1;refGene_gn=GET4;refGene_tx=XM_023246234.1
E3 905222 905308
refGene_gn=GET4;refGene_tx=XM_023246232.1;refGene_gn=GET4;refGene_tx=XM_023246233.1;refGene_gn=GET4;refGene_tx=XM_023246234.1
E3 908399 908428
refGene_gn=ADAP1;refGene_tx=XM_019820317.2;refGene_gn=ADAP1;refGene_tx=XM_023246235.1;refGene_gn=ADAP1;refGene_tx=XM_023246236.1
E3 908513 908742
refGene_gn=ADAP1;refGene_tx=XM_019820317.2;refGene_gn=ADAP1;refGene_tx=XM_023246235.1;refGene_gn=ADAP1;refGene_tx=XM_023246236.1

E3 908920 908992
refGene_gn=ADAP1;refGene_tx=XM_019820317.2;refGene_gn=ADAP1;refGene_tx=XM_023246235.1;refGene_gn=ADAP1;refGene_tx=XM_023246236.1
E3 909597 909660
refGene_gn=ADAP1;refGene_tx=XM_019820317.2;refGene_gn=ADAP1;refGene_tx=XM_023246235.1;refGene_gn=ADAP1;refGene_tx=XM_023246236.1
E3 909885 909969
refGene_gn=ADAP1;refGene_tx=XM_019820317.2;refGene_gn=ADAP1;refGene_tx=XM_023246235.1
E3 916159 916306
refGene_gn=ADAP1;refGene_tx=XM_019820317.2;refGene_gn=ADAP1;refGene_tx=XM_023246235.1;refGene_gn=ADAP1;refGene_tx=XM_023246236.1
E3 916905 917018
refGene_gn=ADAP1;refGene_tx=XM_019820317.2;refGene_gn=ADAP1;refGene_tx=XM_023246235.1;refGene_gn=ADAP1;refGene_tx=XM_023246236.1
E3 923183 923266
refGene_gn=ADAP1;refGene_tx=XM_019820317.2;refGene_gn=ADAP1;refGene_tx=XM_023246235.1;refGene_gn=ADAP1;refGene_tx=XM_023246236.1
E3 926729 926821
refGene_gn=ADAP1;refGene_tx=XM_019820317.2;refGene_gn=ADAP1;refGene_tx=XM_023246235.1;refGene_gn=ADAP1;refGene_tx=XM_023246236.1
E3 931526 931657
refGene_gn=ADAP1;refGene_tx=XM_019820317.2;refGene_gn=ADAP1;refGene_tx=XM_023246235.1;refGene_gn=ADAP1;refGene_tx=XM_023246236.1
E3 943183 943196 refGene_gn=ADAP1;refGene_tx=XM_019820317.2
E3 959084 959166
refGene_gn=ADAP1;refGene_tx=XM_023246235.1;refGene_gn=ADAP1;refGene_tx=XM_023246236.1
E3 977282 977367
refGene_gn=LOC101088538;refGene_tx=XM_011290451.2
E3 981337 981588
refGene_gn=LOC101088538;refGene_tx=XM_011290451.2;refGene_gn=LOC101088538;refGene_tx=XM_019820319.2
E3 982143 982225
refGene_gn=LOC101088538;refGene_tx=XM_011290451.2;refGene_gn=LOC101088538;refGene_tx=XM_019820319.2
E3 989687 989861
refGene_gn=LOC101088802;refGene_tx=XM_003998398.2;refGene_gn=LOC101088802;refGene_tx=XM_019820320.1
E3 991177 991340
refGene_gn=LOC101088802;refGene_tx=XM_003998398.2;refGene_gn=LOC101088802;refGene_tx=XM_019820320.1
E3 991638 991788
refGene_gn=LOC101088802;refGene_tx=XM_003998398.2;refGene_gn=LOC101088802;refGene_tx=XM_019820320.1
E3 991858 992016
refGene_gn=LOC101088802;refGene_tx=XM_003998398.2;refGene_gn=LOC101088802;refGene_tx=XM_019820320.1;refGene_gn=LOC101088802;refGene_tx=XM_019820324.1
E3 993085 993737

refGene_gn=LOC101088802;refGene_tx=XM_003998398.2;refGene_gn=LOC101088802;refGene_tx=XM_019820320.1;refGene_gn=LOC101088802;refGene_tx=XM_019820324.1
E3 993848 994033
refGene_gn=LOC101088802;refGene_tx=XM_003998398.2;refGene_gn=LOC101088802;refGene_tx=XM_019820320.1;refGene_gn=LOC101088802;refGene_tx=XM_019820324.1
E3 994182 994324
refGene_gn=LOC101088802;refGene_tx=XM_003998398.2;refGene_gn=LOC101088802;refGene_tx=XM_019820320.1;refGene_gn=LOC101088802;refGene_tx=XM_019820324.1
E3 994563 994784
refGene_gn=LOC101088802;refGene_tx=XM_003998398.2;refGene_gn=LOC101088802;refGene_tx=XM_019820320.1;refGene_gn=LOC101088802;refGene_tx=XM_019820324.1
E3 1001499 1001682
refGene_gn=CE3H7orf50;refGene_tx=XM_023246546.1;refGene_gn=CE3H7orf50;refGene_tx=XM_023246547.1;refGene_gn=CE3H7orf50;refGene_tx=XM_023246548.1;refGene_gn=CE3H7orf50;refGene_tx=XM_023246549.1
E3 1006268 1006350
refGene_gn=CE3H7orf50;refGene_tx=XM_023246546.1;refGene_gn=CE3H7orf50;refGene_tx=XM_023246547.1;refGene_gn=CE3H7orf50;refGene_tx=XM_023246548.1;refGene_gn=CE3H7orf50;refGene_tx=XM_023246549.1
E3 1011777 1011974
refGene_gn=CE3H7orf50;refGene_tx=XM_023246546.1;refGene_gn=CE3H7orf50;refGene_tx=XM_023246547.1;refGene_gn=CE3H7orf50;refGene_tx=XM_023246548.1;refGene_gn=CE3H7orf50;refGene_tx=XM_023246549.1
E3 1019940 1020375
refGene_gn=CE3H7orf50;refGene_tx=XM_023246546.1
E3 1030440 1030443 refGene_gn=GPR146;refGene_tx=XM_006941852.2
E3 1030451 1030463
refGene_gn=GPR146;refGene_tx=XM_006941851.2;refGene_gn=GPR146;refGene_tx=XM_006941854.3
E3 1033178 1033193
refGene_gn=GPR146;refGene_tx=XM_006941848.2;refGene_gn=GPR146;refGene_tx=XM_006941849.2
E3 1040535 1041546
refGene_gn=GPR146;refGene_tx=XM_006941848.2;refGene_gn=GPR146;refGene_tx=XM_006941849.2;refGene_gn=GPR146;refGene_tx=XM_006941851.2;refGene_gn=GPR146;refGene_tx=XM_006941852.2;refGene_gn=GPR146;refGene_tx=XM_006941853.2;refGene_gn=GPR146;refGene_tx=XM_006941854.3
E3 1070355 1070583
refGene_gn=CE3H7orf50;refGene_tx=XM_023246547.1
E3 1070703 1071783 refGene_gn=GPER1;refGene_tx=XM_003998386.5
E3 1100954 1101078
refGene_gn=CE3H7orf50;refGene_tx=XM_023246548.1;refGene_gn=CE3H7orf50;refGene_tx=XM_023246549.1
E3 1109258 1109341
refGene_gn=CE3H7orf50;refGene_tx=XM_023246548.1
E3 1116510 1116615 refGene_gn=ZFAND2A;refGene_tx=XM_023247066.1

E3	1117805	1117891	refGene_gn=ZFAND2A;refGene_tx=XM_023247066.1
E3	1121704	1121853	refGene_gn=ZFAND2A;refGene_tx=XM_023247066.1
E3	1125009	1125139	refGene_gn=ZFAND2A;refGene_tx=XM_023247066.1
E3	1126952	1127098	refGene_gn=ZFAND2A;refGene_tx=XM_023247066.1
E3	1127974	1128097	refGene_gn=ZFAND2A;refGene_tx=XM_023247066.1
E3	1192053	1192342	refGene_gn=UNCX;refGene_tx=XM_023246223.1
E3	1192577	1192753	refGene_gn=UNCX;refGene_tx=XM_023246223.1
E3	1194323	1194569	refGene_gn=UNCX;refGene_tx=XM_023246224.1
E3	1194854	1196024
refGene_gn=UNCX;refGene_tx=XM_023246223.1;refGene_gn=UNCX;refGene_tx=XM_023246224.1
E3	1339815	1339910
refGene_gn=MICALL2;refGene_tx=XM_011290446.2;refGene_gn=MICALL2;refGene_tx=XM_011290447.2;refGene_gn=MICALL2;refGene_tx=XM_023246474.1
E3	1340248	1340295
refGene_gn=MICALL2;refGene_tx=XM_011290446.2;refGene_gn=MICALL2;refGene_tx=XM_011290447.2;refGene_gn=MICALL2;refGene_tx=XM_023246474.1
E3	1341024	1341139
refGene_gn=MICALL2;refGene_tx=XM_011290446.2;refGene_gn=MICALL2;refGene_tx=XM_011290447.2;refGene_gn=MICALL2;refGene_tx=XM_023246474.1
E3	1341790	1341864
refGene_gn=MICALL2;refGene_tx=XM_011290446.2;refGene_gn=MICALL2;refGene_tx=XM_011290447.2;refGene_gn=MICALL2;refGene_tx=XM_023246474.1
E3	1341987	1342078
refGene_gn=MICALL2;refGene_tx=XM_011290446.2;refGene_gn=MICALL2;refGene_tx=XM_011290447.2;refGene_gn=MICALL2;refGene_tx=XM_023246474.1
E3	1342160	1342281
refGene_gn=MICALL2;refGene_tx=XM_011290446.2;refGene_gn=MICALL2;refGene_tx=XM_011290447.2;refGene_gn=MICALL2;refGene_tx=XM_023246474.1
E3	1342360	1342425
refGene_gn=MICALL2;refGene_tx=XM_011290446.2;refGene_gn=MICALL2;refGene_tx=XM_011290447.2;refGene_gn=MICALL2;refGene_tx=XM_023246474.1
E3	1342906	1343080
refGene_gn=MICALL2;refGene_tx=XM_011290446.2;refGene_gn=MICALL2;refGene_tx=XM_011290447.2;refGene_gn=MICALL2;refGene_tx=XM_023246474.1
E3	1343445	1343606
refGene_gn=MICALL2;refGene_tx=XM_011290446.2;refGene_gn=MICALL2;refGene_tx=XM_011290447.2;refGene_gn=MICALL2;refGene_tx=XM_023246474.1
E3	1344308	1344402
refGene_gn=MICALL2;refGene_tx=XM_011290446.2;refGene_gn=MICALL2;refGene_tx=XM_011290447.2;refGene_gn=MICALL2;refGene_tx=XM_023246474.1
E3	1345432	1345689
refGene_gn=MICALL2;refGene_tx=XM_011290446.2;refGene_gn=MICALL2;refGene_tx=XM_011290447.2;refGene_gn=MICALL2;refGene_tx=XM_023246474.1
E3	1347234	1348011
refGene_gn=MICALL2;refGene_tx=XM_011290446.2;refGene_gn=MICALL2;refGene_tx=XM_011290447.2;refGene_gn=MICALL2;refGene_tx=XM_023246474.1
E3	1348390	1348506
refGene_gn=MICALL2;refGene_tx=XM_011290446.2;refGene_gn=MICALL2;refGene_tx=XM_011290447.2;refGene_gn=MICALL2;refGene_tx=XM_023246474.1

E3	1349290	1349478
refGene_gn=MICALL2;refGene_tx=XM_011290446.2;refGene_gn=MICALL2;refGene_tx=XM_011290447.2;refGene_gn=MICALL2;refGene_tx=XM_023246474.1
E3	1350351	1350493
refGene_gn=MICALL2;refGene_tx=XM_011290446.2;refGene_gn=MICALL2;refGene_tx=XM_011290447.2;refGene_gn=MICALL2;refGene_tx=XM_023246474.1
E3	1352404	1352453
refGene_gn=MICALL2;refGene_tx=XM_011290446.2;refGene_gn=MICALL2;refGene_tx=XM_023246474.1
E3	1358981	1359124
refGene_gn=MICALL2;refGene_tx=XM_011290446.2;refGene_gn=MICALL2;refGene_tx=XM_023246474.1
E3	1368861	1368977	refGene_gn=INTS1;refGene_tx=XM_023247014.1
E3	1369072	1369182	refGene_gn=INTS1;refGene_tx=XM_023247014.1
E3	1369333	1369425	refGene_gn=INTS1;refGene_tx=XM_023247014.1
E3	1369713	1369784	refGene_gn=INTS1;refGene_tx=XM_023247014.1
E3	1370692	1370806	refGene_gn=INTS1;refGene_tx=XM_023247014.1
E3	1371382	1371495	refGene_gn=INTS1;refGene_tx=XM_023247014.1
E3	1371845	1371973	refGene_gn=INTS1;refGene_tx=XM_023247014.1
E3	1372601	1372794	refGene_gn=INTS1;refGene_tx=XM_023247014.1
E3	1373167	1373301	refGene_gn=INTS1;refGene_tx=XM_023247014.1
E3	1374070	1374194	refGene_gn=INTS1;refGene_tx=XM_023247014.1
E3	1374285	1374512	refGene_gn=INTS1;refGene_tx=XM_023247014.1
E3	1374589	1374677	refGene_gn=INTS1;refGene_tx=XM_023247014.1
E3	1375195	1375326	refGene_gn=INTS1;refGene_tx=XM_023247014.1
E3	1375964	1376088	refGene_gn=INTS1;refGene_tx=XM_023247014.1
E3	1376212	1376378	refGene_gn=INTS1;refGene_tx=XM_023247014.1
E3	1376609	1376750	refGene_gn=INTS1;refGene_tx=XM_023247014.1
E3	1377012	1377172	refGene_gn=INTS1;refGene_tx=XM_023247014.1
E3	1377665	1377920	refGene_gn=INTS1;refGene_tx=XM_023247014.1
E3	1378623	1378748	refGene_gn=INTS1;refGene_tx=XM_023247014.1
E3	1378985	1379084	refGene_gn=INTS1;refGene_tx=XM_023247014.1
E3	1379424	1379571	refGene_gn=INTS1;refGene_tx=XM_023247014.1
E3	1380271	1380433	refGene_gn=INTS1;refGene_tx=XM_023247014.1
E3	1381447	1381559	refGene_gn=INTS1;refGene_tx=XM_023247014.1
E3	1381641	1381809	refGene_gn=INTS1;refGene_tx=XM_023247014.1
E3	1382806	1382911	refGene_gn=INTS1;refGene_tx=XM_023247014.1
E3	1383015	1383195	refGene_gn=INTS1;refGene_tx=XM_023247014.1
E3	1383582	1383732	refGene_gn=INTS1;refGene_tx=XM_023247014.1
E3	1383817	1383997	refGene_gn=INTS1;refGene_tx=XM_023247014.1
E3	1384262	1384373	refGene_gn=INTS1;refGene_tx=XM_023247014.1
E3	1384374	1384393	refGene_gn=INTS1;refGene_tx=XM_023247014.1
E3	1384629	1384827	refGene_gn=INTS1;refGene_tx=XM_023247014.1
E3	1385029	1385090	refGene_gn=INTS1;refGene_tx=XM_023247014.1
E3	1385258	1385350	refGene_gn=INTS1;refGene_tx=XM_023247014.1
E3	1388374	1388471	refGene_gn=INTS1;refGene_tx=XM_023247014.1
E3	1389007	1389165	refGene_gn=INTS1;refGene_tx=XM_023247014.1
E3	1389717	1389795	refGene_gn=INTS1;refGene_tx=XM_023247014.1
E3	1390138	1390259	refGene_gn=INTS1;refGene_tx=XM_023247014.1
E3	1390855	1390964	refGene_gn=INTS1;refGene_tx=XM_023247014.1

E3	1391489	1391666	refGene_gn=INTS1;refGene_tx=XM_023247014.1
E3	1392431	1392573	refGene_gn=INTS1;refGene_tx=XM_023247014.1
E3	1392672	1392818	refGene_gn=INTS1;refGene_tx=XM_023247014.1
E3	1392903	1393090	refGene_gn=INTS1;refGene_tx=XM_023247014.1
E3	1393202	1393308	refGene_gn=INTS1;refGene_tx=XM_023247014.1
E3	1393464	1393631	refGene_gn=INTS1;refGene_tx=XM_023247014.1
E3	1393893	1394024	refGene_gn=INTS1;refGene_tx=XM_023247014.1
E3	1394111	1394308	refGene_gn=INTS1;refGene_tx=XM_023247014.1
E3	1396608	1396899	refGene_gn=INTS1;refGene_tx=XM_023247014.1
E3	1397434	1397492	refGene_gn=INTS1;refGene_tx=XM_023247014.1
E3	1422686	1423604
refGene_gn=LOC109495033;refGene_tx=XM_019820233.1
E3	1423902	1424063
refGene_gn=LOC109495033;refGene_tx=XM_019820233.1
E3	1424223	1424281
refGene_gn=LOC109495033;refGene_tx=XM_019820233.1
E3	1424556	1424638
refGene_gn=MAFK;refGene_tx=XM_023246367.1;refGene_gn=MAFK;refGene_tx=XM_023246368.1
E3	1426283	1426363
refGene_gn=MAFK;refGene_tx=XM_023246367.1;refGene_gn=MAFK;refGene_tx=XM_023246368.1;refGene_gn=MAFK;refGene_tx=XM_023246369.1;refGene_gn=MAFK;refGene_tx=XM_023246370.1;refGene_gn=MAFK;refGene_tx=XM_023246371.1;refGene_gn=MAFK;refGene_tx=XM_023246372.1;refGene_gn=MAFK;refGene_tx=XM_023246373.1;refGene_gn=MAFK;refGene_tx=XM_023246374.1
E3	1427276	1427351
refGene_gn=MAFK;refGene_tx=XM_023246367.1;refGene_gn=MAFK;refGene_tx=XM_023246369.1;refGene_gn=MAFK;refGene_tx=XM_023246370.1;refGene_gn=MAFK;refGene_tx=XM_023246371.1
E3	1427615	1428050
refGene_gn=MAFK;refGene_tx=XM_023246367.1;refGene_gn=MAFK;refGene_tx=XM_023246368.1;refGene_gn=MAFK;refGene_tx=XM_023246369.1;refGene_gn=MAFK;refGene_tx=XM_023246370.1;refGene_gn=MAFK;refGene_tx=XM_023246371.1;refGene_gn=MAFK;refGene_tx=XM_023246372.1;refGene_gn=MAFK;refGene_tx=XM_023246373.1;refGene_gn=MAFK;refGene_tx=XM_023246374.1
E3	1433267	1433503
refGene_gn=TMEM184A;refGene_tx=XM_006941856.4;refGene_gn=TMEM184A;refGene_tx=XM_019820331.2
E3	1434147	1434342
refGene_gn=TMEM184A;refGene_tx=XM_006941856.4;refGene_gn=TMEM184A;refGene_tx=XM_019820331.2
E3	1434885	1435055
refGene_gn=TMEM184A;refGene_tx=XM_006941856.4;refGene_gn=TMEM184A;refGene_tx=XM_019820331.2
E3	1435860	1435952
refGene_gn=TMEM184A;refGene_tx=XM_006941856.4;refGene_gn=TMEM184A;refGene_tx=XM_019820331.2
E3	1436033	1436109
refGene_gn=TMEM184A;refGene_tx=XM_006941856.4;refGene_gn=TMEM184A;refGene_tx=XM_019820331.2

E3 1436218 1436309
refGene_gn=TMEM184A;refGene_tx=XM_006941856.4;refGene_gn=TMEM184A;refGene_tx=XM_019820331.2
E3 1436699 1436865
refGene_gn=TMEM184A;refGene_tx=XM_006941856.4;refGene_gn=TMEM184A;refGene_tx=XM_019820331.2
E3 1438971 1439172
refGene_gn=TMEM184A;refGene_tx=XM_006941856.4;refGene_gn=TMEM184A;refGene_tx=XM_019820331.2
E3 1443715 1443721 refGene_gn=TMEM184A;refGene_tx=XM_019820331.2
E3 1447911 1448064
refGene_gn=PSMG3;refGene_tx=XM_011290445.3;refGene_gn=PSMG3;refGene_tx=XM_023246351.1
E3 1448633 1448849
refGene_gn=PSMG3;refGene_tx=XM_011290445.3;refGene_gn=PSMG3;refGene_tx=XM_023246351.1
E3 1535989 1536907
refGene_gn=LOC111558226;refGene_tx=XM_023246429.1
E3 1572922 1575400
refGene_gn=ELFN1;refGene_tx=XM_023246425.1;refGene_gn=ELFN1;refGene_tx=XM_023246426.1;refGene_gn=ELFN1;refGene_tx=XM_023246428.1
E3 1614277 1614436 refGene_gn=MAD1L1;refGene_tx=XM_003998387.4
E3 1652949 1653140 refGene_gn=MAD1L1;refGene_tx=XM_003998387.4
E3 1682969 1683007 refGene_gn=MAD1L1;refGene_tx=XM_003998387.4
E3 1683008 1683178 refGene_gn=MAD1L1;refGene_tx=XM_003998387.4
E3 1707573 1707664 refGene_gn=MAD1L1;refGene_tx=XM_003998387.4
E3 1729349 1729438 refGene_gn=MAD1L1;refGene_tx=XM_003998387.4
E3 1752835 1752892 refGene_gn=MAD1L1;refGene_tx=XM_003998387.4
E3 1765787 1765928 refGene_gn=MAD1L1;refGene_tx=XM_003998387.4
E3 1811188 1811333 refGene_gn=MAD1L1;refGene_tx=XM_003998387.4
E3 1875681 1875768 refGene_gn=MAD1L1;refGene_tx=XM_003998387.4
E3 1919385 1919447 refGene_gn=MAD1L1;refGene_tx=XM_003998387.4
E3 1921307 1921422 refGene_gn=MAD1L1;refGene_tx=XM_003998387.4
E3 1921582 1921713 refGene_gn=MAD1L1;refGene_tx=XM_003998387.4
E3 1922602 1922684 refGene_gn=MAD1L1;refGene_tx=XM_003998387.4
E3 1923499 1923624 refGene_gn=MAD1L1;refGene_tx=XM_003998387.4
E3 1925894 1926074 refGene_gn=MAD1L1;refGene_tx=XM_003998387.4
E3 1928371 1928512 refGene_gn=MAD1L1;refGene_tx=XM_003998387.4
E3 1931773 1931923 refGene_gn=MAD1L1;refGene_tx=XM_003998387.4
E3 1935957 1936400
refGene_gn=MRM2;refGene_tx=XM_003998406.4;refGene_gn=MRM2;refGene_tx=XM_006941859.4
E3 1939057 1939347
refGene_gn=MRM2;refGene_tx=XM_003998406.4;refGene_gn=MRM2;refGene_tx=XM_006941859.4
E3 1940327 1940383 refGene_gn=MRM2;refGene_tx=XM_006941859.4
E3 1940438 1940446 refGene_gn=MRM2;refGene_tx=XM_003998406.4
E3 1942199 1942351 refGene_gn=NUDT1;refGene_tx=XM_003998407.3
E3 1947327 1947473 refGene_gn=NUDT1;refGene_tx=XM_003998407.3
E3 1947706 1947879 refGene_gn=NUDT1;refGene_tx=XM_003998407.3

E3 1951074 1951185
refGene_gn=SNX8;refGene_tx=XM_011290440.3;refGene_gn=SNX8;refGene_tx=XM_023246828.1;refGene_gn=SNX8;refGene_tx=XM_023246829.1;refGene_gn=SNX8;refGene_tx=XM_023246830.1;refGene_gn=SNX8;refGene_tx=XM_023246831.1
E3 1952588 1952738
refGene_gn=SNX8;refGene_tx=XM_011290440.3;refGene_gn=SNX8;refGene_tx=XM_023246828.1;refGene_gn=SNX8;refGene_tx=XM_023246829.1;refGene_gn=SNX8;refGene_tx=XM_023246830.1;refGene_gn=SNX8;refGene_tx=XM_023246831.1
E3 1953125 1953275
refGene_gn=SNX8;refGene_tx=XM_011290440.3;refGene_gn=SNX8;refGene_tx=XM_023246828.1;refGene_gn=SNX8;refGene_tx=XM_023246829.1;refGene_gn=SNX8;refGene_tx=XM_023246830.1;refGene_gn=SNX8;refGene_tx=XM_023246831.1
E3 1954828 1954897
refGene_gn=SNX8;refGene_tx=XM_011290440.3;refGene_gn=SNX8;refGene_tx=XM_023246828.1;refGene_gn=SNX8;refGene_tx=XM_023246829.1;refGene_gn=SNX8;refGene_tx=XM_023246830.1;refGene_gn=SNX8;refGene_tx=XM_023246831.1
E3 1957609 1957742
refGene_gn=SNX8;refGene_tx=XM_011290440.3;refGene_gn=SNX8;refGene_tx=XM_023246828.1;refGene_gn=SNX8;refGene_tx=XM_023246829.1;refGene_gn=SNX8;refGene_tx=XM_023246830.1;refGene_gn=SNX8;refGene_tx=XM_023246831.1
E3 1958246 1958407
refGene_gn=SNX8;refGene_tx=XM_011290440.3;refGene_gn=SNX8;refGene_tx=XM_023246828.1;refGene_gn=SNX8;refGene_tx=XM_023246829.1;refGene_gn=SNX8;refGene_tx=XM_023246830.1;refGene_gn=SNX8;refGene_tx=XM_023246831.1
E3 1959541 1959622
refGene_gn=SNX8;refGene_tx=XM_011290440.3;refGene_gn=SNX8;refGene_tx=XM_023246828.1;refGene_gn=SNX8;refGene_tx=XM_023246829.1;refGene_gn=SNX8;refGene_tx=XM_023246830.1;refGene_gn=SNX8;refGene_tx=XM_023246831.1
E3 1961477 1961599
refGene_gn=SNX8;refGene_tx=XM_011290440.3;refGene_gn=SNX8;refGene_tx=XM_023246828.1;refGene_gn=SNX8;refGene_tx=XM_023246829.1;refGene_gn=SNX8;refGene_tx=XM_023246830.1;refGene_gn=SNX8;refGene_tx=XM_023246831.1
E3 1961863 1961981
refGene_gn=SNX8;refGene_tx=XM_011290440.3;refGene_gn=SNX8;refGene_tx=XM_023246828.1;refGene_gn=SNX8;refGene_tx=XM_023246829.1;refGene_gn=SNX8;refGene_tx=XM_023246830.1;refGene_gn=SNX8;refGene_tx=XM_023246831.1
E3 1964642 1964833
refGene_gn=SNX8;refGene_tx=XM_011290440.3;refGene_gn=SNX8;refGene_tx=XM_023246828.1;refGene_gn=SNX8;refGene_tx=XM_023246829.1;refGene_gn=SNX8;refGene_tx=XM_023246830.1;refGene_gn=SNX8;refGene_tx=XM_023246831.1
E3 1965105 1965256 refGene_gn=SNX8;refGene_tx=XM_023246829.1
E3 1990394 1990500 refGene_gn=SNX8;refGene_tx=XM_023246831.1
E3 1999149 1999189 refGene_gn=SNX8;refGene_tx=XM_023246830.1
E3 2020206 2020423
refGene_gn=SNX8;refGene_tx=XM_011290440.3;refGene_gn=SNX8;refGene_tx=XM_023246828.1
E3 2020879 2021294 refGene_gn=EIF3B;refGene_tx=XM_011290505.3
E3 2025548 2025741 refGene_gn=EIF3B;refGene_tx=XM_011290505.3
E3 2027937 2028057 refGene_gn=EIF3B;refGene_tx=XM_011290505.3
E3 2028314 2028372 refGene_gn=EIF3B;refGene_tx=XM_011290505.3

E3	2028521	2028650	refGene_gn=EIF3B;refGene_tx=XM_011290505.3
E3	2028913	2029071	refGene_gn=EIF3B;refGene_tx=XM_011290505.3
E3	2031646	2031778	refGene_gn=EIF3B;refGene_tx=XM_011290505.3
E3	2031862	2031929	refGene_gn=EIF3B;refGene_tx=XM_011290505.3
E3	2032308	2032355	refGene_gn=EIF3B;refGene_tx=XM_011290505.3
E3	2033307	2033518	refGene_gn=EIF3B;refGene_tx=XM_011290505.3
E3	2034783	2034856	refGene_gn=EIF3B;refGene_tx=XM_011290505.3
E3	2035721	2035844	refGene_gn=EIF3B;refGene_tx=XM_011290505.3
E3	2037276	2037355	refGene_gn=EIF3B;refGene_tx=XM_011290505.3
E3	2038053	2038192	refGene_gn=EIF3B;refGene_tx=XM_011290505.3
E3	2039929	2040055	refGene_gn=EIF3B;refGene_tx=XM_011290505.3
E3	2040630	2040708	refGene_gn=EIF3B;refGene_tx=XM_011290505.3
E3	2040996	2041105	refGene_gn=EIF3B;refGene_tx=XM_011290505.3
E3	2041368	2041472	refGene_gn=EIF3B;refGene_tx=XM_011290505.3
E3	2048405	2048504
refGene_gn=LOC101091832;refGene_tx=XM_023247067.1
E3	2050556	2050644
refGene_gn=LOC101091832;refGene_tx=XM_023247067.1
E3	2051496	2051630
refGene_gn=LOC101091832;refGene_tx=XM_023247067.1
E3	2051832	2051958
refGene_gn=LOC101091832;refGene_tx=XM_023247067.1
E3	2052910	2053105
refGene_gn=LOC101091832;refGene_tx=XM_023247067.1
E3	2055949	2056102
refGene_gn=LOC101091832;refGene_tx=XM_023247067.1
E3	2056650	2056773
refGene_gn=LOC101091832;refGene_tx=XM_023247067.1
E3	2059845	2059996
refGene_gn=LOC101091832;refGene_tx=XM_023247067.1
E3	2066938	2067116
refGene_gn=LOC101091832;refGene_tx=XM_023247067.1
E3	2068285	2068463
refGene_gn=LOC101091832;refGene_tx=XM_023247067.1
E3	2070334	2070680
refGene_gn=LOC101091832;refGene_tx=XM_023247067.1
E3	2071071	2071193
refGene_gn=LOC101091832;refGene_tx=XM_023247067.1
E3	2072746	2072874
refGene_gn=LOC101091832;refGene_tx=XM_023247067.1
E3	2076617	2076726
refGene_gn=LOC101091832;refGene_tx=XM_023247067.1
E3	2077354	2077717
refGene_gn=LOC101091832;refGene_tx=XM_023247067.1
E3	2080200	2080354
refGene_gn=LOC101091832;refGene_tx=XM_023247067.1
E3	2084880	2084988
refGene_gn=LOC101091832;refGene_tx=XM_023247067.1
E3	2086152	2086250
refGene_gn=LOC101091832;refGene_tx=XM_023247067.1

E3 2117091 2118336
refGene_gn=CHST12;refGene_tx=XM_003998388.5;refGene_gn=CHST12;refGene_tx=XM_006941860.4;refGene_gn=CHST12;refGene_tx=XM_006941861.4;refGene_gn=CHST12;refGene_tx=XM_023246472.1
E3 2149884 2149911
refGene_gn=GRIFIN;refGene_tx=XM_006941862.3;refGene_gn=GRIFIN;refGene_tx=XM_011290438.3
E3 2150165 2150332
refGene_gn=GRIFIN;refGene_tx=XM_006941862.3;refGene_gn=GRIFIN;refGene_tx=XM_011290438.3
E3 2150632 2150792
refGene_gn=GRIFIN;refGene_tx=XM_006941862.3;refGene_gn=GRIFIN;refGene_tx=XM_011290438.3
E3 2150889 2150969
refGene_gn=GRIFIN;refGene_tx=XM_006941862.3;refGene_gn=GRIFIN;refGene_tx=XM_011290438.3
E3 2151173 2151185
refGene_gn=GRIFIN;refGene_tx=XM_006941862.3;refGene_gn=GRIFIN;refGene_tx=XM_011290438.3
E3 2185869 2186298 refGene_gn=LFNG;refGene_tx=XM_023246563.1
E3 2190894 2190943 refGene_gn=LFNG;refGene_tx=XM_023246563.1
E3 2191446 2191546 refGene_gn=LFNG;refGene_tx=XM_023246563.1
E3 2191653 2191807 refGene_gn=LFNG;refGene_tx=XM_023246563.1
E3 2191925 2192011 refGene_gn=LFNG;refGene_tx=XM_023246563.1
E3 2192474 2192640 refGene_gn=LFNG;refGene_tx=XM_023246563.1
E3 2193082 2193168 refGene_gn=LFNG;refGene_tx=XM_023246563.1
E3 2193373 2193440 refGene_gn=LFNG;refGene_tx=XM_023246563.1
E3 2204017 2204092
refGene_gn=TTYH3;refGene_tx=XM_023247112.1;refGene_gn=TTYH3;refGene_tx=XM_023247113.1
E3 2205516 2205593 refGene_gn=TTYH3;refGene_tx=XM_023247112.1
E3 2206263 2206339
refGene_gn=TTYH3;refGene_tx=XM_023247112.1;refGene_gn=TTYH3;refGene_tx=XM_023247113.1
E3 2206857 2207031
refGene_gn=TTYH3;refGene_tx=XM_023247112.1;refGene_gn=TTYH3;refGene_tx=XM_023247113.1
E3 2208812 2208949
refGene_gn=TTYH3;refGene_tx=XM_023247112.1;refGene_gn=TTYH3;refGene_tx=XM_023247113.1
E3 2209869 2209962
refGene_gn=TTYH3;refGene_tx=XM_023247112.1;refGene_gn=TTYH3;refGene_tx=XM_023247113.1
E3 2212013 2212106
refGene_gn=TTYH3;refGene_tx=XM_023247112.1;refGene_gn=TTYH3;refGene_tx=XM_023247113.1
E3 2212753 2212809
refGene_gn=TTYH3;refGene_tx=XM_023247112.1;refGene_gn=TTYH3;refGene_tx=XM_023247113.1
E3 2214919 2214995

refGene_gn=TTYH3;refGene_tx=XM_023247112.1;refGene_gn=TTYH3;refGene_tx=XM_023247113.1
E3 2215283 2215356
refGene_gn=TTYH3;refGene_tx=XM_023247112.1;refGene_gn=TTYH3;refGene_tx=XM_023247113.1
E3 2216465 2216561
refGene_gn=TTYH3;refGene_tx=XM_023247112.1;refGene_gn=TTYH3;refGene_tx=XM_023247113.1
E3 2216850 2217071
refGene_gn=TTYH3;refGene_tx=XM_023247112.1;refGene_gn=TTYH3;refGene_tx=XM_023247113.1
E3 2217209 2217321
refGene_gn=TTYH3;refGene_tx=XM_023247112.1;refGene_gn=TTYH3;refGene_tx=XM_023247113.1
E3 2217480 2217650
refGene_gn=TTYH3;refGene_tx=XM_023247112.1;refGene_gn=TTYH3;refGene_tx=XM_023247113.1
E3 2222477 2223440
refGene_gn=LOC109495083;refGene_tx=XM_023247115.1
E3 2231458 2231581
refGene_gn=TTYH3;refGene_tx=XM_023247112.1;refGene_gn=TTYH3;refGene_tx=XM_023247113.1
E3 2250995 2251108 refGene_gn=IQCE;refGene_tx=XM_023246896.1
E3 2252212 2252311 refGene_gn=IQCE;refGene_tx=XM_023246896.1
E3 2253226 2253330 refGene_gn=IQCE;refGene_tx=XM_023246896.1
E3 2254242 2254364 refGene_gn=IQCE;refGene_tx=XM_023246896.1
E3 2254783 2254894 refGene_gn=IQCE;refGene_tx=XM_023246896.1
E3 2257711 2257735 refGene_gn=IQCE;refGene_tx=XM_023246896.1
E3 2261341 2261512 refGene_gn=IQCE;refGene_tx=XM_023246896.1
E3 2262834 2262925 refGene_gn=IQCE;refGene_tx=XM_023246896.1
E3 2264554 2264659 refGene_gn=IQCE;refGene_tx=XM_023246896.1
E3 2266405 2266605 refGene_gn=IQCE;refGene_tx=XM_023246896.1
E3 2267708 2267764 refGene_gn=IQCE;refGene_tx=XM_023246896.1
E3 2269083 2269177 refGene_gn=IQCE;refGene_tx=XM_023246896.1
E3 2269178 2269248 refGene_gn=IQCE;refGene_tx=XM_023246896.1
E3 2270697 2270747 refGene_gn=IQCE;refGene_tx=XM_023246896.1
E3 2270950 2271023 refGene_gn=IQCE;refGene_tx=XM_023246896.1
E3 2271960 2272031 refGene_gn=IQCE;refGene_tx=XM_023246896.1
E3 2275246 2275297 refGene_gn=IQCE;refGene_tx=XM_023246896.1
E3 2275421 2275535 refGene_gn=IQCE;refGene_tx=XM_023246896.1
E3 2279261 2279332 refGene_gn=IQCE;refGene_tx=XM_023246896.1
E3 2280187 2280322 refGene_gn=IQCE;refGene_tx=XM_023246896.1
E3 2281099 2281225 refGene_gn=IQCE;refGene_tx=XM_023246896.1
E3 2285452 2285498 refGene_gn=IQCE;refGene_tx=XM_023246896.1
E3 2286805 2286853 refGene_gn=IQCE;refGene_tx=XM_023246896.1
E3 2292600 2292768 refGene_gn=IQCE;refGene_tx=XM_023246896.1
E3 2296539 2296666
refGene_gn=BRAT1;refGene_tx=XM_006941863.4;refGene_gn=BRAT1;refGene_tx=XM_006941864.4;refGene_gn=BRAT1;refGene_tx=XM_006941865.4;refGene_gn=BRAT1;refGene_tx=XM_006941866.4;refGene_gn=BRAT1;refGene_tx=XM_0198203

50.2
E3 2304606 2304761
refGene_gn=BRAT1;refGene_tx=XM_006941863.4;refGene_gn=BRAT1;refGene_tx=XM_006941864.4;refGene_gn=BRAT1;refGene_tx=XM_006941865.4;refGene_gn=BRAT1;refGene_tx=XM_006941866.4;refGene_gn=BRAT1;refGene_tx=XM_019820350.2;refGene_gn=BRAT1;refGene_tx=XM_023246364.1;refGene_gn=BRAT1;refGene_tx=XM_023246365.1
E3 2307074 2307222
refGene_gn=BRAT1;refGene_tx=XM_006941863.4;refGene_gn=BRAT1;refGene_tx=XM_006941864.4;refGene_gn=BRAT1;refGene_tx=XM_006941865.4;refGene_gn=BRAT1;refGene_tx=XM_006941866.4;refGene_gn=BRAT1;refGene_tx=XM_023246364.1;refGene_gn=BRAT1;refGene_tx=XM_023246365.1
E3 2307689 2308062
refGene_gn=BRAT1;refGene_tx=XM_006941863.4;refGene_gn=BRAT1;refGene_tx=XM_006941864.4;refGene_gn=BRAT1;refGene_tx=XM_006941865.4;refGene_gn=BRAT1;refGene_tx=XM_006941866.4;refGene_gn=BRAT1;refGene_tx=XM_019820350.2;refGene_gn=BRAT1;refGene_tx=XM_023246364.1;refGene_gn=BRAT1;refGene_tx=XM_023246365.1
E3 2308324 2308444
refGene_gn=BRAT1;refGene_tx=XM_006941863.4;refGene_gn=BRAT1;refGene_tx=XM_006941864.4;refGene_gn=BRAT1;refGene_tx=XM_006941865.4;refGene_gn=BRAT1;refGene_tx=XM_006941866.4;refGene_gn=BRAT1;refGene_tx=XM_019820350.2;refGene_gn=BRAT1;refGene_tx=XM_023246364.1;refGene_gn=BRAT1;refGene_tx=XM_023246365.1
E3 2309501 2309593
refGene_gn=BRAT1;refGene_tx=XM_006941863.4;refGene_gn=BRAT1;refGene_tx=XM_006941864.4;refGene_gn=BRAT1;refGene_tx=XM_006941865.4;refGene_gn=BRAT1;refGene_tx=XM_006941866.4;refGene_gn=BRAT1;refGene_tx=XM_019820350.2;refGene_gn=BRAT1;refGene_tx=XM_023246364.1;refGene_gn=BRAT1;refGene_tx=XM_023246365.1
E3 2310284 2310403
refGene_gn=BRAT1;refGene_tx=XM_006941863.4;refGene_gn=BRAT1;refGene_tx=XM_006941864.4;refGene_gn=BRAT1;refGene_tx=XM_006941865.4;refGene_gn=BRAT1;refGene_tx=XM_006941866.4;refGene_gn=BRAT1;refGene_tx=XM_019820350.2;refGene_gn=BRAT1;refGene_tx=XM_023246364.1;refGene_gn=BRAT1;refGene_tx=XM_023246365.1
E3 2310476 2310663
refGene_gn=BRAT1;refGene_tx=XM_006941863.4;refGene_gn=BRAT1;refGene_tx=XM_006941864.4;refGene_gn=BRAT1;refGene_tx=XM_006941865.4;refGene_gn=BRAT1;refGene_tx=XM_006941866.4;refGene_gn=BRAT1;refGene_tx=XM_019820350.2;refGene_gn=BRAT1;refGene_tx=XM_023246364.1;refGene_gn=BRAT1;refGene_tx=XM_023246365.1
E3 2310869 2310943
refGene_gn=BRAT1;refGene_tx=XM_006941863.4;refGene_gn=BRAT1;refGene_tx=XM_006941864.4;refGene_gn=BRAT1;refGene_tx=XM_006941865.4;refGene_gn=BRAT1;refGene_tx=XM_006941866.4;refGene_gn=BRAT1;refGene_tx=XM_019820350.2;refGene_gn=BRAT1;refGene_tx=XM_023246364.1;refGene_gn=BRAT1;refGene_tx=XM_023246365.1
E3 2311587 2311690
refGene_gn=BRAT1;refGene_tx=XM_006941863.4;refGene_gn=BRAT1;refGene_tx

=XM_006941864.4;refGene_gn=BRAT1;refGene_tx=XM_006941865.4;refGene_gn=BRAT1;refGene_tx=XM_006941866.4;refGene_gn=BRAT1;refGene_tx=XM_019820350.2;refGene_gn=BRAT1;refGene_tx=XM_023246364.1;refGene_gn=BRAT1;refGene_tx=XM_023246365.1
E3 2311789 2311888
refGene_gn=BRAT1;refGene_tx=XM_006941863.4;refGene_gn=BRAT1;refGene_tx=XM_006941864.4;refGene_gn=BRAT1;refGene_tx=XM_006941865.4;refGene_gn=BRAT1;refGene_tx=XM_006941866.4;refGene_gn=BRAT1;refGene_tx=XM_019820350.2;refGene_gn=BRAT1;refGene_tx=XM_023246364.1;refGene_gn=BRAT1;refGene_tx=XM_023246365.1
E3 2312062 2312238
refGene_gn=BRAT1;refGene_tx=XM_006941863.4;refGene_gn=BRAT1;refGene_tx=XM_006941864.4;refGene_gn=BRAT1;refGene_tx=XM_006941865.4;refGene_gn=BRAT1;refGene_tx=XM_006941866.4;refGene_gn=BRAT1;refGene_tx=XM_019820350.2;refGene_gn=BRAT1;refGene_tx=XM_023246364.1;refGene_gn=BRAT1;refGene_tx=XM_023246365.1
E3 2312549 2313248
refGene_gn=BRAT1;refGene_tx=XM_006941863.4;refGene_gn=BRAT1;refGene_tx=XM_006941864.4;refGene_gn=BRAT1;refGene_tx=XM_006941865.4;refGene_gn=BRAT1;refGene_tx=XM_006941866.4;refGene_gn=BRAT1;refGene_tx=XM_019820350.2;refGene_gn=BRAT1;refGene_tx=XM_023246364.1;refGene_gn=BRAT1;refGene_tx=XM_023246365.1
E3 2334991 2335295
refGene_gn=AMZ1;refGene_tx=XM_023246311.1;refGene_gn=AMZ1;refGene_tx=XM_023246312.1;refGene_gn=AMZ1;refGene_tx=XM_023246313.1;refGene_gn=AMZ1;refGene_tx=XM_023246314.1;refGene_gn=AMZ1;refGene_tx=XM_023246315.1;refGene_gn=AMZ1;refGene_tx=XM_023246316.1;refGene_gn=AMZ1;refGene_tx=XM_023246317.1;refGene_gn=AMZ1;refGene_tx=XM_023246318.1
E3 2336455 2336623
refGene_gn=AMZ1;refGene_tx=XM_023246311.1;refGene_gn=AMZ1;refGene_tx=XM_023246312.1;refGene_gn=AMZ1;refGene_tx=XM_023246313.1;refGene_gn=AMZ1;refGene_tx=XM_023246314.1;refGene_gn=AMZ1;refGene_tx=XM_023246315.1;refGene_gn=AMZ1;refGene_tx=XM_023246317.1;refGene_gn=AMZ1;refGene_tx=XM_023246318.1
E3 2338466 2338595
refGene_gn=AMZ1;refGene_tx=XM_023246311.1;refGene_gn=AMZ1;refGene_tx=XM_023246312.1;refGene_gn=AMZ1;refGene_tx=XM_023246313.1;refGene_gn=AMZ1;refGene_tx=XM_023246314.1;refGene_gn=AMZ1;refGene_tx=XM_023246315.1;refGene_gn=AMZ1;refGene_tx=XM_023246316.1;refGene_gn=AMZ1;refGene_tx=XM_023246317.1;refGene_gn=AMZ1;refGene_tx=XM_023246318.1
E3 2338854 2339024
refGene_gn=AMZ1;refGene_tx=XM_023246315.1;refGene_gn=AMZ1;refGene_tx=XM_023246317.1
E3 2339440 2339617
refGene_gn=AMZ1;refGene_tx=XM_023246311.1;refGene_gn=AMZ1;refGene_tx=XM_023246312.1;refGene_gn=AMZ1;refGene_tx=XM_023246313.1;refGene_gn=AMZ1;refGene_tx=XM_023246314.1;refGene_gn=AMZ1;refGene_tx=XM_023246315.1;refGene_gn=AMZ1;refGene_tx=XM_023246316.1;refGene_gn=AMZ1;refGene_tx=XM_023246317.1;refGene_gn=AMZ1;refGene_tx=XM_023246318.1
E3 2341147 2341973

refGene_gn=AMZ1;refGene_tx=XM_023246311.1;refGene_gn=AMZ1;refGene_tx=XM_023246312.1;refGene_gn=AMZ1;refGene_tx=XM_023246313.1;refGene_gn=AMZ1;refGene_tx=XM_023246314.1;refGene_gn=AMZ1;refGene_tx=XM_023246315.1;refGene_gn=AMZ1;refGene_tx=XM_023246316.1;refGene_gn=AMZ1;refGene_tx=XM_023246317.1;refGene_gn=AMZ1;refGene_tx=XM_023246318.1
E3 2353486 2354056 refGene_gn=GNA12;refGene_tx=XM_023246889.1
E3 2355311 2355362 refGene_gn=GNA12;refGene_tx=XM_023246889.1
E3 2414339 2414555 refGene_gn=GNA12;refGene_tx=XM_023246889.1
E3 2458415 2458727 refGene_gn=GNA12;refGene_tx=XM_023246889.1
E3 2493808 2494013
refGene_gn=CARD11;refGene_tx=XM_011290436.3;refGene_gn=CARD11;refGene_tx=XM_019820359.2;refGene_gn=CARD11;refGene_tx=XM_019820360.2;refGene_gn=CARD11;refGene_tx=XM_019820361.2;refGene_gn=CARD11;refGene_tx=XM_023246225.1;refGene_gn=CARD11;refGene_tx=XM_023246226.1;refGene_gn=CARD11;refGene_tx=XM_023246227.1
E3 2495983 2496099
refGene_gn=CARD11;refGene_tx=XM_011290436.3;refGene_gn=CARD11;refGene_tx=XM_019820359.2;refGene_gn=CARD11;refGene_tx=XM_019820360.2;refGene_gn=CARD11;refGene_tx=XM_019820361.2;refGene_gn=CARD11;refGene_tx=XM_023246225.1;refGene_gn=CARD11;refGene_tx=XM_023246226.1;refGene_gn=CARD11;refGene_tx=XM_023246227.1
E3 2496403 2496532 refGene_gn=CARD11;refGene_tx=XM_023246228.1
E3 2497519 2497644
refGene_gn=CARD11;refGene_tx=XM_011290436.3;refGene_gn=CARD11;refGene_tx=XM_019820359.2;refGene_gn=CARD11;refGene_tx=XM_019820360.2;refGene_gn=CARD11;refGene_tx=XM_019820361.2;refGene_gn=CARD11;refGene_tx=XM_023246225.1;refGene_gn=CARD11;refGene_tx=XM_023246226.1;refGene_gn=CARD11;refGene_tx=XM_023246227.1;refGene_gn=CARD11;refGene_tx=XM_023246228.1
E3 2502226 2502406
refGene_gn=CARD11;refGene_tx=XM_011290436.3;refGene_gn=CARD11;refGene_tx=XM_019820359.2;refGene_gn=CARD11;refGene_tx=XM_019820360.2;refGene_gn=CARD11;refGene_tx=XM_019820361.2;refGene_gn=CARD11;refGene_tx=XM_023246225.1;refGene_gn=CARD11;refGene_tx=XM_023246226.1;refGene_gn=CARD11;refGene_tx=XM_023246227.1;refGene_gn=CARD11;refGene_tx=XM_023246228.1
E3 2503482 2503618
refGene_gn=CARD11;refGene_tx=XM_011290436.3;refGene_gn=CARD11;refGene_tx=XM_019820359.2;refGene_gn=CARD11;refGene_tx=XM_019820360.2;refGene_gn=CARD11;refGene_tx=XM_019820361.2;refGene_gn=CARD11;refGene_tx=XM_023246225.1;refGene_gn=CARD11;refGene_tx=XM_023246226.1;refGene_gn=CARD11;refGene_tx=XM_023246227.1;refGene_gn=CARD11;refGene_tx=XM_023246228.1
E3 2505119 2505215
refGene_gn=CARD11;refGene_tx=XM_011290436.3;refGene_gn=CARD11;refGene_tx=XM_019820359.2;refGene_gn=CARD11;refGene_tx=XM_019820360.2;refGene_gn=CARD11;refGene_tx=XM_019820361.2;refGene_gn=CARD11;refGene_tx=XM_023246225.1;refGene_gn=CARD11;refGene_tx=XM_023246226.1;refGene_gn=CARD11;refGene_tx=XM_023246227.1;refGene_gn=CARD11;refGene_tx=XM_023246228.1

E3 2505589 2505686
refGene_gn=CARD11;refGene_tx=XM_011290436.3;refGene_gn=CARD11;refGene_tx=XM_019820359.2;refGene_gn=CARD11;refGene_tx=XM_019820360.2;refGene_gn=CARD11;refGene_tx=XM_019820361.2;refGene_gn=CARD11;refGene_tx=XM_023246225.1;refGene_gn=CARD11;refGene_tx=XM_023246226.1;refGene_gn=CARD11;refGene_tx=XM_023246227.1;refGene_gn=CARD11;refGene_tx=XM_023246228.1
E3 2506278 2506519
refGene_gn=CARD11;refGene_tx=XM_011290436.3;refGene_gn=CARD11;refGene_tx=XM_019820359.2;refGene_gn=CARD11;refGene_tx=XM_019820360.2;refGene_gn=CARD11;refGene_tx=XM_019820361.2;refGene_gn=CARD11;refGene_tx=XM_023246225.1;refGene_gn=CARD11;refGene_tx=XM_023246226.1;refGene_gn=CARD11;refGene_tx=XM_023246227.1;refGene_gn=CARD11;refGene_tx=XM_023246228.1
E3 2511278 2511405
refGene_gn=CARD11;refGene_tx=XM_011290436.3;refGene_gn=CARD11;refGene_tx=XM_019820359.2;refGene_gn=CARD11;refGene_tx=XM_019820360.2;refGene_gn=CARD11;refGene_tx=XM_019820361.2;refGene_gn=CARD11;refGene_tx=XM_023246225.1;refGene_gn=CARD11;refGene_tx=XM_023246226.1;refGene_gn=CARD11;refGene_tx=XM_023246227.1;refGene_gn=CARD11;refGene_tx=XM_023246228.1
E3 2511800 2512002
refGene_gn=CARD11;refGene_tx=XM_011290436.3;refGene_gn=CARD11;refGene_tx=XM_019820359.2;refGene_gn=CARD11;refGene_tx=XM_019820360.2;refGene_gn=CARD11;refGene_tx=XM_019820361.2;refGene_gn=CARD11;refGene_tx=XM_023246225.1;refGene_gn=CARD11;refGene_tx=XM_023246226.1;refGene_gn=CARD11;refGene_tx=XM_023246227.1;refGene_gn=CARD11;refGene_tx=XM_023246228.1
E3 2512584 2512717
refGene_gn=CARD11;refGene_tx=XM_011290436.3;refGene_gn=CARD11;refGene_tx=XM_019820359.2;refGene_gn=CARD11;refGene_tx=XM_019820360.2;refGene_gn=CARD11;refGene_tx=XM_019820361.2;refGene_gn=CARD11;refGene_tx=XM_023246225.1;refGene_gn=CARD11;refGene_tx=XM_023246226.1;refGene_gn=CARD11;refGene_tx=XM_023246227.1;refGene_gn=CARD11;refGene_tx=XM_023246228.1
E3 2514325 2514378
refGene_gn=CARD11;refGene_tx=XM_011290436.3;refGene_gn=CARD11;refGene_tx=XM_019820359.2;refGene_gn=CARD11;refGene_tx=XM_019820360.2;refGene_gn=CARD11;refGene_tx=XM_019820361.2;refGene_gn=CARD11;refGene_tx=XM_023246225.1;refGene_gn=CARD11;refGene_tx=XM_023246226.1;refGene_gn=CARD11;refGene_tx=XM_023246227.1;refGene_gn=CARD11;refGene_tx=XM_023246228.1
E3 2515316 2515417
refGene_gn=CARD11;refGene_tx=XM_011290436.3;refGene_gn=CARD11;refGene_tx=XM_019820359.2;refGene_gn=CARD11;refGene_tx=XM_019820360.2;refGene_gn=CARD11;refGene_tx=XM_019820361.2;refGene_gn=CARD11;refGene_tx=XM_023246225.1;refGene_gn=CARD11;refGene_tx=XM_023246226.1;refGene_gn=CARD11;refGene_tx=XM_023246227.1;refGene_gn=CARD11;refGene_tx=XM_023246228.1
E3 2516470 2516553

refGene_gn=CARD11;refGene_tx=XM_011290436.3;refGene_gn=CARD11;refGene_
tx=XM_019820359.2;refGene_gn=CARD11;refGene_tx=XM_019820360.2;refGene_
gn=CARD11;refGene_tx=XM_019820361.2;refGene_gn=CARD11;refGene_tx=XM_02
3246225.1;refGene_gn=CARD11;refGene_tx=XM_023246226.1;refGene_gn=CARD1
1;refGene_tx=XM_023246227.1;refGene_gn=CARD11;refGene_tx=XM_023246228.
1
E3 2517914 2517966
refGene_gn=CARD11;refGene_tx=XM_011290436.3;refGene_gn=CARD11;refGene_
tx=XM_019820359.2;refGene_gn=CARD11;refGene_tx=XM_019820360.2;refGene_
gn=CARD11;refGene_tx=XM_019820361.2;refGene_gn=CARD11;refGene_tx=XM_02
3246225.1;refGene_gn=CARD11;refGene_tx=XM_023246226.1;refGene_gn=CARD1
1;refGene_tx=XM_023246227.1;refGene_gn=CARD11;refGene_tx=XM_023246228.
1
E3 2518887 2519064
refGene_gn=CARD11;refGene_tx=XM_011290436.3;refGene_gn=CARD11;refGene_
tx=XM_019820359.2;refGene_gn=CARD11;refGene_tx=XM_019820360.2;refGene_
gn=CARD11;refGene_tx=XM_019820361.2;refGene_gn=CARD11;refGene_tx=XM_02
3246225.1;refGene_gn=CARD11;refGene_tx=XM_023246226.1;refGene_gn=CARD1
1;refGene_tx=XM_023246227.1;refGene_gn=CARD11;refGene_tx=XM_023246228.
1
E3 2520742 2520940
refGene_gn=CARD11;refGene_tx=XM_011290436.3;refGene_gn=CARD11;refGene_
tx=XM_019820359.2;refGene_gn=CARD11;refGene_tx=XM_019820360.2;refGene_
gn=CARD11;refGene_tx=XM_019820361.2;refGene_gn=CARD11;refGene_tx=XM_02
3246225.1;refGene_gn=CARD11;refGene_tx=XM_023246226.1;refGene_gn=CARD1
1;refGene_tx=XM_023246227.1;refGene_gn=CARD11;refGene_tx=XM_023246228.
1
E3 2521543 2521669
refGene_gn=CARD11;refGene_tx=XM_011290436.3;refGene_gn=CARD11;refGene_
tx=XM_019820359.2;refGene_gn=CARD11;refGene_tx=XM_019820360.2;refGene_
gn=CARD11;refGene_tx=XM_019820361.2;refGene_gn=CARD11;refGene_tx=XM_02
3246225.1;refGene_gn=CARD11;refGene_tx=XM_023246226.1;refGene_gn=CARD1
1;refGene_tx=XM_023246227.1;refGene_gn=CARD11;refGene_tx=XM_023246228.
1
E3 2522178 2522331
refGene_gn=CARD11;refGene_tx=XM_011290436.3;refGene_gn=CARD11;refGene_
tx=XM_019820359.2;refGene_gn=CARD11;refGene_tx=XM_019820360.2;refGene_
gn=CARD11;refGene_tx=XM_019820361.2;refGene_gn=CARD11;refGene_tx=XM_02
3246225.1;refGene_gn=CARD11;refGene_tx=XM_023246226.1;refGene_gn=CARD1
1;refGene_tx=XM_023246227.1;refGene_gn=CARD11;refGene_tx=XM_023246228.
1
E3 2523214 2523394
refGene_gn=CARD11;refGene_tx=XM_011290436.3;refGene_gn=CARD11;refGene_
tx=XM_019820359.2;refGene_gn=CARD11;refGene_tx=XM_019820360.2;refGene_
gn=CARD11;refGene_tx=XM_019820361.2;refGene_gn=CARD11;refGene_tx=XM_02
3246225.1;refGene_gn=CARD11;refGene_tx=XM_023246226.1;refGene_gn=CARD1
1;refGene_tx=XM_023246227.1;refGene_gn=CARD11;refGene_tx=XM_023246228.
1
E3 2526275 2526601
refGene_gn=CARD11;refGene_tx=XM_011290436.3;refGene_gn=CARD11;refGene_

tx=XM_019820359.2;refGene_gn=CARD11;refGene_tx=XM_019820360.2;refGene_
gn=CARD11;refGene_tx=XM_019820361.2;refGene_gn=CARD11;refGene_tx=XM_02
3246225.1;refGene_gn=CARD11;refGene_tx=XM_023246226.1;refGene_gn=CARD1
1;refGene_tx=XM_023246227.1;refGene_gn=CARD11;refGene_tx=XM_023246228.
1
E3 2527638 2527776
refGene_gn=CARD11;refGene_tx=XM_011290436.3;refGene_gn=CARD11;refGene_
tx=XM_019820359.2;refGene_gn=CARD11;refGene_tx=XM_019820360.2;refGene_
gn=CARD11;refGene_tx=XM_019820361.2;refGene_gn=CARD11;refGene_tx=XM_02
3246225.1;refGene_gn=CARD11;refGene_tx=XM_023246226.1;refGene_gn=CARD1
1;refGene_tx=XM_023246227.1;refGene_gn=CARD11;refGene_tx=XM_023246228.
1
E3 2529428 2529641
refGene_gn=CARD11;refGene_tx=XM_011290436.3;refGene_gn=CARD11;refGene_
tx=XM_019820359.2;refGene_gn=CARD11;refGene_tx=XM_019820360.2;refGene_
gn=CARD11;refGene_tx=XM_019820361.2;refGene_gn=CARD11;refGene_tx=XM_02
3246225.1;refGene_gn=CARD11;refGene_tx=XM_023246226.1;refGene_gn=CARD1
1;refGene_tx=XM_023246227.1;refGene_gn=CARD11;refGene_tx=XM_023246228.
1
E3 2540645 2540652
refGene_gn=CARD11;refGene_tx=XM_011290436.3;refGene_gn=CARD11;refGene_
tx=XM_019820359.2;refGene_gn=CARD11;refGene_tx=XM_019820360.2;refGene_
gn=CARD11;refGene_tx=XM_019820361.2;refGene_gn=CARD11;refGene_tx=XM_02
3246225.1;refGene_gn=CARD11;refGene_tx=XM_023246226.1;refGene_gn=CARD1
1;refGene_tx=XM_023246227.1;refGene_gn=CARD11;refGene_tx=XM_023246228.
1
E3 2779096 2779337 refGene_gn=SDK1;refGene_tx=XM_023247068.1
E3 3128409 3128569 refGene_gn=SDK1;refGene_tx=XM_023247068.1
E3 3139542 3139649 refGene_gn=SDK1;refGene_tx=XM_023247068.1
E3 3141849 3141997 refGene_gn=SDK1;refGene_tx=XM_023247068.1
E3 3302049 3302183 refGene_gn=SDK1;refGene_tx=XM_023247068.1
E3 3419112 3419224 refGene_gn=SDK1;refGene_tx=XM_023247068.1
E3 3419943 3420134 refGene_gn=SDK1;refGene_tx=XM_023247068.1
E3 3428450 3428534 refGene_gn=SDK1;refGene_tx=XM_023247068.1
E3 3432241 3432436 refGene_gn=SDK1;refGene_tx=XM_023247068.1
E3 3435928 3436045 refGene_gn=SDK1;refGene_tx=XM_023247068.1
E3 3437565 3437733 refGene_gn=SDK1;refGene_tx=XM_023247068.1
E3 3439594 3439697 refGene_gn=SDK1;refGene_tx=XM_023247068.1
E3 3442054 3442231 refGene_gn=SDK1;refGene_tx=XM_023247068.1
E3 3451662 3451799 refGene_gn=SDK1;refGene_tx=XM_023247068.1
E3 3474634 3474782 refGene_gn=SDK1;refGene_tx=XM_023247068.1
E3 3475740 3475881 refGene_gn=SDK1;refGene_tx=XM_023247068.1
E3 3480261 3480443 refGene_gn=SDK1;refGene_tx=XM_023247068.1
E3 3502299 3502415 refGene_gn=SDK1;refGene_tx=XM_023247068.1
E3 3504168 3504361 refGene_gn=SDK1;refGene_tx=XM_023247068.1
E3 3511118 3511217 refGene_gn=SDK1;refGene_tx=XM_023247068.1
E3 3517031 3517223 refGene_gn=SDK1;refGene_tx=XM_023247068.1
E3 3519979 3520101 refGene_gn=SDK1;refGene_tx=XM_023247068.1
E3 3544156 3544266 refGene_gn=SDK1;refGene_tx=XM_023247068.1
E3 3545911 3546062 refGene_gn=SDK1;refGene_tx=XM_023247068.1

E3 3546661 3546899 refGene_gn=SDK1;refGene_tx=XM_023247068.1
E3 3554230 3554346 refGene_gn=SDK1;refGene_tx=XM_023247068.1
E3 3555642 3555832 refGene_gn=SDK1;refGene_tx=XM_023247068.1
E3 3559027 3559126 refGene_gn=SDK1;refGene_tx=XM_023247068.1
E3 3573091 3573286 refGene_gn=SDK1;refGene_tx=XM_023247068.1
E3 3575710 3575912 refGene_gn=SDK1;refGene_tx=XM_023247068.1
E3 3582473 3582577 refGene_gn=SDK1;refGene_tx=XM_023247068.1
E3 3584776 3584847 refGene_gn=SDK1;refGene_tx=XM_023247068.1
E3 3592315 3592451 refGene_gn=SDK1;refGene_tx=XM_023247068.1
E3 3593958 3594018 refGene_gn=SDK1;refGene_tx=XM_023247068.1
E3 3595910 3596012 refGene_gn=SDK1;refGene_tx=XM_023247068.1
E3 3616975 3617091 refGene_gn=SDK1;refGene_tx=XM_023247068.1
E3 3618630 3618817 refGene_gn=SDK1;refGene_tx=XM_023247068.1
E3 3620236 3620374 refGene_gn=SDK1;refGene_tx=XM_023247068.1
E3 3625924 3626086 refGene_gn=SDK1;refGene_tx=XM_023247068.1
E3 3626988 3627114 refGene_gn=SDK1;refGene_tx=XM_023247068.1
E3 3636154 3636319 refGene_gn=SDK1;refGene_tx=XM_023247068.1
E3 3639218 3639356 refGene_gn=SDK1;refGene_tx=XM_023247068.1
E3 3642155 3642276 refGene_gn=SDK1;refGene_tx=XM_023247068.1
E3 3642682 3642812 refGene_gn=SDK1;refGene_tx=XM_023247068.1
E3 3654966 3655218 refGene_gn=SDK1;refGene_tx=XM_023247068.1
E3 3862648 3863289 refGene_gn=FOXK1;refGene_tx=XM_023246698.1
E3 3909639 3909825 refGene_gn=FOXK1;refGene_tx=XM_023246698.1
E3 3919769 3919926 refGene_gn=FOXK1;refGene_tx=XM_023246698.1
E3 3920467 3920614 refGene_gn=FOXK1;refGene_tx=XM_023246698.1
E3 3921271 3921465 refGene_gn=FOXK1;refGene_tx=XM_023246698.1
E3 3923407 3923574 refGene_gn=FOXK1;refGene_tx=XM_023246698.1
E3 3923661 3923946 refGene_gn=FOXK1;refGene_tx=XM_023246698.1
E3 3924768 3924993 refGene_gn=FOXK1;refGene_tx=XM_023246698.1
E3 3925454 3925723 refGene_gn=FOXK1;refGene_tx=XM_023246698.1
E3 3937465 3937506 refGene_gn=AP5Z1;refGene_tx=XM_023246450.1
E3 3940379 3940517 refGene_gn=AP5Z1;refGene_tx=XM_023246450.1
E3 3940782 3940969 refGene_gn=AP5Z1;refGene_tx=XM_023246450.1
E3 3942498 3942643 refGene_gn=AP5Z1;refGene_tx=XM_023246450.1
E3 3942841 3942954 refGene_gn=AP5Z1;refGene_tx=XM_023246450.1
E3 3943359 3943528 refGene_gn=AP5Z1;refGene_tx=XM_023246450.1
E3 3943869 3944010 refGene_gn=AP5Z1;refGene_tx=XM_023246450.1
E3 3944291 3944329 refGene_gn=AP5Z1;refGene_tx=XM_023246450.1
E3 3944406 3944569 refGene_gn=AP5Z1;refGene_tx=XM_023246450.1
E3 3945048 3945227 refGene_gn=AP5Z1;refGene_tx=XM_023246450.1
E3 3945856 3945999 refGene_gn=AP5Z1;refGene_tx=XM_023246450.1
E3 3946277 3946418 refGene_gn=AP5Z1;refGene_tx=XM_023246450.1
E3 3947026 3947138 refGene_gn=AP5Z1;refGene_tx=XM_023246450.1
E3 3947417 3947515 refGene_gn=AP5Z1;refGene_tx=XM_023246450.1
E3 3947961 3948094 refGene_gn=AP5Z1;refGene_tx=XM_023246450.1
E3 3948172 3948387 refGene_gn=AP5Z1;refGene_tx=XM_023246450.1
E3 3948596 3948867 refGene_gn=AP5Z1;refGene_tx=XM_023246450.1
E3 3952439 3952545
refGene_gn=RADIL;refGene_tx=XM_006941891.3;refGene_gn=RADIL;refGene_tx=XM_006941892.3;refGene_gn=RADIL;refGene_tx=XM_019820372.2;refGene_gn=

RADIL;refGene_tx=XM_019820374.2;refGene_gn=RADIL;refGene_tx=XM_0198203
78.1;refGene_gn=RADIL;refGene_tx=XM_019820379.1;refGene_gn=RADIL;refGe
ne_tx=XM_019820380.1;refGene_gn=RADIL;refGene_tx=XM_019820381.1;refGen
e_gn=RADIL;refGene_tx=XM_019820383.2;refGene_gn=RADIL;refGene_tx=XM_01
9820385.1;refGene_gn=RADIL;refGene_tx=XM_019820387.1;refGene_gn=RADIL;
refGene_tx=XM_023246444.1;refGene_gn=RADIL;refGene_tx=XM_023246445.1;r
efGene_gn=RADIL;refGene_tx=XM_023246446.1;refGene_gn=RADIL;refGene_tx=
XM_023246447.1;refGene_gn=RADIL;refGene_tx=XM_023246449.1
E3 3952813 3952953
refGene_gn=RADIL;refGene_tx=XM_006941891.3;refGene_gn=RADIL;refGene_tx
=XM_006941892.3;refGene_gn=RADIL;refGene_tx=XM_019820372.2;refGene_gn=
RADIL;refGene_tx=XM_019820374.2;refGene_gn=RADIL;refGene_tx=XM_0198203
78.1;refGene_gn=RADIL;refGene_tx=XM_019820379.1;refGene_gn=RADIL;refGe
ne_tx=XM_019820380.1;refGene_gn=RADIL;refGene_tx=XM_019820381.1;refGen
e_gn=RADIL;refGene_tx=XM_019820383.2;refGene_gn=RADIL;refGene_tx=XM_01
9820385.1;refGene_gn=RADIL;refGene_tx=XM_019820387.1;refGene_gn=RADIL;
refGene_tx=XM_023246444.1;refGene_gn=RADIL;refGene_tx=XM_023246445.1;r
efGene_gn=RADIL;refGene_tx=XM_023246446.1;refGene_gn=RADIL;refGene_tx=
XM_023246447.1;refGene_gn=RADIL;refGene_tx=XM_023246449.1
E3 3953298 3953438
refGene_gn=RADIL;refGene_tx=XM_006941891.3;refGene_gn=RADIL;refGene_tx
=XM_006941892.3;refGene_gn=RADIL;refGene_tx=XM_019820372.2;refGene_gn=
RADIL;refGene_tx=XM_019820374.2;refGene_gn=RADIL;refGene_tx=XM_0198203
78.1;refGene_gn=RADIL;refGene_tx=XM_019820379.1;refGene_gn=RADIL;refGe
ne_tx=XM_019820380.1;refGene_gn=RADIL;refGene_tx=XM_019820381.1;refGen
e_gn=RADIL;refGene_tx=XM_019820383.2;refGene_gn=RADIL;refGene_tx=XM_01
9820385.1;refGene_gn=RADIL;refGene_tx=XM_019820387.1;refGene_gn=RADIL;
refGene_tx=XM_023246444.1;refGene_gn=RADIL;refGene_tx=XM_023246445.1;r
efGene_gn=RADIL;refGene_tx=XM_023246446.1;refGene_gn=RADIL;refGene_tx=
XM_023246447.1;refGene_gn=RADIL;refGene_tx=XM_023246449.1
E3 3954435 3954760
refGene_gn=RADIL;refGene_tx=XM_006941891.3;refGene_gn=RADIL;refGene_tx
=XM_006941892.3;refGene_gn=RADIL;refGene_tx=XM_019820372.2;refGene_gn=
RADIL;refGene_tx=XM_019820374.2;refGene_gn=RADIL;refGene_tx=XM_0198203
78.1;refGene_gn=RADIL;refGene_tx=XM_019820379.1;refGene_gn=RADIL;refGe
ne_tx=XM_019820380.1;refGene_gn=RADIL;refGene_tx=XM_019820381.1;refGen
e_gn=RADIL;refGene_tx=XM_019820383.2;refGene_gn=RADIL;refGene_tx=XM_01
9820385.1;refGene_gn=RADIL;refGene_tx=XM_019820387.1;refGene_gn=RADIL;
refGene_tx=XM_023246444.1;refGene_gn=RADIL;refGene_tx=XM_023246445.1;r
efGene_gn=RADIL;refGene_tx=XM_023246446.1;refGene_gn=RADIL;refGene_tx=
XM_023246447.1;refGene_gn=RADIL;refGene_tx=XM_023246449.1
E3 3954935 3955126
refGene_gn=RADIL;refGene_tx=XM_006941891.3;refGene_gn=RADIL;refGene_tx
=XM_006941892.3;refGene_gn=RADIL;refGene_tx=XM_019820372.2;refGene_gn=
RADIL;refGene_tx=XM_019820374.2;refGene_gn=RADIL;refGene_tx=XM_0198203
78.1;refGene_gn=RADIL;refGene_tx=XM_019820379.1;refGene_gn=RADIL;refGe
ne_tx=XM_019820380.1;refGene_gn=RADIL;refGene_tx=XM_019820381.1;refGen
e_gn=RADIL;refGene_tx=XM_019820383.2;refGene_gn=RADIL;refGene_tx=XM_01
9820385.1;refGene_gn=RADIL;refGene_tx=XM_019820387.1;refGene_gn=RADIL;
refGene_tx=XM_023246444.1;refGene_gn=RADIL;refGene_tx=XM_023246445.1;r

efGene_gn=RADIL;refGene_tx=XM_023246446.1;refGene_gn=RADIL;refGene_tx=XM_023246447.1;refGene_gn=RADIL;refGene_tx=XM_023246449.1
E3 3956041 3956192
refGene_gn=RADIL;refGene_tx=XM_006941891.3;refGene_gn=RADIL;refGene_tx=XM_006941892.3;refGene_gn=RADIL;refGene_tx=XM_019820372.2;refGene_gn=RADIL;refGene_tx=XM_019820374.2;refGene_gn=RADIL;refGene_tx=XM_019820378.1;refGene_gn=RADIL;refGene_tx=XM_019820379.1;refGene_gn=RADIL;refGene_tx=XM_019820380.1;refGene_gn=RADIL;refGene_tx=XM_019820381.1;refGene_gn=RADIL;refGene_tx=XM_019820383.2;refGene_gn=RADIL;refGene_tx=XM_019820385.1;refGene_gn=RADIL;refGene_tx=XM_019820387.1;refGene_gn=RADIL;refGene_tx=XM_023246444.1;refGene_gn=RADIL;refGene_tx=XM_023246445.1;refGene_gn=RADIL;refGene_tx=XM_023246446.1;refGene_gn=RADIL;refGene_tx=XM_023246447.1;refGene_gn=RADIL;refGene_tx=XM_023246449.1
E3 3959654 3959827
refGene_gn=RADIL;refGene_tx=XM_006941891.3;refGene_gn=RADIL;refGene_tx=XM_006941892.3;refGene_gn=RADIL;refGene_tx=XM_019820372.2;refGene_gn=RADIL;refGene_tx=XM_019820374.2;refGene_gn=RADIL;refGene_tx=XM_019820378.1;refGene_gn=RADIL;refGene_tx=XM_019820379.1;refGene_gn=RADIL;refGene_tx=XM_019820380.1;refGene_gn=RADIL;refGene_tx=XM_019820381.1;refGene_gn=RADIL;refGene_tx=XM_019820383.2;refGene_gn=RADIL;refGene_tx=XM_019820385.1;refGene_gn=RADIL;refGene_tx=XM_019820387.1;refGene_gn=RADIL;refGene_tx=XM_023246444.1;refGene_gn=RADIL;refGene_tx=XM_023246445.1;refGene_gn=RADIL;refGene_tx=XM_023246446.1;refGene_gn=RADIL;refGene_tx=XM_023246447.1;refGene_gn=RADIL;refGene_tx=XM_023246449.1
E3 3960350 3960588
refGene_gn=RADIL;refGene_tx=XM_006941891.3;refGene_gn=RADIL;refGene_tx=XM_006941892.3;refGene_gn=RADIL;refGene_tx=XM_019820372.2;refGene_gn=RADIL;refGene_tx=XM_019820374.2;refGene_gn=RADIL;refGene_tx=XM_019820378.1;refGene_gn=RADIL;refGene_tx=XM_019820379.1;refGene_gn=RADIL;refGene_tx=XM_019820380.1;refGene_gn=RADIL;refGene_tx=XM_019820381.1;refGene_gn=RADIL;refGene_tx=XM_019820383.2;refGene_gn=RADIL;refGene_tx=XM_019820385.1;refGene_gn=RADIL;refGene_tx=XM_019820387.1;refGene_gn=RADIL;refGene_tx=XM_023246444.1;refGene_gn=RADIL;refGene_tx=XM_023246445.1;refGene_gn=RADIL;refGene_tx=XM_023246446.1;refGene_gn=RADIL;refGene_tx=XM_023246447.1;refGene_gn=RADIL;refGene_tx=XM_023246449.1
E3 3961709 3961822
refGene_gn=RADIL;refGene_tx=XM_006941891.3;refGene_gn=RADIL;refGene_tx=XM_006941892.3;refGene_gn=RADIL;refGene_tx=XM_019820372.2;refGene_gn=RADIL;refGene_tx=XM_019820374.2;refGene_gn=RADIL;refGene_tx=XM_019820378.1;refGene_gn=RADIL;refGene_tx=XM_019820379.1;refGene_gn=RADIL;refGene_tx=XM_019820380.1;refGene_gn=RADIL;refGene_tx=XM_019820381.1;refGene_gn=RADIL;refGene_tx=XM_019820383.2;refGene_gn=RADIL;refGene_tx=XM_019820385.1;refGene_gn=RADIL;refGene_tx=XM_019820387.1;refGene_gn=RADIL;refGene_tx=XM_023246444.1;refGene_gn=RADIL;refGene_tx=XM_023246445.1;refGene_gn=RADIL;refGene_tx=XM_023246446.1;refGene_gn=RADIL;refGene_tx=XM_023246447.1;refGene_gn=RADIL;refGene_tx=XM_023246449.1
E3 3966100 3966261
refGene_gn=RADIL;refGene_tx=XM_006941891.3;refGene_gn=RADIL;refGene_tx=XM_006941892.3;refGene_gn=RADIL;refGene_tx=XM_019820372.2;refGene_gn=RADIL;refGene_tx=XM_019820374.2;refGene_gn=RADIL;refGene_tx=XM_0198203

78.1;refGene_gn=RADIL;refGene_tx=XM_019820379.1;refGene_gn=RADIL;refGene_tx=XM_019820380.1;refGene_gn=RADIL;refGene_tx=XM_019820381.1;refGene_gn=RADIL;refGene_tx=XM_019820383.2;refGene_gn=RADIL;refGene_tx=XM_019820385.1;refGene_gn=RADIL;refGene_tx=XM_019820387.1;refGene_gn=RADIL;refGene_tx=XM_023246444.1;refGene_gn=RADIL;refGene_tx=XM_023246445.1;refGene_gn=RADIL;refGene_tx=XM_023246446.1;refGene_gn=RADIL;refGene_tx=XM_023246447.1;refGene_gn=RADIL;refGene_tx=XM_023246449.1
E3 3972484 3972522
refGene_gn=RADIL;refGene_tx=XM_006941891.3;refGene_gn=RADIL;refGene_tx=XM_006941892.3;refGene_gn=RADIL;refGene_tx=XM_019820372.2;refGene_gn=RADIL;refGene_tx=XM_019820374.2;refGene_gn=RADIL;refGene_tx=XM_019820378.1;refGene_gn=RADIL;refGene_tx=XM_019820379.1;refGene_gn=RADIL;refGene_tx=XM_019820380.1;refGene_gn=RADIL;refGene_tx=XM_019820381.1;refGene_gn=RADIL;refGene_tx=XM_019820383.2;refGene_gn=RADIL;refGene_tx=XM_019820385.1;refGene_gn=RADIL;refGene_tx=XM_019820387.1;refGene_gn=RADIL;refGene_tx=XM_023246444.1;refGene_gn=RADIL;refGene_tx=XM_023246445.1;refGene_gn=RADIL;refGene_tx=XM_023246446.1;refGene_gn=RADIL;refGene_tx=XM_023246447.1;refGene_gn=RADIL;refGene_tx=XM_023246449.1
E3 3974489 3975128
refGene_gn=RADIL;refGene_tx=XM_006941891.3;refGene_gn=RADIL;refGene_tx=XM_006941892.3;refGene_gn=RADIL;refGene_tx=XM_019820372.2;refGene_gn=RADIL;refGene_tx=XM_019820374.2;refGene_gn=RADIL;refGene_tx=XM_019820378.1;refGene_gn=RADIL;refGene_tx=XM_019820379.1;refGene_gn=RADIL;refGene_tx=XM_019820380.1;refGene_gn=RADIL;refGene_tx=XM_019820381.1;refGene_gn=RADIL;refGene_tx=XM_019820383.2;refGene_gn=RADIL;refGene_tx=XM_019820385.1;refGene_gn=RADIL;refGene_tx=XM_019820387.1;refGene_gn=RADIL;refGene_tx=XM_023246444.1;refGene_gn=RADIL;refGene_tx=XM_023246445.1;refGene_gn=RADIL;refGene_tx=XM_023246446.1;refGene_gn=RADIL;refGene_tx=XM_023246447.1;refGene_gn=RADIL;refGene_tx=XM_023246449.1
E3 3976769 3977002
refGene_gn=RADIL;refGene_tx=XM_006941891.3;refGene_gn=RADIL;refGene_tx=XM_006941892.3;refGene_gn=RADIL;refGene_tx=XM_019820372.2;refGene_gn=RADIL;refGene_tx=XM_019820374.2;refGene_gn=RADIL;refGene_tx=XM_019820378.1;refGene_gn=RADIL;refGene_tx=XM_019820379.1;refGene_gn=RADIL;refGene_tx=XM_019820380.1;refGene_gn=RADIL;refGene_tx=XM_019820381.1;refGene_gn=RADIL;refGene_tx=XM_019820383.2;refGene_gn=RADIL;refGene_tx=XM_019820385.1;refGene_gn=RADIL;refGene_tx=XM_019820387.1;refGene_gn=RADIL;refGene_tx=XM_023246444.1;refGene_gn=RADIL;refGene_tx=XM_023246445.1;refGene_gn=RADIL;refGene_tx=XM_023246446.1;refGene_gn=RADIL;refGene_tx=XM_023246447.1;refGene_gn=RADIL;refGene_tx=XM_023246449.1
E3 4002378 4003974 refGene_gn=PAPOLB;refGene_tx=XM_011290434.3
E3 4003975 4004305 refGene_gn=PAPOLB;refGene_tx=XM_011290434.3
E3 4016526 4017125
refGene_gn=RADIL;refGene_tx=XM_006941891.3;refGene_gn=RADIL;refGene_tx=XM_006941892.3;refGene_gn=RADIL;refGene_tx=XM_019820372.2;refGene_gn=RADIL;refGene_tx=XM_019820374.2;refGene_gn=RADIL;refGene_tx=XM_019820378.1;refGene_gn=RADIL;refGene_tx=XM_019820379.1;refGene_gn=RADIL;refGene_tx=XM_019820380.1;refGene_gn=RADIL;refGene_tx=XM_019820381.1;refGene_gn=RADIL;refGene_tx=XM_019820383.2;refGene_gn=RADIL;refGene_tx=XM_019820385.1;refGene_gn=RADIL;refGene_tx=XM_019820387.1;refGene_gn=RADIL;

refGene_tx=XM_023246444.1;refGene_gn=RADIL;refGene_tx=XM_023246445.1;refGene_gn=RADIL;refGene_tx=XM_023246446.1;refGene_gn=RADIL;refGene_tx=XM_023246447.1;refGene_gn=RADIL;refGene_tx=XM_023246449.1
E3 4018264 4018315 refGene_gn=RADIL;refGene_tx=XM_019820385.1
E3 4033207 4033266 refGene_gn=RADIL;refGene_tx=XM_019820380.1;refGene_gn=RADIL;refGene_tx=XM_023246449.1
E3 4040510 4040554 refGene_gn=RADIL;refGene_tx=XM_019820385.1;refGene_gn=RADIL;refGene_tx=XM_019820387.1
E3 4048580 4048655 refGene_gn=RADIL;refGene_tx=XM_019820372.2;refGene_gn=RADIL;refGene_tx=XM_019820374.2;refGene_gn=RADIL;refGene_tx=XM_019820378.1;refGene_gn=RADIL;refGene_tx=XM_019820379.1;refGene_gn=RADIL;refGene_tx=XM_023246444.1;refGene_gn=RADIL;refGene_tx=XM_023246445.1;refGene_gn=RADIL;refGene_tx=XM_023246446.1;refGene_gn=RADIL;refGene_tx=XM_023246447.1
E3 4051515 4051676 refGene_gn=RADIL;refGene_tx=XM_019820372.2;refGene_gn=RADIL;refGene_tx=XM_019820374.2;refGene_gn=RADIL;refGene_tx=XM_023246444.1;refGene_gn=RADIL;refGene_tx=XM_023246445.1;refGene_gn=RADIL;refGene_tx=XM_023246446.1;refGene_gn=RADIL;refGene_tx=XM_023246447.1
E3 4053026 4053108 refGene_gn=RADIL;refGene_tx=XM_019820372.2;refGene_gn=RADIL;refGene_tx=XM_019820374.2;refGene_gn=RADIL;refGene_tx=XM_023246444.1;refGene_gn=RADIL;refGene_tx=XM_023246445.1;refGene_gn=RADIL;refGene_tx=XM_023246446.1;refGene_gn=RADIL;refGene_tx=XM_023246447.1
E3 4070737 4070855 refGene_gn=RADIL;refGene_tx=XM_023246444.1
E3 4081129 4081190 refGene_gn=RADIL;refGene_tx=XM_023246444.1;refGene_gn=RADIL;refGene_tx=XM_023246445.1;refGene_gn=RADIL;refGene_tx=XM_023246446.1
E3 4102600 4102674 refGene_gn=WIPI2;refGene_tx=XM_023247105.1;refGene_gn=WIPI2;refGene_tx=XM_023247106.1
E3 4109686 4109769 refGene_gn=WIPI2;refGene_tx=XM_023247105.1;refGene_gn=WIPI2;refGene_tx=XM_023247106.1
E3 4128349 4128519 refGene_gn=WIPI2;refGene_tx=XM_023247105.1;refGene_gn=WIPI2;refGene_tx=XM_023247106.1
E3 4131486 4131583 refGene_gn=WIPI2;refGene_tx=XM_023247105.1;refGene_gn=WIPI2;refGene_tx=XM_023247106.1;refGene_gn=WIPI2;refGene_tx=XM_023247107.1
E3 4132227 4132325 refGene_gn=WIPI2;refGene_tx=XM_023247105.1;refGene_gn=WIPI2;refGene_tx=XM_023247106.1;refGene_gn=WIPI2;refGene_tx=XM_023247107.1
E3 4132753 4132846 refGene_gn=WIPI2;refGene_tx=XM_023247105.1;refGene_gn=WIPI2;refGene_tx=XM_023247106.1;refGene_gn=WIPI2;refGene_tx=XM_023247107.1
E3 4135512 4135583 refGene_gn=WIPI2;refGene_tx=XM_023247105.1;refGene_gn=WIPI2;refGene_tx

=XM_023247106.1;refGene_gn=WIPI2;refGene_tx=XM_023247107.1
E3	4137872	4137980
refGene_gn=WIPI2;refGene_tx=XM_023247105.1;refGene_gn=WIPI2;refGene_tx=XM_023247106.1;refGene_gn=WIPI2;refGene_tx=XM_023247107.1
E3	4138918	4139083
refGene_gn=WIPI2;refGene_tx=XM_023247105.1;refGene_gn=WIPI2;refGene_tx=XM_023247106.1;refGene_gn=WIPI2;refGene_tx=XM_023247107.1
E3	4139951	4140059
refGene_gn=WIPI2;refGene_tx=XM_023247105.1;refGene_gn=WIPI2;refGene_tx=XM_023247106.1;refGene_gn=WIPI2;refGene_tx=XM_023247107.1
E3	4140882	4141013
refGene_gn=WIPI2;refGene_tx=XM_023247105.1;refGene_gn=WIPI2;refGene_tx=XM_023247106.1;refGene_gn=WIPI2;refGene_tx=XM_023247107.1
E3	4141885	4141998
refGene_gn=WIPI2;refGene_tx=XM_023247105.1;refGene_gn=WIPI2;refGene_tx=XM_023247106.1;refGene_gn=WIPI2;refGene_tx=XM_023247107.1
E3	4181191	4181366
refGene_gn=SLC29A4;refGene_tx=XM_023246820.1;refGene_gn=SLC29A4;refGene_tx=XM_023246821.1;refGene_gn=SLC29A4;refGene_tx=XM_023246822.1;refGene_gn=SLC29A4;refGene_tx=XM_023246823.1
E3	4184388	4184520
refGene_gn=SLC29A4;refGene_tx=XM_023246820.1;refGene_gn=SLC29A4;refGene_tx=XM_023246821.1;refGene_gn=SLC29A4;refGene_tx=XM_023246822.1;refGene_gn=SLC29A4;refGene_tx=XM_023246823.1
E3	4184803	4184917
refGene_gn=SLC29A4;refGene_tx=XM_023246820.1;refGene_gn=SLC29A4;refGene_tx=XM_023246821.1;refGene_gn=SLC29A4;refGene_tx=XM_023246822.1;refGene_gn=SLC29A4;refGene_tx=XM_023246823.1
E3	4185387	4185516
refGene_gn=SLC29A4;refGene_tx=XM_023246820.1;refGene_gn=SLC29A4;refGene_tx=XM_023246821.1;refGene_gn=SLC29A4;refGene_tx=XM_023246822.1;refGene_gn=SLC29A4;refGene_tx=XM_023246823.1
E3	4187209	4187284
refGene_gn=SLC29A4;refGene_tx=XM_023246820.1;refGene_gn=SLC29A4;refGene_tx=XM_023246821.1;refGene_gn=SLC29A4;refGene_tx=XM_023246822.1;refGene_gn=SLC29A4;refGene_tx=XM_023246823.1
E3	4187986	4188055
refGene_gn=SLC29A4;refGene_tx=XM_023246820.1;refGene_gn=SLC29A4;refGene_tx=XM_023246821.1
E3	4188659	4188907
refGene_gn=SLC29A4;refGene_tx=XM_023246820.1;refGene_gn=SLC29A4;refGene_tx=XM_023246821.1;refGene_gn=SLC29A4;refGene_tx=XM_023246822.1;refGene_gn=SLC29A4;refGene_tx=XM_023246823.1
E3	4189473	4189615
refGene_gn=SLC29A4;refGene_tx=XM_023246820.1;refGene_gn=SLC29A4;refGene_tx=XM_023246821.1;refGene_gn=SLC29A4;refGene_tx=XM_023246822.1;refGene_gn=SLC29A4;refGene_tx=XM_023246823.1
E3	4189833	4190021
refGene_gn=SLC29A4;refGene_tx=XM_023246820.1;refGene_gn=SLC29A4;refGene_tx=XM_023246821.1;refGene_gn=SLC29A4;refGene_tx=XM_023246822.1;refGe

ne_gn=SLC29A4;refGene_tx=XM_023246823.1
E3 4190847 4191088
refGene_gn=SLC29A4;refGene_tx=XM_023246820.1;refGene_gn=SLC29A4;refGene_tx=XM_023246821.1;refGene_gn=SLC29A4;refGene_tx=XM_023246822.1;refGene_gn=SLC29A4;refGene_tx=XM_023246823.1
E3 4192291 4192449
refGene_gn=SLC29A4;refGene_tx=XM_023246820.1;refGene_gn=SLC29A4;refGene_tx=XM_023246821.1;refGene_gn=SLC29A4;refGene_tx=XM_023246822.1;refGene_gn=SLC29A4;refGene_tx=XM_023246823.1
E3 4196155 4196362 refGene_gn=TNRC18;refGene_tx=XM_023246817.1
E3 4196671 4196746 refGene_gn=TNRC18;refGene_tx=XM_023246817.1
E3 4196884 4197121 refGene_gn=TNRC18;refGene_tx=XM_023246817.1
E3 4200413 4201475 refGene_gn=TNRC18;refGene_tx=XM_023246817.1
E3 4201476 4201697 refGene_gn=TNRC18;refGene_tx=XM_023246817.1
E3 4201699 4201738 refGene_gn=TNRC18;refGene_tx=XM_023246817.1
E3 4202214 4202379 refGene_gn=TNRC18;refGene_tx=XM_023246817.1
E3 4203074 4203191 refGene_gn=TNRC18;refGene_tx=XM_023246817.1
E3 4205560 4205669 refGene_gn=TNRC18;refGene_tx=XM_023246817.1
E3 4205759 4205835 refGene_gn=TNRC18;refGene_tx=XM_023246817.1
E3 4206287 4206405 refGene_gn=TNRC18;refGene_tx=XM_023246817.1
E3 4208202 4208344 refGene_gn=TNRC18;refGene_tx=XM_023246817.1
E3 4208884 4209040 refGene_gn=TNRC18;refGene_tx=XM_023246817.1
E3 4213237 4213668 refGene_gn=TNRC18;refGene_tx=XM_023246817.1
E3 4225810 4226059 refGene_gn=TNRC18;refGene_tx=XM_023246817.1
E3 4230964 4231240 refGene_gn=TNRC18;refGene_tx=XM_023246817.1
E3 4234381 4234742 refGene_gn=TNRC18;refGene_tx=XM_023246817.1
E3 4236138 4236310 refGene_gn=TNRC18;refGene_tx=XM_023246817.1
E3 4237968 4238097 refGene_gn=TNRC18;refGene_tx=XM_023246817.1
E3 4238333 4238470 refGene_gn=TNRC18;refGene_tx=XM_023246817.1
E3 4239000 4239176 refGene_gn=TNRC18;refGene_tx=XM_023246817.1
E3 4243987 4244986 refGene_gn=TNRC18;refGene_tx=XM_023246817.1
E3 4246210 4246640 refGene_gn=TNRC18;refGene_tx=XM_023246817.1
E3 4247935 4248126 refGene_gn=TNRC18;refGene_tx=XM_023246817.1
E3 4248628 4248775 refGene_gn=TNRC18;refGene_tx=XM_023246817.1
E3 4249195 4249401 refGene_gn=TNRC18;refGene_tx=XM_023246817.1
E3 4249681 4249784 refGene_gn=TNRC18;refGene_tx=XM_023246817.1
E3 4255696 4257418 refGene_gn=TNRC18;refGene_tx=XM_023246817.1
E3 4257874 4258000 refGene_gn=TNRC18;refGene_tx=XM_023246817.1
E3 4260104 4260260 refGene_gn=TNRC18;refGene_tx=XM_023246817.1
E3 4280294 4280481 refGene_gn=TNRC18;refGene_tx=XM_023246817.1
E3 4293537 4293616 refGene_gn=FBXL18;refGene_tx=XM_019820391.2
E3 4330808 4330965
refGene_gn=FBXL18;refGene_tx=XM_011290431.3;refGene_gn=FBXL18;refGene_tx=XM_019820392.2
E3 4340538 4340732 refGene_gn=FBXL18;refGene_tx=XM_023246819.1
E3 4347130 4347351
refGene_gn=FBXL18;refGene_tx=XM_011290431.3;refGene_gn=FBXL18;refGene_tx=XM_019820391.2;refGene_gn=FBXL18;refGene_tx=XM_023246819.1
E3 4360459 4362003
refGene_gn=FBXL18;refGene_tx=XM_011290431.3;refGene_gn=FBXL18;refGene_

tx=XM_019820391.2;refGene_gn=FBXL18;refGene_tx=XM_019820392.2;refGene_gn=FBXL18;refGene_tx=XM_023246819.1
E3 4365323 4365539
refGene_gn=FBXL18;refGene_tx=XM_011290431.3;refGene_gn=FBXL18;refGene_tx=XM_019820391.2;refGene_gn=FBXL18;refGene_tx=XM_019820392.2;refGene_gn=FBXL18;refGene_tx=XM_023246819.1
E3 4371101 4371119
refGene_gn=FBXL18;refGene_tx=XM_011290431.3;refGene_gn=FBXL18;refGene_tx=XM_019820391.2;refGene_gn=FBXL18;refGene_tx=XM_019820392.2;refGene_gn=FBXL18;refGene_tx=XM_023246819.1
E3 4378945 4379089 refGene_gn=ACTB;refGene_tx=XM_006941899.3
E3 4379209 4379391 refGene_gn=ACTB;refGene_tx=XM_006941899.3
E3 4379480 4379919 refGene_gn=ACTB;refGene_tx=XM_006941899.3
E3 4380363 4380603 refGene_gn=ACTB;refGene_tx=XM_006941899.3
E3 4380789 4380912 refGene_gn=ACTB;refGene_tx=XM_006941899.3
E3 4426617 4427449 refGene_gn=FSCN1;refGene_tx=XM_023246455.1
E3 4433594 4433751 refGene_gn=FSCN1;refGene_tx=XM_023246455.1
E3 4433855 4433977 refGene_gn=FSCN1;refGene_tx=XM_023246455.1
E3 4434228 4434396 refGene_gn=FSCN1;refGene_tx=XM_023246455.1
E3 4434751 4434954 refGene_gn=FSCN1;refGene_tx=XM_023246455.1
E3 4451222 4451542 refGene_gn=RNF216;refGene_tx=XM_023246734.1
E3 4452184 4452254 refGene_gn=RNF216;refGene_tx=XM_023246734.1
E3 4467635 4467858 refGene_gn=RNF216;refGene_tx=XM_023246734.1
E3 4475744 4475842 refGene_gn=RNF216;refGene_tx=XM_023246734.1
E3 4539403 4539482 refGene_gn=RNF216;refGene_tx=XM_023246734.1
E3 4539988 4540137 refGene_gn=RNF216;refGene_tx=XM_023246734.1
E3 4541509 4541647 refGene_gn=RNF216;refGene_tx=XM_023246734.1
E3 4542558 4542609 refGene_gn=RNF216;refGene_tx=XM_023246734.1
E3 4546055 4546195 refGene_gn=RNF216;refGene_tx=XM_023246734.1
E3 4549912 4550027 refGene_gn=RNF216;refGene_tx=XM_023246734.1
E3 4557226 4557402 refGene_gn=RNF216;refGene_tx=XM_023246734.1
E3 4559273 4559380 refGene_gn=RNF216;refGene_tx=XM_023246734.1
E3 4567305 4567382 refGene_gn=RNF216;refGene_tx=XM_023246734.1
E3 4568535 4569219 refGene_gn=RNF216;refGene_tx=XM_023246734.1
E3 4577353 4577487 refGene_gn=RNF216;refGene_tx=XM_023246734.1
E3 4582890 4582957 refGene_gn=RNF216;refGene_tx=XM_023246734.1
E3 4651234 4651290
refGene_gn=LOC105261300;refGene_tx=XM_023246250.1;refGene_gn=LOC105261300;refGene_tx=XM_023246251.1;refGene_gn=LOC105261300;refGene_tx=XM_023246252.1;refGene_gn=LOC105261300;refGene_tx=XM_023246253.1
E3 4652362 4652405
refGene_gn=LOC105261300;refGene_tx=XM_023246250.1;refGene_gn=LOC105261300;refGene_tx=XM_023246251.1;refGene_gn=LOC105261300;refGene_tx=XM_023246252.1;refGene_gn=LOC105261300;refGene_tx=XM_023246253.1;refGene_gn=LOC105261300;refGene_tx=XM_023246254.1;refGene_gn=LOC105261300;refGene_tx=XM_023246255.1
E3 4656072 4656183
refGene_gn=LOC105261300;refGene_tx=XM_023246241.1;refGene_gn=LOC105261300;refGene_tx=XM_023246242.1;refGene_gn=LOC105261300;refGene_tx=XM_023246243.1;refGene_gn=LOC105261300;refGene_tx=XM_023246244.1;refGene_gn

=LOC105261300;refGene_tx=XM_023246245.1;refGene_gn=LOC105261300;refGene_tx=XM_023246246.1;refGene_gn=LOC105261300;refGene_tx=XM_023246247.1;refGene_gn=LOC105261300;refGene_tx=XM_023246249.1
E3 4659204 4659387
refGene_gn=LOC105261300;refGene_tx=XM_023246240.1
E3 4659698 4659825
refGene_gn=LOC105261300;refGene_tx=XM_023246240.1;refGene_gn=LOC105261300;refGene_tx=XM_023246241.1;refGene_gn=LOC105261300;refGene_tx=XM_023246242.1;refGene_gn=LOC105261300;refGene_tx=XM_023246243.1;refGene_gn=LOC105261300;refGene_tx=XM_023246244.1;refGene_gn=LOC105261300;refGene_tx=XM_023246245.1;refGene_gn=LOC105261300;refGene_tx=XM_023246246.1;refGene_gn=LOC105261300;refGene_tx=XM_023246247.1;refGene_gn=LOC105261300;refGene_tx=XM_023246249.1;refGene_gn=LOC105261300;refGene_tx=XM_023246250.1;refGene_gn=LOC105261300;refGene_tx=XM_023246251.1;refGene_gn=LOC105261300;refGene_tx=XM_023246252.1;refGene_gn=LOC105261300;refGene_tx=XM_023246253.1;refGene_gn=LOC105261300;refGene_tx=XM_023246254.1;refGene_gn=LOC105261300;refGene_tx=XM_023246255.1;refGene_gn=LOC105261300;refGene_tx=XM_023246256.1;refGene_gn=LOC105261300;refGene_tx=XM_023246257.1
E3 4660073 4660169
refGene_gn=LOC105261300;refGene_tx=XM_023246240.1;refGene_gn=LOC105261300;refGene_tx=XM_023246241.1;refGene_gn=LOC105261300;refGene_tx=XM_023246242.1;refGene_gn=LOC105261300;refGene_tx=XM_023246243.1;refGene_gn=LOC105261300;refGene_tx=XM_023246244.1;refGene_gn=LOC105261300;refGene_tx=XM_023246245.1;refGene_gn=LOC105261300;refGene_tx=XM_023246246.1;refGene_gn=LOC105261300;refGene_tx=XM_023246247.1;refGene_gn=LOC105261300;refGene_tx=XM_023246249.1;refGene_gn=LOC105261300;refGene_tx=XM_023246250.1;refGene_gn=LOC105261300;refGene_tx=XM_023246251.1;refGene_gn=LOC105261300;refGene_tx=XM_023246252.1;refGene_gn=LOC105261300;refGene_tx=XM_023246253.1;refGene_gn=LOC105261300;refGene_tx=XM_023246254.1;refGene_gn=LOC105261300;refGene_tx=XM_023246255.1;refGene_gn=LOC105261300;refGene_tx=XM_023246256.1;refGene_gn=LOC105261300;refGene_tx=XM_023246257.1
E3 4662515 4663480
refGene_gn=LOC105261300;refGene_tx=XM_023246240.1;refGene_gn=LOC105261300;refGene_tx=XM_023246241.1;refGene_gn=LOC105261300;refGene_tx=XM_023246242.1;refGene_gn=LOC105261300;refGene_tx=XM_023246243.1;refGene_gn=LOC105261300;refGene_tx=XM_023246244.1;refGene_gn=LOC105261300;refGene_tx=XM_023246245.1;refGene_gn=LOC105261300;refGene_tx=XM_023246246.1;refGene_gn=LOC105261300;refGene_tx=XM_023246247.1;refGene_gn=LOC105261300;refGene_tx=XM_023246249.1;refGene_gn=LOC105261300;refGene_tx=XM_023246250.1;refGene_gn=LOC105261300;refGene_tx=XM_023246251.1;refGene_gn=LOC105261300;refGene_tx=XM_023246252.1;refGene_gn=LOC105261300;refGene_tx=XM_023246253.1;refGene_gn=LOC105261300;refGene_tx=XM_023246254.1;refGene_gn=LOC105261300;refGene_tx=XM_023246255.1;refGene_gn=LOC105261300;refGene_tx=XM_023246256.1;refGene_gn=LOC105261300;refGene_tx=XM_023246257.1
E3 4669937 4670915
refGene_gn=LOC101099262;refGene_tx=XM_003998438.3
E3 4695338 4697254

refGene_gn=RBAK;refGene_tx=XM_006941900.4;refGene_gn=RBAK;refGene_tx=XM_006941901.4;refGene_gn=RBAK;refGene_tx=XM_006941902.4;refGene_gn=RBAK;refGene_tx=XM_019820394.2
E3 4700447 4700543
refGene_gn=RBAK;refGene_tx=XM_006941900.4;refGene_gn=RBAK;refGene_tx=XM_006941901.4;refGene_gn=RBAK;refGene_tx=XM_006941902.4;refGene_gn=RBAK;refGene_tx=XM_019820394.2
E3 4700829 4700956
refGene_gn=RBAK;refGene_tx=XM_006941900.4;refGene_gn=RBAK;refGene_tx=XM_006941901.4;refGene_gn=RBAK;refGene_tx=XM_006941902.4;refGene_gn=RBAK;refGene_tx=XM_019820394.2
E3 4712332 4712347
refGene_gn=RBAK;refGene_tx=XM_006941900.4;refGene_gn=RBAK;refGene_tx=XM_006941901.4;refGene_gn=RBAK;refGene_tx=XM_006941902.4
E3 4722930 4723039
refGene_gn=LOC101099765;refGene_tx=XM_023246259.1
E3 4723526 4723724
refGene_gn=LOC101099765;refGene_tx=XM_023246259.1
E3 4724346 4724405
refGene_gn=LOC101099765;refGene_tx=XM_023246259.1
E3 4724908 4724994
refGene_gn=LOC101099765;refGene_tx=XM_023246259.1
E3 4727622 4727791
refGene_gn=LOC101099765;refGene_tx=XM_023246259.1
E3 4727951 4728011
refGene_gn=LOC101099765;refGene_tx=XM_023246259.1
E3 4734681 4734720 refGene_gn=ZNF12;refGene_tx=XM_011290426.3
E3 4740471 4740598 refGene_gn=ZNF12;refGene_tx=XM_011290426.3
E3 4740848 4740944 refGene_gn=ZNF12;refGene_tx=XM_011290426.3
E3 4744714 4746564 refGene_gn=ZNF12;refGene_tx=XM_011290426.3
E3 4749437 4749592
refGene_gn=LOC101100018;refGene_tx=XM_023247069.1
E3 4759626 4760772
refGene_gn=LOC102900594;refGene_tx=XM_023246258.1
E3 4765185 4765281
refGene_gn=LOC101100018;refGene_tx=XM_023247069.1
E3 4765568 4765710
refGene_gn=LOC101100018;refGene_tx=XM_023247069.1
E3 4776532 4776590
refGene_gn=LOC101100018;refGene_tx=XM_023247069.1
E3 4777724 4777901
refGene_gn=LOC101100018;refGene_tx=XM_023247069.1
E3 4778575 4778826
refGene_gn=LOC101100018;refGene_tx=XM_023247069.1
E3 4795728 4798052
refGene_gn=ZNF316;refGene_tx=XM_019820245.2;refGene_gn=ZNF316;refGene_tx=XM_023246890.1
E3 4805111 4805225
refGene_gn=ZNF316;refGene_tx=XM_019820245.2;refGene_gn=ZNF316;refGene_tx=XM_023246890.1

E3 4805780 4805907
refGene_gn=ZNF316;refGene_tx=XM_019820245.2;refGene_gn=ZNF316;refGene_tx=XM_023246890.1
E3 4806726 4806851
refGene_gn=ZNF316;refGene_tx=XM_019820245.2;refGene_gn=ZNF316;refGene_tx=XM_023246890.1
E3 4807063 4807412
refGene_gn=ZNF316;refGene_tx=XM_019820245.2;refGene_gn=ZNF316;refGene_tx=XM_023246890.1
E3 4830094 4831543 refGene_gn=ZNF853;refGene_tx=XM_023246891.1
E3 4831684 4832043 refGene_gn=ZNF853;refGene_tx=XM_023246891.1
E3 4834524 4834642 refGene_gn=ZNF853;refGene_tx=XM_023246891.1
E3 4835384 4835396 refGene_gn=ZNF853;refGene_tx=XM_023246891.1
E3 4851943 4852145
refGene_gn=CE3H7orf26;refGene_tx=XM_023246503.1;refGene_gn=CE3H7orf26;refGene_tx=XM_023246504.1
E3 4855978 4856018
refGene_gn=CE3H7orf26;refGene_tx=XM_023246503.1;refGene_gn=CE3H7orf26;refGene_tx=XM_023246504.1
E3 4857015 4857540
refGene_gn=CE3H7orf26;refGene_tx=XM_023246503.1;refGene_gn=CE3H7orf26;refGene_tx=XM_023246504.1
E3 4862168 4862334
refGene_gn=CE3H7orf26;refGene_tx=XM_023246503.1;refGene_gn=CE3H7orf26;refGene_tx=XM_023246504.1
E3 4864446 4864668
refGene_gn=CE3H7orf26;refGene_tx=XM_023246503.1;refGene_gn=CE3H7orf26;refGene_tx=XM_023246504.1
E3 4865183 4865375
refGene_gn=CE3H7orf26;refGene_tx=XM_023246503.1;refGene_gn=CE3H7orf26;refGene_tx=XM_023246504.1
E3 4867136 4867427
refGene_gn=ZDHHC4;refGene_tx=XM_003998427.5;refGene_gn=ZDHHC4;refGene_tx=XM_006941906.4;refGene_gn=ZDHHC4;refGene_tx=XM_006941907.4
E3 4875037 4875282
refGene_gn=ZDHHC4;refGene_tx=XM_003998427.5;refGene_gn=ZDHHC4;refGene_tx=XM_006941906.4;refGene_gn=ZDHHC4;refGene_tx=XM_006941907.4
E3 4877096 4877222
refGene_gn=ZDHHC4;refGene_tx=XM_003998427.5;refGene_gn=ZDHHC4;refGene_tx=XM_006941906.4;refGene_gn=ZDHHC4;refGene_tx=XM_006941907.4
E3 4879245 4879424
refGene_gn=ZDHHC4;refGene_tx=XM_003998427.5;refGene_gn=ZDHHC4;refGene_tx=XM_006941906.4;refGene_gn=ZDHHC4;refGene_tx=XM_006941907.4
E3 4879893 4879967
refGene_gn=ZDHHC4;refGene_tx=XM_003998427.5;refGene_gn=ZDHHC4;refGene_tx=XM_006941906.4;refGene_gn=ZDHHC4;refGene_tx=XM_006941907.4
E3 4880791 4880908
refGene_gn=ZDHHC4;refGene_tx=XM_003998427.5;refGene_gn=ZDHHC4;refGene_tx=XM_006941906.4
E3 4906690 4907104 refGene_gn=GRID2IP;refGene_tx=XM_023246217.1

E3 4914937 4915092 refGene_gn=GRID2IP;refGene_tx=XM_023246217.1
E3 4923048 4923297 refGene_gn=GRID2IP;refGene_tx=XM_023246217.1
E3 4923534 4923620 refGene_gn=GRID2IP;refGene_tx=XM_023246217.1
E3 4926208 4926278 refGene_gn=GRID2IP;refGene_tx=XM_023246217.1
E3 4926776 4926871 refGene_gn=GRID2IP;refGene_tx=XM_023246217.1
E3 4927184 4927368 refGene_gn=GRID2IP;refGene_tx=XM_023246217.1
E3 4930078 4930233 refGene_gn=GRID2IP;refGene_tx=XM_023246217.1
E3 4932571 4932721 refGene_gn=GRID2IP;refGene_tx=XM_023246217.1
E3 4932983 4933081 refGene_gn=GRID2IP;refGene_tx=XM_023246217.1
E3 4933256 4933374 refGene_gn=GRID2IP;refGene_tx=XM_023246217.1
E3 4934120 4934476 refGene_gn=GRID2IP;refGene_tx=XM_023246217.1
E3 4934833 4935250 refGene_gn=GRID2IP;refGene_tx=XM_023246217.1
E3 4935769 4935857 refGene_gn=GRID2IP;refGene_tx=XM_023246217.1
E3 4936670 4936748 refGene_gn=GRID2IP;refGene_tx=XM_023246217.1
E3 4937277 4937474 refGene_gn=GRID2IP;refGene_tx=XM_023246217.1
E3 4937824 4937980 refGene_gn=GRID2IP;refGene_tx=XM_023246217.1
E3 4938118 4938205 refGene_gn=GRID2IP;refGene_tx=XM_023246217.1
E3 4939102 4939232 refGene_gn=GRID2IP;refGene_tx=XM_023246217.1
E3 4939322 4939441 refGene_gn=GRID2IP;refGene_tx=XM_023246217.1
E3 4940058 4940223 refGene_gn=GRID2IP;refGene_tx=XM_023246217.1
E3 4940452 4940524 refGene_gn=GRID2IP;refGene_tx=XM_023246217.1
E3 4959897 4959988 refGene_gn=KDELR2;refGene_tx=XM_011290425.3
E3 4969291 4969392 refGene_gn=KDELR2;refGene_tx=XM_011290425.3
E3 4973165 4973324 refGene_gn=KDELR2;refGene_tx=XM_011290425.3
E3 4974398 4974651 refGene_gn=KDELR2;refGene_tx=XM_011290425.3
E3 4975826 4975861 refGene_gn=KDELR2;refGene_tx=XM_011290425.3
E3 4982289 4982384
refGene_gn=DAGLB;refGene_tx=XM_023246211.1;refGene_gn=DAGLB;refGene_tx
=XM_023246212.1
E3 4984256 4984408
refGene_gn=DAGLB;refGene_tx=XM_023246211.1;refGene_gn=DAGLB;refGene_tx
=XM_023246212.1
E3 4992336 4992508
refGene_gn=DAGLB;refGene_tx=XM_023246211.1;refGene_gn=DAGLB;refGene_tx
=XM_023246212.1;refGene_gn=DAGLB;refGene_tx=XM_023246213.1
E3 4993712 4993971
refGene_gn=DAGLB;refGene_tx=XM_023246211.1;refGene_gn=DAGLB;refGene_tx
=XM_023246212.1;refGene_gn=DAGLB;refGene_tx=XM_023246213.1
E3 4995624 4995747
refGene_gn=DAGLB;refGene_tx=XM_023246211.1;refGene_gn=DAGLB;refGene_tx
=XM_023246212.1;refGene_gn=DAGLB;refGene_tx=XM_023246213.1
E3 4996329 4996457
refGene_gn=DAGLB;refGene_tx=XM_023246211.1;refGene_gn=DAGLB;refGene_tx
=XM_023246212.1;refGene_gn=DAGLB;refGene_tx=XM_023246213.1
E3 4997929 4998056
refGene_gn=DAGLB;refGene_tx=XM_023246211.1;refGene_gn=DAGLB;refGene_tx
=XM_023246212.1;refGene_gn=DAGLB;refGene_tx=XM_023246213.1
E3 4998810 4998894
refGene_gn=DAGLB;refGene_tx=XM_023246211.1;refGene_gn=DAGLB;refGene_tx
=XM_023246212.1;refGene_gn=DAGLB;refGene_tx=XM_023246213.1

E3	5000281	5000359
refGene_gn=DAGLB;refGene_tx=XM_023246211.1;refGene_gn=DAGLB;refGene_tx
=XM_023246212.1;refGene_gn=DAGLB;refGene_tx=XM_023246213.1
E3	5002991	5003072
refGene_gn=DAGLB;refGene_tx=XM_023246211.1;refGene_gn=DAGLB;refGene_tx
=XM_023246212.1;refGene_gn=DAGLB;refGene_tx=XM_023246213.1
E3	5003162	5003290
refGene_gn=DAGLB;refGene_tx=XM_023246211.1;refGene_gn=DAGLB;refGene_tx
=XM_023246212.1;refGene_gn=DAGLB;refGene_tx=XM_023246213.1
E3	5013631	5013700
refGene_gn=DAGLB;refGene_tx=XM_023246211.1;refGene_gn=DAGLB;refGene_tx
=XM_023246212.1;refGene_gn=DAGLB;refGene_tx=XM_023246213.1
E3	5013782	5013855
refGene_gn=DAGLB;refGene_tx=XM_023246211.1;refGene_gn=DAGLB;refGene_tx
=XM_023246212.1;refGene_gn=DAGLB;refGene_tx=XM_023246213.1
E3	5015350	5015601
refGene_gn=DAGLB;refGene_tx=XM_023246211.1;refGene_gn=DAGLB;refGene_tx
=XM_023246212.1;refGene_gn=DAGLB;refGene_tx=XM_023246213.1
E3	5015690	5015882
refGene_gn=DAGLB;refGene_tx=XM_023246211.1;refGene_gn=DAGLB;refGene_tx
=XM_023246212.1;refGene_gn=DAGLB;refGene_tx=XM_023246213.1
E3	5016287	5016333
refGene_gn=DAGLB;refGene_tx=XM_023246211.1;refGene_gn=DAGLB;refGene_tx
=XM_023246213.1
E3	5018760	5018773	refGene_gn=DAGLB;refGene_tx=XM_023246212.1
E3	5020526	5020657
refGene_gn=RAC1;refGene_tx=XM_023246215.1;refGene_gn=RAC1;refGene_tx=X
M_023246216.1
E3	5020910	5021070
refGene_gn=RAC1;refGene_tx=XM_023246215.1;refGene_gn=RAC1;refGene_tx=X
M_023246216.1
E3	5021813	5021876
refGene_gn=RAC1;refGene_tx=XM_023246215.1;refGene_gn=RAC1;refGene_tx=X
M_023246216.1
E3	5023343	5023400	refGene_gn=RAC1;refGene_tx=XM_023246216.1
E3	5029566	5029684
refGene_gn=RAC1;refGene_tx=XM_023246215.1;refGene_gn=RAC1;refGene_tx=X
M_023246216.1
E3	5032356	5032428
refGene_gn=RAC1;refGene_tx=XM_023246215.1;refGene_gn=RAC1;refGene_tx=X
M_023246216.1
E3	5042334	5042369
refGene_gn=RAC1;refGene_tx=XM_023246215.1;refGene_gn=RAC1;refGene_tx=X
M_023246216.1
E3	5074637	5075246
refGene_gn=LOC111558286;refGene_tx=XM_023247071.1
E3	5093662	5093683
refGene_gn=LOC111558286;refGene_tx=XM_023247071.1
E3	5093743	5094523	refGene_gn=FAM220A;refGene_tx=XM_023246214.1
E3	5152276	5152313	refGene_gn=CYTH3;refGene_tx=XM_023246645.1

E3 5224724 5224807 refGene_gn=CYTH3;refGene_tx=XM_023246645.1
E3 5226711 5226776 refGene_gn=CYTH3;refGene_tx=XM_023246645.1
E3 5227637 5227704 refGene_gn=CYTH3;refGene_tx=XM_023246645.1
E3 5236583 5236702 refGene_gn=CYTH3;refGene_tx=XM_023246645.1
E3 5242526 5242607 refGene_gn=CYTH3;refGene_tx=XM_023246645.1
E3 5243449 5243562 refGene_gn=CYTH3;refGene_tx=XM_023246645.1
E3 5243662 5243811 refGene_gn=CYTH3;refGene_tx=XM_023246645.1
E3 5243979 5244091 refGene_gn=CYTH3;refGene_tx=XM_023246645.1
E3 5248321 5248398 refGene_gn=CYTH3;refGene_tx=XM_023246645.1
E3 5248488 5248560 refGene_gn=CYTH3;refGene_tx=XM_023246645.1
E3 5248642 5248797 refGene_gn=CYTH3;refGene_tx=XM_023246645.1
E3 5249077 5249150 refGene_gn=CYTH3;refGene_tx=XM_023246645.1
E3 5254195 5254203
refGene_gn=USP42;refGene_tx=XM_019820404.2;refGene_gn=USP42;refGene_tx=XM_019820405.2;refGene_gn=USP42;refGene_tx=XM_023246644.1
E3 5256109 5256411
refGene_gn=USP42;refGene_tx=XM_019820404.2;refGene_gn=USP42;refGene_tx=XM_019820405.2;refGene_gn=USP42;refGene_tx=XM_023246644.1
E3 5257446 5258931
refGene_gn=USP42;refGene_tx=XM_019820404.2;refGene_gn=USP42;refGene_tx=XM_019820405.2;refGene_gn=USP42;refGene_tx=XM_023246644.1
E3 5260466 5260561
refGene_gn=USP42;refGene_tx=XM_019820404.2;refGene_gn=USP42;refGene_tx=XM_019820405.2;refGene_gn=USP42;refGene_tx=XM_023246644.1
E3 5260700 5261414
refGene_gn=USP42;refGene_tx=XM_019820404.2;refGene_gn=USP42;refGene_tx=XM_019820405.2;refGene_gn=USP42;refGene_tx=XM_023246644.1
E3 5263069 5263223
refGene_gn=USP42;refGene_tx=XM_019820404.2;refGene_gn=USP42;refGene_tx=XM_019820405.2;refGene_gn=USP42;refGene_tx=XM_023246644.1
E3 5264557 5264658
refGene_gn=USP42;refGene_tx=XM_019820404.2;refGene_gn=USP42;refGene_tx=XM_019820405.2;refGene_gn=USP42;refGene_tx=XM_023246644.1
E3 5264831 5264972
refGene_gn=USP42;refGene_tx=XM_019820404.2;refGene_gn=USP42;refGene_tx=XM_019820405.2;refGene_gn=USP42;refGene_tx=XM_023246644.1
E3 5265754 5265866
refGene_gn=USP42;refGene_tx=XM_019820404.2;refGene_gn=USP42;refGene_tx=XM_019820405.2;refGene_gn=USP42;refGene_tx=XM_023246644.1
E3 5266931 5267014
refGene_gn=USP42;refGene_tx=XM_019820404.2;refGene_gn=USP42;refGene_tx=XM_019820405.2;refGene_gn=USP42;refGene_tx=XM_023246644.1
E3 5267855 5267926
refGene_gn=USP42;refGene_tx=XM_019820404.2;refGene_gn=USP42;refGene_tx=XM_019820405.2;refGene_gn=USP42;refGene_tx=XM_023246644.1
E3 5268858 5268926
refGene_gn=USP42;refGene_tx=XM_019820404.2;refGene_gn=USP42;refGene_tx=XM_019820405.2;refGene_gn=USP42;refGene_tx=XM_023246644.1
E3 5270068 5270171
refGene_gn=USP42;refGene_tx=XM_019820404.2;refGene_gn=USP42;refGene_tx

=XM_019820405.2;refGene_gn=USP42;refGene_tx=XM_023246644.1
E3 5271782 5271893
refGene_gn=USP42;refGene_tx=XM_019820404.2;refGene_gn=USP42;refGene_tx=XM_019820405.2;refGene_gn=USP42;refGene_tx=XM_023246644.1
E3 5294382 5294583
refGene_gn=USP42;refGene_tx=XM_019820404.2;refGene_gn=USP42;refGene_tx=XM_019820405.2;refGene_gn=USP42;refGene_tx=XM_023246644.1
E3 5297124 5297365
refGene_gn=USP42;refGene_tx=XM_019820404.2;refGene_gn=USP42;refGene_tx=XM_019820405.2;refGene_gn=USP42;refGene_tx=XM_023246644.1
E3 5333577 5333695 refGene_gn=EIF2AK1;refGene_tx=XM_023246864.1
E3 5339941 5340100
refGene_gn=EIF2AK1;refGene_tx=XM_023246864.1;refGene_gn=EIF2AK1;refGene_tx=XM_023246866.1
E3 5342901 5343035
refGene_gn=EIF2AK1;refGene_tx=XM_023246864.1;refGene_gn=EIF2AK1;refGene_tx=XM_023246865.1
E3 5343707 5343737 refGene_gn=EIF2AK1;refGene_tx=XM_023246867.1
E3 5344407 5344445
refGene_gn=EIF2AK1;refGene_tx=XM_023246864.1;refGene_gn=EIF2AK1;refGene_tx=XM_023246865.1;refGene_gn=EIF2AK1;refGene_tx=XM_023246866.1;refGene_gn=EIF2AK1;refGene_tx=XM_023246867.1
E3 5345168 5345268
refGene_gn=EIF2AK1;refGene_tx=XM_023246864.1;refGene_gn=EIF2AK1;refGene_tx=XM_023246865.1;refGene_gn=EIF2AK1;refGene_tx=XM_023246866.1;refGene_gn=EIF2AK1;refGene_tx=XM_023246867.1
E3 5345792 5345873
refGene_gn=EIF2AK1;refGene_tx=XM_023246864.1;refGene_gn=EIF2AK1;refGene_tx=XM_023246865.1;refGene_gn=EIF2AK1;refGene_tx=XM_023246866.1;refGene_gn=EIF2AK1;refGene_tx=XM_023246867.1
E3 5347016 5347116
refGene_gn=EIF2AK1;refGene_tx=XM_023246864.1;refGene_gn=EIF2AK1;refGene_tx=XM_023246865.1;refGene_gn=EIF2AK1;refGene_tx=XM_023246866.1;refGene_gn=EIF2AK1;refGene_tx=XM_023246867.1
E3 5347928 5347986
refGene_gn=EIF2AK1;refGene_tx=XM_023246864.1;refGene_gn=EIF2AK1;refGene_tx=XM_023246865.1;refGene_gn=EIF2AK1;refGene_tx=XM_023246866.1;refGene_gn=EIF2AK1;refGene_tx=XM_023246867.1
E3 5348267 5348589
refGene_gn=EIF2AK1;refGene_tx=XM_023246864.1;refGene_gn=EIF2AK1;refGene_tx=XM_023246865.1;refGene_gn=EIF2AK1;refGene_tx=XM_023246866.1;refGene_gn=EIF2AK1;refGene_tx=XM_023246867.1
E3 5349794 5349906
refGene_gn=EIF2AK1;refGene_tx=XM_023246864.1;refGene_gn=EIF2AK1;refGene_tx=XM_023246865.1;refGene_gn=EIF2AK1;refGene_tx=XM_023246866.1;refGene_gn=EIF2AK1;refGene_tx=XM_023246867.1
E3 5353075 5353176
refGene_gn=EIF2AK1;refGene_tx=XM_023246864.1;refGene_gn=EIF2AK1;refGene_tx=XM_023246865.1;refGene_gn=EIF2AK1;refGene_tx=XM_023246866.1;refGene_gn=EIF2AK1;refGene_tx=XM_023246867.1

E3	5353870	5354816	refGene_gn=ANKRD61;refGene_tx=XM_006941914.4
E3	5356428	5356526	refGene_gn=ANKRD61;refGene_tx=XM_006941914.4
E3	5357443	5357659	refGene_gn=ANKRD61;refGene_tx=XM_006941914.4
E3	5359345	5359460
refGene_gn=EIF2AK1;refGene_tx=XM_023246864.1;refGene_gn=EIF2AK1;refGene_tx=XM_023246865.1;refGene_gn=EIF2AK1;refGene_tx=XM_023246866.1;refGene_gn=EIF2AK1;refGene_tx=XM_023246867.1
E3	5359923	5360006
refGene_gn=EIF2AK1;refGene_tx=XM_023246864.1;refGene_gn=EIF2AK1;refGene_tx=XM_023246865.1;refGene_gn=EIF2AK1;refGene_tx=XM_023246866.1;refGene_gn=EIF2AK1;refGene_tx=XM_023246867.1
E3	5361072	5361306
refGene_gn=EIF2AK1;refGene_tx=XM_023246864.1;refGene_gn=EIF2AK1;refGene_tx=XM_023246865.1;refGene_gn=EIF2AK1;refGene_tx=XM_023246866.1;refGene_gn=EIF2AK1;refGene_tx=XM_023246867.1
E3	5365098	5365269
refGene_gn=EIF2AK1;refGene_tx=XM_023246864.1;refGene_gn=EIF2AK1;refGene_tx=XM_023246865.1;refGene_gn=EIF2AK1;refGene_tx=XM_023246866.1;refGene_gn=EIF2AK1;refGene_tx=XM_023246867.1
E3	5366258	5366647
refGene_gn=AIMP2;refGene_tx=XM_003998428.3;refGene_gn=AIMP2;refGene_tx=XM_006941913.2;refGene_gn=AIMP2;refGene_tx=XM_019820416.2
E3	5367468	5367700
refGene_gn=AIMP2;refGene_tx=XM_003998428.3;refGene_gn=AIMP2;refGene_tx=XM_006941913.2;refGene_gn=AIMP2;refGene_tx=XM_019820416.2
E3	5369968	5370175
refGene_gn=AIMP2;refGene_tx=XM_003998428.3;refGene_gn=AIMP2;refGene_tx=XM_006941913.2;refGene_gn=AIMP2;refGene_tx=XM_019820416.2
E3	5373006	5373018	refGene_gn=AIMP2;refGene_tx=XM_019820416.2
E3	5373919	5374054
refGene_gn=AIMP2;refGene_tx=XM_003998428.3;refGene_gn=AIMP2;refGene_tx=XM_006941913.2
E3	5374399	5374422
refGene_gn=PMS2;refGene_tx=XM_006941910.4;refGene_gn=PMS2;refGene_tx=XM_019820414.2
E3	5376280	5376420
refGene_gn=PMS2;refGene_tx=XM_006941910.4;refGene_gn=PMS2;refGene_tx=XM_019820414.2
E3	5376916	5376935	refGene_gn=PMS2;refGene_tx=XM_006941912.4
E3	5377040	5377127
refGene_gn=PMS2;refGene_tx=XM_006941910.4;refGene_gn=PMS2;refGene_tx=XM_006941912.4;refGene_gn=PMS2;refGene_tx=XM_019820414.2
E3	5377535	5377638
refGene_gn=PMS2;refGene_tx=XM_006941910.4;refGene_gn=PMS2;refGene_tx=XM_006941912.4;refGene_gn=PMS2;refGene_tx=XM_019820414.2
E3	5378375	5378559
refGene_gn=PMS2;refGene_tx=XM_006941910.4;refGene_gn=PMS2;refGene_tx=XM_006941912.4;refGene_gn=PMS2;refGene_tx=XM_019820414.2
E3	5381320	5381488
refGene_gn=PMS2;refGene_tx=XM_006941910.4;refGene_gn=PMS2;refGene_tx=X

M_006941912.4;refGene_gn=PMS2;refGene_tx=XM_019820414.2
E3 5382274 5382372
refGene_gn=PMS2;refGene_tx=XM_006941910.4;refGene_gn=PMS2;refGene_tx=XM_006941912.4;refGene_gn=PMS2;refGene_tx=XM_019820414.2
E3 5384081 5384181
refGene_gn=PMS2;refGene_tx=XM_006941910.4;refGene_gn=PMS2;refGene_tx=XM_006941912.4;refGene_gn=PMS2;refGene_tx=XM_019820414.2
E3 5385292 5385377
refGene_gn=PMS2;refGene_tx=XM_006941910.4;refGene_gn=PMS2;refGene_tx=XM_006941912.4;refGene_gn=PMS2;refGene_tx=XM_019820414.2;refGene_gn=PMS2;refGene_tx=XM_019820415.2
E3 5386496 5386652
refGene_gn=PMS2;refGene_tx=XM_006941910.4;refGene_gn=PMS2;refGene_tx=XM_006941912.4;refGene_gn=PMS2;refGene_tx=XM_019820414.2;refGene_gn=PMS2;refGene_tx=XM_019820415.2
E3 5388659 5389545
refGene_gn=PMS2;refGene_tx=XM_006941910.4;refGene_gn=PMS2;refGene_tx=XM_006941912.4;refGene_gn=PMS2;refGene_tx=XM_019820414.2;refGene_gn=PMS2;refGene_tx=XM_019820415.2
E3 5393940 5394108
refGene_gn=PMS2;refGene_tx=XM_006941910.4;refGene_gn=PMS2;refGene_tx=XM_006941912.4;refGene_gn=PMS2;refGene_tx=XM_019820415.2
E3 5396601 5396702
refGene_gn=PMS2;refGene_tx=XM_006941910.4;refGene_gn=PMS2;refGene_tx=XM_006941912.4;refGene_gn=PMS2;refGene_tx=XM_019820414.2;refGene_gn=PMS2;refGene_tx=XM_019820415.2
E3 5396909 5397079
refGene_gn=PMS2;refGene_tx=XM_006941910.4;refGene_gn=PMS2;refGene_tx=XM_006941912.4;refGene_gn=PMS2;refGene_tx=XM_019820414.2;refGene_gn=PMS2;refGene_tx=XM_019820415.2
E3 5399814 5399958
refGene_gn=PMS2;refGene_tx=XM_006941910.4;refGene_gn=PMS2;refGene_tx=XM_006941912.4;refGene_gn=PMS2;refGene_tx=XM_019820414.2;refGene_gn=PMS2;refGene_tx=XM_019820415.2
E3 5416433 5416589
refGene_gn=LOC101097033;refGene_tx=XM_023246598.1;refGene_gn=LOC101097033;refGene_tx=XM_023246599.1;refGene_gn=LOC101097033;refGene_tx=XM_023246600.1;refGene_gn=LOC101097033;refGene_tx=XM_023246601.1
E3 5418371 5418457
refGene_gn=LOC101097033;refGene_tx=XM_023246598.1;refGene_gn=LOC101097033;refGene_tx=XM_023246599.1;refGene_gn=LOC101097033;refGene_tx=XM_023246600.1;refGene_gn=LOC101097033;refGene_tx=XM_023246601.1
E3 5419869 5419990
refGene_gn=LOC101097033;refGene_tx=XM_023246598.1;refGene_gn=LOC101097033;refGene_tx=XM_023246599.1;refGene_gn=LOC101097033;refGene_tx=XM_023246600.1;refGene_gn=LOC101097033;refGene_tx=XM_023246601.1
E3 5428185 5428362
refGene_gn=LOC101097033;refGene_tx=XM_023246598.1;refGene_gn=LOC101097033;refGene_tx=XM_023246599.1;refGene_gn=LOC101097033;refGene_tx=XM_023246600.1;refGene_gn=LOC101097033;refGene_tx=XM_023246601.1

E3	5429666	5429826
refGene_gn=LOC101097033;refGene_tx=XM_023246598.1;refGene_gn=LOC101097033;refGene_tx=XM_023246599.1;refGene_gn=LOC101097033;refGene_tx=XM_023246600.1;refGene_gn=LOC101097033;refGene_tx=XM_023246601.1
E3	5430776	5430898
refGene_gn=LOC101097033;refGene_tx=XM_023246598.1;refGene_gn=LOC101097033;refGene_tx=XM_023246599.1;refGene_gn=LOC101097033;refGene_tx=XM_023246600.1;refGene_gn=LOC101097033;refGene_tx=XM_023246601.1
E3	5433001	5433191
refGene_gn=LOC101097033;refGene_tx=XM_023246598.1;refGene_gn=LOC101097033;refGene_tx=XM_023246599.1;refGene_gn=LOC101097033;refGene_tx=XM_023246600.1;refGene_gn=LOC101097033;refGene_tx=XM_023246601.1
E3	5434872	5434987
refGene_gn=LOC101097033;refGene_tx=XM_023246598.1;refGene_gn=LOC101097033;refGene_tx=XM_023246599.1;refGene_gn=LOC101097033;refGene_tx=XM_023246600.1;refGene_gn=LOC101097033;refGene_tx=XM_023246601.1
E3	5435973	5436053
refGene_gn=LOC101097033;refGene_tx=XM_023246598.1;refGene_gn=LOC101097033;refGene_tx=XM_023246599.1;refGene_gn=LOC101097033;refGene_tx=XM_023246600.1;refGene_gn=LOC101097033;refGene_tx=XM_023246601.1
E3	5436156	5436305
refGene_gn=LOC101097033;refGene_tx=XM_023246598.1;refGene_gn=LOC101097033;refGene_tx=XM_023246599.1;refGene_gn=LOC101097033;refGene_tx=XM_023246600.1;refGene_gn=LOC101097033;refGene_tx=XM_023246601.1
E3	5438702	5438810
refGene_gn=LOC101097033;refGene_tx=XM_023246598.1;refGene_gn=LOC101097033;refGene_tx=XM_023246599.1;refGene_gn=LOC101097033;refGene_tx=XM_023246600.1;refGene_gn=LOC101097033;refGene_tx=XM_023246601.1
E3	5443257	5443389
refGene_gn=LOC101097033;refGene_tx=XM_023246598.1;refGene_gn=LOC101097033;refGene_tx=XM_023246599.1;refGene_gn=LOC101097033;refGene_tx=XM_023246600.1;refGene_gn=LOC101097033;refGene_tx=XM_023246601.1
E3	5444878	5445004
refGene_gn=LOC101097033;refGene_tx=XM_023246598.1;refGene_gn=LOC101097033;refGene_tx=XM_023246599.1;refGene_gn=LOC101097033;refGene_tx=XM_023246600.1;refGene_gn=LOC101097033;refGene_tx=XM_023246601.1
E3	5446868	5446962
refGene_gn=LOC101097033;refGene_tx=XM_023246598.1;refGene_gn=LOC101097033;refGene_tx=XM_023246599.1;refGene_gn=LOC101097033;refGene_tx=XM_023246600.1;refGene_gn=LOC101097033;refGene_tx=XM_023246601.1
E3	5459990	5460160
refGene_gn=LOC101097033;refGene_tx=XM_023246601.1
E3	5462780	5462979
refGene_gn=LOC101097033;refGene_tx=XM_023246598.1;refGene_gn=LOC101097033;refGene_tx=XM_023246599.1;refGene_gn=LOC101097033;refGene_tx=XM_023246600.1
E3	5463811	5463986
refGene_gn=LOC101097033;refGene_tx=XM_023246598.1;refGene_gn=LOC101097033;refGene_tx=XM_023246599.1;refGene_gn=LOC101097033;refGene_tx=XM_023246600.1

```
E3	5467209	5467265	refGene_gn=CCZ1;refGene_tx=XM_023246327.1
E3	5468324	5468452	refGene_gn=CCZ1;refGene_tx=XM_023246327.1
E3	5469167	5469326	refGene_gn=CCZ1;refGene_tx=XM_023246327.1
E3	5477830	5477948	refGene_gn=CCZ1;refGene_tx=XM_023246327.1
E3	5478582	5478616	refGene_gn=CCZ1;refGene_tx=XM_023246327.1
E3	5484851	5484963	refGene_gn=CCZ1;refGene_tx=XM_023246327.1
E3	5485221	5485283	refGene_gn=CCZ1;refGene_tx=XM_023246327.1
E3	5487236	5487318	refGene_gn=CCZ1;refGene_tx=XM_023246327.1
E3	5496065	5496241	refGene_gn=CCZ1;refGene_tx=XM_023246327.1
E3	5497952	5498036	refGene_gn=CCZ1;refGene_tx=XM_023246327.1
E3	5498609	5498657	refGene_gn=CCZ1;refGene_tx=XM_023246327.1
E3	5499034	5499112	refGene_gn=CCZ1;refGene_tx=XM_023246327.1
E3	5499654	5499748	refGene_gn=CCZ1;refGene_tx=XM_023246327.1
E3	5499850	5499948	refGene_gn=CCZ1;refGene_tx=XM_023246327.1
E3	5500914	5501034	refGene_gn=CCZ1;refGene_tx=XM_023246327.1
E3	5510451	5510471
refGene_gn=LOC101097531;refGene_tx=XM_023247072.1
E3	5513716	5513826
refGene_gn=LOC101097531;refGene_tx=XM_023247072.1
E3	5514270	5514403
refGene_gn=LOC101097531;refGene_tx=XM_023247072.1
E3	5515098	5515159
refGene_gn=LOC101097531;refGene_tx=XM_023247072.1
E3	5614887	5614975
refGene_gn=LMTK2;refGene_tx=XM_006941918.4;refGene_gn=LMTK2;refGene_tx
=XM_006941919.4
E3	5635916	5636041
refGene_gn=LMTK2;refGene_tx=XM_006941918.4;refGene_gn=LMTK2;refGene_tx
=XM_006941919.4
E3	5639360	5639505
refGene_gn=LMTK2;refGene_tx=XM_006941918.4;refGene_gn=LMTK2;refGene_tx
=XM_006941919.4
E3	5648185	5648259
refGene_gn=LMTK2;refGene_tx=XM_006941918.4;refGene_gn=LMTK2;refGene_tx
=XM_006941919.4
E3	5652333	5652452
refGene_gn=LMTK2;refGene_tx=XM_006941918.4;refGene_gn=LMTK2;refGene_tx
=XM_006941919.4
E3	5652633	5652721
refGene_gn=LMTK2;refGene_tx=XM_006941918.4;refGene_gn=LMTK2;refGene_tx
=XM_006941919.4
E3	5664615	5664749
refGene_gn=LMTK2;refGene_tx=XM_006941918.4;refGene_gn=LMTK2;refGene_tx
=XM_006941919.4
E3	5675940	5676025
refGene_gn=LMTK2;refGene_tx=XM_006941918.4;refGene_gn=LMTK2;refGene_tx
=XM_006941919.4
E3	5677297	5677419
refGene_gn=LMTK2;refGene_tx=XM_006941918.4;refGene_gn=LMTK2;refGene_tx
=XM_006941919.4
```

E3 5682665 5682815
refGene_gn=LMTK2;refGene_tx=XM_006941918.4;refGene_gn=LMTK2;refGene_tx
=XM_006941919.4
E3 5684763 5687746
refGene_gn=LMTK2;refGene_tx=XM_006941918.4;refGene_gn=LMTK2;refGene_tx
=XM_006941919.4
E3 5691262 5691401
refGene_gn=LMTK2;refGene_tx=XM_006941918.4;refGene_gn=LMTK2;refGene_tx
=XM_006941919.4
E3 5691654 5691897
refGene_gn=LMTK2;refGene_tx=XM_006941918.4;refGene_gn=LMTK2;refGene_tx
=XM_006941919.4
E3 5692895 5692924
refGene_gn=LMTK2;refGene_tx=XM_006941918.4;refGene_gn=LMTK2;refGene_tx
=XM_006941919.4
E3 5698873 5699437 refGene_gn=BHLHA15;refGene_tx=XM_023246320.1
E3 5703242 5703404
refGene_gn=TECPR1;refGene_tx=XM_003998453.5;refGene_gn=TECPR1;refGene_
tx=XM_006941920.4;refGene_gn=TECPR1;refGene_tx=XM_006941921.4;refGene_
gn=TECPR1;refGene_tx=XM_019820420.2;refGene_gn=TECPR1;refGene_tx=XM_01
9820421.2
E3 5703566 5703686
refGene_gn=TECPR1;refGene_tx=XM_003998453.5;refGene_gn=TECPR1;refGene_
tx=XM_006941920.4;refGene_gn=TECPR1;refGene_tx=XM_006941921.4;refGene_
gn=TECPR1;refGene_tx=XM_019820420.2;refGene_gn=TECPR1;refGene_tx=XM_01
9820421.2
E3 5703823 5703930
refGene_gn=TECPR1;refGene_tx=XM_003998453.5;refGene_gn=TECPR1;refGene_
tx=XM_006941920.4;refGene_gn=TECPR1;refGene_tx=XM_006941921.4;refGene_
gn=TECPR1;refGene_tx=XM_019820420.2;refGene_gn=TECPR1;refGene_tx=XM_01
9820421.2
E3 5704620 5704713
refGene_gn=TECPR1;refGene_tx=XM_003998453.5;refGene_gn=TECPR1;refGene_
tx=XM_006941920.4;refGene_gn=TECPR1;refGene_tx=XM_006941921.4;refGene_
gn=TECPR1;refGene_tx=XM_019820420.2;refGene_gn=TECPR1;refGene_tx=XM_01
9820421.2
E3 5705576 5705712
refGene_gn=TECPR1;refGene_tx=XM_003998453.5;refGene_gn=TECPR1;refGene_
tx=XM_006941920.4;refGene_gn=TECPR1;refGene_tx=XM_006941921.4;refGene_
gn=TECPR1;refGene_tx=XM_019820420.2;refGene_gn=TECPR1;refGene_tx=XM_01
9820421.2
E3 5706015 5706196
refGene_gn=TECPR1;refGene_tx=XM_003998453.5;refGene_gn=TECPR1;refGene_
tx=XM_006941920.4;refGene_gn=TECPR1;refGene_tx=XM_006941921.4;refGene_
gn=TECPR1;refGene_tx=XM_019820420.2;refGene_gn=TECPR1;refGene_tx=XM_01
9820421.2
E3 5706642 5706699
refGene_gn=TECPR1;refGene_tx=XM_003998453.5;refGene_gn=TECPR1;refGene_
tx=XM_006941920.4;refGene_gn=TECPR1;refGene_tx=XM_006941921.4;refGene_
gn=TECPR1;refGene_tx=XM_019820420.2;refGene_gn=TECPR1;refGene_tx=XM_01

9820421.2
E3 5707623 5707703
refGene_gn=TECPR1;refGene_tx=XM_003998453.5;refGene_gn=TECPR1;refGene_tx=XM_006941920.4;refGene_gn=TECPR1;refGene_tx=XM_006941921.4;refGene_gn=TECPR1;refGene_tx=XM_019820420.2;refGene_gn=TECPR1;refGene_tx=XM_019820421.2
E3 5707782 5707879
refGene_gn=TECPR1;refGene_tx=XM_003998453.5;refGene_gn=TECPR1;refGene_tx=XM_006941920.4;refGene_gn=TECPR1;refGene_tx=XM_006941921.4;refGene_gn=TECPR1;refGene_tx=XM_019820420.2;refGene_gn=TECPR1;refGene_tx=XM_019820421.2
E3 5708818 5708921
refGene_gn=TECPR1;refGene_tx=XM_003998453.5;refGene_gn=TECPR1;refGene_tx=XM_006941920.4;refGene_gn=TECPR1;refGene_tx=XM_006941921.4;refGene_gn=TECPR1;refGene_tx=XM_019820420.2;refGene_gn=TECPR1;refGene_tx=XM_019820421.2
E3 5709912 5710034
refGene_gn=TECPR1;refGene_tx=XM_003998453.5;refGene_gn=TECPR1;refGene_tx=XM_006941920.4;refGene_gn=TECPR1;refGene_tx=XM_006941921.4;refGene_gn=TECPR1;refGene_tx=XM_019820420.2;refGene_gn=TECPR1;refGene_tx=XM_019820421.2
E3 5710895 5711053
refGene_gn=TECPR1;refGene_tx=XM_003998453.5;refGene_gn=TECPR1;refGene_tx=XM_006941920.4;refGene_gn=TECPR1;refGene_tx=XM_006941921.4;refGene_gn=TECPR1;refGene_tx=XM_019820420.2;refGene_gn=TECPR1;refGene_tx=XM_019820421.2
E3 5711169 5711319
refGene_gn=TECPR1;refGene_tx=XM_003998453.5;refGene_gn=TECPR1;refGene_tx=XM_006941920.4;refGene_gn=TECPR1;refGene_tx=XM_006941921.4;refGene_gn=TECPR1;refGene_tx=XM_019820420.2;refGene_gn=TECPR1;refGene_tx=XM_019820421.2
E3 5711847 5712003
refGene_gn=TECPR1;refGene_tx=XM_003998453.5;refGene_gn=TECPR1;refGene_tx=XM_006941920.4;refGene_gn=TECPR1;refGene_tx=XM_006941921.4;refGene_gn=TECPR1;refGene_tx=XM_019820420.2;refGene_gn=TECPR1;refGene_tx=XM_019820421.2
E3 5712313 5712459
refGene_gn=TECPR1;refGene_tx=XM_003998453.5;refGene_gn=TECPR1;refGene_tx=XM_006941920.4;refGene_gn=TECPR1;refGene_tx=XM_006941921.4;refGene_gn=TECPR1;refGene_tx=XM_019820420.2;refGene_gn=TECPR1;refGene_tx=XM_019820421.2
E3 5712824 5713315
refGene_gn=TECPR1;refGene_tx=XM_003998453.5;refGene_gn=TECPR1;refGene_tx=XM_006941920.4;refGene_gn=TECPR1;refGene_tx=XM_006941921.4;refGene_gn=TECPR1;refGene_tx=XM_019820420.2;refGene_gn=TECPR1;refGene_tx=XM_019820421.2
E3 5715406 5715552
refGene_gn=TECPR1;refGene_tx=XM_003998453.5;refGene_gn=TECPR1;refGene_tx=XM_006941920.4;refGene_gn=TECPR1;refGene_tx=XM_006941921.4;refGene_gn=TECPR1;refGene_tx=XM_019820420.2;refGene_gn=TECPR1;refGene_tx=XM_01

9820421.2
E3 5717073 5717175
refGene_gn=TECPR1;refGene_tx=XM_003998453.5;refGene_gn=TECPR1;refGene_tx=XM_006941920.4;refGene_gn=TECPR1;refGene_tx=XM_006941921.4;refGene_gn=TECPR1;refGene_tx=XM_019820420.2;refGene_gn=TECPR1;refGene_tx=XM_019820421.2
E3 5718550 5718651
refGene_gn=TECPR1;refGene_tx=XM_003998453.5;refGene_gn=TECPR1;refGene_tx=XM_006941920.4;refGene_gn=TECPR1;refGene_tx=XM_006941921.4;refGene_gn=TECPR1;refGene_tx=XM_019820420.2;refGene_gn=TECPR1;refGene_tx=XM_019820421.2
E3 5718758 5718933
refGene_gn=TECPR1;refGene_tx=XM_003998453.5;refGene_gn=TECPR1;refGene_tx=XM_006941920.4;refGene_gn=TECPR1;refGene_tx=XM_006941921.4;refGene_gn=TECPR1;refGene_tx=XM_019820420.2;refGene_gn=TECPR1;refGene_tx=XM_019820421.2
E3 5720780 5720906
refGene_gn=TECPR1;refGene_tx=XM_003998453.5;refGene_gn=TECPR1;refGene_tx=XM_006941920.4;refGene_gn=TECPR1;refGene_tx=XM_006941921.4;refGene_gn=TECPR1;refGene_tx=XM_019820421.2
E3 5721875 5721998
refGene_gn=TECPR1;refGene_tx=XM_003998453.5;refGene_gn=TECPR1;refGene_tx=XM_006941920.4;refGene_gn=TECPR1;refGene_tx=XM_006941921.4;refGene_gn=TECPR1;refGene_tx=XM_019820420.2;refGene_gn=TECPR1;refGene_tx=XM_019820421.2
E3 5722256 5722439
refGene_gn=TECPR1;refGene_tx=XM_003998453.5;refGene_gn=TECPR1;refGene_tx=XM_006941920.4;refGene_gn=TECPR1;refGene_tx=XM_006941921.4;refGene_gn=TECPR1;refGene_tx=XM_019820420.2;refGene_gn=TECPR1;refGene_tx=XM_019820421.2
E3 5722940 5723165
refGene_gn=TECPR1;refGene_tx=XM_003998453.5;refGene_gn=TECPR1;refGene_tx=XM_006941920.4;refGene_gn=TECPR1;refGene_tx=XM_006941921.4;refGene_gn=TECPR1;refGene_tx=XM_019820420.2;refGene_gn=TECPR1;refGene_tx=XM_019820421.2
E3 5747878 5748020 refGene_gn=BRI3;refGene_tx=XM_023246539.1
E3 5748555 5748658 refGene_gn=BRI3;refGene_tx=XM_023246539.1
E3 5757080 5757213 refGene_gn=BRI3;refGene_tx=XM_023246539.1
E3 5759453 5759529
refGene_gn=BAIAP2L1;refGene_tx=XM_006941924.3;refGene_gn=BAIAP2L1;refGene_tx=XM_023246537.1;refGene_gn=BAIAP2L1;refGene_tx=XM_023246538.1
E3 5760022 5760060
refGene_gn=BAIAP2L1;refGene_tx=XM_006941924.3;refGene_gn=BAIAP2L1;refGene_tx=XM_023246537.1;refGene_gn=BAIAP2L1;refGene_tx=XM_023246538.1
E3 5773607 5773788
refGene_gn=BAIAP2L1;refGene_tx=XM_006941924.3;refGene_gn=BAIAP2L1;refGene_tx=XM_023246537.1;refGene_gn=BAIAP2L1;refGene_tx=XM_023246538.1
E3 5774837 5774915
refGene_gn=BAIAP2L1;refGene_tx=XM_006941924.3;refGene_gn=BAIAP2L1;refGene_tx=XM_023246537.1;refGene_gn=BAIAP2L1;refGene_tx=XM_023246538.1

E3 5775554 5775771
refGene_gn=BAIAP2L1;refGene_tx=XM_006941924.3;refGene_gn=BAIAP2L1;refGene_tx=XM_023246537.1;refGene_gn=BAIAP2L1;refGene_tx=XM_023246538.1
E3 5777648 5777796
refGene_gn=BAIAP2L1;refGene_tx=XM_006941924.3;refGene_gn=BAIAP2L1;refGene_tx=XM_023246537.1;refGene_gn=BAIAP2L1;refGene_tx=XM_023246538.1
E3 5780280 5780448
refGene_gn=BAIAP2L1;refGene_tx=XM_006941924.3;refGene_gn=BAIAP2L1;refGene_tx=XM_023246537.1;refGene_gn=BAIAP2L1;refGene_tx=XM_023246538.1
E3 5782175 5782328
refGene_gn=BAIAP2L1;refGene_tx=XM_006941924.3;refGene_gn=BAIAP2L1;refGene_tx=XM_023246537.1;refGene_gn=BAIAP2L1;refGene_tx=XM_023246538.1
E3 5782970 5783108
refGene_gn=BAIAP2L1;refGene_tx=XM_006941924.3;refGene_gn=BAIAP2L1;refGene_tx=XM_023246537.1;refGene_gn=BAIAP2L1;refGene_tx=XM_023246538.1
E3 5784327 5784399
refGene_gn=BAIAP2L1;refGene_tx=XM_006941924.3;refGene_gn=BAIAP2L1;refGene_tx=XM_023246537.1;refGene_gn=BAIAP2L1;refGene_tx=XM_023246538.1
E3 5784506 5784568
refGene_gn=BAIAP2L1;refGene_tx=XM_006941924.3;refGene_gn=BAIAP2L1;refGene_tx=XM_023246537.1;refGene_gn=BAIAP2L1;refGene_tx=XM_023246538.1
E3 5822097 5822184
refGene_gn=BAIAP2L1;refGene_tx=XM_023246537.1;refGene_gn=BAIAP2L1;refGene_tx=XM_023246538.1
E3 5822222 5822244 refGene_gn=BAIAP2L1;refGene_tx=XM_006941924.3
E3 5826586 5826662
refGene_gn=BAIAP2L1;refGene_tx=XM_023246537.1;refGene_gn=BAIAP2L1;refGene_tx=XM_023246538.1
E3 5851631 5851682
refGene_gn=BAIAP2L1;refGene_tx=XM_023246537.1;refGene_gn=BAIAP2L1;refGene_tx=XM_023246538.1
E3 6010425 6010851 refGene_gn=NPTX2;refGene_tx=XM_019820247.2
E3 6012428 6012645 refGene_gn=NPTX2;refGene_tx=XM_019820247.2
E3 6016597 6016842 refGene_gn=NPTX2;refGene_tx=XM_019820247.2
E3 6018515 6018695 refGene_gn=NPTX2;refGene_tx=XM_019820247.2
E3 6019604 6019832 refGene_gn=NPTX2;refGene_tx=XM_019820247.2
E3 6176420 6176609
refGene_gn=TMEM130;refGene_tx=XM_003998500.5;refGene_gn=TMEM130;refGene_tx=XM_023246577.1;refGene_gn=TMEM130;refGene_tx=XM_023246579.1
E3 6176969 6177082
refGene_gn=TMEM130;refGene_tx=XM_003998500.5;refGene_gn=TMEM130;refGene_tx=XM_023246577.1;refGene_gn=TMEM130;refGene_tx=XM_023246579.1
E3 6179548 6179751
refGene_gn=TMEM130;refGene_tx=XM_003998500.5;refGene_gn=TMEM130;refGene_tx=XM_023246577.1;refGene_gn=TMEM130;refGene_tx=XM_023246579.1
E3 6180890 6180975
refGene_gn=TMEM130;refGene_tx=XM_003998500.5;refGene_gn=TMEM130;refGene_tx=XM_023246577.1;refGene_gn=TMEM130;refGene_tx=XM_023246579.1
E3 6181603 6181770
refGene_gn=TMEM130;refGene_tx=XM_003998500.5;refGene_gn=TMEM130;refGen

e_tx=XM_023246577.1;refGene_gn=TMEM130;refGene_tx=XM_023246579.1
E3 6184453 6184613
refGene_gn=TMEM130;refGene_tx=XM_003998500.5;refGene_gn=TMEM130;refGen
e_tx=XM_023246577.1;refGene_gn=TMEM130;refGene_tx=XM_023246579.1
E3 6186154 6186460
refGene_gn=TMEM130;refGene_tx=XM_003998500.5;refGene_gn=TMEM130;refGen
e_tx=XM_023246577.1;refGene_gn=TMEM130;refGene_tx=XM_023246579.1
E3 6191171 6191229 refGene_gn=TMEM130;refGene_tx=XM_023246577.1
E3 6192587 6192672
refGene_gn=TMEM130;refGene_tx=XM_003998500.5;refGene_gn=TMEM130;refGen
e_tx=XM_023246579.1
E3 6199128 6199228
refGene_gn=TRRAP;refGene_tx=XM_019820423.2;refGene_gn=TRRAP;refGene_tx
=XM_023246564.1;refGene_gn=TRRAP;refGene_tx=XM_023246565.1;refGene_gn=
TRRAP;refGene_tx=XM_023246566.1;refGene_gn=TRRAP;refGene_tx=XM_0232465
67.1;refGene_gn=TRRAP;refGene_tx=XM_023246568.1;refGene_gn=TRRAP;refGe
ne_tx=XM_023246569.1;refGene_gn=TRRAP;refGene_tx=XM_023246570.1;refGen
e_gn=TRRAP;refGene_tx=XM_023246571.1;refGene_gn=TRRAP;refGene_tx=XM_02
3246572.1;refGene_gn=TRRAP;refGene_tx=XM_023246573.1;refGene_gn=TRRAP;
refGene_tx=XM_023246574.1;refGene_gn=TRRAP;refGene_tx=XM_023246575.1;r
efGene_gn=TRRAP;refGene_tx=XM_023246576.1
E3 6199648 6199698
refGene_gn=TRRAP;refGene_tx=XM_019820423.2;refGene_gn=TRRAP;refGene_tx
=XM_023246564.1;refGene_gn=TRRAP;refGene_tx=XM_023246565.1;refGene_gn=
TRRAP;refGene_tx=XM_023246566.1;refGene_gn=TRRAP;refGene_tx=XM_0232465
67.1;refGene_gn=TRRAP;refGene_tx=XM_023246568.1;refGene_gn=TRRAP;refGe
ne_tx=XM_023246569.1;refGene_gn=TRRAP;refGene_tx=XM_023246570.1;refGen
e_gn=TRRAP;refGene_tx=XM_023246571.1;refGene_gn=TRRAP;refGene_tx=XM_02
3246572.1;refGene_gn=TRRAP;refGene_tx=XM_023246573.1;refGene_gn=TRRAP;
refGene_tx=XM_023246574.1;refGene_gn=TRRAP;refGene_tx=XM_023246575.1;r
efGene_gn=TRRAP;refGene_tx=XM_023246576.1
E3 6205061 6205172
refGene_gn=TRRAP;refGene_tx=XM_019820423.2;refGene_gn=TRRAP;refGene_tx
=XM_023246564.1;refGene_gn=TRRAP;refGene_tx=XM_023246565.1;refGene_gn=
TRRAP;refGene_tx=XM_023246566.1;refGene_gn=TRRAP;refGene_tx=XM_0232465
67.1;refGene_gn=TRRAP;refGene_tx=XM_023246568.1;refGene_gn=TRRAP;refGe
ne_tx=XM_023246569.1;refGene_gn=TRRAP;refGene_tx=XM_023246570.1;refGen
e_gn=TRRAP;refGene_tx=XM_023246571.1;refGene_gn=TRRAP;refGene_tx=XM_02
3246572.1;refGene_gn=TRRAP;refGene_tx=XM_023246573.1;refGene_gn=TRRAP;
refGene_tx=XM_023246574.1;refGene_gn=TRRAP;refGene_tx=XM_023246575.1;r
efGene_gn=TRRAP;refGene_tx=XM_023246576.1
E3 6206400 6206505
refGene_gn=TRRAP;refGene_tx=XM_019820423.2;refGene_gn=TRRAP;refGene_tx
=XM_023246564.1;refGene_gn=TRRAP;refGene_tx=XM_023246565.1;refGene_gn=
TRRAP;refGene_tx=XM_023246566.1;refGene_gn=TRRAP;refGene_tx=XM_0232465
67.1;refGene_gn=TRRAP;refGene_tx=XM_023246568.1;refGene_gn=TRRAP;refGe
ne_tx=XM_023246569.1;refGene_gn=TRRAP;refGene_tx=XM_023246570.1;refGen
e_gn=TRRAP;refGene_tx=XM_023246571.1;refGene_gn=TRRAP;refGene_tx=XM_02
3246572.1;refGene_gn=TRRAP;refGene_tx=XM_023246573.1;refGene_gn=TRRAP;
refGene_tx=XM_023246574.1;refGene_gn=TRRAP;refGene_tx=XM_023246575.1;r

efGene_gn=TRRAP;refGene_tx=XM_023246576.1
E3 6207417 6207501
refGene_gn=TRRAP;refGene_tx=XM_019820423.2;refGene_gn=TRRAP;refGene_tx=XM_023246564.1;refGene_gn=TRRAP;refGene_tx=XM_023246565.1;refGene_gn=TRRAP;refGene_tx=XM_023246566.1;refGene_gn=TRRAP;refGene_tx=XM_023246567.1;refGene_gn=TRRAP;refGene_tx=XM_023246568.1;refGene_gn=TRRAP;refGene_tx=XM_023246569.1;refGene_gn=TRRAP;refGene_tx=XM_023246570.1;refGene_gn=TRRAP;refGene_tx=XM_023246571.1;refGene_gn=TRRAP;refGene_tx=XM_023246572.1;refGene_gn=TRRAP;refGene_tx=XM_023246573.1;refGene_gn=TRRAP;refGene_tx=XM_023246574.1;refGene_gn=TRRAP;refGene_tx=XM_023246575.1;refGene_gn=TRRAP;refGene_tx=XM_023246576.1
E3 6208785 6208842
refGene_gn=TRRAP;refGene_tx=XM_019820423.2;refGene_gn=TRRAP;refGene_tx=XM_023246564.1;refGene_gn=TRRAP;refGene_tx=XM_023246565.1;refGene_gn=TRRAP;refGene_tx=XM_023246566.1;refGene_gn=TRRAP;refGene_tx=XM_023246567.1;refGene_gn=TRRAP;refGene_tx=XM_023246568.1;refGene_gn=TRRAP;refGene_tx=XM_023246569.1;refGene_gn=TRRAP;refGene_tx=XM_023246570.1;refGene_gn=TRRAP;refGene_tx=XM_023246571.1;refGene_gn=TRRAP;refGene_tx=XM_023246572.1;refGene_gn=TRRAP;refGene_tx=XM_023246573.1;refGene_gn=TRRAP;refGene_tx=XM_023246574.1;refGene_gn=TRRAP;refGene_tx=XM_023246575.1;refGene_gn=TRRAP;refGene_tx=XM_023246576.1
E3 6210256 6210382
refGene_gn=TRRAP;refGene_tx=XM_019820423.2;refGene_gn=TRRAP;refGene_tx=XM_023246564.1;refGene_gn=TRRAP;refGene_tx=XM_023246565.1;refGene_gn=TRRAP;refGene_tx=XM_023246566.1;refGene_gn=TRRAP;refGene_tx=XM_023246567.1;refGene_gn=TRRAP;refGene_tx=XM_023246568.1;refGene_gn=TRRAP;refGene_tx=XM_023246569.1;refGene_gn=TRRAP;refGene_tx=XM_023246570.1;refGene_gn=TRRAP;refGene_tx=XM_023246571.1;refGene_gn=TRRAP;refGene_tx=XM_023246572.1;refGene_gn=TRRAP;refGene_tx=XM_023246573.1;refGene_gn=TRRAP;refGene_tx=XM_023246574.1;refGene_gn=TRRAP;refGene_tx=XM_023246575.1;refGene_gn=TRRAP;refGene_tx=XM_023246576.1
E3 6211688 6211766
refGene_gn=TRRAP;refGene_tx=XM_019820423.2;refGene_gn=TRRAP;refGene_tx=XM_023246564.1;refGene_gn=TRRAP;refGene_tx=XM_023246565.1;refGene_gn=TRRAP;refGene_tx=XM_023246566.1;refGene_gn=TRRAP;refGene_tx=XM_023246567.1;refGene_gn=TRRAP;refGene_tx=XM_023246568.1;refGene_gn=TRRAP;refGene_tx=XM_023246569.1;refGene_gn=TRRAP;refGene_tx=XM_023246570.1;refGene_gn=TRRAP;refGene_tx=XM_023246571.1;refGene_gn=TRRAP;refGene_tx=XM_023246572.1;refGene_gn=TRRAP;refGene_tx=XM_023246573.1;refGene_gn=TRRAP;refGene_tx=XM_023246574.1;refGene_gn=TRRAP;refGene_tx=XM_023246575.1;refGene_gn=TRRAP;refGene_tx=XM_023246576.1
E3 6211970 6212059
refGene_gn=TRRAP;refGene_tx=XM_019820423.2;refGene_gn=TRRAP;refGene_tx=XM_023246564.1;refGene_gn=TRRAP;refGene_tx=XM_023246565.1;refGene_gn=TRRAP;refGene_tx=XM_023246566.1;refGene_gn=TRRAP;refGene_tx=XM_023246567.1;refGene_gn=TRRAP;refGene_tx=XM_023246568.1;refGene_gn=TRRAP;refGene_tx=XM_023246569.1;refGene_gn=TRRAP;refGene_tx=XM_023246570.1;refGene_gn=TRRAP;refGene_tx=XM_023246571.1;refGene_gn=TRRAP;refGene_tx=XM_023246572.1;refGene_gn=TRRAP;refGene_tx=XM_023246573.1;refGene_gn=TRRAP;refGene_tx=XM_023246574.1;refGene_gn=TRRAP;refGene_tx=XM_023246575.1;r

efGene_gn=TRRAP;refGene_tx=XM_023246576.1
E3 6212925 6213022
refGene_gn=TRRAP;refGene_tx=XM_019820423.2;refGene_gn=TRRAP;refGene_tx=XM_023246564.1;refGene_gn=TRRAP;refGene_tx=XM_023246565.1;refGene_gn=TRRAP;refGene_tx=XM_023246566.1;refGene_gn=TRRAP;refGene_tx=XM_023246567.1;refGene_gn=TRRAP;refGene_tx=XM_023246568.1;refGene_gn=TRRAP;refGene_tx=XM_023246569.1;refGene_gn=TRRAP;refGene_tx=XM_023246570.1;refGene_gn=TRRAP;refGene_tx=XM_023246571.1;refGene_gn=TRRAP;refGene_tx=XM_023246572.1;refGene_gn=TRRAP;refGene_tx=XM_023246573.1;refGene_gn=TRRAP;refGene_tx=XM_023246574.1;refGene_gn=TRRAP;refGene_tx=XM_023246575.1;refGene_gn=TRRAP;refGene_tx=XM_023246576.1
E3 6216045 6216184
refGene_gn=TRRAP;refGene_tx=XM_019820423.2;refGene_gn=TRRAP;refGene_tx=XM_023246564.1;refGene_gn=TRRAP;refGene_tx=XM_023246565.1;refGene_gn=TRRAP;refGene_tx=XM_023246566.1;refGene_gn=TRRAP;refGene_tx=XM_023246567.1;refGene_gn=TRRAP;refGene_tx=XM_023246568.1;refGene_gn=TRRAP;refGene_tx=XM_023246569.1;refGene_gn=TRRAP;refGene_tx=XM_023246570.1;refGene_gn=TRRAP;refGene_tx=XM_023246571.1;refGene_gn=TRRAP;refGene_tx=XM_023246572.1;refGene_gn=TRRAP;refGene_tx=XM_023246573.1;refGene_gn=TRRAP;refGene_tx=XM_023246574.1;refGene_gn=TRRAP;refGene_tx=XM_023246575.1;refGene_gn=TRRAP;refGene_tx=XM_023246576.1
E3 6219340 6219419
refGene_gn=TRRAP;refGene_tx=XM_019820423.2;refGene_gn=TRRAP;refGene_tx=XM_023246564.1;refGene_gn=TRRAP;refGene_tx=XM_023246565.1;refGene_gn=TRRAP;refGene_tx=XM_023246566.1;refGene_gn=TRRAP;refGene_tx=XM_023246567.1;refGene_gn=TRRAP;refGene_tx=XM_023246568.1;refGene_gn=TRRAP;refGene_tx=XM_023246569.1;refGene_gn=TRRAP;refGene_tx=XM_023246570.1;refGene_gn=TRRAP;refGene_tx=XM_023246571.1;refGene_gn=TRRAP;refGene_tx=XM_023246572.1;refGene_gn=TRRAP;refGene_tx=XM_023246573.1;refGene_gn=TRRAP;refGene_tx=XM_023246574.1;refGene_gn=TRRAP;refGene_tx=XM_023246575.1;refGene_gn=TRRAP;refGene_tx=XM_023246576.1
E3 6220494 6220729
refGene_gn=TRRAP;refGene_tx=XM_019820423.2;refGene_gn=TRRAP;refGene_tx=XM_023246564.1;refGene_gn=TRRAP;refGene_tx=XM_023246565.1;refGene_gn=TRRAP;refGene_tx=XM_023246566.1;refGene_gn=TRRAP;refGene_tx=XM_023246567.1;refGene_gn=TRRAP;refGene_tx=XM_023246568.1;refGene_gn=TRRAP;refGene_tx=XM_023246569.1;refGene_gn=TRRAP;refGene_tx=XM_023246570.1;refGene_gn=TRRAP;refGene_tx=XM_023246571.1;refGene_gn=TRRAP;refGene_tx=XM_023246572.1;refGene_gn=TRRAP;refGene_tx=XM_023246573.1;refGene_gn=TRRAP;refGene_tx=XM_023246574.1;refGene_gn=TRRAP;refGene_tx=XM_023246575.1;refGene_gn=TRRAP;refGene_tx=XM_023246576.1
E3 6221407 6221768
refGene_gn=TRRAP;refGene_tx=XM_019820423.2;refGene_gn=TRRAP;refGene_tx=XM_023246564.1;refGene_gn=TRRAP;refGene_tx=XM_023246565.1;refGene_gn=TRRAP;refGene_tx=XM_023246566.1;refGene_gn=TRRAP;refGene_tx=XM_023246567.1;refGene_gn=TRRAP;refGene_tx=XM_023246568.1;refGene_gn=TRRAP;refGene_tx=XM_023246569.1;refGene_gn=TRRAP;refGene_tx=XM_023246570.1;refGene_gn=TRRAP;refGene_tx=XM_023246571.1;refGene_gn=TRRAP;refGene_tx=XM_023246572.1;refGene_gn=TRRAP;refGene_tx=XM_023246573.1;refGene_gn=TRRAP;refGene_tx=XM_023246574.1;refGene_gn=TRRAP;refGene_tx=XM_023246575.1;r

efGene_gn=TRRAP;refGene_tx=XM_023246576.1
E3 6221856 6221954
refGene_gn=TRRAP;refGene_tx=XM_019820423.2;refGene_gn=TRRAP;refGene_tx=XM_023246564.1;refGene_gn=TRRAP;refGene_tx=XM_023246565.1;refGene_gn=TRRAP;refGene_tx=XM_023246566.1;refGene_gn=TRRAP;refGene_tx=XM_023246567.1;refGene_gn=TRRAP;refGene_tx=XM_023246568.1;refGene_gn=TRRAP;refGene_tx=XM_023246569.1;refGene_gn=TRRAP;refGene_tx=XM_023246570.1;refGene_gn=TRRAP;refGene_tx=XM_023246571.1;refGene_gn=TRRAP;refGene_tx=XM_023246572.1;refGene_gn=TRRAP;refGene_tx=XM_023246573.1;refGene_gn=TRRAP;refGene_tx=XM_023246574.1;refGene_gn=TRRAP;refGene_tx=XM_023246575.1;refGene_gn=TRRAP;refGene_tx=XM_023246576.1
E3 6222432 6222627
refGene_gn=TRRAP;refGene_tx=XM_019820423.2;refGene_gn=TRRAP;refGene_tx=XM_023246564.1;refGene_gn=TRRAP;refGene_tx=XM_023246565.1;refGene_gn=TRRAP;refGene_tx=XM_023246566.1;refGene_gn=TRRAP;refGene_tx=XM_023246567.1;refGene_gn=TRRAP;refGene_tx=XM_023246568.1;refGene_gn=TRRAP;refGene_tx=XM_023246569.1;refGene_gn=TRRAP;refGene_tx=XM_023246570.1;refGene_gn=TRRAP;refGene_tx=XM_023246571.1;refGene_gn=TRRAP;refGene_tx=XM_023246572.1;refGene_gn=TRRAP;refGene_tx=XM_023246573.1;refGene_gn=TRRAP;refGene_tx=XM_023246574.1;refGene_gn=TRRAP;refGene_tx=XM_023246575.1;refGene_gn=TRRAP;refGene_tx=XM_023246576.1
E3 6223077 6223269
refGene_gn=TRRAP;refGene_tx=XM_019820423.2;refGene_gn=TRRAP;refGene_tx=XM_023246564.1;refGene_gn=TRRAP;refGene_tx=XM_023246565.1;refGene_gn=TRRAP;refGene_tx=XM_023246566.1;refGene_gn=TRRAP;refGene_tx=XM_023246567.1;refGene_gn=TRRAP;refGene_tx=XM_023246568.1;refGene_gn=TRRAP;refGene_tx=XM_023246569.1;refGene_gn=TRRAP;refGene_tx=XM_023246570.1;refGene_gn=TRRAP;refGene_tx=XM_023246571.1;refGene_gn=TRRAP;refGene_tx=XM_023246572.1;refGene_gn=TRRAP;refGene_tx=XM_023246573.1;refGene_gn=TRRAP;refGene_tx=XM_023246574.1;refGene_gn=TRRAP;refGene_tx=XM_023246575.1;refGene_gn=TRRAP;refGene_tx=XM_023246576.1
E3 6225121 6225287
refGene_gn=TRRAP;refGene_tx=XM_019820423.2;refGene_gn=TRRAP;refGene_tx=XM_023246564.1;refGene_gn=TRRAP;refGene_tx=XM_023246565.1;refGene_gn=TRRAP;refGene_tx=XM_023246566.1;refGene_gn=TRRAP;refGene_tx=XM_023246567.1;refGene_gn=TRRAP;refGene_tx=XM_023246568.1;refGene_gn=TRRAP;refGene_tx=XM_023246569.1;refGene_gn=TRRAP;refGene_tx=XM_023246570.1;refGene_gn=TRRAP;refGene_tx=XM_023246571.1;refGene_gn=TRRAP;refGene_tx=XM_023246572.1;refGene_gn=TRRAP;refGene_tx=XM_023246573.1;refGene_gn=TRRAP;refGene_tx=XM_023246574.1;refGene_gn=TRRAP;refGene_tx=XM_023246575.1;refGene_gn=TRRAP;refGene_tx=XM_023246576.1
E3 6226787 6227044
refGene_gn=TRRAP;refGene_tx=XM_019820423.2;refGene_gn=TRRAP;refGene_tx=XM_023246564.1;refGene_gn=TRRAP;refGene_tx=XM_023246565.1;refGene_gn=TRRAP;refGene_tx=XM_023246566.1;refGene_gn=TRRAP;refGene_tx=XM_023246567.1;refGene_gn=TRRAP;refGene_tx=XM_023246568.1;refGene_gn=TRRAP;refGene_tx=XM_023246569.1;refGene_gn=TRRAP;refGene_tx=XM_023246570.1;refGene_gn=TRRAP;refGene_tx=XM_023246571.1;refGene_gn=TRRAP;refGene_tx=XM_023246572.1;refGene_gn=TRRAP;refGene_tx=XM_023246573.1;refGene_gn=TRRAP;refGene_tx=XM_023246574.1;refGene_gn=TRRAP;refGene_tx=XM_023246575.1;r

efGene_gn=TRRAP;refGene_tx=XM_023246576.1
E3 6229315 6229516
refGene_gn=TRRAP;refGene_tx=XM_019820423.2;refGene_gn=TRRAP;refGene_tx=XM_023246564.1;refGene_gn=TRRAP;refGene_tx=XM_023246565.1;refGene_gn=TRRAP;refGene_tx=XM_023246566.1;refGene_gn=TRRAP;refGene_tx=XM_023246567.1;refGene_gn=TRRAP;refGene_tx=XM_023246568.1;refGene_gn=TRRAP;refGene_tx=XM_023246569.1;refGene_gn=TRRAP;refGene_tx=XM_023246570.1;refGene_gn=TRRAP;refGene_tx=XM_023246571.1;refGene_gn=TRRAP;refGene_tx=XM_023246572.1;refGene_gn=TRRAP;refGene_tx=XM_023246573.1;refGene_gn=TRRAP;refGene_tx=XM_023246574.1;refGene_gn=TRRAP;refGene_tx=XM_023246575.1;refGene_gn=TRRAP;refGene_tx=XM_023246576.1
E3 6232100 6232252
refGene_gn=TRRAP;refGene_tx=XM_019820423.2;refGene_gn=TRRAP;refGene_tx=XM_023246564.1;refGene_gn=TRRAP;refGene_tx=XM_023246565.1;refGene_gn=TRRAP;refGene_tx=XM_023246566.1;refGene_gn=TRRAP;refGene_tx=XM_023246567.1;refGene_gn=TRRAP;refGene_tx=XM_023246568.1;refGene_gn=TRRAP;refGene_tx=XM_023246569.1;refGene_gn=TRRAP;refGene_tx=XM_023246570.1;refGene_gn=TRRAP;refGene_tx=XM_023246571.1;refGene_gn=TRRAP;refGene_tx=XM_023246572.1;refGene_gn=TRRAP;refGene_tx=XM_023246573.1;refGene_gn=TRRAP;refGene_tx=XM_023246574.1;refGene_gn=TRRAP;refGene_tx=XM_023246575.1;refGene_gn=TRRAP;refGene_tx=XM_023246576.1
E3 6233989 6234189
refGene_gn=TRRAP;refGene_tx=XM_019820423.2;refGene_gn=TRRAP;refGene_tx=XM_023246564.1;refGene_gn=TRRAP;refGene_tx=XM_023246565.1;refGene_gn=TRRAP;refGene_tx=XM_023246566.1;refGene_gn=TRRAP;refGene_tx=XM_023246567.1;refGene_gn=TRRAP;refGene_tx=XM_023246568.1;refGene_gn=TRRAP;refGene_tx=XM_023246569.1;refGene_gn=TRRAP;refGene_tx=XM_023246570.1;refGene_gn=TRRAP;refGene_tx=XM_023246571.1;refGene_gn=TRRAP;refGene_tx=XM_023246572.1;refGene_gn=TRRAP;refGene_tx=XM_023246573.1;refGene_gn=TRRAP;refGene_tx=XM_023246574.1;refGene_gn=TRRAP;refGene_tx=XM_023246575.1;refGene_gn=TRRAP;refGene_tx=XM_023246576.1
E3 6236842 6237060
refGene_gn=TRRAP;refGene_tx=XM_019820423.2;refGene_gn=TRRAP;refGene_tx=XM_023246564.1;refGene_gn=TRRAP;refGene_tx=XM_023246565.1;refGene_gn=TRRAP;refGene_tx=XM_023246566.1;refGene_gn=TRRAP;refGene_tx=XM_023246567.1;refGene_gn=TRRAP;refGene_tx=XM_023246568.1;refGene_gn=TRRAP;refGene_tx=XM_023246569.1;refGene_gn=TRRAP;refGene_tx=XM_023246570.1;refGene_gn=TRRAP;refGene_tx=XM_023246571.1;refGene_gn=TRRAP;refGene_tx=XM_023246572.1;refGene_gn=TRRAP;refGene_tx=XM_023246573.1;refGene_gn=TRRAP;refGene_tx=XM_023246574.1;refGene_gn=TRRAP;refGene_tx=XM_023246575.1;refGene_gn=TRRAP;refGene_tx=XM_023246576.1
E3 6237181 6237379
refGene_gn=TRRAP;refGene_tx=XM_019820423.2;refGene_gn=TRRAP;refGene_tx=XM_023246564.1;refGene_gn=TRRAP;refGene_tx=XM_023246565.1;refGene_gn=TRRAP;refGene_tx=XM_023246566.1;refGene_gn=TRRAP;refGene_tx=XM_023246567.1;refGene_gn=TRRAP;refGene_tx=XM_023246568.1;refGene_gn=TRRAP;refGene_tx=XM_023246569.1;refGene_gn=TRRAP;refGene_tx=XM_023246570.1;refGene_gn=TRRAP;refGene_tx=XM_023246571.1;refGene_gn=TRRAP;refGene_tx=XM_023246572.1;refGene_gn=TRRAP;refGene_tx=XM_023246573.1;refGene_gn=TRRAP;refGene_tx=XM_023246574.1;refGene_gn=TRRAP;refGene_tx=XM_023246575.1;r

efGene_gn=TRRAP;refGene_tx=XM_023246576.1
E3 6237943 6238204
refGene_gn=TRRAP;refGene_tx=XM_019820423.2;refGene_gn=TRRAP;refGene_tx=XM_023246564.1;refGene_gn=TRRAP;refGene_tx=XM_023246565.1;refGene_gn=TRRAP;refGene_tx=XM_023246566.1;refGene_gn=TRRAP;refGene_tx=XM_023246567.1;refGene_gn=TRRAP;refGene_tx=XM_023246568.1;refGene_gn=TRRAP;refGene_tx=XM_023246569.1;refGene_gn=TRRAP;refGene_tx=XM_023246570.1;refGene_gn=TRRAP;refGene_tx=XM_023246571.1;refGene_gn=TRRAP;refGene_tx=XM_023246572.1;refGene_gn=TRRAP;refGene_tx=XM_023246573.1;refGene_gn=TRRAP;refGene_tx=XM_023246574.1;refGene_gn=TRRAP;refGene_tx=XM_023246575.1;refGene_gn=TRRAP;refGene_tx=XM_023246576.1
E3 6238691 6238853
refGene_gn=TRRAP;refGene_tx=XM_019820423.2;refGene_gn=TRRAP;refGene_tx=XM_023246564.1;refGene_gn=TRRAP;refGene_tx=XM_023246565.1;refGene_gn=TRRAP;refGene_tx=XM_023246566.1;refGene_gn=TRRAP;refGene_tx=XM_023246567.1;refGene_gn=TRRAP;refGene_tx=XM_023246568.1;refGene_gn=TRRAP;refGene_tx=XM_023246569.1;refGene_gn=TRRAP;refGene_tx=XM_023246570.1;refGene_gn=TRRAP;refGene_tx=XM_023246571.1;refGene_gn=TRRAP;refGene_tx=XM_023246572.1;refGene_gn=TRRAP;refGene_tx=XM_023246573.1;refGene_gn=TRRAP;refGene_tx=XM_023246574.1;refGene_gn=TRRAP;refGene_tx=XM_023246575.1;refGene_gn=TRRAP;refGene_tx=XM_023246576.1
E3 6240843 6240940
refGene_gn=TRRAP;refGene_tx=XM_019820423.2;refGene_gn=TRRAP;refGene_tx=XM_023246564.1;refGene_gn=TRRAP;refGene_tx=XM_023246565.1;refGene_gn=TRRAP;refGene_tx=XM_023246566.1;refGene_gn=TRRAP;refGene_tx=XM_023246567.1;refGene_gn=TRRAP;refGene_tx=XM_023246568.1;refGene_gn=TRRAP;refGene_tx=XM_023246569.1;refGene_gn=TRRAP;refGene_tx=XM_023246570.1;refGene_gn=TRRAP;refGene_tx=XM_023246571.1;refGene_gn=TRRAP;refGene_tx=XM_023246572.1;refGene_gn=TRRAP;refGene_tx=XM_023246573.1;refGene_gn=TRRAP;refGene_tx=XM_023246574.1;refGene_gn=TRRAP;refGene_tx=XM_023246575.1;refGene_gn=TRRAP;refGene_tx=XM_023246576.1
E3 6242948 6243070
refGene_gn=TRRAP;refGene_tx=XM_019820423.2;refGene_gn=TRRAP;refGene_tx=XM_023246564.1;refGene_gn=TRRAP;refGene_tx=XM_023246565.1;refGene_gn=TRRAP;refGene_tx=XM_023246566.1;refGene_gn=TRRAP;refGene_tx=XM_023246567.1;refGene_gn=TRRAP;refGene_tx=XM_023246568.1;refGene_gn=TRRAP;refGene_tx=XM_023246569.1;refGene_gn=TRRAP;refGene_tx=XM_023246570.1;refGene_gn=TRRAP;refGene_tx=XM_023246571.1;refGene_gn=TRRAP;refGene_tx=XM_023246572.1;refGene_gn=TRRAP;refGene_tx=XM_023246573.1;refGene_gn=TRRAP;refGene_tx=XM_023246574.1;refGene_gn=TRRAP;refGene_tx=XM_023246575.1;refGene_gn=TRRAP;refGene_tx=XM_023246576.1
E3 6243410 6243581
refGene_gn=TRRAP;refGene_tx=XM_019820423.2;refGene_gn=TRRAP;refGene_tx=XM_023246564.1;refGene_gn=TRRAP;refGene_tx=XM_023246565.1;refGene_gn=TRRAP;refGene_tx=XM_023246566.1;refGene_gn=TRRAP;refGene_tx=XM_023246567.1;refGene_gn=TRRAP;refGene_tx=XM_023246568.1;refGene_gn=TRRAP;refGene_tx=XM_023246569.1;refGene_gn=TRRAP;refGene_tx=XM_023246570.1;refGene_gn=TRRAP;refGene_tx=XM_023246571.1;refGene_gn=TRRAP;refGene_tx=XM_023246572.1;refGene_gn=TRRAP;refGene_tx=XM_023246573.1;refGene_gn=TRRAP;refGene_tx=XM_023246574.1;refGene_gn=TRRAP;refGene_tx=XM_023246575.1;r

efGene_gn=TRRAP;refGene_tx=XM_023246576.1
E3 6247409 6247478
refGene_gn=TRRAP;refGene_tx=XM_019820423.2;refGene_gn=TRRAP;refGene_tx=XM_023246564.1;refGene_gn=TRRAP;refGene_tx=XM_023246565.1;refGene_gn=TRRAP;refGene_tx=XM_023246566.1;refGene_gn=TRRAP;refGene_tx=XM_023246567.1;refGene_gn=TRRAP;refGene_tx=XM_023246568.1;refGene_gn=TRRAP;refGene_tx=XM_023246569.1;refGene_gn=TRRAP;refGene_tx=XM_023246570.1;refGene_gn=TRRAP;refGene_tx=XM_023246571.1;refGene_gn=TRRAP;refGene_tx=XM_023246572.1;refGene_gn=TRRAP;refGene_tx=XM_023246573.1;refGene_gn=TRRAP;refGene_tx=XM_023246574.1;refGene_gn=TRRAP;refGene_tx=XM_023246575.1;refGene_gn=TRRAP;refGene_tx=XM_023246576.1
E3 6249790 6249844
refGene_gn=TRRAP;refGene_tx=XM_019820423.2;refGene_gn=TRRAP;refGene_tx=XM_023246564.1;refGene_gn=TRRAP;refGene_tx=XM_023246565.1;refGene_gn=TRRAP;refGene_tx=XM_023246566.1;refGene_gn=TRRAP;refGene_tx=XM_023246567.1;refGene_gn=TRRAP;refGene_tx=XM_023246568.1;refGene_gn=TRRAP;refGene_tx=XM_023246569.1;refGene_gn=TRRAP;refGene_tx=XM_023246570.1;refGene_gn=TRRAP;refGene_tx=XM_023246571.1;refGene_gn=TRRAP;refGene_tx=XM_023246572.1
E3 6249974 6249995
refGene_gn=TRRAP;refGene_tx=XM_019820423.2;refGene_gn=TRRAP;refGene_tx=XM_023246564.1;refGene_gn=TRRAP;refGene_tx=XM_023246565.1;refGene_gn=TRRAP;refGene_tx=XM_023246566.1;refGene_gn=TRRAP;refGene_tx=XM_023246567.1;refGene_gn=TRRAP;refGene_tx=XM_023246568.1;refGene_gn=TRRAP;refGene_tx=XM_023246569.1;refGene_gn=TRRAP;refGene_tx=XM_023246570.1;refGene_gn=TRRAP;refGene_tx=XM_023246571.1;refGene_gn=TRRAP;refGene_tx=XM_023246572.1
E3 6251686 6251806
refGene_gn=TRRAP;refGene_tx=XM_019820423.2;refGene_gn=TRRAP;refGene_tx=XM_023246564.1;refGene_gn=TRRAP;refGene_tx=XM_023246565.1;refGene_gn=TRRAP;refGene_tx=XM_023246566.1;refGene_gn=TRRAP;refGene_tx=XM_023246567.1;refGene_gn=TRRAP;refGene_tx=XM_023246568.1;refGene_gn=TRRAP;refGene_tx=XM_023246569.1;refGene_gn=TRRAP;refGene_tx=XM_023246570.1;refGene_gn=TRRAP;refGene_tx=XM_023246571.1;refGene_gn=TRRAP;refGene_tx=XM_023246572.1;refGene_gn=TRRAP;refGene_tx=XM_023246573.1;refGene_gn=TRRAP;refGene_tx=XM_023246574.1;refGene_gn=TRRAP;refGene_tx=XM_023246575.1;refGene_gn=TRRAP;refGene_tx=XM_023246576.1
E3 6252051 6252171
refGene_gn=TRRAP;refGene_tx=XM_019820423.2;refGene_gn=TRRAP;refGene_tx=XM_023246564.1;refGene_gn=TRRAP;refGene_tx=XM_023246565.1;refGene_gn=TRRAP;refGene_tx=XM_023246566.1;refGene_gn=TRRAP;refGene_tx=XM_023246567.1;refGene_gn=TRRAP;refGene_tx=XM_023246568.1;refGene_gn=TRRAP;refGene_tx=XM_023246569.1;refGene_gn=TRRAP;refGene_tx=XM_023246570.1;refGene_gn=TRRAP;refGene_tx=XM_023246571.1;refGene_gn=TRRAP;refGene_tx=XM_023246572.1;refGene_gn=TRRAP;refGene_tx=XM_023246573.1;refGene_gn=TRRAP;refGene_tx=XM_023246574.1;refGene_gn=TRRAP;refGene_tx=XM_023246575.1;refGene_gn=TRRAP;refGene_tx=XM_023246576.1
E3 6252570 6252735
refGene_gn=TRRAP;refGene_tx=XM_019820423.2;refGene_gn=TRRAP;refGene_tx=XM_023246564.1;refGene_gn=TRRAP;refGene_tx=XM_023246565.1;refGene_gn=

TRRAP;refGene_tx=XM_023246566.1;refGene_gn=TRRAP;refGene_tx=XM_0232465
67.1;refGene_gn=TRRAP;refGene_tx=XM_023246568.1;refGene_gn=TRRAP;refGe
ne_tx=XM_023246569.1;refGene_gn=TRRAP;refGene_tx=XM_023246570.1;refGen
e_gn=TRRAP;refGene_tx=XM_023246571.1;refGene_gn=TRRAP;refGene_tx=XM_02
3246572.1;refGene_gn=TRRAP;refGene_tx=XM_023246573.1;refGene_gn=TRRAP;
refGene_tx=XM_023246574.1;refGene_gn=TRRAP;refGene_tx=XM_023246575.1;r
efGene_gn=TRRAP;refGene_tx=XM_023246576.1
E3 6252829 6253011
refGene_gn=TRRAP;refGene_tx=XM_019820423.2;refGene_gn=TRRAP;refGene_tx
=XM_023246564.1;refGene_gn=TRRAP;refGene_tx=XM_023246565.1;refGene_gn=
TRRAP;refGene_tx=XM_023246566.1;refGene_gn=TRRAP;refGene_tx=XM_0232465
67.1;refGene_gn=TRRAP;refGene_tx=XM_023246568.1;refGene_gn=TRRAP;refGe
ne_tx=XM_023246569.1;refGene_gn=TRRAP;refGene_tx=XM_023246570.1;refGen
e_gn=TRRAP;refGene_tx=XM_023246571.1;refGene_gn=TRRAP;refGene_tx=XM_02
3246572.1;refGene_gn=TRRAP;refGene_tx=XM_023246573.1;refGene_gn=TRRAP;
refGene_tx=XM_023246574.1;refGene_gn=TRRAP;refGene_tx=XM_023246575.1;r
efGene_gn=TRRAP;refGene_tx=XM_023246576.1
E3 6253180 6253379
refGene_gn=TRRAP;refGene_tx=XM_019820423.2;refGene_gn=TRRAP;refGene_tx
=XM_023246564.1;refGene_gn=TRRAP;refGene_tx=XM_023246565.1;refGene_gn=
TRRAP;refGene_tx=XM_023246566.1;refGene_gn=TRRAP;refGene_tx=XM_0232465
67.1;refGene_gn=TRRAP;refGene_tx=XM_023246568.1;refGene_gn=TRRAP;refGe
ne_tx=XM_023246569.1;refGene_gn=TRRAP;refGene_tx=XM_023246570.1;refGen
e_gn=TRRAP;refGene_tx=XM_023246571.1;refGene_gn=TRRAP;refGene_tx=XM_02
3246572.1;refGene_gn=TRRAP;refGene_tx=XM_023246573.1;refGene_gn=TRRAP;
refGene_tx=XM_023246574.1;refGene_gn=TRRAP;refGene_tx=XM_023246575.1;r
efGene_gn=TRRAP;refGene_tx=XM_023246576.1
E3 6253836 6253965
refGene_gn=TRRAP;refGene_tx=XM_019820423.2;refGene_gn=TRRAP;refGene_tx
=XM_023246564.1;refGene_gn=TRRAP;refGene_tx=XM_023246565.1;refGene_gn=
TRRAP;refGene_tx=XM_023246566.1;refGene_gn=TRRAP;refGene_tx=XM_0232465
67.1;refGene_gn=TRRAP;refGene_tx=XM_023246568.1;refGene_gn=TRRAP;refGe
ne_tx=XM_023246569.1;refGene_gn=TRRAP;refGene_tx=XM_023246570.1;refGen
e_gn=TRRAP;refGene_tx=XM_023246571.1;refGene_gn=TRRAP;refGene_tx=XM_02
3246572.1;refGene_gn=TRRAP;refGene_tx=XM_023246573.1;refGene_gn=TRRAP;
refGene_tx=XM_023246574.1;refGene_gn=TRRAP;refGene_tx=XM_023246575.1;r
efGene_gn=TRRAP;refGene_tx=XM_023246576.1
E3 6256138 6256405
refGene_gn=TRRAP;refGene_tx=XM_019820423.2;refGene_gn=TRRAP;refGene_tx
=XM_023246564.1;refGene_gn=TRRAP;refGene_tx=XM_023246565.1;refGene_gn=
TRRAP;refGene_tx=XM_023246566.1;refGene_gn=TRRAP;refGene_tx=XM_0232465
67.1;refGene_gn=TRRAP;refGene_tx=XM_023246568.1;refGene_gn=TRRAP;refGe
ne_tx=XM_023246569.1;refGene_gn=TRRAP;refGene_tx=XM_023246570.1;refGen
e_gn=TRRAP;refGene_tx=XM_023246571.1;refGene_gn=TRRAP;refGene_tx=XM_02
3246572.1;refGene_gn=TRRAP;refGene_tx=XM_023246573.1;refGene_gn=TRRAP;
refGene_tx=XM_023246574.1;refGene_gn=TRRAP;refGene_tx=XM_023246575.1;r
efGene_gn=TRRAP;refGene_tx=XM_023246576.1
E3 6257654 6257861
refGene_gn=TRRAP;refGene_tx=XM_019820423.2;refGene_gn=TRRAP;refGene_tx
=XM_023246564.1;refGene_gn=TRRAP;refGene_tx=XM_023246565.1;refGene_gn=

TRRAP;refGene_tx=XM_023246566.1;refGene_gn=TRRAP;refGene_tx=XM_0232465
67.1;refGene_gn=TRRAP;refGene_tx=XM_023246568.1;refGene_gn=TRRAP;refGe
ne_tx=XM_023246569.1;refGene_gn=TRRAP;refGene_tx=XM_023246570.1;refGen
e_gn=TRRAP;refGene_tx=XM_023246571.1;refGene_gn=TRRAP;refGene_tx=XM_02
3246572.1;refGene_gn=TRRAP;refGene_tx=XM_023246573.1;refGene_gn=TRRAP;
refGene_tx=XM_023246574.1;refGene_gn=TRRAP;refGene_tx=XM_023246575.1;r
efGene_gn=TRRAP;refGene_tx=XM_023246576.1
E3 6258130 6258289
refGene_gn=TRRAP;refGene_tx=XM_019820423.2;refGene_gn=TRRAP;refGene_tx
=XM_023246564.1;refGene_gn=TRRAP;refGene_tx=XM_023246565.1;refGene_gn=
TRRAP;refGene_tx=XM_023246566.1;refGene_gn=TRRAP;refGene_tx=XM_0232465
67.1;refGene_gn=TRRAP;refGene_tx=XM_023246568.1;refGene_gn=TRRAP;refGe
ne_tx=XM_023246569.1;refGene_gn=TRRAP;refGene_tx=XM_023246570.1;refGen
e_gn=TRRAP;refGene_tx=XM_023246571.1;refGene_gn=TRRAP;refGene_tx=XM_02
3246572.1;refGene_gn=TRRAP;refGene_tx=XM_023246573.1;refGene_gn=TRRAP;
refGene_tx=XM_023246574.1;refGene_gn=TRRAP;refGene_tx=XM_023246575.1;r
efGene_gn=TRRAP;refGene_tx=XM_023246576.1
E3 6258385 6258520
refGene_gn=TRRAP;refGene_tx=XM_019820423.2;refGene_gn=TRRAP;refGene_tx
=XM_023246564.1;refGene_gn=TRRAP;refGene_tx=XM_023246565.1;refGene_gn=
TRRAP;refGene_tx=XM_023246566.1;refGene_gn=TRRAP;refGene_tx=XM_0232465
67.1;refGene_gn=TRRAP;refGene_tx=XM_023246568.1;refGene_gn=TRRAP;refGe
ne_tx=XM_023246569.1;refGene_gn=TRRAP;refGene_tx=XM_023246570.1;refGen
e_gn=TRRAP;refGene_tx=XM_023246571.1;refGene_gn=TRRAP;refGene_tx=XM_02
3246572.1;refGene_gn=TRRAP;refGene_tx=XM_023246573.1;refGene_gn=TRRAP;
refGene_tx=XM_023246574.1;refGene_gn=TRRAP;refGene_tx=XM_023246575.1;r
efGene_gn=TRRAP;refGene_tx=XM_023246576.1
E3 6259894 6260005
refGene_gn=TRRAP;refGene_tx=XM_019820423.2;refGene_gn=TRRAP;refGene_tx
=XM_023246564.1;refGene_gn=TRRAP;refGene_tx=XM_023246565.1;refGene_gn=
TRRAP;refGene_tx=XM_023246566.1;refGene_gn=TRRAP;refGene_tx=XM_0232465
67.1;refGene_gn=TRRAP;refGene_tx=XM_023246568.1;refGene_gn=TRRAP;refGe
ne_tx=XM_023246569.1;refGene_gn=TRRAP;refGene_tx=XM_023246570.1;refGen
e_gn=TRRAP;refGene_tx=XM_023246571.1;refGene_gn=TRRAP;refGene_tx=XM_02
3246572.1;refGene_gn=TRRAP;refGene_tx=XM_023246573.1;refGene_gn=TRRAP;
refGene_tx=XM_023246574.1;refGene_gn=TRRAP;refGene_tx=XM_023246575.1;r
efGene_gn=TRRAP;refGene_tx=XM_023246576.1
E3 6260928 6261075
refGene_gn=TRRAP;refGene_tx=XM_019820423.2;refGene_gn=TRRAP;refGene_tx
=XM_023246564.1;refGene_gn=TRRAP;refGene_tx=XM_023246565.1;refGene_gn=
TRRAP;refGene_tx=XM_023246566.1;refGene_gn=TRRAP;refGene_tx=XM_0232465
67.1;refGene_gn=TRRAP;refGene_tx=XM_023246568.1;refGene_gn=TRRAP;refGe
ne_tx=XM_023246569.1;refGene_gn=TRRAP;refGene_tx=XM_023246570.1;refGen
e_gn=TRRAP;refGene_tx=XM_023246571.1;refGene_gn=TRRAP;refGene_tx=XM_02
3246572.1;refGene_gn=TRRAP;refGene_tx=XM_023246573.1;refGene_gn=TRRAP;
refGene_tx=XM_023246574.1;refGene_gn=TRRAP;refGene_tx=XM_023246575.1;r
efGene_gn=TRRAP;refGene_tx=XM_023246576.1
E3 6263166 6263380
refGene_gn=TRRAP;refGene_tx=XM_019820423.2;refGene_gn=TRRAP;refGene_tx
=XM_023246564.1;refGene_gn=TRRAP;refGene_tx=XM_023246565.1;refGene_gn=

TRRAP;refGene_tx=XM_023246566.1;refGene_gn=TRRAP;refGene_tx=XM_0232465
67.1;refGene_gn=TRRAP;refGene_tx=XM_023246568.1;refGene_gn=TRRAP;refGe
ne_tx=XM_023246569.1;refGene_gn=TRRAP;refGene_tx=XM_023246570.1;refGen
e_gn=TRRAP;refGene_tx=XM_023246571.1;refGene_gn=TRRAP;refGene_tx=XM_02
3246572.1;refGene_gn=TRRAP;refGene_tx=XM_023246573.1;refGene_gn=TRRAP;
refGene_tx=XM_023246574.1;refGene_gn=TRRAP;refGene_tx=XM_023246575.1;r
efGene_gn=TRRAP;refGene_tx=XM_023246576.1
E3 6263932 6264058
refGene_gn=TRRAP;refGene_tx=XM_019820423.2;refGene_gn=TRRAP;refGene_tx
=XM_023246564.1;refGene_gn=TRRAP;refGene_tx=XM_023246565.1;refGene_gn=
TRRAP;refGene_tx=XM_023246566.1;refGene_gn=TRRAP;refGene_tx=XM_0232465
67.1;refGene_gn=TRRAP;refGene_tx=XM_023246568.1;refGene_gn=TRRAP;refGe
ne_tx=XM_023246569.1;refGene_gn=TRRAP;refGene_tx=XM_023246570.1;refGen
e_gn=TRRAP;refGene_tx=XM_023246571.1;refGene_gn=TRRAP;refGene_tx=XM_02
3246572.1;refGene_gn=TRRAP;refGene_tx=XM_023246573.1;refGene_gn=TRRAP;
refGene_tx=XM_023246574.1;refGene_gn=TRRAP;refGene_tx=XM_023246575.1;r
efGene_gn=TRRAP;refGene_tx=XM_023246576.1
E3 6266202 6266349
refGene_gn=TRRAP;refGene_tx=XM_019820423.2;refGene_gn=TRRAP;refGene_tx
=XM_023246564.1;refGene_gn=TRRAP;refGene_tx=XM_023246565.1;refGene_gn=
TRRAP;refGene_tx=XM_023246566.1;refGene_gn=TRRAP;refGene_tx=XM_0232465
67.1;refGene_gn=TRRAP;refGene_tx=XM_023246568.1;refGene_gn=TRRAP;refGe
ne_tx=XM_023246569.1;refGene_gn=TRRAP;refGene_tx=XM_023246570.1;refGen
e_gn=TRRAP;refGene_tx=XM_023246571.1;refGene_gn=TRRAP;refGene_tx=XM_02
3246572.1;refGene_gn=TRRAP;refGene_tx=XM_023246573.1;refGene_gn=TRRAP;
refGene_tx=XM_023246574.1;refGene_gn=TRRAP;refGene_tx=XM_023246575.1;r
efGene_gn=TRRAP;refGene_tx=XM_023246576.1
E3 6267238 6267441
refGene_gn=TRRAP;refGene_tx=XM_019820423.2;refGene_gn=TRRAP;refGene_tx
=XM_023246564.1;refGene_gn=TRRAP;refGene_tx=XM_023246565.1;refGene_gn=
TRRAP;refGene_tx=XM_023246566.1;refGene_gn=TRRAP;refGene_tx=XM_0232465
67.1;refGene_gn=TRRAP;refGene_tx=XM_023246568.1;refGene_gn=TRRAP;refGe
ne_tx=XM_023246569.1;refGene_gn=TRRAP;refGene_tx=XM_023246570.1;refGen
e_gn=TRRAP;refGene_tx=XM_023246571.1;refGene_gn=TRRAP;refGene_tx=XM_02
3246572.1;refGene_gn=TRRAP;refGene_tx=XM_023246573.1;refGene_gn=TRRAP;
refGene_tx=XM_023246574.1;refGene_gn=TRRAP;refGene_tx=XM_023246575.1;r
efGene_gn=TRRAP;refGene_tx=XM_023246576.1
E3 6268042 6268164
refGene_gn=TRRAP;refGene_tx=XM_019820423.2;refGene_gn=TRRAP;refGene_tx
=XM_023246564.1;refGene_gn=TRRAP;refGene_tx=XM_023246565.1;refGene_gn=
TRRAP;refGene_tx=XM_023246566.1;refGene_gn=TRRAP;refGene_tx=XM_0232465
67.1;refGene_gn=TRRAP;refGene_tx=XM_023246568.1;refGene_gn=TRRAP;refGe
ne_tx=XM_023246569.1;refGene_gn=TRRAP;refGene_tx=XM_023246570.1;refGen
e_gn=TRRAP;refGene_tx=XM_023246571.1;refGene_gn=TRRAP;refGene_tx=XM_02
3246572.1;refGene_gn=TRRAP;refGene_tx=XM_023246573.1;refGene_gn=TRRAP;
refGene_tx=XM_023246574.1;refGene_gn=TRRAP;refGene_tx=XM_023246575.1;r
efGene_gn=TRRAP;refGene_tx=XM_023246576.1
E3 6268482 6268696
refGene_gn=TRRAP;refGene_tx=XM_019820423.2;refGene_gn=TRRAP;refGene_tx
=XM_023246564.1;refGene_gn=TRRAP;refGene_tx=XM_023246565.1;refGene_gn=

TRRAP;refGene_tx=XM_023246566.1;refGene_gn=TRRAP;refGene_tx=XM_0232465
67.1;refGene_gn=TRRAP;refGene_tx=XM_023246568.1;refGene_gn=TRRAP;refGe
ne_tx=XM_023246569.1;refGene_gn=TRRAP;refGene_tx=XM_023246570.1;refGen
e_gn=TRRAP;refGene_tx=XM_023246571.1;refGene_gn=TRRAP;refGene_tx=XM_02
3246572.1;refGene_gn=TRRAP;refGene_tx=XM_023246573.1;refGene_gn=TRRAP;
refGene_tx=XM_023246574.1;refGene_gn=TRRAP;refGene_tx=XM_023246575.1;r
efGene_gn=TRRAP;refGene_tx=XM_023246576.1
E3 6270486 6270666
refGene_gn=TRRAP;refGene_tx=XM_019820423.2;refGene_gn=TRRAP;refGene_tx
=XM_023246564.1;refGene_gn=TRRAP;refGene_tx=XM_023246565.1;refGene_gn=
TRRAP;refGene_tx=XM_023246566.1;refGene_gn=TRRAP;refGene_tx=XM_0232465
67.1;refGene_gn=TRRAP;refGene_tx=XM_023246568.1;refGene_gn=TRRAP;refGe
ne_tx=XM_023246569.1;refGene_gn=TRRAP;refGene_tx=XM_023246570.1;refGen
e_gn=TRRAP;refGene_tx=XM_023246571.1;refGene_gn=TRRAP;refGene_tx=XM_02
3246572.1;refGene_gn=TRRAP;refGene_tx=XM_023246573.1;refGene_gn=TRRAP;
refGene_tx=XM_023246574.1;refGene_gn=TRRAP;refGene_tx=XM_023246575.1;r
efGene_gn=TRRAP;refGene_tx=XM_023246576.1
E3 6272551 6272698
refGene_gn=TRRAP;refGene_tx=XM_019820423.2;refGene_gn=TRRAP;refGene_tx
=XM_023246564.1;refGene_gn=TRRAP;refGene_tx=XM_023246565.1;refGene_gn=
TRRAP;refGene_tx=XM_023246566.1;refGene_gn=TRRAP;refGene_tx=XM_0232465
67.1;refGene_gn=TRRAP;refGene_tx=XM_023246568.1;refGene_gn=TRRAP;refGe
ne_tx=XM_023246569.1;refGene_gn=TRRAP;refGene_tx=XM_023246570.1;refGen
e_gn=TRRAP;refGene_tx=XM_023246571.1;refGene_gn=TRRAP;refGene_tx=XM_02
3246572.1;refGene_gn=TRRAP;refGene_tx=XM_023246573.1;refGene_gn=TRRAP;
refGene_tx=XM_023246574.1;refGene_gn=TRRAP;refGene_tx=XM_023246575.1;r
efGene_gn=TRRAP;refGene_tx=XM_023246576.1
E3 6275878 6275998
refGene_gn=TRRAP;refGene_tx=XM_019820423.2;refGene_gn=TRRAP;refGene_tx
=XM_023246564.1;refGene_gn=TRRAP;refGene_tx=XM_023246565.1;refGene_gn=
TRRAP;refGene_tx=XM_023246566.1;refGene_gn=TRRAP;refGene_tx=XM_0232465
67.1;refGene_gn=TRRAP;refGene_tx=XM_023246568.1;refGene_gn=TRRAP;refGe
ne_tx=XM_023246569.1;refGene_gn=TRRAP;refGene_tx=XM_023246570.1;refGen
e_gn=TRRAP;refGene_tx=XM_023246571.1;refGene_gn=TRRAP;refGene_tx=XM_02
3246572.1;refGene_gn=TRRAP;refGene_tx=XM_023246573.1;refGene_gn=TRRAP;
refGene_tx=XM_023246574.1;refGene_gn=TRRAP;refGene_tx=XM_023246575.1;r
efGene_gn=TRRAP;refGene_tx=XM_023246576.1
E3 6276194 6276482
refGene_gn=TRRAP;refGene_tx=XM_019820423.2;refGene_gn=TRRAP;refGene_tx
=XM_023246564.1;refGene_gn=TRRAP;refGene_tx=XM_023246565.1;refGene_gn=
TRRAP;refGene_tx=XM_023246566.1;refGene_gn=TRRAP;refGene_tx=XM_0232465
67.1;refGene_gn=TRRAP;refGene_tx=XM_023246568.1;refGene_gn=TRRAP;refGe
ne_tx=XM_023246569.1;refGene_gn=TRRAP;refGene_tx=XM_023246570.1;refGen
e_gn=TRRAP;refGene_tx=XM_023246571.1;refGene_gn=TRRAP;refGene_tx=XM_02
3246572.1;refGene_gn=TRRAP;refGene_tx=XM_023246573.1;refGene_gn=TRRAP;
refGene_tx=XM_023246574.1;refGene_gn=TRRAP;refGene_tx=XM_023246575.1;r
efGene_gn=TRRAP;refGene_tx=XM_023246576.1
E3 6276650 6276788
refGene_gn=TRRAP;refGene_tx=XM_019820423.2;refGene_gn=TRRAP;refGene_tx
=XM_023246564.1;refGene_gn=TRRAP;refGene_tx=XM_023246565.1;refGene_gn=

TRRAP;refGene_tx=XM_023246566.1;refGene_gn=TRRAP;refGene_tx=XM_0232465
67.1;refGene_gn=TRRAP;refGene_tx=XM_023246568.1;refGene_gn=TRRAP;refGe
ne_tx=XM_023246569.1;refGene_gn=TRRAP;refGene_tx=XM_023246570.1;refGen
e_gn=TRRAP;refGene_tx=XM_023246571.1;refGene_gn=TRRAP;refGene_tx=XM_02
3246572.1;refGene_gn=TRRAP;refGene_tx=XM_023246573.1;refGene_gn=TRRAP;
refGene_tx=XM_023246574.1;refGene_gn=TRRAP;refGene_tx=XM_023246575.1;r
efGene_gn=TRRAP;refGene_tx=XM_023246576.1
E3 6277428 6277541
refGene_gn=TRRAP;refGene_tx=XM_019820423.2;refGene_gn=TRRAP;refGene_tx
=XM_023246564.1;refGene_gn=TRRAP;refGene_tx=XM_023246565.1;refGene_gn=
TRRAP;refGene_tx=XM_023246566.1;refGene_gn=TRRAP;refGene_tx=XM_0232465
67.1;refGene_gn=TRRAP;refGene_tx=XM_023246568.1;refGene_gn=TRRAP;refGe
ne_tx=XM_023246569.1;refGene_gn=TRRAP;refGene_tx=XM_023246570.1;refGen
e_gn=TRRAP;refGene_tx=XM_023246571.1;refGene_gn=TRRAP;refGene_tx=XM_02
3246572.1;refGene_gn=TRRAP;refGene_tx=XM_023246573.1;refGene_gn=TRRAP;
refGene_tx=XM_023246574.1;refGene_gn=TRRAP;refGene_tx=XM_023246575.1;r
efGene_gn=TRRAP;refGene_tx=XM_023246576.1
E3 6277926 6278062
refGene_gn=TRRAP;refGene_tx=XM_019820423.2;refGene_gn=TRRAP;refGene_tx
=XM_023246564.1;refGene_gn=TRRAP;refGene_tx=XM_023246565.1;refGene_gn=
TRRAP;refGene_tx=XM_023246566.1;refGene_gn=TRRAP;refGene_tx=XM_0232465
67.1;refGene_gn=TRRAP;refGene_tx=XM_023246568.1;refGene_gn=TRRAP;refGe
ne_tx=XM_023246569.1;refGene_gn=TRRAP;refGene_tx=XM_023246570.1;refGen
e_gn=TRRAP;refGene_tx=XM_023246571.1;refGene_gn=TRRAP;refGene_tx=XM_02
3246572.1;refGene_gn=TRRAP;refGene_tx=XM_023246573.1;refGene_gn=TRRAP;
refGene_tx=XM_023246574.1;refGene_gn=TRRAP;refGene_tx=XM_023246575.1;r
efGene_gn=TRRAP;refGene_tx=XM_023246576.1
E3 6281031 6281223
refGene_gn=TRRAP;refGene_tx=XM_019820423.2;refGene_gn=TRRAP;refGene_tx
=XM_023246564.1;refGene_gn=TRRAP;refGene_tx=XM_023246565.1;refGene_gn=
TRRAP;refGene_tx=XM_023246566.1;refGene_gn=TRRAP;refGene_tx=XM_0232465
67.1;refGene_gn=TRRAP;refGene_tx=XM_023246568.1;refGene_gn=TRRAP;refGe
ne_tx=XM_023246569.1;refGene_gn=TRRAP;refGene_tx=XM_023246570.1;refGen
e_gn=TRRAP;refGene_tx=XM_023246571.1;refGene_gn=TRRAP;refGene_tx=XM_02
3246572.1;refGene_gn=TRRAP;refGene_tx=XM_023246573.1;refGene_gn=TRRAP;
refGene_tx=XM_023246574.1;refGene_gn=TRRAP;refGene_tx=XM_023246575.1;r
efGene_gn=TRRAP;refGene_tx=XM_023246576.1
E3 6283808 6284037
refGene_gn=TRRAP;refGene_tx=XM_019820423.2;refGene_gn=TRRAP;refGene_tx
=XM_023246564.1;refGene_gn=TRRAP;refGene_tx=XM_023246565.1;refGene_gn=
TRRAP;refGene_tx=XM_023246566.1;refGene_gn=TRRAP;refGene_tx=XM_0232465
67.1;refGene_gn=TRRAP;refGene_tx=XM_023246568.1;refGene_gn=TRRAP;refGe
ne_tx=XM_023246569.1;refGene_gn=TRRAP;refGene_tx=XM_023246570.1;refGen
e_gn=TRRAP;refGene_tx=XM_023246571.1;refGene_gn=TRRAP;refGene_tx=XM_02
3246572.1;refGene_gn=TRRAP;refGene_tx=XM_023246573.1;refGene_gn=TRRAP;
refGene_tx=XM_023246574.1;refGene_gn=TRRAP;refGene_tx=XM_023246575.1;r
efGene_gn=TRRAP;refGene_tx=XM_023246576.1
E3 6284630 6284896
refGene_gn=TRRAP;refGene_tx=XM_019820423.2;refGene_gn=TRRAP;refGene_tx
=XM_023246564.1;refGene_gn=TRRAP;refGene_tx=XM_023246565.1;refGene_gn=

TRRAP;refGene_tx=XM_023246566.1;refGene_gn=TRRAP;refGene_tx=XM_023246567.1;refGene_gn=TRRAP;refGene_tx=XM_023246568.1;refGene_gn=TRRAP;refGene_tx=XM_023246569.1;refGene_gn=TRRAP;refGene_tx=XM_023246570.1;refGene_gn=TRRAP;refGene_tx=XM_023246571.1;refGene_gn=TRRAP;refGene_tx=XM_023246572.1;refGene_gn=TRRAP;refGene_tx=XM_023246573.1;refGene_gn=TRRAP;refGene_tx=XM_023246574.1;refGene_gn=TRRAP;refGene_tx=XM_023246575.1;refGene_gn=TRRAP;refGene_tx=XM_023246576.1
E3 6285461 6285562
refGene_gn=TRRAP;refGene_tx=XM_019820423.2;refGene_gn=TRRAP;refGene_tx=XM_023246564.1;refGene_gn=TRRAP;refGene_tx=XM_023246565.1;refGene_gn=TRRAP;refGene_tx=XM_023246566.1;refGene_gn=TRRAP;refGene_tx=XM_023246567.1;refGene_gn=TRRAP;refGene_tx=XM_023246568.1;refGene_gn=TRRAP;refGene_tx=XM_023246569.1;refGene_gn=TRRAP;refGene_tx=XM_023246570.1;refGene_gn=TRRAP;refGene_tx=XM_023246571.1;refGene_gn=TRRAP;refGene_tx=XM_023246572.1;refGene_gn=TRRAP;refGene_tx=XM_023246573.1;refGene_gn=TRRAP;refGene_tx=XM_023246574.1;refGene_gn=TRRAP;refGene_tx=XM_023246575.1;refGene_gn=TRRAP;refGene_tx=XM_023246576.1
E3 6289279 6289481
refGene_gn=TRRAP;refGene_tx=XM_019820423.2;refGene_gn=TRRAP;refGene_tx=XM_023246564.1;refGene_gn=TRRAP;refGene_tx=XM_023246565.1;refGene_gn=TRRAP;refGene_tx=XM_023246566.1;refGene_gn=TRRAP;refGene_tx=XM_023246567.1;refGene_gn=TRRAP;refGene_tx=XM_023246568.1;refGene_gn=TRRAP;refGene_tx=XM_023246569.1;refGene_gn=TRRAP;refGene_tx=XM_023246570.1;refGene_gn=TRRAP;refGene_tx=XM_023246571.1;refGene_gn=TRRAP;refGene_tx=XM_023246572.1;refGene_gn=TRRAP;refGene_tx=XM_023246573.1;refGene_gn=TRRAP;refGene_tx=XM_023246574.1;refGene_gn=TRRAP;refGene_tx=XM_023246575.1;refGene_gn=TRRAP;refGene_tx=XM_023246576.1
E3 6290614 6290779
refGene_gn=TRRAP;refGene_tx=XM_019820423.2;refGene_gn=TRRAP;refGene_tx=XM_023246564.1;refGene_gn=TRRAP;refGene_tx=XM_023246565.1;refGene_gn=TRRAP;refGene_tx=XM_023246566.1;refGene_gn=TRRAP;refGene_tx=XM_023246567.1;refGene_gn=TRRAP;refGene_tx=XM_023246568.1;refGene_gn=TRRAP;refGene_tx=XM_023246569.1;refGene_gn=TRRAP;refGene_tx=XM_023246570.1;refGene_gn=TRRAP;refGene_tx=XM_023246571.1;refGene_gn=TRRAP;refGene_tx=XM_023246572.1;refGene_gn=TRRAP;refGene_tx=XM_023246573.1;refGene_gn=TRRAP;refGene_tx=XM_023246574.1;refGene_gn=TRRAP;refGene_tx=XM_023246575.1;refGene_gn=TRRAP;refGene_tx=XM_023246576.1
E3 6292475 6292566
refGene_gn=TRRAP;refGene_tx=XM_019820423.2;refGene_gn=TRRAP;refGene_tx=XM_023246564.1;refGene_gn=TRRAP;refGene_tx=XM_023246565.1;refGene_gn=TRRAP;refGene_tx=XM_023246566.1;refGene_gn=TRRAP;refGene_tx=XM_023246567.1;refGene_gn=TRRAP;refGene_tx=XM_023246568.1;refGene_gn=TRRAP;refGene_tx=XM_023246569.1;refGene_gn=TRRAP;refGene_tx=XM_023246570.1;refGene_gn=TRRAP;refGene_tx=XM_023246571.1;refGene_gn=TRRAP;refGene_tx=XM_023246572.1;refGene_gn=TRRAP;refGene_tx=XM_023246573.1;refGene_gn=TRRAP;refGene_tx=XM_023246574.1;refGene_gn=TRRAP;refGene_tx=XM_023246575.1;refGene_gn=TRRAP;refGene_tx=XM_023246576.1
E3 6293819 6294019
refGene_gn=TRRAP;refGene_tx=XM_019820423.2;refGene_gn=TRRAP;refGene_tx=XM_023246564.1;refGene_gn=TRRAP;refGene_tx=XM_023246565.1;refGene_gn=

TRRAP;refGene_tx=XM_023246566.1;refGene_gn=TRRAP;refGene_tx=XM_0232465
67.1;refGene_gn=TRRAP;refGene_tx=XM_023246568.1;refGene_gn=TRRAP;refGe
ne_tx=XM_023246569.1;refGene_gn=TRRAP;refGene_tx=XM_023246570.1;refGen
e_gn=TRRAP;refGene_tx=XM_023246571.1;refGene_gn=TRRAP;refGene_tx=XM_02
3246572.1;refGene_gn=TRRAP;refGene_tx=XM_023246573.1;refGene_gn=TRRAP;
refGene_tx=XM_023246574.1;refGene_gn=TRRAP;refGene_tx=XM_023246575.1;r
efGene_gn=TRRAP;refGene_tx=XM_023246576.1
E3 6294807 6295069
refGene_gn=TRRAP;refGene_tx=XM_019820423.2;refGene_gn=TRRAP;refGene_tx
=XM_023246564.1;refGene_gn=TRRAP;refGene_tx=XM_023246565.1;refGene_gn=
TRRAP;refGene_tx=XM_023246566.1;refGene_gn=TRRAP;refGene_tx=XM_0232465
67.1;refGene_gn=TRRAP;refGene_tx=XM_023246568.1;refGene_gn=TRRAP;refGe
ne_tx=XM_023246569.1;refGene_gn=TRRAP;refGene_tx=XM_023246570.1;refGen
e_gn=TRRAP;refGene_tx=XM_023246571.1;refGene_gn=TRRAP;refGene_tx=XM_02
3246572.1;refGene_gn=TRRAP;refGene_tx=XM_023246573.1;refGene_gn=TRRAP;
refGene_tx=XM_023246574.1;refGene_gn=TRRAP;refGene_tx=XM_023246575.1;r
efGene_gn=TRRAP;refGene_tx=XM_023246576.1
E3 6298477 6298703
refGene_gn=TRRAP;refGene_tx=XM_019820423.2;refGene_gn=TRRAP;refGene_tx
=XM_023246564.1;refGene_gn=TRRAP;refGene_tx=XM_023246565.1;refGene_gn=
TRRAP;refGene_tx=XM_023246566.1;refGene_gn=TRRAP;refGene_tx=XM_0232465
67.1;refGene_gn=TRRAP;refGene_tx=XM_023246568.1;refGene_gn=TRRAP;refGe
ne_tx=XM_023246569.1;refGene_gn=TRRAP;refGene_tx=XM_023246570.1;refGen
e_gn=TRRAP;refGene_tx=XM_023246571.1;refGene_gn=TRRAP;refGene_tx=XM_02
3246572.1;refGene_gn=TRRAP;refGene_tx=XM_023246573.1;refGene_gn=TRRAP;
refGene_tx=XM_023246574.1;refGene_gn=TRRAP;refGene_tx=XM_023246575.1;r
efGene_gn=TRRAP;refGene_tx=XM_023246576.1
E3 6299595 6299813
refGene_gn=TRRAP;refGene_tx=XM_019820423.2;refGene_gn=TRRAP;refGene_tx
=XM_023246564.1;refGene_gn=TRRAP;refGene_tx=XM_023246565.1;refGene_gn=
TRRAP;refGene_tx=XM_023246566.1;refGene_gn=TRRAP;refGene_tx=XM_0232465
67.1;refGene_gn=TRRAP;refGene_tx=XM_023246568.1;refGene_gn=TRRAP;refGe
ne_tx=XM_023246569.1;refGene_gn=TRRAP;refGene_tx=XM_023246570.1;refGen
e_gn=TRRAP;refGene_tx=XM_023246571.1;refGene_gn=TRRAP;refGene_tx=XM_02
3246572.1;refGene_gn=TRRAP;refGene_tx=XM_023246573.1;refGene_gn=TRRAP;
refGene_tx=XM_023246574.1;refGene_gn=TRRAP;refGene_tx=XM_023246575.1;r
efGene_gn=TRRAP;refGene_tx=XM_023246576.1
E3 6302258 6302443
refGene_gn=TRRAP;refGene_tx=XM_019820423.2;refGene_gn=TRRAP;refGene_tx
=XM_023246564.1;refGene_gn=TRRAP;refGene_tx=XM_023246565.1;refGene_gn=
TRRAP;refGene_tx=XM_023246566.1;refGene_gn=TRRAP;refGene_tx=XM_0232465
67.1;refGene_gn=TRRAP;refGene_tx=XM_023246568.1;refGene_gn=TRRAP;refGe
ne_tx=XM_023246569.1;refGene_gn=TRRAP;refGene_tx=XM_023246570.1;refGen
e_gn=TRRAP;refGene_tx=XM_023246571.1;refGene_gn=TRRAP;refGene_tx=XM_02
3246572.1;refGene_gn=TRRAP;refGene_tx=XM_023246573.1;refGene_gn=TRRAP;
refGene_tx=XM_023246574.1;refGene_gn=TRRAP;refGene_tx=XM_023246575.1;r
efGene_gn=TRRAP;refGene_tx=XM_023246576.1
E3 6306188 6306392
refGene_gn=TRRAP;refGene_tx=XM_019820423.2;refGene_gn=TRRAP;refGene_tx
=XM_023246564.1;refGene_gn=TRRAP;refGene_tx=XM_023246565.1;refGene_gn=

TRRAP;refGene_tx=XM_023246566.1;refGene_gn=TRRAP;refGene_tx=XM_0232465
67.1;refGene_gn=TRRAP;refGene_tx=XM_023246568.1;refGene_gn=TRRAP;refGe
ne_tx=XM_023246569.1;refGene_gn=TRRAP;refGene_tx=XM_023246570.1;refGen
e_gn=TRRAP;refGene_tx=XM_023246571.1;refGene_gn=TRRAP;refGene_tx=XM_02
3246572.1;refGene_gn=TRRAP;refGene_tx=XM_023246573.1;refGene_gn=TRRAP;
refGene_tx=XM_023246574.1;refGene_gn=TRRAP;refGene_tx=XM_023246575.1;r
efGene_gn=TRRAP;refGene_tx=XM_023246576.1
E3 6306478 6306673
refGene_gn=TRRAP;refGene_tx=XM_019820423.2;refGene_gn=TRRAP;refGene_tx
=XM_023246564.1;refGene_gn=TRRAP;refGene_tx=XM_023246565.1;refGene_gn=
TRRAP;refGene_tx=XM_023246566.1;refGene_gn=TRRAP;refGene_tx=XM_0232465
67.1;refGene_gn=TRRAP;refGene_tx=XM_023246568.1;refGene_gn=TRRAP;refGe
ne_tx=XM_023246569.1;refGene_gn=TRRAP;refGene_tx=XM_023246570.1;refGen
e_gn=TRRAP;refGene_tx=XM_023246571.1;refGene_gn=TRRAP;refGene_tx=XM_02
3246572.1;refGene_gn=TRRAP;refGene_tx=XM_023246573.1;refGene_gn=TRRAP;
refGene_tx=XM_023246574.1;refGene_gn=TRRAP;refGene_tx=XM_023246575.1;r
efGene_gn=TRRAP;refGene_tx=XM_023246576.1
E3 6308655 6308940
refGene_gn=TRRAP;refGene_tx=XM_019820423.2;refGene_gn=TRRAP;refGene_tx
=XM_023246564.1;refGene_gn=TRRAP;refGene_tx=XM_023246565.1;refGene_gn=
TRRAP;refGene_tx=XM_023246566.1;refGene_gn=TRRAP;refGene_tx=XM_0232465
67.1;refGene_gn=TRRAP;refGene_tx=XM_023246568.1;refGene_gn=TRRAP;refGe
ne_tx=XM_023246569.1;refGene_gn=TRRAP;refGene_tx=XM_023246570.1;refGen
e_gn=TRRAP;refGene_tx=XM_023246571.1;refGene_gn=TRRAP;refGene_tx=XM_02
3246572.1;refGene_gn=TRRAP;refGene_tx=XM_023246573.1;refGene_gn=TRRAP;
refGene_tx=XM_023246574.1;refGene_gn=TRRAP;refGene_tx=XM_023246575.1;r
efGene_gn=TRRAP;refGene_tx=XM_023246576.1
E3 6333125 6333225
refGene_gn=SMURF1;refGene_tx=XM_019820434.2;refGene_gn=SMURF1;refGene_
tx=XM_019820435.2;refGene_gn=SMURF1;refGene_tx=XM_019820437.2;refGene_
gn=SMURF1;refGene_tx=XM_023246787.1;refGene_gn=SMURF1;refGene_tx=XM_02
3246788.1;refGene_gn=SMURF1;refGene_tx=XM_023246789.1;refGene_gn=SMURF
1;refGene_tx=XM_023246790.1;refGene_gn=SMURF1;refGene_tx=XM_023246791.
1
E3 6334739 6334824
refGene_gn=SMURF1;refGene_tx=XM_019820434.2;refGene_gn=SMURF1;refGene_
tx=XM_019820435.2;refGene_gn=SMURF1;refGene_tx=XM_019820437.2;refGene_
gn=SMURF1;refGene_tx=XM_023246787.1;refGene_gn=SMURF1;refGene_tx=XM_02
3246788.1;refGene_gn=SMURF1;refGene_tx=XM_023246789.1;refGene_gn=SMURF
1;refGene_tx=XM_023246790.1;refGene_gn=SMURF1;refGene_tx=XM_023246791.
1
E3 6336280 6336482
refGene_gn=SMURF1;refGene_tx=XM_019820434.2;refGene_gn=SMURF1;refGene_
tx=XM_019820435.2;refGene_gn=SMURF1;refGene_tx=XM_019820437.2;refGene_
gn=SMURF1;refGene_tx=XM_023246787.1;refGene_gn=SMURF1;refGene_tx=XM_02
3246788.1;refGene_gn=SMURF1;refGene_tx=XM_023246789.1;refGene_gn=SMURF
1;refGene_tx=XM_023246790.1;refGene_gn=SMURF1;refGene_tx=XM_023246791.
1
E3 6337390 6337511
refGene_gn=SMURF1;refGene_tx=XM_019820434.2;refGene_gn=SMURF1;refGene_

tx=XM_019820435.2;refGene_gn=SMURF1;refGene_tx=XM_019820437.2;refGene_
gn=SMURF1;refGene_tx=XM_023246787.1;refGene_gn=SMURF1;refGene_tx=XM_02
3246788.1;refGene_gn=SMURF1;refGene_tx=XM_023246789.1;refGene_gn=SMURF
1;refGene_tx=XM_023246790.1;refGene_gn=SMURF1;refGene_tx=XM_023246791.
1
E3 6338538 6338676
refGene_gn=SMURF1;refGene_tx=XM_019820434.2;refGene_gn=SMURF1;refGene_
tx=XM_019820435.2;refGene_gn=SMURF1;refGene_tx=XM_019820437.2;refGene_
gn=SMURF1;refGene_tx=XM_023246787.1;refGene_gn=SMURF1;refGene_tx=XM_02
3246788.1;refGene_gn=SMURF1;refGene_tx=XM_023246789.1;refGene_gn=SMURF
1;refGene_tx=XM_023246790.1;refGene_gn=SMURF1;refGene_tx=XM_023246791.
1
E3 6339861 6340040
refGene_gn=SMURF1;refGene_tx=XM_019820434.2;refGene_gn=SMURF1;refGene_
tx=XM_019820435.2;refGene_gn=SMURF1;refGene_tx=XM_019820437.2;refGene_
gn=SMURF1;refGene_tx=XM_023246787.1;refGene_gn=SMURF1;refGene_tx=XM_02
3246788.1;refGene_gn=SMURF1;refGene_tx=XM_023246789.1;refGene_gn=SMURF
1;refGene_tx=XM_023246790.1;refGene_gn=SMURF1;refGene_tx=XM_023246791.
1
E3 6340812 6340927
refGene_gn=SMURF1;refGene_tx=XM_019820434.2;refGene_gn=SMURF1;refGene_
tx=XM_019820435.2;refGene_gn=SMURF1;refGene_tx=XM_019820437.2;refGene_
gn=SMURF1;refGene_tx=XM_023246787.1;refGene_gn=SMURF1;refGene_tx=XM_02
3246788.1;refGene_gn=SMURF1;refGene_tx=XM_023246789.1;refGene_gn=SMURF
1;refGene_tx=XM_023246790.1;refGene_gn=SMURF1;refGene_tx=XM_023246791.
1
E3 6349132 6349236
refGene_gn=SMURF1;refGene_tx=XM_019820434.2;refGene_gn=SMURF1;refGene_
tx=XM_019820435.2;refGene_gn=SMURF1;refGene_tx=XM_019820437.2;refGene_
gn=SMURF1;refGene_tx=XM_023246787.1;refGene_gn=SMURF1;refGene_tx=XM_02
3246788.1;refGene_gn=SMURF1;refGene_tx=XM_023246789.1;refGene_gn=SMURF
1;refGene_tx=XM_023246790.1;refGene_gn=SMURF1;refGene_tx=XM_023246791.
1
E3 6350967 6351166
refGene_gn=SMURF1;refGene_tx=XM_019820434.2;refGene_gn=SMURF1;refGene_
tx=XM_019820435.2;refGene_gn=SMURF1;refGene_tx=XM_019820437.2;refGene_
gn=SMURF1;refGene_tx=XM_023246787.1;refGene_gn=SMURF1;refGene_tx=XM_02
3246788.1;refGene_gn=SMURF1;refGene_tx=XM_023246789.1;refGene_gn=SMURF
1;refGene_tx=XM_023246790.1;refGene_gn=SMURF1;refGene_tx=XM_023246791.
1
E3 6352489 6352636
refGene_gn=SMURF1;refGene_tx=XM_019820434.2;refGene_gn=SMURF1;refGene_
tx=XM_019820435.2;refGene_gn=SMURF1;refGene_tx=XM_019820437.2;refGene_
gn=SMURF1;refGene_tx=XM_023246787.1;refGene_gn=SMURF1;refGene_tx=XM_02
3246788.1;refGene_gn=SMURF1;refGene_tx=XM_023246789.1;refGene_gn=SMURF
1;refGene_tx=XM_023246790.1;refGene_gn=SMURF1;refGene_tx=XM_023246791.
1
E3 6353548 6353626
refGene_gn=SMURF1;refGene_tx=XM_019820434.2;refGene_gn=SMURF1;refGene_
tx=XM_019820435.2;refGene_gn=SMURF1;refGene_tx=XM_019820437.2;refGene_

gn=SMURF1;refGene_tx=XM_023246787.1;refGene_gn=SMURF1;refGene_tx=XM_02
3246788.1;refGene_gn=SMURF1;refGene_tx=XM_023246791.1
E3	6354227	6354312
refGene_gn=SMURF1;refGene_tx=XM_019820434.2;refGene_gn=SMURF1;refGene_
tx=XM_019820435.2;refGene_gn=SMURF1;refGene_tx=XM_019820437.2;refGene_
gn=SMURF1;refGene_tx=XM_023246787.1;refGene_gn=SMURF1;refGene_tx=XM_02
3246788.1;refGene_gn=SMURF1;refGene_tx=XM_023246789.1;refGene_gn=SMURF
1;refGene_tx=XM_023246790.1;refGene_gn=SMURF1;refGene_tx=XM_023246791.
1
E3	6355035	6355277
refGene_gn=SMURF1;refGene_tx=XM_019820434.2;refGene_gn=SMURF1;refGene_
tx=XM_019820435.2;refGene_gn=SMURF1;refGene_tx=XM_019820437.2;refGene_
gn=SMURF1;refGene_tx=XM_023246787.1;refGene_gn=SMURF1;refGene_tx=XM_02
3246788.1;refGene_gn=SMURF1;refGene_tx=XM_023246789.1;refGene_gn=SMURF
1;refGene_tx=XM_023246790.1;refGene_gn=SMURF1;refGene_tx=XM_023246791.
1
E3	6357262	6357338
refGene_gn=SMURF1;refGene_tx=XM_019820434.2;refGene_gn=SMURF1;refGene_
tx=XM_019820435.2;refGene_gn=SMURF1;refGene_tx=XM_019820437.2;refGene_
gn=SMURF1;refGene_tx=XM_023246787.1;refGene_gn=SMURF1;refGene_tx=XM_02
3246788.1;refGene_gn=SMURF1;refGene_tx=XM_023246789.1;refGene_gn=SMURF
1;refGene_tx=XM_023246790.1;refGene_gn=SMURF1;refGene_tx=XM_023246791.
1
E3	6358607	6358673
refGene_gn=SMURF1;refGene_tx=XM_019820434.2;refGene_gn=SMURF1;refGene_
tx=XM_019820435.2;refGene_gn=SMURF1;refGene_tx=XM_019820437.2;refGene_
gn=SMURF1;refGene_tx=XM_023246787.1;refGene_gn=SMURF1;refGene_tx=XM_02
3246789.1;refGene_gn=SMURF1;refGene_tx=XM_023246790.1;refGene_gn=SMURF
1;refGene_tx=XM_023246791.1
E3	6358837	6358971
refGene_gn=SMURF1;refGene_tx=XM_019820434.2;refGene_gn=SMURF1;refGene_
tx=XM_019820435.2;refGene_gn=SMURF1;refGene_tx=XM_019820437.2;refGene_
gn=SMURF1;refGene_tx=XM_023246787.1;refGene_gn=SMURF1;refGene_tx=XM_02
3246788.1;refGene_gn=SMURF1;refGene_tx=XM_023246789.1;refGene_gn=SMURF
1;refGene_tx=XM_023246790.1;refGene_gn=SMURF1;refGene_tx=XM_023246791.
1
E3	6360262	6360371
refGene_gn=SMURF1;refGene_tx=XM_019820434.2;refGene_gn=SMURF1;refGene_
tx=XM_019820435.2;refGene_gn=SMURF1;refGene_tx=XM_019820437.2;refGene_
gn=SMURF1;refGene_tx=XM_023246787.1;refGene_gn=SMURF1;refGene_tx=XM_02
3246788.1;refGene_gn=SMURF1;refGene_tx=XM_023246789.1;refGene_gn=SMURF
1;refGene_tx=XM_023246790.1;refGene_gn=SMURF1;refGene_tx=XM_023246791.
1
E3	6361526	6361565
refGene_gn=SMURF1;refGene_tx=XM_019820434.2;refGene_gn=SMURF1;refGene_
tx=XM_019820435.2;refGene_gn=SMURF1;refGene_tx=XM_019820437.2;refGene_
gn=SMURF1;refGene_tx=XM_023246787.1;refGene_gn=SMURF1;refGene_tx=XM_02
3246788.1;refGene_gn=SMURF1;refGene_tx=XM_023246789.1;refGene_gn=SMURF
1;refGene_tx=XM_023246790.1;refGene_gn=SMURF1;refGene_tx=XM_023246791.
1

E3 6409162 6409250 refGene_gn=SMURF1;refGene_tx=XM_019820435.2
E3 6438095 6438201
refGene_gn=SMURF1;refGene_tx=XM_019820434.2;refGene_gn=SMURF1;refGene_tx=XM_023246787.1;refGene_gn=SMURF1;refGene_tx=XM_023246788.1;refGene_gn=SMURF1;refGene_tx=XM_023246789.1;refGene_gn=SMURF1;refGene_tx=XM_023246790.1
E3 6438584 6438627 refGene_gn=SMURF1;refGene_tx=XM_019820437.2
E3 6439737 6439792 refGene_gn=SMURF1;refGene_tx=XM_023246791.1
E3 6461135 6461249 refGene_gn=KPNA7;refGene_tx=XM_023246792.1
E3 6464199 6464346 refGene_gn=KPNA7;refGene_tx=XM_023246792.1
E3 6470667 6470850 refGene_gn=KPNA7;refGene_tx=XM_023246792.1
E3 6472892 6473126 refGene_gn=KPNA7;refGene_tx=XM_023246792.1
E3 6474539 6474803 refGene_gn=KPNA7;refGene_tx=XM_023246792.1
E3 6476813 6476896 refGene_gn=KPNA7;refGene_tx=XM_023246792.1
E3 6478518 6478787 refGene_gn=KPNA7;refGene_tx=XM_023246792.1
E3 6479217 6479300 refGene_gn=KPNA7;refGene_tx=XM_023246792.1
E3 6483963 6484092 refGene_gn=KPNA7;refGene_tx=XM_023246792.1
E3 6484788 6484854 refGene_gn=KPNA7;refGene_tx=XM_023246792.1
E3 6508793 6509003
refGene_gn=SMYHC;refGene_tx=NM_001009221.1;refGene_gn=SMYHC;refGene_tx=XM_019820223.1
E3 6511666 6511810
refGene_gn=SMYHC;refGene_tx=NM_001009221.1;refGene_gn=SMYHC;refGene_tx=XM_019820223.1
E3 6513299 6513456
refGene_gn=SMYHC;refGene_tx=NM_001009221.1;refGene_gn=SMYHC;refGene_tx=XM_019820223.1
E3 6514077 6514105
refGene_gn=SMYHC;refGene_tx=NM_001009221.1;refGene_gn=SMYHC;refGene_tx=XM_019820223.1
E3 6514424 6514521
refGene_gn=SMYHC;refGene_tx=NM_001009221.1;refGene_gn=SMYHC;refGene_tx=XM_019820223.1
E3 6515337 6515430
refGene_gn=SMYHC;refGene_tx=NM_001009221.1;refGene_gn=SMYHC;refGene_tx=XM_019820223.1
E3 6516435 6516499
refGene_gn=SMYHC;refGene_tx=NM_001009221.1;refGene_gn=SMYHC;refGene_tx=XM_019820223.1
E3 6517012 6517208
refGene_gn=SMYHC;refGene_tx=NM_001009221.1;refGene_gn=SMYHC;refGene_tx=XM_019820223.1;refGene_gn=SMYHC;refGene_tx=XM_023246202.1
E3 6518831 6518935
refGene_gn=SMYHC;refGene_tx=NM_001009221.1;refGene_gn=SMYHC;refGene_tx=XM_019820222.2;refGene_gn=SMYHC;refGene_tx=XM_019820223.1;refGene_gn=SMYHC;refGene_tx=XM_023246202.1
E3 6519732 6519871
refGene_gn=SMYHC;refGene_tx=NM_001009221.1;refGene_gn=SMYHC;refGene_tx=XM_019820222.2;refGene_gn=SMYHC;refGene_tx=XM_019820223.1;refGene_gn=SMYHC;refGene_tx=XM_023246202.1

E3	6520237	6520506
refGene_gn=SMYHC;refGene_tx=NM_001009221.1;refGene_gn=SMYHC;refGene_tx=XM_019820222.2;refGene_gn=SMYHC;refGene_tx=XM_019820223.1;refGene_gn=SMYHC;refGene_tx=XM_023246202.1
E3	6522146	6522317
refGene_gn=SMYHC;refGene_tx=NM_001009221.1;refGene_gn=SMYHC;refGene_tx=XM_019820222.2;refGene_gn=SMYHC;refGene_tx=XM_019820223.1;refGene_gn=SMYHC;refGene_tx=XM_023246202.1
E3	6523986	6524166
refGene_gn=SMYHC;refGene_tx=NM_001009221.1;refGene_gn=SMYHC;refGene_tx=XM_019820222.2;refGene_gn=SMYHC;refGene_tx=XM_019820223.1;refGene_gn=SMYHC;refGene_tx=XM_023246202.1
E3	6525812	6525939
refGene_gn=SMYHC;refGene_tx=NM_001009221.1;refGene_gn=SMYHC;refGene_tx=XM_019820222.2;refGene_gn=SMYHC;refGene_tx=XM_019820223.1;refGene_gn=SMYHC;refGene_tx=XM_023246202.1
E3	6526715	6526774
refGene_gn=SMYHC;refGene_tx=NM_001009221.1;refGene_gn=SMYHC;refGene_tx=XM_019820222.2;refGene_gn=SMYHC;refGene_tx=XM_019820223.1;refGene_gn=SMYHC;refGene_tx=XM_023246202.1
E3	6527901	6527989
refGene_gn=SMYHC;refGene_tx=NM_001009221.1;refGene_gn=SMYHC;refGene_tx=XM_019820222.2;refGene_gn=SMYHC;refGene_tx=XM_019820223.1;refGene_gn=SMYHC;refGene_tx=XM_023246202.1
E3	6528776	6528894
refGene_gn=SMYHC;refGene_tx=NM_001009221.1;refGene_gn=SMYHC;refGene_tx=XM_019820222.2;refGene_gn=SMYHC;refGene_tx=XM_019820223.1;refGene_gn=SMYHC;refGene_tx=XM_023246202.1
E3	6530098	6530219
refGene_gn=SMYHC;refGene_tx=NM_001009221.1;refGene_gn=SMYHC;refGene_tx=XM_019820222.2;refGene_gn=SMYHC;refGene_tx=XM_019820223.1;refGene_gn=SMYHC;refGene_tx=XM_023246202.1
E3	6533822	6533959
refGene_gn=SMYHC;refGene_tx=NM_001009221.1;refGene_gn=SMYHC;refGene_tx=XM_019820222.2;refGene_gn=SMYHC;refGene_tx=XM_019820223.1;refGene_gn=SMYHC;refGene_tx=XM_023246202.1
E3	6535531	6535613
refGene_gn=SMYHC;refGene_tx=NM_001009221.1;refGene_gn=SMYHC;refGene_tx=XM_019820222.2;refGene_gn=SMYHC;refGene_tx=XM_019820223.1;refGene_gn=SMYHC;refGene_tx=XM_023246202.1
E3	6537944	6538118
refGene_gn=SMYHC;refGene_tx=NM_001009221.1;refGene_gn=SMYHC;refGene_tx=XM_019820222.2;refGene_gn=SMYHC;refGene_tx=XM_019820223.1;refGene_gn=SMYHC;refGene_tx=XM_023246202.1
E3	6539017	6539245
refGene_gn=SMYHC;refGene_tx=NM_001009221.1;refGene_gn=SMYHC;refGene_tx=XM_019820222.2;refGene_gn=SMYHC;refGene_tx=XM_019820223.1;refGene_gn=SMYHC;refGene_tx=XM_023246202.1
E3	6540256	6540361
refGene_gn=SMYHC;refGene_tx=NM_001009221.1;refGene_gn=SMYHC;refGene_tx

=XM_019820222.2;refGene_gn=SMYHC;refGene_tx=XM_019820223.1;refGene_gn=SMYHC;refGene_tx=XM_023246202.1
E3 6542723 6542810
refGene_gn=SMYHC;refGene_tx=NM_001009221.1;refGene_gn=SMYHC;refGene_tx=XM_019820222.2;refGene_gn=SMYHC;refGene_tx=XM_019820223.1;refGene_gn=SMYHC;refGene_tx=XM_023246202.1
E3 6543049 6543195
refGene_gn=SMYHC;refGene_tx=NM_001009221.1;refGene_gn=SMYHC;refGene_tx=XM_019820222.2;refGene_gn=SMYHC;refGene_tx=XM_019820223.1;refGene_gn=SMYHC;refGene_tx=XM_023246202.1
E3 6544320 6544411
refGene_gn=SMYHC;refGene_tx=NM_001009221.1;refGene_gn=SMYHC;refGene_tx=XM_019820222.2;refGene_gn=SMYHC;refGene_tx=XM_019820223.1;refGene_gn=SMYHC;refGene_tx=XM_023246202.1
E3 6544869 6544926
refGene_gn=SMYHC;refGene_tx=NM_001009221.1;refGene_gn=SMYHC;refGene_tx=XM_019820222.2;refGene_gn=SMYHC;refGene_tx=XM_019820223.1;refGene_gn=SMYHC;refGene_tx=XM_023246202.1
E3 6545600 6545687
refGene_gn=SMYHC;refGene_tx=NM_001009221.1;refGene_gn=SMYHC;refGene_tx=XM_019820222.2;refGene_gn=SMYHC;refGene_tx=XM_019820223.1;refGene_gn=SMYHC;refGene_tx=XM_023246202.1
E3 6547481 6547727
refGene_gn=SMYHC;refGene_tx=NM_001009221.1;refGene_gn=SMYHC;refGene_tx=XM_019820222.2;refGene_gn=SMYHC;refGene_tx=XM_019820223.1;refGene_gn=SMYHC;refGene_tx=XM_023246202.1
E3 6548605 6548723
refGene_gn=SMYHC;refGene_tx=NM_001009221.1;refGene_gn=SMYHC;refGene_tx=XM_019820222.2;refGene_gn=SMYHC;refGene_tx=XM_019820223.1;refGene_gn=SMYHC;refGene_tx=XM_023246202.1
E3 6549966 6550094
refGene_gn=SMYHC;refGene_tx=NM_001009221.1;refGene_gn=SMYHC;refGene_tx=XM_019820222.2;refGene_gn=SMYHC;refGene_tx=XM_019820223.1;refGene_gn=SMYHC;refGene_tx=XM_023246202.1
E3 6550647 6550844
refGene_gn=SMYHC;refGene_tx=NM_001009221.1;refGene_gn=SMYHC;refGene_tx=XM_019820222.2;refGene_gn=SMYHC;refGene_tx=XM_019820223.1;refGene_gn=SMYHC;refGene_tx=XM_023246202.1
E3 6552553 6552686
refGene_gn=SMYHC;refGene_tx=NM_001009221.1;refGene_gn=SMYHC;refGene_tx=XM_019820222.2;refGene_gn=SMYHC;refGene_tx=XM_019820223.1;refGene_gn=SMYHC;refGene_tx=XM_023246202.1
E3 6554237 6554454
refGene_gn=SMYHC;refGene_tx=NM_001009221.1;refGene_gn=SMYHC;refGene_tx=XM_019820222.2;refGene_gn=SMYHC;refGene_tx=XM_019820223.1;refGene_gn=SMYHC;refGene_tx=XM_023246202.1
E3 6556675 6556812
refGene_gn=SMYHC;refGene_tx=NM_001009221.1;refGene_gn=SMYHC;refGene_tx=XM_019820222.2;refGene_gn=SMYHC;refGene_tx=XM_019820223.1;refGene_gn=SMYHC;refGene_tx=XM_023246202.1

E3 6556905 6557009
refGene_gn=SMYHC;refGene_tx=NM_001009221.1;refGene_gn=SMYHC;refGene_tx=XM_019820222.2;refGene_gn=SMYHC;refGene_tx=XM_019820223.1;refGene_gn=SMYHC;refGene_tx=XM_023246202.1
E3 6561226 6561419
refGene_gn=SMYHC;refGene_tx=NM_001009221.1;refGene_gn=SMYHC;refGene_tx=XM_019820222.2;refGene_gn=SMYHC;refGene_tx=XM_019820223.1;refGene_gn=SMYHC;refGene_tx=XM_023246202.1
E3 6562340 6562544
refGene_gn=SMYHC;refGene_tx=NM_001009221.1;refGene_gn=SMYHC;refGene_tx=XM_019820222.2;refGene_gn=SMYHC;refGene_tx=XM_019820223.1;refGene_gn=SMYHC;refGene_tx=XM_023246202.1
E3 6563718 6563844
refGene_gn=SMYHC;refGene_tx=NM_001009221.1;refGene_gn=SMYHC;refGene_tx=XM_019820222.2;refGene_gn=SMYHC;refGene_tx=XM_019820223.1;refGene_gn=SMYHC;refGene_tx=XM_023246202.1
E3 6565107 6565278
refGene_gn=SMYHC;refGene_tx=NM_001009221.1;refGene_gn=SMYHC;refGene_tx=XM_019820222.2;refGene_gn=SMYHC;refGene_tx=XM_019820223.1;refGene_gn=SMYHC;refGene_tx=XM_023246202.1
E3 6565885 6566086
refGene_gn=SMYHC;refGene_tx=NM_001009221.1;refGene_gn=SMYHC;refGene_tx=XM_019820222.2;refGene_gn=SMYHC;refGene_tx=XM_019820223.1;refGene_gn=SMYHC;refGene_tx=XM_023246202.1
E3 6567074 6567275
refGene_gn=SMYHC;refGene_tx=NM_001009221.1;refGene_gn=SMYHC;refGene_tx=XM_019820222.2;refGene_gn=SMYHC;refGene_tx=XM_019820223.1;refGene_gn=SMYHC;refGene_tx=XM_023246202.1
E3 6589814 6589878 refGene_gn=ARPC1A;refGene_tx=XM_003998457.4
E3 6594581 6594686 refGene_gn=ARPC1A;refGene_tx=XM_003998457.4
E3 6600473 6600696 refGene_gn=ARPC1A;refGene_tx=XM_003998457.4
E3 6603188 6603296 refGene_gn=ARPC1A;refGene_tx=XM_003998457.4
E3 6605931 6606144 refGene_gn=ARPC1A;refGene_tx=XM_003998457.4
E3 6608434 6608510 refGene_gn=ARPC1A;refGene_tx=XM_003998457.4
E3 6608746 6608940 refGene_gn=ARPC1A;refGene_tx=XM_003998457.4
E3 6611198 6611289 refGene_gn=ARPC1A;refGene_tx=XM_003998457.4
E3 6613147 6613186 refGene_gn=ARPC1A;refGene_tx=XM_003998457.4
E3 6626976 6627040
refGene_gn=ARPC1B;refGene_tx=XM_003998458.3;refGene_gn=ARPC1B;refGene_tx=XM_023246536.1
E3 6627788 6627893
refGene_gn=ARPC1B;refGene_tx=XM_003998458.3;refGene_gn=ARPC1B;refGene_tx=XM_023246536.1
E3 6628333 6628556
refGene_gn=ARPC1B;refGene_tx=XM_003998458.3;refGene_gn=ARPC1B;refGene_tx=XM_023246536.1
E3 6629525 6629633
refGene_gn=ARPC1B;refGene_tx=XM_003998458.3;refGene_gn=ARPC1B;refGene_tx=XM_023246536.1
E3 6630597 6630804

refGene_gn=ARPC1B;refGene_tx=XM_003998458.3;refGene_gn=ARPC1B;refGene_tx=XM_023246536.1
E3 6630897 6630973
refGene_gn=ARPC1B;refGene_tx=XM_003998458.3;refGene_gn=ARPC1B;refGene_tx=XM_023246536.1
E3 6631861 6632067
refGene_gn=ARPC1B;refGene_tx=XM_003998458.3;refGene_gn=ARPC1B;refGene_tx=XM_023246536.1
E3 6632789 6632880
refGene_gn=ARPC1B;refGene_tx=XM_003998458.3;refGene_gn=ARPC1B;refGene_tx=XM_023246536.1
E3 6633091 6633130
refGene_gn=ARPC1B;refGene_tx=XM_003998458.3;refGene_gn=ARPC1B;refGene_tx=XM_023246536.1
E3 6637227 6637286 refGene_gn=PDAP1;refGene_tx=XM_003998459.4
E3 6638364 6638516 refGene_gn=PDAP1;refGene_tx=XM_003998459.4
E3 6640588 6640710 refGene_gn=PDAP1;refGene_tx=XM_003998459.4
E3 6642291 6642402 refGene_gn=PDAP1;refGene_tx=XM_003998459.4
E3 6643722 6643814 refGene_gn=PDAP1;refGene_tx=XM_003998459.4
E3 6647031 6647044 refGene_gn=PDAP1;refGene_tx=XM_003998459.4
E3 6649297 6649391 refGene_gn=BUD31;refGene_tx=XM_003998460.4
E3 6654926 6655049 refGene_gn=BUD31;refGene_tx=XM_003998460.4
E3 6655268 6655435 refGene_gn=BUD31;refGene_tx=XM_003998460.4
E3 6656977 6657028 refGene_gn=BUD31;refGene_tx=XM_003998460.4
E3 6657651 6657834
refGene_gn=PTCD1;refGene_tx=XM_006941945.4;refGene_gn=PTCD1;refGene_tx=XM_006941946.4;refGene_gn=PTCD1;refGene_tx=XM_011290410.3
E3 6659503 6659686
refGene_gn=PTCD1;refGene_tx=XM_006941945.4;refGene_gn=PTCD1;refGene_tx=XM_006941946.4;refGene_gn=PTCD1;refGene_tx=XM_011290410.3
E3 6660377 6660494
refGene_gn=PTCD1;refGene_tx=XM_006941945.4;refGene_gn=PTCD1;refGene_tx=XM_006941946.4;refGene_gn=PTCD1;refGene_tx=XM_011290410.3
E3 6661538 6662246
refGene_gn=PTCD1;refGene_tx=XM_006941945.4;refGene_gn=PTCD1;refGene_tx=XM_006941946.4;refGene_gn=PTCD1;refGene_tx=XM_011290410.3
E3 6664169 6664271
refGene_gn=PTCD1;refGene_tx=XM_006941945.4;refGene_gn=PTCD1;refGene_tx=XM_006941946.4;refGene_gn=PTCD1;refGene_tx=XM_011290410.3
E3 6666048 6666267
refGene_gn=PTCD1;refGene_tx=XM_006941945.4;refGene_gn=PTCD1;refGene_tx=XM_006941946.4;refGene_gn=PTCD1;refGene_tx=XM_011290410.3
E3 6669570 6669711
refGene_gn=PTCD1;refGene_tx=XM_006941945.4;refGene_gn=PTCD1;refGene_tx=XM_006941946.4;refGene_gn=PTCD1;refGene_tx=XM_011290410.3
E3 6671967 6672432
refGene_gn=PTCD1;refGene_tx=XM_006941945.4;refGene_gn=PTCD1;refGene_tx=XM_006941946.4;refGene_gn=PTCD1;refGene_tx=XM_011290410.3
E3 6675105 6675208
refGene_gn=CPSF4;refGene_tx=XM_003998461.4;refGene_gn=CPSF4;refGene_tx

=XM_003998462.5;refGene_gn=CPSF4;refGene_tx=XM_006941949.4
E3 6676636 6676687
refGene_gn=CPSF4;refGene_tx=XM_003998461.4;refGene_gn=CPSF4;refGene_tx
=XM_003998462.5;refGene_gn=CPSF4;refGene_tx=XM_006941949.4
E3 6678756 6678909
refGene_gn=CPSF4;refGene_tx=XM_003998461.4;refGene_gn=CPSF4;refGene_tx
=XM_003998462.5;refGene_gn=CPSF4;refGene_tx=XM_006941949.4
E3 6681369 6681465
refGene_gn=CPSF4;refGene_tx=XM_003998461.4;refGene_gn=CPSF4;refGene_tx
=XM_003998462.5;refGene_gn=CPSF4;refGene_tx=XM_006941949.4
E3 6681854 6681948
refGene_gn=CPSF4;refGene_tx=XM_003998461.4;refGene_gn=CPSF4;refGene_tx
=XM_003998462.5;refGene_gn=CPSF4;refGene_tx=XM_006941949.4
E3 6683096 6683169
refGene_gn=CPSF4;refGene_tx=XM_003998461.4;refGene_gn=CPSF4;refGene_tx
=XM_003998462.5;refGene_gn=CPSF4;refGene_tx=XM_006941949.4
E3 6684708 6684879
refGene_gn=CPSF4;refGene_tx=XM_003998461.4;refGene_gn=CPSF4;refGene_tx
=XM_003998462.5;refGene_gn=CPSF4;refGene_tx=XM_006941949.4
E3 6687737 6687806
refGene_gn=CPSF4;refGene_tx=XM_003998461.4;refGene_gn=CPSF4;refGene_tx
=XM_003998462.5;refGene_gn=CPSF4;refGene_tx=XM_006941949.4
E3 6689741 6689770
refGene_gn=ATP5MF;refGene_tx=XM_006941948.3;refGene_gn=ATP5MF;refGene_
tx=XM_019820444.1;refGene_gn=ATP5MF;refGene_tx=XM_019820445.1
E3 6690522 6690639
refGene_gn=ATP5MF;refGene_tx=XM_006941948.3;refGene_gn=ATP5MF;refGene_
tx=XM_019820444.1;refGene_gn=ATP5MF;refGene_tx=XM_019820445.1
E3 6691872 6691980
refGene_gn=ATP5MF;refGene_tx=XM_006941948.3;refGene_gn=ATP5MF;refGene_
tx=XM_019820444.1;refGene_gn=ATP5MF;refGene_tx=XM_019820445.1
E3 6696876 6696889 refGene_gn=ATP5MF;refGene_tx=XM_019820445.1
E3 6699413 6699429 refGene_gn=ATP5MF;refGene_tx=XM_019820444.1
E3 6699871 6699884 refGene_gn=ATP5MF;refGene_tx=XM_006941948.3
E3 6721136 6721170
refGene_gn=LOC102901593;refGene_tx=XM_011290491.3
E3 6721672 6721770
refGene_gn=LOC102901593;refGene_tx=XM_011290491.3
E3 6725833 6725960
refGene_gn=LOC102901593;refGene_tx=XM_011290491.3
E3 6729398 6729994
refGene_gn=LOC102901593;refGene_tx=XM_011290491.3
E3 6729995 6730613
refGene_gn=LOC102901593;refGene_tx=XM_011290491.3
E3 6749985 6750045
refGene_gn=ZNF789;refGene_tx=XM_006941950.4;refGene_gn=ZNF789;refGene_
tx=XM_023246440.1
E3 6753076 6753154
refGene_gn=ZNF789;refGene_tx=XM_003998505.5;refGene_gn=ZNF789;refGene_
tx=XM_006941950.4;refGene_gn=ZNF789;refGene_tx=XM_006941951.4;refGene_

gn=ZNF789;refGene_tx=XM_006941952.4;refGene_gn=ZNF789;refGene_tx=XM_01
1290408.3;refGene_gn=ZNF789;refGene_tx=XM_023246440.1
E3 6758450 6758577
refGene_gn=ZNF789;refGene_tx=XM_003998505.5;refGene_gn=ZNF789;refGene_
tx=XM_006941950.4;refGene_gn=ZNF789;refGene_tx=XM_006941951.4;refGene_
gn=ZNF789;refGene_tx=XM_006941952.4;refGene_gn=ZNF789;refGene_tx=XM_01
1290408.3;refGene_gn=ZNF789;refGene_tx=XM_023246440.1
E3 6760750 6760864
refGene_gn=ZNF789;refGene_tx=XM_003998505.5;refGene_gn=ZNF789;refGene_
tx=XM_006941950.4;refGene_gn=ZNF789;refGene_tx=XM_006941951.4;refGene_
gn=ZNF789;refGene_tx=XM_006941952.4;refGene_gn=ZNF789;refGene_tx=XM_01
1290408.3
E3 6762274 6763287
refGene_gn=ZNF789;refGene_tx=XM_003998505.5;refGene_gn=ZNF789;refGene_
tx=XM_006941950.4;refGene_gn=ZNF789;refGene_tx=XM_006941951.4;refGene_
gn=ZNF789;refGene_tx=XM_006941952.4;refGene_gn=ZNF789;refGene_tx=XM_01
1290408.3;refGene_gn=ZNF789;refGene_tx=XM_023246440.1
E3 6772428 6773504
refGene_gn=ZNF394;refGene_tx=XM_019820462.2;refGene_gn=ZNF394;refGene_
tx=XM_019820463.1;refGene_gn=ZNF394;refGene_tx=XM_023247129.1
E3 6778813 6778949
refGene_gn=ZNF394;refGene_tx=XM_019820462.2;refGene_gn=ZNF394;refGene_
tx=XM_023247129.1
E3 6781106 6781559
refGene_gn=ZNF394;refGene_tx=XM_019820462.2;refGene_gn=ZNF394;refGene_
tx=XM_019820463.1;refGene_gn=ZNF394;refGene_tx=XM_023247129.1
E3 6791725 6792139
refGene_gn=ZKSCAN5;refGene_tx=XM_003998465.4;refGene_gn=ZKSCAN5;refGen
e_tx=XM_006941957.3;refGene_gn=ZKSCAN5;refGene_tx=XM_006941958.3;refGe
ne_gn=ZKSCAN5;refGene_tx=XM_006941959.3;refGene_gn=ZKSCAN5;refGene_tx=
XM_006941960.3
E3 6796746 6796885
refGene_gn=ZKSCAN5;refGene_tx=XM_003998465.4;refGene_gn=ZKSCAN5;refGen
e_tx=XM_006941957.3;refGene_gn=ZKSCAN5;refGene_tx=XM_006941958.3;refGe
ne_gn=ZKSCAN5;refGene_tx=XM_006941959.3;refGene_gn=ZKSCAN5;refGene_tx=
XM_006941960.3;refGene_gn=ZKSCAN5;refGene_tx=XM_006941961.2;refGene_gn
=ZKSCAN5;refGene_tx=XM_023247128.1
E3 6800150 6800245 refGene_gn=ZKSCAN5;refGene_tx=XM_023247128.1
E3 6801239 6801322
refGene_gn=ZKSCAN5;refGene_tx=XM_003998465.4;refGene_gn=ZKSCAN5;refGen
e_tx=XM_006941957.3;refGene_gn=ZKSCAN5;refGene_tx=XM_006941958.3;refGe
ne_gn=ZKSCAN5;refGene_tx=XM_006941959.3;refGene_gn=ZKSCAN5;refGene_tx=
XM_006941961.2;refGene_gn=ZKSCAN5;refGene_tx=XM_023247128.1
E3 6801601 6801737
refGene_gn=ZKSCAN5;refGene_tx=XM_003998465.4;refGene_gn=ZKSCAN5;refGen
e_tx=XM_006941957.3;refGene_gn=ZKSCAN5;refGene_tx=XM_006941958.3;refGe
ne_gn=ZKSCAN5;refGene_tx=XM_006941959.3;refGene_gn=ZKSCAN5;refGene_tx=
XM_006941961.2
E3 6807827 6808442
refGene_gn=ZKSCAN5;refGene_tx=XM_003998465.4;refGene_gn=ZKSCAN5;refGen

e_tx=XM_006941957.3;refGene_gn=ZKSCAN5;refGene_tx=XM_006941958.3;refGene_gn=ZKSCAN5;refGene_tx=XM_006941959.3;refGene_gn=ZKSCAN5;refGene_tx=XM_006941960.3;refGene_gn=ZKSCAN5;refGene_tx=XM_006941961.2;refGene_gn=ZKSCAN5;refGene_tx=XM_023247128.1
E3 6811920 6813044
refGene_gn=ZKSCAN5;refGene_tx=XM_003998465.4;refGene_gn=ZKSCAN5;refGene_tx=XM_006941957.3;refGene_gn=ZKSCAN5;refGene_tx=XM_006941958.3;refGene_gn=ZKSCAN5;refGene_tx=XM_006941959.3;refGene_gn=ZKSCAN5;refGene_tx=XM_006941960.3;refGene_gn=ZKSCAN5;refGene_tx=XM_006941961.2;refGene_gn=ZKSCAN5;refGene_tx=XM_023247128.1
E3 6818921 6820649 refGene_gn=FAM200A;refGene_tx=XM_019820448.1
E3 6829308 6829444
refGene_gn=ZNF655;refGene_tx=XM_019820466.2;refGene_gn=ZNF655;refGene_tx=XM_023247134.1;refGene_gn=ZNF655;refGene_tx=XM_023247135.1;refGene_gn=ZNF655;refGene_tx=XM_023247136.1;refGene_gn=ZNF655;refGene_tx=XM_023247138.1;refGene_gn=ZNF655;refGene_tx=XM_023247139.1
E3 6830615 6830701 refGene_gn=ZNF655;refGene_tx=XM_023247138.1
E3 6831101 6831228 refGene_gn=ZNF655;refGene_tx=XM_023247138.1
E3 6832223 6832420
refGene_gn=ZNF655;refGene_tx=XM_023247138.1;refGene_gn=ZNF655;refGene_tx=XM_023247139.1
E3 6838585 6838690
refGene_gn=ZNF655;refGene_tx=XM_019820466.2;refGene_gn=ZNF655;refGene_tx=XM_023247134.1;refGene_gn=ZNF655;refGene_tx=XM_023247135.1
E3 6839236 6840630
refGene_gn=ZNF655;refGene_tx=XM_019820466.2;refGene_gn=ZNF655;refGene_tx=XM_023247134.1;refGene_gn=ZNF655;refGene_tx=XM_023247135.1;refGene_gn=ZNF655;refGene_tx=XM_023247136.1
E3 6864278 6864290
refGene_gn=TMEM225B;refGene_tx=XM_019820467.2;refGene_gn=TMEM225B;refGene_tx=XM_019820468.2;refGene_gn=TMEM225B;refGene_tx=XM_023247141.1
E3 6865684 6865895
refGene_gn=TMEM225B;refGene_tx=XM_019820467.2;refGene_gn=TMEM225B;refGene_tx=XM_019820468.2;refGene_gn=TMEM225B;refGene_tx=XM_023247141.1
E3 6867330 6867547
refGene_gn=TMEM225B;refGene_tx=XM_019820467.2;refGene_gn=TMEM225B;refGene_tx=XM_019820468.2;refGene_gn=TMEM225B;refGene_tx=XM_023247141.1
E3 6867965 6868108
refGene_gn=TMEM225B;refGene_tx=XM_019820467.2;refGene_gn=TMEM225B;refGene_tx=XM_023247141.1
E3 6868363 6868501
refGene_gn=TMEM225B;refGene_tx=XM_019820467.2;refGene_gn=TMEM225B;refGene_tx=XM_019820468.2;refGene_gn=TMEM225B;refGene_tx=XM_023247141.1
E3 6874132 6874305
refGene_gn=TMEM225B;refGene_tx=XM_019820467.2;refGene_gn=TMEM225B;refGene_tx=XM_019820468.2;refGene_gn=TMEM225B;refGene_tx=XM_023247141.1
E3 6887733 6888124
refGene_gn=ZSCAN25;refGene_tx=XM_006941953.4;refGene_gn=ZSCAN25;refGene_tx=XM_019820465.1;refGene_gn=ZSCAN25;refGene_tx=XM_023247131.1;refGene_gn=ZSCAN25;refGene_tx=XM_023247132.1;refGene_gn=ZSCAN25;refGene_tx=

XM_023247133.1
E3	6889310	6889512
refGene_gn=ZSCAN25;refGene_tx=XM_006941953.4;refGene_gn=ZSCAN25;refGene_tx=XM_019820465.1;refGene_gn=ZSCAN25;refGene_tx=XM_023247131.1;refGene_gn=ZSCAN25;refGene_tx=XM_023247133.1
E3	6890090	6890179
refGene_gn=ZSCAN25;refGene_tx=XM_006941953.4;refGene_gn=ZSCAN25;refGene_tx=XM_019820465.1;refGene_gn=ZSCAN25;refGene_tx=XM_023247131.1;refGene_gn=ZSCAN25;refGene_tx=XM_023247132.1;refGene_gn=ZSCAN25;refGene_tx=XM_023247133.1
E3	6891925	6892052
refGene_gn=ZSCAN25;refGene_tx=XM_006941953.4;refGene_gn=ZSCAN25;refGene_tx=XM_019820465.1;refGene_gn=ZSCAN25;refGene_tx=XM_023247131.1;refGene_gn=ZSCAN25;refGene_tx=XM_023247132.1;refGene_gn=ZSCAN25;refGene_tx=XM_023247133.1
E3	6896335	6897168
refGene_gn=ZSCAN25;refGene_tx=XM_006941953.4;refGene_gn=ZSCAN25;refGene_tx=XM_019820465.1;refGene_gn=ZSCAN25;refGene_tx=XM_023247131.1;refGene_gn=ZSCAN25;refGene_tx=XM_023247132.1
E3	6907756	6907881	refGene_gn=ZSCAN25;refGene_tx=XM_023247133.1
E3	6916938	6917883
refGene_gn=LOC101096691;refGene_tx=XM_003998507.2
E3	6966459	6966530	refGene_gn=CYP3A131;refGene_tx=NM_001246278.2
E3	6968410	6968504	refGene_gn=CYP3A131;refGene_tx=NM_001246278.2
E3	6970425	6970478	refGene_gn=CYP3A131;refGene_tx=NM_001246278.2
E3	6972643	6972713	refGene_gn=CYP3A131;refGene_tx=NM_001246278.2
E3	6972714	6972744	refGene_gn=CYP3A131;refGene_tx=NM_001246278.2
E3	6986801	6986915	refGene_gn=CYP3A131;refGene_tx=NM_001246278.2
E3	6987198	6987287	refGene_gn=CYP3A131;refGene_tx=NM_001246278.2
E3	6988929	6989078	refGene_gn=CYP3A131;refGene_tx=NM_001246278.2
E3	6996108	6996236	refGene_gn=CYP3A131;refGene_tx=NM_001246278.2
E3	6997576	6997643	refGene_gn=CYP3A131;refGene_tx=NM_001246278.2
E3	7001651	7001812	refGene_gn=CYP3A131;refGene_tx=NM_001246278.2
E3	7002879	7003106	refGene_gn=CYP3A131;refGene_tx=NM_001246278.2
E3	7027854	7028017	refGene_gn=CYP3A131;refGene_tx=NM_001246278.2
E3	7029188	7029284	refGene_gn=CYP3A131;refGene_tx=NM_001246278.2
E3	7040885	7040956
refGene_gn=CYP3A132;refGene_tx=NM_001246271.1;refGene_gn=CYP3A132;refGene_tx=XM_019820218.1
E3	7045423	7045517
refGene_gn=CYP3A132;refGene_tx=NM_001246271.1;refGene_gn=CYP3A132;refGene_tx=XM_019820218.1
E3	7047556	7047609
refGene_gn=CYP3A132;refGene_tx=NM_001246271.1;refGene_gn=CYP3A132;refGene_tx=XM_019820218.1
E3	7049856	7049956
refGene_gn=CYP3A132;refGene_tx=NM_001246271.1;refGene_gn=CYP3A132;refGene_tx=XM_019820218.1
E3	7053515	7053629
refGene_gn=CYP3A132;refGene_tx=NM_001246271.1;refGene_gn=CYP3A132;refG

ene_tx=XM_019820218.1
E3 7053886 7053975
refGene_gn=CYP3A132;refGene_tx=NM_001246271.1;refGene_gn=CYP3A132;refG
ene_tx=XM_019820218.1
E3 7055367 7055516
refGene_gn=CYP3A132;refGene_tx=NM_001246271.1;refGene_gn=CYP3A132;refG
ene_tx=XM_019820218.1
E3 7059527 7059655
refGene_gn=CYP3A132;refGene_tx=NM_001246271.1;refGene_gn=CYP3A132;refG
ene_tx=XM_019820218.1
E3 7060692 7060759
refGene_gn=CYP3A132;refGene_tx=NM_001246271.1;refGene_gn=CYP3A132;refG
ene_tx=XM_019820218.1
E3 7063904 7064065
refGene_gn=CYP3A132;refGene_tx=NM_001246271.1;refGene_gn=CYP3A132;refG
ene_tx=XM_019820218.1
E3 7065460 7065687
refGene_gn=CYP3A132;refGene_tx=NM_001246271.1;refGene_gn=CYP3A132;refG
ene_tx=XM_019820218.1
E3 7067898 7068061 refGene_gn=CYP3A132;refGene_tx=NM_001246271.1
E3 7071859 7071955
refGene_gn=CYP3A132;refGene_tx=NM_001246271.1;refGene_gn=CYP3A132;refG
ene_tx=XM_019820218.1
E3 7122750 7122817
refGene_gn=LOC102901745;refGene_tx=XM_019820248.1
E3 7125718 7125879
refGene_gn=LOC102901745;refGene_tx=XM_019820248.1
E3 7127452 7127486
refGene_gn=LOC102901745;refGene_tx=XM_019820248.1
E3 7127488 7127666
refGene_gn=LOC102901745;refGene_tx=XM_019820248.1
E3 7129974 7130137
refGene_gn=LOC102901745;refGene_tx=XM_019820248.1
E3 7134666 7134714
refGene_gn=LOC102901745;refGene_tx=XM_019820248.1
E3 7140500 7141460
refGene_gn=LOC101096939;refGene_tx=XM_003998508.1
E3 7163230 7163817 refGene_gn=TRIM4;refGene_tx=XM_023247140.1
E3 7167163 7167261 refGene_gn=TRIM4;refGene_tx=XM_023247140.1
E3 7167524 7167547 refGene_gn=TRIM4;refGene_tx=XM_023247140.1
E3 7171914 7172145 refGene_gn=TRIM4;refGene_tx=XM_023247140.1
E3 7172813 7172909 refGene_gn=TRIM4;refGene_tx=XM_023247140.1
E3 7177876 7178269 refGene_gn=TRIM4;refGene_tx=XM_023247140.1
E3 7183040 7183048
refGene_gn=GJC3;refGene_tx=XM_006941966.4;refGene_gn=GJC3;refGene_tx=X
M_019820469.1
E3 7183360 7184207
refGene_gn=GJC3;refGene_tx=XM_006941966.4;refGene_gn=GJC3;refGene_tx=X
M_019820469.1
E3 7211043 7211119 refGene_gn=AZGP1;refGene_tx=XM_006941965.3

E3 7212874 7213135 refGene_gn=AZGP1;refGene_tx=XM_006941965.3
E3 7215676 7215952 refGene_gn=AZGP1;refGene_tx=XM_006941965.3
E3 7216557 7216823 refGene_gn=AZGP1;refGene_tx=XM_006941965.3
E3 7226735 7226757
refGene_gn=LOC109491390;refGene_tx=XM_023247073.1
E3 7226813 7227445
refGene_gn=LOC109491390;refGene_tx=XM_023247073.1
E3 7234341 7234353
refGene_gn=LOC109491390;refGene_tx=XM_023247073.1
E3 7264340 7264766 refGene_gn=ZKSCAN1;refGene_tx=XM_023247074.1
E3 7265230 7265384 refGene_gn=ZKSCAN1;refGene_tx=XM_023247074.1
E3 7267346 7267438 refGene_gn=ZKSCAN1;refGene_tx=XM_023247074.1
E3 7267714 7267841 refGene_gn=ZKSCAN1;refGene_tx=XM_023247074.1
E3 7270307 7271197 refGene_gn=ZKSCAN1;refGene_tx=XM_023247074.1
E3 7297976 7298456 refGene_gn=ZKSCAN1;refGene_tx=XM_023247074.1
E3 7299063 7299253 refGene_gn=ZKSCAN1;refGene_tx=XM_023247074.1
E3 7301105 7301929 refGene_gn=ZKSCAN1;refGene_tx=XM_023247074.1
E3 7312166 7312181 refGene_gn=ZKSCAN1;refGene_tx=XM_023247074.1
E3 7319925 7320663 refGene_gn=ZKSCAN1;refGene_tx=XM_023247074.1
E3 7347072 7347130 refGene_gn=COPS6;refGene_tx=XM_003998471.3
E3 7347359 7347485 refGene_gn=COPS6;refGene_tx=XM_003998471.3
E3 7347695 7347827 refGene_gn=COPS6;refGene_tx=XM_003998471.3
E3 7348145 7348234 refGene_gn=COPS6;refGene_tx=XM_003998471.3
E3 7348326 7348389 refGene_gn=COPS6;refGene_tx=XM_003998471.3
E3 7348723 7348771 refGene_gn=COPS6;refGene_tx=XM_003998471.3
E3 7348849 7348964 refGene_gn=COPS6;refGene_tx=XM_003998471.3
E3 7349056 7349149 refGene_gn=COPS6;refGene_tx=XM_003998471.3
E3 7349230 7349331 refGene_gn=COPS6;refGene_tx=XM_003998471.3
E3 7349442 7349583 refGene_gn=COPS6;refGene_tx=XM_003998471.3
E3 7350531 7350733
refGene_gn=MCM7;refGene_tx=XM_006941967.4;refGene_gn=MCM7;refGene_tx=XM_006941968.3;refGene_gn=MCM7;refGene_tx=XM_011290403.2
E3 7350904 7351014
refGene_gn=MCM7;refGene_tx=XM_006941967.4;refGene_gn=MCM7;refGene_tx=XM_006941968.3;refGene_gn=MCM7;refGene_tx=XM_011290403.2
E3 7351771 7351940
refGene_gn=MCM7;refGene_tx=XM_006941967.4;refGene_gn=MCM7;refGene_tx=XM_006941968.3;refGene_gn=MCM7;refGene_tx=XM_011290403.2
E3 7352550 7352634
refGene_gn=MCM7;refGene_tx=XM_006941967.4;refGene_gn=MCM7;refGene_tx=XM_006941968.3;refGene_gn=MCM7;refGene_tx=XM_011290403.2
E3 7352936 7353330
refGene_gn=MCM7;refGene_tx=XM_006941967.4;refGene_gn=MCM7;refGene_tx=XM_006941968.3;refGene_gn=MCM7;refGene_tx=XM_011290403.2
E3 7353592 7353676
refGene_gn=MCM7;refGene_tx=XM_006941967.4;refGene_gn=MCM7;refGene_tx=XM_006941968.3;refGene_gn=MCM7;refGene_tx=XM_011290403.2
E3 7353915 7354047
refGene_gn=MCM7;refGene_tx=XM_006941967.4;refGene_gn=MCM7;refGene_tx=XM_006941968.3;refGene_gn=MCM7;refGene_tx=XM_011290403.2

E3 7354134 7354249
refGene_gn=MCM7;refGene_tx=XM_006941967.4;refGene_gn=MCM7;refGene_tx=XM_006941968.3;refGene_gn=MCM7;refGene_tx=XM_011290403.2
E3 7354437 7354587
refGene_gn=MCM7;refGene_tx=XM_006941967.4;refGene_gn=MCM7;refGene_tx=XM_006941968.3;refGene_gn=MCM7;refGene_tx=XM_011290403.2
E3 7354897 7355035
refGene_gn=MCM7;refGene_tx=XM_006941967.4;refGene_gn=MCM7;refGene_tx=XM_006941968.3;refGene_gn=MCM7;refGene_tx=XM_011290403.2
E3 7355330 7355511
refGene_gn=MCM7;refGene_tx=XM_006941967.4;refGene_gn=MCM7;refGene_tx=XM_006941968.3;refGene_gn=MCM7;refGene_tx=XM_011290403.2
E3 7355586 7355711
refGene_gn=MCM7;refGene_tx=XM_006941967.4;refGene_gn=MCM7;refGene_tx=XM_006941968.3;refGene_gn=MCM7;refGene_tx=XM_011290403.2
E3 7355880 7356045
refGene_gn=MCM7;refGene_tx=XM_006941967.4;refGene_gn=MCM7;refGene_tx=XM_006941968.3;refGene_gn=MCM7;refGene_tx=XM_011290403.2
E3 7356316 7356396 refGene_gn=MCM7;refGene_tx=XM_006941967.4
E3 7357569 7357600 refGene_gn=MCM7;refGene_tx=XM_006941967.4
E3 7358008 7358066
refGene_gn=AP4M1;refGene_tx=XM_003998473.5;refGene_gn=AP4M1;refGene_tx=XM_019820480.2
E3 7358168 7358257
refGene_gn=AP4M1;refGene_tx=XM_003998473.5;refGene_gn=AP4M1;refGene_tx=XM_019820480.2
E3 7358609 7358716
refGene_gn=AP4M1;refGene_tx=XM_003998473.5;refGene_gn=AP4M1;refGene_tx=XM_019820480.2
E3 7358777 7358892
refGene_gn=AP4M1;refGene_tx=XM_003998473.5;refGene_gn=AP4M1;refGene_tx=XM_019820480.2
E3 7359024 7359135
refGene_gn=AP4M1;refGene_tx=XM_003998473.5;refGene_gn=AP4M1;refGene_tx=XM_019820480.2
E3 7359214 7359295
refGene_gn=AP4M1;refGene_tx=XM_003998473.5;refGene_gn=AP4M1;refGene_tx=XM_019820480.2
E3 7359487 7359550
refGene_gn=AP4M1;refGene_tx=XM_003998473.5;refGene_gn=AP4M1;refGene_tx=XM_019820480.2
E3 7359659 7359726
refGene_gn=AP4M1;refGene_tx=XM_003998473.5;refGene_gn=AP4M1;refGene_tx=XM_019820480.2
E3 7359830 7359884
refGene_gn=AP4M1;refGene_tx=XM_003998473.5;refGene_gn=AP4M1;refGene_tx=XM_019820480.2
E3 7360030 7360137
refGene_gn=AP4M1;refGene_tx=XM_003998473.5;refGene_gn=AP4M1;refGene_tx=XM_019820480.2

E3 7360239 7360334
refGene_gn=AP4M1;refGene_tx=XM_003998473.5;refGene_gn=AP4M1;refGene_tx=XM_019820480.2
E3 7360605 7360650
refGene_gn=AP4M1;refGene_tx=XM_003998473.5;refGene_gn=AP4M1;refGene_tx=XM_019820480.2
E3 7360881 7360932
refGene_gn=AP4M1;refGene_tx=XM_003998473.5;refGene_gn=AP4M1;refGene_tx=XM_019820480.2
E3 7361066 7361178
refGene_gn=AP4M1;refGene_tx=XM_003998473.5;refGene_gn=AP4M1;refGene_tx=XM_019820480.2
E3 7361320 7361545
refGene_gn=AP4M1;refGene_tx=XM_003998473.5;refGene_gn=AP4M1;refGene_tx=XM_019820480.2
E3 7361946 7362327
refGene_gn=TAF6;refGene_tx=XM_003998474.4;refGene_gn=TAF6;refGene_tx=XM_006941970.4;refGene_gn=TAF6;refGene_tx=XM_019820471.2
E3 7362597 7362795
refGene_gn=TAF6;refGene_tx=XM_003998474.4;refGene_gn=TAF6;refGene_tx=XM_006941970.4;refGene_gn=TAF6;refGene_tx=XM_019820471.2
E3 7363053 7363227
refGene_gn=TAF6;refGene_tx=XM_003998474.4;refGene_gn=TAF6;refGene_tx=XM_006941970.4;refGene_gn=TAF6;refGene_tx=XM_019820471.2
E3 7363734 7363860
refGene_gn=TAF6;refGene_tx=XM_003998474.4;refGene_gn=TAF6;refGene_tx=XM_006941970.4;refGene_gn=TAF6;refGene_tx=XM_019820471.2
E3 7363986 7364061
refGene_gn=TAF6;refGene_tx=XM_003998474.4;refGene_gn=TAF6;refGene_tx=XM_006941970.4;refGene_gn=TAF6;refGene_tx=XM_019820471.2
E3 7364326 7364509
refGene_gn=TAF6;refGene_tx=XM_003998474.4;refGene_gn=TAF6;refGene_tx=XM_006941970.4;refGene_gn=TAF6;refGene_tx=XM_019820471.2
E3 7364614 7364716
refGene_gn=TAF6;refGene_tx=XM_003998474.4;refGene_gn=TAF6;refGene_tx=XM_006941970.4;refGene_gn=TAF6;refGene_tx=XM_019820471.2
E3 7364826 7364904
refGene_gn=TAF6;refGene_tx=XM_003998474.4;refGene_gn=TAF6;refGene_tx=XM_006941970.4;refGene_gn=TAF6;refGene_tx=XM_019820471.2
E3 7365017 7365163
refGene_gn=TAF6;refGene_tx=XM_003998474.4;refGene_gn=TAF6;refGene_tx=XM_006941970.4;refGene_gn=TAF6;refGene_tx=XM_019820471.2
E3 7365391 7365511
refGene_gn=TAF6;refGene_tx=XM_003998474.4;refGene_gn=TAF6;refGene_tx=XM_006941970.4;refGene_gn=TAF6;refGene_tx=XM_019820471.2
E3 7365620 7365677
refGene_gn=TAF6;refGene_tx=XM_003998474.4;refGene_gn=TAF6;refGene_tx=XM_006941970.4;refGene_gn=TAF6;refGene_tx=XM_019820471.2
E3 7365871 7366025
refGene_gn=TAF6;refGene_tx=XM_003998474.4;refGene_gn=TAF6;refGene_tx=X

M_006941970.4;refGene_gn=TAF6;refGene_tx=XM_019820471.2
E3 7366180 7366267
refGene_gn=TAF6;refGene_tx=XM_003998474.4;refGene_gn=TAF6;refGene_tx=XM_006941970.4;refGene_gn=TAF6;refGene_tx=XM_019820471.2
E3 7366369 7366525
refGene_gn=TAF6;refGene_tx=XM_003998474.4;refGene_gn=TAF6;refGene_tx=XM_006941970.4;refGene_gn=TAF6;refGene_tx=XM_019820471.2
E3 7370831 7370943 refGene_gn=CNPY4;refGene_tx=XM_003998475.3
E3 7372848 7372975 refGene_gn=CNPY4;refGene_tx=XM_003998475.3
E3 7373073 7373170 refGene_gn=CNPY4;refGene_tx=XM_003998475.3
E3 7373404 7373527 refGene_gn=CNPY4;refGene_tx=XM_003998475.3
E3 7374045 7374163 refGene_gn=CNPY4;refGene_tx=XM_003998475.3
E3 7374248 7374412 refGene_gn=CNPY4;refGene_tx=XM_003998475.3
E3 7375742 7376495 refGene_gn=MBLAC1;refGene_tx=XM_003998476.4
E3 7381036 7381041
refGene_gn=LOC111558250;refGene_tx=XM_023246665.1
E3 7382273 7382371
refGene_gn=LOC111558250;refGene_tx=XM_023246664.1
E3 7384942 7385104
refGene_gn=LOC111558250;refGene_tx=XM_023246664.1;refGene_gn=LOC111558250;refGene_tx=XM_023246665.1
E3 7391144 7391534
refGene_gn=LOC111558250;refGene_tx=XM_023246664.1;refGene_gn=LOC111558250;refGene_tx=XM_023246665.1
E3 7391856 7391911
refGene_gn=LOC111558250;refGene_tx=XM_023246664.1;refGene_gn=LOC111558250;refGene_tx=XM_023246665.1
E3 7409566 7410099
refGene_gn=LOC101098422;refGene_tx=XM_019820475.1;refGene_gn=LOC101098422;refGene_tx=XM_019820477.1;refGene_gn=LOC101098422;refGene_tx=XM_023246658.1;refGene_gn=LOC101098422;refGene_tx=XM_023246659.1;refGene_gn=LOC101098422;refGene_tx=XM_023246660.1;refGene_gn=LOC101098422;refGene_tx=XM_023246661.1;refGene_gn=LOC101098422;refGene_tx=XM_023246662.1
E3 7412068 7412281
refGene_gn=LOC101098422;refGene_tx=XM_019820475.1;refGene_gn=LOC101098422;refGene_tx=XM_019820477.1;refGene_gn=LOC101098422;refGene_tx=XM_023246658.1;refGene_gn=LOC101098422;refGene_tx=XM_023246659.1;refGene_gn=LOC101098422;refGene_tx=XM_023246660.1;refGene_gn=LOC101098422;refGene_tx=XM_023246661.1
E3 7412387 7412461
refGene_gn=LOC101098422;refGene_tx=XM_019820475.1;refGene_gn=LOC101098422;refGene_tx=XM_019820477.1;refGene_gn=LOC101098422;refGene_tx=XM_023246658.1;refGene_gn=LOC101098422;refGene_tx=XM_023246659.1;refGene_gn=LOC101098422;refGene_tx=XM_023246660.1;refGene_gn=LOC101098422;refGene_tx=XM_023246661.1;refGene_gn=LOC101098422;refGene_tx=XM_023246662.1
E3 7417801 7418526
refGene_gn=LOC101098422;refGene_tx=XM_019820475.1;refGene_gn=LOC101098422;refGene_tx=XM_019820477.1;refGene_gn=LOC101098422;refGene_tx=XM_023246658.1;refGene_gn=LOC101098422;refGene_tx=XM_023246659.1;refGene_gn=LOC101098422;refGene_tx=XM_023246660.1;refGene_gn=LOC101098422;refGen

e_tx=XM_023246661.1;refGene_gn=LOC101098422;refGene_tx=XM_023246662.1
E3 7432602 7432717
refGene_gn=LOC101098422;refGene_tx=XM_019820475.1;refGene_gn=LOC101098422;refGene_tx=XM_019820477.1;refGene_gn=LOC101098422;refGene_tx=XM_023246658.1;refGene_gn=LOC101098422;refGene_tx=XM_023246659.1;refGene_gn=LOC101098422;refGene_tx=XM_023246660.1;refGene_gn=LOC101098422;refGene_tx=XM_023246661.1;refGene_gn=LOC101098422;refGene_tx=XM_023246662.1
E3 7438984 7438989
refGene_gn=LOC101098422;refGene_tx=XM_019820475.1;refGene_gn=LOC101098422;refGene_tx=XM_023246658.1;refGene_gn=LOC101098422;refGene_tx=XM_023246659.1;refGene_gn=LOC101098422;refGene_tx=XM_023246660.1;refGene_gn=LOC101098422;refGene_tx=XM_023246662.1
E3 7476875 7476878
refGene_gn=LAMTOR4;refGene_tx=XM_003998477.3;refGene_gn=LAMTOR4;refGene_tx=XM_019820481.1;refGene_gn=LAMTOR4;refGene_tx=XM_019820482.1
E3 7477351 7477432
refGene_gn=LAMTOR4;refGene_tx=XM_003998477.3;refGene_gn=LAMTOR4;refGene_tx=XM_019820481.1
E3 7479153 7479271
refGene_gn=LAMTOR4;refGene_tx=XM_003998477.3;refGene_gn=LAMTOR4;refGene_tx=XM_019820481.1;refGene_gn=LAMTOR4;refGene_tx=XM_019820482.1
E3 7479616 7479731
refGene_gn=LAMTOR4;refGene_tx=XM_003998477.3;refGene_gn=LAMTOR4;refGene_tx=XM_019820481.1;refGene_gn=LAMTOR4;refGene_tx=XM_019820482.1
E3 7480869 7481023
refGene_gn=CE3H7orf43;refGene_tx=XM_011290400.3;refGene_gn=CE3H7orf43;refGene_tx=XM_019820490.2
E3 7481156 7481350
refGene_gn=CE3H7orf43;refGene_tx=XM_011290400.3;refGene_gn=CE3H7orf43;refGene_tx=XM_019820490.2
E3 7481578 7481733
refGene_gn=CE3H7orf43;refGene_tx=XM_011290400.3;refGene_gn=CE3H7orf43;refGene_tx=XM_019820490.2
E3 7482258 7482422
refGene_gn=CE3H7orf43;refGene_tx=XM_011290400.3;refGene_gn=CE3H7orf43;refGene_tx=XM_019820490.2
E3 7482510 7482593
refGene_gn=CE3H7orf43;refGene_tx=XM_011290400.3;refGene_gn=CE3H7orf43;refGene_tx=XM_019820490.2
E3 7482723 7482871
refGene_gn=CE3H7orf43;refGene_tx=XM_011290400.3;refGene_gn=CE3H7orf43;refGene_tx=XM_019820490.2
E3 7482974 7483095
refGene_gn=CE3H7orf43;refGene_tx=XM_011290400.3;refGene_gn=CE3H7orf43;refGene_tx=XM_019820490.2
E3 7483258 7483345
refGene_gn=CE3H7orf43;refGene_tx=XM_011290400.3
E3 7483490 7483620
refGene_gn=CE3H7orf43;refGene_tx=XM_011290400.3
E3 7483726 7483822

refGene_gn=CE3H7orf43;refGene_tx=XM_011290400.3
E3 7484022 7484433
refGene_gn=CE3H7orf43;refGene_tx=XM_011290400.3
E3 7485290 7486325 refGene_gn=GAL3ST4;refGene_tx=XM_003998478.3
E3 7488759 7489063 refGene_gn=GAL3ST4;refGene_tx=XM_003998478.3
E3 7489243 7489368 refGene_gn=GAL3ST4;refGene_tx=XM_003998478.3
E3 7492000 7492260 refGene_gn=GPC2;refGene_tx=XM_023247047.1
E3 7493275 7493448 refGene_gn=GPC2;refGene_tx=XM_023247047.1
E3 7493543 7493683 refGene_gn=GPC2;refGene_tx=XM_023247047.1
E3 7493776 7493935 refGene_gn=GPC2;refGene_tx=XM_023247047.1
E3 7494140 7494271 refGene_gn=GPC2;refGene_tx=XM_023247047.1
E3 7494489 7494652 refGene_gn=GPC2;refGene_tx=XM_023247047.1
E3 7495448 7495529 refGene_gn=GPC2;refGene_tx=XM_023247047.1
E3 7496345 7496668 refGene_gn=GPC2;refGene_tx=XM_023247047.1
E3 7496983 7497142 refGene_gn=GPC2;refGene_tx=XM_023247047.1
E3 7497809 7497975 refGene_gn=GPC2;refGene_tx=XM_023247047.1
E3 7499546 7499698
refGene_gn=STAG3;refGene_tx=XM_006941976.3;refGene_gn=STAG3;refGene_tx=XM_019820483.1;refGene_gn=STAG3;refGene_tx=XM_019820484.1;refGene_gn=STAG3;refGene_tx=XM_019820485.1;refGene_gn=STAG3;refGene_tx=XM_019820486.1;refGene_gn=STAG3;refGene_tx=XM_019820487.1;refGene_gn=STAG3;refGene_tx=XM_023247046.1
E3 7501034 7501137
refGene_gn=STAG3;refGene_tx=XM_006941976.3;refGene_gn=STAG3;refGene_tx=XM_019820483.1;refGene_gn=STAG3;refGene_tx=XM_019820484.1;refGene_gn=STAG3;refGene_tx=XM_019820485.1;refGene_gn=STAG3;refGene_tx=XM_019820486.1;refGene_gn=STAG3;refGene_tx=XM_019820487.1;refGene_gn=STAG3;refGene_tx=XM_023247046.1
E3 7501528 7501645
refGene_gn=STAG3;refGene_tx=XM_006941976.3;refGene_gn=STAG3;refGene_tx=XM_019820483.1;refGene_gn=STAG3;refGene_tx=XM_019820484.1;refGene_gn=STAG3;refGene_tx=XM_019820485.1;refGene_gn=STAG3;refGene_tx=XM_019820486.1;refGene_gn=STAG3;refGene_tx=XM_019820487.1;refGene_gn=STAG3;refGene_tx=XM_023247046.1
E3 7505466 7505563
refGene_gn=STAG3;refGene_tx=XM_006941976.3;refGene_gn=STAG3;refGene_tx=XM_019820483.1;refGene_gn=STAG3;refGene_tx=XM_019820484.1;refGene_gn=STAG3;refGene_tx=XM_019820485.1;refGene_gn=STAG3;refGene_tx=XM_019820486.1;refGene_gn=STAG3;refGene_tx=XM_019820487.1;refGene_gn=STAG3;refGene_tx=XM_023247046.1
E3 7506524 7506601
refGene_gn=STAG3;refGene_tx=XM_006941976.3;refGene_gn=STAG3;refGene_tx=XM_019820483.1;refGene_gn=STAG3;refGene_tx=XM_019820484.1;refGene_gn=STAG3;refGene_tx=XM_019820485.1;refGene_gn=STAG3;refGene_tx=XM_019820486.1;refGene_gn=STAG3;refGene_tx=XM_019820487.1;refGene_gn=STAG3;refGene_tx=XM_023247046.1
E3 7506893 7507098
refGene_gn=STAG3;refGene_tx=XM_006941976.3;refGene_gn=STAG3;refGene_tx=XM_019820483.1;refGene_gn=STAG3;refGene_tx=XM_019820484.1;refGene_gn=STAG3;refGene_tx=XM_019820485.1;refGene_gn=STAG3;refGene_tx=XM_0198204

86.1;refGene_gn=STAG3;refGene_tx=XM_019820487.1;refGene_gn=STAG3;refGene_tx=XM_023247046.1
E3 7507610 7507762
refGene_gn=STAG3;refGene_tx=XM_006941976.3;refGene_gn=STAG3;refGene_tx=XM_019820483.1;refGene_gn=STAG3;refGene_tx=XM_019820484.1;refGene_gn=STAG3;refGene_tx=XM_019820485.1;refGene_gn=STAG3;refGene_tx=XM_019820486.1;refGene_gn=STAG3;refGene_tx=XM_019820487.1;refGene_gn=STAG3;refGene_tx=XM_019820488.1;refGene_gn=STAG3;refGene_tx=XM_023247046.1
E3 7511152 7511226
refGene_gn=STAG3;refGene_tx=XM_006941976.3;refGene_gn=STAG3;refGene_tx=XM_019820483.1;refGene_gn=STAG3;refGene_tx=XM_019820484.1;refGene_gn=STAG3;refGene_tx=XM_019820485.1;refGene_gn=STAG3;refGene_tx=XM_019820486.1;refGene_gn=STAG3;refGene_tx=XM_019820487.1;refGene_gn=STAG3;refGene_tx=XM_019820488.1;refGene_gn=STAG3;refGene_tx=XM_023247046.1
E3 7512238 7512362
refGene_gn=STAG3;refGene_tx=XM_006941976.3;refGene_gn=STAG3;refGene_tx=XM_019820483.1;refGene_gn=STAG3;refGene_tx=XM_019820484.1;refGene_gn=STAG3;refGene_tx=XM_019820485.1;refGene_gn=STAG3;refGene_tx=XM_019820486.1;refGene_gn=STAG3;refGene_tx=XM_019820487.1;refGene_gn=STAG3;refGene_tx=XM_019820488.1;refGene_gn=STAG3;refGene_tx=XM_023247046.1
E3 7513078 7513177
refGene_gn=STAG3;refGene_tx=XM_006941976.3;refGene_gn=STAG3;refGene_tx=XM_019820483.1;refGene_gn=STAG3;refGene_tx=XM_019820484.1;refGene_gn=STAG3;refGene_tx=XM_019820485.1;refGene_gn=STAG3;refGene_tx=XM_019820486.1;refGene_gn=STAG3;refGene_tx=XM_019820487.1;refGene_gn=STAG3;refGene_tx=XM_019820488.1;refGene_gn=STAG3;refGene_tx=XM_023247046.1
E3 7513386 7513466
refGene_gn=STAG3;refGene_tx=XM_006941976.3;refGene_gn=STAG3;refGene_tx=XM_019820483.1;refGene_gn=STAG3;refGene_tx=XM_019820484.1;refGene_gn=STAG3;refGene_tx=XM_019820485.1;refGene_gn=STAG3;refGene_tx=XM_019820486.1;refGene_gn=STAG3;refGene_tx=XM_019820487.1;refGene_gn=STAG3;refGene_tx=XM_019820488.1;refGene_gn=STAG3;refGene_tx=XM_023247046.1
E3 7514096 7514204
refGene_gn=STAG3;refGene_tx=XM_006941976.3;refGene_gn=STAG3;refGene_tx=XM_019820483.1;refGene_gn=STAG3;refGene_tx=XM_019820484.1;refGene_gn=STAG3;refGene_tx=XM_019820485.1;refGene_gn=STAG3;refGene_tx=XM_019820486.1;refGene_gn=STAG3;refGene_tx=XM_019820487.1;refGene_gn=STAG3;refGene_tx=XM_019820488.1;refGene_gn=STAG3;refGene_tx=XM_023247046.1
E3 7514470 7514585
refGene_gn=STAG3;refGene_tx=XM_006941976.3;refGene_gn=STAG3;refGene_tx=XM_019820483.1;refGene_gn=STAG3;refGene_tx=XM_019820484.1;refGene_gn=STAG3;refGene_tx=XM_019820485.1;refGene_gn=STAG3;refGene_tx=XM_019820486.1;refGene_gn=STAG3;refGene_tx=XM_019820487.1;refGene_gn=STAG3;refGene_tx=XM_019820488.1;refGene_gn=STAG3;refGene_tx=XM_023247046.1
E3 7514746 7514852
refGene_gn=STAG3;refGene_tx=XM_006941976.3;refGene_gn=STAG3;refGene_tx=XM_019820483.1;refGene_gn=STAG3;refGene_tx=XM_019820484.1;refGene_gn=STAG3;refGene_tx=XM_019820485.1;refGene_gn=STAG3;refGene_tx=XM_019820486.1;refGene_gn=STAG3;refGene_tx=XM_019820487.1;refGene_gn=STAG3;refGene_tx=XM_019820488.1;refGene_gn=STAG3;refGene_tx=XM_023247046.1

E3	7514993	7515097
refGene_gn=STAG3;refGene_tx=XM_006941976.3;refGene_gn=STAG3;refGene_tx
=XM_019820483.1;refGene_gn=STAG3;refGene_tx=XM_019820484.1;refGene_gn=
STAG3;refGene_tx=XM_019820485.1;refGene_gn=STAG3;refGene_tx=XM_0198204
86.1;refGene_gn=STAG3;refGene_tx=XM_019820487.1;refGene_gn=STAG3;refGe
ne_tx=XM_019820488.1;refGene_gn=STAG3;refGene_tx=XM_023247046.1
E3	7515372	7515465
refGene_gn=STAG3;refGene_tx=XM_006941976.3;refGene_gn=STAG3;refGene_tx
=XM_019820483.1;refGene_gn=STAG3;refGene_tx=XM_019820484.1;refGene_gn=
STAG3;refGene_tx=XM_019820485.1;refGene_gn=STAG3;refGene_tx=XM_0198204
87.1;refGene_gn=STAG3;refGene_tx=XM_019820488.1;refGene_gn=STAG3;refGe
ne_tx=XM_023247046.1
E3	7515590	7515680
refGene_gn=STAG3;refGene_tx=XM_006941976.3;refGene_gn=STAG3;refGene_tx
=XM_019820483.1;refGene_gn=STAG3;refGene_tx=XM_019820484.1;refGene_gn=
STAG3;refGene_tx=XM_019820485.1;refGene_gn=STAG3;refGene_tx=XM_0198204
86.1;refGene_gn=STAG3;refGene_tx=XM_019820487.1;refGene_gn=STAG3;refGe
ne_tx=XM_019820488.1;refGene_gn=STAG3;refGene_tx=XM_023247046.1
E3	7515939	7516140
refGene_gn=STAG3;refGene_tx=XM_006941976.3;refGene_gn=STAG3;refGene_tx
=XM_019820483.1;refGene_gn=STAG3;refGene_tx=XM_019820484.1;refGene_gn=
STAG3;refGene_tx=XM_019820485.1;refGene_gn=STAG3;refGene_tx=XM_0198204
86.1;refGene_gn=STAG3;refGene_tx=XM_019820487.1;refGene_gn=STAG3;refGe
ne_tx=XM_019820488.1;refGene_gn=STAG3;refGene_tx=XM_023247046.1
E3	7516260	7516331
refGene_gn=STAG3;refGene_tx=XM_006941976.3;refGene_gn=STAG3;refGene_tx
=XM_019820483.1;refGene_gn=STAG3;refGene_tx=XM_019820484.1;refGene_gn=
STAG3;refGene_tx=XM_019820485.1;refGene_gn=STAG3;refGene_tx=XM_0198204
86.1;refGene_gn=STAG3;refGene_tx=XM_019820487.1;refGene_gn=STAG3;refGe
ne_tx=XM_019820488.1;refGene_gn=STAG3;refGene_tx=XM_023247046.1
E3	7516435	7516523
refGene_gn=STAG3;refGene_tx=XM_006941976.3;refGene_gn=STAG3;refGene_tx
=XM_019820483.1;refGene_gn=STAG3;refGene_tx=XM_019820484.1;refGene_gn=
STAG3;refGene_tx=XM_019820485.1;refGene_gn=STAG3;refGene_tx=XM_0198204
86.1;refGene_gn=STAG3;refGene_tx=XM_019820487.1;refGene_gn=STAG3;refGe
ne_tx=XM_019820488.1;refGene_gn=STAG3;refGene_tx=XM_023247046.1
E3	7517002	7517083
refGene_gn=STAG3;refGene_tx=XM_006941976.3;refGene_gn=STAG3;refGene_tx
=XM_019820483.1;refGene_gn=STAG3;refGene_tx=XM_019820484.1;refGene_gn=
STAG3;refGene_tx=XM_019820485.1;refGene_gn=STAG3;refGene_tx=XM_0198204
86.1;refGene_gn=STAG3;refGene_tx=XM_019820487.1;refGene_gn=STAG3;refGe
ne_tx=XM_019820488.1;refGene_gn=STAG3;refGene_tx=XM_023247046.1
E3	7517165	7517258
refGene_gn=STAG3;refGene_tx=XM_006941976.3;refGene_gn=STAG3;refGene_tx
=XM_019820483.1;refGene_gn=STAG3;refGene_tx=XM_019820484.1;refGene_gn=
STAG3;refGene_tx=XM_019820485.1;refGene_gn=STAG3;refGene_tx=XM_0198204
86.1;refGene_gn=STAG3;refGene_tx=XM_019820487.1;refGene_gn=STAG3;refGe
ne_tx=XM_019820488.1;refGene_gn=STAG3;refGene_tx=XM_023247046.1
E3	7517409	7517578
refGene_gn=STAG3;refGene_tx=XM_006941976.3;refGene_gn=STAG3;refGene_tx

=XM_019820483.1;refGene_gn=STAG3;refGene_tx=XM_019820484.1;refGene_gn=STAG3;refGene_tx=XM_019820485.1;refGene_gn=STAG3;refGene_tx=XM_019820486.1;refGene_gn=STAG3;refGene_tx=XM_019820487.1;refGene_gn=STAG3;refGene_tx=XM_019820488.1;refGene_gn=STAG3;refGene_tx=XM_023247046.1
E3 7517702 7517839
refGene_gn=STAG3;refGene_tx=XM_006941976.3;refGene_gn=STAG3;refGene_tx=XM_019820483.1;refGene_gn=STAG3;refGene_tx=XM_019820484.1;refGene_gn=STAG3;refGene_tx=XM_019820485.1;refGene_gn=STAG3;refGene_tx=XM_019820486.1;refGene_gn=STAG3;refGene_tx=XM_019820487.1;refGene_gn=STAG3;refGene_tx=XM_019820488.1;refGene_gn=STAG3;refGene_tx=XM_023247046.1
E3 7518445 7518547
refGene_gn=STAG3;refGene_tx=XM_006941976.3;refGene_gn=STAG3;refGene_tx=XM_019820483.1;refGene_gn=STAG3;refGene_tx=XM_019820484.1;refGene_gn=STAG3;refGene_tx=XM_019820485.1;refGene_gn=STAG3;refGene_tx=XM_019820486.1;refGene_gn=STAG3;refGene_tx=XM_019820487.1;refGene_gn=STAG3;refGene_tx=XM_019820488.1;refGene_gn=STAG3;refGene_tx=XM_023247046.1
E3 7519062 7519211
refGene_gn=STAG3;refGene_tx=XM_006941976.3;refGene_gn=STAG3;refGene_tx=XM_019820483.1;refGene_gn=STAG3;refGene_tx=XM_019820484.1;refGene_gn=STAG3;refGene_tx=XM_019820485.1;refGene_gn=STAG3;refGene_tx=XM_019820486.1;refGene_gn=STAG3;refGene_tx=XM_019820487.1;refGene_gn=STAG3;refGene_tx=XM_019820488.1;refGene_gn=STAG3;refGene_tx=XM_023247046.1
E3 7519457 7519586
refGene_gn=STAG3;refGene_tx=XM_006941976.3;refGene_gn=STAG3;refGene_tx=XM_019820483.1;refGene_gn=STAG3;refGene_tx=XM_019820484.1;refGene_gn=STAG3;refGene_tx=XM_019820485.1;refGene_gn=STAG3;refGene_tx=XM_019820486.1;refGene_gn=STAG3;refGene_tx=XM_019820487.1;refGene_gn=STAG3;refGene_tx=XM_019820488.1;refGene_gn=STAG3;refGene_tx=XM_023247046.1
E3 7519677 7519835
refGene_gn=STAG3;refGene_tx=XM_006941976.3;refGene_gn=STAG3;refGene_tx=XM_019820483.1;refGene_gn=STAG3;refGene_tx=XM_019820484.1;refGene_gn=STAG3;refGene_tx=XM_019820485.1;refGene_gn=STAG3;refGene_tx=XM_019820486.1;refGene_gn=STAG3;refGene_tx=XM_019820487.1;refGene_gn=STAG3;refGene_tx=XM_019820488.1;refGene_gn=STAG3;refGene_tx=XM_023247046.1
E3 7524284 7524456
refGene_gn=STAG3;refGene_tx=XM_006941976.3;refGene_gn=STAG3;refGene_tx=XM_019820483.1;refGene_gn=STAG3;refGene_tx=XM_019820484.1;refGene_gn=STAG3;refGene_tx=XM_019820485.1;refGene_gn=STAG3;refGene_tx=XM_019820486.1;refGene_gn=STAG3;refGene_tx=XM_019820487.1;refGene_gn=STAG3;refGene_tx=XM_019820488.1;refGene_gn=STAG3;refGene_tx=XM_023247046.1
E3 7524703 7524808
refGene_gn=STAG3;refGene_tx=XM_006941976.3;refGene_gn=STAG3;refGene_tx=XM_019820483.1;refGene_gn=STAG3;refGene_tx=XM_019820484.1;refGene_gn=STAG3;refGene_tx=XM_019820485.1;refGene_gn=STAG3;refGene_tx=XM_019820486.1;refGene_gn=STAG3;refGene_tx=XM_019820487.1;refGene_gn=STAG3;refGene_tx=XM_019820488.1;refGene_gn=STAG3;refGene_tx=XM_023247046.1
E3 7525116 7525189
refGene_gn=STAG3;refGene_tx=XM_006941976.3;refGene_gn=STAG3;refGene_tx=XM_019820483.1;refGene_gn=STAG3;refGene_tx=XM_019820484.1;refGene_gn=STAG3;refGene_tx=XM_019820485.1;refGene_gn=STAG3;refGene_tx=XM_0198204

86.1;refGene_gn=STAG3;refGene_tx=XM_019820487.1;refGene_gn=STAG3;refGene_tx=XM_019820488.1;refGene_gn=STAG3;refGene_tx=XM_023247046.1
E3 7526062 7526224
refGene_gn=STAG3;refGene_tx=XM_006941976.3;refGene_gn=STAG3;refGene_tx=XM_019820483.1;refGene_gn=STAG3;refGene_tx=XM_019820484.1;refGene_gn=STAG3;refGene_tx=XM_019820485.1;refGene_gn=STAG3;refGene_tx=XM_019820486.1;refGene_gn=STAG3;refGene_tx=XM_019820488.1;refGene_gn=STAG3;refGene_tx=XM_023247046.1
E3 7526392 7526401 refGene_gn=STAG3;refGene_tx=XM_006941976.3
E3 7529405 7529429 refGene_gn=STAG3;refGene_tx=XM_019820487.1
E3 7533153 7533214
refGene_gn=PVRIG;refGene_tx=XM_003998518.5;refGene_gn=PVRIG;refGene_tx=XM_019820492.2
E3 7533297 7533651
refGene_gn=PVRIG;refGene_tx=XM_003998518.5;refGene_gn=PVRIG;refGene_tx=XM_019820492.2
E3 7533739 7533863
refGene_gn=PVRIG;refGene_tx=XM_003998518.5;refGene_gn=PVRIG;refGene_tx=XM_019820492.2
E3 7533953 7534011
refGene_gn=PVRIG;refGene_tx=XM_003998518.5;refGene_gn=PVRIG;refGene_tx=XM_019820492.2
E3 7534112 7534442
refGene_gn=PVRIG;refGene_tx=XM_003998518.5;refGene_gn=PVRIG;refGene_tx=XM_019820492.2
E3 7535965 7536215
refGene_gn=PVRIG;refGene_tx=XM_023246506.1;refGene_gn=PVRIG;refGene_tx=XM_023246507.1
E3 7536789 7537688
refGene_gn=PVRIG;refGene_tx=XM_023246506.1;refGene_gn=PVRIG;refGene_tx=XM_023246507.1
E3 7561020 7561075
refGene_gn=PILRA;refGene_tx=XM_011290397.3;refGene_gn=PILRA;refGene_tx=XM_019820493.2
E3 7561397 7561787
refGene_gn=PILRA;refGene_tx=XM_011290397.3;refGene_gn=PILRA;refGene_tx=XM_019820493.2
E3 7572322 7572517
refGene_gn=PILRA;refGene_tx=XM_011290397.3;refGene_gn=PILRA;refGene_tx=XM_019820493.2
E3 7576447 7576478
refGene_gn=PILRA;refGene_tx=XM_011290397.3;refGene_gn=PILRA;refGene_tx=XM_019820493.2
E3 7577004 7577054
refGene_gn=PILRA;refGene_tx=XM_011290397.3;refGene_gn=PILRA;refGene_tx=XM_019820493.2
E3 7577197 7577244 refGene_gn=PILRA;refGene_tx=XM_019820493.2
E3 7577282 7577314 refGene_gn=PILRA;refGene_tx=XM_011290397.3
E3 7577480 7577615 refGene_gn=PILRA;refGene_tx=XM_011290397.3
E3 7578515 7578844

refGene_gn=ZCWPW1;refGene_tx=XM_011290392.2;refGene_gn=ZCWPW1;refGene_
tx=XM_011290395.2;refGene_gn=ZCWPW1;refGene_tx=XM_011290396.2;refGene_
gn=ZCWPW1;refGene_tx=XM_019820494.1;refGene_gn=ZCWPW1;refGene_tx=XM_01
9820495.2;refGene_gn=ZCWPW1;refGene_tx=XM_019820498.1;refGene_gn=ZCWPW
1;refGene_tx=XM_019820499.1;refGene_gn=ZCWPW1;refGene_tx=XM_019820500.
1;refGene_gn=ZCWPW1;refGene_tx=XM_019820501.1;refGene_gn=ZCWPW1;refGen
e_tx=XM_019820502.1
E3 7579372 7579507
refGene_gn=ZCWPW1;refGene_tx=XM_011290392.2;refGene_gn=ZCWPW1;refGene_
tx=XM_011290395.2;refGene_gn=ZCWPW1;refGene_tx=XM_011290396.2;refGene_
gn=ZCWPW1;refGene_tx=XM_019820494.1;refGene_gn=ZCWPW1;refGene_tx=XM_01
9820495.2;refGene_gn=ZCWPW1;refGene_tx=XM_019820498.1;refGene_gn=ZCWPW
1;refGene_tx=XM_019820499.1;refGene_gn=ZCWPW1;refGene_tx=XM_019820500.
1;refGene_gn=ZCWPW1;refGene_tx=XM_019820501.1;refGene_gn=ZCWPW1;refGen
e_tx=XM_019820502.1
E3 7580067 7580128
refGene_gn=ZCWPW1;refGene_tx=XM_011290392.2;refGene_gn=ZCWPW1;refGene_
tx=XM_011290395.2;refGene_gn=ZCWPW1;refGene_tx=XM_011290396.2;refGene_
gn=ZCWPW1;refGene_tx=XM_019820494.1;refGene_gn=ZCWPW1;refGene_tx=XM_01
9820495.2;refGene_gn=ZCWPW1;refGene_tx=XM_019820498.1;refGene_gn=ZCWPW
1;refGene_tx=XM_019820499.1;refGene_gn=ZCWPW1;refGene_tx=XM_019820500.
1;refGene_gn=ZCWPW1;refGene_tx=XM_019820501.1;refGene_gn=ZCWPW1;refGen
e_tx=XM_019820502.1
E3 7587923 7588024
refGene_gn=ZCWPW1;refGene_tx=XM_011290392.2;refGene_gn=ZCWPW1;refGene_
tx=XM_011290395.2;refGene_gn=ZCWPW1;refGene_tx=XM_011290396.2;refGene_
gn=ZCWPW1;refGene_tx=XM_019820494.1;refGene_gn=ZCWPW1;refGene_tx=XM_01
9820495.2;refGene_gn=ZCWPW1;refGene_tx=XM_019820498.1;refGene_gn=ZCWPW
1;refGene_tx=XM_019820499.1;refGene_gn=ZCWPW1;refGene_tx=XM_019820500.
1;refGene_gn=ZCWPW1;refGene_tx=XM_019820501.1;refGene_gn=ZCWPW1;refGen
e_tx=XM_019820502.1
E3 7588461 7588528
refGene_gn=ZCWPW1;refGene_tx=XM_011290392.2;refGene_gn=ZCWPW1;refGene_
tx=XM_011290395.2;refGene_gn=ZCWPW1;refGene_tx=XM_011290396.2;refGene_
gn=ZCWPW1;refGene_tx=XM_019820494.1;refGene_gn=ZCWPW1;refGene_tx=XM_01
9820495.2;refGene_gn=ZCWPW1;refGene_tx=XM_019820498.1;refGene_gn=ZCWPW
1;refGene_tx=XM_019820499.1;refGene_gn=ZCWPW1;refGene_tx=XM_019820500.
1;refGene_gn=ZCWPW1;refGene_tx=XM_019820501.1;refGene_gn=ZCWPW1;refGen
e_tx=XM_019820502.1
E3 7589375 7589459
refGene_gn=ZCWPW1;refGene_tx=XM_011290392.2;refGene_gn=ZCWPW1;refGene_
tx=XM_011290395.2;refGene_gn=ZCWPW1;refGene_tx=XM_011290396.2;refGene_
gn=ZCWPW1;refGene_tx=XM_019820494.1;refGene_gn=ZCWPW1;refGene_tx=XM_01
9820495.2;refGene_gn=ZCWPW1;refGene_tx=XM_019820498.1;refGene_gn=ZCWPW
1;refGene_tx=XM_019820499.1;refGene_gn=ZCWPW1;refGene_tx=XM_019820500.
1;refGene_gn=ZCWPW1;refGene_tx=XM_019820501.1;refGene_gn=ZCWPW1;refGen
e_tx=XM_019820502.1
E3 7590706 7590790
refGene_gn=ZCWPW1;refGene_tx=XM_011290392.2;refGene_gn=ZCWPW1;refGene_
tx=XM_019820494.1;refGene_gn=ZCWPW1;refGene_tx=XM_019820495.2;refGene_

gn=ZCWPW1;refGene_tx=XM_019820498.1;refGene_gn=ZCWPW1;refGene_tx=XM_01
9820499.1;refGene_gn=ZCWPW1;refGene_tx=XM_019820500.1;refGene_gn=ZCWPW
1;refGene_tx=XM_019820501.1
E3 7590872 7590977
refGene_gn=ZCWPW1;refGene_tx=XM_011290392.2;refGene_gn=ZCWPW1;refGene_
tx=XM_011290395.2;refGene_gn=ZCWPW1;refGene_tx=XM_011290396.2;refGene_
gn=ZCWPW1;refGene_tx=XM_019820494.1;refGene_gn=ZCWPW1;refGene_tx=XM_01
9820495.2;refGene_gn=ZCWPW1;refGene_tx=XM_019820498.1;refGene_gn=ZCWPW
1;refGene_tx=XM_019820499.1;refGene_gn=ZCWPW1;refGene_tx=XM_019820500.
1;refGene_gn=ZCWPW1;refGene_tx=XM_019820501.1;refGene_gn=ZCWPW1;refGen
e_tx=XM_019820502.1
E3 7591399 7591475
refGene_gn=ZCWPW1;refGene_tx=XM_011290392.2;refGene_gn=ZCWPW1;refGene_
tx=XM_011290395.2;refGene_gn=ZCWPW1;refGene_tx=XM_011290396.2;refGene_
gn=ZCWPW1;refGene_tx=XM_019820494.1;refGene_gn=ZCWPW1;refGene_tx=XM_01
9820495.2;refGene_gn=ZCWPW1;refGene_tx=XM_019820498.1;refGene_gn=ZCWPW
1;refGene_tx=XM_019820499.1;refGene_gn=ZCWPW1;refGene_tx=XM_019820500.
1;refGene_gn=ZCWPW1;refGene_tx=XM_019820501.1;refGene_gn=ZCWPW1;refGen
e_tx=XM_019820502.1
E3 7592235 7592356
refGene_gn=ZCWPW1;refGene_tx=XM_011290392.2;refGene_gn=ZCWPW1;refGene_
tx=XM_011290395.2;refGene_gn=ZCWPW1;refGene_tx=XM_011290396.2;refGene_
gn=ZCWPW1;refGene_tx=XM_019820494.1;refGene_gn=ZCWPW1;refGene_tx=XM_01
9820495.2;refGene_gn=ZCWPW1;refGene_tx=XM_019820498.1;refGene_gn=ZCWPW
1;refGene_tx=XM_019820499.1;refGene_gn=ZCWPW1;refGene_tx=XM_019820500.
1;refGene_gn=ZCWPW1;refGene_tx=XM_019820501.1;refGene_gn=ZCWPW1;refGen
e_tx=XM_019820502.1
E3 7593098 7593215
refGene_gn=ZCWPW1;refGene_tx=XM_011290392.2;refGene_gn=ZCWPW1;refGene_
tx=XM_011290395.2;refGene_gn=ZCWPW1;refGene_tx=XM_011290396.2;refGene_
gn=ZCWPW1;refGene_tx=XM_019820494.1;refGene_gn=ZCWPW1;refGene_tx=XM_01
9820495.2;refGene_gn=ZCWPW1;refGene_tx=XM_019820498.1;refGene_gn=ZCWPW
1;refGene_tx=XM_019820499.1;refGene_gn=ZCWPW1;refGene_tx=XM_019820500.
1;refGene_gn=ZCWPW1;refGene_tx=XM_019820501.1;refGene_gn=ZCWPW1;refGen
e_tx=XM_019820502.1
E3 7600369 7600492
refGene_gn=ZCWPW1;refGene_tx=XM_011290392.2;refGene_gn=ZCWPW1;refGene_
tx=XM_011290395.2;refGene_gn=ZCWPW1;refGene_tx=XM_011290396.2;refGene_
gn=ZCWPW1;refGene_tx=XM_019820494.1;refGene_gn=ZCWPW1;refGene_tx=XM_01
9820495.2;refGene_gn=ZCWPW1;refGene_tx=XM_019820498.1;refGene_gn=ZCWPW
1;refGene_tx=XM_019820499.1;refGene_gn=ZCWPW1;refGene_tx=XM_019820500.
1;refGene_gn=ZCWPW1;refGene_tx=XM_019820502.1
E3 7600673 7600831
refGene_gn=ZCWPW1;refGene_tx=XM_011290392.2;refGene_gn=ZCWPW1;refGene_
tx=XM_011290395.2;refGene_gn=ZCWPW1;refGene_tx=XM_011290396.2;refGene_
gn=ZCWPW1;refGene_tx=XM_019820494.1;refGene_gn=ZCWPW1;refGene_tx=XM_01
9820495.2;refGene_gn=ZCWPW1;refGene_tx=XM_019820498.1;refGene_gn=ZCWPW
1;refGene_tx=XM_019820499.1;refGene_gn=ZCWPW1;refGene_tx=XM_019820500.
1;refGene_gn=ZCWPW1;refGene_tx=XM_019820501.1;refGene_gn=ZCWPW1;refGen
e_tx=XM_019820502.1

E3 7601339 7601427
refGene_gn=ZCWPW1;refGene_tx=XM_011290392.2;refGene_gn=ZCWPW1;refGene_tx=XM_011290395.2;refGene_gn=ZCWPW1;refGene_tx=XM_011290396.2;refGene_gn=ZCWPW1;refGene_tx=XM_019820494.1;refGene_gn=ZCWPW1;refGene_tx=XM_019820495.2;refGene_gn=ZCWPW1;refGene_tx=XM_019820498.1;refGene_gn=ZCWPW1;refGene_tx=XM_019820499.1;refGene_gn=ZCWPW1;refGene_tx=XM_019820500.1;refGene_gn=ZCWPW1;refGene_tx=XM_019820501.1;refGene_gn=ZCWPW1;refGene_tx=XM_019820502.1
E3 7602277 7602356
refGene_gn=ZCWPW1;refGene_tx=XM_011290392.2;refGene_gn=ZCWPW1;refGene_tx=XM_011290395.2;refGene_gn=ZCWPW1;refGene_tx=XM_011290396.2;refGene_gn=ZCWPW1;refGene_tx=XM_019820494.1;refGene_gn=ZCWPW1;refGene_tx=XM_019820495.2;refGene_gn=ZCWPW1;refGene_tx=XM_019820498.1;refGene_gn=ZCWPW1;refGene_tx=XM_019820499.1;refGene_gn=ZCWPW1;refGene_tx=XM_019820500.1;refGene_gn=ZCWPW1;refGene_tx=XM_019820501.1;refGene_gn=ZCWPW1;refGene_tx=XM_019820502.1
E3 7602775 7602999
refGene_gn=ZCWPW1;refGene_tx=XM_011290392.2;refGene_gn=ZCWPW1;refGene_tx=XM_011290395.2;refGene_gn=ZCWPW1;refGene_tx=XM_011290396.2;refGene_gn=ZCWPW1;refGene_tx=XM_019820494.1;refGene_gn=ZCWPW1;refGene_tx=XM_019820495.2;refGene_gn=ZCWPW1;refGene_tx=XM_019820498.1;refGene_gn=ZCWPW1;refGene_tx=XM_019820499.1;refGene_gn=ZCWPW1;refGene_tx=XM_019820500.1;refGene_gn=ZCWPW1;refGene_tx=XM_019820501.1;refGene_gn=ZCWPW1;refGene_tx=XM_019820502.1
E3 7603738 7603766
refGene_gn=ZCWPW1;refGene_tx=XM_011290392.2;refGene_gn=ZCWPW1;refGene_tx=XM_011290395.2;refGene_gn=ZCWPW1;refGene_tx=XM_011290396.2;refGene_gn=ZCWPW1;refGene_tx=XM_019820494.1;refGene_gn=ZCWPW1;refGene_tx=XM_019820495.2;refGene_gn=ZCWPW1;refGene_tx=XM_019820498.1;refGene_gn=ZCWPW1;refGene_tx=XM_019820499.1;refGene_gn=ZCWPW1;refGene_tx=XM_019820500.1;refGene_gn=ZCWPW1;refGene_tx=XM_019820501.1;refGene_gn=ZCWPW1;refGene_tx=XM_019820502.1
E3 7612006 7613677 refGene_gn=MEPCE;refGene_tx=XM_003998479.5
E3 7615232 7615451 refGene_gn=MEPCE;refGene_tx=XM_003998479.5
E3 7615629 7615756 refGene_gn=MEPCE;refGene_tx=XM_003998479.5
E3 7615879 7615932 refGene_gn=MEPCE;refGene_tx=XM_003998479.5
E3 7617374 7617548 refGene_gn=PPP1R35;refGene_tx=XM_011290390.3
E3 7617634 7617771 refGene_gn=PPP1R35;refGene_tx=XM_011290390.3
E3 7617849 7618102 refGene_gn=PPP1R35;refGene_tx=XM_011290390.3
E3 7618182 7618419 refGene_gn=PPP1R35;refGene_tx=XM_011290390.3
E3 7634983 7635217
refGene_gn=CE3H7orf61;refGene_tx=XM_003998481.5
E3 7638117 7638441
refGene_gn=CE3H7orf61;refGene_tx=XM_003998481.5
E3 7638622 7638676
refGene_gn=CE3H7orf61;refGene_tx=XM_003998481.5
E3 7640413 7640623
refGene_gn=TSC22D4;refGene_tx=XM_003998482.5;refGene_gn=TSC22D4;refGene_tx=XM_019820503.2
E3 7641017 7641066

refGene_gn=TSC22D4;refGene_tx=XM_003998482.5;refGene_gn=TSC22D4;refGene_tx=XM_019820503.2
E3 7646224 7646391
refGene_gn=TSC22D4;refGene_tx=XM_003998482.5;refGene_gn=TSC22D4;refGene_tx=XM_019820503.2
E3 7648492 7649254
refGene_gn=TSC22D4;refGene_tx=XM_003998482.5;refGene_gn=TSC22D4;refGene_tx=XM_019820503.2
E3 7655209 7655277
refGene_gn=NYAP1;refGene_tx=XM_011290386.3;refGene_gn=NYAP1;refGene_tx=XM_011290388.3;refGene_gn=NYAP1;refGene_tx=XM_011290389.3
E3 7656886 7657242
refGene_gn=NYAP1;refGene_tx=XM_011290386.3;refGene_gn=NYAP1;refGene_tx=XM_011290388.3;refGene_gn=NYAP1;refGene_tx=XM_011290389.3
E3 7657717 7659235
refGene_gn=NYAP1;refGene_tx=XM_011290386.3;refGene_gn=NYAP1;refGene_tx=XM_011290388.3;refGene_gn=NYAP1;refGene_tx=XM_011290389.3
E3 7660144 7660363
refGene_gn=NYAP1;refGene_tx=XM_011290386.3;refGene_gn=NYAP1;refGene_tx=XM_011290388.3;refGene_gn=NYAP1;refGene_tx=XM_011290389.3
E3 7660575 7660685
refGene_gn=NYAP1;refGene_tx=XM_011290386.3;refGene_gn=NYAP1;refGene_tx=XM_011290388.3;refGene_gn=NYAP1;refGene_tx=XM_011290389.3
E3 7661262 7661520
refGene_gn=NYAP1;refGene_tx=XM_011290386.3;refGene_gn=NYAP1;refGene_tx=XM_011290388.3;refGene_gn=NYAP1;refGene_tx=XM_011290389.3
E3 7705920 7706141
refGene_gn=AGFG2;refGene_tx=XM_019820505.2;refGene_gn=AGFG2;refGene_tx=XM_023246422.1;refGene_gn=AGFG2;refGene_tx=XM_023246423.1
E3 7714058 7714152
refGene_gn=AGFG2;refGene_tx=XM_019820505.2;refGene_gn=AGFG2;refGene_tx=XM_023246422.1;refGene_gn=AGFG2;refGene_tx=XM_023246423.1
E3 7715021 7715137
refGene_gn=AGFG2;refGene_tx=XM_019820505.2;refGene_gn=AGFG2;refGene_tx=XM_023246422.1;refGene_gn=AGFG2;refGene_tx=XM_023246423.1
E3 7716202 7716356
refGene_gn=AGFG2;refGene_tx=XM_019820505.2;refGene_gn=AGFG2;refGene_tx=XM_023246422.1;refGene_gn=AGFG2;refGene_tx=XM_023246423.1
E3 7716937 7717103
refGene_gn=AGFG2;refGene_tx=XM_019820505.2;refGene_gn=AGFG2;refGene_tx=XM_023246422.1;refGene_gn=AGFG2;refGene_tx=XM_023246423.1
E3 7717783 7717909
refGene_gn=AGFG2;refGene_tx=XM_019820505.2;refGene_gn=AGFG2;refGene_tx=XM_023246422.1;refGene_gn=AGFG2;refGene_tx=XM_023246423.1
E3 7718626 7718659
refGene_gn=AGFG2;refGene_tx=XM_023246422.1;refGene_gn=AGFG2;refGene_tx=XM_023246423.1
E3 7723690 7723811
refGene_gn=AGFG2;refGene_tx=XM_019820505.2;refGene_gn=AGFG2;refGene_tx=XM_023246422.1;refGene_gn=AGFG2;refGene_tx=XM_023246423.1

E3 7723974 7724057
refGene_gn=AGFG2;refGene_tx=XM_019820505.2;refGene_gn=AGFG2;refGene_tx
=XM_023246422.1;refGene_gn=AGFG2;refGene_tx=XM_023246423.1
E3 7724229 7724313
refGene_gn=AGFG2;refGene_tx=XM_019820505.2;refGene_gn=AGFG2;refGene_tx
=XM_023246422.1;refGene_gn=AGFG2;refGene_tx=XM_023246423.1
E3 7725302 7725431
refGene_gn=AGFG2;refGene_tx=XM_019820505.2;refGene_gn=AGFG2;refGene_tx
=XM_023246422.1;refGene_gn=AGFG2;refGene_tx=XM_023246423.1
E3 7725988 7726074
refGene_gn=AGFG2;refGene_tx=XM_019820505.2;refGene_gn=AGFG2;refGene_tx
=XM_023246422.1;refGene_gn=AGFG2;refGene_tx=XM_023246423.1
E3 7726365 7726425
refGene_gn=AGFG2;refGene_tx=XM_019820505.2;refGene_gn=AGFG2;refGene_tx
=XM_023246422.1;refGene_gn=AGFG2;refGene_tx=XM_023246423.1
E3 7731824 7732224
refGene_gn=LOC101100432;refGene_tx=XM_006941985.3
E3 7732435 7733571
refGene_gn=LOC101100432;refGene_tx=XM_006941985.3
E3 7733931 7733940
refGene_gn=LOC101100432;refGene_tx=XM_006941985.3
E3 7734541 7734829
refGene_gn=SAP25;refGene_tx=XM_023246850.1;refGene_gn=SAP25;refGene_tx
=XM_023246851.1;refGene_gn=SAP25;refGene_tx=XM_023246852.1;refGene_gn=
SAP25;refGene_tx=XM_023246853.1;refGene_gn=SAP25;refGene_tx=XM_0232468
54.1;refGene_gn=SAP25;refGene_tx=XM_023246855.1;refGene_gn=SAP25;refGe
ne_tx=XM_023246856.1
E3 7734910 7735008
refGene_gn=SAP25;refGene_tx=XM_023246850.1;refGene_gn=SAP25;refGene_tx
=XM_023246851.1;refGene_gn=SAP25;refGene_tx=XM_023246852.1;refGene_gn=
SAP25;refGene_tx=XM_023246853.1;refGene_gn=SAP25;refGene_tx=XM_0232468
54.1;refGene_gn=SAP25;refGene_tx=XM_023246855.1;refGene_gn=SAP25;refGe
ne_tx=XM_023246856.1
E3 7735127 7735305
refGene_gn=SAP25;refGene_tx=XM_023246850.1;refGene_gn=SAP25;refGene_tx
=XM_023246851.1;refGene_gn=SAP25;refGene_tx=XM_023246852.1;refGene_gn=
SAP25;refGene_tx=XM_023246853.1;refGene_gn=SAP25;refGene_tx=XM_0232468
54.1;refGene_gn=SAP25;refGene_tx=XM_023246855.1;refGene_gn=SAP25;refGe
ne_tx=XM_023246856.1
E3 7735378 7735485
refGene_gn=SAP25;refGene_tx=XM_023246850.1;refGene_gn=SAP25;refGene_tx
=XM_023246851.1;refGene_gn=SAP25;refGene_tx=XM_023246852.1;refGene_gn=
SAP25;refGene_tx=XM_023246853.1;refGene_gn=SAP25;refGene_tx=XM_0232468
54.1;refGene_gn=SAP25;refGene_tx=XM_023246855.1;refGene_gn=SAP25;refGe
ne_tx=XM_023246856.1
E3 7735591 7735692
refGene_gn=SAP25;refGene_tx=XM_023246850.1;refGene_gn=SAP25;refGene_tx
=XM_023246851.1;refGene_gn=SAP25;refGene_tx=XM_023246852.1;refGene_gn=
SAP25;refGene_tx=XM_023246853.1;refGene_gn=SAP25;refGene_tx=XM_0232468
54.1;refGene_gn=SAP25;refGene_tx=XM_023246855.1;refGene_gn=SAP25;refGe

ne_tx=XM_023246856.1
E3 7735867 7736325
refGene_gn=SAP25;refGene_tx=XM_023246850.1;refGene_gn=SAP25;refGene_tx=XM_023246851.1;refGene_gn=SAP25;refGene_tx=XM_023246852.1;refGene_gn=SAP25;refGene_tx=XM_023246853.1;refGene_gn=SAP25;refGene_tx=XM_023246854.1;refGene_gn=SAP25;refGene_tx=XM_023246855.1;refGene_gn=SAP25;refGene_tx=XM_023246856.1
E3 7736691 7736796
refGene_gn=LRCH4;refGene_tx=XM_023246846.1;refGene_gn=LRCH4;refGene_tx=XM_023246847.1
E3 7737616 7737814
refGene_gn=LRCH4;refGene_tx=XM_003998484.5;refGene_gn=LRCH4;refGene_tx=XM_023246844.1;refGene_gn=LRCH4;refGene_tx=XM_023246848.1
E3 7738128 7738206
refGene_gn=LRCH4;refGene_tx=XM_003998484.5;refGene_gn=LRCH4;refGene_tx=XM_023246844.1;refGene_gn=LRCH4;refGene_tx=XM_023246846.1;refGene_gn=LRCH4;refGene_tx=XM_023246847.1;refGene_gn=LRCH4;refGene_tx=XM_023246848.1
E3 7738301 7738439
refGene_gn=LRCH4;refGene_tx=XM_003998484.5;refGene_gn=LRCH4;refGene_tx=XM_023246844.1;refGene_gn=LRCH4;refGene_tx=XM_023246846.1;refGene_gn=LRCH4;refGene_tx=XM_023246847.1;refGene_gn=LRCH4;refGene_tx=XM_023246848.1
E3 7738572 7738658
refGene_gn=LRCH4;refGene_tx=XM_003998484.5;refGene_gn=LRCH4;refGene_tx=XM_023246844.1;refGene_gn=LRCH4;refGene_tx=XM_023246846.1;refGene_gn=LRCH4;refGene_tx=XM_023246847.1;refGene_gn=LRCH4;refGene_tx=XM_023246848.1
E3 7738870 7738954
refGene_gn=LRCH4;refGene_tx=XM_003998484.5;refGene_gn=LRCH4;refGene_tx=XM_023246844.1;refGene_gn=LRCH4;refGene_tx=XM_023246846.1;refGene_gn=LRCH4;refGene_tx=XM_023246847.1;refGene_gn=LRCH4;refGene_tx=XM_023246848.1
E3 7739085 7739198
refGene_gn=LRCH4;refGene_tx=XM_003998484.5;refGene_gn=LRCH4;refGene_tx=XM_023246844.1;refGene_gn=LRCH4;refGene_tx=XM_023246846.1;refGene_gn=LRCH4;refGene_tx=XM_023246847.1;refGene_gn=LRCH4;refGene_tx=XM_023246848.1
E3 7739288 7739402
refGene_gn=LRCH4;refGene_tx=XM_003998484.5;refGene_gn=LRCH4;refGene_tx=XM_023246844.1;refGene_gn=LRCH4;refGene_tx=XM_023246846.1;refGene_gn=LRCH4;refGene_tx=XM_023246847.1;refGene_gn=LRCH4;refGene_tx=XM_023246848.1
E3 7739488 7739611
refGene_gn=LRCH4;refGene_tx=XM_003998484.5;refGene_gn=LRCH4;refGene_tx=XM_023246844.1;refGene_gn=LRCH4;refGene_tx=XM_023246846.1;refGene_gn=LRCH4;refGene_tx=XM_023246847.1;refGene_gn=LRCH4;refGene_tx=XM_023246848.1
E3 7739721 7739783
refGene_gn=LRCH4;refGene_tx=XM_003998484.5;refGene_gn=LRCH4;refGene_tx

=XM_023246844.1;refGene_gn=LRCH4;refGene_tx=XM_023246846.1;refGene_gn=
LRCH4;refGene_tx=XM_023246847.1;refGene_gn=LRCH4;refGene_tx=XM_0232468
48.1
E3 7739915 7739992
refGene_gn=LRCH4;refGene_tx=XM_003998484.5;refGene_gn=LRCH4;refGene_tx
=XM_023246844.1;refGene_gn=LRCH4;refGene_tx=XM_023246846.1;refGene_gn=
LRCH4;refGene_tx=XM_023246847.1;refGene_gn=LRCH4;refGene_tx=XM_0232468
48.1
E3 7740083 7740174
refGene_gn=LRCH4;refGene_tx=XM_003998484.5;refGene_gn=LRCH4;refGene_tx
=XM_023246844.1;refGene_gn=LRCH4;refGene_tx=XM_023246846.1;refGene_gn=
LRCH4;refGene_tx=XM_023246847.1;refGene_gn=LRCH4;refGene_tx=XM_0232468
48.1
E3 7740410 7740510
refGene_gn=LRCH4;refGene_tx=XM_003998484.5;refGene_gn=LRCH4;refGene_tx
=XM_023246844.1;refGene_gn=LRCH4;refGene_tx=XM_023246846.1;refGene_gn=
LRCH4;refGene_tx=XM_023246847.1;refGene_gn=LRCH4;refGene_tx=XM_0232468
48.1
E3 7740652 7740768
refGene_gn=LRCH4;refGene_tx=XM_003998484.5;refGene_gn=LRCH4;refGene_tx
=XM_023246844.1;refGene_gn=LRCH4;refGene_tx=XM_023246846.1;refGene_gn=
LRCH4;refGene_tx=XM_023246847.1;refGene_gn=LRCH4;refGene_tx=XM_0232468
48.1
E3 7740913 7741050
refGene_gn=LRCH4;refGene_tx=XM_003998484.5;refGene_gn=LRCH4;refGene_tx
=XM_023246844.1;refGene_gn=LRCH4;refGene_tx=XM_023246846.1;refGene_gn=
LRCH4;refGene_tx=XM_023246847.1;refGene_gn=LRCH4;refGene_tx=XM_0232468
48.1
E3 7743065 7743171
refGene_gn=LRCH4;refGene_tx=XM_003998484.5;refGene_gn=LRCH4;refGene_tx
=XM_023246844.1;refGene_gn=LRCH4;refGene_tx=XM_023246846.1;refGene_gn=
LRCH4;refGene_tx=XM_023246847.1;refGene_gn=LRCH4;refGene_tx=XM_0232468
48.1
E3 7743553 7743680
refGene_gn=LRCH4;refGene_tx=XM_003998484.5;refGene_gn=LRCH4;refGene_tx
=XM_023246844.1;refGene_gn=LRCH4;refGene_tx=XM_023246846.1;refGene_gn=
LRCH4;refGene_tx=XM_023246847.1;refGene_gn=LRCH4;refGene_tx=XM_0232468
48.1
E3 7743848 7743993
refGene_gn=LRCH4;refGene_tx=XM_003998484.5;refGene_gn=LRCH4;refGene_tx
=XM_023246844.1;refGene_gn=LRCH4;refGene_tx=XM_023246846.1;refGene_gn=
LRCH4;refGene_tx=XM_023246847.1;refGene_gn=LRCH4;refGene_tx=XM_0232468
48.1
E3 7746624 7746631 refGene_gn=LRCH4;refGene_tx=XM_023246848.1
E3 7747536 7747756
refGene_gn=LRCH4;refGene_tx=XM_003998484.5;refGene_gn=LRCH4;refGene_tx
=XM_023246844.1;refGene_gn=LRCH4;refGene_tx=XM_023246846.1;refGene_gn=
LRCH4;refGene_tx=XM_023246847.1
E3 7748281 7748320 refGene_gn=FBXO24;refGene_tx=XM_023246849.1
E3 7750193 7750265

refGene_gn=FBXO24;refGene_tx=XM_006941987.4;refGene_gn=FBXO24;refGene_
tx=XM_006941989.4;refGene_gn=FBXO24;refGene_tx=XM_019820506.2;refGene_
gn=FBXO24;refGene_tx=XM_019820507.2;refGene_gn=FBXO24;refGene_tx=XM_01
9820508.2
E3 7750525 7750624
refGene_gn=FBXO24;refGene_tx=XM_006941987.4;refGene_gn=FBXO24;refGene_
tx=XM_006941989.4;refGene_gn=FBXO24;refGene_tx=XM_019820506.2;refGene_
gn=FBXO24;refGene_tx=XM_019820507.2;refGene_gn=FBXO24;refGene_tx=XM_01
9820508.2;refGene_gn=FBXO24;refGene_tx=XM_023246849.1
E3 7750728 7750912
refGene_gn=FBXO24;refGene_tx=XM_006941987.4;refGene_gn=FBXO24;refGene_
tx=XM_006941989.4;refGene_gn=FBXO24;refGene_tx=XM_019820506.2;refGene_
gn=FBXO24;refGene_tx=XM_019820507.2;refGene_gn=FBXO24;refGene_tx=XM_02
3246849.1
E3 7751941 7752180
refGene_gn=FBXO24;refGene_tx=XM_006941987.4;refGene_gn=FBXO24;refGene_
tx=XM_006941989.4;refGene_gn=FBXO24;refGene_tx=XM_019820506.2;refGene_
gn=FBXO24;refGene_tx=XM_019820507.2;refGene_gn=FBXO24;refGene_tx=XM_01
9820508.2;refGene_gn=FBXO24;refGene_tx=XM_023246849.1
E3 7752422 7752657
refGene_gn=FBXO24;refGene_tx=XM_006941987.4;refGene_gn=FBXO24;refGene_
tx=XM_006941989.4;refGene_gn=FBXO24;refGene_tx=XM_019820506.2;refGene_
gn=FBXO24;refGene_tx=XM_019820507.2;refGene_gn=FBXO24;refGene_tx=XM_01
9820508.2;refGene_gn=FBXO24;refGene_tx=XM_023246849.1
E3 7753701 7753860
refGene_gn=FBXO24;refGene_tx=XM_006941987.4;refGene_gn=FBXO24;refGene_
tx=XM_006941989.4;refGene_gn=FBXO24;refGene_tx=XM_019820506.2;refGene_
gn=FBXO24;refGene_tx=XM_019820507.2;refGene_gn=FBXO24;refGene_tx=XM_01
9820508.2;refGene_gn=FBXO24;refGene_tx=XM_023246849.1
E3 7754121 7754243
refGene_gn=FBXO24;refGene_tx=XM_006941987.4;refGene_gn=FBXO24;refGene_
tx=XM_006941989.4;refGene_gn=FBXO24;refGene_tx=XM_019820506.2;refGene_
gn=FBXO24;refGene_tx=XM_019820507.2;refGene_gn=FBXO24;refGene_tx=XM_01
9820508.2;refGene_gn=FBXO24;refGene_tx=XM_023246849.1
E3 7754507 7754639
refGene_gn=FBXO24;refGene_tx=XM_006941987.4;refGene_gn=FBXO24;refGene_
tx=XM_006941989.4;refGene_gn=FBXO24;refGene_tx=XM_019820506.2;refGene_
gn=FBXO24;refGene_tx=XM_019820507.2;refGene_gn=FBXO24;refGene_tx=XM_01
9820508.2;refGene_gn=FBXO24;refGene_tx=XM_023246849.1
E3 7757417 7757588
refGene_gn=FBXO24;refGene_tx=XM_006941987.4;refGene_gn=FBXO24;refGene_
tx=XM_006941989.4;refGene_gn=FBXO24;refGene_tx=XM_019820506.2;refGene_
gn=FBXO24;refGene_tx=XM_019820508.2;refGene_gn=FBXO24;refGene_tx=XM_02
3246849.1
E3 7757923 7758286
refGene_gn=FBXO24;refGene_tx=XM_006941987.4;refGene_gn=FBXO24;refGene_
tx=XM_006941989.4;refGene_gn=FBXO24;refGene_tx=XM_019820506.2;refGene_
gn=FBXO24;refGene_tx=XM_019820507.2;refGene_gn=FBXO24;refGene_tx=XM_01
9820508.2;refGene_gn=FBXO24;refGene_tx=XM_023246849.1
E3 7759341 7759652

refGene_gn=PCOLCE;refGene_tx=XM_003998487.5;refGene_gn=PCOLCE;refGene_tx=XM_006941986.4
E3	7760474	7760583	refGene_gn=PCOLCE;refGene_tx=XM_003998487.5;refGene_gn=PCOLCE;refGene_tx=XM_006941986.4
E3	7761018	7761277	refGene_gn=PCOLCE;refGene_tx=XM_003998487.5;refGene_gn=PCOLCE;refGene_tx=XM_006941986.4
E3	7761502	7761577	refGene_gn=PCOLCE;refGene_tx=XM_006941986.4
E3	7762179	7762304	refGene_gn=PCOLCE;refGene_tx=XM_003998487.5;refGene_gn=PCOLCE;refGene_tx=XM_006941986.4
E3	7762856	7762993	refGene_gn=PCOLCE;refGene_tx=XM_003998487.5;refGene_gn=PCOLCE;refGene_tx=XM_006941986.4
E3	7763282	7763530	refGene_gn=PCOLCE;refGene_tx=XM_003998487.5;refGene_gn=PCOLCE;refGene_tx=XM_006941986.4
E3	7763780	7763858	refGene_gn=PCOLCE;refGene_tx=XM_003998487.5;refGene_gn=PCOLCE;refGene_tx=XM_006941986.4
E3	7763990	7764161	refGene_gn=PCOLCE;refGene_tx=XM_003998487.5;refGene_gn=PCOLCE;refGene_tx=XM_006941986.4
E3	7764292	7764459	refGene_gn=PCOLCE;refGene_tx=XM_003998487.5;refGene_gn=PCOLCE;refGene_tx=XM_006941986.4
E3	7766923	7767006	refGene_gn=MOSPD3;refGene_tx=XM_019820509.2
E3	7767499	7767764	refGene_gn=MOSPD3;refGene_tx=XM_006941990.3;refGene_gn=MOSPD3;refGene_tx=XM_011290383.3;refGene_gn=MOSPD3;refGene_tx=XM_019820509.2;refGene_gn=MOSPD3;refGene_tx=XM_019820511.2
E3	7767999	7768075	refGene_gn=MOSPD3;refGene_tx=XM_006941990.3;refGene_gn=MOSPD3;refGene_tx=XM_011290383.3;refGene_gn=MOSPD3;refGene_tx=XM_019820509.2;refGene_gn=MOSPD3;refGene_tx=XM_019820511.2
E3	7768304	7768534	refGene_gn=MOSPD3;refGene_tx=XM_006941990.3;refGene_gn=MOSPD3;refGene_tx=XM_011290383.3;refGene_gn=MOSPD3;refGene_tx=XM_019820509.2;refGene_gn=MOSPD3;refGene_tx=XM_019820511.2
E3	7769108	7769273	refGene_gn=MOSPD3;refGene_tx=XM_006941990.3;refGene_gn=MOSPD3;refGene_tx=XM_011290383.3;refGene_gn=MOSPD3;refGene_tx=XM_019820509.2;refGene_gn=MOSPD3;refGene_tx=XM_019820511.2
E3	7769372	7769404	refGene_gn=MOSPD3;refGene_tx=XM_006941990.3;refGene_gn=MOSPD3;refGene_tx=XM_011290383.3;refGene_gn=MOSPD3;refGene_tx=XM_019820509.2;refGene_gn=MOSPD3;refGene_tx=XM_019820511.2
E3	7772868	7773153	refGene_gn=TFR2;refGene_tx=XM_003998490.3;refGene_gn=TFR2;refGene_tx=X

M_019820514.1
E3	7776189	7776330
refGene_gn=TFR2;refGene_tx=XM_003998490.3;refGene_gn=TFR2;refGene_tx=X
M_019820514.1
E3	7776681	7776909
refGene_gn=TFR2;refGene_tx=XM_003998490.3;refGene_gn=TFR2;refGene_tx=X
M_019820514.1
E3	7776997	7777082
refGene_gn=TFR2;refGene_tx=XM_003998490.3;refGene_gn=TFR2;refGene_tx=X
M_019820514.1
E3	7777163	7777240
refGene_gn=TFR2;refGene_tx=XM_003998490.3;refGene_gn=TFR2;refGene_tx=X
M_019820514.1
E3	7777326	7777394
refGene_gn=TFR2;refGene_tx=XM_003998490.3;refGene_gn=TFR2;refGene_tx=X
M_019820514.1
E3	7777492	7777556
refGene_gn=TFR2;refGene_tx=XM_003998490.3;refGene_gn=TFR2;refGene_tx=X
M_019820514.1
E3	7777638	7777721
refGene_gn=TFR2;refGene_tx=XM_003998490.3;refGene_gn=TFR2;refGene_tx=X
M_019820514.1
E3	7778247	7778367
refGene_gn=TFR2;refGene_tx=XM_003998490.3;refGene_gn=TFR2;refGene_tx=X
M_019820514.1
E3	7779372	7779536
refGene_gn=TFR2;refGene_tx=XM_003998490.3;refGene_gn=TFR2;refGene_tx=X
M_019820514.1
E3	7779877	7780017
refGene_gn=TFR2;refGene_tx=XM_003998490.3;refGene_gn=TFR2;refGene_tx=X
M_019820514.1
E3	7780148	7780265
refGene_gn=TFR2;refGene_tx=XM_003998490.3;refGene_gn=TFR2;refGene_tx=X
M_019820514.1
E3	7780642	7780765
refGene_gn=TFR2;refGene_tx=XM_003998490.3;refGene_gn=TFR2;refGene_tx=X
M_019820514.1
E3	7780871	7780983
refGene_gn=TFR2;refGene_tx=XM_003998490.3;refGene_gn=TFR2;refGene_tx=X
M_019820514.1
E3	7781076	7781217
refGene_gn=TFR2;refGene_tx=XM_003998490.3;refGene_gn=TFR2;refGene_tx=X
M_019820514.1
E3	7785041	7785228
refGene_gn=TFR2;refGene_tx=XM_003998490.3;refGene_gn=TFR2;refGene_tx=X
M_019820514.1
E3	7785339	7785589
refGene_gn=TFR2;refGene_tx=XM_003998490.3;refGene_gn=TFR2;refGene_tx=X
M_019820514.1
E3	7785833	7785866

refGene_gn=TFR2;refGene_tx=XM_003998490.3;refGene_gn=TFR2;refGene_tx=XM_019820514.1
E3 7787609 7787690 refGene_gn=ACTL6B;refGene_tx=XM_003998492.5
E3 7790820 7790907 refGene_gn=ACTL6B;refGene_tx=XM_003998492.5
E3 7791124 7791220 refGene_gn=ACTL6B;refGene_tx=XM_003998492.5
E3 7791316 7791397 refGene_gn=ACTL6B;refGene_tx=XM_003998492.5
E3 7791536 7791651 refGene_gn=ACTL6B;refGene_tx=XM_003998492.5
E3 7791763 7791825 refGene_gn=ACTL6B;refGene_tx=XM_003998492.5
E3 7791981 7792071 refGene_gn=ACTL6B;refGene_tx=XM_003998492.5
E3 7792724 7792831 refGene_gn=ACTL6B;refGene_tx=XM_003998492.5
E3 7792897 7792992 refGene_gn=ACTL6B;refGene_tx=XM_003998492.5
E3 7793661 7793759 refGene_gn=ACTL6B;refGene_tx=XM_003998492.5
E3 7798934 7799035 refGene_gn=ACTL6B;refGene_tx=XM_003998492.5
E3 7799314 7799480 refGene_gn=ACTL6B;refGene_tx=XM_003998492.5
E3 7799684 7799761 refGene_gn=ACTL6B;refGene_tx=XM_003998492.5
E3 7800231 7800256 refGene_gn=ACTL6B;refGene_tx=XM_003998492.5
E3 7810178 7810235 refGene_gn=GNB2;refGene_tx=XM_003998493.5
E3 7810435 7810474 refGene_gn=GNB2;refGene_tx=XM_003998493.5
E3 7810592 7810699 refGene_gn=GNB2;refGene_tx=XM_003998493.5
E3 7811029 7811093 refGene_gn=GNB2;refGene_tx=XM_003998493.5
E3 7811176 7811339 refGene_gn=GNB2;refGene_tx=XM_003998493.5
E3 7811433 7811500 refGene_gn=GNB2;refGene_tx=XM_003998493.5
E3 7811782 7811984 refGene_gn=GNB2;refGene_tx=XM_003998493.5
E3 7812074 7812291 refGene_gn=GNB2;refGene_tx=XM_003998493.5
E3 7812358 7812465 refGene_gn=GNB2;refGene_tx=XM_003998493.5
E3 7815555 7815608
refGene_gn=GIGYF1;refGene_tx=XM_019820513.2;refGene_gn=GIGYF1;refGene_tx=XM_023246857.1;refGene_gn=GIGYF1;refGene_tx=XM_023246858.1;refGene_gn=GIGYF1;refGene_tx=XM_023246859.1
E3 7815701 7815831
refGene_gn=GIGYF1;refGene_tx=XM_019820513.2;refGene_gn=GIGYF1;refGene_tx=XM_023246857.1;refGene_gn=GIGYF1;refGene_tx=XM_023246858.1;refGene_gn=GIGYF1;refGene_tx=XM_023246859.1
E3 7815923 7816081
refGene_gn=GIGYF1;refGene_tx=XM_019820513.2;refGene_gn=GIGYF1;refGene_tx=XM_023246857.1;refGene_gn=GIGYF1;refGene_tx=XM_023246858.1;refGene_gn=GIGYF1;refGene_tx=XM_023246859.1
E3 7816175 7816336
refGene_gn=GIGYF1;refGene_tx=XM_019820513.2;refGene_gn=GIGYF1;refGene_tx=XM_023246857.1;refGene_gn=GIGYF1;refGene_tx=XM_023246858.1;refGene_gn=GIGYF1;refGene_tx=XM_023246859.1
E3 7816436 7816624
refGene_gn=GIGYF1;refGene_tx=XM_019820513.2;refGene_gn=GIGYF1;refGene_tx=XM_023246857.1;refGene_gn=GIGYF1;refGene_tx=XM_023246858.1;refGene_gn=GIGYF1;refGene_tx=XM_023246859.1
E3 7816903 7817131
refGene_gn=GIGYF1;refGene_tx=XM_019820513.2;refGene_gn=GIGYF1;refGene_tx=XM_023246857.1;refGene_gn=GIGYF1;refGene_tx=XM_023246858.1;refGene_gn=GIGYF1;refGene_tx=XM_023246859.1
E3 7817265 7817412

refGene_gn=GIGYF1;refGene_tx=XM_019820513.2;refGene_gn=GIGYF1;refGene_tx=XM_023246857.1;refGene_gn=GIGYF1;refGene_tx=XM_023246858.1;refGene_gn=GIGYF1;refGene_tx=XM_023246859.1
E3 7817519 7817602
refGene_gn=GIGYF1;refGene_tx=XM_019820513.2;refGene_gn=GIGYF1;refGene_tx=XM_023246857.1;refGene_gn=GIGYF1;refGene_tx=XM_023246858.1;refGene_gn=GIGYF1;refGene_tx=XM_023246859.1
E3 7817758 7817859
refGene_gn=GIGYF1;refGene_tx=XM_019820513.2;refGene_gn=GIGYF1;refGene_tx=XM_023246857.1;refGene_gn=GIGYF1;refGene_tx=XM_023246858.1;refGene_gn=GIGYF1;refGene_tx=XM_023246859.1
E3 7817942 7818056
refGene_gn=GIGYF1;refGene_tx=XM_019820513.2;refGene_gn=GIGYF1;refGene_tx=XM_023246857.1;refGene_gn=GIGYF1;refGene_tx=XM_023246858.1;refGene_gn=GIGYF1;refGene_tx=XM_023246859.1
E3 7818143 7818244
refGene_gn=GIGYF1;refGene_tx=XM_019820513.2;refGene_gn=GIGYF1;refGene_tx=XM_023246857.1;refGene_gn=GIGYF1;refGene_tx=XM_023246858.1;refGene_gn=GIGYF1;refGene_tx=XM_023246859.1
E3 7818361 7818528
refGene_gn=GIGYF1;refGene_tx=XM_019820513.2;refGene_gn=GIGYF1;refGene_tx=XM_023246857.1;refGene_gn=GIGYF1;refGene_tx=XM_023246858.1;refGene_gn=GIGYF1;refGene_tx=XM_023246859.1
E3 7818663 7818835
refGene_gn=GIGYF1;refGene_tx=XM_019820513.2;refGene_gn=GIGYF1;refGene_tx=XM_023246857.1;refGene_gn=GIGYF1;refGene_tx=XM_023246858.1;refGene_gn=GIGYF1;refGene_tx=XM_023246859.1
E3 7819043 7819141
refGene_gn=GIGYF1;refGene_tx=XM_019820513.2;refGene_gn=GIGYF1;refGene_tx=XM_023246857.1;refGene_gn=GIGYF1;refGene_tx=XM_023246858.1;refGene_gn=GIGYF1;refGene_tx=XM_023246859.1
E3 7819307 7819445
refGene_gn=GIGYF1;refGene_tx=XM_019820513.2;refGene_gn=GIGYF1;refGene_tx=XM_023246857.1;refGene_gn=GIGYF1;refGene_tx=XM_023246858.1;refGene_gn=GIGYF1;refGene_tx=XM_023246859.1
E3 7819930 7820039
refGene_gn=GIGYF1;refGene_tx=XM_019820513.2;refGene_gn=GIGYF1;refGene_tx=XM_023246857.1;refGene_gn=GIGYF1;refGene_tx=XM_023246858.1;refGene_gn=GIGYF1;refGene_tx=XM_023246859.1
E3 7820159 7820413
refGene_gn=GIGYF1;refGene_tx=XM_019820513.2;refGene_gn=GIGYF1;refGene_tx=XM_023246857.1;refGene_gn=GIGYF1;refGene_tx=XM_023246858.1;refGene_gn=GIGYF1;refGene_tx=XM_023246859.1
E3 7820658 7820829
refGene_gn=GIGYF1;refGene_tx=XM_019820513.2;refGene_gn=GIGYF1;refGene_tx=XM_023246857.1;refGene_gn=GIGYF1;refGene_tx=XM_023246858.1;refGene_gn=GIGYF1;refGene_tx=XM_023246859.1
E3 7821052 7821093
refGene_gn=GIGYF1;refGene_tx=XM_019820513.2;refGene_gn=GIGYF1;refGene_tx=XM_023246857.1;refGene_gn=GIGYF1;refGene_tx=XM_023246858.1;refGene_

gn=GIGYF1;refGene_tx=XM_023246859.1
E3 7821458 7821567
refGene_gn=GIGYF1;refGene_tx=XM_019820513.2;refGene_gn=GIGYF1;refGene_tx=XM_023246857.1;refGene_gn=GIGYF1;refGene_tx=XM_023246858.1;refGene_gn=GIGYF1;refGene_tx=XM_023246859.1
E3 7821679 7821791
refGene_gn=GIGYF1;refGene_tx=XM_019820513.2;refGene_gn=GIGYF1;refGene_tx=XM_023246857.1;refGene_gn=GIGYF1;refGene_tx=XM_023246858.1;refGene_gn=GIGYF1;refGene_tx=XM_023246859.1
E3 7821955 7822051
refGene_gn=GIGYF1;refGene_tx=XM_023246857.1;refGene_gn=GIGYF1;refGene_tx=XM_023246858.1;refGene_gn=GIGYF1;refGene_tx=XM_023246859.1
E3 7822159 7822289
refGene_gn=GIGYF1;refGene_tx=XM_023246857.1;refGene_gn=GIGYF1;refGene_tx=XM_023246858.1;refGene_gn=GIGYF1;refGene_tx=XM_023246859.1
E3 7822383 7822418
refGene_gn=GIGYF1;refGene_tx=XM_023246857.1;refGene_gn=GIGYF1;refGene_tx=XM_023246858.1;refGene_gn=GIGYF1;refGene_tx=XM_023246859.1
E3 7838190 7838613
refGene_gn=POP7;refGene_tx=XM_003998496.5;refGene_gn=POP7;refGene_tx=XM_019820515.2;refGene_gn=POP7;refGene_tx=XM_019820516.2
E3 7848429 7848442 refGene_gn=EPO;refGene_tx=NM_001009269.1
E3 7848998 7849141 refGene_gn=EPO;refGene_tx=NM_001009269.1
E3 7849364 7849451 refGene_gn=EPO;refGene_tx=NM_001009269.1
E3 7849701 7849881 refGene_gn=EPO;refGene_tx=NM_001009269.1
E3 7850048 7850204 refGene_gn=EPO;refGene_tx=NM_001009269.1
E3 7858297 7858374 refGene_gn=ZAN;refGene_tx=XM_023246321.1
E3 7859526 7859579 refGene_gn=ZAN;refGene_tx=XM_023246321.1
E3 7860066 7860213 refGene_gn=ZAN;refGene_tx=XM_023246321.1
E3 7860385 7860657 refGene_gn=ZAN;refGene_tx=XM_023246321.1
E3 7860827 7860915 refGene_gn=ZAN;refGene_tx=XM_023246321.1
E3 7861612 7861720 refGene_gn=ZAN;refGene_tx=XM_023246321.1
E3 7861722 7861764 refGene_gn=ZAN;refGene_tx=XM_023246321.1
E3 7864414 7864579 refGene_gn=ZAN;refGene_tx=XM_023246321.1
E3 7865142 7865234 refGene_gn=ZAN;refGene_tx=XM_023246321.1
E3 7865425 7865504 refGene_gn=ZAN;refGene_tx=XM_023246321.1
E3 7865646 7865793 refGene_gn=ZAN;refGene_tx=XM_023246321.1
E3 7866331 7866603 refGene_gn=ZAN;refGene_tx=XM_023246321.1
E3 7866885 7866970 refGene_gn=ZAN;refGene_tx=XM_023246321.1
E3 7867435 7868965 refGene_gn=ZAN;refGene_tx=XM_023246321.1
E3 7869802 7869987 refGene_gn=ZAN;refGene_tx=XM_023246321.1
E3 7872427 7872569 refGene_gn=ZAN;refGene_tx=XM_023246321.1
E3 7872728 7872845 refGene_gn=ZAN;refGene_tx=XM_023246321.1
E3 7873324 7873449 refGene_gn=ZAN;refGene_tx=XM_023246321.1
E3 7873748 7873894 refGene_gn=ZAN;refGene_tx=XM_023246321.1
E3 7875353 7875497 refGene_gn=ZAN;refGene_tx=XM_023246321.1
E3 7875877 7875985 refGene_gn=ZAN;refGene_tx=XM_023246321.1
E3 7876071 7876241 refGene_gn=ZAN;refGene_tx=XM_023246321.1
E3 7876864 7877067 refGene_gn=ZAN;refGene_tx=XM_023246321.1
E3 7877795 7877937 refGene_gn=ZAN;refGene_tx=XM_023246321.1

E3 7878254 7878502 refGene_gn=ZAN;refGene_tx=XM_023246321.1
E3 7878733 7878917 refGene_gn=ZAN;refGene_tx=XM_023246321.1
E3 7879167 7879279 refGene_gn=ZAN;refGene_tx=XM_023246321.1
E3 7879630 7879725 refGene_gn=ZAN;refGene_tx=XM_023246321.1
E3 7880041 7880218 refGene_gn=ZAN;refGene_tx=XM_023246321.1
E3 7881234 7881443 refGene_gn=ZAN;refGene_tx=XM_023246321.1
E3 7881690 7881835 refGene_gn=ZAN;refGene_tx=XM_023246321.1
E3 7882254 7882502 refGene_gn=ZAN;refGene_tx=XM_023246321.1
E3 7882598 7882763 refGene_gn=ZAN;refGene_tx=XM_023246321.1
E3 7883281 7883406 refGene_gn=ZAN;refGene_tx=XM_023246321.1
E3 7883746 7884042 refGene_gn=ZAN;refGene_tx=XM_023246321.1
E3 7889006 7889221 refGene_gn=ZAN;refGene_tx=XM_023246321.1
E3 7889647 7889798 refGene_gn=ZAN;refGene_tx=XM_023246321.1
E3 7890179 7890208 refGene_gn=ZAN;refGene_tx=XM_023246321.1
E3 7890209 7890422 refGene_gn=ZAN;refGene_tx=XM_023246321.1
E3 7890859 7890989 refGene_gn=ZAN;refGene_tx=XM_023246321.1
E3 7891177 7891352 refGene_gn=ZAN;refGene_tx=XM_023246321.1
E3 7891648 7891822 refGene_gn=ZAN;refGene_tx=XM_023246321.1
E3 7892119 7892259 refGene_gn=ZAN;refGene_tx=XM_023246321.1
E3 7892715 7892915 refGene_gn=ZAN;refGene_tx=XM_023246321.1
E3 7893010 7893149 refGene_gn=ZAN;refGene_tx=XM_023246321.1
E3 7893717 7893858 refGene_gn=ZAN;refGene_tx=XM_023246321.1
E3 7895410 7895516 refGene_gn=ZAN;refGene_tx=XM_023246321.1
E3 7895609 7895654 refGene_gn=ZAN;refGene_tx=XM_023246321.1
E3 7895739 7895740 refGene_gn=ZAN;refGene_tx=XM_023246321.1
E3 7898988 7899118
refGene_gn=EPHB4;refGene_tx=XM_003998497.4;refGene_gn=EPHB4;refGene_tx
=XM_019820518.2
E3 7900247 7900403
refGene_gn=EPHB4;refGene_tx=XM_003998497.4;refGene_gn=EPHB4;refGene_tx
=XM_019820518.2
E3 7900606 7900800
refGene_gn=EPHB4;refGene_tx=XM_003998497.4;refGene_gn=EPHB4;refGene_tx
=XM_019820518.2
E3 7901207 7901357
refGene_gn=EPHB4;refGene_tx=XM_003998497.4;refGene_gn=EPHB4;refGene_tx
=XM_019820518.2
E3 7901634 7901850
refGene_gn=EPHB4;refGene_tx=XM_003998497.4;refGene_gn=EPHB4;refGene_tx
=XM_019820518.2
E3 7905232 7905480
refGene_gn=EPHB4;refGene_tx=XM_003998497.4;refGene_gn=EPHB4;refGene_tx
=XM_019820518.2
E3 7905567 7905681
refGene_gn=EPHB4;refGene_tx=XM_003998497.4;refGene_gn=EPHB4;refGene_tx
=XM_019820518.2
E3 7905799 7905864
refGene_gn=EPHB4;refGene_tx=XM_003998497.4;refGene_gn=EPHB4;refGene_tx
=XM_019820518.2
E3 7906075 7906178

refGene_gn=EPHB4;refGene_tx=XM_003998497.4;refGene_gn=EPHB4;refGene_tx=XM_019820518.2
E3 7907959 7908125
refGene_gn=EPHB4;refGene_tx=XM_003998497.4;refGene_gn=EPHB4;refGene_tx=XM_019820518.2
E3 7908514 7908639
refGene_gn=EPHB4;refGene_tx=XM_003998497.4;refGene_gn=EPHB4;refGene_tx=XM_019820518.2
E3 7908945 7909278
refGene_gn=EPHB4;refGene_tx=XM_003998497.4;refGene_gn=EPHB4;refGene_tx=XM_019820518.2
E3 7909463 7909619
refGene_gn=EPHB4;refGene_tx=XM_003998497.4;refGene_gn=EPHB4;refGene_tx=XM_019820518.2
E3 7910588 7910985
refGene_gn=EPHB4;refGene_tx=XM_003998497.4;refGene_gn=EPHB4;refGene_tx=XM_019820518.2
E3 7911689 7911977
refGene_gn=EPHB4;refGene_tx=XM_003998497.4;refGene_gn=EPHB4;refGene_tx=XM_019820518.2
E3 7912162 7912233
refGene_gn=EPHB4;refGene_tx=XM_003998497.4;refGene_gn=EPHB4;refGene_tx=XM_019820518.2
E3 7914972 7915024 refGene_gn=EPHB4;refGene_tx=XM_003998497.4
E3 7928741 7928748 refGene_gn=EPHB4;refGene_tx=XM_019820518.2
E3 7930525 7930706
refGene_gn=SLC12A9;refGene_tx=XM_019820519.1;refGene_gn=SLC12A9;refGene_tx=XM_023246543.1;refGene_gn=SLC12A9;refGene_tx=XM_023246544.1;refGene_gn=SLC12A9;refGene_tx=XM_023246545.1
E3 7930931 7931066
refGene_gn=SLC12A9;refGene_tx=XM_019820519.1;refGene_gn=SLC12A9;refGene_tx=XM_023246543.1;refGene_gn=SLC12A9;refGene_tx=XM_023246544.1
E3 7931515 7931647
refGene_gn=SLC12A9;refGene_tx=XM_019820519.1;refGene_gn=SLC12A9;refGene_tx=XM_023246543.1;refGene_gn=SLC12A9;refGene_tx=XM_023246544.1
E3 7932032 7932341
refGene_gn=SLC12A9;refGene_tx=XM_019820519.1;refGene_gn=SLC12A9;refGene_tx=XM_023246543.1;refGene_gn=SLC12A9;refGene_tx=XM_023246544.1;refGene_gn=SLC12A9;refGene_tx=XM_023246545.1
E3 7932952 7933060
refGene_gn=SLC12A9;refGene_tx=XM_019820519.1;refGene_gn=SLC12A9;refGene_tx=XM_023246543.1;refGene_gn=SLC12A9;refGene_tx=XM_023246544.1;refGene_gn=SLC12A9;refGene_tx=XM_023246545.1
E3 7933192 7933304
refGene_gn=SLC12A9;refGene_tx=XM_019820519.1;refGene_gn=SLC12A9;refGene_tx=XM_023246543.1;refGene_gn=SLC12A9;refGene_tx=XM_023246544.1;refGene_gn=SLC12A9;refGene_tx=XM_023246545.1
E3 7933871 7934029
refGene_gn=SLC12A9;refGene_tx=XM_019820519.1;refGene_gn=SLC12A9;refGene_tx=XM_023246543.1;refGene_gn=SLC12A9;refGene_tx=XM_023246544.1;refGe

ne_gn=SLC12A9;refGene_tx=XM_023246545.1
E3 7934140 7934223
refGene_gn=SLC12A9;refGene_tx=XM_019820519.1;refGene_gn=SLC12A9;refGene_tx=XM_023246543.1;refGene_gn=SLC12A9;refGene_tx=XM_023246544.1;refGene_gn=SLC12A9;refGene_tx=XM_023246545.1
E3 7934400 7934525
refGene_gn=SLC12A9;refGene_tx=XM_019820519.1;refGene_gn=SLC12A9;refGene_tx=XM_023246543.1;refGene_gn=SLC12A9;refGene_tx=XM_023246544.1;refGene_gn=SLC12A9;refGene_tx=XM_023246545.1
E3 7934647 7934840
refGene_gn=SLC12A9;refGene_tx=XM_019820519.1;refGene_gn=SLC12A9;refGene_tx=XM_023246543.1;refGene_gn=SLC12A9;refGene_tx=XM_023246544.1;refGene_gn=SLC12A9;refGene_tx=XM_023246545.1
E3 7935010 7935185
refGene_gn=SLC12A9;refGene_tx=XM_019820519.1;refGene_gn=SLC12A9;refGene_tx=XM_023246543.1;refGene_gn=SLC12A9;refGene_tx=XM_023246544.1;refGene_gn=SLC12A9;refGene_tx=XM_023246545.1
E3 7935401 7935548
refGene_gn=SLC12A9;refGene_tx=XM_019820519.1;refGene_gn=SLC12A9;refGene_tx=XM_023246543.1;refGene_gn=SLC12A9;refGene_tx=XM_023246544.1;refGene_gn=SLC12A9;refGene_tx=XM_023246545.1
E3 7937403 7938293
refGene_gn=SLC12A9;refGene_tx=XM_019820519.1;refGene_gn=SLC12A9;refGene_tx=XM_023246543.1;refGene_gn=SLC12A9;refGene_tx=XM_023246544.1;refGene_gn=SLC12A9;refGene_tx=XM_023246545.1
E3 7939140 7939249
refGene_gn=TRIP6;refGene_tx=XM_011290375.3;refGene_gn=TRIP6;refGene_tx=XM_019820527.2
E3 7939504 7939632
refGene_gn=TRIP6;refGene_tx=XM_011290375.3;refGene_gn=TRIP6;refGene_tx=XM_019820527.2
E3 7939723 7939849
refGene_gn=TRIP6;refGene_tx=XM_011290375.3;refGene_gn=TRIP6;refGene_tx=XM_019820527.2
E3 7940142 7940520
refGene_gn=TRIP6;refGene_tx=XM_011290375.3;refGene_gn=TRIP6;refGene_tx=XM_019820527.2
E3 7940834 7940928
refGene_gn=TRIP6;refGene_tx=XM_011290375.3;refGene_gn=TRIP6;refGene_tx=XM_019820527.2
E3 7941028 7941198
refGene_gn=TRIP6;refGene_tx=XM_011290375.3;refGene_gn=TRIP6;refGene_tx=XM_019820527.2
E3 7941646 7941825
refGene_gn=TRIP6;refGene_tx=XM_011290375.3;refGene_gn=TRIP6;refGene_tx=XM_019820527.2
E3 7942079 7942200 refGene_gn=TRIP6;refGene_tx=XM_011290375.3
E3 7942311 7942540
refGene_gn=TRIP6;refGene_tx=XM_011290375.3;refGene_gn=TRIP6;refGene_tx=XM_019820527.2

E3	7944002	7944124
refGene_gn=SRRT;refGene_tx=XM_011290376.3;refGene_gn=SRRT;refGene_tx=XM_011290377.3;refGene_gn=SRRT;refGene_tx=XM_011290378.3;refGene_gn=SRRT;refGene_tx=XM_019820522.2;refGene_gn=SRRT;refGene_tx=XM_019820523.2;refGene_gn=SRRT;refGene_tx=XM_019820524.2;refGene_gn=SRRT;refGene_tx=XM_019820525.2
E3	7948278	7948407
refGene_gn=SRRT;refGene_tx=XM_011290376.3;refGene_gn=SRRT;refGene_tx=XM_011290377.3;refGene_gn=SRRT;refGene_tx=XM_011290378.3;refGene_gn=SRRT;refGene_tx=XM_019820522.2;refGene_gn=SRRT;refGene_tx=XM_019820523.2;refGene_gn=SRRT;refGene_tx=XM_019820524.2;refGene_gn=SRRT;refGene_tx=XM_019820525.2
E3	7948673	7948820
refGene_gn=SRRT;refGene_tx=XM_011290376.3;refGene_gn=SRRT;refGene_tx=XM_011290377.3;refGene_gn=SRRT;refGene_tx=XM_011290378.3;refGene_gn=SRRT;refGene_tx=XM_019820522.2;refGene_gn=SRRT;refGene_tx=XM_019820523.2;refGene_gn=SRRT;refGene_tx=XM_019820524.2;refGene_gn=SRRT;refGene_tx=XM_019820525.2
E3	7949054	7949267
refGene_gn=SRRT;refGene_tx=XM_011290376.3;refGene_gn=SRRT;refGene_tx=XM_011290377.3;refGene_gn=SRRT;refGene_tx=XM_011290378.3;refGene_gn=SRRT;refGene_tx=XM_019820522.2;refGene_gn=SRRT;refGene_tx=XM_019820523.2;refGene_gn=SRRT;refGene_tx=XM_019820524.2;refGene_gn=SRRT;refGene_tx=XM_019820525.2
E3	7951104	7951274
refGene_gn=SRRT;refGene_tx=XM_011290376.3;refGene_gn=SRRT;refGene_tx=XM_011290377.3;refGene_gn=SRRT;refGene_tx=XM_011290378.3;refGene_gn=SRRT;refGene_tx=XM_019820522.2;refGene_gn=SRRT;refGene_tx=XM_019820523.2;refGene_gn=SRRT;refGene_tx=XM_019820524.2;refGene_gn=SRRT;refGene_tx=XM_019820525.2
E3	7951404	7951589
refGene_gn=SRRT;refGene_tx=XM_011290376.3;refGene_gn=SRRT;refGene_tx=XM_011290377.3;refGene_gn=SRRT;refGene_tx=XM_011290378.3;refGene_gn=SRRT;refGene_tx=XM_019820522.2;refGene_gn=SRRT;refGene_tx=XM_019820523.2;refGene_gn=SRRT;refGene_tx=XM_019820524.2;refGene_gn=SRRT;refGene_tx=XM_019820525.2
E3	7951802	7951901
refGene_gn=SRRT;refGene_tx=XM_011290376.3;refGene_gn=SRRT;refGene_tx=XM_011290377.3;refGene_gn=SRRT;refGene_tx=XM_011290378.3;refGene_gn=SRRT;refGene_tx=XM_019820522.2;refGene_gn=SRRT;refGene_tx=XM_019820523.2;refGene_gn=SRRT;refGene_tx=XM_019820524.2;refGene_gn=SRRT;refGene_tx=XM_019820525.2
E3	7952411	7952529
refGene_gn=SRRT;refGene_tx=XM_011290376.3;refGene_gn=SRRT;refGene_tx=XM_011290377.3;refGene_gn=SRRT;refGene_tx=XM_011290378.3;refGene_gn=SRRT;refGene_tx=XM_019820522.2;refGene_gn=SRRT;refGene_tx=XM_019820523.2;refGene_gn=SRRT;refGene_tx=XM_019820524.2;refGene_gn=SRRT;refGene_tx=XM_019820525.2
E3	7952719	7952877
refGene_gn=SRRT;refGene_tx=XM_011290376.3;refGene_gn=SRRT;refGene_tx=X

M_011290377.3;refGene_gn=SRRT;refGene_tx=XM_011290378.3;refGene_gn=SRRT;refGene_tx=XM_019820522.2;refGene_gn=SRRT;refGene_tx=XM_019820523.2;refGene_gn=SRRT;refGene_tx=XM_019820524.2;refGene_gn=SRRT;refGene_tx=XM_019820525.2
E3 7953184 7953246
refGene_gn=SRRT;refGene_tx=XM_011290376.3;refGene_gn=SRRT;refGene_tx=XM_011290377.3;refGene_gn=SRRT;refGene_tx=XM_011290378.3;refGene_gn=SRRT;refGene_tx=XM_019820522.2;refGene_gn=SRRT;refGene_tx=XM_019820523.2;refGene_gn=SRRT;refGene_tx=XM_019820524.2;refGene_gn=SRRT;refGene_tx=XM_019820525.2
E3 7953360 7953439
refGene_gn=SRRT;refGene_tx=XM_011290376.3;refGene_gn=SRRT;refGene_tx=XM_011290377.3;refGene_gn=SRRT;refGene_tx=XM_011290378.3;refGene_gn=SRRT;refGene_tx=XM_019820522.2;refGene_gn=SRRT;refGene_tx=XM_019820523.2;refGene_gn=SRRT;refGene_tx=XM_019820524.2;refGene_gn=SRRT;refGene_tx=XM_019820525.2
E3 7953955 7954144
refGene_gn=SRRT;refGene_tx=XM_011290376.3;refGene_gn=SRRT;refGene_tx=XM_011290377.3;refGene_gn=SRRT;refGene_tx=XM_011290378.3;refGene_gn=SRRT;refGene_tx=XM_019820522.2;refGene_gn=SRRT;refGene_tx=XM_019820523.2;refGene_gn=SRRT;refGene_tx=XM_019820524.2;refGene_gn=SRRT;refGene_tx=XM_019820525.2
E3 7954447 7954621
refGene_gn=SRRT;refGene_tx=XM_011290376.3;refGene_gn=SRRT;refGene_tx=XM_011290377.3;refGene_gn=SRRT;refGene_tx=XM_011290378.3;refGene_gn=SRRT;refGene_tx=XM_019820522.2;refGene_gn=SRRT;refGene_tx=XM_019820523.2;refGene_gn=SRRT;refGene_tx=XM_019820524.2;refGene_gn=SRRT;refGene_tx=XM_019820525.2
E3 7954729 7954883
refGene_gn=SRRT;refGene_tx=XM_011290376.3;refGene_gn=SRRT;refGene_tx=XM_011290377.3;refGene_gn=SRRT;refGene_tx=XM_011290378.3;refGene_gn=SRRT;refGene_tx=XM_019820522.2;refGene_gn=SRRT;refGene_tx=XM_019820523.2;refGene_gn=SRRT;refGene_tx=XM_019820524.2;refGene_gn=SRRT;refGene_tx=XM_019820525.2
E3 7955154 7955348
refGene_gn=SRRT;refGene_tx=XM_011290376.3;refGene_gn=SRRT;refGene_tx=XM_011290377.3;refGene_gn=SRRT;refGene_tx=XM_011290378.3;refGene_gn=SRRT;refGene_tx=XM_019820522.2;refGene_gn=SRRT;refGene_tx=XM_019820523.2;refGene_gn=SRRT;refGene_tx=XM_019820524.2;refGene_gn=SRRT;refGene_tx=XM_019820525.2
E3 7955523 7955680
refGene_gn=SRRT;refGene_tx=XM_011290376.3;refGene_gn=SRRT;refGene_tx=XM_011290377.3;refGene_gn=SRRT;refGene_tx=XM_011290378.3;refGene_gn=SRRT;refGene_tx=XM_019820522.2;refGene_gn=SRRT;refGene_tx=XM_019820523.2;refGene_gn=SRRT;refGene_tx=XM_019820524.2;refGene_gn=SRRT;refGene_tx=XM_019820525.2
E3 7955870 7955972
refGene_gn=SRRT;refGene_tx=XM_011290376.3;refGene_gn=SRRT;refGene_tx=XM_011290377.3;refGene_gn=SRRT;refGene_tx=XM_011290378.3;refGene_gn=SRRT;refGene_tx=XM_019820522.2;refGene_gn=SRRT;refGene_tx=XM_019820523.2;

refGene_gn=SRRT;refGene_tx=XM_019820524.2;refGene_gn=SRRT;refGene_tx=XM_019820525.2
E3	7956091	7956218
refGene_gn=SRRT;refGene_tx=XM_011290376.3;refGene_gn=SRRT;refGene_tx=XM_011290377.3;refGene_gn=SRRT;refGene_tx=XM_011290378.3;refGene_gn=SRRT;refGene_tx=XM_019820522.2;refGene_gn=SRRT;refGene_tx=XM_019820523.2;refGene_gn=SRRT;refGene_tx=XM_019820524.2;refGene_gn=SRRT;refGene_tx=XM_019820525.2
E3	7956324	7956400
refGene_gn=SRRT;refGene_tx=XM_011290376.3;refGene_gn=SRRT;refGene_tx=XM_011290377.3;refGene_gn=SRRT;refGene_tx=XM_011290378.3;refGene_gn=SRRT;refGene_tx=XM_019820522.2;refGene_gn=SRRT;refGene_tx=XM_019820523.2;refGene_gn=SRRT;refGene_tx=XM_019820524.2;refGene_gn=SRRT;refGene_tx=XM_019820525.2
E3	7956676	7957105	refGene_gn=UFSP1;refGene_tx=XM_011290374.3
E3	7957977	7958099
refGene_gn=ACHE;refGene_tx=NM_001009203.1;refGene_gn=ACHE;refGene_tx=XM_023246200.1
E3	7958705	7958833
refGene_gn=ACHE;refGene_tx=XM_011290322.3;refGene_gn=ACHE;refGene_tx=XM_019820219.2;refGene_gn=ACHE;refGene_tx=XM_019820220.2;refGene_gn=ACHE;refGene_tx=XM_019820221.2
E3	7958946	7959116
refGene_gn=ACHE;refGene_tx=NM_001009203.1;refGene_gn=ACHE;refGene_tx=XM_011290322.3;refGene_gn=ACHE;refGene_tx=XM_019820219.2;refGene_gn=ACHE;refGene_tx=XM_019820220.2;refGene_gn=ACHE;refGene_tx=XM_019820221.2;refGene_gn=ACHE;refGene_tx=XM_023246200.1
E3	7959933	7960418
refGene_gn=ACHE;refGene_tx=NM_001009203.1;refGene_gn=ACHE;refGene_tx=XM_011290322.3;refGene_gn=ACHE;refGene_tx=XM_019820219.2;refGene_gn=ACHE;refGene_tx=XM_019820220.2;refGene_gn=ACHE;refGene_tx=XM_019820221.2;refGene_gn=ACHE;refGene_tx=XM_023246200.1
E3	7960784	7961864
refGene_gn=ACHE;refGene_tx=NM_001009203.1;refGene_gn=ACHE;refGene_tx=XM_011290322.3;refGene_gn=ACHE;refGene_tx=XM_019820219.2;refGene_gn=ACHE;refGene_tx=XM_019820220.2;refGene_gn=ACHE;refGene_tx=XM_019820221.2;refGene_gn=ACHE;refGene_tx=XM_023246200.1
E3	7963917	7963950
refGene_gn=ACHE;refGene_tx=XM_019820219.2;refGene_gn=ACHE;refGene_tx=XM_023246200.1
E3	7980670	7981753
refGene_gn=LOC109495138;refGene_tx=XM_019820528.2
E3	7981784	7982210	refGene_gn=MUC3A;refGene_tx=XM_023247003.1
E3	7982211	7982822	refGene_gn=MUC3A;refGene_tx=XM_023247003.1
E3	7982928	7983086	refGene_gn=MUC3A;refGene_tx=XM_023247003.1
E3	7983088	7983124	refGene_gn=MUC3A;refGene_tx=XM_023247003.1
E3	7984340	7984455	refGene_gn=MUC3A;refGene_tx=XM_023247003.1
E3	7984574	7984634	refGene_gn=MUC3A;refGene_tx=XM_023247003.1
E3	7985257	7985382	refGene_gn=MUC3A;refGene_tx=XM_023247003.1
E3	7985995	7986061	refGene_gn=MUC3A;refGene_tx=XM_023247003.1

E3 7986409 7986572 refGene_gn=MUC3A;refGene_tx=XM_023247003.1
E3 7986755 7987203 refGene_gn=MUC3A;refGene_tx=XM_023247003.1
E3 8000254 8010529 refGene_gn=MUC17;refGene_tx=XM_023247001.1
E3 8033824 8035381
refGene_gn=LOC111558287;refGene_tx=XM_023247075.1
E3 8035454 8035499
refGene_gn=LOC111558287;refGene_tx=XM_023247075.1
E3 8036518 8038516
refGene_gn=LOC111558288;refGene_tx=XM_023247076.1
E3 8087596 8089909
refGene_gn=TRIM56;refGene_tx=XM_019820526.2;refGene_gn=TRIM56;refGene_
tx=XM_023247002.1
E3 8113270 8113541 refGene_gn=SERPINE1;refGene_tx=XM_003998526.5
E3 8114148 8114382 refGene_gn=SERPINE1;refGene_tx=XM_003998526.5
E3 8114889 8115084 refGene_gn=SERPINE1;refGene_tx=XM_003998526.5
E3 8115870 8116069 refGene_gn=SERPINE1;refGene_tx=XM_003998526.5
E3 8117068 8117169 refGene_gn=SERPINE1;refGene_tx=XM_003998526.5
E3 8117288 8117375 refGene_gn=SERPINE1;refGene_tx=XM_003998526.5
E3 8117933 8118017 refGene_gn=SERPINE1;refGene_tx=XM_003998526.5
E3 8118329 8118367 refGene_gn=SERPINE1;refGene_tx=XM_003998526.5
E3 8126805 8126808 refGene_gn=AP1S1;refGene_tx=XM_003998527.3
E3 8128640 8128819 refGene_gn=AP1S1;refGene_tx=XM_003998527.3
E3 8129452 8129561 refGene_gn=AP1S1;refGene_tx=XM_003998527.3
E3 8130720 8130858 refGene_gn=AP1S1;refGene_tx=XM_003998527.3
E3 8132152 8132200 refGene_gn=AP1S1;refGene_tx=XM_003998527.3
E3 8134537 8136388
refGene_gn=VGF;refGene_tx=XM_019820255.2;refGene_gn=VGF;refGene_tx=XM_
023247005.1;refGene_gn=VGF;refGene_tx=XM_023247006.1
E3 8144022 8144407 refGene_gn=NAT16;refGene_tx=XM_019820256.2
E3 8144408 8144524 refGene_gn=NAT16;refGene_tx=XM_019820256.2
E3 8144526 8144575 refGene_gn=NAT16;refGene_tx=XM_019820256.2
E3 8145195 8145809 refGene_gn=NAT16;refGene_tx=XM_019820256.2
E3 8146163 8146355 refGene_gn=NAT16;refGene_tx=XM_019820256.2
E3 8146429 8146498 refGene_gn=NAT16;refGene_tx=XM_019820256.2
E3 8162791 8162946 refGene_gn=MOGAT3;refGene_tx=XM_006942010.4
E3 8163041 8163244 refGene_gn=MOGAT3;refGene_tx=XM_006942010.4
E3 8164581 8164756 refGene_gn=MOGAT3;refGene_tx=XM_006942010.4
E3 8165067 8165272 refGene_gn=MOGAT3;refGene_tx=XM_006942010.4
E3 8165556 8165627 refGene_gn=MOGAT3;refGene_tx=XM_006942010.4
E3 8165712 8165820 refGene_gn=MOGAT3;refGene_tx=XM_006942010.4
E3 8166023 8166096 refGene_gn=MOGAT3;refGene_tx=XM_006942010.4
E3 8170114 8170500
refGene_gn=PLOD3;refGene_tx=XM_023246290.1;refGene_gn=PLOD3;refGene_tx
=XM_023246291.1
E3 8170778 8170908
refGene_gn=PLOD3;refGene_tx=XM_023246290.1;refGene_gn=PLOD3;refGene_tx
=XM_023246291.1
E3 8171174 8171321
refGene_gn=PLOD3;refGene_tx=XM_023246290.1;refGene_gn=PLOD3;refGene_tx
=XM_023246291.1

E3	8171861	8171966
refGene_gn=PLOD3;refGene_tx=XM_023246290.1;refGene_gn=PLOD3;refGene_tx
=XM_023246291.1
E3	8172440	8172509
refGene_gn=PLOD3;refGene_tx=XM_023246290.1;refGene_gn=PLOD3;refGene_tx
=XM_023246291.1
E3	8172725	8172839
refGene_gn=PLOD3;refGene_tx=XM_023246290.1;refGene_gn=PLOD3;refGene_tx
=XM_023246291.1
E3	8172924	8173066
refGene_gn=PLOD3;refGene_tx=XM_023246290.1;refGene_gn=PLOD3;refGene_tx
=XM_023246291.1
E3	8173296	8173422
refGene_gn=PLOD3;refGene_tx=XM_023246290.1;refGene_gn=PLOD3;refGene_tx
=XM_023246291.1
E3	8173518	8173623
refGene_gn=PLOD3;refGene_tx=XM_023246290.1;refGene_gn=PLOD3;refGene_tx
=XM_023246291.1
E3	8173851	8173973
refGene_gn=PLOD3;refGene_tx=XM_023246290.1;refGene_gn=PLOD3;refGene_tx
=XM_023246291.1
E3	8174138	8174264
refGene_gn=PLOD3;refGene_tx=XM_023246290.1;refGene_gn=PLOD3;refGene_tx
=XM_023246291.1
E3	8174427	8174529
refGene_gn=PLOD3;refGene_tx=XM_023246290.1;refGene_gn=PLOD3;refGene_tx
=XM_023246291.1
E3	8174761	8174859
refGene_gn=PLOD3;refGene_tx=XM_023246290.1;refGene_gn=PLOD3;refGene_tx
=XM_023246291.1
E3	8175264	8175328
refGene_gn=PLOD3;refGene_tx=XM_023246290.1;refGene_gn=PLOD3;refGene_tx
=XM_023246291.1
E3	8175451	8175564
refGene_gn=PLOD3;refGene_tx=XM_023246290.1;refGene_gn=PLOD3;refGene_tx
=XM_023246291.1
E3	8175707	8175871
refGene_gn=PLOD3;refGene_tx=XM_023246290.1;refGene_gn=PLOD3;refGene_tx
=XM_023246291.1
E3	8175979	8176116
refGene_gn=PLOD3;refGene_tx=XM_023246290.1;refGene_gn=PLOD3;refGene_tx
=XM_023246291.1
E3	8176280	8176372
refGene_gn=PLOD3;refGene_tx=XM_023246290.1;refGene_gn=PLOD3;refGene_tx
=XM_023246291.1
E3	8176798	8176913
refGene_gn=PLOD3;refGene_tx=XM_023246290.1;refGene_gn=PLOD3;refGene_tx
=XM_023246291.1
E3	8177596	8177654	refGene_gn=ZNHIT1;refGene_tx=XM_006942009.3
E3	8177925	8177947	refGene_gn=ZNHIT1;refGene_tx=XM_003998528.3

E3 8181367 8181538
refGene_gn=ZNHIT1;refGene_tx=XM_003998528.3;refGene_gn=ZNHIT1;refGene_tx=XM_006942009.3
E3 8182605 8182685
refGene_gn=ZNHIT1;refGene_tx=XM_003998528.3;refGene_gn=ZNHIT1;refGene_tx=XM_006942009.3
E3 8182819 8182989
refGene_gn=ZNHIT1;refGene_tx=XM_003998528.3;refGene_gn=ZNHIT1;refGene_tx=XM_006942009.3
E3 8183084 8183106
refGene_gn=ZNHIT1;refGene_tx=XM_003998528.3;refGene_gn=ZNHIT1;refGene_tx=XM_006942009.3
E3 8187501 8187622 refGene_gn=CLDN15;refGene_tx=XM_003998529.4
E3 8187712 8187829 refGene_gn=CLDN15;refGene_tx=XM_003998529.4
E3 8187896 8187978 refGene_gn=CLDN15;refGene_tx=XM_003998529.4
E3 8188920 8189085 refGene_gn=CLDN15;refGene_tx=XM_003998529.4
E3 8191694 8191911 refGene_gn=CLDN15;refGene_tx=XM_003998529.4
E3 8194193 8194291 refGene_gn=FIS1;refGene_tx=XM_003998530.4
E3 8194875 8194981 refGene_gn=FIS1;refGene_tx=XM_003998530.4
E3 8195416 8195493 refGene_gn=FIS1;refGene_tx=XM_003998530.4
E3 8196803 8196936 refGene_gn=FIS1;refGene_tx=XM_003998530.4
E3 8197821 8197866 refGene_gn=FIS1;refGene_tx=XM_003998530.4
E3 8246641 8246790
refGene_gn=IFT22;refGene_tx=XM_023246892.1;refGene_gn=IFT22;refGene_tx=XM_023246893.1;refGene_gn=IFT22;refGene_tx=XM_023246894.1;refGene_gn=IFT22;refGene_tx=XM_023246895.1
E3 8247451 8247714
refGene_gn=IFT22;refGene_tx=XM_023246892.1;refGene_gn=IFT22;refGene_tx=XM_023246893.1;refGene_gn=IFT22;refGene_tx=XM_023246894.1;refGene_gn=IFT22;refGene_tx=XM_023246895.1
E3 8248392 8248482
refGene_gn=IFT22;refGene_tx=XM_023246892.1;refGene_gn=IFT22;refGene_tx=XM_023246893.1;refGene_gn=IFT22;refGene_tx=XM_023246894.1;refGene_gn=IFT22;refGene_tx=XM_023246895.1
E3 8248597 8248674
refGene_gn=IFT22;refGene_tx=XM_023246892.1;refGene_gn=IFT22;refGene_tx=XM_023246893.1
E3 8252324 8252363
refGene_gn=IFT22;refGene_tx=XM_023246892.1;refGene_gn=IFT22;refGene_tx=XM_023246893.1
E3 8284796 8284849 refGene_gn=COL26A1;refGene_tx=XM_011290370.3
E3 8287329 8287487
refGene_gn=COL26A1;refGene_tx=XM_011290368.3;refGene_gn=COL26A1;refGene_tx=XM_011290369.3
E3 8336848 8336971
refGene_gn=COL26A1;refGene_tx=XM_011290368.3;refGene_gn=COL26A1;refGene_tx=XM_011290369.3;refGene_gn=COL26A1;refGene_tx=XM_011290370.3
E3 8363380 8363484
refGene_gn=COL26A1;refGene_tx=XM_011290368.3;refGene_gn=COL26A1;refGene_tx=XM_011290369.3;refGene_gn=COL26A1;refGene_tx=XM_011290370.3

E3	8432379	8432441
refGene_gn=COL26A1;refGene_tx=XM_011290368.3;refGene_gn=COL26A1;refGene_tx=XM_011290369.3;refGene_gn=COL26A1;refGene_tx=XM_011290370.3
E3	8438890	8439047
refGene_gn=COL26A1;refGene_tx=XM_011290368.3;refGene_gn=COL26A1;refGene_tx=XM_011290369.3;refGene_gn=COL26A1;refGene_tx=XM_011290370.3
E3	8441363	8441462
refGene_gn=COL26A1;refGene_tx=XM_011290368.3;refGene_gn=COL26A1;refGene_tx=XM_011290369.3;refGene_gn=COL26A1;refGene_tx=XM_011290370.3
E3	8442078	8442231
refGene_gn=COL26A1;refGene_tx=XM_011290368.3;refGene_gn=COL26A1;refGene_tx=XM_011290369.3;refGene_gn=COL26A1;refGene_tx=XM_011290370.3
E3	8444180	8444264
refGene_gn=COL26A1;refGene_tx=XM_011290368.3;refGene_gn=COL26A1;refGene_tx=XM_011290369.3;refGene_gn=COL26A1;refGene_tx=XM_011290370.3
E3	8445447	8445500
refGene_gn=COL26A1;refGene_tx=XM_011290368.3;refGene_gn=COL26A1;refGene_tx=XM_011290369.3;refGene_gn=COL26A1;refGene_tx=XM_011290370.3
E3	8448920	8448956
refGene_gn=COL26A1;refGene_tx=XM_011290368.3;refGene_gn=COL26A1;refGene_tx=XM_011290369.3;refGene_gn=COL26A1;refGene_tx=XM_011290370.3
E3	8450266	8450314
refGene_gn=COL26A1;refGene_tx=XM_011290368.3;refGene_gn=COL26A1;refGene_tx=XM_011290369.3;refGene_gn=COL26A1;refGene_tx=XM_011290370.3
E3	8451909	8451994
refGene_gn=COL26A1;refGene_tx=XM_011290368.3;refGene_gn=COL26A1;refGene_tx=XM_011290369.3;refGene_gn=COL26A1;refGene_tx=XM_011290370.3
E3	8452734	8452898
refGene_gn=COL26A1;refGene_tx=XM_011290368.3;refGene_gn=COL26A1;refGene_tx=XM_011290369.3;refGene_gn=COL26A1;refGene_tx=XM_011290370.3
E3	8503615	8503714	refGene_gn=MYL10;refGene_tx=XM_023246751.1
E3	8504015	8504064	refGene_gn=MYL10;refGene_tx=XM_023246751.1
E3	8506208	8506287	refGene_gn=MYL10;refGene_tx=XM_023246751.1
E3	8509378	8509483	refGene_gn=MYL10;refGene_tx=XM_023246751.1
E3	8510308	8510402	refGene_gn=MYL10;refGene_tx=XM_023246751.1
E3	8510844	8510937	refGene_gn=MYL10;refGene_tx=XM_023246751.1
E3	8513282	8513285	refGene_gn=MYL10;refGene_tx=XM_023246751.1
E3	8642751	8642799	refGene_gn=CUX1;refGene_tx=XM_023246757.1
E3	8652414	8652477
refGene_gn=CUX1;refGene_tx=XM_023246756.1;refGene_gn=CUX1;refGene_tx=XM_023246758.1;refGene_gn=CUX1;refGene_tx=XM_023246760.1;refGene_gn=CUX1;refGene_tx=XM_023246761.1;refGene_gn=CUX1;refGene_tx=XM_023246762.1
E3	8653711	8654041
refGene_gn=CUX1;refGene_tx=XM_023246752.1;refGene_gn=CUX1;refGene_tx=XM_023246753.1;refGene_gn=CUX1;refGene_tx=XM_023246754.1;refGene_gn=CUX1;refGene_tx=XM_023246759.1
E3	8729492	8729597
refGene_gn=CUX1;refGene_tx=XM_019820541.2;refGene_gn=CUX1;refGene_tx=XM_023246755.1
E3	8731929	8732040

refGene_gn=CUX1;refGene_tx=XM_019820541.2;refGene_gn=CUX1;refGene_tx=X
M_023246752.1;refGene_gn=CUX1;refGene_tx=XM_023246753.1;refGene_gn=CUX
1;refGene_tx=XM_023246754.1;refGene_gn=CUX1;refGene_tx=XM_023246755.1;
refGene_gn=CUX1;refGene_tx=XM_023246756.1;refGene_gn=CUX1;refGene_tx=X
M_023246757.1;refGene_gn=CUX1;refGene_tx=XM_023246758.1;refGene_gn=CUX
1;refGene_tx=XM_023246759.1;refGene_gn=CUX1;refGene_tx=XM_023246760.1;
refGene_gn=CUX1;refGene_tx=XM_023246761.1;refGene_gn=CUX1;refGene_tx=X
M_023246762.1
E3 8820817 8820865
refGene_gn=CUX1;refGene_tx=XM_019820541.2;refGene_gn=CUX1;refGene_tx=X
M_023246752.1;refGene_gn=CUX1;refGene_tx=XM_023246753.1;refGene_gn=CUX
1;refGene_tx=XM_023246754.1;refGene_gn=CUX1;refGene_tx=XM_023246755.1;
refGene_gn=CUX1;refGene_tx=XM_023246756.1;refGene_gn=CUX1;refGene_tx=X
M_023246757.1;refGene_gn=CUX1;refGene_tx=XM_023246758.1;refGene_gn=CUX
1;refGene_tx=XM_023246759.1;refGene_gn=CUX1;refGene_tx=XM_023246760.1;
refGene_gn=CUX1;refGene_tx=XM_023246761.1;refGene_gn=CUX1;refGene_tx=X
M_023246762.1
E3 8849251 8849330
refGene_gn=CUX1;refGene_tx=XM_019820541.2;refGene_gn=CUX1;refGene_tx=X
M_023246752.1;refGene_gn=CUX1;refGene_tx=XM_023246753.1;refGene_gn=CUX
1;refGene_tx=XM_023246754.1;refGene_gn=CUX1;refGene_tx=XM_023246755.1;
refGene_gn=CUX1;refGene_tx=XM_023246756.1;refGene_gn=CUX1;refGene_tx=X
M_023246757.1;refGene_gn=CUX1;refGene_tx=XM_023246758.1;refGene_gn=CUX
1;refGene_tx=XM_023246759.1;refGene_gn=CUX1;refGene_tx=XM_023246760.1;
refGene_gn=CUX1;refGene_tx=XM_023246761.1;refGene_gn=CUX1;refGene_tx=X
M_023246762.1
E3 8869053 8869191
refGene_gn=CUX1;refGene_tx=XM_019820541.2;refGene_gn=CUX1;refGene_tx=X
M_023246752.1;refGene_gn=CUX1;refGene_tx=XM_023246753.1;refGene_gn=CUX
1;refGene_tx=XM_023246754.1;refGene_gn=CUX1;refGene_tx=XM_023246755.1;
refGene_gn=CUX1;refGene_tx=XM_023246756.1;refGene_gn=CUX1;refGene_tx=X
M_023246757.1;refGene_gn=CUX1;refGene_tx=XM_023246758.1;refGene_gn=CUX
1;refGene_tx=XM_023246759.1;refGene_gn=CUX1;refGene_tx=XM_023246760.1;
refGene_gn=CUX1;refGene_tx=XM_023246761.1;refGene_gn=CUX1;refGene_tx=X
M_023246762.1
E3 8873709 8873833
refGene_gn=CUX1;refGene_tx=XM_019820541.2;refGene_gn=CUX1;refGene_tx=X
M_023246752.1;refGene_gn=CUX1;refGene_tx=XM_023246753.1;refGene_gn=CUX
1;refGene_tx=XM_023246754.1;refGene_gn=CUX1;refGene_tx=XM_023246755.1;
refGene_gn=CUX1;refGene_tx=XM_023246756.1;refGene_gn=CUX1;refGene_tx=X
M_023246757.1;refGene_gn=CUX1;refGene_tx=XM_023246758.1;refGene_gn=CUX
1;refGene_tx=XM_023246759.1;refGene_gn=CUX1;refGene_tx=XM_023246760.1;
refGene_gn=CUX1;refGene_tx=XM_023246761.1;refGene_gn=CUX1;refGene_tx=X
M_023246762.1
E3 8877562 8877639
refGene_gn=CUX1;refGene_tx=XM_019820541.2;refGene_gn=CUX1;refGene_tx=X
M_023246752.1;refGene_gn=CUX1;refGene_tx=XM_023246753.1;refGene_gn=CUX
1;refGene_tx=XM_023246754.1;refGene_gn=CUX1;refGene_tx=XM_023246755.1;
refGene_gn=CUX1;refGene_tx=XM_023246756.1;refGene_gn=CUX1;refGene_tx=X
M_023246757.1;refGene_gn=CUX1;refGene_tx=XM_023246758.1;refGene_gn=CUX

1;refGene_tx=XM_023246759.1;refGene_gn=CUX1;refGene_tx=XM_023246760.1;
refGene_gn=CUX1;refGene_tx=XM_023246761.1;refGene_gn=CUX1;refGene_tx=X
M_023246762.1
E3 8879081 8879148
refGene_gn=CUX1;refGene_tx=XM_019820541.2;refGene_gn=CUX1;refGene_tx=X
M_023246752.1;refGene_gn=CUX1;refGene_tx=XM_023246753.1;refGene_gn=CUX
1;refGene_tx=XM_023246754.1;refGene_gn=CUX1;refGene_tx=XM_023246755.1;
refGene_gn=CUX1;refGene_tx=XM_023246756.1;refGene_gn=CUX1;refGene_tx=X
M_023246757.1;refGene_gn=CUX1;refGene_tx=XM_023246758.1;refGene_gn=CUX
1;refGene_tx=XM_023246759.1;refGene_gn=CUX1;refGene_tx=XM_023246760.1;
refGene_gn=CUX1;refGene_tx=XM_023246761.1;refGene_gn=CUX1;refGene_tx=X
M_023246762.1
E3 8910965 8911014
refGene_gn=CUX1;refGene_tx=XM_019820541.2;refGene_gn=CUX1;refGene_tx=X
M_023246752.1;refGene_gn=CUX1;refGene_tx=XM_023246753.1;refGene_gn=CUX
1;refGene_tx=XM_023246754.1;refGene_gn=CUX1;refGene_tx=XM_023246755.1;
refGene_gn=CUX1;refGene_tx=XM_023246756.1;refGene_gn=CUX1;refGene_tx=X
M_023246757.1;refGene_gn=CUX1;refGene_tx=XM_023246758.1;refGene_gn=CUX
1;refGene_tx=XM_023246759.1;refGene_gn=CUX1;refGene_tx=XM_023246760.1;
refGene_gn=CUX1;refGene_tx=XM_023246761.1;refGene_gn=CUX1;refGene_tx=X
M_023246762.1
E3 8925456 8925561
refGene_gn=CUX1;refGene_tx=XM_019820541.2;refGene_gn=CUX1;refGene_tx=X
M_023246752.1;refGene_gn=CUX1;refGene_tx=XM_023246753.1;refGene_gn=CUX
1;refGene_tx=XM_023246754.1;refGene_gn=CUX1;refGene_tx=XM_023246755.1;
refGene_gn=CUX1;refGene_tx=XM_023246756.1;refGene_gn=CUX1;refGene_tx=X
M_023246757.1;refGene_gn=CUX1;refGene_tx=XM_023246758.1;refGene_gn=CUX
1;refGene_tx=XM_023246759.1;refGene_gn=CUX1;refGene_tx=XM_023246760.1;
refGene_gn=CUX1;refGene_tx=XM_023246761.1;refGene_gn=CUX1;refGene_tx=X
M_023246762.1
E3 8932273 8932462
refGene_gn=CUX1;refGene_tx=XM_019820541.2;refGene_gn=CUX1;refGene_tx=X
M_023246752.1;refGene_gn=CUX1;refGene_tx=XM_023246753.1;refGene_gn=CUX
1;refGene_tx=XM_023246754.1;refGene_gn=CUX1;refGene_tx=XM_023246755.1;
refGene_gn=CUX1;refGene_tx=XM_023246756.1;refGene_gn=CUX1;refGene_tx=X
M_023246757.1;refGene_gn=CUX1;refGene_tx=XM_023246758.1;refGene_gn=CUX
1;refGene_tx=XM_023246759.1;refGene_gn=CUX1;refGene_tx=XM_023246760.1;
refGene_gn=CUX1;refGene_tx=XM_023246761.1;refGene_gn=CUX1;refGene_tx=X
M_023246762.1
E3 8941066 8941125
refGene_gn=CUX1;refGene_tx=XM_019820541.2;refGene_gn=CUX1;refGene_tx=X
M_023246752.1;refGene_gn=CUX1;refGene_tx=XM_023246753.1;refGene_gn=CUX
1;refGene_tx=XM_023246754.1;refGene_gn=CUX1;refGene_tx=XM_023246755.1;
refGene_gn=CUX1;refGene_tx=XM_023246756.1;refGene_gn=CUX1;refGene_tx=X
M_023246757.1;refGene_gn=CUX1;refGene_tx=XM_023246758.1;refGene_gn=CUX
1;refGene_tx=XM_023246759.1;refGene_gn=CUX1;refGene_tx=XM_023246760.1;
refGene_gn=CUX1;refGene_tx=XM_023246761.1;refGene_gn=CUX1;refGene_tx=X
M_023246762.1
E3 8944139 8944188
refGene_gn=CUX1;refGene_tx=XM_019820541.2;refGene_gn=CUX1;refGene_tx=X

M_023246752.1;refGene_gn=CUX1;refGene_tx=XM_023246753.1;refGene_gn=CUX
1;refGene_tx=XM_023246754.1;refGene_gn=CUX1;refGene_tx=XM_023246755.1;
refGene_gn=CUX1;refGene_tx=XM_023246756.1;refGene_gn=CUX1;refGene_tx=X
M_023246757.1;refGene_gn=CUX1;refGene_tx=XM_023246758.1;refGene_gn=CUX
1;refGene_tx=XM_023246759.1;refGene_gn=CUX1;refGene_tx=XM_023246760.1;
refGene_gn=CUX1;refGene_tx=XM_023246761.1;refGene_gn=CUX1;refGene_tx=X
M_023246762.1
E3 8944992 8945089
refGene_gn=CUX1;refGene_tx=XM_019820541.2;refGene_gn=CUX1;refGene_tx=X
M_023246752.1;refGene_gn=CUX1;refGene_tx=XM_023246753.1;refGene_gn=CUX
1;refGene_tx=XM_023246754.1;refGene_gn=CUX1;refGene_tx=XM_023246755.1;
refGene_gn=CUX1;refGene_tx=XM_023246756.1;refGene_gn=CUX1;refGene_tx=X
M_023246757.1;refGene_gn=CUX1;refGene_tx=XM_023246758.1;refGene_gn=CUX
1;refGene_tx=XM_023246759.1;refGene_gn=CUX1;refGene_tx=XM_023246760.1;
refGene_gn=CUX1;refGene_tx=XM_023246761.1;refGene_gn=CUX1;refGene_tx=X
M_023246762.1
E3 8946108 8946780
refGene_gn=CUX1;refGene_tx=XM_023246752.1;refGene_gn=CUX1;refGene_tx=X
M_023246753.1;refGene_gn=CUX1;refGene_tx=XM_023246754.1;refGene_gn=CUX
1;refGene_tx=XM_023246755.1;refGene_gn=CUX1;refGene_tx=XM_023246756.1;
refGene_gn=CUX1;refGene_tx=XM_023246757.1;refGene_gn=CUX1;refGene_tx=X
M_023246758.1;refGene_gn=CUX1;refGene_tx=XM_023246762.1
E3 8947873 8947939
refGene_gn=CUX1;refGene_tx=XM_023246752.1;refGene_gn=CUX1;refGene_tx=X
M_023246753.1;refGene_gn=CUX1;refGene_tx=XM_023246755.1;refGene_gn=CUX
1;refGene_tx=XM_023246756.1;refGene_gn=CUX1;refGene_tx=XM_023246757.1;
refGene_gn=CUX1;refGene_tx=XM_023246758.1
E3 8949295 8949397
refGene_gn=CUX1;refGene_tx=XM_023246752.1;refGene_gn=CUX1;refGene_tx=X
M_023246753.1;refGene_gn=CUX1;refGene_tx=XM_023246754.1;refGene_gn=CUX
1;refGene_tx=XM_023246755.1;refGene_gn=CUX1;refGene_tx=XM_023246756.1;
refGene_gn=CUX1;refGene_tx=XM_023246757.1;refGene_gn=CUX1;refGene_tx=X
M_023246758.1;refGene_gn=CUX1;refGene_tx=XM_023246762.1
E3 8951901 8952731
refGene_gn=CUX1;refGene_tx=XM_023246752.1;refGene_gn=CUX1;refGene_tx=X
M_023246753.1;refGene_gn=CUX1;refGene_tx=XM_023246754.1;refGene_gn=CUX
1;refGene_tx=XM_023246755.1;refGene_gn=CUX1;refGene_tx=XM_023246756.1;
refGene_gn=CUX1;refGene_tx=XM_023246757.1;refGene_gn=CUX1;refGene_tx=X
M_023246758.1;refGene_gn=CUX1;refGene_tx=XM_023246762.1
E3 8954671 8954837
refGene_gn=CUX1;refGene_tx=XM_023246752.1;refGene_gn=CUX1;refGene_tx=X
M_023246753.1;refGene_gn=CUX1;refGene_tx=XM_023246754.1;refGene_gn=CUX
1;refGene_tx=XM_023246755.1;refGene_gn=CUX1;refGene_tx=XM_023246756.1;
refGene_gn=CUX1;refGene_tx=XM_023246757.1;refGene_gn=CUX1;refGene_tx=X
M_023246758.1;refGene_gn=CUX1;refGene_tx=XM_023246762.1
E3 8955329 8955386
refGene_gn=CUX1;refGene_tx=XM_023246752.1;refGene_gn=CUX1;refGene_tx=X
M_023246753.1;refGene_gn=CUX1;refGene_tx=XM_023246754.1;refGene_gn=CUX
1;refGene_tx=XM_023246755.1;refGene_gn=CUX1;refGene_tx=XM_023246756.1;
refGene_gn=CUX1;refGene_tx=XM_023246757.1;refGene_gn=CUX1;refGene_tx=X

M_023246758.1;refGene_gn=CUX1;refGene_tx=XM_023246762.1
E3 8976351 8976654
refGene_gn=CUX1;refGene_tx=XM_023246752.1;refGene_gn=CUX1;refGene_tx=XM_023246753.1;refGene_gn=CUX1;refGene_tx=XM_023246754.1;refGene_gn=CUX1;refGene_tx=XM_023246755.1;refGene_gn=CUX1;refGene_tx=XM_023246756.1;refGene_gn=CUX1;refGene_tx=XM_023246757.1;refGene_gn=CUX1;refGene_tx=XM_023246758.1;refGene_gn=CUX1;refGene_tx=XM_023246762.1
E3 8979845 8980034
refGene_gn=CUX1;refGene_tx=XM_023246752.1;refGene_gn=CUX1;refGene_tx=XM_023246753.1;refGene_gn=CUX1;refGene_tx=XM_023246754.1;refGene_gn=CUX1;refGene_tx=XM_023246755.1;refGene_gn=CUX1;refGene_tx=XM_023246756.1;refGene_gn=CUX1;refGene_tx=XM_023246757.1;refGene_gn=CUX1;refGene_tx=XM_023246758.1;refGene_gn=CUX1;refGene_tx=XM_023246762.1
E3 8985679 8985944
refGene_gn=CUX1;refGene_tx=XM_023246752.1;refGene_gn=CUX1;refGene_tx=XM_023246753.1;refGene_gn=CUX1;refGene_tx=XM_023246754.1;refGene_gn=CUX1;refGene_tx=XM_023246755.1;refGene_gn=CUX1;refGene_tx=XM_023246756.1;refGene_gn=CUX1;refGene_tx=XM_023246757.1;refGene_gn=CUX1;refGene_tx=XM_023246758.1;refGene_gn=CUX1;refGene_tx=XM_023246762.1
E3 8991250 8991974
refGene_gn=CUX1;refGene_tx=XM_023246752.1;refGene_gn=CUX1;refGene_tx=XM_023246753.1;refGene_gn=CUX1;refGene_tx=XM_023246754.1;refGene_gn=CUX1;refGene_tx=XM_023246755.1;refGene_gn=CUX1;refGene_tx=XM_023246756.1;refGene_gn=CUX1;refGene_tx=XM_023246757.1;refGene_gn=CUX1;refGene_tx=XM_023246758.1;refGene_gn=CUX1;refGene_tx=XM_023246762.1
E3 9009416 9009544
refGene_gn=CUX1;refGene_tx=XM_019820541.2;refGene_gn=CUX1;refGene_tx=XM_023246759.1;refGene_gn=CUX1;refGene_tx=XM_023246760.1;refGene_gn=CUX1;refGene_tx=XM_023246761.1
E3 9010225 9010292
refGene_gn=CUX1;refGene_tx=XM_019820541.2;refGene_gn=CUX1;refGene_tx=XM_023246759.1;refGene_gn=CUX1;refGene_tx=XM_023246760.1;refGene_gn=CUX1;refGene_tx=XM_023246761.1
E3 9011029 9011142
refGene_gn=CUX1;refGene_tx=XM_019820541.2;refGene_gn=CUX1;refGene_tx=XM_023246759.1;refGene_gn=CUX1;refGene_tx=XM_023246760.1;refGene_gn=CUX1;refGene_tx=XM_023246761.1
E3 9014565 9014682
refGene_gn=CUX1;refGene_tx=XM_019820541.2;refGene_gn=CUX1;refGene_tx=XM_023246759.1;refGene_gn=CUX1;refGene_tx=XM_023246760.1;refGene_gn=CUX1;refGene_tx=XM_023246761.1
E3 9015783 9015867
refGene_gn=CUX1;refGene_tx=XM_019820541.2;refGene_gn=CUX1;refGene_tx=XM_023246759.1;refGene_gn=CUX1;refGene_tx=XM_023246760.1;refGene_gn=CUX1;refGene_tx=XM_023246761.1
E3 9016496 9016553
refGene_gn=CUX1;refGene_tx=XM_019820541.2;refGene_gn=CUX1;refGene_tx=XM_023246759.1;refGene_gn=CUX1;refGene_tx=XM_023246760.1;refGene_gn=CUX1;refGene_tx=XM_023246761.1
E3 9016930 9017011

refGene_gn=CUX1;refGene_tx=XM_019820541.2;refGene_gn=CUX1;refGene_tx=XM_023246759.1;refGene_gn=CUX1;refGene_tx=XM_023246760.1;refGene_gn=CUX1;refGene_tx=XM_023246761.1
E3 9017757 9017822 refGene_gn=CUX1;refGene_tx=XM_019820541.2;refGene_gn=CUX1;refGene_tx=XM_023246759.1;refGene_gn=CUX1;refGene_tx=XM_023246760.1;refGene_gn=CUX1;refGene_tx=XM_023246761.1
E3 9018057 9018127 refGene_gn=CUX1;refGene_tx=XM_019820541.2;refGene_gn=CUX1;refGene_tx=XM_023246759.1;refGene_gn=CUX1;refGene_tx=XM_023246760.1;refGene_gn=CUX1;refGene_tx=XM_023246761.1
E3 9019895 9019941 refGene_gn=SH2B2;refGene_tx=XM_023246766.1
E3 9032398 9033165 refGene_gn=SH2B2;refGene_tx=XM_023246763.1;refGene_gn=SH2B2;refGene_tx=XM_023246764.1;refGene_gn=SH2B2;refGene_tx=XM_023246765.1;refGene_gn=SH2B2;refGene_tx=XM_023246766.1
E3 9036161 9036263 refGene_gn=SH2B2;refGene_tx=XM_023246763.1;refGene_gn=SH2B2;refGene_tx=XM_023246764.1;refGene_gn=SH2B2;refGene_tx=XM_023246765.1;refGene_gn=SH2B2;refGene_tx=XM_023246766.1
E3 9037977 9038069 refGene_gn=SH2B2;refGene_tx=XM_023246763.1;refGene_gn=SH2B2;refGene_tx=XM_023246764.1;refGene_gn=SH2B2;refGene_tx=XM_023246765.1;refGene_gn=SH2B2;refGene_tx=XM_023246766.1
E3 9039365 9039457 refGene_gn=SH2B2;refGene_tx=XM_023246763.1;refGene_gn=SH2B2;refGene_tx=XM_023246764.1;refGene_gn=SH2B2;refGene_tx=XM_023246765.1;refGene_gn=SH2B2;refGene_tx=XM_023246766.1
E3 9039539 9039704 refGene_gn=SH2B2;refGene_tx=XM_023246763.1;refGene_gn=SH2B2;refGene_tx=XM_023246764.1;refGene_gn=SH2B2;refGene_tx=XM_023246765.1;refGene_gn=SH2B2;refGene_tx=XM_023246766.1
E3 9040193 9040402 refGene_gn=SH2B2;refGene_tx=XM_023246763.1;refGene_gn=SH2B2;refGene_tx=XM_023246764.1;refGene_gn=SH2B2;refGene_tx=XM_023246765.1;refGene_gn=SH2B2;refGene_tx=XM_023246766.1
E3 9042892 9043064 refGene_gn=SH2B2;refGene_tx=XM_023246763.1;refGene_gn=SH2B2;refGene_tx=XM_023246764.1;refGene_gn=SH2B2;refGene_tx=XM_023246765.1;refGene_gn=SH2B2;refGene_tx=XM_023246766.1
E3 9043577 9043921 refGene_gn=SH2B2;refGene_tx=XM_023246763.1;refGene_gn=SH2B2;refGene_tx=XM_023246764.1;refGene_gn=SH2B2;refGene_tx=XM_023246765.1;refGene_gn=SH2B2;refGene_tx=XM_023246766.1
E3 9076029 9076305 refGene_gn=PRKRIP1;refGene_tx=XM_023247077.1
E3 9077003 9077134 refGene_gn=PRKRIP1;refGene_tx=XM_023247077.1
E3 9086739 9086841 refGene_gn=PRKRIP1;refGene_tx=XM_023247077.1
E3 9088177 9088245 refGene_gn=PRKRIP1;refGene_tx=XM_023247077.1
E3 9109372 9109482 refGene_gn=PRKRIP1;refGene_tx=XM_023247077.1
E3 9122020 9122245

refGene_gn=ORAI2;refGene_tx=XM_023246675.1;refGene_gn=ORAI2;refGene_tx
=XM_023246676.1
E3	9126979	9127519
refGene_gn=ORAI2;refGene_tx=XM_023246675.1;refGene_gn=ORAI2;refGene_tx
=XM_023246676.1
E3	9133028	9133616	refGene_gn=ALKBH4;refGene_tx=XM_006942050.3
E3	9136372	9136570	refGene_gn=ALKBH4;refGene_tx=XM_006942050.3
E3	9138722	9138845	refGene_gn=ALKBH4;refGene_tx=XM_006942050.3
E3	9139087	9139167
refGene_gn=LRWD1;refGene_tx=XM_023246669.1;refGene_gn=LRWD1;refGene_tx
=XM_023246670.1;refGene_gn=LRWD1;refGene_tx=XM_023246671.1;refGene_gn=
LRWD1;refGene_tx=XM_023246672.1;refGene_gn=LRWD1;refGene_tx=XM_0232466
73.1
E3	9139713	9139948
refGene_gn=LRWD1;refGene_tx=XM_023246669.1;refGene_gn=LRWD1;refGene_tx
=XM_023246670.1;refGene_gn=LRWD1;refGene_tx=XM_023246671.1;refGene_gn=
LRWD1;refGene_tx=XM_023246672.1;refGene_gn=LRWD1;refGene_tx=XM_0232466
73.1
E3	9140063	9140180
refGene_gn=LRWD1;refGene_tx=XM_023246669.1;refGene_gn=LRWD1;refGene_tx
=XM_023246670.1;refGene_gn=LRWD1;refGene_tx=XM_023246671.1;refGene_gn=
LRWD1;refGene_tx=XM_023246672.1;refGene_gn=LRWD1;refGene_tx=XM_0232466
73.1
E3	9140641	9140782
refGene_gn=LRWD1;refGene_tx=XM_023246669.1;refGene_gn=LRWD1;refGene_tx
=XM_023246670.1;refGene_gn=LRWD1;refGene_tx=XM_023246671.1;refGene_gn=
LRWD1;refGene_tx=XM_023246672.1;refGene_gn=LRWD1;refGene_tx=XM_0232466
73.1
E3	9141021	9141126
refGene_gn=LRWD1;refGene_tx=XM_023246669.1;refGene_gn=LRWD1;refGene_tx
=XM_023246670.1;refGene_gn=LRWD1;refGene_tx=XM_023246671.1;refGene_gn=
LRWD1;refGene_tx=XM_023246672.1;refGene_gn=LRWD1;refGene_tx=XM_0232466
73.1
E3	9141387	9141510
refGene_gn=LRWD1;refGene_tx=XM_023246669.1;refGene_gn=LRWD1;refGene_tx
=XM_023246670.1;refGene_gn=LRWD1;refGene_tx=XM_023246671.1;refGene_gn=
LRWD1;refGene_tx=XM_023246672.1;refGene_gn=LRWD1;refGene_tx=XM_0232466
73.1
E3	9141595	9141710
refGene_gn=LRWD1;refGene_tx=XM_023246669.1;refGene_gn=LRWD1;refGene_tx
=XM_023246670.1;refGene_gn=LRWD1;refGene_tx=XM_023246671.1;refGene_gn=
LRWD1;refGene_tx=XM_023246672.1;refGene_gn=LRWD1;refGene_tx=XM_0232466
73.1
E3	9141907	9142017
refGene_gn=LRWD1;refGene_tx=XM_023246669.1;refGene_gn=LRWD1;refGene_tx
=XM_023246670.1;refGene_gn=LRWD1;refGene_tx=XM_023246671.1;refGene_gn=
LRWD1;refGene_tx=XM_023246672.1;refGene_gn=LRWD1;refGene_tx=XM_0232466
73.1
E3	9142203	9142411
refGene_gn=LRWD1;refGene_tx=XM_023246669.1;refGene_gn=LRWD1;refGene_tx

=XM_023246670.1;refGene_gn=LRWD1;refGene_tx=XM_023246671.1;refGene_gn=LRWD1;refGene_tx=XM_023246672.1;refGene_gn=LRWD1;refGene_tx=XM_023246673.1
E3 9142912 9142985 refGene_gn=LRWD1;refGene_tx=XM_023246669.1;refGene_gn=LRWD1;refGene_tx=XM_023246670.1;refGene_gn=LRWD1;refGene_tx=XM_023246671.1;refGene_gn=LRWD1;refGene_tx=XM_023246672.1;refGene_gn=LRWD1;refGene_tx=XM_023246673.1
E3 9143064 9143208 refGene_gn=LRWD1;refGene_tx=XM_023246669.1;refGene_gn=LRWD1;refGene_tx=XM_023246670.1;refGene_gn=LRWD1;refGene_tx=XM_023246671.1;refGene_gn=LRWD1;refGene_tx=XM_023246672.1;refGene_gn=LRWD1;refGene_tx=XM_023246673.1
E3 9144446 9144538 refGene_gn=LRWD1;refGene_tx=XM_023246669.1;refGene_gn=LRWD1;refGene_tx=XM_023246670.1;refGene_gn=LRWD1;refGene_tx=XM_023246671.1;refGene_gn=LRWD1;refGene_tx=XM_023246672.1;refGene_gn=LRWD1;refGene_tx=XM_023246673.1
E3 9144927 9145083 refGene_gn=LRWD1;refGene_tx=XM_023246669.1;refGene_gn=LRWD1;refGene_tx=XM_023246670.1;refGene_gn=LRWD1;refGene_tx=XM_023246671.1;refGene_gn=LRWD1;refGene_tx=XM_023246672.1
E3 9145180 9145293 refGene_gn=LRWD1;refGene_tx=XM_023246669.1;refGene_gn=LRWD1;refGene_tx=XM_023246670.1;refGene_gn=LRWD1;refGene_tx=XM_023246671.1;refGene_gn=LRWD1;refGene_tx=XM_023246672.1;refGene_gn=LRWD1;refGene_tx=XM_023246673.1
E3 9145374 9145515 refGene_gn=LRWD1;refGene_tx=XM_023246671.1;refGene_gn=LRWD1;refGene_tx=XM_023246672.1
E3 9146092 9146128 refGene_gn=POLR2J;refGene_tx=XM_003998535.4
E3 9146721 9146896 refGene_gn=POLR2J;refGene_tx=XM_003998535.4
E3 9147899 9148189 refGene_gn=LRWD1;refGene_tx=XM_023246669.1;refGene_gn=LRWD1;refGene_tx=XM_023246670.1;refGene_gn=LRWD1;refGene_tx=XM_023246673.1;refGene_gn=POLR2J;refGene_tx=XM_003998535.4
E3 9149302 9149355 refGene_gn=POLR2J;refGene_tx=XM_003998535.4
E3 9151810 9151929 refGene_gn=RASA4B;refGene_tx=XM_003998567.5;refGene_gn=RASA4B;refGene_tx=XM_006942015.4;refGene_gn=RASA4B;refGene_tx=XM_006942016.4;refGene_gn=RASA4B;refGene_tx=XM_006942017.4;refGene_gn=RASA4B;refGene_tx=XM_006942019.4
E3 9152077 9152248 refGene_gn=RASA4B;refGene_tx=XM_003998567.5;refGene_gn=RASA4B;refGene_tx=XM_006942015.4;refGene_gn=RASA4B;refGene_tx=XM_006942016.4;refGene_gn=RASA4B;refGene_tx=XM_006942017.4;refGene_gn=RASA4B;refGene_tx=XM_006942019.4
E3 9152907 9152941 refGene_gn=RASA4B;refGene_tx=XM_006942018.4
E3 9153926 9154047 refGene_gn=RASA4B;refGene_tx=XM_003998567.5;refGene_gn=RASA4B;refGene_

tx=XM_006942015.4;refGene_gn=RASA4B;refGene_tx=XM_006942016.4;refGene_
gn=RASA4B;refGene_tx=XM_006942017.4;refGene_gn=RASA4B;refGene_tx=XM_00
6942018.4;refGene_gn=RASA4B;refGene_tx=XM_006942019.4
E3 9155568 9155704
refGene_gn=RASA4B;refGene_tx=XM_003998567.5;refGene_gn=RASA4B;refGene_
tx=XM_006942015.4;refGene_gn=RASA4B;refGene_tx=XM_006942016.4;refGene_
gn=RASA4B;refGene_tx=XM_006942017.4;refGene_gn=RASA4B;refGene_tx=XM_00
6942018.4;refGene_gn=RASA4B;refGene_tx=XM_006942019.4
E3 9156714 9156852
refGene_gn=RASA4B;refGene_tx=XM_003998567.5;refGene_gn=RASA4B;refGene_
tx=XM_006942015.4;refGene_gn=RASA4B;refGene_tx=XM_006942016.4;refGene_
gn=RASA4B;refGene_tx=XM_006942017.4;refGene_gn=RASA4B;refGene_tx=XM_00
6942018.4;refGene_gn=RASA4B;refGene_tx=XM_006942019.4
E3 9159107 9159274
refGene_gn=RASA4B;refGene_tx=XM_003998567.5;refGene_gn=RASA4B;refGene_
tx=XM_006942015.4;refGene_gn=RASA4B;refGene_tx=XM_006942016.4;refGene_
gn=RASA4B;refGene_tx=XM_006942017.4;refGene_gn=RASA4B;refGene_tx=XM_00
6942018.4;refGene_gn=RASA4B;refGene_tx=XM_006942019.4
E3 9159631 9159776
refGene_gn=RASA4B;refGene_tx=XM_003998567.5;refGene_gn=RASA4B;refGene_
tx=XM_006942015.4;refGene_gn=RASA4B;refGene_tx=XM_006942016.4;refGene_
gn=RASA4B;refGene_tx=XM_006942017.4;refGene_gn=RASA4B;refGene_tx=XM_00
6942018.4;refGene_gn=RASA4B;refGene_tx=XM_006942019.4
E3 9160317 9160455
refGene_gn=RASA4B;refGene_tx=XM_003998567.5;refGene_gn=RASA4B;refGene_
tx=XM_006942015.4;refGene_gn=RASA4B;refGene_tx=XM_006942016.4;refGene_
gn=RASA4B;refGene_tx=XM_006942017.4;refGene_gn=RASA4B;refGene_tx=XM_00
6942018.4;refGene_gn=RASA4B;refGene_tx=XM_006942019.4
E3 9160670 9160863
refGene_gn=RASA4B;refGene_tx=XM_003998567.5;refGene_gn=RASA4B;refGene_
tx=XM_006942015.4;refGene_gn=RASA4B;refGene_tx=XM_006942016.4;refGene_
gn=RASA4B;refGene_tx=XM_006942017.4;refGene_gn=RASA4B;refGene_tx=XM_00
6942018.4;refGene_gn=RASA4B;refGene_tx=XM_006942019.4
E3 9161104 9161217
refGene_gn=RASA4B;refGene_tx=XM_003998567.5;refGene_gn=RASA4B;refGene_
tx=XM_006942015.4;refGene_gn=RASA4B;refGene_tx=XM_006942016.4;refGene_
gn=RASA4B;refGene_tx=XM_006942017.4;refGene_gn=RASA4B;refGene_tx=XM_00
6942018.4;refGene_gn=RASA4B;refGene_tx=XM_006942019.4
E3 9161526 9161591
refGene_gn=RASA4B;refGene_tx=XM_003998567.5;refGene_gn=RASA4B;refGene_
tx=XM_006942015.4;refGene_gn=RASA4B;refGene_tx=XM_006942016.4;refGene_
gn=RASA4B;refGene_tx=XM_006942017.4;refGene_gn=RASA4B;refGene_tx=XM_00
6942018.4;refGene_gn=RASA4B;refGene_tx=XM_006942019.4
E3 9161846 9161997
refGene_gn=RASA4B;refGene_tx=XM_003998567.5;refGene_gn=RASA4B;refGene_
tx=XM_006942015.4;refGene_gn=RASA4B;refGene_tx=XM_006942016.4;refGene_
gn=RASA4B;refGene_tx=XM_006942017.4;refGene_gn=RASA4B;refGene_tx=XM_00
6942018.4;refGene_gn=RASA4B;refGene_tx=XM_006942019.4
E3 9162252 9162370
refGene_gn=RASA4B;refGene_tx=XM_003998567.5;refGene_gn=RASA4B;refGene_

tx=XM_006942015.4;refGene_gn=RASA4B;refGene_tx=XM_006942016.4;refGene_
gn=RASA4B;refGene_tx=XM_006942017.4;refGene_gn=RASA4B;refGene_tx=XM_00
6942018.4;refGene_gn=RASA4B;refGene_tx=XM_006942019.4
E3 9163635 9163727
refGene_gn=RASA4B;refGene_tx=XM_003998567.5;refGene_gn=RASA4B;refGene_
tx=XM_006942015.4;refGene_gn=RASA4B;refGene_tx=XM_006942016.4;refGene_
gn=RASA4B;refGene_tx=XM_006942017.4;refGene_gn=RASA4B;refGene_tx=XM_00
6942018.4;refGene_gn=RASA4B;refGene_tx=XM_006942019.4
E3 9163977 9164109
refGene_gn=RASA4B;refGene_tx=XM_003998567.5;refGene_gn=RASA4B;refGene_
tx=XM_006942015.4;refGene_gn=RASA4B;refGene_tx=XM_006942016.4;refGene_
gn=RASA4B;refGene_tx=XM_006942017.4;refGene_gn=RASA4B;refGene_tx=XM_00
6942018.4;refGene_gn=RASA4B;refGene_tx=XM_006942019.4
E3 9164620 9164702
refGene_gn=RASA4B;refGene_tx=XM_003998567.5;refGene_gn=RASA4B;refGene_
tx=XM_006942015.4;refGene_gn=RASA4B;refGene_tx=XM_006942016.4;refGene_
gn=RASA4B;refGene_tx=XM_006942017.4;refGene_gn=RASA4B;refGene_tx=XM_00
6942018.4;refGene_gn=RASA4B;refGene_tx=XM_006942019.4
E3 9166125 9166258
refGene_gn=RASA4B;refGene_tx=XM_003998567.5;refGene_gn=RASA4B;refGene_
tx=XM_006942015.4;refGene_gn=RASA4B;refGene_tx=XM_006942016.4;refGene_
gn=RASA4B;refGene_tx=XM_006942017.4;refGene_gn=RASA4B;refGene_tx=XM_00
6942018.4;refGene_gn=RASA4B;refGene_tx=XM_006942019.4
E3 9167404 9167466
refGene_gn=RASA4B;refGene_tx=XM_003998567.5;refGene_gn=RASA4B;refGene_
tx=XM_006942015.4;refGene_gn=RASA4B;refGene_tx=XM_006942016.4;refGene_
gn=RASA4B;refGene_tx=XM_006942017.4;refGene_gn=RASA4B;refGene_tx=XM_00
6942018.4;refGene_gn=RASA4B;refGene_tx=XM_006942019.4
E3 9167979 9168093
refGene_gn=RASA4B;refGene_tx=XM_003998567.5;refGene_gn=RASA4B;refGene_
tx=XM_006942015.4;refGene_gn=RASA4B;refGene_tx=XM_006942016.4;refGene_
gn=RASA4B;refGene_tx=XM_006942017.4;refGene_gn=RASA4B;refGene_tx=XM_00
6942018.4;refGene_gn=RASA4B;refGene_tx=XM_006942019.4
E3 9170129 9170186
refGene_gn=RASA4B;refGene_tx=XM_003998567.5;refGene_gn=RASA4B;refGene_
tx=XM_006942015.4;refGene_gn=RASA4B;refGene_tx=XM_006942016.4;refGene_
gn=RASA4B;refGene_tx=XM_006942017.4;refGene_gn=RASA4B;refGene_tx=XM_00
6942018.4
E3 9175015 9175080
refGene_gn=RASA4B;refGene_tx=XM_003998567.5;refGene_gn=RASA4B;refGene_
tx=XM_006942015.4;refGene_gn=RASA4B;refGene_tx=XM_006942018.4
E3 9192636 9192745
refGene_gn=UPK3BL2;refGene_tx=XM_023246646.1;refGene_gn=UPK3BL2;refGen
e_tx=XM_023246647.1;refGene_gn=UPK3BL2;refGene_tx=XM_023246648.1;refGe
ne_gn=UPK3BL2;refGene_tx=XM_023246650.1;refGene_gn=UPK3BL2;refGene_tx=
XM_023246651.1;refGene_gn=UPK3BL2;refGene_tx=XM_023246652.1;refGene_gn
=UPK3BL2;refGene_tx=XM_023246653.1;refGene_gn=UPK3BL2;refGene_tx=XM_02
3246654.1
E3 9193601 9193722
refGene_gn=UPK3BL2;refGene_tx=XM_023246646.1;refGene_gn=UPK3BL2;refGen

e_tx=XM_023246647.1;refGene_gn=UPK3BL2;refGene_tx=XM_023246648.1;refGene_gn=UPK3BL2;refGene_tx=XM_023246650.1;refGene_gn=UPK3BL2;refGene_tx=XM_023246651.1;refGene_gn=UPK3BL2;refGene_tx=XM_023246652.1;refGene_gn=UPK3BL2;refGene_tx=XM_023246653.1;refGene_gn=UPK3BL2;refGene_tx=XM_023246654.1
E3 9193985 9194062
refGene_gn=UPK3BL2;refGene_tx=XM_023246646.1;refGene_gn=UPK3BL2;refGene_tx=XM_023246647.1;refGene_gn=UPK3BL2;refGene_tx=XM_023246648.1;refGene_gn=UPK3BL2;refGene_tx=XM_023246650.1;refGene_gn=UPK3BL2;refGene_tx=XM_023246651.1;refGene_gn=UPK3BL2;refGene_tx=XM_023246652.1;refGene_gn=UPK3BL2;refGene_tx=XM_023246653.1;refGene_gn=UPK3BL2;refGene_tx=XM_023246654.1
E3 9194574 9194794
refGene_gn=UPK3BL2;refGene_tx=XM_023246646.1;refGene_gn=UPK3BL2;refGene_tx=XM_023246647.1;refGene_gn=UPK3BL2;refGene_tx=XM_023246648.1;refGene_gn=UPK3BL2;refGene_tx=XM_023246650.1;refGene_gn=UPK3BL2;refGene_tx=XM_023246651.1;refGene_gn=UPK3BL2;refGene_tx=XM_023246652.1;refGene_gn=UPK3BL2;refGene_tx=XM_023246653.1;refGene_gn=UPK3BL2;refGene_tx=XM_023246654.1
E3 9194983 9195136
refGene_gn=UPK3BL2;refGene_tx=XM_023246646.1;refGene_gn=UPK3BL2;refGene_tx=XM_023246647.1;refGene_gn=UPK3BL2;refGene_tx=XM_023246648.1;refGene_gn=UPK3BL2;refGene_tx=XM_023246650.1;refGene_gn=UPK3BL2;refGene_tx=XM_023246651.1;refGene_gn=UPK3BL2;refGene_tx=XM_023246652.1;refGene_gn=UPK3BL2;refGene_tx=XM_023246653.1;refGene_gn=UPK3BL2;refGene_tx=XM_023246654.1
E3 9196712 9196794 refGene_gn=UPK3BL2;refGene_tx=XM_023246652.1
E3 9199095 9199186
refGene_gn=UPK3BL2;refGene_tx=XM_023246646.1;refGene_gn=UPK3BL2;refGene_tx=XM_023246647.1;refGene_gn=UPK3BL2;refGene_tx=XM_023246648.1;refGene_gn=UPK3BL2;refGene_tx=XM_023246650.1;refGene_gn=UPK3BL2;refGene_tx=XM_023246651.1;refGene_gn=UPK3BL2;refGene_tx=XM_023246653.1;refGene_gn=UPK3BL2;refGene_tx=XM_023246654.1
E3 9210355 9210515
refGene_gn=UPK3B;refGene_tx=XM_003998537.3;refGene_gn=UPK3B;refGene_tx=XM_019820557.1
E3 9210969 9211099
refGene_gn=UPK3B;refGene_tx=XM_003998537.3;refGene_gn=UPK3B;refGene_tx=XM_019820557.1
E3 9211953 9212033
refGene_gn=UPK3B;refGene_tx=XM_003998537.3;refGene_gn=UPK3B;refGene_tx=XM_019820557.1
E3 9213314 9213552
refGene_gn=UPK3B;refGene_tx=XM_003998537.3;refGene_gn=UPK3B;refGene_tx=XM_019820557.1
E3 9213771 9213921
refGene_gn=UPK3B;refGene_tx=XM_003998537.3;refGene_gn=UPK3B;refGene_tx=XM_019820557.1
E3 9214038 9214126 refGene_gn=UPK3B;refGene_tx=XM_003998537.3
E3 9214997 9215010 refGene_gn=UPK3B;refGene_tx=XM_019820557.1

E3 9216571 9216799
refGene_gn=DTX2;refGene_tx=XM_006942021.3;refGene_gn=DTX2;refGene_tx=XM_006942022.4;refGene_gn=DTX2;refGene_tx=XM_006942023.3
E3 9217947 9218037
refGene_gn=DTX2;refGene_tx=XM_006942021.3;refGene_gn=DTX2;refGene_tx=XM_006942022.4;refGene_gn=DTX2;refGene_tx=XM_006942023.3
E3 9219282 9219444
refGene_gn=DTX2;refGene_tx=XM_006942021.3;refGene_gn=DTX2;refGene_tx=XM_006942022.4;refGene_gn=DTX2;refGene_tx=XM_006942023.3
E3 9220114 9220273
refGene_gn=DTX2;refGene_tx=XM_006942021.3;refGene_gn=DTX2;refGene_tx=XM_006942022.4;refGene_gn=DTX2;refGene_tx=XM_006942023.3
E3 9221619 9221693
refGene_gn=DTX2;refGene_tx=XM_006942021.3;refGene_gn=DTX2;refGene_tx=XM_006942022.4;refGene_gn=DTX2;refGene_tx=XM_006942023.3
E3 9224759 9224900
refGene_gn=DTX2;refGene_tx=XM_006942021.3;refGene_gn=DTX2;refGene_tx=XM_006942022.4
E3 9229599 9229700
refGene_gn=DTX2;refGene_tx=XM_006942021.3;refGene_gn=DTX2;refGene_tx=XM_006942022.4;refGene_gn=DTX2;refGene_tx=XM_006942023.3
E3 9235535 9236172
refGene_gn=DTX2;refGene_tx=XM_006942021.3;refGene_gn=DTX2;refGene_tx=XM_006942022.4;refGene_gn=DTX2;refGene_tx=XM_006942023.3
E3 9238733 9239001
refGene_gn=DTX2;refGene_tx=XM_006942021.3;refGene_gn=DTX2;refGene_tx=XM_006942022.4;refGene_gn=DTX2;refGene_tx=XM_006942023.3
E3 9260243 9260464 refGene_gn=ZP3;refGene_tx=NM_001009330.2
E3 9260784 9260921 refGene_gn=ZP3;refGene_tx=NM_001009330.2
E3 9261047 9261139 refGene_gn=ZP3;refGene_tx=NM_001009330.2
E3 9262672 9262790 refGene_gn=ZP3;refGene_tx=NM_001009330.2
E3 9262881 9263059 refGene_gn=ZP3;refGene_tx=NM_001009330.2
E3 9263135 9263239 refGene_gn=ZP3;refGene_tx=NM_001009330.2
E3 9265014 9265133 refGene_gn=ZP3;refGene_tx=NM_001009330.2
E3 9266983 9267289 refGene_gn=ZP3;refGene_tx=NM_001009330.2
E3 9283629 9283795 refGene_gn=SSC4D;refGene_tx=XM_023246643.1
E3 9285345 9285381 refGene_gn=SSC4D;refGene_tx=XM_023246643.1
E3 9285651 9285957 refGene_gn=SSC4D;refGene_tx=XM_023246643.1
E3 9286664 9286742 refGene_gn=SSC4D;refGene_tx=XM_023246643.1
E3 9288193 9288508 refGene_gn=SSC4D;refGene_tx=XM_023246643.1
E3 9289617 9289695 refGene_gn=SSC4D;refGene_tx=XM_023246643.1
E3 9290266 9290341 refGene_gn=SSC4D;refGene_tx=XM_023246643.1
E3 9290456 9290768 refGene_gn=SSC4D;refGene_tx=XM_023246643.1
E3 9291602 9291680 refGene_gn=SSC4D;refGene_tx=XM_023246643.1
E3 9292302 9292619 refGene_gn=SSC4D;refGene_tx=XM_023246643.1
E3 9314876 9314963 refGene_gn=YWHAG;refGene_tx=XM_003998539.5
E3 9339038 9339695 refGene_gn=YWHAG;refGene_tx=XM_003998539.5
E3 9356467 9356657 refGene_gn=HSPB1;refGene_tx=XM_003998540.5
E3 9356762 9356826 refGene_gn=HSPB1;refGene_tx=XM_003998540.5
E3 9357346 9357710 refGene_gn=HSPB1;refGene_tx=XM_003998540.5

E3 9364399 9365140 refGene_gn=SRRM3;refGene_tx=XM_023247143.1
E3 9366398 9366437 refGene_gn=SRRM3;refGene_tx=XM_023247143.1
E3 9366438 9366478 refGene_gn=SRRM3;refGene_tx=XM_023247143.1
E3 9366823 9366961 refGene_gn=SRRM3;refGene_tx=XM_023247143.1
E3 9367074 9367299 refGene_gn=SRRM3;refGene_tx=XM_023247143.1
E3 9368164 9368532 refGene_gn=SRRM3;refGene_tx=XM_023247143.1
E3 9378541 9378719 refGene_gn=SRRM3;refGene_tx=XM_023247143.1
E3 9379071 9379176 refGene_gn=SRRM3;refGene_tx=XM_023247143.1
E3 9379643 9379694 refGene_gn=SRRM3;refGene_tx=XM_023247143.1
E3 9381498 9381534 refGene_gn=SRRM3;refGene_tx=XM_023247143.1
E3 9381653 9381716 refGene_gn=SRRM3;refGene_tx=XM_023247143.1
E3 9382165 9382195 refGene_gn=SRRM3;refGene_tx=XM_023247143.1
E3 9382695 9382776 refGene_gn=SRRM3;refGene_tx=XM_023247143.1
E3 9382875 9383004 refGene_gn=SRRM3;refGene_tx=XM_023247143.1
E3 9391892 9391994 refGene_gn=SRRM3;refGene_tx=XM_023247143.1
E3 9398382 9398615 refGene_gn=SRRM3;refGene_tx=XM_023247143.1
E3 9440236 9440368 refGene_gn=MDH2;refGene_tx=NM_001278853.1
E3 9442060 9442212 refGene_gn=MDH2;refGene_tx=NM_001278853.1
E3 9442569 9442669 refGene_gn=MDH2;refGene_tx=NM_001278853.1
E3 9443785 9443863 refGene_gn=MDH2;refGene_tx=NM_001278853.1
E3 9446506 9446632 refGene_gn=MDH2;refGene_tx=NM_001278853.1
E3 9448136 9448246 refGene_gn=MDH2;refGene_tx=NM_001278853.1
E3 9448756 9448840 refGene_gn=MDH2;refGene_tx=NM_001278853.1
E3 9449472 9449489
refGene_gn=STYXL1;refGene_tx=XM_023246749.1;refGene_gn=STYXL1;refGene_tx=XM_023246750.1
E3 9450624 9450793 refGene_gn=MDH2;refGene_tx=NM_001278853.1
E3 9455657 9455723 refGene_gn=MDH2;refGene_tx=NM_001278853.1
E3 9468210 9468317
refGene_gn=STYXL1;refGene_tx=XM_003998542.5;refGene_gn=STYXL1;refGene_tx=XM_006942026.4;refGene_gn=STYXL1;refGene_tx=XM_006942027.4;refGene_gn=STYXL1;refGene_tx=XM_023246749.1;refGene_gn=STYXL1;refGene_tx=XM_023246750.1
E3 9484887 9484949
refGene_gn=STYXL1;refGene_tx=XM_003998542.5;refGene_gn=STYXL1;refGene_tx=XM_006942026.4;refGene_gn=STYXL1;refGene_tx=XM_006942027.4;refGene_gn=STYXL1;refGene_tx=XM_023246749.1;refGene_gn=STYXL1;refGene_tx=XM_023246750.1
E3 9488442 9488578
refGene_gn=STYXL1;refGene_tx=XM_003998542.5;refGene_gn=STYXL1;refGene_tx=XM_006942026.4;refGene_gn=STYXL1;refGene_tx=XM_006942027.4;refGene_gn=STYXL1;refGene_tx=XM_023246749.1;refGene_gn=STYXL1;refGene_tx=XM_023246750.1
E3 9493709 9493855
refGene_gn=STYXL1;refGene_tx=XM_003998542.5;refGene_gn=STYXL1;refGene_tx=XM_006942026.4;refGene_gn=STYXL1;refGene_tx=XM_006942027.4;refGene_gn=STYXL1;refGene_tx=XM_023246749.1;refGene_gn=STYXL1;refGene_tx=XM_023246750.1
E3 9502228 9502374
refGene_gn=STYXL1;refGene_tx=XM_003998542.5;refGene_gn=STYXL1;refGene_

tx=XM_006942026.4;refGene_gn=STYXL1;refGene_tx=XM_006942027.4;refGene_gn=STYXL1;refGene_tx=XM_023246749.1;refGene_gn=STYXL1;refGene_tx=XM_023246750.1
E3 9510443 9510541 refGene_gn=STYXL1;refGene_tx=XM_003998542.5;refGene_gn=STYXL1;refGene_tx=XM_006942026.4;refGene_gn=STYXL1;refGene_tx=XM_006942027.4;refGene_gn=STYXL1;refGene_tx=XM_023246749.1;refGene_gn=STYXL1;refGene_tx=XM_023246750.1
E3 9512216 9512329 refGene_gn=STYXL1;refGene_tx=XM_003998542.5;refGene_gn=STYXL1;refGene_tx=XM_006942026.4;refGene_gn=STYXL1;refGene_tx=XM_006942027.4;refGene_gn=STYXL1;refGene_tx=XM_023246749.1;refGene_gn=STYXL1;refGene_tx=XM_023246750.1
E3 9524612 9524630 refGene_gn=STYXL1;refGene_tx=XM_023246750.1
E3 9528556 9528688 refGene_gn=STYXL1;refGene_tx=XM_003998542.5;refGene_gn=STYXL1;refGene_tx=XM_006942026.4;refGene_gn=STYXL1;refGene_tx=XM_006942027.4;refGene_gn=STYXL1;refGene_tx=XM_023246749.1
E3 9530332 9530413 refGene_gn=TMEM120A;refGene_tx=XM_023246748.1
E3 9532073 9532192 refGene_gn=TMEM120A;refGene_tx=XM_023246748.1
E3 9532426 9532543 refGene_gn=TMEM120A;refGene_tx=XM_023246748.1
E3 9534161 9534221 refGene_gn=TMEM120A;refGene_tx=XM_023246748.1
E3 9534464 9534560 refGene_gn=TMEM120A;refGene_tx=XM_023246748.1
E3 9534634 9534724 refGene_gn=TMEM120A;refGene_tx=XM_023246748.1
E3 9534909 9534975 refGene_gn=TMEM120A;refGene_tx=XM_023246748.1
E3 9535085 9535147 refGene_gn=TMEM120A;refGene_tx=XM_023246748.1
E3 9535248 9535341 refGene_gn=TMEM120A;refGene_tx=XM_023246748.1
E3 9535462 9535527 refGene_gn=TMEM120A;refGene_tx=XM_023246748.1
E3 9535633 9535702 refGene_gn=TMEM120A;refGene_tx=XM_023246748.1
E3 9535795 9535909 refGene_gn=TMEM120A;refGene_tx=XM_023246748.1
E3 9536633 9536777 refGene_gn=LOC101085768;refGene_tx=XM_023246747.1
E3 9536873 9536960 refGene_gn=LOC101085768;refGene_tx=XM_023246747.1
E3 9537058 9537157 refGene_gn=LOC101085768;refGene_tx=XM_023246747.1
E3 9537159 9537206 refGene_gn=LOC101085768;refGene_tx=XM_023246747.1
E3 9537285 9537556 refGene_gn=LOC101085768;refGene_tx=XM_023246747.1
E3 9537843 9537993 refGene_gn=LOC101085768;refGene_tx=XM_023246747.1
E3 9538105 9538287 refGene_gn=LOC101085768;refGene_tx=XM_023246747.1
E3 9538506 9538625 refGene_gn=LOC101085768;refGene_tx=XM_023246747.1
E3 9538728 9538845 refGene_gn=LOC101085768;refGene_tx=XM_023246747.1
E3 9539784 9539883 refGene_gn=LOC101085768;refGene_tx=XM_023246747.1

E3 9540143 9540233
refGene_gn=LOC101085768;refGene_tx=XM_023246747.1
E3 9540540 9540665
refGene_gn=LOC101085768;refGene_tx=XM_023246747.1
E3 9541191 9541341
refGene_gn=LOC101085768;refGene_tx=XM_023246747.1
E3 9542682 9542811
refGene_gn=LOC101085768;refGene_tx=XM_023246747.1
E3 9546895 9546944
refGene_gn=LOC101085768;refGene_tx=XM_023246747.1
E3 9564982 9565161
refGene_gn=LOC101085768;refGene_tx=XM_023246747.1
E3 9624502 9624860 refGene_gn=RHBDD2;refGene_tx=XM_003998546.3
E3 9627612 9627763 refGene_gn=RHBDD2;refGene_tx=XM_003998546.3
E3 9634979 9635387 refGene_gn=RHBDD2;refGene_tx=XM_003998546.3
E3 9637425 9637612 refGene_gn=RHBDD2;refGene_tx=XM_003998546.3
E3 9691731 9691804 refGene_gn=CCL24;refGene_tx=XM_003998547.4
E3 9692004 9692122 refGene_gn=CCL24;refGene_tx=XM_003998547.4
E3 9693106 9693275 refGene_gn=CCL24;refGene_tx=XM_003998547.4
E3 9705981 9706054 refGene_gn=CCL26;refGene_tx=XM_023246860.1
E3 9706142 9706257 refGene_gn=CCL26;refGene_tx=XM_023246860.1
E3 9708529 9708626 refGene_gn=CCL26;refGene_tx=XM_023246860.1
E3 9755855 9755975 refGene_gn=HIP1;refGene_tx=XM_023246656.1
E3 9825825 9825843 refGene_gn=HIP1;refGene_tx=XM_019820566.2
E3 9861263 9861327
refGene_gn=HIP1;refGene_tx=XM_019820566.2;refGene_gn=HIP1;refGene_tx=X
M_023246656.1
E3 9863930 9864073
refGene_gn=HIP1;refGene_tx=XM_019820566.2;refGene_gn=HIP1;refGene_tx=X
M_023246656.1
E3 9864307 9864364
refGene_gn=HIP1;refGene_tx=XM_019820566.2;refGene_gn=HIP1;refGene_tx=X
M_023246656.1
E3 9872192 9872273
refGene_gn=HIP1;refGene_tx=XM_019820566.2;refGene_gn=HIP1;refGene_tx=X
M_023246656.1
E3 9874546 9874623
refGene_gn=HIP1;refGene_tx=XM_019820566.2;refGene_gn=HIP1;refGene_tx=X
M_023246656.1
E3 9875101 9875163
refGene_gn=HIP1;refGene_tx=XM_019820566.2;refGene_gn=HIP1;refGene_tx=X
M_023246656.1
E3 9879939 9880080
refGene_gn=HIP1;refGene_tx=XM_019820566.2;refGene_gn=HIP1;refGene_tx=X
M_023246656.1
E3 9882462 9882520
refGene_gn=HIP1;refGene_tx=XM_019820566.2;refGene_gn=HIP1;refGene_tx=X
M_023246656.1
E3 9885851 9885927
refGene_gn=HIP1;refGene_tx=XM_019820566.2;refGene_gn=HIP1;refGene_tx=X

M_023246656.1
E3	9886036	9886177
refGene_gn=HIP1;refGene_tx=XM_019820566.2;refGene_gn=HIP1;refGene_tx=X
M_023246656.1
E3	9886511	9886609
refGene_gn=HIP1;refGene_tx=XM_019820566.2;refGene_gn=HIP1;refGene_tx=X
M_023246656.1
E3	9886969	9887042
refGene_gn=HIP1;refGene_tx=XM_019820566.2;refGene_gn=HIP1;refGene_tx=X
M_023246656.1
E3	9887697	9887881
refGene_gn=HIP1;refGene_tx=XM_019820566.2;refGene_gn=HIP1;refGene_tx=X
M_023246656.1
E3	9888648	9888737
refGene_gn=HIP1;refGene_tx=XM_019820566.2;refGene_gn=HIP1;refGene_tx=X
M_023246656.1
E3	9889105	9889222
refGene_gn=HIP1;refGene_tx=XM_019820566.2;refGene_gn=HIP1;refGene_tx=X
M_023246656.1
E3	9890749	9890851
refGene_gn=HIP1;refGene_tx=XM_019820566.2;refGene_gn=HIP1;refGene_tx=X
M_023246656.1
E3	9891096	9891240
refGene_gn=HIP1;refGene_tx=XM_019820566.2;refGene_gn=HIP1;refGene_tx=X
M_023246656.1
E3	9891705	9891841
refGene_gn=HIP1;refGene_tx=XM_019820566.2;refGene_gn=HIP1;refGene_tx=X
M_023246656.1
E3	9893597	9893684
refGene_gn=HIP1;refGene_tx=XM_019820566.2;refGene_gn=HIP1;refGene_tx=X
M_023246656.1
E3	9893875	9893983
refGene_gn=HIP1;refGene_tx=XM_019820566.2;refGene_gn=HIP1;refGene_tx=X
M_023246656.1
E3	9894065	9894199
refGene_gn=HIP1;refGene_tx=XM_019820566.2;refGene_gn=HIP1;refGene_tx=X
M_023246656.1
E3	9899374	9899485
refGene_gn=HIP1;refGene_tx=XM_019820566.2;refGene_gn=HIP1;refGene_tx=X
M_023246656.1
E3	9900259	9900318
refGene_gn=HIP1;refGene_tx=XM_019820566.2;refGene_gn=HIP1;refGene_tx=X
M_023246656.1
E3	9900720	9900814
refGene_gn=HIP1;refGene_tx=XM_019820566.2;refGene_gn=HIP1;refGene_tx=X
M_023246656.1
E3	9902243	9902344
refGene_gn=HIP1;refGene_tx=XM_019820566.2;refGene_gn=HIP1;refGene_tx=X
M_023246656.1
E3	9902866	9902972

refGene_gn=HIP1;refGene_tx=XM_019820566.2;refGene_gn=HIP1;refGene_tx=XM_023246656.1
E3	9903687	9903811
refGene_gn=HIP1;refGene_tx=XM_019820566.2;refGene_gn=HIP1;refGene_tx=XM_023246656.1
E3	9904274	9904336
refGene_gn=HIP1;refGene_tx=XM_019820566.2;refGene_gn=HIP1;refGene_tx=XM_023246656.1
E3	9907612	9907721
refGene_gn=HIP1;refGene_tx=XM_019820566.2;refGene_gn=HIP1;refGene_tx=XM_023246656.1
E3	9908458	9908511
refGene_gn=HIP1;refGene_tx=XM_019820566.2;refGene_gn=HIP1;refGene_tx=XM_023246656.1
E3	9948722	9948919	refGene_gn=POM121C;refGene_tx=XM_023246875.1
E3	9949842	9950058
refGene_gn=POM121C;refGene_tx=XM_023246875.1;refGene_gn=POM121C;refGene_tx=XM_023246876.1;refGene_gn=POM121C;refGene_tx=XM_023246878.1;refGene_gn=POM121C;refGene_tx=XM_023246879.1
E3	9950590	9950749
refGene_gn=POM121C;refGene_tx=XM_023246875.1;refGene_gn=POM121C;refGene_tx=XM_023246876.1;refGene_gn=POM121C;refGene_tx=XM_023246878.1;refGene_gn=POM121C;refGene_tx=XM_023246879.1
E3	9952401	9952482
refGene_gn=POM121C;refGene_tx=XM_023246875.1;refGene_gn=POM121C;refGene_tx=XM_023246876.1;refGene_gn=POM121C;refGene_tx=XM_023246878.1;refGene_gn=POM121C;refGene_tx=XM_023246879.1
E3	9953999	9954171
refGene_gn=POM121C;refGene_tx=XM_023246875.1;refGene_gn=POM121C;refGene_tx=XM_023246876.1;refGene_gn=POM121C;refGene_tx=XM_023246878.1;refGene_gn=POM121C;refGene_tx=XM_023246879.1
E3	9966022	9966114
refGene_gn=POM121C;refGene_tx=XM_023246875.1;refGene_gn=POM121C;refGene_tx=XM_023246876.1;refGene_gn=POM121C;refGene_tx=XM_023246878.1;refGene_gn=POM121C;refGene_tx=XM_023246879.1
E3	9966755	9966829
refGene_gn=POM121C;refGene_tx=XM_023246875.1;refGene_gn=POM121C;refGene_tx=XM_023246876.1;refGene_gn=POM121C;refGene_tx=XM_023246878.1;refGene_gn=POM121C;refGene_tx=XM_023246879.1
E3	9967265	9967390
refGene_gn=POM121C;refGene_tx=XM_023246875.1;refGene_gn=POM121C;refGene_tx=XM_023246876.1;refGene_gn=POM121C;refGene_tx=XM_023246878.1;refGene_gn=POM121C;refGene_tx=XM_023246879.1
E3	9967507	9967610
refGene_gn=POM121C;refGene_tx=XM_023246875.1;refGene_gn=POM121C;refGene_tx=XM_023246876.1;refGene_gn=POM121C;refGene_tx=XM_023246878.1;refGene_gn=POM121C;refGene_tx=XM_023246879.1
E3	9968041	9968218
refGene_gn=POM121C;refGene_tx=XM_023246875.1;refGene_gn=POM121C;refGene_tx=XM_023246876.1;refGene_gn=POM121C;refGene_tx=XM_023246878.1;refGe

ne_gn=POM121C;refGene_tx=XM_023246879.1
E3	9969062	9970676
refGene_gn=POM121C;refGene_tx=XM_023246875.1;refGene_gn=POM121C;refGene_tx=XM_023246876.1;refGene_gn=POM121C;refGene_tx=XM_023246878.1;refGene_gn=POM121C;refGene_tx=XM_023246879.1
E3	9971976	9972099
refGene_gn=POM121C;refGene_tx=XM_023246875.1;refGene_gn=POM121C;refGene_tx=XM_023246876.1;refGene_gn=POM121C;refGene_tx=XM_023246878.1;refGene_gn=POM121C;refGene_tx=XM_023246879.1
E3	9972539	9972637
refGene_gn=POM121C;refGene_tx=XM_023246875.1;refGene_gn=POM121C;refGene_tx=XM_023246876.1;refGene_gn=POM121C;refGene_tx=XM_023246878.1;refGene_gn=POM121C;refGene_tx=XM_023246879.1
E3	9974916	9974931
refGene_gn=NSUN5;refGene_tx=XM_003998549.4;refGene_gn=NSUN5;refGene_tx=XM_006942030.3;refGene_gn=NSUN5;refGene_tx=XM_006942031.4;refGene_gn=NSUN5;refGene_tx=XM_006942032.4
E3	9975021	9975127
refGene_gn=NSUN5;refGene_tx=XM_003998549.4;refGene_gn=NSUN5;refGene_tx=XM_006942030.3;refGene_gn=NSUN5;refGene_tx=XM_006942031.4;refGene_gn=NSUN5;refGene_tx=XM_006942032.4
E3	9975218	9975359
refGene_gn=NSUN5;refGene_tx=XM_003998549.4;refGene_gn=NSUN5;refGene_tx=XM_006942030.3;refGene_gn=NSUN5;refGene_tx=XM_006942031.4;refGene_gn=NSUN5;refGene_tx=XM_006942032.4;refGene_gn=NSUN5;refGene_tx=XM_006942033.3
E3	9975432	9975643
refGene_gn=NSUN5;refGene_tx=XM_003998549.4;refGene_gn=NSUN5;refGene_tx=XM_006942030.3;refGene_gn=NSUN5;refGene_tx=XM_006942031.4;refGene_gn=NSUN5;refGene_tx=XM_006942032.4;refGene_gn=NSUN5;refGene_tx=XM_006942033.3
E3	9975826	9976005
refGene_gn=NSUN5;refGene_tx=XM_003998549.4;refGene_gn=NSUN5;refGene_tx=XM_006942030.3;refGene_gn=NSUN5;refGene_tx=XM_006942031.4;refGene_gn=NSUN5;refGene_tx=XM_006942032.4;refGene_gn=NSUN5;refGene_tx=XM_006942033.3
E3	9976360	9976500
refGene_gn=NSUN5;refGene_tx=XM_003998549.4;refGene_gn=NSUN5;refGene_tx=XM_006942030.3;refGene_gn=NSUN5;refGene_tx=XM_006942031.4;refGene_gn=NSUN5;refGene_tx=XM_006942032.4;refGene_gn=NSUN5;refGene_tx=XM_006942033.3
E3	9976558	9976697
refGene_gn=NSUN5;refGene_tx=XM_003998549.4;refGene_gn=NSUN5;refGene_tx=XM_006942030.3;refGene_gn=NSUN5;refGene_tx=XM_006942031.4;refGene_gn=NSUN5;refGene_tx=XM_006942032.4;refGene_gn=NSUN5;refGene_tx=XM_006942033.3
E3	9986737	9986846
refGene_gn=NSUN5;refGene_tx=XM_003998549.4;refGene_gn=NSUN5;refGene_tx=XM_006942030.3;refGene_gn=NSUN5;refGene_tx=XM_006942031.4;refGene_gn=NSUN5;refGene_tx=XM_006942032.4;refGene_gn=NSUN5;refGene_tx=XM_0069420

33.3
E3 9986947 9987125
refGene_gn=NSUN5;refGene_tx=XM_003998549.4;refGene_gn=NSUN5;refGene_tx=XM_006942030.3;refGene_gn=NSUN5;refGene_tx=XM_006942031.4;refGene_gn=NSUN5;refGene_tx=XM_006942032.4;refGene_gn=NSUN5;refGene_tx=XM_006942033.3
E3 9987599 9987722
refGene_gn=NSUN5;refGene_tx=XM_003998549.4;refGene_gn=NSUN5;refGene_tx=XM_006942030.3;refGene_gn=NSUN5;refGene_tx=XM_006942031.4;refGene_gn=NSUN5;refGene_tx=XM_006942032.4;refGene_gn=NSUN5;refGene_tx=XM_006942033.3
E3 9987816 9987909
refGene_gn=NSUN5;refGene_tx=XM_003998549.4;refGene_gn=NSUN5;refGene_tx=XM_006942030.3;refGene_gn=NSUN5;refGene_tx=XM_006942031.4;refGene_gn=NSUN5;refGene_tx=XM_006942032.4;refGene_gn=NSUN5;refGene_tx=XM_006942033.3
E3 9993672 9994262
refGene_gn=TRIM50;refGene_tx=XM_006942034.2;refGene_gn=TRIM50;refGene_tx=XM_019820569.2;refGene_gn=TRIM50;refGene_tx=XM_023246324.1
E3 9995181 9995306
refGene_gn=TRIM50;refGene_tx=XM_006942034.2;refGene_gn=TRIM50;refGene_tx=XM_019820569.2;refGene_gn=TRIM50;refGene_tx=XM_023246324.1
E3 9996405 9996428
refGene_gn=TRIM50;refGene_tx=XM_006942034.2;refGene_gn=TRIM50;refGene_tx=XM_019820569.2;refGene_gn=TRIM50;refGene_tx=XM_023246324.1
E3 9996518 9996749
refGene_gn=TRIM50;refGene_tx=XM_006942034.2;refGene_gn=TRIM50;refGene_tx=XM_019820569.2;refGene_gn=TRIM50;refGene_tx=XM_023246324.1
E3 9997691 9997787
refGene_gn=TRIM50;refGene_tx=XM_006942034.2;refGene_gn=TRIM50;refGene_tx=XM_019820569.2;refGene_gn=TRIM50;refGene_tx=XM_023246324.1
E3 10000855 10001254
refGene_gn=TRIM50;refGene_tx=XM_006942034.2;refGene_gn=TRIM50;refGene_tx=XM_019820569.2;refGene_gn=TRIM50;refGene_tx=XM_023246324.1
E3 10007508 10007634
refGene_gn=FKBP6;refGene_tx=XM_003998552.4;refGene_gn=FKBP6;refGene_tx=XM_006942035.3;refGene_gn=FKBP6;refGene_tx=XM_006942036.3;refGene_gn=FKBP6;refGene_tx=XM_006942037.3;refGene_gn=FKBP6;refGene_tx=XM_006942038.3
E3 10007727 10007845
refGene_gn=FKBP6;refGene_tx=XM_003998552.4;refGene_gn=FKBP6;refGene_tx=XM_006942035.3;refGene_gn=FKBP6;refGene_tx=XM_006942036.3;refGene_gn=FKBP6;refGene_tx=XM_006942037.3;refGene_gn=FKBP6;refGene_tx=XM_006942038.3
E3 10008268 10008358
refGene_gn=FKBP6;refGene_tx=XM_003998552.4;refGene_gn=FKBP6;refGene_tx=XM_006942035.3;refGene_gn=FKBP6;refGene_tx=XM_006942036.3;refGene_gn=FKBP6;refGene_tx=XM_006942037.3;refGene_gn=FKBP6;refGene_tx=XM_006942038.3
E3 10009096 10009299

refGene_gn=FKBP6;refGene_tx=XM_003998552.4;refGene_gn=FKBP6;refGene_tx=XM_006942035.3;refGene_gn=FKBP6;refGene_tx=XM_006942036.3;refGene_gn=FKBP6;refGene_tx=XM_006942037.3;refGene_gn=FKBP6;refGene_tx=XM_006942038.3
E3 10010522 10010642
refGene_gn=FKBP6;refGene_tx=XM_003998552.4;refGene_gn=FKBP6;refGene_tx=XM_006942035.3;refGene_gn=FKBP6;refGene_tx=XM_006942036.3;refGene_gn=FKBP6;refGene_tx=XM_006942037.3;refGene_gn=FKBP6;refGene_tx=XM_006942038.3
E3 10019135 10019330
refGene_gn=FKBP6;refGene_tx=XM_003998552.4;refGene_gn=FKBP6;refGene_tx=XM_006942035.3;refGene_gn=FKBP6;refGene_tx=XM_006942036.3;refGene_gn=FKBP6;refGene_tx=XM_006942037.3;refGene_gn=FKBP6;refGene_tx=XM_006942038.3
E3 10019457 10019567
refGene_gn=FKBP6;refGene_tx=XM_003998552.4;refGene_gn=FKBP6;refGene_tx=XM_006942035.3;refGene_gn=FKBP6;refGene_tx=XM_006942036.3;refGene_gn=FKBP6;refGene_tx=XM_006942037.3;refGene_gn=FKBP6;refGene_tx=XM_006942038.3
E3 10020898 10020989
refGene_gn=FKBP6;refGene_tx=XM_003998552.4;refGene_gn=FKBP6;refGene_tx=XM_006942035.3;refGene_gn=FKBP6;refGene_tx=XM_006942036.3;refGene_gn=FKBP6;refGene_tx=XM_006942037.3;refGene_gn=FKBP6;refGene_tx=XM_006942038.3
E3 10085431 10087207 refGene_gn=FZD9;refGene_tx=XM_003998553.5
E3 10093546 10093904
refGene_gn=BAZ1B;refGene_tx=XM_023246861.1;refGene_gn=BAZ1B;refGene_tx=XM_023246862.1
E3 10094230 10094334
refGene_gn=BAZ1B;refGene_tx=XM_023246861.1;refGene_gn=BAZ1B;refGene_tx=XM_023246862.1
E3 10095443 10095589
refGene_gn=BAZ1B;refGene_tx=XM_023246861.1;refGene_gn=BAZ1B;refGene_tx=XM_023246862.1
E3 10097858 10097974
refGene_gn=BAZ1B;refGene_tx=XM_023246861.1;refGene_gn=BAZ1B;refGene_tx=XM_023246862.1
E3 10100149 10100297
refGene_gn=BAZ1B;refGene_tx=XM_023246861.1;refGene_gn=BAZ1B;refGene_tx=XM_023246862.1
E3 10101325 10101473
refGene_gn=BAZ1B;refGene_tx=XM_023246861.1;refGene_gn=BAZ1B;refGene_tx=XM_023246862.1
E3 10109211 10109394
refGene_gn=BAZ1B;refGene_tx=XM_023246861.1;refGene_gn=BAZ1B;refGene_tx=XM_023246862.1
E3 10111326 10111504
refGene_gn=BAZ1B;refGene_tx=XM_023246861.1;refGene_gn=BAZ1B;refGene_tx=XM_023246862.1
E3 10114333 10114432

refGene_gn=BAZ1B;refGene_tx=XM_023246861.1;refGene_gn=BAZ1B;refGene_tx=XM_023246862.1
E3 10115388 10115494
refGene_gn=BAZ1B;refGene_tx=XM_023246861.1;refGene_gn=BAZ1B;refGene_tx=XM_023246862.1
E3 10117506 10117637
refGene_gn=BAZ1B;refGene_tx=XM_023246861.1;refGene_gn=BAZ1B;refGene_tx=XM_023246862.1
E3 10118127 10118266
refGene_gn=BAZ1B;refGene_tx=XM_023246861.1;refGene_gn=BAZ1B;refGene_tx=XM_023246862.1
E3 10124840 10126542
refGene_gn=BAZ1B;refGene_tx=XM_023246861.1;refGene_gn=BAZ1B;refGene_tx=XM_023246862.1
E3 10134532 10134730
refGene_gn=BAZ1B;refGene_tx=XM_023246861.1;refGene_gn=BAZ1B;refGene_tx=XM_023246862.1
E3 10137068 10137190
refGene_gn=BAZ1B;refGene_tx=XM_023246861.1;refGene_gn=BAZ1B;refGene_tx=XM_023246862.1
E3 10141335 10141562
refGene_gn=BAZ1B;refGene_tx=XM_023246861.1;refGene_gn=BAZ1B;refGene_tx=XM_023246862.1
E3 10150267 10150412 refGene_gn=BAZ1B;refGene_tx=XM_023246861.1
E3 10155583 10155700 refGene_gn=BAZ1B;refGene_tx=XM_023246861.1
E3 10163623 10163730 refGene_gn=BAZ1B;refGene_tx=XM_023246861.1
E3 10174848 10174941 refGene_gn=BCL7B;refGene_tx=XM_003998554.4
E3 10175205 10175285 refGene_gn=BCL7B;refGene_tx=XM_003998554.4
E3 10175870 10176041 refGene_gn=BCL7B;refGene_tx=XM_003998554.4
E3 10178240 10178337 refGene_gn=BCL7B;refGene_tx=XM_003998554.4
E3 10187428 10187504 refGene_gn=BCL7B;refGene_tx=XM_003998554.4
E3 10190843 10190935 refGene_gn=BCL7B;refGene_tx=XM_003998554.4
E3 10213146 10213609 refGene_gn=TBL2;refGene_tx=XM_003998572.5
E3 10213840 10213993 refGene_gn=TBL2;refGene_tx=XM_003998572.5
E3 10214280 10214407 refGene_gn=TBL2;refGene_tx=XM_003998572.5
E3 10214747 10214899 refGene_gn=TBL2;refGene_tx=XM_003998572.5
E3 10215402 10215587 refGene_gn=TBL2;refGene_tx=XM_003998572.5
E3 10215822 10215953 refGene_gn=TBL2;refGene_tx=XM_003998572.5
E3 10217665 10217708 refGene_gn=TBL2;refGene_tx=XM_003998572.5
E3 10217709 10217796 refGene_gn=TBL2;refGene_tx=XM_003998572.5
E3 10224175 10224294
refGene_gn=MLXIPL;refGene_tx=XM_011290353.3;refGene_gn=MLXIPL;refGene_tx=XM_011290354.3;refGene_gn=MLXIPL;refGene_tx=XM_011290355.3;refGene_gn=MLXIPL;refGene_tx=XM_011290356.2;refGene_gn=MLXIPL;refGene_tx=XM_019820575.2
E3 10224603 10224733
refGene_gn=MLXIPL;refGene_tx=XM_011290353.3;refGene_gn=MLXIPL;refGene_tx=XM_011290354.3;refGene_gn=MLXIPL;refGene_tx=XM_011290355.3;refGene_gn=MLXIPL;refGene_tx=XM_011290356.2;refGene_gn=MLXIPL;refGene_tx=XM_019820575.2

E3 10225307 10225431
refGene_gn=MLXIPL;refGene_tx=XM_011290353.3;refGene_gn=MLXIPL;refGene_tx=XM_011290354.3;refGene_gn=MLXIPL;refGene_tx=XM_011290355.3;refGene_gn=MLXIPL;refGene_tx=XM_011290356.2;refGene_gn=MLXIPL;refGene_tx=XM_019820575.2
E3 10225573 10225757
refGene_gn=MLXIPL;refGene_tx=XM_011290353.3;refGene_gn=MLXIPL;refGene_tx=XM_011290354.3;refGene_gn=MLXIPL;refGene_tx=XM_011290355.3;refGene_gn=MLXIPL;refGene_tx=XM_011290356.2;refGene_gn=MLXIPL;refGene_tx=XM_011290357.3;refGene_gn=MLXIPL;refGene_tx=XM_019820575.2
E3 10225865 10225985
refGene_gn=MLXIPL;refGene_tx=XM_011290353.3;refGene_gn=MLXIPL;refGene_tx=XM_011290354.3;refGene_gn=MLXIPL;refGene_tx=XM_011290355.3;refGene_gn=MLXIPL;refGene_tx=XM_011290356.2;refGene_gn=MLXIPL;refGene_tx=XM_011290357.3;refGene_gn=MLXIPL;refGene_tx=XM_019820575.2
E3 10226077 10226193
refGene_gn=MLXIPL;refGene_tx=XM_011290353.3;refGene_gn=MLXIPL;refGene_tx=XM_011290354.3;refGene_gn=MLXIPL;refGene_tx=XM_011290355.3;refGene_gn=MLXIPL;refGene_tx=XM_011290356.2;refGene_gn=MLXIPL;refGene_tx=XM_011290357.3;refGene_gn=MLXIPL;refGene_tx=XM_019820575.2
E3 10226356 10226504
refGene_gn=MLXIPL;refGene_tx=XM_011290353.3;refGene_gn=MLXIPL;refGene_tx=XM_011290354.3;refGene_gn=MLXIPL;refGene_tx=XM_011290355.3;refGene_gn=MLXIPL;refGene_tx=XM_011290356.2;refGene_gn=MLXIPL;refGene_tx=XM_011290357.3;refGene_gn=MLXIPL;refGene_tx=XM_019820575.2
E3 10226581 10226649
refGene_gn=MLXIPL;refGene_tx=XM_011290353.3;refGene_gn=MLXIPL;refGene_tx=XM_011290354.3;refGene_gn=MLXIPL;refGene_tx=XM_011290355.3;refGene_gn=MLXIPL;refGene_tx=XM_011290356.2;refGene_gn=MLXIPL;refGene_tx=XM_011290357.3;refGene_gn=MLXIPL;refGene_tx=XM_019820575.2
E3 10226876 10227423
refGene_gn=MLXIPL;refGene_tx=XM_011290353.3;refGene_gn=MLXIPL;refGene_tx=XM_011290354.3;refGene_gn=MLXIPL;refGene_tx=XM_011290355.3;refGene_gn=MLXIPL;refGene_tx=XM_011290356.2;refGene_gn=MLXIPL;refGene_tx=XM_011290357.3;refGene_gn=MLXIPL;refGene_tx=XM_019820575.2
E3 10228674 10228844
refGene_gn=MLXIPL;refGene_tx=XM_011290353.3;refGene_gn=MLXIPL;refGene_tx=XM_011290354.3;refGene_gn=MLXIPL;refGene_tx=XM_011290355.3;refGene_gn=MLXIPL;refGene_tx=XM_011290356.2;refGene_gn=MLXIPL;refGene_tx=XM_011290357.3;refGene_gn=MLXIPL;refGene_tx=XM_019820575.2
E3 10232402 10232483
refGene_gn=MLXIPL;refGene_tx=XM_011290353.3;refGene_gn=MLXIPL;refGene_tx=XM_011290354.3;refGene_gn=MLXIPL;refGene_tx=XM_011290355.3;refGene_gn=MLXIPL;refGene_tx=XM_011290356.2;refGene_gn=MLXIPL;refGene_tx=XM_011290357.3;refGene_gn=MLXIPL;refGene_tx=XM_019820575.2
E3 10232607 10232809
refGene_gn=MLXIPL;refGene_tx=XM_011290353.3;refGene_gn=MLXIPL;refGene_tx=XM_011290354.3;refGene_gn=MLXIPL;refGene_tx=XM_011290355.3;refGene_gn=MLXIPL;refGene_tx=XM_011290356.2;refGene_gn=MLXIPL;refGene_tx=XM_011290357.3;refGene_gn=MLXIPL;refGene_tx=XM_019820575.2

E3 10233257 10233302
refGene_gn=MLXIPL;refGene_tx=XM_011290353.3;refGene_gn=MLXIPL;refGene_tx=XM_011290354.3;refGene_gn=MLXIPL;refGene_tx=XM_011290356.2;refGene_gn=MLXIPL;refGene_tx=XM_011290357.3;refGene_gn=MLXIPL;refGene_tx=XM_019820575.2
E3 10233583 10233673
refGene_gn=MLXIPL;refGene_tx=XM_011290353.3;refGene_gn=MLXIPL;refGene_tx=XM_011290354.3;refGene_gn=MLXIPL;refGene_tx=XM_011290355.3;refGene_gn=MLXIPL;refGene_tx=XM_011290356.2;refGene_gn=MLXIPL;refGene_tx=XM_011290357.3
E3 10233827 10233910
refGene_gn=MLXIPL;refGene_tx=XM_011290353.3;refGene_gn=MLXIPL;refGene_tx=XM_011290354.3;refGene_gn=MLXIPL;refGene_tx=XM_011290355.3;refGene_gn=MLXIPL;refGene_tx=XM_011290356.2;refGene_gn=MLXIPL;refGene_tx=XM_011290357.3;refGene_gn=MLXIPL;refGene_tx=XM_019820575.2
E3 10237987 10238094
refGene_gn=MLXIPL;refGene_tx=XM_011290353.3;refGene_gn=MLXIPL;refGene_tx=XM_011290354.3;refGene_gn=MLXIPL;refGene_tx=XM_011290355.3;refGene_gn=MLXIPL;refGene_tx=XM_011290356.2;refGene_gn=MLXIPL;refGene_tx=XM_011290357.3;refGene_gn=MLXIPL;refGene_tx=XM_019820575.2
E3 10241819 10242112
refGene_gn=MLXIPL;refGene_tx=XM_011290353.3;refGene_gn=MLXIPL;refGene_tx=XM_011290354.3;refGene_gn=MLXIPL;refGene_tx=XM_011290355.3;refGene_gn=MLXIPL;refGene_tx=XM_011290357.3;refGene_gn=MLXIPL;refGene_tx=XM_019820575.2
E3 10255283 10255309 refGene_gn=MLXIPL;refGene_tx=XM_011290356.2
E3 10268689 10268827 refGene_gn=VPS37D;refGene_tx=XM_023246733.1
E3 10270168 10270340 refGene_gn=VPS37D;refGene_tx=XM_023246733.1
E3 10270810 10270893 refGene_gn=VPS37D;refGene_tx=XM_023246733.1
E3 10271485 10271848 refGene_gn=VPS37D;refGene_tx=XM_023246733.1
E3 10279457 10280138 refGene_gn=DNAJC30;refGene_tx=XM_023247031.1
E3 10280345 10280393
refGene_gn=BUD23;refGene_tx=XM_003998573.4;refGene_gn=BUD23;refGene_tx=XM_006942054.3;refGene_gn=BUD23;refGene_tx=XM_006942055.3;refGene_gn=BUD23;refGene_tx=XM_019820578.2
E3 10280490 10280528
refGene_gn=BUD23;refGene_tx=XM_003998573.4;refGene_gn=BUD23;refGene_tx=XM_006942054.3;refGene_gn=BUD23;refGene_tx=XM_006942055.3;refGene_gn=BUD23;refGene_tx=XM_019820578.2
E3 10281813 10281909
refGene_gn=BUD23;refGene_tx=XM_003998573.4;refGene_gn=BUD23;refGene_tx=XM_006942054.3;refGene_gn=BUD23;refGene_tx=XM_006942055.3;refGene_gn=BUD23;refGene_tx=XM_019820578.2
E3 10282001 10282084
refGene_gn=BUD23;refGene_tx=XM_003998573.4;refGene_gn=BUD23;refGene_tx=XM_006942054.3;refGene_gn=BUD23;refGene_tx=XM_006942055.3;refGene_gn=BUD23;refGene_tx=XM_019820578.2
E3 10282182 10282279
refGene_gn=BUD23;refGene_tx=XM_003998573.4;refGene_gn=BUD23;refGene_tx=XM_006942054.3;refGene_gn=BUD23;refGene_tx=XM_006942055.3;refGene_gn=

BUD23;refGene_tx=XM_019820578.2
E3 10284893 10284990
refGene_gn=BUD23;refGene_tx=XM_003998573.4;refGene_gn=BUD23;refGene_tx=XM_006942054.3;refGene_gn=BUD23;refGene_tx=XM_006942055.3;refGene_gn=BUD23;refGene_tx=XM_019820578.2
E3 10286291 10286342
refGene_gn=BUD23;refGene_tx=XM_003998573.4;refGene_gn=BUD23;refGene_tx=XM_006942054.3;refGene_gn=BUD23;refGene_tx=XM_006942055.3;refGene_gn=BUD23;refGene_tx=XM_019820578.2
E3 10287014 10287100
refGene_gn=BUD23;refGene_tx=XM_003998573.4;refGene_gn=BUD23;refGene_tx=XM_006942054.3;refGene_gn=BUD23;refGene_tx=XM_006942055.3;refGene_gn=BUD23;refGene_tx=XM_019820578.2
E3 10287294 10287340
refGene_gn=BUD23;refGene_tx=XM_003998573.4;refGene_gn=BUD23;refGene_tx=XM_006942054.3;refGene_gn=BUD23;refGene_tx=XM_006942055.3;refGene_gn=BUD23;refGene_tx=XM_019820578.2
E3 10287507 10287566
refGene_gn=BUD23;refGene_tx=XM_003998573.4;refGene_gn=BUD23;refGene_tx=XM_006942054.3;refGene_gn=BUD23;refGene_tx=XM_006942055.3;refGene_gn=BUD23;refGene_tx=XM_019820578.2
E3 10289281 10289366 refGene_gn=BUD23;refGene_tx=XM_019820578.2
E3 10291149 10291246
refGene_gn=BUD23;refGene_tx=XM_006942054.3;refGene_gn=BUD23;refGene_tx=XM_006942055.3
E3 10291895 10291985 refGene_gn=BUD23;refGene_tx=XM_003998573.4
E3 10292118 10292173 refGene_gn=BUD23;refGene_tx=XM_003998573.4
E3 10294450 10294528 refGene_gn=STX1A;refGene_tx=XM_023247028.1
E3 10294779 10294890 refGene_gn=STX1A;refGene_tx=XM_023247028.1
E3 10294907 10294955 refGene_gn=STX1A;refGene_tx=XM_023247029.1
E3 10296333 10296660 refGene_gn=STX1A;refGene_tx=XM_023247027.1
E3 10296814 10296952
refGene_gn=STX1A;refGene_tx=XM_023247027.1;refGene_gn=STX1A;refGene_tx=XM_023247028.1;refGene_gn=STX1A;refGene_tx=XM_023247029.1
E3 10297719 10297793
refGene_gn=STX1A;refGene_tx=XM_023247027.1;refGene_gn=STX1A;refGene_tx=XM_023247028.1;refGene_gn=STX1A;refGene_tx=XM_023247029.1
E3 10297894 10298003
refGene_gn=STX1A;refGene_tx=XM_023247027.1;refGene_gn=STX1A;refGene_tx=XM_023247028.1;refGene_gn=STX1A;refGene_tx=XM_023247029.1
E3 10298102 10298176
refGene_gn=STX1A;refGene_tx=XM_023247027.1;refGene_gn=STX1A;refGene_tx=XM_023247028.1;refGene_gn=STX1A;refGene_tx=XM_023247029.1
E3 10298897 10298972
refGene_gn=STX1A;refGene_tx=XM_023247027.1;refGene_gn=STX1A;refGene_tx=XM_023247028.1;refGene_gn=STX1A;refGene_tx=XM_023247029.1
E3 10301660 10301760
refGene_gn=STX1A;refGene_tx=XM_023247027.1;refGene_gn=STX1A;refGene_tx=XM_023247028.1;refGene_gn=STX1A;refGene_tx=XM_023247029.1
E3 10302134 10302212

refGene_gn=STX1A;refGene_tx=XM_023247027.1;refGene_gn=STX1A;refGene_tx=XM_023247028.1;refGene_gn=STX1A;refGene_tx=XM_023247029.1
E3 10310415 10310445
refGene_gn=STX1A;refGene_tx=XM_023247027.1;refGene_gn=STX1A;refGene_tx=XM_023247028.1;refGene_gn=STX1A;refGene_tx=XM_023247029.1
E3 10322334 10322467
refGene_gn=ABHD11;refGene_tx=XM_003998590.5;refGene_gn=ABHD11;refGene_tx=XM_019820580.1;refGene_gn=ABHD11;refGene_tx=XM_019820581.1
E3 10322678 10322860
refGene_gn=ABHD11;refGene_tx=XM_003998590.5;refGene_gn=ABHD11;refGene_tx=XM_019820580.1;refGene_gn=ABHD11;refGene_tx=XM_019820581.1
E3 10322980 10323151
refGene_gn=ABHD11;refGene_tx=XM_003998590.5;refGene_gn=ABHD11;refGene_tx=XM_019820580.1;refGene_gn=ABHD11;refGene_tx=XM_019820581.1
E3 10323323 10323497
refGene_gn=ABHD11;refGene_tx=XM_003998590.5;refGene_gn=ABHD11;refGene_tx=XM_019820580.1;refGene_gn=ABHD11;refGene_tx=XM_019820581.1
E3 10324012 10324148 refGene_gn=ABHD11;refGene_tx=XM_003998590.5
E3 10324302 10324421 refGene_gn=ABHD11;refGene_tx=XM_003998590.5
E3 10343297 10343954 refGene_gn=CLDN3;refGene_tx=XM_011290351.3
E3 10382244 10382874 refGene_gn=CLDN4;refGene_tx=XM_006942058.2
E3 10384707 10384834 refGene_gn=METTL27;refGene_tx=XM_003998575.2
E3 10384835 10384956 refGene_gn=METTL27;refGene_tx=XM_003998575.2
E3 10386764 10386857 refGene_gn=METTL27;refGene_tx=XM_003998575.2
E3 10387093 10387229 refGene_gn=METTL27;refGene_tx=XM_003998575.2
E3 10387442 10387571 refGene_gn=METTL27;refGene_tx=XM_003998575.2
E3 10387991 10388114 refGene_gn=METTL27;refGene_tx=XM_003998575.2
E3 10400688 10400760 refGene_gn=TMEM270;refGene_tx=XM_023246605.1
E3 10401718 10402114 refGene_gn=TMEM270;refGene_tx=XM_023246605.1
E3 10402216 10402456 refGene_gn=TMEM270;refGene_tx=XM_023246605.1
E3 10484157 10484293
refGene_gn=LOC105261297;refGene_tx=XM_023247078.1
E3 10484388 10484653
refGene_gn=LOC105261297;refGene_tx=XM_023247078.1
E3 10486294 10486350
refGene_gn=LOC105261297;refGene_tx=XM_023247078.1
E3 10486860 10487075
refGene_gn=LOC105261297;refGene_tx=XM_023247078.1
E3 10487403 10487511
refGene_gn=LOC105261297;refGene_tx=XM_023247078.1
E3 10492942 10493019
refGene_gn=LOC105261297;refGene_tx=XM_023247078.1
E3 10493411 10493505
refGene_gn=LOC105261297;refGene_tx=XM_023247078.1
E3 10513940 10514099
refGene_gn=LOC105261297;refGene_tx=XM_023247078.1
E3 10541903 10541985
refGene_gn=ELN;refGene_tx=XM_006942060.4;refGene_gn=ELN;refGene_tx=XM_019820582.2;refGene_gn=ELN;refGene_tx=XM_019820583.2;refGene_gn=ELN;refGene_tx=XM_019820584.2;refGene_gn=ELN;refGene_tx=XM_019820585.2;refGe

ne_gn=ELN;refGene_tx=XM_019820586.2;refGene_gn=ELN;refGene_tx=XM_01982
0587.2;refGene_gn=ELN;refGene_tx=XM_019820588.2;refGene_gn=ELN;refGene
_tx=XM_019820589.2;refGene_gn=ELN;refGene_tx=XM_019820590.2;refGene_gn
=ELN;refGene_tx=XM_019820591.2;refGene_gn=ELN;refGene_tx=XM_019820592.
2;refGene_gn=ELN;refGene_tx=XM_019820593.2;refGene_gn=ELN;refGene_tx=X
M_019820594.2
E3 10547661 10547712
refGene_gn=ELN;refGene_tx=XM_006942060.4;refGene_gn=ELN;refGene_tx=XM_
019820582.2;refGene_gn=ELN;refGene_tx=XM_019820583.2;refGene_gn=ELN;re
fGene_tx=XM_019820584.2;refGene_gn=ELN;refGene_tx=XM_019820585.2;refGe
ne_gn=ELN;refGene_tx=XM_019820586.2;refGene_gn=ELN;refGene_tx=XM_01982
0587.2;refGene_gn=ELN;refGene_tx=XM_019820588.2;refGene_gn=ELN;refGene
_tx=XM_019820589.2;refGene_gn=ELN;refGene_tx=XM_019820590.2;refGene_gn
=ELN;refGene_tx=XM_019820591.2;refGene_gn=ELN;refGene_tx=XM_019820592.
2;refGene_gn=ELN;refGene_tx=XM_019820593.2;refGene_gn=ELN;refGene_tx=X
M_019820594.2
E3 10548827 10548857
refGene_gn=ELN;refGene_tx=XM_006942060.4;refGene_gn=ELN;refGene_tx=XM_
019820582.2;refGene_gn=ELN;refGene_tx=XM_019820583.2;refGene_gn=ELN;re
fGene_tx=XM_019820584.2;refGene_gn=ELN;refGene_tx=XM_019820585.2;refGe
ne_gn=ELN;refGene_tx=XM_019820586.2;refGene_gn=ELN;refGene_tx=XM_01982
0587.2;refGene_gn=ELN;refGene_tx=XM_019820588.2;refGene_gn=ELN;refGene
_tx=XM_019820589.2;refGene_gn=ELN;refGene_tx=XM_019820590.2;refGene_gn
=ELN;refGene_tx=XM_019820591.2;refGene_gn=ELN;refGene_tx=XM_019820592.
2;refGene_gn=ELN;refGene_tx=XM_019820593.2;refGene_gn=ELN;refGene_tx=X
M_019820594.2
E3 10549345 10549378
refGene_gn=ELN;refGene_tx=XM_006942060.4;refGene_gn=ELN;refGene_tx=XM_
019820582.2;refGene_gn=ELN;refGene_tx=XM_019820583.2;refGene_gn=ELN;re
fGene_tx=XM_019820584.2;refGene_gn=ELN;refGene_tx=XM_019820585.2;refGe
ne_gn=ELN;refGene_tx=XM_019820586.2;refGene_gn=ELN;refGene_tx=XM_01982
0587.2;refGene_gn=ELN;refGene_tx=XM_019820588.2;refGene_gn=ELN;refGene
_tx=XM_019820589.2;refGene_gn=ELN;refGene_tx=XM_019820590.2;refGene_gn
=ELN;refGene_tx=XM_019820591.2;refGene_gn=ELN;refGene_tx=XM_019820592.
2;refGene_gn=ELN;refGene_tx=XM_019820593.2;refGene_gn=ELN;refGene_tx=X
M_019820594.2
E3 10551058 10551088
refGene_gn=ELN;refGene_tx=XM_006942060.4;refGene_gn=ELN;refGene_tx=XM_
019820582.2;refGene_gn=ELN;refGene_tx=XM_019820583.2;refGene_gn=ELN;re
fGene_tx=XM_019820584.2;refGene_gn=ELN;refGene_tx=XM_019820585.2;refGe
ne_gn=ELN;refGene_tx=XM_019820586.2;refGene_gn=ELN;refGene_tx=XM_01982
0587.2;refGene_gn=ELN;refGene_tx=XM_019820588.2;refGene_gn=ELN;refGene
_tx=XM_019820589.2;refGene_gn=ELN;refGene_tx=XM_019820590.2;refGene_gn
=ELN;refGene_tx=XM_019820591.2;refGene_gn=ELN;refGene_tx=XM_019820592.
2;refGene_gn=ELN;refGene_tx=XM_019820593.2;refGene_gn=ELN;refGene_tx=X
M_019820594.2
E3 10552998 10553094
refGene_gn=ELN;refGene_tx=XM_006942060.4;refGene_gn=ELN;refGene_tx=XM_
019820582.2;refGene_gn=ELN;refGene_tx=XM_019820583.2;refGene_gn=ELN;re
fGene_tx=XM_019820584.2;refGene_gn=ELN;refGene_tx=XM_019820585.2;refGe

ne_gn=ELN;refGene_tx=XM_019820586.2;refGene_gn=ELN;refGene_tx=XM_01982
0587.2;refGene_gn=ELN;refGene_tx=XM_019820588.2;refGene_gn=ELN;refGene
_tx=XM_019820589.2;refGene_gn=ELN;refGene_tx=XM_019820590.2;refGene_gn
=ELN;refGene_tx=XM_019820591.2;refGene_gn=ELN;refGene_tx=XM_019820592.
2;refGene_gn=ELN;refGene_tx=XM_019820593.2;refGene_gn=ELN;refGene_tx=X
M_019820594.2
E3 10553368 10553428
refGene_gn=ELN;refGene_tx=XM_006942060.4;refGene_gn=ELN;refGene_tx=XM_
019820582.2;refGene_gn=ELN;refGene_tx=XM_019820583.2;refGene_gn=ELN;re
fGene_tx=XM_019820584.2;refGene_gn=ELN;refGene_tx=XM_019820585.2;refGe
ne_gn=ELN;refGene_tx=XM_019820586.2;refGene_gn=ELN;refGene_tx=XM_01982
0587.2;refGene_gn=ELN;refGene_tx=XM_019820588.2;refGene_gn=ELN;refGene
_tx=XM_019820589.2;refGene_gn=ELN;refGene_tx=XM_019820590.2;refGene_gn
=ELN;refGene_tx=XM_019820591.2;refGene_gn=ELN;refGene_tx=XM_019820592.
2;refGene_gn=ELN;refGene_tx=XM_019820593.2;refGene_gn=ELN;refGene_tx=X
M_019820594.2
E3 10553511 10553580
refGene_gn=ELN;refGene_tx=XM_006942060.4;refGene_gn=ELN;refGene_tx=XM_
019820582.2;refGene_gn=ELN;refGene_tx=XM_019820583.2;refGene_gn=ELN;re
fGene_tx=XM_019820584.2;refGene_gn=ELN;refGene_tx=XM_019820585.2;refGe
ne_gn=ELN;refGene_tx=XM_019820586.2;refGene_gn=ELN;refGene_tx=XM_01982
0587.2;refGene_gn=ELN;refGene_tx=XM_019820588.2;refGene_gn=ELN;refGene
_tx=XM_019820589.2;refGene_gn=ELN;refGene_tx=XM_019820590.2;refGene_gn
=ELN;refGene_tx=XM_019820591.2;refGene_gn=ELN;refGene_tx=XM_019820592.
2;refGene_gn=ELN;refGene_tx=XM_019820593.2;refGene_gn=ELN;refGene_tx=X
M_019820594.2
E3 10554248 10554296
refGene_gn=ELN;refGene_tx=XM_006942060.4;refGene_gn=ELN;refGene_tx=XM_
019820582.2;refGene_gn=ELN;refGene_tx=XM_019820583.2;refGene_gn=ELN;re
fGene_tx=XM_019820584.2;refGene_gn=ELN;refGene_tx=XM_019820585.2;refGe
ne_gn=ELN;refGene_tx=XM_019820586.2;refGene_gn=ELN;refGene_tx=XM_01982
0588.2;refGene_gn=ELN;refGene_tx=XM_019820589.2;refGene_gn=ELN;refGene
_tx=XM_019820590.2;refGene_gn=ELN;refGene_tx=XM_019820591.2;refGene_gn
=ELN;refGene_tx=XM_019820594.2
E3 10555244 10555316
refGene_gn=ELN;refGene_tx=XM_006942060.4;refGene_gn=ELN;refGene_tx=XM_
019820582.2;refGene_gn=ELN;refGene_tx=XM_019820583.2;refGene_gn=ELN;re
fGene_tx=XM_019820584.2;refGene_gn=ELN;refGene_tx=XM_019820585.2;refGe
ne_gn=ELN;refGene_tx=XM_019820586.2;refGene_gn=ELN;refGene_tx=XM_01982
0587.2;refGene_gn=ELN;refGene_tx=XM_019820589.2;refGene_gn=ELN;refGene
_tx=XM_019820591.2;refGene_gn=ELN;refGene_tx=XM_019820594.2
E3 10555645 10555675
refGene_gn=ELN;refGene_tx=XM_006942060.4;refGene_gn=ELN;refGene_tx=XM_
019820582.2;refGene_gn=ELN;refGene_tx=XM_019820583.2;refGene_gn=ELN;re
fGene_tx=XM_019820584.2;refGene_gn=ELN;refGene_tx=XM_019820585.2;refGe
ne_gn=ELN;refGene_tx=XM_019820586.2;refGene_gn=ELN;refGene_tx=XM_01982
0587.2;refGene_gn=ELN;refGene_tx=XM_019820588.2;refGene_gn=ELN;refGene
_tx=XM_019820589.2;refGene_gn=ELN;refGene_tx=XM_019820590.2;refGene_gn
=ELN;refGene_tx=XM_019820591.2;refGene_gn=ELN;refGene_tx=XM_019820592.
2;refGene_gn=ELN;refGene_tx=XM_019820593.2;refGene_gn=ELN;refGene_tx=X

M_019820594.2
E3 10556112 10556184
refGene_gn=ELN;refGene_tx=XM_006942060.4;refGene_gn=ELN;refGene_tx=XM_019820582.2;refGene_gn=ELN;refGene_tx=XM_019820583.2;refGene_gn=ELN;refGene_tx=XM_019820584.2;refGene_gn=ELN;refGene_tx=XM_019820585.2;refGene_gn=ELN;refGene_tx=XM_019820586.2;refGene_gn=ELN;refGene_tx=XM_019820587.2;refGene_gn=ELN;refGene_tx=XM_019820588.2;refGene_gn=ELN;refGene_tx=XM_019820589.2;refGene_gn=ELN;refGene_tx=XM_019820590.2;refGene_gn=ELN;refGene_tx=XM_019820591.2;refGene_gn=ELN;refGene_tx=XM_019820592.2;refGene_gn=ELN;refGene_tx=XM_019820593.2;refGene_gn=ELN;refGene_tx=XM_019820594.2
E3 10556690 10556732
refGene_gn=ELN;refGene_tx=XM_006942060.4;refGene_gn=ELN;refGene_tx=XM_019820582.2;refGene_gn=ELN;refGene_tx=XM_019820583.2;refGene_gn=ELN;refGene_tx=XM_019820585.2;refGene_gn=ELN;refGene_tx=XM_019820586.2;refGene_gn=ELN;refGene_tx=XM_019820587.2;refGene_gn=ELN;refGene_tx=XM_019820588.2;refGene_gn=ELN;refGene_tx=XM_019820591.2;refGene_gn=ELN;refGene_tx=XM_019820592.2;refGene_gn=ELN;refGene_tx=XM_019820594.2
E3 10557142 10557202
refGene_gn=ELN;refGene_tx=XM_006942060.4;refGene_gn=ELN;refGene_tx=XM_019820582.2;refGene_gn=ELN;refGene_tx=XM_019820583.2;refGene_gn=ELN;refGene_tx=XM_019820584.2;refGene_gn=ELN;refGene_tx=XM_019820585.2;refGene_gn=ELN;refGene_tx=XM_019820586.2;refGene_gn=ELN;refGene_tx=XM_019820587.2;refGene_gn=ELN;refGene_tx=XM_019820588.2;refGene_gn=ELN;refGene_tx=XM_019820589.2;refGene_gn=ELN;refGene_tx=XM_019820590.2;refGene_gn=ELN;refGene_tx=XM_019820591.2;refGene_gn=ELN;refGene_tx=XM_019820592.2;refGene_gn=ELN;refGene_tx=XM_019820593.2;refGene_gn=ELN;refGene_tx=XM_019820594.2
E3 10557493 10557541
refGene_gn=ELN;refGene_tx=XM_006942060.4;refGene_gn=ELN;refGene_tx=XM_019820582.2;refGene_gn=ELN;refGene_tx=XM_019820583.2;refGene_gn=ELN;refGene_tx=XM_019820584.2;refGene_gn=ELN;refGene_tx=XM_019820585.2;refGene_gn=ELN;refGene_tx=XM_019820587.2;refGene_gn=ELN;refGene_tx=XM_019820588.2;refGene_gn=ELN;refGene_tx=XM_019820590.2;refGene_gn=ELN;refGene_tx=XM_019820591.2;refGene_gn=ELN;refGene_tx=XM_019820592.2;refGene_gn=ELN;refGene_tx=XM_019820593.2;refGene_gn=ELN;refGene_tx=XM_019820594.2
E3 10559090 10559183
refGene_gn=ELN;refGene_tx=XM_006942060.4;refGene_gn=ELN;refGene_tx=XM_019820582.2;refGene_gn=ELN;refGene_tx=XM_019820583.2;refGene_gn=ELN;refGene_tx=XM_019820584.2;refGene_gn=ELN;refGene_tx=XM_019820585.2;refGene_gn=ELN;refGene_tx=XM_019820586.2;refGene_gn=ELN;refGene_tx=XM_019820587.2;refGene_gn=ELN;refGene_tx=XM_019820588.2;refGene_gn=ELN;refGene_tx=XM_019820589.2;refGene_gn=ELN;refGene_tx=XM_019820590.2;refGene_gn=ELN;refGene_tx=XM_019820591.2;refGene_gn=ELN;refGene_tx=XM_019820592.2;refGene_gn=ELN;refGene_tx=XM_019820593.2;refGene_gn=ELN;refGene_tx=XM_019820594.2
E3 10559265 10559328
refGene_gn=ELN;refGene_tx=XM_006942060.4;refGene_gn=ELN;refGene_tx=XM_019820582.2;refGene_gn=ELN;refGene_tx=XM_019820583.2;refGene_gn=ELN;re

fGene_tx=XM_019820584.2;refGene_gn=ELN;refGene_tx=XM_019820585.2;refGene_gn=ELN;refGene_tx=XM_019820586.2;refGene_gn=ELN;refGene_tx=XM_019820587.2;refGene_gn=ELN;refGene_tx=XM_019820588.2;refGene_gn=ELN;refGene_tx=XM_019820589.2;refGene_gn=ELN;refGene_tx=XM_019820590.2;refGene_gn=ELN;refGene_tx=XM_019820591.2;refGene_gn=ELN;refGene_tx=XM_019820592.2;refGene_gn=ELN;refGene_tx=XM_019820593.2;refGene_gn=ELN;refGene_tx=XM_019820594.2
E3 10559892 10560072
refGene_gn=ELN;refGene_tx=XM_006942060.4;refGene_gn=ELN;refGene_tx=XM_019820582.2;refGene_gn=ELN;refGene_tx=XM_019820583.2;refGene_gn=ELN;refGene_tx=XM_019820584.2;refGene_gn=ELN;refGene_tx=XM_019820585.2;refGene_gn=ELN;refGene_tx=XM_019820586.2;refGene_gn=ELN;refGene_tx=XM_019820587.2;refGene_gn=ELN;refGene_tx=XM_019820588.2;refGene_gn=ELN;refGene_tx=XM_019820589.2;refGene_gn=ELN;refGene_tx=XM_019820590.2;refGene_gn=ELN;refGene_tx=XM_019820591.2;refGene_gn=ELN;refGene_tx=XM_019820592.2;refGene_gn=ELN;refGene_tx=XM_019820593.2;refGene_gn=ELN;refGene_tx=XM_019820594.2
E3 10561028 10561082
refGene_gn=ELN;refGene_tx=XM_006942060.4;refGene_gn=ELN;refGene_tx=XM_019820582.2;refGene_gn=ELN;refGene_tx=XM_019820583.2;refGene_gn=ELN;refGene_tx=XM_019820584.2;refGene_gn=ELN;refGene_tx=XM_019820585.2;refGene_gn=ELN;refGene_tx=XM_019820586.2;refGene_gn=ELN;refGene_tx=XM_019820587.2;refGene_gn=ELN;refGene_tx=XM_019820588.2;refGene_gn=ELN;refGene_tx=XM_019820589.2;refGene_gn=ELN;refGene_tx=XM_019820590.2;refGene_gn=ELN;refGene_tx=XM_019820591.2;refGene_gn=ELN;refGene_tx=XM_019820592.2;refGene_gn=ELN;refGene_tx=XM_019820593.2;refGene_gn=ELN;refGene_tx=XM_019820594.2
E3 10561616 10561745
refGene_gn=ELN;refGene_tx=XM_006942060.4;refGene_gn=ELN;refGene_tx=XM_019820582.2;refGene_gn=ELN;refGene_tx=XM_019820583.2;refGene_gn=ELN;refGene_tx=XM_019820584.2;refGene_gn=ELN;refGene_tx=XM_019820585.2;refGene_gn=ELN;refGene_tx=XM_019820586.2;refGene_gn=ELN;refGene_tx=XM_019820587.2;refGene_gn=ELN;refGene_tx=XM_019820588.2;refGene_gn=ELN;refGene_tx=XM_019820589.2;refGene_gn=ELN;refGene_tx=XM_019820590.2;refGene_gn=ELN;refGene_tx=XM_019820591.2;refGene_gn=ELN;refGene_tx=XM_019820592.2;refGene_gn=ELN;refGene_tx=XM_019820593.2;refGene_gn=ELN;refGene_tx=XM_019820594.2
E3 10561929 10561992
refGene_gn=ELN;refGene_tx=XM_006942060.4;refGene_gn=ELN;refGene_tx=XM_019820582.2;refGene_gn=ELN;refGene_tx=XM_019820583.2;refGene_gn=ELN;refGene_tx=XM_019820584.2;refGene_gn=ELN;refGene_tx=XM_019820585.2;refGene_gn=ELN;refGene_tx=XM_019820586.2;refGene_gn=ELN;refGene_tx=XM_019820587.2;refGene_gn=ELN;refGene_tx=XM_019820588.2;refGene_gn=ELN;refGene_tx=XM_019820589.2;refGene_gn=ELN;refGene_tx=XM_019820590.2;refGene_gn=ELN;refGene_tx=XM_019820591.2;refGene_gn=ELN;refGene_tx=XM_019820592.2;refGene_gn=ELN;refGene_tx=XM_019820593.2;refGene_gn=ELN;refGene_tx=XM_019820594.2
E3 10562404 10562491
refGene_gn=ELN;refGene_tx=XM_006942060.4;refGene_gn=ELN;refGene_tx=XM_019820582.2;refGene_gn=ELN;refGene_tx=XM_019820583.2;refGene_gn=ELN;re

fGene_tx=XM_019820584.2;refGene_gn=ELN;refGene_tx=XM_019820585.2;refGene_gn=ELN;refGene_tx=XM_019820586.2;refGene_gn=ELN;refGene_tx=XM_019820587.2;refGene_gn=ELN;refGene_tx=XM_019820588.2;refGene_gn=ELN;refGene_tx=XM_019820589.2;refGene_gn=ELN;refGene_tx=XM_019820590.2;refGene_gn=ELN;refGene_tx=XM_019820591.2;refGene_gn=ELN;refGene_tx=XM_019820592.2;refGene_gn=ELN;refGene_tx=XM_019820593.2;refGene_gn=ELN;refGene_tx=XM_019820594.2
E3 10563351 10563408
refGene_gn=ELN;refGene_tx=XM_006942060.4;refGene_gn=ELN;refGene_tx=XM_019820582.2;refGene_gn=ELN;refGene_tx=XM_019820583.2;refGene_gn=ELN;refGene_tx=XM_019820584.2;refGene_gn=ELN;refGene_tx=XM_019820585.2;refGene_gn=ELN;refGene_tx=XM_019820586.2;refGene_gn=ELN;refGene_tx=XM_019820587.2;refGene_gn=ELN;refGene_tx=XM_019820588.2;refGene_gn=ELN;refGene_tx=XM_019820589.2;refGene_gn=ELN;refGene_tx=XM_019820590.2;refGene_gn=ELN;refGene_tx=XM_019820591.2;refGene_gn=ELN;refGene_tx=XM_019820592.2;refGene_gn=ELN;refGene_tx=XM_019820593.2;refGene_gn=ELN;refGene_tx=XM_019820594.2
E3 10564416 10564524
refGene_gn=ELN;refGene_tx=XM_006942060.4;refGene_gn=ELN;refGene_tx=XM_019820582.2;refGene_gn=ELN;refGene_tx=XM_019820583.2;refGene_gn=ELN;refGene_tx=XM_019820584.2;refGene_gn=ELN;refGene_tx=XM_019820585.2;refGene_gn=ELN;refGene_tx=XM_019820586.2;refGene_gn=ELN;refGene_tx=XM_019820587.2;refGene_gn=ELN;refGene_tx=XM_019820588.2;refGene_gn=ELN;refGene_tx=XM_019820589.2;refGene_gn=ELN;refGene_tx=XM_019820590.2;refGene_gn=ELN;refGene_tx=XM_019820591.2;refGene_gn=ELN;refGene_tx=XM_019820592.2;refGene_gn=ELN;refGene_tx=XM_019820593.2;refGene_gn=ELN;refGene_tx=XM_019820594.2
E3 10564605 10564665
refGene_gn=ELN;refGene_tx=XM_006942060.4;refGene_gn=ELN;refGene_tx=XM_019820582.2;refGene_gn=ELN;refGene_tx=XM_019820583.2;refGene_gn=ELN;refGene_tx=XM_019820584.2;refGene_gn=ELN;refGene_tx=XM_019820585.2;refGene_gn=ELN;refGene_tx=XM_019820586.2;refGene_gn=ELN;refGene_tx=XM_019820587.2;refGene_gn=ELN;refGene_tx=XM_019820588.2;refGene_gn=ELN;refGene_tx=XM_019820589.2;refGene_gn=ELN;refGene_tx=XM_019820590.2;refGene_gn=ELN;refGene_tx=XM_019820591.2;refGene_gn=ELN;refGene_tx=XM_019820592.2;refGene_gn=ELN;refGene_tx=XM_019820593.2;refGene_gn=ELN;refGene_tx=XM_019820594.2
E3 10565201 10565327
refGene_gn=ELN;refGene_tx=XM_006942060.4;refGene_gn=ELN;refGene_tx=XM_019820582.2;refGene_gn=ELN;refGene_tx=XM_019820583.2;refGene_gn=ELN;refGene_tx=XM_019820584.2;refGene_gn=ELN;refGene_tx=XM_019820585.2;refGene_gn=ELN;refGene_tx=XM_019820586.2;refGene_gn=ELN;refGene_tx=XM_019820587.2;refGene_gn=ELN;refGene_tx=XM_019820588.2;refGene_gn=ELN;refGene_tx=XM_019820589.2;refGene_gn=ELN;refGene_tx=XM_019820590.2;refGene_gn=ELN;refGene_tx=XM_019820591.2;refGene_gn=ELN;refGene_tx=XM_019820592.2;refGene_gn=ELN;refGene_tx=XM_019820593.2;refGene_gn=ELN;refGene_tx=XM_019820594.2
E3 10565986 10566025
refGene_gn=ELN;refGene_tx=XM_006942060.4;refGene_gn=ELN;refGene_tx=XM_019820582.2;refGene_gn=ELN;refGene_tx=XM_019820583.2;refGene_gn=ELN;re

fGene_tx=XM_019820584.2;refGene_gn=ELN;refGene_tx=XM_019820585.2;refGene_gn=ELN;refGene_tx=XM_019820586.2;refGene_gn=ELN;refGene_tx=XM_019820587.2;refGene_gn=ELN;refGene_tx=XM_019820588.2;refGene_gn=ELN;refGene_tx=XM_019820589.2;refGene_gn=ELN;refGene_tx=XM_019820590.2;refGene_gn=ELN;refGene_tx=XM_019820591.2;refGene_gn=ELN;refGene_tx=XM_019820592.2;refGene_gn=ELN;refGene_tx=XM_019820593.2;refGene_gn=ELN;refGene_tx=XM_019820594.2
E3 10567444 10567498
refGene_gn=ELN;refGene_tx=XM_006942060.4;refGene_gn=ELN;refGene_tx=XM_019820582.2;refGene_gn=ELN;refGene_tx=XM_019820583.2;refGene_gn=ELN;refGene_tx=XM_019820584.2;refGene_gn=ELN;refGene_tx=XM_019820585.2;refGene_gn=ELN;refGene_tx=XM_019820586.2;refGene_gn=ELN;refGene_tx=XM_019820587.2;refGene_gn=ELN;refGene_tx=XM_019820588.2;refGene_gn=ELN;refGene_tx=XM_019820589.2;refGene_gn=ELN;refGene_tx=XM_019820590.2;refGene_gn=ELN;refGene_tx=XM_019820591.2;refGene_gn=ELN;refGene_tx=XM_019820592.2;refGene_gn=ELN;refGene_tx=XM_019820593.2
E3 10567825 10567885
refGene_gn=ELN;refGene_tx=XM_006942060.4;refGene_gn=ELN;refGene_tx=XM_019820582.2;refGene_gn=ELN;refGene_tx=XM_019820583.2;refGene_gn=ELN;refGene_tx=XM_019820584.2;refGene_gn=ELN;refGene_tx=XM_019820585.2;refGene_gn=ELN;refGene_tx=XM_019820586.2;refGene_gn=ELN;refGene_tx=XM_019820587.2;refGene_gn=ELN;refGene_tx=XM_019820588.2;refGene_gn=ELN;refGene_tx=XM_019820589.2;refGene_gn=ELN;refGene_tx=XM_019820590.2;refGene_gn=ELN;refGene_tx=XM_019820591.2;refGene_gn=ELN;refGene_tx=XM_019820592.2;refGene_gn=ELN;refGene_tx=XM_019820593.2
E3 10568137 10568215
refGene_gn=ELN;refGene_tx=XM_006942060.4;refGene_gn=ELN;refGene_tx=XM_019820582.2;refGene_gn=ELN;refGene_tx=XM_019820583.2;refGene_gn=ELN;refGene_tx=XM_019820584.2;refGene_gn=ELN;refGene_tx=XM_019820585.2;refGene_gn=ELN;refGene_tx=XM_019820586.2;refGene_gn=ELN;refGene_tx=XM_019820587.2;refGene_gn=ELN;refGene_tx=XM_019820588.2;refGene_gn=ELN;refGene_tx=XM_019820589.2;refGene_gn=ELN;refGene_tx=XM_019820590.2;refGene_gn=ELN;refGene_tx=XM_019820591.2;refGene_gn=ELN;refGene_tx=XM_019820592.2;refGene_gn=ELN;refGene_tx=XM_019820593.2
E3 10568809 10568848
refGene_gn=ELN;refGene_tx=XM_006942060.4;refGene_gn=ELN;refGene_tx=XM_019820582.2;refGene_gn=ELN;refGene_tx=XM_019820583.2;refGene_gn=ELN;refGene_tx=XM_019820584.2;refGene_gn=ELN;refGene_tx=XM_019820585.2;refGene_gn=ELN;refGene_tx=XM_019820586.2;refGene_gn=ELN;refGene_tx=XM_019820587.2;refGene_gn=ELN;refGene_tx=XM_019820588.2;refGene_gn=ELN;refGene_tx=XM_019820589.2;refGene_gn=ELN;refGene_tx=XM_019820590.2;refGene_gn=ELN;refGene_tx=XM_019820591.2;refGene_gn=ELN;refGene_tx=XM_019820592.2;refGene_gn=ELN;refGene_tx=XM_019820593.2
E3 10569499 10569544
refGene_gn=ELN;refGene_tx=XM_006942060.4;refGene_gn=ELN;refGene_tx=XM_019820582.2;refGene_gn=ELN;refGene_tx=XM_019820583.2;refGene_gn=ELN;refGene_tx=XM_019820584.2;refGene_gn=ELN;refGene_tx=XM_019820586.2;refGene_gn=ELN;refGene_tx=XM_019820587.2;refGene_gn=ELN;refGene_tx=XM_019820588.2;refGene_gn=ELN;refGene_tx=XM_019820589.2;refGene_gn=ELN;refGene_tx=XM_019820590.2;refGene_gn=ELN;refGene_tx=XM_019820591.2;refGene_gn

=ELN;refGene_tx=XM_019820592.2;refGene_gn=ELN;refGene_tx=XM_019820593.2;refGene_gn=ELN;refGene_tx=XM_019820594.2
E3 10571915 10571959
refGene_gn=ELN;refGene_tx=XM_006942060.4;refGene_gn=ELN;refGene_tx=XM_019820582.2;refGene_gn=ELN;refGene_tx=XM_019820583.2;refGene_gn=ELN;refGene_tx=XM_019820584.2;refGene_gn=ELN;refGene_tx=XM_019820585.2;refGene_gn=ELN;refGene_tx=XM_019820586.2;refGene_gn=ELN;refGene_tx=XM_019820587.2;refGene_gn=ELN;refGene_tx=XM_019820588.2;refGene_gn=ELN;refGene_tx=XM_019820589.2;refGene_gn=ELN;refGene_tx=XM_019820590.2;refGene_gn=ELN;refGene_tx=XM_019820591.2;refGene_gn=ELN;refGene_tx=XM_019820592.2;refGene_gn=ELN;refGene_tx=XM_019820593.2;refGene_gn=ELN;refGene_tx=XM_019820594.2
E3 10582230 10582285 refGene_gn=LIMK1;refGene_tx=XM_023246437.1
E3 10583977 10584074 refGene_gn=LIMK1;refGene_tx=XM_023246437.1
E3 10584967 10585007
refGene_gn=LOC109495166;refGene_tx=XM_023246438.1;refGene_gn=LOC109495166;refGene_tx=XM_023246439.1
E3 10585807 10586113
refGene_gn=LOC109495166;refGene_tx=XM_023246438.1;refGene_gn=LOC109495166;refGene_tx=XM_023246439.1
E3 10586775 10586936
refGene_gn=LOC109495166;refGene_tx=XM_023246438.1;refGene_gn=LOC109495166;refGene_tx=XM_023246439.1
E3 10587482 10587503
refGene_gn=LOC109495166;refGene_tx=XM_023246438.1;refGene_gn=LOC109495166;refGene_tx=XM_023246439.1
E3 10589954 10590093 refGene_gn=LIMK1;refGene_tx=XM_023246437.1
E3 10590375 10590485 refGene_gn=LIMK1;refGene_tx=XM_023246437.1
E3 10591589 10591796 refGene_gn=LIMK1;refGene_tx=XM_023246437.1
E3 10596940 10597046 refGene_gn=LIMK1;refGene_tx=XM_023246437.1
E3 10597162 10597329 refGene_gn=LIMK1;refGene_tx=XM_023246437.1
E3 10598300 10598484 refGene_gn=LIMK1;refGene_tx=XM_023246437.1
E3 10599242 10599329 refGene_gn=LIMK1;refGene_tx=XM_023246437.1
E3 10599632 10599764 refGene_gn=LIMK1;refGene_tx=XM_023246437.1
E3 10601114 10601174 refGene_gn=LIMK1;refGene_tx=XM_023246437.1
E3 10601427 10601493 refGene_gn=LIMK1;refGene_tx=XM_023246437.1
E3 10604190 10604347 refGene_gn=LIMK1;refGene_tx=XM_023246437.1
E3 10605841 10605897 refGene_gn=LIMK1;refGene_tx=XM_023246437.1
E3 10606101 10606259 refGene_gn=LIMK1;refGene_tx=XM_023246437.1
E3 10606364 10606527 refGene_gn=LIMK1;refGene_tx=XM_023246437.1
E3 10648006 10648065
refGene_gn=EIF4H;refGene_tx=XM_023247018.1;refGene_gn=EIF4H;refGene_tx=XM_023247019.1
E3 10662190 10662378
refGene_gn=EIF4H;refGene_tx=XM_023247018.1;refGene_gn=EIF4H;refGene_tx=XM_023247019.1
E3 10664048 10664113
refGene_gn=EIF4H;refGene_tx=XM_023247018.1;refGene_gn=EIF4H;refGene_tx=XM_023247019.1
E3 10664196 10664293

refGene_gn=EIF4H;refGene_tx=XM_023247018.1;refGene_gn=EIF4H;refGene_tx=XM_023247019.1
E3 10664621 10664681 refGene_gn=EIF4H;refGene_tx=XM_023247018.1
E3 10668439 10668577 refGene_gn=EIF4H;refGene_tx=XM_023247018.1;refGene_gn=EIF4H;refGene_tx=XM_023247019.1
E3 10668869 10669009 refGene_gn=EIF4H;refGene_tx=XM_023247018.1;refGene_gn=EIF4H;refGene_tx=XM_023247019.1
E3 10679613 10679674 refGene_gn=LAT2;refGene_tx=XM_019820596.2
E3 10681527 10681650 refGene_gn=LAT2;refGene_tx=XM_006942068.3;refGene_gn=LAT2;refGene_tx=XM_019820596.2
E3 10682057 10682097 refGene_gn=LAT2;refGene_tx=XM_006942068.3;refGene_gn=LAT2;refGene_tx=XM_019820596.2
E3 10684404 10684448 refGene_gn=LAT2;refGene_tx=XM_006942068.3;refGene_gn=LAT2;refGene_tx=XM_019820596.2
E3 10684614 10684675 refGene_gn=LAT2;refGene_tx=XM_006942068.3;refGene_gn=LAT2;refGene_tx=XM_019820596.2
E3 10684920 10684958 refGene_gn=LAT2;refGene_tx=XM_006942068.3;refGene_gn=LAT2;refGene_tx=XM_019820596.2
E3 10685247 10685286 refGene_gn=LAT2;refGene_tx=XM_006942068.3;refGene_gn=LAT2;refGene_tx=XM_019820596.2
E3 10685367 10685398 refGene_gn=LAT2;refGene_tx=XM_006942068.3;refGene_gn=LAT2;refGene_tx=XM_019820596.2
E3 10685889 10685945 refGene_gn=LAT2;refGene_tx=XM_006942068.3;refGene_gn=LAT2;refGene_tx=XM_019820596.2
E3 10687129 10687192 refGene_gn=LAT2;refGene_tx=XM_006942068.3;refGene_gn=LAT2;refGene_tx=XM_019820596.2
E3 10687491 10687563 refGene_gn=LAT2;refGene_tx=XM_006942068.3;refGene_gn=LAT2;refGene_tx=XM_019820596.2
E3 10687963 10688058 refGene_gn=LAT2;refGene_tx=XM_006942068.3;refGene_gn=LAT2;refGene_tx=XM_019820596.2
E3 10693159 10693270 refGene_gn=RFC2;refGene_tx=XM_011290348.3
E3 10695141 10695255 refGene_gn=RFC2;refGene_tx=XM_011290348.3
E3 10696772 10696853 refGene_gn=RFC2;refGene_tx=XM_011290348.3
E3 10698157 10698223 refGene_gn=RFC2;refGene_tx=XM_011290348.3
E3 10698477 10698635 refGene_gn=RFC2;refGene_tx=XM_011290348.3
E3 10699842 10699943 refGene_gn=RFC2;refGene_tx=XM_011290348.3
E3 10701268 10701370 refGene_gn=RFC2;refGene_tx=XM_011290348.3

E3 10705949 10706056 refGene_gn=RFC2;refGene_tx=XM_011290348.3
E3 10706351 10706393 refGene_gn=RFC2;refGene_tx=XM_011290348.3
E3 10709019 10709089 refGene_gn=RFC2;refGene_tx=XM_011290348.3
E3 10712450 10712557 refGene_gn=RFC2;refGene_tx=XM_011290348.3
E3 10761697 10761821
refGene_gn=CLIP2;refGene_tx=XM_023246988.1;refGene_gn=CLIP2;refGene_tx=XM_023246989.1;refGene_gn=CLIP2;refGene_tx=XM_023246990.1
E3 10775739 10776296
refGene_gn=CLIP2;refGene_tx=XM_023246988.1;refGene_gn=CLIP2;refGene_tx=XM_023246989.1;refGene_gn=CLIP2;refGene_tx=XM_023246990.1
E3 10788388 10788513
refGene_gn=CLIP2;refGene_tx=XM_023246988.1;refGene_gn=CLIP2;refGene_tx=XM_023246989.1;refGene_gn=CLIP2;refGene_tx=XM_023246990.1
E3 10790072 10790286
refGene_gn=CLIP2;refGene_tx=XM_023246988.1;refGene_gn=CLIP2;refGene_tx=XM_023246989.1;refGene_gn=CLIP2;refGene_tx=XM_023246990.1
E3 10791323 10791521
refGene_gn=CLIP2;refGene_tx=XM_023246988.1;refGene_gn=CLIP2;refGene_tx=XM_023246989.1;refGene_gn=CLIP2;refGene_tx=XM_023246990.1
E3 10792351 10792455
refGene_gn=CLIP2;refGene_tx=XM_023246988.1;refGene_gn=CLIP2;refGene_tx=XM_023246989.1;refGene_gn=CLIP2;refGene_tx=XM_023246990.1
E3 10794040 10794101
refGene_gn=CLIP2;refGene_tx=XM_023246988.1;refGene_gn=CLIP2;refGene_tx=XM_023246989.1;refGene_gn=CLIP2;refGene_tx=XM_023246990.1
E3 10800028 10800133
refGene_gn=CLIP2;refGene_tx=XM_023246988.1;refGene_gn=CLIP2;refGene_tx=XM_023246989.1
E3 10801869 10802805
refGene_gn=CLIP2;refGene_tx=XM_023246988.1;refGene_gn=CLIP2;refGene_tx=XM_023246989.1;refGene_gn=CLIP2;refGene_tx=XM_023246990.1
E3 10804773 10804831
refGene_gn=CLIP2;refGene_tx=XM_023246988.1;refGene_gn=CLIP2;refGene_tx=XM_023246989.1;refGene_gn=CLIP2;refGene_tx=XM_023246990.1
E3 10808096 10808180
refGene_gn=CLIP2;refGene_tx=XM_023246988.1;refGene_gn=CLIP2;refGene_tx=XM_023246989.1;refGene_gn=CLIP2;refGene_tx=XM_023246990.1
E3 10809149 10809306
refGene_gn=CLIP2;refGene_tx=XM_023246988.1;refGene_gn=CLIP2;refGene_tx=XM_023246989.1;refGene_gn=CLIP2;refGene_tx=XM_023246990.1
E3 10810838 10810998
refGene_gn=CLIP2;refGene_tx=XM_023246988.1;refGene_gn=CLIP2;refGene_tx=XM_023246989.1;refGene_gn=CLIP2;refGene_tx=XM_023246990.1
E3 10812641 10812824
refGene_gn=CLIP2;refGene_tx=XM_023246988.1;refGene_gn=CLIP2;refGene_tx=XM_023246989.1;refGene_gn=CLIP2;refGene_tx=XM_023246990.1
E3 10813113 10813176
refGene_gn=CLIP2;refGene_tx=XM_023246988.1;refGene_gn=CLIP2;refGene_tx=XM_023246989.1;refGene_gn=CLIP2;refGene_tx=XM_023246990.1
E3 10814048 10814060

refGene_gn=CLIP2;refGene_tx=XM_023246988.1;refGene_gn=CLIP2;refGene_tx=XM_023246989.1;refGene_gn=CLIP2;refGene_tx=XM_023246990.1
E3 10879019 10879142
refGene_gn=GTF2IRD1;refGene_tx=XM_006942070.4;refGene_gn=GTF2IRD1;refGene_tx=XM_019820602.2;refGene_gn=GTF2IRD1;refGene_tx=XM_019820604.2;refGene_gn=GTF2IRD1;refGene_tx=XM_019820605.2;refGene_gn=GTF2IRD1;refGene_tx=XM_023247025.1
E3 10882423 10882565
refGene_gn=GTF2IRD1;refGene_tx=XM_006942070.4;refGene_gn=GTF2IRD1;refGene_tx=XM_019820602.2;refGene_gn=GTF2IRD1;refGene_tx=XM_019820604.2;refGene_gn=GTF2IRD1;refGene_tx=XM_019820605.2;refGene_gn=GTF2IRD1;refGene_tx=XM_023247025.1
E3 10884909 10885065
refGene_gn=GTF2IRD1;refGene_tx=XM_006942070.4;refGene_gn=GTF2IRD1;refGene_tx=XM_019820602.2;refGene_gn=GTF2IRD1;refGene_tx=XM_019820603.1;refGene_gn=GTF2IRD1;refGene_tx=XM_019820604.2;refGene_gn=GTF2IRD1;refGene_tx=XM_019820605.2;refGene_gn=GTF2IRD1;refGene_tx=XM_023247025.1
E3 10886815 10886999
refGene_gn=GTF2IRD1;refGene_tx=XM_006942070.4;refGene_gn=GTF2IRD1;refGene_tx=XM_019820602.2;refGene_gn=GTF2IRD1;refGene_tx=XM_019820603.1;refGene_gn=GTF2IRD1;refGene_tx=XM_019820604.2;refGene_gn=GTF2IRD1;refGene_tx=XM_019820605.2;refGene_gn=GTF2IRD1;refGene_tx=XM_023247025.1
E3 10887715 10888026
refGene_gn=GTF2IRD1;refGene_tx=XM_006942070.4;refGene_gn=GTF2IRD1;refGene_tx=XM_019820602.2;refGene_gn=GTF2IRD1;refGene_tx=XM_019820603.1;refGene_gn=GTF2IRD1;refGene_tx=XM_019820604.2;refGene_gn=GTF2IRD1;refGene_tx=XM_019820605.2;refGene_gn=GTF2IRD1;refGene_tx=XM_023247025.1
E3 10889269 10889359
refGene_gn=GTF2IRD1;refGene_tx=XM_006942070.4;refGene_gn=GTF2IRD1;refGene_tx=XM_019820602.2;refGene_gn=GTF2IRD1;refGene_tx=XM_019820603.1;refGene_gn=GTF2IRD1;refGene_tx=XM_019820604.2;refGene_gn=GTF2IRD1;refGene_tx=XM_019820605.2;refGene_gn=GTF2IRD1;refGene_tx=XM_023247025.1
E3 10891414 10891498
refGene_gn=GTF2IRD1;refGene_tx=XM_006942070.4;refGene_gn=GTF2IRD1;refGene_tx=XM_019820602.2;refGene_gn=GTF2IRD1;refGene_tx=XM_019820603.1;refGene_gn=GTF2IRD1;refGene_tx=XM_019820604.2;refGene_gn=GTF2IRD1;refGene_tx=XM_019820605.2;refGene_gn=GTF2IRD1;refGene_tx=XM_023247025.1
E3 10896316 10896500
refGene_gn=GTF2IRD1;refGene_tx=XM_006942070.4;refGene_gn=GTF2IRD1;refGene_tx=XM_019820602.2;refGene_gn=GTF2IRD1;refGene_tx=XM_019820603.1;refGene_gn=GTF2IRD1;refGene_tx=XM_019820604.2;refGene_gn=GTF2IRD1;refGene_tx=XM_019820605.2;refGene_gn=GTF2IRD1;refGene_tx=XM_023247025.1
E3 10901128 10901154
refGene_gn=GTF2IRD1;refGene_tx=XM_006942070.4;refGene_gn=GTF2IRD1;refGene_tx=XM_019820602.2;refGene_gn=GTF2IRD1;refGene_tx=XM_019820603.1;refGene_gn=GTF2IRD1;refGene_tx=XM_019820604.2;refGene_gn=GTF2IRD1;refGene_tx=XM_019820605.2;refGene_gn=GTF2IRD1;refGene_tx=XM_023247025.1
E3 10902045 10902154
refGene_gn=GTF2IRD1;refGene_tx=XM_006942070.4;refGene_gn=GTF2IRD1;refGene_tx=XM_019820602.2;refGene_gn=GTF2IRD1;refGene_tx=XM_019820603.1;re

fGene_gn=GTF2IRD1;refGene_tx=XM_019820604.2;refGene_gn=GTF2IRD1;refGene_tx=XM_019820605.2;refGene_gn=GTF2IRD1;refGene_tx=XM_023247025.1
E3 10903393 10903431
refGene_gn=GTF2IRD1;refGene_tx=XM_006942070.4;refGene_gn=GTF2IRD1;refGene_tx=XM_019820602.2;refGene_gn=GTF2IRD1;refGene_tx=XM_019820603.1;refGene_gn=GTF2IRD1;refGene_tx=XM_019820604.2;refGene_gn=GTF2IRD1;refGene_tx=XM_019820605.2;refGene_gn=GTF2IRD1;refGene_tx=XM_023247025.1
E3 10903919 10904000
refGene_gn=GTF2IRD1;refGene_tx=XM_006942070.4;refGene_gn=GTF2IRD1;refGene_tx=XM_019820602.2;refGene_gn=GTF2IRD1;refGene_tx=XM_019820603.1;refGene_gn=GTF2IRD1;refGene_tx=XM_019820604.2;refGene_gn=GTF2IRD1;refGene_tx=XM_019820605.2;refGene_gn=GTF2IRD1;refGene_tx=XM_023247025.1
E3 10904641 10904731
refGene_gn=GTF2IRD1;refGene_tx=XM_006942070.4;refGene_gn=GTF2IRD1;refGene_tx=XM_019820602.2;refGene_gn=GTF2IRD1;refGene_tx=XM_019820603.1;refGene_gn=GTF2IRD1;refGene_tx=XM_019820604.2;refGene_gn=GTF2IRD1;refGene_tx=XM_019820605.2;refGene_gn=GTF2IRD1;refGene_tx=XM_023247025.1
E3 10912133 10912181
refGene_gn=GTF2IRD1;refGene_tx=XM_006942070.4;refGene_gn=GTF2IRD1;refGene_tx=XM_019820602.2;refGene_gn=GTF2IRD1;refGene_tx=XM_019820603.1;refGene_gn=GTF2IRD1;refGene_tx=XM_019820604.2;refGene_gn=GTF2IRD1;refGene_tx=XM_019820605.2;refGene_gn=GTF2IRD1;refGene_tx=XM_023247025.1
E3 10913099 10913165
refGene_gn=GTF2IRD1;refGene_tx=XM_006942070.4;refGene_gn=GTF2IRD1;refGene_tx=XM_019820602.2;refGene_gn=GTF2IRD1;refGene_tx=XM_019820603.1;refGene_gn=GTF2IRD1;refGene_tx=XM_019820604.2;refGene_gn=GTF2IRD1;refGene_tx=XM_019820605.2;refGene_gn=GTF2IRD1;refGene_tx=XM_023247025.1
E3 10914208 10914392
refGene_gn=GTF2IRD1;refGene_tx=XM_006942070.4;refGene_gn=GTF2IRD1;refGene_tx=XM_019820602.2;refGene_gn=GTF2IRD1;refGene_tx=XM_019820603.1;refGene_gn=GTF2IRD1;refGene_tx=XM_019820604.2;refGene_gn=GTF2IRD1;refGene_tx=XM_019820605.2;refGene_gn=GTF2IRD1;refGene_tx=XM_023247025.1
E3 10922234 10922284
refGene_gn=GTF2IRD1;refGene_tx=XM_006942070.4;refGene_gn=GTF2IRD1;refGene_tx=XM_019820602.2;refGene_gn=GTF2IRD1;refGene_tx=XM_019820603.1;refGene_gn=GTF2IRD1;refGene_tx=XM_019820604.2;refGene_gn=GTF2IRD1;refGene_tx=XM_019820605.2;refGene_gn=GTF2IRD1;refGene_tx=XM_023247025.1
E3 10922509 10922566
refGene_gn=GTF2IRD1;refGene_tx=XM_019820602.2;refGene_gn=GTF2IRD1;refGene_tx=XM_019820603.1;refGene_gn=GTF2IRD1;refGene_tx=XM_019820604.2;refGene_gn=GTF2IRD1;refGene_tx=XM_019820605.2;refGene_gn=GTF2IRD1;refGene_tx=XM_023247025.1
E3 10923774 10923858
refGene_gn=GTF2IRD1;refGene_tx=XM_006942070.4;refGene_gn=GTF2IRD1;refGene_tx=XM_019820602.2;refGene_gn=GTF2IRD1;refGene_tx=XM_019820603.1;refGene_gn=GTF2IRD1;refGene_tx=XM_019820604.2;refGene_gn=GTF2IRD1;refGene_tx=XM_019820605.2;refGene_gn=GTF2IRD1;refGene_tx=XM_023247025.1
E3 10928483 10928667
refGene_gn=GTF2IRD1;refGene_tx=XM_006942070.4;refGene_gn=GTF2IRD1;refGene_tx=XM_019820602.2;refGene_gn=GTF2IRD1;refGene_tx=XM_019820603.1;re

fGene_gn=GTF2IRD1;refGene_tx=XM_019820604.2;refGene_gn=GTF2IRD1;refGene_tx=XM_019820605.2;refGene_gn=GTF2IRD1;refGene_tx=XM_023247025.1
E3 10929190 10929219
refGene_gn=GTF2IRD1;refGene_tx=XM_006942070.4;refGene_gn=GTF2IRD1;refGene_tx=XM_019820602.2;refGene_gn=GTF2IRD1;refGene_tx=XM_019820603.1;refGene_gn=GTF2IRD1;refGene_tx=XM_019820604.2;refGene_gn=GTF2IRD1;refGene_tx=XM_019820605.2;refGene_gn=GTF2IRD1;refGene_tx=XM_023247025.1
E3 10937744 10937764 refGene_gn=GTF2IRD1;refGene_tx=XM_019820605.2
E3 10950562 10950640
refGene_gn=GTF2IRD1;refGene_tx=XM_006942070.4;refGene_gn=GTF2IRD1;refGene_tx=XM_019820602.2;refGene_gn=GTF2IRD1;refGene_tx=XM_019820603.1;refGene_gn=GTF2IRD1;refGene_tx=XM_019820604.2;refGene_gn=GTF2IRD1;refGene_tx=XM_023247025.1
E3 10952212 10952405
refGene_gn=GTF2IRD1;refGene_tx=XM_006942070.4;refGene_gn=GTF2IRD1;refGene_tx=XM_019820602.2;refGene_gn=GTF2IRD1;refGene_tx=XM_019820603.1;refGene_gn=GTF2IRD1;refGene_tx=XM_019820604.2;refGene_gn=GTF2IRD1;refGene_tx=XM_023247025.1
E3 10954753 10954791
refGene_gn=GTF2IRD1;refGene_tx=XM_006942070.4;refGene_gn=GTF2IRD1;refGene_tx=XM_019820602.2;refGene_gn=GTF2IRD1;refGene_tx=XM_019820603.1;refGene_gn=GTF2IRD1;refGene_tx=XM_023247025.1
E3 10956890 10957027
refGene_gn=GTF2IRD1;refGene_tx=XM_006942070.4;refGene_gn=GTF2IRD1;refGene_tx=XM_019820602.2;refGene_gn=GTF2IRD1;refGene_tx=XM_019820603.1;refGene_gn=GTF2IRD1;refGene_tx=XM_023247025.1
E3 10958290 10958359
refGene_gn=GTF2IRD1;refGene_tx=XM_006942070.4;refGene_gn=GTF2IRD1;refGene_tx=XM_019820603.1;refGene_gn=GTF2IRD1;refGene_tx=XM_023247025.1
E3 10965470 10965476 refGene_gn=GTF2IRD1;refGene_tx=XM_019820602.2
E3 11032039 11032138
refGene_gn=GTF2I;refGene_tx=XM_011290342.3;refGene_gn=GTF2I;refGene_tx=XM_011290343.3;refGene_gn=GTF2I;refGene_tx=XM_011290344.3;refGene_gn=GTF2I;refGene_tx=XM_011290345.3;refGene_gn=GTF2I;refGene_tx=XM_011290346.3;refGene_gn=GTF2I;refGene_tx=XM_011290347.3;refGene_gn=GTF2I;refGene_tx=XM_019820600.2;refGene_gn=GTF2I;refGene_tx=XM_019820601.2
E3 11035589 11035728
refGene_gn=GTF2I;refGene_tx=XM_011290342.3;refGene_gn=GTF2I;refGene_tx=XM_011290343.3;refGene_gn=GTF2I;refGene_tx=XM_011290344.3;refGene_gn=GTF2I;refGene_tx=XM_011290345.3;refGene_gn=GTF2I;refGene_tx=XM_011290346.3;refGene_gn=GTF2I;refGene_tx=XM_011290347.3;refGene_gn=GTF2I;refGene_tx=XM_019820600.2;refGene_gn=GTF2I;refGene_tx=XM_019820601.2
E3 11039378 11039513
refGene_gn=GTF2I;refGene_tx=XM_011290342.3;refGene_gn=GTF2I;refGene_tx=XM_011290343.3;refGene_gn=GTF2I;refGene_tx=XM_011290344.3;refGene_gn=GTF2I;refGene_tx=XM_011290345.3;refGene_gn=GTF2I;refGene_tx=XM_011290346.3;refGene_gn=GTF2I;refGene_tx=XM_011290347.3;refGene_gn=GTF2I;refGene_tx=XM_019820600.2;refGene_gn=GTF2I;refGene_tx=XM_019820601.2
E3 11040556 11040740
refGene_gn=GTF2I;refGene_tx=XM_011290342.3;refGene_gn=GTF2I;refGene_tx

=XM_011290343.3;refGene_gn=GTF2I;refGene_tx=XM_011290344.3;refGene_gn=GTF2I;refGene_tx=XM_011290345.3;refGene_gn=GTF2I;refGene_tx=XM_011290346.3;refGene_gn=GTF2I;refGene_tx=XM_011290347.3;refGene_gn=GTF2I;refGene_tx=XM_019820600.2;refGene_gn=GTF2I;refGene_tx=XM_019820601.2
E3 11041136 11041165
refGene_gn=GTF2I;refGene_tx=XM_011290342.3;refGene_gn=GTF2I;refGene_tx=XM_011290343.3;refGene_gn=GTF2I;refGene_tx=XM_011290344.3;refGene_gn=GTF2I;refGene_tx=XM_011290345.3;refGene_gn=GTF2I;refGene_tx=XM_011290346.3;refGene_gn=GTF2I;refGene_tx=XM_011290347.3;refGene_gn=GTF2I;refGene_tx=XM_019820600.2;refGene_gn=GTF2I;refGene_tx=XM_019820601.2
E3 11045586 11045641
refGene_gn=GTF2I;refGene_tx=XM_011290342.3;refGene_gn=GTF2I;refGene_tx=XM_011290343.3;refGene_gn=GTF2I;refGene_tx=XM_011290344.3;refGene_gn=GTF2I;refGene_tx=XM_011290345.3;refGene_gn=GTF2I;refGene_tx=XM_011290346.3;refGene_gn=GTF2I;refGene_tx=XM_011290347.3;refGene_gn=GTF2I;refGene_tx=XM_019820600.2;refGene_gn=GTF2I;refGene_tx=XM_019820601.2
E3 11046083 11046127
refGene_gn=GTF2I;refGene_tx=XM_011290342.3;refGene_gn=GTF2I;refGene_tx=XM_011290343.3;refGene_gn=GTF2I;refGene_tx=XM_011290344.3;refGene_gn=GTF2I;refGene_tx=XM_011290345.3;refGene_gn=GTF2I;refGene_tx=XM_011290346.3;refGene_gn=GTF2I;refGene_tx=XM_011290347.3;refGene_gn=GTF2I;refGene_tx=XM_019820600.2;refGene_gn=GTF2I;refGene_tx=XM_019820601.2
E3 11056140 11056218
refGene_gn=GTF2I;refGene_tx=XM_011290342.3;refGene_gn=GTF2I;refGene_tx=XM_011290343.3;refGene_gn=GTF2I;refGene_tx=XM_011290344.3;refGene_gn=GTF2I;refGene_tx=XM_011290345.3;refGene_gn=GTF2I;refGene_tx=XM_011290346.3;refGene_gn=GTF2I;refGene_tx=XM_011290347.3;refGene_gn=GTF2I;refGene_tx=XM_019820600.2;refGene_gn=GTF2I;refGene_tx=XM_019820601.2
E3 11065118 11065178
refGene_gn=GTF2I;refGene_tx=XM_011290342.3;refGene_gn=GTF2I;refGene_tx=XM_011290343.3;refGene_gn=GTF2I;refGene_tx=XM_011290345.3;refGene_gn=GTF2I;refGene_tx=XM_011290346.3
E3 11067176 11067238
refGene_gn=GTF2I;refGene_tx=XM_011290342.3;refGene_gn=GTF2I;refGene_tx=XM_011290343.3;refGene_gn=GTF2I;refGene_tx=XM_011290344.3;refGene_gn=GTF2I;refGene_tx=XM_011290345.3;refGene_gn=GTF2I;refGene_tx=XM_011290346.3;refGene_gn=GTF2I;refGene_tx=XM_011290347.3;refGene_gn=GTF2I;refGene_tx=XM_019820601.2
E3 11069181 11069244
refGene_gn=GTF2I;refGene_tx=XM_011290342.3;refGene_gn=GTF2I;refGene_tx=XM_011290343.3;refGene_gn=GTF2I;refGene_tx=XM_011290344.3
E3 11076368 11076479
refGene_gn=GTF2I;refGene_tx=XM_011290342.3;refGene_gn=GTF2I;refGene_tx=XM_011290343.3;refGene_gn=GTF2I;refGene_tx=XM_011290344.3;refGene_gn=GTF2I;refGene_tx=XM_011290345.3;refGene_gn=GTF2I;refGene_tx=XM_011290346.3;refGene_gn=GTF2I;refGene_tx=XM_011290347.3;refGene_gn=GTF2I;refGene_tx=XM_019820600.2
E3 11077495 11077561
refGene_gn=GTF2I;refGene_tx=XM_011290342.3;refGene_gn=GTF2I;refGene_tx=XM_011290343.3;refGene_gn=GTF2I;refGene_tx=XM_011290344.3;refGene_gn=

GTF2I;refGene_tx=XM_011290345.3;refGene_gn=GTF2I;refGene_tx=XM_0112903
46.3;refGene_gn=GTF2I;refGene_tx=XM_011290347.3;refGene_gn=GTF2I;refGe
ne_tx=XM_019820600.2
E3 11080655 11080839
refGene_gn=GTF2I;refGene_tx=XM_011290342.3;refGene_gn=GTF2I;refGene_tx
=XM_011290343.3;refGene_gn=GTF2I;refGene_tx=XM_011290344.3;refGene_gn=
GTF2I;refGene_tx=XM_011290345.3;refGene_gn=GTF2I;refGene_tx=XM_0112903
46.3;refGene_gn=GTF2I;refGene_tx=XM_011290347.3;refGene_gn=GTF2I;refGe
ne_tx=XM_019820600.2
E3 11084542 11084601
refGene_gn=GTF2I;refGene_tx=XM_011290342.3;refGene_gn=GTF2I;refGene_tx
=XM_011290343.3;refGene_gn=GTF2I;refGene_tx=XM_011290344.3;refGene_gn=
GTF2I;refGene_tx=XM_011290345.3;refGene_gn=GTF2I;refGene_tx=XM_0112903
46.3;refGene_gn=GTF2I;refGene_tx=XM_011290347.3;refGene_gn=GTF2I;refGe
ne_tx=XM_019820600.2
E3 11085660 11085732
refGene_gn=GTF2I;refGene_tx=XM_011290342.3;refGene_gn=GTF2I;refGene_tx
=XM_011290343.3;refGene_gn=GTF2I;refGene_tx=XM_011290344.3;refGene_gn=
GTF2I;refGene_tx=XM_011290345.3;refGene_gn=GTF2I;refGene_tx=XM_0112903
46.3;refGene_gn=GTF2I;refGene_tx=XM_011290347.3;refGene_gn=GTF2I;refGe
ne_tx=XM_019820600.2
E3 11090926 11091110
refGene_gn=GTF2I;refGene_tx=XM_011290342.3;refGene_gn=GTF2I;refGene_tx
=XM_011290343.3;refGene_gn=GTF2I;refGene_tx=XM_011290344.3;refGene_gn=
GTF2I;refGene_tx=XM_011290345.3;refGene_gn=GTF2I;refGene_tx=XM_0112903
46.3;refGene_gn=GTF2I;refGene_tx=XM_011290347.3;refGene_gn=GTF2I;refGe
ne_tx=XM_019820600.2
E3 11092524 11092583
refGene_gn=GTF2I;refGene_tx=XM_011290342.3;refGene_gn=GTF2I;refGene_tx
=XM_011290343.3;refGene_gn=GTF2I;refGene_tx=XM_011290344.3;refGene_gn=
GTF2I;refGene_tx=XM_011290345.3;refGene_gn=GTF2I;refGene_tx=XM_0112903
46.3;refGene_gn=GTF2I;refGene_tx=XM_011290347.3;refGene_gn=GTF2I;refGe
ne_tx=XM_019820600.2
E3 11096285 11096357
refGene_gn=GTF2I;refGene_tx=XM_011290342.3;refGene_gn=GTF2I;refGene_tx
=XM_011290343.3;refGene_gn=GTF2I;refGene_tx=XM_011290344.3;refGene_gn=
GTF2I;refGene_tx=XM_011290345.3;refGene_gn=GTF2I;refGene_tx=XM_0112903
46.3;refGene_gn=GTF2I;refGene_tx=XM_011290347.3;refGene_gn=GTF2I;refGe
ne_tx=XM_019820600.2
E3 11097375 11097559
refGene_gn=GTF2I;refGene_tx=XM_011290342.3;refGene_gn=GTF2I;refGene_tx
=XM_011290343.3;refGene_gn=GTF2I;refGene_tx=XM_011290344.3;refGene_gn=
GTF2I;refGene_tx=XM_011290345.3;refGene_gn=GTF2I;refGene_tx=XM_0112903
46.3;refGene_gn=GTF2I;refGene_tx=XM_011290347.3;refGene_gn=GTF2I;refGe
ne_tx=XM_019820600.2
E3 11098201 11098260
refGene_gn=GTF2I;refGene_tx=XM_011290342.3;refGene_gn=GTF2I;refGene_tx
=XM_011290343.3;refGene_gn=GTF2I;refGene_tx=XM_011290344.3;refGene_gn=
GTF2I;refGene_tx=XM_011290345.3;refGene_gn=GTF2I;refGene_tx=XM_0112903
46.3;refGene_gn=GTF2I;refGene_tx=XM_011290347.3;refGene_gn=GTF2I;refGe

ne_tx=XM_019820600.2
E3 11098979 11099054
refGene_gn=GTF2I;refGene_tx=XM_011290342.3;refGene_gn=GTF2I;refGene_tx=XM_011290343.3;refGene_gn=GTF2I;refGene_tx=XM_011290344.3;refGene_gn=GTF2I;refGene_tx=XM_011290345.3;refGene_gn=GTF2I;refGene_tx=XM_011290346.3;refGene_gn=GTF2I;refGene_tx=XM_011290347.3;refGene_gn=GTF2I;refGene_tx=XM_019820600.2
E3 11102752 11102854
refGene_gn=GTF2I;refGene_tx=XM_011290342.3;refGene_gn=GTF2I;refGene_tx=XM_011290343.3;refGene_gn=GTF2I;refGene_tx=XM_011290344.3;refGene_gn=GTF2I;refGene_tx=XM_011290345.3;refGene_gn=GTF2I;refGene_tx=XM_011290346.3;refGene_gn=GTF2I;refGene_tx=XM_011290347.3;refGene_gn=GTF2I;refGene_tx=XM_019820600.2
E3 11103613 11103679
refGene_gn=GTF2I;refGene_tx=XM_011290342.3;refGene_gn=GTF2I;refGene_tx=XM_011290343.3;refGene_gn=GTF2I;refGene_tx=XM_011290344.3;refGene_gn=GTF2I;refGene_tx=XM_011290345.3;refGene_gn=GTF2I;refGene_tx=XM_011290346.3;refGene_gn=GTF2I;refGene_tx=XM_011290347.3;refGene_gn=GTF2I;refGene_tx=XM_019820600.2
E3 11104425 11104609
refGene_gn=GTF2I;refGene_tx=XM_011290342.3;refGene_gn=GTF2I;refGene_tx=XM_011290343.3;refGene_gn=GTF2I;refGene_tx=XM_011290344.3;refGene_gn=GTF2I;refGene_tx=XM_011290345.3;refGene_gn=GTF2I;refGene_tx=XM_011290346.3;refGene_gn=GTF2I;refGene_tx=XM_011290347.3;refGene_gn=GTF2I;refGene_tx=XM_019820600.2
E3 11106037 11106096
refGene_gn=GTF2I;refGene_tx=XM_011290342.3;refGene_gn=GTF2I;refGene_tx=XM_011290343.3;refGene_gn=GTF2I;refGene_tx=XM_011290344.3;refGene_gn=GTF2I;refGene_tx=XM_011290345.3;refGene_gn=GTF2I;refGene_tx=XM_011290346.3;refGene_gn=GTF2I;refGene_tx=XM_011290347.3;refGene_gn=GTF2I;refGene_tx=XM_019820600.2
E3 11106605 11106686
refGene_gn=GTF2I;refGene_tx=XM_011290342.3;refGene_gn=GTF2I;refGene_tx=XM_011290343.3;refGene_gn=GTF2I;refGene_tx=XM_011290344.3;refGene_gn=GTF2I;refGene_tx=XM_011290345.3;refGene_gn=GTF2I;refGene_tx=XM_011290346.3;refGene_gn=GTF2I;refGene_tx=XM_011290347.3;refGene_gn=GTF2I;refGene_tx=XM_019820600.2
E3 11107812 11107896
refGene_gn=GTF2I;refGene_tx=XM_011290342.3;refGene_gn=GTF2I;refGene_tx=XM_011290343.3;refGene_gn=GTF2I;refGene_tx=XM_011290344.3;refGene_gn=GTF2I;refGene_tx=XM_011290345.3;refGene_gn=GTF2I;refGene_tx=XM_011290346.3;refGene_gn=GTF2I;refGene_tx=XM_011290347.3;refGene_gn=GTF2I;refGene_tx=XM_019820600.2
E3 11108249 11108433
refGene_gn=GTF2I;refGene_tx=XM_011290342.3;refGene_gn=GTF2I;refGene_tx=XM_011290343.3;refGene_gn=GTF2I;refGene_tx=XM_011290344.3;refGene_gn=GTF2I;refGene_tx=XM_011290345.3;refGene_gn=GTF2I;refGene_tx=XM_011290346.3;refGene_gn=GTF2I;refGene_tx=XM_011290347.3;refGene_gn=GTF2I;refGene_tx=XM_019820600.2
E3 11109627 11109656

refGene_gn=GTF2I;refGene_tx=XM_011290342.3;refGene_gn=GTF2I;refGene_tx=XM_011290343.3;refGene_gn=GTF2I;refGene_tx=XM_011290344.3;refGene_gn=GTF2I;refGene_tx=XM_011290345.3;refGene_gn=GTF2I;refGene_tx=XM_011290346.3;refGene_gn=GTF2I;refGene_tx=XM_011290347.3;refGene_gn=GTF2I;refGene_tx=XM_019820600.2
E3 11111610 11111652
refGene_gn=GTF2I;refGene_tx=XM_011290342.3;refGene_gn=GTF2I;refGene_tx=XM_011290343.3;refGene_gn=GTF2I;refGene_tx=XM_011290344.3;refGene_gn=GTF2I;refGene_tx=XM_011290345.3;refGene_gn=GTF2I;refGene_tx=XM_011290346.3;refGene_gn=GTF2I;refGene_tx=XM_011290347.3;refGene_gn=GTF2I;refGene_tx=XM_019820600.2
E3 11112456 11112498
refGene_gn=GTF2I;refGene_tx=XM_011290342.3;refGene_gn=GTF2I;refGene_tx=XM_011290343.3;refGene_gn=GTF2I;refGene_tx=XM_011290344.3;refGene_gn=GTF2I;refGene_tx=XM_011290345.3;refGene_gn=GTF2I;refGene_tx=XM_011290346.3;refGene_gn=GTF2I;refGene_tx=XM_011290347.3;refGene_gn=GTF2I;refGene_tx=XM_019820600.2
E3 11112970 11113029
refGene_gn=GTF2I;refGene_tx=XM_011290342.3;refGene_gn=GTF2I;refGene_tx=XM_011290343.3;refGene_gn=GTF2I;refGene_tx=XM_011290344.3;refGene_gn=GTF2I;refGene_tx=XM_011290345.3;refGene_gn=GTF2I;refGene_tx=XM_011290346.3;refGene_gn=GTF2I;refGene_tx=XM_011290347.3;refGene_gn=GTF2I;refGene_tx=XM_019820600.2
E3 11123224 11123296
refGene_gn=NCF1;refGene_tx=XM_023246723.1;refGene_gn=NCF1;refGene_tx=XM_023246724.1;refGene_gn=NCF1;refGene_tx=XM_023246725.1
E3 11128772 11128853
refGene_gn=NCF1;refGene_tx=XM_023246723.1;refGene_gn=NCF1;refGene_tx=XM_023246724.1
E3 11131777 11131853
refGene_gn=NCF1;refGene_tx=XM_023246723.1;refGene_gn=NCF1;refGene_tx=XM_023246724.1;refGene_gn=NCF1;refGene_tx=XM_023246725.1
E3 11131951 11132117
refGene_gn=NCF1;refGene_tx=XM_023246723.1;refGene_gn=NCF1;refGene_tx=XM_023246724.1;refGene_gn=NCF1;refGene_tx=XM_023246725.1
E3 11133363 11133422
refGene_gn=NCF1;refGene_tx=XM_023246723.1;refGene_gn=NCF1;refGene_tx=XM_023246724.1;refGene_gn=NCF1;refGene_tx=XM_023246725.1
E3 11134116 11134239
refGene_gn=NCF1;refGene_tx=XM_023246723.1;refGene_gn=NCF1;refGene_tx=XM_023246724.1;refGene_gn=NCF1;refGene_tx=XM_023246725.1
E3 11134660 11134768
refGene_gn=NCF1;refGene_tx=XM_023246723.1;refGene_gn=NCF1;refGene_tx=XM_023246724.1;refGene_gn=NCF1;refGene_tx=XM_023246725.1
E3 11135610 11135728
refGene_gn=NCF1;refGene_tx=XM_023246723.1;refGene_gn=NCF1;refGene_tx=XM_023246724.1;refGene_gn=NCF1;refGene_tx=XM_023246725.1
E3 11137181 11137289
refGene_gn=NCF1;refGene_tx=XM_023246723.1;refGene_gn=NCF1;refGene_tx=XM_023246724.1;refGene_gn=NCF1;refGene_tx=XM_023246725.1

E3 11137824 11137958
refGene_gn=NCF1;refGene_tx=XM_023246723.1;refGene_gn=NCF1;refGene_tx=X
M_023246724.1;refGene_gn=NCF1;refGene_tx=XM_023246725.1
E3 11138193 11138318
refGene_gn=NCF1;refGene_tx=XM_023246723.1;refGene_gn=NCF1;refGene_tx=X
M_023246724.1;refGene_gn=NCF1;refGene_tx=XM_023246725.1
E3 11142949 11144553
refGene_gn=GTF2IRD2B;refGene_tx=XM_023246375.1;refGene_gn=GTF2IRD2B;re
fGene_tx=XM_023246376.1;refGene_gn=GTF2IRD2B;refGene_tx=XM_023246377.1
E3 11145954 11145986
refGene_gn=GTF2IRD2B;refGene_tx=XM_023246378.1;refGene_gn=GTF2IRD2B;re
fGene_tx=XM_023246379.1
E3 11146008 11146019
refGene_gn=GTF2IRD2B;refGene_tx=XM_023246380.1
E3 11146919 11146948
refGene_gn=GTF2IRD2B;refGene_tx=XM_023246375.1;refGene_gn=GTF2IRD2B;re
fGene_tx=XM_023246376.1;refGene_gn=GTF2IRD2B;refGene_tx=XM_023246377.1
;refGene_gn=GTF2IRD2B;refGene_tx=XM_023246378.1;refGene_gn=GTF2IRD2B;r
efGene_tx=XM_023246379.1;refGene_gn=GTF2IRD2B;refGene_tx=XM_023246380.
1
E3 11147268 11147452
refGene_gn=GTF2IRD2B;refGene_tx=XM_023246375.1;refGene_gn=GTF2IRD2B;re
fGene_tx=XM_023246376.1;refGene_gn=GTF2IRD2B;refGene_tx=XM_023246377.1
;refGene_gn=GTF2IRD2B;refGene_tx=XM_023246378.1;refGene_gn=GTF2IRD2B;r
efGene_tx=XM_023246379.1;refGene_gn=GTF2IRD2B;refGene_tx=XM_023246380.
1
E3 11148345 11148429
refGene_gn=GTF2IRD2B;refGene_tx=XM_023246375.1;refGene_gn=GTF2IRD2B;re
fGene_tx=XM_023246376.1;refGene_gn=GTF2IRD2B;refGene_tx=XM_023246377.1
;refGene_gn=GTF2IRD2B;refGene_tx=XM_023246378.1;refGene_gn=GTF2IRD2B;r
efGene_tx=XM_023246379.1;refGene_gn=GTF2IRD2B;refGene_tx=XM_023246380.
1
E3 11149532 11149610
refGene_gn=GTF2IRD2B;refGene_tx=XM_023246375.1;refGene_gn=GTF2IRD2B;re
fGene_tx=XM_023246376.1;refGene_gn=GTF2IRD2B;refGene_tx=XM_023246377.1
;refGene_gn=GTF2IRD2B;refGene_tx=XM_023246378.1;refGene_gn=GTF2IRD2B;r
efGene_tx=XM_023246379.1;refGene_gn=GTF2IRD2B;refGene_tx=XM_023246380.
1
E3 11157678 11157686
refGene_gn=GTF2IRD2B;refGene_tx=XM_023246381.1;refGene_gn=GTF2IRD2B;re
fGene_tx=XM_023246382.1
E3 11157978 11158041
refGene_gn=GTF2IRD2B;refGene_tx=XM_023246375.1;refGene_gn=GTF2IRD2B;re
fGene_tx=XM_023246376.1;refGene_gn=GTF2IRD2B;refGene_tx=XM_023246377.1
;refGene_gn=GTF2IRD2B;refGene_tx=XM_023246378.1;refGene_gn=GTF2IRD2B;r
efGene_tx=XM_023246379.1;refGene_gn=GTF2IRD2B;refGene_tx=XM_023246380.
1;refGene_gn=GTF2IRD2B;refGene_tx=XM_023246381.1;refGene_gn=GTF2IRD2B;
refGene_tx=XM_023246382.1
E3 11161147 11161204
refGene_gn=GTF2IRD2B;refGene_tx=XM_023246375.1;refGene_gn=GTF2IRD2B;re

fGene_tx=XM_023246376.1;refGene_gn=GTF2IRD2B;refGene_tx=XM_023246377.1
;refGene_gn=GTF2IRD2B;refGene_tx=XM_023246378.1;refGene_gn=GTF2IRD2B;r
efGene_tx=XM_023246379.1;refGene_gn=GTF2IRD2B;refGene_tx=XM_023246380.
1;refGene_gn=GTF2IRD2B;refGene_tx=XM_023246381.1;refGene_gn=GTF2IRD2B;
refGene_tx=XM_023246382.1
E3 11164587 11164665
refGene_gn=GTF2IRD2B;refGene_tx=XM_023246375.1;refGene_gn=GTF2IRD2B;re
fGene_tx=XM_023246376.1;refGene_gn=GTF2IRD2B;refGene_tx=XM_023246377.1
;refGene_gn=GTF2IRD2B;refGene_tx=XM_023246378.1;refGene_gn=GTF2IRD2B;r
efGene_tx=XM_023246379.1;refGene_gn=GTF2IRD2B;refGene_tx=XM_023246380.
1;refGene_gn=GTF2IRD2B;refGene_tx=XM_023246381.1;refGene_gn=GTF2IRD2B;
refGene_tx=XM_023246382.1
E3 11167595 11167639
refGene_gn=GTF2IRD2B;refGene_tx=XM_023246375.1;refGene_gn=GTF2IRD2B;re
fGene_tx=XM_023246376.1;refGene_gn=GTF2IRD2B;refGene_tx=XM_023246377.1
;refGene_gn=GTF2IRD2B;refGene_tx=XM_023246378.1;refGene_gn=GTF2IRD2B;r
efGene_tx=XM_023246379.1;refGene_gn=GTF2IRD2B;refGene_tx=XM_023246380.
1;refGene_gn=GTF2IRD2B;refGene_tx=XM_023246381.1;refGene_gn=GTF2IRD2B;
refGene_tx=XM_023246382.1
E3 11167971 11168026
refGene_gn=GTF2IRD2B;refGene_tx=XM_023246375.1;refGene_gn=GTF2IRD2B;re
fGene_tx=XM_023246376.1;refGene_gn=GTF2IRD2B;refGene_tx=XM_023246377.1
;refGene_gn=GTF2IRD2B;refGene_tx=XM_023246378.1;refGene_gn=GTF2IRD2B;r
efGene_tx=XM_023246379.1;refGene_gn=GTF2IRD2B;refGene_tx=XM_023246380.
1;refGene_gn=GTF2IRD2B;refGene_tx=XM_023246381.1;refGene_gn=GTF2IRD2B;
refGene_tx=XM_023246382.1
E3 11169761 11169790
refGene_gn=GTF2IRD2B;refGene_tx=XM_023246375.1;refGene_gn=GTF2IRD2B;re
fGene_tx=XM_023246376.1;refGene_gn=GTF2IRD2B;refGene_tx=XM_023246377.1
;refGene_gn=GTF2IRD2B;refGene_tx=XM_023246378.1;refGene_gn=GTF2IRD2B;r
efGene_tx=XM_023246379.1;refGene_gn=GTF2IRD2B;refGene_tx=XM_023246380.
1;refGene_gn=GTF2IRD2B;refGene_tx=XM_023246381.1;refGene_gn=GTF2IRD2B;
refGene_tx=XM_023246382.1
E3 11169952 11170136
refGene_gn=GTF2IRD2B;refGene_tx=XM_023246375.1;refGene_gn=GTF2IRD2B;re
fGene_tx=XM_023246376.1;refGene_gn=GTF2IRD2B;refGene_tx=XM_023246377.1
;refGene_gn=GTF2IRD2B;refGene_tx=XM_023246378.1;refGene_gn=GTF2IRD2B;r
efGene_tx=XM_023246379.1;refGene_gn=GTF2IRD2B;refGene_tx=XM_023246380.
1;refGene_gn=GTF2IRD2B;refGene_tx=XM_023246381.1;refGene_gn=GTF2IRD2B;
refGene_tx=XM_023246382.1
E3 11172977 11173097
refGene_gn=GTF2IRD2B;refGene_tx=XM_023246375.1;refGene_gn=GTF2IRD2B;re
fGene_tx=XM_023246376.1;refGene_gn=GTF2IRD2B;refGene_tx=XM_023246377.1
;refGene_gn=GTF2IRD2B;refGene_tx=XM_023246378.1;refGene_gn=GTF2IRD2B;r
efGene_tx=XM_023246379.1;refGene_gn=GTF2IRD2B;refGene_tx=XM_023246380.
1;refGene_gn=GTF2IRD2B;refGene_tx=XM_023246381.1;refGene_gn=GTF2IRD2B;
refGene_tx=XM_023246382.1
E3 11175580 11175719
refGene_gn=GTF2IRD2B;refGene_tx=XM_023246375.1;refGene_gn=GTF2IRD2B;re
fGene_tx=XM_023246376.1;refGene_gn=GTF2IRD2B;refGene_tx=XM_023246377.1

;refGene_gn=GTF2IRD2B;refGene_tx=XM_023246378.1;refGene_gn=GTF2IRD2B;refGene_tx=XM_023246379.1;refGene_gn=GTF2IRD2B;refGene_tx=XM_023246380.1;refGene_gn=GTF2IRD2B;refGene_tx=XM_023246381.1;refGene_gn=GTF2IRD2B;refGene_tx=XM_023246382.1
E3 11188238 11188337
refGene_gn=GTF2IRD2B;refGene_tx=XM_023246375.1;refGene_gn=GTF2IRD2B;refGene_tx=XM_023246376.1;refGene_gn=GTF2IRD2B;refGene_tx=XM_023246378.1;refGene_gn=GTF2IRD2B;refGene_tx=XM_023246379.1;refGene_gn=GTF2IRD2B;refGene_tx=XM_023246380.1;refGene_gn=GTF2IRD2B;refGene_tx=XM_023246381.1;refGene_gn=GTF2IRD2B;refGene_tx=XM_023246382.1
E3 11211473 11211794
refGene_gn=RCC1L;refGene_tx=XM_011290469.2;refGene_gn=RCC1L;refGene_tx=XM_023246383.1
E3 11216833 11216963
refGene_gn=RCC1L;refGene_tx=XM_011290469.2;refGene_gn=RCC1L;refGene_tx=XM_023246383.1
E3 11219735 11219864
refGene_gn=RCC1L;refGene_tx=XM_011290469.2;refGene_gn=RCC1L;refGene_tx=XM_023246383.1
E3 11220712 11220779
refGene_gn=RCC1L;refGene_tx=XM_011290469.2;refGene_gn=RCC1L;refGene_tx=XM_023246383.1
E3 11222217 11222269
refGene_gn=RCC1L;refGene_tx=XM_011290469.2;refGene_gn=RCC1L;refGene_tx=XM_023246383.1
E3 11223142 11223227
refGene_gn=RCC1L;refGene_tx=XM_011290469.2;refGene_gn=RCC1L;refGene_tx=XM_023246383.1
E3 11231170 11231352
refGene_gn=RCC1L;refGene_tx=XM_011290469.2;refGene_gn=RCC1L;refGene_tx=XM_023246383.1
E3 11231664 11231823 refGene_gn=RCC1L;refGene_tx=XM_023246383.1
E3 11235635 11235723 refGene_gn=RCC1L;refGene_tx=XM_011290469.2
E3 11236578 11236752 refGene_gn=RCC1L;refGene_tx=XM_011290469.2
E3 11238738 11238824 refGene_gn=RCC1L;refGene_tx=XM_011290469.2
E3 11244789 11244867 refGene_gn=RCC1L;refGene_tx=XM_011290469.2
E3 11254019 11254085 refGene_gn=CASTOR2;refGene_tx=XM_023246384.1
E3 11254164 11254259 refGene_gn=CASTOR2;refGene_tx=XM_023246384.1
E3 11257561 11257644 refGene_gn=CASTOR2;refGene_tx=XM_023246384.1
E3 11258040 11258151 refGene_gn=CASTOR2;refGene_tx=XM_023246384.1
E3 11258923 11259047 refGene_gn=CASTOR2;refGene_tx=XM_023246384.1
E3 11259574 11259707 refGene_gn=CASTOR2;refGene_tx=XM_023246384.1
E3 11260105 11260299 refGene_gn=CASTOR2;refGene_tx=XM_023246384.1
E3 11271143 11271214 refGene_gn=CASTOR2;refGene_tx=XM_023246384.1
E3 11305256 11305369 refGene_gn=CASTOR2;refGene_tx=XM_023246384.1
E3 12909830 12910139
refGene_gn=AUTS2;refGene_tx=XM_023246462.1;refGene_gn=AUTS2;refGene_tx=XM_023246463.1;refGene_gn=AUTS2;refGene_tx=XM_023246464.1;refGene_gn=AUTS2;refGene_tx=XM_023246465.1;refGene_gn=AUTS2;refGene_tx=XM_023246466.1

E3 12983472 12983496 refGene_gn=AUTS2;refGene_tx=XM_023246467.1
E3 13187603 13187816
refGene_gn=AUTS2;refGene_tx=XM_023246462.1;refGene_gn=AUTS2;refGene_tx=XM_023246463.1;refGene_gn=AUTS2;refGene_tx=XM_023246464.1;refGene_gn=AUTS2;refGene_tx=XM_023246465.1;refGene_gn=AUTS2;refGene_tx=XM_023246466.1;refGene_gn=AUTS2;refGene_tx=XM_023246467.1
E3 13410532 13410634
refGene_gn=AUTS2;refGene_tx=XM_023246462.1;refGene_gn=AUTS2;refGene_tx=XM_023246463.1;refGene_gn=AUTS2;refGene_tx=XM_023246464.1;refGene_gn=AUTS2;refGene_tx=XM_023246465.1;refGene_gn=AUTS2;refGene_tx=XM_023246466.1;refGene_gn=AUTS2;refGene_tx=XM_023246467.1
E3 13425535 13425571
refGene_gn=AUTS2;refGene_tx=XM_023246462.1;refGene_gn=AUTS2;refGene_tx=XM_023246463.1;refGene_gn=AUTS2;refGene_tx=XM_023246464.1;refGene_gn=AUTS2;refGene_tx=XM_023246465.1;refGene_gn=AUTS2;refGene_tx=XM_023246466.1;refGene_gn=AUTS2;refGene_tx=XM_023246467.1
E3 13706302 13706332
refGene_gn=AUTS2;refGene_tx=XM_023246462.1;refGene_gn=AUTS2;refGene_tx=XM_023246463.1;refGene_gn=AUTS2;refGene_tx=XM_023246464.1;refGene_gn=AUTS2;refGene_tx=XM_023246465.1;refGene_gn=AUTS2;refGene_tx=XM_023246466.1;refGene_gn=AUTS2;refGene_tx=XM_023246467.1
E3 13941795 13941847
refGene_gn=AUTS2;refGene_tx=XM_023246462.1;refGene_gn=AUTS2;refGene_tx=XM_023246463.1;refGene_gn=AUTS2;refGene_tx=XM_023246464.1;refGene_gn=AUTS2;refGene_tx=XM_023246465.1;refGene_gn=AUTS2;refGene_tx=XM_023246466.1;refGene_gn=AUTS2;refGene_tx=XM_023246467.1
E3 14000127 14000599
refGene_gn=AUTS2;refGene_tx=XM_023246462.1;refGene_gn=AUTS2;refGene_tx=XM_023246463.1;refGene_gn=AUTS2;refGene_tx=XM_023246464.1;refGene_gn=AUTS2;refGene_tx=XM_023246465.1;refGene_gn=AUTS2;refGene_tx=XM_023246466.1;refGene_gn=AUTS2;refGene_tx=XM_023246467.1
E3 14002233 14002487
refGene_gn=AUTS2;refGene_tx=XM_023246462.1;refGene_gn=AUTS2;refGene_tx=XM_023246463.1;refGene_gn=AUTS2;refGene_tx=XM_023246464.1;refGene_gn=AUTS2;refGene_tx=XM_023246465.1;refGene_gn=AUTS2;refGene_tx=XM_023246466.1;refGene_gn=AUTS2;refGene_tx=XM_023246467.1
E3 14003525 14003767
refGene_gn=AUTS2;refGene_tx=XM_023246462.1;refGene_gn=AUTS2;refGene_tx=XM_023246463.1;refGene_gn=AUTS2;refGene_tx=XM_023246464.1;refGene_gn=AUTS2;refGene_tx=XM_023246465.1;refGene_gn=AUTS2;refGene_tx=XM_023246466.1;refGene_gn=AUTS2;refGene_tx=XM_023246467.1
E3 14005493 14005538
refGene_gn=AUTS2;refGene_tx=XM_023246462.1;refGene_gn=AUTS2;refGene_tx=XM_023246463.1;refGene_gn=AUTS2;refGene_tx=XM_023246464.1;refGene_gn=AUTS2;refGene_tx=XM_023246467.1
E3 14008602 14008698
refGene_gn=AUTS2;refGene_tx=XM_023246462.1;refGene_gn=AUTS2;refGene_tx=XM_023246463.1;refGene_gn=AUTS2;refGene_tx=XM_023246464.1;refGene_gn=AUTS2;refGene_tx=XM_023246465.1;refGene_gn=AUTS2;refGene_tx=XM_023246466.1;refGene_gn=AUTS2;refGene_tx=XM_023246467.1

E3 14011339 14011411
refGene_gn=AUTS2;refGene_tx=XM_023246462.1;refGene_gn=AUTS2;refGene_tx=XM_023246463.1;refGene_gn=AUTS2;refGene_tx=XM_023246464.1;refGene_gn=AUTS2;refGene_tx=XM_023246465.1;refGene_gn=AUTS2;refGene_tx=XM_023246466.1;refGene_gn=AUTS2;refGene_tx=XM_023246467.1
E3 14012630 14012660
refGene_gn=AUTS2;refGene_tx=XM_023246462.1;refGene_gn=AUTS2;refGene_tx=XM_023246463.1;refGene_gn=AUTS2;refGene_tx=XM_023246464.1;refGene_gn=AUTS2;refGene_tx=XM_023246465.1;refGene_gn=AUTS2;refGene_tx=XM_023246466.1;refGene_gn=AUTS2;refGene_tx=XM_023246467.1
E3 14014465 14014537
refGene_gn=AUTS2;refGene_tx=XM_023246462.1;refGene_gn=AUTS2;refGene_tx=XM_023246463.1;refGene_gn=AUTS2;refGene_tx=XM_023246464.1;refGene_gn=AUTS2;refGene_tx=XM_023246465.1;refGene_gn=AUTS2;refGene_tx=XM_023246466.1;refGene_gn=AUTS2;refGene_tx=XM_023246467.1
E3 14018895 14019037
refGene_gn=AUTS2;refGene_tx=XM_023246462.1;refGene_gn=AUTS2;refGene_tx=XM_023246463.1;refGene_gn=AUTS2;refGene_tx=XM_023246464.1;refGene_gn=AUTS2;refGene_tx=XM_023246465.1;refGene_gn=AUTS2;refGene_tx=XM_023246466.1;refGene_gn=AUTS2;refGene_tx=XM_023246467.1
E3 14021901 14021979
refGene_gn=AUTS2;refGene_tx=XM_023246462.1;refGene_gn=AUTS2;refGene_tx=XM_023246463.1;refGene_gn=AUTS2;refGene_tx=XM_023246464.1;refGene_gn=AUTS2;refGene_tx=XM_023246465.1;refGene_gn=AUTS2;refGene_tx=XM_023246466.1;refGene_gn=AUTS2;refGene_tx=XM_023246467.1
E3 14022852 14022936
refGene_gn=AUTS2;refGene_tx=XM_023246462.1;refGene_gn=AUTS2;refGene_tx=XM_023246463.1;refGene_gn=AUTS2;refGene_tx=XM_023246464.1;refGene_gn=AUTS2;refGene_tx=XM_023246465.1;refGene_gn=AUTS2;refGene_tx=XM_023246466.1;refGene_gn=AUTS2;refGene_tx=XM_023246467.1
E3 14024086 14024309
refGene_gn=AUTS2;refGene_tx=XM_023246462.1;refGene_gn=AUTS2;refGene_tx=XM_023246463.1;refGene_gn=AUTS2;refGene_tx=XM_023246464.1;refGene_gn=AUTS2;refGene_tx=XM_023246465.1;refGene_gn=AUTS2;refGene_tx=XM_023246466.1;refGene_gn=AUTS2;refGene_tx=XM_023246467.1
E3 14026661 14027907
refGene_gn=AUTS2;refGene_tx=XM_023246462.1;refGene_gn=AUTS2;refGene_tx=XM_023246463.1;refGene_gn=AUTS2;refGene_tx=XM_023246464.1;refGene_gn=AUTS2;refGene_tx=XM_023246465.1;refGene_gn=AUTS2;refGene_tx=XM_023246466.1;refGene_gn=AUTS2;refGene_tx=XM_023246467.1
E3 14230567 14230805 refGene_gn=GALNT17;refGene_tx=XM_011290337.2
E3 14396974 14397158 refGene_gn=GALNT17;refGene_tx=XM_011290337.2
E3 14436901 14436951 refGene_gn=GALNT17;refGene_tx=XM_023246705.1
E3 14442593 14442760
refGene_gn=GALNT17;refGene_tx=XM_011290337.2;refGene_gn=GALNT17;refGene_tx=XM_023246705.1
E3 14469703 14469878
refGene_gn=GALNT17;refGene_tx=XM_011290337.2;refGene_gn=GALNT17;refGene_tx=XM_023246705.1
E3 14475851 14476049

refGene_gn=GALNT17;refGene_tx=XM_011290337.2;refGene_gn=GALNT17;refGene_tx=XM_023246705.1
E3 14568057 14568175
refGene_gn=GALNT17;refGene_tx=XM_011290337.2;refGene_gn=GALNT17;refGene_tx=XM_023246705.1
E3 14644821 14645007
refGene_gn=GALNT17;refGene_tx=XM_011290337.2;refGene_gn=GALNT17;refGene_tx=XM_023246705.1
E3 14647291 14647429
refGene_gn=GALNT17;refGene_tx=XM_011290337.2;refGene_gn=GALNT17;refGene_tx=XM_023246705.1
E3 14655636 14655732
refGene_gn=GALNT17;refGene_tx=XM_011290337.2;refGene_gn=GALNT17;refGene_tx=XM_023246705.1
E3 14679754 14679922
refGene_gn=GALNT17;refGene_tx=XM_011290337.2;refGene_gn=GALNT17;refGene_tx=XM_023246705.1
E3 14680665 14680794
refGene_gn=GALNT17;refGene_tx=XM_011290337.2;refGene_gn=GALNT17;refGene_tx=XM_023246705.1
E3 14744911 14745039
refGene_gn=CALN1;refGene_tx=XM_011290336.3;refGene_gn=CALN1;refGene_tx=XM_023246903.1
E3 14758050 14758280
refGene_gn=CALN1;refGene_tx=XM_023246898.1;refGene_gn=CALN1;refGene_tx=XM_023246899.1;refGene_gn=CALN1;refGene_tx=XM_023246900.1;refGene_gn=CALN1;refGene_tx=XM_023246901.1;refGene_gn=CALN1;refGene_tx=XM_023246902.1
E3 14758876 14759033
refGene_gn=CALN1;refGene_tx=XM_011290336.3;refGene_gn=CALN1;refGene_tx=XM_023246898.1;refGene_gn=CALN1;refGene_tx=XM_023246899.1;refGene_gn=CALN1;refGene_tx=XM_023246900.1;refGene_gn=CALN1;refGene_tx=XM_023246901.1;refGene_gn=CALN1;refGene_tx=XM_023246902.1;refGene_gn=CALN1;refGene_tx=XM_023246903.1
E3 14933258 14933371
refGene_gn=CALN1;refGene_tx=XM_011290336.3;refGene_gn=CALN1;refGene_tx=XM_023246898.1;refGene_gn=CALN1;refGene_tx=XM_023246899.1;refGene_gn=CALN1;refGene_tx=XM_023246900.1;refGene_gn=CALN1;refGene_tx=XM_023246901.1;refGene_gn=CALN1;refGene_tx=XM_023246902.1;refGene_gn=CALN1;refGene_tx=XM_023246903.1
E3 14996269 14996413
refGene_gn=CALN1;refGene_tx=XM_011290336.3;refGene_gn=CALN1;refGene_tx=XM_023246898.1;refGene_gn=CALN1;refGene_tx=XM_023246899.1;refGene_gn=CALN1;refGene_tx=XM_023246900.1;refGene_gn=CALN1;refGene_tx=XM_023246901.1;refGene_gn=CALN1;refGene_tx=XM_023246902.1;refGene_gn=CALN1;refGene_tx=XM_023246903.1
E3 15123280 15123398
refGene_gn=CALN1;refGene_tx=XM_011290336.3;refGene_gn=CALN1;refGene_tx=XM_023246898.1;refGene_gn=CALN1;refGene_tx=XM_023246899.1;refGene_gn=CALN1;refGene_tx=XM_023246900.1;refGene_gn=CALN1;refGene_tx=XM_0232469

01.1;refGene_gn=CALN1;refGene_tx=XM_023246903.1
E3	15359080	15359302
refGene_gn=LOC101083473;refGene_tx=XM_023247038.1;refGene_gn=LOC101083473;refGene_tx=XM_023247039.1;refGene_gn=LOC101083473;refGene_tx=XM_023247040.1
E3	15383713	15383881
refGene_gn=LOC101083473;refGene_tx=XM_023247038.1;refGene_gn=LOC101083473;refGene_tx=XM_023247039.1
E3	15393161	15393272
refGene_gn=LOC101083473;refGene_tx=XM_023247038.1;refGene_gn=LOC101083473;refGene_tx=XM_023247039.1;refGene_gn=LOC101083473;refGene_tx=XM_023247040.1
E3	15441216	15441352
refGene_gn=LOC101083473;refGene_tx=XM_023247038.1;refGene_gn=LOC101083473;refGene_tx=XM_023247039.1;refGene_gn=LOC101083473;refGene_tx=XM_023247040.1
E3	15459064	15459242
refGene_gn=LOC101083473;refGene_tx=XM_023247038.1;refGene_gn=LOC101083473;refGene_tx=XM_023247039.1;refGene_gn=LOC101083473;refGene_tx=XM_023247040.1
E3	15470100	15470210
refGene_gn=LOC101083473;refGene_tx=XM_023247038.1;refGene_gn=LOC101083473;refGene_tx=XM_023247039.1;refGene_gn=LOC101083473;refGene_tx=XM_023247040.1
E3	15495865	15495984
refGene_gn=LOC101083473;refGene_tx=XM_023247038.1;refGene_gn=LOC101083473;refGene_tx=XM_023247039.1;refGene_gn=LOC101083473;refGene_tx=XM_023247040.1
E3	15533154	15533207
refGene_gn=LOC101083473;refGene_tx=XM_023247038.1;refGene_gn=LOC101083473;refGene_tx=XM_023247039.1;refGene_gn=LOC101083473;refGene_tx=XM_023247040.1
E3	15538932	15539050
refGene_gn=LOC101083473;refGene_tx=XM_023247038.1;refGene_gn=LOC101083473;refGene_tx=XM_023247039.1;refGene_gn=LOC101083473;refGene_tx=XM_023247040.1
E3	15554951	15555074
refGene_gn=LOC101083473;refGene_tx=XM_023247038.1;refGene_gn=LOC101083473;refGene_tx=XM_023247039.1;refGene_gn=LOC101083473;refGene_tx=XM_023247040.1
E3	15560836	15561058
refGene_gn=LOC101083473;refGene_tx=XM_023247038.1;refGene_gn=LOC101083473;refGene_tx=XM_023247039.1;refGene_gn=LOC101083473;refGene_tx=XM_023247040.1
E3	15563176	15563371
refGene_gn=LOC101083473;refGene_tx=XM_023247038.1;refGene_gn=LOC101083473;refGene_tx=XM_023247039.1;refGene_gn=LOC101083473;refGene_tx=XM_023247040.1
E3	15565130	15565232
refGene_gn=LOC101083473;refGene_tx=XM_023247038.1;refGene_gn=LOC101083

473;refGene_tx=XM_023247039.1;refGene_gn=LOC101083473;refGene_tx=XM_023247040.1
E3 15570642 15570795
refGene_gn=LOC101083473;refGene_tx=XM_023247038.1;refGene_gn=LOC101083473;refGene_tx=XM_023247039.1;refGene_gn=LOC101083473;refGene_tx=XM_023247040.1
E3 15571406 15571537
refGene_gn=LOC101083473;refGene_tx=XM_023247038.1;refGene_gn=LOC101083473;refGene_tx=XM_023247039.1;refGene_gn=LOC101083473;refGene_tx=XM_023247040.1
E3 15572655 15572659
refGene_gn=LOC101083473;refGene_tx=XM_023247038.1;refGene_gn=LOC101083473;refGene_tx=XM_023247039.1;refGene_gn=LOC101083473;refGene_tx=XM_023247040.1
E3 15573736 15573864 refGene_gn=SBDS;refGene_tx=XM_003998602.5
E3 15574522 15574652 refGene_gn=SBDS;refGene_tx=XM_003998602.5
E3 15576108 15576309 refGene_gn=SBDS;refGene_tx=XM_003998602.5
E3 15577694 15577859 refGene_gn=SBDS;refGene_tx=XM_003998602.5
E3 15578782 15578911 refGene_gn=SBDS;refGene_tx=XM_003998602.5
E3 15631394 15631415
refGene_gn=TMEM248;refGene_tx=XM_003998603.4;refGene_gn=TMEM248;refGene_tx=XM_023246260.1
E3 15632176 15632320
refGene_gn=TMEM248;refGene_tx=XM_003998603.4;refGene_gn=TMEM248;refGene_tx=XM_023246260.1
E3 15633686 15633870
refGene_gn=TMEM248;refGene_tx=XM_003998603.4;refGene_gn=TMEM248;refGene_tx=XM_023246260.1
E3 15634970 15635121
refGene_gn=TMEM248;refGene_tx=XM_003998603.4;refGene_gn=TMEM248;refGene_tx=XM_023246260.1
E3 15636725 15637011
refGene_gn=TMEM248;refGene_tx=XM_003998603.4;refGene_gn=TMEM248;refGene_tx=XM_023246260.1
E3 15640291 15640468
refGene_gn=TMEM248;refGene_tx=XM_003998603.4;refGene_gn=TMEM248;refGene_tx=XM_023246260.1
E3 15666752 15666854 refGene_gn=TMEM248;refGene_tx=XM_003998603.4
E3 15683505 15683904
refGene_gn=RABGEF1;refGene_tx=XM_011290331.2;refGene_gn=RABGEF1;refGene_tx=XM_011290332.2;refGene_gn=RABGEF1;refGene_tx=XM_011290333.2
E3 15687779 15688036
refGene_gn=RABGEF1;refGene_tx=XM_011290331.2;refGene_gn=RABGEF1;refGene_tx=XM_011290332.2;refGene_gn=RABGEF1;refGene_tx=XM_011290333.2
E3 15693142 15693234
refGene_gn=RABGEF1;refGene_tx=XM_011290331.2;refGene_gn=RABGEF1;refGene_tx=XM_011290332.2;refGene_gn=RABGEF1;refGene_tx=XM_011290333.2
E3 15694482 15694615
refGene_gn=RABGEF1;refGene_tx=XM_011290331.2;refGene_gn=RABGEF1;refGene_tx=XM_011290332.2;refGene_gn=RABGEF1;refGene_tx=XM_011290333.2

E3 15696006 15696088
refGene_gn=RABGEF1;refGene_tx=XM_011290331.2;refGene_gn=RABGEF1;refGene_tx=XM_011290332.2;refGene_gn=RABGEF1;refGene_tx=XM_011290333.2
E3 15710004 15710174
refGene_gn=RABGEF1;refGene_tx=XM_011290331.2;refGene_gn=RABGEF1;refGene_tx=XM_011290332.2;refGene_gn=RABGEF1;refGene_tx=XM_011290333.2
E3 15720735 15720902
refGene_gn=RABGEF1;refGene_tx=XM_011290331.2;refGene_gn=RABGEF1;refGene_tx=XM_011290332.2;refGene_gn=RABGEF1;refGene_tx=XM_011290333.2
E3 15724037 15724216
refGene_gn=RABGEF1;refGene_tx=XM_011290331.2;refGene_gn=RABGEF1;refGene_tx=XM_011290332.2;refGene_gn=RABGEF1;refGene_tx=XM_011290333.2
E3 15761208 15761212 refGene_gn=KCTD7;refGene_tx=XM_023246874.1
E3 15764941 15765314 refGene_gn=KCTD7;refGene_tx=XM_023246874.1
E3 15765772 15765951 refGene_gn=KCTD7;refGene_tx=XM_023246874.1
E3 15768848 15769018 refGene_gn=KCTD7;refGene_tx=XM_023246874.1
E3 15772094 15772238 refGene_gn=KCTD7;refGene_tx=XM_023246874.1
E3 15788554 15788572
refGene_gn=TPST1;refGene_tx=XM_019820629.2;refGene_gn=TPST1;refGene_tx=XM_023247101.1;refGene_gn=TPST1;refGene_tx=XM_023247102.1
E3 15792766 15792817
refGene_gn=TPST1;refGene_tx=XM_019820629.2;refGene_gn=TPST1;refGene_tx=XM_023247101.1;refGene_gn=TPST1;refGene_tx=XM_023247102.1
E3 15846049 15846248
refGene_gn=TPST1;refGene_tx=XM_019820629.2;refGene_gn=TPST1;refGene_tx=XM_023247101.1;refGene_gn=TPST1;refGene_tx=XM_023247102.1
E3 15907931 15908776
refGene_gn=TPST1;refGene_tx=XM_019820629.2;refGene_gn=TPST1;refGene_tx=XM_023247101.1;refGene_gn=TPST1;refGene_tx=XM_023247102.1
E3 16014789 16014936
refGene_gn=CRCP;refGene_tx=XM_023246346.1;refGene_gn=CRCP;refGene_tx=XM_023246347.1;refGene_gn=CRCP;refGene_tx=XM_023246348.1;refGene_gn=CRCP;refGene_tx=XM_023246349.1;refGene_gn=CRCP;refGene_tx=XM_023246350.1
E3 16020179 16020237
refGene_gn=CRCP;refGene_tx=XM_023246346.1;refGene_gn=CRCP;refGene_tx=XM_023246347.1;refGene_gn=CRCP;refGene_tx=XM_023246348.1;refGene_gn=CRCP;refGene_tx=XM_023246349.1;refGene_gn=CRCP;refGene_tx=XM_023246350.1
E3 16035834 16035914 refGene_gn=CRCP;refGene_tx=XM_023246347.1
E3 16041801 16041896
refGene_gn=CRCP;refGene_tx=XM_023246346.1;refGene_gn=CRCP;refGene_tx=XM_023246348.1;refGene_gn=CRCP;refGene_tx=XM_023246349.1
E3 16047424 16047523
refGene_gn=CRCP;refGene_tx=XM_023246346.1;refGene_gn=CRCP;refGene_tx=XM_023246350.1
E3 16051917 16051954 refGene_gn=CRCP;refGene_tx=XM_023246346.1
E3 16060129 16060137 refGene_gn=CRCP;refGene_tx=XM_023246346.1
E3 16086013 16086025
refGene_gn=ASL;refGene_tx=XM_003998605.5;refGene_gn=ASL;refGene_tx=XM_011290327.2;refGene_gn=ASL;refGene_tx=XM_019820635.2
E3 16089323 16089518

refGene_gn=ASL;refGene_tx=XM_003998605.5;refGene_gn=ASL;refGene_tx=XM_011290327.2;refGene_gn=ASL;refGene_tx=XM_019820635.2
E3 16089886 16089970 refGene_gn=ASL;refGene_tx=XM_003998605.5;refGene_gn=ASL;refGene_tx=XM_011290327.2;refGene_gn=ASL;refGene_tx=XM_019820635.2
E3 16090213 16090270 refGene_gn=ASL;refGene_tx=XM_003998605.5;refGene_gn=ASL;refGene_tx=XM_011290327.2;refGene_gn=ASL;refGene_tx=XM_019820635.2
E3 16090413 16090511 refGene_gn=ASL;refGene_tx=XM_003998605.5;refGene_gn=ASL;refGene_tx=XM_011290327.2;refGene_gn=ASL;refGene_tx=XM_019820635.2
E3 16091719 16091797 refGene_gn=ASL;refGene_tx=XM_003998605.5;refGene_gn=ASL;refGene_tx=XM_011290327.2;refGene_gn=ASL;refGene_tx=XM_019820635.2
E3 16091891 16091969 refGene_gn=ASL;refGene_tx=XM_003998605.5;refGene_gn=ASL;refGene_tx=XM_011290327.2;refGene_gn=ASL;refGene_tx=XM_019820635.2
E3 16092419 16092472 refGene_gn=ASL;refGene_tx=XM_003998605.5;refGene_gn=ASL;refGene_tx=XM_011290327.2;refGene_gn=ASL;refGene_tx=XM_019820635.2
E3 16092818 16092881 refGene_gn=ASL;refGene_tx=XM_003998605.5;refGene_gn=ASL;refGene_tx=XM_011290327.2;refGene_gn=ASL;refGene_tx=XM_019820635.2
E3 16093059 16093174 refGene_gn=ASL;refGene_tx=XM_003998605.5;refGene_gn=ASL;refGene_tx=XM_011290327.2;refGene_gn=ASL;refGene_tx=XM_019820635.2
E3 16093347 16093432 refGene_gn=ASL;refGene_tx=XM_003998605.5;refGene_gn=ASL;refGene_tx=XM_011290327.2;refGene_gn=ASL;refGene_tx=XM_019820635.2
E3 16093519 16093579 refGene_gn=ASL;refGene_tx=XM_003998605.5;refGene_gn=ASL;refGene_tx=XM_011290327.2;refGene_gn=ASL;refGene_tx=XM_019820635.2
E3 16093850 16093934 refGene_gn=ASL;refGene_tx=XM_003998605.5;refGene_gn=ASL;refGene_tx=XM_011290327.2;refGene_gn=ASL;refGene_tx=XM_019820635.2
E3 16094773 16094854 refGene_gn=ASL;refGene_tx=XM_003998605.5;refGene_gn=ASL;refGene_tx=XM_011290327.2;refGene_gn=ASL;refGene_tx=XM_019820635.2
E3 16095000 16095107 refGene_gn=ASL;refGene_tx=XM_003998605.5;refGene_gn=ASL;refGene_tx=XM_011290327.2;refGene_gn=ASL;refGene_tx=XM_019820635.2
E3 16095210 16095355 refGene_gn=ASL;refGene_tx=XM_003998605.5;refGene_gn=ASL;refGene_tx=XM_011290327.2;refGene_gn=ASL;refGene_tx=XM_019820635.2
E3 16116442 16116652 refGene_gn=GUSB;refGene_tx=NM_001009310.1
E3 16118047 16118233 refGene_gn=GUSB;refGene_tx=NM_001009310.1;refGene_gn=GUSB;refGene_tx=XM_023246203.1
E3 16118565 16118747 refGene_gn=GUSB;refGene_tx=NM_001009310.1
E3 16118927 16119070 refGene_gn=GUSB;refGene_tx=NM_001009310.1

E3 16119522 16119710
refGene_gn=GUSB;refGene_tx=NM_001009310.1;refGene_gn=GUSB;refGene_tx=XM_023246203.1
E3 16120031 16120184
refGene_gn=GUSB;refGene_tx=NM_001009310.1;refGene_gn=GUSB;refGene_tx=XM_023246203.1
E3 16120434 16120613
refGene_gn=GUSB;refGene_tx=NM_001009310.1;refGene_gn=GUSB;refGene_tx=XM_023246203.1
E3 16120691 16120838
refGene_gn=GUSB;refGene_tx=NM_001009310.1;refGene_gn=GUSB;refGene_tx=XM_023246203.1
E3 16123194 16123279
refGene_gn=GUSB;refGene_tx=NM_001009310.1;refGene_gn=GUSB;refGene_tx=XM_023246203.1
E3 16124872 16125049
refGene_gn=GUSB;refGene_tx=NM_001009310.1;refGene_gn=GUSB;refGene_tx=XM_023246203.1
E3 16127865 16128001
refGene_gn=GUSB;refGene_tx=NM_001009310.1;refGene_gn=GUSB;refGene_tx=XM_023246203.1
E3 16129527 16129697
refGene_gn=GUSB;refGene_tx=NM_001009310.1;refGene_gn=GUSB;refGene_tx=XM_023246203.1
E3 16135537 16135764 refGene_gn=VKORC1L1;refGene_tx=XM_023246218.1
E3 16139401 16139511 refGene_gn=VKORC1L1;refGene_tx=XM_023246218.1
E3 16196262 16196456 refGene_gn=VKORC1L1;refGene_tx=XM_023246218.1
E3 16265994 16266306 refGene_gn=NUPR2;refGene_tx=XM_006942088.4
E3 16272092 16272142 refGene_gn=CHCHD2;refGene_tx=XM_003998609.4
E3 16274503 16274753 refGene_gn=CHCHD2;refGene_tx=XM_003998609.4
E3 16275481 16275632 refGene_gn=CHCHD2;refGene_tx=XM_003998609.4
E3 16276393 16276404 refGene_gn=CHCHD2;refGene_tx=XM_003998609.4
E3 16284543 16284626
refGene_gn=PHKG1;refGene_tx=XM_023246262.1;refGene_gn=PHKG1;refGene_tx=XM_023246263.1
E3 16285104 16285283
refGene_gn=PHKG1;refGene_tx=XM_023246262.1;refGene_gn=PHKG1;refGene_tx=XM_023246263.1
E3 16285729 16285784
refGene_gn=PHKG1;refGene_tx=XM_023246262.1;refGene_gn=PHKG1;refGene_tx=XM_023246263.1
E3 16288288 16288354
refGene_gn=PHKG1;refGene_tx=XM_023246262.1;refGene_gn=PHKG1;refGene_tx=XM_023246263.1
E3 16288629 16288793
refGene_gn=PHKG1;refGene_tx=XM_023246262.1;refGene_gn=PHKG1;refGene_tx=XM_023246263.1
E3 16289180 16289271
refGene_gn=PHKG1;refGene_tx=XM_023246262.1;refGene_gn=PHKG1;refGene_tx=XM_023246263.1

E3 16289404 16289558
refGene_gn=PHKG1;refGene_tx=XM_023246262.1;refGene_gn=PHKG1;refGene_tx
=XM_023246263.1
E3 16289734 16289860
refGene_gn=PHKG1;refGene_tx=XM_023246262.1;refGene_gn=PHKG1;refGene_tx
=XM_023246263.1
E3 16290229 16290475
refGene_gn=PHKG1;refGene_tx=XM_023246262.1;refGene_gn=PHKG1;refGene_tx
=XM_023246263.1
E3 16291629 16291714
refGene_gn=SUMF2;refGene_tx=XM_003998624.4;refGene_gn=SUMF2;refGene_tx
=XM_011290326.2
E3 16292930 16293075
refGene_gn=SUMF2;refGene_tx=XM_003998624.4;refGene_gn=SUMF2;refGene_tx
=XM_011290326.2
E3 16293263 16293348
refGene_gn=SUMF2;refGene_tx=XM_003998624.4;refGene_gn=SUMF2;refGene_tx
=XM_011290326.2
E3 16295243 16295299
refGene_gn=SUMF2;refGene_tx=XM_003998624.4;refGene_gn=SUMF2;refGene_tx
=XM_011290326.2
E3 16296159 16296310
refGene_gn=SUMF2;refGene_tx=XM_003998624.4;refGene_gn=SUMF2;refGene_tx
=XM_011290326.2
E3 16296657 16296702
refGene_gn=SUMF2;refGene_tx=XM_003998624.4;refGene_gn=SUMF2;refGene_tx
=XM_011290326.2
E3 16297152 16297267
refGene_gn=SUMF2;refGene_tx=XM_003998624.4;refGene_gn=SUMF2;refGene_tx
=XM_011290326.2
E3 16299914 16300080
refGene_gn=SUMF2;refGene_tx=XM_003998624.4;refGene_gn=SUMF2;refGene_tx
=XM_011290326.2
E3 16301798 16301862
refGene_gn=SUMF2;refGene_tx=XM_003998624.4;refGene_gn=SUMF2;refGene_tx
=XM_011290326.2
E3 16303138 16303211
refGene_gn=CCT6A;refGene_tx=XM_003998612.5;refGene_gn=CCT6A;refGene_tx
=XM_019820636.2
E3 16303423 16303496
refGene_gn=CCT6A;refGene_tx=XM_003998612.5;refGene_gn=CCT6A;refGene_tx
=XM_019820636.2
E3 16303986 16304089
refGene_gn=CCT6A;refGene_tx=XM_003998612.5;refGene_gn=CCT6A;refGene_tx
=XM_019820636.2
E3 16304594 16304728
refGene_gn=CCT6A;refGene_tx=XM_003998612.5;refGene_gn=CCT6A;refGene_tx
=XM_019820636.2
E3 16305381 16305529
refGene_gn=CCT6A;refGene_tx=XM_003998612.5;refGene_gn=CCT6A;refGene_tx

=XM_019820636.2
E3 16305817 16305914
refGene_gn=CCT6A;refGene_tx=XM_003998612.5;refGene_gn=CCT6A;refGene_tx=XM_019820636.2
E3 16306850 16306933
refGene_gn=CCT6A;refGene_tx=XM_003998612.5;refGene_gn=CCT6A;refGene_tx=XM_019820636.2
E3 16307027 16307187
refGene_gn=CCT6A;refGene_tx=XM_003998612.5;refGene_gn=CCT6A;refGene_tx=XM_019820636.2
E3 16307454 16307565
refGene_gn=CCT6A;refGene_tx=XM_003998612.5;refGene_gn=CCT6A;refGene_tx=XM_019820636.2
E3 16308314 16308418
refGene_gn=CCT6A;refGene_tx=XM_003998612.5;refGene_gn=CCT6A;refGene_tx=XM_019820636.2
E3 16309165 16309339
refGene_gn=CCT6A;refGene_tx=XM_003998612.5;refGene_gn=CCT6A;refGene_tx=XM_019820636.2
E3 16310128 16310263
refGene_gn=CCT6A;refGene_tx=XM_003998612.5;refGene_gn=CCT6A;refGene_tx=XM_019820636.2
E3 16311882 16311946
refGene_gn=CCT6A;refGene_tx=XM_003998612.5;refGene_gn=CCT6A;refGene_tx=XM_019820636.2
E3 16312256 16312262
refGene_gn=PSPH;refGene_tx=XM_006942090.3;refGene_gn=PSPH;refGene_tx=XM_023246265.1
E3 16312392 16312529 refGene_gn=CCT6A;refGene_tx=XM_003998612.5
E3 16344264 16344392
refGene_gn=PSPH;refGene_tx=XM_006942090.3;refGene_gn=PSPH;refGene_tx=XM_023246265.1
E3 16349268 16349291
refGene_gn=PSPH;refGene_tx=XM_019820637.2;refGene_gn=PSPH;refGene_tx=XM_019820638.2
E3 16356298 16356457
refGene_gn=PSPH;refGene_tx=XM_003998615.4;refGene_gn=PSPH;refGene_tx=XM_006942090.3;refGene_gn=PSPH;refGene_tx=XM_006942094.3;refGene_gn=PSPH;refGene_tx=XM_019820637.2;refGene_gn=PSPH;refGene_tx=XM_019820638.2;refGene_gn=PSPH;refGene_tx=XM_023246265.1
E3 16357147 16357282
refGene_gn=PSPH;refGene_tx=XM_003998615.4;refGene_gn=PSPH;refGene_tx=XM_006942090.3;refGene_gn=PSPH;refGene_tx=XM_006942094.3;refGene_gn=PSPH;refGene_tx=XM_019820637.2;refGene_gn=PSPH;refGene_tx=XM_019820638.2;refGene_gn=PSPH;refGene_tx=XM_023246265.1
E3 16357752 16357898
refGene_gn=PSPH;refGene_tx=XM_003998615.4;refGene_gn=PSPH;refGene_tx=XM_006942090.3;refGene_gn=PSPH;refGene_tx=XM_006942094.3;refGene_gn=PSPH;refGene_tx=XM_019820637.2;refGene_gn=PSPH;refGene_tx=XM_019820638.2;refGene_gn=PSPH;refGene_tx=XM_023246265.1

E3 16361129 16361282
refGene_gn=PSPH;refGene_tx=XM_003998615.4;refGene_gn=PSPH;refGene_tx=XM_006942090.3;refGene_gn=PSPH;refGene_tx=XM_006942094.3;refGene_gn=PSPH;refGene_tx=XM_019820637.2;refGene_gn=PSPH;refGene_tx=XM_019820638.2;refGene_gn=PSPH;refGene_tx=XM_023246265.1
E3 16363469 16363577
refGene_gn=PSPH;refGene_tx=XM_003998615.4;refGene_gn=PSPH;refGene_tx=XM_006942090.3;refGene_gn=PSPH;refGene_tx=XM_006942094.3;refGene_gn=PSPH;refGene_tx=XM_019820637.2;refGene_gn=PSPH;refGene_tx=XM_019820638.2;refGene_gn=PSPH;refGene_tx=XM_023246265.1
E3 16373214 16373279 refGene_gn=NIPSNAP2;refGene_tx=XM_023246264.1
E3 16374861 16374945 refGene_gn=NIPSNAP2;refGene_tx=XM_023246264.1
E3 16379154 16379249 refGene_gn=NIPSNAP2;refGene_tx=XM_023246264.1
E3 16389373 16389405 refGene_gn=NIPSNAP2;refGene_tx=XM_023246264.1
E3 16390444 16390585 refGene_gn=NIPSNAP2;refGene_tx=XM_023246264.1
E3 16391372 16391443 refGene_gn=NIPSNAP2;refGene_tx=XM_023246264.1
E3 16391778 16391873 refGene_gn=NIPSNAP2;refGene_tx=XM_023246264.1
E3 16395289 16395335 refGene_gn=NIPSNAP2;refGene_tx=XM_023246264.1
E3 16395417 16395557 refGene_gn=NIPSNAP2;refGene_tx=XM_023246264.1
E3 16412318 16412410 refGene_gn=NIPSNAP2;refGene_tx=XM_023246264.1
E3 16415977 16416247
refGene_gn=MRPS17;refGene_tx=XM_006942097.2;refGene_gn=MRPS17;refGene_tx=XM_006942098.2;refGene_gn=MRPS17;refGene_tx=XM_006942099.3;refGene_gn=MRPS17;refGene_tx=XM_006942100.3
E3 16417833 16417973
refGene_gn=MRPS17;refGene_tx=XM_006942097.2;refGene_gn=MRPS17;refGene_tx=XM_006942098.2;refGene_gn=MRPS17;refGene_tx=XM_006942099.3;refGene_gn=MRPS17;refGene_tx=XM_006942100.3
E3 16418515 16418585 refGene_gn=MRPS17;refGene_tx=XM_006942099.3
E3 16459500 16460558
refGene_gn=ZNF713;refGene_tx=XM_019820640.2;refGene_gn=ZNF713;refGene_tx=XM_019820642.2;refGene_gn=ZNF713;refGene_tx=XM_019820644.2;refGene_gn=ZNF713;refGene_tx=XM_019820645.2;refGene_gn=ZNF713;refGene_tx=XM_023246432.1
E3 16469922 16470015
refGene_gn=ZNF713;refGene_tx=XM_019820640.2;refGene_gn=ZNF713;refGene_tx=XM_019820642.2;refGene_gn=ZNF713;refGene_tx=XM_019820644.2;refGene_gn=ZNF713;refGene_tx=XM_019820645.2;refGene_gn=ZNF713;refGene_tx=XM_023246432.1
E3 16470352 16470479
refGene_gn=ZNF713;refGene_tx=XM_019820640.2;refGene_gn=ZNF713;refGene_tx=XM_019820642.2;refGene_gn=ZNF713;refGene_tx=XM_019820644.2;refGene_gn=ZNF713;refGene_tx=XM_019820645.2;refGene_gn=ZNF713;refGene_tx=XM_023246432.1
E3 16491346 16491435
refGene_gn=ZNF713;refGene_tx=XM_019820640.2;refGene_gn=ZNF713;refGene_tx=XM_019820642.2;refGene_gn=ZNF713;refGene_tx=XM_019820644.2;refGene_gn=ZNF713;refGene_tx=XM_019820645.2;refGene_gn=ZNF713;refGene_tx=XM_023246432.1
E3 16504162 16504247 refGene_gn=ZNF713;refGene_tx=XM_019820640.2

E3 16569292 16569346 refGene_gn=SEPT14;refGene_tx=XM_023247079.1
E3 16580570 16580691 refGene_gn=SEPT14;refGene_tx=XM_023247079.1
E3 16581521 16581717 refGene_gn=SEPT14;refGene_tx=XM_023247079.1
E3 16598842 16599029 refGene_gn=SEPT14;refGene_tx=XM_023247079.1
E3 16601447 16601591 refGene_gn=SEPT14;refGene_tx=XM_023247079.1
E3 16634216 16634313 refGene_gn=SEPT14;refGene_tx=XM_023247079.1
E3 16635604 16635773 refGene_gn=SEPT14;refGene_tx=XM_023247079.1
E3 16644636 16644787 refGene_gn=SEPT14;refGene_tx=XM_023247079.1
E3 20007622 20007826 refGene_gn=AHSP;refGene_tx=XM_003998693.4
E3 20008033 20008108 refGene_gn=AHSP;refGene_tx=XM_003998693.4
E3 20016608 20016908
refGene_gn=CE3H16orf58;refGene_tx=XM_003998627.5;refGene_gn=CE3H16orf5
8;refGene_tx=XM_006942110.4;refGene_gn=CE3H16orf58;refGene_tx=XM_00694
2111.4;refGene_gn=CE3H16orf58;refGene_tx=XM_006942112.4;refGene_gn=CE3
H16orf58;refGene_tx=XM_019820650.2;refGene_gn=CE3H16orf58;refGene_tx=X
M_019820651.2
E3 20017126 20017241
refGene_gn=CE3H16orf58;refGene_tx=XM_003998627.5;refGene_gn=CE3H16orf5
8;refGene_tx=XM_006942110.4;refGene_gn=CE3H16orf58;refGene_tx=XM_00694
2111.4;refGene_gn=CE3H16orf58;refGene_tx=XM_006942112.4;refGene_gn=CE3
H16orf58;refGene_tx=XM_019820650.2;refGene_gn=CE3H16orf58;refGene_tx=X
M_019820651.2
E3 20021055 20021101
refGene_gn=CE3H16orf58;refGene_tx=XM_003998627.5;refGene_gn=CE3H16orf5
8;refGene_tx=XM_006942110.4;refGene_gn=CE3H16orf58;refGene_tx=XM_00694
2111.4;refGene_gn=CE3H16orf58;refGene_tx=XM_006942112.4;refGene_gn=CE3
H16orf58;refGene_tx=XM_019820650.2;refGene_gn=CE3H16orf58;refGene_tx=X
M_019820651.2
E3 20022668 20022754
refGene_gn=CE3H16orf58;refGene_tx=XM_003998627.5;refGene_gn=CE3H16orf5
8;refGene_tx=XM_006942110.4;refGene_gn=CE3H16orf58;refGene_tx=XM_00694
2111.4;refGene_gn=CE3H16orf58;refGene_tx=XM_006942112.4;refGene_gn=CE3
H16orf58;refGene_tx=XM_019820650.2;refGene_gn=CE3H16orf58;refGene_tx=X
M_019820651.2
E3 20022843 20022949
refGene_gn=CE3H16orf58;refGene_tx=XM_003998627.5;refGene_gn=CE3H16orf5
8;refGene_tx=XM_006942110.4;refGene_gn=CE3H16orf58;refGene_tx=XM_00694
2111.4;refGene_gn=CE3H16orf58;refGene_tx=XM_006942112.4;refGene_gn=CE3
H16orf58;refGene_tx=XM_019820650.2;refGene_gn=CE3H16orf58;refGene_tx=X
M_019820651.2
E3 20025374 20025476
refGene_gn=CE3H16orf58;refGene_tx=XM_003998627.5;refGene_gn=CE3H16orf5
8;refGene_tx=XM_006942110.4;refGene_gn=CE3H16orf58;refGene_tx=XM_00694
2111.4;refGene_gn=CE3H16orf58;refGene_tx=XM_006942112.4;refGene_gn=CE3
H16orf58;refGene_tx=XM_019820650.2
E3 20027436 20027507
refGene_gn=CE3H16orf58;refGene_tx=XM_003998627.5;refGene_gn=CE3H16orf5
8;refGene_tx=XM_006942110.4;refGene_gn=CE3H16orf58;refGene_tx=XM_00694
2111.4;refGene_gn=CE3H16orf58;refGene_tx=XM_006942112.4;refGene_gn=CE3
H16orf58;refGene_tx=XM_019820650.2;refGene_gn=CE3H16orf58;refGene_tx=X

M_019820651.2
E3	20027586	20027771
refGene_gn=CE3H16orf58;refGene_tx=XM_003998627.5;refGene_gn=CE3H16orf58;refGene_tx=XM_006942110.4;refGene_gn=CE3H16orf58;refGene_tx=XM_006942111.4;refGene_gn=CE3H16orf58;refGene_tx=XM_019820650.2;refGene_gn=CE3H16orf58;refGene_tx=XM_019820651.2
E3	20027858	20027916
refGene_gn=CE3H16orf58;refGene_tx=XM_003998627.5;refGene_gn=CE3H16orf58;refGene_tx=XM_006942110.4;refGene_gn=CE3H16orf58;refGene_tx=XM_006942111.4;refGene_gn=CE3H16orf58;refGene_tx=XM_019820650.2;refGene_gn=CE3H16orf58;refGene_tx=XM_019820651.2
E3	20028281	20028352
refGene_gn=CE3H16orf58;refGene_tx=XM_003998627.5;refGene_gn=CE3H16orf58;refGene_tx=XM_006942110.4;refGene_gn=CE3H16orf58;refGene_tx=XM_006942111.4;refGene_gn=CE3H16orf58;refGene_tx=XM_006942112.4;refGene_gn=CE3H16orf58;refGene_tx=XM_019820650.2;refGene_gn=CE3H16orf58;refGene_tx=XM_019820651.2
E3	20028986	20029130
refGene_gn=CE3H16orf58;refGene_tx=XM_003998627.5;refGene_gn=CE3H16orf58;refGene_tx=XM_006942111.4;refGene_gn=CE3H16orf58;refGene_tx=XM_006942112.4;refGene_gn=CE3H16orf58;refGene_tx=XM_019820650.2;refGene_gn=CE3H16orf58;refGene_tx=XM_019820651.2
E3	20029237	20029315
refGene_gn=CE3H16orf58;refGene_tx=XM_003998627.5;refGene_gn=CE3H16orf58;refGene_tx=XM_006942110.4;refGene_gn=CE3H16orf58;refGene_tx=XM_006942111.4;refGene_gn=CE3H16orf58;refGene_tx=XM_006942112.4;refGene_gn=CE3H16orf58;refGene_tx=XM_019820651.2
E3	20029809	20029907
refGene_gn=CE3H16orf58;refGene_tx=XM_003998627.5;refGene_gn=CE3H16orf58;refGene_tx=XM_006942110.4;refGene_gn=CE3H16orf58;refGene_tx=XM_006942111.4;refGene_gn=CE3H16orf58;refGene_tx=XM_006942112.4;refGene_gn=CE3H16orf58;refGene_tx=XM_019820650.2;refGene_gn=CE3H16orf58;refGene_tx=XM_019820651.2
E3	20030215	20030442	refGene_gn=SLC5A2;refGene_tx=XM_003998628.4
E3	20030527	20030657	refGene_gn=SLC5A2;refGene_tx=XM_003998628.4
E3	20031442	20031658	refGene_gn=SLC5A2;refGene_tx=XM_003998628.4
E3	20031736	20031905	refGene_gn=SLC5A2;refGene_tx=XM_003998628.4
E3	20032030	20032181	refGene_gn=SLC5A2;refGene_tx=XM_003998628.4
E3	20032296	20032404	refGene_gn=SLC5A2;refGene_tx=XM_003998628.4
E3	20032622	20032758	refGene_gn=SLC5A2;refGene_tx=XM_003998628.4
E3	20033075	20033305	refGene_gn=SLC5A2;refGene_tx=XM_003998628.4
E3	20033457	20033538	refGene_gn=SLC5A2;refGene_tx=XM_003998628.4
E3	20033780	20033886	refGene_gn=SLC5A2;refGene_tx=XM_003998628.4
E3	20034187	20034352	refGene_gn=SLC5A2;refGene_tx=XM_003998628.4
E3	20035190	20035295	refGene_gn=SLC5A2;refGene_tx=XM_003998628.4
E3	20035371	20035443	refGene_gn=SLC5A2;refGene_tx=XM_003998628.4
E3	20035986	20036112	refGene_gn=SLC5A2;refGene_tx=XM_003998628.4
E3	20039845	20040112
refGene_gn=TGFB1I1;refGene_tx=XM_003998629.4;refGene_gn=TGFB1I1;refGene_tx=XM_006942108.3;refGene_gn=TGFB1I1;refGene_tx=XM_019820647.2;refGe

ne_gn=TGFB1I1;refGene_tx=XM_019820648.1;refGene_gn=TGFB1I1;refGene_tx=XM_019820649.2
E3 20040351 20040500
refGene_gn=TGFB1I1;refGene_tx=XM_003998629.4;refGene_gn=TGFB1I1;refGene_tx=XM_006942108.3;refGene_gn=TGFB1I1;refGene_tx=XM_019820647.2;refGene_gn=TGFB1I1;refGene_tx=XM_019820648.1;refGene_gn=TGFB1I1;refGene_tx=XM_019820649.2
E3 20040780 20040862
refGene_gn=TGFB1I1;refGene_tx=XM_003998629.4;refGene_gn=TGFB1I1;refGene_tx=XM_006942108.3;refGene_gn=TGFB1I1;refGene_tx=XM_019820647.2;refGene_gn=TGFB1I1;refGene_tx=XM_019820648.1
E3 20041134 20041308
refGene_gn=TGFB1I1;refGene_tx=XM_003998629.4;refGene_gn=TGFB1I1;refGene_tx=XM_006942108.3;refGene_gn=TGFB1I1;refGene_tx=XM_019820647.2;refGene_gn=TGFB1I1;refGene_tx=XM_019820648.1;refGene_gn=TGFB1I1;refGene_tx=XM_019820649.2
E3 20042233 20042428
refGene_gn=TGFB1I1;refGene_tx=XM_003998629.4;refGene_gn=TGFB1I1;refGene_tx=XM_006942108.3;refGene_gn=TGFB1I1;refGene_tx=XM_019820647.2;refGene_gn=TGFB1I1;refGene_tx=XM_019820648.1;refGene_gn=TGFB1I1;refGene_tx=XM_019820649.2
E3 20042530 20042645
refGene_gn=TGFB1I1;refGene_tx=XM_003998629.4;refGene_gn=TGFB1I1;refGene_tx=XM_006942108.3;refGene_gn=TGFB1I1;refGene_tx=XM_019820647.2;refGene_gn=TGFB1I1;refGene_tx=XM_019820648.1;refGene_gn=TGFB1I1;refGene_tx=XM_019820649.2
E3 20042761 20042849
refGene_gn=TGFB1I1;refGene_tx=XM_003998629.4;refGene_gn=TGFB1I1;refGene_tx=XM_006942108.3;refGene_gn=TGFB1I1;refGene_tx=XM_019820647.2;refGene_gn=TGFB1I1;refGene_tx=XM_019820648.1;refGene_gn=TGFB1I1;refGene_tx=XM_019820649.2
E3 20043013 20043156
refGene_gn=TGFB1I1;refGene_tx=XM_003998629.4;refGene_gn=TGFB1I1;refGene_tx=XM_006942108.3;refGene_gn=TGFB1I1;refGene_tx=XM_019820647.2;refGene_gn=TGFB1I1;refGene_tx=XM_019820648.1;refGene_gn=TGFB1I1;refGene_tx=XM_019820649.2
E3 20043258 20043311
refGene_gn=TGFB1I1;refGene_tx=XM_003998629.4;refGene_gn=TGFB1I1;refGene_tx=XM_006942108.3;refGene_gn=TGFB1I1;refGene_tx=XM_019820647.2;refGene_gn=TGFB1I1;refGene_tx=XM_019820648.1;refGene_gn=TGFB1I1;refGene_tx=XM_019820649.2
E3 20043436 20043552
refGene_gn=TGFB1I1;refGene_tx=XM_003998629.4;refGene_gn=TGFB1I1;refGene_tx=XM_006942108.3;refGene_gn=TGFB1I1;refGene_tx=XM_019820647.2;refGene_gn=TGFB1I1;refGene_tx=XM_019820648.1;refGene_gn=TGFB1I1;refGene_tx=XM_019820649.2
E3 20043985 20044079
refGene_gn=TGFB1I1;refGene_tx=XM_019820647.2;refGene_gn=TGFB1I1;refGene_tx=XM_019820648.1;refGene_gn=TGFB1I1;refGene_tx=XM_019820649.2
E3 20044856 20044869

refGene_gn=TGFB1I1;refGene_tx=XM_003998629.4;refGene_gn=TGFB1I1;refGene_tx=XM_006942108.3
E3 20046946 20047766
refGene_gn=ARMC5;refGene_tx=XM_011290549.3;refGene_gn=ARMC5;refGene_tx=XM_023246881.1
E3 20047868 20048001
refGene_gn=ARMC5;refGene_tx=XM_011290549.3;refGene_gn=ARMC5;refGene_tx=XM_023246881.1
E3 20048606 20049112
refGene_gn=ARMC5;refGene_tx=XM_011290549.3;refGene_gn=ARMC5;refGene_tx=XM_023246881.1
E3 20049853 20050640
refGene_gn=ARMC5;refGene_tx=XM_011290549.3;refGene_gn=ARMC5;refGene_tx=XM_023246881.1
E3 20050739 20050847
refGene_gn=ARMC5;refGene_tx=XM_011290549.3;refGene_gn=ARMC5;refGene_tx=XM_023246881.1
E3 20052706 20053184 refGene_gn=ARMC5;refGene_tx=XM_011290549.3
E3 20094604 20094677
refGene_gn=LOC105261306;refGene_tx=XM_011290548.2
E3 20094868 20095005
refGene_gn=LOC105261306;refGene_tx=XM_011290548.2
E3 20095122 20095203
refGene_gn=LOC105261306;refGene_tx=XM_011290548.2
E3 20096447 20096561
refGene_gn=ITGAD;refGene_tx=XM_023246699.1;refGene_gn=ITGAD;refGene_tx=XM_023246701.1
E3 20097023 20097134
refGene_gn=ITGAD;refGene_tx=XM_023246699.1;refGene_gn=ITGAD;refGene_tx=XM_023246701.1
E3 20097392 20097494
refGene_gn=ITGAD;refGene_tx=XM_023246699.1;refGene_gn=ITGAD;refGene_tx=XM_023246701.1
E3 20097646 20097760
refGene_gn=ITGAD;refGene_tx=XM_023246699.1;refGene_gn=ITGAD;refGene_tx=XM_023246701.1
E3 20097963 20098041
refGene_gn=ITGAD;refGene_tx=XM_023246699.1;refGene_gn=ITGAD;refGene_tx=XM_023246701.1
E3 20098149 20098257
refGene_gn=ITGAD;refGene_tx=XM_023246699.1;refGene_gn=ITGAD;refGene_tx=XM_023246700.1;refGene_gn=ITGAD;refGene_tx=XM_023246701.1;refGene_gn=ITGAD;refGene_tx=XM_023246702.1
E3 20098420 20098499
refGene_gn=ITGAD;refGene_tx=XM_023246699.1;refGene_gn=ITGAD;refGene_tx=XM_023246700.1;refGene_gn=ITGAD;refGene_tx=XM_023246701.1;refGene_gn=ITGAD;refGene_tx=XM_023246702.1
E3 20098545 20098576 refGene_gn=ITGAD;refGene_tx=XM_023246703.1
E3 20100801 20100885
refGene_gn=ITGAD;refGene_tx=XM_023246699.1;refGene_gn=ITGAD;refGene_tx

=XM_023246700.1;refGene_gn=ITGAD;refGene_tx=XM_023246701.1;refGene_gn=ITGAD;refGene_tx=XM_023246702.1;refGene_gn=ITGAD;refGene_tx=XM_023246703.1
E3 20100986 20101066 refGene_gn=ITGAD;refGene_tx=XM_023246699.1;refGene_gn=ITGAD;refGene_tx=XM_023246700.1;refGene_gn=ITGAD;refGene_tx=XM_023246701.1;refGene_gn=ITGAD;refGene_tx=XM_023246702.1;refGene_gn=ITGAD;refGene_tx=XM_023246703.1
E3 20101179 20101293 refGene_gn=ITGAD;refGene_tx=XM_023246699.1;refGene_gn=ITGAD;refGene_tx=XM_023246700.1;refGene_gn=ITGAD;refGene_tx=XM_023246701.1;refGene_gn=ITGAD;refGene_tx=XM_023246702.1;refGene_gn=ITGAD;refGene_tx=XM_023246703.1
E3 20102789 20102931 refGene_gn=ITGAD;refGene_tx=XM_023246699.1;refGene_gn=ITGAD;refGene_tx=XM_023246700.1;refGene_gn=ITGAD;refGene_tx=XM_023246701.1;refGene_gn=ITGAD;refGene_tx=XM_023246702.1;refGene_gn=ITGAD;refGene_tx=XM_023246703.1
E3 20103161 20103235 refGene_gn=ITGAD;refGene_tx=XM_023246699.1;refGene_gn=ITGAD;refGene_tx=XM_023246700.1;refGene_gn=ITGAD;refGene_tx=XM_023246701.1;refGene_gn=ITGAD;refGene_tx=XM_023246702.1;refGene_gn=ITGAD;refGene_tx=XM_023246703.1
E3 20104055 20104187 refGene_gn=ITGAD;refGene_tx=XM_023246699.1;refGene_gn=ITGAD;refGene_tx=XM_023246700.1;refGene_gn=ITGAD;refGene_tx=XM_023246701.1;refGene_gn=ITGAD;refGene_tx=XM_023246702.1;refGene_gn=ITGAD;refGene_tx=XM_023246703.1
E3 20104445 20104600 refGene_gn=ITGAD;refGene_tx=XM_023246699.1;refGene_gn=ITGAD;refGene_tx=XM_023246700.1;refGene_gn=ITGAD;refGene_tx=XM_023246701.1;refGene_gn=ITGAD;refGene_tx=XM_023246702.1;refGene_gn=ITGAD;refGene_tx=XM_023246703.1
E3 20105486 20105644 refGene_gn=ITGAD;refGene_tx=XM_023246699.1;refGene_gn=ITGAD;refGene_tx=XM_023246700.1;refGene_gn=ITGAD;refGene_tx=XM_023246701.1;refGene_gn=ITGAD;refGene_tx=XM_023246702.1;refGene_gn=ITGAD;refGene_tx=XM_023246703.1
E3 20105762 20105893 refGene_gn=ITGAD;refGene_tx=XM_023246699.1;refGene_gn=ITGAD;refGene_tx=XM_023246700.1;refGene_gn=ITGAD;refGene_tx=XM_023246701.1;refGene_gn=ITGAD;refGene_tx=XM_023246702.1;refGene_gn=ITGAD;refGene_tx=XM_023246703.1
E3 20106594 20106804 refGene_gn=ITGAD;refGene_tx=XM_023246699.1;refGene_gn=ITGAD;refGene_tx=XM_023246700.1;refGene_gn=ITGAD;refGene_tx=XM_023246702.1;refGene_gn=ITGAD;refGene_tx=XM_023246703.1
E3 20106897 20107038 refGene_gn=ITGAD;refGene_tx=XM_023246699.1;refGene_gn=ITGAD;refGene_tx=XM_023246700.1;refGene_gn=ITGAD;refGene_tx=XM_023246702.1;refGene_gn=

ITGAD;refGene_tx=XM_023246703.1
E3 20107189 20107332
refGene_gn=ITGAD;refGene_tx=XM_023246699.1;refGene_gn=ITGAD;refGene_tx
=XM_023246700.1;refGene_gn=ITGAD;refGene_tx=XM_023246702.1;refGene_gn=
ITGAD;refGene_tx=XM_023246703.1
E3 20107446 20107576
refGene_gn=ITGAD;refGene_tx=XM_023246699.1;refGene_gn=ITGAD;refGene_tx
=XM_023246700.1;refGene_gn=ITGAD;refGene_tx=XM_023246702.1;refGene_gn=
ITGAD;refGene_tx=XM_023246703.1
E3 20108878 20108952
refGene_gn=ITGAD;refGene_tx=XM_023246699.1;refGene_gn=ITGAD;refGene_tx
=XM_023246700.1;refGene_gn=ITGAD;refGene_tx=XM_023246701.1;refGene_gn=
ITGAD;refGene_tx=XM_023246702.1;refGene_gn=ITGAD;refGene_tx=XM_0232467
03.1
E3 20109148 20109299
refGene_gn=ITGAD;refGene_tx=XM_023246699.1;refGene_gn=ITGAD;refGene_tx
=XM_023246700.1;refGene_gn=ITGAD;refGene_tx=XM_023246701.1;refGene_gn=
ITGAD;refGene_tx=XM_023246702.1;refGene_gn=ITGAD;refGene_tx=XM_0232467
03.1
E3 20109395 20109549
refGene_gn=ITGAD;refGene_tx=XM_023246699.1;refGene_gn=ITGAD;refGene_tx
=XM_023246700.1;refGene_gn=ITGAD;refGene_tx=XM_023246701.1;refGene_gn=
ITGAD;refGene_tx=XM_023246702.1;refGene_gn=ITGAD;refGene_tx=XM_0232467
03.1
E3 20110107 20110253
refGene_gn=ITGAD;refGene_tx=XM_023246699.1;refGene_gn=ITGAD;refGene_tx
=XM_023246700.1;refGene_gn=ITGAD;refGene_tx=XM_023246701.1;refGene_gn=
ITGAD;refGene_tx=XM_023246702.1;refGene_gn=ITGAD;refGene_tx=XM_0232467
03.1
E3 20110918 20111049
refGene_gn=ITGAD;refGene_tx=XM_023246699.1;refGene_gn=ITGAD;refGene_tx
=XM_023246700.1;refGene_gn=ITGAD;refGene_tx=XM_023246701.1;refGene_gn=
ITGAD;refGene_tx=XM_023246702.1;refGene_gn=ITGAD;refGene_tx=XM_0232467
03.1
E3 20112598 20112713
refGene_gn=ITGAD;refGene_tx=XM_023246699.1;refGene_gn=ITGAD;refGene_tx
=XM_023246700.1;refGene_gn=ITGAD;refGene_tx=XM_023246701.1;refGene_gn=
ITGAD;refGene_tx=XM_023246702.1;refGene_gn=ITGAD;refGene_tx=XM_0232467
03.1
E3 20112838 20112909
refGene_gn=ITGAD;refGene_tx=XM_023246699.1;refGene_gn=ITGAD;refGene_tx
=XM_023246700.1;refGene_gn=ITGAD;refGene_tx=XM_023246701.1;refGene_gn=
ITGAD;refGene_tx=XM_023246702.1;refGene_gn=ITGAD;refGene_tx=XM_0232467
03.1
E3 20113041 20113145
refGene_gn=ITGAD;refGene_tx=XM_023246699.1;refGene_gn=ITGAD;refGene_tx
=XM_023246700.1;refGene_gn=ITGAD;refGene_tx=XM_023246701.1;refGene_gn=
ITGAD;refGene_tx=XM_023246702.1;refGene_gn=ITGAD;refGene_tx=XM_0232467
03.1
E3 20115762 20115868

refGene_gn=ITGAD;refGene_tx=XM_023246699.1;refGene_gn=ITGAD;refGene_tx=XM_023246700.1;refGene_gn=ITGAD;refGene_tx=XM_023246701.1;refGene_gn=ITGAD;refGene_tx=XM_023246702.1;refGene_gn=ITGAD;refGene_tx=XM_023246703.1
E3 20117620 20117654
refGene_gn=ITGAD;refGene_tx=XM_023246699.1;refGene_gn=ITGAD;refGene_tx=XM_023246700.1;refGene_gn=ITGAD;refGene_tx=XM_023246701.1;refGene_gn=ITGAD;refGene_tx=XM_023246702.1;refGene_gn=ITGAD;refGene_tx=XM_023246703.1
E3 20154053 20154087
refGene_gn=LOC101087216;refGene_tx=XM_023246717.1;refGene_gn=LOC101087216;refGene_tx=XM_023246718.1;refGene_gn=LOC101087216;refGene_tx=XM_023246719.1
E3 20155847 20155953
refGene_gn=LOC101087216;refGene_tx=XM_023246717.1;refGene_gn=LOC101087216;refGene_tx=XM_023246718.1;refGene_gn=LOC101087216;refGene_tx=XM_023246719.1
E3 20158579 20158683
refGene_gn=LOC101087216;refGene_tx=XM_023246717.1;refGene_gn=LOC101087216;refGene_tx=XM_023246718.1;refGene_gn=LOC101087216;refGene_tx=XM_023246719.1
E3 20158814 20158885
refGene_gn=LOC101087216;refGene_tx=XM_023246717.1;refGene_gn=LOC101087216;refGene_tx=XM_023246718.1;refGene_gn=LOC101087216;refGene_tx=XM_023246719.1
E3 20159010 20159125
refGene_gn=LOC101087216;refGene_tx=XM_023246717.1;refGene_gn=LOC101087216;refGene_tx=XM_023246718.1;refGene_gn=LOC101087216;refGene_tx=XM_023246719.1
E3 20162337 20162468
refGene_gn=LOC101087216;refGene_tx=XM_023246717.1;refGene_gn=LOC101087216;refGene_tx=XM_023246718.1;refGene_gn=LOC101087216;refGene_tx=XM_023246719.1
E3 20163126 20163272
refGene_gn=LOC101087216;refGene_tx=XM_023246717.1;refGene_gn=LOC101087216;refGene_tx=XM_023246718.1;refGene_gn=LOC101087216;refGene_tx=XM_023246719.1
E3 20164757 20164911
refGene_gn=LOC101087216;refGene_tx=XM_023246717.1;refGene_gn=LOC101087216;refGene_tx=XM_023246718.1;refGene_gn=LOC101087216;refGene_tx=XM_023246719.1
E3 20165007 20165158
refGene_gn=LOC101087216;refGene_tx=XM_023246717.1;refGene_gn=LOC101087216;refGene_tx=XM_023246718.1;refGene_gn=LOC101087216;refGene_tx=XM_023246719.1
E3 20165354 20165428
refGene_gn=LOC101087216;refGene_tx=XM_023246717.1;refGene_gn=LOC101087216;refGene_tx=XM_023246718.1;refGene_gn=LOC101087216;refGene_tx=XM_023246719.1
E3 20166729 20166859

refGene_gn=LOC101087216;refGene_tx=XM_023246717.1;refGene_gn=LOC101087216;refGene_tx=XM_023246718.1;refGene_gn=LOC101087216;refGene_tx=XM_023246719.1
E3 20166973 20167116
refGene_gn=LOC101087216;refGene_tx=XM_023246717.1;refGene_gn=LOC101087216;refGene_tx=XM_023246718.1;refGene_gn=LOC101087216;refGene_tx=XM_023246719.1
E3 20167267 20167408
refGene_gn=LOC101087216;refGene_tx=XM_023246717.1;refGene_gn=LOC101087216;refGene_tx=XM_023246718.1;refGene_gn=LOC101087216;refGene_tx=XM_023246719.1
E3 20167501 20167711
refGene_gn=LOC101087216;refGene_tx=XM_023246717.1;refGene_gn=LOC101087216;refGene_tx=XM_023246718.1;refGene_gn=LOC101087216;refGene_tx=XM_023246719.1
E3 20168412 20168543
refGene_gn=LOC101087216;refGene_tx=XM_023246717.1;refGene_gn=LOC101087216;refGene_tx=XM_023246718.1;refGene_gn=LOC101087216;refGene_tx=XM_023246719.1
E3 20168661 20168819
refGene_gn=LOC101087216;refGene_tx=XM_023246717.1;refGene_gn=LOC101087216;refGene_tx=XM_023246718.1;refGene_gn=LOC101087216;refGene_tx=XM_023246719.1
E3 20169703 20169858
refGene_gn=LOC101087216;refGene_tx=XM_023246717.1;refGene_gn=LOC101087216;refGene_tx=XM_023246718.1;refGene_gn=LOC101087216;refGene_tx=XM_023246719.1
E3 20170117 20170249
refGene_gn=LOC101087216;refGene_tx=XM_023246717.1;refGene_gn=LOC101087216;refGene_tx=XM_023246718.1;refGene_gn=LOC101087216;refGene_tx=XM_023246719.1
E3 20171067 20171141
refGene_gn=LOC101087216;refGene_tx=XM_023246717.1;refGene_gn=LOC101087216;refGene_tx=XM_023246718.1;refGene_gn=LOC101087216;refGene_tx=XM_023246719.1
E3 20171371 20171513
refGene_gn=LOC101087216;refGene_tx=XM_023246717.1;refGene_gn=LOC101087216;refGene_tx=XM_023246718.1;refGene_gn=LOC101087216;refGene_tx=XM_023246719.1
E3 20173010 20173124
refGene_gn=LOC101087216;refGene_tx=XM_023246717.1;refGene_gn=LOC101087216;refGene_tx=XM_023246718.1;refGene_gn=LOC101087216;refGene_tx=XM_023246719.1
E3 20173237 20173317
refGene_gn=LOC101087216;refGene_tx=XM_023246717.1;refGene_gn=LOC101087216;refGene_tx=XM_023246718.1;refGene_gn=LOC101087216;refGene_tx=XM_023246719.1
E3 20173418 20173502
refGene_gn=LOC101087216;refGene_tx=XM_023246717.1;refGene_gn=LOC101087216;refGene_tx=XM_023246718.1;refGene_gn=LOC101087216;refGene_tx=XM_02

3246719.1
E3 20175805 20175884
refGene_gn=LOC101087216;refGene_tx=XM_023246717.1;refGene_gn=LOC101087
216;refGene_tx=XM_023246718.1;refGene_gn=LOC101087216;refGene_tx=XM_02
3246719.1
E3 20176047 20176155
refGene_gn=LOC101087216;refGene_tx=XM_023246717.1;refGene_gn=LOC101087
216;refGene_tx=XM_023246718.1;refGene_gn=LOC101087216;refGene_tx=XM_02
3246719.1
E3 20176263 20176341
refGene_gn=LOC101087216;refGene_tx=XM_023246717.1
E3 20176544 20176658
refGene_gn=LOC101087216;refGene_tx=XM_023246717.1
E3 20176810 20176912
refGene_gn=LOC101087216;refGene_tx=XM_023246717.1
E3 20177178 20177289
refGene_gn=LOC101087216;refGene_tx=XM_023246717.1
E3 20183425 20183524
refGene_gn=LOC101087216;refGene_tx=XM_023246717.1
E3 20217952 20218051 refGene_gn=ITGAX;refGene_tx=XM_006942113.3
E3 20219796 20219907 refGene_gn=ITGAX;refGene_tx=XM_006942113.3
E3 20220166 20220268 refGene_gn=ITGAX;refGene_tx=XM_006942113.3
E3 20220430 20220544 refGene_gn=ITGAX;refGene_tx=XM_006942113.3
E3 20220751 20220835 refGene_gn=ITGAX;refGene_tx=XM_006942113.3
E3 20220952 20221060 refGene_gn=ITGAX;refGene_tx=XM_006942113.3
E3 20221170 20221249 refGene_gn=ITGAX;refGene_tx=XM_006942113.3
E3 20223203 20223287 refGene_gn=ITGAX;refGene_tx=XM_006942113.3
E3 20223377 20223457 refGene_gn=ITGAX;refGene_tx=XM_006942113.3
E3 20223545 20223662 refGene_gn=ITGAX;refGene_tx=XM_006942113.3
E3 20241245 20241387 refGene_gn=ITGAX;refGene_tx=XM_006942113.3
E3 20241603 20241677 refGene_gn=ITGAX;refGene_tx=XM_006942113.3
E3 20241772 20241904 refGene_gn=ITGAX;refGene_tx=XM_006942113.3
E3 20242820 20242975 refGene_gn=ITGAX;refGene_tx=XM_006942113.3
E3 20243117 20243275 refGene_gn=ITGAX;refGene_tx=XM_006942113.3
E3 20243406 20243537 refGene_gn=ITGAX;refGene_tx=XM_006942113.3
E3 20252670 20252880 refGene_gn=ITGAX;refGene_tx=XM_006942113.3
E3 20252972 20253113 refGene_gn=ITGAX;refGene_tx=XM_006942113.3
E3 20253303 20253446 refGene_gn=ITGAX;refGene_tx=XM_006942113.3
E3 20253555 20253612 refGene_gn=ITGAX;refGene_tx=XM_006942113.3
E3 20253614 20253684 refGene_gn=ITGAX;refGene_tx=XM_006942113.3
E3 20254108 20254182 refGene_gn=ITGAX;refGene_tx=XM_006942113.3
E3 20254389 20254540 refGene_gn=ITGAX;refGene_tx=XM_006942113.3
E3 20254848 20255002 refGene_gn=ITGAX;refGene_tx=XM_006942113.3
E3 20255244 20255390 refGene_gn=ITGAX;refGene_tx=XM_006942113.3
E3 20255507 20255638 refGene_gn=ITGAX;refGene_tx=XM_006942113.3
E3 20257008 20257123 refGene_gn=ITGAX;refGene_tx=XM_006942113.3
E3 20257275 20257346 refGene_gn=ITGAX;refGene_tx=XM_006942113.3
E3 20257642 20257746 refGene_gn=ITGAX;refGene_tx=XM_006942113.3
E3 20260898 20261004 refGene_gn=ITGAX;refGene_tx=XM_006942113.3
E3 20261536 20261570 refGene_gn=ITGAX;refGene_tx=XM_006942113.3

E3 20274834 20274903
refGene_gn=LOC101087957;refGene_tx=XM_003998631.4;refGene_gn=LOC101087957;refGene_tx=XM_006942115.3;refGene_gn=LOC101087957;refGene_tx=XM_006942116.3;refGene_gn=LOC101087957;refGene_tx=XM_006942117.3;refGene_gn=LOC101087957;refGene_tx=XM_006942118.3;refGene_gn=LOC101087957;refGene_tx=XM_006942119.3;refGene_gn=LOC101087957;refGene_tx=XM_006942120.3;refGene_gn=LOC101087957;refGene_tx=XM_006942121.3;refGene_gn=LOC101087957;refGene_tx=XM_019820653.1
E3 20275327 20275438
refGene_gn=LOC101087957;refGene_tx=XM_003998631.4;refGene_gn=LOC101087957;refGene_tx=XM_006942115.3;refGene_gn=LOC101087957;refGene_tx=XM_006942116.3;refGene_gn=LOC101087957;refGene_tx=XM_006942117.3;refGene_gn=LOC101087957;refGene_tx=XM_006942118.3;refGene_gn=LOC101087957;refGene_tx=XM_006942119.3;refGene_gn=LOC101087957;refGene_tx=XM_006942120.3;refGene_gn=LOC101087957;refGene_tx=XM_006942121.3;refGene_gn=LOC101087957;refGene_tx=XM_019820653.1
E3 20275663 20275765
refGene_gn=LOC101087957;refGene_tx=XM_003998631.4;refGene_gn=LOC101087957;refGene_tx=XM_006942115.3;refGene_gn=LOC101087957;refGene_tx=XM_006942116.3;refGene_gn=LOC101087957;refGene_tx=XM_006942117.3;refGene_gn=LOC101087957;refGene_tx=XM_006942118.3;refGene_gn=LOC101087957;refGene_tx=XM_006942119.3;refGene_gn=LOC101087957;refGene_tx=XM_006942120.3;refGene_gn=LOC101087957;refGene_tx=XM_006942121.3;refGene_gn=LOC101087957;refGene_tx=XM_019820653.1
E3 20275851 20275965
refGene_gn=LOC101087957;refGene_tx=XM_003998631.4;refGene_gn=LOC101087957;refGene_tx=XM_006942115.3;refGene_gn=LOC101087957;refGene_tx=XM_006942116.3;refGene_gn=LOC101087957;refGene_tx=XM_006942117.3;refGene_gn=LOC101087957;refGene_tx=XM_006942118.3;refGene_gn=LOC101087957;refGene_tx=XM_006942119.3;refGene_gn=LOC101087957;refGene_tx=XM_006942120.3;refGene_gn=LOC101087957;refGene_tx=XM_006942121.3;refGene_gn=LOC101087957;refGene_tx=XM_019820653.1
E3 20276098 20276182
refGene_gn=LOC101087957;refGene_tx=XM_003998631.4;refGene_gn=LOC101087957;refGene_tx=XM_006942115.3;refGene_gn=LOC101087957;refGene_tx=XM_006942116.3;refGene_gn=LOC101087957;refGene_tx=XM_006942117.3;refGene_gn=LOC101087957;refGene_tx=XM_006942118.3;refGene_gn=LOC101087957;refGene_tx=XM_006942119.3;refGene_gn=LOC101087957;refGene_tx=XM_006942120.3;refGene_gn=LOC101087957;refGene_tx=XM_006942121.3;refGene_gn=LOC101087957;refGene_tx=XM_019820653.1
E3 20276367 20276475
refGene_gn=LOC101087957;refGene_tx=XM_003998631.4;refGene_gn=LOC101087957;refGene_tx=XM_006942115.3;refGene_gn=LOC101087957;refGene_tx=XM_006942116.3;refGene_gn=LOC101087957;refGene_tx=XM_006942117.3;refGene_gn=LOC101087957;refGene_tx=XM_006942118.3;refGene_gn=LOC101087957;refGene_tx=XM_006942119.3;refGene_gn=LOC101087957;refGene_tx=XM_006942120.3;refGene_gn=LOC101087957;refGene_tx=XM_006942121.3;refGene_gn=LOC101087957;refGene_tx=XM_019820653.1
E3 20276989 20277065
refGene_gn=LOC101087957;refGene_tx=XM_003998631.4;refGene_gn=LOC101087

957;refGene_tx=XM_006942115.3;refGene_gn=LOC101087957;refGene_tx=XM_006942116.3;refGene_gn=LOC101087957;refGene_tx=XM_006942117.3;refGene_gn=LOC101087957;refGene_tx=XM_006942118.3;refGene_gn=LOC101087957;refGene_tx=XM_006942119.3;refGene_gn=LOC101087957;refGene_tx=XM_006942120.3;refGene_gn=LOC101087957;refGene_tx=XM_006942121.3;refGene_gn=LOC101087957;refGene_tx=XM_019820653.1
E3 20277407 20277491
refGene_gn=LOC101087957;refGene_tx=XM_003998631.4;refGene_gn=LOC101087957;refGene_tx=XM_006942115.3;refGene_gn=LOC101087957;refGene_tx=XM_006942116.3;refGene_gn=LOC101087957;refGene_tx=XM_006942117.3;refGene_gn=LOC101087957;refGene_tx=XM_006942118.3;refGene_gn=LOC101087957;refGene_tx=XM_006942119.3;refGene_gn=LOC101087957;refGene_tx=XM_006942120.3;refGene_gn=LOC101087957;refGene_tx=XM_006942121.3
E3 20277755 20277835
refGene_gn=LOC101087957;refGene_tx=XM_003998631.4;refGene_gn=LOC101087957;refGene_tx=XM_006942115.3;refGene_gn=LOC101087957;refGene_tx=XM_006942116.3;refGene_gn=LOC101087957;refGene_tx=XM_006942117.3;refGene_gn=LOC101087957;refGene_tx=XM_006942118.3;refGene_gn=LOC101087957;refGene_tx=XM_006942119.3;refGene_gn=LOC101087957;refGene_tx=XM_006942120.3;refGene_gn=LOC101087957;refGene_tx=XM_006942121.3;refGene_gn=LOC101087957;refGene_tx=XM_019820653.1
E3 20278333 20278456
refGene_gn=LOC101087957;refGene_tx=XM_003998631.4;refGene_gn=LOC101087957;refGene_tx=XM_006942115.3;refGene_gn=LOC101087957;refGene_tx=XM_006942116.3;refGene_gn=LOC101087957;refGene_tx=XM_006942117.3;refGene_gn=LOC101087957;refGene_tx=XM_006942118.3;refGene_gn=LOC101087957;refGene_tx=XM_006942119.3;refGene_gn=LOC101087957;refGene_tx=XM_006942120.3;refGene_gn=LOC101087957;refGene_tx=XM_006942121.3;refGene_gn=LOC101087957;refGene_tx=XM_019820653.1
E3 20278547 20278689
refGene_gn=LOC101087957;refGene_tx=XM_003998631.4;refGene_gn=LOC101087957;refGene_tx=XM_006942115.3;refGene_gn=LOC101087957;refGene_tx=XM_006942116.3;refGene_gn=LOC101087957;refGene_tx=XM_006942117.3;refGene_gn=LOC101087957;refGene_tx=XM_006942118.3;refGene_gn=LOC101087957;refGene_tx=XM_006942119.3;refGene_gn=LOC101087957;refGene_tx=XM_006942120.3;refGene_gn=LOC101087957;refGene_tx=XM_006942121.3;refGene_gn=LOC101087957;refGene_tx=XM_019820653.1
E3 20278905 20278979
refGene_gn=LOC101087957;refGene_tx=XM_003998631.4;refGene_gn=LOC101087957;refGene_tx=XM_006942115.3;refGene_gn=LOC101087957;refGene_tx=XM_006942116.3;refGene_gn=LOC101087957;refGene_tx=XM_006942117.3;refGene_gn=LOC101087957;refGene_tx=XM_006942118.3;refGene_gn=LOC101087957;refGene_tx=XM_006942119.3;refGene_gn=LOC101087957;refGene_tx=XM_006942120.3;refGene_gn=LOC101087957;refGene_tx=XM_006942121.3;refGene_gn=LOC101087957;refGene_tx=XM_019820653.1
E3 20279489 20279621
refGene_gn=LOC101087957;refGene_tx=XM_003998631.4;refGene_gn=LOC101087957;refGene_tx=XM_006942115.3;refGene_gn=LOC101087957;refGene_tx=XM_006942116.3;refGene_gn=LOC101087957;refGene_tx=XM_006942117.3;refGene_gn=LOC101087957;refGene_tx=XM_006942118.3;refGene_gn=LOC101087957;refGen

e_tx=XM_006942119.3;refGene_gn=LOC101087957;refGene_tx=XM_006942120.3;
refGene_gn=LOC101087957;refGene_tx=XM_006942121.3;refGene_gn=LOC101087
957;refGene_tx=XM_019820653.1
E3 20279718 20279873
refGene_gn=LOC101087957;refGene_tx=XM_003998631.4;refGene_gn=LOC101087
957;refGene_tx=XM_006942115.3;refGene_gn=LOC101087957;refGene_tx=XM_00
6942116.3;refGene_gn=LOC101087957;refGene_tx=XM_006942117.3;refGene_gn
=LOC101087957;refGene_tx=XM_006942118.3;refGene_gn=LOC101087957;refGen
e_tx=XM_006942119.3;refGene_gn=LOC101087957;refGene_tx=XM_006942120.3;
refGene_gn=LOC101087957;refGene_tx=XM_006942121.3;refGene_gn=LOC101087
957;refGene_tx=XM_019820653.1
E3 20281961 20282125
refGene_gn=LOC101087957;refGene_tx=XM_003998631.4;refGene_gn=LOC101087
957;refGene_tx=XM_006942115.3;refGene_gn=LOC101087957;refGene_tx=XM_00
6942116.3;refGene_gn=LOC101087957;refGene_tx=XM_006942117.3;refGene_gn
=LOC101087957;refGene_tx=XM_006942118.3;refGene_gn=LOC101087957;refGen
e_tx=XM_006942119.3;refGene_gn=LOC101087957;refGene_tx=XM_006942120.3;
refGene_gn=LOC101087957;refGene_tx=XM_006942121.3;refGene_gn=LOC101087
957;refGene_tx=XM_019820653.1
E3 20282219 20282350
refGene_gn=LOC101087957;refGene_tx=XM_003998631.4;refGene_gn=LOC101087
957;refGene_tx=XM_006942115.3;refGene_gn=LOC101087957;refGene_tx=XM_00
6942116.3;refGene_gn=LOC101087957;refGene_tx=XM_006942117.3;refGene_gn
=LOC101087957;refGene_tx=XM_006942118.3;refGene_gn=LOC101087957;refGen
e_tx=XM_006942119.3;refGene_gn=LOC101087957;refGene_tx=XM_006942120.3;
refGene_gn=LOC101087957;refGene_tx=XM_006942121.3;refGene_gn=LOC101087
957;refGene_tx=XM_019820653.1
E3 20286517 20286730
refGene_gn=LOC101087957;refGene_tx=XM_003998631.4;refGene_gn=LOC101087
957;refGene_tx=XM_006942115.3;refGene_gn=LOC101087957;refGene_tx=XM_00
6942116.3;refGene_gn=LOC101087957;refGene_tx=XM_006942117.3;refGene_gn
=LOC101087957;refGene_tx=XM_006942118.3;refGene_gn=LOC101087957;refGen
e_tx=XM_006942119.3;refGene_gn=LOC101087957;refGene_tx=XM_006942120.3;
refGene_gn=LOC101087957;refGene_tx=XM_006942121.3;refGene_gn=LOC101087
957;refGene_tx=XM_019820653.1
E3 20286818 20286959
refGene_gn=LOC101087957;refGene_tx=XM_003998631.4;refGene_gn=LOC101087
957;refGene_tx=XM_006942115.3;refGene_gn=LOC101087957;refGene_tx=XM_00
6942116.3;refGene_gn=LOC101087957;refGene_tx=XM_006942117.3;refGene_gn
=LOC101087957;refGene_tx=XM_006942118.3;refGene_gn=LOC101087957;refGen
e_tx=XM_006942119.3;refGene_gn=LOC101087957;refGene_tx=XM_006942120.3;
refGene_gn=LOC101087957;refGene_tx=XM_006942121.3;refGene_gn=LOC101087
957;refGene_tx=XM_019820653.1
E3 20298361 20298504
refGene_gn=LOC101087957;refGene_tx=XM_003998631.4;refGene_gn=LOC101087
957;refGene_tx=XM_006942115.3;refGene_gn=LOC101087957;refGene_tx=XM_00
6942116.3;refGene_gn=LOC101087957;refGene_tx=XM_006942117.3;refGene_gn
=LOC101087957;refGene_tx=XM_006942118.3;refGene_gn=LOC101087957;refGen
e_tx=XM_006942119.3;refGene_gn=LOC101087957;refGene_tx=XM_006942120.3;
refGene_gn=LOC101087957;refGene_tx=XM_006942121.3;refGene_gn=LOC101087

957;refGene_tx=XM_019820653.1
E3 20298790 20298920
refGene_gn=LOC101087957;refGene_tx=XM_003998631.4;refGene_gn=LOC101087957;refGene_tx=XM_006942115.3;refGene_gn=LOC101087957;refGene_tx=XM_006942116.3;refGene_gn=LOC101087957;refGene_tx=XM_006942117.3;refGene_gn=LOC101087957;refGene_tx=XM_006942118.3;refGene_gn=LOC101087957;refGene_tx=XM_006942119.3;refGene_gn=LOC101087957;refGene_tx=XM_006942120.3;refGene_gn=LOC101087957;refGene_tx=XM_006942121.3;refGene_gn=LOC101087957;refGene_tx=XM_019820653.1
E3 20300638 20300712
refGene_gn=LOC101087957;refGene_tx=XM_003998631.4;refGene_gn=LOC101087957;refGene_tx=XM_006942115.3;refGene_gn=LOC101087957;refGene_tx=XM_006942116.3;refGene_gn=LOC101087957;refGene_tx=XM_006942117.3;refGene_gn=LOC101087957;refGene_tx=XM_006942118.3;refGene_gn=LOC101087957;refGene_tx=XM_006942119.3;refGene_gn=LOC101087957;refGene_tx=XM_006942120.3;refGene_gn=LOC101087957;refGene_tx=XM_006942121.3;refGene_gn=LOC101087957;refGene_tx=XM_019820653.1
E3 20301349 20301500
refGene_gn=LOC101087957;refGene_tx=XM_003998631.4;refGene_gn=LOC101087957;refGene_tx=XM_006942115.3;refGene_gn=LOC101087957;refGene_tx=XM_006942116.3;refGene_gn=LOC101087957;refGene_tx=XM_006942117.3;refGene_gn=LOC101087957;refGene_tx=XM_006942118.3;refGene_gn=LOC101087957;refGene_tx=XM_006942119.3;refGene_gn=LOC101087957;refGene_tx=XM_006942120.3;refGene_gn=LOC101087957;refGene_tx=XM_006942121.3;refGene_gn=LOC101087957;refGene_tx=XM_019820653.1
E3 20303441 20303595
refGene_gn=LOC101087957;refGene_tx=XM_003998631.4;refGene_gn=LOC101087957;refGene_tx=XM_006942115.3;refGene_gn=LOC101087957;refGene_tx=XM_006942116.3;refGene_gn=LOC101087957;refGene_tx=XM_006942117.3;refGene_gn=LOC101087957;refGene_tx=XM_006942118.3;refGene_gn=LOC101087957;refGene_tx=XM_006942119.3;refGene_gn=LOC101087957;refGene_tx=XM_006942120.3;refGene_gn=LOC101087957;refGene_tx=XM_006942121.3;refGene_gn=LOC101087957;refGene_tx=XM_019820653.1
E3 20304340 20304486
refGene_gn=LOC101087957;refGene_tx=XM_003998631.4;refGene_gn=LOC101087957;refGene_tx=XM_006942115.3;refGene_gn=LOC101087957;refGene_tx=XM_006942116.3;refGene_gn=LOC101087957;refGene_tx=XM_006942117.3;refGene_gn=LOC101087957;refGene_tx=XM_006942118.3;refGene_gn=LOC101087957;refGene_tx=XM_006942119.3;refGene_gn=LOC101087957;refGene_tx=XM_006942120.3;refGene_gn=LOC101087957;refGene_tx=XM_006942121.3;refGene_gn=LOC101087957;refGene_tx=XM_019820653.1
E3 20304926 20305057
refGene_gn=LOC101087957;refGene_tx=XM_003998631.4;refGene_gn=LOC101087957;refGene_tx=XM_006942115.3;refGene_gn=LOC101087957;refGene_tx=XM_006942116.3;refGene_gn=LOC101087957;refGene_tx=XM_006942117.3;refGene_gn=LOC101087957;refGene_tx=XM_006942118.3;refGene_gn=LOC101087957;refGene_tx=XM_006942119.3;refGene_gn=LOC101087957;refGene_tx=XM_006942120.3;refGene_gn=LOC101087957;refGene_tx=XM_006942121.3;refGene_gn=LOC101087957;refGene_tx=XM_019820653.1
E3 20309253 20309371

refGene_gn=LOC101087957;refGene_tx=XM_003998631.4;refGene_gn=LOC101087
957;refGene_tx=XM_006942115.3;refGene_gn=LOC101087957;refGene_tx=XM_00
6942116.3;refGene_gn=LOC101087957;refGene_tx=XM_006942117.3;refGene_gn
=LOC101087957;refGene_tx=XM_006942118.3;refGene_gn=LOC101087957;refGen
e_tx=XM_006942119.3;refGene_gn=LOC101087957;refGene_tx=XM_006942120.3;
refGene_gn=LOC101087957;refGene_tx=XM_006942121.3;refGene_gn=LOC101087
957;refGene_tx=XM_019820653.1
E3 20309517 20309588
refGene_gn=LOC101087957;refGene_tx=XM_003998631.4;refGene_gn=LOC101087
957;refGene_tx=XM_006942115.3;refGene_gn=LOC101087957;refGene_tx=XM_00
6942116.3;refGene_gn=LOC101087957;refGene_tx=XM_006942117.3;refGene_gn
=LOC101087957;refGene_tx=XM_006942118.3;refGene_gn=LOC101087957;refGen
e_tx=XM_006942119.3;refGene_gn=LOC101087957;refGene_tx=XM_006942120.3;
refGene_gn=LOC101087957;refGene_tx=XM_006942121.3;refGene_gn=LOC101087
957;refGene_tx=XM_019820653.1
E3 20309903 20310007
refGene_gn=LOC101087957;refGene_tx=XM_003998631.4;refGene_gn=LOC101087
957;refGene_tx=XM_006942115.3;refGene_gn=LOC101087957;refGene_tx=XM_00
6942116.3;refGene_gn=LOC101087957;refGene_tx=XM_006942117.3;refGene_gn
=LOC101087957;refGene_tx=XM_006942118.3;refGene_gn=LOC101087957;refGen
e_tx=XM_006942119.3;refGene_gn=LOC101087957;refGene_tx=XM_006942120.3;
refGene_gn=LOC101087957;refGene_tx=XM_006942121.3;refGene_gn=LOC101087
957;refGene_tx=XM_019820653.1
E3 20313516 20313643
refGene_gn=LOC101087957;refGene_tx=XM_003998631.4;refGene_gn=LOC101087
957;refGene_tx=XM_006942115.3;refGene_gn=LOC101087957;refGene_tx=XM_00
6942116.3;refGene_gn=LOC101087957;refGene_tx=XM_006942117.3;refGene_gn
=LOC101087957;refGene_tx=XM_006942118.3;refGene_gn=LOC101087957;refGen
e_tx=XM_006942119.3;refGene_gn=LOC101087957;refGene_tx=XM_006942120.3;
refGene_gn=LOC101087957;refGene_tx=XM_006942121.3;refGene_gn=LOC101087
957;refGene_tx=XM_019820653.1
E3 20315150 20315178
refGene_gn=LOC101087957;refGene_tx=XM_003998631.4;refGene_gn=LOC101087
957;refGene_tx=XM_006942115.3;refGene_gn=LOC101087957;refGene_tx=XM_00
6942116.3;refGene_gn=LOC101087957;refGene_tx=XM_006942117.3;refGene_gn
=LOC101087957;refGene_tx=XM_006942118.3;refGene_gn=LOC101087957;refGen
e_tx=XM_006942119.3;refGene_gn=LOC101087957;refGene_tx=XM_006942120.3;
refGene_gn=LOC101087957;refGene_tx=XM_006942121.3;refGene_gn=LOC101087
957;refGene_tx=XM_019820653.1
E3 20374955 20375530
refGene_gn=TRIM72;refGene_tx=XM_023246616.1;refGene_gn=TRIM72;refGene_
tx=XM_023246617.1;refGene_gn=TRIM72;refGene_tx=XM_023246618.1
E3 20375662 20375898
refGene_gn=TRIM72;refGene_tx=XM_023246616.1;refGene_gn=TRIM72;refGene_
tx=XM_023246617.1
E3 20376856 20376975
refGene_gn=TRIM72;refGene_tx=XM_023246616.1;refGene_gn=TRIM72;refGene_
tx=XM_023246617.1;refGene_gn=TRIM72;refGene_tx=XM_023246618.1
E3 20377751 20377774
refGene_gn=TRIM72;refGene_tx=XM_023246616.1;refGene_gn=TRIM72;refGene_

tx=XM_023246617.1;refGene_gn=TRIM72;refGene_tx=XM_023246618.1
E3 20378448 20378679
refGene_gn=TRIM72;refGene_tx=XM_023246616.1;refGene_gn=TRIM72;refGene_tx=XM_023246617.1;refGene_gn=TRIM72;refGene_tx=XM_023246618.1
E3 20378766 20378862
refGene_gn=TRIM72;refGene_tx=XM_023246616.1;refGene_gn=TRIM72;refGene_tx=XM_023246617.1;refGene_gn=TRIM72;refGene_tx=XM_023246618.1
E3 20382095 20382485
refGene_gn=TRIM72;refGene_tx=XM_023246616.1;refGene_gn=TRIM72;refGene_tx=XM_023246617.1;refGene_gn=TRIM72;refGene_tx=XM_023246618.1
E3 20393682 20393956
refGene_gn=LOC101088449;refGene_tx=XM_003998633.5
E3 20394216 20394273
refGene_gn=LOC101088449;refGene_tx=XM_003998633.5
E3 20394517 20394774
refGene_gn=LOC101088449;refGene_tx=XM_003998633.5
E3 20404446 20404486
refGene_gn=FUS;refGene_tx=XM_003998635.3;refGene_gn=FUS;refGene_tx=XM_006942124.2;refGene_gn=FUS;refGene_tx=XM_011290547.2
E3 20404771 20404919
refGene_gn=FUS;refGene_tx=XM_003998635.3;refGene_gn=FUS;refGene_tx=XM_006942124.2;refGene_gn=FUS;refGene_tx=XM_011290547.2
E3 20405042 20405143
refGene_gn=FUS;refGene_tx=XM_003998635.3;refGene_gn=FUS;refGene_tx=XM_006942124.2;refGene_gn=FUS;refGene_tx=XM_011290547.2
E3 20405473 20405597
refGene_gn=FUS;refGene_tx=XM_003998635.3;refGene_gn=FUS;refGene_tx=XM_006942124.2;refGene_gn=FUS;refGene_tx=XM_011290547.2
E3 20405731 20405833
refGene_gn=FUS;refGene_tx=XM_003998635.3;refGene_gn=FUS;refGene_tx=XM_006942124.2;refGene_gn=FUS;refGene_tx=XM_011290547.2
E3 20406583 20406713
refGene_gn=FUS;refGene_tx=XM_003998635.3;refGene_gn=FUS;refGene_tx=XM_006942124.2;refGene_gn=FUS;refGene_tx=XM_011290547.2
E3 20407119 20407223
refGene_gn=FUS;refGene_tx=XM_003998635.3;refGene_gn=FUS;refGene_tx=XM_006942124.2;refGene_gn=FUS;refGene_tx=XM_011290547.2
E3 20407993 20408026
refGene_gn=FUS;refGene_tx=XM_003998635.3;refGene_gn=FUS;refGene_tx=XM_006942124.2;refGene_gn=FUS;refGene_tx=XM_011290547.2
E3 20409494 20409529
refGene_gn=FUS;refGene_tx=XM_003998635.3;refGene_gn=FUS;refGene_tx=XM_006942124.2;refGene_gn=FUS;refGene_tx=XM_011290547.2
E3 20411129 20411337
refGene_gn=FUS;refGene_tx=XM_003998635.3;refGene_gn=FUS;refGene_tx=XM_006942124.2;refGene_gn=FUS;refGene_tx=XM_011290547.2
E3 20411533 20411724
refGene_gn=FUS;refGene_tx=XM_003998635.3;refGene_gn=FUS;refGene_tx=XM_006942124.2;refGene_gn=FUS;refGene_tx=XM_011290547.2
E3 20411924 20412069

refGene_gn=FUS;refGene_tx=XM_003998635.3;refGene_gn=FUS;refGene_tx=XM_006942124.2;refGene_gn=FUS;refGene_tx=XM_011290547.2
E3 20412741 20412893
refGene_gn=FUS;refGene_tx=XM_003998635.3;refGene_gn=FUS;refGene_tx=XM_006942124.2;refGene_gn=FUS;refGene_tx=XM_011290547.2
E3 20412986 20413011
refGene_gn=FUS;refGene_tx=XM_003998635.3;refGene_gn=FUS;refGene_tx=XM_006942124.2;refGene_gn=FUS;refGene_tx=XM_011290547.2
E3 20414557 20414570
refGene_gn=FUS;refGene_tx=XM_003998635.3;refGene_gn=FUS;refGene_tx=XM_006942124.2;refGene_gn=FUS;refGene_tx=XM_011290547.2
E3 20444249 20444286
refGene_gn=PRSS36;refGene_tx=XM_011290543.2;refGene_gn=PRSS36;refGene_tx=XM_011290544.2;refGene_gn=PRSS36;refGene_tx=XM_011290545.2
E3 20444575 20444611
refGene_gn=PRSS36;refGene_tx=XM_011290543.2;refGene_gn=PRSS36;refGene_tx=XM_011290544.2;refGene_gn=PRSS36;refGene_tx=XM_011290545.2
E3 20444831 20444867
refGene_gn=PRSS36;refGene_tx=XM_011290543.2;refGene_gn=PRSS36;refGene_tx=XM_011290544.2;refGene_gn=PRSS36;refGene_tx=XM_011290545.2
E3 20445350 20445513
refGene_gn=PRSS36;refGene_tx=XM_011290543.2;refGene_gn=PRSS36;refGene_tx=XM_011290544.2;refGene_gn=PRSS36;refGene_tx=XM_011290545.2
E3 20445955 20446236
refGene_gn=PRSS36;refGene_tx=XM_011290543.2;refGene_gn=PRSS36;refGene_tx=XM_011290544.2;refGene_gn=PRSS36;refGene_tx=XM_011290545.2
E3 20447588 20447755
refGene_gn=PRSS36;refGene_tx=XM_011290543.2;refGene_gn=PRSS36;refGene_tx=XM_011290544.2;refGene_gn=PRSS36;refGene_tx=XM_011290545.2
E3 20449562 20449812
refGene_gn=PRSS36;refGene_tx=XM_011290543.2;refGene_gn=PRSS36;refGene_tx=XM_011290544.2;refGene_gn=PRSS36;refGene_tx=XM_011290545.2
E3 20449920 20450050
refGene_gn=PRSS36;refGene_tx=XM_011290543.2;refGene_gn=PRSS36;refGene_tx=XM_011290544.2;refGene_gn=PRSS36;refGene_tx=XM_011290545.2
E3 20450381 20450638
refGene_gn=PRSS36;refGene_tx=XM_011290543.2;refGene_gn=PRSS36;refGene_tx=XM_011290544.2;refGene_gn=PRSS36;refGene_tx=XM_011290545.2
E3 20450720 20450884
refGene_gn=PRSS36;refGene_tx=XM_011290543.2;refGene_gn=PRSS36;refGene_tx=XM_011290544.2;refGene_gn=PRSS36;refGene_tx=XM_011290545.2
E3 20451287 20451525
refGene_gn=PRSS36;refGene_tx=XM_011290543.2;refGene_gn=PRSS36;refGene_tx=XM_011290544.2;refGene_gn=PRSS36;refGene_tx=XM_011290545.2
E3 20451641 20451783
refGene_gn=PRSS36;refGene_tx=XM_011290543.2;refGene_gn=PRSS36;refGene_tx=XM_011290544.2;refGene_gn=PRSS36;refGene_tx=XM_011290545.2
E3 20452070 20452336
refGene_gn=PRSS36;refGene_tx=XM_011290543.2;refGene_gn=PRSS36;refGene_tx=XM_011290544.2;refGene_gn=PRSS36;refGene_tx=XM_011290545.2

E3 20452415 20452537
refGene_gn=PRSS36;refGene_tx=XM_011290543.2;refGene_gn=PRSS36;refGene_
tx=XM_011290544.2
E3 20452805 20453087
refGene_gn=PRSS36;refGene_tx=XM_011290543.2;refGene_gn=PRSS36;refGene_
tx=XM_011290544.2;refGene_gn=PRSS36;refGene_tx=XM_011290545.2
E3 20455733 20455818
refGene_gn=PRSS8;refGene_tx=XM_003998697.2;refGene_gn=PRSS8;refGene_tx
=XM_019820681.1
E3 20456095 20456113 refGene_gn=PRSS8;refGene_tx=XM_003998697.2
E3 20457554 20457717
refGene_gn=PRSS8;refGene_tx=XM_003998697.2;refGene_gn=PRSS8;refGene_tx
=XM_019820681.1
E3 20457979 20458251
refGene_gn=PRSS8;refGene_tx=XM_003998697.2;refGene_gn=PRSS8;refGene_tx
=XM_019820681.1
E3 20458337 20458504
refGene_gn=PRSS8;refGene_tx=XM_003998697.2;refGene_gn=PRSS8;refGene_tx
=XM_019820681.1
E3 20458627 20458936
refGene_gn=PRSS8;refGene_tx=XM_003998697.2;refGene_gn=PRSS8;refGene_tx
=XM_019820681.1
E3 20459775 20459840 refGene_gn=KAT8;refGene_tx=XM_003998636.3
E3 20460097 20460305
refGene_gn=KAT8;refGene_tx=XM_003998636.3;refGene_gn=KAT8;refGene_tx=X
M_019820666.1;refGene_gn=KAT8;refGene_tx=XM_019820667.1
E3 20460399 20460550
refGene_gn=KAT8;refGene_tx=XM_003998636.3;refGene_gn=KAT8;refGene_tx=X
M_019820666.1;refGene_gn=KAT8;refGene_tx=XM_019820667.1
E3 20460645 20460739
refGene_gn=KAT8;refGene_tx=XM_003998636.3;refGene_gn=KAT8;refGene_tx=X
M_019820666.1;refGene_gn=KAT8;refGene_tx=XM_019820667.1
E3 20460836 20460977
refGene_gn=KAT8;refGene_tx=XM_003998636.3;refGene_gn=KAT8;refGene_tx=X
M_019820666.1;refGene_gn=KAT8;refGene_tx=XM_019820667.1
E3 20462926 20463016
refGene_gn=KAT8;refGene_tx=XM_003998636.3;refGene_gn=KAT8;refGene_tx=X
M_019820666.1;refGene_gn=KAT8;refGene_tx=XM_019820667.1
E3 20463874 20464039
refGene_gn=KAT8;refGene_tx=XM_003998636.3;refGene_gn=KAT8;refGene_tx=X
M_019820666.1;refGene_gn=KAT8;refGene_tx=XM_019820667.1
E3 20464139 20464193
refGene_gn=KAT8;refGene_tx=XM_003998636.3;refGene_gn=KAT8;refGene_tx=X
M_019820666.1;refGene_gn=KAT8;refGene_tx=XM_019820667.1
E3 20468625 20468801
refGene_gn=KAT8;refGene_tx=XM_003998636.3;refGene_gn=KAT8;refGene_tx=X
M_019820666.1;refGene_gn=KAT8;refGene_tx=XM_019820667.1
E3 20468879 20468954
refGene_gn=KAT8;refGene_tx=XM_003998636.3;refGene_gn=KAT8;refGene_tx=X
M_019820666.1;refGene_gn=KAT8;refGene_tx=XM_019820667.1

E3 20470283 20470494
refGene_gn=KAT8;refGene_tx=XM_003998636.3;refGene_gn=KAT8;refGene_tx=XM_019820666.1;refGene_gn=KAT8;refGene_tx=XM_019820667.1
E3 20473760 20473905
refGene_gn=BCKDK;refGene_tx=XM_003998637.4;refGene_gn=BCKDK;refGene_tx=XM_006942125.3
E3 20473995 20474154 refGene_gn=BCKDK;refGene_tx=XM_003998637.4
E3 20474619 20474709
refGene_gn=BCKDK;refGene_tx=XM_003998637.4;refGene_gn=BCKDK;refGene_tx=XM_006942125.3
E3 20474800 20474929
refGene_gn=BCKDK;refGene_tx=XM_003998637.4;refGene_gn=BCKDK;refGene_tx=XM_006942125.3
E3 20475247 20475321
refGene_gn=BCKDK;refGene_tx=XM_003998637.4;refGene_gn=BCKDK;refGene_tx=XM_006942125.3
E3 20475513 20475612
refGene_gn=BCKDK;refGene_tx=XM_003998637.4;refGene_gn=BCKDK;refGene_tx=XM_006942125.3
E3 20475694 20475814
refGene_gn=BCKDK;refGene_tx=XM_003998637.4;refGene_gn=BCKDK;refGene_tx=XM_006942125.3
E3 20475894 20475942
refGene_gn=BCKDK;refGene_tx=XM_003998637.4;refGene_gn=BCKDK;refGene_tx=XM_006942125.3
E3 20476257 20476368
refGene_gn=BCKDK;refGene_tx=XM_003998637.4;refGene_gn=BCKDK;refGene_tx=XM_006942125.3
E3 20476459 20476528
refGene_gn=BCKDK;refGene_tx=XM_003998637.4;refGene_gn=BCKDK;refGene_tx=XM_006942125.3
E3 20476650 20476845
refGene_gn=BCKDK;refGene_tx=XM_003998637.4;refGene_gn=BCKDK;refGene_tx=XM_006942125.3
E3 20484945 20485118
refGene_gn=VKORC1;refGene_tx=XM_003998638.5;refGene_gn=VKORC1;refGene_tx=XM_019820684.2
E3 20485917 20486027 refGene_gn=VKORC1;refGene_tx=XM_003998638.5
E3 20486528 20486737
refGene_gn=VKORC1;refGene_tx=XM_003998638.5;refGene_gn=VKORC1;refGene_tx=XM_019820684.2
E3 20488872 20488930 refGene_gn=PRSS53;refGene_tx=XM_019820665.1
E3 20489856 20489877 refGene_gn=PRSS53;refGene_tx=XM_019820665.1
E3 20489982 20490145 refGene_gn=PRSS53;refGene_tx=XM_019820665.1
E3 20490471 20490737 refGene_gn=PRSS53;refGene_tx=XM_019820665.1
E3 20490875 20491030 refGene_gn=PRSS53;refGene_tx=XM_019820665.1
E3 20491187 20491407 refGene_gn=PRSS53;refGene_tx=XM_019820665.1
E3 20492392 20492543 refGene_gn=PRSS53;refGene_tx=XM_019820665.1
E3 20492629 20492874 refGene_gn=PRSS53;refGene_tx=XM_019820665.1
E3 20492956 20493102 refGene_gn=PRSS53;refGene_tx=XM_019820665.1

E3 20493350 20493567 refGene_gn=PRSS53;refGene_tx=XM_019820665.1
E3 20493854 20493874 refGene_gn=PRSS53;refGene_tx=XM_019820665.1
E3 20494644 20494766
refGene_gn=ZNF646;refGene_tx=XM_003998640.5;refGene_gn=ZNF646;refGene_tx=XM_023246868.1
E3 20496070 20501417
refGene_gn=ZNF646;refGene_tx=XM_003998640.5;refGene_gn=ZNF646;refGene_tx=XM_023246868.1
E3 20509679 20510329
refGene_gn=ZNF668;refGene_tx=XM_003998641.4;refGene_gn=ZNF668;refGene_tx=XM_006942132.4;refGene_gn=ZNF668;refGene_tx=XM_019820660.2;refGene_gn=ZNF668;refGene_tx=XM_019820662.1;refGene_gn=ZNF668;refGene_tx=XM_019820663.1
E3 20510851 20512064
refGene_gn=ZNF668;refGene_tx=XM_003998641.4;refGene_gn=ZNF668;refGene_tx=XM_006942132.4;refGene_gn=ZNF668;refGene_tx=XM_019820660.2;refGene_gn=ZNF668;refGene_tx=XM_019820662.1;refGene_gn=ZNF668;refGene_tx=XM_019820663.1
E3 20527418 20527499
refGene_gn=STX4;refGene_tx=XM_003998642.4;refGene_gn=STX4;refGene_tx=XM_019820682.1
E3 20527593 20527704
refGene_gn=STX4;refGene_tx=XM_003998642.4;refGene_gn=STX4;refGene_tx=XM_019820682.1
E3 20528931 20529069
refGene_gn=STX4;refGene_tx=XM_003998642.4;refGene_gn=STX4;refGene_tx=XM_019820682.1
E3 20529304 20529381
refGene_gn=STX4;refGene_tx=XM_003998642.4;refGene_gn=STX4;refGene_tx=XM_019820682.1
E3 20529453 20529562
refGene_gn=STX4;refGene_tx=XM_003998642.4;refGene_gn=STX4;refGene_tx=XM_019820682.1
E3 20531075 20531146
refGene_gn=STX4;refGene_tx=XM_003998642.4;refGene_gn=STX4;refGene_tx=XM_019820682.1
E3 20531661 20531736
refGene_gn=STX4;refGene_tx=XM_003998642.4;refGene_gn=STX4;refGene_tx=XM_019820682.1
E3 20531871 20531971
refGene_gn=STX4;refGene_tx=XM_003998642.4;refGene_gn=STX4;refGene_tx=XM_019820682.1
E3 20532083 20532185
refGene_gn=STX4;refGene_tx=XM_003998642.4;refGene_gn=STX4;refGene_tx=XM_019820682.1
E3 20532374 20532404
refGene_gn=STX4;refGene_tx=XM_003998642.4;refGene_gn=STX4;refGene_tx=XM_019820682.1
E3 20545865 20545895 refGene_gn=STX1B;refGene_tx=XM_023247009.1
E3 20553853 20553928 refGene_gn=STX1B;refGene_tx=XM_023247009.1

E3 20554129 20554229 refGene_gn=STX1B;refGene_tx=XM_023247009.1
E3 20554343 20554418 refGene_gn=STX1B;refGene_tx=XM_023247009.1
E3 20556937 20557011 refGene_gn=STX1B;refGene_tx=XM_023247009.1
E3 20557376 20557485 refGene_gn=STX1B;refGene_tx=XM_023247009.1
E3 20557666 20557740 refGene_gn=STX1B;refGene_tx=XM_023247009.1
E3 20559891 20560029 refGene_gn=STX1B;refGene_tx=XM_023247009.1
E3 20560129 20560240 refGene_gn=STX1B;refGene_tx=XM_023247009.1
E3 20560496 20560577 refGene_gn=STX1B;refGene_tx=XM_023247009.1
E3 20564317 20564323
refGene_gn=HSD3B7;refGene_tx=XM_019820669.2;refGene_gn=HSD3B7;refGene_tx=XM_019820670.2
E3 20564683 20565099
refGene_gn=HSD3B7;refGene_tx=XM_003998643.3;refGene_gn=HSD3B7;refGene_tx=XM_019820668.2;refGene_gn=HSD3B7;refGene_tx=XM_019820669.2;refGene_gn=HSD3B7;refGene_tx=XM_019820670.2;refGene_gn=HSD3B7;refGene_tx=XM_019820671.2;refGene_gn=HSD3B7;refGene_tx=XM_019820672.2;refGene_gn=HSD3B7;refGene_tx=XM_023247008.1
E3 20565593 20565756
refGene_gn=HSD3B7;refGene_tx=XM_003998643.3;refGene_gn=HSD3B7;refGene_tx=XM_019820668.2;refGene_gn=HSD3B7;refGene_tx=XM_019820669.2;refGene_gn=HSD3B7;refGene_tx=XM_019820670.2;refGene_gn=HSD3B7;refGene_tx=XM_019820671.2;refGene_gn=HSD3B7;refGene_tx=XM_023247008.1
E3 20565916 20566016
refGene_gn=HSD3B7;refGene_tx=XM_003998643.3;refGene_gn=HSD3B7;refGene_tx=XM_019820668.2;refGene_gn=HSD3B7;refGene_tx=XM_019820669.2;refGene_gn=HSD3B7;refGene_tx=XM_019820670.2;refGene_gn=HSD3B7;refGene_tx=XM_019820671.2;refGene_gn=HSD3B7;refGene_tx=XM_019820672.2;refGene_gn=HSD3B7;refGene_tx=XM_023247008.1
E3 20566310 20566419
refGene_gn=HSD3B7;refGene_tx=XM_003998643.3;refGene_gn=HSD3B7;refGene_tx=XM_019820668.2;refGene_gn=HSD3B7;refGene_tx=XM_019820669.2;refGene_gn=HSD3B7;refGene_tx=XM_019820670.2;refGene_gn=HSD3B7;refGene_tx=XM_019820671.2;refGene_gn=HSD3B7;refGene_tx=XM_019820672.2;refGene_gn=HSD3B7;refGene_tx=XM_023247008.1
E3 20566662 20566818
refGene_gn=HSD3B7;refGene_tx=XM_003998643.3;refGene_gn=HSD3B7;refGene_tx=XM_019820668.2;refGene_gn=HSD3B7;refGene_tx=XM_019820669.2;refGene_gn=HSD3B7;refGene_tx=XM_019820670.2;refGene_gn=HSD3B7;refGene_tx=XM_019820672.2;refGene_gn=HSD3B7;refGene_tx=XM_023247008.1
E3 20567028 20567194
refGene_gn=HSD3B7;refGene_tx=XM_003998643.3;refGene_gn=HSD3B7;refGene_tx=XM_019820668.2;refGene_gn=HSD3B7;refGene_tx=XM_019820669.2;refGene_gn=HSD3B7;refGene_tx=XM_019820670.2;refGene_gn=HSD3B7;refGene_tx=XM_019820671.2;refGene_gn=HSD3B7;refGene_tx=XM_019820672.2;refGene_gn=HSD3B7;refGene_tx=XM_023247008.1
E3 20568824 20568998 refGene_gn=SETD1A;refGene_tx=XM_023247007.1
E3 20569079 20569217 refGene_gn=SETD1A;refGene_tx=XM_023247007.1
E3 20570753 20570873 refGene_gn=SETD1A;refGene_tx=XM_023247007.1
E3 20571057 20571168 refGene_gn=SETD1A;refGene_tx=XM_023247007.1
E3 20571257 20571430 refGene_gn=SETD1A;refGene_tx=XM_023247007.1

E3 20571845 20572037 refGene_gn=SETD1A;refGene_tx=XM_023247007.1
E3 20572039 20572888 refGene_gn=SETD1A;refGene_tx=XM_023247007.1
E3 20579339 20579681 refGene_gn=SETD1A;refGene_tx=XM_023247007.1
E3 20581711 20581799 refGene_gn=SETD1A;refGene_tx=XM_023247007.1
E3 20581948 20582106 refGene_gn=SETD1A;refGene_tx=XM_023247007.1
E3 20582846 20582934 refGene_gn=SETD1A;refGene_tx=XM_023247007.1
E3 20583063 20583240 refGene_gn=SETD1A;refGene_tx=XM_023247007.1
E3 20583961 20584747 refGene_gn=SETD1A;refGene_tx=XM_023247007.1
E3 20584904 20585763 refGene_gn=SETD1A;refGene_tx=XM_023247007.1
E3 20586045 20586275 refGene_gn=SETD1A;refGene_tx=XM_023247007.1
E3 20586499 20586621 refGene_gn=SETD1A;refGene_tx=XM_023247007.1
E3 20587762 20588033 refGene_gn=SETD1A;refGene_tx=XM_023247007.1
E3 20589788 20589884 refGene_gn=SETD1A;refGene_tx=XM_023247007.1
E3 20590098 20590248 refGene_gn=SETD1A;refGene_tx=XM_023247007.1
E3 20594248 20594908 refGene_gn=ORAI3;refGene_tx=XM_003998645.5
E3 20598290 20598518 refGene_gn=ORAI3;refGene_tx=XM_003998645.5
E3 20600738 20600917
refGene_gn=FBXL19;refGene_tx=XM_006942138.4;refGene_gn=FBXL19;refGene_tx=XM_011290536.3;refGene_gn=FBXL19;refGene_tx=XM_011290537.3;refGene_gn=FBXL19;refGene_tx=XM_011290538.3
E3 20601016 20601235
refGene_gn=FBXL19;refGene_tx=XM_006942138.4;refGene_gn=FBXL19;refGene_tx=XM_011290536.3;refGene_gn=FBXL19;refGene_tx=XM_011290537.3;refGene_gn=FBXL19;refGene_tx=XM_011290538.3
E3 20606876 20607038
refGene_gn=FBXL19;refGene_tx=XM_006942138.4;refGene_gn=FBXL19;refGene_tx=XM_011290536.3;refGene_gn=FBXL19;refGene_tx=XM_011290537.3;refGene_gn=FBXL19;refGene_tx=XM_011290538.3
E3 20607129 20607293
refGene_gn=FBXL19;refGene_tx=XM_006942138.4;refGene_gn=FBXL19;refGene_tx=XM_011290536.3;refGene_gn=FBXL19;refGene_tx=XM_011290537.3;refGene_gn=FBXL19;refGene_tx=XM_011290538.3
E3 20614169 20614681
refGene_gn=FBXL19;refGene_tx=XM_006942138.4;refGene_gn=FBXL19;refGene_tx=XM_011290536.3;refGene_gn=FBXL19;refGene_tx=XM_011290537.3;refGene_gn=FBXL19;refGene_tx=XM_011290538.3
E3 20615791 20615953
refGene_gn=FBXL19;refGene_tx=XM_006942138.4;refGene_gn=FBXL19;refGene_tx=XM_011290536.3;refGene_gn=FBXL19;refGene_tx=XM_011290537.3;refGene_gn=FBXL19;refGene_tx=XM_011290538.3
E3 20616459 20616678
refGene_gn=FBXL19;refGene_tx=XM_006942138.4;refGene_gn=FBXL19;refGene_tx=XM_011290536.3;refGene_gn=FBXL19;refGene_tx=XM_011290537.3;refGene_gn=FBXL19;refGene_tx=XM_011290538.3
E3 20616760 20616847
refGene_gn=FBXL19;refGene_tx=XM_006942138.4;refGene_gn=FBXL19;refGene_tx=XM_011290536.3;refGene_gn=FBXL19;refGene_tx=XM_011290537.3;refGene_gn=FBXL19;refGene_tx=XM_011290538.3
E3 20616950 20617094
refGene_gn=FBXL19;refGene_tx=XM_006942138.4;refGene_gn=FBXL19;refGene_

tx=XM_011290536.3
E3 20618494 20618671
refGene_gn=FBXL19;refGene_tx=XM_006942138.4;refGene_gn=FBXL19;refGene_
tx=XM_011290536.3
E3 20643130 20643595
refGene_gn=LOC101083389;refGene_tx=XM_019820685.1
E3 20644966 20645434
refGene_gn=LOC101083642;refGene_tx=XM_011290535.3
E3 20646962 20647081
refGene_gn=LOC101083642;refGene_tx=XM_011290535.3
E3 20648258 20648283
refGene_gn=LOC101083642;refGene_tx=XM_011290535.3
E3 20650901 20650993 refGene_gn=BCL7C;refGene_tx=XM_023246433.1
E3 20651506 20651585 refGene_gn=BCL7C;refGene_tx=XM_023246433.1
E3 20651910 20652019 refGene_gn=BCL7C;refGene_tx=XM_023246433.1
E3 20652119 20652281 refGene_gn=BCL7C;refGene_tx=XM_023246433.1
E3 20653687 20653773 refGene_gn=BCL7C;refGene_tx=XM_023246433.1
E3 20654321 20654447 refGene_gn=BCL7C;refGene_tx=XM_023246433.1
E3 20736457 20736530
refGene_gn=ZNF629;refGene_tx=XM_003998647.5;refGene_gn=ZNF629;refGene_
tx=XM_011290534.3;refGene_gn=ZNF629;refGene_tx=XM_023246884.1
E3 20736685 20739219
refGene_gn=ZNF629;refGene_tx=XM_003998647.5;refGene_gn=ZNF629;refGene_
tx=XM_011290534.3;refGene_gn=ZNF629;refGene_tx=XM_023246884.1
E3 20746914 20747091 refGene_gn=RNF40;refGene_tx=XM_003998648.5
E3 20748985 20749087 refGene_gn=RNF40;refGene_tx=XM_003998648.5
E3 20749197 20749338 refGene_gn=RNF40;refGene_tx=XM_003998648.5
E3 20750559 20750685 refGene_gn=RNF40;refGene_tx=XM_003998648.5
E3 20750763 20750976 refGene_gn=RNF40;refGene_tx=XM_003998648.5
E3 20751153 20751303 refGene_gn=RNF40;refGene_tx=XM_003998648.5
E3 20751427 20751545 refGene_gn=RNF40;refGene_tx=XM_003998648.5
E3 20751631 20752059 refGene_gn=RNF40;refGene_tx=XM_003998648.5
E3 20752143 20752265 refGene_gn=RNF40;refGene_tx=XM_003998648.5
E3 20753102 20753238 refGene_gn=RNF40;refGene_tx=XM_003998648.5
E3 20753412 20753592 refGene_gn=RNF40;refGene_tx=XM_003998648.5
E3 20753687 20753807 refGene_gn=RNF40;refGene_tx=XM_003998648.5
E3 20754509 20754584 refGene_gn=RNF40;refGene_tx=XM_003998648.5
E3 20754676 20754823 refGene_gn=RNF40;refGene_tx=XM_003998648.5
E3 20754921 20755043 refGene_gn=RNF40;refGene_tx=XM_003998648.5
E3 20755592 20755799 refGene_gn=RNF40;refGene_tx=XM_003998648.5
E3 20756454 20756596 refGene_gn=RNF40;refGene_tx=XM_003998648.5
E3 20756731 20756899 refGene_gn=RNF40;refGene_tx=XM_003998648.5
E3 20757311 20757515 refGene_gn=RNF40;refGene_tx=XM_003998648.5
E3 20757623 20757731 refGene_gn=RNF40;refGene_tx=XM_003998648.5
E3 20758197 20758401
refGene_gn=CCDC189;refGene_tx=XM_019820674.1;refGene_gn=CCDC189;refGen
e_tx=XM_019820675.1;refGene_gn=CCDC189;refGene_tx=XM_019820676.1;refGe
ne_gn=CCDC189;refGene_tx=XM_019820677.1;refGene_gn=CCDC189;refGene_tx=
XM_019820678.1;refGene_gn=CCDC189;refGene_tx=XM_019820679.1;refGene_gn
=CCDC189;refGene_tx=XM_019820680.1

E3	20758724	20758756
refGene_gn=CCDC189;refGene_tx=XM_019820674.1;refGene_gn=CCDC189;refGene_tx=XM_019820675.1;refGene_gn=CCDC189;refGene_tx=XM_019820676.1;refGene_gn=CCDC189;refGene_tx=XM_019820677.1;refGene_gn=CCDC189;refGene_tx=XM_019820678.1;refGene_gn=CCDC189;refGene_tx=XM_019820679.1
E3	20759264	20759330
refGene_gn=CCDC189;refGene_tx=XM_019820674.1;refGene_gn=CCDC189;refGene_tx=XM_019820675.1;refGene_gn=CCDC189;refGene_tx=XM_019820676.1;refGene_gn=CCDC189;refGene_tx=XM_019820677.1;refGene_gn=CCDC189;refGene_tx=XM_019820678.1;refGene_gn=CCDC189;refGene_tx=XM_019820679.1;refGene_gn=CCDC189;refGene_tx=XM_019820680.1
E3	20759475	20759645
refGene_gn=CCDC189;refGene_tx=XM_019820674.1;refGene_gn=CCDC189;refGene_tx=XM_019820675.1;refGene_gn=CCDC189;refGene_tx=XM_019820676.1;refGene_gn=CCDC189;refGene_tx=XM_019820677.1;refGene_gn=CCDC189;refGene_tx=XM_019820678.1;refGene_gn=CCDC189;refGene_tx=XM_019820679.1;refGene_gn=CCDC189;refGene_tx=XM_019820680.1
E3	20760216	20760287
refGene_gn=CCDC189;refGene_tx=XM_019820674.1;refGene_gn=CCDC189;refGene_tx=XM_019820675.1;refGene_gn=CCDC189;refGene_tx=XM_019820676.1;refGene_gn=CCDC189;refGene_tx=XM_019820677.1;refGene_gn=CCDC189;refGene_tx=XM_019820679.1;refGene_gn=CCDC189;refGene_tx=XM_019820680.1
E3	20760488	20760757
refGene_gn=CCDC189;refGene_tx=XM_019820674.1;refGene_gn=CCDC189;refGene_tx=XM_019820675.1;refGene_gn=CCDC189;refGene_tx=XM_019820676.1;refGene_gn=CCDC189;refGene_tx=XM_019820677.1;refGene_gn=CCDC189;refGene_tx=XM_019820678.1;refGene_gn=CCDC189;refGene_tx=XM_019820679.1;refGene_gn=CCDC189;refGene_tx=XM_019820680.1
E3	20760842	20760929
refGene_gn=CCDC189;refGene_tx=XM_019820674.1;refGene_gn=CCDC189;refGene_tx=XM_019820675.1;refGene_gn=CCDC189;refGene_tx=XM_019820677.1;refGene_gn=CCDC189;refGene_tx=XM_019820678.1;refGene_gn=CCDC189;refGene_tx=XM_019820680.1
E3	20761545	20761724
refGene_gn=CCDC189;refGene_tx=XM_019820674.1;refGene_gn=CCDC189;refGene_tx=XM_019820675.1;refGene_gn=CCDC189;refGene_tx=XM_019820676.1;refGene_gn=CCDC189;refGene_tx=XM_019820677.1;refGene_gn=CCDC189;refGene_tx=XM_019820678.1;refGene_gn=CCDC189;refGene_tx=XM_019820679.1;refGene_gn=CCDC189;refGene_tx=XM_019820680.1
E3	20762121	20762415
refGene_gn=PHKG2;refGene_tx=XM_003998650.5;refGene_gn=PHKG2;refGene_tx=XM_006942139.4;refGene_gn=PHKG2;refGene_tx=XM_019820673.1
E3	20762516	20762642
refGene_gn=PHKG2;refGene_tx=XM_003998650.5;refGene_gn=PHKG2;refGene_tx=XM_006942139.4;refGene_gn=PHKG2;refGene_tx=XM_019820673.1
E3	20762714	20762868
refGene_gn=PHKG2;refGene_tx=XM_003998650.5;refGene_gn=PHKG2;refGene_tx=XM_006942139.4;refGene_gn=PHKG2;refGene_tx=XM_019820673.1
E3	20762962	20763053
refGene_gn=PHKG2;refGene_tx=XM_003998650.5;refGene_gn=PHKG2;refGene_tx

=XM_006942139.4;refGene_gn=PHKG2;refGene_tx=XM_019820673.1
E3 20763961 20764125
refGene_gn=PHKG2;refGene_tx=XM_003998650.5;refGene_gn=PHKG2;refGene_tx
=XM_006942139.4;refGene_gn=PHKG2;refGene_tx=XM_019820673.1
E3 20764216 20764282
refGene_gn=PHKG2;refGene_tx=XM_003998650.5;refGene_gn=PHKG2;refGene_tx
=XM_006942139.4;refGene_gn=PHKG2;refGene_tx=XM_019820673.1
E3 20765338 20765393
refGene_gn=PHKG2;refGene_tx=XM_003998650.5;refGene_gn=PHKG2;refGene_tx
=XM_006942139.4;refGene_gn=PHKG2;refGene_tx=XM_019820673.1
E3 20765668 20765844
refGene_gn=PHKG2;refGene_tx=XM_003998650.5;refGene_gn=PHKG2;refGene_tx
=XM_006942139.4;refGene_gn=PHKG2;refGene_tx=XM_019820673.1
E3 20770223 20770318
refGene_gn=PHKG2;refGene_tx=XM_003998650.5;refGene_gn=PHKG2;refGene_tx
=XM_006942139.4;refGene_gn=PHKG2;refGene_tx=XM_019820673.1
E3 20775811 20775973 refGene_gn=TMEM265;refGene_tx=XM_011290532.2
E3 20777468 20777636 refGene_gn=TMEM265;refGene_tx=XM_011290532.2
E3 20779646 20782337
refGene_gn=SRCAP;refGene_tx=XM_023246885.1;refGene_gn=SRCAP;refGene_tx
=XM_023246886.1;refGene_gn=SRCAP;refGene_tx=XM_023246887.1;refGene_gn=
SRCAP;refGene_tx=XM_023246888.1
E3 20782448 20782532
refGene_gn=SRCAP;refGene_tx=XM_023246885.1;refGene_gn=SRCAP;refGene_tx
=XM_023246886.1;refGene_gn=SRCAP;refGene_tx=XM_023246887.1;refGene_gn=
SRCAP;refGene_tx=XM_023246888.1
E3 20782688 20782883
refGene_gn=SRCAP;refGene_tx=XM_023246885.1;refGene_gn=SRCAP;refGene_tx
=XM_023246886.1;refGene_gn=SRCAP;refGene_tx=XM_023246887.1;refGene_gn=
SRCAP;refGene_tx=XM_023246888.1
E3 20786465 20786585
refGene_gn=SRCAP;refGene_tx=XM_023246885.1;refGene_gn=SRCAP;refGene_tx
=XM_023246886.1;refGene_gn=SRCAP;refGene_tx=XM_023246887.1;refGene_gn=
SRCAP;refGene_tx=XM_023246888.1
E3 20786770 20786885
refGene_gn=SRCAP;refGene_tx=XM_023246885.1;refGene_gn=SRCAP;refGene_tx
=XM_023246886.1;refGene_gn=SRCAP;refGene_tx=XM_023246887.1;refGene_gn=
SRCAP;refGene_tx=XM_023246888.1
E3 20786979 20787176
refGene_gn=SRCAP;refGene_tx=XM_023246885.1;refGene_gn=SRCAP;refGene_tx
=XM_023246886.1;refGene_gn=SRCAP;refGene_tx=XM_023246887.1;refGene_gn=
SRCAP;refGene_tx=XM_023246888.1
E3 20787303 20787473
refGene_gn=SRCAP;refGene_tx=XM_023246885.1;refGene_gn=SRCAP;refGene_tx
=XM_023246886.1;refGene_gn=SRCAP;refGene_tx=XM_023246887.1;refGene_gn=
SRCAP;refGene_tx=XM_023246888.1
E3 20789955 20790158
refGene_gn=SRCAP;refGene_tx=XM_023246885.1;refGene_gn=SRCAP;refGene_tx
=XM_023246886.1;refGene_gn=SRCAP;refGene_tx=XM_023246887.1;refGene_gn=
SRCAP;refGene_tx=XM_023246888.1

E3 20790294 20790560
refGene_gn=SRCAP;refGene_tx=XM_023246885.1;refGene_gn=SRCAP;refGene_tx=XM_023246886.1;refGene_gn=SRCAP;refGene_tx=XM_023246887.1;refGene_gn=SRCAP;refGene_tx=XM_023246888.1
E3 20794714 20796213
refGene_gn=SRCAP;refGene_tx=XM_023246885.1;refGene_gn=SRCAP;refGene_tx=XM_023246886.1;refGene_gn=SRCAP;refGene_tx=XM_023246887.1;refGene_gn=SRCAP;refGene_tx=XM_023246888.1
E3 20796552 20796819
refGene_gn=SRCAP;refGene_tx=XM_023246885.1;refGene_gn=SRCAP;refGene_tx=XM_023246886.1;refGene_gn=SRCAP;refGene_tx=XM_023246887.1;refGene_gn=SRCAP;refGene_tx=XM_023246888.1
E3 20797037 20797223
refGene_gn=SRCAP;refGene_tx=XM_023246885.1;refGene_gn=SRCAP;refGene_tx=XM_023246886.1;refGene_gn=SRCAP;refGene_tx=XM_023246887.1
E3 20797495 20797660
refGene_gn=SRCAP;refGene_tx=XM_023246885.1;refGene_gn=SRCAP;refGene_tx=XM_023246886.1;refGene_gn=SRCAP;refGene_tx=XM_023246887.1;refGene_gn=SRCAP;refGene_tx=XM_023246888.1
E3 20798144 20798429
refGene_gn=SRCAP;refGene_tx=XM_023246885.1;refGene_gn=SRCAP;refGene_tx=XM_023246886.1;refGene_gn=SRCAP;refGene_tx=XM_023246887.1;refGene_gn=SRCAP;refGene_tx=XM_023246888.1
E3 20798637 20798903
refGene_gn=SRCAP;refGene_tx=XM_023246885.1;refGene_gn=SRCAP;refGene_tx=XM_023246886.1;refGene_gn=SRCAP;refGene_tx=XM_023246887.1;refGene_gn=SRCAP;refGene_tx=XM_023246888.1
E3 20799304 20799474
refGene_gn=SRCAP;refGene_tx=XM_023246885.1;refGene_gn=SRCAP;refGene_tx=XM_023246886.1;refGene_gn=SRCAP;refGene_tx=XM_023246887.1;refGene_gn=SRCAP;refGene_tx=XM_023246888.1
E3 20800951 20801138
refGene_gn=SRCAP;refGene_tx=XM_023246885.1;refGene_gn=SRCAP;refGene_tx=XM_023246886.1;refGene_gn=SRCAP;refGene_tx=XM_023246887.1;refGene_gn=SRCAP;refGene_tx=XM_023246888.1
E3 20801229 20801366
refGene_gn=SRCAP;refGene_tx=XM_023246885.1;refGene_gn=SRCAP;refGene_tx=XM_023246886.1;refGene_gn=SRCAP;refGene_tx=XM_023246887.1;refGene_gn=SRCAP;refGene_tx=XM_023246888.1
E3 20802988 20803181
refGene_gn=SRCAP;refGene_tx=XM_023246885.1;refGene_gn=SRCAP;refGene_tx=XM_023246886.1;refGene_gn=SRCAP;refGene_tx=XM_023246887.1;refGene_gn=SRCAP;refGene_tx=XM_023246888.1
E3 20803296 20803466
refGene_gn=SRCAP;refGene_tx=XM_023246885.1;refGene_gn=SRCAP;refGene_tx=XM_023246886.1;refGene_gn=SRCAP;refGene_tx=XM_023246887.1;refGene_gn=SRCAP;refGene_tx=XM_023246888.1
E3 20803693 20803830
refGene_gn=SRCAP;refGene_tx=XM_023246885.1;refGene_gn=SRCAP;refGene_tx=XM_023246886.1;refGene_gn=SRCAP;refGene_tx=XM_023246887.1;refGene_gn=

SRCAP;refGene_tx=XM_023246888.1
E3 20804617 20804795
refGene_gn=SRCAP;refGene_tx=XM_023246885.1;refGene_gn=SRCAP;refGene_tx
=XM_023246886.1;refGene_gn=SRCAP;refGene_tx=XM_023246887.1;refGene_gn=
SRCAP;refGene_tx=XM_023246888.1
E3 20804897 20805226
refGene_gn=SRCAP;refGene_tx=XM_023246885.1;refGene_gn=SRCAP;refGene_tx
=XM_023246886.1;refGene_gn=SRCAP;refGene_tx=XM_023246887.1;refGene_gn=
SRCAP;refGene_tx=XM_023246888.1
E3 20805311 20805500
refGene_gn=SRCAP;refGene_tx=XM_023246885.1;refGene_gn=SRCAP;refGene_tx
=XM_023246886.1;refGene_gn=SRCAP;refGene_tx=XM_023246887.1;refGene_gn=
SRCAP;refGene_tx=XM_023246888.1
E3 20805993 20806083
refGene_gn=SRCAP;refGene_tx=XM_023246885.1;refGene_gn=SRCAP;refGene_tx
=XM_023246886.1;refGene_gn=SRCAP;refGene_tx=XM_023246887.1;refGene_gn=
SRCAP;refGene_tx=XM_023246888.1
E3 20806223 20806314
refGene_gn=SRCAP;refGene_tx=XM_023246885.1;refGene_gn=SRCAP;refGene_tx
=XM_023246886.1;refGene_gn=SRCAP;refGene_tx=XM_023246887.1;refGene_gn=
SRCAP;refGene_tx=XM_023246888.1
E3 20806940 20807221
refGene_gn=SRCAP;refGene_tx=XM_023246885.1;refGene_gn=SRCAP;refGene_tx
=XM_023246886.1;refGene_gn=SRCAP;refGene_tx=XM_023246887.1;refGene_gn=
SRCAP;refGene_tx=XM_023246888.1
E3 20807336 20807559
refGene_gn=SRCAP;refGene_tx=XM_023246885.1;refGene_gn=SRCAP;refGene_tx
=XM_023246886.1;refGene_gn=SRCAP;refGene_tx=XM_023246887.1;refGene_gn=
SRCAP;refGene_tx=XM_023246888.1
E3 20808316 20808457
refGene_gn=SRCAP;refGene_tx=XM_023246885.1;refGene_gn=SRCAP;refGene_tx
=XM_023246886.1;refGene_gn=SRCAP;refGene_tx=XM_023246887.1;refGene_gn=
SRCAP;refGene_tx=XM_023246888.1
E3 20808699 20808885
refGene_gn=SRCAP;refGene_tx=XM_023246885.1;refGene_gn=SRCAP;refGene_tx
=XM_023246886.1;refGene_gn=SRCAP;refGene_tx=XM_023246887.1;refGene_gn=
SRCAP;refGene_tx=XM_023246888.1
E3 20810407 20810659
refGene_gn=SRCAP;refGene_tx=XM_023246885.1;refGene_gn=SRCAP;refGene_tx
=XM_023246886.1;refGene_gn=SRCAP;refGene_tx=XM_023246887.1;refGene_gn=
SRCAP;refGene_tx=XM_023246888.1
E3 20812031 20812085
refGene_gn=SRCAP;refGene_tx=XM_023246885.1;refGene_gn=SRCAP;refGene_tx
=XM_023246886.1;refGene_gn=SRCAP;refGene_tx=XM_023246887.1;refGene_gn=
SRCAP;refGene_tx=XM_023246888.1
E3 20842859 20843442 refGene_gn=FBRS;refGene_tx=XM_023246322.1
E3 20843562 20843770 refGene_gn=FBRS;refGene_tx=XM_023246322.1
E3 20843892 20843976 refGene_gn=FBRS;refGene_tx=XM_023246322.1
E3 20844944 20845025 refGene_gn=FBRS;refGene_tx=XM_023246322.1
E3 20845130 20845251 refGene_gn=FBRS;refGene_tx=XM_023246322.1

E3 20845560 20845632 refGene_gn=FBRS;refGene_tx=XM_023246322.1
E3 20846011 20846041 refGene_gn=FBRS;refGene_tx=XM_023246322.1
E3 20846840 20846909 refGene_gn=FBRS;refGene_tx=XM_023246322.1
E3 20847145 20847241 refGene_gn=FBRS;refGene_tx=XM_023246322.1
E3 20847457 20847502 refGene_gn=FBRS;refGene_tx=XM_023246322.1
E3 20847617 20847823 refGene_gn=FBRS;refGene_tx=XM_023246322.1
E3 20848029 20848146 refGene_gn=FBRS;refGene_tx=XM_023246322.1
E3 20848148 20848330 refGene_gn=FBRS;refGene_tx=XM_023246322.1
E3 20849724 20850025 refGene_gn=FBRS;refGene_tx=XM_023246322.1
E3 20850116 20850165 refGene_gn=FBRS;refGene_tx=XM_023246322.1
E3 20851226 20851256 refGene_gn=FBRS;refGene_tx=XM_023246322.1
E3 20851376 20851412 refGene_gn=FBRS;refGene_tx=XM_023246322.1
E3 20852150 20852330 refGene_gn=FBRS;refGene_tx=XM_023246322.1
E3 20852618 20853029 refGene_gn=FBRS;refGene_tx=XM_023246322.1
E3 20855732 20856012
refGene_gn=PRR14;refGene_tx=XM_006942144.4;refGene_gn=PRR14;refGene_tx=XM_019820686.2;refGene_gn=PRR14;refGene_tx=XM_023246602.1;refGene_gn=PRR14;refGene_tx=XM_023246603.1
E3 20856102 20856174
refGene_gn=PRR14;refGene_tx=XM_006942144.4;refGene_gn=PRR14;refGene_tx=XM_019820686.2;refGene_gn=PRR14;refGene_tx=XM_023246602.1;refGene_gn=PRR14;refGene_tx=XM_023246603.1
E3 20856434 20856526
refGene_gn=PRR14;refGene_tx=XM_006942144.4;refGene_gn=PRR14;refGene_tx=XM_019820686.2;refGene_gn=PRR14;refGene_tx=XM_023246602.1;refGene_gn=PRR14;refGene_tx=XM_023246603.1
E3 20856606 20856676
refGene_gn=PRR14;refGene_tx=XM_006942144.4;refGene_gn=PRR14;refGene_tx=XM_019820686.2;refGene_gn=PRR14;refGene_tx=XM_023246602.1;refGene_gn=PRR14;refGene_tx=XM_023246603.1
E3 20856812 20857401
refGene_gn=PRR14;refGene_tx=XM_006942144.4;refGene_gn=PRR14;refGene_tx=XM_019820686.2;refGene_gn=PRR14;refGene_tx=XM_023246602.1;refGene_gn=PRR14;refGene_tx=XM_023246603.1
E3 20857702 20857812
refGene_gn=PRR14;refGene_tx=XM_006942144.4;refGene_gn=PRR14;refGene_tx=XM_019820686.2;refGene_gn=PRR14;refGene_tx=XM_023246602.1;refGene_gn=PRR14;refGene_tx=XM_023246603.1
E3 20858013 20858057
refGene_gn=PRR14;refGene_tx=XM_006942144.4;refGene_gn=PRR14;refGene_tx=XM_019820686.2;refGene_gn=PRR14;refGene_tx=XM_023246602.1;refGene_gn=PRR14;refGene_tx=XM_023246603.1
E3 20858291 20858481
refGene_gn=PRR14;refGene_tx=XM_006942144.4;refGene_gn=PRR14;refGene_tx=XM_019820686.2;refGene_gn=PRR14;refGene_tx=XM_023246602.1;refGene_gn=PRR14;refGene_tx=XM_023246603.1
E3 20858551 20858673
refGene_gn=PRR14;refGene_tx=XM_006942144.4;refGene_gn=PRR14;refGene_tx=XM_019820686.2;refGene_gn=PRR14;refGene_tx=XM_023246602.1;refGene_gn=PRR14;refGene_tx=XM_023246603.1

E3 20859327 20859496
refGene_gn=PRR14;refGene_tx=XM_006942144.4;refGene_gn=PRR14;refGene_tx
=XM_019820686.2;refGene_gn=PRR14;refGene_tx=XM_023246602.1;refGene_gn=
PRR14;refGene_tx=XM_023246603.1
E3 20859620 20859643
refGene_gn=PRR14;refGene_tx=XM_006942144.4;refGene_gn=PRR14;refGene_tx
=XM_019820686.2;refGene_gn=PRR14;refGene_tx=XM_023246602.1
E3 20884120 20884316
refGene_gn=LOC101085169;refGene_tx=XM_006942146.3;refGene_gn=LOC101085
169;refGene_tx=XM_019820687.1
E3 20884502 20884616
refGene_gn=LOC101085169;refGene_tx=XM_006942146.3
E3 20885541 20886758
refGene_gn=LOC101085169;refGene_tx=XM_006942146.3;refGene_gn=LOC101085
169;refGene_tx=XM_019820687.1
E3 20894450 20894655
refGene_gn=LOC101085424;refGene_tx=XM_006942147.4
E3 20894868 20894982
refGene_gn=LOC101085424;refGene_tx=XM_006942147.4
E3 20896561 20897559
refGene_gn=LOC101085424;refGene_tx=XM_006942147.4
E3 20897560 20897746
refGene_gn=LOC101085424;refGene_tx=XM_006942147.4
E3 20925201 20925397
refGene_gn=ZNF688;refGene_tx=XM_011290527.3;refGene_gn=ZNF688;refGene_
tx=XM_023247117.1
E3 20925707 20925821 refGene_gn=ZNF688;refGene_tx=XM_011290527.3
E3 20926577 20927098
refGene_gn=ZNF688;refGene_tx=XM_011290527.3;refGene_gn=ZNF688;refGene_
tx=XM_023247117.1
E3 20938025 20938221
refGene_gn=LOC111556261;refGene_tx=XM_023247116.1
E3 20938373 20938487
refGene_gn=LOC111556261;refGene_tx=XM_023247116.1
E3 20939269 20940183
refGene_gn=LOC111556261;refGene_tx=XM_023247116.1
E3 20946700 20946788 refGene_gn=ZNF768;refGene_tx=XM_003998653.5
E3 20947079 20948656 refGene_gn=ZNF768;refGene_tx=XM_003998653.5
E3 20955365 20955539
refGene_gn=ITGAL;refGene_tx=XM_006942148.4;refGene_gn=ITGAL;refGene_tx
=XM_011290524.3;refGene_gn=ITGAL;refGene_tx=XM_019820691.2;refGene_gn=
ITGAL;refGene_tx=XM_023246352.1
E3 20957181 20957292
refGene_gn=ITGAL;refGene_tx=XM_006942148.4;refGene_gn=ITGAL;refGene_tx
=XM_011290524.3;refGene_gn=ITGAL;refGene_tx=XM_019820691.2;refGene_gn=
ITGAL;refGene_tx=XM_023246352.1
E3 20958017 20958113
refGene_gn=ITGAL;refGene_tx=XM_006942148.4;refGene_gn=ITGAL;refGene_tx
=XM_011290524.3;refGene_gn=ITGAL;refGene_tx=XM_019820691.2;refGene_gn=
ITGAL;refGene_tx=XM_023246352.1

E3 20958527 20958626
refGene_gn=ITGAL;refGene_tx=XM_006942148.4;refGene_gn=ITGAL;refGene_tx=XM_011290524.3;refGene_gn=ITGAL;refGene_tx=XM_019820691.2;refGene_gn=ITGAL;refGene_tx=XM_023246352.1
E3 20958722 20958779
refGene_gn=ITGAL;refGene_tx=XM_006942148.4;refGene_gn=ITGAL;refGene_tx=XM_011290524.3;refGene_gn=ITGAL;refGene_tx=XM_019820691.2;refGene_gn=ITGAL;refGene_tx=XM_023246352.1
E3 20959017 20959158
refGene_gn=ITGAL;refGene_tx=XM_006942148.4;refGene_gn=ITGAL;refGene_tx=XM_011290524.3;refGene_gn=ITGAL;refGene_tx=XM_019820691.2;refGene_gn=ITGAL;refGene_tx=XM_023246352.1
E3 20963593 20963669
refGene_gn=ITGAL;refGene_tx=XM_006942148.4;refGene_gn=ITGAL;refGene_tx=XM_011290524.3;refGene_gn=ITGAL;refGene_tx=XM_019820691.2;refGene_gn=ITGAL;refGene_tx=XM_023246352.1
E3 20965208 20965295
refGene_gn=ITGAL;refGene_tx=XM_006942148.4;refGene_gn=ITGAL;refGene_tx=XM_011290524.3;refGene_gn=ITGAL;refGene_tx=XM_019820691.2;refGene_gn=ITGAL;refGene_tx=XM_023246352.1
E3 20965412 20965492
refGene_gn=ITGAL;refGene_tx=XM_006942148.4;refGene_gn=ITGAL;refGene_tx=XM_011290524.3;refGene_gn=ITGAL;refGene_tx=XM_019820691.2;refGene_gn=ITGAL;refGene_tx=XM_023246352.1
E3 20965814 20965925
refGene_gn=ITGAL;refGene_tx=XM_006942148.4;refGene_gn=ITGAL;refGene_tx=XM_011290524.3;refGene_gn=ITGAL;refGene_tx=XM_019820691.2;refGene_gn=ITGAL;refGene_tx=XM_023246352.1
E3 20968255 20968397
refGene_gn=ITGAL;refGene_tx=XM_006942148.4;refGene_gn=ITGAL;refGene_tx=XM_011290524.3;refGene_gn=ITGAL;refGene_tx=XM_019820691.2;refGene_gn=ITGAL;refGene_tx=XM_023246352.1
E3 20968736 20968810
refGene_gn=ITGAL;refGene_tx=XM_006942148.4;refGene_gn=ITGAL;refGene_tx=XM_011290524.3;refGene_gn=ITGAL;refGene_tx=XM_019820691.2;refGene_gn=ITGAL;refGene_tx=XM_023246352.1
E3 20968903 20968960
refGene_gn=ITGAL;refGene_tx=XM_006942148.4;refGene_gn=ITGAL;refGene_tx=XM_011290524.3;refGene_gn=ITGAL;refGene_tx=XM_019820691.2;refGene_gn=ITGAL;refGene_tx=XM_023246352.1
E3 20972218 20972308
refGene_gn=ITGAL;refGene_tx=XM_006942148.4;refGene_gn=ITGAL;refGene_tx=XM_011290524.3;refGene_gn=ITGAL;refGene_tx=XM_019820691.2;refGene_gn=ITGAL;refGene_tx=XM_023246352.1
E3 20973645 20973797
refGene_gn=ITGAL;refGene_tx=XM_006942148.4;refGene_gn=ITGAL;refGene_tx=XM_011290524.3;refGene_gn=ITGAL;refGene_tx=XM_019820691.2;refGene_gn=ITGAL;refGene_tx=XM_023246352.1
E3 20973902 20974063
refGene_gn=ITGAL;refGene_tx=XM_006942148.4;refGene_gn=ITGAL;refGene_tx

=XM_011290524.3;refGene_gn=ITGAL;refGene_tx=XM_019820691.2;refGene_gn=ITGAL;refGene_tx=XM_023246352.1
E3 20975583 20975714
refGene_gn=ITGAL;refGene_tx=XM_006942148.4;refGene_gn=ITGAL;refGene_tx=XM_011290524.3;refGene_gn=ITGAL;refGene_tx=XM_019820691.2;refGene_gn=ITGAL;refGene_tx=XM_023246352.1
E3 20975860 20976058
refGene_gn=ITGAL;refGene_tx=XM_006942148.4;refGene_gn=ITGAL;refGene_tx=XM_011290524.3;refGene_gn=ITGAL;refGene_tx=XM_019820691.2;refGene_gn=ITGAL;refGene_tx=XM_023246352.1
E3 20976516 20976654
refGene_gn=ITGAL;refGene_tx=XM_006942148.4;refGene_gn=ITGAL;refGene_tx=XM_011290524.3;refGene_gn=ITGAL;refGene_tx=XM_019820691.2;refGene_gn=ITGAL;refGene_tx=XM_023246352.1
E3 20976976 20977131
refGene_gn=ITGAL;refGene_tx=XM_006942148.4;refGene_gn=ITGAL;refGene_tx=XM_011290524.3;refGene_gn=ITGAL;refGene_tx=XM_019820691.2;refGene_gn=ITGAL;refGene_tx=XM_023246352.1
E3 20980777 20980910
refGene_gn=ITGAL;refGene_tx=XM_006942148.4;refGene_gn=ITGAL;refGene_tx=XM_011290524.3;refGene_gn=ITGAL;refGene_tx=XM_019820691.2;refGene_gn=ITGAL;refGene_tx=XM_023246352.1
E3 20980990 20981064
refGene_gn=ITGAL;refGene_tx=XM_006942148.4;refGene_gn=ITGAL;refGene_tx=XM_011290524.3;refGene_gn=ITGAL;refGene_tx=XM_019820691.2;refGene_gn=ITGAL;refGene_tx=XM_023246352.1
E3 20983236 20983491
refGene_gn=LOC109495204;refGene_tx=XM_023246354.1
E3 20985372 20985523
refGene_gn=ITGAL;refGene_tx=XM_006942148.4;refGene_gn=ITGAL;refGene_tx=XM_011290524.3;refGene_gn=ITGAL;refGene_tx=XM_019820691.2;refGene_gn=ITGAL;refGene_tx=XM_023246352.1
E3 20985677 20985813
refGene_gn=ITGAL;refGene_tx=XM_006942148.4;refGene_gn=ITGAL;refGene_tx=XM_011290524.3;refGene_gn=ITGAL;refGene_tx=XM_019820691.2;refGene_gn=ITGAL;refGene_tx=XM_023246352.1
E3 20991573 20991719
refGene_gn=ITGAL;refGene_tx=XM_006942148.4;refGene_gn=ITGAL;refGene_tx=XM_011290524.3;refGene_gn=ITGAL;refGene_tx=XM_019820691.2;refGene_gn=ITGAL;refGene_tx=XM_023246352.1
E3 20992846 20992977
refGene_gn=ITGAL;refGene_tx=XM_006942148.4;refGene_gn=ITGAL;refGene_tx=XM_011290524.3;refGene_gn=ITGAL;refGene_tx=XM_019820691.2;refGene_gn=ITGAL;refGene_tx=XM_023246352.1
E3 20993102 20993220
refGene_gn=ITGAL;refGene_tx=XM_006942148.4;refGene_gn=ITGAL;refGene_tx=XM_011290524.3;refGene_gn=ITGAL;refGene_tx=XM_019820691.2;refGene_gn=ITGAL;refGene_tx=XM_023246352.1
E3 20993486 20993519 refGene_gn=ITGAL;refGene_tx=XM_019820691.2
E3 20994602 20994670

refGene_gn=ITGAL;refGene_tx=XM_006942148.4;refGene_gn=ITGAL;refGene_tx=XM_011290524.3
E3 20994781 20994876
refGene_gn=ITGAL;refGene_tx=XM_006942148.4;refGene_gn=ITGAL;refGene_tx=XM_011290524.3;refGene_gn=ITGAL;refGene_tx=XM_023246352.1
E3 20995571 20995674
refGene_gn=ITGAL;refGene_tx=XM_006942148.4;refGene_gn=ITGAL;refGene_tx=XM_011290524.3
E3 20996723 20996973
refGene_gn=ITGAL;refGene_tx=XM_006942148.4;refGene_gn=ITGAL;refGene_tx=XM_011290524.3
E3 21014860 21016213 refGene_gn=SEPHS2;refGene_tx=XM_004001417.5
E3 21025013 21025114 refGene_gn=DCTPP1;refGene_tx=XM_003998655.4
E3 21025753 21025864 refGene_gn=DCTPP1;refGene_tx=XM_003998655.4
E3 21027831 21028132 refGene_gn=DCTPP1;refGene_tx=XM_003998655.4
E3 21030450 21031263
refGene_gn=ZNF771;refGene_tx=XM_023247148.1;refGene_gn=ZNF771;refGene_tx=XM_023247149.1;refGene_gn=ZNF771;refGene_tx=XM_023247150.1;refGene_gn=ZNF771;refGene_tx=XM_023247151.1
E3 21038599 21038762
refGene_gn=ZNF771;refGene_tx=XM_023247148.1;refGene_gn=ZNF771;refGene_tx=XM_023247149.1;refGene_gn=ZNF771;refGene_tx=XM_023247150.1;refGene_gn=ZNF771;refGene_tx=XM_023247151.1
E3 21039112 21039228 refGene_gn=ZNF771;refGene_tx=XM_023247148.1
E3 21044304 21046106
refGene_gn=ZNF48;refGene_tx=XM_003998656.5;refGene_gn=ZNF48;refGene_tx=XM_006942151.3;refGene_gn=ZNF48;refGene_tx=XM_023247145.1
E3 21047321 21047400
refGene_gn=ZNF48;refGene_tx=XM_003998656.5;refGene_gn=ZNF48;refGene_tx=XM_006942151.3;refGene_gn=ZNF48;refGene_tx=XM_023247145.1
E3 21056204 21056234 refGene_gn=SEPT1;refGene_tx=XM_019820696.2
E3 21056266 21056482
refGene_gn=SEPT1;refGene_tx=XM_019820696.2;refGene_gn=SEPT1;refGene_tx=XM_023247147.1
E3 21056657 21056748
refGene_gn=SEPT1;refGene_tx=XM_019820696.2;refGene_gn=SEPT1;refGene_tx=XM_023247147.1
E3 21056847 21056934
refGene_gn=SEPT1;refGene_tx=XM_019820696.2;refGene_gn=SEPT1;refGene_tx=XM_023247147.1
E3 21057165 21057289
refGene_gn=SEPT1;refGene_tx=XM_019820696.2;refGene_gn=SEPT1;refGene_tx=XM_023247147.1
E3 21057594 21057729
refGene_gn=SEPT1;refGene_tx=XM_019820696.2;refGene_gn=SEPT1;refGene_tx=XM_023247147.1
E3 21057827 21057945
refGene_gn=SEPT1;refGene_tx=XM_019820696.2;refGene_gn=SEPT1;refGene_tx=XM_023247147.1
E3 21058502 21058604

refGene_gn=SEPT1;refGene_tx=XM_019820696.2;refGene_gn=SEPT1;refGene_tx=XM_023247147.1
E3 21058725 21058825 refGene_gn=SEPT1;refGene_tx=XM_019820696.2;refGene_gn=SEPT1;refGene_tx=XM_023247147.1
E3 21059047 21059213 refGene_gn=SEPT1;refGene_tx=XM_019820696.2;refGene_gn=SEPT1;refGene_tx=XM_023247147.1
E3 21059460 21059551 refGene_gn=SEPT1;refGene_tx=XM_019820696.2;refGene_gn=SEPT1;refGene_tx=XM_023247147.1
E3 21059664 21059751 refGene_gn=SEPT1;refGene_tx=XM_019820696.2;refGene_gn=SEPT1;refGene_tx=XM_023247147.1
E3 21060306 21060411 refGene_gn=MYLPF;refGene_tx=XM_003998657.3
E3 21060500 21060549 refGene_gn=MYLPF;refGene_tx=XM_003998657.3
E3 21060856 21060935 refGene_gn=MYLPF;refGene_tx=XM_003998657.3
E3 21061041 21061146 refGene_gn=MYLPF;refGene_tx=XM_003998657.3
E3 21061344 21061420 refGene_gn=MYLPF;refGene_tx=XM_003998657.3
E3 21061669 21061765 refGene_gn=MYLPF;refGene_tx=XM_003998657.3
E3 21062374 21062377 refGene_gn=MYLPF;refGene_tx=XM_003998657.3
E3 21066378 21067307 refGene_gn=TBC1D10B;refGene_tx=XM_011290552.3
E3 21070510 21070610 refGene_gn=TBC1D10B;refGene_tx=XM_011290552.3
E3 21070905 21071013 refGene_gn=TBC1D10B;refGene_tx=XM_011290552.3
E3 21071104 21071211 refGene_gn=TBC1D10B;refGene_tx=XM_011290552.3
E3 21074151 21074266 refGene_gn=TBC1D10B;refGene_tx=XM_011290552.3
E3 21074391 21074457 refGene_gn=TBC1D10B;refGene_tx=XM_011290552.3
E3 21074679 21074869 refGene_gn=TBC1D10B;refGene_tx=XM_011290552.3
E3 21075213 21075368 refGene_gn=TBC1D10B;refGene_tx=XM_011290552.3
E3 21075455 21076100 refGene_gn=TBC1D10B;refGene_tx=XM_011290552.3
E3 21077147 21077768 refGene_gn=LOC101086689;refGene_tx=XM_003998712.4;refGene_gn=LOC101086689;refGene_tx=XM_006942152.2;refGene_gn=LOC101086689;refGene_tx=XM_019820699.1;refGene_gn=LOC101086689;refGene_tx=XM_019820700.1;refGene_gn=LOC101086689;refGene_tx=XM_019820701.2
E3 21079473 21079551 refGene_gn=CD2BP2;refGene_tx=XM_003998658.4;refGene_gn=CD2BP2;refGene_tx=XM_019820697.1
E3 21079794 21079933 refGene_gn=CD2BP2;refGene_tx=XM_003998658.4;refGene_gn=CD2BP2;refGene_tx=XM_019820697.1
E3 21080083 21080241 refGene_gn=CD2BP2;refGene_tx=XM_003998658.4;refGene_gn=CD2BP2;refGene_tx=XM_019820697.1
E3 21080353 21080789 refGene_gn=CD2BP2;refGene_tx=XM_003998658.4;refGene_gn=CD2BP2;refGene_tx=XM_019820697.1
E3 21080874 21080981 refGene_gn=CD2BP2;refGene_tx=XM_003998658.4;refGene_gn=CD2BP2;refGene_tx=XM_019820697.1

E3 21081049 21081182
refGene_gn=CD2BP2;refGene_tx=XM_003998658.4;refGene_gn=CD2BP2;refGene_tx=XM_019820697.1
E3 21132475 21133900 refGene_gn=SPN;refGene_tx=XM_006942153.3
E3 21138734 21138747 refGene_gn=QPRT;refGene_tx=XM_023246609.1
E3 21141343 21141648
refGene_gn=LOC109495028;refGene_tx=XM_023246736.1
E3 21145032 21145036
refGene_gn=LOC109495028;refGene_tx=XM_023246736.1
E3 21153026 21153562 refGene_gn=QPRT;refGene_tx=XM_023246609.1
E3 21153869 21154001 refGene_gn=QPRT;refGene_tx=XM_023246609.1
E3 21154093 21154312 refGene_gn=QPRT;refGene_tx=XM_023246609.1
E3 21168265 21168940
refGene_gn=CE3H16orf54;refGene_tx=XM_003998659.4;refGene_gn=CE3H16orf54;refGene_tx=XM_006942155.3;refGene_gn=CE3H16orf54;refGene_tx=XM_006942156.4;refGene_gn=CE3H16orf54;refGene_tx=XM_019820704.2
E3 21182310 21182365
refGene_gn=LOC105261304;refGene_tx=XM_019820706.2
E3 21182549 21182682
refGene_gn=LOC105261304;refGene_tx=XM_019820706.2
E3 21183357 21183673
refGene_gn=LOC105261304;refGene_tx=XM_019820706.2
E3 21208824 21208879
refGene_gn=LOC101095538;refGene_tx=XM_023246434.1;refGene_gn=LOC101095538;refGene_tx=XM_023246436.1
E3 21209063 21209196
refGene_gn=LOC101095538;refGene_tx=XM_023246434.1;refGene_gn=LOC101095538;refGene_tx=XM_023246436.1
E3 21209871 21210175
refGene_gn=LOC101095538;refGene_tx=XM_023246434.1;refGene_gn=LOC101095538;refGene_tx=XM_023246436.1
E3 21222843 21222901 refGene_gn=KIF22;refGene_tx=XM_003998661.4
E3 21229197 21229393 refGene_gn=KIF22;refGene_tx=XM_003998661.4
E3 21231174 21231302 refGene_gn=KIF22;refGene_tx=XM_003998661.4
E3 21231392 21231547 refGene_gn=KIF22;refGene_tx=XM_003998661.4
E3 21231780 21231990 refGene_gn=KIF22;refGene_tx=XM_003998661.4
E3 21232070 21232301 refGene_gn=KIF22;refGene_tx=XM_003998661.4
E3 21232640 21232794 refGene_gn=KIF22;refGene_tx=XM_003998661.4
E3 21232928 21233064 refGene_gn=KIF22;refGene_tx=XM_003998661.4
E3 21234841 21235010 refGene_gn=KIF22;refGene_tx=XM_003998661.4
E3 21235530 21235699 refGene_gn=KIF22;refGene_tx=XM_003998661.4
E3 21236064 21236132 refGene_gn=KIF22;refGene_tx=XM_003998661.4
E3 21236845 21237058 refGene_gn=KIF22;refGene_tx=XM_003998661.4
E3 21237162 21237222 refGene_gn=KIF22;refGene_tx=XM_003998661.4
E3 21237306 21237354 refGene_gn=KIF22;refGene_tx=XM_003998661.4
E3 21238217 21238253
refGene_gn=MAZ;refGene_tx=XM_023246740.1;refGene_gn=MAZ;refGene_tx=XM_023246741.1
E3 21238791 21238983
refGene_gn=MAZ;refGene_tx=XM_023246738.1;refGene_gn=MAZ;refGene_tx=XM_

023246739.1;refGene_gn=MAZ;refGene_tx=XM_023246740.1;refGene_gn=MAZ;refGene_tx=XM_023246741.1;refGene_gn=MAZ;refGene_tx=XM_023246742.1
E3 21239066 21239923
refGene_gn=MAZ;refGene_tx=XM_023246738.1;refGene_gn=MAZ;refGene_tx=XM_023246739.1;refGene_gn=MAZ;refGene_tx=XM_023246740.1;refGene_gn=MAZ;refGene_tx=XM_023246741.1;refGene_gn=MAZ;refGene_tx=XM_023246742.1
E3 21240324 21240388
refGene_gn=MAZ;refGene_tx=XM_023246738.1;refGene_gn=MAZ;refGene_tx=XM_023246739.1;refGene_gn=MAZ;refGene_tx=XM_023246740.1;refGene_gn=MAZ;refGene_tx=XM_023246741.1;refGene_gn=MAZ;refGene_tx=XM_023246742.1
E3 21240526 21240625 refGene_gn=MAZ;refGene_tx=XM_023246742.1
E3 21240664 21240836
refGene_gn=MAZ;refGene_tx=XM_023246738.1;refGene_gn=MAZ;refGene_tx=XM_023246739.1;refGene_gn=MAZ;refGene_tx=XM_023246740.1;refGene_gn=MAZ;refGene_tx=XM_023246741.1
E3 21241648 21241851
refGene_gn=MAZ;refGene_tx=XM_023246738.1;refGene_gn=MAZ;refGene_tx=XM_023246740.1
E3 21242184 21242342
refGene_gn=MAZ;refGene_tx=XM_023246739.1;refGene_gn=MAZ;refGene_tx=XM_023246741.1
E3 21244851 21244853 refGene_gn=PRRT2;refGene_tx=XM_019820709.1
E3 21245234 21246201
refGene_gn=PRRT2;refGene_tx=XM_019820709.1;refGene_gn=PRRT2;refGene_tx=XM_019820710.2
E3 21246593 21246726
refGene_gn=PRRT2;refGene_tx=XM_019820709.1;refGene_gn=PRRT2;refGene_tx=XM_019820710.2
E3 21246900 21246911 refGene_gn=PRRT2;refGene_tx=XM_019820710.2
E3 21248353 21248406 refGene_gn=PRRT2;refGene_tx=XM_019820709.1
E3 21248844 21249326 refGene_gn=PAGR1;refGene_tx=XM_003998666.3
E3 21249519 21249602 refGene_gn=PAGR1;refGene_tx=XM_003998666.3
E3 21250714 21250914 refGene_gn=PAGR1;refGene_tx=XM_003998666.3
E3 21257684 21257809
refGene_gn=MVP;refGene_tx=NM_001042562.1;refGene_gn=MVP;refGene_tx=XM_006942106.4;refGene_gn=MVP;refGene_tx=XM_006942107.4;refGene_gn=MVP;refGene_tx=XM_019820224.2
E3 21257992 21258188
refGene_gn=MVP;refGene_tx=NM_001042562.1;refGene_gn=MVP;refGene_tx=XM_006942106.4;refGene_gn=MVP;refGene_tx=XM_006942107.4;refGene_gn=MVP;refGene_tx=XM_019820224.2
E3 21264748 21264872
refGene_gn=MVP;refGene_tx=NM_001042562.1;refGene_gn=MVP;refGene_tx=XM_006942106.4;refGene_gn=MVP;refGene_tx=XM_006942107.4;refGene_gn=MVP;refGene_tx=XM_019820224.2
E3 21264972 21265104
refGene_gn=MVP;refGene_tx=NM_001042562.1;refGene_gn=MVP;refGene_tx=XM_006942106.4;refGene_gn=MVP;refGene_tx=XM_006942107.4;refGene_gn=MVP;refGene_tx=XM_019820224.2
E3 21265339 21265434

refGene_gn=MVP;refGene_tx=NM_001042562.1;refGene_gn=MVP;refGene_tx=XM_006942106.4;refGene_gn=MVP;refGene_tx=XM_006942107.4;refGene_gn=MVP;refGene_tx=XM_019820224.2
E3 21265732 21265969
refGene_gn=MVP;refGene_tx=NM_001042562.1;refGene_gn=MVP;refGene_tx=XM_006942106.4;refGene_gn=MVP;refGene_tx=XM_006942107.4;refGene_gn=MVP;refGene_tx=XM_019820224.2
E3 21268677 21268959
refGene_gn=MVP;refGene_tx=NM_001042562.1;refGene_gn=MVP;refGene_tx=XM_006942106.4;refGene_gn=MVP;refGene_tx=XM_006942107.4;refGene_gn=MVP;refGene_tx=XM_019820224.2
E3 21269787 21270032
refGene_gn=MVP;refGene_tx=NM_001042562.1;refGene_gn=MVP;refGene_tx=XM_006942106.4;refGene_gn=MVP;refGene_tx=XM_006942107.4;refGene_gn=MVP;refGene_tx=XM_019820224.2
E3 21270169 21270367
refGene_gn=MVP;refGene_tx=NM_001042562.1;refGene_gn=MVP;refGene_tx=XM_006942106.4;refGene_gn=MVP;refGene_tx=XM_006942107.4;refGene_gn=MVP;refGene_tx=XM_019820224.2
E3 21271236 21271623
refGene_gn=MVP;refGene_tx=NM_001042562.1;refGene_gn=MVP;refGene_tx=XM_006942106.4;refGene_gn=MVP;refGene_tx=XM_006942107.4;refGene_gn=MVP;refGene_tx=XM_019820224.2
E3 21272681 21272798
refGene_gn=MVP;refGene_tx=NM_001042562.1;refGene_gn=MVP;refGene_tx=XM_006942106.4;refGene_gn=MVP;refGene_tx=XM_006942107.4;refGene_gn=MVP;refGene_tx=XM_019820224.2
E3 21272955 21273082
refGene_gn=MVP;refGene_tx=NM_001042562.1;refGene_gn=MVP;refGene_tx=XM_006942106.4;refGene_gn=MVP;refGene_tx=XM_006942107.4;refGene_gn=MVP;refGene_tx=XM_019820224.2
E3 21273583 21273772
refGene_gn=MVP;refGene_tx=NM_001042562.1;refGene_gn=MVP;refGene_tx=XM_006942106.4;refGene_gn=MVP;refGene_tx=XM_006942107.4;refGene_gn=MVP;refGene_tx=XM_019820224.2
E3 21274456 21274675
refGene_gn=MVP;refGene_tx=NM_001042562.1;refGene_gn=MVP;refGene_tx=XM_006942106.4;refGene_gn=MVP;refGene_tx=XM_006942107.4;refGene_gn=MVP;refGene_tx=XM_019820224.2
E3 21282859 21283005
refGene_gn=CDIPT;refGene_tx=XM_003998667.3;refGene_gn=CDIPT;refGene_tx=XM_006942161.2
E3 21283150 21283232
refGene_gn=CDIPT;refGene_tx=XM_003998667.3;refGene_gn=CDIPT;refGene_tx=XM_006942161.2
E3 21283804 21283886
refGene_gn=CDIPT;refGene_tx=XM_003998667.3;refGene_gn=CDIPT;refGene_tx=XM_006942161.2
E3 21284325 21284479
refGene_gn=CDIPT;refGene_tx=XM_003998667.3;refGene_gn=CDIPT;refGene_tx

=XM_006942161.2
E3	21285193 21285328 refGene_gn=CDIPT;refGene_tx=XM_003998667.3
E3	21285422 21285465 refGene_gn=CDIPT;refGene_tx=XM_003998667.3
E3	21286868 21286961
refGene_gn=LOC102901594;refGene_tx=XM_006942162.4;refGene_gn=LOC102901
594;refGene_tx=XM_006942163.4;refGene_gn=LOC102901594;refGene_tx=XM_02
3246541.1;refGene_gn=LOC102901594;refGene_tx=XM_023246542.1
E3	21287261 21287409
refGene_gn=LOC102901594;refGene_tx=XM_006942162.4;refGene_gn=LOC102901
594;refGene_tx=XM_006942163.4;refGene_gn=LOC102901594;refGene_tx=XM_02
3246541.1;refGene_gn=LOC102901594;refGene_tx=XM_023246542.1
E3	21287674 21287705
refGene_gn=LOC102901594;refGene_tx=XM_006942162.4;refGene_gn=LOC102901
594;refGene_tx=XM_006942163.4;refGene_gn=LOC102901594;refGene_tx=XM_02
3246541.1;refGene_gn=LOC102901594;refGene_tx=XM_023246542.1
E3	21287895 21287953
refGene_gn=LOC102901594;refGene_tx=XM_006942162.4;refGene_gn=LOC102901
594;refGene_tx=XM_006942163.4;refGene_gn=LOC102901594;refGene_tx=XM_02
3246541.1;refGene_gn=LOC102901594;refGene_tx=XM_023246542.1
E3	21290391 21290427
refGene_gn=LOC102901594;refGene_tx=XM_006942162.4;refGene_gn=LOC102901
594;refGene_tx=XM_006942163.4;refGene_gn=LOC102901594;refGene_tx=XM_02
3246541.1;refGene_gn=LOC102901594;refGene_tx=XM_023246542.1
E3	21292811 21292841
refGene_gn=SEZ6L2;refGene_tx=XM_023246613.1;refGene_gn=SEZ6L2;refGene_
tx=XM_023246614.1
E3	21293317 21293414
refGene_gn=SEZ6L2;refGene_tx=XM_023246613.1;refGene_gn=SEZ6L2;refGene_
tx=XM_023246614.1
E3	21293540 21293658
refGene_gn=SEZ6L2;refGene_tx=XM_023246613.1;refGene_gn=SEZ6L2;refGene_
tx=XM_023246614.1
E3	21293836 21293875 refGene_gn=SEZ6L2;refGene_tx=XM_023246613.1
E3	21294363 21294555
refGene_gn=SEZ6L2;refGene_tx=XM_023246613.1;refGene_gn=SEZ6L2;refGene_
tx=XM_023246614.1
E3	21294638 21294830
refGene_gn=SEZ6L2;refGene_tx=XM_023246613.1;refGene_gn=SEZ6L2;refGene_
tx=XM_023246614.1
E3	21296928 21297123
refGene_gn=SEZ6L2;refGene_tx=XM_023246613.1;refGene_gn=SEZ6L2;refGene_
tx=XM_023246614.1
E3	21297263 21297460
refGene_gn=SEZ6L2;refGene_tx=XM_023246613.1;refGene_gn=SEZ6L2;refGene_
tx=XM_023246614.1
E3	21298301 21298440
refGene_gn=SEZ6L2;refGene_tx=XM_023246613.1;refGene_gn=SEZ6L2;refGene_
tx=XM_023246614.1
E3	21298918 21299119
refGene_gn=SEZ6L2;refGene_tx=XM_023246613.1;refGene_gn=SEZ6L2;refGene_

tx=XM_023246614.1
E3 21301109 21301273
refGene_gn=SEZ6L2;refGene_tx=XM_023246613.1;refGene_gn=SEZ6L2;refGene_
tx=XM_023246614.1
E3 21301903 21302072
refGene_gn=SEZ6L2;refGene_tx=XM_023246613.1;refGene_gn=SEZ6L2;refGene_
tx=XM_023246614.1
E3 21302102 21302107
refGene_gn=SEZ6L2;refGene_tx=XM_023246613.1;refGene_gn=SEZ6L2;refGene_
tx=XM_023246614.1
E3 21302725 21302906
refGene_gn=SEZ6L2;refGene_tx=XM_023246613.1;refGene_gn=SEZ6L2;refGene_
tx=XM_023246614.1
E3 21307487 21307689
refGene_gn=SEZ6L2;refGene_tx=XM_023246613.1;refGene_gn=SEZ6L2;refGene_
tx=XM_023246614.1
E3 21307956 21308096
refGene_gn=SEZ6L2;refGene_tx=XM_023246613.1;refGene_gn=SEZ6L2;refGene_
tx=XM_023246614.1
E3 21308463 21308763
refGene_gn=SEZ6L2;refGene_tx=XM_023246613.1;refGene_gn=SEZ6L2;refGene_
tx=XM_023246614.1
E3 21309432 21309564
refGene_gn=SEZ6L2;refGene_tx=XM_023246613.1;refGene_gn=SEZ6L2;refGene_
tx=XM_023246614.1
E3 21310482 21310561
refGene_gn=SEZ6L2;refGene_tx=XM_023246613.1;refGene_gn=SEZ6L2;refGene_
tx=XM_023246614.1
E3 21312674 21313554 refGene_gn=ASPHD1;refGene_tx=XM_003998675.5
E3 21314797 21314911 refGene_gn=ASPHD1;refGene_tx=XM_003998675.5
E3 21315446 21315556 refGene_gn=ASPHD1;refGene_tx=XM_003998675.5
E3 21316444 21316681 refGene_gn=KCTD13;refGene_tx=XM_003998676.5
E3 21318399 21318595 refGene_gn=KCTD13;refGene_tx=XM_003998676.5
E3 21319255 21319308 refGene_gn=KCTD13;refGene_tx=XM_003998676.5
E3 21319397 21319487 refGene_gn=KCTD13;refGene_tx=XM_003998676.5
E3 21327313 21327483 refGene_gn=KCTD13;refGene_tx=XM_003998676.5
E3 21328785 21329029 refGene_gn=KCTD13;refGene_tx=XM_003998676.5
E3 21376899 21377160
refGene_gn=TMEM219;refGene_tx=XM_006942165.4;refGene_gn=TMEM219;refGen
e_tx=XM_011290519.3;refGene_gn=TMEM219;refGene_tx=XM_019820716.2;refGe
ne_gn=TMEM219;refGene_tx=XM_019820717.2
E3 21377237 21377427
refGene_gn=TMEM219;refGene_tx=XM_006942165.4;refGene_gn=TMEM219;refGen
e_tx=XM_011290519.3;refGene_gn=TMEM219;refGene_tx=XM_019820716.2;refGe
ne_gn=TMEM219;refGene_tx=XM_019820717.2
E3 21383768 21383998
refGene_gn=TMEM219;refGene_tx=XM_006942165.4;refGene_gn=TMEM219;refGen
e_tx=XM_011290519.3;refGene_gn=TMEM219;refGene_tx=XM_019820716.2;refGe
ne_gn=TMEM219;refGene_tx=XM_019820717.2
E3 21385403 21385541

refGene_gn=TMEM219;refGene_tx=XM_006942165.4;refGene_gn=TMEM219;refGene_tx=XM_011290519.3;refGene_gn=TMEM219;refGene_tx=XM_019820716.2;refGene_gn=TMEM219;refGene_tx=XM_019820717.2
E3 21391394 21391526
refGene_gn=TAOK2;refGene_tx=XM_003998678.5;refGene_gn=TAOK2;refGene_tx=XM_003998679.4;refGene_gn=TAOK2;refGene_tx=XM_006942168.4;refGene_gn=TAOK2;refGene_tx=XM_006942169.3;refGene_gn=TAOK2;refGene_tx=XM_006942170.4;refGene_gn=TAOK2;refGene_tx=XM_006942171.4;refGene_gn=TAOK2;refGene_tx=XM_019820714.2
E3 21391713 21391785
refGene_gn=TAOK2;refGene_tx=XM_003998678.5;refGene_gn=TAOK2;refGene_tx=XM_003998679.4;refGene_gn=TAOK2;refGene_tx=XM_006942168.4;refGene_gn=TAOK2;refGene_tx=XM_006942169.3;refGene_gn=TAOK2;refGene_tx=XM_006942170.4;refGene_gn=TAOK2;refGene_tx=XM_006942171.4;refGene_gn=TAOK2;refGene_tx=XM_019820714.2
E3 21391883 21391985
refGene_gn=TAOK2;refGene_tx=XM_003998678.5;refGene_gn=TAOK2;refGene_tx=XM_003998679.4;refGene_gn=TAOK2;refGene_tx=XM_006942168.4;refGene_gn=TAOK2;refGene_tx=XM_006942169.3;refGene_gn=TAOK2;refGene_tx=XM_006942170.4;refGene_gn=TAOK2;refGene_tx=XM_006942171.4;refGene_gn=TAOK2;refGene_tx=XM_019820714.2
E3 21392480 21392526
refGene_gn=TAOK2;refGene_tx=XM_003998678.5;refGene_gn=TAOK2;refGene_tx=XM_003998679.4;refGene_gn=TAOK2;refGene_tx=XM_006942168.4;refGene_gn=TAOK2;refGene_tx=XM_006942169.3;refGene_gn=TAOK2;refGene_tx=XM_006942170.4;refGene_gn=TAOK2;refGene_tx=XM_006942171.4;refGene_gn=TAOK2;refGene_tx=XM_019820714.2
E3 21392658 21392755
refGene_gn=TAOK2;refGene_tx=XM_003998678.5;refGene_gn=TAOK2;refGene_tx=XM_003998679.4;refGene_gn=TAOK2;refGene_tx=XM_006942168.4;refGene_gn=TAOK2;refGene_tx=XM_006942169.3;refGene_gn=TAOK2;refGene_tx=XM_006942170.4;refGene_gn=TAOK2;refGene_tx=XM_006942171.4;refGene_gn=TAOK2;refGene_tx=XM_019820714.2
E3 21392904 21393018
refGene_gn=TAOK2;refGene_tx=XM_003998678.5;refGene_gn=TAOK2;refGene_tx=XM_003998679.4;refGene_gn=TAOK2;refGene_tx=XM_006942168.4;refGene_gn=TAOK2;refGene_tx=XM_006942169.3;refGene_gn=TAOK2;refGene_tx=XM_006942170.4;refGene_gn=TAOK2;refGene_tx=XM_006942171.4;refGene_gn=TAOK2;refGene_tx=XM_019820714.2
E3 21393123 21393215
refGene_gn=TAOK2;refGene_tx=XM_003998678.5;refGene_gn=TAOK2;refGene_tx=XM_003998679.4;refGene_gn=TAOK2;refGene_tx=XM_006942168.4;refGene_gn=TAOK2;refGene_tx=XM_006942169.3;refGene_gn=TAOK2;refGene_tx=XM_006942170.4;refGene_gn=TAOK2;refGene_tx=XM_006942171.4;refGene_gn=TAOK2;refGene_tx=XM_019820714.2
E3 21395243 21395337
refGene_gn=TAOK2;refGene_tx=XM_003998678.5;refGene_gn=TAOK2;refGene_tx=XM_003998679.4;refGene_gn=TAOK2;refGene_tx=XM_006942168.4;refGene_gn=TAOK2;refGene_tx=XM_006942169.3;refGene_gn=TAOK2;refGene_tx=XM_006942170.4;refGene_gn=TAOK2;refGene_tx=XM_006942171.4;refGene_gn=TAOK2;refGe

ne_tx=XM_019820714.2
E3 21395435 21395517
refGene_gn=TAOK2;refGene_tx=XM_003998678.5;refGene_gn=TAOK2;refGene_tx=XM_003998679.4;refGene_gn=TAOK2;refGene_tx=XM_006942168.4;refGene_gn=TAOK2;refGene_tx=XM_006942169.3;refGene_gn=TAOK2;refGene_tx=XM_006942170.4;refGene_gn=TAOK2;refGene_tx=XM_006942171.4;refGene_gn=TAOK2;refGene_tx=XM_019820714.2
E3 21396319 21396487
refGene_gn=TAOK2;refGene_tx=XM_003998678.5;refGene_gn=TAOK2;refGene_tx=XM_003998679.4;refGene_gn=TAOK2;refGene_tx=XM_006942168.4;refGene_gn=TAOK2;refGene_tx=XM_006942169.3;refGene_gn=TAOK2;refGene_tx=XM_006942170.4;refGene_gn=TAOK2;refGene_tx=XM_006942171.4;refGene_gn=TAOK2;refGene_tx=XM_019820714.2
E3 21396645 21396924
refGene_gn=TAOK2;refGene_tx=XM_003998678.5;refGene_gn=TAOK2;refGene_tx=XM_003998679.4;refGene_gn=TAOK2;refGene_tx=XM_006942168.4;refGene_gn=TAOK2;refGene_tx=XM_006942169.3;refGene_gn=TAOK2;refGene_tx=XM_006942170.4;refGene_gn=TAOK2;refGene_tx=XM_006942171.4;refGene_gn=TAOK2;refGene_tx=XM_019820714.2
E3 21397091 21397253
refGene_gn=TAOK2;refGene_tx=XM_003998678.5;refGene_gn=TAOK2;refGene_tx=XM_003998679.4;refGene_gn=TAOK2;refGene_tx=XM_006942168.4;refGene_gn=TAOK2;refGene_tx=XM_006942169.3;refGene_gn=TAOK2;refGene_tx=XM_006942170.4;refGene_gn=TAOK2;refGene_tx=XM_006942171.4;refGene_gn=TAOK2;refGene_tx=XM_019820714.2
E3 21398755 21399121
refGene_gn=TAOK2;refGene_tx=XM_003998678.5;refGene_gn=TAOK2;refGene_tx=XM_003998679.4;refGene_gn=TAOK2;refGene_tx=XM_006942168.4;refGene_gn=TAOK2;refGene_tx=XM_006942169.3;refGene_gn=TAOK2;refGene_tx=XM_006942170.4;refGene_gn=TAOK2;refGene_tx=XM_006942171.4;refGene_gn=TAOK2;refGene_tx=XM_019820714.2
E3 21399203 21399407
refGene_gn=TAOK2;refGene_tx=XM_003998678.5;refGene_gn=TAOK2;refGene_tx=XM_003998679.4;refGene_gn=TAOK2;refGene_tx=XM_006942168.4;refGene_gn=TAOK2;refGene_tx=XM_006942169.3;refGene_gn=TAOK2;refGene_tx=XM_006942170.4;refGene_gn=TAOK2;refGene_tx=XM_006942171.4;refGene_gn=TAOK2;refGene_tx=XM_019820714.2
E3 21399695 21401417
refGene_gn=TAOK2;refGene_tx=XM_003998678.5;refGene_gn=TAOK2;refGene_tx=XM_003998679.4;refGene_gn=TAOK2;refGene_tx=XM_006942168.4;refGene_gn=TAOK2;refGene_tx=XM_006942169.3;refGene_gn=TAOK2;refGene_tx=XM_006942170.4;refGene_gn=TAOK2;refGene_tx=XM_006942171.4;refGene_gn=TAOK2;refGene_tx=XM_019820714.2
E3 21403083 21403296
refGene_gn=TAOK2;refGene_tx=XM_003998679.4;refGene_gn=TAOK2;refGene_tx=XM_006942169.3;refGene_gn=TAOK2;refGene_tx=XM_006942170.4;refGene_gn=TAOK2;refGene_tx=XM_006942171.4
E3 21404029 21404122
refGene_gn=TAOK2;refGene_tx=XM_006942170.4;refGene_gn=TAOK2;refGene_tx=XM_006942171.4

E3 21404274 21404457
refGene_gn=TAOK2;refGene_tx=XM_003998679.4;refGene_gn=TAOK2;refGene_tx=XM_006942169.3;refGene_gn=TAOK2;refGene_tx=XM_019820714.2
E3 21404549 21405071
refGene_gn=TAOK2;refGene_tx=XM_003998679.4;refGene_gn=TAOK2;refGene_tx=XM_006942169.3;refGene_gn=TAOK2;refGene_tx=XM_019820714.2
E3 21407071 21407235 refGene_gn=HIRIP3;refGene_tx=XM_006942178.2
E3 21407322 21407421 refGene_gn=HIRIP3;refGene_tx=XM_006942178.2
E3 21407504 21407673 refGene_gn=HIRIP3;refGene_tx=XM_006942178.2
E3 21407773 21408783 refGene_gn=HIRIP3;refGene_tx=XM_006942178.2
E3 21409017 21409132 refGene_gn=HIRIP3;refGene_tx=XM_006942178.2
E3 21409247 21409368 refGene_gn=HIRIP3;refGene_tx=XM_006942178.2
E3 21409445 21409510 refGene_gn=HIRIP3;refGene_tx=XM_006942178.2
E3 21410209 21410290
refGene_gn=INO80E;refGene_tx=XM_003998681.5;refGene_gn=INO80E;refGene_tx=XM_006942172.3;refGene_gn=INO80E;refGene_tx=XM_006942173.3;refGene_gn=INO80E;refGene_tx=XM_006942174.4;refGene_gn=INO80E;refGene_tx=XM_006942175.4;refGene_gn=INO80E;refGene_tx=XM_006942176.4;refGene_gn=INO80E;refGene_tx=XM_006942177.3;refGene_gn=INO80E;refGene_tx=XM_011290517.2;refGene_gn=INO80E;refGene_tx=XM_019820718.1;refGene_gn=INO80E;refGene_tx=XM_019820720.1;refGene_gn=INO80E;refGene_tx=XM_019820721.2
E3 21410447 21410518
refGene_gn=INO80E;refGene_tx=XM_003998681.5;refGene_gn=INO80E;refGene_tx=XM_006942172.3;refGene_gn=INO80E;refGene_tx=XM_006942173.3;refGene_gn=INO80E;refGene_tx=XM_006942174.4;refGene_gn=INO80E;refGene_tx=XM_006942175.4;refGene_gn=INO80E;refGene_tx=XM_006942176.4;refGene_gn=INO80E;refGene_tx=XM_006942177.3;refGene_gn=INO80E;refGene_tx=XM_011290517.2;refGene_gn=INO80E;refGene_tx=XM_019820718.1;refGene_gn=INO80E;refGene_tx=XM_019820720.1;refGene_gn=INO80E;refGene_tx=XM_019820721.2
E3 21410747 21410800
refGene_gn=INO80E;refGene_tx=XM_003998681.5;refGene_gn=INO80E;refGene_tx=XM_006942172.3;refGene_gn=INO80E;refGene_tx=XM_006942173.3;refGene_gn=INO80E;refGene_tx=XM_006942174.4;refGene_gn=INO80E;refGene_tx=XM_006942175.4;refGene_gn=INO80E;refGene_tx=XM_006942176.4;refGene_gn=INO80E;refGene_tx=XM_006942177.3;refGene_gn=INO80E;refGene_tx=XM_011290517.2;refGene_gn=INO80E;refGene_tx=XM_019820718.1;refGene_gn=INO80E;refGene_tx=XM_019820720.1;refGene_gn=INO80E;refGene_tx=XM_019820721.2
E3 21413429 21413508
refGene_gn=INO80E;refGene_tx=XM_003998681.5;refGene_gn=INO80E;refGene_tx=XM_006942172.3;refGene_gn=INO80E;refGene_tx=XM_006942173.3;refGene_gn=INO80E;refGene_tx=XM_006942174.4;refGene_gn=INO80E;refGene_tx=XM_006942175.4;refGene_gn=INO80E;refGene_tx=XM_006942176.4;refGene_gn=INO80E;refGene_tx=XM_006942177.3;refGene_gn=INO80E;refGene_tx=XM_011290517.2;refGene_gn=INO80E;refGene_tx=XM_019820718.1;refGene_gn=INO80E;refGene_tx=XM_019820720.1;refGene_gn=INO80E;refGene_tx=XM_019820721.2
E3 21413585 21413721
refGene_gn=INO80E;refGene_tx=XM_003998681.5;refGene_gn=INO80E;refGene_tx=XM_006942172.3;refGene_gn=INO80E;refGene_tx=XM_006942173.3;refGene_gn=INO80E;refGene_tx=XM_006942174.4;refGene_gn=INO80E;refGene_tx=XM_006942175.4;refGene_gn=INO80E;refGene_tx=XM_006942176.4;refGene_gn=INO80

E;refGene_tx=XM_006942177.3;refGene_gn=INO80E;refGene_tx=XM_011290517.
2;refGene_gn=INO80E;refGene_tx=XM_019820718.1;refGene_gn=INO80E;refGen
e_tx=XM_019820720.1;refGene_gn=INO80E;refGene_tx=XM_019820721.2
E3 21414104 21414221
refGene_gn=INO80E;refGene_tx=XM_003998681.5;refGene_gn=INO80E;refGene_
tx=XM_006942172.3;refGene_gn=INO80E;refGene_tx=XM_006942173.3;refGene_
gn=INO80E;refGene_tx=XM_006942174.4;refGene_gn=INO80E;refGene_tx=XM_00
6942177.3;refGene_gn=INO80E;refGene_tx=XM_011290517.2;refGene_gn=INO80
E;refGene_tx=XM_019820718.1;refGene_gn=INO80E;refGene_tx=XM_019820720.
1;refGene_gn=INO80E;refGene_tx=XM_019820721.2
E3 21416158 21416425
refGene_gn=INO80E;refGene_tx=XM_006942172.3;refGene_gn=INO80E;refGene_
tx=XM_006942173.3;refGene_gn=INO80E;refGene_tx=XM_019820718.1;refGene_
gn=INO80E;refGene_tx=XM_019820720.1
E3 21417327 21417399
refGene_gn=INO80E;refGene_tx=XM_006942177.3;refGene_gn=INO80E;refGene_
tx=XM_011290517.2;refGene_gn=INO80E;refGene_tx=XM_019820720.1
E3 21418015 21418237
refGene_gn=INO80E;refGene_tx=XM_003998681.5;refGene_gn=INO80E;refGene_
tx=XM_006942174.4;refGene_gn=INO80E;refGene_tx=XM_006942175.4;refGene_
gn=INO80E;refGene_tx=XM_006942176.4;refGene_gn=INO80E;refGene_tx=XM_01
9820721.2
E3 21419033 21419179 refGene_gn=DOC2A;refGene_tx=XM_006942179.4
E3 21419279 21419376 refGene_gn=DOC2A;refGene_tx=XM_006942179.4
E3 21419466 21419548 refGene_gn=DOC2A;refGene_tx=XM_006942179.4
E3 21419655 21419819 refGene_gn=DOC2A;refGene_tx=XM_006942179.4
E3 21419892 21419952 refGene_gn=DOC2A;refGene_tx=XM_006942179.4
E3 21420036 21420163 refGene_gn=DOC2A;refGene_tx=XM_006942179.4
E3 21421690 21421800 refGene_gn=DOC2A;refGene_tx=XM_006942179.4
E3 21421922 21421997 refGene_gn=DOC2A;refGene_tx=XM_006942179.4
E3 21422221 21422301 refGene_gn=DOC2A;refGene_tx=XM_006942179.4
E3 21422657 21422922 refGene_gn=DOC2A;refGene_tx=XM_006942179.4
E3 21429294 21429358
refGene_gn=CE3H16orf92;refGene_tx=XM_003998716.4;refGene_gn=CE3H16orf9
2;refGene_tx=XM_019820722.2
E3 21429658 21429892
refGene_gn=CE3H16orf92;refGene_tx=XM_003998716.4;refGene_gn=CE3H16orf9
2;refGene_tx=XM_019820722.2
E3 21429932 21430020
refGene_gn=CE3H16orf92;refGene_tx=XM_003998716.4
E3 21430148 21430240
refGene_gn=CE3H16orf92;refGene_tx=XM_003998716.4;refGene_gn=CE3H16orf9
2;refGene_tx=XM_019820722.2
E3 21431215 21431503 refGene_gn=FAM57B;refGene_tx=XM_003998683.5
E3 21431666 21431762 refGene_gn=FAM57B;refGene_tx=XM_003998683.5
E3 21432194 21432429 refGene_gn=FAM57B;refGene_tx=XM_003998683.5
E3 21434968 21435052 refGene_gn=FAM57B;refGene_tx=XM_003998683.5
E3 21435950 21436075 refGene_gn=FAM57B;refGene_tx=XM_003998683.5
E3 21457222 21457334
refGene_gn=ALDOA;refGene_tx=XM_003998684.4;refGene_gn=ALDOA;refGene_tx

=XM_003998685.4
E3 21457419 21457631
refGene_gn=ALDOA;refGene_tx=XM_003998684.4;refGene_gn=ALDOA;refGene_tx=XM_003998685.4
E3 21458220 21458275
refGene_gn=ALDOA;refGene_tx=XM_003998684.4;refGene_gn=ALDOA;refGene_tx=XM_003998685.4
E3 21458401 21458562
refGene_gn=ALDOA;refGene_tx=XM_003998684.4;refGene_gn=ALDOA;refGene_tx=XM_003998685.4
E3 21458917 21459001
refGene_gn=ALDOA;refGene_tx=XM_003998684.4;refGene_gn=ALDOA;refGene_tx=XM_003998685.4
E3 21459119 21459294
refGene_gn=ALDOA;refGene_tx=XM_003998684.4;refGene_gn=ALDOA;refGene_tx=XM_003998685.4
E3 21459456 21459656
refGene_gn=ALDOA;refGene_tx=XM_003998684.4;refGene_gn=ALDOA;refGene_tx=XM_003998685.4
E3 21459741 21459837
refGene_gn=ALDOA;refGene_tx=XM_003998684.4;refGene_gn=ALDOA;refGene_tx=XM_003998685.4
E3 21462414 21462512 refGene_gn=PPP4C;refGene_tx=XM_003998688.5
E3 21465920 21465972 refGene_gn=PPP4C;refGene_tx=XM_003998688.5
E3 21466548 21466599 refGene_gn=PPP4C;refGene_tx=XM_003998688.5
E3 21466815 21466917 refGene_gn=PPP4C;refGene_tx=XM_003998688.5
E3 21467470 21467644 refGene_gn=PPP4C;refGene_tx=XM_003998688.5
E3 21467737 21467864 refGene_gn=PPP4C;refGene_tx=XM_003998688.5
E3 21468683 21468873 refGene_gn=PPP4C;refGene_tx=XM_003998688.5
E3 21468955 21469085 refGene_gn=PPP4C;refGene_tx=XM_003998688.5
E3 21470181 21470395 refGene_gn=TBX6;refGene_tx=XM_011290514.3
E3 21470478 21470662 refGene_gn=TBX6;refGene_tx=XM_011290514.3
E3 21470746 21470820 refGene_gn=TBX6;refGene_tx=XM_011290514.3
E3 21471970 21472041 refGene_gn=TBX6;refGene_tx=XM_011290514.3
E3 21472117 21472264 refGene_gn=TBX6;refGene_tx=XM_011290514.3
E3 21472342 21472610 refGene_gn=TBX6;refGene_tx=XM_011290514.3
E3 21473504 21473739 refGene_gn=TBX6;refGene_tx=XM_011290514.3
E3 21473821 21473939 refGene_gn=TBX6;refGene_tx=XM_011290514.3
E3 21475491 21475581 refGene_gn=YPEL3;refGene_tx=XM_023247126.1
E3 21477004 21477113 refGene_gn=YPEL3;refGene_tx=XM_023247126.1
E3 21477329 21477373 refGene_gn=YPEL3;refGene_tx=XM_023247126.1
E3 21477465 21477648 refGene_gn=YPEL3;refGene_tx=XM_023247126.1
E3 21478337 21478799 refGene_gn=YPEL3;refGene_tx=XM_023247126.1
E3 21479803 21479941
refGene_gn=GDPD3;refGene_tx=XM_003998691.4;refGene_gn=GDPD3;refGene_tx=XM_019820723.1;refGene_gn=GDPD3;refGene_tx=XM_019820724.2
E3 21481590 21481642
refGene_gn=GDPD3;refGene_tx=XM_003998691.4;refGene_gn=GDPD3;refGene_tx=XM_019820723.1
E3 21481743 21481803 refGene_gn=GDPD3;refGene_tx=XM_003998691.4

E3 21483122 21483256
refGene_gn=GDPD3;refGene_tx=XM_003998691.4;refGene_gn=GDPD3;refGene_tx=XM_019820723.1;refGene_gn=GDPD3;refGene_tx=XM_019820724.2
E3 21483602 21483692
refGene_gn=GDPD3;refGene_tx=XM_003998691.4;refGene_gn=GDPD3;refGene_tx=XM_019820723.1;refGene_gn=GDPD3;refGene_tx=XM_019820724.2
E3 21483781 21483900
refGene_gn=GDPD3;refGene_tx=XM_003998691.4;refGene_gn=GDPD3;refGene_tx=XM_019820723.1;refGene_gn=GDPD3;refGene_tx=XM_019820724.2
E3 21484036 21484082
refGene_gn=GDPD3;refGene_tx=XM_003998691.4;refGene_gn=GDPD3;refGene_tx=XM_019820723.1;refGene_gn=GDPD3;refGene_tx=XM_019820724.2
E3 21484177 21484313
refGene_gn=GDPD3;refGene_tx=XM_003998691.4;refGene_gn=GDPD3;refGene_tx=XM_019820723.1;refGene_gn=GDPD3;refGene_tx=XM_019820724.2
E3 21484524 21484567
refGene_gn=GDPD3;refGene_tx=XM_003998691.4;refGene_gn=GDPD3;refGene_tx=XM_019820723.1;refGene_gn=GDPD3;refGene_tx=XM_019820724.2
E3 21484816 21484955
refGene_gn=GDPD3;refGene_tx=XM_003998691.4;refGene_gn=GDPD3;refGene_tx=XM_019820723.1;refGene_gn=GDPD3;refGene_tx=XM_019820724.2
E3 21487086 21487209 refGene_gn=MAPK3;refGene_tx=XM_023247125.1
E3 21487302 21487412 refGene_gn=MAPK3;refGene_tx=XM_023247125.1
E3 21487579 21487711 refGene_gn=MAPK3;refGene_tx=XM_023247125.1
E3 21488080 21488195 refGene_gn=MAPK3;refGene_tx=XM_023247125.1
E3 21488480 21488597 refGene_gn=MAPK3;refGene_tx=XM_023247125.1
E3 21488779 21488969 refGene_gn=MAPK3;refGene_tx=XM_023247125.1
E3 21490454 21490637 refGene_gn=MAPK3;refGene_tx=XM_023247125.1
E3 21491684 21491854 refGene_gn=MAPK3;refGene_tx=XM_023247125.1
E3 21536622 21536820 refGene_gn=CORO1A;refGene_tx=XM_023246655.1
E3 21537884 21538007 refGene_gn=CORO1A;refGene_tx=XM_023246655.1
E3 21538099 21538229 refGene_gn=CORO1A;refGene_tx=XM_023246655.1
E3 21538308 21538493 refGene_gn=CORO1A;refGene_tx=XM_023246655.1
E3 21538651 21538771 refGene_gn=CORO1A;refGene_tx=XM_023246655.1
E3 21538992 21539097 refGene_gn=CORO1A;refGene_tx=XM_023246655.1
E3 21539229 21539375 refGene_gn=CORO1A;refGene_tx=XM_023246655.1
E3 21539492 21539550 refGene_gn=CORO1A;refGene_tx=XM_023246655.1
E3 21539665 21539881 refGene_gn=CORO1A;refGene_tx=XM_023246655.1
E3 21540215 21540320 refGene_gn=CORO1A;refGene_tx=XM_023246655.1
E3 21546310 21546360
refGene_gn=LOC102900324;refGene_tx=XM_011290550.3
E3 21547010 21547019
refGene_gn=LOC102900324;refGene_tx=XM_011290550.3
E3 21548138 21548230
refGene_gn=LOC102900324;refGene_tx=XM_011290550.3
E3 21551131 21551253
refGene_gn=LOC102900324;refGene_tx=XM_011290550.3
E3 21556178 21556255
refGene_gn=LOC102900324;refGene_tx=XM_011290550.3
E3 21558501 21558563

refGene_gn=LOC102900324;refGene_tx=XM_011290550.3
E3 21562143 21562246
refGene_gn=LOC102900324;refGene_tx=XM_011290550.3
E3 21565735 21565919
refGene_gn=LOC102900324;refGene_tx=XM_011290550.3
E3 21796256 21796356 refGene_gn=BOLA2B;refGene_tx=XM_003998718.4
E3 21796651 21796749 refGene_gn=BOLA2B;refGene_tx=XM_003998718.4
E3 21796838 21796965 refGene_gn=BOLA2B;refGene_tx=XM_003998718.4
E3 21796966 21797052 refGene_gn=BOLA2B;refGene_tx=XM_003998718.4
E3 21797053 21797098 refGene_gn=BOLA2B;refGene_tx=XM_003998718.4
E3 21797362 21797550
refGene_gn=SLX1A;refGene_tx=XM_003998719.4;refGene_gn=SLX1A;refGene_tx
=XM_006942189.3;refGene_gn=SLX1A;refGene_tx=XM_019820731.1
E3 21798109 21798161
refGene_gn=SLX1A;refGene_tx=XM_003998719.4;refGene_gn=SLX1A;refGene_tx
=XM_006942189.3;refGene_gn=SLX1A;refGene_tx=XM_019820731.1
E3 21798266 21798605
refGene_gn=SLX1A;refGene_tx=XM_003998719.4;refGene_gn=SLX1A;refGene_tx
=XM_006942189.3;refGene_gn=SLX1A;refGene_tx=XM_019820731.1
E3 21799659 21799787
refGene_gn=SLX1A;refGene_tx=XM_003998719.4;refGene_gn=SLX1A;refGene_tx
=XM_006942189.3;refGene_gn=SLX1A;refGene_tx=XM_019820731.1
E3 21799890 21799897 refGene_gn=SLX1A;refGene_tx=XM_006942189.3
E3 21799953 21800041
refGene_gn=SLX1A;refGene_tx=XM_003998719.4;refGene_gn=SLX1A;refGene_tx
=XM_019820731.1
E3 21800126 21800156
refGene_gn=SLX1A;refGene_tx=XM_003998719.4;refGene_gn=SLX1A;refGene_tx
=XM_019820731.1
E3 21802155 21802303
refGene_gn=LOC101088902;refGene_tx=XM_019820726.1;refGene_gn=LOC101088
902;refGene_tx=XM_019820727.1;refGene_gn=LOC101088902;refGene_tx=XM_01
9820728.1;refGene_gn=LOC101088902;refGene_tx=XM_019820729.1
E3 21802409 21802535
refGene_gn=LOC101088902;refGene_tx=XM_019820726.1;refGene_gn=LOC101088
902;refGene_tx=XM_019820727.1;refGene_gn=LOC101088902;refGene_tx=XM_01
9820728.1;refGene_gn=LOC101088902;refGene_tx=XM_019820729.1
E3 21802626 21802724
refGene_gn=LOC101088902;refGene_tx=XM_019820726.1;refGene_gn=LOC101088
902;refGene_tx=XM_019820727.1;refGene_gn=LOC101088902;refGene_tx=XM_01
9820728.1;refGene_gn=LOC101088902;refGene_tx=XM_019820729.1
E3 21803146 21803273
refGene_gn=LOC101088902;refGene_tx=XM_019820726.1;refGene_gn=LOC101088
902;refGene_tx=XM_019820727.1;refGene_gn=LOC101088902;refGene_tx=XM_01
9820728.1;refGene_gn=LOC101088902;refGene_tx=XM_019820729.1
E3 21803370 21803465
refGene_gn=LOC101088902;refGene_tx=XM_019820726.1;refGene_gn=LOC101088
902;refGene_tx=XM_019820727.1;refGene_gn=LOC101088902;refGene_tx=XM_01
9820728.1;refGene_gn=LOC101088902;refGene_tx=XM_019820729.1
E3 21803962 21804143

refGene_gn=LOC101088902;refGene_tx=XM_019820726.1;refGene_gn=LOC101088902;refGene_tx=XM_019820727.1;refGene_gn=LOC101088902;refGene_tx=XM_019820728.1;refGene_gn=LOC101088902;refGene_tx=XM_019820729.1
E3 21804273 21804386
refGene_gn=LOC101088902;refGene_tx=XM_019820726.1;refGene_gn=LOC101088902;refGene_tx=XM_019820727.1;refGene_gn=LOC101088902;refGene_tx=XM_019820728.1;refGene_gn=LOC101088902;refGene_tx=XM_019820729.1
E3 21804723 21804840
refGene_gn=SGF29;refGene_tx=XM_006942198.4;refGene_gn=SGF29;refGene_tx=XM_006942199.4
E3 21805551 21805714
refGene_gn=SGF29;refGene_tx=XM_006942198.4;refGene_gn=SGF29;refGene_tx=XM_006942199.4
E3 21805805 21805841
refGene_gn=SGF29;refGene_tx=XM_006942198.4;refGene_gn=SGF29;refGene_tx=XM_006942199.4
E3 21806054 21806201
refGene_gn=SGF29;refGene_tx=XM_006942198.4;refGene_gn=SGF29;refGene_tx=XM_006942199.4
E3 21806275 21806405
refGene_gn=SGF29;refGene_tx=XM_006942198.4;refGene_gn=SGF29;refGene_tx=XM_006942199.4
E3 21807307 21807372
refGene_gn=SGF29;refGene_tx=XM_006942198.4;refGene_gn=SGF29;refGene_tx=XM_006942199.4
E3 21812290 21812363
refGene_gn=SGF29;refGene_tx=XM_006942198.4;refGene_gn=SGF29;refGene_tx=XM_006942199.4
E3 21813058 21813134
refGene_gn=SGF29;refGene_tx=XM_006942198.4;refGene_gn=SGF29;refGene_tx=XM_006942199.4
E3 21815538 21815613
refGene_gn=SGF29;refGene_tx=XM_006942198.4;refGene_gn=SGF29;refGene_tx=XM_006942199.4
E3 21848952 21849064 refGene_gn=NUPR1;refGene_tx=XM_003998723.5
E3 21849468 21849605 refGene_gn=NUPR1;refGene_tx=XM_003998723.5
E3 21867675 21867706 refGene_gn=IL27;refGene_tx=XM_003998744.2
E3 21869470 21869803
refGene_gn=IL27;refGene_tx=XM_003998744.2;refGene_gn=IL27;refGene_tx=XM_019820732.2
E3 21869895 21869994
refGene_gn=IL27;refGene_tx=XM_003998744.2;refGene_gn=IL27;refGene_tx=XM_019820732.2
E3 21870609 21870768
refGene_gn=IL27;refGene_tx=XM_003998744.2;refGene_gn=IL27;refGene_tx=XM_019820732.2
E3 21871467 21871716
refGene_gn=IL27;refGene_tx=XM_003998744.2;refGene_gn=IL27;refGene_tx=XM_019820732.2
E3 21872681 21872751 refGene_gn=APOBR;refGene_tx=XM_023247080.1

E3 21872840 21872967 refGene_gn=APOBR;refGene_tx=XM_023247080.1
E3 21872969 21873105 refGene_gn=APOBR;refGene_tx=XM_023247080.1
E3 21873190 21875220 refGene_gn=APOBR;refGene_tx=XM_023247080.1
E3 21876015 21876201 refGene_gn=APOBR;refGene_tx=XM_023247080.1
E3 21876291 21877171 refGene_gn=APOBR;refGene_tx=XM_023247080.1
E3 21877518 21877824 refGene_gn=APOBR;refGene_tx=XM_023247080.1
E3 21879606 21879652 refGene_gn=CLN3;refGene_tx=XM_003998725.5
E3 21879807 21879886 refGene_gn=CLN3;refGene_tx=XM_003998725.5
E3 21881111 21881208 refGene_gn=CLN3;refGene_tx=XM_003998725.5
E3 21882019 21882091 refGene_gn=CLN3;refGene_tx=XM_003998725.5
E3 21882482 21882562 refGene_gn=CLN3;refGene_tx=XM_003998725.5
E3 21882699 21882785 refGene_gn=CLN3;refGene_tx=XM_003998725.5
E3 21883283 21883356 refGene_gn=CLN3;refGene_tx=XM_003998725.5
E3 21883438 21883582 refGene_gn=CLN3;refGene_tx=XM_003998725.5
E3 21884479 21884592 refGene_gn=CLN3;refGene_tx=XM_003998725.5
E3 21885709 21885756 refGene_gn=CLN3;refGene_tx=XM_003998725.5
E3 21885831 21885900 refGene_gn=CLN3;refGene_tx=XM_003998725.5
E3 21885996 21886052 refGene_gn=CLN3;refGene_tx=XM_003998725.5
E3 21886143 21886237 refGene_gn=CLN3;refGene_tx=XM_003998725.5
E3 21888320 21888461 refGene_gn=CLN3;refGene_tx=XM_003998725.5
E3 21888563 21888683 refGene_gn=CLN3;refGene_tx=XM_003998725.5
E3 21907620 21907721
refGene_gn=LOC101090225;refGene_tx=XM_006942201.2
E3 21909215 21909281
refGene_gn=LOC101090225;refGene_tx=XM_006942201.2
E3 21909383 21909573
refGene_gn=LOC101090225;refGene_tx=XM_006942201.2
E3 21909970 21910096
refGene_gn=LOC101090225;refGene_tx=XM_006942201.2
E3 21910282 21910337
refGene_gn=LOC101090225;refGene_tx=XM_006942201.2
E3 21910465 21910575
refGene_gn=LOC101090225;refGene_tx=XM_006942201.2
E3 21910978 21911106
refGene_gn=LOC101090225;refGene_tx=XM_006942201.2
E3 21917920 21918080
refGene_gn=LOC101090225;refGene_tx=XM_006942201.2
E3 21918203 21918307
refGene_gn=LOC101090225;refGene_tx=XM_006942201.2
E3 21918751 21918893
refGene_gn=LOC101090225;refGene_tx=XM_006942201.2
E3 21919087 21919213
refGene_gn=LOC101090225;refGene_tx=XM_006942201.2
E3 21919346 21919440
refGene_gn=LOC101090225;refGene_tx=XM_006942201.2
E3 21920166 21920357
refGene_gn=LOC101090225;refGene_tx=XM_006942201.2
E3 21920569 21920794
refGene_gn=LOC101090225;refGene_tx=XM_006942201.2
E3 21924754 21925118

refGene_gn=LOC101090225;refGene_tx=XM_006942201.2
E3 21927045 21927205
refGene_gn=LOC101090225;refGene_tx=XM_006942201.2
E3 21927299 21927365
refGene_gn=LOC101090225;refGene_tx=XM_006942201.2
E3 21927454 21927542
refGene_gn=LOC101090225;refGene_tx=XM_006942201.2
E3 21927658 21927821
refGene_gn=LOC101090225;refGene_tx=XM_006942201.2
E3 21927908 21927991
refGene_gn=LOC101090225;refGene_tx=XM_006942201.2
E3 21985349 21985648
refGene_gn=ATXN2L;refGene_tx=XM_006942224.2;refGene_gn=ATXN2L;refGene_tx=XM_023247083.1;refGene_gn=ATXN2L;refGene_tx=XM_023247084.1;refGene_gn=ATXN2L;refGene_tx=XM_023247085.1;refGene_gn=ATXN2L;refGene_tx=XM_023247086.1;refGene_gn=ATXN2L;refGene_tx=XM_023247087.1;refGene_gn=ATXN2L;refGene_tx=XM_023247088.1;refGene_gn=ATXN2L;refGene_tx=XM_023247089.1;refGene_gn=ATXN2L;refGene_tx=XM_023247090.1;refGene_gn=ATXN2L;refGene_tx=XM_023247091.1;refGene_gn=ATXN2L;refGene_tx=XM_023247092.1;refGene_gn=ATXN2L;refGene_tx=XM_023247093.1;refGene_gn=ATXN2L;refGene_tx=XM_023247094.1;refGene_gn=ATXN2L;refGene_tx=XM_023247095.1;refGene_gn=ATXN2L;refGene_tx=XM_023247096.1;refGene_gn=ATXN2L;refGene_tx=XM_023247097.1;refGene_gn=ATXN2L;refGene_tx=XM_023247098.1
E3 21987162 21987199
refGene_gn=ATXN2L;refGene_tx=XM_006942224.2;refGene_gn=ATXN2L;refGene_tx=XM_023247083.1;refGene_gn=ATXN2L;refGene_tx=XM_023247084.1;refGene_gn=ATXN2L;refGene_tx=XM_023247085.1;refGene_gn=ATXN2L;refGene_tx=XM_023247086.1;refGene_gn=ATXN2L;refGene_tx=XM_023247087.1;refGene_gn=ATXN2L;refGene_tx=XM_023247088.1;refGene_gn=ATXN2L;refGene_tx=XM_023247089.1;refGene_gn=ATXN2L;refGene_tx=XM_023247090.1;refGene_gn=ATXN2L;refGene_tx=XM_023247091.1;refGene_gn=ATXN2L;refGene_tx=XM_023247092.1;refGene_gn=ATXN2L;refGene_tx=XM_023247093.1;refGene_gn=ATXN2L;refGene_tx=XM_023247094.1;refGene_gn=ATXN2L;refGene_tx=XM_023247095.1;refGene_gn=ATXN2L;refGene_tx=XM_023247096.1;refGene_gn=ATXN2L;refGene_tx=XM_023247097.1;refGene_gn=ATXN2L;refGene_tx=XM_023247098.1
E3 21987370 21987427
refGene_gn=ATXN2L;refGene_tx=XM_006942224.2;refGene_gn=ATXN2L;refGene_tx=XM_023247083.1;refGene_gn=ATXN2L;refGene_tx=XM_023247084.1;refGene_gn=ATXN2L;refGene_tx=XM_023247085.1;refGene_gn=ATXN2L;refGene_tx=XM_023247086.1;refGene_gn=ATXN2L;refGene_tx=XM_023247087.1;refGene_gn=ATXN2L;refGene_tx=XM_023247088.1;refGene_gn=ATXN2L;refGene_tx=XM_023247089.1;refGene_gn=ATXN2L;refGene_tx=XM_023247090.1;refGene_gn=ATXN2L;refGene_tx=XM_023247091.1;refGene_gn=ATXN2L;refGene_tx=XM_023247092.1;refGene_gn=ATXN2L;refGene_tx=XM_023247093.1;refGene_gn=ATXN2L;refGene_tx=XM_023247094.1;refGene_gn=ATXN2L;refGene_tx=XM_023247095.1;refGene_gn=ATXN2L;refGene_tx=XM_023247096.1;refGene_gn=ATXN2L;refGene_tx=XM_023247097.1;refGene_gn=ATXN2L;refGene_tx=XM_023247098.1
E3 21987512 21987584
refGene_gn=ATXN2L;refGene_tx=XM_006942224.2;refGene_gn=ATXN2L;refGene_tx=XM_023247083.1;refGene_gn=ATXN2L;refGene_tx=XM_023247084.1;refGene_

gn=ATXN2L;refGene_tx=XM_023247085.1;refGene_gn=ATXN2L;refGene_tx=XM_02
3247086.1;refGene_gn=ATXN2L;refGene_tx=XM_023247087.1;refGene_gn=ATXN2
L;refGene_tx=XM_023247088.1;refGene_gn=ATXN2L;refGene_tx=XM_023247089.
1;refGene_gn=ATXN2L;refGene_tx=XM_023247090.1;refGene_gn=ATXN2L;refGen
e_tx=XM_023247091.1;refGene_gn=ATXN2L;refGene_tx=XM_023247092.1;refGen
e_gn=ATXN2L;refGene_tx=XM_023247093.1;refGene_gn=ATXN2L;refGene_tx=XM_
023247094.1;refGene_gn=ATXN2L;refGene_tx=XM_023247095.1;refGene_gn=ATX
N2L;refGene_tx=XM_023247096.1;refGene_gn=ATXN2L;refGene_tx=XM_02324709
7.1;refGene_gn=ATXN2L;refGene_tx=XM_023247098.1
E3 21987933 21988084
refGene_gn=ATXN2L;refGene_tx=XM_006942224.2;refGene_gn=ATXN2L;refGene_
tx=XM_023247083.1;refGene_gn=ATXN2L;refGene_tx=XM_023247084.1;refGene_
gn=ATXN2L;refGene_tx=XM_023247085.1;refGene_gn=ATXN2L;refGene_tx=XM_02
3247086.1;refGene_gn=ATXN2L;refGene_tx=XM_023247087.1;refGene_gn=ATXN2
L;refGene_tx=XM_023247088.1;refGene_gn=ATXN2L;refGene_tx=XM_023247089.
1;refGene_gn=ATXN2L;refGene_tx=XM_023247090.1;refGene_gn=ATXN2L;refGen
e_tx=XM_023247091.1;refGene_gn=ATXN2L;refGene_tx=XM_023247092.1;refGen
e_gn=ATXN2L;refGene_tx=XM_023247093.1;refGene_gn=ATXN2L;refGene_tx=XM_
023247094.1;refGene_gn=ATXN2L;refGene_tx=XM_023247095.1;refGene_gn=ATX
N2L;refGene_tx=XM_023247096.1;refGene_gn=ATXN2L;refGene_tx=XM_02324709
7.1;refGene_gn=ATXN2L;refGene_tx=XM_023247098.1
E3 21988477 21988602
refGene_gn=ATXN2L;refGene_tx=XM_006942224.2;refGene_gn=ATXN2L;refGene_
tx=XM_023247083.1;refGene_gn=ATXN2L;refGene_tx=XM_023247084.1;refGene_
gn=ATXN2L;refGene_tx=XM_023247085.1;refGene_gn=ATXN2L;refGene_tx=XM_02
3247086.1;refGene_gn=ATXN2L;refGene_tx=XM_023247087.1;refGene_gn=ATXN2
L;refGene_tx=XM_023247088.1;refGene_gn=ATXN2L;refGene_tx=XM_023247089.
1;refGene_gn=ATXN2L;refGene_tx=XM_023247090.1;refGene_gn=ATXN2L;refGen
e_tx=XM_023247091.1;refGene_gn=ATXN2L;refGene_tx=XM_023247092.1;refGen
e_gn=ATXN2L;refGene_tx=XM_023247093.1;refGene_gn=ATXN2L;refGene_tx=XM_
023247094.1;refGene_gn=ATXN2L;refGene_tx=XM_023247095.1;refGene_gn=ATX
N2L;refGene_tx=XM_023247096.1;refGene_gn=ATXN2L;refGene_tx=XM_02324709
7.1;refGene_gn=ATXN2L;refGene_tx=XM_023247098.1
E3 21989675 21989767
refGene_gn=ATXN2L;refGene_tx=XM_006942224.2;refGene_gn=ATXN2L;refGene_
tx=XM_023247083.1;refGene_gn=ATXN2L;refGene_tx=XM_023247084.1;refGene_
gn=ATXN2L;refGene_tx=XM_023247085.1;refGene_gn=ATXN2L;refGene_tx=XM_02
3247086.1;refGene_gn=ATXN2L;refGene_tx=XM_023247087.1;refGene_gn=ATXN2
L;refGene_tx=XM_023247088.1;refGene_gn=ATXN2L;refGene_tx=XM_023247089.
1;refGene_gn=ATXN2L;refGene_tx=XM_023247090.1;refGene_gn=ATXN2L;refGen
e_tx=XM_023247091.1;refGene_gn=ATXN2L;refGene_tx=XM_023247092.1;refGen
e_gn=ATXN2L;refGene_tx=XM_023247093.1;refGene_gn=ATXN2L;refGene_tx=XM_
023247094.1;refGene_gn=ATXN2L;refGene_tx=XM_023247095.1;refGene_gn=ATX
N2L;refGene_tx=XM_023247096.1;refGene_gn=ATXN2L;refGene_tx=XM_02324709
7.1;refGene_gn=ATXN2L;refGene_tx=XM_023247098.1
E3 21990188 21990389
refGene_gn=ATXN2L;refGene_tx=XM_006942224.2;refGene_gn=ATXN2L;refGene_
tx=XM_023247083.1;refGene_gn=ATXN2L;refGene_tx=XM_023247084.1;refGene_
gn=ATXN2L;refGene_tx=XM_023247085.1;refGene_gn=ATXN2L;refGene_tx=XM_02
3247086.1;refGene_gn=ATXN2L;refGene_tx=XM_023247087.1;refGene_gn=ATXN2

L;refGene_tx=XM_023247088.1;refGene_gn=ATXN2L;refGene_tx=XM_023247089.
1;refGene_gn=ATXN2L;refGene_tx=XM_023247090.1;refGene_gn=ATXN2L;refGen
e_tx=XM_023247091.1;refGene_gn=ATXN2L;refGene_tx=XM_023247092.1;refGen
e_gn=ATXN2L;refGene_tx=XM_023247093.1;refGene_gn=ATXN2L;refGene_tx=XM_
023247094.1;refGene_gn=ATXN2L;refGene_tx=XM_023247095.1;refGene_gn=ATX
N2L;refGene_tx=XM_023247096.1;refGene_gn=ATXN2L;refGene_tx=XM_02324709
7.1;refGene_gn=ATXN2L;refGene_tx=XM_023247098.1
E3 21991016 21991192
refGene_gn=ATXN2L;refGene_tx=XM_006942224.2;refGene_gn=ATXN2L;refGene_
tx=XM_023247083.1;refGene_gn=ATXN2L;refGene_tx=XM_023247084.1;refGene_
gn=ATXN2L;refGene_tx=XM_023247085.1;refGene_gn=ATXN2L;refGene_tx=XM_02
3247086.1;refGene_gn=ATXN2L;refGene_tx=XM_023247087.1;refGene_gn=ATXN2
L;refGene_tx=XM_023247088.1;refGene_gn=ATXN2L;refGene_tx=XM_023247089.
1;refGene_gn=ATXN2L;refGene_tx=XM_023247090.1;refGene_gn=ATXN2L;refGen
e_tx=XM_023247091.1;refGene_gn=ATXN2L;refGene_tx=XM_023247092.1;refGen
e_gn=ATXN2L;refGene_tx=XM_023247093.1;refGene_gn=ATXN2L;refGene_tx=XM_
023247094.1;refGene_gn=ATXN2L;refGene_tx=XM_023247095.1;refGene_gn=ATX
N2L;refGene_tx=XM_023247096.1;refGene_gn=ATXN2L;refGene_tx=XM_02324709
7.1;refGene_gn=ATXN2L;refGene_tx=XM_023247098.1
E3 21991367 21991481
refGene_gn=ATXN2L;refGene_tx=XM_006942224.2;refGene_gn=ATXN2L;refGene_
tx=XM_023247083.1;refGene_gn=ATXN2L;refGene_tx=XM_023247084.1;refGene_
gn=ATXN2L;refGene_tx=XM_023247085.1;refGene_gn=ATXN2L;refGene_tx=XM_02
3247086.1;refGene_gn=ATXN2L;refGene_tx=XM_023247087.1;refGene_gn=ATXN2
L;refGene_tx=XM_023247088.1;refGene_gn=ATXN2L;refGene_tx=XM_023247089.
1;refGene_gn=ATXN2L;refGene_tx=XM_023247090.1;refGene_gn=ATXN2L;refGen
e_tx=XM_023247091.1;refGene_gn=ATXN2L;refGene_tx=XM_023247092.1;refGen
e_gn=ATXN2L;refGene_tx=XM_023247093.1;refGene_gn=ATXN2L;refGene_tx=XM_
023247094.1;refGene_gn=ATXN2L;refGene_tx=XM_023247095.1;refGene_gn=ATX
N2L;refGene_tx=XM_023247096.1;refGene_gn=ATXN2L;refGene_tx=XM_02324709
7.1;refGene_gn=ATXN2L;refGene_tx=XM_023247098.1
E3 21992563 21992776
refGene_gn=ATXN2L;refGene_tx=XM_006942224.2;refGene_gn=ATXN2L;refGene_
tx=XM_023247083.1;refGene_gn=ATXN2L;refGene_tx=XM_023247084.1;refGene_
gn=ATXN2L;refGene_tx=XM_023247085.1;refGene_gn=ATXN2L;refGene_tx=XM_02
3247086.1;refGene_gn=ATXN2L;refGene_tx=XM_023247087.1;refGene_gn=ATXN2
L;refGene_tx=XM_023247088.1;refGene_gn=ATXN2L;refGene_tx=XM_023247089.
1;refGene_gn=ATXN2L;refGene_tx=XM_023247090.1;refGene_gn=ATXN2L;refGen
e_tx=XM_023247091.1;refGene_gn=ATXN2L;refGene_tx=XM_023247092.1;refGen
e_gn=ATXN2L;refGene_tx=XM_023247093.1;refGene_gn=ATXN2L;refGene_tx=XM_
023247094.1;refGene_gn=ATXN2L;refGene_tx=XM_023247095.1;refGene_gn=ATX
N2L;refGene_tx=XM_023247096.1;refGene_gn=ATXN2L;refGene_tx=XM_02324709
7.1;refGene_gn=ATXN2L;refGene_tx=XM_023247098.1
E3 21992865 21992937
refGene_gn=ATXN2L;refGene_tx=XM_006942224.2;refGene_gn=ATXN2L;refGene_
tx=XM_023247083.1;refGene_gn=ATXN2L;refGene_tx=XM_023247084.1;refGene_
gn=ATXN2L;refGene_tx=XM_023247085.1;refGene_gn=ATXN2L;refGene_tx=XM_02
3247086.1;refGene_gn=ATXN2L;refGene_tx=XM_023247087.1;refGene_gn=ATXN2
L;refGene_tx=XM_023247088.1;refGene_gn=ATXN2L;refGene_tx=XM_023247089.
1;refGene_gn=ATXN2L;refGene_tx=XM_023247090.1;refGene_gn=ATXN2L;refGen

e_tx=XM_023247091.1;refGene_gn=ATXN2L;refGene_tx=XM_023247092.1;refGen
e_gn=ATXN2L;refGene_tx=XM_023247093.1;refGene_gn=ATXN2L;refGene_tx=XM_
023247094.1;refGene_gn=ATXN2L;refGene_tx=XM_023247095.1;refGene_gn=ATX
N2L;refGene_tx=XM_023247096.1;refGene_gn=ATXN2L;refGene_tx=XM_02324709
7.1;refGene_gn=ATXN2L;refGene_tx=XM_023247098.1
E3 21993215 21993286
refGene_gn=ATXN2L;refGene_tx=XM_006942224.2;refGene_gn=ATXN2L;refGene_
tx=XM_023247083.1;refGene_gn=ATXN2L;refGene_tx=XM_023247084.1;refGene_
gn=ATXN2L;refGene_tx=XM_023247085.1;refGene_gn=ATXN2L;refGene_tx=XM_02
3247086.1;refGene_gn=ATXN2L;refGene_tx=XM_023247087.1;refGene_gn=ATXN2
L;refGene_tx=XM_023247088.1;refGene_gn=ATXN2L;refGene_tx=XM_023247089.
1;refGene_gn=ATXN2L;refGene_tx=XM_023247090.1;refGene_gn=ATXN2L;refGen
e_tx=XM_023247091.1;refGene_gn=ATXN2L;refGene_tx=XM_023247092.1;refGen
e_gn=ATXN2L;refGene_tx=XM_023247093.1;refGene_gn=ATXN2L;refGene_tx=XM_
023247094.1;refGene_gn=ATXN2L;refGene_tx=XM_023247095.1;refGene_gn=ATX
N2L;refGene_tx=XM_023247096.1;refGene_gn=ATXN2L;refGene_tx=XM_02324709
7.1;refGene_gn=ATXN2L;refGene_tx=XM_023247098.1
E3 21993472 21993768
refGene_gn=ATXN2L;refGene_tx=XM_006942224.2;refGene_gn=ATXN2L;refGene_
tx=XM_023247083.1;refGene_gn=ATXN2L;refGene_tx=XM_023247084.1;refGene_
gn=ATXN2L;refGene_tx=XM_023247085.1;refGene_gn=ATXN2L;refGene_tx=XM_02
3247086.1;refGene_gn=ATXN2L;refGene_tx=XM_023247087.1;refGene_gn=ATXN2
L;refGene_tx=XM_023247088.1;refGene_gn=ATXN2L;refGene_tx=XM_023247089.
1;refGene_gn=ATXN2L;refGene_tx=XM_023247090.1;refGene_gn=ATXN2L;refGen
e_tx=XM_023247091.1;refGene_gn=ATXN2L;refGene_tx=XM_023247092.1;refGen
e_gn=ATXN2L;refGene_tx=XM_023247093.1;refGene_gn=ATXN2L;refGene_tx=XM_
023247094.1;refGene_gn=ATXN2L;refGene_tx=XM_023247095.1;refGene_gn=ATX
N2L;refGene_tx=XM_023247096.1;refGene_gn=ATXN2L;refGene_tx=XM_02324709
7.1;refGene_gn=ATXN2L;refGene_tx=XM_023247098.1
E3 21993845 21993915
refGene_gn=ATXN2L;refGene_tx=XM_006942224.2;refGene_gn=ATXN2L;refGene_
tx=XM_023247083.1;refGene_gn=ATXN2L;refGene_tx=XM_023247084.1;refGene_
gn=ATXN2L;refGene_tx=XM_023247085.1;refGene_gn=ATXN2L;refGene_tx=XM_02
3247086.1;refGene_gn=ATXN2L;refGene_tx=XM_023247087.1;refGene_gn=ATXN2
L;refGene_tx=XM_023247088.1;refGene_gn=ATXN2L;refGene_tx=XM_023247089.
1;refGene_gn=ATXN2L;refGene_tx=XM_023247090.1;refGene_gn=ATXN2L;refGen
e_tx=XM_023247091.1;refGene_gn=ATXN2L;refGene_tx=XM_023247092.1;refGen
e_gn=ATXN2L;refGene_tx=XM_023247093.1;refGene_gn=ATXN2L;refGene_tx=XM_
023247094.1;refGene_gn=ATXN2L;refGene_tx=XM_023247095.1;refGene_gn=ATX
N2L;refGene_tx=XM_023247096.1;refGene_gn=ATXN2L;refGene_tx=XM_02324709
7.1;refGene_gn=ATXN2L;refGene_tx=XM_023247098.1
E3 21994353 21994500
refGene_gn=ATXN2L;refGene_tx=XM_006942224.2;refGene_gn=ATXN2L;refGene_
tx=XM_023247083.1;refGene_gn=ATXN2L;refGene_tx=XM_023247084.1;refGene_
gn=ATXN2L;refGene_tx=XM_023247085.1;refGene_gn=ATXN2L;refGene_tx=XM_02
3247086.1;refGene_gn=ATXN2L;refGene_tx=XM_023247087.1;refGene_gn=ATXN2
L;refGene_tx=XM_023247088.1;refGene_gn=ATXN2L;refGene_tx=XM_023247089.
1;refGene_gn=ATXN2L;refGene_tx=XM_023247090.1;refGene_gn=ATXN2L;refGen
e_tx=XM_023247091.1;refGene_gn=ATXN2L;refGene_tx=XM_023247092.1;refGen
e_gn=ATXN2L;refGene_tx=XM_023247093.1;refGene_gn=ATXN2L;refGene_tx=XM_

023247094.1;refGene_gn=ATXN2L;refGene_tx=XM_023247095.1;refGene_gn=ATXN2L;refGene_tx=XM_023247096.1;refGene_gn=ATXN2L;refGene_tx=XM_023247097.1;refGene_gn=ATXN2L;refGene_tx=XM_023247098.1
E3 21994624 21994697
refGene_gn=ATXN2L;refGene_tx=XM_006942224.2;refGene_gn=ATXN2L;refGene_tx=XM_023247083.1;refGene_gn=ATXN2L;refGene_tx=XM_023247084.1;refGene_gn=ATXN2L;refGene_tx=XM_023247085.1;refGene_gn=ATXN2L;refGene_tx=XM_023247086.1;refGene_gn=ATXN2L;refGene_tx=XM_023247087.1;refGene_gn=ATXN2L;refGene_tx=XM_023247088.1;refGene_gn=ATXN2L;refGene_tx=XM_023247089.1;refGene_gn=ATXN2L;refGene_tx=XM_023247090.1;refGene_gn=ATXN2L;refGene_tx=XM_023247091.1;refGene_gn=ATXN2L;refGene_tx=XM_023247092.1;refGene_gn=ATXN2L;refGene_tx=XM_023247093.1;refGene_gn=ATXN2L;refGene_tx=XM_023247094.1;refGene_gn=ATXN2L;refGene_tx=XM_023247095.1;refGene_gn=ATXN2L;refGene_tx=XM_023247096.1;refGene_gn=ATXN2L;refGene_tx=XM_023247097.1;refGene_gn=ATXN2L;refGene_tx=XM_023247098.1
E3 21994800 21994988
refGene_gn=ATXN2L;refGene_tx=XM_006942224.2;refGene_gn=ATXN2L;refGene_tx=XM_023247083.1;refGene_gn=ATXN2L;refGene_tx=XM_023247084.1;refGene_gn=ATXN2L;refGene_tx=XM_023247085.1;refGene_gn=ATXN2L;refGene_tx=XM_023247086.1;refGene_gn=ATXN2L;refGene_tx=XM_023247087.1;refGene_gn=ATXN2L;refGene_tx=XM_023247088.1;refGene_gn=ATXN2L;refGene_tx=XM_023247089.1;refGene_gn=ATXN2L;refGene_tx=XM_023247090.1;refGene_gn=ATXN2L;refGene_tx=XM_023247091.1;refGene_gn=ATXN2L;refGene_tx=XM_023247092.1;refGene_gn=ATXN2L;refGene_tx=XM_023247093.1;refGene_gn=ATXN2L;refGene_tx=XM_023247094.1;refGene_gn=ATXN2L;refGene_tx=XM_023247095.1;refGene_gn=ATXN2L;refGene_tx=XM_023247096.1;refGene_gn=ATXN2L;refGene_tx=XM_023247097.1;refGene_gn=ATXN2L;refGene_tx=XM_023247098.1
E3 21995228 21995358
refGene_gn=ATXN2L;refGene_tx=XM_006942224.2;refGene_gn=ATXN2L;refGene_tx=XM_023247083.1;refGene_gn=ATXN2L;refGene_tx=XM_023247084.1;refGene_gn=ATXN2L;refGene_tx=XM_023247085.1;refGene_gn=ATXN2L;refGene_tx=XM_023247086.1;refGene_gn=ATXN2L;refGene_tx=XM_023247087.1;refGene_gn=ATXN2L;refGene_tx=XM_023247088.1;refGene_gn=ATXN2L;refGene_tx=XM_023247089.1;refGene_gn=ATXN2L;refGene_tx=XM_023247090.1;refGene_gn=ATXN2L;refGene_tx=XM_023247091.1;refGene_gn=ATXN2L;refGene_tx=XM_023247092.1;refGene_gn=ATXN2L;refGene_tx=XM_023247093.1;refGene_gn=ATXN2L;refGene_tx=XM_023247094.1;refGene_gn=ATXN2L;refGene_tx=XM_023247095.1;refGene_gn=ATXN2L;refGene_tx=XM_023247096.1;refGene_gn=ATXN2L;refGene_tx=XM_023247097.1;refGene_gn=ATXN2L;refGene_tx=XM_023247098.1
E3 21995443 21995565
refGene_gn=ATXN2L;refGene_tx=XM_006942224.2;refGene_gn=ATXN2L;refGene_tx=XM_023247083.1;refGene_gn=ATXN2L;refGene_tx=XM_023247084.1;refGene_gn=ATXN2L;refGene_tx=XM_023247085.1;refGene_gn=ATXN2L;refGene_tx=XM_023247086.1;refGene_gn=ATXN2L;refGene_tx=XM_023247087.1;refGene_gn=ATXN2L;refGene_tx=XM_023247088.1;refGene_gn=ATXN2L;refGene_tx=XM_023247089.1;refGene_gn=ATXN2L;refGene_tx=XM_023247090.1;refGene_gn=ATXN2L;refGene_tx=XM_023247091.1;refGene_gn=ATXN2L;refGene_tx=XM_023247092.1;refGene_gn=ATXN2L;refGene_tx=XM_023247093.1;refGene_gn=ATXN2L;refGene_tx=XM_023247094.1;refGene_gn=ATXN2L;refGene_tx=XM_023247095.1;refGene_gn=ATXN2L;refGene_tx=XM_023247096.1;refGene_gn=ATXN2L;refGene_tx=XM_02324709

7.1;refGene_gn=ATXN2L;refGene_tx=XM_023247098.1
E3 21995730 21995943
refGene_gn=ATXN2L;refGene_tx=XM_006942224.2;refGene_gn=ATXN2L;refGene_tx=XM_023247083.1;refGene_gn=ATXN2L;refGene_tx=XM_023247084.1;refGene_gn=ATXN2L;refGene_tx=XM_023247085.1;refGene_gn=ATXN2L;refGene_tx=XM_023247086.1;refGene_gn=ATXN2L;refGene_tx=XM_023247087.1;refGene_gn=ATXN2L;refGene_tx=XM_023247088.1;refGene_gn=ATXN2L;refGene_tx=XM_023247089.1;refGene_gn=ATXN2L;refGene_tx=XM_023247090.1;refGene_gn=ATXN2L;refGene_tx=XM_023247091.1;refGene_gn=ATXN2L;refGene_tx=XM_023247092.1;refGene_gn=ATXN2L;refGene_tx=XM_023247093.1;refGene_gn=ATXN2L;refGene_tx=XM_023247094.1;refGene_gn=ATXN2L;refGene_tx=XM_023247095.1;refGene_gn=ATXN2L;refGene_tx=XM_023247096.1;refGene_gn=ATXN2L;refGene_tx=XM_023247097.1;refGene_gn=ATXN2L;refGene_tx=XM_023247098.1
E3 21996114 21996447
refGene_gn=ATXN2L;refGene_tx=XM_006942224.2;refGene_gn=ATXN2L;refGene_tx=XM_023247083.1;refGene_gn=ATXN2L;refGene_tx=XM_023247084.1;refGene_gn=ATXN2L;refGene_tx=XM_023247085.1;refGene_gn=ATXN2L;refGene_tx=XM_023247086.1;refGene_gn=ATXN2L;refGene_tx=XM_023247087.1;refGene_gn=ATXN2L;refGene_tx=XM_023247088.1;refGene_gn=ATXN2L;refGene_tx=XM_023247089.1;refGene_gn=ATXN2L;refGene_tx=XM_023247090.1;refGene_gn=ATXN2L;refGene_tx=XM_023247091.1;refGene_gn=ATXN2L;refGene_tx=XM_023247092.1;refGene_gn=ATXN2L;refGene_tx=XM_023247093.1;refGene_gn=ATXN2L;refGene_tx=XM_023247094.1;refGene_gn=ATXN2L;refGene_tx=XM_023247095.1;refGene_gn=ATXN2L;refGene_tx=XM_023247096.1;refGene_gn=ATXN2L;refGene_tx=XM_023247097.1;refGene_gn=ATXN2L;refGene_tx=XM_023247098.1
E3 21996507 21996662
refGene_gn=ATXN2L;refGene_tx=XM_023247083.1;refGene_gn=ATXN2L;refGene_tx=XM_023247084.1;refGene_gn=ATXN2L;refGene_tx=XM_023247085.1;refGene_gn=ATXN2L;refGene_tx=XM_023247086.1;refGene_gn=ATXN2L;refGene_tx=XM_023247092.1;refGene_gn=ATXN2L;refGene_tx=XM_023247093.1;refGene_gn=ATXN2L;refGene_tx=XM_023247097.1;refGene_gn=ATXN2L;refGene_tx=XM_023247098.1
E3 21996912 21996985
refGene_gn=ATXN2L;refGene_tx=XM_006942224.2;refGene_gn=ATXN2L;refGene_tx=XM_023247088.1;refGene_gn=ATXN2L;refGene_tx=XM_023247089.1;refGene_gn=ATXN2L;refGene_tx=XM_023247090.1;refGene_gn=ATXN2L;refGene_tx=XM_023247091.1;refGene_gn=ATXN2L;refGene_tx=XM_023247094.1;refGene_gn=ATXN2L;refGene_tx=XM_023247095.1;refGene_gn=ATXN2L;refGene_tx=XM_023247096.1
E3 21999344 21999518 refGene_gn=TUFM;refGene_tx=XM_003998729.4
E3 22000042 22000162 refGene_gn=TUFM;refGene_tx=XM_003998729.4
E3 22000268 22000420 refGene_gn=TUFM;refGene_tx=XM_003998729.4
E3 22000534 22000639 refGene_gn=TUFM;refGene_tx=XM_003998729.4
E3 22000724 22000857 refGene_gn=TUFM;refGene_tx=XM_003998729.4
E3 22000999 22001164 refGene_gn=TUFM;refGene_tx=XM_003998729.4
E3 22001252 22001357 refGene_gn=TUFM;refGene_tx=XM_003998729.4
E3 22001604 22001771 refGene_gn=TUFM;refGene_tx=XM_003998729.4
E3 22002218 22002413 refGene_gn=TUFM;refGene_tx=XM_003998729.4
E3 22002516 22003216 refGene_gn=TUFM;refGene_tx=XM_003998729.4
E3 22015903 22016842

refGene_gn=SH2B1;refGene_tx=XM_006942202.3;refGene_gn=SH2B1;refGene_tx=XM_006942205.4;refGene_gn=SH2B1;refGene_tx=XM_019820735.1;refGene_gn=SH2B1;refGene_tx=XM_019820736.1;refGene_gn=SH2B1;refGene_tx=XM_019820737.1;refGene_gn=SH2B1;refGene_tx=XM_019820738.2;refGene_gn=SH2B1;refGene_tx=XM_019820739.2;refGene_gn=SH2B1;refGene_tx=XM_019820740.1;refGene_gn=SH2B1;refGene_tx=XM_019820741.2
E3 22017125 22017227
refGene_gn=SH2B1;refGene_tx=XM_006942202.3;refGene_gn=SH2B1;refGene_tx=XM_006942205.4;refGene_gn=SH2B1;refGene_tx=XM_019820735.1;refGene_gn=SH2B1;refGene_tx=XM_019820736.1;refGene_gn=SH2B1;refGene_tx=XM_019820737.1;refGene_gn=SH2B1;refGene_tx=XM_019820738.2;refGene_gn=SH2B1;refGene_tx=XM_019820739.2;refGene_gn=SH2B1;refGene_tx=XM_019820740.1;refGene_gn=SH2B1;refGene_tx=XM_019820741.2
E3 22017474 22017566
refGene_gn=SH2B1;refGene_tx=XM_006942202.3;refGene_gn=SH2B1;refGene_tx=XM_006942205.4;refGene_gn=SH2B1;refGene_tx=XM_019820735.1;refGene_gn=SH2B1;refGene_tx=XM_019820736.1;refGene_gn=SH2B1;refGene_tx=XM_019820737.1;refGene_gn=SH2B1;refGene_tx=XM_019820738.2;refGene_gn=SH2B1;refGene_tx=XM_019820739.2;refGene_gn=SH2B1;refGene_tx=XM_019820740.1;refGene_gn=SH2B1;refGene_tx=XM_019820741.2
E3 22017678 22017854
refGene_gn=SH2B1;refGene_tx=XM_006942202.3;refGene_gn=SH2B1;refGene_tx=XM_006942205.4;refGene_gn=SH2B1;refGene_tx=XM_019820735.1;refGene_gn=SH2B1;refGene_tx=XM_019820736.1;refGene_gn=SH2B1;refGene_tx=XM_019820737.1;refGene_gn=SH2B1;refGene_tx=XM_019820738.2;refGene_gn=SH2B1;refGene_tx=XM_019820739.2;refGene_gn=SH2B1;refGene_tx=XM_019820740.1;refGene_gn=SH2B1;refGene_tx=XM_019820741.2
E3 22019474 22019687
refGene_gn=SH2B1;refGene_tx=XM_006942202.3;refGene_gn=SH2B1;refGene_tx=XM_006942205.4;refGene_gn=SH2B1;refGene_tx=XM_019820735.1;refGene_gn=SH2B1;refGene_tx=XM_019820736.1;refGene_gn=SH2B1;refGene_tx=XM_019820737.1;refGene_gn=SH2B1;refGene_tx=XM_019820738.2;refGene_gn=SH2B1;refGene_tx=XM_019820739.2;refGene_gn=SH2B1;refGene_tx=XM_019820740.1;refGene_gn=SH2B1;refGene_tx=XM_019820741.2
E3 22019871 22020083
refGene_gn=SH2B1;refGene_tx=XM_006942202.3;refGene_gn=SH2B1;refGene_tx=XM_006942205.4;refGene_gn=SH2B1;refGene_tx=XM_019820735.1;refGene_gn=SH2B1;refGene_tx=XM_019820736.1;refGene_gn=SH2B1;refGene_tx=XM_019820737.1;refGene_gn=SH2B1;refGene_tx=XM_019820738.2;refGene_gn=SH2B1;refGene_tx=XM_019820739.2;refGene_gn=SH2B1;refGene_tx=XM_019820740.1;refGene_gn=SH2B1;refGene_tx=XM_019820741.2
E3 22020194 22020422
refGene_gn=SH2B1;refGene_tx=XM_006942202.3;refGene_gn=SH2B1;refGene_tx=XM_006942205.4;refGene_gn=SH2B1;refGene_tx=XM_019820735.1;refGene_gn=SH2B1;refGene_tx=XM_019820736.1;refGene_gn=SH2B1;refGene_tx=XM_019820737.1;refGene_gn=SH2B1;refGene_tx=XM_019820738.2;refGene_gn=SH2B1;refGene_tx=XM_019820739.2;refGene_gn=SH2B1;refGene_tx=XM_019820740.1;refGene_gn=SH2B1;refGene_tx=XM_019820741.2
E3 22020833 22020933
refGene_gn=SH2B1;refGene_tx=XM_006942202.3;refGene_gn=SH2B1;refGene_tx

=XM_006942205.4;refGene_gn=SH2B1;refGene_tx=XM_019820740.1;refGene_gn=SH2B1;refGene_tx=XM_019820741.2
E3 22021111 22021488
refGene_gn=SH2B1;refGene_tx=XM_006942202.3;refGene_gn=SH2B1;refGene_tx=XM_006942205.4;refGene_gn=SH2B1;refGene_tx=XM_019820735.1;refGene_gn=SH2B1;refGene_tx=XM_019820736.1;refGene_gn=SH2B1;refGene_tx=XM_019820737.1;refGene_gn=SH2B1;refGene_tx=XM_019820738.2;refGene_gn=SH2B1;refGene_tx=XM_019820739.2;refGene_gn=SH2B1;refGene_tx=XM_019820740.1;refGene_gn=SH2B1;refGene_tx=XM_019820741.2
E3 22026042 22026160
refGene_gn=ATP2A1;refGene_tx=XM_003998733.5;refGene_gn=ATP2A1;refGene_tx=XM_003998734.5
E3 22026498 22026516
refGene_gn=ATP2A1;refGene_tx=XM_003998733.5;refGene_gn=ATP2A1;refGene_tx=XM_003998734.5
E3 22026919 22027002
refGene_gn=ATP2A1;refGene_tx=XM_003998733.5;refGene_gn=ATP2A1;refGene_tx=XM_003998734.5
E3 22028435 22028540
refGene_gn=ATP2A1;refGene_tx=XM_003998733.5;refGene_gn=ATP2A1;refGene_tx=XM_003998734.5
E3 22029236 22029375
refGene_gn=ATP2A1;refGene_tx=XM_003998733.5;refGene_gn=ATP2A1;refGene_tx=XM_003998734.5
E3 22031313 22031394
refGene_gn=ATP2A1;refGene_tx=XM_003998733.5;refGene_gn=ATP2A1;refGene_tx=XM_003998734.5
E3 22032248 22032334
refGene_gn=ATP2A1;refGene_tx=XM_003998733.5;refGene_gn=ATP2A1;refGene_tx=XM_003998734.5
E3 22032489 22032787
refGene_gn=ATP2A1;refGene_tx=XM_003998733.5;refGene_gn=ATP2A1;refGene_tx=XM_003998734.5
E3 22032980 22033147
refGene_gn=ATP2A1;refGene_tx=XM_003998733.5;refGene_gn=ATP2A1;refGene_tx=XM_003998734.5
E3 22034840 22034929
refGene_gn=ATP2A1;refGene_tx=XM_003998733.5;refGene_gn=ATP2A1;refGene_tx=XM_003998734.5
E3 22035197 22035300
refGene_gn=ATP2A1;refGene_tx=XM_003998733.5;refGene_gn=ATP2A1;refGene_tx=XM_003998734.5
E3 22035500 22035632
refGene_gn=ATP2A1;refGene_tx=XM_003998733.5;refGene_gn=ATP2A1;refGene_tx=XM_003998734.5
E3 22037185 22037308
refGene_gn=ATP2A1;refGene_tx=XM_003998733.5;refGene_gn=ATP2A1;refGene_tx=XM_003998734.5
E3 22037436 22037655
refGene_gn=ATP2A1;refGene_tx=XM_003998733.5;refGene_gn=ATP2A1;refGene_

tx=XM_003998734.5
E3 22038359 22038695
refGene_gn=ATP2A1;refGene_tx=XM_003998733.5;refGene_gn=ATP2A1;refGene_
tx=XM_003998734.5
E3 22038899 22039120
refGene_gn=ATP2A1;refGene_tx=XM_003998733.5;refGene_gn=ATP2A1;refGene_
tx=XM_003998734.5
E3 22039227 22039430
refGene_gn=ATP2A1;refGene_tx=XM_003998733.5;refGene_gn=ATP2A1;refGene_
tx=XM_003998734.5
E3 22039576 22039662
refGene_gn=ATP2A1;refGene_tx=XM_003998733.5;refGene_gn=ATP2A1;refGene_
tx=XM_003998734.5
E3 22039754 22039888
refGene_gn=ATP2A1;refGene_tx=XM_003998733.5;refGene_gn=ATP2A1;refGene_
tx=XM_003998734.5
E3 22040006 22040124
refGene_gn=ATP2A1;refGene_tx=XM_003998733.5;refGene_gn=ATP2A1;refGene_
tx=XM_003998734.5
E3 22040312 22040430
refGene_gn=ATP2A1;refGene_tx=XM_003998733.5;refGene_gn=ATP2A1;refGene_
tx=XM_003998734.5
E3 22040668 22040673 refGene_gn=ATP2A1;refGene_tx=XM_003998733.5
E3 22041120 22041146 refGene_gn=ATP2A1;refGene_tx=XM_003998734.5
E3 22041927 22042029 refGene_gn=RABEP2;refGene_tx=XM_003998735.5
E3 22042460 22042577 refGene_gn=RABEP2;refGene_tx=XM_003998735.5
E3 22042839 22042895 refGene_gn=RABEP2;refGene_tx=XM_003998735.5
E3 22043002 22043014 refGene_gn=RABEP2;refGene_tx=XM_003998735.5
E3 22043158 22043336 refGene_gn=RABEP2;refGene_tx=XM_003998735.5
E3 22044404 22044560 refGene_gn=RABEP2;refGene_tx=XM_003998735.5
E3 22045832 22045931 refGene_gn=RABEP2;refGene_tx=XM_003998735.5
E3 22046022 22046118 refGene_gn=RABEP2;refGene_tx=XM_003998735.5
E3 22046369 22046717 refGene_gn=RABEP2;refGene_tx=XM_003998735.5
E3 22046805 22046916 refGene_gn=RABEP2;refGene_tx=XM_003998735.5
E3 22048896 22049045 refGene_gn=RABEP2;refGene_tx=XM_003998735.5
E3 22053215 22053425 refGene_gn=RABEP2;refGene_tx=XM_003998735.5
E3 22053884 22053945 refGene_gn=RABEP2;refGene_tx=XM_003998735.5
E3 22059464 22059552
refGene_gn=CD19;refGene_tx=XM_003998738.4;refGene_gn=CD19;refGene_tx=X
M_006942207.3;refGene_gn=CD19;refGene_tx=XM_019820742.2;refGene_gn=CD1
9;refGene_tx=XM_019820743.2;refGene_gn=CD19;refGene_tx=XM_019820744.2
E3 22059810 22060065
refGene_gn=CD19;refGene_tx=XM_003998738.4;refGene_gn=CD19;refGene_tx=X
M_006942207.3;refGene_gn=CD19;refGene_tx=XM_019820742.2;refGene_gn=CD1
9;refGene_tx=XM_019820743.2;refGene_gn=CD19;refGene_tx=XM_019820744.2
E3 22060378 22060582
refGene_gn=CD19;refGene_tx=XM_003998738.4;refGene_gn=CD19;refGene_tx=X
M_006942207.3;refGene_gn=CD19;refGene_tx=XM_019820742.2;refGene_gn=CD1
9;refGene_tx=XM_019820743.2;refGene_gn=CD19;refGene_tx=XM_019820744.2
E3 22060711 22061005

refGene_gn=CD19;refGene_tx=XM_003998738.4;refGene_gn=CD19;refGene_tx=XM_006942207.3;refGene_gn=CD19;refGene_tx=XM_019820742.2;refGene_gn=CD19;refGene_tx=XM_019820743.2;refGene_gn=CD19;refGene_tx=XM_019820744.2
E3 22062020 22062128
refGene_gn=CD19;refGene_tx=XM_003998738.4;refGene_gn=CD19;refGene_tx=XM_006942207.3;refGene_gn=CD19;refGene_tx=XM_019820742.2;refGene_gn=CD19;refGene_tx=XM_019820743.2;refGene_gn=CD19;refGene_tx=XM_019820744.2
E3 22062400 22062449
refGene_gn=CD19;refGene_tx=XM_003998738.4;refGene_gn=CD19;refGene_tx=XM_006942207.3;refGene_gn=CD19;refGene_tx=XM_019820742.2;refGene_gn=CD19;refGene_tx=XM_019820743.2;refGene_gn=CD19;refGene_tx=XM_019820744.2
E3 22062732 22062821
refGene_gn=CD19;refGene_tx=XM_003998738.4;refGene_gn=CD19;refGene_tx=XM_006942207.3;refGene_gn=CD19;refGene_tx=XM_019820742.2;refGene_gn=CD19;refGene_tx=XM_019820743.2;refGene_gn=CD19;refGene_tx=XM_019820744.2
E3 22063213 22063327
refGene_gn=CD19;refGene_tx=XM_003998738.4;refGene_gn=CD19;refGene_tx=XM_006942207.3;refGene_gn=CD19;refGene_tx=XM_019820742.2;refGene_gn=CD19;refGene_tx=XM_019820743.2;refGene_gn=CD19;refGene_tx=XM_019820744.2
E3 22063461 22063566
refGene_gn=CD19;refGene_tx=XM_003998738.4;refGene_gn=CD19;refGene_tx=XM_006942207.3;refGene_gn=CD19;refGene_tx=XM_019820742.2;refGene_gn=CD19;refGene_tx=XM_019820743.2;refGene_gn=CD19;refGene_tx=XM_019820744.2
E3 22063649 22063718
refGene_gn=CD19;refGene_tx=XM_003998738.4;refGene_gn=CD19;refGene_tx=XM_006942207.3;refGene_gn=CD19;refGene_tx=XM_019820742.2;refGene_gn=CD19;refGene_tx=XM_019820743.2;refGene_gn=CD19;refGene_tx=XM_019820744.2
E3 22063824 22063884
refGene_gn=CD19;refGene_tx=XM_003998738.4;refGene_gn=CD19;refGene_tx=XM_006942207.3;refGene_gn=CD19;refGene_tx=XM_019820742.2;refGene_gn=CD19;refGene_tx=XM_019820743.2;refGene_gn=CD19;refGene_tx=XM_019820744.2
E3 22063976 22064299
refGene_gn=CD19;refGene_tx=XM_003998738.4;refGene_gn=CD19;refGene_tx=XM_006942207.3;refGene_gn=CD19;refGene_tx=XM_019820742.2;refGene_gn=CD19;refGene_tx=XM_019820743.2;refGene_gn=CD19;refGene_tx=XM_019820744.2
E3 22064539 22064632
refGene_gn=CD19;refGene_tx=XM_003998738.4;refGene_gn=CD19;refGene_tx=XM_006942207.3;refGene_gn=CD19;refGene_tx=XM_019820743.2
E3 22064731 22064823
refGene_gn=CD19;refGene_tx=XM_003998738.4;refGene_gn=CD19;refGene_tx=XM_006942207.3;refGene_gn=CD19;refGene_tx=XM_019820743.2;refGene_gn=CD19;refGene_tx=XM_019820744.2
E3 22072459 22072846 refGene_gn=NFATC2IP;refGene_tx=XM_023246509.1
E3 22073510 22073583 refGene_gn=NFATC2IP;refGene_tx=XM_023246509.1
E3 22075750 22075868 refGene_gn=NFATC2IP;refGene_tx=XM_023246509.1
E3 22076702 22076783 refGene_gn=NFATC2IP;refGene_tx=XM_023246509.1
E3 22076873 22077060 refGene_gn=NFATC2IP;refGene_tx=XM_023246509.1
E3 22078248 22078393 refGene_gn=NFATC2IP;refGene_tx=XM_023246509.1
E3 22078526 22078636 refGene_gn=NFATC2IP;refGene_tx=XM_023246509.1
E3 22080478 22080637 refGene_gn=NFATC2IP;refGene_tx=XM_023246509.1

E3 22092383 22092624
refGene_gn=SPNS1;refGene_tx=XM_011290558.3;refGene_gn=SPNS1;refGene_tx=XM_011290559.3;refGene_gn=SPNS1;refGene_tx=XM_011290560.3;refGene_gn=SPNS1;refGene_tx=XM_011290561.3;refGene_gn=SPNS1;refGene_tx=XM_019820746.2
E3 22092719 22092785
refGene_gn=SPNS1;refGene_tx=XM_011290558.3;refGene_gn=SPNS1;refGene_tx=XM_011290559.3;refGene_gn=SPNS1;refGene_tx=XM_011290560.3;refGene_gn=SPNS1;refGene_tx=XM_011290561.3;refGene_gn=SPNS1;refGene_tx=XM_019820746.2
E3 22094634 22094771
refGene_gn=SPNS1;refGene_tx=XM_011290558.3;refGene_gn=SPNS1;refGene_tx=XM_011290559.3;refGene_gn=SPNS1;refGene_tx=XM_011290560.3;refGene_gn=SPNS1;refGene_tx=XM_011290561.3
E3 22095883 22096035
refGene_gn=SPNS1;refGene_tx=XM_011290558.3;refGene_gn=SPNS1;refGene_tx=XM_011290559.3;refGene_gn=SPNS1;refGene_tx=XM_011290560.3;refGene_gn=SPNS1;refGene_tx=XM_011290561.3;refGene_gn=SPNS1;refGene_tx=XM_019820746.2
E3 22096141 22096208
refGene_gn=SPNS1;refGene_tx=XM_011290558.3;refGene_gn=SPNS1;refGene_tx=XM_011290559.3;refGene_gn=SPNS1;refGene_tx=XM_011290560.3;refGene_gn=SPNS1;refGene_tx=XM_011290561.3;refGene_gn=SPNS1;refGene_tx=XM_019820746.2
E3 22097033 22097179
refGene_gn=SPNS1;refGene_tx=XM_011290558.3;refGene_gn=SPNS1;refGene_tx=XM_011290559.3;refGene_gn=SPNS1;refGene_tx=XM_011290560.3;refGene_gn=SPNS1;refGene_tx=XM_011290561.3;refGene_gn=SPNS1;refGene_tx=XM_019820746.2
E3 22097479 22097635
refGene_gn=SPNS1;refGene_tx=XM_011290558.3;refGene_gn=SPNS1;refGene_tx=XM_011290559.3;refGene_gn=SPNS1;refGene_tx=XM_011290560.3;refGene_gn=SPNS1;refGene_tx=XM_011290561.3;refGene_gn=SPNS1;refGene_tx=XM_019820746.2
E3 22097926 22098116
refGene_gn=SPNS1;refGene_tx=XM_011290558.3;refGene_gn=SPNS1;refGene_tx=XM_011290559.3;refGene_gn=SPNS1;refGene_tx=XM_011290560.3;refGene_gn=SPNS1;refGene_tx=XM_011290561.3;refGene_gn=SPNS1;refGene_tx=XM_019820746.2
E3 22098497 22098563
refGene_gn=SPNS1;refGene_tx=XM_011290558.3;refGene_gn=SPNS1;refGene_tx=XM_011290559.3;refGene_gn=SPNS1;refGene_tx=XM_011290560.3;refGene_gn=SPNS1;refGene_tx=XM_011290561.3;refGene_gn=SPNS1;refGene_tx=XM_019820746.2
E3 22098828 22098927
refGene_gn=SPNS1;refGene_tx=XM_011290558.3;refGene_gn=SPNS1;refGene_tx=XM_011290559.3;refGene_gn=SPNS1;refGene_tx=XM_011290560.3;refGene_gn=SPNS1;refGene_tx=XM_011290561.3;refGene_gn=SPNS1;refGene_tx=XM_019820746.2
E3 22099261 22099433

refGene_gn=SPNS1;refGene_tx=XM_011290558.3;refGene_gn=SPNS1;refGene_tx=XM_011290559.3;refGene_gn=SPNS1;refGene_tx=XM_011290560.3;refGene_gn=SPNS1;refGene_tx=XM_011290561.3;refGene_gn=SPNS1;refGene_tx=XM_019820746.2
E3 22099723 22099818
refGene_gn=SPNS1;refGene_tx=XM_011290558.3;refGene_gn=SPNS1;refGene_tx=XM_011290559.3;refGene_gn=SPNS1;refGene_tx=XM_011290560.3;refGene_gn=SPNS1;refGene_tx=XM_011290561.3;refGene_gn=SPNS1;refGene_tx=XM_019820746.2
E3 22100558 22100648
refGene_gn=LAT;refGene_tx=XM_006942211.3;refGene_gn=LAT;refGene_tx=XM_019820747.1;refGene_gn=LAT;refGene_tx=XM_019820748.1
E3 22100974 22101068
refGene_gn=LAT;refGene_tx=XM_006942211.3;refGene_gn=LAT;refGene_tx=XM_019820747.1;refGene_gn=LAT;refGene_tx=XM_019820748.1
E3 22101210 22101238
refGene_gn=LAT;refGene_tx=XM_006942211.3;refGene_gn=LAT;refGene_tx=XM_019820747.1;refGene_gn=LAT;refGene_tx=XM_019820748.1
E3 22101366 22101401
refGene_gn=LAT;refGene_tx=XM_006942211.3;refGene_gn=LAT;refGene_tx=XM_019820747.1;refGene_gn=LAT;refGene_tx=XM_019820748.1
E3 22101666 22101754
refGene_gn=LAT;refGene_tx=XM_006942211.3;refGene_gn=LAT;refGene_tx=XM_019820747.1;refGene_gn=LAT;refGene_tx=XM_019820748.1
E3 22101887 22101952
refGene_gn=LAT;refGene_tx=XM_006942211.3;refGene_gn=LAT;refGene_tx=XM_019820747.1;refGene_gn=LAT;refGene_tx=XM_019820748.1
E3 22102045 22102220
refGene_gn=LAT;refGene_tx=XM_006942211.3;refGene_gn=LAT;refGene_tx=XM_019820747.1;refGene_gn=LAT;refGene_tx=XM_019820748.1
E3 22102293 22102388
refGene_gn=LAT;refGene_tx=XM_006942211.3;refGene_gn=LAT;refGene_tx=XM_019820747.1;refGene_gn=LAT;refGene_tx=XM_019820748.1
E3 22103982 22104045
refGene_gn=LAT;refGene_tx=XM_006942211.3;refGene_gn=LAT;refGene_tx=XM_019820747.1;refGene_gn=LAT;refGene_tx=XM_019820748.1
E3 22104172 22104235
refGene_gn=LAT;refGene_tx=XM_006942211.3;refGene_gn=LAT;refGene_tx=XM_019820747.1;refGene_gn=LAT;refGene_tx=XM_019820748.1
E3 22104345 22104419
refGene_gn=LAT;refGene_tx=XM_006942211.3;refGene_gn=LAT;refGene_tx=XM_019820747.1;refGene_gn=LAT;refGene_tx=XM_019820748.1
E3 22133689 22134523
refGene_gn=SBK1;refGene_tx=XM_023246292.1;refGene_gn=SBK1;refGene_tx=XM_023246293.1
E3 22135385 22135588
refGene_gn=SBK1;refGene_tx=XM_023246292.1;refGene_gn=SBK1;refGene_tx=XM_023246293.1
E3 22136990 22137216
refGene_gn=SBK1;refGene_tx=XM_023246292.1;refGene_gn=SBK1;refGene_tx=X

M_023246293.1
E3	22179566	22180394
refGene_gn=LOC109495212;refGene_tx=XM_019820749.1
E3	22181546	22181723
refGene_gn=LOC109495212;refGene_tx=XM_019820749.1
E3	22210056	22210059
refGene_gn=XPO6;refGene_tx=XM_023246726.1;refGene_gn=XPO6;refGene_tx=XM_023246727.1;refGene_gn=XPO6;refGene_tx=XM_023246728.1;refGene_gn=XPO6;refGene_tx=XM_023246729.1;refGene_gn=XPO6;refGene_tx=XM_023246730.1;refGene_gn=XPO6;refGene_tx=XM_023246731.1;refGene_gn=XPO6;refGene_tx=XM_023246732.1
E3	22240081	22240172
refGene_gn=XPO6;refGene_tx=XM_023246726.1;refGene_gn=XPO6;refGene_tx=XM_023246727.1;refGene_gn=XPO6;refGene_tx=XM_023246728.1;refGene_gn=XPO6;refGene_tx=XM_023246729.1;refGene_gn=XPO6;refGene_tx=XM_023246730.1;refGene_gn=XPO6;refGene_tx=XM_023246731.1;refGene_gn=XPO6;refGene_tx=XM_023246732.1
E3	22243432	22243545
refGene_gn=XPO6;refGene_tx=XM_023246726.1;refGene_gn=XPO6;refGene_tx=XM_023246727.1;refGene_gn=XPO6;refGene_tx=XM_023246728.1;refGene_gn=XPO6;refGene_tx=XM_023246729.1;refGene_gn=XPO6;refGene_tx=XM_023246730.1;refGene_gn=XPO6;refGene_tx=XM_023246731.1;refGene_gn=XPO6;refGene_tx=XM_023246732.1
E3	22244346	22244544
refGene_gn=XPO6;refGene_tx=XM_023246726.1;refGene_gn=XPO6;refGene_tx=XM_023246727.1;refGene_gn=XPO6;refGene_tx=XM_023246728.1;refGene_gn=XPO6;refGene_tx=XM_023246729.1;refGene_gn=XPO6;refGene_tx=XM_023246730.1;refGene_gn=XPO6;refGene_tx=XM_023246731.1;refGene_gn=XPO6;refGene_tx=XM_023246732.1
E3	22249572	22249732
refGene_gn=XPO6;refGene_tx=XM_023246726.1;refGene_gn=XPO6;refGene_tx=XM_023246727.1;refGene_gn=XPO6;refGene_tx=XM_023246728.1;refGene_gn=XPO6;refGene_tx=XM_023246729.1;refGene_gn=XPO6;refGene_tx=XM_023246730.1;refGene_gn=XPO6;refGene_tx=XM_023246731.1;refGene_gn=XPO6;refGene_tx=XM_023246732.1
E3	22251934	22252030
refGene_gn=XPO6;refGene_tx=XM_023246726.1;refGene_gn=XPO6;refGene_tx=XM_023246727.1;refGene_gn=XPO6;refGene_tx=XM_023246728.1;refGene_gn=XPO6;refGene_tx=XM_023246729.1;refGene_gn=XPO6;refGene_tx=XM_023246730.1;refGene_gn=XPO6;refGene_tx=XM_023246731.1;refGene_gn=XPO6;refGene_tx=XM_023246732.1
E3	22261811	22262265
refGene_gn=XPO6;refGene_tx=XM_023246726.1;refGene_gn=XPO6;refGene_tx=XM_023246727.1;refGene_gn=XPO6;refGene_tx=XM_023246728.1;refGene_gn=XPO6;refGene_tx=XM_023246729.1;refGene_gn=XPO6;refGene_tx=XM_023246730.1;refGene_gn=XPO6;refGene_tx=XM_023246731.1;refGene_gn=XPO6;refGene_tx=XM_023246732.1
E3	22265578	22265705
refGene_gn=XPO6;refGene_tx=XM_023246726.1;refGene_gn=XPO6;refGene_tx=XM_023246727.1;refGene_gn=XPO6;refGene_tx=XM_023246728.1;refGene_gn=XPO

6;refGene_tx=XM_023246729.1;refGene_gn=XPO6;refGene_tx=XM_023246730.1;
refGene_gn=XPO6;refGene_tx=XM_023246731.1;refGene_gn=XPO6;refGene_tx=X
M_023246732.1
E3 22271452 22271562
refGene_gn=XPO6;refGene_tx=XM_023246726.1;refGene_gn=XPO6;refGene_tx=X
M_023246727.1;refGene_gn=XPO6;refGene_tx=XM_023246728.1;refGene_gn=XPO
6;refGene_tx=XM_023246729.1;refGene_gn=XPO6;refGene_tx=XM_023246730.1;
refGene_gn=XPO6;refGene_tx=XM_023246731.1;refGene_gn=XPO6;refGene_tx=X
M_023246732.1
E3 22280706 22280815
refGene_gn=XPO6;refGene_tx=XM_023246726.1;refGene_gn=XPO6;refGene_tx=X
M_023246727.1;refGene_gn=XPO6;refGene_tx=XM_023246728.1;refGene_gn=XPO
6;refGene_tx=XM_023246729.1;refGene_gn=XPO6;refGene_tx=XM_023246730.1;
refGene_gn=XPO6;refGene_tx=XM_023246731.1;refGene_gn=XPO6;refGene_tx=X
M_023246732.1
E3 22282563 22282719
refGene_gn=XPO6;refGene_tx=XM_023246726.1;refGene_gn=XPO6;refGene_tx=X
M_023246727.1;refGene_gn=XPO6;refGene_tx=XM_023246728.1;refGene_gn=XPO
6;refGene_tx=XM_023246729.1;refGene_gn=XPO6;refGene_tx=XM_023246730.1;
refGene_gn=XPO6;refGene_tx=XM_023246731.1;refGene_gn=XPO6;refGene_tx=X
M_023246732.1
E3 22283812 22283882
refGene_gn=XPO6;refGene_tx=XM_023246726.1;refGene_gn=XPO6;refGene_tx=X
M_023246727.1;refGene_gn=XPO6;refGene_tx=XM_023246728.1;refGene_gn=XPO
6;refGene_tx=XM_023246729.1;refGene_gn=XPO6;refGene_tx=XM_023246730.1;
refGene_gn=XPO6;refGene_tx=XM_023246731.1;refGene_gn=XPO6;refGene_tx=X
M_023246732.1
E3 22289925 22290085
refGene_gn=XPO6;refGene_tx=XM_023246726.1;refGene_gn=XPO6;refGene_tx=X
M_023246727.1;refGene_gn=XPO6;refGene_tx=XM_023246728.1;refGene_gn=XPO
6;refGene_tx=XM_023246729.1;refGene_gn=XPO6;refGene_tx=XM_023246730.1;
refGene_gn=XPO6;refGene_tx=XM_023246731.1;refGene_gn=XPO6;refGene_tx=X
M_023246732.1
E3 22293042 22293135
refGene_gn=XPO6;refGene_tx=XM_023246726.1;refGene_gn=XPO6;refGene_tx=X
M_023246727.1;refGene_gn=XPO6;refGene_tx=XM_023246728.1;refGene_gn=XPO
6;refGene_tx=XM_023246729.1;refGene_gn=XPO6;refGene_tx=XM_023246730.1;
refGene_gn=XPO6;refGene_tx=XM_023246731.1;refGene_gn=XPO6;refGene_tx=X
M_023246732.1
E3 22295700 22295845
refGene_gn=XPO6;refGene_tx=XM_023246726.1;refGene_gn=XPO6;refGene_tx=X
M_023246727.1;refGene_gn=XPO6;refGene_tx=XM_023246728.1;refGene_gn=XPO
6;refGene_tx=XM_023246729.1;refGene_gn=XPO6;refGene_tx=XM_023246730.1;
refGene_gn=XPO6;refGene_tx=XM_023246731.1;refGene_gn=XPO6;refGene_tx=X
M_023246732.1
E3 22297864 22298011
refGene_gn=XPO6;refGene_tx=XM_023246726.1;refGene_gn=XPO6;refGene_tx=X
M_023246727.1;refGene_gn=XPO6;refGene_tx=XM_023246728.1;refGene_gn=XPO
6;refGene_tx=XM_023246729.1;refGene_gn=XPO6;refGene_tx=XM_023246730.1;
refGene_gn=XPO6;refGene_tx=XM_023246731.1;refGene_gn=XPO6;refGene_tx=X

M_023246732.1
E3 22298915 22299105
refGene_gn=XPO6;refGene_tx=XM_023246726.1;refGene_gn=XPO6;refGene_tx=XM_023246727.1;refGene_gn=XPO6;refGene_tx=XM_023246728.1;refGene_gn=XPO6;refGene_tx=XM_023246729.1;refGene_gn=XPO6;refGene_tx=XM_023246730.1;refGene_gn=XPO6;refGene_tx=XM_023246731.1;refGene_gn=XPO6;refGene_tx=XM_023246732.1
E3 22303457 22303613
refGene_gn=XPO6;refGene_tx=XM_023246726.1;refGene_gn=XPO6;refGene_tx=XM_023246727.1;refGene_gn=XPO6;refGene_tx=XM_023246728.1;refGene_gn=XPO6;refGene_tx=XM_023246729.1;refGene_gn=XPO6;refGene_tx=XM_023246730.1;refGene_gn=XPO6;refGene_tx=XM_023246731.1;refGene_gn=XPO6;refGene_tx=XM_023246732.1
E3 22304691 22304806
refGene_gn=XPO6;refGene_tx=XM_023246726.1;refGene_gn=XPO6;refGene_tx=XM_023246727.1;refGene_gn=XPO6;refGene_tx=XM_023246728.1;refGene_gn=XPO6;refGene_tx=XM_023246729.1;refGene_gn=XPO6;refGene_tx=XM_023246730.1;refGene_gn=XPO6;refGene_tx=XM_023246731.1;refGene_gn=XPO6;refGene_tx=XM_023246732.1
E3 22304989 22305161
refGene_gn=XPO6;refGene_tx=XM_023246726.1;refGene_gn=XPO6;refGene_tx=XM_023246727.1;refGene_gn=XPO6;refGene_tx=XM_023246728.1;refGene_gn=XPO6;refGene_tx=XM_023246729.1;refGene_gn=XPO6;refGene_tx=XM_023246730.1;refGene_gn=XPO6;refGene_tx=XM_023246731.1;refGene_gn=XPO6;refGene_tx=XM_023246732.1
E3 22306498 22306660
refGene_gn=XPO6;refGene_tx=XM_023246726.1;refGene_gn=XPO6;refGene_tx=XM_023246727.1;refGene_gn=XPO6;refGene_tx=XM_023246728.1;refGene_gn=XPO6;refGene_tx=XM_023246729.1;refGene_gn=XPO6;refGene_tx=XM_023246730.1;refGene_gn=XPO6;refGene_tx=XM_023246731.1;refGene_gn=XPO6;refGene_tx=XM_023246732.1
E3 22309170 22309269
refGene_gn=XPO6;refGene_tx=XM_023246726.1;refGene_gn=XPO6;refGene_tx=XM_023246727.1;refGene_gn=XPO6;refGene_tx=XM_023246728.1;refGene_gn=XPO6;refGene_tx=XM_023246729.1;refGene_gn=XPO6;refGene_tx=XM_023246730.1;refGene_gn=XPO6;refGene_tx=XM_023246731.1;refGene_gn=XPO6;refGene_tx=XM_023246732.1
E3 22309401 22309632
refGene_gn=XPO6;refGene_tx=XM_023246726.1;refGene_gn=XPO6;refGene_tx=XM_023246727.1;refGene_gn=XPO6;refGene_tx=XM_023246728.1;refGene_gn=XPO6;refGene_tx=XM_023246729.1;refGene_gn=XPO6;refGene_tx=XM_023246730.1;refGene_gn=XPO6;refGene_tx=XM_023246731.1;refGene_gn=XPO6;refGene_tx=XM_023246732.1
E3 22312059 22312161
refGene_gn=XPO6;refGene_tx=XM_023246726.1;refGene_gn=XPO6;refGene_tx=XM_023246727.1;refGene_gn=XPO6;refGene_tx=XM_023246728.1;refGene_gn=XPO6;refGene_tx=XM_023246729.1;refGene_gn=XPO6;refGene_tx=XM_023246730.1;refGene_gn=XPO6;refGene_tx=XM_023246731.1;refGene_gn=XPO6;refGene_tx=XM_023246732.1
E3 22339451 22339809

refGene_gn=GSG1L;refGene_tx=XM_023246295.1;refGene_gn=GSG1L;refGene_tx=XM_023246296.1;refGene_gn=GSG1L;refGene_tx=XM_023246297.1;refGene_gn=GSG1L;refGene_tx=XM_023246298.1
E3 22410637 22410685
refGene_gn=GSG1L;refGene_tx=XM_023246295.1;refGene_gn=GSG1L;refGene_tx=XM_023246296.1;refGene_gn=GSG1L;refGene_tx=XM_023246297.1;refGene_gn=GSG1L;refGene_tx=XM_023246298.1
E3 22469508 22469661
refGene_gn=GSG1L;refGene_tx=XM_023246295.1;refGene_gn=GSG1L;refGene_tx=XM_023246296.1;refGene_gn=GSG1L;refGene_tx=XM_023246297.1;refGene_gn=GSG1L;refGene_tx=XM_023246298.1
E3 22478978 22479301 refGene_gn=GSG1L;refGene_tx=XM_023246298.1
E3 22507837 22507949
refGene_gn=GSG1L;refGene_tx=XM_023246295.1;refGene_gn=GSG1L;refGene_tx=XM_023246296.1;refGene_gn=GSG1L;refGene_tx=XM_023246297.1
E3 22518835 22519003
refGene_gn=GSG1L;refGene_tx=XM_023246295.1;refGene_gn=GSG1L;refGene_tx=XM_023246296.1;refGene_gn=GSG1L;refGene_tx=XM_023246297.1
E3 22523000 22523211 refGene_gn=GSG1L;refGene_tx=XM_023246295.1
E3 22532941 22533012 refGene_gn=GSG1L;refGene_tx=XM_023246297.1
E3 22544957 22545165
refGene_gn=GSG1L;refGene_tx=XM_023246296.1;refGene_gn=GSG1L;refGene_tx=XM_023246297.1
E3 22555423 22555482
refGene_gn=KIAA0556;refGene_tx=XM_006942221.3;refGene_gn=KIAA0556;refGene_tx=XM_006942222.4;refGene_gn=KIAA0556;refGene_tx=XM_019820757.2;refGene_gn=KIAA0556;refGene_tx=XM_019820758.2
E3 22556059 22556148
refGene_gn=KIAA0556;refGene_tx=XM_006942221.3;refGene_gn=KIAA0556;refGene_tx=XM_006942222.4;refGene_gn=KIAA0556;refGene_tx=XM_019820757.2;refGene_gn=KIAA0556;refGene_tx=XM_019820758.2
E3 22556240 22556401
refGene_gn=KIAA0556;refGene_tx=XM_006942221.3;refGene_gn=KIAA0556;refGene_tx=XM_006942222.4;refGene_gn=KIAA0556;refGene_tx=XM_019820757.2;refGene_gn=KIAA0556;refGene_tx=XM_019820758.2
E3 22556963 22557065
refGene_gn=KIAA0556;refGene_tx=XM_006942221.3;refGene_gn=KIAA0556;refGene_tx=XM_006942222.4;refGene_gn=KIAA0556;refGene_tx=XM_019820757.2;refGene_gn=KIAA0556;refGene_tx=XM_019820758.2
E3 22559022 22559162
refGene_gn=KIAA0556;refGene_tx=XM_006942221.3;refGene_gn=KIAA0556;refGene_tx=XM_006942222.4;refGene_gn=KIAA0556;refGene_tx=XM_019820757.2;refGene_gn=KIAA0556;refGene_tx=XM_019820758.2
E3 22561105 22561216
refGene_gn=KIAA0556;refGene_tx=XM_006942221.3;refGene_gn=KIAA0556;refGene_tx=XM_006942222.4;refGene_gn=KIAA0556;refGene_tx=XM_019820757.2;refGene_gn=KIAA0556;refGene_tx=XM_019820758.2
E3 22562444 22562509
refGene_gn=KIAA0556;refGene_tx=XM_006942221.3;refGene_gn=KIAA0556;refGene_tx=XM_006942222.4;refGene_gn=KIAA0556;refGene_tx=XM_019820757.2;re

fGene_gn=KIAA0556;refGene_tx=XM_019820758.2
E3 22563270 22563283 refGene_gn=KIAA0556;refGene_tx=XM_019820760.2
E3 22564033 22564191
refGene_gn=KIAA0556;refGene_tx=XM_006942221.3;refGene_gn=KIAA0556;refGene_tx=XM_006942222.4;refGene_gn=KIAA0556;refGene_tx=XM_019820757.2;refGene_gn=KIAA0556;refGene_tx=XM_019820758.2;refGene_gn=KIAA0556;refGene_tx=XM_019820760.2
E3 22567399 22567565
refGene_gn=KIAA0556;refGene_tx=XM_006942221.3;refGene_gn=KIAA0556;refGene_tx=XM_006942222.4;refGene_gn=KIAA0556;refGene_tx=XM_019820757.2;refGene_gn=KIAA0556;refGene_tx=XM_019820758.2;refGene_gn=KIAA0556;refGene_tx=XM_019820760.2
E3 22571993 22572171
refGene_gn=KIAA0556;refGene_tx=XM_006942221.3;refGene_gn=KIAA0556;refGene_tx=XM_006942222.4;refGene_gn=KIAA0556;refGene_tx=XM_019820757.2;refGene_gn=KIAA0556;refGene_tx=XM_019820758.2;refGene_gn=KIAA0556;refGene_tx=XM_019820760.2
E3 22577867 22577946
refGene_gn=KIAA0556;refGene_tx=XM_006942221.3;refGene_gn=KIAA0556;refGene_tx=XM_006942222.4;refGene_gn=KIAA0556;refGene_tx=XM_019820757.2;refGene_gn=KIAA0556;refGene_tx=XM_019820758.2;refGene_gn=KIAA0556;refGene_tx=XM_019820760.2
E3 22579855 22580061
refGene_gn=KIAA0556;refGene_tx=XM_006942221.3;refGene_gn=KIAA0556;refGene_tx=XM_006942222.4;refGene_gn=KIAA0556;refGene_tx=XM_019820757.2;refGene_gn=KIAA0556;refGene_tx=XM_019820758.2;refGene_gn=KIAA0556;refGene_tx=XM_019820760.2
E3 22580890 22581616
refGene_gn=KIAA0556;refGene_tx=XM_006942221.3;refGene_gn=KIAA0556;refGene_tx=XM_006942222.4;refGene_gn=KIAA0556;refGene_tx=XM_019820757.2;refGene_gn=KIAA0556;refGene_tx=XM_019820758.2;refGene_gn=KIAA0556;refGene_tx=XM_019820760.2
E3 22586880 22587760
refGene_gn=KIAA0556;refGene_tx=XM_006942221.3;refGene_gn=KIAA0556;refGene_tx=XM_006942222.4;refGene_gn=KIAA0556;refGene_tx=XM_019820757.2;refGene_gn=KIAA0556;refGene_tx=XM_019820758.2;refGene_gn=KIAA0556;refGene_tx=XM_019820760.2
E3 22606322 22606460
refGene_gn=KIAA0556;refGene_tx=XM_006942221.3;refGene_gn=KIAA0556;refGene_tx=XM_006942222.4;refGene_gn=KIAA0556;refGene_tx=XM_019820758.2;refGene_gn=KIAA0556;refGene_tx=XM_019820760.2
E3 22618597 22618813
refGene_gn=KIAA0556;refGene_tx=XM_006942221.3;refGene_gn=KIAA0556;refGene_tx=XM_006942222.4;refGene_gn=KIAA0556;refGene_tx=XM_019820757.2;refGene_gn=KIAA0556;refGene_tx=XM_019820758.2;refGene_gn=KIAA0556;refGene_tx=XM_019820760.2
E3 22622810 22622913
refGene_gn=KIAA0556;refGene_tx=XM_006942221.3;refGene_gn=KIAA0556;refGene_tx=XM_006942222.4;refGene_gn=KIAA0556;refGene_tx=XM_019820757.2;refGene_gn=KIAA0556;refGene_tx=XM_019820758.2;refGene_gn=KIAA0556;refGen

e_tx=XM_019820760.2
E3 22624278 22624394
refGene_gn=KIAA0556;refGene_tx=XM_006942221.3;refGene_gn=KIAA0556;refGene_tx=XM_006942222.4;refGene_gn=KIAA0556;refGene_tx=XM_019820757.2;refGene_gn=KIAA0556;refGene_tx=XM_019820758.2;refGene_gn=KIAA0556;refGene_tx=XM_019820760.2
E3 22625809 22625875
refGene_gn=KIAA0556;refGene_tx=XM_006942221.3;refGene_gn=KIAA0556;refGene_tx=XM_006942222.4;refGene_gn=KIAA0556;refGene_tx=XM_019820757.2;refGene_gn=KIAA0556;refGene_tx=XM_019820758.2;refGene_gn=KIAA0556;refGene_tx=XM_019820760.2
E3 22627117 22627290
refGene_gn=KIAA0556;refGene_tx=XM_006942221.3;refGene_gn=KIAA0556;refGene_tx=XM_006942222.4;refGene_gn=KIAA0556;refGene_tx=XM_019820757.2;refGene_gn=KIAA0556;refGene_tx=XM_019820758.2;refGene_gn=KIAA0556;refGene_tx=XM_019820760.2
E3 22642537 22642666
refGene_gn=KIAA0556;refGene_tx=XM_006942221.3;refGene_gn=KIAA0556;refGene_tx=XM_006942222.4;refGene_gn=KIAA0556;refGene_tx=XM_019820757.2;refGene_gn=KIAA0556;refGene_tx=XM_019820758.2;refGene_gn=KIAA0556;refGene_tx=XM_019820760.2
E3 22648901 22649169
refGene_gn=KIAA0556;refGene_tx=XM_006942221.3;refGene_gn=KIAA0556;refGene_tx=XM_006942222.4;refGene_gn=KIAA0556;refGene_tx=XM_019820757.2;refGene_gn=KIAA0556;refGene_tx=XM_019820758.2;refGene_gn=KIAA0556;refGene_tx=XM_019820760.2
E3 22672143 22672272
refGene_gn=KIAA0556;refGene_tx=XM_006942221.3;refGene_gn=KIAA0556;refGene_tx=XM_006942222.4;refGene_gn=KIAA0556;refGene_tx=XM_019820757.2;refGene_gn=KIAA0556;refGene_tx=XM_019820758.2;refGene_gn=KIAA0556;refGene_tx=XM_019820760.2
E3 22677406 22677538
refGene_gn=KIAA0556;refGene_tx=XM_006942221.3;refGene_gn=KIAA0556;refGene_tx=XM_006942222.4;refGene_gn=KIAA0556;refGene_tx=XM_019820757.2;refGene_gn=KIAA0556;refGene_tx=XM_019820758.2;refGene_gn=KIAA0556;refGene_tx=XM_019820760.2
E3 22687375 22687473
refGene_gn=KIAA0556;refGene_tx=XM_006942221.3;refGene_gn=KIAA0556;refGene_tx=XM_006942222.4;refGene_gn=KIAA0556;refGene_tx=XM_019820757.2;refGene_gn=KIAA0556;refGene_tx=XM_019820758.2;refGene_gn=KIAA0556;refGene_tx=XM_019820760.2
E3 22689050 22689220
refGene_gn=KIAA0556;refGene_tx=XM_006942221.3;refGene_gn=KIAA0556;refGene_tx=XM_006942222.4;refGene_gn=KIAA0556;refGene_tx=XM_019820757.2;refGene_gn=KIAA0556;refGene_tx=XM_019820758.2;refGene_gn=KIAA0556;refGene_tx=XM_019820760.2
E3 22697171 22697248
refGene_gn=KIAA0556;refGene_tx=XM_006942221.3;refGene_gn=KIAA0556;refGene_tx=XM_006942222.4;refGene_gn=KIAA0556;refGene_tx=XM_019820757.2;refGene_gn=KIAA0556;refGene_tx=XM_019820758.2;refGene_gn=KIAA0556;refGen

e_tx=XM_019820760.2
E3	22729264	22729320
refGene_gn=KIAA0556;refGene_tx=XM_006942221.3;refGene_gn=KIAA0556;refGene_tx=XM_006942222.4;refGene_gn=KIAA0556;refGene_tx=XM_019820757.2;refGene_gn=KIAA0556;refGene_tx=XM_019820760.2
E3	22735958	22736049
refGene_gn=KIAA0556;refGene_tx=XM_006942221.3;refGene_gn=KIAA0556;refGene_tx=XM_019820757.2;refGene_gn=KIAA0556;refGene_tx=XM_019820760.2
E3	22747757	22747764	refGene_gn=KIAA0556;refGene_tx=XM_006942222.4
E3	22748043	22748264	refGene_gn=GTF3C1;refGene_tx=XM_011290562.3
E3	22750794	22751004	refGene_gn=GTF3C1;refGene_tx=XM_011290562.3
E3	22757168	22757345	refGene_gn=GTF3C1;refGene_tx=XM_011290562.3
E3	22757974	22758118	refGene_gn=GTF3C1;refGene_tx=XM_011290562.3
E3	22761462	22761559	refGene_gn=GTF3C1;refGene_tx=XM_011290562.3
E3	22765839	22765963	refGene_gn=GTF3C1;refGene_tx=XM_011290562.3
E3	22780570	22780738	refGene_gn=GTF3C1;refGene_tx=XM_011290562.3
E3	22782550	22782666	refGene_gn=GTF3C1;refGene_tx=XM_011290562.3
E3	22783729	22784042	refGene_gn=GTF3C1;refGene_tx=XM_011290562.3
E3	22784635	22784850	refGene_gn=GTF3C1;refGene_tx=XM_011290562.3
E3	22787195	22787332	refGene_gn=GTF3C1;refGene_tx=XM_011290562.3
E3	22788420	22788574	refGene_gn=GTF3C1;refGene_tx=XM_011290562.3
E3	22791063	22791167	refGene_gn=GTF3C1;refGene_tx=XM_011290562.3
E3	22791921	22792106	refGene_gn=GTF3C1;refGene_tx=XM_011290562.3
E3	22793555	22793834	refGene_gn=GTF3C1;refGene_tx=XM_011290562.3
E3	22794079	22794225	refGene_gn=GTF3C1;refGene_tx=XM_011290562.3
E3	22794890	22794988	refGene_gn=GTF3C1;refGene_tx=XM_011290562.3
E3	22795494	22795591	refGene_gn=GTF3C1;refGene_tx=XM_011290562.3
E3	22795702	22795880	refGene_gn=GTF3C1;refGene_tx=XM_011290562.3
E3	22799194	22799336	refGene_gn=GTF3C1;refGene_tx=XM_011290562.3
E3	22799716	22799852	refGene_gn=GTF3C1;refGene_tx=XM_011290562.3
E3	22800353	22800435	refGene_gn=GTF3C1;refGene_tx=XM_011290562.3
E3	22800573	22800762	refGene_gn=GTF3C1;refGene_tx=XM_011290562.3
E3	22802455	22802613	refGene_gn=GTF3C1;refGene_tx=XM_011290562.3
E3	22804612	22804755	refGene_gn=GTF3C1;refGene_tx=XM_011290562.3
E3	22806849	22806931	refGene_gn=GTF3C1;refGene_tx=XM_011290562.3
E3	22808690	22808803	refGene_gn=GTF3C1;refGene_tx=XM_011290562.3
E3	22810544	22810607	refGene_gn=GTF3C1;refGene_tx=XM_011290562.3
E3	22811453	22811547	refGene_gn=GTF3C1;refGene_tx=XM_011290562.3
E3	22815537	22815710	refGene_gn=GTF3C1;refGene_tx=XM_011290562.3
E3	22817060	22817348	refGene_gn=GTF3C1;refGene_tx=XM_011290562.3
E3	22817518	22817778	refGene_gn=GTF3C1;refGene_tx=XM_011290562.3
E3	22818757	22819038	refGene_gn=GTF3C1;refGene_tx=XM_011290562.3
E3	22820204	22820694	refGene_gn=GTF3C1;refGene_tx=XM_011290562.3
E3	22821357	22821409	refGene_gn=GTF3C1;refGene_tx=XM_011290562.3
E3	22822210	22822403	refGene_gn=GTF3C1;refGene_tx=XM_011290562.3
E3	22822718	22822931	refGene_gn=GTF3C1;refGene_tx=XM_011290562.3
E3	22832786	22833551
refGene_gn=IL21R;refGene_tx=XM_003998751.4;refGene_gn=IL21R;refGene_tx=XM_023246639.1
E3	22835194	22835276

refGene_gn=IL21R;refGene_tx=XM_003998751.4;refGene_gn=IL21R;refGene_tx=XM_023246639.1
E3 22836280 22836374
refGene_gn=IL21R;refGene_tx=XM_003998751.4;refGene_gn=IL21R;refGene_tx=XM_023246639.1
E3 22836651 22836829
refGene_gn=IL21R;refGene_tx=XM_003998751.4;refGene_gn=IL21R;refGene_tx=XM_023246639.1
E3 22837691 22837837
refGene_gn=IL21R;refGene_tx=XM_003998751.4;refGene_gn=IL21R;refGene_tx=XM_023246639.1
E3 22839624 22839824
refGene_gn=IL21R;refGene_tx=XM_003998751.4;refGene_gn=IL21R;refGene_tx=XM_023246639.1
E3 22842210 22842313
refGene_gn=IL21R;refGene_tx=XM_003998751.4;refGene_gn=IL21R;refGene_tx=XM_023246639.1
E3 22845129 22845178
refGene_gn=IL21R;refGene_tx=XM_003998751.4;refGene_gn=IL21R;refGene_tx=XM_023246639.1
E3 22899370 22900931
refGene_gn=IL4R;refGene_tx=XM_023246308.1;refGene_gn=IL4R;refGene_tx=XM_023246309.1;refGene_gn=IL4R;refGene_tx=XM_023246310.1
E3 22902260 22902310
refGene_gn=IL4R;refGene_tx=XM_023246308.1;refGene_gn=IL4R;refGene_tx=XM_023246309.1;refGene_gn=IL4R;refGene_tx=XM_023246310.1
E3 22903692 22903771
refGene_gn=IL4R;refGene_tx=XM_023246308.1;refGene_gn=IL4R;refGene_tx=XM_023246309.1;refGene_gn=IL4R;refGene_tx=XM_023246310.1
E3 22905644 22905741
refGene_gn=IL4R;refGene_tx=XM_023246308.1;refGene_gn=IL4R;refGene_tx=XM_023246309.1;refGene_gn=IL4R;refGene_tx=XM_023246310.1
E3 22909309 22909466
refGene_gn=IL4R;refGene_tx=XM_023246308.1;refGene_gn=IL4R;refGene_tx=XM_023246309.1;refGene_gn=IL4R;refGene_tx=XM_023246310.1
E3 22914720 22914872
refGene_gn=IL4R;refGene_tx=XM_023246308.1;refGene_gn=IL4R;refGene_tx=XM_023246309.1;refGene_gn=IL4R;refGene_tx=XM_023246310.1
E3 22916011 22916163
refGene_gn=IL4R;refGene_tx=XM_023246308.1;refGene_gn=IL4R;refGene_tx=XM_023246309.1;refGene_gn=IL4R;refGene_tx=XM_023246310.1
E3 22917950 22918089
refGene_gn=IL4R;refGene_tx=XM_023246308.1;refGene_gn=IL4R;refGene_tx=XM_023246309.1;refGene_gn=IL4R;refGene_tx=XM_023246310.1
E3 22919114 22919184
refGene_gn=IL4R;refGene_tx=XM_023246308.1;refGene_gn=IL4R;refGene_tx=XM_023246309.1;refGene_gn=IL4R;refGene_tx=XM_023246310.1
E3 22983934 22983974
refGene_gn=NSMCE1;refGene_tx=XM_023246775.1;refGene_gn=NSMCE1;refGene_tx=XM_023246776.1;refGene_gn=NSMCE1;refGene_tx=XM_023246777.1

E3 22989599 22989746
refGene_gn=NSMCE1;refGene_tx=XM_023246775.1;refGene_gn=NSMCE1;refGene_tx=XM_023246777.1
E3 23008288 23008410
refGene_gn=NSMCE1;refGene_tx=XM_023246775.1;refGene_gn=NSMCE1;refGene_tx=XM_023246776.1;refGene_gn=NSMCE1;refGene_tx=XM_023246777.1
E3 23009276 23009354
refGene_gn=NSMCE1;refGene_tx=XM_023246775.1;refGene_gn=NSMCE1;refGene_tx=XM_023246776.1;refGene_gn=NSMCE1;refGene_tx=XM_023246777.1
E3 23010578 23010725
refGene_gn=NSMCE1;refGene_tx=XM_023246775.1;refGene_gn=NSMCE1;refGene_tx=XM_023246776.1
E3 23013101 23013218
refGene_gn=NSMCE1;refGene_tx=XM_023246775.1;refGene_gn=NSMCE1;refGene_tx=XM_023246776.1;refGene_gn=NSMCE1;refGene_tx=XM_023246777.1
E3 23014197 23014318
refGene_gn=NSMCE1;refGene_tx=XM_023246775.1;refGene_gn=NSMCE1;refGene_tx=XM_023246776.1;refGene_gn=NSMCE1;refGene_tx=XM_023246777.1
E3 23014700 23014780
refGene_gn=NSMCE1;refGene_tx=XM_023246775.1;refGene_gn=NSMCE1;refGene_tx=XM_023246776.1;refGene_gn=NSMCE1;refGene_tx=XM_023246777.1
E3 23017715 23017880 refGene_gn=KDM8;refGene_tx=XM_023246778.1
E3 23017965 23018058 refGene_gn=KDM8;refGene_tx=XM_023246778.1
E3 23018990 23019140 refGene_gn=KDM8;refGene_tx=XM_023246778.1
E3 23023971 23024016 refGene_gn=KDM8;refGene_tx=XM_023246778.1
E3 23026601 23026734 refGene_gn=KDM8;refGene_tx=XM_023246778.1
E3 23027807 23027974 refGene_gn=KDM8;refGene_tx=XM_023246778.1
E3 23029531 23030066 refGene_gn=KDM8;refGene_tx=XM_023246778.1
E3 23039896 23039946 refGene_gn=KDM8;refGene_tx=XM_023246778.1
E3 23147415 23148195
refGene_gn=CE3H16orf82;refGene_tx=XM_019820769.2
E3 23869071 23869708 refGene_gn=HS3ST4;refGene_tx=XM_023247121.1
E3 24262832 24263566 refGene_gn=HS3ST4;refGene_tx=XM_023247121.1
E3 24601446 24601851 refGene_gn=ZKSCAN2;refGene_tx=XM_003998754.5
E3 24602918 24603105 refGene_gn=ZKSCAN2;refGene_tx=XM_003998754.5
E3 24606252 24606344 refGene_gn=ZKSCAN2;refGene_tx=XM_003998754.5
E3 24606696 24606823 refGene_gn=ZKSCAN2;refGene_tx=XM_003998754.5
E3 24609681 24610365 refGene_gn=ZKSCAN2;refGene_tx=XM_003998754.5
E3 24612115 24612607 refGene_gn=ZKSCAN2;refGene_tx=XM_003998754.5
E3 24615184 24616107 refGene_gn=ZKSCAN2;refGene_tx=XM_003998754.5
E3 24626124 24626173
refGene_gn=AQP8;refGene_tx=XM_023246629.1;refGene_gn=AQP8;refGene_tx=XM_023246630.1;refGene_gn=AQP8;refGene_tx=XM_023246631.1;refGene_gn=AQP8;refGene_tx=XM_023246632.1
E3 24627400 24627535
refGene_gn=AQP8;refGene_tx=XM_023246629.1;refGene_gn=AQP8;refGene_tx=XM_023246630.1;refGene_gn=AQP8;refGene_tx=XM_023246631.1;refGene_gn=AQP8;refGene_tx=XM_023246632.1
E3 24629341 24629556
refGene_gn=AQP8;refGene_tx=XM_023246629.1;refGene_gn=AQP8;refGene_tx=X

M_023246630.1;refGene_gn=AQP8;refGene_tx=XM_023246631.1;refGene_gn=AQP8;refGene_tx=XM_023246632.1
E3 24631159 24631286 refGene_gn=AQP8;refGene_tx=XM_023246629.1;refGene_gn=AQP8;refGene_tx=XM_023246630.1;refGene_gn=AQP8;refGene_tx=XM_023246631.1;refGene_gn=AQP8;refGene_tx=XM_023246632.1
E3 24632583 24632837 refGene_gn=AQP8;refGene_tx=XM_023246629.1;refGene_gn=AQP8;refGene_tx=XM_023246630.1;refGene_gn=AQP8;refGene_tx=XM_023246631.1;refGene_gn=AQP8;refGene_tx=XM_023246632.1
E3 24632991 24633003 refGene_gn=AQP8;refGene_tx=XM_023246629.1;refGene_gn=AQP8;refGene_tx=XM_023246630.1;refGene_gn=AQP8;refGene_tx=XM_023246631.1;refGene_gn=AQP8;refGene_tx=XM_023246632.1
E3 24657861 24657884 refGene_gn=LCMT1;refGene_tx=XM_023247057.1;refGene_gn=LCMT1;refGene_tx=XM_023247058.1
E3 24660055 24660153 refGene_gn=LCMT1;refGene_tx=XM_023247057.1;refGene_gn=LCMT1;refGene_tx=XM_023247058.1
E3 24662109 24662201 refGene_gn=LCMT1;refGene_tx=XM_023247057.1;refGene_gn=LCMT1;refGene_tx=XM_023247058.1
E3 24663252 24663354 refGene_gn=LCMT1;refGene_tx=XM_023247057.1;refGene_gn=LCMT1;refGene_tx=XM_023247058.1
E3 24667130 24667251 refGene_gn=LCMT1;refGene_tx=XM_023247057.1;refGene_gn=LCMT1;refGene_tx=XM_023247058.1
E3 24670806 24670909 refGene_gn=LCMT1;refGene_tx=XM_023247057.1;refGene_gn=LCMT1;refGene_tx=XM_023247058.1
E3 24673313 24673375 refGene_gn=LCMT1;refGene_tx=XM_023247057.1;refGene_gn=LCMT1;refGene_tx=XM_023247058.1
E3 24675630 24675707 refGene_gn=LCMT1;refGene_tx=XM_023247057.1;refGene_gn=LCMT1;refGene_tx=XM_023247058.1
E3 24680671 24680793 refGene_gn=LCMT1;refGene_tx=XM_023247057.1;refGene_gn=LCMT1;refGene_tx=XM_023247058.1
E3 24683700 24683792 refGene_gn=LCMT1;refGene_tx=XM_023247057.1;refGene_gn=LCMT1;refGene_tx=XM_023247058.1
E3 24710841 24710926 refGene_gn=LCMT1;refGene_tx=XM_023247058.1
E3 24721156 24721263 refGene_gn=LCMT1;refGene_tx=XM_023247057.1;refGene_gn=LCMT1;refGene_tx=XM_023247058.1
E3 24724033 24724203 refGene_gn=LOC111558278;refGene_tx=XM_023246986.1

E3 24774332 24774373 refGene_gn=ARHGAP17;refGene_tx=XM_023247081.1
E3 24774918 24774958 refGene_gn=ARHGAP17;refGene_tx=XM_023247081.1
E3 24778554 24778656 refGene_gn=ARHGAP17;refGene_tx=XM_023247081.1
E3 24784252 24784326 refGene_gn=ARHGAP17;refGene_tx=XM_023247081.1
E3 24786175 24786287 refGene_gn=ARHGAP17;refGene_tx=XM_023247081.1
E3 24786545 24786622 refGene_gn=ARHGAP17;refGene_tx=XM_023247081.1
E3 24789726 24789838 refGene_gn=ARHGAP17;refGene_tx=XM_023247081.1
E3 24794486 24794555 refGene_gn=ARHGAP17;refGene_tx=XM_023247081.1
E3 24794728 24794810 refGene_gn=ARHGAP17;refGene_tx=XM_023247081.1
E3 24798660 24798788 refGene_gn=ARHGAP17;refGene_tx=XM_023247081.1
E3 24800384 24800496 refGene_gn=ARHGAP17;refGene_tx=XM_023247081.1
E3 24801546 24801628 refGene_gn=ARHGAP17;refGene_tx=XM_023247081.1
E3 24803807 24803888 refGene_gn=ARHGAP17;refGene_tx=XM_023247081.1
E3 24806114 24806228 refGene_gn=ARHGAP17;refGene_tx=XM_023247081.1
E3 24808880 24808972 refGene_gn=ARHGAP17;refGene_tx=XM_023247081.1
E3 24811219 24811376 refGene_gn=ARHGAP17;refGene_tx=XM_023247081.1
E3 24813091 24813322 refGene_gn=ARHGAP17;refGene_tx=XM_023247081.1
E3 24815327 24815497 refGene_gn=ARHGAP17;refGene_tx=XM_023247081.1
E3 24820988 24821615 refGene_gn=ARHGAP17;refGene_tx=XM_023247081.1
E3 24828739 24828870 refGene_gn=ARHGAP17;refGene_tx=XM_023247081.1
E3 24836268 24836474
refGene_gn=SLC5A11;refGene_tx=XM_003998755.4;refGene_gn=SLC5A11;refGen
e_tx=XM_019820776.2;refGene_gn=SLC5A11;refGene_tx=XM_023246560.1;refGe
ne_gn=SLC5A11;refGene_tx=XM_023246562.1
E3 24837992 24838164
refGene_gn=SLC5A11;refGene_tx=XM_003998755.4;refGene_gn=SLC5A11;refGen
e_tx=XM_019820776.2;refGene_gn=SLC5A11;refGene_tx=XM_023246560.1;refGe
ne_gn=SLC5A11;refGene_tx=XM_023246562.1
E3 24838809 24839025
refGene_gn=SLC5A11;refGene_tx=XM_003998755.4;refGene_gn=SLC5A11;refGen
e_tx=XM_019820776.2;refGene_gn=SLC5A11;refGene_tx=XM_023246560.1;refGe
ne_gn=SLC5A11;refGene_tx=XM_023246562.1
E3 24839432 24839601
refGene_gn=SLC5A11;refGene_tx=XM_003998755.4;refGene_gn=SLC5A11;refGen
e_tx=XM_019820776.2;refGene_gn=SLC5A11;refGene_tx=XM_023246560.1;refGe
ne_gn=SLC5A11;refGene_tx=XM_023246562.1
E3 24841564 24841715
refGene_gn=SLC5A11;refGene_tx=XM_003998755.4;refGene_gn=SLC5A11;refGen
e_tx=XM_019820776.2;refGene_gn=SLC5A11;refGene_tx=XM_023246560.1;refGe
ne_gn=SLC5A11;refGene_tx=XM_023246562.1
E3 24842056 24842164
refGene_gn=SLC5A11;refGene_tx=XM_003998755.4;refGene_gn=SLC5A11;refGen
e_tx=XM_019820776.2;refGene_gn=SLC5A11;refGene_tx=XM_023246560.1;refGe
ne_gn=SLC5A11;refGene_tx=XM_023246562.1
E3 24848580 24848716
refGene_gn=SLC5A11;refGene_tx=XM_003998755.4;refGene_gn=SLC5A11;refGen
e_tx=XM_019820776.2;refGene_gn=SLC5A11;refGene_tx=XM_023246560.1;refGe
ne_gn=SLC5A11;refGene_tx=XM_023246562.1
E3 24853333 24853539
refGene_gn=SLC5A11;refGene_tx=XM_003998755.4;refGene_gn=SLC5A11;refGen

e_tx=XM_019820776.2;refGene_gn=SLC5A11;refGene_tx=XM_023246560.1
E3 24857111 24857192
refGene_gn=SLC5A11;refGene_tx=XM_003998755.4;refGene_gn=SLC5A11;refGene_tx=XM_019820776.2;refGene_gn=SLC5A11;refGene_tx=XM_023246560.1;refGene_gn=SLC5A11;refGene_tx=XM_023246562.1
E3 24859790 24859896
refGene_gn=SLC5A11;refGene_tx=XM_003998755.4;refGene_gn=SLC5A11;refGene_tx=XM_019820776.2;refGene_gn=SLC5A11;refGene_tx=XM_023246560.1;refGene_gn=SLC5A11;refGene_tx=XM_023246562.1
E3 24861057 24861162
refGene_gn=SLC5A11;refGene_tx=XM_003998755.4;refGene_gn=SLC5A11;refGene_tx=XM_019820776.2;refGene_gn=SLC5A11;refGene_tx=XM_023246560.1;refGene_gn=SLC5A11;refGene_tx=XM_023246562.1
E3 24867753 24867813
refGene_gn=SLC5A11;refGene_tx=XM_003998755.4;refGene_gn=SLC5A11;refGene_tx=XM_019820776.2;refGene_gn=SLC5A11;refGene_tx=XM_023246560.1;refGene_gn=SLC5A11;refGene_tx=XM_023246562.1
E3 24868978 24869083
refGene_gn=SLC5A11;refGene_tx=XM_003998755.4;refGene_gn=SLC5A11;refGene_tx=XM_019820776.2;refGene_gn=SLC5A11;refGene_tx=XM_023246560.1;refGene_gn=SLC5A11;refGene_tx=XM_023246562.1
E3 24871060 24871132
refGene_gn=SLC5A11;refGene_tx=XM_003998755.4;refGene_gn=SLC5A11;refGene_tx=XM_019820776.2;refGene_gn=SLC5A11;refGene_tx=XM_023246560.1;refGene_gn=SLC5A11;refGene_tx=XM_023246562.1
E3 24874600 24874735
refGene_gn=SLC5A11;refGene_tx=XM_023246560.1;refGene_gn=SLC5A11;refGene_tx=XM_023246562.1
E3 24876209 24876344
refGene_gn=SLC5A11;refGene_tx=XM_003998755.4;refGene_gn=SLC5A11;refGene_tx=XM_019820776.2
E3 24908644 24909020
refGene_gn=TNRC6A;refGene_tx=XM_019820778.2;refGene_gn=TNRC6A;refGene_tx=XM_019820779.2;refGene_gn=TNRC6A;refGene_tx=XM_019820780.2;refGene_gn=TNRC6A;refGene_tx=XM_019820783.2;refGene_gn=TNRC6A;refGene_tx=XM_019820784.2;refGene_gn=TNRC6A;refGene_tx=XM_019820785.2;refGene_gn=TNRC6A;refGene_tx=XM_019820786.2;refGene_gn=TNRC6A;refGene_tx=XM_023246342.1;refGene_gn=TNRC6A;refGene_tx=XM_023246343.1;refGene_gn=TNRC6A;refGene_tx=XM_023246344.1
E3 24909446 24909586
refGene_gn=TNRC6A;refGene_tx=XM_019820778.2;refGene_gn=TNRC6A;refGene_tx=XM_019820779.2;refGene_gn=TNRC6A;refGene_tx=XM_019820780.2;refGene_gn=TNRC6A;refGene_tx=XM_019820783.2;refGene_gn=TNRC6A;refGene_tx=XM_019820784.2;refGene_gn=TNRC6A;refGene_tx=XM_019820785.2;refGene_gn=TNRC6A;refGene_tx=XM_019820786.2;refGene_gn=TNRC6A;refGene_tx=XM_023246342.1;refGene_gn=TNRC6A;refGene_tx=XM_023246343.1;refGene_gn=TNRC6A;refGene_tx=XM_023246344.1
E3 24910314 24910385
refGene_gn=TNRC6A;refGene_tx=XM_019820778.2;refGene_gn=TNRC6A;refGene_tx=XM_019820779.2;refGene_gn=TNRC6A;refGene_tx=XM_019820780.2;refGene_

gn=TNRC6A;refGene_tx=XM_019820783.2;refGene_gn=TNRC6A;refGene_tx=XM_01
9820784.2;refGene_gn=TNRC6A;refGene_tx=XM_019820785.2;refGene_gn=TNRC6
A;refGene_tx=XM_019820786.2;refGene_gn=TNRC6A;refGene_tx=XM_023246342.
1;refGene_gn=TNRC6A;refGene_tx=XM_023246343.1;refGene_gn=TNRC6A;refGen
e_tx=XM_023246344.1
E3 24912744 24912966
refGene_gn=TNRC6A;refGene_tx=XM_019820778.2;refGene_gn=TNRC6A;refGene_
tx=XM_019820779.2;refGene_gn=TNRC6A;refGene_tx=XM_019820780.2;refGene_
gn=TNRC6A;refGene_tx=XM_019820783.2;refGene_gn=TNRC6A;refGene_tx=XM_01
9820784.2;refGene_gn=TNRC6A;refGene_tx=XM_019820785.2;refGene_gn=TNRC6
A;refGene_tx=XM_019820786.2;refGene_gn=TNRC6A;refGene_tx=XM_023246342.
1;refGene_gn=TNRC6A;refGene_tx=XM_023246343.1;refGene_gn=TNRC6A;refGen
e_tx=XM_023246344.1
E3 24914279 24914387
refGene_gn=TNRC6A;refGene_tx=XM_019820778.2;refGene_gn=TNRC6A;refGene_
tx=XM_019820779.2;refGene_gn=TNRC6A;refGene_tx=XM_019820780.2;refGene_
gn=TNRC6A;refGene_tx=XM_019820783.2;refGene_gn=TNRC6A;refGene_tx=XM_01
9820784.2;refGene_gn=TNRC6A;refGene_tx=XM_019820785.2;refGene_gn=TNRC6
A;refGene_tx=XM_019820786.2;refGene_gn=TNRC6A;refGene_tx=XM_023246342.
1;refGene_gn=TNRC6A;refGene_tx=XM_023246343.1;refGene_gn=TNRC6A;refGen
e_tx=XM_023246344.1
E3 24915819 24915960
refGene_gn=TNRC6A;refGene_tx=XM_019820778.2;refGene_gn=TNRC6A;refGene_
tx=XM_019820779.2;refGene_gn=TNRC6A;refGene_tx=XM_019820780.2;refGene_
gn=TNRC6A;refGene_tx=XM_019820783.2;refGene_gn=TNRC6A;refGene_tx=XM_01
9820784.2;refGene_gn=TNRC6A;refGene_tx=XM_019820785.2;refGene_gn=TNRC6
A;refGene_tx=XM_019820786.2;refGene_gn=TNRC6A;refGene_tx=XM_023246342.
1;refGene_gn=TNRC6A;refGene_tx=XM_023246343.1;refGene_gn=TNRC6A;refGen
e_tx=XM_023246344.1
E3 24916559 24916718
refGene_gn=TNRC6A;refGene_tx=XM_019820778.2;refGene_gn=TNRC6A;refGene_
tx=XM_019820779.2;refGene_gn=TNRC6A;refGene_tx=XM_019820780.2;refGene_
gn=TNRC6A;refGene_tx=XM_019820783.2;refGene_gn=TNRC6A;refGene_tx=XM_01
9820784.2;refGene_gn=TNRC6A;refGene_tx=XM_019820785.2;refGene_gn=TNRC6
A;refGene_tx=XM_019820786.2;refGene_gn=TNRC6A;refGene_tx=XM_023246342.
1;refGene_gn=TNRC6A;refGene_tx=XM_023246343.1;refGene_gn=TNRC6A;refGen
e_tx=XM_023246344.1
E3 24925332 24925464
refGene_gn=TNRC6A;refGene_tx=XM_019820778.2;refGene_gn=TNRC6A;refGene_
tx=XM_019820779.2;refGene_gn=TNRC6A;refGene_tx=XM_019820780.2;refGene_
gn=TNRC6A;refGene_tx=XM_019820783.2;refGene_gn=TNRC6A;refGene_tx=XM_01
9820784.2;refGene_gn=TNRC6A;refGene_tx=XM_019820785.2;refGene_gn=TNRC6
A;refGene_tx=XM_019820786.2;refGene_gn=TNRC6A;refGene_tx=XM_023246342.
1;refGene_gn=TNRC6A;refGene_tx=XM_023246343.1;refGene_gn=TNRC6A;refGen
e_tx=XM_023246344.1
E3 24928194 24928405
refGene_gn=TNRC6A;refGene_tx=XM_019820778.2;refGene_gn=TNRC6A;refGene_
tx=XM_019820779.2;refGene_gn=TNRC6A;refGene_tx=XM_019820780.2;refGene_
gn=TNRC6A;refGene_tx=XM_019820783.2;refGene_gn=TNRC6A;refGene_tx=XM_01
9820784.2;refGene_gn=TNRC6A;refGene_tx=XM_019820785.2;refGene_gn=TNRC6

A;refGene_tx=XM_019820786.2;refGene_gn=TNRC6A;refGene_tx=XM_023246342.1;refGene_gn=TNRC6A;refGene_tx=XM_023246343.1;refGene_gn=TNRC6A;refGene_tx=XM_023246344.1
E3 24928678 24928756
refGene_gn=TNRC6A;refGene_tx=XM_019820778.2;refGene_gn=TNRC6A;refGene_tx=XM_019820779.2;refGene_gn=TNRC6A;refGene_tx=XM_019820780.2;refGene_gn=TNRC6A;refGene_tx=XM_019820783.2;refGene_gn=TNRC6A;refGene_tx=XM_019820784.2;refGene_gn=TNRC6A;refGene_tx=XM_019820785.2;refGene_gn=TNRC6A;refGene_tx=XM_019820786.2;refGene_gn=TNRC6A;refGene_tx=XM_023246342.1;refGene_gn=TNRC6A;refGene_tx=XM_023246343.1;refGene_gn=TNRC6A;refGene_tx=XM_023246344.1
E3 24929192 24929321
refGene_gn=TNRC6A;refGene_tx=XM_019820778.2;refGene_gn=TNRC6A;refGene_tx=XM_019820779.2;refGene_gn=TNRC6A;refGene_tx=XM_019820780.2;refGene_gn=TNRC6A;refGene_tx=XM_019820783.2;refGene_gn=TNRC6A;refGene_tx=XM_019820784.2;refGene_gn=TNRC6A;refGene_tx=XM_019820785.2;refGene_gn=TNRC6A;refGene_tx=XM_019820786.2;refGene_gn=TNRC6A;refGene_tx=XM_023246342.1;refGene_gn=TNRC6A;refGene_tx=XM_023246343.1;refGene_gn=TNRC6A;refGene_tx=XM_023246344.1
E3 24929770 24929908
refGene_gn=TNRC6A;refGene_tx=XM_019820778.2;refGene_gn=TNRC6A;refGene_tx=XM_019820779.2;refGene_gn=TNRC6A;refGene_tx=XM_019820780.2;refGene_gn=TNRC6A;refGene_tx=XM_019820783.2;refGene_gn=TNRC6A;refGene_tx=XM_019820784.2;refGene_gn=TNRC6A;refGene_tx=XM_019820785.2;refGene_gn=TNRC6A;refGene_tx=XM_019820786.2;refGene_gn=TNRC6A;refGene_tx=XM_023246342.1;refGene_gn=TNRC6A;refGene_tx=XM_023246343.1;refGene_gn=TNRC6A;refGene_tx=XM_023246344.1
E3 24930075 24930222
refGene_gn=TNRC6A;refGene_tx=XM_019820778.2;refGene_gn=TNRC6A;refGene_tx=XM_019820779.2;refGene_gn=TNRC6A;refGene_tx=XM_019820780.2;refGene_gn=TNRC6A;refGene_tx=XM_019820783.2;refGene_gn=TNRC6A;refGene_tx=XM_019820785.2;refGene_gn=TNRC6A;refGene_tx=XM_019820786.2;refGene_gn=TNRC6A;refGene_tx=XM_023246342.1;refGene_gn=TNRC6A;refGene_tx=XM_023246343.1;refGene_gn=TNRC6A;refGene_tx=XM_023246344.1
E3 24930629 24930772
refGene_gn=TNRC6A;refGene_tx=XM_019820778.2;refGene_gn=TNRC6A;refGene_tx=XM_019820779.2;refGene_gn=TNRC6A;refGene_tx=XM_019820780.2;refGene_gn=TNRC6A;refGene_tx=XM_019820783.2;refGene_gn=TNRC6A;refGene_tx=XM_019820784.2;refGene_gn=TNRC6A;refGene_tx=XM_019820785.2;refGene_gn=TNRC6A;refGene_tx=XM_019820786.2;refGene_gn=TNRC6A;refGene_tx=XM_023246342.1;refGene_gn=TNRC6A;refGene_tx=XM_023246343.1;refGene_gn=TNRC6A;refGene_tx=XM_023246344.1
E3 24934185 24934237
refGene_gn=TNRC6A;refGene_tx=XM_019820778.2;refGene_gn=TNRC6A;refGene_tx=XM_019820779.2;refGene_gn=TNRC6A;refGene_tx=XM_019820780.2;refGene_gn=TNRC6A;refGene_tx=XM_019820783.2;refGene_gn=TNRC6A;refGene_tx=XM_019820784.2;refGene_gn=TNRC6A;refGene_tx=XM_019820785.2;refGene_gn=TNRC6A;refGene_tx=XM_019820786.2;refGene_gn=TNRC6A;refGene_tx=XM_023246342.1;refGene_gn=TNRC6A;refGene_tx=XM_023246343.1;refGene_gn=TNRC6A;refGene_tx=XM_023246344.1

E3	24934581 24934662
refGene_gn=TNRC6A;refGene_tx=XM_019820778.2;refGene_gn=TNRC6A;refGene_tx=XM_019820779.2;refGene_gn=TNRC6A;refGene_tx=XM_019820780.2;refGene_gn=TNRC6A;refGene_tx=XM_019820783.2;refGene_gn=TNRC6A;refGene_tx=XM_019820784.2;refGene_gn=TNRC6A;refGene_tx=XM_019820785.2;refGene_gn=TNRC6A;refGene_tx=XM_019820786.2;refGene_gn=TNRC6A;refGene_tx=XM_023246342.1;refGene_gn=TNRC6A;refGene_tx=XM_023246343.1;refGene_gn=TNRC6A;refGene_tx=XM_023246344.1
E3	24936245 24936278
refGene_gn=TNRC6A;refGene_tx=XM_019820778.2;refGene_gn=TNRC6A;refGene_tx=XM_019820780.2;refGene_gn=TNRC6A;refGene_tx=XM_019820783.2;refGene_gn=TNRC6A;refGene_tx=XM_019820784.2;refGene_gn=TNRC6A;refGene_tx=XM_019820785.2;refGene_gn=TNRC6A;refGene_tx=XM_019820786.2;refGene_gn=TNRC6A;refGene_tx=XM_023246342.1;refGene_gn=TNRC6A;refGene_tx=XM_023246343.1;refGene_gn=TNRC6A;refGene_tx=XM_023246344.1
E3	24937492 24937668
refGene_gn=TNRC6A;refGene_tx=XM_019820778.2;refGene_gn=TNRC6A;refGene_tx=XM_019820779.2;refGene_gn=TNRC6A;refGene_tx=XM_019820780.2;refGene_gn=TNRC6A;refGene_tx=XM_019820783.2;refGene_gn=TNRC6A;refGene_tx=XM_019820784.2;refGene_gn=TNRC6A;refGene_tx=XM_019820785.2;refGene_gn=TNRC6A;refGene_tx=XM_019820786.2;refGene_gn=TNRC6A;refGene_tx=XM_023246342.1;refGene_gn=TNRC6A;refGene_tx=XM_023246343.1;refGene_gn=TNRC6A;refGene_tx=XM_023246344.1
E3	24938685 24938862
refGene_gn=TNRC6A;refGene_tx=XM_019820778.2;refGene_gn=TNRC6A;refGene_tx=XM_019820779.2;refGene_gn=TNRC6A;refGene_tx=XM_019820780.2;refGene_gn=TNRC6A;refGene_tx=XM_019820783.2;refGene_gn=TNRC6A;refGene_tx=XM_019820784.2;refGene_gn=TNRC6A;refGene_tx=XM_019820786.2;refGene_gn=TNRC6A;refGene_tx=XM_023246342.1;refGene_gn=TNRC6A;refGene_tx=XM_023246343.1;refGene_gn=TNRC6A;refGene_tx=XM_023246344.1
E3	24940225 24942808
refGene_gn=TNRC6A;refGene_tx=XM_019820778.2;refGene_gn=TNRC6A;refGene_tx=XM_019820779.2;refGene_gn=TNRC6A;refGene_tx=XM_019820780.2;refGene_gn=TNRC6A;refGene_tx=XM_019820783.2;refGene_gn=TNRC6A;refGene_tx=XM_019820784.2;refGene_gn=TNRC6A;refGene_tx=XM_019820785.2;refGene_gn=TNRC6A;refGene_tx=XM_019820786.2;refGene_gn=TNRC6A;refGene_tx=XM_023246342.1;refGene_gn=TNRC6A;refGene_tx=XM_023246343.1;refGene_gn=TNRC6A;refGene_tx=XM_023246344.1
E3	24952284 24952689
refGene_gn=TNRC6A;refGene_tx=XM_019820778.2;refGene_gn=TNRC6A;refGene_tx=XM_019820779.2;refGene_gn=TNRC6A;refGene_tx=XM_019820780.2;refGene_gn=TNRC6A;refGene_tx=XM_019820783.2;refGene_gn=TNRC6A;refGene_tx=XM_019820784.2;refGene_gn=TNRC6A;refGene_tx=XM_019820785.2;refGene_gn=TNRC6A;refGene_tx=XM_023246342.1;refGene_gn=TNRC6A;refGene_tx=XM_023246343.1
E3	24973953 24973975
refGene_gn=TNRC6A;refGene_tx=XM_019820778.2;refGene_gn=TNRC6A;refGene_tx=XM_019820779.2;refGene_gn=TNRC6A;refGene_tx=XM_019820780.2;refGene_gn=TNRC6A;refGene_tx=XM_019820783.2;refGene_gn=TNRC6A;refGene_tx=XM_019820784.2;refGene_gn=TNRC6A;refGene_tx=XM_019820785.2;refGene_gn=TNRC6

A;refGene_tx=XM_019820786.2;refGene_gn=TNRC6A;refGene_tx=XM_023246342.1;refGene_gn=TNRC6A;refGene_tx=XM_023246343.1
E3 24982510 24982598
refGene_gn=TNRC6A;refGene_tx=XM_019820778.2;refGene_gn=TNRC6A;refGene_tx=XM_019820779.2;refGene_gn=TNRC6A;refGene_tx=XM_019820780.2;refGene_gn=TNRC6A;refGene_tx=XM_019820783.2;refGene_gn=TNRC6A;refGene_tx=XM_019820784.2;refGene_gn=TNRC6A;refGene_tx=XM_019820785.2;refGene_gn=TNRC6A;refGene_tx=XM_019820786.2;refGene_gn=TNRC6A;refGene_tx=XM_023246342.1;refGene_gn=TNRC6A;refGene_tx=XM_023246343.1
E3 24999791 24999799 refGene_gn=TNRC6A;refGene_tx=XM_023246343.1
E3 25004967 25005015
refGene_gn=TNRC6A;refGene_tx=XM_019820778.2;refGene_gn=TNRC6A;refGene_tx=XM_019820779.2;refGene_gn=TNRC6A;refGene_tx=XM_019820780.2;refGene_gn=TNRC6A;refGene_tx=XM_019820783.2;refGene_gn=TNRC6A;refGene_tx=XM_019820784.2;refGene_gn=TNRC6A;refGene_tx=XM_019820785.2;refGene_gn=TNRC6A;refGene_tx=XM_019820786.2;refGene_gn=TNRC6A;refGene_tx=XM_023246342.1
E3 25005414 25005419 refGene_gn=TNRC6A;refGene_tx=XM_023246342.1
E3 25067991 25068071
refGene_gn=TNRC6A;refGene_tx=XM_019820778.2;refGene_gn=TNRC6A;refGene_tx=XM_019820779.2;refGene_gn=TNRC6A;refGene_tx=XM_019820780.2;refGene_gn=TNRC6A;refGene_tx=XM_019820783.2;refGene_gn=TNRC6A;refGene_tx=XM_019820784.2;refGene_gn=TNRC6A;refGene_tx=XM_019820785.2;refGene_gn=TNRC6A;refGene_tx=XM_019820786.2
E3 25130662 25132232
refGene_gn=RBBP6;refGene_tx=XM_023246942.1;refGene_gn=RBBP6;refGene_tx=XM_023246943.1
E3 25133005 25134763
refGene_gn=RBBP6;refGene_tx=XM_023246942.1;refGene_gn=RBBP6;refGene_tx=XM_023246943.1
E3 25135613 25135715 refGene_gn=RBBP6;refGene_tx=XM_023246942.1
E3 25136005 25136368
refGene_gn=RBBP6;refGene_tx=XM_023246942.1;refGene_gn=RBBP6;refGene_tx=XM_023246943.1
E3 25138051 25138120
refGene_gn=RBBP6;refGene_tx=XM_023246942.1;refGene_gn=RBBP6;refGene_tx=XM_023246943.1
E3 25139446 25139501
refGene_gn=RBBP6;refGene_tx=XM_023246942.1;refGene_gn=RBBP6;refGene_tx=XM_023246943.1
E3 25139603 25139682
refGene_gn=RBBP6;refGene_tx=XM_023246942.1;refGene_gn=RBBP6;refGene_tx=XM_023246943.1
E3 25139811 25139908
refGene_gn=RBBP6;refGene_tx=XM_023246942.1;refGene_gn=RBBP6;refGene_tx=XM_023246943.1
E3 25141162 25141497
refGene_gn=RBBP6;refGene_tx=XM_023246942.1;refGene_gn=RBBP6;refGene_tx=XM_023246943.1
E3 25141611 25141715

refGene_gn=RBBP6;refGene_tx=XM_023246942.1;refGene_gn=RBBP6;refGene_tx=XM_023246943.1
E3 25143408 25143581 refGene_gn=RBBP6;refGene_tx=XM_023246942.1;refGene_gn=RBBP6;refGene_tx=XM_023246943.1
E3 25146450 25146590 refGene_gn=RBBP6;refGene_tx=XM_023246942.1;refGene_gn=RBBP6;refGene_tx=XM_023246943.1
E3 25146990 25147087 refGene_gn=RBBP6;refGene_tx=XM_023246942.1;refGene_gn=RBBP6;refGene_tx=XM_023246943.1
E3 25147200 25147289 refGene_gn=RBBP6;refGene_tx=XM_023246942.1;refGene_gn=RBBP6;refGene_tx=XM_023246943.1
E3 25149377 25149422 refGene_gn=RBBP6;refGene_tx=XM_023246942.1;refGene_gn=RBBP6;refGene_tx=XM_023246943.1
E3 25153978 25154015 refGene_gn=RBBP6;refGene_tx=XM_023246942.1;refGene_gn=RBBP6;refGene_tx=XM_023246943.1
E3 25156358 25156458 refGene_gn=RBBP6;refGene_tx=XM_023246942.1;refGene_gn=RBBP6;refGene_tx=XM_023246943.1
E3 25161168 25161334 refGene_gn=RBBP6;refGene_tx=XM_023246942.1;refGene_gn=RBBP6;refGene_tx=XM_023246943.1
E3 25273928 25274440 refGene_gn=CACNG3;refGene_tx=XM_003998761.4;refGene_gn=CACNG3;refGene_tx=XM_019820789.1
E3 25279806 25279947 refGene_gn=CACNG3;refGene_tx=XM_003998761.4;refGene_gn=CACNG3;refGene_tx=XM_019820789.1
E3 25284696 25284780 refGene_gn=CACNG3;refGene_tx=XM_003998761.4;refGene_gn=CACNG3;refGene_tx=XM_019820789.1
E3 25284849 25284892 refGene_gn=CACNG3;refGene_tx=XM_019820789.1
E3 25358565 25358776 refGene_gn=CACNG3;refGene_tx=XM_003998761.4
E3 25395319 25395472 refGene_gn=PRKCB;refGene_tx=XM_023246795.1
E3 25401357 25401516 refGene_gn=PRKCB;refGene_tx=XM_019820790.2;refGene_gn=PRKCB;refGene_tx=XM_023246794.1
E3 25419497 25419638 refGene_gn=PRKCB;refGene_tx=XM_019820790.2;refGene_gn=PRKCB;refGene_tx=XM_023246794.1;refGene_gn=PRKCB;refGene_tx=XM_023246795.1
E3 25424574 25424682 refGene_gn=PRKCB;refGene_tx=XM_019820790.2;refGene_gn=PRKCB;refGene_tx=XM_023246794.1;refGene_gn=PRKCB;refGene_tx=XM_023246795.1
E3 25424969 25425050 refGene_gn=PRKCB;refGene_tx=XM_019820790.2;refGene_gn=PRKCB;refGene_tx=XM_023246794.1;refGene_gn=PRKCB;refGene_tx=XM_023246795.1

E3 25434083 25434222
refGene_gn=PRKCB;refGene_tx=XM_019820790.2;refGene_gn=PRKCB;refGene_tx
=XM_023246794.1;refGene_gn=PRKCB;refGene_tx=XM_023246795.1
E3 25439304 25439367
refGene_gn=PRKCB;refGene_tx=XM_019820790.2;refGene_gn=PRKCB;refGene_tx
=XM_023246794.1;refGene_gn=PRKCB;refGene_tx=XM_023246795.1
E3 25441243 25441335
refGene_gn=PRKCB;refGene_tx=XM_019820790.2;refGene_gn=PRKCB;refGene_tx
=XM_023246794.1;refGene_gn=PRKCB;refGene_tx=XM_023246795.1
E3 25453744 25453918
refGene_gn=PRKCB;refGene_tx=XM_019820790.2;refGene_gn=PRKCB;refGene_tx
=XM_023246794.1;refGene_gn=PRKCB;refGene_tx=XM_023246795.1
E3 25479983 25480130
refGene_gn=PRKCB;refGene_tx=XM_019820790.2;refGene_gn=PRKCB;refGene_tx
=XM_023246794.1;refGene_gn=PRKCB;refGene_tx=XM_023246795.1
E3 25490398 25490495
refGene_gn=PRKCB;refGene_tx=XM_019820790.2;refGene_gn=PRKCB;refGene_tx
=XM_023246794.1;refGene_gn=PRKCB;refGene_tx=XM_023246795.1
E3 25501054 25501189
refGene_gn=PRKCB;refGene_tx=XM_019820790.2;refGene_gn=PRKCB;refGene_tx
=XM_023246794.1;refGene_gn=PRKCB;refGene_tx=XM_023246795.1
E3 25502402 25502559
refGene_gn=PRKCB;refGene_tx=XM_019820790.2;refGene_gn=PRKCB;refGene_tx
=XM_023246794.1;refGene_gn=PRKCB;refGene_tx=XM_023246795.1
E3 25559850 25559979
refGene_gn=PRKCB;refGene_tx=XM_019820790.2;refGene_gn=PRKCB;refGene_tx
=XM_023246794.1;refGene_gn=PRKCB;refGene_tx=XM_023246795.1
E3 25560308 25560420
refGene_gn=PRKCB;refGene_tx=XM_019820790.2;refGene_gn=PRKCB;refGene_tx
=XM_023246794.1;refGene_gn=PRKCB;refGene_tx=XM_023246795.1
E3 25592712 25592795
refGene_gn=PRKCB;refGene_tx=XM_019820790.2;refGene_gn=PRKCB;refGene_tx
=XM_023246794.1;refGene_gn=PRKCB;refGene_tx=XM_023246795.1
E3 25725494 25725612 refGene_gn=PRKCB;refGene_tx=XM_019820790.2
E3 25727630 25727662
refGene_gn=PRKCB;refGene_tx=XM_023246794.1;refGene_gn=PRKCB;refGene_tx
=XM_023246795.1
E3 25728656 25728829
refGene_gn=PRKCB;refGene_tx=XM_023246794.1;refGene_gn=PRKCB;refGene_tx
=XM_023246795.1
E3 25801966 25802020 refGene_gn=CHP2;refGene_tx=XM_023246706.1
E3 25802218 25802341 refGene_gn=CHP2;refGene_tx=XM_023246706.1
E3 25803335 25803397 refGene_gn=CHP2;refGene_tx=XM_023246706.1
E3 25803582 25803713 refGene_gn=CHP2;refGene_tx=XM_023246706.1
E3 25803832 25803913 refGene_gn=CHP2;refGene_tx=XM_023246706.1
E3 25804010 25804083 refGene_gn=CHP2;refGene_tx=XM_023246706.1
E3 25804496 25804563 refGene_gn=CHP2;refGene_tx=XM_023246706.1
E3 25831392 25831488 refGene_gn=ERN2;refGene_tx=XM_019820795.2
E3 25833161 25833267 refGene_gn=ERN2;refGene_tx=XM_019820795.2
E3 25833641 25833675 refGene_gn=ERN2;refGene_tx=XM_019820795.2

E3 25833908 25833981 refGene_gn=ERN2;refGene_tx=XM_019820795.2
E3 25837454 25837527 refGene_gn=ERN2;refGene_tx=XM_019820795.2
E3 25837668 25837776 refGene_gn=ERN2;refGene_tx=XM_019820795.2
E3 25838074 25838176 refGene_gn=ERN2;refGene_tx=XM_019820795.2
E3 25839130 25839395 refGene_gn=ERN2;refGene_tx=XM_019820795.2
E3 25841189 25841268 refGene_gn=ERN2;refGene_tx=XM_019820795.2
E3 25841355 25841503 refGene_gn=ERN2;refGene_tx=XM_019820795.2
E3 25841614 25841736 refGene_gn=ERN2;refGene_tx=XM_019820795.2
E3 25842460 25842616 refGene_gn=ERN2;refGene_tx=XM_019820795.2
E3 25842881 25843044 refGene_gn=ERN2;refGene_tx=XM_019820795.2
E3 25845367 25845452 refGene_gn=ERN2;refGene_tx=XM_019820795.2
E3 25845600 25845790 refGene_gn=ERN2;refGene_tx=XM_019820795.2
E3 25845871 25845971 refGene_gn=ERN2;refGene_tx=XM_019820795.2
E3 25846107 25846310 refGene_gn=ERN2;refGene_tx=XM_019820795.2
E3 25847492 25847640 refGene_gn=ERN2;refGene_tx=XM_019820795.2
E3 25847729 25847857 refGene_gn=ERN2;refGene_tx=XM_019820795.2
E3 25848419 25848543 refGene_gn=ERN2;refGene_tx=XM_019820795.2
E3 25848631 25848699 refGene_gn=ERN2;refGene_tx=XM_019820795.2
E3 25848789 25849002 refGene_gn=ERN2;refGene_tx=XM_019820795.2
E3 25849436 25849643 refGene_gn=PLK1;refGene_tx=XM_003998762.5
E3 25849888 25850071 refGene_gn=PLK1;refGene_tx=XM_003998762.5
E3 25850172 25850327 refGene_gn=PLK1;refGene_tx=XM_003998762.5
E3 25850519 25850597 refGene_gn=PLK1;refGene_tx=XM_003998762.5
E3 25851578 25851734 refGene_gn=PLK1;refGene_tx=XM_003998762.5
E3 25854628 25854848 refGene_gn=PLK1;refGene_tx=XM_003998762.5
E3 25856238 25856332 refGene_gn=PLK1;refGene_tx=XM_003998762.5
E3 25857315 25857460 refGene_gn=PLK1;refGene_tx=XM_003998762.5
E3 25858213 25858382 refGene_gn=PLK1;refGene_tx=XM_003998762.5
E3 25859047 25859455 refGene_gn=PLK1;refGene_tx=XM_003998762.5
E3 25864161 25864175 refGene_gn=DCTN5;refGene_tx=XM_006942234.3
E3 25873393 25873491 refGene_gn=DCTN5;refGene_tx=XM_003998763.5
E3 25876688 25876791
refGene_gn=DCTN5;refGene_tx=XM_003998763.5;refGene_gn=DCTN5;refGene_tx
=XM_006942234.3
E3 25881297 25881409
refGene_gn=DCTN5;refGene_tx=XM_003998763.5;refGene_gn=DCTN5;refGene_tx
=XM_006942234.3
E3 25884461 25884580
refGene_gn=DCTN5;refGene_tx=XM_003998763.5;refGene_gn=DCTN5;refGene_tx
=XM_006942234.3
E3 25890196 25890265
refGene_gn=DCTN5;refGene_tx=XM_003998763.5;refGene_gn=DCTN5;refGene_tx
=XM_006942234.3
E3 25891013 25891061
refGene_gn=DCTN5;refGene_tx=XM_003998763.5;refGene_gn=DCTN5;refGene_tx
=XM_006942234.3
E3 25891445 25891493
refGene_gn=PALB2;refGene_tx=XM_006942235.3;refGene_gn=PALB2;refGene_tx
=XM_019820792.1
E3 25893947 25894007

refGene_gn=PALB2;refGene_tx=XM_006942235.3;refGene_gn=PALB2;refGene_tx=XM_019820792.1
E3 25894143 25894246
refGene_gn=PALB2;refGene_tx=XM_006942235.3;refGene_gn=PALB2;refGene_tx=XM_019820792.1
E3 25896009 25897470
refGene_gn=PALB2;refGene_tx=XM_006942235.3;refGene_gn=PALB2;refGene_tx=XM_019820792.1
E3 25899451 25900266
refGene_gn=PALB2;refGene_tx=XM_006942235.3;refGene_gn=PALB2;refGene_tx=XM_019820792.1
E3 25900618 25900690
refGene_gn=PALB2;refGene_tx=XM_006942235.3;refGene_gn=PALB2;refGene_tx=XM_019820792.1
E3 25904516 25904678
refGene_gn=PALB2;refGene_tx=XM_006942235.3;refGene_gn=PALB2;refGene_tx=XM_019820792.1
E3 25907297 25907383
refGene_gn=PALB2;refGene_tx=XM_006942235.3;refGene_gn=PALB2;refGene_tx=XM_019820792.1
E3 25909900 25910062
refGene_gn=PALB2;refGene_tx=XM_006942235.3;refGene_gn=PALB2;refGene_tx=XM_019820792.1
E3 25911325 25911442
refGene_gn=PALB2;refGene_tx=XM_006942235.3;refGene_gn=PALB2;refGene_tx=XM_019820792.1
E3 25915658 25915746
refGene_gn=PALB2;refGene_tx=XM_006942235.3;refGene_gn=PALB2;refGene_tx=XM_019820792.1
E3 25918331 25918480
refGene_gn=PALB2;refGene_tx=XM_006942235.3;refGene_gn=PALB2;refGene_tx=XM_019820792.1
E3 25920468 25920676
refGene_gn=PALB2;refGene_tx=XM_006942235.3;refGene_gn=PALB2;refGene_tx=XM_019820792.1
E3 25926958 25927126
refGene_gn=NDUFAB1;refGene_tx=XM_011290619.3;refGene_gn=NDUFAB1;refGene_tx=XM_011290620.3
E3 25932840 25932957
refGene_gn=NDUFAB1;refGene_tx=XM_011290619.3;refGene_gn=NDUFAB1;refGene_tx=XM_011290620.3
E3 25934337 25934425
refGene_gn=NDUFAB1;refGene_tx=XM_011290619.3;refGene_gn=NDUFAB1;refGene_tx=XM_011290620.3
E3 25935368 25935460
refGene_gn=NDUFAB1;refGene_tx=XM_011290619.3;refGene_gn=NDUFAB1;refGene_tx=XM_011290620.3
E3 25943978 25944026 refGene_gn=UBFD1;refGene_tx=XM_023246836.1
E3 25946172 25946324
refGene_gn=UBFD1;refGene_tx=XM_011290637.3;refGene_gn=UBFD1;refGene_tx

=XM_023246833.1
E3	25946965 25946974 refGene_gn=UBFD1;refGene_tx=XM_023246837.1
E3	25948708 25948791
refGene_gn=UBFD1;refGene_tx=XM_011290637.3;refGene_gn=UBFD1;refGene_tx
=XM_023246832.1;refGene_gn=UBFD1;refGene_tx=XM_023246834.1;refGene_gn=
UBFD1;refGene_tx=XM_023246835.1;refGene_gn=UBFD1;refGene_tx=XM_0232468
36.1;refGene_gn=UBFD1;refGene_tx=XM_023246837.1
E3	25951340 25951411
refGene_gn=UBFD1;refGene_tx=XM_023246832.1;refGene_gn=UBFD1;refGene_tx
=XM_023246834.1;refGene_gn=UBFD1;refGene_tx=XM_023246835.1
E3	25952689 25952795
refGene_gn=UBFD1;refGene_tx=XM_011290637.3;refGene_gn=UBFD1;refGene_tx
=XM_023246832.1;refGene_gn=UBFD1;refGene_tx=XM_023246833.1;refGene_gn=
UBFD1;refGene_tx=XM_023246834.1;refGene_gn=UBFD1;refGene_tx=XM_0232468
35.1;refGene_gn=UBFD1;refGene_tx=XM_023246836.1;refGene_gn=UBFD1;refGe
ne_tx=XM_023246837.1
E3	25952953 25953019
refGene_gn=UBFD1;refGene_tx=XM_011290637.3;refGene_gn=UBFD1;refGene_tx
=XM_023246832.1;refGene_gn=UBFD1;refGene_tx=XM_023246833.1;refGene_gn=
UBFD1;refGene_tx=XM_023246834.1;refGene_gn=UBFD1;refGene_tx=XM_0232468
35.1;refGene_gn=UBFD1;refGene_tx=XM_023246836.1;refGene_gn=UBFD1;refGe
ne_tx=XM_023246837.1
E3	25955796 25956005
refGene_gn=UBFD1;refGene_tx=XM_011290637.3;refGene_gn=UBFD1;refGene_tx
=XM_023246832.1;refGene_gn=UBFD1;refGene_tx=XM_023246833.1;refGene_gn=
UBFD1;refGene_tx=XM_023246834.1;refGene_gn=UBFD1;refGene_tx=XM_0232468
35.1;refGene_gn=UBFD1;refGene_tx=XM_023246836.1;refGene_gn=UBFD1;refGe
ne_tx=XM_023246837.1
E3	25956917 25957250
refGene_gn=UBFD1;refGene_tx=XM_011290637.3;refGene_gn=UBFD1;refGene_tx
=XM_023246832.1;refGene_gn=UBFD1;refGene_tx=XM_023246833.1;refGene_gn=
UBFD1;refGene_tx=XM_023246834.1;refGene_gn=UBFD1;refGene_tx=XM_0232468
35.1;refGene_gn=UBFD1;refGene_tx=XM_023246836.1;refGene_gn=UBFD1;refGe
ne_tx=XM_023246837.1
E3	25957432 25957457
refGene_gn=UBFD1;refGene_tx=XM_011290637.3;refGene_gn=UBFD1;refGene_tx
=XM_023246832.1;refGene_gn=UBFD1;refGene_tx=XM_023246833.1;refGene_gn=
UBFD1;refGene_tx=XM_023246834.1;refGene_gn=UBFD1;refGene_tx=XM_0232468
35.1;refGene_gn=UBFD1;refGene_tx=XM_023246836.1;refGene_gn=UBFD1;refGe
ne_tx=XM_023246837.1
E3	25957858 25957997 refGene_gn=EARS2;refGene_tx=XM_003998765.5
E3	25960288 25960444 refGene_gn=EARS2;refGene_tx=XM_003998765.5
E3	25963328 25963518 refGene_gn=EARS2;refGene_tx=XM_003998765.5
E3	25967098 25967571 refGene_gn=EARS2;refGene_tx=XM_003998765.5
E3	25968953 25969062 refGene_gn=EARS2;refGene_tx=XM_003998765.5
E3	25971460 25971614 refGene_gn=EARS2;refGene_tx=XM_003998765.5
E3	25971705 25971836 refGene_gn=EARS2;refGene_tx=XM_003998765.5
E3	25977884 25978020 refGene_gn=EARS2;refGene_tx=XM_003998765.5
E3	25978479 25978563 refGene_gn=EARS2;refGene_tx=XM_003998765.5
E3	25990941 25991044 refGene_gn=GGA2;refGene_tx=XM_023246328.1

E3 25999036 25999121 refGene_gn=GGA2;refGene_tx=XM_023246328.1
E3 26000237 26000313 refGene_gn=GGA2;refGene_tx=XM_023246328.1
E3 26001436 26001535 refGene_gn=GGA2;refGene_tx=XM_023246328.1
E3 26003585 26003709 refGene_gn=GGA2;refGene_tx=XM_023246328.1
E3 26005702 26005806 refGene_gn=GGA2;refGene_tx=XM_023246328.1
E3 26007146 26007227 refGene_gn=GGA2;refGene_tx=XM_023246328.1
E3 26007734 26007864 refGene_gn=GGA2;refGene_tx=XM_023246328.1
E3 26011402 26011498 refGene_gn=GGA2;refGene_tx=XM_023246328.1
E3 26015341 26015467 refGene_gn=GGA2;refGene_tx=XM_023246328.1
E3 26016168 26016291 refGene_gn=GGA2;refGene_tx=XM_023246328.1
E3 26017028 26017057 refGene_gn=GGA2;refGene_tx=XM_023246328.1
E3 26017425 26017559 refGene_gn=GGA2;refGene_tx=XM_023246328.1
E3 26019047 26019205 refGene_gn=GGA2;refGene_tx=XM_023246328.1
E3 26022306 26022476 refGene_gn=GGA2;refGene_tx=XM_023246328.1
E3 26023394 26023505 refGene_gn=GGA2;refGene_tx=XM_023246328.1
E3 26024048 26024159 refGene_gn=GGA2;refGene_tx=XM_023246328.1
E3 26036218 26036387 refGene_gn=COG7;refGene_tx=XM_003998766.5
E3 26040936 26041085 refGene_gn=COG7;refGene_tx=XM_003998766.5
E3 26041380 26041497 refGene_gn=COG7;refGene_tx=XM_003998766.5
E3 26044809 26044978 refGene_gn=COG7;refGene_tx=XM_003998766.5
E3 26067811 26067894 refGene_gn=COG7;refGene_tx=XM_003998766.5
E3 26068693 26068816 refGene_gn=COG7;refGene_tx=XM_003998766.5
E3 26074668 26074867 refGene_gn=COG7;refGene_tx=XM_003998766.5
E3 26084022 26084150 refGene_gn=COG7;refGene_tx=XM_003998766.5
E3 26085078 26085233 refGene_gn=COG7;refGene_tx=XM_003998766.5
E3 26090579 26090696 refGene_gn=COG7;refGene_tx=XM_003998766.5
E3 26098113 26098179 refGene_gn=COG7;refGene_tx=XM_003998766.5
E3 26102752 26102939 refGene_gn=COG7;refGene_tx=XM_003998766.5
E3 26104516 26104657 refGene_gn=COG7;refGene_tx=XM_003998766.5
E3 26110768 26110852 refGene_gn=COG7;refGene_tx=XM_003998766.5
E3 26114729 26114844 refGene_gn=COG7;refGene_tx=XM_003998766.5
E3 26116812 26116956 refGene_gn=COG7;refGene_tx=XM_003998766.5
E3 26119429 26119596 refGene_gn=COG7;refGene_tx=XM_003998766.5
E3 26126914 26127295 refGene_gn=SCNN1B;refGene_tx=XM_011290618.3
E3 26127556 26127632 refGene_gn=SCNN1B;refGene_tx=XM_011290618.3
E3 26128984 26129046 refGene_gn=SCNN1B;refGene_tx=XM_011290618.3
E3 26129700 26129758 refGene_gn=SCNN1B;refGene_tx=XM_011290618.3
E3 26129847 26129923 refGene_gn=SCNN1B;refGene_tx=XM_011290618.3
E3 26130794 26130912 refGene_gn=SCNN1B;refGene_tx=XM_011290618.3
E3 26132913 26133021 refGene_gn=SCNN1B;refGene_tx=XM_011290618.3
E3 26133306 26133470 refGene_gn=SCNN1B;refGene_tx=XM_011290618.3
E3 26135995 26136099 refGene_gn=SCNN1B;refGene_tx=XM_011290618.3
E3 26144984 26145175 refGene_gn=SCNN1B;refGene_tx=XM_011290618.3
E3 26147233 26147495 refGene_gn=SCNN1B;refGene_tx=XM_011290618.3
E3 26150168 26150479 refGene_gn=SCNN1B;refGene_tx=XM_011290618.3
E3 26262927 26263290
refGene_gn=SCNN1G;refGene_tx=XM_011290636.3;refGene_gn=SCNN1G;refGene_tx=XM_019820796.2;refGene_gn=SCNN1G;refGene_tx=XM_019820797.2
E3 26263587 26263663
refGene_gn=SCNN1G;refGene_tx=XM_011290636.3;refGene_gn=SCNN1G;refGene_

tx=XM_019820796.2;refGene_gn=SCNN1G;refGene_tx=XM_019820797.2
E3 26265408 26265470
refGene_gn=SCNN1G;refGene_tx=XM_011290636.3;refGene_gn=SCNN1G;refGene_tx=XM_019820796.2;refGene_gn=SCNN1G;refGene_tx=XM_019820797.2
E3 26265661 26265719
refGene_gn=SCNN1G;refGene_tx=XM_011290636.3;refGene_gn=SCNN1G;refGene_tx=XM_019820796.2;refGene_gn=SCNN1G;refGene_tx=XM_019820797.2
E3 26265801 26265880
refGene_gn=SCNN1G;refGene_tx=XM_011290636.3;refGene_gn=SCNN1G;refGene_tx=XM_019820796.2;refGene_gn=SCNN1G;refGene_tx=XM_019820797.2
E3 26266640 26266758
refGene_gn=SCNN1G;refGene_tx=XM_011290636.3;refGene_gn=SCNN1G;refGene_tx=XM_019820796.2;refGene_gn=SCNN1G;refGene_tx=XM_019820797.2
E3 26268498 26268597
refGene_gn=SCNN1G;refGene_tx=XM_011290636.3;refGene_gn=SCNN1G;refGene_tx=XM_019820796.2
E3 26276367 26276531
refGene_gn=SCNN1G;refGene_tx=XM_011290636.3;refGene_gn=SCNN1G;refGene_tx=XM_019820796.2;refGene_gn=SCNN1G;refGene_tx=XM_019820797.2
E3 26279282 26279386
refGene_gn=SCNN1G;refGene_tx=XM_011290636.3;refGene_gn=SCNN1G;refGene_tx=XM_019820796.2;refGene_gn=SCNN1G;refGene_tx=XM_019820797.2
E3 26280367 26280558
refGene_gn=SCNN1G;refGene_tx=XM_011290636.3;refGene_gn=SCNN1G;refGene_tx=XM_019820796.2;refGene_gn=SCNN1G;refGene_tx=XM_019820797.2
E3 26282266 26282576
refGene_gn=SCNN1G;refGene_tx=XM_011290636.3;refGene_gn=SCNN1G;refGene_tx=XM_019820796.2;refGene_gn=SCNN1G;refGene_tx=XM_019820797.2
E3 26284333 26284680
refGene_gn=SCNN1G;refGene_tx=XM_011290636.3;refGene_gn=SCNN1G;refGene_tx=XM_019820796.2;refGene_gn=SCNN1G;refGene_tx=XM_019820797.2
E3 26286531 26286567
refGene_gn=SCNN1G;refGene_tx=XM_019820796.2;refGene_gn=SCNN1G;refGene_tx=XM_019820797.2
E3 26321902 26322529
refGene_gn=USP31;refGene_tx=XM_019820798.2;refGene_gn=USP31;refGene_tx=XM_023247020.1
E3 26349759 26349897
refGene_gn=USP31;refGene_tx=XM_019820798.2;refGene_gn=USP31;refGene_tx=XM_023247020.1
E3 26351125 26351214
refGene_gn=USP31;refGene_tx=XM_019820798.2;refGene_gn=USP31;refGene_tx=XM_023247020.1;refGene_gn=USP31;refGene_tx=XM_023247021.1;refGene_gn=USP31;refGene_tx=XM_023247022.1;refGene_gn=USP31;refGene_tx=XM_023247023.1
E3 26351305 26351398
refGene_gn=USP31;refGene_tx=XM_019820798.2;refGene_gn=USP31;refGene_tx=XM_023247020.1;refGene_gn=USP31;refGene_tx=XM_023247021.1;refGene_gn=USP31;refGene_tx=XM_023247022.1;refGene_gn=USP31;refGene_tx=XM_023247023.1

E3 26351961 26352097
refGene_gn=USP31;refGene_tx=XM_019820798.2;refGene_gn=USP31;refGene_tx=XM_023247020.1;refGene_gn=USP31;refGene_tx=XM_023247021.1;refGene_gn=USP31;refGene_tx=XM_023247022.1;refGene_gn=USP31;refGene_tx=XM_023247023.1
E3 26353237 26353345
refGene_gn=USP31;refGene_tx=XM_019820798.2;refGene_gn=USP31;refGene_tx=XM_023247021.1;refGene_gn=USP31;refGene_tx=XM_023247022.1;refGene_gn=USP31;refGene_tx=XM_023247023.1
E3 26354354 26354499
refGene_gn=USP31;refGene_tx=XM_019820798.2;refGene_gn=USP31;refGene_tx=XM_023247020.1;refGene_gn=USP31;refGene_tx=XM_023247021.1;refGene_gn=USP31;refGene_tx=XM_023247022.1;refGene_gn=USP31;refGene_tx=XM_023247023.1
E3 26360628 26360809
refGene_gn=USP31;refGene_tx=XM_019820798.2;refGene_gn=USP31;refGene_tx=XM_023247020.1;refGene_gn=USP31;refGene_tx=XM_023247021.1;refGene_gn=USP31;refGene_tx=XM_023247022.1;refGene_gn=USP31;refGene_tx=XM_023247023.1
E3 26364125 26364237
refGene_gn=USP31;refGene_tx=XM_019820798.2;refGene_gn=USP31;refGene_tx=XM_023247020.1;refGene_gn=USP31;refGene_tx=XM_023247021.1;refGene_gn=USP31;refGene_tx=XM_023247022.1;refGene_gn=USP31;refGene_tx=XM_023247023.1
E3 26364852 26364947
refGene_gn=USP31;refGene_tx=XM_019820798.2;refGene_gn=USP31;refGene_tx=XM_023247020.1;refGene_gn=USP31;refGene_tx=XM_023247021.1;refGene_gn=USP31;refGene_tx=XM_023247022.1;refGene_gn=USP31;refGene_tx=XM_023247023.1
E3 26366640 26366718
refGene_gn=USP31;refGene_tx=XM_019820798.2;refGene_gn=USP31;refGene_tx=XM_023247020.1;refGene_gn=USP31;refGene_tx=XM_023247021.1;refGene_gn=USP31;refGene_tx=XM_023247022.1;refGene_gn=USP31;refGene_tx=XM_023247023.1
E3 26367299 26367429
refGene_gn=USP31;refGene_tx=XM_019820798.2;refGene_gn=USP31;refGene_tx=XM_023247020.1;refGene_gn=USP31;refGene_tx=XM_023247021.1;refGene_gn=USP31;refGene_tx=XM_023247022.1;refGene_gn=USP31;refGene_tx=XM_023247023.1
E3 26372481 26372601
refGene_gn=USP31;refGene_tx=XM_019820798.2;refGene_gn=USP31;refGene_tx=XM_023247020.1;refGene_gn=USP31;refGene_tx=XM_023247021.1;refGene_gn=USP31;refGene_tx=XM_023247022.1;refGene_gn=USP31;refGene_tx=XM_023247023.1
E3 26374863 26375089
refGene_gn=USP31;refGene_tx=XM_019820798.2;refGene_gn=USP31;refGene_tx=XM_023247020.1;refGene_gn=USP31;refGene_tx=XM_023247021.1;refGene_gn=USP31;refGene_tx=XM_023247022.1;refGene_gn=USP31;refGene_tx=XM_023247023.1
E3 26380837 26380996

refGene_gn=USP31;refGene_tx=XM_019820798.2;refGene_gn=USP31;refGene_tx=XM_023247020.1;refGene_gn=USP31;refGene_tx=XM_023247021.1;refGene_gn=USP31;refGene_tx=XM_023247022.1;refGene_gn=USP31;refGene_tx=XM_023247023.1
E3 26382439 26382592
refGene_gn=USP31;refGene_tx=XM_019820798.2;refGene_gn=USP31;refGene_tx=XM_023247020.1;refGene_gn=USP31;refGene_tx=XM_023247021.1;refGene_gn=USP31;refGene_tx=XM_023247022.1;refGene_gn=USP31;refGene_tx=XM_023247023.1
E3 26384737 26386308
refGene_gn=USP31;refGene_tx=XM_019820798.2;refGene_gn=USP31;refGene_tx=XM_023247020.1;refGene_gn=USP31;refGene_tx=XM_023247021.1;refGene_gn=USP31;refGene_tx=XM_023247022.1;refGene_gn=USP31;refGene_tx=XM_023247023.1
E3 26512448 26513067 refGene_gn=HS3ST2;refGene_tx=XM_003998814.5
E3 26603931 26604422 refGene_gn=HS3ST2;refGene_tx=XM_003998814.5
E3 26804278 26804349
refGene_gn=OTOA;refGene_tx=XM_006942284.3;refGene_gn=OTOA;refGene_tx=XM_023246637.1;refGene_gn=OTOA;refGene_tx=XM_023246638.1
E3 26806712 26806908
refGene_gn=OTOA;refGene_tx=XM_006942284.3;refGene_gn=OTOA;refGene_tx=XM_023246637.1;refGene_gn=OTOA;refGene_tx=XM_023246638.1
E3 26811164 26811262
refGene_gn=OTOA;refGene_tx=XM_006942284.3;refGene_gn=OTOA;refGene_tx=XM_023246637.1;refGene_gn=OTOA;refGene_tx=XM_023246638.1
E3 26811792 26811929
refGene_gn=OTOA;refGene_tx=XM_006942284.3;refGene_gn=OTOA;refGene_tx=XM_023246637.1;refGene_gn=OTOA;refGene_tx=XM_023246638.1
E3 26812358 26812501
refGene_gn=OTOA;refGene_tx=XM_006942284.3;refGene_gn=OTOA;refGene_tx=XM_023246637.1;refGene_gn=OTOA;refGene_tx=XM_023246638.1
E3 26819530 26819686
refGene_gn=OTOA;refGene_tx=XM_006942284.3;refGene_gn=OTOA;refGene_tx=XM_023246637.1;refGene_gn=OTOA;refGene_tx=XM_023246638.1
E3 26821699 26821887
refGene_gn=OTOA;refGene_tx=XM_006942284.3;refGene_gn=OTOA;refGene_tx=XM_023246637.1;refGene_gn=OTOA;refGene_tx=XM_023246638.1
E3 26825050 26825180
refGene_gn=OTOA;refGene_tx=XM_006942284.3;refGene_gn=OTOA;refGene_tx=XM_023246637.1;refGene_gn=OTOA;refGene_tx=XM_023246638.1
E3 26827875 26827969
refGene_gn=OTOA;refGene_tx=XM_006942284.3;refGene_gn=OTOA;refGene_tx=XM_023246637.1
E3 26829842 26830033
refGene_gn=OTOA;refGene_tx=XM_006942284.3;refGene_gn=OTOA;refGene_tx=XM_023246637.1
E3 26831227 26831363
refGene_gn=OTOA;refGene_tx=XM_006942284.3;refGene_gn=OTOA;refGene_tx=XM_023246637.1
E3 26833881 26833955

refGene_gn=OTOA;refGene_tx=XM_006942284.3;refGene_gn=OTOA;refGene_tx=XM_023246637.1
E3 26836802 26836920
refGene_gn=OTOA;refGene_tx=XM_006942284.3;refGene_gn=OTOA;refGene_tx=XM_023246637.1;refGene_gn=OTOA;refGene_tx=XM_023246638.1
E3 26837367 26837426
refGene_gn=OTOA;refGene_tx=XM_006942284.3;refGene_gn=OTOA;refGene_tx=XM_023246637.1;refGene_gn=OTOA;refGene_tx=XM_023246638.1
E3 26841565 26841706
refGene_gn=OTOA;refGene_tx=XM_006942284.3;refGene_gn=OTOA;refGene_tx=XM_023246637.1;refGene_gn=OTOA;refGene_tx=XM_023246638.1
E3 26842484 26842652
refGene_gn=OTOA;refGene_tx=XM_006942284.3;refGene_gn=OTOA;refGene_tx=XM_023246637.1;refGene_gn=OTOA;refGene_tx=XM_023246638.1
E3 26845398 26845614
refGene_gn=OTOA;refGene_tx=XM_006942284.3;refGene_gn=OTOA;refGene_tx=XM_023246637.1;refGene_gn=OTOA;refGene_tx=XM_023246638.1
E3 26848209 26848333
refGene_gn=OTOA;refGene_tx=XM_006942284.3;refGene_gn=OTOA;refGene_tx=XM_023246637.1;refGene_gn=OTOA;refGene_tx=XM_023246638.1
E3 26850343 26850483
refGene_gn=OTOA;refGene_tx=XM_006942284.3;refGene_gn=OTOA;refGene_tx=XM_023246637.1;refGene_gn=OTOA;refGene_tx=XM_023246638.1
E3 26852344 26852445
refGene_gn=OTOA;refGene_tx=XM_006942284.3;refGene_gn=OTOA;refGene_tx=XM_023246637.1;refGene_gn=OTOA;refGene_tx=XM_023246638.1
E3 26858325 26858429
refGene_gn=OTOA;refGene_tx=XM_006942284.3;refGene_gn=OTOA;refGene_tx=XM_023246637.1;refGene_gn=OTOA;refGene_tx=XM_023246638.1
E3 26863049 26863285
refGene_gn=OTOA;refGene_tx=XM_006942284.3;refGene_gn=OTOA;refGene_tx=XM_023246637.1;refGene_gn=OTOA;refGene_tx=XM_023246638.1
E3 26865626 26865758
refGene_gn=OTOA;refGene_tx=XM_006942284.3;refGene_gn=OTOA;refGene_tx=XM_023246637.1;refGene_gn=OTOA;refGene_tx=XM_023246638.1
E3 26869083 26869177
refGene_gn=OTOA;refGene_tx=XM_006942284.3;refGene_gn=OTOA;refGene_tx=XM_023246637.1;refGene_gn=OTOA;refGene_tx=XM_023246638.1
E3 26870977 26871005
refGene_gn=OTOA;refGene_tx=XM_006942284.3;refGene_gn=OTOA;refGene_tx=XM_023246638.1
E3 26871121 26871152
refGene_gn=OTOA;refGene_tx=XM_006942284.3;refGene_gn=OTOA;refGene_tx=XM_023246638.1
E3 26871243 26871272
refGene_gn=OTOA;refGene_tx=XM_006942284.3;refGene_gn=OTOA;refGene_tx=XM_023246638.1
E3 26871853 26871944
refGene_gn=OTOA;refGene_tx=XM_006942284.3;refGene_gn=OTOA;refGene_tx=XM_023246638.1

E3 26888671 26888877
refGene_gn=METTL9;refGene_tx=XM_006942238.3;refGene_gn=METTL9;refGene_tx=XM_023246745.1;refGene_gn=METTL9;refGene_tx=XM_023246746.1
E3 26890817 26890884
refGene_gn=IGSF6;refGene_tx=XM_006942237.4;refGene_gn=IGSF6;refGene_tx=XM_019820804.2
E3 26894557 26894917
refGene_gn=IGSF6;refGene_tx=XM_006942237.4;refGene_gn=IGSF6;refGene_tx=XM_019820804.2
E3 26896804 26896899
refGene_gn=IGSF6;refGene_tx=XM_006942237.4;refGene_gn=IGSF6;refGene_tx=XM_019820804.2
E3 26897550 26897598
refGene_gn=IGSF6;refGene_tx=XM_006942237.4;refGene_gn=IGSF6;refGene_tx=XM_019820804.2
E3 26898391 26898511 refGene_gn=IGSF6;refGene_tx=XM_019820804.2
E3 26900429 26900453 refGene_gn=IGSF6;refGene_tx=XM_006942237.4
E3 26910602 26910787
refGene_gn=METTL9;refGene_tx=XM_006942238.3;refGene_gn=METTL9;refGene_tx=XM_023246745.1;refGene_gn=METTL9;refGene_tx=XM_023246746.1
E3 26918417 26918627
refGene_gn=METTL9;refGene_tx=XM_006942238.3;refGene_gn=METTL9;refGene_tx=XM_023246745.1;refGene_gn=METTL9;refGene_tx=XM_023246746.1
E3 26923877 26924068
refGene_gn=METTL9;refGene_tx=XM_006942238.3;refGene_gn=METTL9;refGene_tx=XM_023246745.1;refGene_gn=METTL9;refGene_tx=XM_023246746.1
E3 26938451 26938616
refGene_gn=METTL9;refGene_tx=XM_023246745.1;refGene_gn=METTL9;refGene_tx=XM_023246746.1
E3 26940768 26940813 refGene_gn=METTL9;refGene_tx=XM_006942238.3
E3 26960280 26960413
refGene_gn=LOC102899571;refGene_tx=XM_006942236.3;refGene_gn=LOC102899571;refGene_tx=XM_023246744.1
E3 26977801 26977924
refGene_gn=LOC102899571;refGene_tx=XM_006942236.3;refGene_gn=LOC102899571;refGene_tx=XM_023246744.1
E3 26982297 26982381
refGene_gn=LOC102899571;refGene_tx=XM_006942236.3;refGene_gn=LOC102899571;refGene_tx=XM_023246744.1
E3 26984190 26984345
refGene_gn=LOC102899571;refGene_tx=XM_006942236.3;refGene_gn=LOC102899571;refGene_tx=XM_023246744.1
E3 26994557 26995280
refGene_gn=LOC102899571;refGene_tx=XM_006942236.3;refGene_gn=LOC102899571;refGene_tx=XM_023246744.1
E3 26996308 26996458
refGene_gn=LOC102899571;refGene_tx=XM_006942236.3;refGene_gn=LOC102899571;refGene_tx=XM_023246744.1
E3 27001449 27001528 refGene_gn=CDR2;refGene_tx=XM_011290634.3
E3 27011051 27011164

refGene_gn=CDR2;refGene_tx=XM_011290634.3;refGene_gn=CDR2;refGene_tx=XM_023246743.1
E3 27024762 27024911
refGene_gn=CDR2;refGene_tx=XM_011290634.3;refGene_gn=CDR2;refGene_tx=XM_023246743.1
E3 27025167 27025332
refGene_gn=CDR2;refGene_tx=XM_011290634.3;refGene_gn=CDR2;refGene_tx=XM_023246743.1
E3 27026604 27027469
refGene_gn=CDR2;refGene_tx=XM_011290634.3;refGene_gn=CDR2;refGene_tx=XM_023246743.1
E3 27041899 27041956
refGene_gn=POLR3E;refGene_tx=XM_019820805.2;refGene_gn=POLR3E;refGene_tx=XM_019820807.2;refGene_gn=POLR3E;refGene_tx=XM_023246302.1;refGene_gn=POLR3E;refGene_tx=XM_023246303.1;refGene_gn=POLR3E;refGene_tx=XM_023246304.1
E3 27042638 27042764
refGene_gn=POLR3E;refGene_tx=XM_019820805.2;refGene_gn=POLR3E;refGene_tx=XM_019820807.2;refGene_gn=POLR3E;refGene_tx=XM_023246302.1;refGene_gn=POLR3E;refGene_tx=XM_023246303.1;refGene_gn=POLR3E;refGene_tx=XM_023246304.1
E3 27043927 27044005
refGene_gn=POLR3E;refGene_tx=XM_019820805.2;refGene_gn=POLR3E;refGene_tx=XM_019820807.2;refGene_gn=POLR3E;refGene_tx=XM_023246302.1;refGene_gn=POLR3E;refGene_tx=XM_023246303.1;refGene_gn=POLR3E;refGene_tx=XM_023246304.1
E3 27044186 27044294
refGene_gn=POLR3E;refGene_tx=XM_023246305.1;refGene_gn=POLR3E;refGene_tx=XM_023246306.1
E3 27044640 27044784
refGene_gn=POLR3E;refGene_tx=XM_019820805.2;refGene_gn=POLR3E;refGene_tx=XM_019820807.2;refGene_gn=POLR3E;refGene_tx=XM_023246302.1;refGene_gn=POLR3E;refGene_tx=XM_023246303.1;refGene_gn=POLR3E;refGene_tx=XM_023246305.1
E3 27045152 27045185 refGene_gn=POLR3E;refGene_tx=XM_023246307.1
E3 27046519 27047043
refGene_gn=POLR3E;refGene_tx=XM_019820805.2;refGene_gn=POLR3E;refGene_tx=XM_019820807.2;refGene_gn=POLR3E;refGene_tx=XM_023246302.1;refGene_gn=POLR3E;refGene_tx=XM_023246303.1;refGene_gn=POLR3E;refGene_tx=XM_023246304.1;refGene_gn=POLR3E;refGene_tx=XM_023246305.1;refGene_gn=POLR3E;refGene_tx=XM_023246306.1;refGene_gn=POLR3E;refGene_tx=XM_023246307.1
E3 27047563 27047625
refGene_gn=POLR3E;refGene_tx=XM_019820805.2;refGene_gn=POLR3E;refGene_tx=XM_019820807.2;refGene_gn=POLR3E;refGene_tx=XM_023246302.1;refGene_gn=POLR3E;refGene_tx=XM_023246303.1;refGene_gn=POLR3E;refGene_tx=XM_023246304.1;refGene_gn=POLR3E;refGene_tx=XM_023246305.1;refGene_gn=POLR3E;refGene_tx=XM_023246306.1;refGene_gn=POLR3E;refGene_tx=XM_023246307.1
E3 27048080 27048238

refGene_gn=POLR3E;refGene_tx=XM_019820805.2;refGene_gn=POLR3E;refGene_tx=XM_019820807.2;refGene_gn=POLR3E;refGene_tx=XM_023246302.1;refGene_gn=POLR3E;refGene_tx=XM_023246303.1;refGene_gn=POLR3E;refGene_tx=XM_023246304.1;refGene_gn=POLR3E;refGene_tx=XM_023246305.1;refGene_gn=POLR3E;refGene_tx=XM_023246306.1;refGene_gn=POLR3E;refGene_tx=XM_023246307.1

E3 27048335 27048395 refGene_gn=POLR3E;refGene_tx=XM_019820805.2;refGene_gn=POLR3E;refGene_tx=XM_019820807.2;refGene_gn=POLR3E;refGene_tx=XM_023246302.1;refGene_gn=POLR3E;refGene_tx=XM_023246303.1;refGene_gn=POLR3E;refGene_tx=XM_023246304.1;refGene_gn=POLR3E;refGene_tx=XM_023246305.1;refGene_gn=POLR3E;refGene_tx=XM_023246306.1;refGene_gn=POLR3E;refGene_tx=XM_023246307.1

E3 27049489 27049571 refGene_gn=POLR3E;refGene_tx=XM_019820805.2;refGene_gn=POLR3E;refGene_tx=XM_019820807.2;refGene_gn=POLR3E;refGene_tx=XM_023246302.1;refGene_gn=POLR3E;refGene_tx=XM_023246303.1;refGene_gn=POLR3E;refGene_tx=XM_023246304.1;refGene_gn=POLR3E;refGene_tx=XM_023246305.1;refGene_gn=POLR3E;refGene_tx=XM_023246306.1;refGene_gn=POLR3E;refGene_tx=XM_023246307.1

E3 27052548 27052669 refGene_gn=POLR3E;refGene_tx=XM_019820805.2;refGene_gn=POLR3E;refGene_tx=XM_019820807.2;refGene_gn=POLR3E;refGene_tx=XM_023246302.1;refGene_gn=POLR3E;refGene_tx=XM_023246303.1;refGene_gn=POLR3E;refGene_tx=XM_023246304.1;refGene_gn=POLR3E;refGene_tx=XM_023246305.1;refGene_gn=POLR3E;refGene_tx=XM_023246306.1;refGene_gn=POLR3E;refGene_tx=XM_023246307.1

E3 27053918 27054010 refGene_gn=POLR3E;refGene_tx=XM_019820805.2;refGene_gn=POLR3E;refGene_tx=XM_019820807.2;refGene_gn=POLR3E;refGene_tx=XM_023246302.1;refGene_gn=POLR3E;refGene_tx=XM_023246303.1;refGene_gn=POLR3E;refGene_tx=XM_023246304.1;refGene_gn=POLR3E;refGene_tx=XM_023246305.1;refGene_gn=POLR3E;refGene_tx=XM_023246306.1;refGene_gn=POLR3E;refGene_tx=XM_023246307.1

E3 27054104 27054149 refGene_gn=POLR3E;refGene_tx=XM_019820805.2;refGene_gn=POLR3E;refGene_tx=XM_019820807.2;refGene_gn=POLR3E;refGene_tx=XM_023246302.1;refGene_gn=POLR3E;refGene_tx=XM_023246303.1;refGene_gn=POLR3E;refGene_tx=XM_023246304.1;refGene_gn=POLR3E;refGene_tx=XM_023246305.1;refGene_gn=POLR3E;refGene_tx=XM_023246306.1;refGene_gn=POLR3E;refGene_tx=XM_023246307.1

E3 27054522 27054608 refGene_gn=POLR3E;refGene_tx=XM_019820805.2;refGene_gn=POLR3E;refGene_tx=XM_019820807.2;refGene_gn=POLR3E;refGene_tx=XM_023246302.1;refGene_gn=POLR3E;refGene_tx=XM_023246303.1;refGene_gn=POLR3E;refGene_tx=XM_023246304.1;refGene_gn=POLR3E;refGene_tx=XM_023246305.1;refGene_gn=POLR3E;refGene_tx=XM_023246306.1;refGene_gn=POLR3E;refGene_tx=XM_023246307.1

E3 27055579 27055699 refGene_gn=POLR3E;refGene_tx=XM_019820805.2;refGene_gn=POLR3E;refGene_

tx=XM_019820807.2;refGene_gn=POLR3E;refGene_tx=XM_023246302.1;refGene_gn=POLR3E;refGene_tx=XM_023246303.1;refGene_gn=POLR3E;refGene_tx=XM_023246304.1;refGene_gn=POLR3E;refGene_tx=XM_023246305.1;refGene_gn=POLR3E;refGene_tx=XM_023246306.1;refGene_gn=POLR3E;refGene_tx=XM_023246307.1
E3 27057379 27057429 refGene_gn=POLR3E;refGene_tx=XM_019820805.2;refGene_gn=POLR3E;refGene_tx=XM_019820807.2;refGene_gn=POLR3E;refGene_tx=XM_023246302.1;refGene_gn=POLR3E;refGene_tx=XM_023246303.1;refGene_gn=POLR3E;refGene_tx=XM_023246304.1;refGene_gn=POLR3E;refGene_tx=XM_023246305.1;refGene_gn=POLR3E;refGene_tx=XM_023246306.1;refGene_gn=POLR3E;refGene_tx=XM_023246307.1
E3 27057774 27057882 refGene_gn=POLR3E;refGene_tx=XM_019820805.2;refGene_gn=POLR3E;refGene_tx=XM_019820807.2;refGene_gn=POLR3E;refGene_tx=XM_023246302.1;refGene_gn=POLR3E;refGene_tx=XM_023246303.1;refGene_gn=POLR3E;refGene_tx=XM_023246304.1;refGene_gn=POLR3E;refGene_tx=XM_023246305.1;refGene_gn=POLR3E;refGene_tx=XM_023246306.1;refGene_gn=POLR3E;refGene_tx=XM_023246307.1
E3 27063543 27063626 refGene_gn=POLR3E;refGene_tx=XM_019820805.2;refGene_gn=POLR3E;refGene_tx=XM_019820807.2;refGene_gn=POLR3E;refGene_tx=XM_023246302.1;refGene_gn=POLR3E;refGene_tx=XM_023246303.1;refGene_gn=POLR3E;refGene_tx=XM_023246304.1;refGene_gn=POLR3E;refGene_tx=XM_023246305.1;refGene_gn=POLR3E;refGene_tx=XM_023246306.1;refGene_gn=POLR3E;refGene_tx=XM_023246307.1
E3 27064021 27064137 refGene_gn=POLR3E;refGene_tx=XM_019820805.2;refGene_gn=POLR3E;refGene_tx=XM_019820807.2;refGene_gn=POLR3E;refGene_tx=XM_023246302.1;refGene_gn=POLR3E;refGene_tx=XM_023246303.1;refGene_gn=POLR3E;refGene_tx=XM_023246304.1;refGene_gn=POLR3E;refGene_tx=XM_023246305.1;refGene_gn=POLR3E;refGene_tx=XM_023246306.1;refGene_gn=POLR3E;refGene_tx=XM_023246307.1
E3 27064886 27064964 refGene_gn=POLR3E;refGene_tx=XM_019820805.2;refGene_gn=POLR3E;refGene_tx=XM_019820807.2;refGene_gn=POLR3E;refGene_tx=XM_023246302.1;refGene_gn=POLR3E;refGene_tx=XM_023246303.1;refGene_gn=POLR3E;refGene_tx=XM_023246304.1;refGene_gn=POLR3E;refGene_tx=XM_023246305.1;refGene_gn=POLR3E;refGene_tx=XM_023246306.1;refGene_gn=POLR3E;refGene_tx=XM_023246307.1
E3 27066394 27066433 refGene_gn=POLR3E;refGene_tx=XM_019820807.2
E3 27067299 27067350 refGene_gn=POLR3E;refGene_tx=XM_019820805.2;refGene_gn=POLR3E;refGene_tx=XM_023246302.1;refGene_gn=POLR3E;refGene_tx=XM_023246303.1;refGene_gn=POLR3E;refGene_tx=XM_023246304.1;refGene_gn=POLR3E;refGene_tx=XM_023246305.1;refGene_gn=POLR3E;refGene_tx=XM_023246306.1;refGene_gn=POLR3E;refGene_tx=XM_023246307.1
E3 27068689 27068725 refGene_gn=POLR3E;refGene_tx=XM_019820805.2;refGene_gn=POLR3E;refGene_tx=XM_023246302.1;refGene_gn=POLR3E;refGene_tx=XM_023246303.1;refGene_

gn=POLR3E;refGene_tx=XM_023246304.1;refGene_gn=POLR3E;refGene_tx=XM_023246305.1;refGene_gn=POLR3E;refGene_tx=XM_023246306.1;refGene_gn=POLR3E;refGene_tx=XM_023246307.1

| | | | |
|---|---|---|---|
| E3 | 27089181 | 27089291 | refGene_gn=EEF2K;refGene_tx=XM_023246209.1 |
| E3 | 27091079 | 27091258 | refGene_gn=EEF2K;refGene_tx=XM_023246209.1 |
| E3 | 27097022 | 27097147 | refGene_gn=EEF2K;refGene_tx=XM_023246209.1 |
| E3 | 27101595 | 27101784 | refGene_gn=EEF2K;refGene_tx=XM_023246209.1 |
| E3 | 27101937 | 27102071 | refGene_gn=EEF2K;refGene_tx=XM_023246209.1 |
| E3 | 27103423 | 27103487 | refGene_gn=EEF2K;refGene_tx=XM_023246209.1 |
| E3 | 27104319 | 27104397 | refGene_gn=EEF2K;refGene_tx=XM_023246209.1 |
| E3 | 27106457 | 27106525 | refGene_gn=EEF2K;refGene_tx=XM_023246209.1 |
| E3 | 27107938 | 27108140 | refGene_gn=EEF2K;refGene_tx=XM_023246209.1 |
| E3 | 27108855 | 27108983 | refGene_gn=EEF2K;refGene_tx=XM_023246209.1 |
| E3 | 27109236 | 27109369 | refGene_gn=EEF2K;refGene_tx=XM_023246209.1 |
| E3 | 27109920 | 27110070 | refGene_gn=EEF2K;refGene_tx=XM_023246209.1 |
| E3 | 27113955 | 27114127 | refGene_gn=EEF2K;refGene_tx=XM_023246209.1 |
| E3 | 27114486 | 27114524 | refGene_gn=EEF2K;refGene_tx=XM_023246209.1 |
| E3 | 27115887 | 27115948 | refGene_gn=EEF2K;refGene_tx=XM_023246209.1 |
| E3 | 27119333 | 27119434 | refGene_gn=EEF2K;refGene_tx=XM_023246209.1 |
| E3 | 27135300 | 27135546 | refGene_gn=EEF2K;refGene_tx=XM_023246209.1 |
| E3 | 27172607 | 27172712 | refGene_gn=SDR42E2;refGene_tx=XM_023246773.1 |
| E3 | 27173240 | 27173378 | refGene_gn=SDR42E2;refGene_tx=XM_023246773.1 |
| E3 | 27174967 | 27175050 | refGene_gn=SDR42E2;refGene_tx=XM_023246773.1 |
| E3 | 27178857 | 27178933 | refGene_gn=SDR42E2;refGene_tx=XM_023246773.1 |
| E3 | 27179564 | 27179683 | refGene_gn=SDR42E2;refGene_tx=XM_023246773.1 |
| E3 | 27180389 | 27180447 | refGene_gn=SDR42E2;refGene_tx=XM_023246773.1 |
| E3 | 27181978 | 27182074 | refGene_gn=SDR42E2;refGene_tx=XM_023246773.1 |
| E3 | 27182555 | 27183341 | refGene_gn=SDR42E2;refGene_tx=XM_023246773.1 |
| E3 | 27190802 | 27190854 | refGene_gn=VWA3A;refGene_tx=XM_023246294.1 |
| E3 | 27191045 | 27191143 | refGene_gn=VWA3A;refGene_tx=XM_023246294.1 |
| E3 | 27192707 | 27192831 | refGene_gn=VWA3A;refGene_tx=XM_023246294.1 |
| E3 | 27193841 | 27193993 | refGene_gn=VWA3A;refGene_tx=XM_023246294.1 |
| E3 | 27194431 | 27194576 | refGene_gn=VWA3A;refGene_tx=XM_023246294.1 |
| E3 | 27195597 | 27195742 | refGene_gn=VWA3A;refGene_tx=XM_023246294.1 |
| E3 | 27198569 | 27198678 | refGene_gn=VWA3A;refGene_tx=XM_023246294.1 |
| E3 | 27199855 | 27199993 | refGene_gn=VWA3A;refGene_tx=XM_023246294.1 |
| E3 | 27200845 | 27200943 | refGene_gn=VWA3A;refGene_tx=XM_023246294.1 |
| E3 | 27201474 | 27201585 | refGene_gn=VWA3A;refGene_tx=XM_023246294.1 |
| E3 | 27202523 | 27202614 | refGene_gn=VWA3A;refGene_tx=XM_023246294.1 |
| E3 | 27204152 | 27204305 | refGene_gn=VWA3A;refGene_tx=XM_023246294.1 |
| E3 | 27205362 | 27205433 | refGene_gn=VWA3A;refGene_tx=XM_023246294.1 |
| E3 | 27208445 | 27208641 | refGene_gn=VWA3A;refGene_tx=XM_023246294.1 |
| E3 | 27209384 | 27209529 | refGene_gn=VWA3A;refGene_tx=XM_023246294.1 |
| E3 | 27209674 | 27209749 | refGene_gn=VWA3A;refGene_tx=XM_023246294.1 |
| E3 | 27215251 | 27215371 | refGene_gn=VWA3A;refGene_tx=XM_023246294.1 |
| E3 | 27216876 | 27216971 | refGene_gn=VWA3A;refGene_tx=XM_023246294.1 |
| E3 | 27217386 | 27217467 | refGene_gn=VWA3A;refGene_tx=XM_023246294.1 |
| E3 | 27219249 | 27219353 | refGene_gn=VWA3A;refGene_tx=XM_023246294.1 |
| E3 | 27219457 | 27219593 | refGene_gn=VWA3A;refGene_tx=XM_023246294.1 |
| E3 | 27222084 | 27222210 | refGene_gn=VWA3A;refGene_tx=XM_023246294.1 |

E3 27225842 27225908 refGene_gn=VWA3A;refGene_tx=XM_023246294.1
E3 27226140 27226249 refGene_gn=VWA3A;refGene_tx=XM_023246294.1
E3 27226570 27226696 refGene_gn=VWA3A;refGene_tx=XM_023246294.1
E3 27229774 27229881 refGene_gn=VWA3A;refGene_tx=XM_023246294.1
E3 27231283 27231382 refGene_gn=VWA3A;refGene_tx=XM_023246294.1
E3 27237477 27237532 refGene_gn=VWA3A;refGene_tx=XM_023246294.1
E3 27240131 27240209 refGene_gn=VWA3A;refGene_tx=XM_023246294.1
E3 27240304 27240429 refGene_gn=VWA3A;refGene_tx=XM_023246294.1
E3 27245045 27245169 refGene_gn=VWA3A;refGene_tx=XM_023246294.1
E3 27245488 27245556 refGene_gn=VWA3A;refGene_tx=XM_023246294.1
E3 27245557 27245576 refGene_gn=VWA3A;refGene_tx=XM_023246294.1
E3 27250467 27250481 refGene_gn=VWA3A;refGene_tx=XM_023246294.1
E3 27261392 27261577
refGene_gn=CE3H16orf52;refGene_tx=XM_023246882.1
E3 27266744 27266957
refGene_gn=CE3H16orf52;refGene_tx=XM_023246882.1
E3 27322553 27322659
refGene_gn=CE3H16orf52;refGene_tx=XM_023246882.1
E3 27329074 27329105 refGene_gn=PDZD9;refGene_tx=XM_019820815.2
E3 27330989 27331169 refGene_gn=PDZD9;refGene_tx=XM_019820815.2
E3 27335648 27335838 refGene_gn=PDZD9;refGene_tx=XM_019820815.2
E3 27340922 27341316 refGene_gn=PDZD9;refGene_tx=XM_019820815.2
E3 27342563 27342647
refGene_gn=LOC101101684;refGene_tx=XM_003998776.4
E3 27343983 27344137
refGene_gn=LOC101101684;refGene_tx=XM_003998776.4
E3 27348373 27348450
refGene_gn=LOC101101684;refGene_tx=XM_003998776.4
E3 27350901 27350982
refGene_gn=LOC101101684;refGene_tx=XM_003998776.4
E3 27359999 27360199
refGene_gn=LOC101101684;refGene_tx=XM_003998776.4
E3 27360526 27360622
refGene_gn=LOC101101684;refGene_tx=XM_003998776.4
E3 27363160 27363218
refGene_gn=LOC101101684;refGene_tx=XM_003998776.4
E3 27365793 27365891
refGene_gn=LOC101101684;refGene_tx=XM_003998776.4
E3 27367719 27367844
refGene_gn=LOC101101684;refGene_tx=XM_003998776.4
E3 27368071 27368128
refGene_gn=LOC101101684;refGene_tx=XM_003998776.4
E3 27369629 27369694
refGene_gn=LOC101101684;refGene_tx=XM_003998776.4
E3 27370076 27370226
refGene_gn=LOC101101684;refGene_tx=XM_003998776.4
E3 27370323 27370407
refGene_gn=LOC101101684;refGene_tx=XM_003998776.4
E3 27373574 27373607
refGene_gn=LOC101101684;refGene_tx=XM_003998776.4

E3 27439833 27439914
refGene_gn=LOC101087463;refGene_tx=XM_023247082.1
E3 27441048 27441180
refGene_gn=LOC101087463;refGene_tx=XM_023247082.1
E3 27441577 27441651
refGene_gn=LOC101087463;refGene_tx=XM_023247082.1
E3 27447862 27448053
refGene_gn=LOC101087463;refGene_tx=XM_023247082.1
E3 27448429 27448600
refGene_gn=LOC101087463;refGene_tx=XM_023247082.1
E3 27449262 27449450
refGene_gn=LOC101087463;refGene_tx=XM_023247082.1
E3 27456583 27457357
refGene_gn=LOC105261313;refGene_tx=XM_011290612.3
E3 27458982 27459177
refGene_gn=LOC101087463;refGene_tx=XM_023247082.1
E3 27459407 27459533
refGene_gn=LOC101087463;refGene_tx=XM_023247082.1
E3 27466994 27467173
refGene_gn=LOC101087463;refGene_tx=XM_023247082.1
E3 27474598 27474769
refGene_gn=LOC101087463;refGene_tx=XM_023247082.1
E3 27480628 27480845
refGene_gn=LOC101087463;refGene_tx=XM_023247082.1
E3 27488347 27488552
refGene_gn=LOC101087463;refGene_tx=XM_023247082.1
E3 27488705 27488955
refGene_gn=LOC101087463;refGene_tx=XM_023247082.1
E3 27496505 27496806
refGene_gn=LOC101087463;refGene_tx=XM_023247082.1
E3 27497794 27497963
refGene_gn=LOC101087463;refGene_tx=XM_023247082.1
E3 27498462 27498561
refGene_gn=LOC101087463;refGene_tx=XM_023247082.1
E3 27504806 27504957
refGene_gn=LOC101087463;refGene_tx=XM_023247082.1
E3 27522729 27522922
refGene_gn=LOC101087463;refGene_tx=XM_023247082.1
E3 27523294 27523450
refGene_gn=LOC101087463;refGene_tx=XM_023247082.1
E3 27523689 27523770
refGene_gn=LOC101087463;refGene_tx=XM_023247082.1
E3 27523772 27523846
refGene_gn=LOC101087463;refGene_tx=XM_023247082.1
E3 27537639 27537769
refGene_gn=LOC101087463;refGene_tx=XM_023247082.1
E3 27554805 27554940
refGene_gn=LOC101087463;refGene_tx=XM_023247082.1
E3 27555723 27555902
refGene_gn=LOC101087463;refGene_tx=XM_023247082.1

E3 27556596 27556770
refGene_gn=LOC101087463;refGene_tx=XM_023247082.1
E3 27579251 27579369
refGene_gn=LOC101087463;refGene_tx=XM_023247082.1
E3 27579943 27580060
refGene_gn=LOC101087463;refGene_tx=XM_023247082.1
E3 27592654 27592914
refGene_gn=LOC101087463;refGene_tx=XM_023247082.1
E3 27598687 27598835
refGene_gn=LOC101087463;refGene_tx=XM_023247082.1
E3 27600631 27600759
refGene_gn=LOC101087463;refGene_tx=XM_023247082.1
E3 27605594 27605696
refGene_gn=LOC101087463;refGene_tx=XM_023247082.1
E3 27605805 27605860
refGene_gn=LOC101087463;refGene_tx=XM_023247082.1
E3 27606210 27606270
refGene_gn=LOC101087463;refGene_tx=XM_023247082.1
E3 27702566 27702736 refGene_gn=CRYM;refGene_tx=XM_003998778.5
E3 27703224 27703378 refGene_gn=CRYM;refGene_tx=XM_003998778.5
E3 27705322 27705385 refGene_gn=CRYM;refGene_tx=XM_003998778.5
E3 27709970 27710072 refGene_gn=CRYM;refGene_tx=XM_003998778.5
E3 27710553 27710737 refGene_gn=CRYM;refGene_tx=XM_003998778.5
E3 27714126 27714143 refGene_gn=CRYM;refGene_tx=XM_003998778.5
E3 27714144 27714249 refGene_gn=CRYM;refGene_tx=XM_003998778.5
E3 27715165 27715250 refGene_gn=CRYM;refGene_tx=XM_003998778.5
E3 27717379 27717444 refGene_gn=CRYM;refGene_tx=XM_003998778.5
E3 27726484 27727574
refGene_gn=ANKS4B;refGene_tx=XM_019820832.2;refGene_gn=TMEM159;refGene
_tx=XM_023247041.1;refGene_gn=TMEM159;refGene_tx=XM_023247042.1
E3 27736058 27736222 refGene_gn=ANKS4B;refGene_tx=XM_019820832.2
E3 27748599 27748735
refGene_gn=TMEM159;refGene_tx=XM_023247041.1;refGene_gn=TMEM159;refGen
e_tx=XM_023247042.1
E3 27750564 27750626 refGene_gn=ZP2;refGene_tx=NM_001009875.1
E3 27750719 27750808 refGene_gn=ZP2;refGene_tx=NM_001009875.1
E3 27751750 27751834 refGene_gn=ZP2;refGene_tx=NM_001009875.1
E3 27752220 27752315 refGene_gn=ZP2;refGene_tx=NM_001009875.1
E3 27753914 27754067 refGene_gn=ZP2;refGene_tx=NM_001009875.1
E3 27754867 27754909 refGene_gn=ZP2;refGene_tx=NM_001009875.1
E3 27755062 27755227 refGene_gn=ZP2;refGene_tx=NM_001009875.1
E3 27755976 27756073 refGene_gn=ZP2;refGene_tx=NM_001009875.1
E3 27756174 27756356 refGene_gn=ZP2;refGene_tx=NM_001009875.1
E3 27757175 27757302 refGene_gn=ZP2;refGene_tx=NM_001009875.1
E3 27757968 27758156 refGene_gn=ZP2;refGene_tx=NM_001009875.1
E3 27758239 27758331 refGene_gn=ZP2;refGene_tx=NM_001009875.1
E3 27758433 27758558 refGene_gn=ZP2;refGene_tx=NM_001009875.1
E3 27758934 27759124 refGene_gn=ZP2;refGene_tx=NM_001009875.1
E3 27760717 27760853 refGene_gn=ZP2;refGene_tx=NM_001009875.1
E3 27760927 27761024 refGene_gn=ZP2;refGene_tx=NM_001009875.1

E3 27761621 27761705 refGene_gn=ZP2;refGene_tx=NM_001009875.1
E3 27762511 27762654 refGene_gn=ZP2;refGene_tx=NM_001009875.1
E3 27763085 27763153
refGene_gn=TMEM159;refGene_tx=XM_023247041.1;refGene_gn=TMEM159;refGene_tx=XM_023247042.1
E3 27773500 27773546 refGene_gn=TMEM159;refGene_tx=XM_019820838.2
E3 27777401 27777483 refGene_gn=TMEM159;refGene_tx=XM_006942241.4
E3 27780194 27780279 refGene_gn=TMEM159;refGene_tx=XM_019820837.2
E3 27782315 27782454
refGene_gn=TMEM159;refGene_tx=XM_006942241.4;refGene_gn=TMEM159;refGene_tx=XM_019820837.2;refGene_gn=TMEM159;refGene_tx=XM_019820838.2;refGene_gn=TMEM159;refGene_tx=XM_023247041.1;refGene_gn=TMEM159;refGene_tx=XM_023247042.1
E3 27784453 27784622
refGene_gn=TMEM159;refGene_tx=XM_006942241.4;refGene_gn=TMEM159;refGene_tx=XM_019820837.2;refGene_gn=TMEM159;refGene_tx=XM_019820838.2;refGene_gn=TMEM159;refGene_tx=XM_023247041.1;refGene_gn=TMEM159;refGene_tx=XM_023247042.1
E3 27791982 27792078
refGene_gn=TMEM159;refGene_tx=XM_006942241.4;refGene_gn=TMEM159;refGene_tx=XM_019820837.2;refGene_gn=TMEM159;refGene_tx=XM_019820838.2;refGene_gn=TMEM159;refGene_tx=XM_023247041.1;refGene_gn=TMEM159;refGene_tx=XM_023247042.1
E3 27800364 27800403
refGene_gn=DNAH3;refGene_tx=XM_019820825.2;refGene_gn=DNAH3;refGene_tx=XM_023246505.1
E3 27804901 27805006
refGene_gn=DNAH3;refGene_tx=XM_019820825.2;refGene_gn=DNAH3;refGene_tx=XM_023246505.1
E3 27805540 27805763
refGene_gn=DNAH3;refGene_tx=XM_019820825.2;refGene_gn=DNAH3;refGene_tx=XM_023246505.1
E3 27808881 27808954
refGene_gn=DNAH3;refGene_tx=XM_019820825.2;refGene_gn=DNAH3;refGene_tx=XM_023246505.1
E3 27810011 27810186
refGene_gn=DNAH3;refGene_tx=XM_019820825.2;refGene_gn=DNAH3;refGene_tx=XM_023246505.1
E3 27811802 27811992
refGene_gn=DNAH3;refGene_tx=XM_019820825.2;refGene_gn=DNAH3;refGene_tx=XM_023246505.1
E3 27813752 27813948
refGene_gn=DNAH3;refGene_tx=XM_019820825.2;refGene_gn=DNAH3;refGene_tx=XM_023246505.1
E3 27818257 27818383
refGene_gn=DNAH3;refGene_tx=XM_019820825.2;refGene_gn=DNAH3;refGene_tx=XM_023246505.1
E3 27819552 27819748
refGene_gn=DNAH3;refGene_tx=XM_019820825.2;refGene_gn=DNAH3;refGene_tx=XM_023246505.1

E3	27821423	27821603
refGene_gn=DNAH3;refGene_tx=XM_019820825.2;refGene_gn=DNAH3;refGene_tx=XM_023246505.1
E3	27827521	27827658
refGene_gn=DNAH3;refGene_tx=XM_019820825.2;refGene_gn=DNAH3;refGene_tx=XM_023246505.1
E3	27829907	27830013
refGene_gn=DNAH3;refGene_tx=XM_019820825.2;refGene_gn=DNAH3;refGene_tx=XM_023246505.1
E3	27830196	27830375
refGene_gn=DNAH3;refGene_tx=XM_019820825.2;refGene_gn=DNAH3;refGene_tx=XM_023246505.1
E3	27833402	27833587
refGene_gn=DNAH3;refGene_tx=XM_019820825.2;refGene_gn=DNAH3;refGene_tx=XM_023246505.1
E3	27834796	27834920
refGene_gn=DNAH3;refGene_tx=XM_019820825.2;refGene_gn=DNAH3;refGene_tx=XM_023246505.1
E3	27837664	27837818
refGene_gn=DNAH3;refGene_tx=XM_019820825.2;refGene_gn=DNAH3;refGene_tx=XM_023246505.1
E3	27838608	27838753
refGene_gn=DNAH3;refGene_tx=XM_019820825.2;refGene_gn=DNAH3;refGene_tx=XM_023246505.1
E3	27844005	27844217
refGene_gn=DNAH3;refGene_tx=XM_019820825.2;refGene_gn=DNAH3;refGene_tx=XM_023246505.1
E3	27848549	27848641
refGene_gn=DNAH3;refGene_tx=XM_019820825.2;refGene_gn=DNAH3;refGene_tx=XM_023246505.1
E3	27853247	27853362
refGene_gn=DNAH3;refGene_tx=XM_019820825.2;refGene_gn=DNAH3;refGene_tx=XM_023246505.1
E3	27855318	27855435
refGene_gn=DNAH3;refGene_tx=XM_019820825.2;refGene_gn=DNAH3;refGene_tx=XM_023246505.1
E3	27856229	27856409
refGene_gn=DNAH3;refGene_tx=XM_019820825.2;refGene_gn=DNAH3;refGene_tx=XM_023246505.1
E3	27857785	27857922
refGene_gn=DNAH3;refGene_tx=XM_019820825.2;refGene_gn=DNAH3;refGene_tx=XM_023246505.1
E3	27861748	27861950
refGene_gn=DNAH3;refGene_tx=XM_019820825.2;refGene_gn=DNAH3;refGene_tx=XM_023246505.1
E3	27863104	27863197
refGene_gn=DNAH3;refGene_tx=XM_019820825.2;refGene_gn=DNAH3;refGene_tx=XM_023246505.1
E3	27864536	27864647
refGene_gn=DNAH3;refGene_tx=XM_019820825.2;refGene_gn=DNAH3;refGene_tx

=XM_023246505.1
E3 27867816 27867931
refGene_gn=DNAH3;refGene_tx=XM_019820825.2;refGene_gn=DNAH3;refGene_tx
=XM_023246505.1
E3 27871164 27871363
refGene_gn=DNAH3;refGene_tx=XM_019820825.2;refGene_gn=DNAH3;refGene_tx
=XM_023246505.1
E3 27872849 27872958
refGene_gn=DNAH3;refGene_tx=XM_019820825.2;refGene_gn=DNAH3;refGene_tx
=XM_023246505.1
E3 27873184 27873298
refGene_gn=DNAH3;refGene_tx=XM_019820825.2;refGene_gn=DNAH3;refGene_tx
=XM_023246505.1
E3 27874060 27874237
refGene_gn=DNAH3;refGene_tx=XM_019820825.2;refGene_gn=DNAH3;refGene_tx
=XM_023246505.1
E3 27876252 27876344
refGene_gn=DNAH3;refGene_tx=XM_019820825.2;refGene_gn=DNAH3;refGene_tx
=XM_023246505.1
E3 27877424 27877644
refGene_gn=DNAH3;refGene_tx=XM_019820825.2;refGene_gn=DNAH3;refGene_tx
=XM_023246505.1
E3 27878438 27878573
refGene_gn=DNAH3;refGene_tx=XM_019820825.2;refGene_gn=DNAH3;refGene_tx
=XM_023246505.1
E3 27880261 27880373
refGene_gn=DNAH3;refGene_tx=XM_019820825.2;refGene_gn=DNAH3;refGene_tx
=XM_023246505.1
E3 27882226 27882468
refGene_gn=DNAH3;refGene_tx=XM_019820825.2;refGene_gn=DNAH3;refGene_tx
=XM_023246505.1
E3 27889154 27889255
refGene_gn=DNAH3;refGene_tx=XM_019820825.2;refGene_gn=DNAH3;refGene_tx
=XM_023246505.1
E3 27890651 27890757
refGene_gn=DNAH3;refGene_tx=XM_019820825.2;refGene_gn=DNAH3;refGene_tx
=XM_023246505.1
E3 27892932 27893062
refGene_gn=DNAH3;refGene_tx=XM_019820825.2;refGene_gn=DNAH3;refGene_tx
=XM_023246505.1
E3 27895516 27895762
refGene_gn=DNAH3;refGene_tx=XM_019820825.2;refGene_gn=DNAH3;refGene_tx
=XM_023246505.1
E3 27913965 27914069
refGene_gn=DNAH3;refGene_tx=XM_019820825.2;refGene_gn=DNAH3;refGene_tx
=XM_023246505.1
E3 27916343 27916638
refGene_gn=DNAH3;refGene_tx=XM_019820825.2;refGene_gn=DNAH3;refGene_tx
=XM_023246505.1
E3 27918655 27918835

refGene_gn=DNAH3;refGene_tx=XM_019820825.2;refGene_gn=DNAH3;refGene_tx=XM_023246505.1
E3 27922423 27922547
refGene_gn=DNAH3;refGene_tx=XM_019820825.2;refGene_gn=DNAH3;refGene_tx=XM_023246505.1
E3 27922637 27922794
refGene_gn=DNAH3;refGene_tx=XM_019820825.2;refGene_gn=DNAH3;refGene_tx=XM_023246505.1
E3 27923024 27923168
refGene_gn=DNAH3;refGene_tx=XM_019820825.2;refGene_gn=DNAH3;refGene_tx=XM_023246505.1
E3 27924119 27924786
refGene_gn=DNAH3;refGene_tx=XM_019820825.2;refGene_gn=DNAH3;refGene_tx=XM_023246505.1
E3 27926030 27926196
refGene_gn=DNAH3;refGene_tx=XM_019820825.2;refGene_gn=DNAH3;refGene_tx=XM_023246505.1
E3 27928156 27928373
refGene_gn=DNAH3;refGene_tx=XM_019820825.2;refGene_gn=DNAH3;refGene_tx=XM_023246505.1
E3 27929920 27930103
refGene_gn=DNAH3;refGene_tx=XM_019820825.2;refGene_gn=DNAH3;refGene_tx=XM_023246505.1
E3 27935670 27935869
refGene_gn=DNAH3;refGene_tx=XM_019820825.2;refGene_gn=DNAH3;refGene_tx=XM_023246505.1
E3 27938907 27941049
refGene_gn=DNAH3;refGene_tx=XM_019820825.2;refGene_gn=DNAH3;refGene_tx=XM_023246505.1
E3 27947898 27948124
refGene_gn=DNAH3;refGene_tx=XM_019820825.2;refGene_gn=DNAH3;refGene_tx=XM_023246505.1
E3 27950604 27950849
refGene_gn=DNAH3;refGene_tx=XM_019820825.2;refGene_gn=DNAH3;refGene_tx=XM_023246505.1
E3 27952429 27952546
refGene_gn=DNAH3;refGene_tx=XM_019820825.2;refGene_gn=DNAH3;refGene_tx=XM_023246505.1
E3 27956660 27956815
refGene_gn=DNAH3;refGene_tx=XM_019820825.2;refGene_gn=DNAH3;refGene_tx=XM_023246505.1
E3 27957643 27957811
refGene_gn=DNAH3;refGene_tx=XM_019820825.2;refGene_gn=DNAH3;refGene_tx=XM_023246505.1
E3 27959463 27959606
refGene_gn=DNAH3;refGene_tx=XM_019820825.2;refGene_gn=DNAH3;refGene_tx=XM_023246505.1
E3 27961936 27962141
refGene_gn=DNAH3;refGene_tx=XM_019820825.2;refGene_gn=DNAH3;refGene_tx=XM_023246505.1

E3 27963003 27963141
refGene_gn=DNAH3;refGene_tx=XM_019820825.2;refGene_gn=DNAH3;refGene_tx=XM_023246505.1
E3 27964532 27964886
refGene_gn=DNAH3;refGene_tx=XM_019820825.2;refGene_gn=DNAH3;refGene_tx=XM_023246505.1
E3 27969255 27969372
refGene_gn=LYRM1;refGene_tx=XM_003998781.4;refGene_gn=LYRM1;refGene_tx=XM_006942244.4;refGene_gn=LYRM1;refGene_tx=XM_006942246.4;refGene_gn=LYRM1;refGene_tx=XM_006942247.4;refGene_gn=LYRM1;refGene_tx=XM_006942248.3;refGene_gn=LYRM1;refGene_tx=XM_006942249.3;refGene_gn=LYRM1;refGene_tx=XM_006942252.4;refGene_gn=LYRM1;refGene_tx=XM_011290601.3;refGene_gn=LYRM1;refGene_tx=XM_019820877.2;refGene_gn=LYRM1;refGene_tx=XM_019820878.2;refGene_gn=LYRM1;refGene_tx=XM_019820879.2;refGene_gn=LYRM1;refGene_tx=XM_019820880.2;refGene_gn=LYRM1;refGene_tx=XM_019820882.2
E3 27970504 27970597
refGene_gn=LYRM1;refGene_tx=XM_003998781.4;refGene_gn=LYRM1;refGene_tx=XM_006942244.4;refGene_gn=LYRM1;refGene_tx=XM_006942246.4;refGene_gn=LYRM1;refGene_tx=XM_006942247.4;refGene_gn=LYRM1;refGene_tx=XM_006942248.3;refGene_gn=LYRM1;refGene_tx=XM_006942249.3;refGene_gn=LYRM1;refGene_tx=XM_006942252.4;refGene_gn=LYRM1;refGene_tx=XM_011290601.3;refGene_gn=LYRM1;refGene_tx=XM_019820877.2;refGene_gn=LYRM1;refGene_tx=XM_019820878.2;refGene_gn=LYRM1;refGene_tx=XM_019820879.2;refGene_gn=LYRM1;refGene_tx=XM_019820880.2;refGene_gn=LYRM1;refGene_tx=XM_019820882.2
E3 27974879 27975049
refGene_gn=LYRM1;refGene_tx=XM_003998781.4;refGene_gn=LYRM1;refGene_tx=XM_006942244.4;refGene_gn=LYRM1;refGene_tx=XM_006942246.4;refGene_gn=LYRM1;refGene_tx=XM_006942247.4;refGene_gn=LYRM1;refGene_tx=XM_006942248.3;refGene_gn=LYRM1;refGene_tx=XM_006942249.3;refGene_gn=LYRM1;refGene_tx=XM_006942252.4;refGene_gn=LYRM1;refGene_tx=XM_011290601.3;refGene_gn=LYRM1;refGene_tx=XM_019820877.2;refGene_gn=LYRM1;refGene_tx=XM_019820878.2;refGene_gn=LYRM1;refGene_tx=XM_019820879.2;refGene_gn=LYRM1;refGene_tx=XM_019820880.2;refGene_gn=LYRM1;refGene_tx=XM_019820882.2
E3 27988503 27988549
refGene_gn=LYRM1;refGene_tx=XM_006942246.4;refGene_gn=LYRM1;refGene_tx=XM_006942247.4;refGene_gn=LYRM1;refGene_tx=XM_019820879.2;refGene_gn=LYRM1;refGene_tx=XM_019820880.2
E3 27989304 27989467
refGene_gn=LYRM1;refGene_tx=XM_006942244.4;refGene_gn=LYRM1;refGene_tx=XM_011290601.3;refGene_gn=LYRM1;refGene_tx=XM_019820878.2
E3 28021126 28021557
refGene_gn=DCUN1D3;refGene_tx=XM_023246919.1;refGene_gn=DCUN1D3;refGene_tx=XM_023246920.1;refGene_gn=DCUN1D3;refGene_tx=XM_023246921.1;refGene_gn=DCUN1D3;refGene_tx=XM_023246923.1;refGene_gn=DCUN1D3;refGene_tx=XM_023246924.1;refGene_gn=DCUN1D3;refGene_tx=XM_023246925.1;refGene_gn=DCUN1D3;refGene_tx=XM_023246926.1;refGene_gn=DCUN1D3;refGene_tx=XM_023246927.1;refGene_gn=DCUN1D3;refGene_tx=XM_023246928.1;refGene_gn=DCUN1D3;refGene_tx=XM_023246929.1;refGene_gn=DCUN1D3;refGene_tx=XM_023246930.1;refGene_gn=DCUN1D3;refGene_tx=XM_023246931.1;refGene_gn=DCUN1D3;refGene_tx=XM_023246932.1;refGene_gn=DCUN1D3;refGene_tx=XM_023246933.1;

refGene_gn=DCUN1D3;refGene_tx=XM_023246934.1;refGene_gn=DCUN1D3;refGene_tx=XM_023246935.1;refGene_gn=DCUN1D3;refGene_tx=XM_023246936.1;refGene_gn=DCUN1D3;refGene_tx=XM_023246937.1;refGene_gn=DCUN1D3;refGene_tx=XM_023246938.1;refGene_gn=DCUN1D3;refGene_tx=XM_023246939.1;refGene_gn=DCUN1D3;refGene_tx=XM_023246940.1
E3 28023311 28023795
refGene_gn=DCUN1D3;refGene_tx=XM_023246919.1;refGene_gn=DCUN1D3;refGene_tx=XM_023246920.1;refGene_gn=DCUN1D3;refGene_tx=XM_023246921.1;refGene_gn=DCUN1D3;refGene_tx=XM_023246923.1;refGene_gn=DCUN1D3;refGene_tx=XM_023246924.1;refGene_gn=DCUN1D3;refGene_tx=XM_023246925.1;refGene_gn=DCUN1D3;refGene_tx=XM_023246926.1;refGene_gn=DCUN1D3;refGene_tx=XM_023246927.1;refGene_gn=DCUN1D3;refGene_tx=XM_023246928.1;refGene_gn=DCUN1D3;refGene_tx=XM_023246929.1;refGene_gn=DCUN1D3;refGene_tx=XM_023246930.1;refGene_gn=DCUN1D3;refGene_tx=XM_023246931.1;refGene_gn=DCUN1D3;refGene_tx=XM_023246932.1;refGene_gn=DCUN1D3;refGene_tx=XM_023246933.1;refGene_gn=DCUN1D3;refGene_tx=XM_023246934.1;refGene_gn=DCUN1D3;refGene_tx=XM_023246935.1;refGene_gn=DCUN1D3;refGene_tx=XM_023246936.1;refGene_gn=DCUN1D3;refGene_tx=XM_023246937.1;refGene_gn=DCUN1D3;refGene_tx=XM_023246938.1;refGene_gn=DCUN1D3;refGene_tx=XM_023246939.1;refGene_gn=DCUN1D3;refGene_tx=XM_023246940.1
E3 28033920 28034002
refGene_gn=REXO5;refGene_tx=XM_003998783.4;refGene_gn=REXO5;refGene_tx=XM_019820843.2;refGene_gn=REXO5;refGene_tx=XM_023246906.1;refGene_gn=REXO5;refGene_tx=XM_023246907.1
E3 28036724 28036843
refGene_gn=REXO5;refGene_tx=XM_003998783.4;refGene_gn=REXO5;refGene_tx=XM_019820843.2;refGene_gn=REXO5;refGene_tx=XM_023246906.1;refGene_gn=REXO5;refGene_tx=XM_023246907.1
E3 28037816 28038001
refGene_gn=REXO5;refGene_tx=XM_003998783.4;refGene_gn=REXO5;refGene_tx=XM_019820843.2;refGene_gn=REXO5;refGene_tx=XM_023246906.1;refGene_gn=REXO5;refGene_tx=XM_023246907.1
E3 28038231 28038448
refGene_gn=REXO5;refGene_tx=XM_003998783.4;refGene_gn=REXO5;refGene_tx=XM_019820843.2;refGene_gn=REXO5;refGene_tx=XM_023246906.1;refGene_gn=REXO5;refGene_tx=XM_023246907.1
E3 28039318 28039411
refGene_gn=REXO5;refGene_tx=XM_003998783.4;refGene_gn=REXO5;refGene_tx=XM_019820843.2;refGene_gn=REXO5;refGene_tx=XM_023246907.1
E3 28042028 28042166
refGene_gn=REXO5;refGene_tx=XM_003998783.4;refGene_gn=REXO5;refGene_tx=XM_019820843.2;refGene_gn=REXO5;refGene_tx=XM_023246906.1;refGene_gn=REXO5;refGene_tx=XM_023246907.1
E3 28042650 28042755
refGene_gn=REXO5;refGene_tx=XM_003998783.4;refGene_gn=REXO5;refGene_tx=XM_019820843.2;refGene_gn=REXO5;refGene_tx=XM_023246906.1
E3 28051628 28051749
refGene_gn=REXO5;refGene_tx=XM_003998783.4;refGene_gn=REXO5;refGene_tx=XM_019820843.2;refGene_gn=REXO5;refGene_tx=XM_023246906.1;refGene_gn=REXO5;refGene_tx=XM_023246907.1

E3 28052473 28052577
refGene_gn=REXO5;refGene_tx=XM_003998783.4;refGene_gn=REXO5;refGene_tx=XM_019820843.2;refGene_gn=REXO5;refGene_tx=XM_023246906.1;refGene_gn=REXO5;refGene_tx=XM_023246907.1
E3 28053958 28054061
refGene_gn=REXO5;refGene_tx=XM_003998783.4;refGene_gn=REXO5;refGene_tx=XM_019820843.2;refGene_gn=REXO5;refGene_tx=XM_023246906.1;refGene_gn=REXO5;refGene_tx=XM_023246907.1
E3 28055062 28055157
refGene_gn=REXO5;refGene_tx=XM_003998783.4;refGene_gn=REXO5;refGene_tx=XM_019820843.2;refGene_gn=REXO5;refGene_tx=XM_023246906.1;refGene_gn=REXO5;refGene_tx=XM_023246907.1
E3 28055307 28055446
refGene_gn=REXO5;refGene_tx=XM_003998783.4;refGene_gn=REXO5;refGene_tx=XM_019820843.2;refGene_gn=REXO5;refGene_tx=XM_023246906.1;refGene_gn=REXO5;refGene_tx=XM_023246907.1
E3 28056538 28056654
refGene_gn=REXO5;refGene_tx=XM_003998783.4;refGene_gn=REXO5;refGene_tx=XM_019820843.2;refGene_gn=REXO5;refGene_tx=XM_023246906.1;refGene_gn=REXO5;refGene_tx=XM_023246907.1
E3 28057891 28057980
refGene_gn=REXO5;refGene_tx=XM_003998783.4;refGene_gn=REXO5;refGene_tx=XM_019820843.2;refGene_gn=REXO5;refGene_tx=XM_023246906.1;refGene_gn=REXO5;refGene_tx=XM_023246907.1
E3 28062150 28062294
refGene_gn=REXO5;refGene_tx=XM_003998783.4;refGene_gn=REXO5;refGene_tx=XM_019820843.2;refGene_gn=REXO5;refGene_tx=XM_023246906.1;refGene_gn=REXO5;refGene_tx=XM_023246907.1
E3 28065493 28065590
refGene_gn=REXO5;refGene_tx=XM_003998783.4;refGene_gn=REXO5;refGene_tx=XM_019820843.2;refGene_gn=REXO5;refGene_tx=XM_023246906.1;refGene_gn=REXO5;refGene_tx=XM_023246907.1
E3 28066874 28067001
refGene_gn=REXO5;refGene_tx=XM_003998783.4;refGene_gn=REXO5;refGene_tx=XM_019820843.2;refGene_gn=REXO5;refGene_tx=XM_023246906.1;refGene_gn=REXO5;refGene_tx=XM_023246907.1
E3 28068296 28068409
refGene_gn=REXO5;refGene_tx=XM_003998783.4;refGene_gn=REXO5;refGene_tx=XM_019820843.2;refGene_gn=REXO5;refGene_tx=XM_023246906.1;refGene_gn=REXO5;refGene_tx=XM_023246907.1
E3 28071402 28071540
refGene_gn=REXO5;refGene_tx=XM_003998783.4;refGene_gn=REXO5;refGene_tx=XM_019820843.2;refGene_gn=REXO5;refGene_tx=XM_023246906.1;refGene_gn=REXO5;refGene_tx=XM_023246907.1
E3 28072078 28072101
refGene_gn=ERI2;refGene_tx=XM_003998786.4;refGene_gn=ERI2;refGene_tx=XM_006942257.3;refGene_gn=ERI2;refGene_tx=XM_006942258.3;refGene_gn=ERI2;refGene_tx=XM_023246908.1;refGene_gn=ERI2;refGene_tx=XM_023246909.1;refGene_gn=ERI2;refGene_tx=XM_023246911.1;refGene_gn=ERI2;refGene_tx=XM_023246912.1

E3	28073383 28073451
refGene_gn=ERI2;refGene_tx=XM_003998786.4;refGene_gn=ERI2;refGene_tx=XM_006942257.3;refGene_gn=ERI2;refGene_tx=XM_006942258.3;refGene_gn=ERI2;refGene_tx=XM_023246908.1;refGene_gn=ERI2;refGene_tx=XM_023246909.1;refGene_gn=ERI2;refGene_tx=XM_023246911.1;refGene_gn=ERI2;refGene_tx=XM_023246912.1
E3	28073536 28073620
refGene_gn=ERI2;refGene_tx=XM_003998786.4;refGene_gn=ERI2;refGene_tx=XM_006942257.3;refGene_gn=ERI2;refGene_tx=XM_006942258.3;refGene_gn=ERI2;refGene_tx=XM_023246908.1;refGene_gn=ERI2;refGene_tx=XM_023246909.1;refGene_gn=ERI2;refGene_tx=XM_023246911.1;refGene_gn=ERI2;refGene_tx=XM_023246912.1
E3	28074827 28074955
refGene_gn=ERI2;refGene_tx=XM_003998786.4;refGene_gn=ERI2;refGene_tx=XM_006942254.4;refGene_gn=ERI2;refGene_tx=XM_006942257.3;refGene_gn=ERI2;refGene_tx=XM_006942258.3;refGene_gn=ERI2;refGene_tx=XM_019820846.2;refGene_gn=ERI2;refGene_tx=XM_023246908.1;refGene_gn=ERI2;refGene_tx=XM_023246909.1;refGene_gn=ERI2;refGene_tx=XM_023246911.1;refGene_gn=ERI2;refGene_tx=XM_023246912.1
E3	28075702 28075859
refGene_gn=ERI2;refGene_tx=XM_003998786.4;refGene_gn=ERI2;refGene_tx=XM_006942254.4;refGene_gn=ERI2;refGene_tx=XM_006942257.3;refGene_gn=ERI2;refGene_tx=XM_006942258.3;refGene_gn=ERI2;refGene_tx=XM_019820846.2;refGene_gn=ERI2;refGene_tx=XM_023246908.1;refGene_gn=ERI2;refGene_tx=XM_023246909.1;refGene_gn=ERI2;refGene_tx=XM_023246911.1;refGene_gn=ERI2;refGene_tx=XM_023246912.1
E3	28076842 28076943
refGene_gn=ERI2;refGene_tx=XM_003998786.4;refGene_gn=ERI2;refGene_tx=XM_006942254.4;refGene_gn=ERI2;refGene_tx=XM_006942257.3;refGene_gn=ERI2;refGene_tx=XM_006942258.3;refGene_gn=ERI2;refGene_tx=XM_019820846.2;refGene_gn=ERI2;refGene_tx=XM_023246908.1;refGene_gn=ERI2;refGene_tx=XM_023246909.1;refGene_gn=ERI2;refGene_tx=XM_023246911.1;refGene_gn=ERI2;refGene_tx=XM_023246912.1
E3	28077213 28077295
refGene_gn=ERI2;refGene_tx=XM_003998786.4;refGene_gn=ERI2;refGene_tx=XM_006942254.4;refGene_gn=ERI2;refGene_tx=XM_006942257.3;refGene_gn=ERI2;refGene_tx=XM_006942258.3;refGene_gn=ERI2;refGene_tx=XM_019820846.2;refGene_gn=ERI2;refGene_tx=XM_023246908.1;refGene_gn=ERI2;refGene_tx=XM_023246909.1;refGene_gn=ERI2;refGene_tx=XM_023246911.1;refGene_gn=ERI2;refGene_tx=XM_023246912.1
E3	28077865 28077954
refGene_gn=ERI2;refGene_tx=XM_003998786.4;refGene_gn=ERI2;refGene_tx=XM_006942254.4;refGene_gn=ERI2;refGene_tx=XM_006942257.3;refGene_gn=ERI2;refGene_tx=XM_006942258.3;refGene_gn=ERI2;refGene_tx=XM_019820846.2;refGene_gn=ERI2;refGene_tx=XM_023246908.1;refGene_gn=ERI2;refGene_tx=XM_023246909.1;refGene_gn=ERI2;refGene_tx=XM_023246911.1;refGene_gn=ERI2;refGene_tx=XM_023246912.1
E3	28078153 28079458
refGene_gn=ACSM3;refGene_tx=XM_019820850.2;refGene_gn=ERI2;refGene_tx=XM_003998786.4;refGene_gn=ERI2;refGene_tx=XM_006942254.4;refGene_gn=ER

I2;refGene_tx=XM_006942257.3;refGene_gn=ERI2;refGene_tx=XM_019820846.2
E3 28080978 28081065 refGene_gn=ACSM3;refGene_tx=XM_023246913.1
E3 28081453 28081573
refGene_gn=ACSM3;refGene_tx=XM_019820850.2;refGene_gn=ACSM3;refGene_tx=XM_023246913.1
E3 28084603 28084703
refGene_gn=ACSM3;refGene_tx=XM_019820850.2;refGene_gn=ACSM3;refGene_tx=XM_023246913.1
E3 28084812 28084940
refGene_gn=ACSM3;refGene_tx=XM_019820850.2;refGene_gn=ACSM3;refGene_tx=XM_023246913.1
E3 28086127 28086229
refGene_gn=ACSM3;refGene_tx=XM_019820850.2;refGene_gn=ACSM3;refGene_tx=XM_023246913.1
E3 28091536 28091617
refGene_gn=ACSM3;refGene_tx=XM_019820850.2;refGene_gn=ACSM3;refGene_tx=XM_023246913.1
E3 28092564 28092688
refGene_gn=ACSM3;refGene_tx=XM_019820850.2;refGene_gn=ACSM3;refGene_tx=XM_023246913.1
E3 28094275 28094355
refGene_gn=ACSM3;refGene_tx=XM_019820850.2;refGene_gn=ACSM3;refGene_tx=XM_023246913.1
E3 28094493 28094502 refGene_gn=ERI2;refGene_tx=XM_006942258.3
E3 28094630 28094787
refGene_gn=ACSM3;refGene_tx=XM_019820850.2;refGene_gn=ACSM3;refGene_tx=XM_023246913.1
E3 28094902 28095188
refGene_gn=ACSM3;refGene_tx=XM_019820850.2;refGene_gn=ACSM3;refGene_tx=XM_023246913.1;refGene_gn=ERI2;refGene_tx=XM_023246908.1;refGene_gn=ERI2;refGene_tx=XM_023246909.1;refGene_gn=ERI2;refGene_tx=XM_023246911.1
E3 28097833 28098041
refGene_gn=ACSM3;refGene_tx=XM_019820850.2;refGene_gn=ACSM3;refGene_tx=XM_023246913.1
E3 28107099 28107219
refGene_gn=ERI2;refGene_tx=XM_023246908.1;refGene_gn=ERI2;refGene_tx=XM_023246909.1
E3 28109302 28109513
refGene_gn=ACSM3;refGene_tx=XM_019820850.2;refGene_gn=ACSM3;refGene_tx=XM_023246913.1
E3 28110124 28110343
refGene_gn=ACSM3;refGene_tx=XM_019820850.2;refGene_gn=ACSM3;refGene_tx=XM_023246913.1
E3 28114241 28114261
refGene_gn=ERI2;refGene_tx=XM_023246908.1;refGene_gn=ERI2;refGene_tx=XM_023246909.1;refGene_gn=ERI2;refGene_tx=XM_023246911.1
E3 28136490 28136721
refGene_gn=THUMPD1;refGene_tx=XM_003998787.5;refGene_gn=THUMPD1;refGene_tx=XM_019820859.2

E3	28138034	28138209
refGene_gn=THUMPD1;refGene_tx=XM_003998787.5;refGene_gn=THUMPD1;refGen
e_tx=XM_019820859.2
E3	28139646	28139895
refGene_gn=THUMPD1;refGene_tx=XM_003998787.5;refGene_gn=THUMPD1;refGen
e_tx=XM_019820859.2
E3	28140441	28140863
refGene_gn=THUMPD1;refGene_tx=XM_003998787.5;refGene_gn=THUMPD1;refGen
e_tx=XM_019820859.2
E3	28145557	28145644
refGene_gn=LOC101083228;refGene_tx=XM_003998788.4
E3	28147242	28147362
refGene_gn=LOC101083228;refGene_tx=XM_003998788.4
E3	28148999	28149099
refGene_gn=LOC101083228;refGene_tx=XM_003998788.4
E3	28149197	28149325
refGene_gn=LOC101083228;refGene_tx=XM_003998788.4
E3	28149790	28149892
refGene_gn=LOC101083228;refGene_tx=XM_003998788.4
E3	28150024	28150105
refGene_gn=LOC101083228;refGene_tx=XM_003998788.4
E3	28150816	28150940
refGene_gn=LOC101083228;refGene_tx=XM_003998788.4
E3	28152380	28152460
refGene_gn=LOC101083228;refGene_tx=XM_003998788.4
E3	28154304	28154461
refGene_gn=LOC101083228;refGene_tx=XM_003998788.4
E3	28155350	28155494
refGene_gn=LOC101083228;refGene_tx=XM_003998788.4
E3	28159780	28159988
refGene_gn=LOC101083228;refGene_tx=XM_003998788.4
E3	28163252	28163463
refGene_gn=LOC101083228;refGene_tx=XM_003998788.4
E3	28165754	28165952
refGene_gn=LOC101083228;refGene_tx=XM_003998788.4
E3	28208718	28208823
refGene_gn=LOC102901082;refGene_tx=XM_023246914.1;refGene_gn=LOC102901
082;refGene_tx=XM_023246915.1;refGene_gn=LOC102901082;refGene_tx=XM_02
3246917.1
E3	28209181	28209244
refGene_gn=LOC102901082;refGene_tx=XM_023246916.1;refGene_gn=LOC102901
082;refGene_tx=XM_023246918.1
E3	28212178	28212310
refGene_gn=LOC102901082;refGene_tx=XM_023246914.1;refGene_gn=LOC102901
082;refGene_tx=XM_023246915.1;refGene_gn=LOC102901082;refGene_tx=XM_02
3246916.1;refGene_gn=LOC102901082;refGene_tx=XM_023246917.1
E3	28215909	28215931
refGene_gn=LOC102901082;refGene_tx=XM_023246914.1;refGene_gn=LOC102901
082;refGene_tx=XM_023246915.1;refGene_gn=LOC102901082;refGene_tx=XM_02
3246916.1;refGene_gn=LOC102901082;refGene_tx=XM_023246917.1;refGene_gn

=LOC102901082;refGene_tx=XM_023246918.1
E3 28216874 28216963
refGene_gn=LOC102901082;refGene_tx=XM_023246914.1;refGene_gn=LOC102901082;refGene_tx=XM_023246915.1;refGene_gn=LOC102901082;refGene_tx=XM_023246916.1;refGene_gn=LOC102901082;refGene_tx=XM_023246917.1;refGene_gn=LOC102901082;refGene_tx=XM_023246918.1
E3 28217199 28217277
refGene_gn=LOC102901082;refGene_tx=XM_023246914.1;refGene_gn=LOC102901082;refGene_tx=XM_023246915.1;refGene_gn=LOC102901082;refGene_tx=XM_023246916.1;refGene_gn=LOC102901082;refGene_tx=XM_023246917.1;refGene_gn=LOC102901082;refGene_tx=XM_023246918.1
E3 28218270 28218343
refGene_gn=LOC102901082;refGene_tx=XM_023246914.1;refGene_gn=LOC102901082;refGene_tx=XM_023246915.1;refGene_gn=LOC102901082;refGene_tx=XM_023246916.1;refGene_gn=LOC102901082;refGene_tx=XM_023246917.1;refGene_gn=LOC102901082;refGene_tx=XM_023246918.1
E3 28218507 28218646
refGene_gn=LOC102901082;refGene_tx=XM_023246914.1;refGene_gn=LOC102901082;refGene_tx=XM_023246915.1;refGene_gn=LOC102901082;refGene_tx=XM_023246916.1;refGene_gn=LOC102901082;refGene_tx=XM_023246917.1;refGene_gn=LOC102901082;refGene_tx=XM_023246918.1
E3 28220359 28220461
refGene_gn=LOC102901082;refGene_tx=XM_023246914.1;refGene_gn=LOC102901082;refGene_tx=XM_023246915.1;refGene_gn=LOC102901082;refGene_tx=XM_023246916.1;refGene_gn=LOC102901082;refGene_tx=XM_023246917.1;refGene_gn=LOC102901082;refGene_tx=XM_023246918.1
E3 28221298 28221334
refGene_gn=LOC102901082;refGene_tx=XM_023246914.1;refGene_gn=LOC102901082;refGene_tx=XM_023246916.1;refGene_gn=LOC102901082;refGene_tx=XM_023246917.1;refGene_gn=LOC102901082;refGene_tx=XM_023246918.1
E3 28221416 28221484
refGene_gn=LOC102901082;refGene_tx=XM_023246914.1;refGene_gn=LOC102901082;refGene_tx=XM_023246915.1;refGene_gn=LOC102901082;refGene_tx=XM_023246916.1;refGene_gn=LOC102901082;refGene_tx=XM_023246917.1;refGene_gn=LOC102901082;refGene_tx=XM_023246918.1
E3 28221606 28221701
refGene_gn=LOC102901082;refGene_tx=XM_023246914.1;refGene_gn=LOC102901082;refGene_tx=XM_023246915.1;refGene_gn=LOC102901082;refGene_tx=XM_023246916.1;refGene_gn=LOC102901082;refGene_tx=XM_023246917.1;refGene_gn=LOC102901082;refGene_tx=XM_023246918.1
E3 28222699 28222782
refGene_gn=LOC102901082;refGene_tx=XM_023246914.1;refGene_gn=LOC102901082;refGene_tx=XM_023246915.1;refGene_gn=LOC102901082;refGene_tx=XM_023246916.1;refGene_gn=LOC102901082;refGene_tx=XM_023246917.1;refGene_gn=LOC102901082;refGene_tx=XM_023246918.1
E3 28224236 28224387
refGene_gn=LOC102901082;refGene_tx=XM_023246914.1;refGene_gn=LOC102901082;refGene_tx=XM_023246915.1;refGene_gn=LOC102901082;refGene_tx=XM_023246916.1;refGene_gn=LOC102901082;refGene_tx=XM_023246917.1;refGene_gn=LOC102901082;refGene_tx=XM_023246918.1

E3 28224988 28225065
refGene_gn=LOC102901082;refGene_tx=XM_023246914.1;refGene_gn=LOC102901082;refGene_tx=XM_023246915.1;refGene_gn=LOC102901082;refGene_tx=XM_023246916.1;refGene_gn=LOC102901082;refGene_tx=XM_023246917.1;refGene_gn=LOC102901082;refGene_tx=XM_023246918.1
E3 28226623 28226762
refGene_gn=LOC102901082;refGene_tx=XM_023246914.1;refGene_gn=LOC102901082;refGene_tx=XM_023246915.1;refGene_gn=LOC102901082;refGene_tx=XM_023246916.1;refGene_gn=LOC102901082;refGene_tx=XM_023246918.1
E3 28228293 28228329
refGene_gn=LOC102901082;refGene_tx=XM_023246914.1;refGene_gn=LOC102901082;refGene_tx=XM_023246915.1;refGene_gn=LOC102901082;refGene_tx=XM_023246916.1;refGene_gn=LOC102901082;refGene_tx=XM_023246917.1;refGene_gn=LOC102901082;refGene_tx=XM_023246918.1
E3 28368856 28368892
refGene_gn=LOC105261315;refGene_tx=XM_023246992.1;refGene_gn=LOC105261315;refGene_tx=XM_023246993.1;refGene_gn=LOC105261315;refGene_tx=XM_023246994.1;refGene_gn=LOC105261315;refGene_tx=XM_023246995.1;refGene_gn=LOC105261315;refGene_tx=XM_023246996.1;refGene_gn=LOC105261315;refGene_tx=XM_023246997.1
E3 28370437 28370575
refGene_gn=LOC105261315;refGene_tx=XM_023246992.1;refGene_gn=LOC105261315;refGene_tx=XM_023246993.1;refGene_gn=LOC105261315;refGene_tx=XM_023246994.1;refGene_gn=LOC105261315;refGene_tx=XM_023246995.1;refGene_gn=LOC105261315;refGene_tx=XM_023246997.1
E3 28372116 28372193
refGene_gn=LOC105261315;refGene_tx=XM_023246992.1;refGene_gn=LOC105261315;refGene_tx=XM_023246993.1;refGene_gn=LOC105261315;refGene_tx=XM_023246994.1;refGene_gn=LOC105261315;refGene_tx=XM_023246995.1;refGene_gn=LOC105261315;refGene_tx=XM_023246996.1;refGene_gn=LOC105261315;refGene_tx=XM_023246997.1
E3 28372794 28372945
refGene_gn=LOC105261315;refGene_tx=XM_023246992.1;refGene_gn=LOC105261315;refGene_tx=XM_023246993.1;refGene_gn=LOC105261315;refGene_tx=XM_023246994.1;refGene_gn=LOC105261315;refGene_tx=XM_023246995.1;refGene_gn=LOC105261315;refGene_tx=XM_023246996.1;refGene_gn=LOC105261315;refGene_tx=XM_023246997.1
E3 28374463 28374543
refGene_gn=LOC105261315;refGene_tx=XM_023246992.1;refGene_gn=LOC105261315;refGene_tx=XM_023246993.1;refGene_gn=LOC105261315;refGene_tx=XM_023246994.1;refGene_gn=LOC105261315;refGene_tx=XM_023246995.1;refGene_gn=LOC105261315;refGene_tx=XM_023246996.1;refGene_gn=LOC105261315;refGene_tx=XM_023246997.1
E3 28375474 28375549
refGene_gn=LOC105261315;refGene_tx=XM_023246992.1;refGene_gn=LOC105261315;refGene_tx=XM_023246993.1;refGene_gn=LOC105261315;refGene_tx=XM_023246995.1;refGene_gn=LOC105261315;refGene_tx=XM_023246996.1;refGene_gn=LOC105261315;refGene_tx=XM_023246997.1
E3 28378644 28378739
refGene_gn=LOC105261315;refGene_tx=XM_023246992.1;refGene_gn=LOC105261

315;refGene_tx=XM_023246993.1;refGene_gn=LOC105261315;refGene_tx=XM_023246994.1;refGene_gn=LOC105261315;refGene_tx=XM_023246995.1;refGene_gn=LOC105261315;refGene_tx=XM_023246996.1;refGene_gn=LOC105261315;refGene_tx=XM_023246997.1
E3 28378866 28378949
refGene_gn=LOC105261315;refGene_tx=XM_023246992.1;refGene_gn=LOC105261315;refGene_tx=XM_023246993.1;refGene_gn=LOC105261315;refGene_tx=XM_023246994.1;refGene_gn=LOC105261315;refGene_tx=XM_023246995.1;refGene_gn=LOC105261315;refGene_tx=XM_023246996.1;refGene_gn=LOC105261315;refGene_tx=XM_023246997.1
E3 28379878 28379980
refGene_gn=LOC105261315;refGene_tx=XM_023246992.1;refGene_gn=LOC105261315;refGene_tx=XM_023246993.1;refGene_gn=LOC105261315;refGene_tx=XM_023246994.1;refGene_gn=LOC105261315;refGene_tx=XM_023246996.1
E3 28381743 28381772
refGene_gn=LOC105261315;refGene_tx=XM_023246993.1
E3 28383752 28383793
refGene_gn=LOC105261315;refGene_tx=XM_023246992.1;refGene_gn=LOC105261315;refGene_tx=XM_023246994.1;refGene_gn=LOC105261315;refGene_tx=XM_023246996.1
E3 28396265 28396352
refGene_gn=LOC101083474;refGene_tx=XM_003998789.2;refGene_gn=LOC101083474;refGene_tx=XM_019820891.2;refGene_gn=LOC101083474;refGene_tx=XM_019820892.1;refGene_gn=LOC101083474;refGene_tx=XM_019820893.1;refGene_gn=LOC101083474;refGene_tx=XM_019820894.1
E3 28397830 28397950
refGene_gn=LOC101083474;refGene_tx=XM_003998789.2;refGene_gn=LOC101083474;refGene_tx=XM_019820891.2;refGene_gn=LOC101083474;refGene_tx=XM_019820892.1;refGene_gn=LOC101083474;refGene_tx=XM_019820893.1;refGene_gn=LOC101083474;refGene_tx=XM_019820894.1
E3 28398825 28398925
refGene_gn=LOC101083474;refGene_tx=XM_003998789.2;refGene_gn=LOC101083474;refGene_tx=XM_019820891.2;refGene_gn=LOC101083474;refGene_tx=XM_019820892.1;refGene_gn=LOC101083474;refGene_tx=XM_019820893.1;refGene_gn=LOC101083474;refGene_tx=XM_019820894.1
E3 28400213 28400341
refGene_gn=LOC101083474;refGene_tx=XM_003998789.2;refGene_gn=LOC101083474;refGene_tx=XM_019820891.2;refGene_gn=LOC101083474;refGene_tx=XM_019820892.1;refGene_gn=LOC101083474;refGene_tx=XM_019820893.1;refGene_gn=LOC101083474;refGene_tx=XM_019820894.1
E3 28405656 28405758
refGene_gn=LOC101083474;refGene_tx=XM_003998789.2;refGene_gn=LOC101083474;refGene_tx=XM_019820891.2;refGene_gn=LOC101083474;refGene_tx=XM_019820892.1;refGene_gn=LOC101083474;refGene_tx=XM_019820893.1;refGene_gn=LOC101083474;refGene_tx=XM_019820894.1
E3 28406308 28406389
refGene_gn=LOC101083474;refGene_tx=XM_003998789.2;refGene_gn=LOC101083474;refGene_tx=XM_019820891.2;refGene_gn=LOC101083474;refGene_tx=XM_019820892.1;refGene_gn=LOC101083474;refGene_tx=XM_019820893.1;refGene_gn=LOC101083474;refGene_tx=XM_019820894.1

E3 28408592 28408716
refGene_gn=LOC101083474;refGene_tx=XM_003998789.2;refGene_gn=LOC101083474;refGene_tx=XM_019820891.2;refGene_gn=LOC101083474;refGene_tx=XM_019820892.1;refGene_gn=LOC101083474;refGene_tx=XM_019820893.1;refGene_gn=LOC101083474;refGene_tx=XM_019820894.1
E3 28416191 28416271
refGene_gn=LOC101083474;refGene_tx=XM_003998789.2;refGene_gn=LOC101083474;refGene_tx=XM_019820891.2;refGene_gn=LOC101083474;refGene_tx=XM_019820892.1;refGene_gn=LOC101083474;refGene_tx=XM_019820893.1;refGene_gn=LOC101083474;refGene_tx=XM_019820894.1
E3 28420045 28420205
refGene_gn=LOC101083474;refGene_tx=XM_003998789.2;refGene_gn=LOC101083474;refGene_tx=XM_019820891.2;refGene_gn=LOC101083474;refGene_tx=XM_019820892.1;refGene_gn=LOC101083474;refGene_tx=XM_019820893.1;refGene_gn=LOC101083474;refGene_tx=XM_019820894.1
E3 28422235 28422376
refGene_gn=LOC101083474;refGene_tx=XM_003998789.2;refGene_gn=LOC101083474;refGene_tx=XM_019820891.2;refGene_gn=LOC101083474;refGene_tx=XM_019820892.1;refGene_gn=LOC101083474;refGene_tx=XM_019820893.1;refGene_gn=LOC101083474;refGene_tx=XM_019820894.1
E3 28424128 28424336
refGene_gn=LOC101083474;refGene_tx=XM_003998789.2;refGene_gn=LOC101083474;refGene_tx=XM_019820891.2;refGene_gn=LOC101083474;refGene_tx=XM_019820892.1;refGene_gn=LOC101083474;refGene_tx=XM_019820893.1;refGene_gn=LOC101083474;refGene_tx=XM_019820894.1
E3 28426471 28426682
refGene_gn=LOC101083474;refGene_tx=XM_003998789.2;refGene_gn=LOC101083474;refGene_tx=XM_019820891.2;refGene_gn=LOC101083474;refGene_tx=XM_019820892.1;refGene_gn=LOC101083474;refGene_tx=XM_019820893.1;refGene_gn=LOC101083474;refGene_tx=XM_019820894.1
E3 28441632 28441824
refGene_gn=LOC101083474;refGene_tx=XM_003998789.2;refGene_gn=LOC101083474;refGene_tx=XM_019820891.2;refGene_gn=LOC101083474;refGene_tx=XM_019820892.1;refGene_gn=LOC101083474;refGene_tx=XM_019820893.1;refGene_gn=LOC101083474;refGene_tx=XM_019820894.1
E3 28481023 28481134
refGene_gn=ACSM5;refGene_tx=XM_003998822.4;refGene_gn=ACSM5;refGene_tx=XM_019820887.2;refGene_gn=ACSM5;refGene_tx=XM_023246991.1
E3 28481527 28481647
refGene_gn=ACSM5;refGene_tx=XM_003998822.4;refGene_gn=ACSM5;refGene_tx=XM_019820887.2;refGene_gn=ACSM5;refGene_tx=XM_023246991.1
E3 28482878 28482950 refGene_gn=ACSM5;refGene_tx=XM_019820889.2
E3 28483889 28483989
refGene_gn=ACSM5;refGene_tx=XM_003998822.4;refGene_gn=ACSM5;refGene_tx=XM_019820887.2;refGene_gn=ACSM5;refGene_tx=XM_019820889.2;refGene_gn=ACSM5;refGene_tx=XM_023246991.1
E3 28484074 28484202
refGene_gn=ACSM5;refGene_tx=XM_003998822.4;refGene_gn=ACSM5;refGene_tx=XM_019820887.2;refGene_gn=ACSM5;refGene_tx=XM_019820889.2;refGene_gn=ACSM5;refGene_tx=XM_023246991.1

E3 28487019 28487121
refGene_gn=ACSM5;refGene_tx=XM_003998822.4;refGene_gn=ACSM5;refGene_tx=XM_019820887.2;refGene_gn=ACSM5;refGene_tx=XM_019820889.2;refGene_gn=ACSM5;refGene_tx=XM_023246991.1
E3 28487262 28487343
refGene_gn=ACSM5;refGene_tx=XM_003998822.4;refGene_gn=ACSM5;refGene_tx=XM_019820887.2;refGene_gn=ACSM5;refGene_tx=XM_019820889.2;refGene_gn=ACSM5;refGene_tx=XM_023246991.1
E3 28487940 28488064
refGene_gn=ACSM5;refGene_tx=XM_003998822.4;refGene_gn=ACSM5;refGene_tx=XM_019820887.2;refGene_gn=ACSM5;refGene_tx=XM_019820889.2;refGene_gn=ACSM5;refGene_tx=XM_023246991.1
E3 28489705 28489785
refGene_gn=ACSM5;refGene_tx=XM_003998822.4;refGene_gn=ACSM5;refGene_tx=XM_019820887.2;refGene_gn=ACSM5;refGene_tx=XM_019820889.2;refGene_gn=ACSM5;refGene_tx=XM_023246991.1
E3 28492126 28492280
refGene_gn=ACSM5;refGene_tx=XM_003998822.4;refGene_gn=ACSM5;refGene_tx=XM_019820887.2;refGene_gn=ACSM5;refGene_tx=XM_019820889.2;refGene_gn=ACSM5;refGene_tx=XM_023246991.1
E3 28494690 28494834
refGene_gn=ACSM5;refGene_tx=XM_003998822.4;refGene_gn=ACSM5;refGene_tx=XM_019820887.2;refGene_gn=ACSM5;refGene_tx=XM_019820889.2;refGene_gn=ACSM5;refGene_tx=XM_023246991.1
E3 28496818 28497026
refGene_gn=ACSM5;refGene_tx=XM_003998822.4;refGene_gn=ACSM5;refGene_tx=XM_019820887.2;refGene_gn=ACSM5;refGene_tx=XM_019820889.2;refGene_gn=ACSM5;refGene_tx=XM_023246991.1
E3 28499572 28499783
refGene_gn=ACSM5;refGene_tx=XM_003998822.4;refGene_gn=ACSM5;refGene_tx=XM_019820887.2;refGene_gn=ACSM5;refGene_tx=XM_019820889.2;refGene_gn=ACSM5;refGene_tx=XM_023246991.1
E3 28502141 28502360
refGene_gn=ACSM5;refGene_tx=XM_003998822.4;refGene_gn=ACSM5;refGene_tx=XM_019820887.2;refGene_gn=ACSM5;refGene_tx=XM_019820889.2;refGene_gn=ACSM5;refGene_tx=XM_023246991.1
E3 28504107 28504125
refGene_gn=ACSM5;refGene_tx=XM_019820887.2;refGene_gn=ACSM5;refGene_tx=XM_019820889.2;refGene_gn=ACSM5;refGene_tx=XM_023246991.1
E3 28516249 28516451 refGene_gn=PDILT;refGene_tx=XM_003998790.4
E3 28528808 28529015 refGene_gn=PDILT;refGene_tx=XM_003998790.4
E3 28534904 28535038 refGene_gn=PDILT;refGene_tx=XM_003998790.4
E3 28535875 28536013 refGene_gn=PDILT;refGene_tx=XM_003998790.4
E3 28538929 28539040 refGene_gn=PDILT;refGene_tx=XM_003998790.4
E3 28539122 28539248 refGene_gn=PDILT;refGene_tx=XM_003998790.4
E3 28541981 28542179 refGene_gn=PDILT;refGene_tx=XM_003998790.4
E3 28545244 28545365 refGene_gn=PDILT;refGene_tx=XM_003998790.4
E3 28547570 28547749 refGene_gn=PDILT;refGene_tx=XM_003998790.4
E3 28549161 28549251 refGene_gn=PDILT;refGene_tx=XM_003998790.4
E3 28550948 28551185 refGene_gn=PDILT;refGene_tx=XM_003998790.4

E3	28558959 28559053
refGene_gn=UMOD;refGene_tx=XM_006942287.2;refGene_gn=UMOD;refGene_tx=XM_019820884.1
E3	28560214 28560988
refGene_gn=UMOD;refGene_tx=XM_006942287.2;refGene_gn=UMOD;refGene_tx=XM_019820884.1
E3	28561095 28561203
refGene_gn=UMOD;refGene_tx=XM_006942287.2;refGene_gn=UMOD;refGene_tx=XM_019820884.1
E3	28562395 28562604
refGene_gn=UMOD;refGene_tx=XM_006942287.2;refGene_gn=UMOD;refGene_tx=XM_019820884.1
E3	28563554 28563703
refGene_gn=UMOD;refGene_tx=XM_006942287.2;refGene_gn=UMOD;refGene_tx=XM_019820884.1
E3	28565582 28565828
refGene_gn=UMOD;refGene_tx=XM_006942287.2;refGene_gn=UMOD;refGene_tx=XM_019820884.1
E3	28566394 28566425 refGene_gn=UMOD;refGene_tx=XM_019820884.1
E3	28568492 28568655 refGene_gn=UMOD;refGene_tx=XM_006942287.2
E3	28569234 28569316 refGene_gn=UMOD;refGene_tx=XM_006942287.2
E3	28570049 28570091 refGene_gn=UMOD;refGene_tx=XM_006942287.2
E3	28571242 28571304 refGene_gn=UMOD;refGene_tx=XM_006942287.2
E3	28577277 28577359 refGene_gn=GP2;refGene_tx=XM_003998792.3
E3	28579691 28580078 refGene_gn=GP2;refGene_tx=XM_003998792.3
E3	28580936 28581047 refGene_gn=GP2;refGene_tx=XM_003998792.3
E3	28583218 28583430 refGene_gn=GP2;refGene_tx=XM_003998792.3
E3	28583928 28584077 refGene_gn=GP2;refGene_tx=XM_003998792.3
E3	28585265 28585511 refGene_gn=GP2;refGene_tx=XM_003998792.3
E3	28586386 28586495 refGene_gn=GP2;refGene_tx=XM_003998792.3
E3	28586496 28586550 refGene_gn=GP2;refGene_tx=XM_003998792.3
E3	28587400 28587485 refGene_gn=GP2;refGene_tx=XM_003998792.3
E3	28588724 28588769 refGene_gn=GP2;refGene_tx=XM_003998792.3
E3	28591434 28591493 refGene_gn=GP2;refGene_tx=XM_003998792.3
E3	28822338 28822462 refGene_gn=GPR139;refGene_tx=XM_003998794.3
E3	28860519 28861454 refGene_gn=GPR139;refGene_tx=XM_003998794.3
E3	29019034 29020040 refGene_gn=GPRC5B;refGene_tx=XM_006942266.3
E3	29028870 29028983 refGene_gn=GPRC5B;refGene_tx=XM_006942266.3
E3	29030335 29030380 refGene_gn=GPRC5B;refGene_tx=XM_006942266.3
E3	29034302 29034337
refGene_gn=IQCK;refGene_tx=XM_006942263.3;refGene_gn=IQCK;refGene_tx=XM_006942264.4;refGene_gn=IQCK;refGene_tx=XM_006942265.3;refGene_gn=IQCK;refGene_tx=XM_019820899.2
E3	29066143 29066170
refGene_gn=IQCK;refGene_tx=XM_019820897.1;refGene_gn=IQCK;refGene_tx=XM_019820898.2
E3	29071988 29072100
refGene_gn=IQCK;refGene_tx=XM_006942263.3;refGene_gn=IQCK;refGene_tx=XM_006942264.4;refGene_gn=IQCK;refGene_tx=XM_006942265.3;refGene_gn=IQCK;refGene_tx=XM_019820897.1;refGene_gn=IQCK;refGene_tx=XM_019820898.2;

refGene_gn=IQCK;refGene_tx=XM_019820899.2
E3	29105033	29105118
refGene_gn=IQCK;refGene_tx=XM_006942263.3;refGene_gn=IQCK;refGene_tx=XM_006942264.4;refGene_gn=IQCK;refGene_tx=XM_006942265.3;refGene_gn=IQCK;refGene_tx=XM_019820897.1
E3	29128617	29128695
refGene_gn=IQCK;refGene_tx=XM_006942263.3;refGene_gn=IQCK;refGene_tx=XM_006942264.4;refGene_gn=IQCK;refGene_tx=XM_006942265.3;refGene_gn=IQCK;refGene_tx=XM_019820897.1;refGene_gn=IQCK;refGene_tx=XM_019820898.2;refGene_gn=IQCK;refGene_tx=XM_019820899.2
E3	29128830	29128883
refGene_gn=IQCK;refGene_tx=XM_006942263.3;refGene_gn=IQCK;refGene_tx=XM_006942264.4;refGene_gn=IQCK;refGene_tx=XM_006942265.3;refGene_gn=IQCK;refGene_tx=XM_019820897.1;refGene_gn=IQCK;refGene_tx=XM_019820898.2;refGene_gn=IQCK;refGene_tx=XM_019820899.2
E3	29148817	29148915
refGene_gn=IQCK;refGene_tx=XM_006942263.3;refGene_gn=IQCK;refGene_tx=XM_006942264.4;refGene_gn=IQCK;refGene_tx=XM_006942265.3;refGene_gn=IQCK;refGene_tx=XM_019820897.1;refGene_gn=IQCK;refGene_tx=XM_019820898.2;refGene_gn=IQCK;refGene_tx=XM_019820899.2
E3	29150375	29150505
refGene_gn=IQCK;refGene_tx=XM_006942263.3;refGene_gn=IQCK;refGene_tx=XM_006942264.4;refGene_gn=IQCK;refGene_tx=XM_006942265.3;refGene_gn=IQCK;refGene_tx=XM_019820897.1;refGene_gn=IQCK;refGene_tx=XM_019820898.2;refGene_gn=IQCK;refGene_tx=XM_019820899.2
E3	29151863	29151928
refGene_gn=IQCK;refGene_tx=XM_006942263.3;refGene_gn=IQCK;refGene_tx=XM_006942264.4;refGene_gn=IQCK;refGene_tx=XM_019820897.1;refGene_gn=IQCK;refGene_tx=XM_019820898.2;refGene_gn=IQCK;refGene_tx=XM_019820899.2
E3	29159851	29160038
refGene_gn=IQCK;refGene_tx=XM_006942263.3;refGene_gn=IQCK;refGene_tx=XM_006942264.4;refGene_gn=IQCK;refGene_tx=XM_006942265.3;refGene_gn=IQCK;refGene_tx=XM_019820897.1;refGene_gn=IQCK;refGene_tx=XM_019820898.2;refGene_gn=IQCK;refGene_tx=XM_019820899.2
E3	29161785	29161837	refGene_gn=KNOP1;refGene_tx=XM_006942267.4
E3	29163170	29164090
refGene_gn=KNOP1;refGene_tx=XM_006942267.4;refGene_gn=KNOP1;refGene_tx=XM_011290596.3;refGene_gn=KNOP1;refGene_tx=XM_011290597.3;refGene_gn=KNOP1;refGene_tx=XM_011290598.3;refGene_gn=KNOP1;refGene_tx=XM_023246636.1
E3	29167680	29167749
refGene_gn=KNOP1;refGene_tx=XM_006942267.4;refGene_gn=KNOP1;refGene_tx=XM_011290596.3;refGene_gn=KNOP1;refGene_tx=XM_011290597.3;refGene_gn=KNOP1;refGene_tx=XM_011290598.3
E3	29174161	29174239
refGene_gn=KNOP1;refGene_tx=XM_006942267.4;refGene_gn=KNOP1;refGene_tx=XM_011290596.3;refGene_gn=KNOP1;refGene_tx=XM_011290597.3;refGene_gn=KNOP1;refGene_tx=XM_011290598.3;refGene_gn=KNOP1;refGene_tx=XM_023246636.1
E3	29176729	29177041

refGene_gn=KNOP1;refGene_tx=XM_006942267.4;refGene_gn=KNOP1;refGene_tx=XM_011290596.3;refGene_gn=KNOP1;refGene_tx=XM_011290597.3;refGene_gn=KNOP1;refGene_tx=XM_011290598.3;refGene_gn=KNOP1;refGene_tx=XM_023246636.1
E3 29193770 29193869
refGene_gn=CE3H16orf62;refGene_tx=XM_011290595.2;refGene_gn=CE3H16orf62;refGene_tx=XM_019820900.1;refGene_gn=CE3H16orf62;refGene_tx=XM_023247154.1;refGene_gn=CE3H16orf62;refGene_tx=XM_023247155.1
E3 29194382 29194529
refGene_gn=CE3H16orf62;refGene_tx=XM_011290595.2;refGene_gn=CE3H16orf62;refGene_tx=XM_019820900.1;refGene_gn=CE3H16orf62;refGene_tx=XM_023247154.1;refGene_gn=CE3H16orf62;refGene_tx=XM_023247155.1
E3 29200379 29200498
refGene_gn=CE3H16orf62;refGene_tx=XM_011290595.2;refGene_gn=CE3H16orf62;refGene_tx=XM_019820900.1;refGene_gn=CE3H16orf62;refGene_tx=XM_023247154.1;refGene_gn=CE3H16orf62;refGene_tx=XM_023247155.1
E3 29206677 29206843
refGene_gn=CE3H16orf62;refGene_tx=XM_011290595.2;refGene_gn=CE3H16orf62;refGene_tx=XM_019820900.1;refGene_gn=CE3H16orf62;refGene_tx=XM_023247154.1;refGene_gn=CE3H16orf62;refGene_tx=XM_023247155.1
E3 29213545 29213685
refGene_gn=CE3H16orf62;refGene_tx=XM_011290595.2;refGene_gn=CE3H16orf62;refGene_tx=XM_019820900.1;refGene_gn=CE3H16orf62;refGene_tx=XM_023247154.1;refGene_gn=CE3H16orf62;refGene_tx=XM_023247155.1
E3 29228619 29228734
refGene_gn=CE3H16orf62;refGene_tx=XM_011290595.2;refGene_gn=CE3H16orf62;refGene_tx=XM_019820900.1;refGene_gn=CE3H16orf62;refGene_tx=XM_023247154.1;refGene_gn=CE3H16orf62;refGene_tx=XM_023247155.1
E3 29229571 29229649
refGene_gn=CE3H16orf62;refGene_tx=XM_011290595.2;refGene_gn=CE3H16orf62;refGene_tx=XM_019820900.1;refGene_gn=CE3H16orf62;refGene_tx=XM_023247154.1;refGene_gn=CE3H16orf62;refGene_tx=XM_023247155.1
E3 29231175 29231274
refGene_gn=CE3H16orf62;refGene_tx=XM_011290595.2;refGene_gn=CE3H16orf62;refGene_tx=XM_019820900.1;refGene_gn=CE3H16orf62;refGene_tx=XM_023247154.1;refGene_gn=CE3H16orf62;refGene_tx=XM_023247155.1
E3 29233728 29233792
refGene_gn=CE3H16orf62;refGene_tx=XM_011290595.2;refGene_gn=CE3H16orf62;refGene_tx=XM_019820900.1;refGene_gn=CE3H16orf62;refGene_tx=XM_023247154.1;refGene_gn=CE3H16orf62;refGene_tx=XM_023247155.1
E3 29235976 29236057
refGene_gn=CE3H16orf62;refGene_tx=XM_011290595.2;refGene_gn=CE3H16orf62;refGene_tx=XM_019820900.1;refGene_gn=CE3H16orf62;refGene_tx=XM_023247154.1;refGene_gn=CE3H16orf62;refGene_tx=XM_023247155.1
E3 29237531 29237617
refGene_gn=CE3H16orf62;refGene_tx=XM_011290595.2;refGene_gn=CE3H16orf62;refGene_tx=XM_019820900.1;refGene_gn=CE3H16orf62;refGene_tx=XM_023247154.1;refGene_gn=CE3H16orf62;refGene_tx=XM_023247155.1
E3 29238728 29238791
refGene_gn=CE3H16orf62;refGene_tx=XM_011290595.2;refGene_gn=CE3H16orf6

2;refGene_tx=XM_019820900.1;refGene_gn=CE3H16orf62;refGene_tx=XM_02324
7154.1;refGene_gn=CE3H16orf62;refGene_tx=XM_023247155.1
E3 29241438 29241519
refGene_gn=CE3H16orf62;refGene_tx=XM_011290595.2;refGene_gn=CE3H16orf6
2;refGene_tx=XM_019820900.1;refGene_gn=CE3H16orf62;refGene_tx=XM_02324
7154.1;refGene_gn=CE3H16orf62;refGene_tx=XM_023247155.1
E3 29245035 29245089
refGene_gn=CE3H16orf62;refGene_tx=XM_011290595.2;refGene_gn=CE3H16orf6
2;refGene_tx=XM_019820900.1;refGene_gn=CE3H16orf62;refGene_tx=XM_02324
7154.1;refGene_gn=CE3H16orf62;refGene_tx=XM_023247155.1
E3 29247728 29247845
refGene_gn=CE3H16orf62;refGene_tx=XM_019820900.1;refGene_gn=CE3H16orf6
2;refGene_tx=XM_023247154.1;refGene_gn=CE3H16orf62;refGene_tx=XM_02324
7155.1
E3 29248377 29248489
refGene_gn=CE3H16orf62;refGene_tx=XM_011290595.2;refGene_gn=CE3H16orf6
2;refGene_tx=XM_019820900.1;refGene_gn=CE3H16orf62;refGene_tx=XM_02324
7154.1;refGene_gn=CE3H16orf62;refGene_tx=XM_023247155.1
E3 29249245 29249292
refGene_gn=CE3H16orf62;refGene_tx=XM_011290595.2;refGene_gn=CE3H16orf6
2;refGene_tx=XM_019820900.1;refGene_gn=CE3H16orf62;refGene_tx=XM_02324
7154.1;refGene_gn=CE3H16orf62;refGene_tx=XM_023247155.1
E3 29258786 29258909
refGene_gn=CE3H16orf62;refGene_tx=XM_011290595.2;refGene_gn=CE3H16orf6
2;refGene_tx=XM_019820900.1;refGene_gn=CE3H16orf62;refGene_tx=XM_02324
7154.1;refGene_gn=CE3H16orf62;refGene_tx=XM_023247155.1
E3 29259401 29259479
refGene_gn=CE3H16orf62;refGene_tx=XM_011290595.2;refGene_gn=CE3H16orf6
2;refGene_tx=XM_019820900.1;refGene_gn=CE3H16orf62;refGene_tx=XM_02324
7154.1;refGene_gn=CE3H16orf62;refGene_tx=XM_023247155.1
E3 29263749 29263843
refGene_gn=CE3H16orf62;refGene_tx=XM_011290595.2;refGene_gn=CE3H16orf6
2;refGene_tx=XM_019820900.1;refGene_gn=CE3H16orf62;refGene_tx=XM_02324
7154.1;refGene_gn=CE3H16orf62;refGene_tx=XM_023247155.1
E3 29265269 29265317
refGene_gn=CE3H16orf62;refGene_tx=XM_011290595.2;refGene_gn=CE3H16orf6
2;refGene_tx=XM_019820900.1;refGene_gn=CE3H16orf62;refGene_tx=XM_02324
7154.1;refGene_gn=CE3H16orf62;refGene_tx=XM_023247155.1
E3 29266016 29266113
refGene_gn=CE3H16orf62;refGene_tx=XM_011290595.2;refGene_gn=CE3H16orf6
2;refGene_tx=XM_019820900.1;refGene_gn=CE3H16orf62;refGene_tx=XM_02324
7154.1;refGene_gn=CE3H16orf62;refGene_tx=XM_023247155.1
E3 29273718 29273778
refGene_gn=CE3H16orf62;refGene_tx=XM_011290595.2;refGene_gn=CE3H16orf6
2;refGene_tx=XM_019820900.1;refGene_gn=CE3H16orf62;refGene_tx=XM_02324
7154.1;refGene_gn=CE3H16orf62;refGene_tx=XM_023247155.1
E3 29280644 29280729
refGene_gn=CE3H16orf62;refGene_tx=XM_011290595.2;refGene_gn=CE3H16orf6
2;refGene_tx=XM_023247154.1;refGene_gn=CE3H16orf62;refGene_tx=XM_02324
7155.1

E3 29287525 29287654
refGene_gn=CE3H16orf62;refGene_tx=XM_011290595.2;refGene_gn=CE3H16orf62;refGene_tx=XM_019820900.1;refGene_gn=CE3H16orf62;refGene_tx=XM_023247154.1;refGene_gn=CE3H16orf62;refGene_tx=XM_023247155.1
E3 29290886 29290960
refGene_gn=CE3H16orf62;refGene_tx=XM_011290595.2;refGene_gn=CE3H16orf62;refGene_tx=XM_023247154.1;refGene_gn=CE3H16orf62;refGene_tx=XM_023247155.1
E3 29293890 29293915
refGene_gn=CE3H16orf62;refGene_tx=XM_011290595.2;refGene_gn=CE3H16orf62;refGene_tx=XM_023247154.1;refGene_gn=CE3H16orf62;refGene_tx=XM_023247155.1
E3 29296225 29296348
refGene_gn=CE3H16orf62;refGene_tx=XM_011290595.2;refGene_gn=CE3H16orf62;refGene_tx=XM_023247154.1
E3 29298169 29298325
refGene_gn=CE3H16orf62;refGene_tx=XM_011290595.2;refGene_gn=CE3H16orf62;refGene_tx=XM_023247154.1
E3 29301056 29301156
refGene_gn=CE3H16orf62;refGene_tx=XM_011290595.2
E3 29311522 29311539
refGene_gn=CE3H16orf62;refGene_tx=XM_011290595.2
E3 29316595 29316671
refGene_gn=CCP110;refGene_tx=XM_003998796.4;refGene_gn=CCP110;refGene_tx=XM_011290592.2;refGene_gn=CCP110;refGene_tx=XM_011290593.2;refGene_gn=CCP110;refGene_tx=XM_011290594.2;refGene_gn=CCP110;refGene_tx=XM_019820901.2;refGene_gn=CCP110;refGene_tx=XM_019820902.1;refGene_gn=CCP110;refGene_tx=XM_019820903.1
E3 29319015 29319101
refGene_gn=CCP110;refGene_tx=XM_003998796.4;refGene_gn=CCP110;refGene_tx=XM_011290592.2;refGene_gn=CCP110;refGene_tx=XM_011290594.2;refGene_gn=CCP110;refGene_tx=XM_019820901.2;refGene_gn=CCP110;refGene_tx=XM_019820902.1
E3 29319594 29319654
refGene_gn=CCP110;refGene_tx=XM_003998796.4;refGene_gn=CCP110;refGene_tx=XM_011290592.2;refGene_gn=CCP110;refGene_tx=XM_011290593.2;refGene_gn=CCP110;refGene_tx=XM_011290594.2;refGene_gn=CCP110;refGene_tx=XM_019820901.2;refGene_gn=CCP110;refGene_tx=XM_019820902.1;refGene_gn=CCP110;refGene_tx=XM_019820903.1
E3 29320346 29320409
refGene_gn=CCP110;refGene_tx=XM_003998796.4;refGene_gn=CCP110;refGene_tx=XM_011290592.2;refGene_gn=CCP110;refGene_tx=XM_011290593.2;refGene_gn=CCP110;refGene_tx=XM_019820901.2;refGene_gn=CCP110;refGene_tx=XM_019820902.1
E3 29321160 29321234
refGene_gn=CCP110;refGene_tx=XM_003998796.4;refGene_gn=CCP110;refGene_tx=XM_011290592.2;refGene_gn=CCP110;refGene_tx=XM_011290593.2;refGene_gn=CCP110;refGene_tx=XM_011290594.2;refGene_gn=CCP110;refGene_tx=XM_019820901.2;refGene_gn=CCP110;refGene_tx=XM_019820902.1;refGene_gn=CCP110;refGene_tx=XM_019820903.1

E3 29323080 29323197
refGene_gn=CCP110;refGene_tx=XM_003998796.4;refGene_gn=CCP110;refGene_tx=XM_011290592.2;refGene_gn=CCP110;refGene_tx=XM_011290593.2;refGene_gn=CCP110;refGene_tx=XM_011290594.2;refGene_gn=CCP110;refGene_tx=XM_019820901.2;refGene_gn=CCP110;refGene_tx=XM_019820902.1;refGene_gn=CCP110;refGene_tx=XM_019820903.1
E3 29323382 29323484
refGene_gn=CCP110;refGene_tx=XM_003998796.4;refGene_gn=CCP110;refGene_tx=XM_011290592.2;refGene_gn=CCP110;refGene_tx=XM_011290593.2;refGene_gn=CCP110;refGene_tx=XM_011290594.2;refGene_gn=CCP110;refGene_tx=XM_019820901.2;refGene_gn=CCP110;refGene_tx=XM_019820902.1;refGene_gn=CCP110;refGene_tx=XM_019820903.1
E3 29323857 29323974
refGene_gn=CCP110;refGene_tx=XM_003998796.4;refGene_gn=CCP110;refGene_tx=XM_011290592.2;refGene_gn=CCP110;refGene_tx=XM_011290593.2;refGene_gn=CCP110;refGene_tx=XM_011290594.2;refGene_gn=CCP110;refGene_tx=XM_019820901.2;refGene_gn=CCP110;refGene_tx=XM_019820902.1;refGene_gn=CCP110;refGene_tx=XM_019820903.1
E3 29324224 29324364
refGene_gn=CCP110;refGene_tx=XM_003998796.4;refGene_gn=CCP110;refGene_tx=XM_011290592.2;refGene_gn=CCP110;refGene_tx=XM_011290593.2;refGene_gn=CCP110;refGene_tx=XM_011290594.2;refGene_gn=CCP110;refGene_tx=XM_019820901.2;refGene_gn=CCP110;refGene_tx=XM_019820902.1;refGene_gn=CCP110;refGene_tx=XM_019820903.1
E3 29325127 29325299
refGene_gn=CCP110;refGene_tx=XM_003998796.4;refGene_gn=CCP110;refGene_tx=XM_011290592.2;refGene_gn=CCP110;refGene_tx=XM_011290593.2;refGene_gn=CCP110;refGene_tx=XM_011290594.2;refGene_gn=CCP110;refGene_tx=XM_019820901.2;refGene_gn=CCP110;refGene_tx=XM_019820902.1;refGene_gn=CCP110;refGene_tx=XM_019820903.1
E3 29325768 29325899
refGene_gn=CCP110;refGene_tx=XM_003998796.4;refGene_gn=CCP110;refGene_tx=XM_011290592.2;refGene_gn=CCP110;refGene_tx=XM_011290593.2;refGene_gn=CCP110;refGene_tx=XM_011290594.2;refGene_gn=CCP110;refGene_tx=XM_019820901.2;refGene_gn=CCP110;refGene_tx=XM_019820902.1;refGene_gn=CCP110;refGene_tx=XM_019820903.1
E3 29327939 29329584
refGene_gn=CCP110;refGene_tx=XM_003998796.4;refGene_gn=CCP110;refGene_tx=XM_011290592.2;refGene_gn=CCP110;refGene_tx=XM_011290593.2;refGene_gn=CCP110;refGene_tx=XM_011290594.2;refGene_gn=CCP110;refGene_tx=XM_019820901.2;refGene_gn=CCP110;refGene_tx=XM_019820902.1;refGene_gn=CCP110;refGene_tx=XM_019820903.1
E3 29332464 29332593
refGene_gn=CCP110;refGene_tx=XM_003998796.4;refGene_gn=CCP110;refGene_tx=XM_011290592.2;refGene_gn=CCP110;refGene_tx=XM_011290593.2;refGene_gn=CCP110;refGene_tx=XM_011290594.2;refGene_gn=CCP110;refGene_tx=XM_019820901.2;refGene_gn=CCP110;refGene_tx=XM_019820902.1;refGene_gn=CCP110;refGene_tx=XM_019820903.1
E3 29335331 29335472
refGene_gn=CCP110;refGene_tx=XM_003998796.4;refGene_gn=CCP110;refGene_

tx=XM_011290592.2;refGene_gn=CCP110;refGene_tx=XM_011290593.2;refGene_gn=CCP110;refGene_tx=XM_011290594.2;refGene_gn=CCP110;refGene_tx=XM_019820901.2;refGene_gn=CCP110;refGene_tx=XM_019820902.1;refGene_gn=CCP110;refGene_tx=XM_019820903.1
E3 29341167 29341428 refGene_gn=GDE1;refGene_tx=XM_003998797.3
E3 29341951 29341967 refGene_gn=GDE1;refGene_tx=XM_023247157.1
E3 29350181 29350357
refGene_gn=GDE1;refGene_tx=XM_003998797.3;refGene_gn=GDE1;refGene_tx=XM_019820904.1;refGene_gn=GDE1;refGene_tx=XM_023247157.1
E3 29353988 29354094
refGene_gn=GDE1;refGene_tx=XM_003998797.3;refGene_gn=GDE1;refGene_tx=XM_019820904.1;refGene_gn=GDE1;refGene_tx=XM_023247157.1
E3 29356962 29357055
refGene_gn=GDE1;refGene_tx=XM_003998797.3;refGene_gn=GDE1;refGene_tx=XM_019820904.1;refGene_gn=GDE1;refGene_tx=XM_023247157.1
E3 29358932 29359144
refGene_gn=GDE1;refGene_tx=XM_003998797.3;refGene_gn=GDE1;refGene_tx=XM_019820904.1;refGene_gn=GDE1;refGene_tx=XM_023247157.1
E3 29360918 29361066
refGene_gn=GDE1;refGene_tx=XM_003998797.3;refGene_gn=GDE1;refGene_tx=XM_019820904.1;refGene_gn=GDE1;refGene_tx=XM_023247157.1
E3 29368695 29368742
refGene_gn=TMC5;refGene_tx=XM_006942270.3;refGene_gn=TMC5;refGene_tx=XM_019820907.2;refGene_gn=TMC5;refGene_tx=XM_023246721.1
E3 29369489 29369532
refGene_gn=TMC5;refGene_tx=XM_006942270.3;refGene_gn=TMC5;refGene_tx=XM_019820907.2;refGene_gn=TMC5;refGene_tx=XM_023246721.1
E3 29371085 29371190
refGene_gn=TMC5;refGene_tx=XM_006942270.3;refGene_gn=TMC5;refGene_tx=XM_019820907.2;refGene_gn=TMC5;refGene_tx=XM_023246721.1
E3 29376061 29376140
refGene_gn=TMC5;refGene_tx=XM_006942270.3;refGene_gn=TMC5;refGene_tx=XM_019820907.2;refGene_gn=TMC5;refGene_tx=XM_023246721.1
E3 29378391 29378565
refGene_gn=TMC5;refGene_tx=XM_006942270.3;refGene_gn=TMC5;refGene_tx=XM_019820907.2;refGene_gn=TMC5;refGene_tx=XM_023246721.1
E3 29380906 29381040
refGene_gn=TMC5;refGene_tx=XM_006942270.3;refGene_gn=TMC5;refGene_tx=XM_019820907.2;refGene_gn=TMC5;refGene_tx=XM_023246721.1
E3 29381227 29381303
refGene_gn=TMC5;refGene_tx=XM_006942270.3;refGene_gn=TMC5;refGene_tx=XM_019820907.2;refGene_gn=TMC5;refGene_tx=XM_023246721.1
E3 29384799 29384895
refGene_gn=TMC5;refGene_tx=XM_006942270.3;refGene_gn=TMC5;refGene_tx=XM_019820907.2;refGene_gn=TMC5;refGene_tx=XM_023246721.1
E3 29386575 29386673
refGene_gn=TMC5;refGene_tx=XM_006942270.3;refGene_gn=TMC5;refGene_tx=XM_019820907.2;refGene_gn=TMC5;refGene_tx=XM_023246721.1
E3 29387432 29387511
refGene_gn=TMC5;refGene_tx=XM_006942270.3;refGene_gn=TMC5;refGene_tx=X

M_019820907.2;refGene_gn=TMC5;refGene_tx=XM_023246721.1
E3 29389879 29390031
refGene_gn=TMC5;refGene_tx=XM_006942270.3;refGene_gn=TMC5;refGene_tx=XM_019820907.2;refGene_gn=TMC5;refGene_tx=XM_023246721.1
E3 29393321 29393477
refGene_gn=TMC5;refGene_tx=XM_006942270.3;refGene_gn=TMC5;refGene_tx=XM_019820907.2;refGene_gn=TMC5;refGene_tx=XM_023246721.1
E3 29394708 29394853
refGene_gn=TMC5;refGene_tx=XM_006942270.3;refGene_gn=TMC5;refGene_tx=XM_019820907.2;refGene_gn=TMC5;refGene_tx=XM_023246721.1
E3 29396582 29396734
refGene_gn=TMC5;refGene_tx=XM_006942270.3;refGene_gn=TMC5;refGene_tx=XM_019820907.2;refGene_gn=TMC5;refGene_tx=XM_023246721.1
E3 29400669 29400918
refGene_gn=TMC5;refGene_tx=XM_006942270.3;refGene_gn=TMC5;refGene_tx=XM_019820907.2;refGene_gn=TMC5;refGene_tx=XM_023246721.1
E3 29401388 29401476
refGene_gn=TMC5;refGene_tx=XM_006942270.3;refGene_gn=TMC5;refGene_tx=XM_019820907.2;refGene_gn=TMC5;refGene_tx=XM_023246721.1
E3 29404319 29404419
refGene_gn=TMC5;refGene_tx=XM_006942270.3;refGene_gn=TMC5;refGene_tx=XM_019820907.2;refGene_gn=TMC5;refGene_tx=XM_023246721.1
E3 29406575 29406855
refGene_gn=TMC5;refGene_tx=XM_006942270.3;refGene_gn=TMC5;refGene_tx=XM_019820907.2
E3 29409610 29409700 refGene_gn=TMC5;refGene_tx=XM_023246721.1
E3 29414565 29414732 refGene_gn=TMC5;refGene_tx=XM_023246721.1
E3 29416964 29417866 refGene_gn=TMC5;refGene_tx=XM_023246721.1
E3 29460816 29460883
refGene_gn=TMC7;refGene_tx=XM_006942271.3;refGene_gn=TMC7;refGene_tx=XM_006942272.3;refGene_gn=TMC7;refGene_tx=XM_006942274.3;refGene_gn=TMC7;refGene_tx=XM_023246456.1
E3 29476760 29477004
refGene_gn=TMC7;refGene_tx=XM_006942271.3;refGene_gn=TMC7;refGene_tx=XM_006942272.3;refGene_gn=TMC7;refGene_tx=XM_006942274.3;refGene_gn=TMC7;refGene_tx=XM_023246456.1
E3 29479830 29479979
refGene_gn=TMC7;refGene_tx=XM_006942271.3;refGene_gn=TMC7;refGene_tx=XM_006942272.3;refGene_gn=TMC7;refGene_tx=XM_006942274.3;refGene_gn=TMC7;refGene_tx=XM_023246456.1
E3 29482139 29482307
refGene_gn=TMC7;refGene_tx=XM_006942271.3;refGene_gn=TMC7;refGene_tx=XM_006942272.3;refGene_gn=TMC7;refGene_tx=XM_006942274.3
E3 29483867 29483950
refGene_gn=TMC7;refGene_tx=XM_006942271.3;refGene_gn=TMC7;refGene_tx=XM_006942272.3;refGene_gn=TMC7;refGene_tx=XM_006942274.3;refGene_gn=TMC7;refGene_tx=XM_023246456.1
E3 29489503 29489649
refGene_gn=TMC7;refGene_tx=XM_006942271.3;refGene_gn=TMC7;refGene_tx=XM_006942272.3;refGene_gn=TMC7;refGene_tx=XM_006942274.3;refGene_gn=TMC

7;refGene_tx=XM_023246456.1
E3 29493290 29493438
refGene_gn=TMC7;refGene_tx=XM_006942271.3;refGene_gn=TMC7;refGene_tx=XM_006942272.3;refGene_gn=TMC7;refGene_tx=XM_006942274.3;refGene_gn=TMC7;refGene_tx=XM_023246456.1
E3 29495612 29495786
refGene_gn=TMC7;refGene_tx=XM_006942271.3;refGene_gn=TMC7;refGene_tx=XM_006942272.3;refGene_gn=TMC7;refGene_tx=XM_006942274.3;refGene_gn=TMC7;refGene_tx=XM_023246456.1
E3 29497424 29497582
refGene_gn=TMC7;refGene_tx=XM_006942271.3;refGene_gn=TMC7;refGene_tx=XM_006942272.3;refGene_gn=TMC7;refGene_tx=XM_006942274.3;refGene_gn=TMC7;refGene_tx=XM_023246456.1
E3 29500040 29500158
refGene_gn=TMC7;refGene_tx=XM_006942271.3;refGene_gn=TMC7;refGene_tx=XM_006942272.3;refGene_gn=TMC7;refGene_tx=XM_006942274.3;refGene_gn=TMC7;refGene_tx=XM_023246456.1
E3 29500588 29500686
refGene_gn=TMC7;refGene_tx=XM_006942271.3;refGene_gn=TMC7;refGene_tx=XM_006942272.3;refGene_gn=TMC7;refGene_tx=XM_006942274.3;refGene_gn=TMC7;refGene_tx=XM_023246456.1
E3 29502952 29503139
refGene_gn=TMC7;refGene_tx=XM_006942271.3;refGene_gn=TMC7;refGene_tx=XM_006942272.3;refGene_gn=TMC7;refGene_tx=XM_006942274.3;refGene_gn=TMC7;refGene_tx=XM_023246456.1
E3 29505378 29505509
refGene_gn=TMC7;refGene_tx=XM_006942271.3;refGene_gn=TMC7;refGene_tx=XM_006942272.3;refGene_gn=TMC7;refGene_tx=XM_023246456.1
E3 29511266 29511422
refGene_gn=TMC7;refGene_tx=XM_006942271.3;refGene_gn=TMC7;refGene_tx=XM_006942274.3;refGene_gn=TMC7;refGene_tx=XM_023246456.1
E3 29514717 29514796
refGene_gn=TMC7;refGene_tx=XM_006942271.3;refGene_gn=TMC7;refGene_tx=XM_006942272.3;refGene_gn=TMC7;refGene_tx=XM_006942274.3;refGene_gn=TMC7;refGene_tx=XM_023246456.1
E3 29516055 29516136
refGene_gn=TMC7;refGene_tx=XM_006942271.3;refGene_gn=TMC7;refGene_tx=XM_006942272.3;refGene_gn=TMC7;refGene_tx=XM_006942274.3;refGene_gn=TMC7;refGene_tx=XM_023246456.1
E3 29522630 29522703
refGene_gn=COQ7;refGene_tx=XM_003998798.5;refGene_gn=COQ7;refGene_tx=XM_019820908.2;refGene_gn=COQ7;refGene_tx=XM_019820909.2
E3 29527166 29527345
refGene_gn=COQ7;refGene_tx=XM_003998798.5;refGene_gn=COQ7;refGene_tx=XM_019820908.2;refGene_gn=COQ7;refGene_tx=XM_019820909.2
E3 29528784 29528899
refGene_gn=COQ7;refGene_tx=XM_003998798.5;refGene_gn=COQ7;refGene_tx=XM_019820908.2;refGene_gn=COQ7;refGene_tx=XM_019820909.2
E3 29530293 29530433
refGene_gn=COQ7;refGene_tx=XM_003998798.5;refGene_gn=COQ7;refGene_tx=X

M_019820908.2;refGene_gn=COQ7;refGene_tx=XM_019820909.2
E3 29531114 29531183
refGene_gn=COQ7;refGene_tx=XM_003998798.5;refGene_gn=COQ7;refGene_tx=XM_019820908.2;refGene_gn=COQ7;refGene_tx=XM_019820909.2
E3 29531598 29531676
refGene_gn=COQ7;refGene_tx=XM_003998798.5;refGene_gn=COQ7;refGene_tx=XM_019820908.2;refGene_gn=COQ7;refGene_tx=XM_019820909.2
E3 29560398 29561997 refGene_gn=ITPRIPL2;refGene_tx=XM_003998828.5
E3 29614505 29614520 refGene_gn=SYT17;refGene_tx=XM_003998799.5
E3 29617900 29618003 refGene_gn=SYT17;refGene_tx=XM_019820910.2
E3 29618170 29618193
refGene_gn=SYT17;refGene_tx=XM_003998799.5;refGene_gn=SYT17;refGene_tx=XM_019820910.2
E3 29618881 29619030
refGene_gn=SYT17;refGene_tx=XM_003998799.5;refGene_gn=SYT17;refGene_tx=XM_019820910.2
E3 29627417 29627566
refGene_gn=SYT17;refGene_tx=XM_003998799.5;refGene_gn=SYT17;refGene_tx=XM_019820910.2
E3 29629945 29630565
refGene_gn=SYT17;refGene_tx=XM_003998799.5;refGene_gn=SYT17;refGene_tx=XM_019820910.2
E3 29663541 29663662
refGene_gn=SYT17;refGene_tx=XM_003998799.5;refGene_gn=SYT17;refGene_tx=XM_019820910.2
E3 29665455 29665611
refGene_gn=SYT17;refGene_tx=XM_003998799.5;refGene_gn=SYT17;refGene_tx=XM_019820910.2
E3 29697320 29697517
refGene_gn=SYT17;refGene_tx=XM_003998799.5;refGene_gn=SYT17;refGene_tx=XM_019820910.2
E3 29705357 29705445 refGene_gn=CLEC19A;refGene_tx=XM_019820911.1
E3 29714453 29714619 refGene_gn=CLEC19A;refGene_tx=XM_019820911.1
E3 29721802 29721896 refGene_gn=CLEC19A;refGene_tx=XM_019820911.1
E3 29725865 29725995 refGene_gn=CLEC19A;refGene_tx=XM_019820911.1
E3 29727277 29727357 refGene_gn=CLEC19A;refGene_tx=XM_019820911.1
E3 29844927 29845016
refGene_gn=SMG1;refGene_tx=XM_023246640.1;refGene_gn=SMG1;refGene_tx=XM_023246641.1
E3 29869446 29869460 refGene_gn=SMG1;refGene_tx=XM_023246642.1
E3 29872529 29872693
refGene_gn=SMG1;refGene_tx=XM_023246640.1;refGene_gn=SMG1;refGene_tx=XM_023246641.1;refGene_gn=SMG1;refGene_tx=XM_023246642.1
E3 29874364 29874520
refGene_gn=SMG1;refGene_tx=XM_023246640.1;refGene_gn=SMG1;refGene_tx=XM_023246641.1;refGene_gn=SMG1;refGene_tx=XM_023246642.1
E3 29876892 29877029
refGene_gn=SMG1;refGene_tx=XM_023246640.1;refGene_gn=SMG1;refGene_tx=XM_023246641.1;refGene_gn=SMG1;refGene_tx=XM_023246642.1
E3 29877972 29878031

refGene_gn=SMG1;refGene_tx=XM_023246640.1;refGene_gn=SMG1;refGene_tx=XM_023246641.1;refGene_gn=SMG1;refGene_tx=XM_023246642.1
E3 29879045 29879259
refGene_gn=SMG1;refGene_tx=XM_023246640.1;refGene_gn=SMG1;refGene_tx=XM_023246641.1;refGene_gn=SMG1;refGene_tx=XM_023246642.1
E3 29882125 29882251
refGene_gn=SMG1;refGene_tx=XM_023246640.1;refGene_gn=SMG1;refGene_tx=XM_023246641.1;refGene_gn=SMG1;refGene_tx=XM_023246642.1
E3 29882663 29882736
refGene_gn=SMG1;refGene_tx=XM_023246640.1;refGene_gn=SMG1;refGene_tx=XM_023246641.1;refGene_gn=SMG1;refGene_tx=XM_023246642.1
E3 29883642 29883740
refGene_gn=SMG1;refGene_tx=XM_023246640.1;refGene_gn=SMG1;refGene_tx=XM_023246641.1;refGene_gn=SMG1;refGene_tx=XM_023246642.1
E3 29885976 29886150
refGene_gn=SMG1;refGene_tx=XM_023246640.1;refGene_gn=SMG1;refGene_tx=XM_023246641.1;refGene_gn=SMG1;refGene_tx=XM_023246642.1
E3 29887848 29888073
refGene_gn=SMG1;refGene_tx=XM_023246640.1;refGene_gn=SMG1;refGene_tx=XM_023246641.1;refGene_gn=SMG1;refGene_tx=XM_023246642.1
E3 29889259 29889361
refGene_gn=SMG1;refGene_tx=XM_023246640.1;refGene_gn=SMG1;refGene_tx=XM_023246641.1;refGene_gn=SMG1;refGene_tx=XM_023246642.1
E3 29890213 29890483
refGene_gn=SMG1;refGene_tx=XM_023246640.1;refGene_gn=SMG1;refGene_tx=XM_023246641.1;refGene_gn=SMG1;refGene_tx=XM_023246642.1
E3 29892056 29892187
refGene_gn=SMG1;refGene_tx=XM_023246640.1;refGene_gn=SMG1;refGene_tx=XM_023246641.1;refGene_gn=SMG1;refGene_tx=XM_023246642.1
E3 29892325 29892487
refGene_gn=SMG1;refGene_tx=XM_023246640.1;refGene_gn=SMG1;refGene_tx=XM_023246641.1;refGene_gn=SMG1;refGene_tx=XM_023246642.1
E3 29892954 29893073
refGene_gn=SMG1;refGene_tx=XM_023246640.1;refGene_gn=SMG1;refGene_tx=XM_023246641.1;refGene_gn=SMG1;refGene_tx=XM_023246642.1
E3 29893798 29893886
refGene_gn=SMG1;refGene_tx=XM_023246640.1;refGene_gn=SMG1;refGene_tx=XM_023246641.1;refGene_gn=SMG1;refGene_tx=XM_023246642.1
E3 29893979 29894081
refGene_gn=SMG1;refGene_tx=XM_023246640.1;refGene_gn=SMG1;refGene_tx=XM_023246641.1;refGene_gn=SMG1;refGene_tx=XM_023246642.1
E3 29894557 29894698
refGene_gn=SMG1;refGene_tx=XM_023246640.1;refGene_gn=SMG1;refGene_tx=XM_023246641.1;refGene_gn=SMG1;refGene_tx=XM_023246642.1
E3 29895273 29895473
refGene_gn=SMG1;refGene_tx=XM_023246640.1;refGene_gn=SMG1;refGene_tx=XM_023246641.1;refGene_gn=SMG1;refGene_tx=XM_023246642.1
E3 29896107 29896304
refGene_gn=SMG1;refGene_tx=XM_023246640.1;refGene_gn=SMG1;refGene_tx=XM_023246641.1;refGene_gn=SMG1;refGene_tx=XM_023246642.1

E3 29896459 29896621
refGene_gn=SMG1;refGene_tx=XM_023246640.1;refGene_gn=SMG1;refGene_tx=XM_023246641.1;refGene_gn=SMG1;refGene_tx=XM_023246642.1
E3 29897366 29897521
refGene_gn=SMG1;refGene_tx=XM_023246640.1;refGene_gn=SMG1;refGene_tx=XM_023246641.1;refGene_gn=SMG1;refGene_tx=XM_023246642.1
E3 29899056 29899199
refGene_gn=SMG1;refGene_tx=XM_023246640.1;refGene_gn=SMG1;refGene_tx=XM_023246641.1;refGene_gn=SMG1;refGene_tx=XM_023246642.1
E3 29899344 29899546
refGene_gn=SMG1;refGene_tx=XM_023246640.1;refGene_gn=SMG1;refGene_tx=XM_023246641.1;refGene_gn=SMG1;refGene_tx=XM_023246642.1
E3 29902711 29902821
refGene_gn=SMG1;refGene_tx=XM_023246640.1;refGene_gn=SMG1;refGene_tx=XM_023246641.1;refGene_gn=SMG1;refGene_tx=XM_023246642.1
E3 29903508 29903656
refGene_gn=SMG1;refGene_tx=XM_023246640.1;refGene_gn=SMG1;refGene_tx=XM_023246641.1;refGene_gn=SMG1;refGene_tx=XM_023246642.1
E3 29904029 29904189
refGene_gn=SMG1;refGene_tx=XM_023246640.1;refGene_gn=SMG1;refGene_tx=XM_023246641.1;refGene_gn=SMG1;refGene_tx=XM_023246642.1
E3 29905287 29905408
refGene_gn=SMG1;refGene_tx=XM_023246640.1;refGene_gn=SMG1;refGene_tx=XM_023246641.1;refGene_gn=SMG1;refGene_tx=XM_023246642.1
E3 29908639 29908888
refGene_gn=SMG1;refGene_tx=XM_023246640.1;refGene_gn=SMG1;refGene_tx=XM_023246641.1;refGene_gn=SMG1;refGene_tx=XM_023246642.1
E3 29909203 29909488
refGene_gn=SMG1;refGene_tx=XM_023246640.1;refGene_gn=SMG1;refGene_tx=XM_023246641.1;refGene_gn=SMG1;refGene_tx=XM_023246642.1
E3 29911305 29911450
refGene_gn=SMG1;refGene_tx=XM_023246640.1;refGene_gn=SMG1;refGene_tx=XM_023246641.1;refGene_gn=SMG1;refGene_tx=XM_023246642.1
E3 29911582 29911721
refGene_gn=SMG1;refGene_tx=XM_023246640.1;refGene_gn=SMG1;refGene_tx=XM_023246641.1;refGene_gn=SMG1;refGene_tx=XM_023246642.1
E3 29912677 29912908
refGene_gn=SMG1;refGene_tx=XM_023246640.1;refGene_gn=SMG1;refGene_tx=XM_023246641.1;refGene_gn=SMG1;refGene_tx=XM_023246642.1
E3 29913010 29913188
refGene_gn=SMG1;refGene_tx=XM_023246640.1;refGene_gn=SMG1;refGene_tx=XM_023246641.1;refGene_gn=SMG1;refGene_tx=XM_023246642.1
E3 29913702 29913864
refGene_gn=SMG1;refGene_tx=XM_023246640.1;refGene_gn=SMG1;refGene_tx=XM_023246641.1;refGene_gn=SMG1;refGene_tx=XM_023246642.1
E3 29914404 29914622
refGene_gn=SMG1;refGene_tx=XM_023246640.1;refGene_gn=SMG1;refGene_tx=XM_023246641.1;refGene_gn=SMG1;refGene_tx=XM_023246642.1
E3 29914951 29915106
refGene_gn=SMG1;refGene_tx=XM_023246640.1;refGene_gn=SMG1;refGene_tx=X

M_023246641.1;refGene_gn=SMG1;refGene_tx=XM_023246642.1
E3 29916530 29916753
refGene_gn=SMG1;refGene_tx=XM_023246640.1;refGene_gn=SMG1;refGene_tx=XM_023246641.1;refGene_gn=SMG1;refGene_tx=XM_023246642.1
E3 29918650 29918897
refGene_gn=SMG1;refGene_tx=XM_023246640.1;refGene_gn=SMG1;refGene_tx=XM_023246641.1;refGene_gn=SMG1;refGene_tx=XM_023246642.1
E3 29919300 29919530
refGene_gn=SMG1;refGene_tx=XM_023246640.1;refGene_gn=SMG1;refGene_tx=XM_023246641.1;refGene_gn=SMG1;refGene_tx=XM_023246642.1
E3 29920853 29921102
refGene_gn=SMG1;refGene_tx=XM_023246640.1;refGene_gn=SMG1;refGene_tx=XM_023246641.1;refGene_gn=SMG1;refGene_tx=XM_023246642.1
E3 29921602 29921741
refGene_gn=SMG1;refGene_tx=XM_023246640.1;refGene_gn=SMG1;refGene_tx=XM_023246641.1;refGene_gn=SMG1;refGene_tx=XM_023246642.1
E3 29921828 29921938
refGene_gn=SMG1;refGene_tx=XM_023246640.1;refGene_gn=SMG1;refGene_tx=XM_023246641.1;refGene_gn=SMG1;refGene_tx=XM_023246642.1
E3 29922058 29922277
refGene_gn=SMG1;refGene_tx=XM_023246640.1;refGene_gn=SMG1;refGene_tx=XM_023246641.1;refGene_gn=SMG1;refGene_tx=XM_023246642.1
E3 29923087 29923278
refGene_gn=SMG1;refGene_tx=XM_023246640.1;refGene_gn=SMG1;refGene_tx=XM_023246641.1;refGene_gn=SMG1;refGene_tx=XM_023246642.1
E3 29924475 29924648
refGene_gn=SMG1;refGene_tx=XM_023246640.1;refGene_gn=SMG1;refGene_tx=XM_023246641.1;refGene_gn=SMG1;refGene_tx=XM_023246642.1
E3 29924796 29925076
refGene_gn=SMG1;refGene_tx=XM_023246640.1;refGene_gn=SMG1;refGene_tx=XM_023246641.1;refGene_gn=SMG1;refGene_tx=XM_023246642.1
E3 29925541 29925814
refGene_gn=SMG1;refGene_tx=XM_023246640.1;refGene_gn=SMG1;refGene_tx=XM_023246641.1;refGene_gn=SMG1;refGene_tx=XM_023246642.1
E3 29926044 29926279
refGene_gn=SMG1;refGene_tx=XM_023246640.1;refGene_gn=SMG1;refGene_tx=XM_023246641.1;refGene_gn=SMG1;refGene_tx=XM_023246642.1
E3 29926543 29926770
refGene_gn=SMG1;refGene_tx=XM_023246640.1;refGene_gn=SMG1;refGene_tx=XM_023246641.1;refGene_gn=SMG1;refGene_tx=XM_023246642.1
E3 29929618 29929769
refGene_gn=SMG1;refGene_tx=XM_023246640.1;refGene_gn=SMG1;refGene_tx=XM_023246641.1;refGene_gn=SMG1;refGene_tx=XM_023246642.1
E3 29929867 29930057
refGene_gn=SMG1;refGene_tx=XM_023246640.1;refGene_gn=SMG1;refGene_tx=XM_023246641.1;refGene_gn=SMG1;refGene_tx=XM_023246642.1
E3 29930433 29930903
refGene_gn=SMG1;refGene_tx=XM_023246640.1;refGene_gn=SMG1;refGene_tx=XM_023246641.1;refGene_gn=SMG1;refGene_tx=XM_023246642.1
E3 29933514 29933652

refGene_gn=SMG1;refGene_tx=XM_023246640.1;refGene_gn=SMG1;refGene_tx=XM_023246641.1;refGene_gn=SMG1;refGene_tx=XM_023246642.1
E3 29935286 29935439
refGene_gn=SMG1;refGene_tx=XM_023246640.1;refGene_gn=SMG1;refGene_tx=XM_023246641.1;refGene_gn=SMG1;refGene_tx=XM_023246642.1
E3 29936412 29936592
refGene_gn=SMG1;refGene_tx=XM_023246640.1;refGene_gn=SMG1;refGene_tx=XM_023246641.1;refGene_gn=SMG1;refGene_tx=XM_023246642.1
E3 29937491 29937719
refGene_gn=SMG1;refGene_tx=XM_023246640.1;refGene_gn=SMG1;refGene_tx=XM_023246641.1;refGene_gn=SMG1;refGene_tx=XM_023246642.1
E3 29938371 29938583
refGene_gn=SMG1;refGene_tx=XM_023246640.1;refGene_gn=SMG1;refGene_tx=XM_023246641.1;refGene_gn=SMG1;refGene_tx=XM_023246642.1
E3 29938743 29938850
refGene_gn=SMG1;refGene_tx=XM_023246640.1;refGene_gn=SMG1;refGene_tx=XM_023246641.1;refGene_gn=SMG1;refGene_tx=XM_023246642.1
E3 29939924 29940104
refGene_gn=SMG1;refGene_tx=XM_023246640.1;refGene_gn=SMG1;refGene_tx=XM_023246641.1;refGene_gn=SMG1;refGene_tx=XM_023246642.1
E3 29940183 29940290
refGene_gn=SMG1;refGene_tx=XM_023246640.1;refGene_gn=SMG1;refGene_tx=XM_023246641.1;refGene_gn=SMG1;refGene_tx=XM_023246642.1
E3 29944513 29944591
refGene_gn=SMG1;refGene_tx=XM_023246640.1;refGene_gn=SMG1;refGene_tx=XM_023246641.1;refGene_gn=SMG1;refGene_tx=XM_023246642.1
E3 29951540 29951576 refGene_gn=ARL6IP1;refGene_tx=XM_003998830.5
E3 29953963 29954097 refGene_gn=ARL6IP1;refGene_tx=XM_003998830.5
E3 29954571 29954691 refGene_gn=ARL6IP1;refGene_tx=XM_003998830.5
E3 29956996 29957114 refGene_gn=ARL6IP1;refGene_tx=XM_003998830.5
E3 29957738 29957823 refGene_gn=ARL6IP1;refGene_tx=XM_003998830.5
E3 29958217 29958336 refGene_gn=ARL6IP1;refGene_tx=XM_003998830.5
E3 29966775 29966908 refGene_gn=RPS15A;refGene_tx=XM_003998831.3
E3 29967495 29967575 refGene_gn=RPS15A;refGene_tx=XM_003998831.3
E3 29970501 29970587 refGene_gn=RPS15A;refGene_tx=XM_003998831.3
E3 29971736 29971830 refGene_gn=RPS15A;refGene_tx=XM_003998831.3
E3 30406634 30406957 refGene_gn=XYLT1;refGene_tx=XM_023246430.1
E3 30413175 30413509 refGene_gn=XYLT1;refGene_tx=XM_023246430.1
E3 30419334 30419530 refGene_gn=XYLT1;refGene_tx=XM_023246430.1
E3 30426019 30426194 refGene_gn=XYLT1;refGene_tx=XM_023246430.1
E3 30426196 30426281 refGene_gn=XYLT1;refGene_tx=XM_023246430.1
E3 30431155 30431332 refGene_gn=XYLT1;refGene_tx=XM_023246430.1
E3 30434390 30434607 refGene_gn=XYLT1;refGene_tx=XM_023246430.1
E3 30449084 30449165 refGene_gn=XYLT1;refGene_tx=XM_023246430.1
E3 30483451 30483654 refGene_gn=XYLT1;refGene_tx=XM_023246430.1
E3 30484660 30484833 refGene_gn=XYLT1;refGene_tx=XM_023246430.1
E3 30542622 30543130 refGene_gn=XYLT1;refGene_tx=XM_023246430.1
E3 30623068 30623107 refGene_gn=XYLT1;refGene_tx=XM_023246430.1
E3 30714383 30714731 refGene_gn=XYLT1;refGene_tx=XM_023246430.1
E3 31416574 31416706

refGene_gn=LOC101097198;refGene_tx=XM_023247123.1;refGene_gn=LOC101097198;refGene_tx=XM_023247124.1
E3 31419627 31419720
refGene_gn=LOC101097198;refGene_tx=XM_023247123.1
E3 31423990 31424110
refGene_gn=LOC101097198;refGene_tx=XM_023247123.1;refGene_gn=LOC101097198;refGene_tx=XM_023247124.1
E3 31425785 31425887
refGene_gn=LOC101097198;refGene_tx=XM_023247123.1;refGene_gn=LOC101097198;refGene_tx=XM_023247124.1
E3 31427541 31427652
refGene_gn=LOC101097198;refGene_tx=XM_023247123.1;refGene_gn=LOC101097198;refGene_tx=XM_023247124.1
E3 31428793 31428877
refGene_gn=LOC101097198;refGene_tx=XM_023247123.1;refGene_gn=LOC101097198;refGene_tx=XM_023247124.1
E3 31432716 31432858
refGene_gn=LOC101097198;refGene_tx=XM_023247123.1;refGene_gn=LOC101097198;refGene_tx=XM_023247124.1
E3 31433817 31433945
refGene_gn=LOC101097198;refGene_tx=XM_023247123.1;refGene_gn=LOC101097198;refGene_tx=XM_023247124.1
E3 31434823 31434937
refGene_gn=LOC101097198;refGene_tx=XM_023247123.1;refGene_gn=LOC101097198;refGene_tx=XM_023247124.1
E3 31436620 31436747
refGene_gn=LOC101097198;refGene_tx=XM_023247123.1;refGene_gn=LOC101097198;refGene_tx=XM_023247124.1
E3 31436860 31437020
refGene_gn=LOC101097198;refGene_tx=XM_023247123.1;refGene_gn=LOC101097198;refGene_tx=XM_023247124.1
E3 31438134 31438217
refGene_gn=LOC101097198;refGene_tx=XM_023247123.1;refGene_gn=LOC101097198;refGene_tx=XM_023247124.1
E3 31438307 31438526
refGene_gn=LOC101097198;refGene_tx=XM_023247123.1;refGene_gn=LOC101097198;refGene_tx=XM_023247124.1
E3 31440960 31441056
refGene_gn=LOC101097198;refGene_tx=XM_023247123.1;refGene_gn=LOC101097198;refGene_tx=XM_023247124.1
E3 31441643 31441707
refGene_gn=LOC101097198;refGene_tx=XM_023247123.1;refGene_gn=LOC101097198;refGene_tx=XM_023247124.1
E3 31443644 31443732
refGene_gn=LOC101097198;refGene_tx=XM_023247123.1;refGene_gn=LOC101097198;refGene_tx=XM_023247124.1
E3 31445445 31445582
refGene_gn=LOC101097198;refGene_tx=XM_023247123.1;refGene_gn=LOC101097198;refGene_tx=XM_023247124.1
E3 31446590 31446722

refGene_gn=LOC101097198;refGene_tx=XM_023247123.1;refGene_gn=LOC101097
198;refGene_tx=XM_023247124.1
E3 31447947 31448089
refGene_gn=LOC101097198;refGene_tx=XM_023247123.1;refGene_gn=LOC101097
198;refGene_tx=XM_023247124.1
E3 31449722 31449897
refGene_gn=LOC101097198;refGene_tx=XM_023247123.1;refGene_gn=LOC101097
198;refGene_tx=XM_023247124.1
E3 31454479 31454630
refGene_gn=LOC101097198;refGene_tx=XM_023247123.1;refGene_gn=LOC101097
198;refGene_tx=XM_023247124.1
E3 31454819 31454925
refGene_gn=LOC101097198;refGene_tx=XM_023247123.1;refGene_gn=LOC101097
198;refGene_tx=XM_023247124.1
E3 31457496 31457586
refGene_gn=LOC101097198;refGene_tx=XM_023247123.1;refGene_gn=LOC101097
198;refGene_tx=XM_023247124.1
E3 31457921 31458059
refGene_gn=LOC101097198;refGene_tx=XM_023247123.1;refGene_gn=LOC101097
198;refGene_tx=XM_023247124.1
E3 31458616 31458769
refGene_gn=LOC101097198;refGene_tx=XM_023247123.1;refGene_gn=LOC101097
198;refGene_tx=XM_023247124.1
E3 31461170 31461243
refGene_gn=LOC101097198;refGene_tx=XM_023247123.1;refGene_gn=LOC101097
198;refGene_tx=XM_023247124.1
E3 31463061 31463168
refGene_gn=LOC101097198;refGene_tx=XM_023247123.1;refGene_gn=LOC101097
198;refGene_tx=XM_023247124.1
E3 31464136 31464237
refGene_gn=LOC101097198;refGene_tx=XM_023247123.1;refGene_gn=LOC101097
198;refGene_tx=XM_023247124.1
E3 31467427 31467473
refGene_gn=LOC101097198;refGene_tx=XM_023247123.1;refGene_gn=LOC101097
198;refGene_tx=XM_023247124.1
E3 31470967 31471057
refGene_gn=LOC101097198;refGene_tx=XM_023247123.1;refGene_gn=LOC101097
198;refGene_tx=XM_023247124.1
E3 31473081 31473246
refGene_gn=LOC101097198;refGene_tx=XM_023247123.1;refGene_gn=LOC101097
198;refGene_tx=XM_023247124.1
E3 31475929 31475965 refGene_gn=ABCC6;refGene_tx=XM_011290585.3
E3 31477456 31477639 refGene_gn=ABCC6;refGene_tx=XM_011290585.3
E3 31480167 31480293 refGene_gn=ABCC6;refGene_tx=XM_011290585.3
E3 31480438 31480567 refGene_gn=ABCC6;refGene_tx=XM_011290585.3
E3 31485814 31485940 refGene_gn=ABCC6;refGene_tx=XM_011290585.3
E3 31487145 31487207 refGene_gn=ABCC6;refGene_tx=XM_011290585.3
E3 31490232 31490364 refGene_gn=ABCC6;refGene_tx=XM_011290585.3
E3 31494774 31494969 refGene_gn=ABCC6;refGene_tx=XM_011290585.3
E3 31496623 31496801 refGene_gn=ABCC6;refGene_tx=XM_011290585.3

E3	31499255	31499417	refGene_gn=ABCC6;refGene_tx=XM_011290585.3
E3	31500908	31501001	refGene_gn=ABCC6;refGene_tx=XM_011290585.3
E3	31502441	31502645	refGene_gn=ABCC6;refGene_tx=XM_011290585.3
E3	31503880	31504027	refGene_gn=ABCC6;refGene_tx=XM_011290585.3
E3	31504937	31505025	refGene_gn=ABCC6;refGene_tx=XM_011290585.3
E3	31505411	31505487	refGene_gn=ABCC6;refGene_tx=XM_011290585.3
E3	31507272	31507399	refGene_gn=ABCC6;refGene_tx=XM_011290585.3
E3	31507587	31507764	refGene_gn=ABCC6;refGene_tx=XM_011290585.3
E3	31509939	31510107	refGene_gn=ABCC6;refGene_tx=XM_011290585.3
E3	31510535	31510710	refGene_gn=ABCC6;refGene_tx=XM_011290585.3
E3	31512044	31512120	refGene_gn=ABCC6;refGene_tx=XM_011290585.3
E3	31513553	31513683	refGene_gn=ABCC6;refGene_tx=XM_011290585.3
E3	31515948	31516156	refGene_gn=ABCC6;refGene_tx=XM_011290585.3
E3	31517101	31517412	refGene_gn=ABCC6;refGene_tx=XM_011290585.3
E3	31519447	31519647	refGene_gn=ABCC6;refGene_tx=XM_011290585.3
E3	31521940	31522067	refGene_gn=ABCC6;refGene_tx=XM_011290585.3
E3	31522776	31522878	refGene_gn=ABCC6;refGene_tx=XM_011290585.3
E3	31523678	31523825	refGene_gn=ABCC6;refGene_tx=XM_011290585.3
E3	31525640	31525799	refGene_gn=ABCC6;refGene_tx=XM_011290585.3
E3	31525878	31526045	refGene_gn=ABCC6;refGene_tx=XM_011290585.3
E3	31528871	31529066	refGene_gn=ABCC6;refGene_tx=XM_011290585.3
E3	31529383	31529492	refGene_gn=ABCC6;refGene_tx=XM_011290585.3
E3	31536973	31537082
refGene_gn=ABCC1;refGene_tx=XM_023246871.1;refGene_gn=ABCC1;refGene_tx=XM_023246872.1
E3	31537607	31537802
refGene_gn=ABCC1;refGene_tx=XM_023246871.1;refGene_gn=ABCC1;refGene_tx=XM_023246872.1
E3	31538874	31539041
refGene_gn=ABCC1;refGene_tx=XM_023246871.1;refGene_gn=ABCC1;refGene_tx=XM_023246872.1
E3	31539689	31539848
refGene_gn=ABCC1;refGene_tx=XM_023246871.1;refGene_gn=ABCC1;refGene_tx=XM_023246872.1
E3	31541911	31542058
refGene_gn=ABCC1;refGene_tx=XM_023246871.1;refGene_gn=ABCC1;refGene_tx=XM_023246872.1
E3	31544343	31544445
refGene_gn=ABCC1;refGene_tx=XM_023246871.1;refGene_gn=ABCC1;refGene_tx=XM_023246872.1
E3	31544971	31545098
refGene_gn=ABCC1;refGene_tx=XM_023246871.1;refGene_gn=ABCC1;refGene_tx=XM_023246872.1
E3	31546957	31547157
refGene_gn=ABCC1;refGene_tx=XM_023246871.1;refGene_gn=ABCC1;refGene_tx=XM_023246872.1
E3	31552178	31552489
refGene_gn=ABCC1;refGene_tx=XM_023246871.1;refGene_gn=ABCC1;refGene_tx=XM_023246872.1
E3	31555731	31555939

refGene_gn=ABCC1;refGene_tx=XM_023246871.1;refGene_gn=ABCC1;refGene_tx=XM_023246872.1
E3 31558484 31558620
refGene_gn=ABCC1;refGene_tx=XM_023246871.1;refGene_gn=ABCC1;refGene_tx=XM_023246872.1
E3 31560861 31560943
refGene_gn=ABCC1;refGene_tx=XM_023246871.1;refGene_gn=ABCC1;refGene_tx=XM_023246872.1
E3 31571552 31571736
refGene_gn=ABCC1;refGene_tx=XM_023246871.1;refGene_gn=ABCC1;refGene_tx=XM_023246872.1
E3 31573729 31573897
refGene_gn=ABCC1;refGene_tx=XM_023246871.1;refGene_gn=ABCC1;refGene_tx=XM_023246872.1
E3 31575613 31575790
refGene_gn=ABCC1;refGene_tx=XM_023246871.1;refGene_gn=ABCC1;refGene_tx=XM_023246872.1
E3 31579552 31579679
refGene_gn=ABCC1;refGene_tx=XM_023246871.1;refGene_gn=ABCC1;refGene_tx=XM_023246872.1
E3 31581779 31581855
refGene_gn=ABCC1;refGene_tx=XM_023246871.1;refGene_gn=ABCC1;refGene_tx=XM_023246872.1
E3 31584763 31584851
refGene_gn=ABCC1;refGene_tx=XM_023246871.1;refGene_gn=ABCC1;refGene_tx=XM_023246872.1
E3 31587198 31587345
refGene_gn=ABCC1;refGene_tx=XM_023246871.1;refGene_gn=ABCC1;refGene_tx=XM_023246872.1
E3 31595951 31596155
refGene_gn=ABCC1;refGene_tx=XM_023246871.1;refGene_gn=ABCC1;refGene_tx=XM_023246872.1
E3 31597668 31597761
refGene_gn=ABCC1;refGene_tx=XM_023246871.1;refGene_gn=ABCC1;refGene_tx=XM_023246872.1
E3 31601677 31601839
refGene_gn=ABCC1;refGene_tx=XM_023246871.1;refGene_gn=ABCC1;refGene_tx=XM_023246872.1
E3 31603587 31603765
refGene_gn=ABCC1;refGene_tx=XM_023246871.1;refGene_gn=ABCC1;refGene_tx=XM_023246872.1
E3 31604404 31604635
refGene_gn=ABCC1;refGene_tx=XM_023246871.1;refGene_gn=ABCC1;refGene_tx=XM_023246872.1
E3 31612913 31613045
refGene_gn=ABCC1;refGene_tx=XM_023246871.1;refGene_gn=ABCC1;refGene_tx=XM_023246872.1
E3 31614989 31615051
refGene_gn=ABCC1;refGene_tx=XM_023246871.1;refGene_gn=ABCC1;refGene_tx=XM_023246872.1

E3 31627413 31627539
refGene_gn=ABCC1;refGene_tx=XM_023246871.1;refGene_gn=ABCC1;refGene_tx=XM_023246872.1
E3 31628473 31628611
refGene_gn=ABCC1;refGene_tx=XM_023246871.1;refGene_gn=ABCC1;refGene_tx=XM_023246872.1
E3 31632075 31632201
refGene_gn=ABCC1;refGene_tx=XM_023246871.1;refGene_gn=ABCC1;refGene_tx=XM_023246872.1
E3 31634235 31634412
refGene_gn=ABCC1;refGene_tx=XM_023246871.1;refGene_gn=ABCC1;refGene_tx=XM_023246872.1
E3 31676006 31676054
refGene_gn=ABCC1;refGene_tx=XM_023246871.1;refGene_gn=ABCC1;refGene_tx=XM_023246872.1
E3 31710136 31710164 refGene_gn=FOPNL;refGene_tx=XM_006942293.4
E3 31710644 31710696
refGene_gn=FOPNL;refGene_tx=XM_023246419.1;refGene_gn=FOPNL;refGene_tx=XM_023246421.1
E3 31713564 31713762
refGene_gn=FOPNL;refGene_tx=XM_006942293.4;refGene_gn=FOPNL;refGene_tx=XM_023246419.1;refGene_gn=FOPNL;refGene_tx=XM_023246421.1
E3 31715452 31715537
refGene_gn=FOPNL;refGene_tx=XM_006942293.4;refGene_gn=FOPNL;refGene_tx=XM_023246419.1;refGene_gn=FOPNL;refGene_tx=XM_023246421.1
E3 31721861 31721998
refGene_gn=FOPNL;refGene_tx=XM_006942293.4;refGene_gn=FOPNL;refGene_tx=XM_023246419.1;refGene_gn=FOPNL;refGene_tx=XM_023246421.1
E3 31726999 31727076
refGene_gn=FOPNL;refGene_tx=XM_006942293.4;refGene_gn=FOPNL;refGene_tx=XM_023246419.1;refGene_gn=FOPNL;refGene_tx=XM_023246421.1
E3 31750971 31751316
refGene_gn=MYH11;refGene_tx=XM_023246707.1;refGene_gn=MYH11;refGene_tx=XM_023246708.1;refGene_gn=MYH11;refGene_tx=XM_023246709.1;refGene_gn=MYH11;refGene_tx=XM_023246710.1;refGene_gn=MYH11;refGene_tx=XM_023246711.1
E3 31760811 31760968
refGene_gn=MYH11;refGene_tx=XM_023246707.1;refGene_gn=MYH11;refGene_tx=XM_023246708.1;refGene_gn=MYH11;refGene_tx=XM_023246709.1;refGene_gn=MYH11;refGene_tx=XM_023246710.1;refGene_gn=MYH11;refGene_tx=XM_023246711.1
E3 31784196 31784224
refGene_gn=MYH11;refGene_tx=XM_023246707.1;refGene_gn=MYH11;refGene_tx=XM_023246708.1;refGene_gn=MYH11;refGene_tx=XM_023246709.1;refGene_gn=MYH11;refGene_tx=XM_023246710.1;refGene_gn=MYH11;refGene_tx=XM_023246711.1
E3 31798461 31798564
refGene_gn=MYH11;refGene_tx=XM_023246707.1;refGene_gn=MYH11;refGene_tx=XM_023246708.1;refGene_gn=MYH11;refGene_tx=XM_023246709.1;refGene_gn=MYH11;refGene_tx=XM_023246710.1;refGene_gn=MYH11;refGene_tx=XM_0232467

11.1
E3	31800074 31800095
refGene_gn=MYH11;refGene_tx=XM_023246707.1;refGene_gn=MYH11;refGene_tx=XM_023246709.1;refGene_gn=MYH11;refGene_tx=XM_023246711.1
E3	31802229 31802322
refGene_gn=MYH11;refGene_tx=XM_023246707.1;refGene_gn=MYH11;refGene_tx=XM_023246708.1;refGene_gn=MYH11;refGene_tx=XM_023246709.1;refGene_gn=MYH11;refGene_tx=XM_023246710.1;refGene_gn=MYH11;refGene_tx=XM_023246711.1
E3	31803749 31803813
refGene_gn=MYH11;refGene_tx=XM_023246707.1;refGene_gn=MYH11;refGene_tx=XM_023246708.1;refGene_gn=MYH11;refGene_tx=XM_023246709.1;refGene_gn=MYH11;refGene_tx=XM_023246710.1;refGene_gn=MYH11;refGene_tx=XM_023246711.1
E3	31806654 31806753
refGene_gn=MYH11;refGene_tx=XM_023246707.1;refGene_gn=MYH11;refGene_tx=XM_023246708.1;refGene_gn=MYH11;refGene_tx=XM_023246709.1;refGene_gn=MYH11;refGene_tx=XM_023246710.1;refGene_gn=MYH11;refGene_tx=XM_023246711.1
E3	31809281 31809425
refGene_gn=MYH11;refGene_tx=XM_023246707.1;refGene_gn=MYH11;refGene_tx=XM_023246708.1;refGene_gn=MYH11;refGene_tx=XM_023246709.1;refGene_gn=MYH11;refGene_tx=XM_023246710.1;refGene_gn=MYH11;refGene_tx=XM_023246711.1
E3	31815627 31815723
refGene_gn=MYH11;refGene_tx=XM_023246707.1;refGene_gn=MYH11;refGene_tx=XM_023246708.1;refGene_gn=MYH11;refGene_tx=XM_023246709.1;refGene_gn=MYH11;refGene_tx=XM_023246710.1;refGene_gn=MYH11;refGene_tx=XM_023246711.1
E3	31817749 31817868
refGene_gn=MYH11;refGene_tx=XM_023246707.1;refGene_gn=MYH11;refGene_tx=XM_023246708.1;refGene_gn=MYH11;refGene_tx=XM_023246709.1;refGene_gn=MYH11;refGene_tx=XM_023246710.1;refGene_gn=MYH11;refGene_tx=XM_023246711.1
E3	31820100 31820253
refGene_gn=MYH11;refGene_tx=XM_023246707.1;refGene_gn=MYH11;refGene_tx=XM_023246708.1;refGene_gn=MYH11;refGene_tx=XM_023246709.1;refGene_gn=MYH11;refGene_tx=XM_023246710.1;refGene_gn=MYH11;refGene_tx=XM_023246711.1
E3	31820676 31820850
refGene_gn=MYH11;refGene_tx=XM_023246707.1;refGene_gn=MYH11;refGene_tx=XM_023246708.1;refGene_gn=MYH11;refGene_tx=XM_023246709.1;refGene_gn=MYH11;refGene_tx=XM_023246710.1;refGene_gn=MYH11;refGene_tx=XM_023246711.1
E3	31822357 31822531
refGene_gn=MYH11;refGene_tx=XM_023246707.1;refGene_gn=MYH11;refGene_tx=XM_023246708.1;refGene_gn=MYH11;refGene_tx=XM_023246709.1;refGene_gn=MYH11;refGene_tx=XM_023246710.1;refGene_gn=MYH11;refGene_tx=XM_023246711.1
E3	31824210 31824325

refGene_gn=MYH11;refGene_tx=XM_023246707.1;refGene_gn=MYH11;refGene_tx=XM_023246708.1;refGene_gn=MYH11;refGene_tx=XM_023246709.1;refGene_gn=MYH11;refGene_tx=XM_023246710.1;refGene_gn=MYH11;refGene_tx=XM_023246711.1
E3 31826729 31826923
refGene_gn=MYH11;refGene_tx=XM_023246707.1;refGene_gn=MYH11;refGene_tx=XM_023246708.1;refGene_gn=MYH11;refGene_tx=XM_023246709.1;refGene_gn=MYH11;refGene_tx=XM_023246710.1;refGene_gn=MYH11;refGene_tx=XM_023246711.1
E3 31828238 31828360
refGene_gn=MYH11;refGene_tx=XM_023246707.1;refGene_gn=MYH11;refGene_tx=XM_023246708.1;refGene_gn=MYH11;refGene_tx=XM_023246709.1;refGene_gn=MYH11;refGene_tx=XM_023246710.1;refGene_gn=MYH11;refGene_tx=XM_023246711.1
E3 31828468 31828538
refGene_gn=MYH11;refGene_tx=XM_023246707.1;refGene_gn=MYH11;refGene_tx=XM_023246708.1;refGene_gn=MYH11;refGene_tx=XM_023246709.1;refGene_gn=MYH11;refGene_tx=XM_023246710.1;refGene_gn=MYH11;refGene_tx=XM_023246711.1
E3 31828703 31828864
refGene_gn=MYH11;refGene_tx=XM_023246707.1;refGene_gn=MYH11;refGene_tx=XM_023246708.1;refGene_gn=MYH11;refGene_tx=XM_023246709.1;refGene_gn=MYH11;refGene_tx=XM_023246710.1;refGene_gn=MYH11;refGene_tx=XM_023246711.1
E3 31830459 31830568
refGene_gn=MYH11;refGene_tx=XM_023246707.1;refGene_gn=MYH11;refGene_tx=XM_023246708.1;refGene_gn=MYH11;refGene_tx=XM_023246709.1;refGene_gn=MYH11;refGene_tx=XM_023246710.1;refGene_gn=MYH11;refGene_tx=XM_023246711.1
E3 31833049 31833181
refGene_gn=MYH11;refGene_tx=XM_023246707.1;refGene_gn=MYH11;refGene_tx=XM_023246708.1;refGene_gn=MYH11;refGene_tx=XM_023246709.1;refGene_gn=MYH11;refGene_tx=XM_023246710.1;refGene_gn=MYH11;refGene_tx=XM_023246711.1
E3 31833272 31833479
refGene_gn=MYH11;refGene_tx=XM_023246707.1;refGene_gn=MYH11;refGene_tx=XM_023246708.1;refGene_gn=MYH11;refGene_tx=XM_023246709.1;refGene_gn=MYH11;refGene_tx=XM_023246710.1;refGene_gn=MYH11;refGene_tx=XM_023246711.1
E3 31834398 31834536
refGene_gn=MYH11;refGene_tx=XM_023246707.1;refGene_gn=MYH11;refGene_tx=XM_023246708.1;refGene_gn=MYH11;refGene_tx=XM_023246709.1;refGene_gn=MYH11;refGene_tx=XM_023246710.1;refGene_gn=MYH11;refGene_tx=XM_023246711.1
E3 31835622 31835746
refGene_gn=MYH11;refGene_tx=XM_023246707.1;refGene_gn=MYH11;refGene_tx=XM_023246708.1;refGene_gn=MYH11;refGene_tx=XM_023246709.1;refGene_gn=MYH11;refGene_tx=XM_023246710.1;refGene_gn=MYH11;refGene_tx=XM_023246711.1
E3 31836093 31836265

refGene_gn=MYH11;refGene_tx=XM_023246707.1;refGene_gn=MYH11;refGene_tx=XM_023246708.1;refGene_gn=MYH11;refGene_tx=XM_023246709.1;refGene_gn=MYH11;refGene_tx=XM_023246710.1;refGene_gn=MYH11;refGene_tx=XM_023246711.1
E3 31837855 31838068
refGene_gn=MYH11;refGene_tx=XM_023246707.1;refGene_gn=MYH11;refGene_tx=XM_023246708.1;refGene_gn=MYH11;refGene_tx=XM_023246709.1;refGene_gn=MYH11;refGene_tx=XM_023246710.1;refGene_gn=MYH11;refGene_tx=XM_023246711.1
E3 31839575 31839720
refGene_gn=MYH11;refGene_tx=XM_023246707.1;refGene_gn=MYH11;refGene_tx=XM_023246708.1;refGene_gn=MYH11;refGene_tx=XM_023246709.1;refGene_gn=MYH11;refGene_tx=XM_023246710.1;refGene_gn=MYH11;refGene_tx=XM_023246711.1
E3 31842375 31842582
refGene_gn=MYH11;refGene_tx=XM_023246707.1;refGene_gn=MYH11;refGene_tx=XM_023246708.1;refGene_gn=MYH11;refGene_tx=XM_023246709.1;refGene_gn=MYH11;refGene_tx=XM_023246710.1;refGene_gn=MYH11;refGene_tx=XM_023246711.1
E3 31843999 31844104
refGene_gn=MYH11;refGene_tx=XM_023246707.1;refGene_gn=MYH11;refGene_tx=XM_023246708.1;refGene_gn=MYH11;refGene_tx=XM_023246709.1;refGene_gn=MYH11;refGene_tx=XM_023246710.1;refGene_gn=MYH11;refGene_tx=XM_023246711.1
E3 31844192 31844345
refGene_gn=MYH11;refGene_tx=XM_023246707.1;refGene_gn=MYH11;refGene_tx=XM_023246708.1;refGene_gn=MYH11;refGene_tx=XM_023246709.1;refGene_gn=MYH11;refGene_tx=XM_023246710.1;refGene_gn=MYH11;refGene_tx=XM_023246711.1
E3 31844571 31844820
refGene_gn=MYH11;refGene_tx=XM_023246707.1;refGene_gn=MYH11;refGene_tx=XM_023246708.1;refGene_gn=MYH11;refGene_tx=XM_023246709.1;refGene_gn=MYH11;refGene_tx=XM_023246710.1;refGene_gn=MYH11;refGene_tx=XM_023246711.1
E3 31846584 31846797
refGene_gn=MYH11;refGene_tx=XM_023246707.1;refGene_gn=MYH11;refGene_tx=XM_023246708.1;refGene_gn=MYH11;refGene_tx=XM_023246709.1;refGene_gn=MYH11;refGene_tx=XM_023246710.1;refGene_gn=MYH11;refGene_tx=XM_023246711.1
E3 31847122 31847335
refGene_gn=MYH11;refGene_tx=XM_023246707.1;refGene_gn=MYH11;refGene_tx=XM_023246708.1;refGene_gn=MYH11;refGene_tx=XM_023246709.1;refGene_gn=MYH11;refGene_tx=XM_023246710.1;refGene_gn=MYH11;refGene_tx=XM_023246711.1
E3 31847690 31847852
refGene_gn=MYH11;refGene_tx=XM_023246707.1;refGene_gn=MYH11;refGene_tx=XM_023246708.1;refGene_gn=MYH11;refGene_tx=XM_023246709.1;refGene_gn=MYH11;refGene_tx=XM_023246710.1;refGene_gn=MYH11;refGene_tx=XM_023246711.1
E3 31848242 31848371

refGene_gn=MYH11;refGene_tx=XM_023246707.1;refGene_gn=MYH11;refGene_tx=XM_023246708.1;refGene_gn=MYH11;refGene_tx=XM_023246709.1;refGene_gn=MYH11;refGene_tx=XM_023246710.1;refGene_gn=MYH11;refGene_tx=XM_023246711.1
E3 31848691 31848780
refGene_gn=MYH11;refGene_tx=XM_023246707.1;refGene_gn=MYH11;refGene_tx=XM_023246708.1;refGene_gn=MYH11;refGene_tx=XM_023246709.1;refGene_gn=MYH11;refGene_tx=XM_023246710.1;refGene_gn=MYH11;refGene_tx=XM_023246711.1
E3 31849224 31849348
refGene_gn=MYH11;refGene_tx=XM_023246707.1;refGene_gn=MYH11;refGene_tx=XM_023246708.1;refGene_gn=MYH11;refGene_tx=XM_023246709.1;refGene_gn=MYH11;refGene_tx=XM_023246710.1;refGene_gn=MYH11;refGene_tx=XM_023246711.1
E3 31849930 31850139
refGene_gn=MYH11;refGene_tx=XM_023246707.1;refGene_gn=MYH11;refGene_tx=XM_023246708.1;refGene_gn=MYH11;refGene_tx=XM_023246709.1;refGene_gn=MYH11;refGene_tx=XM_023246710.1;refGene_gn=MYH11;refGene_tx=XM_023246711.1
E3 31851070 31851179
refGene_gn=MYH11;refGene_tx=XM_023246707.1;refGene_gn=MYH11;refGene_tx=XM_023246708.1;refGene_gn=MYH11;refGene_tx=XM_023246709.1;refGene_gn=MYH11;refGene_tx=XM_023246710.1;refGene_gn=MYH11;refGene_tx=XM_023246711.1
E3 31851260 31851433
refGene_gn=MYH11;refGene_tx=XM_023246707.1;refGene_gn=MYH11;refGene_tx=XM_023246708.1;refGene_gn=MYH11;refGene_tx=XM_023246709.1;refGene_gn=MYH11;refGene_tx=XM_023246710.1;refGene_gn=MYH11;refGene_tx=XM_023246711.1
E3 31851951 31851952 refGene_gn=MYH11;refGene_tx=XM_023246711.1
E3 31855581 31855620
refGene_gn=MYH11;refGene_tx=XM_023246709.1;refGene_gn=MYH11;refGene_tx=XM_023246710.1;refGene_gn=MYH11;refGene_tx=XM_023246711.1
E3 31859068 31859250
refGene_gn=MYH11;refGene_tx=XM_023246707.1;refGene_gn=MYH11;refGene_tx=XM_023246708.1;refGene_gn=MYH11;refGene_tx=XM_023246711.1
E3 31862122 31862284
refGene_gn=NDE1;refGene_tx=XM_023246712.1;refGene_gn=NDE1;refGene_tx=XM_023246714.1
E3 31862325 31862412
refGene_gn=NDE1;refGene_tx=XM_023246713.1;refGene_gn=NDE1;refGene_tx=XM_023246715.1
E3 31862588 31862658
refGene_gn=NDE1;refGene_tx=XM_023246712.1;refGene_gn=NDE1;refGene_tx=XM_023246713.1
E3 31863106 31863261
refGene_gn=NDE1;refGene_tx=XM_023246712.1;refGene_gn=NDE1;refGene_tx=XM_023246713.1;refGene_gn=NDE1;refGene_tx=XM_023246714.1;refGene_gn=NDE1;refGene_tx=XM_023246715.1
E3 31864293 31864385

refGene_gn=NDE1;refGene_tx=XM_023246712.1;refGene_gn=NDE1;refGene_tx=XM_023246713.1;refGene_gn=NDE1;refGene_tx=XM_023246714.1;refGene_gn=NDE1;refGene_tx=XM_023246715.1
E3 31865729 31865909
refGene_gn=NDE1;refGene_tx=XM_023246712.1;refGene_gn=NDE1;refGene_tx=XM_023246713.1;refGene_gn=NDE1;refGene_tx=XM_023246714.1;refGene_gn=NDE1;refGene_tx=XM_023246715.1
E3 31867384 31867521
refGene_gn=NDE1;refGene_tx=XM_023246712.1;refGene_gn=NDE1;refGene_tx=XM_023246713.1;refGene_gn=NDE1;refGene_tx=XM_023246714.1;refGene_gn=NDE1;refGene_tx=XM_023246715.1
E3 31872582 31872731
refGene_gn=NDE1;refGene_tx=XM_023246712.1;refGene_gn=NDE1;refGene_tx=XM_023246713.1;refGene_gn=NDE1;refGene_tx=XM_023246714.1;refGene_gn=NDE1;refGene_tx=XM_023246715.1
E3 31878543 31878697
refGene_gn=NDE1;refGene_tx=XM_023246712.1;refGene_gn=NDE1;refGene_tx=XM_023246713.1;refGene_gn=NDE1;refGene_tx=XM_023246714.1;refGene_gn=NDE1;refGene_tx=XM_023246715.1
E3 31880165 31880248
refGene_gn=NDE1;refGene_tx=XM_023246712.1;refGene_gn=NDE1;refGene_tx=XM_023246713.1;refGene_gn=NDE1;refGene_tx=XM_023246714.1;refGene_gn=NDE1;refGene_tx=XM_023246715.1
E3 31895925 31896069
refGene_gn=MARF1;refGene_tx=XM_023247059.1;refGene_gn=MARF1;refGene_tx=XM_023247060.1;refGene_gn=MARF1;refGene_tx=XM_023247061.1;refGene_gn=MARF1;refGene_tx=XM_023247062.1;refGene_gn=MARF1;refGene_tx=XM_023247063.1;refGene_gn=MARF1;refGene_tx=XM_023247064.1;refGene_gn=MARF1;refGene_tx=XM_023247065.1
E3 31898145 31898838
refGene_gn=MARF1;refGene_tx=XM_023247059.1;refGene_gn=MARF1;refGene_tx=XM_023247060.1;refGene_gn=MARF1;refGene_tx=XM_023247061.1;refGene_gn=MARF1;refGene_tx=XM_023247062.1;refGene_gn=MARF1;refGene_tx=XM_023247063.1;refGene_gn=MARF1;refGene_tx=XM_023247064.1;refGene_gn=MARF1;refGene_tx=XM_023247065.1
E3 31899586 31899761
refGene_gn=MARF1;refGene_tx=XM_023247059.1;refGene_gn=MARF1;refGene_tx=XM_023247060.1;refGene_gn=MARF1;refGene_tx=XM_023247061.1;refGene_gn=MARF1;refGene_tx=XM_023247062.1;refGene_gn=MARF1;refGene_tx=XM_023247063.1;refGene_gn=MARF1;refGene_tx=XM_023247064.1;refGene_gn=MARF1;refGene_tx=XM_023247065.1
E3 31900688 31900915
refGene_gn=MARF1;refGene_tx=XM_023247059.1;refGene_gn=MARF1;refGene_tx=XM_023247060.1;refGene_gn=MARF1;refGene_tx=XM_023247061.1;refGene_gn=MARF1;refGene_tx=XM_023247062.1;refGene_gn=MARF1;refGene_tx=XM_023247063.1;refGene_gn=MARF1;refGene_tx=XM_023247064.1;refGene_gn=MARF1;refGene_tx=XM_023247065.1
E3 31902253 31902371
refGene_gn=MARF1;refGene_tx=XM_023247059.1;refGene_gn=MARF1;refGene_tx=XM_023247060.1;refGene_gn=MARF1;refGene_tx=XM_023247061.1;refGene_gn=

MARF1;refGene_tx=XM_023247062.1;refGene_gn=MARF1;refGene_tx=XM_0232470
63.1;refGene_gn=MARF1;refGene_tx=XM_023247064.1;refGene_gn=MARF1;refGe
ne_tx=XM_023247065.1
E3 31902966 31903139
refGene_gn=MARF1;refGene_tx=XM_023247059.1;refGene_gn=MARF1;refGene_tx
=XM_023247060.1;refGene_gn=MARF1;refGene_tx=XM_023247061.1;refGene_gn=
MARF1;refGene_tx=XM_023247062.1;refGene_gn=MARF1;refGene_tx=XM_0232470
63.1;refGene_gn=MARF1;refGene_tx=XM_023247064.1;refGene_gn=MARF1;refGe
ne_tx=XM_023247065.1
E3 31904150 31904579
refGene_gn=MARF1;refGene_tx=XM_023247059.1;refGene_gn=MARF1;refGene_tx
=XM_023247060.1;refGene_gn=MARF1;refGene_tx=XM_023247061.1;refGene_gn=
MARF1;refGene_tx=XM_023247062.1;refGene_gn=MARF1;refGene_tx=XM_0232470
63.1;refGene_gn=MARF1;refGene_tx=XM_023247065.1
E3 31904780 31904938
refGene_gn=MARF1;refGene_tx=XM_023247059.1;refGene_gn=MARF1;refGene_tx
=XM_023247060.1;refGene_gn=MARF1;refGene_tx=XM_023247061.1;refGene_gn=
MARF1;refGene_tx=XM_023247062.1;refGene_gn=MARF1;refGene_tx=XM_0232470
63.1;refGene_gn=MARF1;refGene_tx=XM_023247064.1;refGene_gn=MARF1;refGe
ne_tx=XM_023247065.1
E3 31905025 31905184
refGene_gn=MARF1;refGene_tx=XM_023247059.1;refGene_gn=MARF1;refGene_tx
=XM_023247060.1;refGene_gn=MARF1;refGene_tx=XM_023247061.1;refGene_gn=
MARF1;refGene_tx=XM_023247062.1;refGene_gn=MARF1;refGene_tx=XM_0232470
63.1;refGene_gn=MARF1;refGene_tx=XM_023247064.1;refGene_gn=MARF1;refGe
ne_tx=XM_023247065.1
E3 31906210 31906403
refGene_gn=MARF1;refGene_tx=XM_023247059.1;refGene_gn=MARF1;refGene_tx
=XM_023247060.1;refGene_gn=MARF1;refGene_tx=XM_023247061.1;refGene_gn=
MARF1;refGene_tx=XM_023247062.1;refGene_gn=MARF1;refGene_tx=XM_0232470
63.1;refGene_gn=MARF1;refGene_tx=XM_023247064.1;refGene_gn=MARF1;refGe
ne_tx=XM_023247065.1
E3 31907120 31907299
refGene_gn=MARF1;refGene_tx=XM_023247059.1;refGene_gn=MARF1;refGene_tx
=XM_023247060.1;refGene_gn=MARF1;refGene_tx=XM_023247061.1;refGene_gn=
MARF1;refGene_tx=XM_023247062.1;refGene_gn=MARF1;refGene_tx=XM_0232470
63.1;refGene_gn=MARF1;refGene_tx=XM_023247064.1;refGene_gn=MARF1;refGe
ne_tx=XM_023247065.1
E3 31908958 31909039
refGene_gn=MARF1;refGene_tx=XM_023247059.1;refGene_gn=MARF1;refGene_tx
=XM_023247060.1;refGene_gn=MARF1;refGene_tx=XM_023247061.1;refGene_gn=
MARF1;refGene_tx=XM_023247062.1;refGene_gn=MARF1;refGene_tx=XM_0232470
63.1;refGene_gn=MARF1;refGene_tx=XM_023247064.1;refGene_gn=MARF1;refGe
ne_tx=XM_023247065.1
E3 31911395 31911632
refGene_gn=MARF1;refGene_tx=XM_023247059.1;refGene_gn=MARF1;refGene_tx
=XM_023247060.1;refGene_gn=MARF1;refGene_tx=XM_023247061.1;refGene_gn=
MARF1;refGene_tx=XM_023247062.1;refGene_gn=MARF1;refGene_tx=XM_0232470
63.1;refGene_gn=MARF1;refGene_tx=XM_023247064.1;refGene_gn=MARF1;refGe
ne_tx=XM_023247065.1

E3	31911758 31911878
refGene_gn=MARF1;refGene_tx=XM_023247059.1;refGene_gn=MARF1;refGene_tx=XM_023247060.1;refGene_gn=MARF1;refGene_tx=XM_023247061.1;refGene_gn=MARF1;refGene_tx=XM_023247062.1;refGene_gn=MARF1;refGene_tx=XM_023247063.1;refGene_gn=MARF1;refGene_tx=XM_023247064.1;refGene_gn=MARF1;refGene_tx=XM_023247065.1
E3	31912519 31912695
refGene_gn=MARF1;refGene_tx=XM_023247059.1;refGene_gn=MARF1;refGene_tx=XM_023247060.1;refGene_gn=MARF1;refGene_tx=XM_023247061.1;refGene_gn=MARF1;refGene_tx=XM_023247062.1;refGene_gn=MARF1;refGene_tx=XM_023247063.1;refGene_gn=MARF1;refGene_tx=XM_023247064.1;refGene_gn=MARF1;refGene_tx=XM_023247065.1
E3	31914468 31914689
refGene_gn=MARF1;refGene_tx=XM_023247059.1;refGene_gn=MARF1;refGene_tx=XM_023247060.1;refGene_gn=MARF1;refGene_tx=XM_023247061.1;refGene_gn=MARF1;refGene_tx=XM_023247062.1;refGene_gn=MARF1;refGene_tx=XM_023247063.1;refGene_gn=MARF1;refGene_tx=XM_023247064.1;refGene_gn=MARF1;refGene_tx=XM_023247065.1
E3	31915406 31915549
refGene_gn=MARF1;refGene_tx=XM_023247059.1;refGene_gn=MARF1;refGene_tx=XM_023247060.1;refGene_gn=MARF1;refGene_tx=XM_023247061.1;refGene_gn=MARF1;refGene_tx=XM_023247062.1;refGene_gn=MARF1;refGene_tx=XM_023247063.1;refGene_gn=MARF1;refGene_tx=XM_023247064.1;refGene_gn=MARF1;refGene_tx=XM_023247065.1
E3	31915727 31915861
refGene_gn=MARF1;refGene_tx=XM_023247059.1;refGene_gn=MARF1;refGene_tx=XM_023247060.1;refGene_gn=MARF1;refGene_tx=XM_023247061.1;refGene_gn=MARF1;refGene_tx=XM_023247062.1;refGene_gn=MARF1;refGene_tx=XM_023247063.1;refGene_gn=MARF1;refGene_tx=XM_023247064.1;refGene_gn=MARF1;refGene_tx=XM_023247065.1
E3	31917631 31917834
refGene_gn=MARF1;refGene_tx=XM_023247059.1;refGene_gn=MARF1;refGene_tx=XM_023247060.1;refGene_gn=MARF1;refGene_tx=XM_023247061.1;refGene_gn=MARF1;refGene_tx=XM_023247062.1;refGene_gn=MARF1;refGene_tx=XM_023247063.1;refGene_gn=MARF1;refGene_tx=XM_023247064.1;refGene_gn=MARF1;refGene_tx=XM_023247065.1
E3	31919209 31919437
refGene_gn=MARF1;refGene_tx=XM_023247059.1;refGene_gn=MARF1;refGene_tx=XM_023247060.1;refGene_gn=MARF1;refGene_tx=XM_023247061.1;refGene_gn=MARF1;refGene_tx=XM_023247062.1;refGene_gn=MARF1;refGene_tx=XM_023247063.1;refGene_gn=MARF1;refGene_tx=XM_023247064.1;refGene_gn=MARF1;refGene_tx=XM_023247065.1
E3	31922609 31922840
refGene_gn=MARF1;refGene_tx=XM_023247059.1;refGene_gn=MARF1;refGene_tx=XM_023247060.1;refGene_gn=MARF1;refGene_tx=XM_023247061.1;refGene_gn=MARF1;refGene_tx=XM_023247062.1;refGene_gn=MARF1;refGene_tx=XM_023247063.1;refGene_gn=MARF1;refGene_tx=XM_023247064.1;refGene_gn=MARF1;refGene_tx=XM_023247065.1
E3	31924518 31924731
refGene_gn=MARF1;refGene_tx=XM_023247059.1;refGene_gn=MARF1;refGene_tx

=XM_023247060.1;refGene_gn=MARF1;refGene_tx=XM_023247061.1;refGene_gn=MARF1;refGene_tx=XM_023247062.1;refGene_gn=MARF1;refGene_tx=XM_023247063.1;refGene_gn=MARF1;refGene_tx=XM_023247064.1;refGene_gn=MARF1;refGene_tx=XM_023247065.1
E3 31925921 31925982
refGene_gn=MARF1;refGene_tx=XM_023247059.1;refGene_gn=MARF1;refGene_tx=XM_023247060.1;refGene_gn=MARF1;refGene_tx=XM_023247061.1;refGene_gn=MARF1;refGene_tx=XM_023247062.1;refGene_gn=MARF1;refGene_tx=XM_023247063.1;refGene_gn=MARF1;refGene_tx=XM_023247064.1;refGene_gn=MARF1;refGene_tx=XM_023247065.1
E3 31926069 31926195
refGene_gn=MARF1;refGene_tx=XM_023247059.1;refGene_gn=MARF1;refGene_tx=XM_023247060.1;refGene_gn=MARF1;refGene_tx=XM_023247061.1;refGene_gn=MARF1;refGene_tx=XM_023247062.1;refGene_gn=MARF1;refGene_tx=XM_023247063.1;refGene_gn=MARF1;refGene_tx=XM_023247064.1;refGene_gn=MARF1;refGene_tx=XM_023247065.1
E3 31927764 31927935
refGene_gn=MARF1;refGene_tx=XM_023247059.1;refGene_gn=MARF1;refGene_tx=XM_023247060.1;refGene_gn=MARF1;refGene_tx=XM_023247061.1;refGene_gn=MARF1;refGene_tx=XM_023247062.1;refGene_gn=MARF1;refGene_tx=XM_023247063.1;refGene_gn=MARF1;refGene_tx=XM_023247064.1;refGene_gn=MARF1;refGene_tx=XM_023247065.1
E3 31929943 31930188
refGene_gn=MARF1;refGene_tx=XM_023247059.1;refGene_gn=MARF1;refGene_tx=XM_023247060.1;refGene_gn=MARF1;refGene_tx=XM_023247061.1;refGene_gn=MARF1;refGene_tx=XM_023247062.1;refGene_gn=MARF1;refGene_tx=XM_023247063.1;refGene_gn=MARF1;refGene_tx=XM_023247064.1;refGene_gn=MARF1;refGene_tx=XM_023247065.1
E3 31943215 31943328
refGene_gn=CE3H16orf45;refGene_tx=XM_003998842.5
E3 31946260 31946343
refGene_gn=CE3H16orf45;refGene_tx=XM_003998842.5
E3 31947610 31947725
refGene_gn=CE3H16orf45;refGene_tx=XM_003998842.5
E3 31954193 31954267
refGene_gn=CE3H16orf45;refGene_tx=XM_003998842.5
E3 31999022 31999146
refGene_gn=CE3H16orf45;refGene_tx=XM_003998842.5
E3 32064385 32064491
refGene_gn=CE3H16orf45;refGene_tx=XM_003998842.5
E3 32080104 32080359
refGene_gn=MPV17L;refGene_tx=XM_023246550.1;refGene_gn=MPV17L;refGene_tx=XM_023246551.1
E3 32110461 32110641
refGene_gn=MPV17L;refGene_tx=XM_003998853.4;refGene_gn=MPV17L;refGene_tx=XM_023246552.1
E3 32110721 32110751
refGene_gn=MPV17L;refGene_tx=XM_003998853.4;refGene_gn=MPV17L;refGene_tx=XM_023246550.1;refGene_gn=MPV17L;refGene_tx=XM_023246551.1;refGene_gn=MPV17L;refGene_tx=XM_023246552.1

E3 32114984 32115055
refGene_gn=MPV17L;refGene_tx=XM_003998853.4;refGene_gn=MPV17L;refGene_tx=XM_023246550.1;refGene_gn=MPV17L;refGene_tx=XM_023246551.1
E3 32118782 32119092
refGene_gn=MPV17L;refGene_tx=XM_003998853.4;refGene_gn=MPV17L;refGene_tx=XM_023246550.1;refGene_gn=MPV17L;refGene_tx=XM_023246552.1
E3 32119269 32119492 refGene_gn=MPV17L;refGene_tx=XM_023246551.1
E3 32139099 32139120 refGene_gn=PDXDC1;refGene_tx=XM_003998843.5
E3 32155707 32155781
refGene_gn=PDXDC1;refGene_tx=XM_003998843.5;refGene_gn=PDXDC1;refGene_tx=XM_006942302.4;refGene_gn=PDXDC1;refGene_tx=XM_006942303.4;refGene_gn=PDXDC1;refGene_tx=XM_019820923.2;refGene_gn=PDXDC1;refGene_tx=XM_023246625.1
E3 32156312 32156378
refGene_gn=PDXDC1;refGene_tx=XM_003998843.5;refGene_gn=PDXDC1;refGene_tx=XM_006942302.4;refGene_gn=PDXDC1;refGene_tx=XM_006942303.4;refGene_gn=PDXDC1;refGene_tx=XM_019820923.2;refGene_gn=PDXDC1;refGene_tx=XM_023246625.1
E3 32158335 32158416
refGene_gn=PDXDC1;refGene_tx=XM_003998843.5;refGene_gn=PDXDC1;refGene_tx=XM_006942302.4;refGene_gn=PDXDC1;refGene_tx=XM_006942303.4;refGene_gn=PDXDC1;refGene_tx=XM_019820923.2;refGene_gn=PDXDC1;refGene_tx=XM_023246625.1
E3 32160295 32160442
refGene_gn=PDXDC1;refGene_tx=XM_003998843.5;refGene_gn=PDXDC1;refGene_tx=XM_006942302.4;refGene_gn=PDXDC1;refGene_tx=XM_006942303.4;refGene_gn=PDXDC1;refGene_tx=XM_019820923.2;refGene_gn=PDXDC1;refGene_tx=XM_023246625.1
E3 32161799 32161989
refGene_gn=PDXDC1;refGene_tx=XM_003998843.5;refGene_gn=PDXDC1;refGene_tx=XM_006942302.4;refGene_gn=PDXDC1;refGene_tx=XM_006942303.4;refGene_gn=PDXDC1;refGene_tx=XM_019820923.2;refGene_gn=PDXDC1;refGene_tx=XM_023246625.1
E3 32164399 32164468
refGene_gn=PDXDC1;refGene_tx=XM_003998843.5;refGene_gn=PDXDC1;refGene_tx=XM_006942302.4;refGene_gn=PDXDC1;refGene_tx=XM_006942303.4;refGene_gn=PDXDC1;refGene_tx=XM_019820923.2;refGene_gn=PDXDC1;refGene_tx=XM_023246625.1
E3 32165292 32165371
refGene_gn=PDXDC1;refGene_tx=XM_003998843.5;refGene_gn=PDXDC1;refGene_tx=XM_006942302.4;refGene_gn=PDXDC1;refGene_tx=XM_006942303.4;refGene_gn=PDXDC1;refGene_tx=XM_019820923.2;refGene_gn=PDXDC1;refGene_tx=XM_023246625.1
E3 32168665 32168750
refGene_gn=PDXDC1;refGene_tx=XM_003998843.5;refGene_gn=PDXDC1;refGene_tx=XM_006942302.4;refGene_gn=PDXDC1;refGene_tx=XM_006942303.4;refGene_gn=PDXDC1;refGene_tx=XM_019820923.2;refGene_gn=PDXDC1;refGene_tx=XM_023246625.1
E3 32169600 32169649
refGene_gn=PDXDC1;refGene_tx=XM_003998843.5;refGene_gn=PDXDC1;refGene_

tx=XM_006942302.4;refGene_gn=PDXDC1;refGene_tx=XM_006942303.4;refGene_
gn=PDXDC1;refGene_tx=XM_019820923.2;refGene_gn=PDXDC1;refGene_tx=XM_02
3246625.1
E3 32169799 32169901
refGene_gn=PDXDC1;refGene_tx=XM_003998843.5;refGene_gn=PDXDC1;refGene_
tx=XM_006942302.4;refGene_gn=PDXDC1;refGene_tx=XM_006942303.4;refGene_
gn=PDXDC1;refGene_tx=XM_019820923.2;refGene_gn=PDXDC1;refGene_tx=XM_02
3246625.1
E3 32170701 32170827
refGene_gn=PDXDC1;refGene_tx=XM_003998843.5;refGene_gn=PDXDC1;refGene_
tx=XM_006942302.4;refGene_gn=PDXDC1;refGene_tx=XM_006942303.4;refGene_
gn=PDXDC1;refGene_tx=XM_019820923.2;refGene_gn=PDXDC1;refGene_tx=XM_02
3246625.1
E3 32174896 32174947
refGene_gn=PDXDC1;refGene_tx=XM_003998843.5;refGene_gn=PDXDC1;refGene_
tx=XM_006942302.4;refGene_gn=PDXDC1;refGene_tx=XM_006942303.4;refGene_
gn=PDXDC1;refGene_tx=XM_019820923.2;refGene_gn=PDXDC1;refGene_tx=XM_02
3246625.1
E3 32179994 32180058
refGene_gn=PDXDC1;refGene_tx=XM_003998843.5;refGene_gn=PDXDC1;refGene_
tx=XM_006942302.4;refGene_gn=PDXDC1;refGene_tx=XM_006942303.4;refGene_
gn=PDXDC1;refGene_tx=XM_019820923.2;refGene_gn=PDXDC1;refGene_tx=XM_02
3246625.1
E3 32182205 32182294
refGene_gn=PDXDC1;refGene_tx=XM_003998843.5;refGene_gn=PDXDC1;refGene_
tx=XM_006942302.4;refGene_gn=PDXDC1;refGene_tx=XM_006942303.4;refGene_
gn=PDXDC1;refGene_tx=XM_019820923.2;refGene_gn=PDXDC1;refGene_tx=XM_02
3246625.1
E3 32183229 32183335
refGene_gn=PDXDC1;refGene_tx=XM_003998843.5;refGene_gn=PDXDC1;refGene_
tx=XM_006942302.4;refGene_gn=PDXDC1;refGene_tx=XM_006942303.4;refGene_
gn=PDXDC1;refGene_tx=XM_019820923.2;refGene_gn=PDXDC1;refGene_tx=XM_02
3246625.1
E3 32185100 32185272
refGene_gn=PDXDC1;refGene_tx=XM_003998843.5;refGene_gn=PDXDC1;refGene_
tx=XM_006942302.4;refGene_gn=PDXDC1;refGene_tx=XM_006942303.4;refGene_
gn=PDXDC1;refGene_tx=XM_019820923.2;refGene_gn=PDXDC1;refGene_tx=XM_02
3246625.1
E3 32186144 32186263
refGene_gn=PDXDC1;refGene_tx=XM_003998843.5;refGene_gn=PDXDC1;refGene_
tx=XM_006942302.4;refGene_gn=PDXDC1;refGene_tx=XM_006942303.4;refGene_
gn=PDXDC1;refGene_tx=XM_019820923.2;refGene_gn=PDXDC1;refGene_tx=XM_02
3246625.1
E3 32186562 32186684
refGene_gn=PDXDC1;refGene_tx=XM_003998843.5;refGene_gn=PDXDC1;refGene_
tx=XM_006942302.4;refGene_gn=PDXDC1;refGene_tx=XM_006942303.4;refGene_
gn=PDXDC1;refGene_tx=XM_019820923.2;refGene_gn=PDXDC1;refGene_tx=XM_02
3246625.1
E3 32187633 32187726
refGene_gn=PDXDC1;refGene_tx=XM_003998843.5;refGene_gn=PDXDC1;refGene_

tx=XM_006942302.4;refGene_gn=PDXDC1;refGene_tx=XM_006942303.4;refGene_gn=PDXDC1;refGene_tx=XM_019820923.2;refGene_gn=PDXDC1;refGene_tx=XM_023246625.1
E3 32187820 32187917
refGene_gn=PDXDC1;refGene_tx=XM_003998843.5;refGene_gn=PDXDC1;refGene_tx=XM_006942302.4;refGene_gn=PDXDC1;refGene_tx=XM_006942303.4;refGene_gn=PDXDC1;refGene_tx=XM_019820923.2;refGene_gn=PDXDC1;refGene_tx=XM_023246625.1
E3 32188848 32188953
refGene_gn=PDXDC1;refGene_tx=XM_003998843.5;refGene_gn=PDXDC1;refGene_tx=XM_006942302.4;refGene_gn=PDXDC1;refGene_tx=XM_006942303.4;refGene_gn=PDXDC1;refGene_tx=XM_019820923.2;refGene_gn=PDXDC1;refGene_tx=XM_023246625.1
E3 32189427 32189687
refGene_gn=PDXDC1;refGene_tx=XM_003998843.5;refGene_gn=PDXDC1;refGene_tx=XM_006942302.4;refGene_gn=PDXDC1;refGene_tx=XM_006942303.4;refGene_gn=PDXDC1;refGene_tx=XM_019820923.2;refGene_gn=PDXDC1;refGene_tx=XM_023246625.1
E3 32191875 32192055
refGene_gn=NTAN1;refGene_tx=XM_023246626.1;refGene_gn=NTAN1;refGene_tx=XM_023246627.1
E3 32192191 32192212 refGene_gn=NTAN1;refGene_tx=XM_023246628.1
E3 32192455 32192569
refGene_gn=NTAN1;refGene_tx=XM_023246626.1;refGene_gn=NTAN1;refGene_tx=XM_023246627.1;refGene_gn=NTAN1;refGene_tx=XM_023246628.1
E3 32193431 32193559
refGene_gn=NTAN1;refGene_tx=XM_023246626.1;refGene_gn=NTAN1;refGene_tx=XM_023246627.1;refGene_gn=NTAN1;refGene_tx=XM_023246628.1
E3 32194366 32194420
refGene_gn=NTAN1;refGene_tx=XM_023246626.1;refGene_gn=NTAN1;refGene_tx=XM_023246627.1;refGene_gn=NTAN1;refGene_tx=XM_023246628.1
E3 32194919 32194973
refGene_gn=NTAN1;refGene_tx=XM_023246626.1;refGene_gn=NTAN1;refGene_tx=XM_023246627.1;refGene_gn=NTAN1;refGene_tx=XM_023246628.1
E3 32196116 32196190
refGene_gn=NTAN1;refGene_tx=XM_023246626.1;refGene_gn=NTAN1;refGene_tx=XM_023246627.1;refGene_gn=NTAN1;refGene_tx=XM_023246628.1
E3 32197856 32197965
refGene_gn=NTAN1;refGene_tx=XM_023246626.1;refGene_gn=NTAN1;refGene_tx=XM_023246627.1;refGene_gn=NTAN1;refGene_tx=XM_023246628.1
E3 32198235 32198301
refGene_gn=NTAN1;refGene_tx=XM_023246626.1;refGene_gn=NTAN1;refGene_tx=XM_023246627.1;refGene_gn=NTAN1;refGene_tx=XM_023246628.1
E3 32198377 32198480
refGene_gn=NTAN1;refGene_tx=XM_023246626.1;refGene_gn=NTAN1;refGene_tx=XM_023246627.1;refGene_gn=NTAN1;refGene_tx=XM_023246628.1
E3 32204557 32204638
refGene_gn=NTAN1;refGene_tx=XM_023246626.1;refGene_gn=NTAN1;refGene_tx=XM_023246627.1;refGene_gn=NTAN1;refGene_tx=XM_023246628.1
E3 32208047 32208209

refGene_gn=RRN3;refGene_tx=XM_011290580.3;refGene_gn=RRN3;refGene_tx=XM_019820926.2
E3 32209718 32209806
refGene_gn=RRN3;refGene_tx=XM_011290580.3;refGene_gn=RRN3;refGene_tx=XM_019820926.2
E3 32210764 32210917
refGene_gn=RRN3;refGene_tx=XM_011290580.3;refGene_gn=RRN3;refGene_tx=XM_019820926.2
E3 32212593 32212702
refGene_gn=RRN3;refGene_tx=XM_011290580.3;refGene_gn=RRN3;refGene_tx=XM_019820926.2
E3 32214814 32214999
refGene_gn=RRN3;refGene_tx=XM_011290580.3;refGene_gn=RRN3;refGene_tx=XM_019820926.2
E3 32215989 32216120
refGene_gn=RRN3;refGene_tx=XM_011290580.3;refGene_gn=RRN3;refGene_tx=XM_019820926.2
E3 32217769 32217900
refGene_gn=RRN3;refGene_tx=XM_011290580.3;refGene_gn=RRN3;refGene_tx=XM_019820926.2
E3 32219546 32219685
refGene_gn=RRN3;refGene_tx=XM_011290580.3;refGene_gn=RRN3;refGene_tx=XM_019820926.2
E3 32220967 32221060
refGene_gn=RRN3;refGene_tx=XM_011290580.3;refGene_gn=RRN3;refGene_tx=XM_019820926.2
E3 32222936 32223035
refGene_gn=RRN3;refGene_tx=XM_011290580.3;refGene_gn=RRN3;refGene_tx=XM_019820926.2
E3 32226851 32226921
refGene_gn=RRN3;refGene_tx=XM_011290580.3;refGene_gn=RRN3;refGene_tx=XM_019820926.2
E3 32227745 32227809
refGene_gn=RRN3;refGene_tx=XM_011290580.3;refGene_gn=RRN3;refGene_tx=XM_019820926.2
E3 32228386 32228446
refGene_gn=RRN3;refGene_tx=XM_011290580.3;refGene_gn=RRN3;refGene_tx=XM_019820926.2
E3 32228857 32228987
refGene_gn=RRN3;refGene_tx=XM_011290580.3;refGene_gn=RRN3;refGene_tx=XM_019820926.2
E3 32229098 32229188
refGene_gn=RRN3;refGene_tx=XM_011290580.3;refGene_gn=RRN3;refGene_tx=XM_019820926.2
E3 32235225 32235282
refGene_gn=RRN3;refGene_tx=XM_011290580.3;refGene_gn=RRN3;refGene_tx=XM_019820926.2
E3 32236066 32236172
refGene_gn=RRN3;refGene_tx=XM_011290580.3;refGene_gn=RRN3;refGene_tx=XM_019820926.2

E3 32237680 32237769
refGene_gn=RRN3;refGene_tx=XM_011290580.3;refGene_gn=RRN3;refGene_tx=XM_019820926.2
E3 32246021 32247578
refGene_gn=LOC102901380;refGene_tx=XM_011290579.3
E3 32283227 32283285 refGene_gn=PLA2G10;refGene_tx=XM_023246870.1
E3 32286385 32286536 refGene_gn=PLA2G10;refGene_tx=XM_023246870.1
E3 32288255 32288362 refGene_gn=PLA2G10;refGene_tx=XM_023246870.1
E3 32299065 32299208 refGene_gn=PLA2G10;refGene_tx=XM_023246870.1
E3 32302846 32303039
refGene_gn=BFAR;refGene_tx=XM_003998845.4;refGene_gn=BFAR;refGene_tx=XM_006942306.4;refGene_gn=BFAR;refGene_tx=XM_006942307.4;refGene_gn=BFAR;refGene_tx=XM_006942309.3;refGene_gn=BFAR;refGene_tx=XM_006942310.4;refGene_gn=BFAR;refGene_tx=XM_019820930.1
E3 32304972 32305175
refGene_gn=BFAR;refGene_tx=XM_003998845.4;refGene_gn=BFAR;refGene_tx=XM_006942306.4;refGene_gn=BFAR;refGene_tx=XM_006942307.4;refGene_gn=BFAR;refGene_tx=XM_006942309.3;refGene_gn=BFAR;refGene_tx=XM_006942310.4;refGene_gn=BFAR;refGene_tx=XM_019820930.1
E3 32307108 32307282
refGene_gn=BFAR;refGene_tx=XM_003998845.4;refGene_gn=BFAR;refGene_tx=XM_006942306.4;refGene_gn=BFAR;refGene_tx=XM_006942307.4;refGene_gn=BFAR;refGene_tx=XM_006942309.3;refGene_gn=BFAR;refGene_tx=XM_006942310.4;refGene_gn=BFAR;refGene_tx=XM_019820930.1
E3 32315165 32315310
refGene_gn=BFAR;refGene_tx=XM_003998845.4;refGene_gn=BFAR;refGene_tx=XM_006942306.4;refGene_gn=BFAR;refGene_tx=XM_006942307.4;refGene_gn=BFAR;refGene_tx=XM_006942309.3;refGene_gn=BFAR;refGene_tx=XM_006942310.4;refGene_gn=BFAR;refGene_tx=XM_019820930.1
E3 32318207 32318377
refGene_gn=BFAR;refGene_tx=XM_003998845.4;refGene_gn=BFAR;refGene_tx=XM_006942306.4;refGene_gn=BFAR;refGene_tx=XM_006942307.4;refGene_gn=BFAR;refGene_tx=XM_006942309.3;refGene_gn=BFAR;refGene_tx=XM_006942310.4;refGene_gn=BFAR;refGene_tx=XM_019820930.1
E3 32319432 32319637
refGene_gn=BFAR;refGene_tx=XM_003998845.4;refGene_gn=BFAR;refGene_tx=XM_006942306.4;refGene_gn=BFAR;refGene_tx=XM_006942307.4;refGene_gn=BFAR;refGene_tx=XM_006942309.3;refGene_gn=BFAR;refGene_tx=XM_006942310.4;refGene_gn=BFAR;refGene_tx=XM_019820930.1
E3 32319931 32319933 refGene_gn=BFAR;refGene_tx=XM_019820930.1
E3 32321185 32321448
refGene_gn=BFAR;refGene_tx=XM_003998845.4;refGene_gn=BFAR;refGene_tx=XM_006942306.4;refGene_gn=BFAR;refGene_tx=XM_006942307.4;refGene_gn=BFAR;refGene_tx=XM_006942309.3;refGene_gn=BFAR;refGene_tx=XM_006942310.4
E3 32340205 32340224
refGene_gn=PARN;refGene_tx=XM_023247035.1;refGene_gn=PARN;refGene_tx=XM_023247036.1;refGene_gn=PARN;refGene_tx=XM_023247037.1
E3 32340673 32340751
refGene_gn=PARN;refGene_tx=XM_023247035.1;refGene_gn=PARN;refGene_tx=XM_023247036.1;refGene_gn=PARN;refGene_tx=XM_023247037.1

E3 32342161 32342241
refGene_gn=PARN;refGene_tx=XM_023247035.1;refGene_gn=PARN;refGene_tx=XM_023247036.1;refGene_gn=PARN;refGene_tx=XM_023247037.1
E3 32342703 32342771
refGene_gn=PARN;refGene_tx=XM_023247035.1;refGene_gn=PARN;refGene_tx=XM_023247036.1;refGene_gn=PARN;refGene_tx=XM_023247037.1
E3 32342855 32342937
refGene_gn=PARN;refGene_tx=XM_023247035.1;refGene_gn=PARN;refGene_tx=XM_023247036.1;refGene_gn=PARN;refGene_tx=XM_023247037.1
E3 32349900 32349961
refGene_gn=PARN;refGene_tx=XM_023247035.1;refGene_gn=PARN;refGene_tx=XM_023247036.1;refGene_gn=PARN;refGene_tx=XM_023247037.1
E3 32354824 32354990
refGene_gn=PARN;refGene_tx=XM_023247035.1;refGene_gn=PARN;refGene_tx=XM_023247036.1;refGene_gn=PARN;refGene_tx=XM_023247037.1
E3 32356434 32356500
refGene_gn=PARN;refGene_tx=XM_023247035.1;refGene_gn=PARN;refGene_tx=XM_023247036.1;refGene_gn=PARN;refGene_tx=XM_023247037.1
E3 32357235 32357274
refGene_gn=PARN;refGene_tx=XM_023247035.1;refGene_gn=PARN;refGene_tx=XM_023247036.1;refGene_gn=PARN;refGene_tx=XM_023247037.1
E3 32358597 32358640
refGene_gn=PARN;refGene_tx=XM_023247035.1;refGene_gn=PARN;refGene_tx=XM_023247036.1;refGene_gn=PARN;refGene_tx=XM_023247037.1
E3 32361472 32361553
refGene_gn=PARN;refGene_tx=XM_023247035.1;refGene_gn=PARN;refGene_tx=XM_023247036.1;refGene_gn=PARN;refGene_tx=XM_023247037.1
E3 32366044 32366101
refGene_gn=PARN;refGene_tx=XM_023247035.1;refGene_gn=PARN;refGene_tx=XM_023247036.1;refGene_gn=PARN;refGene_tx=XM_023247037.1
E3 32370142 32370220
refGene_gn=PARN;refGene_tx=XM_023247035.1;refGene_gn=PARN;refGene_tx=XM_023247036.1
E3 32374523 32374567
refGene_gn=PARN;refGene_tx=XM_023247035.1;refGene_gn=PARN;refGene_tx=XM_023247036.1;refGene_gn=PARN;refGene_tx=XM_023247037.1
E3 32375660 32375703
refGene_gn=PARN;refGene_tx=XM_023247035.1;refGene_gn=PARN;refGene_tx=XM_023247036.1;refGene_gn=PARN;refGene_tx=XM_023247037.1
E3 32377129 32377205
refGene_gn=PARN;refGene_tx=XM_023247035.1;refGene_gn=PARN;refGene_tx=XM_023247036.1;refGene_gn=PARN;refGene_tx=XM_023247037.1
E3 32379647 32379758
refGene_gn=PARN;refGene_tx=XM_023247035.1;refGene_gn=PARN;refGene_tx=XM_023247036.1;refGene_gn=PARN;refGene_tx=XM_023247037.1
E3 32380076 32380146
refGene_gn=PARN;refGene_tx=XM_023247035.1;refGene_gn=PARN;refGene_tx=XM_023247036.1;refGene_gn=PARN;refGene_tx=XM_023247037.1
E3 32401618 32401674
refGene_gn=PARN;refGene_tx=XM_023247035.1;refGene_gn=PARN;refGene_tx=X

M_023247036.1;refGene_gn=PARN;refGene_tx=XM_023247037.1
E3 32403079 32403166
refGene_gn=PARN;refGene_tx=XM_023247035.1;refGene_gn=PARN;refGene_tx=XM_023247036.1;refGene_gn=PARN;refGene_tx=XM_023247037.1
E3 32406626 32406701
refGene_gn=PARN;refGene_tx=XM_023247035.1;refGene_gn=PARN;refGene_tx=XM_023247036.1;refGene_gn=PARN;refGene_tx=XM_023247037.1
E3 32465501 32465691
refGene_gn=PARN;refGene_tx=XM_023247035.1;refGene_gn=PARN;refGene_tx=XM_023247036.1;refGene_gn=PARN;refGene_tx=XM_023247037.1
E3 32495874 32496068
refGene_gn=PARN;refGene_tx=XM_023247035.1;refGene_gn=PARN;refGene_tx=XM_023247036.1;refGene_gn=PARN;refGene_tx=XM_023247037.1
E3 32506031 32506084
refGene_gn=PARN;refGene_tx=XM_023247035.1;refGene_gn=PARN;refGene_tx=XM_023247037.1
E3 32651381 32651914
refGene_gn=MKL2;refGene_tx=XM_006942313.3;refGene_gn=MKL2;refGene_tx=XM_006942314.3;refGene_gn=MKL2;refGene_tx=XM_019820931.2;refGene_gn=MKL2;refGene_tx=XM_019820932.2;refGene_gn=MKL2;refGene_tx=XM_019820933.2;refGene_gn=MKL2;refGene_tx=XM_019820934.2;refGene_gn=MKL2;refGene_tx=XM_019820935.2;refGene_gn=MKL2;refGene_tx=XM_019820936.2;refGene_gn=MKL2;refGene_tx=XM_023246366.1
E3 32654349 32654410
refGene_gn=MKL2;refGene_tx=XM_006942313.3;refGene_gn=MKL2;refGene_tx=XM_006942314.3;refGene_gn=MKL2;refGene_tx=XM_019820931.2;refGene_gn=MKL2;refGene_tx=XM_019820932.2;refGene_gn=MKL2;refGene_tx=XM_019820933.2;refGene_gn=MKL2;refGene_tx=XM_019820934.2;refGene_gn=MKL2;refGene_tx=XM_019820935.2;refGene_gn=MKL2;refGene_tx=XM_019820936.2;refGene_gn=MKL2;refGene_tx=XM_023246366.1
E3 32672517 32672655
refGene_gn=MKL2;refGene_tx=XM_006942313.3;refGene_gn=MKL2;refGene_tx=XM_006942314.3;refGene_gn=MKL2;refGene_tx=XM_019820931.2;refGene_gn=MKL2;refGene_tx=XM_019820932.2;refGene_gn=MKL2;refGene_tx=XM_019820933.2;refGene_gn=MKL2;refGene_tx=XM_019820934.2;refGene_gn=MKL2;refGene_tx=XM_019820935.2;refGene_gn=MKL2;refGene_tx=XM_019820936.2;refGene_gn=MKL2;refGene_tx=XM_023246366.1
E3 32672977 32673136
refGene_gn=MKL2;refGene_tx=XM_006942313.3;refGene_gn=MKL2;refGene_tx=XM_006942314.3;refGene_gn=MKL2;refGene_tx=XM_019820931.2;refGene_gn=MKL2;refGene_tx=XM_019820932.2;refGene_gn=MKL2;refGene_tx=XM_019820933.2;refGene_gn=MKL2;refGene_tx=XM_019820934.2;refGene_gn=MKL2;refGene_tx=XM_019820935.2;refGene_gn=MKL2;refGene_tx=XM_019820936.2;refGene_gn=MKL2;refGene_tx=XM_023246366.1
E3 32675711 32675855
refGene_gn=MKL2;refGene_tx=XM_006942313.3;refGene_gn=MKL2;refGene_tx=XM_006942314.3;refGene_gn=MKL2;refGene_tx=XM_019820931.2;refGene_gn=MKL2;refGene_tx=XM_019820932.2;refGene_gn=MKL2;refGene_tx=XM_019820933.2;refGene_gn=MKL2;refGene_tx=XM_019820934.2;refGene_gn=MKL2;refGene_tx=XM_019820935.2;refGene_gn=MKL2;refGene_tx=XM_019820936.2;refGene_gn=MKL

2;refGene_tx=XM_023246366.1
E3 32677519 32678566
refGene_gn=MKL2;refGene_tx=XM_006942313.3;refGene_gn=MKL2;refGene_tx=X
M_006942314.3;refGene_gn=MKL2;refGene_tx=XM_019820931.2;refGene_gn=MKL
2;refGene_tx=XM_019820932.2;refGene_gn=MKL2;refGene_tx=XM_019820933.2;
refGene_gn=MKL2;refGene_tx=XM_019820934.2;refGene_gn=MKL2;refGene_tx=X
M_019820935.2;refGene_gn=MKL2;refGene_tx=XM_019820936.2;refGene_gn=MKL
2;refGene_tx=XM_023246366.1
E3 32679508 32679641
refGene_gn=MKL2;refGene_tx=XM_006942313.3;refGene_gn=MKL2;refGene_tx=X
M_006942314.3;refGene_gn=MKL2;refGene_tx=XM_019820931.2;refGene_gn=MKL
2;refGene_tx=XM_019820932.2;refGene_gn=MKL2;refGene_tx=XM_019820933.2;
refGene_gn=MKL2;refGene_tx=XM_019820934.2;refGene_gn=MKL2;refGene_tx=X
M_019820935.2;refGene_gn=MKL2;refGene_tx=XM_019820936.2;refGene_gn=MKL
2;refGene_tx=XM_023246366.1
E3 32684825 32685073
refGene_gn=MKL2;refGene_tx=XM_006942313.3;refGene_gn=MKL2;refGene_tx=X
M_006942314.3;refGene_gn=MKL2;refGene_tx=XM_019820931.2;refGene_gn=MKL
2;refGene_tx=XM_019820932.2;refGene_gn=MKL2;refGene_tx=XM_019820933.2;
refGene_gn=MKL2;refGene_tx=XM_019820934.2;refGene_gn=MKL2;refGene_tx=X
M_019820935.2;refGene_gn=MKL2;refGene_tx=XM_019820936.2;refGene_gn=MKL
2;refGene_tx=XM_023246366.1
E3 32687781 32687919
refGene_gn=MKL2;refGene_tx=XM_006942313.3;refGene_gn=MKL2;refGene_tx=X
M_006942314.3;refGene_gn=MKL2;refGene_tx=XM_019820931.2;refGene_gn=MKL
2;refGene_tx=XM_019820932.2;refGene_gn=MKL2;refGene_tx=XM_019820933.2;
refGene_gn=MKL2;refGene_tx=XM_019820934.2;refGene_gn=MKL2;refGene_tx=X
M_019820935.2;refGene_gn=MKL2;refGene_tx=XM_019820936.2;refGene_gn=MKL
2;refGene_tx=XM_023246366.1
E3 32700974 32701153
refGene_gn=MKL2;refGene_tx=XM_006942313.3;refGene_gn=MKL2;refGene_tx=X
M_006942314.3;refGene_gn=MKL2;refGene_tx=XM_019820931.2;refGene_gn=MKL
2;refGene_tx=XM_019820932.2;refGene_gn=MKL2;refGene_tx=XM_019820933.2;
refGene_gn=MKL2;refGene_tx=XM_019820934.2;refGene_gn=MKL2;refGene_tx=X
M_019820935.2;refGene_gn=MKL2;refGene_tx=XM_019820936.2;refGene_gn=MKL
2;refGene_tx=XM_023246366.1
E3 32703139 32703301
refGene_gn=MKL2;refGene_tx=XM_006942313.3;refGene_gn=MKL2;refGene_tx=X
M_006942314.3;refGene_gn=MKL2;refGene_tx=XM_019820931.2;refGene_gn=MKL
2;refGene_tx=XM_019820932.2;refGene_gn=MKL2;refGene_tx=XM_019820933.2;
refGene_gn=MKL2;refGene_tx=XM_019820934.2;refGene_gn=MKL2;refGene_tx=X
M_019820935.2;refGene_gn=MKL2;refGene_tx=XM_019820936.2;refGene_gn=MKL
2;refGene_tx=XM_023246366.1
E3 32706927 32707003
refGene_gn=MKL2;refGene_tx=XM_006942313.3;refGene_gn=MKL2;refGene_tx=X
M_006942314.3;refGene_gn=MKL2;refGene_tx=XM_019820931.2;refGene_gn=MKL
2;refGene_tx=XM_019820932.2;refGene_gn=MKL2;refGene_tx=XM_019820933.2;
refGene_gn=MKL2;refGene_tx=XM_019820934.2;refGene_gn=MKL2;refGene_tx=X
M_019820935.2;refGene_gn=MKL2;refGene_tx=XM_019820936.2
E3 32708174 32708230

refGene_gn=MKL2;refGene_tx=XM_006942313.3;refGene_gn=MKL2;refGene_tx=XM_006942314.3;refGene_gn=MKL2;refGene_tx=XM_019820931.2;refGene_gn=MKL2;refGene_tx=XM_019820932.2;refGene_gn=MKL2;refGene_tx=XM_019820933.2;refGene_gn=MKL2;refGene_tx=XM_019820934.2;refGene_gn=MKL2;refGene_tx=XM_019820935.2;refGene_gn=MKL2;refGene_tx=XM_019820936.2
E3 32710264 32710330
refGene_gn=MKL2;refGene_tx=XM_006942313.3;refGene_gn=MKL2;refGene_tx=XM_006942314.3;refGene_gn=MKL2;refGene_tx=XM_019820931.2;refGene_gn=MKL2;refGene_tx=XM_019820932.2;refGene_gn=MKL2;refGene_tx=XM_019820933.2;refGene_gn=MKL2;refGene_tx=XM_019820934.2;refGene_gn=MKL2;refGene_tx=XM_019820935.2;refGene_gn=MKL2;refGene_tx=XM_019820936.2
E3 32729258 32729379
refGene_gn=MKL2;refGene_tx=XM_006942313.3;refGene_gn=MKL2;refGene_tx=XM_006942314.3
E3 32768643 32768706
refGene_gn=MKL2;refGene_tx=XM_019820931.2;refGene_gn=MKL2;refGene_tx=XM_019820932.2;refGene_gn=MKL2;refGene_tx=XM_019820935.2
E3 32774956 32775110
refGene_gn=MKL2;refGene_tx=XM_019820931.2;refGene_gn=MKL2;refGene_tx=XM_019820932.2;refGene_gn=MKL2;refGene_tx=XM_019820933.2;refGene_gn=MKL2;refGene_tx=XM_019820934.2;refGene_gn=MKL2;refGene_tx=XM_019820935.2;refGene_gn=MKL2;refGene_tx=XM_019820936.2
E3 32940510 32941244
refGene_gn=ERCC4;refGene_tx=XM_003998848.4;refGene_gn=ERCC4;refGene_tx=XM_019820937.2;refGene_gn=ERCC4;refGene_tx=XM_023246355.1
E3 32943554 32943667
refGene_gn=ERCC4;refGene_tx=XM_003998848.4;refGene_gn=ERCC4;refGene_tx=XM_019820937.2;refGene_gn=ERCC4;refGene_tx=XM_023246355.1
E3 32949450 32949543
refGene_gn=ERCC4;refGene_tx=XM_003998848.4;refGene_gn=ERCC4;refGene_tx=XM_019820937.2;refGene_gn=ERCC4;refGene_tx=XM_023246355.1
E3 32951812 32952410
refGene_gn=ERCC4;refGene_tx=XM_003998848.4;refGene_gn=ERCC4;refGene_tx=XM_019820937.2;refGene_gn=ERCC4;refGene_tx=XM_023246355.1
E3 32954822 32954933
refGene_gn=ERCC4;refGene_tx=XM_003998848.4;refGene_gn=ERCC4;refGene_tx=XM_023246355.1
E3 32956101 32956230
refGene_gn=ERCC4;refGene_tx=XM_003998848.4;refGene_gn=ERCC4;refGene_tx=XM_019820937.2;refGene_gn=ERCC4;refGene_tx=XM_023246355.1
E3 32957483 32957664
refGene_gn=ERCC4;refGene_tx=XM_003998848.4;refGene_gn=ERCC4;refGene_tx=XM_019820937.2;refGene_gn=ERCC4;refGene_tx=XM_023246355.1
E3 32960763 32960971
refGene_gn=ERCC4;refGene_tx=XM_003998848.4;refGene_gn=ERCC4;refGene_tx=XM_019820937.2;refGene_gn=ERCC4;refGene_tx=XM_023246355.1
E3 32962350 32962546
refGene_gn=ERCC4;refGene_tx=XM_003998848.4;refGene_gn=ERCC4;refGene_tx=XM_019820937.2;refGene_gn=ERCC4;refGene_tx=XM_023246355.1
E3 32964454 32964650 refGene_gn=ERCC4;refGene_tx=XM_023246355.1

E3 32968285 32968466
refGene_gn=ERCC4;refGene_tx=XM_003998848.4;refGene_gn=ERCC4;refGene_tx=XM_019820937.2
E3 32969987 32970194
refGene_gn=ERCC4;refGene_tx=XM_003998848.4;refGene_gn=ERCC4;refGene_tx=XM_019820937.2
E3 33605081 33605461 refGene_gn=SHISA9;refGene_tx=XM_023246581.1
E3 33622409 33622457 refGene_gn=SHISA9;refGene_tx=XM_023246581.1
E3 33630119 33630275 refGene_gn=SHISA9;refGene_tx=XM_023246581.1
E3 33872491 33872619 refGene_gn=SHISA9;refGene_tx=XM_023246581.1
E3 33875474 33875481 refGene_gn=SHISA9;refGene_tx=XM_023246581.1
E3 33885898 33886124 refGene_gn=SHISA9;refGene_tx=XM_023246581.1
E3 33886125 33886452 refGene_gn=SHISA9;refGene_tx=XM_023246581.1
E3 33967449 33967519 refGene_gn=CPPED1;refGene_tx=XM_011290577.3
E3 33985911 33986130
refGene_gn=CPPED1;refGene_tx=XM_011290577.3;refGene_gn=CPPED1;refGene_tx=XM_019820939.2
E3 34035759 34036182
refGene_gn=CPPED1;refGene_tx=XM_011290577.3;refGene_gn=CPPED1;refGene_tx=XM_019820939.2
E3 34075651 34075881
refGene_gn=CPPED1;refGene_tx=XM_011290577.3;refGene_gn=CPPED1;refGene_tx=XM_019820939.2
E3 34156949 34157079 refGene_gn=SNX29;refGene_tx=XM_023246582.1
E3 34194466 34194606 refGene_gn=SNX29;refGene_tx=XM_023246582.1
E3 34231565 34231706 refGene_gn=SNX29;refGene_tx=XM_023246582.1
E3 34289127 34289209 refGene_gn=SNX29;refGene_tx=XM_023246582.1
E3 34292114 34292170 refGene_gn=SNX29;refGene_tx=XM_023246582.1
E3 34326460 34326577 refGene_gn=SNX29;refGene_tx=XM_023246582.1
E3 34392443 34392550 refGene_gn=SNX29;refGene_tx=XM_023246582.1
E3 34456673 34456756 refGene_gn=SNX29;refGene_tx=XM_023246582.1
E3 34528934 34529063 refGene_gn=SNX29;refGene_tx=XM_023246582.1
E3 34531128 34531192 refGene_gn=SNX29;refGene_tx=XM_023246582.1
E3 34587265 34587427 refGene_gn=SNX29;refGene_tx=XM_023246582.1
E3 34596624 34596743 refGene_gn=SNX29;refGene_tx=XM_023246582.1
E3 34603835 34604211 refGene_gn=SNX29;refGene_tx=XM_023246582.1
E3 34606162 34606411 refGene_gn=SNX29;refGene_tx=XM_023246582.1
E3 34607581 34607652 refGene_gn=SNX29;refGene_tx=XM_023246582.1
E3 34610388 34610569 refGene_gn=SNX29;refGene_tx=XM_023246582.1
E3 34630649 34630713 refGene_gn=SNX29;refGene_tx=XM_023246582.1
E3 34630714 34630775 refGene_gn=SNX29;refGene_tx=XM_023246582.1
E3 34650056 34650109 refGene_gn=SNX29;refGene_tx=XM_023246582.1
E3 34651810 34651875 refGene_gn=SNX29;refGene_tx=XM_023246582.1
E3 34669940 34669947 refGene_gn=SNX29;refGene_tx=XM_023246582.1
E3 34674282 34674563
refGene_gn=TNFRSF17;refGene_tx=XM_006942320.2;refGene_gn=TNFRSF17;refGene_tx=XM_019820946.1;refGene_gn=TNFRSF17;refGene_tx=XM_023246692.1
E3 34676255 34676387
refGene_gn=TNFRSF17;refGene_tx=XM_006942320.2;refGene_gn=TNFRSF17;refGene_tx=XM_019820946.1;refGene_gn=TNFRSF17;refGene_tx=XM_023246692.1

E3 34676743 34676861 refGene_gn=TNFRSF17;refGene_tx=XM_006942320.2
E3 34698176 34698183
refGene_gn=TNFRSF17;refGene_tx=XM_019820946.1;refGene_gn=TNFRSF17;refGene_tx=XM_023246692.1
E3 34747038 34747393 refGene_gn=GSPT1;refGene_tx=XM_023246999.1
E3 34761507 34761552 refGene_gn=GSPT1;refGene_tx=XM_023246999.1
E3 34761666 34761708
refGene_gn=GSPT1;refGene_tx=XM_023246999.1;refGene_gn=GSPT1;refGene_tx=XM_023247000.1
E3 34762439 34762667
refGene_gn=GSPT1;refGene_tx=XM_023246999.1;refGene_gn=GSPT1;refGene_tx=XM_023247000.1
E3 34763666 34763700
refGene_gn=GSPT1;refGene_tx=XM_023246999.1;refGene_gn=GSPT1;refGene_tx=XM_023247000.1
E3 34766891 34766969
refGene_gn=GSPT1;refGene_tx=XM_023246999.1;refGene_gn=GSPT1;refGene_tx=XM_023247000.1
E3 34769198 34769379
refGene_gn=GSPT1;refGene_tx=XM_023246999.1;refGene_gn=GSPT1;refGene_tx=XM_023247000.1
E3 34770007 34770162
refGene_gn=GSPT1;refGene_tx=XM_023246999.1;refGene_gn=GSPT1;refGene_tx=XM_023247000.1
E3 34770559 34770700
refGene_gn=GSPT1;refGene_tx=XM_023246999.1;refGene_gn=GSPT1;refGene_tx=XM_023247000.1
E3 34771791 34771885
refGene_gn=GSPT1;refGene_tx=XM_023246999.1;refGene_gn=GSPT1;refGene_tx=XM_023247000.1
E3 34772888 34772969
refGene_gn=GSPT1;refGene_tx=XM_023246999.1;refGene_gn=GSPT1;refGene_tx=XM_023247000.1
E3 34776736 34776910
refGene_gn=GSPT1;refGene_tx=XM_023246999.1;refGene_gn=GSPT1;refGene_tx=XM_023247000.1
E3 34777785 34777875
refGene_gn=GSPT1;refGene_tx=XM_023246999.1;refGene_gn=GSPT1;refGene_tx=XM_023247000.1
E3 34778054 34778223
refGene_gn=GSPT1;refGene_tx=XM_023246999.1;refGene_gn=GSPT1;refGene_tx=XM_023247000.1
E3 34779916 34779969
refGene_gn=GSPT1;refGene_tx=XM_023246999.1;refGene_gn=GSPT1;refGene_tx=XM_023247000.1
E3 34789241 34789340 refGene_gn=RSL1D1;refGene_tx=XM_023246580.1
E3 34790060 34790200 refGene_gn=RSL1D1;refGene_tx=XM_023246580.1
E3 34791711 34791850 refGene_gn=RSL1D1;refGene_tx=XM_023246580.1
E3 34792497 34792646 refGene_gn=RSL1D1;refGene_tx=XM_023246580.1
E3 34792735 34792837 refGene_gn=RSL1D1;refGene_tx=XM_023246580.1

E3 34794022 34794116 refGene_gn=RSL1D1;refGene_tx=XM_023246580.1
E3 34794203 34794329 refGene_gn=RSL1D1;refGene_tx=XM_023246580.1
E3 34796724 34797021 refGene_gn=RSL1D1;refGene_tx=XM_023246580.1
E3 34797552 34797879 refGene_gn=RSL1D1;refGene_tx=XM_023246580.1
E3 34848736 34848804
refGene_gn=ZC3H7A;refGene_tx=XM_006942322.4;refGene_gn=ZC3H7A;refGene_tx=XM_019820948.2;refGene_gn=ZC3H7A;refGene_tx=XM_023246904.1
E3 34849936 34849976
refGene_gn=ZC3H7A;refGene_tx=XM_006942322.4;refGene_gn=ZC3H7A;refGene_tx=XM_019820948.2;refGene_gn=ZC3H7A;refGene_tx=XM_023246904.1
E3 34851934 34852132
refGene_gn=ZC3H7A;refGene_tx=XM_006942322.4;refGene_gn=ZC3H7A;refGene_tx=XM_019820948.2;refGene_gn=ZC3H7A;refGene_tx=XM_023246904.1
E3 34853532 34853691
refGene_gn=ZC3H7A;refGene_tx=XM_006942322.4;refGene_gn=ZC3H7A;refGene_tx=XM_019820948.2;refGene_gn=ZC3H7A;refGene_tx=XM_023246904.1
E3 34853933 34854014
refGene_gn=ZC3H7A;refGene_tx=XM_006942322.4;refGene_gn=ZC3H7A;refGene_tx=XM_019820948.2;refGene_gn=ZC3H7A;refGene_tx=XM_023246904.1
E3 34854127 34854166
refGene_gn=ZC3H7A;refGene_tx=XM_006942322.4;refGene_gn=ZC3H7A;refGene_tx=XM_019820948.2;refGene_gn=ZC3H7A;refGene_tx=XM_023246904.1
E3 34854819 34854853
refGene_gn=ZC3H7A;refGene_tx=XM_006942322.4;refGene_gn=ZC3H7A;refGene_tx=XM_019820948.2;refGene_gn=ZC3H7A;refGene_tx=XM_023246904.1
E3 34855343 34855627
refGene_gn=ZC3H7A;refGene_tx=XM_006942322.4;refGene_gn=ZC3H7A;refGene_tx=XM_019820948.2;refGene_gn=ZC3H7A;refGene_tx=XM_023246904.1
E3 34858345 34858550
refGene_gn=ZC3H7A;refGene_tx=XM_006942322.4;refGene_gn=ZC3H7A;refGene_tx=XM_019820948.2;refGene_gn=ZC3H7A;refGene_tx=XM_023246904.1
E3 34859639 34859704
refGene_gn=ZC3H7A;refGene_tx=XM_006942322.4;refGene_gn=ZC3H7A;refGene_tx=XM_019820948.2;refGene_gn=ZC3H7A;refGene_tx=XM_023246904.1
E3 34860268 34860455
refGene_gn=ZC3H7A;refGene_tx=XM_006942322.4;refGene_gn=ZC3H7A;refGene_tx=XM_019820948.2;refGene_gn=ZC3H7A;refGene_tx=XM_023246904.1
E3 34861159 34861321
refGene_gn=ZC3H7A;refGene_tx=XM_006942322.4;refGene_gn=ZC3H7A;refGene_tx=XM_019820948.2;refGene_gn=ZC3H7A;refGene_tx=XM_023246904.1
E3 34863783 34863980
refGene_gn=ZC3H7A;refGene_tx=XM_006942322.4;refGene_gn=ZC3H7A;refGene_tx=XM_019820948.2;refGene_gn=ZC3H7A;refGene_tx=XM_023246904.1
E3 34864252 34864353
refGene_gn=ZC3H7A;refGene_tx=XM_006942322.4;refGene_gn=ZC3H7A;refGene_tx=XM_019820948.2;refGene_gn=ZC3H7A;refGene_tx=XM_023246904.1
E3 34864813 34864995
refGene_gn=ZC3H7A;refGene_tx=XM_006942322.4;refGene_gn=ZC3H7A;refGene_tx=XM_019820948.2;refGene_gn=ZC3H7A;refGene_tx=XM_023246904.1
E3 34865882 34865959

refGene_gn=ZC3H7A;refGene_tx=XM_006942322.4;refGene_gn=ZC3H7A;refGene_tx=XM_019820948.2;refGene_gn=ZC3H7A;refGene_tx=XM_023246904.1
E3 34866569 34866703
refGene_gn=ZC3H7A;refGene_tx=XM_006942322.4;refGene_gn=ZC3H7A;refGene_tx=XM_019820948.2;refGene_gn=ZC3H7A;refGene_tx=XM_023246904.1
E3 34867176 34867282
refGene_gn=ZC3H7A;refGene_tx=XM_006942322.4;refGene_gn=ZC3H7A;refGene_tx=XM_019820948.2;refGene_gn=ZC3H7A;refGene_tx=XM_023246904.1
E3 34869657 34869766
refGene_gn=ZC3H7A;refGene_tx=XM_006942322.4;refGene_gn=ZC3H7A;refGene_tx=XM_019820948.2;refGene_gn=ZC3H7A;refGene_tx=XM_023246904.1
E3 34871697 34871831
refGene_gn=ZC3H7A;refGene_tx=XM_006942322.4;refGene_gn=ZC3H7A;refGene_tx=XM_019820948.2;refGene_gn=ZC3H7A;refGene_tx=XM_023246904.1
E3 34873623 34873787
refGene_gn=ZC3H7A;refGene_tx=XM_006942322.4;refGene_gn=ZC3H7A;refGene_tx=XM_019820948.2;refGene_gn=ZC3H7A;refGene_tx=XM_023246904.1
E3 34874600 34874790
refGene_gn=ZC3H7A;refGene_tx=XM_006942322.4;refGene_gn=ZC3H7A;refGene_tx=XM_019820948.2;refGene_gn=ZC3H7A;refGene_tx=XM_023246904.1
E3 34879567 34879830 refGene_gn=TXNDC11;refGene_tx=XM_023246824.1
E3 34886669 34886886 refGene_gn=TXNDC11;refGene_tx=XM_023246824.1
E3 34890381 34890479
refGene_gn=TXNDC11;refGene_tx=XM_019820950.1;refGene_gn=TXNDC11;refGene_tx=XM_023246824.1
E3 34894057 34894187 refGene_gn=TXNDC11;refGene_tx=XM_023246824.1
E3 34903933 34904027
refGene_gn=TXNDC11;refGene_tx=XM_019820950.1;refGene_gn=TXNDC11;refGene_tx=XM_023246824.1
E3 34910206 34910216 refGene_gn=TXNDC11;refGene_tx=XM_019820951.1
E3 34920453 34920566
refGene_gn=TXNDC11;refGene_tx=XM_019820950.1;refGene_gn=TXNDC11;refGene_tx=XM_019820951.1;refGene_gn=TXNDC11;refGene_tx=XM_023246824.1
E3 34922640 34922841
refGene_gn=TXNDC11;refGene_tx=XM_019820950.1;refGene_gn=TXNDC11;refGene_tx=XM_019820951.1;refGene_gn=TXNDC11;refGene_tx=XM_023246824.1
E3 34927040 34927833
refGene_gn=TXNDC11;refGene_tx=XM_019820950.1;refGene_gn=TXNDC11;refGene_tx=XM_019820951.1;refGene_gn=TXNDC11;refGene_tx=XM_023246824.1
E3 34930339 34930482
refGene_gn=TXNDC11;refGene_tx=XM_019820950.1;refGene_gn=TXNDC11;refGene_tx=XM_019820951.1;refGene_gn=TXNDC11;refGene_tx=XM_023246824.1
E3 34930801 34930911
refGene_gn=TXNDC11;refGene_tx=XM_019820950.1;refGene_gn=TXNDC11;refGene_tx=XM_019820951.1;refGene_gn=TXNDC11;refGene_tx=XM_023246824.1
E3 34934079 34934160
refGene_gn=TXNDC11;refGene_tx=XM_019820950.1;refGene_gn=TXNDC11;refGene_tx=XM_019820951.1;refGene_gn=TXNDC11;refGene_tx=XM_023246824.1
E3 34938518 34939161
refGene_gn=TXNDC11;refGene_tx=XM_019820950.1;refGene_gn=TXNDC11;refGen

e_tx=XM_019820951.1;refGene_gn=TXNDC11;refGene_tx=XM_023246824.1
E3 34941930 34942194 refGene_gn=SNN;refGene_tx=XM_023246825.1
E3 35046739 35046959 refGene_gn=LITAF;refGene_tx=XM_023246210.1
E3 35049842 35049941 refGene_gn=LITAF;refGene_tx=XM_023246210.1
E3 35049943 35049998 refGene_gn=LITAF;refGene_tx=XM_023246210.1
E3 35052738 35052847 refGene_gn=LITAF;refGene_tx=XM_023246210.1
E3 35088150 35088175
refGene_gn=LOC111558241;refGene_tx=XM_023246583.1
E3 35088630 35088830
refGene_gn=LOC111558241;refGene_tx=XM_023246583.1
E3 35088904 35089039
refGene_gn=LOC111558241;refGene_tx=XM_023246583.1
E3 35089787 35090009
refGene_gn=LOC111558241;refGene_tx=XM_023246583.1
E3 35092241 35092379
refGene_gn=LOC102899050;refGene_tx=XM_023247099.1;refGene_gn=LOC102899
050;refGene_tx=XM_023247100.1
E3 35097787 35097891
refGene_gn=LOC102899050;refGene_tx=XM_023247099.1;refGene_gn=LOC102899
050;refGene_tx=XM_023247100.1
E3 35099985 35100151
refGene_gn=LOC102899050;refGene_tx=XM_023247099.1;refGene_gn=LOC102899
050;refGene_tx=XM_023247100.1
E3 35106016 35106202
refGene_gn=LOC102899050;refGene_tx=XM_023247099.1;refGene_gn=LOC102899
050;refGene_tx=XM_023247100.1
E3 35106317 35106470
refGene_gn=LOC102899050;refGene_tx=XM_023247099.1;refGene_gn=LOC102899
050;refGene_tx=XM_023247100.1
E3 35108146 35108291
refGene_gn=LOC102899050;refGene_tx=XM_023247099.1;refGene_gn=LOC102899
050;refGene_tx=XM_023247100.1
E3 35112920 35113079
refGene_gn=LOC102899050;refGene_tx=XM_023247099.1;refGene_gn=LOC102899
050;refGene_tx=XM_023247100.1
E3 35113828 35113953
refGene_gn=LOC102899050;refGene_tx=XM_023247099.1;refGene_gn=LOC102899
050;refGene_tx=XM_023247100.1
E3 35116862 35117036
refGene_gn=LOC102899050;refGene_tx=XM_023247099.1;refGene_gn=LOC102899
050;refGene_tx=XM_023247100.1
E3 35120945 35121101
refGene_gn=LOC102899050;refGene_tx=XM_023247099.1;refGene_gn=LOC102899
050;refGene_tx=XM_023247100.1
E3 35121870 35122055
refGene_gn=LOC102899050;refGene_tx=XM_023247099.1;refGene_gn=LOC102899
050;refGene_tx=XM_023247100.1
E3 35123000 35123183
refGene_gn=LOC102899050;refGene_tx=XM_023247099.1;refGene_gn=LOC102899
050;refGene_tx=XM_023247100.1

E3 35124154 35124246
refGene_gn=LOC102899050;refGene_tx=XM_023247099.1;refGene_gn=LOC102899050;refGene_tx=XM_023247100.1
E3 35126028 35126192
refGene_gn=LOC102899050;refGene_tx=XM_023247099.1;refGene_gn=LOC102899050;refGene_tx=XM_023247100.1
E3 35128186 35128346
refGene_gn=LOC102899050;refGene_tx=XM_023247099.1;refGene_gn=LOC102899050;refGene_tx=XM_023247100.1
E3 35128471 35128551
refGene_gn=LOC102899050;refGene_tx=XM_023247099.1;refGene_gn=LOC102899050;refGene_tx=XM_023247100.1
E3 35130797 35130984
refGene_gn=LOC102899050;refGene_tx=XM_023247099.1;refGene_gn=LOC102899050;refGene_tx=XM_023247100.1
E3 35131772 35131928
refGene_gn=LOC102899050;refGene_tx=XM_023247099.1;refGene_gn=LOC102899050;refGene_tx=XM_023247100.1
E3 35132298 35132432
refGene_gn=LOC102899050;refGene_tx=XM_023247099.1;refGene_gn=LOC102899050;refGene_tx=XM_023247100.1
E3 35133507 35133666
refGene_gn=LOC102899050;refGene_tx=XM_023247099.1;refGene_gn=LOC102899050;refGene_tx=XM_023247100.1
E3 35139357 35139538
refGene_gn=LOC102899050;refGene_tx=XM_023247099.1;refGene_gn=LOC102899050;refGene_tx=XM_023247100.1
E3 35142919 35143131
refGene_gn=LOC102899050;refGene_tx=XM_023247099.1;refGene_gn=LOC102899050;refGene_tx=XM_023247100.1
E3 35145063 35145252
refGene_gn=LOC102899050;refGene_tx=XM_023247099.1;refGene_gn=LOC102899050;refGene_tx=XM_023247100.1
E3 35145691 35145831
refGene_gn=LOC102899050;refGene_tx=XM_023247099.1;refGene_gn=LOC102899050;refGene_tx=XM_023247100.1
E3 35146801 35146964
refGene_gn=LOC102899050;refGene_tx=XM_023247099.1;refGene_gn=LOC102899050;refGene_tx=XM_023247100.1
E3 35147089 35147176
refGene_gn=LOC102899050;refGene_tx=XM_023247099.1;refGene_gn=LOC102899050;refGene_tx=XM_023247100.1
E3 35150856 35150994
refGene_gn=LOC102899050;refGene_tx=XM_023247099.1;refGene_gn=LOC102899050;refGene_tx=XM_023247100.1
E3 35151088 35151208
refGene_gn=LOC102899050;refGene_tx=XM_023247099.1;refGene_gn=LOC102899050;refGene_tx=XM_023247100.1
E3 35152132 35152225
refGene_gn=LOC102899050;refGene_tx=XM_023247099.1;refGene_gn=LOC102899

050;refGene_tx=XM_023247100.1
E3	35156885	35157103
refGene_gn=LOC102899050;refGene_tx=XM_023247099.1;refGene_gn=LOC102899050;refGene_tx=XM_023247100.1
E3	35159165	35159359
refGene_gn=LOC102899050;refGene_tx=XM_023247099.1;refGene_gn=LOC102899050;refGene_tx=XM_023247100.1
E3	35170284	35170593
refGene_gn=LOC102899050;refGene_tx=XM_023247099.1
E3	35171362	35171592
refGene_gn=LOC102899050;refGene_tx=XM_023247099.1;refGene_gn=LOC102899050;refGene_tx=XM_023247100.1
E3	35171667	35171829
refGene_gn=LOC102899050;refGene_tx=XM_023247099.1;refGene_gn=LOC102899050;refGene_tx=XM_023247100.1
E3	35172797	35172875
refGene_gn=LOC102899050;refGene_tx=XM_023247099.1;refGene_gn=LOC102899050;refGene_tx=XM_023247100.1
E3	35173796	35174156
refGene_gn=LOC102899050;refGene_tx=XM_023247099.1;refGene_gn=LOC102899050;refGene_tx=XM_023247100.1
E3	35175024	35175381
refGene_gn=LOC102899050;refGene_tx=XM_023247099.1;refGene_gn=LOC102899050;refGene_tx=XM_023247100.1
E3	35176535	35176889
refGene_gn=LOC102899050;refGene_tx=XM_023247099.1;refGene_gn=LOC102899050;refGene_tx=XM_023247100.1
E3	35178836	35178969
refGene_gn=LOC102899050;refGene_tx=XM_023247099.1;refGene_gn=LOC102899050;refGene_tx=XM_023247100.1
E3	35180553	35180703
refGene_gn=LOC102899050;refGene_tx=XM_023247099.1;refGene_gn=LOC102899050;refGene_tx=XM_023247100.1
E3	35180807	35180924
refGene_gn=LOC102899050;refGene_tx=XM_023247099.1;refGene_gn=LOC102899050;refGene_tx=XM_023247100.1
E3	35182984	35183172
refGene_gn=LOC102899050;refGene_tx=XM_023247099.1;refGene_gn=LOC102899050;refGene_tx=XM_023247100.1
E3	35248666	35248815	refGene_gn=RMI2;refGene_tx=XM_023246716.1
E3	35251375	35251688	refGene_gn=RMI2;refGene_tx=XM_023246716.1
E3	35295948	35296242	refGene_gn=PRM3;refGene_tx=XM_003998864.5
E3	35299712	35300055	refGene_gn=TNP2;refGene_tx=XM_003998865.5
E3	35300882	35300899	refGene_gn=TNP2;refGene_tx=XM_003998865.5
E3	35302575	35302606	refGene_gn=SOCS1;refGene_tx=XM_023246585.1
E3	35312111	35312623	refGene_gn=SOCS1;refGene_tx=XM_023246585.1
E3	35363242	35363583	refGene_gn=CLEC16A;refGene_tx=XM_019820956.2
E3	35373345	35373507	refGene_gn=CLEC16A;refGene_tx=XM_019820956.2
E3	35408093	35408139	refGene_gn=CLEC16A;refGene_tx=XM_019820956.2
E3	35408140	35408259	refGene_gn=CLEC16A;refGene_tx=XM_019820956.2

E3 35409959 35410164 refGene_gn=CLEC16A;refGene_tx=XM_019820956.2
E3 35412894 35413046 refGene_gn=CLEC16A;refGene_tx=XM_019820956.2
E3 35461584 35461705 refGene_gn=CLEC16A;refGene_tx=XM_019820956.2
E3 35470056 35470185 refGene_gn=CLEC16A;refGene_tx=XM_019820956.2
E3 35473466 35473517 refGene_gn=CLEC16A;refGene_tx=XM_019820956.2
E3 35476082 35476127 refGene_gn=CLEC16A;refGene_tx=XM_019820956.2
E3 35476874 35476984 refGene_gn=CLEC16A;refGene_tx=XM_019820956.2
E3 35480218 35480341 refGene_gn=CLEC16A;refGene_tx=XM_019820956.2
E3 35492999 35493100 refGene_gn=CLEC16A;refGene_tx=XM_019820956.2
E3 35497170 35497300 refGene_gn=CLEC16A;refGene_tx=XM_019820956.2
E3 35510347 35510579 refGene_gn=CLEC16A;refGene_tx=XM_019820956.2
E3 35526614 35526728 refGene_gn=CLEC16A;refGene_tx=XM_019820956.2
E3 35529794 35529848 refGene_gn=CLEC16A;refGene_tx=XM_019820956.2
E3 35531395 35531570 refGene_gn=CLEC16A;refGene_tx=XM_019820956.2
E3 35534998 35535122 refGene_gn=CLEC16A;refGene_tx=XM_019820956.2
E3 35535515 35535521 refGene_gn=CLEC16A;refGene_tx=XM_019820956.2
E3 35536847 35536953 refGene_gn=CLEC16A;refGene_tx=XM_019820956.2
E3 35540432 35540581 refGene_gn=CLEC16A;refGene_tx=XM_019820956.2
E3 35547713 35547847 refGene_gn=CLEC16A;refGene_tx=XM_019820956.2
E3 35551114 35551243 refGene_gn=CLEC16A;refGene_tx=XM_019820956.2
E3 35563009 35563089 refGene_gn=CLEC16A;refGene_tx=XM_019820956.2
E3 35565950 35566238
refGene_gn=DEXI;refGene_tx=XM_023247033.1;refGene_gn=DEXI;refGene_tx=XM_023247034.1
E3 35582837 35582913
refGene_gn=CIITA;refGene_tx=XM_011290571.3;refGene_gn=CIITA;refGene_tx=XM_011290572.2;refGene_gn=CIITA;refGene_tx=XM_011290573.3;refGene_gn=CIITA;refGene_tx=XM_011290574.2;refGene_gn=CIITA;refGene_tx=XM_011290575.3;refGene_gn=CIITA;refGene_tx=XM_019820963.1;refGene_gn=CIITA;refGene_tx=XM_019820965.1;refGene_gn=CIITA;refGene_tx=XM_019820966.2
E3 35583519 35583603
refGene_gn=CIITA;refGene_tx=XM_011290571.3;refGene_gn=CIITA;refGene_tx=XM_011290572.2;refGene_gn=CIITA;refGene_tx=XM_011290573.3;refGene_gn=CIITA;refGene_tx=XM_011290574.2;refGene_gn=CIITA;refGene_tx=XM_011290575.3;refGene_gn=CIITA;refGene_tx=XM_019820963.1;refGene_gn=CIITA;refGene_tx=XM_019820965.1;refGene_gn=CIITA;refGene_tx=XM_019820966.2
E3 35583783 35583867
refGene_gn=CIITA;refGene_tx=XM_011290571.3;refGene_gn=CIITA;refGene_tx=XM_011290572.2;refGene_gn=CIITA;refGene_tx=XM_011290573.3;refGene_gn=CIITA;refGene_tx=XM_011290574.2;refGene_gn=CIITA;refGene_tx=XM_011290575.3;refGene_gn=CIITA;refGene_tx=XM_019820963.1;refGene_gn=CIITA;refGene_tx=XM_019820965.1;refGene_gn=CIITA;refGene_tx=XM_019820966.2
E3 35586267 35586354
refGene_gn=CIITA;refGene_tx=XM_011290571.3;refGene_gn=CIITA;refGene_tx=XM_011290572.2;refGene_gn=CIITA;refGene_tx=XM_011290573.3;refGene_gn=CIITA;refGene_tx=XM_011290574.2;refGene_gn=CIITA;refGene_tx=XM_011290575.3;refGene_gn=CIITA;refGene_tx=XM_019820963.1;refGene_gn=CIITA;refGene_tx=XM_019820965.1;refGene_gn=CIITA;refGene_tx=XM_019820966.2
E3 35588217 35588310
refGene_gn=CIITA;refGene_tx=XM_011290571.3;refGene_gn=CIITA;refGene_tx

=XM_011290572.2;refGene_gn=CIITA;refGene_tx=XM_011290573.3;refGene_gn=CIITA;refGene_tx=XM_011290574.2;refGene_gn=CIITA;refGene_tx=XM_011290575.3;refGene_gn=CIITA;refGene_tx=XM_019820963.1;refGene_gn=CIITA;refGene_tx=XM_019820965.1;refGene_gn=CIITA;refGene_tx=XM_019820966.2
E3 35588891 35588972
refGene_gn=CIITA;refGene_tx=XM_011290571.3;refGene_gn=CIITA;refGene_tx=XM_011290572.2;refGene_gn=CIITA;refGene_tx=XM_011290573.3;refGene_gn=CIITA;refGene_tx=XM_011290574.2;refGene_gn=CIITA;refGene_tx=XM_011290575.3;refGene_gn=CIITA;refGene_tx=XM_019820963.1;refGene_gn=CIITA;refGene_tx=XM_019820965.1;refGene_gn=CIITA;refGene_tx=XM_019820966.2
E3 35590549 35590641 refGene_gn=CIITA;refGene_tx=XM_023247032.1
E3 35591492 35591564
refGene_gn=CIITA;refGene_tx=XM_011290571.3;refGene_gn=CIITA;refGene_tx=XM_011290572.2;refGene_gn=CIITA;refGene_tx=XM_011290573.3;refGene_gn=CIITA;refGene_tx=XM_011290574.2;refGene_gn=CIITA;refGene_tx=XM_011290575.3;refGene_gn=CIITA;refGene_tx=XM_019820963.1;refGene_gn=CIITA;refGene_tx=XM_019820965.1;refGene_gn=CIITA;refGene_tx=XM_019820966.2;refGene_gn=CIITA;refGene_tx=XM_023247032.1
E3 35592446 35592645
refGene_gn=CIITA;refGene_tx=XM_011290571.3;refGene_gn=CIITA;refGene_tx=XM_011290572.2;refGene_gn=CIITA;refGene_tx=XM_011290573.3;refGene_gn=CIITA;refGene_tx=XM_011290574.2;refGene_gn=CIITA;refGene_tx=XM_011290575.3;refGene_gn=CIITA;refGene_tx=XM_019820963.1;refGene_gn=CIITA;refGene_tx=XM_019820965.1;refGene_gn=CIITA;refGene_tx=XM_019820966.2;refGene_gn=CIITA;refGene_tx=XM_019820967.1;refGene_gn=CIITA;refGene_tx=XM_023247032.1
E3 35593425 35595076
refGene_gn=CIITA;refGene_tx=XM_011290571.3;refGene_gn=CIITA;refGene_tx=XM_011290572.2;refGene_gn=CIITA;refGene_tx=XM_011290573.3;refGene_gn=CIITA;refGene_tx=XM_011290574.2;refGene_gn=CIITA;refGene_tx=XM_011290575.3;refGene_gn=CIITA;refGene_tx=XM_019820963.1;refGene_gn=CIITA;refGene_tx=XM_019820965.1;refGene_gn=CIITA;refGene_tx=XM_019820966.2;refGene_gn=CIITA;refGene_tx=XM_019820967.1;refGene_gn=CIITA;refGene_tx=XM_023247032.1
E3 35595818 35595887
refGene_gn=CIITA;refGene_tx=XM_011290571.3;refGene_gn=CIITA;refGene_tx=XM_011290572.2;refGene_gn=CIITA;refGene_tx=XM_011290573.3;refGene_gn=CIITA;refGene_tx=XM_011290574.2;refGene_gn=CIITA;refGene_tx=XM_011290575.3;refGene_gn=CIITA;refGene_tx=XM_019820963.1;refGene_gn=CIITA;refGene_tx=XM_019820965.1;refGene_gn=CIITA;refGene_tx=XM_019820966.2;refGene_gn=CIITA;refGene_tx=XM_019820967.1;refGene_gn=CIITA;refGene_tx=XM_023247032.1
E3 35596396 35596561
refGene_gn=CIITA;refGene_tx=XM_011290571.3;refGene_gn=CIITA;refGene_tx=XM_011290572.2;refGene_gn=CIITA;refGene_tx=XM_011290573.3;refGene_gn=CIITA;refGene_tx=XM_011290574.2;refGene_gn=CIITA;refGene_tx=XM_011290575.3;refGene_gn=CIITA;refGene_tx=XM_019820963.1;refGene_gn=CIITA;refGene_tx=XM_019820965.1;refGene_gn=CIITA;refGene_tx=XM_019820966.2;refGene_gn=CIITA;refGene_tx=XM_019820967.1;refGene_gn=CIITA;refGene_tx=XM_023247032.1

E3 35597834 35597969
refGene_gn=CIITA;refGene_tx=XM_011290571.3;refGene_gn=CIITA;refGene_tx=XM_011290572.2;refGene_gn=CIITA;refGene_tx=XM_011290573.3;refGene_gn=CIITA;refGene_tx=XM_011290574.2;refGene_gn=CIITA;refGene_tx=XM_011290575.3;refGene_gn=CIITA;refGene_tx=XM_019820963.1;refGene_gn=CIITA;refGene_tx=XM_019820965.1;refGene_gn=CIITA;refGene_tx=XM_019820967.1;refGene_gn=CIITA;refGene_tx=XM_023247032.1
E3 35598391 35598544
refGene_gn=CIITA;refGene_tx=XM_011290571.3;refGene_gn=CIITA;refGene_tx=XM_011290572.2;refGene_gn=CIITA;refGene_tx=XM_011290573.3;refGene_gn=CIITA;refGene_tx=XM_011290574.2;refGene_gn=CIITA;refGene_tx=XM_011290575.3;refGene_gn=CIITA;refGene_tx=XM_019820963.1;refGene_gn=CIITA;refGene_tx=XM_019820965.1;refGene_gn=CIITA;refGene_tx=XM_019820967.1;refGene_gn=CIITA;refGene_tx=XM_023247032.1
E3 35599093 35599138
refGene_gn=CIITA;refGene_tx=XM_011290571.3;refGene_gn=CIITA;refGene_tx=XM_011290572.2;refGene_gn=CIITA;refGene_tx=XM_011290573.3;refGene_gn=CIITA;refGene_tx=XM_011290574.2;refGene_gn=CIITA;refGene_tx=XM_011290575.3;refGene_gn=CIITA;refGene_tx=XM_019820963.1;refGene_gn=CIITA;refGene_tx=XM_019820965.1;refGene_gn=CIITA;refGene_tx=XM_019820967.1;refGene_gn=CIITA;refGene_tx=XM_023247032.1
E3 35601434 35601509
refGene_gn=CIITA;refGene_tx=XM_011290571.3;refGene_gn=CIITA;refGene_tx=XM_011290572.2;refGene_gn=CIITA;refGene_tx=XM_011290573.3;refGene_gn=CIITA;refGene_tx=XM_011290574.2;refGene_gn=CIITA;refGene_tx=XM_011290575.3;refGene_gn=CIITA;refGene_tx=XM_019820965.1;refGene_gn=CIITA;refGene_tx=XM_019820966.2;refGene_gn=CIITA;refGene_tx=XM_019820967.1;refGene_gn=CIITA;refGene_tx=XM_023247032.1
E3 35601690 35601753
refGene_gn=CIITA;refGene_tx=XM_011290571.3;refGene_gn=CIITA;refGene_tx=XM_011290572.2;refGene_gn=CIITA;refGene_tx=XM_011290573.3;refGene_gn=CIITA;refGene_tx=XM_011290574.2;refGene_gn=CIITA;refGene_tx=XM_011290575.3;refGene_gn=CIITA;refGene_tx=XM_019820965.1;refGene_gn=CIITA;refGene_tx=XM_019820966.2;refGene_gn=CIITA;refGene_tx=XM_019820967.1;refGene_gn=CIITA;refGene_tx=XM_023247032.1
E3 35603956 35604052
refGene_gn=CIITA;refGene_tx=XM_011290571.3;refGene_gn=CIITA;refGene_tx=XM_011290572.2;refGene_gn=CIITA;refGene_tx=XM_011290573.3;refGene_gn=CIITA;refGene_tx=XM_011290574.2;refGene_gn=CIITA;refGene_tx=XM_011290575.3;refGene_gn=CIITA;refGene_tx=XM_019820963.1;refGene_gn=CIITA;refGene_tx=XM_019820965.1;refGene_gn=CIITA;refGene_tx=XM_019820966.2;refGene_gn=CIITA;refGene_tx=XM_019820967.1;refGene_gn=CIITA;refGene_tx=XM_023247032.1
E3 35604268 35604412
refGene_gn=CIITA;refGene_tx=XM_011290571.3;refGene_gn=CIITA;refGene_tx=XM_011290572.2;refGene_gn=CIITA;refGene_tx=XM_011290573.3;refGene_gn=CIITA;refGene_tx=XM_011290574.2;refGene_gn=CIITA;refGene_tx=XM_011290575.3;refGene_gn=CIITA;refGene_tx=XM_019820963.1;refGene_gn=CIITA;refGene_tx=XM_019820965.1;refGene_gn=CIITA;refGene_tx=XM_019820966.2;refGene_gn=CIITA;refGene_tx=XM_019820967.1;refGene_gn=CIITA;refGene_tx=XM_02

3247032.1
E3	35617539 35617570 refGene_gn=CIITA;refGene_tx=XM_011290574.2
E3	35619390 35619442 refGene_gn=CIITA;refGene_tx=XM_011290573.3
E3	35627093 35627376
refGene_gn=CIITA;refGene_tx=XM_011290571.3;refGene_gn=CIITA;refGene_tx=XM_011290572.2;refGene_gn=CIITA;refGene_tx=XM_019820963.1;refGene_gn=CIITA;refGene_tx=XM_019820966.2;refGene_gn=CIITA;refGene_tx=XM_019820967.1;refGene_gn=CIITA;refGene_tx=XM_023247032.1
E3	35674862 35674871 refGene_gn=TVP23A;refGene_tx=XM_023247070.1
E3	35675613 35675622
refGene_gn=TVP23A;refGene_tx=XM_011290568.2;refGene_gn=TVP23A;refGene_tx=XM_019820969.1
E3	35675891 35675971
refGene_gn=TVP23A;refGene_tx=XM_011290568.2;refGene_gn=TVP23A;refGene_tx=XM_019820969.1;refGene_gn=TVP23A;refGene_tx=XM_023247070.1
E3	35700909 35701054
refGene_gn=TVP23A;refGene_tx=XM_011290568.2;refGene_gn=TVP23A;refGene_tx=XM_019820969.1;refGene_gn=TVP23A;refGene_tx=XM_023247070.1
E3	35701568 35701658
refGene_gn=TVP23A;refGene_tx=XM_011290568.2;refGene_gn=TVP23A;refGene_tx=XM_019820969.1;refGene_gn=TVP23A;refGene_tx=XM_023247070.1
E3	35701843 35702242
refGene_gn=TVP23A;refGene_tx=XM_011290568.2;refGene_gn=TVP23A;refGene_tx=XM_019820969.1;refGene_gn=TVP23A;refGene_tx=XM_023247070.1
E3	35702936 35703065 refGene_gn=TVP23A;refGene_tx=XM_011290568.2
E3	35703424 35703484 refGene_gn=TVP23A;refGene_tx=XM_011290568.2
E3	35704374 35704433
refGene_gn=NUBP1;refGene_tx=XM_003998868.4;refGene_gn=NUBP1;refGene_tx=XM_011290570.3
E3	35704937 35705033
refGene_gn=NUBP1;refGene_tx=XM_003998868.4;refGene_gn=NUBP1;refGene_tx=XM_011290570.3
E3	35708283 35708386
refGene_gn=NUBP1;refGene_tx=XM_003998868.4;refGene_gn=NUBP1;refGene_tx=XM_011290570.3
E3	35708661 35708772
refGene_gn=NUBP1;refGene_tx=XM_003998868.4;refGene_gn=NUBP1;refGene_tx=XM_011290570.3
E3	35711343 35711498
refGene_gn=NUBP1;refGene_tx=XM_003998868.4;refGene_gn=NUBP1;refGene_tx=XM_011290570.3
E3	35712126 35712217
refGene_gn=NUBP1;refGene_tx=XM_003998868.4;refGene_gn=NUBP1;refGene_tx=XM_011290570.3
E3	35713268 35713301
refGene_gn=NUBP1;refGene_tx=XM_003998868.4;refGene_gn=NUBP1;refGene_tx=XM_011290570.3
E3	35715872 35715941
refGene_gn=NUBP1;refGene_tx=XM_003998868.4;refGene_gn=NUBP1;refGene_tx=XM_011290570.3

E3 35722385 35722519
refGene_gn=NUBP1;refGene_tx=XM_003998868.4;refGene_gn=NUBP1;refGene_tx
=XM_011290570.3
E3 35724244 35724349
refGene_gn=NUBP1;refGene_tx=XM_003998868.4;refGene_gn=NUBP1;refGene_tx
=XM_011290570.3
E3 35724427 35724446
refGene_gn=NUBP1;refGene_tx=XM_003998868.4;refGene_gn=NUBP1;refGene_tx
=XM_011290570.3
E3 35740773 35741349 refGene_gn=TEKT5;refGene_tx=XM_003998869.3
E3 35743988 35744072 refGene_gn=TEKT5;refGene_tx=XM_003998869.3
E3 35744974 35745045 refGene_gn=TEKT5;refGene_tx=XM_003998869.3
E3 35749458 35749602 refGene_gn=TEKT5;refGene_tx=XM_003998869.3
E3 35753568 35753791 refGene_gn=TEKT5;refGene_tx=XM_003998869.3
E3 35775657 35775812 refGene_gn=TEKT5;refGene_tx=XM_003998869.3
E3 35779726 35779943 refGene_gn=TEKT5;refGene_tx=XM_003998869.3
E3 35851591 35851669 refGene_gn=EMP2;refGene_tx=XM_023246206.1
E3 35854017 35854108 refGene_gn=EMP2;refGene_tx=XM_023246206.1
E3 35856188 35856335 refGene_gn=EMP2;refGene_tx=XM_023246206.1
E3 35858298 35858486 refGene_gn=EMP2;refGene_tx=XM_023246206.1
E3 35882293 35882707
refGene_gn=ATF7IP2;refGene_tx=XM_019820970.2;refGene_gn=ATF7IP2;refGen
e_tx=XM_023246204.1;refGene_gn=ATF7IP2;refGene_tx=XM_023246205.1
E3 35883313 35883399
refGene_gn=ATF7IP2;refGene_tx=XM_019820970.2;refGene_gn=ATF7IP2;refGen
e_tx=XM_023246204.1;refGene_gn=ATF7IP2;refGene_tx=XM_023246205.1
E3 35900528 35900598
refGene_gn=ATF7IP2;refGene_tx=XM_019820970.2;refGene_gn=ATF7IP2;refGen
e_tx=XM_023246204.1;refGene_gn=ATF7IP2;refGene_tx=XM_023246205.1
E3 35901278 35901340
refGene_gn=ATF7IP2;refGene_tx=XM_019820970.2;refGene_gn=ATF7IP2;refGen
e_tx=XM_023246204.1;refGene_gn=ATF7IP2;refGene_tx=XM_023246205.1
E3 35902468 35902542
refGene_gn=ATF7IP2;refGene_tx=XM_019820970.2;refGene_gn=ATF7IP2;refGen
e_tx=XM_023246204.1
E3 35920381 35920536
refGene_gn=ATF7IP2;refGene_tx=XM_019820970.2;refGene_gn=ATF7IP2;refGen
e_tx=XM_023246204.1;refGene_gn=ATF7IP2;refGene_tx=XM_023246205.1
E3 35922371 35922470
refGene_gn=ATF7IP2;refGene_tx=XM_019820970.2;refGene_gn=ATF7IP2;refGen
e_tx=XM_023246204.1
E3 35923071 35923206
refGene_gn=ATF7IP2;refGene_tx=XM_019820970.2;refGene_gn=ATF7IP2;refGen
e_tx=XM_023246204.1
E3 35927422 35927547
refGene_gn=ATF7IP2;refGene_tx=XM_019820970.2;refGene_gn=ATF7IP2;refGen
e_tx=XM_023246204.1
E3 35928798 35929651
refGene_gn=ATF7IP2;refGene_tx=XM_019820970.2;refGene_gn=ATF7IP2;refGen
e_tx=XM_023246204.1

E3	36101625	36102039	refGene_gn=GRIN2A;refGene_tx=XM_006942329.4
E3	36326587	36327180	refGene_gn=GRIN2A;refGene_tx=XM_006942329.4
E3	36366067	36366182	refGene_gn=GRIN2A;refGene_tx=XM_006942329.4
E3	36401758	36401964	refGene_gn=GRIN2A;refGene_tx=XM_006942329.4
E3	36408089	36408258	refGene_gn=GRIN2A;refGene_tx=XM_006942329.4
E3	36408367	36408521	refGene_gn=GRIN2A;refGene_tx=XM_006942329.4
E3	36413730	36413856	refGene_gn=GRIN2A;refGene_tx=XM_006942329.4
E3	36417233	36417463	refGene_gn=GRIN2A;refGene_tx=XM_006942329.4
E3	36425139	36425300	refGene_gn=GRIN2A;refGene_tx=XM_006942329.4
E3	36443930	36444118	refGene_gn=GRIN2A;refGene_tx=XM_006942329.4
E3	36467544	36467783	refGene_gn=GRIN2A;refGene_tx=XM_006942329.4
E3	36474248	36476048	refGene_gn=GRIN2A;refGene_tx=XM_006942329.4
E3	37008857	37009102
refGene_gn=CE3H16orf72;refGene_tx=XM_023246780.1
E3	37020818	37021094
refGene_gn=CE3H16orf72;refGene_tx=XM_023246780.1
E3	37032309	37032390
refGene_gn=CE3H16orf72;refGene_tx=XM_023246780.1
E3	37033036	37033262
refGene_gn=CE3H16orf72;refGene_tx=XM_023246780.1
E3	37145860	37145942	refGene_gn=USP7;refGene_tx=XM_023246944.1
E3	37172865	37172959	refGene_gn=USP7;refGene_tx=XM_003998879.5
E3	37177585	37177690
refGene_gn=USP7;refGene_tx=XM_003998879.5;refGene_gn=USP7;refGene_tx=XM_023246944.1
E3	37183477	37183676
refGene_gn=USP7;refGene_tx=XM_003998879.5;refGene_gn=USP7;refGene_tx=XM_019820971.1;refGene_gn=USP7;refGene_tx=XM_023246944.1
E3	37185611	37185750
refGene_gn=USP7;refGene_tx=XM_003998879.5;refGene_gn=USP7;refGene_tx=XM_019820971.1;refGene_gn=USP7;refGene_tx=XM_023246944.1
E3	37186595	37186684
refGene_gn=USP7;refGene_tx=XM_003998879.5;refGene_gn=USP7;refGene_tx=XM_019820971.1;refGene_gn=USP7;refGene_tx=XM_023246944.1
E3	37188648	37188757
refGene_gn=USP7;refGene_tx=XM_003998879.5;refGene_gn=USP7;refGene_tx=XM_019820971.1;refGene_gn=USP7;refGene_tx=XM_023246944.1
E3	37189686	37189817
refGene_gn=USP7;refGene_tx=XM_003998879.5;refGene_gn=USP7;refGene_tx=XM_019820971.1;refGene_gn=USP7;refGene_tx=XM_023246944.1
E3	37190249	37190304
refGene_gn=USP7;refGene_tx=XM_003998879.5;refGene_gn=USP7;refGene_tx=XM_019820971.1;refGene_gn=USP7;refGene_tx=XM_023246944.1
E3	37191309	37191390
refGene_gn=USP7;refGene_tx=XM_003998879.5;refGene_gn=USP7;refGene_tx=XM_019820971.1;refGene_gn=USP7;refGene_tx=XM_023246944.1
E3	37191490	37191581
refGene_gn=USP7;refGene_tx=XM_003998879.5;refGene_gn=USP7;refGene_tx=XM_019820971.1;refGene_gn=USP7;refGene_tx=XM_023246944.1
E3	37194217	37194300

refGene_gn=USP7;refGene_tx=XM_003998879.5;refGene_gn=USP7;refGene_tx=XM_019820971.1;refGene_gn=USP7;refGene_tx=XM_023246944.1
E3 37196762 37196872
refGene_gn=USP7;refGene_tx=XM_003998879.5;refGene_gn=USP7;refGene_tx=XM_019820971.1;refGene_gn=USP7;refGene_tx=XM_023246944.1
E3 37198397 37198554
refGene_gn=USP7;refGene_tx=XM_003998879.5;refGene_gn=USP7;refGene_tx=XM_019820971.1;refGene_gn=USP7;refGene_tx=XM_023246944.1
E3 37200235 37200380
refGene_gn=USP7;refGene_tx=XM_003998879.5;refGene_gn=USP7;refGene_tx=XM_019820971.1;refGene_gn=USP7;refGene_tx=XM_023246944.1
E3 37200655 37200786
refGene_gn=USP7;refGene_tx=XM_003998879.5;refGene_gn=USP7;refGene_tx=XM_019820971.1;refGene_gn=USP7;refGene_tx=XM_023246944.1
E3 37201493 37201628
refGene_gn=USP7;refGene_tx=XM_003998879.5;refGene_gn=USP7;refGene_tx=XM_019820971.1;refGene_gn=USP7;refGene_tx=XM_023246944.1
E3 37202092 37202194
refGene_gn=USP7;refGene_tx=XM_003998879.5;refGene_gn=USP7;refGene_tx=XM_019820971.1;refGene_gn=USP7;refGene_tx=XM_023246944.1
E3 37202347 37202453
refGene_gn=USP7;refGene_tx=XM_003998879.5;refGene_gn=USP7;refGene_tx=XM_019820971.1;refGene_gn=USP7;refGene_tx=XM_023246944.1
E3 37203191 37203284
refGene_gn=USP7;refGene_tx=XM_003998879.5;refGene_gn=USP7;refGene_tx=XM_019820971.1;refGene_gn=USP7;refGene_tx=XM_023246944.1
E3 37203372 37203440
refGene_gn=USP7;refGene_tx=XM_003998879.5;refGene_gn=USP7;refGene_tx=XM_019820971.1;refGene_gn=USP7;refGene_tx=XM_023246944.1
E3 37203767 37203868
refGene_gn=USP7;refGene_tx=XM_003998879.5;refGene_gn=USP7;refGene_tx=XM_019820971.1;refGene_gn=USP7;refGene_tx=XM_023246944.1
E3 37204695 37204849
refGene_gn=USP7;refGene_tx=XM_003998879.5;refGene_gn=USP7;refGene_tx=XM_019820971.1;refGene_gn=USP7;refGene_tx=XM_023246944.1
E3 37205584 37205652
refGene_gn=USP7;refGene_tx=XM_003998879.5;refGene_gn=USP7;refGene_tx=XM_019820971.1;refGene_gn=USP7;refGene_tx=XM_023246944.1
E3 37206098 37206207
refGene_gn=USP7;refGene_tx=XM_003998879.5;refGene_gn=USP7;refGene_tx=XM_019820971.1;refGene_gn=USP7;refGene_tx=XM_023246944.1
E3 37206295 37206373
refGene_gn=USP7;refGene_tx=XM_003998879.5;refGene_gn=USP7;refGene_tx=XM_019820971.1;refGene_gn=USP7;refGene_tx=XM_023246944.1
E3 37207172 37207273
refGene_gn=USP7;refGene_tx=XM_003998879.5;refGene_gn=USP7;refGene_tx=XM_019820971.1;refGene_gn=USP7;refGene_tx=XM_023246944.1
E3 37208294 37208394
refGene_gn=USP7;refGene_tx=XM_003998879.5;refGene_gn=USP7;refGene_tx=XM_019820971.1;refGene_gn=USP7;refGene_tx=XM_023246944.1

E3 37209102 37209222
refGene_gn=USP7;refGene_tx=XM_003998879.5;refGene_gn=USP7;refGene_tx=XM_019820971.1;refGene_gn=USP7;refGene_tx=XM_023246944.1
E3 37209333 37209405
refGene_gn=USP7;refGene_tx=XM_003998879.5;refGene_gn=USP7;refGene_tx=XM_019820971.1;refGene_gn=USP7;refGene_tx=XM_023246944.1
E3 37209526 37209617
refGene_gn=USP7;refGene_tx=XM_003998879.5;refGene_gn=USP7;refGene_tx=XM_019820971.1;refGene_gn=USP7;refGene_tx=XM_023246944.1
E3 37209854 37209961
refGene_gn=USP7;refGene_tx=XM_003998879.5;refGene_gn=USP7;refGene_tx=XM_019820971.1;refGene_gn=USP7;refGene_tx=XM_023246944.1
E3 37216672 37216781
refGene_gn=LOC102899335;refGene_tx=XM_019820977.2;refGene_gn=LOC102899335;refGene_tx=XM_019820978.2;refGene_gn=LOC102899335;refGene_tx=XM_023246813.1;refGene_gn=LOC102899335;refGene_tx=XM_023246814.1
E3 37216905 37217050
refGene_gn=LOC102899335;refGene_tx=XM_019820977.2;refGene_gn=LOC102899335;refGene_tx=XM_019820978.2;refGene_gn=LOC102899335;refGene_tx=XM_023246813.1
E3 37218167 37218269
refGene_gn=LOC102899335;refGene_tx=XM_019820977.2;refGene_gn=LOC102899335;refGene_tx=XM_019820978.2;refGene_gn=LOC102899335;refGene_tx=XM_023246813.1;refGene_gn=LOC102899335;refGene_tx=XM_023246814.1
E3 37219907 37220112
refGene_gn=LOC102899335;refGene_tx=XM_019820977.2;refGene_gn=LOC102899335;refGene_tx=XM_019820978.2;refGene_gn=LOC102899335;refGene_tx=XM_023246813.1;refGene_gn=LOC102899335;refGene_tx=XM_023246814.1
E3 37245576 37245629 refGene_gn=CARHSP1;refGene_tx=XM_019820974.2
E3 37246991 37246993 refGene_gn=CARHSP1;refGene_tx=XM_019820975.2
E3 37247093 37247258
refGene_gn=CARHSP1;refGene_tx=XM_006942337.4;refGene_gn=CARHSP1;refGene_tx=XM_019820974.2;refGene_gn=CARHSP1;refGene_tx=XM_019820975.2;refGene_gn=CARHSP1;refGene_tx=XM_019820976.2;refGene_gn=CARHSP1;refGene_tx=XM_023247104.1
E3 37247938 37248061
refGene_gn=CARHSP1;refGene_tx=XM_006942337.4;refGene_gn=CARHSP1;refGene_tx=XM_019820974.2;refGene_gn=CARHSP1;refGene_tx=XM_019820975.2;refGene_gn=CARHSP1;refGene_tx=XM_019820976.2;refGene_gn=CARHSP1;refGene_tx=XM_023247104.1
E3 37249617 37249780
refGene_gn=CARHSP1;refGene_tx=XM_006942337.4;refGene_gn=CARHSP1;refGene_tx=XM_019820974.2;refGene_gn=CARHSP1;refGene_tx=XM_019820975.2;refGene_gn=CARHSP1;refGene_tx=XM_019820976.2;refGene_gn=CARHSP1;refGene_tx=XM_023247104.1
E3 37256075 37256177
refGene_gn=PMM2;refGene_tx=XM_003998882.4;refGene_gn=PMM2;refGene_tx=XM_006942339.3
E3 37266514 37266630
refGene_gn=PMM2;refGene_tx=XM_003998882.4;refGene_gn=PMM2;refGene_tx=X

M_006942339.3
E3 37267722 37267830
refGene_gn=PMM2;refGene_tx=XM_003998882.4;refGene_gn=PMM2;refGene_tx=XM_006942339.3
E3 37267936 37268036
refGene_gn=PMM2;refGene_tx=XM_003998882.4;refGene_gn=PMM2;refGene_tx=XM_006942339.3
E3 37270777 37270869
refGene_gn=PMM2;refGene_tx=XM_003998882.4;refGene_gn=PMM2;refGene_tx=XM_006942339.3
E3 37271541 37271618
refGene_gn=PMM2;refGene_tx=XM_003998882.4;refGene_gn=PMM2;refGene_tx=XM_006942339.3
E3 37274051 37274163
refGene_gn=PMM2;refGene_tx=XM_003998882.4;refGene_gn=PMM2;refGene_tx=XM_006942339.3
E3 37285806 37285872
refGene_gn=PMM2;refGene_tx=XM_003998882.4;refGene_gn=PMM2;refGene_tx=XM_006942339.3
E3 37286135 37286138 refGene_gn=TMEM186;refGene_tx=XM_006942343.4
E3 37286676 37286679 refGene_gn=TMEM186;refGene_tx=XM_006942344.4
E3 37289679 37290321
refGene_gn=TMEM186;refGene_tx=XM_006942343.4;refGene_gn=TMEM186;refGene_tx=XM_006942344.4
E3 37302753 37302875
refGene_gn=ABAT;refGene_tx=XM_006942340.4;refGene_gn=ABAT;refGene_tx=XM_019820979.2
E3 37303896 37304008
refGene_gn=ABAT;refGene_tx=XM_006942340.4;refGene_gn=ABAT;refGene_tx=XM_019820979.2
E3 37307807 37307954
refGene_gn=ABAT;refGene_tx=XM_006942340.4;refGene_gn=ABAT;refGene_tx=XM_019820979.2
E3 37309254 37309422
refGene_gn=ABAT;refGene_tx=XM_006942340.4;refGene_gn=ABAT;refGene_tx=XM_019820979.2
E3 37310202 37310340
refGene_gn=ABAT;refGene_tx=XM_006942340.4;refGene_gn=ABAT;refGene_tx=XM_019820979.2
E3 37312407 37312556
refGene_gn=ABAT;refGene_tx=XM_006942340.4;refGene_gn=ABAT;refGene_tx=XM_019820979.2
E3 37312979 37313043
refGene_gn=ABAT;refGene_tx=XM_006942340.4;refGene_gn=ABAT;refGene_tx=XM_019820979.2
E3 37315886 37315949
refGene_gn=ABAT;refGene_tx=XM_006942340.4;refGene_gn=ABAT;refGene_tx=XM_019820979.2
E3 37316410 37316503
refGene_gn=ABAT;refGene_tx=XM_006942340.4;refGene_gn=ABAT;refGene_tx=X

M_019820979.2
E3 37317164 37317245
refGene_gn=ABAT;refGene_tx=XM_006942340.4;refGene_gn=ABAT;refGene_tx=XM_019820979.2
E3 37327271 37327321
refGene_gn=ABAT;refGene_tx=XM_006942340.4;refGene_gn=ABAT;refGene_tx=XM_019820979.2
E3 37330515 37330633
refGene_gn=ABAT;refGene_tx=XM_006942340.4;refGene_gn=ABAT;refGene_tx=XM_019820979.2
E3 37332167 37332197
refGene_gn=ABAT;refGene_tx=XM_006942340.4;refGene_gn=ABAT;refGene_tx=XM_019820979.2
E3 37334526 37334624
refGene_gn=ABAT;refGene_tx=XM_006942340.4;refGene_gn=ABAT;refGene_tx=XM_019820979.2
E3 37345517 37345587
refGene_gn=ABAT;refGene_tx=XM_006942340.4;refGene_gn=ABAT;refGene_tx=XM_019820979.2
E3 37409049 37409138
refGene_gn=METTL22;refGene_tx=XM_006942345.2;refGene_gn=METTL22;refGene_tx=XM_011290566.2;refGene_gn=METTL22;refGene_tx=XM_011290567.2
E3 37409886 37409998
refGene_gn=METTL22;refGene_tx=XM_011290566.2;refGene_gn=METTL22;refGene_tx=XM_011290567.2
E3 37410364 37410533
refGene_gn=METTL22;refGene_tx=XM_006942345.2;refGene_gn=METTL22;refGene_tx=XM_011290566.2;refGene_gn=METTL22;refGene_tx=XM_011290567.2
E3 37412382 37412485
refGene_gn=METTL22;refGene_tx=XM_006942345.2;refGene_gn=METTL22;refGene_tx=XM_011290566.2;refGene_gn=METTL22;refGene_tx=XM_011290567.2
E3 37412697 37412778
refGene_gn=METTL22;refGene_tx=XM_006942345.2;refGene_gn=METTL22;refGene_tx=XM_011290566.2;refGene_gn=METTL22;refGene_tx=XM_011290567.2
E3 37413485 37413539
refGene_gn=METTL22;refGene_tx=XM_006942345.2;refGene_gn=METTL22;refGene_tx=XM_011290566.2;refGene_gn=METTL22;refGene_tx=XM_011290567.2
E3 37414847 37414919
refGene_gn=METTL22;refGene_tx=XM_006942345.2;refGene_gn=METTL22;refGene_tx=XM_011290566.2;refGene_gn=METTL22;refGene_tx=XM_011290567.2
E3 37417566 37417711
refGene_gn=METTL22;refGene_tx=XM_006942345.2;refGene_gn=METTL22;refGene_tx=XM_011290566.2
E3 37417800 37417841
refGene_gn=METTL22;refGene_tx=XM_006942345.2;refGene_gn=METTL22;refGene_tx=XM_011290566.2
E3 37422982 37423336
refGene_gn=METTL22;refGene_tx=XM_006942345.2;refGene_gn=METTL22;refGene_tx=XM_011290566.2;refGene_gn=METTL22;refGene_tx=XM_011290567.2
E3 37425984 37426090

refGene_gn=METTL22;refGene_tx=XM_006942345.2;refGene_gn=METTL22;refGene_tx=XM_011290566.2;refGene_gn=METTL22;refGene_tx=XM_011290567.2
E3 37459377 37459597 refGene_gn=TMEM114;refGene_tx=XM_023246586.1
E3 37459894 37459975 refGene_gn=TMEM114;refGene_tx=XM_023246586.1
E3 37469400 37469538 refGene_gn=TMEM114;refGene_tx=XM_023246586.1
E3 37470758 37470839 refGene_gn=TMEM114;refGene_tx=XM_023246586.1
E3 37472468 37472540 refGene_gn=TMEM114;refGene_tx=XM_023246586.1
E3 37473855 37473996 refGene_gn=TMEM114;refGene_tx=XM_023246586.1
E3 37478493 37478573 refGene_gn=TMEM114;refGene_tx=XM_023246586.1
E3 38167196 38167319
refGene_gn=RBFOX1;refGene_tx=XM_023246514.1;refGene_gn=RBFOX1;refGene_tx=XM_023246515.1;refGene_gn=RBFOX1;refGene_tx=XM_023246521.1;refGene_gn=RBFOX1;refGene_tx=XM_023246524.1;refGene_gn=RBFOX1;refGene_tx=XM_023246525.1;refGene_gn=RBFOX1;refGene_tx=XM_023246526.1;refGene_gn=RBFOX1;refGene_tx=XM_023246535.1
E3 38168338 38168486
refGene_gn=RBFOX1;refGene_tx=XM_023246510.1;refGene_gn=RBFOX1;refGene_tx=XM_023246511.1;refGene_gn=RBFOX1;refGene_tx=XM_023246513.1;refGene_gn=RBFOX1;refGene_tx=XM_023246516.1;refGene_gn=RBFOX1;refGene_tx=XM_023246517.1;refGene_gn=RBFOX1;refGene_tx=XM_023246518.1;refGene_gn=RBFOX1;refGene_tx=XM_023246519.1;refGene_gn=RBFOX1;refGene_tx=XM_023246520.1;refGene_gn=RBFOX1;refGene_tx=XM_023246522.1;refGene_gn=RBFOX1;refGene_tx=XM_023246527.1;refGene_gn=RBFOX1;refGene_tx=XM_023246528.1;refGene_gn=RBFOX1;refGene_tx=XM_023246529.1;refGene_gn=RBFOX1;refGene_tx=XM_023246530.1;refGene_gn=RBFOX1;refGene_tx=XM_023246531.1;refGene_gn=RBFOX1;refGene_tx=XM_023246532.1;refGene_gn=RBFOX1;refGene_tx=XM_023246533.1;refGene_gn=RBFOX1;refGene_tx=XM_023246534.1
E3 38168856 38168932
refGene_gn=RBFOX1;refGene_tx=XM_023246510.1;refGene_gn=RBFOX1;refGene_tx=XM_023246511.1;refGene_gn=RBFOX1;refGene_tx=XM_023246513.1;refGene_gn=RBFOX1;refGene_tx=XM_023246514.1;refGene_gn=RBFOX1;refGene_tx=XM_023246515.1;refGene_gn=RBFOX1;refGene_tx=XM_023246516.1;refGene_gn=RBFOX1;refGene_tx=XM_023246517.1;refGene_gn=RBFOX1;refGene_tx=XM_023246518.1;refGene_gn=RBFOX1;refGene_tx=XM_023246519.1;refGene_gn=RBFOX1;refGene_tx=XM_023246520.1;refGene_gn=RBFOX1;refGene_tx=XM_023246521.1;refGene_gn=RBFOX1;refGene_tx=XM_023246522.1;refGene_gn=RBFOX1;refGene_tx=XM_023246524.1;refGene_gn=RBFOX1;refGene_tx=XM_023246525.1;refGene_gn=RBFOX1;refGene_tx=XM_023246526.1;refGene_gn=RBFOX1;refGene_tx=XM_023246527.1;refGene_gn=RBFOX1;refGene_tx=XM_023246528.1;refGene_gn=RBFOX1;refGene_tx=XM_023246529.1;refGene_gn=RBFOX1;refGene_tx=XM_023246530.1;refGene_gn=RBFOX1;refGene_tx=XM_023246531.1;refGene_gn=RBFOX1;refGene_tx=XM_023246532.1;refGene_gn=RBFOX1;refGene_tx=XM_023246533.1;refGene_gn=RBFOX1;refGene_tx=XM_023246534.1;refGene_gn=RBFOX1;refGene_tx=XM_023246535.1
E3 38182267 38182320
refGene_gn=RBFOX1;refGene_tx=XM_023246510.1;refGene_gn=RBFOX1;refGene_tx=XM_023246511.1;refGene_gn=RBFOX1;refGene_tx=XM_023246513.1;refGene_gn=RBFOX1;refGene_tx=XM_023246516.1;refGene_gn=RBFOX1;refGene_tx=XM_023246517.1;refGene_gn=RBFOX1;refGene_tx=XM_023246521.1;refGene_gn=RBFOX1;refGene_tx=XM_023246522.1;refGene_gn=RBFOX1;refGene_tx=XM_023246525.

1;refGene_gn=RBFOX1;refGene_tx=XM_023246527.1;refGene_gn=RBFOX1;refGene_tx=XM_023246528.1;refGene_gn=RBFOX1;refGene_tx=XM_023246529.1;refGene_gn=RBFOX1;refGene_tx=XM_023246530.1;refGene_gn=RBFOX1;refGene_tx=XM_023246531.1;refGene_gn=RBFOX1;refGene_tx=XM_023246532.1;refGene_gn=RBFOX1;refGene_tx=XM_023246533.1;refGene_gn=RBFOX1;refGene_tx=XM_023246534.1
E3 38201163 38201228
refGene_gn=RBFOX1;refGene_tx=XM_023246510.1;refGene_gn=RBFOX1;refGene_tx=XM_023246511.1;refGene_gn=RBFOX1;refGene_tx=XM_023246513.1;refGene_gn=RBFOX1;refGene_tx=XM_023246514.1;refGene_gn=RBFOX1;refGene_tx=XM_023246515.1;refGene_gn=RBFOX1;refGene_tx=XM_023246516.1;refGene_gn=RBFOX1;refGene_tx=XM_023246517.1;refGene_gn=RBFOX1;refGene_tx=XM_023246518.1;refGene_gn=RBFOX1;refGene_tx=XM_023246519.1;refGene_gn=RBFOX1;refGene_tx=XM_023246520.1;refGene_gn=RBFOX1;refGene_tx=XM_023246521.1;refGene_gn=RBFOX1;refGene_tx=XM_023246522.1;refGene_gn=RBFOX1;refGene_tx=XM_023246524.1;refGene_gn=RBFOX1;refGene_tx=XM_023246525.1;refGene_gn=RBFOX1;refGene_tx=XM_023246526.1;refGene_gn=RBFOX1;refGene_tx=XM_023246527.1;refGene_gn=RBFOX1;refGene_tx=XM_023246528.1;refGene_gn=RBFOX1;refGene_tx=XM_023246529.1;refGene_gn=RBFOX1;refGene_tx=XM_023246530.1;refGene_gn=RBFOX1;refGene_tx=XM_023246531.1;refGene_gn=RBFOX1;refGene_tx=XM_023246532.1;refGene_gn=RBFOX1;refGene_tx=XM_023246533.1;refGene_gn=RBFOX1;refGene_tx=XM_023246534.1;refGene_gn=RBFOX1;refGene_tx=XM_023246535.1
E3 38206020 38206063
refGene_gn=RBFOX1;refGene_tx=XM_023246510.1;refGene_gn=RBFOX1;refGene_tx=XM_023246513.1;refGene_gn=RBFOX1;refGene_tx=XM_023246514.1;refGene_gn=RBFOX1;refGene_tx=XM_023246516.1;refGene_gn=RBFOX1;refGene_tx=XM_023246517.1;refGene_gn=RBFOX1;refGene_tx=XM_023246519.1;refGene_gn=RBFOX1;refGene_tx=XM_023246521.1;refGene_gn=RBFOX1;refGene_tx=XM_023246527.1;refGene_gn=RBFOX1;refGene_tx=XM_023246528.1;refGene_gn=RBFOX1;refGene_tx=XM_023246529.1;refGene_gn=RBFOX1;refGene_tx=XM_023246530.1;refGene_gn=RBFOX1;refGene_tx=XM_023246531.1;refGene_gn=RBFOX1;refGene_tx=XM_023246532.1;refGene_gn=RBFOX1;refGene_tx=XM_023246533.1;refGene_gn=RBFOX1;refGene_tx=XM_023246534.1;refGene_gn=RBFOX1;refGene_tx=XM_023246535.1
E3 38211820 38211860
refGene_gn=RBFOX1;refGene_tx=XM_023246511.1;refGene_gn=RBFOX1;refGene_tx=XM_023246515.1;refGene_gn=RBFOX1;refGene_tx=XM_023246520.1;refGene_gn=RBFOX1;refGene_tx=XM_023246524.1
E3 38221242 38221375
refGene_gn=RBFOX1;refGene_tx=XM_023246510.1;refGene_gn=RBFOX1;refGene_tx=XM_023246511.1;refGene_gn=RBFOX1;refGene_tx=XM_023246513.1;refGene_gn=RBFOX1;refGene_tx=XM_023246514.1;refGene_gn=RBFOX1;refGene_tx=XM_023246515.1;refGene_gn=RBFOX1;refGene_tx=XM_023246516.1;refGene_gn=RBFOX1;refGene_tx=XM_023246517.1;refGene_gn=RBFOX1;refGene_tx=XM_023246518.1;refGene_gn=RBFOX1;refGene_tx=XM_023246519.1;refGene_gn=RBFOX1;refGene_tx=XM_023246520.1;refGene_gn=RBFOX1;refGene_tx=XM_023246521.1;refGene_gn=RBFOX1;refGene_tx=XM_023246522.1;refGene_gn=RBFOX1;refGene_tx=XM_023246524.1;refGene_gn=RBFOX1;refGene_tx=XM_023246525.1;refGene_gn=RBFOX1;refGene_tx=XM_023246526.1;refGene_gn=RBFOX1;refGene_tx=XM_02324652

7.1;refGene_gn=RBFOX1;refGene_tx=XM_023246528.1;refGene_gn=RBFOX1;refGene_tx=XM_023246529.1;refGene_gn=RBFOX1;refGene_tx=XM_023246530.1;refGene_gn=RBFOX1;refGene_tx=XM_023246531.1;refGene_gn=RBFOX1;refGene_tx=XM_023246532.1;refGene_gn=RBFOX1;refGene_tx=XM_023246533.1;refGene_gn=RBFOX1;refGene_tx=XM_023246534.1;refGene_gn=RBFOX1;refGene_tx=XM_023246535.1
E3 38249474 38249555
refGene_gn=RBFOX1;refGene_tx=XM_023246510.1;refGene_gn=RBFOX1;refGene_tx=XM_023246511.1;refGene_gn=RBFOX1;refGene_tx=XM_023246513.1;refGene_gn=RBFOX1;refGene_tx=XM_023246514.1;refGene_gn=RBFOX1;refGene_tx=XM_023246515.1;refGene_gn=RBFOX1;refGene_tx=XM_023246517.1;refGene_gn=RBFOX1;refGene_tx=XM_023246518.1;refGene_gn=RBFOX1;refGene_tx=XM_023246519.1;refGene_gn=RBFOX1;refGene_tx=XM_023246520.1;refGene_gn=RBFOX1;refGene_tx=XM_023246521.1;refGene_gn=RBFOX1;refGene_tx=XM_023246522.1;refGene_gn=RBFOX1;refGene_tx=XM_023246525.1;refGene_gn=RBFOX1;refGene_tx=XM_023246526.1;refGene_gn=RBFOX1;refGene_tx=XM_023246527.1;refGene_gn=RBFOX1;refGene_tx=XM_023246528.1;refGene_gn=RBFOX1;refGene_tx=XM_023246529.1;refGene_gn=RBFOX1;refGene_tx=XM_023246530.1;refGene_gn=RBFOX1;refGene_tx=XM_023246531.1;refGene_gn=RBFOX1;refGene_tx=XM_023246532.1;refGene_gn=RBFOX1;refGene_tx=XM_023246533.1;refGene_gn=RBFOX1;refGene_tx=XM_023246534.1;refGene_gn=RBFOX1;refGene_tx=XM_023246535.1
E3 38271017 38271071
refGene_gn=RBFOX1;refGene_tx=XM_023246510.1;refGene_gn=RBFOX1;refGene_tx=XM_023246511.1;refGene_gn=RBFOX1;refGene_tx=XM_023246513.1;refGene_gn=RBFOX1;refGene_tx=XM_023246514.1;refGene_gn=RBFOX1;refGene_tx=XM_023246515.1;refGene_gn=RBFOX1;refGene_tx=XM_023246516.1;refGene_gn=RBFOX1;refGene_tx=XM_023246517.1;refGene_gn=RBFOX1;refGene_tx=XM_023246518.1;refGene_gn=RBFOX1;refGene_tx=XM_023246519.1;refGene_gn=RBFOX1;refGene_tx=XM_023246520.1;refGene_gn=RBFOX1;refGene_tx=XM_023246521.1;refGene_gn=RBFOX1;refGene_tx=XM_023246522.1;refGene_gn=RBFOX1;refGene_tx=XM_023246524.1;refGene_gn=RBFOX1;refGene_tx=XM_023246525.1;refGene_gn=RBFOX1;refGene_tx=XM_023246526.1;refGene_gn=RBFOX1;refGene_tx=XM_023246527.1;refGene_gn=RBFOX1;refGene_tx=XM_023246528.1;refGene_gn=RBFOX1;refGene_tx=XM_023246529.1;refGene_gn=RBFOX1;refGene_tx=XM_023246530.1;refGene_gn=RBFOX1;refGene_tx=XM_023246531.1;refGene_gn=RBFOX1;refGene_tx=XM_023246532.1;refGene_gn=RBFOX1;refGene_tx=XM_023246533.1;refGene_gn=RBFOX1;refGene_tx=XM_023246534.1;refGene_gn=RBFOX1;refGene_tx=XM_023246535.1
E3 38280546 38280607
refGene_gn=RBFOX1;refGene_tx=XM_023246510.1;refGene_gn=RBFOX1;refGene_tx=XM_023246511.1;refGene_gn=RBFOX1;refGene_tx=XM_023246513.1;refGene_gn=RBFOX1;refGene_tx=XM_023246514.1;refGene_gn=RBFOX1;refGene_tx=XM_023246515.1;refGene_gn=RBFOX1;refGene_tx=XM_023246516.1;refGene_gn=RBFOX1;refGene_tx=XM_023246517.1;refGene_gn=RBFOX1;refGene_tx=XM_023246518.1;refGene_gn=RBFOX1;refGene_tx=XM_023246519.1;refGene_gn=RBFOX1;refGene_tx=XM_023246520.1;refGene_gn=RBFOX1;refGene_tx=XM_023246521.1;refGene_gn=RBFOX1;refGene_tx=XM_023246522.1;refGene_gn=RBFOX1;refGene_tx=XM_023246524.1;refGene_gn=RBFOX1;refGene_tx=XM_023246525.1;refGene_gn=RBFOX1;refGene_tx=XM_023246526.1;refGene_gn=RBFOX1;refGene_tx=XM_023246527.1;refGene_gn=RBFOX1;refGene_tx=XM_023246528.1;refGene_gn=RBFOX1;refG

ene_tx=XM_023246529.1;refGene_gn=RBFOX1;refGene_tx=XM_023246530.1;refGene_gn=RBFOX1;refGene_tx=XM_023246531.1;refGene_gn=RBFOX1;refGene_tx=XM_023246532.1;refGene_gn=RBFOX1;refGene_tx=XM_023246533.1;refGene_gn=RBFOX1;refGene_tx=XM_023246534.1;refGene_gn=RBFOX1;refGene_tx=XM_023246535.1

E3 38282407 38282500

refGene_gn=RBFOX1;refGene_tx=XM_023246510.1;refGene_gn=RBFOX1;refGene_tx=XM_023246511.1;refGene_gn=RBFOX1;refGene_tx=XM_023246513.1;refGene_gn=RBFOX1;refGene_tx=XM_023246514.1;refGene_gn=RBFOX1;refGene_tx=XM_023246515.1;refGene_gn=RBFOX1;refGene_tx=XM_023246516.1;refGene_gn=RBFOX1;refGene_tx=XM_023246518.1;refGene_gn=RBFOX1;refGene_tx=XM_023246519.1;refGene_gn=RBFOX1;refGene_tx=XM_023246520.1;refGene_gn=RBFOX1;refGene_tx=XM_023246521.1;refGene_gn=RBFOX1;refGene_tx=XM_023246522.1;refGene_gn=RBFOX1;refGene_tx=XM_023246524.1;refGene_gn=RBFOX1;refGene_tx=XM_023246525.1;refGene_gn=RBFOX1;refGene_tx=XM_023246526.1;refGene_gn=RBFOX1;refGene_tx=XM_023246527.1;refGene_gn=RBFOX1;refGene_tx=XM_023246528.1;refGene_gn=RBFOX1;refGene_tx=XM_023246529.1;refGene_gn=RBFOX1;refGene_tx=XM_023246530.1;refGene_gn=RBFOX1;refGene_tx=XM_023246531.1;refGene_gn=RBFOX1;refGene_tx=XM_023246532.1;refGene_gn=RBFOX1;refGene_tx=XM_023246533.1;refGene_gn=RBFOX1;refGene_tx=XM_023246534.1;refGene_gn=RBFOX1;refGene_tx=XM_023246535.1

E3 38290949 38291003

refGene_gn=RBFOX1;refGene_tx=XM_023246510.1;refGene_gn=RBFOX1;refGene_tx=XM_023246511.1;refGene_gn=RBFOX1;refGene_tx=XM_023246513.1;refGene_gn=RBFOX1;refGene_tx=XM_023246514.1;refGene_gn=RBFOX1;refGene_tx=XM_023246515.1;refGene_gn=RBFOX1;refGene_tx=XM_023246516.1;refGene_gn=RBFOX1;refGene_tx=XM_023246517.1;refGene_gn=RBFOX1;refGene_tx=XM_023246518.1;refGene_gn=RBFOX1;refGene_tx=XM_023246519.1;refGene_gn=RBFOX1;refGene_tx=XM_023246520.1;refGene_gn=RBFOX1;refGene_tx=XM_023246521.1;refGene_gn=RBFOX1;refGene_tx=XM_023246522.1;refGene_gn=RBFOX1;refGene_tx=XM_023246524.1;refGene_gn=RBFOX1;refGene_tx=XM_023246525.1;refGene_gn=RBFOX1;refGene_tx=XM_023246526.1;refGene_gn=RBFOX1;refGene_tx=XM_023246527.1;refGene_gn=RBFOX1;refGene_tx=XM_023246528.1;refGene_gn=RBFOX1;refGene_tx=XM_023246529.1;refGene_gn=RBFOX1;refGene_tx=XM_023246530.1;refGene_gn=RBFOX1;refGene_tx=XM_023246531.1;refGene_gn=RBFOX1;refGene_tx=XM_023246532.1;refGene_gn=RBFOX1;refGene_tx=XM_023246533.1;refGene_gn=RBFOX1;refGene_tx=XM_023246534.1;refGene_gn=RBFOX1;refGene_tx=XM_023246535.1

E3 38297875 38298019

refGene_gn=RBFOX1;refGene_tx=XM_023246510.1;refGene_gn=RBFOX1;refGene_tx=XM_023246511.1;refGene_gn=RBFOX1;refGene_tx=XM_023246513.1;refGene_gn=RBFOX1;refGene_tx=XM_023246514.1;refGene_gn=RBFOX1;refGene_tx=XM_023246515.1;refGene_gn=RBFOX1;refGene_tx=XM_023246516.1;refGene_gn=RBFOX1;refGene_tx=XM_023246517.1;refGene_gn=RBFOX1;refGene_tx=XM_023246518.1;refGene_gn=RBFOX1;refGene_tx=XM_023246519.1;refGene_gn=RBFOX1;refGene_tx=XM_023246520.1;refGene_gn=RBFOX1;refGene_tx=XM_023246521.1;refGene_gn=RBFOX1;refGene_tx=XM_023246522.1;refGene_gn=RBFOX1;refGene_tx=XM_023246524.1;refGene_gn=RBFOX1;refGene_tx=XM_023246525.1;refGene_gn=RBFOX1;refGene_tx=XM_023246526.1;refGene_gn=RBFOX1;refGene_tx=XM_023246527.1;refGene_gn=RBFOX1;refGene_tx=XM_023246528.1;refGene_gn=RBFOX1;refG

ene_tx=XM_023246529.1;refGene_gn=RBFOX1;refGene_tx=XM_023246530.1;refGene_gn=RBFOX1;refGene_tx=XM_023246531.1;refGene_gn=RBFOX1;refGene_tx=XM_023246532.1;refGene_gn=RBFOX1;refGene_tx=XM_023246533.1;refGene_gn=RBFOX1;refGene_tx=XM_023246534.1;refGene_gn=RBFOX1;refGene_tx=XM_023246535.1
E3 38348113 38348356
refGene_gn=RBFOX1;refGene_tx=XM_023246510.1;refGene_gn=RBFOX1;refGene_tx=XM_023246511.1;refGene_gn=RBFOX1;refGene_tx=XM_023246513.1;refGene_gn=RBFOX1;refGene_tx=XM_023246514.1;refGene_gn=RBFOX1;refGene_tx=XM_023246515.1;refGene_gn=RBFOX1;refGene_tx=XM_023246516.1;refGene_gn=RBFOX1;refGene_tx=XM_023246517.1;refGene_gn=RBFOX1;refGene_tx=XM_023246518.1;refGene_gn=RBFOX1;refGene_tx=XM_023246519.1;refGene_gn=RBFOX1;refGene_tx=XM_023246520.1;refGene_gn=RBFOX1;refGene_tx=XM_023246521.1;refGene_gn=RBFOX1;refGene_tx=XM_023246522.1;refGene_gn=RBFOX1;refGene_tx=XM_023246524.1;refGene_gn=RBFOX1;refGene_tx=XM_023246525.1;refGene_gn=RBFOX1;refGene_tx=XM_023246526.1;refGene_gn=RBFOX1;refGene_tx=XM_023246527.1;refGene_gn=RBFOX1;refGene_tx=XM_023246528.1;refGene_gn=RBFOX1;refGene_tx=XM_023246529.1;refGene_gn=RBFOX1;refGene_tx=XM_023246530.1;refGene_gn=RBFOX1;refGene_tx=XM_023246531.1;refGene_gn=RBFOX1;refGene_tx=XM_023246532.1;refGene_gn=RBFOX1;refGene_tx=XM_023246533.1;refGene_gn=RBFOX1;refGene_tx=XM_023246534.1;refGene_gn=RBFOX1;refGene_tx=XM_023246535.1
E3 38516717 38516804
refGene_gn=RBFOX1;refGene_tx=XM_023246534.1;refGene_gn=RBFOX1;refGene_tx=XM_023246535.1
E3 38545255 38545399 refGene_gn=RBFOX1;refGene_tx=XM_023246531.1
E3 38762558 38762600
refGene_gn=RBFOX1;refGene_tx=XM_023246510.1;refGene_gn=RBFOX1;refGene_tx=XM_023246511.1;refGene_gn=RBFOX1;refGene_tx=XM_023246514.1;refGene_gn=RBFOX1;refGene_tx=XM_023246515.1;refGene_gn=RBFOX1;refGene_tx=XM_023246516.1;refGene_gn=RBFOX1;refGene_tx=XM_023246517.1;refGene_gn=RBFOX1;refGene_tx=XM_023246518.1;refGene_gn=RBFOX1;refGene_tx=XM_023246519.1;refGene_gn=RBFOX1;refGene_tx=XM_023246520.1;refGene_gn=RBFOX1;refGene_tx=XM_023246521.1;refGene_gn=RBFOX1;refGene_tx=XM_023246522.1;refGene_gn=RBFOX1;refGene_tx=XM_023246524.1;refGene_gn=RBFOX1;refGene_tx=XM_023246525.1;refGene_gn=RBFOX1;refGene_tx=XM_023246526.1;refGene_gn=RBFOX1;refGene_tx=XM_023246527.1;refGene_gn=RBFOX1;refGene_tx=XM_023246528.1;refGene_gn=RBFOX1;refGene_tx=XM_023246529.1;refGene_gn=RBFOX1;refGene_tx=XM_023246530.1;refGene_gn=RBFOX1;refGene_tx=XM_023246532.1
E3 39088730 39088778
refGene_gn=RBFOX1;refGene_tx=XM_023246510.1;refGene_gn=RBFOX1;refGene_tx=XM_023246511.1;refGene_gn=RBFOX1;refGene_tx=XM_023246513.1;refGene_gn=RBFOX1;refGene_tx=XM_023246514.1;refGene_gn=RBFOX1;refGene_tx=XM_023246515.1;refGene_gn=RBFOX1;refGene_tx=XM_023246516.1;refGene_gn=RBFOX1;refGene_tx=XM_023246517.1;refGene_gn=RBFOX1;refGene_tx=XM_023246518.1;refGene_gn=RBFOX1;refGene_tx=XM_023246519.1;refGene_gn=RBFOX1;refGene_tx=XM_023246520.1;refGene_gn=RBFOX1;refGene_tx=XM_023246521.1;refGene_gn=RBFOX1;refGene_tx=XM_023246522.1;refGene_gn=RBFOX1;refGene_tx=XM_023246524.1;refGene_gn=RBFOX1;refGene_tx=XM_023246525.1;refGene_gn=RBFOX1;refGene_tx=XM_023246526.1;refGene_gn=RBFOX1;refGene_tx=XM_02324652

7.1;refGene_gn=RBFOX1;refGene_tx=XM_023246528.1;refGene_gn=RBFOX1;refGene_tx=XM_023246529.1;refGene_gn=RBFOX1;refGene_tx=XM_023246530.1;refGene_gn=RBFOX1;refGene_tx=XM_023246532.1;refGene_gn=RBFOX1;refGene_tx=XM_023246533.1
E3 39239700 39239745
refGene_gn=RBFOX1;refGene_tx=XM_023246532.1;refGene_gn=RBFOX1;refGene_tx=XM_023246533.1
E3 39375529 39375602
refGene_gn=RBFOX1;refGene_tx=XM_023246527.1;refGene_gn=RBFOX1;refGene_tx=XM_023246528.1;refGene_gn=RBFOX1;refGene_tx=XM_023246529.1
E3 39375687 39375750
refGene_gn=RBFOX1;refGene_tx=XM_023246510.1;refGene_gn=RBFOX1;refGene_tx=XM_023246511.1;refGene_gn=RBFOX1;refGene_tx=XM_023246513.1;refGene_gn=RBFOX1;refGene_tx=XM_023246514.1;refGene_gn=RBFOX1;refGene_tx=XM_023246515.1;refGene_gn=RBFOX1;refGene_tx=XM_023246516.1;refGene_gn=RBFOX1;refGene_tx=XM_023246517.1;refGene_gn=RBFOX1;refGene_tx=XM_023246518.1;refGene_gn=RBFOX1;refGene_tx=XM_023246519.1;refGene_gn=RBFOX1;refGene_tx=XM_023246520.1;refGene_gn=RBFOX1;refGene_tx=XM_023246521.1;refGene_gn=RBFOX1;refGene_tx=XM_023246522.1;refGene_gn=RBFOX1;refGene_tx=XM_023246524.1;refGene_gn=RBFOX1;refGene_tx=XM_023246525.1;refGene_gn=RBFOX1;refGene_tx=XM_023246526.1;refGene_gn=RBFOX1;refGene_tx=XM_023246527.1;refGene_gn=RBFOX1;refGene_tx=XM_023246528.1;refGene_gn=RBFOX1;refGene_tx=XM_023246529.1;refGene_gn=RBFOX1;refGene_tx=XM_023246530.1
E3 39633367 39633370 refGene_gn=RBFOX1;refGene_tx=XM_023246530.1
E3 39743077 39743110
refGene_gn=RBFOX1;refGene_tx=XM_023246510.1;refGene_gn=RBFOX1;refGene_tx=XM_023246511.1;refGene_gn=RBFOX1;refGene_tx=XM_023246513.1;refGene_gn=RBFOX1;refGene_tx=XM_023246514.1;refGene_gn=RBFOX1;refGene_tx=XM_023246515.1;refGene_gn=RBFOX1;refGene_tx=XM_023246516.1;refGene_gn=RBFOX1;refGene_tx=XM_023246517.1;refGene_gn=RBFOX1;refGene_tx=XM_023246518.1;refGene_gn=RBFOX1;refGene_tx=XM_023246519.1;refGene_gn=RBFOX1;refGene_tx=XM_023246520.1;refGene_gn=RBFOX1;refGene_tx=XM_023246521.1;refGene_gn=RBFOX1;refGene_tx=XM_023246522.1;refGene_gn=RBFOX1;refGene_tx=XM_023246524.1;refGene_gn=RBFOX1;refGene_tx=XM_023246525.1;refGene_gn=RBFOX1;refGene_tx=XM_023246526.1
E3 39955982 39956042
refGene_gn=RBFOX1;refGene_tx=XM_023246510.1;refGene_gn=RBFOX1;refGene_tx=XM_023246511.1;refGene_gn=RBFOX1;refGene_tx=XM_023246513.1;refGene_gn=RBFOX1;refGene_tx=XM_023246514.1;refGene_gn=RBFOX1;refGene_tx=XM_023246515.1;refGene_gn=RBFOX1;refGene_tx=XM_023246516.1;refGene_gn=RBFOX1;refGene_tx=XM_023246517.1;refGene_gn=RBFOX1;refGene_tx=XM_023246518.1;refGene_gn=RBFOX1;refGene_tx=XM_023246519.1;refGene_gn=RBFOX1;refGene_tx=XM_023246520.1;refGene_gn=RBFOX1;refGene_tx=XM_023246521.1;refGene_gn=RBFOX1;refGene_tx=XM_023246522.1;refGene_gn=RBFOX1;refGene_tx=XM_023246524.1;refGene_gn=RBFOX1;refGene_tx=XM_023246525.1;refGene_gn=RBFOX1;refGene_tx=XM_023246526.1
E3 40050404 40050443
refGene_gn=RBFOX1;refGene_tx=XM_023246510.1;refGene_gn=RBFOX1;refGene_tx=XM_023246511.1;refGene_gn=RBFOX1;refGene_tx=XM_023246513.1;refGene_gn=RBFOX1;refGene_tx=XM_023246514.1;refGene_gn=RBFOX1;refGene_tx=XM_02

3246515.1;refGene_gn=RBFOX1;refGene_tx=XM_023246516.1;refGene_gn=RBFOX1;refGene_tx=XM_023246517.1;refGene_gn=RBFOX1;refGene_tx=XM_023246518.1;refGene_gn=RBFOX1;refGene_tx=XM_023246519.1;refGene_gn=RBFOX1;refGene_tx=XM_023246520.1;refGene_gn=RBFOX1;refGene_tx=XM_023246521.1;refGene_gn=RBFOX1;refGene_tx=XM_023246522.1;refGene_gn=RBFOX1;refGene_tx=XM_023246524.1;refGene_gn=RBFOX1;refGene_tx=XM_023246525.1;refGene_gn=RBFOX1;refGene_tx=XM_023246526.1
E3 40233137 40233347
refGene_gn=RBFOX1;refGene_tx=XM_023246510.1;refGene_gn=RBFOX1;refGene_tx=XM_023246511.1;refGene_gn=RBFOX1;refGene_tx=XM_023246513.1;refGene_gn=RBFOX1;refGene_tx=XM_023246514.1;refGene_gn=RBFOX1;refGene_tx=XM_023246515.1;refGene_gn=RBFOX1;refGene_tx=XM_023246516.1;refGene_gn=RBFOX1;refGene_tx=XM_023246517.1;refGene_gn=RBFOX1;refGene_tx=XM_023246518.1;refGene_gn=RBFOX1;refGene_tx=XM_023246519.1;refGene_gn=RBFOX1;refGene_tx=XM_023246520.1;refGene_gn=RBFOX1;refGene_tx=XM_023246521.1;refGene_gn=RBFOX1;refGene_tx=XM_023246522.1;refGene_gn=RBFOX1;refGene_tx=XM_023246524.1;refGene_gn=RBFOX1;refGene_tx=XM_023246525.1;refGene_gn=RBFOX1;refGene_tx=XM_023246526.1
E3 40295155 40295405
refGene_gn=LOC101091244;refGene_tx=XM_023246587.1
E3 40312860 40313006
refGene_gn=LOC101091244;refGene_tx=XM_023246587.1
E3 40320171 40320798
refGene_gn=LOC101091244;refGene_tx=XM_023246587.1
E3 40322877 40323124
refGene_gn=LOC101091244;refGene_tx=XM_023246587.1
E3 40323265 40323303
refGene_gn=LOC101091244;refGene_tx=XM_023246587.1
E3 40328071 40328167 refGene_gn=EEF2KMT;refGene_tx=XM_023246604.1
E3 40330419 40330482 refGene_gn=EEF2KMT;refGene_tx=XM_023246604.1
E3 40332377 40332458 refGene_gn=EEF2KMT;refGene_tx=XM_023246604.1
E3 40333602 40333704 refGene_gn=EEF2KMT;refGene_tx=XM_023246604.1
E3 40338271 40338405 refGene_gn=EEF2KMT;refGene_tx=XM_023246604.1
E3 40338491 40338787 refGene_gn=EEF2KMT;refGene_tx=XM_023246604.1
E3 40340224 40340374 refGene_gn=EEF2KMT;refGene_tx=XM_023246604.1
E3 40342101 40342202 refGene_gn=EEF2KMT;refGene_tx=XM_023246604.1
E3 40343197 40343329
refGene_gn=ALG1;refGene_tx=XM_003998892.5;refGene_gn=ALG1;refGene_tx=XM_019820986.1
E3 40344406 40344482
refGene_gn=ALG1;refGene_tx=XM_003998892.5;refGene_gn=ALG1;refGene_tx=XM_019820986.1
E3 40345379 40345494
refGene_gn=ALG1;refGene_tx=XM_003998892.5;refGene_gn=ALG1;refGene_tx=XM_019820986.1
E3 40346697 40346808
refGene_gn=ALG1;refGene_tx=XM_003998892.5;refGene_gn=ALG1;refGene_tx=XM_019820986.1
E3 40347605 40347665
refGene_gn=ALG1;refGene_tx=XM_003998892.5;refGene_gn=ALG1;refGene_tx=X

M_019820986.1
E3	40347977	40348016
refGene_gn=ALG1;refGene_tx=XM_003998892.5;refGene_gn=ALG1;refGene_tx=X
M_019820986.1
E3	40348207	40348329
refGene_gn=ALG1;refGene_tx=XM_003998892.5;refGene_gn=ALG1;refGene_tx=X
M_019820986.1
E3	40348759	40348870
refGene_gn=ALG1;refGene_tx=XM_003998892.5;refGene_gn=ALG1;refGene_tx=X
M_019820986.1
E3	40349175	40349265
refGene_gn=ALG1;refGene_tx=XM_003998892.5;refGene_gn=ALG1;refGene_tx=X
M_019820986.1
E3	40350731	40350880
refGene_gn=ALG1;refGene_tx=XM_003998892.5;refGene_gn=ALG1;refGene_tx=X
M_019820986.1
E3	40351335	40351439	refGene_gn=ALG1;refGene_tx=XM_003998892.5
E3	40351554	40351632	refGene_gn=ALG1;refGene_tx=XM_003998892.5
E3	40352553	40352761	refGene_gn=ALG1;refGene_tx=XM_003998892.5
E3	40357235	40357446
refGene_gn=CE3H16orf89;refGene_tx=XM_003998978.5;refGene_gn=CE3H16orf8
9;refGene_tx=XM_006942357.4
E3	40361789	40361939
refGene_gn=CE3H16orf89;refGene_tx=XM_003998978.5;refGene_gn=CE3H16orf8
9;refGene_tx=XM_006942357.4
E3	40362987	40363138
refGene_gn=CE3H16orf89;refGene_tx=XM_003998978.5;refGene_gn=CE3H16orf8
9;refGene_tx=XM_006942357.4
E3	40364229	40364347
refGene_gn=CE3H16orf89;refGene_tx=XM_003998978.5;refGene_gn=CE3H16orf8
9;refGene_tx=XM_006942357.4
E3	40366497	40366633
refGene_gn=CE3H16orf89;refGene_tx=XM_003998978.5;refGene_gn=CE3H16orf8
9;refGene_tx=XM_006942357.4
E3	40367279	40367384
refGene_gn=CE3H16orf89;refGene_tx=XM_003998978.5;refGene_gn=CE3H16orf8
9;refGene_tx=XM_006942357.4
E3	40368709	40368796
refGene_gn=CE3H16orf89;refGene_tx=XM_003998978.5;refGene_gn=CE3H16orf8
9;refGene_tx=XM_006942357.4
E3	40371390	40371554
refGene_gn=CE3H16orf89;refGene_tx=XM_003998978.5
E3	40376754	40376828
refGene_gn=CE3H16orf89;refGene_tx=XM_006942357.4
E3	40380682	40380768
refGene_gn=NAGPA;refGene_tx=XM_003998893.4;refGene_gn=NAGPA;refGene_tx
=XM_006942358.3
E3	40380854	40381310
refGene_gn=NAGPA;refGene_tx=XM_003998893.4;refGene_gn=NAGPA;refGene_tx
=XM_006942358.3

E3 40381736 40381876
refGene_gn=NAGPA;refGene_tx=XM_003998893.4;refGene_gn=NAGPA;refGene_tx
=XM_006942358.3
E3 40382821 40382930
refGene_gn=NAGPA;refGene_tx=XM_003998893.4;refGene_gn=NAGPA;refGene_tx
=XM_006942358.3
E3 40384008 40384137
refGene_gn=NAGPA;refGene_tx=XM_003998893.4;refGene_gn=NAGPA;refGene_tx
=XM_006942358.3
E3 40384635 40384841
refGene_gn=NAGPA;refGene_tx=XM_003998893.4;refGene_gn=NAGPA;refGene_tx
=XM_006942358.3
E3 40384945 40384993
refGene_gn=NAGPA;refGene_tx=XM_003998893.4;refGene_gn=NAGPA;refGene_tx
=XM_006942358.3
E3 40386005 40386134
refGene_gn=NAGPA;refGene_tx=XM_003998893.4;refGene_gn=NAGPA;refGene_tx
=XM_006942358.3
E3 40386305 40386437
refGene_gn=NAGPA;refGene_tx=XM_003998893.4;refGene_gn=NAGPA;refGene_tx
=XM_006942358.3
E3 40386523 40386591
refGene_gn=NAGPA;refGene_tx=XM_003998893.4;refGene_gn=NAGPA;refGene_tx
=XM_006942358.3
E3 40388090 40388301 refGene_gn=NAGPA;refGene_tx=XM_003998893.4
E3 40398518 40398627 refGene_gn=SEC14L5;refGene_tx=XM_003998895.5
E3 40401165 40401344 refGene_gn=SEC14L5;refGene_tx=XM_003998895.5
E3 40402258 40402428 refGene_gn=SEC14L5;refGene_tx=XM_003998895.5
E3 40402430 40402485 refGene_gn=SEC14L5;refGene_tx=XM_003998895.5
E3 40402845 40402980 refGene_gn=SEC14L5;refGene_tx=XM_003998895.5
E3 40404052 40404187 refGene_gn=SEC14L5;refGene_tx=XM_003998895.5
E3 40408603 40408775 refGene_gn=SEC14L5;refGene_tx=XM_003998895.5
E3 40410184 40410255 refGene_gn=SEC14L5;refGene_tx=XM_003998895.5
E3 40410346 40410435 refGene_gn=SEC14L5;refGene_tx=XM_003998895.5
E3 40412995 40413185 refGene_gn=SEC14L5;refGene_tx=XM_003998895.5
E3 40414993 40415106 refGene_gn=SEC14L5;refGene_tx=XM_003998895.5
E3 40419078 40419271 refGene_gn=SEC14L5;refGene_tx=XM_003998895.5
E3 40419927 40420056 refGene_gn=SEC14L5;refGene_tx=XM_003998895.5
E3 40422742 40422874 refGene_gn=SEC14L5;refGene_tx=XM_003998895.5
E3 40423274 40423424 refGene_gn=SEC14L5;refGene_tx=XM_003998895.5
E3 40438519 40438582 refGene_gn=SEC14L5;refGene_tx=XM_003998895.5
E3 40454874 40454936
refGene_gn=PPL;refGene_tx=XM_003998979.5;refGene_gn=PPL;refGene_tx=XM_
006942360.4
E3 40475991 40476091
refGene_gn=PPL;refGene_tx=XM_003998979.5;refGene_gn=PPL;refGene_tx=XM_
006942360.4
E3 40481349 40481504
refGene_gn=PPL;refGene_tx=XM_003998979.5;refGene_gn=PPL;refGene_tx=XM_
006942360.4

E3 40482236 40482357
refGene_gn=PPL;refGene_tx=XM_003998979.5;refGene_gn=PPL;refGene_tx=XM_006942360.4
E3 40482905 40483031
refGene_gn=PPL;refGene_tx=XM_003998979.5;refGene_gn=PPL;refGene_tx=XM_006942360.4
E3 40483125 40483164
refGene_gn=PPL;refGene_tx=XM_003998979.5;refGene_gn=PPL;refGene_tx=XM_006942360.4
E3 40484521 40484683
refGene_gn=PPL;refGene_tx=XM_003998979.5;refGene_gn=PPL;refGene_tx=XM_006942360.4
E3 40484789 40484897
refGene_gn=PPL;refGene_tx=XM_003998979.5;refGene_gn=PPL;refGene_tx=XM_006942360.4
E3 40486439 40486535
refGene_gn=PPL;refGene_tx=XM_003998979.5;refGene_gn=PPL;refGene_tx=XM_006942360.4
E3 40487995 40488118
refGene_gn=PPL;refGene_tx=XM_003998979.5;refGene_gn=PPL;refGene_tx=XM_006942360.4
E3 40488218 40488365
refGene_gn=PPL;refGene_tx=XM_003998979.5;refGene_gn=PPL;refGene_tx=XM_006942360.4
E3 40488906 40489058
refGene_gn=PPL;refGene_tx=XM_003998979.5;refGene_gn=PPL;refGene_tx=XM_006942360.4
E3 40489965 40490063
refGene_gn=PPL;refGene_tx=XM_003998979.5;refGene_gn=PPL;refGene_tx=XM_006942360.4
E3 40490255 40490413
refGene_gn=PPL;refGene_tx=XM_003998979.5;refGene_gn=PPL;refGene_tx=XM_006942360.4
E3 40491104 40491283
refGene_gn=PPL;refGene_tx=XM_003998979.5;refGene_gn=PPL;refGene_tx=XM_006942360.4
E3 40491366 40491505
refGene_gn=PPL;refGene_tx=XM_003998979.5;refGene_gn=PPL;refGene_tx=XM_006942360.4
E3 40492307 40492501
refGene_gn=PPL;refGene_tx=XM_003998979.5;refGene_gn=PPL;refGene_tx=XM_006942360.4
E3 40493890 40494041
refGene_gn=PPL;refGene_tx=XM_003998979.5;refGene_gn=PPL;refGene_tx=XM_006942360.4
E3 40494791 40494875
refGene_gn=PPL;refGene_tx=XM_003998979.5;refGene_gn=PPL;refGene_tx=XM_006942360.4
E3 40495539 40495650
refGene_gn=PPL;refGene_tx=XM_003998979.5;refGene_gn=PPL;refGene_tx=XM_

006942360.4
E3 40496134 40496227
refGene_gn=PPL;refGene_tx=XM_003998979.5;refGene_gn=PPL;refGene_tx=XM_006942360.4
E3 40496867 40499531
refGene_gn=PPL;refGene_tx=XM_003998979.5;refGene_gn=PPL;refGene_tx=XM_006942360.4
E3 40502679 40502729
refGene_gn=UBN1;refGene_tx=XM_011290643.3;refGene_gn=UBN1;refGene_tx=XM_011290644.3
E3 40505014 40505104
refGene_gn=UBN1;refGene_tx=XM_011290643.3;refGene_gn=UBN1;refGene_tx=XM_011290644.3
E3 40505386 40505630
refGene_gn=UBN1;refGene_tx=XM_011290643.3;refGene_gn=UBN1;refGene_tx=XM_011290644.3
E3 40507026 40508349
refGene_gn=UBN1;refGene_tx=XM_011290643.3;refGene_gn=UBN1;refGene_tx=XM_011290644.3
E3 40509497 40509540
refGene_gn=UBN1;refGene_tx=XM_011290643.3;refGene_gn=UBN1;refGene_tx=XM_011290644.3
E3 40509634 40509685
refGene_gn=UBN1;refGene_tx=XM_011290643.3;refGene_gn=UBN1;refGene_tx=XM_011290644.3
E3 40518732 40518879
refGene_gn=UBN1;refGene_tx=XM_011290643.3;refGene_gn=UBN1;refGene_tx=XM_011290644.3
E3 40519064 40519193
refGene_gn=UBN1;refGene_tx=XM_011290643.3;refGene_gn=UBN1;refGene_tx=XM_011290644.3
E3 40519463 40519582
refGene_gn=UBN1;refGene_tx=XM_011290643.3;refGene_gn=UBN1;refGene_tx=XM_011290644.3
E3 40519749 40519879
refGene_gn=UBN1;refGene_tx=XM_011290643.3;refGene_gn=UBN1;refGene_tx=XM_011290644.3
E3 40521351 40521422
refGene_gn=UBN1;refGene_tx=XM_011290643.3;refGene_gn=UBN1;refGene_tx=XM_011290644.3
E3 40527747 40528186
refGene_gn=UBN1;refGene_tx=XM_011290643.3;refGene_gn=UBN1;refGene_tx=XM_011290644.3
E3 40528899 40529003
refGene_gn=UBN1;refGene_tx=XM_011290643.3;refGene_gn=UBN1;refGene_tx=XM_011290644.3
E3 40529716 40529851
refGene_gn=UBN1;refGene_tx=XM_011290643.3;refGene_gn=UBN1;refGene_tx=XM_011290644.3
E3 40530205 40530301

refGene_gn=UBN1;refGene_tx=XM_011290643.3;refGene_gn=UBN1;refGene_tx=XM_011290644.3
E3 40530850 40530937
refGene_gn=UBN1;refGene_tx=XM_011290643.3;refGene_gn=UBN1;refGene_tx=XM_011290644.3
E3 40534650 40534899
refGene_gn=UBN1;refGene_tx=XM_011290643.3;refGene_gn=UBN1;refGene_tx=XM_011290644.3
E3 40540137 40540175 refGene_gn=GLYR1;refGene_tx=XM_003998897.5
E3 40541214 40541251 refGene_gn=GLYR1;refGene_tx=XM_003998897.5
E3 40542481 40542561 refGene_gn=GLYR1;refGene_tx=XM_003998897.5
E3 40551108 40551247 refGene_gn=GLYR1;refGene_tx=XM_003998897.5
E3 40551789 40552032 refGene_gn=GLYR1;refGene_tx=XM_003998897.5
E3 40556732 40556819 refGene_gn=GLYR1;refGene_tx=XM_003998897.5
E3 40557733 40557790 refGene_gn=GLYR1;refGene_tx=XM_003998897.5
E3 40558544 40558595 refGene_gn=GLYR1;refGene_tx=XM_003998897.5
E3 40558687 40558761 refGene_gn=GLYR1;refGene_tx=XM_003998897.5
E3 40561261 40561361 refGene_gn=GLYR1;refGene_tx=XM_003998897.5
E3 40565046 40565157 refGene_gn=GLYR1;refGene_tx=XM_003998897.5
E3 40565794 40565896 refGene_gn=GLYR1;refGene_tx=XM_003998897.5
E3 40566516 40566679 refGene_gn=GLYR1;refGene_tx=XM_003998897.5
E3 40566996 40567176 refGene_gn=GLYR1;refGene_tx=XM_003998897.5
E3 40567491 40567616 refGene_gn=GLYR1;refGene_tx=XM_003998897.5
E3 40571759 40571834 refGene_gn=GLYR1;refGene_tx=XM_003998897.5
E3 40574565 40574610 refGene_gn=ROGDI;refGene_tx=XM_023246622.1
E3 40574686 40574758 refGene_gn=ROGDI;refGene_tx=XM_023246622.1
E3 40575820 40575909 refGene_gn=ROGDI;refGene_tx=XM_023246622.1
E3 40576100 40576155 refGene_gn=ROGDI;refGene_tx=XM_023246622.1
E3 40576848 40576929 refGene_gn=ROGDI;refGene_tx=XM_023246622.1
E3 40577767 40577863 refGene_gn=ROGDI;refGene_tx=XM_023246622.1
E3 40579116 40579215 refGene_gn=ROGDI;refGene_tx=XM_023246622.1
E3 40579570 40579684 refGene_gn=ROGDI;refGene_tx=XM_023246622.1
E3 40579778 40579828 refGene_gn=ROGDI;refGene_tx=XM_023246622.1
E3 40579943 40580070 refGene_gn=ROGDI;refGene_tx=XM_023246622.1
E3 40580390 40580432 refGene_gn=ROGDI;refGene_tx=XM_023246622.1
E3 40581380 40581422
refGene_gn=SMIM22;refGene_tx=XM_011290647.3;refGene_gn=SMIM22;refGene_tx=XM_019820992.2;refGene_gn=SMIM22;refGene_tx=XM_019820993.2
E3 40581571 40581672
refGene_gn=SMIM22;refGene_tx=XM_011290647.3;refGene_gn=SMIM22;refGene_tx=XM_019820992.2;refGene_gn=SMIM22;refGene_tx=XM_019820993.2
E3 40581782 40581915
refGene_gn=SMIM22;refGene_tx=XM_011290647.3;refGene_gn=SMIM22;refGene_tx=XM_019820992.2;refGene_gn=SMIM22;refGene_tx=XM_019820993.2
E3 40588200 40588363
refGene_gn=SEPT12;refGene_tx=XM_006942361.4;refGene_gn=SEPT12;refGene_tx=XM_019820989.2;refGene_gn=SEPT12;refGene_tx=XM_019820990.2;refGene_gn=SEPT12;refGene_tx=XM_019820991.2;refGene_gn=SEPT12;refGene_tx=XM_023246704.1
E3 40591085 40591211

refGene_gn=SEPT12;refGene_tx=XM_006942361.4;refGene_gn=SEPT12;refGene_tx=XM_019820989.2;refGene_gn=SEPT12;refGene_tx=XM_019820990.2;refGene_gn=SEPT12;refGene_tx=XM_019820991.2;refGene_gn=SEPT12;refGene_tx=XM_023246704.1
E3 40591300 40591382 refGene_gn=SEPT12;refGene_tx=XM_006942361.4;refGene_gn=SEPT12;refGene_tx=XM_019820989.2;refGene_gn=SEPT12;refGene_tx=XM_019820990.2;refGene_gn=SEPT12;refGene_tx=XM_019820991.2;refGene_gn=SEPT12;refGene_tx=XM_023246704.1
E3 40593046 40593184 refGene_gn=SEPT12;refGene_tx=XM_006942361.4;refGene_gn=SEPT12;refGene_tx=XM_019820989.2;refGene_gn=SEPT12;refGene_tx=XM_019820990.2;refGene_gn=SEPT12;refGene_tx=XM_019820991.2;refGene_gn=SEPT12;refGene_tx=XM_023246704.1
E3 40593325 40593443 refGene_gn=SEPT12;refGene_tx=XM_006942361.4;refGene_gn=SEPT12;refGene_tx=XM_019820989.2;refGene_gn=SEPT12;refGene_tx=XM_019820990.2;refGene_gn=SEPT12;refGene_tx=XM_019820991.2;refGene_gn=SEPT12;refGene_tx=XM_023246704.1
E3 40593531 40593627 refGene_gn=SEPT12;refGene_tx=XM_006942361.4;refGene_gn=SEPT12;refGene_tx=XM_019820989.2;refGene_gn=SEPT12;refGene_tx=XM_019820990.2;refGene_gn=SEPT12;refGene_tx=XM_019820991.2;refGene_gn=SEPT12;refGene_tx=XM_023246704.1
E3 40593732 40593894 refGene_gn=SEPT12;refGene_tx=XM_023246704.1
E3 40596437 40596534 refGene_gn=SEPT12;refGene_tx=XM_006942361.4;refGene_gn=SEPT12;refGene_tx=XM_019820989.2;refGene_gn=SEPT12;refGene_tx=XM_019820990.2;refGene_gn=SEPT12;refGene_tx=XM_019820991.2
E3 40596628 40596708 refGene_gn=SEPT12;refGene_tx=XM_019820989.2;refGene_gn=SEPT12;refGene_tx=XM_019820990.2
E3 40597403 40597501 refGene_gn=SEPT12;refGene_tx=XM_006942361.4;refGene_gn=SEPT12;refGene_tx=XM_019820989.2;refGene_gn=SEPT12;refGene_tx=XM_019820990.2;refGene_gn=SEPT12;refGene_tx=XM_019820991.2
E3 40597580 40597791 refGene_gn=SEPT12;refGene_tx=XM_006942361.4;refGene_gn=SEPT12;refGene_tx=XM_019820989.2
E3 40614877 40615279 refGene_gn=ZNF500;refGene_tx=XM_003998982.3;refGene_gn=ZNF500;refGene_tx=XM_006942364.3;refGene_gn=ZNF500;refGene_tx=XM_006942365.4;refGene_gn=ZNF500;refGene_tx=XM_006942366.3;refGene_gn=ZNF500;refGene_tx=XM_019821005.2;refGene_gn=ZNF500;refGene_tx=XM_023246340.1
E3 40616919 40617103 refGene_gn=ZNF500;refGene_tx=XM_003998982.3;refGene_gn=ZNF500;refGene_tx=XM_006942364.3;refGene_gn=ZNF500;refGene_tx=XM_006942365.4;refGene_gn=ZNF500;refGene_tx=XM_006942366.3;refGene_gn=ZNF500;refGene_tx=XM_019821005.2;refGene_gn=ZNF500;refGene_tx=XM_023246340.1
E3 40617319 40617384

refGene_gn=ZNF500;refGene_tx=XM_003998982.3;refGene_gn=ZNF500;refGene_tx=XM_006942364.3;refGene_gn=ZNF500;refGene_tx=XM_006942365.4;refGene_gn=ZNF500;refGene_tx=XM_006942366.3;refGene_gn=ZNF500;refGene_tx=XM_019821005.2;refGene_gn=ZNF500;refGene_tx=XM_023246340.1
E3 40618392 40618504
refGene_gn=ZNF500;refGene_tx=XM_003998982.3;refGene_gn=ZNF500;refGene_tx=XM_006942364.3;refGene_gn=ZNF500;refGene_tx=XM_006942365.4;refGene_gn=ZNF500;refGene_tx=XM_006942366.3;refGene_gn=ZNF500;refGene_tx=XM_019821005.2;refGene_gn=ZNF500;refGene_tx=XM_023246340.1
E3 40620235 40620921
refGene_gn=ZNF500;refGene_tx=XM_003998982.3;refGene_gn=ZNF500;refGene_tx=XM_006942364.3;refGene_gn=ZNF500;refGene_tx=XM_006942365.4;refGene_gn=ZNF500;refGene_tx=XM_006942366.3;refGene_gn=ZNF500;refGene_tx=XM_019821005.2;refGene_gn=ZNF500;refGene_tx=XM_023246340.1
E3 40623646 40623785
refGene_gn=CE3H16orf71;refGene_tx=XM_019820999.2;refGene_gn=CE3H16orf71;refGene_tx=XM_023246335.1;refGene_gn=CE3H16orf71;refGene_tx=XM_023246336.1;refGene_gn=CE3H16orf71;refGene_tx=XM_023246337.1;refGene_gn=CE3H16orf71;refGene_tx=XM_023246338.1
E3 40623980 40624254
refGene_gn=CE3H16orf71;refGene_tx=XM_019820999.2;refGene_gn=CE3H16orf71;refGene_tx=XM_023246335.1;refGene_gn=CE3H16orf71;refGene_tx=XM_023246336.1;refGene_gn=CE3H16orf71;refGene_tx=XM_023246337.1;refGene_gn=CE3H16orf71;refGene_tx=XM_023246338.1;refGene_gn=CE3H16orf71;refGene_tx=XM_023246339.1
E3 40625031 40625130
refGene_gn=CE3H16orf71;refGene_tx=XM_019820999.2;refGene_gn=CE3H16orf71;refGene_tx=XM_023246335.1;refGene_gn=CE3H16orf71;refGene_tx=XM_023246336.1;refGene_gn=CE3H16orf71;refGene_tx=XM_023246337.1;refGene_gn=CE3H16orf71;refGene_tx=XM_023246338.1;refGene_gn=CE3H16orf71;refGene_tx=XM_023246339.1
E3 40625498 40625642
refGene_gn=CE3H16orf71;refGene_tx=XM_019820999.2;refGene_gn=CE3H16orf71;refGene_tx=XM_023246335.1;refGene_gn=CE3H16orf71;refGene_tx=XM_023246336.1;refGene_gn=CE3H16orf71;refGene_tx=XM_023246337.1;refGene_gn=CE3H16orf71;refGene_tx=XM_023246338.1;refGene_gn=CE3H16orf71;refGene_tx=XM_023246339.1
E3 40626888 40627027
refGene_gn=CE3H16orf71;refGene_tx=XM_019820999.2;refGene_gn=CE3H16orf71;refGene_tx=XM_023246335.1;refGene_gn=CE3H16orf71;refGene_tx=XM_023246336.1;refGene_gn=CE3H16orf71;refGene_tx=XM_023246337.1;refGene_gn=CE3H16orf71;refGene_tx=XM_023246338.1;refGene_gn=CE3H16orf71;refGene_tx=XM_023246339.1
E3 40627834 40627843
refGene_gn=CE3H16orf71;refGene_tx=XM_023246335.1;refGene_gn=CE3H16orf71;refGene_tx=XM_023246336.1;refGene_gn=CE3H16orf71;refGene_tx=XM_023246337.1;refGene_gn=CE3H16orf71;refGene_tx=XM_023246338.1;refGene_gn=CE3H16orf71;refGene_tx=XM_023246339.1
E3 40628346 40628461
refGene_gn=CE3H16orf71;refGene_tx=XM_019820999.2;refGene_gn=CE3H16orf7

1;refGene_tx=XM_023246335.1;refGene_gn=CE3H16orf71;refGene_tx=XM_02324
6336.1;refGene_gn=CE3H16orf71;refGene_tx=XM_023246337.1;refGene_gn=CE3
H16orf71;refGene_tx=XM_023246338.1;refGene_gn=CE3H16orf71;refGene_tx=X
M_023246339.1
E3 40630195 40630708
refGene_gn=CE3H16orf71;refGene_tx=XM_019820999.2;refGene_gn=CE3H16orf7
1;refGene_tx=XM_023246335.1;refGene_gn=CE3H16orf71;refGene_tx=XM_02324
6336.1;refGene_gn=CE3H16orf71;refGene_tx=XM_023246337.1;refGene_gn=CE3
H16orf71;refGene_tx=XM_023246338.1;refGene_gn=CE3H16orf71;refGene_tx=X
M_023246339.1
E3 40632754 40632883
refGene_gn=CE3H16orf71;refGene_tx=XM_019820999.2;refGene_gn=CE3H16orf7
1;refGene_tx=XM_023246335.1;refGene_gn=CE3H16orf71;refGene_tx=XM_02324
6336.1;refGene_gn=CE3H16orf71;refGene_tx=XM_023246337.1;refGene_gn=CE3
H16orf71;refGene_tx=XM_023246338.1;refGene_gn=CE3H16orf71;refGene_tx=X
M_023246339.1
E3 40634038 40634161
refGene_gn=CE3H16orf71;refGene_tx=XM_019820999.2;refGene_gn=CE3H16orf7
1;refGene_tx=XM_023246335.1;refGene_gn=CE3H16orf71;refGene_tx=XM_02324
6336.1;refGene_gn=CE3H16orf71;refGene_tx=XM_023246337.1;refGene_gn=CE3
H16orf71;refGene_tx=XM_023246338.1;refGene_gn=CE3H16orf71;refGene_tx=X
M_023246339.1
E3 40637401 40637492
refGene_gn=ANKS3;refGene_tx=XM_003998984.5;refGene_gn=ANKS3;refGene_tx
=XM_006942372.4;refGene_gn=ANKS3;refGene_tx=XM_006942373.4;refGene_gn=
ANKS3;refGene_tx=XM_019820994.2
E3 40640279 40640451
refGene_gn=ANKS3;refGene_tx=XM_003998984.5;refGene_gn=ANKS3;refGene_tx
=XM_006942372.4;refGene_gn=ANKS3;refGene_tx=XM_006942373.4;refGene_gn=
ANKS3;refGene_tx=XM_006942374.3;refGene_gn=ANKS3;refGene_tx=XM_0198209
94.2;refGene_gn=ANKS3;refGene_tx=XM_019820995.1
E3 40642860 40643059
refGene_gn=ANKS3;refGene_tx=XM_003998984.5;refGene_gn=ANKS3;refGene_tx
=XM_006942372.4;refGene_gn=ANKS3;refGene_tx=XM_006942373.4;refGene_gn=
ANKS3;refGene_tx=XM_006942374.3;refGene_gn=ANKS3;refGene_tx=XM_0198209
94.2
E3 40643254 40643376
refGene_gn=ANKS3;refGene_tx=XM_003998984.5;refGene_gn=ANKS3;refGene_tx
=XM_006942372.4;refGene_gn=ANKS3;refGene_tx=XM_006942373.4;refGene_gn=
ANKS3;refGene_tx=XM_006942374.3;refGene_gn=ANKS3;refGene_tx=XM_0198209
94.2;refGene_gn=ANKS3;refGene_tx=XM_019820995.1;refGene_gn=ANKS3;refGe
ne_tx=XM_019820996.1
E3 40645396 40645478
refGene_gn=ANKS3;refGene_tx=XM_003998984.5;refGene_gn=ANKS3;refGene_tx
=XM_006942372.4;refGene_gn=ANKS3;refGene_tx=XM_006942373.4;refGene_gn=
ANKS3;refGene_tx=XM_006942374.3;refGene_gn=ANKS3;refGene_tx=XM_0198209
94.2;refGene_gn=ANKS3;refGene_tx=XM_019820995.1;refGene_gn=ANKS3;refGe
ne_tx=XM_019820996.1
E3 40655917 40656053
refGene_gn=ANKS3;refGene_tx=XM_003998984.5;refGene_gn=ANKS3;refGene_tx

=XM_006942372.4;refGene_gn=ANKS3;refGene_tx=XM_006942373.4;refGene_gn=ANKS3;refGene_tx=XM_006942374.3;refGene_gn=ANKS3;refGene_tx=XM_019820994.2;refGene_gn=ANKS3;refGene_tx=XM_019820995.1;refGene_gn=ANKS3;refGene_tx=XM_019820996.1
E3 40661870 40662029
refGene_gn=ANKS3;refGene_tx=XM_003998984.5;refGene_gn=ANKS3;refGene_tx=XM_006942372.4;refGene_gn=ANKS3;refGene_tx=XM_006942373.4;refGene_gn=ANKS3;refGene_tx=XM_006942374.3;refGene_gn=ANKS3;refGene_tx=XM_019820994.2;refGene_gn=ANKS3;refGene_tx=XM_019820995.1;refGene_gn=ANKS3;refGene_tx=XM_019820996.1;refGene_gn=ANKS3;refGene_tx=XM_019820997.1
E3 40664778 40664916
refGene_gn=ANKS3;refGene_tx=XM_003998984.5;refGene_gn=ANKS3;refGene_tx=XM_006942372.4;refGene_gn=ANKS3;refGene_tx=XM_006942373.4;refGene_gn=ANKS3;refGene_tx=XM_006942374.3;refGene_gn=ANKS3;refGene_tx=XM_019820994.2;refGene_gn=ANKS3;refGene_tx=XM_019820995.1;refGene_gn=ANKS3;refGene_tx=XM_019820996.1;refGene_gn=ANKS3;refGene_tx=XM_019820997.1
E3 40665467 40665577
refGene_gn=ANKS3;refGene_tx=XM_003998984.5;refGene_gn=ANKS3;refGene_tx=XM_006942372.4;refGene_gn=ANKS3;refGene_tx=XM_006942373.4;refGene_gn=ANKS3;refGene_tx=XM_006942374.3;refGene_gn=ANKS3;refGene_tx=XM_019820994.2;refGene_gn=ANKS3;refGene_tx=XM_019820995.1;refGene_gn=ANKS3;refGene_tx=XM_019820996.1;refGene_gn=ANKS3;refGene_tx=XM_019820997.1
E3 40665871 40666048
refGene_gn=ANKS3;refGene_tx=XM_003998984.5;refGene_gn=ANKS3;refGene_tx=XM_006942372.4;refGene_gn=ANKS3;refGene_tx=XM_006942373.4;refGene_gn=ANKS3;refGene_tx=XM_006942374.3;refGene_gn=ANKS3;refGene_tx=XM_019820994.2;refGene_gn=ANKS3;refGene_tx=XM_019820995.1;refGene_gn=ANKS3;refGene_tx=XM_019820996.1;refGene_gn=ANKS3;refGene_tx=XM_019820997.1
E3 40668202 40668327
refGene_gn=ANKS3;refGene_tx=XM_003998984.5;refGene_gn=ANKS3;refGene_tx=XM_006942372.4;refGene_gn=ANKS3;refGene_tx=XM_006942373.4;refGene_gn=ANKS3;refGene_tx=XM_006942374.3;refGene_gn=ANKS3;refGene_tx=XM_019820994.2;refGene_gn=ANKS3;refGene_tx=XM_019820995.1;refGene_gn=ANKS3;refGene_tx=XM_019820996.1;refGene_gn=ANKS3;refGene_tx=XM_019820997.1
E3 40668872 40669014
refGene_gn=ANKS3;refGene_tx=XM_003998984.5;refGene_gn=ANKS3;refGene_tx=XM_006942372.4;refGene_gn=ANKS3;refGene_tx=XM_006942373.4;refGene_gn=ANKS3;refGene_tx=XM_006942374.3;refGene_gn=ANKS3;refGene_tx=XM_019820994.2;refGene_gn=ANKS3;refGene_tx=XM_019820995.1;refGene_gn=ANKS3;refGene_tx=XM_019820996.1;refGene_gn=ANKS3;refGene_tx=XM_019820997.1
E3 40669477 40669650
refGene_gn=ANKS3;refGene_tx=XM_003998984.5;refGene_gn=ANKS3;refGene_tx=XM_006942372.4;refGene_gn=ANKS3;refGene_tx=XM_006942373.4;refGene_gn=ANKS3;refGene_tx=XM_006942374.3;refGene_gn=ANKS3;refGene_tx=XM_019820994.2;refGene_gn=ANKS3;refGene_tx=XM_019820995.1;refGene_gn=ANKS3;refGene_tx=XM_019820996.1;refGene_gn=ANKS3;refGene_tx=XM_019820997.1
E3 40669996 40670082
refGene_gn=ANKS3;refGene_tx=XM_003998984.5;refGene_gn=ANKS3;refGene_tx=XM_006942372.4;refGene_gn=ANKS3;refGene_tx=XM_006942373.4;refGene_gn=ANKS3;refGene_tx=XM_006942374.3;refGene_gn=ANKS3;refGene_tx=XM_0198209

94.2;refGene_gn=ANKS3;refGene_tx=XM_019820995.1;refGene_gn=ANKS3;refGene_tx=XM_019820996.1;refGene_gn=ANKS3;refGene_tx=XM_019820997.1
E3 40670545 40670629
refGene_gn=ANKS3;refGene_tx=XM_003998984.5;refGene_gn=ANKS3;refGene_tx=XM_006942372.4;refGene_gn=ANKS3;refGene_tx=XM_006942373.4;refGene_gn=ANKS3;refGene_tx=XM_006942374.3;refGene_gn=ANKS3;refGene_tx=XM_019820994.2;refGene_gn=ANKS3;refGene_tx=XM_019820995.1;refGene_gn=ANKS3;refGene_tx=XM_019820996.1;refGene_gn=ANKS3;refGene_tx=XM_019820997.1
E3 40670828 40670908
refGene_gn=ANKS3;refGene_tx=XM_003998984.5;refGene_gn=ANKS3;refGene_tx=XM_006942372.4;refGene_gn=ANKS3;refGene_tx=XM_006942374.3;refGene_gn=ANKS3;refGene_tx=XM_019820994.2;refGene_gn=ANKS3;refGene_tx=XM_019820995.1;refGene_gn=ANKS3;refGene_tx=XM_019820996.1;refGene_gn=ANKS3;refGene_tx=XM_019820997.1
E3 40671012 40671029 refGene_gn=ANKS3;refGene_tx=XM_006942373.4
E3 40672730 40672952
refGene_gn=NUDT16L1;refGene_tx=XM_006942375.4;refGene_gn=NUDT16L1;refGene_tx=XM_023246341.1
E3 40673678 40673939
refGene_gn=NUDT16L1;refGene_tx=XM_006942375.4;refGene_gn=NUDT16L1;refGene_tx=XM_023246341.1
E3 40674015 40674165
refGene_gn=NUDT16L1;refGene_tx=XM_006942375.4;refGene_gn=NUDT16L1;refGene_tx=XM_023246341.1
E3 40678563 40678676
refGene_gn=MGRN1;refGene_tx=XM_023246330.1;refGene_gn=MGRN1;refGene_tx=XM_023246331.1
E3 40680409 40680450
refGene_gn=MGRN1;refGene_tx=XM_023246332.1;refGene_gn=MGRN1;refGene_tx=XM_023246333.1
E3 40682560 40682659
refGene_gn=MGRN1;refGene_tx=XM_023246330.1;refGene_gn=MGRN1;refGene_tx=XM_023246331.1;refGene_gn=MGRN1;refGene_tx=XM_023246332.1;refGene_gn=MGRN1;refGene_tx=XM_023246333.1
E3 40684113 40684159
refGene_gn=MGRN1;refGene_tx=XM_023246330.1;refGene_gn=MGRN1;refGene_tx=XM_023246331.1;refGene_gn=MGRN1;refGene_tx=XM_023246332.1;refGene_gn=MGRN1;refGene_tx=XM_023246333.1
E3 40684425 40684543
refGene_gn=MGRN1;refGene_tx=XM_023246330.1;refGene_gn=MGRN1;refGene_tx=XM_023246331.1;refGene_gn=MGRN1;refGene_tx=XM_023246332.1;refGene_gn=MGRN1;refGene_tx=XM_023246333.1
E3 40685453 40685680
refGene_gn=MGRN1;refGene_tx=XM_023246330.1;refGene_gn=MGRN1;refGene_tx=XM_023246331.1;refGene_gn=MGRN1;refGene_tx=XM_023246332.1;refGene_gn=MGRN1;refGene_tx=XM_023246333.1
E3 40687257 40687323
refGene_gn=MGRN1;refGene_tx=XM_023246330.1;refGene_gn=MGRN1;refGene_tx=XM_023246332.1
E3 40689971 40690081

refGene_gn=MGRN1;refGene_tx=XM_023246330.1;refGene_gn=MGRN1;refGene_tx=XM_023246331.1;refGene_gn=MGRN1;refGene_tx=XM_023246332.1;refGene_gn=MGRN1;refGene_tx=XM_023246333.1
E3 40692067 40692227
refGene_gn=MGRN1;refGene_tx=XM_023246330.1;refGene_gn=MGRN1;refGene_tx=XM_023246331.1;refGene_gn=MGRN1;refGene_tx=XM_023246332.1;refGene_gn=MGRN1;refGene_tx=XM_023246333.1
E3 40693675 40693744
refGene_gn=MGRN1;refGene_tx=XM_023246330.1;refGene_gn=MGRN1;refGene_tx=XM_023246331.1;refGene_gn=MGRN1;refGene_tx=XM_023246332.1;refGene_gn=MGRN1;refGene_tx=XM_023246333.1
E3 40694343 40694391
refGene_gn=MGRN1;refGene_tx=XM_023246330.1;refGene_gn=MGRN1;refGene_tx=XM_023246331.1;refGene_gn=MGRN1;refGene_tx=XM_023246332.1;refGene_gn=MGRN1;refGene_tx=XM_023246333.1
E3 40696498 40696548
refGene_gn=MGRN1;refGene_tx=XM_023246330.1;refGene_gn=MGRN1;refGene_tx=XM_023246331.1;refGene_gn=MGRN1;refGene_tx=XM_023246332.1;refGene_gn=MGRN1;refGene_tx=XM_023246333.1
E3 40697200 40697267
refGene_gn=MGRN1;refGene_tx=XM_023246330.1;refGene_gn=MGRN1;refGene_tx=XM_023246331.1;refGene_gn=MGRN1;refGene_tx=XM_023246332.1;refGene_gn=MGRN1;refGene_tx=XM_023246333.1
E3 40703844 40703962
refGene_gn=MGRN1;refGene_tx=XM_023246330.1;refGene_gn=MGRN1;refGene_tx=XM_023246331.1;refGene_gn=MGRN1;refGene_tx=XM_023246332.1;refGene_gn=MGRN1;refGene_tx=XM_023246333.1
E3 40705943 40706090
refGene_gn=MGRN1;refGene_tx=XM_023246330.1;refGene_gn=MGRN1;refGene_tx=XM_023246331.1;refGene_gn=MGRN1;refGene_tx=XM_023246332.1;refGene_gn=MGRN1;refGene_tx=XM_023246333.1
E3 40706670 40706759
refGene_gn=MGRN1;refGene_tx=XM_023246330.1;refGene_gn=MGRN1;refGene_tx=XM_023246331.1;refGene_gn=MGRN1;refGene_tx=XM_023246332.1;refGene_gn=MGRN1;refGene_tx=XM_023246333.1
E3 40708389 40708508
refGene_gn=MGRN1;refGene_tx=XM_023246330.1;refGene_gn=MGRN1;refGene_tx=XM_023246331.1;refGene_gn=MGRN1;refGene_tx=XM_023246332.1;refGene_gn=MGRN1;refGene_tx=XM_023246333.1
E3 40731672 40731760
refGene_gn=MGRN1;refGene_tx=XM_023246330.1;refGene_gn=MGRN1;refGene_tx=XM_023246331.1;refGene_gn=MGRN1;refGene_tx=XM_023246332.1;refGene_gn=MGRN1;refGene_tx=XM_023246333.1
E3 40748919 40748935
refGene_gn=CE3H16orf96;refGene_tx=XM_003998986.5;refGene_gn=CE3H16orf96;refGene_tx=XM_019821007.2;refGene_gn=CE3H16orf96;refGene_tx=XM_019821009.2;refGene_gn=CE3H16orf96;refGene_tx=XM_019821010.2;refGene_gn=CE3H16orf96;refGene_tx=XM_019821013.2
E3 40750287 40750489
refGene_gn=CE3H16orf96;refGene_tx=XM_003998986.5;refGene_gn=CE3H16orf9

6;refGene_tx=XM_019821007.2;refGene_gn=CE3H16orf96;refGene_tx=XM_01982
1009.2;refGene_gn=CE3H16orf96;refGene_tx=XM_019821010.2;refGene_gn=CE3
H16orf96;refGene_tx=XM_019821013.2
E3 40750759 40750872
refGene_gn=CE3H16orf96;refGene_tx=XM_019821014.2
E3 40751259 40751340
refGene_gn=CE3H16orf96;refGene_tx=XM_003998986.5;refGene_gn=CE3H16orf9
6;refGene_tx=XM_019821007.2;refGene_gn=CE3H16orf96;refGene_tx=XM_01982
1009.2;refGene_gn=CE3H16orf96;refGene_tx=XM_019821010.2;refGene_gn=CE3
H16orf96;refGene_tx=XM_019821013.2;refGene_gn=CE3H16orf96;refGene_tx=X
M_019821014.2;refGene_gn=CE3H16orf96;refGene_tx=XM_019821016.2;refGene
_gn=CE3H16orf96;refGene_tx=XM_019821017.2
E3 40757150 40757173
refGene_gn=CE3H16orf96;refGene_tx=XM_023246452.1
E3 40757300 40757403
refGene_gn=CE3H16orf96;refGene_tx=XM_003998986.5;refGene_gn=CE3H16orf9
6;refGene_tx=XM_019821007.2;refGene_gn=CE3H16orf96;refGene_tx=XM_01982
1009.2;refGene_gn=CE3H16orf96;refGene_tx=XM_019821010.2;refGene_gn=CE3
H16orf96;refGene_tx=XM_019821013.2;refGene_gn=CE3H16orf96;refGene_tx=X
M_019821014.2;refGene_gn=CE3H16orf96;refGene_tx=XM_023246452.1
E3 40757528 40757709
refGene_gn=CE3H16orf96;refGene_tx=XM_003998986.5;refGene_gn=CE3H16orf9
6;refGene_tx=XM_019821007.2;refGene_gn=CE3H16orf96;refGene_tx=XM_01982
1009.2;refGene_gn=CE3H16orf96;refGene_tx=XM_019821010.2;refGene_gn=CE3
H16orf96;refGene_tx=XM_019821013.2;refGene_gn=CE3H16orf96;refGene_tx=X
M_019821014.2;refGene_gn=CE3H16orf96;refGene_tx=XM_023246452.1
E3 40758556 40758649
refGene_gn=CE3H16orf96;refGene_tx=XM_003998986.5;refGene_gn=CE3H16orf9
6;refGene_tx=XM_019821007.2;refGene_gn=CE3H16orf96;refGene_tx=XM_01982
1009.2;refGene_gn=CE3H16orf96;refGene_tx=XM_019821010.2;refGene_gn=CE3
H16orf96;refGene_tx=XM_019821013.2;refGene_gn=CE3H16orf96;refGene_tx=X
M_019821014.2;refGene_gn=CE3H16orf96;refGene_tx=XM_019821016.2;refGene
_gn=CE3H16orf96;refGene_tx=XM_019821017.2;refGene_gn=CE3H16orf96;refGe
ne_tx=XM_023246452.1
E3 40760543 40760606
refGene_gn=CE3H16orf96;refGene_tx=XM_003998986.5;refGene_gn=CE3H16orf9
6;refGene_tx=XM_019821007.2;refGene_gn=CE3H16orf96;refGene_tx=XM_01982
1009.2;refGene_gn=CE3H16orf96;refGene_tx=XM_019821010.2;refGene_gn=CE3
H16orf96;refGene_tx=XM_019821013.2;refGene_gn=CE3H16orf96;refGene_tx=X
M_019821014.2;refGene_gn=CE3H16orf96;refGene_tx=XM_019821016.2;refGene
_gn=CE3H16orf96;refGene_tx=XM_019821017.2;refGene_gn=CE3H16orf96;refGe
ne_tx=XM_023246452.1
E3 40761304 40761423
refGene_gn=CE3H16orf96;refGene_tx=XM_003998986.5;refGene_gn=CE3H16orf9
6;refGene_tx=XM_019821007.2;refGene_gn=CE3H16orf96;refGene_tx=XM_01982
1009.2;refGene_gn=CE3H16orf96;refGene_tx=XM_019821010.2;refGene_gn=CE3
H16orf96;refGene_tx=XM_019821013.2;refGene_gn=CE3H16orf96;refGene_tx=X
M_019821014.2;refGene_gn=CE3H16orf96;refGene_tx=XM_019821016.2;refGene
_gn=CE3H16orf96;refGene_tx=XM_019821017.2;refGene_gn=CE3H16orf96;refGe
ne_tx=XM_023246452.1

E3 40764888 40765053
refGene_gn=CE3H16orf96;refGene_tx=XM_003998986.5;refGene_gn=CE3H16orf96;refGene_tx=XM_019821007.2;refGene_gn=CE3H16orf96;refGene_tx=XM_019821009.2;refGene_gn=CE3H16orf96;refGene_tx=XM_019821010.2;refGene_gn=CE3H16orf96;refGene_tx=XM_019821013.2;refGene_gn=CE3H16orf96;refGene_tx=XM_019821014.2;refGene_gn=CE3H16orf96;refGene_tx=XM_019821016.2;refGene_gn=CE3H16orf96;refGene_tx=XM_019821017.2;refGene_gn=CE3H16orf96;refGene_tx=XM_023246452.1
E3 40765497 40765572
refGene_gn=CE3H16orf96;refGene_tx=XM_003998986.5;refGene_gn=CE3H16orf96;refGene_tx=XM_019821007.2;refGene_gn=CE3H16orf96;refGene_tx=XM_019821009.2;refGene_gn=CE3H16orf96;refGene_tx=XM_019821010.2;refGene_gn=CE3H16orf96;refGene_tx=XM_019821013.2;refGene_gn=CE3H16orf96;refGene_tx=XM_019821014.2;refGene_gn=CE3H16orf96;refGene_tx=XM_019821016.2;refGene_gn=CE3H16orf96;refGene_tx=XM_019821017.2;refGene_gn=CE3H16orf96;refGene_tx=XM_023246452.1
E3 40768766 40768877
refGene_gn=CE3H16orf96;refGene_tx=XM_003998986.5;refGene_gn=CE3H16orf96;refGene_tx=XM_019821007.2;refGene_gn=CE3H16orf96;refGene_tx=XM_019821009.2;refGene_gn=CE3H16orf96;refGene_tx=XM_019821010.2;refGene_gn=CE3H16orf96;refGene_tx=XM_019821013.2;refGene_gn=CE3H16orf96;refGene_tx=XM_019821014.2;refGene_gn=CE3H16orf96;refGene_tx=XM_019821016.2;refGene_gn=CE3H16orf96;refGene_tx=XM_019821017.2;refGene_gn=CE3H16orf96;refGene_tx=XM_023246452.1
E3 40769168 40769254
refGene_gn=CE3H16orf96;refGene_tx=XM_003998986.5;refGene_gn=CE3H16orf96;refGene_tx=XM_019821007.2;refGene_gn=CE3H16orf96;refGene_tx=XM_019821009.2;refGene_gn=CE3H16orf96;refGene_tx=XM_019821010.2;refGene_gn=CE3H16orf96;refGene_tx=XM_019821013.2;refGene_gn=CE3H16orf96;refGene_tx=XM_019821014.2;refGene_gn=CE3H16orf96;refGene_tx=XM_019821016.2;refGene_gn=CE3H16orf96;refGene_tx=XM_019821017.2;refGene_gn=CE3H16orf96;refGene_tx=XM_023246452.1
E3 40771686 40773049
refGene_gn=CE3H16orf96;refGene_tx=XM_003998986.5;refGene_gn=CE3H16orf96;refGene_tx=XM_019821007.2;refGene_gn=CE3H16orf96;refGene_tx=XM_019821009.2;refGene_gn=CE3H16orf96;refGene_tx=XM_019821010.2;refGene_gn=CE3H16orf96;refGene_tx=XM_019821013.2;refGene_gn=CE3H16orf96;refGene_tx=XM_019821014.2;refGene_gn=CE3H16orf96;refGene_tx=XM_019821016.2;refGene_gn=CE3H16orf96;refGene_tx=XM_019821017.2;refGene_gn=CE3H16orf96;refGene_tx=XM_023246452.1
E3 40773157 40773244
refGene_gn=CE3H16orf96;refGene_tx=XM_003998986.5;refGene_gn=CE3H16orf96;refGene_tx=XM_019821007.2;refGene_gn=CE3H16orf96;refGene_tx=XM_019821009.2;refGene_gn=CE3H16orf96;refGene_tx=XM_019821010.2;refGene_gn=CE3H16orf96;refGene_tx=XM_019821013.2;refGene_gn=CE3H16orf96;refGene_tx=XM_019821014.2;refGene_gn=CE3H16orf96;refGene_tx=XM_019821016.2;refGene_gn=CE3H16orf96;refGene_tx=XM_019821017.2;refGene_gn=CE3H16orf96;refGene_tx=XM_023246452.1
E3 40773404 40773485
refGene_gn=CE3H16orf96;refGene_tx=XM_003998986.5;refGene_gn=CE3H16orf9

6;refGene_tx=XM_019821007.2;refGene_gn=CE3H16orf96;refGene_tx=XM_01982
1009.2;refGene_gn=CE3H16orf96;refGene_tx=XM_019821010.2;refGene_gn=CE3
H16orf96;refGene_tx=XM_019821014.2;refGene_gn=CE3H16orf96;refGene_tx=X
M_019821016.2;refGene_gn=CE3H16orf96;refGene_tx=XM_019821017.2;refGene
_gn=CE3H16orf96;refGene_tx=XM_023246452.1
E3 40774677 40774782
refGene_gn=CE3H16orf96;refGene_tx=XM_003998986.5;refGene_gn=CE3H16orf9
6;refGene_tx=XM_019821007.2;refGene_gn=CE3H16orf96;refGene_tx=XM_01982
1009.2;refGene_gn=CE3H16orf96;refGene_tx=XM_019821010.2;refGene_gn=CE3
H16orf96;refGene_tx=XM_019821013.2;refGene_gn=CE3H16orf96;refGene_tx=X
M_019821014.2;refGene_gn=CE3H16orf96;refGene_tx=XM_019821016.2;refGene
_gn=CE3H16orf96;refGene_tx=XM_019821017.2;refGene_gn=CE3H16orf96;refGe
ne_tx=XM_023246452.1
E3 40785290 40785710
refGene_gn=CE3H16orf96;refGene_tx=XM_003998986.5;refGene_gn=CE3H16orf9
6;refGene_tx=XM_019821007.2;refGene_gn=CE3H16orf96;refGene_tx=XM_01982
1009.2;refGene_gn=CE3H16orf96;refGene_tx=XM_019821010.2;refGene_gn=CE3
H16orf96;refGene_tx=XM_019821013.2;refGene_gn=CE3H16orf96;refGene_tx=X
M_019821014.2;refGene_gn=CE3H16orf96;refGene_tx=XM_019821016.2;refGene
_gn=CE3H16orf96;refGene_tx=XM_019821017.2;refGene_gn=CE3H16orf96;refGe
ne_tx=XM_023246452.1
E3 40823528 40823613
refGene_gn=CDIP1;refGene_tx=XM_006942377.3;refGene_gn=CDIP1;refGene_tx
=XM_006942378.4;refGene_gn=CDIP1;refGene_tx=XM_006942380.4;refGene_gn=
CDIP1;refGene_tx=XM_019821023.2
E3 40823714 40823870
refGene_gn=CDIP1;refGene_tx=XM_006942377.3;refGene_gn=CDIP1;refGene_tx
=XM_006942378.4;refGene_gn=CDIP1;refGene_tx=XM_006942380.4;refGene_gn=
CDIP1;refGene_tx=XM_019821023.2
E3 40824699 40824973
refGene_gn=CDIP1;refGene_tx=XM_006942377.3;refGene_gn=CDIP1;refGene_tx
=XM_006942378.4;refGene_gn=CDIP1;refGene_tx=XM_006942380.4;refGene_gn=
CDIP1;refGene_tx=XM_019821023.2
E3 40825082 40825194
refGene_gn=CDIP1;refGene_tx=XM_006942377.3;refGene_gn=CDIP1;refGene_tx
=XM_006942378.4;refGene_gn=CDIP1;refGene_tx=XM_006942380.4;refGene_gn=
CDIP1;refGene_tx=XM_019821023.2
E3 40827896 40828024
refGene_gn=HMOX2;refGene_tx=XM_003998903.5;refGene_gn=HMOX2;refGene_tx
=XM_006942384.4;refGene_gn=HMOX2;refGene_tx=XM_019821019.2;refGene_gn=
HMOX2;refGene_tx=XM_019821020.2;refGene_gn=HMOX2;refGene_tx=XM_0198210
21.2;refGene_gn=HMOX2;refGene_tx=XM_019821022.2
E3 40828112 40828239
refGene_gn=HMOX2;refGene_tx=XM_003998903.5;refGene_gn=HMOX2;refGene_tx
=XM_006942384.4;refGene_gn=HMOX2;refGene_tx=XM_019821019.2;refGene_gn=
HMOX2;refGene_tx=XM_019821020.2;refGene_gn=HMOX2;refGene_tx=XM_0198210
21.2;refGene_gn=HMOX2;refGene_tx=XM_019821022.2
E3 40829099 40829591
refGene_gn=HMOX2;refGene_tx=XM_003998903.5;refGene_gn=HMOX2;refGene_tx
=XM_006942384.4;refGene_gn=HMOX2;refGene_tx=XM_019821019.2;refGene_gn=

HMOX2;refGene_tx=XM_019821020.2;refGene_gn=HMOX2;refGene_tx=XM_019821021.2;refGene_gn=HMOX2;refGene_tx=XM_019821022.2
E3 40830005 40830123 refGene_gn=HMOX2;refGene_tx=XM_003998903.5;refGene_gn=HMOX2;refGene_tx=XM_006942384.4;refGene_gn=HMOX2;refGene_tx=XM_019821019.2;refGene_gn=HMOX2;refGene_tx=XM_019821020.2;refGene_gn=HMOX2;refGene_tx=XM_019821021.2;refGene_gn=HMOX2;refGene_tx=XM_019821022.2
E3 40831339 40831425 refGene_gn=HMOX2;refGene_tx=XM_003998903.5;refGene_gn=HMOX2;refGene_tx=XM_006942384.4;refGene_gn=HMOX2;refGene_tx=XM_019821019.2;refGene_gn=HMOX2;refGene_tx=XM_019821020.2;refGene_gn=HMOX2;refGene_tx=XM_019821021.2;refGene_gn=HMOX2;refGene_tx=XM_019821022.2
E3 40862471 40862615 refGene_gn=NMRAL1;refGene_tx=XM_003998905.5
E3 40862904 40862977 refGene_gn=NMRAL1;refGene_tx=XM_003998905.5;refGene_gn=NMRAL1;refGene_tx=XM_019821018.2
E3 40866197 40866436 refGene_gn=NMRAL1;refGene_tx=XM_003998905.5;refGene_gn=NMRAL1;refGene_tx=XM_019821018.2
E3 40868792 40869042 refGene_gn=NMRAL1;refGene_tx=XM_003998905.5;refGene_gn=NMRAL1;refGene_tx=XM_019821018.2
E3 40870091 40870282 refGene_gn=NMRAL1;refGene_tx=XM_003998905.5;refGene_gn=NMRAL1;refGene_tx=XM_019821018.2
E3 40871710 40871890 refGene_gn=NMRAL1;refGene_tx=XM_003998905.5;refGene_gn=NMRAL1;refGene_tx=XM_019821018.2
E3 40878602 40878741 refGene_gn=DNAJA3;refGene_tx=XM_006942387.4;refGene_gn=DNAJA3;refGene_tx=XM_023246363.1
E3 40879408 40879512 refGene_gn=DNAJA3;refGene_tx=XM_006942386.4
E3 40887762 40887860 refGene_gn=DNAJA3;refGene_tx=XM_006942386.4;refGene_gn=DNAJA3;refGene_tx=XM_006942387.4;refGene_gn=DNAJA3;refGene_tx=XM_023246363.1
E3 40889139 40889255 refGene_gn=DNAJA3;refGene_tx=XM_006942386.4;refGene_gn=DNAJA3;refGene_tx=XM_006942387.4
E3 40889792 40889921 refGene_gn=DNAJA3;refGene_tx=XM_006942386.4;refGene_gn=DNAJA3;refGene_tx=XM_006942387.4
E3 40891873 40891938 refGene_gn=DNAJA3;refGene_tx=XM_006942386.4;refGene_gn=DNAJA3;refGene_tx=XM_006942387.4;refGene_gn=DNAJA3;refGene_tx=XM_023246363.1
E3 40892987 40893135 refGene_gn=DNAJA3;refGene_tx=XM_006942386.4;refGene_gn=DNAJA3;refGene_tx=XM_006942387.4;refGene_gn=DNAJA3;refGene_tx=XM_023246363.1
E3 40895986 40896139 refGene_gn=DNAJA3;refGene_tx=XM_006942386.4;refGene_gn=DNAJA3;refGene_tx=XM_006942387.4;refGene_gn=DNAJA3;refGene_tx=XM_023246363.1

E3 40897192 40897387
refGene_gn=DNAJA3;refGene_tx=XM_006942386.4;refGene_gn=DNAJA3;refGene_tx=XM_006942387.4;refGene_gn=DNAJA3;refGene_tx=XM_023246363.1
E3 40898121 40898205
refGene_gn=DNAJA3;refGene_tx=XM_006942386.4;refGene_gn=DNAJA3;refGene_tx=XM_006942387.4;refGene_gn=DNAJA3;refGene_tx=XM_023246363.1
E3 40902738 40902872
refGene_gn=DNAJA3;refGene_tx=XM_006942386.4;refGene_gn=DNAJA3;refGene_tx=XM_006942387.4;refGene_gn=DNAJA3;refGene_tx=XM_023246363.1
E3 40908741 40908952
refGene_gn=DNAJA3;refGene_tx=XM_006942386.4;refGene_gn=DNAJA3;refGene_tx=XM_006942387.4;refGene_gn=DNAJA3;refGene_tx=XM_023246363.1
E3 40920981 40921041
refGene_gn=LOC101093982;refGene_tx=XM_023246553.1
E3 40923616 40923713
refGene_gn=LOC101093982;refGene_tx=XM_023246553.1
E3 40924588 40924663
refGene_gn=LOC101093982;refGene_tx=XM_023246553.1
E3 40928366 40928437
refGene_gn=LOC101093982;refGene_tx=XM_023246553.1
E3 40928879 40929063
refGene_gn=LOC101093982;refGene_tx=XM_023246553.1
E3 40931700 40931777
refGene_gn=LOC101093982;refGene_tx=XM_023246553.1
E3 40942864 40942915
refGene_gn=LOC101093982;refGene_tx=XM_023246553.1
E3 40948842 40948929
refGene_gn=LOC101093982;refGene_tx=XM_023246553.1
E3 40949376 40949459
refGene_gn=LOC101093982;refGene_tx=XM_023246553.1
E3 40954471 40956505 refGene_gn=VASN;refGene_tx=XM_023246554.1
E3 40969596 40969651
refGene_gn=LOC101093982;refGene_tx=XM_023246553.1
E3 40970027 40970085
refGene_gn=LOC101093982;refGene_tx=XM_023246553.1
E3 40970203 40970349
refGene_gn=LOC101093982;refGene_tx=XM_023246553.1
E3 40970431 40970524
refGene_gn=LOC101093982;refGene_tx=XM_023246553.1
E3 40970680 40970812
refGene_gn=LOC101093982;refGene_tx=XM_023246553.1
E3 40971269 40971396
refGene_gn=LOC101093982;refGene_tx=XM_023246553.1
E3 40971612 40971788
refGene_gn=LOC101093982;refGene_tx=XM_023246553.1
E3 40972300 40972409
refGene_gn=LOC101093982;refGene_tx=XM_023246553.1
E3 40972517 40972604
refGene_gn=LOC101093982;refGene_tx=XM_023246553.1
E3 40972684 40972827

refGene_gn=LOC101093982;refGene_tx=XM_023246553.1
E3 40972904 40973009
refGene_gn=LOC101093982;refGene_tx=XM_023246553.1
E3 40973088 40973174
refGene_gn=LOC101093982;refGene_tx=XM_023246553.1
E3 40973919 40974061
refGene_gn=LOC101093982;refGene_tx=XM_023246553.1
E3 40974155 40974245
refGene_gn=LOC101093982;refGene_tx=XM_023246553.1
E3 40975077 40975194
refGene_gn=LOC101093982;refGene_tx=XM_023246553.1
E3 40975698 40975834
refGene_gn=LOC101093982;refGene_tx=XM_023246553.1
E3 40976254 40976346
refGene_gn=LOC101093982;refGene_tx=XM_023246553.1
E3 40976875 40976962
refGene_gn=LOC101093982;refGene_tx=XM_023246553.1
E3 40977074 40977080
refGene_gn=LOC101093982;refGene_tx=XM_023246553.1
E3 40979968 40979971
refGene_gn=PAM16;refGene_tx=XM_006942388.4;refGene_gn=PAM16;refGene_tx
=XM_023246557.1
E3 40982165 40982204 refGene_gn=PAM16;refGene_tx=XM_019821031.2
E3 40985282 40985367
refGene_gn=PAM16;refGene_tx=XM_006942388.4;refGene_gn=PAM16;refGene_tx
=XM_019821031.2;refGene_gn=PAM16;refGene_tx=XM_023246557.1
E3 40986218 40986355
refGene_gn=PAM16;refGene_tx=XM_006942388.4;refGene_gn=PAM16;refGene_tx
=XM_019821031.2;refGene_gn=PAM16;refGene_tx=XM_023246557.1
E3 40986750 40986816
refGene_gn=PAM16;refGene_tx=XM_006942388.4;refGene_gn=PAM16;refGene_tx
=XM_019821031.2;refGene_gn=PAM16;refGene_tx=XM_023246557.1
E3 40987284 40987371
refGene_gn=PAM16;refGene_tx=XM_006942388.4;refGene_gn=PAM16;refGene_tx
=XM_019821031.2;refGene_gn=PAM16;refGene_tx=XM_023246557.1
E3 40990113 40990913
refGene_gn=GLIS2;refGene_tx=XM_019821024.2;refGene_gn=GLIS2;refGene_tx
=XM_019821025.2;refGene_gn=GLIS2;refGene_tx=XM_019821027.2;refGene_gn=
GLIS2;refGene_tx=XM_019821028.2;refGene_gn=GLIS2;refGene_tx=XM_0198210
29.2;refGene_gn=GLIS2;refGene_tx=XM_019821030.2;refGene_gn=GLIS2;refGe
ne_tx=XM_023246555.1;refGene_gn=GLIS2;refGene_tx=XM_023246556.1
E3 40991043 40991244
refGene_gn=GLIS2;refGene_tx=XM_019821024.2;refGene_gn=GLIS2;refGene_tx
=XM_019821025.2;refGene_gn=GLIS2;refGene_tx=XM_019821028.2;refGene_gn=
GLIS2;refGene_tx=XM_019821029.2;refGene_gn=GLIS2;refGene_tx=XM_0198210
30.2;refGene_gn=GLIS2;refGene_tx=XM_023246555.1
E3 40991886 40992008
refGene_gn=GLIS2;refGene_tx=XM_019821024.2;refGene_gn=GLIS2;refGene_tx
=XM_019821025.2;refGene_gn=GLIS2;refGene_tx=XM_019821027.2;refGene_gn=
GLIS2;refGene_tx=XM_019821028.2;refGene_gn=GLIS2;refGene_tx=XM_0198210

29.2;refGene_gn=GLIS2;refGene_tx=XM_019821030.2;refGene_gn=GLIS2;refGene_tx=XM_023246555.1;refGene_gn=GLIS2;refGene_tx=XM_023246556.1
E3 40992109 40992243
refGene_gn=GLIS2;refGene_tx=XM_019821024.2;refGene_gn=GLIS2;refGene_tx=XM_019821025.2;refGene_gn=GLIS2;refGene_tx=XM_019821027.2;refGene_gn=GLIS2;refGene_tx=XM_019821028.2;refGene_gn=GLIS2;refGene_tx=XM_019821029.2;refGene_gn=GLIS2;refGene_tx=XM_019821030.2;refGene_gn=GLIS2;refGene_tx=XM_023246555.1;refGene_gn=GLIS2;refGene_tx=XM_023246556.1
E3 40992323 40992500
refGene_gn=GLIS2;refGene_tx=XM_019821024.2;refGene_gn=GLIS2;refGene_tx=XM_019821025.2;refGene_gn=GLIS2;refGene_tx=XM_019821027.2;refGene_gn=GLIS2;refGene_tx=XM_019821028.2;refGene_gn=GLIS2;refGene_tx=XM_019821029.2;refGene_gn=GLIS2;refGene_tx=XM_019821030.2;refGene_gn=GLIS2;refGene_tx=XM_023246555.1;refGene_gn=GLIS2;refGene_tx=XM_023246556.1
E3 40992628 40992724 refGene_gn=GLIS2;refGene_tx=XM_019821030.2
E3 40993034 40993207
refGene_gn=GLIS2;refGene_tx=XM_019821024.2;refGene_gn=GLIS2;refGene_tx=XM_019821025.2;refGene_gn=GLIS2;refGene_tx=XM_019821027.2;refGene_gn=GLIS2;refGene_tx=XM_019821028.2;refGene_gn=GLIS2;refGene_tx=XM_019821029.2;refGene_gn=GLIS2;refGene_tx=XM_023246555.1;refGene_gn=GLIS2;refGene_tx=XM_023246556.1
E3 40994013 40994256
refGene_gn=GLIS2;refGene_tx=XM_019821024.2;refGene_gn=GLIS2;refGene_tx=XM_019821025.2;refGene_gn=GLIS2;refGene_tx=XM_019821027.2;refGene_gn=GLIS2;refGene_tx=XM_019821028.2;refGene_gn=GLIS2;refGene_tx=XM_019821029.2;refGene_gn=GLIS2;refGene_tx=XM_023246555.1;refGene_gn=GLIS2;refGene_tx=XM_023246556.1
E3 40997842 40997867
refGene_gn=GLIS2;refGene_tx=XM_019821024.2;refGene_gn=GLIS2;refGene_tx=XM_019821025.2;refGene_gn=GLIS2;refGene_tx=XM_019821027.2;refGene_gn=GLIS2;refGene_tx=XM_019821028.2;refGene_gn=GLIS2;refGene_tx=XM_023246556.1
E3 41021605 41021700 refGene_gn=TFAP4;refGene_tx=XM_019821034.2
E3 41037065 41037154 refGene_gn=TFAP4;refGene_tx=XM_006942390.4
E3 41043295 41043414 refGene_gn=TFAP4;refGene_tx=XM_006942389.4
E3 41045445 41045611
refGene_gn=TFAP4;refGene_tx=XM_006942389.4;refGene_gn=TFAP4;refGene_tx=XM_006942390.4;refGene_gn=TFAP4;refGene_tx=XM_019821034.2;refGene_gn=TFAP4;refGene_tx=XM_023246458.1
E3 41045728 41045827
refGene_gn=TFAP4;refGene_tx=XM_006942389.4;refGene_gn=TFAP4;refGene_tx=XM_006942390.4;refGene_gn=TFAP4;refGene_tx=XM_019821034.2;refGene_gn=TFAP4;refGene_tx=XM_023246458.1
E3 41046243 41046414
refGene_gn=TFAP4;refGene_tx=XM_006942389.4;refGene_gn=TFAP4;refGene_tx=XM_006942390.4;refGene_gn=TFAP4;refGene_tx=XM_019821034.2;refGene_gn=TFAP4;refGene_tx=XM_023246458.1
E3 41047021 41047162
refGene_gn=TFAP4;refGene_tx=XM_006942389.4;refGene_gn=TFAP4;refGene_tx=XM_006942390.4;refGene_gn=TFAP4;refGene_tx=XM_019821034.2;refGene_gn=

TFAP4;refGene_tx=XM_023246458.1
E3 41047723 41047876
refGene_gn=TFAP4;refGene_tx=XM_006942389.4;refGene_gn=TFAP4;refGene_tx=XM_006942390.4;refGene_gn=TFAP4;refGene_tx=XM_019821034.2;refGene_gn=TFAP4;refGene_tx=XM_023246458.1
E3 41048786 41048981
refGene_gn=TFAP4;refGene_tx=XM_006942389.4;refGene_gn=TFAP4;refGene_tx=XM_006942390.4;refGene_gn=TFAP4;refGene_tx=XM_019821034.2;refGene_gn=TFAP4;refGene_tx=XM_023246458.1
E3 41062307 41062368
refGene_gn=SRL;refGene_tx=XM_003998911.5;refGene_gn=SRL;refGene_tx=XM_006942392.4;refGene_gn=SRL;refGene_tx=XM_006942393.4;refGene_gn=SRL;refGene_tx=XM_006942395.4
E3 41084973 41084977 refGene_gn=SRL;refGene_tx=XM_023246219.1
E3 41086107 41087178
refGene_gn=SRL;refGene_tx=XM_006942392.4;refGene_gn=SRL;refGene_tx=XM_006942393.4;refGene_gn=SRL;refGene_tx=XM_023246219.1
E3 41088782 41088884
refGene_gn=SRL;refGene_tx=XM_003998911.5;refGene_gn=SRL;refGene_tx=XM_006942392.4;refGene_gn=SRL;refGene_tx=XM_006942393.4;refGene_gn=SRL;refGene_tx=XM_006942395.4;refGene_gn=SRL;refGene_tx=XM_019821035.2;refGene_gn=SRL;refGene_tx=XM_023246219.1
E3 41089662 41089758
refGene_gn=SRL;refGene_tx=XM_003998911.5;refGene_gn=SRL;refGene_tx=XM_006942392.4;refGene_gn=SRL;refGene_tx=XM_006942393.4;refGene_gn=SRL;refGene_tx=XM_006942395.4;refGene_gn=SRL;refGene_tx=XM_019821035.2;refGene_gn=SRL;refGene_tx=XM_023246219.1
E3 41093704 41093821
refGene_gn=SRL;refGene_tx=XM_003998911.5;refGene_gn=SRL;refGene_tx=XM_006942392.4;refGene_gn=SRL;refGene_tx=XM_006942393.4;refGene_gn=SRL;refGene_tx=XM_006942395.4;refGene_gn=SRL;refGene_tx=XM_019821035.2;refGene_gn=SRL;refGene_tx=XM_023246219.1
E3 41094545 41094779
refGene_gn=SRL;refGene_tx=XM_003998911.5;refGene_gn=SRL;refGene_tx=XM_006942392.4;refGene_gn=SRL;refGene_tx=XM_006942393.4;refGene_gn=SRL;refGene_tx=XM_006942395.4;refGene_gn=SRL;refGene_tx=XM_019821035.2;refGene_gn=SRL;refGene_tx=XM_023246219.1
E3 41095999 41096811
refGene_gn=SRL;refGene_tx=XM_003998911.5;refGene_gn=SRL;refGene_tx=XM_006942392.4;refGene_gn=SRL;refGene_tx=XM_006942393.4;refGene_gn=SRL;refGene_tx=XM_006942395.4;refGene_gn=SRL;refGene_tx=XM_019821035.2;refGene_gn=SRL;refGene_tx=XM_023246219.1
E3 41153724 41155417 refGene_gn=ADCY9;refGene_tx=XM_023246667.1
E3 41245379 41245570 refGene_gn=ADCY9;refGene_tx=XM_023246667.1
E3 41256891 41256996 refGene_gn=ADCY9;refGene_tx=XM_023246667.1
E3 41257912 41258130 refGene_gn=ADCY9;refGene_tx=XM_023246667.1
E3 41264299 41264402 refGene_gn=ADCY9;refGene_tx=XM_023246667.1
E3 41265328 41265537 refGene_gn=ADCY9;refGene_tx=XM_023246667.1
E3 41269360 41269520 refGene_gn=ADCY9;refGene_tx=XM_023246667.1
E3 41270422 41270571 refGene_gn=ADCY9;refGene_tx=XM_023246667.1

E3 41274101 41274143 refGene_gn=ADCY9;refGene_tx=XM_023246667.1
E3 41279173 41280362 refGene_gn=ADCY9;refGene_tx=XM_023246667.1
E3 41366493 41366578
refGene_gn=CREBBP;refGene_tx=XM_023246594.1;refGene_gn=CREBBP;refGene_tx=XM_023246595.1;refGene_gn=CREBBP;refGene_tx=XM_023246596.1;refGene_gn=CREBBP;refGene_tx=XM_023246597.1
E3 41389699 41390412
refGene_gn=CREBBP;refGene_tx=XM_023246594.1;refGene_gn=CREBBP;refGene_tx=XM_023246595.1;refGene_gn=CREBBP;refGene_tx=XM_023246596.1
E3 41424893 41425070
refGene_gn=CREBBP;refGene_tx=XM_023246594.1;refGene_gn=CREBBP;refGene_tx=XM_023246595.1;refGene_gn=CREBBP;refGene_tx=XM_023246597.1
E3 41441690 41441931
refGene_gn=CREBBP;refGene_tx=XM_023246594.1;refGene_gn=CREBBP;refGene_tx=XM_023246595.1;refGene_gn=CREBBP;refGene_tx=XM_023246596.1;refGene_gn=CREBBP;refGene_tx=XM_023246597.1
E3 41443254 41443368
refGene_gn=CREBBP;refGene_tx=XM_023246594.1;refGene_gn=CREBBP;refGene_tx=XM_023246595.1;refGene_gn=CREBBP;refGene_tx=XM_023246596.1;refGene_gn=CREBBP;refGene_tx=XM_023246597.1
E3 41450688 41450931
refGene_gn=CREBBP;refGene_tx=XM_023246594.1;refGene_gn=CREBBP;refGene_tx=XM_023246595.1;refGene_gn=CREBBP;refGene_tx=XM_023246596.1;refGene_gn=CREBBP;refGene_tx=XM_023246597.1
E3 41452217 41452320
refGene_gn=CREBBP;refGene_tx=XM_023246594.1;refGene_gn=CREBBP;refGene_tx=XM_023246595.1;refGene_gn=CREBBP;refGene_tx=XM_023246596.1;refGene_gn=CREBBP;refGene_tx=XM_023246597.1
E3 41452545 41452692
refGene_gn=CREBBP;refGene_tx=XM_023246594.1;refGene_gn=CREBBP;refGene_tx=XM_023246595.1;refGene_gn=CREBBP;refGene_tx=XM_023246596.1;refGene_gn=CREBBP;refGene_tx=XM_023246597.1
E3 41453329 41453447
refGene_gn=CREBBP;refGene_tx=XM_023246594.1;refGene_gn=CREBBP;refGene_tx=XM_023246595.1;refGene_gn=CREBBP;refGene_tx=XM_023246596.1;refGene_gn=CREBBP;refGene_tx=XM_023246597.1
E3 41454140 41454312
refGene_gn=CREBBP;refGene_tx=XM_023246594.1;refGene_gn=CREBBP;refGene_tx=XM_023246595.1;refGene_gn=CREBBP;refGene_tx=XM_023246596.1;refGene_gn=CREBBP;refGene_tx=XM_023246597.1
E3 41454641 41454686
refGene_gn=CREBBP;refGene_tx=XM_023246594.1;refGene_gn=CREBBP;refGene_tx=XM_023246595.1;refGene_gn=CREBBP;refGene_tx=XM_023246596.1;refGene_gn=CREBBP;refGene_tx=XM_023246597.1
E3 41456785 41456910
refGene_gn=CREBBP;refGene_tx=XM_023246594.1;refGene_gn=CREBBP;refGene_tx=XM_023246595.1;refGene_gn=CREBBP;refGene_tx=XM_023246596.1;refGene_gn=CREBBP;refGene_tx=XM_023246597.1
E3 41457466 41457655
refGene_gn=CREBBP;refGene_tx=XM_023246594.1;refGene_gn=CREBBP;refGene_

tx=XM_023246595.1;refGene_gn=CREBBP;refGene_tx=XM_023246596.1;refGene_gn=CREBBP;refGene_tx=XM_023246597.1
E3 41459246 41459663 refGene_gn=CREBBP;refGene_tx=XM_023246594.1;refGene_gn=CREBBP;refGene_tx=XM_023246595.1;refGene_gn=CREBBP;refGene_tx=XM_023246596.1;refGene_gn=CREBBP;refGene_tx=XM_023246597.1
E3 41461254 41461434 refGene_gn=CREBBP;refGene_tx=XM_023246594.1;refGene_gn=CREBBP;refGene_tx=XM_023246595.1;refGene_gn=CREBBP;refGene_tx=XM_023246596.1;refGene_gn=CREBBP;refGene_tx=XM_023246597.1
E3 41462396 41462586 refGene_gn=CREBBP;refGene_tx=XM_023246594.1;refGene_gn=CREBBP;refGene_tx=XM_023246595.1;refGene_gn=CREBBP;refGene_tx=XM_023246596.1;refGene_gn=CREBBP;refGene_tx=XM_023246597.1
E3 41466312 41466431 refGene_gn=CREBBP;refGene_tx=XM_023246594.1;refGene_gn=CREBBP;refGene_tx=XM_023246595.1;refGene_gn=CREBBP;refGene_tx=XM_023246596.1;refGene_gn=CREBBP;refGene_tx=XM_023246597.1
E3 41467521 41467761 refGene_gn=CREBBP;refGene_tx=XM_023246594.1;refGene_gn=CREBBP;refGene_tx=XM_023246595.1;refGene_gn=CREBBP;refGene_tx=XM_023246596.1;refGene_gn=CREBBP;refGene_tx=XM_023246597.1
E3 41468216 41468305 refGene_gn=CREBBP;refGene_tx=XM_023246594.1;refGene_gn=CREBBP;refGene_tx=XM_023246595.1;refGene_gn=CREBBP;refGene_tx=XM_023246596.1;refGene_gn=CREBBP;refGene_tx=XM_023246597.1
E3 41473751 41473862 refGene_gn=CREBBP;refGene_tx=XM_023246594.1;refGene_gn=CREBBP;refGene_tx=XM_023246595.1;refGene_gn=CREBBP;refGene_tx=XM_023246596.1;refGene_gn=CREBBP;refGene_tx=XM_023246597.1
E3 41475376 41475433 refGene_gn=CREBBP;refGene_tx=XM_023246594.1;refGene_gn=CREBBP;refGene_tx=XM_023246595.1;refGene_gn=CREBBP;refGene_tx=XM_023246596.1;refGene_gn=CREBBP;refGene_tx=XM_023246597.1
E3 41478679 41478757 refGene_gn=CREBBP;refGene_tx=XM_023246594.1;refGene_gn=CREBBP;refGene_tx=XM_023246595.1;refGene_gn=CREBBP;refGene_tx=XM_023246596.1;refGene_gn=CREBBP;refGene_tx=XM_023246597.1
E3 41479006 41479074 refGene_gn=CREBBP;refGene_tx=XM_023246594.1;refGene_gn=CREBBP;refGene_tx=XM_023246595.1;refGene_gn=CREBBP;refGene_tx=XM_023246596.1;refGene_gn=CREBBP;refGene_tx=XM_023246597.1
E3 41482779 41482930 refGene_gn=CREBBP;refGene_tx=XM_023246594.1;refGene_gn=CREBBP;refGene_tx=XM_023246595.1;refGene_gn=CREBBP;refGene_tx=XM_023246596.1;refGene_gn=CREBBP;refGene_tx=XM_023246597.1
E3 41483404 41483551 refGene_gn=CREBBP;refGene_tx=XM_023246594.1;refGene_gn=CREBBP;refGene_tx=XM_023246595.1;refGene_gn=CREBBP;refGene_tx=XM_023246596.1;refGene_gn=CREBBP;refGene_tx=XM_023246597.1

E3	41485060	41485174
refGene_gn=CREBBP;refGene_tx=XM_023246594.1;refGene_gn=CREBBP;refGene_tx=XM_023246595.1;refGene_gn=CREBBP;refGene_tx=XM_023246596.1;refGene_gn=CREBBP;refGene_tx=XM_023246597.1
E3	41486178	41486344
refGene_gn=CREBBP;refGene_tx=XM_023246594.1;refGene_gn=CREBBP;refGene_tx=XM_023246595.1;refGene_gn=CREBBP;refGene_tx=XM_023246596.1;refGene_gn=CREBBP;refGene_tx=XM_023246597.1
E3	41486859	41487027
refGene_gn=CREBBP;refGene_tx=XM_023246594.1;refGene_gn=CREBBP;refGene_tx=XM_023246595.1;refGene_gn=CREBBP;refGene_tx=XM_023246596.1;refGene_gn=CREBBP;refGene_tx=XM_023246597.1
E3	41490098	41490260
refGene_gn=CREBBP;refGene_tx=XM_023246594.1;refGene_gn=CREBBP;refGene_tx=XM_023246595.1;refGene_gn=CREBBP;refGene_tx=XM_023246596.1;refGene_gn=CREBBP;refGene_tx=XM_023246597.1
E3	41490494	41490776
refGene_gn=CREBBP;refGene_tx=XM_023246594.1;refGene_gn=CREBBP;refGene_tx=XM_023246595.1;refGene_gn=CREBBP;refGene_tx=XM_023246596.1;refGene_gn=CREBBP;refGene_tx=XM_023246597.1
E3	41492584	41494738
refGene_gn=CREBBP;refGene_tx=XM_023246594.1;refGene_gn=CREBBP;refGene_tx=XM_023246595.1;refGene_gn=CREBBP;refGene_tx=XM_023246596.1;refGene_gn=CREBBP;refGene_tx=XM_023246597.1
E3	41503935	41504011	refGene_gn=TRAP1;refGene_tx=XM_023246827.1
E3	41504897	41504930	refGene_gn=TRAP1;refGene_tx=XM_019821039.1
E3	41505637	41505728	refGene_gn=TRAP1;refGene_tx=XM_019821039.1
E3	41527835	41528000
refGene_gn=TRAP1;refGene_tx=XM_019821039.1;refGene_gn=TRAP1;refGene_tx=XM_023246827.1
E3	41528846	41528929
refGene_gn=TRAP1;refGene_tx=XM_019821039.1;refGene_gn=TRAP1;refGene_tx=XM_023246827.1
E3	41529908	41530049
refGene_gn=TRAP1;refGene_tx=XM_019821039.1;refGene_gn=TRAP1;refGene_tx=XM_023246827.1
E3	41532383	41532455
refGene_gn=TRAP1;refGene_tx=XM_019821039.1;refGene_gn=TRAP1;refGene_tx=XM_023246827.1
E3	41534357	41534518
refGene_gn=TRAP1;refGene_tx=XM_019821039.1;refGene_gn=TRAP1;refGene_tx=XM_023246827.1
E3	41535777	41535887
refGene_gn=TRAP1;refGene_tx=XM_019821039.1;refGene_gn=TRAP1;refGene_tx=XM_023246827.1
E3	41536433	41536507
refGene_gn=TRAP1;refGene_tx=XM_019821039.1;refGene_gn=TRAP1;refGene_tx=XM_023246827.1
E3	41537253	41537409
refGene_gn=TRAP1;refGene_tx=XM_019821039.1;refGene_gn=TRAP1;refGene_tx

=XM_023246827.1
E3	41538616	41538737
refGene_gn=TRAP1;refGene_tx=XM_019821039.1;refGene_gn=TRAP1;refGene_tx
=XM_023246827.1
E3	41540313	41540383
refGene_gn=TRAP1;refGene_tx=XM_019821039.1;refGene_gn=TRAP1;refGene_tx
=XM_023246827.1
E3	41542775	41542923
refGene_gn=TRAP1;refGene_tx=XM_019821039.1;refGene_gn=TRAP1;refGene_tx
=XM_023246827.1
E3	41546290	41546476
refGene_gn=TRAP1;refGene_tx=XM_019821039.1;refGene_gn=TRAP1;refGene_tx
=XM_023246827.1
E3	41546858	41546997
refGene_gn=TRAP1;refGene_tx=XM_019821039.1;refGene_gn=TRAP1;refGene_tx
=XM_023246827.1
E3	41547843	41547929
refGene_gn=TRAP1;refGene_tx=XM_019821039.1;refGene_gn=TRAP1;refGene_tx
=XM_023246827.1
E3	41549838	41549984
refGene_gn=TRAP1;refGene_tx=XM_019821039.1;refGene_gn=TRAP1;refGene_tx
=XM_023246827.1
E3	41552683	41552756
refGene_gn=TRAP1;refGene_tx=XM_019821039.1;refGene_gn=TRAP1;refGene_tx
=XM_023246827.1
E3	41552873	41552975
refGene_gn=TRAP1;refGene_tx=XM_019821039.1;refGene_gn=TRAP1;refGene_tx
=XM_023246827.1
E3	41553332	41553386	refGene_gn=DNASE1;refGene_tx=XM_019821040.2
E3	41553522	41553619	refGene_gn=DNASE1;refGene_tx=XM_019821040.2
E3	41553876	41554031	refGene_gn=DNASE1;refGene_tx=XM_019821040.2
E3	41554105	41554218	refGene_gn=DNASE1;refGene_tx=XM_019821040.2
E3	41554339	41554455	refGene_gn=DNASE1;refGene_tx=XM_019821040.2
E3	41554631	41554715	refGene_gn=DNASE1;refGene_tx=XM_019821040.2
E3	41554856	41554945	refGene_gn=DNASE1;refGene_tx=XM_019821040.2
E3	41555254	41555402	refGene_gn=DNASE1;refGene_tx=XM_019821040.2
E3	41555837	41555851	refGene_gn=DNASE1;refGene_tx=XM_019821040.2
E3	41609412	41609785	refGene_gn=SLX4;refGene_tx=XM_023246826.1
E3	41611201	41611426	refGene_gn=SLX4;refGene_tx=XM_023246826.1
E3	41613578	41613768	refGene_gn=SLX4;refGene_tx=XM_023246826.1
E3	41614659	41614872	refGene_gn=SLX4;refGene_tx=XM_023246826.1
E3	41616822	41617031	refGene_gn=SLX4;refGene_tx=XM_023246826.1
E3	41617125	41617442	refGene_gn=SLX4;refGene_tx=XM_023246826.1
E3	41617985	41618226	refGene_gn=SLX4;refGene_tx=XM_023246826.1
E3	41618693	41618782	refGene_gn=SLX4;refGene_tx=XM_023246826.1
E3	41619677	41619824	refGene_gn=SLX4;refGene_tx=XM_023246826.1
E3	41620632	41620802	refGene_gn=SLX4;refGene_tx=XM_023246826.1
E3	41621736	41623931	refGene_gn=SLX4;refGene_tx=XM_023246826.1
E3	41626086	41626186	refGene_gn=SLX4;refGene_tx=XM_023246826.1
E3	41626734	41627055	refGene_gn=SLX4;refGene_tx=XM_023246826.1

E3 41627531 41628000 refGene_gn=SLX4;refGene_tx=XM_023246826.1
E3 41640685 41640863 refGene_gn=NLRC3;refGene_tx=XM_003998914.5
E3 41640959 41642694 refGene_gn=NLRC3;refGene_tx=XM_003998914.5
E3 41643687 41643774 refGene_gn=NLRC3;refGene_tx=XM_003998914.5
E3 41646517 41646601 refGene_gn=NLRC3;refGene_tx=XM_003998914.5
E3 41647058 41647142 refGene_gn=NLRC3;refGene_tx=XM_003998914.5
E3 41648508 41648592 refGene_gn=NLRC3;refGene_tx=XM_003998914.5
E3 41649789 41649873 refGene_gn=NLRC3;refGene_tx=XM_003998914.5
E3 41650317 41650401 refGene_gn=NLRC3;refGene_tx=XM_003998914.5
E3 41651061 41651145 refGene_gn=NLRC3;refGene_tx=XM_003998914.5
E3 41651698 41651782 refGene_gn=NLRC3;refGene_tx=XM_003998914.5
E3 41652268 41652352 refGene_gn=NLRC3;refGene_tx=XM_003998914.5
E3 41652832 41652916 refGene_gn=NLRC3;refGene_tx=XM_003998914.5
E3 41654967 41655051 refGene_gn=NLRC3;refGene_tx=XM_003998914.5
E3 41655463 41655547 refGene_gn=NLRC3;refGene_tx=XM_003998914.5
E3 41656239 41656323 refGene_gn=NLRC3;refGene_tx=XM_003998914.5
E3 41656702 41656786 refGene_gn=NLRC3;refGene_tx=XM_003998914.5
E3 41657099 41657190 refGene_gn=NLRC3;refGene_tx=XM_003998914.5
E3 41664674 41664821
refGene_gn=CLUAP1;refGene_tx=XM_003998915.5;refGene_gn=CLUAP1;refGene_tx=XM_006942398.4;refGene_gn=CLUAP1;refGene_tx=XM_019821042.2;refGene_gn=CLUAP1;refGene_tx=XM_019821043.2
E3 41668363 41668420
refGene_gn=CLUAP1;refGene_tx=XM_019821042.2;refGene_gn=CLUAP1;refGene_tx=XM_019821043.2
E3 41668665 41668721
refGene_gn=CLUAP1;refGene_tx=XM_003998915.5;refGene_gn=CLUAP1;refGene_tx=XM_006942398.4;refGene_gn=CLUAP1;refGene_tx=XM_019821042.2;refGene_gn=CLUAP1;refGene_tx=XM_019821043.2
E3 41671394 41671502
refGene_gn=CLUAP1;refGene_tx=XM_003998915.5;refGene_gn=CLUAP1;refGene_tx=XM_006942398.4;refGene_gn=CLUAP1;refGene_tx=XM_019821042.2;refGene_gn=CLUAP1;refGene_tx=XM_019821043.2
E3 41677015 41677088
refGene_gn=CLUAP1;refGene_tx=XM_003998915.5;refGene_gn=CLUAP1;refGene_tx=XM_006942398.4;refGene_gn=CLUAP1;refGene_tx=XM_019821042.2;refGene_gn=CLUAP1;refGene_tx=XM_019821043.2
E3 41679080 41679222
refGene_gn=CLUAP1;refGene_tx=XM_003998915.5;refGene_gn=CLUAP1;refGene_tx=XM_006942398.4;refGene_gn=CLUAP1;refGene_tx=XM_019821042.2;refGene_gn=CLUAP1;refGene_tx=XM_019821043.2
E3 41683568 41683702
refGene_gn=CLUAP1;refGene_tx=XM_003998915.5;refGene_gn=CLUAP1;refGene_tx=XM_006942398.4;refGene_gn=CLUAP1;refGene_tx=XM_019821042.2;refGene_gn=CLUAP1;refGene_tx=XM_019821043.2
E3 41688084 41688168
refGene_gn=CLUAP1;refGene_tx=XM_003998915.5;refGene_gn=CLUAP1;refGene_tx=XM_006942398.4;refGene_gn=CLUAP1;refGene_tx=XM_019821042.2;refGene_gn=CLUAP1;refGene_tx=XM_019821043.2
E3 41690907 41691003

refGene_gn=CLUAP1;refGene_tx=XM_003998915.5;refGene_gn=CLUAP1;refGene_tx=XM_006942398.4;refGene_gn=CLUAP1;refGene_tx=XM_019821042.2;refGene_gn=CLUAP1;refGene_tx=XM_019821043.2
E3 41693573 41693753
refGene_gn=CLUAP1;refGene_tx=XM_003998915.5;refGene_gn=CLUAP1;refGene_tx=XM_006942398.4;refGene_gn=CLUAP1;refGene_tx=XM_019821042.2;refGene_gn=CLUAP1;refGene_tx=XM_019821043.2
E3 41694595 41694680
refGene_gn=CLUAP1;refGene_tx=XM_003998915.5;refGene_gn=CLUAP1;refGene_tx=XM_006942398.4;refGene_gn=CLUAP1;refGene_tx=XM_019821042.2;refGene_gn=CLUAP1;refGene_tx=XM_019821043.2
E3 41695990 41696102
refGene_gn=CLUAP1;refGene_tx=XM_003998915.5;refGene_gn=CLUAP1;refGene_tx=XM_006942398.4;refGene_gn=CLUAP1;refGene_tx=XM_019821042.2;refGene_gn=CLUAP1;refGene_tx=XM_019821043.2
E3 41697763 41697854
refGene_gn=CLUAP1;refGene_tx=XM_006942398.4;refGene_gn=CLUAP1;refGene_tx=XM_019821042.2
E3 41698714 41698736
refGene_gn=CLUAP1;refGene_tx=XM_003998915.5;refGene_gn=CLUAP1;refGene_tx=XM_019821043.2
E3 41703100 41703303
refGene_gn=CE3H16orf90;refGene_tx=XM_003998916.4;refGene_gn=CE3H16orf90;refGene_tx=XM_006942400.3;refGene_gn=CE3H16orf90;refGene_tx=XM_019821044.2;refGene_gn=CE3H16orf90;refGene_tx=XM_019821045.2
E3 41703797 41704148
refGene_gn=CE3H16orf90;refGene_tx=XM_003998916.4;refGene_gn=CE3H16orf90;refGene_tx=XM_006942400.3;refGene_gn=CE3H16orf90;refGene_tx=XM_019821044.2;refGene_gn=CE3H16orf90;refGene_tx=XM_019821045.2;refGene_gn=CE3H16orf90;refGene_tx=XM_023246288.1;refGene_gn=CE3H16orf90;refGene_tx=XM_023246289.1
E3 41704412 41704453
refGene_gn=CE3H16orf90;refGene_tx=XM_019821045.2
E3 41704553 41704711
refGene_gn=CE3H16orf90;refGene_tx=XM_003998916.4;refGene_gn=CE3H16orf90;refGene_tx=XM_006942400.3;refGene_gn=CE3H16orf90;refGene_tx=XM_019821044.2;refGene_gn=CE3H16orf90;refGene_tx=XM_023246288.1;refGene_gn=CE3H16orf90;refGene_tx=XM_023246289.1
E3 41710386 41710543
refGene_gn=NAA60;refGene_tx=XM_003998917.5;refGene_gn=NAA60;refGene_tx=XM_006942401.4
E3 41711606 41711841
refGene_gn=NAA60;refGene_tx=XM_003998917.5;refGene_gn=NAA60;refGene_tx=XM_006942401.4
E3 41712882 41712979
refGene_gn=NAA60;refGene_tx=XM_003998917.5;refGene_gn=NAA60;refGene_tx=XM_006942401.4
E3 41714321 41714451
refGene_gn=NAA60;refGene_tx=XM_003998917.5;refGene_gn=NAA60;refGene_tx=XM_006942401.4

E3 41716373 41716483
refGene_gn=NAA60;refGene_tx=XM_003998917.5;refGene_gn=NAA60;refGene_tx
=XM_006942401.4
E3 41741730 41741891
refGene_gn=ZNF597;refGene_tx=XM_019821066.2;refGene_gn=ZNF597;refGene_
tx=XM_019821073.2
E3 41742078 41742175
refGene_gn=ZNF597;refGene_tx=XM_019821066.2;refGene_gn=ZNF597;refGene_
tx=XM_019821071.2;refGene_gn=ZNF597;refGene_tx=XM_019821073.2
E3 41742943 41743070
refGene_gn=ZNF597;refGene_tx=XM_019821066.2;refGene_gn=ZNF597;refGene_
tx=XM_019821071.2;refGene_gn=ZNF597;refGene_tx=XM_019821072.2;refGene_
gn=ZNF597;refGene_tx=XM_019821073.2
E3 41744908 41745990
refGene_gn=ZNF597;refGene_tx=XM_019821066.2;refGene_gn=ZNF597;refGene_
tx=XM_019821071.2;refGene_gn=ZNF597;refGene_tx=XM_019821072.2;refGene_
gn=ZNF597;refGene_tx=XM_019821073.2
E3 41764410 41765012
refGene_gn=ZNF174;refGene_tx=XM_003998918.5;refGene_gn=ZNF174;refGene_
tx=XM_011290656.3
E3 41767922 41768002 refGene_gn=ZNF174;refGene_tx=XM_006942402.4
E3 41768262 41768485
refGene_gn=ZNF174;refGene_tx=XM_003998918.5;refGene_gn=ZNF174;refGene_
tx=XM_006942402.4;refGene_gn=ZNF174;refGene_tx=XM_011290656.3
E3 41769497 41769905
refGene_gn=ZNF174;refGene_tx=XM_003998918.5;refGene_gn=ZNF174;refGene_
tx=XM_006942402.4;refGene_gn=ZNF174;refGene_tx=XM_011290656.3
E3 41772879 41772905
refGene_gn=LOC101095709;refGene_tx=XM_019821079.2
E3 41780801 41781100
refGene_gn=LOC109495328;refGene_tx=XM_019821080.2
E3 41781101 41781122
refGene_gn=LOC109495328;refGene_tx=XM_019821080.2
E3 41781123 41781463
refGene_gn=LOC109495328;refGene_tx=XM_019821080.2
E3 41782090 41783111
refGene_gn=LOC101095709;refGene_tx=XM_019821079.2
E3 41793739 41794678
refGene_gn=LOC101095943;refGene_tx=XM_003998993.5
E3 41832640 41833236
refGene_gn=ZNF75A;refGene_tx=XM_023246267.1;refGene_gn=ZNF75A;refGene_
tx=XM_023246268.1;refGene_gn=ZNF75A;refGene_tx=XM_023246269.1;refGene_
gn=ZNF75A;refGene_tx=XM_023246271.1;refGene_gn=ZNF75A;refGene_tx=XM_02
3246273.1;refGene_gn=ZNF75A;refGene_tx=XM_023246278.1;refGene_gn=ZNF75
A;refGene_tx=XM_023246279.1
E3 41833410 41833521
refGene_gn=ZNF75A;refGene_tx=XM_023246267.1;refGene_gn=ZNF75A;refGene_
tx=XM_023246268.1;refGene_gn=ZNF75A;refGene_tx=XM_023246269.1;refGene_
gn=ZNF75A;refGene_tx=XM_023246271.1;refGene_gn=ZNF75A;refGene_tx=XM_02
3246273.1;refGene_gn=ZNF75A;refGene_tx=XM_023246277.1;refGene_gn=ZNF75

A;refGene_tx=XM_023246278.1;refGene_gn=ZNF75A;refGene_tx=XM_023246279.1
E3 41835424 41835551 refGene_gn=ZNF75A;refGene_tx=XM_023246267.1;refGene_gn=ZNF75A;refGene_tx=XM_023246268.1;refGene_gn=ZNF75A;refGene_tx=XM_023246271.1;refGene_gn=ZNF75A;refGene_tx=XM_023246273.1;refGene_gn=ZNF75A;refGene_tx=XM_023246278.1;refGene_gn=ZNF75A;refGene_tx=XM_023246279.1
E3 41835841 41835933 refGene_gn=ZNF75A;refGene_tx=XM_023246267.1;refGene_gn=ZNF75A;refGene_tx=XM_023246268.1;refGene_gn=ZNF75A;refGene_tx=XM_023246273.1;refGene_gn=ZNF75A;refGene_tx=XM_023246277.1;refGene_gn=ZNF75A;refGene_tx=XM_023246278.1;refGene_gn=ZNF75A;refGene_tx=XM_023246279.1
E3 41837363 41837490 refGene_gn=ZNF75A;refGene_tx=XM_023246267.1;refGene_gn=ZNF75A;refGene_tx=XM_023246268.1;refGene_gn=ZNF75A;refGene_tx=XM_023246269.1;refGene_gn=ZNF75A;refGene_tx=XM_023246273.1;refGene_gn=ZNF75A;refGene_tx=XM_023246277.1;refGene_gn=ZNF75A;refGene_tx=XM_023246278.1;refGene_gn=ZNF75A;refGene_tx=XM_023246279.1
E3 41837773 41837865 refGene_gn=ZNF75A;refGene_tx=XM_023246267.1;refGene_gn=ZNF75A;refGene_tx=XM_023246268.1;refGene_gn=ZNF75A;refGene_tx=XM_023246269.1;refGene_gn=ZNF75A;refGene_tx=XM_023246271.1;refGene_gn=ZNF75A;refGene_tx=XM_023246273.1;refGene_gn=ZNF75A;refGene_tx=XM_023246277.1;refGene_gn=ZNF75A;refGene_tx=XM_023246278.1;refGene_gn=ZNF75A;refGene_tx=XM_023246279.1
E3 41838564 41838688 refGene_gn=ZNF75A;refGene_tx=XM_023246278.1;refGene_gn=ZNF75A;refGene_tx=XM_023246279.1
E3 41858780 41858797 refGene_gn=ZNF75A;refGene_tx=XM_023246280.1
E3 41862943 41863623 refGene_gn=ZNF75A;refGene_tx=XM_023246272.1;refGene_gn=ZNF75A;refGene_tx=XM_023246274.1;refGene_gn=ZNF75A;refGene_tx=XM_023246275.1;refGene_gn=ZNF75A;refGene_tx=XM_023246276.1
E3 41863797 41863908 refGene_gn=ZNF75A;refGene_tx=XM_023246272.1;refGene_gn=ZNF75A;refGene_tx=XM_023246275.1
E3 41865809 41865936 refGene_gn=ZNF75A;refGene_tx=XM_023246267.1;refGene_gn=ZNF75A;refGene_tx=XM_023246268.1;refGene_gn=ZNF75A;refGene_tx=XM_023246269.1;refGene_gn=ZNF75A;refGene_tx=XM_023246271.1;refGene_gn=ZNF75A;refGene_tx=XM_023246272.1;refGene_gn=ZNF75A;refGene_tx=XM_023246273.1;refGene_gn=ZNF75A;refGene_tx=XM_023246274.1;refGene_gn=ZNF75A;refGene_tx=XM_023246277.1;refGene_gn=ZNF75A;refGene_tx=XM_023246280.1
E3 41866013 41866025 refGene_gn=ZNF75A;refGene_tx=XM_023246281.1
E3 41866276 41866368 refGene_gn=ZNF75A;refGene_tx=XM_023246267.1;refGene_gn=ZNF75A;refGene_tx=XM_023246268.1;refGene_gn=ZNF75A;refGene_tx=XM_023246269.1;refGene_gn=ZNF75A;refGene_tx=XM_023246271.1;refGene_gn=ZNF75A;refGene_tx=XM_023246272.1;refGene_gn=ZNF75A;refGene_tx=XM_023246273.1;refGene_gn=ZNF75A;refGene_tx=XM_023246274.1;refGene_gn=ZNF75A;refGene_tx=XM_023246277.

1;refGene_gn=ZNF75A;refGene_tx=XM_023246280.1;refGene_gn=ZNF75A;refGen
e_tx=XM_023246281.1
E3 41867433 41867560
refGene_gn=ZNF75A;refGene_tx=XM_023246267.1;refGene_gn=ZNF75A;refGene_
tx=XM_023246268.1;refGene_gn=ZNF75A;refGene_tx=XM_023246269.1;refGene_
gn=ZNF75A;refGene_tx=XM_023246271.1;refGene_gn=ZNF75A;refGene_tx=XM_02
3246272.1;refGene_gn=ZNF75A;refGene_tx=XM_023246273.1;refGene_gn=ZNF75
A;refGene_tx=XM_023246274.1;refGene_gn=ZNF75A;refGene_tx=XM_023246275.
1;refGene_gn=ZNF75A;refGene_tx=XM_023246276.1;refGene_gn=ZNF75A;refGen
e_tx=XM_023246277.1;refGene_gn=ZNF75A;refGene_tx=XM_023246280.1;refGen
e_gn=ZNF75A;refGene_tx=XM_023246281.1
E3 41867834 41867926
refGene_gn=ZNF75A;refGene_tx=XM_023246267.1;refGene_gn=ZNF75A;refGene_
tx=XM_023246268.1;refGene_gn=ZNF75A;refGene_tx=XM_023246269.1;refGene_
gn=ZNF75A;refGene_tx=XM_023246271.1;refGene_gn=ZNF75A;refGene_tx=XM_02
3246272.1;refGene_gn=ZNF75A;refGene_tx=XM_023246273.1;refGene_gn=ZNF75
A;refGene_tx=XM_023246274.1;refGene_gn=ZNF75A;refGene_tx=XM_023246275.
1;refGene_gn=ZNF75A;refGene_tx=XM_023246276.1;refGene_gn=ZNF75A;refGen
e_tx=XM_023246277.1;refGene_gn=ZNF75A;refGene_tx=XM_023246280.1;refGen
e_gn=ZNF75A;refGene_tx=XM_023246281.1
E3 41868557 41868723
refGene_gn=ZNF75A;refGene_tx=XM_023246267.1;refGene_gn=ZNF75A;refGene_
tx=XM_023246268.1;refGene_gn=ZNF75A;refGene_tx=XM_023246269.1;refGene_
gn=ZNF75A;refGene_tx=XM_023246271.1;refGene_gn=ZNF75A;refGene_tx=XM_02
3246272.1;refGene_gn=ZNF75A;refGene_tx=XM_023246273.1;refGene_gn=ZNF75
A;refGene_tx=XM_023246274.1;refGene_gn=ZNF75A;refGene_tx=XM_023246275.
1;refGene_gn=ZNF75A;refGene_tx=XM_023246276.1;refGene_gn=ZNF75A;refGen
e_tx=XM_023246277.1;refGene_gn=ZNF75A;refGene_tx=XM_023246280.1;refGen
e_gn=ZNF75A;refGene_tx=XM_023246281.1
E3 41869212 41869239 refGene_gn=ZNF75A;refGene_tx=XM_023246273.1
E3 41869887 41870295
refGene_gn=ZNF75A;refGene_tx=XM_023246267.1;refGene_gn=ZNF75A;refGene_
tx=XM_023246268.1;refGene_gn=ZNF75A;refGene_tx=XM_023246269.1;refGene_
gn=ZNF75A;refGene_tx=XM_023246271.1;refGene_gn=ZNF75A;refGene_tx=XM_02
3246272.1;refGene_gn=ZNF75A;refGene_tx=XM_023246274.1;refGene_gn=ZNF75
A;refGene_tx=XM_023246275.1;refGene_gn=ZNF75A;refGene_tx=XM_023246276.
1;refGene_gn=ZNF75A;refGene_tx=XM_023246277.1;refGene_gn=ZNF75A;refGen
e_tx=XM_023246280.1;refGene_gn=ZNF75A;refGene_tx=XM_023246281.1
E3 41874734 41874820 refGene_gn=ZNF263;refGene_tx=XM_019821047.2
E3 41874908 41875019 refGene_gn=ZNF263;refGene_tx=XM_019821047.2
E3 41876822 41876827
refGene_gn=ZNF263;refGene_tx=XM_019821049.2;refGene_gn=ZNF263;refGene_
tx=XM_019821050.2
E3 41883659 41884828
refGene_gn=ZNF263;refGene_tx=XM_003998920.5;refGene_gn=ZNF263;refGene_
tx=XM_019821046.2
E3 41885327 41885444
refGene_gn=ZNF263;refGene_tx=XM_003998920.5;refGene_gn=ZNF263;refGene_
tx=XM_019821046.2;refGene_gn=ZNF263;refGene_tx=XM_019821047.2;refGene_
gn=ZNF263;refGene_tx=XM_019821049.2;refGene_gn=ZNF263;refGene_tx=XM_01

9821050.2
E3 41887267 41887394
refGene_gn=ZNF263;refGene_tx=XM_003998920.5;refGene_gn=ZNF263;refGene_tx=XM_019821046.2;refGene_gn=ZNF263;refGene_tx=XM_019821047.2;refGene_gn=ZNF263;refGene_tx=XM_019821049.2;refGene_gn=ZNF263;refGene_tx=XM_019821050.2
E3 41887672 41887746
refGene_gn=ZNF263;refGene_tx=XM_003998920.5;refGene_gn=ZNF263;refGene_tx=XM_019821047.2;refGene_gn=ZNF263;refGene_tx=XM_019821049.2;refGene_gn=ZNF263;refGene_tx=XM_019821050.2
E3 41888207 41888388
refGene_gn=ZNF263;refGene_tx=XM_003998920.5;refGene_gn=ZNF263;refGene_tx=XM_019821047.2;refGene_gn=ZNF263;refGene_tx=XM_019821049.2;refGene_gn=ZNF263;refGene_tx=XM_019821050.2
E3 41889278 41889665
refGene_gn=ZNF263;refGene_tx=XM_003998920.5;refGene_gn=ZNF263;refGene_tx=XM_019821046.2;refGene_gn=ZNF263;refGene_tx=XM_019821047.2;refGene_gn=ZNF263;refGene_tx=XM_019821049.2;refGene_gn=ZNF263;refGene_tx=XM_019821050.2
E3 41914680 41915107
refGene_gn=MEFV;refGene_tx=XM_006942408.4;refGene_gn=MEFV;refGene_tx=XM_006942409.4
E3 41917462 41918299
refGene_gn=MEFV;refGene_tx=XM_006942408.4;refGene_gn=MEFV;refGene_tx=XM_006942409.4
E3 41919909 41920289
refGene_gn=MEFV;refGene_tx=XM_006942408.4;refGene_gn=MEFV;refGene_tx=XM_006942409.4
E3 41920719 41920815
refGene_gn=MEFV;refGene_tx=XM_006942408.4;refGene_gn=MEFV;refGene_tx=XM_006942409.4
E3 41923933 41924164
refGene_gn=MEFV;refGene_tx=XM_006942408.4;refGene_gn=MEFV;refGene_tx=XM_006942409.4
E3 41924531 41924554
refGene_gn=MEFV;refGene_tx=XM_006942408.4;refGene_gn=MEFV;refGene_tx=XM_006942409.4
E3 41926432 41926548
refGene_gn=MEFV;refGene_tx=XM_006942408.4;refGene_gn=MEFV;refGene_tx=XM_006942409.4
E3 41926727 41926760
refGene_gn=MEFV;refGene_tx=XM_006942408.4;refGene_gn=MEFV;refGene_tx=XM_006942409.4
E3 41927104 41927137
refGene_gn=MEFV;refGene_tx=XM_006942408.4;refGene_gn=MEFV;refGene_tx=XM_006942409.4
E3 41927308 41927862
refGene_gn=MEFV;refGene_tx=XM_006942408.4;refGene_gn=MEFV;refGene_tx=XM_006942409.4
E3 41933802 41934049 refGene_gn=ZNF200;refGene_tx=XM_006942410.3

E3 41934655 41934747 refGene_gn=ZNF200;refGene_tx=XM_006942410.3
E3 41935085 41935206 refGene_gn=ZNF200;refGene_tx=XM_006942410.3
E3 41948652 41949404 refGene_gn=ZNF200;refGene_tx=XM_006942410.3
E3 41956231 41957185
refGene_gn=LOC101097200;refGene_tx=XM_023246286.1
E3 41967724 41968660
refGene_gn=LOC102899898;refGene_tx=XM_023246287.1
E3 41974186 41975128
refGene_gn=LOC105261319;refGene_tx=XM_011290658.3
E3 42024113 42024227
refGene_gn=LOC101098676;refGene_tx=XM_003998921.5
E3 42024856 42024960
refGene_gn=LOC101098676;refGene_tx=XM_003998921.5
E3 42025461 42025533
refGene_gn=LOC101098676;refGene_tx=XM_003998921.5
E3 42025965 42026082
refGene_gn=LOC101098676;refGene_tx=XM_003998921.5
E3 42026408 42026631
refGene_gn=LOC101098676;refGene_tx=XM_003998921.5
E3 42027163 42027331
refGene_gn=LOC101098676;refGene_tx=XM_003998921.5
E3 42027467 42027575
refGene_gn=LOC101098676;refGene_tx=XM_003998921.5
E3 42028228 42028317
refGene_gn=LOC101098676;refGene_tx=XM_003998921.5
E3 42028428 42028624
refGene_gn=LOC101098676;refGene_tx=XM_003998921.5
E3 42028728 42028869
refGene_gn=LOC101098676;refGene_tx=XM_003998921.5
E3 42028953 42029028
refGene_gn=LOC101098676;refGene_tx=XM_003998921.5
E3 42031146 42031877
refGene_gn=ZNF213;refGene_tx=XM_023246283.1;refGene_gn=ZNF213;refGene_tx=XM_023246284.1
E3 42032890 42033014
refGene_gn=ZNF213;refGene_tx=XM_023246283.1;refGene_gn=ZNF213;refGene_tx=XM_023246284.1
E3 42033228 42033299
refGene_gn=ZNF213;refGene_tx=XM_023246283.1;refGene_gn=ZNF213;refGene_tx=XM_023246284.1
E3 42033490 42033614
refGene_gn=ZNF213;refGene_tx=XM_023246283.1;refGene_gn=ZNF213;refGene_tx=XM_023246284.1
E3 42034340 42034739
refGene_gn=ZNF213;refGene_tx=XM_023246283.1;refGene_gn=ZNF213;refGene_tx=XM_023246284.1
E3 42044398 42045204 refGene_gn=ZNF205;refGene_tx=XM_023246282.1
E3 42045206 42045464 refGene_gn=ZNF205;refGene_tx=XM_023246282.1
E3 42045716 42045812 refGene_gn=ZNF205;refGene_tx=XM_023246282.1
E3 42047335 42047456 refGene_gn=ZNF205;refGene_tx=XM_023246282.1

E3	42047825	42047917	refGene_gn=ZNF205;refGene_tx=XM_023246282.1
E3	42048213	42048427	refGene_gn=ZNF205;refGene_tx=XM_023246282.1
E3	42051059	42051197	refGene_gn=ZNF205;refGene_tx=XM_023246282.1
E3	42052143	42052198	refGene_gn=ZNF205;refGene_tx=XM_023246282.1
E3	42060671	42061181
refGene_gn=LOC101097447;refGene_tx=XM_019820226.2
E3	42062927	42063007	refGene_gn=ZNF205;refGene_tx=XM_023246282.1
E3	42068637	42068842	refGene_gn=ZSCAN10;refGene_tx=XM_023246266.1
E3	42068844	42069032	refGene_gn=ZSCAN10;refGene_tx=XM_023246266.1
E3	42069248	42069513	refGene_gn=ZSCAN10;refGene_tx=XM_023246266.1
E3	42069718	42069786	refGene_gn=ZSCAN10;refGene_tx=XM_023246266.1
E3	42069960	42070018	refGene_gn=ZSCAN10;refGene_tx=XM_023246266.1
E3	42071229	42071384	refGene_gn=ZSCAN10;refGene_tx=XM_023246266.1
E3	42071385	42071451	refGene_gn=ZSCAN10;refGene_tx=XM_023246266.1
E3	42071452	42071978	refGene_gn=ZSCAN10;refGene_tx=XM_023246266.1
E3	42071980	42072502	refGene_gn=ZSCAN10;refGene_tx=XM_023246266.1
E3	42072504	42072567	refGene_gn=ZSCAN10;refGene_tx=XM_023246266.1
E3	42072569	42072793	refGene_gn=ZSCAN10;refGene_tx=XM_023246266.1
E3	42076564	42076882
refGene_gn=IL32;refGene_tx=XM_023246977.1;refGene_gn=IL32;refGene_tx=XM_023246978.1;refGene_gn=IL32;refGene_tx=XM_023246979.1
E3	42077276	42077339
refGene_gn=IL32;refGene_tx=XM_023246977.1;refGene_gn=IL32;refGene_tx=XM_023246978.1;refGene_gn=IL32;refGene_tx=XM_023246979.1
E3	42077499	42077529
refGene_gn=IL32;refGene_tx=XM_023246977.1;refGene_gn=IL32;refGene_tx=XM_023246978.1;refGene_gn=IL32;refGene_tx=XM_023246979.1
E3	42077937	42077994
refGene_gn=IL32;refGene_tx=XM_023246977.1;refGene_gn=IL32;refGene_tx=XM_023246978.1;refGene_gn=IL32;refGene_tx=XM_023246979.1
E3	42078140	42078179
refGene_gn=IL32;refGene_tx=XM_023246977.1;refGene_gn=IL32;refGene_tx=XM_023246978.1;refGene_gn=IL32;refGene_tx=XM_023246979.1
E3	42079695	42079710	refGene_gn=IL32;refGene_tx=XM_023246977.1
E3	42084367	42084639	refGene_gn=MMP25;refGene_tx=XM_023246956.1
E3	42084787	42085045	refGene_gn=MMP25;refGene_tx=XM_023246956.1
E3	42085133	42085286	refGene_gn=MMP25;refGene_tx=XM_023246956.1
E3	42086048	42086131	refGene_gn=MMP25;refGene_tx=XM_023246956.1
E3	42086281	42086366	refGene_gn=MMP25;refGene_tx=XM_023246956.1
E3	42086458	42086635	refGene_gn=MMP25;refGene_tx=XM_023246956.1
E3	42091587	42091880	refGene_gn=MMP25;refGene_tx=XM_023246956.1
E3	42091958	42092103	refGene_gn=MMP25;refGene_tx=XM_023246956.1
E3	42093458	42093591	refGene_gn=MMP25;refGene_tx=XM_023246956.1
E3	42093989	42094079	refGene_gn=MMP25;refGene_tx=XM_023246956.1
E3	42109269	42109551	refGene_gn=BICDL2;refGene_tx=XM_003998924.5
E3	42111525	42111612	refGene_gn=BICDL2;refGene_tx=XM_019821086.2
E3	42111834	42112050
refGene_gn=BICDL2;refGene_tx=XM_003998924.5;refGene_gn=BICDL2;refGene_tx=XM_019821086.2
E3	42112155	42112272

refGene_gn=BICDL2;refGene_tx=XM_003998924.5;refGene_gn=BICDL2;refGene_tx=XM_019821086.2
E3 42112348 42112495
refGene_gn=BICDL2;refGene_tx=XM_003998924.5;refGene_gn=BICDL2;refGene_tx=XM_019821086.2
E3 42113348 42113540
refGene_gn=BICDL2;refGene_tx=XM_003998924.5;refGene_gn=BICDL2;refGene_tx=XM_019821086.2
E3 42113644 42113794
refGene_gn=BICDL2;refGene_tx=XM_003998924.5;refGene_gn=BICDL2;refGene_tx=XM_019821086.2
E3 42114252 42114383
refGene_gn=BICDL2;refGene_tx=XM_003998924.5;refGene_gn=BICDL2;refGene_tx=XM_019821086.2
E3 42114610 42114731
refGene_gn=BICDL2;refGene_tx=XM_003998924.5;refGene_gn=BICDL2;refGene_tx=XM_019821086.2
E3 42114816 42114978
refGene_gn=BICDL2;refGene_tx=XM_003998924.5;refGene_gn=BICDL2;refGene_tx=XM_019821086.2
E3 42115428 42115509
refGene_gn=THOC6;refGene_tx=XM_003998925.5;refGene_gn=THOC6;refGene_tx=XM_006942419.4
E3 42115592 42115727
refGene_gn=THOC6;refGene_tx=XM_003998925.5;refGene_gn=THOC6;refGene_tx=XM_006942419.4
E3 42115813 42115924
refGene_gn=THOC6;refGene_tx=XM_003998925.5;refGene_gn=THOC6;refGene_tx=XM_006942419.4
E3 42116024 42116087
refGene_gn=THOC6;refGene_tx=XM_003998925.5;refGene_gn=THOC6;refGene_tx=XM_006942419.4
E3 42116175 42116225
refGene_gn=THOC6;refGene_tx=XM_003998925.5;refGene_gn=THOC6;refGene_tx=XM_006942419.4
E3 42116319 42116422
refGene_gn=THOC6;refGene_tx=XM_003998925.5;refGene_gn=THOC6;refGene_tx=XM_006942419.4
E3 42116513 42116585
refGene_gn=THOC6;refGene_tx=XM_003998925.5;refGene_gn=THOC6;refGene_tx=XM_006942419.4
E3 42116708 42116757
refGene_gn=THOC6;refGene_tx=XM_003998925.5;refGene_gn=THOC6;refGene_tx=XM_006942419.4
E3 42116840 42116878
refGene_gn=THOC6;refGene_tx=XM_003998925.5;refGene_gn=THOC6;refGene_tx=XM_006942419.4
E3 42116958 42117062
refGene_gn=THOC6;refGene_tx=XM_003998925.5;refGene_gn=THOC6;refGene_tx=XM_006942419.4

E3 42117154 42117219
refGene_gn=THOC6;refGene_tx=XM_003998925.5;refGene_gn=THOC6;refGene_tx=XM_006942419.4
E3 42117324 42117440
refGene_gn=THOC6;refGene_tx=XM_003998925.5;refGene_gn=THOC6;refGene_tx=XM_006942419.4
E3 42118101 42118155 refGene_gn=THOC6;refGene_tx=XM_006942419.4
E3 42118231 42118270 refGene_gn=THOC6;refGene_tx=XM_003998925.5
E3 42118674 42118769
refGene_gn=HCFC1R1;refGene_tx=XM_003998927.5;refGene_gn=HCFC1R1;refGene_tx=XM_003998928.5;refGene_gn=HCFC1R1;refGene_tx=XM_006942420.4
E3 42119069 42119126 refGene_gn=HCFC1R1;refGene_tx=XM_003998928.5
E3 42119176 42119365
refGene_gn=HCFC1R1;refGene_tx=XM_003998927.5;refGene_gn=HCFC1R1;refGene_tx=XM_003998928.5;refGene_gn=HCFC1R1;refGene_tx=XM_006942420.4
E3 42119595 42119731
refGene_gn=HCFC1R1;refGene_tx=XM_003998927.5;refGene_gn=HCFC1R1;refGene_tx=XM_003998928.5;refGene_gn=HCFC1R1;refGene_tx=XM_006942420.4
E3 42120806 42120862
refGene_gn=TNFRSF12A;refGene_tx=XM_023246982.1;refGene_gn=TNFRSF12A;refGene_tx=XM_023246983.1
E3 42120947 42121083
refGene_gn=TNFRSF12A;refGene_tx=XM_023246982.1;refGene_gn=TNFRSF12A;refGene_tx=XM_023246983.1
E3 42121318 42121432
refGene_gn=TNFRSF12A;refGene_tx=XM_023246982.1;refGene_gn=TNFRSF12A;refGene_tx=XM_023246983.1
E3 42122208 42122302
refGene_gn=TNFRSF12A;refGene_tx=XM_023246982.1;refGene_gn=TNFRSF12A;refGene_tx=XM_023246983.1
E3 42124463 42124527
refGene_gn=CLDN6;refGene_tx=XM_006942421.4;refGene_gn=CLDN6;refGene_tx=XM_006942422.4;refGene_gn=CLDN6;refGene_tx=XM_006942423.4
E3 42124780 42125469
refGene_gn=CLDN6;refGene_tx=XM_003998929.5;refGene_gn=CLDN6;refGene_tx=XM_006942421.4;refGene_gn=CLDN6;refGene_tx=XM_006942422.4;refGene_gn=CLDN6;refGene_tx=XM_006942423.4;refGene_gn=CLDN6;refGene_tx=XM_019821089.2
E3 42126822 42127476
refGene_gn=CLDN9;refGene_tx=XM_003998930.5;refGene_gn=CLDN9;refGene_tx=XM_023246974.1
E3 42138236 42138262
refGene_gn=LOC102900118;refGene_tx=XM_019821091.2;refGene_gn=LOC102900118;refGene_tx=XM_019821092.2;refGene_gn=LOC102900118;refGene_tx=XM_019821093.2;refGene_gn=LOC102900118;refGene_tx=XM_019821094.2;refGene_gn=LOC102900118;refGene_tx=XM_019821095.2;refGene_gn=LOC102900118;refGene_tx=XM_019821096.2;refGene_gn=LOC102900118;refGene_tx=XM_019821097.2;refGene_gn=LOC102900118;refGene_tx=XM_019821098.2;refGene_gn=LOC102900118;refGene_tx=XM_019821099.2;refGene_gn=LOC102900118;refGene_tx=XM_019821100.2;refGene_gn=LOC102900118;refGene_tx=XM_019821101.2;refGene_gn

=LOC102900118;refGene_tx=XM_019821102.2;refGene_gn=LOC102900118;refGen
e_tx=XM_019821103.2;refGene_gn=LOC102900118;refGene_tx=XM_019821104.2;
refGene_gn=LOC102900118;refGene_tx=XM_019821105.2;refGene_gn=LOC102900
118;refGene_tx=XM_019821107.2;refGene_gn=LOC102900118;refGene_tx=XM_01
9821108.2;refGene_gn=LOC102900118;refGene_tx=XM_019821109.2;refGene_gn
=LOC102900118;refGene_tx=XM_019821110.2;refGene_gn=LOC102900118;refGen
e_tx=XM_019821111.2;refGene_gn=LOC102900118;refGene_tx=XM_019821112.2;
refGene_gn=LOC102900118;refGene_tx=XM_019821113.2;refGene_gn=LOC102900
118;refGene_tx=XM_019821114.2;refGene_gn=LOC102900118;refGene_tx=XM_01
9821115.2;refGene_gn=LOC102900118;refGene_tx=XM_019821116.2;refGene_gn
=LOC102900118;refGene_tx=XM_019821117.2;refGene_gn=LOC102900118;refGen
e_tx=XM_019821118.2;refGene_gn=LOC102900118;refGene_tx=XM_023246960.1
E3 42138501 42138555
refGene_gn=LOC102900118;refGene_tx=XM_019821091.2;refGene_gn=LOC102900
118;refGene_tx=XM_019821092.2;refGene_gn=LOC102900118;refGene_tx=XM_01
9821093.2;refGene_gn=LOC102900118;refGene_tx=XM_019821094.2;refGene_gn
=LOC102900118;refGene_tx=XM_019821095.2;refGene_gn=LOC102900118;refGen
e_tx=XM_019821096.2;refGene_gn=LOC102900118;refGene_tx=XM_019821097.2;
refGene_gn=LOC102900118;refGene_tx=XM_019821098.2;refGene_gn=LOC102900
118;refGene_tx=XM_019821099.2;refGene_gn=LOC102900118;refGene_tx=XM_01
9821100.2;refGene_gn=LOC102900118;refGene_tx=XM_019821101.2;refGene_gn
=LOC102900118;refGene_tx=XM_019821102.2;refGene_gn=LOC102900118;refGen
e_tx=XM_019821103.2;refGene_gn=LOC102900118;refGene_tx=XM_019821104.2;
refGene_gn=LOC102900118;refGene_tx=XM_019821105.2;refGene_gn=LOC102900
118;refGene_tx=XM_019821107.2;refGene_gn=LOC102900118;refGene_tx=XM_01
9821108.2;refGene_gn=LOC102900118;refGene_tx=XM_019821109.2;refGene_gn
=LOC102900118;refGene_tx=XM_019821110.2;refGene_gn=LOC102900118;refGen
e_tx=XM_019821111.2;refGene_gn=LOC102900118;refGene_tx=XM_019821112.2;
refGene_gn=LOC102900118;refGene_tx=XM_019821113.2;refGene_gn=LOC102900
118;refGene_tx=XM_019821114.2;refGene_gn=LOC102900118;refGene_tx=XM_01
9821115.2;refGene_gn=LOC102900118;refGene_tx=XM_019821116.2;refGene_gn
=LOC102900118;refGene_tx=XM_019821117.2;refGene_gn=LOC102900118;refGen
e_tx=XM_019821118.2;refGene_gn=LOC102900118;refGene_tx=XM_023246960.1
E3 42138654 42138699
refGene_gn=LOC102900118;refGene_tx=XM_019821091.2;refGene_gn=LOC102900
118;refGene_tx=XM_019821092.2;refGene_gn=LOC102900118;refGene_tx=XM_01
9821093.2;refGene_gn=LOC102900118;refGene_tx=XM_019821094.2;refGene_gn
=LOC102900118;refGene_tx=XM_019821095.2;refGene_gn=LOC102900118;refGen
e_tx=XM_019821096.2;refGene_gn=LOC102900118;refGene_tx=XM_019821097.2;
refGene_gn=LOC102900118;refGene_tx=XM_019821098.2;refGene_gn=LOC102900
118;refGene_tx=XM_019821099.2;refGene_gn=LOC102900118;refGene_tx=XM_01
9821100.2;refGene_gn=LOC102900118;refGene_tx=XM_019821101.2;refGene_gn
=LOC102900118;refGene_tx=XM_019821102.2;refGene_gn=LOC102900118;refGen
e_tx=XM_019821103.2;refGene_gn=LOC102900118;refGene_tx=XM_019821104.2;
refGene_gn=LOC102900118;refGene_tx=XM_019821105.2;refGene_gn=LOC102900
118;refGene_tx=XM_019821107.2;refGene_gn=LOC102900118;refGene_tx=XM_01
9821108.2;refGene_gn=LOC102900118;refGene_tx=XM_019821109.2;refGene_gn
=LOC102900118;refGene_tx=XM_019821110.2;refGene_gn=LOC102900118;refGen
e_tx=XM_019821111.2;refGene_gn=LOC102900118;refGene_tx=XM_019821112.2;
refGene_gn=LOC102900118;refGene_tx=XM_019821113.2;refGene_gn=LOC102900

118;refGene_tx=XM_019821114.2;refGene_gn=LOC102900118;refGene_tx=XM_01
9821115.2;refGene_gn=LOC102900118;refGene_tx=XM_019821116.2;refGene_gn
=LOC102900118;refGene_tx=XM_019821117.2;refGene_gn=LOC102900118;refGen
e_tx=XM_019821118.2;refGene_gn=LOC102900118;refGene_tx=XM_023246960.1
E3 42138965 42139079
refGene_gn=LOC102900118;refGene_tx=XM_019821091.2;refGene_gn=LOC102900
118;refGene_tx=XM_019821092.2;refGene_gn=LOC102900118;refGene_tx=XM_01
9821093.2;refGene_gn=LOC102900118;refGene_tx=XM_019821094.2;refGene_gn
=LOC102900118;refGene_tx=XM_019821095.2;refGene_gn=LOC102900118;refGen
e_tx=XM_019821096.2;refGene_gn=LOC102900118;refGene_tx=XM_019821097.2;
refGene_gn=LOC102900118;refGene_tx=XM_019821098.2;refGene_gn=LOC102900
118;refGene_tx=XM_019821099.2;refGene_gn=LOC102900118;refGene_tx=XM_01
9821100.2;refGene_gn=LOC102900118;refGene_tx=XM_019821101.2;refGene_gn
=LOC102900118;refGene_tx=XM_019821102.2;refGene_gn=LOC102900118;refGen
e_tx=XM_019821103.2;refGene_gn=LOC102900118;refGene_tx=XM_019821104.2;
refGene_gn=LOC102900118;refGene_tx=XM_019821105.2;refGene_gn=LOC102900
118;refGene_tx=XM_019821107.2;refGene_gn=LOC102900118;refGene_tx=XM_01
9821108.2;refGene_gn=LOC102900118;refGene_tx=XM_019821109.2;refGene_gn
=LOC102900118;refGene_tx=XM_019821110.2;refGene_gn=LOC102900118;refGen
e_tx=XM_019821111.2;refGene_gn=LOC102900118;refGene_tx=XM_019821112.2;
refGene_gn=LOC102900118;refGene_tx=XM_019821113.2;refGene_gn=LOC102900
118;refGene_tx=XM_019821114.2;refGene_gn=LOC102900118;refGene_tx=XM_01
9821115.2;refGene_gn=LOC102900118;refGene_tx=XM_019821116.2;refGene_gn
=LOC102900118;refGene_tx=XM_019821117.2;refGene_gn=LOC102900118;refGen
e_tx=XM_019821118.2;refGene_gn=LOC102900118;refGene_tx=XM_023246960.1
E3 42139212 42139272
refGene_gn=LOC102900118;refGene_tx=XM_019821091.2;refGene_gn=LOC102900
118;refGene_tx=XM_019821092.2;refGene_gn=LOC102900118;refGene_tx=XM_01
9821093.2;refGene_gn=LOC102900118;refGene_tx=XM_019821094.2;refGene_gn
=LOC102900118;refGene_tx=XM_019821095.2;refGene_gn=LOC102900118;refGen
e_tx=XM_019821096.2;refGene_gn=LOC102900118;refGene_tx=XM_019821097.2;
refGene_gn=LOC102900118;refGene_tx=XM_019821098.2;refGene_gn=LOC102900
118;refGene_tx=XM_019821099.2;refGene_gn=LOC102900118;refGene_tx=XM_01
9821100.2;refGene_gn=LOC102900118;refGene_tx=XM_019821101.2;refGene_gn
=LOC102900118;refGene_tx=XM_019821102.2;refGene_gn=LOC102900118;refGen
e_tx=XM_019821103.2;refGene_gn=LOC102900118;refGene_tx=XM_019821104.2;
refGene_gn=LOC102900118;refGene_tx=XM_019821105.2;refGene_gn=LOC102900
118;refGene_tx=XM_019821107.2;refGene_gn=LOC102900118;refGene_tx=XM_01
9821108.2;refGene_gn=LOC102900118;refGene_tx=XM_019821109.2;refGene_gn
=LOC102900118;refGene_tx=XM_019821110.2;refGene_gn=LOC102900118;refGen
e_tx=XM_019821111.2;refGene_gn=LOC102900118;refGene_tx=XM_019821112.2;
refGene_gn=LOC102900118;refGene_tx=XM_019821113.2;refGene_gn=LOC102900
118;refGene_tx=XM_019821114.2;refGene_gn=LOC102900118;refGene_tx=XM_01
9821115.2;refGene_gn=LOC102900118;refGene_tx=XM_019821116.2;refGene_gn
=LOC102900118;refGene_tx=XM_019821117.2;refGene_gn=LOC102900118;refGen
e_tx=XM_019821118.2;refGene_gn=LOC102900118;refGene_tx=XM_023246960.1
E3 42139350 42139380
refGene_gn=LOC102900118;refGene_tx=XM_019821091.2;refGene_gn=LOC102900
118;refGene_tx=XM_019821092.2;refGene_gn=LOC102900118;refGene_tx=XM_01
9821093.2;refGene_gn=LOC102900118;refGene_tx=XM_019821094.2;refGene_gn

=LOC102900118;refGene_tx=XM_019821095.2;refGene_gn=LOC102900118;refGene_tx=XM_019821096.2;refGene_gn=LOC102900118;refGene_tx=XM_019821097.2;refGene_gn=LOC102900118;refGene_tx=XM_019821098.2;refGene_gn=LOC102900118;refGene_tx=XM_019821099.2;refGene_gn=LOC102900118;refGene_tx=XM_019821100.2;refGene_gn=LOC102900118;refGene_tx=XM_019821101.2;refGene_gn=LOC102900118;refGene_tx=XM_019821102.2;refGene_gn=LOC102900118;refGene_tx=XM_019821103.2;refGene_gn=LOC102900118;refGene_tx=XM_019821104.2;refGene_gn=LOC102900118;refGene_tx=XM_019821105.2;refGene_gn=LOC102900118;refGene_tx=XM_019821107.2;refGene_gn=LOC102900118;refGene_tx=XM_019821108.2;refGene_gn=LOC102900118;refGene_tx=XM_019821109.2;refGene_gn=LOC102900118;refGene_tx=XM_019821110.2;refGene_gn=LOC102900118;refGene_tx=XM_019821111.2;refGene_gn=LOC102900118;refGene_tx=XM_019821112.2;refGene_gn=LOC102900118;refGene_tx=XM_019821113.2;refGene_gn=LOC102900118;refGene_tx=XM_019821114.2;refGene_gn=LOC102900118;refGene_tx=XM_019821115.2;refGene_gn=LOC102900118;refGene_tx=XM_019821116.2;refGene_gn=LOC102900118;refGene_tx=XM_019821117.2;refGene_gn=LOC102900118;refGene_tx=XM_019821118.2;refGene_gn=LOC102900118;refGene_tx=XM_023246960.1
E3 42139459 42139519
refGene_gn=LOC102900118;refGene_tx=XM_019821091.2;refGene_gn=LOC102900118;refGene_tx=XM_019821092.2;refGene_gn=LOC102900118;refGene_tx=XM_019821093.2;refGene_gn=LOC102900118;refGene_tx=XM_019821094.2;refGene_gn=LOC102900118;refGene_tx=XM_019821095.2;refGene_gn=LOC102900118;refGene_tx=XM_019821096.2;refGene_gn=LOC102900118;refGene_tx=XM_019821097.2;refGene_gn=LOC102900118;refGene_tx=XM_019821098.2;refGene_gn=LOC102900118;refGene_tx=XM_019821099.2;refGene_gn=LOC102900118;refGene_tx=XM_019821100.2;refGene_gn=LOC102900118;refGene_tx=XM_019821101.2;refGene_gn=LOC102900118;refGene_tx=XM_019821102.2;refGene_gn=LOC102900118;refGene_tx=XM_019821103.2;refGene_gn=LOC102900118;refGene_tx=XM_019821104.2;refGene_gn=LOC102900118;refGene_tx=XM_019821105.2;refGene_gn=LOC102900118;refGene_tx=XM_019821107.2;refGene_gn=LOC102900118;refGene_tx=XM_019821108.2;refGene_gn=LOC102900118;refGene_tx=XM_019821109.2;refGene_gn=LOC102900118;refGene_tx=XM_019821110.2;refGene_gn=LOC102900118;refGene_tx=XM_019821111.2;refGene_gn=LOC102900118;refGene_tx=XM_019821112.2;refGene_gn=LOC102900118;refGene_tx=XM_019821113.2;refGene_gn=LOC102900118;refGene_tx=XM_019821114.2;refGene_gn=LOC102900118;refGene_tx=XM_019821115.2;refGene_gn=LOC102900118;refGene_tx=XM_019821116.2;refGene_gn=LOC102900118;refGene_tx=XM_019821117.2;refGene_gn=LOC102900118;refGene_tx=XM_019821118.2;refGene_gn=LOC102900118;refGene_tx=XM_023246960.1
E3 42139946 42139979
refGene_gn=LOC102900118;refGene_tx=XM_019821091.2;refGene_gn=LOC102900118;refGene_tx=XM_019821092.2;refGene_gn=LOC102900118;refGene_tx=XM_019821093.2;refGene_gn=LOC102900118;refGene_tx=XM_019821094.2;refGene_gn=LOC102900118;refGene_tx=XM_019821095.2;refGene_gn=LOC102900118;refGene_tx=XM_019821096.2;refGene_gn=LOC102900118;refGene_tx=XM_019821097.2;refGene_gn=LOC102900118;refGene_tx=XM_019821098.2;refGene_gn=LOC102900118;refGene_tx=XM_019821099.2;refGene_gn=LOC102900118;refGene_tx=XM_019821100.2;refGene_gn=LOC102900118;refGene_tx=XM_019821101.2;refGene_gn=LOC102900118;refGene_tx=XM_019821102.2;refGene_gn=LOC102900118;refGene_tx=XM_019821103.2;refGene_gn=LOC102900118;refGene_tx=XM_019821104.2;refGene_gn=LOC102900118;refGene_tx=XM_019821105.2;refGene_gn=LOC102900

118;refGene_tx=XM_019821107.2;refGene_gn=LOC102900118;refGene_tx=XM_01
9821108.2;refGene_gn=LOC102900118;refGene_tx=XM_019821109.2;refGene_gn
=LOC102900118;refGene_tx=XM_019821110.2;refGene_gn=LOC102900118;refGen
e_tx=XM_019821111.2;refGene_gn=LOC102900118;refGene_tx=XM_019821112.2;
refGene_gn=LOC102900118;refGene_tx=XM_019821113.2;refGene_gn=LOC102900
118;refGene_tx=XM_019821114.2;refGene_gn=LOC102900118;refGene_tx=XM_01
9821115.2;refGene_gn=LOC102900118;refGene_tx=XM_019821116.2;refGene_gn
=LOC102900118;refGene_tx=XM_019821117.2;refGene_gn=LOC102900118;refGen
e_tx=XM_019821118.2;refGene_gn=LOC102900118;refGene_tx=XM_023246960.1
E3 42140093 42140135
refGene_gn=LOC102900118;refGene_tx=XM_019821091.2;refGene_gn=LOC102900
118;refGene_tx=XM_019821092.2;refGene_gn=LOC102900118;refGene_tx=XM_01
9821093.2;refGene_gn=LOC102900118;refGene_tx=XM_019821094.2;refGene_gn
=LOC102900118;refGene_tx=XM_019821095.2;refGene_gn=LOC102900118;refGen
e_tx=XM_019821096.2;refGene_gn=LOC102900118;refGene_tx=XM_019821097.2;
refGene_gn=LOC102900118;refGene_tx=XM_019821098.2;refGene_gn=LOC102900
118;refGene_tx=XM_019821099.2;refGene_gn=LOC102900118;refGene_tx=XM_01
9821100.2;refGene_gn=LOC102900118;refGene_tx=XM_019821101.2;refGene_gn
=LOC102900118;refGene_tx=XM_019821102.2;refGene_gn=LOC102900118;refGen
e_tx=XM_019821103.2;refGene_gn=LOC102900118;refGene_tx=XM_019821104.2;
refGene_gn=LOC102900118;refGene_tx=XM_019821105.2;refGene_gn=LOC102900
118;refGene_tx=XM_019821107.2;refGene_gn=LOC102900118;refGene_tx=XM_01
9821108.2;refGene_gn=LOC102900118;refGene_tx=XM_019821109.2;refGene_gn
=LOC102900118;refGene_tx=XM_019821110.2;refGene_gn=LOC102900118;refGen
e_tx=XM_019821111.2;refGene_gn=LOC102900118;refGene_tx=XM_019821112.2;
refGene_gn=LOC102900118;refGene_tx=XM_019821113.2;refGene_gn=LOC102900
118;refGene_tx=XM_019821114.2;refGene_gn=LOC102900118;refGene_tx=XM_01
9821115.2;refGene_gn=LOC102900118;refGene_tx=XM_019821116.2;refGene_gn
=LOC102900118;refGene_tx=XM_019821117.2;refGene_gn=LOC102900118;refGen
e_tx=XM_019821118.2;refGene_gn=LOC102900118;refGene_tx=XM_023246960.1
E3 42140443 42140488
refGene_gn=LOC102900118;refGene_tx=XM_019821091.2;refGene_gn=LOC102900
118;refGene_tx=XM_019821092.2;refGene_gn=LOC102900118;refGene_tx=XM_01
9821093.2;refGene_gn=LOC102900118;refGene_tx=XM_019821094.2;refGene_gn
=LOC102900118;refGene_tx=XM_019821095.2;refGene_gn=LOC102900118;refGen
e_tx=XM_019821096.2;refGene_gn=LOC102900118;refGene_tx=XM_019821097.2;
refGene_gn=LOC102900118;refGene_tx=XM_019821098.2;refGene_gn=LOC102900
118;refGene_tx=XM_019821099.2;refGene_gn=LOC102900118;refGene_tx=XM_01
9821100.2;refGene_gn=LOC102900118;refGene_tx=XM_019821101.2;refGene_gn
=LOC102900118;refGene_tx=XM_019821102.2;refGene_gn=LOC102900118;refGen
e_tx=XM_019821103.2;refGene_gn=LOC102900118;refGene_tx=XM_019821104.2;
refGene_gn=LOC102900118;refGene_tx=XM_019821105.2;refGene_gn=LOC102900
118;refGene_tx=XM_019821107.2;refGene_gn=LOC102900118;refGene_tx=XM_01
9821108.2;refGene_gn=LOC102900118;refGene_tx=XM_019821109.2;refGene_gn
=LOC102900118;refGene_tx=XM_019821110.2;refGene_gn=LOC102900118;refGen
e_tx=XM_019821111.2;refGene_gn=LOC102900118;refGene_tx=XM_019821112.2;
refGene_gn=LOC102900118;refGene_tx=XM_019821113.2;refGene_gn=LOC102900
118;refGene_tx=XM_019821114.2;refGene_gn=LOC102900118;refGene_tx=XM_01
9821115.2;refGene_gn=LOC102900118;refGene_tx=XM_019821116.2;refGene_gn
=LOC102900118;refGene_tx=XM_019821117.2;refGene_gn=LOC102900118;refGen

e_tx=XM_019821118.2;refGene_gn=LOC102900118;refGene_tx=XM_023246960.1
E3 42140770 42140902
refGene_gn=LOC102900118;refGene_tx=XM_019821091.2;refGene_gn=LOC102900118;refGene_tx=XM_019821092.2;refGene_gn=LOC102900118;refGene_tx=XM_019821093.2;refGene_gn=LOC102900118;refGene_tx=XM_019821094.2;refGene_gn=LOC102900118;refGene_tx=XM_019821095.2;refGene_gn=LOC102900118;refGene_tx=XM_019821096.2;refGene_gn=LOC102900118;refGene_tx=XM_019821097.2;refGene_gn=LOC102900118;refGene_tx=XM_019821098.2;refGene_gn=LOC102900118;refGene_tx=XM_019821099.2;refGene_gn=LOC102900118;refGene_tx=XM_019821100.2;refGene_gn=LOC102900118;refGene_tx=XM_019821101.2;refGene_gn=LOC102900118;refGene_tx=XM_019821102.2;refGene_gn=LOC102900118;refGene_tx=XM_019821104.2;refGene_gn=LOC102900118;refGene_tx=XM_019821105.2;refGene_gn=LOC102900118;refGene_tx=XM_019821107.2;refGene_gn=LOC102900118;refGene_tx=XM_019821108.2;refGene_gn=LOC102900118;refGene_tx=XM_019821109.2;refGene_gn=LOC102900118;refGene_tx=XM_019821110.2;refGene_gn=LOC102900118;refGene_tx=XM_019821111.2;refGene_gn=LOC102900118;refGene_tx=XM_019821112.2;refGene_gn=LOC102900118;refGene_tx=XM_019821113.2;refGene_gn=LOC102900118;refGene_tx=XM_019821114.2;refGene_gn=LOC102900118;refGene_tx=XM_019821115.2;refGene_gn=LOC102900118;refGene_tx=XM_019821116.2;refGene_gn=LOC102900118;refGene_tx=XM_019821117.2
E3 42141221 42141257
refGene_gn=LOC102900118;refGene_tx=XM_019821091.2;refGene_gn=LOC102900118;refGene_tx=XM_019821092.2;refGene_gn=LOC102900118;refGene_tx=XM_019821093.2;refGene_gn=LOC102900118;refGene_tx=XM_019821094.2;refGene_gn=LOC102900118;refGene_tx=XM_019821095.2;refGene_gn=LOC102900118;refGene_tx=XM_019821096.2;refGene_gn=LOC102900118;refGene_tx=XM_019821097.2;refGene_gn=LOC102900118;refGene_tx=XM_019821098.2;refGene_gn=LOC102900118;refGene_tx=XM_019821099.2;refGene_gn=LOC102900118;refGene_tx=XM_019821100.2;refGene_gn=LOC102900118;refGene_tx=XM_019821101.2;refGene_gn=LOC102900118;refGene_tx=XM_019821102.2;refGene_gn=LOC102900118;refGene_tx=XM_019821103.2;refGene_gn=LOC102900118;refGene_tx=XM_019821104.2;refGene_gn=LOC102900118;refGene_tx=XM_019821105.2;refGene_gn=LOC102900118;refGene_tx=XM_019821107.2;refGene_gn=LOC102900118;refGene_tx=XM_019821108.2;refGene_gn=LOC102900118;refGene_tx=XM_019821109.2;refGene_gn=LOC102900118;refGene_tx=XM_019821110.2;refGene_gn=LOC102900118;refGene_tx=XM_019821111.2;refGene_gn=LOC102900118;refGene_tx=XM_019821112.2;refGene_gn=LOC102900118;refGene_tx=XM_019821113.2;refGene_gn=LOC102900118;refGene_tx=XM_019821114.2;refGene_gn=LOC102900118;refGene_tx=XM_019821115.2;refGene_gn=LOC102900118;refGene_tx=XM_019821116.2;refGene_gn=LOC102900118;refGene_tx=XM_019821117.2;refGene_gn=LOC102900118;refGene_tx=XM_019821118.2;refGene_gn=LOC102900118;refGene_tx=XM_023246960.1
E3 42141367 42141400
refGene_gn=LOC102900118;refGene_tx=XM_019821091.2;refGene_gn=LOC102900118;refGene_tx=XM_019821092.2;refGene_gn=LOC102900118;refGene_tx=XM_019821093.2;refGene_gn=LOC102900118;refGene_tx=XM_019821094.2;refGene_gn=LOC102900118;refGene_tx=XM_019821095.2;refGene_gn=LOC102900118;refGene_tx=XM_019821096.2;refGene_gn=LOC102900118;refGene_tx=XM_019821097.2;refGene_gn=LOC102900118;refGene_tx=XM_019821098.2;refGene_gn=LOC102900118;refGene_tx=XM_019821099.2;refGene_gn=LOC102900118;refGene_tx=XM_019821100.2;refGene_gn=LOC102900118;refGene_tx=XM_019821101.2;refGene_gn

=LOC102900118;refGene_tx=XM_019821102.2;refGene_gn=LOC102900118;refGene_tx=XM_019821103.2;refGene_gn=LOC102900118;refGene_tx=XM_019821104.2;refGene_gn=LOC102900118;refGene_tx=XM_019821105.2;refGene_gn=LOC102900118;refGene_tx=XM_019821107.2;refGene_gn=LOC102900118;refGene_tx=XM_019821108.2;refGene_gn=LOC102900118;refGene_tx=XM_019821109.2;refGene_gn=LOC102900118;refGene_tx=XM_019821110.2;refGene_gn=LOC102900118;refGene_tx=XM_019821111.2;refGene_gn=LOC102900118;refGene_tx=XM_019821112.2;refGene_gn=LOC102900118;refGene_tx=XM_019821113.2;refGene_gn=LOC102900118;refGene_tx=XM_019821114.2;refGene_gn=LOC102900118;refGene_tx=XM_019821116.2;refGene_gn=LOC102900118;refGene_tx=XM_019821117.2
E3 42141483 42141519
refGene_gn=LOC102900118;refGene_tx=XM_019821091.2;refGene_gn=LOC102900118;refGene_tx=XM_019821092.2;refGene_gn=LOC102900118;refGene_tx=XM_019821093.2;refGene_gn=LOC102900118;refGene_tx=XM_019821094.2;refGene_gn=LOC102900118;refGene_tx=XM_019821095.2;refGene_gn=LOC102900118;refGene_tx=XM_019821096.2;refGene_gn=LOC102900118;refGene_tx=XM_019821097.2;refGene_gn=LOC102900118;refGene_tx=XM_019821098.2;refGene_gn=LOC102900118;refGene_tx=XM_019821099.2;refGene_gn=LOC102900118;refGene_tx=XM_019821100.2;refGene_gn=LOC102900118;refGene_tx=XM_019821101.2;refGene_gn=LOC102900118;refGene_tx=XM_019821102.2;refGene_gn=LOC102900118;refGene_tx=XM_019821103.2;refGene_gn=LOC102900118;refGene_tx=XM_019821104.2;refGene_gn=LOC102900118;refGene_tx=XM_019821105.2;refGene_gn=LOC102900118;refGene_tx=XM_019821107.2;refGene_gn=LOC102900118;refGene_tx=XM_019821108.2;refGene_gn=LOC102900118;refGene_tx=XM_019821109.2;refGene_gn=LOC102900118;refGene_tx=XM_019821110.2;refGene_gn=LOC102900118;refGene_tx=XM_019821111.2;refGene_gn=LOC102900118;refGene_tx=XM_019821112.2;refGene_gn=LOC102900118;refGene_tx=XM_019821113.2;refGene_gn=LOC102900118;refGene_tx=XM_019821114.2;refGene_gn=LOC102900118;refGene_tx=XM_019821116.2;refGene_gn=LOC102900118;refGene_tx=XM_019821117.2
E3 42141603 42141636
refGene_gn=LOC102900118;refGene_tx=XM_019821091.2;refGene_gn=LOC102900118;refGene_tx=XM_019821092.2;refGene_gn=LOC102900118;refGene_tx=XM_019821093.2;refGene_gn=LOC102900118;refGene_tx=XM_019821095.2;refGene_gn=LOC102900118;refGene_tx=XM_019821096.2;refGene_gn=LOC102900118;refGene_tx=XM_019821097.2;refGene_gn=LOC102900118;refGene_tx=XM_019821098.2;refGene_gn=LOC102900118;refGene_tx=XM_019821099.2;refGene_gn=LOC102900118;refGene_tx=XM_019821100.2;refGene_gn=LOC102900118;refGene_tx=XM_019821101.2;refGene_gn=LOC102900118;refGene_tx=XM_019821102.2;refGene_gn=LOC102900118;refGene_tx=XM_019821103.2;refGene_gn=LOC102900118;refGene_tx=XM_019821104.2;refGene_gn=LOC102900118;refGene_tx=XM_019821105.2;refGene_gn=LOC102900118;refGene_tx=XM_019821107.2;refGene_gn=LOC102900118;refGene_tx=XM_019821108.2;refGene_gn=LOC102900118;refGene_tx=XM_019821109.2;refGene_gn=LOC102900118;refGene_tx=XM_019821110.2;refGene_gn=LOC102900118;refGene_tx=XM_019821111.2;refGene_gn=LOC102900118;refGene_tx=XM_019821112.2;refGene_gn=LOC102900118;refGene_tx=XM_019821113.2;refGene_gn=LOC102900118;refGene_tx=XM_019821114.2;refGene_gn=LOC102900118;refGene_tx=XM_019821116.2;refGene_gn=LOC102900118;refGene_tx=XM_019821117.2
E3 42141806 42141842
refGene_gn=LOC102900118;refGene_tx=XM_019821091.2;refGene_gn=LOC102900

118;refGene_tx=XM_019821092.2;refGene_gn=LOC102900118;refGene_tx=XM_019821093.2;refGene_gn=LOC102900118;refGene_tx=XM_019821094.2;refGene_gn=LOC102900118;refGene_tx=XM_019821095.2;refGene_gn=LOC102900118;refGene_tx=XM_019821096.2;refGene_gn=LOC102900118;refGene_tx=XM_019821097.2;refGene_gn=LOC102900118;refGene_tx=XM_019821098.2;refGene_gn=LOC102900118;refGene_tx=XM_019821099.2;refGene_gn=LOC102900118;refGene_tx=XM_019821100.2;refGene_gn=LOC102900118;refGene_tx=XM_019821101.2;refGene_gn=LOC102900118;refGene_tx=XM_019821102.2;refGene_gn=LOC102900118;refGene_tx=XM_019821103.2;refGene_gn=LOC102900118;refGene_tx=XM_019821104.2;refGene_gn=LOC102900118;refGene_tx=XM_019821105.2;refGene_gn=LOC102900118;refGene_tx=XM_019821107.2;refGene_gn=LOC102900118;refGene_tx=XM_019821108.2;refGene_gn=LOC102900118;refGene_tx=XM_019821109.2;refGene_gn=LOC102900118;refGene_tx=XM_019821110.2;refGene_gn=LOC102900118;refGene_tx=XM_019821111.2;refGene_gn=LOC102900118;refGene_tx=XM_019821112.2;refGene_gn=LOC102900118;refGene_tx=XM_019821113.2;refGene_gn=LOC102900118;refGene_tx=XM_019821114.2;refGene_gn=LOC102900118;refGene_tx=XM_019821116.2;refGene_gn=LOC102900118;refGene_tx=XM_019821117.2

E3 42142674 42142707

refGene_gn=LOC102900118;refGene_tx=XM_019821091.2;refGene_gn=LOC102900118;refGene_tx=XM_019821092.2;refGene_gn=LOC102900118;refGene_tx=XM_019821093.2;refGene_gn=LOC102900118;refGene_tx=XM_019821094.2;refGene_gn=LOC102900118;refGene_tx=XM_019821095.2;refGene_gn=LOC102900118;refGene_tx=XM_019821096.2;refGene_gn=LOC102900118;refGene_tx=XM_019821097.2;refGene_gn=LOC102900118;refGene_tx=XM_019821098.2;refGene_gn=LOC102900118;refGene_tx=XM_019821099.2;refGene_gn=LOC102900118;refGene_tx=XM_019821100.2;refGene_gn=LOC102900118;refGene_tx=XM_019821101.2;refGene_gn=LOC102900118;refGene_tx=XM_019821102.2;refGene_gn=LOC102900118;refGene_tx=XM_019821103.2;refGene_gn=LOC102900118;refGene_tx=XM_019821104.2;refGene_gn=LOC102900118;refGene_tx=XM_019821105.2;refGene_gn=LOC102900118;refGene_tx=XM_019821107.2;refGene_gn=LOC102900118;refGene_tx=XM_019821108.2;refGene_gn=LOC102900118;refGene_tx=XM_019821109.2;refGene_gn=LOC102900118;refGene_tx=XM_019821110.2;refGene_gn=LOC102900118;refGene_tx=XM_019821111.2;refGene_gn=LOC102900118;refGene_tx=XM_019821112.2;refGene_gn=LOC102900118;refGene_tx=XM_019821113.2;refGene_gn=LOC102900118;refGene_tx=XM_019821114.2;refGene_gn=LOC102900118;refGene_tx=XM_019821116.2;refGene_gn=LOC102900118;refGene_tx=XM_019821117.2;refGene_gn=LOC102900118;refGene_tx=XM_023246960.1

E3 42142799 42142829

refGene_gn=LOC102900118;refGene_tx=XM_019821091.2;refGene_gn=LOC102900118;refGene_tx=XM_019821092.2;refGene_gn=LOC102900118;refGene_tx=XM_019821093.2;refGene_gn=LOC102900118;refGene_tx=XM_019821094.2;refGene_gn=LOC102900118;refGene_tx=XM_019821095.2;refGene_gn=LOC102900118;refGene_tx=XM_019821096.2;refGene_gn=LOC102900118;refGene_tx=XM_019821097.2;refGene_gn=LOC102900118;refGene_tx=XM_019821098.2;refGene_gn=LOC102900118;refGene_tx=XM_019821099.2;refGene_gn=LOC102900118;refGene_tx=XM_019821100.2;refGene_gn=LOC102900118;refGene_tx=XM_019821101.2;refGene_gn=LOC102900118;refGene_tx=XM_019821102.2;refGene_gn=LOC102900118;refGene_tx=XM_019821103.2;refGene_gn=LOC102900118;refGene_tx=XM_019821104.2;refGene_gn=LOC102900118;refGene_tx=XM_019821105.2;refGene_gn=LOC102900118;refGene_tx=XM_019821107.2;refGene_gn=LOC102900118;refGene_tx=XM_01

9821108.2;refGene_gn=LOC102900118;refGene_tx=XM_019821109.2;refGene_gn
=LOC102900118;refGene_tx=XM_019821110.2;refGene_gn=LOC102900118;refGen
e_tx=XM_019821111.2;refGene_gn=LOC102900118;refGene_tx=XM_019821112.2;
refGene_gn=LOC102900118;refGene_tx=XM_019821113.2;refGene_gn=LOC102900
118;refGene_tx=XM_019821114.2;refGene_gn=LOC102900118;refGene_tx=XM_01
9821116.2;refGene_gn=LOC102900118;refGene_tx=XM_019821117.2;refGene_gn
=LOC102900118;refGene_tx=XM_023246960.1
E3 42143181 42143217
refGene_gn=LOC102900118;refGene_tx=XM_019821091.2;refGene_gn=LOC102900
118;refGene_tx=XM_019821092.2;refGene_gn=LOC102900118;refGene_tx=XM_01
9821093.2;refGene_gn=LOC102900118;refGene_tx=XM_019821094.2;refGene_gn
=LOC102900118;refGene_tx=XM_019821095.2;refGene_gn=LOC102900118;refGen
e_tx=XM_019821096.2;refGene_gn=LOC102900118;refGene_tx=XM_019821097.2;
refGene_gn=LOC102900118;refGene_tx=XM_019821098.2;refGene_gn=LOC102900
118;refGene_tx=XM_019821099.2;refGene_gn=LOC102900118;refGene_tx=XM_01
9821100.2;refGene_gn=LOC102900118;refGene_tx=XM_019821101.2;refGene_gn
=LOC102900118;refGene_tx=XM_019821102.2;refGene_gn=LOC102900118;refGen
e_tx=XM_019821103.2;refGene_gn=LOC102900118;refGene_tx=XM_019821104.2;
refGene_gn=LOC102900118;refGene_tx=XM_019821105.2;refGene_gn=LOC102900
118;refGene_tx=XM_019821107.2;refGene_gn=LOC102900118;refGene_tx=XM_01
9821108.2;refGene_gn=LOC102900118;refGene_tx=XM_019821109.2;refGene_gn
=LOC102900118;refGene_tx=XM_019821110.2;refGene_gn=LOC102900118;refGen
e_tx=XM_019821111.2;refGene_gn=LOC102900118;refGene_tx=XM_019821112.2;
refGene_gn=LOC102900118;refGene_tx=XM_019821113.2;refGene_gn=LOC102900
118;refGene_tx=XM_019821114.2;refGene_gn=LOC102900118;refGene_tx=XM_01
9821116.2;refGene_gn=LOC102900118;refGene_tx=XM_019821117.2;refGene_gn
=LOC102900118;refGene_tx=XM_023246960.1
E3 42143848 42143881
refGene_gn=LOC102900118;refGene_tx=XM_019821091.2;refGene_gn=LOC102900
118;refGene_tx=XM_019821092.2;refGene_gn=LOC102900118;refGene_tx=XM_01
9821093.2;refGene_gn=LOC102900118;refGene_tx=XM_019821094.2;refGene_gn
=LOC102900118;refGene_tx=XM_019821096.2;refGene_gn=LOC102900118;refGen
e_tx=XM_019821097.2;refGene_gn=LOC102900118;refGene_tx=XM_019821098.2;
refGene_gn=LOC102900118;refGene_tx=XM_019821099.2;refGene_gn=LOC102900
118;refGene_tx=XM_019821100.2;refGene_gn=LOC102900118;refGene_tx=XM_01
9821101.2;refGene_gn=LOC102900118;refGene_tx=XM_019821102.2;refGene_gn
=LOC102900118;refGene_tx=XM_019821103.2;refGene_gn=LOC102900118;refGen
e_tx=XM_019821104.2;refGene_gn=LOC102900118;refGene_tx=XM_019821105.2;
refGene_gn=LOC102900118;refGene_tx=XM_019821107.2;refGene_gn=LOC102900
118;refGene_tx=XM_019821108.2;refGene_gn=LOC102900118;refGene_tx=XM_01
9821109.2;refGene_gn=LOC102900118;refGene_tx=XM_019821110.2;refGene_gn
=LOC102900118;refGene_tx=XM_019821112.2;refGene_gn=LOC102900118;refGen
e_tx=XM_019821113.2;refGene_gn=LOC102900118;refGene_tx=XM_019821114.2;
refGene_gn=LOC102900118;refGene_tx=XM_019821115.2;refGene_gn=LOC102900
118;refGene_tx=XM_019821116.2;refGene_gn=LOC102900118;refGene_tx=XM_01
9821117.2;refGene_gn=LOC102900118;refGene_tx=XM_019821118.2;refGene_gn
=LOC102900118;refGene_tx=XM_023246960.1
E3 42143964 42143994
refGene_gn=LOC102900118;refGene_tx=XM_019821091.2;refGene_gn=LOC102900
118;refGene_tx=XM_019821092.2;refGene_gn=LOC102900118;refGene_tx=XM_01

9821093.2;refGene_gn=LOC102900118;refGene_tx=XM_019821094.2;refGene_gn
=LOC102900118;refGene_tx=XM_019821095.2;refGene_gn=LOC102900118;refGen
e_tx=XM_019821096.2;refGene_gn=LOC102900118;refGene_tx=XM_019821097.2;
refGene_gn=LOC102900118;refGene_tx=XM_019821098.2;refGene_gn=LOC102900
118;refGene_tx=XM_019821099.2;refGene_gn=LOC102900118;refGene_tx=XM_01
9821100.2;refGene_gn=LOC102900118;refGene_tx=XM_019821101.2;refGene_gn
=LOC102900118;refGene_tx=XM_019821102.2;refGene_gn=LOC102900118;refGen
e_tx=XM_019821103.2;refGene_gn=LOC102900118;refGene_tx=XM_019821104.2;
refGene_gn=LOC102900118;refGene_tx=XM_019821105.2;refGene_gn=LOC102900
118;refGene_tx=XM_019821107.2;refGene_gn=LOC102900118;refGene_tx=XM_01
9821108.2;refGene_gn=LOC102900118;refGene_tx=XM_019821109.2;refGene_gn
=LOC102900118;refGene_tx=XM_019821110.2;refGene_gn=LOC102900118;refGen
e_tx=XM_019821111.2;refGene_gn=LOC102900118;refGene_tx=XM_019821112.2;
refGene_gn=LOC102900118;refGene_tx=XM_019821113.2;refGene_gn=LOC102900
118;refGene_tx=XM_019821114.2;refGene_gn=LOC102900118;refGene_tx=XM_01
9821115.2;refGene_gn=LOC102900118;refGene_tx=XM_019821116.2;refGene_gn
=LOC102900118;refGene_tx=XM_019821117.2;refGene_gn=LOC102900118;refGen
e_tx=XM_019821118.2;refGene_gn=LOC102900118;refGene_tx=XM_023246960.1
E3 42144197 42144233
refGene_gn=LOC102900118;refGene_tx=XM_019821091.2;refGene_gn=LOC102900
118;refGene_tx=XM_019821092.2;refGene_gn=LOC102900118;refGene_tx=XM_01
9821093.2;refGene_gn=LOC102900118;refGene_tx=XM_019821094.2;refGene_gn
=LOC102900118;refGene_tx=XM_019821095.2;refGene_gn=LOC102900118;refGen
e_tx=XM_019821096.2;refGene_gn=LOC102900118;refGene_tx=XM_019821097.2;
refGene_gn=LOC102900118;refGene_tx=XM_019821098.2;refGene_gn=LOC102900
118;refGene_tx=XM_019821099.2;refGene_gn=LOC102900118;refGene_tx=XM_01
9821100.2;refGene_gn=LOC102900118;refGene_tx=XM_019821101.2;refGene_gn
=LOC102900118;refGene_tx=XM_019821102.2;refGene_gn=LOC102900118;refGen
e_tx=XM_019821103.2;refGene_gn=LOC102900118;refGene_tx=XM_019821104.2;
refGene_gn=LOC102900118;refGene_tx=XM_019821107.2;refGene_gn=LOC102900
118;refGene_tx=XM_019821108.2;refGene_gn=LOC102900118;refGene_tx=XM_01
9821109.2;refGene_gn=LOC102900118;refGene_tx=XM_019821110.2;refGene_gn
=LOC102900118;refGene_tx=XM_019821112.2;refGene_gn=LOC102900118;refGen
e_tx=XM_019821113.2;refGene_gn=LOC102900118;refGene_tx=XM_019821114.2;
refGene_gn=LOC102900118;refGene_tx=XM_019821115.2;refGene_gn=LOC102900
118;refGene_tx=XM_019821116.2;refGene_gn=LOC102900118;refGene_tx=XM_01
9821117.2;refGene_gn=LOC102900118;refGene_tx=XM_019821118.2;refGene_gn
=LOC102900118;refGene_tx=XM_023246960.1
E3 42144320 42144353
refGene_gn=LOC102900118;refGene_tx=XM_019821091.2;refGene_gn=LOC102900
118;refGene_tx=XM_019821092.2;refGene_gn=LOC102900118;refGene_tx=XM_01
9821093.2;refGene_gn=LOC102900118;refGene_tx=XM_019821094.2;refGene_gn
=LOC102900118;refGene_tx=XM_019821095.2;refGene_gn=LOC102900118;refGen
e_tx=XM_019821096.2;refGene_gn=LOC102900118;refGene_tx=XM_019821097.2;
refGene_gn=LOC102900118;refGene_tx=XM_019821098.2;refGene_gn=LOC102900
118;refGene_tx=XM_019821099.2;refGene_gn=LOC102900118;refGene_tx=XM_01
9821100.2;refGene_gn=LOC102900118;refGene_tx=XM_019821101.2;refGene_gn
=LOC102900118;refGene_tx=XM_019821102.2;refGene_gn=LOC102900118;refGen
e_tx=XM_019821103.2;refGene_gn=LOC102900118;refGene_tx=XM_019821104.2;
refGene_gn=LOC102900118;refGene_tx=XM_019821107.2;refGene_gn=LOC102900

118;refGene_tx=XM_019821108.2;refGene_gn=LOC102900118;refGene_tx=XM_01
9821109.2;refGene_gn=LOC102900118;refGene_tx=XM_019821110.2;refGene_gn
=LOC102900118;refGene_tx=XM_019821112.2;refGene_gn=LOC102900118;refGen
e_tx=XM_019821113.2;refGene_gn=LOC102900118;refGene_tx=XM_019821114.2;
refGene_gn=LOC102900118;refGene_tx=XM_019821115.2;refGene_gn=LOC102900
118;refGene_tx=XM_019821116.2;refGene_gn=LOC102900118;refGene_tx=XM_01
9821117.2;refGene_gn=LOC102900118;refGene_tx=XM_019821118.2;refGene_gn
=LOC102900118;refGene_tx=XM_023246960.1
E3 42144427 42144457
refGene_gn=LOC102900118;refGene_tx=XM_019821091.2;refGene_gn=LOC102900
118;refGene_tx=XM_019821092.2;refGene_gn=LOC102900118;refGene_tx=XM_01
9821093.2;refGene_gn=LOC102900118;refGene_tx=XM_019821094.2;refGene_gn
=LOC102900118;refGene_tx=XM_019821095.2;refGene_gn=LOC102900118;refGen
e_tx=XM_019821096.2;refGene_gn=LOC102900118;refGene_tx=XM_019821097.2;
refGene_gn=LOC102900118;refGene_tx=XM_019821098.2;refGene_gn=LOC102900
118;refGene_tx=XM_019821099.2;refGene_gn=LOC102900118;refGene_tx=XM_01
9821100.2;refGene_gn=LOC102900118;refGene_tx=XM_019821101.2;refGene_gn
=LOC102900118;refGene_tx=XM_019821102.2;refGene_gn=LOC102900118;refGen
e_tx=XM_019821103.2;refGene_gn=LOC102900118;refGene_tx=XM_019821104.2;
refGene_gn=LOC102900118;refGene_tx=XM_019821107.2;refGene_gn=LOC102900
118;refGene_tx=XM_019821108.2;refGene_gn=LOC102900118;refGene_tx=XM_01
9821109.2;refGene_gn=LOC102900118;refGene_tx=XM_019821110.2;refGene_gn
=LOC102900118;refGene_tx=XM_019821112.2;refGene_gn=LOC102900118;refGen
e_tx=XM_019821113.2;refGene_gn=LOC102900118;refGene_tx=XM_019821114.2;
refGene_gn=LOC102900118;refGene_tx=XM_019821115.2;refGene_gn=LOC102900
118;refGene_tx=XM_019821116.2;refGene_gn=LOC102900118;refGene_tx=XM_01
9821117.2;refGene_gn=LOC102900118;refGene_tx=XM_019821118.2;refGene_gn
=LOC102900118;refGene_tx=XM_023246960.1
E3 42145170 42145230
refGene_gn=LOC102900118;refGene_tx=XM_019821091.2;refGene_gn=LOC102900
118;refGene_tx=XM_019821092.2;refGene_gn=LOC102900118;refGene_tx=XM_01
9821093.2;refGene_gn=LOC102900118;refGene_tx=XM_019821094.2;refGene_gn
=LOC102900118;refGene_tx=XM_019821095.2;refGene_gn=LOC102900118;refGen
e_tx=XM_019821096.2;refGene_gn=LOC102900118;refGene_tx=XM_019821097.2;
refGene_gn=LOC102900118;refGene_tx=XM_019821098.2;refGene_gn=LOC102900
118;refGene_tx=XM_019821099.2;refGene_gn=LOC102900118;refGene_tx=XM_01
9821100.2;refGene_gn=LOC102900118;refGene_tx=XM_019821101.2;refGene_gn
=LOC102900118;refGene_tx=XM_019821102.2;refGene_gn=LOC102900118;refGen
e_tx=XM_019821103.2;refGene_gn=LOC102900118;refGene_tx=XM_019821104.2;
refGene_gn=LOC102900118;refGene_tx=XM_019821107.2;refGene_gn=LOC102900
118;refGene_tx=XM_019821108.2;refGene_gn=LOC102900118;refGene_tx=XM_01
9821109.2;refGene_gn=LOC102900118;refGene_tx=XM_019821110.2;refGene_gn
=LOC102900118;refGene_tx=XM_019821112.2;refGene_gn=LOC102900118;refGen
e_tx=XM_019821113.2;refGene_gn=LOC102900118;refGene_tx=XM_019821114.2;
refGene_gn=LOC102900118;refGene_tx=XM_019821115.2;refGene_gn=LOC102900
118;refGene_tx=XM_019821116.2;refGene_gn=LOC102900118;refGene_tx=XM_01
9821117.2;refGene_gn=LOC102900118;refGene_tx=XM_019821118.2;refGene_gn
=LOC102900118;refGene_tx=XM_023246960.1
E3 42145547 42145589
refGene_gn=LOC102900118;refGene_tx=XM_019821091.2;refGene_gn=LOC102900

118;refGene_tx=XM_019821092.2;refGene_gn=LOC102900118;refGene_tx=XM_01
9821093.2;refGene_gn=LOC102900118;refGene_tx=XM_019821094.2;refGene_gn
=LOC102900118;refGene_tx=XM_019821095.2;refGene_gn=LOC102900118;refGen
e_tx=XM_019821096.2;refGene_gn=LOC102900118;refGene_tx=XM_019821097.2;
refGene_gn=LOC102900118;refGene_tx=XM_019821098.2;refGene_gn=LOC102900
118;refGene_tx=XM_019821099.2;refGene_gn=LOC102900118;refGene_tx=XM_01
9821100.2;refGene_gn=LOC102900118;refGene_tx=XM_019821102.2;refGene_gn
=LOC102900118;refGene_tx=XM_019821103.2;refGene_gn=LOC102900118;refGen
e_tx=XM_019821104.2;refGene_gn=LOC102900118;refGene_tx=XM_019821105.2;
refGene_gn=LOC102900118;refGene_tx=XM_019821107.2;refGene_gn=LOC102900
118;refGene_tx=XM_019821108.2;refGene_gn=LOC102900118;refGene_tx=XM_01
9821109.2;refGene_gn=LOC102900118;refGene_tx=XM_019821111.2;refGene_gn
=LOC102900118;refGene_tx=XM_019821112.2;refGene_gn=LOC102900118;refGen
e_tx=XM_019821113.2;refGene_gn=LOC102900118;refGene_tx=XM_019821114.2;
refGene_gn=LOC102900118;refGene_tx=XM_019821115.2;refGene_gn=LOC102900
118;refGene_tx=XM_019821116.2;refGene_gn=LOC102900118;refGene_tx=XM_01
9821118.2;refGene_gn=LOC102900118;refGene_tx=XM_023246960.1
E3 42145673 42145706
refGene_gn=LOC102900118;refGene_tx=XM_019821091.2;refGene_gn=LOC102900
118;refGene_tx=XM_019821092.2;refGene_gn=LOC102900118;refGene_tx=XM_01
9821093.2;refGene_gn=LOC102900118;refGene_tx=XM_019821094.2;refGene_gn
=LOC102900118;refGene_tx=XM_019821095.2;refGene_gn=LOC102900118;refGen
e_tx=XM_019821096.2;refGene_gn=LOC102900118;refGene_tx=XM_019821097.2;
refGene_gn=LOC102900118;refGene_tx=XM_019821098.2;refGene_gn=LOC102900
118;refGene_tx=XM_019821099.2;refGene_gn=LOC102900118;refGene_tx=XM_01
9821100.2;refGene_gn=LOC102900118;refGene_tx=XM_019821102.2;refGene_gn
=LOC102900118;refGene_tx=XM_019821103.2;refGene_gn=LOC102900118;refGen
e_tx=XM_019821104.2;refGene_gn=LOC102900118;refGene_tx=XM_019821105.2;
refGene_gn=LOC102900118;refGene_tx=XM_019821107.2;refGene_gn=LOC102900
118;refGene_tx=XM_019821108.2;refGene_gn=LOC102900118;refGene_tx=XM_01
9821109.2;refGene_gn=LOC102900118;refGene_tx=XM_019821111.2;refGene_gn
=LOC102900118;refGene_tx=XM_019821112.2;refGene_gn=LOC102900118;refGen
e_tx=XM_019821113.2;refGene_gn=LOC102900118;refGene_tx=XM_019821114.2;
refGene_gn=LOC102900118;refGene_tx=XM_019821115.2;refGene_gn=LOC102900
118;refGene_tx=XM_019821116.2;refGene_gn=LOC102900118;refGene_tx=XM_01
9821118.2;refGene_gn=LOC102900118;refGene_tx=XM_023246960.1
E3 42145793 42145814
refGene_gn=LOC102900118;refGene_tx=XM_019821091.2;refGene_gn=LOC102900
118;refGene_tx=XM_019821092.2;refGene_gn=LOC102900118;refGene_tx=XM_01
9821093.2;refGene_gn=LOC102900118;refGene_tx=XM_019821094.2;refGene_gn
=LOC102900118;refGene_tx=XM_019821095.2;refGene_gn=LOC102900118;refGen
e_tx=XM_019821096.2;refGene_gn=LOC102900118;refGene_tx=XM_019821097.2;
refGene_gn=LOC102900118;refGene_tx=XM_019821098.2;refGene_gn=LOC102900
118;refGene_tx=XM_019821099.2;refGene_gn=LOC102900118;refGene_tx=XM_01
9821100.2;refGene_gn=LOC102900118;refGene_tx=XM_019821102.2;refGene_gn
=LOC102900118;refGene_tx=XM_019821103.2;refGene_gn=LOC102900118;refGen
e_tx=XM_019821104.2;refGene_gn=LOC102900118;refGene_tx=XM_019821105.2;
refGene_gn=LOC102900118;refGene_tx=XM_019821107.2;refGene_gn=LOC102900
118;refGene_tx=XM_019821108.2;refGene_gn=LOC102900118;refGene_tx=XM_01
9821109.2;refGene_gn=LOC102900118;refGene_tx=XM_019821111.2;refGene_gn

=LOC102900118;refGene_tx=XM_019821112.2;refGene_gn=LOC102900118;refGene_tx=XM_019821113.2;refGene_gn=LOC102900118;refGene_tx=XM_019821114.2;refGene_gn=LOC102900118;refGene_tx=XM_019821115.2;refGene_gn=LOC102900118;refGene_tx=XM_019821116.2;refGene_gn=LOC102900118;refGene_tx=XM_019821118.2;refGene_gn=LOC102900118;refGene_tx=XM_023246960.1
E3 42146244 42146298
refGene_gn=LOC102900118;refGene_tx=XM_019821091.2;refGene_gn=LOC102900118;refGene_tx=XM_019821092.2;refGene_gn=LOC102900118;refGene_tx=XM_019821093.2;refGene_gn=LOC102900118;refGene_tx=XM_019821094.2;refGene_gn=LOC102900118;refGene_tx=XM_019821095.2;refGene_gn=LOC102900118;refGene_tx=XM_019821096.2;refGene_gn=LOC102900118;refGene_tx=XM_019821097.2;refGene_gn=LOC102900118;refGene_tx=XM_019821098.2;refGene_gn=LOC102900118;refGene_tx=XM_019821099.2;refGene_gn=LOC102900118;refGene_tx=XM_019821100.2;refGene_gn=LOC102900118;refGene_tx=XM_019821101.2;refGene_gn=LOC102900118;refGene_tx=XM_019821102.2;refGene_gn=LOC102900118;refGene_tx=XM_019821103.2;refGene_gn=LOC102900118;refGene_tx=XM_019821104.2;refGene_gn=LOC102900118;refGene_tx=XM_019821105.2;refGene_gn=LOC102900118;refGene_tx=XM_019821107.2;refGene_gn=LOC102900118;refGene_tx=XM_019821108.2;refGene_gn=LOC102900118;refGene_tx=XM_019821109.2;refGene_gn=LOC102900118;refGene_tx=XM_019821110.2;refGene_gn=LOC102900118;refGene_tx=XM_019821111.2;refGene_gn=LOC102900118;refGene_tx=XM_019821112.2;refGene_gn=LOC102900118;refGene_tx=XM_019821113.2;refGene_gn=LOC102900118;refGene_tx=XM_019821114.2;refGene_gn=LOC102900118;refGene_tx=XM_019821115.2;refGene_gn=LOC102900118;refGene_tx=XM_019821118.2;refGene_gn=LOC102900118;refGene_tx=XM_023246960.1
E3 42146404 42146512
refGene_gn=LOC102900118;refGene_tx=XM_019821091.2;refGene_gn=LOC102900118;refGene_tx=XM_019821092.2;refGene_gn=LOC102900118;refGene_tx=XM_019821093.2;refGene_gn=LOC102900118;refGene_tx=XM_019821094.2;refGene_gn=LOC102900118;refGene_tx=XM_019821095.2;refGene_gn=LOC102900118;refGene_tx=XM_019821096.2;refGene_gn=LOC102900118;refGene_tx=XM_019821097.2;refGene_gn=LOC102900118;refGene_tx=XM_019821098.2;refGene_gn=LOC102900118;refGene_tx=XM_019821099.2;refGene_gn=LOC102900118;refGene_tx=XM_019821100.2;refGene_gn=LOC102900118;refGene_tx=XM_019821101.2;refGene_gn=LOC102900118;refGene_tx=XM_019821102.2;refGene_gn=LOC102900118;refGene_tx=XM_019821103.2;refGene_gn=LOC102900118;refGene_tx=XM_019821104.2;refGene_gn=LOC102900118;refGene_tx=XM_019821105.2;refGene_gn=LOC102900118;refGene_tx=XM_019821107.2;refGene_gn=LOC102900118;refGene_tx=XM_019821108.2;refGene_gn=LOC102900118;refGene_tx=XM_019821110.2;refGene_gn=LOC102900118;refGene_tx=XM_019821111.2;refGene_gn=LOC102900118;refGene_tx=XM_019821112.2;refGene_gn=LOC102900118;refGene_tx=XM_019821113.2;refGene_gn=LOC102900118;refGene_tx=XM_019821114.2;refGene_gn=LOC102900118;refGene_tx=XM_019821115.2;refGene_gn=LOC102900118;refGene_tx=XM_019821118.2;refGene_gn=LOC102900118;refGene_tx=XM_023246960.1
E3 42146735 42146795
refGene_gn=LOC102900118;refGene_tx=XM_019821091.2;refGene_gn=LOC102900118;refGene_tx=XM_019821092.2;refGene_gn=LOC102900118;refGene_tx=XM_019821093.2;refGene_gn=LOC102900118;refGene_tx=XM_019821094.2;refGene_gn=LOC102900118;refGene_tx=XM_019821095.2;refGene_gn=LOC102900118;refGene_tx=XM_019821097.2;refGene_gn=LOC102900118;refGene_tx=XM_019821098.2;

refGene_gn=LOC102900118;refGene_tx=XM_019821099.2;refGene_gn=LOC102900118;refGene_tx=XM_019821101.2;refGene_gn=LOC102900118;refGene_tx=XM_019821103.2;refGene_gn=LOC102900118;refGene_tx=XM_019821105.2;refGene_gn=LOC102900118;refGene_tx=XM_019821110.2;refGene_gn=LOC102900118;refGene_tx=XM_019821111.2;refGene_gn=LOC102900118;refGene_tx=XM_019821115.2;refGene_gn=LOC102900118;refGene_tx=XM_019821118.2;refGene_gn=LOC102900118;refGene_tx=XM_023246960.1
E3 42146879 42146912
refGene_gn=LOC102900118;refGene_tx=XM_019821091.2;refGene_gn=LOC102900118;refGene_tx=XM_019821092.2;refGene_gn=LOC102900118;refGene_tx=XM_019821093.2;refGene_gn=LOC102900118;refGene_tx=XM_019821094.2;refGene_gn=LOC102900118;refGene_tx=XM_019821095.2;refGene_gn=LOC102900118;refGene_tx=XM_019821096.2;refGene_gn=LOC102900118;refGene_tx=XM_019821097.2;refGene_gn=LOC102900118;refGene_tx=XM_019821098.2;refGene_gn=LOC102900118;refGene_tx=XM_019821099.2;refGene_gn=LOC102900118;refGene_tx=XM_019821100.2;refGene_gn=LOC102900118;refGene_tx=XM_019821101.2;refGene_gn=LOC102900118;refGene_tx=XM_019821102.2;refGene_gn=LOC102900118;refGene_tx=XM_019821103.2;refGene_gn=LOC102900118;refGene_tx=XM_019821104.2;refGene_gn=LOC102900118;refGene_tx=XM_019821105.2;refGene_gn=LOC102900118;refGene_tx=XM_019821107.2;refGene_gn=LOC102900118;refGene_tx=XM_019821109.2;refGene_gn=LOC102900118;refGene_tx=XM_019821110.2;refGene_gn=LOC102900118;refGene_tx=XM_019821111.2;refGene_gn=LOC102900118;refGene_tx=XM_019821112.2;refGene_gn=LOC102900118;refGene_tx=XM_019821115.2;refGene_gn=LOC102900118;refGene_tx=XM_019821118.2;refGene_gn=LOC102900118;refGene_tx=XM_023246960.1
E3 42147249 42147285
refGene_gn=LOC102900118;refGene_tx=XM_019821091.2;refGene_gn=LOC102900118;refGene_tx=XM_019821092.2;refGene_gn=LOC102900118;refGene_tx=XM_019821093.2;refGene_gn=LOC102900118;refGene_tx=XM_019821094.2;refGene_gn=LOC102900118;refGene_tx=XM_019821095.2;refGene_gn=LOC102900118;refGene_tx=XM_019821096.2;refGene_gn=LOC102900118;refGene_tx=XM_019821097.2;refGene_gn=LOC102900118;refGene_tx=XM_019821098.2;refGene_gn=LOC102900118;refGene_tx=XM_019821099.2;refGene_gn=LOC102900118;refGene_tx=XM_019821100.2;refGene_gn=LOC102900118;refGene_tx=XM_019821101.2;refGene_gn=LOC102900118;refGene_tx=XM_019821102.2;refGene_gn=LOC102900118;refGene_tx=XM_019821103.2;refGene_gn=LOC102900118;refGene_tx=XM_019821104.2;refGene_gn=LOC102900118;refGene_tx=XM_019821105.2;refGene_gn=LOC102900118;refGene_tx=XM_019821107.2;refGene_gn=LOC102900118;refGene_tx=XM_019821108.2;refGene_gn=LOC102900118;refGene_tx=XM_019821109.2;refGene_gn=LOC102900118;refGene_tx=XM_019821110.2;refGene_gn=LOC102900118;refGene_tx=XM_019821111.2;refGene_gn=LOC102900118;refGene_tx=XM_019821112.2;refGene_gn=LOC102900118;refGene_tx=XM_019821113.2;refGene_gn=LOC102900118;refGene_tx=XM_019821114.2;refGene_gn=LOC102900118;refGene_tx=XM_019821115.2;refGene_gn=LOC102900118;refGene_tx=XM_019821116.2;refGene_gn=LOC102900118;refGene_tx=XM_019821117.2;refGene_gn=LOC102900118;refGene_tx=XM_019821118.2;refGene_gn=LOC102900118;refGene_tx=XM_023246960.1
E3 42147942 42147972
refGene_gn=LOC102900118;refGene_tx=XM_019821091.2;refGene_gn=LOC102900118;refGene_tx=XM_019821092.2;refGene_gn=LOC102900118;refGene_tx=XM_019821093.2;refGene_gn=LOC102900118;refGene_tx=XM_019821094.2;refGene_gn

=LOC102900118;refGene_tx=XM_019821095.2;refGene_gn=LOC102900118;refGene_tx=XM_019821096.2;refGene_gn=LOC102900118;refGene_tx=XM_019821098.2;refGene_gn=LOC102900118;refGene_tx=XM_019821099.2;refGene_gn=LOC102900118;refGene_tx=XM_019821100.2;refGene_gn=LOC102900118;refGene_tx=XM_019821101.2;refGene_gn=LOC102900118;refGene_tx=XM_019821103.2;refGene_gn=LOC102900118;refGene_tx=XM_019821104.2;refGene_gn=LOC102900118;refGene_tx=XM_019821105.2;refGene_gn=LOC102900118;refGene_tx=XM_019821107.2;refGene_gn=LOC102900118;refGene_tx=XM_019821108.2;refGene_gn=LOC102900118;refGene_tx=XM_019821109.2;refGene_gn=LOC102900118;refGene_tx=XM_019821110.2;refGene_gn=LOC102900118;refGene_tx=XM_019821111.2;refGene_gn=LOC102900118;refGene_tx=XM_019821113.2;refGene_gn=LOC102900118;refGene_tx=XM_019821115.2;refGene_gn=LOC102900118;refGene_tx=XM_019821116.2;refGene_gn=LOC102900118;refGene_tx=XM_019821117.2;refGene_gn=LOC102900118;refGene_tx=XM_019821118.2;refGene_gn=LOC102900118;refGene_tx=XM_023246960.1

E3 42148924 42148957

refGene_gn=LOC102900118;refGene_tx=XM_019821091.2;refGene_gn=LOC102900118;refGene_tx=XM_019821092.2;refGene_gn=LOC102900118;refGene_tx=XM_019821094.2;refGene_gn=LOC102900118;refGene_tx=XM_019821095.2;refGene_gn=LOC102900118;refGene_tx=XM_019821096.2;refGene_gn=LOC102900118;refGene_tx=XM_019821098.2;refGene_gn=LOC102900118;refGene_tx=XM_019821099.2;refGene_gn=LOC102900118;refGene_tx=XM_019821101.2;refGene_gn=LOC102900118;refGene_tx=XM_019821103.2;refGene_gn=LOC102900118;refGene_tx=XM_019821104.2;refGene_gn=LOC102900118;refGene_tx=XM_019821105.2;refGene_gn=LOC102900118;refGene_tx=XM_019821108.2;refGene_gn=LOC102900118;refGene_tx=XM_019821109.2;refGene_gn=LOC102900118;refGene_tx=XM_019821110.2;refGene_gn=LOC102900118;refGene_tx=XM_019821111.2;refGene_gn=LOC102900118;refGene_tx=XM_019821115.2;refGene_gn=LOC102900118;refGene_tx=XM_019821116.2;refGene_gn=LOC102900118;refGene_tx=XM_019821117.2;refGene_gn=LOC102900118;refGene_tx=XM_019821118.2;refGene_gn=LOC102900118;refGene_tx=XM_023246960.1

E3 42149157 42149224

refGene_gn=LOC102900118;refGene_tx=XM_019821091.2;refGene_gn=LOC102900118;refGene_tx=XM_019821092.2;refGene_gn=LOC102900118;refGene_tx=XM_019821093.2;refGene_gn=LOC102900118;refGene_tx=XM_019821094.2;refGene_gn=LOC102900118;refGene_tx=XM_019821095.2;refGene_gn=LOC102900118;refGene_tx=XM_019821096.2;refGene_gn=LOC102900118;refGene_tx=XM_019821097.2;refGene_gn=LOC102900118;refGene_tx=XM_019821098.2;refGene_gn=LOC102900118;refGene_tx=XM_019821099.2;refGene_gn=LOC102900118;refGene_tx=XM_019821100.2;refGene_gn=LOC102900118;refGene_tx=XM_019821101.2;refGene_gn=LOC102900118;refGene_tx=XM_019821102.2;refGene_gn=LOC102900118;refGene_tx=XM_019821103.2;refGene_gn=LOC102900118;refGene_tx=XM_019821104.2;refGene_gn=LOC102900118;refGene_tx=XM_019821105.2;refGene_gn=LOC102900118;refGene_tx=XM_019821107.2;refGene_gn=LOC102900118;refGene_tx=XM_019821108.2;refGene_gn=LOC102900118;refGene_tx=XM_019821109.2;refGene_gn=LOC102900118;refGene_tx=XM_019821110.2;refGene_gn=LOC102900118;refGene_tx=XM_019821111.2;refGene_gn=LOC102900118;refGene_tx=XM_019821112.2;refGene_gn=LOC102900118;refGene_tx=XM_019821113.2;refGene_gn=LOC102900118;refGene_tx=XM_019821114.2;refGene_gn=LOC102900118;refGene_tx=XM_019821115.2;refGene_gn=LOC102900118;refGene_tx=XM_019821116.2;refGene_gn

=LOC102900118;refGene_tx=XM_019821117.2;refGene_gn=LOC102900118;refGene_tx=XM_019821118.2;refGene_gn=LOC102900118;refGene_tx=XM_023246960.1
E3 42158877 42159423
refGene_gn=PKMYT1;refGene_tx=XM_006942430.4;refGene_gn=PKMYT1;refGene_tx=XM_011290664.3;refGene_gn=PKMYT1;refGene_tx=XM_019821119.2;refGene_gn=PKMYT1;refGene_tx=XM_019821120.2;refGene_gn=PKMYT1;refGene_tx=XM_019821121.2;refGene_gn=PKMYT1;refGene_tx=XM_019821122.2;refGene_gn=PKMYT1;refGene_tx=XM_019821123.2;refGene_gn=PKMYT1;refGene_tx=XM_019821124.2;refGene_gn=PKMYT1;refGene_tx=XM_023246951.1
E3 42160235 42160729
refGene_gn=PKMYT1;refGene_tx=XM_006942430.4;refGene_gn=PKMYT1;refGene_tx=XM_011290664.3;refGene_gn=PKMYT1;refGene_tx=XM_019821119.2;refGene_gn=PKMYT1;refGene_tx=XM_019821120.2;refGene_gn=PKMYT1;refGene_tx=XM_019821121.2;refGene_gn=PKMYT1;refGene_tx=XM_019821122.2;refGene_gn=PKMYT1;refGene_tx=XM_019821123.2;refGene_gn=PKMYT1;refGene_tx=XM_019821124.2;refGene_gn=PKMYT1;refGene_tx=XM_023246951.1
E3 42161390 42161497
refGene_gn=PKMYT1;refGene_tx=XM_006942430.4;refGene_gn=PKMYT1;refGene_tx=XM_011290664.3;refGene_gn=PKMYT1;refGene_tx=XM_019821119.2;refGene_gn=PKMYT1;refGene_tx=XM_019821120.2;refGene_gn=PKMYT1;refGene_tx=XM_019821121.2;refGene_gn=PKMYT1;refGene_tx=XM_019821122.2;refGene_gn=PKMYT1;refGene_tx=XM_019821123.2;refGene_gn=PKMYT1;refGene_tx=XM_019821124.2;refGene_gn=PKMYT1;refGene_tx=XM_023246951.1
E3 42161611 42161784
refGene_gn=PKMYT1;refGene_tx=XM_006942430.4;refGene_gn=PKMYT1;refGene_tx=XM_011290664.3;refGene_gn=PKMYT1;refGene_tx=XM_019821119.2;refGene_gn=PKMYT1;refGene_tx=XM_019821120.2;refGene_gn=PKMYT1;refGene_tx=XM_019821121.2;refGene_gn=PKMYT1;refGene_tx=XM_019821122.2;refGene_gn=PKMYT1;refGene_tx=XM_019821123.2;refGene_gn=PKMYT1;refGene_tx=XM_019821124.2;refGene_gn=PKMYT1;refGene_tx=XM_023246951.1
E3 42161869 42162027
refGene_gn=PKMYT1;refGene_tx=XM_006942430.4;refGene_gn=PKMYT1;refGene_tx=XM_011290664.3;refGene_gn=PKMYT1;refGene_tx=XM_019821119.2;refGene_gn=PKMYT1;refGene_tx=XM_019821120.2;refGene_gn=PKMYT1;refGene_tx=XM_019821121.2;refGene_gn=PKMYT1;refGene_tx=XM_019821122.2;refGene_gn=PKMYT1;refGene_tx=XM_019821123.2;refGene_gn=PKMYT1;refGene_tx=XM_019821124.2;refGene_gn=PKMYT1;refGene_tx=XM_023246951.1
E3 42162238 42162359
refGene_gn=PKMYT1;refGene_tx=XM_006942430.4;refGene_gn=PKMYT1;refGene_tx=XM_011290664.3;refGene_gn=PKMYT1;refGene_tx=XM_019821119.2;refGene_gn=PKMYT1;refGene_tx=XM_019821120.2;refGene_gn=PKMYT1;refGene_tx=XM_019821121.2;refGene_gn=PKMYT1;refGene_tx=XM_019821124.2;refGene_gn=PKMYT1;refGene_tx=XM_023246951.1
E3 42162856 42162955
refGene_gn=PKMYT1;refGene_tx=XM_006942430.4;refGene_gn=PKMYT1;refGene_tx=XM_011290664.3;refGene_gn=PKMYT1;refGene_tx=XM_019821119.2;refGene_gn=PKMYT1;refGene_tx=XM_019821120.2;refGene_gn=PKMYT1;refGene_tx=XM_019821121.2;refGene_gn=PKMYT1;refGene_tx=XM_019821122.2;refGene_gn=PKMYT1;refGene_tx=XM_019821123.2;refGene_gn=PKMYT1;refGene_tx=XM_019821124.2;refGene_gn=PKMYT1;refGene_tx=XM_023246951.1

E3 42163038 42163150
refGene_gn=PKMYT1;refGene_tx=XM_006942430.4;refGene_gn=PKMYT1;refGene_tx=XM_019821119.2;refGene_gn=PKMYT1;refGene_tx=XM_019821120.2;refGene_gn=PKMYT1;refGene_tx=XM_019821121.2;refGene_gn=PKMYT1;refGene_tx=XM_019821122.2;refGene_gn=PKMYT1;refGene_tx=XM_019821123.2;refGene_gn=PKMYT1;refGene_tx=XM_019821124.2;refGene_gn=PKMYT1;refGene_tx=XM_023246951.1
E3 42164134 42164568
refGene_gn=PAQR4;refGene_tx=XM_023246970.1;refGene_gn=PAQR4;refGene_tx=XM_023246971.1
E3 42164720 42164967
refGene_gn=PAQR4;refGene_tx=XM_023246970.1;refGene_gn=PAQR4;refGene_tx=XM_023246971.1
E3 42166326 42166492 refGene_gn=PAQR4;refGene_tx=XM_023246970.1
E3 42168224 42168388 refGene_gn=KREMEN2;refGene_tx=XM_023246984.1
E3 42168788 42169310 refGene_gn=KREMEN2;refGene_tx=XM_023246984.1
E3 42169671 42169825 refGene_gn=KREMEN2;refGene_tx=XM_023246984.1
E3 42170021 42170146 refGene_gn=KREMEN2;refGene_tx=XM_023246984.1
E3 42170246 42170338 refGene_gn=KREMEN2;refGene_tx=XM_023246984.1
E3 42171350 42171391 refGene_gn=KREMEN2;refGene_tx=XM_023246984.1
E3 42171392 42171526 refGene_gn=KREMEN2;refGene_tx=XM_023246984.1
E3 42171767 42171864 refGene_gn=KREMEN2;refGene_tx=XM_023246984.1
E3 42180443 42180473
refGene_gn=FLYWCH1;refGene_tx=XM_023246948.1;refGene_gn=FLYWCH1;refGene_tx=XM_023246949.1
E3 42183961 42184055
refGene_gn=FLYWCH1;refGene_tx=XM_023246947.1;refGene_gn=FLYWCH1;refGene_tx=XM_023246948.1;refGene_gn=FLYWCH1;refGene_tx=XM_023246949.1
E3 42185454 42185752
refGene_gn=FLYWCH1;refGene_tx=XM_023246947.1;refGene_gn=FLYWCH1;refGene_tx=XM_023246948.1;refGene_gn=FLYWCH1;refGene_tx=XM_023246949.1
E3 42186736 42187000
refGene_gn=FLYWCH1;refGene_tx=XM_023246947.1;refGene_gn=FLYWCH1;refGene_tx=XM_023246948.1;refGene_gn=FLYWCH1;refGene_tx=XM_023246949.1
E3 42190803 42191067
refGene_gn=FLYWCH1;refGene_tx=XM_023246947.1;refGene_gn=FLYWCH1;refGene_tx=XM_023246948.1;refGene_gn=FLYWCH1;refGene_tx=XM_023246949.1
E3 42191198 42191624
refGene_gn=FLYWCH1;refGene_tx=XM_023246947.1;refGene_gn=FLYWCH1;refGene_tx=XM_023246948.1;refGene_gn=FLYWCH1;refGene_tx=XM_023246949.1
E3 42192701 42193163
refGene_gn=FLYWCH1;refGene_tx=XM_023246947.1;refGene_gn=FLYWCH1;refGene_tx=XM_023246948.1;refGene_gn=FLYWCH1;refGene_tx=XM_023246949.1
E3 42193435 42193769
refGene_gn=FLYWCH1;refGene_tx=XM_023246947.1;refGene_gn=FLYWCH1;refGene_tx=XM_023246948.1;refGene_gn=FLYWCH1;refGene_tx=XM_023246949.1
E3 42213762 42213866
refGene_gn=FLYWCH2;refGene_tx=XM_006942433.4;refGene_gn=FLYWCH2;refGene_tx=XM_019821128.2;refGene_gn=FLYWCH2;refGene_tx=XM_023246981.1
E3 42215749 42216080

refGene_gn=FLYWCH2;refGene_tx=XM_006942433.4;refGene_gn=FLYWCH2;refGene_tx=XM_019821128.2;refGene_gn=FLYWCH2;refGene_tx=XM_023246981.1
E3 42236687 42236709
refGene_gn=LOC102901314;refGene_tx=XM_023246588.1
E3 42242553 42242691
refGene_gn=LOC102901314;refGene_tx=XM_023246588.1
E3 42243182 42243327
refGene_gn=LOC102901314;refGene_tx=XM_023246588.1
E3 42249239 42249760
refGene_gn=LOC102901314;refGene_tx=XM_023246588.1
E3 42249948 42249992
refGene_gn=LOC102901314;refGene_tx=XM_023246588.1
E3 42250520 42250577
refGene_gn=SRRM2;refGene_tx=XM_006942441.4;refGene_gn=SRRM2;refGene_tx=XM_006942442.4;refGene_gn=SRRM2;refGene_tx=XM_006942444.4;refGene_gn=SRRM2;refGene_tx=XM_011290674.3;refGene_gn=SRRM2;refGene_tx=XM_011290678.3;refGene_gn=SRRM2;refGene_tx=XM_019821130.2;refGene_gn=SRRM2;refGene_tx=XM_019821131.2
E3 42252947 42253219
refGene_gn=SRRM2;refGene_tx=XM_006942441.4;refGene_gn=SRRM2;refGene_tx=XM_006942442.4;refGene_gn=SRRM2;refGene_tx=XM_006942444.4;refGene_gn=SRRM2;refGene_tx=XM_011290674.3;refGene_gn=SRRM2;refGene_tx=XM_011290678.3;refGene_gn=SRRM2;refGene_tx=XM_019821130.2;refGene_gn=SRRM2;refGene_tx=XM_019821131.2
E3 42254099 42254207
refGene_gn=SRRM2;refGene_tx=XM_006942441.4;refGene_gn=SRRM2;refGene_tx=XM_006942442.4;refGene_gn=SRRM2;refGene_tx=XM_006942444.4;refGene_gn=SRRM2;refGene_tx=XM_011290674.3;refGene_gn=SRRM2;refGene_tx=XM_011290678.3;refGene_gn=SRRM2;refGene_tx=XM_019821130.2;refGene_gn=SRRM2;refGene_tx=XM_019821131.2
E3 42254387 42254552
refGene_gn=SRRM2;refGene_tx=XM_006942441.4;refGene_gn=SRRM2;refGene_tx=XM_006942442.4;refGene_gn=SRRM2;refGene_tx=XM_006942444.4;refGene_gn=SRRM2;refGene_tx=XM_011290674.3;refGene_gn=SRRM2;refGene_tx=XM_011290678.3;refGene_gn=SRRM2;refGene_tx=XM_019821130.2;refGene_gn=SRRM2;refGene_tx=XM_019821131.2
E3 42254724 42254802
refGene_gn=SRRM2;refGene_tx=XM_006942441.4;refGene_gn=SRRM2;refGene_tx=XM_006942442.4;refGene_gn=SRRM2;refGene_tx=XM_006942444.4;refGene_gn=SRRM2;refGene_tx=XM_011290674.3;refGene_gn=SRRM2;refGene_tx=XM_011290678.3;refGene_gn=SRRM2;refGene_tx=XM_019821130.2;refGene_gn=SRRM2;refGene_tx=XM_019821131.2
E3 42255251 42255314
refGene_gn=SRRM2;refGene_tx=XM_006942441.4;refGene_gn=SRRM2;refGene_tx=XM_006942442.4;refGene_gn=SRRM2;refGene_tx=XM_006942444.4;refGene_gn=SRRM2;refGene_tx=XM_011290674.3;refGene_gn=SRRM2;refGene_tx=XM_011290678.3;refGene_gn=SRRM2;refGene_tx=XM_019821130.2;refGene_gn=SRRM2;refGene_tx=XM_019821131.2
E3 42255411 42255444
refGene_gn=SRRM2;refGene_tx=XM_006942441.4;refGene_gn=SRRM2;refGene_tx

=XM_006942442.4;refGene_gn=SRRM2;refGene_tx=XM_006942444.4;refGene_gn=SRRM2;refGene_tx=XM_011290674.3;refGene_gn=SRRM2;refGene_tx=XM_011290678.3;refGene_gn=SRRM2;refGene_tx=XM_019821130.2;refGene_gn=SRRM2;refGene_tx=XM_019821131.2
E3 42255628 42255679
refGene_gn=SRRM2;refGene_tx=XM_006942441.4;refGene_gn=SRRM2;refGene_tx=XM_006942442.4;refGene_gn=SRRM2;refGene_tx=XM_006942444.4;refGene_gn=SRRM2;refGene_tx=XM_011290674.3;refGene_gn=SRRM2;refGene_tx=XM_011290678.3;refGene_gn=SRRM2;refGene_tx=XM_019821130.2;refGene_gn=SRRM2;refGene_tx=XM_019821131.2
E3 42255867 42255960
refGene_gn=SRRM2;refGene_tx=XM_006942441.4;refGene_gn=SRRM2;refGene_tx=XM_006942442.4;refGene_gn=SRRM2;refGene_tx=XM_006942444.4;refGene_gn=SRRM2;refGene_tx=XM_011290674.3;refGene_gn=SRRM2;refGene_tx=XM_011290678.3;refGene_gn=SRRM2;refGene_tx=XM_019821130.2;refGene_gn=SRRM2;refGene_tx=XM_019821131.2
E3 42256045 42256088 refGene_gn=SRRM2;refGene_tx=XM_011290678.3
E3 42256615 42256811
refGene_gn=SRRM2;refGene_tx=XM_006942441.4;refGene_gn=SRRM2;refGene_tx=XM_006942442.4;refGene_gn=SRRM2;refGene_tx=XM_006942444.4;refGene_gn=SRRM2;refGene_tx=XM_011290674.3;refGene_gn=SRRM2;refGene_tx=XM_019821130.2;refGene_gn=SRRM2;refGene_tx=XM_019821131.2
E3 42257909 42264598
refGene_gn=SRRM2;refGene_tx=XM_006942441.4;refGene_gn=SRRM2;refGene_tx=XM_006942442.4;refGene_gn=SRRM2;refGene_tx=XM_006942444.4;refGene_gn=SRRM2;refGene_tx=XM_011290674.3;refGene_gn=SRRM2;refGene_tx=XM_019821130.2;refGene_gn=SRRM2;refGene_tx=XM_019821131.2
E3 42264838 42264851 refGene_gn=SRRM2;refGene_tx=XM_006942444.4
E3 42264856 42264933 refGene_gn=SRRM2;refGene_tx=XM_023246945.1
E3 42265331 42265616
refGene_gn=SRRM2;refGene_tx=XM_006942441.4;refGene_gn=SRRM2;refGene_tx=XM_006942442.4;refGene_gn=SRRM2;refGene_tx=XM_011290674.3;refGene_gn=SRRM2;refGene_tx=XM_019821130.2;refGene_gn=SRRM2;refGene_tx=XM_019821131.2;refGene_gn=SRRM2;refGene_tx=XM_023246945.1
E3 42265949 42266082 refGene_gn=SRRM2;refGene_tx=XM_006942441.4
E3 42266186 42266211 refGene_gn=SRRM2;refGene_tx=XM_006942442.4
E3 42266698 42266812
refGene_gn=SRRM2;refGene_tx=XM_011290674.3;refGene_gn=SRRM2;refGene_tx=XM_019821130.2;refGene_gn=SRRM2;refGene_tx=XM_019821131.2;refGene_gn=SRRM2;refGene_tx=XM_023246945.1
E3 42266930 42267041
refGene_gn=SRRM2;refGene_tx=XM_011290674.3;refGene_gn=SRRM2;refGene_tx=XM_019821130.2;refGene_gn=SRRM2;refGene_tx=XM_019821131.2;refGene_gn=SRRM2;refGene_tx=XM_023246945.1
E3 42267125 42267135
refGene_gn=SRRM2;refGene_tx=XM_011290674.3;refGene_gn=SRRM2;refGene_tx=XM_019821130.2;refGene_gn=SRRM2;refGene_tx=XM_019821131.2;refGene_gn=SRRM2;refGene_tx=XM_023246945.1
E3 42268224 42268337
refGene_gn=ELOB;refGene_tx=XM_023246975.1;refGene_gn=ELOB;refGene_tx=X

M_023246976.1
E3 42270923 42271029
refGene_gn=ELOB;refGene_tx=XM_023246975.1;refGene_gn=ELOB;refGene_tx=XM_023246976.1
E3 42272039 42272174
refGene_gn=ELOB;refGene_tx=XM_023246975.1;refGene_gn=ELOB;refGene_tx=XM_023246976.1
E3 42272322 42272325 refGene_gn=ELOB;refGene_tx=XM_023246976.1
E3 42272530 42272785 refGene_gn=ELOB;refGene_tx=XM_023246975.1
E3 42277985 42278144
refGene_gn=LOC101099923;refGene_tx=XM_023246985.1
E3 42278298 42278468
refGene_gn=LOC101099923;refGene_tx=XM_023246985.1
E3 42278695 42278967
refGene_gn=LOC101099923;refGene_tx=XM_023246985.1
E3 42279078 42279241
refGene_gn=LOC101099923;refGene_tx=XM_023246985.1
E3 42279382 42279409
refGene_gn=LOC101099923;refGene_tx=XM_023246985.1
E3 42279780 42280024
refGene_gn=LOC101099923;refGene_tx=XM_023246985.1
E3 42283366 42283603
refGene_gn=PRSS41;refGene_tx=XM_023246963.1;refGene_gn=PRSS41;refGene_tx=XM_023246964.1;refGene_gn=PRSS41;refGene_tx=XM_023246965.1;refGene_gn=PRSS41;refGene_tx=XM_023246966.1
E3 42284027 42284182
refGene_gn=PRSS41;refGene_tx=XM_023246963.1;refGene_gn=PRSS41;refGene_tx=XM_023246964.1;refGene_gn=PRSS41;refGene_tx=XM_023246965.1;refGene_gn=PRSS41;refGene_tx=XM_023246966.1
E3 42292599 42292883
refGene_gn=PRSS41;refGene_tx=XM_023246963.1;refGene_gn=PRSS41;refGene_tx=XM_023246964.1;refGene_gn=PRSS41;refGene_tx=XM_023246965.1;refGene_gn=PRSS41;refGene_tx=XM_023246966.1
E3 42293504 42293639
refGene_gn=PRSS41;refGene_tx=XM_023246963.1;refGene_gn=PRSS41;refGene_tx=XM_023246964.1
E3 42293784 42293950
refGene_gn=PRSS41;refGene_tx=XM_023246963.1;refGene_gn=PRSS41;refGene_tx=XM_023246964.1;refGene_gn=PRSS41;refGene_tx=XM_023246965.1;refGene_gn=PRSS41;refGene_tx=XM_023246966.1
E3 42294147 42294291
refGene_gn=PRSS41;refGene_tx=XM_023246963.1;refGene_gn=PRSS41;refGene_tx=XM_023246964.1;refGene_gn=PRSS41;refGene_tx=XM_023246965.1;refGene_gn=PRSS41;refGene_tx=XM_023246966.1
E3 42294406 42294473
refGene_gn=PRSS41;refGene_tx=XM_023246963.1;refGene_gn=PRSS41;refGene_tx=XM_023246964.1;refGene_gn=PRSS41;refGene_tx=XM_023246965.1;refGene_gn=PRSS41;refGene_tx=XM_023246966.1
E3 42298374 42298495
refGene_gn=LOC101100173;refGene_tx=XM_011290768.3

E3 42298917 42299080
refGene_gn=LOC101100173;refGene_tx=XM_011290768.3
E3 42299640 42299918
refGene_gn=LOC101100173;refGene_tx=XM_011290768.3
E3 42301483 42301650
refGene_gn=LOC101100173;refGene_tx=XM_011290768.3
E3 42302820 42303090
refGene_gn=LOC101100173;refGene_tx=XM_011290768.3
E3 42306692 42306759
refGene_gn=LOC101100434;refGene_tx=XM_023246967.1;refGene_gn=LOC101100434;refGene_tx=XM_023246968.1;refGene_gn=LOC101100434;refGene_tx=XM_023246969.1
E3 42306875 42307019
refGene_gn=LOC101100434;refGene_tx=XM_023246967.1;refGene_gn=LOC101100434;refGene_tx=XM_023246968.1;refGene_gn=LOC101100434;refGene_tx=XM_023246969.1
E3 42307214 42307383
refGene_gn=LOC101100434;refGene_tx=XM_023246967.1;refGene_gn=LOC101100434;refGene_tx=XM_023246968.1;refGene_gn=LOC101100434;refGene_tx=XM_023246969.1
E3 42307896 42308180
refGene_gn=LOC101100434;refGene_tx=XM_023246967.1;refGene_gn=LOC101100434;refGene_tx=XM_023246968.1;refGene_gn=LOC101100434;refGene_tx=XM_023246969.1
E3 42310082 42310237
refGene_gn=LOC101100434;refGene_tx=XM_023246967.1;refGene_gn=LOC101100434;refGene_tx=XM_023246968.1;refGene_gn=LOC101100434;refGene_tx=XM_023246969.1
E3 42310521 42310758
refGene_gn=LOC101100434;refGene_tx=XM_023246967.1;refGene_gn=LOC101100434;refGene_tx=XM_023246968.1;refGene_gn=LOC101100434;refGene_tx=XM_023246969.1
E3 42320687 42320739 refGene_gn=ZG16B;refGene_tx=XM_023246980.1
E3 42320957 42321063 refGene_gn=ZG16B;refGene_tx=XM_023246980.1
E3 42321719 42322068 refGene_gn=ZG16B;refGene_tx=XM_023246980.1
E3 42337342 42337573
refGene_gn=LOC101100937;refGene_tx=XM_011290681.3;refGene_gn=LOC101100937;refGene_tx=XM_011290685.3;refGene_gn=LOC101100937;refGene_tx=XM_019821133.2;refGene_gn=LOC101100937;refGene_tx=XM_019821135.2;refGene_gn=LOC101100937;refGene_tx=XM_019821136.2;refGene_gn=LOC101100937;refGene_tx=XM_019821137.2;refGene_gn=LOC101100937;refGene_tx=XM_019821138.2;refGene_gn=LOC101100937;refGene_tx=XM_019821139.2;refGene_gn=LOC101100937;refGene_tx=XM_019821140.2;refGene_gn=LOC101100937;refGene_tx=XM_019821141.2;refGene_gn=LOC101100937;refGene_tx=XM_019821142.2;refGene_gn=LOC101100937;refGene_tx=XM_019821143.2;refGene_gn=LOC101100937;refGene_tx=XM_019821144.2;refGene_gn=LOC101100937;refGene_tx=XM_019821145.2;refGene_gn=LOC101100937;refGene_tx=XM_019821146.2
E3 42338225 42338383
refGene_gn=LOC101100937;refGene_tx=XM_011290681.3;refGene_gn=LOC101100937;refGene_tx=XM_011290685.3;refGene_gn=LOC101100937;refGene_tx=XM_01

9821133.2;refGene_gn=LOC101100937;refGene_tx=XM_019821135.2;refGene_gn=LOC101100937;refGene_tx=XM_019821136.2;refGene_gn=LOC101100937;refGene_tx=XM_019821137.2;refGene_gn=LOC101100937;refGene_tx=XM_019821138.2;refGene_gn=LOC101100937;refGene_tx=XM_019821139.2;refGene_gn=LOC101100937;refGene_tx=XM_019821140.2;refGene_gn=LOC101100937;refGene_tx=XM_019821141.2;refGene_gn=LOC101100937;refGene_tx=XM_019821142.2;refGene_gn=LOC101100937;refGene_tx=XM_019821143.2;refGene_gn=LOC101100937;refGene_tx=XM_019821144.2;refGene_gn=LOC101100937;refGene_tx=XM_019821145.2;refGene_gn=LOC101100937;refGene_tx=XM_019821146.2
E3 42339537 42339815
refGene_gn=LOC101100937;refGene_tx=XM_011290681.3;refGene_gn=LOC101100937;refGene_tx=XM_011290685.3;refGene_gn=LOC101100937;refGene_tx=XM_019821133.2;refGene_gn=LOC101100937;refGene_tx=XM_019821135.2;refGene_gn=LOC101100937;refGene_tx=XM_019821136.2;refGene_gn=LOC101100937;refGene_tx=XM_019821137.2;refGene_gn=LOC101100937;refGene_tx=XM_019821138.2;refGene_gn=LOC101100937;refGene_tx=XM_019821139.2;refGene_gn=LOC101100937;refGene_tx=XM_019821140.2;refGene_gn=LOC101100937;refGene_tx=XM_019821141.2;refGene_gn=LOC101100937;refGene_tx=XM_019821142.2;refGene_gn=LOC101100937;refGene_tx=XM_019821143.2;refGene_gn=LOC101100937;refGene_tx=XM_019821144.2;refGene_gn=LOC101100937;refGene_tx=XM_019821145.2;refGene_gn=LOC101100937;refGene_tx=XM_019821146.2
E3 42340054 42340226
refGene_gn=LOC101100937;refGene_tx=XM_011290681.3;refGene_gn=LOC101100937;refGene_tx=XM_011290685.3;refGene_gn=LOC101100937;refGene_tx=XM_019821133.2;refGene_gn=LOC101100937;refGene_tx=XM_019821135.2;refGene_gn=LOC101100937;refGene_tx=XM_019821136.2;refGene_gn=LOC101100937;refGene_tx=XM_019821137.2;refGene_gn=LOC101100937;refGene_tx=XM_019821138.2;refGene_gn=LOC101100937;refGene_tx=XM_019821139.2;refGene_gn=LOC101100937;refGene_tx=XM_019821140.2;refGene_gn=LOC101100937;refGene_tx=XM_019821141.2;refGene_gn=LOC101100937;refGene_tx=XM_019821142.2;refGene_gn=LOC101100937;refGene_tx=XM_019821143.2;refGene_gn=LOC101100937;refGene_tx=XM_019821144.2;refGene_gn=LOC101100937;refGene_tx=XM_019821145.2;refGene_gn=LOC101100937;refGene_tx=XM_019821146.2
E3 42340565 42340592
refGene_gn=LOC101100937;refGene_tx=XM_011290681.3;refGene_gn=LOC101100937;refGene_tx=XM_011290685.3;refGene_gn=LOC101100937;refGene_tx=XM_019821133.2;refGene_gn=LOC101100937;refGene_tx=XM_019821135.2;refGene_gn=LOC101100937;refGene_tx=XM_019821136.2;refGene_gn=LOC101100937;refGene_tx=XM_019821137.2;refGene_gn=LOC101100937;refGene_tx=XM_019821138.2;refGene_gn=LOC101100937;refGene_tx=XM_019821139.2;refGene_gn=LOC101100937;refGene_tx=XM_019821140.2;refGene_gn=LOC101100937;refGene_tx=XM_019821141.2;refGene_gn=LOC101100937;refGene_tx=XM_019821142.2;refGene_gn=LOC101100937;refGene_tx=XM_019821143.2;refGene_gn=LOC101100937;refGene_tx=XM_019821144.2;refGene_gn=LOC101100937;refGene_tx=XM_019821145.2;refGene_gn=LOC101100937;refGene_tx=XM_019821146.2
E3 42341112 42341357
refGene_gn=LOC101100937;refGene_tx=XM_011290681.3;refGene_gn=LOC101100937;refGene_tx=XM_011290685.3;refGene_gn=LOC101100937;refGene_tx=XM_019821133.2;refGene_gn=LOC101100937;refGene_tx=XM_019821135.2;refGene_gn=LOC101100937;refGene_tx=XM_019821136.2;refGene_gn=LOC101100937;refGen

e_tx=XM_019821137.2;refGene_gn=LOC101100937;refGene_tx=XM_019821138.2;
refGene_gn=LOC101100937;refGene_tx=XM_019821139.2;refGene_gn=LOC101100
937;refGene_tx=XM_019821140.2;refGene_gn=LOC101100937;refGene_tx=XM_01
9821141.2;refGene_gn=LOC101100937;refGene_tx=XM_019821142.2;refGene_gn
=LOC101100937;refGene_tx=XM_019821143.2;refGene_gn=LOC101100937;refGen
e_tx=XM_019821144.2;refGene_gn=LOC101100937;refGene_tx=XM_019821145.2;
refGene_gn=LOC101100937;refGene_tx=XM_019821146.2
E3 42343871 42343957
refGene_gn=LOC101100937;refGene_tx=XM_011290681.3;refGene_gn=LOC101100
937;refGene_tx=XM_011290685.3;refGene_gn=LOC101100937;refGene_tx=XM_01
9821133.2;refGene_gn=LOC101100937;refGene_tx=XM_019821135.2;refGene_gn
=LOC101100937;refGene_tx=XM_019821136.2;refGene_gn=LOC101100937;refGen
e_tx=XM_019821137.2;refGene_gn=LOC101100937;refGene_tx=XM_019821138.2;
refGene_gn=LOC101100937;refGene_tx=XM_019821139.2;refGene_gn=LOC101100
937;refGene_tx=XM_019821140.2;refGene_gn=LOC101100937;refGene_tx=XM_01
9821141.2;refGene_gn=LOC101100937;refGene_tx=XM_019821142.2;refGene_gn
=LOC101100937;refGene_tx=XM_019821143.2;refGene_gn=LOC101100937;refGen
e_tx=XM_019821144.2;refGene_gn=LOC101100937;refGene_tx=XM_019821145.2
E3 42369735 42369784
refGene_gn=LOC101101183;refGene_tx=XM_003999013.3
E3 42373044 42373071
refGene_gn=LOC101101183;refGene_tx=XM_003999013.3
E3 42373780 42373943
refGene_gn=LOC101101183;refGene_tx=XM_003999013.3
E3 42374841 42375113
refGene_gn=LOC101101183;refGene_tx=XM_003999013.3
E3 42375309 42375479
refGene_gn=LOC101101183;refGene_tx=XM_003999013.3
E3 42375972 42376266
refGene_gn=LOC101101183;refGene_tx=XM_003999013.3
E3 42382058 42382088 refGene_gn=KCTD5;refGene_tx=XM_023246973.1
E3 42385654 42385699 refGene_gn=KCTD5;refGene_tx=XM_023246972.1
E3 42388211 42388337
refGene_gn=KCTD5;refGene_tx=XM_023246972.1;refGene_gn=KCTD5;refGene_tx
=XM_023246973.1
E3 42391332 42391428
refGene_gn=KCTD5;refGene_tx=XM_023246972.1;refGene_gn=KCTD5;refGene_tx
=XM_023246973.1
E3 42393377 42393469
refGene_gn=KCTD5;refGene_tx=XM_023246972.1;refGene_gn=KCTD5;refGene_tx
=XM_023246973.1
E3 42395187 42395296
refGene_gn=KCTD5;refGene_tx=XM_023246972.1;refGene_gn=KCTD5;refGene_tx
=XM_023246973.1
E3 42405196 42405448
refGene_gn=KCTD5;refGene_tx=XM_023246972.1;refGene_gn=KCTD5;refGene_tx
=XM_023246973.1
E3 42411388 42411505
refGene_gn=PDPK1;refGene_tx=XM_011290686.3;refGene_gn=PDPK1;refGene_tx
=XM_011290687.3;refGene_gn=PDPK1;refGene_tx=XM_019821153.2;refGene_gn=

PDPK1;refGene_tx=XM_019821155.2;refGene_gn=PDPK1;refGene_tx=XM_019821156.2;refGene_gn=PDPK1;refGene_tx=XM_019821157.2;refGene_gn=PDPK1;refGene_tx=XM_023246952.1;refGene_gn=PDPK1;refGene_tx=XM_023246953.1;refGene_gn=PDPK1;refGene_tx=XM_023246955.1
E3 42411812 42411965
refGene_gn=PDPK1;refGene_tx=XM_011290686.3;refGene_gn=PDPK1;refGene_tx=XM_011290687.3;refGene_gn=PDPK1;refGene_tx=XM_019821153.2;refGene_gn=PDPK1;refGene_tx=XM_019821155.2;refGene_gn=PDPK1;refGene_tx=XM_019821156.2;refGene_gn=PDPK1;refGene_tx=XM_019821157.2;refGene_gn=PDPK1;refGene_tx=XM_023246952.1;refGene_gn=PDPK1;refGene_tx=XM_023246953.1;refGene_gn=PDPK1;refGene_tx=XM_023246955.1
E3 42412817 42412875
refGene_gn=PDPK1;refGene_tx=XM_011290686.3;refGene_gn=PDPK1;refGene_tx=XM_011290687.3;refGene_gn=PDPK1;refGene_tx=XM_019821153.2;refGene_gn=PDPK1;refGene_tx=XM_019821155.2;refGene_gn=PDPK1;refGene_tx=XM_019821156.2;refGene_gn=PDPK1;refGene_tx=XM_019821157.2;refGene_gn=PDPK1;refGene_tx=XM_023246952.1;refGene_gn=PDPK1;refGene_tx=XM_023246953.1;refGene_gn=PDPK1;refGene_tx=XM_023246955.1
E3 42421468 42421686
refGene_gn=PDPK1;refGene_tx=XM_011290686.3;refGene_gn=PDPK1;refGene_tx=XM_011290687.3;refGene_gn=PDPK1;refGene_tx=XM_019821153.2;refGene_gn=PDPK1;refGene_tx=XM_019821155.2;refGene_gn=PDPK1;refGene_tx=XM_019821156.2;refGene_gn=PDPK1;refGene_tx=XM_019821157.2;refGene_gn=PDPK1;refGene_tx=XM_023246952.1;refGene_gn=PDPK1;refGene_tx=XM_023246953.1;refGene_gn=PDPK1;refGene_tx=XM_023246955.1
E3 42425099 42425273
refGene_gn=PDPK1;refGene_tx=XM_011290686.3;refGene_gn=PDPK1;refGene_tx=XM_011290687.3;refGene_gn=PDPK1;refGene_tx=XM_019821153.2;refGene_gn=PDPK1;refGene_tx=XM_019821155.2;refGene_gn=PDPK1;refGene_tx=XM_019821156.2;refGene_gn=PDPK1;refGene_tx=XM_019821157.2;refGene_gn=PDPK1;refGene_tx=XM_023246952.1;refGene_gn=PDPK1;refGene_tx=XM_023246953.1;refGene_gn=PDPK1;refGene_tx=XM_023246955.1
E3 42426748 42426845
refGene_gn=PDPK1;refGene_tx=XM_011290686.3;refGene_gn=PDPK1;refGene_tx=XM_011290687.3;refGene_gn=PDPK1;refGene_tx=XM_019821153.2;refGene_gn=PDPK1;refGene_tx=XM_019821155.2;refGene_gn=PDPK1;refGene_tx=XM_019821156.2;refGene_gn=PDPK1;refGene_tx=XM_019821157.2;refGene_gn=PDPK1;refGene_tx=XM_023246952.1;refGene_gn=PDPK1;refGene_tx=XM_023246953.1;refGene_gn=PDPK1;refGene_tx=XM_023246955.1
E3 42427055 42427124
refGene_gn=PDPK1;refGene_tx=XM_011290686.3;refGene_gn=PDPK1;refGene_tx=XM_011290687.3;refGene_gn=PDPK1;refGene_tx=XM_019821153.2;refGene_gn=PDPK1;refGene_tx=XM_019821155.2;refGene_gn=PDPK1;refGene_tx=XM_019821156.2;refGene_gn=PDPK1;refGene_tx=XM_019821157.2;refGene_gn=PDPK1;refGene_tx=XM_023246952.1;refGene_gn=PDPK1;refGene_tx=XM_023246953.1;refGene_gn=PDPK1;refGene_tx=XM_023246955.1
E3 42430976 42431052
refGene_gn=PDPK1;refGene_tx=XM_011290686.3;refGene_gn=PDPK1;refGene_tx=XM_011290687.3;refGene_gn=PDPK1;refGene_tx=XM_019821153.2;refGene_gn=PDPK1;refGene_tx=XM_019821155.2;refGene_gn=PDPK1;refGene_tx=XM_0198211

56.2;refGene_gn=PDPK1;refGene_tx=XM_019821157.2;refGene_gn=PDPK1;refGene_tx=XM_023246952.1;refGene_gn=PDPK1;refGene_tx=XM_023246953.1;refGene_gn=PDPK1;refGene_tx=XM_023246955.1
E3 42440937 42441035
refGene_gn=PDPK1;refGene_tx=XM_011290686.3;refGene_gn=PDPK1;refGene_tx=XM_011290687.3;refGene_gn=PDPK1;refGene_tx=XM_019821153.2;refGene_gn=PDPK1;refGene_tx=XM_019821155.2;refGene_gn=PDPK1;refGene_tx=XM_019821156.2;refGene_gn=PDPK1;refGene_tx=XM_019821157.2;refGene_gn=PDPK1;refGene_tx=XM_023246952.1;refGene_gn=PDPK1;refGene_tx=XM_023246953.1;refGene_gn=PDPK1;refGene_tx=XM_023246955.1
E3 42441789 42441934
refGene_gn=PDPK1;refGene_tx=XM_011290686.3;refGene_gn=PDPK1;refGene_tx=XM_011290687.3;refGene_gn=PDPK1;refGene_tx=XM_019821153.2;refGene_gn=PDPK1;refGene_tx=XM_019821155.2;refGene_gn=PDPK1;refGene_tx=XM_019821156.2;refGene_gn=PDPK1;refGene_tx=XM_019821157.2;refGene_gn=PDPK1;refGene_tx=XM_023246952.1;refGene_gn=PDPK1;refGene_tx=XM_023246953.1;refGene_gn=PDPK1;refGene_tx=XM_023246955.1
E3 42444983 42445121
refGene_gn=PDPK1;refGene_tx=XM_011290686.3;refGene_gn=PDPK1;refGene_tx=XM_011290687.3;refGene_gn=PDPK1;refGene_tx=XM_019821153.2;refGene_gn=PDPK1;refGene_tx=XM_019821155.2;refGene_gn=PDPK1;refGene_tx=XM_019821156.2;refGene_gn=PDPK1;refGene_tx=XM_019821157.2;refGene_gn=PDPK1;refGene_tx=XM_023246952.1;refGene_gn=PDPK1;refGene_tx=XM_023246953.1;refGene_gn=PDPK1;refGene_tx=XM_023246955.1
E3 42445390 42445433
refGene_gn=PDPK1;refGene_tx=XM_011290686.3;refGene_gn=PDPK1;refGene_tx=XM_011290687.3;refGene_gn=PDPK1;refGene_tx=XM_019821153.2;refGene_gn=PDPK1;refGene_tx=XM_019821155.2;refGene_gn=PDPK1;refGene_tx=XM_019821156.2;refGene_gn=PDPK1;refGene_tx=XM_019821157.2;refGene_gn=PDPK1;refGene_tx=XM_023246952.1;refGene_gn=PDPK1;refGene_tx=XM_023246953.1;refGene_gn=PDPK1;refGene_tx=XM_023246955.1
E3 42449513 42449780
refGene_gn=PDPK1;refGene_tx=XM_011290686.3;refGene_gn=PDPK1;refGene_tx=XM_011290687.3;refGene_gn=PDPK1;refGene_tx=XM_019821153.2;refGene_gn=PDPK1;refGene_tx=XM_019821155.2;refGene_gn=PDPK1;refGene_tx=XM_019821156.2;refGene_gn=PDPK1;refGene_tx=XM_019821157.2;refGene_gn=PDPK1;refGene_tx=XM_023246952.1;refGene_gn=PDPK1;refGene_tx=XM_023246953.1;refGene_gn=PDPK1;refGene_tx=XM_023246955.1
E3 42452803 42452848 refGene_gn=PDPK1;refGene_tx=XM_019821153.2
E3 42481033 42481057 refGene_gn=PDPK1;refGene_tx=XM_011290686.3
E3 42491660 42491749 refGene_gn=AMDHD2;refGene_tx=XM_023246961.1
E3 42492101 42492203 refGene_gn=AMDHD2;refGene_tx=XM_023246961.1
E3 42493203 42493272 refGene_gn=AMDHD2;refGene_tx=XM_023246961.1
E3 42493432 42493540 refGene_gn=AMDHD2;refGene_tx=XM_023246961.1
E3 42493614 42493759 refGene_gn=AMDHD2;refGene_tx=XM_023246961.1
E3 42493857 42493946 refGene_gn=AMDHD2;refGene_tx=XM_023246961.1
E3 42494027 42494240 refGene_gn=AMDHD2;refGene_tx=XM_023246961.1
E3 42494413 42494468 refGene_gn=AMDHD2;refGene_tx=XM_023246961.1
E3 42500295 42500435 refGene_gn=AMDHD2;refGene_tx=XM_023246961.1
E3 42500514 42500651 refGene_gn=AMDHD2;refGene_tx=XM_023246961.1

E3 42501491 42501574 refGene_gn=AMDHD2;refGene_tx=XM_023246961.1
E3 42502378 42502583 refGene_gn=ATP6V0C;refGene_tx=XM_003998932.4
E3 42502732 42502916 refGene_gn=ATP6V0C;refGene_tx=XM_003998932.4
E3 42506629 42506708 refGene_gn=ATP6V0C;refGene_tx=XM_003998932.4
E3 42517282 42517437
refGene_gn=TBC1D24;refGene_tx=XM_019821158.2;refGene_gn=TBC1D24;refGen
e_tx=XM_019821159.2
E3 42517803 42518026
refGene_gn=TBC1D24;refGene_tx=XM_019821158.2;refGene_gn=TBC1D24;refGen
e_tx=XM_019821159.2
E3 42518245 42518341
refGene_gn=TBC1D24;refGene_tx=XM_019821158.2;refGene_gn=TBC1D24;refGen
e_tx=XM_019821159.2
E3 42519303 42519367
refGene_gn=TBC1D24;refGene_tx=XM_019821158.2;refGene_gn=TBC1D24;refGen
e_tx=XM_019821159.2
E3 42520447 42520606
refGene_gn=TBC1D24;refGene_tx=XM_019821158.2;refGene_gn=TBC1D24;refGen
e_tx=XM_019821159.2
E3 42521100 42521118 refGene_gn=TBC1D24;refGene_tx=XM_019821158.2
E3 42521754 42522719
refGene_gn=TBC1D24;refGene_tx=XM_019821158.2;refGene_gn=TBC1D24;refGen
e_tx=XM_019821159.2
E3 42540364 42540399
refGene_gn=NTN3;refGene_tx=XM_023246238.1;refGene_gn=NTN3;refGene_tx=X
M_023246239.1
E3 42540882 42541223 refGene_gn=NTN3;refGene_tx=XM_023246237.1
E3 42541436 42541529
refGene_gn=NTN3;refGene_tx=XM_023246237.1;refGene_gn=NTN3;refGene_tx=X
M_023246238.1;refGene_gn=NTN3;refGene_tx=XM_023246239.1
E3 42541643 42541694
refGene_gn=NTN3;refGene_tx=XM_023246237.1;refGene_gn=NTN3;refGene_tx=X
M_023246238.1;refGene_gn=NTN3;refGene_tx=XM_023246239.1
E3 42541853 42542003
refGene_gn=NTN3;refGene_tx=XM_023246237.1;refGene_gn=NTN3;refGene_tx=X
M_023246238.1;refGene_gn=NTN3;refGene_tx=XM_023246239.1
E3 42542093 42542282
refGene_gn=NTN3;refGene_tx=XM_023246237.1;refGene_gn=NTN3;refGene_tx=X
M_023246238.1;refGene_gn=NTN3;refGene_tx=XM_023246239.1
E3 42542353 42543281
refGene_gn=NTN3;refGene_tx=XM_023246237.1;refGene_gn=NTN3;refGene_tx=X
M_023246238.1;refGene_gn=NTN3;refGene_tx=XM_023246239.1
E3 42549949 42550087
refGene_gn=TEDC2;refGene_tx=XM_011290690.3;refGene_gn=TEDC2;refGene_tx
=XM_023246496.1;refGene_gn=TEDC2;refGene_tx=XM_023246497.1
E3 42550340 42550531
refGene_gn=TEDC2;refGene_tx=XM_011290690.3;refGene_gn=TEDC2;refGene_tx
=XM_023246496.1;refGene_gn=TEDC2;refGene_tx=XM_023246497.1
E3 42550969 42551071
refGene_gn=TEDC2;refGene_tx=XM_011290690.3;refGene_gn=TEDC2;refGene_tx

=XM_023246496.1;refGene_gn=TEDC2;refGene_tx=XM_023246497.1
E3 42551159 42551271
refGene_gn=TEDC2;refGene_tx=XM_011290690.3;refGene_gn=TEDC2;refGene_tx=XM_023246496.1;refGene_gn=TEDC2;refGene_tx=XM_023246497.1
E3 42551450 42551550
refGene_gn=TEDC2;refGene_tx=XM_011290690.3;refGene_gn=TEDC2;refGene_tx=XM_023246496.1;refGene_gn=TEDC2;refGene_tx=XM_023246497.1
E3 42552169 42552331
refGene_gn=TEDC2;refGene_tx=XM_011290690.3;refGene_gn=TEDC2;refGene_tx=XM_023246496.1;refGene_gn=TEDC2;refGene_tx=XM_023246497.1
E3 42552608 42553014
refGene_gn=TEDC2;refGene_tx=XM_011290690.3;refGene_gn=TEDC2;refGene_tx=XM_023246496.1;refGene_gn=TEDC2;refGene_tx=XM_023246497.1
E3 42553117 42553188
refGene_gn=TEDC2;refGene_tx=XM_011290690.3;refGene_gn=TEDC2;refGene_tx=XM_023246496.1;refGene_gn=TEDC2;refGene_tx=XM_023246497.1
E3 42553356 42553455
refGene_gn=TEDC2;refGene_tx=XM_011290690.3;refGene_gn=TEDC2;refGene_tx=XM_023246496.1;refGene_gn=TEDC2;refGene_tx=XM_023246497.1
E3 42553545 42553571
refGene_gn=TEDC2;refGene_tx=XM_011290690.3;refGene_gn=TEDC2;refGene_tx=XM_023246496.1;refGene_gn=TEDC2;refGene_tx=XM_023246497.1
E3 42556790 42557284 refGene_gn=CCNF;refGene_tx=XM_023246494.1
E3 42557919 42558089 refGene_gn=CCNF;refGene_tx=XM_023246494.1
E3 42558813 42558941 refGene_gn=CCNF;refGene_tx=XM_023246494.1
E3 42559017 42559117 refGene_gn=CCNF;refGene_tx=XM_023246494.1
E3 42560845 42560933 refGene_gn=CCNF;refGene_tx=XM_023246494.1
E3 42561270 42561451 refGene_gn=CCNF;refGene_tx=XM_023246494.1
E3 42563048 42563172 refGene_gn=CCNF;refGene_tx=XM_023246494.1
E3 42564933 42565098 refGene_gn=CCNF;refGene_tx=XM_023246494.1
E3 42567548 42567700 refGene_gn=CCNF;refGene_tx=XM_023246494.1
E3 42569753 42569831 refGene_gn=CCNF;refGene_tx=XM_023246494.1
E3 42570112 42570217 refGene_gn=CCNF;refGene_tx=XM_023246494.1
E3 42570867 42570921 refGene_gn=CCNF;refGene_tx=XM_023246494.1
E3 42571240 42571494 refGene_gn=CCNF;refGene_tx=XM_023246494.1
E3 42573113 42573181 refGene_gn=CCNF;refGene_tx=XM_023246494.1
E3 42574176 42574283 refGene_gn=CCNF;refGene_tx=XM_023246494.1
E3 42575504 42575659 refGene_gn=CCNF;refGene_tx=XM_023246494.1
E3 42576451 42576467 refGene_gn=CCNF;refGene_tx=XM_023246494.1
E3 42579499 42579739
refGene_gn=LOC101081185;refGene_tx=XM_023246589.1
E3 42580249 42580323
refGene_gn=LOC101081185;refGene_tx=XM_023246589.1
E3 42580464 42580655
refGene_gn=LOC101081185;refGene_tx=XM_023246589.1
E3 42580921 42581092
refGene_gn=LOC101081185;refGene_tx=XM_023246589.1
E3 42582114 42582302
refGene_gn=LOC101081185;refGene_tx=XM_023246589.1
E3 42588641 42588836

refGene_gn=LOC101081185;refGene_tx=XM_023246589.1
E3 42589630 42589756
refGene_gn=LOC101081185;refGene_tx=XM_023246589.1
E3 42590356 42590535
refGene_gn=LOC101081185;refGene_tx=XM_023246589.1
E3 42593050 42593209
refGene_gn=LOC101081185;refGene_tx=XM_023246589.1
E3 42598213 42598433
refGene_gn=LOC101081185;refGene_tx=XM_023246589.1
E3 42600229 42600434
refGene_gn=LOC101081185;refGene_tx=XM_023246589.1
E3 42600605 42600879
refGene_gn=LOC101081185;refGene_tx=XM_023246589.1
E3 42602536 42602840
refGene_gn=LOC101081185;refGene_tx=XM_023246589.1
E3 42604636 42604799
refGene_gn=LOC101081185;refGene_tx=XM_023246589.1
E3 42616373 42616472
refGene_gn=LOC101081185;refGene_tx=XM_023246589.1
E3 42622020 42622171
refGene_gn=LOC101081185;refGene_tx=XM_023246589.1
E3 42623825 42624014
refGene_gn=LOC101081185;refGene_tx=XM_023246589.1
E3 42624016 42624035
refGene_gn=LOC101081185;refGene_tx=XM_023246589.1
E3 42624141 42624297
refGene_gn=LOC101081185;refGene_tx=XM_023246589.1
E3 42625677 42625832
refGene_gn=LOC101081185;refGene_tx=XM_023246589.1
E3 42628766 42628908
refGene_gn=LOC101081185;refGene_tx=XM_023246589.1
E3 42629996 42630140
refGene_gn=LOC101081185;refGene_tx=XM_023246589.1
E3 42630525 42630920
refGene_gn=LOC101081185;refGene_tx=XM_023246589.1
E3 42636334 42636508
refGene_gn=LOC101081185;refGene_tx=XM_023246589.1
E3 42644686 42644804
refGene_gn=LOC101081185;refGene_tx=XM_023246589.1
E3 42646165 42646282
refGene_gn=LOC101081185;refGene_tx=XM_023246589.1
E3 42646945 42647205
refGene_gn=LOC101081185;refGene_tx=XM_023246589.1
E3 42648546 42648709
refGene_gn=LOC101081185;refGene_tx=XM_023246589.1
E3 42649764 42649892
refGene_gn=LOC101081185;refGene_tx=XM_023246589.1
E3 42654005 42654264
refGene_gn=LOC101081185;refGene_tx=XM_023246589.1
E3 42654654 42654702

refGene_gn=LOC101081185;refGene_tx=XM_023246589.1
E3 42677613 42677667
refGene_gn=ABCA3;refGene_tx=XM_023247118.1;refGene_gn=ABCA3;refGene_tx
=XM_023247119.1
E3 42677764 42678029
refGene_gn=ABCA3;refGene_tx=XM_023247118.1;refGene_gn=ABCA3;refGene_tx
=XM_023247119.1
E3 42678845 42678973
refGene_gn=ABCA3;refGene_tx=XM_023247118.1;refGene_gn=ABCA3;refGene_tx
=XM_023247119.1
E3 42679653 42679819
refGene_gn=ABCA3;refGene_tx=XM_023247118.1;refGene_gn=ABCA3;refGene_tx
=XM_023247119.1
E3 42683657 42683917
refGene_gn=ABCA3;refGene_tx=XM_023247118.1;refGene_gn=ABCA3;refGene_tx
=XM_023247119.1
E3 42685018 42685135
refGene_gn=ABCA3;refGene_tx=XM_023247118.1;refGene_gn=ABCA3;refGene_tx
=XM_023247119.1
E3 42685380 42685501
refGene_gn=ABCA3;refGene_tx=XM_023247118.1;refGene_gn=ABCA3;refGene_tx
=XM_023247119.1
E3 42688201 42688375
refGene_gn=ABCA3;refGene_tx=XM_023247118.1;refGene_gn=ABCA3;refGene_tx
=XM_023247119.1
E3 42691598 42691780
refGene_gn=ABCA3;refGene_tx=XM_023247118.1;refGene_gn=ABCA3;refGene_tx
=XM_023247119.1
E3 42694416 42694560
refGene_gn=ABCA3;refGene_tx=XM_023247118.1;refGene_gn=ABCA3;refGene_tx
=XM_023247119.1
E3 42695012 42695142
refGene_gn=ABCA3;refGene_tx=XM_023247118.1;refGene_gn=ABCA3;refGene_tx
=XM_023247119.1
E3 42695282 42695437
refGene_gn=ABCA3;refGene_tx=XM_023247118.1;refGene_gn=ABCA3;refGene_tx
=XM_023247119.1
E3 42695675 42695831
refGene_gn=ABCA3;refGene_tx=XM_023247118.1;refGene_gn=ABCA3;refGene_tx
=XM_023247119.1
E3 42695954 42696165
refGene_gn=ABCA3;refGene_tx=XM_023247118.1;refGene_gn=ABCA3;refGene_tx
=XM_023247119.1
E3 42696893 42697044
refGene_gn=ABCA3;refGene_tx=XM_023247118.1;refGene_gn=ABCA3;refGene_tx
=XM_023247119.1
E3 42698720 42698819
refGene_gn=ABCA3;refGene_tx=XM_023247118.1;refGene_gn=ABCA3;refGene_tx
=XM_023247119.1
E3 42699742 42699929

refGene_gn=ABCA3;refGene_tx=XM_023247118.1;refGene_gn=ABCA3;refGene_tx=XM_023247119.1
E3 42700624 42700928
refGene_gn=ABCA3;refGene_tx=XM_023247118.1;refGene_gn=ABCA3;refGene_tx=XM_023247119.1
E3 42702719 42702993
refGene_gn=ABCA3;refGene_tx=XM_023247118.1;refGene_gn=ABCA3;refGene_tx=XM_023247119.1
E3 42706196 42706401
refGene_gn=ABCA3;refGene_tx=XM_023247118.1;refGene_gn=ABCA3;refGene_tx=XM_023247119.1
E3 42706729 42706949
refGene_gn=ABCA3;refGene_tx=XM_023247118.1;refGene_gn=ABCA3;refGene_tx=XM_023247119.1
E3 42707232 42707391
refGene_gn=ABCA3;refGene_tx=XM_023247118.1;refGene_gn=ABCA3;refGene_tx=XM_023247119.1
E3 42708090 42708263
refGene_gn=ABCA3;refGene_tx=XM_023247118.1;refGene_gn=ABCA3;refGene_tx=XM_023247119.1
E3 42708716 42708845
refGene_gn=ABCA3;refGene_tx=XM_023247118.1;refGene_gn=ABCA3;refGene_tx=XM_023247119.1
E3 42708925 42709120
refGene_gn=ABCA3;refGene_tx=XM_023247118.1;refGene_gn=ABCA3;refGene_tx=XM_023247119.1
E3 42710700 42710888
refGene_gn=ABCA3;refGene_tx=XM_023247118.1;refGene_gn=ABCA3;refGene_tx=XM_023247119.1
E3 42711358 42711529
refGene_gn=ABCA3;refGene_tx=XM_023247118.1;refGene_gn=ABCA3;refGene_tx=XM_023247119.1
E3 42711804 42711995
refGene_gn=ABCA3;refGene_tx=XM_023247118.1;refGene_gn=ABCA3;refGene_tx=XM_023247119.1
E3 42712136 42712210
refGene_gn=ABCA3;refGene_tx=XM_023247118.1;refGene_gn=ABCA3;refGene_tx=XM_023247119.1
E3 42712652 42712784
refGene_gn=ABCA3;refGene_tx=XM_023247118.1;refGene_gn=ABCA3;refGene_tx=XM_023247119.1
E3 42720000 42720002 refGene_gn=RNPS1;refGene_tx=XM_011290697.3
E3 42723210 42723281
refGene_gn=RNPS1;refGene_tx=XM_011290692.3;refGene_gn=RNPS1;refGene_tx=XM_011290693.3;refGene_gn=RNPS1;refGene_tx=XM_011290696.3;refGene_gn=RNPS1;refGene_tx=XM_019821167.2
E3 42723518 42723674
refGene_gn=RNPS1;refGene_tx=XM_011290692.3;refGene_gn=RNPS1;refGene_tx=XM_011290693.3;refGene_gn=RNPS1;refGene_tx=XM_011290696.3;refGene_gn=RNPS1;refGene_tx=XM_011290697.3;refGene_gn=RNPS1;refGene_tx=XM_0198211

67.2
E3 42724332 42724524
refGene_gn=RNPS1;refGene_tx=XM_011290692.3;refGene_gn=RNPS1;refGene_tx=XM_011290693.3;refGene_gn=RNPS1;refGene_tx=XM_011290696.3;refGene_gn=RNPS1;refGene_tx=XM_011290697.3;refGene_gn=RNPS1;refGene_tx=XM_019821167.2
E3 42724851 42724954
refGene_gn=RNPS1;refGene_tx=XM_011290692.3;refGene_gn=RNPS1;refGene_tx=XM_011290693.3;refGene_gn=RNPS1;refGene_tx=XM_011290696.3;refGene_gn=RNPS1;refGene_tx=XM_011290697.3;refGene_gn=RNPS1;refGene_tx=XM_019821167.2
E3 42725210 42725364
refGene_gn=RNPS1;refGene_tx=XM_011290692.3;refGene_gn=RNPS1;refGene_tx=XM_011290693.3;refGene_gn=RNPS1;refGene_tx=XM_011290696.3;refGene_gn=RNPS1;refGene_tx=XM_011290697.3;refGene_gn=RNPS1;refGene_tx=XM_019821167.2
E3 42727825 42727967
refGene_gn=RNPS1;refGene_tx=XM_011290692.3;refGene_gn=RNPS1;refGene_tx=XM_011290693.3;refGene_gn=RNPS1;refGene_tx=XM_011290696.3;refGene_gn=RNPS1;refGene_tx=XM_011290697.3;refGene_gn=RNPS1;refGene_tx=XM_019821167.2
E3 42728458 42728558
refGene_gn=RNPS1;refGene_tx=XM_011290692.3;refGene_gn=RNPS1;refGene_tx=XM_011290693.3;refGene_gn=RNPS1;refGene_tx=XM_011290696.3;refGene_gn=RNPS1;refGene_tx=XM_011290697.3;refGene_gn=RNPS1;refGene_tx=XM_019821167.2
E3 42730525 42730577 refGene_gn=ECI1;refGene_tx=XM_003998937.5
E3 42730790 42730904 refGene_gn=ECI1;refGene_tx=XM_003998937.5
E3 42732187 42732244 refGene_gn=DNASE1L2;refGene_tx=XM_019821169.2
E3 42733283 42733411 refGene_gn=ECI1;refGene_tx=XM_003998937.5
E3 42736280 42736427 refGene_gn=ECI1;refGene_tx=XM_003998937.5
E3 42737284 42737406 refGene_gn=ECI1;refGene_tx=XM_003998937.5
E3 42737493 42737672 refGene_gn=ECI1;refGene_tx=XM_003998937.5
E3 42739024 42739191 refGene_gn=ECI1;refGene_tx=XM_003998937.5
E3 42740636 42740690 refGene_gn=DNASE1L2;refGene_tx=XM_019821170.2
E3 42741038 42741290
refGene_gn=DNASE1L2;refGene_tx=XM_019821169.2;refGene_gn=DNASE1L2;refGene_tx=XM_019821170.2
E3 42741385 42741498
refGene_gn=DNASE1L2;refGene_tx=XM_019821169.2;refGene_gn=DNASE1L2;refGene_tx=XM_019821170.2
E3 42741576 42741677
refGene_gn=DNASE1L2;refGene_tx=XM_019821169.2;refGene_gn=DNASE1L2;refGene_tx=XM_019821170.2
E3 42741758 42741842
refGene_gn=DNASE1L2;refGene_tx=XM_019821169.2;refGene_gn=DNASE1L2;refGene_tx=XM_019821170.2
E3 42741935 42742024
refGene_gn=DNASE1L2;refGene_tx=XM_019821169.2;refGene_gn=DNASE1L2;refGene_tx=XM_019821170.2

E3 42742116 42742260
refGene_gn=DNASE1L2;refGene_tx=XM_019821169.2;refGene_gn=DNASE1L2;refGene_tx=XM_019821170.2
E3 42743528 42743885
refGene_gn=E4F1;refGene_tx=XM_011290698.3;refGene_gn=E4F1;refGene_tx=XM_011290699.3;refGene_gn=E4F1;refGene_tx=XM_011290700.3;refGene_gn=E4F1;refGene_tx=XM_011290701.3;refGene_gn=E4F1;refGene_tx=XM_011290702.3
E3 42743974 42744037
refGene_gn=E4F1;refGene_tx=XM_011290698.3;refGene_gn=E4F1;refGene_tx=XM_011290699.3;refGene_gn=E4F1;refGene_tx=XM_011290700.3;refGene_gn=E4F1;refGene_tx=XM_011290701.3;refGene_gn=E4F1;refGene_tx=XM_011290702.3
E3 42744117 42744260
refGene_gn=E4F1;refGene_tx=XM_011290698.3;refGene_gn=E4F1;refGene_tx=XM_011290699.3;refGene_gn=E4F1;refGene_tx=XM_011290700.3;refGene_gn=E4F1;refGene_tx=XM_011290701.3;refGene_gn=E4F1;refGene_tx=XM_011290702.3
E3 42744335 42744534
refGene_gn=E4F1;refGene_tx=XM_011290698.3;refGene_gn=E4F1;refGene_tx=XM_011290699.3;refGene_gn=E4F1;refGene_tx=XM_011290700.3;refGene_gn=E4F1;refGene_tx=XM_011290701.3;refGene_gn=E4F1;refGene_tx=XM_011290702.3
E3 42744692 42744910
refGene_gn=E4F1;refGene_tx=XM_011290698.3;refGene_gn=E4F1;refGene_tx=XM_011290699.3;refGene_gn=E4F1;refGene_tx=XM_011290700.3;refGene_gn=E4F1;refGene_tx=XM_011290701.3;refGene_gn=E4F1;refGene_tx=XM_011290702.3
E3 42745006 42745115
refGene_gn=E4F1;refGene_tx=XM_011290698.3;refGene_gn=E4F1;refGene_tx=XM_011290699.3;refGene_gn=E4F1;refGene_tx=XM_011290700.3;refGene_gn=E4F1;refGene_tx=XM_011290701.3;refGene_gn=E4F1;refGene_tx=XM_011290702.3
E3 42745386 42745596
refGene_gn=E4F1;refGene_tx=XM_011290698.3;refGene_gn=E4F1;refGene_tx=XM_011290699.3;refGene_gn=E4F1;refGene_tx=XM_011290700.3;refGene_gn=E4F1;refGene_tx=XM_011290701.3;refGene_gn=E4F1;refGene_tx=XM_011290702.3
E3 42745918 42746088
refGene_gn=E4F1;refGene_tx=XM_011290698.3;refGene_gn=E4F1;refGene_tx=XM_011290699.3;refGene_gn=E4F1;refGene_tx=XM_011290700.3;refGene_gn=E4F1;refGene_tx=XM_011290701.3;refGene_gn=E4F1;refGene_tx=XM_011290702.3
E3 42746172 42746334
refGene_gn=E4F1;refGene_tx=XM_011290698.3;refGene_gn=E4F1;refGene_tx=XM_011290699.3;refGene_gn=E4F1;refGene_tx=XM_011290700.3;refGene_gn=E4F1;refGene_tx=XM_011290701.3;refGene_gn=E4F1;refGene_tx=XM_011290702.3
E3 42746453 42746574
refGene_gn=E4F1;refGene_tx=XM_011290698.3;refGene_gn=E4F1;refGene_tx=XM_011290699.3;refGene_gn=E4F1;refGene_tx=XM_011290700.3;refGene_gn=E4F1;refGene_tx=XM_011290701.3;refGene_gn=E4F1;refGene_tx=XM_011290702.3
E3 42746680 42746874
refGene_gn=E4F1;refGene_tx=XM_011290698.3;refGene_gn=E4F1;refGene_tx=XM_011290699.3;refGene_gn=E4F1;refGene_tx=XM_011290700.3;refGene_gn=E4F1;refGene_tx=XM_011290701.3;refGene_gn=E4F1;refGene_tx=XM_011290702.3
E3 42749087 42749193
refGene_gn=E4F1;refGene_tx=XM_011290698.3;refGene_gn=E4F1;refGene_tx=XM_011290699.3;refGene_gn=E4F1;refGene_tx=XM_011290700.3;refGene_gn=E4F

1;refGene_tx=XM_011290701.3;refGene_gn=E4F1;refGene_tx=XM_011290702.3
E3 42750460 42750612
refGene_gn=E4F1;refGene_tx=XM_011290698.3;refGene_gn=E4F1;refGene_tx=XM_011290699.3;refGene_gn=E4F1;refGene_tx=XM_011290700.3;refGene_gn=E4F1;refGene_tx=XM_011290701.3;refGene_gn=E4F1;refGene_tx=XM_011290702.3
E3 42752878 42753035
refGene_gn=E4F1;refGene_tx=XM_011290698.3;refGene_gn=E4F1;refGene_tx=XM_011290699.3;refGene_gn=E4F1;refGene_tx=XM_011290700.3;refGene_gn=E4F1;refGene_tx=XM_011290701.3;refGene_gn=E4F1;refGene_tx=XM_011290702.3
E3 42758437 42759080 refGene_gn=PGP;refGene_tx=XM_011290704.2
E3 42759181 42759507 refGene_gn=PGP;refGene_tx=XM_011290704.2
E3 42762429 42762552 refGene_gn=BRICD5;refGene_tx=XM_006942454.4
E3 42762761 42762917 refGene_gn=BRICD5;refGene_tx=XM_006942454.4
E3 42762990 42763092 refGene_gn=BRICD5;refGene_tx=XM_006942454.4
E3 42763358 42763512 refGene_gn=BRICD5;refGene_tx=XM_006942454.4
E3 42763610 42763705 refGene_gn=BRICD5;refGene_tx=XM_006942454.4
E3 42764161 42764280
refGene_gn=MLST8;refGene_tx=XM_011290705.3;refGene_gn=MLST8;refGene_tx=XM_011290706.3
E3 42764382 42764546
refGene_gn=MLST8;refGene_tx=XM_011290705.3;refGene_gn=MLST8;refGene_tx=XM_011290706.3
E3 42764661 42764786
refGene_gn=MLST8;refGene_tx=XM_011290705.3;refGene_gn=MLST8;refGene_tx=XM_011290706.3
E3 42765612 42765765
refGene_gn=MLST8;refGene_tx=XM_011290705.3;refGene_gn=MLST8;refGene_tx=XM_011290706.3
E3 42765849 42765925
refGene_gn=MLST8;refGene_tx=XM_011290705.3;refGene_gn=MLST8;refGene_tx=XM_011290706.3
E3 42766259 42766422
refGene_gn=MLST8;refGene_tx=XM_011290705.3;refGene_gn=MLST8;refGene_tx=XM_011290706.3
E3 42766506 42766558
refGene_gn=MLST8;refGene_tx=XM_011290705.3;refGene_gn=MLST8;refGene_tx=XM_011290706.3
E3 42766716 42766845
refGene_gn=MLST8;refGene_tx=XM_011290705.3;refGene_gn=MLST8;refGene_tx=XM_011290706.3
E3 42773380 42773474 refGene_gn=CASKIN1;refGene_tx=XM_011290707.3
E3 42778535 42778587 refGene_gn=CASKIN1;refGene_tx=XM_011290707.3
E3 42778697 42778795 refGene_gn=CASKIN1;refGene_tx=XM_011290707.3
E3 42779003 42779149 refGene_gn=CASKIN1;refGene_tx=XM_011290707.3
E3 42779248 42779344 refGene_gn=CASKIN1;refGene_tx=XM_011290707.3
E3 42779472 42779603 refGene_gn=CASKIN1;refGene_tx=XM_011290707.3
E3 42780039 42780148 refGene_gn=CASKIN1;refGene_tx=XM_011290707.3
E3 42780227 42780336 refGene_gn=CASKIN1;refGene_tx=XM_011290707.3
E3 42780446 42780541 refGene_gn=CASKIN1;refGene_tx=XM_011290707.3
E3 42780704 42780822 refGene_gn=CASKIN1;refGene_tx=XM_011290707.3

E3 42781759 42781861 refGene_gn=CASKIN1;refGene_tx=XM_011290707.3
E3 42781955 42782044 refGene_gn=CASKIN1;refGene_tx=XM_011290707.3
E3 42782120 42782205 refGene_gn=CASKIN1;refGene_tx=XM_011290707.3
E3 42782291 42782383 refGene_gn=CASKIN1;refGene_tx=XM_011290707.3
E3 42783196 42783307 refGene_gn=CASKIN1;refGene_tx=XM_011290707.3
E3 42783396 42783498 refGene_gn=CASKIN1;refGene_tx=XM_011290707.3
E3 42784874 42785013 refGene_gn=CASKIN1;refGene_tx=XM_011290707.3
E3 42785093 42787160 refGene_gn=CASKIN1;refGene_tx=XM_011290707.3
E3 42787816 42788252 refGene_gn=CASKIN1;refGene_tx=XM_011290707.3
E3 42788518 42788618 refGene_gn=CASKIN1;refGene_tx=XM_011290707.3
E3 42790791 42790806
refGene_gn=TRAF7;refGene_tx=XM_003998941.5;refGene_gn=TRAF7;refGene_tx=XM_006942456.4;refGene_gn=TRAF7;refGene_tx=XM_019821171.2
E3 42790977 42791097
refGene_gn=TRAF7;refGene_tx=XM_003998941.5;refGene_gn=TRAF7;refGene_tx=XM_006942456.4;refGene_gn=TRAF7;refGene_tx=XM_019821171.2
E3 42791183 42791315
refGene_gn=TRAF7;refGene_tx=XM_003998941.5;refGene_gn=TRAF7;refGene_tx=XM_006942456.4;refGene_gn=TRAF7;refGene_tx=XM_019821171.2
E3 42791405 42791525
refGene_gn=TRAF7;refGene_tx=XM_003998941.5;refGene_gn=TRAF7;refGene_tx=XM_006942456.4;refGene_gn=TRAF7;refGene_tx=XM_019821171.2
E3 42791748 42791871
refGene_gn=TRAF7;refGene_tx=XM_003998941.5;refGene_gn=TRAF7;refGene_tx=XM_006942456.4;refGene_gn=TRAF7;refGene_tx=XM_019821171.2
E3 42791957 42792074
refGene_gn=TRAF7;refGene_tx=XM_003998941.5;refGene_gn=TRAF7;refGene_tx=XM_006942456.4;refGene_gn=TRAF7;refGene_tx=XM_019821171.2
E3 42792187 42792227
refGene_gn=TRAF7;refGene_tx=XM_003998941.5;refGene_gn=TRAF7;refGene_tx=XM_006942456.4;refGene_gn=TRAF7;refGene_tx=XM_019821171.2
E3 42792896 42792979
refGene_gn=TRAF7;refGene_tx=XM_003998941.5;refGene_gn=TRAF7;refGene_tx=XM_006942456.4;refGene_gn=TRAF7;refGene_tx=XM_019821171.2
E3 42793171 42793299
refGene_gn=TRAF7;refGene_tx=XM_003998941.5;refGene_gn=TRAF7;refGene_tx=XM_006942456.4;refGene_gn=TRAF7;refGene_tx=XM_019821171.2
E3 42793378 42793427
refGene_gn=TRAF7;refGene_tx=XM_003998941.5;refGene_gn=TRAF7;refGene_tx=XM_006942456.4;refGene_gn=TRAF7;refGene_tx=XM_019821171.2
E3 42793657 42793731
refGene_gn=TRAF7;refGene_tx=XM_003998941.5;refGene_gn=TRAF7;refGene_tx=XM_006942456.4;refGene_gn=TRAF7;refGene_tx=XM_019821171.2
E3 42793805 42794023
refGene_gn=TRAF7;refGene_tx=XM_003998941.5;refGene_gn=TRAF7;refGene_tx=XM_006942456.4;refGene_gn=TRAF7;refGene_tx=XM_019821171.2
E3 42794660 42794795
refGene_gn=TRAF7;refGene_tx=XM_003998941.5;refGene_gn=TRAF7;refGene_tx=XM_006942456.4;refGene_gn=TRAF7;refGene_tx=XM_019821171.2
E3 42794883 42795067

refGene_gn=TRAF7;refGene_tx=XM_003998941.5;refGene_gn=TRAF7;refGene_tx=XM_006942456.4;refGene_gn=TRAF7;refGene_tx=XM_019821171.2
E3 42795723 42795757
refGene_gn=TRAF7;refGene_tx=XM_003998941.5;refGene_gn=TRAF7;refGene_tx=XM_006942456.4;refGene_gn=TRAF7;refGene_tx=XM_019821171.2
E3 42795894 42795987
refGene_gn=TRAF7;refGene_tx=XM_003998941.5;refGene_gn=TRAF7;refGene_tx=XM_006942456.4;refGene_gn=TRAF7;refGene_tx=XM_019821171.2
E3 42796327 42796444
refGene_gn=TRAF7;refGene_tx=XM_003998941.5;refGene_gn=TRAF7;refGene_tx=XM_006942456.4;refGene_gn=TRAF7;refGene_tx=XM_019821171.2
E3 42798212 42798304
refGene_gn=TRAF7;refGene_tx=XM_003998941.5;refGene_gn=TRAF7;refGene_tx=XM_006942456.4;refGene_gn=TRAF7;refGene_tx=XM_019821171.2
E3 42800181 42800239
refGene_gn=TRAF7;refGene_tx=XM_003998941.5;refGene_gn=TRAF7;refGene_tx=XM_006942456.4;refGene_gn=TRAF7;refGene_tx=XM_019821171.2
E3 42800475 42800556
refGene_gn=TRAF7;refGene_tx=XM_003998941.5;refGene_gn=TRAF7;refGene_tx=XM_006942456.4;refGene_gn=TRAF7;refGene_tx=XM_019821171.2
E3 42809880 42809983
refGene_gn=RAB26;refGene_tx=XM_011290709.3;refGene_gn=RAB26;refGene_tx=XM_019821173.2;refGene_gn=RAB26;refGene_tx=XM_019821174.2
E3 42810076 42810185
refGene_gn=RAB26;refGene_tx=XM_011290709.3;refGene_gn=RAB26;refGene_tx=XM_019821173.2;refGene_gn=RAB26;refGene_tx=XM_019821174.2;refGene_gn=RAB26;refGene_tx=XM_019821175.2
E3 42810254 42810311
refGene_gn=RAB26;refGene_tx=XM_011290709.3;refGene_gn=RAB26;refGene_tx=XM_019821173.2;refGene_gn=RAB26;refGene_tx=XM_019821174.2;refGene_gn=RAB26;refGene_tx=XM_019821175.2
E3 42810415 42810481
refGene_gn=RAB26;refGene_tx=XM_011290709.3;refGene_gn=RAB26;refGene_tx=XM_019821173.2;refGene_gn=RAB26;refGene_tx=XM_019821174.2;refGene_gn=RAB26;refGene_tx=XM_019821175.2
E3 42810816 42810869
refGene_gn=RAB26;refGene_tx=XM_011290709.3;refGene_gn=RAB26;refGene_tx=XM_019821173.2;refGene_gn=RAB26;refGene_tx=XM_019821174.2;refGene_gn=RAB26;refGene_tx=XM_019821175.2
E3 42811073 42811140
refGene_gn=RAB26;refGene_tx=XM_011290709.3;refGene_gn=RAB26;refGene_tx=XM_019821173.2;refGene_gn=RAB26;refGene_tx=XM_019821174.2;refGene_gn=RAB26;refGene_tx=XM_019821175.2
E3 42811229 42811271
refGene_gn=RAB26;refGene_tx=XM_011290709.3;refGene_gn=RAB26;refGene_tx=XM_019821173.2;refGene_gn=RAB26;refGene_tx=XM_019821174.2;refGene_gn=RAB26;refGene_tx=XM_019821175.2
E3 42813074 42813185
refGene_gn=RAB26;refGene_tx=XM_011290709.3;refGene_gn=RAB26;refGene_tx=XM_019821173.2;refGene_gn=RAB26;refGene_tx=XM_019821174.2;refGene_gn=

RAB26;refGene_tx=XM_019821175.2
E3 42814243 42814435
refGene_gn=RAB26;refGene_tx=XM_011290709.3;refGene_gn=RAB26;refGene_tx=XM_019821173.2;refGene_gn=RAB26;refGene_tx=XM_019821174.2;refGene_gn=RAB26;refGene_tx=XM_019821175.2
E3 42824331 42824546
refGene_gn=PKD1;refGene_tx=XM_011290710.2;refGene_gn=PKD1;refGene_tx=XM_023247048.1;refGene_gn=PKD1;refGene_tx=XM_023247049.1;refGene_gn=PKD1;refGene_tx=XM_023247050.1;refGene_gn=PKD1;refGene_tx=XM_023247051.1;refGene_gn=PKD1;refGene_tx=XM_023247052.1;refGene_gn=PKD1;refGene_tx=XM_023247053.1
E3 42838778 42838850
refGene_gn=PKD1;refGene_tx=XM_011290710.2;refGene_gn=PKD1;refGene_tx=XM_023247048.1;refGene_gn=PKD1;refGene_tx=XM_023247049.1;refGene_gn=PKD1;refGene_tx=XM_023247050.1;refGene_gn=PKD1;refGene_tx=XM_023247051.1;refGene_gn=PKD1;refGene_tx=XM_023247052.1;refGene_gn=PKD1;refGene_tx=XM_023247053.1
E3 42838957 42839029
refGene_gn=PKD1;refGene_tx=XM_011290710.2;refGene_gn=PKD1;refGene_tx=XM_023247048.1;refGene_gn=PKD1;refGene_tx=XM_023247049.1;refGene_gn=PKD1;refGene_tx=XM_023247050.1;refGene_gn=PKD1;refGene_tx=XM_023247051.1;refGene_gn=PKD1;refGene_tx=XM_023247052.1;refGene_gn=PKD1;refGene_tx=XM_023247053.1
E3 42839309 42839479
refGene_gn=PKD1;refGene_tx=XM_011290710.2;refGene_gn=PKD1;refGene_tx=XM_023247048.1;refGene_gn=PKD1;refGene_tx=XM_023247049.1;refGene_gn=PKD1;refGene_tx=XM_023247050.1;refGene_gn=PKD1;refGene_tx=XM_023247051.1;refGene_gn=PKD1;refGene_tx=XM_023247052.1;refGene_gn=PKD1;refGene_tx=XM_023247053.1
E3 42839839 42840511
refGene_gn=PKD1;refGene_tx=XM_011290710.2;refGene_gn=PKD1;refGene_tx=XM_023247048.1;refGene_gn=PKD1;refGene_tx=XM_023247049.1;refGene_gn=PKD1;refGene_tx=XM_023247050.1;refGene_gn=PKD1;refGene_tx=XM_023247051.1;refGene_gn=PKD1;refGene_tx=XM_023247052.1;refGene_gn=PKD1;refGene_tx=XM_023247053.1
E3 42840605 42840789
refGene_gn=PKD1;refGene_tx=XM_011290710.2;refGene_gn=PKD1;refGene_tx=XM_023247048.1;refGene_gn=PKD1;refGene_tx=XM_023247049.1;refGene_gn=PKD1;refGene_tx=XM_023247050.1;refGene_gn=PKD1;refGene_tx=XM_023247051.1;refGene_gn=PKD1;refGene_tx=XM_023247052.1;refGene_gn=PKD1;refGene_tx=XM_023247053.1
E3 42841881 42842102
refGene_gn=PKD1;refGene_tx=XM_011290710.2;refGene_gn=PKD1;refGene_tx=XM_023247048.1;refGene_gn=PKD1;refGene_tx=XM_023247049.1;refGene_gn=PKD1;refGene_tx=XM_023247050.1;refGene_gn=PKD1;refGene_tx=XM_023247051.1;refGene_gn=PKD1;refGene_tx=XM_023247052.1;refGene_gn=PKD1;refGene_tx=XM_023247053.1
E3 42842511 42842627
refGene_gn=PKD1;refGene_tx=XM_011290710.2;refGene_gn=PKD1;refGene_tx=XM_023247048.1;refGene_gn=PKD1;refGene_tx=XM_023247049.1;refGene_gn=PKD

1;refGene_tx=XM_023247050.1;refGene_gn=PKD1;refGene_tx=XM_023247051.1;
refGene_gn=PKD1;refGene_tx=XM_023247052.1;refGene_gn=PKD1;refGene_tx=X
M_023247053.1
E3 42843040 42843167
refGene_gn=PKD1;refGene_tx=XM_011290710.2;refGene_gn=PKD1;refGene_tx=X
M_023247048.1;refGene_gn=PKD1;refGene_tx=XM_023247049.1;refGene_gn=PKD
1;refGene_tx=XM_023247050.1;refGene_gn=PKD1;refGene_tx=XM_023247051.1;
refGene_gn=PKD1;refGene_tx=XM_023247052.1;refGene_gn=PKD1;refGene_tx=X
M_023247053.1
E3 42843481 42843729
refGene_gn=PKD1;refGene_tx=XM_011290710.2;refGene_gn=PKD1;refGene_tx=X
M_023247048.1;refGene_gn=PKD1;refGene_tx=XM_023247049.1;refGene_gn=PKD
1;refGene_tx=XM_023247050.1;refGene_gn=PKD1;refGene_tx=XM_023247051.1;
refGene_gn=PKD1;refGene_tx=XM_023247052.1;refGene_gn=PKD1;refGene_tx=X
M_023247053.1
E3 42844163 42844919
refGene_gn=PKD1;refGene_tx=XM_011290710.2;refGene_gn=PKD1;refGene_tx=X
M_023247048.1;refGene_gn=PKD1;refGene_tx=XM_023247049.1;refGene_gn=PKD
1;refGene_tx=XM_023247050.1;refGene_gn=PKD1;refGene_tx=XM_023247051.1;
refGene_gn=PKD1;refGene_tx=XM_023247052.1;refGene_gn=PKD1;refGene_tx=X
M_023247053.1
E3 42845719 42845851
refGene_gn=PKD1;refGene_tx=XM_011290710.2;refGene_gn=PKD1;refGene_tx=X
M_023247048.1;refGene_gn=PKD1;refGene_tx=XM_023247049.1;refGene_gn=PKD
1;refGene_tx=XM_023247050.1;refGene_gn=PKD1;refGene_tx=XM_023247051.1;
refGene_gn=PKD1;refGene_tx=XM_023247052.1;refGene_gn=PKD1;refGene_tx=X
M_023247053.1
E3 42846045 42846221
refGene_gn=PKD1;refGene_tx=XM_011290710.2;refGene_gn=PKD1;refGene_tx=X
M_023247048.1;refGene_gn=PKD1;refGene_tx=XM_023247049.1;refGene_gn=PKD
1;refGene_tx=XM_023247050.1;refGene_gn=PKD1;refGene_tx=XM_023247051.1;
refGene_gn=PKD1;refGene_tx=XM_023247052.1;refGene_gn=PKD1;refGene_tx=X
M_023247053.1
E3 42846416 42846550
refGene_gn=PKD1;refGene_tx=XM_011290710.2;refGene_gn=PKD1;refGene_tx=X
M_023247048.1;refGene_gn=PKD1;refGene_tx=XM_023247049.1;refGene_gn=PKD
1;refGene_tx=XM_023247050.1;refGene_gn=PKD1;refGene_tx=XM_023247051.1;
refGene_gn=PKD1;refGene_tx=XM_023247052.1;refGene_gn=PKD1;refGene_tx=X
M_023247053.1
E3 42847097 42850714
refGene_gn=PKD1;refGene_tx=XM_011290710.2;refGene_gn=PKD1;refGene_tx=X
M_023247048.1;refGene_gn=PKD1;refGene_tx=XM_023247049.1;refGene_gn=PKD
1;refGene_tx=XM_023247050.1;refGene_gn=PKD1;refGene_tx=XM_023247051.1;
refGene_gn=PKD1;refGene_tx=XM_023247052.1;refGene_gn=PKD1;refGene_tx=X
M_023247053.1
E3 42850870 42851020
refGene_gn=PKD1;refGene_tx=XM_011290710.2;refGene_gn=PKD1;refGene_tx=X
M_023247048.1;refGene_gn=PKD1;refGene_tx=XM_023247049.1;refGene_gn=PKD
1;refGene_tx=XM_023247050.1;refGene_gn=PKD1;refGene_tx=XM_023247051.1;
refGene_gn=PKD1;refGene_tx=XM_023247052.1;refGene_gn=PKD1;refGene_tx=X

M_023247053.1
E3 42852105 42852249
refGene_gn=PKD1;refGene_tx=XM_011290710.2;refGene_gn=PKD1;refGene_tx=XM_023247048.1;refGene_gn=PKD1;refGene_tx=XM_023247049.1;refGene_gn=PKD1;refGene_tx=XM_023247050.1;refGene_gn=PKD1;refGene_tx=XM_023247051.1;refGene_gn=PKD1;refGene_tx=XM_023247052.1;refGene_gn=PKD1;refGene_tx=XM_023247053.1
E3 42852339 42852622
refGene_gn=PKD1;refGene_tx=XM_011290710.2;refGene_gn=PKD1;refGene_tx=XM_023247048.1;refGene_gn=PKD1;refGene_tx=XM_023247049.1;refGene_gn=PKD1;refGene_tx=XM_023247050.1;refGene_gn=PKD1;refGene_tx=XM_023247051.1;refGene_gn=PKD1;refGene_tx=XM_023247052.1;refGene_gn=PKD1;refGene_tx=XM_023247053.1
E3 42852697 42852911
refGene_gn=PKD1;refGene_tx=XM_011290710.2;refGene_gn=PKD1;refGene_tx=XM_023247048.1;refGene_gn=PKD1;refGene_tx=XM_023247049.1;refGene_gn=PKD1;refGene_tx=XM_023247050.1;refGene_gn=PKD1;refGene_tx=XM_023247051.1;refGene_gn=PKD1;refGene_tx=XM_023247052.1;refGene_gn=PKD1;refGene_tx=XM_023247053.1
E3 42852968 42853167
refGene_gn=PKD1;refGene_tx=XM_011290710.2;refGene_gn=PKD1;refGene_tx=XM_023247048.1;refGene_gn=PKD1;refGene_tx=XM_023247049.1;refGene_gn=PKD1;refGene_tx=XM_023247050.1;refGene_gn=PKD1;refGene_tx=XM_023247051.1;refGene_gn=PKD1;refGene_tx=XM_023247052.1;refGene_gn=PKD1;refGene_tx=XM_023247053.1
E3 42853491 42853644
refGene_gn=PKD1;refGene_tx=XM_011290710.2;refGene_gn=PKD1;refGene_tx=XM_023247048.1;refGene_gn=PKD1;refGene_tx=XM_023247049.1;refGene_gn=PKD1;refGene_tx=XM_023247050.1;refGene_gn=PKD1;refGene_tx=XM_023247051.1;refGene_gn=PKD1;refGene_tx=XM_023247052.1;refGene_gn=PKD1;refGene_tx=XM_023247053.1
E3 42854335 42854480
refGene_gn=PKD1;refGene_tx=XM_011290710.2;refGene_gn=PKD1;refGene_tx=XM_023247048.1;refGene_gn=PKD1;refGene_tx=XM_023247049.1;refGene_gn=PKD1;refGene_tx=XM_023247050.1;refGene_gn=PKD1;refGene_tx=XM_023247051.1;refGene_gn=PKD1;refGene_tx=XM_023247052.1;refGene_gn=PKD1;refGene_tx=XM_023247053.1
E3 42854580 42855201
refGene_gn=PKD1;refGene_tx=XM_011290710.2;refGene_gn=PKD1;refGene_tx=XM_023247048.1;refGene_gn=PKD1;refGene_tx=XM_023247049.1;refGene_gn=PKD1;refGene_tx=XM_023247050.1;refGene_gn=PKD1;refGene_tx=XM_023247051.1;refGene_gn=PKD1;refGene_tx=XM_023247052.1;refGene_gn=PKD1;refGene_tx=XM_023247053.1
E3 42855507 42855664
refGene_gn=PKD1;refGene_tx=XM_011290710.2;refGene_gn=PKD1;refGene_tx=XM_023247048.1;refGene_gn=PKD1;refGene_tx=XM_023247049.1;refGene_gn=PKD1;refGene_tx=XM_023247050.1;refGene_gn=PKD1;refGene_tx=XM_023247051.1;refGene_gn=PKD1;refGene_tx=XM_023247052.1;refGene_gn=PKD1;refGene_tx=XM_023247053.1
E3 42855911 42856164

refGene_gn=PKD1;refGene_tx=XM_011290710.2;refGene_gn=PKD1;refGene_tx=XM_023247048.1;refGene_gn=PKD1;refGene_tx=XM_023247049.1;refGene_gn=PKD1;refGene_tx=XM_023247050.1;refGene_gn=PKD1;refGene_tx=XM_023247051.1;refGene_gn=PKD1;refGene_tx=XM_023247052.1;refGene_gn=PKD1;refGene_tx=XM_023247053.1
E3 42856273 42856469
refGene_gn=PKD1;refGene_tx=XM_011290710.2;refGene_gn=PKD1;refGene_tx=XM_023247048.1;refGene_gn=PKD1;refGene_tx=XM_023247049.1;refGene_gn=PKD1;refGene_tx=XM_023247050.1;refGene_gn=PKD1;refGene_tx=XM_023247051.1;refGene_gn=PKD1;refGene_tx=XM_023247052.1;refGene_gn=PKD1;refGene_tx=XM_023247053.1
E3 42857489 42857660
refGene_gn=PKD1;refGene_tx=XM_011290710.2;refGene_gn=PKD1;refGene_tx=XM_023247048.1;refGene_gn=PKD1;refGene_tx=XM_023247049.1;refGene_gn=PKD1;refGene_tx=XM_023247050.1;refGene_gn=PKD1;refGene_tx=XM_023247051.1;refGene_gn=PKD1;refGene_tx=XM_023247052.1;refGene_gn=PKD1;refGene_tx=XM_023247053.1
E3 42857739 42857883
refGene_gn=PKD1;refGene_tx=XM_011290710.2;refGene_gn=PKD1;refGene_tx=XM_023247048.1;refGene_gn=PKD1;refGene_tx=XM_023247049.1;refGene_gn=PKD1;refGene_tx=XM_023247050.1;refGene_gn=PKD1;refGene_tx=XM_023247051.1;refGene_gn=PKD1;refGene_tx=XM_023247052.1;refGene_gn=PKD1;refGene_tx=XM_023247053.1
E3 42857972 42858183
refGene_gn=PKD1;refGene_tx=XM_011290710.2;refGene_gn=PKD1;refGene_tx=XM_023247048.1;refGene_gn=PKD1;refGene_tx=XM_023247049.1;refGene_gn=PKD1;refGene_tx=XM_023247050.1;refGene_gn=PKD1;refGene_tx=XM_023247051.1;refGene_gn=PKD1;refGene_tx=XM_023247052.1;refGene_gn=PKD1;refGene_tx=XM_023247053.1
E3 42858272 42858399
refGene_gn=PKD1;refGene_tx=XM_011290710.2;refGene_gn=PKD1;refGene_tx=XM_023247048.1;refGene_gn=PKD1;refGene_tx=XM_023247049.1;refGene_gn=PKD1;refGene_tx=XM_023247050.1;refGene_gn=PKD1;refGene_tx=XM_023247051.1;refGene_gn=PKD1;refGene_tx=XM_023247052.1;refGene_gn=PKD1;refGene_tx=XM_023247053.1
E3 42859165 42859282
refGene_gn=PKD1;refGene_tx=XM_011290710.2;refGene_gn=PKD1;refGene_tx=XM_023247048.1;refGene_gn=PKD1;refGene_tx=XM_023247049.1;refGene_gn=PKD1;refGene_tx=XM_023247050.1;refGene_gn=PKD1;refGene_tx=XM_023247051.1;refGene_gn=PKD1;refGene_tx=XM_023247052.1;refGene_gn=PKD1;refGene_tx=XM_023247053.1
E3 42859388 42859438
refGene_gn=PKD1;refGene_tx=XM_011290710.2;refGene_gn=PKD1;refGene_tx=XM_023247048.1;refGene_gn=PKD1;refGene_tx=XM_023247049.1;refGene_gn=PKD1;refGene_tx=XM_023247050.1;refGene_gn=PKD1;refGene_tx=XM_023247051.1;refGene_gn=PKD1;refGene_tx=XM_023247052.1;refGene_gn=PKD1;refGene_tx=XM_023247053.1
E3 42859596 42859769
refGene_gn=PKD1;refGene_tx=XM_011290710.2;refGene_gn=PKD1;refGene_tx=XM_023247048.1;refGene_gn=PKD1;refGene_tx=XM_023247049.1;refGene_gn=PKD

1;refGene_tx=XM_023247050.1;refGene_gn=PKD1;refGene_tx=XM_023247051.1;
refGene_gn=PKD1;refGene_tx=XM_023247052.1;refGene_gn=PKD1;refGene_tx=X
M_023247053.1
E3 42859875 42859969
refGene_gn=PKD1;refGene_tx=XM_011290710.2;refGene_gn=PKD1;refGene_tx=X
M_023247048.1;refGene_gn=PKD1;refGene_tx=XM_023247049.1;refGene_gn=PKD
1;refGene_tx=XM_023247050.1;refGene_gn=PKD1;refGene_tx=XM_023247051.1;
refGene_gn=PKD1;refGene_tx=XM_023247052.1;refGene_gn=PKD1;refGene_tx=X
M_023247053.1
E3 42862534 42862931
refGene_gn=PKD1;refGene_tx=XM_011290710.2;refGene_gn=PKD1;refGene_tx=X
M_023247048.1;refGene_gn=PKD1;refGene_tx=XM_023247049.1;refGene_gn=PKD
1;refGene_tx=XM_023247050.1;refGene_gn=PKD1;refGene_tx=XM_023247051.1;
refGene_gn=PKD1;refGene_tx=XM_023247052.1;refGene_gn=PKD1;refGene_tx=X
M_023247053.1
E3 42863006 42863201
refGene_gn=PKD1;refGene_tx=XM_011290710.2;refGene_gn=PKD1;refGene_tx=X
M_023247048.1;refGene_gn=PKD1;refGene_tx=XM_023247049.1;refGene_gn=PKD
1;refGene_tx=XM_023247050.1;refGene_gn=PKD1;refGene_tx=XM_023247051.1;
refGene_gn=PKD1;refGene_tx=XM_023247052.1;refGene_gn=PKD1;refGene_tx=X
M_023247053.1
E3 42863569 42863709
refGene_gn=PKD1;refGene_tx=XM_011290710.2;refGene_gn=PKD1;refGene_tx=X
M_023247048.1;refGene_gn=PKD1;refGene_tx=XM_023247049.1;refGene_gn=PKD
1;refGene_tx=XM_023247050.1;refGene_gn=PKD1;refGene_tx=XM_023247051.1;
refGene_gn=PKD1;refGene_tx=XM_023247052.1
E3 42864329 42864442
refGene_gn=PKD1;refGene_tx=XM_011290710.2;refGene_gn=PKD1;refGene_tx=X
M_023247048.1;refGene_gn=PKD1;refGene_tx=XM_023247049.1;refGene_gn=PKD
1;refGene_tx=XM_023247050.1;refGene_gn=PKD1;refGene_tx=XM_023247051.1;
refGene_gn=PKD1;refGene_tx=XM_023247052.1
E3 42864647 42864795
refGene_gn=PKD1;refGene_tx=XM_011290710.2;refGene_gn=PKD1;refGene_tx=X
M_023247048.1;refGene_gn=PKD1;refGene_tx=XM_023247049.1;refGene_gn=PKD
1;refGene_tx=XM_023247050.1;refGene_gn=PKD1;refGene_tx=XM_023247051.1;
refGene_gn=PKD1;refGene_tx=XM_023247052.1
E3 42864949 42865075
refGene_gn=PKD1;refGene_tx=XM_011290710.2;refGene_gn=PKD1;refGene_tx=X
M_023247048.1;refGene_gn=PKD1;refGene_tx=XM_023247049.1;refGene_gn=PKD
1;refGene_tx=XM_023247050.1;refGene_gn=PKD1;refGene_tx=XM_023247051.1;
refGene_gn=PKD1;refGene_tx=XM_023247052.1
E3 42865344 42865519
refGene_gn=PKD1;refGene_tx=XM_011290710.2;refGene_gn=PKD1;refGene_tx=X
M_023247048.1;refGene_gn=PKD1;refGene_tx=XM_023247049.1;refGene_gn=PKD
1;refGene_tx=XM_023247050.1;refGene_gn=PKD1;refGene_tx=XM_023247051.1;
refGene_gn=PKD1;refGene_tx=XM_023247052.1
E3 42865654 42865945
refGene_gn=PKD1;refGene_tx=XM_011290710.2;refGene_gn=PKD1;refGene_tx=X
M_023247048.1;refGene_gn=PKD1;refGene_tx=XM_023247049.1;refGene_gn=PKD
1;refGene_tx=XM_023247050.1;refGene_gn=PKD1;refGene_tx=XM_023247051.1;

refGene_gn=PKD1;refGene_tx=XM_023247052.1
E3 42866037 42866172
refGene_gn=PKD1;refGene_tx=XM_011290710.2;refGene_gn=PKD1;refGene_tx=XM_023247048.1;refGene_gn=PKD1;refGene_tx=XM_023247049.1;refGene_gn=PKD1;refGene_tx=XM_023247050.1;refGene_gn=PKD1;refGene_tx=XM_023247051.1;refGene_gn=PKD1;refGene_tx=XM_023247052.1
E3 42866252 42866558
refGene_gn=PKD1;refGene_tx=XM_011290710.2;refGene_gn=PKD1;refGene_tx=XM_023247048.1;refGene_gn=PKD1;refGene_tx=XM_023247049.1;refGene_gn=PKD1;refGene_tx=XM_023247050.1;refGene_gn=PKD1;refGene_tx=XM_023247051.1;refGene_gn=PKD1;refGene_tx=XM_023247052.1
E3 42866653 42867106
refGene_gn=PKD1;refGene_tx=XM_011290710.2;refGene_gn=PKD1;refGene_tx=XM_023247048.1;refGene_gn=PKD1;refGene_tx=XM_023247049.1;refGene_gn=PKD1;refGene_tx=XM_023247050.1;refGene_gn=PKD1;refGene_tx=XM_023247051.1;refGene_gn=PKD1;refGene_tx=XM_023247052.1
E3 42868247 42868406 refGene_gn=TSC2;refGene_tx=XM_023247054.1
E3 42868528 42868627 refGene_gn=TSC2;refGene_tx=XM_023247054.1
E3 42868716 42868808 refGene_gn=TSC2;refGene_tx=XM_023247054.1
E3 42868877 42868956 refGene_gn=TSC2;refGene_tx=XM_023247054.1
E3 42869964 42870104 refGene_gn=TSC2;refGene_tx=XM_023247054.1
E3 42870388 42870575 refGene_gn=TSC2;refGene_tx=XM_023247054.1
E3 42871297 42871384 refGene_gn=TSC2;refGene_tx=XM_023247054.1
E3 42871618 42871694 refGene_gn=TSC2;refGene_tx=XM_023247054.1
E3 42871935 42872435 refGene_gn=TSC2;refGene_tx=XM_023247054.1
E3 42872845 42872976 refGene_gn=TSC2;refGene_tx=XM_023247054.1
E3 42874298 42874367 refGene_gn=TSC2;refGene_tx=XM_023247054.1
E3 42874978 42875182 refGene_gn=TSC2;refGene_tx=XM_023247054.1
E3 42876389 42876584 refGene_gn=TSC2;refGene_tx=XM_023247054.1
E3 42877060 42877173 refGene_gn=TSC2;refGene_tx=XM_023247054.1
E3 42877312 42877465 refGene_gn=TSC2;refGene_tx=XM_023247054.1
E3 42877656 42877821 refGene_gn=TSC2;refGene_tx=XM_023247054.1
E3 42879168 42879297 refGene_gn=TSC2;refGene_tx=XM_023247054.1
E3 42880468 42880563 refGene_gn=TSC2;refGene_tx=XM_023247054.1
E3 42880753 42880856 refGene_gn=TSC2;refGene_tx=XM_023247054.1
E3 42881252 42881346 refGene_gn=TSC2;refGene_tx=XM_023247054.1
E3 42882201 42882391 refGene_gn=TSC2;refGene_tx=XM_023247054.1
E3 42883718 42883853 refGene_gn=TSC2;refGene_tx=XM_023247054.1
E3 42884355 42884478 refGene_gn=TSC2;refGene_tx=XM_023247054.1
E3 42885686 42885837 refGene_gn=TSC2;refGene_tx=XM_023247054.1
E3 42886041 42886148 refGene_gn=TSC2;refGene_tx=XM_023247054.1
E3 42887098 42887221 refGene_gn=TSC2;refGene_tx=XM_023247054.1
E3 42889996 42890017 refGene_gn=TSC2;refGene_tx=XM_023247054.1
E3 42890018 42890114 refGene_gn=TSC2;refGene_tx=XM_023247054.1
E3 42890523 42890679 refGene_gn=TSC2;refGene_tx=XM_023247054.1
E3 42891658 42891740 refGene_gn=TSC2;refGene_tx=XM_023247054.1
E3 42893067 42893171 refGene_gn=TSC2;refGene_tx=XM_023247054.1
E3 42893811 42893949 refGene_gn=TSC2;refGene_tx=XM_023247054.1
E3 42894567 42894711 refGene_gn=TSC2;refGene_tx=XM_023247054.1
E3 42895468 42895595 refGene_gn=TSC2;refGene_tx=XM_023247054.1

E3	42896878	42896952	refGene_gn=TSC2;refGene_tx=XM_023247054.1
E3	42897270	42897396	refGene_gn=TSC2;refGene_tx=XM_023247054.1
E3	42897766	42897815	refGene_gn=TSC2;refGene_tx=XM_023247054.1
E3	42898270	42898388	refGene_gn=TSC2;refGene_tx=XM_023247054.1
E3	42899377	42899522	refGene_gn=TSC2;refGene_tx=XM_023247054.1
E3	42900303	42900414	refGene_gn=TSC2;refGene_tx=XM_023247054.1
E3	42901819	42901906	refGene_gn=TSC2;refGene_tx=XM_023247054.1
E3	42902923	42903061	refGene_gn=TSC2;refGene_tx=XM_023247054.1
E3	42903719	42903903
refGene_gn=NTHL1;refGene_tx=XM_006942460.4;refGene_gn=NTHL1;refGene_tx=XM_023247055.1;refGene_gn=NTHL1;refGene_tx=XM_023247056.1
E3	42904778	42904788	refGene_gn=NTHL1;refGene_tx=XM_019821185.2
E3	42904869	42905096
refGene_gn=NTHL1;refGene_tx=XM_006942460.4;refGene_gn=NTHL1;refGene_tx=XM_019821185.2;refGene_gn=NTHL1;refGene_tx=XM_023247055.1;refGene_gn=NTHL1;refGene_tx=XM_023247056.1
E3	42905820	42905991
refGene_gn=NTHL1;refGene_tx=XM_006942460.4;refGene_gn=NTHL1;refGene_tx=XM_019821185.2;refGene_gn=NTHL1;refGene_tx=XM_023247055.1;refGene_gn=NTHL1;refGene_tx=XM_023247056.1
E3	42906823	42906983
refGene_gn=NTHL1;refGene_tx=XM_006942460.4;refGene_gn=NTHL1;refGene_tx=XM_019821185.2;refGene_gn=NTHL1;refGene_tx=XM_023247055.1;refGene_gn=NTHL1;refGene_tx=XM_023247056.1
E3	42908801	42908924
refGene_gn=NTHL1;refGene_tx=XM_006942460.4;refGene_gn=NTHL1;refGene_tx=XM_019821185.2;refGene_gn=NTHL1;refGene_tx=XM_023247055.1
E3	42908998	42909237
refGene_gn=NTHL1;refGene_tx=XM_006942460.4;refGene_gn=NTHL1;refGene_tx=XM_019821185.2;refGene_gn=NTHL1;refGene_tx=XM_023247055.1;refGene_gn=NTHL1;refGene_tx=XM_023247056.1
E3	42911111	42911270
refGene_gn=SLC9A3R2;refGene_tx=XM_003998943.5;refGene_gn=SLC9A3R2;refGene_tx=XM_006942466.4;refGene_gn=SLC9A3R2;refGene_tx=XM_019821187.2;refGene_gn=SLC9A3R2;refGene_tx=XM_023247109.1;refGene_gn=SLC9A3R2;refGene_tx=XM_023247110.1;refGene_gn=SLC9A3R2;refGene_tx=XM_023247111.1
E3	42911513	42911579
refGene_gn=SLC9A3R2;refGene_tx=XM_003998943.5;refGene_gn=SLC9A3R2;refGene_tx=XM_006942466.4;refGene_gn=SLC9A3R2;refGene_tx=XM_019821187.2;refGene_gn=SLC9A3R2;refGene_tx=XM_023247109.1;refGene_gn=SLC9A3R2;refGene_tx=XM_023247110.1
E3	42912123	42912167
refGene_gn=SLC9A3R2;refGene_tx=XM_003998943.5;refGene_gn=SLC9A3R2;refGene_tx=XM_006942466.4;refGene_gn=SLC9A3R2;refGene_tx=XM_019821187.2;refGene_gn=SLC9A3R2;refGene_tx=XM_023247109.1;refGene_gn=SLC9A3R2;refGene_tx=XM_023247110.1;refGene_gn=SLC9A3R2;refGene_tx=XM_023247111.1
E3	42912236	42912390
refGene_gn=SLC9A3R2;refGene_tx=XM_003998943.5;refGene_gn=SLC9A3R2;refGene_tx=XM_006942466.4;refGene_gn=SLC9A3R2;refGene_tx=XM_019821187.2;refGene_gn=SLC9A3R2;refGene_tx=XM_023247109.1;refGene_gn=SLC9A3R2;refGen

e_tx=XM_023247110.1;refGene_gn=SLC9A3R2;refGene_tx=XM_023247111.1
E3 42912735 42912915
refGene_gn=SLC9A3R2;refGene_tx=XM_003998943.5;refGene_gn=SLC9A3R2;refGene_tx=XM_006942466.4;refGene_gn=SLC9A3R2;refGene_tx=XM_019821187.2;refGene_gn=SLC9A3R2;refGene_tx=XM_023247109.1;refGene_gn=SLC9A3R2;refGene_tx=XM_023247110.1;refGene_gn=SLC9A3R2;refGene_tx=XM_023247111.1
E3 42915322 42915403
refGene_gn=SLC9A3R2;refGene_tx=XM_003998943.5;refGene_gn=SLC9A3R2;refGene_tx=XM_006942466.4
E3 42918014 42918215
refGene_gn=SLC9A3R2;refGene_tx=XM_019821187.2;refGene_gn=SLC9A3R2;refGene_tx=XM_023247109.1;refGene_gn=SLC9A3R2;refGene_tx=XM_023247110.1;refGene_gn=SLC9A3R2;refGene_tx=XM_023247111.1
E3 42920535 42920601 refGene_gn=SLC9A3R2;refGene_tx=XM_019821187.2
E3 42920949 42921162
refGene_gn=SLC9A3R2;refGene_tx=XM_023247109.1;refGene_gn=SLC9A3R2;refGene_tx=XM_023247110.1;refGene_gn=SLC9A3R2;refGene_tx=XM_023247111.1
E3 42926365 42926482
refGene_gn=LOC102902485;refGene_tx=XM_006942462.3
E3 42926699 42926894
refGene_gn=LOC102902485;refGene_tx=XM_006942462.3
E3 42935007 42935503
refGene_gn=LOC102902485;refGene_tx=XM_006942462.3;refGene_gn=ZNF598;refGene_tx=XM_023246469.1;refGene_gn=ZNF598;refGene_tx=XM_023246470.1;refGene_gn=ZNF598;refGene_tx=XM_023246471.1
E3 42940390 42940503
refGene_gn=ZNF598;refGene_tx=XM_023246469.1;refGene_gn=ZNF598;refGene_tx=XM_023246470.1;refGene_gn=ZNF598;refGene_tx=XM_023246471.1
E3 42941020 42941150
refGene_gn=ZNF598;refGene_tx=XM_023246469.1;refGene_gn=ZNF598;refGene_tx=XM_023246470.1;refGene_gn=ZNF598;refGene_tx=XM_023246471.1
E3 42941360 42941572
refGene_gn=ZNF598;refGene_tx=XM_023246469.1;refGene_gn=ZNF598;refGene_tx=XM_023246470.1;refGene_gn=ZNF598;refGene_tx=XM_023246471.1
E3 42941670 42941888
refGene_gn=ZNF598;refGene_tx=XM_023246469.1;refGene_gn=ZNF598;refGene_tx=XM_023246470.1;refGene_gn=ZNF598;refGene_tx=XM_023246471.1
E3 42942260 42942378
refGene_gn=ZNF598;refGene_tx=XM_023246469.1;refGene_gn=ZNF598;refGene_tx=XM_023246470.1;refGene_gn=ZNF598;refGene_tx=XM_023246471.1
E3 42942943 42943224
refGene_gn=ZNF598;refGene_tx=XM_023246469.1;refGene_gn=ZNF598;refGene_tx=XM_023246470.1;refGene_gn=ZNF598;refGene_tx=XM_023246471.1
E3 42943354 42943439
refGene_gn=ZNF598;refGene_tx=XM_023246469.1;refGene_gn=ZNF598;refGene_tx=XM_023246470.1;refGene_gn=ZNF598;refGene_tx=XM_023246471.1
E3 42943547 42944335
refGene_gn=ZNF598;refGene_tx=XM_023246469.1;refGene_gn=ZNF598;refGene_tx=XM_023246470.1;refGene_gn=ZNF598;refGene_tx=XM_023246471.1
E3 42944725 42944876

refGene_gn=ZNF598;refGene_tx=XM_023246469.1;refGene_gn=ZNF598;refGene_tx=XM_023246470.1;refGene_gn=ZNF598;refGene_tx=XM_023246471.1
E3 42944989 42945140
refGene_gn=ZNF598;refGene_tx=XM_023246469.1;refGene_gn=ZNF598;refGene_tx=XM_023246470.1;refGene_gn=ZNF598;refGene_tx=XM_023246471.1
E3 42945231 42945582
refGene_gn=ZNF598;refGene_tx=XM_023246469.1;refGene_gn=ZNF598;refGene_tx=XM_023246470.1;refGene_gn=ZNF598;refGene_tx=XM_023246471.1
E3 42949599 42949809 refGene_gn=SYNGR3;refGene_tx=XM_023246722.1
E3 42949888 42950031 refGene_gn=SYNGR3;refGene_tx=XM_023246722.1
E3 42950134 42950372 refGene_gn=SYNGR3;refGene_tx=XM_023246722.1
E3 42952581 42952680 refGene_gn=SYNGR3;refGene_tx=XM_023246722.1
E3 42956052 42956215 refGene_gn=GFER;refGene_tx=XM_003998945.5
E3 42957114 42957311 refGene_gn=GFER;refGene_tx=XM_003998945.5
E3 42957677 42957941 refGene_gn=GFER;refGene_tx=XM_003998945.5
E3 42961743 42961809 refGene_gn=NOXO1;refGene_tx=XM_003999029.3
E3 42961888 42961969 refGene_gn=NOXO1;refGene_tx=XM_003999029.3
E3 42962146 42962222 refGene_gn=NOXO1;refGene_tx=XM_003999029.3
E3 42962322 42962500 refGene_gn=NOXO1;refGene_tx=XM_003999029.3
E3 42962656 42962838 refGene_gn=NOXO1;refGene_tx=XM_003999029.3
E3 42962932 42963037 refGene_gn=NOXO1;refGene_tx=XM_003999029.3
E3 42963330 42963448 refGene_gn=NOXO1;refGene_tx=XM_003999029.3
E3 42963530 42963670 refGene_gn=NOXO1;refGene_tx=XM_003999029.3
E3 42963672 42963842 refGene_gn=NOXO1;refGene_tx=XM_003999029.3
E3 42964103 42964213 refGene_gn=TBL3;refGene_tx=XM_023247160.1
E3 42964290 42964449 refGene_gn=TBL3;refGene_tx=XM_023247160.1
E3 42964532 42964604 refGene_gn=TBL3;refGene_tx=XM_023247160.1
E3 42964699 42964803 refGene_gn=TBL3;refGene_tx=XM_023247160.1
E3 42965063 42965122 refGene_gn=TBL3;refGene_tx=XM_023247160.1
E3 42965213 42965370 refGene_gn=TBL3;refGene_tx=XM_023247160.1
E3 42965450 42965521 refGene_gn=TBL3;refGene_tx=XM_023247160.1
E3 42965604 42965835 refGene_gn=TBL3;refGene_tx=XM_023247160.1
E3 42965922 42966070 refGene_gn=TBL3;refGene_tx=XM_023247160.1
E3 42967047 42967151 refGene_gn=TBL3;refGene_tx=XM_023247160.1
E3 42967236 42967295 refGene_gn=TBL3;refGene_tx=XM_023247160.1
E3 42967392 42967534 refGene_gn=TBL3;refGene_tx=XM_023247160.1
E3 42967637 42967819 refGene_gn=TBL3;refGene_tx=XM_023247160.1
E3 42967902 42967996 refGene_gn=TBL3;refGene_tx=XM_023247160.1
E3 42968078 42968154 refGene_gn=TBL3;refGene_tx=XM_023247160.1
E3 42968245 42968416 refGene_gn=TBL3;refGene_tx=XM_023247160.1
E3 42968500 42968585 refGene_gn=TBL3;refGene_tx=XM_023247160.1
E3 42968680 42968822 refGene_gn=TBL3;refGene_tx=XM_023247160.1
E3 42968927 42968975 refGene_gn=TBL3;refGene_tx=XM_023247160.1
E3 42969058 42969154 refGene_gn=TBL3;refGene_tx=XM_023247160.1
E3 42969236 42969288 refGene_gn=TBL3;refGene_tx=XM_023247160.1
E3 42970674 42970715 refGene_gn=TBL3;refGene_tx=XM_023247160.1
E3 42971867 42972353
refGene_gn=RNF151;refGene_tx=XM_011290776.3;refGene_gn=RNF151;refGene_tx=XM_019821195.2;refGene_gn=RNF151;refGene_tx=XM_019821196.2
E3 42972891 42972988

refGene_gn=RNF151;refGene_tx=XM_011290776.3;refGene_gn=RNF151;refGene_tx=XM_019821195.2;refGene_gn=RNF151;refGene_tx=XM_019821196.2
E3 42973297 42973443
refGene_gn=RNF151;refGene_tx=XM_011290776.3;refGene_gn=RNF151;refGene_tx=XM_019821195.2;refGene_gn=RNF151;refGene_tx=XM_019821196.2
E3 42973754 42973757
refGene_gn=RNF151;refGene_tx=XM_011290776.3;refGene_gn=RNF151;refGene_tx=XM_019821195.2;refGene_gn=RNF151;refGene_tx=XM_019821196.2
E3 42975893 42976070 refGene_gn=RPS2;refGene_tx=XM_023247162.1
E3 42976156 42976246 refGene_gn=RPS2;refGene_tx=XM_023247162.1
E3 42976443 42976551 refGene_gn=RPS2;refGene_tx=XM_023247162.1
E3 42976805 42976979 refGene_gn=RPS2;refGene_tx=XM_023247162.1
E3 42977062 42977222 refGene_gn=RPS2;refGene_tx=XM_023247162.1
E3 42977442 42977534 refGene_gn=RPS2;refGene_tx=XM_023247162.1
E3 42977623 42977704 refGene_gn=RPS2;refGene_tx=XM_023247162.1
E3 42977922 42978026 refGene_gn=NDUFB10;refGene_tx=XM_003998947.5
E3 42978248 42978388 refGene_gn=NDUFB10;refGene_tx=XM_003998947.5
E3 42978599 42978738 refGene_gn=NDUFB10;refGene_tx=XM_003998947.5
E3 42980618 42980766 refGene_gn=NDUFB10;refGene_tx=XM_003998947.5
E3 42983273 42983276 refGene_gn=RPL3L;refGene_tx=XM_003998948.5
E3 42983730 42983923 refGene_gn=RPL3L;refGene_tx=XM_003998948.5
E3 42985395 42985564 refGene_gn=RPL3L;refGene_tx=XM_003998948.5
E3 42986762 42986898 refGene_gn=RPL3L;refGene_tx=XM_003998948.5
E3 42988133 42988320 refGene_gn=RPL3L;refGene_tx=XM_003998948.5
E3 42988411 42988572 refGene_gn=RPL3L;refGene_tx=XM_003998948.5
E3 42988778 42988880 refGene_gn=RPL3L;refGene_tx=XM_003998948.5
E3 42989337 42989433 refGene_gn=RPL3L;refGene_tx=XM_003998948.5
E3 42989624 42989744 refGene_gn=RPL3L;refGene_tx=XM_003998948.5
E3 42989994 42990051 refGene_gn=RPL3L;refGene_tx=XM_003998948.5
E3 42990587 42990642 refGene_gn=MSRB1;refGene_tx=XM_003999032.5
E3 42992452 42992601 refGene_gn=MSRB1;refGene_tx=XM_003999032.5
E3 42992914 42993029 refGene_gn=MSRB1;refGene_tx=XM_003999032.5
E3 42994583 42994615 refGene_gn=MSRB1;refGene_tx=XM_003999032.5
E3 43003551 43003838
refGene_gn=HS3ST6;refGene_tx=XM_019821200.2;refGene_gn=HS3ST6;refGene_tx=XM_023246500.1;refGene_gn=HS3ST6;refGene_tx=XM_023246501.1
E3 43004088 43004501
refGene_gn=HS3ST6;refGene_tx=XM_003998949.4;refGene_gn=HS3ST6;refGene_tx=XM_019821198.2
E3 43009297 43009913
refGene_gn=HS3ST6;refGene_tx=XM_003998949.4;refGene_gn=HS3ST6;refGene_tx=XM_019821198.2;refGene_gn=HS3ST6;refGene_tx=XM_019821200.2;refGene_gn=HS3ST6;refGene_tx=XM_023246500.1;refGene_gn=HS3ST6;refGene_tx=XM_023246501.1
E3 43062656 43062725
refGene_gn=MEIOB;refGene_tx=XM_023246558.1;refGene_gn=MEIOB;refGene_tx=XM_023246559.1
E3 43064309 43064367
refGene_gn=MEIOB;refGene_tx=XM_023246558.1;refGene_gn=MEIOB;refGene_tx=XM_023246559.1

E3 43066720 43066852 refGene_gn=MEIOB;refGene_tx=XM_023246558.1;refGene_gn=MEIOB;refGene_tx=XM_023246559.1
E3 43068503 43068576 refGene_gn=MEIOB;refGene_tx=XM_023246558.1;refGene_gn=MEIOB;refGene_tx=XM_023246559.1
E3 43070660 43070856 refGene_gn=MEIOB;refGene_tx=XM_023246558.1;refGene_gn=MEIOB;refGene_tx=XM_023246559.1
E3 43074148 43074249 refGene_gn=MEIOB;refGene_tx=XM_023246558.1;refGene_gn=MEIOB;refGene_tx=XM_023246559.1
E3 43075138 43075191 refGene_gn=MEIOB;refGene_tx=XM_023246558.1;refGene_gn=MEIOB;refGene_tx=XM_023246559.1
E3 43075285 43075381 refGene_gn=MEIOB;refGene_tx=XM_023246558.1;refGene_gn=MEIOB;refGene_tx=XM_023246559.1
E3 43078595 43078697 refGene_gn=MEIOB;refGene_tx=XM_023246558.1;refGene_gn=MEIOB;refGene_tx=XM_023246559.1
E3 43079253 43079407 refGene_gn=MEIOB;refGene_tx=XM_023246558.1;refGene_gn=MEIOB;refGene_tx=XM_023246559.1
E3 43082278 43082462 refGene_gn=MEIOB;refGene_tx=XM_023246558.1;refGene_gn=MEIOB;refGene_tx=XM_023246559.1
E3 43083177 43083264 refGene_gn=MEIOB;refGene_tx=XM_023246558.1;refGene_gn=MEIOB;refGene_tx=XM_023246559.1
E3 43086792 43086903 refGene_gn=MEIOB;refGene_tx=XM_023246558.1;refGene_gn=MEIOB;refGene_tx=XM_023246559.1
E3 43096680 43097346 refGene_gn=FAHD1;refGene_tx=XM_006942469.4
E3 43097850 43097926 refGene_gn=HAGH;refGene_tx=XM_023246408.1;refGene_gn=HAGH;refGene_tx=XM_023246409.1;refGene_gn=HAGH;refGene_tx=XM_023246410.1;refGene_gn=HAGH;refGene_tx=XM_023246411.1;refGene_gn=HAGH;refGene_tx=XM_023246412.1;refGene_gn=HAGH;refGene_tx=XM_023246413.1
E3 43100396 43100572 refGene_gn=HAGH;refGene_tx=XM_023246408.1;refGene_gn=HAGH;refGene_tx=XM_023246409.1;refGene_gn=HAGH;refGene_tx=XM_023246410.1;refGene_gn=HAGH;refGene_tx=XM_023246411.1;refGene_gn=HAGH;refGene_tx=XM_023246412.1;refGene_gn=HAGH;refGene_tx=XM_023246413.1
E3 43101128 43101193 refGene_gn=HAGH;refGene_tx=XM_023246408.1;refGene_gn=HAGH;refGene_tx=XM_023246409.1;refGene_gn=HAGH;refGene_tx=XM_023246410.1;refGene_gn=HAGH;refGene_tx=XM_023246411.1;refGene_gn=HAGH;refGene_tx=XM_023246412.1;refGene_gn=HAGH;refGene_tx=XM_023246413.1
E3 43105200 43105254 refGene_gn=HAGH;refGene_tx=XM_023246414.1

E3 43105322 43105460
refGene_gn=HAGH;refGene_tx=XM_023246408.1;refGene_gn=HAGH;refGene_tx=XM_023246409.1;refGene_gn=HAGH;refGene_tx=XM_023246410.1;refGene_gn=HAGH;refGene_tx=XM_023246411.1;refGene_gn=HAGH;refGene_tx=XM_023246412.1;refGene_gn=HAGH;refGene_tx=XM_023246413.1;refGene_gn=HAGH;refGene_tx=XM_023246414.1
E3 43106424 43106533
refGene_gn=HAGH;refGene_tx=XM_023246408.1;refGene_gn=HAGH;refGene_tx=XM_023246409.1;refGene_gn=HAGH;refGene_tx=XM_023246410.1;refGene_gn=HAGH;refGene_tx=XM_023246411.1;refGene_gn=HAGH;refGene_tx=XM_023246412.1;refGene_gn=HAGH;refGene_tx=XM_023246413.1;refGene_gn=HAGH;refGene_tx=XM_023246414.1
E3 43108908 43109012
refGene_gn=HAGH;refGene_tx=XM_023246408.1;refGene_gn=HAGH;refGene_tx=XM_023246409.1;refGene_gn=HAGH;refGene_tx=XM_023246410.1;refGene_gn=HAGH;refGene_tx=XM_023246411.1;refGene_gn=HAGH;refGene_tx=XM_023246412.1;refGene_gn=HAGH;refGene_tx=XM_023246413.1;refGene_gn=HAGH;refGene_tx=XM_023246414.1
E3 43109291 43109393
refGene_gn=HAGH;refGene_tx=XM_023246408.1;refGene_gn=HAGH;refGene_tx=XM_023246409.1;refGene_gn=HAGH;refGene_tx=XM_023246410.1;refGene_gn=HAGH;refGene_tx=XM_023246411.1;refGene_gn=HAGH;refGene_tx=XM_023246412.1;refGene_gn=HAGH;refGene_tx=XM_023246413.1;refGene_gn=HAGH;refGene_tx=XM_023246414.1
E3 43114774 43114872
refGene_gn=HAGH;refGene_tx=XM_023246408.1;refGene_gn=HAGH;refGene_tx=XM_023246409.1;refGene_gn=HAGH;refGene_tx=XM_023246412.1;refGene_gn=HAGH;refGene_tx=XM_023246413.1;refGene_gn=HAGH;refGene_tx=XM_023246414.1
E3 43115656 43115736
refGene_gn=HAGH;refGene_tx=XM_023246408.1;refGene_gn=HAGH;refGene_tx=XM_023246409.1;refGene_gn=HAGH;refGene_tx=XM_023246410.1;refGene_gn=HAGH;refGene_tx=XM_023246411.1;refGene_gn=HAGH;refGene_tx=XM_023246412.1;refGene_gn=HAGH;refGene_tx=XM_023246413.1;refGene_gn=HAGH;refGene_tx=XM_023246414.1
E3 43116200 43116332
refGene_gn=HAGH;refGene_tx=XM_023246408.1;refGene_gn=HAGH;refGene_tx=XM_023246409.1;refGene_gn=HAGH;refGene_tx=XM_023246410.1;refGene_gn=HAGH;refGene_tx=XM_023246411.1;refGene_gn=HAGH;refGene_tx=XM_023246414.1
E3 43121349 43121380 refGene_gn=IGFALS;refGene_tx=XM_023246405.1
E3 43127431 43127603 refGene_gn=IGFALS;refGene_tx=XM_023246404.1
E3 43128546 43130348
refGene_gn=IGFALS;refGene_tx=XM_023246404.1;refGene_gn=IGFALS;refGene_tx=XM_023246405.1
E3 43131612 43131758
refGene_gn=NUBP2;refGene_tx=XM_003999036.5;refGene_gn=NUBP2;refGene_tx=XM_019821219.2;refGene_gn=NUBP2;refGene_tx=XM_019821220.2;refGene_gn=NUBP2;refGene_tx=XM_019821222.2;refGene_gn=NUBP2;refGene_tx=XM_019821223.2
E3 43131865 43131935
refGene_gn=NUBP2;refGene_tx=XM_003999036.5;refGene_gn=NUBP2;refGene_tx

=XM_019821219.2;refGene_gn=NUBP2;refGene_tx=XM_019821220.2;refGene_gn=NUBP2;refGene_tx=XM_019821223.2
E3 43132057 43132168 refGene_gn=NUBP2;refGene_tx=XM_003999036.5;refGene_gn=NUBP2;refGene_tx=XM_019821219.2;refGene_gn=NUBP2;refGene_tx=XM_019821220.2;refGene_gn=NUBP2;refGene_tx=XM_019821222.2;refGene_gn=NUBP2;refGene_tx=XM_019821223.2
E3 43132269 43132424 refGene_gn=NUBP2;refGene_tx=XM_003999036.5;refGene_gn=NUBP2;refGene_tx=XM_019821219.2;refGene_gn=NUBP2;refGene_tx=XM_019821220.2;refGene_gn=NUBP2;refGene_tx=XM_019821222.2;refGene_gn=NUBP2;refGene_tx=XM_019821223.2
E3 43133010 43133209 refGene_gn=NUBP2;refGene_tx=XM_003999036.5;refGene_gn=NUBP2;refGene_tx=XM_019821219.2;refGene_gn=NUBP2;refGene_tx=XM_019821220.2;refGene_gn=NUBP2;refGene_tx=XM_019821222.2;refGene_gn=NUBP2;refGene_tx=XM_019821223.2
E3 43133303 43133422 refGene_gn=NUBP2;refGene_tx=XM_003999036.5;refGene_gn=NUBP2;refGene_tx=XM_019821219.2;refGene_gn=NUBP2;refGene_tx=XM_019821220.2;refGene_gn=NUBP2;refGene_tx=XM_019821222.2
E3 43133694 43133757 refGene_gn=NUBP2;refGene_tx=XM_019821219.2;refGene_gn=NUBP2;refGene_tx=XM_019821220.2;refGene_gn=NUBP2;refGene_tx=XM_019821222.2
E3 43134836 43134852 refGene_gn=NUBP2;refGene_tx=XM_003999036.5;refGene_gn=NUBP2;refGene_tx=XM_019821219.2;refGene_gn=NUBP2;refGene_tx=XM_019821220.2;refGene_gn=NUBP2;refGene_tx=XM_019821222.2
E3 43135135 43135309 refGene_gn=SPSB3;refGene_tx=XM_023246406.1
E3 43136357 43136495 refGene_gn=SPSB3;refGene_tx=XM_023246406.1
E3 43138983 43139161 refGene_gn=SPSB3;refGene_tx=XM_023246406.1
E3 43139254 43139442 refGene_gn=SPSB3;refGene_tx=XM_023246406.1
E3 43139561 43139664 refGene_gn=SPSB3;refGene_tx=XM_023246406.1
E3 43139744 43139870 refGene_gn=SPSB3;refGene_tx=XM_023246406.1
E3 43140083 43140424 refGene_gn=SPSB3;refGene_tx=XM_023246406.1
E3 43141232 43141403 refGene_gn=EME2;refGene_tx=XM_023246407.1
E3 43141489 43141679 refGene_gn=EME2;refGene_tx=XM_023246407.1
E3 43141864 43141980 refGene_gn=EME2;refGene_tx=XM_023246407.1
E3 43142149 43142246 refGene_gn=EME2;refGene_tx=XM_023246407.1
E3 43142430 43142522 refGene_gn=EME2;refGene_tx=XM_023246407.1
E3 43143183 43143276 refGene_gn=EME2;refGene_tx=XM_023246407.1
E3 43143561 43143698 refGene_gn=EME2;refGene_tx=XM_023246407.1
E3 43143906 43144123 refGene_gn=EME2;refGene_tx=XM_023246407.1
E3 43144234 43144555 refGene_gn=MRPS34;refGene_tx=XM_011290727.3
E3 43144644 43144687 refGene_gn=MRPS34;refGene_tx=XM_011290727.3
E3 43144776 43145075 refGene_gn=MRPS34;refGene_tx=XM_011290727.3
E3 43145588 43145634 refGene_gn=NME3;refGene_tx=XM_023246416.1
E3 43145720 43145851 refGene_gn=NME3;refGene_tx=XM_023246416.1
E3 43145943 43146045 refGene_gn=NME3;refGene_tx=XM_023246416.1
E3 43146123 43146236 refGene_gn=NME3;refGene_tx=XM_023246416.1

E3	43146312	43146445	refGene_gn=NME3;refGene_tx=XM_023246416.1
E3	43147865	43147987
refGene_gn=MAPK8IP3;refGene_tx=XM_023246385.1;refGene_gn=MAPK8IP3;refGene_tx=XM_023246386.1;refGene_gn=MAPK8IP3;refGene_tx=XM_023246387.1;refGene_gn=MAPK8IP3;refGene_tx=XM_023246388.1;refGene_gn=MAPK8IP3;refGene_tx=XM_023246389.1;refGene_gn=MAPK8IP3;refGene_tx=XM_023246390.1;refGene_gn=MAPK8IP3;refGene_tx=XM_023246391.1;refGene_gn=MAPK8IP3;refGene_tx=XM_023246392.1;refGene_gn=MAPK8IP3;refGene_tx=XM_023246393.1;refGene_gn=MAPK8IP3;refGene_tx=XM_023246394.1;refGene_gn=MAPK8IP3;refGene_tx=XM_023246395.1;refGene_gn=MAPK8IP3;refGene_tx=XM_023246396.1;refGene_gn=MAPK8IP3;refGene_tx=XM_023246397.1;refGene_gn=MAPK8IP3;refGene_tx=XM_023246398.1;refGene_gn=MAPK8IP3;refGene_tx=XM_023246399.1;refGene_gn=MAPK8IP3;refGene_tx=XM_023246400.1;refGene_gn=MAPK8IP3;refGene_tx=XM_023246401.1
E3	43148059	43148209
refGene_gn=MAPK8IP3;refGene_tx=XM_023246385.1;refGene_gn=MAPK8IP3;refGene_tx=XM_023246386.1;refGene_gn=MAPK8IP3;refGene_tx=XM_023246387.1;refGene_gn=MAPK8IP3;refGene_tx=XM_023246388.1;refGene_gn=MAPK8IP3;refGene_tx=XM_023246389.1;refGene_gn=MAPK8IP3;refGene_tx=XM_023246390.1;refGene_gn=MAPK8IP3;refGene_tx=XM_023246391.1;refGene_gn=MAPK8IP3;refGene_tx=XM_023246392.1;refGene_gn=MAPK8IP3;refGene_tx=XM_023246393.1;refGene_gn=MAPK8IP3;refGene_tx=XM_023246394.1;refGene_gn=MAPK8IP3;refGene_tx=XM_023246395.1;refGene_gn=MAPK8IP3;refGene_tx=XM_023246396.1;refGene_gn=MAPK8IP3;refGene_tx=XM_023246397.1;refGene_gn=MAPK8IP3;refGene_tx=XM_023246398.1;refGene_gn=MAPK8IP3;refGene_tx=XM_023246399.1;refGene_gn=MAPK8IP3;refGene_tx=XM_023246400.1;refGene_gn=MAPK8IP3;refGene_tx=XM_023246401.1
E3	43148295	43148475
refGene_gn=MAPK8IP3;refGene_tx=XM_023246385.1;refGene_gn=MAPK8IP3;refGene_tx=XM_023246386.1;refGene_gn=MAPK8IP3;refGene_tx=XM_023246387.1;refGene_gn=MAPK8IP3;refGene_tx=XM_023246388.1;refGene_gn=MAPK8IP3;refGene_tx=XM_023246389.1;refGene_gn=MAPK8IP3;refGene_tx=XM_023246390.1;refGene_gn=MAPK8IP3;refGene_tx=XM_023246391.1;refGene_gn=MAPK8IP3;refGene_tx=XM_023246392.1;refGene_gn=MAPK8IP3;refGene_tx=XM_023246393.1;refGene_gn=MAPK8IP3;refGene_tx=XM_023246394.1;refGene_gn=MAPK8IP3;refGene_tx=XM_023246395.1;refGene_gn=MAPK8IP3;refGene_tx=XM_023246396.1;refGene_gn=MAPK8IP3;refGene_tx=XM_023246397.1;refGene_gn=MAPK8IP3;refGene_tx=XM_023246398.1;refGene_gn=MAPK8IP3;refGene_tx=XM_023246399.1;refGene_gn=MAPK8IP3;refGene_tx=XM_023246400.1;refGene_gn=MAPK8IP3;refGene_tx=XM_023246401.1
E3	43148649	43148688
refGene_gn=MAPK8IP3;refGene_tx=XM_023246385.1;refGene_gn=MAPK8IP3;refGene_tx=XM_023246386.1;refGene_gn=MAPK8IP3;refGene_tx=XM_023246387.1;refGene_gn=MAPK8IP3;refGene_tx=XM_023246388.1;refGene_gn=MAPK8IP3;refGene_tx=XM_023246389.1;refGene_gn=MAPK8IP3;refGene_tx=XM_023246390.1;refGene_gn=MAPK8IP3;refGene_tx=XM_023246391.1;refGene_gn=MAPK8IP3;refGene_tx=XM_023246392.1;refGene_gn=MAPK8IP3;refGene_tx=XM_023246393.1;refGene_gn=MAPK8IP3;refGene_tx=XM_023246394.1;refGene_gn=MAPK8IP3;refGene_tx=XM_023246395.1;refGene_gn=MAPK8IP3;refGene_tx=XM_023246396.1;refGene_gn=MAPK8IP3;refGene_tx=XM_023246397.1;refGene_gn=MAPK8IP3;refGene_tx=X

M_023246398.1;refGene_gn=MAPK8IP3;refGene_tx=XM_023246399.1;refGene_gn=MAPK8IP3;refGene_tx=XM_023246400.1;refGene_gn=MAPK8IP3;refGene_tx=XM_023246401.1
E3 43148883 43148997
refGene_gn=MAPK8IP3;refGene_tx=XM_023246385.1;refGene_gn=MAPK8IP3;refGene_tx=XM_023246386.1;refGene_gn=MAPK8IP3;refGene_tx=XM_023246387.1;refGene_gn=MAPK8IP3;refGene_tx=XM_023246388.1;refGene_gn=MAPK8IP3;refGene_tx=XM_023246389.1;refGene_gn=MAPK8IP3;refGene_tx=XM_023246390.1;refGene_gn=MAPK8IP3;refGene_tx=XM_023246391.1;refGene_gn=MAPK8IP3;refGene_tx=XM_023246392.1;refGene_gn=MAPK8IP3;refGene_tx=XM_023246393.1;refGene_gn=MAPK8IP3;refGene_tx=XM_023246394.1;refGene_gn=MAPK8IP3;refGene_tx=XM_023246395.1;refGene_gn=MAPK8IP3;refGene_tx=XM_023246396.1;refGene_gn=MAPK8IP3;refGene_tx=XM_023246397.1;refGene_gn=MAPK8IP3;refGene_tx=XM_023246398.1;refGene_gn=MAPK8IP3;refGene_tx=XM_023246399.1;refGene_gn=MAPK8IP3;refGene_tx=XM_023246400.1;refGene_gn=MAPK8IP3;refGene_tx=XM_023246401.1
E3 43149077 43149249
refGene_gn=MAPK8IP3;refGene_tx=XM_023246385.1;refGene_gn=MAPK8IP3;refGene_tx=XM_023246386.1;refGene_gn=MAPK8IP3;refGene_tx=XM_023246387.1;refGene_gn=MAPK8IP3;refGene_tx=XM_023246388.1;refGene_gn=MAPK8IP3;refGene_tx=XM_023246389.1;refGene_gn=MAPK8IP3;refGene_tx=XM_023246390.1;refGene_gn=MAPK8IP3;refGene_tx=XM_023246391.1;refGene_gn=MAPK8IP3;refGene_tx=XM_023246392.1;refGene_gn=MAPK8IP3;refGene_tx=XM_023246393.1;refGene_gn=MAPK8IP3;refGene_tx=XM_023246394.1;refGene_gn=MAPK8IP3;refGene_tx=XM_023246395.1;refGene_gn=MAPK8IP3;refGene_tx=XM_023246396.1;refGene_gn=MAPK8IP3;refGene_tx=XM_023246397.1;refGene_gn=MAPK8IP3;refGene_tx=XM_023246398.1;refGene_gn=MAPK8IP3;refGene_tx=XM_023246399.1;refGene_gn=MAPK8IP3;refGene_tx=XM_023246400.1;refGene_gn=MAPK8IP3;refGene_tx=XM_023246401.1
E3 43149658 43149807
refGene_gn=MAPK8IP3;refGene_tx=XM_023246385.1;refGene_gn=MAPK8IP3;refGene_tx=XM_023246386.1;refGene_gn=MAPK8IP3;refGene_tx=XM_023246387.1;refGene_gn=MAPK8IP3;refGene_tx=XM_023246388.1;refGene_gn=MAPK8IP3;refGene_tx=XM_023246389.1;refGene_gn=MAPK8IP3;refGene_tx=XM_023246390.1;refGene_gn=MAPK8IP3;refGene_tx=XM_023246391.1;refGene_gn=MAPK8IP3;refGene_tx=XM_023246392.1;refGene_gn=MAPK8IP3;refGene_tx=XM_023246393.1;refGene_gn=MAPK8IP3;refGene_tx=XM_023246394.1;refGene_gn=MAPK8IP3;refGene_tx=XM_023246395.1;refGene_gn=MAPK8IP3;refGene_tx=XM_023246396.1;refGene_gn=MAPK8IP3;refGene_tx=XM_023246397.1;refGene_gn=MAPK8IP3;refGene_tx=XM_023246398.1;refGene_gn=MAPK8IP3;refGene_tx=XM_023246399.1;refGene_gn=MAPK8IP3;refGene_tx=XM_023246400.1;refGene_gn=MAPK8IP3;refGene_tx=XM_023246401.1
E3 43149892 43149960
refGene_gn=MAPK8IP3;refGene_tx=XM_023246385.1;refGene_gn=MAPK8IP3;refGene_tx=XM_023246386.1;refGene_gn=MAPK8IP3;refGene_tx=XM_023246387.1;refGene_gn=MAPK8IP3;refGene_tx=XM_023246388.1;refGene_gn=MAPK8IP3;refGene_tx=XM_023246389.1;refGene_gn=MAPK8IP3;refGene_tx=XM_023246390.1;refGene_gn=MAPK8IP3;refGene_tx=XM_023246391.1;refGene_gn=MAPK8IP3;refGene_tx=XM_023246392.1;refGene_gn=MAPK8IP3;refGene_tx=XM_023246393.1;refGene_gn=MAPK8IP3;refGene_tx=XM_023246394.1;refGene_gn=MAPK8IP3;refGene_tx

=XM_023246395.1;refGene_gn=MAPK8IP3;refGene_tx=XM_023246396.1;refGene_gn=MAPK8IP3;refGene_tx=XM_023246397.1;refGene_gn=MAPK8IP3;refGene_tx=XM_023246398.1;refGene_gn=MAPK8IP3;refGene_tx=XM_023246399.1;refGene_gn=MAPK8IP3;refGene_tx=XM_023246400.1;refGene_gn=MAPK8IP3;refGene_tx=XM_023246401.1
E3 43150044 43150125
refGene_gn=MAPK8IP3;refGene_tx=XM_023246385.1;refGene_gn=MAPK8IP3;refGene_tx=XM_023246386.1;refGene_gn=MAPK8IP3;refGene_tx=XM_023246387.1;refGene_gn=MAPK8IP3;refGene_tx=XM_023246388.1;refGene_gn=MAPK8IP3;refGene_tx=XM_023246389.1;refGene_gn=MAPK8IP3;refGene_tx=XM_023246390.1;refGene_gn=MAPK8IP3;refGene_tx=XM_023246391.1;refGene_gn=MAPK8IP3;refGene_tx=XM_023246392.1;refGene_gn=MAPK8IP3;refGene_tx=XM_023246393.1;refGene_gn=MAPK8IP3;refGene_tx=XM_023246394.1;refGene_gn=MAPK8IP3;refGene_tx=XM_023246395.1;refGene_gn=MAPK8IP3;refGene_tx=XM_023246396.1;refGene_gn=MAPK8IP3;refGene_tx=XM_023246397.1;refGene_gn=MAPK8IP3;refGene_tx=XM_023246398.1;refGene_gn=MAPK8IP3;refGene_tx=XM_023246399.1;refGene_gn=MAPK8IP3;refGene_tx=XM_023246400.1;refGene_gn=MAPK8IP3;refGene_tx=XM_023246401.1
E3 43150202 43150322
refGene_gn=MAPK8IP3;refGene_tx=XM_023246385.1;refGene_gn=MAPK8IP3;refGene_tx=XM_023246386.1;refGene_gn=MAPK8IP3;refGene_tx=XM_023246387.1;refGene_gn=MAPK8IP3;refGene_tx=XM_023246388.1;refGene_gn=MAPK8IP3;refGene_tx=XM_023246389.1;refGene_gn=MAPK8IP3;refGene_tx=XM_023246390.1;refGene_gn=MAPK8IP3;refGene_tx=XM_023246391.1;refGene_gn=MAPK8IP3;refGene_tx=XM_023246392.1;refGene_gn=MAPK8IP3;refGene_tx=XM_023246393.1;refGene_gn=MAPK8IP3;refGene_tx=XM_023246394.1;refGene_gn=MAPK8IP3;refGene_tx=XM_023246395.1;refGene_gn=MAPK8IP3;refGene_tx=XM_023246396.1;refGene_gn=MAPK8IP3;refGene_tx=XM_023246397.1;refGene_gn=MAPK8IP3;refGene_tx=XM_023246398.1;refGene_gn=MAPK8IP3;refGene_tx=XM_023246399.1;refGene_gn=MAPK8IP3;refGene_tx=XM_023246400.1;refGene_gn=MAPK8IP3;refGene_tx=XM_023246401.1
E3 43150425 43150615
refGene_gn=MAPK8IP3;refGene_tx=XM_023246385.1;refGene_gn=MAPK8IP3;refGene_tx=XM_023246386.1;refGene_gn=MAPK8IP3;refGene_tx=XM_023246387.1;refGene_gn=MAPK8IP3;refGene_tx=XM_023246388.1;refGene_gn=MAPK8IP3;refGene_tx=XM_023246389.1;refGene_gn=MAPK8IP3;refGene_tx=XM_023246390.1;refGene_gn=MAPK8IP3;refGene_tx=XM_023246391.1;refGene_gn=MAPK8IP3;refGene_tx=XM_023246392.1;refGene_gn=MAPK8IP3;refGene_tx=XM_023246393.1;refGene_gn=MAPK8IP3;refGene_tx=XM_023246394.1;refGene_gn=MAPK8IP3;refGene_tx=XM_023246395.1;refGene_gn=MAPK8IP3;refGene_tx=XM_023246396.1;refGene_gn=MAPK8IP3;refGene_tx=XM_023246397.1;refGene_gn=MAPK8IP3;refGene_tx=XM_023246398.1;refGene_gn=MAPK8IP3;refGene_tx=XM_023246399.1;refGene_gn=MAPK8IP3;refGene_tx=XM_023246400.1;refGene_gn=MAPK8IP3;refGene_tx=XM_023246401.1
E3 43150709 43150892
refGene_gn=MAPK8IP3;refGene_tx=XM_023246385.1;refGene_gn=MAPK8IP3;refGene_tx=XM_023246386.1;refGene_gn=MAPK8IP3;refGene_tx=XM_023246387.1;refGene_gn=MAPK8IP3;refGene_tx=XM_023246388.1;refGene_gn=MAPK8IP3;refGene_tx=XM_023246389.1;refGene_gn=MAPK8IP3;refGene_tx=XM_023246390.1;refGene_gn=MAPK8IP3;refGene_tx=XM_023246391.1;refGene_gn=MAPK8IP3;refGene_

tx=XM_023246392.1;refGene_gn=MAPK8IP3;refGene_tx=XM_023246393.1;refGene_gn=MAPK8IP3;refGene_tx=XM_023246394.1;refGene_gn=MAPK8IP3;refGene_tx=XM_023246395.1;refGene_gn=MAPK8IP3;refGene_tx=XM_023246396.1;refGene_gn=MAPK8IP3;refGene_tx=XM_023246397.1;refGene_gn=MAPK8IP3;refGene_tx=XM_023246398.1;refGene_gn=MAPK8IP3;refGene_tx=XM_023246399.1;refGene_gn=MAPK8IP3;refGene_tx=XM_023246400.1;refGene_gn=MAPK8IP3;refGene_tx=XM_023246401.1
E3 43151546 43151712
refGene_gn=MAPK8IP3;refGene_tx=XM_023246385.1;refGene_gn=MAPK8IP3;refGene_tx=XM_023246386.1;refGene_gn=MAPK8IP3;refGene_tx=XM_023246387.1;refGene_gn=MAPK8IP3;refGene_tx=XM_023246388.1;refGene_gn=MAPK8IP3;refGene_tx=XM_023246389.1;refGene_gn=MAPK8IP3;refGene_tx=XM_023246390.1;refGene_gn=MAPK8IP3;refGene_tx=XM_023246391.1;refGene_gn=MAPK8IP3;refGene_tx=XM_023246392.1;refGene_gn=MAPK8IP3;refGene_tx=XM_023246393.1;refGene_gn=MAPK8IP3;refGene_tx=XM_023246394.1;refGene_gn=MAPK8IP3;refGene_tx=XM_023246395.1;refGene_gn=MAPK8IP3;refGene_tx=XM_023246396.1;refGene_gn=MAPK8IP3;refGene_tx=XM_023246397.1;refGene_gn=MAPK8IP3;refGene_tx=XM_023246398.1;refGene_gn=MAPK8IP3;refGene_tx=XM_023246399.1;refGene_gn=MAPK8IP3;refGene_tx=XM_023246400.1;refGene_gn=MAPK8IP3;refGene_tx=XM_023246401.1
E3 43152296 43152452
refGene_gn=MAPK8IP3;refGene_tx=XM_023246385.1;refGene_gn=MAPK8IP3;refGene_tx=XM_023246386.1;refGene_gn=MAPK8IP3;refGene_tx=XM_023246387.1;refGene_gn=MAPK8IP3;refGene_tx=XM_023246388.1;refGene_gn=MAPK8IP3;refGene_tx=XM_023246389.1;refGene_gn=MAPK8IP3;refGene_tx=XM_023246390.1;refGene_gn=MAPK8IP3;refGene_tx=XM_023246391.1;refGene_gn=MAPK8IP3;refGene_tx=XM_023246392.1;refGene_gn=MAPK8IP3;refGene_tx=XM_023246393.1;refGene_gn=MAPK8IP3;refGene_tx=XM_023246394.1;refGene_gn=MAPK8IP3;refGene_tx=XM_023246395.1;refGene_gn=MAPK8IP3;refGene_tx=XM_023246396.1;refGene_gn=MAPK8IP3;refGene_tx=XM_023246397.1;refGene_gn=MAPK8IP3;refGene_tx=XM_023246398.1;refGene_gn=MAPK8IP3;refGene_tx=XM_023246399.1;refGene_gn=MAPK8IP3;refGene_tx=XM_023246400.1;refGene_gn=MAPK8IP3;refGene_tx=XM_023246401.1
E3 43152571 43152667
refGene_gn=MAPK8IP3;refGene_tx=XM_023246385.1;refGene_gn=MAPK8IP3;refGene_tx=XM_023246386.1;refGene_gn=MAPK8IP3;refGene_tx=XM_023246387.1;refGene_gn=MAPK8IP3;refGene_tx=XM_023246388.1;refGene_gn=MAPK8IP3;refGene_tx=XM_023246389.1;refGene_gn=MAPK8IP3;refGene_tx=XM_023246390.1;refGene_gn=MAPK8IP3;refGene_tx=XM_023246391.1;refGene_gn=MAPK8IP3;refGene_tx=XM_023246392.1;refGene_gn=MAPK8IP3;refGene_tx=XM_023246393.1;refGene_gn=MAPK8IP3;refGene_tx=XM_023246394.1;refGene_gn=MAPK8IP3;refGene_tx=XM_023246395.1;refGene_gn=MAPK8IP3;refGene_tx=XM_023246396.1;refGene_gn=MAPK8IP3;refGene_tx=XM_023246397.1;refGene_gn=MAPK8IP3;refGene_tx=XM_023246398.1;refGene_gn=MAPK8IP3;refGene_tx=XM_023246399.1;refGene_gn=MAPK8IP3;refGene_tx=XM_023246400.1;refGene_gn=MAPK8IP3;refGene_tx=XM_023246401.1
E3 43152810 43152937
refGene_gn=MAPK8IP3;refGene_tx=XM_023246385.1;refGene_gn=MAPK8IP3;refGene_tx=XM_023246386.1;refGene_gn=MAPK8IP3;refGene_tx=XM_023246387.1;refGene_gn=MAPK8IP3;refGene_tx=XM_023246388.1;refGene_gn=MAPK8IP3;refGen

e_tx=XM_023246389.1;refGene_gn=MAPK8IP3;refGene_tx=XM_023246390.1;refGene_gn=MAPK8IP3;refGene_tx=XM_023246391.1;refGene_gn=MAPK8IP3;refGene_tx=XM_023246392.1;refGene_gn=MAPK8IP3;refGene_tx=XM_023246393.1;refGene_gn=MAPK8IP3;refGene_tx=XM_023246394.1;refGene_gn=MAPK8IP3;refGene_tx=XM_023246395.1;refGene_gn=MAPK8IP3;refGene_tx=XM_023246396.1;refGene_gn=MAPK8IP3;refGene_tx=XM_023246397.1;refGene_gn=MAPK8IP3;refGene_tx=XM_023246398.1;refGene_gn=MAPK8IP3;refGene_tx=XM_023246399.1;refGene_gn=MAPK8IP3;refGene_tx=XM_023246400.1;refGene_gn=MAPK8IP3;refGene_tx=XM_023246401.1

E3 43153536 43153707

refGene_gn=MAPK8IP3;refGene_tx=XM_023246385.1;refGene_gn=MAPK8IP3;refGene_tx=XM_023246386.1;refGene_gn=MAPK8IP3;refGene_tx=XM_023246387.1;refGene_gn=MAPK8IP3;refGene_tx=XM_023246388.1;refGene_gn=MAPK8IP3;refGene_tx=XM_023246389.1;refGene_gn=MAPK8IP3;refGene_tx=XM_023246390.1;refGene_gn=MAPK8IP3;refGene_tx=XM_023246391.1;refGene_gn=MAPK8IP3;refGene_tx=XM_023246392.1;refGene_gn=MAPK8IP3;refGene_tx=XM_023246393.1;refGene_gn=MAPK8IP3;refGene_tx=XM_023246394.1;refGene_gn=MAPK8IP3;refGene_tx=XM_023246395.1;refGene_gn=MAPK8IP3;refGene_tx=XM_023246396.1;refGene_gn=MAPK8IP3;refGene_tx=XM_023246397.1;refGene_gn=MAPK8IP3;refGene_tx=XM_023246398.1;refGene_gn=MAPK8IP3;refGene_tx=XM_023246399.1;refGene_gn=MAPK8IP3;refGene_tx=XM_023246400.1;refGene_gn=MAPK8IP3;refGene_tx=XM_023246401.1

E3 43153789 43153846

refGene_gn=MAPK8IP3;refGene_tx=XM_023246385.1;refGene_gn=MAPK8IP3;refGene_tx=XM_023246386.1;refGene_gn=MAPK8IP3;refGene_tx=XM_023246387.1;refGene_gn=MAPK8IP3;refGene_tx=XM_023246388.1;refGene_gn=MAPK8IP3;refGene_tx=XM_023246389.1;refGene_gn=MAPK8IP3;refGene_tx=XM_023246390.1;refGene_gn=MAPK8IP3;refGene_tx=XM_023246391.1;refGene_gn=MAPK8IP3;refGene_tx=XM_023246392.1;refGene_gn=MAPK8IP3;refGene_tx=XM_023246393.1;refGene_gn=MAPK8IP3;refGene_tx=XM_023246394.1;refGene_gn=MAPK8IP3;refGene_tx=XM_023246395.1;refGene_gn=MAPK8IP3;refGene_tx=XM_023246396.1;refGene_gn=MAPK8IP3;refGene_tx=XM_023246397.1;refGene_gn=MAPK8IP3;refGene_tx=XM_023246398.1;refGene_gn=MAPK8IP3;refGene_tx=XM_023246399.1;refGene_gn=MAPK8IP3;refGene_tx=XM_023246400.1;refGene_gn=MAPK8IP3;refGene_tx=XM_023246401.1

E3 43154309 43154440

refGene_gn=MAPK8IP3;refGene_tx=XM_023246385.1;refGene_gn=MAPK8IP3;refGene_tx=XM_023246386.1;refGene_gn=MAPK8IP3;refGene_tx=XM_023246387.1;refGene_gn=MAPK8IP3;refGene_tx=XM_023246388.1;refGene_gn=MAPK8IP3;refGene_tx=XM_023246389.1;refGene_gn=MAPK8IP3;refGene_tx=XM_023246390.1;refGene_gn=MAPK8IP3;refGene_tx=XM_023246391.1;refGene_gn=MAPK8IP3;refGene_tx=XM_023246392.1;refGene_gn=MAPK8IP3;refGene_tx=XM_023246393.1;refGene_gn=MAPK8IP3;refGene_tx=XM_023246394.1;refGene_gn=MAPK8IP3;refGene_tx=XM_023246395.1;refGene_gn=MAPK8IP3;refGene_tx=XM_023246396.1;refGene_gn=MAPK8IP3;refGene_tx=XM_023246397.1;refGene_gn=MAPK8IP3;refGene_tx=XM_023246398.1;refGene_gn=MAPK8IP3;refGene_tx=XM_023246399.1;refGene_gn=MAPK8IP3;refGene_tx=XM_023246400.1;refGene_gn=MAPK8IP3;refGene_tx=XM_023246401.1

E3 43155513 43155595

refGene_gn=MAPK8IP3;refGene_tx=XM_023246385.1;refGene_gn=MAPK8IP3;refG

ene_tx=XM_023246386.1;refGene_gn=MAPK8IP3;refGene_tx=XM_023246387.1;refGene_gn=MAPK8IP3;refGene_tx=XM_023246388.1;refGene_gn=MAPK8IP3;refGene_tx=XM_023246389.1;refGene_gn=MAPK8IP3;refGene_tx=XM_023246390.1;refGene_gn=MAPK8IP3;refGene_tx=XM_023246391.1;refGene_gn=MAPK8IP3;refGene_tx=XM_023246392.1;refGene_gn=MAPK8IP3;refGene_tx=XM_023246393.1;refGene_gn=MAPK8IP3;refGene_tx=XM_023246394.1;refGene_gn=MAPK8IP3;refGene_tx=XM_023246395.1;refGene_gn=MAPK8IP3;refGene_tx=XM_023246396.1;refGene_gn=MAPK8IP3;refGene_tx=XM_023246397.1;refGene_gn=MAPK8IP3;refGene_tx=XM_023246398.1;refGene_gn=MAPK8IP3;refGene_tx=XM_023246399.1;refGene_gn=MAPK8IP3;refGene_tx=XM_023246400.1;refGene_gn=MAPK8IP3;refGene_tx=XM_023246401.1
E3 43156165 43156318
refGene_gn=MAPK8IP3;refGene_tx=XM_023246385.1;refGene_gn=MAPK8IP3;refGene_tx=XM_023246386.1;refGene_gn=MAPK8IP3;refGene_tx=XM_023246387.1;refGene_gn=MAPK8IP3;refGene_tx=XM_023246388.1;refGene_gn=MAPK8IP3;refGene_tx=XM_023246389.1;refGene_gn=MAPK8IP3;refGene_tx=XM_023246390.1;refGene_gn=MAPK8IP3;refGene_tx=XM_023246391.1;refGene_gn=MAPK8IP3;refGene_tx=XM_023246392.1;refGene_gn=MAPK8IP3;refGene_tx=XM_023246393.1;refGene_gn=MAPK8IP3;refGene_tx=XM_023246394.1;refGene_gn=MAPK8IP3;refGene_tx=XM_023246395.1;refGene_gn=MAPK8IP3;refGene_tx=XM_023246396.1;refGene_gn=MAPK8IP3;refGene_tx=XM_023246397.1;refGene_gn=MAPK8IP3;refGene_tx=XM_023246398.1;refGene_gn=MAPK8IP3;refGene_tx=XM_023246399.1;refGene_gn=MAPK8IP3;refGene_tx=XM_023246400.1;refGene_gn=MAPK8IP3;refGene_tx=XM_023246401.1
E3 43156480 43156538
refGene_gn=MAPK8IP3;refGene_tx=XM_023246385.1;refGene_gn=MAPK8IP3;refGene_tx=XM_023246386.1;refGene_gn=MAPK8IP3;refGene_tx=XM_023246387.1;refGene_gn=MAPK8IP3;refGene_tx=XM_023246388.1;refGene_gn=MAPK8IP3;refGene_tx=XM_023246389.1;refGene_gn=MAPK8IP3;refGene_tx=XM_023246390.1;refGene_gn=MAPK8IP3;refGene_tx=XM_023246391.1;refGene_gn=MAPK8IP3;refGene_tx=XM_023246392.1;refGene_gn=MAPK8IP3;refGene_tx=XM_023246393.1;refGene_gn=MAPK8IP3;refGene_tx=XM_023246394.1;refGene_gn=MAPK8IP3;refGene_tx=XM_023246395.1;refGene_gn=MAPK8IP3;refGene_tx=XM_023246396.1;refGene_gn=MAPK8IP3;refGene_tx=XM_023246397.1;refGene_gn=MAPK8IP3;refGene_tx=XM_023246398.1;refGene_gn=MAPK8IP3;refGene_tx=XM_023246399.1;refGene_gn=MAPK8IP3;refGene_tx=XM_023246400.1;refGene_gn=MAPK8IP3;refGene_tx=XM_023246401.1
E3 43157739 43157757
refGene_gn=MAPK8IP3;refGene_tx=XM_023246385.1;refGene_gn=MAPK8IP3;refGene_tx=XM_023246386.1;refGene_gn=MAPK8IP3;refGene_tx=XM_023246388.1;refGene_gn=MAPK8IP3;refGene_tx=XM_023246389.1;refGene_gn=MAPK8IP3;refGene_tx=XM_023246390.1;refGene_gn=MAPK8IP3;refGene_tx=XM_023246395.1;refGene_gn=MAPK8IP3;refGene_tx=XM_023246396.1;refGene_gn=MAPK8IP3;refGene_tx=XM_023246397.1;refGene_gn=MAPK8IP3;refGene_tx=XM_023246398.1;refGene_gn=MAPK8IP3;refGene_tx=XM_023246399.1;refGene_gn=MAPK8IP3;refGene_tx=XM_023246401.1
E3 43159392 43159404
refGene_gn=MAPK8IP3;refGene_tx=XM_023246385.1;refGene_gn=MAPK8IP3;refGene_tx=XM_023246386.1;refGene_gn=MAPK8IP3;refGene_tx=XM_023246387.1;refGene_gn=MAPK8IP3;refGene_tx=XM_023246388.1;refGene_gn=MAPK8IP3;refGen

e_tx=XM_023246389.1;refGene_gn=MAPK8IP3;refGene_tx=XM_023246390.1;refGene_gn=MAPK8IP3;refGene_tx=XM_023246392.1;refGene_gn=MAPK8IP3;refGene_tx=XM_023246394.1;refGene_gn=MAPK8IP3;refGene_tx=XM_023246395.1;refGene_gn=MAPK8IP3;refGene_tx=XM_023246396.1;refGene_gn=MAPK8IP3;refGene_tx=XM_023246397.1;refGene_gn=MAPK8IP3;refGene_tx=XM_023246398.1;refGene_gn=MAPK8IP3;refGene_tx=XM_023246399.1;refGene_gn=MAPK8IP3;refGene_tx=XM_023246400.1;refGene_gn=MAPK8IP3;refGene_tx=XM_023246401.1
E3 43165920 43166039
refGene_gn=MAPK8IP3;refGene_tx=XM_023246385.1;refGene_gn=MAPK8IP3;refGene_tx=XM_023246386.1;refGene_gn=MAPK8IP3;refGene_tx=XM_023246387.1;refGene_gn=MAPK8IP3;refGene_tx=XM_023246388.1;refGene_gn=MAPK8IP3;refGene_tx=XM_023246389.1;refGene_gn=MAPK8IP3;refGene_tx=XM_023246390.1;refGene_gn=MAPK8IP3;refGene_tx=XM_023246391.1;refGene_gn=MAPK8IP3;refGene_tx=XM_023246392.1;refGene_gn=MAPK8IP3;refGene_tx=XM_023246393.1;refGene_gn=MAPK8IP3;refGene_tx=XM_023246394.1;refGene_gn=MAPK8IP3;refGene_tx=XM_023246395.1;refGene_gn=MAPK8IP3;refGene_tx=XM_023246396.1;refGene_gn=MAPK8IP3;refGene_tx=XM_023246397.1;refGene_gn=MAPK8IP3;refGene_tx=XM_023246398.1;refGene_gn=MAPK8IP3;refGene_tx=XM_023246399.1;refGene_gn=MAPK8IP3;refGene_tx=XM_023246400.1;refGene_gn=MAPK8IP3;refGene_tx=XM_023246401.1
E3 43166237 43166340
refGene_gn=MAPK8IP3;refGene_tx=XM_023246385.1;refGene_gn=MAPK8IP3;refGene_tx=XM_023246386.1;refGene_gn=MAPK8IP3;refGene_tx=XM_023246387.1;refGene_gn=MAPK8IP3;refGene_tx=XM_023246388.1;refGene_gn=MAPK8IP3;refGene_tx=XM_023246389.1;refGene_gn=MAPK8IP3;refGene_tx=XM_023246390.1;refGene_gn=MAPK8IP3;refGene_tx=XM_023246391.1;refGene_gn=MAPK8IP3;refGene_tx=XM_023246392.1;refGene_gn=MAPK8IP3;refGene_tx=XM_023246393.1;refGene_gn=MAPK8IP3;refGene_tx=XM_023246394.1;refGene_gn=MAPK8IP3;refGene_tx=XM_023246395.1;refGene_gn=MAPK8IP3;refGene_tx=XM_023246396.1;refGene_gn=MAPK8IP3;refGene_tx=XM_023246397.1;refGene_gn=MAPK8IP3;refGene_tx=XM_023246398.1;refGene_gn=MAPK8IP3;refGene_tx=XM_023246399.1;refGene_gn=MAPK8IP3;refGene_tx=XM_023246400.1;refGene_gn=MAPK8IP3;refGene_tx=XM_023246401.1
E3 43166926 43167173
refGene_gn=MAPK8IP3;refGene_tx=XM_023246385.1;refGene_gn=MAPK8IP3;refGene_tx=XM_023246386.1;refGene_gn=MAPK8IP3;refGene_tx=XM_023246387.1;refGene_gn=MAPK8IP3;refGene_tx=XM_023246388.1;refGene_gn=MAPK8IP3;refGene_tx=XM_023246389.1;refGene_gn=MAPK8IP3;refGene_tx=XM_023246390.1;refGene_gn=MAPK8IP3;refGene_tx=XM_023246391.1;refGene_gn=MAPK8IP3;refGene_tx=XM_023246392.1;refGene_gn=MAPK8IP3;refGene_tx=XM_023246393.1;refGene_gn=MAPK8IP3;refGene_tx=XM_023246394.1;refGene_gn=MAPK8IP3;refGene_tx=XM_023246395.1;refGene_gn=MAPK8IP3;refGene_tx=XM_023246396.1;refGene_gn=MAPK8IP3;refGene_tx=XM_023246397.1;refGene_gn=MAPK8IP3;refGene_tx=XM_023246398.1;refGene_gn=MAPK8IP3;refGene_tx=XM_023246399.1;refGene_gn=MAPK8IP3;refGene_tx=XM_023246400.1;refGene_gn=MAPK8IP3;refGene_tx=XM_023246401.1
E3 43170703 43170848
refGene_gn=MAPK8IP3;refGene_tx=XM_023246385.1;refGene_gn=MAPK8IP3;refGene_tx=XM_023246386.1;refGene_gn=MAPK8IP3;refGene_tx=XM_023246387.1;refGene_gn=MAPK8IP3;refGene_tx=XM_023246388.1;refGene_gn=MAPK8IP3;refGen

e_tx=XM_023246389.1;refGene_gn=MAPK8IP3;refGene_tx=XM_023246390.1;refGene_gn=MAPK8IP3;refGene_tx=XM_023246391.1;refGene_gn=MAPK8IP3;refGene_tx=XM_023246392.1;refGene_gn=MAPK8IP3;refGene_tx=XM_023246393.1;refGene_gn=MAPK8IP3;refGene_tx=XM_023246394.1;refGene_gn=MAPK8IP3;refGene_tx=XM_023246395.1;refGene_gn=MAPK8IP3;refGene_tx=XM_023246396.1;refGene_gn=MAPK8IP3;refGene_tx=XM_023246397.1;refGene_gn=MAPK8IP3;refGene_tx=XM_023246398.1;refGene_gn=MAPK8IP3;refGene_tx=XM_023246399.1;refGene_gn=MAPK8IP3;refGene_tx=XM_023246400.1
E3 43173363 43173387
refGene_gn=MAPK8IP3;refGene_tx=XM_023246385.1;refGene_gn=MAPK8IP3;refGene_tx=XM_023246386.1;refGene_gn=MAPK8IP3;refGene_tx=XM_023246387.1;refGene_gn=MAPK8IP3;refGene_tx=XM_023246390.1;refGene_gn=MAPK8IP3;refGene_tx=XM_023246391.1;refGene_gn=MAPK8IP3;refGene_tx=XM_023246393.1;refGene_gn=MAPK8IP3;refGene_tx=XM_023246395.1;refGene_gn=MAPK8IP3;refGene_tx=XM_023246396.1;refGene_gn=MAPK8IP3;refGene_tx=XM_023246397.1
E3 43177182 43177274
refGene_gn=MAPK8IP3;refGene_tx=XM_023246385.1;refGene_gn=MAPK8IP3;refGene_tx=XM_023246386.1;refGene_gn=MAPK8IP3;refGene_tx=XM_023246387.1;refGene_gn=MAPK8IP3;refGene_tx=XM_023246388.1;refGene_gn=MAPK8IP3;refGene_tx=XM_023246389.1;refGene_gn=MAPK8IP3;refGene_tx=XM_023246390.1;refGene_gn=MAPK8IP3;refGene_tx=XM_023246391.1;refGene_gn=MAPK8IP3;refGene_tx=XM_023246392.1;refGene_gn=MAPK8IP3;refGene_tx=XM_023246393.1;refGene_gn=MAPK8IP3;refGene_tx=XM_023246394.1;refGene_gn=MAPK8IP3;refGene_tx=XM_023246395.1;refGene_gn=MAPK8IP3;refGene_tx=XM_023246396.1;refGene_gn=MAPK8IP3;refGene_tx=XM_023246397.1;refGene_gn=MAPK8IP3;refGene_tx=XM_023246398.1;refGene_gn=MAPK8IP3;refGene_tx=XM_023246399.1;refGene_gn=MAPK8IP3;refGene_tx=XM_023246400.1
E3 43177508 43177579
refGene_gn=MAPK8IP3;refGene_tx=XM_023246385.1;refGene_gn=MAPK8IP3;refGene_tx=XM_023246386.1;refGene_gn=MAPK8IP3;refGene_tx=XM_023246387.1;refGene_gn=MAPK8IP3;refGene_tx=XM_023246388.1;refGene_gn=MAPK8IP3;refGene_tx=XM_023246389.1;refGene_gn=MAPK8IP3;refGene_tx=XM_023246390.1;refGene_gn=MAPK8IP3;refGene_tx=XM_023246391.1;refGene_gn=MAPK8IP3;refGene_tx=XM_023246392.1;refGene_gn=MAPK8IP3;refGene_tx=XM_023246393.1;refGene_gn=MAPK8IP3;refGene_tx=XM_023246394.1;refGene_gn=MAPK8IP3;refGene_tx=XM_023246395.1;refGene_gn=MAPK8IP3;refGene_tx=XM_023246396.1;refGene_gn=MAPK8IP3;refGene_tx=XM_023246397.1;refGene_gn=MAPK8IP3;refGene_tx=XM_023246398.1;refGene_gn=MAPK8IP3;refGene_tx=XM_023246399.1;refGene_gn=MAPK8IP3;refGene_tx=XM_023246400.1
E3 43178847 43178968
refGene_gn=MAPK8IP3;refGene_tx=XM_023246385.1;refGene_gn=MAPK8IP3;refGene_tx=XM_023246386.1;refGene_gn=MAPK8IP3;refGene_tx=XM_023246387.1;refGene_gn=MAPK8IP3;refGene_tx=XM_023246388.1;refGene_gn=MAPK8IP3;refGene_tx=XM_023246389.1;refGene_gn=MAPK8IP3;refGene_tx=XM_023246390.1;refGene_gn=MAPK8IP3;refGene_tx=XM_023246391.1;refGene_gn=MAPK8IP3;refGene_tx=XM_023246392.1;refGene_gn=MAPK8IP3;refGene_tx=XM_023246393.1;refGene_gn=MAPK8IP3;refGene_tx=XM_023246394.1;refGene_gn=MAPK8IP3;refGene_tx=XM_023246395.1;refGene_gn=MAPK8IP3;refGene_tx=XM_023246396.1;refGene_gn=MAPK8IP3;refGene_tx=XM_023246397.1;refGene_gn=MAPK8IP3;refGene_tx=XM_023246398.1;refGene_gn=MAPK8IP3;refGene_tx=XM_023246399.1;refGene_gn

=MAPK8IP3;refGene_tx=XM_023246400.1
E3 43193228 43193546
refGene_gn=MAPK8IP3;refGene_tx=XM_023246385.1;refGene_gn=MAPK8IP3;refGene_tx=XM_023246386.1;refGene_gn=MAPK8IP3;refGene_tx=XM_023246387.1;refGene_gn=MAPK8IP3;refGene_tx=XM_023246388.1;refGene_gn=MAPK8IP3;refGene_tx=XM_023246389.1;refGene_gn=MAPK8IP3;refGene_tx=XM_023246390.1;refGene_gn=MAPK8IP3;refGene_tx=XM_023246391.1;refGene_gn=MAPK8IP3;refGene_tx=XM_023246392.1;refGene_gn=MAPK8IP3;refGene_tx=XM_023246393.1;refGene_gn=MAPK8IP3;refGene_tx=XM_023246394.1;refGene_gn=MAPK8IP3;refGene_tx=XM_023246395.1;refGene_gn=MAPK8IP3;refGene_tx=XM_023246396.1;refGene_gn=MAPK8IP3;refGene_tx=XM_023246397.1;refGene_gn=MAPK8IP3;refGene_tx=XM_023246398.1;refGene_gn=MAPK8IP3;refGene_tx=XM_023246399.1;refGene_gn=MAPK8IP3;refGene_tx=XM_023246400.1
E3 43194410 43194598 refGene_gn=JPT2;refGene_tx=XM_019821229.2
E3 43199772 43199960
refGene_gn=JPT2;refGene_tx=XM_003998951.5;refGene_gn=JPT2;refGene_tx=XM_006942476.4;refGene_gn=JPT2;refGene_tx=XM_019821230.2
E3 43200884 43200933
refGene_gn=JPT2;refGene_tx=XM_003998951.5;refGene_gn=JPT2;refGene_tx=XM_006942476.4;refGene_gn=JPT2;refGene_tx=XM_019821229.2;refGene_gn=JPT2;refGene_tx=XM_019821230.2
E3 43203490 43203633
refGene_gn=JPT2;refGene_tx=XM_003998951.5;refGene_gn=JPT2;refGene_tx=XM_006942476.4;refGene_gn=JPT2;refGene_tx=XM_019821229.2;refGene_gn=JPT2;refGene_tx=XM_019821230.2
E3 43208481 43208630
refGene_gn=JPT2;refGene_tx=XM_003998951.5;refGene_gn=JPT2;refGene_tx=XM_006942476.4;refGene_gn=JPT2;refGene_tx=XM_019821229.2;refGene_gn=JPT2;refGene_tx=XM_019821230.2
E3 43215278 43215322
refGene_gn=JPT2;refGene_tx=XM_003998951.5;refGene_gn=JPT2;refGene_tx=XM_019821229.2
E3 43219506 43219671
refGene_gn=CRAMP1;refGene_tx=XM_023246402.1;refGene_gn=CRAMP1;refGene_tx=XM_023246403.1
E3 43222133 43222279
refGene_gn=CRAMP1;refGene_tx=XM_023246402.1;refGene_gn=CRAMP1;refGene_tx=XM_023246403.1
E3 43223227 43223392
refGene_gn=CRAMP1;refGene_tx=XM_023246402.1;refGene_gn=CRAMP1;refGene_tx=XM_023246403.1
E3 43224036 43224268
refGene_gn=CRAMP1;refGene_tx=XM_023246402.1;refGene_gn=CRAMP1;refGene_tx=XM_023246403.1
E3 43224835 43224904
refGene_gn=CRAMP1;refGene_tx=XM_023246402.1;refGene_gn=CRAMP1;refGene_tx=XM_023246403.1
E3 43225352 43225531
refGene_gn=CRAMP1;refGene_tx=XM_023246402.1;refGene_gn=CRAMP1;refGene_tx=XM_023246403.1

E3 43225715 43225820
refGene_gn=CRAMP1;refGene_tx=XM_023246402.1;refGene_gn=CRAMP1;refGene_tx=XM_023246403.1
E3 43226630 43226712
refGene_gn=CRAMP1;refGene_tx=XM_023246402.1;refGene_gn=CRAMP1;refGene_tx=XM_023246403.1
E3 43228946 43229021
refGene_gn=CRAMP1;refGene_tx=XM_023246402.1;refGene_gn=CRAMP1;refGene_tx=XM_023246403.1
E3 43229115 43229297
refGene_gn=CRAMP1;refGene_tx=XM_023246402.1;refGene_gn=CRAMP1;refGene_tx=XM_023246403.1
E3 43231120 43231298
refGene_gn=CRAMP1;refGene_tx=XM_023246402.1;refGene_gn=CRAMP1;refGene_tx=XM_023246403.1
E3 43234089 43235232
refGene_gn=CRAMP1;refGene_tx=XM_023246402.1;refGene_gn=CRAMP1;refGene_tx=XM_023246403.1
E3 43235656 43235738
refGene_gn=CRAMP1;refGene_tx=XM_023246402.1;refGene_gn=CRAMP1;refGene_tx=XM_023246403.1
E3 43236994 43237118
refGene_gn=CRAMP1;refGene_tx=XM_023246402.1;refGene_gn=CRAMP1;refGene_tx=XM_023246403.1
E3 43237545 43237631
refGene_gn=CRAMP1;refGene_tx=XM_023246402.1;refGene_gn=CRAMP1;refGene_tx=XM_023246403.1
E3 43244053 43244265 refGene_gn=CRAMP1;refGene_tx=XM_023246403.1
E3 43246690 43246739 refGene_gn=CRAMP1;refGene_tx=XM_023246402.1
E3 43249589 43249673 refGene_gn=CRAMP1;refGene_tx=XM_023246402.1
E3 43254280 43254434 refGene_gn=CRAMP1;refGene_tx=XM_023246402.1
E3 43260451 43260654 refGene_gn=CRAMP1;refGene_tx=XM_023246402.1
E3 43270192 43270544 refGene_gn=CRAMP1;refGene_tx=XM_023246402.1
E3 43278770 43278917
refGene_gn=IFT140;refGene_tx=XM_011290728.3;refGene_gn=IFT140;refGene_tx=XM_011290729.3;refGene_gn=IFT140;refGene_tx=XM_011290731.3;refGene_gn=IFT140;refGene_tx=XM_019821231.2;refGene_gn=IFT140;refGene_tx=XM_023246693.1;refGene_gn=IFT140;refGene_tx=XM_023246694.1;refGene_gn=IFT140;refGene_tx=XM_023246695.1;refGene_gn=IFT140;refGene_tx=XM_023246697.1
E3 43280477 43280699
refGene_gn=IFT140;refGene_tx=XM_011290728.3;refGene_gn=IFT140;refGene_tx=XM_011290729.3;refGene_gn=IFT140;refGene_tx=XM_011290731.3;refGene_gn=IFT140;refGene_tx=XM_019821231.2;refGene_gn=IFT140;refGene_tx=XM_023246693.1;refGene_gn=IFT140;refGene_tx=XM_023246694.1;refGene_gn=IFT140;refGene_tx=XM_023246695.1;refGene_gn=IFT140;refGene_tx=XM_023246697.1
E3 43284751 43284873
refGene_gn=IFT140;refGene_tx=XM_011290728.3;refGene_gn=IFT140;refGene_tx=XM_011290729.3;refGene_gn=IFT140;refGene_tx=XM_011290731.3;refGene_

gn=IFT140;refGene_tx=XM_019821231.2;refGene_gn=IFT140;refGene_tx=XM_023246693.1;refGene_gn=IFT140;refGene_tx=XM_023246694.1;refGene_gn=IFT140;refGene_tx=XM_023246695.1;refGene_gn=IFT140;refGene_tx=XM_023246697.1
E3 43284988 43285131
refGene_gn=IFT140;refGene_tx=XM_011290728.3;refGene_gn=IFT140;refGene_tx=XM_011290729.3;refGene_gn=IFT140;refGene_tx=XM_011290731.3;refGene_gn=IFT140;refGene_tx=XM_019821231.2;refGene_gn=IFT140;refGene_tx=XM_023246693.1;refGene_gn=IFT140;refGene_tx=XM_023246694.1;refGene_gn=IFT140;refGene_tx=XM_023246695.1;refGene_gn=IFT140;refGene_tx=XM_023246697.1
E3 43286635 43286811
refGene_gn=IFT140;refGene_tx=XM_011290728.3;refGene_gn=IFT140;refGene_tx=XM_011290729.3;refGene_gn=IFT140;refGene_tx=XM_011290731.3;refGene_gn=IFT140;refGene_tx=XM_019821231.2;refGene_gn=IFT140;refGene_tx=XM_023246693.1;refGene_gn=IFT140;refGene_tx=XM_023246694.1;refGene_gn=IFT140;refGene_tx=XM_023246695.1;refGene_gn=IFT140;refGene_tx=XM_023246697.1
E3 43287779 43287871
refGene_gn=IFT140;refGene_tx=XM_011290728.3;refGene_gn=IFT140;refGene_tx=XM_011290729.3;refGene_gn=IFT140;refGene_tx=XM_011290731.3;refGene_gn=IFT140;refGene_tx=XM_019821231.2;refGene_gn=IFT140;refGene_tx=XM_023246693.1;refGene_gn=IFT140;refGene_tx=XM_023246694.1;refGene_gn=IFT140;refGene_tx=XM_023246695.1;refGene_gn=IFT140;refGene_tx=XM_023246697.1
E3 43288357 43288464
refGene_gn=IFT140;refGene_tx=XM_011290728.3;refGene_gn=IFT140;refGene_tx=XM_011290729.3;refGene_gn=IFT140;refGene_tx=XM_011290731.3;refGene_gn=IFT140;refGene_tx=XM_019821231.2;refGene_gn=IFT140;refGene_tx=XM_023246693.1;refGene_gn=IFT140;refGene_tx=XM_023246694.1;refGene_gn=IFT140;refGene_tx=XM_023246695.1;refGene_gn=IFT140;refGene_tx=XM_023246697.1
E3 43289042 43289188
refGene_gn=IFT140;refGene_tx=XM_011290728.3;refGene_gn=IFT140;refGene_tx=XM_011290729.3;refGene_gn=IFT140;refGene_tx=XM_011290731.3;refGene_gn=IFT140;refGene_tx=XM_019821231.2;refGene_gn=IFT140;refGene_tx=XM_023246693.1;refGene_gn=IFT140;refGene_tx=XM_023246694.1;refGene_gn=IFT140;refGene_tx=XM_023246695.1;refGene_gn=IFT140;refGene_tx=XM_023246697.1
E3 43292172 43292376
refGene_gn=IFT140;refGene_tx=XM_011290728.3;refGene_gn=IFT140;refGene_tx=XM_011290729.3;refGene_gn=IFT140;refGene_tx=XM_011290731.3;refGene_gn=IFT140;refGene_tx=XM_019821231.2;refGene_gn=IFT140;refGene_tx=XM_023246693.1;refGene_gn=IFT140;refGene_tx=XM_023246694.1;refGene_gn=IFT140;refGene_tx=XM_023246695.1
E3 43292510 43292583
refGene_gn=IFT140;refGene_tx=XM_011290728.3;refGene_gn=IFT140;refGene_tx=XM_011290729.3;refGene_gn=IFT140;refGene_tx=XM_011290731.3;refGene_gn=IFT140;refGene_tx=XM_019821231.2;refGene_gn=IFT140;refGene_tx=XM_023246693.1;refGene_gn=IFT140;refGene_tx=XM_023246694.1;refGene_gn=IFT14

0;refGene_tx=XM_023246695.1;refGene_gn=IFT140;refGene_tx=XM_023246697.
1
E3 43294142 43294234
refGene_gn=IFT140;refGene_tx=XM_011290728.3;refGene_gn=IFT140;refGene_
tx=XM_011290729.3;refGene_gn=IFT140;refGene_tx=XM_011290731.3;refGene_
gn=IFT140;refGene_tx=XM_019821231.2;refGene_gn=IFT140;refGene_tx=XM_02
3246693.1;refGene_gn=IFT140;refGene_tx=XM_023246694.1;refGene_gn=IFT14
0;refGene_tx=XM_023246695.1;refGene_gn=IFT140;refGene_tx=XM_023246697.
1
E3 43304034 43304162
refGene_gn=IFT140;refGene_tx=XM_011290728.3;refGene_gn=IFT140;refGene_
tx=XM_011290729.3;refGene_gn=IFT140;refGene_tx=XM_011290731.3;refGene_
gn=IFT140;refGene_tx=XM_019821231.2;refGene_gn=IFT140;refGene_tx=XM_02
3246693.1;refGene_gn=IFT140;refGene_tx=XM_023246694.1;refGene_gn=IFT14
0;refGene_tx=XM_023246695.1;refGene_gn=IFT140;refGene_tx=XM_023246697.
1
E3 43306913 43307031
refGene_gn=IFT140;refGene_tx=XM_011290728.3;refGene_gn=IFT140;refGene_
tx=XM_011290729.3;refGene_gn=IFT140;refGene_tx=XM_011290731.3;refGene_
gn=IFT140;refGene_tx=XM_019821231.2;refGene_gn=IFT140;refGene_tx=XM_02
3246693.1;refGene_gn=IFT140;refGene_tx=XM_023246694.1;refGene_gn=IFT14
0;refGene_tx=XM_023246695.1;refGene_gn=IFT140;refGene_tx=XM_023246697.
1
E3 43308916 43309047
refGene_gn=IFT140;refGene_tx=XM_011290728.3;refGene_gn=IFT140;refGene_
tx=XM_011290729.3;refGene_gn=IFT140;refGene_tx=XM_011290731.3;refGene_
gn=IFT140;refGene_tx=XM_019821231.2;refGene_gn=IFT140;refGene_tx=XM_02
3246693.1;refGene_gn=IFT140;refGene_tx=XM_023246694.1;refGene_gn=IFT14
0;refGene_tx=XM_023246695.1;refGene_gn=IFT140;refGene_tx=XM_023246697.
1
E3 43310381 43310550
refGene_gn=IFT140;refGene_tx=XM_011290728.3;refGene_gn=IFT140;refGene_
tx=XM_011290729.3;refGene_gn=IFT140;refGene_tx=XM_011290731.3;refGene_
gn=IFT140;refGene_tx=XM_019821231.2;refGene_gn=IFT140;refGene_tx=XM_02
3246693.1;refGene_gn=IFT140;refGene_tx=XM_023246694.1;refGene_gn=IFT14
0;refGene_tx=XM_023246695.1;refGene_gn=IFT140;refGene_tx=XM_023246697.
1
E3 43311823 43311955
refGene_gn=IFT140;refGene_tx=XM_011290728.3;refGene_gn=IFT140;refGene_
tx=XM_011290729.3;refGene_gn=IFT140;refGene_tx=XM_011290731.3;refGene_
gn=IFT140;refGene_tx=XM_019821231.2;refGene_gn=IFT140;refGene_tx=XM_02
3246693.1;refGene_gn=IFT140;refGene_tx=XM_023246694.1;refGene_gn=IFT14
0;refGene_tx=XM_023246695.1;refGene_gn=IFT140;refGene_tx=XM_023246697.
1
E3 43315631 43315831
refGene_gn=IFT140;refGene_tx=XM_011290728.3;refGene_gn=IFT140;refGene_
tx=XM_011290729.3;refGene_gn=IFT140;refGene_tx=XM_011290731.3;refGene_
gn=IFT140;refGene_tx=XM_019821231.2;refGene_gn=IFT140;refGene_tx=XM_02
3246693.1;refGene_gn=IFT140;refGene_tx=XM_023246694.1;refGene_gn=IFT14
0;refGene_tx=XM_023246695.1;refGene_gn=IFT140;refGene_tx=XM_023246697.

1
E3	43318674 43318919 refGene_gn=TMEM204;refGene_tx=XM_003998953.5
E3	43327931 43328087 refGene_gn=TMEM204;refGene_tx=XM_003998953.5
E3	43331969 43332249 refGene_gn=TMEM204;refGene_tx=XM_003998953.5
E3	43336551 43336729
refGene_gn=IFT140;refGene_tx=XM_011290728.3;refGene_gn=IFT140;refGene_tx=XM_011290729.3;refGene_gn=IFT140;refGene_tx=XM_011290731.3;refGene_gn=IFT140;refGene_tx=XM_019821231.2;refGene_gn=IFT140;refGene_tx=XM_023246693.1;refGene_gn=IFT140;refGene_tx=XM_023246694.1;refGene_gn=IFT140;refGene_tx=XM_023246695.1;refGene_gn=IFT140;refGene_tx=XM_023246697.1
E3	43337175 43337366
refGene_gn=IFT140;refGene_tx=XM_011290728.3;refGene_gn=IFT140;refGene_tx=XM_011290729.3;refGene_gn=IFT140;refGene_tx=XM_011290731.3;refGene_gn=IFT140;refGene_tx=XM_019821231.2;refGene_gn=IFT140;refGene_tx=XM_023246693.1;refGene_gn=IFT140;refGene_tx=XM_023246694.1;refGene_gn=IFT140;refGene_tx=XM_023246695.1;refGene_gn=IFT140;refGene_tx=XM_023246697.1
E3	43337771 43337867
refGene_gn=IFT140;refGene_tx=XM_011290728.3;refGene_gn=IFT140;refGene_tx=XM_011290729.3;refGene_gn=IFT140;refGene_tx=XM_011290731.3;refGene_gn=IFT140;refGene_tx=XM_019821231.2;refGene_gn=IFT140;refGene_tx=XM_023246693.1;refGene_gn=IFT140;refGene_tx=XM_023246694.1;refGene_gn=IFT140;refGene_tx=XM_023246695.1;refGene_gn=IFT140;refGene_tx=XM_023246697.1
E3	43338086 43338219
refGene_gn=IFT140;refGene_tx=XM_011290728.3;refGene_gn=IFT140;refGene_tx=XM_011290729.3;refGene_gn=IFT140;refGene_tx=XM_011290731.3;refGene_gn=IFT140;refGene_tx=XM_019821231.2;refGene_gn=IFT140;refGene_tx=XM_023246693.1;refGene_gn=IFT140;refGene_tx=XM_023246694.1;refGene_gn=IFT140;refGene_tx=XM_023246695.1;refGene_gn=IFT140;refGene_tx=XM_023246697.1
E3	43338532 43338676
refGene_gn=IFT140;refGene_tx=XM_011290728.3;refGene_gn=IFT140;refGene_tx=XM_011290729.3;refGene_gn=IFT140;refGene_tx=XM_011290731.3;refGene_gn=IFT140;refGene_tx=XM_019821231.2;refGene_gn=IFT140;refGene_tx=XM_023246693.1;refGene_gn=IFT140;refGene_tx=XM_023246694.1;refGene_gn=IFT140;refGene_tx=XM_023246695.1;refGene_gn=IFT140;refGene_tx=XM_023246697.1
E3	43339181 43339310
refGene_gn=IFT140;refGene_tx=XM_011290728.3;refGene_gn=IFT140;refGene_tx=XM_011290729.3;refGene_gn=IFT140;refGene_tx=XM_011290731.3;refGene_gn=IFT140;refGene_tx=XM_019821231.2;refGene_gn=IFT140;refGene_tx=XM_023246693.1;refGene_gn=IFT140;refGene_tx=XM_023246694.1;refGene_gn=IFT140;refGene_tx=XM_023246695.1;refGene_gn=IFT140;refGene_tx=XM_023246697.1
E3	43339421 43339604
refGene_gn=IFT140;refGene_tx=XM_011290728.3;refGene_gn=IFT140;refGene_tx=XM_011290729.3;refGene_gn=IFT140;refGene_tx=XM_011290731.3;refGene_gn=IFT140;refGene_tx=XM_019821231.2;refGene_gn=IFT140;refGene_tx=XM_02

3246693.1;refGene_gn=IFT140;refGene_tx=XM_023246694.1;refGene_gn=IFT14
0;refGene_tx=XM_023246695.1;refGene_gn=IFT140;refGene_tx=XM_023246697.
1
E3 43342574 43342781
refGene_gn=IFT140;refGene_tx=XM_011290728.3;refGene_gn=IFT140;refGene_
tx=XM_011290729.3;refGene_gn=IFT140;refGene_tx=XM_011290731.3;refGene_
gn=IFT140;refGene_tx=XM_019821231.2;refGene_gn=IFT140;refGene_tx=XM_02
3246693.1;refGene_gn=IFT140;refGene_tx=XM_023246694.1;refGene_gn=IFT14
0;refGene_tx=XM_023246695.1;refGene_gn=IFT140;refGene_tx=XM_023246697.
1
E3 43342974 43343187
refGene_gn=IFT140;refGene_tx=XM_011290728.3;refGene_gn=IFT140;refGene_
tx=XM_011290729.3;refGene_gn=IFT140;refGene_tx=XM_011290731.3;refGene_
gn=IFT140;refGene_tx=XM_019821231.2;refGene_gn=IFT140;refGene_tx=XM_02
3246693.1;refGene_gn=IFT140;refGene_tx=XM_023246694.1;refGene_gn=IFT14
0;refGene_tx=XM_023246695.1;refGene_gn=IFT140;refGene_tx=XM_023246697.
1
E3 43343276 43343443
refGene_gn=IFT140;refGene_tx=XM_011290728.3;refGene_gn=IFT140;refGene_
tx=XM_011290729.3;refGene_gn=IFT140;refGene_tx=XM_011290731.3;refGene_
gn=IFT140;refGene_tx=XM_019821231.2;refGene_gn=IFT140;refGene_tx=XM_02
3246693.1;refGene_gn=IFT140;refGene_tx=XM_023246694.1;refGene_gn=IFT14
0;refGene_tx=XM_023246695.1;refGene_gn=IFT140;refGene_tx=XM_023246697.
1
E3 43344646 43344788
refGene_gn=IFT140;refGene_tx=XM_011290728.3;refGene_gn=IFT140;refGene_
tx=XM_011290729.3;refGene_gn=IFT140;refGene_tx=XM_011290731.3;refGene_
gn=IFT140;refGene_tx=XM_019821231.2;refGene_gn=IFT140;refGene_tx=XM_02
3246693.1;refGene_gn=IFT140;refGene_tx=XM_023246694.1;refGene_gn=IFT14
0;refGene_tx=XM_023246695.1;refGene_gn=IFT140;refGene_tx=XM_023246697.
1
E3 43349679 43349889
refGene_gn=IFT140;refGene_tx=XM_011290728.3;refGene_gn=IFT140;refGene_
tx=XM_011290729.3;refGene_gn=IFT140;refGene_tx=XM_011290731.3;refGene_
gn=IFT140;refGene_tx=XM_019821231.2;refGene_gn=IFT140;refGene_tx=XM_02
3246693.1;refGene_gn=IFT140;refGene_tx=XM_023246694.1;refGene_gn=IFT14
0;refGene_tx=XM_023246695.1;refGene_gn=IFT140;refGene_tx=XM_023246697.
1
E3 43350724 43350831 refGene_gn=TELO2;refGene_tx=XM_003998954.5
E3 43351532 43351812
refGene_gn=TELO2;refGene_tx=XM_003998954.5;refGene_gn=TELO2;refGene_tx
=XM_011290733.3;refGene_gn=TELO2;refGene_tx=XM_019821233.2
E3 43352067 43352132
refGene_gn=TELO2;refGene_tx=XM_003998954.5;refGene_gn=TELO2;refGene_tx
=XM_011290733.3;refGene_gn=TELO2;refGene_tx=XM_019821233.2
E3 43352254 43352354
refGene_gn=TELO2;refGene_tx=XM_003998954.5;refGene_gn=TELO2;refGene_tx
=XM_011290733.3;refGene_gn=TELO2;refGene_tx=XM_019821233.2
E3 43352745 43352837
refGene_gn=TELO2;refGene_tx=XM_003998954.5;refGene_gn=TELO2;refGene_tx

=XM_011290733.3;refGene_gn=TELO2;refGene_tx=XM_019821233.2
E3	43353113	43353278
refGene_gn=TELO2;refGene_tx=XM_003998954.5;refGene_gn=TELO2;refGene_tx
=XM_011290733.3;refGene_gn=TELO2;refGene_tx=XM_019821233.2
E3	43354329	43354401
refGene_gn=TELO2;refGene_tx=XM_003998954.5;refGene_gn=TELO2;refGene_tx
=XM_011290733.3;refGene_gn=TELO2;refGene_tx=XM_019821233.2
E3	43354492	43354609
refGene_gn=TELO2;refGene_tx=XM_003998954.5;refGene_gn=TELO2;refGene_tx
=XM_011290733.3;refGene_gn=TELO2;refGene_tx=XM_019821233.2
E3	43354737	43354829
refGene_gn=TELO2;refGene_tx=XM_003998954.5;refGene_gn=TELO2;refGene_tx
=XM_011290733.3;refGene_gn=TELO2;refGene_tx=XM_019821233.2
E3	43355010	43355099
refGene_gn=TELO2;refGene_tx=XM_003998954.5;refGene_gn=TELO2;refGene_tx
=XM_011290733.3;refGene_gn=TELO2;refGene_tx=XM_019821233.2
E3	43355369	43355471
refGene_gn=TELO2;refGene_tx=XM_003998954.5;refGene_gn=TELO2;refGene_tx
=XM_011290733.3;refGene_gn=TELO2;refGene_tx=XM_019821233.2
E3	43355681	43355761
refGene_gn=TELO2;refGene_tx=XM_003998954.5;refGene_gn=TELO2;refGene_tx
=XM_011290733.3;refGene_gn=TELO2;refGene_tx=XM_019821233.2
E3	43356113	43356250
refGene_gn=TELO2;refGene_tx=XM_003998954.5;refGene_gn=TELO2;refGene_tx
=XM_011290733.3;refGene_gn=TELO2;refGene_tx=XM_019821233.2
E3	43356396	43356538
refGene_gn=TELO2;refGene_tx=XM_003998954.5;refGene_gn=TELO2;refGene_tx
=XM_011290733.3;refGene_gn=TELO2;refGene_tx=XM_019821233.2
E3	43356718	43356787
refGene_gn=TELO2;refGene_tx=XM_003998954.5;refGene_gn=TELO2;refGene_tx
=XM_011290733.3;refGene_gn=TELO2;refGene_tx=XM_019821233.2
E3	43356995	43357098
refGene_gn=TELO2;refGene_tx=XM_003998954.5;refGene_gn=TELO2;refGene_tx
=XM_011290733.3;refGene_gn=TELO2;refGene_tx=XM_019821233.2
E3	43358239	43358387
refGene_gn=TELO2;refGene_tx=XM_003998954.5;refGene_gn=TELO2;refGene_tx
=XM_011290733.3;refGene_gn=TELO2;refGene_tx=XM_019821233.2
E3	43358581	43358650
refGene_gn=TELO2;refGene_tx=XM_003998954.5;refGene_gn=TELO2;refGene_tx
=XM_011290733.3;refGene_gn=TELO2;refGene_tx=XM_019821233.2
E3	43359674	43359952
refGene_gn=TELO2;refGene_tx=XM_003998954.5;refGene_gn=TELO2;refGene_tx
=XM_011290733.3;refGene_gn=TELO2;refGene_tx=XM_019821233.2
E3	43360892	43361227
refGene_gn=TELO2;refGene_tx=XM_003998954.5;refGene_gn=TELO2;refGene_tx
=XM_011290733.3;refGene_gn=TELO2;refGene_tx=XM_019821233.2
E3	43367548	43367800	refGene_gn=PTX4;refGene_tx=XM_003999042.4
E3	43368363	43369033	refGene_gn=PTX4;refGene_tx=XM_003999042.4
E3	43370039	43370680	refGene_gn=PTX4;refGene_tx=XM_003999042.4
E3	43371023	43371639

refGene_gn=LOC109495358;refGene_tx=XM_019821235.2;refGene_gn=LOC109495358;refGene_tx=XM_019821236.2
E3 43374445 43374500
refGene_gn=LOC109495358;refGene_tx=XM_019821235.2;refGene_gn=LOC109495358;refGene_tx=XM_019821236.2
E3 43375156 43375309
refGene_gn=CLCN7;refGene_tx=XM_023247015.1;refGene_gn=CLCN7;refGene_tx=XM_023247016.1
E3 43383125 43383197 refGene_gn=CLCN7;refGene_tx=XM_023247015.1
E3 43387609 43387681
refGene_gn=CLCN7;refGene_tx=XM_023247015.1;refGene_gn=CLCN7;refGene_tx=XM_023247016.1
E3 43387799 43387865
refGene_gn=CLCN7;refGene_tx=XM_023247015.1;refGene_gn=CLCN7;refGene_tx=XM_023247016.1
E3 43388194 43388327
refGene_gn=CLCN7;refGene_tx=XM_023247015.1;refGene_gn=CLCN7;refGene_tx=XM_023247016.1
E3 43388548 43388658
refGene_gn=CLCN7;refGene_tx=XM_023247015.1;refGene_gn=CLCN7;refGene_tx=XM_023247016.1
E3 43389250 43389331
refGene_gn=CLCN7;refGene_tx=XM_023247015.1;refGene_gn=CLCN7;refGene_tx=XM_023247016.1
E3 43390289 43390352
refGene_gn=CLCN7;refGene_tx=XM_023247015.1;refGene_gn=CLCN7;refGene_tx=XM_023247016.1
E3 43390471 43390555
refGene_gn=CLCN7;refGene_tx=XM_023247015.1;refGene_gn=CLCN7;refGene_tx=XM_023247016.1
E3 43391182 43391276
refGene_gn=CLCN7;refGene_tx=XM_023247015.1;refGene_gn=CLCN7;refGene_tx=XM_023247016.1
E3 43391510 43391575
refGene_gn=CLCN7;refGene_tx=XM_023247015.1;refGene_gn=CLCN7;refGene_tx=XM_023247016.1
E3 43392023 43392140
refGene_gn=CLCN7;refGene_tx=XM_023247015.1;refGene_gn=CLCN7;refGene_tx=XM_023247016.1
E3 43392757 43392812
refGene_gn=CLCN7;refGene_tx=XM_023247015.1;refGene_gn=CLCN7;refGene_tx=XM_023247016.1
E3 43393272 43393333
refGene_gn=CLCN7;refGene_tx=XM_023247015.1;refGene_gn=CLCN7;refGene_tx=XM_023247016.1
E3 43393893 43394032
refGene_gn=CLCN7;refGene_tx=XM_023247015.1;refGene_gn=CLCN7;refGene_tx=XM_023247016.1
E3 43394902 43394996
refGene_gn=CLCN7;refGene_tx=XM_023247015.1;refGene_gn=CLCN7;refGene_tx

=XM_023247016.1
E3 43395257 43395427
refGene_gn=CLCN7;refGene_tx=XM_023247015.1;refGene_gn=CLCN7;refGene_tx
=XM_023247016.1
E3 43396415 43396467
refGene_gn=CLCN7;refGene_tx=XM_023247015.1;refGene_gn=CLCN7;refGene_tx
=XM_023247016.1
E3 43396747 43396875
refGene_gn=CLCN7;refGene_tx=XM_023247015.1;refGene_gn=CLCN7;refGene_tx
=XM_023247016.1
E3 43397036 43397122
refGene_gn=CLCN7;refGene_tx=XM_023247015.1;refGene_gn=CLCN7;refGene_tx
=XM_023247016.1
E3 43397372 43397502
refGene_gn=CLCN7;refGene_tx=XM_023247015.1;refGene_gn=CLCN7;refGene_tx
=XM_023247016.1
E3 43397979 43398039
refGene_gn=CLCN7;refGene_tx=XM_023247015.1;refGene_gn=CLCN7;refGene_tx
=XM_023247016.1
E3 43398130 43398307
refGene_gn=CLCN7;refGene_tx=XM_023247015.1;refGene_gn=CLCN7;refGene_tx
=XM_023247016.1
E3 43398714 43398795
refGene_gn=CLCN7;refGene_tx=XM_023247015.1;refGene_gn=CLCN7;refGene_tx
=XM_023247016.1
E3 43399060 43399147
refGene_gn=CLCN7;refGene_tx=XM_023247015.1;refGene_gn=CLCN7;refGene_tx
=XM_023247016.1
E3 43400950 43401050 refGene_gn=CCDC154;refGene_tx=XM_019821234.2
E3 43401278 43401353 refGene_gn=CCDC154;refGene_tx=XM_019821234.2
E3 43401655 43401920 refGene_gn=CCDC154;refGene_tx=XM_019821234.2
E3 43402019 43402209 refGene_gn=CCDC154;refGene_tx=XM_019821234.2
E3 43402305 43402352 refGene_gn=CCDC154;refGene_tx=XM_019821234.2
E3 43402621 43402717 refGene_gn=CCDC154;refGene_tx=XM_019821234.2
E3 43403155 43403279 refGene_gn=CCDC154;refGene_tx=XM_019821234.2
E3 43404722 43404824 refGene_gn=CCDC154;refGene_tx=XM_019821234.2
E3 43404903 43405032 refGene_gn=CCDC154;refGene_tx=XM_019821234.2
E3 43405111 43405239 refGene_gn=CCDC154;refGene_tx=XM_019821234.2
E3 43405484 43405611 refGene_gn=CCDC154;refGene_tx=XM_019821234.2
E3 43405693 43405831 refGene_gn=CCDC154;refGene_tx=XM_019821234.2
E3 43406921 43407041 refGene_gn=CCDC154;refGene_tx=XM_019821234.2
E3 43407127 43407204 refGene_gn=CCDC154;refGene_tx=XM_019821234.2
E3 43407470 43407588 refGene_gn=CCDC154;refGene_tx=XM_019821234.2
E3 43407866 43407953 refGene_gn=CCDC154;refGene_tx=XM_019821234.2
E3 43408189 43408377 refGene_gn=CCDC154;refGene_tx=XM_019821234.2
E3 43408472 43408599 refGene_gn=CCDC154;refGene_tx=XM_019821234.2
E3 43409295 43410060
refGene_gn=LOC111558284;refGene_tx=XM_023247017.1
E3 43418621 43418653
refGene_gn=CE3H16orf91;refGene_tx=XM_006942481.4

E3 43418738 43419114
refGene_gn=CE3H16orf91;refGene_tx=XM_006942481.4
E3 43422318 43422395 refGene_gn=UNKL;refGene_tx=XM_023246808.1
E3 43423394 43423604 refGene_gn=UNKL;refGene_tx=XM_023246808.1
E3 43431677 43431828 refGene_gn=UNKL;refGene_tx=XM_023246808.1
E3 43431829 43431855 refGene_gn=UNKL;refGene_tx=XM_023246808.1
E3 43432241 43432375 refGene_gn=UNKL;refGene_tx=XM_023246808.1
E3 43435892 43436028 refGene_gn=UNKL;refGene_tx=XM_023246808.1
E3 43436716 43436834 refGene_gn=UNKL;refGene_tx=XM_023246808.1
E3 43439095 43439180 refGene_gn=UNKL;refGene_tx=XM_023246808.1
E3 43439547 43439627 refGene_gn=UNKL;refGene_tx=XM_023246808.1
E3 43443697 43443760 refGene_gn=UNKL;refGene_tx=XM_023246808.1
E3 43446849 43446991 refGene_gn=UNKL;refGene_tx=XM_023246808.1
E3 43454749 43454842 refGene_gn=UNKL;refGene_tx=XM_023246808.1
E3 43455315 43455543 refGene_gn=UNKL;refGene_tx=XM_023246808.1
E3 43456139 43456342 refGene_gn=UNKL;refGene_tx=XM_023246808.1
E3 43456443 43456701 refGene_gn=UNKL;refGene_tx=XM_023246808.1
E3 43457234 43457342 refGene_gn=UNKL;refGene_tx=XM_023246808.1
E3 43459554 43459652 refGene_gn=GNPTG;refGene_tx=XM_023246811.1
E3 43459730 43459812 refGene_gn=GNPTG;refGene_tx=XM_023246811.1
E3 43459897 43460026 refGene_gn=GNPTG;refGene_tx=XM_023246811.1
E3 43460105 43460188 refGene_gn=GNPTG;refGene_tx=XM_023246811.1
E3 43460278 43460393 refGene_gn=GNPTG;refGene_tx=XM_023246811.1
E3 43460471 43460565 refGene_gn=GNPTG;refGene_tx=XM_023246811.1
E3 43460657 43460741 refGene_gn=GNPTG;refGene_tx=XM_023246811.1
E3 43460806 43460861 refGene_gn=GNPTG;refGene_tx=XM_023246811.1
E3 43472026 43472094 refGene_gn=GNPTG;refGene_tx=XM_023246811.1
E3 43472175 43472233 refGene_gn=GNPTG;refGene_tx=XM_023246811.1
E3 43472319 43472380 refGene_gn=GNPTG;refGene_tx=XM_023246811.1
E3 43472521 43472633
refGene_gn=TSR3;refGene_tx=XM_023246809.1;refGene_gn=TSR3;refGene_tx=XM_023246810.1
E3 43472734 43472954
refGene_gn=TSR3;refGene_tx=XM_023246809.1;refGene_gn=TSR3;refGene_tx=XM_023246810.1
E3 43473258 43473452
refGene_gn=TSR3;refGene_tx=XM_023246809.1;refGene_gn=TSR3;refGene_tx=XM_023246810.1
E3 43473820 43473997
refGene_gn=TSR3;refGene_tx=XM_023246809.1;refGene_gn=TSR3;refGene_tx=XM_023246810.1
E3 43474196 43474260
refGene_gn=TSR3;refGene_tx=XM_023246809.1;refGene_gn=TSR3;refGene_tx=XM_023246810.1
E3 43474525 43474703
refGene_gn=TSR3;refGene_tx=XM_023246809.1;refGene_gn=TSR3;refGene_tx=XM_023246810.1
E3 43475726 43475946
refGene_gn=BAIAP3;refGene_tx=XM_023246797.1;refGene_gn=BAIAP3;refGene_tx=XM_023246798.1;refGene_gn=BAIAP3;refGene_tx=XM_023246799.1;refGene_

gn=BAIAP3;refGene_tx=XM_023246800.1;refGene_gn=BAIAP3;refGene_tx=XM_023246801.1;refGene_gn=BAIAP3;refGene_tx=XM_023246802.1;refGene_gn=BAIAP3;refGene_tx=XM_023246803.1;refGene_gn=BAIAP3;refGene_tx=XM_023246804.1
E3 43476020 43476226
refGene_gn=BAIAP3;refGene_tx=XM_023246797.1;refGene_gn=BAIAP3;refGene_tx=XM_023246798.1;refGene_gn=BAIAP3;refGene_tx=XM_023246799.1;refGene_gn=BAIAP3;refGene_tx=XM_023246801.1;refGene_gn=BAIAP3;refGene_tx=XM_023246802.1;refGene_gn=BAIAP3;refGene_tx=XM_023246803.1;refGene_gn=BAIAP3;refGene_tx=XM_023246804.1
E3 43476315 43476439
refGene_gn=BAIAP3;refGene_tx=XM_023246797.1;refGene_gn=BAIAP3;refGene_tx=XM_023246798.1;refGene_gn=BAIAP3;refGene_tx=XM_023246799.1;refGene_gn=BAIAP3;refGene_tx=XM_023246800.1;refGene_gn=BAIAP3;refGene_tx=XM_023246801.1;refGene_gn=BAIAP3;refGene_tx=XM_023246802.1;refGene_gn=BAIAP3;refGene_tx=XM_023246803.1;refGene_gn=BAIAP3;refGene_tx=XM_023246804.1;refGene_gn=BAIAP3;refGene_tx=XM_023246806.1;refGene_gn=BAIAP3;refGene_tx=XM_023246807.1
E3 43476530 43476841
refGene_gn=BAIAP3;refGene_tx=XM_023246797.1;refGene_gn=BAIAP3;refGene_tx=XM_023246798.1;refGene_gn=BAIAP3;refGene_tx=XM_023246799.1;refGene_gn=BAIAP3;refGene_tx=XM_023246800.1;refGene_gn=BAIAP3;refGene_tx=XM_023246801.1;refGene_gn=BAIAP3;refGene_tx=XM_023246802.1;refGene_gn=BAIAP3;refGene_tx=XM_023246803.1;refGene_gn=BAIAP3;refGene_tx=XM_023246804.1;refGene_gn=BAIAP3;refGene_tx=XM_023246806.1;refGene_gn=BAIAP3;refGene_tx=XM_023246807.1
E3 43476917 43476989
refGene_gn=BAIAP3;refGene_tx=XM_023246797.1;refGene_gn=BAIAP3;refGene_tx=XM_023246798.1;refGene_gn=BAIAP3;refGene_tx=XM_023246799.1;refGene_gn=BAIAP3;refGene_tx=XM_023246800.1;refGene_gn=BAIAP3;refGene_tx=XM_023246801.1;refGene_gn=BAIAP3;refGene_tx=XM_023246802.1;refGene_gn=BAIAP3;refGene_tx=XM_023246803.1;refGene_gn=BAIAP3;refGene_tx=XM_023246804.1;refGene_gn=BAIAP3;refGene_tx=XM_023246806.1;refGene_gn=BAIAP3;refGene_tx=XM_023246807.1
E3 43477373 43477637
refGene_gn=BAIAP3;refGene_tx=XM_023246797.1;refGene_gn=BAIAP3;refGene_tx=XM_023246798.1;refGene_gn=BAIAP3;refGene_tx=XM_023246799.1;refGene_gn=BAIAP3;refGene_tx=XM_023246800.1;refGene_gn=BAIAP3;refGene_tx=XM_023246801.1;refGene_gn=BAIAP3;refGene_tx=XM_023246802.1;refGene_gn=BAIAP3;refGene_tx=XM_023246803.1;refGene_gn=BAIAP3;refGene_tx=XM_023246804.1;refGene_gn=BAIAP3;refGene_tx=XM_023246806.1;refGene_gn=BAIAP3;refGene_tx=XM_023246807.1
E3 43477713 43477782
refGene_gn=BAIAP3;refGene_tx=XM_023246797.1;refGene_gn=BAIAP3;refGene_tx=XM_023246798.1;refGene_gn=BAIAP3;refGene_tx=XM_023246799.1;refGene_gn=BAIAP3;refGene_tx=XM_023246800.1;refGene_gn=BAIAP3;refGene_tx=XM_023246801.1;refGene_gn=BAIAP3;refGene_tx=XM_023246802.1;refGene_gn=BAIAP3;refGene_tx=XM_023246803.1;refGene_gn=BAIAP3;refGene_tx=XM_023246804.1;refGene_gn=BAIAP3;refGene_tx=XM_023246806.1;refGene_gn=BAIAP3;refGene_tx=XM_023246807.1

E3 43477860 43478052
refGene_gn=BAIAP3;refGene_tx=XM_023246797.1;refGene_gn=BAIAP3;refGene_tx=XM_023246798.1;refGene_gn=BAIAP3;refGene_tx=XM_023246799.1;refGene_gn=BAIAP3;refGene_tx=XM_023246800.1;refGene_gn=BAIAP3;refGene_tx=XM_023246801.1;refGene_gn=BAIAP3;refGene_tx=XM_023246802.1;refGene_gn=BAIAP3;refGene_tx=XM_023246803.1;refGene_gn=BAIAP3;refGene_tx=XM_023246804.1;refGene_gn=BAIAP3;refGene_tx=XM_023246806.1;refGene_gn=BAIAP3;refGene_tx=XM_023246807.1
E3 43478148 43478241
refGene_gn=BAIAP3;refGene_tx=XM_023246797.1;refGene_gn=BAIAP3;refGene_tx=XM_023246798.1;refGene_gn=BAIAP3;refGene_tx=XM_023246799.1;refGene_gn=BAIAP3;refGene_tx=XM_023246800.1;refGene_gn=BAIAP3;refGene_tx=XM_023246801.1;refGene_gn=BAIAP3;refGene_tx=XM_023246802.1;refGene_gn=BAIAP3;refGene_tx=XM_023246803.1;refGene_gn=BAIAP3;refGene_tx=XM_023246804.1;refGene_gn=BAIAP3;refGene_tx=XM_023246806.1;refGene_gn=BAIAP3;refGene_tx=XM_023246807.1
E3 43478351 43478495
refGene_gn=BAIAP3;refGene_tx=XM_023246797.1;refGene_gn=BAIAP3;refGene_tx=XM_023246798.1;refGene_gn=BAIAP3;refGene_tx=XM_023246799.1;refGene_gn=BAIAP3;refGene_tx=XM_023246800.1;refGene_gn=BAIAP3;refGene_tx=XM_023246801.1;refGene_gn=BAIAP3;refGene_tx=XM_023246802.1;refGene_gn=BAIAP3;refGene_tx=XM_023246803.1;refGene_gn=BAIAP3;refGene_tx=XM_023246804.1;refGene_gn=BAIAP3;refGene_tx=XM_023246806.1;refGene_gn=BAIAP3;refGene_tx=XM_023246807.1
E3 43478581 43478705
refGene_gn=BAIAP3;refGene_tx=XM_023246797.1;refGene_gn=BAIAP3;refGene_tx=XM_023246798.1;refGene_gn=BAIAP3;refGene_tx=XM_023246799.1;refGene_gn=BAIAP3;refGene_tx=XM_023246800.1;refGene_gn=BAIAP3;refGene_tx=XM_023246801.1;refGene_gn=BAIAP3;refGene_tx=XM_023246802.1;refGene_gn=BAIAP3;refGene_tx=XM_023246803.1;refGene_gn=BAIAP3;refGene_tx=XM_023246804.1;refGene_gn=BAIAP3;refGene_tx=XM_023246806.1;refGene_gn=BAIAP3;refGene_tx=XM_023246807.1
E3 43478948 43479079
refGene_gn=BAIAP3;refGene_tx=XM_023246797.1;refGene_gn=BAIAP3;refGene_tx=XM_023246798.1;refGene_gn=BAIAP3;refGene_tx=XM_023246799.1;refGene_gn=BAIAP3;refGene_tx=XM_023246800.1;refGene_gn=BAIAP3;refGene_tx=XM_023246801.1;refGene_gn=BAIAP3;refGene_tx=XM_023246802.1;refGene_gn=BAIAP3;refGene_tx=XM_023246803.1;refGene_gn=BAIAP3;refGene_tx=XM_023246804.1;refGene_gn=BAIAP3;refGene_tx=XM_023246806.1;refGene_gn=BAIAP3;refGene_tx=XM_023246807.1
E3 43479167 43479219
refGene_gn=BAIAP3;refGene_tx=XM_023246797.1;refGene_gn=BAIAP3;refGene_tx=XM_023246798.1;refGene_gn=BAIAP3;refGene_tx=XM_023246799.1;refGene_gn=BAIAP3;refGene_tx=XM_023246800.1;refGene_gn=BAIAP3;refGene_tx=XM_023246801.1;refGene_gn=BAIAP3;refGene_tx=XM_023246802.1;refGene_gn=BAIAP3;refGene_tx=XM_023246803.1;refGene_gn=BAIAP3;refGene_tx=XM_023246804.1;refGene_gn=BAIAP3;refGene_tx=XM_023246806.1;refGene_gn=BAIAP3;refGene_tx=XM_023246807.1
E3 43479321 43479419
refGene_gn=BAIAP3;refGene_tx=XM_023246797.1;refGene_gn=BAIAP3;refGene_

tx=XM_023246798.1;refGene_gn=BAIAP3;refGene_tx=XM_023246799.1;refGene_gn=BAIAP3;refGene_tx=XM_023246800.1;refGene_gn=BAIAP3;refGene_tx=XM_023246801.1;refGene_gn=BAIAP3;refGene_tx=XM_023246802.1;refGene_gn=BAIAP3;refGene_tx=XM_023246803.1;refGene_gn=BAIAP3;refGene_tx=XM_023246804.1;refGene_gn=BAIAP3;refGene_tx=XM_023246806.1;refGene_gn=BAIAP3;refGene_tx=XM_023246807.1
E3 43479501 43479558
refGene_gn=BAIAP3;refGene_tx=XM_023246797.1;refGene_gn=BAIAP3;refGene_tx=XM_023246798.1;refGene_gn=BAIAP3;refGene_tx=XM_023246799.1;refGene_gn=BAIAP3;refGene_tx=XM_023246800.1;refGene_gn=BAIAP3;refGene_tx=XM_023246801.1;refGene_gn=BAIAP3;refGene_tx=XM_023246802.1;refGene_gn=BAIAP3;refGene_tx=XM_023246803.1;refGene_gn=BAIAP3;refGene_tx=XM_023246804.1;refGene_gn=BAIAP3;refGene_tx=XM_023246806.1;refGene_gn=BAIAP3;refGene_tx=XM_023246807.1
E3 43479721 43479812
refGene_gn=BAIAP3;refGene_tx=XM_023246797.1;refGene_gn=BAIAP3;refGene_tx=XM_023246798.1;refGene_gn=BAIAP3;refGene_tx=XM_023246799.1;refGene_gn=BAIAP3;refGene_tx=XM_023246800.1;refGene_gn=BAIAP3;refGene_tx=XM_023246801.1;refGene_gn=BAIAP3;refGene_tx=XM_023246802.1;refGene_gn=BAIAP3;refGene_tx=XM_023246803.1;refGene_gn=BAIAP3;refGene_tx=XM_023246804.1;refGene_gn=BAIAP3;refGene_tx=XM_023246806.1;refGene_gn=BAIAP3;refGene_tx=XM_023246807.1
E3 43479917 43480042
refGene_gn=BAIAP3;refGene_tx=XM_023246797.1;refGene_gn=BAIAP3;refGene_tx=XM_023246798.1;refGene_gn=BAIAP3;refGene_tx=XM_023246799.1;refGene_gn=BAIAP3;refGene_tx=XM_023246800.1;refGene_gn=BAIAP3;refGene_tx=XM_023246801.1;refGene_gn=BAIAP3;refGene_tx=XM_023246802.1;refGene_gn=BAIAP3;refGene_tx=XM_023246803.1;refGene_gn=BAIAP3;refGene_tx=XM_023246804.1;refGene_gn=BAIAP3;refGene_tx=XM_023246806.1;refGene_gn=BAIAP3;refGene_tx=XM_023246807.1
E3 43480450 43480571
refGene_gn=BAIAP3;refGene_tx=XM_023246797.1;refGene_gn=BAIAP3;refGene_tx=XM_023246798.1;refGene_gn=BAIAP3;refGene_tx=XM_023246799.1;refGene_gn=BAIAP3;refGene_tx=XM_023246800.1;refGene_gn=BAIAP3;refGene_tx=XM_023246801.1;refGene_gn=BAIAP3;refGene_tx=XM_023246802.1;refGene_gn=BAIAP3;refGene_tx=XM_023246803.1;refGene_gn=BAIAP3;refGene_tx=XM_023246804.1;refGene_gn=BAIAP3;refGene_tx=XM_023246806.1;refGene_gn=BAIAP3;refGene_tx=XM_023246807.1
E3 43480827 43480931
refGene_gn=BAIAP3;refGene_tx=XM_023246797.1;refGene_gn=BAIAP3;refGene_tx=XM_023246798.1;refGene_gn=BAIAP3;refGene_tx=XM_023246799.1;refGene_gn=BAIAP3;refGene_tx=XM_023246800.1;refGene_gn=BAIAP3;refGene_tx=XM_023246801.1;refGene_gn=BAIAP3;refGene_tx=XM_023246802.1;refGene_gn=BAIAP3;refGene_tx=XM_023246803.1;refGene_gn=BAIAP3;refGene_tx=XM_023246804.1;refGene_gn=BAIAP3;refGene_tx=XM_023246806.1;refGene_gn=BAIAP3;refGene_tx=XM_023246807.1
E3 43481057 43481153
refGene_gn=BAIAP3;refGene_tx=XM_023246797.1;refGene_gn=BAIAP3;refGene_tx=XM_023246798.1;refGene_gn=BAIAP3;refGene_tx=XM_023246799.1;refGene_gn=BAIAP3;refGene_tx=XM_023246800.1;refGene_gn=BAIAP3;refGene_tx=XM_02

3246801.1;refGene_gn=BAIAP3;refGene_tx=XM_023246803.1;refGene_gn=BAIAP3;refGene_tx=XM_023246804.1;refGene_gn=BAIAP3;refGene_tx=XM_023246806.1;refGene_gn=BAIAP3;refGene_tx=XM_023246807.1
E3 43481250 43481358
refGene_gn=BAIAP3;refGene_tx=XM_023246797.1;refGene_gn=BAIAP3;refGene_tx=XM_023246798.1;refGene_gn=BAIAP3;refGene_tx=XM_023246799.1;refGene_gn=BAIAP3;refGene_tx=XM_023246800.1;refGene_gn=BAIAP3;refGene_tx=XM_023246801.1;refGene_gn=BAIAP3;refGene_tx=XM_023246802.1;refGene_gn=BAIAP3;refGene_tx=XM_023246803.1;refGene_gn=BAIAP3;refGene_tx=XM_023246804.1;refGene_gn=BAIAP3;refGene_tx=XM_023246806.1;refGene_gn=BAIAP3;refGene_tx=XM_023246807.1
E3 43481667 43481770
refGene_gn=BAIAP3;refGene_tx=XM_023246797.1;refGene_gn=BAIAP3;refGene_tx=XM_023246798.1;refGene_gn=BAIAP3;refGene_tx=XM_023246799.1;refGene_gn=BAIAP3;refGene_tx=XM_023246800.1;refGene_gn=BAIAP3;refGene_tx=XM_023246801.1;refGene_gn=BAIAP3;refGene_tx=XM_023246802.1;refGene_gn=BAIAP3;refGene_tx=XM_023246803.1;refGene_gn=BAIAP3;refGene_tx=XM_023246804.1;refGene_gn=BAIAP3;refGene_tx=XM_023246806.1;refGene_gn=BAIAP3;refGene_tx=XM_023246807.1
E3 43481894 43481972
refGene_gn=BAIAP3;refGene_tx=XM_023246797.1;refGene_gn=BAIAP3;refGene_tx=XM_023246798.1;refGene_gn=BAIAP3;refGene_tx=XM_023246799.1;refGene_gn=BAIAP3;refGene_tx=XM_023246800.1;refGene_gn=BAIAP3;refGene_tx=XM_023246801.1;refGene_gn=BAIAP3;refGene_tx=XM_023246802.1;refGene_gn=BAIAP3;refGene_tx=XM_023246803.1;refGene_gn=BAIAP3;refGene_tx=XM_023246804.1;refGene_gn=BAIAP3;refGene_tx=XM_023246806.1;refGene_gn=BAIAP3;refGene_tx=XM_023246807.1
E3 43482104 43482149
refGene_gn=BAIAP3;refGene_tx=XM_023246797.1;refGene_gn=BAIAP3;refGene_tx=XM_023246798.1;refGene_gn=BAIAP3;refGene_tx=XM_023246799.1;refGene_gn=BAIAP3;refGene_tx=XM_023246800.1;refGene_gn=BAIAP3;refGene_tx=XM_023246801.1;refGene_gn=BAIAP3;refGene_tx=XM_023246802.1;refGene_gn=BAIAP3;refGene_tx=XM_023246803.1;refGene_gn=BAIAP3;refGene_tx=XM_023246804.1;refGene_gn=BAIAP3;refGene_tx=XM_023246806.1;refGene_gn=BAIAP3;refGene_tx=XM_023246807.1
E3 43482495 43482691
refGene_gn=BAIAP3;refGene_tx=XM_023246797.1;refGene_gn=BAIAP3;refGene_tx=XM_023246798.1;refGene_gn=BAIAP3;refGene_tx=XM_023246799.1;refGene_gn=BAIAP3;refGene_tx=XM_023246800.1;refGene_gn=BAIAP3;refGene_tx=XM_023246801.1;refGene_gn=BAIAP3;refGene_tx=XM_023246802.1;refGene_gn=BAIAP3;refGene_tx=XM_023246803.1;refGene_gn=BAIAP3;refGene_tx=XM_023246804.1;refGene_gn=BAIAP3;refGene_tx=XM_023246806.1;refGene_gn=BAIAP3;refGene_tx=XM_023246807.1
E3 43482792 43482859
refGene_gn=BAIAP3;refGene_tx=XM_023246797.1;refGene_gn=BAIAP3;refGene_tx=XM_023246798.1;refGene_gn=BAIAP3;refGene_tx=XM_023246799.1;refGene_gn=BAIAP3;refGene_tx=XM_023246800.1;refGene_gn=BAIAP3;refGene_tx=XM_023246801.1;refGene_gn=BAIAP3;refGene_tx=XM_023246802.1;refGene_gn=BAIAP3;refGene_tx=XM_023246803.1;refGene_gn=BAIAP3;refGene_tx=XM_023246804.1;refGene_gn=BAIAP3;refGene_tx=XM_023246806.1;refGene_gn=BAIAP3;refGen

e_tx=XM_023246807.1
E3 43482996 43483056
refGene_gn=BAIAP3;refGene_tx=XM_023246797.1;refGene_gn=BAIAP3;refGene_tx=XM_023246798.1;refGene_gn=BAIAP3;refGene_tx=XM_023246799.1;refGene_gn=BAIAP3;refGene_tx=XM_023246800.1;refGene_gn=BAIAP3;refGene_tx=XM_023246801.1;refGene_gn=BAIAP3;refGene_tx=XM_023246802.1;refGene_gn=BAIAP3;refGene_tx=XM_023246803.1;refGene_gn=BAIAP3;refGene_tx=XM_023246804.1;refGene_gn=BAIAP3;refGene_tx=XM_023246806.1;refGene_gn=BAIAP3;refGene_tx=XM_023246807.1
E3 43483474 43483582
refGene_gn=BAIAP3;refGene_tx=XM_023246797.1;refGene_gn=BAIAP3;refGene_tx=XM_023246798.1;refGene_gn=BAIAP3;refGene_tx=XM_023246799.1;refGene_gn=BAIAP3;refGene_tx=XM_023246800.1;refGene_gn=BAIAP3;refGene_tx=XM_023246801.1;refGene_gn=BAIAP3;refGene_tx=XM_023246802.1;refGene_gn=BAIAP3;refGene_tx=XM_023246803.1;refGene_gn=BAIAP3;refGene_tx=XM_023246804.1;refGene_gn=BAIAP3;refGene_tx=XM_023246806.1;refGene_gn=BAIAP3;refGene_tx=XM_023246807.1
E3 43483955 43484036
refGene_gn=BAIAP3;refGene_tx=XM_023246797.1;refGene_gn=BAIAP3;refGene_tx=XM_023246798.1;refGene_gn=BAIAP3;refGene_tx=XM_023246799.1;refGene_gn=BAIAP3;refGene_tx=XM_023246800.1;refGene_gn=BAIAP3;refGene_tx=XM_023246801.1;refGene_gn=BAIAP3;refGene_tx=XM_023246802.1;refGene_gn=BAIAP3;refGene_tx=XM_023246803.1;refGene_gn=BAIAP3;refGene_tx=XM_023246804.1;refGene_gn=BAIAP3;refGene_tx=XM_023246806.1;refGene_gn=BAIAP3;refGene_tx=XM_023246807.1
E3 43484175 43484263
refGene_gn=BAIAP3;refGene_tx=XM_023246797.1;refGene_gn=BAIAP3;refGene_tx=XM_023246798.1;refGene_gn=BAIAP3;refGene_tx=XM_023246799.1;refGene_gn=BAIAP3;refGene_tx=XM_023246800.1;refGene_gn=BAIAP3;refGene_tx=XM_023246801.1;refGene_gn=BAIAP3;refGene_tx=XM_023246802.1;refGene_gn=BAIAP3;refGene_tx=XM_023246803.1;refGene_gn=BAIAP3;refGene_tx=XM_023246804.1;refGene_gn=BAIAP3;refGene_tx=XM_023246806.1;refGene_gn=BAIAP3;refGene_tx=XM_023246807.1
E3 43484464 43484595
refGene_gn=BAIAP3;refGene_tx=XM_023246797.1;refGene_gn=BAIAP3;refGene_tx=XM_023246798.1;refGene_gn=BAIAP3;refGene_tx=XM_023246799.1;refGene_gn=BAIAP3;refGene_tx=XM_023246800.1;refGene_gn=BAIAP3;refGene_tx=XM_023246801.1;refGene_gn=BAIAP3;refGene_tx=XM_023246802.1;refGene_gn=BAIAP3;refGene_tx=XM_023246803.1;refGene_gn=BAIAP3;refGene_tx=XM_023246804.1;refGene_gn=BAIAP3;refGene_tx=XM_023246806.1;refGene_gn=BAIAP3;refGene_tx=XM_023246807.1
E3 43494958 43495022 refGene_gn=UBE2I;refGene_tx=XM_023246325.1
E3 43497283 43497363 refGene_gn=UBE2I;refGene_tx=XM_023246325.1
E3 43497501 43497611 refGene_gn=UBE2I;refGene_tx=XM_023246325.1
E3 43501343 43501416 refGene_gn=UBE2I;refGene_tx=XM_023246325.1
E3 43502452 43502536 refGene_gn=UBE2I;refGene_tx=XM_023246325.1
E3 43502723 43502789 refGene_gn=UBE2I;refGene_tx=XM_023246325.1
E3 43532998 43533059
refGene_gn=LOC101089067;refGene_tx=XM_003999048.3
E3 43533210 43533391

refGene_gn=LOC101089067;refGene_tx=XM_003999048.3;refGene_gn=LOC101089067;refGene_tx=XM_019821241.2
E3 43533550 43533825
refGene_gn=LOC101089067;refGene_tx=XM_003999048.3;refGene_gn=LOC101089067;refGene_tx=XM_019821241.2
E3 43534428 43534586
refGene_gn=LOC101089067;refGene_tx=XM_003999048.3;refGene_gn=LOC101089067;refGene_tx=XM_019821241.2
E3 43534720 43534891
refGene_gn=LOC101089067;refGene_tx=XM_003999048.3;refGene_gn=LOC101089067;refGene_tx=XM_019821241.2
E3 43537033 43537094
refGene_gn=LOC101089318;refGene_tx=XM_011290780.3
E3 43537274 43537455
refGene_gn=LOC101089318;refGene_tx=XM_011290780.3
E3 43537692 43537958
refGene_gn=LOC101089318;refGene_tx=XM_011290780.3
E3 43538970 43539134
refGene_gn=LOC101089318;refGene_tx=XM_011290780.3
E3 43539230 43539395
refGene_gn=LOC101089318;refGene_tx=XM_011290780.3
E3 43547384 43547445
refGene_gn=LOC101089568;refGene_tx=XM_019821244.2
E3 43547590 43547771
refGene_gn=LOC101089568;refGene_tx=XM_019821244.2
E3 43547899 43548165
refGene_gn=LOC101089568;refGene_tx=XM_019821244.2
E3 43549144 43549308
refGene_gn=LOC101089568;refGene_tx=XM_019821244.2
E3 43549409 43549574
refGene_gn=LOC101089568;refGene_tx=XM_019821244.2
E3 43552213 43552468
refGene_gn=LOC101089824;refGene_tx=XM_023246590.1
E3 43552557 43552618
refGene_gn=LOC101089824;refGene_tx=XM_023246590.1
E3 43552827 43553008
refGene_gn=LOC101089824;refGene_tx=XM_023246590.1
E3 43553149 43553617
refGene_gn=LOC101089824;refGene_tx=XM_023246590.1
E3 43553840 43554117
refGene_gn=LOC101089824;refGene_tx=XM_023246590.1
E3 43554293 43554415
refGene_gn=LOC101089824;refGene_tx=XM_023246590.1
E3 43554519 43554607
refGene_gn=LOC101089824;refGene_tx=XM_023246590.1
E3 43554683 43554848
refGene_gn=LOC101089824;refGene_tx=XM_023246590.1
E3 43579424 43580426
refGene_gn=CACNA1H;refGene_tx=XM_023246207.1;refGene_gn=CACNA1H;refGene_tx=XM_023246208.1

E3 43580982 43581155
refGene_gn=CACNA1H;refGene_tx=XM_023246207.1;refGene_gn=CACNA1H;refGene_tx=XM_023246208.1
E3 43581244 43581647
refGene_gn=CACNA1H;refGene_tx=XM_023246207.1;refGene_gn=CACNA1H;refGene_tx=XM_023246208.1
E3 43581755 43581877
refGene_gn=CACNA1H;refGene_tx=XM_023246207.1;refGene_gn=CACNA1H;refGene_tx=XM_023246208.1
E3 43582970 43583049
refGene_gn=CACNA1H;refGene_tx=XM_023246207.1;refGene_gn=CACNA1H;refGene_tx=XM_023246208.1
E3 43584739 43584810
refGene_gn=CACNA1H;refGene_tx=XM_023246207.1;refGene_gn=CACNA1H;refGene_tx=XM_023246208.1
E3 43585030 43585164
refGene_gn=CACNA1H;refGene_tx=XM_023246207.1;refGene_gn=CACNA1H;refGene_tx=XM_023246208.1
E3 43585342 43585452
refGene_gn=CACNA1H;refGene_tx=XM_023246207.1;refGene_gn=CACNA1H;refGene_tx=XM_023246208.1
E3 43586243 43586395
refGene_gn=CACNA1H;refGene_tx=XM_023246207.1;refGene_gn=CACNA1H;refGene_tx=XM_023246208.1
E3 43587785 43587803 refGene_gn=CACNA1H;refGene_tx=XM_023246207.1
E3 43588295 43588488
refGene_gn=CACNA1H;refGene_tx=XM_023246207.1;refGene_gn=CACNA1H;refGene_tx=XM_023246208.1
E3 43588719 43588809
refGene_gn=CACNA1H;refGene_tx=XM_023246207.1;refGene_gn=CACNA1H;refGene_tx=XM_023246208.1
E3 43589028 43589154
refGene_gn=CACNA1H;refGene_tx=XM_023246207.1;refGene_gn=CACNA1H;refGene_tx=XM_023246208.1
E3 43589398 43589525
refGene_gn=CACNA1H;refGene_tx=XM_023246207.1;refGene_gn=CACNA1H;refGene_tx=XM_023246208.1
E3 43589739 43589924
refGene_gn=CACNA1H;refGene_tx=XM_023246207.1;refGene_gn=CACNA1H;refGene_tx=XM_023246208.1
E3 43590018 43590087
refGene_gn=CACNA1H;refGene_tx=XM_023246207.1;refGene_gn=CACNA1H;refGene_tx=XM_023246208.1
E3 43590164 43590288
refGene_gn=CACNA1H;refGene_tx=XM_023246207.1;refGene_gn=CACNA1H;refGene_tx=XM_023246208.1
E3 43590449 43590550
refGene_gn=CACNA1H;refGene_tx=XM_023246207.1;refGene_gn=CACNA1H;refGene_tx=XM_023246208.1
E3 43591104 43591515

refGene_gn=CACNA1H;refGene_tx=XM_023246207.1;refGene_gn=CACNA1H;refGene_tx=XM_023246208.1
E3 43593343 43593552
refGene_gn=CACNA1H;refGene_tx=XM_023246207.1;refGene_gn=CACNA1H;refGene_tx=XM_023246208.1
E3 43593692 43593783
refGene_gn=CACNA1H;refGene_tx=XM_023246207.1;refGene_gn=CACNA1H;refGene_tx=XM_023246208.1
E3 43593995 43594151
refGene_gn=CACNA1H;refGene_tx=XM_023246207.1;refGene_gn=CACNA1H;refGene_tx=XM_023246208.1
E3 43594323 43594441
refGene_gn=CACNA1H;refGene_tx=XM_023246207.1;refGene_gn=CACNA1H;refGene_tx=XM_023246208.1
E3 43595049 43595235
refGene_gn=CACNA1H;refGene_tx=XM_023246207.1;refGene_gn=CACNA1H;refGene_tx=XM_023246208.1
E3 43596120 43596272
refGene_gn=CACNA1H;refGene_tx=XM_023246207.1;refGene_gn=CACNA1H;refGene_tx=XM_023246208.1
E3 43597512 43597955
refGene_gn=CACNA1H;refGene_tx=XM_023246207.1;refGene_gn=CACNA1H;refGene_tx=XM_023246208.1
E3 43598953 43599734
refGene_gn=CACNA1H;refGene_tx=XM_023246207.1;refGene_gn=CACNA1H;refGene_tx=XM_023246208.1
E3 43600810 43600903
refGene_gn=CACNA1H;refGene_tx=XM_023246207.1;refGene_gn=CACNA1H;refGene_tx=XM_023246208.1
E3 43601097 43601416
refGene_gn=CACNA1H;refGene_tx=XM_023246207.1;refGene_gn=CACNA1H;refGene_tx=XM_023246208.1
E3 43603075 43603235
refGene_gn=CACNA1H;refGene_tx=XM_023246207.1;refGene_gn=CACNA1H;refGene_tx=XM_023246208.1
E3 43605518 43605616
refGene_gn=CACNA1H;refGene_tx=XM_023246207.1;refGene_gn=CACNA1H;refGene_tx=XM_023246208.1
E3 43606037 43606171
refGene_gn=CACNA1H;refGene_tx=XM_023246207.1;refGene_gn=CACNA1H;refGene_tx=XM_023246208.1
E3 43606282 43606394
refGene_gn=CACNA1H;refGene_tx=XM_023246207.1;refGene_gn=CACNA1H;refGene_tx=XM_023246208.1
E3 43640797 43641210
refGene_gn=CACNA1H;refGene_tx=XM_023246207.1;refGene_gn=CACNA1H;refGene_tx=XM_023246208.1
E3 43672819 43673353
refGene_gn=LOC101086027;refGene_tx=XM_003998957.5
E3 43675800 43675871

refGene_gn=LOC101086027;refGene_tx=XM_003998957.5
E3 43676284 43676428
refGene_gn=LOC101086027;refGene_tx=XM_003998957.5
E3 43676936 43677159
refGene_gn=LOC101086027;refGene_tx=XM_003998957.5
E3 43678033 43678188
refGene_gn=LOC101086027;refGene_tx=XM_003998957.5
E3 43678966 43679183
refGene_gn=LOC101086027;refGene_tx=XM_003998957.5
E3 43682716 43682830 refGene_gn=C1QTNF8;refGene_tx=XM_003998958.3
E3 43684104 43684321 refGene_gn=C1QTNF8;refGene_tx=XM_003998958.3
E3 43684809 43685360 refGene_gn=C1QTNF8;refGene_tx=XM_003998958.3
E3 43694067 43695168 refGene_gn=SSTR5;refGene_tx=XM_006942579.4
E3 43734576 43734611
refGene_gn=LOC102901851;refGene_tx=XM_023246591.1
E3 43734906 43735111
refGene_gn=LOC102901851;refGene_tx=XM_023246591.1
E3 43736055 43736180
refGene_gn=LOC102901851;refGene_tx=XM_023246591.1
E3 43736970 43737388
refGene_gn=LOC102901851;refGene_tx=XM_023246591.1
E3 43737899 43738031
refGene_gn=LOC102901851;refGene_tx=XM_023246591.1
E3 43768716 43769453 refGene_gn=SOX8;refGene_tx=XM_023246502.1
E3 43770189 43770419 refGene_gn=SOX8;refGene_tx=XM_023246502.1
E3 43771699 43772121 refGene_gn=SOX8;refGene_tx=XM_023246502.1
E3 43782560 43782732
refGene_gn=LMF1;refGene_tx=XM_003999053.5;refGene_gn=LMF1;refGene_tx=X
M_019821259.2
E3 43797995 43798305
refGene_gn=LMF1;refGene_tx=XM_003999053.5;refGene_gn=LMF1;refGene_tx=X
M_019821259.2
E3 43805868 43805929
refGene_gn=LMF1;refGene_tx=XM_003999053.5;refGene_gn=LMF1;refGene_tx=X
M_019821259.2
E3 43834093 43834242 refGene_gn=LMF1;refGene_tx=XM_003999053.5
E3 43849047 43849113
refGene_gn=LMF1;refGene_tx=XM_003999053.5;refGene_gn=LMF1;refGene_tx=X
M_019821259.2
E3 43859746 43859914
refGene_gn=LMF1;refGene_tx=XM_003999053.5;refGene_gn=LMF1;refGene_tx=X
M_019821259.2
E3 43870689 43870873
refGene_gn=LMF1;refGene_tx=XM_003999053.5;refGene_gn=LMF1;refGene_tx=X
M_019821259.2
E3 43871100 43871254
refGene_gn=LMF1;refGene_tx=XM_003999053.5;refGene_gn=LMF1;refGene_tx=X
M_019821259.2
E3 43871590 43871774
refGene_gn=LMF1;refGene_tx=XM_003999053.5;refGene_gn=LMF1;refGene_tx=X

M_019821259.2
E3 43872303 43872416
refGene_gn=LMF1;refGene_tx=XM_003999053.5;refGene_gn=LMF1;refGene_tx=XM_019821259.2
E3 43872739 43872914
refGene_gn=LMF1;refGene_tx=XM_003999053.5;refGene_gn=LMF1;refGene_tx=XM_019821259.2
E3 43897075 43897463
refGene_gn=LOC109495366;refGene_tx=XM_019821262.2
E3 43897767 43898057
refGene_gn=LOC109495366;refGene_tx=XM_019821262.2
E3 43918287 43918385
refGene_gn=GNG13;refGene_tx=XM_003998959.4;refGene_gn=GNG13;refGene_tx=XM_023246611.1
E3 43918558 43918664
refGene_gn=GNG13;refGene_tx=XM_003998959.4;refGene_gn=GNG13;refGene_tx=XM_023246611.1
E3 43918954 43919091 refGene_gn=CHTF18;refGene_tx=XM_006942484.4
E3 43919172 43919230 refGene_gn=CHTF18;refGene_tx=XM_006942484.4
E3 43919496 43919613 refGene_gn=CHTF18;refGene_tx=XM_006942484.4
E3 43919727 43919872 refGene_gn=CHTF18;refGene_tx=XM_006942484.4
E3 43920542 43920673 refGene_gn=CHTF18;refGene_tx=XM_006942484.4
E3 43920771 43920921 refGene_gn=CHTF18;refGene_tx=XM_006942484.4
E3 43921003 43921228 refGene_gn=CHTF18;refGene_tx=XM_006942484.4
E3 43922014 43922186 refGene_gn=CHTF18;refGene_tx=XM_006942484.4
E3 43922373 43922504 refGene_gn=CHTF18;refGene_tx=XM_006942484.4
E3 43922592 43922691 refGene_gn=CHTF18;refGene_tx=XM_006942484.4
E3 43922994 43923088 refGene_gn=CHTF18;refGene_tx=XM_006942484.4
E3 43923195 43923347 refGene_gn=CHTF18;refGene_tx=XM_006942484.4
E3 43923430 43923554 refGene_gn=CHTF18;refGene_tx=XM_006942484.4
E3 43923797 43923895 refGene_gn=CHTF18;refGene_tx=XM_006942484.4
E3 43924411 43924621 refGene_gn=CHTF18;refGene_tx=XM_006942484.4
E3 43925094 43925239 refGene_gn=CHTF18;refGene_tx=XM_006942484.4
E3 43925368 43925421 refGene_gn=CHTF18;refGene_tx=XM_006942484.4
E3 43925502 43925595 refGene_gn=CHTF18;refGene_tx=XM_006942484.4
E3 43926017 43926186 refGene_gn=CHTF18;refGene_tx=XM_006942484.4
E3 43926357 43926508 refGene_gn=CHTF18;refGene_tx=XM_006942484.4
E3 43926589 43926784 refGene_gn=CHTF18;refGene_tx=XM_006942484.4
E3 43926963 43927054 refGene_gn=CHTF18;refGene_tx=XM_006942484.4
E3 43928053 43928235 refGene_gn=RPUSD1;refGene_tx=XM_023246619.1
E3 43928333 43928457 refGene_gn=RPUSD1;refGene_tx=XM_023246619.1
E3 43928543 43928646
refGene_gn=RPUSD1;refGene_tx=XM_023246619.1;refGene_gn=RPUSD1;refGene_tx=XM_023246620.1
E3 43928828 43928930
refGene_gn=RPUSD1;refGene_tx=XM_023246619.1;refGene_gn=RPUSD1;refGene_tx=XM_023246620.1
E3 43929406 43929804
refGene_gn=RPUSD1;refGene_tx=XM_023246619.1;refGene_gn=RPUSD1;refGene_tx=XM_023246620.1

E3 43934551 43934587
refGene_gn=LOC101091073;refGene_tx=XM_023246838.1
E3 43935442 43935528
refGene_gn=LOC101091073;refGene_tx=XM_023246838.1
E3 43935684 43935811
refGene_gn=LOC101091073;refGene_tx=XM_023246838.1
E3 43935891 43936082
refGene_gn=LOC101091073;refGene_tx=XM_023246838.1;refGene_gn=LOC101091073;refGene_tx=XM_023246839.1
E3 43936413 43936853
refGene_gn=LOC101091073;refGene_tx=XM_023246838.1;refGene_gn=LOC101091073;refGene_tx=XM_023246839.1
E3 43937213 43937407
refGene_gn=LOC101091073;refGene_tx=XM_023246838.1;refGene_gn=LOC101091073;refGene_tx=XM_023246839.1
E3 43937484 43937640
refGene_gn=LOC101091073;refGene_tx=XM_023246838.1;refGene_gn=LOC101091073;refGene_tx=XM_023246839.1
E3 43937712 43937840
refGene_gn=LOC101091073;refGene_tx=XM_023246838.1;refGene_gn=LOC101091073;refGene_tx=XM_023246839.1
E3 43938405 43938710
refGene_gn=LOC101091073;refGene_tx=XM_023246838.1;refGene_gn=LOC101091073;refGene_tx=XM_023246839.1
E3 43938925 43939181
refGene_gn=LOC101091073;refGene_tx=XM_023246838.1;refGene_gn=LOC101091073;refGene_tx=XM_023246839.1
E3 43939607 43939870
refGene_gn=LOC101091073;refGene_tx=XM_023246838.1;refGene_gn=LOC101091073;refGene_tx=XM_023246839.1
E3 43940428 43940517
refGene_gn=LOC101091333;refGene_tx=XM_006942487.3;refGene_gn=LOC101091333;refGene_tx=XM_019821276.2;refGene_gn=LOC101091333;refGene_tx=XM_019821277.2;refGene_gn=LOC101091333;refGene_tx=XM_019821281.2;refGene_gn=LOC101091333;refGene_tx=XM_023246840.1;refGene_gn=LOC101091333;refGene_tx=XM_023246841.1;refGene_gn=LOC101091333;refGene_tx=XM_023246842.1
E3 43940591 43940778
refGene_gn=LOC101091333;refGene_tx=XM_006942487.3;refGene_gn=LOC101091333;refGene_tx=XM_019821276.2;refGene_gn=LOC101091333;refGene_tx=XM_019821277.2;refGene_gn=LOC101091333;refGene_tx=XM_019821281.2;refGene_gn=LOC101091333;refGene_tx=XM_023246840.1;refGene_gn=LOC101091333;refGene_tx=XM_023246841.1;refGene_gn=LOC101091333;refGene_tx=XM_023246842.1
E3 43941026 43941121
refGene_gn=LOC101091333;refGene_tx=XM_006942487.3;refGene_gn=LOC101091333;refGene_tx=XM_019821276.2;refGene_gn=LOC101091333;refGene_tx=XM_019821277.2;refGene_gn=LOC101091333;refGene_tx=XM_019821281.2;refGene_gn=LOC101091333;refGene_tx=XM_023246840.1;refGene_gn=LOC101091333;refGene_tx=XM_023246841.1;refGene_gn=LOC101091333;refGene_tx=XM_023246842.1
E3 43941481 43941609
refGene_gn=LOC101091333;refGene_tx=XM_006942487.3;refGene_gn=LOC101091

333;refGene_tx=XM_019821276.2;refGene_gn=LOC101091333;refGene_tx=XM_019821277.2;refGene_gn=LOC101091333;refGene_tx=XM_019821281.2;refGene_gn=LOC101091333;refGene_tx=XM_023246840.1;refGene_gn=LOC101091333;refGene_tx=XM_023246841.1;refGene_gn=LOC101091333;refGene_tx=XM_023246842.1
E3 43941687 43941830
refGene_gn=LOC101091333;refGene_tx=XM_006942487.3;refGene_gn=LOC101091333;refGene_tx=XM_019821276.2;refGene_gn=LOC101091333;refGene_tx=XM_019821277.2;refGene_gn=LOC101091333;refGene_tx=XM_019821281.2;refGene_gn=LOC101091333;refGene_tx=XM_023246840.1;refGene_gn=LOC101091333;refGene_tx=XM_023246841.1;refGene_gn=LOC101091333;refGene_tx=XM_023246842.1
E3 43941921 43942077
refGene_gn=LOC101091333;refGene_tx=XM_006942487.3;refGene_gn=LOC101091333;refGene_tx=XM_019821276.2;refGene_gn=LOC101091333;refGene_tx=XM_019821277.2;refGene_gn=LOC101091333;refGene_tx=XM_019821281.2;refGene_gn=LOC101091333;refGene_tx=XM_023246840.1;refGene_gn=LOC101091333;refGene_tx=XM_023246841.1;refGene_gn=LOC101091333;refGene_tx=XM_023246842.1
E3 43942154 43942333
refGene_gn=LOC101091333;refGene_tx=XM_006942487.3;refGene_gn=LOC101091333;refGene_tx=XM_019821276.2;refGene_gn=LOC101091333;refGene_tx=XM_019821277.2;refGene_gn=LOC101091333;refGene_tx=XM_019821281.2;refGene_gn=LOC101091333;refGene_tx=XM_023246840.1;refGene_gn=LOC101091333;refGene_tx=XM_023246841.1;refGene_gn=LOC101091333;refGene_tx=XM_023246842.1
E3 43942607 43942707
refGene_gn=LOC101091333;refGene_tx=XM_006942487.3;refGene_gn=LOC101091333;refGene_tx=XM_019821276.2;refGene_gn=LOC101091333;refGene_tx=XM_019821277.2;refGene_gn=LOC101091333;refGene_tx=XM_019821281.2;refGene_gn=LOC101091333;refGene_tx=XM_023246840.1;refGene_gn=LOC101091333;refGene_tx=XM_023246841.1;refGene_gn=LOC101091333;refGene_tx=XM_023246842.1
E3 43942810 43942854
refGene_gn=LOC101091333;refGene_tx=XM_019821276.2;refGene_gn=LOC101091333;refGene_tx=XM_019821277.2;refGene_gn=LOC101091333;refGene_tx=XM_019821281.2;refGene_gn=LOC101091333;refGene_tx=XM_023246840.1;refGene_gn=LOC101091333;refGene_tx=XM_023246841.1
E3 43943508 43943596
refGene_gn=LOC101091333;refGene_tx=XM_019821276.2;refGene_gn=LOC101091333;refGene_tx=XM_019821277.2;refGene_gn=LOC101091333;refGene_tx=XM_019821281.2;refGene_gn=LOC101091333;refGene_tx=XM_023246840.1;refGene_gn=LOC101091333;refGene_tx=XM_023246841.1
E3 43944195 43944355
refGene_gn=LOC101091333;refGene_tx=XM_019821276.2;refGene_gn=LOC101091333;refGene_tx=XM_019821277.2;refGene_gn=LOC101091333;refGene_tx=XM_023246840.1
E3 43944522 43944734
refGene_gn=LOC101091333;refGene_tx=XM_019821276.2
E3 43964240 43964306 refGene_gn=NARFL;refGene_tx=XM_003998962.5
E3 43965485 43965581 refGene_gn=NARFL;refGene_tx=XM_003998962.5
E3 43966280 43966424 refGene_gn=NARFL;refGene_tx=XM_003998962.5
E3 43966703 43966836 refGene_gn=NARFL;refGene_tx=XM_003998962.5
E3 43967340 43967475 refGene_gn=NARFL;refGene_tx=XM_003998962.5
E3 43967766 43967885 refGene_gn=NARFL;refGene_tx=XM_003998962.5

E3 43968107 43968237 refGene_gn=NARFL;refGene_tx=XM_003998962.5
E3 43968915 43968988 refGene_gn=NARFL;refGene_tx=XM_003998962.5
E3 43969483 43969621 refGene_gn=NARFL;refGene_tx=XM_003998962.5
E3 43970047 43970205 refGene_gn=NARFL;refGene_tx=XM_003998962.5
E3 43970439 43970678 refGene_gn=NARFL;refGene_tx=XM_003998962.5
E3 43971735 43971883
refGene_gn=HAGHL;refGene_tx=XM_011290743.2;refGene_gn=HAGHL;refGene_tx=XM_011290744.2;refGene_gn=HAGHL;refGene_tx=XM_011290745.2;refGene_gn=HAGHL;refGene_tx=XM_019821283.1;refGene_gn=HAGHL;refGene_tx=XM_019821284.1
E3 43972146 43972226
refGene_gn=HAGHL;refGene_tx=XM_011290743.2;refGene_gn=HAGHL;refGene_tx=XM_011290744.2;refGene_gn=HAGHL;refGene_tx=XM_011290745.2;refGene_gn=HAGHL;refGene_tx=XM_019821284.1
E3 43972575 43972677
refGene_gn=HAGHL;refGene_tx=XM_011290743.2;refGene_gn=HAGHL;refGene_tx=XM_011290744.2;refGene_gn=HAGHL;refGene_tx=XM_011290745.2;refGene_gn=HAGHL;refGene_tx=XM_019821283.1;refGene_gn=HAGHL;refGene_tx=XM_019821284.1
E3 43972907 43973011
refGene_gn=HAGHL;refGene_tx=XM_011290743.2;refGene_gn=HAGHL;refGene_tx=XM_011290744.2;refGene_gn=HAGHL;refGene_tx=XM_011290745.2;refGene_gn=HAGHL;refGene_tx=XM_019821283.1;refGene_gn=HAGHL;refGene_tx=XM_019821284.1
E3 43973088 43973197
refGene_gn=HAGHL;refGene_tx=XM_011290743.2;refGene_gn=HAGHL;refGene_tx=XM_011290744.2;refGene_gn=HAGHL;refGene_tx=XM_019821283.1
E3 43973284 43973402
refGene_gn=HAGHL;refGene_tx=XM_011290743.2;refGene_gn=HAGHL;refGene_tx=XM_011290744.2;refGene_gn=HAGHL;refGene_tx=XM_011290745.2;refGene_gn=HAGHL;refGene_tx=XM_019821283.1;refGene_gn=HAGHL;refGene_tx=XM_019821284.1
E3 43973492 43973557
refGene_gn=HAGHL;refGene_tx=XM_011290743.2;refGene_gn=HAGHL;refGene_tx=XM_011290744.2;refGene_gn=HAGHL;refGene_tx=XM_011290745.2;refGene_gn=HAGHL;refGene_tx=XM_019821283.1;refGene_gn=HAGHL;refGene_tx=XM_019821284.1
E3 43973905 43974010
refGene_gn=HAGHL;refGene_tx=XM_011290743.2;refGene_gn=HAGHL;refGene_tx=XM_011290744.2;refGene_gn=HAGHL;refGene_tx=XM_011290745.2;refGene_gn=HAGHL;refGene_tx=XM_019821283.1;refGene_gn=HAGHL;refGene_tx=XM_019821284.1
E3 43975084 43975423
refGene_gn=CCDC78;refGene_tx=XM_019821264.2;refGene_gn=CCDC78;refGene_tx=XM_019821265.2;refGene_gn=CCDC78;refGene_tx=XM_019821266.2;refGene_gn=CCDC78;refGene_tx=XM_019821268.2;refGene_gn=CCDC78;refGene_tx=XM_019821269.2;refGene_gn=CCDC78;refGene_tx=XM_019821270.2;refGene_gn=CCDC78;refGene_tx=XM_019821271.2;refGene_gn=CCDC78;refGene_tx=XM_019821272.2;refGene_gn=CCDC78;refGene_tx=XM_019821273.2;refGene_gn=CCDC78;refGene_tx=XM_019821274.2;refGene_gn=CCDC78;refGene_tx=XM_019821275.2

E3 43975691 43975778
refGene_gn=CCDC78;refGene_tx=XM_019821264.2;refGene_gn=CCDC78;refGene_tx=XM_019821265.2;refGene_gn=CCDC78;refGene_tx=XM_019821266.2;refGene_gn=CCDC78;refGene_tx=XM_019821268.2;refGene_gn=CCDC78;refGene_tx=XM_019821269.2;refGene_gn=CCDC78;refGene_tx=XM_019821270.2;refGene_gn=CCDC78;refGene_tx=XM_019821271.2;refGene_gn=CCDC78;refGene_tx=XM_019821272.2;refGene_gn=CCDC78;refGene_tx=XM_019821273.2;refGene_gn=CCDC78;refGene_tx=XM_019821274.2;refGene_gn=CCDC78;refGene_tx=XM_019821275.2
E3 43975852 43975939
refGene_gn=CCDC78;refGene_tx=XM_019821264.2;refGene_gn=CCDC78;refGene_tx=XM_019821265.2;refGene_gn=CCDC78;refGene_tx=XM_019821266.2;refGene_gn=CCDC78;refGene_tx=XM_019821268.2;refGene_gn=CCDC78;refGene_tx=XM_019821269.2;refGene_gn=CCDC78;refGene_tx=XM_019821270.2;refGene_gn=CCDC78;refGene_tx=XM_019821271.2;refGene_gn=CCDC78;refGene_tx=XM_019821272.2;refGene_gn=CCDC78;refGene_tx=XM_019821273.2;refGene_gn=CCDC78;refGene_tx=XM_019821274.2;refGene_gn=CCDC78;refGene_tx=XM_019821275.2
E3 43976174 43976680
refGene_gn=CCDC78;refGene_tx=XM_019821264.2;refGene_gn=CCDC78;refGene_tx=XM_019821265.2;refGene_gn=CCDC78;refGene_tx=XM_019821266.2;refGene_gn=CCDC78;refGene_tx=XM_019821268.2;refGene_gn=CCDC78;refGene_tx=XM_019821269.2;refGene_gn=CCDC78;refGene_tx=XM_019821270.2;refGene_gn=CCDC78;refGene_tx=XM_019821271.2;refGene_gn=CCDC78;refGene_tx=XM_019821272.2;refGene_gn=CCDC78;refGene_tx=XM_019821273.2;refGene_gn=CCDC78;refGene_tx=XM_019821274.2;refGene_gn=CCDC78;refGene_tx=XM_019821275.2
E3 43976805 43976884
refGene_gn=CCDC78;refGene_tx=XM_019821264.2;refGene_gn=CCDC78;refGene_tx=XM_019821265.2;refGene_gn=CCDC78;refGene_tx=XM_019821266.2;refGene_gn=CCDC78;refGene_tx=XM_019821268.2;refGene_gn=CCDC78;refGene_tx=XM_019821269.2;refGene_gn=CCDC78;refGene_tx=XM_019821270.2;refGene_gn=CCDC78;refGene_tx=XM_019821271.2;refGene_gn=CCDC78;refGene_tx=XM_019821272.2;refGene_gn=CCDC78;refGene_tx=XM_019821273.2;refGene_gn=CCDC78;refGene_tx=XM_019821274.2;refGene_gn=CCDC78;refGene_tx=XM_019821275.2
E3 43976988 43977530
refGene_gn=CCDC78;refGene_tx=XM_019821264.2;refGene_gn=CCDC78;refGene_tx=XM_019821265.2;refGene_gn=CCDC78;refGene_tx=XM_019821266.2;refGene_gn=CCDC78;refGene_tx=XM_019821268.2;refGene_gn=CCDC78;refGene_tx=XM_019821269.2;refGene_gn=CCDC78;refGene_tx=XM_019821270.2;refGene_gn=CCDC78;refGene_tx=XM_019821271.2;refGene_gn=CCDC78;refGene_tx=XM_019821272.2;refGene_gn=CCDC78;refGene_tx=XM_019821273.2;refGene_gn=CCDC78;refGene_tx=XM_019821274.2;refGene_gn=CCDC78;refGene_tx=XM_019821275.2
E3 43977597 43977769
refGene_gn=CCDC78;refGene_tx=XM_019821264.2;refGene_gn=CCDC78;refGene_tx=XM_019821265.2;refGene_gn=CCDC78;refGene_tx=XM_019821266.2;refGene_gn=CCDC78;refGene_tx=XM_019821268.2;refGene_gn=CCDC78;refGene_tx=XM_019821269.2;refGene_gn=CCDC78;refGene_tx=XM_019821270.2;refGene_gn=CCDC78;refGene_tx=XM_019821271.2;refGene_gn=CCDC78;refGene_tx=XM_019821272.2;refGene_gn=CCDC78;refGene_tx=XM_019821274.2;refGene_gn=CCDC78;refGene_tx=XM_019821275.2
E3 43977908 43977997
refGene_gn=CCDC78;refGene_tx=XM_019821264.2;refGene_gn=CCDC78;refGene_

tx=XM_019821265.2;refGene_gn=CCDC78;refGene_tx=XM_019821266.2;refGene_gn=CCDC78;refGene_tx=XM_019821268.2;refGene_gn=CCDC78;refGene_tx=XM_019821269.2;refGene_gn=CCDC78;refGene_tx=XM_019821270.2;refGene_gn=CCDC78;refGene_tx=XM_019821271.2;refGene_gn=CCDC78;refGene_tx=XM_019821272.2;refGene_gn=CCDC78;refGene_tx=XM_019821273.2;refGene_gn=CCDC78;refGene_tx=XM_019821274.2;refGene_gn=CCDC78;refGene_tx=XM_019821275.2
E3 43978946 43979013
refGene_gn=CCDC78;refGene_tx=XM_019821264.2;refGene_gn=CCDC78;refGene_tx=XM_019821265.2;refGene_gn=CCDC78;refGene_tx=XM_019821266.2;refGene_gn=CCDC78;refGene_tx=XM_019821268.2;refGene_gn=CCDC78;refGene_tx=XM_019821269.2;refGene_gn=CCDC78;refGene_tx=XM_019821270.2;refGene_gn=CCDC78;refGene_tx=XM_019821271.2;refGene_gn=CCDC78;refGene_tx=XM_019821272.2;refGene_gn=CCDC78;refGene_tx=XM_019821273.2;refGene_gn=CCDC78;refGene_tx=XM_019821274.2;refGene_gn=CCDC78;refGene_tx=XM_019821275.2
E3 43979090 43979191
refGene_gn=CCDC78;refGene_tx=XM_019821265.2;refGene_gn=CCDC78;refGene_tx=XM_019821275.2
E3 43979287 43979503
refGene_gn=CCDC78;refGene_tx=XM_019821264.2;refGene_gn=CCDC78;refGene_tx=XM_019821265.2;refGene_gn=CCDC78;refGene_tx=XM_019821266.2;refGene_gn=CCDC78;refGene_tx=XM_019821268.2;refGene_gn=CCDC78;refGene_tx=XM_019821269.2;refGene_gn=CCDC78;refGene_tx=XM_019821270.2;refGene_gn=CCDC78;refGene_tx=XM_019821271.2;refGene_gn=CCDC78;refGene_tx=XM_019821272.2;refGene_gn=CCDC78;refGene_tx=XM_019821273.2;refGene_gn=CCDC78;refGene_tx=XM_019821274.2;refGene_gn=CCDC78;refGene_tx=XM_019821275.2
E3 43979549 43979771
refGene_gn=FAM173A;refGene_tx=XM_006942488.4;refGene_gn=FAM173A;refGene_tx=XM_006942489.4
E3 43979907 43979958 refGene_gn=FAM173A;refGene_tx=XM_006942488.4
E3 43980168 43980309
refGene_gn=FAM173A;refGene_tx=XM_006942488.4;refGene_gn=FAM173A;refGene_tx=XM_006942489.4
E3 43980403 43980508
refGene_gn=FAM173A;refGene_tx=XM_006942488.4;refGene_gn=FAM173A;refGene_tx=XM_006942489.4
E3 43980705 43980867
refGene_gn=FAM173A;refGene_tx=XM_006942488.4;refGene_gn=FAM173A;refGene_tx=XM_006942489.4
E3 43984271 43984582 refGene_gn=METRN;refGene_tx=XM_023246843.1
E3 43984666 43984726 refGene_gn=METRN;refGene_tx=XM_023246843.1
E3 43985626 43986027 refGene_gn=METRN;refGene_tx=XM_023246843.1
E3 43986175 43986273 refGene_gn=METRN;refGene_tx=XM_023246843.1
E3 43999349 43999994
refGene_gn=FBXL16;refGene_tx=XM_023246477.1;refGene_gn=FBXL16;refGene_tx=XM_023246478.1
E3 44000662 44001171
refGene_gn=FBXL16;refGene_tx=XM_023246477.1;refGene_gn=FBXL16;refGene_tx=XM_023246478.1
E3 44001579 44001664
refGene_gn=FBXL16;refGene_tx=XM_023246477.1;refGene_gn=FBXL16;refGene_

tx=XM_023246478.1
E3 44002019 44002083
refGene_gn=FBXL16;refGene_tx=XM_023246477.1;refGene_gn=FBXL16;refGene_tx=XM_023246478.1
E3 44002244 44002393
refGene_gn=FBXL16;refGene_tx=XM_023246477.1;refGene_gn=FBXL16;refGene_tx=XM_023246478.1
E3 44005980 44006461 refGene_gn=WDR24;refGene_tx=XM_003998964.5
E3 44007445 44007623 refGene_gn=WDR24;refGene_tx=XM_003998964.5
E3 44007760 44008433 refGene_gn=WDR24;refGene_tx=XM_003998964.5
E3 44008989 44009108 refGene_gn=WDR24;refGene_tx=XM_003998964.5
E3 44009186 44009308 refGene_gn=WDR24;refGene_tx=XM_003998964.5
E3 44009402 44009507 refGene_gn=WDR24;refGene_tx=XM_003998964.5
E3 44009586 44009927 refGene_gn=WDR24;refGene_tx=XM_003998964.5
E3 44010081 44010435 refGene_gn=WDR24;refGene_tx=XM_003998964.5
E3 44010732 44010848
refGene_gn=JMJD8;refGene_tx=XM_003999060.5;refGene_gn=JMJD8;refGene_tx=XM_006942493.3
E3 44010907 44011000
refGene_gn=JMJD8;refGene_tx=XM_003999060.5;refGene_gn=JMJD8;refGene_tx=XM_006942493.3
E3 44011155 44011204
refGene_gn=JMJD8;refGene_tx=XM_003999060.5;refGene_gn=JMJD8;refGene_tx=XM_006942493.3
E3 44011281 44011378
refGene_gn=JMJD8;refGene_tx=XM_003999060.5;refGene_gn=JMJD8;refGene_tx=XM_006942493.3
E3 44011452 44011521
refGene_gn=JMJD8;refGene_tx=XM_003999060.5;refGene_gn=JMJD8;refGene_tx=XM_006942493.3
E3 44011625 44011745
refGene_gn=JMJD8;refGene_tx=XM_003999060.5;refGene_gn=JMJD8;refGene_tx=XM_006942493.3
E3 44011838 44011906
refGene_gn=JMJD8;refGene_tx=XM_003999060.5;refGene_gn=JMJD8;refGene_tx=XM_006942493.3
E3 44011981 44012116
refGene_gn=JMJD8;refGene_tx=XM_003999060.5;refGene_gn=JMJD8;refGene_tx=XM_006942493.3
E3 44012202 44012283
refGene_gn=JMJD8;refGene_tx=XM_003999060.5;refGene_gn=JMJD8;refGene_tx=XM_006942493.3
E3 44012576 44012702 refGene_gn=STUB1;refGene_tx=XM_006942583.3
E3 44012780 44012897 refGene_gn=STUB1;refGene_tx=XM_006942583.3
E3 44012986 44013043 refGene_gn=STUB1;refGene_tx=XM_006942583.3
E3 44013167 44013255 refGene_gn=STUB1;refGene_tx=XM_006942583.3
E3 44013465 44013631 refGene_gn=STUB1;refGene_tx=XM_006942583.3
E3 44013717 44013916 refGene_gn=STUB1;refGene_tx=XM_006942583.3
E3 44014444 44014603 refGene_gn=STUB1;refGene_tx=XM_006942583.3
E3 44016631 44016903

refGene_gn=RHBDL1;refGene_tx=XM_023246479.1;refGene_gn=RHBDL1;refGene_tx=XM_023246480.1;refGene_gn=RHBDL1;refGene_tx=XM_023246481.1;refGene_gn=RHBDL1;refGene_tx=XM_023246482.1;refGene_gn=RHBDL1;refGene_tx=XM_023246484.1;refGene_gn=RHBDL1;refGene_tx=XM_023246485.1;refGene_gn=RHBDL1;refGene_tx=XM_023246486.1;refGene_gn=RHBDL1;refGene_tx=XM_023246487.1;refGene_gn=RHBDL1;refGene_tx=XM_023246488.1
E3 44016979 44017040
refGene_gn=RHBDL1;refGene_tx=XM_023246479.1;refGene_gn=RHBDL1;refGene_tx=XM_023246480.1;refGene_gn=RHBDL1;refGene_tx=XM_023246481.1;refGene_gn=RHBDL1;refGene_tx=XM_023246482.1;refGene_gn=RHBDL1;refGene_tx=XM_023246485.1;refGene_gn=RHBDL1;refGene_tx=XM_023246486.1;refGene_gn=RHBDL1;refGene_tx=XM_023246487.1;refGene_gn=RHBDL1;refGene_tx=XM_023246488.1
E3 44017120 44017221
refGene_gn=RHBDL1;refGene_tx=XM_023246479.1;refGene_gn=RHBDL1;refGene_tx=XM_023246481.1;refGene_gn=RHBDL1;refGene_tx=XM_023246485.1;refGene_gn=RHBDL1;refGene_tx=XM_023246486.1;refGene_gn=RHBDL1;refGene_tx=XM_023246487.1;refGene_gn=RHBDL1;refGene_tx=XM_023246488.1
E3 44017260 44017408
refGene_gn=RHBDL1;refGene_tx=XM_023246479.1;refGene_gn=RHBDL1;refGene_tx=XM_023246480.1;refGene_gn=RHBDL1;refGene_tx=XM_023246481.1;refGene_gn=RHBDL1;refGene_tx=XM_023246482.1;refGene_gn=RHBDL1;refGene_tx=XM_023246484.1;refGene_gn=RHBDL1;refGene_tx=XM_023246485.1;refGene_gn=RHBDL1;refGene_tx=XM_023246486.1;refGene_gn=RHBDL1;refGene_tx=XM_023246487.1;refGene_gn=RHBDL1;refGene_tx=XM_023246488.1
E3 44017525 44017674
refGene_gn=RHBDL1;refGene_tx=XM_023246479.1;refGene_gn=RHBDL1;refGene_tx=XM_023246480.1;refGene_gn=RHBDL1;refGene_tx=XM_023246481.1;refGene_gn=RHBDL1;refGene_tx=XM_023246482.1;refGene_gn=RHBDL1;refGene_tx=XM_023246484.1;refGene_gn=RHBDL1;refGene_tx=XM_023246485.1;refGene_gn=RHBDL1;refGene_tx=XM_023246486.1;refGene_gn=RHBDL1;refGene_tx=XM_023246487.1;refGene_gn=RHBDL1;refGene_tx=XM_023246488.1
E3 44017751 44017976
refGene_gn=RHBDL1;refGene_tx=XM_023246479.1;refGene_gn=RHBDL1;refGene_tx=XM_023246480.1;refGene_gn=RHBDL1;refGene_tx=XM_023246481.1;refGene_gn=RHBDL1;refGene_tx=XM_023246482.1;refGene_gn=RHBDL1;refGene_tx=XM_023246484.1;refGene_gn=RHBDL1;refGene_tx=XM_023246485.1;refGene_gn=RHBDL1;refGene_tx=XM_023246486.1
E3 44018217 44018580
refGene_gn=RHBDL1;refGene_tx=XM_023246479.1;refGene_gn=RHBDL1;refGene_tx=XM_023246480.1;refGene_gn=RHBDL1;refGene_tx=XM_023246481.1;refGene_gn=RHBDL1;refGene_tx=XM_023246482.1;refGene_gn=RHBDL1;refGene_tx=XM_023246484.1;refGene_gn=RHBDL1;refGene_tx=XM_023246485.1;refGene_gn=RHBDL1;refGene_tx=XM_023246486.1;refGene_gn=RHBDL1;refGene_tx=XM_023246487.1;refGene_gn=RHBDL1;refGene_tx=XM_023246488.1
E3 44018846 44018885
refGene_gn=RHBDL1;refGene_tx=XM_023246479.1;refGene_gn=RHBDL1;refGene_tx=XM_023246480.1;refGene_gn=RHBDL1;refGene_tx=XM_023246481.1;refGene_gn=RHBDL1;refGene_tx=XM_023246482.1;refGene_gn=RHBDL1;refGene_tx=XM_023246484.1;refGene_gn=RHBDL1;refGene_tx=XM_023246485.1;refGene_gn=RHBDL

1;refGene_tx=XM_023246486.1;refGene_gn=RHBDL1;refGene_tx=XM_023246487.1;refGene_gn=RHBDL1;refGene_tx=XM_023246488.1
E3 44020943 44021070
refGene_gn=RHOT2;refGene_tx=XM_011290746.2;refGene_gn=RHOT2;refGene_tx=XM_011290747.3;refGene_gn=RHOT2;refGene_tx=XM_023246476.1
E3 44021353 44021556
refGene_gn=RHOT2;refGene_tx=XM_011290746.2;refGene_gn=RHOT2;refGene_tx=XM_011290747.3;refGene_gn=RHOT2;refGene_tx=XM_023246476.1
E3 44021644 44021767
refGene_gn=RHOT2;refGene_tx=XM_011290746.2;refGene_gn=RHOT2;refGene_tx=XM_011290747.3;refGene_gn=RHOT2;refGene_tx=XM_023246476.1
E3 44021883 44021964
refGene_gn=RHOT2;refGene_tx=XM_011290746.2;refGene_gn=RHOT2;refGene_tx=XM_011290747.3;refGene_gn=RHOT2;refGene_tx=XM_023246476.1
E3 44022096 44022227
refGene_gn=RHOT2;refGene_tx=XM_011290746.2;refGene_gn=RHOT2;refGene_tx=XM_011290747.3;refGene_gn=RHOT2;refGene_tx=XM_023246476.1
E3 44022306 44022407
refGene_gn=RHOT2;refGene_tx=XM_011290746.2;refGene_gn=RHOT2;refGene_tx=XM_011290747.3;refGene_gn=RHOT2;refGene_tx=XM_023246476.1
E3 44022490 44022633
refGene_gn=RHOT2;refGene_tx=XM_011290746.2;refGene_gn=RHOT2;refGene_tx=XM_011290747.3;refGene_gn=RHOT2;refGene_tx=XM_023246476.1
E3 44022721 44022806
refGene_gn=RHOT2;refGene_tx=XM_011290746.2;refGene_gn=RHOT2;refGene_tx=XM_011290747.3;refGene_gn=RHOT2;refGene_tx=XM_023246476.1
E3 44023042 44023163
refGene_gn=RHOT2;refGene_tx=XM_011290746.2;refGene_gn=RHOT2;refGene_tx=XM_011290747.3;refGene_gn=RHOT2;refGene_tx=XM_023246476.1
E3 44023253 44023362
refGene_gn=RHOT2;refGene_tx=XM_011290746.2;refGene_gn=RHOT2;refGene_tx=XM_011290747.3;refGene_gn=RHOT2;refGene_tx=XM_023246476.1
E3 44023494 44023593
refGene_gn=RHOT2;refGene_tx=XM_011290746.2;refGene_gn=RHOT2;refGene_tx=XM_011290747.3;refGene_gn=RHOT2;refGene_tx=XM_023246476.1
E3 44023679 44023781
refGene_gn=RHOT2;refGene_tx=XM_011290746.2;refGene_gn=RHOT2;refGene_tx=XM_011290747.3;refGene_gn=RHOT2;refGene_tx=XM_023246476.1
E3 44023953 44024062
refGene_gn=RHOT2;refGene_tx=XM_011290746.2;refGene_gn=RHOT2;refGene_tx=XM_011290747.3;refGene_gn=RHOT2;refGene_tx=XM_023246476.1
E3 44024136 44024189
refGene_gn=RHOT2;refGene_tx=XM_011290746.2;refGene_gn=RHOT2;refGene_tx=XM_011290747.3;refGene_gn=RHOT2;refGene_tx=XM_023246476.1
E3 44024674 44024728
refGene_gn=RHOT2;refGene_tx=XM_011290746.2;refGene_gn=RHOT2;refGene_tx=XM_011290747.3;refGene_gn=RHOT2;refGene_tx=XM_023246476.1
E3 44025606 44025650
refGene_gn=RHOT2;refGene_tx=XM_011290746.2;refGene_gn=RHOT2;refGene_tx=XM_011290747.3;refGene_gn=RHOT2;refGene_tx=XM_023246476.1

E3 44025738 44025820
refGene_gn=RHOT2;refGene_tx=XM_011290746.2;refGene_gn=RHOT2;refGene_tx
=XM_011290747.3;refGene_gn=RHOT2;refGene_tx=XM_023246476.1
E3 44025909 44025968
refGene_gn=RHOT2;refGene_tx=XM_011290746.2;refGene_gn=RHOT2;refGene_tx
=XM_011290747.3;refGene_gn=RHOT2;refGene_tx=XM_023246476.1
E3 44026107 44026144
refGene_gn=RHOT2;refGene_tx=XM_011290746.2;refGene_gn=RHOT2;refGene_tx
=XM_011290747.3
E3 44026707 44026919
refGene_gn=RHOT2;refGene_tx=XM_023246476.1;refGene_gn=WDR90;refGene_tx
=XM_006942501.3
E3 44027403 44027488 refGene_gn=WDR90;refGene_tx=XM_006942501.3
E3 44027574 44027694 refGene_gn=WDR90;refGene_tx=XM_006942501.3
E3 44027769 44027913 refGene_gn=WDR90;refGene_tx=XM_006942501.3
E3 44028022 44028153 refGene_gn=WDR90;refGene_tx=XM_006942501.3
E3 44028266 44028441 refGene_gn=WDR90;refGene_tx=XM_006942501.3
E3 44028593 44028716 refGene_gn=WDR90;refGene_tx=XM_006942501.3
E3 44030993 44031165 refGene_gn=WDR90;refGene_tx=XM_006942501.3
E3 44031313 44031425 refGene_gn=WDR90;refGene_tx=XM_006942501.3
E3 44031546 44031715 refGene_gn=WDR90;refGene_tx=XM_006942501.3
E3 44031803 44031994 refGene_gn=WDR90;refGene_tx=XM_006942501.3
E3 44032089 44032249 refGene_gn=WDR90;refGene_tx=XM_006942501.3
E3 44032412 44032534 refGene_gn=WDR90;refGene_tx=XM_006942501.3
E3 44032860 44032963 refGene_gn=WDR90;refGene_tx=XM_006942501.3
E3 44033510 44033608 refGene_gn=WDR90;refGene_tx=XM_006942501.3
E3 44033957 44034140 refGene_gn=WDR90;refGene_tx=XM_006942501.3
E3 44034274 44034406 refGene_gn=WDR90;refGene_tx=XM_006942501.3
E3 44034595 44034636 refGene_gn=WDR90;refGene_tx=XM_006942501.3
E3 44034715 44034846 refGene_gn=WDR90;refGene_tx=XM_006942501.3
E3 44035113 44035242 refGene_gn=WDR90;refGene_tx=XM_006942501.3
E3 44035460 44035622 refGene_gn=WDR90;refGene_tx=XM_006942501.3
E3 44035984 44036115 refGene_gn=WDR90;refGene_tx=XM_006942501.3
E3 44036595 44036726 refGene_gn=WDR90;refGene_tx=XM_006942501.3
E3 44036891 44037031 refGene_gn=WDR90;refGene_tx=XM_006942501.3
E3 44037255 44037491 refGene_gn=WDR90;refGene_tx=XM_006942501.3
E3 44037701 44037818 refGene_gn=WDR90;refGene_tx=XM_006942501.3
E3 44037887 44038018 refGene_gn=WDR90;refGene_tx=XM_006942501.3
E3 44038102 44038246 refGene_gn=WDR90;refGene_tx=XM_006942501.3
E3 44038388 44038507 refGene_gn=WDR90;refGene_tx=XM_006942501.3
E3 44039007 44039065 refGene_gn=WDR90;refGene_tx=XM_006942501.3
E3 44039134 44039280 refGene_gn=WDR90;refGene_tx=XM_006942501.3
E3 44039358 44039469 refGene_gn=WDR90;refGene_tx=XM_006942501.3
E3 44040043 44040112 refGene_gn=WDR90;refGene_tx=XM_006942501.3
E3 44040343 44040406 refGene_gn=WDR90;refGene_tx=XM_006942501.3
E3 44040525 44040735 refGene_gn=WDR90;refGene_tx=XM_006942501.3
E3 44040823 44040927 refGene_gn=WDR90;refGene_tx=XM_006942501.3
E3 44041216 44041284 refGene_gn=WDR90;refGene_tx=XM_006942501.3
E3 44041368 44041483 refGene_gn=WDR90;refGene_tx=XM_006942501.3
E3 44041761 44041932 refGene_gn=WDR90;refGene_tx=XM_006942501.3

E3 44042118 44042227 refGene_gn=WDR90;refGene_tx=XM_006942501.3
E3 44042314 44042491 refGene_gn=WDR90;refGene_tx=XM_006942501.3
E3 44042597 44042689 refGene_gn=WDR90;refGene_tx=XM_006942501.3
E3 44043036 44043046 refGene_gn=WDR90;refGene_tx=XM_006942501.3
E3 44044276 44044347
refGene_gn=MCRIP2;refGene_tx=XM_023246490.1;refGene_gn=MCRIP2;refGene_tx=XM_023246491.1;refGene_gn=MCRIP2;refGene_tx=XM_023246492.1;refGene_gn=MCRIP2;refGene_tx=XM_023246493.1
E3 44044577 44044679
refGene_gn=MCRIP2;refGene_tx=XM_023246490.1;refGene_gn=MCRIP2;refGene_tx=XM_023246491.1
E3 44044909 44045037
refGene_gn=MCRIP2;refGene_tx=XM_023246491.1;refGene_gn=MCRIP2;refGene_tx=XM_023246493.1
E3 44045862 44046002
refGene_gn=MCRIP2;refGene_tx=XM_023246490.1;refGene_gn=MCRIP2;refGene_tx=XM_023246492.1
E3 44048646 44048764
refGene_gn=MCRIP2;refGene_tx=XM_023246490.1;refGene_gn=MCRIP2;refGene_tx=XM_023246491.1;refGene_gn=MCRIP2;refGene_tx=XM_023246492.1;refGene_gn=MCRIP2;refGene_tx=XM_023246493.1
E3 44048866 44048918
refGene_gn=MCRIP2;refGene_tx=XM_023246490.1;refGene_gn=MCRIP2;refGene_tx=XM_023246491.1;refGene_gn=MCRIP2;refGene_tx=XM_023246492.1;refGene_gn=MCRIP2;refGene_tx=XM_023246493.1
E3 44054076 44054273 refGene_gn=METTL26;refGene_tx=XM_003998965.5
E3 44054423 44054586 refGene_gn=METTL26;refGene_tx=XM_003998965.5
E3 44054799 44054859 refGene_gn=METTL26;refGene_tx=XM_003998965.5
E3 44055239 44055307 refGene_gn=METTL26;refGene_tx=XM_003998965.5
E3 44055410 44055489 refGene_gn=METTL26;refGene_tx=XM_003998965.5
E3 44055568 44055616 refGene_gn=METTL26;refGene_tx=XM_003998965.5
E3 44056246 44057725 refGene_gn=WFIKKN1;refGene_tx=XM_023247043.1
E3 44058408 44058579 refGene_gn=WFIKKN1;refGene_tx=XM_023247043.1
E3 44062186 44062467
refGene_gn=RAB40C;refGene_tx=XM_011290748.3;refGene_gn=RAB40C;refGene_tx=XM_011290749.3;refGene_gn=RAB40C;refGene_tx=XM_019821288.2;refGene_gn=RAB40C;refGene_tx=XM_023247044.1
E3 44063047 44063270
refGene_gn=RAB40C;refGene_tx=XM_011290748.3;refGene_gn=RAB40C;refGene_tx=XM_011290749.3;refGene_gn=RAB40C;refGene_tx=XM_019821288.2;refGene_gn=RAB40C;refGene_tx=XM_023247044.1
E3 44063552 44063630
refGene_gn=RAB40C;refGene_tx=XM_011290748.3;refGene_gn=RAB40C;refGene_tx=XM_011290749.3;refGene_gn=RAB40C;refGene_tx=XM_019821288.2;refGene_gn=RAB40C;refGene_tx=XM_023247044.1
E3 44069150 44069211
refGene_gn=RAB40C;refGene_tx=XM_011290748.3;refGene_gn=RAB40C;refGene_tx=XM_011290749.3;refGene_gn=RAB40C;refGene_tx=XM_019821288.2;refGene_gn=RAB40C;refGene_tx=XM_023247044.1
E3 44069696 44069757

refGene_gn=RAB40C;refGene_tx=XM_011290748.3;refGene_gn=RAB40C;refGene_tx=XM_011290749.3;refGene_gn=RAB40C;refGene_tx=XM_019821288.2;refGene_gn=RAB40C;refGene_tx=XM_023247044.1
E3 44069948 44070057 refGene_gn=RAB40C;refGene_tx=XM_019821288.2
E3 44088574 44088716
refGene_gn=RAB40C;refGene_tx=XM_011290748.3;refGene_gn=RAB40C;refGene_tx=XM_011290749.3;refGene_gn=RAB40C;refGene_tx=XM_023247044.1
E3 44096828 44096981 refGene_gn=PIGQ;refGene_tx=XM_019821289.2
E3 44097781 44097843 refGene_gn=PIGQ;refGene_tx=XM_019821289.2
E3 44099395 44099510 refGene_gn=PIGQ;refGene_tx=XM_019821289.2
E3 44099967 44100048 refGene_gn=PIGQ;refGene_tx=XM_019821289.2
E3 44100751 44100863 refGene_gn=PIGQ;refGene_tx=XM_019821289.2
E3 44100972 44101126 refGene_gn=PIGQ;refGene_tx=XM_019821289.2
E3 44101436 44101563 refGene_gn=PIGQ;refGene_tx=XM_019821289.2
E3 44103297 44103418 refGene_gn=PIGQ;refGene_tx=XM_019821289.2
E3 44103922 44104054 refGene_gn=PIGQ;refGene_tx=XM_019821289.2
E3 44105754 44106374 refGene_gn=PIGQ;refGene_tx=XM_019821289.2
E3 44106375 44106444 refGene_gn=PIGQ;refGene_tx=XM_019821289.2
E3 44111160 44111844 refGene_gn=NHLRC4;refGene_tx=XM_006942506.2
E3 44114528 44115162 refGene_gn=PRR35;refGene_tx=XM_023246869.1
E3 44115441 44116207 refGene_gn=PRR35;refGene_tx=XM_023246869.1
E3 44116251 44116561 refGene_gn=PRR35;refGene_tx=XM_023246869.1
E3 44126699 44126877
refGene_gn=CAPN15;refGene_tx=XM_023246220.1;refGene_gn=CAPN15;refGene_tx=XM_023246221.1;refGene_gn=CAPN15;refGene_tx=XM_023246222.1
E3 44127023 44127202
refGene_gn=CAPN15;refGene_tx=XM_023246220.1;refGene_gn=CAPN15;refGene_tx=XM_023246221.1;refGene_gn=CAPN15;refGene_tx=XM_023246222.1
E3 44127282 44127449
refGene_gn=CAPN15;refGene_tx=XM_023246220.1;refGene_gn=CAPN15;refGene_tx=XM_023246221.1;refGene_gn=CAPN15;refGene_tx=XM_023246222.1
E3 44127542 44127775
refGene_gn=CAPN15;refGene_tx=XM_023246220.1;refGene_gn=CAPN15;refGene_tx=XM_023246221.1;refGene_gn=CAPN15;refGene_tx=XM_023246222.1
E3 44127860 44128022
refGene_gn=CAPN15;refGene_tx=XM_023246220.1;refGene_gn=CAPN15;refGene_tx=XM_023246221.1;refGene_gn=CAPN15;refGene_tx=XM_023246222.1
E3 44128437 44128590
refGene_gn=CAPN15;refGene_tx=XM_023246220.1;refGene_gn=CAPN15;refGene_tx=XM_023246221.1;refGene_gn=CAPN15;refGene_tx=XM_023246222.1
E3 44128661 44128787
refGene_gn=CAPN15;refGene_tx=XM_023246220.1;refGene_gn=CAPN15;refGene_tx=XM_023246221.1;refGene_gn=CAPN15;refGene_tx=XM_023246222.1
E3 44129484 44129708
refGene_gn=CAPN15;refGene_tx=XM_023246220.1;refGene_gn=CAPN15;refGene_tx=XM_023246221.1;refGene_gn=CAPN15;refGene_tx=XM_023246222.1
E3 44129795 44129979
refGene_gn=CAPN15;refGene_tx=XM_023246220.1;refGene_gn=CAPN15;refGene_tx=XM_023246221.1;refGene_gn=CAPN15;refGene_tx=XM_023246222.1
E3 44130054 44130263

refGene_gn=CAPN15;refGene_tx=XM_023246220.1;refGene_gn=CAPN15;refGene_tx=XM_023246221.1;refGene_gn=CAPN15;refGene_tx=XM_023246222.1
E3 44131054 44132515
refGene_gn=CAPN15;refGene_tx=XM_023246220.1;refGene_gn=CAPN15;refGene_tx=XM_023246221.1;refGene_gn=CAPN15;refGene_tx=XM_023246222.1
E3 44154270 44154384
refGene_gn=RAB11FIP3;refGene_tx=XM_023246593.1
E3 44154465 44154606
refGene_gn=RAB11FIP3;refGene_tx=XM_023246593.1
E3 44154784 44154940
refGene_gn=RAB11FIP3;refGene_tx=XM_023246593.1
E3 44155195 44155333
refGene_gn=RAB11FIP3;refGene_tx=XM_023246593.1
E3 44156064 44156146
refGene_gn=RAB11FIP3;refGene_tx=XM_023246593.1
E3 44157897 44158038
refGene_gn=RAB11FIP3;refGene_tx=XM_023246593.1
E3 44160907 44161011
refGene_gn=RAB11FIP3;refGene_tx=XM_023246593.1
E3 44161544 44161638
refGene_gn=RAB11FIP3;refGene_tx=XM_023246593.1
E3 44165506 44165542
refGene_gn=RAB11FIP3;refGene_tx=XM_023246593.1
E3 44169509 44169644
refGene_gn=RAB11FIP3;refGene_tx=XM_023246593.1
E3 44171762 44171912
refGene_gn=RAB11FIP3;refGene_tx=XM_023246593.1
E3 44175832 44176044
refGene_gn=RAB11FIP3;refGene_tx=XM_023246593.1
E3 44183284 44183376
refGene_gn=RAB11FIP3;refGene_tx=XM_023246593.1
E3 44193221 44193315
refGene_gn=RAB11FIP3;refGene_tx=XM_023246593.1
E3 44234822 44235379
refGene_gn=RAB11FIP3;refGene_tx=XM_023246593.1
E3 44235429 44235556
refGene_gn=RAB11FIP3;refGene_tx=XM_023246593.1
E3 44240141 44240371
refGene_gn=DECR2;refGene_tx=XM_023246356.1;refGene_gn=DECR2;refGene_tx=XM_023246362.1
E3 44246186 44246251 refGene_gn=DECR2;refGene_tx=XM_023246361.1
E3 44246888 44246981
refGene_gn=DECR2;refGene_tx=XM_023246356.1;refGene_gn=DECR2;refGene_tx=XM_023246361.1;refGene_gn=DECR2;refGene_tx=XM_023246362.1
E3 44248854 44249090 refGene_gn=DECR2;refGene_tx=XM_023246357.1
E3 44250192 44250410
refGene_gn=DECR2;refGene_tx=XM_023246358.1;refGene_gn=DECR2;refGene_tx=XM_023246359.1;refGene_gn=DECR2;refGene_tx=XM_023246360.1
E3 44250884 44250989
refGene_gn=DECR2;refGene_tx=XM_023246356.1;refGene_gn=DECR2;refGene_tx

=XM_023246357.1;refGene_gn=DECR2;refGene_tx=XM_023246358.1;refGene_gn=DECR2;refGene_tx=XM_023246359.1;refGene_gn=DECR2;refGene_tx=XM_023246360.1;refGene_gn=DECR2;refGene_tx=XM_023246361.1;refGene_gn=DECR2;refGene_tx=XM_023246362.1
E3	44251202 44251296
refGene_gn=DECR2;refGene_tx=XM_023246356.1;refGene_gn=DECR2;refGene_tx=XM_023246357.1;refGene_gn=DECR2;refGene_tx=XM_023246358.1;refGene_gn=DECR2;refGene_tx=XM_023246359.1;refGene_gn=DECR2;refGene_tx=XM_023246360.1;refGene_gn=DECR2;refGene_tx=XM_023246361.1;refGene_gn=DECR2;refGene_tx=XM_023246362.1
E3	44251367 44251492
refGene_gn=DECR2;refGene_tx=XM_023246356.1;refGene_gn=DECR2;refGene_tx=XM_023246357.1;refGene_gn=DECR2;refGene_tx=XM_023246358.1;refGene_gn=DECR2;refGene_tx=XM_023246359.1;refGene_gn=DECR2;refGene_tx=XM_023246360.1;refGene_gn=DECR2;refGene_tx=XM_023246361.1;refGene_gn=DECR2;refGene_tx=XM_023246362.1
E3	44252502 44252638
refGene_gn=DECR2;refGene_tx=XM_023246356.1;refGene_gn=DECR2;refGene_tx=XM_023246357.1;refGene_gn=DECR2;refGene_tx=XM_023246358.1;refGene_gn=DECR2;refGene_tx=XM_023246359.1;refGene_gn=DECR2;refGene_tx=XM_023246360.1;refGene_gn=DECR2;refGene_tx=XM_023246361.1
E3	44253091 44253094
refGene_gn=DECR2;refGene_tx=XM_023246356.1;refGene_gn=DECR2;refGene_tx=XM_023246357.1;refGene_gn=DECR2;refGene_tx=XM_023246358.1;refGene_gn=DECR2;refGene_tx=XM_023246359.1;refGene_gn=DECR2;refGene_tx=XM_023246360.1;refGene_gn=DECR2;refGene_tx=XM_023246361.1
E3	44257530 44257654 refGene_gn=NME4;refGene_tx=XM_023246607.1
E3	44258192 44258305 refGene_gn=NME4;refGene_tx=XM_023246607.1
E3	44258416 44258518 refGene_gn=NME4;refGene_tx=XM_023246607.1
E3	44258786 44258920 refGene_gn=NME4;refGene_tx=XM_023246607.1
E3	44260580 44260671 refGene_gn=NME4;refGene_tx=XM_023246607.1
E3	44272443 44272564 refGene_gn=TMEM8A;refGene_tx=XM_023246442.1
E3	44276040 44276218 refGene_gn=TMEM8A;refGene_tx=XM_023246442.1
E3	44276316 44276509
refGene_gn=TMEM8A;refGene_tx=XM_023246442.1;refGene_gn=TMEM8A;refGene_tx=XM_023246443.1
E3	44276729 44276857
refGene_gn=TMEM8A;refGene_tx=XM_023246442.1;refGene_gn=TMEM8A;refGene_tx=XM_023246443.1
E3	44277048 44277317
refGene_gn=TMEM8A;refGene_tx=XM_023246442.1;refGene_gn=TMEM8A;refGene_tx=XM_023246443.1
E3	44277412 44277732
refGene_gn=TMEM8A;refGene_tx=XM_023246442.1;refGene_gn=TMEM8A;refGene_tx=XM_023246443.1
E3	44278117 44278208
refGene_gn=TMEM8A;refGene_tx=XM_023246442.1;refGene_gn=TMEM8A;refGene_tx=XM_023246443.1
E3	44278290 44278414
refGene_gn=TMEM8A;refGene_tx=XM_023246442.1;refGene_gn=TMEM8A;refGene_

tx=XM_023246443.1
E3 44278844 44278981
refGene_gn=TMEM8A;refGene_tx=XM_023246442.1;refGene_gn=TMEM8A;refGene_tx=XM_023246443.1
E3 44279685 44279864
refGene_gn=TMEM8A;refGene_tx=XM_023246442.1;refGene_gn=TMEM8A;refGene_tx=XM_023246443.1
E3 44279951 44280098
refGene_gn=TMEM8A;refGene_tx=XM_023246442.1;refGene_gn=TMEM8A;refGene_tx=XM_023246443.1
E3 44280756 44280873
refGene_gn=TMEM8A;refGene_tx=XM_023246442.1;refGene_gn=TMEM8A;refGene_tx=XM_023246443.1
E3 44281140 44281437
refGene_gn=TMEM8A;refGene_tx=XM_023246442.1;refGene_gn=TMEM8A;refGene_tx=XM_023246443.1
E3 44283392 44283680
refGene_gn=MRPL28;refGene_tx=XM_003999073.3;refGene_gn=MRPL28;refGene_tx=XM_006942515.3
E3 44284406 44284559
refGene_gn=MRPL28;refGene_tx=XM_003999073.3;refGene_gn=MRPL28;refGene_tx=XM_006942515.3
E3 44284777 44284912
refGene_gn=MRPL28;refGene_tx=XM_003999073.3;refGene_gn=MRPL28;refGene_tx=XM_006942515.3
E3 44285002 44285089
refGene_gn=MRPL28;refGene_tx=XM_003999073.3;refGene_gn=MRPL28;refGene_tx=XM_006942515.3
E3 44285942 44286050
refGene_gn=MRPL28;refGene_tx=XM_003999073.3;refGene_gn=MRPL28;refGene_tx=XM_006942515.3
E3 44302133 44303011
refGene_gn=AXIN1;refGene_tx=XM_023247010.1;refGene_gn=AXIN1;refGene_tx=XM_023247011.1
E3 44336814 44336955
refGene_gn=AXIN1;refGene_tx=XM_023247010.1;refGene_gn=AXIN1;refGene_tx=XM_023247011.1
E3 44341195 44341292
refGene_gn=AXIN1;refGene_tx=XM_023247010.1;refGene_gn=AXIN1;refGene_tx=XM_023247011.1
E3 44347640 44347778
refGene_gn=AXIN1;refGene_tx=XM_023247010.1;refGene_gn=AXIN1;refGene_tx=XM_023247011.1
E3 44352328 44352858
refGene_gn=AXIN1;refGene_tx=XM_023247010.1;refGene_gn=AXIN1;refGene_tx=XM_023247011.1
E3 44353467 44353638
refGene_gn=AXIN1;refGene_tx=XM_023247010.1;refGene_gn=AXIN1;refGene_tx=XM_023247011.1
E3 44356216 44356462

refGene_gn=AXIN1;refGene_tx=XM_023247010.1;refGene_gn=AXIN1;refGene_tx=XM_023247011.1
E3 44358289 44358397 refGene_gn=AXIN1;refGene_tx=XM_023247010.1
E3 44360013 44360181
refGene_gn=AXIN1;refGene_tx=XM_023247010.1;refGene_gn=AXIN1;refGene_tx=XM_023247011.1
E3 44360347 44360474
refGene_gn=AXIN1;refGene_tx=XM_023247010.1;refGene_gn=AXIN1;refGene_tx=XM_023247011.1
E3 44361587 44361632
refGene_gn=PDIA2;refGene_tx=XM_003998969.5;refGene_gn=PDIA2;refGene_tx=XM_019821292.2
E3 44361744 44361855
refGene_gn=PDIA2;refGene_tx=XM_003998969.5;refGene_gn=PDIA2;refGene_tx=XM_019821292.2
E3 44362032 44362214
refGene_gn=PDIA2;refGene_tx=XM_003998969.5;refGene_gn=PDIA2;refGene_tx=XM_019821292.2
E3 44362294 44362415
refGene_gn=PDIA2;refGene_tx=XM_003998969.5;refGene_gn=PDIA2;refGene_tx=XM_019821292.2
E3 44362520 44362718
refGene_gn=PDIA2;refGene_tx=XM_003998969.5;refGene_gn=PDIA2;refGene_tx=XM_019821292.2
E3 44362789 44362915 refGene_gn=PDIA2;refGene_tx=XM_003998969.5
E3 44362998 44363115
refGene_gn=PDIA2;refGene_tx=XM_003998969.5;refGene_gn=PDIA2;refGene_tx=XM_019821292.2
E3 44363186 44363324
refGene_gn=PDIA2;refGene_tx=XM_003998969.5;refGene_gn=PDIA2;refGene_tx=XM_019821292.2
E3 44363405 44363539
refGene_gn=PDIA2;refGene_tx=XM_003998969.5;refGene_gn=PDIA2;refGene_tx=XM_019821292.2
E3 44363612 44363819
refGene_gn=PDIA2;refGene_tx=XM_003998969.5;refGene_gn=PDIA2;refGene_tx=XM_019821292.2
E3 44364971 44365164
refGene_gn=PDIA2;refGene_tx=XM_003998969.5;refGene_gn=PDIA2;refGene_tx=XM_019821292.2
E3 44365488 44365688
refGene_gn=ARHGDIG;refGene_tx=XM_023247012.1;refGene_gn=ARHGDIG;refGene_tx=XM_023247013.1
E3 44365781 44365845
refGene_gn=ARHGDIG;refGene_tx=XM_023247012.1;refGene_gn=ARHGDIG;refGene_tx=XM_023247013.1
E3 44365927 44366169
refGene_gn=ARHGDIG;refGene_tx=XM_023247012.1;refGene_gn=ARHGDIG;refGene_tx=XM_023247013.1
E3 44366283 44366463

refGene_gn=ARHGDIG;refGene_tx=XM_023247012.1;refGene_gn=ARHGDIG;refGene_tx=XM_023247013.1
E3 44367572 44367645
refGene_gn=ARHGDIG;refGene_tx=XM_023247012.1;refGene_gn=ARHGDIG;refGene_tx=XM_023247013.1
E3 44370648 44370693
refGene_gn=RGS11;refGene_tx=XM_011290752.3;refGene_gn=RGS11;refGene_tx=XM_011290753.3;refGene_gn=RGS11;refGene_tx=XM_011290754.3;refGene_gn=RGS11;refGene_tx=XM_019821294.2;refGene_gn=RGS11;refGene_tx=XM_019821295.2;refGene_gn=RGS11;refGene_tx=XM_019821297.2;refGene_gn=RGS11;refGene_tx=XM_023246498.1
E3 44371054 44371151
refGene_gn=RGS11;refGene_tx=XM_011290752.3;refGene_gn=RGS11;refGene_tx=XM_011290753.3;refGene_gn=RGS11;refGene_tx=XM_011290754.3;refGene_gn=RGS11;refGene_tx=XM_019821294.2;refGene_gn=RGS11;refGene_tx=XM_019821295.2;refGene_gn=RGS11;refGene_tx=XM_019821297.2;refGene_gn=RGS11;refGene_tx=XM_023246498.1
E3 44371230 44371281
refGene_gn=RGS11;refGene_tx=XM_011290752.3;refGene_gn=RGS11;refGene_tx=XM_011290753.3;refGene_gn=RGS11;refGene_tx=XM_011290754.3;refGene_gn=RGS11;refGene_tx=XM_019821294.2;refGene_gn=RGS11;refGene_tx=XM_019821295.2;refGene_gn=RGS11;refGene_tx=XM_019821297.2;refGene_gn=RGS11;refGene_tx=XM_023246498.1
E3 44371478 44371585
refGene_gn=RGS11;refGene_tx=XM_011290752.3;refGene_gn=RGS11;refGene_tx=XM_011290753.3;refGene_gn=RGS11;refGene_tx=XM_011290754.3;refGene_gn=RGS11;refGene_tx=XM_019821294.2;refGene_gn=RGS11;refGene_tx=XM_019821295.2;refGene_gn=RGS11;refGene_tx=XM_019821297.2;refGene_gn=RGS11;refGene_tx=XM_023246498.1
E3 44372267 44372319
refGene_gn=RGS11;refGene_tx=XM_011290752.3;refGene_gn=RGS11;refGene_tx=XM_011290753.3;refGene_gn=RGS11;refGene_tx=XM_011290754.3;refGene_gn=RGS11;refGene_tx=XM_019821294.2;refGene_gn=RGS11;refGene_tx=XM_019821295.2;refGene_gn=RGS11;refGene_tx=XM_019821297.2;refGene_gn=RGS11;refGene_tx=XM_023246498.1;refGene_gn=RGS11;refGene_tx=XM_023246499.1
E3 44372371 44372787
refGene_gn=RGS11;refGene_tx=XM_011290752.3;refGene_gn=RGS11;refGene_tx=XM_011290753.3;refGene_gn=RGS11;refGene_tx=XM_011290754.3;refGene_gn=RGS11;refGene_tx=XM_019821294.2;refGene_gn=RGS11;refGene_tx=XM_019821295.2;refGene_gn=RGS11;refGene_tx=XM_019821296.2;refGene_gn=RGS11;refGene_tx=XM_019821297.2;refGene_gn=RGS11;refGene_tx=XM_019821298.2;refGene_gn=RGS11;refGene_tx=XM_023246498.1;refGene_gn=RGS11;refGene_tx=XM_023246499.1
E3 44372959 44373041
refGene_gn=RGS11;refGene_tx=XM_011290752.3;refGene_gn=RGS11;refGene_tx=XM_011290753.3;refGene_gn=RGS11;refGene_tx=XM_011290754.3;refGene_gn=RGS11;refGene_tx=XM_019821294.2;refGene_gn=RGS11;refGene_tx=XM_019821295.2;refGene_gn=RGS11;refGene_tx=XM_019821296.2;refGene_gn=RGS11;refGene_tx=XM_019821297.2;refGene_gn=RGS11;refGene_tx=XM_019821298.2;refGene_gn=RGS11;refGene_tx=XM_023246498.1;refGene_gn=RGS11;refGene_tx=XM_02

3246499.1
E3 44374439 44374508
refGene_gn=RGS11;refGene_tx=XM_011290752.3;refGene_gn=RGS11;refGene_tx=XM_011290754.3;refGene_gn=RGS11;refGene_tx=XM_019821294.2;refGene_gn=RGS11;refGene_tx=XM_019821295.2;refGene_gn=RGS11;refGene_tx=XM_019821296.2;refGene_gn=RGS11;refGene_tx=XM_019821297.2;refGene_gn=RGS11;refGene_tx=XM_019821298.2;refGene_gn=RGS11;refGene_tx=XM_023246498.1;refGene_gn=RGS11;refGene_tx=XM_023246499.1
E3 44375520 44375598
refGene_gn=RGS11;refGene_tx=XM_011290752.3;refGene_gn=RGS11;refGene_tx=XM_011290753.3;refGene_gn=RGS11;refGene_tx=XM_011290754.3;refGene_gn=RGS11;refGene_tx=XM_019821294.2;refGene_gn=RGS11;refGene_tx=XM_019821295.2;refGene_gn=RGS11;refGene_tx=XM_019821296.2;refGene_gn=RGS11;refGene_tx=XM_019821297.2;refGene_gn=RGS11;refGene_tx=XM_019821298.2;refGene_gn=RGS11;refGene_tx=XM_023246498.1;refGene_gn=RGS11;refGene_tx=XM_023246499.1
E3 44375683 44375745
refGene_gn=RGS11;refGene_tx=XM_011290752.3;refGene_gn=RGS11;refGene_tx=XM_011290753.3;refGene_gn=RGS11;refGene_tx=XM_011290754.3;refGene_gn=RGS11;refGene_tx=XM_019821294.2;refGene_gn=RGS11;refGene_tx=XM_019821295.2;refGene_gn=RGS11;refGene_tx=XM_019821296.2;refGene_gn=RGS11;refGene_tx=XM_019821297.2;refGene_gn=RGS11;refGene_tx=XM_019821298.2;refGene_gn=RGS11;refGene_tx=XM_023246498.1;refGene_gn=RGS11;refGene_tx=XM_023246499.1
E3 44375867 44375981
refGene_gn=RGS11;refGene_tx=XM_011290752.3;refGene_gn=RGS11;refGene_tx=XM_011290753.3;refGene_gn=RGS11;refGene_tx=XM_011290754.3;refGene_gn=RGS11;refGene_tx=XM_019821294.2;refGene_gn=RGS11;refGene_tx=XM_019821295.2;refGene_gn=RGS11;refGene_tx=XM_019821296.2;refGene_gn=RGS11;refGene_tx=XM_019821297.2;refGene_gn=RGS11;refGene_tx=XM_019821298.2;refGene_gn=RGS11;refGene_tx=XM_023246498.1;refGene_gn=RGS11;refGene_tx=XM_023246499.1
E3 44376088 44376204
refGene_gn=RGS11;refGene_tx=XM_011290752.3;refGene_gn=RGS11;refGene_tx=XM_011290753.3;refGene_gn=RGS11;refGene_tx=XM_011290754.3;refGene_gn=RGS11;refGene_tx=XM_019821294.2;refGene_gn=RGS11;refGene_tx=XM_019821295.2;refGene_gn=RGS11;refGene_tx=XM_019821296.2;refGene_gn=RGS11;refGene_tx=XM_019821297.2;refGene_gn=RGS11;refGene_tx=XM_019821298.2;refGene_gn=RGS11;refGene_tx=XM_023246499.1
E3 44376354 44376442
refGene_gn=RGS11;refGene_tx=XM_011290752.3;refGene_gn=RGS11;refGene_tx=XM_011290753.3;refGene_gn=RGS11;refGene_tx=XM_011290754.3;refGene_gn=RGS11;refGene_tx=XM_019821294.2;refGene_gn=RGS11;refGene_tx=XM_019821295.2;refGene_gn=RGS11;refGene_tx=XM_019821296.2;refGene_gn=RGS11;refGene_tx=XM_019821297.2;refGene_gn=RGS11;refGene_tx=XM_019821298.2;refGene_gn=RGS11;refGene_tx=XM_023246498.1;refGene_gn=RGS11;refGene_tx=XM_023246499.1
E3 44376538 44376677
refGene_gn=RGS11;refGene_tx=XM_011290752.3;refGene_gn=RGS11;refGene_tx=XM_011290753.3;refGene_gn=RGS11;refGene_tx=XM_011290754.3;refGene_gn=

RGS11;refGene_tx=XM_019821294.2;refGene_gn=RGS11;refGene_tx=XM_0198212
95.2;refGene_gn=RGS11;refGene_tx=XM_019821296.2;refGene_gn=RGS11;refGe
ne_tx=XM_019821297.2;refGene_gn=RGS11;refGene_tx=XM_019821298.2;refGen
e_gn=RGS11;refGene_tx=XM_023246499.1
E3 44379655 44379829 refGene_gn=RGS11;refGene_tx=XM_019821294.2
E3 44380403 44380486
refGene_gn=RGS11;refGene_tx=XM_011290752.3;refGene_gn=RGS11;refGene_tx
=XM_011290753.3;refGene_gn=RGS11;refGene_tx=XM_011290754.3;refGene_gn=
RGS11;refGene_tx=XM_019821295.2;refGene_gn=RGS11;refGene_tx=XM_0198212
96.2;refGene_gn=RGS11;refGene_tx=XM_019821297.2;refGene_gn=RGS11;refGe
ne_tx=XM_019821298.2;refGene_gn=RGS11;refGene_tx=XM_023246499.1
E3 44380579 44380682
refGene_gn=RGS11;refGene_tx=XM_011290752.3;refGene_gn=RGS11;refGene_tx
=XM_011290753.3;refGene_gn=RGS11;refGene_tx=XM_011290754.3;refGene_gn=
RGS11;refGene_tx=XM_019821295.2;refGene_gn=RGS11;refGene_tx=XM_0198212
96.2;refGene_gn=RGS11;refGene_tx=XM_019821297.2;refGene_gn=RGS11;refGe
ne_tx=XM_019821298.2;refGene_gn=RGS11;refGene_tx=XM_023246499.1
E3 44384247 44384459
refGene_gn=FAM234A;refGene_tx=XM_006942520.4;refGene_gn=FAM234A;refGen
e_tx=XM_006942521.4;refGene_gn=FAM234A;refGene_tx=XM_006942522.4;refGe
ne_gn=FAM234A;refGene_tx=XM_023246768.1
E3 44384557 44384660
refGene_gn=FAM234A;refGene_tx=XM_006942520.4;refGene_gn=FAM234A;refGen
e_tx=XM_006942521.4;refGene_gn=FAM234A;refGene_tx=XM_006942522.4;refGe
ne_gn=FAM234A;refGene_tx=XM_023246768.1
E3 44385136 44385298
refGene_gn=FAM234A;refGene_tx=XM_006942520.4;refGene_gn=FAM234A;refGen
e_tx=XM_006942521.4;refGene_gn=FAM234A;refGene_tx=XM_006942522.4;refGe
ne_gn=FAM234A;refGene_tx=XM_023246768.1
E3 44385525 44385601
refGene_gn=FAM234A;refGene_tx=XM_006942520.4;refGene_gn=FAM234A;refGen
e_tx=XM_006942521.4;refGene_gn=FAM234A;refGene_tx=XM_006942522.4;refGe
ne_gn=FAM234A;refGene_tx=XM_023246768.1
E3 44385930 44386071
refGene_gn=FAM234A;refGene_tx=XM_006942520.4;refGene_gn=FAM234A;refGen
e_tx=XM_006942521.4;refGene_gn=FAM234A;refGene_tx=XM_006942522.4;refGe
ne_gn=FAM234A;refGene_tx=XM_023246768.1
E3 44386424 44386554
refGene_gn=FAM234A;refGene_tx=XM_006942520.4;refGene_gn=FAM234A;refGen
e_tx=XM_006942521.4;refGene_gn=FAM234A;refGene_tx=XM_006942522.4;refGe
ne_gn=FAM234A;refGene_tx=XM_023246768.1
E3 44386823 44386956
refGene_gn=FAM234A;refGene_tx=XM_006942520.4;refGene_gn=FAM234A;refGen
e_tx=XM_006942521.4;refGene_gn=FAM234A;refGene_tx=XM_006942522.4;refGe
ne_gn=FAM234A;refGene_tx=XM_023246768.1
E3 44387506 44387637
refGene_gn=FAM234A;refGene_tx=XM_006942520.4;refGene_gn=FAM234A;refGen
e_tx=XM_006942521.4;refGene_gn=FAM234A;refGene_tx=XM_006942522.4;refGe
ne_gn=FAM234A;refGene_tx=XM_023246768.1
E3 44388638 44388830

refGene_gn=FAM234A;refGene_tx=XM_006942520.4;refGene_gn=FAM234A;refGene_tx=XM_006942521.4;refGene_gn=FAM234A;refGene_tx=XM_006942522.4;refGene_gn=FAM234A;refGene_tx=XM_023246768.1
E3 44389133 44389250
refGene_gn=FAM234A;refGene_tx=XM_006942520.4;refGene_gn=FAM234A;refGene_tx=XM_006942521.4;refGene_gn=FAM234A;refGene_tx=XM_006942522.4;refGene_gn=FAM234A;refGene_tx=XM_023246768.1
E3 44392448 44392716
refGene_gn=FAM234A;refGene_tx=XM_006942520.4;refGene_gn=FAM234A;refGene_tx=XM_006942521.4;refGene_gn=FAM234A;refGene_tx=XM_006942522.4;refGene_gn=FAM234A;refGene_tx=XM_023246768.1
E3 44414581 44414642
refGene_gn=LUC7L;refGene_tx=XM_023246769.1;refGene_gn=LUC7L;refGene_tx=XM_023246770.1;refGene_gn=LUC7L;refGene_tx=XM_023246771.1
E3 44416558 44416653
refGene_gn=LUC7L;refGene_tx=XM_023246769.1;refGene_gn=LUC7L;refGene_tx=XM_023246770.1;refGene_gn=LUC7L;refGene_tx=XM_023246771.1
E3 44422906 44423005
refGene_gn=LUC7L;refGene_tx=XM_023246769.1;refGene_gn=LUC7L;refGene_tx=XM_023246770.1;refGene_gn=LUC7L;refGene_tx=XM_023246771.1;refGene_gn=LUC7L;refGene_tx=XM_023246772.1
E3 44437357 44437468
refGene_gn=LUC7L;refGene_tx=XM_023246769.1;refGene_gn=LUC7L;refGene_tx=XM_023246770.1;refGene_gn=LUC7L;refGene_tx=XM_023246771.1;refGene_gn=LUC7L;refGene_tx=XM_023246772.1
E3 44438175 44438319
refGene_gn=LUC7L;refGene_tx=XM_023246769.1;refGene_gn=LUC7L;refGene_tx=XM_023246770.1;refGene_gn=LUC7L;refGene_tx=XM_023246771.1;refGene_gn=LUC7L;refGene_tx=XM_023246772.1
E3 44444018 44444195
refGene_gn=LUC7L;refGene_tx=XM_023246769.1;refGene_gn=LUC7L;refGene_tx=XM_023246770.1;refGene_gn=LUC7L;refGene_tx=XM_023246772.1
E3 44447161 44447250
refGene_gn=LUC7L;refGene_tx=XM_023246769.1;refGene_gn=LUC7L;refGene_tx=XM_023246770.1;refGene_gn=LUC7L;refGene_tx=XM_023246771.1;refGene_gn=LUC7L;refGene_tx=XM_023246772.1
E3 44449033 44449063
refGene_gn=LUC7L;refGene_tx=XM_023246769.1;refGene_gn=LUC7L;refGene_tx=XM_023246770.1;refGene_gn=LUC7L;refGene_tx=XM_023246771.1;refGene_gn=LUC7L;refGene_tx=XM_023246772.1
E3 44449506 44449678
refGene_gn=LUC7L;refGene_tx=XM_023246769.1;refGene_gn=LUC7L;refGene_tx=XM_023246770.1;refGene_gn=LUC7L;refGene_tx=XM_023246771.1;refGene_gn=LUC7L;refGene_tx=XM_023246772.1
E3 44450707 44450849
refGene_gn=LUC7L;refGene_tx=XM_023246769.1;refGene_gn=LUC7L;refGene_tx=XM_023246771.1;refGene_gn=LUC7L;refGene_tx=XM_023246772.1
E3 44461892 44462021 refGene_gn=HBQ1;refGene_tx=XM_011290755.3
E3 44462111 44462316 refGene_gn=HBQ1;refGene_tx=XM_011290755.3
E3 44462480 44462575 refGene_gn=HBQ1;refGene_tx=XM_011290755.3

E3 44464567 44464696
refGene_gn=LOC111558266;refGene_tx=XM_023246781.1
E3 44464837 44465042
refGene_gn=LOC111558266;refGene_tx=XM_023246781.1
E3 44465159 44465254
refGene_gn=LOC111558266;refGene_tx=XM_023246781.1
E3 44469038 44469167
refGene_gn=LOC101089822;refGene_tx=XM_003998972.4
E3 44469308 44469513
refGene_gn=LOC101089822;refGene_tx=XM_003998972.4
E3 44469630 44469725
refGene_gn=LOC101089822;refGene_tx=XM_003998972.4
E3 44471439 44471568 refGene_gn=HBM;refGene_tx=XM_006942524.3
E3 44471669 44471874 refGene_gn=HBM;refGene_tx=XM_006942524.3
E3 44472034 44472126 refGene_gn=HBM;refGene_tx=XM_006942524.3
E3 44472929 44473058
refGene_gn=LOC111558267;refGene_tx=XM_023246785.1
E3 44473215 44473420
refGene_gn=LOC111558267;refGene_tx=XM_023246785.1
E3 44474207 44474302
refGene_gn=LOC111558267;refGene_tx=XM_023246785.1
E3 44479882 44480011
refGene_gn=LOC101097290;refGene_tx=XM_023246783.1;refGene_gn=LOC101097
290;refGene_tx=XM_023246784.1
E3 44480170 44480375
refGene_gn=LOC101097290;refGene_tx=XM_023246783.1;refGene_gn=LOC101097
290;refGene_tx=XM_023246784.1
E3 44481161 44481256
refGene_gn=LOC101097290;refGene_tx=XM_023246783.1;refGene_gn=LOC101097
290;refGene_tx=XM_023246784.1
E3 44493767 44493885
refGene_gn=NPRL3;refGene_tx=XM_023246683.1;refGene_gn=NPRL3;refGene_tx
=XM_023246684.1
E3 44494175 44494209
refGene_gn=NPRL3;refGene_tx=XM_023246685.1;refGene_gn=NPRL3;refGene_tx
=XM_023246686.1
E3 44498854 44498924
refGene_gn=NPRL3;refGene_tx=XM_023246683.1;refGene_gn=NPRL3;refGene_tx
=XM_023246684.1;refGene_gn=NPRL3;refGene_tx=XM_023246685.1;refGene_gn=
NPRL3;refGene_tx=XM_023246686.1
E3 44503932 44503971
refGene_gn=NPRL3;refGene_tx=XM_023246683.1;refGene_gn=NPRL3;refGene_tx
=XM_023246685.1
E3 44507310 44507440
refGene_gn=NPRL3;refGene_tx=XM_023246683.1;refGene_gn=NPRL3;refGene_tx
=XM_023246684.1;refGene_gn=NPRL3;refGene_tx=XM_023246685.1;refGene_gn=
NPRL3;refGene_tx=XM_023246686.1
E3 44509128 44509203
refGene_gn=NPRL3;refGene_tx=XM_019821305.2;refGene_gn=NPRL3;refGene_tx
=XM_023246683.1;refGene_gn=NPRL3;refGene_tx=XM_023246684.1;refGene_gn=

NPRL3;refGene_tx=XM_023246685.1;refGene_gn=NPRL3;refGene_tx=XM_023246686.1
E3 44516024 44516178
refGene_gn=NPRL3;refGene_tx=XM_019821305.2;refGene_gn=NPRL3;refGene_tx=XM_023246683.1;refGene_gn=NPRL3;refGene_tx=XM_023246684.1;refGene_gn=NPRL3;refGene_tx=XM_023246685.1;refGene_gn=NPRL3;refGene_tx=XM_023246686.1
E3 44518172 44518254
refGene_gn=NPRL3;refGene_tx=XM_019821305.2;refGene_gn=NPRL3;refGene_tx=XM_023246683.1;refGene_gn=NPRL3;refGene_tx=XM_023246684.1;refGene_gn=NPRL3;refGene_tx=XM_023246685.1;refGene_gn=NPRL3;refGene_tx=XM_023246686.1
E3 44525559 44525697
refGene_gn=NPRL3;refGene_tx=XM_019821305.2;refGene_gn=NPRL3;refGene_tx=XM_023246683.1;refGene_gn=NPRL3;refGene_tx=XM_023246684.1;refGene_gn=NPRL3;refGene_tx=XM_023246685.1;refGene_gn=NPRL3;refGene_tx=XM_023246686.1
E3 44526663 44526820
refGene_gn=NPRL3;refGene_tx=XM_019821305.2;refGene_gn=NPRL3;refGene_tx=XM_023246683.1;refGene_gn=NPRL3;refGene_tx=XM_023246684.1;refGene_gn=NPRL3;refGene_tx=XM_023246685.1;refGene_gn=NPRL3;refGene_tx=XM_023246686.1
E3 44530569 44530676
refGene_gn=NPRL3;refGene_tx=XM_019821305.2;refGene_gn=NPRL3;refGene_tx=XM_023246683.1;refGene_gn=NPRL3;refGene_tx=XM_023246684.1;refGene_gn=NPRL3;refGene_tx=XM_023246685.1;refGene_gn=NPRL3;refGene_tx=XM_023246686.1
E3 44531145 44531275
refGene_gn=NPRL3;refGene_tx=XM_019821305.2;refGene_gn=NPRL3;refGene_tx=XM_023246683.1;refGene_gn=NPRL3;refGene_tx=XM_023246684.1;refGene_gn=NPRL3;refGene_tx=XM_023246685.1;refGene_gn=NPRL3;refGene_tx=XM_023246686.1
E3 44533572 44533762
refGene_gn=NPRL3;refGene_tx=XM_019821305.2;refGene_gn=NPRL3;refGene_tx=XM_023246683.1;refGene_gn=NPRL3;refGene_tx=XM_023246684.1;refGene_gn=NPRL3;refGene_tx=XM_023246685.1;refGene_gn=NPRL3;refGene_tx=XM_023246686.1
E3 44534454 44534647
refGene_gn=NPRL3;refGene_tx=XM_019821305.2;refGene_gn=NPRL3;refGene_tx=XM_023246683.1;refGene_gn=NPRL3;refGene_tx=XM_023246684.1;refGene_gn=NPRL3;refGene_tx=XM_023246685.1;refGene_gn=NPRL3;refGene_tx=XM_023246686.1
E3 44535413 44535579
refGene_gn=NPRL3;refGene_tx=XM_019821305.2;refGene_gn=NPRL3;refGene_tx=XM_023246683.1;refGene_gn=NPRL3;refGene_tx=XM_023246684.1;refGene_gn=NPRL3;refGene_tx=XM_023246685.1;refGene_gn=NPRL3;refGene_tx=XM_023246686.1
E3 44536339 44536734
refGene_gn=MPG;refGene_tx=XM_023246687.1;refGene_gn=MPG;refGene_tx=XM_023246689.1;refGene_gn=MPG;refGene_tx=XM_023246690.1;refGene_gn=MPG;re

fGene_tx=XM_023246691.1
E3 44539447 44539652
refGene_gn=MPG;refGene_tx=XM_023246687.1;refGene_gn=MPG;refGene_tx=XM_023246689.1;refGene_gn=MPG;refGene_tx=XM_023246690.1;refGene_gn=MPG;refGene_tx=XM_023246691.1
E3 44540499 44540733
refGene_gn=MPG;refGene_tx=XM_023246687.1;refGene_gn=MPG;refGene_tx=XM_023246689.1;refGene_gn=MPG;refGene_tx=XM_023246690.1;refGene_gn=MPG;refGene_tx=XM_023246691.1
E3 44543323 44543530 refGene_gn=MPG;refGene_tx=XM_023246687.1
E3 44556044 44556361
refGene_gn=RHBDF1;refGene_tx=XM_023246677.1;refGene_gn=RHBDF1;refGene_tx=XM_023246678.1;refGene_gn=RHBDF1;refGene_tx=XM_023246680.1;refGene_gn=RHBDF1;refGene_tx=XM_023246681.1;refGene_gn=RHBDF1;refGene_tx=XM_023246682.1
E3 44556634 44556739
refGene_gn=RHBDF1;refGene_tx=XM_023246677.1;refGene_gn=RHBDF1;refGene_tx=XM_023246678.1;refGene_gn=RHBDF1;refGene_tx=XM_023246682.1
E3 44557311 44557525
refGene_gn=RHBDF1;refGene_tx=XM_023246677.1;refGene_gn=RHBDF1;refGene_tx=XM_023246678.1;refGene_gn=RHBDF1;refGene_tx=XM_023246680.1;refGene_gn=RHBDF1;refGene_tx=XM_023246681.1;refGene_gn=RHBDF1;refGene_tx=XM_023246682.1
E3 44557954 44558164
refGene_gn=RHBDF1;refGene_tx=XM_023246677.1;refGene_gn=RHBDF1;refGene_tx=XM_023246678.1;refGene_gn=RHBDF1;refGene_tx=XM_023246680.1;refGene_gn=RHBDF1;refGene_tx=XM_023246681.1;refGene_gn=RHBDF1;refGene_tx=XM_023246682.1
E3 44558237 44558360
refGene_gn=RHBDF1;refGene_tx=XM_023246677.1;refGene_gn=RHBDF1;refGene_tx=XM_023246678.1;refGene_gn=RHBDF1;refGene_tx=XM_023246680.1;refGene_gn=RHBDF1;refGene_tx=XM_023246681.1;refGene_gn=RHBDF1;refGene_tx=XM_023246682.1
E3 44558432 44558590
refGene_gn=RHBDF1;refGene_tx=XM_023246677.1;refGene_gn=RHBDF1;refGene_tx=XM_023246678.1;refGene_gn=RHBDF1;refGene_tx=XM_023246680.1;refGene_gn=RHBDF1;refGene_tx=XM_023246681.1;refGene_gn=RHBDF1;refGene_tx=XM_023246682.1
E3 44559084 44559342
refGene_gn=RHBDF1;refGene_tx=XM_023246677.1;refGene_gn=RHBDF1;refGene_tx=XM_023246678.1;refGene_gn=RHBDF1;refGene_tx=XM_023246680.1;refGene_gn=RHBDF1;refGene_tx=XM_023246681.1;refGene_gn=RHBDF1;refGene_tx=XM_023246682.1
E3 44559456 44559568
refGene_gn=RHBDF1;refGene_tx=XM_023246677.1;refGene_gn=RHBDF1;refGene_tx=XM_023246678.1;refGene_gn=RHBDF1;refGene_tx=XM_023246680.1;refGene_gn=RHBDF1;refGene_tx=XM_023246681.1;refGene_gn=RHBDF1;refGene_tx=XM_023246682.1
E3 44559691 44559766
refGene_gn=RHBDF1;refGene_tx=XM_023246677.1;refGene_gn=RHBDF1;refGene_

tx=XM_023246678.1;refGene_gn=RHBDF1;refGene_tx=XM_023246680.1;refGene_gn=RHBDF1;refGene_tx=XM_023246681.1;refGene_gn=RHBDF1;refGene_tx=XM_023246682.1
E3 44559857 44560019 refGene_gn=RHBDF1;refGene_tx=XM_023246677.1;refGene_gn=RHBDF1;refGene_tx=XM_023246678.1;refGene_gn=RHBDF1;refGene_tx=XM_023246680.1;refGene_gn=RHBDF1;refGene_tx=XM_023246681.1;refGene_gn=RHBDF1;refGene_tx=XM_023246682.1
E3 44560624 44560725 refGene_gn=RHBDF1;refGene_tx=XM_023246677.1;refGene_gn=RHBDF1;refGene_tx=XM_023246678.1;refGene_gn=RHBDF1;refGene_tx=XM_023246680.1;refGene_gn=RHBDF1;refGene_tx=XM_023246681.1;refGene_gn=RHBDF1;refGene_tx=XM_023246682.1
E3 44560887 44560951 refGene_gn=RHBDF1;refGene_tx=XM_023246677.1;refGene_gn=RHBDF1;refGene_tx=XM_023246678.1;refGene_gn=RHBDF1;refGene_tx=XM_023246680.1;refGene_gn=RHBDF1;refGene_tx=XM_023246681.1;refGene_gn=RHBDF1;refGene_tx=XM_023246682.1
E3 44561356 44561451 refGene_gn=RHBDF1;refGene_tx=XM_023246677.1;refGene_gn=RHBDF1;refGene_tx=XM_023246678.1;refGene_gn=RHBDF1;refGene_tx=XM_023246680.1;refGene_gn=RHBDF1;refGene_tx=XM_023246681.1;refGene_gn=RHBDF1;refGene_tx=XM_023246682.1
E3 44561706 44561782 refGene_gn=RHBDF1;refGene_tx=XM_023246677.1;refGene_gn=RHBDF1;refGene_tx=XM_023246678.1;refGene_gn=RHBDF1;refGene_tx=XM_023246680.1;refGene_gn=RHBDF1;refGene_tx=XM_023246681.1;refGene_gn=RHBDF1;refGene_tx=XM_023246682.1
E3 44561855 44561956 refGene_gn=RHBDF1;refGene_tx=XM_023246677.1;refGene_gn=RHBDF1;refGene_tx=XM_023246678.1;refGene_gn=RHBDF1;refGene_tx=XM_023246680.1;refGene_gn=RHBDF1;refGene_tx=XM_023246681.1;refGene_gn=RHBDF1;refGene_tx=XM_023246682.1
E3 44562072 44562226 refGene_gn=RHBDF1;refGene_tx=XM_023246677.1;refGene_gn=RHBDF1;refGene_tx=XM_023246678.1;refGene_gn=RHBDF1;refGene_tx=XM_023246680.1;refGene_gn=RHBDF1;refGene_tx=XM_023246681.1;refGene_gn=RHBDF1;refGene_tx=XM_023246682.1
E3 44562463 44562883 refGene_gn=RHBDF1;refGene_tx=XM_023246677.1;refGene_gn=RHBDF1;refGene_tx=XM_023246678.1;refGene_gn=RHBDF1;refGene_tx=XM_023246680.1;refGene_gn=RHBDF1;refGene_tx=XM_023246681.1;refGene_gn=RHBDF1;refGene_tx=XM_023246682.1
E3 44563988 44564046 refGene_gn=SNRNP25;refGene_tx=XM_006942536.3
E3 44564500 44564575 refGene_gn=SNRNP25;refGene_tx=XM_006942536.3
E3 44565124 44565230 refGene_gn=SNRNP25;refGene_tx=XM_006942536.3
E3 44565469 44565560 refGene_gn=SNRNP25;refGene_tx=XM_006942536.3
E3 44566743 44566937 refGene_gn=SNRNP25;refGene_tx=XM_006942536.3
E3 44567211 44567322 refGene_gn=POLR3K;refGene_tx=XM_003998975.5
E3 44567449 44567597 refGene_gn=SNRNP25;refGene_tx=XM_006942536.3

E3 44567707 44567809 refGene_gn=SNRNP25;refGene_tx=XM_006942536.3
E3 44568698 44568786 refGene_gn=POLR3K;refGene_tx=XM_003998975.5
E3 44570916 44571044 refGene_gn=POLR3K;refGene_tx=XM_003998975.5
E3 44576578 44576927
refGene_gn=IL9R;refGene_tx=XM_023246459.1;refGene_gn=IL9R;refGene_tx=XM_023246460.1;refGene_gn=IL9R;refGene_tx=XM_023246461.1
E3 44580616 44580730
refGene_gn=IL9R;refGene_tx=XM_023246459.1;refGene_gn=IL9R;refGene_tx=XM_023246460.1;refGene_gn=IL9R;refGene_tx=XM_023246461.1
E3 44581038 44581150
refGene_gn=IL9R;refGene_tx=XM_023246459.1;refGene_gn=IL9R;refGene_tx=XM_023246460.1;refGene_gn=IL9R;refGene_tx=XM_023246461.1
E3 44581280 44581459
refGene_gn=IL9R;refGene_tx=XM_023246459.1;refGene_gn=IL9R;refGene_tx=XM_023246460.1;refGene_gn=IL9R;refGene_tx=XM_023246461.1
E3 44581971 44582117
refGene_gn=IL9R;refGene_tx=XM_023246459.1;refGene_gn=IL9R;refGene_tx=XM_023246460.1;refGene_gn=IL9R;refGene_tx=XM_023246461.1
E3 44582499 44582707
refGene_gn=IL9R;refGene_tx=XM_023246459.1;refGene_gn=IL9R;refGene_tx=XM_023246460.1;refGene_gn=IL9R;refGene_tx=XM_023246461.1
E3 44583202 44583308
refGene_gn=IL9R;refGene_tx=XM_023246459.1;refGene_gn=IL9R;refGene_tx=XM_023246460.1;refGene_gn=IL9R;refGene_tx=XM_023246461.1
E3 44584350 44584435
refGene_gn=IL9R;refGene_tx=XM_023246459.1;refGene_gn=IL9R;refGene_tx=XM_023246460.1;refGene_gn=IL9R;refGene_tx=XM_023246461.1
E3 44585968 44586469
refGene_gn=IL9R;refGene_tx=XM_023246459.1;refGene_gn=IL9R;refGene_tx=XM_023246460.1
E3 44587442 44587466 refGene_gn=IL9R;refGene_tx=XM_023246461.1
F1 2236333 2236782
refGene_gn=DISP1;refGene_tx=XM_011290802.3;refGene_gn=DISP1;refGene_tx=XM_019821412.2;refGene_gn=DISP1;refGene_tx=XM_023248075.1;refGene_gn=DISP1;refGene_tx=XM_023248076.1;refGene_gn=DISP1;refGene_tx=XM_023248077.1;refGene_gn=DISP1;refGene_tx=XM_023248078.1;refGene_gn=DISP1;refGene_tx=XM_023248079.1;refGene_gn=DISP1;refGene_tx=XM_023248081.1
F1 2268303 2268333
refGene_gn=DISP1;refGene_tx=XM_011290802.3;refGene_gn=DISP1;refGene_tx=XM_019821412.2;refGene_gn=DISP1;refGene_tx=XM_023248075.1;refGene_gn=DISP1;refGene_tx=XM_023248076.1;refGene_gn=DISP1;refGene_tx=XM_023248077.1;refGene_gn=DISP1;refGene_tx=XM_023248078.1;refGene_gn=DISP1;refGene_tx=XM_023248079.1
F1 2274238 2274362
refGene_gn=DISP1;refGene_tx=XM_011290802.3;refGene_gn=DISP1;refGene_tx=XM_019821412.2;refGene_gn=DISP1;refGene_tx=XM_023248075.1;refGene_gn=DISP1;refGene_tx=XM_023248076.1;refGene_gn=DISP1;refGene_tx=XM_023248077.1;refGene_gn=DISP1;refGene_tx=XM_023248078.1;refGene_gn=DISP1;refGene_tx=XM_023248079.1;refGene_gn=DISP1;refGene_tx=XM_023248081.1;refGene_gn=DISP1;refGene_tx=XM_023248082.1

F1 2275099 2275227
refGene_gn=DISP1;refGene_tx=XM_011290802.3;refGene_gn=DISP1;refGene_tx=XM_011290804.2;refGene_gn=DISP1;refGene_tx=XM_019821412.2;refGene_gn=DISP1;refGene_tx=XM_023248075.1;refGene_gn=DISP1;refGene_tx=XM_023248076.1;refGene_gn=DISP1;refGene_tx=XM_023248077.1;refGene_gn=DISP1;refGene_tx=XM_023248078.1;refGene_gn=DISP1;refGene_tx=XM_023248079.1;refGene_gn=DISP1;refGene_tx=XM_023248081.1;refGene_gn=DISP1;refGene_tx=XM_023248082.1
F1 2275880 2275994
refGene_gn=DISP1;refGene_tx=XM_011290802.3;refGene_gn=DISP1;refGene_tx=XM_011290804.2;refGene_gn=DISP1;refGene_tx=XM_019821412.2;refGene_gn=DISP1;refGene_tx=XM_023248075.1;refGene_gn=DISP1;refGene_tx=XM_023248076.1;refGene_gn=DISP1;refGene_tx=XM_023248077.1;refGene_gn=DISP1;refGene_tx=XM_023248078.1;refGene_gn=DISP1;refGene_tx=XM_023248079.1;refGene_gn=DISP1;refGene_tx=XM_023248081.1;refGene_gn=DISP1;refGene_tx=XM_023248082.1
F1 2278079 2278177
refGene_gn=DISP1;refGene_tx=XM_011290802.3;refGene_gn=DISP1;refGene_tx=XM_011290804.2;refGene_gn=DISP1;refGene_tx=XM_019821412.2;refGene_gn=DISP1;refGene_tx=XM_023248075.1;refGene_gn=DISP1;refGene_tx=XM_023248076.1;refGene_gn=DISP1;refGene_tx=XM_023248077.1;refGene_gn=DISP1;refGene_tx=XM_023248078.1;refGene_gn=DISP1;refGene_tx=XM_023248079.1;refGene_gn=DISP1;refGene_tx=XM_023248081.1;refGene_gn=DISP1;refGene_tx=XM_023248082.1
F1 2284619 2288108
refGene_gn=DISP1;refGene_tx=XM_011290802.3;refGene_gn=DISP1;refGene_tx=XM_011290804.2;refGene_gn=DISP1;refGene_tx=XM_019821412.2;refGene_gn=DISP1;refGene_tx=XM_023248075.1;refGene_gn=DISP1;refGene_tx=XM_023248076.1;refGene_gn=DISP1;refGene_tx=XM_023248077.1;refGene_gn=DISP1;refGene_tx=XM_023248078.1;refGene_gn=DISP1;refGene_tx=XM_023248079.1;refGene_gn=DISP1;refGene_tx=XM_023248081.1;refGene_gn=DISP1;refGene_tx=XM_023248082.1
F1 2329671 2332344
refGene_gn=TLR5;refGene_tx=XM_011290806.3;refGene_gn=TLR5;refGene_tx=XM_019821416.2;refGene_gn=TLR5;refGene_tx=XM_019821417.2;refGene_gn=TLR5;refGene_tx=XM_019821419.2;refGene_gn=TLR5;refGene_tx=XM_019821420.2;refGene_gn=TLR5;refGene_tx=XM_023247432.1;refGene_gn=TLR5;refGene_tx=XM_023247433.1;refGene_gn=TLR5;refGene_tx=XM_023247434.1
F1 2378083 2378112
refGene_gn=SUSD4;refGene_tx=XM_023247988.1;refGene_gn=SUSD4;refGene_tx=XM_023247989.1;refGene_gn=SUSD4;refGene_tx=XM_023247990.1;refGene_gn=SUSD4;refGene_tx=XM_023247991.1
F1 2378851 2379228
refGene_gn=SUSD4;refGene_tx=XM_023247988.1;refGene_gn=SUSD4;refGene_tx=XM_023247989.1;refGene_gn=SUSD4;refGene_tx=XM_023247990.1;refGene_gn=SUSD4;refGene_tx=XM_023247991.1
F1 2379658 2379740
refGene_gn=SUSD4;refGene_tx=XM_023247988.1;refGene_gn=SUSD4;refGene_tx=XM_023247989.1;refGene_gn=SUSD4;refGene_tx=XM_023247990.1
F1 2382801 2382831

refGene_gn=SUSD4;refGene_tx=XM_023247988.1;refGene_gn=SUSD4;refGene_tx=XM_023247990.1
F1 2383758 2383903 refGene_gn=SUSD4;refGene_tx=XM_023247988.1;refGene_gn=SUSD4;refGene_tx=XM_023247989.1;refGene_gn=SUSD4;refGene_tx=XM_023247990.1;refGene_gn=SUSD4;refGene_tx=XM_023247991.1
F1 2384867 2385059 refGene_gn=SUSD4;refGene_tx=XM_023247988.1;refGene_gn=SUSD4;refGene_tx=XM_023247989.1;refGene_gn=SUSD4;refGene_tx=XM_023247990.1;refGene_gn=SUSD4;refGene_tx=XM_023247991.1
F1 2411155 2411344 refGene_gn=SUSD4;refGene_tx=XM_023247988.1;refGene_gn=SUSD4;refGene_tx=XM_023247989.1;refGene_gn=SUSD4;refGene_tx=XM_023247990.1;refGene_gn=SUSD4;refGene_tx=XM_023247991.1
F1 2414480 2414654 refGene_gn=SUSD4;refGene_tx=XM_023247988.1;refGene_gn=SUSD4;refGene_tx=XM_023247989.1;refGene_gn=SUSD4;refGene_tx=XM_023247990.1;refGene_gn=SUSD4;refGene_tx=XM_023247991.1
F1 2426448 2426661 refGene_gn=SUSD4;refGene_tx=XM_023247988.1;refGene_gn=SUSD4;refGene_tx=XM_023247989.1;refGene_gn=SUSD4;refGene_tx=XM_023247991.1
F1 2470685 2470836 refGene_gn=SUSD4;refGene_tx=XM_023247988.1;refGene_gn=SUSD4;refGene_tx=XM_023247989.1;refGene_gn=SUSD4;refGene_tx=XM_023247990.1;refGene_gn=SUSD4;refGene_tx=XM_023247991.1
F1 2485526 2487113 refGene_gn=CCDC185;refGene_tx=XM_023247887.1
F1 2553878 2554007 refGene_gn=LOC101082970;refGene_tx=XM_023247998.1
F1 2554561 2554635 refGene_gn=LOC101082970;refGene_tx=XM_023247998.1
F1 2556530 2556599 refGene_gn=LOC101082970;refGene_tx=XM_023247998.1
F1 2556845 2556987 refGene_gn=LOC101082970;refGene_tx=XM_023247998.1
F1 2558669 2558726 refGene_gn=LOC101082970;refGene_tx=XM_023247998.1
F1 2558964 2559059 refGene_gn=LOC101082970;refGene_tx=XM_023247998.1
F1 2560168 2560343 refGene_gn=LOC101082970;refGene_tx=XM_023247998.1
F1 2562982 2563213 refGene_gn=LOC101082970;refGene_tx=XM_023247998.1
F1 2565452 2565542 refGene_gn=LOC101082970;refGene_tx=XM_023247998.1
F1 2570017 2570081 refGene_gn=LOC101082970;refGene_tx=XM_023247998.1
F1 2571030 2571265 refGene_gn=LOC101082970;refGene_tx=XM_023247998.1
F1 2571267 2571327 refGene_gn=LOC101082970;refGene_tx=XM_023247998.1

F1	2573928	2574089
refGene_gn=LOC101082970;refGene_tx=XM_023247998.1
F1	2574645	2574720
refGene_gn=LOC101082970;refGene_tx=XM_023247998.1
F1	2576511	2576597
refGene_gn=LOC101082970;refGene_tx=XM_023247998.1
F1	2577940	2578030
refGene_gn=LOC101082970;refGene_tx=XM_023247998.1
F1	2578785	2578954
refGene_gn=LOC101082970;refGene_tx=XM_023247998.1
F1	2579548	2579682
refGene_gn=LOC101082970;refGene_tx=XM_023247998.1
F1	2580325	2580444
refGene_gn=LOC101082970;refGene_tx=XM_023247998.1
F1	2589751	2589821
refGene_gn=LOC101082970;refGene_tx=XM_023247998.1
F1	2596950	2597187
refGene_gn=LOC101082970;refGene_tx=XM_023247998.1
F1	2621418	2621655	refGene_gn=CAPN2;refGene_tx=XM_023247642.1
F1	2625691	2625761	refGene_gn=CAPN2;refGene_tx=XM_023247642.1
F1	2639305	2639424	refGene_gn=CAPN2;refGene_tx=XM_023247642.1
F1	2640607	2640741	refGene_gn=CAPN2;refGene_tx=XM_023247642.1
F1	2641487	2641656	refGene_gn=CAPN2;refGene_tx=XM_023247642.1
F1	2642664	2642748	refGene_gn=CAPN2;refGene_tx=XM_023247642.1
F1	2644791	2644877	refGene_gn=CAPN2;refGene_tx=XM_023247642.1
F1	2645411	2645486	refGene_gn=CAPN2;refGene_tx=XM_023247642.1
F1	2645940	2646101	refGene_gn=CAPN2;refGene_tx=XM_023247642.1
F1	2647585	2647755	refGene_gn=CAPN2;refGene_tx=XM_023247642.1
F1	2649016	2649028	refGene_gn=CAPN2;refGene_tx=XM_023247642.1
F1	2650272	2650484	refGene_gn=CAPN2;refGene_tx=XM_023247642.1
F1	2651563	2651600	refGene_gn=CAPN2;refGene_tx=XM_023247642.1
F1	2652281	2652347	refGene_gn=CAPN2;refGene_tx=XM_023247642.1
F1	2653233	2653291	refGene_gn=CAPN2;refGene_tx=XM_023247642.1
F1	2654745	2654810	refGene_gn=CAPN2;refGene_tx=XM_023247642.1
F1	2655460	2655529	refGene_gn=CAPN2;refGene_tx=XM_023247642.1
F1	2656040	2656119	refGene_gn=CAPN2;refGene_tx=XM_023247642.1
F1	2657239	2657356	refGene_gn=CAPN2;refGene_tx=XM_023247642.1
F1	2657682	2657741	refGene_gn=CAPN2;refGene_tx=XM_023247642.1
F1	2659069	2659093	refGene_gn=CAPN2;refGene_tx=XM_023247642.1
F1	2662630	2662672
refGene_gn=TP53BP2;refGene_tx=XM_019821423.2;refGene_gn=TP53BP2;refGen
e_tx=XM_019821424.1;refGene_gn=TP53BP2;refGene_tx=XM_023247640.1;refGe
ne_gn=TP53BP2;refGene_tx=XM_023247641.1
F1	2664797	2664997
refGene_gn=TP53BP2;refGene_tx=XM_019821423.2;refGene_gn=TP53BP2;refGen
e_tx=XM_019821424.1;refGene_gn=TP53BP2;refGene_tx=XM_023247640.1;refGe
ne_gn=TP53BP2;refGene_tx=XM_023247641.1
F1	2665355	2665522
refGene_gn=TP53BP2;refGene_tx=XM_019821423.2;refGene_gn=TP53BP2;refGen
e_tx=XM_019821424.1;refGene_gn=TP53BP2;refGene_tx=XM_023247640.1;refGe

ne_gn=TP53BP2;refGene_tx=XM_023247641.1
F1 2667895 2668029
refGene_gn=TP53BP2;refGene_tx=XM_019821423.2;refGene_gn=TP53BP2;refGene_tx=XM_019821424.1;refGene_gn=TP53BP2;refGene_tx=XM_023247640.1;refGene_gn=TP53BP2;refGene_tx=XM_023247641.1
F1 2668711 2668849
refGene_gn=TP53BP2;refGene_tx=XM_019821423.2;refGene_gn=TP53BP2;refGene_tx=XM_019821424.1;refGene_gn=TP53BP2;refGene_tx=XM_023247640.1;refGene_gn=TP53BP2;refGene_tx=XM_023247641.1
F1 2670643 2671419
refGene_gn=TP53BP2;refGene_tx=XM_019821423.2;refGene_gn=TP53BP2;refGene_tx=XM_019821424.1;refGene_gn=TP53BP2;refGene_tx=XM_023247640.1;refGene_gn=TP53BP2;refGene_tx=XM_023247641.1
F1 2672616 2673076
refGene_gn=TP53BP2;refGene_tx=XM_019821423.2;refGene_gn=TP53BP2;refGene_tx=XM_019821424.1;refGene_gn=TP53BP2;refGene_tx=XM_023247640.1;refGene_gn=TP53BP2;refGene_tx=XM_023247641.1
F1 2673781 2673930
refGene_gn=TP53BP2;refGene_tx=XM_019821423.2;refGene_gn=TP53BP2;refGene_tx=XM_019821424.1;refGene_gn=TP53BP2;refGene_tx=XM_023247640.1;refGene_gn=TP53BP2;refGene_tx=XM_023247641.1
F1 2674220 2674343
refGene_gn=TP53BP2;refGene_tx=XM_019821423.2;refGene_gn=TP53BP2;refGene_tx=XM_019821424.1;refGene_gn=TP53BP2;refGene_tx=XM_023247640.1;refGene_gn=TP53BP2;refGene_tx=XM_023247641.1
F1 2675800 2676023
refGene_gn=TP53BP2;refGene_tx=XM_019821423.2;refGene_gn=TP53BP2;refGene_tx=XM_019821424.1;refGene_gn=TP53BP2;refGene_tx=XM_023247640.1;refGene_gn=TP53BP2;refGene_tx=XM_023247641.1
F1 2676207 2676372
refGene_gn=TP53BP2;refGene_tx=XM_019821423.2;refGene_gn=TP53BP2;refGene_tx=XM_019821424.1;refGene_gn=TP53BP2;refGene_tx=XM_023247640.1;refGene_gn=TP53BP2;refGene_tx=XM_023247641.1
F1 2676612 2676794
refGene_gn=TP53BP2;refGene_tx=XM_019821423.2;refGene_gn=TP53BP2;refGene_tx=XM_019821424.1;refGene_gn=TP53BP2;refGene_tx=XM_023247640.1;refGene_gn=TP53BP2;refGene_tx=XM_023247641.1
F1 2677419 2677594
refGene_gn=TP53BP2;refGene_tx=XM_019821423.2;refGene_gn=TP53BP2;refGene_tx=XM_019821424.1;refGene_gn=TP53BP2;refGene_tx=XM_023247640.1;refGene_gn=TP53BP2;refGene_tx=XM_023247641.1
F1 2678785 2678935 refGene_gn=TP53BP2;refGene_tx=XM_019821424.1
F1 2679535 2679637
refGene_gn=TP53BP2;refGene_tx=XM_019821423.2;refGene_gn=TP53BP2;refGene_tx=XM_023247640.1;refGene_gn=TP53BP2;refGene_tx=XM_023247641.1
F1 2683086 2683169
refGene_gn=TP53BP2;refGene_tx=XM_019821423.2;refGene_gn=TP53BP2;refGene_tx=XM_023247640.1;refGene_gn=TP53BP2;refGene_tx=XM_023247641.1
F1 2685898 2686012
refGene_gn=TP53BP2;refGene_tx=XM_019821423.2;refGene_gn=TP53BP2;refGen

e_tx=XM_023247640.1;refGene_gn=TP53BP2;refGene_tx=XM_023247641.1
F1 2691041 2691189
refGene_gn=TP53BP2;refGene_tx=XM_019821423.2;refGene_gn=TP53BP2;refGene_tx=XM_023247640.1;refGene_gn=TP53BP2;refGene_tx=XM_023247641.1
F1 2714856 2714883
refGene_gn=TP53BP2;refGene_tx=XM_023247640.1;refGene_gn=TP53BP2;refGene_tx=XM_023247641.1
F1 2769652 2769919
refGene_gn=FBXO28;refGene_tx=XM_023248109.1;refGene_gn=FBXO28;refGene_tx=XM_023248110.1
F1 2781726 2781836
refGene_gn=FBXO28;refGene_tx=XM_023248109.1;refGene_gn=FBXO28;refGene_tx=XM_023248110.1
F1 2785461 2785600
refGene_gn=FBXO28;refGene_tx=XM_023248109.1;refGene_gn=FBXO28;refGene_tx=XM_023248110.1
F1 2793336 2793532
refGene_gn=FBXO28;refGene_tx=XM_023248109.1;refGene_gn=FBXO28;refGene_tx=XM_023248110.1
F1 2794312 2794395 refGene_gn=FBXO28;refGene_tx=XM_023248110.1
F1 2795780 2796175 refGene_gn=FBXO28;refGene_tx=XM_023248109.1
F1 2812972 2813054 refGene_gn=DEGS1;refGene_tx=XM_023247544.1
F1 2817798 2818541 refGene_gn=DEGS1;refGene_tx=XM_023247544.1
F1 2821329 2821476 refGene_gn=DEGS1;refGene_tx=XM_023247544.1
F1 2840058 2840103
refGene_gn=NVL;refGene_tx=XM_006942592.4;refGene_gn=NVL;refGene_tx=XM_006942593.4;refGene_gn=NVL;refGene_tx=XM_019821429.2;refGene_gn=NVL;refGene_tx=XM_019821430.1
F1 2848229 2848247 refGene_gn=NVL;refGene_tx=XM_019821427.2
F1 2848761 2848832
refGene_gn=NVL;refGene_tx=XM_006942592.4;refGene_gn=NVL;refGene_tx=XM_006942593.4;refGene_gn=NVL;refGene_tx=XM_019821427.2;refGene_gn=NVL;refGene_tx=XM_019821429.2;refGene_gn=NVL;refGene_tx=XM_019821430.1
F1 2851415 2851504
refGene_gn=NVL;refGene_tx=XM_006942592.4;refGene_gn=NVL;refGene_tx=XM_006942593.4;refGene_gn=NVL;refGene_tx=XM_019821427.2;refGene_gn=NVL;refGene_tx=XM_019821429.2;refGene_gn=NVL;refGene_tx=XM_019821430.1
F1 2852944 2853021
refGene_gn=NVL;refGene_tx=XM_006942592.4;refGene_gn=NVL;refGene_tx=XM_006942593.4;refGene_gn=NVL;refGene_tx=XM_019821427.2;refGene_gn=NVL;refGene_tx=XM_019821429.2;refGene_gn=NVL;refGene_tx=XM_019821430.1
F1 2870928 2871035
refGene_gn=NVL;refGene_tx=XM_006942592.4;refGene_gn=NVL;refGene_tx=XM_006942593.4;refGene_gn=NVL;refGene_tx=XM_019821427.2;refGene_gn=NVL;refGene_tx=XM_019821429.2;refGene_gn=NVL;refGene_tx=XM_019821430.1
F1 2883959 2884059
refGene_gn=NVL;refGene_tx=XM_006942592.4;refGene_gn=NVL;refGene_tx=XM_006942593.4;refGene_gn=NVL;refGene_tx=XM_019821427.2;refGene_gn=NVL;refGene_tx=XM_019821429.2;refGene_gn=NVL;refGene_tx=XM_019821430.1
F1 2887874 2887994

refGene_gn=NVL;refGene_tx=XM_006942592.4;refGene_gn=NVL;refGene_tx=XM_006942593.4;refGene_gn=NVL;refGene_tx=XM_019821427.2;refGene_gn=NVL;refGene_tx=XM_019821429.2;refGene_gn=NVL;refGene_tx=XM_019821430.1
F1 2893684 2893747
refGene_gn=NVL;refGene_tx=XM_006942592.4;refGene_gn=NVL;refGene_tx=XM_006942593.4;refGene_gn=NVL;refGene_tx=XM_019821427.2;refGene_gn=NVL;refGene_tx=XM_019821429.2;refGene_gn=NVL;refGene_tx=XM_019821430.1
F1 2897894 2897999
refGene_gn=NVL;refGene_tx=XM_006942592.4;refGene_gn=NVL;refGene_tx=XM_006942593.4;refGene_gn=NVL;refGene_tx=XM_019821427.2;refGene_gn=NVL;refGene_tx=XM_019821430.1
F1 2898750 2898969
refGene_gn=NVL;refGene_tx=XM_006942592.4;refGene_gn=NVL;refGene_tx=XM_006942593.4;refGene_gn=NVL;refGene_tx=XM_019821427.2;refGene_gn=NVL;refGene_tx=XM_019821429.2;refGene_gn=NVL;refGene_tx=XM_019821430.1
F1 2900220 2900524
refGene_gn=NVL;refGene_tx=XM_006942592.4;refGene_gn=NVL;refGene_tx=XM_006942593.4;refGene_gn=NVL;refGene_tx=XM_019821427.2;refGene_gn=NVL;refGene_tx=XM_019821429.2;refGene_gn=NVL;refGene_tx=XM_019821430.1
F1 2908287 2908432
refGene_gn=NVL;refGene_tx=XM_006942592.4;refGene_gn=NVL;refGene_tx=XM_006942593.4;refGene_gn=NVL;refGene_tx=XM_019821427.2;refGene_gn=NVL;refGene_tx=XM_019821429.2;refGene_gn=NVL;refGene_tx=XM_019821430.1
F1 2910980 2911098
refGene_gn=NVL;refGene_tx=XM_006942592.4;refGene_gn=NVL;refGene_tx=XM_006942593.4;refGene_gn=NVL;refGene_tx=XM_019821427.2;refGene_gn=NVL;refGene_tx=XM_019821429.2;refGene_gn=NVL;refGene_tx=XM_019821430.1
F1 2912922 2913024
refGene_gn=NVL;refGene_tx=XM_006942592.4;refGene_gn=NVL;refGene_tx=XM_006942593.4;refGene_gn=NVL;refGene_tx=XM_019821427.2;refGene_gn=NVL;refGene_tx=XM_019821429.2;refGene_gn=NVL;refGene_tx=XM_019821430.1
F1 2915025 2915160
refGene_gn=NVL;refGene_tx=XM_006942592.4;refGene_gn=NVL;refGene_tx=XM_006942593.4;refGene_gn=NVL;refGene_tx=XM_019821427.2;refGene_gn=NVL;refGene_tx=XM_019821429.2;refGene_gn=NVL;refGene_tx=XM_019821430.1
F1 2915724 2915801
refGene_gn=NVL;refGene_tx=XM_006942592.4;refGene_gn=NVL;refGene_tx=XM_006942593.4;refGene_gn=NVL;refGene_tx=XM_019821427.2;refGene_gn=NVL;refGene_tx=XM_019821429.2;refGene_gn=NVL;refGene_tx=XM_019821430.1
F1 2916002 2916132
refGene_gn=NVL;refGene_tx=XM_006942592.4;refGene_gn=NVL;refGene_tx=XM_006942593.4;refGene_gn=NVL;refGene_tx=XM_019821427.2;refGene_gn=NVL;refGene_tx=XM_019821429.2;refGene_gn=NVL;refGene_tx=XM_019821430.1
F1 2919039 2919297
refGene_gn=NVL;refGene_tx=XM_006942592.4;refGene_gn=NVL;refGene_tx=XM_019821427.2;refGene_gn=NVL;refGene_tx=XM_019821429.2;refGene_gn=NVL;refGene_tx=XM_019821430.1
F1 2920694 2920752
refGene_gn=NVL;refGene_tx=XM_006942592.4;refGene_gn=NVL;refGene_tx=XM_006942593.4;refGene_gn=NVL;refGene_tx=XM_019821427.2;refGene_gn=NVL;re

fGene_tx=XM_019821429.2;refGene_gn=NVL;refGene_tx=XM_019821430.1
F1 2924250 2924350
refGene_gn=NVL;refGene_tx=XM_006942592.4;refGene_gn=NVL;refGene_tx=XM_006942593.4;refGene_gn=NVL;refGene_tx=XM_019821427.2;refGene_gn=NVL;refGene_tx=XM_019821429.2
F1 2924431 2924484
refGene_gn=NVL;refGene_tx=XM_006942592.4;refGene_gn=NVL;refGene_tx=XM_006942593.4;refGene_gn=NVL;refGene_tx=XM_019821427.2;refGene_gn=NVL;refGene_tx=XM_019821429.2
F1 2926489 2926563
refGene_gn=NVL;refGene_tx=XM_006942592.4;refGene_gn=NVL;refGene_tx=XM_006942593.4;refGene_gn=NVL;refGene_tx=XM_019821427.2;refGene_gn=NVL;refGene_tx=XM_019821429.2
F1 2930629 2930686
refGene_gn=NVL;refGene_tx=XM_006942592.4;refGene_gn=NVL;refGene_tx=XM_006942593.4;refGene_gn=NVL;refGene_tx=XM_019821427.2;refGene_gn=NVL;refGene_tx=XM_019821429.2
F1 2945544 2945613 refGene_gn=CNIH4;refGene_tx=XM_023247481.1
F1 2949089 2949158 refGene_gn=CNIH4;refGene_tx=XM_023247481.1
F1 2952273 2952386
refGene_gn=CNIH4;refGene_tx=XM_019821432.2;refGene_gn=CNIH4;refGene_tx=XM_023247481.1
F1 2957537 2957678
refGene_gn=CNIH4;refGene_tx=XM_019821432.2;refGene_gn=CNIH4;refGene_tx=XM_023247481.1
F1 2959876 2959904
refGene_gn=CNIH4;refGene_tx=XM_019821432.2;refGene_gn=CNIH4;refGene_tx=XM_023247481.1
F1 2970298 2970324
refGene_gn=WDR26;refGene_tx=XM_023247992.1;refGene_gn=WDR26;refGene_tx=XM_023247993.1
F1 2972579 2972765
refGene_gn=WDR26;refGene_tx=XM_023247992.1;refGene_gn=WDR26;refGene_tx=XM_023247993.1
F1 2975736 2975866
refGene_gn=WDR26;refGene_tx=XM_023247992.1;refGene_gn=WDR26;refGene_tx=XM_023247993.1
F1 2976221 2976300
refGene_gn=WDR26;refGene_tx=XM_023247992.1;refGene_gn=WDR26;refGene_tx=XM_023247993.1
F1 2976598 2976744
refGene_gn=WDR26;refGene_tx=XM_023247992.1;refGene_gn=WDR26;refGene_tx=XM_023247993.1
F1 2978697 2978817
refGene_gn=WDR26;refGene_tx=XM_023247992.1;refGene_gn=WDR26;refGene_tx=XM_023247993.1
F1 2981645 2981786
refGene_gn=WDR26;refGene_tx=XM_023247992.1;refGene_gn=WDR26;refGene_tx=XM_023247993.1
F1 2985240 2985379

refGene_gn=WDR26;refGene_tx=XM_023247992.1;refGene_gn=WDR26;refGene_tx=XM_023247993.1
F1 2990204 2990361
refGene_gn=WDR26;refGene_tx=XM_023247992.1;refGene_gn=WDR26;refGene_tx=XM_023247993.1
F1 2991943 2992041
refGene_gn=WDR26;refGene_tx=XM_023247992.1;refGene_gn=WDR26;refGene_tx=XM_023247993.1
F1 2995294 2995431
refGene_gn=WDR26;refGene_tx=XM_023247992.1;refGene_gn=WDR26;refGene_tx=XM_023247993.1
F1 3001485 3001590
refGene_gn=WDR26;refGene_tx=XM_023247992.1;refGene_gn=WDR26;refGene_tx=XM_023247993.1
F1 3001691 3001791
refGene_gn=WDR26;refGene_tx=XM_023247992.1;refGene_gn=WDR26;refGene_tx=XM_023247993.1
F1 3003391 3003789
refGene_gn=WDR26;refGene_tx=XM_023247992.1;refGene_gn=WDR26;refGene_tx=XM_023247993.1
F1 3116824 3116905
refGene_gn=CNIH3;refGene_tx=XM_023248015.1;refGene_gn=CNIH3;refGene_tx=XM_023248016.1;refGene_gn=CNIH3;refGene_tx=XM_023248017.1;refGene_gn=CNIH3;refGene_tx=XM_023248021.1;refGene_gn=CNIH3;refGene_tx=XM_023248022.1;refGene_gn=CNIH3;refGene_tx=XM_023248023.1;refGene_gn=CNIH3;refGene_tx=XM_023248024.1
F1 3158485 3158554
refGene_gn=CNIH3;refGene_tx=XM_023248015.1;refGene_gn=CNIH3;refGene_tx=XM_023248016.1;refGene_gn=CNIH3;refGene_tx=XM_023248017.1;refGene_gn=CNIH3;refGene_tx=XM_023248021.1;refGene_gn=CNIH3;refGene_tx=XM_023248022.1;refGene_gn=CNIH3;refGene_tx=XM_023248023.1;refGene_gn=CNIH3;refGene_tx=XM_023248024.1
F1 3163419 3163467
refGene_gn=CNIH3;refGene_tx=XM_023248015.1;refGene_gn=CNIH3;refGene_tx=XM_023248017.1;refGene_gn=CNIH3;refGene_tx=XM_023248021.1;refGene_gn=CNIH3;refGene_tx=XM_023248022.1;refGene_gn=CNIH3;refGene_tx=XM_023248023.1
F1 3194118 3194231
refGene_gn=CNIH3;refGene_tx=XM_023248015.1;refGene_gn=CNIH3;refGene_tx=XM_023248016.1;refGene_gn=CNIH3;refGene_tx=XM_023248017.1;refGene_gn=CNIH3;refGene_tx=XM_023248018.1;refGene_gn=CNIH3;refGene_tx=XM_023248019.1;refGene_gn=CNIH3;refGene_tx=XM_023248021.1;refGene_gn=CNIH3;refGene_tx=XM_023248022.1;refGene_gn=CNIH3;refGene_tx=XM_023248023.1;refGene_gn=CNIH3;refGene_tx=XM_023248024.1
F1 3194363 3194526 refGene_gn=CNIH3;refGene_tx=XM_023248023.1
F1 3198477 3198621
refGene_gn=CNIH3;refGene_tx=XM_023248015.1;refGene_gn=CNIH3;refGene_tx=XM_023248016.1;refGene_gn=CNIH3;refGene_tx=XM_023248017.1;refGene_gn=CNIH3;refGene_tx=XM_023248018.1;refGene_gn=CNIH3;refGene_tx=XM_023248019.1;refGene_gn=CNIH3;refGene_tx=XM_023248020.1;refGene_gn=CNIH3;refGe

ne_tx=XM_023248021.1;refGene_gn=CNIH3;refGene_tx=XM_023248022.1;refGene_gn=CNIH3;refGene_tx=XM_023248024.1
F1	3198868	3199048	refGene_gn=CNIH3;refGene_tx=XM_023248015.1;refGene_gn=CNIH3;refGene_tx=XM_023248016.1;refGene_gn=CNIH3;refGene_tx=XM_023248018.1;refGene_gn=CNIH3;refGene_tx=XM_023248019.1;refGene_gn=CNIH3;refGene_tx=XM_023248020.1
F1	3202580	3202608	refGene_gn=CNIH3;refGene_tx=XM_023248022.1;refGene_gn=CNIH3;refGene_tx=XM_023248024.1
F1	3206240	3206802	refGene_gn=CNIH3;refGene_tx=XM_023248015.1;refGene_gn=CNIH3;refGene_tx=XM_023248016.1;refGene_gn=CNIH3;refGene_tx=XM_023248017.1;refGene_gn=CNIH3;refGene_tx=XM_023248018.1;refGene_gn=CNIH3;refGene_tx=XM_023248019.1;refGene_gn=CNIH3;refGene_tx=XM_023248020.1
F1	3251191	3251225	refGene_gn=CNIH3;refGene_tx=XM_023248021.1
F1	3342828	3342875	refGene_gn=DNAH14;refGene_tx=XM_023247999.1
F1	3351822	3351986	refGene_gn=DNAH14;refGene_tx=XM_023247999.1
F1	3359746	3359890	refGene_gn=DNAH14;refGene_tx=XM_023247999.1
F1	3365630	3365761	refGene_gn=DNAH14;refGene_tx=XM_023247999.1
F1	3368361	3368514	refGene_gn=DNAH14;refGene_tx=XM_023247999.1
F1	3369665	3369778	refGene_gn=DNAH14;refGene_tx=XM_023247999.1
F1	3379156	3379219	refGene_gn=DNAH14;refGene_tx=XM_023247999.1
F1	3383913	3384058	refGene_gn=DNAH14;refGene_tx=XM_023247999.1
F1	3388175	3388307	refGene_gn=DNAH14;refGene_tx=XM_023247999.1
F1	3396581	3396832	refGene_gn=DNAH14;refGene_tx=XM_023247999.1
F1	3402593	3402723	refGene_gn=DNAH14;refGene_tx=XM_023247999.1
F1	3407709	3407986	refGene_gn=DNAH14;refGene_tx=XM_023247999.1
F1	3408485	3408531	refGene_gn=DNAH14;refGene_tx=XM_023247999.1
F1	3408620	3408718	refGene_gn=DNAH14;refGene_tx=XM_023247999.1
F1	3412607	3412774	refGene_gn=DNAH14;refGene_tx=XM_023247999.1
F1	3413491	3413836	refGene_gn=DNAH14;refGene_tx=XM_023247999.1
F1	3419999	3420341	refGene_gn=DNAH14;refGene_tx=XM_023247999.1
F1	3422827	3423197	refGene_gn=DNAH14;refGene_tx=XM_023247999.1
F1	3424959	3425147	refGene_gn=DNAH14;refGene_tx=XM_023247999.1
F1	3428078	3428324	refGene_gn=DNAH14;refGene_tx=XM_023247999.1
F1	3434862	3434984	refGene_gn=DNAH14;refGene_tx=XM_023247999.1
F1	3436842	3437014	refGene_gn=DNAH14;refGene_tx=XM_023247999.1
F1	3445539	3445644	refGene_gn=DNAH14;refGene_tx=XM_023247999.1
F1	3445732	3445851	refGene_gn=DNAH14;refGene_tx=XM_023247999.1
F1	3446599	3446674	refGene_gn=DNAH14;refGene_tx=XM_023247999.1
F1	3452299	3452387	refGene_gn=DNAH14;refGene_tx=XM_023247999.1
F1	3455496	3455750	refGene_gn=DNAH14;refGene_tx=XM_023247999.1
F1	3460311	3460543	refGene_gn=DNAH14;refGene_tx=XM_023247999.1
F1	3461217	3461271	refGene_gn=DNAH14;refGene_tx=XM_023247999.1
F1	3462686	3462832	refGene_gn=DNAH14;refGene_tx=XM_023247999.1
F1	3464529	3464598	refGene_gn=DNAH14;refGene_tx=XM_023247999.1
F1	3465404	3465591	refGene_gn=DNAH14;refGene_tx=XM_023247999.1
F1	3466550	3466627	refGene_gn=DNAH14;refGene_tx=XM_023247999.1
F1	3469434	3469606	refGene_gn=DNAH14;refGene_tx=XM_023247999.1

| | | | |
|---|---|---|---|
| F1 | 3480161 | 3480251 | refGene_gn=DNAH14;refGene_tx=XM_023247999.1 |
| F1 | 3481736 | 3481871 | refGene_gn=DNAH14;refGene_tx=XM_023247999.1 |
| F1 | 3490510 | 3490726 | refGene_gn=DNAH14;refGene_tx=XM_023247999.1 |
| F1 | 3493315 | 3493409 | refGene_gn=DNAH14;refGene_tx=XM_023247999.1 |
| F1 | 3494683 | 3494892 | refGene_gn=DNAH14;refGene_tx=XM_023247999.1 |
| F1 | 3495647 | 3495900 | refGene_gn=DNAH14;refGene_tx=XM_023247999.1 |
| F1 | 3507772 | 3507857 | refGene_gn=DNAH14;refGene_tx=XM_023247999.1 |
| F1 | 3509231 | 3509461 | refGene_gn=DNAH14;refGene_tx=XM_023247999.1 |
| F1 | 3512211 | 3512460 | refGene_gn=DNAH14;refGene_tx=XM_023247999.1 |
| F1 | 3517522 | 3517639 | refGene_gn=DNAH14;refGene_tx=XM_023247999.1 |
| F1 | 3521080 | 3521239 | refGene_gn=DNAH14;refGene_tx=XM_023247999.1 |
| F1 | 3522788 | 3522921 | refGene_gn=DNAH14;refGene_tx=XM_023247999.1 |
| F1 | 3524515 | 3524577 | refGene_gn=DNAH14;refGene_tx=XM_023247999.1 |
| F1 | 3525502 | 3525690 | refGene_gn=DNAH14;refGene_tx=XM_023247999.1 |
| F1 | 3526871 | 3527000 | refGene_gn=DNAH14;refGene_tx=XM_023247999.1 |
| F1 | 3528875 | 3529013 | refGene_gn=DNAH14;refGene_tx=XM_023247999.1 |
| F1 | 3530490 | 3530658 | refGene_gn=DNAH14;refGene_tx=XM_023247999.1 |
| F1 | 3531554 | 3531725 | refGene_gn=DNAH14;refGene_tx=XM_023247999.1 |
| F1 | 3535599 | 3535767 | refGene_gn=DNAH14;refGene_tx=XM_023247999.1 |
| F1 | 3537154 | 3537247 | refGene_gn=DNAH14;refGene_tx=XM_023247999.1 |
| F1 | 3546195 | 3546393 | refGene_gn=DNAH14;refGene_tx=XM_023247999.1 |
| F1 | 3548796 | 3548958 | refGene_gn=DNAH14;refGene_tx=XM_023247999.1 |
| F1 | 3551365 | 3551557 | refGene_gn=DNAH14;refGene_tx=XM_023247999.1 |
| F1 | 3552867 | 3553049 | refGene_gn=DNAH14;refGene_tx=XM_023247999.1 |
| F1 | 3553564 | 3553673 | refGene_gn=DNAH14;refGene_tx=XM_023247999.1 |
| F1 | 3554608 | 3554734 | refGene_gn=DNAH14;refGene_tx=XM_023247999.1 |
| F1 | 3557354 | 3557449 | refGene_gn=DNAH14;refGene_tx=XM_023247999.1 |
| F1 | 3560972 | 3561132 | refGene_gn=DNAH14;refGene_tx=XM_023247999.1 |
| F1 | 3568522 | 3568654 | refGene_gn=DNAH14;refGene_tx=XM_023247999.1 |
| F1 | 3568994 | 3569090 | refGene_gn=DNAH14;refGene_tx=XM_023247999.1 |
| F1 | 3578735 | 3578876 | refGene_gn=DNAH14;refGene_tx=XM_023247999.1 |
| F1 | 3579997 | 3580213 | refGene_gn=DNAH14;refGene_tx=XM_023247999.1 |
| F1 | 3583412 | 3583643 | refGene_gn=DNAH14;refGene_tx=XM_023247999.1 |
| F1 | 3583971 | 3584093 | refGene_gn=DNAH14;refGene_tx=XM_023247999.1 |
| F1 | 3585925 | 3586170 | refGene_gn=DNAH14;refGene_tx=XM_023247999.1 |
| F1 | 3589621 | 3590040 | refGene_gn=DNAH14;refGene_tx=XM_023247999.1 |
| F1 | 3590117 | 3590316 | refGene_gn=DNAH14;refGene_tx=XM_023247999.1 |
| F1 | 3596078 | 3596315 | refGene_gn=DNAH14;refGene_tx=XM_023247999.1 |
| F1 | 3598303 | 3598389 | refGene_gn=DNAH14;refGene_tx=XM_023247999.1 |
| F1 | 3600965 | 3601122 | refGene_gn=DNAH14;refGene_tx=XM_023247999.1 |
| F1 | 3602421 | 3602632 | refGene_gn=DNAH14;refGene_tx=XM_023247999.1 |
| F1 | 3605372 | 3605475 | refGene_gn=DNAH14;refGene_tx=XM_023247999.1 |
| F1 | 3607095 | 3607323 | refGene_gn=DNAH14;refGene_tx=XM_023247999.1 |
| F1 | 3616312 | 3616502 | refGene_gn=DNAH14;refGene_tx=XM_023247999.1 |
| F1 | 3617533 | 3617717 | refGene_gn=DNAH14;refGene_tx=XM_023247999.1 |
| F1 | 3621998 | 3622159 | refGene_gn=DNAH14;refGene_tx=XM_023247999.1 |
| F1 | 3623377 | 3623574 | refGene_gn=DNAH14;refGene_tx=XM_023247999.1 |
| F1 | 3639472 | 3639585 | refGene_gn=DNAH14;refGene_tx=XM_023247999.1 |
| F1 | 3640489 | 3640629 | refGene_gn=DNAH14;refGene_tx=XM_023247999.1 |
| F1 | 3643020 | 3643184 | refGene_gn=DNAH14;refGene_tx=XM_023247999.1 |

F1	3644246	3644393	refGene_gn=DNAH14;refGene_tx=XM_023247999.1
F1	3645478	3645694	refGene_gn=DNAH14;refGene_tx=XM_023247999.1
F1	3648105	3648266	refGene_gn=LBR;refGene_tx=XM_023247871.1
F1	3648402	3648525	refGene_gn=LBR;refGene_tx=XM_023247871.1
F1	3648622	3648703	refGene_gn=LBR;refGene_tx=XM_023247871.1
F1	3650195	3650364	refGene_gn=LBR;refGene_tx=XM_023247871.1
F1	3653685	3653811	refGene_gn=LBR;refGene_tx=XM_023247871.1
F1	3654629	3654733	refGene_gn=LBR;refGene_tx=XM_023247871.1
F1	3655363	3655555	refGene_gn=LBR;refGene_tx=XM_023247871.1
F1	3658335	3658390	refGene_gn=LBR;refGene_tx=XM_023247871.1
F1	3659390	3659587	refGene_gn=LBR;refGene_tx=XM_023247871.1
F1	3660286	3660476	refGene_gn=LBR;refGene_tx=XM_023247871.1
F1	3661199	3661283	refGene_gn=LBR;refGene_tx=XM_023247871.1
F1	3663816	3664020	refGene_gn=LBR;refGene_tx=XM_023247871.1
F1	3665490	3665655	refGene_gn=LBR;refGene_tx=XM_023247871.1
F1	3675948	3676467
refGene_gn=LOC109495407;refGene_tx=XM_019821335.1
F1	3689673	3690162
refGene_gn=LOC109495407;refGene_tx=XM_019821335.1
F1	3690711	3690749
refGene_gn=LOC109495407;refGene_tx=XM_019821335.1
F1	3690864	3690913
refGene_gn=LOC109495407;refGene_tx=XM_019821335.1
F1	3705559	3705597
refGene_gn=ENAH;refGene_tx=XM_023247677.1;refGene_gn=ENAH;refGene_tx=XM_023247678.1;refGene_gn=ENAH;refGene_tx=XM_023247679.1;refGene_gn=ENAH;refGene_tx=XM_023247680.1;refGene_gn=ENAH;refGene_tx=XM_023247681.1;refGene_gn=ENAH;refGene_tx=XM_023247682.1
F1	3706131	3706189
refGene_gn=ENAH;refGene_tx=XM_023247677.1;refGene_gn=ENAH;refGene_tx=XM_023247678.1;refGene_gn=ENAH;refGene_tx=XM_023247679.1;refGene_gn=ENAH;refGene_tx=XM_023247680.1;refGene_gn=ENAH;refGene_tx=XM_023247681.1;refGene_gn=ENAH;refGene_tx=XM_023247682.1
F1	3710032	3710111
refGene_gn=ENAH;refGene_tx=XM_023247677.1;refGene_gn=ENAH;refGene_tx=XM_023247678.1;refGene_gn=ENAH;refGene_tx=XM_023247679.1;refGene_gn=ENAH;refGene_tx=XM_023247680.1;refGene_gn=ENAH;refGene_tx=XM_023247681.1;refGene_gn=ENAH;refGene_tx=XM_023247682.1
F1	3713016	3713079	refGene_gn=ENAH;refGene_tx=XM_023247681.1
F1	3715990	3716057
refGene_gn=ENAH;refGene_tx=XM_023247677.1;refGene_gn=ENAH;refGene_tx=XM_023247678.1;refGene_gn=ENAH;refGene_tx=XM_023247679.1;refGene_gn=ENAH;refGene_tx=XM_023247680.1;refGene_gn=ENAH;refGene_tx=XM_023247681.1;refGene_gn=ENAH;refGene_tx=XM_023247682.1
F1	3718195	3718244
refGene_gn=ENAH;refGene_tx=XM_023247677.1;refGene_gn=ENAH;refGene_tx=XM_023247678.1;refGene_gn=ENAH;refGene_tx=XM_023247679.1;refGene_gn=ENAH;refGene_tx=XM_023247680.1;refGene_gn=ENAH;refGene_tx=XM_023247681.1;refGene_gn=ENAH;refGene_tx=XM_023247682.1
F1	3718975	3719033

refGene_gn=ENAH;refGene_tx=XM_023247677.1;refGene_gn=ENAH;refGene_tx=XM_023247678.1;refGene_gn=ENAH;refGene_tx=XM_023247679.1;refGene_gn=ENAH;refGene_tx=XM_023247680.1;refGene_gn=ENAH;refGene_tx=XM_023247681.1;refGene_gn=ENAH;refGene_tx=XM_023247682.1
F1 3719192 3719338
refGene_gn=ENAH;refGene_tx=XM_023247677.1;refGene_gn=ENAH;refGene_tx=XM_023247678.1;refGene_gn=ENAH;refGene_tx=XM_023247679.1;refGene_gn=ENAH;refGene_tx=XM_023247680.1;refGene_gn=ENAH;refGene_tx=XM_023247681.1;refGene_gn=ENAH;refGene_tx=XM_023247682.1
F1 3720248 3720553
refGene_gn=ENAH;refGene_tx=XM_023247677.1;refGene_gn=ENAH;refGene_tx=XM_023247678.1;refGene_gn=ENAH;refGene_tx=XM_023247679.1;refGene_gn=ENAH;refGene_tx=XM_023247680.1;refGene_gn=ENAH;refGene_tx=XM_023247681.1;refGene_gn=ENAH;refGene_tx=XM_023247682.1
F1 3722486 3723281
refGene_gn=ENAH;refGene_tx=XM_023247677.1;refGene_gn=ENAH;refGene_tx=XM_023247678.1;refGene_gn=ENAH;refGene_tx=XM_023247679.1;refGene_gn=ENAH;refGene_tx=XM_023247680.1;refGene_gn=ENAH;refGene_tx=XM_023247681.1;refGene_gn=ENAH;refGene_tx=XM_023247682.1
F1 3724404 3724754
refGene_gn=ENAH;refGene_tx=XM_023247677.1;refGene_gn=ENAH;refGene_tx=XM_023247678.1;refGene_gn=ENAH;refGene_tx=XM_023247679.1;refGene_gn=ENAH;refGene_tx=XM_023247680.1;refGene_gn=ENAH;refGene_tx=XM_023247681.1;refGene_gn=ENAH;refGene_tx=XM_023247682.1
F1 3738734 3738819
refGene_gn=ENAH;refGene_tx=XM_023247677.1;refGene_gn=ENAH;refGene_tx=XM_023247678.1;refGene_gn=ENAH;refGene_tx=XM_023247679.1;refGene_gn=ENAH;refGene_tx=XM_023247680.1;refGene_gn=ENAH;refGene_tx=XM_023247681.1;refGene_gn=ENAH;refGene_tx=XM_023247682.1
F1 3739233 3739245 refGene_gn=ENAH;refGene_tx=XM_023247678.1
F1 3743639 3743696
refGene_gn=ENAH;refGene_tx=XM_023247677.1;refGene_gn=ENAH;refGene_tx=XM_023247680.1
F1 3761715 3761893
refGene_gn=ENAH;refGene_tx=XM_023247677.1;refGene_gn=ENAH;refGene_tx=XM_023247678.1;refGene_gn=ENAH;refGene_tx=XM_023247679.1;refGene_gn=ENAH;refGene_tx=XM_023247680.1;refGene_gn=ENAH;refGene_tx=XM_023247681.1;refGene_gn=ENAH;refGene_tx=XM_023247682.1
F1 3770717 3770883
refGene_gn=ENAH;refGene_tx=XM_023247677.1;refGene_gn=ENAH;refGene_tx=XM_023247678.1;refGene_gn=ENAH;refGene_tx=XM_023247679.1;refGene_gn=ENAH;refGene_tx=XM_023247680.1;refGene_gn=ENAH;refGene_tx=XM_023247681.1;refGene_gn=ENAH;refGene_tx=XM_023247682.1
F1 3811795 3812890
refGene_gn=LOC111558368;refGene_tx=XM_023247683.1;refGene_gn=LOC111558368;refGene_tx=XM_023247684.1
F1 3839510 3839575
refGene_gn=ENAH;refGene_tx=XM_023247677.1;refGene_gn=ENAH;refGene_tx=XM_023247678.1;refGene_gn=ENAH;refGene_tx=XM_023247679.1;refGene_gn=ENAH;refGene_tx=XM_023247681.1;refGene_gn=ENAH;refGene_tx=XM_023247682.1

F1 3839818 3839823 refGene_gn=ENAH;refGene_tx=XM_023247680.1
F1 3921733 3921805
refGene_gn=SRP9;refGene_tx=XM_003999087.5;refGene_gn=SRP9;refGene_tx=XM_006942597.4
F1 3926782 3926851
refGene_gn=SRP9;refGene_tx=XM_003999087.5;refGene_gn=SRP9;refGene_tx=XM_006942597.4
F1 3930794 3930914
refGene_gn=SRP9;refGene_tx=XM_003999087.5;refGene_gn=SRP9;refGene_tx=XM_006942597.4
F1 3965941 3966124
refGene_gn=EPHX1;refGene_tx=XM_011290813.3;refGene_gn=EPHX1;refGene_tx=XM_019821451.2;refGene_gn=EPHX1;refGene_tx=XM_023247923.1
F1 3967908 3968089
refGene_gn=EPHX1;refGene_tx=XM_011290813.3;refGene_gn=EPHX1;refGene_tx=XM_019821451.2;refGene_gn=EPHX1;refGene_tx=XM_023247923.1
F1 3973590 3973818
refGene_gn=EPHX1;refGene_tx=XM_011290813.3;refGene_gn=EPHX1;refGene_tx=XM_019821451.2;refGene_gn=EPHX1;refGene_tx=XM_023247923.1
F1 3974289 3974419
refGene_gn=EPHX1;refGene_tx=XM_011290813.3;refGene_gn=EPHX1;refGene_tx=XM_019821451.2;refGene_gn=EPHX1;refGene_tx=XM_023247923.1
F1 3974839 3975045
refGene_gn=EPHX1;refGene_tx=XM_011290813.3;refGene_gn=EPHX1;refGene_tx=XM_019821451.2;refGene_gn=EPHX1;refGene_tx=XM_023247923.1
F1 3976541 3976650
refGene_gn=EPHX1;refGene_tx=XM_011290813.3;refGene_gn=EPHX1;refGene_tx=XM_019821451.2;refGene_gn=EPHX1;refGene_tx=XM_023247923.1
F1 3979679 3979796
refGene_gn=EPHX1;refGene_tx=XM_011290813.3;refGene_gn=EPHX1;refGene_tx=XM_019821451.2;refGene_gn=EPHX1;refGene_tx=XM_023247923.1
F1 3980240 3980442
refGene_gn=EPHX1;refGene_tx=XM_011290813.3;refGene_gn=EPHX1;refGene_tx=XM_019821451.2;refGene_gn=EPHX1;refGene_tx=XM_023247923.1
F1 3981866 3982046
refGene_gn=TMEM63A;refGene_tx=XM_023247918.1;refGene_gn=TMEM63A;refGene_tx=XM_023247919.1;refGene_gn=TMEM63A;refGene_tx=XM_023247920.1;refGene_gn=TMEM63A;refGene_tx=XM_023247921.1;refGene_gn=TMEM63A;refGene_tx=XM_023247922.1
F1 3982549 3982612
refGene_gn=TMEM63A;refGene_tx=XM_023247918.1;refGene_gn=TMEM63A;refGene_tx=XM_023247919.1;refGene_gn=TMEM63A;refGene_tx=XM_023247920.1;refGene_gn=TMEM63A;refGene_tx=XM_023247921.1;refGene_gn=TMEM63A;refGene_tx=XM_023247922.1
F1 3982843 3982959
refGene_gn=TMEM63A;refGene_tx=XM_023247918.1;refGene_gn=TMEM63A;refGene_tx=XM_023247919.1;refGene_gn=TMEM63A;refGene_tx=XM_023247920.1;refGene_gn=TMEM63A;refGene_tx=XM_023247921.1;refGene_gn=TMEM63A;refGene_tx=XM_023247922.1
F1 3983066 3983154 refGene_gn=TMEM63A;refGene_tx=XM_023247919.1

F1	3983621	3983789
refGene_gn=TMEM63A;refGene_tx=XM_023247918.1;refGene_gn=TMEM63A;refGene_tx=XM_023247919.1;refGene_gn=TMEM63A;refGene_tx=XM_023247920.1;refGene_gn=TMEM63A;refGene_tx=XM_023247921.1;refGene_gn=TMEM63A;refGene_tx=XM_023247922.1
F1	3985150	3985256
refGene_gn=TMEM63A;refGene_tx=XM_023247918.1;refGene_gn=TMEM63A;refGene_tx=XM_023247919.1;refGene_gn=TMEM63A;refGene_tx=XM_023247920.1;refGene_gn=TMEM63A;refGene_tx=XM_023247921.1;refGene_gn=TMEM63A;refGene_tx=XM_023247922.1
F1	3985609	3985772
refGene_gn=TMEM63A;refGene_tx=XM_023247918.1;refGene_gn=TMEM63A;refGene_tx=XM_023247919.1;refGene_gn=TMEM63A;refGene_tx=XM_023247920.1;refGene_gn=TMEM63A;refGene_tx=XM_023247921.1;refGene_gn=TMEM63A;refGene_tx=XM_023247922.1
F1	3987404	3987467
refGene_gn=TMEM63A;refGene_tx=XM_023247918.1;refGene_gn=TMEM63A;refGene_tx=XM_023247919.1;refGene_gn=TMEM63A;refGene_tx=XM_023247920.1;refGene_gn=TMEM63A;refGene_tx=XM_023247921.1;refGene_gn=TMEM63A;refGene_tx=XM_023247922.1
F1	3987878	3987965
refGene_gn=TMEM63A;refGene_tx=XM_023247918.1;refGene_gn=TMEM63A;refGene_tx=XM_023247919.1;refGene_gn=TMEM63A;refGene_tx=XM_023247920.1;refGene_gn=TMEM63A;refGene_tx=XM_023247921.1;refGene_gn=TMEM63A;refGene_tx=XM_023247922.1
F1	3988056	3988163
refGene_gn=TMEM63A;refGene_tx=XM_023247918.1;refGene_gn=TMEM63A;refGene_tx=XM_023247919.1;refGene_gn=TMEM63A;refGene_tx=XM_023247920.1;refGene_gn=TMEM63A;refGene_tx=XM_023247921.1;refGene_gn=TMEM63A;refGene_tx=XM_023247922.1
F1	3990505	3990659
refGene_gn=TMEM63A;refGene_tx=XM_023247918.1;refGene_gn=TMEM63A;refGene_tx=XM_023247919.1;refGene_gn=TMEM63A;refGene_tx=XM_023247920.1;refGene_gn=TMEM63A;refGene_tx=XM_023247921.1;refGene_gn=TMEM63A;refGene_tx=XM_023247922.1
F1	3991794	3991932
refGene_gn=TMEM63A;refGene_tx=XM_023247918.1;refGene_gn=TMEM63A;refGene_tx=XM_023247919.1;refGene_gn=TMEM63A;refGene_tx=XM_023247920.1;refGene_gn=TMEM63A;refGene_tx=XM_023247921.1;refGene_gn=TMEM63A;refGene_tx=XM_023247922.1
F1	3992925	3993059
refGene_gn=TMEM63A;refGene_tx=XM_023247918.1;refGene_gn=TMEM63A;refGene_tx=XM_023247919.1;refGene_gn=TMEM63A;refGene_tx=XM_023247920.1;refGene_gn=TMEM63A;refGene_tx=XM_023247921.1;refGene_gn=TMEM63A;refGene_tx=XM_023247922.1
F1	3993144	3993268
refGene_gn=TMEM63A;refGene_tx=XM_023247918.1;refGene_gn=TMEM63A;refGene_tx=XM_023247919.1;refGene_gn=TMEM63A;refGene_tx=XM_023247920.1;refGene_gn=TMEM63A;refGene_tx=XM_023247921.1;refGene_gn=TMEM63A;refGene_tx=XM_023247922.1

F1 3993713 3993794
refGene_gn=TMEM63A;refGene_tx=XM_023247918.1;refGene_gn=TMEM63A;refGene_tx=XM_023247919.1;refGene_gn=TMEM63A;refGene_tx=XM_023247920.1;refGene_gn=TMEM63A;refGene_tx=XM_023247921.1;refGene_gn=TMEM63A;refGene_tx=XM_023247922.1
F1 3995149 3995220
refGene_gn=TMEM63A;refGene_tx=XM_023247918.1;refGene_gn=TMEM63A;refGene_tx=XM_023247919.1;refGene_gn=TMEM63A;refGene_tx=XM_023247920.1;refGene_gn=TMEM63A;refGene_tx=XM_023247921.1;refGene_gn=TMEM63A;refGene_tx=XM_023247922.1
F1 3997057 3997166
refGene_gn=TMEM63A;refGene_tx=XM_023247918.1;refGene_gn=TMEM63A;refGene_tx=XM_023247919.1;refGene_gn=TMEM63A;refGene_tx=XM_023247920.1;refGene_gn=TMEM63A;refGene_tx=XM_023247921.1;refGene_gn=TMEM63A;refGene_tx=XM_023247922.1
F1 3997335 3997387
refGene_gn=TMEM63A;refGene_tx=XM_023247918.1;refGene_gn=TMEM63A;refGene_tx=XM_023247919.1;refGene_gn=TMEM63A;refGene_tx=XM_023247920.1;refGene_gn=TMEM63A;refGene_tx=XM_023247921.1;refGene_gn=TMEM63A;refGene_tx=XM_023247922.1
F1 3998549 3998692
refGene_gn=TMEM63A;refGene_tx=XM_023247918.1;refGene_gn=TMEM63A;refGene_tx=XM_023247919.1;refGene_gn=TMEM63A;refGene_tx=XM_023247920.1;refGene_gn=TMEM63A;refGene_tx=XM_023247921.1;refGene_gn=TMEM63A;refGene_tx=XM_023247922.1
F1 4001913 4001951
refGene_gn=TMEM63A;refGene_tx=XM_023247918.1;refGene_gn=TMEM63A;refGene_tx=XM_023247919.1;refGene_gn=TMEM63A;refGene_tx=XM_023247920.1;refGene_gn=TMEM63A;refGene_tx=XM_023247921.1
F1 4002800 4002867
refGene_gn=TMEM63A;refGene_tx=XM_023247918.1;refGene_gn=TMEM63A;refGene_tx=XM_023247919.1;refGene_gn=TMEM63A;refGene_tx=XM_023247920.1;refGene_gn=TMEM63A;refGene_tx=XM_023247921.1;refGene_gn=TMEM63A;refGene_tx=XM_023247922.1
F1 4004551 4004631
refGene_gn=TMEM63A;refGene_tx=XM_023247918.1;refGene_gn=TMEM63A;refGene_tx=XM_023247919.1;refGene_gn=TMEM63A;refGene_tx=XM_023247920.1;refGene_gn=TMEM63A;refGene_tx=XM_023247921.1;refGene_gn=TMEM63A;refGene_tx=XM_023247922.1
F1 4007019 4007219
refGene_gn=TMEM63A;refGene_tx=XM_023247918.1;refGene_gn=TMEM63A;refGene_tx=XM_023247919.1;refGene_gn=TMEM63A;refGene_tx=XM_023247920.1;refGene_gn=TMEM63A;refGene_tx=XM_023247921.1
F1 4011433 4011521
refGene_gn=TMEM63A;refGene_tx=XM_023247918.1;refGene_gn=TMEM63A;refGene_tx=XM_023247919.1;refGene_gn=TMEM63A;refGene_tx=XM_023247920.1
F1 4014239 4014603
refGene_gn=LOC101086029;refGene_tx=XM_023247924.1
F1 4014876 4015119
refGene_gn=LOC101086029;refGene_tx=XM_023247924.1

F1 4015537 4015784
refGene_gn=LOC101086029;refGene_tx=XM_023247924.1
F1 4016760 4016862
refGene_gn=LOC101086029;refGene_tx=XM_023247924.1
F1 4016864 4017009
refGene_gn=LOC101086029;refGene_tx=XM_023247924.1
F1 4035218 4035384 refGene_gn=PYCR2;refGene_tx=XM_023247605.1
F1 4035914 4036078 refGene_gn=PYCR2;refGene_tx=XM_023247605.1
F1 4036296 4036389 refGene_gn=PYCR2;refGene_tx=XM_023247605.1
F1 4036566 4036788 refGene_gn=PYCR2;refGene_tx=XM_023247605.1
F1 4036885 4037065 refGene_gn=PYCR2;refGene_tx=XM_023247605.1
F1 4037790 4037861 refGene_gn=PYCR2;refGene_tx=XM_023247605.1
F1 4037941 4038008 refGene_gn=PYCR2;refGene_tx=XM_023247605.1
F1 4044356 4044720
refGene_gn=LOC105261327;refGene_tx=XM_023247870.1
F1 4044993 4045236
refGene_gn=LOC105261327;refGene_tx=XM_023247870.1
F1 4045855 4046102
refGene_gn=LOC105261327;refGene_tx=XM_023247870.1
F1 4047105 4047355
refGene_gn=LOC105261327;refGene_tx=XM_023247870.1
F1 4071604 4071826 refGene_gn=SDE2;refGene_tx=XM_023247502.1
F1 4074521 4075017 refGene_gn=SDE2;refGene_tx=XM_023247502.1
F1 4076839 4076954 refGene_gn=SDE2;refGene_tx=XM_023247502.1
F1 4077677 4077847 refGene_gn=SDE2;refGene_tx=XM_023247502.1
F1 4078090 4078202 refGene_gn=SDE2;refGene_tx=XM_023247502.1
F1 4079861 4079979 refGene_gn=SDE2;refGene_tx=XM_023247502.1
F1 4085631 4085751 refGene_gn=SDE2;refGene_tx=XM_023247502.1
F1 4124912 4125040 refGene_gn=H3F3A;refGene_tx=XM_023247314.1
F1 4126270 4126424 refGene_gn=H3F3A;refGene_tx=XM_023247314.1
F1 4130834 4130963 refGene_gn=H3F3A;refGene_tx=XM_023247314.1
F1 4143225 4143276
refGene_gn=LOC111558411;refGene_tx=XM_023248003.1
F1 4143593 4143743
refGene_gn=LOC111558411;refGene_tx=XM_023248003.1
F1 4148151 4148277
refGene_gn=LOC111558411;refGene_tx=XM_023248003.1
F1 4148627 4148858
refGene_gn=LOC111558411;refGene_tx=XM_023248003.1
F1 4149596 4149694
refGene_gn=LOC111558411;refGene_tx=XM_023248003.1
F1 4157441 4157882
refGene_gn=LOC111558411;refGene_tx=XM_023248003.1
F1 4158444 4158500
refGene_gn=LOC111558411;refGene_tx=XM_023248003.1
F1 4158711 4158776
refGene_gn=LOC111558411;refGene_tx=XM_023248003.1
F1 4184658 4184870
refGene_gn=ACBD3;refGene_tx=XM_023248105.1;refGene_gn=ACBD3;refGene_tx
=XM_023248106.1

F1 4188180 4188465
refGene_gn=ACBD3;refGene_tx=XM_023248105.1;refGene_gn=ACBD3;refGene_tx
=XM_023248106.1
F1 4190771 4190958
refGene_gn=ACBD3;refGene_tx=XM_023248105.1;refGene_gn=ACBD3;refGene_tx
=XM_023248106.1
F1 4193476 4193651
refGene_gn=ACBD3;refGene_tx=XM_023248105.1;refGene_gn=ACBD3;refGene_tx
=XM_023248106.1
F1 4195155 4195314
refGene_gn=ACBD3;refGene_tx=XM_023248105.1;refGene_gn=ACBD3;refGene_tx
=XM_023248106.1
F1 4198611 4198752
refGene_gn=ACBD3;refGene_tx=XM_023248105.1;refGene_gn=ACBD3;refGene_tx
=XM_023248106.1
F1 4199405 4199547
refGene_gn=ACBD3;refGene_tx=XM_023248105.1;refGene_gn=ACBD3;refGene_tx
=XM_023248106.1
F1 4210545 4210642 refGene_gn=ACBD3;refGene_tx=XM_023248106.1
F1 4215666 4215952 refGene_gn=ACBD3;refGene_tx=XM_023248105.1
F1 4238812 4239097
refGene_gn=MIXL1;refGene_tx=XM_023247558.1;refGene_gn=MIXL1;refGene_tx
=XM_023247559.1
F1 4240332 4240662
refGene_gn=MIXL1;refGene_tx=XM_023247558.1;refGene_gn=MIXL1;refGene_tx
=XM_023247559.1
F1 4246741 4246847
refGene_gn=LIN9;refGene_tx=XM_019821456.2;refGene_gn=LIN9;refGene_tx=X
M_023247321.1;refGene_gn=LIN9;refGene_tx=XM_023247322.1;refGene_gn=LIN
9;refGene_tx=XM_023247323.1;refGene_gn=LIN9;refGene_tx=XM_023247324.1;
refGene_gn=LIN9;refGene_tx=XM_023247325.1;refGene_gn=LIN9;refGene_tx=X
M_023247326.1;refGene_gn=LIN9;refGene_tx=XM_023247327.1;refGene_gn=LIN
9;refGene_tx=XM_023247328.1
F1 4247306 4247404
refGene_gn=LIN9;refGene_tx=XM_019821456.2;refGene_gn=LIN9;refGene_tx=X
M_023247321.1;refGene_gn=LIN9;refGene_tx=XM_023247322.1;refGene_gn=LIN
9;refGene_tx=XM_023247323.1;refGene_gn=LIN9;refGene_tx=XM_023247324.1;
refGene_gn=LIN9;refGene_tx=XM_023247325.1;refGene_gn=LIN9;refGene_tx=X
M_023247326.1;refGene_gn=LIN9;refGene_tx=XM_023247327.1;refGene_gn=LIN
9;refGene_tx=XM_023247328.1
F1 4247562 4247742
refGene_gn=LIN9;refGene_tx=XM_019821456.2;refGene_gn=LIN9;refGene_tx=X
M_023247321.1;refGene_gn=LIN9;refGene_tx=XM_023247322.1;refGene_gn=LIN
9;refGene_tx=XM_023247323.1;refGene_gn=LIN9;refGene_tx=XM_023247324.1;
refGene_gn=LIN9;refGene_tx=XM_023247325.1;refGene_gn=LIN9;refGene_tx=X
M_023247326.1;refGene_gn=LIN9;refGene_tx=XM_023247327.1;refGene_gn=LIN
9;refGene_tx=XM_023247328.1
F1 4251923 4252049
refGene_gn=LIN9;refGene_tx=XM_019821456.2;refGene_gn=LIN9;refGene_tx=X
M_023247321.1;refGene_gn=LIN9;refGene_tx=XM_023247322.1;refGene_gn=LIN

9;refGene_tx=XM_023247323.1;refGene_gn=LIN9;refGene_tx=XM_023247324.1;
refGene_gn=LIN9;refGene_tx=XM_023247325.1;refGene_gn=LIN9;refGene_tx=X
M_023247326.1;refGene_gn=LIN9;refGene_tx=XM_023247327.1;refGene_gn=LIN
9;refGene_tx=XM_023247328.1
F1 4256664 4256850 refGene_gn=LIN9;refGene_tx=XM_019821458.2
F1 4268086 4268167
refGene_gn=LIN9;refGene_tx=XM_019821456.2;refGene_gn=LIN9;refGene_tx=X
M_019821458.2;refGene_gn=LIN9;refGene_tx=XM_023247321.1;refGene_gn=LIN
9;refGene_tx=XM_023247322.1;refGene_gn=LIN9;refGene_tx=XM_023247323.1;
refGene_gn=LIN9;refGene_tx=XM_023247324.1;refGene_gn=LIN9;refGene_tx=X
M_023247325.1;refGene_gn=LIN9;refGene_tx=XM_023247326.1;refGene_gn=LIN
9;refGene_tx=XM_023247327.1;refGene_gn=LIN9;refGene_tx=XM_023247328.1
F1 4272169 4272271
refGene_gn=LIN9;refGene_tx=XM_019821456.2;refGene_gn=LIN9;refGene_tx=X
M_019821458.2;refGene_gn=LIN9;refGene_tx=XM_023247321.1;refGene_gn=LIN
9;refGene_tx=XM_023247322.1;refGene_gn=LIN9;refGene_tx=XM_023247323.1;
refGene_gn=LIN9;refGene_tx=XM_023247324.1;refGene_gn=LIN9;refGene_tx=X
M_023247325.1;refGene_gn=LIN9;refGene_tx=XM_023247326.1;refGene_gn=LIN
9;refGene_tx=XM_023247327.1;refGene_gn=LIN9;refGene_tx=XM_023247328.1
F1 4272555 4272675
refGene_gn=LIN9;refGene_tx=XM_019821456.2;refGene_gn=LIN9;refGene_tx=X
M_019821458.2;refGene_gn=LIN9;refGene_tx=XM_023247321.1;refGene_gn=LIN
9;refGene_tx=XM_023247322.1;refGene_gn=LIN9;refGene_tx=XM_023247323.1;
refGene_gn=LIN9;refGene_tx=XM_023247324.1;refGene_gn=LIN9;refGene_tx=X
M_023247325.1;refGene_gn=LIN9;refGene_tx=XM_023247326.1;refGene_gn=LIN
9;refGene_tx=XM_023247327.1;refGene_gn=LIN9;refGene_tx=XM_023247328.1
F1 4274597 4274731
refGene_gn=LIN9;refGene_tx=XM_019821456.2;refGene_gn=LIN9;refGene_tx=X
M_019821458.2;refGene_gn=LIN9;refGene_tx=XM_023247321.1;refGene_gn=LIN
9;refGene_tx=XM_023247322.1;refGene_gn=LIN9;refGene_tx=XM_023247323.1;
refGene_gn=LIN9;refGene_tx=XM_023247324.1;refGene_gn=LIN9;refGene_tx=X
M_023247325.1;refGene_gn=LIN9;refGene_tx=XM_023247326.1;refGene_gn=LIN
9;refGene_tx=XM_023247327.1;refGene_gn=LIN9;refGene_tx=XM_023247328.1
F1 4278935 4279093
refGene_gn=LIN9;refGene_tx=XM_019821456.2;refGene_gn=LIN9;refGene_tx=X
M_019821458.2;refGene_gn=LIN9;refGene_tx=XM_023247321.1;refGene_gn=LIN
9;refGene_tx=XM_023247322.1;refGene_gn=LIN9;refGene_tx=XM_023247323.1;
refGene_gn=LIN9;refGene_tx=XM_023247324.1;refGene_gn=LIN9;refGene_tx=X
M_023247325.1;refGene_gn=LIN9;refGene_tx=XM_023247326.1;refGene_gn=LIN
9;refGene_tx=XM_023247327.1;refGene_gn=LIN9;refGene_tx=XM_023247328.1
F1 4287365 4287491
refGene_gn=LIN9;refGene_tx=XM_019821456.2;refGene_gn=LIN9;refGene_tx=X
M_019821458.2;refGene_gn=LIN9;refGene_tx=XM_023247321.1;refGene_gn=LIN
9;refGene_tx=XM_023247322.1;refGene_gn=LIN9;refGene_tx=XM_023247323.1;
refGene_gn=LIN9;refGene_tx=XM_023247324.1;refGene_gn=LIN9;refGene_tx=X
M_023247325.1;refGene_gn=LIN9;refGene_tx=XM_023247326.1;refGene_gn=LIN
9;refGene_tx=XM_023247327.1;refGene_gn=LIN9;refGene_tx=XM_023247328.1
F1 4289120 4289254
refGene_gn=LIN9;refGene_tx=XM_019821456.2;refGene_gn=LIN9;refGene_tx=X
M_019821458.2;refGene_gn=LIN9;refGene_tx=XM_023247321.1;refGene_gn=LIN

9;refGene_tx=XM_023247322.1;refGene_gn=LIN9;refGene_tx=XM_023247323.1;refGene_gn=LIN9;refGene_tx=XM_023247324.1;refGene_gn=LIN9;refGene_tx=XM_023247325.1;refGene_gn=LIN9;refGene_tx=XM_023247326.1;refGene_gn=LIN9;refGene_tx=XM_023247327.1;refGene_gn=LIN9;refGene_tx=XM_023247328.1
F1 4299096 4299201 refGene_gn=LIN9;refGene_tx=XM_019821456.2;refGene_gn=LIN9;refGene_tx=XM_019821458.2;refGene_gn=LIN9;refGene_tx=XM_023247321.1;refGene_gn=LIN9;refGene_tx=XM_023247322.1;refGene_gn=LIN9;refGene_tx=XM_023247323.1;refGene_gn=LIN9;refGene_tx=XM_023247324.1;refGene_gn=LIN9;refGene_tx=XM_023247325.1;refGene_gn=LIN9;refGene_tx=XM_023247326.1;refGene_gn=LIN9;refGene_tx=XM_023247327.1;refGene_gn=LIN9;refGene_tx=XM_023247328.1
F1 4300444 4300539 refGene_gn=LIN9;refGene_tx=XM_019821456.2;refGene_gn=LIN9;refGene_tx=XM_019821458.2;refGene_gn=LIN9;refGene_tx=XM_023247321.1;refGene_gn=LIN9;refGene_tx=XM_023247322.1;refGene_gn=LIN9;refGene_tx=XM_023247323.1;refGene_gn=LIN9;refGene_tx=XM_023247324.1;refGene_gn=LIN9;refGene_tx=XM_023247328.1
F1 4302469 4302502 refGene_gn=LIN9;refGene_tx=XM_019821456.2;refGene_gn=LIN9;refGene_tx=XM_019821458.2;refGene_gn=LIN9;refGene_tx=XM_023247321.1
F1 4307261 4307538 refGene_gn=LIN9;refGene_tx=XM_019821456.2;refGene_gn=LIN9;refGene_tx=XM_019821458.2
F1 4309859 4309938 refGene_gn=LIN9;refGene_tx=XM_023247321.1
F1 4336735 4336817 refGene_gn=PARP1;refGene_tx=XM_003999089.5
F1 4337060 4337175 refGene_gn=PARP1;refGene_tx=XM_003999089.5
F1 4337732 4337794 refGene_gn=PARP1;refGene_tx=XM_003999089.5
F1 4338441 4338569 refGene_gn=PARP1;refGene_tx=XM_003999089.5
F1 4338923 4339076 refGene_gn=PARP1;refGene_tx=XM_003999089.5
F1 4339471 4339570 refGene_gn=PARP1;refGene_tx=XM_003999089.5
F1 4340296 4340425 refGene_gn=PARP1;refGene_tx=XM_003999089.5
F1 4341245 4341368 refGene_gn=PARP1;refGene_tx=XM_003999089.5
F1 4342754 4342838 refGene_gn=PARP1;refGene_tx=XM_003999089.5
F1 4344969 4345098 refGene_gn=PARP1;refGene_tx=XM_003999089.5
F1 4348341 4348537 refGene_gn=PARP1;refGene_tx=XM_003999089.5
F1 4350173 4350306 refGene_gn=PARP1;refGene_tx=XM_003999089.5
F1 4350526 4350595 refGene_gn=PARP1;refGene_tx=XM_003999089.5
F1 4350749 4350992 refGene_gn=PARP1;refGene_tx=XM_003999089.5
F1 4353094 4353235 refGene_gn=PARP1;refGene_tx=XM_003999089.5
F1 4353821 4353966 refGene_gn=PARP1;refGene_tx=XM_003999089.5
F1 4356544 4356721 refGene_gn=PARP1;refGene_tx=XM_003999089.5
F1 4357135 4357252 refGene_gn=PARP1;refGene_tx=XM_003999089.5
F1 4358906 4359006 refGene_gn=PARP1;refGene_tx=XM_003999089.5
F1 4361246 4361461 refGene_gn=PARP1;refGene_tx=XM_003999089.5
F1 4362492 4362608 refGene_gn=PARP1;refGene_tx=XM_003999089.5
F1 4370886 4371052 refGene_gn=PARP1;refGene_tx=XM_003999089.5
F1 4375337 4375391 refGene_gn=PARP1;refGene_tx=XM_003999089.5
F1 4375392 4375458 refGene_gn=PARP1;refGene_tx=XM_003999089.5
F1 4445486 4445688 refGene_gn=STUM;refGene_tx=XM_023247311.1
F1 4493841 4493886 refGene_gn=STUM;refGene_tx=XM_023247311.1

F1	4493888	4494020	refGene_gn=STUM;refGene_tx=XM_023247311.1
F1	4501087	4501096	refGene_gn=STUM;refGene_tx=XM_023247311.1
F1	4502065	4502100	refGene_gn=STUM;refGene_tx=XM_023247311.1
F1	4519268	4519484	refGene_gn=ITPKB;refGene_tx=XM_023247925.1
F1	4520892	4520964	refGene_gn=ITPKB;refGene_tx=XM_023247925.1
F1	4522423	4522525	refGene_gn=ITPKB;refGene_tx=XM_023247925.1
F1	4524420	4524625	refGene_gn=ITPKB;refGene_tx=XM_023247925.1
F1	4529048	4529262	refGene_gn=ITPKB;refGene_tx=XM_023247925.1
F1	4530506	4530606	refGene_gn=ITPKB;refGene_tx=XM_023247925.1
F1	4603138	4604061	refGene_gn=ITPKB;refGene_tx=XM_023247925.1
F1	4604063	4605033	refGene_gn=ITPKB;refGene_tx=XM_023247925.1
F1	4696586	4696727
refGene_gn=PSEN2;refGene_tx=XM_019821462.2;refGene_gn=PSEN2;refGene_tx
=XM_019821463.2;refGene_gn=PSEN2;refGene_tx=XM_019821464.2;refGene_gn=
PSEN2;refGene_tx=XM_023247308.1;refGene_gn=PSEN2;refGene_tx=XM_0232473
09.1
F1	4698361	4698576
refGene_gn=PSEN2;refGene_tx=XM_019821462.2;refGene_gn=PSEN2;refGene_tx
=XM_019821463.2;refGene_gn=PSEN2;refGene_tx=XM_019821464.2;refGene_gn=
PSEN2;refGene_tx=XM_023247308.1;refGene_gn=PSEN2;refGene_tx=XM_0232473
09.1
F1	4699257	4699450
refGene_gn=PSEN2;refGene_tx=XM_019821462.2;refGene_gn=PSEN2;refGene_tx
=XM_019821463.2;refGene_gn=PSEN2;refGene_tx=XM_019821464.2;refGene_gn=
PSEN2;refGene_tx=XM_023247308.1;refGene_gn=PSEN2;refGene_tx=XM_0232473
09.1
F1	4700267	4700335
refGene_gn=PSEN2;refGene_tx=XM_019821462.2;refGene_gn=PSEN2;refGene_tx
=XM_019821463.2;refGene_gn=PSEN2;refGene_tx=XM_019821464.2;refGene_gn=
PSEN2;refGene_tx=XM_023247308.1;refGene_gn=PSEN2;refGene_tx=XM_0232473
09.1
F1	4701227	4701448
refGene_gn=PSEN2;refGene_tx=XM_019821462.2;refGene_gn=PSEN2;refGene_tx
=XM_019821463.2;refGene_gn=PSEN2;refGene_tx=XM_019821464.2;refGene_gn=
PSEN2;refGene_tx=XM_023247308.1;refGene_gn=PSEN2;refGene_tx=XM_0232473
09.1
F1	4702088	4702187
refGene_gn=PSEN2;refGene_tx=XM_019821462.2;refGene_gn=PSEN2;refGene_tx
=XM_019821463.2;refGene_gn=PSEN2;refGene_tx=XM_019821464.2;refGene_gn=
PSEN2;refGene_tx=XM_023247308.1;refGene_gn=PSEN2;refGene_tx=XM_0232473
09.1
F1	4704521	4704605
refGene_gn=PSEN2;refGene_tx=XM_019821462.2;refGene_gn=PSEN2;refGene_tx
=XM_019821463.2;refGene_gn=PSEN2;refGene_tx=XM_019821464.2;refGene_gn=
PSEN2;refGene_tx=XM_023247308.1;refGene_gn=PSEN2;refGene_tx=XM_0232473
09.1
F1	4704975	4705077
refGene_gn=PSEN2;refGene_tx=XM_019821462.2;refGene_gn=PSEN2;refGene_tx
=XM_019821463.2;refGene_gn=PSEN2;refGene_tx=XM_019821464.2;refGene_gn=
PSEN2;refGene_tx=XM_023247308.1;refGene_gn=PSEN2;refGene_tx=XM_0232473

09.1
F1 4705781 4705900
refGene_gn=PSEN2;refGene_tx=XM_019821462.2;refGene_gn=PSEN2;refGene_tx=XM_019821463.2;refGene_gn=PSEN2;refGene_tx=XM_019821464.2;refGene_gn=PSEN2;refGene_tx=XM_023247308.1;refGene_gn=PSEN2;refGene_tx=XM_023247309.1
F1 4707217 4707373
refGene_gn=PSEN2;refGene_tx=XM_019821462.2;refGene_gn=PSEN2;refGene_tx=XM_019821463.2;refGene_gn=PSEN2;refGene_tx=XM_019821464.2;refGene_gn=PSEN2;refGene_tx=XM_023247308.1;refGene_gn=PSEN2;refGene_tx=XM_023247309.1
F1 4755443 4755620
refGene_gn=COQ8A;refGene_tx=XM_023247952.1;refGene_gn=COQ8A;refGene_tx=XM_023247953.1;refGene_gn=COQ8A;refGene_tx=XM_023247954.1
F1 4757913 4758318
refGene_gn=COQ8A;refGene_tx=XM_023247952.1;refGene_gn=COQ8A;refGene_tx=XM_023247953.1;refGene_gn=COQ8A;refGene_tx=XM_023247954.1
F1 4758566 4758633
refGene_gn=COQ8A;refGene_tx=XM_023247952.1;refGene_gn=COQ8A;refGene_tx=XM_023247953.1;refGene_gn=COQ8A;refGene_tx=XM_023247954.1
F1 4768716 4768791
refGene_gn=COQ8A;refGene_tx=XM_023247952.1;refGene_gn=COQ8A;refGene_tx=XM_023247953.1;refGene_gn=COQ8A;refGene_tx=XM_023247954.1
F1 4771579 4771702
refGene_gn=COQ8A;refGene_tx=XM_023247952.1;refGene_gn=COQ8A;refGene_tx=XM_023247953.1;refGene_gn=COQ8A;refGene_tx=XM_023247954.1
F1 4772087 4772173
refGene_gn=COQ8A;refGene_tx=XM_023247952.1;refGene_gn=COQ8A;refGene_tx=XM_023247953.1;refGene_gn=COQ8A;refGene_tx=XM_023247954.1
F1 4772258 4772399
refGene_gn=COQ8A;refGene_tx=XM_023247952.1;refGene_gn=COQ8A;refGene_tx=XM_023247953.1;refGene_gn=COQ8A;refGene_tx=XM_023247954.1
F1 4772787 4772869
refGene_gn=COQ8A;refGene_tx=XM_023247952.1;refGene_gn=COQ8A;refGene_tx=XM_023247953.1;refGene_gn=COQ8A;refGene_tx=XM_023247954.1
F1 4772966 4773060
refGene_gn=COQ8A;refGene_tx=XM_023247952.1;refGene_gn=COQ8A;refGene_tx=XM_023247953.1;refGene_gn=COQ8A;refGene_tx=XM_023247954.1
F1 4773174 4773316
refGene_gn=COQ8A;refGene_tx=XM_023247952.1;refGene_gn=COQ8A;refGene_tx=XM_023247953.1;refGene_gn=COQ8A;refGene_tx=XM_023247954.1
F1 4773470 4773578
refGene_gn=COQ8A;refGene_tx=XM_023247952.1;refGene_gn=COQ8A;refGene_tx=XM_023247953.1;refGene_gn=COQ8A;refGene_tx=XM_023247954.1
F1 4773665 4773731
refGene_gn=COQ8A;refGene_tx=XM_023247952.1;refGene_gn=COQ8A;refGene_tx=XM_023247953.1;refGene_gn=COQ8A;refGene_tx=XM_023247954.1
F1 4773969 4774077
refGene_gn=COQ8A;refGene_tx=XM_023247952.1;refGene_gn=COQ8A;refGene_tx=XM_023247953.1;refGene_gn=COQ8A;refGene_tx=XM_023247954.1

F1 4774436 4774721
refGene_gn=COQ8A;refGene_tx=XM_023247952.1;refGene_gn=COQ8A;refGene_tx=XM_023247953.1
F1 4780111 4780243
refGene_gn=CDC42BPA;refGene_tx=XM_011290816.3;refGene_gn=CDC42BPA;refGene_tx=XM_011290817.3;refGene_gn=CDC42BPA;refGene_tx=XM_011290818.3;refGene_gn=CDC42BPA;refGene_tx=XM_011290819.3;refGene_gn=CDC42BPA;refGene_tx=XM_011290820.3;refGene_gn=CDC42BPA;refGene_tx=XM_011290821.3;refGene_gn=CDC42BPA;refGene_tx=XM_019821465.2;refGene_gn=CDC42BPA;refGene_tx=XM_019821467.2;refGene_gn=CDC42BPA;refGene_tx=XM_019821468.2;refGene_gn=CDC42BPA;refGene_tx=XM_019821469.2;refGene_gn=CDC42BPA;refGene_tx=XM_019821470.2;refGene_gn=CDC42BPA;refGene_tx=XM_019821471.2;refGene_gn=CDC42BPA;refGene_tx=XM_019821472.2;refGene_gn=CDC42BPA;refGene_tx=XM_019821474.2
F1 4780728 4780886
refGene_gn=CDC42BPA;refGene_tx=XM_011290816.3;refGene_gn=CDC42BPA;refGene_tx=XM_011290817.3;refGene_gn=CDC42BPA;refGene_tx=XM_011290818.3;refGene_gn=CDC42BPA;refGene_tx=XM_011290819.3;refGene_gn=CDC42BPA;refGene_tx=XM_011290820.3;refGene_gn=CDC42BPA;refGene_tx=XM_011290821.3;refGene_gn=CDC42BPA;refGene_tx=XM_019821465.2;refGene_gn=CDC42BPA;refGene_tx=XM_019821467.2;refGene_gn=CDC42BPA;refGene_tx=XM_019821468.2;refGene_gn=CDC42BPA;refGene_tx=XM_019821469.2;refGene_gn=CDC42BPA;refGene_tx=XM_019821470.2;refGene_gn=CDC42BPA;refGene_tx=XM_019821471.2;refGene_gn=CDC42BPA;refGene_tx=XM_019821472.2;refGene_gn=CDC42BPA;refGene_tx=XM_019821474.2
F1 4784946 4785064
refGene_gn=CDC42BPA;refGene_tx=XM_011290816.3;refGene_gn=CDC42BPA;refGene_tx=XM_011290817.3;refGene_gn=CDC42BPA;refGene_tx=XM_011290818.3;refGene_gn=CDC42BPA;refGene_tx=XM_011290819.3;refGene_gn=CDC42BPA;refGene_tx=XM_011290820.3;refGene_gn=CDC42BPA;refGene_tx=XM_011290821.3;refGene_gn=CDC42BPA;refGene_tx=XM_019821465.2;refGene_gn=CDC42BPA;refGene_tx=XM_019821467.2;refGene_gn=CDC42BPA;refGene_tx=XM_019821468.2;refGene_gn=CDC42BPA;refGene_tx=XM_019821469.2;refGene_gn=CDC42BPA;refGene_tx=XM_019821470.2;refGene_gn=CDC42BPA;refGene_tx=XM_019821471.2;refGene_gn=CDC42BPA;refGene_tx=XM_019821472.2;refGene_gn=CDC42BPA;refGene_tx=XM_019821474.2
F1 4787804 4787987
refGene_gn=CDC42BPA;refGene_tx=XM_011290816.3;refGene_gn=CDC42BPA;refGene_tx=XM_011290817.3;refGene_gn=CDC42BPA;refGene_tx=XM_019821465.2;refGene_gn=CDC42BPA;refGene_tx=XM_019821467.2;refGene_gn=CDC42BPA;refGene_tx=XM_019821468.2;refGene_gn=CDC42BPA;refGene_tx=XM_019821469.2;refGene_gn=CDC42BPA;refGene_tx=XM_019821470.2;refGene_gn=CDC42BPA;refGene_tx=XM_019821471.2;refGene_gn=CDC42BPA;refGene_tx=XM_019821472.2
F1 4791182 4791300
refGene_gn=CDC42BPA;refGene_tx=XM_011290816.3;refGene_gn=CDC42BPA;refGene_tx=XM_011290817.3;refGene_gn=CDC42BPA;refGene_tx=XM_011290818.3;refGene_gn=CDC42BPA;refGene_tx=XM_011290819.3;refGene_gn=CDC42BPA;refGene_tx=XM_011290820.3;refGene_gn=CDC42BPA;refGene_tx=XM_011290821.3;refGene_gn=CDC42BPA;refGene_tx=XM_019821465.2;refGene_gn=CDC42BPA;refGene_tx=XM_019821467.2;refGene_gn=CDC42BPA;refGene_tx=XM_019821468.2;refGen

e_gn=CDC42BPA;refGene_tx=XM_019821469.2;refGene_gn=CDC42BPA;refGene_tx
=XM_019821470.2;refGene_gn=CDC42BPA;refGene_tx=XM_019821471.2;refGene_
gn=CDC42BPA;refGene_tx=XM_019821472.2;refGene_gn=CDC42BPA;refGene_tx=X
M_019821474.2
F1 4792018 4792142
refGene_gn=CDC42BPA;refGene_tx=XM_011290816.3;refGene_gn=CDC42BPA;refG
ene_tx=XM_011290817.3;refGene_gn=CDC42BPA;refGene_tx=XM_011290818.3;re
fGene_gn=CDC42BPA;refGene_tx=XM_011290819.3;refGene_gn=CDC42BPA;refGen
e_tx=XM_011290820.3;refGene_gn=CDC42BPA;refGene_tx=XM_011290821.3;refG
ene_gn=CDC42BPA;refGene_tx=XM_019821465.2;refGene_gn=CDC42BPA;refGene_
tx=XM_019821467.2;refGene_gn=CDC42BPA;refGene_tx=XM_019821468.2;refGen
e_gn=CDC42BPA;refGene_tx=XM_019821469.2;refGene_gn=CDC42BPA;refGene_tx
=XM_019821470.2;refGene_gn=CDC42BPA;refGene_tx=XM_019821471.2;refGene_
gn=CDC42BPA;refGene_tx=XM_019821472.2;refGene_gn=CDC42BPA;refGene_tx=X
M_019821474.2
F1 4794163 4794248
refGene_gn=CDC42BPA;refGene_tx=XM_011290816.3;refGene_gn=CDC42BPA;refG
ene_tx=XM_011290817.3;refGene_gn=CDC42BPA;refGene_tx=XM_011290818.3;re
fGene_gn=CDC42BPA;refGene_tx=XM_011290819.3;refGene_gn=CDC42BPA;refGen
e_tx=XM_011290820.3;refGene_gn=CDC42BPA;refGene_tx=XM_011290821.3;refG
ene_gn=CDC42BPA;refGene_tx=XM_019821465.2;refGene_gn=CDC42BPA;refGene_
tx=XM_019821467.2;refGene_gn=CDC42BPA;refGene_tx=XM_019821468.2;refGen
e_gn=CDC42BPA;refGene_tx=XM_019821469.2;refGene_gn=CDC42BPA;refGene_tx
=XM_019821470.2;refGene_gn=CDC42BPA;refGene_tx=XM_019821471.2;refGene_
gn=CDC42BPA;refGene_tx=XM_019821472.2;refGene_gn=CDC42BPA;refGene_tx=X
M_019821474.2
F1 4796889 4796987
refGene_gn=CDC42BPA;refGene_tx=XM_011290816.3;refGene_gn=CDC42BPA;refG
ene_tx=XM_011290817.3;refGene_gn=CDC42BPA;refGene_tx=XM_011290818.3;re
fGene_gn=CDC42BPA;refGene_tx=XM_011290819.3;refGene_gn=CDC42BPA;refGen
e_tx=XM_011290820.3;refGene_gn=CDC42BPA;refGene_tx=XM_011290821.3;refG
ene_gn=CDC42BPA;refGene_tx=XM_019821465.2;refGene_gn=CDC42BPA;refGene_
tx=XM_019821467.2;refGene_gn=CDC42BPA;refGene_tx=XM_019821468.2;refGen
e_gn=CDC42BPA;refGene_tx=XM_019821469.2;refGene_gn=CDC42BPA;refGene_tx
=XM_019821470.2;refGene_gn=CDC42BPA;refGene_tx=XM_019821471.2;refGene_
gn=CDC42BPA;refGene_tx=XM_019821472.2;refGene_gn=CDC42BPA;refGene_tx=X
M_019821474.2
F1 4798837 4799431
refGene_gn=CDC42BPA;refGene_tx=XM_011290816.3;refGene_gn=CDC42BPA;refG
ene_tx=XM_011290817.3;refGene_gn=CDC42BPA;refGene_tx=XM_011290818.3;re
fGene_gn=CDC42BPA;refGene_tx=XM_011290819.3;refGene_gn=CDC42BPA;refGen
e_tx=XM_011290820.3;refGene_gn=CDC42BPA;refGene_tx=XM_011290821.3;refG
ene_gn=CDC42BPA;refGene_tx=XM_019821465.2;refGene_gn=CDC42BPA;refGene_
tx=XM_019821467.2;refGene_gn=CDC42BPA;refGene_tx=XM_019821468.2;refGen
e_gn=CDC42BPA;refGene_tx=XM_019821469.2;refGene_gn=CDC42BPA;refGene_tx
=XM_019821470.2;refGene_gn=CDC42BPA;refGene_tx=XM_019821471.2;refGene_
gn=CDC42BPA;refGene_tx=XM_019821472.2;refGene_gn=CDC42BPA;refGene_tx=X
M_019821474.2
F1 4800277 4800340
refGene_gn=CDC42BPA;refGene_tx=XM_011290816.3;refGene_gn=CDC42BPA;refG

ene_tx=XM_011290817.3;refGene_gn=CDC42BPA;refGene_tx=XM_011290818.3;refGene_gn=CDC42BPA;refGene_tx=XM_011290819.3;refGene_gn=CDC42BPA;refGene_tx=XM_011290820.3;refGene_gn=CDC42BPA;refGene_tx=XM_011290821.3;refGene_gn=CDC42BPA;refGene_tx=XM_019821465.2;refGene_gn=CDC42BPA;refGene_tx=XM_019821467.2;refGene_gn=CDC42BPA;refGene_tx=XM_019821468.2;refGene_gn=CDC42BPA;refGene_tx=XM_019821469.2;refGene_gn=CDC42BPA;refGene_tx=XM_019821470.2;refGene_gn=CDC42BPA;refGene_tx=XM_019821471.2;refGene_gn=CDC42BPA;refGene_tx=XM_019821472.2;refGene_gn=CDC42BPA;refGene_tx=XM_019821474.2

F1 4801160 4801377
refGene_gn=CDC42BPA;refGene_tx=XM_011290816.3;refGene_gn=CDC42BPA;refGene_tx=XM_011290817.3;refGene_gn=CDC42BPA;refGene_tx=XM_011290818.3;refGene_gn=CDC42BPA;refGene_tx=XM_011290819.3;refGene_gn=CDC42BPA;refGene_tx=XM_011290820.3;refGene_gn=CDC42BPA;refGene_tx=XM_011290821.3;refGene_gn=CDC42BPA;refGene_tx=XM_019821465.2;refGene_gn=CDC42BPA;refGene_tx=XM_019821467.2;refGene_gn=CDC42BPA;refGene_tx=XM_019821468.2;refGene_gn=CDC42BPA;refGene_tx=XM_019821469.2;refGene_gn=CDC42BPA;refGene_tx=XM_019821470.2;refGene_gn=CDC42BPA;refGene_tx=XM_019821471.2;refGene_gn=CDC42BPA;refGene_tx=XM_019821472.2;refGene_gn=CDC42BPA;refGene_tx=XM_019821474.2

F1 4803357 4803439
refGene_gn=CDC42BPA;refGene_tx=XM_011290816.3;refGene_gn=CDC42BPA;refGene_tx=XM_011290817.3;refGene_gn=CDC42BPA;refGene_tx=XM_011290818.3;refGene_gn=CDC42BPA;refGene_tx=XM_011290819.3;refGene_gn=CDC42BPA;refGene_tx=XM_011290820.3;refGene_gn=CDC42BPA;refGene_tx=XM_011290821.3;refGene_gn=CDC42BPA;refGene_tx=XM_019821465.2;refGene_gn=CDC42BPA;refGene_tx=XM_019821467.2;refGene_gn=CDC42BPA;refGene_tx=XM_019821468.2;refGene_gn=CDC42BPA;refGene_tx=XM_019821469.2;refGene_gn=CDC42BPA;refGene_tx=XM_019821470.2;refGene_gn=CDC42BPA;refGene_tx=XM_019821471.2;refGene_gn=CDC42BPA;refGene_tx=XM_019821472.2;refGene_gn=CDC42BPA;refGene_tx=XM_019821474.2

F1 4804684 4804824
refGene_gn=CDC42BPA;refGene_tx=XM_011290816.3;refGene_gn=CDC42BPA;refGene_tx=XM_011290817.3;refGene_gn=CDC42BPA;refGene_tx=XM_011290818.3;refGene_gn=CDC42BPA;refGene_tx=XM_011290819.3;refGene_gn=CDC42BPA;refGene_tx=XM_011290820.3;refGene_gn=CDC42BPA;refGene_tx=XM_011290821.3;refGene_gn=CDC42BPA;refGene_tx=XM_019821465.2;refGene_gn=CDC42BPA;refGene_tx=XM_019821467.2;refGene_gn=CDC42BPA;refGene_tx=XM_019821468.2;refGene_gn=CDC42BPA;refGene_tx=XM_019821469.2;refGene_gn=CDC42BPA;refGene_tx=XM_019821470.2;refGene_gn=CDC42BPA;refGene_tx=XM_019821471.2;refGene_gn=CDC42BPA;refGene_tx=XM_019821472.2;refGene_gn=CDC42BPA;refGene_tx=XM_019821474.2

F1 4805871 4806008
refGene_gn=CDC42BPA;refGene_tx=XM_011290816.3;refGene_gn=CDC42BPA;refGene_tx=XM_011290817.3;refGene_gn=CDC42BPA;refGene_tx=XM_011290818.3;refGene_gn=CDC42BPA;refGene_tx=XM_011290819.3;refGene_gn=CDC42BPA;refGene_tx=XM_011290820.3;refGene_gn=CDC42BPA;refGene_tx=XM_011290821.3;refGene_gn=CDC42BPA;refGene_tx=XM_019821465.2;refGene_gn=CDC42BPA;refGene_tx=XM_019821467.2;refGene_gn=CDC42BPA;refGene_tx=XM_019821468.2;refGene_gn=CDC42BPA;refGene_tx=XM_019821469.2;refGene_gn=CDC42BPA;refGene_tx

=XM_019821470.2;refGene_gn=CDC42BPA;refGene_tx=XM_019821471.2;refGene_
gn=CDC42BPA;refGene_tx=XM_019821472.2;refGene_gn=CDC42BPA;refGene_tx=X
M_019821474.2
F1 4808785 4808891
refGene_gn=CDC42BPA;refGene_tx=XM_011290816.3;refGene_gn=CDC42BPA;refG
ene_tx=XM_011290817.3;refGene_gn=CDC42BPA;refGene_tx=XM_011290818.3;re
fGene_gn=CDC42BPA;refGene_tx=XM_011290819.3;refGene_gn=CDC42BPA;refGen
e_tx=XM_011290820.3;refGene_gn=CDC42BPA;refGene_tx=XM_011290821.3;refG
ene_gn=CDC42BPA;refGene_tx=XM_019821465.2;refGene_gn=CDC42BPA;refGene_
tx=XM_019821467.2;refGene_gn=CDC42BPA;refGene_tx=XM_019821468.2;refGen
e_gn=CDC42BPA;refGene_tx=XM_019821469.2;refGene_gn=CDC42BPA;refGene_tx
=XM_019821470.2;refGene_gn=CDC42BPA;refGene_tx=XM_019821471.2;refGene_
gn=CDC42BPA;refGene_tx=XM_019821472.2;refGene_gn=CDC42BPA;refGene_tx=X
M_019821474.2
F1 4823381 4823465
refGene_gn=CDC42BPA;refGene_tx=XM_011290816.3;refGene_gn=CDC42BPA;refG
ene_tx=XM_011290817.3;refGene_gn=CDC42BPA;refGene_tx=XM_011290818.3;re
fGene_gn=CDC42BPA;refGene_tx=XM_011290819.3;refGene_gn=CDC42BPA;refGen
e_tx=XM_011290820.3;refGene_gn=CDC42BPA;refGene_tx=XM_011290821.3;refG
ene_gn=CDC42BPA;refGene_tx=XM_019821465.2;refGene_gn=CDC42BPA;refGene_
tx=XM_019821467.2;refGene_gn=CDC42BPA;refGene_tx=XM_019821468.2;refGen
e_gn=CDC42BPA;refGene_tx=XM_019821469.2;refGene_gn=CDC42BPA;refGene_tx
=XM_019821470.2;refGene_gn=CDC42BPA;refGene_tx=XM_019821471.2;refGene_
gn=CDC42BPA;refGene_tx=XM_019821472.2;refGene_gn=CDC42BPA;refGene_tx=X
M_019821474.2
F1 4834486 4834525
refGene_gn=CDC42BPA;refGene_tx=XM_011290816.3;refGene_gn=CDC42BPA;refG
ene_tx=XM_011290818.3;refGene_gn=CDC42BPA;refGene_tx=XM_011290819.3;re
fGene_gn=CDC42BPA;refGene_tx=XM_019821465.2;refGene_gn=CDC42BPA;refGen
e_tx=XM_019821468.2;refGene_gn=CDC42BPA;refGene_tx=XM_019821470.2;refG
ene_gn=CDC42BPA;refGene_tx=XM_019821471.2;refGene_gn=CDC42BPA;refGene_
tx=XM_019821472.2;refGene_gn=CDC42BPA;refGene_tx=XM_019821474.2
F1 4841013 4841061
refGene_gn=CDC42BPA;refGene_tx=XM_011290818.3;refGene_gn=CDC42BPA;refG
ene_tx=XM_019821465.2;refGene_gn=CDC42BPA;refGene_tx=XM_019821467.2;re
fGene_gn=CDC42BPA;refGene_tx=XM_019821468.2;refGene_gn=CDC42BPA;refGen
e_tx=XM_019821469.2;refGene_gn=CDC42BPA;refGene_tx=XM_019821471.2;refG
ene_gn=CDC42BPA;refGene_tx=XM_019821472.2
F1 4842120 4842197
refGene_gn=CDC42BPA;refGene_tx=XM_011290816.3;refGene_gn=CDC42BPA;refG
ene_tx=XM_011290817.3;refGene_gn=CDC42BPA;refGene_tx=XM_011290818.3;re
fGene_gn=CDC42BPA;refGene_tx=XM_011290819.3;refGene_gn=CDC42BPA;refGen
e_tx=XM_011290820.3;refGene_gn=CDC42BPA;refGene_tx=XM_011290821.3;refG
ene_gn=CDC42BPA;refGene_tx=XM_019821465.2;refGene_gn=CDC42BPA;refGene_
tx=XM_019821467.2;refGene_gn=CDC42BPA;refGene_tx=XM_019821471.2;refGen
e_gn=CDC42BPA;refGene_tx=XM_019821472.2
F1 4844432 4844524
refGene_gn=CDC42BPA;refGene_tx=XM_011290816.3;refGene_gn=CDC42BPA;refG
ene_tx=XM_011290817.3;refGene_gn=CDC42BPA;refGene_tx=XM_011290818.3;re
fGene_gn=CDC42BPA;refGene_tx=XM_011290819.3;refGene_gn=CDC42BPA;refGen

e_tx=XM_011290820.3;refGene_gn=CDC42BPA;refGene_tx=XM_011290821.3;refGene_gn=CDC42BPA;refGene_tx=XM_019821465.2;refGene_gn=CDC42BPA;refGene_tx=XM_019821467.2;refGene_gn=CDC42BPA;refGene_tx=XM_019821468.2;refGene_gn=CDC42BPA;refGene_tx=XM_019821469.2;refGene_gn=CDC42BPA;refGene_tx=XM_019821470.2;refGene_gn=CDC42BPA;refGene_tx=XM_019821471.2;refGene_gn=CDC42BPA;refGene_tx=XM_019821472.2;refGene_gn=CDC42BPA;refGene_tx=XM_019821474.2
F1 4845749 4845898
refGene_gn=CDC42BPA;refGene_tx=XM_011290816.3;refGene_gn=CDC42BPA;refGene_tx=XM_011290817.3;refGene_gn=CDC42BPA;refGene_tx=XM_011290818.3;refGene_gn=CDC42BPA;refGene_tx=XM_011290819.3;refGene_gn=CDC42BPA;refGene_tx=XM_011290820.3;refGene_gn=CDC42BPA;refGene_tx=XM_011290821.3;refGene_gn=CDC42BPA;refGene_tx=XM_019821465.2;refGene_gn=CDC42BPA;refGene_tx=XM_019821467.2;refGene_gn=CDC42BPA;refGene_tx=XM_019821468.2;refGene_gn=CDC42BPA;refGene_tx=XM_019821469.2;refGene_gn=CDC42BPA;refGene_tx=XM_019821470.2;refGene_gn=CDC42BPA;refGene_tx=XM_019821471.2;refGene_gn=CDC42BPA;refGene_tx=XM_019821472.2;refGene_gn=CDC42BPA;refGene_tx=XM_019821474.2
F1 4846283 4846389
refGene_gn=CDC42BPA;refGene_tx=XM_011290816.3;refGene_gn=CDC42BPA;refGene_tx=XM_011290817.3;refGene_gn=CDC42BPA;refGene_tx=XM_011290818.3;refGene_gn=CDC42BPA;refGene_tx=XM_011290819.3;refGene_gn=CDC42BPA;refGene_tx=XM_011290820.3;refGene_gn=CDC42BPA;refGene_tx=XM_011290821.3;refGene_gn=CDC42BPA;refGene_tx=XM_019821465.2;refGene_gn=CDC42BPA;refGene_tx=XM_019821467.2;refGene_gn=CDC42BPA;refGene_tx=XM_019821468.2;refGene_gn=CDC42BPA;refGene_tx=XM_019821469.2;refGene_gn=CDC42BPA;refGene_tx=XM_019821470.2;refGene_gn=CDC42BPA;refGene_tx=XM_019821471.2;refGene_gn=CDC42BPA;refGene_tx=XM_019821472.2;refGene_gn=CDC42BPA;refGene_tx=XM_019821474.2
F1 4863885 4864010
refGene_gn=CDC42BPA;refGene_tx=XM_011290816.3;refGene_gn=CDC42BPA;refGene_tx=XM_011290817.3;refGene_gn=CDC42BPA;refGene_tx=XM_011290818.3;refGene_gn=CDC42BPA;refGene_tx=XM_011290819.3;refGene_gn=CDC42BPA;refGene_tx=XM_011290820.3;refGene_gn=CDC42BPA;refGene_tx=XM_011290821.3;refGene_gn=CDC42BPA;refGene_tx=XM_019821465.2;refGene_gn=CDC42BPA;refGene_tx=XM_019821467.2;refGene_gn=CDC42BPA;refGene_tx=XM_019821468.2;refGene_gn=CDC42BPA;refGene_tx=XM_019821469.2;refGene_gn=CDC42BPA;refGene_tx=XM_019821470.2;refGene_gn=CDC42BPA;refGene_tx=XM_019821471.2;refGene_gn=CDC42BPA;refGene_tx=XM_019821472.2;refGene_gn=CDC42BPA;refGene_tx=XM_019821474.2
F1 4872951 4873057
refGene_gn=CDC42BPA;refGene_tx=XM_011290816.3;refGene_gn=CDC42BPA;refGene_tx=XM_011290817.3;refGene_gn=CDC42BPA;refGene_tx=XM_011290818.3;refGene_gn=CDC42BPA;refGene_tx=XM_011290819.3;refGene_gn=CDC42BPA;refGene_tx=XM_011290820.3;refGene_gn=CDC42BPA;refGene_tx=XM_011290821.3;refGene_gn=CDC42BPA;refGene_tx=XM_019821465.2;refGene_gn=CDC42BPA;refGene_tx=XM_019821467.2;refGene_gn=CDC42BPA;refGene_tx=XM_019821468.2;refGene_gn=CDC42BPA;refGene_tx=XM_019821469.2;refGene_gn=CDC42BPA;refGene_tx=XM_019821470.2;refGene_gn=CDC42BPA;refGene_tx=XM_019821471.2;refGene_gn=CDC42BPA;refGene_tx=XM_019821472.2;refGene_gn=CDC42BPA;refGene_tx=X

M_019821474.2
F1	4881520	4881768
refGene_gn=CDC42BPA;refGene_tx=XM_011290816.3;refGene_gn=CDC42BPA;refGene_tx=XM_011290817.3;refGene_gn=CDC42BPA;refGene_tx=XM_011290818.3;refGene_gn=CDC42BPA;refGene_tx=XM_011290819.3;refGene_gn=CDC42BPA;refGene_tx=XM_011290820.3;refGene_gn=CDC42BPA;refGene_tx=XM_011290821.3;refGene_gn=CDC42BPA;refGene_tx=XM_019821465.2;refGene_gn=CDC42BPA;refGene_tx=XM_019821467.2;refGene_gn=CDC42BPA;refGene_tx=XM_019821468.2;refGene_gn=CDC42BPA;refGene_tx=XM_019821469.2;refGene_gn=CDC42BPA;refGene_tx=XM_019821470.2;refGene_gn=CDC42BPA;refGene_tx=XM_019821471.2;refGene_gn=CDC42BPA;refGene_tx=XM_019821472.2;refGene_gn=CDC42BPA;refGene_tx=XM_019821474.2
F1	4899073	4899184
refGene_gn=CDC42BPA;refGene_tx=XM_011290816.3;refGene_gn=CDC42BPA;refGene_tx=XM_011290817.3;refGene_gn=CDC42BPA;refGene_tx=XM_011290818.3;refGene_gn=CDC42BPA;refGene_tx=XM_011290819.3;refGene_gn=CDC42BPA;refGene_tx=XM_011290820.3;refGene_gn=CDC42BPA;refGene_tx=XM_011290821.3;refGene_gn=CDC42BPA;refGene_tx=XM_019821465.2;refGene_gn=CDC42BPA;refGene_tx=XM_019821467.2;refGene_gn=CDC42BPA;refGene_tx=XM_019821468.2;refGene_gn=CDC42BPA;refGene_tx=XM_019821469.2;refGene_gn=CDC42BPA;refGene_tx=XM_019821470.2;refGene_gn=CDC42BPA;refGene_tx=XM_019821471.2;refGene_gn=CDC42BPA;refGene_tx=XM_019821472.2;refGene_gn=CDC42BPA;refGene_tx=XM_019821474.2
F1	4899436	4899679
refGene_gn=CDC42BPA;refGene_tx=XM_011290816.3;refGene_gn=CDC42BPA;refGene_tx=XM_011290817.3;refGene_gn=CDC42BPA;refGene_tx=XM_011290818.3;refGene_gn=CDC42BPA;refGene_tx=XM_011290819.3;refGene_gn=CDC42BPA;refGene_tx=XM_011290820.3;refGene_gn=CDC42BPA;refGene_tx=XM_019821465.2;refGene_gn=CDC42BPA;refGene_tx=XM_019821467.2;refGene_gn=CDC42BPA;refGene_tx=XM_019821468.2;refGene_gn=CDC42BPA;refGene_tx=XM_019821469.2;refGene_gn=CDC42BPA;refGene_tx=XM_019821470.2;refGene_gn=CDC42BPA;refGene_tx=XM_019821471.2;refGene_gn=CDC42BPA;refGene_tx=XM_019821474.2
F1	4907950	4908084
refGene_gn=CDC42BPA;refGene_tx=XM_011290816.3;refGene_gn=CDC42BPA;refGene_tx=XM_011290817.3;refGene_gn=CDC42BPA;refGene_tx=XM_011290818.3;refGene_gn=CDC42BPA;refGene_tx=XM_011290819.3;refGene_gn=CDC42BPA;refGene_tx=XM_011290820.3;refGene_gn=CDC42BPA;refGene_tx=XM_011290821.3;refGene_gn=CDC42BPA;refGene_tx=XM_019821465.2;refGene_gn=CDC42BPA;refGene_tx=XM_019821467.2;refGene_gn=CDC42BPA;refGene_tx=XM_019821468.2;refGene_gn=CDC42BPA;refGene_tx=XM_019821469.2;refGene_gn=CDC42BPA;refGene_tx=XM_019821470.2;refGene_gn=CDC42BPA;refGene_tx=XM_019821471.2;refGene_gn=CDC42BPA;refGene_tx=XM_019821472.2;refGene_gn=CDC42BPA;refGene_tx=XM_019821474.2
F1	4922647	4922770
refGene_gn=CDC42BPA;refGene_tx=XM_011290816.3;refGene_gn=CDC42BPA;refGene_tx=XM_011290817.3;refGene_gn=CDC42BPA;refGene_tx=XM_011290818.3;refGene_gn=CDC42BPA;refGene_tx=XM_011290819.3;refGene_gn=CDC42BPA;refGene_tx=XM_011290820.3;refGene_gn=CDC42BPA;refGene_tx=XM_011290821.3;refGene_gn=CDC42BPA;refGene_tx=XM_019821465.2;refGene_gn=CDC42BPA;refGene_tx=XM_019821467.2;refGene_gn=CDC42BPA;refGene_tx=XM_019821468.2;refGen

e_gn=CDC42BPA;refGene_tx=XM_019821469.2;refGene_gn=CDC42BPA;refGene_tx=XM_019821470.2;refGene_gn=CDC42BPA;refGene_tx=XM_019821471.2;refGene_gn=CDC42BPA;refGene_tx=XM_019821472.2;refGene_gn=CDC42BPA;refGene_tx=XM_019821474.2
F1 4936945 4937112
refGene_gn=CDC42BPA;refGene_tx=XM_011290816.3;refGene_gn=CDC42BPA;refGene_tx=XM_011290817.3;refGene_gn=CDC42BPA;refGene_tx=XM_011290818.3;refGene_gn=CDC42BPA;refGene_tx=XM_011290819.3;refGene_gn=CDC42BPA;refGene_tx=XM_011290820.3;refGene_gn=CDC42BPA;refGene_tx=XM_011290821.3;refGene_gn=CDC42BPA;refGene_tx=XM_019821465.2;refGene_gn=CDC42BPA;refGene_tx=XM_019821467.2;refGene_gn=CDC42BPA;refGene_tx=XM_019821468.2;refGene_gn=CDC42BPA;refGene_tx=XM_019821469.2;refGene_gn=CDC42BPA;refGene_tx=XM_019821470.2;refGene_gn=CDC42BPA;refGene_tx=XM_019821471.2;refGene_gn=CDC42BPA;refGene_tx=XM_019821472.2;refGene_gn=CDC42BPA;refGene_tx=XM_019821474.2
F1 4939051 4939131
refGene_gn=CDC42BPA;refGene_tx=XM_011290816.3;refGene_gn=CDC42BPA;refGene_tx=XM_011290817.3;refGene_gn=CDC42BPA;refGene_tx=XM_011290818.3;refGene_gn=CDC42BPA;refGene_tx=XM_011290819.3;refGene_gn=CDC42BPA;refGene_tx=XM_011290820.3;refGene_gn=CDC42BPA;refGene_tx=XM_011290821.3;refGene_gn=CDC42BPA;refGene_tx=XM_019821465.2;refGene_gn=CDC42BPA;refGene_tx=XM_019821467.2;refGene_gn=CDC42BPA;refGene_tx=XM_019821468.2;refGene_gn=CDC42BPA;refGene_tx=XM_019821469.2;refGene_gn=CDC42BPA;refGene_tx=XM_019821470.2;refGene_gn=CDC42BPA;refGene_tx=XM_019821471.2;refGene_gn=CDC42BPA;refGene_tx=XM_019821472.2;refGene_gn=CDC42BPA;refGene_tx=XM_019821474.2
F1 4941407 4941656
refGene_gn=CDC42BPA;refGene_tx=XM_011290816.3;refGene_gn=CDC42BPA;refGene_tx=XM_011290817.3;refGene_gn=CDC42BPA;refGene_tx=XM_011290818.3;refGene_gn=CDC42BPA;refGene_tx=XM_011290819.3;refGene_gn=CDC42BPA;refGene_tx=XM_011290820.3;refGene_gn=CDC42BPA;refGene_tx=XM_011290821.3;refGene_gn=CDC42BPA;refGene_tx=XM_019821465.2;refGene_gn=CDC42BPA;refGene_tx=XM_019821467.2;refGene_gn=CDC42BPA;refGene_tx=XM_019821468.2;refGene_gn=CDC42BPA;refGene_tx=XM_019821469.2;refGene_gn=CDC42BPA;refGene_tx=XM_019821470.2;refGene_gn=CDC42BPA;refGene_tx=XM_019821471.2;refGene_gn=CDC42BPA;refGene_tx=XM_019821472.2;refGene_gn=CDC42BPA;refGene_tx=XM_019821474.2
F1 4943743 4943944
refGene_gn=CDC42BPA;refGene_tx=XM_011290816.3;refGene_gn=CDC42BPA;refGene_tx=XM_011290817.3;refGene_gn=CDC42BPA;refGene_tx=XM_011290818.3;refGene_gn=CDC42BPA;refGene_tx=XM_011290819.3;refGene_gn=CDC42BPA;refGene_tx=XM_011290820.3;refGene_gn=CDC42BPA;refGene_tx=XM_011290821.3;refGene_gn=CDC42BPA;refGene_tx=XM_019821465.2;refGene_gn=CDC42BPA;refGene_tx=XM_019821467.2;refGene_gn=CDC42BPA;refGene_tx=XM_019821468.2;refGene_gn=CDC42BPA;refGene_tx=XM_019821469.2;refGene_gn=CDC42BPA;refGene_tx=XM_019821470.2;refGene_gn=CDC42BPA;refGene_tx=XM_019821471.2;refGene_gn=CDC42BPA;refGene_tx=XM_019821472.2;refGene_gn=CDC42BPA;refGene_tx=XM_019821474.2
F1 4953741 4953835
refGene_gn=CDC42BPA;refGene_tx=XM_011290816.3;refGene_gn=CDC42BPA;refG

ene_tx=XM_011290817.3;refGene_gn=CDC42BPA;refGene_tx=XM_011290818.3;re
fGene_gn=CDC42BPA;refGene_tx=XM_011290819.3;refGene_gn=CDC42BPA;refGen
e_tx=XM_011290820.3;refGene_gn=CDC42BPA;refGene_tx=XM_011290821.3;refG
ene_gn=CDC42BPA;refGene_tx=XM_019821465.2;refGene_gn=CDC42BPA;refGene_
tx=XM_019821467.2;refGene_gn=CDC42BPA;refGene_tx=XM_019821468.2;refGen
e_gn=CDC42BPA;refGene_tx=XM_019821469.2;refGene_gn=CDC42BPA;refGene_tx
=XM_019821470.2;refGene_gn=CDC42BPA;refGene_tx=XM_019821471.2;refGene_
gn=CDC42BPA;refGene_tx=XM_019821472.2;refGene_gn=CDC42BPA;refGene_tx=X
M_019821474.2
F1 4976447 4976596
refGene_gn=CDC42BPA;refGene_tx=XM_011290816.3;refGene_gn=CDC42BPA;refG
ene_tx=XM_011290817.3;refGene_gn=CDC42BPA;refGene_tx=XM_011290818.3;re
fGene_gn=CDC42BPA;refGene_tx=XM_011290819.3;refGene_gn=CDC42BPA;refGen
e_tx=XM_011290820.3;refGene_gn=CDC42BPA;refGene_tx=XM_011290821.3;refG
ene_gn=CDC42BPA;refGene_tx=XM_019821465.2;refGene_gn=CDC42BPA;refGene_
tx=XM_019821467.2;refGene_gn=CDC42BPA;refGene_tx=XM_019821468.2;refGen
e_gn=CDC42BPA;refGene_tx=XM_019821469.2;refGene_gn=CDC42BPA;refGene_tx
=XM_019821470.2;refGene_gn=CDC42BPA;refGene_tx=XM_019821471.2;refGene_
gn=CDC42BPA;refGene_tx=XM_019821472.2;refGene_gn=CDC42BPA;refGene_tx=X
M_019821474.2
F1 4989268 4989364
refGene_gn=CDC42BPA;refGene_tx=XM_011290816.3;refGene_gn=CDC42BPA;refG
ene_tx=XM_011290817.3;refGene_gn=CDC42BPA;refGene_tx=XM_011290818.3;re
fGene_gn=CDC42BPA;refGene_tx=XM_011290819.3;refGene_gn=CDC42BPA;refGen
e_tx=XM_011290820.3;refGene_gn=CDC42BPA;refGene_tx=XM_011290821.3;refG
ene_gn=CDC42BPA;refGene_tx=XM_019821465.2;refGene_gn=CDC42BPA;refGene_
tx=XM_019821467.2;refGene_gn=CDC42BPA;refGene_tx=XM_019821468.2;refGen
e_gn=CDC42BPA;refGene_tx=XM_019821469.2;refGene_gn=CDC42BPA;refGene_tx
=XM_019821470.2;refGene_gn=CDC42BPA;refGene_tx=XM_019821471.2;refGene_
gn=CDC42BPA;refGene_tx=XM_019821472.2;refGene_gn=CDC42BPA;refGene_tx=X
M_019821474.2
F1 4996616 4996700
refGene_gn=CDC42BPA;refGene_tx=XM_011290816.3;refGene_gn=CDC42BPA;refG
ene_tx=XM_011290817.3;refGene_gn=CDC42BPA;refGene_tx=XM_011290818.3;re
fGene_gn=CDC42BPA;refGene_tx=XM_011290819.3;refGene_gn=CDC42BPA;refGen
e_tx=XM_011290820.3;refGene_gn=CDC42BPA;refGene_tx=XM_011290821.3;refG
ene_gn=CDC42BPA;refGene_tx=XM_019821465.2;refGene_gn=CDC42BPA;refGene_
tx=XM_019821467.2;refGene_gn=CDC42BPA;refGene_tx=XM_019821468.2;refGen
e_gn=CDC42BPA;refGene_tx=XM_019821469.2;refGene_gn=CDC42BPA;refGene_tx
=XM_019821470.2;refGene_gn=CDC42BPA;refGene_tx=XM_019821471.2;refGene_
gn=CDC42BPA;refGene_tx=XM_019821472.2;refGene_gn=CDC42BPA;refGene_tx=X
M_019821474.2
F1 5031584 5031676
refGene_gn=CDC42BPA;refGene_tx=XM_011290816.3;refGene_gn=CDC42BPA;refG
ene_tx=XM_011290817.3;refGene_gn=CDC42BPA;refGene_tx=XM_011290818.3;re
fGene_gn=CDC42BPA;refGene_tx=XM_011290819.3;refGene_gn=CDC42BPA;refGen
e_tx=XM_011290820.3;refGene_gn=CDC42BPA;refGene_tx=XM_011290821.3;refG
ene_gn=CDC42BPA;refGene_tx=XM_019821465.2;refGene_gn=CDC42BPA;refGene_
tx=XM_019821467.2;refGene_gn=CDC42BPA;refGene_tx=XM_019821468.2;refGen
e_gn=CDC42BPA;refGene_tx=XM_019821469.2;refGene_gn=CDC42BPA;refGene_tx

=XM_019821470.2;refGene_gn=CDC42BPA;refGene_tx=XM_019821472.2;refGene_gn=CDC42BPA;refGene_tx=XM_019821474.2
F1 5100929 5101107
refGene_gn=CDC42BPA;refGene_tx=XM_011290816.3;refGene_gn=CDC42BPA;refGene_tx=XM_011290817.3;refGene_gn=CDC42BPA;refGene_tx=XM_011290818.3;refGene_gn=CDC42BPA;refGene_tx=XM_011290819.3;refGene_gn=CDC42BPA;refGene_tx=XM_011290820.3;refGene_gn=CDC42BPA;refGene_tx=XM_011290821.3;refGene_gn=CDC42BPA;refGene_tx=XM_019821465.2;refGene_gn=CDC42BPA;refGene_tx=XM_019821467.2;refGene_gn=CDC42BPA;refGene_tx=XM_019821468.2;refGene_gn=CDC42BPA;refGene_tx=XM_019821469.2;refGene_gn=CDC42BPA;refGene_tx=XM_019821470.2;refGene_gn=CDC42BPA;refGene_tx=XM_019821471.2;refGene_gn=CDC42BPA;refGene_tx=XM_019821472.2;refGene_gn=CDC42BPA;refGene_tx=XM_019821474.2
F1 5263172 5263290
refGene_gn=AHCTF1;refGene_tx=XM_023247872.1;refGene_gn=AHCTF1;refGene_tx=XM_023247873.1;refGene_gn=AHCTF1;refGene_tx=XM_023247874.1;refGene_gn=AHCTF1;refGene_tx=XM_023247875.1;refGene_gn=AHCTF1;refGene_tx=XM_023247876.1;refGene_gn=AHCTF1;refGene_tx=XM_023247877.1
F1 5265092 5265346
refGene_gn=AHCTF1;refGene_tx=XM_023247872.1;refGene_gn=AHCTF1;refGene_tx=XM_023247873.1;refGene_gn=AHCTF1;refGene_tx=XM_023247874.1;refGene_gn=AHCTF1;refGene_tx=XM_023247875.1;refGene_gn=AHCTF1;refGene_tx=XM_023247876.1;refGene_gn=AHCTF1;refGene_tx=XM_023247877.1
F1 5269397 5269575
refGene_gn=AHCTF1;refGene_tx=XM_023247872.1;refGene_gn=AHCTF1;refGene_tx=XM_023247873.1;refGene_gn=AHCTF1;refGene_tx=XM_023247874.1;refGene_gn=AHCTF1;refGene_tx=XM_023247875.1;refGene_gn=AHCTF1;refGene_tx=XM_023247876.1;refGene_gn=AHCTF1;refGene_tx=XM_023247877.1
F1 5276035 5276243
refGene_gn=AHCTF1;refGene_tx=XM_023247872.1;refGene_gn=AHCTF1;refGene_tx=XM_023247873.1;refGene_gn=AHCTF1;refGene_tx=XM_023247874.1;refGene_gn=AHCTF1;refGene_tx=XM_023247875.1;refGene_gn=AHCTF1;refGene_tx=XM_023247876.1;refGene_gn=AHCTF1;refGene_tx=XM_023247877.1
F1 5277856 5277973
refGene_gn=AHCTF1;refGene_tx=XM_023247872.1;refGene_gn=AHCTF1;refGene_tx=XM_023247873.1;refGene_gn=AHCTF1;refGene_tx=XM_023247874.1;refGene_gn=AHCTF1;refGene_tx=XM_023247875.1;refGene_gn=AHCTF1;refGene_tx=XM_023247876.1;refGene_gn=AHCTF1;refGene_tx=XM_023247877.1
F1 5282903 5282988
refGene_gn=AHCTF1;refGene_tx=XM_023247872.1;refGene_gn=AHCTF1;refGene_tx=XM_023247873.1;refGene_gn=AHCTF1;refGene_tx=XM_023247874.1;refGene_gn=AHCTF1;refGene_tx=XM_023247875.1;refGene_gn=AHCTF1;refGene_tx=XM_023247876.1;refGene_gn=AHCTF1;refGene_tx=XM_023247877.1
F1 5284551 5284702
refGene_gn=AHCTF1;refGene_tx=XM_023247872.1;refGene_gn=AHCTF1;refGene_tx=XM_023247873.1;refGene_gn=AHCTF1;refGene_tx=XM_023247874.1;refGene_gn=AHCTF1;refGene_tx=XM_023247875.1;refGene_gn=AHCTF1;refGene_tx=XM_023247876.1;refGene_gn=AHCTF1;refGene_tx=XM_023247877.1
F1 5286062 5286195
refGene_gn=AHCTF1;refGene_tx=XM_023247872.1;refGene_gn=AHCTF1;refGene_

tx=XM_023247873.1;refGene_gn=AHCTF1;refGene_tx=XM_023247874.1;refGene_gn=AHCTF1;refGene_tx=XM_023247875.1;refGene_gn=AHCTF1;refGene_tx=XM_023247876.1;refGene_gn=AHCTF1;refGene_tx=XM_023247877.1
F1 5286281 5286463
refGene_gn=AHCTF1;refGene_tx=XM_023247872.1;refGene_gn=AHCTF1;refGene_tx=XM_023247873.1;refGene_gn=AHCTF1;refGene_tx=XM_023247874.1;refGene_gn=AHCTF1;refGene_tx=XM_023247875.1;refGene_gn=AHCTF1;refGene_tx=XM_023247876.1;refGene_gn=AHCTF1;refGene_tx=XM_023247877.1
F1 5287119 5287181
refGene_gn=AHCTF1;refGene_tx=XM_023247872.1;refGene_gn=AHCTF1;refGene_tx=XM_023247873.1;refGene_gn=AHCTF1;refGene_tx=XM_023247874.1;refGene_gn=AHCTF1;refGene_tx=XM_023247875.1;refGene_gn=AHCTF1;refGene_tx=XM_023247876.1;refGene_gn=AHCTF1;refGene_tx=XM_023247877.1
F1 5288048 5288177
refGene_gn=AHCTF1;refGene_tx=XM_023247872.1;refGene_gn=AHCTF1;refGene_tx=XM_023247873.1;refGene_gn=AHCTF1;refGene_tx=XM_023247874.1;refGene_gn=AHCTF1;refGene_tx=XM_023247875.1;refGene_gn=AHCTF1;refGene_tx=XM_023247876.1;refGene_gn=AHCTF1;refGene_tx=XM_023247877.1
F1 5291459 5291550
refGene_gn=AHCTF1;refGene_tx=XM_023247872.1;refGene_gn=AHCTF1;refGene_tx=XM_023247873.1;refGene_gn=AHCTF1;refGene_tx=XM_023247874.1;refGene_gn=AHCTF1;refGene_tx=XM_023247875.1;refGene_gn=AHCTF1;refGene_tx=XM_023247876.1;refGene_gn=AHCTF1;refGene_tx=XM_023247877.1
F1 5292818 5292908
refGene_gn=AHCTF1;refGene_tx=XM_023247872.1;refGene_gn=AHCTF1;refGene_tx=XM_023247873.1;refGene_gn=AHCTF1;refGene_tx=XM_023247874.1;refGene_gn=AHCTF1;refGene_tx=XM_023247875.1;refGene_gn=AHCTF1;refGene_tx=XM_023247876.1;refGene_gn=AHCTF1;refGene_tx=XM_023247877.1
F1 5295363 5295504
refGene_gn=AHCTF1;refGene_tx=XM_023247872.1;refGene_gn=AHCTF1;refGene_tx=XM_023247873.1;refGene_gn=AHCTF1;refGene_tx=XM_023247874.1;refGene_gn=AHCTF1;refGene_tx=XM_023247875.1;refGene_gn=AHCTF1;refGene_tx=XM_023247876.1
F1 5296129 5296234
refGene_gn=AHCTF1;refGene_tx=XM_023247872.1;refGene_gn=AHCTF1;refGene_tx=XM_023247873.1;refGene_gn=AHCTF1;refGene_tx=XM_023247874.1;refGene_gn=AHCTF1;refGene_tx=XM_023247875.1;refGene_gn=AHCTF1;refGene_tx=XM_023247876.1
F1 5297115 5297209
refGene_gn=AHCTF1;refGene_tx=XM_023247872.1;refGene_gn=AHCTF1;refGene_tx=XM_023247873.1;refGene_gn=AHCTF1;refGene_tx=XM_023247874.1;refGene_gn=AHCTF1;refGene_tx=XM_023247875.1;refGene_gn=AHCTF1;refGene_tx=XM_023247876.1;refGene_gn=AHCTF1;refGene_tx=XM_023247877.1
F1 5298803 5298927
refGene_gn=AHCTF1;refGene_tx=XM_023247872.1;refGene_gn=AHCTF1;refGene_tx=XM_023247873.1;refGene_gn=AHCTF1;refGene_tx=XM_023247874.1;refGene_gn=AHCTF1;refGene_tx=XM_023247875.1;refGene_gn=AHCTF1;refGene_tx=XM_023247876.1;refGene_gn=AHCTF1;refGene_tx=XM_023247877.1
F1 5299064 5299121
refGene_gn=AHCTF1;refGene_tx=XM_023247872.1;refGene_gn=AHCTF1;refGene_

tx=XM_023247873.1;refGene_gn=AHCTF1;refGene_tx=XM_023247874.1;refGene_gn=AHCTF1;refGene_tx=XM_023247875.1;refGene_gn=AHCTF1;refGene_tx=XM_023247876.1;refGene_gn=AHCTF1;refGene_tx=XM_023247877.1
F1 5299933 5300080
refGene_gn=AHCTF1;refGene_tx=XM_023247872.1;refGene_gn=AHCTF1;refGene_tx=XM_023247873.1;refGene_gn=AHCTF1;refGene_tx=XM_023247874.1;refGene_gn=AHCTF1;refGene_tx=XM_023247875.1;refGene_gn=AHCTF1;refGene_tx=XM_023247876.1;refGene_gn=AHCTF1;refGene_tx=XM_023247877.1
F1 5300625 5300813
refGene_gn=AHCTF1;refGene_tx=XM_023247872.1;refGene_gn=AHCTF1;refGene_tx=XM_023247873.1;refGene_gn=AHCTF1;refGene_tx=XM_023247874.1;refGene_gn=AHCTF1;refGene_tx=XM_023247875.1;refGene_gn=AHCTF1;refGene_tx=XM_023247876.1;refGene_gn=AHCTF1;refGene_tx=XM_023247877.1
F1 5305016 5305161
refGene_gn=AHCTF1;refGene_tx=XM_023247872.1;refGene_gn=AHCTF1;refGene_tx=XM_023247873.1;refGene_gn=AHCTF1;refGene_tx=XM_023247874.1;refGene_gn=AHCTF1;refGene_tx=XM_023247875.1;refGene_gn=AHCTF1;refGene_tx=XM_023247876.1;refGene_gn=AHCTF1;refGene_tx=XM_023247877.1
F1 5305237 5305369
refGene_gn=AHCTF1;refGene_tx=XM_023247872.1;refGene_gn=AHCTF1;refGene_tx=XM_023247873.1;refGene_gn=AHCTF1;refGene_tx=XM_023247874.1;refGene_gn=AHCTF1;refGene_tx=XM_023247875.1;refGene_gn=AHCTF1;refGene_tx=XM_023247876.1;refGene_gn=AHCTF1;refGene_tx=XM_023247877.1
F1 5306279 5306430
refGene_gn=AHCTF1;refGene_tx=XM_023247872.1;refGene_gn=AHCTF1;refGene_tx=XM_023247873.1;refGene_gn=AHCTF1;refGene_tx=XM_023247874.1;refGene_gn=AHCTF1;refGene_tx=XM_023247875.1;refGene_gn=AHCTF1;refGene_tx=XM_023247876.1;refGene_gn=AHCTF1;refGene_tx=XM_023247877.1
F1 5306922 5307073
refGene_gn=AHCTF1;refGene_tx=XM_023247872.1;refGene_gn=AHCTF1;refGene_tx=XM_023247873.1;refGene_gn=AHCTF1;refGene_tx=XM_023247874.1;refGene_gn=AHCTF1;refGene_tx=XM_023247875.1;refGene_gn=AHCTF1;refGene_tx=XM_023247876.1;refGene_gn=AHCTF1;refGene_tx=XM_023247877.1
F1 5307652 5307751
refGene_gn=AHCTF1;refGene_tx=XM_023247872.1;refGene_gn=AHCTF1;refGene_tx=XM_023247873.1;refGene_gn=AHCTF1;refGene_tx=XM_023247874.1;refGene_gn=AHCTF1;refGene_tx=XM_023247875.1;refGene_gn=AHCTF1;refGene_tx=XM_023247876.1;refGene_gn=AHCTF1;refGene_tx=XM_023247877.1
F1 5310212 5310405
refGene_gn=AHCTF1;refGene_tx=XM_023247872.1;refGene_gn=AHCTF1;refGene_tx=XM_023247873.1;refGene_gn=AHCTF1;refGene_tx=XM_023247874.1;refGene_gn=AHCTF1;refGene_tx=XM_023247875.1;refGene_gn=AHCTF1;refGene_tx=XM_023247876.1;refGene_gn=AHCTF1;refGene_tx=XM_023247877.1
F1 5312714 5312918
refGene_gn=AHCTF1;refGene_tx=XM_023247872.1;refGene_gn=AHCTF1;refGene_tx=XM_023247873.1;refGene_gn=AHCTF1;refGene_tx=XM_023247874.1;refGene_gn=AHCTF1;refGene_tx=XM_023247875.1;refGene_gn=AHCTF1;refGene_tx=XM_023247876.1;refGene_gn=AHCTF1;refGene_tx=XM_023247877.1
F1 5313544 5313959
refGene_gn=AHCTF1;refGene_tx=XM_023247872.1;refGene_gn=AHCTF1;refGene_

tx=XM_023247873.1;refGene_gn=AHCTF1;refGene_tx=XM_023247874.1;refGene_gn=AHCTF1;refGene_tx=XM_023247875.1;refGene_gn=AHCTF1;refGene_tx=XM_023247876.1;refGene_gn=AHCTF1;refGene_tx=XM_023247877.1
F1 5315263 5315387
refGene_gn=AHCTF1;refGene_tx=XM_023247872.1;refGene_gn=AHCTF1;refGene_tx=XM_023247873.1;refGene_gn=AHCTF1;refGene_tx=XM_023247874.1;refGene_gn=AHCTF1;refGene_tx=XM_023247875.1;refGene_gn=AHCTF1;refGene_tx=XM_023247876.1;refGene_gn=AHCTF1;refGene_tx=XM_023247877.1
F1 5317291 5317377
refGene_gn=AHCTF1;refGene_tx=XM_023247872.1;refGene_gn=AHCTF1;refGene_tx=XM_023247873.1;refGene_gn=AHCTF1;refGene_tx=XM_023247874.1;refGene_gn=AHCTF1;refGene_tx=XM_023247875.1;refGene_gn=AHCTF1;refGene_tx=XM_023247876.1;refGene_gn=AHCTF1;refGene_tx=XM_023247877.1
F1 5320068 5320277
refGene_gn=AHCTF1;refGene_tx=XM_023247872.1;refGene_gn=AHCTF1;refGene_tx=XM_023247873.1;refGene_gn=AHCTF1;refGene_tx=XM_023247874.1;refGene_gn=AHCTF1;refGene_tx=XM_023247875.1;refGene_gn=AHCTF1;refGene_tx=XM_023247876.1;refGene_gn=AHCTF1;refGene_tx=XM_023247877.1
F1 5322049 5324045
refGene_gn=AHCTF1;refGene_tx=XM_023247872.1;refGene_gn=AHCTF1;refGene_tx=XM_023247873.1;refGene_gn=AHCTF1;refGene_tx=XM_023247874.1;refGene_gn=AHCTF1;refGene_tx=XM_023247875.1;refGene_gn=AHCTF1;refGene_tx=XM_023247876.1;refGene_gn=AHCTF1;refGene_tx=XM_023247877.1
F1 5328527 5328661
refGene_gn=AHCTF1;refGene_tx=XM_023247872.1;refGene_gn=AHCTF1;refGene_tx=XM_023247873.1;refGene_gn=AHCTF1;refGene_tx=XM_023247874.1;refGene_gn=AHCTF1;refGene_tx=XM_023247875.1;refGene_gn=AHCTF1;refGene_tx=XM_023247876.1;refGene_gn=AHCTF1;refGene_tx=XM_023247877.1
F1 5330325 5330408
refGene_gn=AHCTF1;refGene_tx=XM_023247872.1;refGene_gn=AHCTF1;refGene_tx=XM_023247873.1;refGene_gn=AHCTF1;refGene_tx=XM_023247874.1;refGene_gn=AHCTF1;refGene_tx=XM_023247875.1;refGene_gn=AHCTF1;refGene_tx=XM_023247876.1;refGene_gn=AHCTF1;refGene_tx=XM_023247877.1
F1 5331218 5331408
refGene_gn=AHCTF1;refGene_tx=XM_023247872.1;refGene_gn=AHCTF1;refGene_tx=XM_023247873.1;refGene_gn=AHCTF1;refGene_tx=XM_023247874.1;refGene_gn=AHCTF1;refGene_tx=XM_023247875.1;refGene_gn=AHCTF1;refGene_tx=XM_023247876.1;refGene_gn=AHCTF1;refGene_tx=XM_023247877.1
F1 5332659 5332743
refGene_gn=LOC102901264;refGene_tx=XM_023247878.1
F1 5335503 5335698
refGene_gn=LOC102901264;refGene_tx=XM_023247878.1
F1 5348223 5348427
refGene_gn=LOC102901264;refGene_tx=XM_023247878.1
F1 5352833 5352869
refGene_gn=LOC102901264;refGene_tx=XM_023247878.1
F1 5358113 5358338
refGene_gn=LOC102901264;refGene_tx=XM_023247878.1
F1 5373346 5373501
refGene_gn=LOC102901264;refGene_tx=XM_023247878.1

F1 5384085 5384197
refGene_gn=LOC102901264;refGene_tx=XM_023247878.1
F1 5385554 5385622
refGene_gn=LOC102901264;refGene_tx=XM_023247878.1
F1 5388028 5388138
refGene_gn=LOC102901264;refGene_tx=XM_023247878.1
F1 5389577 5389675
refGene_gn=LOC102901264;refGene_tx=XM_023247878.1
F1 5390979 5391070
refGene_gn=LOC102901264;refGene_tx=XM_023247878.1
F1 5392512 5392567
refGene_gn=LOC102901264;refGene_tx=XM_023247878.1
F1 5394014 5394138
refGene_gn=LOC102901264;refGene_tx=XM_023247878.1
F1 5395500 5395645
refGene_gn=LOC102901264;refGene_tx=XM_023247878.1
F1 5396751 5396870
refGene_gn=LOC102901264;refGene_tx=XM_023247878.1
F1 5397366 5397474
refGene_gn=LOC102901264;refGene_tx=XM_023247878.1
F1 5400471 5400746
refGene_gn=LOC102901264;refGene_tx=XM_023247878.1
F1 5401902 5402037
refGene_gn=LOC102901264;refGene_tx=XM_023247878.1
F1 5402980 5403147
refGene_gn=LOC102901264;refGene_tx=XM_023247878.1
F1 5403620 5403727
refGene_gn=LOC102901264;refGene_tx=XM_023247878.1
F1 5403961 5404121
refGene_gn=LOC102901264;refGene_tx=XM_023247878.1
F1 5404604 5404795
refGene_gn=LOC102901264;refGene_tx=XM_023247878.1
F1 5409693 5409822
refGene_gn=LOC102901264;refGene_tx=XM_023247878.1
F1 5411657 5411771
refGene_gn=LOC102901264;refGene_tx=XM_023247878.1
F1 5413316 5413422 refGene_gn=SCCPDH;refGene_tx=XM_011290824.2
F1 5416192 5416274 refGene_gn=SCCPDH;refGene_tx=XM_011290824.2
F1 5417515 5417627 refGene_gn=SCCPDH;refGene_tx=XM_011290824.2
F1 5422771 5422828 refGene_gn=SCCPDH;refGene_tx=XM_011290824.2
F1 5422930 5423050 refGene_gn=SCCPDH;refGene_tx=XM_011290824.2
F1 5423810 5423928 refGene_gn=SCCPDH;refGene_tx=XM_011290824.2
F1 5424533 5424664 refGene_gn=SCCPDH;refGene_tx=XM_011290824.2
F1 5435970 5436020 refGene_gn=SCCPDH;refGene_tx=XM_011290824.2
F1 5440492 5440622 refGene_gn=SCCPDH;refGene_tx=XM_011290824.2
F1 5442246 5442327 refGene_gn=SCCPDH;refGene_tx=XM_011290824.2
F1 5450287 5450400 refGene_gn=SCCPDH;refGene_tx=XM_011290824.2
F1 5452241 5452431 refGene_gn=SCCPDH;refGene_tx=XM_011290824.2
F1 5511166 5511372
refGene_gn=CNST;refGene_tx=XM_011290828.2;refGene_gn=CNST;refGene_tx=X

M_019821479.1;refGene_gn=CNST;refGene_tx=XM_019821480.1
F1 5514889 5515025
refGene_gn=CNST;refGene_tx=XM_011290828.2;refGene_gn=CNST;refGene_tx=X
M_019821479.1;refGene_gn=CNST;refGene_tx=XM_019821480.1
F1 5528768 5529616
refGene_gn=CNST;refGene_tx=XM_011290828.2;refGene_gn=CNST;refGene_tx=X
M_019821479.1;refGene_gn=CNST;refGene_tx=XM_019821480.1
F1 5539195 5539292
refGene_gn=CNST;refGene_tx=XM_011290828.2;refGene_gn=CNST;refGene_tx=X
M_019821479.1;refGene_gn=CNST;refGene_tx=XM_019821480.1
F1 5539469 5539491
refGene_gn=CNST;refGene_tx=XM_011290828.2;refGene_gn=CNST;refGene_tx=X
M_019821479.1;refGene_gn=CNST;refGene_tx=XM_019821480.1
F1 5542134 5542249
refGene_gn=CNST;refGene_tx=XM_011290828.2;refGene_gn=CNST;refGene_tx=X
M_019821479.1;refGene_gn=CNST;refGene_tx=XM_019821480.1
F1 5542708 5542795
refGene_gn=CNST;refGene_tx=XM_011290828.2;refGene_gn=CNST;refGene_tx=X
M_019821479.1;refGene_gn=CNST;refGene_tx=XM_019821480.1
F1 5544653 5544684
refGene_gn=CNST;refGene_tx=XM_011290828.2;refGene_gn=CNST;refGene_tx=X
M_019821479.1;refGene_gn=CNST;refGene_tx=XM_019821480.1
F1 5552447 5552653
refGene_gn=CNST;refGene_tx=XM_011290828.2;refGene_gn=CNST;refGene_tx=X
M_019821479.1;refGene_gn=CNST;refGene_tx=XM_019821480.1
F1 5555011 5555062
refGene_gn=CNST;refGene_tx=XM_019821479.1;refGene_gn=CNST;refGene_tx=X
M_019821480.1
F1 5584253 5584632
refGene_gn=CNST;refGene_tx=XM_011290828.2;refGene_gn=CNST;refGene_tx=X
M_019821479.1;refGene_gn=CNST;refGene_tx=XM_019821480.1
F1 5628202 5628497
refGene_gn=TFB2M;refGene_tx=XM_011290825.3;refGene_gn=TFB2M;refGene_tx
=XM_023247899.1
F1 5629301 5629390
refGene_gn=TFB2M;refGene_tx=XM_011290825.3;refGene_gn=TFB2M;refGene_tx
=XM_023247899.1
F1 5633217 5633371
refGene_gn=TFB2M;refGene_tx=XM_011290825.3;refGene_gn=TFB2M;refGene_tx
=XM_023247899.1
F1 5633676 5633825
refGene_gn=TFB2M;refGene_tx=XM_011290825.3;refGene_gn=TFB2M;refGene_tx
=XM_023247899.1
F1 5636929 5637016
refGene_gn=TFB2M;refGene_tx=XM_011290825.3;refGene_gn=TFB2M;refGene_tx
=XM_023247899.1
F1 5639226 5639289
refGene_gn=TFB2M;refGene_tx=XM_011290825.3;refGene_gn=TFB2M;refGene_tx
=XM_023247899.1
F1 5641782 5641946

refGene_gn=TFB2M;refGene_tx=XM_011290825.3;refGene_gn=TFB2M;refGene_tx=XM_023247899.1
F1 5644253 5644428 refGene_gn=TFB2M;refGene_tx=XM_011290825.3;refGene_gn=TFB2M;refGene_tx=XM_023247899.1
F1 5652160 5652324 refGene_gn=SMYD3;refGene_tx=XM_011290829.3;refGene_gn=SMYD3;refGene_tx=XM_023247957.1
F1 5815942 5816006 refGene_gn=SMYD3;refGene_tx=XM_011290829.3;refGene_gn=SMYD3;refGene_tx=XM_023247957.1
F1 5836457 5836565 refGene_gn=SMYD3;refGene_tx=XM_011290829.3
F1 5838324 5838382 refGene_gn=SMYD3;refGene_tx=XM_011290829.3;refGene_gn=SMYD3;refGene_tx=XM_023247957.1
F1 5840576 5840713 refGene_gn=SMYD3;refGene_tx=XM_011290829.3;refGene_gn=SMYD3;refGene_tx=XM_023247957.1
F1 6188832 6188900 refGene_gn=SMYD3;refGene_tx=XM_011290829.3;refGene_gn=SMYD3;refGene_tx=XM_023247957.1
F1 6190146 6190249 refGene_gn=SMYD3;refGene_tx=XM_011290829.3;refGene_gn=SMYD3;refGene_tx=XM_023247957.1
F1 6199694 6199805 refGene_gn=SMYD3;refGene_tx=XM_011290829.3;refGene_gn=SMYD3;refGene_tx=XM_023247957.1
F1 6239319 6239407 refGene_gn=SMYD3;refGene_tx=XM_011290829.3;refGene_gn=SMYD3;refGene_tx=XM_023247957.1
F1 6244257 6244432 refGene_gn=SMYD3;refGene_tx=XM_011290829.3;refGene_gn=SMYD3;refGene_tx=XM_023247957.1
F1 6328454 6328563 refGene_gn=SMYD3;refGene_tx=XM_011290829.3;refGene_gn=SMYD3;refGene_tx=XM_023247957.1
F1 6340070 6340172 refGene_gn=SMYD3;refGene_tx=XM_011290829.3;refGene_gn=SMYD3;refGene_tx=XM_023247957.1
F1 6394558 6394707 refGene_gn=KIF26B;refGene_tx=XM_023247495.1
F1 6405447 6405598 refGene_gn=KIF26B;refGene_tx=XM_023247495.1
F1 6405989 6406192 refGene_gn=KIF26B;refGene_tx=XM_023247495.1
F1 6411128 6411448 refGene_gn=KIF26B;refGene_tx=XM_023247495.1
F1 6411449 6414547 refGene_gn=KIF26B;refGene_tx=XM_023247495.1
F1 6415484 6415647 refGene_gn=KIF26B;refGene_tx=XM_023247495.1
F1 6455012 6455172 refGene_gn=KIF26B;refGene_tx=XM_023247495.1
F1 6470452 6470636 refGene_gn=KIF26B;refGene_tx=XM_023247495.1
F1 6474020 6474283 refGene_gn=KIF26B;refGene_tx=XM_023247495.1
F1 6475273 6475367 refGene_gn=KIF26B;refGene_tx=XM_023247495.1
F1 6480475 6480682 refGene_gn=KIF26B;refGene_tx=XM_023247495.1

F1 6538331 6538515 refGene_gn=KIF26B;refGene_tx=XM_023247495.1
F1 6648482 6648649 refGene_gn=KIF26B;refGene_tx=XM_023247495.1
F1 6696622 6697159 refGene_gn=KIF26B;refGene_tx=XM_023247495.1
F1 6869521 6869923 refGene_gn=KIF26B;refGene_tx=XM_023247495.1
F1 6870763 6870826 refGene_gn=KIF26B;refGene_tx=XM_023247495.1
F1 6920538 6920654
refGene_gn=EFCAB2;refGene_tx=XM_023248049.1;refGene_gn=EFCAB2;refGene_tx=XM_023248050.1
F1 6925466 6925533
refGene_gn=EFCAB2;refGene_tx=XM_023248049.1;refGene_gn=EFCAB2;refGene_tx=XM_023248050.1
F1 6925631 6925674
refGene_gn=EFCAB2;refGene_tx=XM_023248049.1;refGene_gn=EFCAB2;refGene_tx=XM_023248050.1
F1 6926798 6926878
refGene_gn=EFCAB2;refGene_tx=XM_023248049.1;refGene_gn=EFCAB2;refGene_tx=XM_023248050.1
F1 6934630 6934698 refGene_gn=EFCAB2;refGene_tx=XM_023248049.1
F1 6935201 6935367 refGene_gn=EFCAB2;refGene_tx=XM_023248049.1
F1 6947990 6948063 refGene_gn=EFCAB2;refGene_tx=XM_023248050.1
F1 6989407 6989492 refGene_gn=EFCAB2;refGene_tx=XM_023248050.1
F1 7052889 7052920 refGene_gn=EFCAB2;refGene_tx=XM_023248050.1
F1 7135620 7136308
refGene_gn=HNRNPU;refGene_tx=XM_023247628.1;refGene_gn=HNRNPU;refGene_tx=XM_023247629.1
F1 7137190 7137302
refGene_gn=HNRNPU;refGene_tx=XM_023247628.1;refGene_gn=HNRNPU;refGene_tx=XM_023247629.1
F1 7137387 7137461
refGene_gn=HNRNPU;refGene_tx=XM_023247628.1;refGene_gn=HNRNPU;refGene_tx=XM_023247629.1
F1 7139520 7139660
refGene_gn=HNRNPU;refGene_tx=XM_023247628.1;refGene_gn=HNRNPU;refGene_tx=XM_023247629.1
F1 7140243 7140343
refGene_gn=HNRNPU;refGene_tx=XM_023247628.1;refGene_gn=HNRNPU;refGene_tx=XM_023247629.1
F1 7140732 7140845
refGene_gn=HNRNPU;refGene_tx=XM_023247628.1;refGene_gn=HNRNPU;refGene_tx=XM_023247629.1
F1 7141186 7141450
refGene_gn=HNRNPU;refGene_tx=XM_023247628.1;refGene_gn=HNRNPU;refGene_tx=XM_023247629.1
F1 7141733 7141853
refGene_gn=HNRNPU;refGene_tx=XM_023247628.1;refGene_gn=HNRNPU;refGene_tx=XM_023247629.1
F1 7142277 7142406
refGene_gn=HNRNPU;refGene_tx=XM_023247628.1;refGene_gn=HNRNPU;refGene_tx=XM_023247629.1
F1 7142498 7142667

refGene_gn=HNRNPU;refGene_tx=XM_023247628.1;refGene_gn=HNRNPU;refGene_tx=XM_023247629.1
F1 7142913 7143168
refGene_gn=HNRNPU;refGene_tx=XM_023247628.1;refGene_gn=HNRNPU;refGene_tx=XM_023247629.1
F1 7143670 7143855
refGene_gn=HNRNPU;refGene_tx=XM_023247628.1;refGene_gn=HNRNPU;refGene_tx=XM_023247629.1
F1 7144215 7144287
refGene_gn=HNRNPU;refGene_tx=XM_023247628.1;refGene_gn=HNRNPU;refGene_tx=XM_023247629.1
F1 7144712 7144766
refGene_gn=HNRNPU;refGene_tx=XM_023247628.1;refGene_gn=HNRNPU;refGene_tx=XM_023247629.1
F1 7154613 7154791
refGene_gn=LOC102902396;refGene_tx=XM_023247630.1
F1 7155452 7155516
refGene_gn=LOC102902396;refGene_tx=XM_023247630.1
F1 7155611 7155726
refGene_gn=LOC102902396;refGene_tx=XM_023247630.1
F1 7162202 7162244
refGene_gn=LOC102902396;refGene_tx=XM_023247630.1
F1 7250643 7250877
refGene_gn=DESI2;refGene_tx=XM_011290833.3;refGene_gn=DESI2;refGene_tx=XM_023248031.1;refGene_gn=DESI2;refGene_tx=XM_023248032.1
F1 7258996 7259138
refGene_gn=DESI2;refGene_tx=XM_011290833.3;refGene_gn=DESI2;refGene_tx=XM_023248031.1;refGene_gn=DESI2;refGene_tx=XM_023248032.1
F1 7260792 7260886
refGene_gn=DESI2;refGene_tx=XM_011290833.3;refGene_gn=DESI2;refGene_tx=XM_023248031.1
F1 7263471 7263544
refGene_gn=DESI2;refGene_tx=XM_011290833.3;refGene_gn=DESI2;refGene_tx=XM_023248031.1;refGene_gn=DESI2;refGene_tx=XM_023248032.1
F1 7293951 7293963 refGene_gn=DESI2;refGene_tx=XM_011290833.3
F1 7294702 7294744
refGene_gn=DESI2;refGene_tx=XM_023248031.1;refGene_gn=DESI2;refGene_tx=XM_023248032.1
F1 7508464 7508647
refGene_gn=ADSS;refGene_tx=XM_023247274.1;refGene_gn=ADSS;refGene_tx=XM_023247275.1
F1 7518682 7518736 refGene_gn=ADSS;refGene_tx=XM_019821486.2
F1 7520816 7520919
refGene_gn=ADSS;refGene_tx=XM_019821486.2;refGene_gn=ADSS;refGene_tx=XM_023247274.1;refGene_gn=ADSS;refGene_tx=XM_023247275.1
F1 7521725 7521794
refGene_gn=ADSS;refGene_tx=XM_019821486.2;refGene_gn=ADSS;refGene_tx=XM_023247274.1;refGene_gn=ADSS;refGene_tx=XM_023247275.1
F1 7523473 7523524
refGene_gn=ADSS;refGene_tx=XM_019821486.2;refGene_gn=ADSS;refGene_tx=X

M_023247274.1;refGene_gn=ADSS;refGene_tx=XM_023247275.1
F1	7530061	7530128
refGene_gn=ADSS;refGene_tx=XM_019821486.2;refGene_gn=ADSS;refGene_tx=XM_023247274.1;refGene_gn=ADSS;refGene_tx=XM_023247275.1
F1	7530378	7530486
refGene_gn=ADSS;refGene_tx=XM_019821486.2;refGene_gn=ADSS;refGene_tx=XM_023247274.1;refGene_gn=ADSS;refGene_tx=XM_023247275.1
F1	7531491	7531573
refGene_gn=ADSS;refGene_tx=XM_019821486.2;refGene_gn=ADSS;refGene_tx=XM_023247274.1;refGene_gn=ADSS;refGene_tx=XM_023247275.1
F1	7533859	7533986
refGene_gn=ADSS;refGene_tx=XM_019821486.2;refGene_gn=ADSS;refGene_tx=XM_023247274.1;refGene_gn=ADSS;refGene_tx=XM_023247275.1
F1	7536052	7536207
refGene_gn=ADSS;refGene_tx=XM_019821486.2;refGene_gn=ADSS;refGene_tx=XM_023247274.1;refGene_gn=ADSS;refGene_tx=XM_023247275.1
F1	7538152	7538277
refGene_gn=ADSS;refGene_tx=XM_019821486.2;refGene_gn=ADSS;refGene_tx=XM_023247274.1;refGene_gn=ADSS;refGene_tx=XM_023247275.1
F1	7539505	7539603
refGene_gn=ADSS;refGene_tx=XM_019821486.2;refGene_gn=ADSS;refGene_tx=XM_023247274.1;refGene_gn=ADSS;refGene_tx=XM_023247275.1
F1	7542310	7542360	refGene_gn=ADSS;refGene_tx=XM_023247275.1
F1	7542427	7542577
refGene_gn=ADSS;refGene_tx=XM_019821486.2;refGene_gn=ADSS;refGene_tx=XM_023247274.1
F1	7544001	7544054
refGene_gn=ADSS;refGene_tx=XM_019821486.2;refGene_gn=ADSS;refGene_tx=XM_023247274.1
F1	7553312	7553404
refGene_gn=CF1H1orf100;refGene_tx=XM_006942622.4
F1	7562719	7562804
refGene_gn=CF1H1orf100;refGene_tx=XM_006942622.4
F1	7562805	7562893
refGene_gn=CF1H1orf100;refGene_tx=XM_006942622.4
F1	7567527	7567623
refGene_gn=CF1H1orf100;refGene_tx=XM_006942622.4
F1	7573708	7573764
refGene_gn=CF1H1orf100;refGene_tx=XM_006942622.4
F1	7858559	7860142
refGene_gn=ZBTB18;refGene_tx=XM_011290836.3;refGene_gn=ZBTB18;refGene_tx=XM_023247405.1
F1	7862550	7862563	refGene_gn=ZBTB18;refGene_tx=XM_011290836.3
F1	7864129	7864205
refGene_gn=LOC111558413;refGene_tx=XM_023248007.1
F1	7864328	7865047
refGene_gn=LOC111558413;refGene_tx=XM_023248007.1
F1	8083961	8084007
refGene_gn=AKT3;refGene_tx=XM_023247426.1;refGene_gn=AKT3;refGene_tx=XM_023247427.1;refGene_gn=AKT3;refGene_tx=XM_023247428.1

F1	8208382	8208508
refGene_gn=AKT3;refGene_tx=XM_023247426.1;refGene_gn=AKT3;refGene_tx=X
M_023247428.1
F1	8228013	8228101	refGene_gn=AKT3;refGene_tx=XM_011290838.3
F1	8235652	8235764
refGene_gn=AKT3;refGene_tx=XM_011290838.3;refGene_gn=AKT3;refGene_tx=X
M_023247426.1;refGene_gn=AKT3;refGene_tx=XM_023247427.1;refGene_gn=AKT
3;refGene_tx=XM_023247428.1
F1	8251081	8251226
refGene_gn=AKT3;refGene_tx=XM_011290838.3;refGene_gn=AKT3;refGene_tx=X
M_023247426.1;refGene_gn=AKT3;refGene_tx=XM_023247427.1;refGene_gn=AKT
3;refGene_tx=XM_023247428.1
F1	8259601	8259733
refGene_gn=AKT3;refGene_tx=XM_011290838.3;refGene_gn=AKT3;refGene_tx=X
M_023247426.1;refGene_gn=AKT3;refGene_tx=XM_023247427.1;refGene_gn=AKT
3;refGene_tx=XM_023247428.1
F1	8287240	8287306
refGene_gn=AKT3;refGene_tx=XM_011290838.3;refGene_gn=AKT3;refGene_tx=X
M_023247426.1;refGene_gn=AKT3;refGene_tx=XM_023247427.1
F1	8288612	8288681
refGene_gn=AKT3;refGene_tx=XM_011290838.3;refGene_gn=AKT3;refGene_tx=X
M_023247426.1;refGene_gn=AKT3;refGene_tx=XM_023247427.1
F1	8315710	8315833
refGene_gn=AKT3;refGene_tx=XM_011290838.3;refGene_gn=AKT3;refGene_tx=X
M_023247426.1;refGene_gn=AKT3;refGene_tx=XM_023247427.1;refGene_gn=AKT
3;refGene_tx=XM_023247428.1
F1	8324945	8325074
refGene_gn=AKT3;refGene_tx=XM_011290838.3;refGene_gn=AKT3;refGene_tx=X
M_023247426.1;refGene_gn=AKT3;refGene_tx=XM_023247427.1;refGene_gn=AKT
3;refGene_tx=XM_023247428.1
F1	8336452	8336667
refGene_gn=AKT3;refGene_tx=XM_011290838.3;refGene_gn=AKT3;refGene_tx=X
M_023247426.1;refGene_gn=AKT3;refGene_tx=XM_023247427.1;refGene_gn=AKT
3;refGene_tx=XM_023247428.1
F1	8343384	8343472
refGene_gn=AKT3;refGene_tx=XM_011290838.3;refGene_gn=AKT3;refGene_tx=X
M_023247426.1;refGene_gn=AKT3;refGene_tx=XM_023247427.1;refGene_gn=AKT
3;refGene_tx=XM_023247428.1
F1	8371062	8371165
refGene_gn=AKT3;refGene_tx=XM_011290838.3;refGene_gn=AKT3;refGene_tx=X
M_023247426.1;refGene_gn=AKT3;refGene_tx=XM_023247427.1;refGene_gn=AKT
3;refGene_tx=XM_023247428.1
F1	8378265	8378351
refGene_gn=AKT3;refGene_tx=XM_011290838.3;refGene_gn=AKT3;refGene_tx=X
M_023247426.1;refGene_gn=AKT3;refGene_tx=XM_023247427.1;refGene_gn=AKT
3;refGene_tx=XM_023247428.1
F1	8383713	8383762
refGene_gn=SDCCAG8;refGene_tx=XM_011290841.3;refGene_gn=SDCCAG8;refGen
e_tx=XM_019821498.2;refGene_gn=SDCCAG8;refGene_tx=XM_019821505.2;refGe
ne_gn=SDCCAG8;refGene_tx=XM_023247417.1;refGene_gn=SDCCAG8;refGene_tx=

XM_023247418.1;refGene_gn=SDCCAG8;refGene_tx=XM_023247420.1
F1 8387425 8387541
refGene_gn=SDCCAG8;refGene_tx=XM_019821498.2;refGene_gn=SDCCAG8;refGene_tx=XM_019821505.2;refGene_gn=SDCCAG8;refGene_tx=XM_023247417.1;refGene_gn=SDCCAG8;refGene_tx=XM_023247420.1
F1 8388732 8388744 refGene_gn=SDCCAG8;refGene_tx=XM_023247419.1
F1 8393720 8393847
refGene_gn=SDCCAG8;refGene_tx=XM_011290841.3;refGene_gn=SDCCAG8;refGene_tx=XM_019821498.2;refGene_gn=SDCCAG8;refGene_tx=XM_019821505.2;refGene_gn=SDCCAG8;refGene_tx=XM_023247417.1;refGene_gn=SDCCAG8;refGene_tx=XM_023247418.1;refGene_gn=SDCCAG8;refGene_tx=XM_023247419.1;refGene_gn=SDCCAG8;refGene_tx=XM_023247420.1
F1 8461353 8461485
refGene_gn=SDCCAG8;refGene_tx=XM_011290841.3;refGene_gn=SDCCAG8;refGene_tx=XM_019821498.2;refGene_gn=SDCCAG8;refGene_tx=XM_019821505.2;refGene_gn=SDCCAG8;refGene_tx=XM_023247417.1;refGene_gn=SDCCAG8;refGene_tx=XM_023247418.1;refGene_gn=SDCCAG8;refGene_tx=XM_023247419.1;refGene_gn=SDCCAG8;refGene_tx=XM_023247420.1;refGene_gn=SDCCAG8;refGene_tx=XM_023247424.1
F1 8472065 8472174
refGene_gn=SDCCAG8;refGene_tx=XM_011290841.3;refGene_gn=SDCCAG8;refGene_tx=XM_019821498.2;refGene_gn=SDCCAG8;refGene_tx=XM_023247417.1;refGene_gn=SDCCAG8;refGene_tx=XM_023247418.1;refGene_gn=SDCCAG8;refGene_tx=XM_023247419.1;refGene_gn=SDCCAG8;refGene_tx=XM_023247420.1
F1 8472691 8472765 refGene_gn=SDCCAG8;refGene_tx=XM_023247421.1
F1 8474240 8474368
refGene_gn=SDCCAG8;refGene_tx=XM_011290841.3;refGene_gn=SDCCAG8;refGene_tx=XM_019821498.2;refGene_gn=SDCCAG8;refGene_tx=XM_023247417.1;refGene_gn=SDCCAG8;refGene_tx=XM_023247418.1;refGene_gn=SDCCAG8;refGene_tx=XM_023247419.1;refGene_gn=SDCCAG8;refGene_tx=XM_023247420.1;refGene_gn=SDCCAG8;refGene_tx=XM_023247421.1;refGene_gn=SDCCAG8;refGene_tx=XM_023247422.1;refGene_gn=SDCCAG8;refGene_tx=XM_023247423.1;refGene_gn=SDCCAG8;refGene_tx=XM_023247424.1
F1 8495419 8495487
refGene_gn=SDCCAG8;refGene_tx=XM_023247422.1;refGene_gn=SDCCAG8;refGene_tx=XM_023247423.1
F1 8496281 8496392
refGene_gn=SDCCAG8;refGene_tx=XM_023247417.1;refGene_gn=SDCCAG8;refGene_tx=XM_023247418.1;refGene_gn=SDCCAG8;refGene_tx=XM_023247419.1;refGene_gn=SDCCAG8;refGene_tx=XM_023247420.1;refGene_gn=SDCCAG8;refGene_tx=XM_023247421.1;refGene_gn=SDCCAG8;refGene_tx=XM_023247422.1;refGene_gn=SDCCAG8;refGene_tx=XM_023247423.1;refGene_gn=SDCCAG8;refGene_tx=XM_023247424.1
F1 8510178 8510321
refGene_gn=SDCCAG8;refGene_tx=XM_011290841.3;refGene_gn=SDCCAG8;refGene_tx=XM_019821498.2;refGene_gn=SDCCAG8;refGene_tx=XM_019821505.2;refGene_gn=SDCCAG8;refGene_tx=XM_023247417.1;refGene_gn=SDCCAG8;refGene_tx=XM_023247418.1;refGene_gn=SDCCAG8;refGene_tx=XM_023247419.1;refGene_gn=SDCCAG8;refGene_tx=XM_023247420.1;refGene_gn=SDCCAG8;refGene_tx=XM_023247421.1;refGene_gn=SDCCAG8;refGene_tx=XM_023247422.1;refGene_gn=SDCC

AG8;refGene_tx=XM_023247423.1;refGene_gn=SDCCAG8;refGene_tx=XM_0232474
24.1
F1 8542874 8543168 refGene_gn=SDCCAG8;refGene_tx=XM_023247425.1
F1 8545558 8545675
refGene_gn=SDCCAG8;refGene_tx=XM_011290841.3;refGene_gn=SDCCAG8;refGen
e_tx=XM_019821498.2;refGene_gn=SDCCAG8;refGene_tx=XM_019821505.2;refGe
ne_gn=SDCCAG8;refGene_tx=XM_023247417.1;refGene_gn=SDCCAG8;refGene_tx=
XM_023247418.1;refGene_gn=SDCCAG8;refGene_tx=XM_023247419.1;refGene_gn
=SDCCAG8;refGene_tx=XM_023247420.1;refGene_gn=SDCCAG8;refGene_tx=XM_02
3247421.1;refGene_gn=SDCCAG8;refGene_tx=XM_023247422.1;refGene_gn=SDCC
AG8;refGene_tx=XM_023247423.1;refGene_gn=SDCCAG8;refGene_tx=XM_0232474
24.1;refGene_gn=SDCCAG8;refGene_tx=XM_023247425.1
F1 8548517 8548652
refGene_gn=SDCCAG8;refGene_tx=XM_011290841.3;refGene_gn=SDCCAG8;refGen
e_tx=XM_019821498.2;refGene_gn=SDCCAG8;refGene_tx=XM_019821505.2;refGe
ne_gn=SDCCAG8;refGene_tx=XM_023247417.1;refGene_gn=SDCCAG8;refGene_tx=
XM_023247418.1;refGene_gn=SDCCAG8;refGene_tx=XM_023247419.1;refGene_gn
=SDCCAG8;refGene_tx=XM_023247420.1;refGene_gn=SDCCAG8;refGene_tx=XM_02
3247421.1;refGene_gn=SDCCAG8;refGene_tx=XM_023247422.1;refGene_gn=SDCC
AG8;refGene_tx=XM_023247423.1;refGene_gn=SDCCAG8;refGene_tx=XM_0232474
24.1;refGene_gn=SDCCAG8;refGene_tx=XM_023247425.1
F1 8557034 8557187
refGene_gn=SDCCAG8;refGene_tx=XM_011290841.3;refGene_gn=SDCCAG8;refGen
e_tx=XM_019821498.2;refGene_gn=SDCCAG8;refGene_tx=XM_019821505.2;refGe
ne_gn=SDCCAG8;refGene_tx=XM_023247417.1;refGene_gn=SDCCAG8;refGene_tx=
XM_023247418.1;refGene_gn=SDCCAG8;refGene_tx=XM_023247419.1;refGene_gn
=SDCCAG8;refGene_tx=XM_023247420.1;refGene_gn=SDCCAG8;refGene_tx=XM_02
3247421.1;refGene_gn=SDCCAG8;refGene_tx=XM_023247422.1;refGene_gn=SDCC
AG8;refGene_tx=XM_023247423.1;refGene_gn=SDCCAG8;refGene_tx=XM_0232474
24.1;refGene_gn=SDCCAG8;refGene_tx=XM_023247425.1
F1 8567188 8567327
refGene_gn=SDCCAG8;refGene_tx=XM_011290841.3;refGene_gn=SDCCAG8;refGen
e_tx=XM_019821498.2;refGene_gn=SDCCAG8;refGene_tx=XM_019821505.2;refGe
ne_gn=SDCCAG8;refGene_tx=XM_023247417.1;refGene_gn=SDCCAG8;refGene_tx=
XM_023247418.1;refGene_gn=SDCCAG8;refGene_tx=XM_023247419.1;refGene_gn
=SDCCAG8;refGene_tx=XM_023247420.1;refGene_gn=SDCCAG8;refGene_tx=XM_02
3247421.1;refGene_gn=SDCCAG8;refGene_tx=XM_023247422.1;refGene_gn=SDCC
AG8;refGene_tx=XM_023247423.1;refGene_gn=SDCCAG8;refGene_tx=XM_0232474
24.1;refGene_gn=SDCCAG8;refGene_tx=XM_023247425.1
F1 8576987 8577176
refGene_gn=SDCCAG8;refGene_tx=XM_011290841.3;refGene_gn=SDCCAG8;refGen
e_tx=XM_019821498.2;refGene_gn=SDCCAG8;refGene_tx=XM_019821505.2;refGe
ne_gn=SDCCAG8;refGene_tx=XM_023247417.1;refGene_gn=SDCCAG8;refGene_tx=
XM_023247418.1;refGene_gn=SDCCAG8;refGene_tx=XM_023247419.1;refGene_gn
=SDCCAG8;refGene_tx=XM_023247421.1;refGene_gn=SDCCAG8;refGene_tx=XM_02
3247422.1;refGene_gn=SDCCAG8;refGene_tx=XM_023247423.1;refGene_gn=SDCC
AG8;refGene_tx=XM_023247424.1;refGene_gn=SDCCAG8;refGene_tx=XM_0232474
25.1
F1 8580926 8580991
refGene_gn=SDCCAG8;refGene_tx=XM_011290841.3;refGene_gn=SDCCAG8;refGen

e_tx=XM_019821498.2;refGene_gn=SDCCAG8;refGene_tx=XM_019821505.2;refGene_gn=SDCCAG8;refGene_tx=XM_023247417.1;refGene_gn=SDCCAG8;refGene_tx=XM_023247418.1;refGene_gn=SDCCAG8;refGene_tx=XM_023247419.1;refGene_gn=SDCCAG8;refGene_tx=XM_023247420.1;refGene_gn=SDCCAG8;refGene_tx=XM_023247421.1;refGene_gn=SDCCAG8;refGene_tx=XM_023247422.1;refGene_gn=SDCCAG8;refGene_tx=XM_023247423.1;refGene_gn=SDCCAG8;refGene_tx=XM_023247424.1;refGene_gn=SDCCAG8;refGene_tx=XM_023247425.1
F1 8590694 8590835
refGene_gn=SDCCAG8;refGene_tx=XM_011290841.3;refGene_gn=SDCCAG8;refGene_tx=XM_019821498.2;refGene_gn=SDCCAG8;refGene_tx=XM_019821505.2;refGene_gn=SDCCAG8;refGene_tx=XM_023247417.1;refGene_gn=SDCCAG8;refGene_tx=XM_023247418.1;refGene_gn=SDCCAG8;refGene_tx=XM_023247419.1;refGene_gn=SDCCAG8;refGene_tx=XM_023247420.1;refGene_gn=SDCCAG8;refGene_tx=XM_023247421.1;refGene_gn=SDCCAG8;refGene_tx=XM_023247422.1;refGene_gn=SDCCAG8;refGene_tx=XM_023247423.1;refGene_gn=SDCCAG8;refGene_tx=XM_023247424.1;refGene_gn=SDCCAG8;refGene_tx=XM_023247425.1
F1 8603844 8603982
refGene_gn=SDCCAG8;refGene_tx=XM_011290841.3;refGene_gn=SDCCAG8;refGene_tx=XM_019821498.2;refGene_gn=SDCCAG8;refGene_tx=XM_019821505.2;refGene_gn=SDCCAG8;refGene_tx=XM_023247417.1;refGene_gn=SDCCAG8;refGene_tx=XM_023247418.1;refGene_gn=SDCCAG8;refGene_tx=XM_023247419.1;refGene_gn=SDCCAG8;refGene_tx=XM_023247420.1;refGene_gn=SDCCAG8;refGene_tx=XM_023247421.1;refGene_gn=SDCCAG8;refGene_tx=XM_023247422.1;refGene_gn=SDCCAG8;refGene_tx=XM_023247423.1;refGene_gn=SDCCAG8;refGene_tx=XM_023247424.1;refGene_gn=SDCCAG8;refGene_tx=XM_023247425.1
F1 8615457 8615571
refGene_gn=SDCCAG8;refGene_tx=XM_011290841.3;refGene_gn=SDCCAG8;refGene_tx=XM_019821498.2;refGene_gn=SDCCAG8;refGene_tx=XM_019821505.2;refGene_gn=SDCCAG8;refGene_tx=XM_023247417.1;refGene_gn=SDCCAG8;refGene_tx=XM_023247418.1;refGene_gn=SDCCAG8;refGene_tx=XM_023247419.1;refGene_gn=SDCCAG8;refGene_tx=XM_023247420.1;refGene_gn=SDCCAG8;refGene_tx=XM_023247421.1;refGene_gn=SDCCAG8;refGene_tx=XM_023247422.1;refGene_gn=SDCCAG8;refGene_tx=XM_023247423.1;refGene_gn=SDCCAG8;refGene_tx=XM_023247424.1;refGene_gn=SDCCAG8;refGene_tx=XM_023247425.1
F1 8618426 8618512
refGene_gn=SDCCAG8;refGene_tx=XM_011290841.3;refGene_gn=SDCCAG8;refGene_tx=XM_019821498.2;refGene_gn=SDCCAG8;refGene_tx=XM_019821505.2;refGene_gn=SDCCAG8;refGene_tx=XM_023247417.1;refGene_gn=SDCCAG8;refGene_tx=XM_023247418.1;refGene_gn=SDCCAG8;refGene_tx=XM_023247419.1;refGene_gn=SDCCAG8;refGene_tx=XM_023247420.1;refGene_gn=SDCCAG8;refGene_tx=XM_023247421.1;refGene_gn=SDCCAG8;refGene_tx=XM_023247422.1;refGene_gn=SDCCAG8;refGene_tx=XM_023247423.1;refGene_gn=SDCCAG8;refGene_tx=XM_023247424.1;refGene_gn=SDCCAG8;refGene_tx=XM_023247425.1
F1 8619675 8619834
refGene_gn=SDCCAG8;refGene_tx=XM_011290841.3;refGene_gn=SDCCAG8;refGene_tx=XM_019821498.2;refGene_gn=SDCCAG8;refGene_tx=XM_019821505.2;refGene_gn=SDCCAG8;refGene_tx=XM_023247417.1;refGene_gn=SDCCAG8;refGene_tx=XM_023247418.1;refGene_gn=SDCCAG8;refGene_tx=XM_023247419.1;refGene_gn=SDCCAG8;refGene_tx=XM_023247420.1;refGene_gn=SDCCAG8;refGene_tx=XM_023247421.1;refGene_gn=SDCCAG8;refGene_tx=XM_023247422.1;refGene_gn=SDCC

AG8;refGene_tx=XM_023247423.1;refGene_gn=SDCCAG8;refGene_tx=XM_0232474
24.1;refGene_gn=SDCCAG8;refGene_tx=XM_023247425.1
F1	8632412	8632659
refGene_gn=SDCCAG8;refGene_tx=XM_011290841.3;refGene_gn=SDCCAG8;refGen
e_tx=XM_019821498.2;refGene_gn=SDCCAG8;refGene_tx=XM_019821505.2;refGe
ne_gn=SDCCAG8;refGene_tx=XM_023247417.1;refGene_gn=SDCCAG8;refGene_tx=
XM_023247418.1;refGene_gn=SDCCAG8;refGene_tx=XM_023247419.1;refGene_gn
=SDCCAG8;refGene_tx=XM_023247420.1;refGene_gn=SDCCAG8;refGene_tx=XM_02
3247421.1;refGene_gn=SDCCAG8;refGene_tx=XM_023247422.1;refGene_gn=SDCC
AG8;refGene_tx=XM_023247423.1;refGene_gn=SDCCAG8;refGene_tx=XM_0232474
24.1;refGene_gn=SDCCAG8;refGene_tx=XM_023247425.1
F1	8667762	8667867
refGene_gn=CEP170;refGene_tx=XM_006942628.4;refGene_gn=CEP170;refGene_
tx=XM_006942630.4;refGene_gn=CEP170;refGene_tx=XM_019821494.2;refGene_
gn=CEP170;refGene_tx=XM_023247409.1;refGene_gn=CEP170;refGene_tx=XM_02
3247410.1;refGene_gn=CEP170;refGene_tx=XM_023247411.1;refGene_gn=CEP17
0;refGene_tx=XM_023247412.1;refGene_gn=CEP170;refGene_tx=XM_023247413.
1;refGene_gn=CEP170;refGene_tx=XM_023247414.1;refGene_gn=CEP170;refGen
e_tx=XM_023247415.1;refGene_gn=CEP170;refGene_tx=XM_023247416.1
F1	8682146	8682236
refGene_gn=CEP170;refGene_tx=XM_006942628.4;refGene_gn=CEP170;refGene_
tx=XM_006942630.4;refGene_gn=CEP170;refGene_tx=XM_019821494.2;refGene_
gn=CEP170;refGene_tx=XM_023247409.1;refGene_gn=CEP170;refGene_tx=XM_02
3247410.1;refGene_gn=CEP170;refGene_tx=XM_023247411.1;refGene_gn=CEP17
0;refGene_tx=XM_023247412.1;refGene_gn=CEP170;refGene_tx=XM_023247413.
1;refGene_gn=CEP170;refGene_tx=XM_023247414.1;refGene_gn=CEP170;refGen
e_tx=XM_023247415.1;refGene_gn=CEP170;refGene_tx=XM_023247416.1
F1	8689395	8689474
refGene_gn=CEP170;refGene_tx=XM_006942628.4;refGene_gn=CEP170;refGene_
tx=XM_006942630.4;refGene_gn=CEP170;refGene_tx=XM_019821494.2;refGene_
gn=CEP170;refGene_tx=XM_023247409.1;refGene_gn=CEP170;refGene_tx=XM_02
3247410.1;refGene_gn=CEP170;refGene_tx=XM_023247411.1;refGene_gn=CEP17
0;refGene_tx=XM_023247412.1;refGene_gn=CEP170;refGene_tx=XM_023247413.
1;refGene_gn=CEP170;refGene_tx=XM_023247414.1;refGene_gn=CEP170;refGen
e_tx=XM_023247415.1;refGene_gn=CEP170;refGene_tx=XM_023247416.1
F1	8698921	8698980
refGene_gn=CEP170;refGene_tx=XM_006942628.4;refGene_gn=CEP170;refGene_
tx=XM_006942630.4;refGene_gn=CEP170;refGene_tx=XM_019821494.2;refGene_
gn=CEP170;refGene_tx=XM_023247409.1;refGene_gn=CEP170;refGene_tx=XM_02
3247410.1;refGene_gn=CEP170;refGene_tx=XM_023247411.1;refGene_gn=CEP17
0;refGene_tx=XM_023247412.1;refGene_gn=CEP170;refGene_tx=XM_023247413.
1;refGene_gn=CEP170;refGene_tx=XM_023247414.1;refGene_gn=CEP170;refGen
e_tx=XM_023247415.1;refGene_gn=CEP170;refGene_tx=XM_023247416.1
F1	8699076	8699239
refGene_gn=CEP170;refGene_tx=XM_006942628.4;refGene_gn=CEP170;refGene_
tx=XM_006942630.4;refGene_gn=CEP170;refGene_tx=XM_019821494.2;refGene_
gn=CEP170;refGene_tx=XM_023247409.1;refGene_gn=CEP170;refGene_tx=XM_02
3247410.1;refGene_gn=CEP170;refGene_tx=XM_023247411.1;refGene_gn=CEP17
0;refGene_tx=XM_023247412.1;refGene_gn=CEP170;refGene_tx=XM_023247413.
1;refGene_gn=CEP170;refGene_tx=XM_023247414.1;refGene_gn=CEP170;refGen

e_tx=XM_023247415.1;refGene_gn=CEP170;refGene_tx=XM_023247416.1
F1 8700361 8700496
refGene_gn=CEP170;refGene_tx=XM_006942628.4;refGene_gn=CEP170;refGene_tx=XM_006942630.4;refGene_gn=CEP170;refGene_tx=XM_019821494.2;refGene_gn=CEP170;refGene_tx=XM_023247409.1;refGene_gn=CEP170;refGene_tx=XM_023247410.1;refGene_gn=CEP170;refGene_tx=XM_023247411.1;refGene_gn=CEP170;refGene_tx=XM_023247412.1;refGene_gn=CEP170;refGene_tx=XM_023247413.1;refGene_gn=CEP170;refGene_tx=XM_023247414.1;refGene_gn=CEP170;refGene_tx=XM_023247415.1;refGene_gn=CEP170;refGene_tx=XM_023247416.1
F1 8706234 8706711
refGene_gn=CEP170;refGene_tx=XM_006942628.4;refGene_gn=CEP170;refGene_tx=XM_006942630.4;refGene_gn=CEP170;refGene_tx=XM_019821494.2;refGene_gn=CEP170;refGene_tx=XM_023247409.1;refGene_gn=CEP170;refGene_tx=XM_023247410.1;refGene_gn=CEP170;refGene_tx=XM_023247411.1;refGene_gn=CEP170;refGene_tx=XM_023247412.1;refGene_gn=CEP170;refGene_tx=XM_023247413.1;refGene_gn=CEP170;refGene_tx=XM_023247414.1;refGene_gn=CEP170;refGene_tx=XM_023247415.1;refGene_gn=CEP170;refGene_tx=XM_023247416.1
F1 8711229 8711393
refGene_gn=CEP170;refGene_tx=XM_006942628.4;refGene_gn=CEP170;refGene_tx=XM_006942630.4;refGene_gn=CEP170;refGene_tx=XM_019821494.2;refGene_gn=CEP170;refGene_tx=XM_023247409.1;refGene_gn=CEP170;refGene_tx=XM_023247410.1;refGene_gn=CEP170;refGene_tx=XM_023247411.1;refGene_gn=CEP170;refGene_tx=XM_023247412.1;refGene_gn=CEP170;refGene_tx=XM_023247413.1;refGene_gn=CEP170;refGene_tx=XM_023247414.1;refGene_gn=CEP170;refGene_tx=XM_023247415.1;refGene_gn=CEP170;refGene_tx=XM_023247416.1
F1 8711579 8711873
refGene_gn=CEP170;refGene_tx=XM_006942628.4;refGene_gn=CEP170;refGene_tx=XM_006942630.4;refGene_gn=CEP170;refGene_tx=XM_019821494.2;refGene_gn=CEP170;refGene_tx=XM_023247409.1;refGene_gn=CEP170;refGene_tx=XM_023247410.1;refGene_gn=CEP170;refGene_tx=XM_023247411.1;refGene_gn=CEP170;refGene_tx=XM_023247412.1;refGene_gn=CEP170;refGene_tx=XM_023247413.1
F1 8722632 8722782
refGene_gn=CEP170;refGene_tx=XM_006942628.4;refGene_gn=CEP170;refGene_tx=XM_006942630.4;refGene_gn=CEP170;refGene_tx=XM_019821494.2;refGene_gn=CEP170;refGene_tx=XM_023247409.1;refGene_gn=CEP170;refGene_tx=XM_023247410.1;refGene_gn=CEP170;refGene_tx=XM_023247411.1;refGene_gn=CEP170;refGene_tx=XM_023247412.1;refGene_gn=CEP170;refGene_tx=XM_023247413.1;refGene_gn=CEP170;refGene_tx=XM_023247414.1;refGene_gn=CEP170;refGene_tx=XM_023247415.1;refGene_gn=CEP170;refGene_tx=XM_023247416.1
F1 8725322 8725449
refGene_gn=CEP170;refGene_tx=XM_006942628.4;refGene_gn=CEP170;refGene_tx=XM_006942630.4;refGene_gn=CEP170;refGene_tx=XM_019821494.2;refGene_gn=CEP170;refGene_tx=XM_023247409.1;refGene_gn=CEP170;refGene_tx=XM_023247410.1;refGene_gn=CEP170;refGene_tx=XM_023247411.1;refGene_gn=CEP170;refGene_tx=XM_023247412.1;refGene_gn=CEP170;refGene_tx=XM_023247413.1;refGene_gn=CEP170;refGene_tx=XM_023247414.1;refGene_gn=CEP170;refGene_tx=XM_023247415.1;refGene_gn=CEP170;refGene_tx=XM_023247416.1
F1 8727802 8729617
refGene_gn=CEP170;refGene_tx=XM_006942628.4;refGene_gn=CEP170;refGene_

tx=XM_006942630.4;refGene_gn=CEP170;refGene_tx=XM_019821494.2;refGene_gn=CEP170;refGene_tx=XM_023247409.1;refGene_gn=CEP170;refGene_tx=XM_023247410.1;refGene_gn=CEP170;refGene_tx=XM_023247411.1;refGene_gn=CEP170;refGene_tx=XM_023247412.1;refGene_gn=CEP170;refGene_tx=XM_023247413.1;refGene_gn=CEP170;refGene_tx=XM_023247414.1;refGene_gn=CEP170;refGene_tx=XM_023247415.1;refGene_gn=CEP170;refGene_tx=XM_023247416.1
F1 8736689 8736924
refGene_gn=CEP170;refGene_tx=XM_006942628.4;refGene_gn=CEP170;refGene_tx=XM_006942630.4;refGene_gn=CEP170;refGene_tx=XM_019821494.2;refGene_gn=CEP170;refGene_tx=XM_023247409.1;refGene_gn=CEP170;refGene_tx=XM_023247410.1;refGene_gn=CEP170;refGene_tx=XM_023247411.1;refGene_gn=CEP170;refGene_tx=XM_023247412.1;refGene_gn=CEP170;refGene_tx=XM_023247413.1;refGene_gn=CEP170;refGene_tx=XM_023247414.1;refGene_gn=CEP170;refGene_tx=XM_023247415.1;refGene_gn=CEP170;refGene_tx=XM_023247416.1
F1 8741904 8742082
refGene_gn=CEP170;refGene_tx=XM_006942628.4;refGene_gn=CEP170;refGene_tx=XM_006942630.4;refGene_gn=CEP170;refGene_tx=XM_019821494.2;refGene_gn=CEP170;refGene_tx=XM_023247409.1;refGene_gn=CEP170;refGene_tx=XM_023247410.1;refGene_gn=CEP170;refGene_tx=XM_023247411.1;refGene_gn=CEP170;refGene_tx=XM_023247412.1;refGene_gn=CEP170;refGene_tx=XM_023247413.1;refGene_gn=CEP170;refGene_tx=XM_023247414.1;refGene_gn=CEP170;refGene_tx=XM_023247415.1;refGene_gn=CEP170;refGene_tx=XM_023247416.1
F1 8744107 8744278
refGene_gn=CEP170;refGene_tx=XM_006942628.4;refGene_gn=CEP170;refGene_tx=XM_006942630.4;refGene_gn=CEP170;refGene_tx=XM_019821494.2;refGene_gn=CEP170;refGene_tx=XM_023247409.1;refGene_gn=CEP170;refGene_tx=XM_023247410.1;refGene_gn=CEP170;refGene_tx=XM_023247411.1;refGene_gn=CEP170;refGene_tx=XM_023247412.1;refGene_gn=CEP170;refGene_tx=XM_023247413.1;refGene_gn=CEP170;refGene_tx=XM_023247414.1;refGene_gn=CEP170;refGene_tx=XM_023247415.1;refGene_gn=CEP170;refGene_tx=XM_023247416.1
F1 8748492 8748581
refGene_gn=CEP170;refGene_tx=XM_006942628.4;refGene_gn=CEP170;refGene_tx=XM_006942630.4;refGene_gn=CEP170;refGene_tx=XM_019821494.2;refGene_gn=CEP170;refGene_tx=XM_023247409.1;refGene_gn=CEP170;refGene_tx=XM_023247410.1;refGene_gn=CEP170;refGene_tx=XM_023247411.1;refGene_gn=CEP170;refGene_tx=XM_023247412.1;refGene_gn=CEP170;refGene_tx=XM_023247413.1;refGene_gn=CEP170;refGene_tx=XM_023247414.1;refGene_gn=CEP170;refGene_tx=XM_023247415.1;refGene_gn=CEP170;refGene_tx=XM_023247416.1
F1 8754137 8754231
refGene_gn=CEP170;refGene_tx=XM_006942628.4;refGene_gn=CEP170;refGene_tx=XM_006942630.4;refGene_gn=CEP170;refGene_tx=XM_019821494.2;refGene_gn=CEP170;refGene_tx=XM_023247409.1;refGene_gn=CEP170;refGene_tx=XM_023247410.1;refGene_gn=CEP170;refGene_tx=XM_023247411.1;refGene_gn=CEP170;refGene_tx=XM_023247412.1;refGene_gn=CEP170;refGene_tx=XM_023247413.1;refGene_gn=CEP170;refGene_tx=XM_023247414.1;refGene_gn=CEP170;refGene_tx=XM_023247415.1;refGene_gn=CEP170;refGene_tx=XM_023247416.1
F1 8754965 8755017
refGene_gn=CEP170;refGene_tx=XM_006942628.4;refGene_gn=CEP170;refGene_tx=XM_006942630.4;refGene_gn=CEP170;refGene_tx=XM_019821494.2;refGene_gn=CEP170;refGene_tx=XM_023247409.1;refGene_gn=CEP170;refGene_tx=XM_02

3247410.1;refGene_gn=CEP170;refGene_tx=XM_023247411.1;refGene_gn=CEP170;refGene_tx=XM_023247412.1;refGene_gn=CEP170;refGene_tx=XM_023247413.1;refGene_gn=CEP170;refGene_tx=XM_023247414.1;refGene_gn=CEP170;refGene_tx=XM_023247415.1;refGene_gn=CEP170;refGene_tx=XM_023247416.1
F1 8756356 8756430 refGene_gn=CEP170;refGene_tx=XM_019821494.2
F1 8756630 8756920
refGene_gn=CEP170;refGene_tx=XM_006942628.4;refGene_gn=CEP170;refGene_tx=XM_006942630.4;refGene_gn=CEP170;refGene_tx=XM_023247409.1;refGene_gn=CEP170;refGene_tx=XM_023247410.1;refGene_gn=CEP170;refGene_tx=XM_023247411.1;refGene_gn=CEP170;refGene_tx=XM_023247412.1;refGene_gn=CEP170;refGene_tx=XM_023247413.1;refGene_gn=CEP170;refGene_tx=XM_023247414.1;refGene_gn=CEP170;refGene_tx=XM_023247415.1;refGene_gn=CEP170;refGene_tx=XM_023247416.1
F1 8757236 8757263
refGene_gn=CEP170;refGene_tx=XM_006942628.4;refGene_gn=CEP170;refGene_tx=XM_006942630.4
F1 9228946 9229135
refGene_gn=PLD5;refGene_tx=XM_023247982.1;refGene_gn=PLD5;refGene_tx=XM_023247983.1
F1 9309504 9309507 refGene_gn=PLD5;refGene_tx=XM_023247984.1
F1 9405516 9405660
refGene_gn=PLD5;refGene_tx=XM_019821516.1;refGene_gn=PLD5;refGene_tx=XM_023247982.1;refGene_gn=PLD5;refGene_tx=XM_023247983.1;refGene_gn=PLD5;refGene_tx=XM_023247984.1
F1 9473074 9473243
refGene_gn=PLD5;refGene_tx=XM_019821516.1;refGene_gn=PLD5;refGene_tx=XM_023247982.1;refGene_gn=PLD5;refGene_tx=XM_023247983.1;refGene_gn=PLD5;refGene_tx=XM_023247984.1
F1 9493728 9493840
refGene_gn=PLD5;refGene_tx=XM_019821516.1;refGene_gn=PLD5;refGene_tx=XM_023247982.1;refGene_gn=PLD5;refGene_tx=XM_023247983.1;refGene_gn=PLD5;refGene_tx=XM_023247984.1
F1 9536011 9536108 refGene_gn=PLD5;refGene_tx=XM_019821518.2
F1 9546898 9547026
refGene_gn=PLD5;refGene_tx=XM_019821516.1;refGene_gn=PLD5;refGene_tx=XM_019821518.2;refGene_gn=PLD5;refGene_tx=XM_023247982.1;refGene_gn=PLD5;refGene_tx=XM_023247983.1;refGene_gn=PLD5;refGene_tx=XM_023247984.1
F1 9618768 9618966
refGene_gn=PLD5;refGene_tx=XM_019821516.1;refGene_gn=PLD5;refGene_tx=XM_019821518.2;refGene_gn=PLD5;refGene_tx=XM_023247982.1;refGene_gn=PLD5;refGene_tx=XM_023247983.1;refGene_gn=PLD5;refGene_tx=XM_023247984.1
F1 9629260 9629397
refGene_gn=PLD5;refGene_tx=XM_019821516.1;refGene_gn=PLD5;refGene_tx=XM_019821518.2;refGene_gn=PLD5;refGene_tx=XM_023247982.1;refGene_gn=PLD5;refGene_tx=XM_023247983.1;refGene_gn=PLD5;refGene_tx=XM_023247984.1
F1 9640990 9641159
refGene_gn=PLD5;refGene_tx=XM_019821516.1;refGene_gn=PLD5;refGene_tx=XM_019821518.2;refGene_gn=PLD5;refGene_tx=XM_023247982.1;refGene_gn=PLD5;refGene_tx=XM_023247983.1;refGene_gn=PLD5;refGene_tx=XM_023247984.1
F1 9650948 9651063

refGene_gn=PLD5;refGene_tx=XM_019821516.1;refGene_gn=PLD5;refGene_tx=X
M_019821518.2;refGene_gn=PLD5;refGene_tx=XM_023247982.1;refGene_gn=PLD
5;refGene_tx=XM_023247983.1;refGene_gn=PLD5;refGene_tx=XM_023247984.1
F1	9658032	9658121	refGene_gn=PLD5;refGene_tx=XM_023247983.1
F1	9659207	9659464
refGene_gn=PLD5;refGene_tx=XM_019821516.1;refGene_gn=PLD5;refGene_tx=X
M_019821518.2;refGene_gn=PLD5;refGene_tx=XM_023247982.1;refGene_gn=PLD
5;refGene_tx=XM_023247984.1
F1	9713425	9713483	refGene_gn=MAP1LC3C;refGene_tx=XM_019821523.1
F1	9713584	9713640	refGene_gn=MAP1LC3C;refGene_tx=XM_019821523.1
F1	9713825	9713932	refGene_gn=MAP1LC3C;refGene_tx=XM_019821523.1
F1	9716029	9716234	refGene_gn=MAP1LC3C;refGene_tx=XM_019821523.1
F1	9761943	9763245	refGene_gn=BECN2;refGene_tx=XM_003999135.2
F1	9803654	9803862
refGene_gn=LOC101095070;refGene_tx=XM_003999136.2
F1	9806877	9806993
refGene_gn=LOC101095070;refGene_tx=XM_003999136.2
F1	9807149	9807205
refGene_gn=LOC101095070;refGene_tx=XM_003999136.2
F1	9807315	9807373
refGene_gn=LOC101095070;refGene_tx=XM_003999136.2
F1	9819195	9819337
refGene_gn=EXO1;refGene_tx=XM_019821527.2;refGene_gn=EXO1;refGene_tx=X
M_019821529.2;refGene_gn=EXO1;refGene_tx=XM_019821530.2;refGene_gn=EXO
1;refGene_tx=XM_023247556.1
F1	9821755	9821949
refGene_gn=EXO1;refGene_tx=XM_019821527.2;refGene_gn=EXO1;refGene_tx=X
M_019821529.2;refGene_gn=EXO1;refGene_tx=XM_019821530.2;refGene_gn=EXO
1;refGene_tx=XM_023247556.1
F1	9825696	9825798
refGene_gn=EXO1;refGene_tx=XM_019821527.2;refGene_gn=EXO1;refGene_tx=X
M_019821529.2;refGene_gn=EXO1;refGene_tx=XM_019821530.2;refGene_gn=EXO
1;refGene_tx=XM_023247556.1
F1	9828990	9829882
refGene_gn=EXO1;refGene_tx=XM_019821527.2;refGene_gn=EXO1;refGene_tx=X
M_019821529.2;refGene_gn=EXO1;refGene_tx=XM_019821530.2;refGene_gn=EXO
1;refGene_tx=XM_023247556.1
F1	9835074	9835318
refGene_gn=EXO1;refGene_tx=XM_019821527.2;refGene_gn=EXO1;refGene_tx=X
M_019821529.2;refGene_gn=EXO1;refGene_tx=XM_019821530.2;refGene_gn=EXO
1;refGene_tx=XM_023247556.1
F1	9839699	9839925
refGene_gn=EXO1;refGene_tx=XM_019821527.2;refGene_gn=EXO1;refGene_tx=X
M_019821529.2;refGene_gn=EXO1;refGene_tx=XM_019821530.2;refGene_gn=EXO
1;refGene_tx=XM_023247556.1
F1	9845202	9845299
refGene_gn=EXO1;refGene_tx=XM_019821527.2;refGene_gn=EXO1;refGene_tx=X
M_019821529.2;refGene_gn=EXO1;refGene_tx=XM_019821530.2;refGene_gn=EXO
1;refGene_tx=XM_023247556.1
F1	9845870	9846058

refGene_gn=EXO1;refGene_tx=XM_019821527.2;refGene_gn=EXO1;refGene_tx=XM_019821529.2;refGene_gn=EXO1;refGene_tx=XM_019821530.2;refGene_gn=EXO1;refGene_tx=XM_023247556.1
F1 9847645 9847858
refGene_gn=EXO1;refGene_tx=XM_019821527.2;refGene_gn=EXO1;refGene_tx=XM_019821529.2;refGene_gn=EXO1;refGene_tx=XM_023247556.1
F1 9848789 9848927
refGene_gn=EXO1;refGene_tx=XM_019821527.2;refGene_gn=EXO1;refGene_tx=XM_019821529.2;refGene_gn=EXO1;refGene_tx=XM_019821530.2;refGene_gn=EXO1;refGene_tx=XM_023247556.1
F1 9849922 9850046
refGene_gn=EXO1;refGene_tx=XM_019821527.2;refGene_gn=EXO1;refGene_tx=XM_019821529.2;refGene_gn=EXO1;refGene_tx=XM_019821530.2;refGene_gn=EXO1;refGene_tx=XM_023247556.1
F1 9852786 9852906
refGene_gn=EXO1;refGene_tx=XM_019821527.2;refGene_gn=EXO1;refGene_tx=XM_019821529.2;refGene_gn=EXO1;refGene_tx=XM_019821530.2;refGene_gn=EXO1;refGene_tx=XM_023247556.1
F1 9853773 9853934
refGene_gn=EXO1;refGene_tx=XM_019821527.2;refGene_gn=EXO1;refGene_tx=XM_019821530.2;refGene_gn=EXO1;refGene_tx=XM_023247556.1
F1 9909024 9909108
refGene_gn=WDR64;refGene_tx=XM_003999137.4;refGene_gn=WDR64;refGene_tx=XM_019821525.1
F1 9914524 9914635
refGene_gn=WDR64;refGene_tx=XM_003999137.4;refGene_gn=WDR64;refGene_tx=XM_019821525.1
F1 9915507 9915588
refGene_gn=WDR64;refGene_tx=XM_003999137.4;refGene_gn=WDR64;refGene_tx=XM_019821525.1
F1 9918365 9918471
refGene_gn=WDR64;refGene_tx=XM_003999137.4;refGene_gn=WDR64;refGene_tx=XM_019821525.1
F1 9920503 9920692
refGene_gn=WDR64;refGene_tx=XM_003999137.4;refGene_gn=WDR64;refGene_tx=XM_019821525.1
F1 9923592 9923702
refGene_gn=WDR64;refGene_tx=XM_003999137.4;refGene_gn=WDR64;refGene_tx=XM_019821525.1
F1 9927178 9927237
refGene_gn=WDR64;refGene_tx=XM_003999137.4;refGene_gn=WDR64;refGene_tx=XM_019821525.1
F1 9932216 9932322
refGene_gn=WDR64;refGene_tx=XM_003999137.4;refGene_gn=WDR64;refGene_tx=XM_019821525.1
F1 9934458 9934598
refGene_gn=WDR64;refGene_tx=XM_003999137.4;refGene_gn=WDR64;refGene_tx=XM_019821525.1
F1 9935147 9935184
refGene_gn=WDR64;refGene_tx=XM_003999137.4;refGene_gn=WDR64;refGene_tx

=XM_019821525.1
F1	9936800	9936870
refGene_gn=WDR64;refGene_tx=XM_003999137.4;refGene_gn=WDR64;refGene_tx
=XM_019821525.1
F1	9937914	9938016
refGene_gn=WDR64;refGene_tx=XM_003999137.4;refGene_gn=WDR64;refGene_tx
=XM_019821525.1
F1	9940580	9940711
refGene_gn=WDR64;refGene_tx=XM_003999137.4;refGene_gn=WDR64;refGene_tx
=XM_019821525.1
F1	9946552	9946726
refGene_gn=WDR64;refGene_tx=XM_003999137.4;refGene_gn=WDR64;refGene_tx
=XM_019821525.1
F1	9948335	9948511
refGene_gn=WDR64;refGene_tx=XM_003999137.4;refGene_gn=WDR64;refGene_tx
=XM_019821525.1
F1	9951381	9951505
refGene_gn=WDR64;refGene_tx=XM_003999137.4;refGene_gn=WDR64;refGene_tx
=XM_019821525.1
F1	9953781	9953930
refGene_gn=WDR64;refGene_tx=XM_003999137.4;refGene_gn=WDR64;refGene_tx
=XM_019821525.1
F1	9956901	9957028
refGene_gn=WDR64;refGene_tx=XM_003999137.4;refGene_gn=WDR64;refGene_tx
=XM_019821525.1
F1	9985697	9985837
refGene_gn=WDR64;refGene_tx=XM_003999137.4;refGene_gn=WDR64;refGene_tx
=XM_019821525.1
F1	9997224	9997304
refGene_gn=WDR64;refGene_tx=XM_003999137.4;refGene_gn=WDR64;refGene_tx
=XM_019821525.1
F1	10018332	10018467
refGene_gn=WDR64;refGene_tx=XM_003999137.4;refGene_gn=WDR64;refGene_tx
=XM_019821525.1
F1	10024066	10024281
refGene_gn=WDR64;refGene_tx=XM_003999137.4;refGene_gn=WDR64;refGene_tx
=XM_019821525.1
F1	10026759	10026870
refGene_gn=WDR64;refGene_tx=XM_003999137.4;refGene_gn=WDR64;refGene_tx
=XM_019821525.1
F1	10028078	10028108
refGene_gn=WDR64;refGene_tx=XM_003999137.4;refGene_gn=WDR64;refGene_tx
=XM_019821525.1
F1	10031127	10031231	refGene_gn=WDR64;refGene_tx=XM_003999137.4
F1	10032784	10032887
refGene_gn=WDR64;refGene_tx=XM_003999137.4;refGene_gn=WDR64;refGene_tx
=XM_019821525.1
F1	10050743	10050874	refGene_gn=WDR64;refGene_tx=XM_003999137.4
F1	10056371	10056516	refGene_gn=WDR64;refGene_tx=XM_003999137.4
F1	10063956	10064329	refGene_gn=OPN3;refGene_tx=XM_003999138.5

F1 10068776 10070738 refGene_gn=CHML;refGene_tx=XM_006942644.3
F1 10097780 10098100 refGene_gn=OPN3;refGene_tx=XM_003999138.5
F1 10102897 10103149 refGene_gn=OPN3;refGene_tx=XM_003999138.5
F1 10105702 10105966 refGene_gn=OPN3;refGene_tx=XM_003999138.5
F1 10108469 10108538 refGene_gn=KMO;refGene_tx=XM_023247883.1
F1 10108540 10108672 refGene_gn=KMO;refGene_tx=XM_023247883.1
F1 10109996 10110056 refGene_gn=KMO;refGene_tx=XM_023247883.1
F1 10110145 10110247 refGene_gn=KMO;refGene_tx=XM_023247883.1
F1 10118354 10118437 refGene_gn=KMO;refGene_tx=XM_023247883.1
F1 10120437 10120495 refGene_gn=KMO;refGene_tx=XM_023247883.1
F1 10122898 10123046 refGene_gn=KMO;refGene_tx=XM_023247883.1
F1 10124743 10124865 refGene_gn=KMO;refGene_tx=XM_023247883.1
F1 10125696 10125768 refGene_gn=KMO;refGene_tx=XM_023247883.1
F1 10129524 10129690 refGene_gn=KMO;refGene_tx=XM_023247883.1
F1 10131043 10131131 refGene_gn=KMO;refGene_tx=XM_023247883.1
F1 10134599 10134648 refGene_gn=KMO;refGene_tx=XM_023247883.1
F1 10141490 10141580 refGene_gn=KMO;refGene_tx=XM_023247883.1
F1 10142784 10142882 refGene_gn=KMO;refGene_tx=XM_023247883.1
F1 10143999 10144069 refGene_gn=KMO;refGene_tx=XM_023247883.1
F1 10165029 10165083 refGene_gn=KMO;refGene_tx=XM_023247883.1
F1 10193761 10193887 refGene_gn=FH;refGene_tx=XM_023247450.1
F1 10200733 10200868 refGene_gn=FH;refGene_tx=XM_023247450.1
F1 10203785 10203896 refGene_gn=FH;refGene_tx=XM_023247450.1
F1 10206057 10206234 refGene_gn=FH;refGene_tx=XM_023247450.1
F1 10208785 10208968 refGene_gn=FH;refGene_tx=XM_023247450.1
F1 10211697 10211863 refGene_gn=FH;refGene_tx=XM_023247450.1
F1 10216883 10217087 refGene_gn=FH;refGene_tx=XM_023247450.1
F1 10219022 10219150 refGene_gn=FH;refGene_tx=XM_023247450.1
F1 10221083 10221237 refGene_gn=FH;refGene_tx=XM_023247450.1
F1 10224271 10224414 refGene_gn=FH;refGene_tx=XM_023247450.1
F1 10355608 10355686
refGene_gn=RGS7;refGene_tx=XM_011290846.3;refGene_gn=RGS7;refGene_tx=XM_019821536.1;refGene_gn=RGS7;refGene_tx=XM_023247632.1;refGene_gn=RGS7;refGene_tx=XM_023247634.1;refGene_gn=RGS7;refGene_tx=XM_023247635.1;refGene_gn=RGS7;refGene_tx=XM_023247636.1;refGene_gn=RGS7;refGene_tx=XM_023247637.1;refGene_gn=RGS7;refGene_tx=XM_023247638.1
F1 10601041 10601138
refGene_gn=RGS7;refGene_tx=XM_011290846.3;refGene_gn=RGS7;refGene_tx=XM_019821536.1;refGene_gn=RGS7;refGene_tx=XM_019821537.1;refGene_gn=RGS7;refGene_tx=XM_023247632.1;refGene_gn=RGS7;refGene_tx=XM_023247634.1;refGene_gn=RGS7;refGene_tx=XM_023247635.1;refGene_gn=RGS7;refGene_tx=XM_023247636.1;refGene_gn=RGS7;refGene_tx=XM_023247637.1;refGene_gn=RGS7;refGene_tx=XM_023247638.1
F1 10701429 10701480
refGene_gn=RGS7;refGene_tx=XM_011290846.3;refGene_gn=RGS7;refGene_tx=XM_019821536.1;refGene_gn=RGS7;refGene_tx=XM_019821537.1;refGene_gn=RGS7;refGene_tx=XM_023247632.1;refGene_gn=RGS7;refGene_tx=XM_023247634.1;refGene_gn=RGS7;refGene_tx=XM_023247635.1;refGene_gn=RGS7;refGene_tx=XM_023247636.1;refGene_gn=RGS7;refGene_tx=XM_023247637.1;refGene_gn=RGS7;refGene_tx=XM_023247638.1

F1 10735371 10735478
refGene_gn=RGS7;refGene_tx=XM_011290846.3;refGene_gn=RGS7;refGene_tx=XM_019821536.1;refGene_gn=RGS7;refGene_tx=XM_019821537.1;refGene_gn=RGS7;refGene_tx=XM_023247632.1;refGene_gn=RGS7;refGene_tx=XM_023247634.1;refGene_gn=RGS7;refGene_tx=XM_023247635.1;refGene_gn=RGS7;refGene_tx=XM_023247636.1;refGene_gn=RGS7;refGene_tx=XM_023247637.1;refGene_gn=RGS7;refGene_tx=XM_023247638.1
F1 10740492 10740544
refGene_gn=RGS7;refGene_tx=XM_011290846.3;refGene_gn=RGS7;refGene_tx=XM_019821536.1;refGene_gn=RGS7;refGene_tx=XM_019821537.1;refGene_gn=RGS7;refGene_tx=XM_023247632.1;refGene_gn=RGS7;refGene_tx=XM_023247634.1;refGene_gn=RGS7;refGene_tx=XM_023247635.1;refGene_gn=RGS7;refGene_tx=XM_023247636.1;refGene_gn=RGS7;refGene_tx=XM_023247637.1;refGene_gn=RGS7;refGene_tx=XM_023247638.1
F1 10783940 10784005
refGene_gn=RGS7;refGene_tx=XM_011290846.3;refGene_gn=RGS7;refGene_tx=XM_019821536.1;refGene_gn=RGS7;refGene_tx=XM_019821537.1;refGene_gn=RGS7;refGene_tx=XM_023247632.1;refGene_gn=RGS7;refGene_tx=XM_023247634.1;refGene_gn=RGS7;refGene_tx=XM_023247635.1;refGene_gn=RGS7;refGene_tx=XM_023247636.1;refGene_gn=RGS7;refGene_tx=XM_023247637.1;refGene_gn=RGS7;refGene_tx=XM_023247638.1
F1 10784848 10784925
refGene_gn=RGS7;refGene_tx=XM_011290846.3;refGene_gn=RGS7;refGene_tx=XM_019821536.1;refGene_gn=RGS7;refGene_tx=XM_019821537.1;refGene_gn=RGS7;refGene_tx=XM_023247632.1;refGene_gn=RGS7;refGene_tx=XM_023247634.1;refGene_gn=RGS7;refGene_tx=XM_023247635.1;refGene_gn=RGS7;refGene_tx=XM_023247636.1;refGene_gn=RGS7;refGene_tx=XM_023247637.1;refGene_gn=RGS7;refGene_tx=XM_023247638.1
F1 10785031 10785113
refGene_gn=RGS7;refGene_tx=XM_011290846.3;refGene_gn=RGS7;refGene_tx=XM_019821536.1;refGene_gn=RGS7;refGene_tx=XM_019821537.1;refGene_gn=RGS7;refGene_tx=XM_023247632.1;refGene_gn=RGS7;refGene_tx=XM_023247634.1;refGene_gn=RGS7;refGene_tx=XM_023247635.1;refGene_gn=RGS7;refGene_tx=XM_023247636.1;refGene_gn=RGS7;refGene_tx=XM_023247637.1;refGene_gn=RGS7;refGene_tx=XM_023247638.1
F1 10830251 10830326
refGene_gn=RGS7;refGene_tx=XM_011290846.3;refGene_gn=RGS7;refGene_tx=XM_019821536.1;refGene_gn=RGS7;refGene_tx=XM_019821537.1;refGene_gn=RGS7;refGene_tx=XM_023247632.1;refGene_gn=RGS7;refGene_tx=XM_023247634.1;refGene_gn=RGS7;refGene_tx=XM_023247635.1;refGene_gn=RGS7;refGene_tx=XM_023247636.1;refGene_gn=RGS7;refGene_tx=XM_023247638.1
F1 10837571 10837670
refGene_gn=RGS7;refGene_tx=XM_011290846.3;refGene_gn=RGS7;refGene_tx=XM_019821536.1;refGene_gn=RGS7;refGene_tx=XM_019821537.1;refGene_gn=RGS7;refGene_tx=XM_023247632.1;refGene_gn=RGS7;refGene_tx=XM_023247634.1;refGene_gn=RGS7;refGene_tx=XM_023247635.1;refGene_gn=RGS7;refGene_tx=XM_023247636.1;refGene_gn=RGS7;refGene_tx=XM_023247637.1;refGene_gn=RGS7;refGene_tx=XM_023247638.1
F1 10839347 10839409
refGene_gn=RGS7;refGene_tx=XM_011290846.3;refGene_gn=RGS7;refGene_tx=X

M_019821536.1;refGene_gn=RGS7;refGene_tx=XM_019821537.1;refGene_gn=RGS7;refGene_tx=XM_023247632.1;refGene_gn=RGS7;refGene_tx=XM_023247634.1;refGene_gn=RGS7;refGene_tx=XM_023247635.1;refGene_gn=RGS7;refGene_tx=XM_023247636.1;refGene_gn=RGS7;refGene_tx=XM_023247637.1;refGene_gn=RGS7;refGene_tx=XM_023247638.1
F1 10840743 10840854
refGene_gn=RGS7;refGene_tx=XM_011290846.3;refGene_gn=RGS7;refGene_tx=XM_019821536.1;refGene_gn=RGS7;refGene_tx=XM_019821537.1;refGene_gn=RGS7;refGene_tx=XM_023247632.1;refGene_gn=RGS7;refGene_tx=XM_023247634.1;refGene_gn=RGS7;refGene_tx=XM_023247635.1;refGene_gn=RGS7;refGene_tx=XM_023247636.1;refGene_gn=RGS7;refGene_tx=XM_023247637.1;refGene_gn=RGS7;refGene_tx=XM_023247638.1
F1 10842115 10842241
refGene_gn=RGS7;refGene_tx=XM_011290846.3;refGene_gn=RGS7;refGene_tx=XM_019821536.1;refGene_gn=RGS7;refGene_tx=XM_019821537.1;refGene_gn=RGS7;refGene_tx=XM_023247632.1;refGene_gn=RGS7;refGene_tx=XM_023247634.1;refGene_gn=RGS7;refGene_tx=XM_023247635.1;refGene_gn=RGS7;refGene_tx=XM_023247636.1;refGene_gn=RGS7;refGene_tx=XM_023247637.1;refGene_gn=RGS7;refGene_tx=XM_023247638.1
F1 10847558 10847745
refGene_gn=RGS7;refGene_tx=XM_011290846.3;refGene_gn=RGS7;refGene_tx=XM_019821536.1;refGene_gn=RGS7;refGene_tx=XM_019821537.1;refGene_gn=RGS7;refGene_tx=XM_023247632.1;refGene_gn=RGS7;refGene_tx=XM_023247634.1;refGene_gn=RGS7;refGene_tx=XM_023247635.1;refGene_gn=RGS7;refGene_tx=XM_023247636.1;refGene_gn=RGS7;refGene_tx=XM_023247637.1;refGene_gn=RGS7;refGene_tx=XM_023247638.1
F1 10851022 10851112
refGene_gn=RGS7;refGene_tx=XM_011290846.3;refGene_gn=RGS7;refGene_tx=XM_019821536.1;refGene_gn=RGS7;refGene_tx=XM_019821537.1;refGene_gn=RGS7;refGene_tx=XM_023247632.1;refGene_gn=RGS7;refGene_tx=XM_023247634.1;refGene_gn=RGS7;refGene_tx=XM_023247635.1;refGene_gn=RGS7;refGene_tx=XM_023247636.1;refGene_gn=RGS7;refGene_tx=XM_023247637.1;refGene_gn=RGS7;refGene_tx=XM_023247638.1
F1 10852501 10852555
refGene_gn=RGS7;refGene_tx=XM_011290846.3;refGene_gn=RGS7;refGene_tx=XM_019821536.1;refGene_gn=RGS7;refGene_tx=XM_019821537.1;refGene_gn=RGS7;refGene_tx=XM_023247632.1;refGene_gn=RGS7;refGene_tx=XM_023247634.1;refGene_gn=RGS7;refGene_tx=XM_023247635.1;refGene_gn=RGS7;refGene_tx=XM_023247637.1
F1 10853313 10853388
refGene_gn=RGS7;refGene_tx=XM_011290846.3;refGene_gn=RGS7;refGene_tx=XM_019821536.1;refGene_gn=RGS7;refGene_tx=XM_019821537.1;refGene_gn=RGS7;refGene_tx=XM_023247632.1;refGene_gn=RGS7;refGene_tx=XM_023247634.1;refGene_gn=RGS7;refGene_tx=XM_023247636.1;refGene_gn=RGS7;refGene_tx=XM_023247637.1
F1 10871054 10871105
refGene_gn=RGS7;refGene_tx=XM_023247635.1;refGene_gn=RGS7;refGene_tx=XM_023247638.1
F1 11013226 11013282
refGene_gn=GREM2;refGene_tx=XM_006942647.4;refGene_gn=GREM2;refGene_tx

=XM_023248114.1
F1	11117079	11117587
refGene_gn=GREM2;refGene_tx=XM_006942647.4;refGene_gn=GREM2;refGene_tx=XM_019821538.2;refGene_gn=GREM2;refGene_tx=XM_023248114.1
F1	11151450	11151477	refGene_gn=FMN2;refGene_tx=XM_023247262.1
F1	11153261	11153343	refGene_gn=FMN2;refGene_tx=XM_023247262.1
F1	11181235	11181385	refGene_gn=FMN2;refGene_tx=XM_023247262.1
F1	11224531	11224583	refGene_gn=FMN2;refGene_tx=XM_023247262.1
F1	11254303	11254396	refGene_gn=FMN2;refGene_tx=XM_023247262.1
F1	11281955	11282076	refGene_gn=FMN2;refGene_tx=XM_023247262.1
F1	11282459	11282519	refGene_gn=FMN2;refGene_tx=XM_023247262.1
F1	11285992	11286139	refGene_gn=FMN2;refGene_tx=XM_023247262.1
F1	11287863	11287993	refGene_gn=FMN2;refGene_tx=XM_023247262.1
F1	11288137	11288229	refGene_gn=FMN2;refGene_tx=XM_023247262.1
F1	11324652	11324714	refGene_gn=FMN2;refGene_tx=XM_023247262.1
F1	11355119	11355207	refGene_gn=FMN2;refGene_tx=XM_023247262.1
F1	11403693	11403838	refGene_gn=FMN2;refGene_tx=XM_023247262.1
F1	11405500	11407221	refGene_gn=FMN2;refGene_tx=XM_023247262.1
F1	11428875	11428931	refGene_gn=FMN2;refGene_tx=XM_023247262.1
F1	11437739	11437887	refGene_gn=FMN2;refGene_tx=XM_023247262.1
F1	11506861	11507028	refGene_gn=FMN2;refGene_tx=XM_023247262.1
F1	11549139	11551168	refGene_gn=FMN2;refGene_tx=XM_023247262.1
F1	11667848	11668354
refGene_gn=POGK;refGene_tx=XM_023247263.1;refGene_gn=POGK;refGene_tx=XM_023247264.1
F1	11669480	11669544
refGene_gn=POGK;refGene_tx=XM_023247268.1;refGene_gn=POGK;refGene_tx=XM_023247269.1
F1	11669796	11669930
refGene_gn=POGK;refGene_tx=XM_019821539.2;refGene_gn=POGK;refGene_tx=XM_023247263.1;refGene_gn=POGK;refGene_tx=XM_023247264.1;refGene_gn=POGK;refGene_tx=XM_023247266.1;refGene_gn=POGK;refGene_tx=XM_023247267.1;refGene_gn=POGK;refGene_tx=XM_023247268.1;refGene_gn=POGK;refGene_tx=XM_023247269.1;refGene_gn=POGK;refGene_tx=XM_023247270.1;refGene_gn=POGK;refGene_tx=XM_023247271.1;refGene_gn=POGK;refGene_tx=XM_023247272.1
F1	11675121	11675286
refGene_gn=POGK;refGene_tx=XM_019821539.2;refGene_gn=POGK;refGene_tx=XM_023247266.1;refGene_gn=POGK;refGene_tx=XM_023247267.1
F1	11675990	11676133
refGene_gn=POGK;refGene_tx=XM_019821539.2;refGene_gn=POGK;refGene_tx=XM_019821543.2;refGene_gn=POGK;refGene_tx=XM_023247266.1;refGene_gn=POGK;refGene_tx=XM_023247267.1;refGene_gn=POGK;refGene_tx=XM_023247268.1;refGene_gn=POGK;refGene_tx=XM_023247269.1;refGene_gn=POGK;refGene_tx=XM_023247270.1;refGene_gn=POGK;refGene_tx=XM_023247271.1;refGene_gn=POGK;refGene_tx=XM_023247272.1
F1	11676911	11677010
refGene_gn=POGK;refGene_tx=XM_019821539.2;refGene_gn=POGK;refGene_tx=XM_019821543.2;refGene_gn=POGK;refGene_tx=XM_023247263.1;refGene_gn=POGK;refGene_tx=XM_023247266.1;refGene_gn=POGK;refGene_tx=XM_023247267.1;refGene_gn=POGK;refGene_tx=XM_023247268.1;refGene_gn=POGK;refGene_tx=X

M_023247269.1;refGene_gn=POGK;refGene_tx=XM_023247270.1;refGene_gn=POG
K;refGene_tx=XM_023247271.1;refGene_gn=POGK;refGene_tx=XM_023247272.1
F1 11678279 11679751
refGene_gn=POGK;refGene_tx=XM_019821539.2;refGene_gn=POGK;refGene_tx=X
M_019821543.2;refGene_gn=POGK;refGene_tx=XM_023247263.1;refGene_gn=POG
K;refGene_tx=XM_023247264.1;refGene_gn=POGK;refGene_tx=XM_023247266.1;
refGene_gn=POGK;refGene_tx=XM_023247267.1;refGene_gn=POGK;refGene_tx=X
M_023247268.1;refGene_gn=POGK;refGene_tx=XM_023247269.1;refGene_gn=POG
K;refGene_tx=XM_023247270.1;refGene_gn=POGK;refGene_tx=XM_023247271.1;
refGene_gn=POGK;refGene_tx=XM_023247272.1
F1 11686789 11686942 refGene_gn=TADA1;refGene_tx=XM_023247273.1
F1 11687603 11687766 refGene_gn=TADA1;refGene_tx=XM_023247273.1
F1 11689980 11690132 refGene_gn=TADA1;refGene_tx=XM_023247273.1
F1 11691577 11691787 refGene_gn=TADA1;refGene_tx=XM_023247273.1
F1 11693217 11693382 refGene_gn=TADA1;refGene_tx=XM_023247273.1
F1 11748448 11748484
refGene_gn=ILDR2;refGene_tx=XM_023248047.1;refGene_gn=ILDR2;refGene_tx
=XM_023248048.1
F1 11749783 11750471
refGene_gn=ILDR2;refGene_tx=XM_023248047.1;refGene_gn=ILDR2;refGene_tx
=XM_023248048.1
F1 11751831 11752048
refGene_gn=ILDR2;refGene_tx=XM_023248047.1;refGene_gn=ILDR2;refGene_tx
=XM_023248048.1
F1 11756514 11756628
refGene_gn=ILDR2;refGene_tx=XM_023248047.1;refGene_gn=ILDR2;refGene_tx
=XM_023248048.1
F1 11761402 11761480 refGene_gn=ILDR2;refGene_tx=XM_023248047.1
F1 11765194 11765371
refGene_gn=ILDR2;refGene_tx=XM_023248047.1;refGene_gn=ILDR2;refGene_tx
=XM_023248048.1
F1 11766465 11766612
refGene_gn=ILDR2;refGene_tx=XM_023248047.1;refGene_gn=ILDR2;refGene_tx
=XM_023248048.1
F1 11769386 11769443
refGene_gn=ILDR2;refGene_tx=XM_023248047.1;refGene_gn=ILDR2;refGene_tx
=XM_023248048.1
F1 11790179 11790299
refGene_gn=ILDR2;refGene_tx=XM_023248047.1;refGene_gn=ILDR2;refGene_tx
=XM_023248048.1
F1 11791196 11791529
refGene_gn=ILDR2;refGene_tx=XM_023248047.1;refGene_gn=ILDR2;refGene_tx
=XM_023248048.1
F1 11807459 11807505 refGene_gn=ILDR2;refGene_tx=XM_023248048.1
F1 11821527 11821659 refGene_gn=MAEL;refGene_tx=XM_023247737.1
F1 11821922 11822015 refGene_gn=MAEL;refGene_tx=XM_023247737.1
F1 11823631 11823731 refGene_gn=MAEL;refGene_tx=XM_023247737.1
F1 11824678 11824829 refGene_gn=MAEL;refGene_tx=XM_023247737.1
F1 11826469 11826516 refGene_gn=MAEL;refGene_tx=XM_023247737.1
F1 11836878 11837003 refGene_gn=MAEL;refGene_tx=XM_023247737.1

F1 11837417 11837472 refGene_gn=MAEL;refGene_tx=XM_023247737.1
F1 11837595 11837737 refGene_gn=MAEL;refGene_tx=XM_023247737.1
F1 11853517 11853580 refGene_gn=MAEL;refGene_tx=XM_023247737.1
F1 11854575 11854708 refGene_gn=MAEL;refGene_tx=XM_023247737.1
F1 11858386 11858462 refGene_gn=MAEL;refGene_tx=XM_023247737.1
F1 11859226 11859423 refGene_gn=MAEL;refGene_tx=XM_023247737.1
F1 11894492 11894622 refGene_gn=GPA33;refGene_tx=XM_003999163.5
F1 11895059 11895183 refGene_gn=GPA33;refGene_tx=XM_003999163.5
F1 11895890 11896010 refGene_gn=GPA33;refGene_tx=XM_003999163.5
F1 11901214 11901367 refGene_gn=GPA33;refGene_tx=XM_003999163.5
F1 11910337 11910554 refGene_gn=GPA33;refGene_tx=XM_003999163.5
F1 11916424 11916579 refGene_gn=GPA33;refGene_tx=XM_003999163.5
F1 11931480 11931523 refGene_gn=GPA33;refGene_tx=XM_003999163.5
F1 11936526 11936636 refGene_gn=DUSP27;refGene_tx=XM_023247431.1
F1 11951136 11951231 refGene_gn=DUSP27;refGene_tx=XM_023247431.1
F1 11954480 11954712 refGene_gn=DUSP27;refGene_tx=XM_023247431.1
F1 11957415 11957633 refGene_gn=DUSP27;refGene_tx=XM_023247431.1
F1 11963696 11966102 refGene_gn=DUSP27;refGene_tx=XM_023247431.1
F1 11966104 11966517 refGene_gn=DUSP27;refGene_tx=XM_023247431.1
F1 12054950 12055011
refGene_gn=POU2F1;refGene_tx=XM_023247738.1;refGene_gn=POU2F1;refGene_tx=XM_023247740.1
F1 12160557 12160585 refGene_gn=POU2F1;refGene_tx=XM_019821549.2
F1 12163800 12163866
refGene_gn=POU2F1;refGene_tx=XM_019821549.2;refGene_gn=POU2F1;refGene_tx=XM_023247738.1;refGene_gn=POU2F1;refGene_tx=XM_023247739.1;refGene_gn=POU2F1;refGene_tx=XM_023247740.1
F1 12188940 12189041
refGene_gn=POU2F1;refGene_tx=XM_019821549.2;refGene_gn=POU2F1;refGene_tx=XM_023247738.1;refGene_gn=POU2F1;refGene_tx=XM_023247739.1;refGene_gn=POU2F1;refGene_tx=XM_023247740.1
F1 12191563 12191617
refGene_gn=POU2F1;refGene_tx=XM_019821549.2;refGene_gn=POU2F1;refGene_tx=XM_023247738.1;refGene_gn=POU2F1;refGene_tx=XM_023247739.1;refGene_gn=POU2F1;refGene_tx=XM_023247740.1
F1 12193485 12193605
refGene_gn=POU2F1;refGene_tx=XM_019821549.2;refGene_gn=POU2F1;refGene_tx=XM_023247738.1;refGene_gn=POU2F1;refGene_tx=XM_023247739.1;refGene_gn=POU2F1;refGene_tx=XM_023247740.1
F1 12195949 12196138
refGene_gn=POU2F1;refGene_tx=XM_019821549.2;refGene_gn=POU2F1;refGene_tx=XM_023247738.1;refGene_gn=POU2F1;refGene_tx=XM_023247739.1;refGene_gn=POU2F1;refGene_tx=XM_023247740.1
F1 12198072 12198199
refGene_gn=POU2F1;refGene_tx=XM_019821549.2;refGene_gn=POU2F1;refGene_tx=XM_023247738.1;refGene_gn=POU2F1;refGene_tx=XM_023247739.1;refGene_gn=POU2F1;refGene_tx=XM_023247740.1
F1 12200111 12200206
refGene_gn=POU2F1;refGene_tx=XM_019821549.2;refGene_gn=POU2F1;refGene_tx=XM_023247738.1;refGene_gn=POU2F1;refGene_tx=XM_023247739.1;refGene_

gn=POU2F1;refGene_tx=XM_023247740.1
F1 12205617 12205791
refGene_gn=POU2F1;refGene_tx=XM_019821549.2;refGene_gn=POU2F1;refGene_tx=XM_023247738.1;refGene_gn=POU2F1;refGene_tx=XM_023247739.1;refGene_gn=POU2F1;refGene_tx=XM_023247740.1
F1 12209975 12210117
refGene_gn=POU2F1;refGene_tx=XM_019821549.2;refGene_gn=POU2F1;refGene_tx=XM_023247738.1;refGene_gn=POU2F1;refGene_tx=XM_023247739.1;refGene_gn=POU2F1;refGene_tx=XM_023247740.1
F1 12211501 12211647
refGene_gn=POU2F1;refGene_tx=XM_019821549.2;refGene_gn=POU2F1;refGene_tx=XM_023247738.1;refGene_gn=POU2F1;refGene_tx=XM_023247739.1;refGene_gn=POU2F1;refGene_tx=XM_023247740.1
F1 12213403 12213583
refGene_gn=POU2F1;refGene_tx=XM_019821549.2;refGene_gn=POU2F1;refGene_tx=XM_023247738.1;refGene_gn=POU2F1;refGene_tx=XM_023247739.1;refGene_gn=POU2F1;refGene_tx=XM_023247740.1
F1 12215034 12215140
refGene_gn=POU2F1;refGene_tx=XM_019821549.2;refGene_gn=POU2F1;refGene_tx=XM_023247738.1;refGene_gn=POU2F1;refGene_tx=XM_023247739.1;refGene_gn=POU2F1;refGene_tx=XM_023247740.1
F1 12227174 12227517
refGene_gn=POU2F1;refGene_tx=XM_019821549.2;refGene_gn=POU2F1;refGene_tx=XM_023247738.1;refGene_gn=POU2F1;refGene_tx=XM_023247739.1;refGene_gn=POU2F1;refGene_tx=XM_023247740.1
F1 12228518 12228607
refGene_gn=POU2F1;refGene_tx=XM_019821549.2;refGene_gn=POU2F1;refGene_tx=XM_023247738.1;refGene_gn=POU2F1;refGene_tx=XM_023247739.1;refGene_gn=POU2F1;refGene_tx=XM_023247740.1
F1 12232080 12232391
refGene_gn=POU2F1;refGene_tx=XM_019821549.2;refGene_gn=POU2F1;refGene_tx=XM_023247738.1;refGene_gn=POU2F1;refGene_tx=XM_023247739.1
F1 12248271 12248337
refGene_gn=CD247;refGene_tx=XM_023247741.1;refGene_gn=CD247;refGene_tx=XM_023247742.1
F1 12249564 12249606
refGene_gn=CD247;refGene_tx=XM_023247741.1;refGene_gn=CD247;refGene_tx=XM_023247742.1
F1 12250275 12250529
refGene_gn=CD247;refGene_tx=XM_023247741.1;refGene_gn=CD247;refGene_tx=XM_023247742.1;refGene_gn=POU2F1;refGene_tx=XM_023247740.1
F1 12251377 12251413
refGene_gn=CD247;refGene_tx=XM_023247741.1;refGene_gn=CD247;refGene_tx=XM_023247742.1
F1 12252583 12252664
refGene_gn=CD247;refGene_tx=XM_023247741.1;refGene_gn=CD247;refGene_tx=XM_023247742.1
F1 12253189 12253246
refGene_gn=CD247;refGene_tx=XM_023247741.1;refGene_gn=CD247;refGene_tx=XM_023247742.1

F1	12254221	12254325
refGene_gn=CD247;refGene_tx=XM_023247741.1;refGene_gn=CD247;refGene_tx=XM_023247742.1
F1	12321971	12322080
refGene_gn=CD247;refGene_tx=XM_023247741.1;refGene_gn=CD247;refGene_tx=XM_023247742.1
F1	12341802	12341806	refGene_gn=CREG1;refGene_tx=XM_019821551.2
F1	12343205	12343227	refGene_gn=CREG1;refGene_tx=XM_023247444.1
F1	12345213	12345402
refGene_gn=CREG1;refGene_tx=XM_019821551.2;refGene_gn=CREG1;refGene_tx=XM_023247444.1;refGene_gn=CREG1;refGene_tx=XM_023247445.1
F1	12346560	12346680
refGene_gn=CREG1;refGene_tx=XM_019821551.2;refGene_gn=CREG1;refGene_tx=XM_023247444.1;refGene_gn=CREG1;refGene_tx=XM_023247445.1
F1	12349979	12350333
refGene_gn=CREG1;refGene_tx=XM_019821551.2;refGene_gn=CREG1;refGene_tx=XM_023247444.1;refGene_gn=CREG1;refGene_tx=XM_023247445.1
F1	12389605	12389668	refGene_gn=RCSD1;refGene_tx=XM_011290867.3
F1	12444009	12444111	refGene_gn=RCSD1;refGene_tx=XM_011290867.3
F1	12445176	12445266	refGene_gn=RCSD1;refGene_tx=XM_011290867.3
F1	12449696	12449768	refGene_gn=RCSD1;refGene_tx=XM_011290867.3
F1	12453628	12453832	refGene_gn=RCSD1;refGene_tx=XM_011290867.3
F1	12455922	12456469	refGene_gn=RCSD1;refGene_tx=XM_011290867.3
F1	12456471	12456602	refGene_gn=RCSD1;refGene_tx=XM_011290867.3
F1	12462592	12462625	refGene_gn=RCSD1;refGene_tx=XM_011290867.3
F1	12480426	12480517	refGene_gn=MPZL1;refGene_tx=XM_023247958.1
F1	12523985	12524152	refGene_gn=MPZL1;refGene_tx=XM_023247958.1
F1	12529397	12529611	refGene_gn=MPZL1;refGene_tx=XM_023247958.1
F1	12530360	12530493	refGene_gn=MPZL1;refGene_tx=XM_023247958.1
F1	12532836	12532939	refGene_gn=MPZL1;refGene_tx=XM_023247958.1
F1	12544248	12544350	refGene_gn=MPZL1;refGene_tx=XM_023247958.1
F1	12556485	12556641
refGene_gn=ADCY10;refGene_tx=XM_006942654.3;refGene_gn=ADCY10;refGene_tx=XM_019821558.2;refGene_gn=ADCY10;refGene_tx=XM_019821559.2;refGene_gn=ADCY10;refGene_tx=XM_019821560.2;refGene_gn=ADCY10;refGene_tx=XM_019821561.2;refGene_gn=ADCY10;refGene_tx=XM_019821564.2;refGene_gn=ADCY10;refGene_tx=XM_019821565.2;refGene_gn=ADCY10;refGene_tx=XM_023247221.1
F1	12557408	12557597
refGene_gn=ADCY10;refGene_tx=XM_006942654.3;refGene_gn=ADCY10;refGene_tx=XM_019821558.2;refGene_gn=ADCY10;refGene_tx=XM_019821559.2;refGene_gn=ADCY10;refGene_tx=XM_019821560.2;refGene_gn=ADCY10;refGene_tx=XM_019821561.2;refGene_gn=ADCY10;refGene_tx=XM_019821564.2;refGene_gn=ADCY10;refGene_tx=XM_019821565.2;refGene_gn=ADCY10;refGene_tx=XM_023247221.1
F1	12561972	12562168
refGene_gn=ADCY10;refGene_tx=XM_006942654.3;refGene_gn=ADCY10;refGene_tx=XM_019821558.2;refGene_gn=ADCY10;refGene_tx=XM_019821559.2;refGene_gn=ADCY10;refGene_tx=XM_019821560.2;refGene_gn=ADCY10;refGene_tx=XM_019821561.2;refGene_gn=ADCY10;refGene_tx=XM_019821564.2;refGene_gn=ADCY1

0;refGene_tx=XM_019821565.2;refGene_gn=ADCY10;refGene_tx=XM_023247221.
1
F1 12564280 12564398
refGene_gn=ADCY10;refGene_tx=XM_006942654.3;refGene_gn=ADCY10;refGene_
tx=XM_019821558.2;refGene_gn=ADCY10;refGene_tx=XM_019821559.2;refGene_
gn=ADCY10;refGene_tx=XM_019821560.2;refGene_gn=ADCY10;refGene_tx=XM_01
9821561.2;refGene_gn=ADCY10;refGene_tx=XM_019821564.2;refGene_gn=ADCY1
0;refGene_tx=XM_019821565.2;refGene_gn=ADCY10;refGene_tx=XM_023247221.
1
F1 12565343 12565459
refGene_gn=ADCY10;refGene_tx=XM_006942654.3;refGene_gn=ADCY10;refGene_
tx=XM_019821558.2;refGene_gn=ADCY10;refGene_tx=XM_019821559.2;refGene_
gn=ADCY10;refGene_tx=XM_019821560.2;refGene_gn=ADCY10;refGene_tx=XM_01
9821561.2;refGene_gn=ADCY10;refGene_tx=XM_019821564.2;refGene_gn=ADCY1
0;refGene_tx=XM_019821565.2;refGene_gn=ADCY10;refGene_tx=XM_023247221.
1
F1 12567901 12567998
refGene_gn=ADCY10;refGene_tx=XM_006942654.3;refGene_gn=ADCY10;refGene_
tx=XM_019821558.2;refGene_gn=ADCY10;refGene_tx=XM_019821559.2;refGene_
gn=ADCY10;refGene_tx=XM_019821560.2;refGene_gn=ADCY10;refGene_tx=XM_01
9821561.2;refGene_gn=ADCY10;refGene_tx=XM_019821564.2;refGene_gn=ADCY1
0;refGene_tx=XM_019821565.2;refGene_gn=ADCY10;refGene_tx=XM_023247221.
1
F1 12568077 12568282
refGene_gn=ADCY10;refGene_tx=XM_006942654.3;refGene_gn=ADCY10;refGene_
tx=XM_019821558.2;refGene_gn=ADCY10;refGene_tx=XM_019821559.2;refGene_
gn=ADCY10;refGene_tx=XM_019821560.2;refGene_gn=ADCY10;refGene_tx=XM_01
9821561.2;refGene_gn=ADCY10;refGene_tx=XM_019821564.2;refGene_gn=ADCY1
0;refGene_tx=XM_019821565.2;refGene_gn=ADCY10;refGene_tx=XM_023247221.
1
F1 12572695 12572852
refGene_gn=ADCY10;refGene_tx=XM_006942654.3;refGene_gn=ADCY10;refGene_
tx=XM_019821558.2;refGene_gn=ADCY10;refGene_tx=XM_019821559.2;refGene_
gn=ADCY10;refGene_tx=XM_019821560.2;refGene_gn=ADCY10;refGene_tx=XM_01
9821561.2;refGene_gn=ADCY10;refGene_tx=XM_019821564.2;refGene_gn=ADCY1
0;refGene_tx=XM_019821565.2;refGene_gn=ADCY10;refGene_tx=XM_023247221.
1
F1 12575342 12575518
refGene_gn=ADCY10;refGene_tx=XM_006942654.3;refGene_gn=ADCY10;refGene_
tx=XM_019821558.2;refGene_gn=ADCY10;refGene_tx=XM_019821559.2;refGene_
gn=ADCY10;refGene_tx=XM_019821560.2;refGene_gn=ADCY10;refGene_tx=XM_01
9821561.2;refGene_gn=ADCY10;refGene_tx=XM_019821564.2;refGene_gn=ADCY1
0;refGene_tx=XM_019821565.2;refGene_gn=ADCY10;refGene_tx=XM_023247221.
1
F1 12575966 12576074
refGene_gn=ADCY10;refGene_tx=XM_006942654.3;refGene_gn=ADCY10;refGene_
tx=XM_019821558.2;refGene_gn=ADCY10;refGene_tx=XM_019821559.2;refGene_
gn=ADCY10;refGene_tx=XM_019821560.2;refGene_gn=ADCY10;refGene_tx=XM_01
9821561.2;refGene_gn=ADCY10;refGene_tx=XM_019821564.2;refGene_gn=ADCY1
0;refGene_tx=XM_019821565.2;refGene_gn=ADCY10;refGene_tx=XM_023247221.

1
F1 12578059 12578291
refGene_gn=ADCY10;refGene_tx=XM_006942654.3;refGene_gn=ADCY10;refGene_tx=XM_019821558.2;refGene_gn=ADCY10;refGene_tx=XM_019821559.2;refGene_gn=ADCY10;refGene_tx=XM_019821560.2;refGene_gn=ADCY10;refGene_tx=XM_019821561.2;refGene_gn=ADCY10;refGene_tx=XM_019821564.2;refGene_gn=ADCY10;refGene_tx=XM_019821565.2;refGene_gn=ADCY10;refGene_tx=XM_023247221.1
F1 12578546 12578772
refGene_gn=ADCY10;refGene_tx=XM_006942654.3;refGene_gn=ADCY10;refGene_tx=XM_019821558.2;refGene_gn=ADCY10;refGene_tx=XM_019821559.2;refGene_gn=ADCY10;refGene_tx=XM_019821560.2;refGene_gn=ADCY10;refGene_tx=XM_019821561.2;refGene_gn=ADCY10;refGene_tx=XM_019821564.2;refGene_gn=ADCY10;refGene_tx=XM_019821565.2;refGene_gn=ADCY10;refGene_tx=XM_023247221.1
F1 12590016 12590307
refGene_gn=ADCY10;refGene_tx=XM_006942654.3;refGene_gn=ADCY10;refGene_tx=XM_019821558.2;refGene_gn=ADCY10;refGene_tx=XM_019821559.2;refGene_gn=ADCY10;refGene_tx=XM_019821560.2;refGene_gn=ADCY10;refGene_tx=XM_019821561.2;refGene_gn=ADCY10;refGene_tx=XM_019821564.2;refGene_gn=ADCY10;refGene_tx=XM_019821565.2;refGene_gn=ADCY10;refGene_tx=XM_023247221.1
F1 12590419 12590698
refGene_gn=ADCY10;refGene_tx=XM_006942654.3;refGene_gn=ADCY10;refGene_tx=XM_019821558.2;refGene_gn=ADCY10;refGene_tx=XM_019821559.2;refGene_gn=ADCY10;refGene_tx=XM_019821560.2;refGene_gn=ADCY10;refGene_tx=XM_019821561.2;refGene_gn=ADCY10;refGene_tx=XM_019821564.2;refGene_gn=ADCY10;refGene_tx=XM_019821565.2;refGene_gn=ADCY10;refGene_tx=XM_023247221.1
F1 12591915 12592044
refGene_gn=ADCY10;refGene_tx=XM_006942654.3;refGene_gn=ADCY10;refGene_tx=XM_019821558.2;refGene_gn=ADCY10;refGene_tx=XM_019821559.2;refGene_gn=ADCY10;refGene_tx=XM_019821560.2;refGene_gn=ADCY10;refGene_tx=XM_019821561.2;refGene_gn=ADCY10;refGene_tx=XM_019821564.2;refGene_gn=ADCY10;refGene_tx=XM_019821565.2
F1 12601151 12601288
refGene_gn=ADCY10;refGene_tx=XM_006942654.3;refGene_gn=ADCY10;refGene_tx=XM_019821558.2;refGene_gn=ADCY10;refGene_tx=XM_019821559.2;refGene_gn=ADCY10;refGene_tx=XM_019821560.2;refGene_gn=ADCY10;refGene_tx=XM_019821561.2;refGene_gn=ADCY10;refGene_tx=XM_019821564.2;refGene_gn=ADCY10;refGene_tx=XM_019821565.2;refGene_gn=ADCY10;refGene_tx=XM_023247221.1
F1 12603611 12603886
refGene_gn=ADCY10;refGene_tx=XM_006942654.3;refGene_gn=ADCY10;refGene_tx=XM_019821558.2;refGene_gn=ADCY10;refGene_tx=XM_019821559.2;refGene_gn=ADCY10;refGene_tx=XM_019821560.2;refGene_gn=ADCY10;refGene_tx=XM_019821561.2;refGene_gn=ADCY10;refGene_tx=XM_019821564.2;refGene_gn=ADCY10;refGene_tx=XM_019821565.2;refGene_gn=ADCY10;refGene_tx=XM_023247221.1
F1 12607834 12607921

refGene_gn=ADCY10;refGene_tx=XM_006942654.3;refGene_gn=ADCY10;refGene_tx=XM_019821558.2;refGene_gn=ADCY10;refGene_tx=XM_019821559.2;refGene_gn=ADCY10;refGene_tx=XM_019821560.2;refGene_gn=ADCY10;refGene_tx=XM_019821561.2;refGene_gn=ADCY10;refGene_tx=XM_019821565.2;refGene_gn=ADCY10;refGene_tx=XM_023247221.1
F1 12608425 12608618
refGene_gn=ADCY10;refGene_tx=XM_006942654.3;refGene_gn=ADCY10;refGene_tx=XM_019821558.2;refGene_gn=ADCY10;refGene_tx=XM_019821559.2;refGene_gn=ADCY10;refGene_tx=XM_019821560.2;refGene_gn=ADCY10;refGene_tx=XM_019821561.2;refGene_gn=ADCY10;refGene_tx=XM_019821565.2;refGene_gn=ADCY10;refGene_tx=XM_023247221.1
F1 12610709 12610863
refGene_gn=ADCY10;refGene_tx=XM_006942654.3;refGene_gn=ADCY10;refGene_tx=XM_019821558.2;refGene_gn=ADCY10;refGene_tx=XM_019821559.2;refGene_gn=ADCY10;refGene_tx=XM_019821560.2;refGene_gn=ADCY10;refGene_tx=XM_019821561.2;refGene_gn=ADCY10;refGene_tx=XM_019821565.2;refGene_gn=ADCY10;refGene_tx=XM_023247221.1
F1 12613834 12613890
refGene_gn=ADCY10;refGene_tx=XM_006942654.3;refGene_gn=ADCY10;refGene_tx=XM_019821558.2;refGene_gn=ADCY10;refGene_tx=XM_019821559.2;refGene_gn=ADCY10;refGene_tx=XM_019821560.2;refGene_gn=ADCY10;refGene_tx=XM_019821561.2;refGene_gn=ADCY10;refGene_tx=XM_019821565.2;refGene_gn=ADCY10;refGene_tx=XM_023247221.1
F1 12615638 12615828
refGene_gn=ADCY10;refGene_tx=XM_006942654.3;refGene_gn=ADCY10;refGene_tx=XM_019821558.2;refGene_gn=ADCY10;refGene_tx=XM_019821559.2;refGene_gn=ADCY10;refGene_tx=XM_019821560.2;refGene_gn=ADCY10;refGene_tx=XM_019821561.2;refGene_gn=ADCY10;refGene_tx=XM_019821565.2;refGene_gn=ADCY10;refGene_tx=XM_023247221.1
F1 12616240 12616317
refGene_gn=ADCY10;refGene_tx=XM_006942654.3;refGene_gn=ADCY10;refGene_tx=XM_019821558.2;refGene_gn=ADCY10;refGene_tx=XM_019821559.2;refGene_gn=ADCY10;refGene_tx=XM_019821560.2;refGene_gn=ADCY10;refGene_tx=XM_019821561.2;refGene_gn=ADCY10;refGene_tx=XM_019821565.2;refGene_gn=ADCY10;refGene_tx=XM_023247221.1
F1 12617807 12617926
refGene_gn=ADCY10;refGene_tx=XM_006942654.3;refGene_gn=ADCY10;refGene_tx=XM_019821558.2;refGene_gn=ADCY10;refGene_tx=XM_019821559.2;refGene_gn=ADCY10;refGene_tx=XM_019821560.2;refGene_gn=ADCY10;refGene_tx=XM_019821561.2;refGene_gn=ADCY10;refGene_tx=XM_019821565.2;refGene_gn=ADCY10;refGene_tx=XM_023247221.1
F1 12620681 12620873
refGene_gn=ADCY10;refGene_tx=XM_006942654.3;refGene_gn=ADCY10;refGene_tx=XM_019821558.2;refGene_gn=ADCY10;refGene_tx=XM_019821559.2;refGene_gn=ADCY10;refGene_tx=XM_019821560.2;refGene_gn=ADCY10;refGene_tx=XM_019821561.2;refGene_gn=ADCY10;refGene_tx=XM_019821564.2;refGene_gn=ADCY10;refGene_tx=XM_019821565.2;refGene_gn=ADCY10;refGene_tx=XM_023247221.1
F1 12622204 12622293
refGene_gn=ADCY10;refGene_tx=XM_006942654.3;refGene_gn=ADCY10;refGene_

tx=XM_019821558.2;refGene_gn=ADCY10;refGene_tx=XM_019821559.2;refGene_gn=ADCY10;refGene_tx=XM_019821560.2;refGene_gn=ADCY10;refGene_tx=XM_019821561.2;refGene_gn=ADCY10;refGene_tx=XM_019821564.2;refGene_gn=ADCY10;refGene_tx=XM_023247221.1
F1 12625664 12625761
refGene_gn=ADCY10;refGene_tx=XM_006942654.3;refGene_gn=ADCY10;refGene_tx=XM_019821558.2;refGene_gn=ADCY10;refGene_tx=XM_019821559.2;refGene_gn=ADCY10;refGene_tx=XM_019821560.2;refGene_gn=ADCY10;refGene_tx=XM_019821561.2;refGene_gn=ADCY10;refGene_tx=XM_019821564.2;refGene_gn=ADCY10;refGene_tx=XM_023247221.1
F1 12627678 12627884
refGene_gn=ADCY10;refGene_tx=XM_006942654.3;refGene_gn=ADCY10;refGene_tx=XM_019821558.2;refGene_gn=ADCY10;refGene_tx=XM_019821559.2;refGene_gn=ADCY10;refGene_tx=XM_019821560.2;refGene_gn=ADCY10;refGene_tx=XM_019821561.2;refGene_gn=ADCY10;refGene_tx=XM_019821564.2;refGene_gn=ADCY10;refGene_tx=XM_023247221.1
F1 12629341 12629485
refGene_gn=ADCY10;refGene_tx=XM_006942654.3;refGene_gn=ADCY10;refGene_tx=XM_019821558.2;refGene_gn=ADCY10;refGene_tx=XM_019821559.2;refGene_gn=ADCY10;refGene_tx=XM_019821560.2;refGene_gn=ADCY10;refGene_tx=XM_019821561.2;refGene_gn=ADCY10;refGene_tx=XM_019821564.2;refGene_gn=ADCY10;refGene_tx=XM_023247221.1
F1 12629657 12629696
refGene_gn=ADCY10;refGene_tx=XM_006942654.3;refGene_gn=ADCY10;refGene_tx=XM_019821558.2;refGene_gn=ADCY10;refGene_tx=XM_019821559.2;refGene_gn=ADCY10;refGene_tx=XM_019821560.2;refGene_gn=ADCY10;refGene_tx=XM_019821561.2;refGene_gn=ADCY10;refGene_tx=XM_019821564.2;refGene_gn=ADCY10;refGene_tx=XM_023247221.1
F1 12631056 12631161
refGene_gn=ADCY10;refGene_tx=XM_006942654.3;refGene_gn=ADCY10;refGene_tx=XM_019821558.2;refGene_gn=ADCY10;refGene_tx=XM_019821559.2;refGene_gn=ADCY10;refGene_tx=XM_019821560.2;refGene_gn=ADCY10;refGene_tx=XM_019821561.2;refGene_gn=ADCY10;refGene_tx=XM_019821564.2;refGene_gn=ADCY10;refGene_tx=XM_023247221.1
F1 12631801 12631949
refGene_gn=ADCY10;refGene_tx=XM_006942654.3;refGene_gn=ADCY10;refGene_tx=XM_019821558.2;refGene_gn=ADCY10;refGene_tx=XM_019821559.2;refGene_gn=ADCY10;refGene_tx=XM_019821560.2;refGene_gn=ADCY10;refGene_tx=XM_019821561.2;refGene_gn=ADCY10;refGene_tx=XM_019821564.2;refGene_gn=ADCY10;refGene_tx=XM_023247221.1
F1 12643040 12643077
refGene_gn=MPC2;refGene_tx=XM_003999153.5;refGene_gn=MPC2;refGene_tx=XM_006942662.4;refGene_gn=MPC2;refGene_tx=XM_006942663.3
F1 12647368 12647480
refGene_gn=MPC2;refGene_tx=XM_003999153.5;refGene_gn=MPC2;refGene_tx=XM_006942662.4;refGene_gn=MPC2;refGene_tx=XM_006942663.3
F1 12647956 12648041
refGene_gn=MPC2;refGene_tx=XM_003999153.5;refGene_gn=MPC2;refGene_tx=XM_006942662.4;refGene_gn=MPC2;refGene_tx=XM_006942663.3
F1 12658801 12658842

refGene_gn=MPC2;refGene_tx=XM_003999153.5;refGene_gn=MPC2;refGene_tx=XM_006942662.4;refGene_gn=MPC2;refGene_tx=XM_006942663.3
F1 12673765 12673874
refGene_gn=MPC2;refGene_tx=XM_003999153.5;refGene_gn=MPC2;refGene_tx=XM_006942662.4;refGene_gn=MPC2;refGene_tx=XM_006942663.3
F1 12674943 12675040 refGene_gn=DCAF6;refGene_tx=XM_003999156.5
F1 12691091 12691153 refGene_gn=DCAF6;refGene_tx=XM_003999156.5
F1 12699544 12699637 refGene_gn=DCAF6;refGene_tx=XM_003999156.5
F1 12708804 12708990 refGene_gn=DCAF6;refGene_tx=XM_003999156.5
F1 12720339 12720453 refGene_gn=DCAF6;refGene_tx=XM_003999156.5
F1 12721081 12721217 refGene_gn=DCAF6;refGene_tx=XM_003999156.5
F1 12724082 12724297 refGene_gn=DCAF6;refGene_tx=XM_003999156.5
F1 12730000 12730094 refGene_gn=DCAF6;refGene_tx=XM_003999156.5
F1 12731366 12731486 refGene_gn=DCAF6;refGene_tx=XM_003999156.5
F1 12733934 12734195 refGene_gn=DCAF6;refGene_tx=XM_003999156.5
F1 12745106 12745271 refGene_gn=DCAF6;refGene_tx=XM_003999156.5
F1 12752808 12752868 refGene_gn=DCAF6;refGene_tx=XM_003999156.5
F1 12767152 12767270 refGene_gn=DCAF6;refGene_tx=XM_003999156.5
F1 12771676 12771789 refGene_gn=DCAF6;refGene_tx=XM_003999156.5
F1 12774325 12774412 refGene_gn=DCAF6;refGene_tx=XM_003999156.5
F1 12774632 12774871 refGene_gn=DCAF6;refGene_tx=XM_003999156.5
F1 12774872 12774961 refGene_gn=DCAF6;refGene_tx=XM_003999156.5
F1 12779829 12779871 refGene_gn=DCAF6;refGene_tx=XM_003999156.5
F1 12797132 12797271 refGene_gn=DCAF6;refGene_tx=XM_003999156.5
F1 12798968 12799125 refGene_gn=DCAF6;refGene_tx=XM_003999156.5
F1 12799768 12799857 refGene_gn=DCAF6;refGene_tx=XM_003999156.5
F1 12801959 12802065 refGene_gn=DCAF6;refGene_tx=XM_003999156.5
F1 12809216 12809281 refGene_gn=DCAF6;refGene_tx=XM_003999156.5
F1 12817122 12817388
refGene_gn=GPR161;refGene_tx=XM_006942657.4;refGene_gn=GPR161;refGene_tx=XM_011290866.3;refGene_gn=GPR161;refGene_tx=XM_019821572.2;refGene_gn=GPR161;refGene_tx=XM_023248033.1;refGene_gn=GPR161;refGene_tx=XM_023248034.1;refGene_gn=GPR161;refGene_tx=XM_023248035.1;refGene_gn=GPR161;refGene_tx=XM_023248036.1
F1 12819200 12819320
refGene_gn=GPR161;refGene_tx=XM_006942657.4;refGene_gn=GPR161;refGene_tx=XM_011290866.3;refGene_gn=GPR161;refGene_tx=XM_019821572.2;refGene_gn=GPR161;refGene_tx=XM_023248033.1;refGene_gn=GPR161;refGene_tx=XM_023248034.1;refGene_gn=GPR161;refGene_tx=XM_023248035.1;refGene_gn=GPR161;refGene_tx=XM_023248036.1
F1 12821627 12821732
refGene_gn=GPR161;refGene_tx=XM_006942657.4;refGene_gn=GPR161;refGene_tx=XM_011290866.3;refGene_gn=GPR161;refGene_tx=XM_019821572.2;refGene_gn=GPR161;refGene_tx=XM_023248033.1;refGene_gn=GPR161;refGene_tx=XM_023248034.1;refGene_gn=GPR161;refGene_tx=XM_023248035.1;refGene_gn=GPR161;refGene_tx=XM_023248036.1
F1 12827471 12828196
refGene_gn=GPR161;refGene_tx=XM_006942657.4;refGene_gn=GPR161;refGene_tx=XM_011290866.3;refGene_gn=GPR161;refGene_tx=XM_019821572.2;refGene_gn=GPR161;refGene_tx=XM_023248033.1;refGene_gn=GPR161;refGene_tx=XM_02

3248034.1;refGene_gn=GPR161;refGene_tx=XM_023248035.1;refGene_gn=GPR16
1;refGene_tx=XM_023248036.1
F1	12838028	12838446
refGene_gn=GPR161;refGene_tx=XM_006942657.4;refGene_gn=GPR161;refGene_
tx=XM_011290866.3;refGene_gn=GPR161;refGene_tx=XM_023248033.1;refGene_
gn=GPR161;refGene_tx=XM_023248034.1;refGene_gn=GPR161;refGene_tx=XM_02
3248035.1;refGene_gn=GPR161;refGene_tx=XM_023248036.1
F1	12864912	12864928
refGene_gn=GPR161;refGene_tx=XM_011290866.3;refGene_gn=GPR161;refGene_
tx=XM_023248033.1;refGene_gn=GPR161;refGene_tx=XM_023248034.1;refGene_
gn=GPR161;refGene_tx=XM_023248035.1;refGene_gn=GPR161;refGene_tx=XM_02
3248036.1
F1	12894179	12894283
refGene_gn=TIPRL;refGene_tx=XM_023247350.1;refGene_gn=TIPRL;refGene_tx
=XM_023247351.1
F1	12900989	12901169
refGene_gn=TIPRL;refGene_tx=XM_023247350.1;refGene_gn=TIPRL;refGene_tx
=XM_023247351.1
F1	12902332	12902432
refGene_gn=TIPRL;refGene_tx=XM_023247350.1;refGene_gn=TIPRL;refGene_tx
=XM_023247351.1
F1	12909333	12909465
refGene_gn=TIPRL;refGene_tx=XM_023247350.1;refGene_gn=TIPRL;refGene_tx
=XM_023247351.1
F1	12918897	12918993
refGene_gn=TIPRL;refGene_tx=XM_023247350.1;refGene_gn=TIPRL;refGene_tx
=XM_023247351.1
F1	12922443	12922506
refGene_gn=TIPRL;refGene_tx=XM_023247350.1;refGene_gn=TIPRL;refGene_tx
=XM_023247351.1
F1	12923422	12923566
refGene_gn=TIPRL;refGene_tx=XM_023247350.1;refGene_gn=TIPRL;refGene_tx
=XM_023247351.1
F1	12936935	12936998	refGene_gn=SFT2D2;refGene_tx=XM_023247217.1
F1	12939861	12939948	refGene_gn=SFT2D2;refGene_tx=XM_023247217.1
F1	12940188	12940274	refGene_gn=SFT2D2;refGene_tx=XM_023247217.1
F1	12944036	12944118	refGene_gn=SFT2D2;refGene_tx=XM_023247217.1
F1	12945354	12945390	refGene_gn=SFT2D2;refGene_tx=XM_023247217.1
F1	12945474	12945533	refGene_gn=SFT2D2;refGene_tx=XM_023247217.1
F1	12947331	12947361	refGene_gn=SFT2D2;refGene_tx=XM_023247217.1
F1	12949744	12949784	refGene_gn=SFT2D2;refGene_tx=XM_023247217.1
F1	12975990	12976193	refGene_gn=TBX19;refGene_tx=XM_019821575.2
F1	12984080	12984345	refGene_gn=TBX19;refGene_tx=XM_019821575.2
F1	12985981	12986116	refGene_gn=TBX19;refGene_tx=XM_019821575.2
F1	12989357	12989419	refGene_gn=TBX19;refGene_tx=XM_019821575.2
F1	12990554	12990616	refGene_gn=TBX19;refGene_tx=XM_019821575.2
F1	12992585	12992660	refGene_gn=TBX19;refGene_tx=XM_019821575.2
F1	12992662	12992776	refGene_gn=TBX19;refGene_tx=XM_019821575.2
F1	12994968	12995104	refGene_gn=TBX19;refGene_tx=XM_019821575.2
F1	12998021	12998307	refGene_gn=TBX19;refGene_tx=XM_019821575.2

F1 13210138 13210199
refGene_gn=LOC101081599;refGene_tx=XM_023247600.1
F1 13212057 13212172
refGene_gn=LOC101081599;refGene_tx=XM_023247600.1
F1 13212818 13212981
refGene_gn=LOC101081599;refGene_tx=XM_023247600.1
F1 13330889 13330956 refGene_gn=DPT;refGene_tx=XM_003999161.5
F1 13333876 13333984 refGene_gn=DPT;refGene_tx=XM_003999161.5
F1 13343913 13344039 refGene_gn=DPT;refGene_tx=XM_003999161.5
F1 13355937 13356269 refGene_gn=DPT;refGene_tx=XM_003999161.5
F1 13682575 13682672 refGene_gn=ATP1B1;refGene_tx=XM_011290899.2
F1 13687218 13687347 refGene_gn=ATP1B1;refGene_tx=XM_011290899.2
F1 13699223 13699379 refGene_gn=ATP1B1;refGene_tx=XM_011290899.2
F1 13701111 13701296 refGene_gn=ATP1B1;refGene_tx=XM_011290899.2
F1 13703771 13703852 refGene_gn=ATP1B1;refGene_tx=XM_011290899.2
F1 13704825 13705089 refGene_gn=ATP1B1;refGene_tx=XM_011290899.2
F1 13706329 13706362 refGene_gn=NME7;refGene_tx=XM_019821576.1
F1 13761907 13762024
refGene_gn=NME7;refGene_tx=XM_011290900.2;refGene_gn=NME7;refGene_tx=XM_019821576.1
F1 13831243 13831345
refGene_gn=NME7;refGene_tx=XM_011290900.2;refGene_gn=NME7;refGene_tx=XM_019821576.1
F1 13840721 13840790
refGene_gn=NME7;refGene_tx=XM_011290900.2;refGene_gn=NME7;refGene_tx=XM_019821576.1
F1 13844173 13844238
refGene_gn=NME7;refGene_tx=XM_011290900.2;refGene_gn=NME7;refGene_tx=XM_019821576.1
F1 13882658 13882764
refGene_gn=NME7;refGene_tx=XM_011290900.2;refGene_gn=NME7;refGene_tx=XM_019821576.1
F1 13896035 13896243
refGene_gn=NME7;refGene_tx=XM_011290900.2;refGene_gn=NME7;refGene_tx=XM_019821576.1
F1 13899824 13899875
refGene_gn=NME7;refGene_tx=XM_011290900.2;refGene_gn=NME7;refGene_tx=XM_019821576.1
F1 13904174 13904285
refGene_gn=NME7;refGene_tx=XM_011290900.2;refGene_gn=NME7;refGene_tx=XM_019821576.1
F1 13920810 13920977
refGene_gn=NME7;refGene_tx=XM_011290900.2;refGene_gn=NME7;refGene_tx=XM_019821576.1
F1 13921739 13921847
refGene_gn=NME7;refGene_tx=XM_011290900.2;refGene_gn=NME7;refGene_tx=XM_019821576.1
F1 13969021 13969024
refGene_gn=NME7;refGene_tx=XM_011290900.2;refGene_gn=NME7;refGene_tx=XM_019821576.1

F1 13970924 13970955 refGene_gn=BLZF1;refGene_tx=XM_003999171.5
F1 13981112 13981552
refGene_gn=BLZF1;refGene_tx=XM_003999171.5;refGene_gn=BLZF1;refGene_tx=XM_023247380.1
F1 13983943 13984143
refGene_gn=BLZF1;refGene_tx=XM_003999171.5;refGene_gn=BLZF1;refGene_tx=XM_023247380.1
F1 13986493 13986622
refGene_gn=BLZF1;refGene_tx=XM_003999171.5;refGene_gn=BLZF1;refGene_tx=XM_023247380.1
F1 13989532 13989752
refGene_gn=BLZF1;refGene_tx=XM_003999171.5;refGene_gn=BLZF1;refGene_tx=XM_023247380.1
F1 13998176 13998362
refGene_gn=BLZF1;refGene_tx=XM_003999171.5;refGene_gn=BLZF1;refGene_tx=XM_023247380.1
F1 14003457 14003617
refGene_gn=CCDC181;refGene_tx=XM_019821577.1;refGene_gn=CCDC181;refGene_tx=XM_019821578.1;refGene_gn=CCDC181;refGene_tx=XM_023248040.1;refGene_gn=CCDC181;refGene_tx=XM_023248041.1;refGene_gn=CCDC181;refGene_tx=XM_023248042.1;refGene_gn=CCDC181;refGene_tx=XM_023248043.1;refGene_gn=CCDC181;refGene_tx=XM_023248044.1;refGene_gn=CCDC181;refGene_tx=XM_023248045.1
F1 14005716 14005871
refGene_gn=CCDC181;refGene_tx=XM_019821577.1;refGene_gn=CCDC181;refGene_tx=XM_019821578.1;refGene_gn=CCDC181;refGene_tx=XM_023248040.1;refGene_gn=CCDC181;refGene_tx=XM_023248041.1;refGene_gn=CCDC181;refGene_tx=XM_023248042.1;refGene_gn=CCDC181;refGene_tx=XM_023248043.1;refGene_gn=CCDC181;refGene_tx=XM_023248044.1
F1 14011126 14011273
refGene_gn=CCDC181;refGene_tx=XM_019821577.1;refGene_gn=CCDC181;refGene_tx=XM_019821578.1;refGene_gn=CCDC181;refGene_tx=XM_023248040.1;refGene_gn=CCDC181;refGene_tx=XM_023248041.1;refGene_gn=CCDC181;refGene_tx=XM_023248042.1;refGene_gn=CCDC181;refGene_tx=XM_023248043.1;refGene_gn=CCDC181;refGene_tx=XM_023248044.1;refGene_gn=CCDC181;refGene_tx=XM_023248045.1
F1 14014376 14014474
refGene_gn=CCDC181;refGene_tx=XM_023248040.1;refGene_gn=CCDC181;refGene_tx=XM_023248041.1;refGene_gn=CCDC181;refGene_tx=XM_023248042.1;refGene_gn=CCDC181;refGene_tx=XM_023248043.1;refGene_gn=CCDC181;refGene_tx=XM_023248044.1;refGene_gn=CCDC181;refGene_tx=XM_023248045.1
F1 14022670 14023618
refGene_gn=CCDC181;refGene_tx=XM_019821577.1;refGene_gn=CCDC181;refGene_tx=XM_019821578.1;refGene_gn=CCDC181;refGene_tx=XM_023248040.1;refGene_gn=CCDC181;refGene_tx=XM_023248041.1;refGene_gn=CCDC181;refGene_tx=XM_023248042.1;refGene_gn=CCDC181;refGene_tx=XM_023248043.1;refGene_gn=CCDC181;refGene_tx=XM_023248044.1;refGene_gn=CCDC181;refGene_tx=XM_023248045.1
F1 14026413 14026553
refGene_gn=CCDC181;refGene_tx=XM_019821577.1;refGene_gn=CCDC181;refGen

e_tx=XM_019821578.1;refGene_gn=CCDC181;refGene_tx=XM_023248040.1;refGene_gn=CCDC181;refGene_tx=XM_023248041.1;refGene_gn=CCDC181;refGene_tx=XM_023248042.1;refGene_gn=CCDC181;refGene_tx=XM_023248043.1;refGene_gn=CCDC181;refGene_tx=XM_023248044.1;refGene_gn=CCDC181;refGene_tx=XM_023248045.1
F1 14031854 14031915 refGene_gn=CCDC181;refGene_tx=XM_019821577.1;refGene_gn=CCDC181;refGene_tx=XM_019821578.1;refGene_gn=CCDC181;refGene_tx=XM_023248040.1;refGene_gn=CCDC181;refGene_tx=XM_023248041.1;refGene_gn=CCDC181;refGene_tx=XM_023248045.1
F1 14058717 14058846 refGene_gn=SLC19A2;refGene_tx=NM_001039956.1;refGene_gn=SLC19A2;refGene_tx=XM_019821329.2
F1 14061108 14061250 refGene_gn=SLC19A2;refGene_tx=NM_001039956.1;refGene_gn=SLC19A2;refGene_tx=XM_019821329.2
F1 14061585 14061778 refGene_gn=SLC19A2;refGene_tx=NM_001039956.1;refGene_gn=SLC19A2;refGene_tx=XM_019821329.2
F1 14063298 14063521 refGene_gn=SLC19A2;refGene_tx=NM_001039956.1;refGene_gn=SLC19A2;refGene_tx=XM_019821329.2
F1 14073096 14073684 refGene_gn=SLC19A2;refGene_tx=NM_001039956.1;refGene_gn=SLC19A2;refGene_tx=XM_019821329.2
F1 14077908 14077953 refGene_gn=SLC19A2;refGene_tx=XM_019821329.2
F1 14079749 14079947 refGene_gn=SLC19A2;refGene_tx=NM_001039956.1
F1 14109504 14109651 refGene_gn=F5;refGene_tx=XM_011290903.3
F1 14111005 14111188 refGene_gn=F5;refGene_tx=XM_011290903.3
F1 14114187 14114339 refGene_gn=F5;refGene_tx=XM_011290903.3
F1 14116524 14116669 refGene_gn=F5;refGene_tx=XM_011290903.3
F1 14119109 14119265 refGene_gn=F5;refGene_tx=XM_011290903.3
F1 14119692 14119793 refGene_gn=F5;refGene_tx=XM_011290903.3
F1 14121003 14121075 refGene_gn=F5;refGene_tx=XM_011290903.3
F1 14121936 14122053 refGene_gn=F5;refGene_tx=XM_011290903.3
F1 14124116 14124296 refGene_gn=F5;refGene_tx=XM_011290903.3
F1 14126966 14127183 refGene_gn=F5;refGene_tx=XM_011290903.3
F1 14128032 14128269 refGene_gn=F5;refGene_tx=XM_011290903.3
F1 14129485 14129660 refGene_gn=F5;refGene_tx=XM_011290903.3
F1 14132333 14134788 refGene_gn=F5;refGene_tx=XM_011290903.3
F1 14136326 14136539 refGene_gn=F5;refGene_tx=XM_011290903.3
F1 14138415 14138566 refGene_gn=F5;refGene_tx=XM_011290903.3
F1 14139653 14139865 refGene_gn=F5;refGene_tx=XM_011290903.3
F1 14140242 14140342 refGene_gn=F5;refGene_tx=XM_011290903.3
F1 14141806 14141984 refGene_gn=F5;refGene_tx=XM_011290903.3
F1 14146893 14147059 refGene_gn=F5;refGene_tx=XM_011290903.3
F1 14148806 14149028 refGene_gn=F5;refGene_tx=XM_011290903.3
F1 14151510 14151654 refGene_gn=F5;refGene_tx=XM_011290903.3
F1 14152864 14153077 refGene_gn=F5;refGene_tx=XM_011290903.3
F1 14174639 14174762 refGene_gn=F5;refGene_tx=XM_011290903.3

F1 14180905 14180997 refGene_gn=F5;refGene_tx=XM_011290903.3
F1 14185211 14185369 refGene_gn=F5;refGene_tx=XM_011290903.3
F1 14188812 14188867
refGene_gn=SELP;refGene_tx=XM_019821592.2;refGene_gn=SELP;refGene_tx=XM_023247395.1;refGene_gn=SELP;refGene_tx=XM_023247396.1;refGene_gn=SELP;refGene_tx=XM_023247397.1;refGene_gn=SELP;refGene_tx=XM_023247398.1;refGene_gn=SELP;refGene_tx=XM_023247399.1;refGene_gn=SELP;refGene_tx=XM_023247401.1;refGene_gn=SELP;refGene_tx=XM_023247402.1;refGene_gn=SELP;refGene_tx=XM_023247403.1;refGene_gn=SELP;refGene_tx=XM_023247404.1
F1 14190368 14190399
refGene_gn=SELP;refGene_tx=XM_019821592.2;refGene_gn=SELP;refGene_tx=XM_023247395.1;refGene_gn=SELP;refGene_tx=XM_023247396.1;refGene_gn=SELP;refGene_tx=XM_023247397.1;refGene_gn=SELP;refGene_tx=XM_023247398.1;refGene_gn=SELP;refGene_tx=XM_023247399.1;refGene_gn=SELP;refGene_tx=XM_023247401.1;refGene_gn=SELP;refGene_tx=XM_023247402.1;refGene_gn=SELP;refGene_tx=XM_023247403.1;refGene_gn=SELP;refGene_tx=XM_023247404.1
F1 14193951 14194071
refGene_gn=SELP;refGene_tx=XM_019821592.2;refGene_gn=SELP;refGene_tx=XM_023247395.1;refGene_gn=SELP;refGene_tx=XM_023247396.1;refGene_gn=SELP;refGene_tx=XM_023247397.1;refGene_gn=SELP;refGene_tx=XM_023247398.1;refGene_gn=SELP;refGene_tx=XM_023247399.1;refGene_gn=SELP;refGene_tx=XM_023247401.1;refGene_gn=SELP;refGene_tx=XM_023247402.1;refGene_gn=SELP;refGene_tx=XM_023247403.1;refGene_gn=SELP;refGene_tx=XM_023247404.1
F1 14195782 14195968
refGene_gn=SELP;refGene_tx=XM_019821592.2;refGene_gn=SELP;refGene_tx=XM_023247395.1;refGene_gn=SELP;refGene_tx=XM_023247396.1;refGene_gn=SELP;refGene_tx=XM_023247397.1;refGene_gn=SELP;refGene_tx=XM_023247398.1;refGene_gn=SELP;refGene_tx=XM_023247399.1;refGene_gn=SELP;refGene_tx=XM_023247401.1;refGene_gn=SELP;refGene_tx=XM_023247402.1;refGene_gn=SELP;refGene_tx=XM_023247403.1;refGene_gn=SELP;refGene_tx=XM_023247404.1
F1 14197172 14197385
refGene_gn=SELP;refGene_tx=XM_019821592.2;refGene_gn=SELP;refGene_tx=XM_023247395.1;refGene_gn=SELP;refGene_tx=XM_023247396.1;refGene_gn=SELP;refGene_tx=XM_023247397.1;refGene_gn=SELP;refGene_tx=XM_023247398.1;refGene_gn=SELP;refGene_tx=XM_023247399.1;refGene_gn=SELP;refGene_tx=XM_023247402.1;refGene_gn=SELP;refGene_tx=XM_023247403.1
F1 14198758 14198944
refGene_gn=SELP;refGene_tx=XM_019821592.2;refGene_gn=SELP;refGene_tx=XM_023247395.1;refGene_gn=SELP;refGene_tx=XM_023247396.1;refGene_gn=SELP;refGene_tx=XM_023247397.1;refGene_gn=SELP;refGene_tx=XM_023247399.1;refGene_gn=SELP;refGene_tx=XM_023247401.1;refGene_gn=SELP;refGene_tx=XM_023247403.1
F1 14204789 14204975
refGene_gn=SELP;refGene_tx=XM_019821592.2;refGene_gn=SELP;refGene_tx=XM_023247395.1;refGene_gn=SELP;refGene_tx=XM_023247396.1;refGene_gn=SELP;refGene_tx=XM_023247398.1;refGene_gn=SELP;refGene_tx=XM_023247399.1;refGene_gn=SELP;refGene_tx=XM_023247401.1;refGene_gn=SELP;refGene_tx=XM_023247404.1
F1 14208640 14208826
refGene_gn=SELP;refGene_tx=XM_019821592.2;refGene_gn=SELP;refGene_tx=X

M_023247395.1;refGene_gn=SELP;refGene_tx=XM_023247396.1;refGene_gn=SELP;refGene_tx=XM_023247397.1;refGene_gn=SELP;refGene_tx=XM_023247398.1;refGene_gn=SELP;refGene_tx=XM_023247401.1;refGene_gn=SELP;refGene_tx=XM_023247402.1;refGene_gn=SELP;refGene_tx=XM_023247404.1
F1 14210697 14210883
refGene_gn=SELP;refGene_tx=XM_019821592.2;refGene_gn=SELP;refGene_tx=XM_023247395.1;refGene_gn=SELP;refGene_tx=XM_023247396.1;refGene_gn=SELP;refGene_tx=XM_023247397.1;refGene_gn=SELP;refGene_tx=XM_023247398.1;refGene_gn=SELP;refGene_tx=XM_023247399.1;refGene_gn=SELP;refGene_tx=XM_023247401.1;refGene_gn=SELP;refGene_tx=XM_023247402.1;refGene_gn=SELP;refGene_tx=XM_023247403.1;refGene_gn=SELP;refGene_tx=XM_023247404.1
F1 14212363 14212549
refGene_gn=SELP;refGene_tx=XM_019821592.2;refGene_gn=SELP;refGene_tx=XM_023247395.1;refGene_gn=SELP;refGene_tx=XM_023247397.1;refGene_gn=SELP;refGene_tx=XM_023247398.1;refGene_gn=SELP;refGene_tx=XM_023247399.1;refGene_gn=SELP;refGene_tx=XM_023247401.1;refGene_gn=SELP;refGene_tx=XM_023247402.1;refGene_gn=SELP;refGene_tx=XM_023247403.1;refGene_gn=SELP;refGene_tx=XM_023247404.1
F1 14213152 14213338
refGene_gn=SELP;refGene_tx=XM_019821592.2;refGene_gn=SELP;refGene_tx=XM_023247395.1;refGene_gn=SELP;refGene_tx=XM_023247396.1;refGene_gn=SELP;refGene_tx=XM_023247397.1;refGene_gn=SELP;refGene_tx=XM_023247398.1;refGene_gn=SELP;refGene_tx=XM_023247399.1;refGene_gn=SELP;refGene_tx=XM_023247401.1;refGene_gn=SELP;refGene_tx=XM_023247402.1;refGene_gn=SELP;refGene_tx=XM_023247403.1;refGene_gn=SELP;refGene_tx=XM_023247404.1
F1 14214077 14214263
refGene_gn=SELP;refGene_tx=XM_019821592.2;refGene_gn=SELP;refGene_tx=XM_023247395.1;refGene_gn=SELP;refGene_tx=XM_023247396.1;refGene_gn=SELP;refGene_tx=XM_023247397.1;refGene_gn=SELP;refGene_tx=XM_023247398.1;refGene_gn=SELP;refGene_tx=XM_023247399.1;refGene_gn=SELP;refGene_tx=XM_023247401.1;refGene_gn=SELP;refGene_tx=XM_023247402.1;refGene_gn=SELP;refGene_tx=XM_023247403.1;refGene_gn=SELP;refGene_tx=XM_023247404.1
F1 14214714 14214822
refGene_gn=SELP;refGene_tx=XM_019821592.2;refGene_gn=SELP;refGene_tx=XM_023247395.1;refGene_gn=SELP;refGene_tx=XM_023247396.1;refGene_gn=SELP;refGene_tx=XM_023247397.1;refGene_gn=SELP;refGene_tx=XM_023247398.1;refGene_gn=SELP;refGene_tx=XM_023247399.1;refGene_gn=SELP;refGene_tx=XM_023247401.1;refGene_gn=SELP;refGene_tx=XM_023247402.1;refGene_gn=SELP;refGene_tx=XM_023247403.1;refGene_gn=SELP;refGene_tx=XM_023247404.1
F1 14217046 14217433
refGene_gn=SELP;refGene_tx=XM_019821592.2;refGene_gn=SELP;refGene_tx=XM_023247395.1;refGene_gn=SELP;refGene_tx=XM_023247396.1;refGene_gn=SELP;refGene_tx=XM_023247397.1;refGene_gn=SELP;refGene_tx=XM_023247398.1;refGene_gn=SELP;refGene_tx=XM_023247399.1;refGene_gn=SELP;refGene_tx=XM_023247401.1;refGene_gn=SELP;refGene_tx=XM_023247402.1;refGene_gn=SELP;refGene_tx=XM_023247403.1;refGene_gn=SELP;refGene_tx=XM_023247404.1
F1 14219039 14219130
refGene_gn=SELP;refGene_tx=XM_019821592.2;refGene_gn=SELP;refGene_tx=XM_023247395.1;refGene_gn=SELP;refGene_tx=XM_023247396.1;refGene_gn=SELP;refGene_tx=XM_023247397.1;refGene_gn=SELP;refGene_tx=XM_023247398.1;

refGene_gn=SELP;refGene_tx=XM_023247399.1;refGene_gn=SELP;refGene_tx=XM_023247401.1;refGene_gn=SELP;refGene_tx=XM_023247402.1;refGene_gn=SELP;refGene_tx=XM_023247403.1;refGene_gn=SELP;refGene_tx=XM_023247404.1
F1 14228279 14228282
refGene_gn=SELP;refGene_tx=XM_019821592.2;refGene_gn=SELP;refGene_tx=XM_023247395.1;refGene_gn=SELP;refGene_tx=XM_023247396.1;refGene_gn=SELP;refGene_tx=XM_023247397.1;refGene_gn=SELP;refGene_tx=XM_023247398.1;refGene_gn=SELP;refGene_tx=XM_023247399.1;refGene_gn=SELP;refGene_tx=XM_023247401.1;refGene_gn=SELP;refGene_tx=XM_023247402.1;refGene_gn=SELP;refGene_tx=XM_023247403.1;refGene_gn=SELP;refGene_tx=XM_023247404.1
F1 14259332 14259351 refGene_gn=SELL;refGene_tx=NM_001089310.1
F1 14263899 14263918 refGene_gn=SELL;refGene_tx=NM_001089310.1
F1 14267846 14267975 refGene_gn=SELL;refGene_tx=NM_001089310.1
F1 14269543 14269729 refGene_gn=SELL;refGene_tx=NM_001089310.1
F1 14270919 14271105 refGene_gn=SELL;refGene_tx=NM_001089310.1
F1 14275103 14275211 refGene_gn=SELL;refGene_tx=NM_001089310.1
F1 14276141 14276528 refGene_gn=SELL;refGene_tx=NM_001089310.1
F1 14278722 14278804 refGene_gn=SELL;refGene_tx=NM_001089310.1
F1 14279638 14279653 refGene_gn=SELL;refGene_tx=NM_001089310.1
F1 14288411 14288469 refGene_gn=SELE;refGene_tx=XM_011290902.2
F1 14288574 14288596 refGene_gn=SELE;refGene_tx=XM_011290902.2
F1 14289580 14289691 refGene_gn=SELE;refGene_tx=XM_011290902.2
F1 14290168 14290345 refGene_gn=SELE;refGene_tx=XM_011290902.2
F1 14290555 14290744 refGene_gn=SELE;refGene_tx=XM_011290902.2
F1 14290865 14291054 refGene_gn=SELE;refGene_tx=XM_011290902.2
F1 14291707 14291893 refGene_gn=SELE;refGene_tx=XM_011290902.2
F1 14292843 14293029 refGene_gn=SELE;refGene_tx=XM_011290902.2
F1 14293442 14293550 refGene_gn=SELE;refGene_tx=XM_011290902.2
F1 14293898 14294285 refGene_gn=SELE;refGene_tx=XM_011290902.2
F1 14294884 14294942 refGene_gn=SELE;refGene_tx=XM_011290902.2
F1 14341397 14342504
refGene_gn=METTL18;refGene_tx=XM_006942677.3;refGene_gn=METTL18;refGene_tx=XM_006942678.2;refGene_gn=METTL18;refGene_tx=XM_019821600.2
F1 14344515 14344626
refGene_gn=CF1H1orf112;refGene_tx=XM_003999181.4
F1 14347418 14347458
refGene_gn=CF1H1orf112;refGene_tx=XM_003999181.4
F1 14348773 14348862
refGene_gn=CF1H1orf112;refGene_tx=XM_003999181.4;refGene_gn=CF1H1orf112;refGene_tx=XM_023247889.1;refGene_gn=CF1H1orf112;refGene_tx=XM_023247890.1
F1 14350850 14350955
refGene_gn=CF1H1orf112;refGene_tx=XM_003999181.4;refGene_gn=CF1H1orf112;refGene_tx=XM_023247889.1;refGene_gn=CF1H1orf112;refGene_tx=XM_023247890.1
F1 14351436 14351577
refGene_gn=CF1H1orf112;refGene_tx=XM_003999181.4;refGene_gn=CF1H1orf112;refGene_tx=XM_023247889.1;refGene_gn=CF1H1orf112;refGene_tx=XM_023247890.1
F1 14353239 14353405

refGene_gn=CF1H1orf112;refGene_tx=XM_003999181.4;refGene_gn=CF1H1orf11
2;refGene_tx=XM_023247889.1;refGene_gn=CF1H1orf112;refGene_tx=XM_02324
7890.1
F1 14354839 14354924
refGene_gn=CF1H1orf112;refGene_tx=XM_003999181.4;refGene_gn=CF1H1orf11
2;refGene_tx=XM_023247889.1;refGene_gn=CF1H1orf112;refGene_tx=XM_02324
7890.1
F1 14356806 14356945
refGene_gn=CF1H1orf112;refGene_tx=XM_003999181.4;refGene_gn=CF1H1orf11
2;refGene_tx=XM_023247889.1;refGene_gn=CF1H1orf112;refGene_tx=XM_02324
7890.1
F1 14364188 14364269
refGene_gn=CF1H1orf112;refGene_tx=XM_003999181.4;refGene_gn=CF1H1orf11
2;refGene_tx=XM_023247889.1;refGene_gn=CF1H1orf112;refGene_tx=XM_02324
7890.1
F1 14365697 14365762
refGene_gn=CF1H1orf112;refGene_tx=XM_003999181.4;refGene_gn=CF1H1orf11
2;refGene_tx=XM_023247889.1;refGene_gn=CF1H1orf112;refGene_tx=XM_02324
7890.1
F1 14368721 14368870
refGene_gn=CF1H1orf112;refGene_tx=XM_003999181.4;refGene_gn=CF1H1orf11
2;refGene_tx=XM_023247889.1;refGene_gn=CF1H1orf112;refGene_tx=XM_02324
7890.1
F1 14369139 14369269
refGene_gn=CF1H1orf112;refGene_tx=XM_003999181.4;refGene_gn=CF1H1orf11
2;refGene_tx=XM_023247889.1;refGene_gn=CF1H1orf112;refGene_tx=XM_02324
7890.1
F1 14370796 14370983
refGene_gn=CF1H1orf112;refGene_tx=XM_003999181.4;refGene_gn=CF1H1orf11
2;refGene_tx=XM_023247889.1;refGene_gn=CF1H1orf112;refGene_tx=XM_02324
7890.1
F1 14371711 14371794
refGene_gn=CF1H1orf112;refGene_tx=XM_003999181.4;refGene_gn=CF1H1orf11
2;refGene_tx=XM_023247889.1;refGene_gn=CF1H1orf112;refGene_tx=XM_02324
7890.1
F1 14372136 14372188
refGene_gn=CF1H1orf112;refGene_tx=XM_003999181.4;refGene_gn=CF1H1orf11
2;refGene_tx=XM_023247889.1;refGene_gn=CF1H1orf112;refGene_tx=XM_02324
7890.1
F1 14373592 14373682
refGene_gn=CF1H1orf112;refGene_tx=XM_003999181.4;refGene_gn=CF1H1orf11
2;refGene_tx=XM_023247889.1;refGene_gn=CF1H1orf112;refGene_tx=XM_02324
7890.1
F1 14378547 14378733
refGene_gn=CF1H1orf112;refGene_tx=XM_003999181.4;refGene_gn=CF1H1orf11
2;refGene_tx=XM_023247889.1;refGene_gn=CF1H1orf112;refGene_tx=XM_02324
7890.1
F1 14380687 14380810
refGene_gn=CF1H1orf112;refGene_tx=XM_003999181.4;refGene_gn=CF1H1orf11
2;refGene_tx=XM_023247889.1;refGene_gn=CF1H1orf112;refGene_tx=XM_02324

7890.1
F1 14382082 14382178
refGene_gn=CF1H1orf112;refGene_tx=XM_003999181.4;refGene_gn=CF1H1orf11
2;refGene_tx=XM_023247889.1;refGene_gn=CF1H1orf112;refGene_tx=XM_02324
7890.1
F1 14383979 14384093
refGene_gn=CF1H1orf112;refGene_tx=XM_003999181.4;refGene_gn=CF1H1orf11
2;refGene_tx=XM_023247889.1;refGene_gn=CF1H1orf112;refGene_tx=XM_02324
7890.1
F1 14385906 14386011
refGene_gn=CF1H1orf112;refGene_tx=XM_003999181.4;refGene_gn=CF1H1orf11
2;refGene_tx=XM_023247889.1;refGene_gn=CF1H1orf112;refGene_tx=XM_02324
7890.1
F1 14386652 14386733
refGene_gn=CF1H1orf112;refGene_tx=XM_003999181.4;refGene_gn=CF1H1orf11
2;refGene_tx=XM_023247889.1;refGene_gn=CF1H1orf112;refGene_tx=XM_02324
7890.1
F1 14387658 14387778
refGene_gn=CF1H1orf112;refGene_tx=XM_003999181.4;refGene_gn=CF1H1orf11
2;refGene_tx=XM_023247889.1;refGene_gn=CF1H1orf112;refGene_tx=XM_02324
7890.1
F1 14389438 14389639
refGene_gn=CF1H1orf112;refGene_tx=XM_003999181.4;refGene_gn=CF1H1orf11
2;refGene_tx=XM_023247889.1;refGene_gn=CF1H1orf112;refGene_tx=XM_02324
7890.1
F1 14390357 14390417
refGene_gn=SCYL3;refGene_tx=XM_006942681.4;refGene_gn=SCYL3;refGene_tx
=XM_011290894.3;refGene_gn=SCYL3;refGene_tx=XM_011290895.3;refGene_gn=
SCYL3;refGene_tx=XM_023247891.1;refGene_gn=SCYL3;refGene_tx=XM_0232478
92.1;refGene_gn=SCYL3;refGene_tx=XM_023247893.1;refGene_gn=SCYL3;refGe
ne_tx=XM_023247894.1;refGene_gn=SCYL3;refGene_tx=XM_023247895.1;refGen
e_gn=SCYL3;refGene_tx=XM_023247896.1;refGene_gn=SCYL3;refGene_tx=XM_02
3247897.1;refGene_gn=SCYL3;refGene_tx=XM_023247898.1
F1 14390932 14391627
refGene_gn=SCYL3;refGene_tx=XM_006942681.4;refGene_gn=SCYL3;refGene_tx
=XM_011290894.3;refGene_gn=SCYL3;refGene_tx=XM_011290895.3;refGene_gn=
SCYL3;refGene_tx=XM_023247891.1;refGene_gn=SCYL3;refGene_tx=XM_0232478
92.1;refGene_gn=SCYL3;refGene_tx=XM_023247893.1;refGene_gn=SCYL3;refGe
ne_tx=XM_023247894.1;refGene_gn=SCYL3;refGene_tx=XM_023247895.1;refGen
e_gn=SCYL3;refGene_tx=XM_023247896.1;refGene_gn=SCYL3;refGene_tx=XM_02
3247897.1;refGene_gn=SCYL3;refGene_tx=XM_023247898.1
F1 14397424 14397596
refGene_gn=SCYL3;refGene_tx=XM_006942681.4;refGene_gn=SCYL3;refGene_tx
=XM_011290894.3;refGene_gn=SCYL3;refGene_tx=XM_011290895.3;refGene_gn=
SCYL3;refGene_tx=XM_023247891.1;refGene_gn=SCYL3;refGene_tx=XM_0232478
92.1;refGene_gn=SCYL3;refGene_tx=XM_023247893.1;refGene_gn=SCYL3;refGe
ne_tx=XM_023247894.1;refGene_gn=SCYL3;refGene_tx=XM_023247895.1;refGen
e_gn=SCYL3;refGene_tx=XM_023247896.1;refGene_gn=SCYL3;refGene_tx=XM_02
3247897.1;refGene_gn=SCYL3;refGene_tx=XM_023247898.1
F1 14407125 14407310

refGene_gn=SCYL3;refGene_tx=XM_006942681.4;refGene_gn=SCYL3;refGene_tx=XM_011290894.3;refGene_gn=SCYL3;refGene_tx=XM_011290895.3;refGene_gn=SCYL3;refGene_tx=XM_023247891.1;refGene_gn=SCYL3;refGene_tx=XM_023247892.1;refGene_gn=SCYL3;refGene_tx=XM_023247893.1;refGene_gn=SCYL3;refGene_tx=XM_023247894.1;refGene_gn=SCYL3;refGene_tx=XM_023247895.1;refGene_gn=SCYL3;refGene_tx=XM_023247896.1;refGene_gn=SCYL3;refGene_tx=XM_023247897.1;refGene_gn=SCYL3;refGene_tx=XM_023247898.1
F1 14409025 14409165
refGene_gn=SCYL3;refGene_tx=XM_006942681.4;refGene_gn=SCYL3;refGene_tx=XM_011290894.3;refGene_gn=SCYL3;refGene_tx=XM_011290895.3;refGene_gn=SCYL3;refGene_tx=XM_023247891.1;refGene_gn=SCYL3;refGene_tx=XM_023247892.1;refGene_gn=SCYL3;refGene_tx=XM_023247893.1;refGene_gn=SCYL3;refGene_tx=XM_023247894.1;refGene_gn=SCYL3;refGene_tx=XM_023247895.1;refGene_gn=SCYL3;refGene_tx=XM_023247896.1;refGene_gn=SCYL3;refGene_tx=XM_023247897.1;refGene_gn=SCYL3;refGene_tx=XM_023247898.1
F1 14410953 14411031
refGene_gn=SCYL3;refGene_tx=XM_006942681.4;refGene_gn=SCYL3;refGene_tx=XM_011290894.3;refGene_gn=SCYL3;refGene_tx=XM_011290895.3;refGene_gn=SCYL3;refGene_tx=XM_023247891.1;refGene_gn=SCYL3;refGene_tx=XM_023247892.1;refGene_gn=SCYL3;refGene_tx=XM_023247893.1;refGene_gn=SCYL3;refGene_tx=XM_023247894.1;refGene_gn=SCYL3;refGene_tx=XM_023247895.1;refGene_gn=SCYL3;refGene_tx=XM_023247896.1;refGene_gn=SCYL3;refGene_tx=XM_023247897.1
F1 14413347 14413459
refGene_gn=SCYL3;refGene_tx=XM_006942681.4;refGene_gn=SCYL3;refGene_tx=XM_011290894.3;refGene_gn=SCYL3;refGene_tx=XM_011290895.3;refGene_gn=SCYL3;refGene_tx=XM_023247891.1;refGene_gn=SCYL3;refGene_tx=XM_023247892.1;refGene_gn=SCYL3;refGene_tx=XM_023247893.1;refGene_gn=SCYL3;refGene_tx=XM_023247894.1;refGene_gn=SCYL3;refGene_tx=XM_023247895.1;refGene_gn=SCYL3;refGene_tx=XM_023247896.1;refGene_gn=SCYL3;refGene_tx=XM_023247897.1;refGene_gn=SCYL3;refGene_tx=XM_023247898.1
F1 14414698 14414801
refGene_gn=SCYL3;refGene_tx=XM_006942681.4;refGene_gn=SCYL3;refGene_tx=XM_011290894.3;refGene_gn=SCYL3;refGene_tx=XM_011290895.3;refGene_gn=SCYL3;refGene_tx=XM_023247891.1;refGene_gn=SCYL3;refGene_tx=XM_023247892.1;refGene_gn=SCYL3;refGene_tx=XM_023247893.1;refGene_gn=SCYL3;refGene_tx=XM_023247894.1;refGene_gn=SCYL3;refGene_tx=XM_023247895.1;refGene_gn=SCYL3;refGene_tx=XM_023247896.1;refGene_gn=SCYL3;refGene_tx=XM_023247897.1;refGene_gn=SCYL3;refGene_tx=XM_023247898.1
F1 14415984 14416041
refGene_gn=SCYL3;refGene_tx=XM_006942681.4;refGene_gn=SCYL3;refGene_tx=XM_011290894.3;refGene_gn=SCYL3;refGene_tx=XM_011290895.3;refGene_gn=SCYL3;refGene_tx=XM_023247891.1;refGene_gn=SCYL3;refGene_tx=XM_023247892.1;refGene_gn=SCYL3;refGene_tx=XM_023247893.1;refGene_gn=SCYL3;refGene_tx=XM_023247894.1;refGene_gn=SCYL3;refGene_tx=XM_023247895.1;refGene_gn=SCYL3;refGene_tx=XM_023247896.1;refGene_gn=SCYL3;refGene_tx=XM_023247898.1
F1 14417977 14418091
refGene_gn=SCYL3;refGene_tx=XM_006942681.4;refGene_gn=SCYL3;refGene_tx=XM_011290894.3;refGene_gn=SCYL3;refGene_tx=XM_011290895.3;refGene_gn=

SCYL3;refGene_tx=XM_023247891.1;refGene_gn=SCYL3;refGene_tx=XM_0232478
92.1;refGene_gn=SCYL3;refGene_tx=XM_023247893.1;refGene_gn=SCYL3;refGe
ne_tx=XM_023247894.1;refGene_gn=SCYL3;refGene_tx=XM_023247895.1;refGen
e_gn=SCYL3;refGene_tx=XM_023247896.1;refGene_gn=SCYL3;refGene_tx=XM_02
3247897.1;refGene_gn=SCYL3;refGene_tx=XM_023247898.1
F1 14420112 14420298
refGene_gn=SCYL3;refGene_tx=XM_006942681.4;refGene_gn=SCYL3;refGene_tx
=XM_011290894.3;refGene_gn=SCYL3;refGene_tx=XM_011290895.3;refGene_gn=
SCYL3;refGene_tx=XM_023247891.1;refGene_gn=SCYL3;refGene_tx=XM_0232478
92.1;refGene_gn=SCYL3;refGene_tx=XM_023247893.1;refGene_gn=SCYL3;refGe
ne_tx=XM_023247894.1;refGene_gn=SCYL3;refGene_tx=XM_023247895.1;refGen
e_gn=SCYL3;refGene_tx=XM_023247896.1;refGene_gn=SCYL3;refGene_tx=XM_02
3247897.1;refGene_gn=SCYL3;refGene_tx=XM_023247898.1
F1 14427327 14427492
refGene_gn=SCYL3;refGene_tx=XM_006942681.4;refGene_gn=SCYL3;refGene_tx
=XM_011290894.3;refGene_gn=SCYL3;refGene_tx=XM_011290895.3;refGene_gn=
SCYL3;refGene_tx=XM_023247891.1;refGene_gn=SCYL3;refGene_tx=XM_0232478
92.1;refGene_gn=SCYL3;refGene_tx=XM_023247893.1;refGene_gn=SCYL3;refGe
ne_tx=XM_023247894.1;refGene_gn=SCYL3;refGene_tx=XM_023247895.1;refGen
e_gn=SCYL3;refGene_tx=XM_023247896.1;refGene_gn=SCYL3;refGene_tx=XM_02
3247897.1;refGene_gn=SCYL3;refGene_tx=XM_023247898.1
F1 14445415 14445524
refGene_gn=KIFAP3;refGene_tx=XM_011290890.3;refGene_gn=KIFAP3;refGene_
tx=XM_019821595.2
F1 14476018 14476118
refGene_gn=KIFAP3;refGene_tx=XM_011290890.3;refGene_gn=KIFAP3;refGene_
tx=XM_019821595.2
F1 14481964 14482154
refGene_gn=KIFAP3;refGene_tx=XM_011290890.3;refGene_gn=KIFAP3;refGene_
tx=XM_019821595.2
F1 14495423 14495509
refGene_gn=KIFAP3;refGene_tx=XM_011290890.3;refGene_gn=KIFAP3;refGene_
tx=XM_019821595.2
F1 14500701 14500800
refGene_gn=KIFAP3;refGene_tx=XM_011290890.3;refGene_gn=KIFAP3;refGene_
tx=XM_019821595.2
F1 14505205 14505331
refGene_gn=KIFAP3;refGene_tx=XM_011290890.3;refGene_gn=KIFAP3;refGene_
tx=XM_019821595.2
F1 14505962 14506128
refGene_gn=KIFAP3;refGene_tx=XM_011290890.3;refGene_gn=KIFAP3;refGene_
tx=XM_019821595.2
F1 14506653 14506766
refGene_gn=KIFAP3;refGene_tx=XM_011290890.3;refGene_gn=KIFAP3;refGene_
tx=XM_019821595.2
F1 14508231 14508340
refGene_gn=KIFAP3;refGene_tx=XM_011290890.3;refGene_gn=KIFAP3;refGene_
tx=XM_019821595.2
F1 14515921 14516022
refGene_gn=KIFAP3;refGene_tx=XM_011290890.3;refGene_gn=KIFAP3;refGene_

tx=XM_019821595.2
F1 14540748 14540911
refGene_gn=KIFAP3;refGene_tx=XM_011290890.3;refGene_gn=KIFAP3;refGene_
tx=XM_019821595.2
F1 14553239 14553418
refGene_gn=KIFAP3;refGene_tx=XM_011290890.3;refGene_gn=KIFAP3;refGene_
tx=XM_019821595.2
F1 14577383 14577482
refGene_gn=KIFAP3;refGene_tx=XM_011290890.3;refGene_gn=KIFAP3;refGene_
tx=XM_019821595.2
F1 14579150 14579275
refGene_gn=KIFAP3;refGene_tx=XM_011290890.3;refGene_gn=KIFAP3;refGene_
tx=XM_019821595.2
F1 14580322 14580422
refGene_gn=KIFAP3;refGene_tx=XM_011290890.3;refGene_gn=KIFAP3;refGene_
tx=XM_019821595.2
F1 14589098 14589240
refGene_gn=KIFAP3;refGene_tx=XM_011290890.3;refGene_gn=KIFAP3;refGene_
tx=XM_019821595.2
F1 14589789 14589845
refGene_gn=KIFAP3;refGene_tx=XM_011290890.3;refGene_gn=KIFAP3;refGene_
tx=XM_019821595.2
F1 14591884 14592039
refGene_gn=KIFAP3;refGene_tx=XM_011290890.3;refGene_gn=KIFAP3;refGene_
tx=XM_019821595.2
F1 14594820 14594952
refGene_gn=KIFAP3;refGene_tx=XM_011290890.3;refGene_gn=KIFAP3;refGene_
tx=XM_019821595.2
F1 14608314 14608346 refGene_gn=KIFAP3;refGene_tx=XM_011290890.3
F1 14618959 14619039 refGene_gn=KIFAP3;refGene_tx=XM_019821595.2
F1 14662425 14662579 refGene_gn=METTL11B;refGene_tx=XM_023247955.1
F1 14672908 14673084 refGene_gn=METTL11B;refGene_tx=XM_023247955.1
F1 14678903 14679153 refGene_gn=METTL11B;refGene_tx=XM_023247955.1
F1 14679954 14680226 refGene_gn=METTL11B;refGene_tx=XM_023247955.1
F1 14975039 14975100 refGene_gn=GORAB;refGene_tx=XM_006942686.4
F1 14980623 14980978 refGene_gn=GORAB;refGene_tx=XM_006942686.4
F1 14983593 14983695
refGene_gn=GORAB;refGene_tx=XM_006942686.4;refGene_gn=GORAB;refGene_tx
=XM_019821599.1
F1 14986089 14986230
refGene_gn=GORAB;refGene_tx=XM_006942686.4;refGene_gn=GORAB;refGene_tx
=XM_019821599.1
F1 14993530 14993987
refGene_gn=GORAB;refGene_tx=XM_006942686.4;refGene_gn=GORAB;refGene_tx
=XM_019821599.1
F1 15084885 15085126
refGene_gn=PRRX1;refGene_tx=XM_003999182.5;refGene_gn=PRRX1;refGene_tx
=XM_003999183.5;refGene_gn=PRRX1;refGene_tx=XM_006942687.4;refGene_gn=
PRRX1;refGene_tx=XM_006942688.4
F1 15140182 15140358

refGene_gn=PRRX1;refGene_tx=XM_003999182.5;refGene_gn=PRRX1;refGene_tx=XM_003999183.5;refGene_gn=PRRX1;refGene_tx=XM_006942687.4;refGene_gn=PRRX1;refGene_tx=XM_006942688.4
F1 15145100 15145282 refGene_gn=PRRX1;refGene_tx=XM_003999182.5;refGene_gn=PRRX1;refGene_tx=XM_003999183.5;refGene_gn=PRRX1;refGene_tx=XM_006942687.4;refGene_gn=PRRX1;refGene_tx=XM_006942688.4
F1 15149042 15149097 refGene_gn=PRRX1;refGene_tx=XM_003999182.5;refGene_gn=PRRX1;refGene_tx=XM_006942688.4
F1 15154404 15154543 refGene_gn=PRRX1;refGene_tx=XM_003999183.5
F1 15161721 15161815 refGene_gn=PRRX1;refGene_tx=XM_006942687.4
F1 15315378 15315433 refGene_gn=MROH9;refGene_tx=XM_006942690.2;refGene_gn=MROH9;refGene_tx=XM_019821594.1
F1 15318328 15318375 refGene_gn=MROH9;refGene_tx=XM_006942690.2;refGene_gn=MROH9;refGene_tx=XM_019821594.1
F1 15326221 15326301 refGene_gn=MROH9;refGene_tx=XM_006942690.2;refGene_gn=MROH9;refGene_tx=XM_019821594.1
F1 15340882 15341018 refGene_gn=MROH9;refGene_tx=XM_006942690.2;refGene_gn=MROH9;refGene_tx=XM_019821594.1
F1 15341578 15341665 refGene_gn=MROH9;refGene_tx=XM_006942690.2;refGene_gn=MROH9;refGene_tx=XM_019821594.1
F1 15343834 15343939 refGene_gn=MROH9;refGene_tx=XM_006942690.2;refGene_gn=MROH9;refGene_tx=XM_019821594.1
F1 15358065 15358201 refGene_gn=MROH9;refGene_tx=XM_006942690.2;refGene_gn=MROH9;refGene_tx=XM_019821594.1
F1 15364123 15364248 refGene_gn=MROH9;refGene_tx=XM_006942690.2;refGene_gn=MROH9;refGene_tx=XM_019821594.1
F1 15366610 15366760 refGene_gn=MROH9;refGene_tx=XM_006942690.2;refGene_gn=MROH9;refGene_tx=XM_019821594.1
F1 15370301 15370456 refGene_gn=MROH9;refGene_tx=XM_006942690.2;refGene_gn=MROH9;refGene_tx=XM_019821594.1
F1 15372054 15372220 refGene_gn=MROH9;refGene_tx=XM_006942690.2;refGene_gn=MROH9;refGene_tx=XM_019821594.1
F1 15373382 15373525 refGene_gn=MROH9;refGene_tx=XM_006942690.2;refGene_gn=MROH9;refGene_tx=XM_019821594.1
F1 15374522 15374660 refGene_gn=MROH9;refGene_tx=XM_006942690.2;refGene_gn=MROH9;refGene_tx

=XM_019821594.1
F1 15376447 15376568
refGene_gn=MROH9;refGene_tx=XM_006942690.2;refGene_gn=MROH9;refGene_tx
=XM_019821594.1
F1 15383593 15383731
refGene_gn=MROH9;refGene_tx=XM_006942690.2;refGene_gn=MROH9;refGene_tx
=XM_019821594.1
F1 15386225 15386402
refGene_gn=MROH9;refGene_tx=XM_006942690.2;refGene_gn=MROH9;refGene_tx
=XM_019821594.1
F1 15389048 15389201
refGene_gn=MROH9;refGene_tx=XM_006942690.2;refGene_gn=MROH9;refGene_tx
=XM_019821594.1
F1 15389302 15389419
refGene_gn=MROH9;refGene_tx=XM_006942690.2;refGene_gn=MROH9;refGene_tx
=XM_019821594.1
F1 15389641 15389734 refGene_gn=MROH9;refGene_tx=XM_019821594.1
F1 15390020 15390123 refGene_gn=MROH9;refGene_tx=XM_006942690.2
F1 15396196 15396259 refGene_gn=MROH9;refGene_tx=XM_006942690.2
F1 15397823 15398110 refGene_gn=MROH9;refGene_tx=XM_006942690.2
F1 15456681 15456813 refGene_gn=FMO3;refGene_tx=XM_003999196.2
F1 15464727 15464916 refGene_gn=FMO3;refGene_tx=XM_003999196.2
F1 15468381 15468544 refGene_gn=FMO3;refGene_tx=XM_003999196.2
F1 15468809 15468952 refGene_gn=FMO3;refGene_tx=XM_003999196.2
F1 15472211 15472411 refGene_gn=FMO3;refGene_tx=XM_003999196.2
F1 15475028 15475384 refGene_gn=FMO3;refGene_tx=XM_003999196.2
F1 15478078 15478151 refGene_gn=FMO3;refGene_tx=XM_003999196.2
F1 15478860 15479206 refGene_gn=FMO3;refGene_tx=XM_003999196.2
F1 15513692 15513824
refGene_gn=LOC101086441;refGene_tx=XM_003999184.4
F1 15518750 15518939
refGene_gn=LOC101086441;refGene_tx=XM_003999184.4
F1 15520051 15520214
refGene_gn=LOC101086441;refGene_tx=XM_003999184.4
F1 15521390 15521533
refGene_gn=LOC101086441;refGene_tx=XM_003999184.4
F1 15523416 15523616
refGene_gn=LOC101086441;refGene_tx=XM_003999184.4
F1 15526080 15526436
refGene_gn=LOC101086441;refGene_tx=XM_003999184.4
F1 15528410 15528483
refGene_gn=LOC101086441;refGene_tx=XM_003999184.4
F1 15530362 15530705
refGene_gn=LOC101086441;refGene_tx=XM_003999184.4
F1 15551625 15551757
refGene_gn=FMO2;refGene_tx=XM_003999186.5;refGene_gn=FMO2;refGene_tx=X
M_011290888.2;refGene_gn=FMO2;refGene_tx=XM_019821608.2
F1 15557834 15557861 refGene_gn=FMO2;refGene_tx=XM_019821609.2
F1 15565412 15565601
refGene_gn=FMO2;refGene_tx=XM_003999186.5;refGene_gn=FMO2;refGene_tx=X

M_011290888.2;refGene_gn=FMO2;refGene_tx=XM_019821608.2;refGene_gn=FMO2;refGene_tx=XM_019821609.2;refGene_gn=FMO2;refGene_tx=XM_023247498.1
F1 15567903 15568066 refGene_gn=FMO2;refGene_tx=XM_003999186.5;refGene_gn=FMO2;refGene_tx=XM_011290888.2;refGene_gn=FMO2;refGene_tx=XM_019821608.2;refGene_gn=FMO2;refGene_tx=XM_019821609.2;refGene_gn=FMO2;refGene_tx=XM_023247498.1
F1 15570320 15570463 refGene_gn=FMO2;refGene_tx=XM_003999186.5;refGene_gn=FMO2;refGene_tx=XM_011290888.2;refGene_gn=FMO2;refGene_tx=XM_019821608.2;refGene_gn=FMO2;refGene_tx=XM_019821609.2;refGene_gn=FMO2;refGene_tx=XM_023247498.1
F1 15570781 15570987 refGene_gn=FMO2;refGene_tx=XM_003999186.5;refGene_gn=FMO2;refGene_tx=XM_011290888.2;refGene_gn=FMO2;refGene_tx=XM_019821608.2;refGene_gn=FMO2;refGene_tx=XM_019821609.2;refGene_gn=FMO2;refGene_tx=XM_023247498.1
F1 15574933 15575289 refGene_gn=FMO2;refGene_tx=XM_003999186.5;refGene_gn=FMO2;refGene_tx=XM_011290888.2;refGene_gn=FMO2;refGene_tx=XM_019821608.2;refGene_gn=FMO2;refGene_tx=XM_019821609.2;refGene_gn=FMO2;refGene_tx=XM_023247498.1
F1 15577793 15577866 refGene_gn=FMO2;refGene_tx=XM_003999186.5;refGene_gn=FMO2;refGene_tx=XM_011290888.2;refGene_gn=FMO2;refGene_tx=XM_019821608.2;refGene_gn=FMO2;refGene_tx=XM_019821609.2;refGene_gn=FMO2;refGene_tx=XM_023247498.1
F1 15578175 15578221 refGene_gn=FMO2;refGene_tx=XM_011290888.2
F1 15578881 15579233 refGene_gn=FMO2;refGene_tx=XM_003999186.5;refGene_gn=FMO2;refGene_tx=XM_019821608.2;refGene_gn=FMO2;refGene_tx=XM_019821609.2;refGene_gn=FMO2;refGene_tx=XM_023247498.1
F1 15593497 15593629 refGene_gn=FMO1;refGene_tx=XM_003999187.4
F1 15602743 15602932 refGene_gn=FMO1;refGene_tx=XM_003999187.4
F1 15606285 15606448 refGene_gn=FMO1;refGene_tx=XM_003999187.4
F1 15611189 15611332 refGene_gn=FMO1;refGene_tx=XM_003999187.4
F1 15613564 15613764 refGene_gn=FMO1;refGene_tx=XM_003999187.4
F1 15614607 15614963 refGene_gn=FMO1;refGene_tx=XM_003999187.4
F1 15615482 15615555 refGene_gn=FMO1;refGene_tx=XM_003999187.4
F1 15618293 15618636 refGene_gn=FMO1;refGene_tx=XM_003999187.4
F1 15632608 15632740 refGene_gn=FMO4;refGene_tx=XM_003999197.4;refGene_gn=FMO4;refGene_tx=XM_006942693.3;refGene_gn=FMO4;refGene_tx=XM_023247493.1
F1 15635709 15635898 refGene_gn=FMO4;refGene_tx=XM_003999197.4;refGene_gn=FMO4;refGene_tx=XM_006942693.3;refGene_gn=FMO4;refGene_tx=XM_023247493.1
F1 15636939 15637102 refGene_gn=FMO4;refGene_tx=XM_003999197.4;refGene_gn=FMO4;refGene_tx=XM_006942693.3;refGene_gn=FMO4;refGene_tx=XM_023247493.1
F1 15641208 15641351 refGene_gn=FMO4;refGene_tx=XM_003999197.4;refGene_gn=FMO4;refGene_tx=XM_006942693.3;refGene_gn=FMO4;refGene_tx=XM_023247493.1
F1 15642821 15643021 refGene_gn=FMO4;refGene_tx=XM_003999197.4;refGene_gn=FMO4;refGene_tx=XM_006942693.3;refGene_gn=FMO4;refGene_tx=XM_023247493.1

F1	15644354	15644707
refGene_gn=FMO4;refGene_tx=XM_003999197.4;refGene_gn=FMO4;refGene_tx=XM_006942693.3;refGene_gn=FMO4;refGene_tx=XM_023247493.1
F1	15645042	15645101	refGene_gn=FMO4;refGene_tx=XM_023247493.1
F1	15646835	15646905
refGene_gn=FMO4;refGene_tx=XM_003999197.4;refGene_gn=FMO4;refGene_tx=XM_006942693.3
F1	15648806	15649254
refGene_gn=FMO4;refGene_tx=XM_003999197.4;refGene_gn=FMO4;refGene_tx=XM_006942693.3
F1	15767950	15768062	refGene_gn=PRRC2C;refGene_tx=XM_006942701.4
F1	15769430	15769602	refGene_gn=PRRC2C;refGene_tx=XM_006942701.4
F1	15771335	15771445	refGene_gn=PRRC2C;refGene_tx=XM_006942701.4
F1	15772528	15772654	refGene_gn=PRRC2C;refGene_tx=XM_006942701.4
F1	15776603	15776827	refGene_gn=PRRC2C;refGene_tx=XM_006942701.4
F1	15780762	15780845	refGene_gn=PRRC2C;refGene_tx=XM_006942701.4
F1	15781491	15781625	refGene_gn=PRRC2C;refGene_tx=XM_006942701.4
F1	15781705	15781793	refGene_gn=PRRC2C;refGene_tx=XM_006942701.4
F1	15783080	15783225	refGene_gn=PRRC2C;refGene_tx=XM_006942701.4
F1	15784985	15785039	refGene_gn=PRRC2C;refGene_tx=XM_006942701.4
F1	15789369	15789991	refGene_gn=PRRC2C;refGene_tx=XM_006942701.4
F1	15795381	15795551	refGene_gn=PRRC2C;refGene_tx=XM_006942701.4
F1	15796009	15796271	refGene_gn=PRRC2C;refGene_tx=XM_006942701.4
F1	15797258	15797469	refGene_gn=PRRC2C;refGene_tx=XM_006942701.4
F1	15799666	15801916	refGene_gn=PRRC2C;refGene_tx=XM_006942701.4
F1	15803993	15804111	refGene_gn=PRRC2C;refGene_tx=XM_006942701.4
F1	15804112	15804203	refGene_gn=PRRC2C;refGene_tx=XM_006942701.4
F1	15808491	15808646	refGene_gn=PRRC2C;refGene_tx=XM_006942701.4
F1	15809957	15810732	refGene_gn=PRRC2C;refGene_tx=XM_006942701.4
F1	15811140	15811226	refGene_gn=PRRC2C;refGene_tx=XM_006942701.4
F1	15815937	15816126	refGene_gn=PRRC2C;refGene_tx=XM_006942701.4
F1	15816254	15816506	refGene_gn=PRRC2C;refGene_tx=XM_006942701.4
F1	15817831	15817924	refGene_gn=PRRC2C;refGene_tx=XM_006942701.4
F1	15821614	15821716	refGene_gn=PRRC2C;refGene_tx=XM_006942701.4
F1	15824461	15824663	refGene_gn=PRRC2C;refGene_tx=XM_006942701.4
F1	15827242	15827446	refGene_gn=PRRC2C;refGene_tx=XM_006942701.4
F1	15829121	15829234	refGene_gn=PRRC2C;refGene_tx=XM_006942701.4
F1	15829598	15829735	refGene_gn=PRRC2C;refGene_tx=XM_006942701.4
F1	15831990	15832222	refGene_gn=PRRC2C;refGene_tx=XM_006942701.4
F1	15832441	15832549	refGene_gn=PRRC2C;refGene_tx=XM_006942701.4
F1	15833589	15833808	refGene_gn=PRRC2C;refGene_tx=XM_006942701.4
F1	15834451	15834555	refGene_gn=PRRC2C;refGene_tx=XM_006942701.4
F1	15834941	15835068	refGene_gn=PRRC2C;refGene_tx=XM_006942701.4
F1	15838273	15838531	refGene_gn=PRRC2C;refGene_tx=XM_006942701.4
F1	15870041	15870826	refGene_gn=MYOC;refGene_tx=NM_001278850.1
F1	15872991	15873117	refGene_gn=MYOC;refGene_tx=NM_001278850.1
F1	15883092	15883654	refGene_gn=MYOC;refGene_tx=NM_001278850.1
F1	15917304	15917333
refGene_gn=VAMP4;refGene_tx=XM_003999190.4;refGene_gn=VAMP4;refGene_tx=XM_006942702.4

F1 15919249 15919301
refGene_gn=VAMP4;refGene_tx=XM_003999190.4;refGene_gn=VAMP4;refGene_tx=XM_006942702.4
F1 15921967 15922047
refGene_gn=VAMP4;refGene_tx=XM_003999190.4;refGene_gn=VAMP4;refGene_tx=XM_006942702.4
F1 15922979 15923080
refGene_gn=VAMP4;refGene_tx=XM_003999190.4;refGene_gn=VAMP4;refGene_tx=XM_006942702.4
F1 15929717 15929768
refGene_gn=VAMP4;refGene_tx=XM_003999190.4;refGene_gn=VAMP4;refGene_tx=XM_006942702.4
F1 15933340 15933387
refGene_gn=VAMP4;refGene_tx=XM_003999190.4;refGene_gn=VAMP4;refGene_tx=XM_006942702.4
F1 15940320 15940386
refGene_gn=VAMP4;refGene_tx=XM_003999190.4;refGene_gn=VAMP4;refGene_tx=XM_006942702.4
F1 16016916 16017069
refGene_gn=METTL13;refGene_tx=XM_003999192.5;refGene_gn=METTL13;refGene_tx=XM_023247555.1
F1 16018384 16019144
refGene_gn=METTL13;refGene_tx=XM_003999192.5;refGene_gn=METTL13;refGene_tx=XM_023247555.1
F1 16020452 16020652 refGene_gn=METTL13;refGene_tx=XM_003999192.5
F1 16022533 16022729 refGene_gn=METTL13;refGene_tx=XM_003999192.5
F1 16024332 16024500
refGene_gn=METTL13;refGene_tx=XM_003999192.5;refGene_gn=METTL13;refGene_tx=XM_023247555.1
F1 16025751 16025967
refGene_gn=METTL13;refGene_tx=XM_003999192.5;refGene_gn=METTL13;refGene_tx=XM_023247555.1
F1 16028516 16028648
refGene_gn=METTL13;refGene_tx=XM_003999192.5;refGene_gn=METTL13;refGene_tx=XM_023247555.1
F1 16029965 16030240
refGene_gn=METTL13;refGene_tx=XM_003999192.5;refGene_gn=METTL13;refGene_tx=XM_023247555.1
F1 16073336 16073497
refGene_gn=DNM3;refGene_tx=XM_023247907.1;refGene_gn=DNM3;refGene_tx=XM_023247908.1;refGene_gn=DNM3;refGene_tx=XM_023247909.1;refGene_gn=DNM3;refGene_tx=XM_023247910.1;refGene_gn=DNM3;refGene_tx=XM_023247912.1;refGene_gn=DNM3;refGene_tx=XM_023247913.1;refGene_gn=DNM3;refGene_tx=XM_023247914.1;refGene_gn=DNM3;refGene_tx=XM_023247915.1
F1 16163772 16163846
refGene_gn=DNM3;refGene_tx=XM_023247907.1;refGene_gn=DNM3;refGene_tx=XM_023247908.1;refGene_gn=DNM3;refGene_tx=XM_023247909.1;refGene_gn=DNM3;refGene_tx=XM_023247910.1;refGene_gn=DNM3;refGene_tx=XM_023247912.1;refGene_gn=DNM3;refGene_tx=XM_023247913.1;refGene_gn=DNM3;refGene_tx=XM_023247914.1;refGene_gn=DNM3;refGene_tx=XM_023247915.1

F1 16220564 16220714
refGene_gn=DNM3;refGene_tx=XM_023247907.1;refGene_gn=DNM3;refGene_tx=XM_023247908.1;refGene_gn=DNM3;refGene_tx=XM_023247909.1;refGene_gn=DNM3;refGene_tx=XM_023247910.1;refGene_gn=DNM3;refGene_tx=XM_023247912.1;refGene_gn=DNM3;refGene_tx=XM_023247913.1;refGene_gn=DNM3;refGene_tx=XM_023247914.1;refGene_gn=DNM3;refGene_tx=XM_023247915.1
F1 16221835 16222039
refGene_gn=DNM3;refGene_tx=XM_023247907.1;refGene_gn=DNM3;refGene_tx=XM_023247908.1;refGene_gn=DNM3;refGene_tx=XM_023247909.1;refGene_gn=DNM3;refGene_tx=XM_023247910.1;refGene_gn=DNM3;refGene_tx=XM_023247912.1;refGene_gn=DNM3;refGene_tx=XM_023247913.1;refGene_gn=DNM3;refGene_tx=XM_023247914.1;refGene_gn=DNM3;refGene_tx=XM_023247915.1
F1 16269093 16269192
refGene_gn=DNM3;refGene_tx=XM_023247907.1;refGene_gn=DNM3;refGene_tx=XM_023247908.1;refGene_gn=DNM3;refGene_tx=XM_023247909.1;refGene_gn=DNM3;refGene_tx=XM_023247910.1;refGene_gn=DNM3;refGene_tx=XM_023247912.1;refGene_gn=DNM3;refGene_tx=XM_023247913.1;refGene_gn=DNM3;refGene_tx=XM_023247914.1;refGene_gn=DNM3;refGene_tx=XM_023247915.1
F1 16269779 16269940
refGene_gn=DNM3;refGene_tx=XM_023247907.1;refGene_gn=DNM3;refGene_tx=XM_023247908.1;refGene_gn=DNM3;refGene_tx=XM_023247909.1;refGene_gn=DNM3;refGene_tx=XM_023247910.1;refGene_gn=DNM3;refGene_tx=XM_023247912.1;refGene_gn=DNM3;refGene_tx=XM_023247913.1;refGene_gn=DNM3;refGene_tx=XM_023247914.1;refGene_gn=DNM3;refGene_tx=XM_023247915.1
F1 16273900 16274043
refGene_gn=DNM3;refGene_tx=XM_023247907.1;refGene_gn=DNM3;refGene_tx=XM_023247908.1;refGene_gn=DNM3;refGene_tx=XM_023247909.1;refGene_gn=DNM3;refGene_tx=XM_023247910.1;refGene_gn=DNM3;refGene_tx=XM_023247912.1;refGene_gn=DNM3;refGene_tx=XM_023247913.1;refGene_gn=DNM3;refGene_tx=XM_023247914.1;refGene_gn=DNM3;refGene_tx=XM_023247915.1
F1 16277575 16277711
refGene_gn=DNM3;refGene_tx=XM_023247907.1;refGene_gn=DNM3;refGene_tx=XM_023247908.1;refGene_gn=DNM3;refGene_tx=XM_023247909.1;refGene_gn=DNM3;refGene_tx=XM_023247910.1;refGene_gn=DNM3;refGene_tx=XM_023247912.1;refGene_gn=DNM3;refGene_tx=XM_023247913.1;refGene_gn=DNM3;refGene_tx=XM_023247914.1;refGene_gn=DNM3;refGene_tx=XM_023247915.1
F1 16280001 16280069
refGene_gn=DNM3;refGene_tx=XM_023247907.1;refGene_gn=DNM3;refGene_tx=XM_023247908.1;refGene_gn=DNM3;refGene_tx=XM_023247909.1;refGene_gn=DNM3;refGene_tx=XM_023247910.1;refGene_gn=DNM3;refGene_tx=XM_023247912.1;refGene_gn=DNM3;refGene_tx=XM_023247913.1;refGene_gn=DNM3;refGene_tx=XM_023247914.1;refGene_gn=DNM3;refGene_tx=XM_023247915.1
F1 16283316 16283455
refGene_gn=DNM3;refGene_tx=XM_023247907.1;refGene_gn=DNM3;refGene_tx=XM_023247908.1;refGene_gn=DNM3;refGene_tx=XM_023247909.1;refGene_gn=DNM3;refGene_tx=XM_023247910.1;refGene_gn=DNM3;refGene_tx=XM_023247912.1;refGene_gn=DNM3;refGene_tx=XM_023247913.1;refGene_gn=DNM3;refGene_tx=XM_023247914.1;refGene_gn=DNM3;refGene_tx=XM_023247915.1
F1 16295248 16295335
refGene_gn=DNM3;refGene_tx=XM_023247907.1;refGene_gn=DNM3;refGene_tx=X

M_023247908.1;refGene_gn=DNM3;refGene_tx=XM_023247909.1;refGene_gn=DNM3;refGene_tx=XM_023247910.1;refGene_gn=DNM3;refGene_tx=XM_023247912.1;refGene_gn=DNM3;refGene_tx=XM_023247913.1;refGene_gn=DNM3;refGene_tx=XM_023247914.1;refGene_gn=DNM3;refGene_tx=XM_023247915.1
F1 16301494 16301565
refGene_gn=DNM3;refGene_tx=XM_023247907.1;refGene_gn=DNM3;refGene_tx=XM_023247908.1;refGene_gn=DNM3;refGene_tx=XM_023247909.1;refGene_gn=DNM3;refGene_tx=XM_023247910.1;refGene_gn=DNM3;refGene_tx=XM_023247912.1;refGene_gn=DNM3;refGene_tx=XM_023247913.1;refGene_gn=DNM3;refGene_tx=XM_023247914.1;refGene_gn=DNM3;refGene_tx=XM_023247915.1
F1 16325725 16325777
refGene_gn=DNM3;refGene_tx=XM_023247907.1;refGene_gn=DNM3;refGene_tx=XM_023247908.1;refGene_gn=DNM3;refGene_tx=XM_023247909.1;refGene_gn=DNM3;refGene_tx=XM_023247910.1;refGene_gn=DNM3;refGene_tx=XM_023247912.1;refGene_gn=DNM3;refGene_tx=XM_023247913.1;refGene_gn=DNM3;refGene_tx=XM_023247914.1;refGene_gn=DNM3;refGene_tx=XM_023247915.1
F1 16326603 16326633
refGene_gn=DNM3;refGene_tx=XM_023247907.1;refGene_gn=DNM3;refGene_tx=XM_023247908.1;refGene_gn=DNM3;refGene_tx=XM_023247910.1;refGene_gn=DNM3;refGene_tx=XM_023247912.1;refGene_gn=DNM3;refGene_tx=XM_023247914.1
F1 16369859 16369973
refGene_gn=DNM3;refGene_tx=XM_023247907.1;refGene_gn=DNM3;refGene_tx=XM_023247908.1;refGene_gn=DNM3;refGene_tx=XM_023247909.1;refGene_gn=DNM3;refGene_tx=XM_023247910.1;refGene_gn=DNM3;refGene_tx=XM_023247912.1;refGene_gn=DNM3;refGene_tx=XM_023247913.1;refGene_gn=DNM3;refGene_tx=XM_023247914.1;refGene_gn=DNM3;refGene_tx=XM_023247915.1
F1 16476017 16476127
refGene_gn=DNM3;refGene_tx=XM_023247907.1;refGene_gn=DNM3;refGene_tx=XM_023247908.1;refGene_gn=DNM3;refGene_tx=XM_023247909.1;refGene_gn=DNM3;refGene_tx=XM_023247910.1;refGene_gn=DNM3;refGene_tx=XM_023247912.1;refGene_gn=DNM3;refGene_tx=XM_023247913.1;refGene_gn=DNM3;refGene_tx=XM_023247914.1;refGene_gn=DNM3;refGene_tx=XM_023247915.1
F1 16523529 16523641
refGene_gn=DNM3;refGene_tx=XM_023247907.1;refGene_gn=DNM3;refGene_tx=XM_023247908.1;refGene_gn=DNM3;refGene_tx=XM_023247909.1;refGene_gn=DNM3;refGene_tx=XM_023247910.1;refGene_gn=DNM3;refGene_tx=XM_023247912.1;refGene_gn=DNM3;refGene_tx=XM_023247913.1;refGene_gn=DNM3;refGene_tx=XM_023247914.1;refGene_gn=DNM3;refGene_tx=XM_023247915.1
F1 16531598 16531610
refGene_gn=DNM3;refGene_tx=XM_023247907.1;refGene_gn=DNM3;refGene_tx=XM_023247909.1;refGene_gn=DNM3;refGene_tx=XM_023247910.1;refGene_gn=DNM3;refGene_tx=XM_023247912.1;refGene_gn=DNM3;refGene_tx=XM_023247914.1
F1 16592332 16592497
refGene_gn=DNM3;refGene_tx=XM_023247907.1;refGene_gn=DNM3;refGene_tx=XM_023247908.1;refGene_gn=DNM3;refGene_tx=XM_023247909.1;refGene_gn=DNM3;refGene_tx=XM_023247910.1;refGene_gn=DNM3;refGene_tx=XM_023247912.1;refGene_gn=DNM3;refGene_tx=XM_023247913.1;refGene_gn=DNM3;refGene_tx=XM_023247914.1;refGene_gn=DNM3;refGene_tx=XM_023247915.1
F1 16600886 16601113
refGene_gn=DNM3;refGene_tx=XM_023247907.1;refGene_gn=DNM3;refGene_tx=X

M_023247908.1;refGene_gn=DNM3;refGene_tx=XM_023247909.1;refGene_gn=DNM3;refGene_tx=XM_023247910.1;refGene_gn=DNM3;refGene_tx=XM_023247912.1;refGene_gn=DNM3;refGene_tx=XM_023247913.1;refGene_gn=DNM3;refGene_tx=XM_023247914.1;refGene_gn=DNM3;refGene_tx=XM_023247915.1
F1 16601777 16602017 refGene_gn=DNM3;refGene_tx=XM_023247907.1;refGene_gn=DNM3;refGene_tx=XM_023247908.1;refGene_gn=DNM3;refGene_tx=XM_023247909.1;refGene_gn=DNM3;refGene_tx=XM_023247910.1;refGene_gn=DNM3;refGene_tx=XM_023247912.1;refGene_gn=DNM3;refGene_tx=XM_023247913.1;refGene_gn=DNM3;refGene_tx=XM_023247914.1;refGene_gn=DNM3;refGene_tx=XM_023247915.1
F1 16606647 16606666 refGene_gn=DNM3;refGene_tx=XM_023247914.1
F1 16606720 16606757 refGene_gn=DNM3;refGene_tx=XM_023247912.1;refGene_gn=DNM3;refGene_tx=XM_023247915.1
F1 16622101 16622171 refGene_gn=DNM3;refGene_tx=XM_023247907.1;refGene_gn=DNM3;refGene_tx=XM_023247908.1;refGene_gn=DNM3;refGene_tx=XM_023247909.1;refGene_gn=DNM3;refGene_tx=XM_023247913.1
F1 16631707 16631744 refGene_gn=DNM3;refGene_tx=XM_023247910.1
F1 16634925 16634946 refGene_gn=CF1H1orf105;refGene_tx=XM_006942705.3;refGene_gn=CF1H1orf105;refGene_tx=XM_019821617.2;refGene_gn=CF1H1orf105;refGene_tx=XM_019821618.2;refGene_gn=CF1H1orf105;refGene_tx=XM_019821623.2;refGene_gn=CF1H1orf105;refGene_tx=XM_019821625.2;refGene_gn=CF1H1orf105;refGene_tx=XM_023247916.1;refGene_gn=CF1H1orf105;refGene_tx=XM_023247917.1
F1 16645466 16646360 refGene_gn=PIGC;refGene_tx=XM_006942704.4;refGene_gn=PIGC;refGene_tx=XM_019821613.2
F1 16648777 16648863 refGene_gn=CF1H1orf105;refGene_tx=XM_006942705.3;refGene_gn=CF1H1orf105;refGene_tx=XM_019821617.2;refGene_gn=CF1H1orf105;refGene_tx=XM_019821618.2;refGene_gn=CF1H1orf105;refGene_tx=XM_019821623.2;refGene_gn=CF1H1orf105;refGene_tx=XM_019821625.2;refGene_gn=CF1H1orf105;refGene_tx=XM_023247916.1;refGene_gn=CF1H1orf105;refGene_tx=XM_023247917.1
F1 16652670 16652758 refGene_gn=CF1H1orf105;refGene_tx=XM_006942705.3;refGene_gn=CF1H1orf105;refGene_tx=XM_019821617.2;refGene_gn=CF1H1orf105;refGene_tx=XM_019821618.2;refGene_gn=CF1H1orf105;refGene_tx=XM_019821623.2;refGene_gn=CF1H1orf105;refGene_tx=XM_019821625.2;refGene_gn=CF1H1orf105;refGene_tx=XM_023247916.1;refGene_gn=CF1H1orf105;refGene_tx=XM_023247917.1
F1 16659318 16659372 refGene_gn=CF1H1orf105;refGene_tx=XM_019821617.2;refGene_gn=CF1H1orf105;refGene_tx=XM_019821623.2;refGene_gn=CF1H1orf105;refGene_tx=XM_023247916.1;refGene_gn=CF1H1orf105;refGene_tx=XM_023247917.1
F1 16659886 16659961 refGene_gn=CF1H1orf105;refGene_tx=XM_006942705.3;refGene_gn=CF1H1orf105;refGene_tx=XM_019821617.2;refGene_gn=CF1H1orf105;refGene_tx=XM_019821618.2;refGene_gn=CF1H1orf105;refGene_tx=XM_019821623.2;refGene_gn=CF1H1orf105;refGene_tx=XM_019821625.2;refGene_gn=CF1H1orf105;refGene_tx=XM_019821626.2;refGene_gn=CF1H1orf105;refGene_tx=XM_019821627.2;refGene

_gn=CF1H1orf105;refGene_tx=XM_019821628.2;refGene_gn=CF1H1orf105;refGene_tx=XM_019821629.2;refGene_gn=CF1H1orf105;refGene_tx=XM_023247916.1;refGene_gn=CF1H1orf105;refGene_tx=XM_023247917.1
F1 16660707 16660824
refGene_gn=CF1H1orf105;refGene_tx=XM_019821617.2;refGene_gn=CF1H1orf105;refGene_tx=XM_019821618.2;refGene_gn=CF1H1orf105;refGene_tx=XM_019821626.2;refGene_gn=CF1H1orf105;refGene_tx=XM_019821627.2;refGene_gn=CF1H1orf105;refGene_tx=XM_023247916.1;refGene_gn=CF1H1orf105;refGene_tx=XM_023247917.1
F1 16666320 16666388
refGene_gn=CF1H1orf105;refGene_tx=XM_006942705.3;refGene_gn=CF1H1orf105;refGene_tx=XM_019821617.2;refGene_gn=CF1H1orf105;refGene_tx=XM_019821618.2;refGene_gn=CF1H1orf105;refGene_tx=XM_019821623.2;refGene_gn=CF1H1orf105;refGene_tx=XM_019821625.2;refGene_gn=CF1H1orf105;refGene_tx=XM_019821626.2;refGene_gn=CF1H1orf105;refGene_tx=XM_019821627.2;refGene_gn=CF1H1orf105;refGene_tx=XM_019821628.2;refGene_gn=CF1H1orf105;refGene_tx=XM_019821629.2;refGene_gn=CF1H1orf105;refGene_tx=XM_023247916.1;refGene_gn=CF1H1orf105;refGene_tx=XM_023247917.1
F1 16669248 16669316
refGene_gn=CF1H1orf105;refGene_tx=XM_006942705.3;refGene_gn=CF1H1orf105;refGene_tx=XM_019821617.2;refGene_gn=CF1H1orf105;refGene_tx=XM_019821618.2;refGene_gn=CF1H1orf105;refGene_tx=XM_019821623.2;refGene_gn=CF1H1orf105;refGene_tx=XM_019821625.2;refGene_gn=CF1H1orf105;refGene_tx=XM_019821626.2;refGene_gn=CF1H1orf105;refGene_tx=XM_019821627.2;refGene_gn=CF1H1orf105;refGene_tx=XM_019821628.2;refGene_gn=CF1H1orf105;refGene_tx=XM_019821629.2;refGene_gn=CF1H1orf105;refGene_tx=XM_023247916.1;refGene_gn=CF1H1orf105;refGene_tx=XM_023247917.1
F1 16670792 16670863
refGene_gn=CF1H1orf105;refGene_tx=XM_023247917.1
F1 16672341 16672487
refGene_gn=CF1H1orf105;refGene_tx=XM_006942705.3;refGene_gn=CF1H1orf105;refGene_tx=XM_023247916.1
F1 16690547 16690825
refGene_gn=CF1H1orf105;refGene_tx=XM_019821617.2;refGene_gn=CF1H1orf105;refGene_tx=XM_019821618.2;refGene_gn=CF1H1orf105;refGene_tx=XM_019821623.2;refGene_gn=CF1H1orf105;refGene_tx=XM_019821625.2;refGene_gn=CF1H1orf105;refGene_tx=XM_019821626.2;refGene_gn=CF1H1orf105;refGene_tx=XM_019821627.2;refGene_gn=CF1H1orf105;refGene_tx=XM_019821628.2;refGene_gn=CF1H1orf105;refGene_tx=XM_019821629.2
F1 16735799 16735861
refGene_gn=SUCO;refGene_tx=XM_011290883.3;refGene_gn=SUCO;refGene_tx=XM_019821610.2;refGene_gn=SUCO;refGene_tx=XM_019821611.2;refGene_gn=SUCO;refGene_tx=XM_019821612.2
F1 16760938 16761047
refGene_gn=SUCO;refGene_tx=XM_011290883.3;refGene_gn=SUCO;refGene_tx=XM_019821610.2;refGene_gn=SUCO;refGene_tx=XM_019821611.2;refGene_gn=SUCO;refGene_tx=XM_019821612.2
F1 16764675 16764786
refGene_gn=SUCO;refGene_tx=XM_011290883.3;refGene_gn=SUCO;refGene_tx=XM_019821610.2;refGene_gn=SUCO;refGene_tx=XM_019821611.2;refGene_gn=SUC

O;refGene_tx=XM_019821612.2
F1 16767191 16767349
refGene_gn=SUCO;refGene_tx=XM_011290883.3;refGene_gn=SUCO;refGene_tx=XM_019821610.2;refGene_gn=SUCO;refGene_tx=XM_019821611.2;refGene_gn=SUCO;refGene_tx=XM_019821612.2
F1 16769130 16769268
refGene_gn=SUCO;refGene_tx=XM_011290883.3;refGene_gn=SUCO;refGene_tx=XM_019821610.2;refGene_gn=SUCO;refGene_tx=XM_019821611.2;refGene_gn=SUCO;refGene_tx=XM_019821612.2
F1 16769493 16769644
refGene_gn=SUCO;refGene_tx=XM_011290883.3;refGene_gn=SUCO;refGene_tx=XM_019821610.2;refGene_gn=SUCO;refGene_tx=XM_019821611.2;refGene_gn=SUCO;refGene_tx=XM_019821612.2
F1 16772675 16772799
refGene_gn=SUCO;refGene_tx=XM_011290883.3;refGene_gn=SUCO;refGene_tx=XM_019821610.2;refGene_gn=SUCO;refGene_tx=XM_019821611.2;refGene_gn=SUCO;refGene_tx=XM_019821612.2
F1 16773485 16773622
refGene_gn=SUCO;refGene_tx=XM_011290883.3;refGene_gn=SUCO;refGene_tx=XM_019821610.2;refGene_gn=SUCO;refGene_tx=XM_019821611.2;refGene_gn=SUCO;refGene_tx=XM_019821612.2
F1 16774661 16774729
refGene_gn=SUCO;refGene_tx=XM_011290883.3;refGene_gn=SUCO;refGene_tx=XM_019821610.2;refGene_gn=SUCO;refGene_tx=XM_019821611.2;refGene_gn=SUCO;refGene_tx=XM_019821612.2
F1 16781946 16782054
refGene_gn=SUCO;refGene_tx=XM_011290883.3;refGene_gn=SUCO;refGene_tx=XM_019821610.2;refGene_gn=SUCO;refGene_tx=XM_019821611.2;refGene_gn=SUCO;refGene_tx=XM_019821612.2
F1 16784122 16784228
refGene_gn=SUCO;refGene_tx=XM_011290883.3;refGene_gn=SUCO;refGene_tx=XM_019821610.2;refGene_gn=SUCO;refGene_tx=XM_019821611.2;refGene_gn=SUCO;refGene_tx=XM_019821612.2
F1 16786559 16786580
refGene_gn=SUCO;refGene_tx=XM_011290883.3;refGene_gn=SUCO;refGene_tx=XM_019821610.2;refGene_gn=SUCO;refGene_tx=XM_019821611.2
F1 16786766 16786822
refGene_gn=SUCO;refGene_tx=XM_011290883.3;refGene_gn=SUCO;refGene_tx=XM_019821610.2;refGene_gn=SUCO;refGene_tx=XM_019821611.2;refGene_gn=SUCO;refGene_tx=XM_019821612.2
F1 16787541 16787633
refGene_gn=SUCO;refGene_tx=XM_011290883.3;refGene_gn=SUCO;refGene_tx=XM_019821610.2;refGene_gn=SUCO;refGene_tx=XM_019821611.2;refGene_gn=SUCO;refGene_tx=XM_019821612.2
F1 16788312 16788378
refGene_gn=SUCO;refGene_tx=XM_011290883.3;refGene_gn=SUCO;refGene_tx=XM_019821610.2;refGene_gn=SUCO;refGene_tx=XM_019821611.2;refGene_gn=SUCO;refGene_tx=XM_019821612.2
F1 16794907 16794976
refGene_gn=SUCO;refGene_tx=XM_011290883.3;refGene_gn=SUCO;refGene_tx=X

M_019821610.2;refGene_gn=SUCO;refGene_tx=XM_019821611.2;refGene_gn=SUCO;refGene_tx=XM_019821612.2
F1 16795765 16795856 refGene_gn=SUCO;refGene_tx=XM_011290883.3;refGene_gn=SUCO;refGene_tx=XM_019821610.2;refGene_gn=SUCO;refGene_tx=XM_019821611.2;refGene_gn=SUCO;refGene_tx=XM_019821612.2
F1 16798349 16799510 refGene_gn=SUCO;refGene_tx=XM_011290883.3;refGene_gn=SUCO;refGene_tx=XM_019821610.2;refGene_gn=SUCO;refGene_tx=XM_019821611.2;refGene_gn=SUCO;refGene_tx=XM_019821612.2
F1 16802154 16802242 refGene_gn=SUCO;refGene_tx=XM_011290883.3;refGene_gn=SUCO;refGene_tx=XM_019821610.2;refGene_gn=SUCO;refGene_tx=XM_019821611.2;refGene_gn=SUCO;refGene_tx=XM_019821612.2
F1 16805289 16805394 refGene_gn=SUCO;refGene_tx=XM_011290883.3;refGene_gn=SUCO;refGene_tx=XM_019821610.2;refGene_gn=SUCO;refGene_tx=XM_019821611.2;refGene_gn=SUCO;refGene_tx=XM_019821612.2
F1 16807642 16807797 refGene_gn=SUCO;refGene_tx=XM_011290883.3;refGene_gn=SUCO;refGene_tx=XM_019821610.2;refGene_gn=SUCO;refGene_tx=XM_019821611.2;refGene_gn=SUCO;refGene_tx=XM_019821612.2
F1 16808284 16808376 refGene_gn=SUCO;refGene_tx=XM_011290883.3;refGene_gn=SUCO;refGene_tx=XM_019821610.2;refGene_gn=SUCO;refGene_tx=XM_019821611.2;refGene_gn=SUCO;refGene_tx=XM_019821612.2
F1 16823299 16823355 refGene_gn=SUCO;refGene_tx=XM_011290883.3;refGene_gn=SUCO;refGene_tx=XM_019821610.2;refGene_gn=SUCO;refGene_tx=XM_019821611.2;refGene_gn=SUCO;refGene_tx=XM_019821612.2
F1 16824502 16824946 refGene_gn=SUCO;refGene_tx=XM_011290883.3;refGene_gn=SUCO;refGene_tx=XM_019821610.2;refGene_gn=SUCO;refGene_tx=XM_019821611.2;refGene_gn=SUCO;refGene_tx=XM_019821612.2
F1 16866049 16866394 refGene_gn=FASLG;refGene_tx=NM_001009352.1
F1 16867461 16867507 refGene_gn=FASLG;refGene_tx=NM_001009352.1
F1 16871889 16871946 refGene_gn=FASLG;refGene_tx=NM_001009352.1
F1 16873169 16873564 refGene_gn=FASLG;refGene_tx=NM_001009352.1
F1 17203152 17203499 refGene_gn=TNFSF18;refGene_tx=XM_006942706.4
F1 17206173 17206204 refGene_gn=TNFSF18;refGene_tx=XM_006942706.4
F1 17212510 17212669 refGene_gn=TNFSF18;refGene_tx=XM_006942706.4
F1 17303329 17303671 refGene_gn=TNFSF4;refGene_tx=NM_001079828.1
F1 17305404 17305453 refGene_gn=TNFSF4;refGene_tx=NM_001079828.1
F1 17322216 17322354 refGene_gn=TNFSF4;refGene_tx=NM_001079828.1
F1 17322355 17322370 refGene_gn=TNFSF4;refGene_tx=NM_001079828.1
F1 17496916 17497225 refGene_gn=LOC111558417;refGene_tx=XM_023248028.1
F1 17525913 17526221 refGene_gn=LOC111558417;refGene_tx=XM_023248028.1
F1 17526259 17526410

refGene_gn=LOC111558417;refGene_tx=XM_023248028.1;refGene_gn=PRDX6;refGene_tx=XM_011290881.3
F1 17530532 17530689 refGene_gn=PRDX6;refGene_tx=XM_011290881.3
F1 17533982 17534129 refGene_gn=PRDX6;refGene_tx=XM_011290881.3
F1 17535363 17535510 refGene_gn=PRDX6;refGene_tx=XM_011290881.3
F1 17537175 17537304 refGene_gn=PRDX6;refGene_tx=XM_011290881.3
F1 17542046 17542082 refGene_gn=SLC9C2;refGene_tx=XM_019821631.2
F1 17544235 17544333
refGene_gn=SLC9C2;refGene_tx=XM_019821630.2;refGene_gn=SLC9C2;refGene_tx=XM_019821631.2;refGene_gn=SLC9C2;refGene_tx=XM_019821635.1
F1 17546699 17546784
refGene_gn=SLC9C2;refGene_tx=XM_019821630.2;refGene_gn=SLC9C2;refGene_tx=XM_019821631.2;refGene_gn=SLC9C2;refGene_tx=XM_019821635.1
F1 17548242 17548428
refGene_gn=SLC9C2;refGene_tx=XM_019821630.2;refGene_gn=SLC9C2;refGene_tx=XM_019821631.2;refGene_gn=SLC9C2;refGene_tx=XM_019821635.1
F1 17553652 17553784
refGene_gn=SLC9C2;refGene_tx=XM_019821630.2;refGene_gn=SLC9C2;refGene_tx=XM_019821631.2;refGene_gn=SLC9C2;refGene_tx=XM_019821635.1
F1 17555901 17556069
refGene_gn=SLC9C2;refGene_tx=XM_019821630.2;refGene_gn=SLC9C2;refGene_tx=XM_019821631.2;refGene_gn=SLC9C2;refGene_tx=XM_019821635.1
F1 17559001 17559100
refGene_gn=SLC9C2;refGene_tx=XM_019821630.2;refGene_gn=SLC9C2;refGene_tx=XM_019821631.2;refGene_gn=SLC9C2;refGene_tx=XM_019821635.1
F1 17564377 17564503
refGene_gn=SLC9C2;refGene_tx=XM_019821630.2;refGene_gn=SLC9C2;refGene_tx=XM_019821631.2;refGene_gn=SLC9C2;refGene_tx=XM_019821635.1
F1 17566990 17567139
refGene_gn=SLC9C2;refGene_tx=XM_019821630.2;refGene_gn=SLC9C2;refGene_tx=XM_019821631.2;refGene_gn=SLC9C2;refGene_tx=XM_019821635.1
F1 17569583 17569635
refGene_gn=SLC9C2;refGene_tx=XM_019821630.2;refGene_gn=SLC9C2;refGene_tx=XM_019821631.2;refGene_gn=SLC9C2;refGene_tx=XM_019821635.1
F1 17574610 17574760
refGene_gn=SLC9C2;refGene_tx=XM_019821630.2;refGene_gn=SLC9C2;refGene_tx=XM_019821631.2;refGene_gn=SLC9C2;refGene_tx=XM_019821635.1
F1 17578266 17578455
refGene_gn=SLC9C2;refGene_tx=XM_019821630.2;refGene_gn=SLC9C2;refGene_tx=XM_019821631.2;refGene_gn=SLC9C2;refGene_tx=XM_019821635.1
F1 17578933 17579132
refGene_gn=SLC9C2;refGene_tx=XM_019821630.2;refGene_gn=SLC9C2;refGene_tx=XM_019821631.2;refGene_gn=SLC9C2;refGene_tx=XM_019821635.1
F1 17580314 17580434
refGene_gn=SLC9C2;refGene_tx=XM_019821630.2;refGene_gn=SLC9C2;refGene_tx=XM_019821631.2;refGene_gn=SLC9C2;refGene_tx=XM_019821635.1
F1 17581525 17581623
refGene_gn=SLC9C2;refGene_tx=XM_019821630.2;refGene_gn=SLC9C2;refGene_tx=XM_019821631.2;refGene_gn=SLC9C2;refGene_tx=XM_019821635.1
F1 17588313 17588424

refGene_gn=SLC9C2;refGene_tx=XM_019821630.2;refGene_gn=SLC9C2;refGene_
tx=XM_019821631.2;refGene_gn=SLC9C2;refGene_tx=XM_019821635.1
F1 17589036 17589203
refGene_gn=SLC9C2;refGene_tx=XM_019821630.2;refGene_gn=SLC9C2;refGene_
tx=XM_019821631.2;refGene_gn=SLC9C2;refGene_tx=XM_019821635.1
F1 17593002 17593084
refGene_gn=SLC9C2;refGene_tx=XM_019821630.2;refGene_gn=SLC9C2;refGene_
tx=XM_019821631.2;refGene_gn=SLC9C2;refGene_tx=XM_019821635.1
F1 17596776 17596945
refGene_gn=SLC9C2;refGene_tx=XM_019821630.2;refGene_gn=SLC9C2;refGene_
tx=XM_019821631.2;refGene_gn=SLC9C2;refGene_tx=XM_019821635.1
F1 17599356 17599500
refGene_gn=SLC9C2;refGene_tx=XM_019821630.2;refGene_gn=SLC9C2;refGene_
tx=XM_019821631.2;refGene_gn=SLC9C2;refGene_tx=XM_019821635.1
F1 17603068 17603168
refGene_gn=SLC9C2;refGene_tx=XM_019821630.2;refGene_gn=SLC9C2;refGene_
tx=XM_019821631.2;refGene_gn=SLC9C2;refGene_tx=XM_019821635.1
F1 17608034 17608196
refGene_gn=SLC9C2;refGene_tx=XM_019821630.2;refGene_gn=SLC9C2;refGene_
tx=XM_019821631.2;refGene_gn=SLC9C2;refGene_tx=XM_019821635.1
F1 17609530 17609647
refGene_gn=SLC9C2;refGene_tx=XM_019821630.2;refGene_gn=SLC9C2;refGene_
tx=XM_019821631.2;refGene_gn=SLC9C2;refGene_tx=XM_019821635.1
F1 17617298 17617464
refGene_gn=SLC9C2;refGene_tx=XM_019821630.2;refGene_gn=SLC9C2;refGene_
tx=XM_019821631.2;refGene_gn=SLC9C2;refGene_tx=XM_019821635.1
F1 17621581 17621710
refGene_gn=SLC9C2;refGene_tx=XM_019821630.2;refGene_gn=SLC9C2;refGene_
tx=XM_019821631.2;refGene_gn=SLC9C2;refGene_tx=XM_019821635.1
F1 17623710 17623811
refGene_gn=SLC9C2;refGene_tx=XM_019821630.2;refGene_gn=SLC9C2;refGene_
tx=XM_019821631.2
F1 17626173 17626306
refGene_gn=SLC9C2;refGene_tx=XM_019821630.2;refGene_gn=SLC9C2;refGene_
tx=XM_019821631.2
F1 17635026 17635068 refGene_gn=ANKRD45;refGene_tx=XM_011290879.3
F1 17635070 17635099 refGene_gn=ANKRD45;refGene_tx=XM_011290879.3
F1 17638372 17638511 refGene_gn=ANKRD45;refGene_tx=XM_011290879.3
F1 17640676 17640753 refGene_gn=ANKRD45;refGene_tx=XM_011290879.3
F1 17640755 17640770 refGene_gn=ANKRD45;refGene_tx=XM_011290879.3
F1 17650850 17651174 refGene_gn=TEX50;refGene_tx=XM_011290876.2
F1 17652351 17652567 refGene_gn=TEX50;refGene_tx=XM_011290876.2
F1 17657844 17658012 refGene_gn=ANKRD45;refGene_tx=XM_011290879.3
F1 17673225 17673580 refGene_gn=ANKRD45;refGene_tx=XM_011290879.3
F1 17907991 17908014 refGene_gn=KLHL20;refGene_tx=XM_011290904.3
F1 17924915 17925489 refGene_gn=KLHL20;refGene_tx=XM_011290904.3
F1 17938617 17938776 refGene_gn=KLHL20;refGene_tx=XM_011290904.3
F1 17939781 17939876 refGene_gn=KLHL20;refGene_tx=XM_011290904.3
F1 17942419 17942535 refGene_gn=KLHL20;refGene_tx=XM_011290904.3
F1 17946365 17946549 refGene_gn=KLHL20;refGene_tx=XM_011290904.3

F1 17953610 17953754 refGene_gn=KLHL20;refGene_tx=XM_011290904.3
F1 17961557 17961691 refGene_gn=KLHL20;refGene_tx=XM_011290904.3
F1 17962719 17962779 refGene_gn=KLHL20;refGene_tx=XM_011290904.3
F1 17962781 17962927 refGene_gn=KLHL20;refGene_tx=XM_011290904.3
F1 17966241 17966348 refGene_gn=KLHL20;refGene_tx=XM_011290904.3
F1 17976293 17976378 refGene_gn=KLHL20;refGene_tx=XM_011290904.3
F1 17984993 17985065
refGene_gn=CENPL;refGene_tx=XM_003999206.3;refGene_gn=CENPL;refGene_tx
=XM_019821644.1;refGene_gn=CENPL;refGene_tx=XM_023248046.1
F1 17988089 17988632
refGene_gn=CENPL;refGene_tx=XM_003999206.3;refGene_gn=CENPL;refGene_tx
=XM_019821644.1;refGene_gn=CENPL;refGene_tx=XM_023248046.1
F1 17992915 17993167
refGene_gn=CENPL;refGene_tx=XM_003999206.3;refGene_gn=CENPL;refGene_tx
=XM_019821644.1;refGene_gn=CENPL;refGene_tx=XM_023248046.1
F1 17996250 17996424
refGene_gn=CENPL;refGene_tx=XM_003999206.3;refGene_gn=CENPL;refGene_tx
=XM_019821644.1;refGene_gn=CENPL;refGene_tx=XM_023248046.1
F1 18000757 18000881 refGene_gn=DARS2;refGene_tx=XM_011290905.3
F1 18001546 18001646 refGene_gn=DARS2;refGene_tx=XM_011290905.3
F1 18003314 18003381 refGene_gn=DARS2;refGene_tx=XM_011290905.3
F1 18005611 18005713 refGene_gn=DARS2;refGene_tx=XM_011290905.3
F1 18006109 18006205 refGene_gn=DARS2;refGene_tx=XM_011290905.3
F1 18008184 18008250 refGene_gn=DARS2;refGene_tx=XM_011290905.3
F1 18008251 18008309 refGene_gn=DARS2;refGene_tx=XM_011290905.3
F1 18008982 18009029 refGene_gn=DARS2;refGene_tx=XM_011290905.3
F1 18010967 18011074 refGene_gn=DARS2;refGene_tx=XM_011290905.3
F1 18011867 18011937 refGene_gn=DARS2;refGene_tx=XM_011290905.3
F1 18013352 18013532 refGene_gn=DARS2;refGene_tx=XM_011290905.3
F1 18015318 18015426 refGene_gn=DARS2;refGene_tx=XM_011290905.3
F1 18016764 18016827 refGene_gn=DARS2;refGene_tx=XM_011290905.3
F1 18021401 18021554 refGene_gn=DARS2;refGene_tx=XM_011290905.3
F1 18024145 18024364 refGene_gn=DARS2;refGene_tx=XM_011290905.3
F1 18024580 18024691 refGene_gn=DARS2;refGene_tx=XM_011290905.3
F1 18026447 18026523 refGene_gn=DARS2;refGene_tx=XM_011290905.3
F1 18027965 18028180 refGene_gn=DARS2;refGene_tx=XM_011290905.3
F1 18031860 18032019
refGene_gn=LOC109495442;refGene_tx=XM_019821401.2
F1 18044078 18045001
refGene_gn=ZBTB37;refGene_tx=XM_003999207.5;refGene_gn=ZBTB37;refGene_
tx=XM_019821646.2;refGene_gn=ZBTB37;refGene_tx=XM_019821647.2;refGene_
gn=ZBTB37;refGene_tx=XM_019821648.2
F1 18046491 18046591
refGene_gn=ZBTB37;refGene_tx=XM_003999207.5;refGene_gn=ZBTB37;refGene_
tx=XM_019821646.2;refGene_gn=ZBTB37;refGene_tx=XM_019821647.2;refGene_
gn=ZBTB37;refGene_tx=XM_019821648.2
F1 18051683 18052172
refGene_gn=ZBTB37;refGene_tx=XM_003999207.5;refGene_gn=ZBTB37;refGene_
tx=XM_019821646.2;refGene_gn=ZBTB37;refGene_tx=XM_019821647.2;refGene_
gn=ZBTB37;refGene_tx=XM_019821648.2

F1 18068173 18068350 refGene_gn=SERPINC1;refGene_tx=XM_003999208.3
F1 18071100 18071165 refGene_gn=SERPINC1;refGene_tx=XM_003999208.3
F1 18072210 18072601 refGene_gn=SERPINC1;refGene_tx=XM_003999208.3
F1 18074341 18074479 refGene_gn=SERPINC1;refGene_tx=XM_003999208.3
F1 18075306 18075522 refGene_gn=SERPINC1;refGene_tx=XM_003999208.3
F1 18075802 18076169 refGene_gn=SERPINC1;refGene_tx=XM_003999208.3
F1 18078130 18078171 refGene_gn=SERPINC1;refGene_tx=XM_003999208.3
F1 18094303 18094454
refGene_gn=RC3H1;refGene_tx=XM_023247903.1;refGene_gn=RC3H1;refGene_tx=XM_023247904.1;refGene_gn=RC3H1;refGene_tx=XM_023247906.1
F1 18095786 18095902
refGene_gn=RC3H1;refGene_tx=XM_023247903.1;refGene_gn=RC3H1;refGene_tx=XM_023247904.1;refGene_gn=RC3H1;refGene_tx=XM_023247906.1
F1 18096805 18096979
refGene_gn=RC3H1;refGene_tx=XM_023247903.1;refGene_gn=RC3H1;refGene_tx=XM_023247904.1;refGene_gn=RC3H1;refGene_tx=XM_023247906.1
F1 18099355 18099488
refGene_gn=RC3H1;refGene_tx=XM_023247903.1;refGene_gn=RC3H1;refGene_tx=XM_023247904.1;refGene_gn=RC3H1;refGene_tx=XM_023247906.1
F1 18099982 18100073
refGene_gn=RC3H1;refGene_tx=XM_023247903.1;refGene_gn=RC3H1;refGene_tx=XM_023247904.1;refGene_gn=RC3H1;refGene_tx=XM_023247906.1
F1 18100667 18100881
refGene_gn=RC3H1;refGene_tx=XM_023247903.1;refGene_gn=RC3H1;refGene_tx=XM_023247904.1;refGene_gn=RC3H1;refGene_tx=XM_023247906.1
F1 18103499 18103652
refGene_gn=RC3H1;refGene_tx=XM_023247903.1;refGene_gn=RC3H1;refGene_tx=XM_023247904.1;refGene_gn=RC3H1;refGene_tx=XM_023247906.1
F1 18109851 18110019
refGene_gn=RC3H1;refGene_tx=XM_023247903.1;refGene_gn=RC3H1;refGene_tx=XM_023247904.1;refGene_gn=RC3H1;refGene_tx=XM_023247906.1
F1 18110849 18111220
refGene_gn=RC3H1;refGene_tx=XM_023247903.1;refGene_gn=RC3H1;refGene_tx=XM_023247904.1;refGene_gn=RC3H1;refGene_tx=XM_023247906.1
F1 18113040 18113255
refGene_gn=RC3H1;refGene_tx=XM_023247903.1;refGene_gn=RC3H1;refGene_tx=XM_023247904.1;refGene_gn=RC3H1;refGene_tx=XM_023247906.1
F1 18113788 18114070
refGene_gn=RC3H1;refGene_tx=XM_023247903.1;refGene_gn=RC3H1;refGene_tx=XM_023247904.1;refGene_gn=RC3H1;refGene_tx=XM_023247906.1
F1 18117379 18117492
refGene_gn=RC3H1;refGene_tx=XM_023247903.1;refGene_gn=RC3H1;refGene_tx=XM_023247904.1;refGene_gn=RC3H1;refGene_tx=XM_023247906.1
F1 18119494 18119613
refGene_gn=RC3H1;refGene_tx=XM_023247903.1;refGene_gn=RC3H1;refGene_tx=XM_023247904.1;refGene_gn=RC3H1;refGene_tx=XM_023247906.1
F1 18124228 18124361
refGene_gn=RC3H1;refGene_tx=XM_023247903.1;refGene_gn=RC3H1;refGene_tx=XM_023247904.1;refGene_gn=RC3H1;refGene_tx=XM_023247906.1
F1 18127843 18128044

refGene_gn=RC3H1;refGene_tx=XM_023247903.1;refGene_gn=RC3H1;refGene_tx=XM_023247904.1;refGene_gn=RC3H1;refGene_tx=XM_023247906.1
F1	18131749	18131925
refGene_gn=RC3H1;refGene_tx=XM_023247903.1;refGene_gn=RC3H1;refGene_tx=XM_023247904.1;refGene_gn=RC3H1;refGene_tx=XM_023247906.1
F1	18132371	18132611
refGene_gn=RC3H1;refGene_tx=XM_023247903.1;refGene_gn=RC3H1;refGene_tx=XM_023247904.1;refGene_gn=RC3H1;refGene_tx=XM_023247906.1
F1	18133330	18133451
refGene_gn=RC3H1;refGene_tx=XM_023247903.1;refGene_gn=RC3H1;refGene_tx=XM_023247904.1;refGene_gn=RC3H1;refGene_tx=XM_023247906.1
F1	18136864	18137095
refGene_gn=RC3H1;refGene_tx=XM_023247903.1;refGene_gn=RC3H1;refGene_tx=XM_023247904.1;refGene_gn=RC3H1;refGene_tx=XM_023247906.1
F1	18348958	18349096
refGene_gn=RABGAP1L;refGene_tx=XM_023247520.1;refGene_gn=RABGAP1L;refGene_tx=XM_023247521.1;refGene_gn=RABGAP1L;refGene_tx=XM_023247522.1;refGene_gn=RABGAP1L;refGene_tx=XM_023247523.1
F1	18350651	18350844
refGene_gn=RABGAP1L;refGene_tx=XM_023247520.1;refGene_gn=RABGAP1L;refGene_tx=XM_023247521.1;refGene_gn=RABGAP1L;refGene_tx=XM_023247522.1;refGene_gn=RABGAP1L;refGene_tx=XM_023247523.1
F1	18356313	18356524
refGene_gn=RABGAP1L;refGene_tx=XM_023247520.1;refGene_gn=RABGAP1L;refGene_tx=XM_023247521.1;refGene_gn=RABGAP1L;refGene_tx=XM_023247522.1;refGene_gn=RABGAP1L;refGene_tx=XM_023247523.1
F1	18360144	18360319
refGene_gn=RABGAP1L;refGene_tx=XM_023247520.1;refGene_gn=RABGAP1L;refGene_tx=XM_023247521.1;refGene_gn=RABGAP1L;refGene_tx=XM_023247522.1;refGene_gn=RABGAP1L;refGene_tx=XM_023247523.1
F1	18368280	18368438
refGene_gn=RABGAP1L;refGene_tx=XM_023247520.1;refGene_gn=RABGAP1L;refGene_tx=XM_023247521.1;refGene_gn=RABGAP1L;refGene_tx=XM_023247522.1;refGene_gn=RABGAP1L;refGene_tx=XM_023247523.1
F1	18371106	18371217
refGene_gn=RABGAP1L;refGene_tx=XM_023247520.1;refGene_gn=RABGAP1L;refGene_tx=XM_023247521.1;refGene_gn=RABGAP1L;refGene_tx=XM_023247522.1;refGene_gn=RABGAP1L;refGene_tx=XM_023247523.1
F1	18394903	18394970
refGene_gn=RABGAP1L;refGene_tx=XM_023247520.1;refGene_gn=RABGAP1L;refGene_tx=XM_023247521.1;refGene_gn=RABGAP1L;refGene_tx=XM_023247522.1;refGene_gn=RABGAP1L;refGene_tx=XM_023247523.1
F1	18400841	18400944
refGene_gn=RABGAP1L;refGene_tx=XM_023247520.1;refGene_gn=RABGAP1L;refGene_tx=XM_023247521.1;refGene_gn=RABGAP1L;refGene_tx=XM_023247522.1;refGene_gn=RABGAP1L;refGene_tx=XM_023247523.1
F1	18405071	18405238
refGene_gn=RABGAP1L;refGene_tx=XM_023247520.1;refGene_gn=RABGAP1L;refGene_tx=XM_023247521.1;refGene_gn=RABGAP1L;refGene_tx=XM_023247522.1;refGene_gn=RABGAP1L;refGene_tx=XM_023247523.1

F1 18454096 18454238
refGene_gn=RABGAP1L;refGene_tx=XM_023247520.1;refGene_gn=RABGAP1L;refGene_tx=XM_023247521.1;refGene_gn=RABGAP1L;refGene_tx=XM_023247522.1;refGene_gn=RABGAP1L;refGene_tx=XM_023247523.1
F1 18498217 18498311
refGene_gn=RABGAP1L;refGene_tx=XM_023247520.1;refGene_gn=RABGAP1L;refGene_tx=XM_023247521.1;refGene_gn=RABGAP1L;refGene_tx=XM_023247522.1;refGene_gn=RABGAP1L;refGene_tx=XM_023247523.1
F1 18514430 18514581
refGene_gn=RABGAP1L;refGene_tx=XM_023247520.1;refGene_gn=RABGAP1L;refGene_tx=XM_023247521.1;refGene_gn=RABGAP1L;refGene_tx=XM_023247522.1;refGene_gn=RABGAP1L;refGene_tx=XM_023247523.1
F1 18557321 18558407
refGene_gn=GPR52;refGene_tx=XM_019821654.2;refGene_gn=GPR52;refGene_tx=XM_019821655.2
F1 18719490 18719604
refGene_gn=RABGAP1L;refGene_tx=XM_023247520.1;refGene_gn=RABGAP1L;refGene_tx=XM_023247521.1;refGene_gn=RABGAP1L;refGene_tx=XM_023247522.1;refGene_gn=RABGAP1L;refGene_tx=XM_023247523.1
F1 18751141 18751216
refGene_gn=RABGAP1L;refGene_tx=XM_023247520.1;refGene_gn=RABGAP1L;refGene_tx=XM_023247521.1;refGene_gn=RABGAP1L;refGene_tx=XM_023247522.1;refGene_gn=RABGAP1L;refGene_tx=XM_023247523.1
F1 18770787 18770913
refGene_gn=RABGAP1L;refGene_tx=XM_023247520.1;refGene_gn=RABGAP1L;refGene_tx=XM_023247521.1;refGene_gn=RABGAP1L;refGene_tx=XM_023247522.1;refGene_gn=RABGAP1L;refGene_tx=XM_023247523.1
F1 18773230 18773374
refGene_gn=RABGAP1L;refGene_tx=XM_023247520.1;refGene_gn=RABGAP1L;refGene_tx=XM_023247521.1;refGene_gn=RABGAP1L;refGene_tx=XM_023247522.1;refGene_gn=RABGAP1L;refGene_tx=XM_023247523.1
F1 18826978 18827020
refGene_gn=RABGAP1L;refGene_tx=XM_023247520.1;refGene_gn=RABGAP1L;refGene_tx=XM_023247521.1;refGene_gn=RABGAP1L;refGene_tx=XM_023247522.1;refGene_gn=RABGAP1L;refGene_tx=XM_023247523.1
F1 18882312 18882480 refGene_gn=RABGAP1L;refGene_tx=XM_019821651.2
F1 18891898 18892027
refGene_gn=RABGAP1L;refGene_tx=XM_019821651.2;refGene_gn=RABGAP1L;refGene_tx=XM_023247520.1;refGene_gn=RABGAP1L;refGene_tx=XM_023247521.1;refGene_gn=RABGAP1L;refGene_tx=XM_023247522.1;refGene_gn=RABGAP1L;refGene_tx=XM_023247523.1
F1 18940863 18940974
refGene_gn=RABGAP1L;refGene_tx=XM_019821652.2;refGene_gn=RABGAP1L;refGene_tx=XM_019821653.2
F1 19009878 19009971
refGene_gn=RABGAP1L;refGene_tx=XM_019821651.2;refGene_gn=RABGAP1L;refGene_tx=XM_019821652.2;refGene_gn=RABGAP1L;refGene_tx=XM_019821653.2;refGene_gn=RABGAP1L;refGene_tx=XM_023247520.1;refGene_gn=RABGAP1L;refGene_tx=XM_023247521.1;refGene_gn=RABGAP1L;refGene_tx=XM_023247522.1;refGene_gn=RABGAP1L;refGene_tx=XM_023247523.1

F1	19010164	19010179
refGene_gn=RABGAP1L;refGene_tx=XM_019821653.2;refGene_gn=RABGAP1L;refGene_tx=XM_023247523.1
F1	19016559	19016598	refGene_gn=RABGAP1L;refGene_tx=XM_006942714.4
F1	19020359	19020470
refGene_gn=RABGAP1L;refGene_tx=XM_006942714.4;refGene_gn=RABGAP1L;refGene_tx=XM_019821651.2;refGene_gn=RABGAP1L;refGene_tx=XM_019821652.2;refGene_gn=RABGAP1L;refGene_tx=XM_023247520.1;refGene_gn=RABGAP1L;refGene_tx=XM_023247521.1;refGene_gn=RABGAP1L;refGene_tx=XM_023247522.1
F1	19027566	19027671
refGene_gn=RABGAP1L;refGene_tx=XM_006942714.4;refGene_gn=RABGAP1L;refGene_tx=XM_019821651.2;refGene_gn=RABGAP1L;refGene_tx=XM_019821652.2;refGene_gn=RABGAP1L;refGene_tx=XM_023247520.1;refGene_gn=RABGAP1L;refGene_tx=XM_023247521.1;refGene_gn=RABGAP1L;refGene_tx=XM_023247522.1
F1	19030433	19030517
refGene_gn=RABGAP1L;refGene_tx=XM_006942714.4;refGene_gn=RABGAP1L;refGene_tx=XM_019821651.2;refGene_gn=RABGAP1L;refGene_tx=XM_019821652.2;refGene_gn=RABGAP1L;refGene_tx=XM_023247520.1;refGene_gn=RABGAP1L;refGene_tx=XM_023247521.1;refGene_gn=RABGAP1L;refGene_tx=XM_023247522.1
F1	19033662	19033734
refGene_gn=RABGAP1L;refGene_tx=XM_006942714.4;refGene_gn=RABGAP1L;refGene_tx=XM_019821651.2;refGene_gn=RABGAP1L;refGene_tx=XM_019821652.2;refGene_gn=RABGAP1L;refGene_tx=XM_023247520.1;refGene_gn=RABGAP1L;refGene_tx=XM_023247521.1;refGene_gn=RABGAP1L;refGene_tx=XM_023247522.1
F1	19038088	19038286
refGene_gn=RABGAP1L;refGene_tx=XM_006942714.4;refGene_gn=RABGAP1L;refGene_tx=XM_019821651.2;refGene_gn=RABGAP1L;refGene_tx=XM_019821652.2;refGene_gn=RABGAP1L;refGene_tx=XM_023247520.1;refGene_gn=RABGAP1L;refGene_tx=XM_023247521.1;refGene_gn=RABGAP1L;refGene_tx=XM_023247522.1
F1	19039260	19039413
refGene_gn=RABGAP1L;refGene_tx=XM_006942714.4;refGene_gn=RABGAP1L;refGene_tx=XM_019821651.2;refGene_gn=RABGAP1L;refGene_tx=XM_019821652.2;refGene_gn=RABGAP1L;refGene_tx=XM_023247520.1;refGene_gn=RABGAP1L;refGene_tx=XM_023247521.1;refGene_gn=RABGAP1L;refGene_tx=XM_023247522.1
F1	19049887	19049908	refGene_gn=CACYBP;refGene_tx=XM_011290910.3
F1	19054648	19054868	refGene_gn=CACYBP;refGene_tx=XM_011290910.3
F1	19058080	19058177	refGene_gn=CACYBP;refGene_tx=XM_011290910.3
F1	19058409	19058509	refGene_gn=CACYBP;refGene_tx=XM_011290910.3
F1	19059876	19059974	refGene_gn=CACYBP;refGene_tx=XM_011290910.3
F1	19060779	19060936	refGene_gn=CACYBP;refGene_tx=XM_011290910.3
F1	19064197	19064380
refGene_gn=MRPS14;refGene_tx=XM_003999216.5;refGene_gn=MRPS14;refGene_tx=XM_019821656.2;refGene_gn=MRPS14;refGene_tx=XM_019821657.2
F1	19080106	19080265
refGene_gn=MRPS14;refGene_tx=XM_003999216.5;refGene_gn=MRPS14;refGene_tx=XM_019821656.2;refGene_gn=MRPS14;refGene_tx=XM_019821657.2
F1	19085720	19085738	refGene_gn=MRPS14;refGene_tx=XM_019821656.2
F1	19090401	19090446	refGene_gn=MRPS14;refGene_tx=XM_003999216.5
F1	19142719	19143125	refGene_gn=TNN;refGene_tx=XM_023247526.1
F1	19144623	19144998	refGene_gn=TNN;refGene_tx=XM_023247526.1

F1	19145529	19145793	refGene_gn=TNN;refGene_tx=XM_023247526.1
F1	19149142	19149328	refGene_gn=TNN;refGene_tx=XM_023247526.1
F1	19150561	19150651	refGene_gn=TNN;refGene_tx=XM_023247526.1
F1	19160049	19160313	refGene_gn=TNN;refGene_tx=XM_023247526.1
F1	19163602	19163866	refGene_gn=TNN;refGene_tx=XM_023247526.1
F1	19166525	19166789	refGene_gn=TNN;refGene_tx=XM_023247526.1
F1	19179310	19179574	refGene_gn=TNN;refGene_tx=XM_023247526.1
F1	19180233	19180497	refGene_gn=TNN;refGene_tx=XM_023247526.1
F1	19182253	19182517	refGene_gn=TNN;refGene_tx=XM_023247526.1
F1	19183178	19183442	refGene_gn=TNN;refGene_tx=XM_023247526.1
F1	19184829	19185093	refGene_gn=TNN;refGene_tx=XM_023247526.1
F1	19186228	19186492	refGene_gn=TNN;refGene_tx=XM_023247526.1
F1	19189419	19189683	refGene_gn=TNN;refGene_tx=XM_023247526.1
F1	19192823	19192954	refGene_gn=TNN;refGene_tx=XM_023247526.1
F1	19193992	19194125	refGene_gn=TNN;refGene_tx=XM_023247526.1
F1	19194551	19194703	refGene_gn=TNN;refGene_tx=XM_023247526.1
F1	19200693	19200790	refGene_gn=TNN;refGene_tx=XM_023247526.1
F1	19202152	19202320	refGene_gn=TNN;refGene_tx=XM_023247526.1
F1	19208578	19208742	refGene_gn=TNN;refGene_tx=XM_023247526.1
F1	19211004	19211145	refGene_gn=TNN;refGene_tx=XM_023247526.1
F1	19222712	19223012
refGene_gn=KIAA0040;refGene_tx=XM_011290911.2;refGene_gn=KIAA0040;refG
ene_tx=XM_019821658.2;refGene_gn=KIAA0040;refGene_tx=XM_023247349.1
F1	19347242	19347362	refGene_gn=TNR;refGene_tx=XM_011290913.3
F1	19348283	19348447	refGene_gn=TNR;refGene_tx=XM_011290913.3
F1	19353125	19353287	refGene_gn=TNR;refGene_tx=XM_011290913.3
F1	19357172	19357269	refGene_gn=TNR;refGene_tx=XM_011290913.3
F1	19358684	19358836	refGene_gn=TNR;refGene_tx=XM_011290913.3
F1	19376571	19376704	refGene_gn=TNR;refGene_tx=XM_011290913.3
F1	19377611	19377742	refGene_gn=TNR;refGene_tx=XM_011290913.3
F1	19378417	19378561	refGene_gn=TNR;refGene_tx=XM_011290913.3
F1	19381841	19381961	refGene_gn=TNR;refGene_tx=XM_011290913.3
F1	19384675	19384822	refGene_gn=TNR;refGene_tx=XM_011290913.3
F1	19385749	19385869	refGene_gn=TNR;refGene_tx=XM_011290913.3
F1	19387010	19387280	refGene_gn=TNR;refGene_tx=XM_011290913.3
F1	19387911	19388175	refGene_gn=TNR;refGene_tx=XM_011290913.3
F1	19389278	19389368	refGene_gn=TNR;refGene_tx=XM_011290913.3
F1	19402404	19402579	refGene_gn=TNR;refGene_tx=XM_011290913.3
F1	19402580	19402591	refGene_gn=TNR;refGene_tx=XM_011290913.3
F1	19409855	19410125	refGene_gn=TNR;refGene_tx=XM_011290913.3
F1	19415251	19415402	refGene_gn=TNR;refGene_tx=XM_011290913.3
F1	19417780	19417896	refGene_gn=TNR;refGene_tx=XM_011290913.3
F1	19421193	19421457	refGene_gn=TNR;refGene_tx=XM_011290913.3
F1	19427488	19427965	refGene_gn=TNR;refGene_tx=XM_011290913.3
F1	19430460	19430959	refGene_gn=TNR;refGene_tx=XM_011290913.3
F1	19911420	19911438	refGene_gn=RFWD2;refGene_tx=XM_023247685.1
F1	19913776	19913821	refGene_gn=RFWD2;refGene_tx=XM_023247685.1
F1	19967009	19967170	refGene_gn=RFWD2;refGene_tx=XM_023247685.1
F1	19968276	19968401	refGene_gn=RFWD2;refGene_tx=XM_023247685.1
F1	19971237	19971355	refGene_gn=RFWD2;refGene_tx=XM_023247685.1

F1 20007066 20007183 refGene_gn=RFWD2;refGene_tx=XM_023247685.1
F1 20023509 20023591 refGene_gn=RFWD2;refGene_tx=XM_023247685.1
F1 20023964 20024073 refGene_gn=RFWD2;refGene_tx=XM_023247685.1
F1 20026580 20026724 refGene_gn=RFWD2;refGene_tx=XM_023247685.1
F1 20060312 20060448 refGene_gn=RFWD2;refGene_tx=XM_023247685.1
F1 20064935 20065050 refGene_gn=RFWD2;refGene_tx=XM_023247685.1
F1 20080634 20080692 refGene_gn=RFWD2;refGene_tx=XM_023247685.1
F1 20093172 20093203 refGene_gn=RFWD2;refGene_tx=XM_023247685.1
F1 20093205 20093248 refGene_gn=RFWD2;refGene_tx=XM_023247685.1
F1 20094654 20094714 refGene_gn=RFWD2;refGene_tx=XM_023247685.1
F1 20113397 20113466 refGene_gn=RFWD2;refGene_tx=XM_023247685.1
F1 20121654 20121774 refGene_gn=RFWD2;refGene_tx=XM_023247685.1
F1 20123017 20123094 refGene_gn=RFWD2;refGene_tx=XM_023247685.1
F1 20128599 20128697 refGene_gn=RFWD2;refGene_tx=XM_023247685.1
F1 20140189 20140249 refGene_gn=RFWD2;refGene_tx=XM_023247685.1
F1 20170344 20170745 refGene_gn=RFWD2;refGene_tx=XM_023247685.1
F1 20459798 20460687 refGene_gn=PAPPA2;refGene_tx=XM_023248030.1
F1 20494494 20495566 refGene_gn=PAPPA2;refGene_tx=XM_023248030.1
F1 20559001 20559147 refGene_gn=PAPPA2;refGene_tx=XM_023248030.1
F1 20580235 20580529 refGene_gn=PAPPA2;refGene_tx=XM_023248030.1
F1 20582492 20582685 refGene_gn=PAPPA2;refGene_tx=XM_023248030.1
F1 20586165 20586287 refGene_gn=PAPPA2;refGene_tx=XM_023248030.1
F1 20589342 20589832 refGene_gn=PAPPA2;refGene_tx=XM_023248030.1
F1 20593086 20593215 refGene_gn=PAPPA2;refGene_tx=XM_023248030.1
F1 20596456 20596548 refGene_gn=PAPPA2;refGene_tx=XM_023248030.1
F1 20600420 20600611 refGene_gn=PAPPA2;refGene_tx=XM_023248030.1
F1 20602555 20602702 refGene_gn=PAPPA2;refGene_tx=XM_023248030.1
F1 20623319 20623455 refGene_gn=PAPPA2;refGene_tx=XM_023248030.1
F1 20623653 20623870 refGene_gn=PAPPA2;refGene_tx=XM_023248030.1
F1 20644298 20644470 refGene_gn=PAPPA2;refGene_tx=XM_023248030.1
F1 20648372 20648553 refGene_gn=PAPPA2;refGene_tx=XM_023248030.1
F1 20649761 20649975 refGene_gn=PAPPA2;refGene_tx=XM_023248030.1
F1 20673899 20674068 refGene_gn=PAPPA2;refGene_tx=XM_023248030.1
F1 20675492 20675628 refGene_gn=PAPPA2;refGene_tx=XM_023248030.1
F1 20677332 20677442 refGene_gn=PAPPA2;refGene_tx=XM_023248030.1
F1 20691159 20691230 refGene_gn=PAPPA2;refGene_tx=XM_023248030.1
F1 20691375 20691382 refGene_gn=PAPPA2;refGene_tx=XM_023248030.1
F1 20716609 20716678
refGene_gn=LOC111558335;refGene_tx=XM_023247345.1
F1 20725684 20725783
refGene_gn=LOC111558335;refGene_tx=XM_023247345.1
F1 20727458 20727533
refGene_gn=LOC111558335;refGene_tx=XM_023247345.1
F1 20750210 20750448
refGene_gn=ASTN1;refGene_tx=XM_006942718.4;refGene_gn=ASTN1;refGene_tx
=XM_019821664.2;refGene_gn=ASTN1;refGene_tx=XM_019821665.2;refGene_gn=
ASTN1;refGene_tx=XM_019821666.2
F1 20750724 20750833 refGene_gn=ASTN1;refGene_tx=XM_019821667.2
F1 20753598 20753638 refGene_gn=ASTN1;refGene_tx=XM_019821668.2
F1 20754308 20754492

refGene_gn=ASTN1;refGene_tx=XM_006942718.4;refGene_gn=ASTN1;refGene_tx=XM_019821664.2;refGene_gn=ASTN1;refGene_tx=XM_019821665.2;refGene_gn=ASTN1;refGene_tx=XM_019821666.2;refGene_gn=ASTN1;refGene_tx=XM_019821667.2;refGene_gn=ASTN1;refGene_tx=XM_019821668.2
F1 20763952 20764053
refGene_gn=ASTN1;refGene_tx=XM_006942718.4;refGene_gn=ASTN1;refGene_tx=XM_019821664.2;refGene_gn=ASTN1;refGene_tx=XM_019821665.2;refGene_gn=ASTN1;refGene_tx=XM_019821666.2;refGene_gn=ASTN1;refGene_tx=XM_019821667.2;refGene_gn=ASTN1;refGene_tx=XM_019821668.2
F1 20767840 20767976
refGene_gn=ASTN1;refGene_tx=XM_006942718.4;refGene_gn=ASTN1;refGene_tx=XM_019821664.2;refGene_gn=ASTN1;refGene_tx=XM_019821665.2;refGene_gn=ASTN1;refGene_tx=XM_019821666.2;refGene_gn=ASTN1;refGene_tx=XM_019821667.2;refGene_gn=ASTN1;refGene_tx=XM_019821668.2
F1 20769028 20769180
refGene_gn=ASTN1;refGene_tx=XM_006942718.4;refGene_gn=ASTN1;refGene_tx=XM_019821664.2;refGene_gn=ASTN1;refGene_tx=XM_019821665.2;refGene_gn=ASTN1;refGene_tx=XM_019821666.2;refGene_gn=ASTN1;refGene_tx=XM_019821667.2;refGene_gn=ASTN1;refGene_tx=XM_019821668.2
F1 20772944 20773078
refGene_gn=ASTN1;refGene_tx=XM_006942718.4;refGene_gn=ASTN1;refGene_tx=XM_019821664.2;refGene_gn=ASTN1;refGene_tx=XM_019821665.2;refGene_gn=ASTN1;refGene_tx=XM_019821666.2;refGene_gn=ASTN1;refGene_tx=XM_019821667.2;refGene_gn=ASTN1;refGene_tx=XM_019821668.2
F1 20779390 20779659
refGene_gn=ASTN1;refGene_tx=XM_006942718.4;refGene_gn=ASTN1;refGene_tx=XM_019821664.2;refGene_gn=ASTN1;refGene_tx=XM_019821665.2;refGene_gn=ASTN1;refGene_tx=XM_019821666.2;refGene_gn=ASTN1;refGene_tx=XM_019821667.2;refGene_gn=ASTN1;refGene_tx=XM_019821668.2
F1 20815429 20815618
refGene_gn=ASTN1;refGene_tx=XM_006942718.4;refGene_gn=ASTN1;refGene_tx=XM_019821664.2;refGene_gn=ASTN1;refGene_tx=XM_019821665.2;refGene_gn=ASTN1;refGene_tx=XM_019821666.2;refGene_gn=ASTN1;refGene_tx=XM_019821667.2;refGene_gn=ASTN1;refGene_tx=XM_019821668.2
F1 20817851 20817956
refGene_gn=ASTN1;refGene_tx=XM_006942718.4;refGene_gn=ASTN1;refGene_tx=XM_019821664.2;refGene_gn=ASTN1;refGene_tx=XM_019821665.2;refGene_gn=ASTN1;refGene_tx=XM_019821666.2;refGene_gn=ASTN1;refGene_tx=XM_019821667.2;refGene_gn=ASTN1;refGene_tx=XM_019821668.2
F1 20824876 20825004
refGene_gn=ASTN1;refGene_tx=XM_006942718.4;refGene_gn=ASTN1;refGene_tx=XM_019821664.2;refGene_gn=ASTN1;refGene_tx=XM_019821665.2;refGene_gn=ASTN1;refGene_tx=XM_019821666.2;refGene_gn=ASTN1;refGene_tx=XM_019821667.2;refGene_gn=ASTN1;refGene_tx=XM_019821668.2
F1 20826379 20826574
refGene_gn=ASTN1;refGene_tx=XM_006942718.4;refGene_gn=ASTN1;refGene_tx=XM_019821664.2;refGene_gn=ASTN1;refGene_tx=XM_019821665.2;refGene_gn=ASTN1;refGene_tx=XM_019821666.2;refGene_gn=ASTN1;refGene_tx=XM_019821667.2;refGene_gn=ASTN1;refGene_tx=XM_019821668.2
F1 20829774 20829941

refGene_gn=ASTN1;refGene_tx=XM_006942718.4;refGene_gn=ASTN1;refGene_tx=XM_019821664.2;refGene_gn=ASTN1;refGene_tx=XM_019821665.2;refGene_gn=ASTN1;refGene_tx=XM_019821666.2;refGene_gn=ASTN1;refGene_tx=XM_019821667.2;refGene_gn=ASTN1;refGene_tx=XM_019821668.2
F1 20837954 20838105
refGene_gn=ASTN1;refGene_tx=XM_006942718.4;refGene_gn=ASTN1;refGene_tx=XM_019821664.2;refGene_gn=ASTN1;refGene_tx=XM_019821665.2;refGene_gn=ASTN1;refGene_tx=XM_019821666.2;refGene_gn=ASTN1;refGene_tx=XM_019821667.2;refGene_gn=ASTN1;refGene_tx=XM_019821668.2
F1 20838614 20838752
refGene_gn=ASTN1;refGene_tx=XM_006942718.4;refGene_gn=ASTN1;refGene_tx=XM_019821664.2;refGene_gn=ASTN1;refGene_tx=XM_019821665.2;refGene_gn=ASTN1;refGene_tx=XM_019821666.2;refGene_gn=ASTN1;refGene_tx=XM_019821667.2;refGene_gn=ASTN1;refGene_tx=XM_019821668.2
F1 20845482 20845557
refGene_gn=ASTN1;refGene_tx=XM_006942718.4;refGene_gn=ASTN1;refGene_tx=XM_019821664.2;refGene_gn=ASTN1;refGene_tx=XM_019821667.2;refGene_gn=ASTN1;refGene_tx=XM_019821668.2
F1 20886856 20886941
refGene_gn=ASTN1;refGene_tx=XM_006942718.4;refGene_gn=ASTN1;refGene_tx=XM_019821664.2;refGene_gn=ASTN1;refGene_tx=XM_019821665.2;refGene_gn=ASTN1;refGene_tx=XM_019821666.2;refGene_gn=ASTN1;refGene_tx=XM_019821667.2;refGene_gn=ASTN1;refGene_tx=XM_019821668.2
F1 20894992 20895184
refGene_gn=ASTN1;refGene_tx=XM_006942718.4;refGene_gn=ASTN1;refGene_tx=XM_019821664.2;refGene_gn=ASTN1;refGene_tx=XM_019821665.2;refGene_gn=ASTN1;refGene_tx=XM_019821666.2;refGene_gn=ASTN1;refGene_tx=XM_019821667.2;refGene_gn=ASTN1;refGene_tx=XM_019821668.2
F1 20896204 20896354
refGene_gn=ASTN1;refGene_tx=XM_006942718.4;refGene_gn=ASTN1;refGene_tx=XM_019821664.2;refGene_gn=ASTN1;refGene_tx=XM_019821665.2;refGene_gn=ASTN1;refGene_tx=XM_019821666.2;refGene_gn=ASTN1;refGene_tx=XM_019821667.2;refGene_gn=ASTN1;refGene_tx=XM_019821668.2
F1 20900645 20900753
refGene_gn=ASTN1;refGene_tx=XM_006942718.4;refGene_gn=ASTN1;refGene_tx=XM_019821664.2;refGene_gn=ASTN1;refGene_tx=XM_019821665.2;refGene_gn=ASTN1;refGene_tx=XM_019821666.2;refGene_gn=ASTN1;refGene_tx=XM_019821667.2;refGene_gn=ASTN1;refGene_tx=XM_019821668.2
F1 20901767 20901914
refGene_gn=ASTN1;refGene_tx=XM_006942718.4;refGene_gn=ASTN1;refGene_tx=XM_019821664.2;refGene_gn=ASTN1;refGene_tx=XM_019821665.2;refGene_gn=ASTN1;refGene_tx=XM_019821666.2;refGene_gn=ASTN1;refGene_tx=XM_019821667.2;refGene_gn=ASTN1;refGene_tx=XM_019821668.2
F1 20903400 20903794
refGene_gn=ASTN1;refGene_tx=XM_006942718.4;refGene_gn=ASTN1;refGene_tx=XM_019821664.2;refGene_gn=ASTN1;refGene_tx=XM_019821665.2;refGene_gn=ASTN1;refGene_tx=XM_019821666.2;refGene_gn=ASTN1;refGene_tx=XM_019821667.2;refGene_gn=ASTN1;refGene_tx=XM_019821668.2
F1 20934411 20934599
refGene_gn=ASTN1;refGene_tx=XM_006942718.4;refGene_gn=ASTN1;refGene_tx

=XM_019821664.2;refGene_gn=ASTN1;refGene_tx=XM_019821665.2;refGene_gn=ASTN1;refGene_tx=XM_019821666.2;refGene_gn=ASTN1;refGene_tx=XM_019821667.2;refGene_gn=ASTN1;refGene_tx=XM_019821668.2
F1 21051231 21051514
refGene_gn=ASTN1;refGene_tx=XM_006942718.4;refGene_gn=ASTN1;refGene_tx=XM_019821664.2;refGene_gn=ASTN1;refGene_tx=XM_019821665.2;refGene_gn=ASTN1;refGene_tx=XM_019821666.2;refGene_gn=ASTN1;refGene_tx=XM_019821667.2;refGene_gn=ASTN1;refGene_tx=XM_019821668.2
F1 21113995 21114267
refGene_gn=BRINP2;refGene_tx=XM_023247854.1;refGene_gn=BRINP2;refGene_tx=XM_023247855.1;refGene_gn=BRINP2;refGene_tx=XM_023247856.1
F1 21156409 21156600
refGene_gn=BRINP2;refGene_tx=XM_023247854.1;refGene_gn=BRINP2;refGene_tx=XM_023247855.1;refGene_gn=BRINP2;refGene_tx=XM_023247856.1
F1 21157671 21157880
refGene_gn=BRINP2;refGene_tx=XM_023247854.1;refGene_gn=BRINP2;refGene_tx=XM_023247855.1;refGene_gn=BRINP2;refGene_tx=XM_023247856.1
F1 21169627 21169733
refGene_gn=BRINP2;refGene_tx=XM_023247854.1;refGene_gn=BRINP2;refGene_tx=XM_023247855.1;refGene_gn=BRINP2;refGene_tx=XM_023247856.1
F1 21172456 21172693
refGene_gn=BRINP2;refGene_tx=XM_023247854.1;refGene_gn=BRINP2;refGene_tx=XM_023247855.1;refGene_gn=BRINP2;refGene_tx=XM_023247856.1
F1 21175122 21175345
refGene_gn=BRINP2;refGene_tx=XM_023247854.1;refGene_gn=BRINP2;refGene_tx=XM_023247855.1;refGene_gn=BRINP2;refGene_tx=XM_023247856.1
F1 21176989 21178106
refGene_gn=BRINP2;refGene_tx=XM_023247854.1;refGene_gn=BRINP2;refGene_tx=XM_023247855.1;refGene_gn=BRINP2;refGene_tx=XM_023247856.1
F1 21178585 21178594 refGene_gn=BRINP2;refGene_tx=XM_023247856.1
F1 21778900 21778969
refGene_gn=SEC16B;refGene_tx=XM_019821681.2;refGene_gn=SEC16B;refGene_tx=XM_023247280.1;refGene_gn=SEC16B;refGene_tx=XM_023247281.1;refGene_gn=SEC16B;refGene_tx=XM_023247282.1;refGene_gn=SEC16B;refGene_tx=XM_023247283.1;refGene_gn=SEC16B;refGene_tx=XM_023247284.1;refGene_gn=SEC16B;refGene_tx=XM_023247285.1;refGene_gn=SEC16B;refGene_tx=XM_023247286.1
F1 21779611 21779710
refGene_gn=SEC16B;refGene_tx=XM_019821681.2;refGene_gn=SEC16B;refGene_tx=XM_023247280.1;refGene_gn=SEC16B;refGene_tx=XM_023247281.1;refGene_gn=SEC16B;refGene_tx=XM_023247282.1;refGene_gn=SEC16B;refGene_tx=XM_023247283.1;refGene_gn=SEC16B;refGene_tx=XM_023247284.1;refGene_gn=SEC16B;refGene_tx=XM_023247285.1;refGene_gn=SEC16B;refGene_tx=XM_023247286.1
F1 21781644 21781724
refGene_gn=SEC16B;refGene_tx=XM_019821681.2;refGene_gn=SEC16B;refGene_tx=XM_023247280.1;refGene_gn=SEC16B;refGene_tx=XM_023247281.1;refGene_gn=SEC16B;refGene_tx=XM_023247282.1;refGene_gn=SEC16B;refGene_tx=XM_023247283.1;refGene_gn=SEC16B;refGene_tx=XM_023247284.1;refGene_gn=SEC16B;refGene_tx=XM_023247285.1;refGene_gn=SEC16B;refGene_tx=XM_023247286.

1
F1	21781853 21781959
refGene_gn=SEC16B;refGene_tx=XM_019821681.2;refGene_gn=SEC16B;refGene_tx=XM_023247280.1;refGene_gn=SEC16B;refGene_tx=XM_023247281.1;refGene_gn=SEC16B;refGene_tx=XM_023247282.1;refGene_gn=SEC16B;refGene_tx=XM_023247283.1;refGene_gn=SEC16B;refGene_tx=XM_023247284.1;refGene_gn=SEC16B;refGene_tx=XM_023247285.1;refGene_gn=SEC16B;refGene_tx=XM_023247286.1
F1	21782293 21782389
refGene_gn=SEC16B;refGene_tx=XM_019821681.2;refGene_gn=SEC16B;refGene_tx=XM_023247280.1;refGene_gn=SEC16B;refGene_tx=XM_023247281.1;refGene_gn=SEC16B;refGene_tx=XM_023247282.1;refGene_gn=SEC16B;refGene_tx=XM_023247283.1;refGene_gn=SEC16B;refGene_tx=XM_023247284.1;refGene_gn=SEC16B;refGene_tx=XM_023247285.1;refGene_gn=SEC16B;refGene_tx=XM_023247286.1
F1	21782560 21782713
refGene_gn=SEC16B;refGene_tx=XM_019821681.2;refGene_gn=SEC16B;refGene_tx=XM_023247280.1;refGene_gn=SEC16B;refGene_tx=XM_023247281.1;refGene_gn=SEC16B;refGene_tx=XM_023247282.1;refGene_gn=SEC16B;refGene_tx=XM_023247283.1;refGene_gn=SEC16B;refGene_tx=XM_023247284.1;refGene_gn=SEC16B;refGene_tx=XM_023247285.1;refGene_gn=SEC16B;refGene_tx=XM_023247286.1
F1	21785711 21785776
refGene_gn=SEC16B;refGene_tx=XM_019821681.2;refGene_gn=SEC16B;refGene_tx=XM_023247280.1;refGene_gn=SEC16B;refGene_tx=XM_023247281.1;refGene_gn=SEC16B;refGene_tx=XM_023247282.1;refGene_gn=SEC16B;refGene_tx=XM_023247283.1;refGene_gn=SEC16B;refGene_tx=XM_023247284.1;refGene_gn=SEC16B;refGene_tx=XM_023247285.1;refGene_gn=SEC16B;refGene_tx=XM_023247286.1
F1	21786513 21786786
refGene_gn=SEC16B;refGene_tx=XM_019821681.2;refGene_gn=SEC16B;refGene_tx=XM_023247280.1;refGene_gn=SEC16B;refGene_tx=XM_023247281.1;refGene_gn=SEC16B;refGene_tx=XM_023247282.1;refGene_gn=SEC16B;refGene_tx=XM_023247283.1;refGene_gn=SEC16B;refGene_tx=XM_023247284.1;refGene_gn=SEC16B;refGene_tx=XM_023247285.1;refGene_gn=SEC16B;refGene_tx=XM_023247286.1
F1	21788119 21788192
refGene_gn=SEC16B;refGene_tx=XM_019821681.2;refGene_gn=SEC16B;refGene_tx=XM_023247280.1;refGene_gn=SEC16B;refGene_tx=XM_023247281.1;refGene_gn=SEC16B;refGene_tx=XM_023247282.1;refGene_gn=SEC16B;refGene_tx=XM_023247283.1;refGene_gn=SEC16B;refGene_tx=XM_023247284.1;refGene_gn=SEC16B;refGene_tx=XM_023247285.1;refGene_gn=SEC16B;refGene_tx=XM_023247286.1
F1	21789209 21789320
refGene_gn=SEC16B;refGene_tx=XM_019821681.2;refGene_gn=SEC16B;refGene_tx=XM_023247280.1;refGene_gn=SEC16B;refGene_tx=XM_023247281.1;refGene_gn=SEC16B;refGene_tx=XM_023247282.1;refGene_gn=SEC16B;refGene_tx=XM_023247283.1;refGene_gn=SEC16B;refGene_tx=XM_023247284.1;refGene_gn=SEC16B;refGene_tx=XM_023247285.1;refGene_gn=SEC16B;refGene_tx=XM_023247286.1

F1 21790356 21790497
refGene_gn=SEC16B;refGene_tx=XM_019821681.2;refGene_gn=SEC16B;refGene_tx=XM_023247280.1;refGene_gn=SEC16B;refGene_tx=XM_023247281.1;refGene_gn=SEC16B;refGene_tx=XM_023247282.1;refGene_gn=SEC16B;refGene_tx=XM_023247283.1;refGene_gn=SEC16B;refGene_tx=XM_023247284.1;refGene_gn=SEC16B;refGene_tx=XM_023247285.1;refGene_gn=SEC16B;refGene_tx=XM_023247286.1
F1 21792147 21792253
refGene_gn=SEC16B;refGene_tx=XM_019821681.2;refGene_gn=SEC16B;refGene_tx=XM_023247280.1;refGene_gn=SEC16B;refGene_tx=XM_023247281.1;refGene_gn=SEC16B;refGene_tx=XM_023247282.1;refGene_gn=SEC16B;refGene_tx=XM_023247283.1;refGene_gn=SEC16B;refGene_tx=XM_023247284.1;refGene_gn=SEC16B;refGene_tx=XM_023247285.1;refGene_gn=SEC16B;refGene_tx=XM_023247286.1
F1 21794159 21794271
refGene_gn=SEC16B;refGene_tx=XM_019821681.2;refGene_gn=SEC16B;refGene_tx=XM_023247280.1;refGene_gn=SEC16B;refGene_tx=XM_023247281.1;refGene_gn=SEC16B;refGene_tx=XM_023247282.1;refGene_gn=SEC16B;refGene_tx=XM_023247283.1;refGene_gn=SEC16B;refGene_tx=XM_023247284.1;refGene_gn=SEC16B;refGene_tx=XM_023247285.1;refGene_gn=SEC16B;refGene_tx=XM_023247286.1
F1 21795449 21795567
refGene_gn=SEC16B;refGene_tx=XM_019821681.2;refGene_gn=SEC16B;refGene_tx=XM_023247280.1;refGene_gn=SEC16B;refGene_tx=XM_023247281.1;refGene_gn=SEC16B;refGene_tx=XM_023247282.1;refGene_gn=SEC16B;refGene_tx=XM_023247283.1;refGene_gn=SEC16B;refGene_tx=XM_023247284.1;refGene_gn=SEC16B;refGene_tx=XM_023247285.1;refGene_gn=SEC16B;refGene_tx=XM_023247286.1
F1 21799567 21799649
refGene_gn=SEC16B;refGene_tx=XM_019821681.2;refGene_gn=SEC16B;refGene_tx=XM_023247280.1;refGene_gn=SEC16B;refGene_tx=XM_023247281.1;refGene_gn=SEC16B;refGene_tx=XM_023247282.1;refGene_gn=SEC16B;refGene_tx=XM_023247283.1;refGene_gn=SEC16B;refGene_tx=XM_023247284.1;refGene_gn=SEC16B;refGene_tx=XM_023247285.1;refGene_gn=SEC16B;refGene_tx=XM_023247286.1
F1 21801558 21801656
refGene_gn=SEC16B;refGene_tx=XM_019821681.2;refGene_gn=SEC16B;refGene_tx=XM_023247280.1;refGene_gn=SEC16B;refGene_tx=XM_023247281.1;refGene_gn=SEC16B;refGene_tx=XM_023247282.1;refGene_gn=SEC16B;refGene_tx=XM_023247283.1;refGene_gn=SEC16B;refGene_tx=XM_023247284.1;refGene_gn=SEC16B;refGene_tx=XM_023247285.1;refGene_gn=SEC16B;refGene_tx=XM_023247286.1
F1 21807509 21807740
refGene_gn=SEC16B;refGene_tx=XM_019821681.2;refGene_gn=SEC16B;refGene_tx=XM_023247280.1;refGene_gn=SEC16B;refGene_tx=XM_023247281.1;refGene_gn=SEC16B;refGene_tx=XM_023247282.1;refGene_gn=SEC16B;refGene_tx=XM_023247283.1;refGene_gn=SEC16B;refGene_tx=XM_023247284.1;refGene_gn=SEC16B;refGene_tx=XM_023247285.1;refGene_gn=SEC16B;refGene_tx=XM_023247286.1
F1 21808207 21808343

refGene_gn=SEC16B;refGene_tx=XM_019821681.2;refGene_gn=SEC16B;refGene_tx=XM_023247280.1;refGene_gn=SEC16B;refGene_tx=XM_023247281.1;refGene_gn=SEC16B;refGene_tx=XM_023247282.1;refGene_gn=SEC16B;refGene_tx=XM_023247283.1;refGene_gn=SEC16B;refGene_tx=XM_023247284.1;refGene_gn=SEC16B;refGene_tx=XM_023247285.1;refGene_gn=SEC16B;refGene_tx=XM_023247286.1
F1 21809640 21809702
refGene_gn=SEC16B;refGene_tx=XM_019821681.2;refGene_gn=SEC16B;refGene_tx=XM_023247280.1;refGene_gn=SEC16B;refGene_tx=XM_023247281.1;refGene_gn=SEC16B;refGene_tx=XM_023247282.1;refGene_gn=SEC16B;refGene_tx=XM_023247283.1;refGene_gn=SEC16B;refGene_tx=XM_023247284.1;refGene_gn=SEC16B;refGene_tx=XM_023247285.1;refGene_gn=SEC16B;refGene_tx=XM_023247286.1
F1 21810063 21810215
refGene_gn=SEC16B;refGene_tx=XM_019821681.2;refGene_gn=SEC16B;refGene_tx=XM_023247280.1;refGene_gn=SEC16B;refGene_tx=XM_023247281.1;refGene_gn=SEC16B;refGene_tx=XM_023247282.1;refGene_gn=SEC16B;refGene_tx=XM_023247283.1;refGene_gn=SEC16B;refGene_tx=XM_023247284.1;refGene_gn=SEC16B;refGene_tx=XM_023247285.1;refGene_gn=SEC16B;refGene_tx=XM_023247286.1
F1 21811115 21811260
refGene_gn=SEC16B;refGene_tx=XM_019821681.2;refGene_gn=SEC16B;refGene_tx=XM_023247280.1;refGene_gn=SEC16B;refGene_tx=XM_023247281.1;refGene_gn=SEC16B;refGene_tx=XM_023247282.1;refGene_gn=SEC16B;refGene_tx=XM_023247283.1;refGene_gn=SEC16B;refGene_tx=XM_023247284.1;refGene_gn=SEC16B;refGene_tx=XM_023247285.1;refGene_gn=SEC16B;refGene_tx=XM_023247286.1
F1 21812615 21812724
refGene_gn=SEC16B;refGene_tx=XM_019821681.2;refGene_gn=SEC16B;refGene_tx=XM_023247280.1;refGene_gn=SEC16B;refGene_tx=XM_023247281.1;refGene_gn=SEC16B;refGene_tx=XM_023247282.1;refGene_gn=SEC16B;refGene_tx=XM_023247283.1;refGene_gn=SEC16B;refGene_tx=XM_023247284.1;refGene_gn=SEC16B;refGene_tx=XM_023247285.1;refGene_gn=SEC16B;refGene_tx=XM_023247286.1
F1 21813467 21813565
refGene_gn=SEC16B;refGene_tx=XM_019821681.2;refGene_gn=SEC16B;refGene_tx=XM_023247280.1;refGene_gn=SEC16B;refGene_tx=XM_023247281.1;refGene_gn=SEC16B;refGene_tx=XM_023247282.1;refGene_gn=SEC16B;refGene_tx=XM_023247283.1;refGene_gn=SEC16B;refGene_tx=XM_023247284.1;refGene_gn=SEC16B;refGene_tx=XM_023247285.1;refGene_gn=SEC16B;refGene_tx=XM_023247286.1
F1 21814327 21814440
refGene_gn=SEC16B;refGene_tx=XM_019821681.2;refGene_gn=SEC16B;refGene_tx=XM_023247280.1;refGene_gn=SEC16B;refGene_tx=XM_023247281.1;refGene_gn=SEC16B;refGene_tx=XM_023247282.1;refGene_gn=SEC16B;refGene_tx=XM_023247283.1;refGene_gn=SEC16B;refGene_tx=XM_023247284.1;refGene_gn=SEC16B;refGene_tx=XM_023247285.1;refGene_gn=SEC16B;refGene_tx=XM_023247286.1
F1 21816027 21816323
refGene_gn=SEC16B;refGene_tx=XM_019821681.2;refGene_gn=SEC16B;refGene_

tx=XM_023247280.1;refGene_gn=SEC16B;refGene_tx=XM_023247281.1;refGene_gn=SEC16B;refGene_tx=XM_023247282.1;refGene_gn=SEC16B;refGene_tx=XM_023247283.1;refGene_gn=SEC16B;refGene_tx=XM_023247284.1;refGene_gn=SEC16B;refGene_tx=XM_023247285.1
F1 21848261 21848312
refGene_gn=LOC101083645;refGene_tx=XM_023247218.1
F1 21848565 21848592
refGene_gn=LOC101083645;refGene_tx=XM_011291025.2
F1 21850593 21850650
refGene_gn=LOC101083645;refGene_tx=XM_011291023.2;refGene_gn=LOC101083645;refGene_tx=XM_023247219.1;refGene_gn=LOC101083645;refGene_tx=XM_023247220.1
F1 21851475 21851523
refGene_gn=LOC101083645;refGene_tx=XM_011291023.2;refGene_gn=LOC101083645;refGene_tx=XM_011291025.2;refGene_gn=LOC101083645;refGene_tx=XM_023247218.1;refGene_gn=LOC101083645;refGene_tx=XM_023247219.1;refGene_gn=LOC101083645;refGene_tx=XM_023247220.1
F1 21861837 21862066
refGene_gn=LOC101083645;refGene_tx=XM_011291023.2;refGene_gn=LOC101083645;refGene_tx=XM_011291025.2;refGene_gn=LOC101083645;refGene_tx=XM_023247218.1;refGene_gn=LOC101083645;refGene_tx=XM_023247219.1;refGene_gn=LOC101083645;refGene_tx=XM_023247220.1
F1 21866363 21866545
refGene_gn=LOC101083645;refGene_tx=XM_011291023.2;refGene_gn=LOC101083645;refGene_tx=XM_011291025.2;refGene_gn=LOC101083645;refGene_tx=XM_023247218.1;refGene_gn=LOC101083645;refGene_tx=XM_023247219.1
F1 21871402 21871484
refGene_gn=LOC101083645;refGene_tx=XM_011291023.2;refGene_gn=LOC101083645;refGene_tx=XM_011291025.2;refGene_gn=LOC101083645;refGene_tx=XM_023247218.1
F1 21872265 21872375
refGene_gn=LOC101083645;refGene_tx=XM_011291023.2;refGene_gn=LOC101083645;refGene_tx=XM_011291025.2;refGene_gn=LOC101083645;refGene_tx=XM_023247218.1;refGene_gn=LOC101083645;refGene_tx=XM_023247220.1
F1 21873159 21873225
refGene_gn=LOC101083645;refGene_tx=XM_011291023.2;refGene_gn=LOC101083645;refGene_tx=XM_011291025.2;refGene_gn=LOC101083645;refGene_tx=XM_023247218.1;refGene_gn=LOC101083645;refGene_tx=XM_023247219.1;refGene_gn=LOC101083645;refGene_tx=XM_023247220.1
F1 21873783 21873842
refGene_gn=LOC101083645;refGene_tx=XM_011291023.2;refGene_gn=LOC101083645;refGene_tx=XM_011291025.2;refGene_gn=LOC101083645;refGene_tx=XM_023247218.1;refGene_gn=LOC101083645;refGene_tx=XM_023247219.1;refGene_gn=LOC101083645;refGene_tx=XM_023247220.1
F1 21876318 21876415
refGene_gn=LOC101083645;refGene_tx=XM_011291023.2;refGene_gn=LOC101083645;refGene_tx=XM_011291025.2;refGene_gn=LOC101083645;refGene_tx=XM_023247218.1;refGene_gn=LOC101083645;refGene_tx=XM_023247219.1;refGene_gn=LOC101083645;refGene_tx=XM_023247220.1
F1 21878493 21878602

refGene_gn=LOC101083645;refGene_tx=XM_011291023.2;refGene_gn=LOC101083
645;refGene_tx=XM_011291025.2;refGene_gn=LOC101083645;refGene_tx=XM_02
3247218.1;refGene_gn=LOC101083645;refGene_tx=XM_023247219.1;refGene_gn
=LOC101083645;refGene_tx=XM_023247220.1
F1 21878691 21879020
refGene_gn=LOC101083645;refGene_tx=XM_011291023.2;refGene_gn=LOC101083
645;refGene_tx=XM_011291025.2;refGene_gn=LOC101083645;refGene_tx=XM_02
3247218.1;refGene_gn=LOC101083645;refGene_tx=XM_023247219.1;refGene_gn
=LOC101083645;refGene_tx=XM_023247220.1
F1 21926315 21926517
refGene_gn=RASAL2;refGene_tx=XM_003999225.5;refGene_gn=RASAL2;refGene_
tx=XM_003999226.5;refGene_gn=RASAL2;refGene_tx=XM_019821687.2
F1 22047707 22048305
refGene_gn=RASAL2;refGene_tx=XM_019821682.2;refGene_gn=RASAL2;refGene_
tx=XM_019821683.2;refGene_gn=RASAL2;refGene_tx=XM_019821684.2;refGene_
gn=RASAL2;refGene_tx=XM_019821686.2;refGene_gn=RASAL2;refGene_tx=XM_02
3247310.1
F1 22104243 22104371
refGene_gn=RASAL2;refGene_tx=XM_003999225.5;refGene_gn=RASAL2;refGene_
tx=XM_003999226.5;refGene_gn=RASAL2;refGene_tx=XM_019821682.2;refGene_
gn=RASAL2;refGene_tx=XM_019821683.2;refGene_gn=RASAL2;refGene_tx=XM_01
9821684.2;refGene_gn=RASAL2;refGene_tx=XM_019821686.2;refGene_gn=RASAL
2;refGene_tx=XM_019821687.2;refGene_gn=RASAL2;refGene_tx=XM_023247310.
1
F1 22119738 22119865
refGene_gn=RASAL2;refGene_tx=XM_003999225.5;refGene_gn=RASAL2;refGene_
tx=XM_003999226.5;refGene_gn=RASAL2;refGene_tx=XM_019821682.2;refGene_
gn=RASAL2;refGene_tx=XM_019821683.2;refGene_gn=RASAL2;refGene_tx=XM_01
9821684.2;refGene_gn=RASAL2;refGene_tx=XM_019821686.2;refGene_gn=RASAL
2;refGene_tx=XM_019821687.2;refGene_gn=RASAL2;refGene_tx=XM_023247310.
1
F1 22146042 22146055 refGene_gn=RASAL2;refGene_tx=XM_019821690.2
F1 22176365 22176372 refGene_gn=RASAL2;refGene_tx=XM_019821688.2
F1 22177039 22177159 refGene_gn=RASAL2;refGene_tx=XM_019821688.2
F1 22195960 22195982 refGene_gn=RASAL2;refGene_tx=XM_019821689.2
F1 22196785 22196892
refGene_gn=RASAL2;refGene_tx=XM_003999225.5;refGene_gn=RASAL2;refGene_
tx=XM_003999226.5;refGene_gn=RASAL2;refGene_tx=XM_019821682.2;refGene_
gn=RASAL2;refGene_tx=XM_019821683.2;refGene_gn=RASAL2;refGene_tx=XM_01
9821684.2;refGene_gn=RASAL2;refGene_tx=XM_019821686.2;refGene_gn=RASAL
2;refGene_tx=XM_019821687.2;refGene_gn=RASAL2;refGene_tx=XM_019821688.
2;refGene_gn=RASAL2;refGene_tx=XM_019821689.2;refGene_gn=RASAL2;refGen
e_tx=XM_019821690.2;refGene_gn=RASAL2;refGene_tx=XM_023247310.1
F1 22220870 22220980
refGene_gn=RASAL2;refGene_tx=XM_003999225.5;refGene_gn=RASAL2;refGene_
tx=XM_003999226.5;refGene_gn=RASAL2;refGene_tx=XM_019821682.2;refGene_
gn=RASAL2;refGene_tx=XM_019821683.2;refGene_gn=RASAL2;refGene_tx=XM_01
9821684.2;refGene_gn=RASAL2;refGene_tx=XM_019821686.2;refGene_gn=RASAL
2;refGene_tx=XM_019821687.2;refGene_gn=RASAL2;refGene_tx=XM_019821688.
2;refGene_gn=RASAL2;refGene_tx=XM_019821689.2;refGene_gn=RASAL2;refGen

e_tx=XM_019821690.2;refGene_gn=RASAL2;refGene_tx=XM_023247310.1
F1 22240882 22241036
refGene_gn=RASAL2;refGene_tx=XM_003999225.5;refGene_gn=RASAL2;refGene_tx=XM_003999226.5;refGene_gn=RASAL2;refGene_tx=XM_019821682.2;refGene_gn=RASAL2;refGene_tx=XM_019821683.2;refGene_gn=RASAL2;refGene_tx=XM_019821684.2;refGene_gn=RASAL2;refGene_tx=XM_019821686.2;refGene_gn=RASAL2;refGene_tx=XM_019821687.2;refGene_gn=RASAL2;refGene_tx=XM_019821688.2;refGene_gn=RASAL2;refGene_tx=XM_019821689.2;refGene_gn=RASAL2;refGene_tx=XM_019821690.2;refGene_gn=RASAL2;refGene_tx=XM_023247310.1
F1 22242832 22242931
refGene_gn=RASAL2;refGene_tx=XM_003999225.5;refGene_gn=RASAL2;refGene_tx=XM_003999226.5;refGene_gn=RASAL2;refGene_tx=XM_019821682.2;refGene_gn=RASAL2;refGene_tx=XM_019821683.2;refGene_gn=RASAL2;refGene_tx=XM_019821684.2;refGene_gn=RASAL2;refGene_tx=XM_019821686.2;refGene_gn=RASAL2;refGene_tx=XM_019821687.2;refGene_gn=RASAL2;refGene_tx=XM_019821688.2;refGene_gn=RASAL2;refGene_tx=XM_019821689.2;refGene_gn=RASAL2;refGene_tx=XM_019821690.2;refGene_gn=RASAL2;refGene_tx=XM_023247310.1
F1 22243699 22244254
refGene_gn=RASAL2;refGene_tx=XM_003999225.5;refGene_gn=RASAL2;refGene_tx=XM_003999226.5;refGene_gn=RASAL2;refGene_tx=XM_019821682.2;refGene_gn=RASAL2;refGene_tx=XM_019821683.2;refGene_gn=RASAL2;refGene_tx=XM_019821684.2;refGene_gn=RASAL2;refGene_tx=XM_019821686.2;refGene_gn=RASAL2;refGene_tx=XM_019821687.2;refGene_gn=RASAL2;refGene_tx=XM_019821688.2;refGene_gn=RASAL2;refGene_tx=XM_019821689.2;refGene_gn=RASAL2;refGene_tx=XM_019821690.2;refGene_gn=RASAL2;refGene_tx=XM_023247310.1
F1 22247504 22247649
refGene_gn=RASAL2;refGene_tx=XM_003999225.5;refGene_gn=RASAL2;refGene_tx=XM_003999226.5;refGene_gn=RASAL2;refGene_tx=XM_019821682.2;refGene_gn=RASAL2;refGene_tx=XM_019821683.2;refGene_gn=RASAL2;refGene_tx=XM_019821684.2;refGene_gn=RASAL2;refGene_tx=XM_019821686.2;refGene_gn=RASAL2;refGene_tx=XM_019821687.2;refGene_gn=RASAL2;refGene_tx=XM_019821688.2;refGene_gn=RASAL2;refGene_tx=XM_019821689.2;refGene_gn=RASAL2;refGene_tx=XM_019821690.2;refGene_gn=RASAL2;refGene_tx=XM_023247310.1
F1 22255715 22255860
refGene_gn=RASAL2;refGene_tx=XM_003999225.5;refGene_gn=RASAL2;refGene_tx=XM_003999226.5;refGene_gn=RASAL2;refGene_tx=XM_019821682.2;refGene_gn=RASAL2;refGene_tx=XM_019821683.2;refGene_gn=RASAL2;refGene_tx=XM_019821684.2;refGene_gn=RASAL2;refGene_tx=XM_019821686.2;refGene_gn=RASAL2;refGene_tx=XM_019821687.2;refGene_gn=RASAL2;refGene_tx=XM_019821688.2;refGene_gn=RASAL2;refGene_tx=XM_019821689.2;refGene_gn=RASAL2;refGene_tx=XM_019821690.2;refGene_gn=RASAL2;refGene_tx=XM_023247310.1
F1 22256528 22256765
refGene_gn=RASAL2;refGene_tx=XM_003999225.5;refGene_gn=RASAL2;refGene_tx=XM_003999226.5;refGene_gn=RASAL2;refGene_tx=XM_019821682.2;refGene_gn=RASAL2;refGene_tx=XM_019821683.2;refGene_gn=RASAL2;refGene_tx=XM_019821684.2;refGene_gn=RASAL2;refGene_tx=XM_019821686.2;refGene_gn=RASAL2;refGene_tx=XM_019821687.2;refGene_gn=RASAL2;refGene_tx=XM_019821688.2;refGene_gn=RASAL2;refGene_tx=XM_019821689.2;refGene_gn=RASAL2;refGene_tx=XM_019821690.2;refGene_gn=RASAL2;refGene_tx=XM_023247310.1
F1 22258394 22258596

refGene_gn=RASAL2;refGene_tx=XM_003999225.5;refGene_gn=RASAL2;refGene_tx=XM_003999226.5;refGene_gn=RASAL2;refGene_tx=XM_019821682.2;refGene_gn=RASAL2;refGene_tx=XM_019821683.2;refGene_gn=RASAL2;refGene_tx=XM_019821684.2;refGene_gn=RASAL2;refGene_tx=XM_019821686.2;refGene_gn=RASAL2;refGene_tx=XM_019821687.2;refGene_gn=RASAL2;refGene_tx=XM_019821688.2;refGene_gn=RASAL2;refGene_tx=XM_019821689.2;refGene_gn=RASAL2;refGene_tx=XM_019821690.2;refGene_gn=RASAL2;refGene_tx=XM_023247310.1
F1 22260715 22260915
refGene_gn=RASAL2;refGene_tx=XM_003999225.5;refGene_gn=RASAL2;refGene_tx=XM_003999226.5;refGene_gn=RASAL2;refGene_tx=XM_019821682.2;refGene_gn=RASAL2;refGene_tx=XM_019821683.2;refGene_gn=RASAL2;refGene_tx=XM_019821684.2;refGene_gn=RASAL2;refGene_tx=XM_019821686.2;refGene_gn=RASAL2;refGene_tx=XM_019821687.2;refGene_gn=RASAL2;refGene_tx=XM_019821688.2;refGene_gn=RASAL2;refGene_tx=XM_019821689.2;refGene_gn=RASAL2;refGene_tx=XM_019821690.2;refGene_gn=RASAL2;refGene_tx=XM_023247310.1
F1 22261694 22262556
refGene_gn=RASAL2;refGene_tx=XM_003999225.5;refGene_gn=RASAL2;refGene_tx=XM_003999226.5;refGene_gn=RASAL2;refGene_tx=XM_019821682.2;refGene_gn=RASAL2;refGene_tx=XM_019821683.2;refGene_gn=RASAL2;refGene_tx=XM_019821684.2;refGene_gn=RASAL2;refGene_tx=XM_019821686.2;refGene_gn=RASAL2;refGene_tx=XM_019821687.2;refGene_gn=RASAL2;refGene_tx=XM_019821688.2;refGene_gn=RASAL2;refGene_tx=XM_019821689.2;refGene_gn=RASAL2;refGene_tx=XM_019821690.2;refGene_gn=RASAL2;refGene_tx=XM_023247310.1
F1 22270058 22270193
refGene_gn=RASAL2;refGene_tx=XM_003999225.5;refGene_gn=RASAL2;refGene_tx=XM_003999226.5;refGene_gn=RASAL2;refGene_tx=XM_019821682.2;refGene_gn=RASAL2;refGene_tx=XM_019821683.2;refGene_gn=RASAL2;refGene_tx=XM_019821684.2;refGene_gn=RASAL2;refGene_tx=XM_019821686.2;refGene_gn=RASAL2;refGene_tx=XM_019821687.2;refGene_gn=RASAL2;refGene_tx=XM_019821688.2;refGene_gn=RASAL2;refGene_tx=XM_019821689.2;refGene_gn=RASAL2;refGene_tx=XM_019821690.2;refGene_gn=RASAL2;refGene_tx=XM_023247310.1
F1 22271889 22272092
refGene_gn=RASAL2;refGene_tx=XM_003999225.5;refGene_gn=RASAL2;refGene_tx=XM_003999226.5;refGene_gn=RASAL2;refGene_tx=XM_019821682.2;refGene_gn=RASAL2;refGene_tx=XM_019821683.2;refGene_gn=RASAL2;refGene_tx=XM_019821684.2;refGene_gn=RASAL2;refGene_tx=XM_019821686.2;refGene_gn=RASAL2;refGene_tx=XM_019821687.2;refGene_gn=RASAL2;refGene_tx=XM_019821688.2;refGene_gn=RASAL2;refGene_tx=XM_019821689.2;refGene_gn=RASAL2;refGene_tx=XM_019821690.2;refGene_gn=RASAL2;refGene_tx=XM_023247310.1
F1 22274050 22274138
refGene_gn=RASAL2;refGene_tx=XM_003999225.5;refGene_gn=RASAL2;refGene_tx=XM_003999226.5;refGene_gn=RASAL2;refGene_tx=XM_019821682.2;refGene_gn=RASAL2;refGene_tx=XM_019821683.2;refGene_gn=RASAL2;refGene_tx=XM_019821684.2;refGene_gn=RASAL2;refGene_tx=XM_019821686.2;refGene_gn=RASAL2;refGene_tx=XM_019821687.2;refGene_gn=RASAL2;refGene_tx=XM_019821688.2;refGene_gn=RASAL2;refGene_tx=XM_019821689.2;refGene_gn=RASAL2;refGene_tx=XM_019821690.2;refGene_gn=RASAL2;refGene_tx=XM_023247310.1
F1 22277290 22277321
refGene_gn=RASAL2;refGene_tx=XM_019821682.2;refGene_gn=RASAL2;refGene_tx=XM_019821684.2;refGene_gn=RASAL2;refGene_tx=XM_019821687.2;refGene_

gn=RASAL2;refGene_tx=XM_019821688.2;refGene_gn=RASAL2;refGene_tx=XM_01
9821689.2;refGene_gn=RASAL2;refGene_tx=XM_019821690.2
F1 22279760 22279925
refGene_gn=RASAL2;refGene_tx=XM_003999225.5;refGene_gn=RASAL2;refGene_
tx=XM_003999226.5;refGene_gn=RASAL2;refGene_tx=XM_019821682.2;refGene_
gn=RASAL2;refGene_tx=XM_019821683.2;refGene_gn=RASAL2;refGene_tx=XM_01
9821684.2;refGene_gn=RASAL2;refGene_tx=XM_019821687.2;refGene_gn=RASAL
2;refGene_tx=XM_019821688.2;refGene_gn=RASAL2;refGene_tx=XM_019821689.
2;refGene_gn=RASAL2;refGene_tx=XM_019821690.2;refGene_gn=RASAL2;refGen
e_tx=XM_023247310.1
F1 22285605 22285665 refGene_gn=RASAL2;refGene_tx=XM_019821686.2
F1 22286238 22286370 refGene_gn=CLEC20A;refGene_tx=XM_019821691.2
F1 22288200 22288286 refGene_gn=CLEC20A;refGene_tx=XM_019821691.2
F1 22289067 22289187 refGene_gn=CLEC20A;refGene_tx=XM_019821691.2
F1 22294162 22294255 refGene_gn=CLEC20A;refGene_tx=XM_019821691.2
F1 22295350 22295687 refGene_gn=CLEC20A;refGene_tx=XM_019821691.2
F1 22299340 22299414
refGene_gn=LOC105261341;refGene_tx=XM_019821692.2
F1 22306604 22306995
refGene_gn=LOC105261341;refGene_tx=XM_019821692.2
F1 22317968 22318007
refGene_gn=TEX35;refGene_tx=XM_006942725.4;refGene_gn=TEX35;refGene_tx
=XM_011291021.3;refGene_gn=TEX35;refGene_tx=XM_019821694.1;refGene_gn=
TEX35;refGene_tx=XM_023247609.1
F1 22318765 22318924
refGene_gn=TEX35;refGene_tx=XM_006942725.4;refGene_gn=TEX35;refGene_tx
=XM_011291021.3;refGene_gn=TEX35;refGene_tx=XM_019821694.1;refGene_gn=
TEX35;refGene_tx=XM_023247609.1
F1 22319414 22319483
refGene_gn=TEX35;refGene_tx=XM_006942725.4;refGene_gn=TEX35;refGene_tx
=XM_011291021.3;refGene_gn=TEX35;refGene_tx=XM_019821694.1;refGene_gn=
TEX35;refGene_tx=XM_023247609.1
F1 22320317 22320374
refGene_gn=TEX35;refGene_tx=XM_006942725.4;refGene_gn=TEX35;refGene_tx
=XM_011291021.3;refGene_gn=TEX35;refGene_tx=XM_019821694.1;refGene_gn=
TEX35;refGene_tx=XM_023247609.1
F1 22321067 22321127
refGene_gn=TEX35;refGene_tx=XM_006942725.4;refGene_gn=TEX35;refGene_tx
=XM_011291021.3;refGene_gn=TEX35;refGene_tx=XM_019821694.1;refGene_gn=
TEX35;refGene_tx=XM_023247609.1
F1 22326022 22326084
refGene_gn=TEX35;refGene_tx=XM_006942725.4;refGene_gn=TEX35;refGene_tx
=XM_011291021.3;refGene_gn=TEX35;refGene_tx=XM_019821694.1;refGene_gn=
TEX35;refGene_tx=XM_023247609.1
F1 22326317 22326510
refGene_gn=TEX35;refGene_tx=XM_006942725.4;refGene_gn=TEX35;refGene_tx
=XM_011291021.3;refGene_gn=TEX35;refGene_tx=XM_019821694.1;refGene_gn=
TEX35;refGene_tx=XM_023247609.1
F1 22326871 22326915
refGene_gn=TEX35;refGene_tx=XM_006942725.4;refGene_gn=TEX35;refGene_tx

=XM_011291021.3;refGene_gn=TEX35;refGene_tx=XM_019821694.1;refGene_gn=TEX35;refGene_tx=XM_023247609.1
F1 22327188 22327252 refGene_gn=TEX35;refGene_tx=XM_006942725.4;refGene_gn=TEX35;refGene_tx=XM_011291021.3;refGene_gn=TEX35;refGene_tx=XM_019821694.1
F1 22329490 22329524 refGene_gn=TEX35;refGene_tx=XM_023247609.1
F1 22535593 22535650 refGene_gn=RALGPS2;refGene_tx=XM_019821695.2;refGene_gn=RALGPS2;refGene_tx=XM_023247645.1;refGene_gn=RALGPS2;refGene_tx=XM_023247646.1;refGene_gn=RALGPS2;refGene_tx=XM_023247647.1;refGene_gn=RALGPS2;refGene_tx=XM_023247648.1;refGene_gn=RALGPS2;refGene_tx=XM_023247649.1;refGene_gn=RALGPS2;refGene_tx=XM_023247650.1
F1 22541905 22542010 refGene_gn=RALGPS2;refGene_tx=XM_019821695.2;refGene_gn=RALGPS2;refGene_tx=XM_023247645.1;refGene_gn=RALGPS2;refGene_tx=XM_023247646.1;refGene_gn=RALGPS2;refGene_tx=XM_023247647.1;refGene_gn=RALGPS2;refGene_tx=XM_023247648.1;refGene_gn=RALGPS2;refGene_tx=XM_023247649.1;refGene_gn=RALGPS2;refGene_tx=XM_023247650.1
F1 22544333 22544384 refGene_gn=RALGPS2;refGene_tx=XM_019821695.2;refGene_gn=RALGPS2;refGene_tx=XM_023247645.1;refGene_gn=RALGPS2;refGene_tx=XM_023247646.1;refGene_gn=RALGPS2;refGene_tx=XM_023247647.1;refGene_gn=RALGPS2;refGene_tx=XM_023247648.1;refGene_gn=RALGPS2;refGene_tx=XM_023247649.1;refGene_gn=RALGPS2;refGene_tx=XM_023247650.1
F1 22559834 22559918 refGene_gn=RALGPS2;refGene_tx=XM_019821695.2;refGene_gn=RALGPS2;refGene_tx=XM_023247645.1;refGene_gn=RALGPS2;refGene_tx=XM_023247646.1;refGene_gn=RALGPS2;refGene_tx=XM_023247647.1;refGene_gn=RALGPS2;refGene_tx=XM_023247648.1;refGene_gn=RALGPS2;refGene_tx=XM_023247649.1;refGene_gn=RALGPS2;refGene_tx=XM_023247650.1
F1 22570353 22570443 refGene_gn=RALGPS2;refGene_tx=XM_019821695.2;refGene_gn=RALGPS2;refGene_tx=XM_023247645.1;refGene_gn=RALGPS2;refGene_tx=XM_023247646.1;refGene_gn=RALGPS2;refGene_tx=XM_023247647.1;refGene_gn=RALGPS2;refGene_tx=XM_023247648.1;refGene_gn=RALGPS2;refGene_tx=XM_023247649.1;refGene_gn=RALGPS2;refGene_tx=XM_023247650.1
F1 22576137 22576230 refGene_gn=RALGPS2;refGene_tx=XM_019821695.2;refGene_gn=RALGPS2;refGene_tx=XM_023247645.1;refGene_gn=RALGPS2;refGene_tx=XM_023247646.1;refGene_gn=RALGPS2;refGene_tx=XM_023247647.1;refGene_gn=RALGPS2;refGene_tx=XM_023247648.1;refGene_gn=RALGPS2;refGene_tx=XM_023247649.1;refGene_gn=RALGPS2;refGene_tx=XM_023247650.1
F1 22587719 22587846 refGene_gn=RALGPS2;refGene_tx=XM_019821695.2;refGene_gn=RALGPS2;refGene_tx=XM_023247645.1;refGene_gn=RALGPS2;refGene_tx=XM_023247646.1;refGene_gn=RALGPS2;refGene_tx=XM_023247647.1;refGene_gn=RALGPS2;refGene_tx=XM_023247648.1;refGene_gn=RALGPS2;refGene_tx=XM_023247649.1;refGene_gn=RALGPS2;refGene_tx=XM_023247650.1
F1 22598639 22598827 refGene_gn=ANGPTL1;refGene_tx=XM_003999231.5
F1 22600226 22600497 refGene_gn=ANGPTL1;refGene_tx=XM_003999231.5

F1 22602124 22602318 refGene_gn=ANGPTL1;refGene_tx=XM_003999231.5
F1 22614759 22615605 refGene_gn=ANGPTL1;refGene_tx=XM_003999231.5
F1 22620785 22620795 refGene_gn=ANGPTL1;refGene_tx=XM_003999231.5
F1 22626570 22626708
refGene_gn=RALGPS2;refGene_tx=XM_019821695.2;refGene_gn=RALGPS2;refGene_tx=XM_023247645.1;refGene_gn=RALGPS2;refGene_tx=XM_023247646.1;refGene_gn=RALGPS2;refGene_tx=XM_023247647.1;refGene_gn=RALGPS2;refGene_tx=XM_023247648.1;refGene_gn=RALGPS2;refGene_tx=XM_023247649.1;refGene_gn=RALGPS2;refGene_tx=XM_023247650.1
F1 22627715 22627806
refGene_gn=RALGPS2;refGene_tx=XM_019821695.2;refGene_gn=RALGPS2;refGene_tx=XM_023247645.1;refGene_gn=RALGPS2;refGene_tx=XM_023247646.1;refGene_gn=RALGPS2;refGene_tx=XM_023247647.1;refGene_gn=RALGPS2;refGene_tx=XM_023247648.1;refGene_gn=RALGPS2;refGene_tx=XM_023247649.1;refGene_gn=RALGPS2;refGene_tx=XM_023247650.1
F1 22632437 22632505
refGene_gn=RALGPS2;refGene_tx=XM_019821695.2;refGene_gn=RALGPS2;refGene_tx=XM_023247645.1;refGene_gn=RALGPS2;refGene_tx=XM_023247646.1;refGene_gn=RALGPS2;refGene_tx=XM_023247647.1;refGene_gn=RALGPS2;refGene_tx=XM_023247648.1;refGene_gn=RALGPS2;refGene_tx=XM_023247649.1;refGene_gn=RALGPS2;refGene_tx=XM_023247650.1
F1 22634117 22634253
refGene_gn=RALGPS2;refGene_tx=XM_019821695.2;refGene_gn=RALGPS2;refGene_tx=XM_023247645.1;refGene_gn=RALGPS2;refGene_tx=XM_023247646.1;refGene_gn=RALGPS2;refGene_tx=XM_023247647.1;refGene_gn=RALGPS2;refGene_tx=XM_023247648.1;refGene_gn=RALGPS2;refGene_tx=XM_023247649.1
F1 22635038 22635190
refGene_gn=RALGPS2;refGene_tx=XM_019821695.2;refGene_gn=RALGPS2;refGene_tx=XM_023247645.1;refGene_gn=RALGPS2;refGene_tx=XM_023247646.1;refGene_gn=RALGPS2;refGene_tx=XM_023247647.1;refGene_gn=RALGPS2;refGene_tx=XM_023247648.1;refGene_gn=RALGPS2;refGene_tx=XM_023247649.1;refGene_gn=RALGPS2;refGene_tx=XM_023247650.1
F1 22638489 22638544
refGene_gn=RALGPS2;refGene_tx=XM_019821695.2;refGene_gn=RALGPS2;refGene_tx=XM_023247645.1;refGene_gn=RALGPS2;refGene_tx=XM_023247646.1;refGene_gn=RALGPS2;refGene_tx=XM_023247647.1;refGene_gn=RALGPS2;refGene_tx=XM_023247648.1;refGene_gn=RALGPS2;refGene_tx=XM_023247649.1
F1 22640926 22641004
refGene_gn=RALGPS2;refGene_tx=XM_019821695.2;refGene_gn=RALGPS2;refGene_tx=XM_023247645.1;refGene_gn=RALGPS2;refGene_tx=XM_023247646.1;refGene_gn=RALGPS2;refGene_tx=XM_023247647.1
F1 22642770 22642876
refGene_gn=RALGPS2;refGene_tx=XM_019821695.2;refGene_gn=RALGPS2;refGene_tx=XM_023247645.1;refGene_gn=RALGPS2;refGene_tx=XM_023247646.1;refGene_gn=RALGPS2;refGene_tx=XM_023247647.1;refGene_gn=RALGPS2;refGene_tx=XM_023247648.1;refGene_gn=RALGPS2;refGene_tx=XM_023247649.1
F1 22645910 22646003
refGene_gn=RALGPS2;refGene_tx=XM_019821695.2;refGene_gn=RALGPS2;refGene_tx=XM_023247645.1;refGene_gn=RALGPS2;refGene_tx=XM_023247646.1;refGene_gn=RALGPS2;refGene_tx=XM_023247647.1;refGene_gn=RALGPS2;refGene_tx=

XM_023247648.1;refGene_gn=RALGPS2;refGene_tx=XM_023247649.1
F1 22653337 22653443
refGene_gn=RALGPS2;refGene_tx=XM_019821695.2;refGene_gn=RALGPS2;refGene_tx=XM_023247645.1;refGene_gn=RALGPS2;refGene_tx=XM_023247646.1;refGene_gn=RALGPS2;refGene_tx=XM_023247647.1;refGene_gn=RALGPS2;refGene_tx=XM_023247648.1;refGene_gn=RALGPS2;refGene_tx=XM_023247649.1
F1 22655639 22655731
refGene_gn=RALGPS2;refGene_tx=XM_019821695.2;refGene_gn=RALGPS2;refGene_tx=XM_023247645.1;refGene_gn=RALGPS2;refGene_tx=XM_023247646.1;refGene_gn=RALGPS2;refGene_tx=XM_023247647.1;refGene_gn=RALGPS2;refGene_tx=XM_023247648.1;refGene_gn=RALGPS2;refGene_tx=XM_023247649.1
F1 22661598 22661691
refGene_gn=RALGPS2;refGene_tx=XM_019821695.2;refGene_gn=RALGPS2;refGene_tx=XM_023247645.1;refGene_gn=RALGPS2;refGene_tx=XM_023247646.1;refGene_gn=RALGPS2;refGene_tx=XM_023247648.1
F1 22662124 22662154
refGene_gn=RALGPS2;refGene_tx=XM_023247647.1;refGene_gn=RALGPS2;refGene_tx=XM_023247649.1
F1 22769457 22769834
refGene_gn=FAM20B;refGene_tx=XM_003999232.5;refGene_gn=FAM20B;refGene_tx=XM_006942728.4;refGene_gn=FAM20B;refGene_tx=XM_006942729.4;refGene_gn=FAM20B;refGene_tx=XM_006942730.4;refGene_gn=FAM20B;refGene_tx=XM_006942731.4;refGene_gn=FAM20B;refGene_tx=XM_023247306.1
F1 22775417 22775504
refGene_gn=FAM20B;refGene_tx=XM_003999232.5;refGene_gn=FAM20B;refGene_tx=XM_006942728.4;refGene_gn=FAM20B;refGene_tx=XM_006942729.4;refGene_gn=FAM20B;refGene_tx=XM_006942730.4;refGene_gn=FAM20B;refGene_tx=XM_006942731.4;refGene_gn=FAM20B;refGene_tx=XM_023247306.1
F1 22779348 22779458
refGene_gn=FAM20B;refGene_tx=XM_003999232.5;refGene_gn=FAM20B;refGene_tx=XM_006942728.4;refGene_gn=FAM20B;refGene_tx=XM_006942729.4;refGene_gn=FAM20B;refGene_tx=XM_006942730.4;refGene_gn=FAM20B;refGene_tx=XM_006942731.4;refGene_gn=FAM20B;refGene_tx=XM_023247306.1
F1 22793236 22793408
refGene_gn=FAM20B;refGene_tx=XM_003999232.5;refGene_gn=FAM20B;refGene_tx=XM_006942728.4;refGene_gn=FAM20B;refGene_tx=XM_006942729.4;refGene_gn=FAM20B;refGene_tx=XM_006942730.4;refGene_gn=FAM20B;refGene_tx=XM_006942731.4;refGene_gn=FAM20B;refGene_tx=XM_023247306.1
F1 22793630 22793822
refGene_gn=FAM20B;refGene_tx=XM_003999232.5;refGene_gn=FAM20B;refGene_tx=XM_006942728.4;refGene_gn=FAM20B;refGene_tx=XM_006942729.4;refGene_gn=FAM20B;refGene_tx=XM_006942730.4;refGene_gn=FAM20B;refGene_tx=XM_006942731.4;refGene_gn=FAM20B;refGene_tx=XM_023247306.1
F1 22795164 22795224
refGene_gn=FAM20B;refGene_tx=XM_003999232.5;refGene_gn=FAM20B;refGene_tx=XM_006942728.4;refGene_gn=FAM20B;refGene_tx=XM_006942729.4;refGene_gn=FAM20B;refGene_tx=XM_006942730.4;refGene_gn=FAM20B;refGene_tx=XM_006942731.4;refGene_gn=FAM20B;refGene_tx=XM_023247306.1
F1 22796495 22796727
refGene_gn=FAM20B;refGene_tx=XM_003999232.5;refGene_gn=FAM20B;refGene_

tx=XM_006942728.4;refGene_gn=FAM20B;refGene_tx=XM_006942729.4;refGene_gn=FAM20B;refGene_tx=XM_006942730.4;refGene_gn=FAM20B;refGene_tx=XM_006942731.4;refGene_gn=FAM20B;refGene_tx=XM_023247306.1
F1 22803079 22803782 refGene_gn=TOR3A;refGene_tx=XM_023247305.1
F1 22804752 22804866 refGene_gn=TOR3A;refGene_tx=XM_023247305.1
F1 22805956 22806222 refGene_gn=TOR3A;refGene_tx=XM_023247305.1
F1 22807756 22807935 refGene_gn=TOR3A;refGene_tx=XM_023247305.1
F1 22815082 22815207 refGene_gn=TOR3A;refGene_tx=XM_023247305.1
F1 22815664 22815915 refGene_gn=TOR3A;refGene_tx=XM_023247305.1
F1 22826548 22828272 refGene_gn=ABL2;refGene_tx=XM_023247307.1
F1 22828823 22828997 refGene_gn=ABL2;refGene_tx=XM_023247307.1
F1 22831777 22831867 refGene_gn=ABL2;refGene_tx=XM_023247307.1
F1 22833605 22833758 refGene_gn=ABL2;refGene_tx=XM_023247307.1
F1 22835056 22835241 refGene_gn=ABL2;refGene_tx=XM_023247307.1
F1 22835866 22836044 refGene_gn=ABL2;refGene_tx=XM_023247307.1
F1 22838132 22838217 refGene_gn=ABL2;refGene_tx=XM_023247307.1
F1 22839314 22839587 refGene_gn=ABL2;refGene_tx=XM_023247307.1
F1 22842186 22842494 refGene_gn=ABL2;refGene_tx=XM_023247307.1
F1 22847233 22847404 refGene_gn=ABL2;refGene_tx=XM_023247307.1
F1 22848627 22848690 refGene_gn=ABL2;refGene_tx=XM_023247307.1
F1 22932582 22932739 refGene_gn=ABL2;refGene_tx=XM_023247307.1
F1 23013815 23013933
refGene_gn=SOAT1;refGene_tx=XM_011291017.3;refGene_gn=SOAT1;refGene_tx=XM_011291018.3
F1 23042646 23042705
refGene_gn=SOAT1;refGene_tx=XM_011291017.3;refGene_gn=SOAT1;refGene_tx=XM_011291018.3
F1 23059529 23059678
refGene_gn=SOAT1;refGene_tx=XM_011291017.3;refGene_gn=SOAT1;refGene_tx=XM_011291018.3
F1 23060527 23060587
refGene_gn=SOAT1;refGene_tx=XM_011291017.3;refGene_gn=SOAT1;refGene_tx=XM_011291018.3
F1 23063178 23063286
refGene_gn=SOAT1;refGene_tx=XM_011291017.3;refGene_gn=SOAT1;refGene_tx=XM_011291018.3
F1 23064167 23064450
refGene_gn=SOAT1;refGene_tx=XM_011291017.3;refGene_gn=SOAT1;refGene_tx=XM_011291018.3
F1 23065161 23065240
refGene_gn=SOAT1;refGene_tx=XM_011291017.3;refGene_gn=SOAT1;refGene_tx=XM_011291018.3
F1 23065751 23065833
refGene_gn=SOAT1;refGene_tx=XM_011291017.3;refGene_gn=SOAT1;refGene_tx=XM_011291018.3
F1 23066111 23066157
refGene_gn=SOAT1;refGene_tx=XM_011291017.3;refGene_gn=SOAT1;refGene_tx=XM_011291018.3
F1 23066819 23066949
refGene_gn=SOAT1;refGene_tx=XM_011291017.3;refGene_gn=SOAT1;refGene_tx

=XM_011291018.3
F1	23069452	23069550
refGene_gn=SOAT1;refGene_tx=XM_011291017.3;refGene_gn=SOAT1;refGene_tx
=XM_011291018.3
F1	23070707	23070806
refGene_gn=SOAT1;refGene_tx=XM_011291017.3;refGene_gn=SOAT1;refGene_tx
=XM_011291018.3
F1	23072107	23072243
refGene_gn=SOAT1;refGene_tx=XM_011291017.3;refGene_gn=SOAT1;refGene_tx
=XM_011291018.3
F1	23079557	23079703
refGene_gn=SOAT1;refGene_tx=XM_011291017.3;refGene_gn=SOAT1;refGene_tx
=XM_011291018.3
F1	23080916	23080973
refGene_gn=SOAT1;refGene_tx=XM_011291017.3;refGene_gn=SOAT1;refGene_tx
=XM_011291018.3
F1	23091813	23091907	refGene_gn=AXDND1;refGene_tx=XM_023247451.1
F1	23093668	23093838	refGene_gn=AXDND1;refGene_tx=XM_023247451.1
F1	23094656	23094760	refGene_gn=AXDND1;refGene_tx=XM_023247451.1
F1	23101233	23101354	refGene_gn=AXDND1;refGene_tx=XM_023247451.1
F1	23101929	23102015	refGene_gn=AXDND1;refGene_tx=XM_023247451.1
F1	23103948	23104005	refGene_gn=AXDND1;refGene_tx=XM_023247451.1
F1	23104795	23104898	refGene_gn=AXDND1;refGene_tx=XM_023247451.1
F1	23107255	23107377	refGene_gn=AXDND1;refGene_tx=XM_023247451.1
F1	23129936	23130077	refGene_gn=AXDND1;refGene_tx=XM_023247451.1
F1	23130772	23130877	refGene_gn=AXDND1;refGene_tx=XM_023247451.1
F1	23136696	23136817	refGene_gn=AXDND1;refGene_tx=XM_023247451.1
F1	23139242	23139344	refGene_gn=AXDND1;refGene_tx=XM_023247451.1
F1	23140418	23140582	refGene_gn=AXDND1;refGene_tx=XM_023247451.1
F1	23142987	23143063	refGene_gn=AXDND1;refGene_tx=XM_023247451.1
F1	23145184	23145419	refGene_gn=AXDND1;refGene_tx=XM_023247451.1
F1	23148323	23148525	refGene_gn=AXDND1;refGene_tx=XM_023247451.1
F1	23149338	23149432	refGene_gn=AXDND1;refGene_tx=XM_023247451.1
F1	23153819	23153996	refGene_gn=AXDND1;refGene_tx=XM_023247451.1
F1	23155837	23155948	refGene_gn=AXDND1;refGene_tx=XM_023247451.1
F1	23162622	23162736	refGene_gn=AXDND1;refGene_tx=XM_023247451.1
F1	23164209	23164314	refGene_gn=AXDND1;refGene_tx=XM_023247451.1
F1	23170378	23170461	refGene_gn=AXDND1;refGene_tx=XM_023247451.1
F1	23171536	23171790	refGene_gn=AXDND1;refGene_tx=XM_023247451.1
F1	23174434	23174514	refGene_gn=AXDND1;refGene_tx=XM_023247451.1
F1	23183073	23183352	refGene_gn=NPHS2;refGene_tx=XM_023247332.1
F1	23184456	23184535	refGene_gn=NPHS2;refGene_tx=XM_023247332.1
F1	23186033	23186089	refGene_gn=NPHS2;refGene_tx=XM_023247332.1
F1	23188812	23189016	refGene_gn=NPHS2;refGene_tx=XM_023247332.1
F1	23192058	23192141	refGene_gn=NPHS2;refGene_tx=XM_023247332.1
F1	23194187	23194260	refGene_gn=NPHS2;refGene_tx=XM_023247332.1
F1	23198815	23198919	refGene_gn=NPHS2;refGene_tx=XM_023247332.1
F1	23202413	23202672	refGene_gn=NPHS2;refGene_tx=XM_023247332.1
F1	23213660	23213892
refGene_gn=TDRD5;refGene_tx=XM_006942733.4;refGene_gn=TDRD5;refGene_tx

=XM_019821702.2;refGene_gn=TDRD5;refGene_tx=XM_019821705.2;refGene_gn=TDRD5;refGene_tx=XM_023247330.1;refGene_gn=TDRD5;refGene_tx=XM_023247331.1
F1 23214593 23215001
refGene_gn=TDRD5;refGene_tx=XM_006942733.4;refGene_gn=TDRD5;refGene_tx=XM_019821702.2;refGene_gn=TDRD5;refGene_tx=XM_019821705.2;refGene_gn=TDRD5;refGene_tx=XM_023247330.1;refGene_gn=TDRD5;refGene_tx=XM_023247331.1
F1 23217022 23217219
refGene_gn=TDRD5;refGene_tx=XM_006942733.4;refGene_gn=TDRD5;refGene_tx=XM_019821702.2;refGene_gn=TDRD5;refGene_tx=XM_019821705.2;refGene_gn=TDRD5;refGene_tx=XM_023247330.1;refGene_gn=TDRD5;refGene_tx=XM_023247331.1
F1 23233063 23233147
refGene_gn=TDRD5;refGene_tx=XM_006942733.4;refGene_gn=TDRD5;refGene_tx=XM_019821702.2;refGene_gn=TDRD5;refGene_tx=XM_019821705.2;refGene_gn=TDRD5;refGene_tx=XM_023247330.1;refGene_gn=TDRD5;refGene_tx=XM_023247331.1
F1 23235931 23235988
refGene_gn=TDRD5;refGene_tx=XM_006942733.4;refGene_gn=TDRD5;refGene_tx=XM_019821702.2;refGene_gn=TDRD5;refGene_tx=XM_019821705.2;refGene_gn=TDRD5;refGene_tx=XM_023247330.1;refGene_gn=TDRD5;refGene_tx=XM_023247331.1
F1 23257169 23257323
refGene_gn=TDRD5;refGene_tx=XM_006942733.4;refGene_gn=TDRD5;refGene_tx=XM_019821702.2;refGene_gn=TDRD5;refGene_tx=XM_019821705.2;refGene_gn=TDRD5;refGene_tx=XM_023247330.1;refGene_gn=TDRD5;refGene_tx=XM_023247331.1
F1 23267009 23267218
refGene_gn=TDRD5;refGene_tx=XM_006942733.4;refGene_gn=TDRD5;refGene_tx=XM_019821702.2;refGene_gn=TDRD5;refGene_tx=XM_019821705.2;refGene_gn=TDRD5;refGene_tx=XM_023247330.1;refGene_gn=TDRD5;refGene_tx=XM_023247331.1
F1 23271459 23271680
refGene_gn=TDRD5;refGene_tx=XM_006942733.4;refGene_gn=TDRD5;refGene_tx=XM_019821702.2;refGene_gn=TDRD5;refGene_tx=XM_019821705.2;refGene_gn=TDRD5;refGene_tx=XM_023247330.1;refGene_gn=TDRD5;refGene_tx=XM_023247331.1
F1 23276792 23277005
refGene_gn=TDRD5;refGene_tx=XM_006942733.4;refGene_gn=TDRD5;refGene_tx=XM_019821702.2;refGene_gn=TDRD5;refGene_tx=XM_019821705.2;refGene_gn=TDRD5;refGene_tx=XM_023247330.1;refGene_gn=TDRD5;refGene_tx=XM_023247331.1
F1 23277542 23277609
refGene_gn=TDRD5;refGene_tx=XM_006942733.4;refGene_gn=TDRD5;refGene_tx=XM_019821702.2;refGene_gn=TDRD5;refGene_tx=XM_019821705.2;refGene_gn=TDRD5;refGene_tx=XM_023247330.1;refGene_gn=TDRD5;refGene_tx=XM_023247331.1
F1 23281796 23281997
refGene_gn=TDRD5;refGene_tx=XM_006942733.4;refGene_gn=TDRD5;refGene_tx

=XM_019821702.2;refGene_gn=TDRD5;refGene_tx=XM_019821705.2;refGene_gn=TDRD5;refGene_tx=XM_023247330.1;refGene_gn=TDRD5;refGene_tx=XM_023247331.1
F1 23282743 23282902
refGene_gn=TDRD5;refGene_tx=XM_006942733.4;refGene_gn=TDRD5;refGene_tx=XM_019821702.2;refGene_gn=TDRD5;refGene_tx=XM_019821705.2;refGene_gn=TDRD5;refGene_tx=XM_023247330.1;refGene_gn=TDRD5;refGene_tx=XM_023247331.1
F1 23285933 23286095
refGene_gn=TDRD5;refGene_tx=XM_006942733.4;refGene_gn=TDRD5;refGene_tx=XM_019821702.2;refGene_gn=TDRD5;refGene_tx=XM_019821705.2;refGene_gn=TDRD5;refGene_tx=XM_023247330.1
F1 23298304 23298487
refGene_gn=TDRD5;refGene_tx=XM_006942733.4;refGene_gn=TDRD5;refGene_tx=XM_019821702.2;refGene_gn=TDRD5;refGene_tx=XM_019821705.2;refGene_gn=TDRD5;refGene_tx=XM_023247330.1;refGene_gn=TDRD5;refGene_tx=XM_023247331.1
F1 23299249 23299393
refGene_gn=TDRD5;refGene_tx=XM_006942733.4;refGene_gn=TDRD5;refGene_tx=XM_019821702.2;refGene_gn=TDRD5;refGene_tx=XM_019821705.2;refGene_gn=TDRD5;refGene_tx=XM_023247330.1;refGene_gn=TDRD5;refGene_tx=XM_023247331.1
F1 23302220 23302431
refGene_gn=TDRD5;refGene_tx=XM_006942733.4;refGene_gn=TDRD5;refGene_tx=XM_019821702.2;refGene_gn=TDRD5;refGene_tx=XM_019821705.2;refGene_gn=TDRD5;refGene_tx=XM_023247330.1;refGene_gn=TDRD5;refGene_tx=XM_023247331.1
F1 23313974 23314201
refGene_gn=TDRD5;refGene_tx=XM_006942733.4;refGene_gn=TDRD5;refGene_tx=XM_019821702.2;refGene_gn=TDRD5;refGene_tx=XM_019821705.2;refGene_gn=TDRD5;refGene_tx=XM_023247330.1;refGene_gn=TDRD5;refGene_tx=XM_023247331.1
F1 23431059 23431152
refGene_gn=FAM163A;refGene_tx=XM_003999235.5;refGene_gn=FAM163A;refGene_tx=XM_019821709.2;refGene_gn=FAM163A;refGene_tx=XM_023247587.1;refGene_gn=FAM163A;refGene_tx=XM_023247588.1
F1 23431724 23432132
refGene_gn=FAM163A;refGene_tx=XM_003999235.5;refGene_gn=FAM163A;refGene_tx=XM_019821709.2;refGene_gn=FAM163A;refGene_tx=XM_023247587.1;refGene_gn=FAM163A;refGene_tx=XM_023247588.1
F1 23506770 23506828
refGene_gn=LOC101085428;refGene_tx=XM_019821711.2;refGene_gn=LOC101085428;refGene_tx=XM_023247799.1
F1 23509534 23509729
refGene_gn=LOC101085428;refGene_tx=XM_019821711.2;refGene_gn=LOC101085428;refGene_tx=XM_023247799.1
F1 23511213 23511390
refGene_gn=LOC101085428;refGene_tx=XM_019821711.2
F1 23528178 23528361
refGene_gn=LOC101085428;refGene_tx=XM_019821711.2;refGene_gn=LOC101085

428;refGene_tx=XM_023247799.1
F1 23528814 23529023
refGene_gn=LOC101085428;refGene_tx=XM_019821711.2;refGene_gn=LOC101085428;refGene_tx=XM_023247799.1
F1 23539577 23539705
refGene_gn=LOC102899478;refGene_tx=XM_019821712.2
F1 23560012 23560141
refGene_gn=LOC102899478;refGene_tx=XM_019821712.2
F1 23561640 23561986
refGene_gn=LOC102899478;refGene_tx=XM_019821712.2
F1 23604167 23604299
refGene_gn=PLA2G4A;refGene_tx=XM_006942736.3;refGene_gn=PLA2G4A;refGene_tx=XM_006942737.2;refGene_gn=PLA2G4A;refGene_tx=XM_019821715.2;refGene_gn=PLA2G4A;refGene_tx=XM_019821716.1;refGene_gn=PLA2G4A;refGene_tx=XM_019821718.2
F1 23612692 23612850
refGene_gn=PLA2G4A;refGene_tx=XM_006942736.3;refGene_gn=PLA2G4A;refGene_tx=XM_006942737.2;refGene_gn=PLA2G4A;refGene_tx=XM_019821715.2;refGene_gn=PLA2G4A;refGene_tx=XM_019821716.1;refGene_gn=PLA2G4A;refGene_tx=XM_019821718.2
F1 23614390 23614586
refGene_gn=PLA2G4A;refGene_tx=XM_006942736.3;refGene_gn=PLA2G4A;refGene_tx=XM_006942737.2;refGene_gn=PLA2G4A;refGene_tx=XM_019821715.2;refGene_gn=PLA2G4A;refGene_tx=XM_019821716.1;refGene_gn=PLA2G4A;refGene_tx=XM_019821718.2
F1 23625850 23626035
refGene_gn=PLA2G4A;refGene_tx=XM_006942736.3;refGene_gn=PLA2G4A;refGene_tx=XM_006942737.2;refGene_gn=PLA2G4A;refGene_tx=XM_019821715.2;refGene_gn=PLA2G4A;refGene_tx=XM_019821716.1;refGene_gn=PLA2G4A;refGene_tx=XM_019821718.2
F1 23633743 23633986
refGene_gn=PLA2G4A;refGene_tx=XM_006942736.3;refGene_gn=PLA2G4A;refGene_tx=XM_006942737.2;refGene_gn=PLA2G4A;refGene_tx=XM_019821715.2;refGene_gn=PLA2G4A;refGene_tx=XM_019821716.1;refGene_gn=PLA2G4A;refGene_tx=XM_019821718.2
F1 23640984 23641056
refGene_gn=PLA2G4A;refGene_tx=XM_006942736.3;refGene_gn=PLA2G4A;refGene_tx=XM_006942737.2;refGene_gn=PLA2G4A;refGene_tx=XM_019821715.2;refGene_gn=PLA2G4A;refGene_tx=XM_019821716.1;refGene_gn=PLA2G4A;refGene_tx=XM_019821718.2
F1 23644498 23644591
refGene_gn=PLA2G4A;refGene_tx=XM_006942736.3;refGene_gn=PLA2G4A;refGene_tx=XM_006942737.2;refGene_gn=PLA2G4A;refGene_tx=XM_019821715.2;refGene_gn=PLA2G4A;refGene_tx=XM_019821716.1;refGene_gn=PLA2G4A;refGene_tx=XM_019821718.2
F1 23644679 23644817
refGene_gn=PLA2G4A;refGene_tx=XM_006942736.3;refGene_gn=PLA2G4A;refGene_tx=XM_006942737.2;refGene_gn=PLA2G4A;refGene_tx=XM_019821715.2;refGene_gn=PLA2G4A;refGene_tx=XM_019821716.1;refGene_gn=PLA2G4A;refGene_tx=XM_019821718.2

F1 23650796 23650911
refGene_gn=PLA2G4A;refGene_tx=XM_006942736.3;refGene_gn=PLA2G4A;refGene_tx=XM_006942737.2;refGene_gn=PLA2G4A;refGene_tx=XM_019821715.2;refGene_gn=PLA2G4A;refGene_tx=XM_019821716.1;refGene_gn=PLA2G4A;refGene_tx=XM_019821718.2
F1 23651769 23651992
refGene_gn=PLA2G4A;refGene_tx=XM_006942736.3;refGene_gn=PLA2G4A;refGene_tx=XM_006942737.2;refGene_gn=PLA2G4A;refGene_tx=XM_019821715.2;refGene_gn=PLA2G4A;refGene_tx=XM_019821716.1;refGene_gn=PLA2G4A;refGene_tx=XM_019821718.2
F1 23659382 23659519
refGene_gn=PLA2G4A;refGene_tx=XM_006942736.3;refGene_gn=PLA2G4A;refGene_tx=XM_006942737.2;refGene_gn=PLA2G4A;refGene_tx=XM_019821715.2;refGene_gn=PLA2G4A;refGene_tx=XM_019821716.1;refGene_gn=PLA2G4A;refGene_tx=XM_019821718.2
F1 23676024 23676166
refGene_gn=PLA2G4A;refGene_tx=XM_006942736.3;refGene_gn=PLA2G4A;refGene_tx=XM_006942737.2;refGene_gn=PLA2G4A;refGene_tx=XM_019821715.2;refGene_gn=PLA2G4A;refGene_tx=XM_019821716.1;refGene_gn=PLA2G4A;refGene_tx=XM_019821718.2
F1 23681299 23681337
refGene_gn=PLA2G4A;refGene_tx=XM_006942736.3;refGene_gn=PLA2G4A;refGene_tx=XM_006942737.2;refGene_gn=PLA2G4A;refGene_tx=XM_019821715.2;refGene_gn=PLA2G4A;refGene_tx=XM_019821716.1;refGene_gn=PLA2G4A;refGene_tx=XM_019821718.2
F1 23694335 23694449
refGene_gn=PLA2G4A;refGene_tx=XM_006942736.3;refGene_gn=PLA2G4A;refGene_tx=XM_019821715.2;refGene_gn=PLA2G4A;refGene_tx=XM_019821716.1;refGene_gn=PLA2G4A;refGene_tx=XM_019821718.2
F1 23695161 23695310
refGene_gn=PLA2G4A;refGene_tx=XM_006942736.3;refGene_gn=PLA2G4A;refGene_tx=XM_019821715.2;refGene_gn=PLA2G4A;refGene_tx=XM_019821716.1;refGene_gn=PLA2G4A;refGene_tx=XM_019821718.2
F1 23717834 23717916
refGene_gn=PLA2G4A;refGene_tx=XM_006942736.3;refGene_gn=PLA2G4A;refGene_tx=XM_019821715.2;refGene_gn=PLA2G4A;refGene_tx=XM_019821716.1;refGene_gn=PLA2G4A;refGene_tx=XM_019821718.2
F1 23718037 23718046 refGene_gn=PLA2G4A;refGene_tx=XM_019821718.2
F1 23730831 23730864
refGene_gn=PLA2G4A;refGene_tx=XM_006942736.3;refGene_gn=PLA2G4A;refGene_tx=XM_019821715.2;refGene_gn=PLA2G4A;refGene_tx=XM_019821716.1
F1 23931221 23931273 refGene_gn=PTGS2;refGene_tx=NM_001110449.1
F1 23931977 23932094 refGene_gn=PTGS2;refGene_tx=NM_001110449.1
F1 23932218 23932362 refGene_gn=PTGS2;refGene_tx=NM_001110449.1
F1 23933261 23933405 refGene_gn=PTGS2;refGene_tx=NM_001110449.1
F1 23933871 23934053 refGene_gn=PTGS2;refGene_tx=NM_001110449.1
F1 23934730 23934814 refGene_gn=PTGS2;refGene_tx=NM_001110449.1
F1 23934921 23935168 refGene_gn=PTGS2;refGene_tx=NM_001110449.1
F1 23935452 23935739 refGene_gn=PTGS2;refGene_tx=NM_001110449.1
F1 23936308 23936456 refGene_gn=PTGS2;refGene_tx=NM_001110449.1

F1 23936601 23937011 refGene_gn=PTGS2;refGene_tx=NM_001110449.1
F1 24126220 24126281
refGene_gn=PDC;refGene_tx=NM_001043340.1;refGene_gn=PDC;refGene_tx=XM_
019821330.1;refGene_gn=PDC;refGene_tx=XM_019821331.1
F1 24132006 24132158
refGene_gn=PDC;refGene_tx=NM_001043340.1;refGene_gn=PDC;refGene_tx=XM_
019821330.1;refGene_gn=PDC;refGene_tx=XM_019821331.1
F1 24133593 24134118
refGene_gn=PDC;refGene_tx=NM_001043340.1;refGene_gn=PDC;refGene_tx=XM_
019821330.1;refGene_gn=PDC;refGene_tx=XM_019821331.1
F1 24151618 24151686 refGene_gn=ODR4;refGene_tx=XM_023247797.1
F1 24154441 24154552 refGene_gn=ODR4;refGene_tx=XM_023247797.1
F1 24161156 24161342 refGene_gn=ODR4;refGene_tx=XM_023247797.1
F1 24164584 24164675 refGene_gn=ODR4;refGene_tx=XM_023247797.1
F1 24165563 24165692 refGene_gn=ODR4;refGene_tx=XM_023247797.1
F1 24168286 24168355 refGene_gn=ODR4;refGene_tx=XM_023247797.1
F1 24169460 24169556 refGene_gn=ODR4;refGene_tx=XM_023247797.1
F1 24170548 24170689 refGene_gn=ODR4;refGene_tx=XM_023247797.1
F1 24171497 24171534 refGene_gn=ODR4;refGene_tx=XM_023247797.1
F1 24172602 24172709 refGene_gn=ODR4;refGene_tx=XM_023247797.1
F1 24174780 24174876 refGene_gn=ODR4;refGene_tx=XM_023247797.1
F1 24178353 24178491 refGene_gn=ODR4;refGene_tx=XM_023247797.1
F1 24181336 24181435 refGene_gn=ODR4;refGene_tx=XM_023247797.1
F1 24185085 24185236
refGene_gn=TPR;refGene_tx=XM_011291010.3;refGene_gn=TPR;refGene_tx=XM_
019821720.2
F1 24186990 24187095
refGene_gn=TPR;refGene_tx=XM_011291010.3;refGene_gn=TPR;refGene_tx=XM_
019821720.2
F1 24189403 24189477
refGene_gn=TPR;refGene_tx=XM_011291010.3;refGene_gn=TPR;refGene_tx=XM_
019821720.2
F1 24190975 24191072
refGene_gn=TPR;refGene_tx=XM_011291010.3;refGene_gn=TPR;refGene_tx=XM_
019821720.2
F1 24192593 24192697
refGene_gn=TPR;refGene_tx=XM_011291010.3;refGene_gn=TPR;refGene_tx=XM_
019821720.2
F1 24193059 24193224
refGene_gn=TPR;refGene_tx=XM_011291010.3;refGene_gn=TPR;refGene_tx=XM_
019821720.2
F1 24193700 24193793
refGene_gn=TPR;refGene_tx=XM_011291010.3;refGene_gn=TPR;refGene_tx=XM_
019821720.2
F1 24194429 24194510
refGene_gn=TPR;refGene_tx=XM_011291010.3;refGene_gn=TPR;refGene_tx=XM_
019821720.2
F1 24194589 24194677
refGene_gn=TPR;refGene_tx=XM_011291010.3;refGene_gn=TPR;refGene_tx=XM_
019821720.2

F1 24195376 24195517
refGene_gn=TPR;refGene_tx=XM_011291010.3;refGene_gn=TPR;refGene_tx=XM_019821720.2
F1 24195948 24196040
refGene_gn=TPR;refGene_tx=XM_011291010.3;refGene_gn=TPR;refGene_tx=XM_019821720.2
F1 24196312 24196510
refGene_gn=TPR;refGene_tx=XM_011291010.3;refGene_gn=TPR;refGene_tx=XM_019821720.2
F1 24198596 24198704
refGene_gn=TPR;refGene_tx=XM_011291010.3;refGene_gn=TPR;refGene_tx=XM_019821720.2
F1 24199842 24200069
refGene_gn=TPR;refGene_tx=XM_011291010.3;refGene_gn=TPR;refGene_tx=XM_019821720.2
F1 24200753 24200917
refGene_gn=TPR;refGene_tx=XM_011291010.3;refGene_gn=TPR;refGene_tx=XM_019821720.2
F1 24201568 24201702
refGene_gn=TPR;refGene_tx=XM_011291010.3;refGene_gn=TPR;refGene_tx=XM_019821720.2
F1 24201788 24201937
refGene_gn=TPR;refGene_tx=XM_011291010.3;refGene_gn=TPR;refGene_tx=XM_019821720.2
F1 24203185 24203348
refGene_gn=TPR;refGene_tx=XM_011291010.3;refGene_gn=TPR;refGene_tx=XM_019821720.2
F1 24204980 24205115
refGene_gn=TPR;refGene_tx=XM_011291010.3;refGene_gn=TPR;refGene_tx=XM_019821720.2
F1 24205589 24205730
refGene_gn=TPR;refGene_tx=XM_011291010.3;refGene_gn=TPR;refGene_tx=XM_019821720.2
F1 24206650 24206816
refGene_gn=TPR;refGene_tx=XM_011291010.3;refGene_gn=TPR;refGene_tx=XM_019821720.2
F1 24207793 24207960
refGene_gn=TPR;refGene_tx=XM_011291010.3;refGene_gn=TPR;refGene_tx=XM_019821720.2
F1 24208725 24208878
refGene_gn=TPR;refGene_tx=XM_011291010.3;refGene_gn=TPR;refGene_tx=XM_019821720.2
F1 24209334 24209451
refGene_gn=TPR;refGene_tx=XM_011291010.3;refGene_gn=TPR;refGene_tx=XM_019821720.2
F1 24214644 24214848
refGene_gn=TPR;refGene_tx=XM_011291010.3;refGene_gn=TPR;refGene_tx=XM_019821720.2
F1 24215167 24215352
refGene_gn=TPR;refGene_tx=XM_011291010.3;refGene_gn=TPR;refGene_tx=XM_

019821720.2
F1	24216027	24216175
refGene_gn=TPR;refGene_tx=XM_011291010.3;refGene_gn=TPR;refGene_tx=XM_019821720.2
F1	24220029	24220167
refGene_gn=TPR;refGene_tx=XM_011291010.3;refGene_gn=TPR;refGene_tx=XM_019821720.2
F1	24220264	24220396
refGene_gn=TPR;refGene_tx=XM_011291010.3;refGene_gn=TPR;refGene_tx=XM_019821720.2
F1	24221432	24221563
refGene_gn=TPR;refGene_tx=XM_011291010.3;refGene_gn=TPR;refGene_tx=XM_019821720.2
F1	24222507	24222718
refGene_gn=TPR;refGene_tx=XM_011291010.3;refGene_gn=TPR;refGene_tx=XM_019821720.2
F1	24223840	24223984
refGene_gn=TPR;refGene_tx=XM_011291010.3;refGene_gn=TPR;refGene_tx=XM_019821720.2
F1	24224319	24224518
refGene_gn=TPR;refGene_tx=XM_011291010.3;refGene_gn=TPR;refGene_tx=XM_019821720.2
F1	24225417	24225623
refGene_gn=TPR;refGene_tx=XM_011291010.3;refGene_gn=TPR;refGene_tx=XM_019821720.2
F1	24225829	24225891
refGene_gn=TPR;refGene_tx=XM_011291010.3;refGene_gn=TPR;refGene_tx=XM_019821720.2
F1	24228783	24228992
refGene_gn=TPR;refGene_tx=XM_011291010.3;refGene_gn=TPR;refGene_tx=XM_019821720.2
F1	24229774	24230047
refGene_gn=TPR;refGene_tx=XM_011291010.3;refGene_gn=TPR;refGene_tx=XM_019821720.2
F1	24230910	24231059
refGene_gn=TPR;refGene_tx=XM_011291010.3;refGene_gn=TPR;refGene_tx=XM_019821720.2
F1	24231692	24231776
refGene_gn=TPR;refGene_tx=XM_011291010.3;refGene_gn=TPR;refGene_tx=XM_019821720.2
F1	24234751	24234949
refGene_gn=TPR;refGene_tx=XM_011291010.3;refGene_gn=TPR;refGene_tx=XM_019821720.2
F1	24236026	24236158
refGene_gn=TPR;refGene_tx=XM_011291010.3;refGene_gn=TPR;refGene_tx=XM_019821720.2
F1	24236349	24236440
refGene_gn=TPR;refGene_tx=XM_011291010.3;refGene_gn=TPR;refGene_tx=XM_019821720.2
F1	24238770	24238955

refGene_gn=TPR;refGene_tx=XM_011291010.3;refGene_gn=TPR;refGene_tx=XM_019821720.2
F1 24240383 24240452
refGene_gn=TPR;refGene_tx=XM_011291010.3;refGene_gn=TPR;refGene_tx=XM_019821720.2
F1 24240565 24240660
refGene_gn=TPR;refGene_tx=XM_011291010.3;refGene_gn=TPR;refGene_tx=XM_019821720.2
F1 24242513 24242620
refGene_gn=TPR;refGene_tx=XM_011291010.3;refGene_gn=TPR;refGene_tx=XM_019821720.2
F1 24243793 24243889
refGene_gn=TPR;refGene_tx=XM_011291010.3;refGene_gn=TPR;refGene_tx=XM_019821720.2
F1 24244015 24244172
refGene_gn=TPR;refGene_tx=XM_011291010.3;refGene_gn=TPR;refGene_tx=XM_019821720.2
F1 24246377 24246496
refGene_gn=TPR;refGene_tx=XM_011291010.3;refGene_gn=TPR;refGene_tx=XM_019821720.2
F1 24249053 24249149
refGene_gn=TPR;refGene_tx=XM_011291010.3;refGene_gn=TPR;refGene_tx=XM_019821720.2
F1 24249760 24249816
refGene_gn=TPR;refGene_tx=XM_011291010.3;refGene_gn=TPR;refGene_tx=XM_019821720.2
F1 24250015 24250113 refGene_gn=PRG4;refGene_tx=XM_023247798.1
F1 24250869 24250995 refGene_gn=PRG4;refGene_tx=XM_023247798.1
F1 24251455 24251653 refGene_gn=PRG4;refGene_tx=XM_023247798.1
F1 24252125 24252282 refGene_gn=PRG4;refGene_tx=XM_023247798.1
F1 24252560 24252697 refGene_gn=PRG4;refGene_tx=XM_023247798.1
F1 24253530 24253608 refGene_gn=PRG4;refGene_tx=XM_023247798.1
F1 24254287 24255129 refGene_gn=PRG4;refGene_tx=XM_023247798.1
F1 24255321 24255914 refGene_gn=PRG4;refGene_tx=XM_023247798.1
F1 24256468 24256995 refGene_gn=PRG4;refGene_tx=XM_023247798.1
F1 24257172 24257945 refGene_gn=PRG4;refGene_tx=XM_023247798.1
F1 24259427 24259550 refGene_gn=PRG4;refGene_tx=XM_023247798.1
F1 24260062 24260209 refGene_gn=PRG4;refGene_tx=XM_023247798.1
F1 24262774 24262894 refGene_gn=PRG4;refGene_tx=XM_023247798.1
F1 24264149 24264272 refGene_gn=PRG4;refGene_tx=XM_023247798.1
F1 24266830 24266903 refGene_gn=PRG4;refGene_tx=XM_023247798.1
F1 24383075 24383442
refGene_gn=HMCN1;refGene_tx=XM_003999241.4;refGene_gn=HMCN1;refGene_tx=XM_019821722.2;refGene_gn=HMCN1;refGene_tx=XM_019821723.2;refGene_gn=HMCN1;refGene_tx=XM_023247312.1;refGene_gn=HMCN1;refGene_tx=XM_023247313.1
F1 24385183 24385310
refGene_gn=HMCN1;refGene_tx=XM_003999241.4;refGene_gn=HMCN1;refGene_tx=XM_019821722.2;refGene_gn=HMCN1;refGene_tx=XM_019821723.2;refGene_gn=HMCN1;refGene_tx=XM_023247312.1;refGene_gn=HMCN1;refGene_tx=XM_0232473

13.1
F1 24391749 24391869
refGene_gn=HMCN1;refGene_tx=XM_003999241.4;refGene_gn=HMCN1;refGene_tx=XM_019821722.2;refGene_gn=HMCN1;refGene_tx=XM_019821723.2;refGene_gn=HMCN1;refGene_tx=XM_023247312.1;refGene_gn=HMCN1;refGene_tx=XM_023247313.1
F1 24395720 24396071
refGene_gn=HMCN1;refGene_tx=XM_003999241.4;refGene_gn=HMCN1;refGene_tx=XM_019821723.2;refGene_gn=HMCN1;refGene_tx=XM_023247312.1;refGene_gn=HMCN1;refGene_tx=XM_023247313.1
F1 24397942 24398071
refGene_gn=HMCN1;refGene_tx=XM_003999241.4;refGene_gn=HMCN1;refGene_tx=XM_019821722.2;refGene_gn=HMCN1;refGene_tx=XM_019821723.2;refGene_gn=HMCN1;refGene_tx=XM_023247312.1;refGene_gn=HMCN1;refGene_tx=XM_023247313.1
F1 24401154 24401280
refGene_gn=HMCN1;refGene_tx=XM_003999241.4;refGene_gn=HMCN1;refGene_tx=XM_019821722.2;refGene_gn=HMCN1;refGene_tx=XM_019821723.2;refGene_gn=HMCN1;refGene_tx=XM_023247312.1;refGene_gn=HMCN1;refGene_tx=XM_023247313.1
F1 24401783 24401897
refGene_gn=HMCN1;refGene_tx=XM_003999241.4;refGene_gn=HMCN1;refGene_tx=XM_019821722.2;refGene_gn=HMCN1;refGene_tx=XM_019821723.2;refGene_gn=HMCN1;refGene_tx=XM_023247312.1;refGene_gn=HMCN1;refGene_tx=XM_023247313.1
F1 24404351 24404486
refGene_gn=HMCN1;refGene_tx=XM_003999241.4;refGene_gn=HMCN1;refGene_tx=XM_019821722.2;refGene_gn=HMCN1;refGene_tx=XM_019821723.2;refGene_gn=HMCN1;refGene_tx=XM_023247312.1;refGene_gn=HMCN1;refGene_tx=XM_023247313.1
F1 24405655 24405775
refGene_gn=HMCN1;refGene_tx=XM_003999241.4;refGene_gn=HMCN1;refGene_tx=XM_019821722.2;refGene_gn=HMCN1;refGene_tx=XM_019821723.2;refGene_gn=HMCN1;refGene_tx=XM_023247312.1;refGene_gn=HMCN1;refGene_tx=XM_023247313.1
F1 24406810 24406873
refGene_gn=HMCN1;refGene_tx=XM_003999241.4;refGene_gn=HMCN1;refGene_tx=XM_019821722.2;refGene_gn=HMCN1;refGene_tx=XM_019821723.2;refGene_gn=HMCN1;refGene_tx=XM_023247312.1;refGene_gn=HMCN1;refGene_tx=XM_023247313.1
F1 24411900 24412138
refGene_gn=HMCN1;refGene_tx=XM_003999241.4;refGene_gn=HMCN1;refGene_tx=XM_019821722.2;refGene_gn=HMCN1;refGene_tx=XM_019821723.2;refGene_gn=HMCN1;refGene_tx=XM_023247312.1;refGene_gn=HMCN1;refGene_tx=XM_023247313.1
F1 24412817 24412939
refGene_gn=HMCN1;refGene_tx=XM_003999241.4;refGene_gn=HMCN1;refGene_tx=XM_019821722.2;refGene_gn=HMCN1;refGene_tx=XM_019821723.2;refGene_gn=HMCN1;refGene_tx=XM_023247312.1;refGene_gn=HMCN1;refGene_tx=XM_023247313.1

F1 24413975 24414113
refGene_gn=HMCN1;refGene_tx=XM_003999241.4;refGene_gn=HMCN1;refGene_tx=XM_019821722.2;refGene_gn=HMCN1;refGene_tx=XM_019821723.2;refGene_gn=HMCN1;refGene_tx=XM_023247312.1;refGene_gn=HMCN1;refGene_tx=XM_023247313.1
F1 24414386 24414536
refGene_gn=HMCN1;refGene_tx=XM_003999241.4;refGene_gn=HMCN1;refGene_tx=XM_019821722.2;refGene_gn=HMCN1;refGene_tx=XM_019821723.2;refGene_gn=HMCN1;refGene_tx=XM_023247312.1;refGene_gn=HMCN1;refGene_tx=XM_023247313.1
F1 24418189 24418360
refGene_gn=HMCN1;refGene_tx=XM_003999241.4;refGene_gn=HMCN1;refGene_tx=XM_019821722.2;refGene_gn=HMCN1;refGene_tx=XM_019821723.2;refGene_gn=HMCN1;refGene_tx=XM_023247312.1;refGene_gn=HMCN1;refGene_tx=XM_023247313.1
F1 24418531 24418702
refGene_gn=HMCN1;refGene_tx=XM_003999241.4;refGene_gn=HMCN1;refGene_tx=XM_019821722.2;refGene_gn=HMCN1;refGene_tx=XM_019821723.2;refGene_gn=HMCN1;refGene_tx=XM_019821724.2;refGene_gn=HMCN1;refGene_tx=XM_023247312.1;refGene_gn=HMCN1;refGene_tx=XM_023247313.1
F1 24419563 24419750
refGene_gn=HMCN1;refGene_tx=XM_003999241.4;refGene_gn=HMCN1;refGene_tx=XM_019821722.2;refGene_gn=HMCN1;refGene_tx=XM_019821723.2;refGene_gn=HMCN1;refGene_tx=XM_019821724.2;refGene_gn=HMCN1;refGene_tx=XM_023247312.1;refGene_gn=HMCN1;refGene_tx=XM_023247313.1
F1 24420521 24420692
refGene_gn=HMCN1;refGene_tx=XM_003999241.4;refGene_gn=HMCN1;refGene_tx=XM_019821722.2;refGene_gn=HMCN1;refGene_tx=XM_019821723.2;refGene_gn=HMCN1;refGene_tx=XM_019821724.2;refGene_gn=HMCN1;refGene_tx=XM_023247312.1;refGene_gn=HMCN1;refGene_tx=XM_023247313.1
F1 24423950 24424121
refGene_gn=HMCN1;refGene_tx=XM_003999241.4;refGene_gn=HMCN1;refGene_tx=XM_019821722.2;refGene_gn=HMCN1;refGene_tx=XM_019821723.2;refGene_gn=HMCN1;refGene_tx=XM_019821724.2;refGene_gn=HMCN1;refGene_tx=XM_023247312.1;refGene_gn=HMCN1;refGene_tx=XM_023247313.1
F1 24424225 24424396
refGene_gn=HMCN1;refGene_tx=XM_003999241.4;refGene_gn=HMCN1;refGene_tx=XM_019821722.2;refGene_gn=HMCN1;refGene_tx=XM_019821723.2;refGene_gn=HMCN1;refGene_tx=XM_019821724.2;refGene_gn=HMCN1;refGene_tx=XM_023247312.1;refGene_gn=HMCN1;refGene_tx=XM_023247313.1
F1 24424957 24425227
refGene_gn=HMCN1;refGene_tx=XM_003999241.4;refGene_gn=HMCN1;refGene_tx=XM_019821722.2;refGene_gn=HMCN1;refGene_tx=XM_019821723.2;refGene_gn=HMCN1;refGene_tx=XM_019821724.2;refGene_gn=HMCN1;refGene_tx=XM_023247312.1;refGene_gn=HMCN1;refGene_tx=XM_023247313.1
F1 24429424 24429506
refGene_gn=HMCN1;refGene_tx=XM_003999241.4;refGene_gn=HMCN1;refGene_tx=XM_019821722.2;refGene_gn=HMCN1;refGene_tx=XM_019821723.2;refGene_gn=HMCN1;refGene_tx=XM_019821724.2;refGene_gn=HMCN1;refGene_tx=XM_023247312.1;refGene_gn=HMCN1;refGene_tx=XM_023247313.1

F1 24431195 24431386
refGene_gn=HMCN1;refGene_tx=XM_003999241.4;refGene_gn=HMCN1;refGene_tx=XM_019821722.2;refGene_gn=HMCN1;refGene_tx=XM_019821723.2;refGene_gn=HMCN1;refGene_tx=XM_019821724.2;refGene_gn=HMCN1;refGene_tx=XM_023247312.1;refGene_gn=HMCN1;refGene_tx=XM_023247313.1
F1 24431775 24431910
refGene_gn=HMCN1;refGene_tx=XM_003999241.4;refGene_gn=HMCN1;refGene_tx=XM_019821722.2;refGene_gn=HMCN1;refGene_tx=XM_019821723.2;refGene_gn=HMCN1;refGene_tx=XM_019821724.2;refGene_gn=HMCN1;refGene_tx=XM_023247312.1;refGene_gn=HMCN1;refGene_tx=XM_023247313.1
F1 24434035 24434249
refGene_gn=HMCN1;refGene_tx=XM_003999241.4;refGene_gn=HMCN1;refGene_tx=XM_019821722.2;refGene_gn=HMCN1;refGene_tx=XM_019821723.2;refGene_gn=HMCN1;refGene_tx=XM_019821724.2;refGene_gn=HMCN1;refGene_tx=XM_023247312.1;refGene_gn=HMCN1;refGene_tx=XM_023247313.1
F1 24438267 24438458
refGene_gn=HMCN1;refGene_tx=XM_003999241.4;refGene_gn=HMCN1;refGene_tx=XM_019821722.2;refGene_gn=HMCN1;refGene_tx=XM_019821723.2;refGene_gn=HMCN1;refGene_tx=XM_019821724.2;refGene_gn=HMCN1;refGene_tx=XM_023247312.1;refGene_gn=HMCN1;refGene_tx=XM_023247313.1
F1 24440517 24440787
refGene_gn=HMCN1;refGene_tx=XM_003999241.4;refGene_gn=HMCN1;refGene_tx=XM_019821722.2;refGene_gn=HMCN1;refGene_tx=XM_019821723.2;refGene_gn=HMCN1;refGene_tx=XM_019821724.2;refGene_gn=HMCN1;refGene_tx=XM_023247312.1;refGene_gn=HMCN1;refGene_tx=XM_023247313.1
F1 24444100 24444235
refGene_gn=HMCN1;refGene_tx=XM_003999241.4;refGene_gn=HMCN1;refGene_tx=XM_019821722.2;refGene_gn=HMCN1;refGene_tx=XM_019821723.2;refGene_gn=HMCN1;refGene_tx=XM_019821724.2;refGene_gn=HMCN1;refGene_tx=XM_023247312.1;refGene_gn=HMCN1;refGene_tx=XM_023247313.1
F1 24444368 24444506
refGene_gn=HMCN1;refGene_tx=XM_003999241.4;refGene_gn=HMCN1;refGene_tx=XM_019821722.2;refGene_gn=HMCN1;refGene_tx=XM_019821723.2;refGene_gn=HMCN1;refGene_tx=XM_019821724.2;refGene_gn=HMCN1;refGene_tx=XM_023247312.1;refGene_gn=HMCN1;refGene_tx=XM_023247313.1
F1 24445170 24445278
refGene_gn=HMCN1;refGene_tx=XM_003999241.4;refGene_gn=HMCN1;refGene_tx=XM_019821722.2;refGene_gn=HMCN1;refGene_tx=XM_019821723.2;refGene_gn=HMCN1;refGene_tx=XM_019821724.2;refGene_gn=HMCN1;refGene_tx=XM_023247312.1;refGene_gn=HMCN1;refGene_tx=XM_023247313.1
F1 24446878 24447043
refGene_gn=HMCN1;refGene_tx=XM_003999241.4;refGene_gn=HMCN1;refGene_tx=XM_019821722.2;refGene_gn=HMCN1;refGene_tx=XM_019821723.2;refGene_gn=HMCN1;refGene_tx=XM_019821724.2;refGene_gn=HMCN1;refGene_tx=XM_023247312.1;refGene_gn=HMCN1;refGene_tx=XM_023247313.1
F1 24447241 24447363
refGene_gn=HMCN1;refGene_tx=XM_003999241.4;refGene_gn=HMCN1;refGene_tx=XM_019821722.2;refGene_gn=HMCN1;refGene_tx=XM_019821723.2;refGene_gn=HMCN1;refGene_tx=XM_019821724.2;refGene_gn=HMCN1;refGene_tx=XM_023247312.1;refGene_gn=HMCN1;refGene_tx=XM_023247313.1

F1 24449446 24449603
refGene_gn=HMCN1;refGene_tx=XM_003999241.4;refGene_gn=HMCN1;refGene_tx=XM_019821722.2;refGene_gn=HMCN1;refGene_tx=XM_019821723.2;refGene_gn=HMCN1;refGene_tx=XM_019821724.2;refGene_gn=HMCN1;refGene_tx=XM_023247312.1;refGene_gn=HMCN1;refGene_tx=XM_023247313.1
F1 24449916 24450044
refGene_gn=HMCN1;refGene_tx=XM_003999241.4;refGene_gn=HMCN1;refGene_tx=XM_019821722.2;refGene_gn=HMCN1;refGene_tx=XM_019821723.2;refGene_gn=HMCN1;refGene_tx=XM_019821724.2;refGene_gn=HMCN1;refGene_tx=XM_023247312.1;refGene_gn=HMCN1;refGene_tx=XM_023247313.1
F1 24450440 24450585
refGene_gn=HMCN1;refGene_tx=XM_003999241.4;refGene_gn=HMCN1;refGene_tx=XM_019821722.2;refGene_gn=HMCN1;refGene_tx=XM_019821723.2;refGene_gn=HMCN1;refGene_tx=XM_019821724.2;refGene_gn=HMCN1;refGene_tx=XM_023247312.1;refGene_gn=HMCN1;refGene_tx=XM_023247313.1
F1 24451599 24451741
refGene_gn=HMCN1;refGene_tx=XM_003999241.4;refGene_gn=HMCN1;refGene_tx=XM_019821722.2;refGene_gn=HMCN1;refGene_tx=XM_019821723.2;refGene_gn=HMCN1;refGene_tx=XM_019821724.2;refGene_gn=HMCN1;refGene_tx=XM_023247312.1;refGene_gn=HMCN1;refGene_tx=XM_023247313.1
F1 24455706 24455843
refGene_gn=HMCN1;refGene_tx=XM_003999241.4;refGene_gn=HMCN1;refGene_tx=XM_019821722.2;refGene_gn=HMCN1;refGene_tx=XM_019821723.2;refGene_gn=HMCN1;refGene_tx=XM_019821724.2;refGene_gn=HMCN1;refGene_tx=XM_023247312.1;refGene_gn=HMCN1;refGene_tx=XM_023247313.1
F1 24457377 24457459
refGene_gn=HMCN1;refGene_tx=XM_003999241.4;refGene_gn=HMCN1;refGene_tx=XM_019821722.2;refGene_gn=HMCN1;refGene_tx=XM_019821723.2;refGene_gn=HMCN1;refGene_tx=XM_019821724.2;refGene_gn=HMCN1;refGene_tx=XM_023247312.1;refGene_gn=HMCN1;refGene_tx=XM_023247313.1
F1 24460753 24460950
refGene_gn=HMCN1;refGene_tx=XM_003999241.4;refGene_gn=HMCN1;refGene_tx=XM_019821722.2;refGene_gn=HMCN1;refGene_tx=XM_019821723.2;refGene_gn=HMCN1;refGene_tx=XM_019821724.2;refGene_gn=HMCN1;refGene_tx=XM_023247312.1;refGene_gn=HMCN1;refGene_tx=XM_023247313.1
F1 24466290 24466569
refGene_gn=HMCN1;refGene_tx=XM_003999241.4;refGene_gn=HMCN1;refGene_tx=XM_019821722.2;refGene_gn=HMCN1;refGene_tx=XM_019821723.2;refGene_gn=HMCN1;refGene_tx=XM_019821724.2;refGene_gn=HMCN1;refGene_tx=XM_023247312.1;refGene_gn=HMCN1;refGene_tx=XM_023247313.1
F1 24467466 24467564
refGene_gn=HMCN1;refGene_tx=XM_003999241.4;refGene_gn=HMCN1;refGene_tx=XM_019821722.2;refGene_gn=HMCN1;refGene_tx=XM_019821723.2;refGene_gn=HMCN1;refGene_tx=XM_019821724.2;refGene_gn=HMCN1;refGene_tx=XM_023247312.1;refGene_gn=HMCN1;refGene_tx=XM_023247313.1
F1 24468153 24468337
refGene_gn=HMCN1;refGene_tx=XM_003999241.4;refGene_gn=HMCN1;refGene_tx=XM_019821722.2;refGene_gn=HMCN1;refGene_tx=XM_019821723.2;refGene_gn=HMCN1;refGene_tx=XM_019821724.2;refGene_gn=HMCN1;refGene_tx=XM_023247312.1;refGene_gn=HMCN1;refGene_tx=XM_023247313.1

F1	24468565 24468690
refGene_gn=HMCN1;refGene_tx=XM_003999241.4;refGene_gn=HMCN1;refGene_tx=XM_019821722.2;refGene_gn=HMCN1;refGene_tx=XM_019821723.2;refGene_gn=HMCN1;refGene_tx=XM_019821724.2;refGene_gn=HMCN1;refGene_tx=XM_023247312.1;refGene_gn=HMCN1;refGene_tx=XM_023247313.1
F1	24470751 24470911
refGene_gn=HMCN1;refGene_tx=XM_003999241.4;refGene_gn=HMCN1;refGene_tx=XM_019821722.2;refGene_gn=HMCN1;refGene_tx=XM_019821723.2;refGene_gn=HMCN1;refGene_tx=XM_019821724.2;refGene_gn=HMCN1;refGene_tx=XM_023247312.1;refGene_gn=HMCN1;refGene_tx=XM_023247313.1
F1	24473302 24473452
refGene_gn=HMCN1;refGene_tx=XM_003999241.4;refGene_gn=HMCN1;refGene_tx=XM_019821722.2;refGene_gn=HMCN1;refGene_tx=XM_019821723.2;refGene_gn=HMCN1;refGene_tx=XM_019821724.2;refGene_gn=HMCN1;refGene_tx=XM_023247312.1;refGene_gn=HMCN1;refGene_tx=XM_023247313.1
F1	24473791 24473923
refGene_gn=HMCN1;refGene_tx=XM_003999241.4;refGene_gn=HMCN1;refGene_tx=XM_019821722.2;refGene_gn=HMCN1;refGene_tx=XM_019821723.2;refGene_gn=HMCN1;refGene_tx=XM_019821724.2;refGene_gn=HMCN1;refGene_tx=XM_023247312.1;refGene_gn=HMCN1;refGene_tx=XM_023247313.1
F1	24474030 24474112
refGene_gn=HMCN1;refGene_tx=XM_003999241.4;refGene_gn=HMCN1;refGene_tx=XM_019821722.2;refGene_gn=HMCN1;refGene_tx=XM_019821723.2;refGene_gn=HMCN1;refGene_tx=XM_019821724.2;refGene_gn=HMCN1;refGene_tx=XM_023247312.1;refGene_gn=HMCN1;refGene_tx=XM_023247313.1
F1	24474750 24474953
refGene_gn=HMCN1;refGene_tx=XM_003999241.4;refGene_gn=HMCN1;refGene_tx=XM_019821722.2;refGene_gn=HMCN1;refGene_tx=XM_019821723.2;refGene_gn=HMCN1;refGene_tx=XM_019821724.2;refGene_gn=HMCN1;refGene_tx=XM_023247312.1;refGene_gn=HMCN1;refGene_tx=XM_023247313.1
F1	24475074 24475188
refGene_gn=HMCN1;refGene_tx=XM_003999241.4;refGene_gn=HMCN1;refGene_tx=XM_019821722.2;refGene_gn=HMCN1;refGene_tx=XM_019821723.2;refGene_gn=HMCN1;refGene_tx=XM_019821724.2;refGene_gn=HMCN1;refGene_tx=XM_023247312.1;refGene_gn=HMCN1;refGene_tx=XM_023247313.1
F1	24476323 24476485
refGene_gn=HMCN1;refGene_tx=XM_003999241.4;refGene_gn=HMCN1;refGene_tx=XM_019821722.2;refGene_gn=HMCN1;refGene_tx=XM_019821723.2;refGene_gn=HMCN1;refGene_tx=XM_019821724.2;refGene_gn=HMCN1;refGene_tx=XM_023247312.1;refGene_gn=HMCN1;refGene_tx=XM_023247313.1
F1	24482389 24482486
refGene_gn=HMCN1;refGene_tx=XM_003999241.4;refGene_gn=HMCN1;refGene_tx=XM_019821722.2;refGene_gn=HMCN1;refGene_tx=XM_019821723.2;refGene_gn=HMCN1;refGene_tx=XM_019821724.2;refGene_gn=HMCN1;refGene_tx=XM_023247312.1;refGene_gn=HMCN1;refGene_tx=XM_023247313.1
F1	24483916 24484104
refGene_gn=HMCN1;refGene_tx=XM_003999241.4;refGene_gn=HMCN1;refGene_tx=XM_019821722.2;refGene_gn=HMCN1;refGene_tx=XM_019821723.2;refGene_gn=HMCN1;refGene_tx=XM_019821724.2;refGene_gn=HMCN1;refGene_tx=XM_023247312.1;refGene_gn=HMCN1;refGene_tx=XM_023247313.1

F1 24488291 24488405
refGene_gn=HMCN1;refGene_tx=XM_003999241.4;refGene_gn=HMCN1;refGene_tx=XM_019821722.2;refGene_gn=HMCN1;refGene_tx=XM_019821723.2;refGene_gn=HMCN1;refGene_tx=XM_019821724.2;refGene_gn=HMCN1;refGene_tx=XM_023247312.1;refGene_gn=HMCN1;refGene_tx=XM_023247313.1
F1 24489902 24490097
refGene_gn=HMCN1;refGene_tx=XM_003999241.4;refGene_gn=HMCN1;refGene_tx=XM_019821722.2;refGene_gn=HMCN1;refGene_tx=XM_019821723.2;refGene_gn=HMCN1;refGene_tx=XM_019821724.2;refGene_gn=HMCN1;refGene_tx=XM_023247312.1;refGene_gn=HMCN1;refGene_tx=XM_023247313.1
F1 24492105 24492256
refGene_gn=HMCN1;refGene_tx=XM_003999241.4;refGene_gn=HMCN1;refGene_tx=XM_019821722.2;refGene_gn=HMCN1;refGene_tx=XM_019821723.2;refGene_gn=HMCN1;refGene_tx=XM_019821724.2;refGene_gn=HMCN1;refGene_tx=XM_023247312.1;refGene_gn=HMCN1;refGene_tx=XM_023247313.1
F1 24496101 24496247
refGene_gn=HMCN1;refGene_tx=XM_003999241.4;refGene_gn=HMCN1;refGene_tx=XM_019821722.2;refGene_gn=HMCN1;refGene_tx=XM_019821723.2;refGene_gn=HMCN1;refGene_tx=XM_019821724.2;refGene_gn=HMCN1;refGene_tx=XM_023247312.1;refGene_gn=HMCN1;refGene_tx=XM_023247313.1
F1 24497354 24497468
refGene_gn=HMCN1;refGene_tx=XM_003999241.4;refGene_gn=HMCN1;refGene_tx=XM_019821722.2;refGene_gn=HMCN1;refGene_tx=XM_019821723.2;refGene_gn=HMCN1;refGene_tx=XM_019821724.2;refGene_gn=HMCN1;refGene_tx=XM_023247312.1;refGene_gn=HMCN1;refGene_tx=XM_023247313.1
F1 24498795 24498969
refGene_gn=HMCN1;refGene_tx=XM_003999241.4;refGene_gn=HMCN1;refGene_tx=XM_019821722.2;refGene_gn=HMCN1;refGene_tx=XM_019821723.2;refGene_gn=HMCN1;refGene_tx=XM_019821724.2;refGene_gn=HMCN1;refGene_tx=XM_023247312.1;refGene_gn=HMCN1;refGene_tx=XM_023247313.1
F1 24502071 24502263
refGene_gn=HMCN1;refGene_tx=XM_003999241.4;refGene_gn=HMCN1;refGene_tx=XM_019821722.2;refGene_gn=HMCN1;refGene_tx=XM_019821723.2;refGene_gn=HMCN1;refGene_tx=XM_019821724.2;refGene_gn=HMCN1;refGene_tx=XM_023247312.1;refGene_gn=HMCN1;refGene_tx=XM_023247313.1
F1 24513899 24513986
refGene_gn=HMCN1;refGene_tx=XM_003999241.4;refGene_gn=HMCN1;refGene_tx=XM_019821722.2;refGene_gn=HMCN1;refGene_tx=XM_019821723.2;refGene_gn=HMCN1;refGene_tx=XM_019821724.2;refGene_gn=HMCN1;refGene_tx=XM_023247312.1;refGene_gn=HMCN1;refGene_tx=XM_023247313.1
F1 24514565 24514679
refGene_gn=HMCN1;refGene_tx=XM_003999241.4;refGene_gn=HMCN1;refGene_tx=XM_019821722.2;refGene_gn=HMCN1;refGene_tx=XM_019821723.2;refGene_gn=HMCN1;refGene_tx=XM_019821724.2;refGene_gn=HMCN1;refGene_tx=XM_023247312.1;refGene_gn=HMCN1;refGene_tx=XM_023247313.1
F1 24519089 24519257
refGene_gn=HMCN1;refGene_tx=XM_003999241.4;refGene_gn=HMCN1;refGene_tx=XM_019821722.2;refGene_gn=HMCN1;refGene_tx=XM_019821723.2;refGene_gn=HMCN1;refGene_tx=XM_019821724.2;refGene_gn=HMCN1;refGene_tx=XM_023247312.1;refGene_gn=HMCN1;refGene_tx=XM_023247313.1

F1 24522342 24522624
refGene_gn=HMCN1;refGene_tx=XM_003999241.4;refGene_gn=HMCN1;refGene_tx=XM_019821722.2;refGene_gn=HMCN1;refGene_tx=XM_019821723.2;refGene_gn=HMCN1;refGene_tx=XM_019821724.2;refGene_gn=HMCN1;refGene_tx=XM_023247312.1;refGene_gn=HMCN1;refGene_tx=XM_023247313.1
F1 24524016 24524178
refGene_gn=HMCN1;refGene_tx=XM_003999241.4;refGene_gn=HMCN1;refGene_tx=XM_019821722.2;refGene_gn=HMCN1;refGene_tx=XM_019821723.2;refGene_gn=HMCN1;refGene_tx=XM_019821724.2;refGene_gn=HMCN1;refGene_tx=XM_023247312.1;refGene_gn=HMCN1;refGene_tx=XM_023247313.1
F1 24525241 24525364
refGene_gn=HMCN1;refGene_tx=XM_003999241.4;refGene_gn=HMCN1;refGene_tx=XM_019821722.2;refGene_gn=HMCN1;refGene_tx=XM_019821723.2;refGene_gn=HMCN1;refGene_tx=XM_019821724.2;refGene_gn=HMCN1;refGene_tx=XM_023247312.1;refGene_gn=HMCN1;refGene_tx=XM_023247313.1
F1 24529241 24529338
refGene_gn=HMCN1;refGene_tx=XM_003999241.4;refGene_gn=HMCN1;refGene_tx=XM_019821722.2;refGene_gn=HMCN1;refGene_tx=XM_019821723.2;refGene_gn=HMCN1;refGene_tx=XM_019821724.2;refGene_gn=HMCN1;refGene_tx=XM_023247312.1;refGene_gn=HMCN1;refGene_tx=XM_023247313.1
F1 24533834 24534010
refGene_gn=HMCN1;refGene_tx=XM_003999241.4;refGene_gn=HMCN1;refGene_tx=XM_019821722.2;refGene_gn=HMCN1;refGene_tx=XM_019821723.2;refGene_gn=HMCN1;refGene_tx=XM_019821724.2;refGene_gn=HMCN1;refGene_tx=XM_023247312.1;refGene_gn=HMCN1;refGene_tx=XM_023247313.1
F1 24538665 24538789
refGene_gn=HMCN1;refGene_tx=XM_003999241.4;refGene_gn=HMCN1;refGene_tx=XM_019821722.2;refGene_gn=HMCN1;refGene_tx=XM_019821723.2;refGene_gn=HMCN1;refGene_tx=XM_019821724.2;refGene_gn=HMCN1;refGene_tx=XM_023247312.1;refGene_gn=HMCN1;refGene_tx=XM_023247313.1
F1 24540870 24541022
refGene_gn=HMCN1;refGene_tx=XM_003999241.4;refGene_gn=HMCN1;refGene_tx=XM_019821722.2;refGene_gn=HMCN1;refGene_tx=XM_019821723.2;refGene_gn=HMCN1;refGene_tx=XM_019821724.2;refGene_gn=HMCN1;refGene_tx=XM_023247312.1;refGene_gn=HMCN1;refGene_tx=XM_023247313.1
F1 24543631 24543808
refGene_gn=HMCN1;refGene_tx=XM_003999241.4;refGene_gn=HMCN1;refGene_tx=XM_019821722.2;refGene_gn=HMCN1;refGene_tx=XM_019821723.2;refGene_gn=HMCN1;refGene_tx=XM_019821724.2;refGene_gn=HMCN1;refGene_tx=XM_023247312.1;refGene_gn=HMCN1;refGene_tx=XM_023247313.1
F1 24544601 24544703
refGene_gn=HMCN1;refGene_tx=XM_003999241.4;refGene_gn=HMCN1;refGene_tx=XM_019821722.2;refGene_gn=HMCN1;refGene_tx=XM_019821723.2;refGene_gn=HMCN1;refGene_tx=XM_019821724.2;refGene_gn=HMCN1;refGene_tx=XM_023247312.1;refGene_gn=HMCN1;refGene_tx=XM_023247313.1
F1 24559666 24559790
refGene_gn=HMCN1;refGene_tx=XM_003999241.4;refGene_gn=HMCN1;refGene_tx=XM_019821722.2;refGene_gn=HMCN1;refGene_tx=XM_019821723.2;refGene_gn=HMCN1;refGene_tx=XM_019821724.2;refGene_gn=HMCN1;refGene_tx=XM_023247312.1;refGene_gn=HMCN1;refGene_tx=XM_023247313.1

F1 24572204 24572359
refGene_gn=HMCN1;refGene_tx=XM_003999241.4;refGene_gn=HMCN1;refGene_tx=XM_019821722.2;refGene_gn=HMCN1;refGene_tx=XM_019821723.2;refGene_gn=HMCN1;refGene_tx=XM_019821724.2;refGene_gn=HMCN1;refGene_tx=XM_023247312.1;refGene_gn=HMCN1;refGene_tx=XM_023247313.1
F1 24573676 24573846
refGene_gn=HMCN1;refGene_tx=XM_003999241.4;refGene_gn=HMCN1;refGene_tx=XM_019821722.2;refGene_gn=HMCN1;refGene_tx=XM_019821723.2;refGene_gn=HMCN1;refGene_tx=XM_019821724.2;refGene_gn=HMCN1;refGene_tx=XM_023247312.1;refGene_gn=HMCN1;refGene_tx=XM_023247313.1
F1 24574840 24574949
refGene_gn=HMCN1;refGene_tx=XM_003999241.4;refGene_gn=HMCN1;refGene_tx=XM_019821722.2;refGene_gn=HMCN1;refGene_tx=XM_019821723.2;refGene_gn=HMCN1;refGene_tx=XM_019821724.2;refGene_gn=HMCN1;refGene_tx=XM_023247312.1;refGene_gn=HMCN1;refGene_tx=XM_023247313.1
F1 24575673 24575955
refGene_gn=HMCN1;refGene_tx=XM_003999241.4;refGene_gn=HMCN1;refGene_tx=XM_019821722.2;refGene_gn=HMCN1;refGene_tx=XM_019821723.2;refGene_gn=HMCN1;refGene_tx=XM_019821724.2;refGene_gn=HMCN1;refGene_tx=XM_023247312.1;refGene_gn=HMCN1;refGene_tx=XM_023247313.1
F1 24576461 24576740
refGene_gn=HMCN1;refGene_tx=XM_003999241.4;refGene_gn=HMCN1;refGene_tx=XM_019821722.2;refGene_gn=HMCN1;refGene_tx=XM_019821723.2;refGene_gn=HMCN1;refGene_tx=XM_019821724.2;refGene_gn=HMCN1;refGene_tx=XM_023247312.1;refGene_gn=HMCN1;refGene_tx=XM_023247313.1
F1 24584046 24584201
refGene_gn=HMCN1;refGene_tx=XM_003999241.4;refGene_gn=HMCN1;refGene_tx=XM_019821722.2;refGene_gn=HMCN1;refGene_tx=XM_019821723.2;refGene_gn=HMCN1;refGene_tx=XM_019821724.2;refGene_gn=HMCN1;refGene_tx=XM_023247312.1;refGene_gn=HMCN1;refGene_tx=XM_023247313.1
F1 24587477 24587604
refGene_gn=HMCN1;refGene_tx=XM_003999241.4;refGene_gn=HMCN1;refGene_tx=XM_019821722.2;refGene_gn=HMCN1;refGene_tx=XM_019821723.2;refGene_gn=HMCN1;refGene_tx=XM_019821724.2;refGene_gn=HMCN1;refGene_tx=XM_023247312.1;refGene_gn=HMCN1;refGene_tx=XM_023247313.1
F1 24589832 24589980
refGene_gn=HMCN1;refGene_tx=XM_003999241.4;refGene_gn=HMCN1;refGene_tx=XM_019821722.2;refGene_gn=HMCN1;refGene_tx=XM_019821723.2;refGene_gn=HMCN1;refGene_tx=XM_019821724.2;refGene_gn=HMCN1;refGene_tx=XM_023247312.1;refGene_gn=HMCN1;refGene_tx=XM_023247313.1
F1 24590145 24590276
refGene_gn=HMCN1;refGene_tx=XM_003999241.4;refGene_gn=HMCN1;refGene_tx=XM_019821722.2;refGene_gn=HMCN1;refGene_tx=XM_019821723.2;refGene_gn=HMCN1;refGene_tx=XM_019821724.2;refGene_gn=HMCN1;refGene_tx=XM_023247312.1;refGene_gn=HMCN1;refGene_tx=XM_023247313.1
F1 24591586 24591781
refGene_gn=HMCN1;refGene_tx=XM_003999241.4;refGene_gn=HMCN1;refGene_tx=XM_019821722.2;refGene_gn=HMCN1;refGene_tx=XM_019821723.2;refGene_gn=HMCN1;refGene_tx=XM_019821724.2;refGene_gn=HMCN1;refGene_tx=XM_023247312.1;refGene_gn=HMCN1;refGene_tx=XM_023247313.1

F1 24594263 24594359
refGene_gn=HMCN1;refGene_tx=XM_003999241.4;refGene_gn=HMCN1;refGene_tx=XM_019821722.2;refGene_gn=HMCN1;refGene_tx=XM_019821723.2;refGene_gn=HMCN1;refGene_tx=XM_019821724.2;refGene_gn=HMCN1;refGene_tx=XM_023247312.1;refGene_gn=HMCN1;refGene_tx=XM_023247313.1
F1 24596668 24596941
refGene_gn=HMCN1;refGene_tx=XM_003999241.4;refGene_gn=HMCN1;refGene_tx=XM_019821722.2;refGene_gn=HMCN1;refGene_tx=XM_019821723.2;refGene_gn=HMCN1;refGene_tx=XM_019821724.2;refGene_gn=HMCN1;refGene_tx=XM_023247312.1;refGene_gn=HMCN1;refGene_tx=XM_023247313.1
F1 24599112 24599240
refGene_gn=HMCN1;refGene_tx=XM_003999241.4;refGene_gn=HMCN1;refGene_tx=XM_019821722.2;refGene_gn=HMCN1;refGene_tx=XM_019821723.2;refGene_gn=HMCN1;refGene_tx=XM_019821724.2;refGene_gn=HMCN1;refGene_tx=XM_023247312.1;refGene_gn=HMCN1;refGene_tx=XM_023247313.1
F1 24602772 24602941
refGene_gn=HMCN1;refGene_tx=XM_003999241.4;refGene_gn=HMCN1;refGene_tx=XM_019821722.2;refGene_gn=HMCN1;refGene_tx=XM_019821723.2;refGene_gn=HMCN1;refGene_tx=XM_019821724.2;refGene_gn=HMCN1;refGene_tx=XM_023247312.1;refGene_gn=HMCN1;refGene_tx=XM_023247313.1
F1 24603544 24603704
refGene_gn=HMCN1;refGene_tx=XM_003999241.4;refGene_gn=HMCN1;refGene_tx=XM_019821722.2;refGene_gn=HMCN1;refGene_tx=XM_019821723.2;refGene_gn=HMCN1;refGene_tx=XM_019821724.2;refGene_gn=HMCN1;refGene_tx=XM_023247312.1;refGene_gn=HMCN1;refGene_tx=XM_023247313.1
F1 24605393 24605506
refGene_gn=HMCN1;refGene_tx=XM_003999241.4;refGene_gn=HMCN1;refGene_tx=XM_019821722.2;refGene_gn=HMCN1;refGene_tx=XM_019821723.2;refGene_gn=HMCN1;refGene_tx=XM_019821724.2;refGene_gn=HMCN1;refGene_tx=XM_023247312.1;refGene_gn=HMCN1;refGene_tx=XM_023247313.1
F1 24608572 24608717
refGene_gn=HMCN1;refGene_tx=XM_003999241.4;refGene_gn=HMCN1;refGene_tx=XM_019821722.2;refGene_gn=HMCN1;refGene_tx=XM_019821723.2;refGene_gn=HMCN1;refGene_tx=XM_019821724.2;refGene_gn=HMCN1;refGene_tx=XM_023247312.1;refGene_gn=HMCN1;refGene_tx=XM_023247313.1
F1 24610023 24610151
refGene_gn=HMCN1;refGene_tx=XM_003999241.4;refGene_gn=HMCN1;refGene_tx=XM_019821722.2;refGene_gn=HMCN1;refGene_tx=XM_019821723.2;refGene_gn=HMCN1;refGene_tx=XM_019821724.2;refGene_gn=HMCN1;refGene_tx=XM_023247312.1;refGene_gn=HMCN1;refGene_tx=XM_023247313.1
F1 24611650 24611746
refGene_gn=HMCN1;refGene_tx=XM_003999241.4;refGene_gn=HMCN1;refGene_tx=XM_019821722.2;refGene_gn=HMCN1;refGene_tx=XM_019821723.2;refGene_gn=HMCN1;refGene_tx=XM_019821724.2;refGene_gn=HMCN1;refGene_tx=XM_023247312.1;refGene_gn=HMCN1;refGene_tx=XM_023247313.1
F1 24614332 24614527
refGene_gn=HMCN1;refGene_tx=XM_003999241.4;refGene_gn=HMCN1;refGene_tx=XM_019821722.2;refGene_gn=HMCN1;refGene_tx=XM_019821723.2;refGene_gn=HMCN1;refGene_tx=XM_019821724.2;refGene_gn=HMCN1;refGene_tx=XM_023247312.1;refGene_gn=HMCN1;refGene_tx=XM_023247313.1

F1	24621446	24621605
refGene_gn=HMCN1;refGene_tx=XM_003999241.4;refGene_gn=HMCN1;refGene_tx=XM_019821722.2;refGene_gn=HMCN1;refGene_tx=XM_019821723.2;refGene_gn=HMCN1;refGene_tx=XM_019821724.2;refGene_gn=HMCN1;refGene_tx=XM_023247312.1;refGene_gn=HMCN1;refGene_tx=XM_023247313.1
F1	24627002	24627116
refGene_gn=HMCN1;refGene_tx=XM_003999241.4;refGene_gn=HMCN1;refGene_tx=XM_019821722.2;refGene_gn=HMCN1;refGene_tx=XM_019821723.2;refGene_gn=HMCN1;refGene_tx=XM_019821724.2;refGene_gn=HMCN1;refGene_tx=XM_023247312.1;refGene_gn=HMCN1;refGene_tx=XM_023247313.1
F1	24629330	24629458
refGene_gn=HMCN1;refGene_tx=XM_003999241.4;refGene_gn=HMCN1;refGene_tx=XM_019821722.2;refGene_gn=HMCN1;refGene_tx=XM_019821723.2;refGene_gn=HMCN1;refGene_tx=XM_019821724.2;refGene_gn=HMCN1;refGene_tx=XM_023247312.1;refGene_gn=HMCN1;refGene_tx=XM_023247313.1
F1	24630552	24630701
refGene_gn=HMCN1;refGene_tx=XM_003999241.4;refGene_gn=HMCN1;refGene_tx=XM_019821722.2;refGene_gn=HMCN1;refGene_tx=XM_019821723.2;refGene_gn=HMCN1;refGene_tx=XM_019821724.2;refGene_gn=HMCN1;refGene_tx=XM_023247312.1;refGene_gn=HMCN1;refGene_tx=XM_023247313.1
F1	24650555	24650831
refGene_gn=HMCN1;refGene_tx=XM_003999241.4;refGene_gn=HMCN1;refGene_tx=XM_019821722.2;refGene_gn=HMCN1;refGene_tx=XM_019821724.2;refGene_gn=HMCN1;refGene_tx=XM_023247312.1;refGene_gn=HMCN1;refGene_tx=XM_023247313.1
F1	24656134	24656256
refGene_gn=HMCN1;refGene_tx=XM_003999241.4;refGene_gn=HMCN1;refGene_tx=XM_019821722.2;refGene_gn=HMCN1;refGene_tx=XM_019821724.2;refGene_gn=HMCN1;refGene_tx=XM_023247312.1;refGene_gn=HMCN1;refGene_tx=XM_023247313.1
F1	24659676	24659821
refGene_gn=HMCN1;refGene_tx=XM_003999241.4;refGene_gn=HMCN1;refGene_tx=XM_019821722.2;refGene_gn=HMCN1;refGene_tx=XM_019821724.2;refGene_gn=HMCN1;refGene_tx=XM_023247312.1;refGene_gn=HMCN1;refGene_tx=XM_023247313.1
F1	24661518	24661782
refGene_gn=HMCN1;refGene_tx=XM_003999241.4;refGene_gn=HMCN1;refGene_tx=XM_019821722.2;refGene_gn=HMCN1;refGene_tx=XM_019821724.2;refGene_gn=HMCN1;refGene_tx=XM_023247312.1;refGene_gn=HMCN1;refGene_tx=XM_023247313.1
F1	24662676	24662797
refGene_gn=HMCN1;refGene_tx=XM_003999241.4;refGene_gn=HMCN1;refGene_tx=XM_019821722.2;refGene_gn=HMCN1;refGene_tx=XM_019821724.2;refGene_gn=HMCN1;refGene_tx=XM_023247312.1;refGene_gn=HMCN1;refGene_tx=XM_023247313.1
F1	24674591	24674698
refGene_gn=HMCN1;refGene_tx=XM_003999241.4;refGene_gn=HMCN1;refGene_tx=XM_019821722.2;refGene_gn=HMCN1;refGene_tx=XM_019821724.2;refGene_gn=HMCN1;refGene_tx=XM_023247312.1;refGene_gn=HMCN1;refGene_tx=XM_023247313.1

F1 24676502 24676674
refGene_gn=HMCN1;refGene_tx=XM_003999241.4;refGene_gn=HMCN1;refGene_tx=XM_019821722.2;refGene_gn=HMCN1;refGene_tx=XM_019821724.2;refGene_gn=HMCN1;refGene_tx=XM_023247312.1;refGene_gn=HMCN1;refGene_tx=XM_023247313.1
F1 24724931 24725054
refGene_gn=HMCN1;refGene_tx=XM_003999241.4;refGene_gn=HMCN1;refGene_tx=XM_019821722.2;refGene_gn=HMCN1;refGene_tx=XM_019821724.2;refGene_gn=HMCN1;refGene_tx=XM_023247312.1;refGene_gn=HMCN1;refGene_tx=XM_023247313.1
F1 24726782 24726941
refGene_gn=HMCN1;refGene_tx=XM_003999241.4;refGene_gn=HMCN1;refGene_tx=XM_019821722.2;refGene_gn=HMCN1;refGene_tx=XM_019821724.2;refGene_gn=HMCN1;refGene_tx=XM_023247312.1;refGene_gn=HMCN1;refGene_tx=XM_023247313.1
F1 24744442 24744513
refGene_gn=HMCN1;refGene_tx=XM_003999241.4;refGene_gn=HMCN1;refGene_tx=XM_019821722.2;refGene_gn=HMCN1;refGene_tx=XM_019821724.2;refGene_gn=HMCN1;refGene_tx=XM_023247312.1;refGene_gn=HMCN1;refGene_tx=XM_023247313.1
F1 24845282 24845550
refGene_gn=HMCN1;refGene_tx=XM_003999241.4;refGene_gn=HMCN1;refGene_tx=XM_019821722.2;refGene_gn=HMCN1;refGene_tx=XM_019821724.2;refGene_gn=HMCN1;refGene_tx=XM_023247312.1;refGene_gn=HMCN1;refGene_tx=XM_023247313.1
F1 24849059 24849078 refGene_gn=HMCN1;refGene_tx=XM_023247312.1
F1 25198602 25198789
refGene_gn=LOC109495616;refGene_tx=XM_023247452.1
F1 25199270 25199758
refGene_gn=LOC109495616;refGene_tx=XM_023247452.1
F1 25207142 25207253
refGene_gn=IVNS1ABP;refGene_tx=XM_003999242.4;refGene_gn=IVNS1ABP;refGene_tx=XM_011291008.3
F1 25207476 25207646
refGene_gn=IVNS1ABP;refGene_tx=XM_003999242.4;refGene_gn=IVNS1ABP;refGene_tx=XM_011291008.3
F1 25207776 25207852
refGene_gn=IVNS1ABP;refGene_tx=XM_003999242.4;refGene_gn=IVNS1ABP;refGene_tx=XM_011291008.3
F1 25209233 25209407
refGene_gn=IVNS1ABP;refGene_tx=XM_003999242.4;refGene_gn=IVNS1ABP;refGene_tx=XM_011291008.3
F1 25209708 25209834
refGene_gn=IVNS1ABP;refGene_tx=XM_003999242.4;refGene_gn=IVNS1ABP;refGene_tx=XM_011291008.3
F1 25211153 25211261
refGene_gn=IVNS1ABP;refGene_tx=XM_003999242.4;refGene_gn=IVNS1ABP;refGene_tx=XM_011291008.3
F1 25215644 25215774
refGene_gn=IVNS1ABP;refGene_tx=XM_003999242.4;refGene_gn=IVNS1ABP;refG

ene_tx=XM_011291008.3
F1 25216007 25216232
refGene_gn=IVNS1ABP;refGene_tx=XM_003999242.4;refGene_gn=IVNS1ABP;refGene_tx=XM_011291008.3
F1 25216700 25216822
refGene_gn=IVNS1ABP;refGene_tx=XM_003999242.4;refGene_gn=IVNS1ABP;refGene_tx=XM_011291008.3
F1 25216903 25217030
refGene_gn=IVNS1ABP;refGene_tx=XM_003999242.4;refGene_gn=IVNS1ABP;refGene_tx=XM_011291008.3
F1 25217116 25217248
refGene_gn=IVNS1ABP;refGene_tx=XM_003999242.4;refGene_gn=IVNS1ABP;refGene_tx=XM_011291008.3
F1 25217376 25217550
refGene_gn=IVNS1ABP;refGene_tx=XM_003999242.4;refGene_gn=IVNS1ABP;refGene_tx=XM_011291008.3
F1 25218453 25218707
refGene_gn=IVNS1ABP;refGene_tx=XM_003999242.4;refGene_gn=IVNS1ABP;refGene_tx=XM_011291008.3
F1 25222716 25222720 refGene_gn=SWT1;refGene_tx=XM_019821726.2
F1 25226945 25227075
refGene_gn=SWT1;refGene_tx=XM_011291009.3;refGene_gn=SWT1;refGene_tx=XM_019821725.2
F1 25239247 25239312
refGene_gn=SWT1;refGene_tx=XM_011291009.3;refGene_gn=SWT1;refGene_tx=XM_019821725.2;refGene_gn=SWT1;refGene_tx=XM_019821726.2
F1 25242489 25242556
refGene_gn=SWT1;refGene_tx=XM_011291009.3;refGene_gn=SWT1;refGene_tx=XM_019821725.2;refGene_gn=SWT1;refGene_tx=XM_019821726.2
F1 25267194 25267326
refGene_gn=SWT1;refGene_tx=XM_011291009.3;refGene_gn=SWT1;refGene_tx=XM_019821725.2;refGene_gn=SWT1;refGene_tx=XM_019821726.2
F1 25272965 25273153
refGene_gn=SWT1;refGene_tx=XM_011291009.3;refGene_gn=SWT1;refGene_tx=XM_019821725.2;refGene_gn=SWT1;refGene_tx=XM_019821726.2
F1 25279750 25279899
refGene_gn=SWT1;refGene_tx=XM_011291009.3;refGene_gn=SWT1;refGene_tx=XM_019821725.2;refGene_gn=SWT1;refGene_tx=XM_019821726.2
F1 25284264 25284403
refGene_gn=SWT1;refGene_tx=XM_011291009.3;refGene_gn=SWT1;refGene_tx=XM_019821725.2;refGene_gn=SWT1;refGene_tx=XM_019821726.2
F1 25285629 25285793
refGene_gn=SWT1;refGene_tx=XM_011291009.3;refGene_gn=SWT1;refGene_tx=XM_019821725.2;refGene_gn=SWT1;refGene_tx=XM_019821726.2
F1 25287251 25287397
refGene_gn=SWT1;refGene_tx=XM_011291009.3;refGene_gn=SWT1;refGene_tx=XM_019821725.2;refGene_gn=SWT1;refGene_tx=XM_019821726.2
F1 25294740 25294834
refGene_gn=SWT1;refGene_tx=XM_011291009.3;refGene_gn=SWT1;refGene_tx=XM_019821725.2;refGene_gn=SWT1;refGene_tx=XM_019821726.2

F1 25301054 25301243
refGene_gn=SWT1;refGene_tx=XM_011291009.3;refGene_gn=SWT1;refGene_tx=XM_019821725.2;refGene_gn=SWT1;refGene_tx=XM_019821726.2
F1 25301841 25301943
refGene_gn=SWT1;refGene_tx=XM_011291009.3;refGene_gn=SWT1;refGene_tx=XM_019821725.2;refGene_gn=SWT1;refGene_tx=XM_019821726.2
F1 25302898 25303028
refGene_gn=SWT1;refGene_tx=XM_011291009.3;refGene_gn=SWT1;refGene_tx=XM_019821725.2;refGene_gn=SWT1;refGene_tx=XM_019821726.2
F1 25304256 25304316
refGene_gn=SWT1;refGene_tx=XM_011291009.3;refGene_gn=SWT1;refGene_tx=XM_019821726.2
F1 25308584 25309323
refGene_gn=SWT1;refGene_tx=XM_011291009.3;refGene_gn=SWT1;refGene_tx=XM_019821725.2;refGene_gn=SWT1;refGene_tx=XM_019821726.2
F1 25319665 25319721
refGene_gn=SWT1;refGene_tx=XM_011291009.3;refGene_gn=SWT1;refGene_tx=XM_019821725.2;refGene_gn=SWT1;refGene_tx=XM_019821726.2
F1 25321614 25321695
refGene_gn=SWT1;refGene_tx=XM_011291009.3;refGene_gn=SWT1;refGene_tx=XM_019821725.2;refGene_gn=SWT1;refGene_tx=XM_019821726.2
F1 25324572 25324653
refGene_gn=SWT1;refGene_tx=XM_011291009.3;refGene_gn=SWT1;refGene_tx=XM_019821725.2;refGene_gn=SWT1;refGene_tx=XM_019821726.2
F1 25328207 25328424
refGene_gn=TRMT1L;refGene_tx=XM_003999251.5;refGene_gn=TRMT1L;refGene_tx=XM_011291006.3;refGene_gn=TRMT1L;refGene_tx=XM_019821728.2
F1 25332793 25332904
refGene_gn=TRMT1L;refGene_tx=XM_003999251.5;refGene_gn=TRMT1L;refGene_tx=XM_011291006.3;refGene_gn=TRMT1L;refGene_tx=XM_011291007.2;refGene_gn=TRMT1L;refGene_tx=XM_019821728.2
F1 25334179 25334293
refGene_gn=TRMT1L;refGene_tx=XM_003999251.5;refGene_gn=TRMT1L;refGene_tx=XM_011291006.3;refGene_gn=TRMT1L;refGene_tx=XM_011291007.2;refGene_gn=TRMT1L;refGene_tx=XM_019821728.2
F1 25337614 25337679
refGene_gn=TRMT1L;refGene_tx=XM_003999251.5;refGene_gn=TRMT1L;refGene_tx=XM_019821728.2
F1 25339388 25339518
refGene_gn=TRMT1L;refGene_tx=XM_003999251.5;refGene_gn=TRMT1L;refGene_tx=XM_011291006.3;refGene_gn=TRMT1L;refGene_tx=XM_011291007.2;refGene_gn=TRMT1L;refGene_tx=XM_019821728.2
F1 25341567 25341691
refGene_gn=TRMT1L;refGene_tx=XM_003999251.5;refGene_gn=TRMT1L;refGene_tx=XM_011291006.3;refGene_gn=TRMT1L;refGene_tx=XM_011291007.2;refGene_gn=TRMT1L;refGene_tx=XM_019821728.2
F1 25342165 25342245
refGene_gn=TRMT1L;refGene_tx=XM_003999251.5;refGene_gn=TRMT1L;refGene_tx=XM_011291006.3;refGene_gn=TRMT1L;refGene_tx=XM_011291007.2;refGene_gn=TRMT1L;refGene_tx=XM_019821728.2

F1 25345644 25345894
refGene_gn=TRMT1L;refGene_tx=XM_003999251.5;refGene_gn=TRMT1L;refGene_tx=XM_011291006.3;refGene_gn=TRMT1L;refGene_tx=XM_011291007.2;refGene_gn=TRMT1L;refGene_tx=XM_019821728.2
F1 25346296 25346509
refGene_gn=TRMT1L;refGene_tx=XM_003999251.5;refGene_gn=TRMT1L;refGene_tx=XM_011291006.3;refGene_gn=TRMT1L;refGene_tx=XM_011291007.2;refGene_gn=TRMT1L;refGene_tx=XM_019821728.2
F1 25348171 25348362
refGene_gn=TRMT1L;refGene_tx=XM_003999251.5;refGene_gn=TRMT1L;refGene_tx=XM_011291006.3;refGene_gn=TRMT1L;refGene_tx=XM_011291007.2;refGene_gn=TRMT1L;refGene_tx=XM_019821728.2
F1 25363575 25363654
refGene_gn=TRMT1L;refGene_tx=XM_003999251.5;refGene_gn=TRMT1L;refGene_tx=XM_011291006.3;refGene_gn=TRMT1L;refGene_tx=XM_011291007.2;refGene_gn=TRMT1L;refGene_tx=XM_019821728.2
F1 25365931 25366095
refGene_gn=TRMT1L;refGene_tx=XM_003999251.5;refGene_gn=TRMT1L;refGene_tx=XM_011291006.3;refGene_gn=TRMT1L;refGene_tx=XM_011291007.2;refGene_gn=TRMT1L;refGene_tx=XM_019821728.2
F1 25366604 25366667
refGene_gn=TRMT1L;refGene_tx=XM_003999251.5;refGene_gn=TRMT1L;refGene_tx=XM_011291006.3;refGene_gn=TRMT1L;refGene_tx=XM_011291007.2;refGene_gn=TRMT1L;refGene_tx=XM_019821728.2
F1 25368340 25368467
refGene_gn=TRMT1L;refGene_tx=XM_003999251.5;refGene_gn=TRMT1L;refGene_tx=XM_011291006.3;refGene_gn=TRMT1L;refGene_tx=XM_011291007.2;refGene_gn=TRMT1L;refGene_tx=XM_019821728.2
F1 25368568 25368821
refGene_gn=TRMT1L;refGene_tx=XM_003999251.5;refGene_gn=TRMT1L;refGene_tx=XM_011291006.3;refGene_gn=TRMT1L;refGene_tx=XM_011291007.2;refGene_gn=TRMT1L;refGene_tx=XM_019821728.2
F1 25384274 25384376
refGene_gn=RNF2;refGene_tx=XM_004001340.4;refGene_gn=RNF2;refGene_tx=XM_011291005.3;refGene_gn=RNF2;refGene_tx=XM_023247243.1
F1 25384665 25384837
refGene_gn=RNF2;refGene_tx=XM_004001340.4;refGene_gn=RNF2;refGene_tx=XM_011291005.3;refGene_gn=RNF2;refGene_tx=XM_023247243.1
F1 25385664 25385937
refGene_gn=RNF2;refGene_tx=XM_004001340.4;refGene_gn=RNF2;refGene_tx=XM_011291005.3;refGene_gn=RNF2;refGene_tx=XM_023247243.1
F1 25390557 25390773
refGene_gn=RNF2;refGene_tx=XM_004001340.4;refGene_gn=RNF2;refGene_tx=XM_011291005.3;refGene_gn=RNF2;refGene_tx=XM_023247243.1
F1 25391394 25391555
refGene_gn=RNF2;refGene_tx=XM_004001340.4;refGene_gn=RNF2;refGene_tx=XM_011291005.3;refGene_gn=RNF2;refGene_tx=XM_023247243.1
F1 25395240 25395327
refGene_gn=RNF2;refGene_tx=XM_004001340.4;refGene_gn=RNF2;refGene_tx=XM_011291005.3;refGene_gn=RNF2;refGene_tx=XM_023247243.1

F1	25480140	25480195	refGene_gn=FAM129A;refGene_tx=XM_023247242.1
F1	25543081	25543212	refGene_gn=FAM129A;refGene_tx=XM_023247242.1
F1	25546227	25546359	refGene_gn=FAM129A;refGene_tx=XM_023247242.1
F1	25550346	25550461	refGene_gn=FAM129A;refGene_tx=XM_023247242.1
F1	25555342	25555510	refGene_gn=FAM129A;refGene_tx=XM_023247242.1
F1	25595681	25595797	refGene_gn=FAM129A;refGene_tx=XM_023247242.1
F1	25602621	25602726	refGene_gn=FAM129A;refGene_tx=XM_023247242.1
F1	25603039	25603202	refGene_gn=FAM129A;refGene_tx=XM_023247242.1
F1	25610484	25610672	refGene_gn=FAM129A;refGene_tx=XM_023247242.1
F1	25613537	25613699	refGene_gn=FAM129A;refGene_tx=XM_023247242.1
F1	25615277	25615388	refGene_gn=FAM129A;refGene_tx=XM_023247242.1
F1	25617277	25617385	refGene_gn=FAM129A;refGene_tx=XM_023247242.1
F1	25621230	25621342	refGene_gn=FAM129A;refGene_tx=XM_023247242.1
F1	25622959	25624083	refGene_gn=FAM129A;refGene_tx=XM_023247242.1
F1	25653865	25654026
refGene_gn=EDEM3;refGene_tx=XM_011291002.3;refGene_gn=EDEM3;refGene_tx=XM_011291003.3;refGene_gn=EDEM3;refGene_tx=XM_011291004.3;refGene_gn=EDEM3;refGene_tx=XM_019821730.2;refGene_gn=EDEM3;refGene_tx=XM_019821731.2;refGene_gn=EDEM3;refGene_tx=XM_019821732.2;refGene_gn=EDEM3;refGene_tx=XM_019821733.2;refGene_gn=EDEM3;refGene_tx=XM_019821734.2;refGene_gn=EDEM3;refGene_tx=XM_019821735.2
F1	25659223	25659269
refGene_gn=EDEM3;refGene_tx=XM_011291002.3;refGene_gn=EDEM3;refGene_tx=XM_011291003.3;refGene_gn=EDEM3;refGene_tx=XM_011291004.3;refGene_gn=EDEM3;refGene_tx=XM_019821730.2;refGene_gn=EDEM3;refGene_tx=XM_019821731.2;refGene_gn=EDEM3;refGene_tx=XM_019821732.2;refGene_gn=EDEM3;refGene_tx=XM_019821733.2;refGene_gn=EDEM3;refGene_tx=XM_019821734.2;refGene_gn=EDEM3;refGene_tx=XM_019821735.2
F1	25673526	25673627
refGene_gn=EDEM3;refGene_tx=XM_011291002.3;refGene_gn=EDEM3;refGene_tx=XM_011291003.3;refGene_gn=EDEM3;refGene_tx=XM_011291004.3;refGene_gn=EDEM3;refGene_tx=XM_019821730.2;refGene_gn=EDEM3;refGene_tx=XM_019821731.2;refGene_gn=EDEM3;refGene_tx=XM_019821732.2;refGene_gn=EDEM3;refGene_tx=XM_019821733.2;refGene_gn=EDEM3;refGene_tx=XM_019821734.2;refGene_gn=EDEM3;refGene_tx=XM_019821735.2;refGene_gn=EDEM3;refGene_tx=XM_023247241.1
F1	25674151	25674191
refGene_gn=EDEM3;refGene_tx=XM_011291002.3;refGene_gn=EDEM3;refGene_tx=XM_011291003.3;refGene_gn=EDEM3;refGene_tx=XM_011291004.3;refGene_gn=EDEM3;refGene_tx=XM_019821730.2;refGene_gn=EDEM3;refGene_tx=XM_019821731.2;refGene_gn=EDEM3;refGene_tx=XM_019821732.2;refGene_gn=EDEM3;refGene_tx=XM_019821733.2;refGene_gn=EDEM3;refGene_tx=XM_019821734.2;refGene_gn=EDEM3;refGene_tx=XM_019821735.2;refGene_gn=EDEM3;refGene_tx=XM_023247241.1
F1	25679086	25679199
refGene_gn=EDEM3;refGene_tx=XM_011291002.3;refGene_gn=EDEM3;refGene_tx=XM_011291003.3;refGene_gn=EDEM3;refGene_tx=XM_011291004.3;refGene_gn=EDEM3;refGene_tx=XM_019821730.2;refGene_gn=EDEM3;refGene_tx=XM_019821731.2;refGene_gn=EDEM3;refGene_tx=XM_019821732.2;refGene_gn=EDEM3;refGene_tx=XM_019821733.2;refGene_gn=EDEM3;refGene_tx=XM_019821734.2;refGen

e_gn=EDEM3;refGene_tx=XM_019821735.2;refGene_gn=EDEM3;refGene_tx=XM_023247241.1
F1 25680171 25680325
refGene_gn=EDEM3;refGene_tx=XM_011291002.3;refGene_gn=EDEM3;refGene_tx=XM_011291003.3;refGene_gn=EDEM3;refGene_tx=XM_011291004.3;refGene_gn=EDEM3;refGene_tx=XM_019821730.2;refGene_gn=EDEM3;refGene_tx=XM_019821731.2;refGene_gn=EDEM3;refGene_tx=XM_019821732.2;refGene_gn=EDEM3;refGene_tx=XM_019821733.2;refGene_gn=EDEM3;refGene_tx=XM_019821734.2;refGene_gn=EDEM3;refGene_tx=XM_019821735.2;refGene_gn=EDEM3;refGene_tx=XM_023247241.1
F1 25685429 25685564
refGene_gn=EDEM3;refGene_tx=XM_011291002.3;refGene_gn=EDEM3;refGene_tx=XM_011291003.3;refGene_gn=EDEM3;refGene_tx=XM_011291004.3;refGene_gn=EDEM3;refGene_tx=XM_019821730.2;refGene_gn=EDEM3;refGene_tx=XM_019821731.2;refGene_gn=EDEM3;refGene_tx=XM_019821732.2;refGene_gn=EDEM3;refGene_tx=XM_019821733.2;refGene_gn=EDEM3;refGene_tx=XM_019821734.2;refGene_gn=EDEM3;refGene_tx=XM_019821735.2;refGene_gn=EDEM3;refGene_tx=XM_023247241.1
F1 25687824 25687930
refGene_gn=EDEM3;refGene_tx=XM_011291002.3;refGene_gn=EDEM3;refGene_tx=XM_011291003.3;refGene_gn=EDEM3;refGene_tx=XM_011291004.3;refGene_gn=EDEM3;refGene_tx=XM_019821730.2;refGene_gn=EDEM3;refGene_tx=XM_019821731.2;refGene_gn=EDEM3;refGene_tx=XM_019821732.2;refGene_gn=EDEM3;refGene_tx=XM_019821733.2;refGene_gn=EDEM3;refGene_tx=XM_019821734.2;refGene_gn=EDEM3;refGene_tx=XM_019821735.2;refGene_gn=EDEM3;refGene_tx=XM_023247241.1
F1 25695136 25695234
refGene_gn=EDEM3;refGene_tx=XM_011291002.3;refGene_gn=EDEM3;refGene_tx=XM_011291003.3;refGene_gn=EDEM3;refGene_tx=XM_011291004.3;refGene_gn=EDEM3;refGene_tx=XM_019821730.2;refGene_gn=EDEM3;refGene_tx=XM_019821731.2;refGene_gn=EDEM3;refGene_tx=XM_019821732.2;refGene_gn=EDEM3;refGene_tx=XM_019821733.2;refGene_gn=EDEM3;refGene_tx=XM_019821734.2;refGene_gn=EDEM3;refGene_tx=XM_019821735.2;refGene_gn=EDEM3;refGene_tx=XM_023247241.1
F1 25696750 25696876
refGene_gn=EDEM3;refGene_tx=XM_011291002.3;refGene_gn=EDEM3;refGene_tx=XM_011291003.3;refGene_gn=EDEM3;refGene_tx=XM_011291004.3;refGene_gn=EDEM3;refGene_tx=XM_019821730.2;refGene_gn=EDEM3;refGene_tx=XM_019821731.2;refGene_gn=EDEM3;refGene_tx=XM_019821732.2;refGene_gn=EDEM3;refGene_tx=XM_019821733.2;refGene_gn=EDEM3;refGene_tx=XM_019821734.2;refGene_gn=EDEM3;refGene_tx=XM_019821735.2;refGene_gn=EDEM3;refGene_tx=XM_023247241.1
F1 25697081 25697165
refGene_gn=EDEM3;refGene_tx=XM_011291002.3;refGene_gn=EDEM3;refGene_tx=XM_011291003.3;refGene_gn=EDEM3;refGene_tx=XM_011291004.3;refGene_gn=EDEM3;refGene_tx=XM_019821730.2;refGene_gn=EDEM3;refGene_tx=XM_019821731.2;refGene_gn=EDEM3;refGene_tx=XM_019821732.2;refGene_gn=EDEM3;refGene_tx=XM_019821733.2;refGene_gn=EDEM3;refGene_tx=XM_019821734.2;refGene_gn=EDEM3;refGene_tx=XM_019821735.2;refGene_gn=EDEM3;refGene_tx=XM_023247241.1

F1 25699052 25699136
refGene_gn=EDEM3;refGene_tx=XM_011291002.3;refGene_gn=EDEM3;refGene_tx=XM_011291003.3;refGene_gn=EDEM3;refGene_tx=XM_011291004.3;refGene_gn=EDEM3;refGene_tx=XM_019821730.2;refGene_gn=EDEM3;refGene_tx=XM_019821731.2;refGene_gn=EDEM3;refGene_tx=XM_019821732.2;refGene_gn=EDEM3;refGene_tx=XM_019821733.2;refGene_gn=EDEM3;refGene_tx=XM_019821734.2;refGene_gn=EDEM3;refGene_tx=XM_019821735.2;refGene_gn=EDEM3;refGene_tx=XM_023247241.1
F1 25699676 25699801
refGene_gn=EDEM3;refGene_tx=XM_011291002.3;refGene_gn=EDEM3;refGene_tx=XM_011291003.3;refGene_gn=EDEM3;refGene_tx=XM_011291004.3;refGene_gn=EDEM3;refGene_tx=XM_019821730.2;refGene_gn=EDEM3;refGene_tx=XM_019821731.2;refGene_gn=EDEM3;refGene_tx=XM_019821732.2;refGene_gn=EDEM3;refGene_tx=XM_019821733.2;refGene_gn=EDEM3;refGene_tx=XM_019821734.2;refGene_gn=EDEM3;refGene_tx=XM_019821735.2;refGene_gn=EDEM3;refGene_tx=XM_023247241.1
F1 25704807 25704973
refGene_gn=EDEM3;refGene_tx=XM_011291002.3;refGene_gn=EDEM3;refGene_tx=XM_011291003.3;refGene_gn=EDEM3;refGene_tx=XM_011291004.3;refGene_gn=EDEM3;refGene_tx=XM_019821730.2;refGene_gn=EDEM3;refGene_tx=XM_019821731.2;refGene_gn=EDEM3;refGene_tx=XM_019821732.2;refGene_gn=EDEM3;refGene_tx=XM_019821733.2;refGene_gn=EDEM3;refGene_tx=XM_019821734.2;refGene_gn=EDEM3;refGene_tx=XM_019821735.2;refGene_gn=EDEM3;refGene_tx=XM_023247241.1
F1 25705221 25705376
refGene_gn=EDEM3;refGene_tx=XM_011291002.3;refGene_gn=EDEM3;refGene_tx=XM_011291003.3;refGene_gn=EDEM3;refGene_tx=XM_011291004.3;refGene_gn=EDEM3;refGene_tx=XM_019821730.2;refGene_gn=EDEM3;refGene_tx=XM_019821731.2;refGene_gn=EDEM3;refGene_tx=XM_019821732.2;refGene_gn=EDEM3;refGene_tx=XM_019821733.2;refGene_gn=EDEM3;refGene_tx=XM_019821734.2;refGene_gn=EDEM3;refGene_tx=XM_019821735.2;refGene_gn=EDEM3;refGene_tx=XM_023247241.1
F1 25706298 25706455
refGene_gn=EDEM3;refGene_tx=XM_011291002.3;refGene_gn=EDEM3;refGene_tx=XM_011291003.3;refGene_gn=EDEM3;refGene_tx=XM_011291004.3;refGene_gn=EDEM3;refGene_tx=XM_019821730.2;refGene_gn=EDEM3;refGene_tx=XM_019821731.2;refGene_gn=EDEM3;refGene_tx=XM_019821732.2;refGene_gn=EDEM3;refGene_tx=XM_019821733.2;refGene_gn=EDEM3;refGene_tx=XM_019821734.2;refGene_gn=EDEM3;refGene_tx=XM_019821735.2;refGene_gn=EDEM3;refGene_tx=XM_023247241.1
F1 25708477 25708669
refGene_gn=EDEM3;refGene_tx=XM_011291002.3;refGene_gn=EDEM3;refGene_tx=XM_011291003.3;refGene_gn=EDEM3;refGene_tx=XM_011291004.3;refGene_gn=EDEM3;refGene_tx=XM_019821730.2;refGene_gn=EDEM3;refGene_tx=XM_019821731.2;refGene_gn=EDEM3;refGene_tx=XM_019821732.2;refGene_gn=EDEM3;refGene_tx=XM_019821733.2;refGene_gn=EDEM3;refGene_tx=XM_019821734.2;refGene_gn=EDEM3;refGene_tx=XM_019821735.2;refGene_gn=EDEM3;refGene_tx=XM_023247241.1
F1 25710004 25710170
refGene_gn=EDEM3;refGene_tx=XM_011291002.3;refGene_gn=EDEM3;refGene_tx

=XM_011291003.3;refGene_gn=EDEM3;refGene_tx=XM_011291004.3;refGene_gn=EDEM3;refGene_tx=XM_019821730.2;refGene_gn=EDEM3;refGene_tx=XM_019821731.2;refGene_gn=EDEM3;refGene_tx=XM_019821732.2;refGene_gn=EDEM3;refGene_tx=XM_019821733.2;refGene_gn=EDEM3;refGene_tx=XM_019821734.2;refGene_gn=EDEM3;refGene_tx=XM_019821735.2;refGene_gn=EDEM3;refGene_tx=XM_023247241.1
F1 25713356 25713542
refGene_gn=EDEM3;refGene_tx=XM_011291002.3;refGene_gn=EDEM3;refGene_tx=XM_011291003.3;refGene_gn=EDEM3;refGene_tx=XM_011291004.3;refGene_gn=EDEM3;refGene_tx=XM_019821730.2;refGene_gn=EDEM3;refGene_tx=XM_019821731.2;refGene_gn=EDEM3;refGene_tx=XM_019821732.2;refGene_gn=EDEM3;refGene_tx=XM_019821733.2;refGene_gn=EDEM3;refGene_tx=XM_019821734.2;refGene_gn=EDEM3;refGene_tx=XM_019821735.2;refGene_gn=EDEM3;refGene_tx=XM_023247241.1
F1 25714813 25714861
refGene_gn=EDEM3;refGene_tx=XM_011291002.3;refGene_gn=EDEM3;refGene_tx=XM_019821730.2;refGene_gn=EDEM3;refGene_tx=XM_019821731.2;refGene_gn=EDEM3;refGene_tx=XM_019821732.2;refGene_gn=EDEM3;refGene_tx=XM_019821733.2;refGene_gn=EDEM3;refGene_tx=XM_023247241.1
F1 25720409 25720813
refGene_gn=EDEM3;refGene_tx=XM_011291002.3;refGene_gn=EDEM3;refGene_tx=XM_011291003.3;refGene_gn=EDEM3;refGene_tx=XM_011291004.3;refGene_gn=EDEM3;refGene_tx=XM_019821730.2;refGene_gn=EDEM3;refGene_tx=XM_019821731.2;refGene_gn=EDEM3;refGene_tx=XM_019821732.2;refGene_gn=EDEM3;refGene_tx=XM_019821733.2;refGene_gn=EDEM3;refGene_tx=XM_019821734.2;refGene_gn=EDEM3;refGene_tx=XM_023247241.1
F1 25734652 25734657 refGene_gn=EDEM3;refGene_tx=XM_019821735.2
F1 25779161 25779200
refGene_gn=CF1H1orf21;refGene_tx=XM_003999255.5
F1 25802281 25802342
refGene_gn=CF1H1orf21;refGene_tx=XM_003999255.5
F1 25809496 25809573
refGene_gn=CF1H1orf21;refGene_tx=XM_003999255.5
F1 25886341 25886436
refGene_gn=CF1H1orf21;refGene_tx=XM_003999255.5
F1 25925799 25925893
refGene_gn=CF1H1orf21;refGene_tx=XM_003999255.5
F1 26258892 26258913 refGene_gn=TSEN15;refGene_tx=XM_023247552.1
F1 26259435 26259577 refGene_gn=TSEN15;refGene_tx=XM_023247552.1
F1 26285426 26285562 refGene_gn=TSEN15;refGene_tx=XM_023247552.1
F1 26285805 26285887 refGene_gn=TSEN15;refGene_tx=XM_023247552.1
F1 26288698 26288821 refGene_gn=TSEN15;refGene_tx=XM_023247552.1
F1 26300359 26300622
refGene_gn=COLGALT2;refGene_tx=XM_023247550.1;refGene_gn=COLGALT2;refGene_tx=XM_023247551.1
F1 26366867 26366978
refGene_gn=COLGALT2;refGene_tx=XM_023247550.1;refGene_gn=COLGALT2;refGene_tx=XM_023247551.1
F1 26369738 26369856
refGene_gn=COLGALT2;refGene_tx=XM_023247550.1;refGene_gn=COLGALT2;refG

ene_tx=XM_023247551.1
F1 26372551 26372686
refGene_gn=COLGALT2;refGene_tx=XM_023247550.1;refGene_gn=COLGALT2;refGene_tx=XM_023247551.1
F1 26377083 26377288
refGene_gn=COLGALT2;refGene_tx=XM_023247550.1;refGene_gn=COLGALT2;refGene_tx=XM_023247551.1
F1 26382870 26382990
refGene_gn=COLGALT2;refGene_tx=XM_023247550.1;refGene_gn=COLGALT2;refGene_tx=XM_023247551.1
F1 26393193 26393270
refGene_gn=COLGALT2;refGene_tx=XM_023247550.1;refGene_gn=COLGALT2;refGene_tx=XM_023247551.1
F1 26397363 26397470
refGene_gn=COLGALT2;refGene_tx=XM_023247550.1;refGene_gn=COLGALT2;refGene_tx=XM_023247551.1
F1 26403559 26403692
refGene_gn=COLGALT2;refGene_tx=XM_023247550.1;refGene_gn=COLGALT2;refGene_tx=XM_023247551.1
F1 26405055 26405183
refGene_gn=COLGALT2;refGene_tx=XM_023247550.1;refGene_gn=COLGALT2;refGene_tx=XM_023247551.1
F1 26408214 26408421
refGene_gn=COLGALT2;refGene_tx=XM_023247550.1;refGene_gn=COLGALT2;refGene_tx=XM_023247551.1
F1 26409841 26409860 refGene_gn=COLGALT2;refGene_tx=XM_023247551.1
F1 26410032 26410309 refGene_gn=COLGALT2;refGene_tx=XM_023247550.1
F1 26422338 26422526
refGene_gn=RGL1;refGene_tx=XM_011290999.3;refGene_gn=RGL1;refGene_tx=XM_011291000.3;refGene_gn=RGL1;refGene_tx=XM_019821742.1
F1 26425744 26425859
refGene_gn=RGL1;refGene_tx=XM_011290999.3;refGene_gn=RGL1;refGene_tx=XM_011291000.3;refGene_gn=RGL1;refGene_tx=XM_019821742.1
F1 26430042 26430297
refGene_gn=RGL1;refGene_tx=XM_011290999.3;refGene_gn=RGL1;refGene_tx=XM_011291000.3;refGene_gn=RGL1;refGene_tx=XM_019821742.1
F1 26434336 26434523
refGene_gn=RGL1;refGene_tx=XM_011290999.3;refGene_gn=RGL1;refGene_tx=XM_011291000.3;refGene_gn=RGL1;refGene_tx=XM_019821742.1
F1 26440846 26440936
refGene_gn=RGL1;refGene_tx=XM_011290999.3;refGene_gn=RGL1;refGene_tx=XM_011291000.3;refGene_gn=RGL1;refGene_tx=XM_019821742.1
F1 26443407 26443529
refGene_gn=RGL1;refGene_tx=XM_011290999.3;refGene_gn=RGL1;refGene_tx=XM_011291000.3;refGene_gn=RGL1;refGene_tx=XM_019821742.1
F1 26445775 26445808
refGene_gn=RGL1;refGene_tx=XM_011290999.3;refGene_gn=RGL1;refGene_tx=XM_011291000.3;refGene_gn=RGL1;refGene_tx=XM_019821742.1
F1 26447374 26447461
refGene_gn=RGL1;refGene_tx=XM_011290999.3;refGene_gn=RGL1;refGene_tx=X

M_011291000.3;refGene_gn=RGL1;refGene_tx=XM_019821742.1
F1 26449839 26449929
refGene_gn=RGL1;refGene_tx=XM_011290999.3;refGene_gn=RGL1;refGene_tx=X
M_011291000.3;refGene_gn=RGL1;refGene_tx=XM_019821742.1
F1 26455076 26455161
refGene_gn=RGL1;refGene_tx=XM_011290999.3;refGene_gn=RGL1;refGene_tx=X
M_011291000.3;refGene_gn=RGL1;refGene_tx=XM_019821742.1
F1 26458589 26458693
refGene_gn=RGL1;refGene_tx=XM_011290999.3;refGene_gn=RGL1;refGene_tx=X
M_011291000.3;refGene_gn=RGL1;refGene_tx=XM_019821742.1
F1 26462423 26462639
refGene_gn=RGL1;refGene_tx=XM_011290999.3;refGene_gn=RGL1;refGene_tx=X
M_011291000.3;refGene_gn=RGL1;refGene_tx=XM_019821742.1
F1 26464043 26464168
refGene_gn=RGL1;refGene_tx=XM_011290999.3;refGene_gn=RGL1;refGene_tx=X
M_011291000.3;refGene_gn=RGL1;refGene_tx=XM_019821742.1
F1 26467597 26467782
refGene_gn=RGL1;refGene_tx=XM_011290999.3;refGene_gn=RGL1;refGene_tx=X
M_011291000.3;refGene_gn=RGL1;refGene_tx=XM_019821742.1
F1 26484882 26484960
refGene_gn=RGL1;refGene_tx=XM_011290999.3;refGene_gn=RGL1;refGene_tx=X
M_011291000.3;refGene_gn=RGL1;refGene_tx=XM_019821742.1
F1 26507500 26507709
refGene_gn=RGL1;refGene_tx=XM_011290999.3;refGene_gn=RGL1;refGene_tx=X
M_011291000.3;refGene_gn=RGL1;refGene_tx=XM_019821742.1
F1 26546091 26546202
refGene_gn=RGL1;refGene_tx=XM_011290999.3;refGene_gn=RGL1;refGene_tx=X
M_011291000.3
F1 26547200 26547227 refGene_gn=RGL1;refGene_tx=XM_011290999.3
F1 26583123 26583255
refGene_gn=RGL1;refGene_tx=XM_011291000.3;refGene_gn=RGL1;refGene_tx=X
M_019821742.1
F1 26667600 26668707 refGene_gn=APOBEC4;refGene_tx=XM_003999257.5
F1 26681641 26681784 refGene_gn=ARPC5;refGene_tx=XM_003999258.5
F1 26684124 26684197 refGene_gn=ARPC5;refGene_tx=XM_003999258.5
F1 26686186 26686363 refGene_gn=ARPC5;refGene_tx=XM_003999258.5
F1 26688993 26689056 refGene_gn=ARPC5;refGene_tx=XM_003999258.5
F1 26722081 26722255 refGene_gn=NCF2;refGene_tx=XM_003999260.5
F1 26724442 26724525 refGene_gn=NCF2;refGene_tx=XM_003999260.5
F1 26732382 26732491 refGene_gn=NCF2;refGene_tx=XM_003999260.5
F1 26737074 26737209 refGene_gn=NCF2;refGene_tx=XM_003999260.5
F1 26738110 26738218 refGene_gn=NCF2;refGene_tx=XM_003999260.5
F1 26739758 26739818 refGene_gn=NCF2;refGene_tx=XM_003999260.5
F1 26741223 26741267 refGene_gn=NCF2;refGene_tx=XM_003999260.5
F1 26742988 26743130 refGene_gn=NCF2;refGene_tx=XM_003999260.5
F1 26743352 26743421 refGene_gn=NCF2;refGene_tx=XM_003999260.5
F1 26744427 26744503 refGene_gn=NCF2;refGene_tx=XM_003999260.5
F1 26746411 26746437 refGene_gn=NCF2;refGene_tx=XM_003999260.5
F1 26746829 26746981 refGene_gn=NCF2;refGene_tx=XM_003999260.5
F1 26747091 26747203 refGene_gn=NCF2;refGene_tx=XM_003999260.5

F1 26751728 26751906 refGene_gn=NCF2;refGene_tx=XM_003999260.5
F1 26755053 26755166 refGene_gn=NCF2;refGene_tx=XM_003999260.5
F1 26758740 26758902
refGene_gn=SMG7;refGene_tx=XM_006942748.3;refGene_gn=SMG7;refGene_tx=XM_006942749.3;refGene_gn=SMG7;refGene_tx=XM_006942751.3;refGene_gn=SMG7;refGene_tx=XM_006942752.4;refGene_gn=SMG7;refGene_tx=XM_006942754.4;refGene_gn=SMG7;refGene_tx=XM_019821744.2;refGene_gn=SMG7;refGene_tx=XM_023247825.1;refGene_gn=SMG7;refGene_tx=XM_023247826.1;refGene_gn=SMG7;refGene_tx=XM_023247828.1
F1 26759484 26759630
refGene_gn=SMG7;refGene_tx=XM_006942748.3;refGene_gn=SMG7;refGene_tx=XM_006942749.3;refGene_gn=SMG7;refGene_tx=XM_006942751.3;refGene_gn=SMG7;refGene_tx=XM_006942752.4;refGene_gn=SMG7;refGene_tx=XM_006942754.4;refGene_gn=SMG7;refGene_tx=XM_019821744.2;refGene_gn=SMG7;refGene_tx=XM_023247825.1;refGene_gn=SMG7;refGene_tx=XM_023247826.1;refGene_gn=SMG7;refGene_tx=XM_023247828.1
F1 26759746 26759917
refGene_gn=SMG7;refGene_tx=XM_006942748.3;refGene_gn=SMG7;refGene_tx=XM_006942749.3;refGene_gn=SMG7;refGene_tx=XM_006942751.3;refGene_gn=SMG7;refGene_tx=XM_006942752.4;refGene_gn=SMG7;refGene_tx=XM_006942754.4;refGene_gn=SMG7;refGene_tx=XM_019821744.2;refGene_gn=SMG7;refGene_tx=XM_023247825.1;refGene_gn=SMG7;refGene_tx=XM_023247826.1;refGene_gn=SMG7;refGene_tx=XM_023247828.1
F1 26760592 26760752
refGene_gn=SMG7;refGene_tx=XM_006942748.3;refGene_gn=SMG7;refGene_tx=XM_006942749.3;refGene_gn=SMG7;refGene_tx=XM_006942751.3;refGene_gn=SMG7;refGene_tx=XM_006942752.4;refGene_gn=SMG7;refGene_tx=XM_006942754.4;refGene_gn=SMG7;refGene_tx=XM_019821744.2;refGene_gn=SMG7;refGene_tx=XM_023247825.1;refGene_gn=SMG7;refGene_tx=XM_023247826.1;refGene_gn=SMG7;refGene_tx=XM_023247828.1
F1 26761478 26761559
refGene_gn=SMG7;refGene_tx=XM_006942748.3;refGene_gn=SMG7;refGene_tx=XM_006942749.3;refGene_gn=SMG7;refGene_tx=XM_006942751.3;refGene_gn=SMG7;refGene_tx=XM_006942752.4;refGene_gn=SMG7;refGene_tx=XM_006942754.4;refGene_gn=SMG7;refGene_tx=XM_019821744.2;refGene_gn=SMG7;refGene_tx=XM_023247825.1;refGene_gn=SMG7;refGene_tx=XM_023247826.1;refGene_gn=SMG7;refGene_tx=XM_023247828.1
F1 26763507 26763657
refGene_gn=SMG7;refGene_tx=XM_006942748.3;refGene_gn=SMG7;refGene_tx=XM_006942749.3;refGene_gn=SMG7;refGene_tx=XM_006942751.3;refGene_gn=SMG7;refGene_tx=XM_019821744.2;refGene_gn=SMG7;refGene_tx=XM_023247825.1;refGene_gn=SMG7;refGene_tx=XM_023247826.1;refGene_gn=SMG7;refGene_tx=XM_023247828.1
F1 26764423 26764795
refGene_gn=SMG7;refGene_tx=XM_006942748.3;refGene_gn=SMG7;refGene_tx=XM_006942749.3;refGene_gn=SMG7;refGene_tx=XM_006942751.3;refGene_gn=SMG7;refGene_tx=XM_006942752.4;refGene_gn=SMG7;refGene_tx=XM_006942754.4;refGene_gn=SMG7;refGene_tx=XM_019821744.2;refGene_gn=SMG7;refGene_tx=XM_023247825.1;refGene_gn=SMG7;refGene_tx=XM_023247826.1;refGene_gn=SMG7;refGene_tx=XM_023247828.1

F1 26765479 26765859
refGene_gn=SMG7;refGene_tx=XM_006942748.3;refGene_gn=SMG7;refGene_tx=XM_006942749.3;refGene_gn=SMG7;refGene_tx=XM_006942751.3;refGene_gn=SMG7;refGene_tx=XM_006942752.4;refGene_gn=SMG7;refGene_tx=XM_006942754.4;refGene_gn=SMG7;refGene_tx=XM_019821744.2;refGene_gn=SMG7;refGene_tx=XM_023247825.1;refGene_gn=SMG7;refGene_tx=XM_023247826.1;refGene_gn=SMG7;refGene_tx=XM_023247828.1
F1 26766343 26766488
refGene_gn=SMG7;refGene_tx=XM_006942748.3;refGene_gn=SMG7;refGene_tx=XM_006942749.3;refGene_gn=SMG7;refGene_tx=XM_006942751.3;refGene_gn=SMG7;refGene_tx=XM_006942752.4;refGene_gn=SMG7;refGene_tx=XM_006942754.4;refGene_gn=SMG7;refGene_tx=XM_019821744.2;refGene_gn=SMG7;refGene_tx=XM_023247825.1;refGene_gn=SMG7;refGene_tx=XM_023247826.1;refGene_gn=SMG7;refGene_tx=XM_023247828.1
F1 26768295 26768722
refGene_gn=SMG7;refGene_tx=XM_006942748.3;refGene_gn=SMG7;refGene_tx=XM_006942749.3;refGene_gn=SMG7;refGene_tx=XM_006942751.3;refGene_gn=SMG7;refGene_tx=XM_006942752.4;refGene_gn=SMG7;refGene_tx=XM_006942754.4;refGene_gn=SMG7;refGene_tx=XM_019821744.2;refGene_gn=SMG7;refGene_tx=XM_023247825.1;refGene_gn=SMG7;refGene_tx=XM_023247826.1;refGene_gn=SMG7;refGene_tx=XM_023247828.1
F1 26769566 26769686
refGene_gn=SMG7;refGene_tx=XM_006942748.3;refGene_gn=SMG7;refGene_tx=XM_006942749.3;refGene_gn=SMG7;refGene_tx=XM_006942751.3;refGene_gn=SMG7;refGene_tx=XM_006942752.4;refGene_gn=SMG7;refGene_tx=XM_006942754.4;refGene_gn=SMG7;refGene_tx=XM_019821744.2;refGene_gn=SMG7;refGene_tx=XM_023247825.1;refGene_gn=SMG7;refGene_tx=XM_023247826.1;refGene_gn=SMG7;refGene_tx=XM_023247828.1
F1 26772834 26772895
refGene_gn=SMG7;refGene_tx=XM_006942748.3;refGene_gn=SMG7;refGene_tx=XM_006942749.3;refGene_gn=SMG7;refGene_tx=XM_006942751.3;refGene_gn=SMG7;refGene_tx=XM_006942752.4;refGene_gn=SMG7;refGene_tx=XM_006942754.4;refGene_gn=SMG7;refGene_tx=XM_019821744.2;refGene_gn=SMG7;refGene_tx=XM_023247825.1;refGene_gn=SMG7;refGene_tx=XM_023247826.1;refGene_gn=SMG7;refGene_tx=XM_023247828.1
F1 26773994 26774065
refGene_gn=SMG7;refGene_tx=XM_006942748.3;refGene_gn=SMG7;refGene_tx=XM_006942749.3;refGene_gn=SMG7;refGene_tx=XM_006942751.3;refGene_gn=SMG7;refGene_tx=XM_006942752.4;refGene_gn=SMG7;refGene_tx=XM_006942754.4;refGene_gn=SMG7;refGene_tx=XM_019821744.2;refGene_gn=SMG7;refGene_tx=XM_023247825.1;refGene_gn=SMG7;refGene_tx=XM_023247826.1;refGene_gn=SMG7;refGene_tx=XM_023247828.1
F1 26775156 26775313
refGene_gn=SMG7;refGene_tx=XM_006942748.3;refGene_gn=SMG7;refGene_tx=XM_006942749.3;refGene_gn=SMG7;refGene_tx=XM_006942751.3;refGene_gn=SMG7;refGene_tx=XM_006942752.4;refGene_gn=SMG7;refGene_tx=XM_006942754.4;refGene_gn=SMG7;refGene_tx=XM_019821744.2;refGene_gn=SMG7;refGene_tx=XM_023247825.1;refGene_gn=SMG7;refGene_tx=XM_023247826.1;refGene_gn=SMG7;refGene_tx=XM_023247828.1
F1 26776233 26776396

refGene_gn=SMG7;refGene_tx=XM_006942748.3;refGene_gn=SMG7;refGene_tx=XM_006942749.3;refGene_gn=SMG7;refGene_tx=XM_006942751.3;refGene_gn=SMG7;refGene_tx=XM_006942752.4;refGene_gn=SMG7;refGene_tx=XM_006942754.4;refGene_gn=SMG7;refGene_tx=XM_019821744.2;refGene_gn=SMG7;refGene_tx=XM_023247825.1;refGene_gn=SMG7;refGene_tx=XM_023247826.1;refGene_gn=SMG7;refGene_tx=XM_023247828.1
F1 26780623 26780759
refGene_gn=SMG7;refGene_tx=XM_006942748.3;refGene_gn=SMG7;refGene_tx=XM_006942749.3;refGene_gn=SMG7;refGene_tx=XM_006942751.3;refGene_gn=SMG7;refGene_tx=XM_006942752.4;refGene_gn=SMG7;refGene_tx=XM_006942754.4;refGene_gn=SMG7;refGene_tx=XM_019821744.2;refGene_gn=SMG7;refGene_tx=XM_023247825.1;refGene_gn=SMG7;refGene_tx=XM_023247826.1;refGene_gn=SMG7;refGene_tx=XM_023247828.1
F1 26781137 26781288
refGene_gn=SMG7;refGene_tx=XM_006942748.3;refGene_gn=SMG7;refGene_tx=XM_006942749.3;refGene_gn=SMG7;refGene_tx=XM_006942751.3;refGene_gn=SMG7;refGene_tx=XM_006942752.4;refGene_gn=SMG7;refGene_tx=XM_006942754.4;refGene_gn=SMG7;refGene_tx=XM_019821744.2;refGene_gn=SMG7;refGene_tx=XM_023247825.1;refGene_gn=SMG7;refGene_tx=XM_023247826.1;refGene_gn=SMG7;refGene_tx=XM_023247828.1
F1 26782116 26782188
refGene_gn=SMG7;refGene_tx=XM_006942748.3;refGene_gn=SMG7;refGene_tx=XM_006942749.3;refGene_gn=SMG7;refGene_tx=XM_006942751.3;refGene_gn=SMG7;refGene_tx=XM_006942752.4;refGene_gn=SMG7;refGene_tx=XM_006942754.4;refGene_gn=SMG7;refGene_tx=XM_019821744.2;refGene_gn=SMG7;refGene_tx=XM_023247825.1;refGene_gn=SMG7;refGene_tx=XM_023247826.1;refGene_gn=SMG7;refGene_tx=XM_023247828.1
F1 26783390 26783562
refGene_gn=SMG7;refGene_tx=XM_006942748.3;refGene_gn=SMG7;refGene_tx=XM_006942749.3;refGene_gn=SMG7;refGene_tx=XM_006942751.3;refGene_gn=SMG7;refGene_tx=XM_006942752.4;refGene_gn=SMG7;refGene_tx=XM_006942754.4;refGene_gn=SMG7;refGene_tx=XM_019821744.2;refGene_gn=SMG7;refGene_tx=XM_023247825.1;refGene_gn=SMG7;refGene_tx=XM_023247826.1;refGene_gn=SMG7;refGene_tx=XM_023247828.1
F1 26791557 26791690
refGene_gn=SMG7;refGene_tx=XM_006942748.3;refGene_gn=SMG7;refGene_tx=XM_006942749.3;refGene_gn=SMG7;refGene_tx=XM_006942751.3;refGene_gn=SMG7;refGene_tx=XM_006942752.4;refGene_gn=SMG7;refGene_tx=XM_006942754.4;refGene_gn=SMG7;refGene_tx=XM_019821744.2;refGene_gn=SMG7;refGene_tx=XM_023247825.1;refGene_gn=SMG7;refGene_tx=XM_023247826.1;refGene_gn=SMG7;refGene_tx=XM_023247828.1
F1 26792868 26792903 refGene_gn=SMG7;refGene_tx=XM_023247828.1
F1 26793438 26793556
refGene_gn=SMG7;refGene_tx=XM_006942748.3;refGene_gn=SMG7;refGene_tx=XM_006942749.3;refGene_gn=SMG7;refGene_tx=XM_006942751.3;refGene_gn=SMG7;refGene_tx=XM_006942752.4;refGene_gn=SMG7;refGene_tx=XM_006942754.4;refGene_gn=SMG7;refGene_tx=XM_019821744.2;refGene_gn=SMG7;refGene_tx=XM_023247825.1;refGene_gn=SMG7;refGene_tx=XM_023247826.1
F1 26796085 26796117
refGene_gn=SMG7;refGene_tx=XM_006942748.3;refGene_gn=SMG7;refGene_tx=X

M_006942749.3;refGene_gn=SMG7;refGene_tx=XM_006942751.3;refGene_gn=SMG7;refGene_tx=XM_006942752.4;refGene_gn=SMG7;refGene_tx=XM_006942754.4;refGene_gn=SMG7;refGene_tx=XM_019821744.2;refGene_gn=SMG7;refGene_tx=XM_023247825.1;refGene_gn=SMG7;refGene_tx=XM_023247826.1
F1	26811317	26811421	refGene_gn=SMG7;refGene_tx=XM_006942748.3;refGene_gn=SMG7;refGene_tx=XM_006942751.3;refGene_gn=SMG7;refGene_tx=XM_006942752.4;refGene_gn=SMG7;refGene_tx=XM_006942754.4;refGene_gn=SMG7;refGene_tx=XM_023247825.1;refGene_gn=SMG7;refGene_tx=XM_023247826.1
F1	26848201	26848230	refGene_gn=SMG7;refGene_tx=XM_006942749.3
F1	26893795	26893880	refGene_gn=NMNAT2;refGene_tx=XM_011290996.3
F1	27006027	27006116	refGene_gn=NMNAT2;refGene_tx=XM_011290996.3
F1	27006928	27006996	refGene_gn=NMNAT2;refGene_tx=XM_011290996.3
F1	27009048	27009127	refGene_gn=NMNAT2;refGene_tx=XM_011290996.3
F1	27012167	27012294	refGene_gn=NMNAT2;refGene_tx=XM_011290996.3
F1	27013959	27014040	refGene_gn=NMNAT2;refGene_tx=XM_011290996.3
F1	27015221	27015266	refGene_gn=NMNAT2;refGene_tx=XM_011290996.3
F1	27019406	27019483	refGene_gn=NMNAT2;refGene_tx=XM_011290996.3
F1	27033756	27033858	refGene_gn=NMNAT2;refGene_tx=XM_011290996.3
F1	27033995	27034063	refGene_gn=NMNAT2;refGene_tx=XM_011290996.3
F1	27036015	27036118	refGene_gn=NMNAT2;refGene_tx=XM_011290996.3
F1	27044823	27045077	refGene_gn=LAMC2;refGene_tx=XM_023248065.1;refGene_gn=LAMC2;refGene_tx=XM_023248066.1
F1	27047478	27047578	refGene_gn=LAMC2;refGene_tx=XM_023248065.1;refGene_gn=LAMC2;refGene_tx=XM_023248066.1
F1	27047671	27047830	refGene_gn=LAMC2;refGene_tx=XM_023248065.1;refGene_gn=LAMC2;refGene_tx=XM_023248066.1
F1	27050896	27051096	refGene_gn=LAMC2;refGene_tx=XM_023248065.1;refGene_gn=LAMC2;refGene_tx=XM_023248066.1
F1	27051623	27051738	refGene_gn=LAMC2;refGene_tx=XM_023248065.1;refGene_gn=LAMC2;refGene_tx=XM_023248066.1
F1	27052247	27052397	refGene_gn=LAMC2;refGene_tx=XM_023248065.1;refGene_gn=LAMC2;refGene_tx=XM_023248066.1
F1	27053038	27053183	refGene_gn=LAMC2;refGene_tx=XM_023248065.1;refGene_gn=LAMC2;refGene_tx=XM_023248066.1
F1	27053662	27053812	refGene_gn=LAMC2;refGene_tx=XM_023248065.1;refGene_gn=LAMC2;refGene_tx=XM_023248066.1
F1	27054893	27054973	refGene_gn=LAMC2;refGene_tx=XM_023248065.1;refGene_gn=LAMC2;refGene_tx=XM_023248066.1
F1	27056149	27056355	refGene_gn=LAMC2;refGene_tx=XM_023248065.1;refGene_gn=LAMC2;refGene_tx

=XM_023248066.1
F1 27056612 27056769
refGene_gn=LAMC2;refGene_tx=XM_023248065.1;refGene_gn=LAMC2;refGene_tx
=XM_023248066.1
F1 27057698 27057844
refGene_gn=LAMC2;refGene_tx=XM_023248065.1;refGene_gn=LAMC2;refGene_tx
=XM_023248066.1
F1 27059762 27060008
refGene_gn=LAMC2;refGene_tx=XM_023248065.1;refGene_gn=LAMC2;refGene_tx
=XM_023248066.1
F1 27060737 27060920
refGene_gn=LAMC2;refGene_tx=XM_023248065.1;refGene_gn=LAMC2;refGene_tx
=XM_023248066.1
F1 27061543 27061762
refGene_gn=LAMC2;refGene_tx=XM_023248065.1;refGene_gn=LAMC2;refGene_tx
=XM_023248066.1
F1 27062817 27062930
refGene_gn=LAMC2;refGene_tx=XM_023248065.1;refGene_gn=LAMC2;refGene_tx
=XM_023248066.1
F1 27063912 27064102
refGene_gn=LAMC2;refGene_tx=XM_023248065.1;refGene_gn=LAMC2;refGene_tx
=XM_023248066.1
F1 27065531 27065654
refGene_gn=LAMC2;refGene_tx=XM_023248065.1;refGene_gn=LAMC2;refGene_tx
=XM_023248066.1
F1 27066386 27066523
refGene_gn=LAMC2;refGene_tx=XM_023248065.1;refGene_gn=LAMC2;refGene_tx
=XM_023248066.1
F1 27068691 27068790
refGene_gn=LAMC2;refGene_tx=XM_023248065.1;refGene_gn=LAMC2;refGene_tx
=XM_023248066.1
F1 27071781 27071917
refGene_gn=LAMC2;refGene_tx=XM_023248065.1;refGene_gn=LAMC2;refGene_tx
=XM_023248066.1
F1 27080161 27080350
refGene_gn=LAMC2;refGene_tx=XM_023248065.1;refGene_gn=LAMC2;refGene_tx
=XM_023248066.1
F1 27100124 27100203
refGene_gn=LAMC2;refGene_tx=XM_023248065.1;refGene_gn=LAMC2;refGene_tx
=XM_023248066.1
F1 27134364 27134621 refGene_gn=LAMC1;refGene_tx=XM_011291050.3
F1 27136201 27136301 refGene_gn=LAMC1;refGene_tx=XM_011291050.3
F1 27138542 27138701 refGene_gn=LAMC1;refGene_tx=XM_011291050.3
F1 27139808 27140008 refGene_gn=LAMC1;refGene_tx=XM_011291050.3
F1 27141227 27141342 refGene_gn=LAMC1;refGene_tx=XM_011291050.3
F1 27141599 27141749 refGene_gn=LAMC1;refGene_tx=XM_011291050.3
F1 27143698 27143843 refGene_gn=LAMC1;refGene_tx=XM_011291050.3
F1 27144576 27144714 refGene_gn=LAMC1;refGene_tx=XM_011291050.3
F1 27146275 27146355 refGene_gn=LAMC1;refGene_tx=XM_011291050.3
F1 27146784 27146990 refGene_gn=LAMC1;refGene_tx=XM_011291050.3

F1 27151021 27151178 refGene_gn=LAMC1;refGene_tx=XM_011291050.3
F1 27152232 27152411 refGene_gn=LAMC1;refGene_tx=XM_011291050.3
F1 27153368 27153511 refGene_gn=LAMC1;refGene_tx=XM_011291050.3
F1 27154114 27154268 refGene_gn=LAMC1;refGene_tx=XM_011291050.3
F1 27154741 27154987 refGene_gn=LAMC1;refGene_tx=XM_011291050.3
F1 27160413 27160602 refGene_gn=LAMC1;refGene_tx=XM_011291050.3
F1 27160721 27160943 refGene_gn=LAMC1;refGene_tx=XM_011291050.3
F1 27163092 27163205 refGene_gn=LAMC1;refGene_tx=XM_011291050.3
F1 27163515 27163705 refGene_gn=LAMC1;refGene_tx=XM_011291050.3
F1 27163796 27163919 refGene_gn=LAMC1;refGene_tx=XM_011291050.3
F1 27164532 27164669 refGene_gn=LAMC1;refGene_tx=XM_011291050.3
F1 27164759 27164858 refGene_gn=LAMC1;refGene_tx=XM_011291050.3
F1 27166250 27166368 refGene_gn=LAMC1;refGene_tx=XM_011291050.3
F1 27167135 27167324 refGene_gn=LAMC1;refGene_tx=XM_011291050.3
F1 27170329 27170496 refGene_gn=LAMC1;refGene_tx=XM_011291050.3
F1 27172825 27172956 refGene_gn=LAMC1;refGene_tx=XM_011291050.3
F1 27177526 27177831 refGene_gn=LAMC1;refGene_tx=XM_011291050.3
F1 27260687 27261102 refGene_gn=LAMC1;refGene_tx=XM_011291050.3
F1 27314562 27314985 refGene_gn=SHCBP1L;refGene_tx=XM_023247661.1
F1 27316421 27316571 refGene_gn=SHCBP1L;refGene_tx=XM_023247661.1
F1 27326014 27326229 refGene_gn=SHCBP1L;refGene_tx=XM_023247661.1
F1 27327192 27327279 refGene_gn=SHCBP1L;refGene_tx=XM_023247661.1
F1 27327587 27327806 refGene_gn=SHCBP1L;refGene_tx=XM_023247661.1
F1 27337306 27337412 refGene_gn=SHCBP1L;refGene_tx=XM_023247661.1
F1 27347036 27347190 refGene_gn=SHCBP1L;refGene_tx=XM_023247661.1
F1 27348844 27349095 refGene_gn=SHCBP1L;refGene_tx=XM_023247661.1
F1 27349853 27349976 refGene_gn=SHCBP1L;refGene_tx=XM_023247661.1
F1 27350793 27351042 refGene_gn=SHCBP1L;refGene_tx=XM_023247661.1
F1 27359060 27359427
refGene_gn=DHX9;refGene_tx=XM_011290994.3;refGene_gn=DHX9;refGene_tx=X
M_019821747.2
F1 27361638 27361839
refGene_gn=DHX9;refGene_tx=XM_011290994.3;refGene_gn=DHX9;refGene_tx=X
M_019821747.2
F1 27362543 27362659
refGene_gn=DHX9;refGene_tx=XM_011290994.3;refGene_gn=DHX9;refGene_tx=X
M_019821747.2
F1 27363062 27363292
refGene_gn=DHX9;refGene_tx=XM_011290994.3;refGene_gn=DHX9;refGene_tx=X
M_019821747.2
F1 27364226 27364354
refGene_gn=DHX9;refGene_tx=XM_011290994.3;refGene_gn=DHX9;refGene_tx=X
M_019821747.2
F1 27364450 27364612
refGene_gn=DHX9;refGene_tx=XM_011290994.3;refGene_gn=DHX9;refGene_tx=X
M_019821747.2
F1 27365182 27365294
refGene_gn=DHX9;refGene_tx=XM_011290994.3;refGene_gn=DHX9;refGene_tx=X
M_019821747.2
F1 27366046 27366207

refGene_gn=DHX9;refGene_tx=XM_011290994.3;refGene_gn=DHX9;refGene_tx=XM_019821747.2
F1 27367540 27367693
refGene_gn=DHX9;refGene_tx=XM_011290994.3;refGene_gn=DHX9;refGene_tx=XM_019821747.2
F1 27368898 27368972
refGene_gn=DHX9;refGene_tx=XM_011290994.3;refGene_gn=DHX9;refGene_tx=XM_019821747.2
F1 27369262 27369357
refGene_gn=DHX9;refGene_tx=XM_011290994.3;refGene_gn=DHX9;refGene_tx=XM_019821747.2
F1 27369551 27369765
refGene_gn=DHX9;refGene_tx=XM_011290994.3;refGene_gn=DHX9;refGene_tx=XM_019821747.2
F1 27371276 27371377
refGene_gn=DHX9;refGene_tx=XM_011290994.3;refGene_gn=DHX9;refGene_tx=XM_019821747.2
F1 27373347 27373504
refGene_gn=DHX9;refGene_tx=XM_011290994.3;refGene_gn=DHX9;refGene_tx=XM_019821747.2
F1 27375944 27376027
refGene_gn=DHX9;refGene_tx=XM_011290994.3;refGene_gn=DHX9;refGene_tx=XM_019821747.2
F1 27376459 27376601
refGene_gn=DHX9;refGene_tx=XM_011290994.3;refGene_gn=DHX9;refGene_tx=XM_019821747.2
F1 27380957 27381149
refGene_gn=DHX9;refGene_tx=XM_011290994.3;refGene_gn=DHX9;refGene_tx=XM_019821747.2
F1 27382138 27382216
refGene_gn=DHX9;refGene_tx=XM_011290994.3;refGene_gn=DHX9;refGene_tx=XM_019821747.2
F1 27382357 27382519
refGene_gn=DHX9;refGene_tx=XM_011290994.3;refGene_gn=DHX9;refGene_tx=XM_019821747.2
F1 27382615 27382705
refGene_gn=DHX9;refGene_tx=XM_011290994.3;refGene_gn=DHX9;refGene_tx=XM_019821747.2
F1 27383067 27383204
refGene_gn=DHX9;refGene_tx=XM_011290994.3;refGene_gn=DHX9;refGene_tx=XM_019821747.2
F1 27384813 27384860
refGene_gn=DHX9;refGene_tx=XM_011290994.3;refGene_gn=DHX9;refGene_tx=XM_019821747.2
F1 27387179 27387328 refGene_gn=DHX9;refGene_tx=XM_019821747.2
F1 27388099 27388212 refGene_gn=DHX9;refGene_tx=XM_019821747.2
F1 27389684 27389799 refGene_gn=DHX9;refGene_tx=XM_019821747.2
F1 27406124 27406265 refGene_gn=DHX9;refGene_tx=XM_019821747.2
F1 27407548 27407659 refGene_gn=DHX9;refGene_tx=XM_019821747.2
F1 27425848 27426033

refGene_gn=NPL;refGene_tx=XM_003999270.5;refGene_gn=NPL;refGene_tx=XM_006942759.4;refGene_gn=NPL;refGene_tx=XM_011290993.3;refGene_gn=NPL;refGene_tx=XM_019821757.2;refGene_gn=NPL;refGene_tx=XM_023247373.1
F1 27429345 27429385
refGene_gn=NPL;refGene_tx=XM_003999270.5;refGene_gn=NPL;refGene_tx=XM_006942759.4;refGene_gn=NPL;refGene_tx=XM_011290993.3;refGene_gn=NPL;refGene_tx=XM_019821757.2;refGene_gn=NPL;refGene_tx=XM_023247373.1
F1 27432523 27432608
refGene_gn=NPL;refGene_tx=XM_003999270.5;refGene_gn=NPL;refGene_tx=XM_006942759.4;refGene_gn=NPL;refGene_tx=XM_019821757.2;refGene_gn=NPL;refGene_tx=XM_023247373.1
F1 27435187 27435234
refGene_gn=NPL;refGene_tx=XM_003999270.5;refGene_gn=NPL;refGene_tx=XM_006942759.4;refGene_gn=NPL;refGene_tx=XM_011290993.3;refGene_gn=NPL;refGene_tx=XM_019821757.2;refGene_gn=NPL;refGene_tx=XM_023247373.1
F1 27435353 27435502
refGene_gn=NPL;refGene_tx=XM_003999270.5;refGene_gn=NPL;refGene_tx=XM_006942759.4;refGene_gn=NPL;refGene_tx=XM_011290993.3;refGene_gn=NPL;refGene_tx=XM_019821757.2;refGene_gn=NPL;refGene_tx=XM_023247373.1
F1 27436224 27436317
refGene_gn=NPL;refGene_tx=XM_003999270.5;refGene_gn=NPL;refGene_tx=XM_006942759.4;refGene_gn=NPL;refGene_tx=XM_011290993.3;refGene_gn=NPL;refGene_tx=XM_023247373.1
F1 27437861 27437937
refGene_gn=NPL;refGene_tx=XM_003999270.5;refGene_gn=NPL;refGene_tx=XM_006942759.4;refGene_gn=NPL;refGene_tx=XM_011290993.3;refGene_gn=NPL;refGene_tx=XM_019821757.2;refGene_gn=NPL;refGene_tx=XM_023247373.1
F1 27443060 27443118
refGene_gn=NPL;refGene_tx=XM_003999270.5;refGene_gn=NPL;refGene_tx=XM_006942759.4;refGene_gn=NPL;refGene_tx=XM_011290993.3;refGene_gn=NPL;refGene_tx=XM_019821757.2;refGene_gn=NPL;refGene_tx=XM_023247373.1
F1 27447114 27447202
refGene_gn=NPL;refGene_tx=XM_003999270.5;refGene_gn=NPL;refGene_tx=XM_006942759.4;refGene_gn=NPL;refGene_tx=XM_011290993.3;refGene_gn=NPL;refGene_tx=XM_019821757.2
F1 27448906 27448980
refGene_gn=NPL;refGene_tx=XM_003999270.5;refGene_gn=NPL;refGene_tx=XM_006942759.4;refGene_gn=NPL;refGene_tx=XM_011290993.3;refGene_gn=NPL;refGene_tx=XM_019821757.2;refGene_gn=NPL;refGene_tx=XM_023247373.1
F1 27455487 27455490 refGene_gn=NPL;refGene_tx=XM_023247373.1
F1 27459004 27459072
refGene_gn=NPL;refGene_tx=XM_003999270.5;refGene_gn=NPL;refGene_tx=XM_006942759.4;refGene_gn=NPL;refGene_tx=XM_011290993.3;refGene_gn=NPL;refGene_tx=XM_019821757.2
F1 27528155 27528158 refGene_gn=RGS8;refGene_tx=XM_019821754.2
F1 27546351 27546450 refGene_gn=RGS8;refGene_tx=XM_023247246.1
F1 27554233 27554629
refGene_gn=RGS8;refGene_tx=XM_019821748.2;refGene_gn=RGS8;refGene_tx=XM_019821749.2
F1 27567729 27567735 refGene_gn=RGS8;refGene_tx=XM_019821753.2

F1	27568852	27568926
refGene_gn=RGS8;refGene_tx=XM_019821748.2;refGene_gn=RGS8;refGene_tx=XM_019821749.2;refGene_gn=RGS8;refGene_tx=XM_019821753.2;refGene_gn=RGS8;refGene_tx=XM_019821754.2;refGene_gn=RGS8;refGene_tx=XM_023247246.1
F1	27570910	27570936
refGene_gn=RGS8;refGene_tx=XM_006942760.4;refGene_gn=RGS8;refGene_tx=XM_019821755.2;refGene_gn=RGS8;refGene_tx=XM_019821756.2
F1	27573398	27573500
refGene_gn=RGS8;refGene_tx=XM_006942760.4;refGene_gn=RGS8;refGene_tx=XM_019821748.2;refGene_gn=RGS8;refGene_tx=XM_019821753.2;refGene_gn=RGS8;refGene_tx=XM_019821754.2;refGene_gn=RGS8;refGene_tx=XM_019821755.2;refGene_gn=RGS8;refGene_tx=XM_019821756.2;refGene_gn=RGS8;refGene_tx=XM_023247246.1
F1	27574562	27574627
refGene_gn=RGS8;refGene_tx=XM_006942760.4;refGene_gn=RGS8;refGene_tx=XM_019821748.2;refGene_gn=RGS8;refGene_tx=XM_019821749.2;refGene_gn=RGS8;refGene_tx=XM_019821753.2;refGene_gn=RGS8;refGene_tx=XM_019821754.2;refGene_gn=RGS8;refGene_tx=XM_019821755.2;refGene_gn=RGS8;refGene_tx=XM_019821756.2;refGene_gn=RGS8;refGene_tx=XM_023247246.1
F1	27591068	27591235
refGene_gn=RGS8;refGene_tx=XM_006942760.4;refGene_gn=RGS8;refGene_tx=XM_019821748.2;refGene_gn=RGS8;refGene_tx=XM_019821749.2;refGene_gn=RGS8;refGene_tx=XM_019821753.2;refGene_gn=RGS8;refGene_tx=XM_019821754.2;refGene_gn=RGS8;refGene_tx=XM_019821755.2;refGene_gn=RGS8;refGene_tx=XM_019821756.2;refGene_gn=RGS8;refGene_tx=XM_023247246.1
F1	27592464	27592647
refGene_gn=RGS8;refGene_tx=XM_006942760.4;refGene_gn=RGS8;refGene_tx=XM_019821748.2;refGene_gn=RGS8;refGene_tx=XM_019821749.2;refGene_gn=RGS8;refGene_tx=XM_019821753.2;refGene_gn=RGS8;refGene_tx=XM_019821754.2;refGene_gn=RGS8;refGene_tx=XM_019821755.2;refGene_gn=RGS8;refGene_tx=XM_019821756.2;refGene_gn=RGS8;refGene_tx=XM_023247246.1
F1	27633720	27633764
refGene_gn=RGS16;refGene_tx=XM_003999274.5;refGene_gn=RGS16;refGene_tx=XM_006942761.4
F1	27634665	27634776
refGene_gn=RGS16;refGene_tx=XM_003999274.5;refGene_gn=RGS16;refGene_tx=XM_006942761.4
F1	27635528	27635593
refGene_gn=RGS16;refGene_tx=XM_003999274.5;refGene_gn=RGS16;refGene_tx=XM_006942761.4
F1	27635881	27636048
refGene_gn=RGS16;refGene_tx=XM_003999274.5;refGene_gn=RGS16;refGene_tx=XM_006942761.4
F1	27637929	27638151
refGene_gn=RGS16;refGene_tx=XM_003999274.5;refGene_gn=RGS16;refGene_tx=XM_006942761.4
F1	27650682	27652159	refGene_gn=RNASEL;refGene_tx=XM_006942762.3
F1	27661868	27661954	refGene_gn=RNASEL;refGene_tx=XM_006942762.3
F1	27663176	27663382	refGene_gn=RNASEL;refGene_tx=XM_006942762.3
F1	27664267	27664400	refGene_gn=RNASEL;refGene_tx=XM_006942762.3

F1 27667128 27667262 refGene_gn=RNASEL;refGene_tx=XM_006942762.3
F1 27668350 27668489 refGene_gn=RNASEL;refGene_tx=XM_006942762.3
F1 27782400 27782471 refGene_gn=RGSL1;refGene_tx=XM_006942763.4
F1 27783487 27783606
refGene_gn=RGSL1;refGene_tx=XM_006942763.4;refGene_gn=RGSL1;refGene_tx=XM_019821761.2
F1 27785367 27785477
refGene_gn=RGSL1;refGene_tx=XM_006942763.4;refGene_gn=RGSL1;refGene_tx=XM_019821761.2
F1 27787780 27787905
refGene_gn=RGSL1;refGene_tx=XM_006942763.4;refGene_gn=RGSL1;refGene_tx=XM_019821761.2
F1 27788158 27788297
refGene_gn=RGSL1;refGene_tx=XM_006942763.4;refGene_gn=RGSL1;refGene_tx=XM_019821761.2
F1 27799567 27799742
refGene_gn=RGSL1;refGene_tx=XM_006942763.4;refGene_gn=RGSL1;refGene_tx=XM_019821761.2
F1 27806607 27806737
refGene_gn=RGSL1;refGene_tx=XM_006942763.4;refGene_gn=RGSL1;refGene_tx=XM_019821761.2
F1 27808971 27809083
refGene_gn=RGSL1;refGene_tx=XM_006942763.4;refGene_gn=RGSL1;refGene_tx=XM_019821761.2
F1 27809510 27809628
refGene_gn=RGSL1;refGene_tx=XM_006942763.4;refGene_gn=RGSL1;refGene_tx=XM_019821761.2
F1 27810938 27811132
refGene_gn=RGSL1;refGene_tx=XM_006942763.4;refGene_gn=RGSL1;refGene_tx=XM_019821761.2
F1 27814255 27814361
refGene_gn=RGSL1;refGene_tx=XM_006942763.4;refGene_gn=RGSL1;refGene_tx=XM_019821761.2
F1 27855874 27855982
refGene_gn=RGSL1;refGene_tx=XM_006942763.4;refGene_gn=RGSL1;refGene_tx=XM_019821761.2
F1 27860012 27860235
refGene_gn=RGSL1;refGene_tx=XM_006942763.4;refGene_gn=RGSL1;refGene_tx=XM_019821761.2
F1 27860629 27860692
refGene_gn=RGSL1;refGene_tx=XM_006942763.4;refGene_gn=RGSL1;refGene_tx=XM_019821761.2
F1 27871182 27872162
refGene_gn=RGSL1;refGene_tx=XM_006942763.4;refGene_gn=RGSL1;refGene_tx=XM_019821761.2
F1 27873187 27873349
refGene_gn=RGSL1;refGene_tx=XM_006942763.4;refGene_gn=RGSL1;refGene_tx=XM_019821761.2
F1 27874332 27874437
refGene_gn=RGSL1;refGene_tx=XM_006942763.4;refGene_gn=RGSL1;refGene_tx

=XM_019821761.2
F1 27876436 27876566
refGene_gn=RGSL1;refGene_tx=XM_006942763.4;refGene_gn=RGSL1;refGene_tx
=XM_019821761.2
F1 27881631 27881706 refGene_gn=RGSL1;refGene_tx=XM_006942763.4
F1 27884808 27884891
refGene_gn=RGSL1;refGene_tx=XM_006942763.4;refGene_gn=RGSL1;refGene_tx
=XM_019821761.2
F1 27888952 27888965
refGene_gn=RGSL1;refGene_tx=XM_006942763.4;refGene_gn=RGSL1;refGene_tx
=XM_019821761.2
F1 27899419 27900193
refGene_gn=LOC101092577;refGene_tx=XM_003999275.4
F1 27931198 27932104
refGene_gn=LOC101092820;refGene_tx=XM_003999276.5
F1 27945493 27946399 refGene_gn=TEDDM1;refGene_tx=XM_003999277.4
F1 27946407 27946566
refGene_gn=LOC111558341;refGene_tx=XM_023247453.1
F1 27953042 27953480
refGene_gn=LOC111558341;refGene_tx=XM_023247453.1
F1 27954582 27954689 refGene_gn=GLUL;refGene_tx=XM_003999279.5
F1 27956087 27956266
refGene_gn=GLUL;refGene_tx=XM_003999278.5;refGene_gn=GLUL;refGene_tx=X
M_003999279.5;refGene_gn=GLUL;refGene_tx=XM_006942764.4
F1 27958256 27958418
refGene_gn=GLUL;refGene_tx=XM_003999278.5;refGene_gn=GLUL;refGene_tx=X
M_003999279.5;refGene_gn=GLUL;refGene_tx=XM_006942764.4
F1 27960028 27960175
refGene_gn=GLUL;refGene_tx=XM_003999278.5;refGene_gn=GLUL;refGene_tx=X
M_003999279.5;refGene_gn=GLUL;refGene_tx=XM_006942764.4
F1 27960565 27960693
refGene_gn=GLUL;refGene_tx=XM_003999278.5;refGene_gn=GLUL;refGene_tx=X
M_003999279.5;refGene_gn=GLUL;refGene_tx=XM_006942764.4
F1 27960864 27961064
refGene_gn=GLUL;refGene_tx=XM_003999278.5;refGene_gn=GLUL;refGene_tx=X
M_003999279.5;refGene_gn=GLUL;refGene_tx=XM_006942764.4
F1 27961630 27961949
refGene_gn=GLUL;refGene_tx=XM_003999278.5;refGene_gn=GLUL;refGene_tx=X
M_003999279.5;refGene_gn=GLUL;refGene_tx=XM_006942764.4
F1 28214443 28216150
refGene_gn=ZNF648;refGene_tx=XM_003999280.3;refGene_gn=ZNF648;refGene_
tx=XM_019821763.2;refGene_gn=ZNF648;refGene_tx=XM_023247844.1
F1 28435834 28436377
refGene_gn=CACNA1E;refGene_tx=XM_023247743.1;refGene_gn=CACNA1E;refGen
e_tx=XM_023247744.1;refGene_gn=CACNA1E;refGene_tx=XM_023247745.1;refGe
ne_gn=CACNA1E;refGene_tx=XM_023247746.1;refGene_gn=CACNA1E;refGene_tx=
XM_023247747.1;refGene_gn=CACNA1E;refGene_tx=XM_023247748.1;refGene_gn
=CACNA1E;refGene_tx=XM_023247749.1;refGene_gn=CACNA1E;refGene_tx=XM_02
3247750.1
F1 28437841 28438032

refGene_gn=CACNA1E;refGene_tx=XM_023247743.1;refGene_gn=CACNA1E;refGene_tx=XM_023247744.1;refGene_gn=CACNA1E;refGene_tx=XM_023247745.1;refGene_gn=CACNA1E;refGene_tx=XM_023247746.1;refGene_gn=CACNA1E;refGene_tx=XM_023247747.1;refGene_gn=CACNA1E;refGene_tx=XM_023247748.1;refGene_gn=CACNA1E;refGene_tx=XM_023247749.1;refGene_gn=CACNA1E;refGene_tx=XM_023247750.1
F1 28439051 28439232
refGene_gn=CACNA1E;refGene_tx=XM_023247743.1;refGene_gn=CACNA1E;refGene_tx=XM_023247744.1;refGene_gn=CACNA1E;refGene_tx=XM_023247745.1;refGene_gn=CACNA1E;refGene_tx=XM_023247746.1;refGene_gn=CACNA1E;refGene_tx=XM_023247747.1;refGene_gn=CACNA1E;refGene_tx=XM_023247748.1;refGene_gn=CACNA1E;refGene_tx=XM_023247749.1;refGene_gn=CACNA1E;refGene_tx=XM_023247750.1
F1 28440254 28440383
refGene_gn=CACNA1E;refGene_tx=XM_023247743.1;refGene_gn=CACNA1E;refGene_tx=XM_023247744.1;refGene_gn=CACNA1E;refGene_tx=XM_023247745.1;refGene_gn=CACNA1E;refGene_tx=XM_023247746.1;refGene_gn=CACNA1E;refGene_tx=XM_023247747.1
F1 28443698 28443810
refGene_gn=CACNA1E;refGene_tx=XM_023247743.1;refGene_gn=CACNA1E;refGene_tx=XM_023247744.1;refGene_gn=CACNA1E;refGene_tx=XM_023247745.1;refGene_gn=CACNA1E;refGene_tx=XM_023247746.1;refGene_gn=CACNA1E;refGene_tx=XM_023247747.1;refGene_gn=CACNA1E;refGene_tx=XM_023247748.1;refGene_gn=CACNA1E;refGene_tx=XM_023247749.1;refGene_gn=CACNA1E;refGene_tx=XM_023247750.1
F1 28447697 28447804
refGene_gn=CACNA1E;refGene_tx=XM_023247743.1;refGene_gn=CACNA1E;refGene_tx=XM_023247744.1;refGene_gn=CACNA1E;refGene_tx=XM_023247745.1;refGene_gn=CACNA1E;refGene_tx=XM_023247746.1;refGene_gn=CACNA1E;refGene_tx=XM_023247747.1;refGene_gn=CACNA1E;refGene_tx=XM_023247748.1;refGene_gn=CACNA1E;refGene_tx=XM_023247749.1;refGene_gn=CACNA1E;refGene_tx=XM_023247750.1
F1 28448088 28448189
refGene_gn=CACNA1E;refGene_tx=XM_023247743.1;refGene_gn=CACNA1E;refGene_tx=XM_023247744.1;refGene_gn=CACNA1E;refGene_tx=XM_023247745.1;refGene_gn=CACNA1E;refGene_tx=XM_023247746.1;refGene_gn=CACNA1E;refGene_tx=XM_023247747.1;refGene_gn=CACNA1E;refGene_tx=XM_023247748.1;refGene_gn=CACNA1E;refGene_tx=XM_023247749.1;refGene_gn=CACNA1E;refGene_tx=XM_023247750.1
F1 28448756 28448864
refGene_gn=CACNA1E;refGene_tx=XM_023247743.1;refGene_gn=CACNA1E;refGene_tx=XM_023247744.1;refGene_gn=CACNA1E;refGene_tx=XM_023247745.1;refGene_gn=CACNA1E;refGene_tx=XM_023247746.1;refGene_gn=CACNA1E;refGene_tx=XM_023247747.1;refGene_gn=CACNA1E;refGene_tx=XM_023247748.1;refGene_gn=CACNA1E;refGene_tx=XM_023247749.1;refGene_gn=CACNA1E;refGene_tx=XM_023247750.1
F1 28449728 28449834
refGene_gn=CACNA1E;refGene_tx=XM_023247743.1;refGene_gn=CACNA1E;refGene_tx=XM_023247744.1;refGene_gn=CACNA1E;refGene_tx=XM_023247745.1;refGene_gn=CACNA1E;refGene_tx=XM_023247746.1;refGene_gn=CACNA1E;refGene_tx=

XM_023247747.1;refGene_gn=CACNA1E;refGene_tx=XM_023247748.1;refGene_gn=CACNA1E;refGene_tx=XM_023247749.1;refGene_gn=CACNA1E;refGene_tx=XM_023247750.1
F1 28451892 28451989
refGene_gn=CACNA1E;refGene_tx=XM_023247743.1;refGene_gn=CACNA1E;refGene_tx=XM_023247745.1;refGene_gn=CACNA1E;refGene_tx=XM_023247747.1;refGene_gn=CACNA1E;refGene_tx=XM_023247748.1;refGene_gn=CACNA1E;refGene_tx=XM_023247749.1
F1 28453876 28453973
refGene_gn=CACNA1E;refGene_tx=XM_023247744.1;refGene_gn=CACNA1E;refGene_tx=XM_023247746.1;refGene_gn=CACNA1E;refGene_tx=XM_023247750.1
F1 28457146 28457274
refGene_gn=CACNA1E;refGene_tx=XM_023247743.1;refGene_gn=CACNA1E;refGene_tx=XM_023247744.1;refGene_gn=CACNA1E;refGene_tx=XM_023247745.1;refGene_gn=CACNA1E;refGene_tx=XM_023247746.1;refGene_gn=CACNA1E;refGene_tx=XM_023247747.1;refGene_gn=CACNA1E;refGene_tx=XM_023247748.1;refGene_gn=CACNA1E;refGene_tx=XM_023247749.1;refGene_gn=CACNA1E;refGene_tx=XM_023247750.1
F1 28461443 28461609
refGene_gn=CACNA1E;refGene_tx=XM_023247743.1;refGene_gn=CACNA1E;refGene_tx=XM_023247744.1;refGene_gn=CACNA1E;refGene_tx=XM_023247745.1;refGene_gn=CACNA1E;refGene_tx=XM_023247746.1;refGene_gn=CACNA1E;refGene_tx=XM_023247747.1;refGene_gn=CACNA1E;refGene_tx=XM_023247748.1;refGene_gn=CACNA1E;refGene_tx=XM_023247749.1;refGene_gn=CACNA1E;refGene_tx=XM_023247750.1
F1 28462204 28462296
refGene_gn=CACNA1E;refGene_tx=XM_023247743.1;refGene_gn=CACNA1E;refGene_tx=XM_023247744.1;refGene_gn=CACNA1E;refGene_tx=XM_023247745.1;refGene_gn=CACNA1E;refGene_tx=XM_023247746.1;refGene_gn=CACNA1E;refGene_tx=XM_023247747.1;refGene_gn=CACNA1E;refGene_tx=XM_023247748.1;refGene_gn=CACNA1E;refGene_tx=XM_023247749.1;refGene_gn=CACNA1E;refGene_tx=XM_023247750.1
F1 28466400 28466466
refGene_gn=CACNA1E;refGene_tx=XM_023247743.1;refGene_gn=CACNA1E;refGene_tx=XM_023247744.1;refGene_gn=CACNA1E;refGene_tx=XM_023247745.1;refGene_gn=CACNA1E;refGene_tx=XM_023247746.1;refGene_gn=CACNA1E;refGene_tx=XM_023247747.1;refGene_gn=CACNA1E;refGene_tx=XM_023247748.1;refGene_gn=CACNA1E;refGene_tx=XM_023247749.1;refGene_gn=CACNA1E;refGene_tx=XM_023247750.1
F1 28469967 28470093
refGene_gn=CACNA1E;refGene_tx=XM_023247743.1;refGene_gn=CACNA1E;refGene_tx=XM_023247744.1;refGene_gn=CACNA1E;refGene_tx=XM_023247745.1;refGene_gn=CACNA1E;refGene_tx=XM_023247746.1;refGene_gn=CACNA1E;refGene_tx=XM_023247747.1;refGene_gn=CACNA1E;refGene_tx=XM_023247748.1;refGene_gn=CACNA1E;refGene_tx=XM_023247749.1;refGene_gn=CACNA1E;refGene_tx=XM_023247750.1
F1 28470876 28470960
refGene_gn=CACNA1E;refGene_tx=XM_023247743.1;refGene_gn=CACNA1E;refGene_tx=XM_023247744.1;refGene_gn=CACNA1E;refGene_tx=XM_023247745.1;refGene_gn=CACNA1E;refGene_tx=XM_023247746.1;refGene_gn=CACNA1E;refGene_tx=

XM_023247747.1;refGene_gn=CACNA1E;refGene_tx=XM_023247748.1;refGene_gn=CACNA1E;refGene_tx=XM_023247749.1;refGene_gn=CACNA1E;refGene_tx=XM_023247750.1
F1 28474607 28474718
refGene_gn=CACNA1E;refGene_tx=XM_023247743.1;refGene_gn=CACNA1E;refGene_tx=XM_023247744.1;refGene_gn=CACNA1E;refGene_tx=XM_023247745.1;refGene_gn=CACNA1E;refGene_tx=XM_023247746.1;refGene_gn=CACNA1E;refGene_tx=XM_023247747.1;refGene_gn=CACNA1E;refGene_tx=XM_023247748.1;refGene_gn=CACNA1E;refGene_tx=XM_023247749.1;refGene_gn=CACNA1E;refGene_tx=XM_023247750.1
F1 28475382 28475547
refGene_gn=CACNA1E;refGene_tx=XM_023247743.1;refGene_gn=CACNA1E;refGene_tx=XM_023247744.1;refGene_gn=CACNA1E;refGene_tx=XM_023247745.1;refGene_gn=CACNA1E;refGene_tx=XM_023247746.1;refGene_gn=CACNA1E;refGene_tx=XM_023247747.1;refGene_gn=CACNA1E;refGene_tx=XM_023247748.1;refGene_gn=CACNA1E;refGene_tx=XM_023247749.1;refGene_gn=CACNA1E;refGene_tx=XM_023247750.1
F1 28476359 28476561
refGene_gn=CACNA1E;refGene_tx=XM_023247743.1;refGene_gn=CACNA1E;refGene_tx=XM_023247744.1;refGene_gn=CACNA1E;refGene_tx=XM_023247745.1;refGene_gn=CACNA1E;refGene_tx=XM_023247746.1;refGene_gn=CACNA1E;refGene_tx=XM_023247747.1;refGene_gn=CACNA1E;refGene_tx=XM_023247748.1;refGene_gn=CACNA1E;refGene_tx=XM_023247749.1;refGene_gn=CACNA1E;refGene_tx=XM_023247750.1
F1 28477389 28477527
refGene_gn=CACNA1E;refGene_tx=XM_023247743.1;refGene_gn=CACNA1E;refGene_tx=XM_023247744.1;refGene_gn=CACNA1E;refGene_tx=XM_023247745.1;refGene_gn=CACNA1E;refGene_tx=XM_023247746.1;refGene_gn=CACNA1E;refGene_tx=XM_023247747.1;refGene_gn=CACNA1E;refGene_tx=XM_023247748.1;refGene_gn=CACNA1E;refGene_tx=XM_023247749.1;refGene_gn=CACNA1E;refGene_tx=XM_023247750.1
F1 28478042 28478203
refGene_gn=CACNA1E;refGene_tx=XM_023247743.1;refGene_gn=CACNA1E;refGene_tx=XM_023247744.1;refGene_gn=CACNA1E;refGene_tx=XM_023247745.1;refGene_gn=CACNA1E;refGene_tx=XM_023247746.1;refGene_gn=CACNA1E;refGene_tx=XM_023247747.1;refGene_gn=CACNA1E;refGene_tx=XM_023247748.1;refGene_gn=CACNA1E;refGene_tx=XM_023247749.1;refGene_gn=CACNA1E;refGene_tx=XM_023247750.1
F1 28480378 28480475
refGene_gn=CACNA1E;refGene_tx=XM_023247743.1;refGene_gn=CACNA1E;refGene_tx=XM_023247744.1;refGene_gn=CACNA1E;refGene_tx=XM_023247745.1;refGene_gn=CACNA1E;refGene_tx=XM_023247746.1;refGene_gn=CACNA1E;refGene_tx=XM_023247747.1;refGene_gn=CACNA1E;refGene_tx=XM_023247748.1;refGene_gn=CACNA1E;refGene_tx=XM_023247749.1;refGene_gn=CACNA1E;refGene_tx=XM_023247750.1
F1 28482149 28482161
refGene_gn=CACNA1E;refGene_tx=XM_023247743.1;refGene_gn=CACNA1E;refGene_tx=XM_023247744.1;refGene_gn=CACNA1E;refGene_tx=XM_023247747.1;refGene_gn=CACNA1E;refGene_tx=XM_023247748.1;refGene_gn=CACNA1E;refGene_tx=XM_023247749.1;refGene_gn=CACNA1E;refGene_tx=XM_023247750.1

F1 28493250 28493357
refGene_gn=CACNA1E;refGene_tx=XM_023247743.1;refGene_gn=CACNA1E;refGene_tx=XM_023247744.1;refGene_gn=CACNA1E;refGene_tx=XM_023247745.1;refGene_gn=CACNA1E;refGene_tx=XM_023247746.1;refGene_gn=CACNA1E;refGene_tx=XM_023247747.1;refGene_gn=CACNA1E;refGene_tx=XM_023247748.1;refGene_gn=CACNA1E;refGene_tx=XM_023247749.1;refGene_gn=CACNA1E;refGene_tx=XM_023247750.1
F1 28494141 28494201
refGene_gn=CACNA1E;refGene_tx=XM_023247743.1;refGene_gn=CACNA1E;refGene_tx=XM_023247744.1;refGene_gn=CACNA1E;refGene_tx=XM_023247745.1;refGene_gn=CACNA1E;refGene_tx=XM_023247746.1;refGene_gn=CACNA1E;refGene_tx=XM_023247747.1;refGene_gn=CACNA1E;refGene_tx=XM_023247748.1;refGene_gn=CACNA1E;refGene_tx=XM_023247749.1;refGene_gn=CACNA1E;refGene_tx=XM_023247750.1
F1 28494900 28495030
refGene_gn=CACNA1E;refGene_tx=XM_023247743.1;refGene_gn=CACNA1E;refGene_tx=XM_023247744.1;refGene_gn=CACNA1E;refGene_tx=XM_023247745.1;refGene_gn=CACNA1E;refGene_tx=XM_023247746.1;refGene_gn=CACNA1E;refGene_tx=XM_023247747.1;refGene_gn=CACNA1E;refGene_tx=XM_023247748.1;refGene_gn=CACNA1E;refGene_tx=XM_023247749.1;refGene_gn=CACNA1E;refGene_tx=XM_023247750.1
F1 28496174 28496334
refGene_gn=CACNA1E;refGene_tx=XM_023247743.1;refGene_gn=CACNA1E;refGene_tx=XM_023247744.1;refGene_gn=CACNA1E;refGene_tx=XM_023247745.1;refGene_gn=CACNA1E;refGene_tx=XM_023247746.1;refGene_gn=CACNA1E;refGene_tx=XM_023247747.1;refGene_gn=CACNA1E;refGene_tx=XM_023247748.1;refGene_gn=CACNA1E;refGene_tx=XM_023247749.1;refGene_gn=CACNA1E;refGene_tx=XM_023247750.1
F1 28498617 28498931
refGene_gn=CACNA1E;refGene_tx=XM_023247743.1;refGene_gn=CACNA1E;refGene_tx=XM_023247744.1;refGene_gn=CACNA1E;refGene_tx=XM_023247745.1;refGene_gn=CACNA1E;refGene_tx=XM_023247746.1;refGene_gn=CACNA1E;refGene_tx=XM_023247747.1;refGene_gn=CACNA1E;refGene_tx=XM_023247748.1;refGene_gn=CACNA1E;refGene_tx=XM_023247749.1;refGene_gn=CACNA1E;refGene_tx=XM_023247750.1
F1 28499361 28500009
refGene_gn=CACNA1E;refGene_tx=XM_023247743.1;refGene_gn=CACNA1E;refGene_tx=XM_023247744.1;refGene_gn=CACNA1E;refGene_tx=XM_023247745.1;refGene_gn=CACNA1E;refGene_tx=XM_023247746.1;refGene_gn=CACNA1E;refGene_tx=XM_023247747.1;refGene_gn=CACNA1E;refGene_tx=XM_023247748.1;refGene_gn=CACNA1E;refGene_tx=XM_023247749.1;refGene_gn=CACNA1E;refGene_tx=XM_023247750.1
F1 28501135 28501192
refGene_gn=CACNA1E;refGene_tx=XM_023247743.1;refGene_gn=CACNA1E;refGene_tx=XM_023247744.1;refGene_gn=CACNA1E;refGene_tx=XM_023247745.1;refGene_gn=CACNA1E;refGene_tx=XM_023247746.1;refGene_gn=CACNA1E;refGene_tx=XM_023247748.1
F1 28505387 28505485
refGene_gn=CACNA1E;refGene_tx=XM_023247743.1;refGene_gn=CACNA1E;refGene_tx=XM_023247744.1;refGene_gn=CACNA1E;refGene_tx=XM_023247745.1;refGe

ne_gn=CACNA1E;refGene_tx=XM_023247746.1;refGene_gn=CACNA1E;refGene_tx=XM_023247747.1;refGene_gn=CACNA1E;refGene_tx=XM_023247748.1;refGene_gn=CACNA1E;refGene_tx=XM_023247749.1;refGene_gn=CACNA1E;refGene_tx=XM_023247750.1
F1 28506247 28506315
refGene_gn=CACNA1E;refGene_tx=XM_023247743.1;refGene_gn=CACNA1E;refGene_tx=XM_023247744.1;refGene_gn=CACNA1E;refGene_tx=XM_023247745.1;refGene_gn=CACNA1E;refGene_tx=XM_023247746.1;refGene_gn=CACNA1E;refGene_tx=XM_023247747.1;refGene_gn=CACNA1E;refGene_tx=XM_023247748.1;refGene_gn=CACNA1E;refGene_tx=XM_023247749.1;refGene_gn=CACNA1E;refGene_tx=XM_023247750.1
F1 28508108 28508226
refGene_gn=CACNA1E;refGene_tx=XM_023247743.1;refGene_gn=CACNA1E;refGene_tx=XM_023247744.1;refGene_gn=CACNA1E;refGene_tx=XM_023247745.1;refGene_gn=CACNA1E;refGene_tx=XM_023247746.1;refGene_gn=CACNA1E;refGene_tx=XM_023247747.1;refGene_gn=CACNA1E;refGene_tx=XM_023247748.1;refGene_gn=CACNA1E;refGene_tx=XM_023247749.1;refGene_gn=CACNA1E;refGene_tx=XM_023247750.1
F1 28510047 28510120
refGene_gn=CACNA1E;refGene_tx=XM_023247743.1;refGene_gn=CACNA1E;refGene_tx=XM_023247744.1;refGene_gn=CACNA1E;refGene_tx=XM_023247745.1;refGene_gn=CACNA1E;refGene_tx=XM_023247746.1;refGene_gn=CACNA1E;refGene_tx=XM_023247747.1;refGene_gn=CACNA1E;refGene_tx=XM_023247748.1;refGene_gn=CACNA1E;refGene_tx=XM_023247749.1;refGene_gn=CACNA1E;refGene_tx=XM_023247750.1
F1 28510599 28510731
refGene_gn=CACNA1E;refGene_tx=XM_023247743.1;refGene_gn=CACNA1E;refGene_tx=XM_023247744.1;refGene_gn=CACNA1E;refGene_tx=XM_023247745.1;refGene_gn=CACNA1E;refGene_tx=XM_023247746.1;refGene_gn=CACNA1E;refGene_tx=XM_023247747.1;refGene_gn=CACNA1E;refGene_tx=XM_023247748.1;refGene_gn=CACNA1E;refGene_tx=XM_023247749.1;refGene_gn=CACNA1E;refGene_tx=XM_023247750.1
F1 28511086 28511199
refGene_gn=CACNA1E;refGene_tx=XM_023247743.1;refGene_gn=CACNA1E;refGene_tx=XM_023247744.1;refGene_gn=CACNA1E;refGene_tx=XM_023247745.1;refGene_gn=CACNA1E;refGene_tx=XM_023247746.1;refGene_gn=CACNA1E;refGene_tx=XM_023247747.1;refGene_gn=CACNA1E;refGene_tx=XM_023247748.1;refGene_gn=CACNA1E;refGene_tx=XM_023247749.1;refGene_gn=CACNA1E;refGene_tx=XM_023247750.1
F1 28512902 28513015
refGene_gn=CACNA1E;refGene_tx=XM_023247743.1;refGene_gn=CACNA1E;refGene_tx=XM_023247744.1;refGene_gn=CACNA1E;refGene_tx=XM_023247745.1;refGene_gn=CACNA1E;refGene_tx=XM_023247746.1;refGene_gn=CACNA1E;refGene_tx=XM_023247747.1;refGene_gn=CACNA1E;refGene_tx=XM_023247748.1;refGene_gn=CACNA1E;refGene_tx=XM_023247749.1;refGene_gn=CACNA1E;refGene_tx=XM_023247750.1
F1 28513751 28513961
refGene_gn=CACNA1E;refGene_tx=XM_023247743.1;refGene_gn=CACNA1E;refGene_tx=XM_023247744.1;refGene_gn=CACNA1E;refGene_tx=XM_023247745.1;refGene_gn=CACNA1E;refGene_tx=XM_023247746.1;refGene_gn=CACNA1E;refGene_tx=

XM_023247747.1;refGene_gn=CACNA1E;refGene_tx=XM_023247748.1;refGene_gn=CACNA1E;refGene_tx=XM_023247749.1;refGene_gn=CACNA1E;refGene_tx=XM_023247750.1
F1 28515450 28515540
refGene_gn=CACNA1E;refGene_tx=XM_023247743.1;refGene_gn=CACNA1E;refGene_tx=XM_023247744.1;refGene_gn=CACNA1E;refGene_tx=XM_023247745.1;refGene_gn=CACNA1E;refGene_tx=XM_023247746.1;refGene_gn=CACNA1E;refGene_tx=XM_023247747.1;refGene_gn=CACNA1E;refGene_tx=XM_023247748.1;refGene_gn=CACNA1E;refGene_tx=XM_023247749.1;refGene_gn=CACNA1E;refGene_tx=XM_023247750.1
F1 28516392 28516446
refGene_gn=CACNA1E;refGene_tx=XM_023247743.1;refGene_gn=CACNA1E;refGene_tx=XM_023247744.1;refGene_gn=CACNA1E;refGene_tx=XM_023247745.1;refGene_gn=CACNA1E;refGene_tx=XM_023247746.1;refGene_gn=CACNA1E;refGene_tx=XM_023247747.1;refGene_gn=CACNA1E;refGene_tx=XM_023247748.1;refGene_gn=CACNA1E;refGene_tx=XM_023247749.1;refGene_gn=CACNA1E;refGene_tx=XM_023247750.1
F1 28520776 28520892
refGene_gn=CACNA1E;refGene_tx=XM_023247743.1;refGene_gn=CACNA1E;refGene_tx=XM_023247744.1;refGene_gn=CACNA1E;refGene_tx=XM_023247745.1;refGene_gn=CACNA1E;refGene_tx=XM_023247746.1;refGene_gn=CACNA1E;refGene_tx=XM_023247747.1;refGene_gn=CACNA1E;refGene_tx=XM_023247748.1;refGene_gn=CACNA1E;refGene_tx=XM_023247749.1;refGene_gn=CACNA1E;refGene_tx=XM_023247750.1
F1 28582520 28582624
refGene_gn=CACNA1E;refGene_tx=XM_023247743.1;refGene_gn=CACNA1E;refGene_tx=XM_023247744.1;refGene_gn=CACNA1E;refGene_tx=XM_023247745.1;refGene_gn=CACNA1E;refGene_tx=XM_023247746.1;refGene_gn=CACNA1E;refGene_tx=XM_023247747.1;refGene_gn=CACNA1E;refGene_tx=XM_023247748.1;refGene_gn=CACNA1E;refGene_tx=XM_023247749.1;refGene_gn=CACNA1E;refGene_tx=XM_023247750.1
F1 28653856 28654038
refGene_gn=CACNA1E;refGene_tx=XM_023247743.1;refGene_gn=CACNA1E;refGene_tx=XM_023247744.1;refGene_gn=CACNA1E;refGene_tx=XM_023247745.1;refGene_gn=CACNA1E;refGene_tx=XM_023247746.1;refGene_gn=CACNA1E;refGene_tx=XM_023247747.1;refGene_gn=CACNA1E;refGene_tx=XM_023247748.1;refGene_gn=CACNA1E;refGene_tx=XM_023247749.1;refGene_gn=CACNA1E;refGene_tx=XM_023247750.1
F1 28655227 28655380
refGene_gn=CACNA1E;refGene_tx=XM_023247743.1;refGene_gn=CACNA1E;refGene_tx=XM_023247744.1;refGene_gn=CACNA1E;refGene_tx=XM_023247745.1;refGene_gn=CACNA1E;refGene_tx=XM_023247746.1;refGene_gn=CACNA1E;refGene_tx=XM_023247747.1;refGene_gn=CACNA1E;refGene_tx=XM_023247748.1;refGene_gn=CACNA1E;refGene_tx=XM_023247749.1;refGene_gn=CACNA1E;refGene_tx=XM_023247750.1
F1 28656418 28656522
refGene_gn=CACNA1E;refGene_tx=XM_023247743.1;refGene_gn=CACNA1E;refGene_tx=XM_023247744.1;refGene_gn=CACNA1E;refGene_tx=XM_023247745.1;refGene_gn=CACNA1E;refGene_tx=XM_023247746.1;refGene_gn=CACNA1E;refGene_tx=XM_023247747.1;refGene_gn=CACNA1E;refGene_tx=XM_023247748.1;refGene_gn

=CACNA1E;refGene_tx=XM_023247749.1;refGene_gn=CACNA1E;refGene_tx=XM_023247750.1
F1 28714453 28714593 refGene_gn=CACNA1E;refGene_tx=XM_023247743.1;refGene_gn=CACNA1E;refGene_tx=XM_023247744.1;refGene_gn=CACNA1E;refGene_tx=XM_023247745.1;refGene_gn=CACNA1E;refGene_tx=XM_023247746.1;refGene_gn=CACNA1E;refGene_tx=XM_023247747.1;refGene_gn=CACNA1E;refGene_tx=XM_023247748.1;refGene_gn=CACNA1E;refGene_tx=XM_023247749.1;refGene_gn=CACNA1E;refGene_tx=XM_023247750.1
F1 28715471 28715577 refGene_gn=CACNA1E;refGene_tx=XM_023247743.1;refGene_gn=CACNA1E;refGene_tx=XM_023247744.1;refGene_gn=CACNA1E;refGene_tx=XM_023247745.1;refGene_gn=CACNA1E;refGene_tx=XM_023247746.1;refGene_gn=CACNA1E;refGene_tx=XM_023247747.1;refGene_gn=CACNA1E;refGene_tx=XM_023247748.1;refGene_gn=CACNA1E;refGene_tx=XM_023247749.1;refGene_gn=CACNA1E;refGene_tx=XM_023247750.1
F1 28742413 28742680 refGene_gn=CACNA1E;refGene_tx=XM_023247743.1;refGene_gn=CACNA1E;refGene_tx=XM_023247744.1;refGene_gn=CACNA1E;refGene_tx=XM_023247745.1;refGene_gn=CACNA1E;refGene_tx=XM_023247746.1;refGene_gn=CACNA1E;refGene_tx=XM_023247747.1;refGene_gn=CACNA1E;refGene_tx=XM_023247748.1;refGene_gn=CACNA1E;refGene_tx=XM_023247749.1;refGene_gn=CACNA1E;refGene_tx=XM_023247750.1
F1 28817209 28817649 refGene_gn=CACNA1E;refGene_tx=XM_023247743.1;refGene_gn=CACNA1E;refGene_tx=XM_023247744.1;refGene_gn=CACNA1E;refGene_tx=XM_023247745.1;refGene_gn=CACNA1E;refGene_tx=XM_023247746.1;refGene_gn=CACNA1E;refGene_tx=XM_023247747.1;refGene_gn=CACNA1E;refGene_tx=XM_023247748.1;refGene_gn=CACNA1E;refGene_tx=XM_023247749.1;refGene_gn=CACNA1E;refGene_tx=XM_023247750.1
F1 28849184 28849304 refGene_gn=LOC101101608;refGene_tx=XM_023247454.1
F1 28849305 28849422 refGene_gn=LOC101101608;refGene_tx=XM_023247454.1
F1 28849672 28849801 refGene_gn=LOC101101608;refGene_tx=XM_023247454.1
F1 28850826 28850939 refGene_gn=LOC101101608;refGene_tx=XM_023247454.1
F1 28871289 28871488 refGene_gn=LOC101101608;refGene_tx=XM_023247454.1
F1 29126897 29127857 refGene_gn=IER5;refGene_tx=XM_023247660.1
F1 29162750 29162961 refGene_gn=LOC105261339;refGene_tx=XM_019821358.1
F1 29162962 29163146 refGene_gn=LOC105261339;refGene_tx=XM_019821358.1
F1 29163792 29163907 refGene_gn=LOC105261339;refGene_tx=XM_019821358.1
F1 29185351 29185386 refGene_gn=STX6;refGene_tx=XM_003999284.5
F1 29206473 29206643 refGene_gn=STX6;refGene_tx=XM_003999284.5
F1 29209444 29209539 refGene_gn=STX6;refGene_tx=XM_003999284.5

F1 29217425 29217488 refGene_gn=STX6;refGene_tx=XM_003999284.5
F1 29221082 29221208 refGene_gn=STX6;refGene_tx=XM_003999284.5
F1 29222407 29222514 refGene_gn=STX6;refGene_tx=XM_003999284.5
F1 29226852 29226947 refGene_gn=STX6;refGene_tx=XM_003999284.5
F1 29231207 29231284 refGene_gn=STX6;refGene_tx=XM_003999284.5
F1 29312223 29312284 refGene_gn=XPR1;refGene_tx=XM_011290988.3
F1 29315451 29315673 refGene_gn=XPR1;refGene_tx=XM_011290988.3
F1 29321518 29321658 refGene_gn=XPR1;refGene_tx=XM_011290988.3
F1 29325736 29325903 refGene_gn=XPR1;refGene_tx=XM_011290988.3
F1 29353684 29353879 refGene_gn=XPR1;refGene_tx=XM_011290988.3
F1 29355032 29355204 refGene_gn=XPR1;refGene_tx=XM_011290988.3
F1 29360809 29360989 refGene_gn=XPR1;refGene_tx=XM_011290988.3
F1 29361205 29361396 refGene_gn=XPR1;refGene_tx=XM_011290988.3
F1 29370530 29370612 refGene_gn=XPR1;refGene_tx=XM_011290988.3
F1 29375349 29375433 refGene_gn=XPR1;refGene_tx=XM_011290988.3
F1 29375702 29375852 refGene_gn=XPR1;refGene_tx=XM_011290988.3
F1 29378709 29378933 refGene_gn=XPR1;refGene_tx=XM_011290988.3
F1 29390177 29390279 refGene_gn=XPR1;refGene_tx=XM_011290988.3
F1 29496731 29496783 refGene_gn=XPR1;refGene_tx=XM_011290988.3
F1 29524473 29524542 refGene_gn=XPR1;refGene_tx=XM_011290988.3
F1 29640902 29641124
refGene_gn=ACBD6;refGene_tx=XM_003999286.5;refGene_gn=ACBD6;refGene_tx
=XM_006942765.4;refGene_gn=ACBD6;refGene_tx=XM_019821774.2;refGene_gn=
ACBD6;refGene_tx=XM_019821776.2;refGene_gn=ACBD6;refGene_tx=XM_0198217
77.2;refGene_gn=ACBD6;refGene_tx=XM_019821778.2;refGene_gn=ACBD6;refGe
ne_tx=XM_019821779.2;refGene_gn=ACBD6;refGene_tx=XM_019821780.2
F1 29647861 29647926
refGene_gn=ACBD6;refGene_tx=XM_003999286.5;refGene_gn=ACBD6;refGene_tx
=XM_006942765.4;refGene_gn=ACBD6;refGene_tx=XM_019821774.2;refGene_gn=
ACBD6;refGene_tx=XM_019821776.2;refGene_gn=ACBD6;refGene_tx=XM_0198217
77.2;refGene_gn=ACBD6;refGene_tx=XM_019821778.2;refGene_gn=ACBD6;refGe
ne_tx=XM_019821779.2
F1 29650588 29650685
refGene_gn=ACBD6;refGene_tx=XM_003999286.5;refGene_gn=ACBD6;refGene_tx
=XM_006942765.4;refGene_gn=ACBD6;refGene_tx=XM_019821774.2;refGene_gn=
ACBD6;refGene_tx=XM_019821776.2;refGene_gn=ACBD6;refGene_tx=XM_0198217
77.2;refGene_gn=ACBD6;refGene_tx=XM_019821778.2;refGene_gn=ACBD6;refGe
ne_tx=XM_019821779.2
F1 29711298 29711381
refGene_gn=ACBD6;refGene_tx=XM_003999286.5;refGene_gn=ACBD6;refGene_tx
=XM_006942765.4;refGene_gn=ACBD6;refGene_tx=XM_019821774.2;refGene_gn=
ACBD6;refGene_tx=XM_019821776.2;refGene_gn=ACBD6;refGene_tx=XM_0198217
77.2;refGene_gn=ACBD6;refGene_tx=XM_019821778.2;refGene_gn=ACBD6;refGe
ne_tx=XM_019821779.2;refGene_gn=ACBD6;refGene_tx=XM_019821780.2
F1 29728352 29728458
refGene_gn=ACBD6;refGene_tx=XM_003999286.5;refGene_gn=ACBD6;refGene_tx
=XM_006942765.4;refGene_gn=ACBD6;refGene_tx=XM_019821774.2;refGene_gn=
ACBD6;refGene_tx=XM_019821776.2;refGene_gn=ACBD6;refGene_tx=XM_0198217
77.2;refGene_gn=ACBD6;refGene_tx=XM_019821778.2;refGene_gn=ACBD6;refGe
ne_tx=XM_019821779.2;refGene_gn=ACBD6;refGene_tx=XM_019821780.2

F1 29746205 29746295
refGene_gn=ACBD6;refGene_tx=XM_003999286.5;refGene_gn=ACBD6;refGene_tx=XM_006942765.4;refGene_gn=ACBD6;refGene_tx=XM_019821774.2;refGene_gn=ACBD6;refGene_tx=XM_019821776.2;refGene_gn=ACBD6;refGene_tx=XM_019821777.2;refGene_gn=ACBD6;refGene_tx=XM_019821778.2;refGene_gn=ACBD6;refGene_tx=XM_019821779.2;refGene_gn=ACBD6;refGene_tx=XM_019821780.2
F1 29770677 29770752 refGene_gn=ACBD6;refGene_tx=XM_019821778.2
F1 29818188 29818219
refGene_gn=ACBD6;refGene_tx=XM_003999286.5;refGene_gn=ACBD6;refGene_tx=XM_006942765.4;refGene_gn=ACBD6;refGene_tx=XM_019821774.2;refGene_gn=ACBD6;refGene_tx=XM_019821776.2;refGene_gn=ACBD6;refGene_tx=XM_019821777.2;refGene_gn=ACBD6;refGene_tx=XM_019821779.2;refGene_gn=ACBD6;refGene_tx=XM_019821780.2
F1 29828831 29828866
refGene_gn=ACBD6;refGene_tx=XM_006942765.4;refGene_gn=ACBD6;refGene_tx=XM_019821776.2
F1 29830406 29830439 refGene_gn=ACBD6;refGene_tx=XM_019821776.2
F1 29839702 29839749 refGene_gn=ACBD6;refGene_tx=XM_019821777.2
F1 29843064 29843219
refGene_gn=ACBD6;refGene_tx=XM_003999286.5;refGene_gn=ACBD6;refGene_tx=XM_006942765.4;refGene_gn=ACBD6;refGene_tx=XM_019821780.2
F1 29850940 29850951 refGene_gn=ACBD6;refGene_tx=XM_019821779.2
F1 29854094 29854224 refGene_gn=ACBD6;refGene_tx=XM_019821774.2
F1 29854448 29854452 refGene_gn=ACBD6;refGene_tx=XM_019821774.2
F1 29861650 29862045 refGene_gn=LHX4;refGene_tx=XM_003999287.4
F1 29864223 29864395 refGene_gn=LHX4;refGene_tx=XM_003999287.4
F1 29864742 29864897 refGene_gn=LHX4;refGene_tx=XM_003999287.4
F1 29869373 29869576 refGene_gn=LHX4;refGene_tx=XM_003999287.4
F1 29886842 29887014 refGene_gn=LHX4;refGene_tx=XM_003999287.4
F1 29903023 29903099 refGene_gn=LHX4;refGene_tx=XM_003999287.4
F1 29927372 29928112 refGene_gn=QSOX1;refGene_tx=XM_023248115.1
F1 29930040 29930220 refGene_gn=QSOX1;refGene_tx=XM_023248115.1
F1 29933704 29933852 refGene_gn=QSOX1;refGene_tx=XM_023248115.1
F1 29934653 29934776 refGene_gn=QSOX1;refGene_tx=XM_023248115.1
F1 29936350 29936480 refGene_gn=QSOX1;refGene_tx=XM_023248115.1
F1 29938536 29938671 refGene_gn=QSOX1;refGene_tx=XM_023248115.1
F1 29940276 29940422 refGene_gn=QSOX1;refGene_tx=XM_023248115.1
F1 29942684 29942775 refGene_gn=QSOX1;refGene_tx=XM_023248115.1
F1 29945280 29945383 refGene_gn=QSOX1;refGene_tx=XM_023248115.1
F1 29945936 29945982 refGene_gn=QSOX1;refGene_tx=XM_023248115.1
F1 29954131 29954232 refGene_gn=QSOX1;refGene_tx=XM_023248115.1
F1 29960922 29961487 refGene_gn=QSOX1;refGene_tx=XM_023248115.1
F1 29999928 30000093
refGene_gn=CEP350;refGene_tx=XM_023247839.1;refGene_gn=CEP350;refGene_tx=XM_023247840.1;refGene_gn=CEP350;refGene_tx=XM_023247842.1
F1 30004825 30004873 refGene_gn=CEP350;refGene_tx=XM_023247841.1
F1 30012066 30012189
refGene_gn=CEP350;refGene_tx=XM_023247839.1;refGene_gn=CEP350;refGene_tx=XM_023247840.1;refGene_gn=CEP350;refGene_tx=XM_023247841.1;refGene_gn=CEP350;refGene_tx=XM_023247842.1

F1 30015192 30015339
refGene_gn=CEP350;refGene_tx=XM_023247839.1;refGene_gn=CEP350;refGene_tx=XM_023247840.1;refGene_gn=CEP350;refGene_tx=XM_023247841.1;refGene_gn=CEP350;refGene_tx=XM_023247842.1
F1 30015441 30015849
refGene_gn=CEP350;refGene_tx=XM_023247839.1;refGene_gn=CEP350;refGene_tx=XM_023247840.1;refGene_gn=CEP350;refGene_tx=XM_023247841.1;refGene_gn=CEP350;refGene_tx=XM_023247842.1
F1 30016703 30018697
refGene_gn=CEP350;refGene_tx=XM_023247839.1;refGene_gn=CEP350;refGene_tx=XM_023247840.1;refGene_gn=CEP350;refGene_tx=XM_023247841.1;refGene_gn=CEP350;refGene_tx=XM_023247842.1
F1 30019693 30019776
refGene_gn=CEP350;refGene_tx=XM_023247839.1;refGene_gn=CEP350;refGene_tx=XM_023247840.1;refGene_gn=CEP350;refGene_tx=XM_023247841.1;refGene_gn=CEP350;refGene_tx=XM_023247842.1
F1 30022367 30022504
refGene_gn=CEP350;refGene_tx=XM_023247839.1;refGene_gn=CEP350;refGene_tx=XM_023247840.1;refGene_gn=CEP350;refGene_tx=XM_023247841.1;refGene_gn=CEP350;refGene_tx=XM_023247842.1
F1 30025507 30025668
refGene_gn=CEP350;refGene_tx=XM_023247839.1;refGene_gn=CEP350;refGene_tx=XM_023247840.1;refGene_gn=CEP350;refGene_tx=XM_023247841.1;refGene_gn=CEP350;refGene_tx=XM_023247842.1
F1 30032728 30032873
refGene_gn=CEP350;refGene_tx=XM_023247839.1;refGene_gn=CEP350;refGene_tx=XM_023247840.1;refGene_gn=CEP350;refGene_tx=XM_023247841.1;refGene_gn=CEP350;refGene_tx=XM_023247842.1
F1 30036714 30036926
refGene_gn=CEP350;refGene_tx=XM_023247839.1;refGene_gn=CEP350;refGene_tx=XM_023247840.1;refGene_gn=CEP350;refGene_tx=XM_023247841.1;refGene_gn=CEP350;refGene_tx=XM_023247842.1
F1 30042254 30042454
refGene_gn=CEP350;refGene_tx=XM_023247839.1;refGene_gn=CEP350;refGene_tx=XM_023247840.1;refGene_gn=CEP350;refGene_tx=XM_023247841.1;refGene_gn=CEP350;refGene_tx=XM_023247842.1
F1 30043561 30043684
refGene_gn=CEP350;refGene_tx=XM_023247839.1;refGene_gn=CEP350;refGene_tx=XM_023247840.1;refGene_gn=CEP350;refGene_tx=XM_023247841.1
F1 30051124 30051282
refGene_gn=CEP350;refGene_tx=XM_023247839.1;refGene_gn=CEP350;refGene_tx=XM_023247840.1;refGene_gn=CEP350;refGene_tx=XM_023247841.1;refGene_gn=CEP350;refGene_tx=XM_023247842.1
F1 30055116 30055263
refGene_gn=CEP350;refGene_tx=XM_023247839.1;refGene_gn=CEP350;refGene_tx=XM_023247840.1;refGene_gn=CEP350;refGene_tx=XM_023247841.1;refGene_gn=CEP350;refGene_tx=XM_023247842.1
F1 30062561 30062649
refGene_gn=CEP350;refGene_tx=XM_023247839.1;refGene_gn=CEP350;refGene_tx=XM_023247840.1;refGene_gn=CEP350;refGene_tx=XM_023247841.1;refGene_

gn=CEP350;refGene_tx=XM_023247842.1
F1 30063122 30063307
refGene_gn=CEP350;refGene_tx=XM_023247839.1;refGene_gn=CEP350;refGene_tx=XM_023247840.1;refGene_gn=CEP350;refGene_tx=XM_023247841.1;refGene_gn=CEP350;refGene_tx=XM_023247842.1
F1 30063907 30064104
refGene_gn=CEP350;refGene_tx=XM_023247839.1;refGene_gn=CEP350;refGene_tx=XM_023247840.1;refGene_gn=CEP350;refGene_tx=XM_023247841.1;refGene_gn=CEP350;refGene_tx=XM_023247842.1
F1 30071280 30071450
refGene_gn=CEP350;refGene_tx=XM_023247839.1;refGene_gn=CEP350;refGene_tx=XM_023247840.1;refGene_gn=CEP350;refGene_tx=XM_023247841.1;refGene_gn=CEP350;refGene_tx=XM_023247842.1
F1 30077230 30077353
refGene_gn=CEP350;refGene_tx=XM_023247839.1;refGene_gn=CEP350;refGene_tx=XM_023247840.1;refGene_gn=CEP350;refGene_tx=XM_023247841.1;refGene_gn=CEP350;refGene_tx=XM_023247842.1
F1 30078171 30078308
refGene_gn=CEP350;refGene_tx=XM_023247839.1;refGene_gn=CEP350;refGene_tx=XM_023247840.1;refGene_gn=CEP350;refGene_tx=XM_023247841.1;refGene_gn=CEP350;refGene_tx=XM_023247842.1
F1 30080642 30080783
refGene_gn=CEP350;refGene_tx=XM_023247839.1;refGene_gn=CEP350;refGene_tx=XM_023247840.1;refGene_gn=CEP350;refGene_tx=XM_023247841.1;refGene_gn=CEP350;refGene_tx=XM_023247842.1
F1 30081241 30081352
refGene_gn=CEP350;refGene_tx=XM_023247839.1;refGene_gn=CEP350;refGene_tx=XM_023247840.1;refGene_gn=CEP350;refGene_tx=XM_023247841.1;refGene_gn=CEP350;refGene_tx=XM_023247842.1
F1 30085575 30085739
refGene_gn=CEP350;refGene_tx=XM_023247839.1;refGene_gn=CEP350;refGene_tx=XM_023247840.1;refGene_gn=CEP350;refGene_tx=XM_023247841.1;refGene_gn=CEP350;refGene_tx=XM_023247842.1
F1 30086312 30086533
refGene_gn=CEP350;refGene_tx=XM_023247839.1;refGene_gn=CEP350;refGene_tx=XM_023247840.1;refGene_gn=CEP350;refGene_tx=XM_023247841.1;refGene_gn=CEP350;refGene_tx=XM_023247842.1
F1 30089593 30089780
refGene_gn=CEP350;refGene_tx=XM_023247839.1;refGene_gn=CEP350;refGene_tx=XM_023247840.1;refGene_gn=CEP350;refGene_tx=XM_023247841.1;refGene_gn=CEP350;refGene_tx=XM_023247842.1
F1 30094154 30094318
refGene_gn=CEP350;refGene_tx=XM_023247839.1;refGene_gn=CEP350;refGene_tx=XM_023247840.1;refGene_gn=CEP350;refGene_tx=XM_023247841.1;refGene_gn=CEP350;refGene_tx=XM_023247842.1
F1 30095887 30096038
refGene_gn=CEP350;refGene_tx=XM_023247839.1;refGene_gn=CEP350;refGene_tx=XM_023247840.1;refGene_gn=CEP350;refGene_tx=XM_023247841.1;refGene_gn=CEP350;refGene_tx=XM_023247842.1
F1 30097635 30098696

refGene_gn=CEP350;refGene_tx=XM_023247839.1;refGene_gn=CEP350;refGene_tx=XM_023247840.1;refGene_gn=CEP350;refGene_tx=XM_023247841.1;refGene_gn=CEP350;refGene_tx=XM_023247842.1
F1 30103232 30103354
refGene_gn=CEP350;refGene_tx=XM_023247839.1;refGene_gn=CEP350;refGene_tx=XM_023247840.1;refGene_gn=CEP350;refGene_tx=XM_023247841.1;refGene_gn=CEP350;refGene_tx=XM_023247842.1
F1 30104130 30104789
refGene_gn=CEP350;refGene_tx=XM_023247839.1;refGene_gn=CEP350;refGene_tx=XM_023247840.1;refGene_gn=CEP350;refGene_tx=XM_023247841.1;refGene_gn=CEP350;refGene_tx=XM_023247842.1
F1 30106290 30106437
refGene_gn=CEP350;refGene_tx=XM_023247839.1;refGene_gn=CEP350;refGene_tx=XM_023247840.1;refGene_gn=CEP350;refGene_tx=XM_023247841.1;refGene_gn=CEP350;refGene_tx=XM_023247842.1
F1 30109979 30110093
refGene_gn=CEP350;refGene_tx=XM_023247839.1;refGene_gn=CEP350;refGene_tx=XM_023247840.1;refGene_gn=CEP350;refGene_tx=XM_023247841.1;refGene_gn=CEP350;refGene_tx=XM_023247842.1
F1 30113757 30113871
refGene_gn=CEP350;refGene_tx=XM_023247839.1;refGene_gn=CEP350;refGene_tx=XM_023247840.1;refGene_gn=CEP350;refGene_tx=XM_023247841.1;refGene_gn=CEP350;refGene_tx=XM_023247842.1
F1 30119572 30120186
refGene_gn=CEP350;refGene_tx=XM_023247839.1;refGene_gn=CEP350;refGene_tx=XM_023247840.1;refGene_gn=CEP350;refGene_tx=XM_023247841.1;refGene_gn=CEP350;refGene_tx=XM_023247842.1
F1 30123658 30123818
refGene_gn=CEP350;refGene_tx=XM_023247839.1;refGene_gn=CEP350;refGene_tx=XM_023247840.1;refGene_gn=CEP350;refGene_tx=XM_023247841.1;refGene_gn=CEP350;refGene_tx=XM_023247842.1
F1 30124652 30124767
refGene_gn=CEP350;refGene_tx=XM_023247839.1;refGene_gn=CEP350;refGene_tx=XM_023247840.1;refGene_gn=CEP350;refGene_tx=XM_023247841.1;refGene_gn=CEP350;refGene_tx=XM_023247842.1
F1 30127013 30127060
refGene_gn=CEP350;refGene_tx=XM_023247839.1;refGene_gn=CEP350;refGene_tx=XM_023247840.1;refGene_gn=CEP350;refGene_tx=XM_023247841.1;refGene_gn=CEP350;refGene_tx=XM_023247842.1
F1 30128083 30128156
refGene_gn=CEP350;refGene_tx=XM_023247839.1;refGene_gn=CEP350;refGene_tx=XM_023247840.1;refGene_gn=CEP350;refGene_tx=XM_023247841.1;refGene_gn=CEP350;refGene_tx=XM_023247842.1
F1 30174140 30174928
refGene_gn=TOR1AIP1;refGene_tx=XM_011290985.3;refGene_gn=TOR1AIP1;refGene_tx=XM_011290986.3;refGene_gn=TOR1AIP1;refGene_tx=XM_019821789.2;refGene_gn=TOR1AIP1;refGene_tx=XM_019821791.2
F1 30177686 30177743
refGene_gn=TOR1AIP1;refGene_tx=XM_011290985.3;refGene_gn=TOR1AIP1;refGene_tx=XM_011290986.3;refGene_gn=TOR1AIP1;refGene_tx=XM_019821789.2;re

fGene_gn=TOR1AIP1;refGene_tx=XM_019821791.2
F1 30178107 30178152
refGene_gn=TOR1AIP1;refGene_tx=XM_011290985.3;refGene_gn=TOR1AIP1;refGene_tx=XM_011290986.3
F1 30188605 30188674
refGene_gn=TOR1AIP1;refGene_tx=XM_011290985.3;refGene_gn=TOR1AIP1;refGene_tx=XM_011290986.3;refGene_gn=TOR1AIP1;refGene_tx=XM_019821789.2;refGene_gn=TOR1AIP1;refGene_tx=XM_019821791.2
F1 30189214 30189256
refGene_gn=TOR1AIP1;refGene_tx=XM_011290985.3;refGene_gn=TOR1AIP1;refGene_tx=XM_011290986.3;refGene_gn=TOR1AIP1;refGene_tx=XM_019821789.2;refGene_gn=TOR1AIP1;refGene_tx=XM_019821791.2
F1 30192472 30192526
refGene_gn=TOR1AIP1;refGene_tx=XM_011290985.3;refGene_gn=TOR1AIP1;refGene_tx=XM_011290986.3;refGene_gn=TOR1AIP1;refGene_tx=XM_019821789.2;refGene_gn=TOR1AIP1;refGene_tx=XM_019821791.2
F1 30195330 30195417
refGene_gn=TOR1AIP1;refGene_tx=XM_011290985.3;refGene_gn=TOR1AIP1;refGene_tx=XM_011290986.3;refGene_gn=TOR1AIP1;refGene_tx=XM_019821789.2;refGene_gn=TOR1AIP1;refGene_tx=XM_019821791.2
F1 30196208 30196250
refGene_gn=TOR1AIP1;refGene_tx=XM_011290985.3;refGene_gn=TOR1AIP1;refGene_tx=XM_011290986.3;refGene_gn=TOR1AIP1;refGene_tx=XM_019821789.2;refGene_gn=TOR1AIP1;refGene_tx=XM_019821791.2
F1 30206820 30206865
refGene_gn=TOR1AIP1;refGene_tx=XM_011290985.3;refGene_gn=TOR1AIP1;refGene_tx=XM_019821789.2
F1 30207835 30207895
refGene_gn=TOR1AIP1;refGene_tx=XM_011290985.3;refGene_gn=TOR1AIP1;refGene_tx=XM_011290986.3;refGene_gn=TOR1AIP1;refGene_tx=XM_019821789.2;refGene_gn=TOR1AIP1;refGene_tx=XM_019821791.2
F1 30213977 30214055
refGene_gn=TOR1AIP1;refGene_tx=XM_011290985.3;refGene_gn=TOR1AIP1;refGene_tx=XM_011290986.3;refGene_gn=TOR1AIP1;refGene_tx=XM_019821789.2;refGene_gn=TOR1AIP1;refGene_tx=XM_019821791.2
F1 30215951 30216429
refGene_gn=TOR1AIP1;refGene_tx=XM_011290985.3;refGene_gn=TOR1AIP1;refGene_tx=XM_011290986.3;refGene_gn=TOR1AIP1;refGene_tx=XM_019821789.2;refGene_gn=TOR1AIP1;refGene_tx=XM_019821791.2
F1 30235005 30235401
refGene_gn=LOC102902503;refGene_tx=XM_006942767.4;refGene_gn=LOC102902503;refGene_tx=XM_006942768.4;refGene_gn=LOC102902503;refGene_tx=XM_006942769.4;refGene_gn=LOC102902503;refGene_tx=XM_006942770.4;refGene_gn=LOC102902503;refGene_tx=XM_019821795.2
F1 30244224 30244258
refGene_gn=LOC101096439;refGene_tx=XM_019821793.2;refGene_gn=LOC101096439;refGene_tx=XM_019821794.1
F1 30246793 30247315
refGene_gn=LOC101096439;refGene_tx=XM_019821793.2;refGene_gn=LOC101096439;refGene_tx=XM_019821794.1

F1	30250509	30250614
refGene_gn=LOC101096439;refGene_tx=XM_019821793.2;refGene_gn=LOC101096439;refGene_tx=XM_019821794.1
F1	30251386	30252147
refGene_gn=LOC101096439;refGene_tx=XM_019821793.2;refGene_gn=LOC101096439;refGene_tx=XM_019821794.1
F1	30557761	30558079
refGene_gn=KCNK2;refGene_tx=XM_003999292.4;refGene_gn=KCNK2;refGene_tx=XM_019821797.2;refGene_gn=KCNK2;refGene_tx=XM_019821798.2
F1	30599074	30599214
refGene_gn=KCNK2;refGene_tx=XM_003999292.4;refGene_gn=KCNK2;refGene_tx=XM_019821797.2;refGene_gn=KCNK2;refGene_tx=XM_019821798.2
F1	30612213	30612400
refGene_gn=KCNK2;refGene_tx=XM_003999292.4;refGene_gn=KCNK2;refGene_tx=XM_019821797.2;refGene_gn=KCNK2;refGene_tx=XM_019821798.2
F1	30615463	30615624
refGene_gn=KCNK2;refGene_tx=XM_003999292.4;refGene_gn=KCNK2;refGene_tx=XM_019821797.2;refGene_gn=KCNK2;refGene_tx=XM_019821798.2
F1	30645217	30645335
refGene_gn=KCNK2;refGene_tx=XM_003999292.4;refGene_gn=KCNK2;refGene_tx=XM_019821797.2;refGene_gn=KCNK2;refGene_tx=XM_019821798.2
F1	30679483	30679794
refGene_gn=KCNK2;refGene_tx=XM_003999292.4;refGene_gn=KCNK2;refGene_tx=XM_019821797.2;refGene_gn=KCNK2;refGene_tx=XM_019821798.2
F1	30682838	30682884	refGene_gn=KCNK2;refGene_tx=XM_003999292.4
F1	30682958	30682959	refGene_gn=KCNK2;refGene_tx=XM_019821798.2
F1	30753338	30753372	refGene_gn=KCNK2;refGene_tx=XM_019821797.2
F1	31051551	31051755
refGene_gn=CENPF;refGene_tx=XM_006942776.4;refGene_gn=CENPF;refGene_tx=XM_019821799.2;refGene_gn=CENPF;refGene_tx=XM_019821800.2
F1	31056842	31057021
refGene_gn=CENPF;refGene_tx=XM_006942776.4;refGene_gn=CENPF;refGene_tx=XM_019821799.2;refGene_gn=CENPF;refGene_tx=XM_019821800.2
F1	31058524	31058998
refGene_gn=CENPF;refGene_tx=XM_006942776.4;refGene_gn=CENPF;refGene_tx=XM_019821799.2;refGene_gn=CENPF;refGene_tx=XM_019821800.2
F1	31060233	31060396
refGene_gn=CENPF;refGene_tx=XM_006942776.4;refGene_gn=CENPF;refGene_tx=XM_019821799.2;refGene_gn=CENPF;refGene_tx=XM_019821800.2
F1	31063504	31063666
refGene_gn=CENPF;refGene_tx=XM_006942776.4;refGene_gn=CENPF;refGene_tx=XM_019821799.2;refGene_gn=CENPF;refGene_tx=XM_019821800.2
F1	31064009	31064186
refGene_gn=CENPF;refGene_tx=XM_006942776.4;refGene_gn=CENPF;refGene_tx=XM_019821799.2;refGene_gn=CENPF;refGene_tx=XM_019821800.2
F1	31066956	31067109
refGene_gn=CENPF;refGene_tx=XM_006942776.4;refGene_gn=CENPF;refGene_tx=XM_019821799.2;refGene_gn=CENPF;refGene_tx=XM_019821800.2
F1	31068604	31071457
refGene_gn=CENPF;refGene_tx=XM_006942776.4;refGene_gn=CENPF;refGene_tx

=XM_019821799.2;refGene_gn=CENPF;refGene_tx=XM_019821800.2
F1 31072952 31076341
refGene_gn=CENPF;refGene_tx=XM_006942776.4;refGene_gn=CENPF;refGene_tx=XM_019821799.2;refGene_gn=CENPF;refGene_tx=XM_019821800.2
F1 31079638 31079774
refGene_gn=CENPF;refGene_tx=XM_006942776.4;refGene_gn=CENPF;refGene_tx=XM_019821799.2;refGene_gn=CENPF;refGene_tx=XM_019821800.2
F1 31085554 31085677
refGene_gn=CENPF;refGene_tx=XM_006942776.4;refGene_gn=CENPF;refGene_tx=XM_019821799.2;refGene_gn=CENPF;refGene_tx=XM_019821800.2
F1 31089224 31089353
refGene_gn=CENPF;refGene_tx=XM_006942776.4;refGene_gn=CENPF;refGene_tx=XM_019821799.2;refGene_gn=CENPF;refGene_tx=XM_019821800.2
F1 31090317 31090443
refGene_gn=CENPF;refGene_tx=XM_006942776.4;refGene_gn=CENPF;refGene_tx=XM_019821799.2;refGene_gn=CENPF;refGene_tx=XM_019821800.2
F1 31098992 31099195
refGene_gn=CENPF;refGene_tx=XM_006942776.4;refGene_gn=CENPF;refGene_tx=XM_019821799.2;refGene_gn=CENPF;refGene_tx=XM_019821800.2
F1 31100316 31100608
refGene_gn=CENPF;refGene_tx=XM_006942776.4;refGene_gn=CENPF;refGene_tx=XM_019821799.2;refGene_gn=CENPF;refGene_tx=XM_019821800.2
F1 31102437 31102529
refGene_gn=CENPF;refGene_tx=XM_006942776.4;refGene_gn=CENPF;refGene_tx=XM_019821799.2;refGene_gn=CENPF;refGene_tx=XM_019821800.2
F1 31102967 31103089
refGene_gn=CENPF;refGene_tx=XM_006942776.4;refGene_gn=CENPF;refGene_tx=XM_019821799.2;refGene_gn=CENPF;refGene_tx=XM_019821800.2
F1 31108025 31108222
refGene_gn=CENPF;refGene_tx=XM_006942776.4;refGene_gn=CENPF;refGene_tx=XM_019821799.2;refGene_gn=CENPF;refGene_tx=XM_019821800.2
F1 31109515 31109677
refGene_gn=CENPF;refGene_tx=XM_006942776.4;refGene_gn=CENPF;refGene_tx=XM_019821799.2;refGene_gn=CENPF;refGene_tx=XM_019821800.2
F1 31225570 31225744
refGene_gn=PTPN14;refGene_tx=XM_003999294.5;refGene_gn=PTPN14;refGene_tx=XM_006942777.4;refGene_gn=PTPN14;refGene_tx=XM_019821801.2
F1 31237648 31237818
refGene_gn=PTPN14;refGene_tx=XM_003999294.5;refGene_gn=PTPN14;refGene_tx=XM_006942777.4;refGene_gn=PTPN14;refGene_tx=XM_019821801.2
F1 31276316 31276414
refGene_gn=PTPN14;refGene_tx=XM_003999294.5;refGene_gn=PTPN14;refGene_tx=XM_006942777.4;refGene_gn=PTPN14;refGene_tx=XM_019821801.2
F1 31278767 31278835
refGene_gn=PTPN14;refGene_tx=XM_003999294.5;refGene_gn=PTPN14;refGene_tx=XM_006942777.4;refGene_gn=PTPN14;refGene_tx=XM_019821801.2
F1 31284699 31284770
refGene_gn=PTPN14;refGene_tx=XM_003999294.5;refGene_gn=PTPN14;refGene_tx=XM_006942777.4;refGene_gn=PTPN14;refGene_tx=XM_019821801.2
F1 31285638 31285726

refGene_gn=PTPN14;refGene_tx=XM_003999294.5;refGene_gn=PTPN14;refGene_tx=XM_006942777.4;refGene_gn=PTPN14;refGene_tx=XM_019821801.2
F1 31289318 31289407 refGene_gn=PTPN14;refGene_tx=XM_003999294.5;refGene_gn=PTPN14;refGene_tx=XM_006942777.4;refGene_gn=PTPN14;refGene_tx=XM_019821801.2
F1 31292292 31292380 refGene_gn=PTPN14;refGene_tx=XM_003999294.5;refGene_gn=PTPN14;refGene_tx=XM_006942777.4;refGene_gn=PTPN14;refGene_tx=XM_019821801.2
F1 31293196 31293279 refGene_gn=PTPN14;refGene_tx=XM_003999294.5;refGene_gn=PTPN14;refGene_tx=XM_006942777.4;refGene_gn=PTPN14;refGene_tx=XM_019821801.2
F1 31296373 31296431 refGene_gn=PTPN14;refGene_tx=XM_003999294.5;refGene_gn=PTPN14;refGene_tx=XM_006942777.4;refGene_gn=PTPN14;refGene_tx=XM_019821801.2
F1 31299887 31299966 refGene_gn=PTPN14;refGene_tx=XM_003999294.5;refGene_gn=PTPN14;refGene_tx=XM_006942777.4;refGene_gn=PTPN14;refGene_tx=XM_019821801.2
F1 31301985 31303463 refGene_gn=PTPN14;refGene_tx=XM_003999294.5;refGene_gn=PTPN14;refGene_tx=XM_006942777.4;refGene_gn=PTPN14;refGene_tx=XM_019821801.2
F1 31307619 31307763 refGene_gn=PTPN14;refGene_tx=XM_003999294.5;refGene_gn=PTPN14;refGene_tx=XM_006942777.4;refGene_gn=PTPN14;refGene_tx=XM_019821801.2
F1 31309321 31309540 refGene_gn=PTPN14;refGene_tx=XM_003999294.5;refGene_gn=PTPN14;refGene_tx=XM_006942777.4;refGene_gn=PTPN14;refGene_tx=XM_019821801.2
F1 31309716 31309848 refGene_gn=PTPN14;refGene_tx=XM_006942777.4
F1 31312840 31312969 refGene_gn=PTPN14;refGene_tx=XM_003999294.5;refGene_gn=PTPN14;refGene_tx=XM_019821801.2
F1 31315117 31315352 refGene_gn=PTPN14;refGene_tx=XM_003999294.5;refGene_gn=PTPN14;refGene_tx=XM_019821801.2
F1 31319103 31319267 refGene_gn=PTPN14;refGene_tx=XM_003999294.5;refGene_gn=PTPN14;refGene_tx=XM_019821801.2
F1 31324993 31325122 refGene_gn=PTPN14;refGene_tx=XM_003999294.5;refGene_gn=PTPN14;refGene_tx=XM_019821801.2
F1 31336904 31336985 refGene_gn=SMYD2;refGene_tx=XM_023248067.1
F1 31339148 31339257 refGene_gn=SMYD2;refGene_tx=XM_023248067.1
F1 31341517 31341692 refGene_gn=SMYD2;refGene_tx=XM_023248067.1
F1 31342783 31342904 refGene_gn=SMYD2;refGene_tx=XM_023248067.1
F1 31343889 31344000 refGene_gn=SMYD2;refGene_tx=XM_023248067.1
F1 31346851 31346954 refGene_gn=SMYD2;refGene_tx=XM_023248067.1
F1 31349533 31349601 refGene_gn=SMYD2;refGene_tx=XM_023248067.1
F1 31355540 31355665 refGene_gn=SMYD2;refGene_tx=XM_023248067.1
F1 31356372 31356433 refGene_gn=SMYD2;refGene_tx=XM_023248067.1
F1 31358877 31358988 refGene_gn=SMYD2;refGene_tx=XM_023248067.1
F1 31368143 31368207 refGene_gn=SMYD2;refGene_tx=XM_023248067.1

F1 31386995 31387168 refGene_gn=SMYD2;refGene_tx=XM_023248067.1
F1 31640389 31640578
refGene_gn=PROX1;refGene_tx=XM_003999296.5;refGene_gn=PROX1;refGene_tx=XM_019821804.2;refGene_gn=PROX1;refGene_tx=XM_019821805.2;refGene_gn=PROX1;refGene_tx=XM_019821806.2;refGene_gn=PROX1;refGene_tx=XM_019821807.2;refGene_gn=PROX1;refGene_tx=XM_019821808.2
F1 31663479 31663674
refGene_gn=PROX1;refGene_tx=XM_003999296.5;refGene_gn=PROX1;refGene_tx=XM_019821804.2;refGene_gn=PROX1;refGene_tx=XM_019821805.2;refGene_gn=PROX1;refGene_tx=XM_019821806.2;refGene_gn=PROX1;refGene_tx=XM_019821807.2;refGene_gn=PROX1;refGene_tx=XM_019821808.2
F1 31670049 31670157
refGene_gn=PROX1;refGene_tx=XM_003999296.5;refGene_gn=PROX1;refGene_tx=XM_019821804.2;refGene_gn=PROX1;refGene_tx=XM_019821805.2;refGene_gn=PROX1;refGene_tx=XM_019821806.2;refGene_gn=PROX1;refGene_tx=XM_019821807.2;refGene_gn=PROX1;refGene_tx=XM_019821808.2
F1 31677157 31678882
refGene_gn=PROX1;refGene_tx=XM_003999296.5;refGene_gn=PROX1;refGene_tx=XM_019821804.2;refGene_gn=PROX1;refGene_tx=XM_019821805.2;refGene_gn=PROX1;refGene_tx=XM_019821806.2;refGene_gn=PROX1;refGene_tx=XM_019821807.2;refGene_gn=PROX1;refGene_tx=XM_019821808.2
F1 31688804 31689111
refGene_gn=LOC109495638;refGene_tx=XM_023247852.1
F1 31690622 31690930
refGene_gn=LOC109495638;refGene_tx=XM_023247852.1
F1 32250579 32250653 refGene_gn=RPS6KC1;refGene_tx=XM_023247755.1
F1 32346522 32346599 refGene_gn=RPS6KC1;refGene_tx=XM_023247754.1
F1 32381971 32381988 refGene_gn=RPS6KC1;refGene_tx=XM_023247752.1
F1 32385231 32385464
refGene_gn=RPS6KC1;refGene_tx=XM_023247751.1;refGene_gn=RPS6KC1;refGene_tx=XM_023247752.1;refGene_gn=RPS6KC1;refGene_tx=XM_023247753.1;refGene_gn=RPS6KC1;refGene_tx=XM_023247759.1;refGene_gn=RPS6KC1;refGene_tx=XM_023247761.1;refGene_gn=RPS6KC1;refGene_tx=XM_023247762.1;refGene_gn=RPS6KC1;refGene_tx=XM_023247763.1;refGene_gn=RPS6KC1;refGene_tx=XM_023247764.1
F1 32389849 32389979
refGene_gn=RPS6KC1;refGene_tx=XM_023247751.1;refGene_gn=RPS6KC1;refGene_tx=XM_023247752.1;refGene_gn=RPS6KC1;refGene_tx=XM_023247753.1;refGene_gn=RPS6KC1;refGene_tx=XM_023247754.1;refGene_gn=RPS6KC1;refGene_tx=XM_023247755.1;refGene_gn=RPS6KC1;refGene_tx=XM_023247759.1;refGene_gn=RPS6KC1;refGene_tx=XM_023247761.1;refGene_gn=RPS6KC1;refGene_tx=XM_023247762.1;refGene_gn=RPS6KC1;refGene_tx=XM_023247763.1;refGene_gn=RPS6KC1;refGene_tx=XM_023247764.1
F1 32391509 32391620
refGene_gn=RPS6KC1;refGene_tx=XM_023247756.1;refGene_gn=RPS6KC1;refGene_tx=XM_023247760.1
F1 32396100 32396133 refGene_gn=RPS6KC1;refGene_tx=XM_023247757.1
F1 32400864 32400960
refGene_gn=RPS6KC1;refGene_tx=XM_023247751.1;refGene_gn=RPS6KC1;refGene_tx=XM_023247752.1;refGene_gn=RPS6KC1;refGene_tx=XM_023247753.1;refGe

ne_gn=RPS6KC1;refGene_tx=XM_023247754.1;refGene_gn=RPS6KC1;refGene_tx=XM_023247755.1;refGene_gn=RPS6KC1;refGene_tx=XM_023247756.1;refGene_gn=RPS6KC1;refGene_tx=XM_023247757.1;refGene_gn=RPS6KC1;refGene_tx=XM_023247759.1;refGene_gn=RPS6KC1;refGene_tx=XM_023247760.1;refGene_gn=RPS6KC1;refGene_tx=XM_023247761.1;refGene_gn=RPS6KC1;refGene_tx=XM_023247762.1;refGene_gn=RPS6KC1;refGene_tx=XM_023247763.1;refGene_gn=RPS6KC1;refGene_tx=XM_023247764.1
F1 32402033 32402116
refGene_gn=RPS6KC1;refGene_tx=XM_023247751.1;refGene_gn=RPS6KC1;refGene_tx=XM_023247752.1;refGene_gn=RPS6KC1;refGene_tx=XM_023247753.1;refGene_gn=RPS6KC1;refGene_tx=XM_023247754.1;refGene_gn=RPS6KC1;refGene_tx=XM_023247755.1;refGene_gn=RPS6KC1;refGene_tx=XM_023247756.1;refGene_gn=RPS6KC1;refGene_tx=XM_023247757.1;refGene_gn=RPS6KC1;refGene_tx=XM_023247759.1;refGene_gn=RPS6KC1;refGene_tx=XM_023247760.1;refGene_gn=RPS6KC1;refGene_tx=XM_023247761.1;refGene_gn=RPS6KC1;refGene_tx=XM_023247762.1;refGene_gn=RPS6KC1;refGene_tx=XM_023247763.1;refGene_gn=RPS6KC1;refGene_tx=XM_023247764.1
F1 32419028 32419118
refGene_gn=RPS6KC1;refGene_tx=XM_023247751.1;refGene_gn=RPS6KC1;refGene_tx=XM_023247752.1;refGene_gn=RPS6KC1;refGene_tx=XM_023247753.1;refGene_gn=RPS6KC1;refGene_tx=XM_023247754.1;refGene_gn=RPS6KC1;refGene_tx=XM_023247755.1;refGene_gn=RPS6KC1;refGene_tx=XM_023247756.1;refGene_gn=RPS6KC1;refGene_tx=XM_023247757.1;refGene_gn=RPS6KC1;refGene_tx=XM_023247759.1;refGene_gn=RPS6KC1;refGene_tx=XM_023247760.1;refGene_gn=RPS6KC1;refGene_tx=XM_023247761.1;refGene_gn=RPS6KC1;refGene_tx=XM_023247762.1;refGene_gn=RPS6KC1;refGene_tx=XM_023247763.1;refGene_gn=RPS6KC1;refGene_tx=XM_023247764.1
F1 32419389 32420985
refGene_gn=RPS6KC1;refGene_tx=XM_023247751.1;refGene_gn=RPS6KC1;refGene_tx=XM_023247752.1;refGene_gn=RPS6KC1;refGene_tx=XM_023247753.1;refGene_gn=RPS6KC1;refGene_tx=XM_023247754.1;refGene_gn=RPS6KC1;refGene_tx=XM_023247755.1;refGene_gn=RPS6KC1;refGene_tx=XM_023247756.1;refGene_gn=RPS6KC1;refGene_tx=XM_023247757.1;refGene_gn=RPS6KC1;refGene_tx=XM_023247759.1;refGene_gn=RPS6KC1;refGene_tx=XM_023247760.1;refGene_gn=RPS6KC1;refGene_tx=XM_023247761.1;refGene_gn=RPS6KC1;refGene_tx=XM_023247762.1;refGene_gn=RPS6KC1;refGene_tx=XM_023247763.1;refGene_gn=RPS6KC1;refGene_tx=XM_023247764.1
F1 32430103 32430236
refGene_gn=RPS6KC1;refGene_tx=XM_023247751.1;refGene_gn=RPS6KC1;refGene_tx=XM_023247752.1;refGene_gn=RPS6KC1;refGene_tx=XM_023247753.1;refGene_gn=RPS6KC1;refGene_tx=XM_023247754.1;refGene_gn=RPS6KC1;refGene_tx=XM_023247755.1;refGene_gn=RPS6KC1;refGene_tx=XM_023247756.1;refGene_gn=RPS6KC1;refGene_tx=XM_023247757.1;refGene_gn=RPS6KC1;refGene_tx=XM_023247759.1;refGene_gn=RPS6KC1;refGene_tx=XM_023247760.1;refGene_gn=RPS6KC1;refGene_tx=XM_023247761.1;refGene_gn=RPS6KC1;refGene_tx=XM_023247762.1;refGene_gn=RPS6KC1;refGene_tx=XM_023247763.1;refGene_gn=RPS6KC1;refGene_tx=XM_023247764.1
F1 32431392 32431440
refGene_gn=RPS6KC1;refGene_tx=XM_023247751.1;refGene_gn=RPS6KC1;refGene_tx=XM_023247752.1;refGene_gn=RPS6KC1;refGene_tx=XM_023247753.1;refGe

ne_gn=RPS6KC1;refGene_tx=XM_023247754.1;refGene_gn=RPS6KC1;refGene_tx=XM_023247755.1;refGene_gn=RPS6KC1;refGene_tx=XM_023247756.1;refGene_gn=RPS6KC1;refGene_tx=XM_023247757.1;refGene_gn=RPS6KC1;refGene_tx=XM_023247759.1;refGene_gn=RPS6KC1;refGene_tx=XM_023247760.1;refGene_gn=RPS6KC1;refGene_tx=XM_023247761.1;refGene_gn=RPS6KC1;refGene_tx=XM_023247762.1;refGene_gn=RPS6KC1;refGene_tx=XM_023247763.1;refGene_gn=RPS6KC1;refGene_tx=XM_023247764.1
F1 32449316 32449478 refGene_gn=RPS6KC1;refGene_tx=XM_023247763.1
F1 32471514 32471607
refGene_gn=RPS6KC1;refGene_tx=XM_023247751.1;refGene_gn=RPS6KC1;refGene_tx=XM_023247752.1;refGene_gn=RPS6KC1;refGene_tx=XM_023247754.1;refGene_gn=RPS6KC1;refGene_tx=XM_023247755.1;refGene_gn=RPS6KC1;refGene_tx=XM_023247756.1;refGene_gn=RPS6KC1;refGene_tx=XM_023247757.1;refGene_gn=RPS6KC1;refGene_tx=XM_023247759.1;refGene_gn=RPS6KC1;refGene_tx=XM_023247761.1;refGene_gn=RPS6KC1;refGene_tx=XM_023247762.1;refGene_gn=RPS6KC1;refGene_tx=XM_023247764.1
F1 32478890 32479006
refGene_gn=RPS6KC1;refGene_tx=XM_023247751.1;refGene_gn=RPS6KC1;refGene_tx=XM_023247752.1;refGene_gn=RPS6KC1;refGene_tx=XM_023247753.1;refGene_gn=RPS6KC1;refGene_tx=XM_023247754.1;refGene_gn=RPS6KC1;refGene_tx=XM_023247755.1;refGene_gn=RPS6KC1;refGene_tx=XM_023247756.1;refGene_gn=RPS6KC1;refGene_tx=XM_023247757.1;refGene_gn=RPS6KC1;refGene_tx=XM_023247759.1;refGene_gn=RPS6KC1;refGene_tx=XM_023247760.1;refGene_gn=RPS6KC1;refGene_tx=XM_023247761.1;refGene_gn=RPS6KC1;refGene_tx=XM_023247762.1;refGene_gn=RPS6KC1;refGene_tx=XM_023247764.1
F1 32515329 32515692
refGene_gn=RPS6KC1;refGene_tx=XM_023247751.1;refGene_gn=RPS6KC1;refGene_tx=XM_023247752.1;refGene_gn=RPS6KC1;refGene_tx=XM_023247753.1;refGene_gn=RPS6KC1;refGene_tx=XM_023247754.1;refGene_gn=RPS6KC1;refGene_tx=XM_023247755.1;refGene_gn=RPS6KC1;refGene_tx=XM_023247756.1;refGene_gn=RPS6KC1;refGene_tx=XM_023247757.1;refGene_gn=RPS6KC1;refGene_tx=XM_023247759.1;refGene_gn=RPS6KC1;refGene_tx=XM_023247760.1;refGene_gn=RPS6KC1;refGene_tx=XM_023247761.1;refGene_gn=RPS6KC1;refGene_tx=XM_023247762.1
F1 32527073 32527167
refGene_gn=RPS6KC1;refGene_tx=XM_023247751.1;refGene_gn=RPS6KC1;refGene_tx=XM_023247752.1;refGene_gn=RPS6KC1;refGene_tx=XM_023247753.1;refGene_gn=RPS6KC1;refGene_tx=XM_023247754.1;refGene_gn=RPS6KC1;refGene_tx=XM_023247755.1;refGene_gn=RPS6KC1;refGene_tx=XM_023247756.1;refGene_gn=RPS6KC1;refGene_tx=XM_023247757.1;refGene_gn=RPS6KC1;refGene_tx=XM_023247759.1;refGene_gn=RPS6KC1;refGene_tx=XM_023247760.1
F1 32535793 32535909
refGene_gn=RPS6KC1;refGene_tx=XM_023247751.1;refGene_gn=RPS6KC1;refGene_tx=XM_023247752.1;refGene_gn=RPS6KC1;refGene_tx=XM_023247753.1;refGene_gn=RPS6KC1;refGene_tx=XM_023247754.1;refGene_gn=RPS6KC1;refGene_tx=XM_023247755.1;refGene_gn=RPS6KC1;refGene_tx=XM_023247756.1;refGene_gn=RPS6KC1;refGene_tx=XM_023247757.1;refGene_gn=RPS6KC1;refGene_tx=XM_023247759.1;refGene_gn=RPS6KC1;refGene_tx=XM_023247760.1
F1 32550385 32550506
refGene_gn=RPS6KC1;refGene_tx=XM_023247751.1;refGene_gn=RPS6KC1;refGen

e_tx=XM_023247752.1;refGene_gn=RPS6KC1;refGene_tx=XM_023247753.1;refGene_gn=RPS6KC1;refGene_tx=XM_023247754.1;refGene_gn=RPS6KC1;refGene_tx=XM_023247755.1;refGene_gn=RPS6KC1;refGene_tx=XM_023247756.1;refGene_gn=RPS6KC1;refGene_tx=XM_023247757.1;refGene_gn=RPS6KC1;refGene_tx=XM_023247759.1;refGene_gn=RPS6KC1;refGene_tx=XM_023247760.1
F1 32557364 32557400
refGene_gn=RPS6KC1;refGene_tx=XM_023247751.1;refGene_gn=RPS6KC1;refGene_tx=XM_023247752.1;refGene_gn=RPS6KC1;refGene_tx=XM_023247753.1;refGene_gn=RPS6KC1;refGene_tx=XM_023247754.1;refGene_gn=RPS6KC1;refGene_tx=XM_023247755.1;refGene_gn=RPS6KC1;refGene_tx=XM_023247756.1;refGene_gn=RPS6KC1;refGene_tx=XM_023247757.1;refGene_gn=RPS6KC1;refGene_tx=XM_023247759.1;refGene_gn=RPS6KC1;refGene_tx=XM_023247760.1
F1 32569627 32569732
refGene_gn=RPS6KC1;refGene_tx=XM_023247751.1;refGene_gn=RPS6KC1;refGene_tx=XM_023247752.1;refGene_gn=RPS6KC1;refGene_tx=XM_023247753.1;refGene_gn=RPS6KC1;refGene_tx=XM_023247754.1;refGene_gn=RPS6KC1;refGene_tx=XM_023247755.1;refGene_gn=RPS6KC1;refGene_tx=XM_023247756.1;refGene_gn=RPS6KC1;refGene_tx=XM_023247757.1;refGene_gn=RPS6KC1;refGene_tx=XM_023247759.1;refGene_gn=RPS6KC1;refGene_tx=XM_023247760.1
F1 32591040 32591099 refGene_gn=ANGEL2;refGene_tx=XM_011290977.3
F1 32593534 32593860
refGene_gn=ANGEL2;refGene_tx=XM_011290977.3;refGene_gn=ANGEL2;refGene_tx=XM_011290978.3;refGene_gn=ANGEL2;refGene_tx=XM_011290979.3;refGene_gn=ANGEL2;refGene_tx=XM_019821832.2;refGene_gn=ANGEL2;refGene_tx=XM_023247960.1
F1 32595789 32595796 refGene_gn=ANGEL2;refGene_tx=XM_011290981.3
F1 32598418 32598675
refGene_gn=ANGEL2;refGene_tx=XM_011290977.3;refGene_gn=ANGEL2;refGene_tx=XM_011290978.3;refGene_gn=ANGEL2;refGene_tx=XM_011290979.3;refGene_gn=ANGEL2;refGene_tx=XM_011290981.3;refGene_gn=ANGEL2;refGene_tx=XM_011290982.3;refGene_gn=ANGEL2;refGene_tx=XM_019821832.2;refGene_gn=ANGEL2;refGene_tx=XM_023247960.1
F1 32599369 32599439
refGene_gn=ANGEL2;refGene_tx=XM_011290977.3;refGene_gn=ANGEL2;refGene_tx=XM_011290978.3;refGene_gn=ANGEL2;refGene_tx=XM_011290979.3;refGene_gn=ANGEL2;refGene_tx=XM_011290981.3;refGene_gn=ANGEL2;refGene_tx=XM_011290982.3;refGene_gn=ANGEL2;refGene_tx=XM_019821832.2;refGene_gn=ANGEL2;refGene_tx=XM_023247960.1
F1 32600561 32600983
refGene_gn=ANGEL2;refGene_tx=XM_011290977.3;refGene_gn=ANGEL2;refGene_tx=XM_011290978.3;refGene_gn=ANGEL2;refGene_tx=XM_011290979.3;refGene_gn=ANGEL2;refGene_tx=XM_011290981.3;refGene_gn=ANGEL2;refGene_tx=XM_011290982.3;refGene_gn=ANGEL2;refGene_tx=XM_019821832.2;refGene_gn=ANGEL2;refGene_tx=XM_023247960.1
F1 32601998 32602125
refGene_gn=ANGEL2;refGene_tx=XM_011290977.3;refGene_gn=ANGEL2;refGene_tx=XM_011290978.3;refGene_gn=ANGEL2;refGene_tx=XM_011290979.3;refGene_gn=ANGEL2;refGene_tx=XM_011290981.3;refGene_gn=ANGEL2;refGene_tx=XM_011290982.3;refGene_gn=ANGEL2;refGene_tx=XM_019821832.2;refGene_gn=ANGEL2;refGene_tx=XM_023247960.1

F1 32602530 32602588
refGene_gn=ANGEL2;refGene_tx=XM_011290977.3;refGene_gn=ANGEL2;refGene_tx=XM_011290978.3;refGene_gn=ANGEL2;refGene_tx=XM_011290979.3;refGene_gn=ANGEL2;refGene_tx=XM_011290981.3;refGene_gn=ANGEL2;refGene_tx=XM_011290982.3;refGene_gn=ANGEL2;refGene_tx=XM_019821832.2;refGene_gn=ANGEL2;refGene_tx=XM_023247960.1
F1 32605451 32605615
refGene_gn=ANGEL2;refGene_tx=XM_011290977.3;refGene_gn=ANGEL2;refGene_tx=XM_011290978.3;refGene_gn=ANGEL2;refGene_tx=XM_011290979.3;refGene_gn=ANGEL2;refGene_tx=XM_011290981.3;refGene_gn=ANGEL2;refGene_tx=XM_011290982.3;refGene_gn=ANGEL2;refGene_tx=XM_019821832.2;refGene_gn=ANGEL2;refGene_tx=XM_023247960.1
F1 32607465 32607617
refGene_gn=ANGEL2;refGene_tx=XM_011290977.3;refGene_gn=ANGEL2;refGene_tx=XM_011290978.3;refGene_gn=ANGEL2;refGene_tx=XM_011290979.3;refGene_gn=ANGEL2;refGene_tx=XM_011290981.3;refGene_gn=ANGEL2;refGene_tx=XM_011290982.3;refGene_gn=ANGEL2;refGene_tx=XM_019821832.2;refGene_gn=ANGEL2;refGene_tx=XM_023247960.1
F1 32613247 32613320
refGene_gn=VASH2;refGene_tx=XM_003999298.5;refGene_gn=VASH2;refGene_tx=XM_003999299.5;refGene_gn=VASH2;refGene_tx=XM_019821833.2
F1 32626940 32627056
refGene_gn=VASH2;refGene_tx=XM_003999298.5;refGene_gn=VASH2;refGene_tx=XM_003999299.5;refGene_gn=VASH2;refGene_tx=XM_019821833.2
F1 32628471 32628853
refGene_gn=VASH2;refGene_tx=XM_003999298.5;refGene_gn=VASH2;refGene_tx=XM_003999299.5;refGene_gn=VASH2;refGene_tx=XM_019821833.2
F1 32635074 32635149
refGene_gn=VASH2;refGene_tx=XM_003999298.5;refGene_gn=VASH2;refGene_tx=XM_019821833.2
F1 32635709 32635766
refGene_gn=VASH2;refGene_tx=XM_003999298.5;refGene_gn=VASH2;refGene_tx=XM_019821833.2
F1 32640830 32640919
refGene_gn=VASH2;refGene_tx=XM_003999298.5;refGene_gn=VASH2;refGene_tx=XM_003999299.5;refGene_gn=VASH2;refGene_tx=XM_019821833.2
F1 32650525 32650801
refGene_gn=VASH2;refGene_tx=XM_003999298.5;refGene_gn=VASH2;refGene_tx=XM_003999299.5;refGene_gn=VASH2;refGene_tx=XM_019821833.2
F1 32685704 32685794 refGene_gn=FLVCR1;refGene_tx=NM_001009302.1
F1 32685950 32686018 refGene_gn=FLVCR1;refGene_tx=NM_001009302.1
F1 32687407 32687519 refGene_gn=FLVCR1;refGene_tx=NM_001009302.1
F1 32687946 32688052 refGene_gn=FLVCR1;refGene_tx=NM_001009302.1
F1 32688551 32688662 refGene_gn=FLVCR1;refGene_tx=NM_001009302.1
F1 32698422 32698526 refGene_gn=FLVCR1;refGene_tx=NM_001009302.1
F1 32700021 32700089 refGene_gn=FLVCR1;refGene_tx=NM_001009302.1
F1 32708162 32708303 refGene_gn=FLVCR1;refGene_tx=NM_001009302.1
F1 32714135 32714280 refGene_gn=FLVCR1;refGene_tx=NM_001009302.1
F1 32719637 32720375 refGene_gn=FLVCR1;refGene_tx=NM_001009302.1
F1 32720574 32720771 refGene_gn=SPATA45;refGene_tx=XM_019821834.1

F1 32725171 32725337 refGene_gn=SPATA45;refGene_tx=XM_019821834.1
F1 32745358 32745635
refGene_gn=SPATA45;refGene_tx=XM_023247257.1;refGene_gn=SPATA45;refGene_tx=XM_023247259.1;refGene_gn=SPATA45;refGene_tx=XM_023247260.1
F1 32751449 32751509
refGene_gn=SPATA45;refGene_tx=XM_019821834.1;refGene_gn=SPATA45;refGene_tx=XM_023247257.1;refGene_gn=SPATA45;refGene_tx=XM_023247259.1;refGene_gn=SPATA45;refGene_tx=XM_023247260.1
F1 32763960 32764104
refGene_gn=TATDN3;refGene_tx=XM_003999301.4;refGene_gn=TATDN3;refGene_tx=XM_006942780.3;refGene_gn=TATDN3;refGene_tx=XM_019821835.2
F1 32766656 32766737
refGene_gn=TATDN3;refGene_tx=XM_003999301.4;refGene_gn=TATDN3;refGene_tx=XM_006942780.3;refGene_gn=TATDN3;refGene_tx=XM_019821835.2
F1 32774517 32774630
refGene_gn=TATDN3;refGene_tx=XM_003999301.4;refGene_gn=TATDN3;refGene_tx=XM_006942780.3;refGene_gn=TATDN3;refGene_tx=XM_019821835.2
F1 32775948 32776004
refGene_gn=TATDN3;refGene_tx=XM_003999301.4;refGene_gn=TATDN3;refGene_tx=XM_006942780.3;refGene_gn=TATDN3;refGene_tx=XM_019821835.2
F1 32776167 32776277
refGene_gn=TATDN3;refGene_tx=XM_003999301.4;refGene_gn=TATDN3;refGene_tx=XM_006942780.3;refGene_gn=TATDN3;refGene_tx=XM_019821835.2
F1 32777407 32777470
refGene_gn=TATDN3;refGene_tx=XM_003999301.4;refGene_gn=TATDN3;refGene_tx=XM_006942780.3;refGene_gn=TATDN3;refGene_tx=XM_019821835.2
F1 32781614 32781699
refGene_gn=TATDN3;refGene_tx=XM_003999301.4;refGene_gn=TATDN3;refGene_tx=XM_006942780.3;refGene_gn=TATDN3;refGene_tx=XM_019821835.2
F1 32781916 32781990
refGene_gn=TATDN3;refGene_tx=XM_003999301.4;refGene_gn=TATDN3;refGene_tx=XM_006942780.3;refGene_gn=TATDN3;refGene_tx=XM_019821835.2
F1 32782904 32782937
refGene_gn=TATDN3;refGene_tx=XM_003999301.4;refGene_gn=TATDN3;refGene_tx=XM_006942780.3;refGene_gn=TATDN3;refGene_tx=XM_019821835.2
F1 32784451 32784517
refGene_gn=TATDN3;refGene_tx=XM_003999301.4;refGene_gn=TATDN3;refGene_tx=XM_006942780.3;refGene_gn=TATDN3;refGene_tx=XM_019821835.2
F1 32784674 32784908 refGene_gn=NSL1;refGene_tx=XM_003999316.5
F1 32788332 32788408 refGene_gn=NSL1;refGene_tx=XM_003999316.5
F1 32792124 32792255 refGene_gn=NSL1;refGene_tx=XM_003999316.5
F1 32793824 32793879 refGene_gn=NSL1;refGene_tx=XM_003999316.5
F1 32816585 32816653 refGene_gn=NSL1;refGene_tx=XM_003999316.5
F1 32818186 32818468 refGene_gn=NSL1;refGene_tx=XM_003999316.5
F1 32852106 32852205
refGene_gn=BATF3;refGene_tx=XM_011291046.3;refGene_gn=BATF3;refGene_tx=XM_023247790.1
F1 32852900 32852978
refGene_gn=BATF3;refGene_tx=XM_023247791.1;refGene_gn=BATF3;refGene_tx=XM_023247792.1

F1 32855623 32855728
refGene_gn=BATF3;refGene_tx=XM_011291046.3;refGene_gn=BATF3;refGene_tx
=XM_023247790.1;refGene_gn=BATF3;refGene_tx=XM_023247791.1;refGene_gn=
BATF3;refGene_tx=XM_023247792.1
F1 32864483 32864669
refGene_gn=BATF3;refGene_tx=XM_011291046.3;refGene_gn=BATF3;refGene_tx
=XM_023247792.1
F1 32877058 32877274
refGene_gn=BATF3;refGene_tx=XM_023247790.1;refGene_gn=BATF3;refGene_tx
=XM_023247791.1
F1 32910983 32912771 refGene_gn=FAM71A;refGene_tx=XM_003999318.4
F1 32918899 32919097
refGene_gn=ATF3;refGene_tx=XM_019821845.2;refGene_gn=ATF3;refGene_tx=X
M_023247488.1;refGene_gn=ATF3;refGene_tx=XM_023247489.1
F1 32920292 32920400
refGene_gn=ATF3;refGene_tx=XM_019821845.2;refGene_gn=ATF3;refGene_tx=X
M_023247488.1;refGene_gn=ATF3;refGene_tx=XM_023247489.1
F1 32923504 32923748
refGene_gn=ATF3;refGene_tx=XM_019821845.2;refGene_gn=ATF3;refGene_tx=X
M_023247488.1;refGene_gn=ATF3;refGene_tx=XM_023247489.1
F1 32929732 32929962 refGene_gn=ATF3;refGene_tx=XM_023247488.1
F1 33025232 33025369
refGene_gn=TMEM206;refGene_tx=XM_023248084.1;refGene_gn=TMEM206;refGen
e_tx=XM_023248085.1;refGene_gn=TMEM206;refGene_tx=XM_023248086.1;refGe
ne_gn=TMEM206;refGene_tx=XM_023248087.1
F1 33033772 33033859
refGene_gn=TMEM206;refGene_tx=XM_023248084.1;refGene_gn=TMEM206;refGen
e_tx=XM_023248085.1;refGene_gn=TMEM206;refGene_tx=XM_023248086.1;refGe
ne_gn=TMEM206;refGene_tx=XM_023248087.1
F1 33072460 33072510 refGene_gn=TMEM206;refGene_tx=XM_023248088.1
F1 33097571 33097748
refGene_gn=NENF;refGene_tx=XM_023248089.1;refGene_gn=NENF;refGene_tx=X
M_023248090.1
F1 33099153 33099266
refGene_gn=NENF;refGene_tx=XM_023248089.1;refGene_gn=NENF;refGene_tx=X
M_023248090.1
F1 33099993 33100054
refGene_gn=NENF;refGene_tx=XM_023248089.1;refGene_gn=NENF;refGene_tx=X
M_023248090.1
F1 33103355 33103458
refGene_gn=TMEM206;refGene_tx=XM_023248084.1;refGene_gn=TMEM206;refGen
e_tx=XM_023248085.1;refGene_gn=TMEM206;refGene_tx=XM_023248086.1;refGe
ne_gn=TMEM206;refGene_tx=XM_023248087.1;refGene_gn=TMEM206;refGene_tx=
XM_023248088.1
F1 33108357 33108540
refGene_gn=NENF;refGene_tx=XM_023248089.1;refGene_gn=NENF;refGene_tx=X
M_023248090.1
F1 33114861 33114897 refGene_gn=TMEM206;refGene_tx=XM_003999307.5
F1 33119192 33119289
refGene_gn=TMEM206;refGene_tx=XM_003999307.5;refGene_gn=TMEM206;refGen

e_tx=XM_023248084.1;refGene_gn=TMEM206;refGene_tx=XM_023248085.1;refGene_gn=TMEM206;refGene_tx=XM_023248086.1;refGene_gn=TMEM206;refGene_tx=XM_023248087.1;refGene_gn=TMEM206;refGene_tx=XM_023248088.1
F1	33136854	33137064	refGene_gn=TMEM206;refGene_tx=XM_003999307.5;refGene_gn=TMEM206;refGene_tx=XM_023248084.1;refGene_gn=TMEM206;refGene_tx=XM_023248085.1;refGene_gn=TMEM206;refGene_tx=XM_023248086.1;refGene_gn=TMEM206;refGene_tx=XM_023248087.1;refGene_gn=TMEM206;refGene_tx=XM_023248088.1
F1	33138443	33138595	refGene_gn=TMEM206;refGene_tx=XM_003999307.5;refGene_gn=TMEM206;refGene_tx=XM_023248084.1;refGene_gn=TMEM206;refGene_tx=XM_023248085.1;refGene_gn=TMEM206;refGene_tx=XM_023248086.1;refGene_gn=TMEM206;refGene_tx=XM_023248087.1;refGene_gn=TMEM206;refGene_tx=XM_023248088.1
F1	33143513	33143656	refGene_gn=TMEM206;refGene_tx=XM_003999307.5;refGene_gn=TMEM206;refGene_tx=XM_023248084.1;refGene_gn=TMEM206;refGene_tx=XM_023248085.1;refGene_gn=TMEM206;refGene_tx=XM_023248086.1;refGene_gn=TMEM206;refGene_tx=XM_023248087.1;refGene_gn=TMEM206;refGene_tx=XM_023248088.1
F1	33144594	33144739	refGene_gn=TMEM206;refGene_tx=XM_003999307.5;refGene_gn=TMEM206;refGene_tx=XM_023248084.1;refGene_gn=TMEM206;refGene_tx=XM_023248085.1;refGene_gn=TMEM206;refGene_tx=XM_023248086.1;refGene_gn=TMEM206;refGene_tx=XM_023248087.1;refGene_gn=TMEM206;refGene_tx=XM_023248088.1
F1	33146852	33146960	refGene_gn=TMEM206;refGene_tx=XM_003999307.5;refGene_gn=TMEM206;refGene_tx=XM_023248084.1;refGene_gn=TMEM206;refGene_tx=XM_023248085.1;refGene_gn=TMEM206;refGene_tx=XM_023248086.1;refGene_gn=TMEM206;refGene_tx=XM_023248087.1;refGene_gn=TMEM206;refGene_tx=XM_023248088.1
F1	33148637	33148799	refGene_gn=TMEM206;refGene_tx=XM_003999307.5;refGene_gn=TMEM206;refGene_tx=XM_023248084.1;refGene_gn=TMEM206;refGene_tx=XM_023248085.1;refGene_gn=TMEM206;refGene_tx=XM_023248086.1;refGene_gn=TMEM206;refGene_tx=XM_023248087.1;refGene_gn=TMEM206;refGene_tx=XM_023248088.1
F1	33152073	33152206	refGene_gn=PPP2R5A;refGene_tx=XM_003999320.5
F1	33153900	33154002	refGene_gn=PPP2R5A;refGene_tx=XM_003999320.5
F1	33154720	33154848	refGene_gn=PPP2R5A;refGene_tx=XM_003999320.5
F1	33154931	33155051	refGene_gn=PPP2R5A;refGene_tx=XM_003999320.5
F1	33155350	33155401	refGene_gn=PPP2R5A;refGene_tx=XM_003999320.5
F1	33161286	33161340	refGene_gn=PPP2R5A;refGene_tx=XM_003999320.5
F1	33162016	33162125	refGene_gn=PPP2R5A;refGene_tx=XM_003999320.5
F1	33163033	33163093	refGene_gn=PPP2R5A;refGene_tx=XM_003999320.5
F1	33164174	33164305	refGene_gn=PPP2R5A;refGene_tx=XM_003999320.5
F1	33167907	33168000	refGene_gn=PPP2R5A;refGene_tx=XM_003999320.5
F1	33180447	33180549	refGene_gn=PPP2R5A;refGene_tx=XM_003999320.5
F1	33183609	33183806	refGene_gn=PPP2R5A;refGene_tx=XM_003999320.5
F1	33216282	33216466	refGene_gn=PPP2R5A;refGene_tx=XM_003999320.5
F1	33360813	33360912	refGene_gn=DTL;refGene_tx=XM_006942785.4;refGene_gn=DTL;refGene_tx=XM_006942786.3;refGene_gn=DTL;refGene_tx=XM_006942787.4;refGene_gn=DTL;refGene_tx=XM_011290974.3;refGene_gn=DTL;refGene_tx=XM_019821837.2

F1 33362818 33363651
refGene_gn=DTL;refGene_tx=XM_006942785.4;refGene_gn=DTL;refGene_tx=XM_006942786.3;refGene_gn=DTL;refGene_tx=XM_006942787.4;refGene_gn=DTL;refGene_tx=XM_011290974.3;refGene_gn=DTL;refGene_tx=XM_019821837.2
F1 33373068 33373204
refGene_gn=DTL;refGene_tx=XM_006942785.4;refGene_gn=DTL;refGene_tx=XM_006942786.3;refGene_gn=DTL;refGene_tx=XM_006942787.4;refGene_gn=DTL;refGene_tx=XM_011290974.3;refGene_gn=DTL;refGene_tx=XM_019821837.2
F1 33375775 33375865
refGene_gn=DTL;refGene_tx=XM_006942785.4;refGene_gn=DTL;refGene_tx=XM_006942786.3;refGene_gn=DTL;refGene_tx=XM_006942787.4;refGene_gn=DTL;refGene_tx=XM_011290974.3;refGene_gn=DTL;refGene_tx=XM_019821837.2
F1 33382566 33382679
refGene_gn=DTL;refGene_tx=XM_006942785.4;refGene_gn=DTL;refGene_tx=XM_006942786.3;refGene_gn=DTL;refGene_tx=XM_006942787.4;refGene_gn=DTL;refGene_tx=XM_011290974.3;refGene_gn=DTL;refGene_tx=XM_019821837.2
F1 33385833 33385938
refGene_gn=DTL;refGene_tx=XM_006942785.4;refGene_gn=DTL;refGene_tx=XM_006942786.3;refGene_gn=DTL;refGene_tx=XM_006942787.4;refGene_gn=DTL;refGene_tx=XM_011290974.3;refGene_gn=DTL;refGene_tx=XM_019821837.2
F1 33386227 33386331
refGene_gn=DTL;refGene_tx=XM_006942785.4;refGene_gn=DTL;refGene_tx=XM_006942786.3;refGene_gn=DTL;refGene_tx=XM_006942787.4;refGene_gn=DTL;refGene_tx=XM_011290974.3;refGene_gn=DTL;refGene_tx=XM_019821837.2
F1 33387638 33387712
refGene_gn=DTL;refGene_tx=XM_006942785.4;refGene_gn=DTL;refGene_tx=XM_006942786.3;refGene_gn=DTL;refGene_tx=XM_006942787.4;refGene_gn=DTL;refGene_tx=XM_011290974.3;refGene_gn=DTL;refGene_tx=XM_019821837.2
F1 33388937 33389050
refGene_gn=DTL;refGene_tx=XM_006942785.4;refGene_gn=DTL;refGene_tx=XM_006942786.3;refGene_gn=DTL;refGene_tx=XM_006942787.4;refGene_gn=DTL;refGene_tx=XM_011290974.3;refGene_gn=DTL;refGene_tx=XM_019821837.2
F1 33390234 33390300
refGene_gn=DTL;refGene_tx=XM_006942785.4;refGene_gn=DTL;refGene_tx=XM_006942786.3;refGene_gn=DTL;refGene_tx=XM_006942787.4;refGene_gn=DTL;refGene_tx=XM_011290974.3;refGene_gn=DTL;refGene_tx=XM_019821837.2
F1 33393190 33393311
refGene_gn=DTL;refGene_tx=XM_006942785.4;refGene_gn=DTL;refGene_tx=XM_006942786.3;refGene_gn=DTL;refGene_tx=XM_006942787.4;refGene_gn=DTL;refGene_tx=XM_011290974.3;refGene_gn=DTL;refGene_tx=XM_019821837.2
F1 33393395 33393457
refGene_gn=DTL;refGene_tx=XM_006942785.4;refGene_gn=DTL;refGene_tx=XM_006942786.3;refGene_gn=DTL;refGene_tx=XM_006942787.4;refGene_gn=DTL;refGene_tx=XM_011290974.3;refGene_gn=DTL;refGene_tx=XM_019821837.2
F1 33394717 33394816
refGene_gn=DTL;refGene_tx=XM_006942785.4;refGene_gn=DTL;refGene_tx=XM_006942786.3;refGene_gn=DTL;refGene_tx=XM_006942787.4;refGene_gn=DTL;refGene_tx=XM_011290974.3;refGene_gn=DTL;refGene_tx=XM_019821837.2
F1 33395842 33395968
refGene_gn=DTL;refGene_tx=XM_006942785.4;refGene_gn=DTL;refGene_tx=XM_

006942786.3;refGene_gn=DTL;refGene_tx=XM_011290974.3;refGene_gn=DTL;refGene_tx=XM_019821837.2
F1 33401011 33401063
refGene_gn=DTL;refGene_tx=XM_006942785.4;refGene_gn=DTL;refGene_tx=XM_006942786.3;refGene_gn=DTL;refGene_tx=XM_011290974.3;refGene_gn=DTL;refGene_tx=XM_019821837.2
F1 33401514 33401608 refGene_gn=INTS7;refGene_tx=XM_011290973.3
F1 33413312 33413442 refGene_gn=INTS7;refGene_tx=XM_011290973.3
F1 33414258 33414405 refGene_gn=INTS7;refGene_tx=XM_011290973.3
F1 33421520 33421658 refGene_gn=INTS7;refGene_tx=XM_011290973.3
F1 33428944 33428991 refGene_gn=INTS7;refGene_tx=XM_011290973.3
F1 33432777 33432977 refGene_gn=INTS7;refGene_tx=XM_011290973.3
F1 33433446 33433569 refGene_gn=INTS7;refGene_tx=XM_011290973.3
F1 33450784 33450902 refGene_gn=INTS7;refGene_tx=XM_011290973.3
F1 33455553 33455688 refGene_gn=INTS7;refGene_tx=XM_011290973.3
F1 33458795 33458893 refGene_gn=INTS7;refGene_tx=XM_011290973.3
F1 33461419 33461659 refGene_gn=INTS7;refGene_tx=XM_011290973.3
F1 33463227 33463365 refGene_gn=INTS7;refGene_tx=XM_011290973.3
F1 33464436 33464643 refGene_gn=INTS7;refGene_tx=XM_011290973.3
F1 33474749 33474944 refGene_gn=INTS7;refGene_tx=XM_011290973.3
F1 33475430 33475534 refGene_gn=INTS7;refGene_tx=XM_011290973.3
F1 33476581 33476650 refGene_gn=INTS7;refGene_tx=XM_011290973.3
F1 33484299 33484432 refGene_gn=INTS7;refGene_tx=XM_011290973.3
F1 33488095 33488194 refGene_gn=INTS7;refGene_tx=XM_011290973.3
F1 33489474 33489660 refGene_gn=INTS7;refGene_tx=XM_011290973.3
F1 33492755 33493043 refGene_gn=INTS7;refGene_tx=XM_011290973.3
F1 33571686 33571924
refGene_gn=LPGAT1;refGene_tx=XM_003999322.4;refGene_gn=LPGAT1;refGene_tx=XM_019821846.2;refGene_gn=LPGAT1;refGene_tx=XM_023247836.1
F1 33603464 33603583
refGene_gn=LPGAT1;refGene_tx=XM_003999322.4;refGene_gn=LPGAT1;refGene_tx=XM_019821846.2;refGene_gn=LPGAT1;refGene_tx=XM_023247836.1
F1 33608516 33608612
refGene_gn=LPGAT1;refGene_tx=XM_003999322.4;refGene_gn=LPGAT1;refGene_tx=XM_019821846.2;refGene_gn=LPGAT1;refGene_tx=XM_023247836.1
F1 33610492 33610766
refGene_gn=LPGAT1;refGene_tx=XM_003999322.4;refGene_gn=LPGAT1;refGene_tx=XM_019821846.2;refGene_gn=LPGAT1;refGene_tx=XM_023247836.1
F1 33614464 33614591
refGene_gn=LPGAT1;refGene_tx=XM_003999322.4;refGene_gn=LPGAT1;refGene_tx=XM_019821846.2;refGene_gn=LPGAT1;refGene_tx=XM_023247836.1
F1 33638750 33638857
refGene_gn=LPGAT1;refGene_tx=XM_003999322.4;refGene_gn=LPGAT1;refGene_tx=XM_019821846.2;refGene_gn=LPGAT1;refGene_tx=XM_023247836.1
F1 33641256 33641408
refGene_gn=LPGAT1;refGene_tx=XM_003999322.4;refGene_gn=LPGAT1;refGene_tx=XM_019821846.2;refGene_gn=LPGAT1;refGene_tx=XM_023247836.1
F1 33684105 33684201 refGene_gn=NEK2;refGene_tx=XM_003999323.5
F1 33685313 33685531 refGene_gn=NEK2;refGene_tx=XM_003999323.5
F1 33685809 33686050 refGene_gn=NEK2;refGene_tx=XM_003999323.5

F1	33687870	33687953	refGene_gn=NEK2;refGene_tx=XM_003999323.5
F1	33688679	33688806	refGene_gn=NEK2;refGene_tx=XM_003999323.5
F1	33689648	33689865	refGene_gn=NEK2;refGene_tx=XM_003999323.5
F1	33691322	33691448	refGene_gn=NEK2;refGene_tx=XM_003999323.5
F1	33695490	33695717	refGene_gn=NEK2;refGene_tx=XM_003999323.5
F1	33769945	33770185	refGene_gn=SLC30A1;refGene_tx=XM_006942941.4
F1	33770186	33770556	refGene_gn=SLC30A1;refGene_tx=XM_006942941.4
F1	33772450	33773352	refGene_gn=SLC30A1;refGene_tx=XM_006942941.4
F1	33854420	33854427
refGene_gn=RD3;refGene_tx=XM_011291045.3;refGene_gn=RD3;refGene_tx=XM_019821847.2;refGene_gn=RD3;refGene_tx=XM_019821850.2
F1	33854909	33855259
refGene_gn=RD3;refGene_tx=XM_011291045.3;refGene_gn=RD3;refGene_tx=XM_019821847.2;refGene_gn=RD3;refGene_tx=XM_019821850.2
F1	33857080	33857378
refGene_gn=RD3;refGene_tx=XM_011291045.3;refGene_gn=RD3;refGene_tx=XM_019821847.2;refGene_gn=RD3;refGene_tx=XM_019821850.2
F1	33891087	33891665
refGene_gn=TRAF5;refGene_tx=XM_003999324.5;refGene_gn=TRAF5;refGene_tx=XM_006942799.4;refGene_gn=TRAF5;refGene_tx=XM_006942800.4
F1	33892597	33892766
refGene_gn=TRAF5;refGene_tx=XM_003999324.5;refGene_gn=TRAF5;refGene_tx=XM_006942799.4;refGene_gn=TRAF5;refGene_tx=XM_006942800.4
F1	33894891	33895032
refGene_gn=TRAF5;refGene_tx=XM_003999324.5;refGene_gn=TRAF5;refGene_tx=XM_006942799.4;refGene_gn=TRAF5;refGene_tx=XM_006942800.4
F1	33898978	33899071
refGene_gn=TRAF5;refGene_tx=XM_003999324.5;refGene_gn=TRAF5;refGene_tx=XM_006942799.4;refGene_gn=TRAF5;refGene_tx=XM_006942800.4
F1	33902874	33902949
refGene_gn=TRAF5;refGene_tx=XM_003999324.5;refGene_gn=TRAF5;refGene_tx=XM_006942799.4;refGene_gn=TRAF5;refGene_tx=XM_006942800.4
F1	33903279	33903357
refGene_gn=TRAF5;refGene_tx=XM_003999324.5;refGene_gn=TRAF5;refGene_tx=XM_006942799.4;refGene_gn=TRAF5;refGene_tx=XM_006942800.4
F1	33904158	33904323
refGene_gn=TRAF5;refGene_tx=XM_003999324.5;refGene_gn=TRAF5;refGene_tx=XM_006942799.4
F1	33906854	33906956
refGene_gn=TRAF5;refGene_tx=XM_003999324.5;refGene_gn=TRAF5;refGene_tx=XM_006942799.4;refGene_gn=TRAF5;refGene_tx=XM_006942800.4
F1	33909766	33909824
refGene_gn=TRAF5;refGene_tx=XM_003999324.5;refGene_gn=TRAF5;refGene_tx=XM_006942799.4;refGene_gn=TRAF5;refGene_tx=XM_006942800.4
F1	33911415	33911633
refGene_gn=TRAF5;refGene_tx=XM_003999324.5;refGene_gn=TRAF5;refGene_tx=XM_006942799.4;refGene_gn=TRAF5;refGene_tx=XM_006942800.4
F1	33953237	33953582
refGene_gn=RCOR3;refGene_tx=XM_003999326.5;refGene_gn=RCOR3;refGene_tx=XM_003999327.5;refGene_gn=RCOR3;refGene_tx=XM_006942801.3;refGene_gn=

RCOR3;refGene_tx=XM_019821851.2;refGene_gn=RCOR3;refGene_tx=XM_0232480
68.1
F1 33954047 33954289
refGene_gn=RCOR3;refGene_tx=XM_003999326.5;refGene_gn=RCOR3;refGene_tx
=XM_006942801.3;refGene_gn=RCOR3;refGene_tx=XM_019821851.2;refGene_gn=
RCOR3;refGene_tx=XM_023248068.1
F1 33962325 33962383
refGene_gn=RCOR3;refGene_tx=XM_003999326.5;refGene_gn=RCOR3;refGene_tx
=XM_003999327.5;refGene_gn=RCOR3;refGene_tx=XM_006942801.3;refGene_gn=
RCOR3;refGene_tx=XM_019821851.2;refGene_gn=RCOR3;refGene_tx=XM_0232480
68.1
F1 33970441 33970519
refGene_gn=RCOR3;refGene_tx=XM_003999326.5;refGene_gn=RCOR3;refGene_tx
=XM_003999327.5;refGene_gn=RCOR3;refGene_tx=XM_006942801.3;refGene_gn=
RCOR3;refGene_tx=XM_019821851.2;refGene_gn=RCOR3;refGene_tx=XM_0232480
68.1
F1 33976414 33976633
refGene_gn=RCOR3;refGene_tx=XM_003999326.5;refGene_gn=RCOR3;refGene_tx
=XM_003999327.5;refGene_gn=RCOR3;refGene_tx=XM_006942801.3;refGene_gn=
RCOR3;refGene_tx=XM_019821851.2;refGene_gn=RCOR3;refGene_tx=XM_0232480
68.1
F1 33987213 33987292
refGene_gn=RCOR3;refGene_tx=XM_003999326.5;refGene_gn=RCOR3;refGene_tx
=XM_003999327.5;refGene_gn=RCOR3;refGene_tx=XM_006942801.3;refGene_gn=
RCOR3;refGene_tx=XM_019821851.2;refGene_gn=RCOR3;refGene_tx=XM_0232480
68.1
F1 33988047 33988172
refGene_gn=RCOR3;refGene_tx=XM_003999326.5;refGene_gn=RCOR3;refGene_tx
=XM_003999327.5;refGene_gn=RCOR3;refGene_tx=XM_006942801.3;refGene_gn=
RCOR3;refGene_tx=XM_019821851.2;refGene_gn=RCOR3;refGene_tx=XM_0232480
68.1
F1 33989835 33989997
refGene_gn=RCOR3;refGene_tx=XM_003999326.5;refGene_gn=RCOR3;refGene_tx
=XM_003999327.5;refGene_gn=RCOR3;refGene_tx=XM_006942801.3;refGene_gn=
RCOR3;refGene_tx=XM_023248068.1
F1 33992074 33992127
refGene_gn=RCOR3;refGene_tx=XM_003999326.5;refGene_gn=RCOR3;refGene_tx
=XM_003999327.5;refGene_gn=RCOR3;refGene_tx=XM_006942801.3;refGene_gn=
RCOR3;refGene_tx=XM_019821851.2
F1 33994753 33994831
refGene_gn=RCOR3;refGene_tx=XM_003999326.5;refGene_gn=RCOR3;refGene_tx
=XM_003999327.5;refGene_gn=RCOR3;refGene_tx=XM_006942801.3;refGene_gn=
RCOR3;refGene_tx=XM_019821851.2
F1 34003898 34003955
refGene_gn=RCOR3;refGene_tx=XM_003999326.5;refGene_gn=RCOR3;refGene_tx
=XM_003999327.5;refGene_gn=RCOR3;refGene_tx=XM_006942801.3;refGene_gn=
RCOR3;refGene_tx=XM_019821851.2
F1 34004353 34004528
refGene_gn=RCOR3;refGene_tx=XM_003999326.5;refGene_gn=RCOR3;refGene_tx
=XM_003999327.5;refGene_gn=RCOR3;refGene_tx=XM_006942801.3;refGene_gn=

RCOR3;refGene_tx=XM_019821851.2
F1 34094527 34094606
refGene_gn=KCNH1;refGene_tx=XM_011290971.3;refGene_gn=KCNH1;refGene_tx=XM_023248008.1
F1 34123298 34123422
refGene_gn=KCNH1;refGene_tx=XM_011290971.3;refGene_gn=KCNH1;refGene_tx=XM_023248008.1;refGene_gn=KCNH1;refGene_tx=XM_023248009.1
F1 34127118 34127225
refGene_gn=KCNH1;refGene_tx=XM_011290971.3;refGene_gn=KCNH1;refGene_tx=XM_023248008.1;refGene_gn=KCNH1;refGene_tx=XM_023248009.1
F1 34140228 34140357
refGene_gn=KCNH1;refGene_tx=XM_011290971.3;refGene_gn=KCNH1;refGene_tx=XM_023248008.1;refGene_gn=KCNH1;refGene_tx=XM_023248009.1
F1 34148298 34148417
refGene_gn=KCNH1;refGene_tx=XM_011290971.3;refGene_gn=KCNH1;refGene_tx=XM_023248008.1;refGene_gn=KCNH1;refGene_tx=XM_023248009.1
F1 34170454 34170928
refGene_gn=KCNH1;refGene_tx=XM_011290971.3;refGene_gn=KCNH1;refGene_tx=XM_023248008.1;refGene_gn=KCNH1;refGene_tx=XM_023248009.1
F1 34237782 34238212
refGene_gn=KCNH1;refGene_tx=XM_011290971.3;refGene_gn=KCNH1;refGene_tx=XM_023248008.1;refGene_gn=KCNH1;refGene_tx=XM_023248009.1
F1 34357271 34357471
refGene_gn=KCNH1;refGene_tx=XM_011290971.3;refGene_gn=KCNH1;refGene_tx=XM_023248008.1;refGene_gn=KCNH1;refGene_tx=XM_023248009.1
F1 34363882 34364135
refGene_gn=KCNH1;refGene_tx=XM_011290971.3;refGene_gn=KCNH1;refGene_tx=XM_023248008.1;refGene_gn=KCNH1;refGene_tx=XM_023248009.1
F1 34386113 34386310
refGene_gn=KCNH1;refGene_tx=XM_011290971.3;refGene_gn=KCNH1;refGene_tx=XM_023248008.1;refGene_gn=KCNH1;refGene_tx=XM_023248009.1
F1 34468162 34469017
refGene_gn=KCNH1;refGene_tx=XM_011290971.3;refGene_gn=KCNH1;refGene_tx=XM_023248008.1;refGene_gn=KCNH1;refGene_tx=XM_023248009.1
F1 34478180 34478272 refGene_gn=HHAT;refGene_tx=XM_023247996.1
F1 34535128 34535313
refGene_gn=HHAT;refGene_tx=XM_019821853.2;refGene_gn=HHAT;refGene_tx=XM_019821854.2;refGene_gn=HHAT;refGene_tx=XM_019821855.2;refGene_gn=HHAT;refGene_tx=XM_019821858.2;refGene_gn=HHAT;refGene_tx=XM_023247994.1;refGene_gn=HHAT;refGene_tx=XM_023247995.1
F1 34535668 34535813
refGene_gn=HHAT;refGene_tx=XM_019821853.2;refGene_gn=HHAT;refGene_tx=XM_019821854.2;refGene_gn=HHAT;refGene_tx=XM_019821855.2;refGene_gn=HHAT;refGene_tx=XM_019821858.2;refGene_gn=HHAT;refGene_tx=XM_023247994.1;refGene_gn=HHAT;refGene_tx=XM_023247995.1;refGene_gn=HHAT;refGene_tx=XM_023247996.1
F1 34558870 34559072
refGene_gn=HHAT;refGene_tx=XM_019821853.2;refGene_gn=HHAT;refGene_tx=XM_019821854.2;refGene_gn=HHAT;refGene_tx=XM_019821855.2;refGene_gn=HHAT;refGene_tx=XM_019821858.2;refGene_gn=HHAT;refGene_tx=XM_023247994.1;

refGene_gn=HHAT;refGene_tx=XM_023247995.1;refGene_gn=HHAT;refGene_tx=XM_023247996.1
F1 34585362 34585528 refGene_gn=HHAT;refGene_tx=XM_023247997.1
F1 34621431 34621467
refGene_gn=HHAT;refGene_tx=XM_019821853.2;refGene_gn=HHAT;refGene_tx=XM_019821854.2;refGene_gn=HHAT;refGene_tx=XM_019821855.2;refGene_gn=HHAT;refGene_tx=XM_019821858.2;refGene_gn=HHAT;refGene_tx=XM_023247994.1;refGene_gn=HHAT;refGene_tx=XM_023247995.1;refGene_gn=HHAT;refGene_tx=XM_023247996.1;refGene_gn=HHAT;refGene_tx=XM_023247997.1
F1 34662926 34663077
refGene_gn=HHAT;refGene_tx=XM_019821853.2;refGene_gn=HHAT;refGene_tx=XM_019821854.2;refGene_gn=HHAT;refGene_tx=XM_019821855.2;refGene_gn=HHAT;refGene_tx=XM_019821858.2;refGene_gn=HHAT;refGene_tx=XM_023247994.1;refGene_gn=HHAT;refGene_tx=XM_023247995.1;refGene_gn=HHAT;refGene_tx=XM_023247996.1;refGene_gn=HHAT;refGene_tx=XM_023247997.1
F1 34709195 34709367
refGene_gn=HHAT;refGene_tx=XM_019821853.2;refGene_gn=HHAT;refGene_tx=XM_019821854.2;refGene_gn=HHAT;refGene_tx=XM_019821855.2;refGene_gn=HHAT;refGene_tx=XM_019821858.2;refGene_gn=HHAT;refGene_tx=XM_023247994.1;refGene_gn=HHAT;refGene_tx=XM_023247995.1;refGene_gn=HHAT;refGene_tx=XM_023247996.1;refGene_gn=HHAT;refGene_tx=XM_023247997.1
F1 34717303 34717513
refGene_gn=HHAT;refGene_tx=XM_019821853.2;refGene_gn=HHAT;refGene_tx=XM_019821854.2;refGene_gn=HHAT;refGene_tx=XM_019821855.2;refGene_gn=HHAT;refGene_tx=XM_019821858.2;refGene_gn=HHAT;refGene_tx=XM_023247994.1;refGene_gn=HHAT;refGene_tx=XM_023247995.1;refGene_gn=HHAT;refGene_tx=XM_023247996.1;refGene_gn=HHAT;refGene_tx=XM_023247997.1
F1 34719860 34720055
refGene_gn=HHAT;refGene_tx=XM_019821853.2;refGene_gn=HHAT;refGene_tx=XM_019821854.2;refGene_gn=HHAT;refGene_tx=XM_019821855.2;refGene_gn=HHAT;refGene_tx=XM_019821858.2;refGene_gn=HHAT;refGene_tx=XM_023247994.1;refGene_gn=HHAT;refGene_tx=XM_023247995.1;refGene_gn=HHAT;refGene_tx=XM_023247996.1;refGene_gn=HHAT;refGene_tx=XM_023247997.1
F1 34734348 34734462
refGene_gn=HHAT;refGene_tx=XM_019821853.2;refGene_gn=HHAT;refGene_tx=XM_019821854.2;refGene_gn=HHAT;refGene_tx=XM_019821855.2;refGene_gn=HHAT;refGene_tx=XM_019821858.2;refGene_gn=HHAT;refGene_tx=XM_023247994.1;refGene_gn=HHAT;refGene_tx=XM_023247995.1;refGene_gn=HHAT;refGene_tx=XM_023247996.1;refGene_gn=HHAT;refGene_tx=XM_023247997.1
F1 34758144 34758212
refGene_gn=HHAT;refGene_tx=XM_019821853.2;refGene_gn=HHAT;refGene_tx=XM_019821854.2;refGene_gn=HHAT;refGene_tx=XM_019821855.2;refGene_gn=HHAT;refGene_tx=XM_023247994.1;refGene_gn=HHAT;refGene_tx=XM_023247995.1;refGene_gn=HHAT;refGene_tx=XM_023247996.1;refGene_gn=HHAT;refGene_tx=XM_023247997.1
F1 34771336 34771427
refGene_gn=HHAT;refGene_tx=XM_019821853.2;refGene_gn=HHAT;refGene_tx=XM_019821854.2;refGene_gn=HHAT;refGene_tx=XM_019821855.2;refGene_gn=HHAT;refGene_tx=XM_019821858.2;refGene_gn=HHAT;refGene_tx=XM_023247994.1;refGene_gn=HHAT;refGene_tx=XM_023247995.1;refGene_gn=HHAT;refGene_tx=X

M_023247996.1;refGene_gn=HHAT;refGene_tx=XM_023247997.1
F1 34859373 34860135 refGene_gn=SERTAD4;refGene_tx=XM_023247485.1
F1 34862221 34862337 refGene_gn=SERTAD4;refGene_tx=XM_023247485.1
F1 34863531 34863706 refGene_gn=SERTAD4;refGene_tx=XM_023247485.1
F1 34867841 34868143
refGene_gn=LOC109495656;refGene_tx=XM_023247486.1
F1 34870000 34870049
refGene_gn=LOC109495656;refGene_tx=XM_023247486.1
F1 34920031 34920039 refGene_gn=SYT14;refGene_tx=XM_019821862.2
F1 34932613 34932927
refGene_gn=SYT14;refGene_tx=XM_019821860.2;refGene_gn=SYT14;refGene_tx
=XM_019821861.2;refGene_gn=SYT14;refGene_tx=XM_023248111.1;refGene_gn=
SYT14;refGene_tx=XM_023248112.1;refGene_gn=SYT14;refGene_tx=XM_0232481
13.1
F1 34934214 34934271
refGene_gn=SYT14;refGene_tx=XM_019821860.2;refGene_gn=SYT14;refGene_tx
=XM_023248111.1;refGene_gn=SYT14;refGene_tx=XM_023248113.1
F1 34936803 34936993
refGene_gn=SYT14;refGene_tx=XM_019821860.2;refGene_gn=SYT14;refGene_tx
=XM_019821861.2;refGene_gn=SYT14;refGene_tx=XM_019821862.2;refGene_gn=
SYT14;refGene_tx=XM_023248111.1;refGene_gn=SYT14;refGene_tx=XM_0232481
12.1;refGene_gn=SYT14;refGene_tx=XM_023248113.1
F1 34993325 34993775
refGene_gn=SYT14;refGene_tx=XM_019821860.2;refGene_gn=SYT14;refGene_tx
=XM_019821861.2;refGene_gn=SYT14;refGene_tx=XM_019821862.2;refGene_gn=
SYT14;refGene_tx=XM_023248111.1;refGene_gn=SYT14;refGene_tx=XM_0232481
12.1;refGene_gn=SYT14;refGene_tx=XM_023248113.1
F1 34998819 34999091
refGene_gn=SYT14;refGene_tx=XM_019821860.2;refGene_gn=SYT14;refGene_tx
=XM_019821861.2;refGene_gn=SYT14;refGene_tx=XM_019821862.2;refGene_gn=
SYT14;refGene_tx=XM_023248111.1;refGene_gn=SYT14;refGene_tx=XM_0232481
12.1;refGene_gn=SYT14;refGene_tx=XM_023248113.1
F1 35067790 35068006
refGene_gn=SYT14;refGene_tx=XM_019821860.2;refGene_gn=SYT14;refGene_tx
=XM_019821861.2;refGene_gn=SYT14;refGene_tx=XM_019821862.2;refGene_gn=
SYT14;refGene_tx=XM_023248111.1;refGene_gn=SYT14;refGene_tx=XM_0232481
12.1;refGene_gn=SYT14;refGene_tx=XM_023248113.1
F1 35071942 35072873
refGene_gn=SYT14;refGene_tx=XM_019821860.2;refGene_gn=SYT14;refGene_tx
=XM_019821861.2;refGene_gn=SYT14;refGene_tx=XM_019821862.2
F1 35075068 35075233
refGene_gn=SYT14;refGene_tx=XM_023248111.1;refGene_gn=SYT14;refGene_tx
=XM_023248112.1;refGene_gn=SYT14;refGene_tx=XM_023248113.1
F1 35130453 35130501
refGene_gn=SYT14;refGene_tx=XM_023248111.1;refGene_gn=SYT14;refGene_tx
=XM_023248112.1
F1 35144347 35144360
refGene_gn=SYT14;refGene_tx=XM_023248111.1;refGene_gn=SYT14;refGene_tx
=XM_023248112.1
F1 35223615 35223859

refGene_gn=DIEXF;refGene_tx=XM_003999442.4;refGene_gn=DIEXF;refGene_tx=XM_019821867.1
F1 35231148 35231394 refGene_gn=DIEXF;refGene_tx=XM_003999442.4;refGene_gn=DIEXF;refGene_tx=XM_019821867.1
F1 35232160 35232273 refGene_gn=DIEXF;refGene_tx=XM_003999442.4;refGene_gn=DIEXF;refGene_tx=XM_019821867.1
F1 35232409 35232592 refGene_gn=DIEXF;refGene_tx=XM_003999442.4;refGene_gn=DIEXF;refGene_tx=XM_019821867.1
F1 35233967 35234170 refGene_gn=DIEXF;refGene_tx=XM_003999442.4;refGene_gn=DIEXF;refGene_tx=XM_019821867.1
F1 35236009 35236229 refGene_gn=DIEXF;refGene_tx=XM_003999442.4;refGene_gn=DIEXF;refGene_tx=XM_019821867.1
F1 35238005 35238416 refGene_gn=DIEXF;refGene_tx=XM_003999442.4;refGene_gn=DIEXF;refGene_tx=XM_019821867.1
F1 35240362 35240451 refGene_gn=DIEXF;refGene_tx=XM_003999442.4;refGene_gn=DIEXF;refGene_tx=XM_019821867.1
F1 35242381 35242543 refGene_gn=DIEXF;refGene_tx=XM_003999442.4;refGene_gn=DIEXF;refGene_tx=XM_019821867.1
F1 35244271 35244506 refGene_gn=DIEXF;refGene_tx=XM_003999442.4;refGene_gn=DIEXF;refGene_tx=XM_019821867.1
F1 35245763 35245803 refGene_gn=DIEXF;refGene_tx=XM_003999442.4
F1 35246766 35246873 refGene_gn=DIEXF;refGene_tx=XM_003999442.4
F1 35273378 35273552 refGene_gn=IRF6;refGene_tx=XM_003999329.4;refGene_gn=IRF6;refGene_tx=XM_019821870.1;refGene_gn=IRF6;refGene_tx=XM_019821871.2
F1 35277640 35277845 refGene_gn=IRF6;refGene_tx=XM_003999329.4;refGene_gn=IRF6;refGene_tx=XM_019821870.1;refGene_gn=IRF6;refGene_tx=XM_019821871.2
F1 35279095 35279224 refGene_gn=IRF6;refGene_tx=XM_003999329.4;refGene_gn=IRF6;refGene_tx=XM_019821870.1;refGene_gn=IRF6;refGene_tx=XM_019821871.2
F1 35281225 35281384 refGene_gn=IRF6;refGene_tx=XM_003999329.4;refGene_gn=IRF6;refGene_tx=XM_019821870.1;refGene_gn=IRF6;refGene_tx=XM_019821871.2
F1 35282455 35282848 refGene_gn=IRF6;refGene_tx=XM_003999329.4;refGene_gn=IRF6;refGene_tx=XM_019821870.1;refGene_gn=IRF6;refGene_tx=XM_019821871.2
F1 35283529 35283648 refGene_gn=IRF6;refGene_tx=XM_003999329.4;refGene_gn=IRF6;refGene_tx=XM_019821870.1;refGene_gn=IRF6;refGene_tx=XM_019821871.2
F1 35284750 35284975

refGene_gn=IRF6;refGene_tx=XM_003999329.4;refGene_gn=IRF6;refGene_tx=XM_019821870.1;refGene_gn=IRF6;refGene_tx=XM_019821871.2
F1	35289862 35290672
refGene_gn=CF1H1orf74;refGene_tx=XM_019821881.2
F1	35291321 35291414
refGene_gn=TRAF3IP3;refGene_tx=XM_003999331.4;refGene_gn=TRAF3IP3;refGene_tx=XM_019821868.2
F1	35292056 35292170
refGene_gn=TRAF3IP3;refGene_tx=XM_003999331.4;refGene_gn=TRAF3IP3;refGene_tx=XM_019821868.2
F1	35292922 35293059
refGene_gn=TRAF3IP3;refGene_tx=XM_003999331.4;refGene_gn=TRAF3IP3;refGene_tx=XM_019821868.2
F1	35294146 35294206
refGene_gn=TRAF3IP3;refGene_tx=XM_003999331.4;refGene_gn=TRAF3IP3;refGene_tx=XM_019821868.2
F1	35295101 35295304 refGene_gn=TRAF3IP3;refGene_tx=XM_019821869.1
F1	35295657 35295720
refGene_gn=TRAF3IP3;refGene_tx=XM_003999331.4;refGene_gn=TRAF3IP3;refGene_tx=XM_019821868.2;refGene_gn=TRAF3IP3;refGene_tx=XM_019821869.1
F1	35296292 35296428
refGene_gn=TRAF3IP3;refGene_tx=XM_003999331.4;refGene_gn=TRAF3IP3;refGene_tx=XM_019821868.2;refGene_gn=TRAF3IP3;refGene_tx=XM_019821869.1
F1	35298100 35298238
refGene_gn=TRAF3IP3;refGene_tx=XM_003999331.4;refGene_gn=TRAF3IP3;refGene_tx=XM_019821868.2;refGene_gn=TRAF3IP3;refGene_tx=XM_019821869.1
F1	35298389 35298530
refGene_gn=TRAF3IP3;refGene_tx=XM_003999331.4;refGene_gn=TRAF3IP3;refGene_tx=XM_019821868.2;refGene_gn=TRAF3IP3;refGene_tx=XM_019821869.1
F1	35300931 35301003
refGene_gn=TRAF3IP3;refGene_tx=XM_003999331.4;refGene_gn=TRAF3IP3;refGene_tx=XM_019821868.2;refGene_gn=TRAF3IP3;refGene_tx=XM_019821869.1
F1	35302275 35302306
refGene_gn=IRF6;refGene_tx=XM_019821870.1;refGene_gn=IRF6;refGene_tx=XM_019821871.2
F1	35306963 35307059
refGene_gn=TRAF3IP3;refGene_tx=XM_003999331.4;refGene_gn=TRAF3IP3;refGene_tx=XM_019821868.2;refGene_gn=TRAF3IP3;refGene_tx=XM_019821869.1
F1	35307153 35307183
refGene_gn=TRAF3IP3;refGene_tx=XM_003999331.4;refGene_gn=TRAF3IP3;refGene_tx=XM_019821868.2;refGene_gn=TRAF3IP3;refGene_tx=XM_019821869.1
F1	35307829 35307854
refGene_gn=TRAF3IP3;refGene_tx=XM_003999331.4;refGene_gn=TRAF3IP3;refGene_tx=XM_019821868.2;refGene_gn=TRAF3IP3;refGene_tx=XM_019821869.1
F1	35308105 35308169
refGene_gn=TRAF3IP3;refGene_tx=XM_003999331.4;refGene_gn=TRAF3IP3;refGene_tx=XM_019821868.2;refGene_gn=TRAF3IP3;refGene_tx=XM_019821869.1
F1	35308310 35308455
refGene_gn=TRAF3IP3;refGene_tx=XM_003999331.4;refGene_gn=TRAF3IP3;refGene_tx=XM_019821868.2;refGene_gn=TRAF3IP3;refGene_tx=XM_019821869.1

F1	35310342	35310690
refGene_gn=TRAF3IP3;refGene_tx=XM_003999331.4;refGene_gn=TRAF3IP3;refGene_tx=XM_019821868.2;refGene_gn=TRAF3IP3;refGene_tx=XM_019821869.1
F1	35326248	35326466
refGene_gn=HSD11B1;refGene_tx=XM_003999332.5;refGene_gn=HSD11B1;refGene_tx=XM_006942808.3;refGene_gn=HSD11B1;refGene_tx=XM_006942809.3;refGene_gn=HSD11B1;refGene_tx=XM_019821872.2;refGene_gn=HSD11B1;refGene_tx=XM_019821874.2;refGene_gn=HSD11B1;refGene_tx=XM_019821875.2;refGene_gn=HSD11B1;refGene_tx=XM_019821876.2;refGene_gn=HSD11B1;refGene_tx=XM_019821878.2
F1	35326579	35326629
refGene_gn=HSD11B1;refGene_tx=XM_019821879.2;refGene_gn=HSD11B1;refGene_tx=XM_019821880.2
F1	35328821	35328965
refGene_gn=HSD11B1;refGene_tx=XM_003999332.5;refGene_gn=HSD11B1;refGene_tx=XM_006942808.3;refGene_gn=HSD11B1;refGene_tx=XM_006942809.3;refGene_gn=HSD11B1;refGene_tx=XM_019821872.2;refGene_gn=HSD11B1;refGene_tx=XM_019821874.2;refGene_gn=HSD11B1;refGene_tx=XM_019821875.2;refGene_gn=HSD11B1;refGene_tx=XM_019821876.2;refGene_gn=HSD11B1;refGene_tx=XM_019821878.2;refGene_gn=HSD11B1;refGene_tx=XM_019821879.2;refGene_gn=HSD11B1;refGene_tx=XM_019821880.2
F1	35352196	35352382
refGene_gn=HSD11B1;refGene_tx=XM_003999332.5;refGene_gn=HSD11B1;refGene_tx=XM_006942808.3;refGene_gn=HSD11B1;refGene_tx=XM_006942809.3;refGene_gn=HSD11B1;refGene_tx=XM_019821872.2;refGene_gn=HSD11B1;refGene_tx=XM_019821874.2;refGene_gn=HSD11B1;refGene_tx=XM_019821875.2;refGene_gn=HSD11B1;refGene_tx=XM_019821876.2;refGene_gn=HSD11B1;refGene_tx=XM_019821878.2;refGene_gn=HSD11B1;refGene_tx=XM_019821879.2;refGene_gn=HSD11B1;refGene_tx=XM_019821880.2
F1	35352513	35352625
refGene_gn=HSD11B1;refGene_tx=XM_003999332.5;refGene_gn=HSD11B1;refGene_tx=XM_006942808.3;refGene_gn=HSD11B1;refGene_tx=XM_006942809.3;refGene_gn=HSD11B1;refGene_tx=XM_019821872.2;refGene_gn=HSD11B1;refGene_tx=XM_019821874.2;refGene_gn=HSD11B1;refGene_tx=XM_019821875.2;refGene_gn=HSD11B1;refGene_tx=XM_019821876.2;refGene_gn=HSD11B1;refGene_tx=XM_019821878.2;refGene_gn=HSD11B1;refGene_tx=XM_019821879.2;refGene_gn=HSD11B1;refGene_tx=XM_019821880.2
F1	35353347	35353478
refGene_gn=HSD11B1;refGene_tx=XM_003999332.5;refGene_gn=HSD11B1;refGene_tx=XM_006942808.3;refGene_gn=HSD11B1;refGene_tx=XM_006942809.3;refGene_gn=HSD11B1;refGene_tx=XM_019821872.2;refGene_gn=HSD11B1;refGene_tx=XM_019821874.2;refGene_gn=HSD11B1;refGene_tx=XM_019821875.2;refGene_gn=HSD11B1;refGene_tx=XM_019821876.2;refGene_gn=HSD11B1;refGene_tx=XM_019821878.2;refGene_gn=HSD11B1;refGene_tx=XM_019821879.2;refGene_gn=HSD11B1;refGene_tx=XM_019821880.2
F1	35354217	35354305
refGene_gn=HSD11B1;refGene_tx=XM_003999332.5;refGene_gn=HSD11B1;refGene_tx=XM_006942808.3;refGene_gn=HSD11B1;refGene_tx=XM_019821872.2;refGene_gn=HSD11B1;refGene_tx=XM_019821874.2;refGene_gn=HSD11B1;refGene_tx=XM_019821875.2;refGene_gn=HSD11B1;refGene_tx=XM_019821876.2;refGene_gn

=HSD11B1;refGene_tx=XM_019821879.2;refGene_gn=HSD11B1;refGene_tx=XM_019821880.2
F1 35388864 35389176 refGene_gn=G0S2;refGene_tx=XM_011290970.3
F1 35415192 35415220 refGene_gn=LAMB3;refGene_tx=XM_019821882.1
F1 35416013 35416168 refGene_gn=LAMB3;refGene_tx=XM_019821882.1
F1 35428197 35428312 refGene_gn=LAMB3;refGene_tx=XM_019821882.1
F1 35428846 35428920 refGene_gn=LAMB3;refGene_tx=XM_019821882.1
F1 35432718 35432910 refGene_gn=LAMB3;refGene_tx=XM_019821882.1
F1 35434516 35434580 refGene_gn=LAMB3;refGene_tx=XM_019821882.1
F1 35434982 35435176 refGene_gn=LAMB3;refGene_tx=XM_019821882.1
F1 35437064 35437185 refGene_gn=LAMB3;refGene_tx=XM_019821882.1
F1 35437944 35438133 refGene_gn=LAMB3;refGene_tx=XM_019821882.1
F1 35439538 35439694 refGene_gn=LAMB3;refGene_tx=XM_019821882.1
F1 35440049 35440246 refGene_gn=LAMB3;refGene_tx=XM_019821882.1
F1 35440589 35440701 refGene_gn=LAMB3;refGene_tx=XM_019821882.1
F1 35441568 35441947 refGene_gn=LAMB3;refGene_tx=XM_019821882.1
F1 35443164 35443325 refGene_gn=LAMB3;refGene_tx=XM_019821882.1
F1 35443432 35443653 refGene_gn=LAMB3;refGene_tx=XM_019821882.1
F1 35443934 35444132 refGene_gn=LAMB3;refGene_tx=XM_019821882.1
F1 35444490 35444635 refGene_gn=LAMB3;refGene_tx=XM_019821882.1
F1 35448068 35448276 refGene_gn=LAMB3;refGene_tx=XM_019821882.1
F1 35448709 35448851 refGene_gn=LAMB3;refGene_tx=XM_019821882.1
F1 35449241 35449418 refGene_gn=LAMB3;refGene_tx=XM_019821882.1
F1 35450151 35450305 refGene_gn=LAMB3;refGene_tx=XM_019821882.1
F1 35452174 35452311 refGene_gn=LAMB3;refGene_tx=XM_019821882.1
F1 35454709 35454800
refGene_gn=CAMK1G;refGene_tx=XM_006942811.4;refGene_gn=CAMK1G;refGene_tx=XM_011290967.3
F1 35455313 35455738
refGene_gn=CAMK1G;refGene_tx=XM_006942811.4;refGene_gn=CAMK1G;refGene_tx=XM_011290967.3
F1 35455974 35456062
refGene_gn=CAMK1G;refGene_tx=XM_006942811.4;refGene_gn=CAMK1G;refGene_tx=XM_011290967.3
F1 35457566 35457645
refGene_gn=CAMK1G;refGene_tx=XM_006942811.4;refGene_gn=CAMK1G;refGene_tx=XM_011290967.3
F1 35458409 35458522
refGene_gn=CAMK1G;refGene_tx=XM_006942811.4;refGene_gn=CAMK1G;refGene_tx=XM_011290967.3
F1 35459580 35459656
refGene_gn=CAMK1G;refGene_tx=XM_006942811.4;refGene_gn=CAMK1G;refGene_tx=XM_011290967.3
F1 35461019 35461143
refGene_gn=CAMK1G;refGene_tx=XM_006942811.4;refGene_gn=CAMK1G;refGene_tx=XM_011290967.3
F1 35461786 35461925
refGene_gn=CAMK1G;refGene_tx=XM_006942811.4;refGene_gn=CAMK1G;refGene_tx=XM_011290967.3
F1 35464352 35464427

refGene_gn=CAMK1G;refGene_tx=XM_006942811.4;refGene_gn=CAMK1G;refGene_tx=XM_011290967.3
F1 35467118 35467247
refGene_gn=CAMK1G;refGene_tx=XM_006942811.4;refGene_gn=CAMK1G;refGene_tx=XM_011290967.3
F1 35472558 35472676
refGene_gn=CAMK1G;refGene_tx=XM_006942811.4;refGene_gn=CAMK1G;refGene_tx=XM_011290967.3
F1 35476935 35476963 refGene_gn=CAMK1G;refGene_tx=XM_011290967.3
F1 36711574 36712807
refGene_gn=PLXNA2;refGene_tx=XM_003999337.4;refGene_gn=PLXNA2;refGene_tx=XM_006942812.4;refGene_gn=PLXNA2;refGene_tx=XM_019821884.2;refGene_gn=PLXNA2;refGene_tx=XM_019821886.2
F1 36719079 36719262
refGene_gn=PLXNA2;refGene_tx=XM_003999337.4;refGene_gn=PLXNA2;refGene_tx=XM_006942812.4;refGene_gn=PLXNA2;refGene_tx=XM_019821884.2;refGene_gn=PLXNA2;refGene_tx=XM_019821886.2
F1 36725900 36726032 refGene_gn=PLXNA2;refGene_tx=XM_019821886.2
F1 36782345 36782480
refGene_gn=PLXNA2;refGene_tx=XM_003999337.4;refGene_gn=PLXNA2;refGene_tx=XM_006942812.4;refGene_gn=PLXNA2;refGene_tx=XM_019821884.2
F1 36820746 36820847
refGene_gn=PLXNA2;refGene_tx=XM_003999337.4;refGene_gn=PLXNA2;refGene_tx=XM_006942812.4;refGene_gn=PLXNA2;refGene_tx=XM_019821884.2
F1 36824101 36824225
refGene_gn=PLXNA2;refGene_tx=XM_003999337.4;refGene_gn=PLXNA2;refGene_tx=XM_006942812.4;refGene_gn=PLXNA2;refGene_tx=XM_019821884.2
F1 36825926 36826080
refGene_gn=PLXNA2;refGene_tx=XM_003999337.4;refGene_gn=PLXNA2;refGene_tx=XM_006942812.4;refGene_gn=PLXNA2;refGene_tx=XM_019821884.2
F1 36826709 36826806
refGene_gn=PLXNA2;refGene_tx=XM_003999337.4;refGene_gn=PLXNA2;refGene_tx=XM_006942812.4;refGene_gn=PLXNA2;refGene_tx=XM_019821884.2
F1 36830345 36830460
refGene_gn=PLXNA2;refGene_tx=XM_003999337.4;refGene_gn=PLXNA2;refGene_tx=XM_006942812.4;refGene_gn=PLXNA2;refGene_tx=XM_019821884.2
F1 36838927 36839128
refGene_gn=PLXNA2;refGene_tx=XM_003999337.4;refGene_gn=PLXNA2;refGene_tx=XM_006942812.4;refGene_gn=PLXNA2;refGene_tx=XM_019821884.2
F1 36840883 36840980
refGene_gn=PLXNA2;refGene_tx=XM_003999337.4;refGene_gn=PLXNA2;refGene_tx=XM_006942812.4;refGene_gn=PLXNA2;refGene_tx=XM_019821884.2
F1 36844273 36844464
refGene_gn=PLXNA2;refGene_tx=XM_003999337.4;refGene_gn=PLXNA2;refGene_tx=XM_006942812.4;refGene_gn=PLXNA2;refGene_tx=XM_019821884.2
F1 36851572 36851584 refGene_gn=PLXNA2;refGene_tx=XM_006942812.4
F1 36860461 36860613
refGene_gn=PLXNA2;refGene_tx=XM_003999337.4;refGene_gn=PLXNA2;refGene_tx=XM_019821884.2
F1 36866593 36866711

refGene_gn=PLXNA2;refGene_tx=XM_003999337.4;refGene_gn=PLXNA2;refGene_tx=XM_019821884.2
F1 36868555 36868692
refGene_gn=PLXNA2;refGene_tx=XM_003999337.4;refGene_gn=PLXNA2;refGene_tx=XM_019821884.2
F1 36869629 36869797
refGene_gn=PLXNA2;refGene_tx=XM_003999337.4;refGene_gn=PLXNA2;refGene_tx=XM_019821884.2
F1 36869928 36870022
refGene_gn=PLXNA2;refGene_tx=XM_003999337.4;refGene_gn=PLXNA2;refGene_tx=XM_019821884.2
F1 36874098 36874338
refGene_gn=PLXNA2;refGene_tx=XM_003999337.4;refGene_gn=PLXNA2;refGene_tx=XM_019821884.2
F1 36874984 36875128
refGene_gn=PLXNA2;refGene_tx=XM_003999337.4;refGene_gn=PLXNA2;refGene_tx=XM_019821884.2
F1 36875412 36875647
refGene_gn=PLXNA2;refGene_tx=XM_003999337.4;refGene_gn=PLXNA2;refGene_tx=XM_019821884.2
F1 36877580 36877723
refGene_gn=PLXNA2;refGene_tx=XM_003999337.4;refGene_gn=PLXNA2;refGene_tx=XM_019821884.2
F1 36878418 36878687 refGene_gn=PLXNA2;refGene_tx=XM_003999337.4
F1 36880659 36880726 refGene_gn=PLXNA2;refGene_tx=XM_003999337.4
F1 36880943 36881090 refGene_gn=PLXNA2;refGene_tx=XM_003999337.4
F1 36881729 36881889 refGene_gn=PLXNA2;refGene_tx=XM_003999337.4
F1 36882263 36882367 refGene_gn=PLXNA2;refGene_tx=XM_003999337.4
F1 36886136 36886236 refGene_gn=PLXNA2;refGene_tx=XM_003999337.4
F1 36887504 36887695 refGene_gn=PLXNA2;refGene_tx=XM_003999337.4
F1 36889117 36889287 refGene_gn=PLXNA2;refGene_tx=XM_003999337.4
F1 36891816 36892029 refGene_gn=PLXNA2;refGene_tx=XM_003999337.4
F1 36892719 36892870 refGene_gn=PLXNA2;refGene_tx=XM_003999337.4
F1 36894054 36894156 refGene_gn=PLXNA2;refGene_tx=XM_003999337.4
F1 37008457 37008536
refGene_gn=CD34;refGene_tx=NM_001009318.1;refGene_gn=CD34;refGene_tx=XM_023247164.1;refGene_gn=CD34;refGene_tx=XM_023247165.1
F1 37010502 37010571
refGene_gn=CD34;refGene_tx=NM_001009318.1;refGene_gn=CD34;refGene_tx=XM_023247165.1
F1 37018426 37018609
refGene_gn=CD34;refGene_tx=NM_001009318.1;refGene_gn=CD34;refGene_tx=XM_023247164.1;refGene_gn=CD34;refGene_tx=XM_023247165.1
F1 37019271 37019525
refGene_gn=CD34;refGene_tx=NM_001009318.1;refGene_gn=CD34;refGene_tx=XM_023247164.1;refGene_gn=CD34;refGene_tx=XM_023247165.1
F1 37020559 37020640
refGene_gn=CD34;refGene_tx=NM_001009318.1;refGene_gn=CD34;refGene_tx=XM_023247164.1;refGene_gn=CD34;refGene_tx=XM_023247165.1
F1 37028865 37029016

refGene_gn=CD34;refGene_tx=NM_001009318.1;refGene_gn=CD34;refGene_tx=XM_023247164.1;refGene_gn=CD34;refGene_tx=XM_023247165.1
F1 37029252 37029305
refGene_gn=CD34;refGene_tx=NM_001009318.1;refGene_gn=CD34;refGene_tx=XM_023247164.1;refGene_gn=CD34;refGene_tx=XM_023247165.1
F1 37029622 37029787
refGene_gn=CD34;refGene_tx=NM_001009318.1;refGene_gn=CD34;refGene_tx=XM_023247164.1;refGene_gn=CD34;refGene_tx=XM_023247165.1
F1 37030350 37030365 refGene_gn=CD34;refGene_tx=XM_023247165.1
F1 37030497 37030683
refGene_gn=CD34;refGene_tx=NM_001009318.1;refGene_gn=CD34;refGene_tx=XM_023247164.1
F1 37048972 37049181
refGene_gn=LOC109495671;refGene_tx=XM_023248014.1
F1 37050546 37050643
refGene_gn=LOC109495671;refGene_tx=XM_023248014.1
F1 37051140 37051221
refGene_gn=LOC109495671;refGene_tx=XM_023248014.1
F1 37051345 37051512
refGene_gn=LOC109495671;refGene_tx=XM_023248014.1
F1 37051621 37051691
refGene_gn=LOC109495671;refGene_tx=XM_023248014.1
F1 37124245 37124318
refGene_gn=LOC102899273;refGene_tx=XM_011290966.3
F1 37130683 37130744
refGene_gn=LOC102899273;refGene_tx=XM_011290966.3
F1 37131250 37131286
refGene_gn=LOC102899273;refGene_tx=XM_011290966.3
F1 37132785 37132833
refGene_gn=LOC102899273;refGene_tx=XM_011290966.3
F1 37137318 37137381
refGene_gn=LOC102899273;refGene_tx=XM_011290966.3
F1 37139127 37139172
refGene_gn=LOC102899273;refGene_tx=XM_011290966.3
F1 37139862 37140045
refGene_gn=LOC102899273;refGene_tx=XM_011290966.3
F1 37142290 37142488
refGene_gn=LOC102899273;refGene_tx=XM_011290966.3
F1 37145041 37145136
refGene_gn=LOC102899273;refGene_tx=XM_011290966.3
F1 37151288 37151391
refGene_gn=LOC102899273;refGene_tx=XM_011290966.3
F1 37151705 37151888
refGene_gn=LOC102899273;refGene_tx=XM_011290966.3
F1 37159485 37159603
refGene_gn=LOC102899273;refGene_tx=XM_011290966.3
F1 37188751 37188813
refGene_gn=LOC101094152;refGene_tx=XM_023247354.1;refGene_gn=LOC101094152;refGene_tx=XM_023247355.1;refGene_gn=LOC101094152;refGene_tx=XM_023247356.1

F1 37189309 37189345
refGene_gn=LOC101094152;refGene_tx=XM_023247354.1;refGene_gn=LOC101094152;refGene_tx=XM_023247355.1;refGene_gn=LOC101094152;refGene_tx=XM_023247356.1
F1 37189856 37189907
refGene_gn=LOC101094152;refGene_tx=XM_023247354.1;refGene_gn=LOC101094152;refGene_tx=XM_023247355.1;refGene_gn=LOC101094152;refGene_tx=XM_023247356.1
F1 37195929 37195977
refGene_gn=LOC101094152;refGene_tx=XM_023247354.1;refGene_gn=LOC101094152;refGene_tx=XM_023247355.1
F1 37198573 37198756
refGene_gn=LOC101094152;refGene_tx=XM_023247354.1;refGene_gn=LOC101094152;refGene_tx=XM_023247355.1
F1 37207210 37207408
refGene_gn=LOC101094152;refGene_tx=XM_023247354.1;refGene_gn=LOC101094152;refGene_tx=XM_023247355.1;refGene_gn=LOC101094152;refGene_tx=XM_023247356.1
F1 37208219 37208305
refGene_gn=LOC101094152;refGene_tx=XM_023247354.1;refGene_gn=LOC101094152;refGene_tx=XM_023247355.1;refGene_gn=LOC101094152;refGene_tx=XM_023247356.1
F1 37212771 37212871
refGene_gn=LOC101094152;refGene_tx=XM_023247354.1;refGene_gn=LOC101094152;refGene_tx=XM_023247355.1;refGene_gn=LOC101094152;refGene_tx=XM_023247356.1
F1 37213200 37213380
refGene_gn=LOC101094152;refGene_tx=XM_023247354.1;refGene_gn=LOC101094152;refGene_tx=XM_023247355.1;refGene_gn=LOC101094152;refGene_tx=XM_023247356.1
F1 37215223 37215333
refGene_gn=LOC101094152;refGene_tx=XM_023247354.1;refGene_gn=LOC101094152;refGene_tx=XM_023247356.1
F1 37216120 37216134
refGene_gn=LOC101094152;refGene_tx=XM_023247354.1;refGene_gn=LOC101094152;refGene_tx=XM_023247356.1
F1 37220712 37220830
refGene_gn=LOC101094152;refGene_tx=XM_023247355.1
F1 37227091 37227104 refGene_gn=CR1L;refGene_tx=XM_011290964.3
F1 37230752 37230854 refGene_gn=CR1L;refGene_tx=XM_011290964.3
F1 37235527 37235603 refGene_gn=CR1L;refGene_tx=XM_011290964.3
F1 37236396 37236420 refGene_gn=CR1L;refGene_tx=XM_011290964.3
F1 37236688 37236874 refGene_gn=CR1L;refGene_tx=XM_011290964.3
F1 37239740 37239917 refGene_gn=CR1L;refGene_tx=XM_011290964.3
F1 37240905 37241133 refGene_gn=CR1L;refGene_tx=XM_011290964.3
F1 37249653 37249739 refGene_gn=CR1L;refGene_tx=XM_011290964.3
F1 37255062 37255162 refGene_gn=CR1L;refGene_tx=XM_011290964.3
F1 37255778 37255958 refGene_gn=CR1L;refGene_tx=XM_011290964.3
F1 37258226 37258454 refGene_gn=CR1L;refGene_tx=XM_011290964.3
F1 37263384 37263470 refGene_gn=CR1L;refGene_tx=XM_011290964.3

F1 37264749 37264852 refGene_gn=CR1L;refGene_tx=XM_011290964.3
F1 37264942 37265119 refGene_gn=CR1L;refGene_tx=XM_011290964.3
F1 37266820 37267219 refGene_gn=CR1L;refGene_tx=XM_011290964.3
F1 37276982 37277068 refGene_gn=CR1L;refGene_tx=XM_011290964.3
F1 37283451 37283551 refGene_gn=CR1L;refGene_tx=XM_011290964.3
F1 37284198 37284378 refGene_gn=CR1L;refGene_tx=XM_011290964.3
F1 37289509 37289627 refGene_gn=CR1L;refGene_tx=XM_011290964.3
F1 37296003 37296097
refGene_gn=CR2;refGene_tx=XM_006942818.2;refGene_gn=CR2;refGene_tx=XM_019821898.1;refGene_gn=CR2;refGene_tx=XM_019821899.1;refGene_gn=CR2;refGene_tx=XM_019821901.1;refGene_gn=CR2;refGene_tx=XM_019821902.1
F1 37300363 37300439
refGene_gn=CR2;refGene_tx=XM_006942818.2;refGene_gn=CR2;refGene_tx=XM_019821898.1;refGene_gn=CR2;refGene_tx=XM_019821899.1;refGene_gn=CR2;refGene_tx=XM_019821901.1;refGene_gn=CR2;refGene_tx=XM_019821902.1
F1 37301529 37301553
refGene_gn=CR2;refGene_tx=XM_006942818.2;refGene_gn=CR2;refGene_tx=XM_019821898.1;refGene_gn=CR2;refGene_tx=XM_019821899.1;refGene_gn=CR2;refGene_tx=XM_019821901.1;refGene_gn=CR2;refGene_tx=XM_019821902.1
F1 37301728 37301736 refGene_gn=CR2;refGene_tx=XM_019821900.1
F1 37302289 37302478
refGene_gn=CR2;refGene_tx=XM_006942818.2;refGene_gn=CR2;refGene_tx=XM_019821898.1;refGene_gn=CR2;refGene_tx=XM_019821899.1;refGene_gn=CR2;refGene_tx=XM_019821900.1;refGene_gn=CR2;refGene_tx=XM_019821901.1;refGene_gn=CR2;refGene_tx=XM_019821902.1
F1 37303528 37303714
refGene_gn=CR2;refGene_tx=XM_006942818.2;refGene_gn=CR2;refGene_tx=XM_019821898.1;refGene_gn=CR2;refGene_tx=XM_019821899.1;refGene_gn=CR2;refGene_tx=XM_019821900.1;refGene_gn=CR2;refGene_tx=XM_019821901.1;refGene_gn=CR2;refGene_tx=XM_019821902.1
F1 37304939 37305332
refGene_gn=CR2;refGene_tx=XM_006942818.2;refGene_gn=CR2;refGene_tx=XM_019821898.1;refGene_gn=CR2;refGene_tx=XM_019821899.1;refGene_gn=CR2;refGene_tx=XM_019821900.1;refGene_gn=CR2;refGene_tx=XM_019821901.1;refGene_gn=CR2;refGene_tx=XM_019821902.1
F1 37306308 37306391
refGene_gn=CR2;refGene_tx=XM_006942818.2;refGene_gn=CR2;refGene_tx=XM_019821898.1;refGene_gn=CR2;refGene_tx=XM_019821899.1;refGene_gn=CR2;refGene_tx=XM_019821900.1;refGene_gn=CR2;refGene_tx=XM_019821901.1;refGene_gn=CR2;refGene_tx=XM_019821902.1
F1 37306744 37306832
refGene_gn=CR2;refGene_tx=XM_006942818.2;refGene_gn=CR2;refGene_tx=XM_019821898.1;refGene_gn=CR2;refGene_tx=XM_019821899.1;refGene_gn=CR2;refGene_tx=XM_019821900.1;refGene_gn=CR2;refGene_tx=XM_019821901.1;refGene_gn=CR2;refGene_tx=XM_019821902.1
F1 37306906 37307083
refGene_gn=CR2;refGene_tx=XM_006942818.2;refGene_gn=CR2;refGene_tx=XM_019821898.1;refGene_gn=CR2;refGene_tx=XM_019821900.1;refGene_gn=CR2;refGene_tx=XM_019821901.1;refGene_gn=CR2;refGene_tx=XM_019821902.1
F1 37307676 37308084

refGene_gn=CR2;refGene_tx=XM_006942818.2;refGene_gn=CR2;refGene_tx=XM_019821898.1;refGene_gn=CR2;refGene_tx=XM_019821899.1;refGene_gn=CR2;refGene_tx=XM_019821900.1;refGene_gn=CR2;refGene_tx=XM_019821901.1;refGene_gn=CR2;refGene_tx=XM_019821902.1
F1 37309132 37309209
refGene_gn=CR2;refGene_tx=XM_006942818.2;refGene_gn=CR2;refGene_tx=XM_019821898.1;refGene_gn=CR2;refGene_tx=XM_019821899.1;refGene_gn=CR2;refGene_tx=XM_019821900.1;refGene_gn=CR2;refGene_tx=XM_019821901.1;refGene_gn=CR2;refGene_tx=XM_019821902.1
F1 37309505 37309596
refGene_gn=CR2;refGene_tx=XM_006942818.2;refGene_gn=CR2;refGene_tx=XM_019821898.1;refGene_gn=CR2;refGene_tx=XM_019821899.1;refGene_gn=CR2;refGene_tx=XM_019821900.1;refGene_gn=CR2;refGene_tx=XM_019821901.1;refGene_gn=CR2;refGene_tx=XM_019821902.1
F1 37309675 37309852
refGene_gn=CR2;refGene_tx=XM_006942818.2;refGene_gn=CR2;refGene_tx=XM_019821898.1;refGene_gn=CR2;refGene_tx=XM_019821899.1;refGene_gn=CR2;refGene_tx=XM_019821900.1;refGene_gn=CR2;refGene_tx=XM_019821901.1;refGene_gn=CR2;refGene_tx=XM_019821902.1
F1 37310430 37310841
refGene_gn=CR2;refGene_tx=XM_006942818.2;refGene_gn=CR2;refGene_tx=XM_019821898.1;refGene_gn=CR2;refGene_tx=XM_019821899.1;refGene_gn=CR2;refGene_tx=XM_019821900.1;refGene_gn=CR2;refGene_tx=XM_019821901.1;refGene_gn=CR2;refGene_tx=XM_019821902.1
F1 37311346 37311429
refGene_gn=CR2;refGene_tx=XM_006942818.2;refGene_gn=CR2;refGene_tx=XM_019821898.1;refGene_gn=CR2;refGene_tx=XM_019821899.1;refGene_gn=CR2;refGene_tx=XM_019821900.1;refGene_gn=CR2;refGene_tx=XM_019821901.1;refGene_gn=CR2;refGene_tx=XM_019821902.1
F1 37311681 37311781
refGene_gn=CR2;refGene_tx=XM_006942818.2;refGene_gn=CR2;refGene_tx=XM_019821898.1;refGene_gn=CR2;refGene_tx=XM_019821899.1;refGene_gn=CR2;refGene_tx=XM_019821900.1;refGene_gn=CR2;refGene_tx=XM_019821901.1;refGene_gn=CR2;refGene_tx=XM_019821902.1
F1 37311865 37312054
refGene_gn=CR2;refGene_tx=XM_006942818.2;refGene_gn=CR2;refGene_tx=XM_019821898.1;refGene_gn=CR2;refGene_tx=XM_019821899.1;refGene_gn=CR2;refGene_tx=XM_019821900.1;refGene_gn=CR2;refGene_tx=XM_019821901.1;refGene_gn=CR2;refGene_tx=XM_019821902.1
F1 37313388 37313778
refGene_gn=CR2;refGene_tx=XM_006942818.2;refGene_gn=CR2;refGene_tx=XM_019821898.1;refGene_gn=CR2;refGene_tx=XM_019821899.1;refGene_gn=CR2;refGene_tx=XM_019821900.1;refGene_gn=CR2;refGene_tx=XM_019821901.1;refGene_gn=CR2;refGene_tx=XM_019821902.1
F1 37314955 37315041
refGene_gn=CR2;refGene_tx=XM_006942818.2;refGene_gn=CR2;refGene_tx=XM_019821898.1;refGene_gn=CR2;refGene_tx=XM_019821899.1;refGene_gn=CR2;refGene_tx=XM_019821900.1;refGene_gn=CR2;refGene_tx=XM_019821901.1
F1 37315954 37316057
refGene_gn=CR2;refGene_tx=XM_006942818.2;refGene_gn=CR2;refGene_tx=XM_

019821898.1;refGene_gn=CR2;refGene_tx=XM_019821899.1;refGene_gn=CR2;refGene_tx=XM_019821900.1;refGene_gn=CR2;refGene_tx=XM_019821901.1
F1 37316138 37316315
refGene_gn=CR2;refGene_tx=XM_006942818.2;refGene_gn=CR2;refGene_tx=XM_019821898.1;refGene_gn=CR2;refGene_tx=XM_019821899.1;refGene_gn=CR2;refGene_tx=XM_019821900.1;refGene_gn=CR2;refGene_tx=XM_019821901.1
F1 37317315 37317714
refGene_gn=CR2;refGene_tx=XM_006942818.2;refGene_gn=CR2;refGene_tx=XM_019821898.1;refGene_gn=CR2;refGene_tx=XM_019821899.1;refGene_gn=CR2;refGene_tx=XM_019821900.1
F1 37318286 37318372
refGene_gn=CR2;refGene_tx=XM_006942818.2;refGene_gn=CR2;refGene_tx=XM_019821898.1;refGene_gn=CR2;refGene_tx=XM_019821899.1;refGene_gn=CR2;refGene_tx=XM_019821900.1;refGene_gn=CR2;refGene_tx=XM_019821901.1
F1 37318789 37318889
refGene_gn=CR2;refGene_tx=XM_006942818.2;refGene_gn=CR2;refGene_tx=XM_019821898.1;refGene_gn=CR2;refGene_tx=XM_019821899.1;refGene_gn=CR2;refGene_tx=XM_019821900.1;refGene_gn=CR2;refGene_tx=XM_019821901.1
F1 37320198 37320378
refGene_gn=CR2;refGene_tx=XM_006942818.2;refGene_gn=CR2;refGene_tx=XM_019821898.1;refGene_gn=CR2;refGene_tx=XM_019821899.1;refGene_gn=CR2;refGene_tx=XM_019821900.1;refGene_gn=CR2;refGene_tx=XM_019821901.1
F1 37328319 37328377
refGene_gn=CR2;refGene_tx=XM_006942818.2;refGene_gn=CR2;refGene_tx=XM_019821898.1;refGene_gn=CR2;refGene_tx=XM_019821899.1;refGene_gn=CR2;refGene_tx=XM_019821900.1;refGene_gn=CR2;refGene_tx=XM_019821901.1;refGene_gn=CR2;refGene_tx=XM_019821902.1
F1 37378863 37378928 refGene_gn=CD55;refGene_tx=XM_023247291.1
F1 37395263 37395387
refGene_gn=CD55;refGene_tx=XM_023247288.1;refGene_gn=CD55;refGene_tx=XM_023247293.1
F1 37396244 37396347 refGene_gn=CD55;refGene_tx=XM_023247288.1
F1 37399601 37399675 refGene_gn=CD55;refGene_tx=XM_023247290.1
F1 37402225 37402305 refGene_gn=CD55;refGene_tx=XM_023247289.1
F1 37402861 37402917 refGene_gn=CD55;refGene_tx=XM_023247294.1
F1 37404732 37405067
refGene_gn=CD55;refGene_tx=XM_023247287.1;refGene_gn=CD55;refGene_tx=XM_023247288.1;refGene_gn=CD55;refGene_tx=XM_023247289.1;refGene_gn=CD55;refGene_tx=XM_023247290.1;refGene_gn=CD55;refGene_tx=XM_023247291.1;refGene_gn=CD55;refGene_tx=XM_023247293.1;refGene_gn=CD55;refGene_tx=XM_023247294.1
F1 37409226 37409313
refGene_gn=CD55;refGene_tx=XM_023247287.1;refGene_gn=CD55;refGene_tx=XM_023247288.1;refGene_gn=CD55;refGene_tx=XM_023247289.1;refGene_gn=CD55;refGene_tx=XM_023247290.1;refGene_gn=CD55;refGene_tx=XM_023247291.1;refGene_gn=CD55;refGene_tx=XM_023247293.1;refGene_gn=CD55;refGene_tx=XM_023247294.1
F1 37410217 37410484
refGene_gn=CD55;refGene_tx=XM_023247287.1;refGene_gn=CD55;refGene_tx=XM_023247288.1;refGene_gn=CD55;refGene_tx=XM_023247289.1;refGene_gn=CD5

5;refGene_tx=XM_023247290.1;refGene_gn=CD55;refGene_tx=XM_023247291.1;
refGene_gn=CD55;refGene_tx=XM_023247293.1;refGene_gn=CD55;refGene_tx=X
M_023247294.1
F1 37416199 37416388
refGene_gn=CD55;refGene_tx=XM_023247287.1;refGene_gn=CD55;refGene_tx=X
M_023247288.1;refGene_gn=CD55;refGene_tx=XM_023247289.1;refGene_gn=CD5
5;refGene_tx=XM_023247290.1;refGene_gn=CD55;refGene_tx=XM_023247291.1;
refGene_gn=CD55;refGene_tx=XM_023247293.1;refGene_gn=CD55;refGene_tx=X
M_023247294.1
F1 37422540 37422626
refGene_gn=CD55;refGene_tx=XM_023247287.1;refGene_gn=CD55;refGene_tx=X
M_023247288.1;refGene_gn=CD55;refGene_tx=XM_023247289.1;refGene_gn=CD5
5;refGene_tx=XM_023247290.1;refGene_gn=CD55;refGene_tx=XM_023247291.1;
refGene_gn=CD55;refGene_tx=XM_023247293.1;refGene_gn=CD55;refGene_tx=X
M_023247294.1
F1 37423937 37424037
refGene_gn=CD55;refGene_tx=XM_023247287.1;refGene_gn=CD55;refGene_tx=X
M_023247288.1;refGene_gn=CD55;refGene_tx=XM_023247289.1;refGene_gn=CD5
5;refGene_tx=XM_023247290.1;refGene_gn=CD55;refGene_tx=XM_023247291.1;
refGene_gn=CD55;refGene_tx=XM_023247293.1;refGene_gn=CD55;refGene_tx=X
M_023247294.1
F1 37424643 37424835
refGene_gn=CD55;refGene_tx=XM_023247287.1;refGene_gn=CD55;refGene_tx=X
M_023247288.1;refGene_gn=CD55;refGene_tx=XM_023247289.1;refGene_gn=CD5
5;refGene_tx=XM_023247290.1;refGene_gn=CD55;refGene_tx=XM_023247291.1;
refGene_gn=CD55;refGene_tx=XM_023247293.1;refGene_gn=CD55;refGene_tx=X
M_023247294.1
F1 37426261 37426453
refGene_gn=CD55;refGene_tx=XM_023247287.1;refGene_gn=CD55;refGene_tx=X
M_023247288.1;refGene_gn=CD55;refGene_tx=XM_023247289.1;refGene_gn=CD5
5;refGene_tx=XM_023247290.1;refGene_gn=CD55;refGene_tx=XM_023247291.1;
refGene_gn=CD55;refGene_tx=XM_023247293.1;refGene_gn=CD55;refGene_tx=X
M_023247294.1
F1 37426525 37426610
refGene_gn=CD55;refGene_tx=XM_023247287.1;refGene_gn=CD55;refGene_tx=X
M_023247288.1;refGene_gn=CD55;refGene_tx=XM_023247289.1;refGene_gn=CD5
5;refGene_tx=XM_023247290.1;refGene_gn=CD55;refGene_tx=XM_023247291.1;
refGene_gn=CD55;refGene_tx=XM_023247293.1;refGene_gn=CD55;refGene_tx=X
M_023247294.1
F1 37502374 37502515
refGene_gn=LOC102899713;refGene_tx=XM_006942825.4
F1 37503070 37503246
refGene_gn=LOC102899713;refGene_tx=XM_006942825.4
F1 37505932 37506103
refGene_gn=LOC102899713;refGene_tx=XM_006942825.4
F1 37509982 37510067
refGene_gn=LOC102899713;refGene_tx=XM_006942825.4
F1 37523967 37524141
refGene_gn=LOC101094897;refGene_tx=XM_023247334.1
F1 37524697 37524873

refGene_gn=LOC101094897;refGene_tx=XM_023247334.1
F1 37526074 37526248
refGene_gn=LOC101094897;refGene_tx=XM_023247334.1
F1 37548501 37548675
refGene_gn=LOC101094897;refGene_tx=XM_006942826.4
F1 37549197 37549373
refGene_gn=LOC101094897;refGene_tx=XM_006942826.4
F1 37549851 37550025
refGene_gn=LOC101094897;refGene_tx=XM_006942826.4
F1 37553089 37553275
refGene_gn=LOC101094897;refGene_tx=XM_006942826.4;refGene_gn=LOC101094897;refGene_tx=XM_023247334.1
F1 37557766 37557967
refGene_gn=LOC101094897;refGene_tx=XM_006942826.4;refGene_gn=LOC101094897;refGene_tx=XM_023247334.1
F1 37558636 37558819
refGene_gn=LOC101094897;refGene_tx=XM_006942826.4;refGene_gn=LOC101094897;refGene_tx=XM_023247334.1
F1 37559814 37560009
refGene_gn=LOC101094897;refGene_tx=XM_006942826.4;refGene_gn=LOC101094897;refGene_tx=XM_023247334.1
F1 37560184 37560270
refGene_gn=LOC101094897;refGene_tx=XM_006942826.4;refGene_gn=LOC101094897;refGene_tx=XM_023247334.1
F1 37564677 37564777
refGene_gn=LOC101094897;refGene_tx=XM_006942826.4;refGene_gn=LOC101094897;refGene_tx=XM_023247334.1
F1 37565842 37566025
refGene_gn=LOC101094897;refGene_tx=XM_006942826.4;refGene_gn=LOC101094897;refGene_tx=XM_023247334.1
F1 37566830 37566972
refGene_gn=LOC101094897;refGene_tx=XM_006942826.4;refGene_gn=LOC101094897;refGene_tx=XM_023247334.1
F1 37581885 37582038
refGene_gn=C4BPB;refGene_tx=XM_003999447.4;refGene_gn=C4BPB;refGene_tx=XM_006942827.3;refGene_gn=C4BPB;refGene_tx=XM_019821908.1
F1 37582591 37582718
refGene_gn=C4BPB;refGene_tx=XM_003999447.4;refGene_gn=C4BPB;refGene_tx=XM_006942827.3;refGene_gn=C4BPB;refGene_tx=XM_019821908.1
F1 37584411 37584505
refGene_gn=C4BPB;refGene_tx=XM_003999447.4;refGene_gn=C4BPB;refGene_tx=XM_006942827.3
F1 37589863 37590040
refGene_gn=C4BPB;refGene_tx=XM_003999447.4;refGene_gn=C4BPB;refGene_tx=XM_006942827.3;refGene_gn=C4BPB;refGene_tx=XM_019821908.1
F1 37591240 37591411
refGene_gn=C4BPB;refGene_tx=XM_003999447.4;refGene_gn=C4BPB;refGene_tx=XM_006942827.3;refGene_gn=C4BPB;refGene_tx=XM_019821908.1
F1 37591933 37591991
refGene_gn=C4BPB;refGene_tx=XM_003999447.4;refGene_gn=C4BPB;refGene_tx

=XM_006942827.3;refGene_gn=C4BPB;refGene_tx=XM_019821908.1
F1 37601420 37601486
refGene_gn=PFKFB2;refGene_tx=XM_006942833.4;refGene_gn=PFKFB2;refGene_tx=XM_023247866.1;refGene_gn=PFKFB2;refGene_tx=XM_023247867.1;refGene_gn=PFKFB2;refGene_tx=XM_023247868.1;refGene_gn=PFKFB2;refGene_tx=XM_023247869.1
F1 37607840 37608020
refGene_gn=PFKFB2;refGene_tx=XM_023247866.1;refGene_gn=PFKFB2;refGene_tx=XM_023247867.1;refGene_gn=PFKFB2;refGene_tx=XM_023247868.1;refGene_gn=PFKFB2;refGene_tx=XM_023247869.1
F1 37608641 37608706
refGene_gn=PFKFB2;refGene_tx=XM_006942833.4;refGene_gn=PFKFB2;refGene_tx=XM_023247866.1;refGene_gn=PFKFB2;refGene_tx=XM_023247867.1;refGene_gn=PFKFB2;refGene_tx=XM_023247868.1;refGene_gn=PFKFB2;refGene_tx=XM_023247869.1
F1 37608979 37609042
refGene_gn=PFKFB2;refGene_tx=XM_006942833.4;refGene_gn=PFKFB2;refGene_tx=XM_023247866.1;refGene_gn=PFKFB2;refGene_tx=XM_023247867.1;refGene_gn=PFKFB2;refGene_tx=XM_023247868.1;refGene_gn=PFKFB2;refGene_tx=XM_023247869.1
F1 37609801 37609931
refGene_gn=PFKFB2;refGene_tx=XM_006942833.4;refGene_gn=PFKFB2;refGene_tx=XM_023247866.1;refGene_gn=PFKFB2;refGene_tx=XM_023247867.1;refGene_gn=PFKFB2;refGene_tx=XM_023247868.1;refGene_gn=PFKFB2;refGene_tx=XM_023247869.1
F1 37610658 37610763
refGene_gn=PFKFB2;refGene_tx=XM_006942833.4;refGene_gn=PFKFB2;refGene_tx=XM_023247866.1;refGene_gn=PFKFB2;refGene_tx=XM_023247867.1;refGene_gn=PFKFB2;refGene_tx=XM_023247868.1;refGene_gn=PFKFB2;refGene_tx=XM_023247869.1
F1 37612140 37612287
refGene_gn=PFKFB2;refGene_tx=XM_006942833.4;refGene_gn=PFKFB2;refGene_tx=XM_023247866.1;refGene_gn=PFKFB2;refGene_tx=XM_023247867.1;refGene_gn=PFKFB2;refGene_tx=XM_023247868.1;refGene_gn=PFKFB2;refGene_tx=XM_023247869.1
F1 37612692 37612900
refGene_gn=PFKFB2;refGene_tx=XM_006942833.4;refGene_gn=PFKFB2;refGene_tx=XM_023247866.1;refGene_gn=PFKFB2;refGene_tx=XM_023247867.1;refGene_gn=PFKFB2;refGene_tx=XM_023247868.1;refGene_gn=PFKFB2;refGene_tx=XM_023247869.1
F1 37614804 37614929
refGene_gn=PFKFB2;refGene_tx=XM_006942833.4;refGene_gn=PFKFB2;refGene_tx=XM_023247866.1;refGene_gn=PFKFB2;refGene_tx=XM_023247867.1;refGene_gn=PFKFB2;refGene_tx=XM_023247868.1;refGene_gn=PFKFB2;refGene_tx=XM_023247869.1
F1 37615278 37615326 refGene_gn=PFKFB2;refGene_tx=XM_023247869.1
F1 37615833 37615890
refGene_gn=PFKFB2;refGene_tx=XM_006942833.4;refGene_gn=PFKFB2;refGene_tx=XM_023247866.1;refGene_gn=PFKFB2;refGene_tx=XM_023247867.1;refGene_gn=PFKFB2;refGene_tx=XM_023247868.1

F1 37616247 37616322
refGene_gn=PFKFB2;refGene_tx=XM_006942833.4;refGene_gn=PFKFB2;refGene_tx=XM_023247866.1;refGene_gn=PFKFB2;refGene_tx=XM_023247867.1;refGene_gn=PFKFB2;refGene_tx=XM_023247868.1
F1 37616459 37616526
refGene_gn=PFKFB2;refGene_tx=XM_006942833.4;refGene_gn=PFKFB2;refGene_tx=XM_023247866.1;refGene_gn=PFKFB2;refGene_tx=XM_023247867.1;refGene_gn=PFKFB2;refGene_tx=XM_023247868.1
F1 37616940 37617037
refGene_gn=PFKFB2;refGene_tx=XM_006942833.4;refGene_gn=PFKFB2;refGene_tx=XM_023247866.1;refGene_gn=PFKFB2;refGene_tx=XM_023247867.1;refGene_gn=PFKFB2;refGene_tx=XM_023247868.1
F1 37617549 37617675
refGene_gn=PFKFB2;refGene_tx=XM_006942833.4;refGene_gn=PFKFB2;refGene_tx=XM_023247866.1
F1 37622736 37622827
refGene_gn=PFKFB2;refGene_tx=XM_006942833.4;refGene_gn=PFKFB2;refGene_tx=XM_023247866.1
F1 37626272 37626615
refGene_gn=YOD1;refGene_tx=XM_003999339.4;refGene_gn=YOD1;refGene_tx=XM_019821909.2
F1 37627666 37628370 refGene_gn=YOD1;refGene_tx=XM_003999339.4
F1 37640932 37640960
refGene_gn=YOD1;refGene_tx=XM_023247859.1;refGene_gn=YOD1;refGene_tx=XM_023247860.1;refGene_gn=YOD1;refGene_tx=XM_023247861.1
F1 37649925 37650110
refGene_gn=YOD1;refGene_tx=XM_019821909.2;refGene_gn=YOD1;refGene_tx=XM_019821912.2;refGene_gn=YOD1;refGene_tx=XM_019821913.2;refGene_gn=YOD1;refGene_tx=XM_023247859.1;refGene_gn=YOD1;refGene_tx=XM_023247860.1;refGene_gn=YOD1;refGene_tx=XM_023247861.1;refGene_gn=YOD1;refGene_tx=XM_023247862.1;refGene_gn=YOD1;refGene_tx=XM_023247863.1;refGene_gn=YOD1;refGene_tx=XM_023247864.1;refGene_gn=YOD1;refGene_tx=XM_023247865.1
F1 37652570 37652748
refGene_gn=YOD1;refGene_tx=XM_019821909.2;refGene_gn=YOD1;refGene_tx=XM_019821912.2;refGene_gn=YOD1;refGene_tx=XM_019821913.2;refGene_gn=YOD1;refGene_tx=XM_023247859.1;refGene_gn=YOD1;refGene_tx=XM_023247860.1;refGene_gn=YOD1;refGene_tx=XM_023247861.1;refGene_gn=YOD1;refGene_tx=XM_023247862.1;refGene_gn=YOD1;refGene_tx=XM_023247863.1;refGene_gn=YOD1;refGene_tx=XM_023247864.1;refGene_gn=YOD1;refGene_tx=XM_023247865.1
F1 37653965 37655548
refGene_gn=YOD1;refGene_tx=XM_019821909.2;refGene_gn=YOD1;refGene_tx=XM_019821912.2;refGene_gn=YOD1;refGene_tx=XM_019821913.2;refGene_gn=YOD1;refGene_tx=XM_023247859.1;refGene_gn=YOD1;refGene_tx=XM_023247860.1;refGene_gn=YOD1;refGene_tx=XM_023247861.1;refGene_gn=YOD1;refGene_tx=XM_023247862.1;refGene_gn=YOD1;refGene_tx=XM_023247863.1;refGene_gn=YOD1;refGene_tx=XM_023247864.1;refGene_gn=YOD1;refGene_tx=XM_023247865.1
F1 37706048 37706087
refGene_gn=FCAMR;refGene_tx=XM_019821918.2;refGene_gn=FCAMR;refGene_tx=XM_019821919.2;refGene_gn=FCAMR;refGene_tx=XM_019821920.2
F1 37708932 37709005

refGene_gn=FCAMR;refGene_tx=XM_019821918.2;refGene_gn=FCAMR;refGene_tx=XM_019821919.2;refGene_gn=FCAMR;refGene_tx=XM_019821920.2
F1 37710492 37710636 refGene_gn=FCAMR;refGene_tx=XM_019821918.2;refGene_gn=FCAMR;refGene_tx=XM_019821919.2;refGene_gn=FCAMR;refGene_tx=XM_019821920.2
F1 37713587 37713926 refGene_gn=FCAMR;refGene_tx=XM_019821918.2;refGene_gn=FCAMR;refGene_tx=XM_019821919.2;refGene_gn=FCAMR;refGene_tx=XM_019821920.2
F1 37715055 37715815 refGene_gn=FCAMR;refGene_tx=XM_019821918.2;refGene_gn=FCAMR;refGene_tx=XM_019821919.2;refGene_gn=FCAMR;refGene_tx=XM_019821920.2
F1 37716448 37716567 refGene_gn=FCAMR;refGene_tx=XM_019821918.2;refGene_gn=FCAMR;refGene_tx=XM_019821919.2;refGene_gn=FCAMR;refGene_tx=XM_019821920.2
F1 37717178 37717360 refGene_gn=FCAMR;refGene_tx=XM_019821918.2;refGene_gn=FCAMR;refGene_tx=XM_019821919.2
F1 37718338 37718376 refGene_gn=FCAMR;refGene_tx=XM_019821920.2
F1 37738717 37738760 refGene_gn=PIGR;refGene_tx=XM_006942944.3
F1 37739706 37740051 refGene_gn=PIGR;refGene_tx=XM_006942944.3
F1 37741419 37742073 refGene_gn=PIGR;refGene_tx=XM_006942944.3
F1 37743352 37743685 refGene_gn=PIGR;refGene_tx=XM_006942944.3
F1 37744250 37744574 refGene_gn=PIGR;refGene_tx=XM_006942944.3
F1 37745501 37745676 refGene_gn=PIGR;refGene_tx=XM_006942944.3
F1 37746023 37746145 refGene_gn=PIGR;refGene_tx=XM_006942944.3
F1 37746785 37746917 refGene_gn=PIGR;refGene_tx=XM_006942944.3
F1 37747040 37747099 refGene_gn=PIGR;refGene_tx=XM_006942944.3
F1 37748126 37748222 refGene_gn=PIGR;refGene_tx=XM_006942944.3
F1 37754703 37754740 refGene_gn=FCMR;refGene_tx=XM_023247654.1;refGene_gn=FCMR;refGene_tx=XM_023247655.1
F1 37761938 37762274 refGene_gn=FCMR;refGene_tx=XM_023247654.1
F1 37763032 37763164 refGene_gn=FCMR;refGene_tx=XM_023247654.1;refGene_gn=FCMR;refGene_tx=XM_023247655.1
F1 37764114 37764340 refGene_gn=FCMR;refGene_tx=XM_023247654.1;refGene_gn=FCMR;refGene_tx=XM_023247655.1
F1 37765770 37765901 refGene_gn=FCMR;refGene_tx=XM_023247654.1;refGene_gn=FCMR;refGene_tx=XM_023247655.1
F1 37766276 37766432 refGene_gn=FCMR;refGene_tx=XM_023247654.1;refGene_gn=FCMR;refGene_tx=XM_023247655.1
F1 37766680 37766739 refGene_gn=FCMR;refGene_tx=XM_023247654.1;refGene_gn=FCMR;refGene_tx=XM_023247655.1
F1 37769051 37769228 refGene_gn=FCMR;refGene_tx=XM_023247654.1;refGene_gn=FCMR;refGene_tx=XM_023247655.1

F1 37771443 37771527 refGene_gn=IL24;refGene_tx=XM_003999450.3
F1 37772445 37772520 refGene_gn=IL24;refGene_tx=XM_003999450.3
F1 37773213 37773372 refGene_gn=IL24;refGene_tx=XM_003999450.3
F1 37774025 37774088 refGene_gn=IL24;refGene_tx=XM_003999450.3
F1 37775208 37775379 refGene_gn=IL24;refGene_tx=XM_003999450.3
F1 37796537 37796615 refGene_gn=IL20;refGene_tx=XM_023247455.1
F1 37797627 37797702 refGene_gn=IL20;refGene_tx=XM_023247455.1
F1 37798287 37798440 refGene_gn=IL20;refGene_tx=XM_023247455.1
F1 37798534 37798600 refGene_gn=IL20;refGene_tx=XM_023247455.1
F1 37798884 37799043 refGene_gn=IL20;refGene_tx=XM_023247455.1
F1 37810848 37810944 refGene_gn=IL19;refGene_tx=XM_003999342.2
F1 37812416 37812491 refGene_gn=IL19;refGene_tx=XM_003999342.2
F1 37813434 37813587 refGene_gn=IL19;refGene_tx=XM_003999342.2
F1 37816301 37816367 refGene_gn=IL19;refGene_tx=XM_003999342.2
F1 37816524 37816668 refGene_gn=IL19;refGene_tx=XM_003999342.2
F1 37878796 37878961 refGene_gn=IL10;refGene_tx=NM_001009209.1
F1 37879759 37879819 refGene_gn=IL10;refGene_tx=NM_001009209.1
F1 37880085 37880238 refGene_gn=IL10;refGene_tx=NM_001009209.1
F1 37881255 37881321 refGene_gn=IL10;refGene_tx=NM_001009209.1
F1 37882394 37882487 refGene_gn=IL10;refGene_tx=NM_001009209.1
F1 37911707 37911851
refGene_gn=MAPKAPK2;refGene_tx=XM_023248055.1;refGene_gn=MAPKAPK2;refG
ene_tx=XM_023248056.1
F1 37912343 37912424
refGene_gn=MAPKAPK2;refGene_tx=XM_023248055.1;refGene_gn=MAPKAPK2;refG
ene_tx=XM_023248056.1
F1 37912543 37912629
refGene_gn=MAPKAPK2;refGene_tx=XM_023248055.1;refGene_gn=MAPKAPK2;refG
ene_tx=XM_023248056.1
F1 37912953 37913078
refGene_gn=MAPKAPK2;refGene_tx=XM_023248055.1;refGene_gn=MAPKAPK2;refG
ene_tx=XM_023248056.1
F1 37913484 37913560
refGene_gn=MAPKAPK2;refGene_tx=XM_023248055.1;refGene_gn=MAPKAPK2;refG
ene_tx=XM_023248056.1
F1 37914176 37914303
refGene_gn=MAPKAPK2;refGene_tx=XM_023248055.1;refGene_gn=MAPKAPK2;refG
ene_tx=XM_023248056.1
F1 37914785 37914865
refGene_gn=MAPKAPK2;refGene_tx=XM_023248055.1;refGene_gn=MAPKAPK2;refG
ene_tx=XM_023248056.1
F1 37915143 37915217
refGene_gn=MAPKAPK2;refGene_tx=XM_023248055.1;refGene_gn=MAPKAPK2;refG
ene_tx=XM_023248056.1
F1 37915407 37915547
refGene_gn=MAPKAPK2;refGene_tx=XM_023248055.1;refGene_gn=MAPKAPK2;refG
ene_tx=XM_023248056.1
F1 37955009 37955294
refGene_gn=MAPKAPK2;refGene_tx=XM_023248055.1;refGene_gn=MAPKAPK2;refG
ene_tx=XM_023248056.1

F1 37989455 37991030
refGene_gn=DYRK3;refGene_tx=XM_003999344.5;refGene_gn=DYRK3;refGene_tx
=XM_019821923.2;refGene_gn=DYRK3;refGene_tx=XM_019821924.2;refGene_gn=
DYRK3;refGene_tx=XM_023247347.1
F1 37996639 37996751
refGene_gn=DYRK3;refGene_tx=XM_003999344.5;refGene_gn=DYRK3;refGene_tx
=XM_019821923.2;refGene_gn=DYRK3;refGene_tx=XM_019821924.2;refGene_gn=
DYRK3;refGene_tx=XM_023247347.1
F1 37998217 37998234 refGene_gn=DYRK3;refGene_tx=XM_023247347.1
F1 37998786 37998863 refGene_gn=DYRK3;refGene_tx=XM_003999344.5
F1 38135158 38135216 refGene_gn=CFHR5;refGene_tx=XM_019821926.2
F1 38138511 38138706 refGene_gn=CFHR5;refGene_tx=XM_019821926.2
F1 38139855 38140032 refGene_gn=CFHR5;refGene_tx=XM_019821926.2
F1 38148124 38148301 refGene_gn=CFHR5;refGene_tx=XM_019821926.2
F1 38149382 38149565 refGene_gn=CFHR5;refGene_tx=XM_019821926.2
F1 38149946 38150126 refGene_gn=CFHR5;refGene_tx=XM_019821926.2
F1 38154611 38154788 refGene_gn=CFHR5;refGene_tx=XM_019821926.2
F1 38161932 38162115 refGene_gn=CFHR5;refGene_tx=XM_019821926.2
F1 38164626 38164809 refGene_gn=CFHR5;refGene_tx=XM_019821926.2
F1 38166135 38166332 refGene_gn=CFHR5;refGene_tx=XM_019821926.2
F1 38184825 38184859
refGene_gn=F13B;refGene_tx=XM_019821927.2;refGene_gn=F13B;refGene_tx=X
M_019821928.2;refGene_gn=F13B;refGene_tx=XM_019821929.2
F1 38187659 38187873
refGene_gn=F13B;refGene_tx=XM_019821927.2;refGene_gn=F13B;refGene_tx=X
M_019821928.2;refGene_gn=F13B;refGene_tx=XM_019821929.2
F1 38193832 38194015
refGene_gn=F13B;refGene_tx=XM_019821927.2;refGene_gn=F13B;refGene_tx=X
M_019821928.2;refGene_gn=F13B;refGene_tx=XM_019821929.2
F1 38194750 38194951
refGene_gn=F13B;refGene_tx=XM_019821927.2;refGene_gn=F13B;refGene_tx=X
M_019821928.2;refGene_gn=F13B;refGene_tx=XM_019821929.2
F1 38203284 38203467
refGene_gn=F13B;refGene_tx=XM_019821927.2;refGene_gn=F13B;refGene_tx=X
M_019821928.2;refGene_gn=F13B;refGene_tx=XM_019821929.2
F1 38204528 38204714
refGene_gn=F13B;refGene_tx=XM_019821927.2;refGene_gn=F13B;refGene_tx=X
M_019821928.2;refGene_gn=F13B;refGene_tx=XM_019821929.2
F1 38204802 38204982
refGene_gn=F13B;refGene_tx=XM_019821927.2;refGene_gn=F13B;refGene_tx=X
M_019821928.2;refGene_gn=F13B;refGene_tx=XM_019821929.2
F1 38207537 38207714
refGene_gn=F13B;refGene_tx=XM_019821927.2;refGene_gn=F13B;refGene_tx=X
M_019821929.2
F1 38208195 38208372
refGene_gn=F13B;refGene_tx=XM_019821927.2;refGene_gn=F13B;refGene_tx=X
M_019821928.2;refGene_gn=F13B;refGene_tx=XM_019821929.2
F1 38210439 38210625
refGene_gn=F13B;refGene_tx=XM_019821927.2;refGene_gn=F13B;refGene_tx=X
M_019821928.2;refGene_gn=F13B;refGene_tx=XM_019821929.2

F1 38211927 38212128
refGene_gn=F13B;refGene_tx=XM_019821927.2;refGene_gn=F13B;refGene_tx=XM_019821928.2
F1 38215658 38215722
refGene_gn=F13B;refGene_tx=XM_019821927.2;refGene_gn=F13B;refGene_tx=XM_019821928.2;refGene_gn=F13B;refGene_tx=XM_019821929.2
F1 38230881 38230984
refGene_gn=ASPM;refGene_tx=XM_003999347.5;refGene_gn=ASPM;refGene_tx=XM_023247980.1;refGene_gn=ASPM;refGene_tx=XM_023247981.1
F1 38235593 38235763
refGene_gn=ASPM;refGene_tx=XM_003999347.5;refGene_gn=ASPM;refGene_tx=XM_023247980.1;refGene_gn=ASPM;refGene_tx=XM_023247981.1
F1 38236971 38237148
refGene_gn=ASPM;refGene_tx=XM_003999347.5;refGene_gn=ASPM;refGene_tx=XM_023247980.1;refGene_gn=ASPM;refGene_tx=XM_023247981.1
F1 38238636 38238791
refGene_gn=ASPM;refGene_tx=XM_003999347.5;refGene_gn=ASPM;refGene_tx=XM_023247980.1;refGene_gn=ASPM;refGene_tx=XM_023247981.1
F1 38238941 38239134
refGene_gn=ASPM;refGene_tx=XM_003999347.5;refGene_gn=ASPM;refGene_tx=XM_023247980.1;refGene_gn=ASPM;refGene_tx=XM_023247981.1
F1 38239600 38239792
refGene_gn=ASPM;refGene_tx=XM_003999347.5;refGene_gn=ASPM;refGene_tx=XM_023247980.1;refGene_gn=ASPM;refGene_tx=XM_023247981.1
F1 38240647 38240797
refGene_gn=ASPM;refGene_tx=XM_003999347.5;refGene_gn=ASPM;refGene_tx=XM_023247980.1;refGene_gn=ASPM;refGene_tx=XM_023247981.1
F1 38241790 38242000
refGene_gn=ASPM;refGene_tx=XM_003999347.5;refGene_gn=ASPM;refGene_tx=XM_023247980.1;refGene_gn=ASPM;refGene_tx=XM_023247981.1
F1 38243127 38243224
refGene_gn=ASPM;refGene_tx=XM_003999347.5;refGene_gn=ASPM;refGene_tx=XM_023247980.1;refGene_gn=ASPM;refGene_tx=XM_023247981.1
F1 38244487 38244654
refGene_gn=ASPM;refGene_tx=XM_003999347.5;refGene_gn=ASPM;refGene_tx=XM_023247980.1;refGene_gn=ASPM;refGene_tx=XM_023247981.1
F1 38247396 38252151
refGene_gn=ASPM;refGene_tx=XM_003999347.5;refGene_gn=ASPM;refGene_tx=XM_023247980.1;refGene_gn=ASPM;refGene_tx=XM_023247981.1
F1 38254211 38254406
refGene_gn=ASPM;refGene_tx=XM_003999347.5;refGene_gn=ASPM;refGene_tx=XM_023247980.1;refGene_gn=ASPM;refGene_tx=XM_023247981.1
F1 38259634 38259763
refGene_gn=ASPM;refGene_tx=XM_003999347.5;refGene_gn=ASPM;refGene_tx=XM_023247980.1;refGene_gn=ASPM;refGene_tx=XM_023247981.1
F1 38259872 38260018
refGene_gn=ASPM;refGene_tx=XM_003999347.5;refGene_gn=ASPM;refGene_tx=XM_023247980.1;refGene_gn=ASPM;refGene_tx=XM_023247981.1
F1 38260112 38260320
refGene_gn=ASPM;refGene_tx=XM_003999347.5;refGene_gn=ASPM;refGene_tx=X

M_023247980.1;refGene_gn=ASPM;refGene_tx=XM_023247981.1
F1 38261627 38261849
refGene_gn=ASPM;refGene_tx=XM_003999347.5;refGene_gn=ASPM;refGene_tx=XM_023247980.1;refGene_gn=ASPM;refGene_tx=XM_023247981.1
F1 38262275 38262361
refGene_gn=ASPM;refGene_tx=XM_003999347.5;refGene_gn=ASPM;refGene_tx=XM_023247980.1;refGene_gn=ASPM;refGene_tx=XM_023247981.1
F1 38262449 38262595
refGene_gn=ASPM;refGene_tx=XM_003999347.5;refGene_gn=ASPM;refGene_tx=XM_023247980.1;refGene_gn=ASPM;refGene_tx=XM_023247981.1
F1 38265107 38265283
refGene_gn=ASPM;refGene_tx=XM_003999347.5;refGene_gn=ASPM;refGene_tx=XM_023247980.1;refGene_gn=ASPM;refGene_tx=XM_023247981.1
F1 38266262 38266393
refGene_gn=ASPM;refGene_tx=XM_003999347.5;refGene_gn=ASPM;refGene_tx=XM_023247980.1;refGene_gn=ASPM;refGene_tx=XM_023247981.1
F1 38266980 38267122
refGene_gn=ASPM;refGene_tx=XM_003999347.5;refGene_gn=ASPM;refGene_tx=XM_023247980.1;refGene_gn=ASPM;refGene_tx=XM_023247981.1
F1 38269368 38269436
refGene_gn=ASPM;refGene_tx=XM_003999347.5;refGene_gn=ASPM;refGene_tx=XM_023247980.1;refGene_gn=ASPM;refGene_tx=XM_023247981.1
F1 38269710 38269956
refGene_gn=ASPM;refGene_tx=XM_003999347.5;refGene_gn=ASPM;refGene_tx=XM_023247980.1;refGene_gn=ASPM;refGene_tx=XM_023247981.1
F1 38271308 38271455
refGene_gn=ASPM;refGene_tx=XM_003999347.5;refGene_gn=ASPM;refGene_tx=XM_023247980.1;refGene_gn=ASPM;refGene_tx=XM_023247981.1
F1 38271862 38271967
refGene_gn=ASPM;refGene_tx=XM_003999347.5;refGene_gn=ASPM;refGene_tx=XM_023247980.1;refGene_gn=ASPM;refGene_tx=XM_023247981.1
F1 38278707 38280193
refGene_gn=ASPM;refGene_tx=XM_003999347.5;refGene_gn=ASPM;refGene_tx=XM_023247980.1;refGene_gn=ASPM;refGene_tx=XM_023247981.1
F1 38280369 38280513
refGene_gn=ASPM;refGene_tx=XM_003999347.5;refGene_gn=ASPM;refGene_tx=XM_023247980.1;refGene_gn=ASPM;refGene_tx=XM_023247981.1
F1 38282223 38282517
refGene_gn=ASPM;refGene_tx=XM_003999347.5;refGene_gn=ASPM;refGene_tx=XM_023247980.1;refGene_gn=ASPM;refGene_tx=XM_023247981.1
F1 38290807 38291463 refGene_gn=ZBTB41;refGene_tx=XM_003999348.5
F1 38309531 38309620 refGene_gn=ZBTB41;refGene_tx=XM_003999348.5
F1 38315578 38315684 refGene_gn=ZBTB41;refGene_tx=XM_003999348.5
F1 38317367 38317474 refGene_gn=ZBTB41;refGene_tx=XM_003999348.5
F1 38319068 38319164 refGene_gn=ZBTB41;refGene_tx=XM_003999348.5
F1 38323334 38323464 refGene_gn=ZBTB41;refGene_tx=XM_003999348.5
F1 38327207 38327355 refGene_gn=ZBTB41;refGene_tx=XM_003999348.5
F1 38329419 38329489 refGene_gn=ZBTB41;refGene_tx=XM_003999348.5
F1 38330168 38330376 refGene_gn=ZBTB41;refGene_tx=XM_003999348.5
F1 38338627 38339750 refGene_gn=ZBTB41;refGene_tx=XM_003999348.5

F1 38426112 38426182 refGene_gn=CRB1;refGene_tx=XM_019821930.2
F1 38492366 38492948 refGene_gn=CRB1;refGene_tx=XM_019821930.2
F1 38512325 38512457 refGene_gn=CRB1;refGene_tx=XM_019821930.2
F1 38512459 38512520 refGene_gn=CRB1;refGene_tx=XM_019821930.2
F1 38515862 38516002 refGene_gn=CRB1;refGene_tx=XM_019821930.2
F1 38523247 38523430 refGene_gn=CRB1;refGene_tx=XM_019821930.2
F1 38582262 38583219 refGene_gn=CRB1;refGene_tx=XM_019821930.2
F1 38587074 38587622 refGene_gn=CRB1;refGene_tx=XM_019821930.2
F1 38589047 38589213 refGene_gn=CRB1;refGene_tx=XM_019821930.2
F1 38594503 38595410 refGene_gn=CRB1;refGene_tx=XM_019821930.2
F1 38598014 38598143 refGene_gn=CRB1;refGene_tx=XM_019821930.2
F1 38602484 38602611 refGene_gn=CRB1;refGene_tx=XM_019821930.2
F1 38636619 38636835 refGene_gn=CRB1;refGene_tx=XM_019821930.2
F1 38652942 38653479
refGene_gn=DENND1B;refGene_tx=XM_019821934.2;refGene_gn=DENND1B;refGene_tx=XM_019821935.2;refGene_gn=DENND1B;refGene_tx=XM_019821936.2;refGene_gn=DENND1B;refGene_tx=XM_023247768.1;refGene_gn=DENND1B;refGene_tx=XM_023247769.1
F1 38654516 38654733
refGene_gn=DENND1B;refGene_tx=XM_019821934.2;refGene_gn=DENND1B;refGene_tx=XM_019821935.2;refGene_gn=DENND1B;refGene_tx=XM_019821936.2;refGene_gn=DENND1B;refGene_tx=XM_023247768.1;refGene_gn=DENND1B;refGene_tx=XM_023247769.1
F1 38655701 38655775
refGene_gn=DENND1B;refGene_tx=XM_019821934.2;refGene_gn=DENND1B;refGene_tx=XM_019821935.2;refGene_gn=DENND1B;refGene_tx=XM_019821936.2;refGene_gn=DENND1B;refGene_tx=XM_023247768.1;refGene_gn=DENND1B;refGene_tx=XM_023247769.1
F1 38680924 38681032
refGene_gn=DENND1B;refGene_tx=XM_019821934.2;refGene_gn=DENND1B;refGene_tx=XM_019821935.2;refGene_gn=DENND1B;refGene_tx=XM_019821936.2;refGene_gn=DENND1B;refGene_tx=XM_023247768.1;refGene_gn=DENND1B;refGene_tx=XM_023247769.1
F1 38681855 38681912
refGene_gn=DENND1B;refGene_tx=XM_019821934.2;refGene_gn=DENND1B;refGene_tx=XM_019821935.2;refGene_gn=DENND1B;refGene_tx=XM_019821936.2;refGene_gn=DENND1B;refGene_tx=XM_023247768.1;refGene_gn=DENND1B;refGene_tx=XM_023247769.1
F1 38682405 38682477
refGene_gn=DENND1B;refGene_tx=XM_019821934.2;refGene_gn=DENND1B;refGene_tx=XM_019821935.2;refGene_gn=DENND1B;refGene_tx=XM_023247768.1
F1 38686427 38686496
refGene_gn=DENND1B;refGene_tx=XM_019821934.2;refGene_gn=DENND1B;refGene_tx=XM_019821935.2;refGene_gn=DENND1B;refGene_tx=XM_019821936.2;refGene_gn=DENND1B;refGene_tx=XM_023247768.1;refGene_gn=DENND1B;refGene_tx=XM_023247769.1
F1 38687429 38687470
refGene_gn=DENND1B;refGene_tx=XM_019821934.2;refGene_gn=DENND1B;refGene_tx=XM_019821935.2;refGene_gn=DENND1B;refGene_tx=XM_019821936.2;refGene_gn=DENND1B;refGene_tx=XM_023247768.1;refGene_gn=DENND1B;refGene_tx=

XM_023247769.1
F1 38694319 38694410
refGene_gn=DENND1B;refGene_tx=XM_019821934.2;refGene_gn=DENND1B;refGene_tx=XM_019821935.2;refGene_gn=DENND1B;refGene_tx=XM_019821936.2;refGene_gn=DENND1B;refGene_tx=XM_023247768.1;refGene_gn=DENND1B;refGene_tx=XM_023247769.1
F1 38717253 38717355
refGene_gn=DENND1B;refGene_tx=XM_019821934.2;refGene_gn=DENND1B;refGene_tx=XM_019821935.2;refGene_gn=DENND1B;refGene_tx=XM_019821936.2;refGene_gn=DENND1B;refGene_tx=XM_023247768.1;refGene_gn=DENND1B;refGene_tx=XM_023247769.1
F1 38729451 38729577
refGene_gn=DENND1B;refGene_tx=XM_019821934.2;refGene_gn=DENND1B;refGene_tx=XM_019821935.2;refGene_gn=DENND1B;refGene_tx=XM_019821936.2;refGene_gn=DENND1B;refGene_tx=XM_023247768.1;refGene_gn=DENND1B;refGene_tx=XM_023247769.1
F1 38741760 38741862
refGene_gn=DENND1B;refGene_tx=XM_019821934.2;refGene_gn=DENND1B;refGene_tx=XM_019821935.2;refGene_gn=DENND1B;refGene_tx=XM_019821936.2;refGene_gn=DENND1B;refGene_tx=XM_023247768.1;refGene_gn=DENND1B;refGene_tx=XM_023247769.1
F1 38746746 38746792
refGene_gn=DENND1B;refGene_tx=XM_019821934.2;refGene_gn=DENND1B;refGene_tx=XM_019821935.2;refGene_gn=DENND1B;refGene_tx=XM_019821936.2;refGene_gn=DENND1B;refGene_tx=XM_023247768.1;refGene_gn=DENND1B;refGene_tx=XM_023247769.1
F1 38753955 38754056
refGene_gn=DENND1B;refGene_tx=XM_019821934.2;refGene_gn=DENND1B;refGene_tx=XM_019821935.2;refGene_gn=DENND1B;refGene_tx=XM_019821936.2;refGene_gn=DENND1B;refGene_tx=XM_023247768.1;refGene_gn=DENND1B;refGene_tx=XM_023247769.1
F1 38771526 38771637
refGene_gn=DENND1B;refGene_tx=XM_019821934.2;refGene_gn=DENND1B;refGene_tx=XM_019821935.2;refGene_gn=DENND1B;refGene_tx=XM_019821936.2;refGene_gn=DENND1B;refGene_tx=XM_023247768.1;refGene_gn=DENND1B;refGene_tx=XM_023247769.1
F1 38773829 38773883
refGene_gn=DENND1B;refGene_tx=XM_019821934.2;refGene_gn=DENND1B;refGene_tx=XM_019821935.2;refGene_gn=DENND1B;refGene_tx=XM_019821936.2;refGene_gn=DENND1B;refGene_tx=XM_023247768.1;refGene_gn=DENND1B;refGene_tx=XM_023247769.1
F1 38775121 38775181
refGene_gn=DENND1B;refGene_tx=XM_019821934.2;refGene_gn=DENND1B;refGene_tx=XM_019821936.2;refGene_gn=DENND1B;refGene_tx=XM_023247768.1;refGene_gn=DENND1B;refGene_tx=XM_023247769.1
F1 38778282 38778363
refGene_gn=DENND1B;refGene_tx=XM_019821934.2;refGene_gn=DENND1B;refGene_tx=XM_019821935.2;refGene_gn=DENND1B;refGene_tx=XM_019821936.2;refGene_gn=DENND1B;refGene_tx=XM_023247768.1;refGene_gn=DENND1B;refGene_tx=XM_023247769.1

F1 38787428 38787498
refGene_gn=DENND1B;refGene_tx=XM_019821934.2;refGene_gn=DENND1B;refGene_tx=XM_019821935.2;refGene_gn=DENND1B;refGene_tx=XM_019821936.2;refGene_gn=DENND1B;refGene_tx=XM_023247768.1;refGene_gn=DENND1B;refGene_tx=XM_023247769.1
F1 38804499 38804619
refGene_gn=DENND1B;refGene_tx=XM_019821934.2;refGene_gn=DENND1B;refGene_tx=XM_019821935.2;refGene_gn=DENND1B;refGene_tx=XM_019821936.2;refGene_gn=DENND1B;refGene_tx=XM_023247768.1;refGene_gn=DENND1B;refGene_tx=XM_023247769.1
F1 38806586 38806636
refGene_gn=DENND1B;refGene_tx=XM_019821934.2;refGene_gn=DENND1B;refGene_tx=XM_019821935.2;refGene_gn=DENND1B;refGene_tx=XM_019821936.2;refGene_gn=DENND1B;refGene_tx=XM_023247768.1;refGene_gn=DENND1B;refGene_tx=XM_023247769.1
F1 38837215 38837259
refGene_gn=DENND1B;refGene_tx=XM_019821934.2;refGene_gn=DENND1B;refGene_tx=XM_019821935.2;refGene_gn=DENND1B;refGene_tx=XM_019821936.2;refGene_gn=DENND1B;refGene_tx=XM_023247768.1;refGene_gn=DENND1B;refGene_tx=XM_023247769.1
F1 38907456 38907521
refGene_gn=DENND1B;refGene_tx=XM_019821934.2;refGene_gn=DENND1B;refGene_tx=XM_019821935.2;refGene_gn=DENND1B;refGene_tx=XM_019821936.2;refGene_gn=DENND1B;refGene_tx=XM_023247768.1;refGene_gn=DENND1B;refGene_tx=XM_023247769.1
F1 38909653 38909670
refGene_gn=DENND1B;refGene_tx=XM_023247768.1;refGene_gn=DENND1B;refGene_tx=XM_023247769.1
F1 38911137 38911171
refGene_gn=CF1H1orf53;refGene_tx=XM_023247772.1
F1 38912600 38912728
refGene_gn=CF1H1orf53;refGene_tx=XM_023247772.1
F1 38916229 38916285
refGene_gn=DENND1B;refGene_tx=XM_019821934.2;refGene_gn=DENND1B;refGene_tx=XM_019821935.2;refGene_gn=DENND1B;refGene_tx=XM_019821936.2
F1 39018106 39018370
refGene_gn=CF1H1orf53;refGene_tx=XM_023247770.1;refGene_gn=CF1H1orf53;refGene_tx=XM_023247771.1
F1 39021275 39021377
refGene_gn=CF1H1orf53;refGene_tx=XM_023247770.1;refGene_gn=CF1H1orf53;refGene_tx=XM_023247771.1;refGene_gn=CF1H1orf53;refGene_tx=XM_023247772.1
F1 39022829 39022922
refGene_gn=CF1H1orf53;refGene_tx=XM_023247770.1;refGene_gn=CF1H1orf53;refGene_tx=XM_023247771.1;refGene_gn=CF1H1orf53;refGene_tx=XM_023247772.1
F1 39028418 39028446
refGene_gn=LHX9;refGene_tx=XM_003999352.2;refGene_gn=LHX9;refGene_tx=XM_023247652.1
F1 39033747 39033921

refGene_gn=LHX9;refGene_tx=XM_003999352.2;refGene_gn=LHX9;refGene_tx=XM_006942839.3;refGene_gn=LHX9;refGene_tx=XM_019821939.2;refGene_gn=LHX9;refGene_tx=XM_019821940.2;refGene_gn=LHX9;refGene_tx=XM_023247652.1
F1 39035879 39036082
refGene_gn=LHX9;refGene_tx=XM_003999352.2;refGene_gn=LHX9;refGene_tx=XM_006942839.3;refGene_gn=LHX9;refGene_tx=XM_019821939.2;refGene_gn=LHX9;refGene_tx=XM_019821940.2;refGene_gn=LHX9;refGene_tx=XM_023247652.1
F1 39037272 39037628
refGene_gn=LHX9;refGene_tx=XM_003999352.2;refGene_gn=LHX9;refGene_tx=XM_006942839.3;refGene_gn=LHX9;refGene_tx=XM_019821939.2;refGene_gn=LHX9;refGene_tx=XM_019821940.2;refGene_gn=LHX9;refGene_tx=XM_023247652.1
F1 39043548 39043751
refGene_gn=LHX9;refGene_tx=XM_003999352.2;refGene_gn=LHX9;refGene_tx=XM_006942839.3;refGene_gn=LHX9;refGene_tx=XM_019821939.2;refGene_gn=LHX9;refGene_tx=XM_019821940.2;refGene_gn=LHX9;refGene_tx=XM_023247652.1
F1 39045072 39045330
refGene_gn=LHX9;refGene_tx=XM_003999352.2;refGene_gn=LHX9;refGene_tx=XM_006942839.3;refGene_gn=LHX9;refGene_tx=XM_019821939.2
F1 39048040 39048097
refGene_gn=LHX9;refGene_tx=XM_019821940.2;refGene_gn=LHX9;refGene_tx=XM_023247652.1
F1 39257765 39257773 refGene_gn=NEK7;refGene_tx=XM_023247961.1
F1 39286796 39286881
refGene_gn=NEK7;refGene_tx=XM_023247961.1;refGene_gn=NEK7;refGene_tx=XM_023247962.1;refGene_gn=NEK7;refGene_tx=XM_023247963.1
F1 39306918 39306954 refGene_gn=NEK7;refGene_tx=XM_023247964.1
F1 39311158 39311299
refGene_gn=NEK7;refGene_tx=XM_023247961.1;refGene_gn=NEK7;refGene_tx=XM_023247962.1;refGene_gn=NEK7;refGene_tx=XM_023247963.1;refGene_gn=NEK7;refGene_tx=XM_023247964.1
F1 39325815 39325878
refGene_gn=NEK7;refGene_tx=XM_023247961.1;refGene_gn=NEK7;refGene_tx=XM_023247962.1;refGene_gn=NEK7;refGene_tx=XM_023247963.1;refGene_gn=NEK7;refGene_tx=XM_023247964.1
F1 39327586 39327697
refGene_gn=NEK7;refGene_tx=XM_023247961.1;refGene_gn=NEK7;refGene_tx=XM_023247962.1;refGene_gn=NEK7;refGene_tx=XM_023247963.1;refGene_gn=NEK7;refGene_tx=XM_023247964.1
F1 39344306 39344415
refGene_gn=NEK7;refGene_tx=XM_023247961.1;refGene_gn=NEK7;refGene_tx=XM_023247962.1;refGene_gn=NEK7;refGene_tx=XM_023247963.1;refGene_gn=NEK7;refGene_tx=XM_023247964.1
F1 39345560 39345668
refGene_gn=NEK7;refGene_tx=XM_023247961.1;refGene_gn=NEK7;refGene_tx=XM_023247962.1;refGene_gn=NEK7;refGene_tx=XM_023247963.1;refGene_gn=NEK7;refGene_tx=XM_023247964.1
F1 39359819 39359914
refGene_gn=NEK7;refGene_tx=XM_023247961.1;refGene_gn=NEK7;refGene_tx=XM_023247962.1;refGene_gn=NEK7;refGene_tx=XM_023247963.1;refGene_gn=NEK7;refGene_tx=XM_023247964.1

F1 39364123 39364237
refGene_gn=NEK7;refGene_tx=XM_023247961.1;refGene_gn=NEK7;refGene_tx=XM_023247962.1;refGene_gn=NEK7;refGene_tx=XM_023247963.1;refGene_gn=NEK7;refGene_tx=XM_023247964.1
F1 39379798 39379824 refGene_gn=NEK7;refGene_tx=XM_023247962.1
F1 39382367 39382478
refGene_gn=NEK7;refGene_tx=XM_023247961.1;refGene_gn=NEK7;refGene_tx=XM_023247962.1;refGene_gn=NEK7;refGene_tx=XM_023247963.1;refGene_gn=NEK7;refGene_tx=XM_023247964.1
F1 39559124 39559298
refGene_gn=ATP6V1G3;refGene_tx=XM_003999355.4;refGene_gn=ATP6V1G3;refGene_tx=XM_019821941.2
F1 39566563 39566664
refGene_gn=ATP6V1G3;refGene_tx=XM_003999355.4;refGene_gn=ATP6V1G3;refGene_tx=XM_019821941.2
F1 39579020 39579102
refGene_gn=ATP6V1G3;refGene_tx=XM_003999355.4;refGene_gn=ATP6V1G3;refGene_tx=XM_019821941.2
F1 39670494 39670567
refGene_gn=PTPRC;refGene_tx=XM_006942841.3;refGene_gn=PTPRC;refGene_tx=XM_011290957.2;refGene_gn=PTPRC;refGene_tx=XM_019821942.1;refGene_gn=PTPRC;refGene_tx=XM_019821943.1;refGene_gn=PTPRC;refGene_tx=XM_019821944.1;refGene_gn=PTPRC;refGene_tx=XM_019821945.1
F1 39727014 39727041
refGene_gn=PTPRC;refGene_tx=XM_006942841.3;refGene_gn=PTPRC;refGene_tx=XM_011290957.2;refGene_gn=PTPRC;refGene_tx=XM_019821942.1;refGene_gn=PTPRC;refGene_tx=XM_019821943.1;refGene_gn=PTPRC;refGene_tx=XM_019821944.1;refGene_gn=PTPRC;refGene_tx=XM_019821945.1
F1 39730742 39730940
refGene_gn=PTPRC;refGene_tx=XM_019821942.1;refGene_gn=PTPRC;refGene_tx=XM_019821943.1;refGene_gn=PTPRC;refGene_tx=XM_019821944.1;refGene_gn=PTPRC;refGene_tx=XM_019821945.1
F1 39733434 39733569
refGene_gn=PTPRC;refGene_tx=XM_006942841.3;refGene_gn=PTPRC;refGene_tx=XM_019821942.1;refGene_gn=PTPRC;refGene_tx=XM_019821943.1;refGene_gn=PTPRC;refGene_tx=XM_019821944.1;refGene_gn=PTPRC;refGene_tx=XM_019821945.1
F1 39736331 39736475
refGene_gn=PTPRC;refGene_tx=XM_006942841.3;refGene_gn=PTPRC;refGene_tx=XM_019821942.1;refGene_gn=PTPRC;refGene_tx=XM_019821943.1;refGene_gn=PTPRC;refGene_tx=XM_019821944.1
F1 39737206 39737263
refGene_gn=PTPRC;refGene_tx=XM_006942841.3;refGene_gn=PTPRC;refGene_tx=XM_011290957.2;refGene_gn=PTPRC;refGene_tx=XM_019821942.1;refGene_gn=PTPRC;refGene_tx=XM_019821943.1;refGene_gn=PTPRC;refGene_tx=XM_019821944.1;refGene_gn=PTPRC;refGene_tx=XM_019821945.1
F1 39744634 39744661
refGene_gn=PTPRC;refGene_tx=XM_006942841.3;refGene_gn=PTPRC;refGene_tx=XM_011290957.2;refGene_gn=PTPRC;refGene_tx=XM_019821942.1;refGene_gn=PTPRC;refGene_tx=XM_019821943.1;refGene_gn=PTPRC;refGene_tx=XM_0198219

44.1;refGene_gn=PTPRC;refGene_tx=XM_019821945.1
F1	39746409	39746628
refGene_gn=PTPRC;refGene_tx=XM_006942841.3;refGene_gn=PTPRC;refGene_tx=XM_011290957.2;refGene_gn=PTPRC;refGene_tx=XM_019821942.1;refGene_gn=PTPRC;refGene_tx=XM_019821943.1;refGene_gn=PTPRC;refGene_tx=XM_019821944.1;refGene_gn=PTPRC;refGene_tx=XM_019821945.1
F1	39748622	39748751
refGene_gn=PTPRC;refGene_tx=XM_006942841.3;refGene_gn=PTPRC;refGene_tx=XM_011290957.2;refGene_gn=PTPRC;refGene_tx=XM_019821942.1;refGene_gn=PTPRC;refGene_tx=XM_019821943.1;refGene_gn=PTPRC;refGene_tx=XM_019821944.1;refGene_gn=PTPRC;refGene_tx=XM_019821945.1
F1	39750104	39750245
refGene_gn=PTPRC;refGene_tx=XM_006942841.3;refGene_gn=PTPRC;refGene_tx=XM_011290957.2;refGene_gn=PTPRC;refGene_tx=XM_019821942.1;refGene_gn=PTPRC;refGene_tx=XM_019821943.1;refGene_gn=PTPRC;refGene_tx=XM_019821944.1;refGene_gn=PTPRC;refGene_tx=XM_019821945.1
F1	39757207	39757327
refGene_gn=PTPRC;refGene_tx=XM_006942841.3;refGene_gn=PTPRC;refGene_tx=XM_011290957.2;refGene_gn=PTPRC;refGene_tx=XM_019821942.1;refGene_gn=PTPRC;refGene_tx=XM_019821943.1;refGene_gn=PTPRC;refGene_tx=XM_019821944.1;refGene_gn=PTPRC;refGene_tx=XM_019821945.1
F1	39757704	39757863
refGene_gn=PTPRC;refGene_tx=XM_006942841.3;refGene_gn=PTPRC;refGene_tx=XM_011290957.2;refGene_gn=PTPRC;refGene_tx=XM_019821942.1;refGene_gn=PTPRC;refGene_tx=XM_019821943.1;refGene_gn=PTPRC;refGene_tx=XM_019821944.1;refGene_gn=PTPRC;refGene_tx=XM_019821945.1
F1	39759736	39759948
refGene_gn=PTPRC;refGene_tx=XM_006942841.3;refGene_gn=PTPRC;refGene_tx=XM_011290957.2;refGene_gn=PTPRC;refGene_tx=XM_019821942.1;refGene_gn=PTPRC;refGene_tx=XM_019821943.1;refGene_gn=PTPRC;refGene_tx=XM_019821944.1;refGene_gn=PTPRC;refGene_tx=XM_019821945.1
F1	39765409	39765470
refGene_gn=PTPRC;refGene_tx=XM_006942841.3;refGene_gn=PTPRC;refGene_tx=XM_011290957.2;refGene_gn=PTPRC;refGene_tx=XM_019821942.1;refGene_gn=PTPRC;refGene_tx=XM_019821943.1;refGene_gn=PTPRC;refGene_tx=XM_019821944.1;refGene_gn=PTPRC;refGene_tx=XM_019821945.1
F1	39773781	39773890
refGene_gn=PTPRC;refGene_tx=XM_006942841.3;refGene_gn=PTPRC;refGene_tx=XM_011290957.2;refGene_gn=PTPRC;refGene_tx=XM_019821942.1;refGene_gn=PTPRC;refGene_tx=XM_019821943.1;refGene_gn=PTPRC;refGene_tx=XM_019821944.1;refGene_gn=PTPRC;refGene_tx=XM_019821945.1
F1	39774599	39774634
refGene_gn=PTPRC;refGene_tx=XM_006942841.3;refGene_gn=PTPRC;refGene_tx=XM_011290957.2;refGene_gn=PTPRC;refGene_tx=XM_019821942.1;refGene_gn=PTPRC;refGene_tx=XM_019821943.1;refGene_gn=PTPRC;refGene_tx=XM_019821944.1;refGene_gn=PTPRC;refGene_tx=XM_019821945.1
F1	39777060	39777170
refGene_gn=PTPRC;refGene_tx=XM_006942841.3;refGene_gn=PTPRC;refGene_tx=XM_011290957.2;refGene_gn=PTPRC;refGene_tx=XM_019821942.1;refGene_gn=PTPRC;refGene_tx=XM_019821943.1;refGene_gn=PTPRC;refGene_tx=XM_0198219

44.1;refGene_gn=PTPRC;refGene_tx=XM_019821945.1
F1	39777618	39777709
refGene_gn=PTPRC;refGene_tx=XM_006942841.3;refGene_gn=PTPRC;refGene_tx=XM_011290957.2;refGene_gn=PTPRC;refGene_tx=XM_019821942.1;refGene_gn=PTPRC;refGene_tx=XM_019821943.1;refGene_gn=PTPRC;refGene_tx=XM_019821944.1;refGene_gn=PTPRC;refGene_tx=XM_019821945.1
F1	39777799	39777876
refGene_gn=PTPRC;refGene_tx=XM_006942841.3;refGene_gn=PTPRC;refGene_tx=XM_011290957.2;refGene_gn=PTPRC;refGene_tx=XM_019821942.1;refGene_gn=PTPRC;refGene_tx=XM_019821943.1;refGene_gn=PTPRC;refGene_tx=XM_019821944.1;refGene_gn=PTPRC;refGene_tx=XM_019821945.1
F1	39781841	39781878
refGene_gn=PTPRC;refGene_tx=XM_006942841.3;refGene_gn=PTPRC;refGene_tx=XM_011290957.2;refGene_gn=PTPRC;refGene_tx=XM_019821942.1;refGene_gn=PTPRC;refGene_tx=XM_019821943.1;refGene_gn=PTPRC;refGene_tx=XM_019821944.1;refGene_gn=PTPRC;refGene_tx=XM_019821945.1
F1	39781971	39782069
refGene_gn=PTPRC;refGene_tx=XM_006942841.3;refGene_gn=PTPRC;refGene_tx=XM_011290957.2;refGene_gn=PTPRC;refGene_tx=XM_019821942.1;refGene_gn=PTPRC;refGene_tx=XM_019821943.1;refGene_gn=PTPRC;refGene_tx=XM_019821944.1;refGene_gn=PTPRC;refGene_tx=XM_019821945.1
F1	39782799	39782925
refGene_gn=PTPRC;refGene_tx=XM_006942841.3;refGene_gn=PTPRC;refGene_tx=XM_011290957.2;refGene_gn=PTPRC;refGene_tx=XM_019821942.1;refGene_gn=PTPRC;refGene_tx=XM_019821943.1;refGene_gn=PTPRC;refGene_tx=XM_019821944.1;refGene_gn=PTPRC;refGene_tx=XM_019821945.1
F1	39785019	39785177
refGene_gn=PTPRC;refGene_tx=XM_006942841.3;refGene_gn=PTPRC;refGene_tx=XM_011290957.2;refGene_gn=PTPRC;refGene_tx=XM_019821942.1;refGene_gn=PTPRC;refGene_tx=XM_019821943.1;refGene_gn=PTPRC;refGene_tx=XM_019821944.1;refGene_gn=PTPRC;refGene_tx=XM_019821945.1
F1	39785401	39785537
refGene_gn=PTPRC;refGene_tx=XM_006942841.3;refGene_gn=PTPRC;refGene_tx=XM_011290957.2;refGene_gn=PTPRC;refGene_tx=XM_019821942.1;refGene_gn=PTPRC;refGene_tx=XM_019821943.1;refGene_gn=PTPRC;refGene_tx=XM_019821944.1;refGene_gn=PTPRC;refGene_tx=XM_019821945.1
F1	39787644	39787794
refGene_gn=PTPRC;refGene_tx=XM_006942841.3;refGene_gn=PTPRC;refGene_tx=XM_011290957.2;refGene_gn=PTPRC;refGene_tx=XM_019821942.1;refGene_gn=PTPRC;refGene_tx=XM_019821943.1;refGene_gn=PTPRC;refGene_tx=XM_019821944.1;refGene_gn=PTPRC;refGene_tx=XM_019821945.1
F1	39792157	39792248
refGene_gn=PTPRC;refGene_tx=XM_006942841.3;refGene_gn=PTPRC;refGene_tx=XM_011290957.2;refGene_gn=PTPRC;refGene_tx=XM_019821942.1;refGene_gn=PTPRC;refGene_tx=XM_019821943.1;refGene_gn=PTPRC;refGene_tx=XM_019821944.1;refGene_gn=PTPRC;refGene_tx=XM_019821945.1
F1	39793508	39793642
refGene_gn=PTPRC;refGene_tx=XM_006942841.3;refGene_gn=PTPRC;refGene_tx=XM_011290957.2;refGene_gn=PTPRC;refGene_tx=XM_019821942.1;refGene_gn=PTPRC;refGene_tx=XM_019821943.1;refGene_gn=PTPRC;refGene_tx=XM_0198219

44.1;refGene_gn=PTPRC;refGene_tx=XM_019821945.1
F1 39794320 39794455
refGene_gn=PTPRC;refGene_tx=XM_006942841.3;refGene_gn=PTPRC;refGene_tx=XM_011290957.2;refGene_gn=PTPRC;refGene_tx=XM_019821942.1;refGene_gn=PTPRC;refGene_tx=XM_019821943.1;refGene_gn=PTPRC;refGene_tx=XM_019821944.1;refGene_gn=PTPRC;refGene_tx=XM_019821945.1
F1 39796870 39796993
refGene_gn=PTPRC;refGene_tx=XM_006942841.3;refGene_gn=PTPRC;refGene_tx=XM_011290957.2;refGene_gn=PTPRC;refGene_tx=XM_019821942.1;refGene_gn=PTPRC;refGene_tx=XM_019821943.1;refGene_gn=PTPRC;refGene_tx=XM_019821944.1;refGene_gn=PTPRC;refGene_tx=XM_019821945.1
F1 39797215 39797394
refGene_gn=PTPRC;refGene_tx=XM_006942841.3;refGene_gn=PTPRC;refGene_tx=XM_011290957.2;refGene_gn=PTPRC;refGene_tx=XM_019821942.1;refGene_gn=PTPRC;refGene_tx=XM_019821943.1;refGene_gn=PTPRC;refGene_tx=XM_019821944.1;refGene_gn=PTPRC;refGene_tx=XM_019821945.1
F1 39798489 39798625
refGene_gn=PTPRC;refGene_tx=XM_006942841.3;refGene_gn=PTPRC;refGene_tx=XM_011290957.2;refGene_gn=PTPRC;refGene_tx=XM_019821942.1;refGene_gn=PTPRC;refGene_tx=XM_019821943.1;refGene_gn=PTPRC;refGene_tx=XM_019821944.1;refGene_gn=PTPRC;refGene_tx=XM_019821945.1
F1 39799966 39800242
refGene_gn=PTPRC;refGene_tx=XM_006942841.3;refGene_gn=PTPRC;refGene_tx=XM_011290957.2;refGene_gn=PTPRC;refGene_tx=XM_019821942.1;refGene_gn=PTPRC;refGene_tx=XM_019821943.1;refGene_gn=PTPRC;refGene_tx=XM_019821944.1;refGene_gn=PTPRC;refGene_tx=XM_019821945.1
F1 40637672 40637927
refGene_gn=LOC101093986;refGene_tx=XM_011290956.2
F1 40988950 40989014
refGene_gn=NR5A2;refGene_tx=XM_019821946.2;refGene_gn=NR5A2;refGene_tx=XM_023247519.1
F1 41000496 41000634
refGene_gn=NR5A2;refGene_tx=XM_023247518.1;refGene_gn=NR5A2;refGene_tx=XM_023247519.1
F1 41004761 41004880
refGene_gn=NR5A2;refGene_tx=XM_006942842.4;refGene_gn=NR5A2;refGene_tx=XM_019821946.2;refGene_gn=NR5A2;refGene_tx=XM_023247518.1;refGene_gn=NR5A2;refGene_tx=XM_023247519.1
F1 41006422 41006564
refGene_gn=NR5A2;refGene_tx=XM_006942842.4;refGene_gn=NR5A2;refGene_tx=XM_019821946.2;refGene_gn=NR5A2;refGene_tx=XM_023247518.1;refGene_gn=NR5A2;refGene_tx=XM_023247519.1
F1 41008832 41009479
refGene_gn=NR5A2;refGene_tx=XM_006942842.4;refGene_gn=NR5A2;refGene_tx=XM_019821946.2;refGene_gn=NR5A2;refGene_tx=XM_023247518.1;refGene_gn=NR5A2;refGene_tx=XM_023247519.1
F1 41062766 41062886
refGene_gn=NR5A2;refGene_tx=XM_006942842.4;refGene_gn=NR5A2;refGene_tx=XM_019821946.2;refGene_gn=NR5A2;refGene_tx=XM_023247518.1;refGene_gn=NR5A2;refGene_tx=XM_023247519.1

F1 41072475 41072623
refGene_gn=NR5A2;refGene_tx=XM_006942842.4;refGene_gn=NR5A2;refGene_tx=XM_019821946.2;refGene_gn=NR5A2;refGene_tx=XM_023247518.1;refGene_gn=NR5A2;refGene_tx=XM_023247519.1
F1 41122877 41123125
refGene_gn=NR5A2;refGene_tx=XM_006942842.4;refGene_gn=NR5A2;refGene_tx=XM_019821946.2;refGene_gn=NR5A2;refGene_tx=XM_023247518.1;refGene_gn=NR5A2;refGene_tx=XM_023247519.1
F1 41327573 41330285
refGene_gn=ZNF281;refGene_tx=XM_006942844.3;refGene_gn=ZNF281;refGene_tx=XM_023247447.1;refGene_gn=ZNF281;refGene_tx=XM_023247448.1;refGene_gn=ZNF281;refGene_tx=XM_023247449.1
F1 41330507 41330916 refGene_gn=ZNF281;refGene_tx=XM_023247447.1
F1 41331225 41331274
refGene_gn=ZNF281;refGene_tx=XM_006942844.3;refGene_gn=ZNF281;refGene_tx=XM_023247447.1
F1 41332258 41332278
refGene_gn=ZNF281;refGene_tx=XM_006942844.3;refGene_gn=ZNF281;refGene_tx=XM_023247447.1
F1 41456183 41456563 refGene_gn=KIF14;refGene_tx=XM_023247456.1
F1 41456935 41457074 refGene_gn=KIF14;refGene_tx=XM_023247456.1
F1 41461878 41461953 refGene_gn=KIF14;refGene_tx=XM_023247456.1
F1 41465690 41465813 refGene_gn=KIF14;refGene_tx=XM_023247456.1
F1 41466393 41466552 refGene_gn=KIF14;refGene_tx=XM_023247456.1
F1 41469387 41469572 refGene_gn=KIF14;refGene_tx=XM_023247456.1
F1 41470052 41470277 refGene_gn=KIF14;refGene_tx=XM_023247456.1
F1 41473885 41473980 refGene_gn=KIF14;refGene_tx=XM_023247456.1
F1 41479455 41479556 refGene_gn=KIF14;refGene_tx=XM_023247456.1
F1 41480908 41481041 refGene_gn=KIF14;refGene_tx=XM_023247456.1
F1 41481629 41481723 refGene_gn=KIF14;refGene_tx=XM_023247456.1
F1 41484140 41484267 refGene_gn=KIF14;refGene_tx=XM_023247456.1
F1 41485916 41486069 refGene_gn=KIF14;refGene_tx=XM_023247456.1
F1 41486611 41486759 refGene_gn=KIF14;refGene_tx=XM_023247456.1
F1 41488767 41488928 refGene_gn=KIF14;refGene_tx=XM_023247456.1
F1 41490199 41490302 refGene_gn=KIF14;refGene_tx=XM_023247456.1
F1 41500056 41500241 refGene_gn=KIF14;refGene_tx=XM_023247456.1
F1 41502683 41502747 refGene_gn=KIF14;refGene_tx=XM_023247456.1
F1 41503008 41503156 refGene_gn=KIF14;refGene_tx=XM_023247456.1
F1 41504737 41504910 refGene_gn=KIF14;refGene_tx=XM_023247456.1
F1 41506234 41506371 refGene_gn=KIF14;refGene_tx=XM_023247456.1
F1 41507911 41507942 refGene_gn=KIF14;refGene_tx=XM_023247456.1
F1 41509402 41509455 refGene_gn=KIF14;refGene_tx=XM_023247456.1
F1 41510238 41510337 refGene_gn=KIF14;refGene_tx=XM_023247456.1
F1 41511380 41511468 refGene_gn=KIF14;refGene_tx=XM_023247456.1
F1 41512193 41512448 refGene_gn=KIF14;refGene_tx=XM_023247456.1
F1 41514392 41515507 refGene_gn=KIF14;refGene_tx=XM_023247456.1
F1 41543129 41543393
refGene_gn=DDX59;refGene_tx=XM_003999359.5;refGene_gn=DDX59;refGene_tx=XM_011290955.3;refGene_gn=DDX59;refGene_tx=XM_023247358.1
F1 41548738 41548867

refGene_gn=DDX59;refGene_tx=XM_003999359.5;refGene_gn=DDX59;refGene_tx=XM_011290955.3;refGene_gn=DDX59;refGene_tx=XM_023247358.1
F1 41549358 41549511
refGene_gn=DDX59;refGene_tx=XM_003999359.5;refGene_gn=DDX59;refGene_tx=XM_011290955.3;refGene_gn=DDX59;refGene_tx=XM_023247358.1
F1 41550208 41550460
refGene_gn=DDX59;refGene_tx=XM_003999359.5;refGene_gn=DDX59;refGene_tx=XM_011290955.3;refGene_gn=DDX59;refGene_tx=XM_023247358.1
F1 41556365 41556455
refGene_gn=DDX59;refGene_tx=XM_003999359.5;refGene_gn=DDX59;refGene_tx=XM_011290955.3;refGene_gn=DDX59;refGene_tx=XM_023247358.1
F1 41560723 41560891
refGene_gn=DDX59;refGene_tx=XM_003999359.5;refGene_gn=DDX59;refGene_tx=XM_011290955.3;refGene_gn=DDX59;refGene_tx=XM_023247358.1
F1 41562062 41562869
refGene_gn=DDX59;refGene_tx=XM_003999359.5;refGene_gn=DDX59;refGene_tx=XM_011290955.3;refGene_gn=DDX59;refGene_tx=XM_023247358.1
F1 41635414 41635553
refGene_gn=CAMSAP2;refGene_tx=XM_003999361.4;refGene_gn=CAMSAP2;refGene_tx=XM_011290950.2;refGene_gn=CAMSAP2;refGene_tx=XM_011290951.2;refGene_gn=CAMSAP2;refGene_tx=XM_011290952.2;refGene_gn=CAMSAP2;refGene_tx=XM_011290953.2;refGene_gn=CAMSAP2;refGene_tx=XM_011290954.2;refGene_gn=CAMSAP2;refGene_tx=XM_019821952.2
F1 41657243 41657503
refGene_gn=CAMSAP2;refGene_tx=XM_003999361.4;refGene_gn=CAMSAP2;refGene_tx=XM_011290950.2;refGene_gn=CAMSAP2;refGene_tx=XM_011290951.2;refGene_gn=CAMSAP2;refGene_tx=XM_011290952.2;refGene_gn=CAMSAP2;refGene_tx=XM_011290953.2;refGene_gn=CAMSAP2;refGene_tx=XM_011290954.2;refGene_gn=CAMSAP2;refGene_tx=XM_019821952.2
F1 41696093 41696255
refGene_gn=CAMSAP2;refGene_tx=XM_003999361.4;refGene_gn=CAMSAP2;refGene_tx=XM_011290950.2;refGene_gn=CAMSAP2;refGene_tx=XM_011290951.2;refGene_gn=CAMSAP2;refGene_tx=XM_011290952.2;refGene_gn=CAMSAP2;refGene_tx=XM_011290953.2;refGene_gn=CAMSAP2;refGene_tx=XM_011290954.2;refGene_gn=CAMSAP2;refGene_tx=XM_019821952.2
F1 41708379 41708463
refGene_gn=CAMSAP2;refGene_tx=XM_003999361.4;refGene_gn=CAMSAP2;refGene_tx=XM_011290950.2;refGene_gn=CAMSAP2;refGene_tx=XM_011290951.2;refGene_gn=CAMSAP2;refGene_tx=XM_011290952.2;refGene_gn=CAMSAP2;refGene_tx=XM_011290953.2;refGene_gn=CAMSAP2;refGene_tx=XM_011290954.2;refGene_gn=CAMSAP2;refGene_tx=XM_019821952.2
F1 41718233 41718266
refGene_gn=CAMSAP2;refGene_tx=XM_011290950.2;refGene_gn=CAMSAP2;refGene_tx=XM_011290951.2;refGene_gn=CAMSAP2;refGene_tx=XM_011290952.2;refGene_gn=CAMSAP2;refGene_tx=XM_019821952.2
F1 41721061 41721203
refGene_gn=CAMSAP2;refGene_tx=XM_003999361.4;refGene_gn=CAMSAP2;refGene_tx=XM_011290950.2;refGene_gn=CAMSAP2;refGene_tx=XM_011290951.2;refGene_gn=CAMSAP2;refGene_tx=XM_011290952.2;refGene_gn=CAMSAP2;refGene_tx=XM_011290953.2;refGene_gn=CAMSAP2;refGene_tx=XM_011290954.2;refGene_gn

=CAMSAP2;refGene_tx=XM_019821952.2
F1 41721512 41721652
refGene_gn=CAMSAP2;refGene_tx=XM_003999361.4;refGene_gn=CAMSAP2;refGene_tx=XM_011290950.2;refGene_gn=CAMSAP2;refGene_tx=XM_011290951.2;refGene_gn=CAMSAP2;refGene_tx=XM_011290952.2;refGene_gn=CAMSAP2;refGene_tx=XM_011290953.2;refGene_gn=CAMSAP2;refGene_tx=XM_011290954.2;refGene_gn=CAMSAP2;refGene_tx=XM_019821952.2
F1 41729302 41729399
refGene_gn=CAMSAP2;refGene_tx=XM_003999361.4;refGene_gn=CAMSAP2;refGene_tx=XM_011290950.2;refGene_gn=CAMSAP2;refGene_tx=XM_011290951.2;refGene_gn=CAMSAP2;refGene_tx=XM_011290952.2;refGene_gn=CAMSAP2;refGene_tx=XM_011290953.2;refGene_gn=CAMSAP2;refGene_tx=XM_011290954.2;refGene_gn=CAMSAP2;refGene_tx=XM_019821952.2
F1 41731542 41731636
refGene_gn=CAMSAP2;refGene_tx=XM_003999361.4;refGene_gn=CAMSAP2;refGene_tx=XM_011290950.2;refGene_gn=CAMSAP2;refGene_tx=XM_011290951.2;refGene_gn=CAMSAP2;refGene_tx=XM_011290952.2;refGene_gn=CAMSAP2;refGene_tx=XM_011290953.2;refGene_gn=CAMSAP2;refGene_tx=XM_011290954.2;refGene_gn=CAMSAP2;refGene_tx=XM_019821952.2
F1 41735312 41735395
refGene_gn=CAMSAP2;refGene_tx=XM_003999361.4;refGene_gn=CAMSAP2;refGene_tx=XM_011290950.2;refGene_gn=CAMSAP2;refGene_tx=XM_011290951.2;refGene_gn=CAMSAP2;refGene_tx=XM_011290952.2;refGene_gn=CAMSAP2;refGene_tx=XM_011290953.2;refGene_gn=CAMSAP2;refGene_tx=XM_011290954.2;refGene_gn=CAMSAP2;refGene_tx=XM_019821952.2
F1 41736232 41736302
refGene_gn=CAMSAP2;refGene_tx=XM_003999361.4;refGene_gn=CAMSAP2;refGene_tx=XM_011290950.2;refGene_gn=CAMSAP2;refGene_tx=XM_011290951.2;refGene_gn=CAMSAP2;refGene_tx=XM_011290952.2;refGene_gn=CAMSAP2;refGene_tx=XM_011290953.2;refGene_gn=CAMSAP2;refGene_tx=XM_011290954.2;refGene_gn=CAMSAP2;refGene_tx=XM_019821952.2
F1 41736694 41738897
refGene_gn=CAMSAP2;refGene_tx=XM_003999361.4;refGene_gn=CAMSAP2;refGene_tx=XM_011290950.2;refGene_gn=CAMSAP2;refGene_tx=XM_011290951.2;refGene_gn=CAMSAP2;refGene_tx=XM_011290952.2;refGene_gn=CAMSAP2;refGene_tx=XM_011290953.2;refGene_gn=CAMSAP2;refGene_tx=XM_011290954.2;refGene_gn=CAMSAP2;refGene_tx=XM_019821952.2
F1 41740151 41740288
refGene_gn=CAMSAP2;refGene_tx=XM_003999361.4;refGene_gn=CAMSAP2;refGene_tx=XM_011290950.2;refGene_gn=CAMSAP2;refGene_tx=XM_011290951.2;refGene_gn=CAMSAP2;refGene_tx=XM_011290952.2;refGene_gn=CAMSAP2;refGene_tx=XM_011290953.2;refGene_gn=CAMSAP2;refGene_tx=XM_011290954.2;refGene_gn=CAMSAP2;refGene_tx=XM_019821952.2
F1 41740643 41740864
refGene_gn=CAMSAP2;refGene_tx=XM_003999361.4;refGene_gn=CAMSAP2;refGene_tx=XM_011290950.2;refGene_gn=CAMSAP2;refGene_tx=XM_011290951.2;refGene_gn=CAMSAP2;refGene_tx=XM_011290952.2;refGene_gn=CAMSAP2;refGene_tx=XM_011290953.2;refGene_gn=CAMSAP2;refGene_tx=XM_011290954.2;refGene_gn=CAMSAP2;refGene_tx=XM_019821952.2
F1 41741991 41742091

refGene_gn=CAMSAP2;refGene_tx=XM_003999361.4;refGene_gn=CAMSAP2;refGene_tx=XM_011290950.2;refGene_gn=CAMSAP2;refGene_tx=XM_011290951.2;refGene_gn=CAMSAP2;refGene_tx=XM_011290952.2;refGene_gn=CAMSAP2;refGene_tx=XM_011290953.2;refGene_gn=CAMSAP2;refGene_tx=XM_011290954.2;refGene_gn=CAMSAP2;refGene_tx=XM_019821952.2
F1 41742861 41742977
refGene_gn=CAMSAP2;refGene_tx=XM_003999361.4;refGene_gn=CAMSAP2;refGene_tx=XM_011290950.2;refGene_gn=CAMSAP2;refGene_tx=XM_011290951.2;refGene_gn=CAMSAP2;refGene_tx=XM_011290952.2;refGene_gn=CAMSAP2;refGene_tx=XM_011290953.2;refGene_gn=CAMSAP2;refGene_tx=XM_011290954.2;refGene_gn=CAMSAP2;refGene_tx=XM_019821952.2
F1 41744795 41744836
refGene_gn=CAMSAP2;refGene_tx=XM_003999361.4;refGene_gn=CAMSAP2;refGene_tx=XM_011290950.2;refGene_gn=CAMSAP2;refGene_tx=XM_011290951.2;refGene_gn=CAMSAP2;refGene_tx=XM_011290952.2;refGene_gn=CAMSAP2;refGene_tx=XM_011290953.2;refGene_gn=CAMSAP2;refGene_tx=XM_011290954.2;refGene_gn=CAMSAP2;refGene_tx=XM_019821952.2
F1 41761870 41762974 refGene_gn=GPR25;refGene_tx=XM_023247599.1
F1 41784861 41785064
refGene_gn=CF1H1orf106;refGene_tx=XM_023247625.1
F1 41791338 41791487
refGene_gn=CF1H1orf106;refGene_tx=XM_023247625.1;refGene_gn=CF1H1orf106;refGene_tx=XM_023247626.1
F1 41792239 41792364
refGene_gn=CF1H1orf106;refGene_tx=XM_023247625.1;refGene_gn=CF1H1orf106;refGene_tx=XM_023247626.1
F1 41792875 41792992
refGene_gn=CF1H1orf106;refGene_tx=XM_023247625.1;refGene_gn=CF1H1orf106;refGene_tx=XM_023247626.1
F1 41793714 41793937
refGene_gn=CF1H1orf106;refGene_tx=XM_023247625.1;refGene_gn=CF1H1orf106;refGene_tx=XM_023247626.1
F1 41798380 41798434
refGene_gn=CF1H1orf106;refGene_tx=XM_023247625.1;refGene_gn=CF1H1orf106;refGene_tx=XM_023247626.1
F1 41799266 41799477
refGene_gn=CF1H1orf106;refGene_tx=XM_023247625.1;refGene_gn=CF1H1orf106;refGene_tx=XM_023247626.1
F1 41799701 41799875
refGene_gn=CF1H1orf106;refGene_tx=XM_023247625.1;refGene_gn=CF1H1orf106;refGene_tx=XM_023247626.1
F1 41801742 41802400
refGene_gn=CF1H1orf106;refGene_tx=XM_023247625.1;refGene_gn=CF1H1orf106;refGene_tx=XM_023247626.1
F1 41803722 41803815
refGene_gn=CF1H1orf106;refGene_tx=XM_023247625.1;refGene_gn=CF1H1orf106;refGene_tx=XM_023247626.1
F1 41845528 41845583
refGene_gn=KIF21B;refGene_tx=XM_023247203.1;refGene_gn=KIF21B;refGene_tx=XM_023247204.1

F1	41846102	41846202
refGene_gn=KIF21B;refGene_tx=XM_023247199.1;refGene_gn=KIF21B;refGene_tx=XM_023247200.1;refGene_gn=KIF21B;refGene_tx=XM_023247202.1
F1	41846722	41846922
refGene_gn=KIF21B;refGene_tx=XM_023247199.1;refGene_gn=KIF21B;refGene_tx=XM_023247200.1;refGene_gn=KIF21B;refGene_tx=XM_023247202.1;refGene_gn=KIF21B;refGene_tx=XM_023247203.1;refGene_gn=KIF21B;refGene_tx=XM_023247204.1
F1	41847461	41847632
refGene_gn=KIF21B;refGene_tx=XM_023247199.1;refGene_gn=KIF21B;refGene_tx=XM_023247200.1;refGene_gn=KIF21B;refGene_tx=XM_023247202.1;refGene_gn=KIF21B;refGene_tx=XM_023247203.1;refGene_gn=KIF21B;refGene_tx=XM_023247204.1
F1	41848479	41848597
refGene_gn=KIF21B;refGene_tx=XM_023247199.1;refGene_gn=KIF21B;refGene_tx=XM_023247200.1;refGene_gn=KIF21B;refGene_tx=XM_023247202.1;refGene_gn=KIF21B;refGene_tx=XM_023247203.1;refGene_gn=KIF21B;refGene_tx=XM_023247204.1
F1	41848932	41849097
refGene_gn=KIF21B;refGene_tx=XM_023247199.1;refGene_gn=KIF21B;refGene_tx=XM_023247200.1;refGene_gn=KIF21B;refGene_tx=XM_023247202.1;refGene_gn=KIF21B;refGene_tx=XM_023247203.1;refGene_gn=KIF21B;refGene_tx=XM_023247204.1
F1	41850276	41850457
refGene_gn=KIF21B;refGene_tx=XM_023247199.1;refGene_gn=KIF21B;refGene_tx=XM_023247200.1;refGene_gn=KIF21B;refGene_tx=XM_023247202.1;refGene_gn=KIF21B;refGene_tx=XM_023247203.1;refGene_gn=KIF21B;refGene_tx=XM_023247204.1
F1	41851396	41851533
refGene_gn=KIF21B;refGene_tx=XM_023247199.1;refGene_gn=KIF21B;refGene_tx=XM_023247200.1;refGene_gn=KIF21B;refGene_tx=XM_023247202.1;refGene_gn=KIF21B;refGene_tx=XM_023247203.1;refGene_gn=KIF21B;refGene_tx=XM_023247204.1
F1	41853503	41853542
refGene_gn=KIF21B;refGene_tx=XM_023247199.1;refGene_gn=KIF21B;refGene_tx=XM_023247200.1;refGene_gn=KIF21B;refGene_tx=XM_023247203.1
F1	41855263	41855377
refGene_gn=KIF21B;refGene_tx=XM_023247199.1;refGene_gn=KIF21B;refGene_tx=XM_023247200.1;refGene_gn=KIF21B;refGene_tx=XM_023247202.1;refGene_gn=KIF21B;refGene_tx=XM_023247203.1;refGene_gn=KIF21B;refGene_tx=XM_023247204.1
F1	41857142	41857217
refGene_gn=KIF21B;refGene_tx=XM_023247199.1;refGene_gn=KIF21B;refGene_tx=XM_023247200.1;refGene_gn=KIF21B;refGene_tx=XM_023247202.1;refGene_gn=KIF21B;refGene_tx=XM_023247203.1;refGene_gn=KIF21B;refGene_tx=XM_023247204.1
F1	41857295	41857501
refGene_gn=KIF21B;refGene_tx=XM_023247199.1;refGene_gn=KIF21B;refGene_tx=XM_023247200.1;refGene_gn=KIF21B;refGene_tx=XM_023247202.1;refGene_gn=KIF21B;refGene_tx=XM_023247203.1;refGene_gn=KIF21B;refGene_tx=XM_02

3247204.1
F1 41858266 41858324
refGene_gn=KIF21B;refGene_tx=XM_023247199.1;refGene_gn=KIF21B;refGene_tx=XM_023247200.1;refGene_gn=KIF21B;refGene_tx=XM_023247202.1;refGene_gn=KIF21B;refGene_tx=XM_023247203.1;refGene_gn=KIF21B;refGene_tx=XM_023247204.1
F1 41858509 41858561
refGene_gn=KIF21B;refGene_tx=XM_023247199.1;refGene_gn=KIF21B;refGene_tx=XM_023247200.1;refGene_gn=KIF21B;refGene_tx=XM_023247202.1;refGene_gn=KIF21B;refGene_tx=XM_023247203.1;refGene_gn=KIF21B;refGene_tx=XM_023247204.1
F1 41858771 41858937
refGene_gn=KIF21B;refGene_tx=XM_023247199.1;refGene_gn=KIF21B;refGene_tx=XM_023247200.1;refGene_gn=KIF21B;refGene_tx=XM_023247202.1;refGene_gn=KIF21B;refGene_tx=XM_023247203.1;refGene_gn=KIF21B;refGene_tx=XM_023247204.1
F1 41860175 41860277
refGene_gn=KIF21B;refGene_tx=XM_023247199.1;refGene_gn=KIF21B;refGene_tx=XM_023247200.1;refGene_gn=KIF21B;refGene_tx=XM_023247202.1;refGene_gn=KIF21B;refGene_tx=XM_023247203.1;refGene_gn=KIF21B;refGene_tx=XM_023247204.1
F1 41860375 41860570
refGene_gn=KIF21B;refGene_tx=XM_023247199.1;refGene_gn=KIF21B;refGene_tx=XM_023247200.1;refGene_gn=KIF21B;refGene_tx=XM_023247202.1;refGene_gn=KIF21B;refGene_tx=XM_023247203.1;refGene_gn=KIF21B;refGene_tx=XM_023247204.1
F1 41860815 41860963
refGene_gn=KIF21B;refGene_tx=XM_023247199.1;refGene_gn=KIF21B;refGene_tx=XM_023247200.1;refGene_gn=KIF21B;refGene_tx=XM_023247202.1;refGene_gn=KIF21B;refGene_tx=XM_023247203.1;refGene_gn=KIF21B;refGene_tx=XM_023247204.1
F1 41861332 41861565
refGene_gn=KIF21B;refGene_tx=XM_023247199.1;refGene_gn=KIF21B;refGene_tx=XM_023247200.1;refGene_gn=KIF21B;refGene_tx=XM_023247202.1;refGene_gn=KIF21B;refGene_tx=XM_023247203.1;refGene_gn=KIF21B;refGene_tx=XM_023247204.1
F1 41861966 41862035
refGene_gn=KIF21B;refGene_tx=XM_023247199.1;refGene_gn=KIF21B;refGene_tx=XM_023247200.1;refGene_gn=KIF21B;refGene_tx=XM_023247202.1;refGene_gn=KIF21B;refGene_tx=XM_023247203.1;refGene_gn=KIF21B;refGene_tx=XM_023247204.1
F1 41862500 41862608
refGene_gn=KIF21B;refGene_tx=XM_023247199.1;refGene_gn=KIF21B;refGene_tx=XM_023247200.1;refGene_gn=KIF21B;refGene_tx=XM_023247202.1;refGene_gn=KIF21B;refGene_tx=XM_023247203.1;refGene_gn=KIF21B;refGene_tx=XM_023247204.1
F1 41865779 41865979
refGene_gn=KIF21B;refGene_tx=XM_023247199.1;refGene_gn=KIF21B;refGene_tx=XM_023247200.1;refGene_gn=KIF21B;refGene_tx=XM_023247202.1;refGene_gn=KIF21B;refGene_tx=XM_023247203.1;refGene_gn=KIF21B;refGene_tx=XM_02

3247204.1
F1 41867763 41867955
refGene_gn=KIF21B;refGene_tx=XM_023247199.1;refGene_gn=KIF21B;refGene_tx=XM_023247200.1;refGene_gn=KIF21B;refGene_tx=XM_023247202.1;refGene_gn=KIF21B;refGene_tx=XM_023247203.1;refGene_gn=KIF21B;refGene_tx=XM_023247204.1
F1 41868787 41868905
refGene_gn=KIF21B;refGene_tx=XM_023247199.1;refGene_gn=KIF21B;refGene_tx=XM_023247200.1;refGene_gn=KIF21B;refGene_tx=XM_023247202.1;refGene_gn=KIF21B;refGene_tx=XM_023247203.1;refGene_gn=KIF21B;refGene_tx=XM_023247204.1
F1 41869356 41869438
refGene_gn=KIF21B;refGene_tx=XM_023247199.1;refGene_gn=KIF21B;refGene_tx=XM_023247200.1;refGene_gn=KIF21B;refGene_tx=XM_023247202.1;refGene_gn=KIF21B;refGene_tx=XM_023247203.1;refGene_gn=KIF21B;refGene_tx=XM_023247204.1
F1 41869774 41869993
refGene_gn=KIF21B;refGene_tx=XM_023247199.1;refGene_gn=KIF21B;refGene_tx=XM_023247200.1;refGene_gn=KIF21B;refGene_tx=XM_023247202.1;refGene_gn=KIF21B;refGene_tx=XM_023247203.1;refGene_gn=KIF21B;refGene_tx=XM_023247204.1
F1 41870099 41870163
refGene_gn=KIF21B;refGene_tx=XM_023247199.1;refGene_gn=KIF21B;refGene_tx=XM_023247200.1;refGene_gn=KIF21B;refGene_tx=XM_023247202.1;refGene_gn=KIF21B;refGene_tx=XM_023247203.1;refGene_gn=KIF21B;refGene_tx=XM_023247204.1
F1 41871161 41871351
refGene_gn=KIF21B;refGene_tx=XM_023247199.1;refGene_gn=KIF21B;refGene_tx=XM_023247200.1;refGene_gn=KIF21B;refGene_tx=XM_023247202.1;refGene_gn=KIF21B;refGene_tx=XM_023247203.1;refGene_gn=KIF21B;refGene_tx=XM_023247204.1
F1 41872481 41872677
refGene_gn=KIF21B;refGene_tx=XM_023247199.1;refGene_gn=KIF21B;refGene_tx=XM_023247200.1;refGene_gn=KIF21B;refGene_tx=XM_023247202.1;refGene_gn=KIF21B;refGene_tx=XM_023247203.1;refGene_gn=KIF21B;refGene_tx=XM_023247204.1
F1 41873188 41873304
refGene_gn=KIF21B;refGene_tx=XM_023247199.1;refGene_gn=KIF21B;refGene_tx=XM_023247200.1;refGene_gn=KIF21B;refGene_tx=XM_023247202.1;refGene_gn=KIF21B;refGene_tx=XM_023247203.1;refGene_gn=KIF21B;refGene_tx=XM_023247204.1
F1 41873643 41873811
refGene_gn=KIF21B;refGene_tx=XM_023247199.1;refGene_gn=KIF21B;refGene_tx=XM_023247200.1;refGene_gn=KIF21B;refGene_tx=XM_023247202.1;refGene_gn=KIF21B;refGene_tx=XM_023247203.1;refGene_gn=KIF21B;refGene_tx=XM_023247204.1
F1 41874209 41874344
refGene_gn=KIF21B;refGene_tx=XM_023247199.1;refGene_gn=KIF21B;refGene_tx=XM_023247200.1;refGene_gn=KIF21B;refGene_tx=XM_023247202.1;refGene_gn=KIF21B;refGene_tx=XM_023247203.1;refGene_gn=KIF21B;refGene_tx=XM_02

3247204.1
F1	41874427	41874577
refGene_gn=KIF21B;refGene_tx=XM_023247199.1;refGene_gn=KIF21B;refGene_tx=XM_023247200.1;refGene_gn=KIF21B;refGene_tx=XM_023247202.1;refGene_gn=KIF21B;refGene_tx=XM_023247203.1;refGene_gn=KIF21B;refGene_tx=XM_023247204.1
F1	41876938	41877121
refGene_gn=KIF21B;refGene_tx=XM_023247199.1;refGene_gn=KIF21B;refGene_tx=XM_023247200.1;refGene_gn=KIF21B;refGene_tx=XM_023247202.1;refGene_gn=KIF21B;refGene_tx=XM_023247203.1;refGene_gn=KIF21B;refGene_tx=XM_023247204.1
F1	41877451	41877674
refGene_gn=KIF21B;refGene_tx=XM_023247199.1;refGene_gn=KIF21B;refGene_tx=XM_023247200.1;refGene_gn=KIF21B;refGene_tx=XM_023247202.1;refGene_gn=KIF21B;refGene_tx=XM_023247203.1;refGene_gn=KIF21B;refGene_tx=XM_023247204.1
F1	41890955	41890996
refGene_gn=KIF21B;refGene_tx=XM_023247199.1;refGene_gn=KIF21B;refGene_tx=XM_023247200.1;refGene_gn=KIF21B;refGene_tx=XM_023247202.1;refGene_gn=KIF21B;refGene_tx=XM_023247203.1;refGene_gn=KIF21B;refGene_tx=XM_023247204.1
F1	41904361	41904613
refGene_gn=CACNA1S;refGene_tx=NM_001038605.2;refGene_gn=CACNA1S;refGene_tx=XM_006942798.3
F1	41904760	41904898
refGene_gn=CACNA1S;refGene_tx=NM_001038605.2;refGene_gn=CACNA1S;refGene_tx=XM_006942798.3
F1	41905103	41905195
refGene_gn=CACNA1S;refGene_tx=NM_001038605.2;refGene_gn=CACNA1S;refGene_tx=XM_006942798.3
F1	41905982	41906068
refGene_gn=CACNA1S;refGene_tx=NM_001038605.2;refGene_gn=CACNA1S;refGene_tx=XM_006942798.3
F1	41907445	41907696
refGene_gn=CACNA1S;refGene_tx=NM_001038605.2;refGene_gn=CACNA1S;refGene_tx=XM_006942798.3
F1	41908485	41908614
refGene_gn=CACNA1S;refGene_tx=NM_001038605.2;refGene_gn=CACNA1S;refGene_tx=XM_006942798.3
F1	41909802	41909927
refGene_gn=CACNA1S;refGene_tx=NM_001038605.2;refGene_gn=CACNA1S;refGene_tx=XM_006942798.3
F1	41910219	41910321
refGene_gn=CACNA1S;refGene_tx=NM_001038605.2;refGene_gn=CACNA1S;refGene_tx=XM_006942798.3
F1	41911250	41911353
refGene_gn=CACNA1S;refGene_tx=NM_001038605.2;refGene_gn=CACNA1S;refGene_tx=XM_006942798.3
F1	41911411	41911426	refGene_gn=CACNA1S;refGene_tx=XM_019821327.1
F1	41911706	41911803

refGene_gn=CACNA1S;refGene_tx=NM_001038605.2;refGene_gn=CACNA1S;refGene_tx=XM_006942798.3;refGene_gn=CACNA1S;refGene_tx=XM_019821327.1
F1 41912920 41913048
refGene_gn=CACNA1S;refGene_tx=NM_001038605.2;refGene_gn=CACNA1S;refGene_tx=XM_006942798.3;refGene_gn=CACNA1S;refGene_tx=XM_019821327.1
F1 41914038 41914198
refGene_gn=CACNA1S;refGene_tx=NM_001038605.2;refGene_gn=CACNA1S;refGene_tx=XM_006942798.3;refGene_gn=CACNA1S;refGene_tx=XM_019821327.1
F1 41915357 41915449
refGene_gn=CACNA1S;refGene_tx=NM_001038605.2;refGene_gn=CACNA1S;refGene_tx=XM_006942798.3;refGene_gn=CACNA1S;refGene_tx=XM_019821327.1
F1 41916041 41916107
refGene_gn=CACNA1S;refGene_tx=NM_001038605.2;refGene_gn=CACNA1S;refGene_tx=XM_006942798.3;refGene_gn=CACNA1S;refGene_tx=XM_019821327.1
F1 41916307 41916436
refGene_gn=CACNA1S;refGene_tx=NM_001038605.2;refGene_gn=CACNA1S;refGene_tx=XM_006942798.3;refGene_gn=CACNA1S;refGene_tx=XM_019821327.1
F1 41917281 41917338
refGene_gn=CACNA1S;refGene_tx=NM_001038605.2;refGene_gn=CACNA1S;refGene_tx=XM_019821327.1
F1 41922986 41923070
refGene_gn=CACNA1S;refGene_tx=NM_001038605.2;refGene_gn=CACNA1S;refGene_tx=XM_006942798.3;refGene_gn=CACNA1S;refGene_tx=XM_019821327.1
F1 41923785 41923896
refGene_gn=CACNA1S;refGene_tx=NM_001038605.2;refGene_gn=CACNA1S;refGene_tx=XM_006942798.3;refGene_gn=CACNA1S;refGene_tx=XM_019821327.1
F1 41925236 41925395
refGene_gn=CACNA1S;refGene_tx=NM_001038605.2;refGene_gn=CACNA1S;refGene_tx=XM_006942798.3;refGene_gn=CACNA1S;refGene_tx=XM_019821327.1
F1 41925874 41926076
refGene_gn=CACNA1S;refGene_tx=NM_001038605.2;refGene_gn=CACNA1S;refGene_tx=XM_006942798.3;refGene_gn=CACNA1S;refGene_tx=XM_019821327.1
F1 41926549 41926696
refGene_gn=CACNA1S;refGene_tx=NM_001038605.2;refGene_gn=CACNA1S;refGene_tx=XM_006942798.3;refGene_gn=CACNA1S;refGene_tx=XM_019821327.1
F1 41926953 41927006
refGene_gn=CACNA1S;refGene_tx=NM_001038605.2;refGene_gn=CACNA1S;refGene_tx=XM_006942798.3;refGene_gn=CACNA1S;refGene_tx=XM_019821327.1
F1 41929343 41929451
refGene_gn=CACNA1S;refGene_tx=NM_001038605.2;refGene_gn=CACNA1S;refGene_tx=XM_006942798.3;refGene_gn=CACNA1S;refGene_tx=XM_019821327.1
F1 41929687 41929775
refGene_gn=CACNA1S;refGene_tx=NM_001038605.2;refGene_gn=CACNA1S;refGene_tx=XM_006942798.3;refGene_gn=CACNA1S;refGene_tx=XM_019821327.1
F1 41930222 41930329
refGene_gn=CACNA1S;refGene_tx=NM_001038605.2;refGene_gn=CACNA1S;refGene_tx=XM_006942798.3;refGene_gn=CACNA1S;refGene_tx=XM_019821327.1
F1 41931885 41931945
refGene_gn=CACNA1S;refGene_tx=NM_001038605.2;refGene_gn=CACNA1S;refGene_tx=XM_006942798.3;refGene_gn=CACNA1S;refGene_tx=XM_019821327.1

F1	41932254	41932384
refGene_gn=CACNA1S;refGene_tx=NM_001038605.2;refGene_gn=CACNA1S;refGene_tx=XM_006942798.3;refGene_gn=CACNA1S;refGene_tx=XM_019821327.1
F1	41932967	41933100
refGene_gn=CACNA1S;refGene_tx=NM_001038605.2;refGene_gn=CACNA1S;refGene_tx=XM_006942798.3;refGene_gn=CACNA1S;refGene_tx=XM_019821327.1
F1	41935660	41935730
refGene_gn=CACNA1S;refGene_tx=NM_001038605.2;refGene_gn=CACNA1S;refGene_tx=XM_006942798.3;refGene_gn=CACNA1S;refGene_tx=XM_019821327.1
F1	41936390	41936484
refGene_gn=CACNA1S;refGene_tx=NM_001038605.2;refGene_gn=CACNA1S;refGene_tx=XM_006942798.3;refGene_gn=CACNA1S;refGene_tx=XM_019821327.1
F1	41937581	41937696
refGene_gn=CACNA1S;refGene_tx=NM_001038605.2;refGene_gn=CACNA1S;refGene_tx=XM_006942798.3;refGene_gn=CACNA1S;refGene_tx=XM_019821327.1
F1	41938541	41938662
refGene_gn=CACNA1S;refGene_tx=NM_001038605.2;refGene_gn=CACNA1S;refGene_tx=XM_006942798.3;refGene_gn=CACNA1S;refGene_tx=XM_019821327.1
F1	41939435	41939643
refGene_gn=CACNA1S;refGene_tx=NM_001038605.2;refGene_gn=CACNA1S;refGene_tx=XM_006942798.3;refGene_gn=CACNA1S;refGene_tx=XM_019821327.1
F1	41939740	41939966
refGene_gn=CACNA1S;refGene_tx=NM_001038605.2;refGene_gn=CACNA1S;refGene_tx=XM_006942798.3;refGene_gn=CACNA1S;refGene_tx=XM_019821327.1
F1	41941547	41941708
refGene_gn=CACNA1S;refGene_tx=NM_001038605.2;refGene_gn=CACNA1S;refGene_tx=XM_006942798.3;refGene_gn=CACNA1S;refGene_tx=XM_019821327.1
F1	41943678	41943760
refGene_gn=CACNA1S;refGene_tx=NM_001038605.2;refGene_gn=CACNA1S;refGene_tx=XM_006942798.3;refGene_gn=CACNA1S;refGene_tx=XM_019821327.1
F1	41944185	41944331
refGene_gn=CACNA1S;refGene_tx=NM_001038605.2;refGene_gn=CACNA1S;refGene_tx=XM_006942798.3;refGene_gn=CACNA1S;refGene_tx=XM_019821327.1
F1	41946184	41946288
refGene_gn=CACNA1S;refGene_tx=NM_001038605.2;refGene_gn=CACNA1S;refGene_tx=XM_006942798.3;refGene_gn=CACNA1S;refGene_tx=XM_019821327.1
F1	41947881	41948087
refGene_gn=CACNA1S;refGene_tx=NM_001038605.2;refGene_gn=CACNA1S;refGene_tx=XM_006942798.3;refGene_gn=CACNA1S;refGene_tx=XM_019821327.1
F1	41950337	41950490
refGene_gn=CACNA1S;refGene_tx=NM_001038605.2;refGene_gn=CACNA1S;refGene_tx=XM_006942798.3;refGene_gn=CACNA1S;refGene_tx=XM_019821327.1
F1	41950661	41950804
refGene_gn=CACNA1S;refGene_tx=NM_001038605.2;refGene_gn=CACNA1S;refGene_tx=XM_006942798.3;refGene_gn=CACNA1S;refGene_tx=XM_019821327.1
F1	41953271	41953411
refGene_gn=CACNA1S;refGene_tx=NM_001038605.2;refGene_gn=CACNA1S;refGene_tx=XM_006942798.3;refGene_gn=CACNA1S;refGene_tx=XM_019821327.1
F1	41967273	41967379
refGene_gn=CACNA1S;refGene_tx=NM_001038605.2;refGene_gn=CACNA1S;refGen

e_tx=XM_006942798.3;refGene_gn=CACNA1S;refGene_tx=XM_019821327.1
F1	41969132	41969284
refGene_gn=CACNA1S;refGene_tx=NM_001038605.2;refGene_gn=CACNA1S;refGen
e_tx=XM_006942798.3;refGene_gn=CACNA1S;refGene_tx=XM_019821327.1
F1	41970899	41971472	refGene_gn=ASCL5;refGene_tx=XM_023247362.1
F1	41991575	41991728
refGene_gn=TMEM9;refGene_tx=XM_003999364.4;refGene_gn=TMEM9;refGene_tx
=XM_011290948.3;refGene_gn=TMEM9;refGene_tx=XM_011290949.3
F1	42000464	42000596
refGene_gn=TMEM9;refGene_tx=XM_003999364.4;refGene_gn=TMEM9;refGene_tx
=XM_011290948.3;refGene_gn=TMEM9;refGene_tx=XM_011290949.3
F1	42002909	42003018
refGene_gn=TMEM9;refGene_tx=XM_003999364.4;refGene_gn=TMEM9;refGene_tx
=XM_011290948.3;refGene_gn=TMEM9;refGene_tx=XM_011290949.3
F1	42008509	42008601
refGene_gn=TMEM9;refGene_tx=XM_003999364.4;refGene_gn=TMEM9;refGene_tx
=XM_011290948.3;refGene_gn=TMEM9;refGene_tx=XM_011290949.3
F1	42009703	42009778
refGene_gn=TMEM9;refGene_tx=XM_003999364.4;refGene_gn=TMEM9;refGene_tx
=XM_011290948.3;refGene_gn=TMEM9;refGene_tx=XM_011290949.3
F1	42046245	42046252	refGene_gn=IGFN1;refGene_tx=XM_023247443.1
F1	42046731	42046854	refGene_gn=IGFN1;refGene_tx=XM_023247443.1
F1	42048175	42048315	refGene_gn=IGFN1;refGene_tx=XM_023247443.1
F1	42049498	42049598	refGene_gn=IGFN1;refGene_tx=XM_023247443.1
F1	42050987	42051032	refGene_gn=IGFN1;refGene_tx=XM_023247443.1
F1	42051296	42051342	refGene_gn=IGFN1;refGene_tx=XM_023247443.1
F1	42051915	42052090	refGene_gn=IGFN1;refGene_tx=XM_023247443.1
F1	42052959	42053073	refGene_gn=IGFN1;refGene_tx=XM_023247443.1
F1	42054573	42054742	refGene_gn=IGFN1;refGene_tx=XM_023247443.1
F1	42055937	42056210	refGene_gn=IGFN1;refGene_tx=XM_023247443.1
F1	42056949	42056999	refGene_gn=IGFN1;refGene_tx=XM_023247443.1
F1	42065339	42067278
refGene_gn=IGFN1;refGene_tx=XM_023247442.1;refGene_gn=IGFN1;refGene_tx
=XM_023247443.1
F1	42067791	42067916
refGene_gn=IGFN1;refGene_tx=XM_023247442.1;refGene_gn=IGFN1;refGene_tx
=XM_023247443.1
F1	42068038	42068180
refGene_gn=IGFN1;refGene_tx=XM_023247442.1;refGene_gn=IGFN1;refGene_tx
=XM_023247443.1
F1	42068528	42068831
refGene_gn=IGFN1;refGene_tx=XM_023247442.1;refGene_gn=IGFN1;refGene_tx
=XM_023247443.1
F1	42069350	42069650
refGene_gn=IGFN1;refGene_tx=XM_023247442.1;refGene_gn=IGFN1;refGene_tx
=XM_023247443.1
F1	42070124	42070298
refGene_gn=IGFN1;refGene_tx=XM_023247442.1;refGene_gn=IGFN1;refGene_tx
=XM_023247443.1
F1	42071328	42071457

refGene_gn=IGFN1;refGene_tx=XM_023247442.1;refGene_gn=IGFN1;refGene_tx=XM_023247443.1
F1 42071734 42071778 refGene_gn=IGFN1;refGene_tx=XM_023247442.1
F1 42073506 42073809 refGene_gn=IGFN1;refGene_tx=XM_023247443.1
F1 42074794 42074883 refGene_gn=IGFN1;refGene_tx=XM_023247443.1
F1 42076613 42076809 refGene_gn=IGFN1;refGene_tx=XM_023247443.1
F1 42077537 42077837 refGene_gn=IGFN1;refGene_tx=XM_023247443.1
F1 42078601 42078928 refGene_gn=IGFN1;refGene_tx=XM_023247443.1
F1 42079822 42079836 refGene_gn=IGFN1;refGene_tx=XM_023247443.1
F1 42122029 42122231 refGene_gn=PKP1;refGene_tx=XM_019821959.2
F1 42132741 42132848 refGene_gn=PKP1;refGene_tx=XM_019821959.2
F1 42155063 42155458 refGene_gn=PKP1;refGene_tx=XM_019821959.2
F1 42158104 42158249 refGene_gn=PKP1;refGene_tx=XM_019821959.2
F1 42159219 42159427 refGene_gn=PKP1;refGene_tx=XM_019821959.2
F1 42160204 42160382 refGene_gn=PKP1;refGene_tx=XM_019821959.2
F1 42161794 42161909 refGene_gn=PKP1;refGene_tx=XM_019821959.2
F1 42163829 42163985 refGene_gn=PKP1;refGene_tx=XM_019821959.2
F1 42164842 42165019 refGene_gn=PKP1;refGene_tx=XM_019821959.2
F1 42166311 42166465 refGene_gn=PKP1;refGene_tx=XM_019821959.2
F1 42166819 42167006 refGene_gn=PKP1;refGene_tx=XM_019821959.2
F1 42167608 42167693 refGene_gn=PKP1;refGene_tx=XM_019821959.2
F1 42170732 42170807 refGene_gn=PKP1;refGene_tx=XM_019821959.2
F1 42194973 42195019
refGene_gn=TNNT2;refGene_tx=NM_001009347.1;refGene_gn=TNNT2;refGene_tx=XM_006942793.4;refGene_gn=TNNT2;refGene_tx=XM_006942794.4;refGene_gn=TNNT2;refGene_tx=XM_006942796.4;refGene_gn=TNNT2;refGene_tx=XM_006942797.4;refGene_gn=TNNT2;refGene_tx=XM_019821319.2;refGene_gn=TNNT2;refGene_tx=XM_019821320.2;refGene_gn=TNNT2;refGene_tx=XM_019821321.2;refGene_gn=TNNT2;refGene_tx=XM_019821322.2;refGene_gn=TNNT2;refGene_tx=XM_019821323.2;refGene_gn=TNNT2;refGene_tx=XM_019821324.2;refGene_gn=TNNT2;refGene_tx=XM_019821325.2;refGene_gn=TNNT2;refGene_tx=XM_019821326.2
F1 42195351 42195392
refGene_gn=TNNT2;refGene_tx=NM_001009347.1;refGene_gn=TNNT2;refGene_tx=XM_006942793.4;refGene_gn=TNNT2;refGene_tx=XM_006942794.4;refGene_gn=TNNT2;refGene_tx=XM_006942796.4;refGene_gn=TNNT2;refGene_tx=XM_006942797.4;refGene_gn=TNNT2;refGene_tx=XM_019821319.2;refGene_gn=TNNT2;refGene_tx=XM_019821320.2;refGene_gn=TNNT2;refGene_tx=XM_019821321.2;refGene_gn=TNNT2;refGene_tx=XM_019821322.2;refGene_gn=TNNT2;refGene_tx=XM_019821323.2;refGene_gn=TNNT2;refGene_tx=XM_019821324.2;refGene_gn=TNNT2;refGene_tx=XM_019821325.2;refGene_gn=TNNT2;refGene_tx=XM_019821326.2
F1 42196928 42197019
refGene_gn=TNNT2;refGene_tx=NM_001009347.1;refGene_gn=TNNT2;refGene_tx=XM_006942793.4;refGene_gn=TNNT2;refGene_tx=XM_006942794.4;refGene_gn=TNNT2;refGene_tx=XM_006942796.4;refGene_gn=TNNT2;refGene_tx=XM_006942797.4;refGene_gn=TNNT2;refGene_tx=XM_019821319.2;refGene_gn=TNNT2;refGene_tx=XM_019821320.2;refGene_gn=TNNT2;refGene_tx=XM_019821321.2;refGene_gn=TNNT2;refGene_tx=XM_019821322.2;refGene_gn=TNNT2;refGene_tx=XM_019821323.2;refGene_gn=TNNT2;refGene_tx=XM_019821324.2;refGene_gn=TNNT2;refGene_tx=XM_019821325.2;refGene_gn=TNNT2;refGene_tx=XM_019821326.2
F1 42197530 42197640

refGene_gn=TNNT2;refGene_tx=NM_001009347.1;refGene_gn=TNNT2;refGene_tx=XM_006942793.4;refGene_gn=TNNT2;refGene_tx=XM_006942794.4;refGene_gn=TNNT2;refGene_tx=XM_006942796.4;refGene_gn=TNNT2;refGene_tx=XM_006942797.4;refGene_gn=TNNT2;refGene_tx=XM_019821319.2;refGene_gn=TNNT2;refGene_tx=XM_019821320.2;refGene_gn=TNNT2;refGene_tx=XM_019821321.2;refGene_gn=TNNT2;refGene_tx=XM_019821322.2;refGene_gn=TNNT2;refGene_tx=XM_019821323.2;refGene_gn=TNNT2;refGene_tx=XM_019821324.2;refGene_gn=TNNT2;refGene_tx=XM_019821325.2;refGene_gn=TNNT2;refGene_tx=XM_019821326.2
F1 42198003 42198009
refGene_gn=TNNT2;refGene_tx=XM_006942793.4;refGene_gn=TNNT2;refGene_tx=XM_006942794.4;refGene_gn=TNNT2;refGene_tx=XM_006942796.4;refGene_gn=TNNT2;refGene_tx=XM_006942797.4;refGene_gn=TNNT2;refGene_tx=XM_019821319.2;refGene_gn=TNNT2;refGene_tx=XM_019821320.2;refGene_gn=TNNT2;refGene_tx=XM_019821322.2;refGene_gn=TNNT2;refGene_tx=XM_019821323.2;refGene_gn=TNNT2;refGene_tx=XM_019821324.2;refGene_gn=TNNT2;refGene_tx=XM_019821326.2
F1 42198859 42198970
refGene_gn=TNNT2;refGene_tx=NM_001009347.1;refGene_gn=TNNT2;refGene_tx=XM_006942793.4;refGene_gn=TNNT2;refGene_tx=XM_006942794.4;refGene_gn=TNNT2;refGene_tx=XM_006942796.4;refGene_gn=TNNT2;refGene_tx=XM_006942797.4;refGene_gn=TNNT2;refGene_tx=XM_019821319.2;refGene_gn=TNNT2;refGene_tx=XM_019821320.2;refGene_gn=TNNT2;refGene_tx=XM_019821321.2;refGene_gn=TNNT2;refGene_tx=XM_019821322.2;refGene_gn=TNNT2;refGene_tx=XM_019821323.2;refGene_gn=TNNT2;refGene_tx=XM_019821324.2;refGene_gn=TNNT2;refGene_tx=XM_019821325.2;refGene_gn=TNNT2;refGene_tx=XM_019821326.2
F1 42200035 42200113
refGene_gn=TNNT2;refGene_tx=NM_001009347.1;refGene_gn=TNNT2;refGene_tx=XM_006942793.4;refGene_gn=TNNT2;refGene_tx=XM_006942794.4;refGene_gn=TNNT2;refGene_tx=XM_006942796.4;refGene_gn=TNNT2;refGene_tx=XM_006942797.4;refGene_gn=TNNT2;refGene_tx=XM_019821319.2;refGene_gn=TNNT2;refGene_tx=XM_019821320.2;refGene_gn=TNNT2;refGene_tx=XM_019821321.2;refGene_gn=TNNT2;refGene_tx=XM_019821322.2;refGene_gn=TNNT2;refGene_tx=XM_019821323.2;refGene_gn=TNNT2;refGene_tx=XM_019821324.2;refGene_gn=TNNT2;refGene_tx=XM_019821325.2;refGene_gn=TNNT2;refGene_tx=XM_019821326.2
F1 42201007 42201124
refGene_gn=TNNT2;refGene_tx=NM_001009347.1;refGene_gn=TNNT2;refGene_tx=XM_006942793.4;refGene_gn=TNNT2;refGene_tx=XM_006942794.4;refGene_gn=TNNT2;refGene_tx=XM_006942796.4;refGene_gn=TNNT2;refGene_tx=XM_006942797.4;refGene_gn=TNNT2;refGene_tx=XM_019821319.2;refGene_gn=TNNT2;refGene_tx=XM_019821320.2;refGene_gn=TNNT2;refGene_tx=XM_019821321.2;refGene_gn=TNNT2;refGene_tx=XM_019821322.2;refGene_gn=TNNT2;refGene_tx=XM_019821323.2;refGene_gn=TNNT2;refGene_tx=XM_019821324.2;refGene_gn=TNNT2;refGene_tx=XM_019821325.2;refGene_gn=TNNT2;refGene_tx=XM_019821326.2
F1 42201392 42201453
refGene_gn=TNNT2;refGene_tx=NM_001009347.1;refGene_gn=TNNT2;refGene_tx=XM_006942793.4;refGene_gn=TNNT2;refGene_tx=XM_006942794.4;refGene_gn=TNNT2;refGene_tx=XM_006942796.4;refGene_gn=TNNT2;refGene_tx=XM_006942797.4;refGene_gn=TNNT2;refGene_tx=XM_019821319.2;refGene_gn=TNNT2;refGene_tx=XM_019821320.2;refGene_gn=TNNT2;refGene_tx=XM_019821321.2;refGene_gn=TNNT2;refGene_tx=XM_019821322.2;refGene_gn=TNNT2;refGene_tx=XM_01

9821323.2;refGene_gn=TNNT2;refGene_tx=XM_019821324.2;refGene_gn=TNNT2;
refGene_tx=XM_019821325.2;refGene_gn=TNNT2;refGene_tx=XM_019821326.2
F1 42202573 42202607
refGene_gn=TNNT2;refGene_tx=NM_001009347.1;refGene_gn=TNNT2;refGene_tx
=XM_006942793.4;refGene_gn=TNNT2;refGene_tx=XM_006942794.4;refGene_gn=
TNNT2;refGene_tx=XM_006942796.4;refGene_gn=TNNT2;refGene_tx=XM_0069427
97.4;refGene_gn=TNNT2;refGene_tx=XM_019821319.2;refGene_gn=TNNT2;refGe
ne_tx=XM_019821320.2;refGene_gn=TNNT2;refGene_tx=XM_019821321.2;refGen
e_gn=TNNT2;refGene_tx=XM_019821322.2;refGene_gn=TNNT2;refGene_tx=XM_01
9821323.2;refGene_gn=TNNT2;refGene_tx=XM_019821324.2;refGene_gn=TNNT2;
refGene_tx=XM_019821325.2;refGene_gn=TNNT2;refGene_tx=XM_019821326.2
F1 42203475 42203511
refGene_gn=TNNT2;refGene_tx=NM_001009347.1;refGene_gn=TNNT2;refGene_tx
=XM_006942793.4;refGene_gn=TNNT2;refGene_tx=XM_006942794.4;refGene_gn=
TNNT2;refGene_tx=XM_006942796.4;refGene_gn=TNNT2;refGene_tx=XM_0069427
97.4;refGene_gn=TNNT2;refGene_tx=XM_019821319.2;refGene_gn=TNNT2;refGe
ne_tx=XM_019821320.2;refGene_gn=TNNT2;refGene_tx=XM_019821321.2;refGen
e_gn=TNNT2;refGene_tx=XM_019821322.2;refGene_gn=TNNT2;refGene_tx=XM_01
9821323.2;refGene_gn=TNNT2;refGene_tx=XM_019821324.2;refGene_gn=TNNT2;
refGene_tx=XM_019821325.2;refGene_gn=TNNT2;refGene_tx=XM_019821326.2
F1 42203878 42203944
refGene_gn=TNNT2;refGene_tx=NM_001009347.1;refGene_gn=TNNT2;refGene_tx
=XM_006942793.4;refGene_gn=TNNT2;refGene_tx=XM_006942794.4;refGene_gn=
TNNT2;refGene_tx=XM_006942796.4;refGene_gn=TNNT2;refGene_tx=XM_0069427
97.4;refGene_gn=TNNT2;refGene_tx=XM_019821319.2;refGene_gn=TNNT2;refGe
ne_tx=XM_019821320.2;refGene_gn=TNNT2;refGene_tx=XM_019821321.2;refGen
e_gn=TNNT2;refGene_tx=XM_019821322.2;refGene_gn=TNNT2;refGene_tx=XM_01
9821323.2;refGene_gn=TNNT2;refGene_tx=XM_019821324.2;refGene_gn=TNNT2;
refGene_tx=XM_019821325.2;refGene_gn=TNNT2;refGene_tx=XM_019821326.2
F1 42205885 42205915
refGene_gn=TNNT2;refGene_tx=XM_006942793.4;refGene_gn=TNNT2;refGene_tx
=XM_006942794.4;refGene_gn=TNNT2;refGene_tx=XM_019821319.2;refGene_gn=
TNNT2;refGene_tx=XM_019821320.2;refGene_gn=TNNT2;refGene_tx=XM_0198213
21.2;refGene_gn=TNNT2;refGene_tx=XM_019821322.2;refGene_gn=TNNT2;refGe
ne_tx=XM_019821323.2
F1 42208440 42208455
refGene_gn=TNNT2;refGene_tx=NM_001009347.1;refGene_gn=TNNT2;refGene_tx
=XM_006942793.4;refGene_gn=TNNT2;refGene_tx=XM_006942794.4;refGene_gn=
TNNT2;refGene_tx=XM_006942796.4;refGene_gn=TNNT2;refGene_tx=XM_0198213
19.2;refGene_gn=TNNT2;refGene_tx=XM_019821320.2;refGene_gn=TNNT2;refGe
ne_tx=XM_019821321.2;refGene_gn=TNNT2;refGene_tx=XM_019821324.2;refGen
e_gn=TNNT2;refGene_tx=XM_019821325.2
F1 42208559 42208570
refGene_gn=TNNT2;refGene_tx=NM_001009347.1;refGene_gn=TNNT2;refGene_tx
=XM_006942793.4;refGene_gn=TNNT2;refGene_tx=XM_006942794.4;refGene_gn=
TNNT2;refGene_tx=XM_006942796.4;refGene_gn=TNNT2;refGene_tx=XM_0069427
97.4;refGene_gn=TNNT2;refGene_tx=XM_019821319.2;refGene_gn=TNNT2;refGe
ne_tx=XM_019821320.2;refGene_gn=TNNT2;refGene_tx=XM_019821321.2;refGen
e_gn=TNNT2;refGene_tx=XM_019821322.2;refGene_gn=TNNT2;refGene_tx=XM_01
9821323.2;refGene_gn=TNNT2;refGene_tx=XM_019821324.2;refGene_gn=TNNT2;

refGene_tx=XM_019821325.2;refGene_gn=TNNT2;refGene_tx=XM_019821326.2
F1	42209489	42209527
refGene_gn=TNNT2;refGene_tx=NM_001009347.1;refGene_gn=TNNT2;refGene_tx=XM_006942793.4;refGene_gn=TNNT2;refGene_tx=XM_006942794.4;refGene_gn=TNNT2;refGene_tx=XM_006942796.4;refGene_gn=TNNT2;refGene_tx=XM_006942797.4;refGene_gn=TNNT2;refGene_tx=XM_019821319.2;refGene_gn=TNNT2;refGene_tx=XM_019821320.2;refGene_gn=TNNT2;refGene_tx=XM_019821321.2;refGene_gn=TNNT2;refGene_tx=XM_019821322.2;refGene_gn=TNNT2;refGene_tx=XM_019821323.2;refGene_gn=TNNT2;refGene_tx=XM_019821324.2;refGene_gn=TNNT2;refGene_tx=XM_019821325.2;refGene_gn=TNNT2;refGene_tx=XM_019821326.2
F1	42218542	42218548	refGene_gn=LAD1;refGene_tx=XM_023247261.1
F1	42218892	42218967	refGene_gn=LAD1;refGene_tx=XM_023247261.1
F1	42219240	42219327	refGene_gn=LAD1;refGene_tx=XM_023247261.1
F1	42219849	42219987	refGene_gn=LAD1;refGene_tx=XM_023247261.1
F1	42220090	42220163	refGene_gn=LAD1;refGene_tx=XM_023247261.1
F1	42221561	42221605	refGene_gn=LAD1;refGene_tx=XM_023247261.1
F1	42222430	42222535	refGene_gn=LAD1;refGene_tx=XM_023247261.1
F1	42223047	42223873	refGene_gn=LAD1;refGene_tx=XM_023247261.1
F1	42225163	42225301	refGene_gn=LAD1;refGene_tx=XM_023247261.1
F1	42234466	42234504	refGene_gn=LAD1;refGene_tx=XM_023247261.1
F1	42244983	42245091	refGene_gn=TNNI1;refGene_tx=XM_006942848.3
F1	42246470	42246647	refGene_gn=TNNI1;refGene_tx=XM_006942848.3
F1	42247862	42247952	refGene_gn=TNNI1;refGene_tx=XM_006942848.3
F1	42248924	42249056	refGene_gn=TNNI1;refGene_tx=XM_006942848.3
F1	42249715	42249757	refGene_gn=TNNI1;refGene_tx=XM_006942848.3
F1	42252306	42252310	refGene_gn=TNNI1;refGene_tx=XM_006942848.3
F1	42252954	42252990	refGene_gn=TNNI1;refGene_tx=XM_006942848.3
F1	42256467	42256520	refGene_gn=TNNI1;refGene_tx=XM_006942848.3
F1	42262884	42262890	refGene_gn=TNNI1;refGene_tx=XM_006942848.3
F1	42293161	42293539	refGene_gn=PHLDA3;refGene_tx=XM_003999461.5
F1	42310423	42310500	refGene_gn=CSRP1;refGene_tx=XM_023247215.1
F1	42310834	42310928	refGene_gn=CSRP1;refGene_tx=XM_023247215.1
F1	42314182	42314312	refGene_gn=CSRP1;refGene_tx=XM_023247215.1
F1	42315489	42315658	refGene_gn=CSRP1;refGene_tx=XM_023247215.1
F1	42320915	42321027	refGene_gn=CSRP1;refGene_tx=XM_023247215.1
F1	42429349	42430053	refGene_gn=NAV1;refGene_tx=XM_023247662.1
F1	42436762	42436866	refGene_gn=NAV1;refGene_tx=XM_023247662.1
F1	42454861	42455656	refGene_gn=NAV1;refGene_tx=XM_023247662.1
F1	42509921	42510024	refGene_gn=NAV1;refGene_tx=XM_023247662.1
F1	42515181	42515365	refGene_gn=NAV1;refGene_tx=XM_023247662.1
F1	42515366	42515548	refGene_gn=NAV1;refGene_tx=XM_023247662.1
F1	42571261	42571400	refGene_gn=NAV1;refGene_tx=XM_023247662.1
F1	42571843	42572141	refGene_gn=NAV1;refGene_tx=XM_023247662.1
F1	42573488	42574182	refGene_gn=NAV1;refGene_tx=XM_023247662.1
F1	42574579	42575023	refGene_gn=NAV1;refGene_tx=XM_023247662.1
F1	42575621	42575663	refGene_gn=NAV1;refGene_tx=XM_023247662.1
F1	42576341	42576490	refGene_gn=NAV1;refGene_tx=XM_023247662.1
F1	42578334	42578505	refGene_gn=NAV1;refGene_tx=XM_023247662.1
F1	42579640	42579693	refGene_gn=NAV1;refGene_tx=XM_023247662.1
F1	42580477	42580501	refGene_gn=NAV1;refGene_tx=XM_023247662.1

F1 42580612 42580690 refGene_gn=NAV1;refGene_tx=XM_023247662.1
F1 42583602 42583686 refGene_gn=NAV1;refGene_tx=XM_023247662.1
F1 42583904 42584016 refGene_gn=NAV1;refGene_tx=XM_023247662.1
F1 42588450 42588572 refGene_gn=NAV1;refGene_tx=XM_023247662.1
F1 42589340 42589349 refGene_gn=NAV1;refGene_tx=XM_023247662.1
F1 42592276 42592473 refGene_gn=NAV1;refGene_tx=XM_023247662.1
F1 42592730 42592923 refGene_gn=NAV1;refGene_tx=XM_023247662.1
F1 42593015 42593184 refGene_gn=NAV1;refGene_tx=XM_023247662.1
F1 42593713 42593811 refGene_gn=NAV1;refGene_tx=XM_023247662.1
F1 42593979 42594075 refGene_gn=NAV1;refGene_tx=XM_023247662.1
F1 42594314 42594474 refGene_gn=NAV1;refGene_tx=XM_023247662.1
F1 42595157 42595393 refGene_gn=NAV1;refGene_tx=XM_023247662.1
F1 42596210 42596365 refGene_gn=NAV1;refGene_tx=XM_023247662.1
F1 42596507 42596579 refGene_gn=NAV1;refGene_tx=XM_023247662.1
F1 42597079 42597276 refGene_gn=NAV1;refGene_tx=XM_023247662.1
F1 42598016 42598135 refGene_gn=NAV1;refGene_tx=XM_023247662.1
F1 42600133 42600331 refGene_gn=NAV1;refGene_tx=XM_023247662.1
F1 42602229 42602325 refGene_gn=NAV1;refGene_tx=XM_023247662.1
F1 42611187 42611350 refGene_gn=IPO9;refGene_tx=XM_003999369.4
F1 42631060 42631122 refGene_gn=IPO9;refGene_tx=XM_003999369.4
F1 42631351 42631438 refGene_gn=IPO9;refGene_tx=XM_003999369.4
F1 42632209 42632411 refGene_gn=IPO9;refGene_tx=XM_003999369.4
F1 42636704 42636793 refGene_gn=IPO9;refGene_tx=XM_003999369.4
F1 42637601 42637688 refGene_gn=IPO9;refGene_tx=XM_003999369.4
F1 42638239 42638359 refGene_gn=IPO9;refGene_tx=XM_003999369.4
F1 42638473 42638574 refGene_gn=IPO9;refGene_tx=XM_003999369.4
F1 42638772 42638831 refGene_gn=IPO9;refGene_tx=XM_003999369.4
F1 42639412 42639564 refGene_gn=IPO9;refGene_tx=XM_003999369.4
F1 42640616 42640715 refGene_gn=IPO9;refGene_tx=XM_003999369.4
F1 42641606 42641713 refGene_gn=IPO9;refGene_tx=XM_003999369.4
F1 42642076 42642216 refGene_gn=IPO9;refGene_tx=XM_003999369.4
F1 42644641 42644801 refGene_gn=IPO9;refGene_tx=XM_003999369.4
F1 42655915 42656142 refGene_gn=IPO9;refGene_tx=XM_003999369.4
F1 42658406 42658555 refGene_gn=IPO9;refGene_tx=XM_003999369.4
F1 42659075 42659204 refGene_gn=IPO9;refGene_tx=XM_003999369.4
F1 42659913 42660189 refGene_gn=IPO9;refGene_tx=XM_003999369.4
F1 42660700 42660867 refGene_gn=IPO9;refGene_tx=XM_003999369.4
F1 42661483 42661617 refGene_gn=IPO9;refGene_tx=XM_003999369.4
F1 42662283 42662406 refGene_gn=IPO9;refGene_tx=XM_003999369.4
F1 42662842 42662947 refGene_gn=IPO9;refGene_tx=XM_003999369.4
F1 42663161 42663238 refGene_gn=IPO9;refGene_tx=XM_003999369.4
F1 42663683 42663794 refGene_gn=IPO9;refGene_tx=XM_003999369.4
F1 42674696 42674769 refGene_gn=SHISA4;refGene_tx=XM_023248026.1
F1 42675068 42675237 refGene_gn=SHISA4;refGene_tx=XM_023248026.1
F1 42676045 42676179 refGene_gn=SHISA4;refGene_tx=XM_023248026.1
F1 42677004 42677169 refGene_gn=SHISA4;refGene_tx=XM_023248026.1
F1 42677393 42677440 refGene_gn=SHISA4;refGene_tx=XM_023248026.1
F1 42680423 42680763
refGene_gn=LOC109495696;refGene_tx=XM_019821964.2;refGene_gn=LOC109495
696;refGene_tx=XM_023248027.1

F1 42683436 42683463 refGene_gn=LMOD1;refGene_tx=XM_006942858.4
F1 42684465 42685929 refGene_gn=LMOD1;refGene_tx=XM_006942858.4
F1 42699414 42699551
refGene_gn=LOC109495696;refGene_tx=XM_023248027.1
F1 42720863 42720984
refGene_gn=LOC109495696;refGene_tx=XM_019821964.2
F1 42723776 42723842
refGene_gn=LMOD1;refGene_tx=XM_006942858.4;refGene_gn=LOC109495696;refGene_tx=XM_019821964.2
F1 42723843 42724038 refGene_gn=LMOD1;refGene_tx=XM_006942858.4
F1 42732683 42732709 refGene_gn=TIMM17A;refGene_tx=XM_003999370.4
F1 42737175 42737275 refGene_gn=TIMM17A;refGene_tx=XM_003999370.4
F1 42737393 42737457 refGene_gn=TIMM17A;refGene_tx=XM_003999370.4
F1 42742586 42742715 refGene_gn=TIMM17A;refGene_tx=XM_003999370.4
F1 42743508 42743619 refGene_gn=TIMM17A;refGene_tx=XM_003999370.4
F1 42746400 42746486 refGene_gn=TIMM17A;refGene_tx=XM_003999370.4
F1 42760874 42761321 refGene_gn=RNPEP;refGene_tx=XM_011290946.3
F1 42765422 42765563 refGene_gn=RNPEP;refGene_tx=XM_011290946.3
F1 42765879 42766028 refGene_gn=RNPEP;refGene_tx=XM_011290946.3
F1 42770506 42770623 refGene_gn=RNPEP;refGene_tx=XM_011290946.3
F1 42771320 42771556 refGene_gn=RNPEP;refGene_tx=XM_011290946.3
F1 42774171 42774285 refGene_gn=RNPEP;refGene_tx=XM_011290946.3
F1 42775172 42775285 refGene_gn=RNPEP;refGene_tx=XM_011290946.3
F1 42775464 42775573 refGene_gn=RNPEP;refGene_tx=XM_011290946.3
F1 42776709 42776934 refGene_gn=RNPEP;refGene_tx=XM_011290946.3
F1 42777742 42777885 refGene_gn=RNPEP;refGene_tx=XM_011290946.3
F1 42778880 42779039 refGene_gn=RNPEP;refGene_tx=XM_011290946.3
F1 42783486 42783649 refGene_gn=ELF3;refGene_tx=XM_003999371.5
F1 42784000 42784222 refGene_gn=ELF3;refGene_tx=XM_003999371.5
F1 42784433 42784520 refGene_gn=ELF3;refGene_tx=XM_003999371.5
F1 42784718 42784841 refGene_gn=ELF3;refGene_tx=XM_003999371.5
F1 42785038 42785128 refGene_gn=ELF3;refGene_tx=XM_003999371.5
F1 42785298 42785415 refGene_gn=ELF3;refGene_tx=XM_003999371.5
F1 42786559 42786755 refGene_gn=ELF3;refGene_tx=XM_003999371.5
F1 42787442 42787557 refGene_gn=ELF3;refGene_tx=XM_003999371.5
F1 42865746 42866376 refGene_gn=GPR37L1;refGene_tx=XM_003999372.5
F1 42869142 42869955 refGene_gn=GPR37L1;refGene_tx=XM_003999372.5
F1 42875190 42875418
refGene_gn=ARL8A;refGene_tx=XM_019821967.2;refGene_gn=ARL8A;refGene_tx=XM_023247429.1;refGene_gn=ARL8A;refGene_tx=XM_023247430.1
F1 42875963 42876057
refGene_gn=ARL8A;refGene_tx=XM_019821967.2;refGene_gn=ARL8A;refGene_tx=XM_023247429.1;refGene_gn=ARL8A;refGene_tx=XM_023247430.1
F1 42876271 42876339
refGene_gn=ARL8A;refGene_tx=XM_019821967.2;refGene_gn=ARL8A;refGene_tx=XM_023247429.1;refGene_gn=ARL8A;refGene_tx=XM_023247430.1
F1 42876503 42876633
refGene_gn=ARL8A;refGene_tx=XM_019821967.2;refGene_gn=ARL8A;refGene_tx=XM_023247429.1;refGene_gn=ARL8A;refGene_tx=XM_023247430.1
F1 42876705 42876788 refGene_gn=ARL8A;refGene_tx=XM_019821967.2

F1 42878081 42878155
refGene_gn=ARL8A;refGene_tx=XM_023247429.1;refGene_gn=ARL8A;refGene_tx
=XM_023247430.1
F1 42878524 42878605
refGene_gn=ARL8A;refGene_tx=XM_023247429.1;refGene_gn=ARL8A;refGene_tx
=XM_023247430.1
F1 42883111 42883234
refGene_gn=ARL8A;refGene_tx=XM_023247429.1;refGene_gn=ARL8A;refGene_tx
=XM_023247430.1
F1 42886294 42886388 refGene_gn=PTPN7;refGene_tx=XM_019821968.2
F1 42887304 42887418 refGene_gn=PTPN7;refGene_tx=XM_019821968.2
F1 42889309 42889467 refGene_gn=PTPN7;refGene_tx=XM_019821968.2
F1 42890182 42890293 refGene_gn=PTPN7;refGene_tx=XM_019821968.2
F1 42890673 42890811 refGene_gn=PTPN7;refGene_tx=XM_019821968.2
F1 42891978 42892055 refGene_gn=PTPN7;refGene_tx=XM_019821968.2
F1 42893744 42893829 refGene_gn=PTPN7;refGene_tx=XM_019821968.2
F1 42894127 42894311 refGene_gn=PTPN7;refGene_tx=XM_019821968.2
F1 42894850 42894979 refGene_gn=PTPN7;refGene_tx=XM_019821968.2
F1 42894980 42895041 refGene_gn=PTPN7;refGene_tx=XM_019821968.2
F1 42896854 42896987 refGene_gn=PTPN7;refGene_tx=XM_019821968.2
F1 42896989 42897052 refGene_gn=PTPN7;refGene_tx=XM_019821968.2
F1 42902449 42902506
refGene_gn=LOC101101103;refGene_tx=XM_006942864.4;refGene_gn=LOC101101
103;refGene_tx=XM_019821969.2;refGene_gn=LOC101101103;refGene_tx=XM_01
9821972.2;refGene_gn=LOC101101103;refGene_tx=XM_019821973.2;refGene_gn
=LOC101101103;refGene_tx=XM_019821974.2;refGene_gn=LOC101101103;refGen
e_tx=XM_019821975.2
F1 42902855 42902904
refGene_gn=LOC101101103;refGene_tx=XM_006942864.4;refGene_gn=LOC101101
103;refGene_tx=XM_019821969.2;refGene_gn=LOC101101103;refGene_tx=XM_01
9821972.2;refGene_gn=LOC101101103;refGene_tx=XM_019821973.2;refGene_gn
=LOC101101103;refGene_tx=XM_019821974.2;refGene_gn=LOC101101103;refGen
e_tx=XM_019821975.2
F1 42903082 42903355
refGene_gn=LOC101101103;refGene_tx=XM_006942864.4;refGene_gn=LOC101101
103;refGene_tx=XM_019821969.2;refGene_gn=LOC101101103;refGene_tx=XM_01
9821972.2;refGene_gn=LOC101101103;refGene_tx=XM_019821973.2;refGene_gn
=LOC101101103;refGene_tx=XM_019821974.2;refGene_gn=LOC101101103;refGen
e_tx=XM_019821975.2
F1 42903548 42903818
refGene_gn=LOC101101103;refGene_tx=XM_006942864.4;refGene_gn=LOC101101
103;refGene_tx=XM_019821969.2;refGene_gn=LOC101101103;refGene_tx=XM_01
9821972.2;refGene_gn=LOC101101103;refGene_tx=XM_019821973.2;refGene_gn
=LOC101101103;refGene_tx=XM_019821974.2;refGene_gn=LOC101101103;refGen
e_tx=XM_019821975.2
F1 42904013 42904277
refGene_gn=LOC101101103;refGene_tx=XM_006942864.4;refGene_gn=LOC101101
103;refGene_tx=XM_019821969.2;refGene_gn=LOC101101103;refGene_tx=XM_01
9821972.2;refGene_gn=LOC101101103;refGene_tx=XM_019821973.2;refGene_gn
=LOC101101103;refGene_tx=XM_019821974.2;refGene_gn=LOC101101103;refGen

e_tx=XM_019821975.2;refGene_gn=LOC101101103;refGene_tx=XM_023247492.1
F1 42905136 42905391
refGene_gn=LOC101101103;refGene_tx=XM_006942864.4;refGene_gn=LOC101101103;refGene_tx=XM_019821969.2;refGene_gn=LOC101101103;refGene_tx=XM_019821972.2;refGene_gn=LOC101101103;refGene_tx=XM_019821973.2;refGene_gn=LOC101101103;refGene_tx=XM_019821974.2;refGene_gn=LOC101101103;refGene_tx=XM_019821975.2;refGene_gn=LOC101101103;refGene_tx=XM_023247492.1
F1 42905492 42905738
refGene_gn=LOC101101103;refGene_tx=XM_006942864.4;refGene_gn=LOC101101103;refGene_tx=XM_019821969.2;refGene_gn=LOC101101103;refGene_tx=XM_019821972.2;refGene_gn=LOC101101103;refGene_tx=XM_019821973.2;refGene_gn=LOC101101103;refGene_tx=XM_019821974.2;refGene_gn=LOC101101103;refGene_tx=XM_019821975.2;refGene_gn=LOC101101103;refGene_tx=XM_023247492.1
F1 42905905 42906181
refGene_gn=LOC101101103;refGene_tx=XM_006942864.4;refGene_gn=LOC101101103;refGene_tx=XM_019821969.2;refGene_gn=LOC101101103;refGene_tx=XM_019821972.2;refGene_gn=LOC101101103;refGene_tx=XM_019821973.2;refGene_gn=LOC101101103;refGene_tx=XM_019821974.2;refGene_gn=LOC101101103;refGene_tx=XM_019821975.2;refGene_gn=LOC101101103;refGene_tx=XM_023247492.1
F1 42907181 42907454
refGene_gn=LOC101101103;refGene_tx=XM_006942864.4;refGene_gn=LOC101101103;refGene_tx=XM_019821969.2;refGene_gn=LOC101101103;refGene_tx=XM_019821972.2;refGene_gn=LOC101101103;refGene_tx=XM_019821973.2;refGene_gn=LOC101101103;refGene_tx=XM_019821974.2;refGene_gn=LOC101101103;refGene_tx=XM_019821975.2;refGene_gn=LOC101101103;refGene_tx=XM_023247492.1
F1 42909276 42909540
refGene_gn=LOC101101103;refGene_tx=XM_006942864.4;refGene_gn=LOC101101103;refGene_tx=XM_019821969.2;refGene_gn=LOC101101103;refGene_tx=XM_019821972.2;refGene_gn=LOC101101103;refGene_tx=XM_019821973.2;refGene_gn=LOC101101103;refGene_tx=XM_019821974.2;refGene_gn=LOC101101103;refGene_tx=XM_019821975.2;refGene_gn=LOC101101103;refGene_tx=XM_023247492.1
F1 42909738 42910005
refGene_gn=LOC101101103;refGene_tx=XM_006942864.4;refGene_gn=LOC101101103;refGene_tx=XM_019821969.2;refGene_gn=LOC101101103;refGene_tx=XM_019821972.2;refGene_gn=LOC101101103;refGene_tx=XM_019821973.2;refGene_gn=LOC101101103;refGene_tx=XM_019821974.2;refGene_gn=LOC101101103;refGene_tx=XM_019821975.2;refGene_gn=LOC101101103;refGene_tx=XM_023247492.1
F1 42910324 42910606
refGene_gn=LOC101101103;refGene_tx=XM_006942864.4;refGene_gn=LOC101101103;refGene_tx=XM_019821969.2;refGene_gn=LOC101101103;refGene_tx=XM_019821972.2;refGene_gn=LOC101101103;refGene_tx=XM_019821973.2;refGene_gn=LOC101101103;refGene_tx=XM_019821974.2;refGene_gn=LOC101101103;refGene_tx=XM_019821975.2;refGene_gn=LOC101101103;refGene_tx=XM_023247492.1
F1 42910938 42911079
refGene_gn=LOC101101103;refGene_tx=XM_006942864.4;refGene_gn=LOC101101103;refGene_tx=XM_019821969.2;refGene_gn=LOC101101103;refGene_tx=XM_019821972.2;refGene_gn=LOC101101103;refGene_tx=XM_019821973.2;refGene_gn=LOC101101103;refGene_tx=XM_019821974.2;refGene_gn=LOC101101103;refGene_tx=XM_019821975.2;refGene_gn=LOC101101103;refGene_tx=XM_023247492.1
F1 42911360 42911571

refGene_gn=LOC101101103;refGene_tx=XM_006942864.4;refGene_gn=LOC101101103;refGene_tx=XM_019821969.2;refGene_gn=LOC101101103;refGene_tx=XM_019821972.2;refGene_gn=LOC101101103;refGene_tx=XM_019821973.2;refGene_gn=LOC101101103;refGene_tx=XM_019821974.2;refGene_gn=LOC101101103;refGene_tx=XM_019821975.2;refGene_gn=LOC101101103;refGene_tx=XM_023247492.1
F1 42912038 42912106
refGene_gn=LOC101101103;refGene_tx=XM_006942864.4;refGene_gn=LOC101101103;refGene_tx=XM_019821969.2;refGene_gn=LOC101101103;refGene_tx=XM_019821972.2;refGene_gn=LOC101101103;refGene_tx=XM_019821973.2;refGene_gn=LOC101101103;refGene_tx=XM_019821974.2;refGene_gn=LOC101101103;refGene_tx=XM_019821975.2;refGene_gn=LOC101101103;refGene_tx=XM_023247492.1
F1 42913063 42913190
refGene_gn=LOC101101103;refGene_tx=XM_006942864.4;refGene_gn=LOC101101103;refGene_tx=XM_019821969.2;refGene_gn=LOC101101103;refGene_tx=XM_019821972.2;refGene_gn=LOC101101103;refGene_tx=XM_019821973.2;refGene_gn=LOC101101103;refGene_tx=XM_019821974.2;refGene_gn=LOC101101103;refGene_tx=XM_019821975.2;refGene_gn=LOC101101103;refGene_tx=XM_023247492.1
F1 42913309 42913357
refGene_gn=LOC101101103;refGene_tx=XM_006942864.4;refGene_gn=LOC101101103;refGene_tx=XM_019821969.2;refGene_gn=LOC101101103;refGene_tx=XM_019821972.2;refGene_gn=LOC101101103;refGene_tx=XM_019821973.2;refGene_gn=LOC101101103;refGene_tx=XM_019821974.2;refGene_gn=LOC101101103;refGene_tx=XM_019821975.2;refGene_gn=LOC101101103;refGene_tx=XM_023247492.1
F1 42913984 42914078
refGene_gn=LOC101101103;refGene_tx=XM_006942864.4;refGene_gn=LOC101101103;refGene_tx=XM_019821969.2;refGene_gn=LOC101101103;refGene_tx=XM_019821972.2;refGene_gn=LOC101101103;refGene_tx=XM_019821973.2;refGene_gn=LOC101101103;refGene_tx=XM_019821974.2;refGene_gn=LOC101101103;refGene_tx=XM_019821975.2;refGene_gn=LOC101101103;refGene_tx=XM_023247492.1
F1 42914258 42914349
refGene_gn=LOC101101103;refGene_tx=XM_006942864.4;refGene_gn=LOC101101103;refGene_tx=XM_019821969.2;refGene_gn=LOC101101103;refGene_tx=XM_019821972.2;refGene_gn=LOC101101103;refGene_tx=XM_019821973.2;refGene_gn=LOC101101103;refGene_tx=XM_019821974.2;refGene_gn=LOC101101103;refGene_tx=XM_019821975.2;refGene_gn=LOC101101103;refGene_tx=XM_023247492.1
F1 42914457 42914534
refGene_gn=LOC101101103;refGene_tx=XM_006942864.4;refGene_gn=LOC101101103;refGene_tx=XM_019821969.2;refGene_gn=LOC101101103;refGene_tx=XM_019821972.2;refGene_gn=LOC101101103;refGene_tx=XM_019821973.2;refGene_gn=LOC101101103;refGene_tx=XM_019821974.2;refGene_gn=LOC101101103;refGene_tx=XM_019821975.2;refGene_gn=LOC101101103;refGene_tx=XM_023247492.1
F1 42915438 42915573
refGene_gn=LOC101101103;refGene_tx=XM_006942864.4;refGene_gn=LOC101101103;refGene_tx=XM_019821969.2;refGene_gn=LOC101101103;refGene_tx=XM_019821972.2;refGene_gn=LOC101101103;refGene_tx=XM_019821973.2;refGene_gn=LOC101101103;refGene_tx=XM_019821974.2;refGene_gn=LOC101101103;refGene_tx=XM_019821975.2;refGene_gn=LOC101101103;refGene_tx=XM_023247492.1
F1 42916591 42916714
refGene_gn=LOC101101103;refGene_tx=XM_006942864.4;refGene_gn=LOC101101103;refGene_tx=XM_019821969.2;refGene_gn=LOC101101103;refGene_tx=XM_01

9821972.2;refGene_gn=LOC101101103;refGene_tx=XM_019821973.2;refGene_gn=LOC101101103;refGene_tx=XM_019821974.2;refGene_gn=LOC101101103;refGene_tx=XM_019821975.2;refGene_gn=LOC101101103;refGene_tx=XM_023247492.1
F1 42917279 42917440
refGene_gn=LOC101101103;refGene_tx=XM_006942864.4;refGene_gn=LOC101101103;refGene_tx=XM_019821969.2;refGene_gn=LOC101101103;refGene_tx=XM_019821972.2;refGene_gn=LOC101101103;refGene_tx=XM_019821973.2;refGene_gn=LOC101101103;refGene_tx=XM_019821974.2;refGene_gn=LOC101101103;refGene_tx=XM_019821975.2;refGene_gn=LOC101101103;refGene_tx=XM_023247492.1
F1 42917529 42917665
refGene_gn=LOC101101103;refGene_tx=XM_006942864.4;refGene_gn=LOC101101103;refGene_tx=XM_019821969.2;refGene_gn=LOC101101103;refGene_tx=XM_019821972.2;refGene_gn=LOC101101103;refGene_tx=XM_019821973.2;refGene_gn=LOC101101103;refGene_tx=XM_019821974.2;refGene_gn=LOC101101103;refGene_tx=XM_019821975.2;refGene_gn=LOC101101103;refGene_tx=XM_023247492.1
F1 42918040 42918108
refGene_gn=LOC101101103;refGene_tx=XM_006942864.4;refGene_gn=LOC101101103;refGene_tx=XM_019821969.2;refGene_gn=LOC101101103;refGene_tx=XM_019821972.2;refGene_gn=LOC101101103;refGene_tx=XM_019821973.2;refGene_gn=LOC101101103;refGene_tx=XM_019821974.2;refGene_gn=LOC101101103;refGene_tx=XM_019821975.2;refGene_gn=LOC101101103;refGene_tx=XM_023247492.1
F1 42918201 42918358
refGene_gn=LOC101101103;refGene_tx=XM_006942864.4;refGene_gn=LOC101101103;refGene_tx=XM_019821969.2;refGene_gn=LOC101101103;refGene_tx=XM_019821972.2;refGene_gn=LOC101101103;refGene_tx=XM_019821973.2;refGene_gn=LOC101101103;refGene_tx=XM_019821974.2;refGene_gn=LOC101101103;refGene_tx=XM_019821975.2;refGene_gn=LOC101101103;refGene_tx=XM_023247492.1
F1 42918474 42918553
refGene_gn=LOC101101103;refGene_tx=XM_006942864.4;refGene_gn=LOC101101103;refGene_tx=XM_019821969.2;refGene_gn=LOC101101103;refGene_tx=XM_019821972.2;refGene_gn=LOC101101103;refGene_tx=XM_019821973.2;refGene_gn=LOC101101103;refGene_tx=XM_019821974.2;refGene_gn=LOC101101103;refGene_tx=XM_019821975.2;refGene_gn=LOC101101103;refGene_tx=XM_023247492.1
F1 42918644 42918721
refGene_gn=LOC101101103;refGene_tx=XM_006942864.4;refGene_gn=LOC101101103;refGene_tx=XM_019821969.2;refGene_gn=LOC101101103;refGene_tx=XM_019821972.2;refGene_gn=LOC101101103;refGene_tx=XM_019821973.2;refGene_gn=LOC101101103;refGene_tx=XM_019821974.2;refGene_gn=LOC101101103;refGene_tx=XM_019821975.2;refGene_gn=LOC101101103;refGene_tx=XM_023247492.1
F1 42918859 42918988
refGene_gn=LOC101101103;refGene_tx=XM_006942864.4;refGene_gn=LOC101101103;refGene_tx=XM_019821969.2;refGene_gn=LOC101101103;refGene_tx=XM_019821972.2;refGene_gn=LOC101101103;refGene_tx=XM_019821973.2;refGene_gn=LOC101101103;refGene_tx=XM_019821974.2;refGene_gn=LOC101101103;refGene_tx=XM_023247492.1
F1 42919205 42919316
refGene_gn=LOC101101103;refGene_tx=XM_006942864.4;refGene_gn=LOC101101103;refGene_tx=XM_019821969.2;refGene_gn=LOC101101103;refGene_tx=XM_019821972.2;refGene_gn=LOC101101103;refGene_tx=XM_019821973.2;refGene_gn=LOC101101103;refGene_tx=XM_019821975.2;refGene_gn=LOC101101103;refGen

e_tx=XM_023247492.1
F1 42919422 42919583
refGene_gn=LOC101101103;refGene_tx=XM_006942864.4;refGene_gn=LOC101101
103;refGene_tx=XM_019821969.2;refGene_gn=LOC101101103;refGene_tx=XM_01
9821972.2;refGene_gn=LOC101101103;refGene_tx=XM_019821973.2;refGene_gn
=LOC101101103;refGene_tx=XM_019821974.2;refGene_gn=LOC101101103;refGen
e_tx=XM_019821975.2;refGene_gn=LOC101101103;refGene_tx=XM_023247492.1
F1 42919948 42920084
refGene_gn=LOC101101103;refGene_tx=XM_006942864.4;refGene_gn=LOC101101
103;refGene_tx=XM_019821969.2;refGene_gn=LOC101101103;refGene_tx=XM_01
9821972.2;refGene_gn=LOC101101103;refGene_tx=XM_019821973.2;refGene_gn
=LOC101101103;refGene_tx=XM_019821974.2;refGene_gn=LOC101101103;refGen
e_tx=XM_019821975.2;refGene_gn=LOC101101103;refGene_tx=XM_023247492.1
F1 42920307 42920367
refGene_gn=LOC101101103;refGene_tx=XM_006942864.4;refGene_gn=LOC101101
103;refGene_tx=XM_019821969.2;refGene_gn=LOC101101103;refGene_tx=XM_01
9821972.2;refGene_gn=LOC101101103;refGene_tx=XM_019821973.2;refGene_gn
=LOC101101103;refGene_tx=XM_019821974.2;refGene_gn=LOC101101103;refGen
e_tx=XM_019821975.2;refGene_gn=LOC101101103;refGene_tx=XM_023247492.1
F1 42923340 42923552
refGene_gn=LGR6;refGene_tx=XM_011290944.3;refGene_gn=LGR6;refGene_tx=X
M_023247376.1
F1 42940474 42940503 refGene_gn=LGR6;refGene_tx=XM_023247375.1
F1 42950561 42950633
refGene_gn=LGR6;refGene_tx=XM_011290944.3;refGene_gn=LGR6;refGene_tx=X
M_023247375.1;refGene_gn=LGR6;refGene_tx=XM_023247376.1;refGene_gn=LGR
6;refGene_tx=XM_023247377.1;refGene_gn=LGR6;refGene_tx=XM_023247378.1
F1 42954047 42954119
refGene_gn=LGR6;refGene_tx=XM_011290944.3;refGene_gn=LGR6;refGene_tx=X
M_023247375.1;refGene_gn=LGR6;refGene_tx=XM_023247376.1;refGene_gn=LGR
6;refGene_tx=XM_023247377.1;refGene_gn=LGR6;refGene_tx=XM_023247378.1
F1 42962151 42962223
refGene_gn=LGR6;refGene_tx=XM_011290944.3;refGene_gn=LGR6;refGene_tx=X
M_023247375.1;refGene_gn=LGR6;refGene_tx=XM_023247377.1;refGene_gn=LGR
6;refGene_tx=XM_023247378.1
F1 42999393 42999609
refGene_gn=LGR6;refGene_tx=XM_011290944.3;refGene_gn=LGR6;refGene_tx=X
M_023247375.1;refGene_gn=LGR6;refGene_tx=XM_023247376.1;refGene_gn=LGR
6;refGene_tx=XM_023247377.1;refGene_gn=LGR6;refGene_tx=XM_023247378.1
F1 43004729 43004801
refGene_gn=LGR6;refGene_tx=XM_011290944.3;refGene_gn=LGR6;refGene_tx=X
M_023247375.1;refGene_gn=LGR6;refGene_tx=XM_023247376.1;refGene_gn=LGR
6;refGene_tx=XM_023247377.1;refGene_gn=LGR6;refGene_tx=XM_023247378.1
F1 43022778 43022847
refGene_gn=LGR6;refGene_tx=XM_011290944.3;refGene_gn=LGR6;refGene_tx=X
M_023247375.1;refGene_gn=LGR6;refGene_tx=XM_023247376.1;refGene_gn=LGR
6;refGene_tx=XM_023247377.1;refGene_gn=LGR6;refGene_tx=XM_023247378.1
F1 43024885 43024957
refGene_gn=LGR6;refGene_tx=XM_011290944.3;refGene_gn=LGR6;refGene_tx=X
M_023247375.1;refGene_gn=LGR6;refGene_tx=XM_023247376.1;refGene_gn=LGR

6;refGene_tx=XM_023247377.1;refGene_gn=LGR6;refGene_tx=XM_023247378.1
F1 43025153 43025225
refGene_gn=LGR6;refGene_tx=XM_011290944.3;refGene_gn=LGR6;refGene_tx=XM_023247375.1;refGene_gn=LGR6;refGene_tx=XM_023247376.1;refGene_gn=LGR6;refGene_tx=XM_023247377.1;refGene_gn=LGR6;refGene_tx=XM_023247378.1
F1 43026526 43026595
refGene_gn=LGR6;refGene_tx=XM_011290944.3;refGene_gn=LGR6;refGene_tx=XM_023247375.1;refGene_gn=LGR6;refGene_tx=XM_023247376.1;refGene_gn=LGR6;refGene_tx=XM_023247377.1;refGene_gn=LGR6;refGene_tx=XM_023247378.1
F1 43027851 43027923
refGene_gn=LGR6;refGene_tx=XM_011290944.3;refGene_gn=LGR6;refGene_tx=XM_023247375.1;refGene_gn=LGR6;refGene_tx=XM_023247376.1;refGene_gn=LGR6;refGene_tx=XM_023247377.1;refGene_gn=LGR6;refGene_tx=XM_023247378.1
F1 43029083 43029149
refGene_gn=LGR6;refGene_tx=XM_011290944.3;refGene_gn=LGR6;refGene_tx=XM_023247375.1;refGene_gn=LGR6;refGene_tx=XM_023247376.1;refGene_gn=LGR6;refGene_tx=XM_023247377.1;refGene_gn=LGR6;refGene_tx=XM_023247378.1
F1 43030255 43030327
refGene_gn=LGR6;refGene_tx=XM_011290944.3;refGene_gn=LGR6;refGene_tx=XM_023247375.1;refGene_gn=LGR6;refGene_tx=XM_023247376.1;refGene_gn=LGR6;refGene_tx=XM_023247377.1;refGene_gn=LGR6;refGene_tx=XM_023247378.1
F1 43030712 43030784
refGene_gn=LGR6;refGene_tx=XM_011290944.3;refGene_gn=LGR6;refGene_tx=XM_023247375.1;refGene_gn=LGR6;refGene_tx=XM_023247376.1;refGene_gn=LGR6;refGene_tx=XM_023247377.1;refGene_gn=LGR6;refGene_tx=XM_023247378.1
F1 43032383 43032509
refGene_gn=LGR6;refGene_tx=XM_011290944.3;refGene_gn=LGR6;refGene_tx=XM_023247375.1;refGene_gn=LGR6;refGene_tx=XM_023247376.1;refGene_gn=LGR6;refGene_tx=XM_023247377.1;refGene_gn=LGR6;refGene_tx=XM_023247378.1
F1 43033512 43033673
refGene_gn=LGR6;refGene_tx=XM_011290944.3;refGene_gn=LGR6;refGene_tx=XM_023247375.1;refGene_gn=LGR6;refGene_tx=XM_023247376.1;refGene_gn=LGR6;refGene_tx=XM_023247377.1;refGene_gn=LGR6;refGene_tx=XM_023247378.1
F1 43037247 43037328
refGene_gn=LGR6;refGene_tx=XM_011290944.3;refGene_gn=LGR6;refGene_tx=XM_023247375.1;refGene_gn=LGR6;refGene_tx=XM_023247376.1;refGene_gn=LGR6;refGene_tx=XM_023247377.1;refGene_gn=LGR6;refGene_tx=XM_023247378.1
F1 43039858 43041114
refGene_gn=LGR6;refGene_tx=XM_011290944.3;refGene_gn=LGR6;refGene_tx=XM_023247375.1;refGene_gn=LGR6;refGene_tx=XM_023247376.1;refGene_gn=LGR6;refGene_tx=XM_023247377.1;refGene_gn=LGR6;refGene_tx=XM_023247378.1
F1 43053622 43053748
refGene_gn=UBE2T;refGene_tx=XM_006942866.3;refGene_gn=UBE2T;refGene_tx=XM_019821976.2;refGene_gn=UBE2T;refGene_tx=XM_019821977.2;refGene_gn=UBE2T;refGene_tx=XM_019821978.2
F1 43056066 43056150
refGene_gn=UBE2T;refGene_tx=XM_006942866.3;refGene_gn=UBE2T;refGene_tx=XM_019821976.2;refGene_gn=UBE2T;refGene_tx=XM_019821977.2;refGene_gn=UBE2T;refGene_tx=XM_019821978.2
F1 43056294 43056393

refGene_gn=UBE2T;refGene_tx=XM_006942866.3;refGene_gn=UBE2T;refGene_tx=XM_019821976.2;refGene_gn=UBE2T;refGene_tx=XM_019821977.2;refGene_gn=UBE2T;refGene_tx=XM_019821978.2
F1 43056511 43056617
refGene_gn=UBE2T;refGene_tx=XM_006942866.3;refGene_gn=UBE2T;refGene_tx=XM_019821976.2;refGene_gn=UBE2T;refGene_tx=XM_019821977.2;refGene_gn=UBE2T;refGene_tx=XM_019821978.2
F1 43057714 43057784
refGene_gn=UBE2T;refGene_tx=XM_006942866.3;refGene_gn=UBE2T;refGene_tx=XM_019821976.2;refGene_gn=UBE2T;refGene_tx=XM_019821977.2;refGene_gn=UBE2T;refGene_tx=XM_019821978.2
F1 43058361 43058470
refGene_gn=UBE2T;refGene_tx=XM_006942866.3;refGene_gn=UBE2T;refGene_tx=XM_019821976.2;refGene_gn=UBE2T;refGene_tx=XM_019821977.2;refGene_gn=UBE2T;refGene_tx=XM_019821978.2
F1 43067278 43067854
refGene_gn=PPP1R12B;refGene_tx=XM_003999376.4;refGene_gn=PPP1R12B;refGene_tx=XM_011290936.3;refGene_gn=PPP1R12B;refGene_tx=XM_019821983.2;refGene_gn=PPP1R12B;refGene_tx=XM_019821984.2;refGene_gn=PPP1R12B;refGene_tx=XM_019821985.2;refGene_gn=PPP1R12B;refGene_tx=XM_019821986.2;refGene_gn=PPP1R12B;refGene_tx=XM_019821987.2
F1 43124602 43124733
refGene_gn=PPP1R12B;refGene_tx=XM_003999376.4;refGene_gn=PPP1R12B;refGene_tx=XM_011290936.3;refGene_gn=PPP1R12B;refGene_tx=XM_019821983.2;refGene_gn=PPP1R12B;refGene_tx=XM_019821984.2;refGene_gn=PPP1R12B;refGene_tx=XM_019821985.2;refGene_gn=PPP1R12B;refGene_tx=XM_019821986.2;refGene_gn=PPP1R12B;refGene_tx=XM_019821987.2
F1 43130271 43130390
refGene_gn=PPP1R12B;refGene_tx=XM_003999376.4;refGene_gn=PPP1R12B;refGene_tx=XM_011290936.3;refGene_gn=PPP1R12B;refGene_tx=XM_019821983.2;refGene_gn=PPP1R12B;refGene_tx=XM_019821984.2;refGene_gn=PPP1R12B;refGene_tx=XM_019821985.2;refGene_gn=PPP1R12B;refGene_tx=XM_019821986.2;refGene_gn=PPP1R12B;refGene_tx=XM_019821987.2
F1 43136782 43136942
refGene_gn=PPP1R12B;refGene_tx=XM_003999376.4;refGene_gn=PPP1R12B;refGene_tx=XM_011290936.3;refGene_gn=PPP1R12B;refGene_tx=XM_019821983.2;refGene_gn=PPP1R12B;refGene_tx=XM_019821984.2;refGene_gn=PPP1R12B;refGene_tx=XM_019821985.2;refGene_gn=PPP1R12B;refGene_tx=XM_019821986.2;refGene_gn=PPP1R12B;refGene_tx=XM_019821987.2
F1 43138220 43138365
refGene_gn=PPP1R12B;refGene_tx=XM_003999376.4;refGene_gn=PPP1R12B;refGene_tx=XM_011290936.3;refGene_gn=PPP1R12B;refGene_tx=XM_019821983.2;refGene_gn=PPP1R12B;refGene_tx=XM_019821984.2;refGene_gn=PPP1R12B;refGene_tx=XM_019821985.2;refGene_gn=PPP1R12B;refGene_tx=XM_019821986.2;refGene_gn=PPP1R12B;refGene_tx=XM_019821987.2
F1 43139419 43139494
refGene_gn=PPP1R12B;refGene_tx=XM_003999376.4;refGene_gn=PPP1R12B;refGene_tx=XM_011290936.3;refGene_gn=PPP1R12B;refGene_tx=XM_019821983.2;refGene_gn=PPP1R12B;refGene_tx=XM_019821984.2;refGene_gn=PPP1R12B;refGene_tx=XM_019821985.2;refGene_gn=PPP1R12B;refGene_tx=XM_019821986.2;refG

ene_gn=PPP1R12B;refGene_tx=XM_019821987.2
F1 43142125 43142205
refGene_gn=PPP1R12B;refGene_tx=XM_003999376.4;refGene_gn=PPP1R12B;refGene_tx=XM_011290936.3;refGene_gn=PPP1R12B;refGene_tx=XM_019821983.2;refGene_gn=PPP1R12B;refGene_tx=XM_019821984.2;refGene_gn=PPP1R12B;refGene_tx=XM_019821985.2;refGene_gn=PPP1R12B;refGene_tx=XM_019821986.2;refGene_gn=PPP1R12B;refGene_tx=XM_019821987.2
F1 43142859 43142999
refGene_gn=PPP1R12B;refGene_tx=XM_003999376.4;refGene_gn=PPP1R12B;refGene_tx=XM_011290936.3;refGene_gn=PPP1R12B;refGene_tx=XM_019821983.2;refGene_gn=PPP1R12B;refGene_tx=XM_019821984.2;refGene_gn=PPP1R12B;refGene_tx=XM_019821985.2;refGene_gn=PPP1R12B;refGene_tx=XM_019821986.2;refGene_gn=PPP1R12B;refGene_tx=XM_019821987.2
F1 43149158 43149271
refGene_gn=PPP1R12B;refGene_tx=XM_003999376.4;refGene_gn=PPP1R12B;refGene_tx=XM_011290936.3;refGene_gn=PPP1R12B;refGene_tx=XM_019821983.2;refGene_gn=PPP1R12B;refGene_tx=XM_019821984.2;refGene_gn=PPP1R12B;refGene_tx=XM_019821985.2;refGene_gn=PPP1R12B;refGene_tx=XM_019821986.2;refGene_gn=PPP1R12B;refGene_tx=XM_019821987.2
F1 43151963 43152167
refGene_gn=PPP1R12B;refGene_tx=XM_003999376.4;refGene_gn=PPP1R12B;refGene_tx=XM_011290936.3;refGene_gn=PPP1R12B;refGene_tx=XM_019821983.2;refGene_gn=PPP1R12B;refGene_tx=XM_019821984.2;refGene_gn=PPP1R12B;refGene_tx=XM_019821985.2;refGene_gn=PPP1R12B;refGene_tx=XM_019821986.2;refGene_gn=PPP1R12B;refGene_tx=XM_019821987.2
F1 43153231 43153314
refGene_gn=PPP1R12B;refGene_tx=XM_003999376.4;refGene_gn=PPP1R12B;refGene_tx=XM_011290936.3;refGene_gn=PPP1R12B;refGene_tx=XM_019821983.2;refGene_gn=PPP1R12B;refGene_tx=XM_019821984.2;refGene_gn=PPP1R12B;refGene_tx=XM_019821985.2;refGene_gn=PPP1R12B;refGene_tx=XM_019821986.2;refGene_gn=PPP1R12B;refGene_tx=XM_019821987.2
F1 43154535 43154661
refGene_gn=PPP1R12B;refGene_tx=XM_003999376.4;refGene_gn=PPP1R12B;refGene_tx=XM_011290936.3;refGene_gn=PPP1R12B;refGene_tx=XM_019821983.2;refGene_gn=PPP1R12B;refGene_tx=XM_019821984.2;refGene_gn=PPP1R12B;refGene_tx=XM_019821985.2;refGene_gn=PPP1R12B;refGene_tx=XM_019821986.2;refGene_gn=PPP1R12B;refGene_tx=XM_019821987.2
F1 43155200 43155241 refGene_gn=PPP1R12B;refGene_tx=XM_019821987.2
F1 43157359 43157542
refGene_gn=PPP1R12B;refGene_tx=XM_019821983.2;refGene_gn=PPP1R12B;refGene_tx=XM_019821984.2;refGene_gn=PPP1R12B;refGene_tx=XM_019821985.2;refGene_gn=PPP1R12B;refGene_tx=XM_019821986.2;refGene_gn=PPP1R12B;refGene_tx=XM_019821987.2
F1 43160871 43161054
refGene_gn=PPP1R12B;refGene_tx=XM_003999376.4;refGene_gn=PPP1R12B;refGene_tx=XM_011290936.3;refGene_gn=PPP1R12B;refGene_tx=XM_019821983.2;refGene_gn=PPP1R12B;refGene_tx=XM_019821984.2;refGene_gn=PPP1R12B;refGene_tx=XM_019821985.2;refGene_gn=PPP1R12B;refGene_tx=XM_019821986.2;refGene_gn=PPP1R12B;refGene_tx=XM_019821987.2
F1 43180551 43180571 refGene_gn=PPP1R12B;refGene_tx=XM_011290939.3

F1 43190198 43190289
refGene_gn=PPP1R12B;refGene_tx=XM_003999376.4;refGene_gn=PPP1R12B;refGene_tx=XM_011290936.3;refGene_gn=PPP1R12B;refGene_tx=XM_011290939.3;refGene_gn=PPP1R12B;refGene_tx=XM_019821983.2;refGene_gn=PPP1R12B;refGene_tx=XM_019821984.2;refGene_gn=PPP1R12B;refGene_tx=XM_019821985.2;refGene_gn=PPP1R12B;refGene_tx=XM_019821986.2
F1 43194833 43195037
refGene_gn=PPP1R12B;refGene_tx=XM_003999376.4;refGene_gn=PPP1R12B;refGene_tx=XM_011290936.3;refGene_gn=PPP1R12B;refGene_tx=XM_011290939.3;refGene_gn=PPP1R12B;refGene_tx=XM_019821983.2;refGene_gn=PPP1R12B;refGene_tx=XM_019821984.2;refGene_gn=PPP1R12B;refGene_tx=XM_019821985.2;refGene_gn=PPP1R12B;refGene_tx=XM_019821986.2
F1 43196905 43197095
refGene_gn=PPP1R12B;refGene_tx=XM_003999376.4;refGene_gn=PPP1R12B;refGene_tx=XM_011290936.3;refGene_gn=PPP1R12B;refGene_tx=XM_011290939.3;refGene_gn=PPP1R12B;refGene_tx=XM_011290940.2;refGene_gn=PPP1R12B;refGene_tx=XM_019821983.2;refGene_gn=PPP1R12B;refGene_tx=XM_019821984.2;refGene_gn=PPP1R12B;refGene_tx=XM_019821985.2;refGene_gn=PPP1R12B;refGene_tx=XM_019821986.2;refGene_gn=PPP1R12B;refGene_tx=XM_019821988.1;refGene_gn=PPP1R12B;refGene_tx=XM_019821989.1
F1 43197182 43197295
refGene_gn=PPP1R12B;refGene_tx=XM_003999376.4;refGene_gn=PPP1R12B;refGene_tx=XM_011290936.3;refGene_gn=PPP1R12B;refGene_tx=XM_011290939.3;refGene_gn=PPP1R12B;refGene_tx=XM_011290940.2;refGene_gn=PPP1R12B;refGene_tx=XM_019821983.2;refGene_gn=PPP1R12B;refGene_tx=XM_019821984.2;refGene_gn=PPP1R12B;refGene_tx=XM_019821985.2;refGene_gn=PPP1R12B;refGene_tx=XM_019821986.2;refGene_gn=PPP1R12B;refGene_tx=XM_019821988.1;refGene_gn=PPP1R12B;refGene_tx=XM_019821989.1
F1 43199641 43199683
refGene_gn=PPP1R12B;refGene_tx=XM_003999376.4;refGene_gn=PPP1R12B;refGene_tx=XM_011290936.3;refGene_gn=PPP1R12B;refGene_tx=XM_011290939.3;refGene_gn=PPP1R12B;refGene_tx=XM_011290940.2;refGene_gn=PPP1R12B;refGene_tx=XM_019821983.2;refGene_gn=PPP1R12B;refGene_tx=XM_019821984.2;refGene_gn=PPP1R12B;refGene_tx=XM_019821985.2;refGene_gn=PPP1R12B;refGene_tx=XM_019821986.2;refGene_gn=PPP1R12B;refGene_tx=XM_019821988.1;refGene_gn=PPP1R12B;refGene_tx=XM_019821989.1
F1 43217547 43217583 refGene_gn=PPP1R12B;refGene_tx=XM_019821986.2
F1 43240077 43240179 refGene_gn=PPP1R12B;refGene_tx=XM_011290942.3
F1 43260240 43260257
refGene_gn=PPP1R12B;refGene_tx=XM_003999376.4;refGene_gn=PPP1R12B;refGene_tx=XM_011290936.3;refGene_gn=PPP1R12B;refGene_tx=XM_011290939.3;refGene_gn=PPP1R12B;refGene_tx=XM_011290940.2;refGene_gn=PPP1R12B;refGene_tx=XM_011290942.3;refGene_gn=PPP1R12B;refGene_tx=XM_019821983.2;refGene_gn=PPP1R12B;refGene_tx=XM_019821984.2;refGene_gn=PPP1R12B;refGene_tx=XM_019821985.2;refGene_gn=PPP1R12B;refGene_tx=XM_019821988.1;refGene_gn=PPP1R12B;refGene_tx=XM_019821989.1
F1 43263248 43263399
refGene_gn=PPP1R12B;refGene_tx=XM_003999376.4;refGene_gn=PPP1R12B;refGene_tx=XM_011290936.3;refGene_gn=PPP1R12B;refGene_tx=XM_011290939.3;refGene_gn=PPP1R12B;refGene_tx=XM_011290940.2;refGene_gn=PPP1R12B;refGen

e_tx=XM_011290942.3;refGene_gn=PPP1R12B;refGene_tx=XM_019821983.2;refGene_gn=PPP1R12B;refGene_tx=XM_019821984.2;refGene_gn=PPP1R12B;refGene_tx=XM_019821985.2;refGene_gn=PPP1R12B;refGene_tx=XM_019821988.1;refGene_gn=PPP1R12B;refGene_tx=XM_019821989.1
F1 43264298 43264403
refGene_gn=PPP1R12B;refGene_tx=XM_003999376.4;refGene_gn=PPP1R12B;refGene_tx=XM_011290936.3;refGene_gn=PPP1R12B;refGene_tx=XM_011290939.3;refGene_gn=PPP1R12B;refGene_tx=XM_011290940.2;refGene_gn=PPP1R12B;refGene_tx=XM_011290942.3;refGene_gn=PPP1R12B;refGene_tx=XM_019821983.2;refGene_gn=PPP1R12B;refGene_tx=XM_019821984.2;refGene_gn=PPP1R12B;refGene_tx=XM_019821985.2;refGene_gn=PPP1R12B;refGene_tx=XM_019821988.1;refGene_gn=PPP1R12B;refGene_tx=XM_019821989.1
F1 43267847 43267901
refGene_gn=PPP1R12B;refGene_tx=XM_003999376.4;refGene_gn=PPP1R12B;refGene_tx=XM_011290936.3;refGene_gn=PPP1R12B;refGene_tx=XM_011290939.3;refGene_gn=PPP1R12B;refGene_tx=XM_011290940.2;refGene_gn=PPP1R12B;refGene_tx=XM_011290942.3;refGene_gn=PPP1R12B;refGene_tx=XM_019821983.2;refGene_gn=PPP1R12B;refGene_tx=XM_019821984.2;refGene_gn=PPP1R12B;refGene_tx=XM_019821985.2;refGene_gn=PPP1R12B;refGene_tx=XM_019821988.1;refGene_gn=PPP1R12B;refGene_tx=XM_019821989.1
F1 43269128 43269179
refGene_gn=PPP1R12B;refGene_tx=XM_003999376.4;refGene_gn=PPP1R12B;refGene_tx=XM_011290936.3;refGene_gn=PPP1R12B;refGene_tx=XM_011290939.3;refGene_gn=PPP1R12B;refGene_tx=XM_011290940.2;refGene_gn=PPP1R12B;refGene_tx=XM_011290942.3;refGene_gn=PPP1R12B;refGene_tx=XM_019821983.2;refGene_gn=PPP1R12B;refGene_tx=XM_019821984.2;refGene_gn=PPP1R12B;refGene_tx=XM_019821985.2;refGene_gn=PPP1R12B;refGene_tx=XM_019821988.1;refGene_gn=PPP1R12B;refGene_tx=XM_019821989.1
F1 43274368 43274503
refGene_gn=PPP1R12B;refGene_tx=XM_011290936.3;refGene_gn=PPP1R12B;refGene_tx=XM_011290939.3;refGene_gn=PPP1R12B;refGene_tx=XM_011290940.2;refGene_gn=PPP1R12B;refGene_tx=XM_019821983.2;refGene_gn=PPP1R12B;refGene_tx=XM_019821984.2;refGene_gn=PPP1R12B;refGene_tx=XM_019821988.1;refGene_gn=PPP1R12B;refGene_tx=XM_019821989.1
F1 43280525 43280612
refGene_gn=PPP1R12B;refGene_tx=XM_003999376.4;refGene_gn=PPP1R12B;refGene_tx=XM_011290942.3;refGene_gn=PPP1R12B;refGene_tx=XM_019821985.2
F1 43296341 43296548
refGene_gn=SYT2;refGene_tx=XM_003999377.5;refGene_gn=SYT2;refGene_tx=XM_019821995.2;refGene_gn=SYT2;refGene_tx=XM_019821996.2;refGene_gn=SYT2;refGene_tx=XM_019821997.2;refGene_gn=SYT2;refGene_tx=XM_019821998.2
F1 43298672 43298806
refGene_gn=SYT2;refGene_tx=XM_003999377.5;refGene_gn=SYT2;refGene_tx=XM_019821995.2;refGene_gn=SYT2;refGene_tx=XM_019821996.2;refGene_gn=SYT2;refGene_tx=XM_019821997.2;refGene_gn=SYT2;refGene_tx=XM_019821998.2
F1 43299731 43299849
refGene_gn=SYT2;refGene_tx=XM_003999377.5;refGene_gn=SYT2;refGene_tx=XM_019821995.2;refGene_gn=SYT2;refGene_tx=XM_019821996.2;refGene_gn=SYT2;refGene_tx=XM_019821997.2;refGene_gn=SYT2;refGene_tx=XM_019821998.2
F1 43301171 43301339

refGene_gn=SYT2;refGene_tx=XM_003999377.5;refGene_gn=SYT2;refGene_tx=XM_019821995.2;refGene_gn=SYT2;refGene_tx=XM_019821996.2;refGene_gn=SYT2;refGene_tx=XM_019821997.2;refGene_gn=SYT2;refGene_tx=XM_019821998.2
F1 43301709 43301877
refGene_gn=SYT2;refGene_tx=XM_003999377.5;refGene_gn=SYT2;refGene_tx=XM_019821995.2;refGene_gn=SYT2;refGene_tx=XM_019821996.2;refGene_gn=SYT2;refGene_tx=XM_019821997.2;refGene_gn=SYT2;refGene_tx=XM_019821998.2
F1 43302345 43302465
refGene_gn=SYT2;refGene_tx=XM_003999377.5;refGene_gn=SYT2;refGene_tx=XM_019821995.2;refGene_gn=SYT2;refGene_tx=XM_019821996.2;refGene_gn=SYT2;refGene_tx=XM_019821997.2;refGene_gn=SYT2;refGene_tx=XM_019821998.2
F1 43303811 43303978
refGene_gn=SYT2;refGene_tx=XM_003999377.5;refGene_gn=SYT2;refGene_tx=XM_019821995.2;refGene_gn=SYT2;refGene_tx=XM_019821996.2;refGene_gn=SYT2;refGene_tx=XM_019821997.2;refGene_gn=SYT2;refGene_tx=XM_019821998.2
F1 43304912 43305090
refGene_gn=SYT2;refGene_tx=XM_003999377.5;refGene_gn=SYT2;refGene_tx=XM_019821995.2;refGene_gn=SYT2;refGene_tx=XM_019821996.2;refGene_gn=SYT2;refGene_tx=XM_019821997.2;refGene_gn=SYT2;refGene_tx=XM_019821998.2
F1 43411509 43411548 refGene_gn=KDM5B;refGene_tx=XM_023247927.1
F1 43415491 43415629
refGene_gn=KDM5B;refGene_tx=XM_023247926.1;refGene_gn=KDM5B;refGene_tx=XM_023247928.1
F1 43416162 43416483
refGene_gn=KDM5B;refGene_tx=XM_023247926.1;refGene_gn=KDM5B;refGene_tx=XM_023247927.1;refGene_gn=KDM5B;refGene_tx=XM_023247928.1
F1 43418026 43418181
refGene_gn=KDM5B;refGene_tx=XM_023247926.1;refGene_gn=KDM5B;refGene_tx=XM_023247927.1;refGene_gn=KDM5B;refGene_tx=XM_023247928.1
F1 43419294 43419406
refGene_gn=KDM5B;refGene_tx=XM_023247926.1;refGene_gn=KDM5B;refGene_tx=XM_023247927.1;refGene_gn=KDM5B;refGene_tx=XM_023247928.1
F1 43420846 43421332
refGene_gn=KDM5B;refGene_tx=XM_023247926.1;refGene_gn=KDM5B;refGene_tx=XM_023247927.1;refGene_gn=KDM5B;refGene_tx=XM_023247928.1
F1 43425595 43425754
refGene_gn=KDM5B;refGene_tx=XM_023247926.1;refGene_gn=KDM5B;refGene_tx=XM_023247927.1;refGene_gn=KDM5B;refGene_tx=XM_023247928.1
F1 43426399 43426579
refGene_gn=KDM5B;refGene_tx=XM_023247926.1;refGene_gn=KDM5B;refGene_tx=XM_023247927.1;refGene_gn=KDM5B;refGene_tx=XM_023247928.1
F1 43430971 43431110
refGene_gn=KDM5B;refGene_tx=XM_023247926.1;refGene_gn=KDM5B;refGene_tx=XM_023247927.1;refGene_gn=KDM5B;refGene_tx=XM_023247928.1
F1 43432175 43432531
refGene_gn=KDM5B;refGene_tx=XM_023247926.1;refGene_gn=KDM5B;refGene_tx=XM_023247927.1;refGene_gn=KDM5B;refGene_tx=XM_023247928.1
F1 43433231 43433346
refGene_gn=KDM5B;refGene_tx=XM_023247926.1;refGene_gn=KDM5B;refGene_tx=XM_023247927.1;refGene_gn=KDM5B;refGene_tx=XM_023247928.1

F1	43433503	43433654
refGene_gn=KDM5B;refGene_tx=XM_023247926.1;refGene_gn=KDM5B;refGene_tx=XM_023247927.1;refGene_gn=KDM5B;refGene_tx=XM_023247928.1
F1	43435289	43435414
refGene_gn=KDM5B;refGene_tx=XM_023247926.1;refGene_gn=KDM5B;refGene_tx=XM_023247927.1;refGene_gn=KDM5B;refGene_tx=XM_023247928.1
F1	43435569	43435751
refGene_gn=KDM5B;refGene_tx=XM_023247926.1;refGene_gn=KDM5B;refGene_tx=XM_023247927.1;refGene_gn=KDM5B;refGene_tx=XM_023247928.1
F1	43438603	43438798
refGene_gn=KDM5B;refGene_tx=XM_023247926.1;refGene_gn=KDM5B;refGene_tx=XM_023247927.1;refGene_gn=KDM5B;refGene_tx=XM_023247928.1
F1	43440191	43440311
refGene_gn=KDM5B;refGene_tx=XM_023247926.1;refGene_gn=KDM5B;refGene_tx=XM_023247927.1;refGene_gn=KDM5B;refGene_tx=XM_023247928.1
F1	43443370	43443533
refGene_gn=KDM5B;refGene_tx=XM_023247926.1;refGene_gn=KDM5B;refGene_tx=XM_023247927.1;refGene_gn=KDM5B;refGene_tx=XM_023247928.1
F1	43444822	43445004
refGene_gn=KDM5B;refGene_tx=XM_023247926.1;refGene_gn=KDM5B;refGene_tx=XM_023247927.1;refGene_gn=KDM5B;refGene_tx=XM_023247928.1
F1	43445997	43446156
refGene_gn=KDM5B;refGene_tx=XM_023247926.1;refGene_gn=KDM5B;refGene_tx=XM_023247927.1;refGene_gn=KDM5B;refGene_tx=XM_023247928.1
F1	43447905	43448025
refGene_gn=KDM5B;refGene_tx=XM_023247926.1;refGene_gn=KDM5B;refGene_tx=XM_023247927.1;refGene_gn=KDM5B;refGene_tx=XM_023247928.1
F1	43448492	43448651
refGene_gn=KDM5B;refGene_tx=XM_023247926.1;refGene_gn=KDM5B;refGene_tx=XM_023247927.1;refGene_gn=KDM5B;refGene_tx=XM_023247928.1
F1	43450495	43450605
refGene_gn=KDM5B;refGene_tx=XM_023247926.1;refGene_gn=KDM5B;refGene_tx=XM_023247927.1;refGene_gn=KDM5B;refGene_tx=XM_023247928.1
F1	43451854	43451951
refGene_gn=KDM5B;refGene_tx=XM_023247926.1;refGene_gn=KDM5B;refGene_tx=XM_023247927.1;refGene_gn=KDM5B;refGene_tx=XM_023247928.1
F1	43453572	43453707
refGene_gn=KDM5B;refGene_tx=XM_023247926.1;refGene_gn=KDM5B;refGene_tx=XM_023247927.1;refGene_gn=KDM5B;refGene_tx=XM_023247928.1
F1	43460214	43460385
refGene_gn=KDM5B;refGene_tx=XM_023247926.1;refGene_gn=KDM5B;refGene_tx=XM_023247927.1;refGene_gn=KDM5B;refGene_tx=XM_023247928.1
F1	43461984	43462107
refGene_gn=KDM5B;refGene_tx=XM_023247926.1;refGene_gn=KDM5B;refGene_tx=XM_023247927.1;refGene_gn=KDM5B;refGene_tx=XM_023247928.1
F1	43463735	43463813
refGene_gn=KDM5B;refGene_tx=XM_023247926.1;refGene_gn=KDM5B;refGene_tx=XM_023247927.1;refGene_gn=KDM5B;refGene_tx=XM_023247928.1
F1	43490659	43491187
refGene_gn=KDM5B;refGene_tx=XM_023247926.1;refGene_gn=KDM5B;refGene_tx

=XM_023247927.1
F1 43491375 43491579 refGene_gn=KDM5B;refGene_tx=XM_023247928.1
F1 43502787 43502913
refGene_gn=LOC109495700;refGene_tx=XM_019821990.2;refGene_gn=LOC109495700;refGene_tx=XM_019821991.2;refGene_gn=LOC109495700;refGene_tx=XM_019821992.2
F1 43503288 43503385
refGene_gn=LOC109495700;refGene_tx=XM_019821990.2;refGene_gn=LOC109495700;refGene_tx=XM_019821991.2;refGene_gn=LOC109495700;refGene_tx=XM_019821992.2
F1 43504221 43505369
refGene_gn=LOC109495700;refGene_tx=XM_019821990.2;refGene_gn=LOC109495700;refGene_tx=XM_019821991.2;refGene_gn=LOC109495700;refGene_tx=XM_019821992.2
F1 43527832 43528126
refGene_gn=LOC101099268;refGene_tx=XM_006942867.4;refGene_gn=LOC101099268;refGene_tx=XM_023248011.1
F1 43530049 43530730
refGene_gn=LOC101099268;refGene_tx=XM_006942867.4;refGene_gn=LOC101099268;refGene_tx=XM_023248011.1
F1 43532210 43532359
refGene_gn=LOC101099268;refGene_tx=XM_006942867.4;refGene_gn=LOC101099268;refGene_tx=XM_023248011.1
F1 43533709 43533932
refGene_gn=LOC101099268;refGene_tx=XM_006942867.4;refGene_gn=LOC101099268;refGene_tx=XM_023248011.1
F1 43535583 43535784
refGene_gn=LOC101099268;refGene_tx=XM_006942867.4
F1 43546609 43546855
refGene_gn=RABIF;refGene_tx=XM_023248012.1;refGene_gn=RABIF;refGene_tx=XM_023248013.1
F1 43552830 43553023
refGene_gn=RABIF;refGene_tx=XM_023248012.1;refGene_gn=RABIF;refGene_tx=XM_023248013.1
F1 43553230 43553283 refGene_gn=RABIF;refGene_tx=XM_023248012.1
F1 43556926 43557053
refGene_gn=KLHL12;refGene_tx=XM_003999378.4;refGene_gn=KLHL12;refGene_tx=XM_019822002.2
F1 43557234 43557421
refGene_gn=KLHL12;refGene_tx=XM_003999378.4;refGene_gn=KLHL12;refGene_tx=XM_019822002.2
F1 43557771 43557870
refGene_gn=KLHL12;refGene_tx=XM_003999378.4;refGene_gn=KLHL12;refGene_tx=XM_019822002.2
F1 43558176 43558335
refGene_gn=KLHL12;refGene_tx=XM_003999378.4;refGene_gn=KLHL12;refGene_tx=XM_019822002.2
F1 43559005 43559201
refGene_gn=KLHL12;refGene_tx=XM_003999378.4;refGene_gn=KLHL12;refGene_tx=XM_019822002.2

F1 43560049 43560156
refGene_gn=KLHL12;refGene_tx=XM_003999378.4;refGene_gn=KLHL12;refGene_tx=XM_019822002.2
F1 43572448 43572563
refGene_gn=KLHL12;refGene_tx=XM_003999378.4;refGene_gn=KLHL12;refGene_tx=XM_019822002.2
F1 43574086 43574236
refGene_gn=KLHL12;refGene_tx=XM_003999378.4;refGene_gn=KLHL12;refGene_tx=XM_019822002.2
F1 43582903 43583121
refGene_gn=KLHL12;refGene_tx=XM_003999378.4;refGene_gn=KLHL12;refGene_tx=XM_019822002.2
F1 43584199 43584353
refGene_gn=KLHL12;refGene_tx=XM_003999378.4;refGene_gn=KLHL12;refGene_tx=XM_019822002.2
F1 43589249 43589483
refGene_gn=KLHL12;refGene_tx=XM_003999378.4;refGene_gn=KLHL12;refGene_tx=XM_019822002.2
F1 43592227 43592530 refGene_gn=KLHL12;refGene_tx=XM_019822002.2
F1 43600240 43600369 refGene_gn=ADIPOR1;refGene_tx=NM_001134681.1
F1 43600683 43600877 refGene_gn=ADIPOR1;refGene_tx=NM_001134681.1
F1 43602127 43602315 refGene_gn=ADIPOR1;refGene_tx=NM_001134681.1
F1 43603601 43603788 refGene_gn=ADIPOR1;refGene_tx=NM_001134681.1
F1 43604266 43604438 refGene_gn=ADIPOR1;refGene_tx=NM_001134681.1
F1 43605454 43605508 refGene_gn=ADIPOR1;refGene_tx=NM_001134681.1
F1 43605509 43605572 refGene_gn=ADIPOR1;refGene_tx=NM_001134681.1
F1 43607243 43607384 refGene_gn=ADIPOR1;refGene_tx=NM_001134681.1
F1 43617504 43617677
refGene_gn=LOC101099773;refGene_tx=XM_003999379.4
F1 43618025 43618125
refGene_gn=LOC101099773;refGene_tx=XM_003999379.4
F1 43618596 43618682
refGene_gn=LOC101099773;refGene_tx=XM_003999379.4
F1 43619774 43619858
refGene_gn=LOC101099773;refGene_tx=XM_003999379.4
F1 43620550 43620680
refGene_gn=LOC101099773;refGene_tx=XM_003999379.4
F1 43620898 43621005
refGene_gn=LOC101099773;refGene_tx=XM_003999379.4
F1 43621455 43621528
refGene_gn=LOC101099773;refGene_tx=XM_003999379.4
F1 43621646 43621796
refGene_gn=LOC101099773;refGene_tx=XM_003999379.4
F1 43622105 43622120
refGene_gn=LOC101099773;refGene_tx=XM_003999379.4
F1 43631985 43633190
refGene_gn=LOC101080681;refGene_tx=XM_011290934.3;refGene_gn=LOC101080681;refGene_tx=XM_019822003.2;refGene_gn=LOC101080681;refGene_tx=XM_019822005.2;refGene_gn=LOC101080681;refGene_tx=XM_023247335.1;refGene_gn=LOC101080681;refGene_tx=XM_023247336.1;refGene_gn=LOC101080681;refGen

e_tx=XM_023247338.1;refGene_gn=LOC101080681;refGene_tx=XM_023247339.1;
refGene_gn=LOC101080681;refGene_tx=XM_023247340.1;refGene_gn=LOC101080
681;refGene_tx=XM_023247341.1;refGene_gn=LOC101080681;refGene_tx=XM_02
3247342.1
F1 43634488 43634648
refGene_gn=LOC101080681;refGene_tx=XM_011290934.3;refGene_gn=LOC101080
681;refGene_tx=XM_019822003.2;refGene_gn=LOC101080681;refGene_tx=XM_01
9822005.2;refGene_gn=LOC101080681;refGene_tx=XM_023247335.1;refGene_gn
=LOC101080681;refGene_tx=XM_023247336.1;refGene_gn=LOC101080681;refGen
e_tx=XM_023247338.1;refGene_gn=LOC101080681;refGene_tx=XM_023247339.1;
refGene_gn=LOC101080681;refGene_tx=XM_023247340.1;refGene_gn=LOC101080
681;refGene_tx=XM_023247341.1;refGene_gn=LOC101080681;refGene_tx=XM_02
3247342.1
F1 43635125 43635335
refGene_gn=LOC101080681;refGene_tx=XM_019822003.2
F1 43635615 43635723
refGene_gn=LOC101080681;refGene_tx=XM_011290934.3;refGene_gn=LOC101080
681;refGene_tx=XM_019822005.2;refGene_gn=LOC101080681;refGene_tx=XM_02
3247335.1;refGene_gn=LOC101080681;refGene_tx=XM_023247336.1;refGene_gn
=LOC101080681;refGene_tx=XM_023247338.1;refGene_gn=LOC101080681;refGen
e_tx=XM_023247339.1;refGene_gn=LOC101080681;refGene_tx=XM_023247340.1
F1 43636366 43636369
refGene_gn=LOC101080681;refGene_tx=XM_023247341.1;refGene_gn=LOC101080
681;refGene_tx=XM_023247342.1
F1 43664005 43664114
refGene_gn=TMEM183A;refGene_tx=XM_003999380.3;refGene_gn=TMEM183A;refG
ene_tx=XM_006942870.3;refGene_gn=TMEM183A;refGene_tx=XM_006942871.3;re
fGene_gn=TMEM183A;refGene_tx=XM_019822006.2
F1 43664312 43664402
refGene_gn=TMEM183A;refGene_tx=XM_003999380.3;refGene_gn=TMEM183A;refG
ene_tx=XM_006942870.3;refGene_gn=TMEM183A;refGene_tx=XM_006942871.3;re
fGene_gn=TMEM183A;refGene_tx=XM_019822006.2
F1 43665259 43665427
refGene_gn=TMEM183A;refGene_tx=XM_003999380.3;refGene_gn=TMEM183A;refG
ene_tx=XM_006942870.3
F1 43670474 43670634
refGene_gn=TMEM183A;refGene_tx=XM_003999380.3;refGene_gn=TMEM183A;refG
ene_tx=XM_006942870.3;refGene_gn=TMEM183A;refGene_tx=XM_006942871.3;re
fGene_gn=TMEM183A;refGene_tx=XM_019822006.2
F1 43671559 43671740
refGene_gn=TMEM183A;refGene_tx=XM_003999380.3;refGene_gn=TMEM183A;refG
ene_tx=XM_006942870.3;refGene_gn=TMEM183A;refGene_tx=XM_006942871.3;re
fGene_gn=TMEM183A;refGene_tx=XM_019822006.2
F1 43674202 43674283
refGene_gn=TMEM183A;refGene_tx=XM_003999380.3;refGene_gn=TMEM183A;refG
ene_tx=XM_006942870.3;refGene_gn=TMEM183A;refGene_tx=XM_006942871.3;re
fGene_gn=TMEM183A;refGene_tx=XM_019822006.2
F1 43675870 43676026
refGene_gn=TMEM183A;refGene_tx=XM_003999380.3;refGene_gn=TMEM183A;refG
ene_tx=XM_006942870.3;refGene_gn=TMEM183A;refGene_tx=XM_006942871.3;re

fGene_gn=TMEM183A;refGene_tx=XM_019822006.2
F1 43677269 43677455
refGene_gn=TMEM183A;refGene_tx=XM_003999380.3;refGene_gn=TMEM183A;refGene_tx=XM_006942870.3;refGene_gn=TMEM183A;refGene_tx=XM_006942871.3;refGene_gn=TMEM183A;refGene_tx=XM_019822006.2
F1 43693910 43694144
refGene_gn=PPFIA4;refGene_tx=XM_023247297.1;refGene_gn=PPFIA4;refGene_tx=XM_023247298.1;refGene_gn=PPFIA4;refGene_tx=XM_023247299.1;refGene_gn=PPFIA4;refGene_tx=XM_023247300.1;refGene_gn=PPFIA4;refGene_tx=XM_023247301.1;refGene_gn=PPFIA4;refGene_tx=XM_023247302.1;refGene_gn=PPFIA4;refGene_tx=XM_023247303.1;refGene_gn=PPFIA4;refGene_tx=XM_023247304.1
F1 43697460 43697562
refGene_gn=PPFIA4;refGene_tx=XM_023247297.1;refGene_gn=PPFIA4;refGene_tx=XM_023247298.1;refGene_gn=PPFIA4;refGene_tx=XM_023247299.1;refGene_gn=PPFIA4;refGene_tx=XM_023247300.1;refGene_gn=PPFIA4;refGene_tx=XM_023247301.1;refGene_gn=PPFIA4;refGene_tx=XM_023247302.1;refGene_gn=PPFIA4;refGene_tx=XM_023247303.1;refGene_gn=PPFIA4;refGene_tx=XM_023247304.1
F1 43697956 43698121
refGene_gn=PPFIA4;refGene_tx=XM_023247297.1;refGene_gn=PPFIA4;refGene_tx=XM_023247298.1;refGene_gn=PPFIA4;refGene_tx=XM_023247299.1;refGene_gn=PPFIA4;refGene_tx=XM_023247300.1;refGene_gn=PPFIA4;refGene_tx=XM_023247301.1;refGene_gn=PPFIA4;refGene_tx=XM_023247302.1;refGene_gn=PPFIA4;refGene_tx=XM_023247303.1;refGene_gn=PPFIA4;refGene_tx=XM_023247304.1
F1 43698420 43698495
refGene_gn=PPFIA4;refGene_tx=XM_023247297.1;refGene_gn=PPFIA4;refGene_tx=XM_023247298.1;refGene_gn=PPFIA4;refGene_tx=XM_023247299.1;refGene_gn=PPFIA4;refGene_tx=XM_023247300.1;refGene_gn=PPFIA4;refGene_tx=XM_023247301.1;refGene_gn=PPFIA4;refGene_tx=XM_023247302.1;refGene_gn=PPFIA4;refGene_tx=XM_023247303.1;refGene_gn=PPFIA4;refGene_tx=XM_023247304.1
F1 43698713 43698803
refGene_gn=PPFIA4;refGene_tx=XM_023247297.1;refGene_gn=PPFIA4;refGene_tx=XM_023247298.1;refGene_gn=PPFIA4;refGene_tx=XM_023247299.1;refGene_gn=PPFIA4;refGene_tx=XM_023247300.1;refGene_gn=PPFIA4;refGene_tx=XM_023247301.1;refGene_gn=PPFIA4;refGene_tx=XM_023247302.1;refGene_gn=PPFIA4;refGene_tx=XM_023247303.1;refGene_gn=PPFIA4;refGene_tx=XM_023247304.1
F1 43699277 43699469
refGene_gn=PPFIA4;refGene_tx=XM_023247297.1;refGene_gn=PPFIA4;refGene_tx=XM_023247298.1;refGene_gn=PPFIA4;refGene_tx=XM_023247299.1;refGene_gn=PPFIA4;refGene_tx=XM_023247300.1;refGene_gn=PPFIA4;refGene_tx=XM_023247301.1;refGene_gn=PPFIA4;refGene_tx=XM_023247302.1;refGene_gn=PPFIA4;refGene_tx=XM_023247303.1;refGene_gn=PPFIA4;refGene_tx=XM_023247304.1
F1 43699783 43699930
refGene_gn=PPFIA4;refGene_tx=XM_023247297.1;refGene_gn=PPFIA4;refGene_tx=XM_023247298.1;refGene_gn=PPFIA4;refGene_tx=XM_023247299.1;refGene_

gn=PPFIA4;refGene_tx=XM_023247300.1;refGene_gn=PPFIA4;refGene_tx=XM_02
3247301.1;refGene_gn=PPFIA4;refGene_tx=XM_023247302.1;refGene_gn=PPFIA
4;refGene_tx=XM_023247303.1;refGene_gn=PPFIA4;refGene_tx=XM_023247304.
1
F1 43700215 43700350
refGene_gn=PPFIA4;refGene_tx=XM_023247297.1;refGene_gn=PPFIA4;refGene_
tx=XM_023247298.1;refGene_gn=PPFIA4;refGene_tx=XM_023247299.1;refGene_
gn=PPFIA4;refGene_tx=XM_023247300.1;refGene_gn=PPFIA4;refGene_tx=XM_02
3247301.1;refGene_gn=PPFIA4;refGene_tx=XM_023247302.1;refGene_gn=PPFIA
4;refGene_tx=XM_023247303.1;refGene_gn=PPFIA4;refGene_tx=XM_023247304.
1
F1 43702939 43703023
refGene_gn=PPFIA4;refGene_tx=XM_023247297.1;refGene_gn=PPFIA4;refGene_
tx=XM_023247298.1;refGene_gn=PPFIA4;refGene_tx=XM_023247299.1;refGene_
gn=PPFIA4;refGene_tx=XM_023247300.1;refGene_gn=PPFIA4;refGene_tx=XM_02
3247301.1;refGene_gn=PPFIA4;refGene_tx=XM_023247302.1;refGene_gn=PPFIA
4;refGene_tx=XM_023247303.1;refGene_gn=PPFIA4;refGene_tx=XM_023247304.
1
F1 43703238 43703370
refGene_gn=PPFIA4;refGene_tx=XM_023247297.1;refGene_gn=PPFIA4;refGene_
tx=XM_023247298.1;refGene_gn=PPFIA4;refGene_tx=XM_023247299.1;refGene_
gn=PPFIA4;refGene_tx=XM_023247300.1;refGene_gn=PPFIA4;refGene_tx=XM_02
3247301.1;refGene_gn=PPFIA4;refGene_tx=XM_023247302.1;refGene_gn=PPFIA
4;refGene_tx=XM_023247303.1;refGene_gn=PPFIA4;refGene_tx=XM_023247304.
1
F1 43703625 43703688
refGene_gn=PPFIA4;refGene_tx=XM_023247297.1;refGene_gn=PPFIA4;refGene_
tx=XM_023247298.1;refGene_gn=PPFIA4;refGene_tx=XM_023247299.1;refGene_
gn=PPFIA4;refGene_tx=XM_023247300.1;refGene_gn=PPFIA4;refGene_tx=XM_02
3247301.1;refGene_gn=PPFIA4;refGene_tx=XM_023247302.1;refGene_gn=PPFIA
4;refGene_tx=XM_023247303.1;refGene_gn=PPFIA4;refGene_tx=XM_023247304.
1
F1 43704355 43704447
refGene_gn=PPFIA4;refGene_tx=XM_023247297.1;refGene_gn=PPFIA4;refGene_
tx=XM_023247298.1;refGene_gn=PPFIA4;refGene_tx=XM_023247299.1;refGene_
gn=PPFIA4;refGene_tx=XM_023247300.1;refGene_gn=PPFIA4;refGene_tx=XM_02
3247301.1;refGene_gn=PPFIA4;refGene_tx=XM_023247302.1;refGene_gn=PPFIA
4;refGene_tx=XM_023247303.1
F1 43707024 43707133
refGene_gn=PPFIA4;refGene_tx=XM_023247297.1;refGene_gn=PPFIA4;refGene_
tx=XM_023247298.1;refGene_gn=PPFIA4;refGene_tx=XM_023247299.1;refGene_
gn=PPFIA4;refGene_tx=XM_023247300.1;refGene_gn=PPFIA4;refGene_tx=XM_02
3247301.1;refGene_gn=PPFIA4;refGene_tx=XM_023247302.1;refGene_gn=PPFIA
4;refGene_tx=XM_023247303.1
F1 43708704 43708913
refGene_gn=PPFIA4;refGene_tx=XM_023247297.1;refGene_gn=PPFIA4;refGene_
tx=XM_023247298.1;refGene_gn=PPFIA4;refGene_tx=XM_023247299.1;refGene_
gn=PPFIA4;refGene_tx=XM_023247300.1;refGene_gn=PPFIA4;refGene_tx=XM_02
3247301.1;refGene_gn=PPFIA4;refGene_tx=XM_023247302.1;refGene_gn=PPFIA
4;refGene_tx=XM_023247303.1;refGene_gn=PPFIA4;refGene_tx=XM_023247304.

1
F1 43710417 43710658
refGene_gn=PPFIA4;refGene_tx=XM_023247297.1;refGene_gn=PPFIA4;refGene_tx=XM_023247298.1;refGene_gn=PPFIA4;refGene_tx=XM_023247299.1;refGene_gn=PPFIA4;refGene_tx=XM_023247300.1;refGene_gn=PPFIA4;refGene_tx=XM_023247301.1;refGene_gn=PPFIA4;refGene_tx=XM_023247302.1;refGene_gn=PPFIA4;refGene_tx=XM_023247303.1;refGene_gn=PPFIA4;refGene_tx=XM_023247304.1
F1 43711114 43711150
refGene_gn=PPFIA4;refGene_tx=XM_023247297.1;refGene_gn=PPFIA4;refGene_tx=XM_023247298.1;refGene_gn=PPFIA4;refGene_tx=XM_023247299.1;refGene_gn=PPFIA4;refGene_tx=XM_023247300.1;refGene_gn=PPFIA4;refGene_tx=XM_023247301.1;refGene_gn=PPFIA4;refGene_tx=XM_023247302.1;refGene_gn=PPFIA4;refGene_tx=XM_023247303.1;refGene_gn=PPFIA4;refGene_tx=XM_023247304.1
F1 43711366 43711500
refGene_gn=PPFIA4;refGene_tx=XM_023247297.1;refGene_gn=PPFIA4;refGene_tx=XM_023247298.1;refGene_gn=PPFIA4;refGene_tx=XM_023247299.1;refGene_gn=PPFIA4;refGene_tx=XM_023247300.1;refGene_gn=PPFIA4;refGene_tx=XM_023247301.1;refGene_gn=PPFIA4;refGene_tx=XM_023247302.1;refGene_gn=PPFIA4;refGene_tx=XM_023247303.1;refGene_gn=PPFIA4;refGene_tx=XM_023247304.1
F1 43711763 43711930
refGene_gn=PPFIA4;refGene_tx=XM_023247297.1;refGene_gn=PPFIA4;refGene_tx=XM_023247298.1;refGene_gn=PPFIA4;refGene_tx=XM_023247299.1;refGene_gn=PPFIA4;refGene_tx=XM_023247300.1;refGene_gn=PPFIA4;refGene_tx=XM_023247301.1;refGene_gn=PPFIA4;refGene_tx=XM_023247302.1;refGene_gn=PPFIA4;refGene_tx=XM_023247303.1;refGene_gn=PPFIA4;refGene_tx=XM_023247304.1
F1 43714036 43714130
refGene_gn=PPFIA4;refGene_tx=XM_023247297.1;refGene_gn=PPFIA4;refGene_tx=XM_023247298.1;refGene_gn=PPFIA4;refGene_tx=XM_023247299.1;refGene_gn=PPFIA4;refGene_tx=XM_023247300.1;refGene_gn=PPFIA4;refGene_tx=XM_023247301.1;refGene_gn=PPFIA4;refGene_tx=XM_023247302.1;refGene_gn=PPFIA4;refGene_tx=XM_023247303.1;refGene_gn=PPFIA4;refGene_tx=XM_023247304.1
F1 43714623 43714705
refGene_gn=PPFIA4;refGene_tx=XM_023247297.1;refGene_gn=PPFIA4;refGene_tx=XM_023247298.1;refGene_gn=PPFIA4;refGene_tx=XM_023247299.1;refGene_gn=PPFIA4;refGene_tx=XM_023247300.1;refGene_gn=PPFIA4;refGene_tx=XM_023247301.1;refGene_gn=PPFIA4;refGene_tx=XM_023247302.1;refGene_gn=PPFIA4;refGene_tx=XM_023247303.1;refGene_gn=PPFIA4;refGene_tx=XM_023247304.1
F1 43715262 43715463
refGene_gn=PPFIA4;refGene_tx=XM_023247297.1;refGene_gn=PPFIA4;refGene_tx=XM_023247298.1;refGene_gn=PPFIA4;refGene_tx=XM_023247299.1;refGene_gn=PPFIA4;refGene_tx=XM_023247300.1;refGene_gn=PPFIA4;refGene_tx=XM_023247301.1;refGene_gn=PPFIA4;refGene_tx=XM_023247302.1;refGene_gn=PPFIA4;refGene_tx=XM_023247303.1;refGene_gn=PPFIA4;refGene_tx=XM_023247304.1

F1 43716117 43716180
refGene_gn=PPFIA4;refGene_tx=XM_023247297.1;refGene_gn=PPFIA4;refGene_tx=XM_023247298.1;refGene_gn=PPFIA4;refGene_tx=XM_023247299.1;refGene_gn=PPFIA4;refGene_tx=XM_023247300.1;refGene_gn=PPFIA4;refGene_tx=XM_023247301.1;refGene_gn=PPFIA4;refGene_tx=XM_023247302.1;refGene_gn=PPFIA4;refGene_tx=XM_023247303.1;refGene_gn=PPFIA4;refGene_tx=XM_023247304.1
F1 43716779 43716806
refGene_gn=PPFIA4;refGene_tx=XM_023247297.1;refGene_gn=PPFIA4;refGene_tx=XM_023247298.1;refGene_gn=PPFIA4;refGene_tx=XM_023247299.1;refGene_gn=PPFIA4;refGene_tx=XM_023247300.1;refGene_gn=PPFIA4;refGene_tx=XM_023247302.1;refGene_gn=PPFIA4;refGene_tx=XM_023247304.1
F1 43718480 43718656
refGene_gn=PPFIA4;refGene_tx=XM_023247297.1;refGene_gn=PPFIA4;refGene_tx=XM_023247298.1;refGene_gn=PPFIA4;refGene_tx=XM_023247299.1;refGene_gn=PPFIA4;refGene_tx=XM_023247300.1;refGene_gn=PPFIA4;refGene_tx=XM_023247301.1;refGene_gn=PPFIA4;refGene_tx=XM_023247302.1;refGene_gn=PPFIA4;refGene_tx=XM_023247303.1;refGene_gn=PPFIA4;refGene_tx=XM_023247304.1
F1 43722306 43722404
refGene_gn=PPFIA4;refGene_tx=XM_023247297.1;refGene_gn=PPFIA4;refGene_tx=XM_023247298.1;refGene_gn=PPFIA4;refGene_tx=XM_023247299.1;refGene_gn=PPFIA4;refGene_tx=XM_023247300.1;refGene_gn=PPFIA4;refGene_tx=XM_023247301.1;refGene_gn=PPFIA4;refGene_tx=XM_023247302.1;refGene_gn=PPFIA4;refGene_tx=XM_023247303.1;refGene_gn=PPFIA4;refGene_tx=XM_023247304.1
F1 43723065 43723241
refGene_gn=PPFIA4;refGene_tx=XM_023247297.1;refGene_gn=PPFIA4;refGene_tx=XM_023247298.1;refGene_gn=PPFIA4;refGene_tx=XM_023247299.1;refGene_gn=PPFIA4;refGene_tx=XM_023247300.1;refGene_gn=PPFIA4;refGene_tx=XM_023247301.1;refGene_gn=PPFIA4;refGene_tx=XM_023247302.1;refGene_gn=PPFIA4;refGene_tx=XM_023247303.1;refGene_gn=PPFIA4;refGene_tx=XM_023247304.1
F1 43725599 43725668
refGene_gn=PPFIA4;refGene_tx=XM_023247297.1;refGene_gn=PPFIA4;refGene_tx=XM_023247298.1;refGene_gn=PPFIA4;refGene_tx=XM_023247299.1;refGene_gn=PPFIA4;refGene_tx=XM_023247300.1;refGene_gn=PPFIA4;refGene_tx=XM_023247301.1;refGene_gn=PPFIA4;refGene_tx=XM_023247302.1;refGene_gn=PPFIA4;refGene_tx=XM_023247303.1;refGene_gn=PPFIA4;refGene_tx=XM_023247304.1
F1 43729176 43729351
refGene_gn=PPFIA4;refGene_tx=XM_023247297.1;refGene_gn=PPFIA4;refGene_tx=XM_023247298.1;refGene_gn=PPFIA4;refGene_tx=XM_023247299.1;refGene_gn=PPFIA4;refGene_tx=XM_023247300.1;refGene_gn=PPFIA4;refGene_tx=XM_023247301.1;refGene_gn=PPFIA4;refGene_tx=XM_023247302.1;refGene_gn=PPFIA4;refGene_tx=XM_023247303.1;refGene_gn=PPFIA4;refGene_tx=XM_023247304.1
F1 43729488 43729547
refGene_gn=PPFIA4;refGene_tx=XM_023247297.1;refGene_gn=PPFIA4;refGene_tx=XM_023247298.1;refGene_gn=PPFIA4;refGene_tx=XM_023247299.1;refGene_

gn=PPFIA4;refGene_tx=XM_023247301.1;refGene_gn=PPFIA4;refGene_tx=XM_023247302.1;refGene_gn=PPFIA4;refGene_tx=XM_023247304.1
F1 43729881 43729931 refGene_gn=PPFIA4;refGene_tx=XM_023247300.1;refGene_gn=PPFIA4;refGene_tx=XM_023247303.1
F1 43737672 43737794 refGene_gn=MYOG;refGene_tx=XM_003999381.5
F1 43738440 43738522 refGene_gn=MYOG;refGene_tx=XM_003999381.5
F1 43739288 43739834 refGene_gn=MYOG;refGene_tx=XM_003999381.5
F1 43781227 43781568 refGene_gn=ADORA1;refGene_tx=XM_023247833.1
F1 43795903 43796049 refGene_gn=ADORA1;refGene_tx=XM_019822016.2
F1 43812283 43812923 refGene_gn=ADORA1;refGene_tx=XM_019822016.2;refGene_gn=ADORA1;refGene_tx=XM_023247833.1
F1 43815653 43815797 refGene_gn=MYBPH;refGene_tx=XM_006942872.4
F1 43815798 43815912 refGene_gn=MYBPH;refGene_tx=XM_006942872.4
F1 43816037 43816174 refGene_gn=MYBPH;refGene_tx=XM_006942872.4
F1 43817415 43817575 refGene_gn=MYBPH;refGene_tx=XM_006942872.4
F1 43818184 43818324 refGene_gn=MYBPH;refGene_tx=XM_006942872.4
F1 43818515 43818607 refGene_gn=MYBPH;refGene_tx=XM_006942872.4
F1 43818608 43818695 refGene_gn=MYBPH;refGene_tx=XM_006942872.4
F1 43818894 43818899 refGene_gn=MYBPH;refGene_tx=XM_006942872.4
F1 43819081 43819170 refGene_gn=MYBPH;refGene_tx=XM_006942872.4
F1 43821621 43821727 refGene_gn=MYBPH;refGene_tx=XM_006942872.4
F1 43821728 43821778 refGene_gn=MYBPH;refGene_tx=XM_006942872.4
F1 43822528 43822663 refGene_gn=MYBPH;refGene_tx=XM_006942872.4
F1 43822780 43822991 refGene_gn=MYBPH;refGene_tx=XM_006942872.4
F1 43826574 43826715 refGene_gn=CHI3L1;refGene_tx=XM_003999383.5;refGene_gn=CHI3L1;refGene_tx=XM_019822014.2;refGene_gn=CHI3L1;refGene_tx=XM_019822015.2
F1 43826956 43827073 refGene_gn=CHI3L1;refGene_tx=XM_003999383.5;refGene_gn=CHI3L1;refGene_tx=XM_019822014.2;refGene_gn=CHI3L1;refGene_tx=XM_019822015.2
F1 43828364 43828547 refGene_gn=CHI3L1;refGene_tx=XM_003999383.5;refGene_gn=CHI3L1;refGene_tx=XM_019822014.2;refGene_gn=CHI3L1;refGene_tx=XM_019822015.2
F1 43829260 43829384 refGene_gn=CHI3L1;refGene_tx=XM_003999383.5;refGene_gn=CHI3L1;refGene_tx=XM_019822014.2;refGene_gn=CHI3L1;refGene_tx=XM_019822015.2
F1 43830679 43830801 refGene_gn=CHI3L1;refGene_tx=XM_003999383.5;refGene_gn=CHI3L1;refGene_tx=XM_019822014.2;refGene_gn=CHI3L1;refGene_tx=XM_019822015.2
F1 43831887 43832038 refGene_gn=CHI3L1;refGene_tx=XM_003999383.5;refGene_gn=CHI3L1;refGene_tx=XM_019822014.2;refGene_gn=CHI3L1;refGene_tx=XM_019822015.2
F1 43832659 43832716 refGene_gn=CHI3L1;refGene_tx=XM_003999383.5;refGene_gn=CHI3L1;refGene_tx=XM_019822014.2;refGene_gn=CHI3L1;refGene_tx=XM_019822015.2
F1 43833213 43833415 refGene_gn=CHI3L1;refGene_tx=XM_003999383.5;refGene_gn=CHI3L1;refGene_tx=XM_019822014.2;refGene_gn=CHI3L1;refGene_tx=XM_019822015.2

F1 43834352 43834382
refGene_gn=CHI3L1;refGene_tx=XM_003999383.5;refGene_gn=CHI3L1;refGene_tx=XM_019822014.2;refGene_gn=CHI3L1;refGene_tx=XM_019822015.2
F1 43834619 43834674
refGene_gn=CHI3L1;refGene_tx=XM_003999383.5;refGene_gn=CHI3L1;refGene_tx=XM_019822014.2;refGene_gn=CHI3L1;refGene_tx=XM_019822015.2
F1 43862182 43862418
refGene_gn=CHIT1;refGene_tx=XM_003999472.5;refGene_gn=CHIT1;refGene_tx=XM_019822008.2;refGene_gn=CHIT1;refGene_tx=XM_019822009.2;refGene_gn=CHIT1;refGene_tx=XM_019822010.2;refGene_gn=CHIT1;refGene_tx=XM_019822011.2;refGene_gn=CHIT1;refGene_tx=XM_019822012.2;refGene_gn=CHIT1;refGene_tx=XM_023247831.1
F1 43862976 43863103
refGene_gn=CHIT1;refGene_tx=XM_003999472.5;refGene_gn=CHIT1;refGene_tx=XM_019822008.2;refGene_gn=CHIT1;refGene_tx=XM_019822009.2;refGene_gn=CHIT1;refGene_tx=XM_019822010.2;refGene_gn=CHIT1;refGene_tx=XM_019822011.2;refGene_gn=CHIT1;refGene_tx=XM_019822012.2;refGene_gn=CHIT1;refGene_tx=XM_023247831.1
F1 43864433 43864547
refGene_gn=CHIT1;refGene_tx=XM_003999472.5;refGene_gn=CHIT1;refGene_tx=XM_019822008.2;refGene_gn=CHIT1;refGene_tx=XM_019822009.2;refGene_gn=CHIT1;refGene_tx=XM_019822010.2;refGene_gn=CHIT1;refGene_tx=XM_019822011.2;refGene_gn=CHIT1;refGene_tx=XM_019822012.2;refGene_gn=CHIT1;refGene_tx=XM_023247831.1
F1 43864864 43865050
refGene_gn=CHIT1;refGene_tx=XM_003999472.5;refGene_gn=CHIT1;refGene_tx=XM_019822008.2;refGene_gn=CHIT1;refGene_tx=XM_019822009.2;refGene_gn=CHIT1;refGene_tx=XM_019822010.2;refGene_gn=CHIT1;refGene_tx=XM_019822011.2;refGene_gn=CHIT1;refGene_tx=XM_019822012.2;refGene_gn=CHIT1;refGene_tx=XM_023247831.1
F1 43866875 43866999
refGene_gn=CHIT1;refGene_tx=XM_003999472.5;refGene_gn=CHIT1;refGene_tx=XM_019822008.2;refGene_gn=CHIT1;refGene_tx=XM_019822009.2;refGene_gn=CHIT1;refGene_tx=XM_019822010.2;refGene_gn=CHIT1;refGene_tx=XM_019822011.2;refGene_gn=CHIT1;refGene_tx=XM_019822012.2;refGene_gn=CHIT1;refGene_tx=XM_023247831.1
F1 43867704 43867829
refGene_gn=CHIT1;refGene_tx=XM_003999472.5;refGene_gn=CHIT1;refGene_tx=XM_019822008.2;refGene_gn=CHIT1;refGene_tx=XM_019822009.2;refGene_gn=CHIT1;refGene_tx=XM_019822010.2;refGene_gn=CHIT1;refGene_tx=XM_019822011.2;refGene_gn=CHIT1;refGene_tx=XM_019822012.2;refGene_gn=CHIT1;refGene_tx=XM_023247831.1
F1 43868057 43868223
refGene_gn=CHIT1;refGene_tx=XM_003999472.5;refGene_gn=CHIT1;refGene_tx=XM_019822008.2;refGene_gn=CHIT1;refGene_tx=XM_019822009.2;refGene_gn=CHIT1;refGene_tx=XM_019822010.2;refGene_gn=CHIT1;refGene_tx=XM_019822011.2;refGene_gn=CHIT1;refGene_tx=XM_019822012.2;refGene_gn=CHIT1;refGene_tx=XM_023247831.1
F1 43869560 43869617
refGene_gn=CHIT1;refGene_tx=XM_003999472.5;refGene_gn=CHIT1;refGene_tx

=XM_019822008.2;refGene_gn=CHIT1;refGene_tx=XM_019822009.2;refGene_gn=CHIT1;refGene_tx=XM_019822010.2;refGene_gn=CHIT1;refGene_tx=XM_019822011.2;refGene_gn=CHIT1;refGene_tx=XM_019822012.2;refGene_gn=CHIT1;refGene_tx=XM_023247831.1
F1 43870217 43870419
refGene_gn=CHIT1;refGene_tx=XM_003999472.5;refGene_gn=CHIT1;refGene_tx=XM_019822008.2;refGene_gn=CHIT1;refGene_tx=XM_019822009.2;refGene_gn=CHIT1;refGene_tx=XM_019822010.2;refGene_gn=CHIT1;refGene_tx=XM_019822011.2;refGene_gn=CHIT1;refGene_tx=XM_019822012.2;refGene_gn=CHIT1;refGene_tx=XM_023247831.1
F1 43872952 43872982
refGene_gn=CHIT1;refGene_tx=XM_003999472.5;refGene_gn=CHIT1;refGene_tx=XM_019822008.2;refGene_gn=CHIT1;refGene_tx=XM_019822009.2;refGene_gn=CHIT1;refGene_tx=XM_019822010.2;refGene_gn=CHIT1;refGene_tx=XM_019822011.2;refGene_gn=CHIT1;refGene_tx=XM_019822012.2;refGene_gn=CHIT1;refGene_tx=XM_023247831.1
F1 43874088 43874113 refGene_gn=CHIT1;refGene_tx=XM_003999472.5
F1 43874516 43874593 refGene_gn=CHIT1;refGene_tx=XM_019822008.2
F1 43897505 43897516 refGene_gn=CHIT1;refGene_tx=XM_019822008.2
F1 43929584 43929738 refGene_gn=BTG2;refGene_tx=XM_003999473.5
F1 43931003 43931338 refGene_gn=BTG2;refGene_tx=XM_003999473.5
F1 43959333 43959485 refGene_gn=FMOD;refGene_tx=XM_006942878.4
F1 43964738 43965717 refGene_gn=FMOD;refGene_tx=XM_006942878.4
F1 44068098 44069062
refGene_gn=PRELP;refGene_tx=XM_003999385.5;refGene_gn=PRELP;refGene_tx=XM_006942879.4
F1 44071606 44071782
refGene_gn=PRELP;refGene_tx=XM_003999385.5;refGene_gn=PRELP;refGene_tx=XM_006942879.4
F1 44077096 44077290
refGene_gn=OPTC;refGene_tx=XM_006942880.2;refGene_gn=OPTC;refGene_tx=XM_011290930.3;refGene_gn=OPTC;refGene_tx=XM_019822017.2;refGene_gn=OPTC;refGene_tx=XM_019822018.2
F1 44078942 44079174
refGene_gn=OPTC;refGene_tx=XM_006942880.2;refGene_gn=OPTC;refGene_tx=XM_011290930.3;refGene_gn=OPTC;refGene_tx=XM_019822017.2;refGene_gn=OPTC;refGene_tx=XM_019822018.2
F1 44079904 44080043
refGene_gn=OPTC;refGene_tx=XM_006942880.2;refGene_gn=OPTC;refGene_tx=XM_011290930.3;refGene_gn=OPTC;refGene_tx=XM_019822017.2;refGene_gn=OPTC;refGene_tx=XM_019822018.2
F1 44081555 44081723
refGene_gn=OPTC;refGene_tx=XM_006942880.2;refGene_gn=OPTC;refGene_tx=XM_011290930.3;refGene_gn=OPTC;refGene_tx=XM_019822017.2;refGene_gn=OPTC;refGene_tx=XM_019822018.2
F1 44082596 44082799
refGene_gn=OPTC;refGene_tx=XM_006942880.2;refGene_gn=OPTC;refGene_tx=XM_011290930.3;refGene_gn=OPTC;refGene_tx=XM_019822017.2;refGene_gn=OPTC;refGene_tx=XM_019822018.2
F1 44083001 44083229 refGene_gn=OPTC;refGene_tx=XM_019822018.2

F1 44083972 44084068
refGene_gn=OPTC;refGene_tx=XM_006942880.2;refGene_gn=OPTC;refGene_tx=XM_011290930.3
F1 44084510 44084681
refGene_gn=OPTC;refGene_tx=XM_006942880.2;refGene_gn=OPTC;refGene_tx=XM_011290930.3
F1 44085184 44085403 refGene_gn=OPTC;refGene_tx=XM_019822017.2
F1 44225536 44225729
refGene_gn=ATP2B4;refGene_tx=XM_003999386.5;refGene_gn=ATP2B4;refGene_tx=XM_003999387.5;refGene_gn=ATP2B4;refGene_tx=XM_003999388.5;refGene_gn=ATP2B4;refGene_tx=XM_006942881.4;refGene_gn=ATP2B4;refGene_tx=XM_006942883.4;refGene_gn=ATP2B4;refGene_tx=XM_006942884.4;refGene_gn=ATP2B4;refGene_tx=XM_011290928.3;refGene_gn=ATP2B4;refGene_tx=XM_019822019.2
F1 44236867 44237065
refGene_gn=ATP2B4;refGene_tx=XM_003999386.5;refGene_gn=ATP2B4;refGene_tx=XM_003999387.5;refGene_gn=ATP2B4;refGene_tx=XM_003999388.5;refGene_gn=ATP2B4;refGene_tx=XM_006942881.4;refGene_gn=ATP2B4;refGene_tx=XM_006942883.4;refGene_gn=ATP2B4;refGene_tx=XM_006942884.4;refGene_gn=ATP2B4;refGene_tx=XM_011290928.3;refGene_gn=ATP2B4;refGene_tx=XM_019822019.2
F1 44237677 44237935
refGene_gn=ATP2B4;refGene_tx=XM_003999386.5;refGene_gn=ATP2B4;refGene_tx=XM_003999387.5;refGene_gn=ATP2B4;refGene_tx=XM_003999388.5;refGene_gn=ATP2B4;refGene_tx=XM_006942881.4;refGene_gn=ATP2B4;refGene_tx=XM_006942883.4;refGene_gn=ATP2B4;refGene_tx=XM_006942884.4;refGene_gn=ATP2B4;refGene_tx=XM_011290928.3;refGene_gn=ATP2B4;refGene_tx=XM_019822019.2
F1 44238379 44238505
refGene_gn=ATP2B4;refGene_tx=XM_003999386.5;refGene_gn=ATP2B4;refGene_tx=XM_003999387.5;refGene_gn=ATP2B4;refGene_tx=XM_003999388.5;refGene_gn=ATP2B4;refGene_tx=XM_006942881.4;refGene_gn=ATP2B4;refGene_tx=XM_006942883.4;refGene_gn=ATP2B4;refGene_tx=XM_006942884.4;refGene_gn=ATP2B4;refGene_tx=XM_011290928.3;refGene_gn=ATP2B4;refGene_tx=XM_019822019.2;refGene_gn=ATP2B4;refGene_tx=XM_019822020.2
F1 44239418 44239544
refGene_gn=ATP2B4;refGene_tx=XM_003999386.5;refGene_gn=ATP2B4;refGene_tx=XM_003999387.5;refGene_gn=ATP2B4;refGene_tx=XM_003999388.5;refGene_gn=ATP2B4;refGene_tx=XM_006942881.4;refGene_gn=ATP2B4;refGene_tx=XM_006942883.4;refGene_gn=ATP2B4;refGene_tx=XM_006942884.4;refGene_gn=ATP2B4;refGene_tx=XM_011290928.3;refGene_gn=ATP2B4;refGene_tx=XM_019822019.2;refGene_gn=ATP2B4;refGene_tx=XM_019822020.2
F1 44240661 44240697
refGene_gn=ATP2B4;refGene_tx=XM_003999386.5;refGene_gn=ATP2B4;refGene_tx=XM_003999387.5;refGene_gn=ATP2B4;refGene_tx=XM_003999388.5;refGene_gn=ATP2B4;refGene_tx=XM_006942881.4;refGene_gn=ATP2B4;refGene_tx=XM_011290928.3;refGene_gn=ATP2B4;refGene_tx=XM_019822019.2;refGene_gn=ATP2B4;refGene_tx=XM_019822020.2
F1 44242280 44242442
refGene_gn=ATP2B4;refGene_tx=XM_003999386.5;refGene_gn=ATP2B4;refGene_

tx=XM_003999387.5;refGene_gn=ATP2B4;refGene_tx=XM_003999388.5;refGene_gn=ATP2B4;refGene_tx=XM_006942881.4;refGene_gn=ATP2B4;refGene_tx=XM_006942883.4;refGene_gn=ATP2B4;refGene_tx=XM_006942884.4;refGene_gn=ATP2B4;refGene_tx=XM_011290928.3;refGene_gn=ATP2B4;refGene_tx=XM_019822019.2;refGene_gn=ATP2B4;refGene_tx=XM_019822020.2
F1 44245695 44245910
refGene_gn=ATP2B4;refGene_tx=XM_003999386.5;refGene_gn=ATP2B4;refGene_tx=XM_003999387.5;refGene_gn=ATP2B4;refGene_tx=XM_003999388.5;refGene_gn=ATP2B4;refGene_tx=XM_006942881.4;refGene_gn=ATP2B4;refGene_tx=XM_006942883.4;refGene_gn=ATP2B4;refGene_tx=XM_006942884.4;refGene_gn=ATP2B4;refGene_tx=XM_011290928.3;refGene_gn=ATP2B4;refGene_tx=XM_019822019.2;refGene_gn=ATP2B4;refGene_tx=XM_019822020.2
F1 44246272 44246515
refGene_gn=ATP2B4;refGene_tx=XM_003999386.5;refGene_gn=ATP2B4;refGene_tx=XM_003999387.5;refGene_gn=ATP2B4;refGene_tx=XM_003999388.5;refGene_gn=ATP2B4;refGene_tx=XM_006942881.4;refGene_gn=ATP2B4;refGene_tx=XM_006942883.4;refGene_gn=ATP2B4;refGene_tx=XM_006942884.4;refGene_gn=ATP2B4;refGene_tx=XM_011290928.3;refGene_gn=ATP2B4;refGene_tx=XM_019822019.2;refGene_gn=ATP2B4;refGene_tx=XM_019822020.2
F1 44247048 44247290
refGene_gn=ATP2B4;refGene_tx=XM_003999386.5;refGene_gn=ATP2B4;refGene_tx=XM_003999387.5;refGene_gn=ATP2B4;refGene_tx=XM_003999388.5;refGene_gn=ATP2B4;refGene_tx=XM_006942881.4;refGene_gn=ATP2B4;refGene_tx=XM_006942883.4;refGene_gn=ATP2B4;refGene_tx=XM_006942884.4;refGene_gn=ATP2B4;refGene_tx=XM_011290928.3;refGene_gn=ATP2B4;refGene_tx=XM_019822019.2;refGene_gn=ATP2B4;refGene_tx=XM_019822020.2
F1 44251096 44251328
refGene_gn=ATP2B4;refGene_tx=XM_003999386.5;refGene_gn=ATP2B4;refGene_tx=XM_003999387.5;refGene_gn=ATP2B4;refGene_tx=XM_003999388.5;refGene_gn=ATP2B4;refGene_tx=XM_006942881.4;refGene_gn=ATP2B4;refGene_tx=XM_006942883.4;refGene_gn=ATP2B4;refGene_tx=XM_006942884.4;refGene_gn=ATP2B4;refGene_tx=XM_011290928.3;refGene_gn=ATP2B4;refGene_tx=XM_019822019.2;refGene_gn=ATP2B4;refGene_tx=XM_019822020.2
F1 44252251 44252431
refGene_gn=ATP2B4;refGene_tx=XM_003999386.5;refGene_gn=ATP2B4;refGene_tx=XM_003999387.5;refGene_gn=ATP2B4;refGene_tx=XM_003999388.5;refGene_gn=ATP2B4;refGene_tx=XM_006942881.4;refGene_gn=ATP2B4;refGene_tx=XM_006942883.4;refGene_gn=ATP2B4;refGene_tx=XM_006942884.4;refGene_gn=ATP2B4;refGene_tx=XM_011290928.3;refGene_gn=ATP2B4;refGene_tx=XM_019822019.2;refGene_gn=ATP2B4;refGene_tx=XM_019822020.2
F1 44253272 44253360
refGene_gn=ATP2B4;refGene_tx=XM_003999386.5;refGene_gn=ATP2B4;refGene_tx=XM_003999387.5;refGene_gn=ATP2B4;refGene_tx=XM_003999388.5;refGene_gn=ATP2B4;refGene_tx=XM_006942881.4;refGene_gn=ATP2B4;refGene_tx=XM_006942883.4;refGene_gn=ATP2B4;refGene_tx=XM_006942884.4;refGene_gn=ATP2B4;refGene_tx=XM_011290928.3;refGene_gn=ATP2B4;refGene_tx=XM_019822019.2;refGene_gn=ATP2B4;refGene_tx=XM_019822020.2
F1 44255146 44255253
refGene_gn=ATP2B4;refGene_tx=XM_003999386.5;refGene_gn=ATP2B4;refGene_tx=XM_003999387.5;refGene_gn=ATP2B4;refGene_tx=XM_003999388.5;refGene_

gn=ATP2B4;refGene_tx=XM_006942881.4;refGene_gn=ATP2B4;refGene_tx=XM_006942883.4;refGene_gn=ATP2B4;refGene_tx=XM_006942884.4;refGene_gn=ATP2B4;refGene_tx=XM_011290928.3;refGene_gn=ATP2B4;refGene_tx=XM_019822019.2;refGene_gn=ATP2B4;refGene_tx=XM_019822020.2
F1 44259029 44259221
refGene_gn=ATP2B4;refGene_tx=XM_003999386.5;refGene_gn=ATP2B4;refGene_tx=XM_003999387.5;refGene_gn=ATP2B4;refGene_tx=XM_003999388.5;refGene_gn=ATP2B4;refGene_tx=XM_006942881.4;refGene_gn=ATP2B4;refGene_tx=XM_006942883.4;refGene_gn=ATP2B4;refGene_tx=XM_006942884.4;refGene_gn=ATP2B4;refGene_tx=XM_011290928.3;refGene_gn=ATP2B4;refGene_tx=XM_019822019.2;refGene_gn=ATP2B4;refGene_tx=XM_019822020.2
F1 44259694 44259908
refGene_gn=ATP2B4;refGene_tx=XM_003999386.5;refGene_gn=ATP2B4;refGene_tx=XM_003999387.5;refGene_gn=ATP2B4;refGene_tx=XM_003999388.5;refGene_gn=ATP2B4;refGene_tx=XM_006942881.4;refGene_gn=ATP2B4;refGene_tx=XM_006942883.4;refGene_gn=ATP2B4;refGene_tx=XM_006942884.4;refGene_gn=ATP2B4;refGene_tx=XM_011290928.3;refGene_gn=ATP2B4;refGene_tx=XM_019822019.2;refGene_gn=ATP2B4;refGene_tx=XM_019822020.2
F1 44260735 44260947
refGene_gn=ATP2B4;refGene_tx=XM_003999386.5;refGene_gn=ATP2B4;refGene_tx=XM_003999387.5;refGene_gn=ATP2B4;refGene_tx=XM_003999388.5;refGene_gn=ATP2B4;refGene_tx=XM_006942881.4;refGene_gn=ATP2B4;refGene_tx=XM_006942883.4;refGene_gn=ATP2B4;refGene_tx=XM_006942884.4;refGene_gn=ATP2B4;refGene_tx=XM_011290928.3;refGene_gn=ATP2B4;refGene_tx=XM_019822019.2;refGene_gn=ATP2B4;refGene_tx=XM_019822020.2
F1 44261960 44262068
refGene_gn=ATP2B4;refGene_tx=XM_003999386.5;refGene_gn=ATP2B4;refGene_tx=XM_003999387.5;refGene_gn=ATP2B4;refGene_tx=XM_003999388.5;refGene_gn=ATP2B4;refGene_tx=XM_006942881.4;refGene_gn=ATP2B4;refGene_tx=XM_006942883.4;refGene_gn=ATP2B4;refGene_tx=XM_006942884.4;refGene_gn=ATP2B4;refGene_tx=XM_011290928.3;refGene_gn=ATP2B4;refGene_tx=XM_019822020.2
F1 44264482 44264659
refGene_gn=ATP2B4;refGene_tx=XM_003999386.5;refGene_gn=ATP2B4;refGene_tx=XM_003999387.5;refGene_gn=ATP2B4;refGene_tx=XM_003999388.5;refGene_gn=ATP2B4;refGene_tx=XM_006942881.4;refGene_gn=ATP2B4;refGene_tx=XM_006942883.4;refGene_gn=ATP2B4;refGene_tx=XM_006942884.4;refGene_gn=ATP2B4;refGene_tx=XM_011290928.3;refGene_gn=ATP2B4;refGene_tx=XM_019822019.2;refGene_gn=ATP2B4;refGene_tx=XM_019822020.2
F1 44267902 44268083
refGene_gn=ATP2B4;refGene_tx=XM_003999387.5;refGene_gn=ATP2B4;refGene_tx=XM_003999388.5;refGene_gn=ATP2B4;refGene_tx=XM_006942884.4
F1 44272815 44273130
refGene_gn=ATP2B4;refGene_tx=XM_003999386.5;refGene_gn=ATP2B4;refGene_tx=XM_003999387.5;refGene_gn=ATP2B4;refGene_tx=XM_003999388.5;refGene_gn=ATP2B4;refGene_tx=XM_006942881.4;refGene_gn=ATP2B4;refGene_tx=XM_006942883.4;refGene_gn=ATP2B4;refGene_tx=XM_006942884.4;refGene_gn=ATP2B4;refGene_tx=XM_011290928.3;refGene_gn=ATP2B4;refGene_tx=XM_019822019.2;refGene_gn=ATP2B4;refGene_tx=XM_019822020.2
F1 44287310 44287405

refGene_gn=LAX1;refGene_tx=XM_006942885.4;refGene_gn=LAX1;refGene_tx=XM_006942886.4;refGene_gn=LAX1;refGene_tx=XM_019822021.2;refGene_gn=LAX1;refGene_tx=XM_019822022.2;refGene_gn=LAX1;refGene_tx=XM_019822023.2
F1 44290034 44290144
refGene_gn=LAX1;refGene_tx=XM_006942885.4;refGene_gn=LAX1;refGene_tx=XM_006942886.4;refGene_gn=LAX1;refGene_tx=XM_019822021.2;refGene_gn=LAX1;refGene_tx=XM_019822022.2;refGene_gn=LAX1;refGene_tx=XM_019822023.2;refGene_gn=LAX1;refGene_tx=XM_019822025.2;refGene_gn=LAX1;refGene_tx=XM_019822026.2;refGene_gn=LAX1;refGene_tx=XM_019822027.2;refGene_gn=LAX1;refGene_tx=XM_019822028.2
F1 44290575 44290686
refGene_gn=LAX1;refGene_tx=XM_006942885.4;refGene_gn=LAX1;refGene_tx=XM_006942886.4;refGene_gn=LAX1;refGene_tx=XM_019822021.2;refGene_gn=LAX1;refGene_tx=XM_019822022.2;refGene_gn=LAX1;refGene_tx=XM_019822023.2;refGene_gn=LAX1;refGene_tx=XM_019822025.2;refGene_gn=LAX1;refGene_tx=XM_019822026.2;refGene_gn=LAX1;refGene_tx=XM_019822027.2;refGene_gn=LAX1;refGene_tx=XM_019822028.2
F1 44291257 44291343
refGene_gn=LAX1;refGene_tx=XM_006942885.4;refGene_gn=LAX1;refGene_tx=XM_006942886.4;refGene_gn=LAX1;refGene_tx=XM_019822021.2;refGene_gn=LAX1;refGene_tx=XM_019822022.2;refGene_gn=LAX1;refGene_tx=XM_019822023.2;refGene_gn=LAX1;refGene_tx=XM_019822025.2;refGene_gn=LAX1;refGene_tx=XM_019822026.2;refGene_gn=LAX1;refGene_tx=XM_019822027.2;refGene_gn=LAX1;refGene_tx=XM_019822028.2
F1 44293641 44294436
refGene_gn=LAX1;refGene_tx=XM_006942885.4;refGene_gn=LAX1;refGene_tx=XM_006942886.4;refGene_gn=LAX1;refGene_tx=XM_019822021.2;refGene_gn=LAX1;refGene_tx=XM_019822022.2;refGene_gn=LAX1;refGene_tx=XM_019822023.2;refGene_gn=LAX1;refGene_tx=XM_019822025.2;refGene_gn=LAX1;refGene_tx=XM_019822026.2;refGene_gn=LAX1;refGene_tx=XM_019822027.2;refGene_gn=LAX1;refGene_tx=XM_019822028.2
F1 44314804 44317747 refGene_gn=ZBED6;refGene_tx=XM_019822029.2
F1 44327757 44327811
refGene_gn=ZC3H11A;refGene_tx=XM_006942887.4;refGene_gn=ZC3H11A;refGene_tx=XM_006942888.4;refGene_gn=ZC3H11A;refGene_tx=XM_006942889.4;refGene_gn=ZC3H11A;refGene_tx=XM_006942890.4;refGene_gn=ZC3H11A;refGene_tx=XM_006942891.4;refGene_gn=ZC3H11A;refGene_tx=XM_006942892.4;refGene_gn=ZC3H11A;refGene_tx=XM_006942893.4;refGene_gn=ZC3H11A;refGene_tx=XM_006942896.4;refGene_gn=ZC3H11A;refGene_tx=XM_011290924.3;refGene_gn=ZC3H11A;refGene_tx=XM_019822030.2;refGene_gn=ZC3H11A;refGene_tx=XM_019822031.2;refGene_gn=ZC3H11A;refGene_tx=XM_019822032.2;refGene_gn=ZC3H11A;refGene_tx=XM_019822033.2;refGene_gn=ZC3H11A;refGene_tx=XM_019822035.2;refGene_gn=ZC3H11A;refGene_tx=XM_019822037.2;refGene_gn=ZC3H11A;refGene_tx=XM_019822038.2;refGene_gn=ZC3H11A;refGene_tx=XM_023247369.1
F1 44328917 44329037
refGene_gn=ZC3H11A;refGene_tx=XM_006942887.4;refGene_gn=ZC3H11A;refGene_tx=XM_006942888.4;refGene_gn=ZC3H11A;refGene_tx=XM_006942889.4;refGene_gn=ZC3H11A;refGene_tx=XM_006942890.4;refGene_gn=ZC3H11A;refGene_tx=XM_006942891.4;refGene_gn=ZC3H11A;refGene_tx=XM_006942892.4;refGene_gn=ZC3H11A;refGene_tx=XM_006942893.4;refGene_gn=ZC3H11A;refGene_tx=XM_00

6942896.4;refGene_gn=ZC3H11A;refGene_tx=XM_011290924.3;refGene_gn=ZC3H11A;refGene_tx=XM_019822030.2;refGene_gn=ZC3H11A;refGene_tx=XM_019822031.2;refGene_gn=ZC3H11A;refGene_tx=XM_019822032.2;refGene_gn=ZC3H11A;refGene_tx=XM_019822033.2;refGene_gn=ZC3H11A;refGene_tx=XM_019822035.2;refGene_gn=ZC3H11A;refGene_tx=XM_019822037.2;refGene_gn=ZC3H11A;refGene_tx=XM_019822038.2;refGene_gn=ZC3H11A;refGene_tx=XM_023247369.1
F1 44337217 44337341
refGene_gn=ZC3H11A;refGene_tx=XM_006942887.4;refGene_gn=ZC3H11A;refGene_tx=XM_006942888.4;refGene_gn=ZC3H11A;refGene_tx=XM_006942889.4;refGene_gn=ZC3H11A;refGene_tx=XM_006942890.4;refGene_gn=ZC3H11A;refGene_tx=XM_006942891.4;refGene_gn=ZC3H11A;refGene_tx=XM_006942892.4;refGene_gn=ZC3H11A;refGene_tx=XM_006942893.4;refGene_gn=ZC3H11A;refGene_tx=XM_006942896.4;refGene_gn=ZC3H11A;refGene_tx=XM_011290924.3;refGene_gn=ZC3H11A;refGene_tx=XM_019822030.2;refGene_gn=ZC3H11A;refGene_tx=XM_019822031.2;refGene_gn=ZC3H11A;refGene_tx=XM_019822032.2;refGene_gn=ZC3H11A;refGene_tx=XM_019822033.2;refGene_gn=ZC3H11A;refGene_tx=XM_019822035.2;refGene_gn=ZC3H11A;refGene_tx=XM_019822037.2;refGene_gn=ZC3H11A;refGene_tx=XM_019822038.2;refGene_gn=ZC3H11A;refGene_tx=XM_023247369.1
F1 44338316 44338520
refGene_gn=ZC3H11A;refGene_tx=XM_006942887.4;refGene_gn=ZC3H11A;refGene_tx=XM_006942888.4;refGene_gn=ZC3H11A;refGene_tx=XM_006942889.4;refGene_gn=ZC3H11A;refGene_tx=XM_006942890.4;refGene_gn=ZC3H11A;refGene_tx=XM_006942891.4;refGene_gn=ZC3H11A;refGene_tx=XM_006942892.4;refGene_gn=ZC3H11A;refGene_tx=XM_006942893.4;refGene_gn=ZC3H11A;refGene_tx=XM_006942896.4;refGene_gn=ZC3H11A;refGene_tx=XM_011290924.3;refGene_gn=ZC3H11A;refGene_tx=XM_019822030.2;refGene_gn=ZC3H11A;refGene_tx=XM_019822031.2;refGene_gn=ZC3H11A;refGene_tx=XM_019822032.2;refGene_gn=ZC3H11A;refGene_tx=XM_019822033.2;refGene_gn=ZC3H11A;refGene_tx=XM_019822035.2;refGene_gn=ZC3H11A;refGene_tx=XM_019822037.2;refGene_gn=ZC3H11A;refGene_tx=XM_019822038.2;refGene_gn=ZC3H11A;refGene_tx=XM_023247369.1
F1 44338637 44338754
refGene_gn=ZC3H11A;refGene_tx=XM_006942887.4;refGene_gn=ZC3H11A;refGene_tx=XM_006942888.4;refGene_gn=ZC3H11A;refGene_tx=XM_006942889.4;refGene_gn=ZC3H11A;refGene_tx=XM_006942890.4;refGene_gn=ZC3H11A;refGene_tx=XM_006942891.4;refGene_gn=ZC3H11A;refGene_tx=XM_006942892.4;refGene_gn=ZC3H11A;refGene_tx=XM_006942893.4;refGene_gn=ZC3H11A;refGene_tx=XM_006942896.4;refGene_gn=ZC3H11A;refGene_tx=XM_011290924.3;refGene_gn=ZC3H11A;refGene_tx=XM_019822030.2;refGene_gn=ZC3H11A;refGene_tx=XM_019822031.2;refGene_gn=ZC3H11A;refGene_tx=XM_019822032.2;refGene_gn=ZC3H11A;refGene_tx=XM_019822033.2;refGene_gn=ZC3H11A;refGene_tx=XM_019822035.2;refGene_gn=ZC3H11A;refGene_tx=XM_019822037.2;refGene_gn=ZC3H11A;refGene_tx=XM_019822038.2;refGene_gn=ZC3H11A;refGene_tx=XM_023247369.1
F1 44339017 44339098
refGene_gn=ZC3H11A;refGene_tx=XM_006942887.4;refGene_gn=ZC3H11A;refGene_tx=XM_006942888.4;refGene_gn=ZC3H11A;refGene_tx=XM_006942889.4;refGene_gn=ZC3H11A;refGene_tx=XM_006942890.4;refGene_gn=ZC3H11A;refGene_tx=XM_006942891.4;refGene_gn=ZC3H11A;refGene_tx=XM_006942892.4;refGene_gn=ZC3H11A;refGene_tx=XM_006942893.4;refGene_gn=ZC3H11A;refGene_tx=XM_006942896.4;refGene_gn=ZC3H11A;refGene_tx=XM_011290924.3;refGene_gn=ZC3H11A;refGene_tx=XM_019822030.2;refGene_gn=ZC3H11A;refGene_tx=XM_0198220

31.2;refGene_gn=ZC3H11A;refGene_tx=XM_019822032.2;refGene_gn=ZC3H11A;refGene_tx=XM_019822033.2;refGene_gn=ZC3H11A;refGene_tx=XM_019822035.2;refGene_gn=ZC3H11A;refGene_tx=XM_019822037.2;refGene_gn=ZC3H11A;refGene_tx=XM_019822038.2;refGene_gn=ZC3H11A;refGene_tx=XM_023247369.1
F1 44339906 44340017
refGene_gn=ZC3H11A;refGene_tx=XM_006942887.4;refGene_gn=ZC3H11A;refGene_tx=XM_006942888.4;refGene_gn=ZC3H11A;refGene_tx=XM_006942889.4;refGene_gn=ZC3H11A;refGene_tx=XM_006942890.4;refGene_gn=ZC3H11A;refGene_tx=XM_006942891.4;refGene_gn=ZC3H11A;refGene_tx=XM_006942892.4;refGene_gn=ZC3H11A;refGene_tx=XM_006942893.4;refGene_gn=ZC3H11A;refGene_tx=XM_006942896.4;refGene_gn=ZC3H11A;refGene_tx=XM_011290924.3;refGene_gn=ZC3H11A;refGene_tx=XM_019822030.2;refGene_gn=ZC3H11A;refGene_tx=XM_019822031.2;refGene_gn=ZC3H11A;refGene_tx=XM_019822032.2;refGene_gn=ZC3H11A;refGene_tx=XM_019822033.2;refGene_gn=ZC3H11A;refGene_tx=XM_019822035.2;refGene_gn=ZC3H11A;refGene_tx=XM_019822037.2;refGene_gn=ZC3H11A;refGene_tx=XM_019822038.2;refGene_gn=ZC3H11A;refGene_tx=XM_023247369.1
F1 44341090 44341153
refGene_gn=ZC3H11A;refGene_tx=XM_006942887.4;refGene_gn=ZC3H11A;refGene_tx=XM_006942888.4;refGene_gn=ZC3H11A;refGene_tx=XM_006942889.4;refGene_gn=ZC3H11A;refGene_tx=XM_006942890.4;refGene_gn=ZC3H11A;refGene_tx=XM_006942891.4;refGene_gn=ZC3H11A;refGene_tx=XM_006942892.4;refGene_gn=ZC3H11A;refGene_tx=XM_006942893.4;refGene_gn=ZC3H11A;refGene_tx=XM_006942896.4;refGene_gn=ZC3H11A;refGene_tx=XM_011290924.3;refGene_gn=ZC3H11A;refGene_tx=XM_019822030.2;refGene_gn=ZC3H11A;refGene_tx=XM_019822031.2;refGene_gn=ZC3H11A;refGene_tx=XM_019822032.2;refGene_gn=ZC3H11A;refGene_tx=XM_019822033.2;refGene_gn=ZC3H11A;refGene_tx=XM_019822035.2;refGene_gn=ZC3H11A;refGene_tx=XM_019822037.2;refGene_gn=ZC3H11A;refGene_tx=XM_019822038.2;refGene_gn=ZC3H11A;refGene_tx=XM_023247369.1
F1 44342236 44342335
refGene_gn=ZC3H11A;refGene_tx=XM_006942887.4;refGene_gn=ZC3H11A;refGene_tx=XM_006942888.4;refGene_gn=ZC3H11A;refGene_tx=XM_006942889.4;refGene_gn=ZC3H11A;refGene_tx=XM_006942890.4;refGene_gn=ZC3H11A;refGene_tx=XM_006942891.4;refGene_gn=ZC3H11A;refGene_tx=XM_006942892.4;refGene_gn=ZC3H11A;refGene_tx=XM_006942893.4;refGene_gn=ZC3H11A;refGene_tx=XM_006942896.4;refGene_gn=ZC3H11A;refGene_tx=XM_011290924.3;refGene_gn=ZC3H11A;refGene_tx=XM_019822030.2;refGene_gn=ZC3H11A;refGene_tx=XM_019822031.2;refGene_gn=ZC3H11A;refGene_tx=XM_019822032.2;refGene_gn=ZC3H11A;refGene_tx=XM_019822033.2;refGene_gn=ZC3H11A;refGene_tx=XM_019822035.2;refGene_gn=ZC3H11A;refGene_tx=XM_019822037.2;refGene_gn=ZC3H11A;refGene_tx=XM_019822038.2;refGene_gn=ZC3H11A;refGene_tx=XM_023247369.1
F1 44344948 44345017
refGene_gn=ZC3H11A;refGene_tx=XM_006942887.4;refGene_gn=ZC3H11A;refGene_tx=XM_006942888.4;refGene_gn=ZC3H11A;refGene_tx=XM_006942889.4;refGene_gn=ZC3H11A;refGene_tx=XM_006942890.4;refGene_gn=ZC3H11A;refGene_tx=XM_006942891.4;refGene_gn=ZC3H11A;refGene_tx=XM_006942892.4;refGene_gn=ZC3H11A;refGene_tx=XM_006942893.4;refGene_gn=ZC3H11A;refGene_tx=XM_006942896.4;refGene_gn=ZC3H11A;refGene_tx=XM_011290924.3;refGene_gn=ZC3H11A;refGene_tx=XM_019822030.2;refGene_gn=ZC3H11A;refGene_tx=XM_019822031.2;refGene_gn=ZC3H11A;refGene_tx=XM_019822032.2;refGene_gn=ZC3H11A;refGene_tx=XM_019822033.2;refGene_gn=ZC3H11A;refGene_tx=XM_019822035.2;

refGene_gn=ZC3H11A;refGene_tx=XM_019822037.2;refGene_gn=ZC3H11A;refGene_tx=XM_019822038.2;refGene_gn=ZC3H11A;refGene_tx=XM_023247369.1
F1 44349437 44349941
refGene_gn=ZC3H11A;refGene_tx=XM_006942887.4;refGene_gn=ZC3H11A;refGene_tx=XM_006942888.4;refGene_gn=ZC3H11A;refGene_tx=XM_006942889.4;refGene_gn=ZC3H11A;refGene_tx=XM_006942890.4;refGene_gn=ZC3H11A;refGene_tx=XM_006942891.4;refGene_gn=ZC3H11A;refGene_tx=XM_006942892.4;refGene_gn=ZC3H11A;refGene_tx=XM_006942893.4;refGene_gn=ZC3H11A;refGene_tx=XM_006942896.4;refGene_gn=ZC3H11A;refGene_tx=XM_011290924.3;refGene_gn=ZC3H11A;refGene_tx=XM_019822030.2;refGene_gn=ZC3H11A;refGene_tx=XM_019822031.2;refGene_gn=ZC3H11A;refGene_tx=XM_019822032.2;refGene_gn=ZC3H11A;refGene_tx=XM_019822033.2;refGene_gn=ZC3H11A;refGene_tx=XM_019822035.2;refGene_gn=ZC3H11A;refGene_tx=XM_019822037.2;refGene_gn=ZC3H11A;refGene_tx=XM_019822038.2;refGene_gn=ZC3H11A;refGene_tx=XM_023247369.1
F1 44350275 44350355
refGene_gn=ZC3H11A;refGene_tx=XM_006942887.4;refGene_gn=ZC3H11A;refGene_tx=XM_006942888.4;refGene_gn=ZC3H11A;refGene_tx=XM_006942889.4;refGene_gn=ZC3H11A;refGene_tx=XM_006942890.4;refGene_gn=ZC3H11A;refGene_tx=XM_006942891.4;refGene_gn=ZC3H11A;refGene_tx=XM_006942892.4;refGene_gn=ZC3H11A;refGene_tx=XM_006942893.4;refGene_gn=ZC3H11A;refGene_tx=XM_006942896.4;refGene_gn=ZC3H11A;refGene_tx=XM_011290924.3;refGene_gn=ZC3H11A;refGene_tx=XM_019822030.2;refGene_gn=ZC3H11A;refGene_tx=XM_019822031.2;refGene_gn=ZC3H11A;refGene_tx=XM_019822032.2;refGene_gn=ZC3H11A;refGene_tx=XM_019822033.2;refGene_gn=ZC3H11A;refGene_tx=XM_019822035.2;refGene_gn=ZC3H11A;refGene_tx=XM_019822037.2;refGene_gn=ZC3H11A;refGene_tx=XM_019822038.2;refGene_gn=ZC3H11A;refGene_tx=XM_023247369.1
F1 44352515 44352846
refGene_gn=ZC3H11A;refGene_tx=XM_006942887.4;refGene_gn=ZC3H11A;refGene_tx=XM_006942888.4;refGene_gn=ZC3H11A;refGene_tx=XM_006942889.4;refGene_gn=ZC3H11A;refGene_tx=XM_006942890.4;refGene_gn=ZC3H11A;refGene_tx=XM_006942891.4;refGene_gn=ZC3H11A;refGene_tx=XM_006942892.4;refGene_gn=ZC3H11A;refGene_tx=XM_006942893.4;refGene_gn=ZC3H11A;refGene_tx=XM_006942896.4;refGene_gn=ZC3H11A;refGene_tx=XM_011290924.3;refGene_gn=ZC3H11A;refGene_tx=XM_019822030.2;refGene_gn=ZC3H11A;refGene_tx=XM_019822031.2;refGene_gn=ZC3H11A;refGene_tx=XM_019822032.2;refGene_gn=ZC3H11A;refGene_tx=XM_019822033.2;refGene_gn=ZC3H11A;refGene_tx=XM_019822035.2;refGene_gn=ZC3H11A;refGene_tx=XM_019822037.2;refGene_gn=ZC3H11A;refGene_tx=XM_019822038.2;refGene_gn=ZC3H11A;refGene_tx=XM_023247369.1
F1 44354068 44354235
refGene_gn=ZC3H11A;refGene_tx=XM_006942887.4;refGene_gn=ZC3H11A;refGene_tx=XM_006942888.4;refGene_gn=ZC3H11A;refGene_tx=XM_006942889.4;refGene_gn=ZC3H11A;refGene_tx=XM_006942890.4;refGene_gn=ZC3H11A;refGene_tx=XM_006942891.4;refGene_gn=ZC3H11A;refGene_tx=XM_006942892.4;refGene_gn=ZC3H11A;refGene_tx=XM_006942893.4;refGene_gn=ZC3H11A;refGene_tx=XM_006942896.4;refGene_gn=ZC3H11A;refGene_tx=XM_011290924.3;refGene_gn=ZC3H11A;refGene_tx=XM_019822030.2;refGene_gn=ZC3H11A;refGene_tx=XM_019822031.2;refGene_gn=ZC3H11A;refGene_tx=XM_019822032.2;refGene_gn=ZC3H11A;refGene_tx=XM_019822033.2;refGene_gn=ZC3H11A;refGene_tx=XM_019822035.2;refGene_gn=ZC3H11A;refGene_tx=XM_019822037.2;refGene_gn=ZC3H11A;refGene_tx=XM_019822038.2;refGene_gn=ZC3H11A;refGene_tx=XM_023247369.1

F1 44354599 44354667
refGene_gn=ZC3H11A;refGene_tx=XM_006942887.4;refGene_gn=ZC3H11A;refGene_tx=XM_006942888.4;refGene_gn=ZC3H11A;refGene_tx=XM_006942889.4;refGene_gn=ZC3H11A;refGene_tx=XM_006942890.4;refGene_gn=ZC3H11A;refGene_tx=XM_006942891.4;refGene_gn=ZC3H11A;refGene_tx=XM_006942892.4;refGene_gn=ZC3H11A;refGene_tx=XM_006942893.4;refGene_gn=ZC3H11A;refGene_tx=XM_006942896.4;refGene_gn=ZC3H11A;refGene_tx=XM_011290924.3;refGene_gn=ZC3H11A;refGene_tx=XM_019822030.2;refGene_gn=ZC3H11A;refGene_tx=XM_019822031.2;refGene_gn=ZC3H11A;refGene_tx=XM_019822032.2;refGene_gn=ZC3H11A;refGene_tx=XM_019822033.2;refGene_gn=ZC3H11A;refGene_tx=XM_019822035.2;refGene_gn=ZC3H11A;refGene_tx=XM_019822037.2;refGene_gn=ZC3H11A;refGene_tx=XM_019822038.2;refGene_gn=ZC3H11A;refGene_tx=XM_023247369.1
F1 44355106 44355365
refGene_gn=ZC3H11A;refGene_tx=XM_006942887.4;refGene_gn=ZC3H11A;refGene_tx=XM_006942888.4;refGene_gn=ZC3H11A;refGene_tx=XM_006942889.4;refGene_gn=ZC3H11A;refGene_tx=XM_006942890.4;refGene_gn=ZC3H11A;refGene_tx=XM_006942891.4;refGene_gn=ZC3H11A;refGene_tx=XM_006942892.4;refGene_gn=ZC3H11A;refGene_tx=XM_006942893.4;refGene_gn=ZC3H11A;refGene_tx=XM_006942896.4;refGene_gn=ZC3H11A;refGene_tx=XM_011290924.3;refGene_gn=ZC3H11A;refGene_tx=XM_019822030.2;refGene_gn=ZC3H11A;refGene_tx=XM_019822031.2;refGene_gn=ZC3H11A;refGene_tx=XM_019822032.2;refGene_gn=ZC3H11A;refGene_tx=XM_019822033.2;refGene_gn=ZC3H11A;refGene_tx=XM_019822035.2;refGene_gn=ZC3H11A;refGene_tx=XM_019822037.2;refGene_gn=ZC3H11A;refGene_tx=XM_019822038.2;refGene_gn=ZC3H11A;refGene_tx=XM_023247369.1
F1 44364163 44364217
refGene_gn=SNRPE;refGene_tx=XM_003999391.4;refGene_gn=SNRPE;refGene_tx=XM_019822043.2;refGene_gn=SNRPE;refGene_tx=XM_019822044.2
F1 44364724 44364751
refGene_gn=SNRPE;refGene_tx=XM_003999391.4;refGene_gn=SNRPE;refGene_tx=XM_019822043.2;refGene_gn=SNRPE;refGene_tx=XM_019822044.2
F1 44366192 44366255
refGene_gn=SNRPE;refGene_tx=XM_003999391.4;refGene_gn=SNRPE;refGene_tx=XM_019822043.2;refGene_gn=SNRPE;refGene_tx=XM_019822044.2
F1 44367143 44367222
refGene_gn=SNRPE;refGene_tx=XM_003999391.4;refGene_gn=SNRPE;refGene_tx=XM_019822043.2;refGene_gn=SNRPE;refGene_tx=XM_019822044.2
F1 44369475 44369531 refGene_gn=SNRPE;refGene_tx=XM_003999391.4
F1 44410176 44410247
refGene_gn=SNRPE;refGene_tx=XM_019822043.2;refGene_gn=SNRPE;refGene_tx=XM_019822044.2
F1 44482365 44482635
refGene_gn=LOC101082973;refGene_tx=XM_023247457.1
F1 44563801 44563999
refGene_gn=SOX13;refGene_tx=XM_023247975.1;refGene_gn=SOX13;refGene_tx=XM_023247976.1;refGene_gn=SOX13;refGene_tx=XM_023247977.1
F1 44565141 44565253
refGene_gn=SOX13;refGene_tx=XM_023247975.1;refGene_gn=SOX13;refGene_tx=XM_023247976.1;refGene_gn=SOX13;refGene_tx=XM_023247977.1
F1 44565335 44565422
refGene_gn=SOX13;refGene_tx=XM_023247975.1;refGene_gn=SOX13;refGene_tx

=XM_023247976.1;refGene_gn=SOX13;refGene_tx=XM_023247977.1
F1	44566868	44567041
refGene_gn=SOX13;refGene_tx=XM_023247975.1;refGene_gn=SOX13;refGene_tx=XM_023247976.1;refGene_gn=SOX13;refGene_tx=XM_023247977.1
F1	44567425	44567494
refGene_gn=SOX13;refGene_tx=XM_023247975.1;refGene_gn=SOX13;refGene_tx=XM_023247976.1;refGene_gn=SOX13;refGene_tx=XM_023247977.1
F1	44567868	44567983
refGene_gn=SOX13;refGene_tx=XM_023247975.1;refGene_gn=SOX13;refGene_tx=XM_023247976.1;refGene_gn=SOX13;refGene_tx=XM_023247977.1
F1	44570173	44570259
refGene_gn=SOX13;refGene_tx=XM_023247975.1;refGene_gn=SOX13;refGene_tx=XM_023247976.1;refGene_gn=SOX13;refGene_tx=XM_023247977.1
F1	44570478	44570638
refGene_gn=SOX13;refGene_tx=XM_023247975.1;refGene_gn=SOX13;refGene_tx=XM_023247976.1;refGene_gn=SOX13;refGene_tx=XM_023247977.1
F1	44571068	44571178
refGene_gn=SOX13;refGene_tx=XM_023247975.1;refGene_gn=SOX13;refGene_tx=XM_023247976.1;refGene_gn=SOX13;refGene_tx=XM_023247977.1
F1	44571320	44571417
refGene_gn=SOX13;refGene_tx=XM_023247975.1;refGene_gn=SOX13;refGene_tx=XM_023247976.1;refGene_gn=SOX13;refGene_tx=XM_023247977.1
F1	44571728	44571872
refGene_gn=SOX13;refGene_tx=XM_023247975.1;refGene_gn=SOX13;refGene_tx=XM_023247976.1;refGene_gn=SOX13;refGene_tx=XM_023247977.1
F1	44573066	44573283
refGene_gn=SOX13;refGene_tx=XM_023247975.1;refGene_gn=SOX13;refGene_tx=XM_023247976.1;refGene_gn=SOX13;refGene_tx=XM_023247977.1
F1	44573998	44574275
refGene_gn=SOX13;refGene_tx=XM_023247975.1;refGene_gn=SOX13;refGene_tx=XM_023247976.1;refGene_gn=SOX13;refGene_tx=XM_023247977.1
F1	44579709	44579782	refGene_gn=ETNK2;refGene_tx=XM_006942899.4
F1	44581356	44581430	refGene_gn=ETNK2;refGene_tx=XM_006942899.4
F1	44582874	44583020	refGene_gn=ETNK2;refGene_tx=XM_006942899.4
F1	44585602	44585686	refGene_gn=ETNK2;refGene_tx=XM_006942899.4
F1	44586505	44586648	refGene_gn=ETNK2;refGene_tx=XM_006942899.4
F1	44591382	44591505	refGene_gn=ETNK2;refGene_tx=XM_006942899.4
F1	44594475	44594735	refGene_gn=ETNK2;refGene_tx=XM_006942899.4
F1	44596420	44596678	refGene_gn=ETNK2;refGene_tx=XM_006942899.4
F1	44599257	44599419
refGene_gn=REN;refGene_tx=XM_003999393.4;refGene_gn=REN;refGene_tx=XM_019822047.2
F1	44600472	44600571
refGene_gn=REN;refGene_tx=XM_003999393.4;refGene_gn=REN;refGene_tx=XM_019822047.2
F1	44600846	44600991
refGene_gn=REN;refGene_tx=XM_003999393.4;refGene_gn=REN;refGene_tx=XM_019822047.2
F1	44601367	44601487
refGene_gn=REN;refGene_tx=XM_003999393.4;refGene_gn=REN;refGene_tx=XM_

019822047.2
F1 44602067 44602076 refGene_gn=REN;refGene_tx=XM_003999393.4
F1 44603793 44603990
refGene_gn=REN;refGene_tx=XM_003999393.4;refGene_gn=REN;refGene_tx=XM_
019822047.2
F1 44604889 44605008
refGene_gn=REN;refGene_tx=XM_003999393.4;refGene_gn=REN;refGene_tx=XM_
019822047.2
F1 44605603 44605727
refGene_gn=REN;refGene_tx=XM_003999393.4;refGene_gn=REN;refGene_tx=XM_
019822047.2
F1 44606259 44606410
refGene_gn=REN;refGene_tx=XM_003999393.4;refGene_gn=REN;refGene_tx=XM_
019822047.2
F1 44611278 44611376
refGene_gn=REN;refGene_tx=XM_003999393.4;refGene_gn=REN;refGene_tx=XM_
019822047.2
F1 44625397 44625744 refGene_gn=KISS1;refGene_tx=XM_023247458.1
F1 44627704 44627807 refGene_gn=KISS1;refGene_tx=XM_023247458.1
F1 44632348 44632387
refGene_gn=GOLT1A;refGene_tx=XM_003999394.4;refGene_gn=GOLT1A;refGene_
tx=XM_019822048.2
F1 44633014 44633078
refGene_gn=GOLT1A;refGene_tx=XM_003999394.4;refGene_gn=GOLT1A;refGene_
tx=XM_019822048.2
F1 44635215 44635394
refGene_gn=GOLT1A;refGene_tx=XM_003999394.4;refGene_gn=GOLT1A;refGene_
tx=XM_019822048.2
F1 44636333 44636425
refGene_gn=GOLT1A;refGene_tx=XM_003999394.4;refGene_gn=GOLT1A;refGene_
tx=XM_019822048.2
F1 44648763 44648788
refGene_gn=GOLT1A;refGene_tx=XM_003999394.4;refGene_gn=GOLT1A;refGene_
tx=XM_019822048.2
F1 44657497 44657550 refGene_gn=PLEKHA6;refGene_tx=XM_023247806.1
F1 44658179 44658295
refGene_gn=PLEKHA6;refGene_tx=XM_019822049.2;refGene_gn=PLEKHA6;refGen
e_tx=XM_019822052.2;refGene_gn=PLEKHA6;refGene_tx=XM_019822063.2;refGe
ne_gn=PLEKHA6;refGene_tx=XM_019822064.2;refGene_gn=PLEKHA6;refGene_tx=
XM_019822065.2;refGene_gn=PLEKHA6;refGene_tx=XM_019822066.2;refGene_gn
=PLEKHA6;refGene_tx=XM_019822067.2;refGene_gn=PLEKHA6;refGene_tx=XM_01
9822068.2;refGene_gn=PLEKHA6;refGene_tx=XM_019822069.2;refGene_gn=PLEK
HA6;refGene_tx=XM_019822075.2;refGene_gn=PLEKHA6;refGene_tx=XM_0232478
02.1;refGene_gn=PLEKHA6;refGene_tx=XM_023247803.1;refGene_gn=PLEKHA6;r
efGene_tx=XM_023247804.1;refGene_gn=PLEKHA6;refGene_tx=XM_023247805.1;
refGene_gn=PLEKHA6;refGene_tx=XM_023247807.1;refGene_gn=PLEKHA6;refGen
e_tx=XM_023247808.1;refGene_gn=PLEKHA6;refGene_tx=XM_023247809.1;refGe
ne_gn=PLEKHA6;refGene_tx=XM_023247810.1;refGene_gn=PLEKHA6;refGene_tx=
XM_023247811.1;refGene_gn=PLEKHA6;refGene_tx=XM_023247812.1;refGene_gn
=PLEKHA6;refGene_tx=XM_023247813.1;refGene_gn=PLEKHA6;refGene_tx=XM_02

3247814.1;refGene_gn=PLEKHA6;refGene_tx=XM_023247815.1;refGene_gn=PLEKHA6;refGene_tx=XM_023247817.1;refGene_gn=PLEKHA6;refGene_tx=XM_023247818.1
F1 44663048 44663194
refGene_gn=PLEKHA6;refGene_tx=XM_019822049.2;refGene_gn=PLEKHA6;refGene_tx=XM_019822052.2;refGene_gn=PLEKHA6;refGene_tx=XM_019822063.2;refGene_gn=PLEKHA6;refGene_tx=XM_019822064.2;refGene_gn=PLEKHA6;refGene_tx=XM_019822065.2;refGene_gn=PLEKHA6;refGene_tx=XM_019822066.2;refGene_gn=PLEKHA6;refGene_tx=XM_019822067.2;refGene_gn=PLEKHA6;refGene_tx=XM_019822068.2;refGene_gn=PLEKHA6;refGene_tx=XM_019822069.2;refGene_gn=PLEKHA6;refGene_tx=XM_019822075.2;refGene_gn=PLEKHA6;refGene_tx=XM_023247802.1;refGene_gn=PLEKHA6;refGene_tx=XM_023247803.1;refGene_gn=PLEKHA6;refGene_tx=XM_023247804.1;refGene_gn=PLEKHA6;refGene_tx=XM_023247805.1;refGene_gn=PLEKHA6;refGene_tx=XM_023247806.1;refGene_gn=PLEKHA6;refGene_tx=XM_023247807.1;refGene_gn=PLEKHA6;refGene_tx=XM_023247808.1;refGene_gn=PLEKHA6;refGene_tx=XM_023247809.1;refGene_gn=PLEKHA6;refGene_tx=XM_023247810.1;refGene_gn=PLEKHA6;refGene_tx=XM_023247811.1;refGene_gn=PLEKHA6;refGene_tx=XM_023247812.1;refGene_gn=PLEKHA6;refGene_tx=XM_023247813.1;refGene_gn=PLEKHA6;refGene_tx=XM_023247814.1;refGene_gn=PLEKHA6;refGene_tx=XM_023247815.1;refGene_gn=PLEKHA6;refGene_tx=XM_023247817.1;refGene_gn=PLEKHA6;refGene_tx=XM_023247818.1
F1 44663693 44663827
refGene_gn=PLEKHA6;refGene_tx=XM_019822049.2;refGene_gn=PLEKHA6;refGene_tx=XM_019822052.2;refGene_gn=PLEKHA6;refGene_tx=XM_019822063.2;refGene_gn=PLEKHA6;refGene_tx=XM_019822064.2;refGene_gn=PLEKHA6;refGene_tx=XM_019822065.2;refGene_gn=PLEKHA6;refGene_tx=XM_019822066.2;refGene_gn=PLEKHA6;refGene_tx=XM_019822067.2;refGene_gn=PLEKHA6;refGene_tx=XM_019822068.2;refGene_gn=PLEKHA6;refGene_tx=XM_019822069.2;refGene_gn=PLEKHA6;refGene_tx=XM_019822075.2;refGene_gn=PLEKHA6;refGene_tx=XM_023247802.1;refGene_gn=PLEKHA6;refGene_tx=XM_023247803.1;refGene_gn=PLEKHA6;refGene_tx=XM_023247804.1;refGene_gn=PLEKHA6;refGene_tx=XM_023247805.1;refGene_gn=PLEKHA6;refGene_tx=XM_023247806.1;refGene_gn=PLEKHA6;refGene_tx=XM_023247807.1;refGene_gn=PLEKHA6;refGene_tx=XM_023247808.1;refGene_gn=PLEKHA6;refGene_tx=XM_023247809.1;refGene_gn=PLEKHA6;refGene_tx=XM_023247810.1;refGene_gn=PLEKHA6;refGene_tx=XM_023247811.1;refGene_gn=PLEKHA6;refGene_tx=XM_023247812.1;refGene_gn=PLEKHA6;refGene_tx=XM_023247813.1;refGene_gn=PLEKHA6;refGene_tx=XM_023247814.1;refGene_gn=PLEKHA6;refGene_tx=XM_023247815.1;refGene_gn=PLEKHA6;refGene_tx=XM_023247817.1;refGene_gn=PLEKHA6;refGene_tx=XM_023247818.1
F1 44663908 44664076
refGene_gn=PLEKHA6;refGene_tx=XM_019822049.2;refGene_gn=PLEKHA6;refGene_tx=XM_019822052.2;refGene_gn=PLEKHA6;refGene_tx=XM_019822063.2;refGene_gn=PLEKHA6;refGene_tx=XM_019822064.2;refGene_gn=PLEKHA6;refGene_tx=XM_019822065.2;refGene_gn=PLEKHA6;refGene_tx=XM_019822066.2;refGene_gn=PLEKHA6;refGene_tx=XM_019822067.2;refGene_gn=PLEKHA6;refGene_tx=XM_019822068.2;refGene_gn=PLEKHA6;refGene_tx=XM_019822069.2;refGene_gn=PLEKHA6;refGene_tx=XM_019822075.2;refGene_gn=PLEKHA6;refGene_tx=XM_023247802.1;refGene_gn=PLEKHA6;refGene_tx=XM_023247803.1;refGene_gn=PLEKHA6;refGene_tx=XM_023247804.1;refGene_gn=PLEKHA6;refGene_tx=XM_023247805.1;refGene_gn=PLEKHA6;refGene_tx=XM_023247806.1;refGene_gn=PLEKHA6;refGen

e_tx=XM_023247807.1;refGene_gn=PLEKHA6;refGene_tx=XM_023247808.1;refGene_gn=PLEKHA6;refGene_tx=XM_023247809.1;refGene_gn=PLEKHA6;refGene_tx=XM_023247810.1;refGene_gn=PLEKHA6;refGene_tx=XM_023247811.1;refGene_gn=PLEKHA6;refGene_tx=XM_023247812.1;refGene_gn=PLEKHA6;refGene_tx=XM_023247813.1;refGene_gn=PLEKHA6;refGene_tx=XM_023247814.1;refGene_gn=PLEKHA6;refGene_tx=XM_023247815.1;refGene_gn=PLEKHA6;refGene_tx=XM_023247817.1;refGene_gn=PLEKHA6;refGene_tx=XM_023247818.1
F1 44665119 44665296
refGene_gn=PLEKHA6;refGene_tx=XM_019822049.2;refGene_gn=PLEKHA6;refGene_tx=XM_019822052.2;refGene_gn=PLEKHA6;refGene_tx=XM_019822063.2;refGene_gn=PLEKHA6;refGene_tx=XM_019822064.2;refGene_gn=PLEKHA6;refGene_tx=XM_019822065.2;refGene_gn=PLEKHA6;refGene_tx=XM_019822066.2;refGene_gn=PLEKHA6;refGene_tx=XM_019822067.2;refGene_gn=PLEKHA6;refGene_tx=XM_019822068.2;refGene_gn=PLEKHA6;refGene_tx=XM_019822069.2;refGene_gn=PLEKHA6;refGene_tx=XM_019822075.2;refGene_gn=PLEKHA6;refGene_tx=XM_023247802.1;refGene_gn=PLEKHA6;refGene_tx=XM_023247803.1;refGene_gn=PLEKHA6;refGene_tx=XM_023247804.1;refGene_gn=PLEKHA6;refGene_tx=XM_023247805.1;refGene_gn=PLEKHA6;refGene_tx=XM_023247806.1;refGene_gn=PLEKHA6;refGene_tx=XM_023247807.1;refGene_gn=PLEKHA6;refGene_tx=XM_023247808.1;refGene_gn=PLEKHA6;refGene_tx=XM_023247809.1;refGene_gn=PLEKHA6;refGene_tx=XM_023247810.1;refGene_gn=PLEKHA6;refGene_tx=XM_023247811.1;refGene_gn=PLEKHA6;refGene_tx=XM_023247812.1;refGene_gn=PLEKHA6;refGene_tx=XM_023247813.1;refGene_gn=PLEKHA6;refGene_tx=XM_023247814.1;refGene_gn=PLEKHA6;refGene_tx=XM_023247815.1;refGene_gn=PLEKHA6;refGene_tx=XM_023247817.1;refGene_gn=PLEKHA6;refGene_tx=XM_023247818.1
F1 44666224 44666331
refGene_gn=PLEKHA6;refGene_tx=XM_019822049.2;refGene_gn=PLEKHA6;refGene_tx=XM_019822052.2;refGene_gn=PLEKHA6;refGene_tx=XM_019822063.2;refGene_gn=PLEKHA6;refGene_tx=XM_019822064.2;refGene_gn=PLEKHA6;refGene_tx=XM_019822065.2;refGene_gn=PLEKHA6;refGene_tx=XM_019822066.2;refGene_gn=PLEKHA6;refGene_tx=XM_019822067.2;refGene_gn=PLEKHA6;refGene_tx=XM_019822068.2;refGene_gn=PLEKHA6;refGene_tx=XM_019822069.2;refGene_gn=PLEKHA6;refGene_tx=XM_019822075.2;refGene_gn=PLEKHA6;refGene_tx=XM_023247802.1;refGene_gn=PLEKHA6;refGene_tx=XM_023247803.1;refGene_gn=PLEKHA6;refGene_tx=XM_023247804.1;refGene_gn=PLEKHA6;refGene_tx=XM_023247805.1;refGene_gn=PLEKHA6;refGene_tx=XM_023247806.1;refGene_gn=PLEKHA6;refGene_tx=XM_023247807.1;refGene_gn=PLEKHA6;refGene_tx=XM_023247808.1;refGene_gn=PLEKHA6;refGene_tx=XM_023247809.1;refGene_gn=PLEKHA6;refGene_tx=XM_023247810.1;refGene_gn=PLEKHA6;refGene_tx=XM_023247811.1;refGene_gn=PLEKHA6;refGene_tx=XM_023247812.1;refGene_gn=PLEKHA6;refGene_tx=XM_023247813.1;refGene_gn=PLEKHA6;refGene_tx=XM_023247814.1;refGene_gn=PLEKHA6;refGene_tx=XM_023247815.1;refGene_gn=PLEKHA6;refGene_tx=XM_023247817.1;refGene_gn=PLEKHA6;refGene_tx=XM_023247818.1
F1 44666547 44666677
refGene_gn=PLEKHA6;refGene_tx=XM_019822049.2;refGene_gn=PLEKHA6;refGene_tx=XM_019822052.2;refGene_gn=PLEKHA6;refGene_tx=XM_019822063.2;refGene_gn=PLEKHA6;refGene_tx=XM_019822064.2;refGene_gn=PLEKHA6;refGene_tx=XM_019822065.2;refGene_gn=PLEKHA6;refGene_tx=XM_019822066.2;refGene_gn=PLEKHA6;refGene_tx=XM_019822067.2;refGene_gn=PLEKHA6;refGene_tx=XM_019822068.2;refGene_gn=PLEKHA6;refGene_tx=XM_019822069.2;refGene_gn=PLEK

HA6;refGene_tx=XM_019822075.2;refGene_gn=PLEKHA6;refGene_tx=XM_0232478
02.1;refGene_gn=PLEKHA6;refGene_tx=XM_023247803.1;refGene_gn=PLEKHA6;r
efGene_tx=XM_023247804.1;refGene_gn=PLEKHA6;refGene_tx=XM_023247805.1;
refGene_gn=PLEKHA6;refGene_tx=XM_023247806.1;refGene_gn=PLEKHA6;refGen
e_tx=XM_023247807.1;refGene_gn=PLEKHA6;refGene_tx=XM_023247808.1;refGe
ne_gn=PLEKHA6;refGene_tx=XM_023247809.1;refGene_gn=PLEKHA6;refGene_tx=
XM_023247810.1;refGene_gn=PLEKHA6;refGene_tx=XM_023247811.1;refGene_gn
=PLEKHA6;refGene_tx=XM_023247812.1;refGene_gn=PLEKHA6;refGene_tx=XM_02
3247813.1;refGene_gn=PLEKHA6;refGene_tx=XM_023247814.1;refGene_gn=PLEK
HA6;refGene_tx=XM_023247815.1;refGene_gn=PLEKHA6;refGene_tx=XM_0232478
17.1;refGene_gn=PLEKHA6;refGene_tx=XM_023247818.1
F1 44667949 44668099
refGene_gn=PLEKHA6;refGene_tx=XM_019822049.2;refGene_gn=PLEKHA6;refGen
e_tx=XM_019822052.2;refGene_gn=PLEKHA6;refGene_tx=XM_019822063.2;refGe
ne_gn=PLEKHA6;refGene_tx=XM_019822064.2;refGene_gn=PLEKHA6;refGene_tx=
XM_019822065.2;refGene_gn=PLEKHA6;refGene_tx=XM_019822066.2;refGene_gn
=PLEKHA6;refGene_tx=XM_019822067.2;refGene_gn=PLEKHA6;refGene_tx=XM_01
9822068.2;refGene_gn=PLEKHA6;refGene_tx=XM_019822069.2;refGene_gn=PLEK
HA6;refGene_tx=XM_019822075.2;refGene_gn=PLEKHA6;refGene_tx=XM_0232478
02.1;refGene_gn=PLEKHA6;refGene_tx=XM_023247803.1;refGene_gn=PLEKHA6;r
efGene_tx=XM_023247804.1;refGene_gn=PLEKHA6;refGene_tx=XM_023247805.1;
refGene_gn=PLEKHA6;refGene_tx=XM_023247806.1;refGene_gn=PLEKHA6;refGen
e_tx=XM_023247807.1;refGene_gn=PLEKHA6;refGene_tx=XM_023247808.1;refGe
ne_gn=PLEKHA6;refGene_tx=XM_023247809.1;refGene_gn=PLEKHA6;refGene_tx=
XM_023247810.1;refGene_gn=PLEKHA6;refGene_tx=XM_023247811.1;refGene_gn
=PLEKHA6;refGene_tx=XM_023247812.1;refGene_gn=PLEKHA6;refGene_tx=XM_02
3247814.1;refGene_gn=PLEKHA6;refGene_tx=XM_023247815.1
F1 44669064 44669204
refGene_gn=PLEKHA6;refGene_tx=XM_019822049.2;refGene_gn=PLEKHA6;refGen
e_tx=XM_019822052.2;refGene_gn=PLEKHA6;refGene_tx=XM_019822063.2;refGe
ne_gn=PLEKHA6;refGene_tx=XM_019822064.2;refGene_gn=PLEKHA6;refGene_tx=
XM_019822065.2;refGene_gn=PLEKHA6;refGene_tx=XM_019822066.2;refGene_gn
=PLEKHA6;refGene_tx=XM_019822067.2;refGene_gn=PLEKHA6;refGene_tx=XM_01
9822068.2;refGene_gn=PLEKHA6;refGene_tx=XM_019822069.2;refGene_gn=PLEK
HA6;refGene_tx=XM_019822075.2;refGene_gn=PLEKHA6;refGene_tx=XM_0232478
02.1;refGene_gn=PLEKHA6;refGene_tx=XM_023247803.1;refGene_gn=PLEKHA6;r
efGene_tx=XM_023247804.1;refGene_gn=PLEKHA6;refGene_tx=XM_023247805.1;
refGene_gn=PLEKHA6;refGene_tx=XM_023247806.1;refGene_gn=PLEKHA6;refGen
e_tx=XM_023247807.1;refGene_gn=PLEKHA6;refGene_tx=XM_023247808.1;refGe
ne_gn=PLEKHA6;refGene_tx=XM_023247809.1;refGene_gn=PLEKHA6;refGene_tx=
XM_023247810.1;refGene_gn=PLEKHA6;refGene_tx=XM_023247811.1;refGene_gn
=PLEKHA6;refGene_tx=XM_023247812.1;refGene_gn=PLEKHA6;refGene_tx=XM_02
3247813.1;refGene_gn=PLEKHA6;refGene_tx=XM_023247814.1;refGene_gn=PLEK
HA6;refGene_tx=XM_023247815.1;refGene_gn=PLEKHA6;refGene_tx=XM_0232478
17.1;refGene_gn=PLEKHA6;refGene_tx=XM_023247818.1
F1 44669810 44669922
refGene_gn=PLEKHA6;refGene_tx=XM_019822049.2;refGene_gn=PLEKHA6;refGen
e_tx=XM_019822052.2;refGene_gn=PLEKHA6;refGene_tx=XM_019822063.2;refGe
ne_gn=PLEKHA6;refGene_tx=XM_019822064.2;refGene_gn=PLEKHA6;refGene_tx=
XM_019822065.2;refGene_gn=PLEKHA6;refGene_tx=XM_019822066.2;refGene_gn

=PLEKHA6;refGene_tx=XM_019822067.2;refGene_gn=PLEKHA6;refGene_tx=XM_01
9822068.2;refGene_gn=PLEKHA6;refGene_tx=XM_019822069.2;refGene_gn=PLEK
HA6;refGene_tx=XM_019822075.2;refGene_gn=PLEKHA6;refGene_tx=XM_0232478
02.1;refGene_gn=PLEKHA6;refGene_tx=XM_023247803.1;refGene_gn=PLEKHA6;r
efGene_tx=XM_023247804.1;refGene_gn=PLEKHA6;refGene_tx=XM_023247805.1;
refGene_gn=PLEKHA6;refGene_tx=XM_023247806.1;refGene_gn=PLEKHA6;refGen
e_tx=XM_023247807.1;refGene_gn=PLEKHA6;refGene_tx=XM_023247808.1;refGe
ne_gn=PLEKHA6;refGene_tx=XM_023247809.1;refGene_gn=PLEKHA6;refGene_tx=
XM_023247810.1;refGene_gn=PLEKHA6;refGene_tx=XM_023247811.1;refGene_gn
=PLEKHA6;refGene_tx=XM_023247812.1;refGene_gn=PLEKHA6;refGene_tx=XM_02
3247813.1;refGene_gn=PLEKHA6;refGene_tx=XM_023247814.1;refGene_gn=PLEK
HA6;refGene_tx=XM_023247815.1;refGene_gn=PLEKHA6;refGene_tx=XM_0232478
17.1;refGene_gn=PLEKHA6;refGene_tx=XM_023247818.1
F1 44671388 44671484
refGene_gn=PLEKHA6;refGene_tx=XM_019822049.2;refGene_gn=PLEKHA6;refGen
e_tx=XM_019822052.2;refGene_gn=PLEKHA6;refGene_tx=XM_019822063.2;refGe
ne_gn=PLEKHA6;refGene_tx=XM_019822064.2;refGene_gn=PLEKHA6;refGene_tx=
XM_019822065.2;refGene_gn=PLEKHA6;refGene_tx=XM_019822066.2;refGene_gn
=PLEKHA6;refGene_tx=XM_019822067.2;refGene_gn=PLEKHA6;refGene_tx=XM_01
9822068.2;refGene_gn=PLEKHA6;refGene_tx=XM_019822069.2;refGene_gn=PLEK
HA6;refGene_tx=XM_019822075.2;refGene_gn=PLEKHA6;refGene_tx=XM_0232478
02.1;refGene_gn=PLEKHA6;refGene_tx=XM_023247803.1;refGene_gn=PLEKHA6;r
efGene_tx=XM_023247804.1;refGene_gn=PLEKHA6;refGene_tx=XM_023247805.1;
refGene_gn=PLEKHA6;refGene_tx=XM_023247806.1;refGene_gn=PLEKHA6;refGen
e_tx=XM_023247807.1;refGene_gn=PLEKHA6;refGene_tx=XM_023247808.1;refGe
ne_gn=PLEKHA6;refGene_tx=XM_023247809.1;refGene_gn=PLEKHA6;refGene_tx=
XM_023247810.1;refGene_gn=PLEKHA6;refGene_tx=XM_023247811.1;refGene_gn
=PLEKHA6;refGene_tx=XM_023247812.1;refGene_gn=PLEKHA6;refGene_tx=XM_02
3247813.1;refGene_gn=PLEKHA6;refGene_tx=XM_023247814.1;refGene_gn=PLEK
HA6;refGene_tx=XM_023247815.1;refGene_gn=PLEKHA6;refGene_tx=XM_0232478
17.1;refGene_gn=PLEKHA6;refGene_tx=XM_023247818.1
F1 44672323 44672473
refGene_gn=PLEKHA6;refGene_tx=XM_019822049.2;refGene_gn=PLEKHA6;refGen
e_tx=XM_019822052.2;refGene_gn=PLEKHA6;refGene_tx=XM_019822063.2;refGe
ne_gn=PLEKHA6;refGene_tx=XM_019822064.2;refGene_gn=PLEKHA6;refGene_tx=
XM_019822065.2;refGene_gn=PLEKHA6;refGene_tx=XM_019822066.2;refGene_gn
=PLEKHA6;refGene_tx=XM_019822067.2;refGene_gn=PLEKHA6;refGene_tx=XM_01
9822068.2;refGene_gn=PLEKHA6;refGene_tx=XM_019822069.2;refGene_gn=PLEK
HA6;refGene_tx=XM_019822075.2;refGene_gn=PLEKHA6;refGene_tx=XM_0232478
02.1;refGene_gn=PLEKHA6;refGene_tx=XM_023247803.1;refGene_gn=PLEKHA6;r
efGene_tx=XM_023247804.1;refGene_gn=PLEKHA6;refGene_tx=XM_023247805.1;
refGene_gn=PLEKHA6;refGene_tx=XM_023247806.1;refGene_gn=PLEKHA6;refGen
e_tx=XM_023247807.1;refGene_gn=PLEKHA6;refGene_tx=XM_023247808.1;refGe
ne_gn=PLEKHA6;refGene_tx=XM_023247809.1;refGene_gn=PLEKHA6;refGene_tx=
XM_023247810.1;refGene_gn=PLEKHA6;refGene_tx=XM_023247811.1;refGene_gn
=PLEKHA6;refGene_tx=XM_023247812.1;refGene_gn=PLEKHA6;refGene_tx=XM_02
3247813.1;refGene_gn=PLEKHA6;refGene_tx=XM_023247814.1;refGene_gn=PLEK
HA6;refGene_tx=XM_023247815.1;refGene_gn=PLEKHA6;refGene_tx=XM_0232478
17.1;refGene_gn=PLEKHA6;refGene_tx=XM_023247818.1
F1 44672766 44672847

refGene_gn=PLEKHA6;refGene_tx=XM_019822049.2;refGene_gn=PLEKHA6;refGen
e_tx=XM_019822052.2;refGene_gn=PLEKHA6;refGene_tx=XM_019822063.2;refGe
ne_gn=PLEKHA6;refGene_tx=XM_019822064.2;refGene_gn=PLEKHA6;refGene_tx=
XM_019822065.2;refGene_gn=PLEKHA6;refGene_tx=XM_019822066.2;refGene_gn
=PLEKHA6;refGene_tx=XM_019822067.2;refGene_gn=PLEKHA6;refGene_tx=XM_01
9822068.2;refGene_gn=PLEKHA6;refGene_tx=XM_019822069.2;refGene_gn=PLEK
HA6;refGene_tx=XM_019822075.2;refGene_gn=PLEKHA6;refGene_tx=XM_0232478
02.1;refGene_gn=PLEKHA6;refGene_tx=XM_023247803.1;refGene_gn=PLEKHA6;r
efGene_tx=XM_023247804.1;refGene_gn=PLEKHA6;refGene_tx=XM_023247805.1;
refGene_gn=PLEKHA6;refGene_tx=XM_023247806.1;refGene_gn=PLEKHA6;refGen
e_tx=XM_023247807.1;refGene_gn=PLEKHA6;refGene_tx=XM_023247808.1;refGe
ne_gn=PLEKHA6;refGene_tx=XM_023247809.1;refGene_gn=PLEKHA6;refGene_tx=
XM_023247810.1;refGene_gn=PLEKHA6;refGene_tx=XM_023247811.1;refGene_gn
=PLEKHA6;refGene_tx=XM_023247812.1;refGene_gn=PLEKHA6;refGene_tx=XM_02
3247813.1;refGene_gn=PLEKHA6;refGene_tx=XM_023247814.1;refGene_gn=PLEK
HA6;refGene_tx=XM_023247815.1;refGene_gn=PLEKHA6;refGene_tx=XM_0232478
17.1;refGene_gn=PLEKHA6;refGene_tx=XM_023247818.1
F1 44674100 44674169
refGene_gn=PLEKHA6;refGene_tx=XM_019822049.2;refGene_gn=PLEKHA6;refGen
e_tx=XM_019822052.2;refGene_gn=PLEKHA6;refGene_tx=XM_019822063.2;refGe
ne_gn=PLEKHA6;refGene_tx=XM_019822064.2;refGene_gn=PLEKHA6;refGene_tx=
XM_019822065.2;refGene_gn=PLEKHA6;refGene_tx=XM_019822066.2;refGene_gn
=PLEKHA6;refGene_tx=XM_019822067.2;refGene_gn=PLEKHA6;refGene_tx=XM_01
9822068.2;refGene_gn=PLEKHA6;refGene_tx=XM_019822069.2;refGene_gn=PLEK
HA6;refGene_tx=XM_019822075.2;refGene_gn=PLEKHA6;refGene_tx=XM_0232478
02.1;refGene_gn=PLEKHA6;refGene_tx=XM_023247803.1;refGene_gn=PLEKHA6;r
efGene_tx=XM_023247804.1;refGene_gn=PLEKHA6;refGene_tx=XM_023247805.1;
refGene_gn=PLEKHA6;refGene_tx=XM_023247806.1;refGene_gn=PLEKHA6;refGen
e_tx=XM_023247807.1;refGene_gn=PLEKHA6;refGene_tx=XM_023247808.1;refGe
ne_gn=PLEKHA6;refGene_tx=XM_023247809.1;refGene_gn=PLEKHA6;refGene_tx=
XM_023247810.1;refGene_gn=PLEKHA6;refGene_tx=XM_023247811.1;refGene_gn
=PLEKHA6;refGene_tx=XM_023247813.1;refGene_gn=PLEKHA6;refGene_tx=XM_02
3247815.1;refGene_gn=PLEKHA6;refGene_tx=XM_023247817.1;refGene_gn=PLEK
HA6;refGene_tx=XM_023247818.1
F1 44675055 44675136
refGene_gn=PLEKHA6;refGene_tx=XM_019822049.2;refGene_gn=PLEKHA6;refGen
e_tx=XM_019822052.2;refGene_gn=PLEKHA6;refGene_tx=XM_019822063.2;refGe
ne_gn=PLEKHA6;refGene_tx=XM_019822064.2;refGene_gn=PLEKHA6;refGene_tx=
XM_019822065.2;refGene_gn=PLEKHA6;refGene_tx=XM_019822066.2;refGene_gn
=PLEKHA6;refGene_tx=XM_019822067.2;refGene_gn=PLEKHA6;refGene_tx=XM_01
9822068.2;refGene_gn=PLEKHA6;refGene_tx=XM_019822069.2;refGene_gn=PLEK
HA6;refGene_tx=XM_019822075.2;refGene_gn=PLEKHA6;refGene_tx=XM_0232478
02.1;refGene_gn=PLEKHA6;refGene_tx=XM_023247803.1;refGene_gn=PLEKHA6;r
efGene_tx=XM_023247804.1;refGene_gn=PLEKHA6;refGene_tx=XM_023247805.1;
refGene_gn=PLEKHA6;refGene_tx=XM_023247806.1;refGene_gn=PLEKHA6;refGen
e_tx=XM_023247807.1;refGene_gn=PLEKHA6;refGene_tx=XM_023247809.1;refGe
ne_gn=PLEKHA6;refGene_tx=XM_023247811.1;refGene_gn=PLEKHA6;refGene_tx=
XM_023247813.1
F1 44680651 44680705
refGene_gn=PLEKHA6;refGene_tx=XM_019822049.2;refGene_gn=PLEKHA6;refGen

e_tx=XM_019822052.2;refGene_gn=PLEKHA6;refGene_tx=XM_019822063.2;refGene_gn=PLEKHA6;refGene_tx=XM_019822064.2;refGene_gn=PLEKHA6;refGene_tx=XM_019822065.2;refGene_gn=PLEKHA6;refGene_tx=XM_019822066.2;refGene_gn=PLEKHA6;refGene_tx=XM_019822067.2;refGene_gn=PLEKHA6;refGene_tx=XM_019822068.2;refGene_gn=PLEKHA6;refGene_tx=XM_019822069.2;refGene_gn=PLEKHA6;refGene_tx=XM_019822075.2;refGene_gn=PLEKHA6;refGene_tx=XM_023247802.1;refGene_gn=PLEKHA6;refGene_tx=XM_023247803.1;refGene_gn=PLEKHA6;refGene_tx=XM_023247805.1;refGene_gn=PLEKHA6;refGene_tx=XM_023247806.1;refGene_gn=PLEKHA6;refGene_tx=XM_023247807.1;refGene_gn=PLEKHA6;refGene_tx=XM_023247808.1;refGene_gn=PLEKHA6;refGene_tx=XM_023247809.1;refGene_gn=PLEKHA6;refGene_tx=XM_023247812.1;refGene_gn=PLEKHA6;refGene_tx=XM_023247813.1

F1 44681835 44681922

refGene_gn=PLEKHA6;refGene_tx=XM_019822049.2;refGene_gn=PLEKHA6;refGene_tx=XM_019822052.2;refGene_gn=PLEKHA6;refGene_tx=XM_019822063.2;refGene_gn=PLEKHA6;refGene_tx=XM_019822064.2;refGene_gn=PLEKHA6;refGene_tx=XM_019822065.2;refGene_gn=PLEKHA6;refGene_tx=XM_019822066.2;refGene_gn=PLEKHA6;refGene_tx=XM_019822067.2;refGene_gn=PLEKHA6;refGene_tx=XM_019822068.2;refGene_gn=PLEKHA6;refGene_tx=XM_019822069.2;refGene_gn=PLEKHA6;refGene_tx=XM_019822075.2;refGene_gn=PLEKHA6;refGene_tx=XM_023247802.1;refGene_gn=PLEKHA6;refGene_tx=XM_023247803.1;refGene_gn=PLEKHA6;refGene_tx=XM_023247804.1;refGene_gn=PLEKHA6;refGene_tx=XM_023247805.1;refGene_gn=PLEKHA6;refGene_tx=XM_023247806.1;refGene_gn=PLEKHA6;refGene_tx=XM_023247807.1;refGene_gn=PLEKHA6;refGene_tx=XM_023247808.1;refGene_gn=PLEKHA6;refGene_tx=XM_023247809.1;refGene_gn=PLEKHA6;refGene_tx=XM_023247810.1;refGene_gn=PLEKHA6;refGene_tx=XM_023247812.1;refGene_gn=PLEKHA6;refGene_tx=XM_023247813.1;refGene_gn=PLEKHA6;refGene_tx=XM_023247814.1

F1 44682410 44682933

refGene_gn=PLEKHA6;refGene_tx=XM_019822049.2;refGene_gn=PLEKHA6;refGene_tx=XM_019822052.2;refGene_gn=PLEKHA6;refGene_tx=XM_019822063.2;refGene_gn=PLEKHA6;refGene_tx=XM_019822064.2;refGene_gn=PLEKHA6;refGene_tx=XM_019822065.2;refGene_gn=PLEKHA6;refGene_tx=XM_019822066.2;refGene_gn=PLEKHA6;refGene_tx=XM_019822067.2;refGene_gn=PLEKHA6;refGene_tx=XM_019822068.2;refGene_gn=PLEKHA6;refGene_tx=XM_019822069.2;refGene_gn=PLEKHA6;refGene_tx=XM_019822075.2;refGene_gn=PLEKHA6;refGene_tx=XM_023247802.1;refGene_gn=PLEKHA6;refGene_tx=XM_023247803.1;refGene_gn=PLEKHA6;refGene_tx=XM_023247804.1;refGene_gn=PLEKHA6;refGene_tx=XM_023247805.1;refGene_gn=PLEKHA6;refGene_tx=XM_023247806.1;refGene_gn=PLEKHA6;refGene_tx=XM_023247807.1;refGene_gn=PLEKHA6;refGene_tx=XM_023247808.1;refGene_gn=PLEKHA6;refGene_tx=XM_023247809.1;refGene_gn=PLEKHA6;refGene_tx=XM_023247810.1;refGene_gn=PLEKHA6;refGene_tx=XM_023247811.1;refGene_gn=PLEKHA6;refGene_tx=XM_023247812.1;refGene_gn=PLEKHA6;refGene_tx=XM_023247813.1;refGene_gn=PLEKHA6;refGene_tx=XM_023247814.1;refGene_gn=PLEKHA6;refGene_tx=XM_023247815.1;refGene_gn=PLEKHA6;refGene_tx=XM_023247817.1;refGene_gn=PLEKHA6;refGene_tx=XM_023247818.1

F1 44684625 44685180

refGene_gn=PLEKHA6;refGene_tx=XM_019822049.2;refGene_gn=PLEKHA6;refGene_tx=XM_019822052.2;refGene_gn=PLEKHA6;refGene_tx=XM_019822063.2;refGene_gn=PLEKHA6;refGene_tx=XM_019822064.2;refGene_gn=PLEKHA6;refGene_tx=

XM_019822065.2;refGene_gn=PLEKHA6;refGene_tx=XM_019822066.2;refGene_gn=PLEKHA6;refGene_tx=XM_019822067.2;refGene_gn=PLEKHA6;refGene_tx=XM_019822068.2;refGene_gn=PLEKHA6;refGene_tx=XM_019822069.2;refGene_gn=PLEKHA6;refGene_tx=XM_019822075.2;refGene_gn=PLEKHA6;refGene_tx=XM_023247802.1;refGene_gn=PLEKHA6;refGene_tx=XM_023247803.1;refGene_gn=PLEKHA6;refGene_tx=XM_023247804.1;refGene_gn=PLEKHA6;refGene_tx=XM_023247805.1;refGene_gn=PLEKHA6;refGene_tx=XM_023247806.1;refGene_gn=PLEKHA6;refGene_tx=XM_023247807.1;refGene_gn=PLEKHA6;refGene_tx=XM_023247808.1;refGene_gn=PLEKHA6;refGene_tx=XM_023247809.1;refGene_gn=PLEKHA6;refGene_tx=XM_023247810.1;refGene_gn=PLEKHA6;refGene_tx=XM_023247811.1;refGene_gn=PLEKHA6;refGene_tx=XM_023247812.1;refGene_gn=PLEKHA6;refGene_tx=XM_023247813.1;refGene_gn=PLEKHA6;refGene_tx=XM_023247814.1;refGene_gn=PLEKHA6;refGene_tx=XM_023247815.1;refGene_gn=PLEKHA6;refGene_tx=XM_023247817.1;refGene_gn=PLEKHA6;refGene_tx=XM_023247818.1

F1 44686672 44686875

refGene_gn=PLEKHA6;refGene_tx=XM_019822049.2;refGene_gn=PLEKHA6;refGene_tx=XM_019822052.2;refGene_gn=PLEKHA6;refGene_tx=XM_019822063.2;refGene_gn=PLEKHA6;refGene_tx=XM_019822064.2;refGene_gn=PLEKHA6;refGene_tx=XM_019822065.2;refGene_gn=PLEKHA6;refGene_tx=XM_019822066.2;refGene_gn=PLEKHA6;refGene_tx=XM_019822067.2;refGene_gn=PLEKHA6;refGene_tx=XM_019822068.2;refGene_gn=PLEKHA6;refGene_tx=XM_019822069.2;refGene_gn=PLEKHA6;refGene_tx=XM_019822075.2;refGene_gn=PLEKHA6;refGene_tx=XM_023247802.1;refGene_gn=PLEKHA6;refGene_tx=XM_023247803.1;refGene_gn=PLEKHA6;refGene_tx=XM_023247804.1;refGene_gn=PLEKHA6;refGene_tx=XM_023247805.1;refGene_gn=PLEKHA6;refGene_tx=XM_023247806.1;refGene_gn=PLEKHA6;refGene_tx=XM_023247807.1;refGene_gn=PLEKHA6;refGene_tx=XM_023247808.1;refGene_gn=PLEKHA6;refGene_tx=XM_023247809.1;refGene_gn=PLEKHA6;refGene_tx=XM_023247810.1;refGene_gn=PLEKHA6;refGene_tx=XM_023247811.1;refGene_gn=PLEKHA6;refGene_tx=XM_023247812.1;refGene_gn=PLEKHA6;refGene_tx=XM_023247813.1;refGene_gn=PLEKHA6;refGene_tx=XM_023247814.1;refGene_gn=PLEKHA6;refGene_tx=XM_023247815.1;refGene_gn=PLEKHA6;refGene_tx=XM_023247817.1;refGene_gn=PLEKHA6;refGene_tx=XM_023247818.1

F1 44690365 44690466

refGene_gn=PLEKHA6;refGene_tx=XM_019822049.2;refGene_gn=PLEKHA6;refGene_tx=XM_019822052.2;refGene_gn=PLEKHA6;refGene_tx=XM_019822063.2;refGene_gn=PLEKHA6;refGene_tx=XM_019822064.2;refGene_gn=PLEKHA6;refGene_tx=XM_019822065.2;refGene_gn=PLEKHA6;refGene_tx=XM_019822066.2;refGene_gn=PLEKHA6;refGene_tx=XM_019822067.2;refGene_gn=PLEKHA6;refGene_tx=XM_019822068.2;refGene_gn=PLEKHA6;refGene_tx=XM_019822069.2;refGene_gn=PLEKHA6;refGene_tx=XM_023247802.1;refGene_gn=PLEKHA6;refGene_tx=XM_023247803.1;refGene_gn=PLEKHA6;refGene_tx=XM_023247804.1;refGene_gn=PLEKHA6;refGene_tx=XM_023247805.1;refGene_gn=PLEKHA6;refGene_tx=XM_023247806.1;refGene_gn=PLEKHA6;refGene_tx=XM_023247807.1;refGene_gn=PLEKHA6;refGene_tx=XM_023247808.1;refGene_gn=PLEKHA6;refGene_tx=XM_023247809.1;refGene_gn=PLEKHA6;refGene_tx=XM_023247810.1;refGene_gn=PLEKHA6;refGene_tx=XM_023247811.1;refGene_gn=PLEKHA6;refGene_tx=XM_023247812.1;refGene_gn=PLEKHA6;refGene_tx=XM_023247813.1;refGene_gn=PLEKHA6;refGene_tx=XM_023247814.1;refGene_gn=PLEKHA6;refGene_tx=XM_023247815.1;refGene_gn=PLEKHA6;refGene_tx=XM_023247817.1;refGene_gn=PLEKHA6;refGene_tx=XM_023247818.1

F1 44693258 44693331
refGene_gn=PLEKHA6;refGene_tx=XM_019822049.2;refGene_gn=PLEKHA6;refGene_tx=XM_019822052.2;refGene_gn=PLEKHA6;refGene_tx=XM_019822063.2;refGene_gn=PLEKHA6;refGene_tx=XM_019822064.2;refGene_gn=PLEKHA6;refGene_tx=XM_019822065.2;refGene_gn=PLEKHA6;refGene_tx=XM_019822066.2;refGene_gn=PLEKHA6;refGene_tx=XM_019822067.2;refGene_gn=PLEKHA6;refGene_tx=XM_019822068.2;refGene_gn=PLEKHA6;refGene_tx=XM_019822069.2;refGene_gn=PLEKHA6;refGene_tx=XM_023247802.1;refGene_gn=PLEKHA6;refGene_tx=XM_023247803.1;refGene_gn=PLEKHA6;refGene_tx=XM_023247804.1;refGene_gn=PLEKHA6;refGene_tx=XM_023247805.1;refGene_gn=PLEKHA6;refGene_tx=XM_023247806.1;refGene_gn=PLEKHA6;refGene_tx=XM_023247807.1;refGene_gn=PLEKHA6;refGene_tx=XM_023247808.1;refGene_gn=PLEKHA6;refGene_tx=XM_023247809.1;refGene_gn=PLEKHA6;refGene_tx=XM_023247810.1;refGene_gn=PLEKHA6;refGene_tx=XM_023247811.1;refGene_gn=PLEKHA6;refGene_tx=XM_023247812.1;refGene_gn=PLEKHA6;refGene_tx=XM_023247813.1;refGene_gn=PLEKHA6;refGene_tx=XM_023247814.1;refGene_gn=PLEKHA6;refGene_tx=XM_023247815.1;refGene_gn=PLEKHA6;refGene_tx=XM_023247817.1;refGene_gn=PLEKHA6;refGene_tx=XM_023247818.1
F1 44693986 44694091
refGene_gn=PLEKHA6;refGene_tx=XM_019822049.2;refGene_gn=PLEKHA6;refGene_tx=XM_019822052.2;refGene_gn=PLEKHA6;refGene_tx=XM_019822063.2;refGene_gn=PLEKHA6;refGene_tx=XM_019822064.2;refGene_gn=PLEKHA6;refGene_tx=XM_019822065.2;refGene_gn=PLEKHA6;refGene_tx=XM_019822066.2;refGene_gn=PLEKHA6;refGene_tx=XM_019822067.2;refGene_gn=PLEKHA6;refGene_tx=XM_019822068.2;refGene_gn=PLEKHA6;refGene_tx=XM_019822069.2;refGene_gn=PLEKHA6;refGene_tx=XM_023247802.1;refGene_gn=PLEKHA6;refGene_tx=XM_023247803.1;refGene_gn=PLEKHA6;refGene_tx=XM_023247804.1;refGene_gn=PLEKHA6;refGene_tx=XM_023247805.1;refGene_gn=PLEKHA6;refGene_tx=XM_023247806.1;refGene_gn=PLEKHA6;refGene_tx=XM_023247807.1;refGene_gn=PLEKHA6;refGene_tx=XM_023247808.1;refGene_gn=PLEKHA6;refGene_tx=XM_023247809.1;refGene_gn=PLEKHA6;refGene_tx=XM_023247810.1;refGene_gn=PLEKHA6;refGene_tx=XM_023247811.1;refGene_gn=PLEKHA6;refGene_tx=XM_023247812.1;refGene_gn=PLEKHA6;refGene_tx=XM_023247813.1;refGene_gn=PLEKHA6;refGene_tx=XM_023247814.1;refGene_gn=PLEKHA6;refGene_tx=XM_023247815.1;refGene_gn=PLEKHA6;refGene_tx=XM_023247817.1;refGene_gn=PLEKHA6;refGene_tx=XM_023247818.1
F1 44699129 44699244
refGene_gn=PLEKHA6;refGene_tx=XM_019822049.2;refGene_gn=PLEKHA6;refGene_tx=XM_019822052.2;refGene_gn=PLEKHA6;refGene_tx=XM_019822063.2;refGene_gn=PLEKHA6;refGene_tx=XM_019822064.2;refGene_gn=PLEKHA6;refGene_tx=XM_019822065.2;refGene_gn=PLEKHA6;refGene_tx=XM_019822066.2;refGene_gn=PLEKHA6;refGene_tx=XM_019822067.2;refGene_gn=PLEKHA6;refGene_tx=XM_019822068.2;refGene_gn=PLEKHA6;refGene_tx=XM_019822069.2;refGene_gn=PLEKHA6;refGene_tx=XM_023247802.1;refGene_gn=PLEKHA6;refGene_tx=XM_023247803.1;refGene_gn=PLEKHA6;refGene_tx=XM_023247804.1;refGene_gn=PLEKHA6;refGene_tx=XM_023247805.1;refGene_gn=PLEKHA6;refGene_tx=XM_023247806.1;refGene_gn=PLEKHA6;refGene_tx=XM_023247807.1;refGene_gn=PLEKHA6;refGene_tx=XM_023247808.1;refGene_gn=PLEKHA6;refGene_tx=XM_023247809.1;refGene_gn=PLEKHA6;refGene_tx=XM_023247810.1;refGene_gn=PLEKHA6;refGene_tx=XM_023247811.1;refGene_gn=PLEKHA6;refGene_tx=XM_023247812.1;refGene_gn

=PLEKHA6;refGene_tx=XM_023247813.1;refGene_gn=PLEKHA6;refGene_tx=XM_02
3247814.1;refGene_gn=PLEKHA6;refGene_tx=XM_023247815.1;refGene_gn=PLEK
HA6;refGene_tx=XM_023247817.1;refGene_gn=PLEKHA6;refGene_tx=XM_0232478
18.1
F1 44699925 44700006
refGene_gn=PLEKHA6;refGene_tx=XM_019822049.2;refGene_gn=PLEKHA6;refGen
e_tx=XM_019822052.2;refGene_gn=PLEKHA6;refGene_tx=XM_023247802.1;refGe
ne_gn=PLEKHA6;refGene_tx=XM_023247803.1;refGene_gn=PLEKHA6;refGene_tx=
XM_023247804.1;refGene_gn=PLEKHA6;refGene_tx=XM_023247805.1;refGene_gn
=PLEKHA6;refGene_tx=XM_023247806.1;refGene_gn=PLEKHA6;refGene_tx=XM_02
3247807.1;refGene_gn=PLEKHA6;refGene_tx=XM_023247808.1;refGene_gn=PLEK
HA6;refGene_tx=XM_023247809.1;refGene_gn=PLEKHA6;refGene_tx=XM_0232478
10.1;refGene_gn=PLEKHA6;refGene_tx=XM_023247811.1;refGene_gn=PLEKHA6;r
efGene_tx=XM_023247812.1;refGene_gn=PLEKHA6;refGene_tx=XM_023247813.1;
refGene_gn=PLEKHA6;refGene_tx=XM_023247814.1;refGene_gn=PLEKHA6;refGen
e_tx=XM_023247815.1;refGene_gn=PLEKHA6;refGene_tx=XM_023247817.1;refGe
ne_gn=PLEKHA6;refGene_tx=XM_023247818.1
F1 44702152 44702184 refGene_gn=PLEKHA6;refGene_tx=XM_019822052.2
F1 44768552 44768614 refGene_gn=PLEKHA6;refGene_tx=XM_019822049.2
F1 44788274 44788332
refGene_gn=PLEKHA6;refGene_tx=XM_023247802.1;refGene_gn=PLEKHA6;refGen
e_tx=XM_023247803.1;refGene_gn=PLEKHA6;refGene_tx=XM_023247804.1;refGe
ne_gn=PLEKHA6;refGene_tx=XM_023247805.1;refGene_gn=PLEKHA6;refGene_tx=
XM_023247806.1;refGene_gn=PLEKHA6;refGene_tx=XM_023247807.1;refGene_gn
=PLEKHA6;refGene_tx=XM_023247808.1;refGene_gn=PLEKHA6;refGene_tx=XM_02
3247809.1;refGene_gn=PLEKHA6;refGene_tx=XM_023247810.1;refGene_gn=PLEK
HA6;refGene_tx=XM_023247811.1;refGene_gn=PLEKHA6;refGene_tx=XM_0232478
12.1;refGene_gn=PLEKHA6;refGene_tx=XM_023247813.1;refGene_gn=PLEKHA6;r
efGene_tx=XM_023247814.1;refGene_gn=PLEKHA6;refGene_tx=XM_023247815.1;
refGene_gn=PLEKHA6;refGene_tx=XM_023247817.1;refGene_gn=PLEKHA6;refGen
e_tx=XM_023247818.1
F1 44794233 44794310
refGene_gn=PLEKHA6;refGene_tx=XM_023247802.1;refGene_gn=PLEKHA6;refGen
e_tx=XM_023247803.1;refGene_gn=PLEKHA6;refGene_tx=XM_023247804.1;refGe
ne_gn=PLEKHA6;refGene_tx=XM_023247805.1;refGene_gn=PLEKHA6;refGene_tx=
XM_023247806.1;refGene_gn=PLEKHA6;refGene_tx=XM_023247807.1;refGene_gn
=PLEKHA6;refGene_tx=XM_023247808.1;refGene_gn=PLEKHA6;refGene_tx=XM_02
3247809.1;refGene_gn=PLEKHA6;refGene_tx=XM_023247810.1;refGene_gn=PLEK
HA6;refGene_tx=XM_023247811.1;refGene_gn=PLEKHA6;refGene_tx=XM_0232478
12.1;refGene_gn=PLEKHA6;refGene_tx=XM_023247813.1;refGene_gn=PLEKHA6;r
efGene_tx=XM_023247814.1;refGene_gn=PLEKHA6;refGene_tx=XM_023247815.1;
refGene_gn=PLEKHA6;refGene_tx=XM_023247817.1;refGene_gn=PLEKHA6;refGen
e_tx=XM_023247818.1
F1 44800760 44800843
refGene_gn=PLEKHA6;refGene_tx=XM_023247802.1;refGene_gn=PLEKHA6;refGen
e_tx=XM_023247803.1;refGene_gn=PLEKHA6;refGene_tx=XM_023247804.1;refGe
ne_gn=PLEKHA6;refGene_tx=XM_023247805.1;refGene_gn=PLEKHA6;refGene_tx=
XM_023247806.1;refGene_gn=PLEKHA6;refGene_tx=XM_023247807.1;refGene_gn
=PLEKHA6;refGene_tx=XM_023247808.1;refGene_gn=PLEKHA6;refGene_tx=XM_02
3247809.1;refGene_gn=PLEKHA6;refGene_tx=XM_023247810.1;refGene_gn=PLEK

HA6;refGene_tx=XM_023247811.1;refGene_gn=PLEKHA6;refGene_tx=XM_023247812.1;refGene_gn=PLEKHA6;refGene_tx=XM_023247813.1;refGene_gn=PLEKHA6;refGene_tx=XM_023247814.1;refGene_gn=PLEKHA6;refGene_tx=XM_023247815.1;refGene_gn=PLEKHA6;refGene_tx=XM_023247817.1;refGene_gn=PLEKHA6;refGene_tx=XM_023247818.1
F1 44813712 44813735 refGene_gn=PPP1R15B;refGene_tx=XM_023248108.1
F1 44818825 44818950
refGene_gn=PPP1R15B;refGene_tx=XM_019822077.2;refGene_gn=PPP1R15B;refGene_tx=XM_019822080.2;refGene_gn=PPP1R15B;refGene_tx=XM_023248107.1;refGene_gn=PPP1R15B;refGene_tx=XM_023248108.1
F1 44825078 44825300
refGene_gn=PPP1R15B;refGene_tx=XM_006942902.4;refGene_gn=PPP1R15B;refGene_tx=XM_011291066.3
F1 44829146 44831066
refGene_gn=PPP1R15B;refGene_tx=XM_006942902.4;refGene_gn=PPP1R15B;refGene_tx=XM_011291066.3;refGene_gn=PPP1R15B;refGene_tx=XM_019822077.2;refGene_gn=PPP1R15B;refGene_tx=XM_019822080.2;refGene_gn=PPP1R15B;refGene_tx=XM_023248107.1;refGene_gn=PPP1R15B;refGene_tx=XM_023248108.1
F1 44842532 44842721
refGene_gn=PIK3C2B;refGene_tx=XM_019822082.2;refGene_gn=PIK3C2B;refGene_tx=XM_023248051.1;refGene_gn=PIK3C2B;refGene_tx=XM_023248052.1;refGene_gn=PIK3C2B;refGene_tx=XM_023248054.1
F1 44843895 44844024
refGene_gn=PIK3C2B;refGene_tx=XM_019822082.2;refGene_gn=PIK3C2B;refGene_tx=XM_023248051.1;refGene_gn=PIK3C2B;refGene_tx=XM_023248052.1;refGene_gn=PIK3C2B;refGene_tx=XM_023248053.1;refGene_gn=PIK3C2B;refGene_tx=XM_023248054.1
F1 44845667 44845774
refGene_gn=PIK3C2B;refGene_tx=XM_019822082.2;refGene_gn=PIK3C2B;refGene_tx=XM_023248051.1;refGene_gn=PIK3C2B;refGene_tx=XM_023248052.1;refGene_gn=PIK3C2B;refGene_tx=XM_023248054.1
F1 44846200 44846282
refGene_gn=PIK3C2B;refGene_tx=XM_019822082.2;refGene_gn=PIK3C2B;refGene_tx=XM_023248051.1;refGene_gn=PIK3C2B;refGene_tx=XM_023248052.1;refGene_gn=PIK3C2B;refGene_tx=XM_023248053.1;refGene_gn=PIK3C2B;refGene_tx=XM_023248054.1
F1 44847555 44847673
refGene_gn=PIK3C2B;refGene_tx=XM_019822082.2;refGene_gn=PIK3C2B;refGene_tx=XM_023248051.1;refGene_gn=PIK3C2B;refGene_tx=XM_023248052.1;refGene_gn=PIK3C2B;refGene_tx=XM_023248053.1;refGene_gn=PIK3C2B;refGene_tx=XM_023248054.1
F1 44849275 44849400
refGene_gn=PIK3C2B;refGene_tx=XM_019822082.2;refGene_gn=PIK3C2B;refGene_tx=XM_023248051.1;refGene_gn=PIK3C2B;refGene_tx=XM_023248052.1;refGene_gn=PIK3C2B;refGene_tx=XM_023248053.1;refGene_gn=PIK3C2B;refGene_tx=XM_023248054.1
F1 44849824 44850026
refGene_gn=PIK3C2B;refGene_tx=XM_019822082.2;refGene_gn=PIK3C2B;refGene_tx=XM_023248051.1;refGene_gn=PIK3C2B;refGene_tx=XM_023248052.1;refGene_gn=PIK3C2B;refGene_tx=XM_023248053.1;refGene_gn=PIK3C2B;refGene_tx=

XM_023248054.1
F1 44850809 44850919
refGene_gn=PIK3C2B;refGene_tx=XM_019822082.2;refGene_gn=PIK3C2B;refGene_tx=XM_023248051.1;refGene_gn=PIK3C2B;refGene_tx=XM_023248052.1;refGene_gn=PIK3C2B;refGene_tx=XM_023248053.1;refGene_gn=PIK3C2B;refGene_tx=XM_023248054.1
F1 44851257 44851414
refGene_gn=PIK3C2B;refGene_tx=XM_019822082.2;refGene_gn=PIK3C2B;refGene_tx=XM_023248051.1;refGene_gn=PIK3C2B;refGene_tx=XM_023248052.1;refGene_gn=PIK3C2B;refGene_tx=XM_023248053.1;refGene_gn=PIK3C2B;refGene_tx=XM_023248054.1
F1 44852137 44852307
refGene_gn=PIK3C2B;refGene_tx=XM_019822082.2;refGene_gn=PIK3C2B;refGene_tx=XM_023248051.1;refGene_gn=PIK3C2B;refGene_tx=XM_023248052.1;refGene_gn=PIK3C2B;refGene_tx=XM_023248053.1;refGene_gn=PIK3C2B;refGene_tx=XM_023248054.1
F1 44854901 44855038
refGene_gn=PIK3C2B;refGene_tx=XM_019822082.2;refGene_gn=PIK3C2B;refGene_tx=XM_023248051.1;refGene_gn=PIK3C2B;refGene_tx=XM_023248052.1;refGene_gn=PIK3C2B;refGene_tx=XM_023248053.1;refGene_gn=PIK3C2B;refGene_tx=XM_023248054.1
F1 44856019 44856149
refGene_gn=PIK3C2B;refGene_tx=XM_019822082.2;refGene_gn=PIK3C2B;refGene_tx=XM_023248051.1;refGene_gn=PIK3C2B;refGene_tx=XM_023248052.1;refGene_gn=PIK3C2B;refGene_tx=XM_023248053.1;refGene_gn=PIK3C2B;refGene_tx=XM_023248054.1
F1 44856949 44857042
refGene_gn=PIK3C2B;refGene_tx=XM_019822082.2;refGene_gn=PIK3C2B;refGene_tx=XM_023248051.1;refGene_gn=PIK3C2B;refGene_tx=XM_023248052.1;refGene_gn=PIK3C2B;refGene_tx=XM_023248053.1;refGene_gn=PIK3C2B;refGene_tx=XM_023248054.1
F1 44857656 44857764
refGene_gn=PIK3C2B;refGene_tx=XM_019822082.2;refGene_gn=PIK3C2B;refGene_tx=XM_023248051.1;refGene_gn=PIK3C2B;refGene_tx=XM_023248052.1;refGene_gn=PIK3C2B;refGene_tx=XM_023248053.1;refGene_gn=PIK3C2B;refGene_tx=XM_023248054.1
F1 44858799 44858980
refGene_gn=PIK3C2B;refGene_tx=XM_019822082.2;refGene_gn=PIK3C2B;refGene_tx=XM_023248051.1;refGene_gn=PIK3C2B;refGene_tx=XM_023248052.1;refGene_gn=PIK3C2B;refGene_tx=XM_023248053.1;refGene_gn=PIK3C2B;refGene_tx=XM_023248054.1
F1 44859870 44859965
refGene_gn=PIK3C2B;refGene_tx=XM_019822082.2;refGene_gn=PIK3C2B;refGene_tx=XM_023248051.1;refGene_gn=PIK3C2B;refGene_tx=XM_023248052.1;refGene_gn=PIK3C2B;refGene_tx=XM_023248053.1;refGene_gn=PIK3C2B;refGene_tx=XM_023248054.1
F1 44860120 44860214
refGene_gn=PIK3C2B;refGene_tx=XM_019822082.2;refGene_gn=PIK3C2B;refGene_tx=XM_023248051.1;refGene_gn=PIK3C2B;refGene_tx=XM_023248052.1;refGene_gn=PIK3C2B;refGene_tx=XM_023248053.1;refGene_gn=PIK3C2B;refGene_tx=

XM_023248054.1
F1 44861403 44861592
refGene_gn=PIK3C2B;refGene_tx=XM_019822082.2;refGene_gn=PIK3C2B;refGene_tx=XM_023248051.1;refGene_gn=PIK3C2B;refGene_tx=XM_023248052.1;refGene_gn=PIK3C2B;refGene_tx=XM_023248053.1;refGene_gn=PIK3C2B;refGene_tx=XM_023248054.1
F1 44863139 44863282
refGene_gn=PIK3C2B;refGene_tx=XM_019822082.2;refGene_gn=PIK3C2B;refGene_tx=XM_023248051.1;refGene_gn=PIK3C2B;refGene_tx=XM_023248052.1;refGene_gn=PIK3C2B;refGene_tx=XM_023248053.1;refGene_gn=PIK3C2B;refGene_tx=XM_023248054.1
F1 44864469 44864581
refGene_gn=PIK3C2B;refGene_tx=XM_019822082.2;refGene_gn=PIK3C2B;refGene_tx=XM_023248051.1;refGene_gn=PIK3C2B;refGene_tx=XM_023248052.1;refGene_gn=PIK3C2B;refGene_tx=XM_023248053.1;refGene_gn=PIK3C2B;refGene_tx=XM_023248054.1
F1 44865095 44865263
refGene_gn=PIK3C2B;refGene_tx=XM_019822082.2;refGene_gn=PIK3C2B;refGene_tx=XM_023248051.1;refGene_gn=PIK3C2B;refGene_tx=XM_023248052.1;refGene_gn=PIK3C2B;refGene_tx=XM_023248053.1;refGene_gn=PIK3C2B;refGene_tx=XM_023248054.1
F1 44868916 44869039
refGene_gn=PIK3C2B;refGene_tx=XM_019822082.2;refGene_gn=PIK3C2B;refGene_tx=XM_023248051.1;refGene_gn=PIK3C2B;refGene_tx=XM_023248052.1;refGene_gn=PIK3C2B;refGene_tx=XM_023248053.1;refGene_gn=PIK3C2B;refGene_tx=XM_023248054.1
F1 44870077 44870273
refGene_gn=PIK3C2B;refGene_tx=XM_019822082.2;refGene_gn=PIK3C2B;refGene_tx=XM_023248051.1;refGene_gn=PIK3C2B;refGene_tx=XM_023248052.1;refGene_gn=PIK3C2B;refGene_tx=XM_023248053.1;refGene_gn=PIK3C2B;refGene_tx=XM_023248054.1
F1 44871113 44871147
refGene_gn=PIK3C2B;refGene_tx=XM_019822082.2;refGene_gn=PIK3C2B;refGene_tx=XM_023248051.1;refGene_gn=PIK3C2B;refGene_tx=XM_023248052.1;refGene_gn=PIK3C2B;refGene_tx=XM_023248053.1;refGene_gn=PIK3C2B;refGene_tx=XM_023248054.1
F1 44871742 44871889
refGene_gn=PIK3C2B;refGene_tx=XM_019822082.2;refGene_gn=PIK3C2B;refGene_tx=XM_023248051.1;refGene_gn=PIK3C2B;refGene_tx=XM_023248052.1;refGene_gn=PIK3C2B;refGene_tx=XM_023248053.1;refGene_gn=PIK3C2B;refGene_tx=XM_023248054.1
F1 44874068 44874132
refGene_gn=PIK3C2B;refGene_tx=XM_019822082.2;refGene_gn=PIK3C2B;refGene_tx=XM_023248051.1;refGene_gn=PIK3C2B;refGene_tx=XM_023248052.1;refGene_gn=PIK3C2B;refGene_tx=XM_023248053.1;refGene_gn=PIK3C2B;refGene_tx=XM_023248054.1
F1 44874515 44874595
refGene_gn=PIK3C2B;refGene_tx=XM_019822082.2;refGene_gn=PIK3C2B;refGene_tx=XM_023248051.1;refGene_gn=PIK3C2B;refGene_tx=XM_023248052.1;refGene_gn=PIK3C2B;refGene_tx=XM_023248053.1;refGene_gn=PIK3C2B;refGene_tx=

XM_023248054.1
F1 44874850 44874962
refGene_gn=PIK3C2B;refGene_tx=XM_019822082.2;refGene_gn=PIK3C2B;refGene_tx=XM_023248051.1;refGene_gn=PIK3C2B;refGene_tx=XM_023248052.1;refGene_gn=PIK3C2B;refGene_tx=XM_023248053.1;refGene_gn=PIK3C2B;refGene_tx=XM_023248054.1
F1 44877879 44878000
refGene_gn=PIK3C2B;refGene_tx=XM_019822082.2;refGene_gn=PIK3C2B;refGene_tx=XM_023248051.1;refGene_gn=PIK3C2B;refGene_tx=XM_023248052.1;refGene_gn=PIK3C2B;refGene_tx=XM_023248053.1;refGene_gn=PIK3C2B;refGene_tx=XM_023248054.1
F1 44878349 44878541
refGene_gn=PIK3C2B;refGene_tx=XM_019822082.2;refGene_gn=PIK3C2B;refGene_tx=XM_023248051.1;refGene_gn=PIK3C2B;refGene_tx=XM_023248052.1;refGene_gn=PIK3C2B;refGene_tx=XM_023248053.1;refGene_gn=PIK3C2B;refGene_tx=XM_023248054.1
F1 44879106 44879209
refGene_gn=PIK3C2B;refGene_tx=XM_019822082.2;refGene_gn=PIK3C2B;refGene_tx=XM_023248051.1;refGene_gn=PIK3C2B;refGene_tx=XM_023248052.1;refGene_gn=PIK3C2B;refGene_tx=XM_023248053.1;refGene_gn=PIK3C2B;refGene_tx=XM_023248054.1
F1 44881061 44881155 refGene_gn=PIK3C2B;refGene_tx=XM_019822082.2
F1 44883098 44884047
refGene_gn=PIK3C2B;refGene_tx=XM_023248051.1;refGene_gn=PIK3C2B;refGene_tx=XM_023248052.1;refGene_gn=PIK3C2B;refGene_tx=XM_023248053.1;refGene_gn=PIK3C2B;refGene_tx=XM_023248054.1
F1 44923726 44923780
refGene_gn=MDM4;refGene_tx=XM_023248004.1;refGene_gn=MDM4;refGene_tx=XM_023248006.1
F1 44938167 44938281
refGene_gn=MDM4;refGene_tx=XM_003999396.5;refGene_gn=MDM4;refGene_tx=XM_019822086.2;refGene_gn=MDM4;refGene_tx=XM_019822087.2;refGene_gn=MDM4;refGene_tx=XM_023248004.1;refGene_gn=MDM4;refGene_tx=XM_023248006.1
F1 44939766 44939841 refGene_gn=MDM4;refGene_tx=XM_023248006.1
F1 44941281 44941356
refGene_gn=MDM4;refGene_tx=XM_003999396.5;refGene_gn=MDM4;refGene_tx=XM_019822086.2;refGene_gn=MDM4;refGene_tx=XM_019822087.2;refGene_gn=MDM4;refGene_tx=XM_023248004.1
F1 44946275 44946409
refGene_gn=MDM4;refGene_tx=XM_003999396.5;refGene_gn=MDM4;refGene_tx=XM_019822086.2;refGene_gn=MDM4;refGene_tx=XM_019822087.2;refGene_gn=MDM4;refGene_tx=XM_023248004.1;refGene_gn=MDM4;refGene_tx=XM_023248006.1
F1 44947477 44947533
refGene_gn=MDM4;refGene_tx=XM_003999396.5;refGene_gn=MDM4;refGene_tx=XM_019822086.2;refGene_gn=MDM4;refGene_tx=XM_019822087.2;refGene_gn=MDM4;refGene_tx=XM_023248004.1;refGene_gn=MDM4;refGene_tx=XM_023248006.1
F1 44951331 44951399
refGene_gn=MDM4;refGene_tx=XM_003999396.5;refGene_gn=MDM4;refGene_tx=XM_019822086.2;refGene_gn=MDM4;refGene_tx=XM_019822087.2;refGene_gn=MDM4;refGene_tx=XM_023248004.1;refGene_gn=MDM4;refGene_tx=XM_023248006.1

F1	44952128	44952228
refGene_gn=MDM4;refGene_tx=XM_003999396.5;refGene_gn=MDM4;refGene_tx=XM_019822086.2;refGene_gn=MDM4;refGene_tx=XM_019822087.2;refGene_gn=MDM4;refGene_tx=XM_023248004.1;refGene_gn=MDM4;refGene_tx=XM_023248006.1
F1	44954397	44954558
refGene_gn=MDM4;refGene_tx=XM_003999396.5;refGene_gn=MDM4;refGene_tx=XM_019822086.2;refGene_gn=MDM4;refGene_tx=XM_019822087.2;refGene_gn=MDM4;refGene_tx=XM_023248004.1;refGene_gn=MDM4;refGene_tx=XM_023248006.1
F1	44955814	44955964
refGene_gn=MDM4;refGene_tx=XM_003999396.5;refGene_gn=MDM4;refGene_tx=XM_019822086.2;refGene_gn=MDM4;refGene_tx=XM_019822087.2;refGene_gn=MDM4;refGene_tx=XM_023248004.1;refGene_gn=MDM4;refGene_tx=XM_023248006.1
F1	44957647	44957728
refGene_gn=MDM4;refGene_tx=XM_003999396.5;refGene_gn=MDM4;refGene_tx=XM_019822086.2;refGene_gn=MDM4;refGene_tx=XM_019822087.2;refGene_gn=MDM4;refGene_tx=XM_023248004.1;refGene_gn=MDM4;refGene_tx=XM_023248006.1
F1	44959667	44960240
refGene_gn=MDM4;refGene_tx=XM_003999396.5;refGene_gn=MDM4;refGene_tx=XM_019822086.2;refGene_gn=MDM4;refGene_tx=XM_019822087.2;refGene_gn=MDM4;refGene_tx=XM_023248004.1;refGene_gn=MDM4;refGene_tx=XM_023248006.1
F1	45019419	45021561
refGene_gn=LRRN2;refGene_tx=XM_006942904.4;refGene_gn=LRRN2;refGene_tx=XM_019822089.2;refGene_gn=LRRN2;refGene_tx=XM_023248116.1;refGene_gn=LRRN2;refGene_tx=XM_023248117.1
F1	45211604	45211711
refGene_gn=NFASC;refGene_tx=XM_023247774.1;refGene_gn=NFASC;refGene_tx=XM_023247775.1;refGene_gn=NFASC;refGene_tx=XM_023247776.1;refGene_gn=NFASC;refGene_tx=XM_023247777.1;refGene_gn=NFASC;refGene_tx=XM_023247778.1;refGene_gn=NFASC;refGene_tx=XM_023247779.1;refGene_gn=NFASC;refGene_tx=XM_023247780.1;refGene_gn=NFASC;refGene_tx=XM_023247782.1;refGene_gn=NFASC;refGene_tx=XM_023247783.1;refGene_gn=NFASC;refGene_tx=XM_023247784.1;refGene_gn=NFASC;refGene_tx=XM_023247785.1;refGene_gn=NFASC;refGene_tx=XM_023247786.1;refGene_gn=NFASC;refGene_tx=XM_023247787.1;refGene_gn=NFASC;refGene_tx=XM_023247788.1;refGene_gn=NFASC;refGene_tx=XM_023247789.1
F1	45250667	45250759	refGene_gn=NFASC;refGene_tx=XM_011291070.3
F1	45300357	45300463
refGene_gn=NFASC;refGene_tx=XM_011291070.3;refGene_gn=NFASC;refGene_tx=XM_023247774.1;refGene_gn=NFASC;refGene_tx=XM_023247775.1;refGene_gn=NFASC;refGene_tx=XM_023247776.1;refGene_gn=NFASC;refGene_tx=XM_023247777.1;refGene_gn=NFASC;refGene_tx=XM_023247778.1;refGene_gn=NFASC;refGene_tx=XM_023247779.1;refGene_gn=NFASC;refGene_tx=XM_023247780.1;refGene_gn=NFASC;refGene_tx=XM_023247782.1;refGene_gn=NFASC;refGene_tx=XM_023247783.1;refGene_gn=NFASC;refGene_tx=XM_023247784.1;refGene_gn=NFASC;refGene_tx=XM_023247785.1;refGene_gn=NFASC;refGene_tx=XM_023247786.1;refGene_gn=NFASC;refGene_tx=XM_023247787.1;refGene_gn=NFASC;refGene_tx=XM_023247788.1;refGene_gn=NFASC;refGene_tx=XM_023247789.1
F1	45322750	45322934
refGene_gn=NFASC;refGene_tx=XM_011291070.3;refGene_gn=NFASC;refGene_tx=XM_023247774.1;refGene_gn=NFASC;refGene_tx=XM_023247775.1;refGene_gn=

NFASC;refGene_tx=XM_023247776.1;refGene_gn=NFASC;refGene_tx=XM_0232477
77.1;refGene_gn=NFASC;refGene_tx=XM_023247778.1;refGene_gn=NFASC;refGe
ne_tx=XM_023247779.1;refGene_gn=NFASC;refGene_tx=XM_023247780.1;refGen
e_gn=NFASC;refGene_tx=XM_023247781.1;refGene_gn=NFASC;refGene_tx=XM_02
3247782.1;refGene_gn=NFASC;refGene_tx=XM_023247783.1;refGene_gn=NFASC;
refGene_tx=XM_023247784.1;refGene_gn=NFASC;refGene_tx=XM_023247785.1;r
efGene_gn=NFASC;refGene_tx=XM_023247786.1;refGene_gn=NFASC;refGene_tx=
XM_023247787.1;refGene_gn=NFASC;refGene_tx=XM_023247788.1;refGene_gn=N
FASC;refGene_tx=XM_023247789.1
F1 45328943 45328961
refGene_gn=NFASC;refGene_tx=XM_011291070.3;refGene_gn=NFASC;refGene_tx
=XM_023247774.1;refGene_gn=NFASC;refGene_tx=XM_023247776.1;refGene_gn=
NFASC;refGene_tx=XM_023247777.1;refGene_gn=NFASC;refGene_tx=XM_0232477
79.1;refGene_gn=NFASC;refGene_tx=XM_023247780.1;refGene_gn=NFASC;refGe
ne_tx=XM_023247781.1;refGene_gn=NFASC;refGene_tx=XM_023247782.1;refGen
e_gn=NFASC;refGene_tx=XM_023247783.1;refGene_gn=NFASC;refGene_tx=XM_02
3247784.1;refGene_gn=NFASC;refGene_tx=XM_023247785.1
F1 45329740 45329846
refGene_gn=NFASC;refGene_tx=XM_011291070.3;refGene_gn=NFASC;refGene_tx
=XM_023247774.1;refGene_gn=NFASC;refGene_tx=XM_023247775.1;refGene_gn=
NFASC;refGene_tx=XM_023247776.1;refGene_gn=NFASC;refGene_tx=XM_0232477
77.1;refGene_gn=NFASC;refGene_tx=XM_023247778.1;refGene_gn=NFASC;refGe
ne_tx=XM_023247779.1;refGene_gn=NFASC;refGene_tx=XM_023247780.1;refGen
e_gn=NFASC;refGene_tx=XM_023247781.1;refGene_gn=NFASC;refGene_tx=XM_02
3247782.1;refGene_gn=NFASC;refGene_tx=XM_023247783.1;refGene_gn=NFASC;
refGene_tx=XM_023247784.1;refGene_gn=NFASC;refGene_tx=XM_023247785.1;r
efGene_gn=NFASC;refGene_tx=XM_023247786.1;refGene_gn=NFASC;refGene_tx=
XM_023247787.1;refGene_gn=NFASC;refGene_tx=XM_023247788.1;refGene_gn=N
FASC;refGene_tx=XM_023247789.1
F1 45331362 45331559
refGene_gn=NFASC;refGene_tx=XM_011291070.3;refGene_gn=NFASC;refGene_tx
=XM_023247774.1;refGene_gn=NFASC;refGene_tx=XM_023247775.1;refGene_gn=
NFASC;refGene_tx=XM_023247776.1;refGene_gn=NFASC;refGene_tx=XM_0232477
77.1;refGene_gn=NFASC;refGene_tx=XM_023247778.1;refGene_gn=NFASC;refGe
ne_tx=XM_023247779.1;refGene_gn=NFASC;refGene_tx=XM_023247780.1;refGen
e_gn=NFASC;refGene_tx=XM_023247781.1;refGene_gn=NFASC;refGene_tx=XM_02
3247782.1;refGene_gn=NFASC;refGene_tx=XM_023247783.1;refGene_gn=NFASC;
refGene_tx=XM_023247784.1;refGene_gn=NFASC;refGene_tx=XM_023247785.1;r
efGene_gn=NFASC;refGene_tx=XM_023247786.1;refGene_gn=NFASC;refGene_tx=
XM_023247787.1;refGene_gn=NFASC;refGene_tx=XM_023247788.1;refGene_gn=N
FASC;refGene_tx=XM_023247789.1
F1 45332018 45332141
refGene_gn=NFASC;refGene_tx=XM_011291070.3;refGene_gn=NFASC;refGene_tx
=XM_023247774.1;refGene_gn=NFASC;refGene_tx=XM_023247775.1;refGene_gn=
NFASC;refGene_tx=XM_023247776.1;refGene_gn=NFASC;refGene_tx=XM_0232477
77.1;refGene_gn=NFASC;refGene_tx=XM_023247778.1;refGene_gn=NFASC;refGe
ne_tx=XM_023247779.1;refGene_gn=NFASC;refGene_tx=XM_023247780.1;refGen
e_gn=NFASC;refGene_tx=XM_023247781.1;refGene_gn=NFASC;refGene_tx=XM_02
3247782.1;refGene_gn=NFASC;refGene_tx=XM_023247783.1;refGene_gn=NFASC;
refGene_tx=XM_023247784.1;refGene_gn=NFASC;refGene_tx=XM_023247785.1;r

efGene_gn=NFASC;refGene_tx=XM_023247786.1;refGene_gn=NFASC;refGene_tx=XM_023247787.1;refGene_gn=NFASC;refGene_tx=XM_023247788.1;refGene_gn=NFASC;refGene_tx=XM_023247789.1
F1 45334645 45334816
refGene_gn=NFASC;refGene_tx=XM_011291070.3;refGene_gn=NFASC;refGene_tx=XM_023247774.1;refGene_gn=NFASC;refGene_tx=XM_023247775.1;refGene_gn=NFASC;refGene_tx=XM_023247776.1;refGene_gn=NFASC;refGene_tx=XM_023247777.1;refGene_gn=NFASC;refGene_tx=XM_023247778.1;refGene_gn=NFASC;refGene_tx=XM_023247779.1;refGene_gn=NFASC;refGene_tx=XM_023247780.1;refGene_gn=NFASC;refGene_tx=XM_023247781.1;refGene_gn=NFASC;refGene_tx=XM_023247782.1;refGene_gn=NFASC;refGene_tx=XM_023247783.1;refGene_gn=NFASC;refGene_tx=XM_023247784.1;refGene_gn=NFASC;refGene_tx=XM_023247785.1;refGene_gn=NFASC;refGene_tx=XM_023247786.1;refGene_gn=NFASC;refGene_tx=XM_023247787.1;refGene_gn=NFASC;refGene_tx=XM_023247788.1;refGene_gn=NFASC;refGene_tx=XM_023247789.1
F1 45339043 45339094
refGene_gn=NFASC;refGene_tx=XM_011291070.3;refGene_gn=NFASC;refGene_tx=XM_023247774.1;refGene_gn=NFASC;refGene_tx=XM_023247775.1;refGene_gn=NFASC;refGene_tx=XM_023247776.1;refGene_gn=NFASC;refGene_tx=XM_023247779.1;refGene_gn=NFASC;refGene_tx=XM_023247780.1;refGene_gn=NFASC;refGene_tx=XM_023247781.1;refGene_gn=NFASC;refGene_tx=XM_023247782.1;refGene_gn=NFASC;refGene_tx=XM_023247784.1;refGene_gn=NFASC;refGene_tx=XM_023247785.1;refGene_gn=NFASC;refGene_tx=XM_023247786.1;refGene_gn=NFASC;refGene_tx=XM_023247787.1;refGene_gn=NFASC;refGene_tx=XM_023247788.1
F1 45345030 45345142
refGene_gn=NFASC;refGene_tx=XM_011291070.3;refGene_gn=NFASC;refGene_tx=XM_023247774.1;refGene_gn=NFASC;refGene_tx=XM_023247775.1;refGene_gn=NFASC;refGene_tx=XM_023247776.1;refGene_gn=NFASC;refGene_tx=XM_023247777.1;refGene_gn=NFASC;refGene_tx=XM_023247778.1;refGene_gn=NFASC;refGene_tx=XM_023247779.1;refGene_gn=NFASC;refGene_tx=XM_023247780.1;refGene_gn=NFASC;refGene_tx=XM_023247781.1;refGene_gn=NFASC;refGene_tx=XM_023247782.1;refGene_gn=NFASC;refGene_tx=XM_023247783.1;refGene_gn=NFASC;refGene_tx=XM_023247784.1;refGene_gn=NFASC;refGene_tx=XM_023247785.1;refGene_gn=NFASC;refGene_tx=XM_023247786.1;refGene_gn=NFASC;refGene_tx=XM_023247787.1;refGene_gn=NFASC;refGene_tx=XM_023247788.1;refGene_gn=NFASC;refGene_tx=XM_023247789.1
F1 45345527 45345712
refGene_gn=NFASC;refGene_tx=XM_011291070.3;refGene_gn=NFASC;refGene_tx=XM_023247774.1;refGene_gn=NFASC;refGene_tx=XM_023247775.1;refGene_gn=NFASC;refGene_tx=XM_023247776.1;refGene_gn=NFASC;refGene_tx=XM_023247777.1;refGene_gn=NFASC;refGene_tx=XM_023247778.1;refGene_gn=NFASC;refGene_tx=XM_023247779.1;refGene_gn=NFASC;refGene_tx=XM_023247780.1;refGene_gn=NFASC;refGene_tx=XM_023247781.1;refGene_gn=NFASC;refGene_tx=XM_023247782.1;refGene_gn=NFASC;refGene_tx=XM_023247783.1;refGene_gn=NFASC;refGene_tx=XM_023247784.1;refGene_gn=NFASC;refGene_tx=XM_023247785.1;refGene_gn=NFASC;refGene_tx=XM_023247786.1;refGene_gn=NFASC;refGene_tx=XM_023247787.1;refGene_gn=NFASC;refGene_tx=XM_023247788.1;refGene_gn=NFASC;refGene_tx=XM_023247789.1
F1 45347026 45347158
refGene_gn=NFASC;refGene_tx=XM_011291070.3;refGene_gn=NFASC;refGene_tx

=XM_023247774.1;refGene_gn=NFASC;refGene_tx=XM_023247775.1;refGene_gn=NFASC;refGene_tx=XM_023247776.1;refGene_gn=NFASC;refGene_tx=XM_023247777.1;refGene_gn=NFASC;refGene_tx=XM_023247778.1;refGene_gn=NFASC;refGene_tx=XM_023247779.1;refGene_gn=NFASC;refGene_tx=XM_023247780.1;refGene_gn=NFASC;refGene_tx=XM_023247781.1;refGene_gn=NFASC;refGene_tx=XM_023247782.1;refGene_gn=NFASC;refGene_tx=XM_023247783.1;refGene_gn=NFASC;refGene_tx=XM_023247784.1;refGene_gn=NFASC;refGene_tx=XM_023247785.1;refGene_gn=NFASC;refGene_tx=XM_023247786.1;refGene_gn=NFASC;refGene_tx=XM_023247787.1;refGene_gn=NFASC;refGene_tx=XM_023247788.1;refGene_gn=NFASC;refGene_tx=XM_023247789.1
F1 45349641 45349785
refGene_gn=NFASC;refGene_tx=XM_011291070.3;refGene_gn=NFASC;refGene_tx=XM_023247774.1;refGene_gn=NFASC;refGene_tx=XM_023247775.1;refGene_gn=NFASC;refGene_tx=XM_023247776.1;refGene_gn=NFASC;refGene_tx=XM_023247777.1;refGene_gn=NFASC;refGene_tx=XM_023247778.1;refGene_gn=NFASC;refGene_tx=XM_023247779.1;refGene_gn=NFASC;refGene_tx=XM_023247780.1;refGene_gn=NFASC;refGene_tx=XM_023247781.1;refGene_gn=NFASC;refGene_tx=XM_023247782.1;refGene_gn=NFASC;refGene_tx=XM_023247783.1;refGene_gn=NFASC;refGene_tx=XM_023247784.1;refGene_gn=NFASC;refGene_tx=XM_023247785.1;refGene_gn=NFASC;refGene_tx=XM_023247786.1;refGene_gn=NFASC;refGene_tx=XM_023247787.1;refGene_gn=NFASC;refGene_tx=XM_023247788.1;refGene_gn=NFASC;refGene_tx=XM_023247789.1
F1 45350427 45350539
refGene_gn=NFASC;refGene_tx=XM_011291070.3;refGene_gn=NFASC;refGene_tx=XM_023247774.1;refGene_gn=NFASC;refGene_tx=XM_023247775.1;refGene_gn=NFASC;refGene_tx=XM_023247776.1;refGene_gn=NFASC;refGene_tx=XM_023247777.1;refGene_gn=NFASC;refGene_tx=XM_023247778.1;refGene_gn=NFASC;refGene_tx=XM_023247779.1;refGene_gn=NFASC;refGene_tx=XM_023247780.1;refGene_gn=NFASC;refGene_tx=XM_023247781.1;refGene_gn=NFASC;refGene_tx=XM_023247782.1;refGene_gn=NFASC;refGene_tx=XM_023247783.1;refGene_gn=NFASC;refGene_tx=XM_023247784.1;refGene_gn=NFASC;refGene_tx=XM_023247785.1;refGene_gn=NFASC;refGene_tx=XM_023247786.1;refGene_gn=NFASC;refGene_tx=XM_023247787.1;refGene_gn=NFASC;refGene_tx=XM_023247788.1;refGene_gn=NFASC;refGene_tx=XM_023247789.1
F1 45351298 45351465
refGene_gn=NFASC;refGene_tx=XM_011291070.3;refGene_gn=NFASC;refGene_tx=XM_023247774.1;refGene_gn=NFASC;refGene_tx=XM_023247775.1;refGene_gn=NFASC;refGene_tx=XM_023247776.1;refGene_gn=NFASC;refGene_tx=XM_023247777.1;refGene_gn=NFASC;refGene_tx=XM_023247778.1;refGene_gn=NFASC;refGene_tx=XM_023247779.1;refGene_gn=NFASC;refGene_tx=XM_023247780.1;refGene_gn=NFASC;refGene_tx=XM_023247781.1;refGene_gn=NFASC;refGene_tx=XM_023247782.1;refGene_gn=NFASC;refGene_tx=XM_023247783.1;refGene_gn=NFASC;refGene_tx=XM_023247784.1;refGene_gn=NFASC;refGene_tx=XM_023247785.1;refGene_gn=NFASC;refGene_tx=XM_023247786.1;refGene_gn=NFASC;refGene_tx=XM_023247787.1;refGene_gn=NFASC;refGene_tx=XM_023247788.1;refGene_gn=NFASC;refGene_tx=XM_023247789.1
F1 45351935 45352083
refGene_gn=NFASC;refGene_tx=XM_011291070.3;refGene_gn=NFASC;refGene_tx=XM_023247774.1;refGene_gn=NFASC;refGene_tx=XM_023247775.1;refGene_gn=NFASC;refGene_tx=XM_023247776.1;refGene_gn=NFASC;refGene_tx=XM_0232477

77.1;refGene_gn=NFASC;refGene_tx=XM_023247778.1;refGene_gn=NFASC;refGene_tx=XM_023247779.1;refGene_gn=NFASC;refGene_tx=XM_023247780.1;refGene_gn=NFASC;refGene_tx=XM_023247781.1;refGene_gn=NFASC;refGene_tx=XM_023247782.1;refGene_gn=NFASC;refGene_tx=XM_023247783.1;refGene_gn=NFASC;refGene_tx=XM_023247784.1;refGene_gn=NFASC;refGene_tx=XM_023247785.1;refGene_gn=NFASC;refGene_tx=XM_023247786.1;refGene_gn=NFASC;refGene_tx=XM_023247787.1;refGene_gn=NFASC;refGene_tx=XM_023247788.1;refGene_gn=NFASC;refGene_tx=XM_023247789.1
F1 45353255 45353380
refGene_gn=NFASC;refGene_tx=XM_011291070.3;refGene_gn=NFASC;refGene_tx=XM_023247774.1;refGene_gn=NFASC;refGene_tx=XM_023247775.1;refGene_gn=NFASC;refGene_tx=XM_023247776.1;refGene_gn=NFASC;refGene_tx=XM_023247777.1;refGene_gn=NFASC;refGene_tx=XM_023247778.1;refGene_gn=NFASC;refGene_tx=XM_023247779.1;refGene_gn=NFASC;refGene_tx=XM_023247780.1;refGene_gn=NFASC;refGene_tx=XM_023247781.1;refGene_gn=NFASC;refGene_tx=XM_023247782.1;refGene_gn=NFASC;refGene_tx=XM_023247783.1;refGene_gn=NFASC;refGene_tx=XM_023247784.1;refGene_gn=NFASC;refGene_tx=XM_023247785.1;refGene_gn=NFASC;refGene_tx=XM_023247786.1;refGene_gn=NFASC;refGene_tx=XM_023247787.1;refGene_gn=NFASC;refGene_tx=XM_023247788.1;refGene_gn=NFASC;refGene_tx=XM_023247789.1
F1 45354060 45354105
refGene_gn=NFASC;refGene_tx=XM_011291070.3;refGene_gn=NFASC;refGene_tx=XM_023247774.1;refGene_gn=NFASC;refGene_tx=XM_023247775.1;refGene_gn=NFASC;refGene_tx=XM_023247777.1;refGene_gn=NFASC;refGene_tx=XM_023247778.1;refGene_gn=NFASC;refGene_tx=XM_023247779.1;refGene_gn=NFASC;refGene_tx=XM_023247780.1;refGene_gn=NFASC;refGene_tx=XM_023247781.1;refGene_gn=NFASC;refGene_tx=XM_023247782.1;refGene_gn=NFASC;refGene_tx=XM_023247783.1;refGene_gn=NFASC;refGene_tx=XM_023247784.1;refGene_gn=NFASC;refGene_tx=XM_023247785.1;refGene_gn=NFASC;refGene_tx=XM_023247786.1
F1 45355309 45355411
refGene_gn=NFASC;refGene_tx=XM_011291070.3;refGene_gn=NFASC;refGene_tx=XM_023247774.1;refGene_gn=NFASC;refGene_tx=XM_023247775.1;refGene_gn=NFASC;refGene_tx=XM_023247776.1;refGene_gn=NFASC;refGene_tx=XM_023247777.1;refGene_gn=NFASC;refGene_tx=XM_023247778.1;refGene_gn=NFASC;refGene_tx=XM_023247779.1;refGene_gn=NFASC;refGene_tx=XM_023247780.1;refGene_gn=NFASC;refGene_tx=XM_023247781.1;refGene_gn=NFASC;refGene_tx=XM_023247782.1;refGene_gn=NFASC;refGene_tx=XM_023247783.1;refGene_gn=NFASC;refGene_tx=XM_023247784.1;refGene_gn=NFASC;refGene_tx=XM_023247785.1;refGene_gn=NFASC;refGene_tx=XM_023247786.1;refGene_gn=NFASC;refGene_tx=XM_023247787.1;refGene_gn=NFASC;refGene_tx=XM_023247788.1;refGene_gn=NFASC;refGene_tx=XM_023247789.1
F1 45355716 45355914
refGene_gn=NFASC;refGene_tx=XM_011291070.3;refGene_gn=NFASC;refGene_tx=XM_023247774.1;refGene_gn=NFASC;refGene_tx=XM_023247775.1;refGene_gn=NFASC;refGene_tx=XM_023247776.1;refGene_gn=NFASC;refGene_tx=XM_023247777.1;refGene_gn=NFASC;refGene_tx=XM_023247778.1;refGene_gn=NFASC;refGene_tx=XM_023247779.1;refGene_gn=NFASC;refGene_tx=XM_023247780.1;refGene_gn=NFASC;refGene_tx=XM_023247781.1;refGene_gn=NFASC;refGene_tx=XM_023247782.1;refGene_gn=NFASC;refGene_tx=XM_023247783.1;refGene_gn=NFASC;refGene_tx=XM_023247784.1;refGene_gn=NFASC;refGene_tx=XM_023247785.1;r

efGene_gn=NFASC;refGene_tx=XM_023247786.1;refGene_gn=NFASC;refGene_tx=
XM_023247787.1;refGene_gn=NFASC;refGene_tx=XM_023247788.1;refGene_gn=N
FASC;refGene_tx=XM_023247789.1
F1 45356730 45356801
refGene_gn=NFASC;refGene_tx=XM_011291070.3;refGene_gn=NFASC;refGene_tx
=XM_023247774.1;refGene_gn=NFASC;refGene_tx=XM_023247775.1;refGene_gn=
NFASC;refGene_tx=XM_023247776.1;refGene_gn=NFASC;refGene_tx=XM_0232477
77.1;refGene_gn=NFASC;refGene_tx=XM_023247778.1;refGene_gn=NFASC;refGe
ne_tx=XM_023247779.1;refGene_gn=NFASC;refGene_tx=XM_023247780.1;refGen
e_gn=NFASC;refGene_tx=XM_023247781.1;refGene_gn=NFASC;refGene_tx=XM_02
3247782.1;refGene_gn=NFASC;refGene_tx=XM_023247783.1;refGene_gn=NFASC;
refGene_tx=XM_023247784.1;refGene_gn=NFASC;refGene_tx=XM_023247785.1;r
efGene_gn=NFASC;refGene_tx=XM_023247786.1;refGene_gn=NFASC;refGene_tx=
XM_023247787.1;refGene_gn=NFASC;refGene_tx=XM_023247788.1;refGene_gn=N
FASC;refGene_tx=XM_023247789.1
F1 45358341 45358564
refGene_gn=NFASC;refGene_tx=XM_011291070.3;refGene_gn=NFASC;refGene_tx
=XM_023247774.1;refGene_gn=NFASC;refGene_tx=XM_023247775.1;refGene_gn=
NFASC;refGene_tx=XM_023247776.1;refGene_gn=NFASC;refGene_tx=XM_0232477
77.1;refGene_gn=NFASC;refGene_tx=XM_023247778.1;refGene_gn=NFASC;refGe
ne_tx=XM_023247779.1;refGene_gn=NFASC;refGene_tx=XM_023247780.1;refGen
e_gn=NFASC;refGene_tx=XM_023247781.1;refGene_gn=NFASC;refGene_tx=XM_02
3247782.1;refGene_gn=NFASC;refGene_tx=XM_023247783.1;refGene_gn=NFASC;
refGene_tx=XM_023247784.1;refGene_gn=NFASC;refGene_tx=XM_023247785.1;r
efGene_gn=NFASC;refGene_tx=XM_023247786.1;refGene_gn=NFASC;refGene_tx=
XM_023247787.1;refGene_gn=NFASC;refGene_tx=XM_023247788.1;refGene_gn=N
FASC;refGene_tx=XM_023247789.1
F1 45360562 45360678
refGene_gn=NFASC;refGene_tx=XM_011291070.3;refGene_gn=NFASC;refGene_tx
=XM_023247774.1;refGene_gn=NFASC;refGene_tx=XM_023247775.1;refGene_gn=
NFASC;refGene_tx=XM_023247776.1;refGene_gn=NFASC;refGene_tx=XM_0232477
77.1;refGene_gn=NFASC;refGene_tx=XM_023247778.1;refGene_gn=NFASC;refGe
ne_tx=XM_023247779.1;refGene_gn=NFASC;refGene_tx=XM_023247780.1;refGen
e_gn=NFASC;refGene_tx=XM_023247781.1;refGene_gn=NFASC;refGene_tx=XM_02
3247784.1;refGene_gn=NFASC;refGene_tx=XM_023247785.1;refGene_gn=NFASC;
refGene_tx=XM_023247786.1;refGene_gn=NFASC;refGene_tx=XM_023247787.1;r
efGene_gn=NFASC;refGene_tx=XM_023247788.1;refGene_gn=NFASC;refGene_tx=
XM_023247789.1
F1 45362322 45362527
refGene_gn=NFASC;refGene_tx=XM_011291070.3;refGene_gn=NFASC;refGene_tx
=XM_023247774.1;refGene_gn=NFASC;refGene_tx=XM_023247775.1;refGene_gn=
NFASC;refGene_tx=XM_023247776.1;refGene_gn=NFASC;refGene_tx=XM_0232477
77.1;refGene_gn=NFASC;refGene_tx=XM_023247778.1;refGene_gn=NFASC;refGe
ne_tx=XM_023247779.1;refGene_gn=NFASC;refGene_tx=XM_023247780.1;refGen
e_gn=NFASC;refGene_tx=XM_023247781.1;refGene_gn=NFASC;refGene_tx=XM_02
3247784.1;refGene_gn=NFASC;refGene_tx=XM_023247785.1;refGene_gn=NFASC;
refGene_tx=XM_023247786.1;refGene_gn=NFASC;refGene_tx=XM_023247787.1;r
efGene_gn=NFASC;refGene_tx=XM_023247788.1;refGene_gn=NFASC;refGene_tx=
XM_023247789.1
F1 45363684 45363807

refGene_gn=NFASC;refGene_tx=XM_011291070.3;refGene_gn=NFASC;refGene_tx=XM_023247774.1;refGene_gn=NFASC;refGene_tx=XM_023247775.1;refGene_gn=NFASC;refGene_tx=XM_023247776.1;refGene_gn=NFASC;refGene_tx=XM_023247777.1;refGene_gn=NFASC;refGene_tx=XM_023247778.1;refGene_gn=NFASC;refGene_tx=XM_023247779.1;refGene_gn=NFASC;refGene_tx=XM_023247780.1;refGene_gn=NFASC;refGene_tx=XM_023247781.1;refGene_gn=NFASC;refGene_tx=XM_023247782.1;refGene_gn=NFASC;refGene_tx=XM_023247783.1;refGene_gn=NFASC;refGene_tx=XM_023247784.1;refGene_gn=NFASC;refGene_tx=XM_023247785.1;refGene_gn=NFASC;refGene_tx=XM_023247786.1;refGene_gn=NFASC;refGene_tx=XM_023247787.1;refGene_gn=NFASC;refGene_tx=XM_023247788.1;refGene_gn=NFASC;refGene_tx=XM_023247789.1

F1 45364822 45364996

refGene_gn=NFASC;refGene_tx=XM_011291070.3;refGene_gn=NFASC;refGene_tx=XM_023247774.1;refGene_gn=NFASC;refGene_tx=XM_023247775.1;refGene_gn=NFASC;refGene_tx=XM_023247776.1;refGene_gn=NFASC;refGene_tx=XM_023247777.1;refGene_gn=NFASC;refGene_tx=XM_023247778.1;refGene_gn=NFASC;refGene_tx=XM_023247779.1;refGene_gn=NFASC;refGene_tx=XM_023247780.1;refGene_gn=NFASC;refGene_tx=XM_023247781.1;refGene_gn=NFASC;refGene_tx=XM_023247782.1;refGene_gn=NFASC;refGene_tx=XM_023247783.1;refGene_gn=NFASC;refGene_tx=XM_023247784.1;refGene_gn=NFASC;refGene_tx=XM_023247785.1;refGene_gn=NFASC;refGene_tx=XM_023247786.1;refGene_gn=NFASC;refGene_tx=XM_023247787.1;refGene_gn=NFASC;refGene_tx=XM_023247788.1;refGene_gn=NFASC;refGene_tx=XM_023247789.1

F1 45367157 45367172

refGene_gn=NFASC;refGene_tx=XM_011291070.3;refGene_gn=NFASC;refGene_tx=XM_023247774.1;refGene_gn=NFASC;refGene_tx=XM_023247775.1;refGene_gn=NFASC;refGene_tx=XM_023247776.1;refGene_gn=NFASC;refGene_tx=XM_023247777.1;refGene_gn=NFASC;refGene_tx=XM_023247778.1;refGene_gn=NFASC;refGene_tx=XM_023247779.1;refGene_gn=NFASC;refGene_tx=XM_023247780.1;refGene_gn=NFASC;refGene_tx=XM_023247781.1;refGene_gn=NFASC;refGene_tx=XM_023247782.1;refGene_gn=NFASC;refGene_tx=XM_023247783.1;refGene_gn=NFASC;refGene_tx=XM_023247784.1;refGene_gn=NFASC;refGene_tx=XM_023247786.1;refGene_gn=NFASC;refGene_tx=XM_023247787.1;refGene_gn=NFASC;refGene_tx=XM_023247789.1

F1 45372322 45372568

refGene_gn=NFASC;refGene_tx=XM_011291070.3;refGene_gn=NFASC;refGene_tx=XM_023247774.1;refGene_gn=NFASC;refGene_tx=XM_023247775.1;refGene_gn=NFASC;refGene_tx=XM_023247776.1;refGene_gn=NFASC;refGene_tx=XM_023247777.1;refGene_gn=NFASC;refGene_tx=XM_023247778.1;refGene_gn=NFASC;refGene_tx=XM_023247780.1;refGene_gn=NFASC;refGene_tx=XM_023247781.1;refGene_gn=NFASC;refGene_tx=XM_023247782.1;refGene_gn=NFASC;refGene_tx=XM_023247783.1

F1 45375997 45376114

refGene_gn=NFASC;refGene_tx=XM_011291070.3;refGene_gn=NFASC;refGene_tx=XM_023247774.1;refGene_gn=NFASC;refGene_tx=XM_023247775.1;refGene_gn=NFASC;refGene_tx=XM_023247776.1;refGene_gn=NFASC;refGene_tx=XM_023247777.1;refGene_gn=NFASC;refGene_tx=XM_023247778.1;refGene_gn=NFASC;refGene_tx=XM_023247779.1;refGene_gn=NFASC;refGene_tx=XM_023247781.1;refGene_gn=NFASC;refGene_tx=XM_023247782.1;refGene_gn=NFASC;refGene_tx=XM_023247783.1

F1 45377376 45377529
refGene_gn=NFASC;refGene_tx=XM_011291070.3;refGene_gn=NFASC;refGene_tx=XM_023247774.1;refGene_gn=NFASC;refGene_tx=XM_023247775.1;refGene_gn=NFASC;refGene_tx=XM_023247776.1;refGene_gn=NFASC;refGene_tx=XM_023247777.1;refGene_gn=NFASC;refGene_tx=XM_023247778.1;refGene_gn=NFASC;refGene_tx=XM_023247779.1;refGene_gn=NFASC;refGene_tx=XM_023247781.1;refGene_gn=NFASC;refGene_tx=XM_023247782.1;refGene_gn=NFASC;refGene_tx=XM_023247783.1
F1 45384165 45384297
refGene_gn=NFASC;refGene_tx=XM_011291070.3;refGene_gn=NFASC;refGene_tx=XM_023247774.1;refGene_gn=NFASC;refGene_tx=XM_023247775.1;refGene_gn=NFASC;refGene_tx=XM_023247776.1;refGene_gn=NFASC;refGene_tx=XM_023247777.1;refGene_gn=NFASC;refGene_tx=XM_023247778.1;refGene_gn=NFASC;refGene_tx=XM_023247779.1;refGene_gn=NFASC;refGene_tx=XM_023247780.1;refGene_gn=NFASC;refGene_tx=XM_023247781.1;refGene_gn=NFASC;refGene_tx=XM_023247782.1;refGene_gn=NFASC;refGene_tx=XM_023247783.1;refGene_gn=NFASC;refGene_tx=XM_023247784.1;refGene_gn=NFASC;refGene_tx=XM_023247785.1;refGene_gn=NFASC;refGene_tx=XM_023247786.1;refGene_gn=NFASC;refGene_tx=XM_023247787.1;refGene_gn=NFASC;refGene_tx=XM_023247788.1;refGene_gn=NFASC;refGene_tx=XM_023247789.1
F1 45386349 45386419
refGene_gn=NFASC;refGene_tx=XM_011291070.3;refGene_gn=NFASC;refGene_tx=XM_023247774.1;refGene_gn=NFASC;refGene_tx=XM_023247775.1;refGene_gn=NFASC;refGene_tx=XM_023247776.1;refGene_gn=NFASC;refGene_tx=XM_023247777.1;refGene_gn=NFASC;refGene_tx=XM_023247778.1;refGene_gn=NFASC;refGene_tx=XM_023247779.1;refGene_gn=NFASC;refGene_tx=XM_023247780.1;refGene_gn=NFASC;refGene_tx=XM_023247781.1;refGene_gn=NFASC;refGene_tx=XM_023247782.1;refGene_gn=NFASC;refGene_tx=XM_023247783.1;refGene_gn=NFASC;refGene_tx=XM_023247784.1;refGene_gn=NFASC;refGene_tx=XM_023247785.1;refGene_gn=NFASC;refGene_tx=XM_023247786.1;refGene_gn=NFASC;refGene_tx=XM_023247787.1;refGene_gn=NFASC;refGene_tx=XM_023247788.1;refGene_gn=NFASC;refGene_tx=XM_023247789.1
F1 45389942 45390174
refGene_gn=NFASC;refGene_tx=XM_011291070.3;refGene_gn=NFASC;refGene_tx=XM_023247774.1;refGene_gn=NFASC;refGene_tx=XM_023247775.1;refGene_gn=NFASC;refGene_tx=XM_023247776.1;refGene_gn=NFASC;refGene_tx=XM_023247777.1;refGene_gn=NFASC;refGene_tx=XM_023247778.1;refGene_gn=NFASC;refGene_tx=XM_023247779.1;refGene_gn=NFASC;refGene_tx=XM_023247780.1;refGene_gn=NFASC;refGene_tx=XM_023247781.1;refGene_gn=NFASC;refGene_tx=XM_023247782.1;refGene_gn=NFASC;refGene_tx=XM_023247783.1;refGene_gn=NFASC;refGene_tx=XM_023247784.1;refGene_gn=NFASC;refGene_tx=XM_023247785.1;refGene_gn=NFASC;refGene_tx=XM_023247786.1;refGene_gn=NFASC;refGene_tx=XM_023247787.1;refGene_gn=NFASC;refGene_tx=XM_023247788.1;refGene_gn=NFASC;refGene_tx=XM_023247789.1
F1 45425011 45425020 refGene_gn=CNTN2;refGene_tx=XM_019822106.1
F1 45425351 45425439 refGene_gn=CNTN2;refGene_tx=XM_019822106.1
F1 45428478 45428640
refGene_gn=CNTN2;refGene_tx=XM_011291071.3;refGene_gn=CNTN2;refGene_tx=XM_019822105.2;refGene_gn=CNTN2;refGene_tx=XM_019822106.1;refGene_gn=CNTN2;refGene_tx=XM_023247440.1

F1 45432801 45432946
refGene_gn=CNTN2;refGene_tx=XM_011291071.3;refGene_gn=CNTN2;refGene_tx=XM_019822105.2;refGene_gn=CNTN2;refGene_tx=XM_019822106.1;refGene_gn=CNTN2;refGene_tx=XM_023247440.1
F1 45433076 45433252
refGene_gn=CNTN2;refGene_tx=XM_011291071.3;refGene_gn=CNTN2;refGene_tx=XM_019822105.2;refGene_gn=CNTN2;refGene_tx=XM_019822106.1;refGene_gn=CNTN2;refGene_tx=XM_023247440.1
F1 45433411 45433507
refGene_gn=CNTN2;refGene_tx=XM_011291071.3;refGene_gn=CNTN2;refGene_tx=XM_019822105.2;refGene_gn=CNTN2;refGene_tx=XM_019822106.1;refGene_gn=CNTN2;refGene_tx=XM_023247440.1
F1 45433897 45434107
refGene_gn=CNTN2;refGene_tx=XM_011291071.3;refGene_gn=CNTN2;refGene_tx=XM_019822105.2;refGene_gn=CNTN2;refGene_tx=XM_019822106.1;refGene_gn=CNTN2;refGene_tx=XM_023247440.1
F1 45434390 45434490
refGene_gn=CNTN2;refGene_tx=XM_011291071.3;refGene_gn=CNTN2;refGene_tx=XM_019822105.2;refGene_gn=CNTN2;refGene_tx=XM_019822106.1;refGene_gn=CNTN2;refGene_tx=XM_023247440.1
F1 45435941 45436117
refGene_gn=CNTN2;refGene_tx=XM_011291071.3;refGene_gn=CNTN2;refGene_tx=XM_019822105.2;refGene_gn=CNTN2;refGene_tx=XM_019822106.1;refGene_gn=CNTN2;refGene_tx=XM_023247440.1
F1 45436524 45436661
refGene_gn=CNTN2;refGene_tx=XM_011291071.3;refGene_gn=CNTN2;refGene_tx=XM_019822105.2;refGene_gn=CNTN2;refGene_tx=XM_019822106.1;refGene_gn=CNTN2;refGene_tx=XM_023247440.1
F1 45437142 45437272
refGene_gn=CNTN2;refGene_tx=XM_011291071.3;refGene_gn=CNTN2;refGene_tx=XM_019822105.2;refGene_gn=CNTN2;refGene_tx=XM_019822106.1;refGene_gn=CNTN2;refGene_tx=XM_023247440.1
F1 45439077 45439228
refGene_gn=CNTN2;refGene_tx=XM_011291071.3;refGene_gn=CNTN2;refGene_tx=XM_019822105.2;refGene_gn=CNTN2;refGene_tx=XM_019822106.1;refGene_gn=CNTN2;refGene_tx=XM_023247440.1
F1 45439353 45439481
refGene_gn=CNTN2;refGene_tx=XM_011291071.3;refGene_gn=CNTN2;refGene_tx=XM_019822105.2;refGene_gn=CNTN2;refGene_tx=XM_019822106.1;refGene_gn=CNTN2;refGene_tx=XM_023247440.1
F1 45439827 45440003
refGene_gn=CNTN2;refGene_tx=XM_011291071.3;refGene_gn=CNTN2;refGene_tx=XM_019822105.2;refGene_gn=CNTN2;refGene_tx=XM_019822106.1;refGene_gn=CNTN2;refGene_tx=XM_023247440.1
F1 45440535 45440656
refGene_gn=CNTN2;refGene_tx=XM_011291071.3;refGene_gn=CNTN2;refGene_tx=XM_019822105.2;refGene_gn=CNTN2;refGene_tx=XM_019822106.1;refGene_gn=CNTN2;refGene_tx=XM_023247440.1
F1 45441147 45441306
refGene_gn=CNTN2;refGene_tx=XM_011291071.3;refGene_gn=CNTN2;refGene_tx

=XM_019822105.2;refGene_gn=CNTN2;refGene_tx=XM_019822106.1;refGene_gn=CNTN2;refGene_tx=XM_023247440.1
F1 45441713 45441863 refGene_gn=CNTN2;refGene_tx=XM_011291071.3;refGene_gn=CNTN2;refGene_tx=XM_019822105.2;refGene_gn=CNTN2;refGene_tx=XM_019822106.1;refGene_gn=CNTN2;refGene_tx=XM_023247440.1
F1 45444190 45444261 refGene_gn=CNTN2;refGene_tx=XM_011291071.3;refGene_gn=CNTN2;refGene_tx=XM_019822105.2;refGene_gn=CNTN2;refGene_tx=XM_019822106.1;refGene_gn=CNTN2;refGene_tx=XM_023247440.1
F1 45444550 45444785 refGene_gn=CNTN2;refGene_tx=XM_011291071.3;refGene_gn=CNTN2;refGene_tx=XM_019822105.2;refGene_gn=CNTN2;refGene_tx=XM_019822106.1;refGene_gn=CNTN2;refGene_tx=XM_023247440.1
F1 45445112 45445225 refGene_gn=CNTN2;refGene_tx=XM_011291071.3;refGene_gn=CNTN2;refGene_tx=XM_019822105.2;refGene_gn=CNTN2;refGene_tx=XM_019822106.1;refGene_gn=CNTN2;refGene_tx=XM_023247440.1
F1 45446254 45446441 refGene_gn=CNTN2;refGene_tx=XM_011291071.3;refGene_gn=CNTN2;refGene_tx=XM_019822105.2;refGene_gn=CNTN2;refGene_tx=XM_019822106.1;refGene_gn=CNTN2;refGene_tx=XM_023247440.1
F1 45446772 45446885 refGene_gn=CNTN2;refGene_tx=XM_011291071.3;refGene_gn=CNTN2;refGene_tx=XM_019822105.2;refGene_gn=CNTN2;refGene_tx=XM_019822106.1;refGene_gn=CNTN2;refGene_tx=XM_023247440.1
F1 45447328 45447497 refGene_gn=CNTN2;refGene_tx=XM_011291071.3;refGene_gn=CNTN2;refGene_tx=XM_019822105.2;refGene_gn=CNTN2;refGene_tx=XM_019822106.1;refGene_gn=CNTN2;refGene_tx=XM_023247440.1
F1 45447918 45448028 refGene_gn=CNTN2;refGene_tx=XM_011291071.3;refGene_gn=CNTN2;refGene_tx=XM_019822105.2;refGene_gn=CNTN2;refGene_tx=XM_019822106.1;refGene_gn=CNTN2;refGene_tx=XM_023247440.1
F1 45456355 45457201 refGene_gn=TMEM81;refGene_tx=XM_003999480.3
F1 45461023 45461052 refGene_gn=RBBP5;refGene_tx=XM_003999407.5;refGene_gn=RBBP5;refGene_tx=XM_003999408.5
F1 45467389 45467581 refGene_gn=RBBP5;refGene_tx=XM_003999407.5;refGene_gn=RBBP5;refGene_tx=XM_003999408.5
F1 45468662 45468892 refGene_gn=RBBP5;refGene_tx=XM_003999407.5;refGene_gn=RBBP5;refGene_tx=XM_003999408.5
F1 45469264 45469334 refGene_gn=RBBP5;refGene_tx=XM_003999407.5;refGene_gn=RBBP5;refGene_tx=XM_003999408.5
F1 45469955 45470073 refGene_gn=RBBP5;refGene_tx=XM_003999407.5;refGene_gn=RBBP5;refGene_tx=XM_003999408.5

F1 45470684 45470756
refGene_gn=RBBP5;refGene_tx=XM_003999407.5;refGene_gn=RBBP5;refGene_tx=XM_003999408.5
F1 45470852 45471006
refGene_gn=RBBP5;refGene_tx=XM_003999407.5;refGene_gn=RBBP5;refGene_tx=XM_003999408.5
F1 45471094 45471214
refGene_gn=RBBP5;refGene_tx=XM_003999407.5;refGene_gn=RBBP5;refGene_tx=XM_003999408.5
F1 45472243 45472353
refGene_gn=RBBP5;refGene_tx=XM_003999407.5;refGene_gn=RBBP5;refGene_tx=XM_003999408.5
F1 45473894 45474057
refGene_gn=RBBP5;refGene_tx=XM_003999407.5;refGene_gn=RBBP5;refGene_tx=XM_003999408.5
F1 45474849 45474990
refGene_gn=RBBP5;refGene_tx=XM_003999407.5;refGene_gn=RBBP5;refGene_tx=XM_003999408.5
F1 45483791 45483964
refGene_gn=RBBP5;refGene_tx=XM_003999407.5;refGene_gn=RBBP5;refGene_tx=XM_003999408.5
F1 45484881 45484907
refGene_gn=RBBP5;refGene_tx=XM_003999407.5;refGene_gn=RBBP5;refGene_tx=XM_003999408.5
F1 45495161 45495180
refGene_gn=RBBP5;refGene_tx=XM_003999407.5;refGene_gn=RBBP5;refGene_tx=XM_003999408.5
F1 45514027 45514215 refGene_gn=DSTYK;refGene_tx=XM_003999411.5
F1 45514630 45514765 refGene_gn=DSTYK;refGene_tx=XM_003999411.5
F1 45517407 45517522 refGene_gn=DSTYK;refGene_tx=XM_003999411.5
F1 45518957 45519071 refGene_gn=DSTYK;refGene_tx=XM_003999411.5
F1 45521078 45521211 refGene_gn=DSTYK;refGene_tx=XM_003999411.5
F1 45521667 45521824 refGene_gn=DSTYK;refGene_tx=XM_003999411.5
F1 45523038 45523168 refGene_gn=DSTYK;refGene_tx=XM_003999411.5
F1 45523782 45523959 refGene_gn=DSTYK;refGene_tx=XM_003999411.5
F1 45524590 45524674 refGene_gn=DSTYK;refGene_tx=XM_003999411.5
F1 45525077 45525310 refGene_gn=DSTYK;refGene_tx=XM_003999411.5
F1 45529260 45529930 refGene_gn=DSTYK;refGene_tx=XM_003999411.5
F1 45540699 45541085 refGene_gn=DSTYK;refGene_tx=XM_003999411.5
F1 45559457 45559725 refGene_gn=DSTYK;refGene_tx=XM_003999411.5
F1 45577330 45577537 refGene_gn=TMCC2;refGene_tx=XM_011291073.3
F1 45589645 45590185
refGene_gn=TMCC2;refGene_tx=XM_011291073.3;refGene_gn=TMCC2;refGene_tx=XM_011291074.2;refGene_gn=TMCC2;refGene_tx=XM_019822108.1;refGene_gn=TMCC2;refGene_tx=XM_019822109.2
F1 45603050 45603077 refGene_gn=TMCC2;refGene_tx=XM_019822112.2
F1 45603842 45604025 refGene_gn=TMCC2;refGene_tx=XM_019822110.2
F1 45611897 45611939 refGene_gn=TMCC2;refGene_tx=XM_011291076.3
F1 45612913 45613854
refGene_gn=TMCC2;refGene_tx=XM_011291073.3;refGene_gn=TMCC2;refGene_tx

=XM_011291074.2;refGene_gn=TMCC2;refGene_tx=XM_011291076.3;refGene_gn=TMCC2;refGene_tx=XM_019822108.1;refGene_gn=TMCC2;refGene_tx=XM_019822109.2;refGene_gn=TMCC2;refGene_tx=XM_019822110.2;refGene_gn=TMCC2;refGene_tx=XM_019822112.2
F1 45615068 45615204
refGene_gn=TMCC2;refGene_tx=XM_011291073.3;refGene_gn=TMCC2;refGene_tx=XM_011291074.2;refGene_gn=TMCC2;refGene_tx=XM_011291076.3;refGene_gn=TMCC2;refGene_tx=XM_019822108.1;refGene_gn=TMCC2;refGene_tx=XM_019822109.2;refGene_gn=TMCC2;refGene_tx=XM_019822110.2;refGene_gn=TMCC2;refGene_tx=XM_019822112.2
F1 45615754 45616066
refGene_gn=TMCC2;refGene_tx=XM_011291073.3;refGene_gn=TMCC2;refGene_tx=XM_011291074.2;refGene_gn=TMCC2;refGene_tx=XM_011291076.3;refGene_gn=TMCC2;refGene_tx=XM_019822108.1;refGene_gn=TMCC2;refGene_tx=XM_019822109.2;refGene_gn=TMCC2;refGene_tx=XM_019822110.2;refGene_gn=TMCC2;refGene_tx=XM_019822112.2
F1 45639818 45639976 refGene_gn=NUAK2;refGene_tx=XM_019822114.1
F1 45640523 45641587
refGene_gn=NUAK2;refGene_tx=XM_011291077.3;refGene_gn=NUAK2;refGene_tx=XM_019822114.1;refGene_gn=NUAK2;refGene_tx=XM_019822115.2
F1 45642527 45642660
refGene_gn=NUAK2;refGene_tx=XM_011291077.3;refGene_gn=NUAK2;refGene_tx=XM_019822114.1;refGene_gn=NUAK2;refGene_tx=XM_019822115.2
F1 45643234 45643354
refGene_gn=NUAK2;refGene_tx=XM_011291077.3;refGene_gn=NUAK2;refGene_tx=XM_019822114.1;refGene_gn=NUAK2;refGene_tx=XM_019822115.2
F1 45645237 45645303
refGene_gn=NUAK2;refGene_tx=XM_011291077.3;refGene_gn=NUAK2;refGene_tx=XM_019822114.1;refGene_gn=NUAK2;refGene_tx=XM_019822115.2
F1 45645635 45645787
refGene_gn=NUAK2;refGene_tx=XM_011291077.3;refGene_gn=NUAK2;refGene_tx=XM_019822114.1;refGene_gn=NUAK2;refGene_tx=XM_019822115.2
F1 45648281 45648402
refGene_gn=NUAK2;refGene_tx=XM_011291077.3;refGene_gn=NUAK2;refGene_tx=XM_019822114.1;refGene_gn=NUAK2;refGene_tx=XM_019822115.2
F1 45657509 45657755
refGene_gn=NUAK2;refGene_tx=XM_011291077.3;refGene_gn=NUAK2;refGene_tx=XM_019822114.1
F1 45675215 45675409
refGene_gn=KLHDC8A;refGene_tx=XM_023247478.1;refGene_gn=KLHDC8A;refGene_tx=XM_023247479.1;refGene_gn=KLHDC8A;refGene_tx=XM_023247480.1
F1 45675840 45675942
refGene_gn=KLHDC8A;refGene_tx=XM_023247478.1;refGene_gn=KLHDC8A;refGene_tx=XM_023247479.1;refGene_gn=KLHDC8A;refGene_tx=XM_023247480.1
F1 45676544 45676760
refGene_gn=KLHDC8A;refGene_tx=XM_023247478.1;refGene_gn=KLHDC8A;refGene_tx=XM_023247479.1;refGene_gn=KLHDC8A;refGene_tx=XM_023247480.1
F1 45677011 45677176
refGene_gn=KLHDC8A;refGene_tx=XM_023247478.1;refGene_gn=KLHDC8A;refGene_tx=XM_023247479.1;refGene_gn=KLHDC8A;refGene_tx=XM_023247480.1

F1 45679927 45680303
refGene_gn=KLHDC8A;refGene_tx=XM_023247478.1;refGene_gn=KLHDC8A;refGen
e_tx=XM_023247479.1;refGene_gn=KLHDC8A;refGene_tx=XM_023247480.1
F1 45712597 45712796
refGene_gn=LEMD1;refGene_tx=XM_023247820.1;refGene_gn=LEMD1;refGene_tx
=XM_023247821.1;refGene_gn=LEMD1;refGene_tx=XM_023247822.1
F1 45713735 45713812
refGene_gn=LEMD1;refGene_tx=XM_023247820.1;refGene_gn=LEMD1;refGene_tx
=XM_023247821.1;refGene_gn=LEMD1;refGene_tx=XM_023247822.1
F1 45732428 45732482
refGene_gn=LEMD1;refGene_tx=XM_023247820.1;refGene_gn=LEMD1;refGene_tx
=XM_023247822.1
F1 45733322 45733393
refGene_gn=LEMD1;refGene_tx=XM_023247820.1;refGene_gn=LEMD1;refGene_tx
=XM_023247821.1;refGene_gn=LEMD1;refGene_tx=XM_023247822.1
F1 45736367 45736487
refGene_gn=LEMD1;refGene_tx=XM_023247820.1;refGene_gn=LEMD1;refGene_tx
=XM_023247821.1
F1 45737108 45737190
refGene_gn=LEMD1;refGene_tx=XM_023247820.1;refGene_gn=LEMD1;refGene_tx
=XM_023247821.1;refGene_gn=LEMD1;refGene_tx=XM_023247822.1
F1 45816289 45816320 refGene_gn=CDK18;refGene_tx=XM_019822119.1
F1 45833606 45833756
refGene_gn=CDK18;refGene_tx=XM_019822119.1;refGene_gn=CDK18;refGene_tx
=XM_023247364.1;refGene_gn=CDK18;refGene_tx=XM_023247365.1;refGene_gn=
CDK18;refGene_tx=XM_023247366.1;refGene_gn=CDK18;refGene_tx=XM_0232473
67.1;refGene_gn=CDK18;refGene_tx=XM_023247368.1
F1 45833941 45834081
refGene_gn=CDK18;refGene_tx=XM_019822119.1;refGene_gn=CDK18;refGene_tx
=XM_023247364.1;refGene_gn=CDK18;refGene_tx=XM_023247365.1;refGene_gn=
CDK18;refGene_tx=XM_023247366.1;refGene_gn=CDK18;refGene_tx=XM_0232473
67.1;refGene_gn=CDK18;refGene_tx=XM_023247368.1
F1 45834677 45834803
refGene_gn=CDK18;refGene_tx=XM_019822119.1;refGene_gn=CDK18;refGene_tx
=XM_023247364.1;refGene_gn=CDK18;refGene_tx=XM_023247365.1;refGene_gn=
CDK18;refGene_tx=XM_023247366.1;refGene_gn=CDK18;refGene_tx=XM_0232473
67.1;refGene_gn=CDK18;refGene_tx=XM_023247368.1
F1 45835626 45835683
refGene_gn=CDK18;refGene_tx=XM_019822119.1;refGene_gn=CDK18;refGene_tx
=XM_023247364.1;refGene_gn=CDK18;refGene_tx=XM_023247365.1;refGene_gn=
CDK18;refGene_tx=XM_023247366.1;refGene_gn=CDK18;refGene_tx=XM_0232473
67.1;refGene_gn=CDK18;refGene_tx=XM_023247368.1
F1 45836499 45836614
refGene_gn=CDK18;refGene_tx=XM_019822119.1;refGene_gn=CDK18;refGene_tx
=XM_023247364.1;refGene_gn=CDK18;refGene_tx=XM_023247365.1;refGene_gn=
CDK18;refGene_tx=XM_023247366.1;refGene_gn=CDK18;refGene_tx=XM_0232473
67.1;refGene_gn=CDK18;refGene_tx=XM_023247368.1
F1 45836806 45836901
refGene_gn=CDK18;refGene_tx=XM_019822119.1;refGene_gn=CDK18;refGene_tx
=XM_023247364.1;refGene_gn=CDK18;refGene_tx=XM_023247365.1;refGene_gn=

CDK18;refGene_tx=XM_023247366.1;refGene_gn=CDK18;refGene_tx=XM_0232473
67.1;refGene_gn=CDK18;refGene_tx=XM_023247368.1
F1 45837124 45837187
refGene_gn=CDK18;refGene_tx=XM_019822119.1;refGene_gn=CDK18;refGene_tx
=XM_023247364.1;refGene_gn=CDK18;refGene_tx=XM_023247365.1;refGene_gn=
CDK18;refGene_tx=XM_023247366.1;refGene_gn=CDK18;refGene_tx=XM_0232473
67.1;refGene_gn=CDK18;refGene_tx=XM_023247368.1
F1 45838084 45838208
refGene_gn=CDK18;refGene_tx=XM_019822119.1;refGene_gn=CDK18;refGene_tx
=XM_023247364.1;refGene_gn=CDK18;refGene_tx=XM_023247365.1;refGene_gn=
CDK18;refGene_tx=XM_023247366.1;refGene_gn=CDK18;refGene_tx=XM_0232473
67.1;refGene_gn=CDK18;refGene_tx=XM_023247368.1
F1 45838309 45838430
refGene_gn=CDK18;refGene_tx=XM_019822119.1;refGene_gn=CDK18;refGene_tx
=XM_023247364.1;refGene_gn=CDK18;refGene_tx=XM_023247365.1;refGene_gn=
CDK18;refGene_tx=XM_023247366.1;refGene_gn=CDK18;refGene_tx=XM_0232473
67.1;refGene_gn=CDK18;refGene_tx=XM_023247368.1
F1 45838867 45839084 refGene_gn=CDK18;refGene_tx=XM_023247368.1
F1 45839200 45839298
refGene_gn=CDK18;refGene_tx=XM_019822119.1;refGene_gn=CDK18;refGene_tx
=XM_023247364.1;refGene_gn=CDK18;refGene_tx=XM_023247365.1;refGene_gn=
CDK18;refGene_tx=XM_023247366.1;refGene_gn=CDK18;refGene_tx=XM_0232473
67.1
F1 45839526 45839632
refGene_gn=CDK18;refGene_tx=XM_019822119.1;refGene_gn=CDK18;refGene_tx
=XM_023247364.1;refGene_gn=CDK18;refGene_tx=XM_023247365.1;refGene_gn=
CDK18;refGene_tx=XM_023247366.1;refGene_gn=CDK18;refGene_tx=XM_0232473
67.1
F1 45839760 45839803
refGene_gn=CDK18;refGene_tx=XM_019822119.1;refGene_gn=CDK18;refGene_tx
=XM_023247364.1;refGene_gn=CDK18;refGene_tx=XM_023247365.1;refGene_gn=
CDK18;refGene_tx=XM_023247366.1;refGene_gn=CDK18;refGene_tx=XM_0232473
67.1
F1 45840465 45840556
refGene_gn=CDK18;refGene_tx=XM_019822119.1;refGene_gn=CDK18;refGene_tx
=XM_023247364.1;refGene_gn=CDK18;refGene_tx=XM_023247365.1;refGene_gn=
CDK18;refGene_tx=XM_023247366.1;refGene_gn=CDK18;refGene_tx=XM_0232473
67.1
F1 45840874 45840952
refGene_gn=CDK18;refGene_tx=XM_019822119.1;refGene_gn=CDK18;refGene_tx
=XM_023247364.1;refGene_gn=CDK18;refGene_tx=XM_023247365.1;refGene_gn=
CDK18;refGene_tx=XM_023247366.1;refGene_gn=CDK18;refGene_tx=XM_0232473
67.1
F1 45841704 45841846
refGene_gn=CDK18;refGene_tx=XM_019822119.1;refGene_gn=CDK18;refGene_tx
=XM_023247364.1;refGene_gn=CDK18;refGene_tx=XM_023247365.1;refGene_gn=
CDK18;refGene_tx=XM_023247366.1;refGene_gn=CDK18;refGene_tx=XM_0232473
67.1
F1 45870448 45870678
refGene_gn=MFSD4A;refGene_tx=XM_023247210.1;refGene_gn=MFSD4A;refGene_

tx=XM_023247211.1;refGene_gn=MFSD4A;refGene_tx=XM_023247212.1;refGene_gn=MFSD4A;refGene_tx=XM_023247213.1;refGene_gn=MFSD4A;refGene_tx=XM_023247214.1
F1 45881249 45881448
refGene_gn=MFSD4A;refGene_tx=XM_019822121.2;refGene_gn=MFSD4A;refGene_tx=XM_023247210.1;refGene_gn=MFSD4A;refGene_tx=XM_023247211.1;refGene_gn=MFSD4A;refGene_tx=XM_023247212.1;refGene_gn=MFSD4A;refGene_tx=XM_023247213.1;refGene_gn=MFSD4A;refGene_tx=XM_023247214.1
F1 45882120 45882381
refGene_gn=MFSD4A;refGene_tx=XM_019822121.2;refGene_gn=MFSD4A;refGene_tx=XM_023247210.1;refGene_gn=MFSD4A;refGene_tx=XM_023247211.1;refGene_gn=MFSD4A;refGene_tx=XM_023247212.1;refGene_gn=MFSD4A;refGene_tx=XM_023247213.1;refGene_gn=MFSD4A;refGene_tx=XM_023247214.1
F1 45885408 45885577
refGene_gn=MFSD4A;refGene_tx=XM_019822121.2;refGene_gn=MFSD4A;refGene_tx=XM_023247210.1;refGene_gn=MFSD4A;refGene_tx=XM_023247211.1;refGene_gn=MFSD4A;refGene_tx=XM_023247212.1;refGene_gn=MFSD4A;refGene_tx=XM_023247213.1;refGene_gn=MFSD4A;refGene_tx=XM_023247214.1
F1 45886013 45886129
refGene_gn=MFSD4A;refGene_tx=XM_019822121.2;refGene_gn=MFSD4A;refGene_tx=XM_023247210.1;refGene_gn=MFSD4A;refGene_tx=XM_023247211.1;refGene_gn=MFSD4A;refGene_tx=XM_023247212.1;refGene_gn=MFSD4A;refGene_tx=XM_023247213.1;refGene_gn=MFSD4A;refGene_tx=XM_023247214.1
F1 45887192 45887366
refGene_gn=MFSD4A;refGene_tx=XM_019822121.2;refGene_gn=MFSD4A;refGene_tx=XM_023247210.1;refGene_gn=MFSD4A;refGene_tx=XM_023247211.1;refGene_gn=MFSD4A;refGene_tx=XM_023247212.1;refGene_gn=MFSD4A;refGene_tx=XM_023247214.1
F1 45892676 45892824
refGene_gn=MFSD4A;refGene_tx=XM_019822121.2;refGene_gn=MFSD4A;refGene_tx=XM_023247210.1;refGene_gn=MFSD4A;refGene_tx=XM_023247211.1;refGene_gn=MFSD4A;refGene_tx=XM_023247212.1;refGene_gn=MFSD4A;refGene_tx=XM_023247213.1;refGene_gn=MFSD4A;refGene_tx=XM_023247214.1
F1 45896341 45896412
refGene_gn=MFSD4A;refGene_tx=XM_019822121.2;refGene_gn=MFSD4A;refGene_tx=XM_023247210.1;refGene_gn=MFSD4A;refGene_tx=XM_023247211.1;refGene_gn=MFSD4A;refGene_tx=XM_023247212.1;refGene_gn=MFSD4A;refGene_tx=XM_023247213.1
F1 45897598 45897719
refGene_gn=MFSD4A;refGene_tx=XM_019822121.2;refGene_gn=MFSD4A;refGene_tx=XM_023247210.1;refGene_gn=MFSD4A;refGene_tx=XM_023247212.1;refGene_gn=MFSD4A;refGene_tx=XM_023247213.1
F1 45898878 45899077
refGene_gn=MFSD4A;refGene_tx=XM_019822121.2;refGene_gn=MFSD4A;refGene_tx=XM_023247210.1;refGene_gn=MFSD4A;refGene_tx=XM_023247211.1;refGene_gn=MFSD4A;refGene_tx=XM_023247212.1;refGene_gn=MFSD4A;refGene_tx=XM_023247213.1;refGene_gn=MFSD4A;refGene_tx=XM_023247214.1
F1 45911951 45912050
refGene_gn=ELK4;refGene_tx=XM_003999416.5;refGene_gn=ELK4;refGene_tx=XM_023247236.1

F1 45915211 45915328
refGene_gn=ELK4;refGene_tx=XM_003999416.5;refGene_gn=ELK4;refGene_tx=X
M_023247236.1
F1 45916074 45916947
refGene_gn=ELK4;refGene_tx=XM_003999416.5;refGene_gn=ELK4;refGene_tx=X
M_023247236.1
F1 45918037 45918244
refGene_gn=ELK4;refGene_tx=XM_003999416.5;refGene_gn=ELK4;refGene_tx=X
M_023247236.1
F1 45944477 45944915
refGene_gn=SLC45A3;refGene_tx=XM_006942912.4;refGene_gn=SLC45A3;refGen
e_tx=XM_023247589.1
F1 45947032 45947298
refGene_gn=SLC45A3;refGene_tx=XM_006942912.4;refGene_gn=SLC45A3;refGen
e_tx=XM_023247589.1
F1 45948001 45948787
refGene_gn=SLC45A3;refGene_tx=XM_006942912.4;refGene_gn=SLC45A3;refGen
e_tx=XM_023247589.1
F1 45949654 45949826
refGene_gn=SLC45A3;refGene_tx=XM_006942912.4;refGene_gn=SLC45A3;refGen
e_tx=XM_023247589.1
F1 45993526 45993726 refGene_gn=NUCKS1;refGene_tx=XM_023248039.1
F1 45995419 45995569 refGene_gn=NUCKS1;refGene_tx=XM_023248039.1
F1 45996347 45996500 refGene_gn=NUCKS1;refGene_tx=XM_023248039.1
F1 45998450 45998506 refGene_gn=NUCKS1;refGene_tx=XM_023248039.1
F1 46000931 46001037 refGene_gn=NUCKS1;refGene_tx=XM_023248039.1
F1 46002891 46002941 refGene_gn=NUCKS1;refGene_tx=XM_023248039.1
F1 46019754 46019771 refGene_gn=NUCKS1;refGene_tx=XM_023248039.1
F1 46030005 46030036 refGene_gn=RAB29;refGene_tx=XM_006942918.3
F1 46030208 46030320
refGene_gn=RAB29;refGene_tx=XM_003999417.4;refGene_gn=RAB29;refGene_tx
=XM_006942915.3;refGene_gn=RAB29;refGene_tx=XM_006942916.3;refGene_gn=
RAB29;refGene_tx=XM_006942917.3
F1 46030617 46030739
refGene_gn=RAB29;refGene_tx=XM_003999417.4;refGene_gn=RAB29;refGene_tx
=XM_006942915.3;refGene_gn=RAB29;refGene_tx=XM_006942916.3;refGene_gn=
RAB29;refGene_tx=XM_006942917.3;refGene_gn=RAB29;refGene_tx=XM_0069429
18.3
F1 46031027 46031209
refGene_gn=RAB29;refGene_tx=XM_003999417.4;refGene_gn=RAB29;refGene_tx
=XM_006942915.3;refGene_gn=RAB29;refGene_tx=XM_006942916.3;refGene_gn=
RAB29;refGene_tx=XM_006942917.3;refGene_gn=RAB29;refGene_tx=XM_0069429
18.3
F1 46031892 46031964
refGene_gn=RAB29;refGene_tx=XM_003999417.4;refGene_gn=RAB29;refGene_tx
=XM_006942915.3;refGene_gn=RAB29;refGene_tx=XM_006942916.3;refGene_gn=
RAB29;refGene_tx=XM_006942917.3;refGene_gn=RAB29;refGene_tx=XM_0069429
18.3
F1 46034952 46035076
refGene_gn=RAB29;refGene_tx=XM_003999417.4;refGene_gn=RAB29;refGene_tx

=XM_006942915.3;refGene_gn=RAB29;refGene_tx=XM_006942916.3;refGene_gn=RAB29;refGene_tx=XM_006942917.3;refGene_gn=RAB29;refGene_tx=XM_006942918.3
F1 46049764 46049950 refGene_gn=SLC41A1;refGene_tx=XM_003999419.5
F1 46052891 46053040 refGene_gn=SLC41A1;refGene_tx=XM_003999419.5
F1 46053350 46053485 refGene_gn=SLC41A1;refGene_tx=XM_003999419.5
F1 46054396 46054476 refGene_gn=SLC41A1;refGene_tx=XM_003999419.5
F1 46055361 46055509 refGene_gn=SLC41A1;refGene_tx=XM_003999419.5
F1 46056113 46056260 refGene_gn=SLC41A1;refGene_tx=XM_003999419.5
F1 46056375 46056520 refGene_gn=SLC41A1;refGene_tx=XM_003999419.5
F1 46057160 46057232 refGene_gn=SLC41A1;refGene_tx=XM_003999419.5
F1 46058335 46058443 refGene_gn=SLC41A1;refGene_tx=XM_003999419.5
F1 46066667 46067042 refGene_gn=SLC41A1;refGene_tx=XM_003999419.5
F1 46082238 46082362 refGene_gn=PM20D1;refGene_tx=XM_003999420.5;refGene_gn=PM20D1;refGene_tx=XM_019822127.2
F1 46083778 46083878 refGene_gn=PM20D1;refGene_tx=XM_003999420.5;refGene_gn=PM20D1;refGene_tx=XM_019822127.2
F1 46086028 46086197 refGene_gn=PM20D1;refGene_tx=XM_003999420.5;refGene_gn=PM20D1;refGene_tx=XM_019822127.2
F1 46092451 46092523 refGene_gn=PM20D1;refGene_tx=XM_003999420.5;refGene_gn=PM20D1;refGene_tx=XM_019822127.2
F1 46094562 46094641 refGene_gn=PM20D1;refGene_tx=XM_003999420.5;refGene_gn=PM20D1;refGene_tx=XM_019822127.2
F1 46094888 46094950 refGene_gn=PM20D1;refGene_tx=XM_003999420.5;refGene_gn=PM20D1;refGene_tx=XM_019822127.2
F1 46095558 46095634 refGene_gn=PM20D1;refGene_tx=XM_003999420.5;refGene_gn=PM20D1;refGene_tx=XM_019822127.2
F1 46096338 46096458 refGene_gn=PM20D1;refGene_tx=XM_003999420.5;refGene_gn=PM20D1;refGene_tx=XM_019822127.2
F1 46096743 46096877 refGene_gn=PM20D1;refGene_tx=XM_003999420.5;refGene_gn=PM20D1;refGene_tx=XM_019822127.2
F1 46097964 46098051 refGene_gn=PM20D1;refGene_tx=XM_003999420.5;refGene_gn=PM20D1;refGene_tx=XM_019822127.2
F1 46098717 46098950 refGene_gn=PM20D1;refGene_tx=XM_003999420.5;refGene_gn=PM20D1;refGene_tx=XM_019822127.2
F1 46101604 46101691 refGene_gn=PM20D1;refGene_tx=XM_003999420.5;refGene_gn=PM20D1;refGene_tx=XM_019822127.2
F1 46113152 46113321

refGene_gn=PM20D1;refGene_tx=XM_003999420.5;refGene_gn=PM20D1;refGene_tx=XM_019822127.2
F1 46120326 46120374 refGene_gn=SLC26A9;refGene_tx=XM_023247800.1
F1 46121778 46121850 refGene_gn=SLC26A9;refGene_tx=XM_023247800.1
F1 46123317 46123463 refGene_gn=SLC26A9;refGene_tx=XM_023247800.1
F1 46124372 46124427 refGene_gn=SLC26A9;refGene_tx=XM_023247800.1
F1 46125518 46125800 refGene_gn=SLC26A9;refGene_tx=XM_023247800.1
F1 46126437 46126551 refGene_gn=SLC26A9;refGene_tx=XM_023247800.1
F1 46126682 46126775 refGene_gn=SLC26A9;refGene_tx=XM_023247800.1
F1 46127108 46127178 refGene_gn=SLC26A9;refGene_tx=XM_023247800.1
F1 46127858 46127965 refGene_gn=SLC26A9;refGene_tx=XM_023247800.1
F1 46129857 46129953 refGene_gn=SLC26A9;refGene_tx=XM_023247800.1
F1 46130495 46130573 refGene_gn=SLC26A9;refGene_tx=XM_023247800.1
F1 46130786 46130900 refGene_gn=SLC26A9;refGene_tx=XM_023247800.1
F1 46131194 46131342 refGene_gn=SLC26A9;refGene_tx=XM_023247800.1
F1 46132090 46132173 refGene_gn=SLC26A9;refGene_tx=XM_023247800.1
F1 46132488 46132641 refGene_gn=SLC26A9;refGene_tx=XM_023247800.1
F1 46133211 46133376 refGene_gn=SLC26A9;refGene_tx=XM_023247800.1
F1 46134867 46135043 refGene_gn=SLC26A9;refGene_tx=XM_023247800.1
F1 46135685 46135796 refGene_gn=SLC26A9;refGene_tx=XM_023247800.1
F1 46135916 46136056 refGene_gn=SLC26A9;refGene_tx=XM_023247800.1
F1 46138764 46138889 refGene_gn=SLC26A9;refGene_tx=XM_023247800.1
F1 46178314 46178395
refGene_gn=RAB7B;refGene_tx=XM_019822131.2;refGene_gn=RAB7B;refGene_tx=XM_023247496.1;refGene_gn=RAB7B;refGene_tx=XM_023247497.1
F1 46185732 46185944
refGene_gn=RAB7B;refGene_tx=XM_019822131.2;refGene_gn=RAB7B;refGene_tx=XM_023247496.1;refGene_gn=RAB7B;refGene_tx=XM_023247497.1
F1 46191470 46191686
refGene_gn=RAB7B;refGene_tx=XM_019822131.2;refGene_gn=RAB7B;refGene_tx=XM_023247496.1;refGene_gn=RAB7B;refGene_tx=XM_023247497.1
F1 46192439 46192568
refGene_gn=RAB7B;refGene_tx=XM_019822131.2;refGene_gn=RAB7B;refGene_tx=XM_023247496.1;refGene_gn=RAB7B;refGene_tx=XM_023247497.1
F1 46193323 46193392
refGene_gn=RAB7B;refGene_tx=XM_019822131.2;refGene_gn=RAB7B;refGene_tx=XM_023247496.1;refGene_gn=RAB7B;refGene_tx=XM_023247497.1
F1 46193651 46193786
refGene_gn=RAB7B;refGene_tx=XM_019822131.2;refGene_gn=RAB7B;refGene_tx=XM_023247496.1
F1 46210776 46211074
refGene_gn=CTSE;refGene_tx=XM_003999424.5;refGene_gn=CTSE;refGene_tx=XM_006942920.4
F1 46214109 46214208
refGene_gn=CTSE;refGene_tx=XM_003999424.5;refGene_gn=CTSE;refGene_tx=XM_006942920.4
F1 46214309 46214451 refGene_gn=CTSE;refGene_tx=XM_003999424.5
F1 46215650 46215773
refGene_gn=CTSE;refGene_tx=XM_003999424.5;refGene_gn=CTSE;refGene_tx=XM_006942920.4

F1 46217759 46217959
refGene_gn=CTSE;refGene_tx=XM_003999424.5;refGene_gn=CTSE;refGene_tx=XM_006942920.4
F1 46220721 46220840
refGene_gn=CTSE;refGene_tx=XM_003999424.5;refGene_gn=CTSE;refGene_tx=XM_006942920.4
F1 46221797 46221915
refGene_gn=CTSE;refGene_tx=XM_003999424.5;refGene_gn=CTSE;refGene_tx=XM_006942920.4
F1 46222502 46222659
refGene_gn=CTSE;refGene_tx=XM_003999424.5;refGene_gn=CTSE;refGene_tx=XM_006942920.4
F1 46223281 46223349
refGene_gn=CTSE;refGene_tx=XM_003999424.5;refGene_gn=CTSE;refGene_tx=XM_006942920.4
F1 46249102 46249130
refGene_gn=LOC111558370;refGene_tx=XM_023247728.1
F1 46249590 46249643
refGene_gn=LOC111558370;refGene_tx=XM_023247728.1
F1 46249915 46250074
refGene_gn=LOC111558370;refGene_tx=XM_023247728.1
F1 46288196 46288525 refGene_gn=AVPR1B;refGene_tx=XM_003999425.4
F1 46292784 46293724 refGene_gn=AVPR1B;refGene_tx=XM_003999425.4
F1 46345555 46345650
refGene_gn=FAM72A;refGene_tx=XM_003999426.5;refGene_gn=FAM72A;refGene_tx=XM_019822146.2
F1 46349402 46349527 refGene_gn=FAM72A;refGene_tx=XM_003999426.5
F1 46352255 46352333
refGene_gn=FAM72A;refGene_tx=XM_003999426.5;refGene_gn=FAM72A;refGene_tx=XM_019822146.2
F1 46354731 46354883
refGene_gn=FAM72A;refGene_tx=XM_003999426.5;refGene_gn=FAM72A;refGene_tx=XM_019822146.2
F1 46358857 46358924
refGene_gn=SRGAP2;refGene_tx=XM_003999427.5;refGene_gn=SRGAP2;refGene_tx=XM_006942921.4;refGene_gn=SRGAP2;refGene_tx=XM_006942922.4;refGene_gn=SRGAP2;refGene_tx=XM_006942923.4;refGene_gn=SRGAP2;refGene_tx=XM_019822139.1;refGene_gn=SRGAP2;refGene_tx=XM_019822140.1;refGene_gn=SRGAP2;refGene_tx=XM_019822141.1
F1 46446846 46447039
refGene_gn=SRGAP2;refGene_tx=XM_003999427.5;refGene_gn=SRGAP2;refGene_tx=XM_006942921.4;refGene_gn=SRGAP2;refGene_tx=XM_006942922.4;refGene_gn=SRGAP2;refGene_tx=XM_006942923.4;refGene_gn=SRGAP2;refGene_tx=XM_019822139.1;refGene_gn=SRGAP2;refGene_tx=XM_019822140.1;refGene_gn=SRGAP2;refGene_tx=XM_019822141.1
F1 46473484 46473647
refGene_gn=SRGAP2;refGene_tx=XM_003999427.5;refGene_gn=SRGAP2;refGene_tx=XM_006942921.4;refGene_gn=SRGAP2;refGene_tx=XM_006942922.4;refGene_gn=SRGAP2;refGene_tx=XM_006942923.4;refGene_gn=SRGAP2;refGene_tx=XM_019822139.1;refGene_gn=SRGAP2;refGene_tx=XM_019822140.1;refGene_gn=SRGAP

2;refGene_tx=XM_019822141.1
F1 46510677 46510740
refGene_gn=SRGAP2;refGene_tx=XM_003999427.5;refGene_gn=SRGAP2;refGene_tx=XM_006942921.4;refGene_gn=SRGAP2;refGene_tx=XM_006942922.4;refGene_gn=SRGAP2;refGene_tx=XM_006942923.4;refGene_gn=SRGAP2;refGene_tx=XM_019822139.1;refGene_gn=SRGAP2;refGene_tx=XM_019822140.1;refGene_gn=SRGAP2;refGene_tx=XM_019822141.1
F1 46519113 46519329
refGene_gn=SRGAP2;refGene_tx=XM_003999427.5;refGene_gn=SRGAP2;refGene_tx=XM_006942921.4;refGene_gn=SRGAP2;refGene_tx=XM_006942922.4;refGene_gn=SRGAP2;refGene_tx=XM_006942923.4;refGene_gn=SRGAP2;refGene_tx=XM_019822139.1;refGene_gn=SRGAP2;refGene_tx=XM_019822140.1;refGene_gn=SRGAP2;refGene_tx=XM_019822141.1
F1 46519955 46520084
refGene_gn=SRGAP2;refGene_tx=XM_003999427.5;refGene_gn=SRGAP2;refGene_tx=XM_006942921.4;refGene_gn=SRGAP2;refGene_tx=XM_006942922.4;refGene_gn=SRGAP2;refGene_tx=XM_006942923.4;refGene_gn=SRGAP2;refGene_tx=XM_019822139.1;refGene_gn=SRGAP2;refGene_tx=XM_019822140.1;refGene_gn=SRGAP2;refGene_tx=XM_019822141.1
F1 46528272 46528497
refGene_gn=SRGAP2;refGene_tx=XM_003999427.5;refGene_gn=SRGAP2;refGene_tx=XM_006942921.4;refGene_gn=SRGAP2;refGene_tx=XM_006942922.4;refGene_gn=SRGAP2;refGene_tx=XM_006942923.4;refGene_gn=SRGAP2;refGene_tx=XM_019822139.1;refGene_gn=SRGAP2;refGene_tx=XM_019822140.1;refGene_gn=SRGAP2;refGene_tx=XM_019822141.1
F1 46532114 46532216
refGene_gn=SRGAP2;refGene_tx=XM_003999427.5;refGene_gn=SRGAP2;refGene_tx=XM_006942921.4;refGene_gn=SRGAP2;refGene_tx=XM_006942922.4;refGene_gn=SRGAP2;refGene_tx=XM_006942923.4;refGene_gn=SRGAP2;refGene_tx=XM_019822139.1;refGene_gn=SRGAP2;refGene_tx=XM_019822140.1;refGene_gn=SRGAP2;refGene_tx=XM_019822141.1
F1 46533225 46533423
refGene_gn=SRGAP2;refGene_tx=XM_003999427.5;refGene_gn=SRGAP2;refGene_tx=XM_006942921.4;refGene_gn=SRGAP2;refGene_tx=XM_006942922.4;refGene_gn=SRGAP2;refGene_tx=XM_006942923.4;refGene_gn=SRGAP2;refGene_tx=XM_019822139.1;refGene_gn=SRGAP2;refGene_tx=XM_019822140.1;refGene_gn=SRGAP2;refGene_tx=XM_019822141.1
F1 46542577 46542662
refGene_gn=SRGAP2;refGene_tx=XM_003999427.5;refGene_gn=SRGAP2;refGene_tx=XM_006942921.4;refGene_gn=SRGAP2;refGene_tx=XM_006942922.4;refGene_gn=SRGAP2;refGene_tx=XM_006942923.4;refGene_gn=SRGAP2;refGene_tx=XM_019822139.1;refGene_gn=SRGAP2;refGene_tx=XM_019822140.1;refGene_gn=SRGAP2;refGene_tx=XM_019822141.1
F1 46545394 46545422
refGene_gn=SRGAP2;refGene_tx=XM_003999427.5;refGene_gn=SRGAP2;refGene_tx=XM_006942921.4;refGene_gn=SRGAP2;refGene_tx=XM_006942922.4;refGene_gn=SRGAP2;refGene_tx=XM_006942923.4;refGene_gn=SRGAP2;refGene_tx=XM_019822139.1;refGene_gn=SRGAP2;refGene_tx=XM_019822140.1;refGene_gn=SRGAP2;refGene_tx=XM_019822141.1
F1 46547099 46547124

refGene_gn=SRGAP2;refGene_tx=XM_003999427.5;refGene_gn=SRGAP2;refGene_tx=XM_006942921.4;refGene_gn=SRGAP2;refGene_tx=XM_006942922.4;refGene_gn=SRGAP2;refGene_tx=XM_006942923.4;refGene_gn=SRGAP2;refGene_tx=XM_019822139.1;refGene_gn=SRGAP2;refGene_tx=XM_019822140.1;refGene_gn=SRGAP2;refGene_tx=XM_019822141.1
F1 46553406 46553467
refGene_gn=SRGAP2;refGene_tx=XM_003999427.5;refGene_gn=SRGAP2;refGene_tx=XM_006942921.4;refGene_gn=SRGAP2;refGene_tx=XM_006942922.4;refGene_gn=SRGAP2;refGene_tx=XM_006942923.4;refGene_gn=SRGAP2;refGene_tx=XM_019822139.1;refGene_gn=SRGAP2;refGene_tx=XM_019822140.1;refGene_gn=SRGAP2;refGene_tx=XM_019822141.1
F1 46559681 46559759
refGene_gn=SRGAP2;refGene_tx=XM_003999427.5;refGene_gn=SRGAP2;refGene_tx=XM_006942921.4;refGene_gn=SRGAP2;refGene_tx=XM_006942922.4;refGene_gn=SRGAP2;refGene_tx=XM_006942923.4;refGene_gn=SRGAP2;refGene_tx=XM_019822139.1;refGene_gn=SRGAP2;refGene_tx=XM_019822140.1;refGene_gn=SRGAP2;refGene_tx=XM_019822141.1
F1 46560671 46560806
refGene_gn=SRGAP2;refGene_tx=XM_003999427.5;refGene_gn=SRGAP2;refGene_tx=XM_006942921.4;refGene_gn=SRGAP2;refGene_tx=XM_006942922.4;refGene_gn=SRGAP2;refGene_tx=XM_006942923.4;refGene_gn=SRGAP2;refGene_tx=XM_019822139.1;refGene_gn=SRGAP2;refGene_tx=XM_019822140.1;refGene_gn=SRGAP2;refGene_tx=XM_019822141.1
F1 46562824 46562930
refGene_gn=SRGAP2;refGene_tx=XM_003999427.5;refGene_gn=SRGAP2;refGene_tx=XM_006942921.4;refGene_gn=SRGAP2;refGene_tx=XM_006942922.4;refGene_gn=SRGAP2;refGene_tx=XM_006942923.4;refGene_gn=SRGAP2;refGene_tx=XM_019822139.1;refGene_gn=SRGAP2;refGene_tx=XM_019822140.1;refGene_gn=SRGAP2;refGene_tx=XM_019822141.1
F1 46572263 46572488
refGene_gn=SRGAP2;refGene_tx=XM_003999427.5;refGene_gn=SRGAP2;refGene_tx=XM_006942921.4;refGene_gn=SRGAP2;refGene_tx=XM_006942922.4;refGene_gn=SRGAP2;refGene_tx=XM_006942923.4;refGene_gn=SRGAP2;refGene_tx=XM_019822139.1;refGene_gn=SRGAP2;refGene_tx=XM_019822140.1;refGene_gn=SRGAP2;refGene_tx=XM_019822141.1
F1 46576151 46576231
refGene_gn=SRGAP2;refGene_tx=XM_003999427.5;refGene_gn=SRGAP2;refGene_tx=XM_006942921.4;refGene_gn=SRGAP2;refGene_tx=XM_006942922.4;refGene_gn=SRGAP2;refGene_tx=XM_006942923.4;refGene_gn=SRGAP2;refGene_tx=XM_019822139.1;refGene_gn=SRGAP2;refGene_tx=XM_019822140.1;refGene_gn=SRGAP2;refGene_tx=XM_019822141.1
F1 46578420 46578601
refGene_gn=SRGAP2;refGene_tx=XM_003999427.5;refGene_gn=SRGAP2;refGene_tx=XM_006942921.4;refGene_gn=SRGAP2;refGene_tx=XM_006942922.4;refGene_gn=SRGAP2;refGene_tx=XM_006942923.4;refGene_gn=SRGAP2;refGene_tx=XM_019822139.1;refGene_gn=SRGAP2;refGene_tx=XM_019822140.1;refGene_gn=SRGAP2;refGene_tx=XM_019822141.1
F1 46579373 46579386 refGene_gn=SRGAP2;refGene_tx=XM_019822141.1
F1 46580104 46580251
refGene_gn=SRGAP2;refGene_tx=XM_003999427.5;refGene_gn=SRGAP2;refGene_

tx=XM_006942921.4;refGene_gn=SRGAP2;refGene_tx=XM_006942922.4;refGene_
gn=SRGAP2;refGene_tx=XM_006942923.4;refGene_gn=SRGAP2;refGene_tx=XM_01
9822139.1;refGene_gn=SRGAP2;refGene_tx=XM_019822140.1
F1 46580698 46580708 refGene_gn=SRGAP2;refGene_tx=XM_019822140.1
F1 46583491 46583530
refGene_gn=SRGAP2;refGene_tx=XM_006942921.4;refGene_gn=SRGAP2;refGene_
tx=XM_006942922.4;refGene_gn=SRGAP2;refGene_tx=XM_019822139.1
F1 46583744 46584069
refGene_gn=SRGAP2;refGene_tx=XM_003999427.5;refGene_gn=SRGAP2;refGene_
tx=XM_006942921.4;refGene_gn=SRGAP2;refGene_tx=XM_006942922.4;refGene_
gn=SRGAP2;refGene_tx=XM_006942923.4;refGene_gn=SRGAP2;refGene_tx=XM_01
9822139.1
F1 46584717 46584807 refGene_gn=SRGAP2;refGene_tx=XM_019822139.1
F1 46586165 46586549
refGene_gn=SRGAP2;refGene_tx=XM_003999427.5;refGene_gn=SRGAP2;refGene_
tx=XM_006942921.4;refGene_gn=SRGAP2;refGene_tx=XM_006942922.4;refGene_
gn=SRGAP2;refGene_tx=XM_006942923.4
F1 46597234 46597321
refGene_gn=IKBKE;refGene_tx=XM_006942924.4;refGene_gn=IKBKE;refGene_tx
=XM_006942925.4;refGene_gn=IKBKE;refGene_tx=XM_006942926.4;refGene_gn=
IKBKE;refGene_tx=XM_006942927.4;refGene_gn=IKBKE;refGene_tx=XM_0069429
30.4;refGene_gn=IKBKE;refGene_tx=XM_019822142.2;refGene_gn=IKBKE;refGe
ne_tx=XM_019822144.2;refGene_gn=IKBKE;refGene_tx=XM_019822145.2
F1 46598021 46598270
refGene_gn=IKBKE;refGene_tx=XM_006942924.4;refGene_gn=IKBKE;refGene_tx
=XM_006942925.4;refGene_gn=IKBKE;refGene_tx=XM_006942926.4;refGene_gn=
IKBKE;refGene_tx=XM_006942927.4;refGene_gn=IKBKE;refGene_tx=XM_0069429
30.4;refGene_gn=IKBKE;refGene_tx=XM_019822142.2;refGene_gn=IKBKE;refGe
ne_tx=XM_019822144.2;refGene_gn=IKBKE;refGene_tx=XM_019822145.2
F1 46598576 46598706
refGene_gn=IKBKE;refGene_tx=XM_006942924.4;refGene_gn=IKBKE;refGene_tx
=XM_006942925.4;refGene_gn=IKBKE;refGene_tx=XM_006942926.4;refGene_gn=
IKBKE;refGene_tx=XM_006942927.4;refGene_gn=IKBKE;refGene_tx=XM_0069429
30.4;refGene_gn=IKBKE;refGene_tx=XM_019822142.2;refGene_gn=IKBKE;refGe
ne_tx=XM_019822144.2;refGene_gn=IKBKE;refGene_tx=XM_019822145.2
F1 46600392 46600574
refGene_gn=IKBKE;refGene_tx=XM_006942924.4;refGene_gn=IKBKE;refGene_tx
=XM_006942925.4;refGene_gn=IKBKE;refGene_tx=XM_006942926.4;refGene_gn=
IKBKE;refGene_tx=XM_006942927.4;refGene_gn=IKBKE;refGene_tx=XM_0069429
30.4;refGene_gn=IKBKE;refGene_tx=XM_019822142.2;refGene_gn=IKBKE;refGe
ne_tx=XM_019822144.2;refGene_gn=IKBKE;refGene_tx=XM_019822145.2
F1 46604204 46604365
refGene_gn=IKBKE;refGene_tx=XM_006942924.4;refGene_gn=IKBKE;refGene_tx
=XM_006942925.4;refGene_gn=IKBKE;refGene_tx=XM_006942926.4;refGene_gn=
IKBKE;refGene_tx=XM_006942927.4;refGene_gn=IKBKE;refGene_tx=XM_0069429
30.4;refGene_gn=IKBKE;refGene_tx=XM_019822142.2;refGene_gn=IKBKE;refGe
ne_tx=XM_019822144.2;refGene_gn=IKBKE;refGene_tx=XM_019822145.2
F1 46605209 46605320
refGene_gn=IKBKE;refGene_tx=XM_006942924.4;refGene_gn=IKBKE;refGene_tx
=XM_006942925.4;refGene_gn=IKBKE;refGene_tx=XM_006942926.4;refGene_gn=

IKBKE;refGene_tx=XM_006942927.4;refGene_gn=IKBKE;refGene_tx=XM_0069429
30.4;refGene_gn=IKBKE;refGene_tx=XM_019822142.2;refGene_gn=IKBKE;refGe
ne_tx=XM_019822144.2;refGene_gn=IKBKE;refGene_tx=XM_019822145.2
F1 46605635 46605815
refGene_gn=IKBKE;refGene_tx=XM_006942924.4;refGene_gn=IKBKE;refGene_tx
=XM_006942925.4;refGene_gn=IKBKE;refGene_tx=XM_006942926.4;refGene_gn=
IKBKE;refGene_tx=XM_006942927.4;refGene_gn=IKBKE;refGene_tx=XM_0069429
30.4;refGene_gn=IKBKE;refGene_tx=XM_019822142.2;refGene_gn=IKBKE;refGe
ne_tx=XM_019822144.2;refGene_gn=IKBKE;refGene_tx=XM_019822145.2
F1 46606424 46606621
refGene_gn=IKBKE;refGene_tx=XM_006942924.4;refGene_gn=IKBKE;refGene_tx
=XM_006942925.4;refGene_gn=IKBKE;refGene_tx=XM_006942926.4;refGene_gn=
IKBKE;refGene_tx=XM_006942927.4;refGene_gn=IKBKE;refGene_tx=XM_0069429
30.4;refGene_gn=IKBKE;refGene_tx=XM_019822142.2;refGene_gn=IKBKE;refGe
ne_tx=XM_019822144.2;refGene_gn=IKBKE;refGene_tx=XM_019822145.2
F1 46607371 46607436
refGene_gn=IKBKE;refGene_tx=XM_006942924.4;refGene_gn=IKBKE;refGene_tx
=XM_006942925.4;refGene_gn=IKBKE;refGene_tx=XM_006942926.4;refGene_gn=
IKBKE;refGene_tx=XM_006942927.4;refGene_gn=IKBKE;refGene_tx=XM_0069429
30.4;refGene_gn=IKBKE;refGene_tx=XM_019822142.2;refGene_gn=IKBKE;refGe
ne_tx=XM_019822144.2;refGene_gn=IKBKE;refGene_tx=XM_019822145.2
F1 46607517 46607609
refGene_gn=IKBKE;refGene_tx=XM_006942924.4;refGene_gn=IKBKE;refGene_tx
=XM_006942925.4;refGene_gn=IKBKE;refGene_tx=XM_006942926.4;refGene_gn=
IKBKE;refGene_tx=XM_006942927.4;refGene_gn=IKBKE;refGene_tx=XM_0069429
30.4;refGene_gn=IKBKE;refGene_tx=XM_019822142.2;refGene_gn=IKBKE;refGe
ne_tx=XM_019822144.2;refGene_gn=IKBKE;refGene_tx=XM_019822145.2
F1 46607736 46607823
refGene_gn=IKBKE;refGene_tx=XM_006942924.4;refGene_gn=IKBKE;refGene_tx
=XM_006942925.4;refGene_gn=IKBKE;refGene_tx=XM_006942926.4;refGene_gn=
IKBKE;refGene_tx=XM_006942927.4;refGene_gn=IKBKE;refGene_tx=XM_0069429
30.4;refGene_gn=IKBKE;refGene_tx=XM_019822142.2;refGene_gn=IKBKE;refGe
ne_tx=XM_019822144.2;refGene_gn=IKBKE;refGene_tx=XM_019822145.2
F1 46611618 46611694
refGene_gn=IKBKE;refGene_tx=XM_006942924.4;refGene_gn=IKBKE;refGene_tx
=XM_006942925.4;refGene_gn=IKBKE;refGene_tx=XM_006942926.4;refGene_gn=
IKBKE;refGene_tx=XM_006942927.4;refGene_gn=IKBKE;refGene_tx=XM_0069429
30.4;refGene_gn=IKBKE;refGene_tx=XM_019822142.2;refGene_gn=IKBKE;refGe
ne_tx=XM_019822144.2;refGene_gn=IKBKE;refGene_tx=XM_019822145.2
F1 46611816 46611929
refGene_gn=IKBKE;refGene_tx=XM_006942924.4;refGene_gn=IKBKE;refGene_tx
=XM_006942925.4;refGene_gn=IKBKE;refGene_tx=XM_006942926.4;refGene_gn=
IKBKE;refGene_tx=XM_006942927.4;refGene_gn=IKBKE;refGene_tx=XM_0069429
30.4;refGene_gn=IKBKE;refGene_tx=XM_019822142.2;refGene_gn=IKBKE;refGe
ne_tx=XM_019822144.2;refGene_gn=IKBKE;refGene_tx=XM_019822145.2
F1 46613837 46613914
refGene_gn=IKBKE;refGene_tx=XM_006942924.4;refGene_gn=IKBKE;refGene_tx
=XM_006942925.4;refGene_gn=IKBKE;refGene_tx=XM_006942926.4;refGene_gn=
IKBKE;refGene_tx=XM_006942927.4;refGene_gn=IKBKE;refGene_tx=XM_0069429
30.4;refGene_gn=IKBKE;refGene_tx=XM_019822142.2;refGene_gn=IKBKE;refGe

ne_tx=XM_019822144.2;refGene_gn=IKBKE;refGene_tx=XM_019822145.2
F1	46615308	46615408
refGene_gn=IKBKE;refGene_tx=XM_019822144.2;refGene_gn=IKBKE;refGene_tx=XM_019822145.2
F1	46616476	46616516
refGene_gn=IKBKE;refGene_tx=XM_006942924.4;refGene_gn=IKBKE;refGene_tx=XM_006942925.4;refGene_gn=IKBKE;refGene_tx=XM_006942926.4;refGene_gn=IKBKE;refGene_tx=XM_006942927.4;refGene_gn=IKBKE;refGene_tx=XM_006942930.4;refGene_gn=IKBKE;refGene_tx=XM_019822142.2
F1	46617304	46617406
refGene_gn=IKBKE;refGene_tx=XM_006942924.4;refGene_gn=IKBKE;refGene_tx=XM_006942925.4;refGene_gn=IKBKE;refGene_tx=XM_006942926.4;refGene_gn=IKBKE;refGene_tx=XM_006942927.4;refGene_gn=IKBKE;refGene_tx=XM_019822142.2
F1	46618606	46618703
refGene_gn=IKBKE;refGene_tx=XM_006942924.4;refGene_gn=IKBKE;refGene_tx=XM_006942925.4;refGene_gn=IKBKE;refGene_tx=XM_006942926.4;refGene_gn=IKBKE;refGene_tx=XM_006942927.4;refGene_gn=IKBKE;refGene_tx=XM_019822142.2
F1	46619446	46619559
refGene_gn=IKBKE;refGene_tx=XM_006942924.4;refGene_gn=IKBKE;refGene_tx=XM_006942925.4;refGene_gn=IKBKE;refGene_tx=XM_006942926.4;refGene_gn=IKBKE;refGene_tx=XM_006942927.4;refGene_gn=IKBKE;refGene_tx=XM_019822142.2
F1	46619764	46619836
refGene_gn=IKBKE;refGene_tx=XM_006942924.4;refGene_gn=IKBKE;refGene_tx=XM_006942925.4;refGene_gn=IKBKE;refGene_tx=XM_006942926.4;refGene_gn=IKBKE;refGene_tx=XM_006942927.4;refGene_gn=IKBKE;refGene_tx=XM_019822142.2
F1	46621530	46621564
refGene_gn=IKBKE;refGene_tx=XM_006942924.4;refGene_gn=IKBKE;refGene_tx=XM_006942925.4;refGene_gn=IKBKE;refGene_tx=XM_006942926.4;refGene_gn=IKBKE;refGene_tx=XM_006942927.4;refGene_gn=IKBKE;refGene_tx=XM_019822142.2
F1	46630910	46631388	refGene_gn=RASSF5;refGene_tx=XM_003999428.5
F1	46654735	46654857	refGene_gn=RASSF5;refGene_tx=XM_003999428.5
F1	46668556	46668676	refGene_gn=RASSF5;refGene_tx=XM_011291080.2
F1	46691810	46691921
refGene_gn=RASSF5;refGene_tx=XM_003999428.5;refGene_gn=RASSF5;refGene_tx=XM_011291080.2
F1	46693060	46693358
refGene_gn=RASSF5;refGene_tx=XM_003999428.5;refGene_gn=RASSF5;refGene_tx=XM_011291080.2
F1	46693829	46693945
refGene_gn=RASSF5;refGene_tx=XM_003999428.5;refGene_gn=RASSF5;refGene_tx=XM_011291080.2
F1	46695558	46695711
refGene_gn=RASSF5;refGene_tx=XM_003999428.5;refGene_gn=RASSF5;refGene_tx=XM_011291080.2
F1	46701153	46701224	refGene_gn=EIF2D;refGene_tx=XM_023247459.1

F1 46702753 46702928 refGene_gn=EIF2D;refGene_tx=XM_023247459.1
F1 46707445 46707566 refGene_gn=EIF2D;refGene_tx=XM_023247459.1
F1 46708159 46708267 refGene_gn=EIF2D;refGene_tx=XM_023247459.1
F1 46708834 46708930 refGene_gn=EIF2D;refGene_tx=XM_023247459.1
F1 46709621 46709711 refGene_gn=EIF2D;refGene_tx=XM_023247459.1
F1 46710081 46710231 refGene_gn=EIF2D;refGene_tx=XM_023247459.1
F1 46710328 46710432 refGene_gn=EIF2D;refGene_tx=XM_023247459.1
F1 46710863 46710909 refGene_gn=EIF2D;refGene_tx=XM_023247459.1
F1 46712570 46712688 refGene_gn=EIF2D;refGene_tx=XM_023247459.1
F1 46713160 46713414 refGene_gn=EIF2D;refGene_tx=XM_023247459.1
F1 46715339 46715447 refGene_gn=EIF2D;refGene_tx=XM_023247459.1
F1 46725017 46725108 refGene_gn=EIF2D;refGene_tx=XM_023247459.1
F1 46725869 46725953 refGene_gn=EIF2D;refGene_tx=XM_023247459.1
F1 46739726 46739932
refGene_gn=LOC109497067;refGene_tx=XM_023247987.1
F1 46741818 46742028
refGene_gn=LOC109497067;refGene_tx=XM_023247987.1
F1 46742483 46742658
refGene_gn=LOC109497067;refGene_tx=XM_023247987.1
F1 46843821 46844030
refGene_gn=LOC101089505;refGene_tx=XM_023247985.1;refGene_gn=LOC101089
505;refGene_tx=XM_023247986.1
F1 46844487 46844670
refGene_gn=LOC101089505;refGene_tx=XM_023247985.1;refGene_gn=LOC101089
505;refGene_tx=XM_023247986.1
F1 46851050 46851227
refGene_gn=LOC101089505;refGene_tx=XM_023247985.1;refGene_gn=LOC101089
505;refGene_tx=XM_023247986.1
F1 46851932 46852106
refGene_gn=LOC101089505;refGene_tx=XM_023247985.1;refGene_gn=LOC101089
505;refGene_tx=XM_023247986.1
F1 46853955 46854165
refGene_gn=LOC101089505;refGene_tx=XM_023247985.1;refGene_gn=LOC101089
505;refGene_tx=XM_023247986.1
F1 46854621 46854804
refGene_gn=LOC101089505;refGene_tx=XM_023247985.1;refGene_gn=LOC101089
505;refGene_tx=XM_023247986.1
F1 46858883 46859057
refGene_gn=LOC101089505;refGene_tx=XM_023247985.1;refGene_gn=LOC101089
505;refGene_tx=XM_023247986.1
F1 46861552 46861735
refGene_gn=LOC101089505;refGene_tx=XM_023247985.1;refGene_gn=LOC101089
505;refGene_tx=XM_023247986.1
F1 46861841 46862024
refGene_gn=LOC101089505;refGene_tx=XM_023247985.1;refGene_gn=LOC101089
505;refGene_tx=XM_023247986.1
F1 46863249 46863426
refGene_gn=LOC101089505;refGene_tx=XM_023247985.1;refGene_gn=LOC101089
505;refGene_tx=XM_023247986.1
F1 46868511 46868685

refGene_gn=LOC101089505;refGene_tx=XM_023247985.1;refGene_gn=LOC101089505;refGene_tx=XM_023247986.1
F1 46870537 46870720
refGene_gn=LOC101089505;refGene_tx=XM_023247985.1;refGene_gn=LOC101089505;refGene_tx=XM_023247986.1
F1 46888588 46888765
refGene_gn=LOC101089505;refGene_tx=XM_023247985.1;refGene_gn=LOC101089505;refGene_tx=XM_023247986.1
F1 46889404 46889599
refGene_gn=LOC101089505;refGene_tx=XM_023247985.1;refGene_gn=LOC101089505;refGene_tx=XM_023247986.1
F1 46907257 46907431
refGene_gn=LOC101089505;refGene_tx=XM_023247985.1;refGene_gn=LOC101089505;refGene_tx=XM_023247986.1
F1 46914708 46914879
refGene_gn=LOC101089505;refGene_tx=XM_023247985.1;refGene_gn=LOC101089505;refGene_tx=XM_023247986.1
F1 46917319 46917511
refGene_gn=LOC101089505;refGene_tx=XM_023247985.1
F1 46919264 46919341
refGene_gn=LOC101089505;refGene_tx=XM_023247985.1;refGene_gn=LOC101089505;refGene_tx=XM_023247986.1
F1 46929752 46929858
refGene_gn=LOC101089505;refGene_tx=XM_023247985.1;refGene_gn=LOC101089505;refGene_tx=XM_023247986.1
F1 46930414 46930600
refGene_gn=LOC101089505;refGene_tx=XM_023247985.1;refGene_gn=LOC101089505;refGene_tx=XM_023247986.1
F1 46943237 46943295
refGene_gn=LOC101089505;refGene_tx=XM_023247985.1;refGene_gn=LOC101089505;refGene_tx=XM_023247986.1
F1 46952879 46953070 refGene_gn=EIF2D;refGene_tx=XM_023247459.1
F1 46954038 46954428 refGene_gn=EIF2D;refGene_tx=XM_023247459.1
F1 46954968 46954973 refGene_gn=EIF2D;refGene_tx=XM_023247459.1
F1 46959626 46959823
refGene_gn=LOC102900754;refGene_tx=XM_023247460.1
F1 46960285 46960468
refGene_gn=LOC102900754;refGene_tx=XM_023247460.1
F1 46964228 46964402
refGene_gn=LOC102900754;refGene_tx=XM_023247460.1
F1 46965405 46965582
refGene_gn=LOC102900754;refGene_tx=XM_023247460.1
F1 46966289 46966503
refGene_gn=LOC102900754;refGene_tx=XM_023247460.1
F1 46970787 46970846
refGene_gn=LOC102900754;refGene_tx=XM_023247460.1
F1 46972725 46972899
refGene_gn=LOC102900754;refGene_tx=XM_023247460.1
F1 46975357 46975540
refGene_gn=LOC102900754;refGene_tx=XM_023247460.1

F1 46975647 46975830
refGene_gn=LOC102900754;refGene_tx=XM_023247460.1
F1 46976831 46977008
refGene_gn=LOC102900754;refGene_tx=XM_023247460.1
F1 46978413 46978587
refGene_gn=LOC102900754;refGene_tx=XM_023247460.1
F1 46980170 46980353
refGene_gn=LOC102900754;refGene_tx=XM_023247460.1
F1 46997971 46998148
refGene_gn=LOC102900754;refGene_tx=XM_023247460.1
F1 46998478 46998673
refGene_gn=LOC102900754;refGene_tx=XM_023247460.1
F1 47006419 47006596
refGene_gn=LOC102900754;refGene_tx=XM_023247460.1
F1 47007035 47007230
refGene_gn=LOC102900754;refGene_tx=XM_023247460.1
F1 47029319 47029490
refGene_gn=LOC102900754;refGene_tx=XM_023247460.1
F1 47037257 47037428
refGene_gn=LOC102900754;refGene_tx=XM_023247460.1
F1 47041791 47041868
refGene_gn=LOC102900754;refGene_tx=XM_023247460.1
F1 47047472 47047578
refGene_gn=LOC102900754;refGene_tx=XM_023247460.1
F1 47047916 47048102
refGene_gn=LOC102900754;refGene_tx=XM_023247460.1
F1 47070888 47071209
refGene_gn=LOC101090165;refGene_tx=XM_011291081.3
F1 47078305 47078363
refGene_gn=LOC102900754;refGene_tx=XM_023247460.1
F1 47129715 47129810
refGene_gn=KCNT2;refGene_tx=XM_011291082.3;refGene_gn=KCNT2;refGene_tx=XM_019822157.2;refGene_gn=KCNT2;refGene_tx=XM_019822158.2;refGene_gn=KCNT2;refGene_tx=XM_019822159.2;refGene_gn=KCNT2;refGene_tx=XM_019822160.2;refGene_gn=KCNT2;refGene_tx=XM_019822161.2;refGene_gn=KCNT2;refGene_tx=XM_019822162.2;refGene_gn=KCNT2;refGene_tx=XM_019822163.2;refGene_gn=KCNT2;refGene_tx=XM_019822164.2
F1 47233399 47233479
refGene_gn=KCNT2;refGene_tx=XM_011291082.3;refGene_gn=KCNT2;refGene_tx=XM_019822157.2;refGene_gn=KCNT2;refGene_tx=XM_019822158.2;refGene_gn=KCNT2;refGene_tx=XM_019822159.2;refGene_gn=KCNT2;refGene_tx=XM_019822160.2;refGene_gn=KCNT2;refGene_tx=XM_019822161.2;refGene_gn=KCNT2;refGene_tx=XM_019822162.2;refGene_gn=KCNT2;refGene_tx=XM_019822163.2;refGene_gn=KCNT2;refGene_tx=XM_019822164.2
F1 47235634 47235734
refGene_gn=KCNT2;refGene_tx=XM_011291082.3;refGene_gn=KCNT2;refGene_tx=XM_019822157.2;refGene_gn=KCNT2;refGene_tx=XM_019822158.2;refGene_gn=KCNT2;refGene_tx=XM_019822159.2;refGene_gn=KCNT2;refGene_tx=XM_019822160.2;refGene_gn=KCNT2;refGene_tx=XM_019822161.2;refGene_gn=KCNT2;refGene_tx=XM_019822162.2;refGene_gn=KCNT2;refGene_tx=XM_019822163.2;refGen

e_gn=KCNT2;refGene_tx=XM_019822164.2
F1 47245257 47245306
refGene_gn=KCNT2;refGene_tx=XM_011291082.3;refGene_gn=KCNT2;refGene_tx=XM_019822157.2;refGene_gn=KCNT2;refGene_tx=XM_019822158.2;refGene_gn=KCNT2;refGene_tx=XM_019822159.2;refGene_gn=KCNT2;refGene_tx=XM_019822160.2;refGene_gn=KCNT2;refGene_tx=XM_019822161.2;refGene_gn=KCNT2;refGene_tx=XM_019822162.2;refGene_gn=KCNT2;refGene_tx=XM_019822163.2;refGene_gn=KCNT2;refGene_tx=XM_019822164.2
F1 47247755 47247815
refGene_gn=KCNT2;refGene_tx=XM_011291082.3;refGene_gn=KCNT2;refGene_tx=XM_019822157.2;refGene_gn=KCNT2;refGene_tx=XM_019822158.2;refGene_gn=KCNT2;refGene_tx=XM_019822159.2;refGene_gn=KCNT2;refGene_tx=XM_019822160.2;refGene_gn=KCNT2;refGene_tx=XM_019822161.2;refGene_gn=KCNT2;refGene_tx=XM_019822162.2;refGene_gn=KCNT2;refGene_tx=XM_019822163.2;refGene_gn=KCNT2;refGene_tx=XM_019822164.2
F1 47257644 47257719
refGene_gn=KCNT2;refGene_tx=XM_011291082.3;refGene_gn=KCNT2;refGene_tx=XM_019822157.2;refGene_gn=KCNT2;refGene_tx=XM_019822158.2;refGene_gn=KCNT2;refGene_tx=XM_019822159.2;refGene_gn=KCNT2;refGene_tx=XM_019822160.2;refGene_gn=KCNT2;refGene_tx=XM_019822161.2;refGene_gn=KCNT2;refGene_tx=XM_019822162.2;refGene_gn=KCNT2;refGene_tx=XM_019822163.2;refGene_gn=KCNT2;refGene_tx=XM_019822164.2
F1 47259232 47259334
refGene_gn=KCNT2;refGene_tx=XM_011291082.3;refGene_gn=KCNT2;refGene_tx=XM_019822157.2;refGene_gn=KCNT2;refGene_tx=XM_019822158.2;refGene_gn=KCNT2;refGene_tx=XM_019822159.2;refGene_gn=KCNT2;refGene_tx=XM_019822160.2;refGene_gn=KCNT2;refGene_tx=XM_019822161.2;refGene_gn=KCNT2;refGene_tx=XM_019822162.2;refGene_gn=KCNT2;refGene_tx=XM_019822163.2;refGene_gn=KCNT2;refGene_tx=XM_019822164.2;refGene_gn=KCNT2;refGene_tx=XM_023248070.1
F1 47263177 47263272
refGene_gn=KCNT2;refGene_tx=XM_011291082.3;refGene_gn=KCNT2;refGene_tx=XM_019822157.2;refGene_gn=KCNT2;refGene_tx=XM_019822158.2;refGene_gn=KCNT2;refGene_tx=XM_019822159.2;refGene_gn=KCNT2;refGene_tx=XM_019822160.2;refGene_gn=KCNT2;refGene_tx=XM_019822161.2;refGene_gn=KCNT2;refGene_tx=XM_019822162.2;refGene_gn=KCNT2;refGene_tx=XM_019822163.2;refGene_gn=KCNT2;refGene_tx=XM_019822164.2;refGene_gn=KCNT2;refGene_tx=XM_023248070.1
F1 47299049 47299230
refGene_gn=KCNT2;refGene_tx=XM_011291082.3;refGene_gn=KCNT2;refGene_tx=XM_019822157.2;refGene_gn=KCNT2;refGene_tx=XM_019822158.2;refGene_gn=KCNT2;refGene_tx=XM_019822159.2;refGene_gn=KCNT2;refGene_tx=XM_019822160.2;refGene_gn=KCNT2;refGene_tx=XM_019822161.2;refGene_gn=KCNT2;refGene_tx=XM_019822162.2;refGene_gn=KCNT2;refGene_tx=XM_019822163.2;refGene_gn=KCNT2;refGene_tx=XM_019822164.2;refGene_gn=KCNT2;refGene_tx=XM_023248070.1
F1 47300409 47300574
refGene_gn=KCNT2;refGene_tx=XM_011291082.3;refGene_gn=KCNT2;refGene_tx=XM_019822157.2;refGene_gn=KCNT2;refGene_tx=XM_019822158.2;refGene_gn=KCNT2;refGene_tx=XM_019822159.2;refGene_gn=KCNT2;refGene_tx=XM_0198221

60.2;refGene_gn=KCNT2;refGene_tx=XM_019822161.2;refGene_gn=KCNT2;refGene_tx=XM_019822162.2;refGene_gn=KCNT2;refGene_tx=XM_019822163.2;refGene_gn=KCNT2;refGene_tx=XM_019822164.2;refGene_gn=KCNT2;refGene_tx=XM_023248070.1
F1 47302550 47302687
refGene_gn=KCNT2;refGene_tx=XM_011291082.3;refGene_gn=KCNT2;refGene_tx=XM_019822157.2;refGene_gn=KCNT2;refGene_tx=XM_019822158.2;refGene_gn=KCNT2;refGene_tx=XM_019822159.2;refGene_gn=KCNT2;refGene_tx=XM_019822160.2;refGene_gn=KCNT2;refGene_tx=XM_019822161.2;refGene_gn=KCNT2;refGene_tx=XM_019822162.2;refGene_gn=KCNT2;refGene_tx=XM_019822163.2;refGene_gn=KCNT2;refGene_tx=XM_019822164.2;refGene_gn=KCNT2;refGene_tx=XM_023248070.1
F1 47306564 47306628
refGene_gn=KCNT2;refGene_tx=XM_011291082.3;refGene_gn=KCNT2;refGene_tx=XM_019822157.2;refGene_gn=KCNT2;refGene_tx=XM_019822158.2;refGene_gn=KCNT2;refGene_tx=XM_019822159.2;refGene_gn=KCNT2;refGene_tx=XM_019822160.2;refGene_gn=KCNT2;refGene_tx=XM_019822161.2;refGene_gn=KCNT2;refGene_tx=XM_019822162.2;refGene_gn=KCNT2;refGene_tx=XM_019822163.2;refGene_gn=KCNT2;refGene_tx=XM_019822164.2;refGene_gn=KCNT2;refGene_tx=XM_023248070.1
F1 47331164 47331273
refGene_gn=KCNT2;refGene_tx=XM_011291082.3;refGene_gn=KCNT2;refGene_tx=XM_019822157.2;refGene_gn=KCNT2;refGene_tx=XM_019822158.2;refGene_gn=KCNT2;refGene_tx=XM_019822159.2;refGene_gn=KCNT2;refGene_tx=XM_019822160.2;refGene_gn=KCNT2;refGene_tx=XM_019822161.2;refGene_gn=KCNT2;refGene_tx=XM_019822162.2;refGene_gn=KCNT2;refGene_tx=XM_019822163.2;refGene_gn=KCNT2;refGene_tx=XM_019822164.2;refGene_gn=KCNT2;refGene_tx=XM_023248070.1
F1 47356317 47356426
refGene_gn=KCNT2;refGene_tx=XM_011291082.3;refGene_gn=KCNT2;refGene_tx=XM_019822157.2;refGene_gn=KCNT2;refGene_tx=XM_019822158.2;refGene_gn=KCNT2;refGene_tx=XM_019822159.2;refGene_gn=KCNT2;refGene_tx=XM_019822160.2;refGene_gn=KCNT2;refGene_tx=XM_019822161.2;refGene_gn=KCNT2;refGene_tx=XM_019822162.2;refGene_gn=KCNT2;refGene_tx=XM_019822163.2;refGene_gn=KCNT2;refGene_tx=XM_019822164.2;refGene_gn=KCNT2;refGene_tx=XM_023248070.1
F1 47384112 47384262
refGene_gn=KCNT2;refGene_tx=XM_011291082.3;refGene_gn=KCNT2;refGene_tx=XM_019822157.2;refGene_gn=KCNT2;refGene_tx=XM_019822158.2;refGene_gn=KCNT2;refGene_tx=XM_019822159.2;refGene_gn=KCNT2;refGene_tx=XM_019822160.2;refGene_gn=KCNT2;refGene_tx=XM_019822161.2;refGene_gn=KCNT2;refGene_tx=XM_019822162.2;refGene_gn=KCNT2;refGene_tx=XM_019822163.2;refGene_gn=KCNT2;refGene_tx=XM_019822164.2;refGene_gn=KCNT2;refGene_tx=XM_023248070.1
F1 47386594 47386824
refGene_gn=KCNT2;refGene_tx=XM_011291082.3;refGene_gn=KCNT2;refGene_tx=XM_019822157.2;refGene_gn=KCNT2;refGene_tx=XM_019822158.2;refGene_gn=KCNT2;refGene_tx=XM_019822159.2;refGene_gn=KCNT2;refGene_tx=XM_019822160.2;refGene_gn=KCNT2;refGene_tx=XM_019822161.2;refGene_gn=KCNT2;refGene_tx=XM_019822162.2;refGene_gn=KCNT2;refGene_tx=XM_019822163.2;refGen

e_gn=KCNT2;refGene_tx=XM_019822164.2;refGene_gn=KCNT2;refGene_tx=XM_02
3248070.1
F1 47394701 47394915
refGene_gn=KCNT2;refGene_tx=XM_011291082.3;refGene_gn=KCNT2;refGene_tx
=XM_019822157.2;refGene_gn=KCNT2;refGene_tx=XM_019822158.2;refGene_gn=
KCNT2;refGene_tx=XM_019822159.2;refGene_gn=KCNT2;refGene_tx=XM_0198221
60.2;refGene_gn=KCNT2;refGene_tx=XM_019822161.2;refGene_gn=KCNT2;refGe
ne_tx=XM_019822162.2;refGene_gn=KCNT2;refGene_tx=XM_019822163.2;refGen
e_gn=KCNT2;refGene_tx=XM_019822164.2;refGene_gn=KCNT2;refGene_tx=XM_02
3248070.1
F1 47397415 47397521
refGene_gn=KCNT2;refGene_tx=XM_011291082.3;refGene_gn=KCNT2;refGene_tx
=XM_019822157.2;refGene_gn=KCNT2;refGene_tx=XM_019822158.2;refGene_gn=
KCNT2;refGene_tx=XM_019822159.2;refGene_gn=KCNT2;refGene_tx=XM_0198221
60.2;refGene_gn=KCNT2;refGene_tx=XM_019822161.2;refGene_gn=KCNT2;refGe
ne_tx=XM_019822162.2;refGene_gn=KCNT2;refGene_tx=XM_019822163.2;refGen
e_gn=KCNT2;refGene_tx=XM_019822164.2;refGene_gn=KCNT2;refGene_tx=XM_02
3248070.1
F1 47401900 47402073
refGene_gn=KCNT2;refGene_tx=XM_011291082.3;refGene_gn=KCNT2;refGene_tx
=XM_019822157.2;refGene_gn=KCNT2;refGene_tx=XM_019822158.2;refGene_gn=
KCNT2;refGene_tx=XM_019822159.2;refGene_gn=KCNT2;refGene_tx=XM_0198221
60.2;refGene_gn=KCNT2;refGene_tx=XM_019822161.2;refGene_gn=KCNT2;refGe
ne_tx=XM_019822162.2;refGene_gn=KCNT2;refGene_tx=XM_019822163.2;refGen
e_gn=KCNT2;refGene_tx=XM_019822164.2;refGene_gn=KCNT2;refGene_tx=XM_02
3248070.1
F1 47411367 47411439
refGene_gn=KCNT2;refGene_tx=XM_019822157.2;refGene_gn=KCNT2;refGene_tx
=XM_019822158.2;refGene_gn=KCNT2;refGene_tx=XM_019822159.2;refGene_gn=
KCNT2;refGene_tx=XM_019822160.2;refGene_gn=KCNT2;refGene_tx=XM_0198221
62.2;refGene_gn=KCNT2;refGene_tx=XM_019822163.2;refGene_gn=KCNT2;refGe
ne_tx=XM_023248070.1
F1 47414823 47414958
refGene_gn=KCNT2;refGene_tx=XM_011291082.3;refGene_gn=KCNT2;refGene_tx
=XM_019822157.2;refGene_gn=KCNT2;refGene_tx=XM_019822158.2;refGene_gn=
KCNT2;refGene_tx=XM_019822159.2;refGene_gn=KCNT2;refGene_tx=XM_0198221
60.2;refGene_gn=KCNT2;refGene_tx=XM_019822161.2;refGene_gn=KCNT2;refGe
ne_tx=XM_019822162.2;refGene_gn=KCNT2;refGene_tx=XM_019822163.2;refGen
e_gn=KCNT2;refGene_tx=XM_019822164.2;refGene_gn=KCNT2;refGene_tx=XM_02
3248070.1
F1 47424965 47425077
refGene_gn=KCNT2;refGene_tx=XM_011291082.3;refGene_gn=KCNT2;refGene_tx
=XM_019822157.2;refGene_gn=KCNT2;refGene_tx=XM_019822158.2;refGene_gn=
KCNT2;refGene_tx=XM_019822159.2;refGene_gn=KCNT2;refGene_tx=XM_0198221
60.2;refGene_gn=KCNT2;refGene_tx=XM_019822161.2;refGene_gn=KCNT2;refGe
ne_tx=XM_019822162.2;refGene_gn=KCNT2;refGene_tx=XM_019822163.2;refGen
e_gn=KCNT2;refGene_tx=XM_019822164.2;refGene_gn=KCNT2;refGene_tx=XM_02
3248070.1
F1 47435572 47435674
refGene_gn=KCNT2;refGene_tx=XM_011291082.3;refGene_gn=KCNT2;refGene_tx

=XM_019822157.2;refGene_gn=KCNT2;refGene_tx=XM_019822158.2;refGene_gn=KCNT2;refGene_tx=XM_019822159.2;refGene_gn=KCNT2;refGene_tx=XM_019822160.2;refGene_gn=KCNT2;refGene_tx=XM_019822161.2;refGene_gn=KCNT2;refGene_tx=XM_019822162.2;refGene_gn=KCNT2;refGene_tx=XM_019822163.2;refGene_gn=KCNT2;refGene_tx=XM_019822164.2;refGene_gn=KCNT2;refGene_tx=XM_023248070.1
F1 47440072 47440156
refGene_gn=KCNT2;refGene_tx=XM_011291082.3;refGene_gn=KCNT2;refGene_tx=XM_019822157.2;refGene_gn=KCNT2;refGene_tx=XM_019822158.2;refGene_gn=KCNT2;refGene_tx=XM_019822159.2;refGene_gn=KCNT2;refGene_tx=XM_019822160.2;refGene_gn=KCNT2;refGene_tx=XM_019822161.2;refGene_gn=KCNT2;refGene_tx=XM_019822162.2;refGene_gn=KCNT2;refGene_tx=XM_019822163.2;refGene_gn=KCNT2;refGene_tx=XM_019822164.2;refGene_gn=KCNT2;refGene_tx=XM_023248070.1
F1 47441051 47441180
refGene_gn=KCNT2;refGene_tx=XM_011291082.3;refGene_gn=KCNT2;refGene_tx=XM_019822157.2;refGene_gn=KCNT2;refGene_tx=XM_019822158.2;refGene_gn=KCNT2;refGene_tx=XM_019822159.2;refGene_gn=KCNT2;refGene_tx=XM_019822160.2;refGene_gn=KCNT2;refGene_tx=XM_019822161.2;refGene_gn=KCNT2;refGene_tx=XM_019822162.2;refGene_gn=KCNT2;refGene_tx=XM_019822163.2;refGene_gn=KCNT2;refGene_tx=XM_019822164.2;refGene_gn=KCNT2;refGene_tx=XM_023248070.1
F1 47449161 47449182
refGene_gn=KCNT2;refGene_tx=XM_019822157.2;refGene_gn=KCNT2;refGene_tx=XM_019822158.2;refGene_gn=KCNT2;refGene_tx=XM_019822161.2;refGene_gn=KCNT2;refGene_tx=XM_019822162.2;refGene_gn=KCNT2;refGene_tx=XM_019822163.2;refGene_gn=KCNT2;refGene_tx=XM_019822164.2;refGene_gn=KCNT2;refGene_tx=XM_023248070.1
F1 47466354 47466655
refGene_gn=KCNT2;refGene_tx=XM_011291082.3;refGene_gn=KCNT2;refGene_tx=XM_019822157.2;refGene_gn=KCNT2;refGene_tx=XM_019822158.2;refGene_gn=KCNT2;refGene_tx=XM_019822159.2;refGene_gn=KCNT2;refGene_tx=XM_019822160.2;refGene_gn=KCNT2;refGene_tx=XM_019822161.2;refGene_gn=KCNT2;refGene_tx=XM_019822162.2;refGene_gn=KCNT2;refGene_tx=XM_019822163.2;refGene_gn=KCNT2;refGene_tx=XM_019822164.2;refGene_gn=KCNT2;refGene_tx=XM_023248070.1
F1 47482166 47482251
refGene_gn=KCNT2;refGene_tx=XM_011291082.3;refGene_gn=KCNT2;refGene_tx=XM_019822157.2;refGene_gn=KCNT2;refGene_tx=XM_019822158.2;refGene_gn=KCNT2;refGene_tx=XM_019822159.2;refGene_gn=KCNT2;refGene_tx=XM_019822160.2;refGene_gn=KCNT2;refGene_tx=XM_019822161.2;refGene_gn=KCNT2;refGene_tx=XM_019822162.2;refGene_gn=KCNT2;refGene_tx=XM_019822163.2;refGene_gn=KCNT2;refGene_tx=XM_019822164.2;refGene_gn=KCNT2;refGene_tx=XM_023248070.1
F1 47491941 47492053
refGene_gn=KCNT2;refGene_tx=XM_011291082.3;refGene_gn=KCNT2;refGene_tx=XM_019822157.2;refGene_gn=KCNT2;refGene_tx=XM_019822158.2;refGene_gn=KCNT2;refGene_tx=XM_019822159.2;refGene_gn=KCNT2;refGene_tx=XM_019822160.2;refGene_gn=KCNT2;refGene_tx=XM_019822161.2;refGene_gn=KCNT2;refGene_tx=XM_019822162.2;refGene_gn=KCNT2;refGene_tx=XM_019822163.2;refGen

e_gn=KCNT2;refGene_tx=XM_019822164.2;refGene_gn=KCNT2;refGene_tx=XM_023248070.1
F1 50008434 50008471 refGene_gn=CDC73;refGene_tx=XM_003999432.5
F1 50009417 50009559 refGene_gn=CDC73;refGene_tx=XM_003999432.5
F1 50015061 50015162 refGene_gn=CDC73;refGene_tx=XM_003999432.5
F1 50018147 50018309 refGene_gn=CDC73;refGene_tx=XM_003999432.5
F1 50035373 50035461 refGene_gn=CDC73;refGene_tx=XM_003999432.5
F1 50035756 50035792 refGene_gn=CDC73;refGene_tx=XM_003999432.5
F1 50044182 50044240 refGene_gn=CDC73;refGene_tx=XM_003999432.5
F1 50069149 50070418 refGene_gn=B3GALT2;refGene_tx=XM_003999433.5
F1 50099036 50099101 refGene_gn=CDC73;refGene_tx=XM_003999432.5
F1 50100776 50100855 refGene_gn=CDC73;refGene_tx=XM_003999432.5
F1 50103867 50103966 refGene_gn=CDC73;refGene_tx=XM_003999432.5
F1 50107555 50107772 refGene_gn=CDC73;refGene_tx=XM_003999432.5
F1 50117136 50117225 refGene_gn=CDC73;refGene_tx=XM_003999432.5
F1 50121512 50121565 refGene_gn=CDC73;refGene_tx=XM_003999432.5
F1 50121647 50121710 refGene_gn=CDC73;refGene_tx=XM_003999432.5
F1 50127407 50127477 refGene_gn=CDC73;refGene_tx=XM_003999432.5
F1 50130367 50130473 refGene_gn=CDC73;refGene_tx=XM_003999432.5
F1 50133109 50133240 refGene_gn=CDC73;refGene_tx=XM_003999432.5
F1 50153901 50154002 refGene_gn=GLRX2;refGene_tx=XM_023247603.1
F1 50157260 50157321 refGene_gn=GLRX2;refGene_tx=XM_003999434.5;refGene_gn=GLRX2;refGene_tx=XM_006942936.4;refGene_gn=GLRX2;refGene_tx=XM_023247603.1
F1 50159950 50160127 refGene_gn=GLRX2;refGene_tx=XM_003999434.5;refGene_gn=GLRX2;refGene_tx=XM_006942936.4;refGene_gn=GLRX2;refGene_tx=XM_023247603.1
F1 50161062 50161197 refGene_gn=GLRX2;refGene_tx=XM_003999434.5;refGene_gn=GLRX2;refGene_tx=XM_006942936.4;refGene_gn=GLRX2;refGene_tx=XM_023247603.1
F1 50171677 50171830 refGene_gn=TROVE2;refGene_tx=XM_003999435.4;refGene_gn=TROVE2;refGene_tx=XM_006942938.3;refGene_gn=TROVE2;refGene_tx=XM_023247959.1
F1 50173396 50173543 refGene_gn=TROVE2;refGene_tx=XM_003999435.4;refGene_gn=TROVE2;refGene_tx=XM_006942938.3;refGene_gn=TROVE2;refGene_tx=XM_023247959.1
F1 50173820 50173934 refGene_gn=TROVE2;refGene_tx=XM_003999435.4;refGene_gn=TROVE2;refGene_tx=XM_006942938.3;refGene_gn=TROVE2;refGene_tx=XM_023247959.1
F1 50174250 50174367 refGene_gn=TROVE2;refGene_tx=XM_003999435.4;refGene_gn=TROVE2;refGene_tx=XM_006942938.3;refGene_gn=TROVE2;refGene_tx=XM_023247959.1
F1 50175137 50175275 refGene_gn=TROVE2;refGene_tx=XM_003999435.4;refGene_gn=TROVE2;refGene_tx=XM_006942938.3;refGene_gn=TROVE2;refGene_tx=XM_023247959.1
F1 50175528 50175675 refGene_gn=TROVE2;refGene_tx=XM_003999435.4;refGene_gn=TROVE2;refGene_tx=XM_006942938.3;refGene_gn=TROVE2;refGene_tx=XM_023247959.1
F1 50176174 50176395 refGene_gn=TROVE2;refGene_tx=XM_003999435.4;refGene_gn=TROVE2;refGene_

tx=XM_006942938.3;refGene_gn=TROVE2;refGene_tx=XM_023247959.1
F1 50180366 50180403 refGene_gn=TROVE2;refGene_tx=XM_023247959.1
F1 50180846 50181446
refGene_gn=TROVE2;refGene_tx=XM_003999435.4;refGene_gn=TROVE2;refGene_tx=XM_006942938.3
F1 50186532 50186539 refGene_gn=TROVE2;refGene_tx=XM_006942938.3
F1 50191398 50191474
refGene_gn=UCHL5;refGene_tx=XM_011291085.3;refGene_gn=UCHL5;refGene_tx=XM_011291086.3
F1 50201124 50201188
refGene_gn=UCHL5;refGene_tx=XM_011291085.3;refGene_gn=UCHL5;refGene_tx=XM_011291086.3
F1 50202134 50202240
refGene_gn=UCHL5;refGene_tx=XM_011291085.3;refGene_gn=UCHL5;refGene_tx=XM_011291086.3
F1 50215852 50215978
refGene_gn=UCHL5;refGene_tx=XM_011291085.3;refGene_gn=UCHL5;refGene_tx=XM_011291086.3
F1 50216069 50216131
refGene_gn=UCHL5;refGene_tx=XM_011291085.3;refGene_gn=UCHL5;refGene_tx=XM_011291086.3
F1 50216206 50216337
refGene_gn=UCHL5;refGene_tx=XM_011291085.3;refGene_gn=UCHL5;refGene_tx=XM_011291086.3
F1 50217143 50217207
refGene_gn=UCHL5;refGene_tx=XM_011291085.3;refGene_gn=UCHL5;refGene_tx=XM_011291086.3
F1 50222011 50222114
refGene_gn=UCHL5;refGene_tx=XM_011291085.3;refGene_gn=UCHL5;refGene_tx=XM_011291086.3
F1 50223204 50223318
refGene_gn=UCHL5;refGene_tx=XM_011291085.3;refGene_gn=UCHL5;refGene_tx=XM_011291086.3
F1 50226072 50226171
refGene_gn=UCHL5;refGene_tx=XM_011291085.3;refGene_gn=UCHL5;refGene_tx=XM_011291086.3
F1 50231928 50231973
refGene_gn=UCHL5;refGene_tx=XM_011291085.3;refGene_gn=UCHL5;refGene_tx=XM_011291086.3
F1 50395576 50395771 refGene_gn=RGS2;refGene_tx=XM_003999491.5
F1 50396036 50396203 refGene_gn=RGS2;refGene_tx=XM_003999491.5
F1 50396764 50396826 refGene_gn=RGS2;refGene_tx=XM_003999491.5
F1 50396926 50397028 refGene_gn=RGS2;refGene_tx=XM_003999491.5
F1 50398071 50398181 refGene_gn=RGS2;refGene_tx=XM_003999491.5
F1 50494426 50494609
refGene_gn=RGS13;refGene_tx=XM_003999492.5;refGene_gn=RGS13;refGene_tx=XM_019822167.2
F1 50495763 50495930
refGene_gn=RGS13;refGene_tx=XM_003999492.5;refGene_gn=RGS13;refGene_tx=XM_019822167.2

F1 50509765 50509827
refGene_gn=RGS13;refGene_tx=XM_003999492.5;refGene_gn=RGS13;refGene_tx
=XM_019822167.2
F1 50513497 50513566
refGene_gn=RGS13;refGene_tx=XM_003999492.5;refGene_gn=RGS13;refGene_tx
=XM_019822167.2
F1 50515588 50515650 refGene_gn=RGS13;refGene_tx=XM_019822167.2
F1 50580519 50580705
refGene_gn=RGS1;refGene_tx=XM_003999493.4;refGene_gn=RGS1;refGene_tx=X
M_006942967.2
F1 50581469 50581633
refGene_gn=RGS1;refGene_tx=XM_003999493.4;refGene_gn=RGS1;refGene_tx=X
M_006942967.2
F1 50582943 50583008
refGene_gn=RGS1;refGene_tx=XM_003999493.4;refGene_gn=RGS1;refGene_tx=X
M_006942967.2
F1 50583428 50583509
refGene_gn=RGS1;refGene_tx=XM_003999493.4;refGene_gn=RGS1;refGene_tx=X
M_006942967.2
F1 50583868 50584005
refGene_gn=RGS1;refGene_tx=XM_003999493.4;refGene_gn=RGS1;refGene_tx=X
M_006942967.2
F1 50778533 50778737 refGene_gn=RGS21;refGene_tx=XM_003999494.1
F1 50792713 50792880 refGene_gn=RGS21;refGene_tx=XM_003999494.1
F1 50797083 50797160 refGene_gn=RGS21;refGene_tx=XM_003999494.1
F1 50801594 50801605 refGene_gn=RGS21;refGene_tx=XM_003999494.1
F1 50965480 50965738 refGene_gn=RGS18;refGene_tx=XM_003999495.4
F1 50967202 50967369 refGene_gn=RGS18;refGene_tx=XM_003999495.4
F1 50984927 50984989 refGene_gn=RGS18;refGene_tx=XM_003999495.4
F1 50986061 50986163 refGene_gn=RGS18;refGene_tx=XM_003999495.4
F1 50986627 50986746 refGene_gn=RGS18;refGene_tx=XM_003999495.4
F1 52478234 52478470
refGene_gn=BRINP3;refGene_tx=XM_019822169.1;refGene_gn=BRINP3;refGene_
tx=XM_019822170.1;refGene_gn=BRINP3;refGene_tx=XM_019822171.2
F1 52650805 52650996
refGene_gn=BRINP3;refGene_tx=XM_019822169.1;refGene_gn=BRINP3;refGene_
tx=XM_019822170.1;refGene_gn=BRINP3;refGene_tx=XM_019822171.2
F1 52674007 52674198
refGene_gn=BRINP3;refGene_tx=XM_019822169.1;refGene_gn=BRINP3;refGene_
tx=XM_019822170.1;refGene_gn=BRINP3;refGene_tx=XM_019822171.2
F1 52731232 52731338
refGene_gn=BRINP3;refGene_tx=XM_019822169.1;refGene_gn=BRINP3;refGene_
tx=XM_019822170.1;refGene_gn=BRINP3;refGene_tx=XM_019822171.2
F1 52739232 52739469
refGene_gn=BRINP3;refGene_tx=XM_019822169.1;refGene_gn=BRINP3;refGene_
tx=XM_019822170.1;refGene_gn=BRINP3;refGene_tx=XM_019822171.2
F1 52790129 52790352
refGene_gn=BRINP3;refGene_tx=XM_019822169.1;refGene_gn=BRINP3;refGene_
tx=XM_019822170.1;refGene_gn=BRINP3;refGene_tx=XM_019822171.2
F1 52862306 52863423

refGene_gn=BRINP3;refGene_tx=XM_019822169.1;refGene_gn=BRINP3;refGene_tx=XM_019822170.1;refGene_gn=BRINP3;refGene_tx=XM_019822171.2
F1 53670698 53670989
refGene_gn=LOC101090736;refGene_tx=XM_011291088.2
F1 53671370 53671403
refGene_gn=LOC101090736;refGene_tx=XM_011291088.2
F1 55368333 55368416 refGene_gn=KCTD3;refGene_tx=XM_023248104.1
F1 55372736 55372790 refGene_gn=KCTD3;refGene_tx=XM_023248104.1
F1 55373054 55373100 refGene_gn=KCTD3;refGene_tx=XM_023248104.1
F1 55374718 55374792 refGene_gn=KCTD3;refGene_tx=XM_023248104.1
F1 55379148 55379207 refGene_gn=KCTD3;refGene_tx=XM_023248104.1
F1 55379484 55379565 refGene_gn=KCTD3;refGene_tx=XM_023248104.1
F1 55380494 55380635 refGene_gn=KCTD3;refGene_tx=XM_023248104.1
F1 55381298 55381389 refGene_gn=KCTD3;refGene_tx=XM_023248104.1
F1 55386316 55386507 refGene_gn=KCTD3;refGene_tx=XM_023248104.1
F1 55397275 55397391 refGene_gn=KCTD3;refGene_tx=XM_023248104.1
F1 55403816 55403904 refGene_gn=KCTD3;refGene_tx=XM_023248104.1
F1 55404029 55404146 refGene_gn=KCTD3;refGene_tx=XM_023248104.1
F1 55406818 55406989 refGene_gn=KCTD3;refGene_tx=XM_023248104.1
F1 55411009 55411165 refGene_gn=KCTD3;refGene_tx=XM_023248104.1
F1 55415189 55415286 refGene_gn=KCTD3;refGene_tx=XM_023248104.1
F1 55418787 55418972 refGene_gn=KCTD3;refGene_tx=XM_023248104.1
F1 55419058 55419197 refGene_gn=KCTD3;refGene_tx=XM_023248104.1
F1 55420225 55420784 refGene_gn=KCTD3;refGene_tx=XM_023248104.1
F1 55426303 55426393
refGene_gn=USH2A;refGene_tx=XM_019822176.1;refGene_gn=USH2A;refGene_tx=XM_023248102.1
F1 55428418 55428640
refGene_gn=USH2A;refGene_tx=XM_019822176.1;refGene_gn=USH2A;refGene_tx=XM_023248103.1
F1 55434706 55434951
refGene_gn=USH2A;refGene_tx=XM_019822176.1;refGene_gn=USH2A;refGene_tx=XM_023248102.1
F1 55439545 55439629
refGene_gn=USH2A;refGene_tx=XM_019822176.1;refGene_gn=USH2A;refGene_tx=XM_023248102.1;refGene_gn=USH2A;refGene_tx=XM_023248103.1
F1 55440948 55441125
refGene_gn=USH2A;refGene_tx=XM_019822176.1;refGene_gn=USH2A;refGene_tx=XM_023248102.1;refGene_gn=USH2A;refGene_tx=XM_023248103.1
F1 55448758 55448967
refGene_gn=USH2A;refGene_tx=XM_019822176.1;refGene_gn=USH2A;refGene_tx=XM_023248102.1;refGene_gn=USH2A;refGene_tx=XM_023248103.1
F1 55449749 55449988
refGene_gn=USH2A;refGene_tx=XM_019822176.1;refGene_gn=USH2A;refGene_tx=XM_023248102.1;refGene_gn=USH2A;refGene_tx=XM_023248103.1
F1 55452889 55453096
refGene_gn=USH2A;refGene_tx=XM_019822176.1;refGene_gn=USH2A;refGene_tx=XM_023248102.1;refGene_gn=USH2A;refGene_tx=XM_023248103.1
F1 55473706 55474028
refGene_gn=USH2A;refGene_tx=XM_019822176.1;refGene_gn=USH2A;refGene_tx

=XM_023248102.1;refGene_gn=USH2A;refGene_tx=XM_023248103.1
F1 55476773 55478290
refGene_gn=USH2A;refGene_tx=XM_019822176.1;refGene_gn=USH2A;refGene_tx=XM_023248102.1;refGene_gn=USH2A;refGene_tx=XM_023248103.1
F1 55482769 55482997
refGene_gn=USH2A;refGene_tx=XM_019822176.1;refGene_gn=USH2A;refGene_tx=XM_023248102.1;refGene_gn=USH2A;refGene_tx=XM_023248103.1
F1 55523678 55524033
refGene_gn=USH2A;refGene_tx=XM_019822176.1;refGene_gn=USH2A;refGene_tx=XM_023248102.1;refGene_gn=USH2A;refGene_tx=XM_023248103.1
F1 55537773 55537936
refGene_gn=USH2A;refGene_tx=XM_019822176.1;refGene_gn=USH2A;refGene_tx=XM_023248102.1;refGene_gn=USH2A;refGene_tx=XM_023248103.1
F1 55539387 55539546
refGene_gn=USH2A;refGene_tx=XM_019822176.1;refGene_gn=USH2A;refGene_tx=XM_023248102.1;refGene_gn=USH2A;refGene_tx=XM_023248103.1
F1 55555308 55555466
refGene_gn=USH2A;refGene_tx=XM_019822176.1;refGene_gn=USH2A;refGene_tx=XM_023248102.1;refGene_gn=USH2A;refGene_tx=XM_023248103.1
F1 55556924 55557108
refGene_gn=USH2A;refGene_tx=XM_019822176.1;refGene_gn=USH2A;refGene_tx=XM_023248102.1;refGene_gn=USH2A;refGene_tx=XM_023248103.1
F1 55564377 55564485
refGene_gn=USH2A;refGene_tx=XM_019822176.1;refGene_gn=USH2A;refGene_tx=XM_023248102.1;refGene_gn=USH2A;refGene_tx=XM_023248103.1
F1 55577709 55577908
refGene_gn=USH2A;refGene_tx=XM_019822176.1;refGene_gn=USH2A;refGene_tx=XM_023248102.1;refGene_gn=USH2A;refGene_tx=XM_023248103.1
F1 55579951 55580106
refGene_gn=USH2A;refGene_tx=XM_019822176.1;refGene_gn=USH2A;refGene_tx=XM_023248102.1;refGene_gn=USH2A;refGene_tx=XM_023248103.1
F1 55580621 55580819
refGene_gn=USH2A;refGene_tx=XM_019822176.1;refGene_gn=USH2A;refGene_tx=XM_023248102.1;refGene_gn=USH2A;refGene_tx=XM_023248103.1
F1 55586470 55586675
refGene_gn=USH2A;refGene_tx=XM_019822176.1;refGene_gn=USH2A;refGene_tx=XM_023248102.1;refGene_gn=USH2A;refGene_tx=XM_023248103.1
F1 55591451 55591675
refGene_gn=USH2A;refGene_tx=XM_019822176.1;refGene_gn=USH2A;refGene_tx=XM_023248102.1;refGene_gn=USH2A;refGene_tx=XM_023248103.1
F1 55603909 55604131
refGene_gn=USH2A;refGene_tx=XM_019822176.1;refGene_gn=USH2A;refGene_tx=XM_023248102.1;refGene_gn=USH2A;refGene_tx=XM_023248103.1
F1 55618276 55618445
refGene_gn=USH2A;refGene_tx=XM_019822176.1;refGene_gn=USH2A;refGene_tx=XM_023248102.1;refGene_gn=USH2A;refGene_tx=XM_023248103.1
F1 55621930 55622129
refGene_gn=USH2A;refGene_tx=XM_019822176.1;refGene_gn=USH2A;refGene_tx=XM_023248102.1;refGene_gn=USH2A;refGene_tx=XM_023248103.1
F1 55635328 55635441

refGene_gn=USH2A;refGene_tx=XM_019822176.1;refGene_gn=USH2A;refGene_tx=XM_023248102.1;refGene_gn=USH2A;refGene_tx=XM_023248103.1
F1 55639844 55640047
refGene_gn=USH2A;refGene_tx=XM_019822176.1;refGene_gn=USH2A;refGene_tx=XM_023248102.1;refGene_gn=USH2A;refGene_tx=XM_023248103.1
F1 55641374 55641584
refGene_gn=USH2A;refGene_tx=XM_019822176.1;refGene_gn=USH2A;refGene_tx=XM_023248102.1;refGene_gn=USH2A;refGene_tx=XM_023248103.1
F1 55652716 55652880
refGene_gn=USH2A;refGene_tx=XM_019822176.1;refGene_gn=USH2A;refGene_tx=XM_023248102.1;refGene_gn=USH2A;refGene_tx=XM_023248103.1
F1 55662947 55663070
refGene_gn=USH2A;refGene_tx=XM_019822176.1;refGene_gn=USH2A;refGene_tx=XM_023248102.1;refGene_gn=USH2A;refGene_tx=XM_023248103.1
F1 55663960 55664295
refGene_gn=USH2A;refGene_tx=XM_019822176.1;refGene_gn=USH2A;refGene_tx=XM_023248102.1;refGene_gn=USH2A;refGene_tx=XM_023248103.1
F1 55675118 55675747
refGene_gn=USH2A;refGene_tx=XM_019822176.1;refGene_gn=USH2A;refGene_tx=XM_023248102.1;refGene_gn=USH2A;refGene_tx=XM_023248103.1
F1 55686809 55686952
refGene_gn=USH2A;refGene_tx=XM_019822176.1;refGene_gn=USH2A;refGene_tx=XM_023248102.1;refGene_gn=USH2A;refGene_tx=XM_023248103.1
F1 55687545 55687696
refGene_gn=USH2A;refGene_tx=XM_019822176.1;refGene_gn=USH2A;refGene_tx=XM_023248102.1;refGene_gn=USH2A;refGene_tx=XM_023248103.1
F1 55721697 55721877
refGene_gn=USH2A;refGene_tx=XM_019822176.1;refGene_gn=USH2A;refGene_tx=XM_023248102.1;refGene_gn=USH2A;refGene_tx=XM_023248103.1
F1 55748999 55749162
refGene_gn=USH2A;refGene_tx=XM_019822176.1;refGene_gn=USH2A;refGene_tx=XM_023248102.1;refGene_gn=USH2A;refGene_tx=XM_023248103.1
F1 55753878 55754030
refGene_gn=USH2A;refGene_tx=XM_019822176.1;refGene_gn=USH2A;refGene_tx=XM_023248102.1;refGene_gn=USH2A;refGene_tx=XM_023248103.1
F1 55775071 55775219
refGene_gn=USH2A;refGene_tx=XM_019822176.1;refGene_gn=USH2A;refGene_tx=XM_023248102.1;refGene_gn=USH2A;refGene_tx=XM_023248103.1
F1 55780707 55780879
refGene_gn=USH2A;refGene_tx=XM_019822176.1;refGene_gn=USH2A;refGene_tx=XM_023248102.1;refGene_gn=USH2A;refGene_tx=XM_023248103.1
F1 55782266 55782426
refGene_gn=USH2A;refGene_tx=XM_019822176.1;refGene_gn=USH2A;refGene_tx=XM_023248102.1;refGene_gn=USH2A;refGene_tx=XM_023248103.1
F1 55822121 55822283
refGene_gn=USH2A;refGene_tx=XM_019822176.1;refGene_gn=USH2A;refGene_tx=XM_023248102.1;refGene_gn=USH2A;refGene_tx=XM_023248103.1
F1 55824248 55824362
refGene_gn=USH2A;refGene_tx=XM_019822176.1;refGene_gn=USH2A;refGene_tx=XM_023248102.1;refGene_gn=USH2A;refGene_tx=XM_023248103.1

F1	55844124	55844319
refGene_gn=USH2A;refGene_tx=XM_019822176.1;refGene_gn=USH2A;refGene_tx=XM_023248102.1;refGene_gn=USH2A;refGene_tx=XM_023248103.1
F1	55847172	55847253
refGene_gn=USH2A;refGene_tx=XM_019822176.1;refGene_gn=USH2A;refGene_tx=XM_023248102.1;refGene_gn=USH2A;refGene_tx=XM_023248103.1
F1	55847380	55847584
refGene_gn=USH2A;refGene_tx=XM_019822176.1;refGene_gn=USH2A;refGene_tx=XM_023248102.1;refGene_gn=USH2A;refGene_tx=XM_023248103.1
F1	55852795	55853069
refGene_gn=USH2A;refGene_tx=XM_019822176.1;refGene_gn=USH2A;refGene_tx=XM_023248102.1;refGene_gn=USH2A;refGene_tx=XM_023248103.1
F1	55858709	55858840
refGene_gn=USH2A;refGene_tx=XM_019822176.1;refGene_gn=USH2A;refGene_tx=XM_023248102.1;refGene_gn=USH2A;refGene_tx=XM_023248103.1
F1	55859861	55860041
refGene_gn=USH2A;refGene_tx=XM_019822176.1;refGene_gn=USH2A;refGene_tx=XM_023248102.1;refGene_gn=USH2A;refGene_tx=XM_023248103.1
F1	55862471	55862573
refGene_gn=USH2A;refGene_tx=XM_019822176.1;refGene_gn=USH2A;refGene_tx=XM_023248102.1;refGene_gn=USH2A;refGene_tx=XM_023248103.1
F1	55864910	55865037
refGene_gn=USH2A;refGene_tx=XM_019822176.1;refGene_gn=USH2A;refGene_tx=XM_023248102.1;refGene_gn=USH2A;refGene_tx=XM_023248103.1
F1	55873327	55873458
refGene_gn=USH2A;refGene_tx=XM_019822176.1;refGene_gn=USH2A;refGene_tx=XM_023248102.1;refGene_gn=USH2A;refGene_tx=XM_023248103.1
F1	55934589	55934820
refGene_gn=USH2A;refGene_tx=XM_019822176.1;refGene_gn=USH2A;refGene_tx=XM_023248102.1;refGene_gn=USH2A;refGene_tx=XM_023248103.1
F1	55947784	55947929
refGene_gn=USH2A;refGene_tx=XM_019822176.1;refGene_gn=USH2A;refGene_tx=XM_023248102.1;refGene_gn=USH2A;refGene_tx=XM_023248103.1
F1	55953901	55954071
refGene_gn=USH2A;refGene_tx=XM_019822176.1;refGene_gn=USH2A;refGene_tx=XM_023248102.1;refGene_gn=USH2A;refGene_tx=XM_023248103.1
F1	55955674	55955944
refGene_gn=USH2A;refGene_tx=XM_019822176.1;refGene_gn=USH2A;refGene_tx=XM_023248102.1;refGene_gn=USH2A;refGene_tx=XM_023248103.1
F1	55956981	55957476
refGene_gn=USH2A;refGene_tx=XM_019822176.1;refGene_gn=USH2A;refGene_tx=XM_023248102.1;refGene_gn=USH2A;refGene_tx=XM_023248103.1
F1	55965142	55965301
refGene_gn=USH2A;refGene_tx=XM_019822176.1;refGene_gn=USH2A;refGene_tx=XM_023248102.1;refGene_gn=USH2A;refGene_tx=XM_023248103.1
F1	55975915	55976079
refGene_gn=USH2A;refGene_tx=XM_019822176.1;refGene_gn=USH2A;refGene_tx=XM_023248102.1;refGene_gn=USH2A;refGene_tx=XM_023248103.1
F1	55990250	55990434
refGene_gn=USH2A;refGene_tx=XM_019822176.1;refGene_gn=USH2A;refGene_tx

=XM_023248102.1;refGene_gn=USH2A;refGene_tx=XM_023248103.1
F1 56002491 56003133
refGene_gn=USH2A;refGene_tx=XM_019822176.1;refGene_gn=USH2A;refGene_tx
=XM_023248102.1;refGene_gn=USH2A;refGene_tx=XM_023248103.1
F1 56006590 56006786
refGene_gn=USH2A;refGene_tx=XM_019822176.1;refGene_gn=USH2A;refGene_tx
=XM_023248102.1;refGene_gn=USH2A;refGene_tx=XM_023248103.1
F1 56043492 56043623
refGene_gn=USH2A;refGene_tx=XM_019822176.1;refGene_gn=USH2A;refGene_tx
=XM_023248102.1;refGene_gn=USH2A;refGene_tx=XM_023248103.1
F1 56047081 56047277
refGene_gn=USH2A;refGene_tx=XM_019822176.1;refGene_gn=USH2A;refGene_tx
=XM_023248102.1;refGene_gn=USH2A;refGene_tx=XM_023248103.1
F1 56071320 56071414
refGene_gn=USH2A;refGene_tx=XM_019822176.1;refGene_gn=USH2A;refGene_tx
=XM_023248102.1;refGene_gn=USH2A;refGene_tx=XM_023248103.1
F1 56072543 56072765
refGene_gn=USH2A;refGene_tx=XM_019822176.1;refGene_gn=USH2A;refGene_tx
=XM_023248102.1;refGene_gn=USH2A;refGene_tx=XM_023248103.1
F1 56073248 56073433
refGene_gn=USH2A;refGene_tx=XM_019822176.1;refGene_gn=USH2A;refGene_tx
=XM_023248102.1;refGene_gn=USH2A;refGene_tx=XM_023248103.1
F1 56074611 56074906
refGene_gn=USH2A;refGene_tx=XM_019822176.1;refGene_gn=USH2A;refGene_tx
=XM_023248102.1;refGene_gn=USH2A;refGene_tx=XM_023248103.1
F1 56077701 56077765
refGene_gn=USH2A;refGene_tx=XM_019822176.1;refGene_gn=USH2A;refGene_tx
=XM_023248102.1;refGene_gn=USH2A;refGene_tx=XM_023248103.1
F1 56102411 56102544
refGene_gn=USH2A;refGene_tx=XM_019822176.1;refGene_gn=USH2A;refGene_tx
=XM_023248102.1;refGene_gn=USH2A;refGene_tx=XM_023248103.1
F1 56155368 56155534
refGene_gn=USH2A;refGene_tx=XM_019822176.1;refGene_gn=USH2A;refGene_tx
=XM_023248102.1;refGene_gn=USH2A;refGene_tx=XM_023248103.1
F1 56159305 56159790
refGene_gn=USH2A;refGene_tx=XM_019822176.1;refGene_gn=USH2A;refGene_tx
=XM_023248102.1;refGene_gn=USH2A;refGene_tx=XM_023248103.1
F1 56245919 56246164
refGene_gn=ESRRG;refGene_tx=XM_011291097.3;refGene_gn=ESRRG;refGene_tx
=XM_019822188.2;refGene_gn=ESRRG;refGene_tx=XM_019822189.2;refGene_gn=
ESRRG;refGene_tx=XM_019822191.2;refGene_gn=ESRRG;refGene_tx=XM_0198221
93.2;refGene_gn=ESRRG;refGene_tx=XM_019822195.2;refGene_gn=ESRRG;refGe
ne_tx=XM_019822196.2;refGene_gn=ESRRG;refGene_tx=XM_019822197.2;refGen
e_gn=ESRRG;refGene_tx=XM_019822198.2;refGene_gn=ESRRG;refGene_tx=XM_01
9822199.2;refGene_gn=ESRRG;refGene_tx=XM_019822200.2
F1 56259741 56260011
refGene_gn=ESRRG;refGene_tx=XM_011291097.3;refGene_gn=ESRRG;refGene_tx
=XM_019822188.2;refGene_gn=ESRRG;refGene_tx=XM_019822189.2;refGene_gn=
ESRRG;refGene_tx=XM_019822191.2;refGene_gn=ESRRG;refGene_tx=XM_0198221
93.2;refGene_gn=ESRRG;refGene_tx=XM_019822195.2;refGene_gn=ESRRG;refGe

ne_tx=XM_019822196.2;refGene_gn=ESRRG;refGene_tx=XM_019822197.2;refGene_gn=ESRRG;refGene_tx=XM_019822198.2;refGene_gn=ESRRG;refGene_tx=XM_019822199.2;refGene_gn=ESRRG;refGene_tx=XM_019822200.2
F1 56304924 56305107
refGene_gn=ESRRG;refGene_tx=XM_011291097.3;refGene_gn=ESRRG;refGene_tx=XM_019822188.2;refGene_gn=ESRRG;refGene_tx=XM_019822189.2;refGene_gn=ESRRG;refGene_tx=XM_019822191.2;refGene_gn=ESRRG;refGene_tx=XM_019822193.2;refGene_gn=ESRRG;refGene_tx=XM_019822195.2;refGene_gn=ESRRG;refGene_tx=XM_019822196.2;refGene_gn=ESRRG;refGene_tx=XM_019822197.2;refGene_gn=ESRRG;refGene_tx=XM_019822198.2;refGene_gn=ESRRG;refGene_tx=XM_019822199.2;refGene_gn=ESRRG;refGene_tx=XM_019822200.2
F1 56308725 56308836
refGene_gn=ESRRG;refGene_tx=XM_011291097.3;refGene_gn=ESRRG;refGene_tx=XM_019822188.2;refGene_gn=ESRRG;refGene_tx=XM_019822189.2;refGene_gn=ESRRG;refGene_tx=XM_019822191.2;refGene_gn=ESRRG;refGene_tx=XM_019822193.2;refGene_gn=ESRRG;refGene_tx=XM_019822195.2;refGene_gn=ESRRG;refGene_tx=XM_019822196.2;refGene_gn=ESRRG;refGene_tx=XM_019822197.2;refGene_gn=ESRRG;refGene_tx=XM_019822198.2;refGene_gn=ESRRG;refGene_tx=XM_019822199.2;refGene_gn=ESRRG;refGene_tx=XM_019822200.2
F1 56399480 56399597
refGene_gn=ESRRG;refGene_tx=XM_011291097.3;refGene_gn=ESRRG;refGene_tx=XM_019822188.2;refGene_gn=ESRRG;refGene_tx=XM_019822189.2;refGene_gn=ESRRG;refGene_tx=XM_019822191.2;refGene_gn=ESRRG;refGene_tx=XM_019822193.2;refGene_gn=ESRRG;refGene_tx=XM_019822195.2;refGene_gn=ESRRG;refGene_tx=XM_019822196.2;refGene_gn=ESRRG;refGene_tx=XM_019822197.2;refGene_gn=ESRRG;refGene_tx=XM_019822198.2;refGene_gn=ESRRG;refGene_tx=XM_019822199.2;refGene_gn=ESRRG;refGene_tx=XM_019822200.2
F1 56424742 56425158
refGene_gn=ESRRG;refGene_tx=XM_011291097.3;refGene_gn=ESRRG;refGene_tx=XM_019822188.2;refGene_gn=ESRRG;refGene_tx=XM_019822189.2;refGene_gn=ESRRG;refGene_tx=XM_019822191.2;refGene_gn=ESRRG;refGene_tx=XM_019822193.2;refGene_gn=ESRRG;refGene_tx=XM_019822195.2;refGene_gn=ESRRG;refGene_tx=XM_019822196.2;refGene_gn=ESRRG;refGene_tx=XM_019822197.2;refGene_gn=ESRRG;refGene_tx=XM_019822198.2;refGene_gn=ESRRG;refGene_tx=XM_019822199.2;refGene_gn=ESRRG;refGene_tx=XM_019822200.2
F1 56461645 56461650 refGene_gn=ESRRG;refGene_tx=XM_019822189.2
F1 56470280 56470336
refGene_gn=ESRRG;refGene_tx=XM_011291097.3;refGene_gn=ESRRG;refGene_tx=XM_019822188.2
F1 57125273 57125494 refGene_gn=GPATCH2;refGene_tx=XM_006942970.3
F1 57126159 57126173 refGene_gn=GPATCH2;refGene_tx=XM_019822201.2
F1 57137450 57137539
refGene_gn=GPATCH2;refGene_tx=XM_006942970.3;refGene_gn=GPATCH2;refGene_tx=XM_019822201.2
F1 57147469 57147554 refGene_gn=GPATCH2;refGene_tx=XM_019822202.2
F1 57167107 57167178
refGene_gn=GPATCH2;refGene_tx=XM_006942970.3;refGene_gn=GPATCH2;refGene_tx=XM_019822201.2;refGene_gn=GPATCH2;refGene_tx=XM_019822202.2
F1 57174673 57174713
refGene_gn=GPATCH2;refGene_tx=XM_006942970.3;refGene_gn=GPATCH2;refGen

e_tx=XM_019822201.2;refGene_gn=GPATCH2;refGene_tx=XM_019822202.2
F1 57193230 57193287
refGene_gn=GPATCH2;refGene_tx=XM_006942970.3;refGene_gn=GPATCH2;refGene_tx=XM_019822201.2;refGene_gn=GPATCH2;refGene_tx=XM_019822202.2
F1 57226738 57226745
refGene_gn=GPATCH2;refGene_tx=XM_006942970.3;refGene_gn=GPATCH2;refGene_tx=XM_019822201.2;refGene_gn=GPATCH2;refGene_tx=XM_019822202.2
F1 57280716 57280749 refGene_gn=GPATCH2;refGene_tx=XM_011291145.2
F1 57280872 57280956
refGene_gn=GPATCH2;refGene_tx=XM_006942970.3;refGene_gn=GPATCH2;refGene_tx=XM_011291145.2;refGene_gn=GPATCH2;refGene_tx=XM_019822201.2;refGene_gn=GPATCH2;refGene_tx=XM_019822202.2
F1 57281472 57281655
refGene_gn=GPATCH2;refGene_tx=XM_006942970.3;refGene_gn=GPATCH2;refGene_tx=XM_011291145.2;refGene_gn=GPATCH2;refGene_tx=XM_019822201.2;refGene_gn=GPATCH2;refGene_tx=XM_019822202.2
F1 57284682 57284744
refGene_gn=GPATCH2;refGene_tx=XM_006942970.3;refGene_gn=GPATCH2;refGene_tx=XM_011291145.2;refGene_gn=GPATCH2;refGene_tx=XM_019822201.2;refGene_gn=GPATCH2;refGene_tx=XM_019822202.2
F1 57289365 57290082
refGene_gn=GPATCH2;refGene_tx=XM_006942970.3;refGene_gn=GPATCH2;refGene_tx=XM_011291145.2;refGene_gn=GPATCH2;refGene_tx=XM_019822201.2;refGene_gn=GPATCH2;refGene_tx=XM_019822202.2
F1 57300846 57300902
refGene_gn=GPATCH2;refGene_tx=XM_006942970.3;refGene_gn=GPATCH2;refGene_tx=XM_011291145.2;refGene_gn=GPATCH2;refGene_tx=XM_019822201.2;refGene_gn=GPATCH2;refGene_tx=XM_019822202.2
F1 57301312 57301380
refGene_gn=SPATA17;refGene_tx=XM_003999504.5;refGene_gn=SPATA17;refGene_tx=XM_019822203.2;refGene_gn=SPATA17;refGene_tx=XM_019822204.2;refGene_gn=SPATA17;refGene_tx=XM_019822205.2
F1 57313971 57314061
refGene_gn=SPATA17;refGene_tx=XM_003999504.5;refGene_gn=SPATA17;refGene_tx=XM_019822203.2;refGene_gn=SPATA17;refGene_tx=XM_019822204.2;refGene_gn=SPATA17;refGene_tx=XM_019822205.2
F1 57315185 57315267
refGene_gn=SPATA17;refGene_tx=XM_003999504.5;refGene_gn=SPATA17;refGene_tx=XM_019822203.2;refGene_gn=SPATA17;refGene_tx=XM_019822204.2;refGene_gn=SPATA17;refGene_tx=XM_019822205.2
F1 57330158 57330209
refGene_gn=SPATA17;refGene_tx=XM_003999504.5;refGene_gn=SPATA17;refGene_tx=XM_019822203.2;refGene_gn=SPATA17;refGene_tx=XM_019822204.2
F1 57337482 57337494 refGene_gn=SPATA17;refGene_tx=XM_019822207.1
F1 57340701 57340805
refGene_gn=SPATA17;refGene_tx=XM_003999504.5;refGene_gn=SPATA17;refGene_tx=XM_019822203.2;refGene_gn=SPATA17;refGene_tx=XM_019822204.2;refGene_gn=SPATA17;refGene_tx=XM_019822205.2;refGene_gn=SPATA17;refGene_tx=XM_019822207.1
F1 57343876 57344035

refGene_gn=SPATA17;refGene_tx=XM_019822203.2;refGene_gn=SPATA17;refGene_tx=XM_019822204.2;refGene_gn=SPATA17;refGene_tx=XM_019822205.2;refGene_gn=SPATA17;refGene_tx=XM_019822207.1
F1 57380318 57380442
refGene_gn=SPATA17;refGene_tx=XM_003999504.5;refGene_gn=SPATA17;refGene_tx=XM_019822203.2;refGene_gn=SPATA17;refGene_tx=XM_019822204.2;refGene_gn=SPATA17;refGene_tx=XM_019822205.2;refGene_gn=SPATA17;refGene_tx=XM_019822207.1
F1 57406870 57407074
refGene_gn=SPATA17;refGene_tx=XM_003999504.5;refGene_gn=SPATA17;refGene_tx=XM_019822203.2;refGene_gn=SPATA17;refGene_tx=XM_019822204.2;refGene_gn=SPATA17;refGene_tx=XM_019822205.2;refGene_gn=SPATA17;refGene_tx=XM_019822207.1
F1 57414176 57414325
refGene_gn=SPATA17;refGene_tx=XM_003999504.5;refGene_gn=SPATA17;refGene_tx=XM_019822203.2;refGene_gn=SPATA17;refGene_tx=XM_019822204.2;refGene_gn=SPATA17;refGene_tx=XM_019822205.2;refGene_gn=SPATA17;refGene_tx=XM_019822207.1
F1 57441110 57441246
refGene_gn=SPATA17;refGene_tx=XM_003999504.5;refGene_gn=SPATA17;refGene_tx=XM_019822203.2;refGene_gn=SPATA17;refGene_tx=XM_019822204.2;refGene_gn=SPATA17;refGene_tx=XM_019822205.2;refGene_gn=SPATA17;refGene_tx=XM_019822207.1
F1 57491981 57492062
refGene_gn=SPATA17;refGene_tx=XM_003999504.5;refGene_gn=SPATA17;refGene_tx=XM_019822203.2;refGene_gn=SPATA17;refGene_tx=XM_019822204.2;refGene_gn=SPATA17;refGene_tx=XM_019822205.2;refGene_gn=SPATA17;refGene_tx=XM_019822207.1
F1 57847794 57847939 refGene_gn=RRP15;refGene_tx=XM_003999505.4
F1 57860223 57860498 refGene_gn=RRP15;refGene_tx=XM_003999505.4
F1 57863227 57863325 refGene_gn=RRP15;refGene_tx=XM_003999505.4
F1 57865704 57865906 refGene_gn=RRP15;refGene_tx=XM_003999505.4
F1 57887545 57887689 refGene_gn=RRP15;refGene_tx=XM_003999505.4
F1 57903508 57903854
refGene_gn=TGFB2;refGene_tx=XM_003999506.5;refGene_gn=TGFB2;refGene_tx=XM_003999507.5;refGene_gn=TGFB2;refGene_tx=XM_006942972.4
F1 57918190 57918274
refGene_gn=TGFB2;refGene_tx=XM_003999507.5;refGene_gn=TGFB2;refGene_tx=XM_006942972.4
F1 57953462 57953626
refGene_gn=TGFB2;refGene_tx=XM_003999506.5;refGene_gn=TGFB2;refGene_tx=XM_003999507.5;refGene_gn=TGFB2;refGene_tx=XM_006942972.4
F1 57976529 57976662
refGene_gn=TGFB2;refGene_tx=XM_003999506.5;refGene_gn=TGFB2;refGene_tx=XM_003999507.5;refGene_gn=TGFB2;refGene_tx=XM_006942972.4
F1 57976785 57976896
refGene_gn=TGFB2;refGene_tx=XM_003999506.5;refGene_gn=TGFB2;refGene_tx=XM_003999507.5;refGene_gn=TGFB2;refGene_tx=XM_006942972.4
F1 57978397 57978575
refGene_gn=TGFB2;refGene_tx=XM_003999506.5;refGene_gn=TGFB2;refGene_tx

=XM_003999507.5;refGene_gn=TGFB2;refGene_tx=XM_006942972.4
F1	57979774	57979928
refGene_gn=TGFB2;refGene_tx=XM_003999506.5;refGene_gn=TGFB2;refGene_tx
=XM_003999507.5;refGene_gn=TGFB2;refGene_tx=XM_006942972.4
F1	57983277	57983436
refGene_gn=TGFB2;refGene_tx=XM_003999506.5;refGene_gn=TGFB2;refGene_tx
=XM_003999507.5;refGene_gn=TGFB2;refGene_tx=XM_006942972.4
F1	58556677	58556771	refGene_gn=LYPLAL1;refGene_tx=XM_003999509.4
F1	58562206	58562306	refGene_gn=LYPLAL1;refGene_tx=XM_003999509.4
F1	58580051	58580221	refGene_gn=LYPLAL1;refGene_tx=XM_003999509.4
F1	58593002	58593118	refGene_gn=LYPLAL1;refGene_tx=XM_003999509.4
F1	58593897	58594137	refGene_gn=LYPLAL1;refGene_tx=XM_003999509.4
F1	59190569	59191069	refGene_gn=SLC30A10;refGene_tx=XM_023247956.1
F1	59194178	59194418	refGene_gn=SLC30A10;refGene_tx=XM_023247956.1
F1	59198641	59198719	refGene_gn=SLC30A10;refGene_tx=XM_023247956.1
F1	59199441	59199572	refGene_gn=SLC30A10;refGene_tx=XM_023247956.1
F1	59199573	59200037	refGene_gn=SLC30A10;refGene_tx=XM_023247956.1
F1	59233677	59233828
refGene_gn=EPRS;refGene_tx=XM_011291144.3;refGene_gn=EPRS;refGene_tx=X
M_019822211.2;refGene_gn=EPRS;refGene_tx=XM_019822212.2
F1	59233925	59233990
refGene_gn=EPRS;refGene_tx=XM_011291144.3;refGene_gn=EPRS;refGene_tx=X
M_019822211.2;refGene_gn=EPRS;refGene_tx=XM_019822212.2
F1	59239700	59239779
refGene_gn=EPRS;refGene_tx=XM_011291144.3;refGene_gn=EPRS;refGene_tx=X
M_019822211.2;refGene_gn=EPRS;refGene_tx=XM_019822212.2
F1	59242147	59242308
refGene_gn=EPRS;refGene_tx=XM_011291144.3;refGene_gn=EPRS;refGene_tx=X
M_019822211.2;refGene_gn=EPRS;refGene_tx=XM_019822212.2
F1	59246918	59247092
refGene_gn=EPRS;refGene_tx=XM_011291144.3;refGene_gn=EPRS;refGene_tx=X
M_019822211.2;refGene_gn=EPRS;refGene_tx=XM_019822212.2
F1	59247569	59247767
refGene_gn=EPRS;refGene_tx=XM_011291144.3;refGene_gn=EPRS;refGene_tx=X
M_019822211.2;refGene_gn=EPRS;refGene_tx=XM_019822212.2
F1	59248251	59248407
refGene_gn=EPRS;refGene_tx=XM_011291144.3;refGene_gn=EPRS;refGene_tx=X
M_019822211.2;refGene_gn=EPRS;refGene_tx=XM_019822212.2
F1	59249428	59249530
refGene_gn=EPRS;refGene_tx=XM_011291144.3;refGene_gn=EPRS;refGene_tx=X
M_019822211.2;refGene_gn=EPRS;refGene_tx=XM_019822212.2
F1	59249760	59249840
refGene_gn=EPRS;refGene_tx=XM_011291144.3;refGene_gn=EPRS;refGene_tx=X
M_019822211.2;refGene_gn=EPRS;refGene_tx=XM_019822212.2
F1	59251333	59251406
refGene_gn=EPRS;refGene_tx=XM_011291144.3;refGene_gn=EPRS;refGene_tx=X
M_019822211.2;refGene_gn=EPRS;refGene_tx=XM_019822212.2
F1	59251793	59252003
refGene_gn=EPRS;refGene_tx=XM_011291144.3;refGene_gn=EPRS;refGene_tx=X
M_019822211.2;refGene_gn=EPRS;refGene_tx=XM_019822212.2

F1 59252907 59252959
refGene_gn=EPRS;refGene_tx=XM_011291144.3;refGene_gn=EPRS;refGene_tx=XM_019822211.2;refGene_gn=EPRS;refGene_tx=XM_019822212.2
F1 59255279 59255542
refGene_gn=EPRS;refGene_tx=XM_011291144.3;refGene_gn=EPRS;refGene_tx=XM_019822211.2;refGene_gn=EPRS;refGene_tx=XM_019822212.2
F1 59256580 59256814
refGene_gn=EPRS;refGene_tx=XM_011291144.3;refGene_gn=EPRS;refGene_tx=XM_019822211.2;refGene_gn=EPRS;refGene_tx=XM_019822212.2
F1 59264205 59264562
refGene_gn=EPRS;refGene_tx=XM_011291144.3;refGene_gn=EPRS;refGene_tx=XM_019822211.2;refGene_gn=EPRS;refGene_tx=XM_019822212.2
F1 59269780 59269898
refGene_gn=EPRS;refGene_tx=XM_011291144.3;refGene_gn=EPRS;refGene_tx=XM_019822211.2;refGene_gn=EPRS;refGene_tx=XM_019822212.2
F1 59273932 59274045
refGene_gn=EPRS;refGene_tx=XM_011291144.3;refGene_gn=EPRS;refGene_tx=XM_019822211.2;refGene_gn=EPRS;refGene_tx=XM_019822212.2
F1 59274627 59274835
refGene_gn=EPRS;refGene_tx=XM_011291144.3;refGene_gn=EPRS;refGene_tx=XM_019822211.2;refGene_gn=EPRS;refGene_tx=XM_019822212.2
F1 59275583 59275720
refGene_gn=EPRS;refGene_tx=XM_011291144.3;refGene_gn=EPRS;refGene_tx=XM_019822211.2;refGene_gn=EPRS;refGene_tx=XM_019822212.2
F1 59277743 59277854
refGene_gn=EPRS;refGene_tx=XM_011291144.3;refGene_gn=EPRS;refGene_tx=XM_019822211.2;refGene_gn=EPRS;refGene_tx=XM_019822212.2
F1 59282941 59282962
refGene_gn=EPRS;refGene_tx=XM_019822211.2;refGene_gn=EPRS;refGene_tx=XM_019822212.2
F1 59283262 59283322
refGene_gn=EPRS;refGene_tx=XM_011291144.3;refGene_gn=EPRS;refGene_tx=XM_019822211.2;refGene_gn=EPRS;refGene_tx=XM_019822212.2
F1 59283747 59283832
refGene_gn=EPRS;refGene_tx=XM_011291144.3;refGene_gn=EPRS;refGene_tx=XM_019822211.2;refGene_gn=EPRS;refGene_tx=XM_019822212.2
F1 59285122 59285356
refGene_gn=EPRS;refGene_tx=XM_011291144.3;refGene_gn=EPRS;refGene_tx=XM_019822211.2;refGene_gn=EPRS;refGene_tx=XM_019822212.2
F1 59287102 59287274
refGene_gn=EPRS;refGene_tx=XM_011291144.3;refGene_gn=EPRS;refGene_tx=XM_019822211.2;refGene_gn=EPRS;refGene_tx=XM_019822212.2
F1 59289039 59289232
refGene_gn=EPRS;refGene_tx=XM_011291144.3;refGene_gn=EPRS;refGene_tx=XM_019822211.2;refGene_gn=EPRS;refGene_tx=XM_019822212.2
F1 59289927 59290054
refGene_gn=EPRS;refGene_tx=XM_011291144.3;refGene_gn=EPRS;refGene_tx=XM_019822211.2
F1 59294082 59294177
refGene_gn=EPRS;refGene_tx=XM_011291144.3;refGene_gn=EPRS;refGene_tx=X

M_019822211.2
F1 59295462 59295602
refGene_gn=EPRS;refGene_tx=XM_011291144.3;refGene_gn=EPRS;refGene_tx=X
M_019822211.2
F1 59297119 59297276
refGene_gn=EPRS;refGene_tx=XM_011291144.3;refGene_gn=EPRS;refGene_tx=X
M_019822211.2
F1 59298153 59298253
refGene_gn=EPRS;refGene_tx=XM_011291144.3;refGene_gn=EPRS;refGene_tx=X
M_019822211.2
F1 59301804 59301889
refGene_gn=EPRS;refGene_tx=XM_011291144.3;refGene_gn=EPRS;refGene_tx=X
M_019822211.2
F1 59305809 59305855
refGene_gn=EPRS;refGene_tx=XM_011291144.3;refGene_gn=EPRS;refGene_tx=X
M_019822211.2
F1 59318542 59318691
refGene_gn=BPNT1;refGene_tx=XM_006942973.4;refGene_gn=BPNT1;refGene_tx
=XM_006942974.4;refGene_gn=BPNT1;refGene_tx=XM_006942975.4;refGene_gn=
BPNT1;refGene_tx=XM_006942976.4;refGene_gn=BPNT1;refGene_tx=XM_0198222
14.2
F1 59320392 59320498
refGene_gn=BPNT1;refGene_tx=XM_006942973.4;refGene_gn=BPNT1;refGene_tx
=XM_006942974.4;refGene_gn=BPNT1;refGene_tx=XM_006942975.4;refGene_gn=
BPNT1;refGene_tx=XM_006942976.4;refGene_gn=BPNT1;refGene_tx=XM_0198222
14.2
F1 59322826 59323024
refGene_gn=BPNT1;refGene_tx=XM_006942973.4;refGene_gn=BPNT1;refGene_tx
=XM_006942974.4;refGene_gn=BPNT1;refGene_tx=XM_006942975.4;refGene_gn=
BPNT1;refGene_tx=XM_006942976.4;refGene_gn=BPNT1;refGene_tx=XM_0198222
14.2
F1 59325601 59325646
refGene_gn=BPNT1;refGene_tx=XM_006942973.4;refGene_gn=BPNT1;refGene_tx
=XM_006942974.4;refGene_gn=BPNT1;refGene_tx=XM_006942975.4;refGene_gn=
BPNT1;refGene_tx=XM_006942976.4
F1 59326757 59326849
refGene_gn=BPNT1;refGene_tx=XM_006942973.4;refGene_gn=BPNT1;refGene_tx
=XM_006942974.4;refGene_gn=BPNT1;refGene_tx=XM_006942975.4;refGene_gn=
BPNT1;refGene_tx=XM_006942976.4;refGene_gn=BPNT1;refGene_tx=XM_0198222
14.2
F1 59328819 59328868
refGene_gn=BPNT1;refGene_tx=XM_006942973.4;refGene_gn=BPNT1;refGene_tx
=XM_006942974.4;refGene_gn=BPNT1;refGene_tx=XM_006942975.4;refGene_gn=
BPNT1;refGene_tx=XM_006942976.4;refGene_gn=BPNT1;refGene_tx=XM_0198222
14.2
F1 59330314 59330422
refGene_gn=BPNT1;refGene_tx=XM_006942973.4;refGene_gn=BPNT1;refGene_tx
=XM_006942974.4;refGene_gn=BPNT1;refGene_tx=XM_006942975.4;refGene_gn=
BPNT1;refGene_tx=XM_006942976.4;refGene_gn=BPNT1;refGene_tx=XM_0198222
14.2

F1 59331164 59331269
refGene_gn=BPNT1;refGene_tx=XM_006942973.4;refGene_gn=BPNT1;refGene_tx=XM_006942974.4;refGene_gn=BPNT1;refGene_tx=XM_006942975.4;refGene_gn=BPNT1;refGene_tx=XM_006942976.4;refGene_gn=BPNT1;refGene_tx=XM_019822214.2
F1 59333875 59333995
refGene_gn=BPNT1;refGene_tx=XM_006942973.4;refGene_gn=BPNT1;refGene_tx=XM_006942974.4;refGene_gn=BPNT1;refGene_tx=XM_006942975.4;refGene_gn=BPNT1;refGene_tx=XM_006942976.4;refGene_gn=BPNT1;refGene_tx=XM_019822214.2
F1 59343314 59343581 refGene_gn=IARS2;refGene_tx=XM_023247462.1
F1 59345030 59345153 refGene_gn=IARS2;refGene_tx=XM_023247462.1
F1 59362955 59363115 refGene_gn=IARS2;refGene_tx=XM_023247462.1
F1 59364162 59364311 refGene_gn=IARS2;refGene_tx=XM_023247462.1
F1 59364393 59364443 refGene_gn=IARS2;refGene_tx=XM_023247462.1
F1 59364524 59364634 refGene_gn=IARS2;refGene_tx=XM_023247462.1
F1 59364959 59365050 refGene_gn=IARS2;refGene_tx=XM_023247462.1
F1 59365377 59365493 refGene_gn=IARS2;refGene_tx=XM_023247462.1
F1 59369437 59369673 refGene_gn=IARS2;refGene_tx=XM_023247462.1
F1 59374014 59374170 refGene_gn=IARS2;refGene_tx=XM_023247462.1
F1 59376251 59376412 refGene_gn=IARS2;refGene_tx=XM_023247462.1
F1 59386198 59386301 refGene_gn=IARS2;refGene_tx=XM_023247462.1
F1 59387143 59387237 refGene_gn=IARS2;refGene_tx=XM_023247462.1
F1 59397181 59397290 refGene_gn=IARS2;refGene_tx=XM_023247462.1
F1 59399779 59399882 refGene_gn=IARS2;refGene_tx=XM_023247462.1
F1 59400878 59401004 refGene_gn=IARS2;refGene_tx=XM_023247462.1
F1 59401578 59401710 refGene_gn=IARS2;refGene_tx=XM_023247462.1
F1 59402829 59402936 refGene_gn=IARS2;refGene_tx=XM_023247462.1
F1 59403721 59403867 refGene_gn=IARS2;refGene_tx=XM_023247462.1
F1 59404576 59404767 refGene_gn=IARS2;refGene_tx=XM_023247462.1
F1 59406676 59406821 refGene_gn=IARS2;refGene_tx=XM_023247462.1
F1 59408661 59408807 refGene_gn=IARS2;refGene_tx=XM_023247462.1
F1 59412239 59412395 refGene_gn=RAB3GAP2;refGene_tx=XM_011291141.3
F1 59412815 59412854 refGene_gn=RAB3GAP2;refGene_tx=XM_011291141.3
F1 59412855 59412975 refGene_gn=RAB3GAP2;refGene_tx=XM_011291141.3
F1 59413869 59414091 refGene_gn=RAB3GAP2;refGene_tx=XM_011291141.3
F1 59414611 59414701 refGene_gn=RAB3GAP2;refGene_tx=XM_011291141.3
F1 59417748 59417967 refGene_gn=RAB3GAP2;refGene_tx=XM_011291141.3
F1 59418464 59418539 refGene_gn=RAB3GAP2;refGene_tx=XM_011291141.3
F1 59419754 59419790 refGene_gn=RAB3GAP2;refGene_tx=XM_011291141.3
F1 59422550 59422621 refGene_gn=RAB3GAP2;refGene_tx=XM_011291141.3
F1 59424805 59424872 refGene_gn=RAB3GAP2;refGene_tx=XM_011291141.3
F1 59427859 59427966 refGene_gn=RAB3GAP2;refGene_tx=XM_011291141.3
F1 59428070 59428244 refGene_gn=RAB3GAP2;refGene_tx=XM_011291141.3
F1 59433056 59433285 refGene_gn=RAB3GAP2;refGene_tx=XM_011291141.3
F1 59434356 59434517 refGene_gn=RAB3GAP2;refGene_tx=XM_011291141.3
F1 59435370 59435476 refGene_gn=RAB3GAP2;refGene_tx=XM_011291141.3
F1 59442589 59442687 refGene_gn=RAB3GAP2;refGene_tx=XM_011291141.3
F1 59443069 59443286 refGene_gn=RAB3GAP2;refGene_tx=XM_011291141.3
F1 59445561 59445689 refGene_gn=RAB3GAP2;refGene_tx=XM_011291141.3

F1 59448361 59448452 refGene_gn=RAB3GAP2;refGene_tx=XM_011291141.3
F1 59451161 59451217 refGene_gn=RAB3GAP2;refGene_tx=XM_011291141.3
F1 59451219 59451225 refGene_gn=RAB3GAP2;refGene_tx=XM_011291141.3
F1 59451773 59451856 refGene_gn=RAB3GAP2;refGene_tx=XM_011291141.3
F1 59452307 59452451 refGene_gn=RAB3GAP2;refGene_tx=XM_011291141.3
F1 59453016 59453233 refGene_gn=RAB3GAP2;refGene_tx=XM_011291141.3
F1 59455012 59455152 refGene_gn=RAB3GAP2;refGene_tx=XM_011291141.3
F1 59457516 59457606 refGene_gn=RAB3GAP2;refGene_tx=XM_011291141.3
F1 59457735 59457815 refGene_gn=RAB3GAP2;refGene_tx=XM_011291141.3
F1 59458657 59458806 refGene_gn=RAB3GAP2;refGene_tx=XM_011291141.3
F1 59465548 59465647 refGene_gn=RAB3GAP2;refGene_tx=XM_011291141.3
F1 59467616 59467716 refGene_gn=RAB3GAP2;refGene_tx=XM_011291141.3
F1 59471752 59471854 refGene_gn=RAB3GAP2;refGene_tx=XM_011291141.3
F1 59472181 59472257 refGene_gn=RAB3GAP2;refGene_tx=XM_011291141.3
F1 59472341 59472389 refGene_gn=RAB3GAP2;refGene_tx=XM_011291141.3
F1 59474184 59474266 refGene_gn=RAB3GAP2;refGene_tx=XM_011291141.3
F1 59475348 59475472 refGene_gn=RAB3GAP2;refGene_tx=XM_011291141.3
F1 59491465 59491530 refGene_gn=RAB3GAP2;refGene_tx=XM_011291141.3
F1 59511739 59511854 refGene_gn=RAB3GAP2;refGene_tx=XM_011291141.3
F1 59732859 59732910
refGene_gn=MARK1;refGene_tx=XM_023247971.1;refGene_gn=MARK1;refGene_tx
=XM_023247972.1;refGene_gn=MARK1;refGene_tx=XM_023247973.1
F1 59781563 59781764
refGene_gn=MARK1;refGene_tx=XM_023247971.1;refGene_gn=MARK1;refGene_tx
=XM_023247972.1;refGene_gn=MARK1;refGene_tx=XM_023247973.1
F1 59782917 59782971
refGene_gn=MARK1;refGene_tx=XM_023247971.1;refGene_gn=MARK1;refGene_tx
=XM_023247972.1;refGene_gn=MARK1;refGene_tx=XM_023247973.1
F1 59801039 59801060 refGene_gn=MARK1;refGene_tx=XM_023247973.1
F1 59801133 59801182
refGene_gn=MARK1;refGene_tx=XM_023247971.1;refGene_gn=MARK1;refGene_tx
=XM_023247972.1
F1 59802181 59802247
refGene_gn=MARK1;refGene_tx=XM_023247971.1;refGene_gn=MARK1;refGene_tx
=XM_023247972.1
F1 59806000 59806071
refGene_gn=MARK1;refGene_tx=XM_023247971.1;refGene_gn=MARK1;refGene_tx
=XM_023247972.1
F1 59811126 59811183
refGene_gn=MARK1;refGene_tx=XM_023247971.1;refGene_gn=MARK1;refGene_tx
=XM_023247972.1
F1 59814802 59815039
refGene_gn=MARK1;refGene_tx=XM_023247971.1;refGene_gn=MARK1;refGene_tx
=XM_023247972.1
F1 59815125 59815245
refGene_gn=MARK1;refGene_tx=XM_023247971.1;refGene_gn=MARK1;refGene_tx
=XM_023247972.1
F1 59825752 59825852
refGene_gn=MARK1;refGene_tx=XM_023247971.1;refGene_gn=MARK1;refGene_tx
=XM_023247972.1

F1 59826889 59827002
refGene_gn=MARK1;refGene_tx=XM_023247971.1;refGene_gn=MARK1;refGene_tx=XM_023247972.1
F1 59829512 59829666
refGene_gn=MARK1;refGene_tx=XM_023247971.1;refGene_gn=MARK1;refGene_tx=XM_023247972.1
F1 59829965 59830159
refGene_gn=MARK1;refGene_tx=XM_023247971.1;refGene_gn=MARK1;refGene_tx=XM_023247972.1
F1 59836813 59836914
refGene_gn=MARK1;refGene_tx=XM_023247971.1;refGene_gn=MARK1;refGene_tx=XM_023247972.1
F1 59838208 59838373
refGene_gn=MARK1;refGene_tx=XM_023247971.1;refGene_gn=MARK1;refGene_tx=XM_023247972.1
F1 59839525 59839777
refGene_gn=MARK1;refGene_tx=XM_023247971.1;refGene_gn=MARK1;refGene_tx=XM_023247972.1
F1 59846231 59846276 refGene_gn=MARK1;refGene_tx=XM_023247971.1
F1 59849828 59850183
refGene_gn=MARK1;refGene_tx=XM_023247971.1;refGene_gn=MARK1;refGene_tx=XM_023247972.1
F1 59869271 59869559
refGene_gn=CF1H1orf115;refGene_tx=XM_023247490.1
F1 59875413 59875533
refGene_gn=CF1H1orf115;refGene_tx=XM_023247490.1
F1 59917887 59918174
refGene_gn=MARC2;refGene_tx=XM_023247231.1;refGene_gn=MARC2;refGene_tx=XM_023247232.1;refGene_gn=MARC2;refGene_tx=XM_023247233.1
F1 59923829 59924003
refGene_gn=MARC2;refGene_tx=XM_023247231.1;refGene_gn=MARC2;refGene_tx=XM_023247232.1;refGene_gn=MARC2;refGene_tx=XM_023247233.1
F1 59926672 59926835
refGene_gn=MARC2;refGene_tx=XM_023247231.1;refGene_gn=MARC2;refGene_tx=XM_023247232.1;refGene_gn=MARC2;refGene_tx=XM_023247233.1
F1 59927806 59927947
refGene_gn=MARC2;refGene_tx=XM_023247231.1;refGene_gn=MARC2;refGene_tx=XM_023247232.1;refGene_gn=MARC2;refGene_tx=XM_023247233.1
F1 59944796 59944858
refGene_gn=MARC2;refGene_tx=XM_023247231.1;refGene_gn=MARC2;refGene_tx=XM_023247232.1
F1 59944945 59945017 refGene_gn=MARC2;refGene_tx=XM_023247231.1
F1 59946762 59946889
refGene_gn=MARC2;refGene_tx=XM_023247231.1;refGene_gn=MARC2;refGene_tx=XM_023247232.1
F1 59956555 59956579 refGene_gn=MARC2;refGene_tx=XM_023247233.1
F1 59963361 59963648 refGene_gn=MARC1;refGene_tx=XM_023247482.1
F1 59966872 59967046 refGene_gn=MARC1;refGene_tx=XM_023247482.1
F1 59973021 59973184 refGene_gn=MARC1;refGene_tx=XM_023247482.1
F1 59974137 59974278 refGene_gn=MARC1;refGene_tx=XM_023247482.1

F1 59978483 59978545 refGene_gn=MARC1;refGene_tx=XM_023247482.1
F1 59978644 59978716 refGene_gn=MARC1;refGene_tx=XM_023247482.1
F1 59986313 59986440 refGene_gn=MARC1;refGene_tx=XM_023247482.1
F1 60041376 60041971 refGene_gn=HLX;refGene_tx=XM_011291138.3
F1 60042711 60042891 refGene_gn=HLX;refGene_tx=XM_011291138.3
F1 60043772 60043957 refGene_gn=HLX;refGene_tx=XM_011291138.3
F1 60045711 60046093 refGene_gn=HLX;refGene_tx=XM_011291138.3
F1 60046095 60046199 refGene_gn=HLX;refGene_tx=XM_011291138.3
F1 60192842 60193994
refGene_gn=LOC111558343;refGene_tx=XM_023247463.1
F1 60193995 60194334
refGene_gn=LOC111558343;refGene_tx=XM_023247463.1
F1 60711326 60711592 refGene_gn=DUSP10;refGene_tx=XM_003999516.5
F1 60715135 60715507 refGene_gn=DUSP10;refGene_tx=XM_003999516.5
F1 60746375 60747171 refGene_gn=DUSP10;refGene_tx=XM_003999516.5
F1 61318078 61318443
refGene_gn=HHIPL2;refGene_tx=XM_019822233.2;refGene_gn=HHIPL2;refGene_tx=XM_019822234.2
F1 61319607 61319690 refGene_gn=HHIPL2;refGene_tx=XM_019822233.2
F1 61322148 61322230
refGene_gn=HHIPL2;refGene_tx=XM_019822233.2;refGene_gn=HHIPL2;refGene_tx=XM_019822234.2
F1 61326411 61326557
refGene_gn=HHIPL2;refGene_tx=XM_019822233.2;refGene_gn=HHIPL2;refGene_tx=XM_019822234.2
F1 61332347 61332474
refGene_gn=HHIPL2;refGene_tx=XM_019822233.2;refGene_gn=HHIPL2;refGene_tx=XM_019822234.2
F1 61333268 61333600
refGene_gn=HHIPL2;refGene_tx=XM_019822233.2;refGene_gn=HHIPL2;refGene_tx=XM_019822234.2
F1 61336416 61336560
refGene_gn=HHIPL2;refGene_tx=XM_019822233.2;refGene_gn=HHIPL2;refGene_tx=XM_019822234.2
F1 61337433 61338086
refGene_gn=HHIPL2;refGene_tx=XM_019822233.2;refGene_gn=HHIPL2;refGene_tx=XM_019822234.2
F1 61342363 61342666
refGene_gn=HHIPL2;refGene_tx=XM_019822233.2;refGene_gn=HHIPL2;refGene_tx=XM_019822234.2
F1 61354968 61355081
refGene_gn=TAF1A;refGene_tx=XM_003999519.5;refGene_gn=TAF1A;refGene_tx=XM_019822235.2;refGene_gn=TAF1A;refGene_tx=XM_019822237.2;refGene_gn=TAF1A;refGene_tx=XM_019822238.2;refGene_gn=TAF1A;refGene_tx=XM_019822239.1
F1 61357906 61358061
refGene_gn=TAF1A;refGene_tx=XM_003999519.5;refGene_gn=TAF1A;refGene_tx=XM_019822235.2;refGene_gn=TAF1A;refGene_tx=XM_019822237.2;refGene_gn=TAF1A;refGene_tx=XM_019822238.2;refGene_gn=TAF1A;refGene_tx=XM_019822239.1

F1 61359117 61359241
refGene_gn=TAF1A;refGene_tx=XM_003999519.5;refGene_gn=TAF1A;refGene_tx=XM_019822235.2;refGene_gn=TAF1A;refGene_tx=XM_019822237.2;refGene_gn=TAF1A;refGene_tx=XM_019822238.2;refGene_gn=TAF1A;refGene_tx=XM_019822239.1
F1 61360056 61360123
refGene_gn=TAF1A;refGene_tx=XM_003999519.5;refGene_gn=TAF1A;refGene_tx=XM_019822235.2;refGene_gn=TAF1A;refGene_tx=XM_019822237.2;refGene_gn=TAF1A;refGene_tx=XM_019822238.2;refGene_gn=TAF1A;refGene_tx=XM_019822239.1
F1 61363395 61363554
refGene_gn=TAF1A;refGene_tx=XM_003999519.5;refGene_gn=TAF1A;refGene_tx=XM_019822235.2;refGene_gn=TAF1A;refGene_tx=XM_019822237.2;refGene_gn=TAF1A;refGene_tx=XM_019822239.1
F1 61365092 61365223
refGene_gn=TAF1A;refGene_tx=XM_003999519.5;refGene_gn=TAF1A;refGene_tx=XM_019822235.2;refGene_gn=TAF1A;refGene_tx=XM_019822237.2;refGene_gn=TAF1A;refGene_tx=XM_019822238.2;refGene_gn=TAF1A;refGene_tx=XM_019822239.1
F1 61368575 61368774
refGene_gn=TAF1A;refGene_tx=XM_003999519.5;refGene_gn=TAF1A;refGene_tx=XM_019822235.2;refGene_gn=TAF1A;refGene_tx=XM_019822237.2;refGene_gn=TAF1A;refGene_tx=XM_019822238.2;refGene_gn=TAF1A;refGene_tx=XM_019822239.1
F1 61374672 61374786
refGene_gn=TAF1A;refGene_tx=XM_003999519.5;refGene_gn=TAF1A;refGene_tx=XM_019822235.2;refGene_gn=TAF1A;refGene_tx=XM_019822238.2;refGene_gn=TAF1A;refGene_tx=XM_019822239.1
F1 61379271 61379441
refGene_gn=TAF1A;refGene_tx=XM_003999519.5;refGene_gn=TAF1A;refGene_tx=XM_019822235.2;refGene_gn=TAF1A;refGene_tx=XM_019822237.2;refGene_gn=TAF1A;refGene_tx=XM_019822238.2
F1 61382849 61382970
refGene_gn=TAF1A;refGene_tx=XM_003999519.5;refGene_gn=TAF1A;refGene_tx=XM_019822235.2;refGene_gn=TAF1A;refGene_tx=XM_019822237.2;refGene_gn=TAF1A;refGene_tx=XM_019822238.2
F1 61413456 61413589
refGene_gn=MIA3;refGene_tx=XM_023247248.1;refGene_gn=MIA3;refGene_tx=XM_023247249.1;refGene_gn=MIA3;refGene_tx=XM_023247250.1;refGene_gn=MIA3;refGene_tx=XM_023247251.1;refGene_gn=MIA3;refGene_tx=XM_023247252.1;refGene_gn=MIA3;refGene_tx=XM_023247253.1;refGene_gn=MIA3;refGene_tx=XM_023247254.1
F1 61416507 61416641
refGene_gn=MIA3;refGene_tx=XM_023247248.1;refGene_gn=MIA3;refGene_tx=XM_023247249.1;refGene_gn=MIA3;refGene_tx=XM_023247250.1;refGene_gn=MIA3;refGene_tx=XM_023247251.1;refGene_gn=MIA3;refGene_tx=XM_023247252.1;refGene_gn=MIA3;refGene_tx=XM_023247253.1;refGene_gn=MIA3;refGene_tx=XM_023247254.1
F1 61420190 61420277
refGene_gn=MIA3;refGene_tx=XM_023247248.1;refGene_gn=MIA3;refGene_tx=X

M_023247249.1;refGene_gn=MIA3;refGene_tx=XM_023247250.1;refGene_gn=MIA3;refGene_tx=XM_023247251.1;refGene_gn=MIA3;refGene_tx=XM_023247252.1;refGene_gn=MIA3;refGene_tx=XM_023247253.1;refGene_gn=MIA3;refGene_tx=XM_023247254.1
F1 61422929 61425756
refGene_gn=MIA3;refGene_tx=XM_023247248.1;refGene_gn=MIA3;refGene_tx=XM_023247249.1;refGene_gn=MIA3;refGene_tx=XM_023247250.1;refGene_gn=MIA3;refGene_tx=XM_023247251.1;refGene_gn=MIA3;refGene_tx=XM_023247252.1;refGene_gn=MIA3;refGene_tx=XM_023247253.1;refGene_gn=MIA3;refGene_tx=XM_023247254.1
F1 61428242 61428404
refGene_gn=MIA3;refGene_tx=XM_023247248.1;refGene_gn=MIA3;refGene_tx=XM_023247249.1;refGene_gn=MIA3;refGene_tx=XM_023247250.1;refGene_gn=MIA3;refGene_tx=XM_023247251.1;refGene_gn=MIA3;refGene_tx=XM_023247252.1;refGene_gn=MIA3;refGene_tx=XM_023247253.1;refGene_gn=MIA3;refGene_tx=XM_023247254.1
F1 61430965 61431111
refGene_gn=MIA3;refGene_tx=XM_023247248.1;refGene_gn=MIA3;refGene_tx=XM_023247249.1;refGene_gn=MIA3;refGene_tx=XM_023247250.1;refGene_gn=MIA3;refGene_tx=XM_023247251.1;refGene_gn=MIA3;refGene_tx=XM_023247252.1;refGene_gn=MIA3;refGene_tx=XM_023247253.1;refGene_gn=MIA3;refGene_tx=XM_023247254.1
F1 61447691 61447802
refGene_gn=MIA3;refGene_tx=XM_011291137.3;refGene_gn=MIA3;refGene_tx=XM_019822232.2
F1 61448757 61448889
refGene_gn=MIA3;refGene_tx=XM_011291137.3;refGene_gn=MIA3;refGene_tx=XM_019822232.2;refGene_gn=MIA3;refGene_tx=XM_023247248.1;refGene_gn=MIA3;refGene_tx=XM_023247249.1;refGene_gn=MIA3;refGene_tx=XM_023247250.1;refGene_gn=MIA3;refGene_tx=XM_023247251.1;refGene_gn=MIA3;refGene_tx=XM_023247252.1;refGene_gn=MIA3;refGene_tx=XM_023247253.1;refGene_gn=MIA3;refGene_tx=XM_023247254.1
F1 61451807 61451829
refGene_gn=MIA3;refGene_tx=XM_011291137.3;refGene_gn=MIA3;refGene_tx=XM_019822232.2;refGene_gn=MIA3;refGene_tx=XM_023247248.1;refGene_gn=MIA3;refGene_tx=XM_023247249.1;refGene_gn=MIA3;refGene_tx=XM_023247250.1;refGene_gn=MIA3;refGene_tx=XM_023247251.1;refGene_gn=MIA3;refGene_tx=XM_023247252.1;refGene_gn=MIA3;refGene_tx=XM_023247253.1;refGene_gn=MIA3;refGene_tx=XM_023247254.1
F1 61452552 61452641
refGene_gn=MIA3;refGene_tx=XM_011291137.3;refGene_gn=MIA3;refGene_tx=XM_019822232.2;refGene_gn=MIA3;refGene_tx=XM_023247248.1;refGene_gn=MIA3;refGene_tx=XM_023247249.1;refGene_gn=MIA3;refGene_tx=XM_023247250.1;refGene_gn=MIA3;refGene_tx=XM_023247251.1;refGene_gn=MIA3;refGene_tx=XM_023247252.1;refGene_gn=MIA3;refGene_tx=XM_023247253.1;refGene_gn=MIA3;refGene_tx=XM_023247254.1
F1 61452895 61452973
refGene_gn=MIA3;refGene_tx=XM_011291137.3;refGene_gn=MIA3;refGene_tx=XM_019822232.2;refGene_gn=MIA3;refGene_tx=XM_023247248.1;refGene_gn=MIA3;refGene_tx=XM_023247249.1;refGene_gn=MIA3;refGene_tx=XM_023247250.1;

refGene_gn=MIA3;refGene_tx=XM_023247251.1;refGene_gn=MIA3;refGene_tx=XM_023247252.1;refGene_gn=MIA3;refGene_tx=XM_023247253.1;refGene_gn=MIA3;refGene_tx=XM_023247254.1
F1 61453056 61453167
refGene_gn=MIA3;refGene_tx=XM_011291137.3;refGene_gn=MIA3;refGene_tx=XM_019822232.2;refGene_gn=MIA3;refGene_tx=XM_023247248.1;refGene_gn=MIA3;refGene_tx=XM_023247249.1;refGene_gn=MIA3;refGene_tx=XM_023247250.1;refGene_gn=MIA3;refGene_tx=XM_023247251.1;refGene_gn=MIA3;refGene_tx=XM_023247252.1;refGene_gn=MIA3;refGene_tx=XM_023247253.1;refGene_gn=MIA3;refGene_tx=XM_023247254.1
F1 61454372 61454444
refGene_gn=MIA3;refGene_tx=XM_011291137.3;refGene_gn=MIA3;refGene_tx=XM_019822232.2;refGene_gn=MIA3;refGene_tx=XM_023247248.1;refGene_gn=MIA3;refGene_tx=XM_023247249.1;refGene_gn=MIA3;refGene_tx=XM_023247250.1;refGene_gn=MIA3;refGene_tx=XM_023247252.1;refGene_gn=MIA3;refGene_tx=XM_023247253.1;refGene_gn=MIA3;refGene_tx=XM_023247254.1
F1 61454623 61454728
refGene_gn=MIA3;refGene_tx=XM_011291137.3;refGene_gn=MIA3;refGene_tx=XM_019822232.2;refGene_gn=MIA3;refGene_tx=XM_023247248.1;refGene_gn=MIA3;refGene_tx=XM_023247249.1;refGene_gn=MIA3;refGene_tx=XM_023247250.1;refGene_gn=MIA3;refGene_tx=XM_023247251.1;refGene_gn=MIA3;refGene_tx=XM_023247252.1;refGene_gn=MIA3;refGene_tx=XM_023247253.1;refGene_gn=MIA3;refGene_tx=XM_023247254.1
F1 61455407 61455530
refGene_gn=MIA3;refGene_tx=XM_011291137.3;refGene_gn=MIA3;refGene_tx=XM_019822232.2;refGene_gn=MIA3;refGene_tx=XM_023247248.1;refGene_gn=MIA3;refGene_tx=XM_023247249.1;refGene_gn=MIA3;refGene_tx=XM_023247250.1;refGene_gn=MIA3;refGene_tx=XM_023247251.1;refGene_gn=MIA3;refGene_tx=XM_023247252.1;refGene_gn=MIA3;refGene_tx=XM_023247253.1;refGene_gn=MIA3;refGene_tx=XM_023247254.1
F1 61455626 61455738
refGene_gn=MIA3;refGene_tx=XM_011291137.3;refGene_gn=MIA3;refGene_tx=XM_019822232.2;refGene_gn=MIA3;refGene_tx=XM_023247248.1;refGene_gn=MIA3;refGene_tx=XM_023247249.1;refGene_gn=MIA3;refGene_tx=XM_023247250.1;refGene_gn=MIA3;refGene_tx=XM_023247251.1;refGene_gn=MIA3;refGene_tx=XM_023247252.1;refGene_gn=MIA3;refGene_tx=XM_023247253.1;refGene_gn=MIA3;refGene_tx=XM_023247254.1
F1 61457104 61457160
refGene_gn=MIA3;refGene_tx=XM_011291137.3;refGene_gn=MIA3;refGene_tx=XM_019822232.2;refGene_gn=MIA3;refGene_tx=XM_023247248.1;refGene_gn=MIA3;refGene_tx=XM_023247249.1;refGene_gn=MIA3;refGene_tx=XM_023247250.1;refGene_gn=MIA3;refGene_tx=XM_023247251.1;refGene_gn=MIA3;refGene_tx=XM_023247252.1;refGene_gn=MIA3;refGene_tx=XM_023247253.1;refGene_gn=MIA3;refGene_tx=XM_023247254.1
F1 61457251 61457342
refGene_gn=MIA3;refGene_tx=XM_011291137.3;refGene_gn=MIA3;refGene_tx=XM_019822232.2;refGene_gn=MIA3;refGene_tx=XM_023247248.1;refGene_gn=MIA3;refGene_tx=XM_023247249.1;refGene_gn=MIA3;refGene_tx=XM_023247250.1;refGene_gn=MIA3;refGene_tx=XM_023247251.1;refGene_gn=MIA3;refGene_tx=XM_023247252.1;refGene_gn=MIA3;refGene_tx=XM_023247253.1;refGene_gn=MIA

3;refGene_tx=XM_023247254.1
F1 61457456 61457595
refGene_gn=MIA3;refGene_tx=XM_011291137.3;refGene_gn=MIA3;refGene_tx=XM_019822232.2;refGene_gn=MIA3;refGene_tx=XM_023247248.1;refGene_gn=MIA3;refGene_tx=XM_023247249.1;refGene_gn=MIA3;refGene_tx=XM_023247250.1;refGene_gn=MIA3;refGene_tx=XM_023247251.1;refGene_gn=MIA3;refGene_tx=XM_023247252.1;refGene_gn=MIA3;refGene_tx=XM_023247253.1;refGene_gn=MIA3;refGene_tx=XM_023247254.1
F1 61462167 61462269
refGene_gn=MIA3;refGene_tx=XM_011291137.3;refGene_gn=MIA3;refGene_tx=XM_019822232.2;refGene_gn=MIA3;refGene_tx=XM_023247248.1;refGene_gn=MIA3;refGene_tx=XM_023247249.1;refGene_gn=MIA3;refGene_tx=XM_023247250.1;refGene_gn=MIA3;refGene_tx=XM_023247251.1;refGene_gn=MIA3;refGene_tx=XM_023247252.1;refGene_gn=MIA3;refGene_tx=XM_023247253.1;refGene_gn=MIA3;refGene_tx=XM_023247254.1
F1 61462929 61462990
refGene_gn=MIA3;refGene_tx=XM_011291137.3;refGene_gn=MIA3;refGene_tx=XM_019822232.2;refGene_gn=MIA3;refGene_tx=XM_023247248.1;refGene_gn=MIA3;refGene_tx=XM_023247249.1;refGene_gn=MIA3;refGene_tx=XM_023247250.1;refGene_gn=MIA3;refGene_tx=XM_023247251.1;refGene_gn=MIA3;refGene_tx=XM_023247252.1;refGene_gn=MIA3;refGene_tx=XM_023247253.1;refGene_gn=MIA3;refGene_tx=XM_023247254.1
F1 61463101 61463137
refGene_gn=MIA3;refGene_tx=XM_011291137.3;refGene_gn=MIA3;refGene_tx=XM_019822232.2;refGene_gn=MIA3;refGene_tx=XM_023247248.1;refGene_gn=MIA3;refGene_tx=XM_023247249.1;refGene_gn=MIA3;refGene_tx=XM_023247250.1;refGene_gn=MIA3;refGene_tx=XM_023247251.1;refGene_gn=MIA3;refGene_tx=XM_023247252.1;refGene_gn=MIA3;refGene_tx=XM_023247253.1;refGene_gn=MIA3;refGene_tx=XM_023247254.1
F1 61463213 61463281
refGene_gn=MIA3;refGene_tx=XM_011291137.3;refGene_gn=MIA3;refGene_tx=XM_019822232.2;refGene_gn=MIA3;refGene_tx=XM_023247248.1;refGene_gn=MIA3;refGene_tx=XM_023247249.1;refGene_gn=MIA3;refGene_tx=XM_023247250.1;refGene_gn=MIA3;refGene_tx=XM_023247251.1;refGene_gn=MIA3;refGene_tx=XM_023247252.1;refGene_gn=MIA3;refGene_tx=XM_023247253.1;refGene_gn=MIA3;refGene_tx=XM_023247254.1
F1 61463400 61463501
refGene_gn=MIA3;refGene_tx=XM_011291137.3;refGene_gn=MIA3;refGene_tx=XM_019822232.2;refGene_gn=MIA3;refGene_tx=XM_023247248.1;refGene_gn=MIA3;refGene_tx=XM_023247249.1;refGene_gn=MIA3;refGene_tx=XM_023247250.1;refGene_gn=MIA3;refGene_tx=XM_023247251.1;refGene_gn=MIA3;refGene_tx=XM_023247252.1;refGene_gn=MIA3;refGene_tx=XM_023247253.1;refGene_gn=MIA3;refGene_tx=XM_023247254.1
F1 61463661 61463799
refGene_gn=MIA3;refGene_tx=XM_011291137.3;refGene_gn=MIA3;refGene_tx=XM_019822232.2;refGene_gn=MIA3;refGene_tx=XM_023247248.1;refGene_gn=MIA3;refGene_tx=XM_023247249.1;refGene_gn=MIA3;refGene_tx=XM_023247250.1;refGene_gn=MIA3;refGene_tx=XM_023247251.1;refGene_gn=MIA3;refGene_tx=XM_023247252.1
F1 61464517 61464586

refGene_gn=MIA3;refGene_tx=XM_011291137.3;refGene_gn=MIA3;refGene_tx=XM_019822232.2;refGene_gn=MIA3;refGene_tx=XM_023247248.1;refGene_gn=MIA3;refGene_tx=XM_023247249.1;refGene_gn=MIA3;refGene_tx=XM_023247250.1;refGene_gn=MIA3;refGene_tx=XM_023247251.1;refGene_gn=MIA3;refGene_tx=XM_023247252.1;refGene_gn=MIA3;refGene_tx=XM_023247253.1;refGene_gn=MIA3;refGene_tx=XM_023247254.1
F1 61464691 61464774
refGene_gn=MIA3;refGene_tx=XM_011291137.3;refGene_gn=MIA3;refGene_tx=XM_019822232.2;refGene_gn=MIA3;refGene_tx=XM_023247248.1;refGene_gn=MIA3;refGene_tx=XM_023247249.1;refGene_gn=MIA3;refGene_tx=XM_023247250.1;refGene_gn=MIA3;refGene_tx=XM_023247251.1;refGene_gn=MIA3;refGene_tx=XM_023247252.1;refGene_gn=MIA3;refGene_tx=XM_023247253.1;refGene_gn=MIA3;refGene_tx=XM_023247254.1
F1 61464906 61464978
refGene_gn=MIA3;refGene_tx=XM_019822232.2;refGene_gn=MIA3;refGene_tx=XM_023247248.1;refGene_gn=MIA3;refGene_tx=XM_023247249.1;refGene_gn=MIA3;refGene_tx=XM_023247251.1;refGene_gn=MIA3;refGene_tx=XM_023247253.1;refGene_gn=MIA3;refGene_tx=XM_023247254.1
F1 61466446 61466597
refGene_gn=MIA3;refGene_tx=XM_011291137.3;refGene_gn=MIA3;refGene_tx=XM_019822232.2;refGene_gn=MIA3;refGene_tx=XM_023247248.1;refGene_gn=MIA3;refGene_tx=XM_023247249.1;refGene_gn=MIA3;refGene_tx=XM_023247250.1;refGene_gn=MIA3;refGene_tx=XM_023247251.1;refGene_gn=MIA3;refGene_tx=XM_023247252.1;refGene_gn=MIA3;refGene_tx=XM_023247253.1;refGene_gn=MIA3;refGene_tx=XM_023247254.1
F1 61467479 61467790
refGene_gn=MIA3;refGene_tx=XM_011291137.3;refGene_gn=MIA3;refGene_tx=XM_019822232.2;refGene_gn=MIA3;refGene_tx=XM_023247248.1;refGene_gn=MIA3;refGene_tx=XM_023247249.1;refGene_gn=MIA3;refGene_tx=XM_023247250.1;refGene_gn=MIA3;refGene_tx=XM_023247251.1;refGene_gn=MIA3;refGene_tx=XM_023247252.1;refGene_gn=MIA3;refGene_tx=XM_023247253.1;refGene_gn=MIA3;refGene_tx=XM_023247254.1
F1 61471806 61471903
refGene_gn=AIDA;refGene_tx=XM_003999521.5;refGene_gn=AIDA;refGene_tx=XM_019822240.2
F1 61472029 61472147
refGene_gn=AIDA;refGene_tx=XM_003999521.5;refGene_gn=AIDA;refGene_tx=XM_019822240.2
F1 61475010 61475133 refGene_gn=AIDA;refGene_tx=XM_003999521.5
F1 61477330 61477453
refGene_gn=AIDA;refGene_tx=XM_003999521.5;refGene_gn=AIDA;refGene_tx=XM_019822240.2
F1 61486626 61486733
refGene_gn=AIDA;refGene_tx=XM_003999521.5;refGene_gn=AIDA;refGene_tx=XM_019822240.2
F1 61486960 61487024
refGene_gn=AIDA;refGene_tx=XM_003999521.5;refGene_gn=AIDA;refGene_tx=XM_019822240.2
F1 61493229 61493284
refGene_gn=AIDA;refGene_tx=XM_003999521.5;refGene_gn=AIDA;refGene_tx=X

M_019822240.2
F1 61493682 61493736
refGene_gn=AIDA;refGene_tx=XM_003999521.5;refGene_gn=AIDA;refGene_tx=X
M_019822240.2
F1 61500134 61500204
refGene_gn=AIDA;refGene_tx=XM_003999521.5;refGene_gn=AIDA;refGene_tx=X
M_019822240.2
F1 61507914 61508024
refGene_gn=AIDA;refGene_tx=XM_003999521.5;refGene_gn=AIDA;refGene_tx=X
M_019822240.2
F1 61511150 61511251
refGene_gn=BROX;refGene_tx=XM_003999522.5;refGene_gn=BROX;refGene_tx=X
M_006942981.3
F1 61514960 61515067
refGene_gn=BROX;refGene_tx=XM_003999522.5;refGene_gn=BROX;refGene_tx=X
M_006942981.3
F1 61515649 61515746
refGene_gn=BROX;refGene_tx=XM_003999522.5;refGene_gn=BROX;refGene_tx=X
M_006942981.3
F1 61518035 61518131
refGene_gn=BROX;refGene_tx=XM_003999522.5;refGene_gn=BROX;refGene_tx=X
M_006942981.3
F1 61519216 61519289
refGene_gn=BROX;refGene_tx=XM_003999522.5;refGene_gn=BROX;refGene_tx=X
M_006942981.3
F1 61521114 61521220
refGene_gn=BROX;refGene_tx=XM_003999522.5;refGene_gn=BROX;refGene_tx=X
M_006942981.3
F1 61521919 61522009
refGene_gn=BROX;refGene_tx=XM_003999522.5;refGene_gn=BROX;refGene_tx=X
M_006942981.3
F1 61523959 61524045
refGene_gn=BROX;refGene_tx=XM_003999522.5;refGene_gn=BROX;refGene_tx=X
M_006942981.3
F1 61526165 61526247
refGene_gn=BROX;refGene_tx=XM_003999522.5;refGene_gn=BROX;refGene_tx=X
M_006942981.3
F1 61526555 61526706
refGene_gn=BROX;refGene_tx=XM_003999522.5;refGene_gn=BROX;refGene_tx=X
M_006942981.3
F1 61527559 61527719
refGene_gn=BROX;refGene_tx=XM_003999522.5;refGene_gn=BROX;refGene_tx=X
M_006942981.3
F1 61528973 61529060
refGene_gn=BROX;refGene_tx=XM_003999522.5;refGene_gn=BROX;refGene_tx=X
M_006942981.3
F1 61542433 61542607 refGene_gn=FAM177B;refGene_tx=XM_003999530.3
F1 61543291 61543358 refGene_gn=FAM177B;refGene_tx=XM_003999530.3
F1 61545848 61545946 refGene_gn=FAM177B;refGene_tx=XM_003999530.3
F1 61546250 61546445 refGene_gn=FAM177B;refGene_tx=XM_003999530.3

F1 61852843 61853106
refGene_gn=FAM78B;refGene_tx=XM_023248057.1;refGene_gn=FAM78B;refGene_tx=XM_023248058.1
F1 61934472 61934995
refGene_gn=FAM78B;refGene_tx=XM_023248057.1;refGene_gn=FAM78B;refGene_tx=XM_023248058.1
F1 62057179 62057319
refGene_gn=UCK2;refGene_tx=XM_019822244.2;refGene_gn=UCK2;refGene_tx=XM_023248101.1
F1 62058937 62058986
refGene_gn=UCK2;refGene_tx=XM_019822244.2;refGene_gn=UCK2;refGene_tx=XM_023248101.1
F1 62061085 62061183
refGene_gn=UCK2;refGene_tx=XM_019822244.2;refGene_gn=UCK2;refGene_tx=XM_023248101.1
F1 62067028 62067171
refGene_gn=UCK2;refGene_tx=XM_019822244.2;refGene_gn=UCK2;refGene_tx=XM_023248101.1
F1 62070713 62070810
refGene_gn=UCK2;refGene_tx=XM_019822244.2;refGene_gn=UCK2;refGene_tx=XM_023248101.1
F1 62071598 62071758
refGene_gn=UCK2;refGene_tx=XM_019822244.2;refGene_gn=UCK2;refGene_tx=XM_023248101.1
F1 62106625 62106691 refGene_gn=UCK2;refGene_tx=XM_019822244.2
F1 62140114 62140213 refGene_gn=UCK2;refGene_tx=XM_023248101.1
F1 62170023 62170093 refGene_gn=TMCO1;refGene_tx=XM_023247216.1
F1 62170485 62170563 refGene_gn=TMCO1;refGene_tx=XM_023247216.1
F1 62175297 62175357 refGene_gn=TMCO1;refGene_tx=XM_023247216.1
F1 62181082 62181129 refGene_gn=TMCO1;refGene_tx=XM_023247216.1
F1 62182902 62182970 refGene_gn=TMCO1;refGene_tx=XM_023247216.1
F1 62193659 62193804 refGene_gn=TMCO1;refGene_tx=XM_023247216.1
F1 62209560 62209659 refGene_gn=TMCO1;refGene_tx=XM_023247216.1
F1 62225745 62225926
refGene_gn=ALDH9A1;refGene_tx=XM_003999532.3;refGene_gn=ALDH9A1;refGene_tx=XM_019822246.2;refGene_gn=ALDH9A1;refGene_tx=XM_019822247.2
F1 62228322 62228468
refGene_gn=ALDH9A1;refGene_tx=XM_003999532.3;refGene_gn=ALDH9A1;refGene_tx=XM_019822246.2;refGene_gn=ALDH9A1;refGene_tx=XM_019822247.2
F1 62234128 62234258
refGene_gn=ALDH9A1;refGene_tx=XM_003999532.3;refGene_gn=ALDH9A1;refGene_tx=XM_019822246.2;refGene_gn=ALDH9A1;refGene_tx=XM_019822247.2
F1 62235208 62235343
refGene_gn=ALDH9A1;refGene_tx=XM_003999532.3;refGene_gn=ALDH9A1;refGene_tx=XM_019822246.2;refGene_gn=ALDH9A1;refGene_tx=XM_019822247.2
F1 62237105 62237302
refGene_gn=ALDH9A1;refGene_tx=XM_003999532.3;refGene_gn=ALDH9A1;refGene_tx=XM_019822246.2;refGene_gn=ALDH9A1;refGene_tx=XM_019822247.2
F1 62238201 62238342
refGene_gn=ALDH9A1;refGene_tx=XM_003999532.3;refGene_gn=ALDH9A1;refGen

e_tx=XM_019822246.2;refGene_gn=ALDH9A1;refGene_tx=XM_019822247.2
F1 62240003 62240192
refGene_gn=ALDH9A1;refGene_tx=XM_003999532.3;refGene_gn=ALDH9A1;refGene_tx=XM_019822246.2;refGene_gn=ALDH9A1;refGene_tx=XM_019822247.2
F1 62240637 62240725
refGene_gn=ALDH9A1;refGene_tx=XM_003999532.3;refGene_gn=ALDH9A1;refGene_tx=XM_019822246.2;refGene_gn=ALDH9A1;refGene_tx=XM_019822247.2
F1 62241184 62241318 refGene_gn=ALDH9A1;refGene_tx=XM_019822247.2
F1 62243430 62243572
refGene_gn=ALDH9A1;refGene_tx=XM_003999532.3;refGene_gn=ALDH9A1;refGene_tx=XM_019822246.2
F1 62246305 62246418
refGene_gn=ALDH9A1;refGene_tx=XM_003999532.3;refGene_gn=ALDH9A1;refGene_tx=XM_019822246.2
F1 62248178 62248290
refGene_gn=ALDH9A1;refGene_tx=XM_003999532.3;refGene_gn=ALDH9A1;refGene_tx=XM_019822246.2
F1 62256337 62256474
refGene_gn=MGST3;refGene_tx=XM_003999533.5;refGene_gn=MGST3;refGene_tx=XM_006942984.4
F1 62257396 62257469
refGene_gn=MGST3;refGene_tx=XM_003999533.5;refGene_gn=MGST3;refGene_tx=XM_006942984.4
F1 62259758 62259816
refGene_gn=MGST3;refGene_tx=XM_003999533.5;refGene_gn=MGST3;refGene_tx=XM_006942984.4
F1 62260592 62260666
refGene_gn=MGST3;refGene_tx=XM_003999533.5;refGene_gn=MGST3;refGene_tx=XM_006942984.4
F1 62262985 62263102
refGene_gn=MGST3;refGene_tx=XM_003999533.5;refGene_gn=MGST3;refGene_tx=XM_006942984.4
F1 62350654 62350980 refGene_gn=LRRC52;refGene_tx=XM_003999534.4
F1 62369090 62369712 refGene_gn=LRRC52;refGene_tx=XM_003999534.4
F1 62459820 62459869 refGene_gn=RXRG;refGene_tx=XM_003999535.5
F1 62475061 62475080 refGene_gn=RXRG;refGene_tx=XM_019822249.1
F1 62477471 62477719
refGene_gn=RXRG;refGene_tx=XM_003999535.5;refGene_gn=RXRG;refGene_tx=XM_019822249.1
F1 62487362 62487507
refGene_gn=RXRG;refGene_tx=XM_003999535.5;refGene_gn=RXRG;refGene_tx=XM_006942985.4;refGene_gn=RXRG;refGene_tx=XM_006942986.4;refGene_gn=RXRG;refGene_tx=XM_006942987.3;refGene_gn=RXRG;refGene_tx=XM_019822249.1
F1 62489056 62489236
refGene_gn=RXRG;refGene_tx=XM_003999535.5;refGene_gn=RXRG;refGene_tx=XM_006942985.4;refGene_gn=RXRG;refGene_tx=XM_006942986.4;refGene_gn=RXRG;refGene_tx=XM_006942987.3;refGene_gn=RXRG;refGene_tx=XM_019822249.1
F1 62495195 62495356
refGene_gn=RXRG;refGene_tx=XM_003999535.5;refGene_gn=RXRG;refGene_tx=XM_006942985.4;refGene_gn=RXRG;refGene_tx=XM_006942986.4;refGene_gn=RXR

G;refGene_tx=XM_006942987.3;refGene_gn=RXRG;refGene_tx=XM_019822249.1
F1 62495472 62495602
refGene_gn=RXRG;refGene_tx=XM_003999535.5;refGene_gn=RXRG;refGene_tx=XM_006942985.4;refGene_gn=RXRG;refGene_tx=XM_006942986.4;refGene_gn=RXRG;refGene_tx=XM_006942987.3;refGene_gn=RXRG;refGene_tx=XM_019822249.1
F1 62496920 62497053
refGene_gn=RXRG;refGene_tx=XM_003999535.5;refGene_gn=RXRG;refGene_tx=XM_006942985.4;refGene_gn=RXRG;refGene_tx=XM_006942986.4;refGene_gn=RXRG;refGene_tx=XM_006942987.3;refGene_gn=RXRG;refGene_tx=XM_019822249.1
F1 62498444 62498536
refGene_gn=RXRG;refGene_tx=XM_003999535.5;refGene_gn=RXRG;refGene_tx=XM_006942985.4;refGene_gn=RXRG;refGene_tx=XM_006942986.4;refGene_gn=RXRG;refGene_tx=XM_006942987.3;refGene_gn=RXRG;refGene_tx=XM_019822249.1
F1 62500372 62500478
refGene_gn=RXRG;refGene_tx=XM_003999535.5;refGene_gn=RXRG;refGene_tx=XM_006942985.4;refGene_gn=RXRG;refGene_tx=XM_006942986.4;refGene_gn=RXRG;refGene_tx=XM_006942987.3;refGene_gn=RXRG;refGene_tx=XM_019822249.1
F1 62503989 62504137
refGene_gn=RXRG;refGene_tx=XM_003999535.5;refGene_gn=RXRG;refGene_tx=XM_006942985.4;refGene_gn=RXRG;refGene_tx=XM_006942986.4;refGene_gn=RXRG;refGene_tx=XM_006942987.3;refGene_gn=RXRG;refGene_tx=XM_019822249.1
F1 62548414 62548608 refGene_gn=LMX1A;refGene_tx=XM_023247553.1
F1 62549102 62549200
refGene_gn=LMX1A;refGene_tx=XM_023247553.1;refGene_gn=LMX1A;refGene_tx=XM_023247554.1
F1 62551542 62551729
refGene_gn=LMX1A;refGene_tx=XM_023247553.1;refGene_gn=LMX1A;refGene_tx=XM_023247554.1
F1 62660564 62660797
refGene_gn=LMX1A;refGene_tx=XM_023247553.1;refGene_gn=LMX1A;refGene_tx=XM_023247554.1
F1 62699080 62699253
refGene_gn=LMX1A;refGene_tx=XM_023247553.1;refGene_gn=LMX1A;refGene_tx=XM_023247554.1
F1 62702033 62702111
refGene_gn=LMX1A;refGene_tx=XM_023247553.1;refGene_gn=LMX1A;refGene_tx=XM_023247554.1
F1 62703701 62703771
refGene_gn=LMX1A;refGene_tx=XM_023247553.1;refGene_gn=LMX1A;refGene_tx=XM_023247554.1
F1 62706210 62706381
refGene_gn=LMX1A;refGene_tx=XM_023247553.1;refGene_gn=LMX1A;refGene_tx=XM_023247554.1
F1 62708009 62708170
refGene_gn=LMX1A;refGene_tx=XM_023247553.1;refGene_gn=LMX1A;refGene_tx=XM_023247554.1
F1 63013402 63013495 refGene_gn=PBX1;refGene_tx=XM_006942988.4
F1 63040027 63040117
refGene_gn=PBX1;refGene_tx=XM_006942988.4;refGene_gn=PBX1;refGene_tx=XM_006942990.3

F1	63041520	63041633	refGene_gn=PBX1;refGene_tx=XM_006942988.4
F1	63048447	63048607	refGene_gn=PBX1;refGene_tx=XM_006942988.4;refGene_gn=PBX1;refGene_tx=XM_006942990.3
F1	63053732	63053868	refGene_gn=PBX1;refGene_tx=XM_006942988.4;refGene_gn=PBX1;refGene_tx=XM_006942990.3
F1	63060554	63060745	refGene_gn=PBX1;refGene_tx=XM_006942988.4;refGene_gn=PBX1;refGene_tx=XM_006942990.3
F1	63066812	63067057	refGene_gn=PBX1;refGene_tx=XM_006942988.4;refGene_gn=PBX1;refGene_tx=XM_006942990.3
F1	63305532	63305606	refGene_gn=PBX1;refGene_tx=XM_006942988.4;refGene_gn=PBX1;refGene_tx=XM_006942990.3
F1	63308742	63308933	refGene_gn=PBX1;refGene_tx=XM_006942988.4;refGene_gn=PBX1;refGene_tx=XM_006942990.3
F1	64210777	64210912	refGene_gn=NUF2;refGene_tx=XM_006942992.3;refGene_gn=NUF2;refGene_tx=XM_019822253.2;refGene_gn=NUF2;refGene_tx=XM_019822254.2
F1	64216721	64216857	refGene_gn=NUF2;refGene_tx=XM_006942992.3;refGene_gn=NUF2;refGene_tx=XM_019822253.2;refGene_gn=NUF2;refGene_tx=XM_019822254.2;refGene_gn=NUF2;refGene_tx=XM_019822255.2
F1	64217953	64218129	refGene_gn=NUF2;refGene_tx=XM_006942992.3;refGene_gn=NUF2;refGene_tx=XM_019822253.2;refGene_gn=NUF2;refGene_tx=XM_019822254.2;refGene_gn=NUF2;refGene_tx=XM_019822255.2
F1	64219781	64219922	refGene_gn=NUF2;refGene_tx=XM_006942992.3;refGene_gn=NUF2;refGene_tx=XM_019822253.2;refGene_gn=NUF2;refGene_tx=XM_019822254.2;refGene_gn=NUF2;refGene_tx=XM_019822255.2
F1	64221927	64222065	refGene_gn=NUF2;refGene_tx=XM_006942992.3;refGene_gn=NUF2;refGene_tx=XM_019822253.2;refGene_gn=NUF2;refGene_tx=XM_019822254.2;refGene_gn=NUF2;refGene_tx=XM_019822255.2
F1	64225531	64225594	refGene_gn=NUF2;refGene_tx=XM_006942992.3;refGene_gn=NUF2;refGene_tx=XM_019822253.2;refGene_gn=NUF2;refGene_tx=XM_019822254.2;refGene_gn=NUF2;refGene_tx=XM_019822255.2
F1	64226876	64226973	refGene_gn=NUF2;refGene_tx=XM_006942992.3;refGene_gn=NUF2;refGene_tx=XM_019822253.2;refGene_gn=NUF2;refGene_tx=XM_019822254.2;refGene_gn=NUF2;refGene_tx=XM_019822255.2
F1	64228279	64228353	refGene_gn=NUF2;refGene_tx=XM_006942992.3;refGene_gn=NUF2;refGene_tx=XM_019822253.2;refGene_gn=NUF2;refGene_tx=XM_019822254.2;refGene_gn=NUF2;refGene_tx=XM_019822255.2

F1 64229495 64229593
refGene_gn=NUF2;refGene_tx=XM_006942992.3;refGene_gn=NUF2;refGene_tx=XM_019822253.2;refGene_gn=NUF2;refGene_tx=XM_019822254.2;refGene_gn=NUF2;refGene_tx=XM_019822255.2
F1 64234335 64234397
refGene_gn=NUF2;refGene_tx=XM_006942992.3;refGene_gn=NUF2;refGene_tx=XM_019822253.2;refGene_gn=NUF2;refGene_tx=XM_019822254.2;refGene_gn=NUF2;refGene_tx=XM_019822255.2
F1 64234908 64234985
refGene_gn=NUF2;refGene_tx=XM_006942992.3;refGene_gn=NUF2;refGene_tx=XM_019822253.2;refGene_gn=NUF2;refGene_tx=XM_019822254.2;refGene_gn=NUF2;refGene_tx=XM_019822255.2
F1 64235949 64236024
refGene_gn=NUF2;refGene_tx=XM_006942992.3;refGene_gn=NUF2;refGene_tx=XM_019822253.2;refGene_gn=NUF2;refGene_tx=XM_019822254.2;refGene_gn=NUF2;refGene_tx=XM_019822255.2
F1 64237269 64237392
refGene_gn=NUF2;refGene_tx=XM_006942992.3;refGene_gn=NUF2;refGene_tx=XM_019822253.2;refGene_gn=NUF2;refGene_tx=XM_019822254.2;refGene_gn=NUF2;refGene_tx=XM_019822255.2
F1 64324840 64324884 refGene_gn=RGS5;refGene_tx=XM_003999541.5
F1 64351878 64351989 refGene_gn=RGS5;refGene_tx=XM_003999541.5
F1 64358854 64358916 refGene_gn=RGS5;refGene_tx=XM_003999541.5
F1 64368328 64368495
refGene_gn=RGS5;refGene_tx=XM_003999541.5;refGene_gn=RGS5;refGene_tx=XM_023248083.1
F1 64371687 64371849
refGene_gn=RGS5;refGene_tx=XM_003999541.5;refGene_gn=RGS5;refGene_tx=XM_023248083.1
F1 64424045 64424285
refGene_gn=RGS4;refGene_tx=XM_003999542.5;refGene_gn=RGS4;refGene_tx=XM_003999543.4
F1 64424726 64424893 refGene_gn=RGS4;refGene_tx=XM_003999542.5
F1 64425475 64425537
refGene_gn=RGS4;refGene_tx=XM_003999542.5;refGene_gn=RGS4;refGene_tx=XM_003999543.4
F1 64425825 64425930
refGene_gn=RGS4;refGene_tx=XM_003999542.5;refGene_gn=RGS4;refGene_tx=XM_003999543.4
F1 64428854 64428898
refGene_gn=RGS4;refGene_tx=XM_003999542.5;refGene_gn=RGS4;refGene_tx=XM_003999543.4
F1 64595183 64595851
refGene_gn=LOC102901893;refGene_tx=XM_023247438.1
F1 64610899 64611086
refGene_gn=CCDC190;refGene_tx=XM_019822256.2;refGene_gn=CCDC190;refGene_tx=XM_019822257.2
F1 64616028 64616152
refGene_gn=CCDC190;refGene_tx=XM_019822256.2;refGene_gn=CCDC190;refGene_tx=XM_019822257.2

F1 64616422 64616990
refGene_gn=CCDC190;refGene_tx=XM_019822256.2;refGene_gn=CCDC190;refGene_tx=XM_019822257.2
F1 64617190 64617307
refGene_gn=CCDC190;refGene_tx=XM_019822256.2;refGene_gn=CCDC190;refGene_tx=XM_019822257.2
F1 64630890 64631296
refGene_gn=LOC102901893;refGene_tx=XM_023247438.1
F1 64644663 64644765 refGene_gn=HSD17B7;refGene_tx=XM_019822263.2
F1 64646175 64646208 refGene_gn=HSD17B7;refGene_tx=XM_019822263.2
F1 64647397 64647496 refGene_gn=HSD17B7;refGene_tx=XM_019822263.2
F1 64650882 64650939 refGene_gn=HSD17B7;refGene_tx=XM_019822263.2
F1 64651650 64651755 refGene_gn=HSD17B7;refGene_tx=XM_019822263.2
F1 64655277 64655315 refGene_gn=HSD17B7;refGene_tx=XM_019822263.2
F1 64655317 64655471 refGene_gn=HSD17B7;refGene_tx=XM_019822263.2
F1 64658968 64659083 refGene_gn=HSD17B7;refGene_tx=XM_019822263.2
F1 64660861 64660954 refGene_gn=HSD17B7;refGene_tx=XM_019822263.2
F1 64665138 64665342 refGene_gn=HSD17B7;refGene_tx=XM_019822263.2
F1 64667047 64667082 refGene_gn=HSD17B7;refGene_tx=XM_019822263.2
F1 64678490 64678625
refGene_gn=DDR2;refGene_tx=XM_006942994.4;refGene_gn=DDR2;refGene_tx=XM_006942995.4;refGene_gn=DDR2;refGene_tx=XM_019822258.2;refGene_gn=DDR2;refGene_tx=XM_019822259.2;refGene_gn=DDR2;refGene_tx=XM_019822260.2;refGene_gn=DDR2;refGene_tx=XM_019822262.2
F1 64680700 64680850
refGene_gn=DDR2;refGene_tx=XM_006942994.4;refGene_gn=DDR2;refGene_tx=XM_006942995.4;refGene_gn=DDR2;refGene_tx=XM_019822258.2;refGene_gn=DDR2;refGene_tx=XM_019822259.2;refGene_gn=DDR2;refGene_tx=XM_019822262.2
F1 64683589 64683824
refGene_gn=DDR2;refGene_tx=XM_006942994.4;refGene_gn=DDR2;refGene_tx=XM_006942995.4;refGene_gn=DDR2;refGene_tx=XM_019822258.2;refGene_gn=DDR2;refGene_tx=XM_019822259.2;refGene_gn=DDR2;refGene_tx=XM_019822260.2;refGene_gn=DDR2;refGene_tx=XM_019822262.2
F1 64684118 64684310
refGene_gn=DDR2;refGene_tx=XM_006942994.4;refGene_gn=DDR2;refGene_tx=XM_006942995.4;refGene_gn=DDR2;refGene_tx=XM_019822258.2;refGene_gn=DDR2;refGene_tx=XM_019822259.2;refGene_gn=DDR2;refGene_tx=XM_019822260.2;refGene_gn=DDR2;refGene_tx=XM_019822262.2
F1 64686587 64686715
refGene_gn=DDR2;refGene_tx=XM_006942994.4;refGene_gn=DDR2;refGene_tx=XM_006942995.4;refGene_gn=DDR2;refGene_tx=XM_019822258.2;refGene_gn=DDR2;refGene_tx=XM_019822259.2;refGene_gn=DDR2;refGene_tx=XM_019822260.2;refGene_gn=DDR2;refGene_tx=XM_019822262.2
F1 64688094 64688318
refGene_gn=DDR2;refGene_tx=XM_006942994.4;refGene_gn=DDR2;refGene_tx=XM_006942995.4;refGene_gn=DDR2;refGene_tx=XM_019822258.2;refGene_gn=DDR2;refGene_tx=XM_019822259.2;refGene_gn=DDR2;refGene_tx=XM_019822260.2;refGene_gn=DDR2;refGene_tx=XM_019822262.2
F1 64689929 64690140
refGene_gn=DDR2;refGene_tx=XM_006942994.4;refGene_gn=DDR2;refGene_tx=X

M_006942995.4;refGene_gn=DDR2;refGene_tx=XM_019822258.2;refGene_gn=DDR2;refGene_tx=XM_019822259.2;refGene_gn=DDR2;refGene_tx=XM_019822260.2;refGene_gn=DDR2;refGene_tx=XM_019822262.2
F1 64692686 64692817
refGene_gn=DDR2;refGene_tx=XM_006942994.4;refGene_gn=DDR2;refGene_tx=XM_006942995.4;refGene_gn=DDR2;refGene_tx=XM_019822258.2;refGene_gn=DDR2;refGene_tx=XM_019822259.2;refGene_gn=DDR2;refGene_tx=XM_019822260.2;refGene_gn=DDR2;refGene_tx=XM_019822262.2
F1 64693663 64693726
refGene_gn=DDR2;refGene_tx=XM_006942994.4;refGene_gn=DDR2;refGene_tx=XM_006942995.4;refGene_gn=DDR2;refGene_tx=XM_019822258.2;refGene_gn=DDR2;refGene_tx=XM_019822259.2;refGene_gn=DDR2;refGene_tx=XM_019822260.2;refGene_gn=DDR2;refGene_tx=XM_019822262.2
F1 64695402 64695646
refGene_gn=DDR2;refGene_tx=XM_006942994.4;refGene_gn=DDR2;refGene_tx=XM_006942995.4;refGene_gn=DDR2;refGene_tx=XM_019822258.2;refGene_gn=DDR2;refGene_tx=XM_019822259.2;refGene_gn=DDR2;refGene_tx=XM_019822260.2;refGene_gn=DDR2;refGene_tx=XM_019822262.2
F1 64700150 64700334
refGene_gn=DDR2;refGene_tx=XM_006942994.4;refGene_gn=DDR2;refGene_tx=XM_006942995.4;refGene_gn=DDR2;refGene_tx=XM_019822258.2;refGene_gn=DDR2;refGene_tx=XM_019822259.2;refGene_gn=DDR2;refGene_tx=XM_019822260.2;refGene_gn=DDR2;refGene_tx=XM_019822262.2
F1 64704200 64704306
refGene_gn=DDR2;refGene_tx=XM_006942994.4;refGene_gn=DDR2;refGene_tx=XM_006942995.4;refGene_gn=DDR2;refGene_tx=XM_019822258.2;refGene_gn=DDR2;refGene_tx=XM_019822259.2;refGene_gn=DDR2;refGene_tx=XM_019822260.2;refGene_gn=DDR2;refGene_tx=XM_019822262.2
F1 64704686 64704834
refGene_gn=DDR2;refGene_tx=XM_006942994.4;refGene_gn=DDR2;refGene_tx=XM_006942995.4;refGene_gn=DDR2;refGene_tx=XM_019822258.2;refGene_gn=DDR2;refGene_tx=XM_019822259.2;refGene_gn=DDR2;refGene_tx=XM_019822260.2;refGene_gn=DDR2;refGene_tx=XM_019822262.2
F1 64705142 64705374
refGene_gn=DDR2;refGene_tx=XM_006942994.4;refGene_gn=DDR2;refGene_tx=XM_006942995.4;refGene_gn=DDR2;refGene_tx=XM_019822258.2;refGene_gn=DDR2;refGene_tx=XM_019822259.2;refGene_gn=DDR2;refGene_tx=XM_019822260.2;refGene_gn=DDR2;refGene_tx=XM_019822262.2
F1 64706778 64706881
refGene_gn=DDR2;refGene_tx=XM_006942994.4;refGene_gn=DDR2;refGene_tx=XM_006942995.4;refGene_gn=DDR2;refGene_tx=XM_019822258.2;refGene_gn=DDR2;refGene_tx=XM_019822259.2;refGene_gn=DDR2;refGene_tx=XM_019822260.2
F1 64734976 64735085
refGene_gn=DDR2;refGene_tx=XM_006942994.4;refGene_gn=DDR2;refGene_tx=XM_006942995.4;refGene_gn=DDR2;refGene_tx=XM_019822258.2;refGene_gn=DDR2;refGene_tx=XM_019822260.2
F1 64799308 64799327
refGene_gn=DDR2;refGene_tx=XM_006942994.4;refGene_gn=DDR2;refGene_tx=XM_019822258.2;refGene_gn=DDR2;refGene_tx=XM_019822259.2;refGene_gn=DDR2;refGene_tx=XM_019822260.2

F1 64865814 64865907 refGene_gn=UAP1;refGene_tx=XM_023247656.1
F1 64867162 64867229 refGene_gn=UAP1;refGene_tx=XM_023247656.1
F1 64871726 64871777 refGene_gn=UAP1;refGene_tx=XM_023247656.1
F1 64872882 64873071 refGene_gn=UAP1;refGene_tx=XM_023247656.1
F1 64874128 64874205 refGene_gn=UAP1;refGene_tx=XM_023247656.1
F1 64874206 64874270 refGene_gn=UAP1;refGene_tx=XM_023247656.1
F1 64875181 64875375 refGene_gn=UAP1;refGene_tx=XM_023247656.1
F1 64879861 64880034 refGene_gn=UAP1;refGene_tx=XM_023247656.1
F1 64881857 64882033 refGene_gn=UAP1;refGene_tx=XM_023247656.1
F1 64884336 64884541 refGene_gn=UAP1;refGene_tx=XM_023247656.1
F1 64894605 64894885 refGene_gn=UAP1;refGene_tx=XM_023247656.1
F1 64899054 64899150
refGene_gn=LOC111558344;refGene_tx=XM_023247465.1
F1 64902549 64903209
refGene_gn=LOC111558344;refGene_tx=XM_023247465.1
F1 64937048 64937195
refGene_gn=UHMK1;refGene_tx=XM_023247657.1;refGene_gn=UHMK1;refGene_tx
=XM_023247658.1
F1 64941189 64941278 refGene_gn=UHMK1;refGene_tx=XM_023247657.1
F1 64943570 64943669
refGene_gn=UHMK1;refGene_tx=XM_023247657.1;refGene_gn=UHMK1;refGene_tx
=XM_023247658.1
F1 64943819 64943896
refGene_gn=UHMK1;refGene_tx=XM_023247657.1;refGene_gn=UHMK1;refGene_tx
=XM_023247658.1
F1 64948029 64948124
refGene_gn=UHMK1;refGene_tx=XM_023247657.1;refGene_gn=UHMK1;refGene_tx
=XM_023247658.1
F1 64950149 64950341
refGene_gn=UHMK1;refGene_tx=XM_023247657.1;refGene_gn=UHMK1;refGene_tx
=XM_023247658.1
F1 64951350 64951643
refGene_gn=UHMK1;refGene_tx=XM_023247657.1;refGene_gn=UHMK1;refGene_tx
=XM_023247658.1
F1 64953053 64953321
refGene_gn=UHMK1;refGene_tx=XM_023247657.1;refGene_gn=UHMK1;refGene_tx
=XM_023247658.1
F1 64975050 64975214 refGene_gn=SH2D1B;refGene_tx=XM_003999546.4
F1 64993073 64993137 refGene_gn=SH2D1B;refGene_tx=XM_003999546.4
F1 64995835 64996001 refGene_gn=SH2D1B;refGene_tx=XM_003999546.4
F1 64997765 64997803 refGene_gn=SH2D1B;refGene_tx=XM_003999546.4
F1 65014580 65015055
refGene_gn=CF1H1orf226;refGene_tx=XM_019822278.2;refGene_gn=CF1H1orf22
6;refGene_tx=XM_023247344.1
F1 65016060 65016367
refGene_gn=CF1H1orf226;refGene_tx=XM_019822278.2;refGene_gn=CF1H1orf22
6;refGene_tx=XM_023247344.1
F1 65019104 65019300
refGene_gn=CF1H1orf226;refGene_tx=XM_019822278.2
F1 65020977 65021080 refGene_gn=SPATA46;refGene_tx=XM_003999553.5

F1 65022199 65022440
refGene_gn=SPATA46;refGene_tx=XM_003999553.5;refGene_gn=SPATA46;refGene_tx=XM_011291125.3;refGene_gn=SPATA46;refGene_tx=XM_011291126.3
F1 65023292 65023813
refGene_gn=SPATA46;refGene_tx=XM_003999553.5;refGene_gn=SPATA46;refGene_tx=XM_011291125.3;refGene_gn=SPATA46;refGene_tx=XM_011291126.3
F1 65029391 65029685 refGene_gn=NOS1AP;refGene_tx=XM_023247343.1
F1 65029686 65029787 refGene_gn=NOS1AP;refGene_tx=XM_023247343.1
F1 65030994 65031160 refGene_gn=NOS1AP;refGene_tx=XM_023247343.1
F1 65038231 65038411 refGene_gn=NOS1AP;refGene_tx=XM_023247343.1
F1 65040045 65040212 refGene_gn=NOS1AP;refGene_tx=XM_023247343.1
F1 65046901 65047043 refGene_gn=NOS1AP;refGene_tx=XM_023247343.1
F1 65056369 65056478 refGene_gn=NOS1AP;refGene_tx=XM_023247343.1
F1 65087770 65087829 refGene_gn=NOS1AP;refGene_tx=XM_023247343.1
F1 65099010 65099103 refGene_gn=NOS1AP;refGene_tx=XM_023247343.1
F1 65220673 65220745 refGene_gn=NOS1AP;refGene_tx=XM_023247343.1
F1 65296672 65296777 refGene_gn=NOS1AP;refGene_tx=XM_023247343.1
F1 65339878 65340055 refGene_gn=OLFML2B;refGene_tx=XM_011291123.3
F1 65342418 65342682 refGene_gn=OLFML2B;refGene_tx=XM_011291123.3
F1 65344924 65345032 refGene_gn=OLFML2B;refGene_tx=XM_011291123.3
F1 65360059 65360236 refGene_gn=OLFML2B;refGene_tx=XM_011291123.3
F1 65365280 65365506 refGene_gn=OLFML2B;refGene_tx=XM_011291123.3
F1 65367448 65367729 refGene_gn=OLFML2B;refGene_tx=XM_011291123.3
F1 65367730 65367962 refGene_gn=OLFML2B;refGene_tx=XM_011291123.3
F1 65376556 65376733 refGene_gn=OLFML2B;refGene_tx=XM_011291123.3
F1 65377344 65377946 refGene_gn=OLFML2B;refGene_tx=XM_011291123.3
F1 65399375 65399584
refGene_gn=ATF6;refGene_tx=XM_023247596.1;refGene_gn=ATF6;refGene_tx=XM_023247597.1
F1 65408980 65409057 refGene_gn=ATF6;refGene_tx=XM_023247598.1
F1 65449524 65449609
refGene_gn=ATF6;refGene_tx=XM_023247596.1;refGene_gn=ATF6;refGene_tx=XM_023247597.1;refGene_gn=ATF6;refGene_tx=XM_023247598.1
F1 65514399 65514514
refGene_gn=ATF6;refGene_tx=XM_023247596.1;refGene_gn=ATF6;refGene_tx=XM_023247597.1;refGene_gn=ATF6;refGene_tx=XM_023247598.1
F1 65518189 65518257
refGene_gn=ATF6;refGene_tx=XM_023247596.1;refGene_gn=ATF6;refGene_tx=XM_023247597.1;refGene_gn=ATF6;refGene_tx=XM_023247598.1
F1 65526381 65526481
refGene_gn=ATF6;refGene_tx=XM_023247596.1;refGene_gn=ATF6;refGene_tx=XM_023247597.1;refGene_gn=ATF6;refGene_tx=XM_023247598.1
F1 65527368 65527482
refGene_gn=ATF6;refGene_tx=XM_023247596.1;refGene_gn=ATF6;refGene_tx=XM_023247597.1;refGene_gn=ATF6;refGene_tx=XM_023247598.1
F1 65532159 65532291
refGene_gn=ATF6;refGene_tx=XM_023247596.1;refGene_gn=ATF6;refGene_tx=XM_023247597.1;refGene_gn=ATF6;refGene_tx=XM_023247598.1
F1 65553574 65553666
refGene_gn=ATF6;refGene_tx=XM_023247596.1;refGene_gn=ATF6;refGene_tx=X

M_023247597.1;refGene_gn=ATF6;refGene_tx=XM_023247598.1
F1 65554786 65554972
refGene_gn=ATF6;refGene_tx=XM_023247596.1;refGene_gn=ATF6;refGene_tx=X
M_023247597.1;refGene_gn=ATF6;refGene_tx=XM_023247598.1
F1 65572026 65572247
refGene_gn=ATF6;refGene_tx=XM_023247596.1;refGene_gn=ATF6;refGene_tx=X
M_023247597.1;refGene_gn=ATF6;refGene_tx=XM_023247598.1
F1 65579866 65580070
refGene_gn=ATF6;refGene_tx=XM_023247596.1;refGene_gn=ATF6;refGene_tx=X
M_023247597.1;refGene_gn=ATF6;refGene_tx=XM_023247598.1
F1 65581377 65581507
refGene_gn=ATF6;refGene_tx=XM_023247596.1;refGene_gn=ATF6;refGene_tx=X
M_023247597.1;refGene_gn=ATF6;refGene_tx=XM_023247598.1
F1 65584422 65584523
refGene_gn=ATF6;refGene_tx=XM_023247596.1;refGene_gn=ATF6;refGene_tx=X
M_023247597.1;refGene_gn=ATF6;refGene_tx=XM_023247598.1
F1 65586472 65586560
refGene_gn=ATF6;refGene_tx=XM_023247596.1;refGene_gn=ATF6;refGene_tx=X
M_023247597.1;refGene_gn=ATF6;refGene_tx=XM_023247598.1
F1 65590097 65590174
refGene_gn=ATF6;refGene_tx=XM_023247596.1;refGene_gn=ATF6;refGene_tx=X
M_023247597.1;refGene_gn=ATF6;refGene_tx=XM_023247598.1
F1 65600260 65600312
refGene_gn=ATF6;refGene_tx=XM_023247596.1;refGene_gn=ATF6;refGene_tx=X
M_023247597.1;refGene_gn=ATF6;refGene_tx=XM_023247598.1
F1 65609311 65609473 refGene_gn=DUSP12;refGene_tx=XM_003999556.4
F1 65611137 65611324 refGene_gn=DUSP12;refGene_tx=XM_003999556.4
F1 65612001 65612098 refGene_gn=DUSP12;refGene_tx=XM_003999556.4
F1 65612512 65612631 refGene_gn=DUSP12;refGene_tx=XM_003999556.4
F1 65612721 65612835 refGene_gn=DUSP12;refGene_tx=XM_003999556.4
F1 65614163 65614546 refGene_gn=DUSP12;refGene_tx=XM_003999556.4
F1 65634409 65634828
refGene_gn=FCRLB;refGene_tx=XM_023247794.1;refGene_gn=FCRLB;refGene_tx
=XM_023247795.1;refGene_gn=FCRLB;refGene_tx=XM_023247796.1
F1 65635838 65636117
refGene_gn=FCRLB;refGene_tx=XM_023247794.1;refGene_gn=FCRLB;refGene_tx
=XM_023247795.1;refGene_gn=FCRLB;refGene_tx=XM_023247796.1
F1 65636726 65636993
refGene_gn=FCRLB;refGene_tx=XM_023247794.1;refGene_gn=FCRLB;refGene_tx
=XM_023247795.1;refGene_gn=FCRLB;refGene_tx=XM_023247796.1
F1 65638188 65638443
refGene_gn=FCRLB;refGene_tx=XM_023247794.1;refGene_gn=FCRLB;refGene_tx
=XM_023247795.1;refGene_gn=FCRLB;refGene_tx=XM_023247796.1
F1 65638791 65638812
refGene_gn=FCRLB;refGene_tx=XM_023247794.1;refGene_gn=FCRLB;refGene_tx
=XM_023247795.1;refGene_gn=FCRLB;refGene_tx=XM_023247796.1
F1 65639136 65639284
refGene_gn=FCRLB;refGene_tx=XM_023247794.1;refGene_gn=FCRLB;refGene_tx
=XM_023247795.1;refGene_gn=FCRLB;refGene_tx=XM_023247796.1
F1 65647722 65648015 refGene_gn=FCRLA;refGene_tx=XM_006943003.3

F1 65648879 65649164 refGene_gn=FCRLA;refGene_tx=XM_006943003.3
F1 65649745 65650012 refGene_gn=FCRLA;refGene_tx=XM_006943003.3
F1 65650786 65650933 refGene_gn=FCRLA;refGene_tx=XM_006943003.3
F1 65651188 65651236 refGene_gn=FCRLA;refGene_tx=XM_006943003.3
F1 65654060 65654139 refGene_gn=FCRLA;refGene_tx=XM_006943003.3
F1 65669843 65669933
refGene_gn=FCGR2B;refGene_tx=NM_001290240.1;refGene_gn=FCGR2B;refGene_tx=XM_006943005.2;refGene_gn=FCGR2B;refGene_tx=XM_019821318.1
F1 65670011 65670049
refGene_gn=FCGR2B;refGene_tx=NM_001290240.1;refGene_gn=FCGR2B;refGene_tx=XM_006943005.2;refGene_gn=FCGR2B;refGene_tx=XM_019821318.1
F1 65671875 65672006
refGene_gn=FCGR2B;refGene_tx=XM_006943005.2;refGene_gn=FCGR2B;refGene_tx=XM_019821318.1
F1 65673888 65674011
refGene_gn=FCGR2B;refGene_tx=NM_001290240.1;refGene_gn=FCGR2B;refGene_tx=XM_006943005.2;refGene_gn=FCGR2B;refGene_tx=XM_019821318.1
F1 65674715 65674970
refGene_gn=FCGR2B;refGene_tx=NM_001290240.1;refGene_gn=FCGR2B;refGene_tx=XM_006943005.2;refGene_gn=FCGR2B;refGene_tx=XM_019821318.1
F1 65678222 65678477
refGene_gn=FCGR2B;refGene_tx=NM_001290240.1;refGene_gn=FCGR2B;refGene_tx=XM_006943005.2;refGene_gn=FCGR2B;refGene_tx=XM_019821318.1
F1 65679943 65679964
refGene_gn=FCGR2B;refGene_tx=NM_001290240.1;refGene_gn=FCGR2B;refGene_tx=XM_006943005.2;refGene_gn=FCGR2B;refGene_tx=XM_019821318.1
F1 65684663 65684775
refGene_gn=FCGR2B;refGene_tx=NM_001290240.1;refGene_gn=FCGR2B;refGene_tx=XM_006943005.2;refGene_gn=FCGR2B;refGene_tx=XM_019821318.1
F1 65743379 65743419
refGene_gn=FCGR3A;refGene_tx=NM_001009205.1;refGene_gn=FCGR3A;refGene_tx=XM_006942999.4
F1 65744113 65744134
refGene_gn=FCGR3A;refGene_tx=NM_001009205.1;refGene_gn=FCGR3A;refGene_tx=XM_006942999.4
F1 65744501 65744759
refGene_gn=FCGR3A;refGene_tx=NM_001009205.1;refGene_gn=FCGR3A;refGene_tx=XM_006942999.4;refGene_gn=FCGR3A;refGene_tx=XM_006943000.4;refGene_gn=FCGR3A;refGene_tx=XM_006943001.4
F1 65748330 65748585
refGene_gn=FCGR3A;refGene_tx=NM_001009205.1;refGene_gn=FCGR3A;refGene_tx=XM_006942999.4;refGene_gn=FCGR3A;refGene_tx=XM_006943000.4;refGene_gn=FCGR3A;refGene_tx=XM_006943001.4
F1 65749370 65749549
refGene_gn=FCGR3A;refGene_tx=NM_001009205.1;refGene_gn=FCGR3A;refGene_tx=XM_006942999.4;refGene_gn=FCGR3A;refGene_tx=XM_006943000.4;refGene_gn=FCGR3A;refGene_tx=XM_006943001.4
F1 65901366 65901393 refGene_gn=CFAP126;refGene_tx=XM_003999641.5
F1 65902875 65902938 refGene_gn=CFAP126;refGene_tx=XM_003999641.5
F1 65903140 65903221 refGene_gn=CFAP126;refGene_tx=XM_003999641.5

F1 65903699 65903876 refGene_gn=CFAP126;refGene_tx=XM_003999641.5
F1 65904190 65904529 refGene_gn=CFAP126;refGene_tx=XM_003999641.5
F1 65906961 65907066 refGene_gn=SDHC;refGene_tx=XM_003999558.4
F1 65910128 65910292 refGene_gn=SDHC;refGene_tx=XM_003999558.4
F1 65925353 65925415 refGene_gn=SDHC;refGene_tx=XM_003999558.4
F1 65935321 65935423 refGene_gn=SDHC;refGene_tx=XM_003999558.4
F1 65937068 65937125 refGene_gn=SDHC;refGene_tx=XM_003999558.4
F1 65942710 65942730 refGene_gn=SDHC;refGene_tx=XM_003999558.4
F1 65946835 65946902 refGene_gn=MPZ;refGene_tx=XM_003999560.3
F1 65949393 65949560 refGene_gn=MPZ;refGene_tx=XM_003999560.3
F1 65949798 65950012 refGene_gn=MPZ;refGene_tx=XM_003999560.3
F1 65950264 65950400 refGene_gn=MPZ;refGene_tx=XM_003999560.3
F1 65950831 65950889 refGene_gn=MPZ;refGene_tx=XM_003999560.3
F1 65951013 65951307 refGene_gn=MPZ;refGene_tx=XM_003999560.3
F1 65966905 65967000 refGene_gn=PCP4L1;refGene_tx=XM_023247606.1
F1 65967130 65967273
refGene_gn=PCP4L1;refGene_tx=XM_023247606.1;refGene_gn=PCP4L1;refGene_
tx=XM_023247607.1
F1 65967870 65967925
refGene_gn=PCP4L1;refGene_tx=XM_023247606.1;refGene_gn=PCP4L1;refGene_
tx=XM_023247607.1
F1 65987840 65987849
refGene_gn=PCP4L1;refGene_tx=XM_023247606.1;refGene_gn=PCP4L1;refGene_
tx=XM_023247607.1
F1 65997863 65997970
refGene_gn=LOC101088457;refGene_tx=XM_006943014.4;refGene_gn=LOC101088
457;refGene_tx=XM_019822293.1
F1 65998392 65998523
refGene_gn=LOC101088457;refGene_tx=XM_006943014.4;refGene_gn=LOC101088
457;refGene_tx=XM_019822293.1
F1 65999637 65999807
refGene_gn=LOC101088457;refGene_tx=XM_006943014.4;refGene_gn=LOC101088
457;refGene_tx=XM_019822293.1
F1 66000019 66000159
refGene_gn=LOC101088457;refGene_tx=XM_006943014.4;refGene_gn=LOC101088
457;refGene_tx=XM_019822293.1
F1 66000651 66000797
refGene_gn=LOC101088457;refGene_tx=XM_006943014.4;refGene_gn=LOC101088
457;refGene_tx=XM_019822293.1
F1 66000893 66001010
refGene_gn=LOC101088457;refGene_tx=XM_006943014.4;refGene_gn=LOC101088
457;refGene_tx=XM_019822293.1
F1 66001181 66001287
refGene_gn=LOC101088457;refGene_tx=XM_006943014.4;refGene_gn=LOC101088
457;refGene_tx=XM_019822293.1
F1 66001875 66001957
refGene_gn=TOMM40L;refGene_tx=XM_006943009.3;refGene_gn=TOMM40L;refGen
e_tx=XM_006943010.3
F1 66002182 66002312
refGene_gn=LOC101088457;refGene_tx=XM_006943014.4

F1 66003035 66003175
refGene_gn=TOMM40L;refGene_tx=XM_006943009.3;refGene_gn=TOMM40L;refGene_tx=XM_006943010.3;refGene_gn=TOMM40L;refGene_tx=XM_006943011.3
F1 66003541 66003644
refGene_gn=TOMM40L;refGene_tx=XM_006943009.3;refGene_gn=TOMM40L;refGene_tx=XM_006943010.3;refGene_gn=TOMM40L;refGene_tx=XM_006943011.3
F1 66003855 66003932
refGene_gn=TOMM40L;refGene_tx=XM_006943009.3;refGene_gn=TOMM40L;refGene_tx=XM_006943010.3;refGene_gn=TOMM40L;refGene_tx=XM_006943011.3
F1 66004048 66004171
refGene_gn=TOMM40L;refGene_tx=XM_006943009.3;refGene_gn=TOMM40L;refGene_tx=XM_006943010.3;refGene_gn=TOMM40L;refGene_tx=XM_006943011.3
F1 66004350 66004456
refGene_gn=TOMM40L;refGene_tx=XM_006943009.3;refGene_gn=TOMM40L;refGene_tx=XM_006943010.3;refGene_gn=TOMM40L;refGene_tx=XM_006943011.3
F1 66004599 66004701
refGene_gn=TOMM40L;refGene_tx=XM_006943009.3;refGene_gn=TOMM40L;refGene_tx=XM_006943010.3;refGene_gn=TOMM40L;refGene_tx=XM_006943011.3
F1 66004980 66005073
refGene_gn=TOMM40L;refGene_tx=XM_006943009.3;refGene_gn=TOMM40L;refGene_tx=XM_006943010.3;refGene_gn=TOMM40L;refGene_tx=XM_006943011.3
F1 66005360 66005428
refGene_gn=TOMM40L;refGene_tx=XM_006943009.3;refGene_gn=TOMM40L;refGene_tx=XM_006943010.3;refGene_gn=TOMM40L;refGene_tx=XM_006943011.3
F1 66005718 66005833
refGene_gn=TOMM40L;refGene_tx=XM_006943009.3;refGene_gn=TOMM40L;refGene_tx=XM_006943010.3;refGene_gn=TOMM40L;refGene_tx=XM_006943011.3
F1 66008654 66008758
refGene_gn=LOC101088457;refGene_tx=XM_003999562.3;refGene_gn=LOC101088457;refGene_tx=XM_019822293.1;refGene_gn=LOC101088457;refGene_tx=XM_019822294.1
F1 66008991 66009124
refGene_gn=LOC101088457;refGene_tx=XM_003999562.3;refGene_gn=LOC101088457;refGene_tx=XM_019822293.1;refGene_gn=LOC101088457;refGene_tx=XM_019822294.1
F1 66009956 66010074
refGene_gn=LOC101088457;refGene_tx=XM_003999562.3;refGene_gn=LOC101088457;refGene_tx=XM_019822293.1;refGene_gn=LOC101088457;refGene_tx=XM_019822294.1
F1 66012453 66012516 refGene_gn=FCER1G;refGene_tx=XM_003999564.5
F1 66012652 66012673 refGene_gn=FCER1G;refGene_tx=XM_003999564.5
F1 66013066 66013102 refGene_gn=FCER1G;refGene_tx=XM_003999564.5
F1 66013273 66013365 refGene_gn=FCER1G;refGene_tx=XM_003999564.5
F1 66015987 66016036 refGene_gn=FCER1G;refGene_tx=XM_003999564.5
F1 66016854 66016892 refGene_gn=NDUFS2;refGene_tx=XM_023247974.1
F1 66017084 66017142 refGene_gn=NDUFS2;refGene_tx=XM_023247974.1
F1 66017284 66017368 refGene_gn=NDUFS2;refGene_tx=XM_023247974.1
F1 66017546 66017642 refGene_gn=NDUFS2;refGene_tx=XM_023247974.1
F1 66017924 66018054 refGene_gn=NDUFS2;refGene_tx=XM_023247974.1
F1 66018712 66018832 refGene_gn=NDUFS2;refGene_tx=XM_023247974.1

F1 66019238 66019324 refGene_gn=NDUFS2;refGene_tx=XM_023247974.1
F1 66019500 66019578 refGene_gn=NDUFS2;refGene_tx=XM_023247974.1
F1 66019733 66019808 refGene_gn=NDUFS2;refGene_tx=XM_023247974.1
F1 66020071 66020184 refGene_gn=NDUFS2;refGene_tx=XM_023247974.1
F1 66020356 66020477 refGene_gn=NDUFS2;refGene_tx=XM_023247974.1
F1 66022440 66022631 refGene_gn=NDUFS2;refGene_tx=XM_023247974.1
F1 66025411 66025518 refGene_gn=NDUFS2;refGene_tx=XM_023247974.1
F1 66026336 66026431 refGene_gn=NDUFS2;refGene_tx=XM_023247974.1
F1 66030151 66030775 refGene_gn=ADAMTS4;refGene_tx=XM_003999565.5
F1 66031880 66032204 refGene_gn=ADAMTS4;refGene_tx=XM_003999565.5
F1 66032445 66032578 refGene_gn=ADAMTS4;refGene_tx=XM_003999565.5
F1 66033173 66033344 refGene_gn=ADAMTS4;refGene_tx=XM_003999565.5
F1 66034065 66034352 refGene_gn=ADAMTS4;refGene_tx=XM_003999565.5
F1 66034459 66034646 refGene_gn=ADAMTS4;refGene_tx=XM_003999565.5
F1 66034890 66035066 refGene_gn=ADAMTS4;refGene_tx=XM_003999565.5
F1 66035955 66036131 refGene_gn=ADAMTS4;refGene_tx=XM_003999565.5
F1 66036645 66037072 refGene_gn=ADAMTS4;refGene_tx=XM_003999565.5
F1 66047632 66047894
refGene_gn=B4GALT3;refGene_tx=XM_003999566.5;refGene_gn=B4GALT3;refGen
e_tx=XM_006943019.4;refGene_gn=B4GALT3;refGene_tx=XM_011291120.3;refGe
ne_gn=B4GALT3;refGene_tx=XM_019822295.2
F1 66048436 66048672
refGene_gn=B4GALT3;refGene_tx=XM_003999566.5;refGene_gn=B4GALT3;refGen
e_tx=XM_006943019.4;refGene_gn=B4GALT3;refGene_tx=XM_011291120.3;refGe
ne_gn=B4GALT3;refGene_tx=XM_019822295.2
F1 66049262 66049453
refGene_gn=B4GALT3;refGene_tx=XM_003999566.5;refGene_gn=B4GALT3;refGen
e_tx=XM_006943019.4;refGene_gn=B4GALT3;refGene_tx=XM_011291120.3;refGe
ne_gn=B4GALT3;refGene_tx=XM_019822295.2
F1 66049582 66049705
refGene_gn=B4GALT3;refGene_tx=XM_003999566.5;refGene_gn=B4GALT3;refGen
e_tx=XM_006943019.4;refGene_gn=B4GALT3;refGene_tx=XM_011291120.3;refGe
ne_gn=B4GALT3;refGene_tx=XM_019822295.2
F1 66050881 66050986
refGene_gn=B4GALT3;refGene_tx=XM_003999566.5;refGene_gn=B4GALT3;refGen
e_tx=XM_006943019.4;refGene_gn=B4GALT3;refGene_tx=XM_011291120.3;refGe
ne_gn=B4GALT3;refGene_tx=XM_019822295.2
F1 66051116 66051390
refGene_gn=B4GALT3;refGene_tx=XM_003999566.5;refGene_gn=B4GALT3;refGen
e_tx=XM_006943019.4;refGene_gn=B4GALT3;refGene_tx=XM_011291120.3;refGe
ne_gn=B4GALT3;refGene_tx=XM_019822295.2
F1 66052073 66052216
refGene_gn=PPOX;refGene_tx=XM_003999569.4;refGene_gn=PPOX;refGene_tx=X
M_011291117.3;refGene_gn=PPOX;refGene_tx=XM_011291118.3;refGene_gn=PPO
X;refGene_tx=XM_011291119.3;refGene_gn=PPOX;refGene_tx=XM_019822292.1
F1 66052299 66052342
refGene_gn=PPOX;refGene_tx=XM_003999569.4;refGene_gn=PPOX;refGene_tx=X
M_011291117.3;refGene_gn=PPOX;refGene_tx=XM_011291119.3;refGene_gn=PPO
X;refGene_tx=XM_019822292.1
F1 66052463 66052613

refGene_gn=PPOX;refGene_tx=XM_003999569.4;refGene_gn=PPOX;refGene_tx=XM_011291117.3;refGene_gn=PPOX;refGene_tx=XM_011291118.3;refGene_gn=PPOX;refGene_tx=XM_011291119.3;refGene_gn=PPOX;refGene_tx=XM_019822292.1
F1 66052712 66052823
refGene_gn=PPOX;refGene_tx=XM_003999569.4;refGene_gn=PPOX;refGene_tx=XM_011291117.3;refGene_gn=PPOX;refGene_tx=XM_011291118.3;refGene_gn=PPOX;refGene_tx=XM_011291119.3;refGene_gn=PPOX;refGene_tx=XM_019822292.1
F1 66052903 66053022
refGene_gn=PPOX;refGene_tx=XM_003999569.4;refGene_gn=PPOX;refGene_tx=XM_011291117.3;refGene_gn=PPOX;refGene_tx=XM_011291118.3;refGene_gn=PPOX;refGene_tx=XM_011291119.3;refGene_gn=PPOX;refGene_tx=XM_019822292.1
F1 66053203 66053264
refGene_gn=PPOX;refGene_tx=XM_003999569.4;refGene_gn=PPOX;refGene_tx=XM_011291117.3;refGene_gn=PPOX;refGene_tx=XM_011291118.3;refGene_gn=PPOX;refGene_tx=XM_011291119.3;refGene_gn=PPOX;refGene_tx=XM_019822292.1
F1 66053713 66053904
refGene_gn=PPOX;refGene_tx=XM_003999569.4;refGene_gn=PPOX;refGene_tx=XM_011291117.3;refGene_gn=PPOX;refGene_tx=XM_011291118.3;refGene_gn=PPOX;refGene_tx=XM_011291119.3;refGene_gn=PPOX;refGene_tx=XM_019822292.1
F1 66054011 66054156
refGene_gn=PPOX;refGene_tx=XM_003999569.4;refGene_gn=PPOX;refGene_tx=XM_011291117.3;refGene_gn=PPOX;refGene_tx=XM_011291118.3;refGene_gn=PPOX;refGene_tx=XM_011291119.3;refGene_gn=PPOX;refGene_tx=XM_019822292.1
F1 66054458 66054591
refGene_gn=PPOX;refGene_tx=XM_003999569.4;refGene_gn=PPOX;refGene_tx=XM_011291117.3;refGene_gn=PPOX;refGene_tx=XM_011291118.3;refGene_gn=PPOX;refGene_tx=XM_011291119.3;refGene_gn=PPOX;refGene_tx=XM_019822292.1
F1 66054727 66054876
refGene_gn=PPOX;refGene_tx=XM_003999569.4;refGene_gn=PPOX;refGene_tx=XM_011291117.3;refGene_gn=PPOX;refGene_tx=XM_011291118.3;refGene_gn=PPOX;refGene_tx=XM_011291119.3;refGene_gn=PPOX;refGene_tx=XM_019822292.1
F1 66054973 66055108
refGene_gn=PPOX;refGene_tx=XM_003999569.4;refGene_gn=PPOX;refGene_tx=XM_011291117.3;refGene_gn=PPOX;refGene_tx=XM_011291118.3;refGene_gn=PPOX;refGene_tx=XM_019822292.1
F1 66055246 66055333
refGene_gn=PPOX;refGene_tx=XM_003999569.4;refGene_gn=PPOX;refGene_tx=XM_011291117.3;refGene_gn=PPOX;refGene_tx=XM_011291118.3;refGene_gn=PPOX;refGene_tx=XM_019822292.1
F1 66056743 66056834
refGene_gn=USP21;refGene_tx=XM_003999570.4;refGene_gn=USP21;refGene_tx=XM_006943023.3;refGene_gn=USP21;refGene_tx=XM_019822290.2
F1 66057011 66057126
refGene_gn=USP21;refGene_tx=XM_003999570.4;refGene_gn=USP21;refGene_tx=XM_006943023.3;refGene_gn=USP21;refGene_tx=XM_019822290.2
F1 66057233 66057341
refGene_gn=USP21;refGene_tx=XM_003999570.4;refGene_gn=USP21;refGene_tx=XM_006943023.3;refGene_gn=USP21;refGene_tx=XM_019822290.2
F1 66057573 66057652
refGene_gn=USP21;refGene_tx=XM_003999570.4;refGene_gn=USP21;refGene_tx

=XM_006943023.3;refGene_gn=USP21;refGene_tx=XM_019822290.2;refGene_gn=USP21;refGene_tx=XM_019822291.2
F1 66057932 66058019
refGene_gn=USP21;refGene_tx=XM_003999570.4;refGene_gn=USP21;refGene_tx=XM_006943023.3;refGene_gn=USP21;refGene_tx=XM_019822290.2;refGene_gn=USP21;refGene_tx=XM_019822291.2
F1 66058207 66058311
refGene_gn=USP21;refGene_tx=XM_003999570.4;refGene_gn=USP21;refGene_tx=XM_006943023.3;refGene_gn=USP21;refGene_tx=XM_019822290.2;refGene_gn=USP21;refGene_tx=XM_019822291.2
F1 66058550 66058615
refGene_gn=USP21;refGene_tx=XM_003999570.4;refGene_gn=USP21;refGene_tx=XM_006943023.3;refGene_gn=USP21;refGene_tx=XM_019822290.2;refGene_gn=USP21;refGene_tx=XM_019822291.2
F1 66059128 66059284
refGene_gn=USP21;refGene_tx=XM_003999570.4;refGene_gn=USP21;refGene_tx=XM_006943023.3;refGene_gn=USP21;refGene_tx=XM_019822290.2;refGene_gn=USP21;refGene_tx=XM_019822291.2
F1 66059458 66059570
refGene_gn=USP21;refGene_tx=XM_003999570.4;refGene_gn=USP21;refGene_tx=XM_006943023.3;refGene_gn=USP21;refGene_tx=XM_019822290.2;refGene_gn=USP21;refGene_tx=XM_019822291.2
F1 66059777 66059898
refGene_gn=USP21;refGene_tx=XM_003999570.4;refGene_gn=USP21;refGene_tx=XM_006943023.3;refGene_gn=USP21;refGene_tx=XM_019822290.2;refGene_gn=USP21;refGene_tx=XM_019822291.2
F1 66060068 66060128
refGene_gn=USP21;refGene_tx=XM_003999570.4;refGene_gn=USP21;refGene_tx=XM_006943023.3;refGene_gn=USP21;refGene_tx=XM_019822290.2;refGene_gn=USP21;refGene_tx=XM_019822291.2
F1 66060865 66061540
refGene_gn=USP21;refGene_tx=XM_003999570.4;refGene_gn=USP21;refGene_tx=XM_006943023.3;refGene_gn=USP21;refGene_tx=XM_019822290.2;refGene_gn=USP21;refGene_tx=XM_019822291.2
F1 66063552 66063633 refGene_gn=UFC1;refGene_tx=XM_003999573.5
F1 66063837 66063928 refGene_gn=UFC1;refGene_tx=XM_003999573.5
F1 66064169 66064246 refGene_gn=UFC1;refGene_tx=XM_003999573.5
F1 66064541 66064605 refGene_gn=UFC1;refGene_tx=XM_003999573.5
F1 66064844 66064912 refGene_gn=UFC1;refGene_tx=XM_003999573.5
F1 66066722 66066845 refGene_gn=UFC1;refGene_tx=XM_003999573.5
F1 66094939 66095264
refGene_gn=DEDD;refGene_tx=XM_023247881.1;refGene_gn=DEDD;refGene_tx=XM_023247882.1
F1 66095460 66095568
refGene_gn=DEDD;refGene_tx=XM_023247881.1;refGene_gn=DEDD;refGene_tx=XM_023247882.1
F1 66095973 66096120
refGene_gn=DEDD;refGene_tx=XM_023247881.1;refGene_gn=DEDD;refGene_tx=XM_023247882.1
F1 66096969 66097346

refGene_gn=DEDD;refGene_tx=XM_023247881.1;refGene_gn=DEDD;refGene_tx=XM_023247882.1
F1 66098709 66098976 refGene_gn=NIT1;refGene_tx=XM_003999574.4;refGene_gn=NIT1;refGene_tx=XM_003999576.4;refGene_gn=NIT1;refGene_tx=XM_006943024.3;refGene_gn=NIT1;refGene_tx=XM_019822300.1;refGene_gn=NIT1;refGene_tx=XM_019822301.2;refGene_gn=NIT1;refGene_tx=XM_023247880.1
F1 66099260 66099386 refGene_gn=NIT1;refGene_tx=XM_003999574.4;refGene_gn=NIT1;refGene_tx=XM_003999576.4;refGene_gn=NIT1;refGene_tx=XM_006943024.3;refGene_gn=NIT1;refGene_tx=XM_019822301.2;refGene_gn=NIT1;refGene_tx=XM_023247880.1
F1 66099552 66099686 refGene_gn=NIT1;refGene_tx=XM_003999574.4;refGene_gn=NIT1;refGene_tx=XM_003999576.4;refGene_gn=NIT1;refGene_tx=XM_006943024.3;refGene_gn=NIT1;refGene_tx=XM_019822300.1;refGene_gn=NIT1;refGene_tx=XM_019822301.2;refGene_gn=NIT1;refGene_tx=XM_023247880.1
F1 66099890 66099994 refGene_gn=NIT1;refGene_tx=XM_003999574.4;refGene_gn=NIT1;refGene_tx=XM_003999576.4;refGene_gn=NIT1;refGene_tx=XM_006943024.3;refGene_gn=NIT1;refGene_tx=XM_019822300.1;refGene_gn=NIT1;refGene_tx=XM_019822301.2;refGene_gn=NIT1;refGene_tx=XM_023247880.1
F1 66100125 66100382 refGene_gn=NIT1;refGene_tx=XM_003999574.4;refGene_gn=NIT1;refGene_tx=XM_003999576.4;refGene_gn=NIT1;refGene_tx=XM_006943024.3;refGene_gn=NIT1;refGene_tx=XM_019822300.1;refGene_gn=NIT1;refGene_tx=XM_019822301.2;refGene_gn=NIT1;refGene_tx=XM_023247880.1
F1 66100629 66100728 refGene_gn=NIT1;refGene_tx=XM_003999574.4;refGene_gn=NIT1;refGene_tx=XM_006943024.3;refGene_gn=NIT1;refGene_tx=XM_019822300.1
F1 66100965 66101022 refGene_gn=NIT1;refGene_tx=XM_003999576.4
F1 66101299 66101352 refGene_gn=NIT1;refGene_tx=XM_003999574.4;refGene_gn=NIT1;refGene_tx=XM_006943024.3;refGene_gn=NIT1;refGene_tx=XM_019822300.1
F1 66101481 66101556 refGene_gn=PFDN2;refGene_tx=XM_003999578.3
F1 66111107 66111196 refGene_gn=PFDN2;refGene_tx=XM_003999578.3
F1 66111307 66111431 refGene_gn=PFDN2;refGene_tx=XM_003999578.3
F1 66112161 66112338 refGene_gn=PFDN2;refGene_tx=XM_003999578.3
F1 66112801 66112968 refGene_gn=KLHDC9;refGene_tx=XM_003999644.5;refGene_gn=KLHDC9;refGene_tx=XM_023247879.1
F1 66113088 66113287 refGene_gn=KLHDC9;refGene_tx=XM_003999644.5;refGene_gn=KLHDC9;refGene_tx=XM_023247879.1
F1 66113380 66113540 refGene_gn=KLHDC9;refGene_tx=XM_003999644.5;refGene_gn=KLHDC9;refGene_tx=XM_023247879.1
F1 66113849 66114367 refGene_gn=KLHDC9;refGene_tx=XM_003999644.5;refGene_gn=KLHDC9;refGene_tx=XM_023247879.1
F1 66123289 66123311 refGene_gn=NECTIN4;refGene_tx=XM_019822298.2

F1 66124483 66124562
refGene_gn=NECTIN4;refGene_tx=XM_019822296.2;refGene_gn=NECTIN4;refGene_tx=XM_019822297.2;refGene_gn=NECTIN4;refGene_tx=XM_019822299.2
F1 66132978 66133338
refGene_gn=NECTIN4;refGene_tx=XM_019822296.2;refGene_gn=NECTIN4;refGene_tx=XM_019822297.2;refGene_gn=NECTIN4;refGene_tx=XM_019822298.2;refGene_gn=NECTIN4;refGene_tx=XM_019822299.2
F1 66135111 66135402
refGene_gn=NECTIN4;refGene_tx=XM_019822296.2;refGene_gn=NECTIN4;refGene_tx=XM_019822297.2;refGene_gn=NECTIN4;refGene_tx=XM_019822298.2;refGene_gn=NECTIN4;refGene_tx=XM_019822299.2
F1 66136331 66136452
refGene_gn=NECTIN4;refGene_tx=XM_019822296.2;refGene_gn=NECTIN4;refGene_tx=XM_019822297.2;refGene_gn=NECTIN4;refGene_tx=XM_019822298.2;refGene_gn=NECTIN4;refGene_tx=XM_019822299.2
F1 66137316 66137465
refGene_gn=NECTIN4;refGene_tx=XM_019822296.2;refGene_gn=NECTIN4;refGene_tx=XM_019822297.2;refGene_gn=NECTIN4;refGene_tx=XM_019822298.2;refGene_gn=NECTIN4;refGene_tx=XM_019822299.2
F1 66137699 66137856
refGene_gn=NECTIN4;refGene_tx=XM_019822296.2;refGene_gn=NECTIN4;refGene_tx=XM_019822297.2;refGene_gn=NECTIN4;refGene_tx=XM_019822298.2;refGene_gn=NECTIN4;refGene_tx=XM_019822299.2
F1 66138272 66138360
refGene_gn=NECTIN4;refGene_tx=XM_019822296.2;refGene_gn=NECTIN4;refGene_tx=XM_019822297.2;refGene_gn=NECTIN4;refGene_tx=XM_019822298.2;refGene_gn=NECTIN4;refGene_tx=XM_019822299.2
F1 66138747 66138822
refGene_gn=NECTIN4;refGene_tx=XM_019822296.2;refGene_gn=NECTIN4;refGene_tx=XM_019822297.2;refGene_gn=NECTIN4;refGene_tx=XM_019822298.2
F1 66139112 66139379
refGene_gn=NECTIN4;refGene_tx=XM_019822296.2;refGene_gn=NECTIN4;refGene_tx=XM_019822297.2;refGene_gn=NECTIN4;refGene_tx=XM_019822298.2;refGene_gn=NECTIN4;refGene_tx=XM_019822299.2
F1 66141893 66141990
refGene_gn=ARHGAP30;refGene_tx=XM_003999580.4;refGene_gn=ARHGAP30;refGene_tx=XM_006943029.3
F1 66147575 66147678
refGene_gn=ARHGAP30;refGene_tx=XM_003999580.4;refGene_gn=ARHGAP30;refGene_tx=XM_006943029.3
F1 66150114 66150259
refGene_gn=ARHGAP30;refGene_tx=XM_003999580.4;refGene_gn=ARHGAP30;refGene_tx=XM_006943029.3
F1 66151143 66151226
refGene_gn=ARHGAP30;refGene_tx=XM_003999580.4;refGene_gn=ARHGAP30;refGene_tx=XM_006943029.3
F1 66151384 66151492
refGene_gn=ARHGAP30;refGene_tx=XM_003999580.4;refGene_gn=ARHGAP30;refGene_tx=XM_006943029.3
F1 66152043 66152171

refGene_gn=ARHGAP30;refGene_tx=XM_003999580.4;refGene_gn=ARHGAP30;refGene_tx=XM_006943029.3
F1 66152906 66153078
refGene_gn=ARHGAP30;refGene_tx=XM_003999580.4;refGene_gn=ARHGAP30;refGene_tx=XM_006943029.3
F1 66153156 66153260
refGene_gn=ARHGAP30;refGene_tx=XM_003999580.4;refGene_gn=ARHGAP30;refGene_tx=XM_006943029.3
F1 66153343 66153421
refGene_gn=ARHGAP30;refGene_tx=XM_003999580.4;refGene_gn=ARHGAP30;refGene_tx=XM_006943029.3
F1 66153970 66154372
refGene_gn=ARHGAP30;refGene_tx=XM_003999580.4;refGene_gn=ARHGAP30;refGene_tx=XM_006943029.3
F1 66155321 66155587
refGene_gn=ARHGAP30;refGene_tx=XM_003999580.4;refGene_gn=ARHGAP30;refGene_tx=XM_006943029.3
F1 66155679 66157308
refGene_gn=ARHGAP30;refGene_tx=XM_003999580.4;refGene_gn=ARHGAP30;refGene_tx=XM_006943029.3
F1 66161246 66161254
refGene_gn=USF1;refGene_tx=XM_006943030.2;refGene_gn=USF1;refGene_tx=XM_019822303.2;refGene_gn=USF1;refGene_tx=XM_019822304.2;refGene_gn=USF1;refGene_tx=XM_019822305.2;refGene_gn=USF1;refGene_tx=XM_019822306.2
F1 66161669 66161719
refGene_gn=USF1;refGene_tx=XM_006943030.2;refGene_gn=USF1;refGene_tx=XM_019822303.2;refGene_gn=USF1;refGene_tx=XM_019822304.2;refGene_gn=USF1;refGene_tx=XM_019822305.2;refGene_gn=USF1;refGene_tx=XM_019822306.2
F1 66161874 66161990
refGene_gn=USF1;refGene_tx=XM_006943030.2;refGene_gn=USF1;refGene_tx=XM_019822303.2;refGene_gn=USF1;refGene_tx=XM_019822304.2;refGene_gn=USF1;refGene_tx=XM_019822305.2;refGene_gn=USF1;refGene_tx=XM_019822306.2
F1 66162342 66162444
refGene_gn=USF1;refGene_tx=XM_006943030.2;refGene_gn=USF1;refGene_tx=XM_019822303.2;refGene_gn=USF1;refGene_tx=XM_019822304.2;refGene_gn=USF1;refGene_tx=XM_019822305.2;refGene_gn=USF1;refGene_tx=XM_019822306.2;refGene_gn=USF1;refGene_tx=XM_019822307.2
F1 66162695 66162891
refGene_gn=USF1;refGene_tx=XM_006943030.2;refGene_gn=USF1;refGene_tx=XM_019822303.2;refGene_gn=USF1;refGene_tx=XM_019822304.2;refGene_gn=USF1;refGene_tx=XM_019822305.2;refGene_gn=USF1;refGene_tx=XM_019822306.2;refGene_gn=USF1;refGene_tx=XM_019822307.2
F1 66163114 66163202
refGene_gn=USF1;refGene_tx=XM_006943030.2;refGene_gn=USF1;refGene_tx=XM_019822303.2;refGene_gn=USF1;refGene_tx=XM_019822304.2;refGene_gn=USF1;refGene_tx=XM_019822305.2;refGene_gn=USF1;refGene_tx=XM_019822306.2;refGene_gn=USF1;refGene_tx=XM_019822307.2
F1 66163337 66163396
refGene_gn=USF1;refGene_tx=XM_006943030.2;refGene_gn=USF1;refGene_tx=XM_019822303.2;refGene_gn=USF1;refGene_tx=XM_019822304.2;refGene_gn=USF

1;refGene_tx=XM_019822305.2;refGene_gn=USF1;refGene_tx=XM_019822306.2;
refGene_gn=USF1;refGene_tx=XM_019822307.2
F1 66163521 66163616
refGene_gn=USF1;refGene_tx=XM_006943030.2;refGene_gn=USF1;refGene_tx=XM_019822303.2;refGene_gn=USF1;refGene_tx=XM_019822304.2;refGene_gn=USF1;refGene_tx=XM_019822305.2;refGene_gn=USF1;refGene_tx=XM_019822306.2;refGene_gn=USF1;refGene_tx=XM_019822307.2
F1 66163860 66163989
refGene_gn=USF1;refGene_tx=XM_006943030.2;refGene_gn=USF1;refGene_tx=XM_019822303.2;refGene_gn=USF1;refGene_tx=XM_019822304.2;refGene_gn=USF1;refGene_tx=XM_019822305.2;refGene_gn=USF1;refGene_tx=XM_019822306.2;refGene_gn=USF1;refGene_tx=XM_019822307.2
F1 66164072 66164264
refGene_gn=USF1;refGene_tx=XM_006943030.2;refGene_gn=USF1;refGene_tx=XM_019822303.2;refGene_gn=USF1;refGene_tx=XM_019822304.2;refGene_gn=USF1;refGene_tx=XM_019822305.2;refGene_gn=USF1;refGene_tx=XM_019822306.2;refGene_gn=USF1;refGene_tx=XM_019822307.2
F1 66165070 66165320
refGene_gn=TSTD1;refGene_tx=XM_023247884.1;refGene_gn=TSTD1;refGene_tx=XM_023247885.1;refGene_gn=TSTD1;refGene_tx=XM_023247886.1
F1 66165513 66165636
refGene_gn=TSTD1;refGene_tx=XM_023247884.1;refGene_gn=TSTD1;refGene_tx=XM_023247886.1
F1 66166102 66166265
refGene_gn=TSTD1;refGene_tx=XM_023247884.1;refGene_gn=TSTD1;refGene_tx=XM_023247885.1
F1 66166359 66166415
refGene_gn=TSTD1;refGene_tx=XM_023247884.1;refGene_gn=TSTD1;refGene_tx=XM_023247885.1;refGene_gn=TSTD1;refGene_tx=XM_023247886.1
F1 66184040 66184101
refGene_gn=F11R;refGene_tx=NM_001037864.2;refGene_gn=F11R;refGene_tx=XM_023247166.1
F1 66203839 66203908
refGene_gn=F11R;refGene_tx=NM_001037864.2;refGene_gn=F11R;refGene_tx=XM_023247166.1
F1 66204074 66204182
refGene_gn=F11R;refGene_tx=NM_001037864.2;refGene_gn=F11R;refGene_tx=XM_023247166.1
F1 66204369 66204516
refGene_gn=F11R;refGene_tx=NM_001037864.2;refGene_gn=F11R;refGene_tx=XM_023247166.1
F1 66204827 66205030
refGene_gn=F11R;refGene_tx=NM_001037864.2;refGene_gn=F11R;refGene_tx=XM_023247166.1
F1 66205427 66205530
refGene_gn=F11R;refGene_tx=NM_001037864.2;refGene_gn=F11R;refGene_tx=XM_023247166.1
F1 66205667 66205775 refGene_gn=F11R;refGene_tx=NM_001037864.2
F1 66205823 66205855 refGene_gn=F11R;refGene_tx=XM_023247166.1
F1 66205994 66206007 refGene_gn=F11R;refGene_tx=NM_001037864.2

F1	66206337	66206386	refGene_gn=F11R;refGene_tx=NM_001037864.2
F1	66206519	66206555	refGene_gn=F11R;refGene_tx=NM_001037864.2
F1	66256473	66256531
refGene_gn=CD244;refGene_tx=XM_006943031.4;refGene_gn=CD244;refGene_tx
=XM_019822309.2
F1	66274936	66275269
refGene_gn=CD244;refGene_tx=XM_006943031.4;refGene_gn=CD244;refGene_tx
=XM_019822309.2
F1	66275358	66275628
refGene_gn=CD244;refGene_tx=XM_006943031.4;refGene_gn=CD244;refGene_tx
=XM_019822309.2
F1	66277481	66277589	refGene_gn=CD244;refGene_tx=XM_006943031.4
F1	66278043	66278111
refGene_gn=CD244;refGene_tx=XM_006943031.4;refGene_gn=CD244;refGene_tx
=XM_019822309.2
F1	66280758	66280818
refGene_gn=CD244;refGene_tx=XM_006943031.4;refGene_gn=CD244;refGene_tx
=XM_019822309.2
F1	66283008	66283071
refGene_gn=CD244;refGene_tx=XM_006943031.4;refGene_gn=CD244;refGene_tx
=XM_019822309.2
F1	66284297	66284354
refGene_gn=CD244;refGene_tx=XM_006943031.4;refGene_gn=CD244;refGene_tx
=XM_019822309.2
F1	66285432	66285513
refGene_gn=CD244;refGene_tx=XM_006943031.4;refGene_gn=CD244;refGene_tx
=XM_019822309.2
F1	66289755	66289821
refGene_gn=LY9;refGene_tx=XM_006943032.2;refGene_gn=LY9;refGene_tx=XM_
006943033.2;refGene_gn=LY9;refGene_tx=XM_006943034.2
F1	66292798	66292870
refGene_gn=LY9;refGene_tx=XM_006943032.2;refGene_gn=LY9;refGene_tx=XM_
006943033.2;refGene_gn=LY9;refGene_tx=XM_006943034.2
F1	66293079	66293498
refGene_gn=LY9;refGene_tx=XM_006943032.2;refGene_gn=LY9;refGene_tx=XM_
006943033.2;refGene_gn=LY9;refGene_tx=XM_006943034.2;refGene_gn=LY9;re
fGene_tx=XM_019822310.1;refGene_gn=LY9;refGene_tx=XM_019822311.1;refGe
ne_gn=LY9;refGene_tx=XM_019822313.1;refGene_gn=LY9;refGene_tx=XM_01982
2314.1
F1	66295759	66295813
refGene_gn=LY9;refGene_tx=XM_006943032.2;refGene_gn=LY9;refGene_tx=XM_
006943033.2;refGene_gn=LY9;refGene_tx=XM_006943034.2;refGene_gn=LY9;re
fGene_tx=XM_019822310.1;refGene_gn=LY9;refGene_tx=XM_019822311.1;refGe
ne_gn=LY9;refGene_tx=XM_019822313.1;refGene_gn=LY9;refGene_tx=XM_01982
2314.1
F1	66296642	66296810
refGene_gn=LY9;refGene_tx=XM_006943032.2;refGene_gn=LY9;refGene_tx=XM_
006943033.2;refGene_gn=LY9;refGene_tx=XM_006943034.2;refGene_gn=LY9;re
fGene_tx=XM_019822310.1;refGene_gn=LY9;refGene_tx=XM_019822311.1;refGe
ne_gn=LY9;refGene_tx=XM_019822313.1;refGene_gn=LY9;refGene_tx=XM_01982

2314.1
F1	66298068 66298338
refGene_gn=LY9;refGene_tx=XM_006943032.2;refGene_gn=LY9;refGene_tx=XM_006943033.2;refGene_gn=LY9;refGene_tx=XM_019822310.1;refGene_gn=LY9;refGene_tx=XM_019822311.1
F1	66299447 66299753
refGene_gn=LY9;refGene_tx=XM_006943032.2;refGene_gn=LY9;refGene_tx=XM_006943033.2;refGene_gn=LY9;refGene_tx=XM_006943034.2;refGene_gn=LY9;refGene_tx=XM_019822310.1;refGene_gn=LY9;refGene_tx=XM_019822311.1;refGene_gn=LY9;refGene_tx=XM_019822313.1;refGene_gn=LY9;refGene_tx=XM_019822314.1
F1	66300242 66300518
refGene_gn=LY9;refGene_tx=XM_006943032.2;refGene_gn=LY9;refGene_tx=XM_006943033.2;refGene_gn=LY9;refGene_tx=XM_006943034.2;refGene_gn=LY9;refGene_tx=XM_019822310.1;refGene_gn=LY9;refGene_tx=XM_019822311.1;refGene_gn=LY9;refGene_tx=XM_019822313.1;refGene_gn=LY9;refGene_tx=XM_019822314.1
F1	66313009 66313336
refGene_gn=LY9;refGene_tx=XM_006943032.2;refGene_gn=LY9;refGene_tx=XM_006943033.2;refGene_gn=LY9;refGene_tx=XM_006943034.2;refGene_gn=LY9;refGene_tx=XM_019822310.1;refGene_gn=LY9;refGene_tx=XM_019822311.1;refGene_gn=LY9;refGene_tx=XM_019822313.1;refGene_gn=LY9;refGene_tx=XM_019822314.1
F1	66316895 66317019
refGene_gn=LY9;refGene_tx=XM_006943032.2;refGene_gn=LY9;refGene_tx=XM_006943033.2;refGene_gn=LY9;refGene_tx=XM_006943034.2;refGene_gn=LY9;refGene_tx=XM_019822310.1;refGene_gn=LY9;refGene_tx=XM_019822311.1;refGene_gn=LY9;refGene_tx=XM_019822313.1;refGene_gn=LY9;refGene_tx=XM_019822314.1
F1	66344576 66344648 refGene_gn=SLAMF7;refGene_tx=XM_006943035.3
F1	66345404 66345467 refGene_gn=SLAMF7;refGene_tx=XM_006943035.3
F1	66346198 66346302 refGene_gn=SLAMF7;refGene_tx=XM_006943035.3
F1	66347242 66347359 refGene_gn=SLAMF7;refGene_tx=XM_006943035.3
F1	66347573 66347846 refGene_gn=SLAMF7;refGene_tx=XM_006943035.3
F1	66349124 66349457 refGene_gn=SLAMF7;refGene_tx=XM_006943035.3
F1	66358854 66358906 refGene_gn=SLAMF7;refGene_tx=XM_006943035.3
F1	66376133 66376212
refGene_gn=CD48;refGene_tx=XM_006943036.4;refGene_gn=CD48;refGene_tx=XM_019822315.2;refGene_gn=CD48;refGene_tx=XM_019822316.2
F1	66380949 66381246
refGene_gn=CD48;refGene_tx=XM_006943036.4;refGene_gn=CD48;refGene_tx=XM_019822315.2;refGene_gn=CD48;refGene_tx=XM_019822316.2
F1	66384889 66385150
refGene_gn=CD48;refGene_tx=XM_006943036.4;refGene_gn=CD48;refGene_tx=XM_019822315.2;refGene_gn=CD48;refGene_tx=XM_019822316.2
F1	66386551 66386730
refGene_gn=CD48;refGene_tx=XM_019822315.2;refGene_gn=CD48;refGene_tx=XM_019822316.2
F1	66387072 66387152 refGene_gn=CD48;refGene_tx=XM_006943036.4
F1	66414863 66414939

refGene_gn=SLAMF1;refGene_tx=NM_001278826.1;refGene_gn=SLAMF1;refGene_tx=XM_019821312.2;refGene_gn=SLAMF1;refGene_tx=XM_019821314.2
F1 66422678 66423017
refGene_gn=SLAMF1;refGene_tx=NM_001278826.1;refGene_gn=SLAMF1;refGene_tx=XM_019821312.2;refGene_gn=SLAMF1;refGene_tx=XM_019821314.2
F1 66424721 66425003
refGene_gn=SLAMF1;refGene_tx=NM_001278826.1;refGene_gn=SLAMF1;refGene_tx=XM_019821312.2;refGene_gn=SLAMF1;refGene_tx=XM_019821314.2
F1 66434481 66434580
refGene_gn=SLAMF1;refGene_tx=NM_001278826.1;refGene_gn=SLAMF1;refGene_tx=XM_019821312.2
F1 66438507 66438581
refGene_gn=SLAMF1;refGene_tx=NM_001278826.1;refGene_gn=SLAMF1;refGene_tx=XM_019821312.2;refGene_gn=SLAMF1;refGene_tx=XM_019821314.2
F1 66446380 66446473
refGene_gn=SLAMF1;refGene_tx=NM_001278826.1;refGene_gn=SLAMF1;refGene_tx=XM_019821312.2;refGene_gn=SLAMF1;refGene_tx=XM_019821314.2
F1 66448074 66448172
refGene_gn=SLAMF1;refGene_tx=NM_001278826.1;refGene_gn=SLAMF1;refGene_tx=XM_019821312.2;refGene_gn=SLAMF1;refGene_tx=XM_019821314.2
F1 66457874 66457941 refGene_gn=CD84;refGene_tx=XM_006943037.4
F1 66469034 66469382 refGene_gn=CD84;refGene_tx=XM_006943037.4
F1 66475487 66475739 refGene_gn=CD84;refGene_tx=XM_006943037.4
F1 66476109 66476229 refGene_gn=CD84;refGene_tx=XM_006943037.4
F1 66476879 66476912 refGene_gn=CD84;refGene_tx=XM_006943037.4
F1 66477944 66478042 refGene_gn=CD84;refGene_tx=XM_006943037.4
F1 66478684 66478744 refGene_gn=CD84;refGene_tx=XM_006943037.4
F1 66480678 66480744 refGene_gn=CD84;refGene_tx=XM_006943037.4
F1 66499570 66499685
refGene_gn=SLAMF6;refGene_tx=XM_006943038.4;refGene_gn=SLAMF6;refGene_tx=XM_019822317.2
F1 66512115 66512448
refGene_gn=SLAMF6;refGene_tx=XM_006943038.4;refGene_gn=SLAMF6;refGene_tx=XM_019822317.2
F1 66516508 66516772
refGene_gn=SLAMF6;refGene_tx=XM_006943038.4;refGene_gn=SLAMF6;refGene_tx=XM_019822317.2
F1 66517200 66517320
refGene_gn=SLAMF6;refGene_tx=XM_006943038.4;refGene_gn=SLAMF6;refGene_tx=XM_019822317.2
F1 66517546 66517585 refGene_gn=SLAMF6;refGene_tx=XM_006943038.4
F1 66518613 66518693
refGene_gn=SLAMF6;refGene_tx=XM_006943038.4;refGene_gn=SLAMF6;refGene_tx=XM_019822317.2
F1 66520100 66520175
refGene_gn=SLAMF6;refGene_tx=XM_006943038.4;refGene_gn=SLAMF6;refGene_tx=XM_019822317.2
F1 66520538 66520685
refGene_gn=SLAMF6;refGene_tx=XM_006943038.4;refGene_gn=SLAMF6;refGene_tx=XM_019822317.2

| | | | |
|---|---|---|---|
| F1 | 66582336 | 66582597 | refGene_gn=VANGL2;refGene_tx=XM_019822318.2 |
| F1 | 66583253 | 66583485 | refGene_gn=VANGL2;refGene_tx=XM_019822318.2 |
| F1 | 66585809 | 66585945 | refGene_gn=VANGL2;refGene_tx=XM_019822318.2 |
| F1 | 66586401 | 66586538 | refGene_gn=VANGL2;refGene_tx=XM_019822318.2 |
| F1 | 66587238 | 66587798 | refGene_gn=VANGL2;refGene_tx=XM_019822318.2 |
| F1 | 66587799 | 66587856 | refGene_gn=VANGL2;refGene_tx=XM_019822318.2 |
| F1 | 66591051 | 66591172 | refGene_gn=VANGL2;refGene_tx=XM_019822318.2 |
| F1 | 66591321 | 66591392 | refGene_gn=VANGL2;refGene_tx=XM_019822318.2 |
| F1 | 66631402 | 66631804 | refGene_gn=NHLH1;refGene_tx=XM_023247823.1 |
| F1 | 66642500 | 66642623 | refGene_gn=NCSTN;refGene_tx=XM_011291113.3 |
| F1 | 66643427 | 66643640 | refGene_gn=NCSTN;refGene_tx=XM_011291113.3 |
| F1 | 66643872 | 66644027 | refGene_gn=NCSTN;refGene_tx=XM_011291113.3 |
| F1 | 66644287 | 66644375 | refGene_gn=NCSTN;refGene_tx=XM_011291113.3 |
| F1 | 66644678 | 66644774 | refGene_gn=NCSTN;refGene_tx=XM_011291113.3 |
| F1 | 66644912 | 66645015 | refGene_gn=NCSTN;refGene_tx=XM_011291113.3 |
| F1 | 66646345 | 66646518 | refGene_gn=NCSTN;refGene_tx=XM_011291113.3 |
| F1 | 66647227 | 66647305 | refGene_gn=NCSTN;refGene_tx=XM_011291113.3 |
| F1 | 66647470 | 66647575 | refGene_gn=NCSTN;refGene_tx=XM_011291113.3 |
| F1 | 66648269 | 66648422 | refGene_gn=NCSTN;refGene_tx=XM_011291113.3 |
| F1 | 66648680 | 66648790 | refGene_gn=NCSTN;refGene_tx=XM_011291113.3 |
| F1 | 66649056 | 66649207 | refGene_gn=NCSTN;refGene_tx=XM_011291113.3 |
| F1 | 66649905 | 66650051 | refGene_gn=NCSTN;refGene_tx=XM_011291113.3 |
| F1 | 66650474 | 66650596 | refGene_gn=NCSTN;refGene_tx=XM_011291113.3 |
| F1 | 66651243 | 66651367 | refGene_gn=NCSTN;refGene_tx=XM_011291113.3 |
| F1 | 66654988 | 66655093 | refGene_gn=NCSTN;refGene_tx=XM_011291113.3 |
| F1 | 66656595 | 66656680 | refGene_gn=NCSTN;refGene_tx=XM_011291113.3 |
| F1 | 66656905 | 66656945 | refGene_gn=COPA;refGene_tx=XM_011291111.3 |
| F1 | 66660140 | 66660254 | refGene_gn=COPA;refGene_tx=XM_011291111.3 |
| F1 | 66660439 | 66660513 | refGene_gn=COPA;refGene_tx=XM_011291111.3 |
| F1 | 66664966 | 66665047 | refGene_gn=COPA;refGene_tx=XM_011291111.3 |
| F1 | 66669344 | 66669421 | refGene_gn=COPA;refGene_tx=XM_011291111.3 |
| F1 | 66670139 | 66670249 | refGene_gn=COPA;refGene_tx=XM_011291111.3 |
| F1 | 66675180 | 66675290 | refGene_gn=COPA;refGene_tx=XM_011291111.3 |
| F1 | 66676897 | 66676997 | refGene_gn=COPA;refGene_tx=XM_011291111.3 |
| F1 | 66677783 | 66677919 | refGene_gn=COPA;refGene_tx=XM_011291111.3 |
| F1 | 66678361 | 66678444 | refGene_gn=COPA;refGene_tx=XM_011291111.3 |
| F1 | 66680996 | 66681147 | refGene_gn=COPA;refGene_tx=XM_011291111.3 |
| F1 | 66682103 | 66682170 | refGene_gn=COPA;refGene_tx=XM_011291111.3 |
| F1 | 66683074 | 66683150 | refGene_gn=COPA;refGene_tx=XM_011291111.3 |
| F1 | 66685027 | 66685110 | refGene_gn=COPA;refGene_tx=XM_011291111.3 |
| F1 | 66686022 | 66686162 | refGene_gn=COPA;refGene_tx=XM_011291111.3 |
| F1 | 66686966 | 66687052 | refGene_gn=COPA;refGene_tx=XM_011291111.3 |
| F1 | 66687166 | 66687305 | refGene_gn=COPA;refGene_tx=XM_011291111.3 |
| F1 | 66692478 | 66692641 | refGene_gn=COPA;refGene_tx=XM_011291111.3 |
| F1 | 66692739 | 66692886 | refGene_gn=COPA;refGene_tx=XM_011291111.3 |
| F1 | 66693660 | 66693850 | refGene_gn=COPA;refGene_tx=XM_011291111.3 |
| F1 | 66693973 | 66694069 | refGene_gn=COPA;refGene_tx=XM_011291111.3 |
| F1 | 66695490 | 66695579 | refGene_gn=COPA;refGene_tx=XM_011291111.3 |
| F1 | 66695768 | 66695892 | refGene_gn=COPA;refGene_tx=XM_011291111.3 |
| F1 | 66697149 | 66697239 | refGene_gn=COPA;refGene_tx=XM_011291111.3 |

F1 66697428 66697538 refGene_gn=COPA;refGene_tx=XM_011291111.3
F1 66697918 66697996 refGene_gn=COPA;refGene_tx=XM_011291111.3
F1 66698162 66698231 refGene_gn=COPA;refGene_tx=XM_011291111.3
F1 66698741 66698878 refGene_gn=COPA;refGene_tx=XM_011291111.3
F1 66699118 66699305 refGene_gn=COPA;refGene_tx=XM_011291111.3
F1 66699389 66699500 refGene_gn=COPA;refGene_tx=XM_011291111.3
F1 66699803 66699965 refGene_gn=COPA;refGene_tx=XM_011291111.3
F1 66700592 66700787 refGene_gn=COPA;refGene_tx=XM_011291111.3
F1 66701072 66701132 refGene_gn=COPA;refGene_tx=XM_011291111.3
F1 66704895 66704965
refGene_gn=PEX19;refGene_tx=XM_003999588.4;refGene_gn=PEX19;refGene_tx
=XM_006943040.3
F1 66706399 66706509
refGene_gn=PEX19;refGene_tx=XM_003999588.4;refGene_gn=PEX19;refGene_tx
=XM_006943040.3
F1 66706937 66707103
refGene_gn=PEX19;refGene_tx=XM_003999588.4;refGene_gn=PEX19;refGene_tx
=XM_006943040.3
F1 66707535 66707621
refGene_gn=PEX19;refGene_tx=XM_003999588.4;refGene_gn=PEX19;refGene_tx
=XM_006943040.3
F1 66708285 66708447
refGene_gn=PEX19;refGene_tx=XM_003999588.4;refGene_gn=PEX19;refGene_tx
=XM_006943040.3
F1 66709286 66709463
refGene_gn=PEX19;refGene_tx=XM_003999588.4;refGene_gn=PEX19;refGene_tx
=XM_006943040.3
F1 66709690 66709735 refGene_gn=PEX19;refGene_tx=XM_003999588.4
F1 66709900 66709984
refGene_gn=PEX19;refGene_tx=XM_003999588.4;refGene_gn=PEX19;refGene_tx
=XM_006943040.3
F1 66737439 66737488
refGene_gn=DCAF8;refGene_tx=XM_003999590.5;refGene_gn=DCAF8;refGene_tx
=XM_003999591.5;refGene_gn=DCAF8;refGene_tx=XM_006943041.4
F1 66740076 66740750
refGene_gn=DCAF8;refGene_tx=XM_003999590.5;refGene_gn=DCAF8;refGene_tx
=XM_003999591.5;refGene_gn=DCAF8;refGene_tx=XM_006943041.4
F1 66741683 66741824
refGene_gn=DCAF8;refGene_tx=XM_003999590.5;refGene_gn=DCAF8;refGene_tx
=XM_003999591.5;refGene_gn=DCAF8;refGene_tx=XM_006943041.4
F1 66743351 66743446
refGene_gn=DCAF8;refGene_tx=XM_003999590.5;refGene_gn=DCAF8;refGene_tx
=XM_003999591.5;refGene_gn=DCAF8;refGene_tx=XM_006943041.4
F1 66748671 66748782
refGene_gn=DCAF8;refGene_tx=XM_003999590.5;refGene_gn=DCAF8;refGene_tx
=XM_003999591.5;refGene_gn=DCAF8;refGene_tx=XM_006943041.4
F1 66756194 66756267
refGene_gn=DCAF8;refGene_tx=XM_003999590.5;refGene_gn=DCAF8;refGene_tx
=XM_003999591.5;refGene_gn=DCAF8;refGene_tx=XM_006943041.4
F1 66756741 66756799

refGene_gn=DCAF8;refGene_tx=XM_003999590.5;refGene_gn=DCAF8;refGene_tx=XM_003999591.5;refGene_gn=DCAF8;refGene_tx=XM_006943041.4
F1 66757310 66757418
refGene_gn=DCAF8;refGene_tx=XM_003999590.5;refGene_gn=DCAF8;refGene_tx=XM_003999591.5;refGene_gn=DCAF8;refGene_tx=XM_006943041.4
F1 66758716 66758847
refGene_gn=DCAF8;refGene_tx=XM_003999590.5;refGene_gn=DCAF8;refGene_tx=XM_003999591.5;refGene_gn=DCAF8;refGene_tx=XM_006943041.4
F1 66762523 66762643
refGene_gn=DCAF8;refGene_tx=XM_003999590.5;refGene_gn=DCAF8;refGene_tx=XM_003999591.5;refGene_gn=DCAF8;refGene_tx=XM_006943041.4
F1 66763009 66763126
refGene_gn=DCAF8;refGene_tx=XM_003999590.5;refGene_gn=DCAF8;refGene_tx=XM_003999591.5;refGene_gn=DCAF8;refGene_tx=XM_006943041.4
F1 66763745 66763862
refGene_gn=DCAF8;refGene_tx=XM_003999590.5;refGene_gn=DCAF8;refGene_tx=XM_003999591.5;refGene_gn=DCAF8;refGene_tx=XM_006943041.4
F1 66767953 66768018 refGene_gn=PEA15;refGene_tx=XM_023247767.1
F1 66768171 66768327 refGene_gn=PEA15;refGene_tx=XM_023247767.1
F1 66769490 66769662 refGene_gn=PEA15;refGene_tx=XM_023247767.1
F1 66778647 66778791 refGene_gn=CASQ1;refGene_tx=XM_011291109.3
F1 66779529 66779604 refGene_gn=CASQ1;refGene_tx=XM_011291109.3
F1 66780689 66780790 refGene_gn=CASQ1;refGene_tx=XM_011291109.3
F1 66780993 66781048 refGene_gn=CASQ1;refGene_tx=XM_011291109.3
F1 66781459 66781505 refGene_gn=CASQ1;refGene_tx=XM_011291109.3
F1 66782257 66782388 refGene_gn=CASQ1;refGene_tx=XM_011291109.3
F1 66782725 66782799 refGene_gn=CASQ1;refGene_tx=XM_011291109.3
F1 66783108 66783220 refGene_gn=CASQ1;refGene_tx=XM_011291109.3
F1 66783457 66783558 refGene_gn=CASQ1;refGene_tx=XM_011291109.3
F1 66784312 66784397 refGene_gn=CASQ1;refGene_tx=XM_011291109.3
F1 66785964 66786246 refGene_gn=CASQ1;refGene_tx=XM_011291109.3
F1 66790032 66790061 refGene_gn=ATP1A4;refGene_tx=XM_023247850.1
F1 66790333 66790425
refGene_gn=ATP1A4;refGene_tx=XM_023247850.1;refGene_gn=ATP1A4;refGene_tx=XM_023247851.1
F1 66791972 66792230
refGene_gn=ATP1A4;refGene_tx=XM_023247846.1;refGene_gn=ATP1A4;refGene_tx=XM_023247847.1;refGene_gn=ATP1A4;refGene_tx=XM_023247848.1;refGene_gn=ATP1A4;refGene_tx=XM_023247849.1
F1 66792684 66792786
refGene_gn=ATP1A4;refGene_tx=XM_023247846.1;refGene_gn=ATP1A4;refGene_tx=XM_023247847.1;refGene_gn=ATP1A4;refGene_tx=XM_023247848.1;refGene_gn=ATP1A4;refGene_tx=XM_023247849.1;refGene_gn=ATP1A4;refGene_tx=XM_023247850.1;refGene_gn=ATP1A4;refGene_tx=XM_023247851.1
F1 66792892 66793023
refGene_gn=ATP1A4;refGene_tx=XM_023247846.1;refGene_gn=ATP1A4;refGene_tx=XM_023247847.1;refGene_gn=ATP1A4;refGene_tx=XM_023247848.1;refGene_gn=ATP1A4;refGene_tx=XM_023247849.1;refGene_gn=ATP1A4;refGene_tx=XM_023247850.1;refGene_gn=ATP1A4;refGene_tx=XM_023247851.1
F1 66795229 66795375

refGene_gn=ATP1A4;refGene_tx=XM_023247846.1;refGene_gn=ATP1A4;refGene_tx=XM_023247847.1;refGene_gn=ATP1A4;refGene_tx=XM_023247848.1;refGene_gn=ATP1A4;refGene_tx=XM_023247849.1;refGene_gn=ATP1A4;refGene_tx=XM_023247850.1;refGene_gn=ATP1A4;refGene_tx=XM_023247851.1
F1 66796604 66796728
refGene_gn=ATP1A4;refGene_tx=XM_023247846.1;refGene_gn=ATP1A4;refGene_tx=XM_023247847.1;refGene_gn=ATP1A4;refGene_tx=XM_023247848.1;refGene_gn=ATP1A4;refGene_tx=XM_023247849.1;refGene_gn=ATP1A4;refGene_tx=XM_023247850.1;refGene_gn=ATP1A4;refGene_tx=XM_023247851.1
F1 66796880 66797035
refGene_gn=ATP1A4;refGene_tx=XM_023247846.1;refGene_gn=ATP1A4;refGene_tx=XM_023247847.1;refGene_gn=ATP1A4;refGene_tx=XM_023247848.1;refGene_gn=ATP1A4;refGene_tx=XM_023247849.1;refGene_gn=ATP1A4;refGene_tx=XM_023247850.1;refGene_gn=ATP1A4;refGene_tx=XM_023247851.1
F1 66797268 66797437
refGene_gn=ATP1A4;refGene_tx=XM_023247846.1;refGene_gn=ATP1A4;refGene_tx=XM_023247847.1;refGene_gn=ATP1A4;refGene_tx=XM_023247848.1;refGene_gn=ATP1A4;refGene_tx=XM_023247849.1;refGene_gn=ATP1A4;refGene_tx=XM_023247850.1;refGene_gn=ATP1A4;refGene_tx=XM_023247851.1
F1 66797710 66797861
refGene_gn=ATP1A4;refGene_tx=XM_023247846.1;refGene_gn=ATP1A4;refGene_tx=XM_023247847.1;refGene_gn=ATP1A4;refGene_tx=XM_023247848.1;refGene_gn=ATP1A4;refGene_tx=XM_023247849.1;refGene_gn=ATP1A4;refGene_tx=XM_023247850.1;refGene_gn=ATP1A4;refGene_tx=XM_023247851.1
F1 66798181 66798321
refGene_gn=ATP1A4;refGene_tx=XM_023247846.1;refGene_gn=ATP1A4;refGene_tx=XM_023247847.1;refGene_gn=ATP1A4;refGene_tx=XM_023247848.1;refGene_gn=ATP1A4;refGene_tx=XM_023247849.1;refGene_gn=ATP1A4;refGene_tx=XM_023247850.1;refGene_gn=ATP1A4;refGene_tx=XM_023247851.1
F1 66801197 66801370
refGene_gn=ATP1A4;refGene_tx=XM_023247846.1;refGene_gn=ATP1A4;refGene_tx=XM_023247847.1;refGene_gn=ATP1A4;refGene_tx=XM_023247848.1;refGene_gn=ATP1A4;refGene_tx=XM_023247849.1;refGene_gn=ATP1A4;refGene_tx=XM_023247850.1;refGene_gn=ATP1A4;refGene_tx=XM_023247851.1
F1 66801508 66801698
refGene_gn=ATP1A4;refGene_tx=XM_023247846.1;refGene_gn=ATP1A4;refGene_tx=XM_023247847.1;refGene_gn=ATP1A4;refGene_tx=XM_023247848.1;refGene_gn=ATP1A4;refGene_tx=XM_023247849.1;refGene_gn=ATP1A4;refGene_tx=XM_023247850.1;refGene_gn=ATP1A4;refGene_tx=XM_023247851.1
F1 66805065 66805200
refGene_gn=ATP1A4;refGene_tx=XM_023247846.1;refGene_gn=ATP1A4;refGene_tx=XM_023247847.1;refGene_gn=ATP1A4;refGene_tx=XM_023247848.1;refGene_gn=ATP1A4;refGene_tx=XM_023247849.1;refGene_gn=ATP1A4;refGene_tx=XM_023247850.1;refGene_gn=ATP1A4;refGene_tx=XM_023247851.1
F1 66805368 66805478
refGene_gn=ATP1A4;refGene_tx=XM_023247846.1;refGene_gn=ATP1A4;refGene_tx=XM_023247847.1;refGene_gn=ATP1A4;refGene_tx=XM_023247848.1;refGene_gn=ATP1A4;refGene_tx=XM_023247849.1;refGene_gn=ATP1A4;refGene_tx=XM_023247850.1;refGene_gn=ATP1A4;refGene_tx=XM_023247851.1
F1 66805716 66805915

refGene_gn=ATP1A4;refGene_tx=XM_023247846.1;refGene_gn=ATP1A4;refGene_tx=XM_023247847.1;refGene_gn=ATP1A4;refGene_tx=XM_023247848.1;refGene_gn=ATP1A4;refGene_tx=XM_023247849.1;refGene_gn=ATP1A4;refGene_tx=XM_023247850.1;refGene_gn=ATP1A4;refGene_tx=XM_023247851.1
F1 66807684 66807953
refGene_gn=ATP1A4;refGene_tx=XM_023247846.1;refGene_gn=ATP1A4;refGene_tx=XM_023247847.1;refGene_gn=ATP1A4;refGene_tx=XM_023247848.1;refGene_gn=ATP1A4;refGene_tx=XM_023247849.1;refGene_gn=ATP1A4;refGene_tx=XM_023247850.1;refGene_gn=ATP1A4;refGene_tx=XM_023247851.1
F1 66811661 66811779
refGene_gn=ATP1A4;refGene_tx=XM_023247846.1;refGene_gn=ATP1A4;refGene_tx=XM_023247847.1;refGene_gn=ATP1A4;refGene_tx=XM_023247848.1;refGene_gn=ATP1A4;refGene_tx=XM_023247849.1;refGene_gn=ATP1A4;refGene_tx=XM_023247850.1;refGene_gn=ATP1A4;refGene_tx=XM_023247851.1
F1 66812062 66812197
refGene_gn=ATP1A4;refGene_tx=XM_023247846.1;refGene_gn=ATP1A4;refGene_tx=XM_023247847.1;refGene_gn=ATP1A4;refGene_tx=XM_023247848.1;refGene_gn=ATP1A4;refGene_tx=XM_023247849.1;refGene_gn=ATP1A4;refGene_tx=XM_023247850.1;refGene_gn=ATP1A4;refGene_tx=XM_023247851.1
F1 66813685 66813799
refGene_gn=ATP1A4;refGene_tx=XM_023247846.1;refGene_gn=ATP1A4;refGene_tx=XM_023247847.1;refGene_gn=ATP1A4;refGene_tx=XM_023247848.1;refGene_gn=ATP1A4;refGene_tx=XM_023247849.1;refGene_gn=ATP1A4;refGene_tx=XM_023247850.1;refGene_gn=ATP1A4;refGene_tx=XM_023247851.1
F1 66814320 66814524
refGene_gn=ATP1A4;refGene_tx=XM_023247846.1;refGene_gn=ATP1A4;refGene_tx=XM_023247847.1;refGene_gn=ATP1A4;refGene_tx=XM_023247848.1;refGene_gn=ATP1A4;refGene_tx=XM_023247849.1;refGene_gn=ATP1A4;refGene_tx=XM_023247850.1;refGene_gn=ATP1A4;refGene_tx=XM_023247851.1
F1 66816246 66816306
refGene_gn=ATP1A4;refGene_tx=XM_023247846.1;refGene_gn=ATP1A4;refGene_tx=XM_023247847.1;refGene_gn=ATP1A4;refGene_tx=XM_023247848.1;refGene_gn=ATP1A4;refGene_tx=XM_023247849.1;refGene_gn=ATP1A4;refGene_tx=XM_023247850.1;refGene_gn=ATP1A4;refGene_tx=XM_023247851.1
F1 66818040 66818193
refGene_gn=ATP1A4;refGene_tx=XM_023247846.1;refGene_gn=ATP1A4;refGene_tx=XM_023247847.1;refGene_gn=ATP1A4;refGene_tx=XM_023247848.1;refGene_gn=ATP1A4;refGene_tx=XM_023247849.1;refGene_gn=ATP1A4;refGene_tx=XM_023247850.1;refGene_gn=ATP1A4;refGene_tx=XM_023247851.1
F1 66826310 66826339
refGene_gn=LOC101096785;refGene_tx=XM_019822324.2
F1 66827271 66827363
refGene_gn=LOC101096785;refGene_tx=XM_019822324.2
F1 66827471 66827573
refGene_gn=LOC101096785;refGene_tx=XM_019822324.2
F1 66829689 66829820
refGene_gn=LOC101096785;refGene_tx=XM_019822324.2
F1 66830014 66830160
refGene_gn=LOC101096785;refGene_tx=XM_019822324.2
F1 66830347 66830471

refGene_gn=LOC101096785;refGene_tx=XM_019822324.2
F1 66830715 66830870
refGene_gn=LOC101096785;refGene_tx=XM_019822324.2
F1 66831103 66831272
refGene_gn=LOC101096785;refGene_tx=XM_019822324.2
F1 66831545 66831696
refGene_gn=LOC101096785;refGene_tx=XM_019822324.2
F1 66832214 66832351
refGene_gn=LOC101096785;refGene_tx=XM_019822324.2
F1 66835709 66835885
refGene_gn=LOC101096785;refGene_tx=XM_019822324.2
F1 66836019 66836209
refGene_gn=LOC101096785;refGene_tx=XM_019822324.2
F1 66836813 66836948
refGene_gn=LOC101096785;refGene_tx=XM_019822324.2
F1 66837082 66837192
refGene_gn=LOC101096785;refGene_tx=XM_019822324.2
F1 66837319 66837518
refGene_gn=LOC101096785;refGene_tx=XM_019822324.2
F1 66838271 66838540
refGene_gn=LOC101096785;refGene_tx=XM_019822324.2
F1 66840066 66840184
refGene_gn=LOC101096785;refGene_tx=XM_019822324.2
F1 66841073 66841208
refGene_gn=LOC101096785;refGene_tx=XM_019822324.2
F1 66841465 66841579
refGene_gn=LOC101096785;refGene_tx=XM_019822324.2
F1 66842042 66842246
refGene_gn=LOC101096785;refGene_tx=XM_019822324.2
F1 66844265 66844325
refGene_gn=LOC101096785;refGene_tx=XM_019822324.2
F1 66844498 66844603
refGene_gn=LOC101096785;refGene_tx=XM_019822324.2
F1 66848822 66848834
refGene_gn=LOC101096785;refGene_tx=XM_019822324.2
F1 66862433 66862497 refGene_gn=IGSF8;refGene_tx=XM_003999597.5
F1 66865374 66865489 refGene_gn=IGSF8;refGene_tx=XM_019822325.2
F1 66865589 66865967
refGene_gn=IGSF8;refGene_tx=XM_003999597.5;refGene_gn=IGSF8;refGene_tx
=XM_019822325.2
F1 66866651 66867113
refGene_gn=IGSF8;refGene_tx=XM_003999597.5;refGene_gn=IGSF8;refGene_tx
=XM_019822325.2
F1 66867459 66867867
refGene_gn=IGSF8;refGene_tx=XM_003999597.5;refGene_gn=IGSF8;refGene_tx
=XM_019822325.2
F1 66868062 66868476
refGene_gn=IGSF8;refGene_tx=XM_003999597.5;refGene_gn=IGSF8;refGene_tx
=XM_019822325.2
F1 66868746 66868862

refGene_gn=IGSF8;refGene_tx=XM_003999597.5;refGene_gn=IGSF8;refGene_tx
=XM_019822325.2
F1 66873728 66874060 refGene_gn=KCNJ9;refGene_tx=XM_019821370.2
F1 66876369 66877216 refGene_gn=KCNJ9;refGene_tx=XM_019821370.2
F1 66895285 66895318 refGene_gn=KCNJ10;refGene_tx=XM_019822326.2
F1 66896574 66896772
refGene_gn=KCNJ10;refGene_tx=XM_019822326.2;refGene_gn=KCNJ10;refGene_
tx=XM_019822328.2
F1 66911931 66911934
refGene_gn=KCNJ10;refGene_tx=XM_019822329.2;refGene_gn=KCNJ10;refGene_
tx=XM_019822330.2
F1 66912046 66913186
refGene_gn=KCNJ10;refGene_tx=XM_011291108.3;refGene_gn=KCNJ10;refGene_
tx=XM_019822326.2;refGene_gn=KCNJ10;refGene_tx=XM_019822328.2;refGene_
gn=KCNJ10;refGene_tx=XM_019822329.2;refGene_gn=KCNJ10;refGene_tx=XM_01
9822330.2;refGene_gn=KCNJ10;refGene_tx=XM_019822331.2
F1 66923334 66924003 refGene_gn=PIGM;refGene_tx=XM_003999599.5
F1 66924004 66924610 refGene_gn=PIGM;refGene_tx=XM_003999599.5
F1 66976678 66976762 refGene_gn=SLAMF9;refGene_tx=XM_023247466.1
F1 66977667 66977698 refGene_gn=SLAMF9;refGene_tx=XM_023247466.1
F1 66977967 66978268 refGene_gn=SLAMF9;refGene_tx=XM_023247466.1
F1 66984046 66984547 refGene_gn=SLAMF9;refGene_tx=XM_023247466.1
F1 66984677 66984682 refGene_gn=SLAMF9;refGene_tx=XM_023247466.1
F1 66985340 66985516 refGene_gn=SLAMF9;refGene_tx=XM_023247466.1
F1 66985518 66985651 refGene_gn=SLAMF9;refGene_tx=XM_023247466.1
F1 66986019 66986231 refGene_gn=SLAMF9;refGene_tx=XM_023247466.1
F1 66993814 66993872
refGene_gn=IGSF9;refGene_tx=XM_003999600.5;refGene_gn=IGSF9;refGene_tx
=XM_019822333.2;refGene_gn=IGSF9;refGene_tx=XM_019822334.2;refGene_gn=
IGSF9;refGene_tx=XM_023247167.1;refGene_gn=IGSF9;refGene_tx=XM_0232471
68.1
F1 66994114 66994303
refGene_gn=IGSF9;refGene_tx=XM_003999600.5;refGene_gn=IGSF9;refGene_tx
=XM_019822333.2;refGene_gn=IGSF9;refGene_tx=XM_019822334.2;refGene_gn=
IGSF9;refGene_tx=XM_023247167.1;refGene_gn=IGSF9;refGene_tx=XM_0232471
68.1
F1 66998985 66999138
refGene_gn=IGSF9;refGene_tx=XM_003999600.5;refGene_gn=IGSF9;refGene_tx
=XM_019822333.2;refGene_gn=IGSF9;refGene_tx=XM_019822334.2;refGene_gn=
IGSF9;refGene_tx=XM_023247167.1
F1 66999820 66999975
refGene_gn=IGSF9;refGene_tx=XM_003999600.5;refGene_gn=IGSF9;refGene_tx
=XM_019822333.2;refGene_gn=IGSF9;refGene_tx=XM_019822334.2;refGene_gn=
IGSF9;refGene_tx=XM_023247167.1;refGene_gn=IGSF9;refGene_tx=XM_0232471
68.1
F1 67000249 67000367
refGene_gn=IGSF9;refGene_tx=XM_003999600.5;refGene_gn=IGSF9;refGene_tx
=XM_019822333.2;refGene_gn=IGSF9;refGene_tx=XM_019822334.2;refGene_gn=
IGSF9;refGene_tx=XM_023247167.1;refGene_gn=IGSF9;refGene_tx=XM_0232471
68.1

F1 67002287 67002429
refGene_gn=IGSF9;refGene_tx=XM_003999600.5;refGene_gn=IGSF9;refGene_tx=XM_019822333.2;refGene_gn=IGSF9;refGene_tx=XM_019822334.2;refGene_gn=IGSF9;refGene_tx=XM_023247167.1;refGene_gn=IGSF9;refGene_tx=XM_023247168.1
F1 67002526 67002672
refGene_gn=IGSF9;refGene_tx=XM_003999600.5;refGene_gn=IGSF9;refGene_tx=XM_019822333.2;refGene_gn=IGSF9;refGene_tx=XM_019822334.2;refGene_gn=IGSF9;refGene_tx=XM_023247167.1;refGene_gn=IGSF9;refGene_tx=XM_023247168.1
F1 67002766 67002909
refGene_gn=IGSF9;refGene_tx=XM_003999600.5;refGene_gn=IGSF9;refGene_tx=XM_019822333.2;refGene_gn=IGSF9;refGene_tx=XM_019822334.2;refGene_gn=IGSF9;refGene_tx=XM_023247167.1;refGene_gn=IGSF9;refGene_tx=XM_023247168.1
F1 67003995 67004136
refGene_gn=IGSF9;refGene_tx=XM_003999600.5;refGene_gn=IGSF9;refGene_tx=XM_019822333.2;refGene_gn=IGSF9;refGene_tx=XM_019822334.2;refGene_gn=IGSF9;refGene_tx=XM_023247167.1;refGene_gn=IGSF9;refGene_tx=XM_023247168.1
F1 67004552 67004669
refGene_gn=IGSF9;refGene_tx=XM_003999600.5;refGene_gn=IGSF9;refGene_tx=XM_019822333.2;refGene_gn=IGSF9;refGene_tx=XM_019822334.2;refGene_gn=IGSF9;refGene_tx=XM_023247167.1;refGene_gn=IGSF9;refGene_tx=XM_023247168.1
F1 67004847 67004998
refGene_gn=IGSF9;refGene_tx=XM_003999600.5;refGene_gn=IGSF9;refGene_tx=XM_019822333.2;refGene_gn=IGSF9;refGene_tx=XM_019822334.2;refGene_gn=IGSF9;refGene_tx=XM_023247167.1;refGene_gn=IGSF9;refGene_tx=XM_023247168.1
F1 67005200 67005324
refGene_gn=IGSF9;refGene_tx=XM_003999600.5;refGene_gn=IGSF9;refGene_tx=XM_019822333.2;refGene_gn=IGSF9;refGene_tx=XM_019822334.2;refGene_gn=IGSF9;refGene_tx=XM_023247167.1;refGene_gn=IGSF9;refGene_tx=XM_023247168.1
F1 67005566 67005742
refGene_gn=IGSF9;refGene_tx=XM_003999600.5;refGene_gn=IGSF9;refGene_tx=XM_019822333.2;refGene_gn=IGSF9;refGene_tx=XM_019822334.2;refGene_gn=IGSF9;refGene_tx=XM_023247167.1;refGene_gn=IGSF9;refGene_tx=XM_023247168.1
F1 67006012 67006263
refGene_gn=IGSF9;refGene_tx=XM_003999600.5;refGene_gn=IGSF9;refGene_tx=XM_019822333.2;refGene_gn=IGSF9;refGene_tx=XM_019822334.2;refGene_gn=IGSF9;refGene_tx=XM_023247167.1;refGene_gn=IGSF9;refGene_tx=XM_023247168.1
F1 67006435 67006520
refGene_gn=IGSF9;refGene_tx=XM_003999600.5;refGene_gn=IGSF9;refGene_tx=XM_019822333.2;refGene_gn=IGSF9;refGene_tx=XM_019822334.2;refGene_gn=IGSF9;refGene_tx=XM_023247167.1;refGene_gn=IGSF9;refGene_tx=XM_023247168.1

F1 67006617 67006794
refGene_gn=IGSF9;refGene_tx=XM_003999600.5;refGene_gn=IGSF9;refGene_tx=XM_019822333.2;refGene_gn=IGSF9;refGene_tx=XM_019822334.2;refGene_gn=IGSF9;refGene_tx=XM_023247167.1;refGene_gn=IGSF9;refGene_tx=XM_023247168.1
F1 67007017 67007060
refGene_gn=IGSF9;refGene_tx=XM_003999600.5;refGene_gn=IGSF9;refGene_tx=XM_019822333.2;refGene_gn=IGSF9;refGene_tx=XM_019822334.2;refGene_gn=IGSF9;refGene_tx=XM_023247167.1;refGene_gn=IGSF9;refGene_tx=XM_023247168.1
F1 67007373 67008234
refGene_gn=IGSF9;refGene_tx=XM_003999600.5;refGene_gn=IGSF9;refGene_tx=XM_019822333.2;refGene_gn=IGSF9;refGene_tx=XM_019822334.2;refGene_gn=IGSF9;refGene_tx=XM_023247167.1;refGene_gn=IGSF9;refGene_tx=XM_023247168.1
F1 67008496 67008624
refGene_gn=IGSF9;refGene_tx=XM_003999600.5;refGene_gn=IGSF9;refGene_tx=XM_019822333.2;refGene_gn=IGSF9;refGene_tx=XM_019822334.2;refGene_gn=IGSF9;refGene_tx=XM_023247167.1;refGene_gn=IGSF9;refGene_tx=XM_023247168.1
F1 67008787 67008969
refGene_gn=IGSF9;refGene_tx=XM_003999600.5;refGene_gn=IGSF9;refGene_tx=XM_019822333.2;refGene_gn=IGSF9;refGene_tx=XM_019822334.2;refGene_gn=IGSF9;refGene_tx=XM_023247167.1;refGene_gn=IGSF9;refGene_tx=XM_023247168.1
F1 67015513 67015693 refGene_gn=TAGLN2;refGene_tx=XM_023247169.1
F1 67016098 67016273 refGene_gn=TAGLN2;refGene_tx=XM_023247169.1
F1 67016576 67016679 refGene_gn=TAGLN2;refGene_tx=XM_023247169.1
F1 67016921 67017063 refGene_gn=TAGLN2;refGene_tx=XM_023247169.1
F1 67034000 67034003
refGene_gn=CFAP45;refGene_tx=XM_023247499.1;refGene_gn=CFAP45;refGene_tx=XM_023247500.1
F1 67040248 67040374
refGene_gn=CFAP45;refGene_tx=XM_023247499.1;refGene_gn=CFAP45;refGene_tx=XM_023247500.1
F1 67042705 67042842
refGene_gn=CFAP45;refGene_tx=XM_023247499.1;refGene_gn=CFAP45;refGene_tx=XM_023247500.1;refGene_gn=CFAP45;refGene_tx=XM_023247501.1
F1 67045139 67045284
refGene_gn=CFAP45;refGene_tx=XM_023247499.1;refGene_gn=CFAP45;refGene_tx=XM_023247500.1;refGene_gn=CFAP45;refGene_tx=XM_023247501.1
F1 67045573 67045744
refGene_gn=CFAP45;refGene_tx=XM_023247499.1;refGene_gn=CFAP45;refGene_tx=XM_023247500.1;refGene_gn=CFAP45;refGene_tx=XM_023247501.1
F1 67047090 67047269
refGene_gn=CFAP45;refGene_tx=XM_023247499.1;refGene_gn=CFAP45;refGene_tx=XM_023247501.1
F1 67049272 67049402
refGene_gn=CFAP45;refGene_tx=XM_023247499.1;refGene_gn=CFAP45;refGene_tx=XM_023247501.1

F1 67053820 67053967
refGene_gn=CFAP45;refGene_tx=XM_023247499.1;refGene_gn=CFAP45;refGene_tx=XM_023247500.1;refGene_gn=CFAP45;refGene_tx=XM_023247501.1
F1 67056364 67056478
refGene_gn=CFAP45;refGene_tx=XM_023247499.1;refGene_gn=CFAP45;refGene_tx=XM_023247500.1;refGene_gn=CFAP45;refGene_tx=XM_023247501.1
F1 67057065 67057259
refGene_gn=CFAP45;refGene_tx=XM_023247499.1;refGene_gn=CFAP45;refGene_tx=XM_023247500.1;refGene_gn=CFAP45;refGene_tx=XM_023247501.1
F1 67060974 67061199
refGene_gn=CFAP45;refGene_tx=XM_023247499.1;refGene_gn=CFAP45;refGene_tx=XM_023247500.1;refGene_gn=CFAP45;refGene_tx=XM_023247501.1
F1 67061488 67061567
refGene_gn=CFAP45;refGene_tx=XM_023247499.1;refGene_gn=CFAP45;refGene_tx=XM_023247500.1;refGene_gn=CFAP45;refGene_tx=XM_023247501.1
F1 67071544 67071593 refGene_gn=VSIG8;refGene_tx=XM_006943048.3
F1 67072738 67072784 refGene_gn=VSIG8;refGene_tx=XM_019822336.2
F1 67074585 67074761
refGene_gn=VSIG8;refGene_tx=XM_006943048.3;refGene_gn=VSIG8;refGene_tx=XM_019822336.2
F1 67075190 67075392
refGene_gn=VSIG8;refGene_tx=XM_006943048.3;refGene_gn=VSIG8;refGene_tx=XM_019822336.2
F1 67075490 67075712
refGene_gn=VSIG8;refGene_tx=XM_006943048.3;refGene_gn=VSIG8;refGene_tx=XM_019822336.2
F1 67075907 67076033
refGene_gn=VSIG8;refGene_tx=XM_006943048.3;refGene_gn=VSIG8;refGene_tx=XM_019822336.2
F1 67076323 67076522
refGene_gn=VSIG8;refGene_tx=XM_006943048.3;refGene_gn=VSIG8;refGene_tx=XM_019822336.2
F1 67077393 67077670
refGene_gn=VSIG8;refGene_tx=XM_006943048.3;refGene_gn=VSIG8;refGene_tx=XM_019822336.2
F1 67094800 67094865 refGene_gn=SLAMF8;refGene_tx=XM_003999659.4
F1 67095716 67095821 refGene_gn=SLAMF8;refGene_tx=XM_003999659.4
F1 67095902 67096208 refGene_gn=SLAMF8;refGene_tx=XM_003999659.4
F1 67098908 67099235 refGene_gn=SLAMF8;refGene_tx=XM_003999659.4
F1 67101068 67101159 refGene_gn=SLAMF8;refGene_tx=XM_003999659.4
F1 67114612 67114761 refGene_gn=FCRL6;refGene_tx=XM_023247467.1
F1 67115097 67115385 refGene_gn=FCRL6;refGene_tx=XM_023247467.1
F1 67115531 67115816 refGene_gn=FCRL6;refGene_tx=XM_023247467.1
F1 67116214 67116481 refGene_gn=FCRL6;refGene_tx=XM_023247467.1
F1 67117874 67117938 refGene_gn=FCRL6;refGene_tx=XM_023247467.1
F1 67123018 67123065 refGene_gn=FCRL6;refGene_tx=XM_023247467.1
F1 67123415 67123475 refGene_gn=FCRL6;refGene_tx=XM_023247467.1
F1 67137541 67137707 refGene_gn=FCRL6;refGene_tx=XM_023247467.1
F1 67139357 67139402 refGene_gn=FCRL6;refGene_tx=XM_023247467.1
F1 67139621 67139687 refGene_gn=FCRL6;refGene_tx=XM_023247467.1

F1 67146409 67146595 refGene_gn=DUSP23;refGene_tx=XM_023247560.1
F1 67147639 67147906 refGene_gn=DUSP23;refGene_tx=XM_023247560.1
F1 67206091 67206155 refGene_gn=CRP;refGene_tx=XM_003999662.4
F1 67206739 67207215 refGene_gn=CRP;refGene_tx=XM_003999662.4
F1 67207216 67207348 refGene_gn=CRP;refGene_tx=XM_003999662.4
F1 67305353 67305951 refGene_gn=APCS;refGene_tx=XM_003999663.1
F1 67306062 67306121 refGene_gn=APCS;refGene_tx=XM_003999663.1
F1 67356343 67357258
refGene_gn=LOC101099173;refGene_tx=XM_011291104.2
F1 67413763 67413895
refGene_gn=LOC101099425;refGene_tx=XM_003999604.4
F1 67413896 67414709
refGene_gn=LOC101099425;refGene_tx=XM_003999604.4
F1 67423575 67424511
refGene_gn=LOC101099681;refGene_tx=XM_003999605.3
F1 67444203 67444620
refGene_gn=LOC101093579;refGene_tx=XM_023247468.1
F1 67444621 67444921
refGene_gn=LOC101093579;refGene_tx=XM_023247468.1
F1 67466452 67467399
refGene_gn=LOC101099928;refGene_tx=XM_003999606.3
F1 67491566 67492071
refGene_gn=LOC101099928;refGene_tx=XM_003999606.3
F1 67512915 67513054
refGene_gn=LOC101094078;refGene_tx=XM_019822340.2
F1 67513056 67513240
refGene_gn=LOC101094078;refGene_tx=XM_019822340.2
F1 67513272 67513975
refGene_gn=LOC101094078;refGene_tx=XM_019822340.2
F1 67518998 67519198 refGene_gn=FCER1A;refGene_tx=XM_006943049.3
F1 67520651 67520909 refGene_gn=FCER1A;refGene_tx=XM_006943049.3
F1 67522403 67522658 refGene_gn=FCER1A;refGene_tx=XM_006943049.3
F1 67523694 67523715 refGene_gn=FCER1A;refGene_tx=XM_006943049.3
F1 67524320 67524378 refGene_gn=FCER1A;refGene_tx=XM_006943049.3
F1 67550536 67551469
refGene_gn=LOC101100178;refGene_tx=XM_023247659.1
F1 67603832 67604828 refGene_gn=ACKR1;refGene_tx=XM_003999667.4
F1 67605236 67605260 refGene_gn=ACKR1;refGene_tx=XM_003999667.4
F1 67609057 67609176
refGene_gn=CADM3;refGene_tx=XM_023248000.1;refGene_gn=CADM3;refGene_tx
=XM_023248001.1
F1 67611445 67611571
refGene_gn=CADM3;refGene_tx=XM_023248000.1;refGene_gn=CADM3;refGene_tx
=XM_023248001.1
F1 67614010 67614180
refGene_gn=CADM3;refGene_tx=XM_023248000.1;refGene_gn=CADM3;refGene_tx
=XM_023248001.1
F1 67614575 67614666
refGene_gn=CADM3;refGene_tx=XM_023248000.1;refGene_gn=CADM3;refGene_tx
=XM_023248001.1

F1 67616794 67616965
refGene_gn=CADM3;refGene_tx=XM_023248000.1;refGene_gn=CADM3;refGene_tx=XM_023248001.1
F1 67617282 67617420
refGene_gn=CADM3;refGene_tx=XM_023248000.1;refGene_gn=CADM3;refGene_tx=XM_023248001.1
F1 67618081 67618234
refGene_gn=CADM3;refGene_tx=XM_023248000.1;refGene_gn=CADM3;refGene_tx=XM_023248001.1
F1 67618734 67618875
refGene_gn=CADM3;refGene_tx=XM_023248000.1;refGene_gn=CADM3;refGene_tx=XM_023248001.1
F1 67620913 67621015 refGene_gn=CADM3;refGene_tx=XM_023248000.1
F1 67638523 67638605
refGene_gn=CADM3;refGene_tx=XM_023248000.1;refGene_gn=CADM3;refGene_tx=XM_023248001.1
F1 67737193 67737262
refGene_gn=IFI16;refGene_tx=XM_023247223.1;refGene_gn=IFI16;refGene_tx=XM_023247224.1;refGene_gn=IFI16;refGene_tx=XM_023247225.1;refGene_gn=IFI16;refGene_tx=XM_023247226.1;refGene_gn=IFI16;refGene_tx=XM_023247228.1;refGene_gn=IFI16;refGene_tx=XM_023247229.1
F1 67737466 67737582
refGene_gn=IFI16;refGene_tx=XM_023247223.1;refGene_gn=IFI16;refGene_tx=XM_023247224.1;refGene_gn=IFI16;refGene_tx=XM_023247226.1;refGene_gn=IFI16;refGene_tx=XM_023247229.1
F1 67738207 67738393
refGene_gn=IFI16;refGene_tx=XM_023247223.1;refGene_gn=IFI16;refGene_tx=XM_023247224.1;refGene_gn=IFI16;refGene_tx=XM_023247225.1;refGene_gn=IFI16;refGene_tx=XM_023247226.1;refGene_gn=IFI16;refGene_tx=XM_023247228.1;refGene_gn=IFI16;refGene_tx=XM_023247229.1
F1 67739263 67739683
refGene_gn=IFI16;refGene_tx=XM_023247223.1;refGene_gn=IFI16;refGene_tx=XM_023247224.1;refGene_gn=IFI16;refGene_tx=XM_023247225.1;refGene_gn=IFI16;refGene_tx=XM_023247226.1;refGene_gn=IFI16;refGene_tx=XM_023247228.1;refGene_gn=IFI16;refGene_tx=XM_023247229.1
F1 67742321 67742489
refGene_gn=IFI16;refGene_tx=XM_023247223.1;refGene_gn=IFI16;refGene_tx=XM_023247224.1;refGene_gn=IFI16;refGene_tx=XM_023247225.1;refGene_gn=IFI16;refGene_tx=XM_023247226.1;refGene_gn=IFI16;refGene_tx=XM_023247228.1;refGene_gn=IFI16;refGene_tx=XM_023247229.1
F1 67744994 67745162
refGene_gn=IFI16;refGene_tx=XM_023247223.1;refGene_gn=IFI16;refGene_tx=XM_023247224.1;refGene_gn=IFI16;refGene_tx=XM_023247225.1;refGene_gn=IFI16;refGene_tx=XM_023247226.1;refGene_gn=IFI16;refGene_tx=XM_023247228.1;refGene_gn=IFI16;refGene_tx=XM_023247229.1
F1 67747746 67747914
refGene_gn=IFI16;refGene_tx=XM_023247223.1;refGene_gn=IFI16;refGene_tx=XM_023247224.1;refGene_gn=IFI16;refGene_tx=XM_023247225.1;refGene_gn=IFI16;refGene_tx=XM_023247228.1
F1 67750980 67751148

refGene_gn=IFI16;refGene_tx=XM_023247223.1;refGene_gn=IFI16;refGene_tx=XM_023247224.1;refGene_gn=IFI16;refGene_tx=XM_023247225.1;refGene_gn=IFI16;refGene_tx=XM_023247226.1;refGene_gn=IFI16;refGene_tx=XM_023247228.1
F1 67762736 67762904
refGene_gn=IFI16;refGene_tx=XM_023247223.1;refGene_gn=IFI16;refGene_tx=XM_023247224.1;refGene_gn=IFI16;refGene_tx=XM_023247225.1;refGene_gn=IFI16;refGene_tx=XM_023247226.1;refGene_gn=IFI16;refGene_tx=XM_023247228.1;refGene_gn=IFI16;refGene_tx=XM_023247229.1
F1 67763515 67763646
refGene_gn=IFI16;refGene_tx=XM_023247223.1;refGene_gn=IFI16;refGene_tx=XM_023247224.1;refGene_gn=IFI16;refGene_tx=XM_023247225.1;refGene_gn=IFI16;refGene_tx=XM_023247226.1;refGene_gn=IFI16;refGene_tx=XM_023247228.1;refGene_gn=IFI16;refGene_tx=XM_023247229.1
F1 67764530 67764795
refGene_gn=IFI16;refGene_tx=XM_023247223.1;refGene_gn=IFI16;refGene_tx=XM_023247224.1;refGene_gn=IFI16;refGene_tx=XM_023247225.1;refGene_gn=IFI16;refGene_tx=XM_023247226.1;refGene_gn=IFI16;refGene_tx=XM_023247228.1;refGene_gn=IFI16;refGene_tx=XM_023247229.1
F1 67786764 67786902
refGene_gn=LOC102902635;refGene_tx=XM_023247230.1
F1 67789033 67789222
refGene_gn=LOC102902635;refGene_tx=XM_023247230.1
F1 67790821 67791244
refGene_gn=LOC102902635;refGene_tx=XM_023247230.1
F1 67821299 67821419
refGene_gn=LOC111558418;refGene_tx=XM_023248029.1
F1 67821420 67822245
refGene_gn=LOC111558418;refGene_tx=XM_023248029.1
F1 67845998 67846943
refGene_gn=LOC101095793;refGene_tx=XM_011291156.2
F1 67857487 67858501
refGene_gn=LOC101096032;refGene_tx=XM_003999671.3
F1 67870272 67871244
refGene_gn=LOC101096285;refGene_tx=XM_003999672.2
F1 67894754 67894763
refGene_gn=LOC101096536;refGene_tx=XM_023247469.1
F1 67895105 67895156
refGene_gn=LOC101096536;refGene_tx=XM_023247469.1
F1 67895213 67896104
refGene_gn=LOC101096536;refGene_tx=XM_023247469.1
F1 67900325 67900737
refGene_gn=LOC101096786;refGene_tx=XM_019821376.2
F1 67900739 67901266
refGene_gn=LOC101096786;refGene_tx=XM_019821376.2
F1 67905686 67905766
refGene_gn=LOC101096786;refGene_tx=XM_019821376.2
F1 67907270 67908236
refGene_gn=LOC101096786;refGene_tx=XM_019821376.2;refGene_gn=LOC101097042;refGene_tx=XM_006943103.2

F1 67918275 67919238
refGene_gn=LOC101100439;refGene_tx=XM_011291102.1
F1 67930823 67931060
refGene_gn=SPTA1;refGene_tx=XM_019822344.2;refGene_gn=SPTA1;refGene_tx=XM_019822345.2
F1 67932840 67932966
refGene_gn=SPTA1;refGene_tx=XM_019822344.2;refGene_gn=SPTA1;refGene_tx=XM_019822345.2
F1 67934883 67935024
refGene_gn=SPTA1;refGene_tx=XM_019822344.2;refGene_gn=SPTA1;refGene_tx=XM_019822345.2
F1 67935743 67935890
refGene_gn=SPTA1;refGene_tx=XM_019822344.2;refGene_gn=SPTA1;refGene_tx=XM_019822345.2
F1 67937923 67938057
refGene_gn=SPTA1;refGene_tx=XM_019822344.2;refGene_gn=SPTA1;refGene_tx=XM_019822345.2
F1 67938610 67938755
refGene_gn=SPTA1;refGene_tx=XM_019822344.2;refGene_gn=SPTA1;refGene_tx=XM_019822345.2
F1 67940900 67941055
refGene_gn=SPTA1;refGene_tx=XM_019822344.2;refGene_gn=SPTA1;refGene_tx=XM_019822345.2
F1 67942020 67942156
refGene_gn=SPTA1;refGene_tx=XM_019822344.2;refGene_gn=SPTA1;refGene_tx=XM_019822345.2
F1 67942256 67942358
refGene_gn=SPTA1;refGene_tx=XM_019822344.2;refGene_gn=SPTA1;refGene_tx=XM_019822345.2
F1 67943809 67943947
refGene_gn=SPTA1;refGene_tx=XM_019822344.2;refGene_gn=SPTA1;refGene_tx=XM_019822345.2
F1 67944540 67944651
refGene_gn=SPTA1;refGene_tx=XM_019822344.2;refGene_gn=SPTA1;refGene_tx=XM_019822345.2
F1 67946144 67946222
refGene_gn=SPTA1;refGene_tx=XM_019822344.2;refGene_gn=SPTA1;refGene_tx=XM_019822345.2
F1 67946369 67946525
refGene_gn=SPTA1;refGene_tx=XM_019822344.2;refGene_gn=SPTA1;refGene_tx=XM_019822345.2
F1 67947892 67948097
refGene_gn=SPTA1;refGene_tx=XM_019822344.2;refGene_gn=SPTA1;refGene_tx=XM_019822345.2
F1 67949780 67949962
refGene_gn=SPTA1;refGene_tx=XM_019822344.2;refGene_gn=SPTA1;refGene_tx=XM_019822345.2
F1 67951348 67951592
refGene_gn=SPTA1;refGene_tx=XM_019822344.2;refGene_gn=SPTA1;refGene_tx=XM_019822345.2

F1 67953069 67953192
refGene_gn=SPTA1;refGene_tx=XM_019822344.2;refGene_gn=SPTA1;refGene_tx
=XM_019822345.2
F1 67956003 67956221
refGene_gn=SPTA1;refGene_tx=XM_019822344.2;refGene_gn=SPTA1;refGene_tx
=XM_019822345.2
F1 67957005 67957098
refGene_gn=SPTA1;refGene_tx=XM_019822344.2;refGene_gn=SPTA1;refGene_tx
=XM_019822345.2
F1 67959207 67959345
refGene_gn=SPTA1;refGene_tx=XM_019822344.2;refGene_gn=SPTA1;refGene_tx
=XM_019822345.2
F1 67960113 67960265
refGene_gn=SPTA1;refGene_tx=XM_019822344.2;refGene_gn=SPTA1;refGene_tx
=XM_019822345.2
F1 67960382 67960569
refGene_gn=SPTA1;refGene_tx=XM_019822344.2;refGene_gn=SPTA1;refGene_tx
=XM_019822345.2
F1 67962386 67962488
refGene_gn=SPTA1;refGene_tx=XM_019822344.2;refGene_gn=SPTA1;refGene_tx
=XM_019822345.2
F1 67963463 67963555
refGene_gn=SPTA1;refGene_tx=XM_019822344.2;refGene_gn=SPTA1;refGene_tx
=XM_019822345.2
F1 67964333 67964478
refGene_gn=SPTA1;refGene_tx=XM_019822344.2;refGene_gn=SPTA1;refGene_tx
=XM_019822345.2
F1 67965241 67965423
refGene_gn=SPTA1;refGene_tx=XM_019822344.2;refGene_gn=SPTA1;refGene_tx
=XM_019822345.2
F1 67967641 67967741
refGene_gn=SPTA1;refGene_tx=XM_019822344.2;refGene_gn=SPTA1;refGene_tx
=XM_019822345.2
F1 67967855 67968053
refGene_gn=SPTA1;refGene_tx=XM_019822344.2;refGene_gn=SPTA1;refGene_tx
=XM_019822345.2
F1 67968847 67968991
refGene_gn=SPTA1;refGene_tx=XM_019822344.2;refGene_gn=SPTA1;refGene_tx
=XM_019822345.2
F1 67969831 67969935
refGene_gn=SPTA1;refGene_tx=XM_019822344.2;refGene_gn=SPTA1;refGene_tx
=XM_019822345.2
F1 67970543 67970706
refGene_gn=SPTA1;refGene_tx=XM_019822344.2;refGene_gn=SPTA1;refGene_tx
=XM_019822345.2
F1 67970967 67971099
refGene_gn=SPTA1;refGene_tx=XM_019822344.2;refGene_gn=SPTA1;refGene_tx
=XM_019822345.2
F1 67971599 67971737
refGene_gn=SPTA1;refGene_tx=XM_019822344.2;refGene_gn=SPTA1;refGene_tx

=XM_019822345.2
F1 67971904 67972009
refGene_gn=SPTA1;refGene_tx=XM_019822344.2;refGene_gn=SPTA1;refGene_tx
=XM_019822345.2
F1 67973303 67973512
refGene_gn=SPTA1;refGene_tx=XM_019822344.2;refGene_gn=SPTA1;refGene_tx
=XM_019822345.2
F1 67974863 67974984
refGene_gn=SPTA1;refGene_tx=XM_019822344.2;refGene_gn=SPTA1;refGene_tx
=XM_019822345.2
F1 67975733 67975855
refGene_gn=SPTA1;refGene_tx=XM_019822344.2;refGene_gn=SPTA1;refGene_tx
=XM_019822345.2
F1 67976602 67976735
refGene_gn=SPTA1;refGene_tx=XM_019822344.2;refGene_gn=SPTA1;refGene_tx
=XM_019822345.2
F1 67980407 67980506
refGene_gn=SPTA1;refGene_tx=XM_019822344.2;refGene_gn=SPTA1;refGene_tx
=XM_019822345.2
F1 67981445 67981608
refGene_gn=SPTA1;refGene_tx=XM_019822344.2;refGene_gn=SPTA1;refGene_tx
=XM_019822345.2
F1 67982237 67982314
refGene_gn=SPTA1;refGene_tx=XM_019822344.2;refGene_gn=SPTA1;refGene_tx
=XM_019822345.2
F1 67985484 67985694
refGene_gn=SPTA1;refGene_tx=XM_019822344.2;refGene_gn=SPTA1;refGene_tx
=XM_019822345.2
F1 67988158 67988455 refGene_gn=SPTA1;refGene_tx=XM_019822344.2
F1 67989489 67989602
refGene_gn=SPTA1;refGene_tx=XM_019822344.2;refGene_gn=SPTA1;refGene_tx
=XM_019822345.2
F1 67990769 67990787
refGene_gn=SPTA1;refGene_tx=XM_019822344.2;refGene_gn=SPTA1;refGene_tx
=XM_019822345.2
F1 67991275 67991327
refGene_gn=SPTA1;refGene_tx=XM_019822344.2;refGene_gn=SPTA1;refGene_tx
=XM_019822345.2
F1 67992301 67992489
refGene_gn=SPTA1;refGene_tx=XM_019822344.2;refGene_gn=SPTA1;refGene_tx
=XM_019822345.2
F1 67993397 67993451
refGene_gn=SPTA1;refGene_tx=XM_019822344.2;refGene_gn=SPTA1;refGene_tx
=XM_019822345.2
F1 67993835 67993982
refGene_gn=SPTA1;refGene_tx=XM_019822344.2;refGene_gn=SPTA1;refGene_tx
=XM_019822345.2
F1 67994591 67994736
refGene_gn=SPTA1;refGene_tx=XM_019822344.2;refGene_gn=SPTA1;refGene_tx
=XM_019822345.2

F1 67995246 67995372
refGene_gn=SPTA1;refGene_tx=XM_019822344.2;refGene_gn=SPTA1;refGene_tx=XM_019822345.2
F1 68001129 68002071
refGene_gn=LOC101100692;refGene_tx=XM_003999608.1
F1 68014740 68015673
refGene_gn=LOC101089999;refGene_tx=XM_023247470.1
F1 68022125 68022632
refGene_gn=LOC101090249;refGene_tx=XM_023247471.1
F1 68022633 68023050
refGene_gn=LOC101090249;refGene_tx=XM_023247471.1
F1 68040395 68041328
refGene_gn=LOC105261351;refGene_tx=XM_011291151.2
F1 68063522 68064476
refGene_gn=LOC101100940;refGene_tx=XM_003999609.2
F1 68073409 68074387
refGene_gn=LOC101101187;refGene_tx=XM_023248025.1
F1 68081949 68082120
refGene_gn=LOC111558345;refGene_tx=XM_023247472.1
F1 68085485 68085583
refGene_gn=LOC111558345;refGene_tx=XM_023247472.1
F1 68086750 68087206
refGene_gn=LOC111558345;refGene_tx=XM_023247472.1
F1 68087208 68087593
refGene_gn=LOC111558345;refGene_tx=XM_023247472.1
F1 68087595 68087683
refGene_gn=LOC111558345;refGene_tx=XM_023247472.1
F1 68090788 68090915
refGene_gn=LOC111558345;refGene_tx=XM_023247472.1
F1 68101899 68101951
refGene_gn=LOC111558345;refGene_tx=XM_023247472.1
F1 68102815 68103014
refGene_gn=LOC111558345;refGene_tx=XM_023247472.1
F1 68103097 68103249
refGene_gn=LOC111558345;refGene_tx=XM_023247472.1
F1 68105890 68106658
refGene_gn=LOC101098033;refGene_tx=XM_011291149.1
F1 68106659 68106737
refGene_gn=LOC101098033;refGene_tx=XM_011291149.1
F1 68106738 68106855
refGene_gn=LOC101098033;refGene_tx=XM_011291149.1
F1 68113299 68114241
refGene_gn=LOC101101439;refGene_tx=XM_023247829.1
F1 68131185 68132130
refGene_gn=LOC101080258;refGene_tx=XM_003999612.1
F1 68145706 68146651
refGene_gn=LOC101080507;refGene_tx=XM_003999613.3
F1 68152071 68152301
refGene_gn=LOC101098280;refGene_tx=XM_019821377.2
F1 68163294 68163367

refGene_gn=LOC101098280;refGene_tx=XM_019821377.2
F1 68167158 68167555
refGene_gn=LOC101098280;refGene_tx=XM_019821377.2
F1 68167557 68168192
refGene_gn=LOC101098280;refGene_tx=XM_019821377.2
F1 68178523 68178698
refGene_gn=LOC101098516;refGene_tx=XM_006943058.4
F1 68178822 68178916
refGene_gn=LOC101098516;refGene_tx=XM_006943058.4
F1 68179313 68179592
refGene_gn=LOC101098516;refGene_tx=XM_006943058.4
F1 68179809 68180088
refGene_gn=LOC101098516;refGene_tx=XM_006943058.4
F1 68180731 68181028
refGene_gn=LOC101098516;refGene_tx=XM_006943058.4
F1 68181296 68181432
refGene_gn=LOC101098516;refGene_tx=XM_006943058.4
F1 68195000 68195019
refGene_gn=LOC101098769;refGene_tx=XM_003999680.5
F1 68195924 68196018
refGene_gn=LOC101098769;refGene_tx=XM_003999680.5
F1 68196259 68196538
refGene_gn=LOC101098769;refGene_tx=XM_003999680.5
F1 68196743 68197022
refGene_gn=LOC101098769;refGene_tx=XM_003999680.5
F1 68197597 68197864
refGene_gn=LOC101098769;refGene_tx=XM_003999680.5
F1 68198112 68198173
refGene_gn=LOC101098769;refGene_tx=XM_003999680.5
F1 68207278 68207300
refGene_gn=LOC101080762;refGene_tx=XM_023247843.1
F1 68207430 68207521
refGene_gn=LOC101080762;refGene_tx=XM_023247843.1
F1 68207866 68208145
refGene_gn=LOC101080762;refGene_tx=XM_023247843.1
F1 68208352 68208630
refGene_gn=LOC101080762;refGene_tx=XM_023247843.1
F1 68209250 68209521
refGene_gn=LOC101080762;refGene_tx=XM_023247843.1
F1 68209902 68209963
refGene_gn=LOC101080762;refGene_tx=XM_023247843.1
F1 68233788 68233834
refGene_gn=LOC101099272;refGene_tx=XM_019822350.2
F1 68233981 68234072
refGene_gn=LOC101099272;refGene_tx=XM_019822350.2
F1 68234400 68234679
refGene_gn=LOC101099272;refGene_tx=XM_019822350.2
F1 68234878 68235157
refGene_gn=LOC101099272;refGene_tx=XM_019822350.2
F1 68235830 68236097

refGene_gn=LOC101099272;refGene_tx=XM_019822350.2
F1 68236643 68236704
refGene_gn=LOC101099272;refGene_tx=XM_019822350.2
F1 68253103 68253149
refGene_gn=LOC111558346;refGene_tx=XM_023247473.1
F1 68253293 68253384
refGene_gn=LOC111558346;refGene_tx=XM_023247473.1
F1 68253716 68253995
refGene_gn=LOC111558346;refGene_tx=XM_023247473.1
F1 68254081 68254091
refGene_gn=LOC111558346;refGene_tx=XM_023247473.1
F1 68254261 68254458
refGene_gn=LOC109495813;refGene_tx=XM_023247529.1
F1 68255143 68255410
refGene_gn=LOC109495813;refGene_tx=XM_023247529.1
F1 68255957 68256018
refGene_gn=LOC109495813;refGene_tx=XM_023247529.1
F1 68282731 68282883
refGene_gn=LOC105259666;refGene_tx=XM_011291099.3;refGene_gn=LOC105259
666;refGene_tx=XM_023247528.1
F1 68282947 68283038
refGene_gn=LOC105259666;refGene_tx=XM_011291099.3
F1 68283369 68283648
refGene_gn=LOC105259666;refGene_tx=XM_011291099.3;refGene_gn=LOC105259
666;refGene_tx=XM_023247528.1
F1 68283845 68284124
refGene_gn=LOC105259666;refGene_tx=XM_011291099.3;refGene_gn=LOC105259
666;refGene_tx=XM_023247528.1
F1 68284802 68285069
refGene_gn=LOC105259666;refGene_tx=XM_011291099.3;refGene_gn=LOC105259
666;refGene_tx=XM_023247528.1
F1 68333641 68333720
refGene_gn=LOC101100279;refGene_tx=XM_011291202.3
F1 68333865 68333956
refGene_gn=LOC101100279;refGene_tx=XM_011291202.3
F1 68334283 68334562
refGene_gn=LOC101100279;refGene_tx=XM_011291202.3
F1 68334758 68335037
refGene_gn=LOC101100279;refGene_tx=XM_011291202.3
F1 68335720 68335987
refGene_gn=LOC101100279;refGene_tx=XM_011291202.3
F1 68336825 68336886
refGene_gn=LOC101100279;refGene_tx=XM_011291202.3
F1 68370734 68370813
refGene_gn=LOC111556282;refGene_tx=XM_023247686.1
F1 68370958 68371049
refGene_gn=LOC111556282;refGene_tx=XM_023247686.1
F1 68371377 68371656
refGene_gn=LOC111556282;refGene_tx=XM_023247686.1
F1 68371850 68372129

refGene_gn=LOC111556282;refGene_tx=XM_023247686.1
F1 68372818 68373085
refGene_gn=LOC111556282;refGene_tx=XM_023247686.1
F1 68373932 68373993
refGene_gn=LOC111556282;refGene_tx=XM_023247686.1
F1 68459851 68459873
refGene_gn=LOC101100784;refGene_tx=XM_023247474.1
F1 68459984 68460072
refGene_gn=LOC101100784;refGene_tx=XM_023247474.1
F1 68460794 68461076
refGene_gn=LOC101100784;refGene_tx=XM_023247474.1
F1 68461546 68462129
refGene_gn=LOC101100784;refGene_tx=XM_023247474.1
F1 68462519 68462928
refGene_gn=LOC101100784;refGene_tx=XM_023247474.1
F1 68463072 68463116
refGene_gn=LOC101100784;refGene_tx=XM_023247474.1
F1 68511304 68511781
refGene_gn=KIRREL1;refGene_tx=XM_023247946.1;refGene_gn=KIRREL1;refGen
e_tx=XM_023247947.1;refGene_gn=KIRREL1;refGene_tx=XM_023247948.1;refGe
ne_gn=KIRREL1;refGene_tx=XM_023247949.1
F1 68511890 68511968
refGene_gn=KIRREL1;refGene_tx=XM_023247946.1;refGene_gn=KIRREL1;refGen
e_tx=XM_023247947.1;refGene_gn=KIRREL1;refGene_tx=XM_023247948.1;refGe
ne_gn=KIRREL1;refGene_tx=XM_023247949.1
F1 68512499 68512639
refGene_gn=KIRREL1;refGene_tx=XM_023247946.1;refGene_gn=KIRREL1;refGen
e_tx=XM_023247947.1;refGene_gn=KIRREL1;refGene_tx=XM_023247948.1;refGe
ne_gn=KIRREL1;refGene_tx=XM_023247949.1
F1 68512773 68512881
refGene_gn=KIRREL1;refGene_tx=XM_023247946.1;refGene_gn=KIRREL1;refGen
e_tx=XM_023247947.1;refGene_gn=KIRREL1;refGene_tx=XM_023247948.1;refGe
ne_gn=KIRREL1;refGene_tx=XM_023247949.1
F1 68513999 68514198
refGene_gn=KIRREL1;refGene_tx=XM_023247946.1;refGene_gn=KIRREL1;refGen
e_tx=XM_023247947.1;refGene_gn=KIRREL1;refGene_tx=XM_023247948.1;refGe
ne_gn=KIRREL1;refGene_tx=XM_023247949.1
F1 68515311 68515412
refGene_gn=KIRREL1;refGene_tx=XM_023247946.1;refGene_gn=KIRREL1;refGen
e_tx=XM_023247947.1;refGene_gn=KIRREL1;refGene_tx=XM_023247948.1;refGe
ne_gn=KIRREL1;refGene_tx=XM_023247949.1
F1 68515499 68515674
refGene_gn=KIRREL1;refGene_tx=XM_023247946.1;refGene_gn=KIRREL1;refGen
e_tx=XM_023247947.1;refGene_gn=KIRREL1;refGene_tx=XM_023247948.1;refGe
ne_gn=KIRREL1;refGene_tx=XM_023247949.1
F1 68516232 68516360
refGene_gn=KIRREL1;refGene_tx=XM_023247946.1;refGene_gn=KIRREL1;refGen
e_tx=XM_023247947.1;refGene_gn=KIRREL1;refGene_tx=XM_023247948.1;refGe
ne_gn=KIRREL1;refGene_tx=XM_023247949.1
F1 68516536 68516685

refGene_gn=KIRREL1;refGene_tx=XM_023247946.1;refGene_gn=KIRREL1;refGene_tx=XM_023247947.1;refGene_gn=KIRREL1;refGene_tx=XM_023247948.1;refGene_gn=KIRREL1;refGene_tx=XM_023247949.1
F1 68516817 68516923
refGene_gn=KIRREL1;refGene_tx=XM_023247946.1;refGene_gn=KIRREL1;refGene_tx=XM_023247947.1;refGene_gn=KIRREL1;refGene_tx=XM_023247948.1;refGene_gn=KIRREL1;refGene_tx=XM_023247949.1
F1 68517880 68518031
refGene_gn=KIRREL1;refGene_tx=XM_023247946.1;refGene_gn=KIRREL1;refGene_tx=XM_023247947.1;refGene_gn=KIRREL1;refGene_tx=XM_023247948.1;refGene_gn=KIRREL1;refGene_tx=XM_023247949.1
F1 68519781 68519939
refGene_gn=KIRREL1;refGene_tx=XM_023247946.1;refGene_gn=KIRREL1;refGene_tx=XM_023247947.1;refGene_gn=KIRREL1;refGene_tx=XM_023247948.1;refGene_gn=KIRREL1;refGene_tx=XM_023247949.1
F1 68520918 68521893
refGene_gn=LOC109495817;refGene_tx=XM_019822357.2;refGene_gn=LOC109495817;refGene_tx=XM_019822358.2
F1 68524148 68524298
refGene_gn=KIRREL1;refGene_tx=XM_023247946.1;refGene_gn=KIRREL1;refGene_tx=XM_023247947.1;refGene_gn=KIRREL1;refGene_tx=XM_023247948.1;refGene_gn=KIRREL1;refGene_tx=XM_023247949.1
F1 68525654 68525804
refGene_gn=KIRREL1;refGene_tx=XM_023247946.1;refGene_gn=KIRREL1;refGene_tx=XM_023247947.1;refGene_gn=KIRREL1;refGene_tx=XM_023247948.1;refGene_gn=KIRREL1;refGene_tx=XM_023247949.1
F1 68605331 68605383
refGene_gn=KIRREL1;refGene_tx=XM_023247946.1;refGene_gn=KIRREL1;refGene_tx=XM_023247947.1;refGene_gn=KIRREL1;refGene_tx=XM_023247949.1
F1 68680429 68680580
refGene_gn=CD5L;refGene_tx=XM_023247601.1;refGene_gn=CD5L;refGene_tx=XM_023247602.1
F1 68682910 68682937
refGene_gn=CD5L;refGene_tx=XM_023247601.1;refGene_gn=CD5L;refGene_tx=XM_023247602.1
F1 68685478 68685799
refGene_gn=CD5L;refGene_tx=XM_023247601.1;refGene_gn=CD5L;refGene_tx=XM_023247602.1
F1 68693511 68693826
refGene_gn=CD5L;refGene_tx=XM_023247601.1;refGene_gn=CD5L;refGene_tx=XM_023247602.1
F1 68694768 68695089
refGene_gn=CD5L;refGene_tx=XM_023247601.1;refGene_gn=CD5L;refGene_tx=XM_023247602.1
F1 68695201 68695233 refGene_gn=CD5L;refGene_tx=XM_023247601.1
F1 68695910 68695939 refGene_gn=CD5L;refGene_tx=XM_023247602.1
F1 68705739 68705770
refGene_gn=FCRL1;refGene_tx=XM_006943062.4;refGene_gn=FCRL1;refGene_tx=XM_019822360.2;refGene_gn=FCRL1;refGene_tx=XM_019822362.2;refGene_gn=FCRL1;refGene_tx=XM_019822363.2

F1 68709204 68709225
refGene_gn=FCRL1;refGene_tx=XM_006943062.4;refGene_gn=FCRL1;refGene_tx=XM_019822360.2;refGene_gn=FCRL1;refGene_tx=XM_019822363.2
F1 68711335 68711605
refGene_gn=FCRL1;refGene_tx=XM_006943062.4;refGene_gn=FCRL1;refGene_tx=XM_019822360.2;refGene_gn=FCRL1;refGene_tx=XM_019822362.2;refGene_gn=FCRL1;refGene_tx=XM_019822363.2
F1 68712648 68712936
refGene_gn=FCRL1;refGene_tx=XM_006943062.4;refGene_gn=FCRL1;refGene_tx=XM_019822360.2;refGene_gn=FCRL1;refGene_tx=XM_019822362.2;refGene_gn=FCRL1;refGene_tx=XM_019822363.2
F1 68713091 68713370
refGene_gn=FCRL1;refGene_tx=XM_006943062.4;refGene_gn=FCRL1;refGene_tx=XM_019822360.2;refGene_gn=FCRL1;refGene_tx=XM_019822362.2;refGene_gn=FCRL1;refGene_tx=XM_019822363.2
F1 68714200 68714314
refGene_gn=FCRL1;refGene_tx=XM_006943062.4;refGene_gn=FCRL1;refGene_tx=XM_019822360.2;refGene_gn=FCRL1;refGene_tx=XM_019822362.2;refGene_gn=FCRL1;refGene_tx=XM_019822363.2
F1 68715377 68715405
refGene_gn=FCRL1;refGene_tx=XM_006943062.4;refGene_gn=FCRL1;refGene_tx=XM_019822360.2;refGene_gn=FCRL1;refGene_tx=XM_019822362.2;refGene_gn=FCRL1;refGene_tx=XM_019822363.2
F1 68716399 68716482
refGene_gn=FCRL1;refGene_tx=XM_006943062.4;refGene_gn=FCRL1;refGene_tx=XM_019822360.2;refGene_gn=FCRL1;refGene_tx=XM_019822362.2;refGene_gn=FCRL1;refGene_tx=XM_019822363.2
F1 68716689 68716758
refGene_gn=FCRL1;refGene_tx=XM_006943062.4;refGene_gn=FCRL1;refGene_tx=XM_019822360.2;refGene_gn=FCRL1;refGene_tx=XM_019822362.2;refGene_gn=FCRL1;refGene_tx=XM_019822363.2
F1 68717271 68717378
refGene_gn=FCRL1;refGene_tx=XM_006943062.4;refGene_gn=FCRL1;refGene_tx=XM_019822360.2;refGene_gn=FCRL1;refGene_tx=XM_019822362.2;refGene_gn=FCRL1;refGene_tx=XM_019822363.2
F1 68718238 68718310 refGene_gn=FCRL1;refGene_tx=XM_006943062.4
F1 68726902 68726933
refGene_gn=LOC102900134;refGene_tx=XM_019822375.2;refGene_gn=LOC102900134;refGene_tx=XM_019822376.2;refGene_gn=LOC102900134;refGene_tx=XM_019822377.2;refGene_gn=LOC102900134;refGene_tx=XM_023247516.1;refGene_gn=LOC102900134;refGene_tx=XM_023247517.1
F1 68728309 68728330
refGene_gn=LOC102900134;refGene_tx=XM_019822375.2;refGene_gn=LOC102900134;refGene_tx=XM_019822376.2;refGene_gn=LOC102900134;refGene_tx=XM_019822377.2;refGene_gn=LOC102900134;refGene_tx=XM_023247516.1;refGene_gn=LOC102900134;refGene_tx=XM_023247517.1
F1 68730265 68730535
refGene_gn=LOC102900134;refGene_tx=XM_019822375.2;refGene_gn=LOC102900134;refGene_tx=XM_019822376.2;refGene_gn=LOC102900134;refGene_tx=XM_019822377.2;refGene_gn=LOC102900134;refGene_tx=XM_023247516.1;refGene_gn

=LOC102900134;refGene_tx=XM_023247517.1
F1 68730769 68731054
refGene_gn=LOC102900134;refGene_tx=XM_019822375.2;refGene_gn=LOC102900134;refGene_tx=XM_019822376.2;refGene_gn=LOC102900134;refGene_tx=XM_019822377.2;refGene_gn=LOC102900134;refGene_tx=XM_023247516.1;refGene_gn=LOC102900134;refGene_tx=XM_023247517.1
F1 68732489 68732777
refGene_gn=LOC102900134;refGene_tx=XM_019822375.2;refGene_gn=LOC102900134;refGene_tx=XM_019822376.2;refGene_gn=LOC102900134;refGene_tx=XM_019822377.2;refGene_gn=LOC102900134;refGene_tx=XM_023247516.1;refGene_gn=LOC102900134;refGene_tx=XM_023247517.1
F1 68733161 68733440
refGene_gn=LOC102900134;refGene_tx=XM_019822375.2;refGene_gn=LOC102900134;refGene_tx=XM_019822376.2;refGene_gn=LOC102900134;refGene_tx=XM_019822377.2;refGene_gn=LOC102900134;refGene_tx=XM_023247516.1;refGene_gn=LOC102900134;refGene_tx=XM_023247517.1
F1 68733720 68733834
refGene_gn=LOC102900134;refGene_tx=XM_019822375.2;refGene_gn=LOC102900134;refGene_tx=XM_019822376.2;refGene_gn=LOC102900134;refGene_tx=XM_019822377.2;refGene_gn=LOC102900134;refGene_tx=XM_023247516.1;refGene_gn=LOC102900134;refGene_tx=XM_023247517.1
F1 68734309 68734404
refGene_gn=LOC102900134;refGene_tx=XM_019822377.2
F1 68738304 68738332
refGene_gn=LOC102900134;refGene_tx=XM_019822375.2;refGene_gn=LOC102900134;refGene_tx=XM_023247516.1;refGene_gn=LOC102900134;refGene_tx=XM_023247517.1
F1 68739334 68739420
refGene_gn=LOC102900134;refGene_tx=XM_019822375.2;refGene_gn=LOC102900134;refGene_tx=XM_019822376.2;refGene_gn=LOC102900134;refGene_tx=XM_023247516.1;refGene_gn=LOC102900134;refGene_tx=XM_023247517.1
F1 68739660 68739726
refGene_gn=LOC102900134;refGene_tx=XM_019822375.2;refGene_gn=LOC102900134;refGene_tx=XM_019822376.2;refGene_gn=LOC102900134;refGene_tx=XM_023247516.1;refGene_gn=LOC102900134;refGene_tx=XM_023247517.1
F1 68741350 68741382
refGene_gn=LOC102900134;refGene_tx=XM_019822375.2;refGene_gn=LOC102900134;refGene_tx=XM_019822376.2;refGene_gn=LOC102900134;refGene_tx=XM_023247516.1;refGene_gn=LOC102900134;refGene_tx=XM_023247517.1
F1 68741465 68741504
refGene_gn=LOC102900134;refGene_tx=XM_019822375.2;refGene_gn=LOC102900134;refGene_tx=XM_019822376.2;refGene_gn=LOC102900134;refGene_tx=XM_023247516.1;refGene_gn=LOC102900134;refGene_tx=XM_023247517.1
F1 68755026 68755054
refGene_gn=FCRL3;refGene_tx=XM_019822364.2;refGene_gn=FCRL3;refGene_tx=XM_019822368.2;refGene_gn=FCRL3;refGene_tx=XM_019822369.2;refGene_gn=FCRL3;refGene_tx=XM_019822370.2;refGene_gn=FCRL3;refGene_tx=XM_023247510.1;refGene_gn=FCRL3;refGene_tx=XM_023247511.1;refGene_gn=FCRL3;refGene_tx=XM_023247512.1;refGene_gn=FCRL3;refGene_tx=XM_023247513.1;refGene_gn=FCRL3;refGene_tx=XM_023247514.1

F1 68755648 68755669
refGene_gn=FCRL3;refGene_tx=XM_019822364.2;refGene_gn=FCRL3;refGene_tx=XM_019822368.2;refGene_gn=FCRL3;refGene_tx=XM_019822369.2;refGene_gn=FCRL3;refGene_tx=XM_019822370.2;refGene_gn=FCRL3;refGene_tx=XM_023247510.1;refGene_gn=FCRL3;refGene_tx=XM_023247511.1;refGene_gn=FCRL3;refGene_tx=XM_023247512.1;refGene_gn=FCRL3;refGene_tx=XM_023247513.1;refGene_gn=FCRL3;refGene_tx=XM_023247514.1
F1 68756590 68756833
refGene_gn=FCRL3;refGene_tx=XM_019822364.2;refGene_gn=FCRL3;refGene_tx=XM_019822368.2;refGene_gn=FCRL3;refGene_tx=XM_019822369.2;refGene_gn=FCRL3;refGene_tx=XM_019822370.2;refGene_gn=FCRL3;refGene_tx=XM_023247510.1;refGene_gn=FCRL3;refGene_tx=XM_023247511.1;refGene_gn=FCRL3;refGene_tx=XM_023247512.1;refGene_gn=FCRL3;refGene_tx=XM_023247513.1;refGene_gn=FCRL3;refGene_tx=XM_023247514.1
F1 68757304 68757562
refGene_gn=FCRL3;refGene_tx=XM_019822364.2;refGene_gn=FCRL3;refGene_tx=XM_019822368.2;refGene_gn=FCRL3;refGene_tx=XM_019822369.2;refGene_gn=FCRL3;refGene_tx=XM_019822370.2;refGene_gn=FCRL3;refGene_tx=XM_023247510.1;refGene_gn=FCRL3;refGene_tx=XM_023247511.1;refGene_gn=FCRL3;refGene_tx=XM_023247512.1;refGene_gn=FCRL3;refGene_tx=XM_023247513.1;refGene_gn=FCRL3;refGene_tx=XM_023247514.1
F1 68757799 68758084
refGene_gn=FCRL3;refGene_tx=XM_019822364.2;refGene_gn=FCRL3;refGene_tx=XM_019822368.2;refGene_gn=FCRL3;refGene_tx=XM_019822369.2;refGene_gn=FCRL3;refGene_tx=XM_019822370.2;refGene_gn=FCRL3;refGene_tx=XM_023247510.1;refGene_gn=FCRL3;refGene_tx=XM_023247511.1;refGene_gn=FCRL3;refGene_tx=XM_023247512.1;refGene_gn=FCRL3;refGene_tx=XM_023247513.1;refGene_gn=FCRL3;refGene_tx=XM_023247514.1
F1 68759100 68759388
refGene_gn=FCRL3;refGene_tx=XM_019822364.2;refGene_gn=FCRL3;refGene_tx=XM_019822368.2;refGene_gn=FCRL3;refGene_tx=XM_019822369.2;refGene_gn=FCRL3;refGene_tx=XM_019822370.2;refGene_gn=FCRL3;refGene_tx=XM_023247510.1;refGene_gn=FCRL3;refGene_tx=XM_023247511.1;refGene_gn=FCRL3;refGene_tx=XM_023247512.1;refGene_gn=FCRL3;refGene_tx=XM_023247513.1;refGene_gn=FCRL3;refGene_tx=XM_023247514.1
F1 68759716 68759995
refGene_gn=FCRL3;refGene_tx=XM_019822364.2;refGene_gn=FCRL3;refGene_tx=XM_019822368.2;refGene_gn=FCRL3;refGene_tx=XM_019822369.2;refGene_gn=FCRL3;refGene_tx=XM_019822370.2;refGene_gn=FCRL3;refGene_tx=XM_023247510.1;refGene_gn=FCRL3;refGene_tx=XM_023247511.1;refGene_gn=FCRL3;refGene_tx=XM_023247512.1;refGene_gn=FCRL3;refGene_tx=XM_023247513.1;refGene_gn=FCRL3;refGene_tx=XM_023247514.1
F1 68761057 68761336
refGene_gn=FCRL3;refGene_tx=XM_019822364.2;refGene_gn=FCRL3;refGene_tx=XM_019822368.2;refGene_gn=FCRL3;refGene_tx=XM_019822369.2;refGene_gn=FCRL3;refGene_tx=XM_019822370.2;refGene_gn=FCRL3;refGene_tx=XM_023247510.1;refGene_gn=FCRL3;refGene_tx=XM_023247511.1;refGene_gn=FCRL3;refGene_tx=XM_023247512.1;refGene_gn=FCRL3;refGene_tx=XM_023247513.1;refGene_gn=FCRL3;refGene_tx=XM_023247514.1
F1 68761693 68761813

refGene_gn=FCRL3;refGene_tx=XM_019822364.2;refGene_gn=FCRL3;refGene_tx=XM_019822369.2;refGene_gn=FCRL3;refGene_tx=XM_019822370.2;refGene_gn=FCRL3;refGene_tx=XM_023247510.1;refGene_gn=FCRL3;refGene_tx=XM_023247511.1;refGene_gn=FCRL3;refGene_tx=XM_023247512.1;refGene_gn=FCRL3;refGene_tx=XM_023247513.1;refGene_gn=FCRL3;refGene_tx=XM_023247514.1
F1 68767528 68767585
refGene_gn=FCRL3;refGene_tx=XM_023247510.1;refGene_gn=FCRL3;refGene_tx=XM_023247511.1;refGene_gn=FCRL3;refGene_tx=XM_023247512.1;refGene_gn=FCRL3;refGene_tx=XM_023247513.1;refGene_gn=FCRL3;refGene_tx=XM_023247514.1
F1 68767772 68767792 refGene_gn=FCRL3;refGene_tx=XM_023247514.1
F1 68768679 68768799
refGene_gn=FCRL3;refGene_tx=XM_019822364.2;refGene_gn=FCRL3;refGene_tx=XM_019822368.2;refGene_gn=FCRL3;refGene_tx=XM_023247510.1;refGene_gn=FCRL3;refGene_tx=XM_023247511.1;refGene_gn=FCRL3;refGene_tx=XM_023247512.1;refGene_gn=FCRL3;refGene_tx=XM_023247513.1
F1 68769245 68769314
refGene_gn=FCRL3;refGene_tx=XM_019822364.2;refGene_gn=FCRL3;refGene_tx=XM_019822368.2;refGene_gn=FCRL3;refGene_tx=XM_023247510.1;refGene_gn=FCRL3;refGene_tx=XM_023247511.1
F1 68770916 68770948
refGene_gn=FCRL3;refGene_tx=XM_019822364.2;refGene_gn=FCRL3;refGene_tx=XM_019822368.2;refGene_gn=FCRL3;refGene_tx=XM_019822369.2;refGene_gn=FCRL3;refGene_tx=XM_019822370.2;refGene_gn=FCRL3;refGene_tx=XM_023247510.1;refGene_gn=FCRL3;refGene_tx=XM_023247511.1;refGene_gn=FCRL3;refGene_tx=XM_023247512.1;refGene_gn=FCRL3;refGene_tx=XM_023247513.1
F1 68771020 68771158
refGene_gn=FCRL3;refGene_tx=XM_019822364.2;refGene_gn=FCRL3;refGene_tx=XM_019822368.2;refGene_gn=FCRL3;refGene_tx=XM_019822369.2;refGene_gn=FCRL3;refGene_tx=XM_019822370.2;refGene_gn=FCRL3;refGene_tx=XM_023247510.1;refGene_gn=FCRL3;refGene_tx=XM_023247511.1;refGene_gn=FCRL3;refGene_tx=XM_023247512.1;refGene_gn=FCRL3;refGene_tx=XM_023247513.1
F1 68798142 68798173 refGene_gn=FCRL4;refGene_tx=XM_006943109.4
F1 68800514 68800535 refGene_gn=FCRL4;refGene_tx=XM_006943109.4
F1 68804217 68804472 refGene_gn=FCRL4;refGene_tx=XM_006943109.4
F1 68805527 68805782 refGene_gn=FCRL4;refGene_tx=XM_006943109.4
F1 68806058 68806340 refGene_gn=FCRL4;refGene_tx=XM_006943109.4
F1 68807347 68807635 refGene_gn=FCRL4;refGene_tx=XM_006943109.4
F1 68816327 68816441 refGene_gn=FCRL4;refGene_tx=XM_006943109.4
F1 68817536 68817564 refGene_gn=FCRL4;refGene_tx=XM_006943109.4
F1 68818467 68818562 refGene_gn=FCRL4;refGene_tx=XM_006943109.4
F1 68818791 68818860 refGene_gn=FCRL4;refGene_tx=XM_006943109.4
F1 68822671 68822703 refGene_gn=FCRL4;refGene_tx=XM_006943109.4
F1 68822801 68822918 refGene_gn=FCRL4;refGene_tx=XM_006943109.4
F1 68841860 68842002
refGene_gn=FCRL5;refGene_tx=XM_023247237.1;refGene_gn=FCRL5;refGene_tx=XM_023247238.1;refGene_gn=FCRL5;refGene_tx=XM_023247239.1
F1 68852359 68852380
refGene_gn=FCRL5;refGene_tx=XM_023247237.1;refGene_gn=FCRL5;refGene_tx=XM_023247239.1

F1 68853606 68853861
refGene_gn=FCRL5;refGene_tx=XM_023247237.1;refGene_gn=FCRL5;refGene_tx=XM_023247238.1;refGene_gn=FCRL5;refGene_tx=XM_023247239.1
F1 68855740 68855995
refGene_gn=FCRL5;refGene_tx=XM_023247237.1;refGene_gn=FCRL5;refGene_tx=XM_023247238.1;refGene_gn=FCRL5;refGene_tx=XM_023247239.1
F1 68856276 68856561
refGene_gn=FCRL5;refGene_tx=XM_023247237.1;refGene_gn=FCRL5;refGene_tx=XM_023247238.1;refGene_gn=FCRL5;refGene_tx=XM_023247239.1
F1 68857979 68858252
refGene_gn=FCRL5;refGene_tx=XM_023247237.1;refGene_gn=FCRL5;refGene_tx=XM_023247238.1;refGene_gn=FCRL5;refGene_tx=XM_023247239.1
F1 68860916 68861195
refGene_gn=FCRL5;refGene_tx=XM_023247237.1;refGene_gn=FCRL5;refGene_tx=XM_023247238.1
F1 68867522 68867801
refGene_gn=FCRL5;refGene_tx=XM_023247237.1;refGene_gn=FCRL5;refGene_tx=XM_023247238.1;refGene_gn=FCRL5;refGene_tx=XM_023247239.1
F1 68868174 68868453
refGene_gn=FCRL5;refGene_tx=XM_023247237.1;refGene_gn=FCRL5;refGene_tx=XM_023247238.1;refGene_gn=FCRL5;refGene_tx=XM_023247239.1
F1 68868826 68869105
refGene_gn=FCRL5;refGene_tx=XM_023247237.1;refGene_gn=FCRL5;refGene_tx=XM_023247238.1;refGene_gn=FCRL5;refGene_tx=XM_023247239.1
F1 68871090 68871369
refGene_gn=FCRL5;refGene_tx=XM_023247237.1;refGene_gn=FCRL5;refGene_tx=XM_023247238.1;refGene_gn=FCRL5;refGene_tx=XM_023247239.1
F1 68874069 68874348
refGene_gn=FCRL5;refGene_tx=XM_023247237.1;refGene_gn=FCRL5;refGene_tx=XM_023247238.1;refGene_gn=FCRL5;refGene_tx=XM_023247239.1
F1 68874778 68874895
refGene_gn=FCRL5;refGene_tx=XM_023247237.1;refGene_gn=FCRL5;refGene_tx=XM_023247238.1;refGene_gn=FCRL5;refGene_tx=XM_023247239.1
F1 68875476 68875504
refGene_gn=FCRL5;refGene_tx=XM_023247237.1;refGene_gn=FCRL5;refGene_tx=XM_023247238.1;refGene_gn=FCRL5;refGene_tx=XM_023247239.1
F1 68876456 68876539
refGene_gn=FCRL5;refGene_tx=XM_023247237.1;refGene_gn=FCRL5;refGene_tx=XM_023247238.1;refGene_gn=FCRL5;refGene_tx=XM_023247239.1
F1 68876745 68876814
refGene_gn=FCRL5;refGene_tx=XM_023247237.1;refGene_gn=FCRL5;refGene_tx=XM_023247238.1;refGene_gn=FCRL5;refGene_tx=XM_023247239.1
F1 68878869 68879024
refGene_gn=FCRL5;refGene_tx=XM_023247237.1;refGene_gn=FCRL5;refGene_tx=XM_023247238.1;refGene_gn=FCRL5;refGene_tx=XM_023247239.1
F1 68900017 68900090
refGene_gn=LOC102902050;refGene_tx=XM_019822386.2;refGene_gn=LOC102902050;refGene_tx=XM_019822387.2;refGene_gn=LOC102902050;refGene_tx=XM_019822388.2;refGene_gn=LOC102902050;refGene_tx=XM_019822389.2
F1 68904768 68905161

refGene_gn=LOC102902050;refGene_tx=XM_019822386.2;refGene_gn=LOC102902050;refGene_tx=XM_019822387.2;refGene_gn=LOC102902050;refGene_tx=XM_019822388.2;refGene_gn=LOC102902050;refGene_tx=XM_019822389.2;refGene_gn=LOC102902050;refGene_tx=XM_019822390.2
F1 68905529 68905826
refGene_gn=LOC102902050;refGene_tx=XM_019822386.2;refGene_gn=LOC102902050;refGene_tx=XM_019822387.2;refGene_gn=LOC102902050;refGene_tx=XM_019822388.2;refGene_gn=LOC102902050;refGene_tx=XM_019822389.2;refGene_gn=LOC102902050;refGene_tx=XM_019822390.2
F1 68907161 68907300
refGene_gn=LOC102902050;refGene_tx=XM_019822386.2;refGene_gn=LOC102902050;refGene_tx=XM_019822387.2;refGene_gn=LOC102902050;refGene_tx=XM_019822388.2;refGene_gn=LOC102902050;refGene_tx=XM_019822390.2
F1 68907498 68907584
refGene_gn=LOC102902050;refGene_tx=XM_019822386.2;refGene_gn=LOC102902050;refGene_tx=XM_019822387.2;refGene_gn=LOC102902050;refGene_tx=XM_019822388.2;refGene_gn=LOC102902050;refGene_tx=XM_019822389.2;refGene_gn=LOC102902050;refGene_tx=XM_019822390.2
F1 68907840 68907897
refGene_gn=LOC102902050;refGene_tx=XM_019822386.2;refGene_gn=LOC102902050;refGene_tx=XM_019822387.2;refGene_gn=LOC102902050;refGene_tx=XM_019822388.2;refGene_gn=LOC102902050;refGene_tx=XM_019822389.2;refGene_gn=LOC102902050;refGene_tx=XM_019822390.2
F1 68908040 68908145
refGene_gn=LOC102902050;refGene_tx=XM_019822386.2;refGene_gn=LOC102902050;refGene_tx=XM_019822389.2;refGene_gn=LOC102902050;refGene_tx=XM_019822390.2
F1 68956248 68956341
refGene_gn=LOC105259618;refGene_tx=XM_011291198.3;refGene_gn=LOC105259618;refGene_tx=XM_023247435.1;refGene_gn=LOC105259618;refGene_tx=XM_023247436.1;refGene_gn=LOC105259618;refGene_tx=XM_023247437.1
F1 68956581 68956658
refGene_gn=LOC105259618;refGene_tx=XM_011291198.3;refGene_gn=LOC105259618;refGene_tx=XM_023247435.1
F1 68956858 68957005
refGene_gn=LOC105259618;refGene_tx=XM_011291198.3;refGene_gn=LOC105259618;refGene_tx=XM_019822396.2;refGene_gn=LOC105259618;refGene_tx=XM_023247435.1;refGene_gn=LOC105259618;refGene_tx=XM_023247436.1;refGene_gn=LOC105259618;refGene_tx=XM_023247437.1
F1 68957789 68958086
refGene_gn=LOC105259618;refGene_tx=XM_011291198.3;refGene_gn=LOC105259618;refGene_tx=XM_019822396.2;refGene_gn=LOC105259618;refGene_tx=XM_023247435.1;refGene_gn=LOC105259618;refGene_tx=XM_023247436.1;refGene_gn=LOC105259618;refGene_tx=XM_023247437.1
F1 68958455 68958848
refGene_gn=LOC105259618;refGene_tx=XM_011291198.3;refGene_gn=LOC105259618;refGene_tx=XM_019822396.2;refGene_gn=LOC105259618;refGene_tx=XM_023247435.1;refGene_gn=LOC105259618;refGene_tx=XM_023247436.1;refGene_gn=LOC105259618;refGene_tx=XM_023247437.1
F1 68962427 68962500

refGene_gn=LOC105259618;refGene_tx=XM_011291198.3;refGene_gn=LOC105259618;refGene_tx=XM_019822396.2;refGene_gn=LOC105259618;refGene_tx=XM_023247435.1;refGene_gn=LOC105259618;refGene_tx=XM_023247436.1;refGene_gn=LOC105259618;refGene_tx=XM_023247437.1
F1 68971495 68971544
refGene_gn=LOC111556210;refGene_tx=XM_019822392.2
F1 68971740 68971882
refGene_gn=LOC111556210;refGene_tx=XM_019822392.2
F1 68972675 68972972
refGene_gn=LOC111556210;refGene_tx=XM_019822392.2
F1 68973338 68973737
refGene_gn=LOC111556210;refGene_tx=XM_019822392.2
F1 68974951 68975024
refGene_gn=LOC111556210;refGene_tx=XM_019822392.2
F1 69001073 69001099 refGene_gn=ETV3;refGene_tx=XM_023247361.1
F1 69001239 69001298
refGene_gn=ETV3;refGene_tx=XM_023247359.1;refGene_gn=ETV3;refGene_tx=XM_023247360.1;refGene_gn=ETV3;refGene_tx=XM_023247361.1
F1 69001902 69002140
refGene_gn=ETV3;refGene_tx=XM_023247359.1;refGene_gn=ETV3;refGene_tx=XM_023247360.1;refGene_gn=ETV3;refGene_tx=XM_023247361.1
F1 69003381 69003497
refGene_gn=ETV3;refGene_tx=XM_023247359.1;refGene_gn=ETV3;refGene_tx=XM_023247360.1;refGene_gn=ETV3;refGene_tx=XM_023247361.1
F1 69004196 69004411 refGene_gn=ETV3;refGene_tx=XM_023247361.1
F1 69009428 69010558
refGene_gn=ETV3;refGene_tx=XM_023247359.1;refGene_gn=ETV3;refGene_tx=XM_023247360.1
F1 69031506 69031564 refGene_gn=ETV3L;refGene_tx=XM_023247503.1
F1 69031683 69031921 refGene_gn=ETV3L;refGene_tx=XM_023247503.1
F1 69032096 69032283 refGene_gn=ETV3L;refGene_tx=XM_023247503.1
F1 69032609 69032730 refGene_gn=ETV3L;refGene_tx=XM_023247503.1
F1 69035084 69035563 refGene_gn=ETV3L;refGene_tx=XM_023247503.1
F1 69057648 69057680
refGene_gn=ARHGEF11;refGene_tx=XM_023247561.1;refGene_gn=ARHGEF11;refGene_tx=XM_023247562.1;refGene_gn=ARHGEF11;refGene_tx=XM_023247563.1;refGene_gn=ARHGEF11;refGene_tx=XM_023247564.1;refGene_gn=ARHGEF11;refGene_tx=XM_023247565.1;refGene_gn=ARHGEF11;refGene_tx=XM_023247566.1;refGene_gn=ARHGEF11;refGene_tx=XM_023247567.1;refGene_gn=ARHGEF11;refGene_tx=XM_023247568.1;refGene_gn=ARHGEF11;refGene_tx=XM_023247569.1
F1 69101268 69101283
refGene_gn=ARHGEF11;refGene_tx=XM_023247561.1;refGene_gn=ARHGEF11;refGene_tx=XM_023247562.1;refGene_gn=ARHGEF11;refGene_tx=XM_023247563.1;refGene_gn=ARHGEF11;refGene_tx=XM_023247564.1;refGene_gn=ARHGEF11;refGene_tx=XM_023247566.1;refGene_gn=ARHGEF11;refGene_tx=XM_023247567.1;refGene_gn=ARHGEF11;refGene_tx=XM_023247568.1;refGene_gn=ARHGEF11;refGene_tx=XM_023247569.1
F1 69101594 69101692
refGene_gn=ARHGEF11;refGene_tx=XM_023247561.1;refGene_gn=ARHGEF11;refGene_tx=XM_023247562.1;refGene_gn=ARHGEF11;refGene_tx=XM_023247563.1;re

fGene_gn=ARHGEF11;refGene_tx=XM_023247564.1;refGene_gn=ARHGEF11;refGene_tx=XM_023247565.1;refGene_gn=ARHGEF11;refGene_tx=XM_023247566.1;refGene_gn=ARHGEF11;refGene_tx=XM_023247567.1;refGene_gn=ARHGEF11;refGene_tx=XM_023247568.1;refGene_gn=ARHGEF11;refGene_tx=XM_023247569.1
F1 69102606 69102705
refGene_gn=ARHGEF11;refGene_tx=XM_023247561.1;refGene_gn=ARHGEF11;refGene_tx=XM_023247562.1;refGene_gn=ARHGEF11;refGene_tx=XM_023247563.1;refGene_gn=ARHGEF11;refGene_tx=XM_023247564.1;refGene_gn=ARHGEF11;refGene_tx=XM_023247565.1;refGene_gn=ARHGEF11;refGene_tx=XM_023247566.1;refGene_gn=ARHGEF11;refGene_tx=XM_023247567.1;refGene_gn=ARHGEF11;refGene_tx=XM_023247568.1;refGene_gn=ARHGEF11;refGene_tx=XM_023247569.1
F1 69108578 69108628
refGene_gn=ARHGEF11;refGene_tx=XM_023247561.1;refGene_gn=ARHGEF11;refGene_tx=XM_023247562.1;refGene_gn=ARHGEF11;refGene_tx=XM_023247563.1;refGene_gn=ARHGEF11;refGene_tx=XM_023247564.1;refGene_gn=ARHGEF11;refGene_tx=XM_023247565.1;refGene_gn=ARHGEF11;refGene_tx=XM_023247566.1;refGene_gn=ARHGEF11;refGene_tx=XM_023247567.1;refGene_gn=ARHGEF11;refGene_tx=XM_023247568.1;refGene_gn=ARHGEF11;refGene_tx=XM_023247569.1
F1 69109301 69109359
refGene_gn=ARHGEF11;refGene_tx=XM_023247561.1;refGene_gn=ARHGEF11;refGene_tx=XM_023247562.1;refGene_gn=ARHGEF11;refGene_tx=XM_023247563.1;refGene_gn=ARHGEF11;refGene_tx=XM_023247564.1;refGene_gn=ARHGEF11;refGene_tx=XM_023247565.1;refGene_gn=ARHGEF11;refGene_tx=XM_023247566.1;refGene_gn=ARHGEF11;refGene_tx=XM_023247567.1;refGene_gn=ARHGEF11;refGene_tx=XM_023247568.1;refGene_gn=ARHGEF11;refGene_tx=XM_023247569.1
F1 69110071 69110250
refGene_gn=ARHGEF11;refGene_tx=XM_023247561.1;refGene_gn=ARHGEF11;refGene_tx=XM_023247562.1;refGene_gn=ARHGEF11;refGene_tx=XM_023247563.1;refGene_gn=ARHGEF11;refGene_tx=XM_023247564.1;refGene_gn=ARHGEF11;refGene_tx=XM_023247565.1;refGene_gn=ARHGEF11;refGene_tx=XM_023247566.1;refGene_gn=ARHGEF11;refGene_tx=XM_023247567.1;refGene_gn=ARHGEF11;refGene_tx=XM_023247568.1;refGene_gn=ARHGEF11;refGene_tx=XM_023247569.1
F1 69110934 69111006
refGene_gn=ARHGEF11;refGene_tx=XM_023247561.1;refGene_gn=ARHGEF11;refGene_tx=XM_023247562.1;refGene_gn=ARHGEF11;refGene_tx=XM_023247563.1;refGene_gn=ARHGEF11;refGene_tx=XM_023247564.1;refGene_gn=ARHGEF11;refGene_tx=XM_023247565.1;refGene_gn=ARHGEF11;refGene_tx=XM_023247566.1;refGene_gn=ARHGEF11;refGene_tx=XM_023247567.1;refGene_gn=ARHGEF11;refGene_tx=XM_023247568.1;refGene_gn=ARHGEF11;refGene_tx=XM_023247569.1
F1 69115197 69115317
refGene_gn=ARHGEF11;refGene_tx=XM_023247561.1;refGene_gn=ARHGEF11;refGene_tx=XM_023247562.1;refGene_gn=ARHGEF11;refGene_tx=XM_023247563.1;refGene_gn=ARHGEF11;refGene_tx=XM_023247564.1;refGene_gn=ARHGEF11;refGene_tx=XM_023247565.1;refGene_gn=ARHGEF11;refGene_tx=XM_023247566.1;refGene_gn=ARHGEF11;refGene_tx=XM_023247567.1;refGene_gn=ARHGEF11;refGene_tx=XM_023247568.1;refGene_gn=ARHGEF11;refGene_tx=XM_023247569.1
F1 69116724 69116770
refGene_gn=ARHGEF11;refGene_tx=XM_023247561.1;refGene_gn=ARHGEF11;refGene_tx=XM_023247562.1;refGene_gn=ARHGEF11;refGene_tx=XM_023247563.1;refGene_gn=ARHGEF11;refGene_tx=XM_023247564.1;refGene_gn=ARHGEF11;refGen

e_tx=XM_023247565.1;refGene_gn=ARHGEF11;refGene_tx=XM_023247566.1;refGene_gn=ARHGEF11;refGene_tx=XM_023247567.1;refGene_gn=ARHGEF11;refGene_tx=XM_023247568.1;refGene_gn=ARHGEF11;refGene_tx=XM_023247569.1
F1 69117247 69117324
refGene_gn=ARHGEF11;refGene_tx=XM_023247561.1;refGene_gn=ARHGEF11;refGene_tx=XM_023247562.1;refGene_gn=ARHGEF11;refGene_tx=XM_023247563.1;refGene_gn=ARHGEF11;refGene_tx=XM_023247564.1;refGene_gn=ARHGEF11;refGene_tx=XM_023247565.1;refGene_gn=ARHGEF11;refGene_tx=XM_023247566.1;refGene_gn=ARHGEF11;refGene_tx=XM_023247567.1;refGene_gn=ARHGEF11;refGene_tx=XM_023247568.1;refGene_gn=ARHGEF11;refGene_tx=XM_023247569.1
F1 69118273 69118411
refGene_gn=ARHGEF11;refGene_tx=XM_023247561.1;refGene_gn=ARHGEF11;refGene_tx=XM_023247562.1;refGene_gn=ARHGEF11;refGene_tx=XM_023247563.1;refGene_gn=ARHGEF11;refGene_tx=XM_023247564.1;refGene_gn=ARHGEF11;refGene_tx=XM_023247565.1;refGene_gn=ARHGEF11;refGene_tx=XM_023247566.1;refGene_gn=ARHGEF11;refGene_tx=XM_023247567.1;refGene_gn=ARHGEF11;refGene_tx=XM_023247568.1;refGene_gn=ARHGEF11;refGene_tx=XM_023247569.1
F1 69120079 69120154
refGene_gn=ARHGEF11;refGene_tx=XM_023247561.1;refGene_gn=ARHGEF11;refGene_tx=XM_023247562.1;refGene_gn=ARHGEF11;refGene_tx=XM_023247563.1;refGene_gn=ARHGEF11;refGene_tx=XM_023247564.1;refGene_gn=ARHGEF11;refGene_tx=XM_023247565.1;refGene_gn=ARHGEF11;refGene_tx=XM_023247566.1;refGene_gn=ARHGEF11;refGene_tx=XM_023247567.1;refGene_gn=ARHGEF11;refGene_tx=XM_023247568.1;refGene_gn=ARHGEF11;refGene_tx=XM_023247569.1
F1 69120363 69120465
refGene_gn=ARHGEF11;refGene_tx=XM_023247561.1;refGene_gn=ARHGEF11;refGene_tx=XM_023247562.1;refGene_gn=ARHGEF11;refGene_tx=XM_023247563.1;refGene_gn=ARHGEF11;refGene_tx=XM_023247564.1;refGene_gn=ARHGEF11;refGene_tx=XM_023247565.1;refGene_gn=ARHGEF11;refGene_tx=XM_023247566.1;refGene_gn=ARHGEF11;refGene_tx=XM_023247567.1;refGene_gn=ARHGEF11;refGene_tx=XM_023247568.1;refGene_gn=ARHGEF11;refGene_tx=XM_023247569.1
F1 69121359 69121458
refGene_gn=ARHGEF11;refGene_tx=XM_023247561.1;refGene_gn=ARHGEF11;refGene_tx=XM_023247562.1;refGene_gn=ARHGEF11;refGene_tx=XM_023247563.1;refGene_gn=ARHGEF11;refGene_tx=XM_023247564.1;refGene_gn=ARHGEF11;refGene_tx=XM_023247565.1;refGene_gn=ARHGEF11;refGene_tx=XM_023247566.1;refGene_gn=ARHGEF11;refGene_tx=XM_023247567.1;refGene_gn=ARHGEF11;refGene_tx=XM_023247568.1;refGene_gn=ARHGEF11;refGene_tx=XM_023247569.1
F1 69122440 69122483
refGene_gn=ARHGEF11;refGene_tx=XM_023247561.1;refGene_gn=ARHGEF11;refGene_tx=XM_023247562.1;refGene_gn=ARHGEF11;refGene_tx=XM_023247563.1;refGene_gn=ARHGEF11;refGene_tx=XM_023247564.1;refGene_gn=ARHGEF11;refGene_tx=XM_023247565.1;refGene_gn=ARHGEF11;refGene_tx=XM_023247566.1;refGene_gn=ARHGEF11;refGene_tx=XM_023247567.1;refGene_gn=ARHGEF11;refGene_tx=XM_023247568.1;refGene_gn=ARHGEF11;refGene_tx=XM_023247569.1
F1 69123488 69123585
refGene_gn=ARHGEF11;refGene_tx=XM_023247561.1;refGene_gn=ARHGEF11;refGene_tx=XM_023247562.1;refGene_gn=ARHGEF11;refGene_tx=XM_023247563.1;refGene_gn=ARHGEF11;refGene_tx=XM_023247564.1;refGene_gn=ARHGEF11;refGene_tx=XM_023247565.1;refGene_gn=ARHGEF11;refGene_tx=XM_023247566.1;refG

ene_gn=ARHGEF11;refGene_tx=XM_023247567.1;refGene_gn=ARHGEF11;refGene_tx=XM_023247568.1;refGene_gn=ARHGEF11;refGene_tx=XM_023247569.1
F1 69123719 69123842
refGene_gn=ARHGEF11;refGene_tx=XM_023247561.1;refGene_gn=ARHGEF11;refGene_tx=XM_023247562.1;refGene_gn=ARHGEF11;refGene_tx=XM_023247563.1;refGene_gn=ARHGEF11;refGene_tx=XM_023247564.1;refGene_gn=ARHGEF11;refGene_tx=XM_023247565.1;refGene_gn=ARHGEF11;refGene_tx=XM_023247566.1;refGene_gn=ARHGEF11;refGene_tx=XM_023247567.1;refGene_gn=ARHGEF11;refGene_tx=XM_023247568.1;refGene_gn=ARHGEF11;refGene_tx=XM_023247569.1
F1 69125827 69125851
refGene_gn=ARHGEF11;refGene_tx=XM_023247561.1;refGene_gn=ARHGEF11;refGene_tx=XM_023247562.1;refGene_gn=ARHGEF11;refGene_tx=XM_023247563.1;refGene_gn=ARHGEF11;refGene_tx=XM_023247564.1;refGene_gn=ARHGEF11;refGene_tx=XM_023247565.1;refGene_gn=ARHGEF11;refGene_tx=XM_023247566.1;refGene_gn=ARHGEF11;refGene_tx=XM_023247567.1;refGene_gn=ARHGEF11;refGene_tx=XM_023247568.1;refGene_gn=ARHGEF11;refGene_tx=XM_023247569.1
F1 69126860 69127005
refGene_gn=ARHGEF11;refGene_tx=XM_023247561.1;refGene_gn=ARHGEF11;refGene_tx=XM_023247562.1;refGene_gn=ARHGEF11;refGene_tx=XM_023247563.1;refGene_gn=ARHGEF11;refGene_tx=XM_023247564.1;refGene_gn=ARHGEF11;refGene_tx=XM_023247565.1;refGene_gn=ARHGEF11;refGene_tx=XM_023247566.1;refGene_gn=ARHGEF11;refGene_tx=XM_023247567.1;refGene_gn=ARHGEF11;refGene_tx=XM_023247568.1;refGene_gn=ARHGEF11;refGene_tx=XM_023247569.1
F1 69127293 69127393
refGene_gn=ARHGEF11;refGene_tx=XM_023247561.1;refGene_gn=ARHGEF11;refGene_tx=XM_023247562.1;refGene_gn=ARHGEF11;refGene_tx=XM_023247563.1;refGene_gn=ARHGEF11;refGene_tx=XM_023247564.1;refGene_gn=ARHGEF11;refGene_tx=XM_023247565.1;refGene_gn=ARHGEF11;refGene_tx=XM_023247566.1;refGene_gn=ARHGEF11;refGene_tx=XM_023247567.1;refGene_gn=ARHGEF11;refGene_tx=XM_023247568.1;refGene_gn=ARHGEF11;refGene_tx=XM_023247569.1
F1 69128174 69128204
refGene_gn=ARHGEF11;refGene_tx=XM_023247561.1;refGene_gn=ARHGEF11;refGene_tx=XM_023247562.1;refGene_gn=ARHGEF11;refGene_tx=XM_023247563.1;refGene_gn=ARHGEF11;refGene_tx=XM_023247564.1;refGene_gn=ARHGEF11;refGene_tx=XM_023247565.1;refGene_gn=ARHGEF11;refGene_tx=XM_023247568.1;refGene_gn=ARHGEF11;refGene_tx=XM_023247569.1
F1 69130980 69131107
refGene_gn=ARHGEF11;refGene_tx=XM_023247561.1;refGene_gn=ARHGEF11;refGene_tx=XM_023247562.1;refGene_gn=ARHGEF11;refGene_tx=XM_023247563.1;refGene_gn=ARHGEF11;refGene_tx=XM_023247564.1;refGene_gn=ARHGEF11;refGene_tx=XM_023247565.1;refGene_gn=ARHGEF11;refGene_tx=XM_023247566.1;refGene_gn=ARHGEF11;refGene_tx=XM_023247567.1;refGene_gn=ARHGEF11;refGene_tx=XM_023247568.1;refGene_gn=ARHGEF11;refGene_tx=XM_023247569.1
F1 69133502 69133682
refGene_gn=ARHGEF11;refGene_tx=XM_023247561.1;refGene_gn=ARHGEF11;refGene_tx=XM_023247562.1;refGene_gn=ARHGEF11;refGene_tx=XM_023247563.1;refGene_gn=ARHGEF11;refGene_tx=XM_023247564.1;refGene_gn=ARHGEF11;refGene_tx=XM_023247565.1;refGene_gn=ARHGEF11;refGene_tx=XM_023247566.1;refGene_gn=ARHGEF11;refGene_tx=XM_023247567.1;refGene_gn=ARHGEF11;refGene_tx=XM_023247568.1;refGene_gn=ARHGEF11;refGene_tx=XM_023247569.1

F1 69133761 69133809
refGene_gn=ARHGEF11;refGene_tx=XM_023247561.1;refGene_gn=ARHGEF11;refGene_tx=XM_023247562.1;refGene_gn=ARHGEF11;refGene_tx=XM_023247563.1;refGene_gn=ARHGEF11;refGene_tx=XM_023247564.1;refGene_gn=ARHGEF11;refGene_tx=XM_023247565.1;refGene_gn=ARHGEF11;refGene_tx=XM_023247566.1;refGene_gn=ARHGEF11;refGene_tx=XM_023247567.1;refGene_gn=ARHGEF11;refGene_tx=XM_023247568.1;refGene_gn=ARHGEF11;refGene_tx=XM_023247569.1
F1 69133939 69134127
refGene_gn=ARHGEF11;refGene_tx=XM_023247561.1;refGene_gn=ARHGEF11;refGene_tx=XM_023247562.1;refGene_gn=ARHGEF11;refGene_tx=XM_023247563.1;refGene_gn=ARHGEF11;refGene_tx=XM_023247564.1;refGene_gn=ARHGEF11;refGene_tx=XM_023247565.1;refGene_gn=ARHGEF11;refGene_tx=XM_023247566.1;refGene_gn=ARHGEF11;refGene_tx=XM_023247567.1;refGene_gn=ARHGEF11;refGene_tx=XM_023247568.1;refGene_gn=ARHGEF11;refGene_tx=XM_023247569.1
F1 69134387 69134534
refGene_gn=ARHGEF11;refGene_tx=XM_023247561.1;refGene_gn=ARHGEF11;refGene_tx=XM_023247562.1;refGene_gn=ARHGEF11;refGene_tx=XM_023247563.1;refGene_gn=ARHGEF11;refGene_tx=XM_023247564.1;refGene_gn=ARHGEF11;refGene_tx=XM_023247565.1;refGene_gn=ARHGEF11;refGene_tx=XM_023247566.1;refGene_gn=ARHGEF11;refGene_tx=XM_023247567.1;refGene_gn=ARHGEF11;refGene_tx=XM_023247568.1;refGene_gn=ARHGEF11;refGene_tx=XM_023247569.1
F1 69134837 69134917
refGene_gn=ARHGEF11;refGene_tx=XM_023247561.1;refGene_gn=ARHGEF11;refGene_tx=XM_023247562.1;refGene_gn=ARHGEF11;refGene_tx=XM_023247563.1;refGene_gn=ARHGEF11;refGene_tx=XM_023247564.1;refGene_gn=ARHGEF11;refGene_tx=XM_023247565.1;refGene_gn=ARHGEF11;refGene_tx=XM_023247566.1;refGene_gn=ARHGEF11;refGene_tx=XM_023247567.1;refGene_gn=ARHGEF11;refGene_tx=XM_023247568.1;refGene_gn=ARHGEF11;refGene_tx=XM_023247569.1
F1 69135024 69135150
refGene_gn=ARHGEF11;refGene_tx=XM_023247561.1;refGene_gn=ARHGEF11;refGene_tx=XM_023247562.1;refGene_gn=ARHGEF11;refGene_tx=XM_023247563.1;refGene_gn=ARHGEF11;refGene_tx=XM_023247564.1;refGene_gn=ARHGEF11;refGene_tx=XM_023247565.1;refGene_gn=ARHGEF11;refGene_tx=XM_023247566.1;refGene_gn=ARHGEF11;refGene_tx=XM_023247567.1;refGene_gn=ARHGEF11;refGene_tx=XM_023247568.1;refGene_gn=ARHGEF11;refGene_tx=XM_023247569.1
F1 69135680 69135798
refGene_gn=ARHGEF11;refGene_tx=XM_023247561.1;refGene_gn=ARHGEF11;refGene_tx=XM_023247562.1;refGene_gn=ARHGEF11;refGene_tx=XM_023247563.1;refGene_gn=ARHGEF11;refGene_tx=XM_023247564.1;refGene_gn=ARHGEF11;refGene_tx=XM_023247565.1;refGene_gn=ARHGEF11;refGene_tx=XM_023247566.1;refGene_gn=ARHGEF11;refGene_tx=XM_023247567.1;refGene_gn=ARHGEF11;refGene_tx=XM_023247568.1;refGene_gn=ARHGEF11;refGene_tx=XM_023247569.1
F1 69136594 69136773
refGene_gn=ARHGEF11;refGene_tx=XM_023247561.1;refGene_gn=ARHGEF11;refGene_tx=XM_023247562.1;refGene_gn=ARHGEF11;refGene_tx=XM_023247563.1;refGene_gn=ARHGEF11;refGene_tx=XM_023247564.1;refGene_gn=ARHGEF11;refGene_tx=XM_023247565.1;refGene_gn=ARHGEF11;refGene_tx=XM_023247566.1;refGene_gn=ARHGEF11;refGene_tx=XM_023247567.1;refGene_gn=ARHGEF11;refGene_tx=XM_023247568.1;refGene_gn=ARHGEF11;refGene_tx=XM_023247569.1
F1 69137192 69137268

refGene_gn=ARHGEF11;refGene_tx=XM_023247561.1;refGene_gn=ARHGEF11;refGene_tx=XM_023247562.1;refGene_gn=ARHGEF11;refGene_tx=XM_023247563.1;refGene_gn=ARHGEF11;refGene_tx=XM_023247564.1;refGene_gn=ARHGEF11;refGene_tx=XM_023247565.1;refGene_gn=ARHGEF11;refGene_tx=XM_023247566.1;refGene_gn=ARHGEF11;refGene_tx=XM_023247567.1;refGene_gn=ARHGEF11;refGene_tx=XM_023247568.1;refGene_gn=ARHGEF11;refGene_tx=XM_023247569.1
F1 69137416 69137584
refGene_gn=ARHGEF11;refGene_tx=XM_023247561.1;refGene_gn=ARHGEF11;refGene_tx=XM_023247562.1;refGene_gn=ARHGEF11;refGene_tx=XM_023247563.1;refGene_gn=ARHGEF11;refGene_tx=XM_023247564.1;refGene_gn=ARHGEF11;refGene_tx=XM_023247565.1;refGene_gn=ARHGEF11;refGene_tx=XM_023247566.1;refGene_gn=ARHGEF11;refGene_tx=XM_023247567.1;refGene_gn=ARHGEF11;refGene_tx=XM_023247568.1;refGene_gn=ARHGEF11;refGene_tx=XM_023247569.1
F1 69138030 69138121
refGene_gn=ARHGEF11;refGene_tx=XM_023247561.1;refGene_gn=ARHGEF11;refGene_tx=XM_023247562.1;refGene_gn=ARHGEF11;refGene_tx=XM_023247563.1;refGene_gn=ARHGEF11;refGene_tx=XM_023247564.1;refGene_gn=ARHGEF11;refGene_tx=XM_023247565.1;refGene_gn=ARHGEF11;refGene_tx=XM_023247566.1;refGene_gn=ARHGEF11;refGene_tx=XM_023247567.1;refGene_gn=ARHGEF11;refGene_tx=XM_023247568.1;refGene_gn=ARHGEF11;refGene_tx=XM_023247569.1
F1 69138597 69138723
refGene_gn=ARHGEF11;refGene_tx=XM_023247561.1;refGene_gn=ARHGEF11;refGene_tx=XM_023247562.1;refGene_gn=ARHGEF11;refGene_tx=XM_023247563.1;refGene_gn=ARHGEF11;refGene_tx=XM_023247564.1;refGene_gn=ARHGEF11;refGene_tx=XM_023247565.1;refGene_gn=ARHGEF11;refGene_tx=XM_023247566.1;refGene_gn=ARHGEF11;refGene_tx=XM_023247567.1;refGene_gn=ARHGEF11;refGene_tx=XM_023247568.1;refGene_gn=ARHGEF11;refGene_tx=XM_023247569.1
F1 69139043 69139105
refGene_gn=ARHGEF11;refGene_tx=XM_023247561.1;refGene_gn=ARHGEF11;refGene_tx=XM_023247562.1;refGene_gn=ARHGEF11;refGene_tx=XM_023247563.1;refGene_gn=ARHGEF11;refGene_tx=XM_023247564.1;refGene_gn=ARHGEF11;refGene_tx=XM_023247565.1;refGene_gn=ARHGEF11;refGene_tx=XM_023247566.1;refGene_gn=ARHGEF11;refGene_tx=XM_023247567.1;refGene_gn=ARHGEF11;refGene_tx=XM_023247568.1;refGene_gn=ARHGEF11;refGene_tx=XM_023247569.1
F1 69139859 69140078
refGene_gn=ARHGEF11;refGene_tx=XM_023247561.1;refGene_gn=ARHGEF11;refGene_tx=XM_023247562.1;refGene_gn=ARHGEF11;refGene_tx=XM_023247563.1;refGene_gn=ARHGEF11;refGene_tx=XM_023247564.1;refGene_gn=ARHGEF11;refGene_tx=XM_023247565.1;refGene_gn=ARHGEF11;refGene_tx=XM_023247566.1;refGene_gn=ARHGEF11;refGene_tx=XM_023247567.1;refGene_gn=ARHGEF11;refGene_tx=XM_023247568.1;refGene_gn=ARHGEF11;refGene_tx=XM_023247569.1
F1 69140250 69140568
refGene_gn=ARHGEF11;refGene_tx=XM_023247561.1;refGene_gn=ARHGEF11;refGene_tx=XM_023247562.1;refGene_gn=ARHGEF11;refGene_tx=XM_023247563.1;refGene_gn=ARHGEF11;refGene_tx=XM_023247564.1;refGene_gn=ARHGEF11;refGene_tx=XM_023247565.1;refGene_gn=ARHGEF11;refGene_tx=XM_023247566.1;refGene_gn=ARHGEF11;refGene_tx=XM_023247567.1;refGene_gn=ARHGEF11;refGene_tx=XM_023247568.1;refGene_gn=ARHGEF11;refGene_tx=XM_023247569.1
F1 69141484 69141580
refGene_gn=ARHGEF11;refGene_tx=XM_023247561.1;refGene_gn=ARHGEF11;refG

ene_tx=XM_023247562.1;refGene_gn=ARHGEF11;refGene_tx=XM_023247563.1;refGene_gn=ARHGEF11;refGene_tx=XM_023247564.1;refGene_gn=ARHGEF11;refGene_tx=XM_023247565.1;refGene_gn=ARHGEF11;refGene_tx=XM_023247566.1;refGene_gn=ARHGEF11;refGene_tx=XM_023247567.1;refGene_gn=ARHGEF11;refGene_tx=XM_023247569.1
F1 69142350 69142589
refGene_gn=ARHGEF11;refGene_tx=XM_023247561.1;refGene_gn=ARHGEF11;refGene_tx=XM_023247562.1;refGene_gn=ARHGEF11;refGene_tx=XM_023247563.1;refGene_gn=ARHGEF11;refGene_tx=XM_023247564.1;refGene_gn=ARHGEF11;refGene_tx=XM_023247565.1;refGene_gn=ARHGEF11;refGene_tx=XM_023247566.1;refGene_gn=ARHGEF11;refGene_tx=XM_023247567.1;refGene_gn=ARHGEF11;refGene_tx=XM_023247568.1;refGene_gn=ARHGEF11;refGene_tx=XM_023247569.1
F1 69142671 69142861
refGene_gn=ARHGEF11;refGene_tx=XM_023247561.1;refGene_gn=ARHGEF11;refGene_tx=XM_023247562.1;refGene_gn=ARHGEF11;refGene_tx=XM_023247563.1;refGene_gn=ARHGEF11;refGene_tx=XM_023247564.1;refGene_gn=ARHGEF11;refGene_tx=XM_023247565.1;refGene_gn=ARHGEF11;refGene_tx=XM_023247566.1;refGene_gn=ARHGEF11;refGene_tx=XM_023247567.1;refGene_gn=ARHGEF11;refGene_tx=XM_023247568.1;refGene_gn=ARHGEF11;refGene_tx=XM_023247569.1
F1 69143299 69143358
refGene_gn=ARHGEF11;refGene_tx=XM_023247561.1;refGene_gn=ARHGEF11;refGene_tx=XM_023247562.1;refGene_gn=ARHGEF11;refGene_tx=XM_023247563.1;refGene_gn=ARHGEF11;refGene_tx=XM_023247564.1;refGene_gn=ARHGEF11;refGene_tx=XM_023247565.1;refGene_gn=ARHGEF11;refGene_tx=XM_023247566.1;refGene_gn=ARHGEF11;refGene_tx=XM_023247567.1;refGene_gn=ARHGEF11;refGene_tx=XM_023247568.1;refGene_gn=ARHGEF11;refGene_tx=XM_023247569.1
F1 69145517 69145625
refGene_gn=LRRC71;refGene_tx=XM_023247570.1;refGene_gn=LRRC71;refGene_tx=XM_023247572.1;refGene_gn=LRRC71;refGene_tx=XM_023247573.1;refGene_gn=LRRC71;refGene_tx=XM_023247574.1;refGene_gn=LRRC71;refGene_tx=XM_023247575.1;refGene_gn=LRRC71;refGene_tx=XM_023247576.1;refGene_gn=LRRC71;refGene_tx=XM_023247577.1;refGene_gn=LRRC71;refGene_tx=XM_023247578.1;refGene_gn=LRRC71;refGene_tx=XM_023247579.1;refGene_gn=LRRC71;refGene_tx=XM_023247580.1;refGene_gn=LRRC71;refGene_tx=XM_023247581.1;refGene_gn=LRRC71;refGene_tx=XM_023247583.1;refGene_gn=LRRC71;refGene_tx=XM_023247584.1
F1 69145731 69145853
refGene_gn=LRRC71;refGene_tx=XM_023247570.1;refGene_gn=LRRC71;refGene_tx=XM_023247572.1;refGene_gn=LRRC71;refGene_tx=XM_023247573.1;refGene_gn=LRRC71;refGene_tx=XM_023247574.1;refGene_gn=LRRC71;refGene_tx=XM_023247575.1;refGene_gn=LRRC71;refGene_tx=XM_023247576.1;refGene_gn=LRRC71;refGene_tx=XM_023247577.1;refGene_gn=LRRC71;refGene_tx=XM_023247578.1;refGene_gn=LRRC71;refGene_tx=XM_023247579.1;refGene_gn=LRRC71;refGene_tx=XM_023247580.1;refGene_gn=LRRC71;refGene_tx=XM_023247581.1;refGene_gn=LRRC71;refGene_tx=XM_023247583.1;refGene_gn=LRRC71;refGene_tx=XM_023247584.1
F1 69146236 69146351
refGene_gn=LRRC71;refGene_tx=XM_023247570.1;refGene_gn=LRRC71;refGene_tx=XM_023247572.1;refGene_gn=LRRC71;refGene_tx=XM_023247573.1;refGene_gn=LRRC71;refGene_tx=XM_023247574.1;refGene_gn=LRRC71;refGene_tx=XM_02

3247575.1;refGene_gn=LRRC71;refGene_tx=XM_023247576.1;refGene_gn=LRRC7
1;refGene_tx=XM_023247577.1;refGene_gn=LRRC71;refGene_tx=XM_023247578.
1;refGene_gn=LRRC71;refGene_tx=XM_023247579.1;refGene_gn=LRRC71;refGen
e_tx=XM_023247580.1;refGene_gn=LRRC71;refGene_tx=XM_023247581.1;refGen
e_gn=LRRC71;refGene_tx=XM_023247583.1;refGene_gn=LRRC71;refGene_tx=XM_
023247584.1
F1 69146411 69146462
refGene_gn=LRRC71;refGene_tx=XM_023247570.1;refGene_gn=LRRC71;refGene_
tx=XM_023247572.1;refGene_gn=LRRC71;refGene_tx=XM_023247575.1;refGene_
gn=LRRC71;refGene_tx=XM_023247576.1;refGene_gn=LRRC71;refGene_tx=XM_02
3247577.1;refGene_gn=LRRC71;refGene_tx=XM_023247578.1;refGene_gn=LRRC7
1;refGene_tx=XM_023247579.1;refGene_gn=LRRC71;refGene_tx=XM_023247583.
1;refGene_gn=LRRC71;refGene_tx=XM_023247584.1;refGene_gn=LRRC71;refGen
e_tx=XM_023247585.1
F1 69147498 69147587
refGene_gn=LRRC71;refGene_tx=XM_023247570.1;refGene_gn=LRRC71;refGene_
tx=XM_023247572.1;refGene_gn=LRRC71;refGene_tx=XM_023247573.1;refGene_
gn=LRRC71;refGene_tx=XM_023247574.1;refGene_gn=LRRC71;refGene_tx=XM_02
3247575.1;refGene_gn=LRRC71;refGene_tx=XM_023247576.1;refGene_gn=LRRC7
1;refGene_tx=XM_023247577.1;refGene_gn=LRRC71;refGene_tx=XM_023247578.
1;refGene_gn=LRRC71;refGene_tx=XM_023247579.1;refGene_gn=LRRC71;refGen
e_tx=XM_023247580.1;refGene_gn=LRRC71;refGene_tx=XM_023247581.1;refGen
e_gn=LRRC71;refGene_tx=XM_023247583.1;refGene_gn=LRRC71;refGene_tx=XM_
023247584.1;refGene_gn=LRRC71;refGene_tx=XM_023247585.1
F1 69147971 69148065
refGene_gn=LRRC71;refGene_tx=XM_023247570.1;refGene_gn=LRRC71;refGene_
tx=XM_023247572.1;refGene_gn=LRRC71;refGene_tx=XM_023247573.1;refGene_
gn=LRRC71;refGene_tx=XM_023247574.1;refGene_gn=LRRC71;refGene_tx=XM_02
3247575.1;refGene_gn=LRRC71;refGene_tx=XM_023247576.1;refGene_gn=LRRC7
1;refGene_tx=XM_023247577.1;refGene_gn=LRRC71;refGene_tx=XM_023247578.
1;refGene_gn=LRRC71;refGene_tx=XM_023247579.1;refGene_gn=LRRC71;refGen
e_tx=XM_023247580.1;refGene_gn=LRRC71;refGene_tx=XM_023247581.1;refGen
e_gn=LRRC71;refGene_tx=XM_023247583.1;refGene_gn=LRRC71;refGene_tx=XM_
023247584.1;refGene_gn=LRRC71;refGene_tx=XM_023247585.1
F1 69148207 69148360
refGene_gn=LRRC71;refGene_tx=XM_023247570.1;refGene_gn=LRRC71;refGene_
tx=XM_023247572.1;refGene_gn=LRRC71;refGene_tx=XM_023247573.1;refGene_
gn=LRRC71;refGene_tx=XM_023247574.1;refGene_gn=LRRC71;refGene_tx=XM_02
3247575.1;refGene_gn=LRRC71;refGene_tx=XM_023247576.1;refGene_gn=LRRC7
1;refGene_tx=XM_023247577.1;refGene_gn=LRRC71;refGene_tx=XM_023247578.
1;refGene_gn=LRRC71;refGene_tx=XM_023247579.1;refGene_gn=LRRC71;refGen
e_tx=XM_023247580.1;refGene_gn=LRRC71;refGene_tx=XM_023247581.1;refGen
e_gn=LRRC71;refGene_tx=XM_023247583.1;refGene_gn=LRRC71;refGene_tx=XM_
023247584.1;refGene_gn=LRRC71;refGene_tx=XM_023247585.1
F1 69148990 69149080
refGene_gn=LRRC71;refGene_tx=XM_023247570.1;refGene_gn=LRRC71;refGene_
tx=XM_023247572.1;refGene_gn=LRRC71;refGene_tx=XM_023247573.1;refGene_
gn=LRRC71;refGene_tx=XM_023247574.1;refGene_gn=LRRC71;refGene_tx=XM_02
3247575.1;refGene_gn=LRRC71;refGene_tx=XM_023247576.1;refGene_gn=LRRC7
1;refGene_tx=XM_023247577.1;refGene_gn=LRRC71;refGene_tx=XM_023247578.

1;refGene_gn=LRRC71;refGene_tx=XM_023247579.1;refGene_gn=LRRC71;refGene_tx=XM_023247580.1;refGene_gn=LRRC71;refGene_tx=XM_023247581.1;refGene_gn=LRRC71;refGene_tx=XM_023247583.1;refGene_gn=LRRC71;refGene_tx=XM_023247584.1;refGene_gn=LRRC71;refGene_tx=XM_023247585.1
F1 69149151 69149235
refGene_gn=LRRC71;refGene_tx=XM_023247570.1;refGene_gn=LRRC71;refGene_tx=XM_023247572.1;refGene_gn=LRRC71;refGene_tx=XM_023247573.1;refGene_gn=LRRC71;refGene_tx=XM_023247574.1;refGene_gn=LRRC71;refGene_tx=XM_023247575.1;refGene_gn=LRRC71;refGene_tx=XM_023247576.1;refGene_gn=LRRC71;refGene_tx=XM_023247577.1;refGene_gn=LRRC71;refGene_tx=XM_023247578.1;refGene_gn=LRRC71;refGene_tx=XM_023247579.1;refGene_gn=LRRC71;refGene_tx=XM_023247580.1;refGene_gn=LRRC71;refGene_tx=XM_023247581.1;refGene_gn=LRRC71;refGene_tx=XM_023247583.1;refGene_gn=LRRC71;refGene_tx=XM_023247584.1;refGene_gn=LRRC71;refGene_tx=XM_023247585.1
F1 69149320 69149480
refGene_gn=LRRC71;refGene_tx=XM_023247570.1;refGene_gn=LRRC71;refGene_tx=XM_023247572.1;refGene_gn=LRRC71;refGene_tx=XM_023247573.1;refGene_gn=LRRC71;refGene_tx=XM_023247574.1;refGene_gn=LRRC71;refGene_tx=XM_023247575.1;refGene_gn=LRRC71;refGene_tx=XM_023247576.1;refGene_gn=LRRC71;refGene_tx=XM_023247577.1;refGene_gn=LRRC71;refGene_tx=XM_023247578.1;refGene_gn=LRRC71;refGene_tx=XM_023247579.1;refGene_gn=LRRC71;refGene_tx=XM_023247580.1;refGene_gn=LRRC71;refGene_tx=XM_023247581.1;refGene_gn=LRRC71;refGene_tx=XM_023247583.1;refGene_gn=LRRC71;refGene_tx=XM_023247584.1;refGene_gn=LRRC71;refGene_tx=XM_023247585.1
F1 69149701 69149770
refGene_gn=LRRC71;refGene_tx=XM_023247570.1;refGene_gn=LRRC71;refGene_tx=XM_023247572.1;refGene_gn=LRRC71;refGene_tx=XM_023247573.1;refGene_gn=LRRC71;refGene_tx=XM_023247574.1;refGene_gn=LRRC71;refGene_tx=XM_023247575.1;refGene_gn=LRRC71;refGene_tx=XM_023247577.1;refGene_gn=LRRC71;refGene_tx=XM_023247578.1;refGene_gn=LRRC71;refGene_tx=XM_023247579.1;refGene_gn=LRRC71;refGene_tx=XM_023247580.1;refGene_gn=LRRC71;refGene_tx=XM_023247581.1;refGene_gn=LRRC71;refGene_tx=XM_023247584.1;refGene_gn=LRRC71;refGene_tx=XM_023247585.1
F1 69151701 69151779
refGene_gn=LRRC71;refGene_tx=XM_023247570.1;refGene_gn=LRRC71;refGene_tx=XM_023247572.1;refGene_gn=LRRC71;refGene_tx=XM_023247573.1;refGene_gn=LRRC71;refGene_tx=XM_023247574.1;refGene_gn=LRRC71;refGene_tx=XM_023247575.1;refGene_gn=LRRC71;refGene_tx=XM_023247576.1;refGene_gn=LRRC71;refGene_tx=XM_023247578.1;refGene_gn=LRRC71;refGene_tx=XM_023247579.1;refGene_gn=LRRC71;refGene_tx=XM_023247580.1;refGene_gn=LRRC71;refGene_tx=XM_023247581.1;refGene_gn=LRRC71;refGene_tx=XM_023247584.1;refGene_gn=LRRC71;refGene_tx=XM_023247585.1
F1 69151942 69152018
refGene_gn=LRRC71;refGene_tx=XM_023247570.1;refGene_gn=LRRC71;refGene_tx=XM_023247572.1;refGene_gn=LRRC71;refGene_tx=XM_023247573.1;refGene_gn=LRRC71;refGene_tx=XM_023247574.1;refGene_gn=LRRC71;refGene_tx=XM_023247575.1;refGene_gn=LRRC71;refGene_tx=XM_023247576.1;refGene_gn=LRRC71;refGene_tx=XM_023247577.1;refGene_gn=LRRC71;refGene_tx=XM_023247578.1;refGene_gn=LRRC71;refGene_tx=XM_023247579.1;refGene_gn=LRRC71;refGene_tx=XM_023247580.1;refGene_gn=LRRC71;refGene_tx=XM_023247581.1;refGen

e_gn=LRRC71;refGene_tx=XM_023247583.1;refGene_gn=LRRC71;refGene_tx=XM_023247584.1;refGene_gn=LRRC71;refGene_tx=XM_023247585.1
F1 69152353 69152479
refGene_gn=LRRC71;refGene_tx=XM_023247570.1;refGene_gn=LRRC71;refGene_tx=XM_023247572.1;refGene_gn=LRRC71;refGene_tx=XM_023247573.1;refGene_gn=LRRC71;refGene_tx=XM_023247574.1;refGene_gn=LRRC71;refGene_tx=XM_023247575.1;refGene_gn=LRRC71;refGene_tx=XM_023247576.1;refGene_gn=LRRC71;refGene_tx=XM_023247577.1;refGene_gn=LRRC71;refGene_tx=XM_023247578.1;refGene_gn=LRRC71;refGene_tx=XM_023247579.1;refGene_gn=LRRC71;refGene_tx=XM_023247580.1;refGene_gn=LRRC71;refGene_tx=XM_023247581.1;refGene_gn=LRRC71;refGene_tx=XM_023247583.1;refGene_gn=LRRC71;refGene_tx=XM_023247584.1;refGene_gn=LRRC71;refGene_tx=XM_023247585.1
F1 69152729 69152879
refGene_gn=LRRC71;refGene_tx=XM_023247570.1;refGene_gn=LRRC71;refGene_tx=XM_023247572.1;refGene_gn=LRRC71;refGene_tx=XM_023247573.1;refGene_gn=LRRC71;refGene_tx=XM_023247574.1;refGene_gn=LRRC71;refGene_tx=XM_023247575.1;refGene_gn=LRRC71;refGene_tx=XM_023247576.1;refGene_gn=LRRC71;refGene_tx=XM_023247577.1;refGene_gn=LRRC71;refGene_tx=XM_023247578.1;refGene_gn=LRRC71;refGene_tx=XM_023247579.1;refGene_gn=LRRC71;refGene_tx=XM_023247580.1;refGene_gn=LRRC71;refGene_tx=XM_023247581.1;refGene_gn=LRRC71;refGene_tx=XM_023247583.1;refGene_gn=LRRC71;refGene_tx=XM_023247584.1;refGene_gn=LRRC71;refGene_tx=XM_023247585.1
F1 69156165 69156325
refGene_gn=LRRC71;refGene_tx=XM_023247570.1;refGene_gn=LRRC71;refGene_tx=XM_023247572.1;refGene_gn=LRRC71;refGene_tx=XM_023247573.1;refGene_gn=LRRC71;refGene_tx=XM_023247574.1;refGene_gn=LRRC71;refGene_tx=XM_023247575.1;refGene_gn=LRRC71;refGene_tx=XM_023247576.1;refGene_gn=LRRC71;refGene_tx=XM_023247577.1;refGene_gn=LRRC71;refGene_tx=XM_023247578.1;refGene_gn=LRRC71;refGene_tx=XM_023247579.1;refGene_gn=LRRC71;refGene_tx=XM_023247580.1;refGene_gn=LRRC71;refGene_tx=XM_023247581.1;refGene_gn=LRRC71;refGene_tx=XM_023247583.1;refGene_gn=LRRC71;refGene_tx=XM_023247585.1
F1 69158361 69158513
refGene_gn=PEAR1;refGene_tx=XM_023247315.1;refGene_gn=PEAR1;refGene_tx=XM_023247316.1;refGene_gn=PEAR1;refGene_tx=XM_023247317.1;refGene_gn=PEAR1;refGene_tx=XM_023247318.1;refGene_gn=PEAR1;refGene_tx=XM_023247319.1;refGene_gn=PEAR1;refGene_tx=XM_023247320.1
F1 69158977 69159211
refGene_gn=PEAR1;refGene_tx=XM_023247315.1;refGene_gn=PEAR1;refGene_tx=XM_023247316.1;refGene_gn=PEAR1;refGene_tx=XM_023247317.1;refGene_gn=PEAR1;refGene_tx=XM_023247318.1;refGene_gn=PEAR1;refGene_tx=XM_023247319.1;refGene_gn=PEAR1;refGene_tx=XM_023247320.1
F1 69159314 69159383
refGene_gn=PEAR1;refGene_tx=XM_023247315.1;refGene_gn=PEAR1;refGene_tx=XM_023247316.1;refGene_gn=PEAR1;refGene_tx=XM_023247317.1;refGene_gn=PEAR1;refGene_tx=XM_023247318.1;refGene_gn=PEAR1;refGene_tx=XM_023247319.1;refGene_gn=PEAR1;refGene_tx=XM_023247320.1
F1 69159569 69159705
refGene_gn=PEAR1;refGene_tx=XM_023247315.1;refGene_gn=PEAR1;refGene_tx=XM_023247316.1;refGene_gn=PEAR1;refGene_tx=XM_023247317.1;refGene_gn=

PEAR1;refGene_tx=XM_023247318.1;refGene_gn=PEAR1;refGene_tx=XM_023247319.1;refGene_gn=PEAR1;refGene_tx=XM_023247320.1
F1 69159817 69159906
refGene_gn=PEAR1;refGene_tx=XM_023247315.1;refGene_gn=PEAR1;refGene_tx=XM_023247316.1;refGene_gn=PEAR1;refGene_tx=XM_023247317.1;refGene_gn=PEAR1;refGene_tx=XM_023247318.1;refGene_gn=PEAR1;refGene_tx=XM_023247319.1;refGene_gn=PEAR1;refGene_tx=XM_023247320.1
F1 69160126 69160339
refGene_gn=PEAR1;refGene_tx=XM_023247315.1;refGene_gn=PEAR1;refGene_tx=XM_023247316.1;refGene_gn=PEAR1;refGene_tx=XM_023247317.1;refGene_gn=PEAR1;refGene_tx=XM_023247318.1;refGene_gn=PEAR1;refGene_tx=XM_023247319.1;refGene_gn=PEAR1;refGene_tx=XM_023247320.1
F1 69160491 69160620
refGene_gn=PEAR1;refGene_tx=XM_023247315.1;refGene_gn=PEAR1;refGene_tx=XM_023247316.1;refGene_gn=PEAR1;refGene_tx=XM_023247317.1;refGene_gn=PEAR1;refGene_tx=XM_023247318.1;refGene_gn=PEAR1;refGene_tx=XM_023247319.1;refGene_gn=PEAR1;refGene_tx=XM_023247320.1
F1 69160719 69160848
refGene_gn=PEAR1;refGene_tx=XM_023247315.1;refGene_gn=PEAR1;refGene_tx=XM_023247316.1;refGene_gn=PEAR1;refGene_tx=XM_023247317.1;refGene_gn=PEAR1;refGene_tx=XM_023247318.1;refGene_gn=PEAR1;refGene_tx=XM_023247319.1;refGene_gn=PEAR1;refGene_tx=XM_023247320.1
F1 69161723 69161849
refGene_gn=PEAR1;refGene_tx=XM_023247315.1;refGene_gn=PEAR1;refGene_tx=XM_023247316.1;refGene_gn=PEAR1;refGene_tx=XM_023247317.1;refGene_gn=PEAR1;refGene_tx=XM_023247318.1;refGene_gn=PEAR1;refGene_tx=XM_023247319.1;refGene_gn=PEAR1;refGene_tx=XM_023247320.1
F1 69162054 69162201
refGene_gn=PEAR1;refGene_tx=XM_023247315.1;refGene_gn=PEAR1;refGene_tx=XM_023247316.1;refGene_gn=PEAR1;refGene_tx=XM_023247317.1;refGene_gn=PEAR1;refGene_tx=XM_023247318.1;refGene_gn=PEAR1;refGene_tx=XM_023247319.1;refGene_gn=PEAR1;refGene_tx=XM_023247320.1
F1 69162334 69162437
refGene_gn=PEAR1;refGene_tx=XM_023247315.1;refGene_gn=PEAR1;refGene_tx=XM_023247316.1;refGene_gn=PEAR1;refGene_tx=XM_023247317.1;refGene_gn=PEAR1;refGene_tx=XM_023247318.1;refGene_gn=PEAR1;refGene_tx=XM_023247319.1;refGene_gn=PEAR1;refGene_tx=XM_023247320.1
F1 69162533 69162697
refGene_gn=PEAR1;refGene_tx=XM_023247315.1;refGene_gn=PEAR1;refGene_tx=XM_023247316.1;refGene_gn=PEAR1;refGene_tx=XM_023247317.1;refGene_gn=PEAR1;refGene_tx=XM_023247318.1;refGene_gn=PEAR1;refGene_tx=XM_023247319.1;refGene_gn=PEAR1;refGene_tx=XM_023247320.1
F1 69163431 69163552
refGene_gn=PEAR1;refGene_tx=XM_023247315.1;refGene_gn=PEAR1;refGene_tx=XM_023247316.1;refGene_gn=PEAR1;refGene_tx=XM_023247317.1;refGene_gn=PEAR1;refGene_tx=XM_023247318.1;refGene_gn=PEAR1;refGene_tx=XM_023247319.1;refGene_gn=PEAR1;refGene_tx=XM_023247320.1
F1 69163633 69163808
refGene_gn=PEAR1;refGene_tx=XM_023247315.1;refGene_gn=PEAR1;refGene_tx=XM_023247316.1;refGene_gn=PEAR1;refGene_tx=XM_023247317.1;refGene_gn=

PEAR1;refGene_tx=XM_023247318.1;refGene_gn=PEAR1;refGene_tx=XM_023247319.1;refGene_gn=PEAR1;refGene_tx=XM_023247320.1
F1 69164044 69164257 refGene_gn=PEAR1;refGene_tx=XM_023247315.1;refGene_gn=PEAR1;refGene_tx=XM_023247316.1;refGene_gn=PEAR1;refGene_tx=XM_023247317.1;refGene_gn=PEAR1;refGene_tx=XM_023247318.1;refGene_gn=PEAR1;refGene_tx=XM_023247319.1;refGene_gn=PEAR1;refGene_tx=XM_023247320.1
F1 69164331 69164468 refGene_gn=PEAR1;refGene_tx=XM_023247315.1;refGene_gn=PEAR1;refGene_tx=XM_023247316.1;refGene_gn=PEAR1;refGene_tx=XM_023247317.1;refGene_gn=PEAR1;refGene_tx=XM_023247318.1;refGene_gn=PEAR1;refGene_tx=XM_023247319.1;refGene_gn=PEAR1;refGene_tx=XM_023247320.1
F1 69164647 69164771 refGene_gn=PEAR1;refGene_tx=XM_023247315.1;refGene_gn=PEAR1;refGene_tx=XM_023247316.1;refGene_gn=PEAR1;refGene_tx=XM_023247317.1;refGene_gn=PEAR1;refGene_tx=XM_023247318.1;refGene_gn=PEAR1;refGene_tx=XM_023247319.1;refGene_gn=PEAR1;refGene_tx=XM_023247320.1
F1 69165416 69165660 refGene_gn=PEAR1;refGene_tx=XM_023247315.1;refGene_gn=PEAR1;refGene_tx=XM_023247316.1;refGene_gn=PEAR1;refGene_tx=XM_023247317.1;refGene_gn=PEAR1;refGene_tx=XM_023247318.1;refGene_gn=PEAR1;refGene_tx=XM_023247319.1;refGene_gn=PEAR1;refGene_tx=XM_023247320.1
F1 69165897 69165990 refGene_gn=PEAR1;refGene_tx=XM_023247315.1;refGene_gn=PEAR1;refGene_tx=XM_023247316.1;refGene_gn=PEAR1;refGene_tx=XM_023247317.1;refGene_gn=PEAR1;refGene_tx=XM_023247318.1;refGene_gn=PEAR1;refGene_tx=XM_023247319.1;refGene_gn=PEAR1;refGene_tx=XM_023247320.1
F1 69166598 69166699 refGene_gn=PEAR1;refGene_tx=XM_023247315.1;refGene_gn=PEAR1;refGene_tx=XM_023247316.1;refGene_gn=PEAR1;refGene_tx=XM_023247317.1;refGene_gn=PEAR1;refGene_tx=XM_023247318.1;refGene_gn=PEAR1;refGene_tx=XM_023247319.1;refGene_gn=PEAR1;refGene_tx=XM_023247320.1
F1 69166944 69167049 refGene_gn=PEAR1;refGene_tx=XM_023247315.1;refGene_gn=PEAR1;refGene_tx=XM_023247316.1;refGene_gn=PEAR1;refGene_tx=XM_023247317.1;refGene_gn=PEAR1;refGene_tx=XM_023247318.1;refGene_gn=PEAR1;refGene_tx=XM_023247319.1;refGene_gn=PEAR1;refGene_tx=XM_023247320.1
F1 69167230 69167331 refGene_gn=PEAR1;refGene_tx=XM_023247315.1;refGene_gn=PEAR1;refGene_tx=XM_023247316.1;refGene_gn=PEAR1;refGene_tx=XM_023247317.1;refGene_gn=PEAR1;refGene_tx=XM_023247318.1;refGene_gn=PEAR1;refGene_tx=XM_023247319.1;refGene_gn=PEAR1;refGene_tx=XM_023247320.1
F1 69186515 69186701 refGene_gn=NTRK1;refGene_tx=XM_023247543.1
F1 69187876 69188035 refGene_gn=NTRK1;refGene_tx=XM_023247543.1
F1 69188311 69188552 refGene_gn=NTRK1;refGene_tx=XM_023247543.1
F1 69191733 69191906 refGene_gn=NTRK1;refGene_tx=XM_023247543.1
F1 69192026 69192157 refGene_gn=NTRK1;refGene_tx=XM_023247543.1
F1 69192457 69192604 refGene_gn=NTRK1;refGene_tx=XM_023247543.1
F1 69193115 69193218 refGene_gn=NTRK1;refGene_tx=XM_023247543.1
F1 69193498 69193554 refGene_gn=NTRK1;refGene_tx=XM_023247543.1

F1 69193714 69193732 refGene_gn=NTRK1;refGene_tx=XM_023247543.1
F1 69194208 69194535 refGene_gn=NTRK1;refGene_tx=XM_023247543.1
F1 69196124 69196257 refGene_gn=NTRK1;refGene_tx=XM_023247543.1
F1 69198506 69198649 refGene_gn=NTRK1;refGene_tx=XM_023247543.1
F1 69198857 69199003 refGene_gn=NTRK1;refGene_tx=XM_023247543.1
F1 69199630 69199699 refGene_gn=NTRK1;refGene_tx=XM_023247543.1
F1 69201290 69201362 refGene_gn=NTRK1;refGene_tx=XM_023247543.1
F1 69201594 69201669 refGene_gn=NTRK1;refGene_tx=XM_023247543.1
F1 69205000 69205209 refGene_gn=NTRK1;refGene_tx=XM_023247543.1
F1 69207367 69207452 refGene_gn=INSRR;refGene_tx=XM_023247540.1
F1 69210365 69211062
refGene_gn=INSRR;refGene_tx=XM_023247539.1;refGene_gn=INSRR;refGene_tx=XM_023247540.1;refGene_gn=INSRR;refGene_tx=XM_023247541.1;refGene_gn=INSRR;refGene_tx=XM_023247542.1
F1 69212309 69212613
refGene_gn=INSRR;refGene_tx=XM_023247539.1;refGene_gn=INSRR;refGene_tx=XM_023247540.1;refGene_gn=INSRR;refGene_tx=XM_023247541.1;refGene_gn=INSRR;refGene_tx=XM_023247542.1
F1 69212708 69212851
refGene_gn=INSRR;refGene_tx=XM_023247539.1;refGene_gn=INSRR;refGene_tx=XM_023247540.1;refGene_gn=INSRR;refGene_tx=XM_023247541.1;refGene_gn=INSRR;refGene_tx=XM_023247542.1
F1 69213056 69213201
refGene_gn=INSRR;refGene_tx=XM_023247539.1;refGene_gn=INSRR;refGene_tx=XM_023247540.1;refGene_gn=INSRR;refGene_tx=XM_023247541.1;refGene_gn=INSRR;refGene_tx=XM_023247542.1
F1 69213867 69214082
refGene_gn=INSRR;refGene_tx=XM_023247539.1;refGene_gn=INSRR;refGene_tx=XM_023247540.1;refGene_gn=INSRR;refGene_tx=XM_023247541.1;refGene_gn=INSRR;refGene_tx=XM_023247542.1
F1 69214330 69214457
refGene_gn=INSRR;refGene_tx=XM_023247539.1;refGene_gn=INSRR;refGene_tx=XM_023247540.1;refGene_gn=INSRR;refGene_tx=XM_023247541.1;refGene_gn=INSRR;refGene_tx=XM_023247542.1
F1 69215834 69216073
refGene_gn=INSRR;refGene_tx=XM_023247539.1;refGene_gn=INSRR;refGene_tx=XM_023247540.1;refGene_gn=INSRR;refGene_tx=XM_023247541.1;refGene_gn=INSRR;refGene_tx=XM_023247542.1
F1 69216434 69216602
refGene_gn=INSRR;refGene_tx=XM_023247539.1;refGene_gn=INSRR;refGene_tx=XM_023247540.1;refGene_gn=INSRR;refGene_tx=XM_023247541.1;refGene_gn=INSRR;refGene_tx=XM_023247542.1
F1 69216689 69216885
refGene_gn=INSRR;refGene_tx=XM_023247539.1;refGene_gn=INSRR;refGene_tx=XM_023247540.1;refGene_gn=INSRR;refGene_tx=XM_023247541.1;refGene_gn=INSRR;refGene_tx=XM_023247542.1
F1 69217082 69217124
refGene_gn=INSRR;refGene_tx=XM_023247539.1;refGene_gn=INSRR;refGene_tx=XM_023247540.1;refGene_gn=INSRR;refGene_tx=XM_023247541.1;refGene_gn=INSRR;refGene_tx=XM_023247542.1

F1 69217194 69217415
refGene_gn=INSRR;refGene_tx=XM_023247539.1;refGene_gn=INSRR;refGene_tx=XM_023247540.1;refGene_gn=INSRR;refGene_tx=XM_023247541.1;refGene_gn=INSRR;refGene_tx=XM_023247542.1
F1 69217658 69217795
refGene_gn=INSRR;refGene_tx=XM_023247539.1;refGene_gn=INSRR;refGene_tx=XM_023247540.1;refGene_gn=INSRR;refGene_tx=XM_023247541.1;refGene_gn=INSRR;refGene_tx=XM_023247542.1
F1 69217878 69218041
refGene_gn=INSRR;refGene_tx=XM_023247539.1;refGene_gn=INSRR;refGene_tx=XM_023247540.1;refGene_gn=INSRR;refGene_tx=XM_023247541.1;refGene_gn=INSRR;refGene_tx=XM_023247542.1
F1 69218240 69218346
refGene_gn=INSRR;refGene_tx=XM_023247539.1;refGene_gn=INSRR;refGene_tx=XM_023247540.1;refGene_gn=INSRR;refGene_tx=XM_023247541.1;refGene_gn=INSRR;refGene_tx=XM_023247542.1
F1 69218958 69219011
refGene_gn=INSRR;refGene_tx=XM_023247539.1;refGene_gn=INSRR;refGene_tx=XM_023247540.1;refGene_gn=INSRR;refGene_tx=XM_023247541.1;refGene_gn=INSRR;refGene_tx=XM_023247542.1
F1 69219195 69219425
refGene_gn=INSRR;refGene_tx=XM_023247539.1;refGene_gn=INSRR;refGene_tx=XM_023247540.1;refGene_gn=INSRR;refGene_tx=XM_023247541.1;refGene_gn=INSRR;refGene_tx=XM_023247542.1
F1 69219915 69220026
refGene_gn=INSRR;refGene_tx=XM_023247539.1;refGene_gn=INSRR;refGene_tx=XM_023247540.1;refGene_gn=INSRR;refGene_tx=XM_023247542.1
F1 69220158 69220318
refGene_gn=INSRR;refGene_tx=XM_023247539.1;refGene_gn=INSRR;refGene_tx=XM_023247540.1;refGene_gn=INSRR;refGene_tx=XM_023247541.1;refGene_gn=INSRR;refGene_tx=XM_023247542.1
F1 69220605 69220735
refGene_gn=INSRR;refGene_tx=XM_023247539.1;refGene_gn=INSRR;refGene_tx=XM_023247540.1;refGene_gn=INSRR;refGene_tx=XM_023247541.1;refGene_gn=INSRR;refGene_tx=XM_023247542.1
F1 69220852 69220987
refGene_gn=INSRR;refGene_tx=XM_023247539.1;refGene_gn=INSRR;refGene_tx=XM_023247540.1;refGene_gn=INSRR;refGene_tx=XM_023247541.1;refGene_gn=INSRR;refGene_tx=XM_023247542.1
F1 69221198 69221439
refGene_gn=INSRR;refGene_tx=XM_023247539.1;refGene_gn=INSRR;refGene_tx=XM_023247540.1;refGene_gn=INSRR;refGene_tx=XM_023247541.1;refGene_gn=INSRR;refGene_tx=XM_023247542.1
F1 69232438 69232472
refGene_gn=SH2D2A;refGene_tx=XM_023247205.1;refGene_gn=SH2D2A;refGene_tx=XM_023247206.1;refGene_gn=SH2D2A;refGene_tx=XM_023247207.1;refGene_gn=SH2D2A;refGene_tx=XM_023247209.1
F1 69232978 69233139
refGene_gn=SH2D2A;refGene_tx=XM_023247205.1;refGene_gn=SH2D2A;refGene_tx=XM_023247206.1;refGene_gn=SH2D2A;refGene_tx=XM_023247207.1;refGene_

gn=SH2D2A;refGene_tx=XM_023247209.1
F1 69233850 69234032
refGene_gn=SH2D2A;refGene_tx=XM_023247205.1;refGene_gn=SH2D2A;refGene_tx=XM_023247206.1;refGene_gn=SH2D2A;refGene_tx=XM_023247207.1;refGene_gn=SH2D2A;refGene_tx=XM_023247209.1
F1 69234920 69235010
refGene_gn=SH2D2A;refGene_tx=XM_023247205.1;refGene_gn=SH2D2A;refGene_tx=XM_023247206.1;refGene_gn=SH2D2A;refGene_tx=XM_023247207.1;refGene_gn=SH2D2A;refGene_tx=XM_023247209.1
F1 69235150 69235319
refGene_gn=SH2D2A;refGene_tx=XM_023247205.1;refGene_gn=SH2D2A;refGene_tx=XM_023247206.1;refGene_gn=SH2D2A;refGene_tx=XM_023247207.1;refGene_gn=SH2D2A;refGene_tx=XM_023247209.1
F1 69238379 69238523
refGene_gn=SH2D2A;refGene_tx=XM_023247205.1;refGene_gn=SH2D2A;refGene_tx=XM_023247206.1;refGene_gn=SH2D2A;refGene_tx=XM_023247207.1;refGene_gn=SH2D2A;refGene_tx=XM_023247209.1
F1 69238733 69239039
refGene_gn=SH2D2A;refGene_tx=XM_023247205.1;refGene_gn=SH2D2A;refGene_tx=XM_023247206.1;refGene_gn=SH2D2A;refGene_tx=XM_023247207.1;refGene_gn=SH2D2A;refGene_tx=XM_023247209.1
F1 69240565 69240733
refGene_gn=SH2D2A;refGene_tx=XM_023247205.1;refGene_gn=SH2D2A;refGene_tx=XM_023247206.1;refGene_gn=SH2D2A;refGene_tx=XM_023247207.1;refGene_gn=SH2D2A;refGene_tx=XM_023247209.1
F1 69246843 69246930 refGene_gn=PRCC;refGene_tx=XM_019822399.2
F1 69248992 69249058 refGene_gn=PRCC;refGene_tx=XM_019822399.2
F1 69251036 69251180 refGene_gn=PRCC;refGene_tx=XM_019822399.2
F1 69253497 69253593 refGene_gn=PRCC;refGene_tx=XM_019822399.2
F1 69255710 69256277 refGene_gn=PRCC;refGene_tx=XM_019822399.2
F1 69258439 69258487 refGene_gn=PRCC;refGene_tx=XM_019822399.2
F1 69271449 69271917 refGene_gn=PRCC;refGene_tx=XM_019822399.2
F1 69283112 69283199 refGene_gn=HDGF;refGene_tx=XM_023247932.1
F1 69289070 69289147
refGene_gn=HDGF;refGene_tx=XM_023247932.1;refGene_gn=HDGF;refGene_tx=XM_023247933.1;refGene_gn=HDGF;refGene_tx=XM_023247934.1;refGene_gn=HDGF;refGene_tx=XM_023247935.1
F1 69289274 69289413
refGene_gn=HDGF;refGene_tx=XM_023247932.1;refGene_gn=HDGF;refGene_tx=XM_023247933.1;refGene_gn=HDGF;refGene_tx=XM_023247934.1;refGene_gn=HDGF;refGene_tx=XM_023247935.1
F1 69290077 69290263
refGene_gn=HDGF;refGene_tx=XM_023247932.1;refGene_gn=HDGF;refGene_tx=XM_023247933.1;refGene_gn=HDGF;refGene_tx=XM_023247934.1;refGene_gn=HDGF;refGene_tx=XM_023247935.1
F1 69290520 69290744
refGene_gn=HDGF;refGene_tx=XM_023247932.1;refGene_gn=HDGF;refGene_tx=XM_023247933.1;refGene_gn=HDGF;refGene_tx=XM_023247934.1;refGene_gn=HDGF;refGene_tx=XM_023247935.1
F1 69290926 69290933

refGene_gn=HDGF;refGene_tx=XM_023247932.1;refGene_gn=HDGF;refGene_tx=XM_023247933.1;refGene_gn=HDGF;refGene_tx=XM_023247934.1;refGene_gn=HDGF;refGene_tx=XM_023247935.1
F1	69295604	69295787	refGene_gn=MRPL24;refGene_tx=XM_023247937.1;refGene_gn=MRPL24;refGene_tx=XM_023247938.1;refGene_gn=MRPL24;refGene_tx=XM_023247939.1
F1	69295863	69295959	refGene_gn=MRPL24;refGene_tx=XM_023247937.1;refGene_gn=MRPL24;refGene_tx=XM_023247938.1;refGene_gn=MRPL24;refGene_tx=XM_023247939.1
F1	69296164	69296268	refGene_gn=MRPL24;refGene_tx=XM_023247937.1;refGene_gn=MRPL24;refGene_tx=XM_023247938.1;refGene_gn=MRPL24;refGene_tx=XM_023247939.1
F1	69296539	69296670	refGene_gn=MRPL24;refGene_tx=XM_023247937.1;refGene_gn=MRPL24;refGene_tx=XM_023247938.1;refGene_gn=MRPL24;refGene_tx=XM_023247939.1
F1	69296767	69296904	refGene_gn=MRPL24;refGene_tx=XM_023247937.1;refGene_gn=MRPL24;refGene_tx=XM_023247938.1;refGene_gn=MRPL24;refGene_tx=XM_023247939.1
F1	69297351	69297473	refGene_gn=RRNAD1;refGene_tx=XM_003999618.5;refGene_gn=RRNAD1;refGene_tx=XM_006943068.4;refGene_gn=RRNAD1;refGene_tx=XM_023247929.1
F1	69297846	69298031	refGene_gn=RRNAD1;refGene_tx=XM_003999618.5;refGene_gn=RRNAD1;refGene_tx=XM_006943068.4;refGene_gn=RRNAD1;refGene_tx=XM_023247929.1
F1	69298604	69299086	refGene_gn=RRNAD1;refGene_tx=XM_003999618.5;refGene_gn=RRNAD1;refGene_tx=XM_006943068.4;refGene_gn=RRNAD1;refGene_tx=XM_023247929.1
F1	69299535	69299679	refGene_gn=RRNAD1;refGene_tx=XM_003999618.5;refGene_gn=RRNAD1;refGene_tx=XM_006943068.4;refGene_gn=RRNAD1;refGene_tx=XM_023247929.1
F1	69299934	69299997	refGene_gn=RRNAD1;refGene_tx=XM_003999618.5;refGene_gn=RRNAD1;refGene_tx=XM_006943068.4;refGene_gn=RRNAD1;refGene_tx=XM_023247929.1
F1	69300431	69300624	refGene_gn=RRNAD1;refGene_tx=XM_003999618.5;refGene_gn=RRNAD1;refGene_tx=XM_006943068.4;refGene_gn=RRNAD1;refGene_tx=XM_023247929.1
F1	69300720	69300845	refGene_gn=RRNAD1;refGene_tx=XM_003999618.5;refGene_gn=RRNAD1;refGene_tx=XM_006943068.4;refGene_gn=RRNAD1;refGene_tx=XM_023247929.1
F1	69304001	69304112	refGene_gn=RRNAD1;refGene_tx=XM_003999618.5;refGene_gn=RRNAD1;refGene_tx=XM_023247929.1
F1	69305309	69306041	refGene_gn=ISG20L2;refGene_tx=XM_023247930.1;refGene_gn=ISG20L2;refGene_tx=XM_023247931.1
F1	69307833	69308034	refGene_gn=ISG20L2;refGene_tx=XM_023247930.1;refGene_gn=ISG20L2;refGene_tx=XM_023247931.1
F1	69308732	69308846	refGene_gn=ISG20L2;refGene_tx=XM_023247930.1;refGene_gn=ISG20L2;refGen

e_tx=XM_023247931.1
F1 69320330 69320454
refGene_gn=CRABP2;refGene_tx=XM_011291195.3;refGene_gn=CRABP2;refGene_tx=XM_023247546.1;refGene_gn=CRABP2;refGene_tx=XM_023247547.1;refGene_gn=CRABP2;refGene_tx=XM_023247548.1
F1 69321786 69321841
refGene_gn=CRABP2;refGene_tx=XM_011291195.3;refGene_gn=CRABP2;refGene_tx=XM_023247546.1;refGene_gn=CRABP2;refGene_tx=XM_023247547.1;refGene_gn=CRABP2;refGene_tx=XM_023247548.1
F1 69322036 69322192
refGene_gn=CRABP2;refGene_tx=XM_011291195.3;refGene_gn=CRABP2;refGene_tx=XM_023247546.1;refGene_gn=CRABP2;refGene_tx=XM_023247547.1
F1 69322507 69322738
refGene_gn=CRABP2;refGene_tx=XM_011291195.3;refGene_gn=CRABP2;refGene_tx=XM_019822401.2;refGene_gn=CRABP2;refGene_tx=XM_023247546.1;refGene_gn=CRABP2;refGene_tx=XM_023247547.1;refGene_gn=CRABP2;refGene_tx=XM_023247548.1
F1 69324810 69324889 refGene_gn=CRABP2;refGene_tx=XM_011291195.3
F1 69325100 69325131 refGene_gn=CRABP2;refGene_tx=XM_006943072.4
F1 69325341 69325520
refGene_gn=CRABP2;refGene_tx=XM_006943072.4;refGene_gn=CRABP2;refGene_tx=XM_019822401.2;refGene_gn=CRABP2;refGene_tx=XM_023247546.1;refGene_gn=CRABP2;refGene_tx=XM_023247547.1;refGene_gn=CRABP2;refGene_tx=XM_023247548.1
F1 69325665 69325782
refGene_gn=CRABP2;refGene_tx=XM_006943072.4;refGene_gn=CRABP2;refGene_tx=XM_019822401.2
F1 69326190 69326241
refGene_gn=CRABP2;refGene_tx=XM_006943072.4;refGene_gn=CRABP2;refGene_tx=XM_019822401.2;refGene_gn=CRABP2;refGene_tx=XM_023247546.1;refGene_gn=CRABP2;refGene_tx=XM_023247547.1;refGene_gn=CRABP2;refGene_tx=XM_023247548.1
F1 69341991 69342771 refGene_gn=NES;refGene_tx=XM_023247371.1
F1 69343746 69343871 refGene_gn=NES;refGene_tx=XM_023247371.1
F1 69345597 69345671 refGene_gn=NES;refGene_tx=XM_023247371.1
F1 69345909 69347248 refGene_gn=NES;refGene_tx=XM_023247371.1
F1 69348073 69350699 refGene_gn=NES;refGene_tx=XM_023247371.1
F1 69358796 69358901
refGene_gn=BCAN;refGene_tx=XM_023247530.1;refGene_gn=BCAN;refGene_tx=XM_023247531.1;refGene_gn=BCAN;refGene_tx=XM_023247532.1;refGene_gn=BCAN;refGene_tx=XM_023247533.1;refGene_gn=BCAN;refGene_tx=XM_023247534.1;refGene_gn=BCAN;refGene_tx=XM_023247535.1;refGene_gn=BCAN;refGene_tx=XM_023247537.1;refGene_gn=BCAN;refGene_tx=XM_023247538.1
F1 69359204 69359395
refGene_gn=BCAN;refGene_tx=XM_023247530.1;refGene_gn=BCAN;refGene_tx=XM_023247531.1;refGene_gn=BCAN;refGene_tx=XM_023247532.1;refGene_gn=BCAN;refGene_tx=XM_023247533.1;refGene_gn=BCAN;refGene_tx=XM_023247534.1;refGene_gn=BCAN;refGene_tx=XM_023247535.1;refGene_gn=BCAN;refGene_tx=XM_023247537.1;refGene_gn=BCAN;refGene_tx=XM_023247538.1
F1 69359860 69360005

refGene_gn=BCAN;refGene_tx=XM_023247530.1;refGene_gn=BCAN;refGene_tx=XM_023247531.1;refGene_gn=BCAN;refGene_tx=XM_023247532.1;refGene_gn=BCAN;refGene_tx=XM_023247533.1;refGene_gn=BCAN;refGene_tx=XM_023247534.1;refGene_gn=BCAN;refGene_tx=XM_023247535.1;refGene_gn=BCAN;refGene_tx=XM_023247537.1;refGene_gn=BCAN;refGene_tx=XM_023247538.1
F1 69360390 69360473
refGene_gn=BCAN;refGene_tx=XM_023247530.1;refGene_gn=BCAN;refGene_tx=XM_023247531.1;refGene_gn=BCAN;refGene_tx=XM_023247532.1;refGene_gn=BCAN;refGene_tx=XM_023247533.1;refGene_gn=BCAN;refGene_tx=XM_023247534.1;refGene_gn=BCAN;refGene_tx=XM_023247535.1;refGene_gn=BCAN;refGene_tx=XM_023247537.1;refGene_gn=BCAN;refGene_tx=XM_023247538.1
F1 69361032 69361191
refGene_gn=BCAN;refGene_tx=XM_023247530.1;refGene_gn=BCAN;refGene_tx=XM_023247531.1;refGene_gn=BCAN;refGene_tx=XM_023247532.1;refGene_gn=BCAN;refGene_tx=XM_023247533.1;refGene_gn=BCAN;refGene_tx=XM_023247534.1;refGene_gn=BCAN;refGene_tx=XM_023247535.1;refGene_gn=BCAN;refGene_tx=XM_023247537.1;refGene_gn=BCAN;refGene_tx=XM_023247538.1
F1 69361678 69361786
refGene_gn=BCAN;refGene_tx=XM_023247530.1;refGene_gn=BCAN;refGene_tx=XM_023247531.1;refGene_gn=BCAN;refGene_tx=XM_023247532.1;refGene_gn=BCAN;refGene_tx=XM_023247533.1;refGene_gn=BCAN;refGene_tx=XM_023247534.1;refGene_gn=BCAN;refGene_tx=XM_023247535.1;refGene_gn=BCAN;refGene_tx=XM_023247537.1;refGene_gn=BCAN;refGene_tx=XM_023247538.1
F1 69364925 69365549
refGene_gn=BCAN;refGene_tx=XM_023247530.1;refGene_gn=BCAN;refGene_tx=XM_023247531.1;refGene_gn=BCAN;refGene_tx=XM_023247532.1;refGene_gn=BCAN;refGene_tx=XM_023247533.1;refGene_gn=BCAN;refGene_tx=XM_023247534.1;refGene_gn=BCAN;refGene_tx=XM_023247535.1;refGene_gn=BCAN;refGene_tx=XM_023247537.1;refGene_gn=BCAN;refGene_tx=XM_023247538.1
F1 69366030 69366252
refGene_gn=BCAN;refGene_tx=XM_023247530.1;refGene_gn=BCAN;refGene_tx=XM_023247531.1;refGene_gn=BCAN;refGene_tx=XM_023247532.1;refGene_gn=BCAN;refGene_tx=XM_023247533.1;refGene_gn=BCAN;refGene_tx=XM_023247534.1;refGene_gn=BCAN;refGene_tx=XM_023247535.1;refGene_gn=BCAN;refGene_tx=XM_023247537.1;refGene_gn=BCAN;refGene_tx=XM_023247538.1
F1 69368162 69368456
refGene_gn=BCAN;refGene_tx=XM_023247530.1;refGene_gn=BCAN;refGene_tx=XM_023247531.1;refGene_gn=BCAN;refGene_tx=XM_023247532.1;refGene_gn=BCAN;refGene_tx=XM_023247533.1;refGene_gn=BCAN;refGene_tx=XM_023247534.1;refGene_gn=BCAN;refGene_tx=XM_023247535.1;refGene_gn=BCAN;refGene_tx=XM_023247537.1;refGene_gn=BCAN;refGene_tx=XM_023247538.1
F1 69368955 69369083
refGene_gn=BCAN;refGene_tx=XM_023247530.1;refGene_gn=BCAN;refGene_tx=XM_023247531.1;refGene_gn=BCAN;refGene_tx=XM_023247532.1;refGene_gn=BCAN;refGene_tx=XM_023247533.1;refGene_gn=BCAN;refGene_tx=XM_023247534.1;refGene_gn=BCAN;refGene_tx=XM_023247535.1;refGene_gn=BCAN;refGene_tx=XM_023247537.1;refGene_gn=BCAN;refGene_tx=XM_023247538.1
F1 69369354 69369529
refGene_gn=BCAN;refGene_tx=XM_023247530.1;refGene_gn=BCAN;refGene_tx=XM_023247531.1;refGene_gn=BCAN;refGene_tx=XM_023247532.1;refGene_gn=BCA

N;refGene_tx=XM_023247533.1;refGene_gn=BCAN;refGene_tx=XM_023247534.1;refGene_gn=BCAN;refGene_tx=XM_023247535.1;refGene_gn=BCAN;refGene_tx=XM_023247537.1;refGene_gn=BCAN;refGene_tx=XM_023247538.1
F1 69369848 69370223
refGene_gn=BCAN;refGene_tx=XM_023247530.1;refGene_gn=BCAN;refGene_tx=XM_023247531.1;refGene_gn=BCAN;refGene_tx=XM_023247532.1;refGene_gn=BCAN;refGene_tx=XM_023247533.1;refGene_gn=BCAN;refGene_tx=XM_023247534.1;refGene_gn=BCAN;refGene_tx=XM_023247535.1;refGene_gn=BCAN;refGene_tx=XM_023247537.1;refGene_gn=BCAN;refGene_tx=XM_023247538.1
F1 69370802 69370912
refGene_gn=BCAN;refGene_tx=XM_023247530.1;refGene_gn=BCAN;refGene_tx=XM_023247531.1;refGene_gn=BCAN;refGene_tx=XM_023247532.1;refGene_gn=BCAN;refGene_tx=XM_023247533.1;refGene_gn=BCAN;refGene_tx=XM_023247534.1;refGene_gn=BCAN;refGene_tx=XM_023247535.1;refGene_gn=BCAN;refGene_tx=XM_023247537.1;refGene_gn=BCAN;refGene_tx=XM_023247538.1
F1 69371828 69371836
refGene_gn=BCAN;refGene_tx=XM_023247530.1;refGene_gn=BCAN;refGene_tx=XM_023247531.1;refGene_gn=BCAN;refGene_tx=XM_023247532.1;refGene_gn=BCAN;refGene_tx=XM_023247533.1
F1 69386750 69387031
refGene_gn=HAPLN2;refGene_tx=XM_023247664.1;refGene_gn=HAPLN2;refGene_tx=XM_023247665.1;refGene_gn=HAPLN2;refGene_tx=XM_023247666.1;refGene_gn=HAPLN2;refGene_tx=XM_023247667.1;refGene_gn=HAPLN2;refGene_tx=XM_023247669.1;refGene_gn=HAPLN2;refGene_tx=XM_023247670.1;refGene_gn=HAPLN2;refGene_tx=XM_023247671.1;refGene_gn=HAPLN2;refGene_tx=XM_023247672.1
F1 69387231 69387414
refGene_gn=HAPLN2;refGene_tx=XM_023247664.1;refGene_gn=HAPLN2;refGene_tx=XM_023247665.1;refGene_gn=HAPLN2;refGene_tx=XM_023247666.1;refGene_gn=HAPLN2;refGene_tx=XM_023247667.1;refGene_gn=HAPLN2;refGene_tx=XM_023247669.1;refGene_gn=HAPLN2;refGene_tx=XM_023247670.1;refGene_gn=HAPLN2;refGene_tx=XM_023247671.1;refGene_gn=HAPLN2;refGene_tx=XM_023247672.1
F1 69387541 69387658
refGene_gn=HAPLN2;refGene_tx=XM_023247664.1;refGene_gn=HAPLN2;refGene_tx=XM_023247665.1;refGene_gn=HAPLN2;refGene_tx=XM_023247666.1;refGene_gn=HAPLN2;refGene_tx=XM_023247667.1;refGene_gn=HAPLN2;refGene_tx=XM_023247669.1;refGene_gn=HAPLN2;refGene_tx=XM_023247670.1;refGene_gn=HAPLN2;refGene_tx=XM_023247671.1;refGene_gn=HAPLN2;refGene_tx=XM_023247672.1
F1 69387795 69388149
refGene_gn=HAPLN2;refGene_tx=XM_023247664.1;refGene_gn=HAPLN2;refGene_tx=XM_023247665.1;refGene_gn=HAPLN2;refGene_tx=XM_023247666.1;refGene_gn=HAPLN2;refGene_tx=XM_023247667.1;refGene_gn=HAPLN2;refGene_tx=XM_023247669.1;refGene_gn=HAPLN2;refGene_tx=XM_023247670.1;refGene_gn=HAPLN2;refGene_tx=XM_023247671.1;refGene_gn=HAPLN2;refGene_tx=XM_023247672.1
F1 69388369 69388457
refGene_gn=HAPLN2;refGene_tx=XM_023247664.1;refGene_gn=HAPLN2;refGene_tx=XM_023247665.1;refGene_gn=HAPLN2;refGene_tx=XM_023247666.1;refGene_

gn=HAPLN2;refGene_tx=XM_023247667.1;refGene_gn=HAPLN2;refGene_tx=XM_023247669.1;refGene_gn=HAPLN2;refGene_tx=XM_023247670.1;refGene_gn=HAPLN2;refGene_tx=XM_023247671.1;refGene_gn=HAPLN2;refGene_tx=XM_023247672.1
F1	69405723	69405802	refGene_gn=GPATCH4;refGene_tx=XM_006943074.4
F1	69406249	69406323	refGene_gn=GPATCH4;refGene_tx=XM_006943074.4
F1	69406426	69406519	refGene_gn=GPATCH4;refGene_tx=XM_006943074.4
F1	69406601	69406688	refGene_gn=GPATCH4;refGene_tx=XM_006943074.4
F1	69408054	69408146	refGene_gn=GPATCH4;refGene_tx=XM_006943074.4
F1	69408388	69408439	refGene_gn=GPATCH4;refGene_tx=XM_006943074.4
F1	69408852	69409588	refGene_gn=GPATCH4;refGene_tx=XM_006943074.4
F1	69410272	69410475	refGene_gn=NAXE;refGene_tx=XM_023247279.1
F1	69410832	69410980	refGene_gn=NAXE;refGene_tx=XM_023247279.1
F1	69411607	69411721	refGene_gn=NAXE;refGene_tx=XM_023247279.1
F1	69411796	69411907	refGene_gn=NAXE;refGene_tx=XM_023247279.1
F1	69412007	69412116	refGene_gn=NAXE;refGene_tx=XM_023247279.1
F1	69412226	69412408	refGene_gn=NAXE;refGene_tx=XM_023247279.1
F1	69419673	69419781	refGene_gn=TTC24;refGene_tx=XM_023247276.1
F1	69420097	69420210	refGene_gn=TTC24;refGene_tx=XM_023247276.1
F1	69420306	69420435	refGene_gn=TTC24;refGene_tx=XM_023247276.1
F1	69421041	69421245	refGene_gn=TTC24;refGene_tx=XM_023247276.1
F1	69421436	69422145	refGene_gn=TTC24;refGene_tx=XM_023247276.1
F1	69427335	69427372	refGene_gn=IQGAP3;refGene_tx=XM_006943075.4
F1	69429789	69429877	refGene_gn=IQGAP3;refGene_tx=XM_006943075.4
F1	69434245	69434402	refGene_gn=IQGAP3;refGene_tx=XM_006943075.4
F1	69434920	69434998	refGene_gn=IQGAP3;refGene_tx=XM_006943075.4
F1	69436280	69436357	refGene_gn=IQGAP3;refGene_tx=XM_006943075.4
F1	69437019	69437087	refGene_gn=IQGAP3;refGene_tx=XM_006943075.4
F1	69437175	69437289	refGene_gn=IQGAP3;refGene_tx=XM_006943075.4
F1	69437507	69437686	refGene_gn=IQGAP3;refGene_tx=XM_006943075.4
F1	69438340	69438419	refGene_gn=IQGAP3;refGene_tx=XM_006943075.4
F1	69438969	69439133	refGene_gn=IQGAP3;refGene_tx=XM_006943075.4
F1	69439553	69439641	refGene_gn=IQGAP3;refGene_tx=XM_006943075.4
F1	69441201	69441365	refGene_gn=IQGAP3;refGene_tx=XM_006943075.4
F1	69442743	69442901	refGene_gn=IQGAP3;refGene_tx=XM_006943075.4
F1	69444166	69444285	refGene_gn=IQGAP3;refGene_tx=XM_006943075.4
F1	69444363	69444527	refGene_gn=IQGAP3;refGene_tx=XM_006943075.4
F1	69445997	69446088	refGene_gn=IQGAP3;refGene_tx=XM_006943075.4
F1	69446881	69447049	refGene_gn=IQGAP3;refGene_tx=XM_006943075.4
F1	69447133	69447273	refGene_gn=IQGAP3;refGene_tx=XM_006943075.4
F1	69447371	69447542	refGene_gn=IQGAP3;refGene_tx=XM_006943075.4
F1	69450344	69450428	refGene_gn=IQGAP3;refGene_tx=XM_006943075.4
F1	69450582	69450654	refGene_gn=IQGAP3;refGene_tx=XM_006943075.4
F1	69450748	69450818	refGene_gn=IQGAP3;refGene_tx=XM_006943075.4
F1	69452052	69452261	refGene_gn=IQGAP3;refGene_tx=XM_006943075.4
F1	69452838	69452991	refGene_gn=IQGAP3;refGene_tx=XM_006943075.4
F1	69453247	69453411	refGene_gn=IQGAP3;refGene_tx=XM_006943075.4
F1	69453740	69453965	refGene_gn=IQGAP3;refGene_tx=XM_006943075.4
F1	69454789	69454930	refGene_gn=IQGAP3;refGene_tx=XM_006943075.4
F1	69455748	69455833	refGene_gn=IQGAP3;refGene_tx=XM_006943075.4

F1 69456176 69456409 refGene_gn=IQGAP3;refGene_tx=XM_006943075.4
F1 69456699 69456832 refGene_gn=IQGAP3;refGene_tx=XM_006943075.4
F1 69456942 69457045 refGene_gn=IQGAP3;refGene_tx=XM_006943075.4
F1 69457656 69457783 refGene_gn=IQGAP3;refGene_tx=XM_006943075.4
F1 69458686 69458774 refGene_gn=IQGAP3;refGene_tx=XM_006943075.4
F1 69459435 69459648 refGene_gn=IQGAP3;refGene_tx=XM_006943075.4
F1 69460027 69460194 refGene_gn=IQGAP3;refGene_tx=XM_006943075.4
F1 69460408 69460507 refGene_gn=IQGAP3;refGene_tx=XM_006943075.4
F1 69460839 69460948 refGene_gn=IQGAP3;refGene_tx=XM_006943075.4
F1 69461809 69461923 refGene_gn=IQGAP3;refGene_tx=XM_006943075.4
F1 69500738 69500792
refGene_gn=MEF2D;refGene_tx=XM_023247965.1;refGene_gn=MEF2D;refGene_tx=XM_023247966.1;refGene_gn=MEF2D;refGene_tx=XM_023247967.1;refGene_gn=MEF2D;refGene_tx=XM_023247968.1;refGene_gn=MEF2D;refGene_tx=XM_023247969.1;refGene_gn=MEF2D;refGene_tx=XM_023247970.1
F1 69501365 69501569
refGene_gn=MEF2D;refGene_tx=XM_023247965.1;refGene_gn=MEF2D;refGene_tx=XM_023247966.1;refGene_gn=MEF2D;refGene_tx=XM_023247967.1;refGene_gn=MEF2D;refGene_tx=XM_023247968.1;refGene_gn=MEF2D;refGene_tx=XM_023247969.1;refGene_gn=MEF2D;refGene_tx=XM_023247970.1
F1 69502655 69502793
refGene_gn=MEF2D;refGene_tx=XM_023247965.1;refGene_gn=MEF2D;refGene_tx=XM_023247966.1;refGene_gn=MEF2D;refGene_tx=XM_023247967.1;refGene_gn=MEF2D;refGene_tx=XM_023247969.1
F1 69502842 69502977
refGene_gn=MEF2D;refGene_tx=XM_023247968.1;refGene_gn=MEF2D;refGene_tx=XM_023247970.1
F1 69503706 69503917
refGene_gn=MEF2D;refGene_tx=XM_023247965.1;refGene_gn=MEF2D;refGene_tx=XM_023247966.1;refGene_gn=MEF2D;refGene_tx=XM_023247967.1;refGene_gn=MEF2D;refGene_tx=XM_023247968.1;refGene_gn=MEF2D;refGene_tx=XM_023247969.1;refGene_gn=MEF2D;refGene_tx=XM_023247970.1
F1 69504095 69504152
refGene_gn=MEF2D;refGene_tx=XM_023247965.1;refGene_gn=MEF2D;refGene_tx=XM_023247966.1;refGene_gn=MEF2D;refGene_tx=XM_023247967.1;refGene_gn=MEF2D;refGene_tx=XM_023247968.1;refGene_gn=MEF2D;refGene_tx=XM_023247969.1;refGene_gn=MEF2D;refGene_tx=XM_023247970.1
F1 69505625 69505816
refGene_gn=MEF2D;refGene_tx=XM_023247965.1;refGene_gn=MEF2D;refGene_tx=XM_023247966.1;refGene_gn=MEF2D;refGene_tx=XM_023247967.1;refGene_gn=MEF2D;refGene_tx=XM_023247968.1;refGene_gn=MEF2D;refGene_tx=XM_023247969.1;refGene_gn=MEF2D;refGene_tx=XM_023247970.1
F1 69506244 69506265
refGene_gn=MEF2D;refGene_tx=XM_023247965.1;refGene_gn=MEF2D;refGene_tx=XM_023247966.1;refGene_gn=MEF2D;refGene_tx=XM_023247967.1;refGene_gn=MEF2D;refGene_tx=XM_023247968.1
F1 69507284 69507414
refGene_gn=MEF2D;refGene_tx=XM_023247965.1;refGene_gn=MEF2D;refGene_tx=XM_023247966.1;refGene_gn=MEF2D;refGene_tx=XM_023247967.1;refGene_gn=MEF2D;refGene_tx=XM_023247968.1;refGene_gn=MEF2D;refGene_tx=XM_0232479

69.1;refGene_gn=MEF2D;refGene_tx=XM_023247970.1
F1 69510766 69511001
refGene_gn=MEF2D;refGene_tx=XM_023247965.1;refGene_gn=MEF2D;refGene_tx
=XM_023247966.1;refGene_gn=MEF2D;refGene_tx=XM_023247967.1;refGene_gn=
MEF2D;refGene_tx=XM_023247968.1;refGene_gn=MEF2D;refGene_tx=XM_0232479
69.1;refGene_gn=MEF2D;refGene_tx=XM_023247970.1
F1 69511373 69511680
refGene_gn=MEF2D;refGene_tx=XM_023247965.1;refGene_gn=MEF2D;refGene_tx
=XM_023247966.1;refGene_gn=MEF2D;refGene_tx=XM_023247967.1;refGene_gn=
MEF2D;refGene_tx=XM_023247968.1;refGene_gn=MEF2D;refGene_tx=XM_0232479
69.1;refGene_gn=MEF2D;refGene_tx=XM_023247970.1
F1 69511954 69511966
refGene_gn=MEF2D;refGene_tx=XM_023247965.1;refGene_gn=MEF2D;refGene_tx
=XM_023247966.1;refGene_gn=MEF2D;refGene_tx=XM_023247967.1;refGene_gn=
MEF2D;refGene_tx=XM_023247968.1;refGene_gn=MEF2D;refGene_tx=XM_0232479
69.1;refGene_gn=MEF2D;refGene_tx=XM_023247970.1
F1 69541504 69541558
refGene_gn=LOC105261353;refGene_tx=XM_019822407.2;refGene_gn=LOC105261
353;refGene_tx=XM_019822408.2;refGene_gn=LOC105261353;refGene_tx=XM_01
9822409.1
F1 69541723 69541900
refGene_gn=LOC105261353;refGene_tx=XM_019822407.2;refGene_gn=LOC105261
353;refGene_tx=XM_019822408.2;refGene_gn=LOC105261353;refGene_tx=XM_01
9822409.1
F1 69544152 69544224
refGene_gn=LOC105261353;refGene_tx=XM_019822407.2;refGene_gn=LOC105261
353;refGene_tx=XM_019822408.2;refGene_gn=LOC105261353;refGene_tx=XM_01
9822409.1
F1 69555407 69555506
refGene_gn=LOC105261353;refGene_tx=XM_019822407.2;refGene_gn=LOC105261
353;refGene_tx=XM_019822408.2;refGene_gn=LOC105261353;refGene_tx=XM_01
9822409.1
F1 69555827 69555966
refGene_gn=LOC105261353;refGene_tx=XM_019822407.2;refGene_gn=LOC105261
353;refGene_tx=XM_019822408.2;refGene_gn=LOC105261353;refGene_tx=XM_01
9822409.1
F1 69557800 69558030
refGene_gn=LOC105261353;refGene_tx=XM_019822407.2
F1 69560213 69560296
refGene_gn=LOC105261353;refGene_tx=XM_019822408.2
F1 69560746 69560869
refGene_gn=LOC105261353;refGene_tx=XM_019822407.2
F1 69561509 69561562
refGene_gn=LOC105261353;refGene_tx=XM_019822409.1
F1 69578363 69578432 refGene_gn=RHBG;refGene_tx=XM_023247170.1
F1 69578627 69578701 refGene_gn=RHBG;refGene_tx=XM_023247170.1
F1 69579484 69579606 refGene_gn=RHBG;refGene_tx=XM_023247170.1
F1 69579928 69580062 refGene_gn=RHBG;refGene_tx=XM_023247170.1
F1 69580184 69580322 refGene_gn=RHBG;refGene_tx=XM_023247170.1
F1 69580584 69580751 refGene_gn=RHBG;refGene_tx=XM_023247170.1

F1 69582162 69582310 refGene_gn=RHBG;refGene_tx=XM_023247170.1
F1 69582441 69582592 refGene_gn=RHBG;refGene_tx=XM_023247170.1
F1 69582896 69583083 refGene_gn=RHBG;refGene_tx=XM_023247170.1
F1 69587919 69588106 refGene_gn=RHBG;refGene_tx=XM_023247170.1
F1 69597026 69597287
refGene_gn=TSACC;refGene_tx=XM_019822415.2;refGene_gn=TSACC;refGene_tx
=XM_019822417.2;refGene_gn=TSACC;refGene_tx=XM_019822418.2
F1 69598533 69598656
refGene_gn=TSACC;refGene_tx=XM_019822415.2;refGene_gn=TSACC;refGene_tx
=XM_019822417.2;refGene_gn=TSACC;refGene_tx=XM_019822418.2
F1 69603081 69603132
refGene_gn=TSACC;refGene_tx=XM_019822415.2;refGene_gn=TSACC;refGene_tx
=XM_019822417.2;refGene_gn=TSACC;refGene_tx=XM_019822418.2
F1 69603806 69603912
refGene_gn=TSACC;refGene_tx=XM_019822415.2;refGene_gn=TSACC;refGene_tx
=XM_019822417.2;refGene_gn=TSACC;refGene_tx=XM_019822418.2
F1 69604276 69604307 refGene_gn=CCT3;refGene_tx=XM_003999621.5
F1 69606093 69606155 refGene_gn=CCT3;refGene_tx=XM_003999621.5
F1 69607218 69607269
refGene_gn=CCT3;refGene_tx=XM_003999621.5;refGene_gn=CCT3;refGene_tx=X
M_019822414.2
F1 69607356 69607419
refGene_gn=CCT3;refGene_tx=XM_003999621.5;refGene_gn=CCT3;refGene_tx=X
M_019822414.2
F1 69608180 69608277
refGene_gn=CCT3;refGene_tx=XM_003999621.5;refGene_gn=CCT3;refGene_tx=X
M_019822414.2
F1 69610971 69611089
refGene_gn=CCT3;refGene_tx=XM_003999621.5;refGene_gn=CCT3;refGene_tx=X
M_019822414.2
F1 69612637 69612824
refGene_gn=CCT3;refGene_tx=XM_003999621.5;refGene_gn=CCT3;refGene_tx=X
M_019822414.2
F1 69613698 69613848
refGene_gn=CCT3;refGene_tx=XM_003999621.5;refGene_gn=CCT3;refGene_tx=X
M_019822414.2
F1 69614829 69614962
refGene_gn=CCT3;refGene_tx=XM_003999621.5;refGene_gn=CCT3;refGene_tx=X
M_019822414.2
F1 69615128 69615210
refGene_gn=CCT3;refGene_tx=XM_003999621.5;refGene_gn=CCT3;refGene_tx=X
M_019822414.2
F1 69616038 69616219
refGene_gn=CCT3;refGene_tx=XM_003999621.5;refGene_gn=CCT3;refGene_tx=X
M_019822414.2
F1 69617494 69617740
refGene_gn=CCT3;refGene_tx=XM_003999621.5;refGene_gn=CCT3;refGene_tx=X
M_019822414.2
F1 69618026 69618158
refGene_gn=CCT3;refGene_tx=XM_003999621.5;refGene_gn=CCT3;refGene_tx=X

M_019822414.2
F1 69618465 69618570
refGene_gn=CCT3;refGene_tx=XM_003999621.5;refGene_gn=CCT3;refGene_tx=X
M_019822414.2
F1 69622479 69622599 refGene_gn=GLMP;refGene_tx=XM_003999707.5
F1 69623014 69623272 refGene_gn=GLMP;refGene_tx=XM_003999707.5
F1 69623467 69623665 refGene_gn=GLMP;refGene_tx=XM_003999707.5
F1 69623780 69623999 refGene_gn=GLMP;refGene_tx=XM_003999707.5
F1 69624410 69624664 refGene_gn=GLMP;refGene_tx=XM_003999707.5
F1 69624788 69624954 refGene_gn=GLMP;refGene_tx=XM_003999707.5
F1 69626281 69626323 refGene_gn=TMEM79;refGene_tx=XM_023247591.1
F1 69626324 69626496 refGene_gn=TMEM79;refGene_tx=XM_023247591.1
F1 69628786 69629000 refGene_gn=TMEM79;refGene_tx=XM_023247591.1
F1 69629196 69629445 refGene_gn=TMEM79;refGene_tx=XM_023247591.1
F1 69629447 69629955 refGene_gn=TMEM79;refGene_tx=XM_023247591.1
F1 69632428 69632502
refGene_gn=SMG5;refGene_tx=XM_023247592.1;refGene_gn=SMG5;refGene_tx=X
M_023247593.1
F1 69636136 69636235
refGene_gn=SMG5;refGene_tx=XM_023247592.1;refGene_gn=SMG5;refGene_tx=X
M_023247593.1
F1 69636746 69636870
refGene_gn=SMG5;refGene_tx=XM_023247592.1;refGene_gn=SMG5;refGene_tx=X
M_023247593.1
F1 69637234 69637391
refGene_gn=SMG5;refGene_tx=XM_019822413.2;refGene_gn=SMG5;refGene_tx=X
M_023247592.1;refGene_gn=SMG5;refGene_tx=XM_023247593.1
F1 69639477 69639567
refGene_gn=SMG5;refGene_tx=XM_019822413.2;refGene_gn=SMG5;refGene_tx=X
M_023247592.1;refGene_gn=SMG5;refGene_tx=XM_023247593.1
F1 69640407 69640592
refGene_gn=SMG5;refGene_tx=XM_019822413.2;refGene_gn=SMG5;refGene_tx=X
M_023247592.1;refGene_gn=SMG5;refGene_tx=XM_023247593.1
F1 69641233 69641312
refGene_gn=SMG5;refGene_tx=XM_019822413.2;refGene_gn=SMG5;refGene_tx=X
M_023247592.1;refGene_gn=SMG5;refGene_tx=XM_023247593.1
F1 69643374 69643500
refGene_gn=SMG5;refGene_tx=XM_019822413.2;refGene_gn=SMG5;refGene_tx=X
M_023247592.1;refGene_gn=SMG5;refGene_tx=XM_023247593.1
F1 69643571 69643640
refGene_gn=SMG5;refGene_tx=XM_019822413.2;refGene_gn=SMG5;refGene_tx=X
M_023247592.1;refGene_gn=SMG5;refGene_tx=XM_023247593.1
F1 69643786 69643995
refGene_gn=SMG5;refGene_tx=XM_019822413.2;refGene_gn=SMG5;refGene_tx=X
M_023247592.1;refGene_gn=SMG5;refGene_tx=XM_023247593.1
F1 69644651 69644789
refGene_gn=SMG5;refGene_tx=XM_019822413.2;refGene_gn=SMG5;refGene_tx=X
M_023247592.1;refGene_gn=SMG5;refGene_tx=XM_023247593.1
F1 69644916 69645501
refGene_gn=SMG5;refGene_tx=XM_019822413.2;refGene_gn=SMG5;refGene_tx=X

M_023247592.1;refGene_gn=SMG5;refGene_tx=XM_023247593.1
F1 69647160 69647336
refGene_gn=SMG5;refGene_tx=XM_019822413.2;refGene_gn=SMG5;refGene_tx=XM_023247592.1;refGene_gn=SMG5;refGene_tx=XM_023247593.1
F1 69648329 69648405
refGene_gn=SMG5;refGene_tx=XM_019822413.2;refGene_gn=SMG5;refGene_tx=XM_023247592.1;refGene_gn=SMG5;refGene_tx=XM_023247593.1
F1 69648981 69649157
refGene_gn=SMG5;refGene_tx=XM_019822413.2;refGene_gn=SMG5;refGene_tx=XM_023247592.1;refGene_gn=SMG5;refGene_tx=XM_023247593.1
F1 69650191 69650350
refGene_gn=SMG5;refGene_tx=XM_019822413.2;refGene_gn=SMG5;refGene_tx=XM_023247592.1;refGene_gn=SMG5;refGene_tx=XM_023247593.1
F1 69653511 69653571
refGene_gn=SMG5;refGene_tx=XM_019822413.2;refGene_gn=SMG5;refGene_tx=XM_023247592.1;refGene_gn=SMG5;refGene_tx=XM_023247593.1
F1 69654091 69654251
refGene_gn=SMG5;refGene_tx=XM_019822413.2;refGene_gn=SMG5;refGene_tx=XM_023247592.1;refGene_gn=SMG5;refGene_tx=XM_023247593.1
F1 69654668 69654759
refGene_gn=SMG5;refGene_tx=XM_019822413.2;refGene_gn=SMG5;refGene_tx=XM_023247592.1;refGene_gn=SMG5;refGene_tx=XM_023247593.1
F1 69655605 69655680
refGene_gn=SMG5;refGene_tx=XM_019822413.2;refGene_gn=SMG5;refGene_tx=XM_023247592.1;refGene_gn=SMG5;refGene_tx=XM_023247593.1
F1 69656001 69656140
refGene_gn=SMG5;refGene_tx=XM_019822413.2;refGene_gn=SMG5;refGene_tx=XM_023247592.1;refGene_gn=SMG5;refGene_tx=XM_023247593.1
F1 69656333 69656417
refGene_gn=SMG5;refGene_tx=XM_019822413.2;refGene_gn=SMG5;refGene_tx=XM_023247592.1;refGene_gn=SMG5;refGene_tx=XM_023247593.1
F1 69659284 69659344 refGene_gn=PAQR6;refGene_tx=XM_023247595.1
F1 69659947 69660023
refGene_gn=PAQR6;refGene_tx=XM_023247594.1;refGene_gn=PAQR6;refGene_tx=XM_023247595.1
F1 69660447 69660575
refGene_gn=PAQR6;refGene_tx=XM_023247594.1;refGene_gn=PAQR6;refGene_tx=XM_023247595.1
F1 69660650 69660856
refGene_gn=PAQR6;refGene_tx=XM_023247594.1;refGene_gn=PAQR6;refGene_tx=XM_023247595.1
F1 69660977 69661104
refGene_gn=PAQR6;refGene_tx=XM_023247594.1;refGene_gn=PAQR6;refGene_tx=XM_023247595.1
F1 69661496 69661593 refGene_gn=PAQR6;refGene_tx=XM_023247594.1
F1 69661671 69661822
refGene_gn=PAQR6;refGene_tx=XM_023247594.1;refGene_gn=PAQR6;refGene_tx=XM_023247595.1
F1 69662132 69662407
refGene_gn=PAQR6;refGene_tx=XM_023247594.1;refGene_gn=PAQR6;refGene_tx

=XM_023247595.1
F1	69663328	69663458	refGene_gn=BGLAP;refGene_tx=XM_003999711.5
F1	69663581	69663651	refGene_gn=BGLAP;refGene_tx=XM_003999711.5
F1	69663747	69663786	refGene_gn=BGLAP;refGene_tx=XM_003999711.5
F1	69663926	69663996	refGene_gn=BGLAP;refGene_tx=XM_003999711.5
F1	69665835	69665889	refGene_gn=PMF1;refGene_tx=XM_003999624.5
F1	69667016	69667212	refGene_gn=PMF1;refGene_tx=XM_003999624.5
F1	69668089	69668190	refGene_gn=PMF1;refGene_tx=XM_003999624.5
F1	69668816	69668922	refGene_gn=PMF1;refGene_tx=XM_003999624.5
F1	69682823	69682984	refGene_gn=PMF1;refGene_tx=XM_003999624.5
F1	69685438	69685630	refGene_gn=SLC25A44;refGene_tx=XM_011291189.3
F1	69686756	69686942	refGene_gn=SLC25A44;refGene_tx=XM_019822419.2
F1	69687187	69687315
refGene_gn=SLC25A44;refGene_tx=XM_011291189.3;refGene_gn=SLC25A44;refG
ene_tx=XM_019822419.2
F1	69693183	69693808
refGene_gn=SLC25A44;refGene_tx=XM_011291189.3;refGene_gn=SLC25A44;refG
ene_tx=XM_019822419.2
F1	69703914	69704507
refGene_gn=SEMA4A;refGene_tx=XM_003999626.5;refGene_gn=SEMA4A;refGene_
tx=XM_019822421.2;refGene_gn=SEMA4A;refGene_tx=XM_023247484.1
F1	69704894	69704995
refGene_gn=SEMA4A;refGene_tx=XM_003999626.5;refGene_gn=SEMA4A;refGene_
tx=XM_019822421.2;refGene_gn=SEMA4A;refGene_tx=XM_023247484.1
F1	69705210	69705368
refGene_gn=SEMA4A;refGene_tx=XM_003999626.5;refGene_gn=SEMA4A;refGene_
tx=XM_019822421.2;refGene_gn=SEMA4A;refGene_tx=XM_023247484.1
F1	69705457	69705576
refGene_gn=SEMA4A;refGene_tx=XM_003999626.5;refGene_gn=SEMA4A;refGene_
tx=XM_019822421.2;refGene_gn=SEMA4A;refGene_tx=XM_023247484.1
F1	69706949	69707130
refGene_gn=SEMA4A;refGene_tx=XM_003999626.5;refGene_gn=SEMA4A;refGene_
tx=XM_019822421.2;refGene_gn=SEMA4A;refGene_tx=XM_023247484.1
F1	69711519	69711670
refGene_gn=SEMA4A;refGene_tx=XM_003999626.5;refGene_gn=SEMA4A;refGene_
tx=XM_019822421.2;refGene_gn=SEMA4A;refGene_tx=XM_023247484.1
F1	69712702	69712875
refGene_gn=SEMA4A;refGene_tx=XM_003999626.5;refGene_gn=SEMA4A;refGene_
tx=XM_019822421.2;refGene_gn=SEMA4A;refGene_tx=XM_023247484.1
F1	69713048	69713173
refGene_gn=SEMA4A;refGene_tx=XM_003999626.5;refGene_gn=SEMA4A;refGene_
tx=XM_019822421.2;refGene_gn=SEMA4A;refGene_tx=XM_023247484.1
F1	69713489	69713606
refGene_gn=SEMA4A;refGene_tx=XM_003999626.5;refGene_gn=SEMA4A;refGene_
tx=XM_019822421.2;refGene_gn=SEMA4A;refGene_tx=XM_023247484.1
F1	69714378	69714484
refGene_gn=SEMA4A;refGene_tx=XM_003999626.5;refGene_gn=SEMA4A;refGene_
tx=XM_019822421.2;refGene_gn=SEMA4A;refGene_tx=XM_023247484.1
F1	69714705	69714804
refGene_gn=SEMA4A;refGene_tx=XM_003999626.5;refGene_gn=SEMA4A;refGene_

tx=XM_019822421.2;refGene_gn=SEMA4A;refGene_tx=XM_023247484.1
F1 69715011 69715074
refGene_gn=SEMA4A;refGene_tx=XM_003999626.5;refGene_gn=SEMA4A;refGene_
tx=XM_019822421.2;refGene_gn=SEMA4A;refGene_tx=XM_023247484.1
F1 69715986 69716147
refGene_gn=SEMA4A;refGene_tx=XM_003999626.5;refGene_gn=SEMA4A;refGene_
tx=XM_019822421.2;refGene_gn=SEMA4A;refGene_tx=XM_023247484.1
F1 69717657 69717822
refGene_gn=SEMA4A;refGene_tx=XM_003999626.5;refGene_gn=SEMA4A;refGene_
tx=XM_019822421.2;refGene_gn=SEMA4A;refGene_tx=XM_023247484.1
F1 69718524 69718582 refGene_gn=SEMA4A;refGene_tx=XM_019822421.2
F1 69733952 69733979 refGene_gn=LMNA;refGene_tx=XM_023247978.1
F1 69734267 69734537 refGene_gn=LMNA;refGene_tx=XM_023247978.1
F1 69735196 69735286 refGene_gn=LMNA;refGene_tx=XM_023247978.1
F1 69735713 69735833 refGene_gn=LMNA;refGene_tx=XM_023247978.1
F1 69735940 69736048 refGene_gn=LMNA;refGene_tx=XM_023247978.1
F1 69736449 69736672 refGene_gn=LMNA;refGene_tx=XM_023247978.1
F1 69736776 69736997 refGene_gn=LMNA;refGene_tx=XM_023247978.1
F1 69737702 69737828 refGene_gn=LMNA;refGene_tx=XM_023247978.1
F1 69737920 69738091 refGene_gn=LMNA;refGene_tx=XM_023247978.1
F1 69738358 69738484 refGene_gn=LMNA;refGene_tx=XM_023247978.1
F1 69740018 69740175 refGene_gn=LMNA;refGene_tx=XM_023247978.1
F1 69750154 69750510 refGene_gn=LMNA;refGene_tx=XM_023247978.1
F1 69773821 69774275 refGene_gn=MEX3A;refGene_tx=XM_019821388.2
F1 69777866 69778975 refGene_gn=MEX3A;refGene_tx=XM_019821388.2
F1 69784510 69784638 refGene_gn=RAB25;refGene_tx=XM_003999627.5
F1 69785065 69785146 refGene_gn=RAB25;refGene_tx=XM_003999627.5
F1 69785966 69786160 refGene_gn=RAB25;refGene_tx=XM_003999627.5
F1 69787345 69787541 refGene_gn=RAB25;refGene_tx=XM_003999627.5
F1 69790166 69790209 refGene_gn=RAB25;refGene_tx=XM_003999627.5
F1 69791710 69791767 refGene_gn=LAMTOR2;refGene_tx=XM_003999628.3
F1 69791968 69792058 refGene_gn=LAMTOR2;refGene_tx=XM_003999628.3
F1 69793877 69794040 refGene_gn=LAMTOR2;refGene_tx=XM_003999628.3
F1 69794287 69794355 refGene_gn=LAMTOR2;refGene_tx=XM_003999628.3
F1 69795320 69795428
refGene_gn=UBQLN4;refGene_tx=XM_023248092.1;refGene_gn=UBQLN4;refGene_
tx=XM_023248093.1
F1 69796851 69797003
refGene_gn=UBQLN4;refGene_tx=XM_023248092.1;refGene_gn=UBQLN4;refGene_
tx=XM_023248093.1
F1 69797356 69797574
refGene_gn=UBQLN4;refGene_tx=XM_023248092.1;refGene_gn=UBQLN4;refGene_
tx=XM_023248093.1
F1 69798149 69798412
refGene_gn=UBQLN4;refGene_tx=XM_023248092.1;refGene_gn=UBQLN4;refGene_
tx=XM_023248093.1
F1 69799980 69800139
refGene_gn=UBQLN4;refGene_tx=XM_023248092.1;refGene_gn=UBQLN4;refGene_
tx=XM_023248093.1
F1 69803564 69803790

refGene_gn=UBQLN4;refGene_tx=XM_023248092.1;refGene_gn=UBQLN4;refGene_tx=XM_023248093.1
F1 69804802 69804942
refGene_gn=UBQLN4;refGene_tx=XM_023248092.1;refGene_gn=UBQLN4;refGene_tx=XM_023248093.1
F1 69805404 69805488
refGene_gn=UBQLN4;refGene_tx=XM_023248092.1;refGene_gn=UBQLN4;refGene_tx=XM_023248093.1
F1 69805599 69805715
refGene_gn=UBQLN4;refGene_tx=XM_023248092.1;refGene_gn=UBQLN4;refGene_tx=XM_023248093.1
F1 69805859 69806083
refGene_gn=UBQLN4;refGene_tx=XM_023248092.1;refGene_gn=UBQLN4;refGene_tx=XM_023248093.1
F1 69807482 69807638
refGene_gn=UBQLN4;refGene_tx=XM_023248092.1;refGene_gn=UBQLN4;refGene_tx=XM_023248093.1
F1 69817110 69817510
refGene_gn=LOC102901020;refGene_tx=XM_023247189.1;refGene_gn=LOC102901020;refGene_tx=XM_023247190.1;refGene_gn=LOC102901020;refGene_tx=XM_023247191.1
F1 69817612 69817695
refGene_gn=LOC102901020;refGene_tx=XM_023247189.1;refGene_gn=LOC102901020;refGene_tx=XM_023247190.1
F1 69818557 69818686
refGene_gn=LOC102901020;refGene_tx=XM_023247189.1;refGene_gn=LOC102901020;refGene_tx=XM_023247190.1;refGene_gn=LOC102901020;refGene_tx=XM_023247191.1
F1 69818931 69819042
refGene_gn=LOC102901020;refGene_tx=XM_023247189.1;refGene_gn=LOC102901020;refGene_tx=XM_023247190.1;refGene_gn=LOC102901020;refGene_tx=XM_023247191.1
F1 69825234 69825389 refGene_gn=SSR2;refGene_tx=XM_003999630.5
F1 69826508 69826607 refGene_gn=SSR2;refGene_tx=XM_003999630.5
F1 69827238 69827347 refGene_gn=SSR2;refGene_tx=XM_003999630.5
F1 69829396 69829474 refGene_gn=SSR2;refGene_tx=XM_003999630.5
F1 69830517 69830628 refGene_gn=SSR2;refGene_tx=XM_003999630.5
F1 69849403 69849446
refGene_gn=ARHGEF2;refGene_tx=XM_023247171.1;refGene_gn=ARHGEF2;refGene_tx=XM_023247172.1
F1 69850016 69850056
refGene_gn=ARHGEF2;refGene_tx=XM_023247171.1;refGene_gn=ARHGEF2;refGene_tx=XM_023247172.1
F1 69851225 69851319
refGene_gn=ARHGEF2;refGene_tx=XM_023247171.1;refGene_gn=ARHGEF2;refGene_tx=XM_023247172.1;refGene_gn=ARHGEF2;refGene_tx=XM_023247177.1
F1 69855291 69855354
refGene_gn=ARHGEF2;refGene_tx=XM_023247173.1;refGene_gn=ARHGEF2;refGene_tx=XM_023247179.1
F1 69859842 69859905 refGene_gn=ARHGEF2;refGene_tx=XM_023247175.1

F1 69861632 69861650 refGene_gn=ARHGEF2;refGene_tx=XM_023247178.1
F1 69861789 69861828 refGene_gn=ARHGEF2;refGene_tx=XM_023247176.1
F1 69862031 69862176
refGene_gn=ARHGEF2;refGene_tx=XM_023247171.1;refGene_gn=ARHGEF2;refGen
e_tx=XM_023247172.1;refGene_gn=ARHGEF2;refGene_tx=XM_023247173.1;refGe
ne_gn=ARHGEF2;refGene_tx=XM_023247175.1;refGene_gn=ARHGEF2;refGene_tx=
XM_023247176.1;refGene_gn=ARHGEF2;refGene_tx=XM_023247177.1;refGene_gn
=ARHGEF2;refGene_tx=XM_023247178.1;refGene_gn=ARHGEF2;refGene_tx=XM_02
3247179.1;refGene_gn=ARHGEF2;refGene_tx=XM_023247180.1;refGene_gn=ARHG
EF2;refGene_tx=XM_023247181.1;refGene_gn=ARHGEF2;refGene_tx=XM_0232471
82.1;refGene_gn=ARHGEF2;refGene_tx=XM_023247183.1;refGene_gn=ARHGEF2;r
efGene_tx=XM_023247184.1;refGene_gn=ARHGEF2;refGene_tx=XM_023247185.1;
refGene_gn=ARHGEF2;refGene_tx=XM_023247186.1
F1 69864224 69864292
refGene_gn=ARHGEF2;refGene_tx=XM_023247171.1;refGene_gn=ARHGEF2;refGen
e_tx=XM_023247172.1;refGene_gn=ARHGEF2;refGene_tx=XM_023247173.1;refGe
ne_gn=ARHGEF2;refGene_tx=XM_023247175.1;refGene_gn=ARHGEF2;refGene_tx=
XM_023247176.1;refGene_gn=ARHGEF2;refGene_tx=XM_023247177.1;refGene_gn
=ARHGEF2;refGene_tx=XM_023247178.1;refGene_gn=ARHGEF2;refGene_tx=XM_02
3247179.1;refGene_gn=ARHGEF2;refGene_tx=XM_023247180.1
F1 69864608 69864672
refGene_gn=ARHGEF2;refGene_tx=XM_023247171.1;refGene_gn=ARHGEF2;refGen
e_tx=XM_023247172.1;refGene_gn=ARHGEF2;refGene_tx=XM_023247173.1;refGe
ne_gn=ARHGEF2;refGene_tx=XM_023247175.1;refGene_gn=ARHGEF2;refGene_tx=
XM_023247176.1;refGene_gn=ARHGEF2;refGene_tx=XM_023247177.1;refGene_gn
=ARHGEF2;refGene_tx=XM_023247178.1;refGene_gn=ARHGEF2;refGene_tx=XM_02
3247179.1;refGene_gn=ARHGEF2;refGene_tx=XM_023247180.1;refGene_gn=ARHG
EF2;refGene_tx=XM_023247181.1;refGene_gn=ARHGEF2;refGene_tx=XM_0232471
82.1;refGene_gn=ARHGEF2;refGene_tx=XM_023247183.1;refGene_gn=ARHGEF2;r
efGene_tx=XM_023247184.1;refGene_gn=ARHGEF2;refGene_tx=XM_023247185.1;
refGene_gn=ARHGEF2;refGene_tx=XM_023247186.1
F1 69865157 69865287
refGene_gn=ARHGEF2;refGene_tx=XM_023247171.1;refGene_gn=ARHGEF2;refGen
e_tx=XM_023247172.1;refGene_gn=ARHGEF2;refGene_tx=XM_023247173.1;refGe
ne_gn=ARHGEF2;refGene_tx=XM_023247175.1;refGene_gn=ARHGEF2;refGene_tx=
XM_023247176.1;refGene_gn=ARHGEF2;refGene_tx=XM_023247177.1;refGene_gn
=ARHGEF2;refGene_tx=XM_023247178.1;refGene_gn=ARHGEF2;refGene_tx=XM_02
3247179.1;refGene_gn=ARHGEF2;refGene_tx=XM_023247180.1;refGene_gn=ARHG
EF2;refGene_tx=XM_023247181.1;refGene_gn=ARHGEF2;refGene_tx=XM_0232471
82.1;refGene_gn=ARHGEF2;refGene_tx=XM_023247183.1;refGene_gn=ARHGEF2;r
efGene_tx=XM_023247184.1;refGene_gn=ARHGEF2;refGene_tx=XM_023247185.1;
refGene_gn=ARHGEF2;refGene_tx=XM_023247186.1
F1 69865504 69865614
refGene_gn=ARHGEF2;refGene_tx=XM_023247171.1;refGene_gn=ARHGEF2;refGen
e_tx=XM_023247172.1;refGene_gn=ARHGEF2;refGene_tx=XM_023247173.1;refGe
ne_gn=ARHGEF2;refGene_tx=XM_023247175.1;refGene_gn=ARHGEF2;refGene_tx=
XM_023247176.1;refGene_gn=ARHGEF2;refGene_tx=XM_023247177.1;refGene_gn
=ARHGEF2;refGene_tx=XM_023247178.1;refGene_gn=ARHGEF2;refGene_tx=XM_02
3247179.1;refGene_gn=ARHGEF2;refGene_tx=XM_023247180.1;refGene_gn=ARHG
EF2;refGene_tx=XM_023247181.1;refGene_gn=ARHGEF2;refGene_tx=XM_0232471

82.1;refGene_gn=ARHGEF2;refGene_tx=XM_023247183.1;refGene_gn=ARHGEF2;refGene_tx=XM_023247184.1;refGene_gn=ARHGEF2;refGene_tx=XM_023247185.1;refGene_gn=ARHGEF2;refGene_tx=XM_023247186.1
F1	69865728 69865875
refGene_gn=ARHGEF2;refGene_tx=XM_023247171.1;refGene_gn=ARHGEF2;refGene_tx=XM_023247172.1;refGene_gn=ARHGEF2;refGene_tx=XM_023247173.1;refGene_gn=ARHGEF2;refGene_tx=XM_023247175.1;refGene_gn=ARHGEF2;refGene_tx=XM_023247176.1;refGene_gn=ARHGEF2;refGene_tx=XM_023247177.1;refGene_gn=ARHGEF2;refGene_tx=XM_023247178.1;refGene_gn=ARHGEF2;refGene_tx=XM_023247179.1;refGene_gn=ARHGEF2;refGene_tx=XM_023247180.1;refGene_gn=ARHGEF2;refGene_tx=XM_023247181.1;refGene_gn=ARHGEF2;refGene_tx=XM_023247182.1;refGene_gn=ARHGEF2;refGene_tx=XM_023247183.1;refGene_gn=ARHGEF2;refGene_tx=XM_023247184.1;refGene_gn=ARHGEF2;refGene_tx=XM_023247185.1;refGene_gn=ARHGEF2;refGene_tx=XM_023247186.1
F1	69867274 69867525
refGene_gn=ARHGEF2;refGene_tx=XM_023247171.1;refGene_gn=ARHGEF2;refGene_tx=XM_023247172.1;refGene_gn=ARHGEF2;refGene_tx=XM_023247173.1;refGene_gn=ARHGEF2;refGene_tx=XM_023247175.1;refGene_gn=ARHGEF2;refGene_tx=XM_023247176.1;refGene_gn=ARHGEF2;refGene_tx=XM_023247177.1;refGene_gn=ARHGEF2;refGene_tx=XM_023247178.1;refGene_gn=ARHGEF2;refGene_tx=XM_023247179.1;refGene_gn=ARHGEF2;refGene_tx=XM_023247180.1;refGene_gn=ARHGEF2;refGene_tx=XM_023247181.1;refGene_gn=ARHGEF2;refGene_tx=XM_023247182.1;refGene_gn=ARHGEF2;refGene_tx=XM_023247183.1;refGene_gn=ARHGEF2;refGene_tx=XM_023247184.1;refGene_gn=ARHGEF2;refGene_tx=XM_023247185.1;refGene_gn=ARHGEF2;refGene_tx=XM_023247186.1
F1	69867656 69867782
refGene_gn=ARHGEF2;refGene_tx=XM_023247171.1;refGene_gn=ARHGEF2;refGene_tx=XM_023247172.1;refGene_gn=ARHGEF2;refGene_tx=XM_023247173.1;refGene_gn=ARHGEF2;refGene_tx=XM_023247175.1;refGene_gn=ARHGEF2;refGene_tx=XM_023247176.1;refGene_gn=ARHGEF2;refGene_tx=XM_023247177.1;refGene_gn=ARHGEF2;refGene_tx=XM_023247178.1;refGene_gn=ARHGEF2;refGene_tx=XM_023247179.1;refGene_gn=ARHGEF2;refGene_tx=XM_023247180.1;refGene_gn=ARHGEF2;refGene_tx=XM_023247181.1;refGene_gn=ARHGEF2;refGene_tx=XM_023247182.1;refGene_gn=ARHGEF2;refGene_tx=XM_023247183.1;refGene_gn=ARHGEF2;refGene_tx=XM_023247184.1;refGene_gn=ARHGEF2;refGene_tx=XM_023247185.1;refGene_gn=ARHGEF2;refGene_tx=XM_023247186.1
F1	69868181 69868299
refGene_gn=ARHGEF2;refGene_tx=XM_023247171.1;refGene_gn=ARHGEF2;refGene_tx=XM_023247172.1;refGene_gn=ARHGEF2;refGene_tx=XM_023247173.1;refGene_gn=ARHGEF2;refGene_tx=XM_023247175.1;refGene_gn=ARHGEF2;refGene_tx=XM_023247176.1;refGene_gn=ARHGEF2;refGene_tx=XM_023247177.1;refGene_gn=ARHGEF2;refGene_tx=XM_023247178.1;refGene_gn=ARHGEF2;refGene_tx=XM_023247179.1;refGene_gn=ARHGEF2;refGene_tx=XM_023247180.1;refGene_gn=ARHGEF2;refGene_tx=XM_023247181.1;refGene_gn=ARHGEF2;refGene_tx=XM_023247182.1;refGene_gn=ARHGEF2;refGene_tx=XM_023247183.1;refGene_gn=ARHGEF2;refGene_tx=XM_023247184.1;refGene_gn=ARHGEF2;refGene_tx=XM_023247185.1;refGene_gn=ARHGEF2;refGene_tx=XM_023247186.1
F1	69868487 69868736
refGene_gn=ARHGEF2;refGene_tx=XM_023247171.1;refGene_gn=ARHGEF2;refGene_tx=XM_023247172.1;refGene_gn=ARHGEF2;refGene_tx=XM_023247173.1;refGe

ne_gn=ARHGEF2;refGene_tx=XM_023247175.1;refGene_gn=ARHGEF2;refGene_tx=XM_023247176.1;refGene_gn=ARHGEF2;refGene_tx=XM_023247177.1;refGene_gn=ARHGEF2;refGene_tx=XM_023247178.1;refGene_gn=ARHGEF2;refGene_tx=XM_023247179.1;refGene_gn=ARHGEF2;refGene_tx=XM_023247180.1;refGene_gn=ARHGEF2;refGene_tx=XM_023247181.1;refGene_gn=ARHGEF2;refGene_tx=XM_023247182.1;refGene_gn=ARHGEF2;refGene_tx=XM_023247183.1;refGene_gn=ARHGEF2;refGene_tx=XM_023247184.1;refGene_gn=ARHGEF2;refGene_tx=XM_023247185.1;refGene_gn=ARHGEF2;refGene_tx=XM_023247186.1
F1 69870757 69870834
refGene_gn=ARHGEF2;refGene_tx=XM_023247171.1;refGene_gn=ARHGEF2;refGene_tx=XM_023247172.1;refGene_gn=ARHGEF2;refGene_tx=XM_023247173.1;refGene_gn=ARHGEF2;refGene_tx=XM_023247175.1;refGene_gn=ARHGEF2;refGene_tx=XM_023247176.1;refGene_gn=ARHGEF2;refGene_tx=XM_023247177.1;refGene_gn=ARHGEF2;refGene_tx=XM_023247178.1;refGene_gn=ARHGEF2;refGene_tx=XM_023247179.1;refGene_gn=ARHGEF2;refGene_tx=XM_023247180.1;refGene_gn=ARHGEF2;refGene_tx=XM_023247181.1;refGene_gn=ARHGEF2;refGene_tx=XM_023247182.1;refGene_gn=ARHGEF2;refGene_tx=XM_023247183.1;refGene_gn=ARHGEF2;refGene_tx=XM_023247184.1;refGene_gn=ARHGEF2;refGene_tx=XM_023247185.1;refGene_gn=ARHGEF2;refGene_tx=XM_023247186.1
F1 69871387 69871557
refGene_gn=ARHGEF2;refGene_tx=XM_023247171.1;refGene_gn=ARHGEF2;refGene_tx=XM_023247172.1;refGene_gn=ARHGEF2;refGene_tx=XM_023247173.1;refGene_gn=ARHGEF2;refGene_tx=XM_023247175.1;refGene_gn=ARHGEF2;refGene_tx=XM_023247176.1;refGene_gn=ARHGEF2;refGene_tx=XM_023247177.1;refGene_gn=ARHGEF2;refGene_tx=XM_023247178.1;refGene_gn=ARHGEF2;refGene_tx=XM_023247179.1;refGene_gn=ARHGEF2;refGene_tx=XM_023247180.1;refGene_gn=ARHGEF2;refGene_tx=XM_023247181.1;refGene_gn=ARHGEF2;refGene_tx=XM_023247182.1;refGene_gn=ARHGEF2;refGene_tx=XM_023247183.1;refGene_gn=ARHGEF2;refGene_tx=XM_023247184.1;refGene_gn=ARHGEF2;refGene_tx=XM_023247185.1;refGene_gn=ARHGEF2;refGene_tx=XM_023247186.1
F1 69873133 69873201
refGene_gn=ARHGEF2;refGene_tx=XM_023247171.1;refGene_gn=ARHGEF2;refGene_tx=XM_023247172.1;refGene_gn=ARHGEF2;refGene_tx=XM_023247173.1;refGene_gn=ARHGEF2;refGene_tx=XM_023247175.1;refGene_gn=ARHGEF2;refGene_tx=XM_023247176.1;refGene_gn=ARHGEF2;refGene_tx=XM_023247177.1;refGene_gn=ARHGEF2;refGene_tx=XM_023247178.1;refGene_gn=ARHGEF2;refGene_tx=XM_023247179.1;refGene_gn=ARHGEF2;refGene_tx=XM_023247180.1;refGene_gn=ARHGEF2;refGene_tx=XM_023247181.1;refGene_gn=ARHGEF2;refGene_tx=XM_023247182.1;refGene_gn=ARHGEF2;refGene_tx=XM_023247183.1;refGene_gn=ARHGEF2;refGene_tx=XM_023247184.1;refGene_gn=ARHGEF2;refGene_tx=XM_023247185.1;refGene_gn=ARHGEF2;refGene_tx=XM_023247186.1
F1 69874946 69875144
refGene_gn=ARHGEF2;refGene_tx=XM_023247171.1;refGene_gn=ARHGEF2;refGene_tx=XM_023247172.1;refGene_gn=ARHGEF2;refGene_tx=XM_023247173.1;refGene_gn=ARHGEF2;refGene_tx=XM_023247175.1;refGene_gn=ARHGEF2;refGene_tx=XM_023247176.1;refGene_gn=ARHGEF2;refGene_tx=XM_023247177.1;refGene_gn=ARHGEF2;refGene_tx=XM_023247178.1;refGene_gn=ARHGEF2;refGene_tx=XM_023247179.1;refGene_gn=ARHGEF2;refGene_tx=XM_023247180.1;refGene_gn=ARHGEF2;refGene_tx=XM_023247181.1;refGene_gn=ARHGEF2;refGene_tx=XM_023247182.1;refGene_gn=ARHGEF2;refGene_tx=XM_023247183.1;refGene_gn=ARHGEF2;r

efGene_tx=XM_023247184.1;refGene_gn=ARHGEF2;refGene_tx=XM_023247185.1;
refGene_gn=ARHGEF2;refGene_tx=XM_023247186.1
F1 69875436 69875556
refGene_gn=ARHGEF2;refGene_tx=XM_023247171.1;refGene_gn=ARHGEF2;refGen
e_tx=XM_023247172.1;refGene_gn=ARHGEF2;refGene_tx=XM_023247173.1;refGe
ne_gn=ARHGEF2;refGene_tx=XM_023247175.1;refGene_gn=ARHGEF2;refGene_tx=
XM_023247176.1;refGene_gn=ARHGEF2;refGene_tx=XM_023247177.1;refGene_gn
=ARHGEF2;refGene_tx=XM_023247178.1;refGene_gn=ARHGEF2;refGene_tx=XM_02
3247179.1;refGene_gn=ARHGEF2;refGene_tx=XM_023247180.1;refGene_gn=ARHG
EF2;refGene_tx=XM_023247181.1;refGene_gn=ARHGEF2;refGene_tx=XM_0232471
82.1;refGene_gn=ARHGEF2;refGene_tx=XM_023247183.1;refGene_gn=ARHGEF2;r
efGene_tx=XM_023247184.1;refGene_gn=ARHGEF2;refGene_tx=XM_023247185.1;
refGene_gn=ARHGEF2;refGene_tx=XM_023247186.1
F1 69875664 69875732
refGene_gn=ARHGEF2;refGene_tx=XM_023247171.1;refGene_gn=ARHGEF2;refGen
e_tx=XM_023247172.1;refGene_gn=ARHGEF2;refGene_tx=XM_023247173.1;refGe
ne_gn=ARHGEF2;refGene_tx=XM_023247175.1;refGene_gn=ARHGEF2;refGene_tx=
XM_023247176.1;refGene_gn=ARHGEF2;refGene_tx=XM_023247177.1;refGene_gn
=ARHGEF2;refGene_tx=XM_023247178.1;refGene_gn=ARHGEF2;refGene_tx=XM_02
3247179.1;refGene_gn=ARHGEF2;refGene_tx=XM_023247180.1;refGene_gn=ARHG
EF2;refGene_tx=XM_023247181.1;refGene_gn=ARHGEF2;refGene_tx=XM_0232471
82.1;refGene_gn=ARHGEF2;refGene_tx=XM_023247183.1;refGene_gn=ARHGEF2;r
efGene_tx=XM_023247184.1;refGene_gn=ARHGEF2;refGene_tx=XM_023247185.1;
refGene_gn=ARHGEF2;refGene_tx=XM_023247186.1
F1 69875870 69875906
refGene_gn=ARHGEF2;refGene_tx=XM_023247171.1;refGene_gn=ARHGEF2;refGen
e_tx=XM_023247172.1;refGene_gn=ARHGEF2;refGene_tx=XM_023247173.1;refGe
ne_gn=ARHGEF2;refGene_tx=XM_023247175.1;refGene_gn=ARHGEF2;refGene_tx=
XM_023247176.1;refGene_gn=ARHGEF2;refGene_tx=XM_023247177.1;refGene_gn
=ARHGEF2;refGene_tx=XM_023247178.1;refGene_gn=ARHGEF2;refGene_tx=XM_02
3247179.1;refGene_gn=ARHGEF2;refGene_tx=XM_023247180.1;refGene_gn=ARHG
EF2;refGene_tx=XM_023247181.1;refGene_gn=ARHGEF2;refGene_tx=XM_0232471
82.1;refGene_gn=ARHGEF2;refGene_tx=XM_023247183.1;refGene_gn=ARHGEF2;r
efGene_tx=XM_023247184.1;refGene_gn=ARHGEF2;refGene_tx=XM_023247185.1;
refGene_gn=ARHGEF2;refGene_tx=XM_023247186.1
F1 69876113 69876164
refGene_gn=ARHGEF2;refGene_tx=XM_023247171.1;refGene_gn=ARHGEF2;refGen
e_tx=XM_023247172.1;refGene_gn=ARHGEF2;refGene_tx=XM_023247173.1;refGe
ne_gn=ARHGEF2;refGene_tx=XM_023247175.1;refGene_gn=ARHGEF2;refGene_tx=
XM_023247176.1;refGene_gn=ARHGEF2;refGene_tx=XM_023247177.1;refGene_gn
=ARHGEF2;refGene_tx=XM_023247178.1;refGene_gn=ARHGEF2;refGene_tx=XM_02
3247179.1;refGene_gn=ARHGEF2;refGene_tx=XM_023247180.1;refGene_gn=ARHG
EF2;refGene_tx=XM_023247181.1;refGene_gn=ARHGEF2;refGene_tx=XM_0232471
82.1;refGene_gn=ARHGEF2;refGene_tx=XM_023247183.1;refGene_gn=ARHGEF2;r
efGene_tx=XM_023247184.1;refGene_gn=ARHGEF2;refGene_tx=XM_023247185.1;
refGene_gn=ARHGEF2;refGene_tx=XM_023247186.1
F1 69876305 69876749
refGene_gn=ARHGEF2;refGene_tx=XM_023247171.1;refGene_gn=ARHGEF2;refGen
e_tx=XM_023247172.1;refGene_gn=ARHGEF2;refGene_tx=XM_023247173.1;refGe
ne_gn=ARHGEF2;refGene_tx=XM_023247175.1;refGene_gn=ARHGEF2;refGene_tx=

XM_023247176.1;refGene_gn=ARHGEF2;refGene_tx=XM_023247177.1;refGene_gn=ARHGEF2;refGene_tx=XM_023247178.1;refGene_gn=ARHGEF2;refGene_tx=XM_023247179.1;refGene_gn=ARHGEF2;refGene_tx=XM_023247180.1;refGene_gn=ARHGEF2;refGene_tx=XM_023247181.1;refGene_gn=ARHGEF2;refGene_tx=XM_023247182.1;refGene_gn=ARHGEF2;refGene_tx=XM_023247183.1;refGene_gn=ARHGEF2;refGene_tx=XM_023247184.1;refGene_gn=ARHGEF2;refGene_tx=XM_023247185.1;refGene_gn=ARHGEF2;refGene_tx=XM_023247186.1
F1 69876981 69877165 refGene_gn=ARHGEF2;refGene_tx=XM_023247171.1;refGene_gn=ARHGEF2;refGene_tx=XM_023247172.1;refGene_gn=ARHGEF2;refGene_tx=XM_023247173.1;refGene_gn=ARHGEF2;refGene_tx=XM_023247175.1;refGene_gn=ARHGEF2;refGene_tx=XM_023247176.1;refGene_gn=ARHGEF2;refGene_tx=XM_023247177.1;refGene_gn=ARHGEF2;refGene_tx=XM_023247178.1;refGene_gn=ARHGEF2;refGene_tx=XM_023247179.1;refGene_gn=ARHGEF2;refGene_tx=XM_023247180.1;refGene_gn=ARHGEF2;refGene_tx=XM_023247181.1;refGene_gn=ARHGEF2;refGene_tx=XM_023247182.1;refGene_gn=ARHGEF2;refGene_tx=XM_023247183.1;refGene_gn=ARHGEF2;refGene_tx=XM_023247184.1;refGene_gn=ARHGEF2;refGene_tx=XM_023247185.1;refGene_gn=ARHGEF2;refGene_tx=XM_023247186.1
F1 69878169 69878243 refGene_gn=ARHGEF2;refGene_tx=XM_023247171.1;refGene_gn=ARHGEF2;refGene_tx=XM_023247172.1;refGene_gn=ARHGEF2;refGene_tx=XM_023247173.1;refGene_gn=ARHGEF2;refGene_tx=XM_023247175.1;refGene_gn=ARHGEF2;refGene_tx=XM_023247176.1;refGene_gn=ARHGEF2;refGene_tx=XM_023247177.1;refGene_gn=ARHGEF2;refGene_tx=XM_023247178.1;refGene_gn=ARHGEF2;refGene_tx=XM_023247179.1;refGene_gn=ARHGEF2;refGene_tx=XM_023247180.1;refGene_gn=ARHGEF2;refGene_tx=XM_023247181.1;refGene_gn=ARHGEF2;refGene_tx=XM_023247182.1;refGene_gn=ARHGEF2;refGene_tx=XM_023247183.1;refGene_gn=ARHGEF2;refGene_tx=XM_023247184.1;refGene_gn=ARHGEF2;refGene_tx=XM_023247185.1;refGene_gn=ARHGEF2;refGene_tx=XM_023247186.1
F1 69894945 69894984 refGene_gn=KHDC4;refGene_tx=XM_023247187.1;refGene_gn=KHDC4;refGene_tx=XM_023247188.1
F1 69895184 69895327 refGene_gn=KHDC4;refGene_tx=XM_023247187.1;refGene_gn=KHDC4;refGene_tx=XM_023247188.1
F1 69895408 69895446 refGene_gn=KHDC4;refGene_tx=XM_003999631.4
F1 69895950 69896167 refGene_gn=KHDC4;refGene_tx=XM_003999631.4;refGene_gn=KHDC4;refGene_tx=XM_019822422.2;refGene_gn=KHDC4;refGene_tx=XM_023247187.1;refGene_gn=KHDC4;refGene_tx=XM_023247188.1
F1 69896398 69896492 refGene_gn=KHDC4;refGene_tx=XM_019822422.2
F1 69899674 69899803 refGene_gn=KHDC4;refGene_tx=XM_003999631.4;refGene_gn=KHDC4;refGene_tx=XM_019822422.2;refGene_gn=KHDC4;refGene_tx=XM_023247187.1;refGene_gn=KHDC4;refGene_tx=XM_023247188.1
F1 69900152 69900232 refGene_gn=KHDC4;refGene_tx=XM_003999631.4;refGene_gn=KHDC4;refGene_tx=XM_019822422.2;refGene_gn=KHDC4;refGene_tx=XM_023247187.1;refGene_gn=KHDC4;refGene_tx=XM_023247188.1
F1 69901337 69901390

refGene_gn=KHDC4;refGene_tx=XM_003999631.4;refGene_gn=KHDC4;refGene_tx=XM_019822422.2;refGene_gn=KHDC4;refGene_tx=XM_023247187.1;refGene_gn=KHDC4;refGene_tx=XM_023247188.1
F1 69901661 69901825
refGene_gn=KHDC4;refGene_tx=XM_003999631.4;refGene_gn=KHDC4;refGene_tx=XM_019822422.2;refGene_gn=KHDC4;refGene_tx=XM_023247187.1;refGene_gn=KHDC4;refGene_tx=XM_023247188.1
F1 69902767 69902979
refGene_gn=KHDC4;refGene_tx=XM_003999631.4;refGene_gn=KHDC4;refGene_tx=XM_019822422.2;refGene_gn=KHDC4;refGene_tx=XM_023247187.1;refGene_gn=KHDC4;refGene_tx=XM_023247188.1
F1 69903527 69903588
refGene_gn=KHDC4;refGene_tx=XM_003999631.4;refGene_gn=KHDC4;refGene_tx=XM_019822422.2;refGene_gn=KHDC4;refGene_tx=XM_023247187.1;refGene_gn=KHDC4;refGene_tx=XM_023247188.1
F1 69905087 69905145
refGene_gn=KHDC4;refGene_tx=XM_003999631.4;refGene_gn=KHDC4;refGene_tx=XM_019822422.2;refGene_gn=KHDC4;refGene_tx=XM_023247187.1;refGene_gn=KHDC4;refGene_tx=XM_023247188.1
F1 69905372 69905626
refGene_gn=KHDC4;refGene_tx=XM_003999631.4;refGene_gn=KHDC4;refGene_tx=XM_019822422.2;refGene_gn=KHDC4;refGene_tx=XM_023247187.1;refGene_gn=KHDC4;refGene_tx=XM_023247188.1
F1 69907734 69907914
refGene_gn=KHDC4;refGene_tx=XM_003999631.4;refGene_gn=KHDC4;refGene_tx=XM_019822422.2;refGene_gn=KHDC4;refGene_tx=XM_023247187.1;refGene_gn=KHDC4;refGene_tx=XM_023247188.1
F1 69908800 69908913
refGene_gn=KHDC4;refGene_tx=XM_003999631.4;refGene_gn=KHDC4;refGene_tx=XM_019822422.2;refGene_gn=KHDC4;refGene_tx=XM_023247187.1;refGene_gn=KHDC4;refGene_tx=XM_023247188.1
F1 69909554 69909646
refGene_gn=KHDC4;refGene_tx=XM_003999631.4;refGene_gn=KHDC4;refGene_tx=XM_019822422.2;refGene_gn=KHDC4;refGene_tx=XM_023247187.1;refGene_gn=KHDC4;refGene_tx=XM_023247188.1
F1 69910650 69910850
refGene_gn=KHDC4;refGene_tx=XM_003999631.4;refGene_gn=KHDC4;refGene_tx=XM_019822422.2;refGene_gn=KHDC4;refGene_tx=XM_023247187.1
F1 69913985 69914091
refGene_gn=RIT1;refGene_tx=XM_023247192.1;refGene_gn=RIT1;refGene_tx=XM_023247193.1;refGene_gn=RIT1;refGene_tx=XM_023247194.1;refGene_gn=RIT1;refGene_tx=XM_023247195.1;refGene_gn=RIT1;refGene_tx=XM_023247196.1
F1 69914242 69914299
refGene_gn=RIT1;refGene_tx=XM_023247192.1;refGene_gn=RIT1;refGene_tx=XM_023247193.1;refGene_gn=RIT1;refGene_tx=XM_023247194.1;refGene_gn=RIT1;refGene_tx=XM_023247195.1;refGene_gn=RIT1;refGene_tx=XM_023247196.1;refGene_gn=RIT1;refGene_tx=XM_023247197.1;refGene_gn=RIT1;refGene_tx=XM_023247198.1
F1 69917784 69917858
refGene_gn=RIT1;refGene_tx=XM_023247192.1;refGene_gn=RIT1;refGene_tx=X

M_023247193.1;refGene_gn=RIT1;refGene_tx=XM_023247194.1;refGene_gn=RIT1;refGene_tx=XM_023247195.1;refGene_gn=RIT1;refGene_tx=XM_023247196.1;refGene_gn=RIT1;refGene_tx=XM_023247197.1;refGene_gn=RIT1;refGene_tx=XM_023247198.1
F1 69917983 69917986 refGene_gn=RIT1;refGene_tx=XM_023247192.1;refGene_gn=RIT1;refGene_tx=XM_023247193.1;refGene_gn=RIT1;refGene_tx=XM_023247194.1;refGene_gn=RIT1;refGene_tx=XM_023247195.1;refGene_gn=RIT1;refGene_tx=XM_023247197.1
F1 69918081 69918273 refGene_gn=RIT1;refGene_tx=XM_023247192.1;refGene_gn=RIT1;refGene_tx=XM_023247193.1;refGene_gn=RIT1;refGene_tx=XM_023247194.1;refGene_gn=RIT1;refGene_tx=XM_023247195.1;refGene_gn=RIT1;refGene_tx=XM_023247196.1;refGene_gn=RIT1;refGene_tx=XM_023247197.1;refGene_gn=RIT1;refGene_tx=XM_023247198.1
F1 69919478 69919709 refGene_gn=RIT1;refGene_tx=XM_023247192.1;refGene_gn=RIT1;refGene_tx=XM_023247193.1;refGene_gn=RIT1;refGene_tx=XM_023247194.1;refGene_gn=RIT1;refGene_tx=XM_023247195.1;refGene_gn=RIT1;refGene_tx=XM_023247196.1;refGene_gn=RIT1;refGene_tx=XM_023247197.1;refGene_gn=RIT1;refGene_tx=XM_023247198.1
F1 69925441 69925618 refGene_gn=LOC111558352;refGene_tx=XM_023247524.1
F1 69926823 69927054 refGene_gn=LOC111558352;refGene_tx=XM_023247524.1
F1 69940375 69940686 refGene_gn=SYT11;refGene_tx=XM_011291183.3
F1 69941233 69941357 refGene_gn=SYT11;refGene_tx=XM_011291183.3
F1 69949238 69950065 refGene_gn=SYT11;refGene_tx=XM_011291183.3
F1 69955855 69955889 refGene_gn=SYT11;refGene_tx=XM_011291183.3
F1 69960482 69960987 refGene_gn=LOC101087446;refGene_tx=XM_011291184.3;refGene_gn=LOC101087446;refGene_tx=XM_011291185.3;refGene_gn=LOC101087446;refGene_tx=XM_011291186.3;refGene_gn=LOC101087446;refGene_tx=XM_011291187.3;refGene_gn=LOC101087446;refGene_tx=XM_023247673.1;refGene_gn=LOC101087446;refGene_tx=XM_023247674.1
F1 69966765 69966951 refGene_gn=LOC101087446;refGene_tx=XM_011291184.3;refGene_gn=LOC101087446;refGene_tx=XM_011291185.3;refGene_gn=LOC101087446;refGene_tx=XM_011291186.3;refGene_gn=LOC101087446;refGene_tx=XM_011291187.3;refGene_gn=LOC101087446;refGene_tx=XM_023247673.1
F1 69983115 69983309 refGene_gn=LOC101087446;refGene_tx=XM_011291184.3;refGene_gn=LOC101087446;refGene_tx=XM_011291185.3;refGene_gn=LOC101087446;refGene_tx=XM_011291186.3;refGene_gn=LOC101087446;refGene_tx=XM_011291187.3;refGene_gn=LOC101087446;refGene_tx=XM_023247673.1;refGene_gn=LOC101087446;refGene_tx=XM_023247674.1
F1 69984003 69984078 refGene_gn=LOC101087446;refGene_tx=XM_011291184.3;refGene_gn=LOC101087446;refGene_tx=XM_011291185.3;refGene_gn=LOC101087446;refGene_tx=XM_011291186.3;refGene_gn=LOC101087446;refGene_tx=XM_011291187.3;refGene_gn=LOC101087446;refGene_tx=XM_023247673.1;refGene_gn=LOC101087446;refGen

e_tx=XM_023247674.1
F1 69984260 69984311
refGene_gn=LOC101087446;refGene_tx=XM_011291184.3;refGene_gn=LOC101087446;refGene_tx=XM_011291185.3;refGene_gn=LOC101087446;refGene_tx=XM_011291186.3;refGene_gn=LOC101087446;refGene_tx=XM_011291187.3;refGene_gn=LOC101087446;refGene_tx=XM_023247673.1;refGene_gn=LOC101087446;refGene_tx=XM_023247674.1
F1 69987004 69987055
refGene_gn=LOC101087446;refGene_tx=XM_011291184.3;refGene_gn=LOC101087446;refGene_tx=XM_011291185.3;refGene_gn=LOC101087446;refGene_tx=XM_011291186.3;refGene_gn=LOC101087446;refGene_tx=XM_011291187.3;refGene_gn=LOC101087446;refGene_tx=XM_023247673.1;refGene_gn=LOC101087446;refGene_tx=XM_023247674.1
F1 69987377 69987473
refGene_gn=LOC101087446;refGene_tx=XM_011291184.3;refGene_gn=LOC101087446;refGene_tx=XM_011291185.3;refGene_gn=LOC101087446;refGene_tx=XM_011291186.3;refGene_gn=LOC101087446;refGene_tx=XM_011291187.3;refGene_gn=LOC101087446;refGene_tx=XM_023247673.1;refGene_gn=LOC101087446;refGene_tx=XM_023247674.1
F1 69988446 69988566
refGene_gn=LOC101087446;refGene_tx=XM_011291184.3;refGene_gn=LOC101087446;refGene_tx=XM_011291185.3;refGene_gn=LOC101087446;refGene_tx=XM_011291186.3;refGene_gn=LOC101087446;refGene_tx=XM_011291187.3;refGene_gn=LOC101087446;refGene_tx=XM_023247673.1;refGene_gn=LOC101087446;refGene_tx=XM_023247674.1
F1 69989137 69989308
refGene_gn=LOC101087446;refGene_tx=XM_011291184.3;refGene_gn=LOC101087446;refGene_tx=XM_011291185.3;refGene_gn=LOC101087446;refGene_tx=XM_011291186.3;refGene_gn=LOC101087446;refGene_tx=XM_011291187.3;refGene_gn=LOC101087446;refGene_tx=XM_023247673.1;refGene_gn=LOC101087446;refGene_tx=XM_023247674.1
F1 69992836 69993029
refGene_gn=LOC101087446;refGene_tx=XM_011291184.3;refGene_gn=LOC101087446;refGene_tx=XM_011291185.3;refGene_gn=LOC101087446;refGene_tx=XM_011291186.3;refGene_gn=LOC101087446;refGene_tx=XM_011291187.3;refGene_gn=LOC101087446;refGene_tx=XM_023247673.1;refGene_gn=LOC101087446;refGene_tx=XM_023247674.1
F1 70000988 70001090
refGene_gn=LOC101087446;refGene_tx=XM_011291184.3;refGene_gn=LOC101087446;refGene_tx=XM_011291185.3;refGene_gn=LOC101087446;refGene_tx=XM_011291186.3;refGene_gn=LOC101087446;refGene_tx=XM_011291187.3;refGene_gn=LOC101087446;refGene_tx=XM_023247673.1;refGene_gn=LOC101087446;refGene_tx=XM_023247674.1
F1 70006391 70006432
refGene_gn=LOC101087446;refGene_tx=XM_011291184.3;refGene_gn=LOC101087446;refGene_tx=XM_011291185.3;refGene_gn=LOC101087446;refGene_tx=XM_011291186.3;refGene_gn=LOC101087446;refGene_tx=XM_011291187.3;refGene_gn=LOC101087446;refGene_tx=XM_023247673.1;refGene_gn=LOC101087446;refGene_tx=XM_023247674.1
F1 70006943 70007047

refGene_gn=LOC101087446;refGene_tx=XM_011291184.3;refGene_gn=LOC101087446;refGene_tx=XM_011291185.3;refGene_gn=LOC101087446;refGene_tx=XM_011291186.3;refGene_gn=LOC101087446;refGene_tx=XM_011291187.3;refGene_gn=LOC101087446;refGene_tx=XM_023247673.1;refGene_gn=LOC101087446;refGene_tx=XM_023247674.1
F1 70009354 70009553
refGene_gn=LOC101087446;refGene_tx=XM_011291184.3;refGene_gn=LOC101087446;refGene_tx=XM_011291185.3;refGene_gn=LOC101087446;refGene_tx=XM_011291186.3;refGene_gn=LOC101087446;refGene_tx=XM_011291187.3;refGene_gn=LOC101087446;refGene_tx=XM_023247673.1;refGene_gn=LOC101087446;refGene_tx=XM_023247674.1
F1 70010054 70010141
refGene_gn=LOC101087446;refGene_tx=XM_011291184.3;refGene_gn=LOC101087446;refGene_tx=XM_011291185.3;refGene_gn=LOC101087446;refGene_tx=XM_011291186.3;refGene_gn=LOC101087446;refGene_tx=XM_011291187.3;refGene_gn=LOC101087446;refGene_tx=XM_023247673.1;refGene_gn=LOC101087446;refGene_tx=XM_023247674.1
F1 70011086 70011258
refGene_gn=LOC101087446;refGene_tx=XM_011291184.3;refGene_gn=LOC101087446;refGene_tx=XM_011291185.3;refGene_gn=LOC101087446;refGene_tx=XM_011291186.3;refGene_gn=LOC101087446;refGene_tx=XM_011291187.3;refGene_gn=LOC101087446;refGene_tx=XM_023247673.1;refGene_gn=LOC101087446;refGene_tx=XM_023247674.1
F1 70012654 70012799
refGene_gn=LOC101087446;refGene_tx=XM_011291184.3;refGene_gn=LOC101087446;refGene_tx=XM_011291185.3;refGene_gn=LOC101087446;refGene_tx=XM_011291186.3;refGene_gn=LOC101087446;refGene_tx=XM_011291187.3;refGene_gn=LOC101087446;refGene_tx=XM_023247673.1;refGene_gn=LOC101087446;refGene_tx=XM_023247674.1
F1 70014153 70014304
refGene_gn=LOC101087446;refGene_tx=XM_011291184.3;refGene_gn=LOC101087446;refGene_tx=XM_011291185.3;refGene_gn=LOC101087446;refGene_tx=XM_011291186.3;refGene_gn=LOC101087446;refGene_tx=XM_011291187.3;refGene_gn=LOC101087446;refGene_tx=XM_023247673.1;refGene_gn=LOC101087446;refGene_tx=XM_023247674.1
F1 70018599 70018716
refGene_gn=LOC101087446;refGene_tx=XM_011291184.3;refGene_gn=LOC101087446;refGene_tx=XM_011291185.3;refGene_gn=LOC101087446;refGene_tx=XM_011291186.3;refGene_gn=LOC101087446;refGene_tx=XM_011291187.3;refGene_gn=LOC101087446;refGene_tx=XM_023247673.1;refGene_gn=LOC101087446;refGene_tx=XM_023247674.1
F1 70019089 70020748
refGene_gn=LOC101087446;refGene_tx=XM_011291184.3;refGene_gn=LOC101087446;refGene_tx=XM_011291185.3;refGene_gn=LOC101087446;refGene_tx=XM_011291186.3;refGene_gn=LOC101087446;refGene_tx=XM_011291187.3;refGene_gn=LOC101087446;refGene_tx=XM_023247673.1;refGene_gn=LOC101087446;refGene_tx=XM_023247674.1
F1 70021254 70021510
refGene_gn=LOC101087446;refGene_tx=XM_011291184.3;refGene_gn=LOC101087446;refGene_tx=XM_011291185.3;refGene_gn=LOC101087446;refGene_tx=XM_01

1291186.3;refGene_gn=LOC101087446;refGene_tx=XM_011291187.3;refGene_gn=LOC101087446;refGene_tx=XM_023247673.1;refGene_gn=LOC101087446;refGene_tx=XM_023247674.1
F1 70022071 70022256
refGene_gn=LOC101087446;refGene_tx=XM_011291184.3;refGene_gn=LOC101087446;refGene_tx=XM_011291185.3;refGene_gn=LOC101087446;refGene_tx=XM_011291186.3;refGene_gn=LOC101087446;refGene_tx=XM_011291187.3;refGene_gn=LOC101087446;refGene_tx=XM_023247673.1;refGene_gn=LOC101087446;refGene_tx=XM_023247674.1
F1 70022999 70023197
refGene_gn=LOC101087446;refGene_tx=XM_011291184.3;refGene_gn=LOC101087446;refGene_tx=XM_011291185.3;refGene_gn=LOC101087446;refGene_tx=XM_011291186.3;refGene_gn=LOC101087446;refGene_tx=XM_011291187.3;refGene_gn=LOC101087446;refGene_tx=XM_023247673.1;refGene_gn=LOC101087446;refGene_tx=XM_023247674.1
F1 70024749 70024893
refGene_gn=LOC101087446;refGene_tx=XM_011291184.3;refGene_gn=LOC101087446;refGene_tx=XM_011291185.3;refGene_gn=LOC101087446;refGene_tx=XM_011291186.3;refGene_gn=LOC101087446;refGene_tx=XM_011291187.3;refGene_gn=LOC101087446;refGene_tx=XM_023247673.1;refGene_gn=LOC101087446;refGene_tx=XM_023247674.1
F1 70025382 70025526
refGene_gn=LOC101087446;refGene_tx=XM_011291184.3;refGene_gn=LOC101087446;refGene_tx=XM_011291185.3;refGene_gn=LOC101087446;refGene_tx=XM_011291186.3;refGene_gn=LOC101087446;refGene_tx=XM_011291187.3;refGene_gn=LOC101087446;refGene_tx=XM_023247673.1;refGene_gn=LOC101087446;refGene_tx=XM_023247674.1
F1 70025628 70025748
refGene_gn=LOC101087446;refGene_tx=XM_011291184.3;refGene_gn=LOC101087446;refGene_tx=XM_011291185.3;refGene_gn=LOC101087446;refGene_tx=XM_011291186.3;refGene_gn=LOC101087446;refGene_tx=XM_011291187.3;refGene_gn=LOC101087446;refGene_tx=XM_023247673.1;refGene_gn=LOC101087446;refGene_tx=XM_023247674.1
F1 70026794 70026908
refGene_gn=LOC101087446;refGene_tx=XM_011291184.3;refGene_gn=LOC101087446;refGene_tx=XM_011291185.3;refGene_gn=LOC101087446;refGene_tx=XM_011291186.3;refGene_gn=LOC101087446;refGene_tx=XM_011291187.3;refGene_gn=LOC101087446;refGene_tx=XM_023247673.1;refGene_gn=LOC101087446;refGene_tx=XM_023247674.1
F1 70027880 70028079
refGene_gn=LOC101087446;refGene_tx=XM_011291184.3;refGene_gn=LOC101087446;refGene_tx=XM_011291185.3;refGene_gn=LOC101087446;refGene_tx=XM_011291186.3;refGene_gn=LOC101087446;refGene_tx=XM_011291187.3;refGene_gn=LOC101087446;refGene_tx=XM_023247673.1;refGene_gn=LOC101087446;refGene_tx=XM_023247674.1
F1 70028525 70029144
refGene_gn=LOC101087446;refGene_tx=XM_011291184.3;refGene_gn=LOC101087446;refGene_tx=XM_011291185.3;refGene_gn=LOC101087446;refGene_tx=XM_011291186.3;refGene_gn=LOC101087446;refGene_tx=XM_011291187.3;refGene_gn=LOC101087446;refGene_tx=XM_023247673.1;refGene_gn=LOC101087446;refGen

e_tx=XM_023247674.1
F1 70029239 70029342
refGene_gn=LOC101087446;refGene_tx=XM_011291184.3;refGene_gn=LOC101087446;refGene_tx=XM_011291185.3;refGene_gn=LOC101087446;refGene_tx=XM_011291186.3;refGene_gn=LOC101087446;refGene_tx=XM_011291187.3;refGene_gn=LOC101087446;refGene_tx=XM_023247673.1;refGene_gn=LOC101087446;refGene_tx=XM_023247674.1
F1 70029722 70029872
refGene_gn=LOC101087446;refGene_tx=XM_011291184.3;refGene_gn=LOC101087446;refGene_tx=XM_011291185.3;refGene_gn=LOC101087446;refGene_tx=XM_011291186.3;refGene_gn=LOC101087446;refGene_tx=XM_011291187.3;refGene_gn=LOC101087446;refGene_tx=XM_023247673.1;refGene_gn=LOC101087446;refGene_tx=XM_023247674.1
F1 70030270 70030485 refGene_gn=MSTO1;refGene_tx=XM_023247675.1
F1 70030734 70030847 refGene_gn=MSTO1;refGene_tx=XM_023247675.1
F1 70030920 70031025 refGene_gn=MSTO1;refGene_tx=XM_023247675.1
F1 70031248 70031433 refGene_gn=MSTO1;refGene_tx=XM_023247675.1
F1 70031504 70031636 refGene_gn=MSTO1;refGene_tx=XM_023247675.1
F1 70031859 70032012 refGene_gn=MSTO1;refGene_tx=XM_023247675.1
F1 70032100 70032235 refGene_gn=MSTO1;refGene_tx=XM_023247675.1
F1 70032307 70032425 refGene_gn=MSTO1;refGene_tx=XM_023247675.1
F1 70032596 70032735 refGene_gn=MSTO1;refGene_tx=XM_023247675.1
F1 70032817 70032872 refGene_gn=MSTO1;refGene_tx=XM_023247675.1
F1 70033106 70033182 refGene_gn=MSTO1;refGene_tx=XM_023247675.1
F1 70033279 70033349 refGene_gn=MSTO1;refGene_tx=XM_023247675.1
F1 70033657 70033793 refGene_gn=MSTO1;refGene_tx=XM_023247675.1
F1 70033881 70033965 refGene_gn=MSTO1;refGene_tx=XM_023247675.1
F1 70039029 70039052 refGene_gn=DAP3;refGene_tx=XM_023247622.1
F1 70039550 70039636
refGene_gn=DAP3;refGene_tx=XM_023247616.1;refGene_gn=DAP3;refGene_tx=XM_023247617.1;refGene_gn=DAP3;refGene_tx=XM_023247618.1;refGene_gn=DAP3;refGene_tx=XM_023247620.1;refGene_gn=DAP3;refGene_tx=XM_023247621.1;refGene_gn=DAP3;refGene_tx=XM_023247623.1;refGene_gn=DAP3;refGene_tx=XM_023247624.1
F1 70040400 70040518
refGene_gn=DAP3;refGene_tx=XM_023247616.1;refGene_gn=DAP3;refGene_tx=XM_023247617.1;refGene_gn=DAP3;refGene_tx=XM_023247618.1;refGene_gn=DAP3;refGene_tx=XM_023247620.1;refGene_gn=DAP3;refGene_tx=XM_023247621.1;refGene_gn=DAP3;refGene_tx=XM_023247622.1;refGene_gn=DAP3;refGene_tx=XM_023247623.1;refGene_gn=DAP3;refGene_tx=XM_023247624.1
F1 70042717 70042807
refGene_gn=DAP3;refGene_tx=XM_023247616.1;refGene_gn=DAP3;refGene_tx=XM_023247617.1;refGene_gn=DAP3;refGene_tx=XM_023247618.1;refGene_gn=DAP3;refGene_tx=XM_023247620.1;refGene_gn=DAP3;refGene_tx=XM_023247621.1;refGene_gn=DAP3;refGene_tx=XM_023247622.1;refGene_gn=DAP3;refGene_tx=XM_023247623.1;refGene_gn=DAP3;refGene_tx=XM_023247624.1
F1 70043600 70043660
refGene_gn=DAP3;refGene_tx=XM_023247616.1;refGene_gn=DAP3;refGene_tx=XM_023247617.1;refGene_gn=DAP3;refGene_tx=XM_023247618.1;refGene_gn=DAP3;refGene_tx=XM_023247620.1;refGene_gn=DAP3;refGene_tx=XM_023247621.1;

refGene_gn=DAP3;refGene_tx=XM_023247622.1;refGene_gn=DAP3;refGene_tx=XM_023247623.1;refGene_gn=DAP3;refGene_tx=XM_023247624.1
F1 70044152 70044311
refGene_gn=DAP3;refGene_tx=XM_023247616.1;refGene_gn=DAP3;refGene_tx=XM_023247617.1;refGene_gn=DAP3;refGene_tx=XM_023247618.1;refGene_gn=DAP3;refGene_tx=XM_023247620.1;refGene_gn=DAP3;refGene_tx=XM_023247621.1;refGene_gn=DAP3;refGene_tx=XM_023247622.1;refGene_gn=DAP3;refGene_tx=XM_023247623.1
F1 70044403 70044484
refGene_gn=DAP3;refGene_tx=XM_023247616.1;refGene_gn=DAP3;refGene_tx=XM_023247617.1;refGene_gn=DAP3;refGene_tx=XM_023247618.1;refGene_gn=DAP3;refGene_tx=XM_023247620.1;refGene_gn=DAP3;refGene_tx=XM_023247621.1;refGene_gn=DAP3;refGene_tx=XM_023247622.1;refGene_gn=DAP3;refGene_tx=XM_023247623.1;refGene_gn=DAP3;refGene_tx=XM_023247624.1
F1 70045670 70045801
refGene_gn=DAP3;refGene_tx=XM_023247616.1;refGene_gn=DAP3;refGene_tx=XM_023247617.1;refGene_gn=DAP3;refGene_tx=XM_023247618.1;refGene_gn=DAP3;refGene_tx=XM_023247620.1;refGene_gn=DAP3;refGene_tx=XM_023247621.1;refGene_gn=DAP3;refGene_tx=XM_023247622.1;refGene_gn=DAP3;refGene_tx=XM_023247623.1;refGene_gn=DAP3;refGene_tx=XM_023247624.1
F1 70046776 70046869
refGene_gn=DAP3;refGene_tx=XM_023247616.1;refGene_gn=DAP3;refGene_tx=XM_023247617.1;refGene_gn=DAP3;refGene_tx=XM_023247618.1;refGene_gn=DAP3;refGene_tx=XM_023247620.1;refGene_gn=DAP3;refGene_tx=XM_023247621.1;refGene_gn=DAP3;refGene_tx=XM_023247622.1;refGene_gn=DAP3;refGene_tx=XM_023247623.1;refGene_gn=DAP3;refGene_tx=XM_023247624.1
F1 70047185 70047294
refGene_gn=DAP3;refGene_tx=XM_023247616.1;refGene_gn=DAP3;refGene_tx=XM_023247617.1;refGene_gn=DAP3;refGene_tx=XM_023247618.1;refGene_gn=DAP3;refGene_tx=XM_023247620.1;refGene_gn=DAP3;refGene_tx=XM_023247621.1;refGene_gn=DAP3;refGene_tx=XM_023247622.1;refGene_gn=DAP3;refGene_tx=XM_023247623.1;refGene_gn=DAP3;refGene_tx=XM_023247624.1
F1 70049319 70049421
refGene_gn=DAP3;refGene_tx=XM_023247616.1;refGene_gn=DAP3;refGene_tx=XM_023247617.1;refGene_gn=DAP3;refGene_tx=XM_023247618.1;refGene_gn=DAP3;refGene_tx=XM_023247620.1;refGene_gn=DAP3;refGene_tx=XM_023247621.1;refGene_gn=DAP3;refGene_tx=XM_023247622.1;refGene_gn=DAP3;refGene_tx=XM_023247623.1;refGene_gn=DAP3;refGene_tx=XM_023247624.1
F1 70051277 70051400
refGene_gn=DAP3;refGene_tx=XM_023247616.1;refGene_gn=DAP3;refGene_tx=XM_023247617.1;refGene_gn=DAP3;refGene_tx=XM_023247618.1;refGene_gn=DAP3;refGene_tx=XM_023247620.1;refGene_gn=DAP3;refGene_tx=XM_023247621.1;refGene_gn=DAP3;refGene_tx=XM_023247622.1;refGene_gn=DAP3;refGene_tx=XM_023247623.1;refGene_gn=DAP3;refGene_tx=XM_023247624.1
F1 70054831 70054885
refGene_gn=DAP3;refGene_tx=XM_023247616.1;refGene_gn=DAP3;refGene_tx=XM_023247617.1;refGene_gn=DAP3;refGene_tx=XM_023247618.1;refGene_gn=DAP3;refGene_tx=XM_023247620.1;refGene_gn=DAP3;refGene_tx=XM_023247621.1;refGene_gn=DAP3;refGene_tx=XM_023247622.1;refGene_gn=DAP3;refGene_tx=XM_023247623.1;refGene_gn=DAP3;refGene_tx=XM_023247624.1

F1 70055005 70055084
refGene_gn=DAP3;refGene_tx=XM_023247620.1;refGene_gn=DAP3;refGene_tx=XM_023247621.1
F1 70062364 70062408 refGene_gn=DAP3;refGene_tx=XM_023247620.1
F1 70101341 70101761
refGene_gn=ASH1L;refGene_tx=XM_023247610.1;refGene_gn=ASH1L;refGene_tx=XM_023247611.1
F1 70135654 70140212
refGene_gn=ASH1L;refGene_tx=XM_023247610.1;refGene_gn=ASH1L;refGene_tx=XM_023247611.1
F1 70155910 70156012
refGene_gn=ASH1L;refGene_tx=XM_023247610.1;refGene_gn=ASH1L;refGene_tx=XM_023247611.1
F1 70163517 70164268
refGene_gn=ASH1L;refGene_tx=XM_023247610.1;refGene_gn=ASH1L;refGene_tx=XM_023247611.1
F1 70174794 70174965 refGene_gn=ASH1L;refGene_tx=XM_023247610.1
F1 70187296 70187306 refGene_gn=ASH1L;refGene_tx=XM_023247610.1
F1 70205857 70205952
refGene_gn=ASH1L;refGene_tx=XM_023247610.1;refGene_gn=ASH1L;refGene_tx=XM_023247611.1
F1 70221980 70222085 refGene_gn=ASH1L;refGene_tx=XM_023247611.1
F1 70229398 70229472
refGene_gn=ASH1L;refGene_tx=XM_023247610.1;refGene_gn=ASH1L;refGene_tx=XM_023247611.1
F1 70231065 70231111
refGene_gn=ASH1L;refGene_tx=XM_023247610.1;refGene_gn=ASH1L;refGene_tx=XM_023247611.1
F1 70231227 70231336
refGene_gn=ASH1L;refGene_tx=XM_023247610.1;refGene_gn=ASH1L;refGene_tx=XM_023247611.1
F1 70235326 70235533
refGene_gn=ASH1L;refGene_tx=XM_023247610.1;refGene_gn=ASH1L;refGene_tx=XM_023247611.1
F1 70235661 70235808
refGene_gn=ASH1L;refGene_tx=XM_023247610.1;refGene_gn=ASH1L;refGene_tx=XM_023247611.1
F1 70248726 70248835
refGene_gn=ASH1L;refGene_tx=XM_023247610.1;refGene_gn=ASH1L;refGene_tx=XM_023247611.1
F1 70249242 70249407
refGene_gn=ASH1L;refGene_tx=XM_023247610.1;refGene_gn=ASH1L;refGene_tx=XM_023247611.1
F1 70249565 70249660
refGene_gn=ASH1L;refGene_tx=XM_023247610.1;refGene_gn=ASH1L;refGene_tx=XM_023247611.1
F1 70251526 70251684
refGene_gn=ASH1L;refGene_tx=XM_023247610.1;refGene_gn=ASH1L;refGene_tx=XM_023247611.1
F1 70252774 70252927

refGene_gn=ASH1L;refGene_tx=XM_023247610.1;refGene_gn=ASH1L;refGene_tx=XM_023247611.1
F1 70256040 70256095
refGene_gn=ASH1L;refGene_tx=XM_023247610.1;refGene_gn=ASH1L;refGene_tx=XM_023247611.1
F1 70256177 70256310
refGene_gn=ASH1L;refGene_tx=XM_023247610.1;refGene_gn=ASH1L;refGene_tx=XM_023247611.1
F1 70257614 70257863
refGene_gn=ASH1L;refGene_tx=XM_023247610.1;refGene_gn=ASH1L;refGene_tx=XM_023247611.1
F1 70258736 70258823
refGene_gn=ASH1L;refGene_tx=XM_023247610.1;refGene_gn=ASH1L;refGene_tx=XM_023247611.1
F1 70259629 70259720
refGene_gn=ASH1L;refGene_tx=XM_023247610.1;refGene_gn=ASH1L;refGene_tx=XM_023247611.1
F1 70260721 70260860
refGene_gn=ASH1L;refGene_tx=XM_023247610.1;refGene_gn=ASH1L;refGene_tx=XM_023247611.1
F1 70260957 70261130
refGene_gn=ASH1L;refGene_tx=XM_023247610.1;refGene_gn=ASH1L;refGene_tx=XM_023247611.1
F1 70262280 70262447
refGene_gn=ASH1L;refGene_tx=XM_023247610.1;refGene_gn=ASH1L;refGene_tx=XM_023247611.1
F1 70263611 70263652
refGene_gn=ASH1L;refGene_tx=XM_023247610.1;refGene_gn=ASH1L;refGene_tx=XM_023247611.1
F1 70264578 70264880
refGene_gn=ASH1L;refGene_tx=XM_023247610.1;refGene_gn=ASH1L;refGene_tx=XM_023247611.1
F1 70265213 70265305
refGene_gn=ASH1L;refGene_tx=XM_023247610.1;refGene_gn=ASH1L;refGene_tx=XM_023247611.1
F1 70265625 70265662 refGene_gn=ASH1L;refGene_tx=XM_023247611.1
F1 70274302 70274471
refGene_gn=RUSC1;refGene_tx=XM_023247612.1;refGene_gn=RUSC1;refGene_tx=XM_023247613.1;refGene_gn=RUSC1;refGene_tx=XM_023247615.1
F1 70275660 70275789
refGene_gn=RUSC1;refGene_tx=XM_023247612.1;refGene_gn=RUSC1;refGene_tx=XM_023247613.1;refGene_gn=RUSC1;refGene_tx=XM_023247615.1
F1 70276275 70276825
refGene_gn=RUSC1;refGene_tx=XM_023247612.1;refGene_gn=RUSC1;refGene_tx=XM_023247613.1;refGene_gn=RUSC1;refGene_tx=XM_023247615.1
F1 70277327 70277374
refGene_gn=RUSC1;refGene_tx=XM_023247612.1;refGene_gn=RUSC1;refGene_tx=XM_023247613.1;refGene_gn=RUSC1;refGene_tx=XM_023247615.1
F1 70277525 70277631
refGene_gn=RUSC1;refGene_tx=XM_023247612.1;refGene_gn=RUSC1;refGene_tx

=XM_023247613.1;refGene_gn=RUSC1;refGene_tx=XM_023247615.1
F1 70277705 70277880
refGene_gn=RUSC1;refGene_tx=XM_023247612.1;refGene_gn=RUSC1;refGene_tx
=XM_023247613.1;refGene_gn=RUSC1;refGene_tx=XM_023247615.1
F1 70277984 70278061
refGene_gn=RUSC1;refGene_tx=XM_023247612.1;refGene_gn=RUSC1;refGene_tx
=XM_023247613.1;refGene_gn=RUSC1;refGene_tx=XM_023247615.1
F1 70278206 70278305
refGene_gn=RUSC1;refGene_tx=XM_023247612.1;refGene_gn=RUSC1;refGene_tx
=XM_023247613.1;refGene_gn=RUSC1;refGene_tx=XM_023247615.1
F1 70279554 70280926
refGene_gn=RUSC1;refGene_tx=XM_023247612.1;refGene_gn=RUSC1;refGene_tx
=XM_023247613.1
F1 70282101 70282302
refGene_gn=FDPS;refGene_tx=XM_006943088.4;refGene_gn=FDPS;refGene_tx=X
M_006943089.4;refGene_gn=FDPS;refGene_tx=XM_019822435.2;refGene_gn=FDP
S;refGene_tx=XM_019822436.2
F1 70282551 70282569 refGene_gn=FDPS;refGene_tx=XM_006943087.3
F1 70282763 70282898
refGene_gn=FDPS;refGene_tx=XM_006943087.3;refGene_gn=FDPS;refGene_tx=X
M_006943088.4;refGene_gn=FDPS;refGene_tx=XM_006943089.4;refGene_gn=FDP
S;refGene_tx=XM_019822435.2;refGene_gn=FDPS;refGene_tx=XM_019822436.2
F1 70282977 70283055
refGene_gn=FDPS;refGene_tx=XM_006943087.3;refGene_gn=FDPS;refGene_tx=X
M_006943088.4;refGene_gn=FDPS;refGene_tx=XM_006943089.4;refGene_gn=FDP
S;refGene_tx=XM_019822435.2;refGene_gn=FDPS;refGene_tx=XM_019822436.2
F1 70283673 70283746
refGene_gn=FDPS;refGene_tx=XM_006943087.3;refGene_gn=FDPS;refGene_tx=X
M_006943088.4;refGene_gn=FDPS;refGene_tx=XM_006943089.4;refGene_gn=FDP
S;refGene_tx=XM_019822435.2;refGene_gn=FDPS;refGene_tx=XM_019822436.2
F1 70283836 70283925
refGene_gn=FDPS;refGene_tx=XM_006943087.3;refGene_gn=FDPS;refGene_tx=X
M_006943088.4;refGene_gn=FDPS;refGene_tx=XM_006943089.4;refGene_gn=FDP
S;refGene_tx=XM_019822435.2;refGene_gn=FDPS;refGene_tx=XM_019822436.2
F1 70284372 70284495
refGene_gn=FDPS;refGene_tx=XM_006943087.3;refGene_gn=FDPS;refGene_tx=X
M_006943088.4;refGene_gn=FDPS;refGene_tx=XM_006943089.4;refGene_gn=FDP
S;refGene_tx=XM_019822435.2;refGene_gn=FDPS;refGene_tx=XM_019822436.2
F1 70284625 70284706
refGene_gn=FDPS;refGene_tx=XM_006943087.3;refGene_gn=FDPS;refGene_tx=X
M_006943088.4;refGene_gn=FDPS;refGene_tx=XM_006943089.4;refGene_gn=FDP
S;refGene_tx=XM_019822435.2;refGene_gn=FDPS;refGene_tx=XM_019822436.2
F1 70289244 70289385
refGene_gn=FDPS;refGene_tx=XM_006943087.3;refGene_gn=FDPS;refGene_tx=X
M_006943088.4;refGene_gn=FDPS;refGene_tx=XM_006943089.4;refGene_gn=FDP
S;refGene_tx=XM_019822435.2;refGene_gn=FDPS;refGene_tx=XM_019822436.2
F1 70291385 70291548
refGene_gn=FDPS;refGene_tx=XM_006943087.3;refGene_gn=FDPS;refGene_tx=X
M_006943088.4;refGene_gn=FDPS;refGene_tx=XM_006943089.4;refGene_gn=FDP
S;refGene_tx=XM_019822435.2;refGene_gn=FDPS;refGene_tx=XM_019822436.2

F1 70291623 70291802
refGene_gn=FDPS;refGene_tx=XM_006943087.3;refGene_gn=FDPS;refGene_tx=XM_019822435.2;refGene_gn=FDPS;refGene_tx=XM_019822436.2
F1 70292636 70292713
refGene_gn=FDPS;refGene_tx=XM_006943088.4;refGene_gn=FDPS;refGene_tx=XM_006943089.4
F1 70305381 70305481 refGene_gn=PKLR;refGene_tx=XM_023247387.1
F1 70305861 70305868 refGene_gn=PKLR;refGene_tx=XM_023247388.1
F1 70306184 70306296 refGene_gn=PKLR;refGene_tx=XM_023247386.1
F1 70306432 70306615
refGene_gn=PKLR;refGene_tx=XM_023247386.1;refGene_gn=PKLR;refGene_tx=XM_023247387.1;refGene_gn=PKLR;refGene_tx=XM_023247388.1
F1 70309199 70309291
refGene_gn=PKLR;refGene_tx=XM_023247386.1;refGene_gn=PKLR;refGene_tx=XM_023247387.1;refGene_gn=PKLR;refGene_tx=XM_023247388.1
F1 70309398 70309530
refGene_gn=PKLR;refGene_tx=XM_023247386.1;refGene_gn=PKLR;refGene_tx=XM_023247387.1;refGene_gn=PKLR;refGene_tx=XM_023247388.1
F1 70309619 70309806
refGene_gn=PKLR;refGene_tx=XM_023247386.1;refGene_gn=PKLR;refGene_tx=XM_023247387.1;refGene_gn=PKLR;refGene_tx=XM_023247388.1
F1 70310162 70310433
refGene_gn=PKLR;refGene_tx=XM_023247386.1;refGene_gn=PKLR;refGene_tx=XM_023247387.1;refGene_gn=PKLR;refGene_tx=XM_023247388.1
F1 70310554 70310705
refGene_gn=PKLR;refGene_tx=XM_023247386.1;refGene_gn=PKLR;refGene_tx=XM_023247387.1;refGene_gn=PKLR;refGene_tx=XM_023247388.1
F1 70311001 70311154
refGene_gn=PKLR;refGene_tx=XM_023247386.1;refGene_gn=PKLR;refGene_tx=XM_023247387.1;refGene_gn=PKLR;refGene_tx=XM_023247388.1
F1 70311246 70311413
refGene_gn=PKLR;refGene_tx=XM_023247386.1;refGene_gn=PKLR;refGene_tx=XM_023247387.1;refGene_gn=PKLR;refGene_tx=XM_023247388.1
F1 70312129 70312311
refGene_gn=PKLR;refGene_tx=XM_023247386.1;refGene_gn=PKLR;refGene_tx=XM_023247387.1;refGene_gn=PKLR;refGene_tx=XM_023247388.1
F1 70312798 70312905
refGene_gn=PKLR;refGene_tx=XM_023247386.1;refGene_gn=PKLR;refGene_tx=XM_023247387.1;refGene_gn=PKLR;refGene_tx=XM_023247388.1
F1 70314890 70315591
refGene_gn=HCN3;refGene_tx=XM_011291181.3;refGene_gn=HCN3;refGene_tx=XM_023247382.1;refGene_gn=HCN3;refGene_tx=XM_023247383.1;refGene_gn=HCN3;refGene_tx=XM_023247384.1
F1 70316023 70316188
refGene_gn=HCN3;refGene_tx=XM_011291181.3;refGene_gn=HCN3;refGene_tx=XM_023247382.1;refGene_gn=HCN3;refGene_tx=XM_023247383.1;refGene_gn=HCN3;refGene_tx=XM_023247384.1
F1 70316996 70317237
refGene_gn=HCN3;refGene_tx=XM_011291181.3;refGene_gn=HCN3;refGene_tx=XM_023247382.1;refGene_gn=HCN3;refGene_tx=XM_023247383.1;refGene_gn=HCN

3;refGene_tx=XM_023247384.1
F1 70317650 70317797
refGene_gn=HCN3;refGene_tx=XM_011291181.3;refGene_gn=HCN3;refGene_tx=XM_023247382.1;refGene_gn=HCN3;refGene_tx=XM_023247383.1;refGene_gn=HCN3;refGene_tx=XM_023247384.1
F1 70318209 70318428
refGene_gn=HCN3;refGene_tx=XM_011291181.3;refGene_gn=HCN3;refGene_tx=XM_023247382.1;refGene_gn=HCN3;refGene_tx=XM_023247383.1;refGene_gn=HCN3;refGene_tx=XM_023247384.1
F1 70318799 70318961
refGene_gn=HCN3;refGene_tx=XM_011291181.3;refGene_gn=HCN3;refGene_tx=XM_023247382.1;refGene_gn=HCN3;refGene_tx=XM_023247383.1;refGene_gn=HCN3;refGene_tx=XM_023247384.1
F1 70319867 70320297
refGene_gn=HCN3;refGene_tx=XM_011291181.3;refGene_gn=HCN3;refGene_tx=XM_023247382.1;refGene_gn=HCN3;refGene_tx=XM_023247383.1;refGene_gn=HCN3;refGene_tx=XM_023247384.1
F1 70323233 70323511
refGene_gn=HCN3;refGene_tx=XM_011291181.3;refGene_gn=HCN3;refGene_tx=XM_023247382.1;refGene_gn=HCN3;refGene_tx=XM_023247383.1;refGene_gn=HCN3;refGene_tx=XM_023247384.1
F1 70327919 70328089
refGene_gn=CLK2;refGene_tx=XM_006944267.3;refGene_gn=CLK2;refGene_tx=XM_011291179.2;refGene_gn=CLK2;refGene_tx=XM_011291180.2;refGene_gn=CLK2;refGene_tx=XM_023247389.1;refGene_gn=CLK2;refGene_tx=XM_023247391.1;refGene_gn=CLK2;refGene_tx=XM_023247392.1
F1 70328802 70329031
refGene_gn=CLK2;refGene_tx=XM_006944267.3;refGene_gn=CLK2;refGene_tx=XM_011291179.2;refGene_gn=CLK2;refGene_tx=XM_011291180.2;refGene_gn=CLK2;refGene_tx=XM_023247389.1;refGene_gn=CLK2;refGene_tx=XM_023247391.1;refGene_gn=CLK2;refGene_tx=XM_023247392.1
F1 70329718 70329806
refGene_gn=CLK2;refGene_tx=XM_006944267.3;refGene_gn=CLK2;refGene_tx=XM_011291179.2;refGene_gn=CLK2;refGene_tx=XM_011291180.2;refGene_gn=CLK2;refGene_tx=XM_023247389.1;refGene_gn=CLK2;refGene_tx=XM_023247391.1;refGene_gn=CLK2;refGene_tx=XM_023247392.1
F1 70330154 70330221
refGene_gn=CLK2;refGene_tx=XM_006944267.3;refGene_gn=CLK2;refGene_tx=XM_011291179.2;refGene_gn=CLK2;refGene_tx=XM_011291180.2;refGene_gn=CLK2;refGene_tx=XM_023247389.1;refGene_gn=CLK2;refGene_tx=XM_023247391.1;refGene_gn=CLK2;refGene_tx=XM_023247392.1
F1 70330393 70330510
refGene_gn=CLK2;refGene_tx=XM_006944267.3;refGene_gn=CLK2;refGene_tx=XM_011291179.2;refGene_gn=CLK2;refGene_tx=XM_011291180.2;refGene_gn=CLK2;refGene_tx=XM_023247389.1;refGene_gn=CLK2;refGene_tx=XM_023247391.1;refGene_gn=CLK2;refGene_tx=XM_023247392.1
F1 70331296 70331463
refGene_gn=CLK2;refGene_tx=XM_006944267.3;refGene_gn=CLK2;refGene_tx=XM_011291179.2;refGene_gn=CLK2;refGene_tx=XM_011291180.2;refGene_gn=CLK2;refGene_tx=XM_023247389.1;refGene_gn=CLK2;refGene_tx=XM_023247391.1;

refGene_gn=CLK2;refGene_tx=XM_023247392.1
F1 70332229 70332324
refGene_gn=CLK2;refGene_tx=XM_006944267.3;refGene_gn=CLK2;refGene_tx=XM_011291179.2;refGene_gn=CLK2;refGene_tx=XM_011291180.2;refGene_gn=CLK2;refGene_tx=XM_023247389.1;refGene_gn=CLK2;refGene_tx=XM_023247391.1;refGene_gn=CLK2;refGene_tx=XM_023247392.1
F1 70333023 70333153
refGene_gn=CLK2;refGene_tx=XM_006944267.3;refGene_gn=CLK2;refGene_tx=XM_011291179.2;refGene_gn=CLK2;refGene_tx=XM_011291180.2;refGene_gn=CLK2;refGene_tx=XM_023247389.1;refGene_gn=CLK2;refGene_tx=XM_023247391.1;refGene_gn=CLK2;refGene_tx=XM_023247392.1
F1 70333258 70333341
refGene_gn=CLK2;refGene_tx=XM_006944267.3;refGene_gn=CLK2;refGene_tx=XM_011291179.2;refGene_gn=CLK2;refGene_tx=XM_011291180.2;refGene_gn=CLK2;refGene_tx=XM_023247389.1;refGene_gn=CLK2;refGene_tx=XM_023247391.1;refGene_gn=CLK2;refGene_tx=XM_023247392.1
F1 70333505 70333585
refGene_gn=CLK2;refGene_tx=XM_006944267.3;refGene_gn=CLK2;refGene_tx=XM_011291179.2;refGene_gn=CLK2;refGene_tx=XM_011291180.2;refGene_gn=CLK2;refGene_tx=XM_023247389.1;refGene_gn=CLK2;refGene_tx=XM_023247391.1;refGene_gn=CLK2;refGene_tx=XM_023247392.1
F1 70333771 70333862
refGene_gn=CLK2;refGene_tx=XM_006944267.3;refGene_gn=CLK2;refGene_tx=XM_011291179.2;refGene_gn=CLK2;refGene_tx=XM_011291180.2;refGene_gn=CLK2;refGene_tx=XM_023247389.1;refGene_gn=CLK2;refGene_tx=XM_023247391.1;refGene_gn=CLK2;refGene_tx=XM_023247392.1
F1 70334097 70334127 refGene_gn=CLK2;refGene_tx=XM_023247392.1
F1 70334398 70334584
refGene_gn=CLK2;refGene_tx=XM_006944267.3;refGene_gn=CLK2;refGene_tx=XM_011291179.2;refGene_gn=CLK2;refGene_tx=XM_011291180.2;refGene_gn=CLK2;refGene_tx=XM_023247389.1;refGene_gn=CLK2;refGene_tx=XM_023247391.1
F1 70335494 70335560 refGene_gn=SCAMP3;refGene_tx=XM_023247393.1
F1 70335886 70335964 refGene_gn=SCAMP3;refGene_tx=XM_023247393.1
F1 70336823 70336946 refGene_gn=SCAMP3;refGene_tx=XM_023247393.1
F1 70337040 70337161 refGene_gn=SCAMP3;refGene_tx=XM_023247393.1
F1 70337957 70338086 refGene_gn=SCAMP3;refGene_tx=XM_023247393.1
F1 70338997 70339157 refGene_gn=SCAMP3;refGene_tx=XM_023247393.1
F1 70339261 70339363 refGene_gn=SCAMP3;refGene_tx=XM_023247393.1
F1 70339923 70340041 refGene_gn=SCAMP3;refGene_tx=XM_023247393.1
F1 70340376 70340523 refGene_gn=SCAMP3;refGene_tx=XM_023247393.1
F1 70358772 70358799
refGene_gn=GBA;refGene_tx=XM_003999723.5;refGene_gn=GBA;refGene_tx=XM_019822442.2
F1 70359169 70359257
refGene_gn=GBA;refGene_tx=XM_003999723.5;refGene_gn=GBA;refGene_tx=XM_019822442.2
F1 70359418 70359443 refGene_gn=GBA;refGene_tx=XM_019822444.2
F1 70359504 70359696
refGene_gn=GBA;refGene_tx=XM_003999723.5;refGene_gn=GBA;refGene_tx=XM_019822442.2;refGene_gn=GBA;refGene_tx=XM_019822443.2

F1 70359830 70359977
refGene_gn=GBA;refGene_tx=XM_003999723.5;refGene_gn=GBA;refGene_tx=XM_019822442.2;refGene_gn=GBA;refGene_tx=XM_019822443.2;refGene_gn=GBA;refGene_tx=XM_019822444.2
F1 70360260 70360394
refGene_gn=GBA;refGene_tx=XM_003999723.5;refGene_gn=GBA;refGene_tx=XM_019822442.2;refGene_gn=GBA;refGene_tx=XM_019822443.2;refGene_gn=GBA;refGene_tx=XM_019822444.2
F1 70360620 70360793
refGene_gn=GBA;refGene_tx=XM_003999723.5;refGene_gn=GBA;refGene_tx=XM_019822442.2;refGene_gn=GBA;refGene_tx=XM_019822443.2;refGene_gn=GBA;refGene_tx=XM_019822444.2
F1 70360999 70361237
refGene_gn=GBA;refGene_tx=XM_003999723.5;refGene_gn=GBA;refGene_tx=XM_019822442.2;refGene_gn=GBA;refGene_tx=XM_019822443.2;refGene_gn=GBA;refGene_tx=XM_019822444.2
F1 70361554 70361779
refGene_gn=GBA;refGene_tx=XM_003999723.5;refGene_gn=GBA;refGene_tx=XM_019822442.2;refGene_gn=GBA;refGene_tx=XM_019822443.2;refGene_gn=GBA;refGene_tx=XM_019822444.2
F1 70362146 70362310
refGene_gn=GBA;refGene_tx=XM_003999723.5;refGene_gn=GBA;refGene_tx=XM_019822442.2;refGene_gn=GBA;refGene_tx=XM_019822443.2;refGene_gn=GBA;refGene_tx=XM_019822444.2
F1 70362671 70362788
refGene_gn=GBA;refGene_tx=XM_003999723.5;refGene_gn=GBA;refGene_tx=XM_019822442.2;refGene_gn=GBA;refGene_tx=XM_019822443.2;refGene_gn=GBA;refGene_tx=XM_019822444.2
F1 70362856 70363094
refGene_gn=GBA;refGene_tx=XM_003999723.5;refGene_gn=GBA;refGene_tx=XM_019822442.2;refGene_gn=GBA;refGene_tx=XM_019822443.2;refGene_gn=GBA;refGene_tx=XM_019822444.2
F1 70363662 70363877 refGene_gn=MTX1;refGene_tx=XM_023247394.1
F1 70363975 70364130 refGene_gn=MTX1;refGene_tx=XM_023247394.1
F1 70364490 70364567 refGene_gn=MTX1;refGene_tx=XM_023247394.1
F1 70364685 70364868 refGene_gn=MTX1;refGene_tx=XM_023247394.1
F1 70365008 70365101 refGene_gn=MTX1;refGene_tx=XM_023247394.1
F1 70366536 70366616 refGene_gn=MTX1;refGene_tx=XM_023247394.1
F1 70366735 70366805 refGene_gn=MTX1;refGene_tx=XM_023247394.1
F1 70369221 70369236 refGene_gn=MTX1;refGene_tx=XM_023247394.1
F1 70369555 70369634 refGene_gn=THBS3;refGene_tx=XM_023247381.1
F1 70370670 70370877 refGene_gn=THBS3;refGene_tx=XM_023247381.1
F1 70371427 70371684 refGene_gn=THBS3;refGene_tx=XM_023247381.1
F1 70371784 70371887 refGene_gn=THBS3;refGene_tx=XM_023247381.1
F1 70372693 70372720 refGene_gn=THBS3;refGene_tx=XM_023247381.1
F1 70372922 70373005 refGene_gn=THBS3;refGene_tx=XM_023247381.1
F1 70373106 70373158 refGene_gn=THBS3;refGene_tx=XM_023247381.1
F1 70373283 70373432 refGene_gn=THBS3;refGene_tx=XM_023247381.1
F1 70373607 70373748 refGene_gn=THBS3;refGene_tx=XM_023247381.1
F1 70374151 70374229 refGene_gn=THBS3;refGene_tx=XM_023247381.1

F1 70374592 70374745 refGene_gn=THBS3;refGene_tx=XM_023247381.1
F1 70374965 70375076 refGene_gn=THBS3;refGene_tx=XM_023247381.1
F1 70375178 70375286 refGene_gn=THBS3;refGene_tx=XM_023247381.1
F1 70375543 70375703 refGene_gn=THBS3;refGene_tx=XM_023247381.1
F1 70375956 70376075 refGene_gn=THBS3;refGene_tx=XM_023247381.1
F1 70376188 70376241 refGene_gn=THBS3;refGene_tx=XM_023247381.1
F1 70376556 70376750 refGene_gn=THBS3;refGene_tx=XM_023247381.1
F1 70376868 70377047 refGene_gn=THBS3;refGene_tx=XM_023247381.1
F1 70377170 70377219 refGene_gn=THBS3;refGene_tx=XM_023247381.1
F1 70377406 70377559 refGene_gn=THBS3;refGene_tx=XM_023247381.1
F1 70377561 70377602 refGene_gn=THBS3;refGene_tx=XM_023247381.1
F1 70377840 70378013 refGene_gn=THBS3;refGene_tx=XM_023247381.1
F1 70378951 70379091 refGene_gn=THBS3;refGene_tx=XM_023247381.1
F1 70379185 70379244 refGene_gn=THBS3;refGene_tx=XM_023247381.1
F1 70386144 70386237 refGene_gn=MUC1;refGene_tx=XM_023247487.1
F1 70386320 70386470 refGene_gn=MUC1;refGene_tx=XM_023247487.1
F1 70387358 70387427 refGene_gn=MUC1;refGene_tx=XM_023247487.1
F1 70389302 70389696
refGene_gn=TRIM46;refGene_tx=XM_023247505.1;refGene_gn=TRIM46;refGene_tx=XM_023247506.1;refGene_gn=TRIM46;refGene_tx=XM_023247507.1
F1 70390633 70390931
refGene_gn=TRIM46;refGene_tx=XM_023247505.1;refGene_gn=TRIM46;refGene_tx=XM_023247506.1;refGene_gn=TRIM46;refGene_tx=XM_023247507.1
F1 70391689 70391992
refGene_gn=TRIM46;refGene_tx=XM_023247505.1;refGene_gn=TRIM46;refGene_tx=XM_023247506.1;refGene_gn=TRIM46;refGene_tx=XM_023247507.1;refGene_gn=TRIM46;refGene_tx=XM_023247508.1
F1 70392726 70392764 refGene_gn=TRIM46;refGene_tx=XM_023247508.1
F1 70392966 70393088
refGene_gn=TRIM46;refGene_tx=XM_023247505.1;refGene_gn=TRIM46;refGene_tx=XM_023247506.1;refGene_gn=TRIM46;refGene_tx=XM_023247507.1;refGene_gn=TRIM46;refGene_tx=XM_023247508.1
F1 70393398 70393652
refGene_gn=TRIM46;refGene_tx=XM_023247505.1;refGene_gn=TRIM46;refGene_tx=XM_023247506.1;refGene_gn=TRIM46;refGene_tx=XM_023247507.1;refGene_gn=TRIM46;refGene_tx=XM_023247508.1
F1 70394302 70394398
refGene_gn=TRIM46;refGene_tx=XM_023247505.1;refGene_gn=TRIM46;refGene_tx=XM_023247506.1;refGene_gn=TRIM46;refGene_tx=XM_023247507.1;refGene_gn=TRIM46;refGene_tx=XM_023247508.1
F1 70394506 70394650
refGene_gn=TRIM46;refGene_tx=XM_023247505.1;refGene_gn=TRIM46;refGene_tx=XM_023247506.1;refGene_gn=TRIM46;refGene_tx=XM_023247507.1;refGene_gn=TRIM46;refGene_tx=XM_023247508.1
F1 70395320 70395664
refGene_gn=TRIM46;refGene_tx=XM_023247505.1;refGene_gn=TRIM46;refGene_tx=XM_023247506.1;refGene_gn=TRIM46;refGene_tx=XM_023247507.1;refGene_gn=TRIM46;refGene_tx=XM_023247508.1
F1 70395913 70396175
refGene_gn=TRIM46;refGene_tx=XM_023247505.1;refGene_gn=TRIM46;refGene_

tx=XM_023247506.1;refGene_gn=TRIM46;refGene_tx=XM_023247507.1;refGene_gn=TRIM46;refGene_tx=XM_023247508.1
F1 70396655 70396667 refGene_gn=TRIM46;refGene_tx=XM_023247507.1
F1 70396942 70396966 refGene_gn=TRIM46;refGene_tx=XM_023247506.1
F1 70397557 70398010
refGene_gn=TRIM46;refGene_tx=XM_023247505.1;refGene_gn=TRIM46;refGene_tx=XM_023247508.1
F1 70398287 70398375 refGene_gn=KRTCAP2;refGene_tx=XM_023247509.1
F1 70398631 70398786 refGene_gn=KRTCAP2;refGene_tx=XM_023247509.1
F1 70398873 70398937 refGene_gn=KRTCAP2;refGene_tx=XM_023247509.1
F1 70400319 70400386 refGene_gn=KRTCAP2;refGene_tx=XM_023247509.1
F1 70400566 70400687 refGene_gn=KRTCAP2;refGene_tx=XM_023247509.1
F1 70411562 70411841 refGene_gn=DPM3;refGene_tx=XM_003999727.3
F1 70412627 70412729
refGene_gn=SLC50A1;refGene_tx=XM_003999737.5;refGene_gn=SLC50A1;refGene_tx=XM_006943130.4
F1 70412813 70412933
refGene_gn=SLC50A1;refGene_tx=XM_003999737.5;refGene_gn=SLC50A1;refGene_tx=XM_006943130.4
F1 70413172 70413334
refGene_gn=SLC50A1;refGene_tx=XM_003999737.5;refGene_gn=SLC50A1;refGene_tx=XM_006943130.4
F1 70413944 70414068
refGene_gn=SLC50A1;refGene_tx=XM_003999737.5;refGene_gn=SLC50A1;refGene_tx=XM_006943130.4
F1 70414203 70414281
refGene_gn=SLC50A1;refGene_tx=XM_003999737.5;refGene_gn=SLC50A1;refGene_tx=XM_006943130.4
F1 70414579 70414659
refGene_gn=SLC50A1;refGene_tx=XM_003999737.5;refGene_gn=SLC50A1;refGene_tx=XM_006943130.4
F1 70416243 70416356
refGene_gn=EFNA1;refGene_tx=XM_011291176.2;refGene_gn=EFNA1;refGene_tx=XM_023247549.1
F1 70416529 70416580
refGene_gn=EFNA1;refGene_tx=XM_011291176.2;refGene_gn=EFNA1;refGene_tx=XM_023247549.1
F1 70416749 70416815
refGene_gn=EFNA1;refGene_tx=XM_011291176.2;refGene_gn=EFNA1;refGene_tx=XM_023247549.1
F1 70417772 70418068
refGene_gn=EFNA1;refGene_tx=XM_011291176.2;refGene_gn=EFNA1;refGene_tx=XM_023247549.1
F1 70418373 70418447 refGene_gn=EFNA1;refGene_tx=XM_023247549.1
F1 70421092 70421184 refGene_gn=EFNA1;refGene_tx=XM_011291176.2
F1 70446536 70446753
refGene_gn=LOC105261357;refGene_tx=XM_011291207.2
F1 70447257 70447337
refGene_gn=LOC105261357;refGene_tx=XM_011291207.2
F1 70447970 70449980

refGene_gn=LOC105261357;refGene_tx=XM_011291207.2
F1 70454642 70454773 refGene_gn=EFNA3;refGene_tx=XM_023247475.1
F1 70454984 70455062 refGene_gn=EFNA3;refGene_tx=XM_023247475.1
F1 70455240 70455306 refGene_gn=EFNA3;refGene_tx=XM_023247475.1
F1 70457476 70457790 refGene_gn=EFNA3;refGene_tx=XM_023247475.1
F1 70463683 70463808 refGene_gn=EFNA3;refGene_tx=XM_023247475.1
F1 70474060 70474295
refGene_gn=EFNA4;refGene_tx=XM_023247724.1;refGene_gn=EFNA4;refGene_tx=XM_023247725.1
F1 70474762 70474835
refGene_gn=EFNA4;refGene_tx=XM_023247724.1;refGene_gn=EFNA4;refGene_tx=XM_023247725.1
F1 70474900 70474932 refGene_gn=EFNA4;refGene_tx=XM_023247726.1
F1 70475140 70475436
refGene_gn=EFNA4;refGene_tx=XM_023247724.1;refGene_gn=EFNA4;refGene_tx=XM_023247725.1;refGene_gn=EFNA4;refGene_tx=XM_023247726.1
F1 70477474 70477920
refGene_gn=EFNA4;refGene_tx=XM_023247724.1;refGene_gn=EFNA4;refGene_tx=XM_023247725.1;refGene_gn=EFNA4;refGene_tx=XM_023247726.1
F1 70479003 70479046
refGene_gn=ADAM15;refGene_tx=XM_019822455.2;refGene_gn=ADAM15;refGene_tx=XM_023247695.1;refGene_gn=ADAM15;refGene_tx=XM_023247696.1;refGene_gn=ADAM15;refGene_tx=XM_023247697.1;refGene_gn=ADAM15;refGene_tx=XM_023247698.1;refGene_gn=ADAM15;refGene_tx=XM_023247699.1;refGene_gn=ADAM15;refGene_tx=XM_023247700.1;refGene_gn=ADAM15;refGene_tx=XM_023247701.1;refGene_gn=ADAM15;refGene_tx=XM_023247702.1;refGene_gn=ADAM15;refGene_tx=XM_023247703.1;refGene_gn=ADAM15;refGene_tx=XM_023247705.1;refGene_gn=ADAM15;refGene_tx=XM_023247706.1
F1 70479139 70479380
refGene_gn=ADAM15;refGene_tx=XM_019822455.2;refGene_gn=ADAM15;refGene_tx=XM_023247695.1;refGene_gn=ADAM15;refGene_tx=XM_023247696.1;refGene_gn=ADAM15;refGene_tx=XM_023247697.1;refGene_gn=ADAM15;refGene_tx=XM_023247698.1;refGene_gn=ADAM15;refGene_tx=XM_023247699.1;refGene_gn=ADAM15;refGene_tx=XM_023247700.1;refGene_gn=ADAM15;refGene_tx=XM_023247701.1;refGene_gn=ADAM15;refGene_tx=XM_023247702.1;refGene_gn=ADAM15;refGene_tx=XM_023247703.1;refGene_gn=ADAM15;refGene_tx=XM_023247704.1;refGene_gn=ADAM15;refGene_tx=XM_023247705.1;refGene_gn=ADAM15;refGene_tx=XM_023247706.1;refGene_gn=ADAM15;refGene_tx=XM_023247708.1;refGene_gn=ADAM15;refGene_tx=XM_023247710.1
F1 70479519 70479591
refGene_gn=ADAM15;refGene_tx=XM_019822455.2;refGene_gn=ADAM15;refGene_tx=XM_023247695.1;refGene_gn=ADAM15;refGene_tx=XM_023247696.1;refGene_gn=ADAM15;refGene_tx=XM_023247697.1;refGene_gn=ADAM15;refGene_tx=XM_023247700.1;refGene_gn=ADAM15;refGene_tx=XM_023247703.1;refGene_gn=ADAM15;refGene_tx=XM_023247704.1
F1 70480003 70480078
refGene_gn=ADAM15;refGene_tx=XM_019822455.2;refGene_gn=ADAM15;refGene_tx=XM_023247695.1;refGene_gn=ADAM15;refGene_tx=XM_023247696.1;refGene_gn=ADAM15;refGene_tx=XM_023247697.1;refGene_gn=ADAM15;refGene_tx=XM_023247698.1;refGene_gn=ADAM15;refGene_tx=XM_023247699.1;refGene_gn=ADAM1

5;refGene_tx=XM_023247701.1;refGene_gn=ADAM15;refGene_tx=XM_023247702.1;refGene_gn=ADAM15;refGene_tx=XM_023247704.1;refGene_gn=ADAM15;refGene_tx=XM_023247710.1
F1	70480430	70480523	refGene_gn=ADAM15;refGene_tx=XM_023247707.1
F1	70480687	70480757	refGene_gn=ADAM15;refGene_tx=XM_019822455.2;refGene_gn=ADAM15;refGene_tx=XM_023247695.1;refGene_gn=ADAM15;refGene_tx=XM_023247696.1;refGene_gn=ADAM15;refGene_tx=XM_023247697.1;refGene_gn=ADAM15;refGene_tx=XM_023247698.1;refGene_gn=ADAM15;refGene_tx=XM_023247699.1;refGene_gn=ADAM15;refGene_tx=XM_023247700.1;refGene_gn=ADAM15;refGene_tx=XM_023247701.1;refGene_gn=ADAM15;refGene_tx=XM_023247702.1;refGene_gn=ADAM15;refGene_tx=XM_023247703.1;refGene_gn=ADAM15;refGene_tx=XM_023247704.1;refGene_gn=ADAM15;refGene_tx=XM_023247705.1;refGene_gn=ADAM15;refGene_tx=XM_023247706.1;refGene_gn=ADAM15;refGene_tx=XM_023247707.1;refGene_gn=ADAM15;refGene_tx=XM_023247708.1
F1	70481636	70481775	refGene_gn=ADAM15;refGene_tx=XM_019822455.2;refGene_gn=ADAM15;refGene_tx=XM_023247695.1;refGene_gn=ADAM15;refGene_tx=XM_023247696.1;refGene_gn=ADAM15;refGene_tx=XM_023247697.1;refGene_gn=ADAM15;refGene_tx=XM_023247698.1;refGene_gn=ADAM15;refGene_tx=XM_023247699.1;refGene_gn=ADAM15;refGene_tx=XM_023247700.1;refGene_gn=ADAM15;refGene_tx=XM_023247701.1;refGene_gn=ADAM15;refGene_tx=XM_023247702.1;refGene_gn=ADAM15;refGene_tx=XM_023247703.1;refGene_gn=ADAM15;refGene_tx=XM_023247704.1;refGene_gn=ADAM15;refGene_tx=XM_023247705.1;refGene_gn=ADAM15;refGene_tx=XM_023247706.1;refGene_gn=ADAM15;refGene_tx=XM_023247707.1;refGene_gn=ADAM15;refGene_tx=XM_023247708.1;refGene_gn=ADAM15;refGene_tx=XM_023247710.1
F1	70481938	70482071	refGene_gn=ADAM15;refGene_tx=XM_019822455.2;refGene_gn=ADAM15;refGene_tx=XM_023247695.1;refGene_gn=ADAM15;refGene_tx=XM_023247696.1;refGene_gn=ADAM15;refGene_tx=XM_023247697.1;refGene_gn=ADAM15;refGene_tx=XM_023247698.1;refGene_gn=ADAM15;refGene_tx=XM_023247699.1;refGene_gn=ADAM15;refGene_tx=XM_023247700.1;refGene_gn=ADAM15;refGene_tx=XM_023247701.1;refGene_gn=ADAM15;refGene_tx=XM_023247702.1;refGene_gn=ADAM15;refGene_tx=XM_023247703.1;refGene_gn=ADAM15;refGene_tx=XM_023247704.1;refGene_gn=ADAM15;refGene_tx=XM_023247705.1;refGene_gn=ADAM15;refGene_tx=XM_023247706.1;refGene_gn=ADAM15;refGene_tx=XM_023247707.1;refGene_gn=ADAM15;refGene_tx=XM_023247708.1;refGene_gn=ADAM15;refGene_tx=XM_023247709.1;refGene_gn=ADAM15;refGene_tx=XM_023247710.1
F1	70482430	70482508	refGene_gn=ADAM15;refGene_tx=XM_019822455.2;refGene_gn=ADAM15;refGene_tx=XM_023247695.1;refGene_gn=ADAM15;refGene_tx=XM_023247696.1;refGene_gn=ADAM15;refGene_tx=XM_023247697.1;refGene_gn=ADAM15;refGene_tx=XM_023247698.1;refGene_gn=ADAM15;refGene_tx=XM_023247699.1;refGene_gn=ADAM15;refGene_tx=XM_023247700.1;refGene_gn=ADAM15;refGene_tx=XM_023247701.1;refGene_gn=ADAM15;refGene_tx=XM_023247702.1;refGene_gn=ADAM15;refGene_tx=XM_023247703.1;refGene_gn=ADAM15;refGene_tx=XM_023247704.1;refGene_gn=ADAM15;refGene_tx=XM_023247705.1;refGene_gn=ADAM15;refGene_tx=XM_023247706.1;refGene_gn=ADAM15;refGene_tx=XM_023247707.1;refGene_gn=ADAM15;refGene_tx=XM_023247708.1;refGene_gn=ADAM15;refGene_tx=XM_02324770

9.1;refGene_gn=ADAM15;refGene_tx=XM_023247710.1
F1 70482759 70482961
refGene_gn=ADAM15;refGene_tx=XM_019822455.2;refGene_gn=ADAM15;refGene_tx=XM_023247695.1;refGene_gn=ADAM15;refGene_tx=XM_023247696.1;refGene_gn=ADAM15;refGene_tx=XM_023247697.1;refGene_gn=ADAM15;refGene_tx=XM_023247698.1;refGene_gn=ADAM15;refGene_tx=XM_023247699.1;refGene_gn=ADAM15;refGene_tx=XM_023247700.1;refGene_gn=ADAM15;refGene_tx=XM_023247701.1;refGene_gn=ADAM15;refGene_tx=XM_023247702.1;refGene_gn=ADAM15;refGene_tx=XM_023247703.1;refGene_gn=ADAM15;refGene_tx=XM_023247704.1;refGene_gn=ADAM15;refGene_tx=XM_023247705.1;refGene_gn=ADAM15;refGene_tx=XM_023247706.1;refGene_gn=ADAM15;refGene_tx=XM_023247707.1;refGene_gn=ADAM15;refGene_tx=XM_023247708.1;refGene_gn=ADAM15;refGene_tx=XM_023247709.1;refGene_gn=ADAM15;refGene_tx=XM_023247710.1
F1 70483038 70483343
refGene_gn=ADAM15;refGene_tx=XM_019822455.2;refGene_gn=ADAM15;refGene_tx=XM_023247695.1;refGene_gn=ADAM15;refGene_tx=XM_023247696.1;refGene_gn=ADAM15;refGene_tx=XM_023247697.1;refGene_gn=ADAM15;refGene_tx=XM_023247698.1;refGene_gn=ADAM15;refGene_tx=XM_023247699.1;refGene_gn=ADAM15;refGene_tx=XM_023247700.1;refGene_gn=ADAM15;refGene_tx=XM_023247701.1;refGene_gn=ADAM15;refGene_tx=XM_023247702.1;refGene_gn=ADAM15;refGene_tx=XM_023247703.1;refGene_gn=ADAM15;refGene_tx=XM_023247704.1;refGene_gn=ADAM15;refGene_tx=XM_023247705.1;refGene_gn=ADAM15;refGene_tx=XM_023247706.1;refGene_gn=ADAM15;refGene_tx=XM_023247707.1;refGene_gn=ADAM15;refGene_tx=XM_023247708.1;refGene_gn=ADAM15;refGene_tx=XM_023247709.1;refGene_gn=ADAM15;refGene_tx=XM_023247710.1
F1 70483470 70483563
refGene_gn=ADAM15;refGene_tx=XM_019822455.2;refGene_gn=ADAM15;refGene_tx=XM_023247695.1;refGene_gn=ADAM15;refGene_tx=XM_023247696.1;refGene_gn=ADAM15;refGene_tx=XM_023247697.1;refGene_gn=ADAM15;refGene_tx=XM_023247698.1;refGene_gn=ADAM15;refGene_tx=XM_023247699.1;refGene_gn=ADAM15;refGene_tx=XM_023247700.1;refGene_gn=ADAM15;refGene_tx=XM_023247701.1;refGene_gn=ADAM15;refGene_tx=XM_023247702.1;refGene_gn=ADAM15;refGene_tx=XM_023247703.1;refGene_gn=ADAM15;refGene_tx=XM_023247704.1;refGene_gn=ADAM15;refGene_tx=XM_023247705.1;refGene_gn=ADAM15;refGene_tx=XM_023247706.1;refGene_gn=ADAM15;refGene_tx=XM_023247707.1;refGene_gn=ADAM15;refGene_tx=XM_023247708.1;refGene_gn=ADAM15;refGene_tx=XM_023247709.1;refGene_gn=ADAM15;refGene_tx=XM_023247710.1
F1 70483839 70484014
refGene_gn=ADAM15;refGene_tx=XM_019822455.2;refGene_gn=ADAM15;refGene_tx=XM_023247695.1;refGene_gn=ADAM15;refGene_tx=XM_023247696.1;refGene_gn=ADAM15;refGene_tx=XM_023247697.1;refGene_gn=ADAM15;refGene_tx=XM_023247698.1;refGene_gn=ADAM15;refGene_tx=XM_023247699.1;refGene_gn=ADAM15;refGene_tx=XM_023247700.1;refGene_gn=ADAM15;refGene_tx=XM_023247701.1;refGene_gn=ADAM15;refGene_tx=XM_023247702.1;refGene_gn=ADAM15;refGene_tx=XM_023247703.1;refGene_gn=ADAM15;refGene_tx=XM_023247704.1;refGene_gn=ADAM15;refGene_tx=XM_023247705.1;refGene_gn=ADAM15;refGene_tx=XM_023247706.1;refGene_gn=ADAM15;refGene_tx=XM_023247707.1;refGene_gn=ADAM15;refGene_tx=XM_023247708.1;refGene_gn=ADAM15;refGene_tx=XM_023247709.1;refGene_gn=ADAM15;refGene_tx=XM_023247710.1
F1 70484120 70484269

refGene_gn=ADAM15;refGene_tx=XM_019822455.2;refGene_gn=ADAM15;refGene_tx=XM_023247695.1;refGene_gn=ADAM15;refGene_tx=XM_023247696.1;refGene_gn=ADAM15;refGene_tx=XM_023247697.1;refGene_gn=ADAM15;refGene_tx=XM_023247698.1;refGene_gn=ADAM15;refGene_tx=XM_023247699.1;refGene_gn=ADAM15;refGene_tx=XM_023247700.1;refGene_gn=ADAM15;refGene_tx=XM_023247701.1;refGene_gn=ADAM15;refGene_tx=XM_023247702.1;refGene_gn=ADAM15;refGene_tx=XM_023247703.1;refGene_gn=ADAM15;refGene_tx=XM_023247704.1;refGene_gn=ADAM15;refGene_tx=XM_023247705.1;refGene_gn=ADAM15;refGene_tx=XM_023247706.1;refGene_gn=ADAM15;refGene_tx=XM_023247707.1;refGene_gn=ADAM15;refGene_tx=XM_023247708.1;refGene_gn=ADAM15;refGene_tx=XM_023247709.1;refGene_gn=ADAM15;refGene_tx=XM_023247710.1
F1 70484691 70484776
refGene_gn=ADAM15;refGene_tx=XM_019822455.2;refGene_gn=ADAM15;refGene_tx=XM_023247695.1;refGene_gn=ADAM15;refGene_tx=XM_023247696.1;refGene_gn=ADAM15;refGene_tx=XM_023247697.1;refGene_gn=ADAM15;refGene_tx=XM_023247698.1;refGene_gn=ADAM15;refGene_tx=XM_023247699.1;refGene_gn=ADAM15;refGene_tx=XM_023247700.1;refGene_gn=ADAM15;refGene_tx=XM_023247701.1;refGene_gn=ADAM15;refGene_tx=XM_023247702.1;refGene_gn=ADAM15;refGene_tx=XM_023247703.1;refGene_gn=ADAM15;refGene_tx=XM_023247704.1;refGene_gn=ADAM15;refGene_tx=XM_023247705.1;refGene_gn=ADAM15;refGene_tx=XM_023247706.1;refGene_gn=ADAM15;refGene_tx=XM_023247707.1;refGene_gn=ADAM15;refGene_tx=XM_023247708.1;refGene_gn=ADAM15;refGene_tx=XM_023247709.1;refGene_gn=ADAM15;refGene_tx=XM_023247710.1
F1 70484856 70485026
refGene_gn=ADAM15;refGene_tx=XM_019822455.2;refGene_gn=ADAM15;refGene_tx=XM_023247695.1;refGene_gn=ADAM15;refGene_tx=XM_023247696.1;refGene_gn=ADAM15;refGene_tx=XM_023247697.1;refGene_gn=ADAM15;refGene_tx=XM_023247698.1;refGene_gn=ADAM15;refGene_tx=XM_023247699.1;refGene_gn=ADAM15;refGene_tx=XM_023247700.1;refGene_gn=ADAM15;refGene_tx=XM_023247701.1;refGene_gn=ADAM15;refGene_tx=XM_023247702.1;refGene_gn=ADAM15;refGene_tx=XM_023247703.1;refGene_gn=ADAM15;refGene_tx=XM_023247704.1;refGene_gn=ADAM15;refGene_tx=XM_023247705.1;refGene_gn=ADAM15;refGene_tx=XM_023247706.1;refGene_gn=ADAM15;refGene_tx=XM_023247707.1;refGene_gn=ADAM15;refGene_tx=XM_023247708.1;refGene_gn=ADAM15;refGene_tx=XM_023247709.1;refGene_gn=ADAM15;refGene_tx=XM_023247710.1
F1 70485104 70485173
refGene_gn=ADAM15;refGene_tx=XM_019822455.2;refGene_gn=ADAM15;refGene_tx=XM_023247695.1;refGene_gn=ADAM15;refGene_tx=XM_023247696.1;refGene_gn=ADAM15;refGene_tx=XM_023247697.1;refGene_gn=ADAM15;refGene_tx=XM_023247698.1;refGene_gn=ADAM15;refGene_tx=XM_023247699.1;refGene_gn=ADAM15;refGene_tx=XM_023247700.1;refGene_gn=ADAM15;refGene_tx=XM_023247701.1;refGene_gn=ADAM15;refGene_tx=XM_023247702.1;refGene_gn=ADAM15;refGene_tx=XM_023247703.1;refGene_gn=ADAM15;refGene_tx=XM_023247704.1;refGene_gn=ADAM15;refGene_tx=XM_023247705.1;refGene_gn=ADAM15;refGene_tx=XM_023247706.1;refGene_gn=ADAM15;refGene_tx=XM_023247707.1;refGene_gn=ADAM15;refGene_tx=XM_023247708.1;refGene_gn=ADAM15;refGene_tx=XM_023247709.1;refGene_gn=ADAM15;refGene_tx=XM_023247710.1
F1 70485263 70485329
refGene_gn=ADAM15;refGene_tx=XM_019822455.2;refGene_gn=ADAM15;refGene_tx=XM_023247695.1;refGene_gn=ADAM15;refGene_tx=XM_023247696.1;refGene_

gn=ADAM15;refGene_tx=XM_023247697.1;refGene_gn=ADAM15;refGene_tx=XM_023247698.1;refGene_gn=ADAM15;refGene_tx=XM_023247699.1;refGene_gn=ADAM15;refGene_tx=XM_023247700.1;refGene_gn=ADAM15;refGene_tx=XM_023247701.1;refGene_gn=ADAM15;refGene_tx=XM_023247702.1;refGene_gn=ADAM15;refGene_tx=XM_023247703.1;refGene_gn=ADAM15;refGene_tx=XM_023247704.1;refGene_gn=ADAM15;refGene_tx=XM_023247705.1;refGene_gn=ADAM15;refGene_tx=XM_023247706.1;refGene_gn=ADAM15;refGene_tx=XM_023247707.1;refGene_gn=ADAM15;refGene_tx=XM_023247708.1;refGene_gn=ADAM15;refGene_tx=XM_023247709.1;refGene_gn=ADAM15;refGene_tx=XM_023247710.1
F1 70486461 70486666
refGene_gn=ADAM15;refGene_tx=XM_019822455.2;refGene_gn=ADAM15;refGene_tx=XM_023247695.1;refGene_gn=ADAM15;refGene_tx=XM_023247696.1;refGene_gn=ADAM15;refGene_tx=XM_023247697.1;refGene_gn=ADAM15;refGene_tx=XM_023247698.1;refGene_gn=ADAM15;refGene_tx=XM_023247699.1;refGene_gn=ADAM15;refGene_tx=XM_023247700.1;refGene_gn=ADAM15;refGene_tx=XM_023247701.1;refGene_gn=ADAM15;refGene_tx=XM_023247702.1;refGene_gn=ADAM15;refGene_tx=XM_023247703.1;refGene_gn=ADAM15;refGene_tx=XM_023247704.1;refGene_gn=ADAM15;refGene_tx=XM_023247705.1;refGene_gn=ADAM15;refGene_tx=XM_023247706.1;refGene_gn=ADAM15;refGene_tx=XM_023247707.1;refGene_gn=ADAM15;refGene_tx=XM_023247708.1;refGene_gn=ADAM15;refGene_tx=XM_023247709.1;refGene_gn=ADAM15;refGene_tx=XM_023247710.1
F1 70486755 70486832
refGene_gn=ADAM15;refGene_tx=XM_019822455.2;refGene_gn=ADAM15;refGene_tx=XM_023247695.1;refGene_gn=ADAM15;refGene_tx=XM_023247696.1;refGene_gn=ADAM15;refGene_tx=XM_023247697.1;refGene_gn=ADAM15;refGene_tx=XM_023247698.1;refGene_gn=ADAM15;refGene_tx=XM_023247699.1;refGene_gn=ADAM15;refGene_tx=XM_023247700.1;refGene_gn=ADAM15;refGene_tx=XM_023247701.1;refGene_gn=ADAM15;refGene_tx=XM_023247702.1;refGene_gn=ADAM15;refGene_tx=XM_023247703.1;refGene_gn=ADAM15;refGene_tx=XM_023247704.1;refGene_gn=ADAM15;refGene_tx=XM_023247705.1;refGene_gn=ADAM15;refGene_tx=XM_023247706.1;refGene_gn=ADAM15;refGene_tx=XM_023247707.1;refGene_gn=ADAM15;refGene_tx=XM_023247708.1;refGene_gn=ADAM15;refGene_tx=XM_023247709.1;refGene_gn=ADAM15;refGene_tx=XM_023247710.1
F1 70487002 70487081
refGene_gn=ADAM15;refGene_tx=XM_019822455.2;refGene_gn=ADAM15;refGene_tx=XM_023247695.1;refGene_gn=ADAM15;refGene_tx=XM_023247696.1;refGene_gn=ADAM15;refGene_tx=XM_023247697.1;refGene_gn=ADAM15;refGene_tx=XM_023247698.1;refGene_gn=ADAM15;refGene_tx=XM_023247699.1;refGene_gn=ADAM15;refGene_tx=XM_023247700.1;refGene_gn=ADAM15;refGene_tx=XM_023247701.1;refGene_gn=ADAM15;refGene_tx=XM_023247702.1;refGene_gn=ADAM15;refGene_tx=XM_023247703.1;refGene_gn=ADAM15;refGene_tx=XM_023247704.1;refGene_gn=ADAM15;refGene_tx=XM_023247705.1;refGene_gn=ADAM15;refGene_tx=XM_023247706.1;refGene_gn=ADAM15;refGene_tx=XM_023247707.1;refGene_gn=ADAM15;refGene_tx=XM_023247708.1;refGene_gn=ADAM15;refGene_tx=XM_023247709.1;refGene_gn=ADAM15;refGene_tx=XM_023247710.1
F1 70487453 70487530
refGene_gn=ADAM15;refGene_tx=XM_023247695.1;refGene_gn=ADAM15;refGene_tx=XM_023247696.1;refGene_gn=ADAM15;refGene_tx=XM_023247697.1;refGene_gn=ADAM15;refGene_tx=XM_023247698.1;refGene_gn=ADAM15;refGene_tx=XM_023247699.1;refGene_gn=ADAM15;refGene_tx=XM_023247700.1;refGene_gn=ADAM1

5;refGene_tx=XM_023247701.1;refGene_gn=ADAM15;refGene_tx=XM_023247702.
1;refGene_gn=ADAM15;refGene_tx=XM_023247703.1;refGene_gn=ADAM15;refGen
e_tx=XM_023247704.1;refGene_gn=ADAM15;refGene_tx=XM_023247705.1;refGen
e_gn=ADAM15;refGene_tx=XM_023247706.1;refGene_gn=ADAM15;refGene_tx=XM_
023247707.1;refGene_gn=ADAM15;refGene_tx=XM_023247708.1;refGene_gn=ADA
M15;refGene_tx=XM_023247709.1;refGene_gn=ADAM15;refGene_tx=XM_02324771
0.1
F1 70488133 70488240
refGene_gn=ADAM15;refGene_tx=XM_023247695.1;refGene_gn=ADAM15;refGene_
tx=XM_023247696.1;refGene_gn=ADAM15;refGene_tx=XM_023247697.1;refGene_
gn=ADAM15;refGene_tx=XM_023247698.1;refGene_gn=ADAM15;refGene_tx=XM_02
3247699.1;refGene_gn=ADAM15;refGene_tx=XM_023247700.1;refGene_gn=ADAM1
5;refGene_tx=XM_023247701.1;refGene_gn=ADAM15;refGene_tx=XM_023247702.
1;refGene_gn=ADAM15;refGene_tx=XM_023247703.1;refGene_gn=ADAM15;refGen
e_tx=XM_023247704.1;refGene_gn=ADAM15;refGene_tx=XM_023247705.1;refGen
e_gn=ADAM15;refGene_tx=XM_023247706.1;refGene_gn=ADAM15;refGene_tx=XM_
023247707.1;refGene_gn=ADAM15;refGene_tx=XM_023247708.1;refGene_gn=ADA
M15;refGene_tx=XM_023247709.1;refGene_gn=ADAM15;refGene_tx=XM_02324771
0.1
F1 70489555 70489634
refGene_gn=ADAM15;refGene_tx=XM_023247695.1;refGene_gn=ADAM15;refGene_
tx=XM_023247696.1;refGene_gn=ADAM15;refGene_tx=XM_023247697.1;refGene_
gn=ADAM15;refGene_tx=XM_023247698.1;refGene_gn=ADAM15;refGene_tx=XM_02
3247699.1;refGene_gn=ADAM15;refGene_tx=XM_023247700.1;refGene_gn=ADAM1
5;refGene_tx=XM_023247701.1;refGene_gn=ADAM15;refGene_tx=XM_023247702.
1;refGene_gn=ADAM15;refGene_tx=XM_023247703.1;refGene_gn=ADAM15;refGen
e_tx=XM_023247704.1;refGene_gn=ADAM15;refGene_tx=XM_023247705.1;refGen
e_gn=ADAM15;refGene_tx=XM_023247706.1;refGene_gn=ADAM15;refGene_tx=XM_
023247707.1;refGene_gn=ADAM15;refGene_tx=XM_023247708.1;refGene_gn=ADA
M15;refGene_tx=XM_023247709.1;refGene_gn=ADAM15;refGene_tx=XM_02324771
0.1
F1 70490132 70490384
refGene_gn=DCST1;refGene_tx=XM_023247715.1;refGene_gn=DCST1;refGene_tx
=XM_023247717.1
F1 70491784 70492177
refGene_gn=DCST1;refGene_tx=XM_023247713.1;refGene_gn=DCST1;refGene_tx
=XM_023247714.1;refGene_gn=DCST1;refGene_tx=XM_023247715.1;refGene_gn=
DCST1;refGene_tx=XM_023247716.1;refGene_gn=DCST1;refGene_tx=XM_0232477
17.1;refGene_gn=DCST1;refGene_tx=XM_023247718.1
F1 70492278 70492421
refGene_gn=DCST1;refGene_tx=XM_023247713.1;refGene_gn=DCST1;refGene_tx
=XM_023247714.1;refGene_gn=DCST1;refGene_tx=XM_023247715.1;refGene_gn=
DCST1;refGene_tx=XM_023247716.1;refGene_gn=DCST1;refGene_tx=XM_0232477
17.1;refGene_gn=DCST1;refGene_tx=XM_023247718.1
F1 70492795 70492912
refGene_gn=DCST1;refGene_tx=XM_023247713.1;refGene_gn=DCST1;refGene_tx
=XM_023247714.1;refGene_gn=DCST1;refGene_tx=XM_023247715.1;refGene_gn=
DCST1;refGene_tx=XM_023247716.1;refGene_gn=DCST1;refGene_tx=XM_0232477
17.1;refGene_gn=DCST1;refGene_tx=XM_023247718.1
F1 70493232 70493359

refGene_gn=DCST1;refGene_tx=XM_023247713.1;refGene_gn=DCST1;refGene_tx
=XM_023247714.1;refGene_gn=DCST1;refGene_tx=XM_023247715.1;refGene_gn=
DCST1;refGene_tx=XM_023247716.1;refGene_gn=DCST1;refGene_tx=XM_0232477
17.1;refGene_gn=DCST1;refGene_tx=XM_023247718.1
F1 70493471 70493567
refGene_gn=DCST1;refGene_tx=XM_023247713.1;refGene_gn=DCST1;refGene_tx
=XM_023247714.1;refGene_gn=DCST1;refGene_tx=XM_023247715.1;refGene_gn=
DCST1;refGene_tx=XM_023247716.1;refGene_gn=DCST1;refGene_tx=XM_0232477
17.1;refGene_gn=DCST1;refGene_tx=XM_023247718.1
F1 70493694 70493794
refGene_gn=DCST1;refGene_tx=XM_023247713.1;refGene_gn=DCST1;refGene_tx
=XM_023247714.1;refGene_gn=DCST1;refGene_tx=XM_023247715.1;refGene_gn=
DCST1;refGene_tx=XM_023247716.1;refGene_gn=DCST1;refGene_tx=XM_0232477
17.1;refGene_gn=DCST1;refGene_tx=XM_023247718.1
F1 70495511 70495669
refGene_gn=DCST1;refGene_tx=XM_023247713.1;refGene_gn=DCST1;refGene_tx
=XM_023247714.1;refGene_gn=DCST1;refGene_tx=XM_023247715.1;refGene_gn=
DCST1;refGene_tx=XM_023247716.1;refGene_gn=DCST1;refGene_tx=XM_0232477
17.1;refGene_gn=DCST1;refGene_tx=XM_023247718.1
F1 70495998 70496120
refGene_gn=DCST1;refGene_tx=XM_023247713.1;refGene_gn=DCST1;refGene_tx
=XM_023247714.1;refGene_gn=DCST1;refGene_tx=XM_023247715.1;refGene_gn=
DCST1;refGene_tx=XM_023247716.1;refGene_gn=DCST1;refGene_tx=XM_0232477
17.1;refGene_gn=DCST1;refGene_tx=XM_023247718.1
F1 70496641 70496785
refGene_gn=DCST1;refGene_tx=XM_023247713.1;refGene_gn=DCST1;refGene_tx
=XM_023247714.1;refGene_gn=DCST1;refGene_tx=XM_023247715.1;refGene_gn=
DCST1;refGene_tx=XM_023247716.1;refGene_gn=DCST1;refGene_tx=XM_0232477
17.1;refGene_gn=DCST1;refGene_tx=XM_023247718.1
F1 70496877 70497094
refGene_gn=DCST1;refGene_tx=XM_023247713.1;refGene_gn=DCST1;refGene_tx
=XM_023247714.1;refGene_gn=DCST1;refGene_tx=XM_023247715.1;refGene_gn=
DCST1;refGene_tx=XM_023247716.1;refGene_gn=DCST1;refGene_tx=XM_0232477
17.1;refGene_gn=DCST1;refGene_tx=XM_023247718.1
F1 70497717 70497857
refGene_gn=DCST1;refGene_tx=XM_023247713.1;refGene_gn=DCST1;refGene_tx
=XM_023247714.1;refGene_gn=DCST1;refGene_tx=XM_023247715.1;refGene_gn=
DCST1;refGene_tx=XM_023247716.1;refGene_gn=DCST1;refGene_tx=XM_0232477
17.1;refGene_gn=DCST1;refGene_tx=XM_023247718.1
F1 70498076 70498205
refGene_gn=DCST1;refGene_tx=XM_023247713.1;refGene_gn=DCST1;refGene_tx
=XM_023247714.1;refGene_gn=DCST1;refGene_tx=XM_023247715.1;refGene_gn=
DCST1;refGene_tx=XM_023247716.1;refGene_gn=DCST1;refGene_tx=XM_0232477
17.1;refGene_gn=DCST1;refGene_tx=XM_023247718.1
F1 70502639 70502912
refGene_gn=DCST1;refGene_tx=XM_023247713.1;refGene_gn=DCST1;refGene_tx
=XM_023247714.1;refGene_gn=DCST1;refGene_tx=XM_023247715.1;refGene_gn=
DCST1;refGene_tx=XM_023247716.1;refGene_gn=DCST1;refGene_tx=XM_0232477
17.1
F1 70503243 70503846

refGene_gn=DCST1;refGene_tx=XM_023247713.1;refGene_gn=DCST1;refGene_tx=XM_023247714.1;refGene_gn=DCST1;refGene_tx=XM_023247715.1;refGene_gn=DCST1;refGene_tx=XM_023247716.1;refGene_gn=DCST1;refGene_tx=XM_023247717.1;refGene_gn=DCST2;refGene_tx=XM_023247711.1;refGene_gn=DCST2;refGene_tx=XM_023247712.1
F1 70503995 70504166
refGene_gn=DCST2;refGene_tx=XM_023247711.1;refGene_gn=DCST2;refGene_tx=XM_023247712.1
F1 70504368 70504470
refGene_gn=DCST2;refGene_tx=XM_023247711.1;refGene_gn=DCST2;refGene_tx=XM_023247712.1
F1 70504732 70504930
refGene_gn=DCST2;refGene_tx=XM_023247711.1;refGene_gn=DCST2;refGene_tx=XM_023247712.1
F1 70505114 70505180
refGene_gn=DCST2;refGene_tx=XM_023247711.1;refGene_gn=DCST2;refGene_tx=XM_023247712.1
F1 70505575 70505789
refGene_gn=DCST2;refGene_tx=XM_023247711.1;refGene_gn=DCST2;refGene_tx=XM_023247712.1
F1 70505941 70506099
refGene_gn=DCST2;refGene_tx=XM_023247711.1;refGene_gn=DCST2;refGene_tx=XM_023247712.1
F1 70506838 70507003
refGene_gn=DCST2;refGene_tx=XM_023247711.1;refGene_gn=DCST2;refGene_tx=XM_023247712.1
F1 70507801 70507969
refGene_gn=DCST2;refGene_tx=XM_023247711.1;refGene_gn=DCST2;refGene_tx=XM_023247712.1
F1 70508089 70508190
refGene_gn=DCST2;refGene_tx=XM_023247711.1;refGene_gn=DCST2;refGene_tx=XM_023247712.1
F1 70509582 70509713
refGene_gn=DCST2;refGene_tx=XM_023247711.1;refGene_gn=DCST2;refGene_tx=XM_023247712.1
F1 70510034 70510162
refGene_gn=DCST2;refGene_tx=XM_023247711.1;refGene_gn=DCST2;refGene_tx=XM_023247712.1
F1 70510538 70510632
refGene_gn=DCST2;refGene_tx=XM_023247711.1;refGene_gn=DCST2;refGene_tx=XM_023247712.1
F1 70510733 70510874
refGene_gn=DCST2;refGene_tx=XM_023247711.1;refGene_gn=DCST2;refGene_tx=XM_023247712.1
F1 70513472 70513503
refGene_gn=DCST2;refGene_tx=XM_023247711.1;refGene_gn=DCST2;refGene_tx=XM_023247712.1
F1 70515434 70515900
refGene_gn=ZBTB7B;refGene_tx=XM_023247719.1;refGene_gn=ZBTB7B;refGene_tx=XM_023247720.1;refGene_gn=ZBTB7B;refGene_tx=XM_023247722.1;refGene_

gn=ZBTB7B;refGene_tx=XM_023247723.1
F1	70516247	70517404
refGene_gn=ZBTB7B;refGene_tx=XM_023247719.1;refGene_gn=ZBTB7B;refGene_tx=XM_023247720.1;refGene_gn=ZBTB7B;refGene_tx=XM_023247722.1;refGene_gn=ZBTB7B;refGene_tx=XM_023247723.1
F1	70537029	70537293	refGene_gn=LENEP;refGene_tx=XM_006943131.4
F1	70537783	70537919
refGene_gn=FLAD1;refGene_tx=XM_023247406.1;refGene_gn=FLAD1;refGene_tx=XM_023247407.1
F1	70538024	70538098
refGene_gn=FLAD1;refGene_tx=XM_023247406.1;refGene_gn=FLAD1;refGene_tx=XM_023247407.1
F1	70538578	70538608	refGene_gn=FLAD1;refGene_tx=XM_019822464.2
F1	70538816	70539006
refGene_gn=FLAD1;refGene_tx=XM_019822464.2;refGene_gn=FLAD1;refGene_tx=XM_019822465.2;refGene_gn=FLAD1;refGene_tx=XM_023247406.1;refGene_gn=FLAD1;refGene_tx=XM_023247407.1
F1	70539085	70539266
refGene_gn=FLAD1;refGene_tx=XM_019822464.2;refGene_gn=FLAD1;refGene_tx=XM_019822465.2;refGene_gn=FLAD1;refGene_tx=XM_023247406.1;refGene_gn=FLAD1;refGene_tx=XM_023247407.1
F1	70539449	70539597
refGene_gn=FLAD1;refGene_tx=XM_019822464.2;refGene_gn=FLAD1;refGene_tx=XM_019822465.2;refGene_gn=FLAD1;refGene_tx=XM_023247406.1;refGene_gn=FLAD1;refGene_tx=XM_023247407.1
F1	70539916	70540113	refGene_gn=FLAD1;refGene_tx=XM_006943132.4
F1	70540228	70540973
refGene_gn=FLAD1;refGene_tx=XM_006943132.4;refGene_gn=FLAD1;refGene_tx=XM_019822464.2;refGene_gn=FLAD1;refGene_tx=XM_019822465.2;refGene_gn=FLAD1;refGene_tx=XM_023247406.1;refGene_gn=FLAD1;refGene_tx=XM_023247407.1
F1	70542399	70542480
refGene_gn=FLAD1;refGene_tx=XM_006943132.4;refGene_gn=FLAD1;refGene_tx=XM_019822464.2;refGene_gn=FLAD1;refGene_tx=XM_019822465.2;refGene_gn=FLAD1;refGene_tx=XM_023247406.1;refGene_gn=FLAD1;refGene_tx=XM_023247407.1
F1	70547677	70547730
refGene_gn=CKS1B;refGene_tx=XM_003999730.5;refGene_gn=CKS1B;refGene_tx=XM_006943133.4
F1	70548345	70548473
refGene_gn=CKS1B;refGene_tx=XM_003999730.5;refGene_gn=CKS1B;refGene_tx=XM_006943133.4
F1	70549959	70550105	refGene_gn=CKS1B;refGene_tx=XM_006943133.4
F1	70550307	70550366	refGene_gn=CKS1B;refGene_tx=XM_003999730.5
F1	70553778	70554273	refGene_gn=SHC1;refGene_tx=XM_003999731.5
F1	70554810	70554881	refGene_gn=SHC1;refGene_tx=XM_003999731.5
F1	70555383	70555447	refGene_gn=SHC1;refGene_tx=XM_003999731.5
F1	70555561	70555681	refGene_gn=SHC1;refGene_tx=XM_003999731.5
F1	70555913	70555967	refGene_gn=SHC1;refGene_tx=XM_003999731.5
F1	70556142	70556194	refGene_gn=SHC1;refGene_tx=XM_003999731.5

F1 70556376 70556503 refGene_gn=SHC1;refGene_tx=XM_003999731.5
F1 70557236 70557333 refGene_gn=SHC1;refGene_tx=XM_003999731.5
F1 70557334 70557436 refGene_gn=SHC1;refGene_tx=XM_003999731.5
F1 70557523 70557593 refGene_gn=SHC1;refGene_tx=XM_003999731.5
F1 70557685 70557817 refGene_gn=SHC1;refGene_tx=XM_003999731.5
F1 70557959 70558198 refGene_gn=SHC1;refGene_tx=XM_003999731.5
F1 70558863 70558992 refGene_gn=SHC1;refGene_tx=XM_003999731.5
F1 70560830 70560933 refGene_gn=PYGO2;refGene_tx=XM_023247693.1
F1 70561333 70561383 refGene_gn=PYGO2;refGene_tx=XM_023247693.1
F1 70561915 70562983 refGene_gn=PYGO2;refGene_tx=XM_023247693.1
F1 70568103 70568154
refGene_gn=PBXIP1;refGene_tx=XM_023247687.1;refGene_gn=PBXIP1;refGene_tx=XM_023247688.1;refGene_gn=PBXIP1;refGene_tx=XM_023247689.1;refGene_gn=PBXIP1;refGene_tx=XM_023247690.1;refGene_gn=PBXIP1;refGene_tx=XM_023247691.1;refGene_gn=PBXIP1;refGene_tx=XM_023247692.1
F1 70569582 70569724
refGene_gn=PBXIP1;refGene_tx=XM_023247687.1;refGene_gn=PBXIP1;refGene_tx=XM_023247688.1;refGene_gn=PBXIP1;refGene_tx=XM_023247689.1;refGene_gn=PBXIP1;refGene_tx=XM_023247690.1;refGene_gn=PBXIP1;refGene_tx=XM_023247691.1;refGene_gn=PBXIP1;refGene_tx=XM_023247692.1
F1 70569869 70569934
refGene_gn=PBXIP1;refGene_tx=XM_023247687.1;refGene_gn=PBXIP1;refGene_tx=XM_023247688.1;refGene_gn=PBXIP1;refGene_tx=XM_023247689.1;refGene_gn=PBXIP1;refGene_tx=XM_023247690.1;refGene_gn=PBXIP1;refGene_tx=XM_023247691.1;refGene_gn=PBXIP1;refGene_tx=XM_023247692.1
F1 70570012 70570184
refGene_gn=PBXIP1;refGene_tx=XM_023247687.1;refGene_gn=PBXIP1;refGene_tx=XM_023247688.1;refGene_gn=PBXIP1;refGene_tx=XM_023247689.1;refGene_gn=PBXIP1;refGene_tx=XM_023247690.1;refGene_gn=PBXIP1;refGene_tx=XM_023247691.1;refGene_gn=PBXIP1;refGene_tx=XM_023247692.1
F1 70571496 70571721
refGene_gn=PBXIP1;refGene_tx=XM_023247687.1;refGene_gn=PBXIP1;refGene_tx=XM_023247688.1;refGene_gn=PBXIP1;refGene_tx=XM_023247689.1;refGene_gn=PBXIP1;refGene_tx=XM_023247690.1;refGene_gn=PBXIP1;refGene_tx=XM_023247691.1;refGene_gn=PBXIP1;refGene_tx=XM_023247692.1
F1 70572106 70572162
refGene_gn=PBXIP1;refGene_tx=XM_023247687.1;refGene_gn=PBXIP1;refGene_tx=XM_023247688.1;refGene_gn=PBXIP1;refGene_tx=XM_023247689.1;refGene_gn=PBXIP1;refGene_tx=XM_023247690.1;refGene_gn=PBXIP1;refGene_tx=XM_023247691.1;refGene_gn=PBXIP1;refGene_tx=XM_023247692.1
F1 70572256 70572388
refGene_gn=PBXIP1;refGene_tx=XM_023247687.1;refGene_gn=PBXIP1;refGene_tx=XM_023247688.1;refGene_gn=PBXIP1;refGene_tx=XM_023247689.1;refGene_gn=PBXIP1;refGene_tx=XM_023247690.1;refGene_gn=PBXIP1;refGene_tx=XM_023247691.1;refGene_gn=PBXIP1;refGene_tx=XM_023247692.1
F1 70572927 70574108
refGene_gn=PBXIP1;refGene_tx=XM_023247687.1;refGene_gn=PBXIP1;refGene_tx=XM_023247688.1;refGene_gn=PBXIP1;refGene_tx=XM_023247689.1;refGene_gn=PBXIP1;refGene_tx=XM_023247690.1;refGene_gn=PBXIP1;refGene_tx=XM_023247691.1;refGene_gn=PBXIP1;refGene_tx=XM_023247692.1

F1 70574515 70574561
refGene_gn=PBXIP1;refGene_tx=XM_023247687.1;refGene_gn=PBXIP1;refGene_tx=XM_023247688.1;refGene_gn=PBXIP1;refGene_tx=XM_023247689.1;refGene_gn=PBXIP1;refGene_tx=XM_023247690.1;refGene_gn=PBXIP1;refGene_tx=XM_023247691.1;refGene_gn=PBXIP1;refGene_tx=XM_023247692.1
F1 70574943 70575022
refGene_gn=LOC109495842;refGene_tx=XM_023247476.1
F1 70575090 70575131
refGene_gn=LOC109495842;refGene_tx=XM_023247476.1
F1 70577345 70577682
refGene_gn=LOC109495842;refGene_tx=XM_023247476.1
F1 70578956 70579043
refGene_gn=LOC109495842;refGene_tx=XM_023247476.1
F1 70579834 70579953
refGene_gn=LOC109495842;refGene_tx=XM_023247476.1
F1 70581313 70581544
refGene_gn=LOC109495842;refGene_tx=XM_023247476.1
F1 70585732 70585827 refGene_gn=PMVK;refGene_tx=XM_023247694.1
F1 70587240 70587304 refGene_gn=PMVK;refGene_tx=XM_023247694.1
F1 70589905 70590058
refGene_gn=PMVK;refGene_tx=XM_019822470.2;refGene_gn=PMVK;refGene_tx=XM_023247694.1
F1 70592191 70592321
refGene_gn=PMVK;refGene_tx=XM_019822470.2;refGene_gn=PMVK;refGene_tx=XM_023247694.1
F1 70593111 70593248
refGene_gn=PMVK;refGene_tx=XM_019822470.2;refGene_gn=PMVK;refGene_tx=XM_023247694.1
F1 70593706 70594157
refGene_gn=LOC102902091;refGene_tx=XM_006943136.4;refGene_gn=LOC102902091;refGene_tx=XM_006943137.4
F1 70594269 70594811
refGene_gn=LOC102902091;refGene_tx=XM_006943136.4;refGene_gn=LOC102902091;refGene_tx=XM_006943137.4
F1 70599618 70599658
refGene_gn=LOC109495425;refGene_tx=XM_019821397.2
F1 70600751 70601165
refGene_gn=LOC109495425;refGene_tx=XM_019821397.2
F1 70603438 70603590
refGene_gn=LOC109495425;refGene_tx=XM_019821397.2
F1 70603741 70603897
refGene_gn=LOC109495425;refGene_tx=XM_019821397.2
F1 70630133 70630200
refGene_gn=KCNN3;refGene_tx=XM_011291173.3;refGene_gn=KCNN3;refGene_tx=XM_023248059.1
F1 70630877 70631974
refGene_gn=KCNN3;refGene_tx=XM_011291173.3;refGene_gn=KCNN3;refGene_tx=XM_023248059.1;refGene_gn=KCNN3;refGene_tx=XM_023248060.1
F1 70668378 70668474
refGene_gn=KCNN3;refGene_tx=XM_011291173.3;refGene_gn=KCNN3;refGene_tx

=XM_023248059.1;refGene_gn=KCNN3;refGene_tx=XM_023248060.1;refGene_gn=KCNN3;refGene_tx=XM_023248061.1;refGene_gn=KCNN3;refGene_tx=XM_023248062.1;refGene_gn=KCNN3;refGene_tx=XM_023248063.1
F1 70699192 70699611
refGene_gn=KCNN3;refGene_tx=XM_011291173.3;refGene_gn=KCNN3;refGene_tx=XM_023248059.1;refGene_gn=KCNN3;refGene_tx=XM_023248060.1;refGene_gn=KCNN3;refGene_tx=XM_023248061.1;refGene_gn=KCNN3;refGene_tx=XM_023248062.1;refGene_gn=KCNN3;refGene_tx=XM_023248063.1
F1 70701740 70702763
refGene_gn=LOC111558424;refGene_tx=XM_023248064.1
F1 70725599 70725741
refGene_gn=KCNN3;refGene_tx=XM_011291173.3;refGene_gn=KCNN3;refGene_tx=XM_023248059.1;refGene_gn=KCNN3;refGene_tx=XM_023248060.1;refGene_gn=KCNN3;refGene_tx=XM_023248061.1;refGene_gn=KCNN3;refGene_tx=XM_023248062.1;refGene_gn=KCNN3;refGene_tx=XM_023248063.1
F1 70731632 70731743
refGene_gn=KCNN3;refGene_tx=XM_011291173.3;refGene_gn=KCNN3;refGene_tx=XM_023248059.1;refGene_gn=KCNN3;refGene_tx=XM_023248060.1;refGene_gn=KCNN3;refGene_tx=XM_023248061.1;refGene_gn=KCNN3;refGene_tx=XM_023248062.1;refGene_gn=KCNN3;refGene_tx=XM_023248063.1
F1 70740079 70740207
refGene_gn=KCNN3;refGene_tx=XM_011291173.3;refGene_gn=KCNN3;refGene_tx=XM_023248059.1;refGene_gn=KCNN3;refGene_tx=XM_023248060.1;refGene_gn=KCNN3;refGene_tx=XM_023248061.1;refGene_gn=KCNN3;refGene_tx=XM_023248062.1;refGene_gn=KCNN3;refGene_tx=XM_023248063.1
F1 70742405 70742475
refGene_gn=KCNN3;refGene_tx=XM_011291173.3;refGene_gn=KCNN3;refGene_tx=XM_023248059.1;refGene_gn=KCNN3;refGene_tx=XM_023248060.1;refGene_gn=KCNN3;refGene_tx=XM_023248061.1;refGene_gn=KCNN3;refGene_tx=XM_023248062.1;refGene_gn=KCNN3;refGene_tx=XM_023248063.1
F1 70743433 70743451 refGene_gn=KCNN3;refGene_tx=XM_011291173.3
F1 70745919 70746210
refGene_gn=KCNN3;refGene_tx=XM_023248059.1;refGene_gn=KCNN3;refGene_tx=XM_023248060.1;refGene_gn=KCNN3;refGene_tx=XM_023248061.1;refGene_gn=KCNN3;refGene_tx=XM_023248062.1;refGene_gn=KCNN3;refGene_tx=XM_023248063.1
F1 70800656 70800671
refGene_gn=ADAR;refGene_tx=XM_023247940.1;refGene_gn=ADAR;refGene_tx=XM_023247941.1;refGene_gn=ADAR;refGene_tx=XM_023247942.1;refGene_gn=ADAR;refGene_tx=XM_023247943.1
F1 70804930 70806360
refGene_gn=ADAR;refGene_tx=XM_023247940.1;refGene_gn=ADAR;refGene_tx=XM_023247941.1;refGene_gn=ADAR;refGene_tx=XM_023247942.1;refGene_gn=ADAR;refGene_tx=XM_023247943.1;refGene_gn=ADAR;refGene_tx=XM_023247944.1
F1 70807909 70808093
refGene_gn=ADAR;refGene_tx=XM_023247940.1;refGene_gn=ADAR;refGene_tx=XM_023247941.1;refGene_gn=ADAR;refGene_tx=XM_023247942.1;refGene_gn=ADAR;refGene_tx=XM_023247943.1;refGene_gn=ADAR;refGene_tx=XM_023247944.1
F1 70808721 70808870
refGene_gn=ADAR;refGene_tx=XM_023247940.1;refGene_gn=ADAR;refGene_tx=X

M_023247941.1;refGene_gn=ADAR;refGene_tx=XM_023247942.1;refGene_gn=ADAR;refGene_tx=XM_023247943.1;refGene_gn=ADAR;refGene_tx=XM_023247944.1
F1 70809358 70809503 refGene_gn=ADAR;refGene_tx=XM_023247940.1;refGene_gn=ADAR;refGene_tx=XM_023247941.1;refGene_gn=ADAR;refGene_tx=XM_023247942.1;refGene_gn=ADAR;refGene_tx=XM_023247943.1;refGene_gn=ADAR;refGene_tx=XM_023247944.1
F1 70809596 70809787 refGene_gn=ADAR;refGene_tx=XM_023247940.1;refGene_gn=ADAR;refGene_tx=XM_023247941.1;refGene_gn=ADAR;refGene_tx=XM_023247942.1;refGene_gn=ADAR;refGene_tx=XM_023247943.1;refGene_gn=ADAR;refGene_tx=XM_023247944.1
F1 70817859 70818085 refGene_gn=ADAR;refGene_tx=XM_023247940.1;refGene_gn=ADAR;refGene_tx=XM_023247941.1;refGene_gn=ADAR;refGene_tx=XM_023247942.1;refGene_gn=ADAR;refGene_tx=XM_023247943.1;refGene_gn=ADAR;refGene_tx=XM_023247944.1
F1 70818252 70818424 refGene_gn=ADAR;refGene_tx=XM_023247940.1;refGene_gn=ADAR;refGene_tx=XM_023247941.1;refGene_gn=ADAR;refGene_tx=XM_023247942.1;refGene_gn=ADAR;refGene_tx=XM_023247943.1;refGene_gn=ADAR;refGene_tx=XM_023247944.1
F1 70818564 70818658 refGene_gn=ADAR;refGene_tx=XM_023247940.1;refGene_gn=ADAR;refGene_tx=XM_023247941.1;refGene_gn=ADAR;refGene_tx=XM_023247942.1;refGene_gn=ADAR;refGene_tx=XM_023247943.1;refGene_gn=ADAR;refGene_tx=XM_023247944.1
F1 70818740 70818863 refGene_gn=ADAR;refGene_tx=XM_023247940.1;refGene_gn=ADAR;refGene_tx=XM_023247941.1;refGene_gn=ADAR;refGene_tx=XM_023247942.1;refGene_gn=ADAR;refGene_tx=XM_023247943.1;refGene_gn=ADAR;refGene_tx=XM_023247944.1
F1 70819148 70819282 refGene_gn=ADAR;refGene_tx=XM_023247940.1;refGene_gn=ADAR;refGene_tx=XM_023247941.1;refGene_gn=ADAR;refGene_tx=XM_023247942.1;refGene_gn=ADAR;refGene_tx=XM_023247943.1;refGene_gn=ADAR;refGene_tx=XM_023247944.1
F1 70820703 70820886 refGene_gn=ADAR;refGene_tx=XM_023247940.1;refGene_gn=ADAR;refGene_tx=XM_023247941.1;refGene_gn=ADAR;refGene_tx=XM_023247942.1;refGene_gn=ADAR;refGene_tx=XM_023247943.1;refGene_gn=ADAR;refGene_tx=XM_023247944.1
F1 70821180 70821293 refGene_gn=ADAR;refGene_tx=XM_023247940.1;refGene_gn=ADAR;refGene_tx=XM_023247941.1;refGene_gn=ADAR;refGene_tx=XM_023247942.1;refGene_gn=ADAR;refGene_tx=XM_023247943.1;refGene_gn=ADAR;refGene_tx=XM_023247944.1
F1 70821594 70821722 refGene_gn=ADAR;refGene_tx=XM_023247940.1;refGene_gn=ADAR;refGene_tx=XM_023247941.1;refGene_gn=ADAR;refGene_tx=XM_023247942.1;refGene_gn=ADAR;refGene_tx=XM_023247943.1;refGene_gn=ADAR;refGene_tx=XM_023247944.1
F1 70821858 70822096 refGene_gn=ADAR;refGene_tx=XM_023247940.1;refGene_gn=ADAR;refGene_tx=XM_023247941.1;refGene_gn=ADAR;refGene_tx=XM_023247942.1;refGene_gn=ADAR;refGene_tx=XM_023247943.1;refGene_gn=ADAR;refGene_tx=XM_023247944.1
F1 70828880 70829051 refGene_gn=CHRNB2;refGene_tx=XM_023247557.1
F1 70832095 70832975 refGene_gn=CHRNB2;refGene_tx=XM_023247557.1
F1 70833742 70833852 refGene_gn=CHRNB2;refGene_tx=XM_023247557.1
F1 70834290 70834335 refGene_gn=CHRNB2;refGene_tx=XM_023247557.1

F1 70834515 70834661 refGene_gn=CHRNB2;refGene_tx=XM_023247557.1
F1 70835853 70835917 refGene_gn=CHRNB2;refGene_tx=XM_023247557.1
F1 70843737 70844064 refGene_gn=UBE2Q1;refGene_tx=XM_023247296.1
F1 70846170 70846275 refGene_gn=UBE2Q1;refGene_tx=XM_023247296.1
F1 70846606 70846711 refGene_gn=UBE2Q1;refGene_tx=XM_023247296.1
F1 70847248 70847299 refGene_gn=UBE2Q1;refGene_tx=XM_023247296.1
F1 70848703 70848844 refGene_gn=UBE2Q1;refGene_tx=XM_023247296.1
F1 70850169 70850254 refGene_gn=UBE2Q1;refGene_tx=XM_023247296.1
F1 70850680 70850741 refGene_gn=UBE2Q1;refGene_tx=XM_023247296.1
F1 70850878 70850969 refGene_gn=UBE2Q1;refGene_tx=XM_023247296.1
F1 70851075 70851134 refGene_gn=UBE2Q1;refGene_tx=XM_023247296.1
F1 70851221 70851270 refGene_gn=UBE2Q1;refGene_tx=XM_023247296.1
F1 70851483 70851579 refGene_gn=UBE2Q1;refGene_tx=XM_023247296.1
F1 70851784 70851851 refGene_gn=UBE2Q1;refGene_tx=XM_023247296.1
F1 70852287 70852319 refGene_gn=UBE2Q1;refGene_tx=XM_023247296.1
F1 70896387 70896879 refGene_gn=SHE;refGene_tx=XM_006943205.3
F1 70899072 70899199 refGene_gn=SHE;refGene_tx=XM_006943205.3
F1 70906259 70906553 refGene_gn=SHE;refGene_tx=XM_006943205.3
F1 70907415 70907572 refGene_gn=SHE;refGene_tx=XM_006943205.3
F1 70908267 70908387 refGene_gn=SHE;refGene_tx=XM_006943205.3
F1 70909655 70909842 refGene_gn=SHE;refGene_tx=XM_006943205.3
F1 70922038 70922285
refGene_gn=IL6R;refGene_tx=XM_023248073.1;refGene_gn=IL6R;refGene_tx=XM_023248074.1
F1 70930027 70930118 refGene_gn=IL6R;refGene_tx=XM_023248073.1
F1 70932784 70932854 refGene_gn=IL6R;refGene_tx=XM_023248073.1
F1 70933980 70934027
refGene_gn=IL6R;refGene_tx=XM_023248073.1;refGene_gn=IL6R;refGene_tx=XM_023248074.1
F1 70945905 70946047
refGene_gn=IL6R;refGene_tx=XM_023248073.1;refGene_gn=IL6R;refGene_tx=XM_023248074.1
F1 70946498 70946665
refGene_gn=IL6R;refGene_tx=XM_023248073.1;refGene_gn=IL6R;refGene_tx=XM_023248074.1
F1 70946903 70947085
refGene_gn=IL6R;refGene_tx=XM_023248073.1;refGene_gn=IL6R;refGene_tx=XM_023248074.1
F1 70948699 70948823
refGene_gn=IL6R;refGene_tx=XM_023248073.1;refGene_gn=IL6R;refGene_tx=XM_023248074.1
F1 70949652 70949904
refGene_gn=IL6R;refGene_tx=XM_023248073.1;refGene_gn=IL6R;refGene_tx=XM_023248074.1
F1 70964562 70964647
refGene_gn=IL6R;refGene_tx=XM_023248073.1;refGene_gn=IL6R;refGene_tx=XM_023248074.1
F1 70991813 70992080
refGene_gn=ATP8B2;refGene_tx=XM_011291237.2;refGene_gn=ATP8B2;refGene_tx=XM_023248097.1;refGene_gn=ATP8B2;refGene_tx=XM_023248099.1;refGene_

gn=ATP8B2;refGene_tx=XM_023248100.1
F1 70992263 70992394
refGene_gn=ATP8B2;refGene_tx=XM_011291237.2;refGene_gn=ATP8B2;refGene_tx=XM_023248097.1;refGene_gn=ATP8B2;refGene_tx=XM_023248099.1;refGene_gn=ATP8B2;refGene_tx=XM_023248100.1
F1 70993467 70993724 refGene_gn=ATP8B2;refGene_tx=XM_023248098.1
F1 70994231 70994370
refGene_gn=ATP8B2;refGene_tx=XM_011291237.2;refGene_gn=ATP8B2;refGene_tx=XM_023248097.1;refGene_gn=ATP8B2;refGene_tx=XM_023248098.1;refGene_gn=ATP8B2;refGene_tx=XM_023248099.1;refGene_gn=ATP8B2;refGene_tx=XM_023248100.1
F1 70994480 70994726
refGene_gn=ATP8B2;refGene_tx=XM_011291237.2;refGene_gn=ATP8B2;refGene_tx=XM_023248097.1;refGene_gn=ATP8B2;refGene_tx=XM_023248098.1;refGene_gn=ATP8B2;refGene_tx=XM_023248099.1;refGene_gn=ATP8B2;refGene_tx=XM_023248100.1
F1 70994993 70995077
refGene_gn=ATP8B2;refGene_tx=XM_011291237.2;refGene_gn=ATP8B2;refGene_tx=XM_023248097.1;refGene_gn=ATP8B2;refGene_tx=XM_023248098.1;refGene_gn=ATP8B2;refGene_tx=XM_023248099.1;refGene_gn=ATP8B2;refGene_tx=XM_023248100.1
F1 70995446 70995670
refGene_gn=ATP8B2;refGene_tx=XM_011291237.2;refGene_gn=ATP8B2;refGene_tx=XM_023248097.1;refGene_gn=ATP8B2;refGene_tx=XM_023248098.1;refGene_gn=ATP8B2;refGene_tx=XM_023248099.1;refGene_gn=ATP8B2;refGene_tx=XM_023248100.1
F1 70995778 70995962
refGene_gn=ATP8B2;refGene_tx=XM_011291237.2;refGene_gn=ATP8B2;refGene_tx=XM_023248097.1;refGene_gn=ATP8B2;refGene_tx=XM_023248098.1;refGene_gn=ATP8B2;refGene_tx=XM_023248099.1;refGene_gn=ATP8B2;refGene_tx=XM_023248100.1
F1 70996161 70996306
refGene_gn=ATP8B2;refGene_tx=XM_011291237.2;refGene_gn=ATP8B2;refGene_tx=XM_023248097.1;refGene_gn=ATP8B2;refGene_tx=XM_023248098.1;refGene_gn=ATP8B2;refGene_tx=XM_023248099.1;refGene_gn=ATP8B2;refGene_tx=XM_023248100.1
F1 70996396 70996502
refGene_gn=ATP8B2;refGene_tx=XM_011291237.2;refGene_gn=ATP8B2;refGene_tx=XM_023248097.1;refGene_gn=ATP8B2;refGene_tx=XM_023248098.1;refGene_gn=ATP8B2;refGene_tx=XM_023248099.1;refGene_gn=ATP8B2;refGene_tx=XM_023248100.1
F1 70996601 70996713
refGene_gn=ATP8B2;refGene_tx=XM_011291237.2;refGene_gn=ATP8B2;refGene_tx=XM_023248097.1;refGene_gn=ATP8B2;refGene_tx=XM_023248098.1;refGene_gn=ATP8B2;refGene_tx=XM_023248099.1;refGene_gn=ATP8B2;refGene_tx=XM_023248100.1
F1 70996820 70996985
refGene_gn=ATP8B2;refGene_tx=XM_011291237.2;refGene_gn=ATP8B2;refGene_tx=XM_023248097.1;refGene_gn=ATP8B2;refGene_tx=XM_023248098.1;refGene_gn=ATP8B2;refGene_tx=XM_023248099.1;refGene_gn=ATP8B2;refGene_tx=XM_02

3248100.1
F1 70997068 70997184
refGene_gn=ATP8B2;refGene_tx=XM_011291237.2;refGene_gn=ATP8B2;refGene_tx=XM_023248097.1;refGene_gn=ATP8B2;refGene_tx=XM_023248098.1;refGene_gn=ATP8B2;refGene_tx=XM_023248099.1;refGene_gn=ATP8B2;refGene_tx=XM_023248100.1
F1 70997357 70997546
refGene_gn=ATP8B2;refGene_tx=XM_011291237.2;refGene_gn=ATP8B2;refGene_tx=XM_023248097.1;refGene_gn=ATP8B2;refGene_tx=XM_023248098.1;refGene_gn=ATP8B2;refGene_tx=XM_023248099.1;refGene_gn=ATP8B2;refGene_tx=XM_023248100.1
F1 70997668 70997834
refGene_gn=ATP8B2;refGene_tx=XM_011291237.2;refGene_gn=ATP8B2;refGene_tx=XM_023248097.1;refGene_gn=ATP8B2;refGene_tx=XM_023248098.1;refGene_gn=ATP8B2;refGene_tx=XM_023248099.1;refGene_gn=ATP8B2;refGene_tx=XM_023248100.1
F1 70998112 70998156
refGene_gn=ATP8B2;refGene_tx=XM_011291237.2;refGene_gn=ATP8B2;refGene_tx=XM_023248097.1;refGene_gn=ATP8B2;refGene_tx=XM_023248098.1;refGene_gn=ATP8B2;refGene_tx=XM_023248099.1;refGene_gn=ATP8B2;refGene_tx=XM_023248100.1
F1 70998804 70999013
refGene_gn=ATP8B2;refGene_tx=XM_011291237.2;refGene_gn=ATP8B2;refGene_tx=XM_023248097.1;refGene_gn=ATP8B2;refGene_tx=XM_023248098.1;refGene_gn=ATP8B2;refGene_tx=XM_023248099.1;refGene_gn=ATP8B2;refGene_tx=XM_023248100.1
F1 71001057 71001254
refGene_gn=ATP8B2;refGene_tx=XM_011291237.2;refGene_gn=ATP8B2;refGene_tx=XM_023248097.1;refGene_gn=ATP8B2;refGene_tx=XM_023248098.1;refGene_gn=ATP8B2;refGene_tx=XM_023248099.1;refGene_gn=ATP8B2;refGene_tx=XM_023248100.1
F1 71002742 71002831
refGene_gn=ATP8B2;refGene_tx=XM_011291237.2;refGene_gn=ATP8B2;refGene_tx=XM_023248097.1;refGene_gn=ATP8B2;refGene_tx=XM_023248098.1;refGene_gn=ATP8B2;refGene_tx=XM_023248099.1;refGene_gn=ATP8B2;refGene_tx=XM_023248100.1
F1 71003015 71003174
refGene_gn=ATP8B2;refGene_tx=XM_011291237.2;refGene_gn=ATP8B2;refGene_tx=XM_023248097.1;refGene_gn=ATP8B2;refGene_tx=XM_023248098.1;refGene_gn=ATP8B2;refGene_tx=XM_023248099.1;refGene_gn=ATP8B2;refGene_tx=XM_023248100.1
F1 71004002 71004082
refGene_gn=ATP8B2;refGene_tx=XM_011291237.2;refGene_gn=ATP8B2;refGene_tx=XM_023248097.1;refGene_gn=ATP8B2;refGene_tx=XM_023248098.1;refGene_gn=ATP8B2;refGene_tx=XM_023248099.1;refGene_gn=ATP8B2;refGene_tx=XM_023248100.1
F1 71004311 71004382
refGene_gn=ATP8B2;refGene_tx=XM_011291237.2;refGene_gn=ATP8B2;refGene_tx=XM_023248097.1;refGene_gn=ATP8B2;refGene_tx=XM_023248098.1;refGene_gn=ATP8B2;refGene_tx=XM_023248099.1;refGene_gn=ATP8B2;refGene_tx=XM_02

3248100.1
F1 71004682 71004755
refGene_gn=ATP8B2;refGene_tx=XM_011291237.2;refGene_gn=ATP8B2;refGene_tx=XM_023248097.1;refGene_gn=ATP8B2;refGene_tx=XM_023248098.1;refGene_gn=ATP8B2;refGene_tx=XM_023248099.1;refGene_gn=ATP8B2;refGene_tx=XM_023248100.1
F1 71004853 71004915
refGene_gn=ATP8B2;refGene_tx=XM_011291237.2;refGene_gn=ATP8B2;refGene_tx=XM_023248097.1;refGene_gn=ATP8B2;refGene_tx=XM_023248098.1;refGene_gn=ATP8B2;refGene_tx=XM_023248099.1;refGene_gn=ATP8B2;refGene_tx=XM_023248100.1
F1 71005136 71005235
refGene_gn=ATP8B2;refGene_tx=XM_011291237.2;refGene_gn=ATP8B2;refGene_tx=XM_023248097.1;refGene_gn=ATP8B2;refGene_tx=XM_023248098.1;refGene_gn=ATP8B2;refGene_tx=XM_023248099.1;refGene_gn=ATP8B2;refGene_tx=XM_023248100.1
F1 71005351 71005465
refGene_gn=ATP8B2;refGene_tx=XM_011291237.2;refGene_gn=ATP8B2;refGene_tx=XM_023248097.1;refGene_gn=ATP8B2;refGene_tx=XM_023248098.1;refGene_gn=ATP8B2;refGene_tx=XM_023248099.1;refGene_gn=ATP8B2;refGene_tx=XM_023248100.1
F1 71005788 71005847
refGene_gn=ATP8B2;refGene_tx=XM_011291237.2;refGene_gn=ATP8B2;refGene_tx=XM_023248097.1;refGene_gn=ATP8B2;refGene_tx=XM_023248098.1;refGene_gn=ATP8B2;refGene_tx=XM_023248099.1;refGene_gn=ATP8B2;refGene_tx=XM_023248100.1
F1 71007098 71007186 refGene_gn=ATP8B2;refGene_tx=XM_011291237.2
F1 71007810 71007841
refGene_gn=ATP8B2;refGene_tx=XM_023248097.1;refGene_gn=ATP8B2;refGene_tx=XM_023248098.1;refGene_gn=ATP8B2;refGene_tx=XM_023248099.1;refGene_gn=ATP8B2;refGene_tx=XM_023248100.1
F1 71011663 71011817
refGene_gn=AQP10;refGene_tx=XM_003999749.5;refGene_gn=AQP10;refGene_tx=XM_019822497.2;refGene_gn=AQP10;refGene_tx=XM_019822498.2;refGene_gn=AQP10;refGene_tx=XM_019822500.2
F1 71012247 71012465
refGene_gn=AQP10;refGene_tx=XM_003999749.5;refGene_gn=AQP10;refGene_tx=XM_019822497.2;refGene_gn=AQP10;refGene_tx=XM_019822500.2
F1 71012714 71012833
refGene_gn=AQP10;refGene_tx=XM_003999749.5;refGene_gn=AQP10;refGene_tx=XM_019822497.2;refGene_gn=AQP10;refGene_tx=XM_019822498.2;refGene_gn=AQP10;refGene_tx=XM_019822500.2
F1 71012931 71013069
refGene_gn=AQP10;refGene_tx=XM_003999749.5;refGene_gn=AQP10;refGene_tx=XM_019822497.2;refGene_gn=AQP10;refGene_tx=XM_019822498.2;refGene_gn=AQP10;refGene_tx=XM_019822500.2
F1 71013588 71014210
refGene_gn=AQP10;refGene_tx=XM_003999749.5;refGene_gn=AQP10;refGene_tx=XM_019822497.2;refGene_gn=AQP10;refGene_tx=XM_019822498.2;refGene_gn=AQP10;refGene_tx=XM_019822500.2

F1 71014573 71014720
refGene_gn=AQP10;refGene_tx=XM_003999749.5;refGene_gn=AQP10;refGene_tx=XM_019822497.2;refGene_gn=AQP10;refGene_tx=XM_019822498.2
F1 71018260 71018346
refGene_gn=HAX1;refGene_tx=XM_011291240.3;refGene_gn=HAX1;refGene_tx=XM_019822501.2
F1 71018498 71018589
refGene_gn=HAX1;refGene_tx=XM_011291240.3;refGene_gn=HAX1;refGene_tx=XM_019822501.2
F1 71018717 71018824
refGene_gn=HAX1;refGene_tx=XM_011291240.3;refGene_gn=HAX1;refGene_tx=XM_019822501.2
F1 71018936 71019000
refGene_gn=HAX1;refGene_tx=XM_011291240.3;refGene_gn=HAX1;refGene_tx=XM_019822501.2
F1 71019469 71019663
refGene_gn=HAX1;refGene_tx=XM_011291240.3;refGene_gn=HAX1;refGene_tx=XM_019822501.2
F1 71019841 71020104
refGene_gn=HAX1;refGene_tx=XM_011291240.3;refGene_gn=HAX1;refGene_tx=XM_019822501.2
F1 71020564 71020617
refGene_gn=HAX1;refGene_tx=XM_011291240.3;refGene_gn=HAX1;refGene_tx=XM_019822501.2
F1 71021651 71021690
refGene_gn=UBAP2L;refGene_tx=XM_011291249.3;refGene_gn=UBAP2L;refGene_tx=XM_019822489.2;refGene_gn=UBAP2L;refGene_tx=XM_019822491.2;refGene_gn=UBAP2L;refGene_tx=XM_019822492.2;refGene_gn=UBAP2L;refGene_tx=XM_019822493.1;refGene_gn=UBAP2L;refGene_tx=XM_019822496.2
F1 71022263 71022359
refGene_gn=UBAP2L;refGene_tx=XM_011291248.3;refGene_gn=UBAP2L;refGene_tx=XM_019822486.2;refGene_gn=UBAP2L;refGene_tx=XM_019822487.2;refGene_gn=UBAP2L;refGene_tx=XM_019822488.2;refGene_gn=UBAP2L;refGene_tx=XM_019822490.2
F1 71022797 71023244
refGene_gn=UBAP2L;refGene_tx=XM_011291242.3;refGene_gn=UBAP2L;refGene_tx=XM_019822480.2;refGene_gn=UBAP2L;refGene_tx=XM_019822481.2;refGene_gn=UBAP2L;refGene_tx=XM_019822482.2;refGene_gn=UBAP2L;refGene_tx=XM_019822483.2;refGene_gn=UBAP2L;refGene_tx=XM_019822484.2;refGene_gn=UBAP2L;refGene_tx=XM_019822485.2;refGene_gn=UBAP2L;refGene_tx=XM_019822486.2;refGene_gn=UBAP2L;refGene_tx=XM_019822489.2;refGene_gn=UBAP2L;refGene_tx=XM_019822496.2
F1 71023649 71023847
refGene_gn=UBAP2L;refGene_tx=XM_011291242.3;refGene_gn=UBAP2L;refGene_tx=XM_011291248.3;refGene_gn=UBAP2L;refGene_tx=XM_011291249.3;refGene_gn=UBAP2L;refGene_tx=XM_019822480.2;refGene_gn=UBAP2L;refGene_tx=XM_019822481.2;refGene_gn=UBAP2L;refGene_tx=XM_019822482.2;refGene_gn=UBAP2L;refGene_tx=XM_019822483.2;refGene_gn=UBAP2L;refGene_tx=XM_019822484.2;refGene_gn=UBAP2L;refGene_tx=XM_019822485.2;refGene_gn=UBAP2L;refGene_tx=XM_019822486.2;refGene_gn=UBAP2L;refGene_tx=XM_019822487.2;refGen

e_gn=UBAP2L;refGene_tx=XM_019822488.2;refGene_gn=UBAP2L;refGene_tx=XM_019822489.2;refGene_gn=UBAP2L;refGene_tx=XM_019822490.2;refGene_gn=UBAP2L;refGene_tx=XM_019822491.2;refGene_gn=UBAP2L;refGene_tx=XM_019822492.2;refGene_gn=UBAP2L;refGene_tx=XM_019822493.1;refGene_gn=UBAP2L;refGene_tx=XM_019822496.2
F1 71024575 71024643
refGene_gn=UBAP2L;refGene_tx=XM_011291242.3;refGene_gn=UBAP2L;refGene_tx=XM_011291248.3;refGene_gn=UBAP2L;refGene_tx=XM_011291249.3;refGene_gn=UBAP2L;refGene_tx=XM_019822480.2;refGene_gn=UBAP2L;refGene_tx=XM_019822481.2;refGene_gn=UBAP2L;refGene_tx=XM_019822482.2;refGene_gn=UBAP2L;refGene_tx=XM_019822483.2;refGene_gn=UBAP2L;refGene_tx=XM_019822484.2;refGene_gn=UBAP2L;refGene_tx=XM_019822485.2;refGene_gn=UBAP2L;refGene_tx=XM_019822486.2;refGene_gn=UBAP2L;refGene_tx=XM_019822487.2;refGene_gn=UBAP2L;refGene_tx=XM_019822488.2;refGene_gn=UBAP2L;refGene_tx=XM_019822489.2;refGene_gn=UBAP2L;refGene_tx=XM_019822490.2;refGene_gn=UBAP2L;refGene_tx=XM_019822491.2;refGene_gn=UBAP2L;refGene_tx=XM_019822492.2;refGene_gn=UBAP2L;refGene_tx=XM_019822493.1;refGene_gn=UBAP2L;refGene_tx=XM_019822496.2
F1 71027798 71027848
refGene_gn=UBAP2L;refGene_tx=XM_011291251.3;refGene_gn=UBAP2L;refGene_tx=XM_019822494.1;refGene_gn=UBAP2L;refGene_tx=XM_019822495.2
F1 71029176 71029282
refGene_gn=UBAP2L;refGene_tx=XM_011291242.3;refGene_gn=UBAP2L;refGene_tx=XM_011291248.3;refGene_gn=UBAP2L;refGene_tx=XM_011291249.3;refGene_gn=UBAP2L;refGene_tx=XM_011291251.3;refGene_gn=UBAP2L;refGene_tx=XM_019822480.2;refGene_gn=UBAP2L;refGene_tx=XM_019822481.2;refGene_gn=UBAP2L;refGene_tx=XM_019822482.2;refGene_gn=UBAP2L;refGene_tx=XM_019822483.2;refGene_gn=UBAP2L;refGene_tx=XM_019822484.2;refGene_gn=UBAP2L;refGene_tx=XM_019822485.2;refGene_gn=UBAP2L;refGene_tx=XM_019822486.2;refGene_gn=UBAP2L;refGene_tx=XM_019822487.2;refGene_gn=UBAP2L;refGene_tx=XM_019822488.2;refGene_gn=UBAP2L;refGene_tx=XM_019822489.2;refGene_gn=UBAP2L;refGene_tx=XM_019822490.2;refGene_gn=UBAP2L;refGene_tx=XM_019822491.2;refGene_gn=UBAP2L;refGene_tx=XM_019822492.2;refGene_gn=UBAP2L;refGene_tx=XM_019822493.1;refGene_gn=UBAP2L;refGene_tx=XM_019822494.1;refGene_gn=UBAP2L;refGene_tx=XM_019822495.2;refGene_gn=UBAP2L;refGene_tx=XM_019822496.2
F1 71029735 71029953
refGene_gn=UBAP2L;refGene_tx=XM_011291242.3;refGene_gn=UBAP2L;refGene_tx=XM_011291248.3;refGene_gn=UBAP2L;refGene_tx=XM_011291249.3;refGene_gn=UBAP2L;refGene_tx=XM_011291251.3;refGene_gn=UBAP2L;refGene_tx=XM_019822480.2;refGene_gn=UBAP2L;refGene_tx=XM_019822481.2;refGene_gn=UBAP2L;refGene_tx=XM_019822482.2;refGene_gn=UBAP2L;refGene_tx=XM_019822483.2;refGene_gn=UBAP2L;refGene_tx=XM_019822484.2;refGene_gn=UBAP2L;refGene_tx=XM_019822485.2;refGene_gn=UBAP2L;refGene_tx=XM_019822486.2;refGene_gn=UBAP2L;refGene_tx=XM_019822487.2;refGene_gn=UBAP2L;refGene_tx=XM_019822488.2;refGene_gn=UBAP2L;refGene_tx=XM_019822489.2;refGene_gn=UBAP2L;refGene_tx=XM_019822490.2;refGene_gn=UBAP2L;refGene_tx=XM_019822491.2;refGene_gn=UBAP2L;refGene_tx=XM_019822492.2;refGene_gn=UBAP2L;refGene_tx=XM_019822493.1;refGene_gn=UBAP2L;refGene_tx=XM_019822494.1;refGene_gn=UBAP2L;refGene_tx=XM_019822495.2;refGene_gn=UBAP2L;refGene_tx=X

M_019822496.2
F1 71030685 71030767
refGene_gn=UBAP2L;refGene_tx=XM_011291242.3;refGene_gn=UBAP2L;refGene_tx=XM_011291248.3;refGene_gn=UBAP2L;refGene_tx=XM_011291249.3;refGene_gn=UBAP2L;refGene_tx=XM_011291251.3;refGene_gn=UBAP2L;refGene_tx=XM_019822480.2;refGene_gn=UBAP2L;refGene_tx=XM_019822481.2;refGene_gn=UBAP2L;refGene_tx=XM_019822482.2;refGene_gn=UBAP2L;refGene_tx=XM_019822483.2;refGene_gn=UBAP2L;refGene_tx=XM_019822484.2;refGene_gn=UBAP2L;refGene_tx=XM_019822485.2;refGene_gn=UBAP2L;refGene_tx=XM_019822486.2;refGene_gn=UBAP2L;refGene_tx=XM_019822487.2;refGene_gn=UBAP2L;refGene_tx=XM_019822488.2;refGene_gn=UBAP2L;refGene_tx=XM_019822489.2;refGene_gn=UBAP2L;refGene_tx=XM_019822490.2;refGene_gn=UBAP2L;refGene_tx=XM_019822491.2;refGene_gn=UBAP2L;refGene_tx=XM_019822492.2;refGene_gn=UBAP2L;refGene_tx=XM_019822493.1;refGene_gn=UBAP2L;refGene_tx=XM_019822494.1;refGene_gn=UBAP2L;refGene_tx=XM_019822495.2;refGene_gn=UBAP2L;refGene_tx=XM_019822496.2
F1 71031355 71031409
refGene_gn=UBAP2L;refGene_tx=XM_011291242.3;refGene_gn=UBAP2L;refGene_tx=XM_011291248.3;refGene_gn=UBAP2L;refGene_tx=XM_011291249.3;refGene_gn=UBAP2L;refGene_tx=XM_011291251.3;refGene_gn=UBAP2L;refGene_tx=XM_019822480.2;refGene_gn=UBAP2L;refGene_tx=XM_019822481.2;refGene_gn=UBAP2L;refGene_tx=XM_019822482.2;refGene_gn=UBAP2L;refGene_tx=XM_019822483.2;refGene_gn=UBAP2L;refGene_tx=XM_019822484.2;refGene_gn=UBAP2L;refGene_tx=XM_019822485.2;refGene_gn=UBAP2L;refGene_tx=XM_019822486.2;refGene_gn=UBAP2L;refGene_tx=XM_019822487.2;refGene_gn=UBAP2L;refGene_tx=XM_019822488.2;refGene_gn=UBAP2L;refGene_tx=XM_019822489.2;refGene_gn=UBAP2L;refGene_tx=XM_019822490.2;refGene_gn=UBAP2L;refGene_tx=XM_019822491.2;refGene_gn=UBAP2L;refGene_tx=XM_019822492.2;refGene_gn=UBAP2L;refGene_tx=XM_019822493.1;refGene_gn=UBAP2L;refGene_tx=XM_019822494.1;refGene_gn=UBAP2L;refGene_tx=XM_019822495.2;refGene_gn=UBAP2L;refGene_tx=XM_019822496.2
F1 71032364 71032453
refGene_gn=UBAP2L;refGene_tx=XM_011291242.3;refGene_gn=UBAP2L;refGene_tx=XM_011291248.3;refGene_gn=UBAP2L;refGene_tx=XM_011291249.3;refGene_gn=UBAP2L;refGene_tx=XM_011291251.3;refGene_gn=UBAP2L;refGene_tx=XM_019822480.2;refGene_gn=UBAP2L;refGene_tx=XM_019822481.2;refGene_gn=UBAP2L;refGene_tx=XM_019822482.2;refGene_gn=UBAP2L;refGene_tx=XM_019822483.2;refGene_gn=UBAP2L;refGene_tx=XM_019822484.2;refGene_gn=UBAP2L;refGene_tx=XM_019822485.2;refGene_gn=UBAP2L;refGene_tx=XM_019822486.2;refGene_gn=UBAP2L;refGene_tx=XM_019822487.2;refGene_gn=UBAP2L;refGene_tx=XM_019822488.2;refGene_gn=UBAP2L;refGene_tx=XM_019822489.2;refGene_gn=UBAP2L;refGene_tx=XM_019822490.2;refGene_gn=UBAP2L;refGene_tx=XM_019822491.2;refGene_gn=UBAP2L;refGene_tx=XM_019822492.2;refGene_gn=UBAP2L;refGene_tx=XM_019822493.1;refGene_gn=UBAP2L;refGene_tx=XM_019822494.1;refGene_gn=UBAP2L;refGene_tx=XM_019822495.2;refGene_gn=UBAP2L;refGene_tx=XM_019822496.2
F1 71032533 71032729
refGene_gn=UBAP2L;refGene_tx=XM_011291242.3;refGene_gn=UBAP2L;refGene_tx=XM_011291248.3;refGene_gn=UBAP2L;refGene_tx=XM_011291249.3;refGene_gn=UBAP2L;refGene_tx=XM_011291251.3;refGene_gn=UBAP2L;refGene_tx=XM_01

9822480.2;refGene_gn=UBAP2L;refGene_tx=XM_019822481.2;refGene_gn=UBAP2L;refGene_tx=XM_019822482.2;refGene_gn=UBAP2L;refGene_tx=XM_019822483.2;refGene_gn=UBAP2L;refGene_tx=XM_019822484.2;refGene_gn=UBAP2L;refGene_tx=XM_019822485.2;refGene_gn=UBAP2L;refGene_tx=XM_019822486.2;refGene_gn=UBAP2L;refGene_tx=XM_019822487.2;refGene_gn=UBAP2L;refGene_tx=XM_019822488.2;refGene_gn=UBAP2L;refGene_tx=XM_019822489.2;refGene_gn=UBAP2L;refGene_tx=XM_019822490.2;refGene_gn=UBAP2L;refGene_tx=XM_019822491.2;refGene_gn=UBAP2L;refGene_tx=XM_019822492.2;refGene_gn=UBAP2L;refGene_tx=XM_019822493.1;refGene_gn=UBAP2L;refGene_tx=XM_019822494.1;refGene_gn=UBAP2L;refGene_tx=XM_019822495.2;refGene_gn=UBAP2L;refGene_tx=XM_019822496.2

F1 71033700 71033773
refGene_gn=UBAP2L;refGene_tx=XM_011291242.3;refGene_gn=UBAP2L;refGene_tx=XM_011291248.3;refGene_gn=UBAP2L;refGene_tx=XM_011291249.3;refGene_gn=UBAP2L;refGene_tx=XM_011291251.3;refGene_gn=UBAP2L;refGene_tx=XM_019822480.2;refGene_gn=UBAP2L;refGene_tx=XM_019822481.2;refGene_gn=UBAP2L;refGene_tx=XM_019822482.2;refGene_gn=UBAP2L;refGene_tx=XM_019822483.2;refGene_gn=UBAP2L;refGene_tx=XM_019822484.2;refGene_gn=UBAP2L;refGene_tx=XM_019822485.2;refGene_gn=UBAP2L;refGene_tx=XM_019822486.2;refGene_gn=UBAP2L;refGene_tx=XM_019822487.2;refGene_gn=UBAP2L;refGene_tx=XM_019822488.2;refGene_gn=UBAP2L;refGene_tx=XM_019822489.2;refGene_gn=UBAP2L;refGene_tx=XM_019822490.2;refGene_gn=UBAP2L;refGene_tx=XM_019822491.2;refGene_gn=UBAP2L;refGene_tx=XM_019822492.2;refGene_gn=UBAP2L;refGene_tx=XM_019822493.1;refGene_gn=UBAP2L;refGene_tx=XM_019822494.1;refGene_gn=UBAP2L;refGene_tx=XM_019822495.2;refGene_gn=UBAP2L;refGene_tx=XM_019822496.2

F1 71034091 71034266
refGene_gn=UBAP2L;refGene_tx=XM_011291242.3;refGene_gn=UBAP2L;refGene_tx=XM_011291248.3;refGene_gn=UBAP2L;refGene_tx=XM_011291249.3;refGene_gn=UBAP2L;refGene_tx=XM_011291251.3;refGene_gn=UBAP2L;refGene_tx=XM_019822480.2;refGene_gn=UBAP2L;refGene_tx=XM_019822481.2;refGene_gn=UBAP2L;refGene_tx=XM_019822482.2;refGene_gn=UBAP2L;refGene_tx=XM_019822483.2;refGene_gn=UBAP2L;refGene_tx=XM_019822484.2;refGene_gn=UBAP2L;refGene_tx=XM_019822485.2;refGene_gn=UBAP2L;refGene_tx=XM_019822486.2;refGene_gn=UBAP2L;refGene_tx=XM_019822487.2;refGene_gn=UBAP2L;refGene_tx=XM_019822488.2;refGene_gn=UBAP2L;refGene_tx=XM_019822489.2;refGene_gn=UBAP2L;refGene_tx=XM_019822490.2;refGene_gn=UBAP2L;refGene_tx=XM_019822491.2;refGene_gn=UBAP2L;refGene_tx=XM_019822492.2;refGene_gn=UBAP2L;refGene_tx=XM_019822493.1;refGene_gn=UBAP2L;refGene_tx=XM_019822494.1;refGene_gn=UBAP2L;refGene_tx=XM_019822495.2;refGene_gn=UBAP2L;refGene_tx=XM_019822496.2

F1 71034878 71034933
refGene_gn=UBAP2L;refGene_tx=XM_011291242.3;refGene_gn=UBAP2L;refGene_tx=XM_011291248.3;refGene_gn=UBAP2L;refGene_tx=XM_011291249.3;refGene_gn=UBAP2L;refGene_tx=XM_011291251.3;refGene_gn=UBAP2L;refGene_tx=XM_019822480.2;refGene_gn=UBAP2L;refGene_tx=XM_019822481.2;refGene_gn=UBAP2L;refGene_tx=XM_019822482.2;refGene_gn=UBAP2L;refGene_tx=XM_019822483.2;refGene_gn=UBAP2L;refGene_tx=XM_019822484.2;refGene_gn=UBAP2L;refGene_tx=XM_019822485.2;refGene_gn=UBAP2L;refGene_tx=XM_019822486.2;refGene_gn=UBAP2L;refGene_tx=XM_019822487.2;refGene_gn=UBAP2L;refGene_tx=XM_

019822488.2;refGene_gn=UBAP2L;refGene_tx=XM_019822489.2;refGene_gn=UBAP2L;refGene_tx=XM_019822490.2;refGene_gn=UBAP2L;refGene_tx=XM_019822491.2;refGene_gn=UBAP2L;refGene_tx=XM_019822492.2;refGene_gn=UBAP2L;refGene_tx=XM_019822493.1;refGene_gn=UBAP2L;refGene_tx=XM_019822494.1;refGene_gn=UBAP2L;refGene_tx=XM_019822495.2;refGene_gn=UBAP2L;refGene_tx=XM_019822496.2

F1 71035641 71035831

refGene_gn=UBAP2L;refGene_tx=XM_011291242.3;refGene_gn=UBAP2L;refGene_tx=XM_011291248.3;refGene_gn=UBAP2L;refGene_tx=XM_011291249.3;refGene_gn=UBAP2L;refGene_tx=XM_011291251.3;refGene_gn=UBAP2L;refGene_tx=XM_019822480.2;refGene_gn=UBAP2L;refGene_tx=XM_019822481.2;refGene_gn=UBAP2L;refGene_tx=XM_019822482.2;refGene_gn=UBAP2L;refGene_tx=XM_019822483.2;refGene_gn=UBAP2L;refGene_tx=XM_019822484.2;refGene_gn=UBAP2L;refGene_tx=XM_019822485.2;refGene_gn=UBAP2L;refGene_tx=XM_019822486.2;refGene_gn=UBAP2L;refGene_tx=XM_019822487.2;refGene_gn=UBAP2L;refGene_tx=XM_019822488.2;refGene_gn=UBAP2L;refGene_tx=XM_019822489.2;refGene_gn=UBAP2L;refGene_tx=XM_019822490.2;refGene_gn=UBAP2L;refGene_tx=XM_019822491.2;refGene_gn=UBAP2L;refGene_tx=XM_019822492.2;refGene_gn=UBAP2L;refGene_tx=XM_019822493.1;refGene_gn=UBAP2L;refGene_tx=XM_019822494.1;refGene_gn=UBAP2L;refGene_tx=XM_019822495.2;refGene_gn=UBAP2L;refGene_tx=XM_019822496.2

F1 71037285 71037458

refGene_gn=UBAP2L;refGene_tx=XM_011291242.3;refGene_gn=UBAP2L;refGene_tx=XM_011291248.3;refGene_gn=UBAP2L;refGene_tx=XM_011291249.3;refGene_gn=UBAP2L;refGene_tx=XM_011291251.3;refGene_gn=UBAP2L;refGene_tx=XM_019822480.2;refGene_gn=UBAP2L;refGene_tx=XM_019822481.2;refGene_gn=UBAP2L;refGene_tx=XM_019822482.2;refGene_gn=UBAP2L;refGene_tx=XM_019822483.2;refGene_gn=UBAP2L;refGene_tx=XM_019822484.2;refGene_gn=UBAP2L;refGene_tx=XM_019822485.2;refGene_gn=UBAP2L;refGene_tx=XM_019822486.2;refGene_gn=UBAP2L;refGene_tx=XM_019822487.2;refGene_gn=UBAP2L;refGene_tx=XM_019822488.2;refGene_gn=UBAP2L;refGene_tx=XM_019822489.2;refGene_gn=UBAP2L;refGene_tx=XM_019822490.2;refGene_gn=UBAP2L;refGene_tx=XM_019822491.2;refGene_gn=UBAP2L;refGene_tx=XM_019822492.2;refGene_gn=UBAP2L;refGene_tx=XM_019822493.1;refGene_gn=UBAP2L;refGene_tx=XM_019822494.1;refGene_gn=UBAP2L;refGene_tx=XM_019822495.2;refGene_gn=UBAP2L;refGene_tx=XM_019822496.2

F1 71037615 71037893

refGene_gn=UBAP2L;refGene_tx=XM_011291242.3;refGene_gn=UBAP2L;refGene_tx=XM_011291248.3;refGene_gn=UBAP2L;refGene_tx=XM_011291249.3;refGene_gn=UBAP2L;refGene_tx=XM_011291251.3;refGene_gn=UBAP2L;refGene_tx=XM_019822480.2;refGene_gn=UBAP2L;refGene_tx=XM_019822481.2;refGene_gn=UBAP2L;refGene_tx=XM_019822482.2;refGene_gn=UBAP2L;refGene_tx=XM_019822483.2;refGene_gn=UBAP2L;refGene_tx=XM_019822484.2;refGene_gn=UBAP2L;refGene_tx=XM_019822485.2;refGene_gn=UBAP2L;refGene_tx=XM_019822486.2;refGene_gn=UBAP2L;refGene_tx=XM_019822487.2;refGene_gn=UBAP2L;refGene_tx=XM_019822488.2;refGene_gn=UBAP2L;refGene_tx=XM_019822489.2;refGene_gn=UBAP2L;refGene_tx=XM_019822490.2;refGene_gn=UBAP2L;refGene_tx=XM_019822491.2;refGene_gn=UBAP2L;refGene_tx=XM_019822492.2;refGene_gn=UBAP2L;refGene_tx=XM_019822493.1;refGene_gn=UBAP2L;refGene_tx=XM_019822494.1;refGene_gn=UBAP2L;refGene_tx=XM_019822495.2;refGene_gn=UBAP2L;refGene_tx=X

M_019822496.2
F1 71038480 71038679
refGene_gn=UBAP2L;refGene_tx=XM_011291242.3;refGene_gn=UBAP2L;refGene_tx=XM_011291248.3;refGene_gn=UBAP2L;refGene_tx=XM_011291249.3;refGene_gn=UBAP2L;refGene_tx=XM_011291251.3;refGene_gn=UBAP2L;refGene_tx=XM_019822480.2;refGene_gn=UBAP2L;refGene_tx=XM_019822481.2;refGene_gn=UBAP2L;refGene_tx=XM_019822482.2;refGene_gn=UBAP2L;refGene_tx=XM_019822483.2;refGene_gn=UBAP2L;refGene_tx=XM_019822484.2;refGene_gn=UBAP2L;refGene_tx=XM_019822485.2;refGene_gn=UBAP2L;refGene_tx=XM_019822486.2;refGene_gn=UBAP2L;refGene_tx=XM_019822487.2;refGene_gn=UBAP2L;refGene_tx=XM_019822488.2;refGene_gn=UBAP2L;refGene_tx=XM_019822489.2;refGene_gn=UBAP2L;refGene_tx=XM_019822490.2;refGene_gn=UBAP2L;refGene_tx=XM_019822491.2;refGene_gn=UBAP2L;refGene_tx=XM_019822492.2;refGene_gn=UBAP2L;refGene_tx=XM_019822493.1;refGene_gn=UBAP2L;refGene_tx=XM_019822494.1;refGene_gn=UBAP2L;refGene_tx=XM_019822495.2;refGene_gn=UBAP2L;refGene_tx=XM_019822496.2
F1 71041693 71041865
refGene_gn=UBAP2L;refGene_tx=XM_011291242.3;refGene_gn=UBAP2L;refGene_tx=XM_011291248.3;refGene_gn=UBAP2L;refGene_tx=XM_011291249.3;refGene_gn=UBAP2L;refGene_tx=XM_011291251.3;refGene_gn=UBAP2L;refGene_tx=XM_019822480.2;refGene_gn=UBAP2L;refGene_tx=XM_019822481.2;refGene_gn=UBAP2L;refGene_tx=XM_019822482.2;refGene_gn=UBAP2L;refGene_tx=XM_019822483.2;refGene_gn=UBAP2L;refGene_tx=XM_019822484.2;refGene_gn=UBAP2L;refGene_tx=XM_019822485.2;refGene_gn=UBAP2L;refGene_tx=XM_019822486.2;refGene_gn=UBAP2L;refGene_tx=XM_019822487.2;refGene_gn=UBAP2L;refGene_tx=XM_019822488.2;refGene_gn=UBAP2L;refGene_tx=XM_019822489.2;refGene_gn=UBAP2L;refGene_tx=XM_019822490.2;refGene_gn=UBAP2L;refGene_tx=XM_019822491.2;refGene_gn=UBAP2L;refGene_tx=XM_019822492.2;refGene_gn=UBAP2L;refGene_tx=XM_019822493.1;refGene_gn=UBAP2L;refGene_tx=XM_019822494.1;refGene_gn=UBAP2L;refGene_tx=XM_019822495.2;refGene_gn=UBAP2L;refGene_tx=XM_019822496.2
F1 71043699 71043785
refGene_gn=UBAP2L;refGene_tx=XM_011291242.3;refGene_gn=UBAP2L;refGene_tx=XM_011291248.3;refGene_gn=UBAP2L;refGene_tx=XM_011291249.3;refGene_gn=UBAP2L;refGene_tx=XM_011291251.3;refGene_gn=UBAP2L;refGene_tx=XM_019822480.2;refGene_gn=UBAP2L;refGene_tx=XM_019822481.2;refGene_gn=UBAP2L;refGene_tx=XM_019822482.2;refGene_gn=UBAP2L;refGene_tx=XM_019822483.2;refGene_gn=UBAP2L;refGene_tx=XM_019822484.2;refGene_gn=UBAP2L;refGene_tx=XM_019822485.2;refGene_gn=UBAP2L;refGene_tx=XM_019822486.2;refGene_gn=UBAP2L;refGene_tx=XM_019822487.2;refGene_gn=UBAP2L;refGene_tx=XM_019822488.2;refGene_gn=UBAP2L;refGene_tx=XM_019822489.2;refGene_gn=UBAP2L;refGene_tx=XM_019822490.2;refGene_gn=UBAP2L;refGene_tx=XM_019822491.2;refGene_gn=UBAP2L;refGene_tx=XM_019822492.2;refGene_gn=UBAP2L;refGene_tx=XM_019822493.1;refGene_gn=UBAP2L;refGene_tx=XM_019822494.1;refGene_gn=UBAP2L;refGene_tx=XM_019822495.2;refGene_gn=UBAP2L;refGene_tx=XM_019822496.2
F1 71045354 71045407
refGene_gn=UBAP2L;refGene_tx=XM_011291242.3;refGene_gn=UBAP2L;refGene_tx=XM_011291248.3;refGene_gn=UBAP2L;refGene_tx=XM_011291249.3;refGene_gn=UBAP2L;refGene_tx=XM_011291251.3;refGene_gn=UBAP2L;refGene_tx=XM_01

9822480.2;refGene_gn=UBAP2L;refGene_tx=XM_019822481.2;refGene_gn=UBAP2
L;refGene_tx=XM_019822482.2;refGene_gn=UBAP2L;refGene_tx=XM_019822483.
2;refGene_gn=UBAP2L;refGene_tx=XM_019822484.2;refGene_gn=UBAP2L;refGen
e_tx=XM_019822485.2;refGene_gn=UBAP2L;refGene_tx=XM_019822486.2;refGen
e_gn=UBAP2L;refGene_tx=XM_019822487.2;refGene_gn=UBAP2L;refGene_tx=XM_
019822488.2;refGene_gn=UBAP2L;refGene_tx=XM_019822489.2;refGene_gn=UBA
P2L;refGene_tx=XM_019822490.2;refGene_gn=UBAP2L;refGene_tx=XM_01982249
1.2;refGene_gn=UBAP2L;refGene_tx=XM_019822492.2;refGene_gn=UBAP2L;refG
ene_tx=XM_019822493.1;refGene_gn=UBAP2L;refGene_tx=XM_019822494.1;refG
ene_gn=UBAP2L;refGene_tx=XM_019822495.2;refGene_gn=UBAP2L;refGene_tx=X
M_019822496.2
F1 71047541 71047616
refGene_gn=UBAP2L;refGene_tx=XM_019822480.2;refGene_gn=UBAP2L;refGene_
tx=XM_019822481.2;refGene_gn=UBAP2L;refGene_tx=XM_019822482.2;refGene_
gn=UBAP2L;refGene_tx=XM_019822483.2;refGene_gn=UBAP2L;refGene_tx=XM_01
9822484.2;refGene_gn=UBAP2L;refGene_tx=XM_019822486.2;refGene_gn=UBAP2
L;refGene_tx=XM_019822487.2;refGene_gn=UBAP2L;refGene_tx=XM_019822488.
2;refGene_gn=UBAP2L;refGene_tx=XM_019822489.2;refGene_gn=UBAP2L;refGen
e_tx=XM_019822490.2;refGene_gn=UBAP2L;refGene_tx=XM_019822491.2;refGen
e_gn=UBAP2L;refGene_tx=XM_019822494.1;refGene_gn=UBAP2L;refGene_tx=XM_
019822496.2
F1 71049403 71049516
refGene_gn=UBAP2L;refGene_tx=XM_011291242.3;refGene_gn=UBAP2L;refGene_
tx=XM_011291248.3;refGene_gn=UBAP2L;refGene_tx=XM_011291249.3;refGene_
gn=UBAP2L;refGene_tx=XM_011291251.3;refGene_gn=UBAP2L;refGene_tx=XM_01
9822480.2;refGene_gn=UBAP2L;refGene_tx=XM_019822481.2;refGene_gn=UBAP2
L;refGene_tx=XM_019822482.2;refGene_gn=UBAP2L;refGene_tx=XM_019822483.
2;refGene_gn=UBAP2L;refGene_tx=XM_019822484.2;refGene_gn=UBAP2L;refGen
e_tx=XM_019822485.2;refGene_gn=UBAP2L;refGene_tx=XM_019822486.2;refGen
e_gn=UBAP2L;refGene_tx=XM_019822487.2;refGene_gn=UBAP2L;refGene_tx=XM_
019822488.2;refGene_gn=UBAP2L;refGene_tx=XM_019822489.2;refGene_gn=UBA
P2L;refGene_tx=XM_019822490.2;refGene_gn=UBAP2L;refGene_tx=XM_01982249
1.2;refGene_gn=UBAP2L;refGene_tx=XM_019822492.2;refGene_gn=UBAP2L;refG
ene_tx=XM_019822493.1;refGene_gn=UBAP2L;refGene_tx=XM_019822494.1;refG
ene_gn=UBAP2L;refGene_tx=XM_019822495.2;refGene_gn=UBAP2L;refGene_tx=X
M_019822496.2
F1 71049914 71049960
refGene_gn=UBAP2L;refGene_tx=XM_011291242.3;refGene_gn=UBAP2L;refGene_
tx=XM_011291248.3;refGene_gn=UBAP2L;refGene_tx=XM_011291249.3;refGene_
gn=UBAP2L;refGene_tx=XM_011291251.3;refGene_gn=UBAP2L;refGene_tx=XM_01
9822480.2;refGene_gn=UBAP2L;refGene_tx=XM_019822481.2;refGene_gn=UBAP2
L;refGene_tx=XM_019822482.2;refGene_gn=UBAP2L;refGene_tx=XM_019822483.
2;refGene_gn=UBAP2L;refGene_tx=XM_019822484.2;refGene_gn=UBAP2L;refGen
e_tx=XM_019822485.2;refGene_gn=UBAP2L;refGene_tx=XM_019822486.2;refGen
e_gn=UBAP2L;refGene_tx=XM_019822487.2;refGene_gn=UBAP2L;refGene_tx=XM_
019822488.2;refGene_gn=UBAP2L;refGene_tx=XM_019822489.2;refGene_gn=UBA
P2L;refGene_tx=XM_019822490.2;refGene_gn=UBAP2L;refGene_tx=XM_01982249
1.2;refGene_gn=UBAP2L;refGene_tx=XM_019822492.2;refGene_gn=UBAP2L;refG
ene_tx=XM_019822493.1;refGene_gn=UBAP2L;refGene_tx=XM_019822494.1;refG
ene_gn=UBAP2L;refGene_tx=XM_019822495.2;refGene_gn=UBAP2L;refGene_tx=X

M_019822496.2
F1 71050364 71050460
refGene_gn=UBAP2L;refGene_tx=XM_011291242.3;refGene_gn=UBAP2L;refGene_tx=XM_011291248.3;refGene_gn=UBAP2L;refGene_tx=XM_011291249.3;refGene_gn=UBAP2L;refGene_tx=XM_011291251.3;refGene_gn=UBAP2L;refGene_tx=XM_019822480.2;refGene_gn=UBAP2L;refGene_tx=XM_019822481.2;refGene_gn=UBAP2L;refGene_tx=XM_019822482.2;refGene_gn=UBAP2L;refGene_tx=XM_019822483.2;refGene_gn=UBAP2L;refGene_tx=XM_019822484.2;refGene_gn=UBAP2L;refGene_tx=XM_019822485.2;refGene_gn=UBAP2L;refGene_tx=XM_019822486.2;refGene_gn=UBAP2L;refGene_tx=XM_019822487.2;refGene_gn=UBAP2L;refGene_tx=XM_019822488.2;refGene_gn=UBAP2L;refGene_tx=XM_019822489.2;refGene_gn=UBAP2L;refGene_tx=XM_019822490.2;refGene_gn=UBAP2L;refGene_tx=XM_019822491.2;refGene_gn=UBAP2L;refGene_tx=XM_019822492.2;refGene_gn=UBAP2L;refGene_tx=XM_019822493.1;refGene_gn=UBAP2L;refGene_tx=XM_019822494.1;refGene_gn=UBAP2L;refGene_tx=XM_019822495.2;refGene_gn=UBAP2L;refGene_tx=XM_019822496.2
F1 71050827 71051029
refGene_gn=UBAP2L;refGene_tx=XM_011291242.3;refGene_gn=UBAP2L;refGene_tx=XM_011291248.3;refGene_gn=UBAP2L;refGene_tx=XM_011291249.3;refGene_gn=UBAP2L;refGene_tx=XM_011291251.3;refGene_gn=UBAP2L;refGene_tx=XM_019822480.2;refGene_gn=UBAP2L;refGene_tx=XM_019822481.2;refGene_gn=UBAP2L;refGene_tx=XM_019822482.2;refGene_gn=UBAP2L;refGene_tx=XM_019822483.2;refGene_gn=UBAP2L;refGene_tx=XM_019822484.2;refGene_gn=UBAP2L;refGene_tx=XM_019822485.2;refGene_gn=UBAP2L;refGene_tx=XM_019822486.2;refGene_gn=UBAP2L;refGene_tx=XM_019822487.2;refGene_gn=UBAP2L;refGene_tx=XM_019822488.2;refGene_gn=UBAP2L;refGene_tx=XM_019822489.2;refGene_gn=UBAP2L;refGene_tx=XM_019822490.2;refGene_gn=UBAP2L;refGene_tx=XM_019822491.2;refGene_gn=UBAP2L;refGene_tx=XM_019822492.2;refGene_gn=UBAP2L;refGene_tx=XM_019822493.1;refGene_gn=UBAP2L;refGene_tx=XM_019822494.1;refGene_gn=UBAP2L;refGene_tx=XM_019822495.2;refGene_gn=UBAP2L;refGene_tx=XM_019822496.2
F1 71055451 71055562
refGene_gn=UBAP2L;refGene_tx=XM_011291242.3;refGene_gn=UBAP2L;refGene_tx=XM_011291248.3;refGene_gn=UBAP2L;refGene_tx=XM_011291249.3;refGene_gn=UBAP2L;refGene_tx=XM_011291251.3;refGene_gn=UBAP2L;refGene_tx=XM_019822480.2;refGene_gn=UBAP2L;refGene_tx=XM_019822481.2;refGene_gn=UBAP2L;refGene_tx=XM_019822482.2;refGene_gn=UBAP2L;refGene_tx=XM_019822483.2;refGene_gn=UBAP2L;refGene_tx=XM_019822484.2;refGene_gn=UBAP2L;refGene_tx=XM_019822485.2;refGene_gn=UBAP2L;refGene_tx=XM_019822486.2;refGene_gn=UBAP2L;refGene_tx=XM_019822487.2;refGene_gn=UBAP2L;refGene_tx=XM_019822488.2;refGene_gn=UBAP2L;refGene_tx=XM_019822489.2;refGene_gn=UBAP2L;refGene_tx=XM_019822490.2;refGene_gn=UBAP2L;refGene_tx=XM_019822491.2;refGene_gn=UBAP2L;refGene_tx=XM_019822492.2;refGene_gn=UBAP2L;refGene_tx=XM_019822493.1;refGene_gn=UBAP2L;refGene_tx=XM_019822494.1;refGene_gn=UBAP2L;refGene_tx=XM_019822495.2;refGene_gn=UBAP2L;refGene_tx=XM_019822496.2
F1 71056120 71056198
refGene_gn=UBAP2L;refGene_tx=XM_011291242.3;refGene_gn=UBAP2L;refGene_tx=XM_011291248.3;refGene_gn=UBAP2L;refGene_tx=XM_011291249.3;refGene_gn=UBAP2L;refGene_tx=XM_011291251.3;refGene_gn=UBAP2L;refGene_tx=XM_01

9822480.2;refGene_gn=UBAP2L;refGene_tx=XM_019822481.2;refGene_gn=UBAP2L;refGene_tx=XM_019822482.2;refGene_gn=UBAP2L;refGene_tx=XM_019822483.2;refGene_gn=UBAP2L;refGene_tx=XM_019822484.2;refGene_gn=UBAP2L;refGene_tx=XM_019822485.2;refGene_gn=UBAP2L;refGene_tx=XM_019822486.2;refGene_gn=UBAP2L;refGene_tx=XM_019822487.2;refGene_gn=UBAP2L;refGene_tx=XM_019822488.2;refGene_gn=UBAP2L;refGene_tx=XM_019822489.2;refGene_gn=UBAP2L;refGene_tx=XM_019822490.2;refGene_gn=UBAP2L;refGene_tx=XM_019822491.2;refGene_gn=UBAP2L;refGene_tx=XM_019822492.2;refGene_gn=UBAP2L;refGene_tx=XM_019822493.1;refGene_gn=UBAP2L;refGene_tx=XM_019822494.1;refGene_gn=UBAP2L;refGene_tx=XM_019822495.2;refGene_gn=UBAP2L;refGene_tx=XM_019822496.2
F1 71056882 71056972
refGene_gn=UBAP2L;refGene_tx=XM_011291242.3;refGene_gn=UBAP2L;refGene_tx=XM_011291248.3;refGene_gn=UBAP2L;refGene_tx=XM_011291249.3;refGene_gn=UBAP2L;refGene_tx=XM_011291251.3;refGene_gn=UBAP2L;refGene_tx=XM_019822480.2;refGene_gn=UBAP2L;refGene_tx=XM_019822481.2;refGene_gn=UBAP2L;refGene_tx=XM_019822482.2;refGene_gn=UBAP2L;refGene_tx=XM_019822483.2;refGene_gn=UBAP2L;refGene_tx=XM_019822484.2;refGene_gn=UBAP2L;refGene_tx=XM_019822485.2;refGene_gn=UBAP2L;refGene_tx=XM_019822486.2;refGene_gn=UBAP2L;refGene_tx=XM_019822487.2;refGene_gn=UBAP2L;refGene_tx=XM_019822488.2;refGene_gn=UBAP2L;refGene_tx=XM_019822489.2;refGene_gn=UBAP2L;refGene_tx=XM_019822490.2;refGene_gn=UBAP2L;refGene_tx=XM_019822491.2;refGene_gn=UBAP2L;refGene_tx=XM_019822492.2;refGene_gn=UBAP2L;refGene_tx=XM_019822493.1;refGene_gn=UBAP2L;refGene_tx=XM_019822494.1;refGene_gn=UBAP2L;refGene_tx=XM_019822495.2;refGene_gn=UBAP2L;refGene_tx=XM_019822496.2
F1 71061559 71061625
refGene_gn=CF1H1orf43;refGene_tx=XM_003999753.5;refGene_gn=CF1H1orf43;refGene_tx=XM_019822502.2;refGene_gn=CF1H1orf43;refGene_tx=XM_019822503.2
F1 71061887 71061989
refGene_gn=CF1H1orf43;refGene_tx=XM_003999753.5;refGene_gn=CF1H1orf43;refGene_tx=XM_019822502.2;refGene_gn=CF1H1orf43;refGene_tx=XM_019822503.2
F1 71066498 71066582
refGene_gn=CF1H1orf43;refGene_tx=XM_019822502.2;refGene_gn=CF1H1orf43;refGene_tx=XM_019822503.2
F1 71067325 71067443
refGene_gn=CF1H1orf43;refGene_tx=XM_003999753.5;refGene_gn=CF1H1orf43;refGene_tx=XM_019822502.2;refGene_gn=CF1H1orf43;refGene_tx=XM_019822503.2
F1 71067648 71067702
refGene_gn=CF1H1orf43;refGene_tx=XM_003999753.5;refGene_gn=CF1H1orf43;refGene_tx=XM_019822502.2;refGene_gn=CF1H1orf43;refGene_tx=XM_019822503.2
F1 71068313 71068480
refGene_gn=CF1H1orf43;refGene_tx=XM_003999753.5;refGene_gn=CF1H1orf43;refGene_tx=XM_019822502.2;refGene_gn=CF1H1orf43;refGene_tx=XM_019822503.2
F1 71068567 71068626

refGene_gn=CF1H1orf43;refGene_tx=XM_003999753.5;refGene_gn=CF1H1orf43;
refGene_tx=XM_019822502.2
F1 71072126 71072322
refGene_gn=CF1H1orf43;refGene_tx=XM_003999753.5;refGene_gn=CF1H1orf43;
refGene_tx=XM_019822502.2;refGene_gn=CF1H1orf43;refGene_tx=XM_01982250
3.2
F1 71073369 71073402
refGene_gn=CF1H1orf189;refGene_tx=XM_003999790.4
F1 71073689 71073730
refGene_gn=CF1H1orf189;refGene_tx=XM_003999790.4
F1 71077420 71077550
refGene_gn=CF1H1orf189;refGene_tx=XM_003999790.4
F1 71078110 71078212
refGene_gn=CF1H1orf189;refGene_tx=XM_003999790.4
F1 71083405 71083522
refGene_gn=TPM3;refGene_tx=XM_003999756.4;refGene_gn=TPM3;refGene_tx=X
M_003999758.5;refGene_gn=TPM3;refGene_tx=XM_011291252.3;refGene_gn=TPM
3;refGene_tx=XM_011291254.3;refGene_gn=TPM3;refGene_tx=XM_019822522.2;
refGene_gn=TPM3;refGene_tx=XM_019822523.2;refGene_gn=TPM3;refGene_tx=X
M_019822524.2
F1 71084192 71084318
refGene_gn=TPM3;refGene_tx=XM_003999756.4;refGene_gn=TPM3;refGene_tx=X
M_003999758.5;refGene_gn=TPM3;refGene_tx=XM_011291252.3;refGene_gn=TPM
3;refGene_tx=XM_011291254.3;refGene_gn=TPM3;refGene_tx=XM_019822522.2;
refGene_gn=TPM3;refGene_tx=XM_019822523.2;refGene_gn=TPM3;refGene_tx=X
M_019822524.2
F1 71088846 71088978
refGene_gn=TPM3;refGene_tx=XM_003999757.5;refGene_gn=TPM3;refGene_tx=X
M_006943170.3;refGene_gn=TPM3;refGene_tx=XM_011291257.3;refGene_gn=TPM
3;refGene_tx=XM_019822525.2;refGene_gn=TPM3;refGene_tx=XM_019822526.2;
refGene_gn=TPM3;refGene_tx=XM_019822529.2;refGene_gn=TPM3;refGene_tx=X
M_019822530.2
F1 71095650 71095784
refGene_gn=TPM3;refGene_tx=XM_003999756.4;refGene_gn=TPM3;refGene_tx=X
M_003999757.5;refGene_gn=TPM3;refGene_tx=XM_003999758.5;refGene_gn=TPM
3;refGene_tx=XM_011291252.3;refGene_gn=TPM3;refGene_tx=XM_011291254.3;
refGene_gn=TPM3;refGene_tx=XM_011291257.3;refGene_gn=TPM3;refGene_tx=X
M_019822522.2;refGene_gn=TPM3;refGene_tx=XM_019822523.2;refGene_gn=TPM
3;refGene_tx=XM_019822524.2;refGene_gn=TPM3;refGene_tx=XM_019822525.2;
refGene_gn=TPM3;refGene_tx=XM_019822526.2;refGene_gn=TPM3;refGene_tx=X
M_019822529.2;refGene_gn=TPM3;refGene_tx=XM_019822530.2
F1 71097338 71097456
refGene_gn=TPM3;refGene_tx=XM_003999756.4;refGene_gn=TPM3;refGene_tx=X
M_003999757.5;refGene_gn=TPM3;refGene_tx=XM_003999758.5;refGene_gn=TPM
3;refGene_tx=XM_006943170.3;refGene_gn=TPM3;refGene_tx=XM_011291252.3;
refGene_gn=TPM3;refGene_tx=XM_011291254.3;refGene_gn=TPM3;refGene_tx=X
M_011291257.3;refGene_gn=TPM3;refGene_tx=XM_019822522.2;refGene_gn=TPM
3;refGene_tx=XM_019822523.2;refGene_gn=TPM3;refGene_tx=XM_019822524.2;
refGene_gn=TPM3;refGene_tx=XM_019822525.2;refGene_gn=TPM3;refGene_tx=X
M_019822526.2;refGene_gn=TPM3;refGene_tx=XM_019822529.2;refGene_gn=TPM

3;refGene_tx=XM_019822530.2
F1 71097572 71097643
refGene_gn=TPM3;refGene_tx=XM_003999756.4;refGene_gn=TPM3;refGene_tx=XM_003999757.5;refGene_gn=TPM3;refGene_tx=XM_003999758.5;refGene_gn=TPM3;refGene_tx=XM_006943170.3;refGene_gn=TPM3;refGene_tx=XM_011291252.3;refGene_gn=TPM3;refGene_tx=XM_011291254.3;refGene_gn=TPM3;refGene_tx=XM_011291257.3;refGene_gn=TPM3;refGene_tx=XM_019822522.2;refGene_gn=TPM3;refGene_tx=XM_019822523.2;refGene_gn=TPM3;refGene_tx=XM_019822524.2;refGene_gn=TPM3;refGene_tx=XM_019822525.2;refGene_gn=TPM3;refGene_tx=XM_019822526.2;refGene_gn=TPM3;refGene_tx=XM_019822529.2;refGene_gn=TPM3;refGene_tx=XM_019822530.2
F1 71098288 71098364
refGene_gn=TPM3;refGene_tx=XM_003999756.4;refGene_gn=TPM3;refGene_tx=XM_003999757.5;refGene_gn=TPM3;refGene_tx=XM_003999758.5;refGene_gn=TPM3;refGene_tx=XM_006943170.3;refGene_gn=TPM3;refGene_tx=XM_019822523.2;refGene_gn=TPM3;refGene_tx=XM_019822526.2;refGene_gn=TPM3;refGene_tx=XM_019822530.2
F1 71098919 71098995
refGene_gn=TPM3;refGene_tx=XM_011291252.3;refGene_gn=TPM3;refGene_tx=XM_011291254.3;refGene_gn=TPM3;refGene_tx=XM_011291257.3;refGene_gn=TPM3;refGene_tx=XM_019822522.2;refGene_gn=TPM3;refGene_tx=XM_019822524.2;refGene_gn=TPM3;refGene_tx=XM_019822525.2;refGene_gn=TPM3;refGene_tx=XM_019822529.2
F1 71099699 71099762
refGene_gn=TPM3;refGene_tx=XM_003999756.4;refGene_gn=TPM3;refGene_tx=XM_003999757.5;refGene_gn=TPM3;refGene_tx=XM_003999758.5;refGene_gn=TPM3;refGene_tx=XM_006943170.3;refGene_gn=TPM3;refGene_tx=XM_011291252.3;refGene_gn=TPM3;refGene_tx=XM_011291254.3;refGene_gn=TPM3;refGene_tx=XM_011291257.3;refGene_gn=TPM3;refGene_tx=XM_019822522.2;refGene_gn=TPM3;refGene_tx=XM_019822523.2;refGene_gn=TPM3;refGene_tx=XM_019822524.2;refGene_gn=TPM3;refGene_tx=XM_019822525.2;refGene_gn=TPM3;refGene_tx=XM_019822526.2;refGene_gn=TPM3;refGene_tx=XM_019822529.2;refGene_gn=TPM3;refGene_tx=XM_019822530.2
F1 71099931 71100001
refGene_gn=TPM3;refGene_tx=XM_003999756.4;refGene_gn=TPM3;refGene_tx=XM_003999757.5;refGene_gn=TPM3;refGene_tx=XM_003999758.5;refGene_gn=TPM3;refGene_tx=XM_006943170.3;refGene_gn=TPM3;refGene_tx=XM_011291252.3;refGene_gn=TPM3;refGene_tx=XM_011291254.3;refGene_gn=TPM3;refGene_tx=XM_011291257.3;refGene_gn=TPM3;refGene_tx=XM_019822522.2;refGene_gn=TPM3;refGene_tx=XM_019822523.2;refGene_gn=TPM3;refGene_tx=XM_019822524.2;refGene_gn=TPM3;refGene_tx=XM_019822525.2;refGene_gn=TPM3;refGene_tx=XM_019822526.2;refGene_gn=TPM3;refGene_tx=XM_019822529.2;refGene_gn=TPM3;refGene_tx=XM_019822530.2
F1 71101007 71101090
refGene_gn=TPM3;refGene_tx=XM_003999756.4;refGene_gn=TPM3;refGene_tx=XM_011291252.3;refGene_gn=TPM3;refGene_tx=XM_019822522.2;refGene_gn=TPM3;refGene_tx=XM_019822523.2;refGene_gn=TPM3;refGene_tx=XM_019822524.2;refGene_gn=TPM3;refGene_tx=XM_019822525.2;refGene_gn=TPM3;refGene_tx=XM_019822526.2
F1 71101838 71101842 refGene_gn=TPM3;refGene_tx=XM_011291252.3

F1 71109634 71109714
refGene_gn=TPM3;refGene_tx=XM_019822522.2;refGene_gn=TPM3;refGene_tx=XM_019822523.2;refGene_gn=TPM3;refGene_tx=XM_019822525.2;refGene_gn=TPM3;refGene_tx=XM_019822526.2;refGene_gn=TPM3;refGene_tx=XM_019822529.2;refGene_gn=TPM3;refGene_tx=XM_019822530.2
F1 71110864 71110947
refGene_gn=TPM3;refGene_tx=XM_003999757.5;refGene_gn=TPM3;refGene_tx=XM_003999758.5;refGene_gn=TPM3;refGene_tx=XM_006943170.3;refGene_gn=TPM3;refGene_tx=XM_011291254.3;refGene_gn=TPM3;refGene_tx=XM_011291257.3;refGene_gn=TPM3;refGene_tx=XM_019822524.2
F1 71112870 71112938 refGene_gn=NUP210L;refGene_tx=XM_019822511.2
F1 71113201 71113401
refGene_gn=NUP210L;refGene_tx=XM_003999791.5;refGene_gn=NUP210L;refGene_tx=XM_019822505.2;refGene_gn=NUP210L;refGene_tx=XM_019822506.2;refGene_gn=NUP210L;refGene_tx=XM_019822507.2;refGene_gn=NUP210L;refGene_tx=XM_019822508.2;refGene_gn=NUP210L;refGene_tx=XM_019822509.2;refGene_gn=NUP210L;refGene_tx=XM_019822510.2;refGene_gn=NUP210L;refGene_tx=XM_019822512.2;refGene_gn=NUP210L;refGene_tx=XM_019822513.2;refGene_gn=NUP210L;refGene_tx=XM_019822514.2
F1 71114535 71114672
refGene_gn=NUP210L;refGene_tx=XM_003999791.5;refGene_gn=NUP210L;refGene_tx=XM_019822505.2;refGene_gn=NUP210L;refGene_tx=XM_019822506.2;refGene_gn=NUP210L;refGene_tx=XM_019822507.2;refGene_gn=NUP210L;refGene_tx=XM_019822508.2;refGene_gn=NUP210L;refGene_tx=XM_019822509.2;refGene_gn=NUP210L;refGene_tx=XM_019822510.2;refGene_gn=NUP210L;refGene_tx=XM_019822511.2;refGene_gn=NUP210L;refGene_tx=XM_019822512.2;refGene_gn=NUP210L;refGene_tx=XM_019822513.2;refGene_gn=NUP210L;refGene_tx=XM_019822514.2
F1 71123638 71123788
refGene_gn=NUP210L;refGene_tx=XM_003999791.5;refGene_gn=NUP210L;refGene_tx=XM_019822505.2;refGene_gn=NUP210L;refGene_tx=XM_019822506.2;refGene_gn=NUP210L;refGene_tx=XM_019822507.2;refGene_gn=NUP210L;refGene_tx=XM_019822508.2;refGene_gn=NUP210L;refGene_tx=XM_019822509.2;refGene_gn=NUP210L;refGene_tx=XM_019822510.2;refGene_gn=NUP210L;refGene_tx=XM_019822511.2;refGene_gn=NUP210L;refGene_tx=XM_019822512.2;refGene_gn=NUP210L;refGene_tx=XM_019822513.2;refGene_gn=NUP210L;refGene_tx=XM_019822514.2;refGene_gn=NUP210L;refGene_tx=XM_019822517.2;refGene_gn=NUP210L;refGene_tx=XM_019822518.2
F1 71124915 71125009
refGene_gn=NUP210L;refGene_tx=XM_003999791.5;refGene_gn=NUP210L;refGene_tx=XM_019822505.2;refGene_gn=NUP210L;refGene_tx=XM_019822506.2;refGene_gn=NUP210L;refGene_tx=XM_019822507.2;refGene_gn=NUP210L;refGene_tx=XM_019822508.2;refGene_gn=NUP210L;refGene_tx=XM_019822509.2;refGene_gn=NUP210L;refGene_tx=XM_019822510.2;refGene_gn=NUP210L;refGene_tx=XM_019822511.2;refGene_gn=NUP210L;refGene_tx=XM_019822512.2;refGene_gn=NUP210L;refGene_tx=XM_019822513.2;refGene_gn=NUP210L;refGene_tx=XM_019822514.2;refGene_gn=NUP210L;refGene_tx=XM_019822517.2;refGene_gn=NUP210L;refGene_tx=XM_019822518.2;refGene_gn=NUP210L;refGene_tx=XM_023248094.1
F1 71125578 71125729
refGene_gn=NUP210L;refGene_tx=XM_003999791.5;refGene_gn=NUP210L;refGen

e_tx=XM_019822505.2;refGene_gn=NUP210L;refGene_tx=XM_019822506.2;refGene_gn=NUP210L;refGene_tx=XM_019822507.2;refGene_gn=NUP210L;refGene_tx=XM_019822508.2;refGene_gn=NUP210L;refGene_tx=XM_019822509.2;refGene_gn=NUP210L;refGene_tx=XM_019822510.2;refGene_gn=NUP210L;refGene_tx=XM_019822511.2;refGene_gn=NUP210L;refGene_tx=XM_019822512.2;refGene_gn=NUP210L;refGene_tx=XM_019822513.2;refGene_gn=NUP210L;refGene_tx=XM_019822514.2;refGene_gn=NUP210L;refGene_tx=XM_019822517.2;refGene_gn=NUP210L;refGene_tx=XM_019822518.2;refGene_gn=NUP210L;refGene_tx=XM_019822521.2;refGene_gn=NUP210L;refGene_tx=XM_023248094.1
F1 71126655 71126788
refGene_gn=NUP210L;refGene_tx=XM_003999791.5;refGene_gn=NUP210L;refGene_tx=XM_019822505.2;refGene_gn=NUP210L;refGene_tx=XM_019822506.2;refGene_gn=NUP210L;refGene_tx=XM_019822507.2;refGene_gn=NUP210L;refGene_tx=XM_019822508.2;refGene_gn=NUP210L;refGene_tx=XM_019822509.2;refGene_gn=NUP210L;refGene_tx=XM_019822510.2;refGene_gn=NUP210L;refGene_tx=XM_019822511.2;refGene_gn=NUP210L;refGene_tx=XM_019822512.2;refGene_gn=NUP210L;refGene_tx=XM_019822513.2;refGene_gn=NUP210L;refGene_tx=XM_019822514.2;refGene_gn=NUP210L;refGene_tx=XM_019822517.2;refGene_gn=NUP210L;refGene_tx=XM_019822518.2;refGene_gn=NUP210L;refGene_tx=XM_019822521.2;refGene_gn=NUP210L;refGene_tx=XM_023248094.1
F1 71127852 71128011
refGene_gn=NUP210L;refGene_tx=XM_003999791.5;refGene_gn=NUP210L;refGene_tx=XM_019822505.2;refGene_gn=NUP210L;refGene_tx=XM_019822506.2;refGene_gn=NUP210L;refGene_tx=XM_019822507.2;refGene_gn=NUP210L;refGene_tx=XM_019822508.2;refGene_gn=NUP210L;refGene_tx=XM_019822509.2;refGene_gn=NUP210L;refGene_tx=XM_019822510.2;refGene_gn=NUP210L;refGene_tx=XM_019822511.2;refGene_gn=NUP210L;refGene_tx=XM_019822512.2;refGene_gn=NUP210L;refGene_tx=XM_019822513.2;refGene_gn=NUP210L;refGene_tx=XM_019822514.2;refGene_gn=NUP210L;refGene_tx=XM_019822517.2;refGene_gn=NUP210L;refGene_tx=XM_019822518.2;refGene_gn=NUP210L;refGene_tx=XM_019822521.2;refGene_gn=NUP210L;refGene_tx=XM_023248094.1
F1 71128880 71128949
refGene_gn=NUP210L;refGene_tx=XM_003999791.5;refGene_gn=NUP210L;refGene_tx=XM_019822505.2;refGene_gn=NUP210L;refGene_tx=XM_019822506.2;refGene_gn=NUP210L;refGene_tx=XM_019822507.2;refGene_gn=NUP210L;refGene_tx=XM_019822508.2;refGene_gn=NUP210L;refGene_tx=XM_019822509.2;refGene_gn=NUP210L;refGene_tx=XM_019822510.2;refGene_gn=NUP210L;refGene_tx=XM_019822511.2;refGene_gn=NUP210L;refGene_tx=XM_019822512.2;refGene_gn=NUP210L;refGene_tx=XM_019822513.2;refGene_gn=NUP210L;refGene_tx=XM_019822514.2;refGene_gn=NUP210L;refGene_tx=XM_019822517.2;refGene_gn=NUP210L;refGene_tx=XM_019822518.2;refGene_gn=NUP210L;refGene_tx=XM_019822521.2;refGene_gn=NUP210L;refGene_tx=XM_023248094.1
F1 71129759 71129866
refGene_gn=NUP210L;refGene_tx=XM_003999791.5;refGene_gn=NUP210L;refGene_tx=XM_019822505.2;refGene_gn=NUP210L;refGene_tx=XM_019822506.2;refGene_gn=NUP210L;refGene_tx=XM_019822507.2;refGene_gn=NUP210L;refGene_tx=XM_019822508.2;refGene_gn=NUP210L;refGene_tx=XM_019822509.2;refGene_gn=NUP210L;refGene_tx=XM_019822510.2;refGene_gn=NUP210L;refGene_tx=XM_019822511.2;refGene_gn=NUP210L;refGene_tx=XM_019822512.2;refGene_gn=NUP210L;refGene_tx=XM_019822513.2;refGene_gn=NUP210L;refGene_tx=XM_0198225

14.2;refGene_gn=NUP210L;refGene_tx=XM_019822517.2;refGene_gn=NUP210L;refGene_tx=XM_019822518.2;refGene_gn=NUP210L;refGene_tx=XM_019822521.2;refGene_gn=NUP210L;refGene_tx=XM_023248094.1
F1 71130486 71130627
refGene_gn=NUP210L;refGene_tx=XM_003999791.5;refGene_gn=NUP210L;refGene_tx=XM_019822505.2;refGene_gn=NUP210L;refGene_tx=XM_019822506.2;refGene_gn=NUP210L;refGene_tx=XM_019822507.2;refGene_gn=NUP210L;refGene_tx=XM_019822508.2;refGene_gn=NUP210L;refGene_tx=XM_019822509.2;refGene_gn=NUP210L;refGene_tx=XM_019822510.2;refGene_gn=NUP210L;refGene_tx=XM_019822511.2;refGene_gn=NUP210L;refGene_tx=XM_019822512.2;refGene_gn=NUP210L;refGene_tx=XM_019822513.2;refGene_gn=NUP210L;refGene_tx=XM_019822514.2;refGene_gn=NUP210L;refGene_tx=XM_019822517.2;refGene_gn=NUP210L;refGene_tx=XM_019822518.2;refGene_gn=NUP210L;refGene_tx=XM_019822521.2;refGene_gn=NUP210L;refGene_tx=XM_023248094.1
F1 71144235 71144373
refGene_gn=NUP210L;refGene_tx=XM_003999791.5;refGene_gn=NUP210L;refGene_tx=XM_019822505.2;refGene_gn=NUP210L;refGene_tx=XM_019822506.2;refGene_gn=NUP210L;refGene_tx=XM_019822507.2;refGene_gn=NUP210L;refGene_tx=XM_019822508.2;refGene_gn=NUP210L;refGene_tx=XM_019822509.2;refGene_gn=NUP210L;refGene_tx=XM_019822510.2;refGene_gn=NUP210L;refGene_tx=XM_019822511.2;refGene_gn=NUP210L;refGene_tx=XM_019822513.2;refGene_gn=NUP210L;refGene_tx=XM_019822514.2;refGene_gn=NUP210L;refGene_tx=XM_019822517.2;refGene_gn=NUP210L;refGene_tx=XM_019822518.2;refGene_gn=NUP210L;refGene_tx=XM_019822521.2;refGene_gn=NUP210L;refGene_tx=XM_023248094.1
F1 71145073 71145229
refGene_gn=NUP210L;refGene_tx=XM_003999791.5;refGene_gn=NUP210L;refGene_tx=XM_019822505.2;refGene_gn=NUP210L;refGene_tx=XM_019822506.2;refGene_gn=NUP210L;refGene_tx=XM_019822507.2;refGene_gn=NUP210L;refGene_tx=XM_019822508.2;refGene_gn=NUP210L;refGene_tx=XM_019822509.2;refGene_gn=NUP210L;refGene_tx=XM_019822510.2;refGene_gn=NUP210L;refGene_tx=XM_019822511.2;refGene_gn=NUP210L;refGene_tx=XM_019822512.2;refGene_gn=NUP210L;refGene_tx=XM_019822513.2;refGene_gn=NUP210L;refGene_tx=XM_019822514.2;refGene_gn=NUP210L;refGene_tx=XM_019822517.2;refGene_gn=NUP210L;refGene_tx=XM_019822518.2;refGene_gn=NUP210L;refGene_tx=XM_019822521.2;refGene_gn=NUP210L;refGene_tx=XM_023248094.1
F1 71148508 71148707
refGene_gn=NUP210L;refGene_tx=XM_003999791.5;refGene_gn=NUP210L;refGene_tx=XM_019822505.2;refGene_gn=NUP210L;refGene_tx=XM_019822506.2;refGene_gn=NUP210L;refGene_tx=XM_019822507.2;refGene_gn=NUP210L;refGene_tx=XM_019822508.2;refGene_gn=NUP210L;refGene_tx=XM_019822509.2;refGene_gn=NUP210L;refGene_tx=XM_019822510.2;refGene_gn=NUP210L;refGene_tx=XM_019822511.2;refGene_gn=NUP210L;refGene_tx=XM_019822512.2;refGene_gn=NUP210L;refGene_tx=XM_019822513.2;refGene_gn=NUP210L;refGene_tx=XM_019822514.2;refGene_gn=NUP210L;refGene_tx=XM_019822517.2;refGene_gn=NUP210L;refGene_tx=XM_019822518.2;refGene_gn=NUP210L;refGene_tx=XM_019822521.2;refGene_gn=NUP210L;refGene_tx=XM_023248094.1
F1 71150890 71151036
refGene_gn=NUP210L;refGene_tx=XM_003999791.5;refGene_gn=NUP210L;refGene_tx=XM_019822505.2;refGene_gn=NUP210L;refGene_tx=XM_019822506.2;refGene_gn=NUP210L;refGene_tx=XM_019822507.2;refGene_gn=NUP210L;refGene_tx=

XM_019822508.2;refGene_gn=NUP210L;refGene_tx=XM_019822509.2;refGene_gn=NUP210L;refGene_tx=XM_019822510.2;refGene_gn=NUP210L;refGene_tx=XM_019822511.2;refGene_gn=NUP210L;refGene_tx=XM_019822512.2;refGene_gn=NUP210L;refGene_tx=XM_019822513.2;refGene_gn=NUP210L;refGene_tx=XM_019822514.2;refGene_gn=NUP210L;refGene_tx=XM_019822517.2;refGene_gn=NUP210L;refGene_tx=XM_019822518.2;refGene_gn=NUP210L;refGene_tx=XM_019822521.2;refGene_gn=NUP210L;refGene_tx=XM_023248094.1
F1 71158187 71158409
refGene_gn=NUP210L;refGene_tx=XM_003999791.5;refGene_gn=NUP210L;refGene_tx=XM_019822505.2;refGene_gn=NUP210L;refGene_tx=XM_019822506.2;refGene_gn=NUP210L;refGene_tx=XM_019822507.2;refGene_gn=NUP210L;refGene_tx=XM_019822508.2;refGene_gn=NUP210L;refGene_tx=XM_019822509.2;refGene_gn=NUP210L;refGene_tx=XM_019822510.2;refGene_gn=NUP210L;refGene_tx=XM_019822511.2;refGene_gn=NUP210L;refGene_tx=XM_019822512.2;refGene_gn=NUP210L;refGene_tx=XM_019822513.2;refGene_gn=NUP210L;refGene_tx=XM_019822514.2;refGene_gn=NUP210L;refGene_tx=XM_019822517.2;refGene_gn=NUP210L;refGene_tx=XM_019822518.2;refGene_gn=NUP210L;refGene_tx=XM_019822521.2;refGene_gn=NUP210L;refGene_tx=XM_023248094.1
F1 71159697 71159871
refGene_gn=NUP210L;refGene_tx=XM_003999791.5;refGene_gn=NUP210L;refGene_tx=XM_019822505.2;refGene_gn=NUP210L;refGene_tx=XM_019822506.2;refGene_gn=NUP210L;refGene_tx=XM_019822507.2;refGene_gn=NUP210L;refGene_tx=XM_019822508.2;refGene_gn=NUP210L;refGene_tx=XM_019822509.2;refGene_gn=NUP210L;refGene_tx=XM_019822510.2;refGene_gn=NUP210L;refGene_tx=XM_019822511.2;refGene_gn=NUP210L;refGene_tx=XM_019822512.2;refGene_gn=NUP210L;refGene_tx=XM_019822513.2;refGene_gn=NUP210L;refGene_tx=XM_019822514.2;refGene_gn=NUP210L;refGene_tx=XM_019822517.2;refGene_gn=NUP210L;refGene_tx=XM_019822518.2;refGene_gn=NUP210L;refGene_tx=XM_019822521.2;refGene_gn=NUP210L;refGene_tx=XM_023248094.1
F1 71164150 71164343
refGene_gn=NUP210L;refGene_tx=XM_003999791.5;refGene_gn=NUP210L;refGene_tx=XM_019822505.2;refGene_gn=NUP210L;refGene_tx=XM_019822506.2;refGene_gn=NUP210L;refGene_tx=XM_019822507.2;refGene_gn=NUP210L;refGene_tx=XM_019822508.2;refGene_gn=NUP210L;refGene_tx=XM_019822509.2;refGene_gn=NUP210L;refGene_tx=XM_019822510.2;refGene_gn=NUP210L;refGene_tx=XM_019822511.2;refGene_gn=NUP210L;refGene_tx=XM_019822512.2;refGene_gn=NUP210L;refGene_tx=XM_019822513.2;refGene_gn=NUP210L;refGene_tx=XM_019822514.2;refGene_gn=NUP210L;refGene_tx=XM_019822517.2;refGene_gn=NUP210L;refGene_tx=XM_019822518.2;refGene_gn=NUP210L;refGene_tx=XM_019822521.2;refGene_gn=NUP210L;refGene_tx=XM_023248094.1
F1 71166100 71166189
refGene_gn=NUP210L;refGene_tx=XM_003999791.5;refGene_gn=NUP210L;refGene_tx=XM_019822505.2;refGene_gn=NUP210L;refGene_tx=XM_019822506.2;refGene_gn=NUP210L;refGene_tx=XM_019822507.2;refGene_gn=NUP210L;refGene_tx=XM_019822508.2;refGene_gn=NUP210L;refGene_tx=XM_019822509.2;refGene_gn=NUP210L;refGene_tx=XM_019822510.2;refGene_gn=NUP210L;refGene_tx=XM_019822511.2;refGene_gn=NUP210L;refGene_tx=XM_019822512.2;refGene_gn=NUP210L;refGene_tx=XM_019822513.2;refGene_gn=NUP210L;refGene_tx=XM_019822514.2;refGene_gn=NUP210L;refGene_tx=XM_019822517.2;refGene_gn=NUP210L;refGene_tx=XM_019822518.2;refGene_gn=NUP210L;refGene_tx=XM_019822521.2;

refGene_gn=NUP210L;refGene_tx=XM_023248094.1
F1	71166625 71166730
refGene_gn=NUP210L;refGene_tx=XM_003999791.5;refGene_gn=NUP210L;refGene_tx=XM_019822505.2;refGene_gn=NUP210L;refGene_tx=XM_019822506.2;refGene_gn=NUP210L;refGene_tx=XM_019822507.2;refGene_gn=NUP210L;refGene_tx=XM_019822508.2;refGene_gn=NUP210L;refGene_tx=XM_019822509.2;refGene_gn=NUP210L;refGene_tx=XM_019822510.2;refGene_gn=NUP210L;refGene_tx=XM_019822511.2;refGene_gn=NUP210L;refGene_tx=XM_019822512.2;refGene_gn=NUP210L;refGene_tx=XM_019822513.2;refGene_gn=NUP210L;refGene_tx=XM_019822514.2;refGene_gn=NUP210L;refGene_tx=XM_019822517.2;refGene_gn=NUP210L;refGene_tx=XM_019822518.2;refGene_gn=NUP210L;refGene_tx=XM_019822521.2;refGene_gn=NUP210L;refGene_tx=XM_023248094.1
F1	71167034 71167136
refGene_gn=NUP210L;refGene_tx=XM_003999791.5;refGene_gn=NUP210L;refGene_tx=XM_019822505.2;refGene_gn=NUP210L;refGene_tx=XM_019822506.2;refGene_gn=NUP210L;refGene_tx=XM_019822507.2;refGene_gn=NUP210L;refGene_tx=XM_019822508.2;refGene_gn=NUP210L;refGene_tx=XM_019822509.2;refGene_gn=NUP210L;refGene_tx=XM_019822510.2;refGene_gn=NUP210L;refGene_tx=XM_019822511.2;refGene_gn=NUP210L;refGene_tx=XM_019822512.2;refGene_gn=NUP210L;refGene_tx=XM_019822513.2;refGene_gn=NUP210L;refGene_tx=XM_019822514.2;refGene_gn=NUP210L;refGene_tx=XM_019822517.2;refGene_gn=NUP210L;refGene_tx=XM_019822518.2;refGene_gn=NUP210L;refGene_tx=XM_019822521.2;refGene_gn=NUP210L;refGene_tx=XM_023248094.1
F1	71168076 71168205
refGene_gn=NUP210L;refGene_tx=XM_003999791.5;refGene_gn=NUP210L;refGene_tx=XM_019822505.2;refGene_gn=NUP210L;refGene_tx=XM_019822506.2;refGene_gn=NUP210L;refGene_tx=XM_019822507.2;refGene_gn=NUP210L;refGene_tx=XM_019822508.2;refGene_gn=NUP210L;refGene_tx=XM_019822509.2;refGene_gn=NUP210L;refGene_tx=XM_019822510.2;refGene_gn=NUP210L;refGene_tx=XM_019822511.2;refGene_gn=NUP210L;refGene_tx=XM_019822512.2;refGene_gn=NUP210L;refGene_tx=XM_019822513.2;refGene_gn=NUP210L;refGene_tx=XM_019822514.2;refGene_gn=NUP210L;refGene_tx=XM_019822517.2;refGene_gn=NUP210L;refGene_tx=XM_019822518.2;refGene_gn=NUP210L;refGene_tx=XM_019822521.2;refGene_gn=NUP210L;refGene_tx=XM_023248094.1
F1	71168551 71168679
refGene_gn=NUP210L;refGene_tx=XM_003999791.5;refGene_gn=NUP210L;refGene_tx=XM_019822505.2;refGene_gn=NUP210L;refGene_tx=XM_019822506.2;refGene_gn=NUP210L;refGene_tx=XM_019822507.2;refGene_gn=NUP210L;refGene_tx=XM_019822508.2;refGene_gn=NUP210L;refGene_tx=XM_019822509.2;refGene_gn=NUP210L;refGene_tx=XM_019822510.2;refGene_gn=NUP210L;refGene_tx=XM_019822511.2;refGene_gn=NUP210L;refGene_tx=XM_019822512.2;refGene_gn=NUP210L;refGene_tx=XM_019822513.2;refGene_gn=NUP210L;refGene_tx=XM_019822514.2;refGene_gn=NUP210L;refGene_tx=XM_019822517.2;refGene_gn=NUP210L;refGene_tx=XM_019822518.2;refGene_gn=NUP210L;refGene_tx=XM_019822521.2;refGene_gn=NUP210L;refGene_tx=XM_023248094.1
F1	71169470 71169603
refGene_gn=NUP210L;refGene_tx=XM_003999791.5;refGene_gn=NUP210L;refGene_tx=XM_019822505.2;refGene_gn=NUP210L;refGene_tx=XM_019822506.2;refGene_gn=NUP210L;refGene_tx=XM_019822507.2;refGene_gn=NUP210L;refGene_tx=XM_019822508.2;refGene_gn=NUP210L;refGene_tx=XM_019822509.2;refGene_gn

=NUP210L;refGene_tx=XM_019822510.2;refGene_gn=NUP210L;refGene_tx=XM_019822511.2;refGene_gn=NUP210L;refGene_tx=XM_019822512.2;refGene_gn=NUP210L;refGene_tx=XM_019822513.2;refGene_gn=NUP210L;refGene_tx=XM_019822514.2;refGene_gn=NUP210L;refGene_tx=XM_019822517.2;refGene_gn=NUP210L;refGene_tx=XM_019822518.2;refGene_gn=NUP210L;refGene_tx=XM_019822521.2;refGene_gn=NUP210L;refGene_tx=XM_023248094.1
F1 71170256 71170319
refGene_gn=NUP210L;refGene_tx=XM_003999791.5;refGene_gn=NUP210L;refGene_tx=XM_019822505.2;refGene_gn=NUP210L;refGene_tx=XM_019822506.2;refGene_gn=NUP210L;refGene_tx=XM_019822507.2;refGene_gn=NUP210L;refGene_tx=XM_019822508.2;refGene_gn=NUP210L;refGene_tx=XM_019822509.2;refGene_gn=NUP210L;refGene_tx=XM_019822510.2;refGene_gn=NUP210L;refGene_tx=XM_019822511.2;refGene_gn=NUP210L;refGene_tx=XM_019822512.2;refGene_gn=NUP210L;refGene_tx=XM_019822513.2;refGene_gn=NUP210L;refGene_tx=XM_019822514.2;refGene_gn=NUP210L;refGene_tx=XM_019822517.2;refGene_gn=NUP210L;refGene_tx=XM_019822518.2;refGene_gn=NUP210L;refGene_tx=XM_019822521.2;refGene_gn=NUP210L;refGene_tx=XM_023248094.1
F1 71170684 71170864
refGene_gn=NUP210L;refGene_tx=XM_003999791.5;refGene_gn=NUP210L;refGene_tx=XM_019822505.2;refGene_gn=NUP210L;refGene_tx=XM_019822506.2;refGene_gn=NUP210L;refGene_tx=XM_019822507.2;refGene_gn=NUP210L;refGene_tx=XM_019822508.2;refGene_gn=NUP210L;refGene_tx=XM_019822509.2;refGene_gn=NUP210L;refGene_tx=XM_019822510.2;refGene_gn=NUP210L;refGene_tx=XM_019822511.2;refGene_gn=NUP210L;refGene_tx=XM_019822512.2;refGene_gn=NUP210L;refGene_tx=XM_019822517.2;refGene_gn=NUP210L;refGene_tx=XM_019822518.2;refGene_gn=NUP210L;refGene_tx=XM_019822521.2;refGene_gn=NUP210L;refGene_tx=XM_023248094.1
F1 71171479 71171560
refGene_gn=NUP210L;refGene_tx=XM_003999791.5;refGene_gn=NUP210L;refGene_tx=XM_019822505.2;refGene_gn=NUP210L;refGene_tx=XM_019822506.2;refGene_gn=NUP210L;refGene_tx=XM_019822507.2;refGene_gn=NUP210L;refGene_tx=XM_019822508.2;refGene_gn=NUP210L;refGene_tx=XM_019822509.2;refGene_gn=NUP210L;refGene_tx=XM_019822510.2;refGene_gn=NUP210L;refGene_tx=XM_019822511.2;refGene_gn=NUP210L;refGene_tx=XM_019822512.2;refGene_gn=NUP210L;refGene_tx=XM_019822513.2;refGene_gn=NUP210L;refGene_tx=XM_019822514.2;refGene_gn=NUP210L;refGene_tx=XM_019822517.2;refGene_gn=NUP210L;refGene_tx=XM_019822518.2;refGene_gn=NUP210L;refGene_tx=XM_019822521.2;refGene_gn=NUP210L;refGene_tx=XM_023248094.1
F1 71171633 71171786
refGene_gn=NUP210L;refGene_tx=XM_003999791.5;refGene_gn=NUP210L;refGene_tx=XM_019822505.2;refGene_gn=NUP210L;refGene_tx=XM_019822506.2;refGene_gn=NUP210L;refGene_tx=XM_019822507.2;refGene_gn=NUP210L;refGene_tx=XM_019822508.2;refGene_gn=NUP210L;refGene_tx=XM_019822509.2;refGene_gn=NUP210L;refGene_tx=XM_019822510.2;refGene_gn=NUP210L;refGene_tx=XM_019822511.2;refGene_gn=NUP210L;refGene_tx=XM_019822512.2;refGene_gn=NUP210L;refGene_tx=XM_019822513.2;refGene_gn=NUP210L;refGene_tx=XM_019822514.2;refGene_gn=NUP210L;refGene_tx=XM_019822517.2;refGene_gn=NUP210L;refGene_tx=XM_019822518.2;refGene_gn=NUP210L;refGene_tx=XM_019822521.2;refGene_gn=NUP210L;refGene_tx=XM_023248094.1
F1 71174414 71174573

refGene_gn=NUP210L;refGene_tx=XM_003999791.5;refGene_gn=NUP210L;refGene_tx=XM_019822505.2;refGene_gn=NUP210L;refGene_tx=XM_019822506.2;refGene_gn=NUP210L;refGene_tx=XM_019822507.2;refGene_gn=NUP210L;refGene_tx=XM_019822508.2;refGene_gn=NUP210L;refGene_tx=XM_019822509.2;refGene_gn=NUP210L;refGene_tx=XM_019822510.2;refGene_gn=NUP210L;refGene_tx=XM_019822511.2;refGene_gn=NUP210L;refGene_tx=XM_019822512.2;refGene_gn=NUP210L;refGene_tx=XM_019822513.2;refGene_gn=NUP210L;refGene_tx=XM_019822514.2;refGene_gn=NUP210L;refGene_tx=XM_019822517.2;refGene_gn=NUP210L;refGene_tx=XM_019822518.2;refGene_gn=NUP210L;refGene_tx=XM_019822521.2;refGene_gn=NUP210L;refGene_tx=XM_023248094.1
F1 71176129 71176221
refGene_gn=NUP210L;refGene_tx=XM_003999791.5;refGene_gn=NUP210L;refGene_tx=XM_019822505.2;refGene_gn=NUP210L;refGene_tx=XM_019822506.2;refGene_gn=NUP210L;refGene_tx=XM_019822507.2;refGene_gn=NUP210L;refGene_tx=XM_019822508.2;refGene_gn=NUP210L;refGene_tx=XM_019822509.2;refGene_gn=NUP210L;refGene_tx=XM_019822510.2;refGene_gn=NUP210L;refGene_tx=XM_019822511.2;refGene_gn=NUP210L;refGene_tx=XM_019822512.2;refGene_gn=NUP210L;refGene_tx=XM_019822513.2;refGene_gn=NUP210L;refGene_tx=XM_019822514.2;refGene_gn=NUP210L;refGene_tx=XM_019822517.2;refGene_gn=NUP210L;refGene_tx=XM_019822518.2;refGene_gn=NUP210L;refGene_tx=XM_019822521.2;refGene_gn=NUP210L;refGene_tx=XM_023248094.1
F1 71178326 71178501
refGene_gn=NUP210L;refGene_tx=XM_003999791.5;refGene_gn=NUP210L;refGene_tx=XM_019822505.2;refGene_gn=NUP210L;refGene_tx=XM_019822506.2;refGene_gn=NUP210L;refGene_tx=XM_019822507.2;refGene_gn=NUP210L;refGene_tx=XM_019822508.2;refGene_gn=NUP210L;refGene_tx=XM_019822509.2;refGene_gn=NUP210L;refGene_tx=XM_019822510.2;refGene_gn=NUP210L;refGene_tx=XM_019822511.2;refGene_gn=NUP210L;refGene_tx=XM_019822512.2;refGene_gn=NUP210L;refGene_tx=XM_019822513.2;refGene_gn=NUP210L;refGene_tx=XM_019822514.2;refGene_gn=NUP210L;refGene_tx=XM_019822517.2;refGene_gn=NUP210L;refGene_tx=XM_019822518.2;refGene_gn=NUP210L;refGene_tx=XM_019822521.2;refGene_gn=NUP210L;refGene_tx=XM_023248094.1
F1 71179330 71179506
refGene_gn=NUP210L;refGene_tx=XM_003999791.5;refGene_gn=NUP210L;refGene_tx=XM_019822505.2;refGene_gn=NUP210L;refGene_tx=XM_019822506.2;refGene_gn=NUP210L;refGene_tx=XM_019822507.2;refGene_gn=NUP210L;refGene_tx=XM_019822508.2;refGene_gn=NUP210L;refGene_tx=XM_019822509.2;refGene_gn=NUP210L;refGene_tx=XM_019822510.2;refGene_gn=NUP210L;refGene_tx=XM_019822511.2;refGene_gn=NUP210L;refGene_tx=XM_019822512.2;refGene_gn=NUP210L;refGene_tx=XM_019822513.2;refGene_gn=NUP210L;refGene_tx=XM_019822514.2;refGene_gn=NUP210L;refGene_tx=XM_019822517.2;refGene_gn=NUP210L;refGene_tx=XM_019822518.2;refGene_gn=NUP210L;refGene_tx=XM_019822521.2;refGene_gn=NUP210L;refGene_tx=XM_023248094.1
F1 71180322 71180540
refGene_gn=NUP210L;refGene_tx=XM_003999791.5;refGene_gn=NUP210L;refGene_tx=XM_019822505.2;refGene_gn=NUP210L;refGene_tx=XM_019822506.2;refGene_gn=NUP210L;refGene_tx=XM_019822507.2;refGene_gn=NUP210L;refGene_tx=XM_019822508.2;refGene_gn=NUP210L;refGene_tx=XM_019822509.2;refGene_gn=NUP210L;refGene_tx=XM_019822510.2;refGene_gn=NUP210L;refGene_tx=XM_019822511.2;refGene_gn=NUP210L;refGene_tx=XM_019822512.2;refGene_gn=NUP2

10L;refGene_tx=XM_019822513.2;refGene_gn=NUP210L;refGene_tx=XM_0198225
14.2;refGene_gn=NUP210L;refGene_tx=XM_019822517.2;refGene_gn=NUP210L;r
efGene_tx=XM_019822518.2;refGene_gn=NUP210L;refGene_tx=XM_019822521.2;
refGene_gn=NUP210L;refGene_tx=XM_023248094.1
F1 71182340 71182477
refGene_gn=NUP210L;refGene_tx=XM_003999791.5;refGene_gn=NUP210L;refGen
e_tx=XM_019822505.2;refGene_gn=NUP210L;refGene_tx=XM_019822506.2;refGe
ne_gn=NUP210L;refGene_tx=XM_019822507.2;refGene_gn=NUP210L;refGene_tx=
XM_019822508.2;refGene_gn=NUP210L;refGene_tx=XM_019822509.2;refGene_gn
=NUP210L;refGene_tx=XM_019822510.2;refGene_gn=NUP210L;refGene_tx=XM_01
9822511.2;refGene_gn=NUP210L;refGene_tx=XM_019822512.2;refGene_gn=NUP2
10L;refGene_tx=XM_019822513.2;refGene_gn=NUP210L;refGene_tx=XM_0198225
14.2;refGene_gn=NUP210L;refGene_tx=XM_019822517.2;refGene_gn=NUP210L;r
efGene_tx=XM_019822518.2;refGene_gn=NUP210L;refGene_tx=XM_019822521.2;
refGene_gn=NUP210L;refGene_tx=XM_023248094.1
F1 71187266 71187393
refGene_gn=NUP210L;refGene_tx=XM_003999791.5;refGene_gn=NUP210L;refGen
e_tx=XM_019822505.2;refGene_gn=NUP210L;refGene_tx=XM_019822506.2;refGe
ne_gn=NUP210L;refGene_tx=XM_019822507.2;refGene_gn=NUP210L;refGene_tx=
XM_019822508.2;refGene_gn=NUP210L;refGene_tx=XM_019822509.2;refGene_gn
=NUP210L;refGene_tx=XM_019822510.2;refGene_gn=NUP210L;refGene_tx=XM_01
9822511.2;refGene_gn=NUP210L;refGene_tx=XM_019822512.2;refGene_gn=NUP2
10L;refGene_tx=XM_019822513.2;refGene_gn=NUP210L;refGene_tx=XM_0198225
14.2;refGene_gn=NUP210L;refGene_tx=XM_019822517.2;refGene_gn=NUP210L;r
efGene_tx=XM_019822518.2;refGene_gn=NUP210L;refGene_tx=XM_019822521.2;
refGene_gn=NUP210L;refGene_tx=XM_023248094.1
F1 71190516 71190666
refGene_gn=NUP210L;refGene_tx=XM_003999791.5;refGene_gn=NUP210L;refGen
e_tx=XM_019822505.2;refGene_gn=NUP210L;refGene_tx=XM_019822506.2;refGe
ne_gn=NUP210L;refGene_tx=XM_019822507.2;refGene_gn=NUP210L;refGene_tx=
XM_019822508.2;refGene_gn=NUP210L;refGene_tx=XM_019822509.2;refGene_gn
=NUP210L;refGene_tx=XM_019822510.2;refGene_gn=NUP210L;refGene_tx=XM_01
9822511.2;refGene_gn=NUP210L;refGene_tx=XM_019822512.2;refGene_gn=NUP2
10L;refGene_tx=XM_019822513.2;refGene_gn=NUP210L;refGene_tx=XM_0198225
14.2;refGene_gn=NUP210L;refGene_tx=XM_019822517.2;refGene_gn=NUP210L;r
efGene_tx=XM_019822518.2;refGene_gn=NUP210L;refGene_tx=XM_019822521.2;
refGene_gn=NUP210L;refGene_tx=XM_023248094.1
F1 71192079 71192330
refGene_gn=NUP210L;refGene_tx=XM_003999791.5;refGene_gn=NUP210L;refGen
e_tx=XM_019822505.2;refGene_gn=NUP210L;refGene_tx=XM_019822506.2;refGe
ne_gn=NUP210L;refGene_tx=XM_019822507.2;refGene_gn=NUP210L;refGene_tx=
XM_019822508.2;refGene_gn=NUP210L;refGene_tx=XM_019822509.2;refGene_gn
=NUP210L;refGene_tx=XM_019822510.2;refGene_gn=NUP210L;refGene_tx=XM_01
9822511.2;refGene_gn=NUP210L;refGene_tx=XM_019822512.2;refGene_gn=NUP2
10L;refGene_tx=XM_019822513.2;refGene_gn=NUP210L;refGene_tx=XM_0198225
14.2;refGene_gn=NUP210L;refGene_tx=XM_019822517.2;refGene_gn=NUP210L;r
efGene_tx=XM_019822518.2;refGene_gn=NUP210L;refGene_tx=XM_019822521.2;
refGene_gn=NUP210L;refGene_tx=XM_023248094.1
F1 71193017 71193219
refGene_gn=NUP210L;refGene_tx=XM_003999791.5;refGene_gn=NUP210L;refGen

e_tx=XM_019822505.2;refGene_gn=NUP210L;refGene_tx=XM_019822506.2;refGene_gn=NUP210L;refGene_tx=XM_019822507.2;refGene_gn=NUP210L;refGene_tx=XM_019822508.2;refGene_gn=NUP210L;refGene_tx=XM_019822509.2;refGene_gn=NUP210L;refGene_tx=XM_019822510.2;refGene_gn=NUP210L;refGene_tx=XM_019822511.2;refGene_gn=NUP210L;refGene_tx=XM_019822512.2;refGene_gn=NUP210L;refGene_tx=XM_019822513.2;refGene_gn=NUP210L;refGene_tx=XM_019822514.2;refGene_gn=NUP210L;refGene_tx=XM_019822517.2;refGene_gn=NUP210L;refGene_tx=XM_019822518.2;refGene_gn=NUP210L;refGene_tx=XM_019822521.2;refGene_gn=NUP210L;refGene_tx=XM_023248094.1
F1 71195727 71195818
refGene_gn=NUP210L;refGene_tx=XM_019822505.2;refGene_gn=NUP210L;refGene_tx=XM_019822506.2;refGene_gn=NUP210L;refGene_tx=XM_019822507.2;refGene_gn=NUP210L;refGene_tx=XM_019822508.2;refGene_gn=NUP210L;refGene_tx=XM_019822509.2;refGene_gn=NUP210L;refGene_tx=XM_019822511.2;refGene_gn=NUP210L;refGene_tx=XM_019822512.2;refGene_gn=NUP210L;refGene_tx=XM_019822513.2;refGene_gn=NUP210L;refGene_tx=XM_019822514.2;refGene_gn=NUP210L;refGene_tx=XM_019822517.2;refGene_gn=NUP210L;refGene_tx=XM_019822518.2;refGene_gn=NUP210L;refGene_tx=XM_019822521.2;refGene_gn=NUP210L;refGene_tx=XM_023248094.1
F1 71195884 71195989
refGene_gn=NUP210L;refGene_tx=XM_003999791.5;refGene_gn=NUP210L;refGene_tx=XM_019822510.2
F1 71196728 71196803
refGene_gn=NUP210L;refGene_tx=XM_003999791.5;refGene_gn=NUP210L;refGene_tx=XM_019822505.2;refGene_gn=NUP210L;refGene_tx=XM_019822506.2;refGene_gn=NUP210L;refGene_tx=XM_019822507.2;refGene_gn=NUP210L;refGene_tx=XM_019822508.2;refGene_gn=NUP210L;refGene_tx=XM_019822509.2;refGene_gn=NUP210L;refGene_tx=XM_019822510.2;refGene_gn=NUP210L;refGene_tx=XM_019822511.2;refGene_gn=NUP210L;refGene_tx=XM_019822512.2;refGene_gn=NUP210L;refGene_tx=XM_019822513.2;refGene_gn=NUP210L;refGene_tx=XM_019822514.2;refGene_gn=NUP210L;refGene_tx=XM_019822517.2;refGene_gn=NUP210L;refGene_tx=XM_019822518.2;refGene_gn=NUP210L;refGene_tx=XM_019822521.2;refGene_gn=NUP210L;refGene_tx=XM_023248094.1
F1 71196880 71197130
refGene_gn=NUP210L;refGene_tx=XM_003999791.5;refGene_gn=NUP210L;refGene_tx=XM_019822505.2;refGene_gn=NUP210L;refGene_tx=XM_019822506.2;refGene_gn=NUP210L;refGene_tx=XM_019822507.2;refGene_gn=NUP210L;refGene_tx=XM_019822508.2;refGene_gn=NUP210L;refGene_tx=XM_019822509.2;refGene_gn=NUP210L;refGene_tx=XM_019822510.2;refGene_gn=NUP210L;refGene_tx=XM_019822511.2;refGene_gn=NUP210L;refGene_tx=XM_019822512.2;refGene_gn=NUP210L;refGene_tx=XM_019822513.2;refGene_gn=NUP210L;refGene_tx=XM_019822514.2;refGene_gn=NUP210L;refGene_tx=XM_019822517.2;refGene_gn=NUP210L;refGene_tx=XM_019822518.2;refGene_gn=NUP210L;refGene_tx=XM_019822521.2;refGene_gn=NUP210L;refGene_tx=XM_023248094.1
F1 71197641 71197670 refGene_gn=RPS27;refGene_tx=XM_023248096.1
F1 71197957 71198068 refGene_gn=RPS27;refGene_tx=XM_023248096.1
F1 71198170 71198279 refGene_gn=RPS27;refGene_tx=XM_023248096.1
F1 71198542 71198548 refGene_gn=RPS27;refGene_tx=XM_023248096.1
F1 71200598 71200722
refGene_gn=RAB13;refGene_tx=XM_003999761.5;refGene_gn=RAB13;refGene_tx

=XM_006943173.4
F1 71202020 71202081
refGene_gn=RAB13;refGene_tx=XM_003999761.5;refGene_gn=RAB13;refGene_tx
=XM_006943173.4
F1 71202865 71202926
refGene_gn=RAB13;refGene_tx=XM_003999761.5;refGene_gn=RAB13;refGene_tx
=XM_006943173.4;refGene_gn=RAB13;refGene_tx=XM_019822532.2
F1 71203161 71203239
refGene_gn=RAB13;refGene_tx=XM_003999761.5;refGene_gn=RAB13;refGene_tx
=XM_006943173.4;refGene_gn=RAB13;refGene_tx=XM_019822532.2
F1 71203346 71203436
refGene_gn=RAB13;refGene_tx=XM_003999761.5;refGene_gn=RAB13;refGene_tx
=XM_006943173.4;refGene_gn=RAB13;refGene_tx=XM_019822532.2
F1 71203524 71203590
refGene_gn=RAB13;refGene_tx=XM_003999761.5;refGene_gn=RAB13;refGene_tx
=XM_006943173.4;refGene_gn=RAB13;refGene_tx=XM_019822532.2
F1 71203691 71203745
refGene_gn=RAB13;refGene_tx=XM_003999761.5;refGene_gn=RAB13;refGene_tx
=XM_006943173.4;refGene_gn=RAB13;refGene_tx=XM_019822532.2
F1 71203922 71204000
refGene_gn=RAB13;refGene_tx=XM_003999761.5;refGene_gn=RAB13;refGene_tx
=XM_006943173.4;refGene_gn=RAB13;refGene_tx=XM_019822532.2
F1 71207552 71207635 refGene_gn=JTB;refGene_tx=XM_003999762.5
F1 71207808 71207846 refGene_gn=JTB;refGene_tx=XM_003999762.5
F1 71208042 71208125 refGene_gn=JTB;refGene_tx=XM_003999762.5
F1 71208665 71208745 refGene_gn=JTB;refGene_tx=XM_003999762.5
F1 71209127 71209284 refGene_gn=JTB;refGene_tx=XM_003999762.5
F1 71209815 71210009
refGene_gn=CREB3L4;refGene_tx=XM_003999763.5;refGene_gn=CREB3L4;refGen
e_tx=XM_006943177.4
F1 71210132 71210229
refGene_gn=CREB3L4;refGene_tx=XM_003999763.5;refGene_gn=CREB3L4;refGen
e_tx=XM_006943177.4
F1 71210405 71210490
refGene_gn=CREB3L4;refGene_tx=XM_003999763.5;refGene_gn=CREB3L4;refGen
e_tx=XM_006943177.4
F1 71210673 71210742
refGene_gn=CREB3L4;refGene_tx=XM_003999763.5;refGene_gn=CREB3L4;refGen
e_tx=XM_006943177.4
F1 71210861 71210968
refGene_gn=CREB3L4;refGene_tx=XM_003999763.5;refGene_gn=CREB3L4;refGen
e_tx=XM_006943177.4
F1 71211098 71211275
refGene_gn=CREB3L4;refGene_tx=XM_003999763.5;refGene_gn=CREB3L4;refGen
e_tx=XM_006943177.4
F1 71212616 71212735
refGene_gn=CREB3L4;refGene_tx=XM_003999763.5;refGene_gn=CREB3L4;refGen
e_tx=XM_006943177.4
F1 71212891 71213141
refGene_gn=CREB3L4;refGene_tx=XM_003999763.5;refGene_gn=CREB3L4;refGen

e_tx=XM_006943177.4
F1 71213371 71213545
refGene_gn=CREB3L4;refGene_tx=XM_003999763.5;refGene_gn=CREB3L4;refGen
e_tx=XM_006943177.4
F1 71218389 71218576
refGene_gn=SLC39A1;refGene_tx=XM_003999765.3;refGene_gn=SLC39A1;refGen
e_tx=XM_023248095.1
F1 71218742 71218873
refGene_gn=SLC39A1;refGene_tx=XM_003999765.3;refGene_gn=SLC39A1;refGen
e_tx=XM_023248095.1
F1 71219965 71220619
refGene_gn=SLC39A1;refGene_tx=XM_003999765.3;refGene_gn=SLC39A1;refGen
e_tx=XM_023248095.1
F1 71222199 71222352 refGene_gn=CRTC2;refGene_tx=XM_003999766.5
F1 71225021 71225123 refGene_gn=CRTC2;refGene_tx=XM_003999766.5
F1 71225211 71225328 refGene_gn=CRTC2;refGene_tx=XM_003999766.5
F1 71225862 71225924 refGene_gn=CRTC2;refGene_tx=XM_003999766.5
F1 71226501 71226570 refGene_gn=CRTC2;refGene_tx=XM_003999766.5
F1 71226721 71226825 refGene_gn=CRTC2;refGene_tx=XM_003999766.5
F1 71226928 71226958 refGene_gn=CRTC2;refGene_tx=XM_003999766.5
F1 71227098 71227163 refGene_gn=CRTC2;refGene_tx=XM_003999766.5
F1 71227245 71227295 refGene_gn=CRTC2;refGene_tx=XM_003999766.5
F1 71227414 71227665 refGene_gn=CRTC2;refGene_tx=XM_003999766.5
F1 71227969 71228400 refGene_gn=CRTC2;refGene_tx=XM_003999766.5
F1 71229511 71229766 refGene_gn=CRTC2;refGene_tx=XM_003999766.5
F1 71230212 71230399 refGene_gn=CRTC2;refGene_tx=XM_003999766.5
F1 71230526 71230747 refGene_gn=CRTC2;refGene_tx=XM_003999766.5
F1 71232773 71232783
refGene_gn=DENND4B;refGene_tx=XM_011291261.2;refGene_gn=DENND4B;refGen
e_tx=XM_019822533.2;refGene_gn=DENND4B;refGene_tx=XM_019822538.2;refGe
ne_gn=DENND4B;refGene_tx=XM_019822539.2;refGene_gn=DENND4B;refGene_tx=
XM_019822540.2;refGene_gn=DENND4B;refGene_tx=XM_019822541.2
F1 71233447 71233787
refGene_gn=DENND4B;refGene_tx=XM_011291261.2;refGene_gn=DENND4B;refGen
e_tx=XM_019822533.2;refGene_gn=DENND4B;refGene_tx=XM_019822534.2;refGe
ne_gn=DENND4B;refGene_tx=XM_019822535.2;refGene_gn=DENND4B;refGene_tx=
XM_019822536.2;refGene_gn=DENND4B;refGene_tx=XM_019822538.2;refGene_gn
=DENND4B;refGene_tx=XM_019822539.2;refGene_gn=DENND4B;refGene_tx=XM_01
9822540.2;refGene_gn=DENND4B;refGene_tx=XM_019822541.2;refGene_gn=DENN
D4B;refGene_tx=XM_023247278.1
F1 71234207 71234460
refGene_gn=DENND4B;refGene_tx=XM_011291261.2;refGene_gn=DENND4B;refGen
e_tx=XM_019822533.2;refGene_gn=DENND4B;refGene_tx=XM_019822534.2;refGe
ne_gn=DENND4B;refGene_tx=XM_019822535.2;refGene_gn=DENND4B;refGene_tx=
XM_019822536.2;refGene_gn=DENND4B;refGene_tx=XM_019822538.2;refGene_gn
=DENND4B;refGene_tx=XM_019822539.2;refGene_gn=DENND4B;refGene_tx=XM_01
9822540.2;refGene_gn=DENND4B;refGene_tx=XM_019822541.2;refGene_gn=DENN
D4B;refGene_tx=XM_023247278.1
F1 71234655 71234725
refGene_gn=DENND4B;refGene_tx=XM_011291261.2;refGene_gn=DENND4B;refGen

e_tx=XM_019822533.2;refGene_gn=DENND4B;refGene_tx=XM_019822534.2;refGene_gn=DENND4B;refGene_tx=XM_019822535.2;refGene_gn=DENND4B;refGene_tx=XM_019822536.2;refGene_gn=DENND4B;refGene_tx=XM_019822538.2;refGene_gn=DENND4B;refGene_tx=XM_019822539.2;refGene_gn=DENND4B;refGene_tx=XM_019822540.2;refGene_gn=DENND4B;refGene_tx=XM_019822541.2;refGene_gn=DENND4B;refGene_tx=XM_023247278.1
F1 71234923 71235093
refGene_gn=DENND4B;refGene_tx=XM_011291261.2;refGene_gn=DENND4B;refGene_tx=XM_019822533.2;refGene_gn=DENND4B;refGene_tx=XM_019822534.2;refGene_gn=DENND4B;refGene_tx=XM_019822535.2;refGene_gn=DENND4B;refGene_tx=XM_019822536.2;refGene_gn=DENND4B;refGene_tx=XM_019822538.2;refGene_gn=DENND4B;refGene_tx=XM_019822539.2;refGene_gn=DENND4B;refGene_tx=XM_019822540.2;refGene_gn=DENND4B;refGene_tx=XM_019822541.2;refGene_gn=DENND4B;refGene_tx=XM_023247278.1
F1 71235167 71235412
refGene_gn=DENND4B;refGene_tx=XM_011291261.2;refGene_gn=DENND4B;refGene_tx=XM_019822533.2;refGene_gn=DENND4B;refGene_tx=XM_019822534.2;refGene_gn=DENND4B;refGene_tx=XM_019822535.2;refGene_gn=DENND4B;refGene_tx=XM_019822536.2;refGene_gn=DENND4B;refGene_tx=XM_019822538.2;refGene_gn=DENND4B;refGene_tx=XM_019822539.2;refGene_gn=DENND4B;refGene_tx=XM_019822540.2;refGene_gn=DENND4B;refGene_tx=XM_019822541.2;refGene_gn=DENND4B;refGene_tx=XM_023247278.1
F1 71235707 71235774
refGene_gn=DENND4B;refGene_tx=XM_011291261.2;refGene_gn=DENND4B;refGene_tx=XM_019822533.2;refGene_gn=DENND4B;refGene_tx=XM_019822534.2;refGene_gn=DENND4B;refGene_tx=XM_019822535.2;refGene_gn=DENND4B;refGene_tx=XM_019822536.2;refGene_gn=DENND4B;refGene_tx=XM_019822538.2;refGene_gn=DENND4B;refGene_tx=XM_019822539.2;refGene_gn=DENND4B;refGene_tx=XM_019822540.2;refGene_gn=DENND4B;refGene_tx=XM_019822541.2;refGene_gn=DENND4B;refGene_tx=XM_023247278.1
F1 71235875 71235934
refGene_gn=DENND4B;refGene_tx=XM_011291261.2;refGene_gn=DENND4B;refGene_tx=XM_019822533.2;refGene_gn=DENND4B;refGene_tx=XM_019822534.2;refGene_gn=DENND4B;refGene_tx=XM_019822535.2;refGene_gn=DENND4B;refGene_tx=XM_019822536.2;refGene_gn=DENND4B;refGene_tx=XM_019822538.2;refGene_gn=DENND4B;refGene_tx=XM_019822539.2;refGene_gn=DENND4B;refGene_tx=XM_019822540.2;refGene_gn=DENND4B;refGene_tx=XM_019822541.2;refGene_gn=DENND4B;refGene_tx=XM_023247278.1
F1 71236126 71236271
refGene_gn=DENND4B;refGene_tx=XM_011291261.2;refGene_gn=DENND4B;refGene_tx=XM_019822533.2;refGene_gn=DENND4B;refGene_tx=XM_019822534.2;refGene_gn=DENND4B;refGene_tx=XM_019822535.2;refGene_gn=DENND4B;refGene_tx=XM_019822536.2;refGene_gn=DENND4B;refGene_tx=XM_019822538.2;refGene_gn=DENND4B;refGene_tx=XM_019822539.2;refGene_gn=DENND4B;refGene_tx=XM_019822540.2;refGene_gn=DENND4B;refGene_tx=XM_019822541.2;refGene_gn=DENND4B;refGene_tx=XM_023247278.1
F1 71236476 71236652
refGene_gn=DENND4B;refGene_tx=XM_011291261.2;refGene_gn=DENND4B;refGene_tx=XM_019822533.2;refGene_gn=DENND4B;refGene_tx=XM_019822534.2;refGene_gn=DENND4B;refGene_tx=XM_019822535.2;refGene_gn=DENND4B;refGene_tx=

XM_019822536.2;refGene_gn=DENND4B;refGene_tx=XM_019822538.2;refGene_gn=DENND4B;refGene_tx=XM_019822539.2;refGene_gn=DENND4B;refGene_tx=XM_019822540.2;refGene_gn=DENND4B;refGene_tx=XM_019822541.2;refGene_gn=DENND4B;refGene_tx=XM_023247278.1
F1 71236793 71236894
refGene_gn=DENND4B;refGene_tx=XM_011291261.2;refGene_gn=DENND4B;refGene_tx=XM_019822533.2;refGene_gn=DENND4B;refGene_tx=XM_019822534.2;refGene_gn=DENND4B;refGene_tx=XM_019822535.2;refGene_gn=DENND4B;refGene_tx=XM_019822536.2;refGene_gn=DENND4B;refGene_tx=XM_019822538.2;refGene_gn=DENND4B;refGene_tx=XM_019822539.2;refGene_gn=DENND4B;refGene_tx=XM_019822540.2;refGene_gn=DENND4B;refGene_tx=XM_019822541.2;refGene_gn=DENND4B;refGene_tx=XM_023247278.1
F1 71237166 71237480
refGene_gn=DENND4B;refGene_tx=XM_011291261.2;refGene_gn=DENND4B;refGene_tx=XM_019822533.2;refGene_gn=DENND4B;refGene_tx=XM_019822534.2;refGene_gn=DENND4B;refGene_tx=XM_019822535.2;refGene_gn=DENND4B;refGene_tx=XM_019822536.2;refGene_gn=DENND4B;refGene_tx=XM_019822538.2;refGene_gn=DENND4B;refGene_tx=XM_019822539.2;refGene_gn=DENND4B;refGene_tx=XM_019822540.2;refGene_gn=DENND4B;refGene_tx=XM_019822541.2;refGene_gn=DENND4B;refGene_tx=XM_023247278.1
F1 71237675 71237821
refGene_gn=DENND4B;refGene_tx=XM_011291261.2;refGene_gn=DENND4B;refGene_tx=XM_019822533.2;refGene_gn=DENND4B;refGene_tx=XM_019822534.2;refGene_gn=DENND4B;refGene_tx=XM_019822535.2;refGene_gn=DENND4B;refGene_tx=XM_019822536.2;refGene_gn=DENND4B;refGene_tx=XM_019822538.2;refGene_gn=DENND4B;refGene_tx=XM_019822539.2;refGene_gn=DENND4B;refGene_tx=XM_019822540.2;refGene_gn=DENND4B;refGene_tx=XM_019822541.2;refGene_gn=DENND4B;refGene_tx=XM_023247278.1
F1 71238149 71238289
refGene_gn=DENND4B;refGene_tx=XM_011291261.2;refGene_gn=DENND4B;refGene_tx=XM_019822533.2;refGene_gn=DENND4B;refGene_tx=XM_019822534.2;refGene_gn=DENND4B;refGene_tx=XM_019822535.2;refGene_gn=DENND4B;refGene_tx=XM_019822536.2;refGene_gn=DENND4B;refGene_tx=XM_019822538.2;refGene_gn=DENND4B;refGene_tx=XM_019822539.2;refGene_gn=DENND4B;refGene_tx=XM_019822540.2;refGene_gn=DENND4B;refGene_tx=XM_019822541.2;refGene_gn=DENND4B;refGene_tx=XM_023247278.1
F1 71238368 71238495
refGene_gn=DENND4B;refGene_tx=XM_011291261.2;refGene_gn=DENND4B;refGene_tx=XM_019822533.2;refGene_gn=DENND4B;refGene_tx=XM_019822534.2;refGene_gn=DENND4B;refGene_tx=XM_019822535.2;refGene_gn=DENND4B;refGene_tx=XM_019822536.2;refGene_gn=DENND4B;refGene_tx=XM_019822538.2;refGene_gn=DENND4B;refGene_tx=XM_019822539.2;refGene_gn=DENND4B;refGene_tx=XM_019822540.2;refGene_gn=DENND4B;refGene_tx=XM_019822541.2;refGene_gn=DENND4B;refGene_tx=XM_023247278.1
F1 71238866 71239092
refGene_gn=DENND4B;refGene_tx=XM_011291261.2;refGene_gn=DENND4B;refGene_tx=XM_019822533.2;refGene_gn=DENND4B;refGene_tx=XM_019822534.2;refGene_gn=DENND4B;refGene_tx=XM_019822535.2;refGene_gn=DENND4B;refGene_tx=XM_019822536.2;refGene_gn=DENND4B;refGene_tx=XM_019822538.2;refGene_gn=DENND4B;refGene_tx=XM_019822539.2;refGene_gn=DENND4B;refGene_tx=XM_01

9822540.2;refGene_gn=DENND4B;refGene_tx=XM_019822541.2;refGene_gn=DENND4B;refGene_tx=XM_023247278.1
F1 71239358 71239487
refGene_gn=DENND4B;refGene_tx=XM_011291261.2;refGene_gn=DENND4B;refGene_tx=XM_019822533.2;refGene_gn=DENND4B;refGene_tx=XM_019822534.2;refGene_gn=DENND4B;refGene_tx=XM_019822535.2;refGene_gn=DENND4B;refGene_tx=XM_019822536.2;refGene_gn=DENND4B;refGene_tx=XM_019822538.2;refGene_gn=DENND4B;refGene_tx=XM_019822539.2;refGene_gn=DENND4B;refGene_tx=XM_019822540.2;refGene_gn=DENND4B;refGene_tx=XM_019822541.2;refGene_gn=DENND4B;refGene_tx=XM_023247278.1
F1 71240268 71240434
refGene_gn=DENND4B;refGene_tx=XM_011291261.2;refGene_gn=DENND4B;refGene_tx=XM_019822533.2;refGene_gn=DENND4B;refGene_tx=XM_019822534.2;refGene_gn=DENND4B;refGene_tx=XM_019822535.2;refGene_gn=DENND4B;refGene_tx=XM_019822536.2;refGene_gn=DENND4B;refGene_tx=XM_019822538.2;refGene_gn=DENND4B;refGene_tx=XM_019822539.2;refGene_gn=DENND4B;refGene_tx=XM_019822540.2;refGene_gn=DENND4B;refGene_tx=XM_019822541.2;refGene_gn=DENND4B;refGene_tx=XM_023247278.1
F1 71240688 71240856
refGene_gn=DENND4B;refGene_tx=XM_011291261.2;refGene_gn=DENND4B;refGene_tx=XM_019822533.2;refGene_gn=DENND4B;refGene_tx=XM_019822534.2;refGene_gn=DENND4B;refGene_tx=XM_019822535.2;refGene_gn=DENND4B;refGene_tx=XM_019822536.2;refGene_gn=DENND4B;refGene_tx=XM_019822538.2;refGene_gn=DENND4B;refGene_tx=XM_019822539.2;refGene_gn=DENND4B;refGene_tx=XM_019822540.2;refGene_gn=DENND4B;refGene_tx=XM_019822541.2;refGene_gn=DENND4B;refGene_tx=XM_023247278.1
F1 71241047 71241436
refGene_gn=DENND4B;refGene_tx=XM_011291261.2;refGene_gn=DENND4B;refGene_tx=XM_019822533.2;refGene_gn=DENND4B;refGene_tx=XM_019822534.2;refGene_gn=DENND4B;refGene_tx=XM_019822535.2;refGene_gn=DENND4B;refGene_tx=XM_019822536.2;refGene_gn=DENND4B;refGene_tx=XM_019822538.2;refGene_gn=DENND4B;refGene_tx=XM_019822539.2;refGene_gn=DENND4B;refGene_tx=XM_019822540.2;refGene_gn=DENND4B;refGene_tx=XM_019822541.2;refGene_gn=DENND4B;refGene_tx=XM_023247278.1
F1 71241548 71241671
refGene_gn=DENND4B;refGene_tx=XM_011291261.2;refGene_gn=DENND4B;refGene_tx=XM_019822533.2;refGene_gn=DENND4B;refGene_tx=XM_019822534.2;refGene_gn=DENND4B;refGene_tx=XM_019822535.2;refGene_gn=DENND4B;refGene_tx=XM_019822536.2;refGene_gn=DENND4B;refGene_tx=XM_019822538.2;refGene_gn=DENND4B;refGene_tx=XM_019822539.2;refGene_gn=DENND4B;refGene_tx=XM_019822540.2;refGene_gn=DENND4B;refGene_tx=XM_019822541.2;refGene_gn=DENND4B;refGene_tx=XM_023247278.1
F1 71241760 71241930
refGene_gn=DENND4B;refGene_tx=XM_011291261.2;refGene_gn=DENND4B;refGene_tx=XM_019822533.2;refGene_gn=DENND4B;refGene_tx=XM_019822534.2;refGene_gn=DENND4B;refGene_tx=XM_019822535.2;refGene_gn=DENND4B;refGene_tx=XM_019822536.2;refGene_gn=DENND4B;refGene_tx=XM_019822538.2;refGene_gn=DENND4B;refGene_tx=XM_019822539.2;refGene_gn=DENND4B;refGene_tx=XM_019822540.2;refGene_gn=DENND4B;refGene_tx=XM_019822541.2;refGene_gn=DENND4B;refGene_tx=XM_023247278.1

F1 71242015 71242151
refGene_gn=DENND4B;refGene_tx=XM_011291261.2;refGene_gn=DENND4B;refGene_tx=XM_019822533.2;refGene_gn=DENND4B;refGene_tx=XM_019822534.2;refGene_gn=DENND4B;refGene_tx=XM_019822535.2;refGene_gn=DENND4B;refGene_tx=XM_019822536.2;refGene_gn=DENND4B;refGene_tx=XM_019822538.2;refGene_gn=DENND4B;refGene_tx=XM_019822539.2;refGene_gn=DENND4B;refGene_tx=XM_019822540.2;refGene_gn=DENND4B;refGene_tx=XM_019822541.2;refGene_gn=DENND4B;refGene_tx=XM_023247278.1
F1 71242341 71242578
refGene_gn=DENND4B;refGene_tx=XM_011291261.2;refGene_gn=DENND4B;refGene_tx=XM_019822533.2;refGene_gn=DENND4B;refGene_tx=XM_019822534.2;refGene_gn=DENND4B;refGene_tx=XM_019822535.2;refGene_gn=DENND4B;refGene_tx=XM_019822536.2;refGene_gn=DENND4B;refGene_tx=XM_019822538.2;refGene_gn=DENND4B;refGene_tx=XM_019822539.2;refGene_gn=DENND4B;refGene_tx=XM_019822540.2;refGene_gn=DENND4B;refGene_tx=XM_019822541.2;refGene_gn=DENND4B;refGene_tx=XM_023247278.1
F1 71242977 71243152
refGene_gn=DENND4B;refGene_tx=XM_011291261.2;refGene_gn=DENND4B;refGene_tx=XM_019822533.2;refGene_gn=DENND4B;refGene_tx=XM_019822534.2;refGene_gn=DENND4B;refGene_tx=XM_019822535.2;refGene_gn=DENND4B;refGene_tx=XM_019822536.2;refGene_gn=DENND4B;refGene_tx=XM_019822538.2;refGene_gn=DENND4B;refGene_tx=XM_019822539.2;refGene_gn=DENND4B;refGene_tx=XM_019822540.2;refGene_gn=DENND4B;refGene_tx=XM_019822541.2;refGene_gn=DENND4B;refGene_tx=XM_023247278.1
F1 71243243 71243409
refGene_gn=DENND4B;refGene_tx=XM_011291261.2;refGene_gn=DENND4B;refGene_tx=XM_019822533.2;refGene_gn=DENND4B;refGene_tx=XM_019822534.2;refGene_gn=DENND4B;refGene_tx=XM_019822535.2;refGene_gn=DENND4B;refGene_tx=XM_019822536.2;refGene_gn=DENND4B;refGene_tx=XM_019822538.2;refGene_gn=DENND4B;refGene_tx=XM_019822539.2;refGene_gn=DENND4B;refGene_tx=XM_019822540.2;refGene_gn=DENND4B;refGene_tx=XM_019822541.2;refGene_gn=DENND4B;refGene_tx=XM_023247278.1
F1 71243499 71243564
refGene_gn=DENND4B;refGene_tx=XM_011291261.2;refGene_gn=DENND4B;refGene_tx=XM_019822533.2;refGene_gn=DENND4B;refGene_tx=XM_019822534.2;refGene_gn=DENND4B;refGene_tx=XM_019822535.2;refGene_gn=DENND4B;refGene_tx=XM_019822536.2;refGene_gn=DENND4B;refGene_tx=XM_019822538.2;refGene_gn=DENND4B;refGene_tx=XM_019822539.2;refGene_gn=DENND4B;refGene_tx=XM_019822540.2;refGene_gn=DENND4B;refGene_tx=XM_019822541.2;refGene_gn=DENND4B;refGene_tx=XM_023247278.1
F1 71243683 71243829
refGene_gn=DENND4B;refGene_tx=XM_011291261.2;refGene_gn=DENND4B;refGene_tx=XM_019822533.2;refGene_gn=DENND4B;refGene_tx=XM_019822534.2;refGene_gn=DENND4B;refGene_tx=XM_019822535.2;refGene_gn=DENND4B;refGene_tx=XM_019822536.2;refGene_gn=DENND4B;refGene_tx=XM_019822538.2;refGene_gn=DENND4B;refGene_tx=XM_019822539.2;refGene_gn=DENND4B;refGene_tx=XM_023247278.1
F1 71290482 71290535 refGene_gn=GATAD2B;refGene_tx=XM_019822542.2
F1 71320004 71320340
refGene_gn=GATAD2B;refGene_tx=XM_019822542.2;refGene_gn=GATAD2B;refGen

e_tx=XM_019822543.2;refGene_gn=GATAD2B;refGene_tx=XM_023247900.1;refGene_gn=GATAD2B;refGene_tx=XM_023247901.1
F1 71325601 71325731
refGene_gn=GATAD2B;refGene_tx=XM_019822542.2;refGene_gn=GATAD2B;refGene_tx=XM_019822543.2;refGene_gn=GATAD2B;refGene_tx=XM_023247900.1;refGene_gn=GATAD2B;refGene_tx=XM_023247901.1
F1 71327652 71327784
refGene_gn=GATAD2B;refGene_tx=XM_019822542.2;refGene_gn=GATAD2B;refGene_tx=XM_019822543.2;refGene_gn=GATAD2B;refGene_tx=XM_023247900.1;refGene_gn=GATAD2B;refGene_tx=XM_023247901.1
F1 71328089 71328221
refGene_gn=GATAD2B;refGene_tx=XM_019822542.2;refGene_gn=GATAD2B;refGene_tx=XM_019822543.2;refGene_gn=GATAD2B;refGene_tx=XM_023247900.1;refGene_gn=GATAD2B;refGene_tx=XM_023247901.1
F1 71328592 71328763
refGene_gn=GATAD2B;refGene_tx=XM_019822542.2;refGene_gn=GATAD2B;refGene_tx=XM_019822543.2;refGene_gn=GATAD2B;refGene_tx=XM_023247900.1;refGene_gn=GATAD2B;refGene_tx=XM_023247901.1
F1 71329251 71329567
refGene_gn=GATAD2B;refGene_tx=XM_019822542.2;refGene_gn=GATAD2B;refGene_tx=XM_019822543.2;refGene_gn=GATAD2B;refGene_tx=XM_023247900.1;refGene_gn=GATAD2B;refGene_tx=XM_023247901.1
F1 71330540 71330743
refGene_gn=GATAD2B;refGene_tx=XM_019822542.2;refGene_gn=GATAD2B;refGene_tx=XM_019822543.2;refGene_gn=GATAD2B;refGene_tx=XM_023247900.1;refGene_gn=GATAD2B;refGene_tx=XM_023247901.1
F1 71331606 71331717
refGene_gn=GATAD2B;refGene_tx=XM_019822542.2;refGene_gn=GATAD2B;refGene_tx=XM_019822543.2;refGene_gn=GATAD2B;refGene_tx=XM_023247900.1;refGene_gn=GATAD2B;refGene_tx=XM_023247901.1
F1 71331862 71331980
refGene_gn=GATAD2B;refGene_tx=XM_019822542.2;refGene_gn=GATAD2B;refGene_tx=XM_019822543.2;refGene_gn=GATAD2B;refGene_tx=XM_023247900.1;refGene_gn=GATAD2B;refGene_tx=XM_023247901.1
F1 71332641 71332775
refGene_gn=GATAD2B;refGene_tx=XM_019822542.2;refGene_gn=GATAD2B;refGene_tx=XM_019822543.2;refGene_gn=GATAD2B;refGene_tx=XM_023247900.1;refGene_gn=GATAD2B;refGene_tx=XM_023247901.1
F1 71357307 71357484 refGene_gn=SLC27A3;refGene_tx=XM_023247477.1
F1 71357779 71357910 refGene_gn=SLC27A3;refGene_tx=XM_023247477.1
F1 71358036 71358134 refGene_gn=SLC27A3;refGene_tx=XM_023247477.1
F1 71358338 71358537 refGene_gn=SLC27A3;refGene_tx=XM_023247477.1
F1 71358656 71358747 refGene_gn=SLC27A3;refGene_tx=XM_023247477.1
F1 71358851 71359046 refGene_gn=SLC27A3;refGene_tx=XM_023247477.1
F1 71359244 71359369 refGene_gn=SLC27A3;refGene_tx=XM_023247477.1
F1 71359915 71360074 refGene_gn=SLC27A3;refGene_tx=XM_023247477.1
F1 71360370 71360580 refGene_gn=SLC27A3;refGene_tx=XM_023247477.1
F1 71360831 71361897 refGene_gn=SLC27A3;refGene_tx=XM_023247477.1
F1 71363300 71363378 refGene_gn=INTS3;refGene_tx=XM_003999768.5
F1 71363485 71363642 refGene_gn=INTS3;refGene_tx=XM_003999768.5

| | | | | |
|---|---|---|---|---|
| F1 | 71363807 | 71363880 | refGene_gn=INTS3; | refGene_tx=XM_003999768.5 |
| F1 | 71364029 | 71364130 | refGene_gn=INTS3; | refGene_tx=XM_003999768.5 |
| F1 | 71364350 | 71364518 | refGene_gn=INTS3; | refGene_tx=XM_003999768.5 |
| F1 | 71365317 | 71365366 | refGene_gn=INTS3; | refGene_tx=XM_003999768.5 |
| F1 | 71365687 | 71365801 | refGene_gn=INTS3; | refGene_tx=XM_003999768.5 |
| F1 | 71366157 | 71366233 | refGene_gn=INTS3; | refGene_tx=XM_003999768.5 |
| F1 | 71367019 | 71367088 | refGene_gn=INTS3; | refGene_tx=XM_003999768.5 |
| F1 | 71368098 | 71368252 | refGene_gn=INTS3; | refGene_tx=XM_003999768.5 |
| F1 | 71369908 | 71370028 | refGene_gn=INTS3; | refGene_tx=XM_003999768.5 |
| F1 | 71370225 | 71370270 | refGene_gn=INTS3; | refGene_tx=XM_003999768.5 |
| F1 | 71371090 | 71371194 | refGene_gn=INTS3; | refGene_tx=XM_003999768.5 |
| F1 | 71371443 | 71371498 | refGene_gn=INTS3; | refGene_tx=XM_003999768.5 |
| F1 | 71372139 | 71372269 | refGene_gn=INTS3; | refGene_tx=XM_003999768.5 |
| F1 | 71372549 | 71372669 | refGene_gn=INTS3; | refGene_tx=XM_003999768.5 |
| F1 | 71373355 | 71373462 | refGene_gn=INTS3; | refGene_tx=XM_003999768.5 |
| F1 | 71373786 | 71373878 | refGene_gn=INTS3; | refGene_tx=XM_003999768.5 |
| F1 | 71374260 | 71374340 | refGene_gn=INTS3; | refGene_tx=XM_003999768.5 |
| F1 | 71374820 | 71374908 | refGene_gn=INTS3; | refGene_tx=XM_003999768.5 |
| F1 | 71375848 | 71376040 | refGene_gn=INTS3; | refGene_tx=XM_003999768.5 |
| F1 | 71377552 | 71377650 | refGene_gn=INTS3; | refGene_tx=XM_003999768.5 |
| F1 | 71378925 | 71379055 | refGene_gn=INTS3; | refGene_tx=XM_003999768.5 |
| F1 | 71379964 | 71380109 | refGene_gn=INTS3; | refGene_tx=XM_003999768.5 |
| F1 | 71382467 | 71382534 | refGene_gn=INTS3; | refGene_tx=XM_003999768.5 |
| F1 | 71384759 | 71384844 | refGene_gn=INTS3; | refGene_tx=XM_003999768.5 |
| F1 | 71385038 | 71385152 | refGene_gn=INTS3; | refGene_tx=XM_003999768.5 |
| F1 | 71390261 | 71390345 | refGene_gn=INTS3; | refGene_tx=XM_003999768.5 |
| F1 | 71391736 | 71391820 | refGene_gn=INTS3; | refGene_tx=XM_003999768.5 |
| F1 | 71400163 | 71400307 | refGene_gn=INTS3; | refGene_tx=XM_003999768.5 |
| F1 | 71421457 | 71421520 | refGene_gn=NPR1; | refGene_tx=XM_023247379.1 |
| F1 | 71421690 | 71421782 | refGene_gn=NPR1; | refGene_tx=XM_023247379.1 |
| F1 | 71423613 | 71423712 | refGene_gn=NPR1; | refGene_tx=XM_023247379.1 |
| F1 | 71423838 | 71424013 | refGene_gn=NPR1; | refGene_tx=XM_023247379.1 |
| F1 | 71424263 | 71424332 | refGene_gn=NPR1; | refGene_tx=XM_023247379.1 |
| F1 | 71424485 | 71424609 | refGene_gn=NPR1; | refGene_tx=XM_023247379.1 |
| F1 | 71424695 | 71424842 | refGene_gn=NPR1; | refGene_tx=XM_023247379.1 |
| F1 | 71425361 | 71425498 | refGene_gn=NPR1; | refGene_tx=XM_023247379.1 |
| F1 | 71425499 | 71425531 | refGene_gn=NPR1; | refGene_tx=XM_023247379.1 |
| F1 | 71425770 | 71425926 | refGene_gn=NPR1; | refGene_tx=XM_023247379.1 |
| F1 | 71426229 | 71426386 | refGene_gn=NPR1; | refGene_tx=XM_023247379.1 |
| F1 | 71426500 | 71426572 | refGene_gn=NPR1; | refGene_tx=XM_023247379.1 |
| F1 | 71426823 | 71426928 | refGene_gn=NPR1; | refGene_tx=XM_023247379.1 |
| F1 | 71427216 | 71427294 | refGene_gn=NPR1; | refGene_tx=XM_023247379.1 |
| F1 | 71427436 | 71427511 | refGene_gn=NPR1; | refGene_tx=XM_023247379.1 |
| F1 | 71428120 | 71428241 | refGene_gn=NPR1; | refGene_tx=XM_023247379.1 |
| F1 | 71428926 | 71429011 | refGene_gn=NPR1; | refGene_tx=XM_023247379.1 |
| F1 | 71429227 | 71429363 | refGene_gn=NPR1; | refGene_tx=XM_023247379.1 |
| F1 | 71430068 | 71430160 | refGene_gn=NPR1; | refGene_tx=XM_023247379.1 |
| F1 | 71430520 | 71430656 | refGene_gn=NPR1; | refGene_tx=XM_023247379.1 |
| F1 | 71431074 | 71431188 | refGene_gn=NPR1; | refGene_tx=XM_023247379.1 |
| F1 | 71431614 | 71431814 | refGene_gn=NPR1; | refGene_tx=XM_023247379.1 |

F1 71432388 71433115 refGene_gn=NPR1;refGene_tx=XM_023247379.1
F1 71439307 71439312 refGene_gn=ILF2;refGene_tx=XM_003999770.4
F1 71439797 71439857 refGene_gn=ILF2;refGene_tx=XM_003999770.4
F1 71440032 71440075 refGene_gn=ILF2;refGene_tx=XM_003999770.4
F1 71440348 71440453
refGene_gn=ILF2;refGene_tx=XM_003999770.4;refGene_gn=ILF2;refGene_tx=XM_023247732.1
F1 71440836 71440914 refGene_gn=ILF2;refGene_tx=XM_003999770.4
F1 71441230 71441333
refGene_gn=ILF2;refGene_tx=XM_003999770.4;refGene_gn=ILF2;refGene_tx=XM_023247732.1
F1 71441927 71441993
refGene_gn=ILF2;refGene_tx=XM_003999770.4;refGene_gn=ILF2;refGene_tx=XM_023247732.1
F1 71442650 71442767
refGene_gn=ILF2;refGene_tx=XM_003999770.4;refGene_gn=ILF2;refGene_tx=XM_023247732.1
F1 71443198 71443277
refGene_gn=ILF2;refGene_tx=XM_003999770.4;refGene_gn=ILF2;refGene_tx=XM_023247732.1
F1 71443762 71443850
refGene_gn=ILF2;refGene_tx=XM_003999770.4;refGene_gn=ILF2;refGene_tx=XM_023247732.1
F1 71444239 71444301
refGene_gn=ILF2;refGene_tx=XM_003999770.4;refGene_gn=ILF2;refGene_tx=XM_023247732.1
F1 71444387 71444502
refGene_gn=ILF2;refGene_tx=XM_003999770.4;refGene_gn=ILF2;refGene_tx=XM_023247732.1
F1 71444718 71444809
refGene_gn=ILF2;refGene_tx=XM_003999770.4;refGene_gn=ILF2;refGene_tx=XM_023247732.1
F1 71444952 71445113
refGene_gn=ILF2;refGene_tx=XM_003999770.4;refGene_gn=ILF2;refGene_tx=XM_023247732.1
F1 71446142 71446244 refGene_gn=SNAPIN;refGene_tx=XM_011291270.3
F1 71446546 71446665 refGene_gn=SNAPIN;refGene_tx=XM_011291270.3
F1 71446697 71446816 refGene_gn=SNAPIN;refGene_tx=XM_019822546.2
F1 71446909 71446956
refGene_gn=SNAPIN;refGene_tx=XM_011291270.3;refGene_gn=SNAPIN;refGene_tx=XM_019822546.2
F1 71447167 71447569
refGene_gn=SNAPIN;refGene_tx=XM_011291270.3;refGene_gn=SNAPIN;refGene_tx=XM_019822546.2
F1 71449446 71449611
refGene_gn=SNAPIN;refGene_tx=XM_011291270.3;refGene_gn=SNAPIN;refGene_tx=XM_019822546.2
F1 71452357 71452497 refGene_gn=SNAPIN;refGene_tx=XM_019822546.2
F1 71456367 71456573
refGene_gn=CHTOP;refGene_tx=XM_006943184.4;refGene_gn=CHTOP;refGene_tx

=XM_006943185.3;refGene_gn=CHTOP;refGene_tx=XM_006943186.4;refGene_gn=CHTOP;refGene_tx=XM_011291272.3;refGene_gn=CHTOP;refGene_tx=XM_019822547.2;refGene_gn=CHTOP;refGene_tx=XM_019822548.2;refGene_gn=CHTOP;refGene_tx=XM_019822550.1;refGene_gn=CHTOP;refGene_tx=XM_019822551.2;refGene_gn=CHTOP;refGene_tx=XM_019822552.2;refGene_gn=CHTOP;refGene_tx=XM_019822553.2;refGene_gn=CHTOP;refGene_tx=XM_019822554.2;refGene_gn=CHTOP;refGene_tx=XM_023247734.1
F1 71458232 71458370
refGene_gn=CHTOP;refGene_tx=XM_006943184.4;refGene_gn=CHTOP;refGene_tx=XM_006943185.3;refGene_gn=CHTOP;refGene_tx=XM_006943186.4;refGene_gn=CHTOP;refGene_tx=XM_011291272.3;refGene_gn=CHTOP;refGene_tx=XM_019822547.2;refGene_gn=CHTOP;refGene_tx=XM_019822548.2;refGene_gn=CHTOP;refGene_tx=XM_019822550.1;refGene_gn=CHTOP;refGene_tx=XM_019822551.2;refGene_gn=CHTOP;refGene_tx=XM_019822552.2;refGene_gn=CHTOP;refGene_tx=XM_023247734.1
F1 71459186 71459373
refGene_gn=CHTOP;refGene_tx=XM_006943184.4;refGene_gn=CHTOP;refGene_tx=XM_006943185.3;refGene_gn=CHTOP;refGene_tx=XM_006943186.4;refGene_gn=CHTOP;refGene_tx=XM_011291272.3;refGene_gn=CHTOP;refGene_tx=XM_019822547.2;refGene_gn=CHTOP;refGene_tx=XM_019822548.2;refGene_gn=CHTOP;refGene_tx=XM_019822550.1;refGene_gn=CHTOP;refGene_tx=XM_019822551.2;refGene_gn=CHTOP;refGene_tx=XM_019822552.2;refGene_gn=CHTOP;refGene_tx=XM_019822553.2;refGene_gn=CHTOP;refGene_tx=XM_019822554.2;refGene_gn=CHTOP;refGene_tx=XM_023247734.1
F1 71461456 71461540
refGene_gn=CHTOP;refGene_tx=XM_006943184.4;refGene_gn=CHTOP;refGene_tx=XM_006943185.3;refGene_gn=CHTOP;refGene_tx=XM_006943186.4;refGene_gn=CHTOP;refGene_tx=XM_019822547.2;refGene_gn=CHTOP;refGene_tx=XM_019822548.2;refGene_gn=CHTOP;refGene_tx=XM_019822553.2;refGene_gn=CHTOP;refGene_tx=XM_023247734.1
F1 71462222 71462376
refGene_gn=CHTOP;refGene_tx=XM_006943184.4;refGene_gn=CHTOP;refGene_tx=XM_006943185.3;refGene_gn=CHTOP;refGene_tx=XM_006943186.4;refGene_gn=CHTOP;refGene_tx=XM_011291272.3;refGene_gn=CHTOP;refGene_tx=XM_019822547.2;refGene_gn=CHTOP;refGene_tx=XM_019822548.2;refGene_gn=CHTOP;refGene_tx=XM_019822550.1;refGene_gn=CHTOP;refGene_tx=XM_019822551.2;refGene_gn=CHTOP;refGene_tx=XM_019822552.2;refGene_gn=CHTOP;refGene_tx=XM_019822553.2;refGene_gn=CHTOP;refGene_tx=XM_019822554.2;refGene_gn=CHTOP;refGene_tx=XM_023247734.1
F1 71463911 71463976
refGene_gn=CHTOP;refGene_tx=XM_006943184.4;refGene_gn=CHTOP;refGene_tx=XM_006943185.3;refGene_gn=CHTOP;refGene_tx=XM_006943186.4;refGene_gn=CHTOP;refGene_tx=XM_011291272.3;refGene_gn=CHTOP;refGene_tx=XM_019822552.2;refGene_gn=CHTOP;refGene_tx=XM_019822554.2
F1 71464161 71464262
refGene_gn=CHTOP;refGene_tx=XM_019822547.2;refGene_gn=CHTOP;refGene_tx=XM_019822548.2;refGene_gn=CHTOP;refGene_tx=XM_019822550.1;refGene_gn=CHTOP;refGene_tx=XM_019822551.2;refGene_gn=CHTOP;refGene_tx=XM_019822553.2
F1 71466315 71466459

refGene_gn=S100A1;refGene_tx=XM_003999773.4;refGene_gn=S100A1;refGene_tx=XM_011291273.2
F1 71467549 71467690
refGene_gn=S100A1;refGene_tx=XM_003999773.4;refGene_gn=S100A1;refGene_tx=XM_011291273.2
F1 71471141 71471294 refGene_gn=S100A13;refGene_tx=XM_023247736.1
F1 71477085 71477229 refGene_gn=S100A13;refGene_tx=XM_023247736.1
F1 71480233 71480263
refGene_gn=S100A14;refGene_tx=XM_003999776.4;refGene_gn=S100A14;refGene_tx=XM_019822557.1
F1 71480687 71480834
refGene_gn=S100A14;refGene_tx=XM_003999776.4;refGene_gn=S100A14;refGene_tx=XM_019822557.1
F1 71481048 71481186
refGene_gn=S100A14;refGene_tx=XM_003999776.4;refGene_gn=S100A14;refGene_tx=XM_019822557.1
F1 71487061 71487214
refGene_gn=S100A16;refGene_tx=XM_003999777.4;refGene_gn=S100A16;refGene_tx=XM_006943196.4;refGene_gn=S100A16;refGene_tx=XM_006943197.4
F1 71487458 71487617
refGene_gn=S100A16;refGene_tx=XM_003999777.4;refGene_gn=S100A16;refGene_tx=XM_006943196.4;refGene_gn=S100A16;refGene_tx=XM_006943197.4
F1 71517861 71518002
refGene_gn=S100A2;refGene_tx=XM_003999778.3;refGene_gn=S100A2;refGene_tx=XM_006943199.2
F1 71518815 71518968
refGene_gn=S100A2;refGene_tx=XM_003999778.3;refGene_gn=S100A2;refGene_tx=XM_006943199.2
F1 71532754 71532895 refGene_gn=S100A3;refGene_tx=XM_019821405.1
F1 71533258 71533441 refGene_gn=S100A3;refGene_tx=XM_019821405.1
F1 71535885 71536122
refGene_gn=S100A4;refGene_tx=XM_003999779.3;refGene_gn=S100A4;refGene_tx=XM_019822558.1
F1 71537099 71537264
refGene_gn=S100A4;refGene_tx=XM_003999779.3;refGene_gn=S100A4;refGene_tx=XM_019822558.1
F1 71540509 71540692 refGene_gn=S100A5;refGene_tx=XM_023247801.1
F1 71542370 71542514 refGene_gn=S100A5;refGene_tx=XM_023247801.1
F1 71544163 71544301 refGene_gn=S100A6;refGene_tx=XM_023247824.1
F1 71544600 71544735 refGene_gn=S100A6;refGene_tx=XM_023247824.1
F1 71612616 71612766
refGene_gn=LOC101087705;refGene_tx=XM_019821408.2;refGene_gn=LOC101087705;refGene_tx=XM_019821409.2
F1 71614721 71614898
refGene_gn=LOC101087705;refGene_tx=XM_019821408.2;refGene_gn=LOC101087705;refGene_tx=XM_019821409.2
F1 71622550 71622691 refGene_gn=S100A8;refGene_tx=XM_003999782.5
F1 71622837 71622966 refGene_gn=S100A8;refGene_tx=XM_003999782.5
F1 71639923 71640061
refGene_gn=S100A12;refGene_tx=NM_001198929.1;refGene_gn=S100A12;refGen

e_tx=XM_006943148.4
F1 71640456 71640597
refGene_gn=S100A12;refGene_tx=NM_001198929.1;refGene_gn=S100A12;refGene_tx=XM_006943148.4
F1 71653849 71654113
refGene_gn=S100A9;refGene_tx=XM_003999783.5;refGene_gn=S100A9;refGene_tx=XM_023247845.1
F1 71655919 71656085
refGene_gn=S100A9;refGene_tx=XM_003999783.5;refGene_gn=S100A9;refGene_tx=XM_023247845.1
F1 71656463 71656513
refGene_gn=S100A9;refGene_tx=XM_003999783.5;refGene_gn=S100A9;refGene_tx=XM_023247845.1
F2 2056564 2056589
refGene_gn=LOC102899846;refGene_tx=XM_023248723.1;refGene_gn=LOC102899846;refGene_tx=XM_023248724.1
F2 2091931 2092094
refGene_gn=LOC102899846;refGene_tx=XM_023248723.1;refGene_gn=LOC102899846;refGene_tx=XM_023248724.1
F2 2092564 2092575
refGene_gn=LOC102899846;refGene_tx=XM_023248725.1
F2 2093653 2093806
refGene_gn=LOC102899846;refGene_tx=XM_023248723.1;refGene_gn=LOC102899846;refGene_tx=XM_023248724.1;refGene_gn=LOC102899846;refGene_tx=XM_023248725.1
F2 2094768 2094864
refGene_gn=LOC102899846;refGene_tx=XM_023248723.1;refGene_gn=LOC102899846;refGene_tx=XM_023248725.1
F2 2097111 2097268
refGene_gn=LOC102899846;refGene_tx=XM_023248723.1;refGene_gn=LOC102899846;refGene_tx=XM_023248724.1;refGene_gn=LOC102899846;refGene_tx=XM_023248725.1
F2 2150040 2150129
refGene_gn=LOC109496973;refGene_tx=XM_023248462.1
F2 2150359 2150486
refGene_gn=LOC109496973;refGene_tx=XM_023248462.1
F2 2153789 2153901
refGene_gn=LOC109496973;refGene_tx=XM_023248462.1
F2 2155257 2155259
refGene_gn=LOC109496973;refGene_tx=XM_023248462.1
F2 2211871 2212168
refGene_gn=LOC111558517;refGene_tx=XM_023248305.1
F2 2212701 2212719
refGene_gn=LOC111558517;refGene_tx=XM_023248305.1
F2 2294479 2294512 refGene_gn=SPIDR;refGene_tx=XM_019822628.2
F2 2323581 2323737 refGene_gn=SPIDR;refGene_tx=XM_019822628.2
F2 2324386 2324453 refGene_gn=SPIDR;refGene_tx=XM_019822628.2
F2 2328317 2328503 refGene_gn=SPIDR;refGene_tx=XM_019822628.2
F2 2338679 2338843 refGene_gn=SPIDR;refGene_tx=XM_019822628.2
F2 2471507 2471752 refGene_gn=SPIDR;refGene_tx=XM_019822628.2

F2	2487825	2487926	refGene_gn=SPIDR;refGene_tx=XM_019822628.2
F2	2520153	2520367	refGene_gn=SPIDR;refGene_tx=XM_019822628.2
F2	2652797	2652993	refGene_gn=SPIDR;refGene_tx=XM_019822628.2
F2	2654434	2654682	refGene_gn=SPIDR;refGene_tx=XM_019822628.2
F2	2725877	2726018	refGene_gn=SPIDR;refGene_tx=XM_019822628.2
F2	2761344	2761432	refGene_gn=SPIDR;refGene_tx=XM_019822628.2
F2	2762843	2762987	refGene_gn=SPIDR;refGene_tx=XM_019822628.2
F2	2763079	2763139	refGene_gn=SPIDR;refGene_tx=XM_019822628.2
F2	2770979	2771166	refGene_gn=SPIDR;refGene_tx=XM_019822628.2
F2	2771870	2772023	refGene_gn=SPIDR;refGene_tx=XM_019822628.2
F2	2788331	2788425	refGene_gn=SPIDR;refGene_tx=XM_019822628.2
F2	2789481	2789596	refGene_gn=SPIDR;refGene_tx=XM_019822628.2
F2	2789869	2789923	refGene_gn=SPIDR;refGene_tx=XM_019822628.2
F2	2795105	2795270	refGene_gn=SPIDR;refGene_tx=XM_019822628.2
F2	2796972	2797794	refGene_gn=CEBPD;refGene_tx=XM_023248495.1
F2	2849627	2849832
refGene_gn=PRKDC;refGene_tx=XM_011291282.3;refGene_gn=PRKDC;refGene_tx
=XM_019822629.1
F2	2851732	2851872
refGene_gn=PRKDC;refGene_tx=XM_011291282.3;refGene_gn=PRKDC;refGene_tx
=XM_019822629.1
F2	2852433	2852622
refGene_gn=PRKDC;refGene_tx=XM_011291282.3;refGene_gn=PRKDC;refGene_tx
=XM_019822629.1
F2	2853473	2853675
refGene_gn=PRKDC;refGene_tx=XM_011291282.3;refGene_gn=PRKDC;refGene_tx
=XM_019822629.1
F2	2853750	2853822
refGene_gn=PRKDC;refGene_tx=XM_011291282.3;refGene_gn=PRKDC;refGene_tx
=XM_019822629.1
F2	2854032	2854122
refGene_gn=PRKDC;refGene_tx=XM_011291282.3;refGene_gn=PRKDC;refGene_tx
=XM_019822629.1
F2	2857328	2857421
refGene_gn=PRKDC;refGene_tx=XM_011291282.3;refGene_gn=PRKDC;refGene_tx
=XM_019822629.1
F2	2857721	2857942
refGene_gn=PRKDC;refGene_tx=XM_011291282.3;refGene_gn=PRKDC;refGene_tx
=XM_019822629.1
F2	2863936	2864004
refGene_gn=PRKDC;refGene_tx=XM_011291282.3;refGene_gn=PRKDC;refGene_tx
=XM_019822629.1
F2	2864406	2864611
refGene_gn=PRKDC;refGene_tx=XM_011291282.3;refGene_gn=PRKDC;refGene_tx
=XM_019822629.1
F2	2869182	2869326
refGene_gn=PRKDC;refGene_tx=XM_011291282.3;refGene_gn=PRKDC;refGene_tx
=XM_019822629.1
F2	2869434	2869522
refGene_gn=PRKDC;refGene_tx=XM_011291282.3;refGene_gn=PRKDC;refGene_tx

=XM_019822629.1
F2 2873068 2873280
refGene_gn=PRKDC;refGene_tx=XM_011291282.3;refGene_gn=PRKDC;refGene_tx
=XM_019822629.1
F2 2880253 2880414
refGene_gn=PRKDC;refGene_tx=XM_011291282.3;refGene_gn=PRKDC;refGene_tx
=XM_019822629.1
F2 2881222 2881403
refGene_gn=PRKDC;refGene_tx=XM_011291282.3;refGene_gn=PRKDC;refGene_tx
=XM_019822629.1
F2 2882174 2882365
refGene_gn=PRKDC;refGene_tx=XM_011291282.3;refGene_gn=PRKDC;refGene_tx
=XM_019822629.1
F2 2883793 2883968
refGene_gn=PRKDC;refGene_tx=XM_011291282.3;refGene_gn=PRKDC;refGene_tx
=XM_019822629.1
F2 2889005 2889195
refGene_gn=PRKDC;refGene_tx=XM_011291282.3;refGene_gn=PRKDC;refGene_tx
=XM_019822629.1
F2 2904439 2904551
refGene_gn=PRKDC;refGene_tx=XM_011291282.3;refGene_gn=PRKDC;refGene_tx
=XM_019822629.1
F2 2906141 2906250
refGene_gn=PRKDC;refGene_tx=XM_011291282.3;refGene_gn=PRKDC;refGene_tx
=XM_019822629.1
F2 2907475 2907700
refGene_gn=PRKDC;refGene_tx=XM_011291282.3;refGene_gn=PRKDC;refGene_tx
=XM_019822629.1
F2 2908397 2908586
refGene_gn=PRKDC;refGene_tx=XM_011291282.3;refGene_gn=PRKDC;refGene_tx
=XM_019822629.1
F2 2909717 2909856
refGene_gn=PRKDC;refGene_tx=XM_011291282.3;refGene_gn=PRKDC;refGene_tx
=XM_019822629.1
F2 2911991 2912197
refGene_gn=PRKDC;refGene_tx=XM_011291282.3;refGene_gn=PRKDC;refGene_tx
=XM_019822629.1
F2 2916972 2917152
refGene_gn=PRKDC;refGene_tx=XM_011291282.3;refGene_gn=PRKDC;refGene_tx
=XM_019822629.1
F2 2920073 2920205
refGene_gn=PRKDC;refGene_tx=XM_011291282.3;refGene_gn=PRKDC;refGene_tx
=XM_019822629.1
F2 2922409 2922522
refGene_gn=PRKDC;refGene_tx=XM_011291282.3;refGene_gn=PRKDC;refGene_tx
=XM_019822629.1
F2 2927610 2927823
refGene_gn=PRKDC;refGene_tx=XM_011291282.3;refGene_gn=PRKDC;refGene_tx
=XM_019822629.1
F2 2929412 2929602

refGene_gn=PRKDC;refGene_tx=XM_011291282.3;refGene_gn=PRKDC;refGene_tx=XM_019822629.1
F2 2931598 2931806
refGene_gn=PRKDC;refGene_tx=XM_011291282.3;refGene_gn=PRKDC;refGene_tx=XM_019822629.1
F2 2932812 2932911
refGene_gn=PRKDC;refGene_tx=XM_011291282.3;refGene_gn=PRKDC;refGene_tx=XM_019822629.1
F2 2933578 2933752
refGene_gn=PRKDC;refGene_tx=XM_011291282.3;refGene_gn=PRKDC;refGene_tx=XM_019822629.1
F2 2935401 2935551
refGene_gn=PRKDC;refGene_tx=XM_011291282.3;refGene_gn=PRKDC;refGene_tx=XM_019822629.1
F2 2935636 2935761
refGene_gn=PRKDC;refGene_tx=XM_011291282.3;refGene_gn=PRKDC;refGene_tx=XM_019822629.1
F2 2938435 2938547
refGene_gn=PRKDC;refGene_tx=XM_011291282.3;refGene_gn=PRKDC;refGene_tx=XM_019822629.1
F2 2940040 2940172
refGene_gn=PRKDC;refGene_tx=XM_011291282.3;refGene_gn=PRKDC;refGene_tx=XM_019822629.1
F2 2940731 2940883
refGene_gn=PRKDC;refGene_tx=XM_011291282.3;refGene_gn=PRKDC;refGene_tx=XM_019822629.1
F2 2942402 2942546
refGene_gn=PRKDC;refGene_tx=XM_011291282.3;refGene_gn=PRKDC;refGene_tx=XM_019822629.1
F2 2954461 2954581
refGene_gn=PRKDC;refGene_tx=XM_011291282.3;refGene_gn=PRKDC;refGene_tx=XM_019822629.1
F2 2955013 2955151
refGene_gn=PRKDC;refGene_tx=XM_011291282.3;refGene_gn=PRKDC;refGene_tx=XM_019822629.1
F2 2957701 2957850
refGene_gn=PRKDC;refGene_tx=XM_011291282.3;refGene_gn=PRKDC;refGene_tx=XM_019822629.1
F2 2958949 2959022
refGene_gn=PRKDC;refGene_tx=XM_011291282.3;refGene_gn=PRKDC;refGene_tx=XM_019822629.1
F2 2960362 2960428
refGene_gn=PRKDC;refGene_tx=XM_011291282.3;refGene_gn=PRKDC;refGene_tx=XM_019822629.1
F2 2960676 2960845
refGene_gn=PRKDC;refGene_tx=XM_011291282.3;refGene_gn=PRKDC;refGene_tx=XM_019822629.1
F2 2962491 2962670
refGene_gn=PRKDC;refGene_tx=XM_011291282.3;refGene_gn=PRKDC;refGene_tx=XM_019822629.1

F2	2965441	2965649
refGene_gn=PRKDC;refGene_tx=XM_011291282.3;refGene_gn=PRKDC;refGene_tx
=XM_019822629.1
F2	2971493	2971621
refGene_gn=PRKDC;refGene_tx=XM_011291282.3;refGene_gn=PRKDC;refGene_tx
=XM_019822629.1
F2	2974700	2974868
refGene_gn=PRKDC;refGene_tx=XM_011291282.3;refGene_gn=PRKDC;refGene_tx
=XM_019822629.1
F2	2975949	2976048
refGene_gn=PRKDC;refGene_tx=XM_011291282.3;refGene_gn=PRKDC;refGene_tx
=XM_019822629.1
F2	2976428	2976614
refGene_gn=PRKDC;refGene_tx=XM_011291282.3;refGene_gn=PRKDC;refGene_tx
=XM_019822629.1
F2	2984055	2984259
refGene_gn=PRKDC;refGene_tx=XM_011291282.3;refGene_gn=PRKDC;refGene_tx
=XM_019822629.1
F2	2985103	2985259
refGene_gn=PRKDC;refGene_tx=XM_011291282.3;refGene_gn=PRKDC;refGene_tx
=XM_019822629.1
F2	2985696	2985829
refGene_gn=PRKDC;refGene_tx=XM_011291282.3;refGene_gn=PRKDC;refGene_tx
=XM_019822629.1
F2	2986253	2986465
refGene_gn=PRKDC;refGene_tx=XM_011291282.3;refGene_gn=PRKDC;refGene_tx
=XM_019822629.1
F2	2987189	2987413
refGene_gn=PRKDC;refGene_tx=XM_011291282.3;refGene_gn=PRKDC;refGene_tx
=XM_019822629.1
F2	2991618	2991873
refGene_gn=PRKDC;refGene_tx=XM_011291282.3;refGene_gn=PRKDC;refGene_tx
=XM_019822629.1
F2	2995587	2995721
refGene_gn=PRKDC;refGene_tx=XM_011291282.3;refGene_gn=PRKDC;refGene_tx
=XM_019822629.1
F2	2996843	2996943
refGene_gn=PRKDC;refGene_tx=XM_011291282.3;refGene_gn=PRKDC;refGene_tx
=XM_019822629.1
F2	2998654	2998749
refGene_gn=PRKDC;refGene_tx=XM_011291282.3;refGene_gn=PRKDC;refGene_tx
=XM_019822629.1
F2	3006887	3007114
refGene_gn=PRKDC;refGene_tx=XM_011291282.3;refGene_gn=PRKDC;refGene_tx
=XM_019822629.1
F2	3010112	3010220
refGene_gn=PRKDC;refGene_tx=XM_011291282.3;refGene_gn=PRKDC;refGene_tx
=XM_019822629.1
F2	3014689	3014842
refGene_gn=PRKDC;refGene_tx=XM_011291282.3;refGene_gn=PRKDC;refGene_tx

=XM_019822629.1
F2 3015544 3015708
refGene_gn=PRKDC;refGene_tx=XM_011291282.3;refGene_gn=PRKDC;refGene_tx=XM_019822629.1
F2 3016498 3016589
refGene_gn=PRKDC;refGene_tx=XM_011291282.3;refGene_gn=PRKDC;refGene_tx=XM_019822629.1
F2 3019135 3019239
refGene_gn=PRKDC;refGene_tx=XM_011291282.3;refGene_gn=PRKDC;refGene_tx=XM_019822629.1
F2 3031884 3032044
refGene_gn=PRKDC;refGene_tx=XM_011291282.3;refGene_gn=PRKDC;refGene_tx=XM_019822629.1
F2 3032479 3032599
refGene_gn=PRKDC;refGene_tx=XM_011291282.3;refGene_gn=PRKDC;refGene_tx=XM_019822629.1
F2 3033049 3033136
refGene_gn=PRKDC;refGene_tx=XM_011291282.3;refGene_gn=PRKDC;refGene_tx=XM_019822629.1
F2 3034110 3034270
refGene_gn=PRKDC;refGene_tx=XM_011291282.3;refGene_gn=PRKDC;refGene_tx=XM_019822629.1
F2 3034838 3034954
refGene_gn=PRKDC;refGene_tx=XM_011291282.3;refGene_gn=PRKDC;refGene_tx=XM_019822629.1
F2 3036433 3036586
refGene_gn=PRKDC;refGene_tx=XM_011291282.3;refGene_gn=PRKDC;refGene_tx=XM_019822629.1
F2 3038125 3038251
refGene_gn=PRKDC;refGene_tx=XM_011291282.3;refGene_gn=PRKDC;refGene_tx=XM_019822629.1
F2 3038882 3038932
refGene_gn=PRKDC;refGene_tx=XM_011291282.3;refGene_gn=PRKDC;refGene_tx=XM_019822629.1
F2 3039271 3039440
refGene_gn=PRKDC;refGene_tx=XM_011291282.3;refGene_gn=PRKDC;refGene_tx=XM_019822629.1
F2 3039889 3040054
refGene_gn=PRKDC;refGene_tx=XM_011291282.3;refGene_gn=PRKDC;refGene_tx=XM_019822629.1
F2 3041609 3041756
refGene_gn=PRKDC;refGene_tx=XM_011291282.3;refGene_gn=PRKDC;refGene_tx=XM_019822629.1
F2 3046820 3046978
refGene_gn=PRKDC;refGene_tx=XM_011291282.3;refGene_gn=PRKDC;refGene_tx=XM_019822629.1
F2 3047469 3047500
refGene_gn=PRKDC;refGene_tx=XM_011291282.3;refGene_gn=PRKDC;refGene_tx=XM_019822629.1
F2 3047581 3047637

refGene_gn=PRKDC;refGene_tx=XM_011291282.3;refGene_gn=PRKDC;refGene_tx
=XM_019822629.1
F2 3054596 3054696
refGene_gn=PRKDC;refGene_tx=XM_011291282.3;refGene_gn=PRKDC;refGene_tx
=XM_019822629.1
F2 3054785 3054898 refGene_gn=PRKDC;refGene_tx=XM_011291282.3
F2 3055446 3055555
refGene_gn=PRKDC;refGene_tx=XM_011291282.3;refGene_gn=PRKDC;refGene_tx
=XM_019822629.1
F2 3056300 3056375 refGene_gn=PRKDC;refGene_tx=XM_011291282.3
F2 3058280 3058373 refGene_gn=PRKDC;refGene_tx=XM_011291282.3
F2 3058465 3058542 refGene_gn=PRKDC;refGene_tx=XM_011291282.3
F2 3060033 3060187 refGene_gn=PRKDC;refGene_tx=XM_011291282.3
F2 3061143 3061213
refGene_gn=MCM4;refGene_tx=XM_006943208.4;refGene_gn=MCM4;refGene_tx=X
M_019822630.2;refGene_gn=MCM4;refGene_tx=XM_023248352.1
F2 3061601 3061766
refGene_gn=MCM4;refGene_tx=XM_006943208.4;refGene_gn=MCM4;refGene_tx=X
M_019822630.2;refGene_gn=MCM4;refGene_tx=XM_023248352.1
F2 3062101 3062265
refGene_gn=MCM4;refGene_tx=XM_006943208.4;refGene_gn=MCM4;refGene_tx=X
M_019822630.2;refGene_gn=MCM4;refGene_tx=XM_023248352.1
F2 3062359 3062461
refGene_gn=MCM4;refGene_tx=XM_006943208.4;refGene_gn=MCM4;refGene_tx=X
M_019822630.2;refGene_gn=MCM4;refGene_tx=XM_023248352.1
F2 3062650 3062746
refGene_gn=MCM4;refGene_tx=XM_006943208.4;refGene_gn=MCM4;refGene_tx=X
M_019822630.2;refGene_gn=MCM4;refGene_tx=XM_023248352.1
F2 3062818 3062914
refGene_gn=MCM4;refGene_tx=XM_006943208.4;refGene_gn=MCM4;refGene_tx=X
M_019822630.2;refGene_gn=MCM4;refGene_tx=XM_023248352.1
F2 3064038 3064177
refGene_gn=MCM4;refGene_tx=XM_006943208.4;refGene_gn=MCM4;refGene_tx=X
M_019822630.2;refGene_gn=MCM4;refGene_tx=XM_023248352.1
F2 3066529 3066750
refGene_gn=MCM4;refGene_tx=XM_006943208.4;refGene_gn=MCM4;refGene_tx=X
M_019822630.2;refGene_gn=MCM4;refGene_tx=XM_023248352.1
F2 3068820 3068941
refGene_gn=MCM4;refGene_tx=XM_006943208.4;refGene_gn=MCM4;refGene_tx=X
M_019822630.2;refGene_gn=MCM4;refGene_tx=XM_023248352.1
F2 3071203 3071463
refGene_gn=MCM4;refGene_tx=XM_006943208.4;refGene_gn=MCM4;refGene_tx=X
M_019822630.2;refGene_gn=MCM4;refGene_tx=XM_023248352.1
F2 3071906 3072272
refGene_gn=MCM4;refGene_tx=XM_006943208.4;refGene_gn=MCM4;refGene_tx=X
M_019822630.2;refGene_gn=MCM4;refGene_tx=XM_023248352.1
F2 3072721 3072849
refGene_gn=MCM4;refGene_tx=XM_006943208.4;refGene_gn=MCM4;refGene_tx=X
M_019822630.2;refGene_gn=MCM4;refGene_tx=XM_023248352.1
F2 3074056 3074264

refGene_gn=MCM4;refGene_tx=XM_006943208.4;refGene_gn=MCM4;refGene_tx=XM_019822630.2;refGene_gn=MCM4;refGene_tx=XM_023248352.1
F2 3075275 3075504
refGene_gn=MCM4;refGene_tx=XM_006943208.4;refGene_gn=MCM4;refGene_tx=XM_019822630.2;refGene_gn=MCM4;refGene_tx=XM_023248352.1
F2 3076007 3076141
refGene_gn=MCM4;refGene_tx=XM_006943208.4;refGene_gn=MCM4;refGene_tx=XM_019822630.2;refGene_gn=MCM4;refGene_tx=XM_023248352.1
F2 3077406 3077499
refGene_gn=MCM4;refGene_tx=XM_006943208.4;refGene_gn=MCM4;refGene_tx=XM_019822630.2;refGene_gn=MCM4;refGene_tx=XM_023248352.1
F2 3144761 3144789 refGene_gn=UBE2V2;refGene_tx=XM_019822631.2
F2 3151243 3151392
refGene_gn=UBE2V2;refGene_tx=XM_019822631.2;refGene_gn=UBE2V2;refGene_tx=XM_019822632.2
F2 3156206 3156332
refGene_gn=UBE2V2;refGene_tx=XM_019822631.2;refGene_gn=UBE2V2;refGene_tx=XM_019822632.2
F2 3172573 3172720
refGene_gn=UBE2V2;refGene_tx=XM_019822631.2;refGene_gn=UBE2V2;refGene_tx=XM_019822632.2
F2 3749226 3749261 refGene_gn=EFCAB1;refGene_tx=XM_011291285.3
F2 3749263 3749285 refGene_gn=EFCAB1;refGene_tx=XM_011291285.3
F2 3752158 3752263 refGene_gn=EFCAB1;refGene_tx=XM_011291285.3
F2 3752548 3752700 refGene_gn=EFCAB1;refGene_tx=XM_011291285.3
F2 3753365 3753470 refGene_gn=EFCAB1;refGene_tx=XM_011291285.3
F2 3754193 3754352 refGene_gn=EFCAB1;refGene_tx=XM_011291285.3
F2 3759367 3759425 refGene_gn=EFCAB1;refGene_tx=XM_011291285.3
F2 3759559 3759933
refGene_gn=LOC109495881;refGene_tx=XM_019822569.1
F2 3761250 3761355
refGene_gn=LOC109495881;refGene_tx=XM_019822569.1
F2 3764061 3764260
refGene_gn=LOC109495881;refGene_tx=XM_019822569.1
F2 3952301 3952483
refGene_gn=SNAI2;refGene_tx=XM_023248318.1;refGene_gn=SNAI2;refGene_tx=XM_023248319.1;refGene_gn=SNAI2;refGene_tx=XM_023248320.1
F2 3953385 3953931
refGene_gn=SNAI2;refGene_tx=XM_023248318.1;refGene_gn=SNAI2;refGene_tx=XM_023248319.1;refGene_gn=SNAI2;refGene_tx=XM_023248320.1
F2 3954745 3954824 refGene_gn=SNAI2;refGene_tx=XM_023248318.1
F2 4092185 4092240
refGene_gn=PPDPFL;refGene_tx=XM_006943210.3;refGene_gn=PPDPFL;refGene_tx=XM_023248585.1
F2 4094541 4094619
refGene_gn=PPDPFL;refGene_tx=XM_006943210.3;refGene_gn=PPDPFL;refGene_tx=XM_023248585.1
F2 4094721 4094821
refGene_gn=PPDPFL;refGene_tx=XM_006943210.3;refGene_gn=PPDPFL;refGene_tx=XM_023248585.1

```
F2	4096307	4096320	refGene_gn=PPDPFL;refGene_tx=XM_023248585.1
F2	4096417	4096604	refGene_gn=PPDPFL;refGene_tx=XM_006943210.3
F2	4258331	4258374	
refGene_gn=LOC109495894;refGene_tx=XM_019822619.2
F2	4258606	4258658	
refGene_gn=LOC109495894;refGene_tx=XM_019822618.2
F2	4264606	4264851	
refGene_gn=LOC109495894;refGene_tx=XM_019822618.2;refGene_gn=LOC109495
894;refGene_tx=XM_019822619.2
F2	5401371	5401398	
refGene_gn=SNTG1;refGene_tx=XM_019822636.2;refGene_gn=SNTG1;refGene_tx
=XM_019822637.2;refGene_gn=SNTG1;refGene_tx=XM_019822638.1
F2	5408983	5409118	
refGene_gn=SNTG1;refGene_tx=XM_019822636.2;refGene_gn=SNTG1;refGene_tx
=XM_019822637.2;refGene_gn=SNTG1;refGene_tx=XM_019822638.1
F2	5461259	5461316	
refGene_gn=SNTG1;refGene_tx=XM_019822636.2;refGene_gn=SNTG1;refGene_tx
=XM_019822637.2;refGene_gn=SNTG1;refGene_tx=XM_019822638.1
F2	5471590	5471648	
refGene_gn=SNTG1;refGene_tx=XM_019822636.2;refGene_gn=SNTG1;refGene_tx
=XM_019822637.2;refGene_gn=SNTG1;refGene_tx=XM_019822638.1
F2	5472458	5472502	
refGene_gn=SNTG1;refGene_tx=XM_019822636.2;refGene_gn=SNTG1;refGene_tx
=XM_019822637.2;refGene_gn=SNTG1;refGene_tx=XM_019822638.1
F2	5472593	5472635	
refGene_gn=SNTG1;refGene_tx=XM_019822636.2;refGene_gn=SNTG1;refGene_tx
=XM_019822637.2;refGene_gn=SNTG1;refGene_tx=XM_019822638.1
F2	5514217	5514320	
refGene_gn=SNTG1;refGene_tx=XM_019822636.2;refGene_gn=SNTG1;refGene_tx
=XM_019822637.2;refGene_gn=SNTG1;refGene_tx=XM_019822638.1
F2	5526066	5526149	
refGene_gn=SNTG1;refGene_tx=XM_019822636.2;refGene_gn=SNTG1;refGene_tx
=XM_019822637.2;refGene_gn=SNTG1;refGene_tx=XM_019822638.1
F2	5532842	5532973	
refGene_gn=SNTG1;refGene_tx=XM_019822636.2;refGene_gn=SNTG1;refGene_tx
=XM_019822637.2;refGene_gn=SNTG1;refGene_tx=XM_019822638.1
F2	5549788	5549918	
refGene_gn=SNTG1;refGene_tx=XM_019822636.2;refGene_gn=SNTG1;refGene_tx
=XM_019822637.2;refGene_gn=SNTG1;refGene_tx=XM_019822638.1
F2	5577439	5577478	
refGene_gn=SNTG1;refGene_tx=XM_019822636.2;refGene_gn=SNTG1;refGene_tx
=XM_019822638.1
F2	5648079	5648196	
refGene_gn=SNTG1;refGene_tx=XM_019822636.2;refGene_gn=SNTG1;refGene_tx
=XM_019822637.2;refGene_gn=SNTG1;refGene_tx=XM_019822638.1
F2	5649574	5649646	
refGene_gn=SNTG1;refGene_tx=XM_019822636.2;refGene_gn=SNTG1;refGene_tx
=XM_019822637.2;refGene_gn=SNTG1;refGene_tx=XM_019822638.1
F2	5677096	5677249	
refGene_gn=SNTG1;refGene_tx=XM_019822636.2;refGene_gn=SNTG1;refGene_tx
```

=XM_019822637.2;refGene_gn=SNTG1;refGene_tx=XM_019822638.1
F2	5681150	5681243
refGene_gn=SNTG1;refGene_tx=XM_019822636.2;refGene_gn=SNTG1;refGene_tx=XM_019822637.2;refGene_gn=SNTG1;refGene_tx=XM_019822638.1
F2	5686932	5686980	refGene_gn=SNTG1;refGene_tx=XM_019822638.1
F2	5703314	5703425
refGene_gn=SNTG1;refGene_tx=XM_019822636.2;refGene_gn=SNTG1;refGene_tx=XM_019822637.2
F2	5733051	5733210
refGene_gn=SNTG1;refGene_tx=XM_019822636.2;refGene_gn=SNTG1;refGene_tx=XM_019822637.2
F2	6214269	6214338	refGene_gn=PXDNL;refGene_tx=XM_023248448.1
F2	6235320	6235443
refGene_gn=PXDNL;refGene_tx=XM_023248447.1;refGene_gn=PXDNL;refGene_tx=XM_023248449.1;refGene_gn=PXDNL;refGene_tx=XM_023248450.1;refGene_gn=PXDNL;refGene_tx=XM_023248452.1;refGene_gn=PXDNL;refGene_tx=XM_023248453.1
F2	6236210	6236324
refGene_gn=PXDNL;refGene_tx=XM_023248447.1;refGene_gn=PXDNL;refGene_tx=XM_023248448.1;refGene_gn=PXDNL;refGene_tx=XM_023248449.1;refGene_gn=PXDNL;refGene_tx=XM_023248450.1;refGene_gn=PXDNL;refGene_tx=XM_023248452.1;refGene_gn=PXDNL;refGene_tx=XM_023248453.1
F2	6249269	6249305	refGene_gn=PXDNL;refGene_tx=XM_023248451.1
F2	6250921	6251051
refGene_gn=PXDNL;refGene_tx=XM_023248447.1;refGene_gn=PXDNL;refGene_tx=XM_023248448.1;refGene_gn=PXDNL;refGene_tx=XM_023248449.1;refGene_gn=PXDNL;refGene_tx=XM_023248450.1;refGene_gn=PXDNL;refGene_tx=XM_023248451.1;refGene_gn=PXDNL;refGene_tx=XM_023248452.1;refGene_gn=PXDNL;refGene_tx=XM_023248453.1
F2	6255809	6255924
refGene_gn=PXDNL;refGene_tx=XM_023248447.1;refGene_gn=PXDNL;refGene_tx=XM_023248448.1;refGene_gn=PXDNL;refGene_tx=XM_023248449.1;refGene_gn=PXDNL;refGene_tx=XM_023248450.1;refGene_gn=PXDNL;refGene_tx=XM_023248451.1;refGene_gn=PXDNL;refGene_tx=XM_023248452.1;refGene_gn=PXDNL;refGene_tx=XM_023248453.1
F2	6273105	6273404
refGene_gn=PXDNL;refGene_tx=XM_023248447.1;refGene_gn=PXDNL;refGene_tx=XM_023248448.1;refGene_gn=PXDNL;refGene_tx=XM_023248449.1;refGene_gn=PXDNL;refGene_tx=XM_023248450.1;refGene_gn=PXDNL;refGene_tx=XM_023248451.1;refGene_gn=PXDNL;refGene_tx=XM_023248452.1;refGene_gn=PXDNL;refGene_tx=XM_023248453.1;refGene_gn=PXDNL;refGene_tx=XM_023248454.1
F2	6276095	6276230
refGene_gn=PXDNL;refGene_tx=XM_023248447.1;refGene_gn=PXDNL;refGene_tx=XM_023248448.1;refGene_gn=PXDNL;refGene_tx=XM_023248449.1;refGene_gn=PXDNL;refGene_tx=XM_023248450.1;refGene_gn=PXDNL;refGene_tx=XM_023248451.1;refGene_gn=PXDNL;refGene_tx=XM_023248452.1;refGene_gn=PXDNL;refGene_tx=XM_023248453.1;refGene_gn=PXDNL;refGene_tx=XM_023248454.1
F2	6305318	6306804
refGene_gn=PXDNL;refGene_tx=XM_023248447.1;refGene_gn=PXDNL;refGene_tx=XM_023248448.1;refGene_gn=PXDNL;refGene_tx=XM_023248449.1;refGene_gn=

PXDNL;refGene_tx=XM_023248450.1;refGene_gn=PXDNL;refGene_tx=XM_0232484
51.1;refGene_gn=PXDNL;refGene_tx=XM_023248452.1;refGene_gn=PXDNL;refGe
ne_tx=XM_023248453.1;refGene_gn=PXDNL;refGene_tx=XM_023248454.1
F2 6309024 6309182
refGene_gn=PXDNL;refGene_tx=XM_023248447.1;refGene_gn=PXDNL;refGene_tx
=XM_023248448.1;refGene_gn=PXDNL;refGene_tx=XM_023248449.1;refGene_gn=
PXDNL;refGene_tx=XM_023248450.1;refGene_gn=PXDNL;refGene_tx=XM_0232484
51.1;refGene_gn=PXDNL;refGene_tx=XM_023248452.1;refGene_gn=PXDNL;refGe
ne_tx=XM_023248453.1;refGene_gn=PXDNL;refGene_tx=XM_023248454.1
F2 6310897 6311006
refGene_gn=PXDNL;refGene_tx=XM_023248447.1;refGene_gn=PXDNL;refGene_tx
=XM_023248448.1;refGene_gn=PXDNL;refGene_tx=XM_023248449.1;refGene_gn=
PXDNL;refGene_tx=XM_023248450.1;refGene_gn=PXDNL;refGene_tx=XM_0232484
51.1;refGene_gn=PXDNL;refGene_tx=XM_023248452.1;refGene_gn=PXDNL;refGe
ne_tx=XM_023248453.1;refGene_gn=PXDNL;refGene_tx=XM_023248454.1
F2 6320676 6320833
refGene_gn=PXDNL;refGene_tx=XM_023248447.1;refGene_gn=PXDNL;refGene_tx
=XM_023248448.1;refGene_gn=PXDNL;refGene_tx=XM_023248449.1;refGene_gn=
PXDNL;refGene_tx=XM_023248450.1;refGene_gn=PXDNL;refGene_tx=XM_0232484
51.1;refGene_gn=PXDNL;refGene_tx=XM_023248452.1;refGene_gn=PXDNL;refGe
ne_tx=XM_023248453.1;refGene_gn=PXDNL;refGene_tx=XM_023248454.1
F2 6323583 6323696
refGene_gn=PXDNL;refGene_tx=XM_023248447.1;refGene_gn=PXDNL;refGene_tx
=XM_023248448.1;refGene_gn=PXDNL;refGene_tx=XM_023248449.1;refGene_gn=
PXDNL;refGene_tx=XM_023248450.1;refGene_gn=PXDNL;refGene_tx=XM_0232484
51.1;refGene_gn=PXDNL;refGene_tx=XM_023248452.1;refGene_gn=PXDNL;refGe
ne_tx=XM_023248453.1;refGene_gn=PXDNL;refGene_tx=XM_023248454.1
F2 6337519 6337678
refGene_gn=PXDNL;refGene_tx=XM_023248447.1;refGene_gn=PXDNL;refGene_tx
=XM_023248448.1;refGene_gn=PXDNL;refGene_tx=XM_023248449.1;refGene_gn=
PXDNL;refGene_tx=XM_023248450.1;refGene_gn=PXDNL;refGene_tx=XM_0232484
51.1;refGene_gn=PXDNL;refGene_tx=XM_023248452.1;refGene_gn=PXDNL;refGe
ne_tx=XM_023248453.1;refGene_gn=PXDNL;refGene_tx=XM_023248454.1
F2 6339412 6339529
refGene_gn=PXDNL;refGene_tx=XM_023248447.1;refGene_gn=PXDNL;refGene_tx
=XM_023248448.1;refGene_gn=PXDNL;refGene_tx=XM_023248449.1;refGene_gn=
PXDNL;refGene_tx=XM_023248450.1;refGene_gn=PXDNL;refGene_tx=XM_0232484
51.1;refGene_gn=PXDNL;refGene_tx=XM_023248452.1;refGene_gn=PXDNL;refGe
ne_tx=XM_023248453.1;refGene_gn=PXDNL;refGene_tx=XM_023248454.1
F2 6343322 6343589
refGene_gn=PXDNL;refGene_tx=XM_023248447.1;refGene_gn=PXDNL;refGene_tx
=XM_023248448.1;refGene_gn=PXDNL;refGene_tx=XM_023248449.1;refGene_gn=
PXDNL;refGene_tx=XM_023248450.1;refGene_gn=PXDNL;refGene_tx=XM_0232484
51.1;refGene_gn=PXDNL;refGene_tx=XM_023248452.1;refGene_gn=PXDNL;refGe
ne_tx=XM_023248453.1;refGene_gn=PXDNL;refGene_tx=XM_023248454.1
F2 6347189 6347359
refGene_gn=PXDNL;refGene_tx=XM_023248447.1;refGene_gn=PXDNL;refGene_tx
=XM_023248448.1;refGene_gn=PXDNL;refGene_tx=XM_023248449.1;refGene_gn=
PXDNL;refGene_tx=XM_023248450.1;refGene_gn=PXDNL;refGene_tx=XM_0232484
51.1;refGene_gn=PXDNL;refGene_tx=XM_023248452.1;refGene_gn=PXDNL;refGe

ne_tx=XM_023248453.1;refGene_gn=PXDNL;refGene_tx=XM_023248454.1
F2 6354751 6354869
refGene_gn=PXDNL;refGene_tx=XM_023248447.1;refGene_gn=PXDNL;refGene_tx=XM_023248448.1;refGene_gn=PXDNL;refGene_tx=XM_023248449.1;refGene_gn=PXDNL;refGene_tx=XM_023248450.1;refGene_gn=PXDNL;refGene_tx=XM_023248451.1;refGene_gn=PXDNL;refGene_tx=XM_023248452.1;refGene_gn=PXDNL;refGene_tx=XM_023248453.1;refGene_gn=PXDNL;refGene_tx=XM_023248454.1
F2 6357726 6357896
refGene_gn=PXDNL;refGene_tx=XM_023248447.1;refGene_gn=PXDNL;refGene_tx=XM_023248448.1;refGene_gn=PXDNL;refGene_tx=XM_023248449.1;refGene_gn=PXDNL;refGene_tx=XM_023248450.1;refGene_gn=PXDNL;refGene_tx=XM_023248451.1;refGene_gn=PXDNL;refGene_tx=XM_023248452.1;refGene_gn=PXDNL;refGene_tx=XM_023248453.1;refGene_gn=PXDNL;refGene_tx=XM_023248454.1
F2 6370189 6370261
refGene_gn=PXDNL;refGene_tx=XM_023248447.1;refGene_gn=PXDNL;refGene_tx=XM_023248448.1;refGene_gn=PXDNL;refGene_tx=XM_023248449.1;refGene_gn=PXDNL;refGene_tx=XM_023248450.1;refGene_gn=PXDNL;refGene_tx=XM_023248451.1;refGene_gn=PXDNL;refGene_tx=XM_023248452.1;refGene_gn=PXDNL;refGene_tx=XM_023248453.1;refGene_gn=PXDNL;refGene_tx=XM_023248454.1
F2 6370410 6370412 refGene_gn=PXDNL;refGene_tx=XM_023248453.1
F2 6375491 6375563
refGene_gn=PXDNL;refGene_tx=XM_023248447.1;refGene_gn=PXDNL;refGene_tx=XM_023248448.1;refGene_gn=PXDNL;refGene_tx=XM_023248449.1;refGene_gn=PXDNL;refGene_tx=XM_023248450.1;refGene_gn=PXDNL;refGene_tx=XM_023248451.1;refGene_gn=PXDNL;refGene_tx=XM_023248452.1;refGene_gn=PXDNL;refGene_tx=XM_023248454.1
F2 6391275 6391283 refGene_gn=PXDNL;refGene_tx=XM_023248452.1
F2 6425416 6425488
refGene_gn=PXDNL;refGene_tx=XM_023248447.1;refGene_gn=PXDNL;refGene_tx=XM_023248448.1;refGene_gn=PXDNL;refGene_tx=XM_023248449.1;refGene_gn=PXDNL;refGene_tx=XM_023248450.1;refGene_gn=PXDNL;refGene_tx=XM_023248451.1;refGene_gn=PXDNL;refGene_tx=XM_023248454.1
F2 6450433 6450505
refGene_gn=PXDNL;refGene_tx=XM_023248447.1;refGene_gn=PXDNL;refGene_tx=XM_023248448.1;refGene_gn=PXDNL;refGene_tx=XM_023248450.1;refGene_gn=PXDNL;refGene_tx=XM_023248451.1;refGene_gn=PXDNL;refGene_tx=XM_023248454.1
F2 6503134 6503206
refGene_gn=PXDNL;refGene_tx=XM_023248447.1;refGene_gn=PXDNL;refGene_tx=XM_023248448.1;refGene_gn=PXDNL;refGene_tx=XM_023248449.1;refGene_gn=PXDNL;refGene_tx=XM_023248450.1;refGene_gn=PXDNL;refGene_tx=XM_023248451.1;refGene_gn=PXDNL;refGene_tx=XM_023248454.1
F2 6668762 6668926
refGene_gn=PXDNL;refGene_tx=XM_023248447.1;refGene_gn=PXDNL;refGene_tx=XM_023248448.1;refGene_gn=PXDNL;refGene_tx=XM_023248449.1;refGene_gn=PXDNL;refGene_tx=XM_023248450.1;refGene_gn=PXDNL;refGene_tx=XM_023248451.1;refGene_gn=PXDNL;refGene_tx=XM_023248454.1
F2 6678700 6679068
refGene_gn=PCMTD1;refGene_tx=XM_003999806.5;refGene_gn=PCMTD1;refGene_tx=XM_023248502.1;refGene_gn=PCMTD1;refGene_tx=XM_023248503.1

F2 6686798 6686922
refGene_gn=PCMTD1;refGene_tx=XM_003999806.5;refGene_gn=PCMTD1;refGene_tx=XM_023248502.1;refGene_gn=PCMTD1;refGene_tx=XM_023248503.1
F2 6688747 6688919
refGene_gn=PCMTD1;refGene_tx=XM_003999806.5;refGene_gn=PCMTD1;refGene_tx=XM_023248502.1;refGene_gn=PCMTD1;refGene_tx=XM_023248503.1
F2 6700472 6700575
refGene_gn=PCMTD1;refGene_tx=XM_003999806.5;refGene_gn=PCMTD1;refGene_tx=XM_023248502.1
F2 6712425 6712732
refGene_gn=PCMTD1;refGene_tx=XM_003999806.5;refGene_gn=PCMTD1;refGene_tx=XM_023248502.1
F2 6970951 6971092
refGene_gn=ST18;refGene_tx=XM_023248496.1;refGene_gn=ST18;refGene_tx=XM_023248497.1;refGene_gn=ST18;refGene_tx=XM_023248498.1;refGene_gn=ST18;refGene_tx=XM_023248499.1;refGene_gn=ST18;refGene_tx=XM_023248500.1;refGene_gn=ST18;refGene_tx=XM_023248501.1
F2 6974047 6974191
refGene_gn=ST18;refGene_tx=XM_023248496.1;refGene_gn=ST18;refGene_tx=XM_023248497.1;refGene_gn=ST18;refGene_tx=XM_023248498.1;refGene_gn=ST18;refGene_tx=XM_023248499.1;refGene_gn=ST18;refGene_tx=XM_023248500.1;refGene_gn=ST18;refGene_tx=XM_023248501.1
F2 6976382 6976486
refGene_gn=ST18;refGene_tx=XM_023248496.1;refGene_gn=ST18;refGene_tx=XM_023248497.1;refGene_gn=ST18;refGene_tx=XM_023248498.1;refGene_gn=ST18;refGene_tx=XM_023248499.1;refGene_gn=ST18;refGene_tx=XM_023248500.1;refGene_gn=ST18;refGene_tx=XM_023248501.1
F2 6981864 6981953
refGene_gn=ST18;refGene_tx=XM_023248496.1;refGene_gn=ST18;refGene_tx=XM_023248497.1;refGene_gn=ST18;refGene_tx=XM_023248498.1;refGene_gn=ST18;refGene_tx=XM_023248499.1;refGene_gn=ST18;refGene_tx=XM_023248500.1;refGene_gn=ST18;refGene_tx=XM_023248501.1
F2 6986794 6987016
refGene_gn=ST18;refGene_tx=XM_023248496.1;refGene_gn=ST18;refGene_tx=XM_023248497.1;refGene_gn=ST18;refGene_tx=XM_023248498.1;refGene_gn=ST18;refGene_tx=XM_023248499.1;refGene_gn=ST18;refGene_tx=XM_023248500.1
F2 6987986 6988067
refGene_gn=ST18;refGene_tx=XM_023248496.1;refGene_gn=ST18;refGene_tx=XM_023248497.1;refGene_gn=ST18;refGene_tx=XM_023248498.1;refGene_gn=ST18;refGene_tx=XM_023248499.1;refGene_gn=ST18;refGene_tx=XM_023248500.1
F2 6988169 6988232
refGene_gn=ST18;refGene_tx=XM_023248496.1;refGene_gn=ST18;refGene_tx=XM_023248497.1;refGene_gn=ST18;refGene_tx=XM_023248498.1;refGene_gn=ST18;refGene_tx=XM_023248499.1;refGene_gn=ST18;refGene_tx=XM_023248500.1;refGene_gn=ST18;refGene_tx=XM_023248501.1
F2 6991352 6991421
refGene_gn=ST18;refGene_tx=XM_023248496.1;refGene_gn=ST18;refGene_tx=XM_023248497.1;refGene_gn=ST18;refGene_tx=XM_023248498.1;refGene_gn=ST18;refGene_tx=XM_023248499.1;refGene_gn=ST18;refGene_tx=XM_023248500.1;refGene_gn=ST18;refGene_tx=XM_023248501.1

F2 6992282 6992345
refGene_gn=ST18;refGene_tx=XM_023248496.1;refGene_gn=ST18;refGene_tx=XM_023248497.1;refGene_gn=ST18;refGene_tx=XM_023248498.1;refGene_gn=ST18;refGene_tx=XM_023248499.1;refGene_gn=ST18;refGene_tx=XM_023248500.1;refGene_gn=ST18;refGene_tx=XM_023248501.1
F2 7000860 7000976
refGene_gn=ST18;refGene_tx=XM_023248496.1;refGene_gn=ST18;refGene_tx=XM_023248497.1;refGene_gn=ST18;refGene_tx=XM_023248498.1;refGene_gn=ST18;refGene_tx=XM_023248499.1;refGene_gn=ST18;refGene_tx=XM_023248500.1;refGene_gn=ST18;refGene_tx=XM_023248501.1
F2 7006843 7007089
refGene_gn=ST18;refGene_tx=XM_023248496.1;refGene_gn=ST18;refGene_tx=XM_023248497.1;refGene_gn=ST18;refGene_tx=XM_023248498.1;refGene_gn=ST18;refGene_tx=XM_023248499.1;refGene_gn=ST18;refGene_tx=XM_023248500.1;refGene_gn=ST18;refGene_tx=XM_023248501.1
F2 7015492 7015704
refGene_gn=ST18;refGene_tx=XM_023248496.1;refGene_gn=ST18;refGene_tx=XM_023248497.1;refGene_gn=ST18;refGene_tx=XM_023248498.1;refGene_gn=ST18;refGene_tx=XM_023248499.1;refGene_gn=ST18;refGene_tx=XM_023248500.1;refGene_gn=ST18;refGene_tx=XM_023248501.1
F2 7017831 7018025
refGene_gn=ST18;refGene_tx=XM_023248496.1;refGene_gn=ST18;refGene_tx=XM_023248497.1;refGene_gn=ST18;refGene_tx=XM_023248498.1;refGene_gn=ST18;refGene_tx=XM_023248499.1;refGene_gn=ST18;refGene_tx=XM_023248500.1;refGene_gn=ST18;refGene_tx=XM_023248501.1
F2 7020520 7020625
refGene_gn=ST18;refGene_tx=XM_023248496.1;refGene_gn=ST18;refGene_tx=XM_023248497.1;refGene_gn=ST18;refGene_tx=XM_023248498.1;refGene_gn=ST18;refGene_tx=XM_023248499.1;refGene_gn=ST18;refGene_tx=XM_023248500.1;refGene_gn=ST18;refGene_tx=XM_023248501.1
F2 7023285 7023376
refGene_gn=ST18;refGene_tx=XM_023248496.1;refGene_gn=ST18;refGene_tx=XM_023248497.1;refGene_gn=ST18;refGene_tx=XM_023248498.1;refGene_gn=ST18;refGene_tx=XM_023248499.1;refGene_gn=ST18;refGene_tx=XM_023248500.1;refGene_gn=ST18;refGene_tx=XM_023248501.1
F2 7025144 7025279
refGene_gn=ST18;refGene_tx=XM_023248496.1;refGene_gn=ST18;refGene_tx=XM_023248497.1;refGene_gn=ST18;refGene_tx=XM_023248498.1;refGene_gn=ST18;refGene_tx=XM_023248499.1;refGene_gn=ST18;refGene_tx=XM_023248500.1;refGene_gn=ST18;refGene_tx=XM_023248501.1
F2 7029834 7030626
refGene_gn=ST18;refGene_tx=XM_023248496.1;refGene_gn=ST18;refGene_tx=XM_023248497.1;refGene_gn=ST18;refGene_tx=XM_023248498.1;refGene_gn=ST18;refGene_tx=XM_023248499.1;refGene_gn=ST18;refGene_tx=XM_023248500.1;refGene_gn=ST18;refGene_tx=XM_023248501.1
F2 7040142 7040342
refGene_gn=ST18;refGene_tx=XM_023248496.1;refGene_gn=ST18;refGene_tx=XM_023248497.1;refGene_gn=ST18;refGene_tx=XM_023248498.1;refGene_gn=ST18;refGene_tx=XM_023248499.1;refGene_gn=ST18;refGene_tx=XM_023248500.1;refGene_gn=ST18;refGene_tx=XM_023248501.1

F2 7046966 7047053
refGene_gn=ST18;refGene_tx=XM_023248496.1;refGene_gn=ST18;refGene_tx=XM_023248497.1;refGene_gn=ST18;refGene_tx=XM_023248498.1;refGene_gn=ST18;refGene_tx=XM_023248501.1
F2 7073469 7073500
refGene_gn=ST18;refGene_tx=XM_023248496.1;refGene_gn=ST18;refGene_tx=XM_023248497.1;refGene_gn=ST18;refGene_tx=XM_023248498.1;refGene_gn=ST18;refGene_tx=XM_023248499.1;refGene_gn=ST18;refGene_tx=XM_023248500.1;refGene_gn=ST18;refGene_tx=XM_023248501.1
F2 7075832 7075887
refGene_gn=ST18;refGene_tx=XM_023248496.1;refGene_gn=ST18;refGene_tx=XM_023248497.1;refGene_gn=ST18;refGene_tx=XM_023248498.1;refGene_gn=ST18;refGene_tx=XM_023248499.1;refGene_gn=ST18;refGene_tx=XM_023248500.1;refGene_gn=ST18;refGene_tx=XM_023248501.1
F2 7382085 7382115 refGene_gn=ALKAL1;refGene_tx=XM_023248729.1
F2 7384588 7384665 refGene_gn=ALKAL1;refGene_tx=XM_023248729.1
F2 7385747 7385828 refGene_gn=ALKAL1;refGene_tx=XM_023248729.1
F2 7387572 7387626 refGene_gn=ALKAL1;refGene_tx=XM_023248729.1
F2 7395118 7395714 refGene_gn=ALKAL1;refGene_tx=XM_023248729.1
F2 7395808 7395888 refGene_gn=ALKAL1;refGene_tx=XM_023248729.1
F2 7434828 7434906
refGene_gn=RB1CC1;refGene_tx=XM_023248504.1;refGene_gn=RB1CC1;refGene_tx=XM_023248505.1
F2 7435692 7435763
refGene_gn=RB1CC1;refGene_tx=XM_023248504.1;refGene_gn=RB1CC1;refGene_tx=XM_023248505.1
F2 7438106 7438243
refGene_gn=RB1CC1;refGene_tx=XM_023248504.1;refGene_gn=RB1CC1;refGene_tx=XM_023248505.1
F2 7440368 7440427
refGene_gn=RB1CC1;refGene_tx=XM_023248504.1;refGene_gn=RB1CC1;refGene_tx=XM_023248505.1
F2 7442722 7442770
refGene_gn=RB1CC1;refGene_tx=XM_023248504.1;refGene_gn=RB1CC1;refGene_tx=XM_023248505.1
F2 7443886 7443941
refGene_gn=RB1CC1;refGene_tx=XM_023248504.1;refGene_gn=RB1CC1;refGene_tx=XM_023248505.1
F2 7450908 7451149
refGene_gn=RB1CC1;refGene_tx=XM_023248504.1;refGene_gn=RB1CC1;refGene_tx=XM_023248505.1
F2 7451249 7451358
refGene_gn=RB1CC1;refGene_tx=XM_023248504.1;refGene_gn=RB1CC1;refGene_tx=XM_023248505.1
F2 7454316 7454482
refGene_gn=RB1CC1;refGene_tx=XM_023248504.1;refGene_gn=RB1CC1;refGene_tx=XM_023248505.1
F2 7464138 7464154 refGene_gn=RB1CC1;refGene_tx=XM_023248506.1
F2 7471138 7473039
refGene_gn=RB1CC1;refGene_tx=XM_023248504.1;refGene_gn=RB1CC1;refGene_

tx=XM_023248505.1;refGene_gn=RB1CC1;refGene_tx=XM_023248506.1
F2	7473129	7473256
refGene_gn=RB1CC1;refGene_tx=XM_023248504.1;refGene_gn=RB1CC1;refGene_tx=XM_023248505.1;refGene_gn=RB1CC1;refGene_tx=XM_023248506.1
F2	7473701	7473805
refGene_gn=RB1CC1;refGene_tx=XM_023248504.1;refGene_gn=RB1CC1;refGene_tx=XM_023248505.1;refGene_gn=RB1CC1;refGene_tx=XM_023248506.1
F2	7475645	7475707
refGene_gn=RB1CC1;refGene_tx=XM_023248504.1;refGene_gn=RB1CC1;refGene_tx=XM_023248505.1;refGene_gn=RB1CC1;refGene_tx=XM_023248506.1
F2	7475972	7476054
refGene_gn=RB1CC1;refGene_tx=XM_023248504.1;refGene_gn=RB1CC1;refGene_tx=XM_023248505.1;refGene_gn=RB1CC1;refGene_tx=XM_023248506.1
F2	7476138	7476325
refGene_gn=RB1CC1;refGene_tx=XM_023248504.1;refGene_gn=RB1CC1;refGene_tx=XM_023248505.1;refGene_gn=RB1CC1;refGene_tx=XM_023248506.1
F2	7476495	7476680
refGene_gn=RB1CC1;refGene_tx=XM_023248504.1;refGene_gn=RB1CC1;refGene_tx=XM_023248505.1;refGene_gn=RB1CC1;refGene_tx=XM_023248506.1
F2	7485511	7485682
refGene_gn=RB1CC1;refGene_tx=XM_023248504.1;refGene_gn=RB1CC1;refGene_tx=XM_023248505.1;refGene_gn=RB1CC1;refGene_tx=XM_023248506.1
F2	7490531	7490958
refGene_gn=RB1CC1;refGene_tx=XM_023248504.1;refGene_gn=RB1CC1;refGene_tx=XM_023248505.1;refGene_gn=RB1CC1;refGene_tx=XM_023248506.1
F2	7492320	7492523
refGene_gn=RB1CC1;refGene_tx=XM_023248504.1;refGene_gn=RB1CC1;refGene_tx=XM_023248505.1;refGene_gn=RB1CC1;refGene_tx=XM_023248506.1
F2	7498610	7498781
refGene_gn=RB1CC1;refGene_tx=XM_023248504.1;refGene_gn=RB1CC1;refGene_tx=XM_023248505.1;refGene_gn=RB1CC1;refGene_tx=XM_023248506.1
F2	7498955	7499082
refGene_gn=RB1CC1;refGene_tx=XM_023248504.1;refGene_gn=RB1CC1;refGene_tx=XM_023248505.1;refGene_gn=RB1CC1;refGene_tx=XM_023248506.1
F2	7500316	7500387
refGene_gn=RB1CC1;refGene_tx=XM_023248504.1;refGene_gn=RB1CC1;refGene_tx=XM_023248505.1;refGene_gn=RB1CC1;refGene_tx=XM_023248506.1
F2	7525362	7526403
refGene_gn=LOC109495967;refGene_tx=XM_023248730.1
F2	7731960	7732950	refGene_gn=NPBWR1;refGene_tx=XM_023248443.1
F2	7944964	7945497
refGene_gn=OPRK1;refGene_tx=XM_019822660.2;refGene_gn=OPRK1;refGene_tx=XM_023248439.1
F2	7956293	7956646
refGene_gn=OPRK1;refGene_tx=XM_019822660.2;refGene_gn=OPRK1;refGene_tx=XM_023248439.1
F2	7971629	7971886	refGene_gn=OPRK1;refGene_tx=XM_023248439.1
F2	8309870	8309931	refGene_gn=ATP6V1H;refGene_tx=XM_003999812.5
F2	8336770	8336884	refGene_gn=ATP6V1H;refGene_tx=XM_003999812.5
F2	8353709	8353811	refGene_gn=ATP6V1H;refGene_tx=XM_003999812.5

F2	8357685	8357811	refGene_gn=ATP6V1H;refGene_tx=XM_003999812.5
F2	8360185	8360364	refGene_gn=ATP6V1H;refGene_tx=XM_003999812.5
F2	8375621	8375814	refGene_gn=ATP6V1H;refGene_tx=XM_003999812.5
F2	8383582	8383680	refGene_gn=ATP6V1H;refGene_tx=XM_003999812.5
F2	8397190	8397244	refGene_gn=ATP6V1H;refGene_tx=XM_003999812.5
F2	8400343	8400448	refGene_gn=ATP6V1H;refGene_tx=XM_003999812.5
F2	8401138	8401228	refGene_gn=ATP6V1H;refGene_tx=XM_003999812.5
F2	8401229	8401253	refGene_gn=ATP6V1H;refGene_tx=XM_003999812.5
F2	8405829	8405919	refGene_gn=ATP6V1H;refGene_tx=XM_003999812.5
F2	8411073	8411176	refGene_gn=ATP6V1H;refGene_tx=XM_003999812.5
F2	8417438	8417551	refGene_gn=ATP6V1H;refGene_tx=XM_003999812.5
F2	8433026	8433191	refGene_gn=RGS20;refGene_tx=XM_023248731.1
F2	8461346	8461805	refGene_gn=RGS20;refGene_tx=XM_023248731.1
F2	8462281	8462480	refGene_gn=RGS20;refGene_tx=XM_023248731.1
F2	8462812	8462962	refGene_gn=RGS20;refGene_tx=XM_023248731.1
F2	8463085	8463176	refGene_gn=RGS20;refGene_tx=XM_023248731.1
F2	8513618	8513705	refGene_gn=RGS20;refGene_tx=XM_023248731.1
F2	8518381	8518616	refGene_gn=RGS20;refGene_tx=XM_023248731.1
F2	8521879	8522068	refGene_gn=RGS20;refGene_tx=XM_023248731.1
F2	8529475	8529484	refGene_gn=TCEA1;refGene_tx=XM_023248658.1
F2	8531136	8531208	refGene_gn=TCEA1;refGene_tx=XM_023248658.1
F2	8537757	8537904	refGene_gn=TCEA1;refGene_tx=XM_023248658.1
F2	8540024	8540179	refGene_gn=TCEA1;refGene_tx=XM_023248658.1
F2	8541149	8541206	refGene_gn=TCEA1;refGene_tx=XM_023248658.1
F2	8542647	8542793	refGene_gn=TCEA1;refGene_tx=XM_023248658.1
F2	8546382	8546470	refGene_gn=TCEA1;refGene_tx=XM_023248658.1
F2	8552096	8552202	refGene_gn=TCEA1;refGene_tx=XM_023248658.1
F2	8557151	8557214	refGene_gn=TCEA1;refGene_tx=XM_023248658.1
F2	8570183	8570246	refGene_gn=TCEA1;refGene_tx=XM_023248658.1
F2	8586486	8586540	refGene_gn=LYPLA1;refGene_tx=XM_006943218.4;refGene_gn=LYPLA1;refGene_tx=XM_023248760.1;refGene_gn=LYPLA1;refGene_tx=XM_023248761.1
F2	8590263	8590440	refGene_gn=LYPLA1;refGene_tx=XM_006943218.4;refGene_gn=LYPLA1;refGene_tx=XM_023248760.1;refGene_gn=LYPLA1;refGene_tx=XM_023248761.1
F2	8591824	8591926	refGene_gn=LYPLA1;refGene_tx=XM_006943218.4;refGene_gn=LYPLA1;refGene_tx=XM_023248760.1;refGene_gn=LYPLA1;refGene_tx=XM_023248761.1
F2	8594083	8594157	refGene_gn=LYPLA1;refGene_tx=XM_006943218.4;refGene_gn=LYPLA1;refGene_tx=XM_023248760.1;refGene_gn=LYPLA1;refGene_tx=XM_023248761.1
F2	8596888	8596959	refGene_gn=LYPLA1;refGene_tx=XM_006943218.4;refGene_gn=LYPLA1;refGene_tx=XM_023248760.1;refGene_gn=LYPLA1;refGene_tx=XM_023248761.1
F2	8598209	8598257	refGene_gn=LYPLA1;refGene_tx=XM_006943218.4;refGene_gn=LYPLA1;refGene_tx=XM_023248760.1;refGene_gn=LYPLA1;refGene_tx=XM_023248761.1
F2	8600028	8600094	refGene_gn=LYPLA1;refGene_tx=XM_023248760.1
F2	8614901	8614933	refGene_gn=LYPLA1;refGene_tx=XM_023248760.1
F2	8615883	8615952	refGene_gn=LYPLA1;refGene_tx=XM_023248760.1

F2 8628558 8628666 refGene_gn=MRPL15;refGene_tx=XM_003999814.5
F2 8629546 8629701 refGene_gn=MRPL15;refGene_tx=XM_003999814.5
F2 8630582 8630748 refGene_gn=MRPL15;refGene_tx=XM_003999814.5
F2 8635666 8635790 refGene_gn=MRPL15;refGene_tx=XM_003999814.5
F2 8636467 8636805 refGene_gn=MRPL15;refGene_tx=XM_003999814.5
F2 8914628 8914935 refGene_gn=SOX17;refGene_tx=XM_023248516.1
F2 8915542 8916456 refGene_gn=SOX17;refGene_tx=XM_023248516.1
F2 9055132 9055135 refGene_gn=RP1;refGene_tx=XM_023248465.1
F2 9057578 9058214
refGene_gn=RP1;refGene_tx=XM_019822681.2;refGene_gn=RP1;refGene_tx=XM_
023248465.1
F2 9059034 9059206
refGene_gn=RP1;refGene_tx=XM_019822681.2;refGene_gn=RP1;refGene_tx=XM_
023248465.1
F2 9061629 9067247
refGene_gn=RP1;refGene_tx=XM_019822681.2;refGene_gn=RP1;refGene_tx=XM_
023248465.1
F2 9078664 9078944
refGene_gn=LOC109495978;refGene_tx=XM_019822683.2
F2 9086870 9086954
refGene_gn=LOC109495978;refGene_tx=XM_019822683.2
F2 9092768 9092932
refGene_gn=LOC109495978;refGene_tx=XM_019822683.2
F2 9110835 9110922
refGene_gn=LOC109495978;refGene_tx=XM_019822683.2
F2 9113694 9113827
refGene_gn=LOC109495978;refGene_tx=XM_019822683.2
F2 9131588 9131740
refGene_gn=LOC109495978;refGene_tx=XM_019822683.2
F2 9135033 9135112
refGene_gn=LOC109495978;refGene_tx=XM_019822683.2
F2 9155221 9155297
refGene_gn=LOC109495978;refGene_tx=XM_019822683.2
F2 9156189 9156266
refGene_gn=LOC109495978;refGene_tx=XM_019822683.2
F2 9156370 9156402
refGene_gn=LOC109495978;refGene_tx=XM_019822683.2
F2 9156549 9156679
refGene_gn=LOC109495978;refGene_tx=XM_019822683.2
F2 9181218 9181322
refGene_gn=LOC109495978;refGene_tx=XM_019822683.2
F2 9183101 9183278
refGene_gn=LOC109495978;refGene_tx=XM_019822683.2
F2 9186064 9186277
refGene_gn=LOC109495978;refGene_tx=XM_019822683.2
F2 9197970 9198148
refGene_gn=LOC109495978;refGene_tx=XM_019822683.2
F2 9212578 9212710
refGene_gn=LOC109495978;refGene_tx=XM_019822683.2
F2 9213383 9213400

refGene_gn=LOC109495978;refGene_tx=XM_019822683.2
F2	9560195	9561016	refGene_gn=XKR4;refGene_tx=XM_023248669.1
F2	9825003	9825203	refGene_gn=XKR4;refGene_tx=XM_023248669.1
F2	9982711	9983658	refGene_gn=XKR4;refGene_tx=XM_023248669.1
F2	10082430	10082469	refGene_gn=TMEM68;refGene_tx=XM_019822685.2
F2	10099497	10099614
refGene_gn=TMEM68;refGene_tx=XM_006943220.3;refGene_gn=TMEM68;refGene_tx=XM_006943221.3;refGene_gn=TMEM68;refGene_tx=XM_019822687.2;refGene_gn=TMEM68;refGene_tx=XM_019822689.2;refGene_gn=TMEM68;refGene_tx=XM_019822690.2
F2	10111275	10111415
refGene_gn=TMEM68;refGene_tx=XM_006943220.3;refGene_gn=TMEM68;refGene_tx=XM_006943221.3;refGene_gn=TMEM68;refGene_tx=XM_019822685.2;refGene_gn=TMEM68;refGene_tx=XM_019822687.2;refGene_gn=TMEM68;refGene_tx=XM_019822689.2;refGene_gn=TMEM68;refGene_tx=XM_019822690.2
F2	10113078	10113139
refGene_gn=TMEM68;refGene_tx=XM_006943220.3;refGene_gn=TMEM68;refGene_tx=XM_006943221.3;refGene_gn=TMEM68;refGene_tx=XM_019822685.2;refGene_gn=TMEM68;refGene_tx=XM_019822687.2;refGene_gn=TMEM68;refGene_tx=XM_019822689.2
F2	10118078	10118272
refGene_gn=TMEM68;refGene_tx=XM_006943220.3;refGene_gn=TMEM68;refGene_tx=XM_006943221.3;refGene_gn=TMEM68;refGene_tx=XM_019822685.2;refGene_gn=TMEM68;refGene_tx=XM_019822687.2;refGene_gn=TMEM68;refGene_tx=XM_019822689.2
F2	10120579	10120747
refGene_gn=TMEM68;refGene_tx=XM_006943220.3;refGene_gn=TMEM68;refGene_tx=XM_006943221.3;refGene_gn=TMEM68;refGene_tx=XM_019822685.2;refGene_gn=TMEM68;refGene_tx=XM_019822687.2;refGene_gn=TMEM68;refGene_tx=XM_019822689.2;refGene_gn=TMEM68;refGene_tx=XM_019822690.2
F2	10131019	10131344
refGene_gn=TMEM68;refGene_tx=XM_006943220.3;refGene_gn=TMEM68;refGene_tx=XM_006943221.3;refGene_gn=TMEM68;refGene_tx=XM_019822685.2;refGene_gn=TMEM68;refGene_tx=XM_019822687.2;refGene_gn=TMEM68;refGene_tx=XM_019822689.2;refGene_gn=TMEM68;refGene_tx=XM_019822690.2
F2	10142116	10142220	refGene_gn=TGS1;refGene_tx=XM_003999816.5
F2	10149464	10149529	refGene_gn=TGS1;refGene_tx=XM_003999816.5
F2	10151854	10152015	refGene_gn=TGS1;refGene_tx=XM_003999816.5
F2	10152577	10153403	refGene_gn=TGS1;refGene_tx=XM_003999816.5
F2	10155727	10155845	refGene_gn=TGS1;refGene_tx=XM_003999816.5
F2	10157321	10157408	refGene_gn=TGS1;refGene_tx=XM_003999816.5
F2	10159733	10159908	refGene_gn=TGS1;refGene_tx=XM_003999816.5
F2	10162057	10162364	refGene_gn=TGS1;refGene_tx=XM_003999816.5
F2	10164723	10164879	refGene_gn=TGS1;refGene_tx=XM_003999816.5
F2	10166499	10166643	refGene_gn=TGS1;refGene_tx=XM_003999816.5
F2	10170748	10170965	refGene_gn=TGS1;refGene_tx=XM_003999816.5
F2	10171914	10171993	refGene_gn=TGS1;refGene_tx=XM_003999816.5
F2	10184179	10184302	refGene_gn=TGS1;refGene_tx=XM_003999816.5
F2	10302931	10303063
refGene_gn=LYN;refGene_tx=XM_003999817.4;refGene_gn=LYN;refGene_tx=XM_

003999818.4;refGene_gn=LYN;refGene_tx=XM_019822692.2;refGene_gn=LYN;refGene_tx=XM_019822694.2;refGene_gn=LYN;refGene_tx=XM_023248683.1;refGene_gn=LYN;refGene_tx=XM_023248684.1
F2 10306252 10306298
refGene_gn=LYN;refGene_tx=XM_003999817.4;refGene_gn=LYN;refGene_tx=XM_003999818.4;refGene_gn=LYN;refGene_tx=XM_019822692.2;refGene_gn=LYN;refGene_tx=XM_019822694.2;refGene_gn=LYN;refGene_tx=XM_023248683.1;refGene_gn=LYN;refGene_tx=XM_023248684.1
F2 10307539 10307645
refGene_gn=LYN;refGene_tx=XM_003999817.4;refGene_gn=LYN;refGene_tx=XM_003999818.4;refGene_gn=LYN;refGene_tx=XM_019822692.2;refGene_gn=LYN;refGene_tx=XM_019822694.2;refGene_gn=LYN;refGene_tx=XM_023248683.1;refGene_gn=LYN;refGene_tx=XM_023248684.1
F2 10309486 10309585
refGene_gn=LYN;refGene_tx=XM_003999817.4;refGene_gn=LYN;refGene_tx=XM_003999818.4;refGene_gn=LYN;refGene_tx=XM_019822692.2;refGene_gn=LYN;refGene_tx=XM_019822694.2;refGene_gn=LYN;refGene_tx=XM_023248683.1;refGene_gn=LYN;refGene_tx=XM_023248684.1
F2 10309960 10310064
refGene_gn=LYN;refGene_tx=XM_003999817.4;refGene_gn=LYN;refGene_tx=XM_003999818.4;refGene_gn=LYN;refGene_tx=XM_019822692.2;refGene_gn=LYN;refGene_tx=XM_019822694.2;refGene_gn=LYN;refGene_tx=XM_023248683.1;refGene_gn=LYN;refGene_tx=XM_023248684.1
F2 10310599 10310749
refGene_gn=LYN;refGene_tx=XM_003999817.4;refGene_gn=LYN;refGene_tx=XM_003999818.4;refGene_gn=LYN;refGene_tx=XM_019822692.2;refGene_gn=LYN;refGene_tx=XM_019822694.2;refGene_gn=LYN;refGene_tx=XM_023248683.1;refGene_gn=LYN;refGene_tx=XM_023248684.1
F2 10311590 10311743
refGene_gn=LYN;refGene_tx=XM_003999817.4;refGene_gn=LYN;refGene_tx=XM_003999818.4;refGene_gn=LYN;refGene_tx=XM_019822692.2;refGene_gn=LYN;refGene_tx=XM_019822694.2;refGene_gn=LYN;refGene_tx=XM_023248683.1;refGene_gn=LYN;refGene_tx=XM_023248684.1
F2 10325644 10325827
refGene_gn=LYN;refGene_tx=XM_003999817.4;refGene_gn=LYN;refGene_tx=XM_003999818.4;refGene_gn=LYN;refGene_tx=XM_019822692.2;refGene_gn=LYN;refGene_tx=XM_019822694.2;refGene_gn=LYN;refGene_tx=XM_023248683.1;refGene_gn=LYN;refGene_tx=XM_023248684.1
F2 10328140 10328217
refGene_gn=LYN;refGene_tx=XM_003999817.4;refGene_gn=LYN;refGene_tx=XM_003999818.4;refGene_gn=LYN;refGene_tx=XM_019822692.2;refGene_gn=LYN;refGene_tx=XM_019822694.2;refGene_gn=LYN;refGene_tx=XM_023248683.1;refGene_gn=LYN;refGene_tx=XM_023248684.1
F2 10356457 10356611
refGene_gn=LYN;refGene_tx=XM_003999817.4;refGene_gn=LYN;refGene_tx=XM_003999818.4;refGene_gn=LYN;refGene_tx=XM_019822692.2;refGene_gn=LYN;refGene_tx=XM_019822694.2;refGene_gn=LYN;refGene_tx=XM_023248683.1;refGene_gn=LYN;refGene_tx=XM_023248684.1
F2 10359503 10359635
refGene_gn=LYN;refGene_tx=XM_003999817.4;refGene_gn=LYN;refGene_tx=XM_

003999818.4;refGene_gn=LYN;refGene_tx=XM_019822692.2;refGene_gn=LYN;refGene_tx=XM_019822694.2;refGene_gn=LYN;refGene_tx=XM_023248683.1;refGene_gn=LYN;refGene_tx=XM_023248684.1
F2 10369430 10369633
refGene_gn=LYN;refGene_tx=XM_003999817.4;refGene_gn=LYN;refGene_tx=XM_003999818.4;refGene_gn=LYN;refGene_tx=XM_019822692.2;refGene_gn=LYN;refGene_tx=XM_019822694.2;refGene_gn=LYN;refGene_tx=XM_023248683.1;refGene_gn=LYN;refGene_tx=XM_023248684.1
F2 10404199 10404682
refGene_gn=LOC111558513;refGene_tx=XM_023248299.1
F2 10413175 10413358 refGene_gn=RPS20;refGene_tx=XM_003999819.5
F2 10413666 10413740 refGene_gn=RPS20;refGene_tx=XM_003999819.5
F2 10414052 10414152 refGene_gn=RPS20;refGene_tx=XM_003999819.5
F2 10414360 10414363 refGene_gn=RPS20;refGene_tx=XM_003999819.5
F2 10450430 10451477 refGene_gn=MOS;refGene_tx=XM_003999842.3
F2 10504598 10505856
refGene_gn=PLAG1;refGene_tx=XM_006943223.4;refGene_gn=PLAG1;refGene_tx=XM_019822695.2;refGene_gn=PLAG1;refGene_tx=XM_023248247.1;refGene_gn=PLAG1;refGene_tx=XM_023248248.1;refGene_gn=PLAG1;refGene_tx=XM_023248249.1
F2 10506376 10506618
refGene_gn=PLAG1;refGene_tx=XM_006943223.4;refGene_gn=PLAG1;refGene_tx=XM_023248247.1;refGene_gn=PLAG1;refGene_tx=XM_023248248.1
F2 10553506 10553560
refGene_gn=CHCHD7;refGene_tx=XM_023248414.1;refGene_gn=CHCHD7;refGene_tx=XM_023248415.1
F2 10554994 10555137
refGene_gn=CHCHD7;refGene_tx=XM_023248414.1;refGene_gn=CHCHD7;refGene_tx=XM_023248415.1
F2 10556477 10556582 refGene_gn=CHCHD7;refGene_tx=XM_023248414.1
F2 10608724 10608818 refGene_gn=SDR16C5;refGene_tx=XM_003999843.5
F2 10610451 10610577 refGene_gn=SDR16C5;refGene_tx=XM_003999843.5
F2 10611438 10611583 refGene_gn=SDR16C5;refGene_tx=XM_003999843.5
F2 10613969 10614069 refGene_gn=SDR16C5;refGene_tx=XM_003999843.5
F2 10616171 10616303 refGene_gn=SDR16C5;refGene_tx=XM_003999843.5
F2 10620178 10620511 refGene_gn=SDR16C5;refGene_tx=XM_003999843.5
F2 10701536 10702199 refGene_gn=PENK;refGene_tx=XM_003999824.4
F2 10705403 10705544 refGene_gn=PENK;refGene_tx=XM_003999824.4
F2 10706039 10706078 refGene_gn=PENK;refGene_tx=XM_003999824.4
F2 10737196 10737402
refGene_gn=LOC109495884;refGene_tx=XM_019822572.2
F2 10737404 10737507
refGene_gn=LOC109495884;refGene_tx=XM_019822572.2
F2 10737739 10737865
refGene_gn=LOC109495884;refGene_tx=XM_019822572.2
F2 10743478 10743655
refGene_gn=LOC109495884;refGene_tx=XM_019822572.2
F2 10747060 10747078
refGene_gn=LOC109495884;refGene_tx=XM_019822572.2
F2 11135731 11136003

refGene_gn=IMPAD1;refGene_tx=XM_006943232.3;refGene_gn=IMPAD1;refGene_tx=XM_023248681.1
F2 11137474 11137636 refGene_gn=IMPAD1;refGene_tx=XM_006943232.3;refGene_gn=IMPAD1;refGene_tx=XM_023248681.1
F2 11148009 11148105 refGene_gn=IMPAD1;refGene_tx=XM_006943232.3
F2 11149326 11149489 refGene_gn=IMPAD1;refGene_tx=XM_006943232.3;refGene_gn=IMPAD1;refGene_tx=XM_023248681.1
F2 11165820 11166210 refGene_gn=IMPAD1;refGene_tx=XM_006943232.3;refGene_gn=IMPAD1;refGene_tx=XM_023248681.1
F2 11910260 11911373 refGene_gn=FAM110B;refGene_tx=XM_023248663.1;refGene_gn=FAM110B;refGene_tx=XM_023248664.1;refGene_gn=FAM110B;refGene_tx=XM_023248665.1;refGene_gn=FAM110B;refGene_tx=XM_023248666.1;refGene_gn=FAM110B;refGene_tx=XM_023248667.1;refGene_gn=FAM110B;refGene_tx=XM_023248668.1
F2 12128770 12128851 refGene_gn=UBXN2B;refGene_tx=XM_003999827.5
F2 12132306 12132402 refGene_gn=UBXN2B;refGene_tx=XM_019822698.2;refGene_gn=UBXN2B;refGene_tx=XM_019822699.2
F2 12133471 12133575 refGene_gn=UBXN2B;refGene_tx=XM_003999827.5;refGene_gn=UBXN2B;refGene_tx=XM_019822698.2;refGene_gn=UBXN2B;refGene_tx=XM_019822699.2
F2 12139440 12139591 refGene_gn=UBXN2B;refGene_tx=XM_003999827.5;refGene_gn=UBXN2B;refGene_tx=XM_019822698.2;refGene_gn=UBXN2B;refGene_tx=XM_019822699.2
F2 12142368 12142452 refGene_gn=UBXN2B;refGene_tx=XM_003999827.5;refGene_gn=UBXN2B;refGene_tx=XM_019822698.2;refGene_gn=UBXN2B;refGene_tx=XM_019822699.2
F2 12143244 12143354 refGene_gn=UBXN2B;refGene_tx=XM_003999827.5;refGene_gn=UBXN2B;refGene_tx=XM_019822698.2;refGene_gn=UBXN2B;refGene_tx=XM_019822699.2
F2 12146698 12146836 refGene_gn=UBXN2B;refGene_tx=XM_003999827.5;refGene_gn=UBXN2B;refGene_tx=XM_019822698.2;refGene_gn=UBXN2B;refGene_tx=XM_019822699.2
F2 12151742 12151904 refGene_gn=UBXN2B;refGene_tx=XM_003999827.5;refGene_gn=UBXN2B;refGene_tx=XM_019822698.2
F2 12152863 12153026 refGene_gn=UBXN2B;refGene_tx=XM_003999827.5;refGene_gn=UBXN2B;refGene_tx=XM_019822698.2
F2 12156173 12156480 refGene_gn=UBXN2B;refGene_tx=XM_019822699.2
F2 12186829 12187129 refGene_gn=LOC101099612;refGene_tx=XM_003999828.3
F2 12187720 12187896 refGene_gn=LOC101099612;refGene_tx=XM_003999828.3
F2 12189244 12189375 refGene_gn=LOC101099612;refGene_tx=XM_003999828.3
F2 12190873 12191451

refGene_gn=LOC101099612;refGene_tx=XM_003999828.3
F2 12192343 12192584
refGene_gn=LOC101099612;refGene_tx=XM_003999828.3
F2 12194365 12194445
refGene_gn=LOC101099612;refGene_tx=XM_003999828.3
F2 12245298 12245349
refGene_gn=SDCBP;refGene_tx=XM_006943225.3;refGene_gn=SDCBP;refGene_tx=XM_011291377.3;refGene_gn=SDCBP;refGene_tx=XM_019822702.2
F2 12251244 12251323
refGene_gn=SDCBP;refGene_tx=XM_006943225.3;refGene_gn=SDCBP;refGene_tx=XM_011291377.3;refGene_gn=SDCBP;refGene_tx=XM_019822702.2
F2 12252530 12252640
refGene_gn=SDCBP;refGene_tx=XM_006943225.3;refGene_gn=SDCBP;refGene_tx=XM_011291377.3;refGene_gn=SDCBP;refGene_tx=XM_019822702.2
F2 12258346 12258508
refGene_gn=SDCBP;refGene_tx=XM_006943225.3;refGene_gn=SDCBP;refGene_tx=XM_011291377.3;refGene_gn=SDCBP;refGene_tx=XM_019822702.2
F2 12260851 12261027
refGene_gn=SDCBP;refGene_tx=XM_006943225.3;refGene_gn=SDCBP;refGene_tx=XM_011291377.3;refGene_gn=SDCBP;refGene_tx=XM_019822702.2
F2 12261750 12261922
refGene_gn=SDCBP;refGene_tx=XM_006943225.3;refGene_gn=SDCBP;refGene_tx=XM_011291377.3;refGene_gn=SDCBP;refGene_tx=XM_019822702.2
F2 12262655 12262747
refGene_gn=SDCBP;refGene_tx=XM_006943225.3;refGene_gn=SDCBP;refGene_tx=XM_011291377.3;refGene_gn=SDCBP;refGene_tx=XM_019822702.2
F2 12263757 12263812
refGene_gn=SDCBP;refGene_tx=XM_006943225.3;refGene_gn=SDCBP;refGene_tx=XM_011291377.3;refGene_gn=SDCBP;refGene_tx=XM_019822702.2
F2 12266093 12266188 refGene_gn=NSMAF;refGene_tx=XM_023248601.1
F2 12267178 12267288 refGene_gn=NSMAF;refGene_tx=XM_023248601.1
F2 12267432 12267535 refGene_gn=NSMAF;refGene_tx=XM_023248601.1
F2 12267946 12268097 refGene_gn=NSMAF;refGene_tx=XM_023248601.1
F2 12269248 12269332 refGene_gn=NSMAF;refGene_tx=XM_023248601.1
F2 12270823 12270947 refGene_gn=NSMAF;refGene_tx=XM_023248601.1
F2 12271364 12271432 refGene_gn=NSMAF;refGene_tx=XM_023248601.1
F2 12272242 12272310 refGene_gn=NSMAF;refGene_tx=XM_023248601.1
F2 12275127 12275186 refGene_gn=NSMAF;refGene_tx=XM_023248601.1
F2 12276517 12276626 refGene_gn=NSMAF;refGene_tx=XM_023248601.1
F2 12278116 12278280 refGene_gn=NSMAF;refGene_tx=XM_023248601.1
F2 12278803 12278846 refGene_gn=NSMAF;refGene_tx=XM_023248601.1
F2 12279608 12279740 refGene_gn=NSMAF;refGene_tx=XM_023248601.1
F2 12280127 12280248 refGene_gn=NSMAF;refGene_tx=XM_023248601.1
F2 12280366 12280418 refGene_gn=NSMAF;refGene_tx=XM_023248601.1
F2 12281610 12281674 refGene_gn=NSMAF;refGene_tx=XM_023248601.1
F2 12281769 12281860 refGene_gn=NSMAF;refGene_tx=XM_023248601.1
F2 12282418 12282498 refGene_gn=NSMAF;refGene_tx=XM_023248601.1
F2 12284238 12284415 refGene_gn=NSMAF;refGene_tx=XM_023248601.1
F2 12287128 12287237 refGene_gn=NSMAF;refGene_tx=XM_023248601.1
F2 12287599 12287671 refGene_gn=NSMAF;refGene_tx=XM_023248601.1

F2 12290034 12290164 refGene_gn=NSMAF;refGene_tx=XM_023248601.1
F2 12298384 12298437 refGene_gn=NSMAF;refGene_tx=XM_023248601.1
F2 12298533 12298581 refGene_gn=NSMAF;refGene_tx=XM_023248601.1
F2 12298899 12298971 refGene_gn=NSMAF;refGene_tx=XM_023248601.1
F2 12301922 12301973 refGene_gn=NSMAF;refGene_tx=XM_023248601.1
F2 12305769 12305806 refGene_gn=NSMAF;refGene_tx=XM_023248601.1
F2 12305886 12305954 refGene_gn=NSMAF;refGene_tx=XM_023248601.1
F2 12306051 12306130 refGene_gn=NSMAF;refGene_tx=XM_023248601.1
F2 12309904 12309994 refGene_gn=NSMAF;refGene_tx=XM_023248601.1
F2 12327825 12327884 refGene_gn=NSMAF;refGene_tx=XM_023248601.1
F2 12451363 12451400
refGene_gn=TOX;refGene_tx=XM_019822707.2;refGene_gn=TOX;refGene_tx=XM_
023248541.1;refGene_gn=TOX;refGene_tx=XM_023248542.1
F2 12451742 12451894
refGene_gn=TOX;refGene_tx=XM_019822707.2;refGene_gn=TOX;refGene_tx=XM_
023248541.1;refGene_gn=TOX;refGene_tx=XM_023248542.1
F2 12459117 12459495
refGene_gn=TOX;refGene_tx=XM_019822707.2;refGene_gn=TOX;refGene_tx=XM_
023248541.1;refGene_gn=TOX;refGene_tx=XM_023248542.1
F2 12472989 12473070
refGene_gn=TOX;refGene_tx=XM_019822707.2;refGene_gn=TOX;refGene_tx=XM_
023248541.1;refGene_gn=TOX;refGene_tx=XM_023248542.1
F2 12484634 12484865
refGene_gn=TOX;refGene_tx=XM_019822707.2;refGene_gn=TOX;refGene_tx=XM_
023248541.1;refGene_gn=TOX;refGene_tx=XM_023248542.1
F2 12499310 12499592
refGene_gn=TOX;refGene_tx=XM_019822707.2;refGene_gn=TOX;refGene_tx=XM_
023248541.1;refGene_gn=TOX;refGene_tx=XM_023248542.1
F2 12579404 12579647
refGene_gn=TOX;refGene_tx=XM_019822707.2;refGene_gn=TOX;refGene_tx=XM_
023248541.1;refGene_gn=TOX;refGene_tx=XM_023248542.1
F2 12598597 12598663
refGene_gn=TOX;refGene_tx=XM_019822707.2;refGene_gn=TOX;refGene_tx=XM_
023248541.1
F2 12600182 12600200 refGene_gn=TOX;refGene_tx=XM_019822707.2
F2 12756785 12756887
refGene_gn=TOX;refGene_tx=XM_023248541.1;refGene_gn=TOX;refGene_tx=XM_
023248542.1
F2 13634958 13635093 refGene_gn=CA8;refGene_tx=XM_023248778.1
F2 13647978 13648091 refGene_gn=CA8;refGene_tx=XM_023248778.1
F2 13649738 13649787 refGene_gn=CA8;refGene_tx=XM_023248778.1
F2 13653118 13653181 refGene_gn=CA8;refGene_tx=XM_023248778.1
F2 13658759 13658855 refGene_gn=CA8;refGene_tx=XM_023248778.1
F2 13690308 13690433 refGene_gn=CA8;refGene_tx=XM_023248778.1
F2 13701635 13701827 refGene_gn=CA8;refGene_tx=XM_023248778.1
F2 13702907 13703007 refGene_gn=CA8;refGene_tx=XM_023248778.1
F2 13882075 13882121 refGene_gn=RAB2A;refGene_tx=XM_023248779.1
F2 13900328 13900368 refGene_gn=RAB2A;refGene_tx=XM_019822712.2
F2 13913875 13913947
refGene_gn=RAB2A;refGene_tx=XM_019822712.2;refGene_gn=RAB2A;refGene_tx

=XM_023248779.1
F2 13924203 13924271
refGene_gn=RAB2A;refGene_tx=XM_019822712.2;refGene_gn=RAB2A;refGene_tx
=XM_023248779.1
F2 13935489 13935572
refGene_gn=RAB2A;refGene_tx=XM_019822712.2;refGene_gn=RAB2A;refGene_tx
=XM_023248779.1
F2 13935960 13936053
refGene_gn=RAB2A;refGene_tx=XM_019822712.2;refGene_gn=RAB2A;refGene_tx
=XM_023248779.1
F2 13944309 13944421
refGene_gn=RAB2A;refGene_tx=XM_019822712.2;refGene_gn=RAB2A;refGene_tx
=XM_023248779.1
F2 13964815 13964884
refGene_gn=RAB2A;refGene_tx=XM_019822712.2;refGene_gn=RAB2A;refGene_tx
=XM_023248779.1
F2 13966968 13967064
refGene_gn=RAB2A;refGene_tx=XM_019822712.2;refGene_gn=RAB2A;refGene_tx
=XM_023248779.1
F2 14083668 14085330
refGene_gn=CHD7;refGene_tx=XM_023248436.1;refGene_gn=CHD7;refGene_tx=X
M_023248437.1;refGene_gn=CHD7;refGene_tx=XM_023248438.1
F2 14124377 14124808
refGene_gn=CHD7;refGene_tx=XM_023248436.1;refGene_gn=CHD7;refGene_tx=X
M_023248437.1;refGene_gn=CHD7;refGene_tx=XM_023248438.1
F2 14138930 14139072
refGene_gn=CHD7;refGene_tx=XM_023248436.1;refGene_gn=CHD7;refGene_tx=X
M_023248437.1;refGene_gn=CHD7;refGene_tx=XM_023248438.1
F2 14143655 14143793
refGene_gn=CHD7;refGene_tx=XM_023248436.1;refGene_gn=CHD7;refGene_tx=X
M_023248437.1;refGene_gn=CHD7;refGene_tx=XM_023248438.1
F2 14144875 14144941
refGene_gn=CHD7;refGene_tx=XM_023248436.1;refGene_gn=CHD7;refGene_tx=X
M_023248437.1;refGene_gn=CHD7;refGene_tx=XM_023248438.1
F2 14151576 14151632
refGene_gn=CHD7;refGene_tx=XM_023248436.1;refGene_gn=CHD7;refGene_tx=X
M_023248437.1;refGene_gn=CHD7;refGene_tx=XM_023248438.1
F2 14158082 14158197
refGene_gn=CHD7;refGene_tx=XM_023248436.1;refGene_gn=CHD7;refGene_tx=X
M_023248437.1;refGene_gn=CHD7;refGene_tx=XM_023248438.1
F2 14161433 14161517
refGene_gn=CHD7;refGene_tx=XM_023248436.1;refGene_gn=CHD7;refGene_tx=X
M_023248437.1;refGene_gn=CHD7;refGene_tx=XM_023248438.1
F2 14163219 14163357
refGene_gn=CHD7;refGene_tx=XM_023248436.1;refGene_gn=CHD7;refGene_tx=X
M_023248437.1;refGene_gn=CHD7;refGene_tx=XM_023248438.1
F2 14163453 14163575
refGene_gn=CHD7;refGene_tx=XM_023248436.1;refGene_gn=CHD7;refGene_tx=X
M_023248437.1;refGene_gn=CHD7;refGene_tx=XM_023248438.1
F2 14164172 14164416

refGene_gn=CHD7;refGene_tx=XM_023248436.1;refGene_gn=CHD7;refGene_tx=XM_023248437.1;refGene_gn=CHD7;refGene_tx=XM_023248438.1
F2 14166096 14166273
refGene_gn=CHD7;refGene_tx=XM_023248436.1;refGene_gn=CHD7;refGene_tx=XM_023248437.1;refGene_gn=CHD7;refGene_tx=XM_023248438.1
F2 14170423 14170567
refGene_gn=CHD7;refGene_tx=XM_023248436.1;refGene_gn=CHD7;refGene_tx=XM_023248437.1;refGene_gn=CHD7;refGene_tx=XM_023248438.1
F2 14171728 14171984
refGene_gn=CHD7;refGene_tx=XM_023248436.1;refGene_gn=CHD7;refGene_tx=XM_023248437.1;refGene_gn=CHD7;refGene_tx=XM_023248438.1
F2 14177629 14177840
refGene_gn=CHD7;refGene_tx=XM_023248436.1;refGene_gn=CHD7;refGene_tx=XM_023248437.1;refGene_gn=CHD7;refGene_tx=XM_023248438.1
F2 14178591 14178787
refGene_gn=CHD7;refGene_tx=XM_023248436.1;refGene_gn=CHD7;refGene_tx=XM_023248437.1;refGene_gn=CHD7;refGene_tx=XM_023248438.1
F2 14179117 14179285
refGene_gn=CHD7;refGene_tx=XM_023248436.1;refGene_gn=CHD7;refGene_tx=XM_023248437.1;refGene_gn=CHD7;refGene_tx=XM_023248438.1
F2 14179802 14179982
refGene_gn=CHD7;refGene_tx=XM_023248436.1;refGene_gn=CHD7;refGene_tx=XM_023248437.1;refGene_gn=CHD7;refGene_tx=XM_023248438.1
F2 14182921 14183032
refGene_gn=CHD7;refGene_tx=XM_023248436.1;refGene_gn=CHD7;refGene_tx=XM_023248437.1;refGene_gn=CHD7;refGene_tx=XM_023248438.1
F2 14183121 14183327
refGene_gn=CHD7;refGene_tx=XM_023248436.1;refGene_gn=CHD7;refGene_tx=XM_023248437.1;refGene_gn=CHD7;refGene_tx=XM_023248438.1
F2 14185300 14185502
refGene_gn=CHD7;refGene_tx=XM_023248436.1;refGene_gn=CHD7;refGene_tx=XM_023248437.1;refGene_gn=CHD7;refGene_tx=XM_023248438.1
F2 14185698 14185856
refGene_gn=CHD7;refGene_tx=XM_023248436.1;refGene_gn=CHD7;refGene_tx=XM_023248437.1;refGene_gn=CHD7;refGene_tx=XM_023248438.1
F2 14190749 14190839
refGene_gn=CHD7;refGene_tx=XM_023248436.1;refGene_gn=CHD7;refGene_tx=XM_023248437.1;refGene_gn=CHD7;refGene_tx=XM_023248438.1
F2 14191300 14191404
refGene_gn=CHD7;refGene_tx=XM_023248436.1;refGene_gn=CHD7;refGene_tx=XM_023248437.1;refGene_gn=CHD7;refGene_tx=XM_023248438.1
F2 14192759 14192889
refGene_gn=CHD7;refGene_tx=XM_023248436.1;refGene_gn=CHD7;refGene_tx=XM_023248437.1;refGene_gn=CHD7;refGene_tx=XM_023248438.1
F2 14193281 14193354
refGene_gn=CHD7;refGene_tx=XM_023248436.1;refGene_gn=CHD7;refGene_tx=XM_023248437.1;refGene_gn=CHD7;refGene_tx=XM_023248438.1
F2 14193511 14193569
refGene_gn=CHD7;refGene_tx=XM_023248436.1;refGene_gn=CHD7;refGene_tx=XM_023248437.1;refGene_gn=CHD7;refGene_tx=XM_023248438.1

F2 14194248 14194477
refGene_gn=CHD7;refGene_tx=XM_023248436.1;refGene_gn=CHD7;refGene_tx=XM_023248437.1;refGene_gn=CHD7;refGene_tx=XM_023248438.1
F2 14194729 14194938
refGene_gn=CHD7;refGene_tx=XM_023248436.1;refGene_gn=CHD7;refGene_tx=XM_023248437.1;refGene_gn=CHD7;refGene_tx=XM_023248438.1
F2 14195063 14195735
refGene_gn=CHD7;refGene_tx=XM_023248436.1;refGene_gn=CHD7;refGene_tx=XM_023248437.1;refGene_gn=CHD7;refGene_tx=XM_023248438.1
F2 14196641 14196802
refGene_gn=CHD7;refGene_tx=XM_023248436.1;refGene_gn=CHD7;refGene_tx=XM_023248437.1;refGene_gn=CHD7;refGene_tx=XM_023248438.1
F2 14199441 14199669
refGene_gn=CHD7;refGene_tx=XM_023248436.1;refGene_gn=CHD7;refGene_tx=XM_023248437.1;refGene_gn=CHD7;refGene_tx=XM_023248438.1
F2 14199893 14200337
refGene_gn=CHD7;refGene_tx=XM_023248436.1;refGene_gn=CHD7;refGene_tx=XM_023248437.1;refGene_gn=CHD7;refGene_tx=XM_023248438.1
F2 14203719 14203941
refGene_gn=CHD7;refGene_tx=XM_023248436.1;refGene_gn=CHD7;refGene_tx=XM_023248437.1;refGene_gn=CHD7;refGene_tx=XM_023248438.1
F2 14205253 14205394
refGene_gn=CHD7;refGene_tx=XM_023248436.1;refGene_gn=CHD7;refGene_tx=XM_023248437.1;refGene_gn=CHD7;refGene_tx=XM_023248438.1
F2 14205586 14205691
refGene_gn=CHD7;refGene_tx=XM_023248436.1;refGene_gn=CHD7;refGene_tx=XM_023248437.1;refGene_gn=CHD7;refGene_tx=XM_023248438.1
F2 14206972 14207911
refGene_gn=CHD7;refGene_tx=XM_023248436.1;refGene_gn=CHD7;refGene_tx=XM_023248437.1;refGene_gn=CHD7;refGene_tx=XM_023248438.1
F2 14571086 14571541 refGene_gn=CLVS1;refGene_tx=XM_006943226.4
F2 14644638 14644813 refGene_gn=CLVS1;refGene_tx=XM_006943226.4
F2 14697593 14697704 refGene_gn=CLVS1;refGene_tx=XM_006943226.4
F2 14701674 14701910 refGene_gn=CLVS1;refGene_tx=XM_006943226.4
F2 14731108 14731196 refGene_gn=CLVS1;refGene_tx=XM_006943226.4
F2 14734896 14735047 refGene_gn=ASPH;refGene_tx=XM_023248216.1
F2 14747160 14747294 refGene_gn=ASPH;refGene_tx=XM_023248216.1
F2 14747681 14747773 refGene_gn=ASPH;refGene_tx=XM_023248216.1
F2 14753186 14753322 refGene_gn=ASPH;refGene_tx=XM_023248216.1
F2 14775977 14776115 refGene_gn=ASPH;refGene_tx=XM_023248216.1
F2 14781799 14781889 refGene_gn=ASPH;refGene_tx=XM_023248216.1
F2 14783989 14784088 refGene_gn=ASPH;refGene_tx=XM_023248216.1
F2 14791397 14791534 refGene_gn=ASPH;refGene_tx=XM_023248216.1
F2 14793930 14794081 refGene_gn=ASPH;refGene_tx=XM_023248216.1
F2 14800446 14800533 refGene_gn=ASPH;refGene_tx=XM_023248216.1
F2 14807276 14807362 refGene_gn=ASPH;refGene_tx=XM_023248216.1
F2 14848292 14848334 refGene_gn=ASPH;refGene_tx=XM_023248216.1
F2 14861336 14861381 refGene_gn=ASPH;refGene_tx=XM_023248216.1
F2 14865623 14865659 refGene_gn=ASPH;refGene_tx=XM_023248216.1
F2 14866272 14866311 refGene_gn=ASPH;refGene_tx=XM_023248216.1

F2 14871150 14871183 refGene_gn=ASPH;refGene_tx=XM_023248216.1
F2 14871637 14871688 refGene_gn=ASPH;refGene_tx=XM_023248216.1
F2 14872164 14872221 refGene_gn=ASPH;refGene_tx=XM_023248216.1
F2 14873263 14873296 refGene_gn=ASPH;refGene_tx=XM_023248216.1
F2 14877162 14877288 refGene_gn=ASPH;refGene_tx=XM_023248216.1
F2 14881439 14881514 refGene_gn=ASPH;refGene_tx=XM_023248216.1
F2 14884562 14884655 refGene_gn=ASPH;refGene_tx=XM_023248216.1
F2 14895445 14895807
refGene_gn=ASPH;refGene_tx=XM_023248213.1;refGene_gn=ASPH;refGene_tx=XM_023248214.1;refGene_gn=ASPH;refGene_tx=XM_023248215.1
F2 14898026 14898068
refGene_gn=ASPH;refGene_tx=XM_023248213.1;refGene_gn=ASPH;refGene_tx=XM_023248214.1;refGene_gn=ASPH;refGene_tx=XM_023248215.1
F2 14911613 14911682
refGene_gn=ASPH;refGene_tx=XM_023248213.1;refGene_gn=ASPH;refGene_tx=XM_023248214.1;refGene_gn=ASPH;refGene_tx=XM_023248215.1;refGene_gn=ASPH;refGene_tx=XM_023248216.1
F2 14913786 14913831 refGene_gn=ASPH;refGene_tx=XM_023248213.1
F2 14915503 14915653
refGene_gn=ASPH;refGene_tx=XM_023248213.1;refGene_gn=ASPH;refGene_tx=XM_023248214.1;refGene_gn=ASPH;refGene_tx=XM_023248215.1;refGene_gn=ASPH;refGene_tx=XM_023248216.1
F2 14921504 14921514 refGene_gn=ASPH;refGene_tx=XM_023248215.1
F2 14954790 14954926
refGene_gn=ASPH;refGene_tx=XM_023248213.1;refGene_gn=ASPH;refGene_tx=XM_023248214.1;refGene_gn=ASPH;refGene_tx=XM_023248216.1
F2 15384460 15384514
refGene_gn=NKAIN3;refGene_tx=XM_023248645.1;refGene_gn=NKAIN3;refGene_tx=XM_023248646.1
F2 15699350 15699488
refGene_gn=NKAIN3;refGene_tx=XM_023248645.1;refGene_gn=NKAIN3;refGene_tx=XM_023248646.1
F2 15710031 15710112
refGene_gn=NKAIN3;refGene_tx=XM_023248645.1;refGene_gn=NKAIN3;refGene_tx=XM_023248646.1
F2 15841265 15841463
refGene_gn=NKAIN3;refGene_tx=XM_023248645.1;refGene_gn=NKAIN3;refGene_tx=XM_023248646.1
F2 15952327 15952388
refGene_gn=NKAIN3;refGene_tx=XM_023248645.1;refGene_gn=NKAIN3;refGene_tx=XM_023248646.1
F2 15978274 15978344 refGene_gn=NKAIN3;refGene_tx=XM_023248646.1
F2 15987093 15987107
refGene_gn=NKAIN3;refGene_tx=XM_023248645.1;refGene_gn=NKAIN3;refGene_tx=XM_023248646.1
F2 16026293 16026418 refGene_gn=GGH;refGene_tx=XM_003999834.4
F2 16028597 16028735 refGene_gn=GGH;refGene_tx=XM_003999834.4
F2 16030968 16031059 refGene_gn=GGH;refGene_tx=XM_003999834.4
F2 16031140 16031247 refGene_gn=GGH;refGene_tx=XM_003999834.4
F2 16033541 16033680 refGene_gn=GGH;refGene_tx=XM_003999834.4

F2 16036831 16036916 refGene_gn=GGH;refGene_tx=XM_003999834.4
F2 16038190 16038241 refGene_gn=GGH;refGene_tx=XM_003999834.4
F2 16042388 16042500 refGene_gn=GGH;refGene_tx=XM_003999834.4
F2 16045627 16045736 refGene_gn=GGH;refGene_tx=XM_003999834.4
F2 16064511 16064685
refGene_gn=TTPA;refGene_tx=XM_011291375.3;refGene_gn=TTPA;refGene_tx=XM_023248562.1;refGene_gn=TTPA;refGene_tx=XM_023248563.1;refGene_gn=TTPA;refGene_tx=XM_023248564.1;refGene_gn=TTPA;refGene_tx=XM_023248565.1;refGene_gn=TTPA;refGene_tx=XM_023248566.1
F2 16067057 16067168
refGene_gn=TTPA;refGene_tx=XM_011291375.3;refGene_gn=TTPA;refGene_tx=XM_023248562.1;refGene_gn=TTPA;refGene_tx=XM_023248563.1;refGene_gn=TTPA;refGene_tx=XM_023248564.1;refGene_gn=TTPA;refGene_tx=XM_023248565.1;refGene_gn=TTPA;refGene_tx=XM_023248566.1
F2 16068666 16068860
refGene_gn=TTPA;refGene_tx=XM_011291375.3;refGene_gn=TTPA;refGene_tx=XM_023248562.1;refGene_gn=TTPA;refGene_tx=XM_023248563.1;refGene_gn=TTPA;refGene_tx=XM_023248564.1;refGene_gn=TTPA;refGene_tx=XM_023248565.1;refGene_gn=TTPA;refGene_tx=XM_023248566.1
F2 16074222 16074376
refGene_gn=TTPA;refGene_tx=XM_011291375.3;refGene_gn=TTPA;refGene_tx=XM_023248562.1;refGene_gn=TTPA;refGene_tx=XM_023248563.1;refGene_gn=TTPA;refGene_tx=XM_023248564.1;refGene_gn=TTPA;refGene_tx=XM_023248565.1;refGene_gn=TTPA;refGene_tx=XM_023248566.1
F2 16079839 16079890
refGene_gn=TTPA;refGene_tx=XM_023248565.1;refGene_gn=TTPA;refGene_tx=XM_023248566.1
F2 16080454 16080517 refGene_gn=TTPA;refGene_tx=XM_023248564.1
F2 16081340 16081544 refGene_gn=TTPA;refGene_tx=XM_023248563.1
F2 16082692 16082743 refGene_gn=TTPA;refGene_tx=XM_023248569.1
F2 16083851 16083941
refGene_gn=TTPA;refGene_tx=XM_011291375.3;refGene_gn=TTPA;refGene_tx=XM_023248562.1;refGene_gn=TTPA;refGene_tx=XM_023248567.1;refGene_gn=TTPA;refGene_tx=XM_023248568.1;refGene_gn=TTPA;refGene_tx=XM_023248569.1
F2 16121493 16122097
refGene_gn=TTPA;refGene_tx=XM_011291375.3;refGene_gn=TTPA;refGene_tx=XM_023248562.1;refGene_gn=TTPA;refGene_tx=XM_023248567.1;refGene_gn=TTPA;refGene_tx=XM_023248568.1;refGene_gn=TTPA;refGene_tx=XM_023248569.1
F2 16122348 16122372 refGene_gn=YTHDF3;refGene_tx=XM_003999835.5
F2 16122855 16122880 refGene_gn=YTHDF3;refGene_tx=XM_003999835.5
F2 16126291 16126349 refGene_gn=YTHDF3;refGene_tx=XM_011291374.3
F2 16128654 16128740
refGene_gn=YTHDF3;refGene_tx=XM_003999835.5;refGene_gn=YTHDF3;refGene_tx=XM_011291374.3
F2 16138310 16139909
refGene_gn=YTHDF3;refGene_tx=XM_003999835.5;refGene_gn=YTHDF3;refGene_tx=XM_011291374.3;refGene_gn=YTHDF3;refGene_tx=XM_019822721.2;refGene_gn=YTHDF3;refGene_tx=XM_023248560.1;refGene_gn=YTHDF3;refGene_tx=XM_023248561.1
F2 16160053 16160077

refGene_gn=YTHDF3;refGene_tx=XM_003999835.5;refGene_gn=YTHDF3;refGene_tx=XM_011291374.3;refGene_gn=YTHDF3;refGene_tx=XM_019822721.2;refGene_gn=YTHDF3;refGene_tx=XM_023248560.1;refGene_gn=YTHDF3;refGene_tx=XM_023248561.1
F2 17324599 17325724 refGene_gn=BHLHE22;refGene_tx=XM_023248517.1
F2 17340200 17340488
refGene_gn=LOC101086873;refGene_tx=XM_019822725.1;refGene_gn=LOC101086873;refGene_tx=XM_023248780.1
F2 17347420 17347596
refGene_gn=LOC101086873;refGene_tx=XM_019822725.1;refGene_gn=LOC101086873;refGene_tx=XM_023248780.1
F2 17361002 17361209
refGene_gn=LOC101086873;refGene_tx=XM_019822725.1;refGene_gn=LOC101086873;refGene_tx=XM_023248780.1
F2 17361675 17362263
refGene_gn=LOC101086873;refGene_tx=XM_019822725.1;refGene_gn=LOC101086873;refGene_tx=XM_023248780.1
F2 17365807 17365838
refGene_gn=LOC101086873;refGene_tx=XM_019822725.1
F2 17370291 17370428
refGene_gn=LOC101086873;refGene_tx=XM_023248780.1
F2 17512318 17512431
refGene_gn=LOC101086873;refGene_tx=XM_023248780.1
F2 18175956 18176148 refGene_gn=ARMC1;refGene_tx=XM_004001415.5
F2 18177116 18177191 refGene_gn=ARMC1;refGene_tx=XM_004001415.5
F2 18177274 18177391 refGene_gn=ARMC1;refGene_tx=XM_004001415.5
F2 18190580 18190770 refGene_gn=ARMC1;refGene_tx=XM_004001415.5
F2 18205938 18206030 refGene_gn=ARMC1;refGene_tx=XM_004001415.5
F2 18212311 18212494 refGene_gn=ARMC1;refGene_tx=XM_004001415.5
F2 18250212 18250278 refGene_gn=MTFR1;refGene_tx=XM_023248552.1
F2 18256304 18256403 refGene_gn=MTFR1;refGene_tx=XM_023248552.1
F2 18261423 18261462 refGene_gn=MTFR1;refGene_tx=XM_023248553.1
F2 18267502 18267618
refGene_gn=MTFR1;refGene_tx=XM_023248552.1;refGene_gn=MTFR1;refGene_tx=XM_023248553.1
F2 18285006 18285239
refGene_gn=MTFR1;refGene_tx=XM_023248552.1;refGene_gn=MTFR1;refGene_tx=XM_023248553.1
F2 18286900 18287159
refGene_gn=MTFR1;refGene_tx=XM_023248552.1;refGene_gn=MTFR1;refGene_tx=XM_023248553.1
F2 18287955 18288124
refGene_gn=MTFR1;refGene_tx=XM_023248552.1;refGene_gn=MTFR1;refGene_tx=XM_023248553.1
F2 18289025 18289091
refGene_gn=MTFR1;refGene_tx=XM_023248552.1;refGene_gn=MTFR1;refGene_tx=XM_023248553.1
F2 18301428 18301637
refGene_gn=PDE7A;refGene_tx=XM_011291371.3;refGene_gn=PDE7A;refGene_tx=XM_011291372.3;refGene_gn=PDE7A;refGene_tx=XM_023248551.1

F2 18305184 18305265
refGene_gn=PDE7A;refGene_tx=XM_011291371.3;refGene_gn=PDE7A;refGene_tx
=XM_011291372.3;refGene_gn=PDE7A;refGene_tx=XM_023248551.1
F2 18305946 18306043
refGene_gn=PDE7A;refGene_tx=XM_011291371.3;refGene_gn=PDE7A;refGene_tx
=XM_011291372.3;refGene_gn=PDE7A;refGene_tx=XM_023248551.1
F2 18307744 18307889
refGene_gn=PDE7A;refGene_tx=XM_011291371.3;refGene_gn=PDE7A;refGene_tx
=XM_011291372.3;refGene_gn=PDE7A;refGene_tx=XM_023248551.1
F2 18309820 18309912
refGene_gn=PDE7A;refGene_tx=XM_011291371.3;refGene_gn=PDE7A;refGene_tx
=XM_011291372.3;refGene_gn=PDE7A;refGene_tx=XM_023248551.1
F2 18310099 18310231
refGene_gn=PDE7A;refGene_tx=XM_011291371.3;refGene_gn=PDE7A;refGene_tx
=XM_011291372.3;refGene_gn=PDE7A;refGene_tx=XM_023248551.1
F2 18314692 18314793
refGene_gn=PDE7A;refGene_tx=XM_011291371.3;refGene_gn=PDE7A;refGene_tx
=XM_011291372.3;refGene_gn=PDE7A;refGene_tx=XM_023248551.1
F2 18316865 18316961
refGene_gn=PDE7A;refGene_tx=XM_011291371.3;refGene_gn=PDE7A;refGene_tx
=XM_011291372.3;refGene_gn=PDE7A;refGene_tx=XM_023248551.1
F2 18333568 18333632
refGene_gn=PDE7A;refGene_tx=XM_011291371.3;refGene_gn=PDE7A;refGene_tx
=XM_011291372.3;refGene_gn=PDE7A;refGene_tx=XM_023248551.1
F2 18335590 18335742
refGene_gn=PDE7A;refGene_tx=XM_011291371.3;refGene_gn=PDE7A;refGene_tx
=XM_011291372.3;refGene_gn=PDE7A;refGene_tx=XM_023248551.1
F2 18361897 18361981
refGene_gn=PDE7A;refGene_tx=XM_011291371.3;refGene_gn=PDE7A;refGene_tx
=XM_011291372.3;refGene_gn=PDE7A;refGene_tx=XM_023248551.1
F2 18365498 18365559
refGene_gn=PDE7A;refGene_tx=XM_011291371.3;refGene_gn=PDE7A;refGene_tx
=XM_011291372.3;refGene_gn=PDE7A;refGene_tx=XM_023248551.1
F2 18373639 18373699 refGene_gn=PDE7A;refGene_tx=XM_011291372.3
F2 18428881 18429019 refGene_gn=PDE7A;refGene_tx=XM_011291371.3
F2 18620858 18620977
refGene_gn=DNAJC5B;refGene_tx=XM_019822728.2;refGene_gn=DNAJC5B;refGen
e_tx=XM_019822729.2;refGene_gn=DNAJC5B;refGene_tx=XM_019822730.2
F2 18627877 18627913
refGene_gn=DNAJC5B;refGene_tx=XM_019822728.2;refGene_gn=DNAJC5B;refGen
e_tx=XM_019822729.2
F2 18654214 18654428
refGene_gn=DNAJC5B;refGene_tx=XM_019822728.2;refGene_gn=DNAJC5B;refGen
e_tx=XM_019822729.2;refGene_gn=DNAJC5B;refGene_tx=XM_019822730.2
F2 18658325 18658497
refGene_gn=DNAJC5B;refGene_tx=XM_019822728.2;refGene_gn=DNAJC5B;refGen
e_tx=XM_019822729.2;refGene_gn=DNAJC5B;refGene_tx=XM_019822730.2
F2 18675259 18675354
refGene_gn=DNAJC5B;refGene_tx=XM_019822728.2;refGene_gn=DNAJC5B;refGen
e_tx=XM_019822729.2;refGene_gn=DNAJC5B;refGene_tx=XM_019822730.2

F2 18700612 18700780
refGene_gn=TRIM55;refGene_tx=XM_003999853.4;refGene_gn=TRIM55;refGene_tx=XM_003999854.4;refGene_gn=TRIM55;refGene_tx=XM_006943243.3;refGene_gn=TRIM55;refGene_tx=XM_006943244.3;refGene_gn=TRIM55;refGene_tx=XM_019822732.1
F2 18701343 18701516
refGene_gn=TRIM55;refGene_tx=XM_003999853.4;refGene_gn=TRIM55;refGene_tx=XM_003999854.4;refGene_gn=TRIM55;refGene_tx=XM_006943243.3;refGene_gn=TRIM55;refGene_tx=XM_006943244.3;refGene_gn=TRIM55;refGene_tx=XM_019822732.1
F2 18707902 18708068
refGene_gn=TRIM55;refGene_tx=XM_003999853.4;refGene_gn=TRIM55;refGene_tx=XM_003999854.4;refGene_gn=TRIM55;refGene_tx=XM_006943243.3;refGene_gn=TRIM55;refGene_tx=XM_006943244.3;refGene_gn=TRIM55;refGene_tx=XM_019822732.1
F2 18709965 18710061
refGene_gn=TRIM55;refGene_tx=XM_003999853.4;refGene_gn=TRIM55;refGene_tx=XM_003999854.4;refGene_gn=TRIM55;refGene_tx=XM_006943243.3;refGene_gn=TRIM55;refGene_tx=XM_006943244.3;refGene_gn=TRIM55;refGene_tx=XM_019822732.1
F2 18721734 18721968
refGene_gn=TRIM55;refGene_tx=XM_003999853.4;refGene_gn=TRIM55;refGene_tx=XM_003999854.4;refGene_gn=TRIM55;refGene_tx=XM_006943243.3;refGene_gn=TRIM55;refGene_tx=XM_006943244.3;refGene_gn=TRIM55;refGene_tx=XM_019822732.1
F2 18722313 18722336
refGene_gn=TRIM55;refGene_tx=XM_003999853.4;refGene_gn=TRIM55;refGene_tx=XM_003999854.4;refGene_gn=TRIM55;refGene_tx=XM_006943243.3;refGene_gn=TRIM55;refGene_tx=XM_006943244.3;refGene_gn=TRIM55;refGene_tx=XM_019822732.1
F2 18722439 18722564
refGene_gn=TRIM55;refGene_tx=XM_003999853.4;refGene_gn=TRIM55;refGene_tx=XM_003999854.4;refGene_gn=TRIM55;refGene_tx=XM_006943243.3;refGene_gn=TRIM55;refGene_tx=XM_006943244.3;refGene_gn=TRIM55;refGene_tx=XM_019822732.1
F2 18723837 18724106
refGene_gn=TRIM55;refGene_tx=XM_003999853.4;refGene_gn=TRIM55;refGene_tx=XM_003999854.4;refGene_gn=TRIM55;refGene_tx=XM_006943243.3;refGene_gn=TRIM55;refGene_tx=XM_006943244.3;refGene_gn=TRIM55;refGene_tx=XM_019822732.1
F2 18725459 18725963
refGene_gn=TRIM55;refGene_tx=XM_003999853.4;refGene_gn=TRIM55;refGene_tx=XM_006943243.3;refGene_gn=TRIM55;refGene_tx=XM_006943244.3;refGene_gn=TRIM55;refGene_tx=XM_019822732.1
F2 18727050 18727138 refGene_gn=TRIM55;refGene_tx=XM_006943243.3
F2 18743653 18743776
refGene_gn=TRIM55;refGene_tx=XM_003999853.4;refGene_gn=TRIM55;refGene_tx=XM_003999854.4;refGene_gn=TRIM55;refGene_tx=XM_006943243.3;refGene_gn=TRIM55;refGene_tx=XM_006943244.3
F2 18746089 18746686

refGene_gn=CRH;refGene_tx=NM_001040679.1;refGene_gn=CRH;refGene_tx=XM_006943242.4;refGene_gn=CRH;refGene_tx=XM_019822559.1;refGene_gn=CRH;refGene_tx=XM_019822560.2;refGene_gn=CRH;refGene_tx=XM_019822562.2;refGene_gn=CRH;refGene_tx=XM_019822565.1;refGene_gn=CRH;refGene_tx=XM_023248120.1;refGene_gn=CRH;refGene_tx=XM_023248121.1
F2 18972438 18973536 refGene_gn=RRS1;refGene_tx=XM_003999856.5
F2 18975837 18975896 refGene_gn=ADHFE1;refGene_tx=XM_003999857.3
F2 18982798 18982836 refGene_gn=ADHFE1;refGene_tx=XM_003999857.3
F2 18984974 18985021 refGene_gn=ADHFE1;refGene_tx=XM_003999857.3
F2 18985830 18985884
refGene_gn=ADHFE1;refGene_tx=XM_003999857.3;refGene_gn=ADHFE1;refGene_tx=XM_006943245.2
F2 18986076 18986231
refGene_gn=ADHFE1;refGene_tx=XM_003999857.3;refGene_gn=ADHFE1;refGene_tx=XM_006943245.2
F2 18986374 18986571
refGene_gn=ADHFE1;refGene_tx=XM_003999857.3;refGene_gn=ADHFE1;refGene_tx=XM_006943245.2
F2 18988168 18988246
refGene_gn=ADHFE1;refGene_tx=XM_003999857.3;refGene_gn=ADHFE1;refGene_tx=XM_006943245.2
F2 18989568 18989674
refGene_gn=ADHFE1;refGene_tx=XM_003999857.3;refGene_gn=ADHFE1;refGene_tx=XM_006943245.2
F2 18991789 18991942
refGene_gn=ADHFE1;refGene_tx=XM_003999857.3;refGene_gn=ADHFE1;refGene_tx=XM_006943245.2
F2 18993883 18993982
refGene_gn=ADHFE1;refGene_tx=XM_003999857.3;refGene_gn=ADHFE1;refGene_tx=XM_006943245.2
F2 18995769 18995848
refGene_gn=ADHFE1;refGene_tx=XM_003999857.3;refGene_gn=ADHFE1;refGene_tx=XM_006943245.2
F2 18996259 18996356
refGene_gn=ADHFE1;refGene_tx=XM_003999857.3;refGene_gn=ADHFE1;refGene_tx=XM_006943245.2
F2 18999204 18999362
refGene_gn=ADHFE1;refGene_tx=XM_003999857.3;refGene_gn=ADHFE1;refGene_tx=XM_006943245.2
F2 19006662 19006746
refGene_gn=ADHFE1;refGene_tx=XM_003999857.3;refGene_gn=ADHFE1;refGene_tx=XM_006943245.2
F2 19036111 19036178
refGene_gn=CF2H8orf46;refGene_tx=XM_003999858.4
F2 19040574 19040630
refGene_gn=CF2H8orf46;refGene_tx=XM_003999858.4
F2 19048372 19048523
refGene_gn=CF2H8orf46;refGene_tx=XM_003999858.4
F2 19052462 19052524
refGene_gn=CF2H8orf46;refGene_tx=XM_003999858.4

F2 19055492 19055584
refGene_gn=CF2H8orf46;refGene_tx=XM_003999858.4
F2 19056283 19056467
refGene_gn=CF2H8orf46;refGene_tx=XM_003999858.4
F2 19109848 19109977
refGene_gn=MYBL1;refGene_tx=XM_003999859.5;refGene_gn=MYBL1;refGene_tx=XM_003999860.5
F2 19111504 19111684 refGene_gn=MYBL1;refGene_tx=XM_003999859.5
F2 19112098 19112203
refGene_gn=MYBL1;refGene_tx=XM_003999859.5;refGene_gn=MYBL1;refGene_tx=XM_003999860.5
F2 19112296 19112413
refGene_gn=MYBL1;refGene_tx=XM_003999859.5;refGene_gn=MYBL1;refGene_tx=XM_003999860.5
F2 19116212 19116327
refGene_gn=MYBL1;refGene_tx=XM_003999859.5;refGene_gn=MYBL1;refGene_tx=XM_003999860.5
F2 19116889 19117032
refGene_gn=MYBL1;refGene_tx=XM_003999859.5;refGene_gn=MYBL1;refGene_tx=XM_003999860.5
F2 19118940 19119309
refGene_gn=MYBL1;refGene_tx=XM_003999859.5;refGene_gn=MYBL1;refGene_tx=XM_003999860.5
F2 19120338 19120572
refGene_gn=MYBL1;refGene_tx=XM_003999859.5;refGene_gn=MYBL1;refGene_tx=XM_003999860.5
F2 19124398 19124503
refGene_gn=MYBL1;refGene_tx=XM_003999859.5;refGene_gn=MYBL1;refGene_tx=XM_003999860.5
F2 19126996 19127071
refGene_gn=MYBL1;refGene_tx=XM_003999859.5;refGene_gn=MYBL1;refGene_tx=XM_003999860.5
F2 19129256 19129431
refGene_gn=MYBL1;refGene_tx=XM_003999859.5;refGene_gn=MYBL1;refGene_tx=XM_003999860.5
F2 19139162 19139383
refGene_gn=MYBL1;refGene_tx=XM_003999859.5;refGene_gn=MYBL1;refGene_tx=XM_003999860.5
F2 19141016 19141109
refGene_gn=MYBL1;refGene_tx=XM_003999859.5;refGene_gn=MYBL1;refGene_tx=XM_003999860.5
F2 19143256 19143328
refGene_gn=MYBL1;refGene_tx=XM_003999859.5;refGene_gn=MYBL1;refGene_tx=XM_003999860.5
F2 19143665 19143771
refGene_gn=MYBL1;refGene_tx=XM_003999859.5;refGene_gn=MYBL1;refGene_tx=XM_003999860.5
F2 19153661 19153681
refGene_gn=MYBL1;refGene_tx=XM_003999859.5;refGene_gn=MYBL1;refGene_tx=XM_003999860.5

F2 19153914 19154418
refGene_gn=LOC109496024;refGene_tx=XM_019822737.2
F2 19154798 19154834
refGene_gn=LOC109496024;refGene_tx=XM_019822737.2
F2 19171746 19172618 refGene_gn=VCPIP1;refGene_tx=XM_003999861.4
F2 19184638 19184725 refGene_gn=VCPIP1;refGene_tx=XM_003999861.4
F2 19192857 19195570 refGene_gn=VCPIP1;refGene_tx=XM_003999861.4
F2 19316815 19316893
refGene_gn=SGK3;refGene_tx=XM_011291369.2;refGene_gn=SGK3;refGene_tx=XM_019822738.1;refGene_gn=SGK3;refGene_tx=XM_023248737.1
F2 19320705 19320789
refGene_gn=SGK3;refGene_tx=XM_011291369.2;refGene_gn=SGK3;refGene_tx=XM_019822738.1;refGene_gn=SGK3;refGene_tx=XM_023248737.1
F2 19325207 19325280
refGene_gn=SGK3;refGene_tx=XM_011291369.2;refGene_gn=SGK3;refGene_tx=XM_019822738.1;refGene_gn=SGK3;refGene_tx=XM_023248737.1
F2 19337532 19337608
refGene_gn=SGK3;refGene_tx=XM_011291369.2;refGene_gn=SGK3;refGene_tx=XM_019822738.1;refGene_gn=SGK3;refGene_tx=XM_023248737.1
F2 19345647 19345735
refGene_gn=SGK3;refGene_tx=XM_011291369.2;refGene_gn=SGK3;refGene_tx=XM_019822738.1;refGene_gn=SGK3;refGene_tx=XM_023248737.1
F2 19350171 19350221
refGene_gn=SGK3;refGene_tx=XM_011291369.2;refGene_gn=SGK3;refGene_tx=XM_019822738.1;refGene_gn=SGK3;refGene_tx=XM_023248737.1
F2 19352616 19352674
refGene_gn=SGK3;refGene_tx=XM_011291369.2;refGene_gn=SGK3;refGene_tx=XM_019822738.1;refGene_gn=SGK3;refGene_tx=XM_023248737.1
F2 19359696 19359780
refGene_gn=SGK3;refGene_tx=XM_011291369.2;refGene_gn=SGK3;refGene_tx=XM_019822738.1;refGene_gn=SGK3;refGene_tx=XM_023248737.1
F2 19359894 19360026
refGene_gn=SGK3;refGene_tx=XM_011291369.2;refGene_gn=SGK3;refGene_tx=XM_019822738.1;refGene_gn=SGK3;refGene_tx=XM_023248737.1
F2 19362284 19362299
refGene_gn=SGK3;refGene_tx=XM_019822738.1;refGene_gn=SGK3;refGene_tx=XM_023248737.1
F2 19364020 19364133
refGene_gn=SGK3;refGene_tx=XM_011291369.2;refGene_gn=SGK3;refGene_tx=XM_019822738.1;refGene_gn=SGK3;refGene_tx=XM_023248737.1
F2 19364217 19364254
refGene_gn=SGK3;refGene_tx=XM_011291369.2;refGene_gn=SGK3;refGene_tx=XM_019822738.1;refGene_gn=SGK3;refGene_tx=XM_023248737.1
F2 19364821 19364908
refGene_gn=SGK3;refGene_tx=XM_011291369.2;refGene_gn=SGK3;refGene_tx=XM_019822738.1;refGene_gn=SGK3;refGene_tx=XM_023248737.1
F2 19366696 19366792
refGene_gn=SGK3;refGene_tx=XM_011291369.2;refGene_gn=SGK3;refGene_tx=XM_019822738.1;refGene_gn=SGK3;refGene_tx=XM_023248737.1
F2 19370229 19370385

refGene_gn=SGK3;refGene_tx=XM_011291369.2;refGene_gn=SGK3;refGene_tx=XM_019822738.1;refGene_gn=SGK3;refGene_tx=XM_023248737.1
F2 19374067 19374157
refGene_gn=SGK3;refGene_tx=XM_011291369.2;refGene_gn=SGK3;refGene_tx=XM_019822738.1;refGene_gn=SGK3;refGene_tx=XM_023248737.1
F2 19378808 19378979
refGene_gn=SGK3;refGene_tx=XM_011291369.2;refGene_gn=SGK3;refGene_tx=XM_019822738.1;refGene_gn=SGK3;refGene_tx=XM_023248737.1
F2 19401000 19401094 refGene_gn=MCMDC2;refGene_tx=XM_023248515.1
F2 19401182 19401313 refGene_gn=MCMDC2;refGene_tx=XM_023248515.1
F2 19401383 19401443 refGene_gn=MCMDC2;refGene_tx=XM_023248515.1
F2 19403875 19404071 refGene_gn=MCMDC2;refGene_tx=XM_023248515.1
F2 19404711 19404834 refGene_gn=MCMDC2;refGene_tx=XM_023248515.1
F2 19404955 19405060 refGene_gn=MCMDC2;refGene_tx=XM_023248515.1
F2 19407559 19407685 refGene_gn=MCMDC2;refGene_tx=XM_023248515.1
F2 19410312 19410550 refGene_gn=MCMDC2;refGene_tx=XM_023248515.1
F2 19415295 19415501 refGene_gn=MCMDC2;refGene_tx=XM_023248515.1
F2 19418869 19419036 refGene_gn=MCMDC2;refGene_tx=XM_023248515.1
F2 19419527 19419707 refGene_gn=MCMDC2;refGene_tx=XM_023248515.1
F2 19422538 19422681 refGene_gn=MCMDC2;refGene_tx=XM_023248515.1
F2 19427299 19427409 refGene_gn=MCMDC2;refGene_tx=XM_023248515.1
F2 19436571 19436738 refGene_gn=MCMDC2;refGene_tx=XM_023248515.1
F2 19465603 19465717 refGene_gn=TCF24;refGene_tx=XM_023248306.1
F2 19473080 19473470 refGene_gn=TCF24;refGene_tx=XM_023248306.1
F2 19475776 19475822 refGene_gn=PPP1R42;refGene_tx=XM_011291368.3
F2 19482202 19482362 refGene_gn=PPP1R42;refGene_tx=XM_011291368.3
F2 19489713 19489845 refGene_gn=PPP1R42;refGene_tx=XM_011291368.3
F2 19493588 19493706 refGene_gn=PPP1R42;refGene_tx=XM_011291368.3
F2 19514640 19514757 refGene_gn=PPP1R42;refGene_tx=XM_011291368.3
F2 19516194 19516333 refGene_gn=PPP1R42;refGene_tx=XM_011291368.3
F2 19518450 19518521 refGene_gn=PPP1R42;refGene_tx=XM_011291368.3
F2 19518522 19518618 refGene_gn=PPP1R42;refGene_tx=XM_011291368.3
F2 19520974 19521103 refGene_gn=PPP1R42;refGene_tx=XM_011291368.3
F2 19538003 19538088
refGene_gn=COPS5;refGene_tx=XM_003999866.4;refGene_gn=COPS5;refGene_tx=XM_006943252.3;refGene_gn=COPS5;refGene_tx=XM_011291365.2;refGene_gn=COPS5;refGene_tx=XM_019822753.2;refGene_gn=COPS5;refGene_tx=XM_019822754.1
F2 19540279 19540428
refGene_gn=COPS5;refGene_tx=XM_003999866.4;refGene_gn=COPS5;refGene_tx=XM_006943252.3;refGene_gn=COPS5;refGene_tx=XM_011291365.2;refGene_gn=COPS5;refGene_tx=XM_019822753.2;refGene_gn=COPS5;refGene_tx=XM_019822754.1
F2 19544736 19544848
refGene_gn=COPS5;refGene_tx=XM_003999866.4;refGene_gn=COPS5;refGene_tx=XM_006943252.3;refGene_gn=COPS5;refGene_tx=XM_011291365.2;refGene_gn=COPS5;refGene_tx=XM_019822753.2;refGene_gn=COPS5;refGene_tx=XM_019822754.1
F2 19549687 19549773
refGene_gn=COPS5;refGene_tx=XM_003999866.4;refGene_gn=COPS5;refGene_tx

=XM_006943252.3;refGene_gn=COPS5;refGene_tx=XM_011291365.2;refGene_gn=COPS5;refGene_tx=XM_019822753.2;refGene_gn=COPS5;refGene_tx=XM_019822754.1
F2 19550415 19550481 refGene_gn=COPS5;refGene_tx=XM_003999866.4;refGene_gn=COPS5;refGene_tx=XM_006943252.3;refGene_gn=COPS5;refGene_tx=XM_011291365.2;refGene_gn=COPS5;refGene_tx=XM_019822753.2;refGene_gn=COPS5;refGene_tx=XM_019822754.1
F2 19551117 19551246 refGene_gn=COPS5;refGene_tx=XM_003999866.4;refGene_gn=COPS5;refGene_tx=XM_006943252.3;refGene_gn=COPS5;refGene_tx=XM_011291365.2;refGene_gn=COPS5;refGene_tx=XM_019822753.2;refGene_gn=COPS5;refGene_tx=XM_019822754.1
F2 19552199 19552434 refGene_gn=COPS5;refGene_tx=XM_003999866.4;refGene_gn=COPS5;refGene_tx=XM_006943252.3;refGene_gn=COPS5;refGene_tx=XM_011291365.2;refGene_gn=COPS5;refGene_tx=XM_019822753.2;refGene_gn=COPS5;refGene_tx=XM_019822754.1
F2 19554797 19554940 refGene_gn=COPS5;refGene_tx=XM_003999866.4
F2 19554990 19555106 refGene_gn=COPS5;refGene_tx=XM_006943252.3;refGene_gn=COPS5;refGene_tx=XM_019822753.2;refGene_gn=COPS5;refGene_tx=XM_019822754.1
F2 19567698 19567797 refGene_gn=CSPP1;refGene_tx=XM_006943251.3;refGene_gn=CSPP1;refGene_tx=XM_011291366.2;refGene_gn=CSPP1;refGene_tx=XM_011291367.2;refGene_gn=CSPP1;refGene_tx=XM_019822740.1;refGene_gn=CSPP1;refGene_tx=XM_019822742.1
F2 19574860 19574960 refGene_gn=CSPP1;refGene_tx=XM_006943251.3;refGene_gn=CSPP1;refGene_tx=XM_011291366.2;refGene_gn=CSPP1;refGene_tx=XM_011291367.2;refGene_gn=CSPP1;refGene_tx=XM_019822740.1;refGene_gn=CSPP1;refGene_tx=XM_019822741.2;refGene_gn=CSPP1;refGene_tx=XM_019822742.1
F2 19601942 19602046 refGene_gn=CSPP1;refGene_tx=XM_006943251.3;refGene_gn=CSPP1;refGene_tx=XM_011291366.2;refGene_gn=CSPP1;refGene_tx=XM_011291367.2;refGene_gn=CSPP1;refGene_tx=XM_019822740.1;refGene_gn=CSPP1;refGene_tx=XM_019822741.2;refGene_gn=CSPP1;refGene_tx=XM_019822742.1
F2 19602768 19602792 refGene_gn=CSPP1;refGene_tx=XM_006943251.3;refGene_gn=CSPP1;refGene_tx=XM_011291367.2;refGene_gn=CSPP1;refGene_tx=XM_019822740.1;refGene_gn=CSPP1;refGene_tx=XM_019822741.2;refGene_gn=CSPP1;refGene_tx=XM_019822742.1
F2 19606868 19606949 refGene_gn=CSPP1;refGene_tx=XM_006943251.3;refGene_gn=CSPP1;refGene_tx=XM_011291366.2;refGene_gn=CSPP1;refGene_tx=XM_011291367.2;refGene_gn=CSPP1;refGene_tx=XM_019822740.1;refGene_gn=CSPP1;refGene_tx=XM_019822741.2;refGene_gn=CSPP1;refGene_tx=XM_019822742.1
F2 19608249 19608351 refGene_gn=CSPP1;refGene_tx=XM_006943251.3;refGene_gn=CSPP1;refGene_tx=XM_011291366.2;refGene_gn=CSPP1;refGene_tx=XM_011291367.2;refGene_gn=

CSPP1;refGene_tx=XM_019822740.1;refGene_gn=CSPP1;refGene_tx=XM_019822741.2;refGene_gn=CSPP1;refGene_tx=XM_019822742.1
F2 19609516 19609956
refGene_gn=CSPP1;refGene_tx=XM_006943251.3;refGene_gn=CSPP1;refGene_tx=XM_011291366.2;refGene_gn=CSPP1;refGene_tx=XM_011291367.2;refGene_gn=CSPP1;refGene_tx=XM_019822740.1;refGene_gn=CSPP1;refGene_tx=XM_019822741.2;refGene_gn=CSPP1;refGene_tx=XM_019822742.1
F2 19620232 19620331
refGene_gn=CSPP1;refGene_tx=XM_006943251.3;refGene_gn=CSPP1;refGene_tx=XM_011291366.2;refGene_gn=CSPP1;refGene_tx=XM_011291367.2;refGene_gn=CSPP1;refGene_tx=XM_019822741.2
F2 19621489 19621560
refGene_gn=CSPP1;refGene_tx=XM_006943251.3;refGene_gn=CSPP1;refGene_tx=XM_011291366.2;refGene_gn=CSPP1;refGene_tx=XM_011291367.2;refGene_gn=CSPP1;refGene_tx=XM_019822740.1;refGene_gn=CSPP1;refGene_tx=XM_019822741.2
F2 19629942 19630036
refGene_gn=CSPP1;refGene_tx=XM_006943251.3;refGene_gn=CSPP1;refGene_tx=XM_011291366.2;refGene_gn=CSPP1;refGene_tx=XM_011291367.2;refGene_gn=CSPP1;refGene_tx=XM_019822740.1;refGene_gn=CSPP1;refGene_tx=XM_019822741.2
F2 19631074 19631132
refGene_gn=CSPP1;refGene_tx=XM_006943251.3;refGene_gn=CSPP1;refGene_tx=XM_011291366.2;refGene_gn=CSPP1;refGene_tx=XM_011291367.2;refGene_gn=CSPP1;refGene_tx=XM_019822740.1;refGene_gn=CSPP1;refGene_tx=XM_019822741.2
F2 19632827 19633036
refGene_gn=CSPP1;refGene_tx=XM_006943251.3;refGene_gn=CSPP1;refGene_tx=XM_011291366.2;refGene_gn=CSPP1;refGene_tx=XM_011291367.2;refGene_gn=CSPP1;refGene_tx=XM_019822740.1;refGene_gn=CSPP1;refGene_tx=XM_019822741.2
F2 19635395 19635517
refGene_gn=CSPP1;refGene_tx=XM_006943251.3;refGene_gn=CSPP1;refGene_tx=XM_011291366.2;refGene_gn=CSPP1;refGene_tx=XM_011291367.2;refGene_gn=CSPP1;refGene_tx=XM_019822740.1;refGene_gn=CSPP1;refGene_tx=XM_019822741.2;refGene_gn=CSPP1;refGene_tx=XM_019822742.1
F2 19635879 19635958
refGene_gn=CSPP1;refGene_tx=XM_006943251.3;refGene_gn=CSPP1;refGene_tx=XM_011291366.2;refGene_gn=CSPP1;refGene_tx=XM_011291367.2;refGene_gn=CSPP1;refGene_tx=XM_019822740.1;refGene_gn=CSPP1;refGene_tx=XM_019822741.2;refGene_gn=CSPP1;refGene_tx=XM_019822742.1
F2 19666492 19666622
refGene_gn=CSPP1;refGene_tx=XM_006943251.3;refGene_gn=CSPP1;refGene_tx=XM_011291366.2;refGene_gn=CSPP1;refGene_tx=XM_011291367.2;refGene_gn=CSPP1;refGene_tx=XM_019822740.1;refGene_gn=CSPP1;refGene_tx=XM_019822741.2;refGene_gn=CSPP1;refGene_tx=XM_019822742.1
F2 19668606 19668754
refGene_gn=CSPP1;refGene_tx=XM_006943251.3;refGene_gn=CSPP1;refGene_tx=XM_011291366.2;refGene_gn=CSPP1;refGene_tx=XM_011291367.2;refGene_gn=CSPP1;refGene_tx=XM_019822740.1;refGene_gn=CSPP1;refGene_tx=XM_0198227

41.2;refGene_gn=CSPP1;refGene_tx=XM_019822742.1
F2 19680084 19680237
refGene_gn=CSPP1;refGene_tx=XM_006943251.3;refGene_gn=CSPP1;refGene_tx=XM_011291366.2;refGene_gn=CSPP1;refGene_tx=XM_019822740.1;refGene_gn=CSPP1;refGene_tx=XM_019822741.2;refGene_gn=CSPP1;refGene_tx=XM_019822742.1
F2 19684377 19684490
refGene_gn=CSPP1;refGene_tx=XM_006943251.3;refGene_gn=CSPP1;refGene_tx=XM_011291366.2;refGene_gn=CSPP1;refGene_tx=XM_011291367.2;refGene_gn=CSPP1;refGene_tx=XM_019822740.1;refGene_gn=CSPP1;refGene_tx=XM_019822741.2;refGene_gn=CSPP1;refGene_tx=XM_019822742.1
F2 19688431 19688581
refGene_gn=CSPP1;refGene_tx=XM_006943251.3;refGene_gn=CSPP1;refGene_tx=XM_011291366.2;refGene_gn=CSPP1;refGene_tx=XM_011291367.2;refGene_gn=CSPP1;refGene_tx=XM_019822740.1;refGene_gn=CSPP1;refGene_tx=XM_019822741.2;refGene_gn=CSPP1;refGene_tx=XM_019822742.1
F2 19688978 19689125
refGene_gn=CSPP1;refGene_tx=XM_006943251.3;refGene_gn=CSPP1;refGene_tx=XM_011291366.2;refGene_gn=CSPP1;refGene_tx=XM_011291367.2;refGene_gn=CSPP1;refGene_tx=XM_019822740.1;refGene_gn=CSPP1;refGene_tx=XM_019822741.2;refGene_gn=CSPP1;refGene_tx=XM_019822742.1
F2 19692024 19692129
refGene_gn=CSPP1;refGene_tx=XM_006943251.3;refGene_gn=CSPP1;refGene_tx=XM_011291366.2;refGene_gn=CSPP1;refGene_tx=XM_011291367.2;refGene_gn=CSPP1;refGene_tx=XM_019822740.1;refGene_gn=CSPP1;refGene_tx=XM_019822741.2;refGene_gn=CSPP1;refGene_tx=XM_019822742.1
F2 19694301 19694368
refGene_gn=CSPP1;refGene_tx=XM_006943251.3;refGene_gn=CSPP1;refGene_tx=XM_011291366.2;refGene_gn=CSPP1;refGene_tx=XM_011291367.2;refGene_gn=CSPP1;refGene_tx=XM_019822740.1;refGene_gn=CSPP1;refGene_tx=XM_019822741.2;refGene_gn=CSPP1;refGene_tx=XM_019822742.1
F2 19694954 19695072
refGene_gn=CSPP1;refGene_tx=XM_006943251.3;refGene_gn=CSPP1;refGene_tx=XM_011291366.2;refGene_gn=CSPP1;refGene_tx=XM_011291367.2;refGene_gn=CSPP1;refGene_tx=XM_019822740.1;refGene_gn=CSPP1;refGene_tx=XM_019822741.2;refGene_gn=CSPP1;refGene_tx=XM_019822742.1
F2 19699846 19699986
refGene_gn=CSPP1;refGene_tx=XM_006943251.3;refGene_gn=CSPP1;refGene_tx=XM_011291366.2;refGene_gn=CSPP1;refGene_tx=XM_011291367.2;refGene_gn=CSPP1;refGene_tx=XM_019822740.1;refGene_gn=CSPP1;refGene_tx=XM_019822741.2;refGene_gn=CSPP1;refGene_tx=XM_019822742.1
F2 19703215 19703356
refGene_gn=CSPP1;refGene_tx=XM_006943251.3;refGene_gn=CSPP1;refGene_tx=XM_011291366.2;refGene_gn=CSPP1;refGene_tx=XM_011291367.2;refGene_gn=CSPP1;refGene_tx=XM_019822740.1;refGene_gn=CSPP1;refGene_tx=XM_019822741.2;refGene_gn=CSPP1;refGene_tx=XM_019822742.1
F2 19705670 19705717
refGene_gn=CSPP1;refGene_tx=XM_006943251.3;refGene_gn=CSPP1;refGene_tx=XM_011291366.2;refGene_gn=CSPP1;refGene_tx=XM_011291367.2;refGene_gn=CSPP1;refGene_tx=XM_019822740.1;refGene_gn=CSPP1;refGene_tx=XM_0198227

41.2;refGene_gn=CSPP1;refGene_tx=XM_019822742.1
F2 19709786 19709850
refGene_gn=CSPP1;refGene_tx=XM_006943251.3;refGene_gn=CSPP1;refGene_tx=XM_011291366.2;refGene_gn=CSPP1;refGene_tx=XM_011291367.2;refGene_gn=CSPP1;refGene_tx=XM_019822740.1;refGene_gn=CSPP1;refGene_tx=XM_019822741.2;refGene_gn=CSPP1;refGene_tx=XM_019822742.1
F2 19713812 19713922
refGene_gn=CSPP1;refGene_tx=XM_006943251.3;refGene_gn=CSPP1;refGene_tx=XM_011291366.2;refGene_gn=CSPP1;refGene_tx=XM_011291367.2;refGene_gn=CSPP1;refGene_tx=XM_019822740.1;refGene_gn=CSPP1;refGene_tx=XM_019822741.2;refGene_gn=CSPP1;refGene_tx=XM_019822742.1
F2 19716277 19716416
refGene_gn=CSPP1;refGene_tx=XM_006943251.3;refGene_gn=CSPP1;refGene_tx=XM_011291366.2;refGene_gn=CSPP1;refGene_tx=XM_011291367.2;refGene_gn=CSPP1;refGene_tx=XM_019822740.1;refGene_gn=CSPP1;refGene_tx=XM_019822741.2;refGene_gn=CSPP1;refGene_tx=XM_019822742.1
F2 19718218 19718430
refGene_gn=CSPP1;refGene_tx=XM_006943251.3;refGene_gn=CSPP1;refGene_tx=XM_011291366.2;refGene_gn=CSPP1;refGene_tx=XM_011291367.2;refGene_gn=CSPP1;refGene_tx=XM_019822740.1;refGene_gn=CSPP1;refGene_tx=XM_019822741.2;refGene_gn=CSPP1;refGene_tx=XM_019822742.1
F2 19721204 19721369
refGene_gn=ARFGEF1;refGene_tx=XM_011291364.3;refGene_gn=ARFGEF1;refGene_tx=XM_019822739.2
F2 19722864 19722982
refGene_gn=ARFGEF1;refGene_tx=XM_011291364.3;refGene_gn=ARFGEF1;refGene_tx=XM_019822739.2
F2 19723942 19724081
refGene_gn=ARFGEF1;refGene_tx=XM_011291364.3;refGene_gn=ARFGEF1;refGene_tx=XM_019822739.2
F2 19725379 19725548
refGene_gn=ARFGEF1;refGene_tx=XM_011291364.3;refGene_gn=ARFGEF1;refGene_tx=XM_019822739.2
F2 19727074 19727214
refGene_gn=ARFGEF1;refGene_tx=XM_011291364.3;refGene_gn=ARFGEF1;refGene_tx=XM_019822739.2
F2 19731271 19731404
refGene_gn=ARFGEF1;refGene_tx=XM_011291364.3;refGene_gn=ARFGEF1;refGene_tx=XM_019822739.2
F2 19739453 19739526
refGene_gn=ARFGEF1;refGene_tx=XM_011291364.3;refGene_gn=ARFGEF1;refGene_tx=XM_019822739.2
F2 19741683 19741822
refGene_gn=ARFGEF1;refGene_tx=XM_011291364.3;refGene_gn=ARFGEF1;refGene_tx=XM_019822739.2
F2 19741903 19742039
refGene_gn=ARFGEF1;refGene_tx=XM_011291364.3;refGene_gn=ARFGEF1;refGene_tx=XM_019822739.2
F2 19743355 19743485
refGene_gn=ARFGEF1;refGene_tx=XM_011291364.3;refGene_gn=ARFGEF1;refGen

e_tx=XM_019822739.2
F2 19746790 19746921
refGene_gn=ARFGEF1;refGene_tx=XM_011291364.3;refGene_gn=ARFGEF1;refGene_tx=XM_019822739.2
F2 19748121 19748282
refGene_gn=ARFGEF1;refGene_tx=XM_011291364.3;refGene_gn=ARFGEF1;refGene_tx=XM_019822739.2
F2 19749054 19749227
refGene_gn=ARFGEF1;refGene_tx=XM_011291364.3;refGene_gn=ARFGEF1;refGene_tx=XM_019822739.2
F2 19749366 19749518
refGene_gn=ARFGEF1;refGene_tx=XM_011291364.3;refGene_gn=ARFGEF1;refGene_tx=XM_019822739.2
F2 19749878 19750048
refGene_gn=ARFGEF1;refGene_tx=XM_011291364.3;refGene_gn=ARFGEF1;refGene_tx=XM_019822739.2
F2 19750139 19750180
refGene_gn=ARFGEF1;refGene_tx=XM_011291364.3;refGene_gn=ARFGEF1;refGene_tx=XM_019822739.2
F2 19755348 19755439
refGene_gn=ARFGEF1;refGene_tx=XM_011291364.3;refGene_gn=ARFGEF1;refGene_tx=XM_019822739.2
F2 19760488 19760639
refGene_gn=ARFGEF1;refGene_tx=XM_011291364.3;refGene_gn=ARFGEF1;refGene_tx=XM_019822739.2
F2 19760880 19761039
refGene_gn=ARFGEF1;refGene_tx=XM_011291364.3;refGene_gn=ARFGEF1;refGene_tx=XM_019822739.2
F2 19761887 19762016
refGene_gn=ARFGEF1;refGene_tx=XM_011291364.3;refGene_gn=ARFGEF1;refGene_tx=XM_019822739.2
F2 19765988 19766140
refGene_gn=ARFGEF1;refGene_tx=XM_011291364.3;refGene_gn=ARFGEF1;refGene_tx=XM_019822739.2
F2 19767055 19767227
refGene_gn=ARFGEF1;refGene_tx=XM_011291364.3;refGene_gn=ARFGEF1;refGene_tx=XM_019822739.2
F2 19771400 19771485
refGene_gn=ARFGEF1;refGene_tx=XM_011291364.3;refGene_gn=ARFGEF1;refGene_tx=XM_019822739.2
F2 19771740 19771946
refGene_gn=ARFGEF1;refGene_tx=XM_011291364.3;refGene_gn=ARFGEF1;refGene_tx=XM_019822739.2
F2 19772283 19772395
refGene_gn=ARFGEF1;refGene_tx=XM_011291364.3;refGene_gn=ARFGEF1;refGene_tx=XM_019822739.2
F2 19776598 19776800
refGene_gn=ARFGEF1;refGene_tx=XM_011291364.3;refGene_gn=ARFGEF1;refGene_tx=XM_019822739.2
F2 19777485 19777594

refGene_gn=ARFGEF1;refGene_tx=XM_011291364.3;refGene_gn=ARFGEF1;refGene_tx=XM_019822739.2
F2 19777700 19777840
refGene_gn=ARFGEF1;refGene_tx=XM_011291364.3;refGene_gn=ARFGEF1;refGene_tx=XM_019822739.2
F2 19777945 19778045
refGene_gn=ARFGEF1;refGene_tx=XM_011291364.3;refGene_gn=ARFGEF1;refGene_tx=XM_019822739.2
F2 19784924 19785159
refGene_gn=ARFGEF1;refGene_tx=XM_011291364.3;refGene_gn=ARFGEF1;refGene_tx=XM_019822739.2
F2 19788748 19788882
refGene_gn=ARFGEF1;refGene_tx=XM_011291364.3;refGene_gn=ARFGEF1;refGene_tx=XM_019822739.2
F2 19790051 19790227
refGene_gn=ARFGEF1;refGene_tx=XM_011291364.3;refGene_gn=ARFGEF1;refGene_tx=XM_019822739.2
F2 19801151 19801179 refGene_gn=ARFGEF1;refGene_tx=XM_019822739.2
F2 19802045 19802156 refGene_gn=ARFGEF1;refGene_tx=XM_011291364.3
F2 19805551 19805828 refGene_gn=ARFGEF1;refGene_tx=XM_011291364.3
F2 19812401 19812581 refGene_gn=ARFGEF1;refGene_tx=XM_011291364.3
F2 19814508 19814655 refGene_gn=ARFGEF1;refGene_tx=XM_011291364.3
F2 19815784 19815941 refGene_gn=ARFGEF1;refGene_tx=XM_011291364.3
F2 19816461 19816492 refGene_gn=ARFGEF1;refGene_tx=XM_011291364.3
F2 19867074 19867198 refGene_gn=ARFGEF1;refGene_tx=XM_011291364.3
F2 19937925 19938113
refGene_gn=CPA6;refGene_tx=XM_003999869.5;refGene_gn=CPA6;refGene_tx=XM_019822757.2
F2 19941312 19941397
refGene_gn=CPA6;refGene_tx=XM_003999869.5;refGene_gn=CPA6;refGene_tx=XM_019822757.2
F2 19945423 19945626
refGene_gn=CPA6;refGene_tx=XM_003999869.5;refGene_gn=CPA6;refGene_tx=XM_019822757.2
F2 19983606 19983697
refGene_gn=CPA6;refGene_tx=XM_003999869.5;refGene_gn=CPA6;refGene_tx=XM_019822757.2
F2 19984172 19984283
refGene_gn=CPA6;refGene_tx=XM_003999869.5;refGene_gn=CPA6;refGene_tx=XM_019822757.2
F2 20010041 20010146
refGene_gn=CPA6;refGene_tx=XM_003999869.5;refGene_gn=CPA6;refGene_tx=XM_019822757.2
F2 20012426 20012528
refGene_gn=CPA6;refGene_tx=XM_003999869.5;refGene_gn=CPA6;refGene_tx=XM_019822757.2
F2 20014742 20014857
refGene_gn=CPA6;refGene_tx=XM_003999869.5;refGene_gn=CPA6;refGene_tx=XM_019822757.2
F2 20021877 20022002

refGene_gn=CPA6;refGene_tx=XM_003999869.5;refGene_gn=CPA6;refGene_tx=XM_019822757.2
F2 20112156 20112232
refGene_gn=CPA6;refGene_tx=XM_003999869.5;refGene_gn=CPA6;refGene_tx=XM_019822757.2
F2 20255560 20255676 refGene_gn=CPA6;refGene_tx=XM_003999869.5
F2 20328517 20328564 refGene_gn=CPA6;refGene_tx=XM_019822757.2
F2 20463282 20463423
refGene_gn=PREX2;refGene_tx=XM_023248570.1;refGene_gn=PREX2;refGene_tx=XM_023248571.1;refGene_gn=PREX2;refGene_tx=XM_023248572.1
F2 20538839 20538911
refGene_gn=PREX2;refGene_tx=XM_023248570.1;refGene_gn=PREX2;refGene_tx=XM_023248571.1;refGene_gn=PREX2;refGene_tx=XM_023248572.1
F2 20540304 20540427
refGene_gn=PREX2;refGene_tx=XM_023248570.1;refGene_gn=PREX2;refGene_tx=XM_023248571.1;refGene_gn=PREX2;refGene_tx=XM_023248572.1
F2 20543550 20543655
refGene_gn=PREX2;refGene_tx=XM_023248570.1;refGene_gn=PREX2;refGene_tx=XM_023248571.1;refGene_gn=PREX2;refGene_tx=XM_023248572.1
F2 20550126 20550228
refGene_gn=PREX2;refGene_tx=XM_023248570.1;refGene_gn=PREX2;refGene_tx=XM_023248571.1;refGene_gn=PREX2;refGene_tx=XM_023248572.1
F2 20553854 20554016
refGene_gn=PREX2;refGene_tx=XM_023248570.1;refGene_gn=PREX2;refGene_tx=XM_023248571.1;refGene_gn=PREX2;refGene_tx=XM_023248572.1
F2 20568065 20568199
refGene_gn=PREX2;refGene_tx=XM_023248570.1;refGene_gn=PREX2;refGene_tx=XM_023248571.1;refGene_gn=PREX2;refGene_tx=XM_023248572.1
F2 20579829 20579933
refGene_gn=PREX2;refGene_tx=XM_023248570.1;refGene_gn=PREX2;refGene_tx=XM_023248571.1;refGene_gn=PREX2;refGene_tx=XM_023248572.1
F2 20590714 20590864
refGene_gn=PREX2;refGene_tx=XM_023248570.1;refGene_gn=PREX2;refGene_tx=XM_023248571.1;refGene_gn=PREX2;refGene_tx=XM_023248572.1
F2 20593754 20593899
refGene_gn=PREX2;refGene_tx=XM_023248570.1;refGene_gn=PREX2;refGene_tx=XM_023248571.1;refGene_gn=PREX2;refGene_tx=XM_023248572.1
F2 20598316 20598417
refGene_gn=PREX2;refGene_tx=XM_023248570.1;refGene_gn=PREX2;refGene_tx=XM_023248571.1;refGene_gn=PREX2;refGene_tx=XM_023248572.1
F2 20605501 20605605
refGene_gn=PREX2;refGene_tx=XM_023248570.1;refGene_gn=PREX2;refGene_tx=XM_023248571.1;refGene_gn=PREX2;refGene_tx=XM_023248572.1
F2 20606237 20606287
refGene_gn=PREX2;refGene_tx=XM_023248570.1;refGene_gn=PREX2;refGene_tx=XM_023248571.1;refGene_gn=PREX2;refGene_tx=XM_023248572.1
F2 20609080 20609156
refGene_gn=PREX2;refGene_tx=XM_023248570.1;refGene_gn=PREX2;refGene_tx=XM_023248571.1;refGene_gn=PREX2;refGene_tx=XM_023248572.1
F2 20611085 20611158

refGene_gn=PREX2;refGene_tx=XM_023248570.1;refGene_gn=PREX2;refGene_tx=XM_023248571.1;refGene_gn=PREX2;refGene_tx=XM_023248572.1
F2 20615009 20615152
refGene_gn=PREX2;refGene_tx=XM_023248570.1;refGene_gn=PREX2;refGene_tx=XM_023248571.1;refGene_gn=PREX2;refGene_tx=XM_023248572.1
F2 20615253 20615346
refGene_gn=PREX2;refGene_tx=XM_023248570.1;refGene_gn=PREX2;refGene_tx=XM_023248571.1
F2 20617022 20617171
refGene_gn=PREX2;refGene_tx=XM_023248570.1;refGene_gn=PREX2;refGene_tx=XM_023248571.1;refGene_gn=PREX2;refGene_tx=XM_023248572.1
F2 20621535 20621621
refGene_gn=PREX2;refGene_tx=XM_023248570.1;refGene_gn=PREX2;refGene_tx=XM_023248571.1;refGene_gn=PREX2;refGene_tx=XM_023248572.1
F2 20624741 20624878
refGene_gn=PREX2;refGene_tx=XM_023248570.1;refGene_gn=PREX2;refGene_tx=XM_023248571.1;refGene_gn=PREX2;refGene_tx=XM_023248572.1
F2 20627378 20627496
refGene_gn=PREX2;refGene_tx=XM_023248570.1;refGene_gn=PREX2;refGene_tx=XM_023248571.1;refGene_gn=PREX2;refGene_tx=XM_023248572.1
F2 20632338 20632523
refGene_gn=PREX2;refGene_tx=XM_023248570.1;refGene_gn=PREX2;refGene_tx=XM_023248571.1;refGene_gn=PREX2;refGene_tx=XM_023248572.1
F2 20634765 20634927
refGene_gn=PREX2;refGene_tx=XM_023248570.1;refGene_gn=PREX2;refGene_tx=XM_023248571.1;refGene_gn=PREX2;refGene_tx=XM_023248572.1
F2 20650425 20650645
refGene_gn=PREX2;refGene_tx=XM_023248570.1;refGene_gn=PREX2;refGene_tx=XM_023248571.1;refGene_gn=PREX2;refGene_tx=XM_023248572.1
F2 20651188 20651396
refGene_gn=PREX2;refGene_tx=XM_023248570.1;refGene_gn=PREX2;refGene_tx=XM_023248571.1;refGene_gn=PREX2;refGene_tx=XM_023248572.1
F2 20658475 20658655
refGene_gn=PREX2;refGene_tx=XM_023248570.1;refGene_gn=PREX2;refGene_tx=XM_023248571.1;refGene_gn=PREX2;refGene_tx=XM_023248572.1
F2 20660964 20661059
refGene_gn=PREX2;refGene_tx=XM_023248570.1;refGene_gn=PREX2;refGene_tx=XM_023248571.1;refGene_gn=PREX2;refGene_tx=XM_023248572.1
F2 20661841 20661924
refGene_gn=PREX2;refGene_tx=XM_023248570.1;refGene_gn=PREX2;refGene_tx=XM_023248571.1;refGene_gn=PREX2;refGene_tx=XM_023248572.1
F2 20662934 20663025
refGene_gn=PREX2;refGene_tx=XM_023248570.1;refGene_gn=PREX2;refGene_tx=XM_023248571.1;refGene_gn=PREX2;refGene_tx=XM_023248572.1
F2 20663675 20663804
refGene_gn=PREX2;refGene_tx=XM_023248570.1;refGene_gn=PREX2;refGene_tx=XM_023248571.1;refGene_gn=PREX2;refGene_tx=XM_023248572.1
F2 20670450 20670492
refGene_gn=PREX2;refGene_tx=XM_023248570.1;refGene_gn=PREX2;refGene_tx=XM_023248571.1;refGene_gn=PREX2;refGene_tx=XM_023248572.1

F2 20677008 20677226
refGene_gn=PREX2;refGene_tx=XM_023248570.1;refGene_gn=PREX2;refGene_tx=XM_023248571.1;refGene_gn=PREX2;refGene_tx=XM_023248572.1
F2 20681881 20681984
refGene_gn=PREX2;refGene_tx=XM_023248570.1;refGene_gn=PREX2;refGene_tx=XM_023248571.1;refGene_gn=PREX2;refGene_tx=XM_023248572.1
F2 20686956 20687100
refGene_gn=PREX2;refGene_tx=XM_023248570.1;refGene_gn=PREX2;refGene_tx=XM_023248571.1;refGene_gn=PREX2;refGene_tx=XM_023248572.1
F2 20697058 20697173
refGene_gn=PREX2;refGene_tx=XM_023248570.1;refGene_gn=PREX2;refGene_tx=XM_023248571.1;refGene_gn=PREX2;refGene_tx=XM_023248572.1
F2 20728779 20728846
refGene_gn=PREX2;refGene_tx=XM_023248570.1;refGene_gn=PREX2;refGene_tx=XM_023248571.1;refGene_gn=PREX2;refGene_tx=XM_023248572.1
F2 20729115 20729306
refGene_gn=PREX2;refGene_tx=XM_023248570.1;refGene_gn=PREX2;refGene_tx=XM_023248571.1;refGene_gn=PREX2;refGene_tx=XM_023248572.1
F2 20754724 20754827
refGene_gn=PREX2;refGene_tx=XM_023248570.1;refGene_gn=PREX2;refGene_tx=XM_023248571.1;refGene_gn=PREX2;refGene_tx=XM_023248572.1
F2 20760786 20760854
refGene_gn=PREX2;refGene_tx=XM_023248570.1;refGene_gn=PREX2;refGene_tx=XM_023248571.1;refGene_gn=PREX2;refGene_tx=XM_023248572.1
F2 20766189 20766235
refGene_gn=PREX2;refGene_tx=XM_023248570.1;refGene_gn=PREX2;refGene_tx=XM_023248571.1;refGene_gn=PREX2;refGene_tx=XM_023248572.1
F2 20853210 20853345
refGene_gn=CF2H8orf34;refGene_tx=XM_019822764.2
F2 20853346 20853463
refGene_gn=CF2H8orf34;refGene_tx=XM_019822764.2
F2 20853464 20853539
refGene_gn=CF2H8orf34;refGene_tx=XM_019822764.2
F2 20933275 20933423
refGene_gn=CF2H8orf34;refGene_tx=XM_019822764.2
F2 20947235 20947367
refGene_gn=CF2H8orf34;refGene_tx=XM_019822764.2
F2 20950090 20950219
refGene_gn=CF2H8orf34;refGene_tx=XM_019822764.2
F2 20967159 20967188
refGene_gn=CF2H8orf34;refGene_tx=XM_019822764.2
F2 20999045 20999218
refGene_gn=CF2H8orf34;refGene_tx=XM_019822764.2
F2 21015780 21015947
refGene_gn=CF2H8orf34;refGene_tx=XM_019822764.2
F2 21119303 21119439
refGene_gn=CF2H8orf34;refGene_tx=XM_019822764.2
F2 21161235 21161321
refGene_gn=CF2H8orf34;refGene_tx=XM_019822764.2
F2 21168697 21168774

refGene_gn=CF2H8orf34;refGene_tx=XM_019822764.2
F2 21216385 21216433
refGene_gn=CF2H8orf34;refGene_tx=XM_019822764.2
F2 21228561 21228655
refGene_gn=CF2H8orf34;refGene_tx=XM_019822764.2
F2 21254537 21254597
refGene_gn=CF2H8orf34;refGene_tx=XM_019822764.2
F2 21257682 21257690
refGene_gn=CF2H8orf34;refGene_tx=XM_019822764.2
F2 21879556 21879728
refGene_gn=SULF1;refGene_tx=XM_006943254.4;refGene_gn=SULF1;refGene_tx=XM_011291362.2;refGene_gn=SULF1;refGene_tx=XM_019822760.1;refGene_gn=SULF1;refGene_tx=XM_019822761.2;refGene_gn=SULF1;refGene_tx=XM_019822762.1
F2 21891425 21891665
refGene_gn=SULF1;refGene_tx=XM_006943254.4;refGene_gn=SULF1;refGene_tx=XM_011291362.2;refGene_gn=SULF1;refGene_tx=XM_019822760.1;refGene_gn=SULF1;refGene_tx=XM_019822761.2;refGene_gn=SULF1;refGene_tx=XM_019822762.1
F2 21900327 21900479
refGene_gn=SULF1;refGene_tx=XM_006943254.4;refGene_gn=SULF1;refGene_tx=XM_011291362.2;refGene_gn=SULF1;refGene_tx=XM_019822760.1;refGene_gn=SULF1;refGene_tx=XM_019822761.2;refGene_gn=SULF1;refGene_tx=XM_019822762.1
F2 21902834 21903004
refGene_gn=SULF1;refGene_tx=XM_006943254.4;refGene_gn=SULF1;refGene_tx=XM_011291362.2;refGene_gn=SULF1;refGene_tx=XM_019822760.1;refGene_gn=SULF1;refGene_tx=XM_019822761.2;refGene_gn=SULF1;refGene_tx=XM_019822762.1
F2 21914191 21914342
refGene_gn=SULF1;refGene_tx=XM_006943254.4;refGene_gn=SULF1;refGene_tx=XM_011291362.2;refGene_gn=SULF1;refGene_tx=XM_019822760.1;refGene_gn=SULF1;refGene_tx=XM_019822761.2;refGene_gn=SULF1;refGene_tx=XM_019822762.1
F2 21915252 21915428
refGene_gn=SULF1;refGene_tx=XM_006943254.4;refGene_gn=SULF1;refGene_tx=XM_011291362.2;refGene_gn=SULF1;refGene_tx=XM_019822760.1;refGene_gn=SULF1;refGene_tx=XM_019822761.2;refGene_gn=SULF1;refGene_tx=XM_019822762.1
F2 21917161 21917290
refGene_gn=SULF1;refGene_tx=XM_006943254.4;refGene_gn=SULF1;refGene_tx=XM_011291362.2;refGene_gn=SULF1;refGene_tx=XM_019822760.1;refGene_gn=SULF1;refGene_tx=XM_019822761.2;refGene_gn=SULF1;refGene_tx=XM_019822762.1
F2 21917560 21917617
refGene_gn=SULF1;refGene_tx=XM_006943254.4;refGene_gn=SULF1;refGene_tx=XM_011291362.2;refGene_gn=SULF1;refGene_tx=XM_019822760.1;refGene_gn=SULF1;refGene_tx=XM_019822761.2;refGene_gn=SULF1;refGene_tx=XM_019822762.1
F2 21918337 21918467

refGene_gn=SULF1;refGene_tx=XM_006943254.4;refGene_gn=SULF1;refGene_tx=XM_011291362.2;refGene_gn=SULF1;refGene_tx=XM_019822760.1;refGene_gn=SULF1;refGene_tx=XM_019822761.2;refGene_gn=SULF1;refGene_tx=XM_019822762.1
F2 21930541 21930752
refGene_gn=SULF1;refGene_tx=XM_006943254.4;refGene_gn=SULF1;refGene_tx=XM_011291362.2;refGene_gn=SULF1;refGene_tx=XM_019822760.1;refGene_gn=SULF1;refGene_tx=XM_019822761.2;refGene_gn=SULF1;refGene_tx=XM_019822762.1
F2 21933381 21933628
refGene_gn=SULF1;refGene_tx=XM_006943254.4;refGene_gn=SULF1;refGene_tx=XM_011291362.2;refGene_gn=SULF1;refGene_tx=XM_019822760.1;refGene_gn=SULF1;refGene_tx=XM_019822761.2;refGene_gn=SULF1;refGene_tx=XM_019822762.1
F2 21935433 21935530
refGene_gn=SULF1;refGene_tx=XM_006943254.4;refGene_gn=SULF1;refGene_tx=XM_011291362.2;refGene_gn=SULF1;refGene_tx=XM_019822760.1;refGene_gn=SULF1;refGene_tx=XM_019822761.2;refGene_gn=SULF1;refGene_tx=XM_019822762.1
F2 21936000 21936095
refGene_gn=SULF1;refGene_tx=XM_006943254.4;refGene_gn=SULF1;refGene_tx=XM_011291362.2;refGene_gn=SULF1;refGene_tx=XM_019822760.1;refGene_gn=SULF1;refGene_tx=XM_019822761.2;refGene_gn=SULF1;refGene_tx=XM_019822762.1
F2 21936413 21936479
refGene_gn=SULF1;refGene_tx=XM_006943254.4;refGene_gn=SULF1;refGene_tx=XM_011291362.2;refGene_gn=SULF1;refGene_tx=XM_019822760.1;refGene_gn=SULF1;refGene_tx=XM_019822761.2;refGene_gn=SULF1;refGene_tx=XM_019822762.1
F2 21937567 21937740
refGene_gn=SULF1;refGene_tx=XM_006943254.4;refGene_gn=SULF1;refGene_tx=XM_011291362.2;refGene_gn=SULF1;refGene_tx=XM_019822760.1;refGene_gn=SULF1;refGene_tx=XM_019822761.2;refGene_gn=SULF1;refGene_tx=XM_019822762.1
F2 21941084 21941227
refGene_gn=SULF1;refGene_tx=XM_006943254.4;refGene_gn=SULF1;refGene_tx=XM_011291362.2;refGene_gn=SULF1;refGene_tx=XM_019822760.1;refGene_gn=SULF1;refGene_tx=XM_019822761.2;refGene_gn=SULF1;refGene_tx=XM_019822762.1
F2 21941321 21941445
refGene_gn=SULF1;refGene_tx=XM_006943254.4;refGene_gn=SULF1;refGene_tx=XM_011291362.2;refGene_gn=SULF1;refGene_tx=XM_019822760.1;refGene_gn=SULF1;refGene_tx=XM_019822761.2;refGene_gn=SULF1;refGene_tx=XM_019822762.1
F2 21943293 21943327
refGene_gn=SULF1;refGene_tx=XM_006943254.4;refGene_gn=SULF1;refGene_tx=XM_011291362.2;refGene_gn=SULF1;refGene_tx=XM_019822760.1;refGene_gn=SULF1;refGene_tx=XM_019822761.2;refGene_gn=SULF1;refGene_tx=XM_019822762.1
F2 21944589 21944655

refGene_gn=SULF1;refGene_tx=XM_006943254.4;refGene_gn=SULF1;refGene_tx=XM_011291362.2;refGene_gn=SULF1;refGene_tx=XM_019822760.1;refGene_gn=SULF1;refGene_tx=XM_019822761.2
F2 21958585 21958616 refGene_gn=SULF1;refGene_tx=XM_006943254.4;refGene_gn=SULF1;refGene_tx=XM_011291362.2;refGene_gn=SULF1;refGene_tx=XM_019822760.1;refGene_gn=SULF1;refGene_tx=XM_019822761.2;refGene_gn=SULF1;refGene_tx=XM_019822762.1
F2 21971217 21971687 refGene_gn=SLCO5A1;refGene_tx=XM_019822769.2;refGene_gn=SLCO5A1;refGene_tx=XM_023248210.1;refGene_gn=SLCO5A1;refGene_tx=XM_023248211.1;refGene_gn=SLCO5A1;refGene_tx=XM_023248212.1
F2 21974303 21974368 refGene_gn=SLCO5A1;refGene_tx=XM_019822769.2;refGene_gn=SLCO5A1;refGene_tx=XM_023248210.1;refGene_gn=SLCO5A1;refGene_tx=XM_023248211.1;refGene_gn=SLCO5A1;refGene_tx=XM_023248212.1
F2 21976564 21976806 refGene_gn=SLCO5A1;refGene_tx=XM_019822769.2;refGene_gn=SLCO5A1;refGene_tx=XM_023248210.1;refGene_gn=SLCO5A1;refGene_tx=XM_023248211.1;refGene_gn=SLCO5A1;refGene_tx=XM_023248212.1
F2 21978899 21979059 refGene_gn=SLCO5A1;refGene_tx=XM_019822769.2;refGene_gn=SLCO5A1;refGene_tx=XM_023248210.1;refGene_gn=SLCO5A1;refGene_tx=XM_023248211.1;refGene_gn=SLCO5A1;refGene_tx=XM_023248212.1
F2 21997848 21998047 refGene_gn=SLCO5A1;refGene_tx=XM_019822769.2;refGene_gn=SLCO5A1;refGene_tx=XM_023248210.1;refGene_gn=SLCO5A1;refGene_tx=XM_023248211.1;refGene_gn=SLCO5A1;refGene_tx=XM_023248212.1
F2 22029628 22029793 refGene_gn=SLCO5A1;refGene_tx=XM_019822769.2;refGene_gn=SLCO5A1;refGene_tx=XM_023248210.1;refGene_gn=SLCO5A1;refGene_tx=XM_023248211.1
F2 22046834 22047052 refGene_gn=SLCO5A1;refGene_tx=XM_019822769.2;refGene_gn=SLCO5A1;refGene_tx=XM_023248210.1;refGene_gn=SLCO5A1;refGene_tx=XM_023248211.1;refGene_gn=SLCO5A1;refGene_tx=XM_023248212.1
F2 22054349 22054482 refGene_gn=SLCO5A1;refGene_tx=XM_019822769.2;refGene_gn=SLCO5A1;refGene_tx=XM_023248210.1;refGene_gn=SLCO5A1;refGene_tx=XM_023248211.1;refGene_gn=SLCO5A1;refGene_tx=XM_023248212.1
F2 22114016 22114932 refGene_gn=SLCO5A1;refGene_tx=XM_019822769.2;refGene_gn=SLCO5A1;refGene_tx=XM_023248210.1;refGene_gn=SLCO5A1;refGene_tx=XM_023248211.1;refGene_gn=SLCO5A1;refGene_tx=XM_023248212.1
F2 22296463 22296691 refGene_gn=PRDM14;refGene_tx=XM_003999875.2
F2 22299264 22299366 refGene_gn=PRDM14;refGene_tx=XM_003999875.2
F2 22305488 22305691 refGene_gn=PRDM14;refGene_tx=XM_003999875.2
F2 22311072 22311343 refGene_gn=PRDM14;refGene_tx=XM_003999875.2
F2 22312497 22312655 refGene_gn=PRDM14;refGene_tx=XM_003999875.2
F2 22312743 22312797 refGene_gn=PRDM14;refGene_tx=XM_003999875.2
F2 22313372 22314075 refGene_gn=PRDM14;refGene_tx=XM_003999875.2

F2 22353364 22353376
refGene_gn=NCOA2;refGene_tx=XM_023248763.1;refGene_gn=NCOA2;refGene_tx=XM_023248764.1;refGene_gn=NCOA2;refGene_tx=XM_023248765.1;refGene_gn=NCOA2;refGene_tx=XM_023248766.1;refGene_gn=NCOA2;refGene_tx=XM_023248767.1;refGene_gn=NCOA2;refGene_tx=XM_023248768.1;refGene_gn=NCOA2;refGene_tx=XM_023248769.1;refGene_gn=NCOA2;refGene_tx=XM_023248770.1
F2 22359183 22359273
refGene_gn=NCOA2;refGene_tx=XM_023248763.1;refGene_gn=NCOA2;refGene_tx=XM_023248764.1;refGene_gn=NCOA2;refGene_tx=XM_023248765.1;refGene_gn=NCOA2;refGene_tx=XM_023248766.1;refGene_gn=NCOA2;refGene_tx=XM_023248767.1;refGene_gn=NCOA2;refGene_tx=XM_023248768.1;refGene_gn=NCOA2;refGene_tx=XM_023248770.1
F2 22362201 22362400
refGene_gn=NCOA2;refGene_tx=XM_023248763.1;refGene_gn=NCOA2;refGene_tx=XM_023248764.1;refGene_gn=NCOA2;refGene_tx=XM_023248765.1;refGene_gn=NCOA2;refGene_tx=XM_023248766.1;refGene_gn=NCOA2;refGene_tx=XM_023248767.1;refGene_gn=NCOA2;refGene_tx=XM_023248768.1;refGene_gn=NCOA2;refGene_tx=XM_023248769.1;refGene_gn=NCOA2;refGene_tx=XM_023248770.1
F2 22363113 22363291
refGene_gn=NCOA2;refGene_tx=XM_023248763.1;refGene_gn=NCOA2;refGene_tx=XM_023248764.1;refGene_gn=NCOA2;refGene_tx=XM_023248765.1;refGene_gn=NCOA2;refGene_tx=XM_023248766.1;refGene_gn=NCOA2;refGene_tx=XM_023248767.1;refGene_gn=NCOA2;refGene_tx=XM_023248768.1;refGene_gn=NCOA2;refGene_tx=XM_023248769.1;refGene_gn=NCOA2;refGene_tx=XM_023248770.1
F2 22364861 22365096
refGene_gn=NCOA2;refGene_tx=XM_023248763.1;refGene_gn=NCOA2;refGene_tx=XM_023248764.1;refGene_gn=NCOA2;refGene_tx=XM_023248765.1;refGene_gn=NCOA2;refGene_tx=XM_023248766.1;refGene_gn=NCOA2;refGene_tx=XM_023248767.1;refGene_gn=NCOA2;refGene_tx=XM_023248768.1;refGene_gn=NCOA2;refGene_tx=XM_023248769.1;refGene_gn=NCOA2;refGene_tx=XM_023248770.1
F2 22366728 22366806
refGene_gn=NCOA2;refGene_tx=XM_023248763.1;refGene_gn=NCOA2;refGene_tx=XM_023248764.1;refGene_gn=NCOA2;refGene_tx=XM_023248765.1;refGene_gn=NCOA2;refGene_tx=XM_023248766.1;refGene_gn=NCOA2;refGene_tx=XM_023248767.1;refGene_gn=NCOA2;refGene_tx=XM_023248768.1;refGene_gn=NCOA2;refGene_tx=XM_023248769.1;refGene_gn=NCOA2;refGene_tx=XM_023248770.1
F2 22367505 22367784
refGene_gn=NCOA2;refGene_tx=XM_023248763.1;refGene_gn=NCOA2;refGene_tx=XM_023248764.1;refGene_gn=NCOA2;refGene_tx=XM_023248765.1;refGene_gn=NCOA2;refGene_tx=XM_023248766.1;refGene_gn=NCOA2;refGene_tx=XM_023248767.1;refGene_gn=NCOA2;refGene_tx=XM_023248768.1;refGene_gn=NCOA2;refGene_tx=XM_023248769.1;refGene_gn=NCOA2;refGene_tx=XM_023248770.1
F2 22370442 22370608
refGene_gn=NCOA2;refGene_tx=XM_023248763.1;refGene_gn=NCOA2;refGene_tx=XM_023248764.1;refGene_gn=NCOA2;refGene_tx=XM_023248765.1;refGene_gn=NCOA2;refGene_tx=XM_023248766.1;refGene_gn=NCOA2;refGene_tx=XM_023248767.1;refGene_gn=NCOA2;refGene_tx=XM_023248768.1;refGene_gn=NCOA2;refGene_tx=XM_023248769.1;refGene_gn=NCOA2;refGene_tx=XM_023248770.1
F2 22378730 22378860
refGene_gn=NCOA2;refGene_tx=XM_023248763.1;refGene_gn=NCOA2;refGene_tx

=XM_023248764.1;refGene_gn=NCOA2;refGene_tx=XM_023248765.1;refGene_gn=NCOA2;refGene_tx=XM_023248766.1;refGene_gn=NCOA2;refGene_tx=XM_023248767.1;refGene_gn=NCOA2;refGene_tx=XM_023248768.1;refGene_gn=NCOA2;refGene_tx=XM_023248769.1;refGene_gn=NCOA2;refGene_tx=XM_023248770.1
F2 22381827 22382043
refGene_gn=NCOA2;refGene_tx=XM_023248763.1;refGene_gn=NCOA2;refGene_tx=XM_023248764.1;refGene_gn=NCOA2;refGene_tx=XM_023248765.1;refGene_gn=NCOA2;refGene_tx=XM_023248766.1;refGene_gn=NCOA2;refGene_tx=XM_023248767.1;refGene_gn=NCOA2;refGene_tx=XM_023248768.1;refGene_gn=NCOA2;refGene_tx=XM_023248769.1;refGene_gn=NCOA2;refGene_tx=XM_023248770.1
F2 22384720 22384927
refGene_gn=NCOA2;refGene_tx=XM_023248763.1;refGene_gn=NCOA2;refGene_tx=XM_023248764.1;refGene_gn=NCOA2;refGene_tx=XM_023248765.1;refGene_gn=NCOA2;refGene_tx=XM_023248766.1;refGene_gn=NCOA2;refGene_tx=XM_023248767.1;refGene_gn=NCOA2;refGene_tx=XM_023248768.1;refGene_gn=NCOA2;refGene_tx=XM_023248769.1
F2 22386964 22387175
refGene_gn=NCOA2;refGene_tx=XM_023248763.1;refGene_gn=NCOA2;refGene_tx=XM_023248764.1;refGene_gn=NCOA2;refGene_tx=XM_023248765.1;refGene_gn=NCOA2;refGene_tx=XM_023248766.1;refGene_gn=NCOA2;refGene_tx=XM_023248767.1;refGene_gn=NCOA2;refGene_tx=XM_023248768.1;refGene_gn=NCOA2;refGene_tx=XM_023248769.1;refGene_gn=NCOA2;refGene_tx=XM_023248770.1
F2 22394726 22395996
refGene_gn=NCOA2;refGene_tx=XM_023248763.1;refGene_gn=NCOA2;refGene_tx=XM_023248764.1;refGene_gn=NCOA2;refGene_tx=XM_023248765.1;refGene_gn=NCOA2;refGene_tx=XM_023248766.1;refGene_gn=NCOA2;refGene_tx=XM_023248767.1;refGene_gn=NCOA2;refGene_tx=XM_023248768.1;refGene_gn=NCOA2;refGene_tx=XM_023248769.1;refGene_gn=NCOA2;refGene_tx=XM_023248770.1
F2 22397718 22397866
refGene_gn=NCOA2;refGene_tx=XM_023248763.1;refGene_gn=NCOA2;refGene_tx=XM_023248764.1;refGene_gn=NCOA2;refGene_tx=XM_023248765.1;refGene_gn=NCOA2;refGene_tx=XM_023248766.1;refGene_gn=NCOA2;refGene_tx=XM_023248767.1;refGene_gn=NCOA2;refGene_tx=XM_023248768.1;refGene_gn=NCOA2;refGene_tx=XM_023248769.1;refGene_gn=NCOA2;refGene_tx=XM_023248770.1
F2 22400989 22401133
refGene_gn=NCOA2;refGene_tx=XM_023248763.1;refGene_gn=NCOA2;refGene_tx=XM_023248764.1;refGene_gn=NCOA2;refGene_tx=XM_023248765.1;refGene_gn=NCOA2;refGene_tx=XM_023248766.1;refGene_gn=NCOA2;refGene_tx=XM_023248767.1;refGene_gn=NCOA2;refGene_tx=XM_023248768.1;refGene_gn=NCOA2;refGene_tx=XM_023248769.1;refGene_gn=NCOA2;refGene_tx=XM_023248770.1
F2 22401642 22401744
refGene_gn=NCOA2;refGene_tx=XM_023248763.1;refGene_gn=NCOA2;refGene_tx=XM_023248764.1;refGene_gn=NCOA2;refGene_tx=XM_023248765.1;refGene_gn=NCOA2;refGene_tx=XM_023248766.1;refGene_gn=NCOA2;refGene_tx=XM_023248767.1;refGene_gn=NCOA2;refGene_tx=XM_023248768.1;refGene_gn=NCOA2;refGene_tx=XM_023248769.1;refGene_gn=NCOA2;refGene_tx=XM_023248770.1
F2 22404632 22404821
refGene_gn=NCOA2;refGene_tx=XM_023248763.1;refGene_gn=NCOA2;refGene_tx=XM_023248764.1;refGene_gn=NCOA2;refGene_tx=XM_023248765.1;refGene_gn=NCOA2;refGene_tx=XM_023248766.1;refGene_gn=NCOA2;refGene_tx=XM_0232487

67.1;refGene_gn=NCOA2;refGene_tx=XM_023248768.1;refGene_gn=NCOA2;refGene_tx=XM_023248769.1;refGene_gn=NCOA2;refGene_tx=XM_023248770.1
F2 22407939 22408117
refGene_gn=NCOA2;refGene_tx=XM_023248763.1;refGene_gn=NCOA2;refGene_tx=XM_023248764.1;refGene_gn=NCOA2;refGene_tx=XM_023248765.1;refGene_gn=NCOA2;refGene_tx=XM_023248766.1;refGene_gn=NCOA2;refGene_tx=XM_023248767.1;refGene_gn=NCOA2;refGene_tx=XM_023248768.1;refGene_gn=NCOA2;refGene_tx=XM_023248769.1;refGene_gn=NCOA2;refGene_tx=XM_023248770.1
F2 22412016 22412120
refGene_gn=NCOA2;refGene_tx=XM_023248763.1;refGene_gn=NCOA2;refGene_tx=XM_023248764.1;refGene_gn=NCOA2;refGene_tx=XM_023248765.1;refGene_gn=NCOA2;refGene_tx=XM_023248766.1;refGene_gn=NCOA2;refGene_tx=XM_023248767.1;refGene_gn=NCOA2;refGene_tx=XM_023248768.1;refGene_gn=NCOA2;refGene_tx=XM_023248769.1;refGene_gn=NCOA2;refGene_tx=XM_023248770.1
F2 22455180 22455353
refGene_gn=NCOA2;refGene_tx=XM_023248763.1;refGene_gn=NCOA2;refGene_tx=XM_023248764.1;refGene_gn=NCOA2;refGene_tx=XM_023248765.1;refGene_gn=NCOA2;refGene_tx=XM_023248766.1;refGene_gn=NCOA2;refGene_tx=XM_023248767.1;refGene_gn=NCOA2;refGene_tx=XM_023248768.1;refGene_gn=NCOA2;refGene_tx=XM_023248769.1;refGene_gn=NCOA2;refGene_tx=XM_023248770.1
F2 22457614 22457700
refGene_gn=NCOA2;refGene_tx=XM_023248763.1;refGene_gn=NCOA2;refGene_tx=XM_023248764.1;refGene_gn=NCOA2;refGene_tx=XM_023248765.1;refGene_gn=NCOA2;refGene_tx=XM_023248766.1;refGene_gn=NCOA2;refGene_tx=XM_023248767.1;refGene_gn=NCOA2;refGene_tx=XM_023248768.1;refGene_gn=NCOA2;refGene_tx=XM_023248769.1;refGene_gn=NCOA2;refGene_tx=XM_023248770.1
F2 22767996 22768070 refGene_gn=TRAM1;refGene_tx=XM_023248676.1
F2 22775848 22776009 refGene_gn=TRAM1;refGene_tx=XM_023248676.1
F2 22776355 22776499 refGene_gn=TRAM1;refGene_tx=XM_023248676.1
F2 22778088 22778193 refGene_gn=TRAM1;refGene_tx=XM_023248676.1
F2 22778294 22778365 refGene_gn=TRAM1;refGene_tx=XM_023248676.1
F2 22785628 22785713 refGene_gn=TRAM1;refGene_tx=XM_023248676.1
F2 22787375 22787434 refGene_gn=TRAM1;refGene_tx=XM_023248676.1
F2 22788895 22789012 refGene_gn=TRAM1;refGene_tx=XM_023248676.1
F2 22789135 22789257 refGene_gn=TRAM1;refGene_tx=XM_023248676.1
F2 22791282 22791346 refGene_gn=TRAM1;refGene_tx=XM_023248676.1
F2 22799213 22799336 refGene_gn=TRAM1;refGene_tx=XM_023248676.1
F2 22825160 22825204 refGene_gn=LACTB2;refGene_tx=XM_003999877.5
F2 22825819 22825901 refGene_gn=LACTB2;refGene_tx=XM_003999877.5
F2 22829488 22829637 refGene_gn=LACTB2;refGene_tx=XM_003999877.5
F2 22831884 22832063 refGene_gn=LACTB2;refGene_tx=XM_003999877.5
F2 22838313 22838440 refGene_gn=LACTB2;refGene_tx=XM_003999877.5
F2 22844143 22844307 refGene_gn=LACTB2;refGene_tx=XM_003999877.5
F2 22856925 22857047 refGene_gn=LACTB2;refGene_tx=XM_003999877.5
F2 22866184 22866456 refGene_gn=XKR9;refGene_tx=XM_023248677.1
F2 22887176 22887397
refGene_gn=XKR9;refGene_tx=XM_019822776.2;refGene_gn=XKR9;refGene_tx=XM_023248677.1
F2 22896699 22897328
refGene_gn=XKR9;refGene_tx=XM_019822776.2;refGene_gn=XKR9;refGene_tx=X

M_023248677.1
F2 23288661 23288742
refGene_gn=EYA1;refGene_tx=XM_011291357.2;refGene_gn=EYA1;refGene_tx=XM_019822779.2;refGene_gn=EYA1;refGene_tx=XM_019822781.1;refGene_gn=EYA1;refGene_tx=XM_023248256.1;refGene_gn=EYA1;refGene_tx=XM_023248257.1;refGene_gn=EYA1;refGene_tx=XM_023248259.1;refGene_gn=EYA1;refGene_tx=XM_023248260.1;refGene_gn=EYA1;refGene_tx=XM_023248261.1;refGene_gn=EYA1;refGene_tx=XM_023248262.1;refGene_gn=EYA1;refGene_tx=XM_023248263.1;refGene_gn=EYA1;refGene_tx=XM_023248264.1;refGene_gn=EYA1;refGene_tx=XM_023248265.1;refGene_gn=EYA1;refGene_tx=XM_023248266.1;refGene_gn=EYA1;refGene_tx=XM_023248267.1
F2 23307252 23307353
refGene_gn=EYA1;refGene_tx=XM_011291357.2;refGene_gn=EYA1;refGene_tx=XM_019822779.2;refGene_gn=EYA1;refGene_tx=XM_019822781.1;refGene_gn=EYA1;refGene_tx=XM_023248256.1;refGene_gn=EYA1;refGene_tx=XM_023248257.1;refGene_gn=EYA1;refGene_tx=XM_023248259.1;refGene_gn=EYA1;refGene_tx=XM_023248260.1;refGene_gn=EYA1;refGene_tx=XM_023248261.1;refGene_gn=EYA1;refGene_tx=XM_023248262.1;refGene_gn=EYA1;refGene_tx=XM_023248263.1;refGene_gn=EYA1;refGene_tx=XM_023248264.1;refGene_gn=EYA1;refGene_tx=XM_023248265.1;refGene_gn=EYA1;refGene_tx=XM_023248266.1;refGene_gn=EYA1;refGene_tx=XM_023248267.1
F2 23311668 23311790
refGene_gn=EYA1;refGene_tx=XM_011291357.2;refGene_gn=EYA1;refGene_tx=XM_019822779.2;refGene_gn=EYA1;refGene_tx=XM_019822781.1;refGene_gn=EYA1;refGene_tx=XM_023248256.1;refGene_gn=EYA1;refGene_tx=XM_023248257.1;refGene_gn=EYA1;refGene_tx=XM_023248259.1;refGene_gn=EYA1;refGene_tx=XM_023248260.1;refGene_gn=EYA1;refGene_tx=XM_023248261.1;refGene_gn=EYA1;refGene_tx=XM_023248262.1;refGene_gn=EYA1;refGene_tx=XM_023248263.1;refGene_gn=EYA1;refGene_tx=XM_023248264.1;refGene_gn=EYA1;refGene_tx=XM_023248265.1;refGene_gn=EYA1;refGene_tx=XM_023248266.1;refGene_gn=EYA1;refGene_tx=XM_023248267.1
F2 23311953 23312068
refGene_gn=EYA1;refGene_tx=XM_011291357.2;refGene_gn=EYA1;refGene_tx=XM_019822779.2;refGene_gn=EYA1;refGene_tx=XM_019822781.1;refGene_gn=EYA1;refGene_tx=XM_023248256.1;refGene_gn=EYA1;refGene_tx=XM_023248257.1;refGene_gn=EYA1;refGene_tx=XM_023248259.1;refGene_gn=EYA1;refGene_tx=XM_023248260.1;refGene_gn=EYA1;refGene_tx=XM_023248261.1;refGene_gn=EYA1;refGene_tx=XM_023248262.1;refGene_gn=EYA1;refGene_tx=XM_023248263.1;refGene_gn=EYA1;refGene_tx=XM_023248264.1;refGene_gn=EYA1;refGene_tx=XM_023248265.1;refGene_gn=EYA1;refGene_tx=XM_023248266.1;refGene_gn=EYA1;refGene_tx=XM_023248267.1
F2 23312893 23313054
refGene_gn=EYA1;refGene_tx=XM_011291357.2;refGene_gn=EYA1;refGene_tx=XM_019822779.2;refGene_gn=EYA1;refGene_tx=XM_019822781.1;refGene_gn=EYA1;refGene_tx=XM_023248256.1;refGene_gn=EYA1;refGene_tx=XM_023248257.1;refGene_gn=EYA1;refGene_tx=XM_023248259.1;refGene_gn=EYA1;refGene_tx=XM_023248260.1;refGene_gn=EYA1;refGene_tx=XM_023248261.1;refGene_gn=EYA1;refGene_tx=XM_023248262.1;refGene_gn=EYA1;refGene_tx=XM_023248263.1;refGene_gn=EYA1;refGene_tx=XM_023248264.1;refGene_gn=EYA1;refGene_tx=XM_023248265.1;refGene_gn=EYA1;refGene_tx=XM_023248266.1;refGene_gn=EYA

1;refGene_tx=XM_023248267.1
F2 23313166 23313225
refGene_gn=EYA1;refGene_tx=XM_011291357.2;refGene_gn=EYA1;refGene_tx=XM_019822779.2;refGene_gn=EYA1;refGene_tx=XM_019822781.1;refGene_gn=EYA1;refGene_tx=XM_023248256.1;refGene_gn=EYA1;refGene_tx=XM_023248257.1;refGene_gn=EYA1;refGene_tx=XM_023248259.1;refGene_gn=EYA1;refGene_tx=XM_023248260.1;refGene_gn=EYA1;refGene_tx=XM_023248261.1;refGene_gn=EYA1;refGene_tx=XM_023248262.1;refGene_gn=EYA1;refGene_tx=XM_023248263.1;refGene_gn=EYA1;refGene_tx=XM_023248264.1;refGene_gn=EYA1;refGene_tx=XM_023248265.1;refGene_gn=EYA1;refGene_tx=XM_023248266.1;refGene_gn=EYA1;refGene_tx=XM_023248267.1
F2 23342345 23342435
refGene_gn=EYA1;refGene_tx=XM_011291357.2;refGene_gn=EYA1;refGene_tx=XM_019822779.2;refGene_gn=EYA1;refGene_tx=XM_019822781.1;refGene_gn=EYA1;refGene_tx=XM_023248256.1;refGene_gn=EYA1;refGene_tx=XM_023248257.1;refGene_gn=EYA1;refGene_tx=XM_023248259.1;refGene_gn=EYA1;refGene_tx=XM_023248260.1;refGene_gn=EYA1;refGene_tx=XM_023248261.1;refGene_gn=EYA1;refGene_tx=XM_023248262.1;refGene_gn=EYA1;refGene_tx=XM_023248263.1;refGene_gn=EYA1;refGene_tx=XM_023248264.1;refGene_gn=EYA1;refGene_tx=XM_023248265.1;refGene_gn=EYA1;refGene_tx=XM_023248266.1;refGene_gn=EYA1;refGene_tx=XM_023248267.1
F2 23367223 23367307
refGene_gn=EYA1;refGene_tx=XM_011291357.2;refGene_gn=EYA1;refGene_tx=XM_019822779.2;refGene_gn=EYA1;refGene_tx=XM_019822781.1;refGene_gn=EYA1;refGene_tx=XM_023248256.1;refGene_gn=EYA1;refGene_tx=XM_023248257.1;refGene_gn=EYA1;refGene_tx=XM_023248259.1;refGene_gn=EYA1;refGene_tx=XM_023248260.1;refGene_gn=EYA1;refGene_tx=XM_023248261.1;refGene_gn=EYA1;refGene_tx=XM_023248262.1;refGene_gn=EYA1;refGene_tx=XM_023248263.1;refGene_gn=EYA1;refGene_tx=XM_023248264.1;refGene_gn=EYA1;refGene_tx=XM_023248265.1;refGene_gn=EYA1;refGene_tx=XM_023248266.1;refGene_gn=EYA1;refGene_tx=XM_023248267.1
F2 23368657 23368797
refGene_gn=EYA1;refGene_tx=XM_011291357.2;refGene_gn=EYA1;refGene_tx=XM_019822779.2;refGene_gn=EYA1;refGene_tx=XM_019822781.1;refGene_gn=EYA1;refGene_tx=XM_023248256.1;refGene_gn=EYA1;refGene_tx=XM_023248257.1;refGene_gn=EYA1;refGene_tx=XM_023248259.1;refGene_gn=EYA1;refGene_tx=XM_023248260.1;refGene_gn=EYA1;refGene_tx=XM_023248261.1;refGene_gn=EYA1;refGene_tx=XM_023248262.1;refGene_gn=EYA1;refGene_tx=XM_023248263.1;refGene_gn=EYA1;refGene_tx=XM_023248264.1;refGene_gn=EYA1;refGene_tx=XM_023248265.1;refGene_gn=EYA1;refGene_tx=XM_023248266.1;refGene_gn=EYA1;refGene_tx=XM_023248267.1
F2 23396359 23396546
refGene_gn=EYA1;refGene_tx=XM_011291357.2;refGene_gn=EYA1;refGene_tx=XM_019822779.2;refGene_gn=EYA1;refGene_tx=XM_019822781.1;refGene_gn=EYA1;refGene_tx=XM_023248256.1;refGene_gn=EYA1;refGene_tx=XM_023248257.1;refGene_gn=EYA1;refGene_tx=XM_023248259.1;refGene_gn=EYA1;refGene_tx=XM_023248260.1;refGene_gn=EYA1;refGene_tx=XM_023248261.1;refGene_gn=EYA1;refGene_tx=XM_023248262.1;refGene_gn=EYA1;refGene_tx=XM_023248263.1;refGene_gn=EYA1;refGene_tx=XM_023248264.1;refGene_gn=EYA1;refGene_tx=XM_023248265.1;refGene_gn=EYA1;refGene_tx=XM_023248266.1;refGene_gn=EYA

1;refGene_tx=XM_023248267.1
F2 23396965 23397048
refGene_gn=EYA1;refGene_tx=XM_011291357.2;refGene_gn=EYA1;refGene_tx=XM_019822779.2;refGene_gn=EYA1;refGene_tx=XM_019822781.1;refGene_gn=EYA1;refGene_tx=XM_023248256.1;refGene_gn=EYA1;refGene_tx=XM_023248257.1;refGene_gn=EYA1;refGene_tx=XM_023248259.1;refGene_gn=EYA1;refGene_tx=XM_023248260.1;refGene_gn=EYA1;refGene_tx=XM_023248261.1;refGene_gn=EYA1;refGene_tx=XM_023248262.1;refGene_gn=EYA1;refGene_tx=XM_023248263.1;refGene_gn=EYA1;refGene_tx=XM_023248264.1;refGene_gn=EYA1;refGene_tx=XM_023248265.1;refGene_gn=EYA1;refGene_tx=XM_023248266.1;refGene_gn=EYA1;refGene_tx=XM_023248267.1
F2 23414669 23414807
refGene_gn=EYA1;refGene_tx=XM_011291357.2;refGene_gn=EYA1;refGene_tx=XM_019822779.2;refGene_gn=EYA1;refGene_tx=XM_019822781.1;refGene_gn=EYA1;refGene_tx=XM_023248256.1;refGene_gn=EYA1;refGene_tx=XM_023248257.1;refGene_gn=EYA1;refGene_tx=XM_023248259.1;refGene_gn=EYA1;refGene_tx=XM_023248260.1;refGene_gn=EYA1;refGene_tx=XM_023248261.1;refGene_gn=EYA1;refGene_tx=XM_023248262.1;refGene_gn=EYA1;refGene_tx=XM_023248263.1;refGene_gn=EYA1;refGene_tx=XM_023248264.1;refGene_gn=EYA1;refGene_tx=XM_023248265.1;refGene_gn=EYA1;refGene_tx=XM_023248266.1;refGene_gn=EYA1;refGene_tx=XM_023248267.1
F2 23418469 23418615
refGene_gn=EYA1;refGene_tx=XM_011291357.2;refGene_gn=EYA1;refGene_tx=XM_019822779.2;refGene_gn=EYA1;refGene_tx=XM_019822781.1;refGene_gn=EYA1;refGene_tx=XM_023248256.1;refGene_gn=EYA1;refGene_tx=XM_023248257.1;refGene_gn=EYA1;refGene_tx=XM_023248259.1;refGene_gn=EYA1;refGene_tx=XM_023248260.1;refGene_gn=EYA1;refGene_tx=XM_023248261.1;refGene_gn=EYA1;refGene_tx=XM_023248262.1;refGene_gn=EYA1;refGene_tx=XM_023248263.1;refGene_gn=EYA1;refGene_tx=XM_023248264.1;refGene_gn=EYA1;refGene_tx=XM_023248265.1;refGene_gn=EYA1;refGene_tx=XM_023248266.1;refGene_gn=EYA1;refGene_tx=XM_023248267.1
F2 23418931 23419001
refGene_gn=EYA1;refGene_tx=XM_019822779.2;refGene_gn=EYA1;refGene_tx=XM_023248256.1;refGene_gn=EYA1;refGene_tx=XM_023248257.1;refGene_gn=EYA1;refGene_tx=XM_023248259.1;refGene_gn=EYA1;refGene_tx=XM_023248260.1;refGene_gn=EYA1;refGene_tx=XM_023248261.1;refGene_gn=EYA1;refGene_tx=XM_023248262.1;refGene_gn=EYA1;refGene_tx=XM_023248263.1;refGene_gn=EYA1;refGene_tx=XM_023248264.1;refGene_gn=EYA1;refGene_tx=XM_023248265.1;refGene_gn=EYA1;refGene_tx=XM_023248266.1;refGene_gn=EYA1;refGene_tx=XM_023248267.1
F2 23430163 23430241
refGene_gn=EYA1;refGene_tx=XM_019822779.2;refGene_gn=EYA1;refGene_tx=XM_023248256.1;refGene_gn=EYA1;refGene_tx=XM_023248257.1;refGene_gn=EYA1;refGene_tx=XM_023248259.1;refGene_gn=EYA1;refGene_tx=XM_023248260.1;refGene_gn=EYA1;refGene_tx=XM_023248261.1;refGene_gn=EYA1;refGene_tx=XM_023248262.1;refGene_gn=EYA1;refGene_tx=XM_023248265.1;refGene_gn=EYA1;refGene_tx=XM_023248266.1
F2 23440474 23440565 refGene_gn=EYA1;refGene_tx=XM_019822779.2
F2 23451733 23451857
refGene_gn=EYA1;refGene_tx=XM_023248256.1;refGene_gn=EYA1;refGene_tx=X

M_023248257.1;refGene_gn=EYA1;refGene_tx=XM_023248259.1;refGene_gn=EYA1;refGene_tx=XM_023248260.1;refGene_gn=EYA1;refGene_tx=XM_023248261.1;refGene_gn=EYA1;refGene_tx=XM_023248262.1;refGene_gn=EYA1;refGene_tx=XM_023248263.1;refGene_gn=EYA1;refGene_tx=XM_023248264.1
F2 23453397 23453422
refGene_gn=EYA1;refGene_tx=XM_023248265.1;refGene_gn=EYA1;refGene_tx=XM_023248266.1;refGene_gn=EYA1;refGene_tx=XM_023248267.1
F2 23916982 23917069 refGene_gn=MSC;refGene_tx=XM_023248413.1
F2 23918016 23918526 refGene_gn=MSC;refGene_tx=XM_023248413.1
F2 24072210 24072415 refGene_gn=TRPA1;refGene_tx=XM_011291355.3
F2 24073245 24073343 refGene_gn=TRPA1;refGene_tx=XM_011291355.3
F2 24074971 24075082 refGene_gn=TRPA1;refGene_tx=XM_011291355.3
F2 24077695 24077764 refGene_gn=TRPA1;refGene_tx=XM_011291355.3
F2 24080967 24081150 refGene_gn=TRPA1;refGene_tx=XM_011291355.3
F2 24081543 24081673 refGene_gn=TRPA1;refGene_tx=XM_011291355.3
F2 24083212 24083382 refGene_gn=TRPA1;refGene_tx=XM_011291355.3
F2 24085345 24085435 refGene_gn=TRPA1;refGene_tx=XM_011291355.3
F2 24086215 24086378 refGene_gn=TRPA1;refGene_tx=XM_011291355.3
F2 24087291 24087362 refGene_gn=TRPA1;refGene_tx=XM_011291355.3
F2 24089856 24089955 refGene_gn=TRPA1;refGene_tx=XM_011291355.3
F2 24090706 24090766 refGene_gn=TRPA1;refGene_tx=XM_011291355.3
F2 24095179 24095273 refGene_gn=TRPA1;refGene_tx=XM_011291355.3
F2 24096883 24097050 refGene_gn=TRPA1;refGene_tx=XM_011291355.3
F2 24097970 24098085 refGene_gn=TRPA1;refGene_tx=XM_011291355.3
F2 24099547 24099712 refGene_gn=TRPA1;refGene_tx=XM_011291355.3
F2 24099799 24099969 refGene_gn=TRPA1;refGene_tx=XM_011291355.3
F2 24101489 24101590 refGene_gn=TRPA1;refGene_tx=XM_011291355.3
F2 24102239 24102339 refGene_gn=TRPA1;refGene_tx=XM_011291355.3
F2 24104052 24104101 refGene_gn=TRPA1;refGene_tx=XM_011291355.3
F2 24106896 24107033 refGene_gn=TRPA1;refGene_tx=XM_011291355.3
F2 24107446 24107592 refGene_gn=TRPA1;refGene_tx=XM_011291355.3
F2 24107777 24107886 refGene_gn=TRPA1;refGene_tx=XM_011291355.3
F2 24110769 24110877 refGene_gn=TRPA1;refGene_tx=XM_011291355.3
F2 24115774 24115950 refGene_gn=TRPA1;refGene_tx=XM_011291355.3
F2 24119231 24119388 refGene_gn=TRPA1;refGene_tx=XM_011291355.3
F2 24125032 24125143 refGene_gn=TRPA1;refGene_tx=XM_011291355.3
F2 24391231 24391810 refGene_gn=KCNB2;refGene_tx=XM_011291354.3
F2 24745475 24747632 refGene_gn=KCNB2;refGene_tx=XM_011291354.3
F2 24809856 24810163
refGene_gn=TERF1;refGene_tx=XM_003999887.4;refGene_gn=TERF1;refGene_tx=XM_003999889.4;refGene_gn=TERF1;refGene_tx=XM_006943262.3;refGene_gn=TERF1;refGene_tx=XM_019822782.2
F2 24817180 24817276
refGene_gn=TERF1;refGene_tx=XM_003999887.4;refGene_gn=TERF1;refGene_tx=XM_003999889.4;refGene_gn=TERF1;refGene_tx=XM_006943262.3;refGene_gn=TERF1;refGene_tx=XM_019822782.2
F2 24821346 24821468
refGene_gn=TERF1;refGene_tx=XM_003999887.4;refGene_gn=TERF1;refGene_tx=XM_003999889.4;refGene_gn=TERF1;refGene_tx=XM_006943262.3;refGene_gn=TERF1;refGene_tx=XM_019822782.2

F2 24823167 24823254
refGene_gn=TERF1;refGene_tx=XM_003999887.4;refGene_gn=TERF1;refGene_tx
=XM_003999889.4;refGene_gn=TERF1;refGene_tx=XM_019822782.2
F2 24826353 24826503
refGene_gn=TERF1;refGene_tx=XM_003999887.4;refGene_gn=TERF1;refGene_tx
=XM_003999889.4;refGene_gn=TERF1;refGene_tx=XM_006943262.3;refGene_gn=
TERF1;refGene_tx=XM_019822782.2
F2 24827179 24827289
refGene_gn=TERF1;refGene_tx=XM_003999887.4;refGene_gn=TERF1;refGene_tx
=XM_003999889.4;refGene_gn=TERF1;refGene_tx=XM_006943262.3;refGene_gn=
TERF1;refGene_tx=XM_019822782.2
F2 24830963 24831026
refGene_gn=TERF1;refGene_tx=XM_003999889.4;refGene_gn=TERF1;refGene_tx
=XM_006943262.3;refGene_gn=TERF1;refGene_tx=XM_019822782.2
F2 24832532 24832624
refGene_gn=TERF1;refGene_tx=XM_003999887.4;refGene_gn=TERF1;refGene_tx
=XM_003999889.4;refGene_gn=TERF1;refGene_tx=XM_006943262.3;refGene_gn=
TERF1;refGene_tx=XM_019822782.2
F2 24839102 24839206
refGene_gn=TERF1;refGene_tx=XM_003999887.4;refGene_gn=TERF1;refGene_tx
=XM_003999889.4;refGene_gn=TERF1;refGene_tx=XM_006943262.3;refGene_gn=
TERF1;refGene_tx=XM_019822782.2
F2 24844822 24844999
refGene_gn=TERF1;refGene_tx=XM_003999887.4;refGene_gn=TERF1;refGene_tx
=XM_003999889.4;refGene_gn=TERF1;refGene_tx=XM_006943262.3
F2 24855393 24855412 refGene_gn=SBSPON;refGene_tx=XM_006943264.2
F2 24856056 24856174
refGene_gn=SBSPON;refGene_tx=XM_003999891.3;refGene_gn=SBSPON;refGene_
tx=XM_006943264.2
F2 24858158 24858335
refGene_gn=SBSPON;refGene_tx=XM_003999891.3;refGene_gn=SBSPON;refGene_
tx=XM_006943264.2
F2 24859940 24860031
refGene_gn=SBSPON;refGene_tx=XM_003999891.3;refGene_gn=SBSPON;refGene_
tx=XM_006943264.2
F2 24869892 24870087
refGene_gn=SBSPON;refGene_tx=XM_003999891.3;refGene_gn=SBSPON;refGene_
tx=XM_006943264.2
F2 24881074 24881288
refGene_gn=SBSPON;refGene_tx=XM_003999891.3;refGene_gn=SBSPON;refGene_
tx=XM_006943264.2
F2 25014161 25014310
refGene_gn=CF2H8orf89;refGene_tx=XM_006943265.3;refGene_gn=CF2H8orf89;
refGene_tx=XM_011291351.3;refGene_gn=CF2H8orf89;refGene_tx=XM_01129135
2.3;refGene_gn=CF2H8orf89;refGene_tx=XM_011291353.3;refGene_gn=CF2H8or
f89;refGene_tx=XM_023248287.1;refGene_gn=CF2H8orf89;refGene_tx=XM_0232
48288.1
F2 25016705 25016741
refGene_gn=CF2H8orf89;refGene_tx=XM_006943265.3;refGene_gn=CF2H8orf89;
refGene_tx=XM_011291352.3;refGene_gn=CF2H8orf89;refGene_tx=XM_02324828

7.1;refGene_gn=CF2H8orf89;refGene_tx=XM_023248288.1
F2 25019324 25019380
refGene_gn=CF2H8orf89;refGene_tx=XM_006943265.3;refGene_gn=CF2H8orf89;refGene_tx=XM_011291351.3;refGene_gn=CF2H8orf89;refGene_tx=XM_011291352.3;refGene_gn=CF2H8orf89;refGene_tx=XM_011291353.3;refGene_gn=CF2H8orf89;refGene_tx=XM_023248287.1;refGene_gn=CF2H8orf89;refGene_tx=XM_023248288.1;refGene_gn=CF2H8orf89;refGene_tx=XM_023248289.1
F2 25020607 25020703
refGene_gn=CF2H8orf89;refGene_tx=XM_006943265.3;refGene_gn=CF2H8orf89;refGene_tx=XM_011291351.3;refGene_gn=CF2H8orf89;refGene_tx=XM_023248287.1;refGene_gn=CF2H8orf89;refGene_tx=XM_023248288.1;refGene_gn=CF2H8orf89;refGene_tx=XM_023248289.1
F2 25022690 25022832
refGene_gn=CF2H8orf89;refGene_tx=XM_006943265.3;refGene_gn=CF2H8orf89;refGene_tx=XM_011291351.3;refGene_gn=CF2H8orf89;refGene_tx=XM_011291352.3;refGene_gn=CF2H8orf89;refGene_tx=XM_011291353.3;refGene_gn=CF2H8orf89;refGene_tx=XM_023248287.1;refGene_gn=CF2H8orf89;refGene_tx=XM_023248288.1;refGene_gn=CF2H8orf89;refGene_tx=XM_023248289.1
F2 25024613 25024737
refGene_gn=CF2H8orf89;refGene_tx=XM_006943265.3;refGene_gn=CF2H8orf89;refGene_tx=XM_011291351.3;refGene_gn=CF2H8orf89;refGene_tx=XM_011291352.3;refGene_gn=CF2H8orf89;refGene_tx=XM_011291353.3;refGene_gn=CF2H8orf89;refGene_tx=XM_023248287.1;refGene_gn=CF2H8orf89;refGene_tx=XM_023248289.1
F2 25040297 25040422
refGene_gn=LOC111558523;refGene_tx=XM_023248329.1
F2 25040574 25041121
refGene_gn=LOC111558523;refGene_tx=XM_023248329.1
F2 25045999 25046208
refGene_gn=RPL7;refGene_tx=XM_023248285.1;refGene_gn=RPL7;refGene_tx=XM_023248286.1
F2 25046479 25046589
refGene_gn=RPL7;refGene_tx=XM_023248285.1;refGene_gn=RPL7;refGene_tx=XM_023248286.1
F2 25046715 25046853
refGene_gn=RPL7;refGene_tx=XM_023248285.1;refGene_gn=RPL7;refGene_tx=XM_023248286.1
F2 25047362 25047529
refGene_gn=RPL7;refGene_tx=XM_023248285.1;refGene_gn=RPL7;refGene_tx=XM_023248286.1
F2 25047828 25047937
refGene_gn=RPL7;refGene_tx=XM_023248285.1;refGene_gn=RPL7;refGene_tx=XM_023248286.1
F2 25048315 25048341 refGene_gn=RPL7;refGene_tx=XM_023248285.1
F2 25048684 25048698 refGene_gn=RPL7;refGene_tx=XM_023248286.1
F2 25050409 25050698 refGene_gn=RDH10;refGene_tx=XM_003999902.4
F2 25052304 25052540 refGene_gn=RDH10;refGene_tx=XM_003999902.4
F2 25072186 25072285 refGene_gn=RDH10;refGene_tx=XM_003999902.4
F2 25073475 25073621 refGene_gn=RDH10;refGene_tx=XM_003999902.4
F2 25075370 25075502 refGene_gn=RDH10;refGene_tx=XM_003999902.4

F2 25075614 25075738 refGene_gn=RDH10;refGene_tx=XM_003999902.4
F2 25163181 25163275
refGene_gn=STAU2;refGene_tx=XM_006943271.3;refGene_gn=STAU2;refGene_tx=XM_006943274.4;refGene_gn=STAU2;refGene_tx=XM_019822787.1;refGene_gn=STAU2;refGene_tx=XM_019822791.1;refGene_gn=STAU2;refGene_tx=XM_023248475.1;refGene_gn=STAU2;refGene_tx=XM_023248476.1;refGene_gn=STAU2;refGene_tx=XM_023248477.1;refGene_gn=STAU2;refGene_tx=XM_023248478.1;refGene_gn=STAU2;refGene_tx=XM_023248479.1;refGene_gn=STAU2;refGene_tx=XM_023248480.1;refGene_gn=STAU2;refGene_tx=XM_023248481.1;refGene_gn=STAU2;refGene_tx=XM_023248482.1;refGene_gn=STAU2;refGene_tx=XM_023248483.1
F2 25164407 25164496
refGene_gn=STAU2;refGene_tx=XM_006943271.3;refGene_gn=STAU2;refGene_tx=XM_006943274.4;refGene_gn=STAU2;refGene_tx=XM_019822787.1;refGene_gn=STAU2;refGene_tx=XM_019822791.1;refGene_gn=STAU2;refGene_tx=XM_023248475.1;refGene_gn=STAU2;refGene_tx=XM_023248476.1;refGene_gn=STAU2;refGene_tx=XM_023248477.1;refGene_gn=STAU2;refGene_tx=XM_023248478.1;refGene_gn=STAU2;refGene_tx=XM_023248479.1;refGene_gn=STAU2;refGene_tx=XM_023248480.1;refGene_gn=STAU2;refGene_tx=XM_023248481.1;refGene_gn=STAU2;refGene_tx=XM_023248482.1;refGene_gn=STAU2;refGene_tx=XM_023248483.1
F2 25258847 25258913 refGene_gn=STAU2;refGene_tx=XM_019822788.2
F2 25265682 25265796
refGene_gn=STAU2;refGene_tx=XM_006943271.3;refGene_gn=STAU2;refGene_tx=XM_006943274.4;refGene_gn=STAU2;refGene_tx=XM_019822788.2;refGene_gn=STAU2;refGene_tx=XM_019822791.1;refGene_gn=STAU2;refGene_tx=XM_023248475.1;refGene_gn=STAU2;refGene_tx=XM_023248476.1;refGene_gn=STAU2;refGene_tx=XM_023248477.1;refGene_gn=STAU2;refGene_tx=XM_023248478.1;refGene_gn=STAU2;refGene_tx=XM_023248479.1;refGene_gn=STAU2;refGene_tx=XM_023248480.1;refGene_gn=STAU2;refGene_tx=XM_023248481.1;refGene_gn=STAU2;refGene_tx=XM_023248482.1;refGene_gn=STAU2;refGene_tx=XM_023248483.1
F2 25287857 25288171
refGene_gn=STAU2;refGene_tx=XM_006943271.3;refGene_gn=STAU2;refGene_tx=XM_006943274.4;refGene_gn=STAU2;refGene_tx=XM_019822787.1;refGene_gn=STAU2;refGene_tx=XM_019822788.2;refGene_gn=STAU2;refGene_tx=XM_019822791.1;refGene_gn=STAU2;refGene_tx=XM_023248475.1;refGene_gn=STAU2;refGene_tx=XM_023248476.1;refGene_gn=STAU2;refGene_tx=XM_023248477.1;refGene_gn=STAU2;refGene_tx=XM_023248478.1;refGene_gn=STAU2;refGene_tx=XM_023248479.1;refGene_gn=STAU2;refGene_tx=XM_023248480.1;refGene_gn=STAU2;refGene_tx=XM_023248481.1;refGene_gn=STAU2;refGene_tx=XM_023248482.1;refGene_gn=STAU2;refGene_tx=XM_023248483.1;refGene_gn=STAU2;refGene_tx=XM_023248484.1
F2 25319897 25319958
refGene_gn=STAU2;refGene_tx=XM_006943271.3;refGene_gn=STAU2;refGene_tx=XM_006943274.4;refGene_gn=STAU2;refGene_tx=XM_019822787.1;refGene_gn=STAU2;refGene_tx=XM_019822788.2;refGene_gn=STAU2;refGene_tx=XM_019822791.1;refGene_gn=STAU2;refGene_tx=XM_023248475.1;refGene_gn=STAU2;refGene_tx=XM_023248476.1;refGene_gn=STAU2;refGene_tx=XM_023248477.1;refGene_gn=STAU2;refGene_tx=XM_023248478.1;refGene_gn=STAU2;refGene_tx=XM_023248479.1;refGene_gn=STAU2;refGene_tx=XM_023248480.1;refGene_gn=STAU2;refGene_tx=XM_023248481.1;refGene_gn=STAU2;refGene_tx=XM_023248482.1;refGene_gn=STAU2;refGene_tx=XM_023248483.1;refGene_gn=STAU2;refGene_tx=

XM_023248484.1
F2	25334356 25334488
refGene_gn=STAU2;refGene_tx=XM_006943271.3;refGene_gn=STAU2;refGene_tx=XM_006943274.4;refGene_gn=STAU2;refGene_tx=XM_019822787.1;refGene_gn=STAU2;refGene_tx=XM_019822788.2;refGene_gn=STAU2;refGene_tx=XM_019822791.1;refGene_gn=STAU2;refGene_tx=XM_023248475.1;refGene_gn=STAU2;refGene_tx=XM_023248476.1;refGene_gn=STAU2;refGene_tx=XM_023248477.1;refGene_gn=STAU2;refGene_tx=XM_023248478.1;refGene_gn=STAU2;refGene_tx=XM_023248479.1;refGene_gn=STAU2;refGene_tx=XM_023248480.1;refGene_gn=STAU2;refGene_tx=XM_023248481.1;refGene_gn=STAU2;refGene_tx=XM_023248482.1;refGene_gn=STAU2;refGene_tx=XM_023248483.1;refGene_gn=STAU2;refGene_tx=XM_023248484.1
F2	25342983 25343121
refGene_gn=STAU2;refGene_tx=XM_006943271.3;refGene_gn=STAU2;refGene_tx=XM_006943274.4;refGene_gn=STAU2;refGene_tx=XM_019822787.1;refGene_gn=STAU2;refGene_tx=XM_019822788.2;refGene_gn=STAU2;refGene_tx=XM_019822791.1;refGene_gn=STAU2;refGene_tx=XM_023248475.1;refGene_gn=STAU2;refGene_tx=XM_023248476.1;refGene_gn=STAU2;refGene_tx=XM_023248477.1;refGene_gn=STAU2;refGene_tx=XM_023248478.1;refGene_gn=STAU2;refGene_tx=XM_023248479.1;refGene_gn=STAU2;refGene_tx=XM_023248480.1;refGene_gn=STAU2;refGene_tx=XM_023248481.1;refGene_gn=STAU2;refGene_tx=XM_023248482.1;refGene_gn=STAU2;refGene_tx=XM_023248483.1;refGene_gn=STAU2;refGene_tx=XM_023248484.1
F2	25354013 25354226
refGene_gn=STAU2;refGene_tx=XM_006943271.3;refGene_gn=STAU2;refGene_tx=XM_006943274.4;refGene_gn=STAU2;refGene_tx=XM_019822787.1;refGene_gn=STAU2;refGene_tx=XM_019822788.2;refGene_gn=STAU2;refGene_tx=XM_023248475.1;refGene_gn=STAU2;refGene_tx=XM_023248476.1;refGene_gn=STAU2;refGene_tx=XM_023248477.1;refGene_gn=STAU2;refGene_tx=XM_023248478.1;refGene_gn=STAU2;refGene_tx=XM_023248479.1;refGene_gn=STAU2;refGene_tx=XM_023248480.1;refGene_gn=STAU2;refGene_tx=XM_023248481.1;refGene_gn=STAU2;refGene_tx=XM_023248482.1;refGene_gn=STAU2;refGene_tx=XM_023248483.1;refGene_gn=STAU2;refGene_tx=XM_023248484.1
F2	25356412 25356520
refGene_gn=STAU2;refGene_tx=XM_006943271.3;refGene_gn=STAU2;refGene_tx=XM_006943274.4;refGene_gn=STAU2;refGene_tx=XM_019822787.1;refGene_gn=STAU2;refGene_tx=XM_019822788.2;refGene_gn=STAU2;refGene_tx=XM_019822791.1;refGene_gn=STAU2;refGene_tx=XM_023248475.1;refGene_gn=STAU2;refGene_tx=XM_023248476.1;refGene_gn=STAU2;refGene_tx=XM_023248477.1;refGene_gn=STAU2;refGene_tx=XM_023248478.1;refGene_gn=STAU2;refGene_tx=XM_023248479.1;refGene_gn=STAU2;refGene_tx=XM_023248480.1;refGene_gn=STAU2;refGene_tx=XM_023248481.1;refGene_gn=STAU2;refGene_tx=XM_023248482.1;refGene_gn=STAU2;refGene_tx=XM_023248483.1;refGene_gn=STAU2;refGene_tx=XM_023248484.1
F2	25357776 25357936
refGene_gn=STAU2;refGene_tx=XM_006943271.3;refGene_gn=STAU2;refGene_tx=XM_006943274.4;refGene_gn=STAU2;refGene_tx=XM_019822787.1;refGene_gn=STAU2;refGene_tx=XM_019822788.2;refGene_gn=STAU2;refGene_tx=XM_019822791.1;refGene_gn=STAU2;refGene_tx=XM_023248475.1;refGene_gn=STAU2;refGene_tx=XM_023248476.1;refGene_gn=STAU2;refGene_tx=XM_023248477.1;refGen

e_gn=STAU2;refGene_tx=XM_023248478.1;refGene_gn=STAU2;refGene_tx=XM_023248479.1;refGene_gn=STAU2;refGene_tx=XM_023248480.1;refGene_gn=STAU2;refGene_tx=XM_023248481.1;refGene_gn=STAU2;refGene_tx=XM_023248482.1;refGene_gn=STAU2;refGene_tx=XM_023248483.1;refGene_gn=STAU2;refGene_tx=XM_023248484.1
F2 25413136 25413272
refGene_gn=STAU2;refGene_tx=XM_006943271.3;refGene_gn=STAU2;refGene_tx=XM_006943274.4;refGene_gn=STAU2;refGene_tx=XM_019822787.1;refGene_gn=STAU2;refGene_tx=XM_019822788.2;refGene_gn=STAU2;refGene_tx=XM_019822791.1;refGene_gn=STAU2;refGene_tx=XM_023248475.1;refGene_gn=STAU2;refGene_tx=XM_023248476.1;refGene_gn=STAU2;refGene_tx=XM_023248477.1;refGene_gn=STAU2;refGene_tx=XM_023248478.1;refGene_gn=STAU2;refGene_tx=XM_023248479.1;refGene_gn=STAU2;refGene_tx=XM_023248480.1;refGene_gn=STAU2;refGene_tx=XM_023248481.1;refGene_gn=STAU2;refGene_tx=XM_023248482.1;refGene_gn=STAU2;refGene_tx=XM_023248483.1;refGene_gn=STAU2;refGene_tx=XM_023248484.1
F2 25421159 25421319
refGene_gn=STAU2;refGene_tx=XM_006943271.3;refGene_gn=STAU2;refGene_tx=XM_006943274.4;refGene_gn=STAU2;refGene_tx=XM_019822787.1;refGene_gn=STAU2;refGene_tx=XM_019822788.2;refGene_gn=STAU2;refGene_tx=XM_019822791.1;refGene_gn=STAU2;refGene_tx=XM_023248475.1;refGene_gn=STAU2;refGene_tx=XM_023248476.1;refGene_gn=STAU2;refGene_tx=XM_023248477.1;refGene_gn=STAU2;refGene_tx=XM_023248478.1;refGene_gn=STAU2;refGene_tx=XM_023248479.1;refGene_gn=STAU2;refGene_tx=XM_023248480.1;refGene_gn=STAU2;refGene_tx=XM_023248481.1;refGene_gn=STAU2;refGene_tx=XM_023248482.1;refGene_gn=STAU2;refGene_tx=XM_023248483.1;refGene_gn=STAU2;refGene_tx=XM_023248484.1
F2 25437728 25437859
refGene_gn=STAU2;refGene_tx=XM_006943271.3;refGene_gn=STAU2;refGene_tx=XM_019822787.1;refGene_gn=STAU2;refGene_tx=XM_019822788.2;refGene_gn=STAU2;refGene_tx=XM_019822791.1;refGene_gn=STAU2;refGene_tx=XM_023248475.1;refGene_gn=STAU2;refGene_tx=XM_023248476.1;refGene_gn=STAU2;refGene_tx=XM_023248477.1;refGene_gn=STAU2;refGene_tx=XM_023248478.1
F2 25443574 25443614 refGene_gn=STAU2;refGene_tx=XM_023248475.1
F2 25462931 25462937 refGene_gn=STAU2;refGene_tx=XM_023248483.1
F2 25465473 25465491
refGene_gn=STAU2;refGene_tx=XM_006943274.4;refGene_gn=STAU2;refGene_tx=XM_023248480.1;refGene_gn=STAU2;refGene_tx=XM_023248481.1;refGene_gn=STAU2;refGene_tx=XM_023248482.1
F2 25473823 25473877 refGene_gn=STAU2;refGene_tx=XM_023248479.1
F2 25519277 25519291 refGene_gn=UBE2W;refGene_tx=XM_003999895.5
F2 25528499 25528575 refGene_gn=UBE2W;refGene_tx=XM_003999895.5
F2 25532227 25532383 refGene_gn=UBE2W;refGene_tx=XM_003999895.5
F2 25542301 25542404 refGene_gn=UBE2W;refGene_tx=XM_003999895.5
F2 25547667 25547759 refGene_gn=UBE2W;refGene_tx=XM_003999895.5
F2 25587448 25587511 refGene_gn=UBE2W;refGene_tx=XM_003999895.5
F2 25587512 25587551 refGene_gn=UBE2W;refGene_tx=XM_003999895.5
F2 25630581 25630772 refGene_gn=ELOC;refGene_tx=NM_001246227.1
F2 25637581 25637726 refGene_gn=ELOC;refGene_tx=NM_001246227.1
F2 25657665 25657875 refGene_gn=TMEM70;refGene_tx=XM_003999896.4

F2 25659070 25659176 refGene_gn=TMEM70;refGene_tx=XM_003999896.4
F2 25664182 25664640 refGene_gn=TMEM70;refGene_tx=XM_003999896.4
F2 25670523 25670635 refGene_gn=LY96;refGene_tx=XM_003999897.4
F2 25682133 25682223 refGene_gn=LY96;refGene_tx=XM_003999897.4
F2 25689262 25689391 refGene_gn=LY96;refGene_tx=XM_003999897.4
F2 25698604 25698657 refGene_gn=LY96;refGene_tx=XM_003999897.4
F2 25699635 25699734 refGene_gn=LY96;refGene_tx=XM_003999897.4
F2 25894567 25894770 refGene_gn=JPH1;refGene_tx=XM_023248222.1
F2 25896062 25896121
refGene_gn=JPH1;refGene_tx=XM_023248223.1;refGene_gn=JPH1;refGene_tx=XM_023248224.1
F2 25896386 25896397 refGene_gn=JPH1;refGene_tx=XM_023248225.1
F2 25896564 25896661
refGene_gn=JPH1;refGene_tx=XM_023248222.1;refGene_gn=JPH1;refGene_tx=XM_023248223.1;refGene_gn=JPH1;refGene_tx=XM_023248224.1;refGene_gn=JPH1;refGene_tx=XM_023248225.1
F2 25902859 25903509
refGene_gn=JPH1;refGene_tx=XM_023248222.1;refGene_gn=JPH1;refGene_tx=XM_023248223.1;refGene_gn=JPH1;refGene_tx=XM_023248224.1;refGene_gn=JPH1;refGene_tx=XM_023248225.1
F2 25918931 25919050
refGene_gn=JPH1;refGene_tx=XM_023248222.1;refGene_gn=JPH1;refGene_tx=XM_023248223.1;refGene_gn=JPH1;refGene_tx=XM_023248224.1;refGene_gn=JPH1;refGene_tx=XM_023248225.1
F2 25970716 25971476
refGene_gn=JPH1;refGene_tx=XM_023248222.1;refGene_gn=JPH1;refGene_tx=XM_023248223.1;refGene_gn=JPH1;refGene_tx=XM_023248224.1;refGene_gn=JPH1;refGene_tx=XM_023248225.1
F2 25977060 25977439
refGene_gn=JPH1;refGene_tx=XM_023248222.1;refGene_gn=JPH1;refGene_tx=XM_023248223.1;refGene_gn=JPH1;refGene_tx=XM_023248224.1;refGene_gn=JPH1;refGene_tx=XM_023248225.1
F2 25985270 25985312
refGene_gn=GDAP1;refGene_tx=XM_011291345.3;refGene_gn=GDAP1;refGene_tx=XM_011291346.3
F2 26005389 26005506 refGene_gn=GDAP1;refGene_tx=XM_003999898.4
F2 26006219 26006412
refGene_gn=GDAP1;refGene_tx=XM_003999898.4;refGene_gn=GDAP1;refGene_tx=XM_011291345.3;refGene_gn=GDAP1;refGene_tx=XM_011291346.3
F2 26016246 26016420
refGene_gn=GDAP1;refGene_tx=XM_003999898.4;refGene_gn=GDAP1;refGene_tx=XM_011291345.3;refGene_gn=GDAP1;refGene_tx=XM_011291346.3
F2 26017616 26017711
refGene_gn=GDAP1;refGene_tx=XM_003999898.4;refGene_gn=GDAP1;refGene_tx=XM_011291345.3;refGene_gn=GDAP1;refGene_tx=XM_011291346.3
F2 26020403 26020518
refGene_gn=GDAP1;refGene_tx=XM_003999898.4;refGene_gn=GDAP1;refGene_tx=XM_011291345.3;refGene_gn=GDAP1;refGene_tx=XM_011291346.3
F2 26021524 26021907
refGene_gn=GDAP1;refGene_tx=XM_003999898.4;refGene_gn=GDAP1;refGene_tx

=XM_011291345.3;refGene_gn=GDAP1;refGene_tx=XM_011291346.3
F2 26461835 26462108 refGene_gn=PI15;refGene_tx=XM_003999904.4
F2 26481633 26481752 refGene_gn=PI15;refGene_tx=XM_003999904.4
F2 26482778 26482911 refGene_gn=PI15;refGene_tx=XM_003999904.4
F2 26483031 26483147 refGene_gn=PI15;refGene_tx=XM_003999904.4
F2 26486320 26486456 refGene_gn=PI15;refGene_tx=XM_003999904.4
F2 26605336 26605594 refGene_gn=CRISPLD1;refGene_tx=XM_011291343.2
F2 26630708 26630827 refGene_gn=CRISPLD1;refGene_tx=XM_011291343.2
F2 26631162 26631295
refGene_gn=CRISPLD1;refGene_tx=XM_011291343.2;refGene_gn=CRISPLD1;refGene_tx=XM_011291344.2
F2 26632961 26633077 refGene_gn=CRISPLD1;refGene_tx=XM_011291343.2
F2 26633977 26634078
refGene_gn=CRISPLD1;refGene_tx=XM_011291343.2;refGene_gn=CRISPLD1;refGene_tx=XM_011291344.2
F2 26635734 26635875
refGene_gn=CRISPLD1;refGene_tx=XM_011291343.2;refGene_gn=CRISPLD1;refGene_tx=XM_011291344.2
F2 26636099 26636160
refGene_gn=CRISPLD1;refGene_tx=XM_011291343.2;refGene_gn=CRISPLD1;refGene_tx=XM_011291344.2
F2 26636269 26636336
refGene_gn=CRISPLD1;refGene_tx=XM_011291343.2;refGene_gn=CRISPLD1;refGene_tx=XM_011291344.2
F2 26636543 26636674
refGene_gn=CRISPLD1;refGene_tx=XM_011291343.2;refGene_gn=CRISPLD1;refGene_tx=XM_011291344.2
F2 26638390 26638434
refGene_gn=CRISPLD1;refGene_tx=XM_011291343.2;refGene_gn=CRISPLD1;refGene_tx=XM_011291344.2
F2 26638527 26638600
refGene_gn=CRISPLD1;refGene_tx=XM_011291343.2;refGene_gn=CRISPLD1;refGene_tx=XM_011291344.2
F2 26644040 26644116
refGene_gn=CRISPLD1;refGene_tx=XM_011291343.2;refGene_gn=CRISPLD1;refGene_tx=XM_011291344.2
F2 26648734 26648865
refGene_gn=CRISPLD1;refGene_tx=XM_011291343.2;refGene_gn=CRISPLD1;refGene_tx=XM_011291344.2
F2 26652024 26652076
refGene_gn=CRISPLD1;refGene_tx=XM_011291343.2;refGene_gn=CRISPLD1;refGene_tx=XM_011291344.2
F2 27053536 27053672 refGene_gn=HNF4G;refGene_tx=XM_003999905.3
F2 27056567 27056739
refGene_gn=HNF4G;refGene_tx=XM_003999905.3;refGene_gn=HNF4G;refGene_tx=XM_023248474.1
F2 27062496 27062591
refGene_gn=HNF4G;refGene_tx=XM_003999905.3;refGene_gn=HNF4G;refGene_tx=XM_023248474.1
F2 27065384 27065491

refGene_gn=HNF4G;refGene_tx=XM_003999905.3;refGene_gn=HNF4G;refGene_tx=XM_023248474.1
F2 27067045 27067201
refGene_gn=HNF4G;refGene_tx=XM_003999905.3;refGene_gn=HNF4G;refGene_tx=XM_023248474.1
F2 27070225 27070313
refGene_gn=HNF4G;refGene_tx=XM_003999905.3;refGene_gn=HNF4G;refGene_tx=XM_023248474.1
F2 27074673 27074826
refGene_gn=HNF4G;refGene_tx=XM_003999905.3;refGene_gn=HNF4G;refGene_tx=XM_023248474.1
F2 27074953 27075190
refGene_gn=HNF4G;refGene_tx=XM_003999905.3;refGene_gn=HNF4G;refGene_tx=XM_023248474.1
F2 27076248 27076371
refGene_gn=HNF4G;refGene_tx=XM_003999905.3;refGene_gn=HNF4G;refGene_tx=XM_023248474.1
F2 27080514 27080636
refGene_gn=HNF4G;refGene_tx=XM_003999905.3;refGene_gn=HNF4G;refGene_tx=XM_023248474.1
F2 27986241 27988831
refGene_gn=ZFHX4;refGene_tx=XM_011291342.3;refGene_gn=ZFHX4;refGene_tx=XM_019822795.2;refGene_gn=ZFHX4;refGene_tx=XM_023248241.1;refGene_gn=ZFHX4;refGene_tx=XM_023248243.1;refGene_gn=ZFHX4;refGene_tx=XM_023248244.1
F2 27989693 27990196
refGene_gn=ZFHX4;refGene_tx=XM_011291342.3;refGene_gn=ZFHX4;refGene_tx=XM_019822795.2;refGene_gn=ZFHX4;refGene_tx=XM_023248241.1;refGene_gn=ZFHX4;refGene_tx=XM_023248243.1;refGene_gn=ZFHX4;refGene_tx=XM_023248244.1
F2 28059413 28059645
refGene_gn=ZFHX4;refGene_tx=XM_011291342.3;refGene_gn=ZFHX4;refGene_tx=XM_019822795.2;refGene_gn=ZFHX4;refGene_tx=XM_023248241.1;refGene_gn=ZFHX4;refGene_tx=XM_023248243.1;refGene_gn=ZFHX4;refGene_tx=XM_023248244.1
F2 28112865 28112934
refGene_gn=ZFHX4;refGene_tx=XM_011291342.3;refGene_gn=ZFHX4;refGene_tx=XM_019822795.2;refGene_gn=ZFHX4;refGene_tx=XM_023248241.1;refGene_gn=ZFHX4;refGene_tx=XM_023248243.1;refGene_gn=ZFHX4;refGene_tx=XM_023248244.1
F2 28121769 28121886
refGene_gn=ZFHX4;refGene_tx=XM_011291342.3;refGene_gn=ZFHX4;refGene_tx=XM_019822795.2;refGene_gn=ZFHX4;refGene_tx=XM_023248241.1;refGene_gn=ZFHX4;refGene_tx=XM_023248243.1;refGene_gn=ZFHX4;refGene_tx=XM_023248244.1
F2 28128131 28128265
refGene_gn=ZFHX4;refGene_tx=XM_011291342.3;refGene_gn=ZFHX4;refGene_tx=XM_019822795.2;refGene_gn=ZFHX4;refGene_tx=XM_023248241.1;refGene_gn=ZFHX4;refGene_tx=XM_023248243.1;refGene_gn=ZFHX4;refGene_tx=XM_023248244.1

F2 28128547 28128748
refGene_gn=ZFHX4;refGene_tx=XM_011291342.3;refGene_gn=ZFHX4;refGene_tx=XM_019822795.2;refGene_gn=ZFHX4;refGene_tx=XM_023248241.1;refGene_gn=ZFHX4;refGene_tx=XM_023248243.1;refGene_gn=ZFHX4;refGene_tx=XM_023248244.1
F2 28129515 28129633
refGene_gn=ZFHX4;refGene_tx=XM_011291342.3;refGene_gn=ZFHX4;refGene_tx=XM_019822795.2;refGene_gn=ZFHX4;refGene_tx=XM_023248241.1;refGene_gn=ZFHX4;refGene_tx=XM_023248243.1;refGene_gn=ZFHX4;refGene_tx=XM_023248244.1
F2 28130173 28135618
refGene_gn=ZFHX4;refGene_tx=XM_011291342.3;refGene_gn=ZFHX4;refGene_tx=XM_019822795.2;refGene_gn=ZFHX4;refGene_tx=XM_023248241.1;refGene_gn=ZFHX4;refGene_tx=XM_023248243.1;refGene_gn=ZFHX4;refGene_tx=XM_023248244.1
F2 28143625 28145097
refGene_gn=ZFHX4;refGene_tx=XM_011291342.3;refGene_gn=ZFHX4;refGene_tx=XM_019822795.2;refGene_gn=ZFHX4;refGene_tx=XM_023248241.1;refGene_gn=ZFHX4;refGene_tx=XM_023248243.1;refGene_gn=ZFHX4;refGene_tx=XM_023248244.1
F2 28250011 28250946
refGene_gn=PEX2;refGene_tx=XM_003999908.4;refGene_gn=PEX2;refGene_tx=XM_003999910.4;refGene_gn=PEX2;refGene_tx=XM_006943281.3;refGene_gn=PEX2;refGene_tx=XM_019822798.1;refGene_gn=PEX2;refGene_tx=XM_019822799.1
F2 28262118 28262137 refGene_gn=PEX2;refGene_tx=XM_019822798.1
F2 29631576 29631681 refGene_gn=PKIA;refGene_tx=XM_011291339.2
F2 29638312 29638490
refGene_gn=PKIA;refGene_tx=XM_011291339.2;refGene_gn=PKIA;refGene_tx=XM_011291340.3;refGene_gn=PKIA;refGene_tx=XM_011291341.2
F2 29641828 29641908
refGene_gn=PKIA;refGene_tx=XM_011291339.2;refGene_gn=PKIA;refGene_tx=XM_011291340.3;refGene_gn=PKIA;refGene_tx=XM_011291341.2
F2 29695340 29695485 refGene_gn=ZC2HC1A;refGene_tx=XM_006943283.4
F2 29706848 29706922 refGene_gn=ZC2HC1A;refGene_tx=XM_006943283.4
F2 29709031 29709148 refGene_gn=ZC2HC1A;refGene_tx=XM_006943283.4
F2 29716520 29716662 refGene_gn=ZC2HC1A;refGene_tx=XM_006943283.4
F2 29718474 29718623 refGene_gn=ZC2HC1A;refGene_tx=XM_006943283.4
F2 29725892 29725992 refGene_gn=ZC2HC1A;refGene_tx=XM_006943283.4
F2 29726627 29726727 refGene_gn=ZC2HC1A;refGene_tx=XM_006943283.4
F2 29739567 29739675 refGene_gn=ZC2HC1A;refGene_tx=XM_006943283.4
F2 29744339 29744505 refGene_gn=ZC2HC1A;refGene_tx=XM_006943283.4
F2 29753571 29753691
refGene_gn=IL7;refGene_tx=XM_003999913.4;refGene_gn=IL7;refGene_tx=XM_019822801.1
F2 29764853 29764985
refGene_gn=IL7;refGene_tx=XM_003999913.4;refGene_gn=IL7;refGene_tx=XM_019822801.1
F2 29766080 29766161 refGene_gn=IL7;refGene_tx=XM_003999913.4
F2 29806432 29806566
refGene_gn=IL7;refGene_tx=XM_003999913.4;refGene_gn=IL7;refGene_tx=XM_

019822801.1
F2	29812555	29812565
refGene_gn=IL7;refGene_tx=XM_003999913.4;refGene_gn=IL7;refGene_tx=XM_
019822801.1
F2	30457172	30457191	refGene_gn=STMN2;refGene_tx=XM_003999916.5
F2	30483522	30483618	refGene_gn=STMN2;refGene_tx=XM_003999916.5
F2	30485747	30485920	refGene_gn=STMN2;refGene_tx=XM_003999916.5
F2	30496100	30496292	refGene_gn=STMN2;refGene_tx=XM_003999916.5
F2	30502721	30502781	refGene_gn=STMN2;refGene_tx=XM_003999916.5
F2	30583250	30583834	refGene_gn=HEY1;refGene_tx=XM_003999917.5
F2	30584684	30584766	refGene_gn=HEY1;refGene_tx=XM_003999917.5
F2	30585055	30585139	refGene_gn=HEY1;refGene_tx=XM_003999917.5
F2	30585266	30585342	refGene_gn=HEY1;refGene_tx=XM_003999917.5
F2	30585604	30585693	refGene_gn=HEY1;refGene_tx=XM_003999917.5
F2	30714111	30714277	refGene_gn=MRPS28;refGene_tx=XM_003999933.5
F2	30799033	30799215	refGene_gn=MRPS28;refGene_tx=XM_003999933.5
F2	30820556	30820772	refGene_gn=MRPS28;refGene_tx=XM_003999933.5
F2	30826424	30826541
refGene_gn=TPD52;refGene_tx=XM_011291336.3;refGene_gn=TPD52;refGene_tx
=XM_019822803.2;refGene_gn=TPD52;refGene_tx=XM_019822804.2;refGene_gn=
TPD52;refGene_tx=XM_019822805.2;refGene_gn=TPD52;refGene_tx=XM_0198228
07.2
F2	30831013	30831062
refGene_gn=TPD52;refGene_tx=XM_011291336.3;refGene_gn=TPD52;refGene_tx
=XM_019822803.2;refGene_gn=TPD52;refGene_tx=XM_019822804.2;refGene_gn=
TPD52;refGene_tx=XM_019822805.2;refGene_gn=TPD52;refGene_tx=XM_0198228
07.2
F2	30832560	30832602
refGene_gn=TPD52;refGene_tx=XM_011291336.3;refGene_gn=TPD52;refGene_tx
=XM_019822803.2;refGene_gn=TPD52;refGene_tx=XM_019822804.2;refGene_gn=
TPD52;refGene_tx=XM_019822807.2
F2	30839134	30839161
refGene_gn=TPD52;refGene_tx=XM_011291336.3;refGene_gn=TPD52;refGene_tx
=XM_019822803.2;refGene_gn=TPD52;refGene_tx=XM_019822807.2
F2	30840263	30840365
refGene_gn=TPD52;refGene_tx=XM_011291336.3;refGene_gn=TPD52;refGene_tx
=XM_019822803.2;refGene_gn=TPD52;refGene_tx=XM_019822804.2;refGene_gn=
TPD52;refGene_tx=XM_019822805.2;refGene_gn=TPD52;refGene_tx=XM_0198228
07.2
F2	30842112	30842261
refGene_gn=TPD52;refGene_tx=XM_011291336.3;refGene_gn=TPD52;refGene_tx
=XM_019822803.2;refGene_gn=TPD52;refGene_tx=XM_019822804.2;refGene_gn=
TPD52;refGene_tx=XM_019822805.2;refGene_gn=TPD52;refGene_tx=XM_0198228
07.2
F2	30847005	30847121
refGene_gn=TPD52;refGene_tx=XM_011291336.3;refGene_gn=TPD52;refGene_tx
=XM_019822803.2;refGene_gn=TPD52;refGene_tx=XM_019822804.2;refGene_gn=
TPD52;refGene_tx=XM_019822805.2;refGene_gn=TPD52;refGene_tx=XM_0198228
07.2
F2	30863461	30863585	refGene_gn=TPD52;refGene_tx=XM_019822807.2

F2 30933714 30933820
refGene_gn=TPD52;refGene_tx=XM_019822803.2;refGene_gn=TPD52;refGene_tx
=XM_019822804.2;refGene_gn=TPD52;refGene_tx=XM_019822805.2
F2 30935981 30936000 refGene_gn=TPD52;refGene_tx=XM_011291336.3
F2 31176940 31177036 refGene_gn=ZBTB10;refGene_tx=XM_011291335.3
F2 31177967 31178951 refGene_gn=ZBTB10;refGene_tx=XM_006943284.4
F2 31187939 31188825
refGene_gn=ZBTB10;refGene_tx=XM_006943284.4;refGene_gn=ZBTB10;refGene_
tx=XM_011291335.3
F2 31204677 31204776
refGene_gn=ZBTB10;refGene_tx=XM_006943284.4;refGene_gn=ZBTB10;refGene_
tx=XM_011291335.3
F2 31208877 31209054
refGene_gn=ZBTB10;refGene_tx=XM_006943284.4;refGene_gn=ZBTB10;refGene_
tx=XM_011291335.3
F2 31209281 31209454
refGene_gn=ZBTB10;refGene_tx=XM_006943284.4;refGene_gn=ZBTB10;refGene_
tx=XM_011291335.3
F2 31209718 31210024
refGene_gn=ZBTB10;refGene_tx=XM_006943284.4;refGene_gn=ZBTB10;refGene_
tx=XM_011291335.3
F2 31340334 31340446 refGene_gn=ZNF704;refGene_tx=XM_023248442.1
F2 31341438 31341536 refGene_gn=ZNF704;refGene_tx=XM_023248442.1
F2 31350737 31350842 refGene_gn=ZNF704;refGene_tx=XM_023248442.1
F2 31353929 31354197 refGene_gn=ZNF704;refGene_tx=XM_023248442.1
F2 31358451 31358552 refGene_gn=ZNF704;refGene_tx=XM_023248442.1
F2 31372779 31373027 refGene_gn=ZNF704;refGene_tx=XM_023248442.1
F2 31377503 31377607 refGene_gn=ZNF704;refGene_tx=XM_023248442.1
F2 31521406 31521648 refGene_gn=ZNF704;refGene_tx=XM_023248442.1
F2 31575472 31575985 refGene_gn=ZNF704;refGene_tx=XM_023248442.1
F2 31672018 31672387 refGene_gn=PAG1;refGene_tx=XM_003999919.5
F2 31676823 31676883 refGene_gn=PAG1;refGene_tx=XM_003999919.5
F2 31680610 31681212 refGene_gn=PAG1;refGene_tx=XM_003999919.5
F2 31683515 31683612 refGene_gn=PAG1;refGene_tx=XM_003999919.5
F2 31688119 31688171 refGene_gn=PAG1;refGene_tx=XM_003999919.5
F2 31689964 31690089 refGene_gn=PAG1;refGene_tx=XM_003999919.5
F2 31951220 31951299 refGene_gn=FABP5;refGene_tx=XM_003999920.5
F2 31953758 31953931 refGene_gn=FABP5;refGene_tx=XM_003999920.5
F2 31954328 31954430 refGene_gn=FABP5;refGene_tx=XM_003999920.5
F2 31954930 31954984 refGene_gn=FABP5;refGene_tx=XM_003999920.5
F2 32090502 32090553
refGene_gn=LOC101086947;refGene_tx=XM_003999921.4
F2 32091577 32091679
refGene_gn=LOC101086947;refGene_tx=XM_003999921.4
F2 32092012 32092185
refGene_gn=LOC101086947;refGene_tx=XM_003999921.4
F2 32094624 32094724
refGene_gn=LOC101086947;refGene_tx=XM_003999921.4
F2 32101048 32101054
refGene_gn=LOC101086947;refGene_tx=XM_003999921.4

F2 32105300 32105351
refGene_gn=LOC101087200;refGene_tx=XM_003999922.5
F2 32105509 32105611
refGene_gn=LOC101087200;refGene_tx=XM_003999922.5
F2 32106083 32106256
refGene_gn=LOC101087200;refGene_tx=XM_003999922.5
F2 32108523 32108596
refGene_gn=LOC101087200;refGene_tx=XM_003999922.5
F2 32121283 32121334
refGene_gn=LOC101087449;refGene_tx=XM_003999923.5
F2 32121788 32121890
refGene_gn=LOC101087449;refGene_tx=XM_003999923.5
F2 32122671 32122844
refGene_gn=LOC101087449;refGene_tx=XM_003999923.5
F2 32125270 32125343
refGene_gn=LOC101087449;refGene_tx=XM_003999923.5
F2 32169796 32169871
refGene_gn=FABP12;refGene_tx=XM_006943286.3;refGene_gn=FABP12;refGene_tx=XM_019822813.2;refGene_gn=FABP12;refGene_tx=XM_019822820.2;refGene_gn=FABP12;refGene_tx=XM_019822821.2;refGene_gn=FABP12;refGene_tx=XM_019822822.2;refGene_gn=FABP12;refGene_tx=XM_019822823.2
F2 32171313 32171346 refGene_gn=FABP12;refGene_tx=XM_019822829.2
F2 32171416 32171578
refGene_gn=FABP12;refGene_tx=XM_006943286.3;refGene_gn=FABP12;refGene_tx=XM_019822813.2;refGene_gn=FABP12;refGene_tx=XM_019822820.2;refGene_gn=FABP12;refGene_tx=XM_019822821.2;refGene_gn=FABP12;refGene_tx=XM_019822822.2;refGene_gn=FABP12;refGene_tx=XM_019822823.2;refGene_gn=FABP12;refGene_tx=XM_019822829.2;refGene_gn=FABP12;refGene_tx=XM_023248671.1
F2 32172411 32172584
refGene_gn=FABP12;refGene_tx=XM_006943286.3;refGene_gn=FABP12;refGene_tx=XM_019822813.2;refGene_gn=FABP12;refGene_tx=XM_019822820.2;refGene_gn=FABP12;refGene_tx=XM_019822821.2;refGene_gn=FABP12;refGene_tx=XM_019822822.2;refGene_gn=FABP12;refGene_tx=XM_019822823.2;refGene_gn=FABP12;refGene_tx=XM_019822829.2;refGene_gn=FABP12;refGene_tx=XM_023248671.1
F2 32174327 32174400
refGene_gn=FABP12;refGene_tx=XM_006943286.3;refGene_gn=FABP12;refGene_tx=XM_019822813.2;refGene_gn=FABP12;refGene_tx=XM_019822820.2;refGene_gn=FABP12;refGene_tx=XM_019822821.2;refGene_gn=FABP12;refGene_tx=XM_019822822.2;refGene_gn=FABP12;refGene_tx=XM_019822823.2;refGene_gn=FABP12;refGene_tx=XM_019822829.2;refGene_gn=FABP12;refGene_tx=XM_023248671.1
F2 32253360 32253476
refGene_gn=IMPA1;refGene_tx=XM_003999926.5;refGene_gn=IMPA1;refGene_tx=XM_006943288.4;refGene_gn=IMPA1;refGene_tx=XM_019822809.2
F2 32254363 32254515
refGene_gn=IMPA1;refGene_tx=XM_003999926.5;refGene_gn=IMPA1;refGene_tx=XM_006943288.4;refGene_gn=IMPA1;refGene_tx=XM_019822809.2
F2 32274266 32274375

refGene_gn=IMPA1;refGene_tx=XM_003999926.5;refGene_gn=IMPA1;refGene_tx=XM_006943288.4;refGene_gn=IMPA1;refGene_tx=XM_019822809.2
F2 32275707 32275816
refGene_gn=IMPA1;refGene_tx=XM_003999926.5;refGene_gn=IMPA1;refGene_tx=XM_006943288.4;refGene_gn=IMPA1;refGene_tx=XM_019822809.2
F2 32278050 32278096
refGene_gn=IMPA1;refGene_tx=XM_003999926.5;refGene_gn=IMPA1;refGene_tx=XM_006943288.4;refGene_gn=IMPA1;refGene_tx=XM_019822809.2
F2 32279869 32279974
refGene_gn=IMPA1;refGene_tx=XM_003999926.5;refGene_gn=IMPA1;refGene_tx=XM_006943288.4;refGene_gn=IMPA1;refGene_tx=XM_019822809.2
F2 32281968 32282102
refGene_gn=IMPA1;refGene_tx=XM_003999926.5;refGene_gn=IMPA1;refGene_tx=XM_006943288.4;refGene_gn=IMPA1;refGene_tx=XM_019822809.2
F2 32282479 32282542
refGene_gn=IMPA1;refGene_tx=XM_003999926.5;refGene_gn=IMPA1;refGene_tx=XM_006943288.4;refGene_gn=IMPA1;refGene_tx=XM_019822809.2
F2 32291421 32291611 refGene_gn=SLC10A5;refGene_tx=XM_003999936.5
F2 32291612 32291766 refGene_gn=SLC10A5;refGene_tx=XM_003999936.5
F2 32291768 32291821 refGene_gn=SLC10A5;refGene_tx=XM_003999936.5
F2 32291822 32291953 refGene_gn=SLC10A5;refGene_tx=XM_003999936.5
F2 32291955 32292123 refGene_gn=SLC10A5;refGene_tx=XM_003999936.5
F2 32292124 32292290 refGene_gn=SLC10A5;refGene_tx=XM_003999936.5
F2 32292291 32292460 refGene_gn=SLC10A5;refGene_tx=XM_003999936.5
F2 32292462 32292543 refGene_gn=SLC10A5;refGene_tx=XM_003999936.5
F2 32292545 32292699 refGene_gn=SLC10A5;refGene_tx=XM_003999936.5
F2 32294696 32294741 refGene_gn=SLC10A5;refGene_tx=XM_003999936.5
F2 32295156 32295327
refGene_gn=ZFAND1;refGene_tx=XM_019822831.2;refGene_gn=ZFAND1;refGene_tx=XM_019822832.2
F2 32295420 32295576
refGene_gn=ZFAND1;refGene_tx=XM_019822831.2;refGene_gn=ZFAND1;refGene_tx=XM_019822832.2
F2 32306061 32306183
refGene_gn=ZFAND1;refGene_tx=XM_019822831.2;refGene_gn=ZFAND1;refGene_tx=XM_019822832.2
F2 32307377 32307469
refGene_gn=ZFAND1;refGene_tx=XM_019822831.2;refGene_gn=ZFAND1;refGene_tx=XM_019822832.2
F2 32307554 32307682
refGene_gn=ZFAND1;refGene_tx=XM_019822831.2;refGene_gn=ZFAND1;refGene_tx=XM_019822832.2
F2 32309914 32309954
refGene_gn=ZFAND1;refGene_tx=XM_019822831.2;refGene_gn=ZFAND1;refGene_tx=XM_019822832.2
F2 32310854 32310897
refGene_gn=ZFAND1;refGene_tx=XM_019822831.2;refGene_gn=ZFAND1;refGene_tx=XM_019822832.2
F2 32318372 32318427
refGene_gn=ZFAND1;refGene_tx=XM_019822831.2;refGene_gn=ZFAND1;refGene_

tx=XM_019822832.2
F2 32329747 32329937 refGene_gn=CHMP4C;refGene_tx=XM_003999930.5
F2 32349283 32349461 refGene_gn=CHMP4C;refGene_tx=XM_003999930.5
F2 32352315 32352430 refGene_gn=CHMP4C;refGene_tx=XM_003999930.5
F2 32355340 32355506 refGene_gn=CHMP4C;refGene_tx=XM_003999930.5
F2 32355693 32355758 refGene_gn=CHMP4C;refGene_tx=XM_003999930.5
F2 32376238 32376335
refGene_gn=SNX16;refGene_tx=XM_019822833.2;refGene_gn=SNX16;refGene_tx=XM_019822834.2;refGene_gn=SNX16;refGene_tx=XM_023248654.1;refGene_gn=SNX16;refGene_tx=XM_023248655.1;refGene_gn=SNX16;refGene_tx=XM_023248656.1
F2 32377032 32377152
refGene_gn=SNX16;refGene_tx=XM_019822833.2;refGene_gn=SNX16;refGene_tx=XM_019822834.2;refGene_gn=SNX16;refGene_tx=XM_023248654.1;refGene_gn=SNX16;refGene_tx=XM_023248655.1;refGene_gn=SNX16;refGene_tx=XM_023248656.1
F2 32377897 32378034
refGene_gn=SNX16;refGene_tx=XM_019822833.2;refGene_gn=SNX16;refGene_tx=XM_019822834.2;refGene_gn=SNX16;refGene_tx=XM_023248654.1;refGene_gn=SNX16;refGene_tx=XM_023248655.1;refGene_gn=SNX16;refGene_tx=XM_023248656.1
F2 32390546 32390640
refGene_gn=SNX16;refGene_tx=XM_019822833.2;refGene_gn=SNX16;refGene_tx=XM_019822834.2;refGene_gn=SNX16;refGene_tx=XM_023248654.1;refGene_gn=SNX16;refGene_tx=XM_023248655.1;refGene_gn=SNX16;refGene_tx=XM_023248656.1
F2 32407435 32407584
refGene_gn=SNX16;refGene_tx=XM_019822833.2;refGene_gn=SNX16;refGene_tx=XM_019822834.2;refGene_gn=SNX16;refGene_tx=XM_023248654.1;refGene_gn=SNX16;refGene_tx=XM_023248655.1;refGene_gn=SNX16;refGene_tx=XM_023248656.1
F2 32410795 32410882
refGene_gn=SNX16;refGene_tx=XM_019822833.2;refGene_gn=SNX16;refGene_tx=XM_019822834.2;refGene_gn=SNX16;refGene_tx=XM_023248654.1;refGene_gn=SNX16;refGene_tx=XM_023248655.1;refGene_gn=SNX16;refGene_tx=XM_023248656.1
F2 32419279 32419654
refGene_gn=SNX16;refGene_tx=XM_019822833.2;refGene_gn=SNX16;refGene_tx=XM_019822834.2;refGene_gn=SNX16;refGene_tx=XM_023248654.1;refGene_gn=SNX16;refGene_tx=XM_023248655.1;refGene_gn=SNX16;refGene_tx=XM_023248656.1
F2 34721214 34721470
refGene_gn=RALYL;refGene_tx=XM_011291329.3;refGene_gn=RALYL;refGene_tx=XM_019822839.2;refGene_gn=RALYL;refGene_tx=XM_019822840.2;refGene_gn=RALYL;refGene_tx=XM_019822841.2;refGene_gn=RALYL;refGene_tx=XM_019822842.2;refGene_gn=RALYL;refGene_tx=XM_019822844.2;refGene_gn=RALYL;refGene_tx=XM_023248459.1;refGene_gn=RALYL;refGene_tx=XM_023248460.1
F2 34896948 34896985 refGene_gn=RALYL;refGene_tx=XM_006943292.4
F2 34956883 34956959
refGene_gn=RALYL;refGene_tx=XM_006943292.4;refGene_gn=RALYL;refGene_tx

=XM_011291329.3;refGene_gn=RALYL;refGene_tx=XM_019822839.2;refGene_gn=RALYL;refGene_tx=XM_019822840.2;refGene_gn=RALYL;refGene_tx=XM_019822841.2;refGene_gn=RALYL;refGene_tx=XM_019822842.2;refGene_gn=RALYL;refGene_tx=XM_019822844.2;refGene_gn=RALYL;refGene_tx=XM_023248459.1;refGene_gn=RALYL;refGene_tx=XM_023248460.1
F2 34983627 34983660
refGene_gn=RALYL;refGene_tx=XM_006943292.4;refGene_gn=RALYL;refGene_tx=XM_011291329.3;refGene_gn=RALYL;refGene_tx=XM_019822839.2;refGene_gn=RALYL;refGene_tx=XM_019822840.2;refGene_gn=RALYL;refGene_tx=XM_019822841.2;refGene_gn=RALYL;refGene_tx=XM_019822842.2;refGene_gn=RALYL;refGene_tx=XM_019822844.2;refGene_gn=RALYL;refGene_tx=XM_023248459.1;refGene_gn=RALYL;refGene_tx=XM_023248460.1
F2 35013407 35013455
refGene_gn=RALYL;refGene_tx=XM_006943292.4;refGene_gn=RALYL;refGene_tx=XM_011291329.3;refGene_gn=RALYL;refGene_tx=XM_019822839.2;refGene_gn=RALYL;refGene_tx=XM_019822840.2;refGene_gn=RALYL;refGene_tx=XM_019822841.2;refGene_gn=RALYL;refGene_tx=XM_019822842.2;refGene_gn=RALYL;refGene_tx=XM_019822844.2;refGene_gn=RALYL;refGene_tx=XM_023248459.1;refGene_gn=RALYL;refGene_tx=XM_023248460.1
F2 35024551 35024715
refGene_gn=RALYL;refGene_tx=XM_006943292.4;refGene_gn=RALYL;refGene_tx=XM_011291329.3;refGene_gn=RALYL;refGene_tx=XM_019822839.2;refGene_gn=RALYL;refGene_tx=XM_019822840.2;refGene_gn=RALYL;refGene_tx=XM_019822841.2;refGene_gn=RALYL;refGene_tx=XM_019822842.2;refGene_gn=RALYL;refGene_tx=XM_019822844.2;refGene_gn=RALYL;refGene_tx=XM_023248459.1;refGene_gn=RALYL;refGene_tx=XM_023248460.1
F2 35057051 35057165
refGene_gn=RALYL;refGene_tx=XM_006943292.4;refGene_gn=RALYL;refGene_tx=XM_011291329.3;refGene_gn=RALYL;refGene_tx=XM_019822839.2;refGene_gn=RALYL;refGene_tx=XM_019822840.2;refGene_gn=RALYL;refGene_tx=XM_019822841.2;refGene_gn=RALYL;refGene_tx=XM_019822842.2;refGene_gn=RALYL;refGene_tx=XM_019822844.2;refGene_gn=RALYL;refGene_tx=XM_023248459.1;refGene_gn=RALYL;refGene_tx=XM_023248460.1
F2 35072980 35073153
refGene_gn=RALYL;refGene_tx=XM_006943292.4;refGene_gn=RALYL;refGene_tx=XM_011291329.3;refGene_gn=RALYL;refGene_tx=XM_019822839.2;refGene_gn=RALYL;refGene_tx=XM_019822840.2;refGene_gn=RALYL;refGene_tx=XM_019822841.2;refGene_gn=RALYL;refGene_tx=XM_019822842.2;refGene_gn=RALYL;refGene_tx=XM_023248459.1;refGene_gn=RALYL;refGene_tx=XM_023248460.1
F2 35109964 35109987
refGene_gn=RALYL;refGene_tx=XM_006943292.4;refGene_gn=RALYL;refGene_tx=XM_011291329.3;refGene_gn=RALYL;refGene_tx=XM_019822839.2;refGene_gn=RALYL;refGene_tx=XM_019822840.2;refGene_gn=RALYL;refGene_tx=XM_019822841.2;refGene_gn=RALYL;refGene_tx=XM_019822842.2;refGene_gn=RALYL;refGene_tx=XM_019822844.2;refGene_gn=RALYL;refGene_tx=XM_023248459.1;refGene_gn=RALYL;refGene_tx=XM_023248460.1
F2 35247576 35248243
refGene_gn=LOC101096635;refGene_tx=XM_019822838.1
F2 35252907 35253039
refGene_gn=LOC101096635;refGene_tx=XM_019822838.1

F2 35258053 35258415
refGene_gn=LOC101096635;refGene_tx=XM_019822838.1
F2 35261465 35261741
refGene_gn=LOC101096635;refGene_tx=XM_019822838.1
F2 35281901 35281990
refGene_gn=LRRCC1;refGene_tx=XM_011291327.3;refGene_gn=LRRCC1;refGene_tx=XM_011291328.3;refGene_gn=LRRCC1;refGene_tx=XM_023248234.1
F2 35284320 35284526
refGene_gn=LRRCC1;refGene_tx=XM_011291327.3;refGene_gn=LRRCC1;refGene_tx=XM_011291328.3;refGene_gn=LRRCC1;refGene_tx=XM_019822837.1;refGene_gn=LRRCC1;refGene_tx=XM_023248234.1
F2 35284821 35284887
refGene_gn=LRRCC1;refGene_tx=XM_011291327.3;refGene_gn=LRRCC1;refGene_tx=XM_011291328.3;refGene_gn=LRRCC1;refGene_tx=XM_019822837.1;refGene_gn=LRRCC1;refGene_tx=XM_023248234.1
F2 35289273 35289441
refGene_gn=LRRCC1;refGene_tx=XM_011291327.3;refGene_gn=LRRCC1;refGene_tx=XM_011291328.3;refGene_gn=LRRCC1;refGene_tx=XM_019822837.1;refGene_gn=LRRCC1;refGene_tx=XM_023248234.1
F2 35292014 35292190
refGene_gn=LRRCC1;refGene_tx=XM_011291327.3;refGene_gn=LRRCC1;refGene_tx=XM_011291328.3;refGene_gn=LRRCC1;refGene_tx=XM_019822837.1;refGene_gn=LRRCC1;refGene_tx=XM_023248234.1
F2 35292288 35292495
refGene_gn=LRRCC1;refGene_tx=XM_011291327.3;refGene_gn=LRRCC1;refGene_tx=XM_019822837.1;refGene_gn=LRRCC1;refGene_tx=XM_023248234.1
F2 35306556 35306750
refGene_gn=LRRCC1;refGene_tx=XM_011291327.3;refGene_gn=LRRCC1;refGene_tx=XM_011291328.3;refGene_gn=LRRCC1;refGene_tx=XM_019822837.1;refGene_gn=LRRCC1;refGene_tx=XM_023248234.1;refGene_gn=LRRCC1;refGene_tx=XM_023248235.1
F2 35307021 35307169
refGene_gn=LRRCC1;refGene_tx=XM_011291327.3;refGene_gn=LRRCC1;refGene_tx=XM_011291328.3;refGene_gn=LRRCC1;refGene_tx=XM_019822837.1;refGene_gn=LRRCC1;refGene_tx=XM_023248234.1;refGene_gn=LRRCC1;refGene_tx=XM_023248235.1
F2 35308218 35308367
refGene_gn=LRRCC1;refGene_tx=XM_011291327.3;refGene_gn=LRRCC1;refGene_tx=XM_011291328.3;refGene_gn=LRRCC1;refGene_tx=XM_019822837.1;refGene_gn=LRRCC1;refGene_tx=XM_023248234.1;refGene_gn=LRRCC1;refGene_tx=XM_023248235.1
F2 35312002 35312207
refGene_gn=LRRCC1;refGene_tx=XM_011291327.3;refGene_gn=LRRCC1;refGene_tx=XM_011291328.3;refGene_gn=LRRCC1;refGene_tx=XM_019822837.1;refGene_gn=LRRCC1;refGene_tx=XM_023248234.1;refGene_gn=LRRCC1;refGene_tx=XM_023248235.1
F2 35313054 35313194
refGene_gn=LRRCC1;refGene_tx=XM_011291327.3;refGene_gn=LRRCC1;refGene_tx=XM_011291328.3;refGene_gn=LRRCC1;refGene_tx=XM_019822837.1;refGene_gn=LRRCC1;refGene_tx=XM_023248234.1;refGene_gn=LRRCC1;refGene_tx=XM_02

3248235.1
F2	35315068	35315270
refGene_gn=LRRCC1;refGene_tx=XM_011291327.3;refGene_gn=LRRCC1;refGene_tx=XM_011291328.3;refGene_gn=LRRCC1;refGene_tx=XM_019822837.1;refGene_gn=LRRCC1;refGene_tx=XM_023248234.1;refGene_gn=LRRCC1;refGene_tx=XM_023248235.1
F2	35315662	35315848
refGene_gn=LRRCC1;refGene_tx=XM_011291327.3;refGene_gn=LRRCC1;refGene_tx=XM_011291328.3;refGene_gn=LRRCC1;refGene_tx=XM_019822837.1;refGene_gn=LRRCC1;refGene_tx=XM_023248234.1;refGene_gn=LRRCC1;refGene_tx=XM_023248235.1
F2	35316586	35316761
refGene_gn=LRRCC1;refGene_tx=XM_011291327.3;refGene_gn=LRRCC1;refGene_tx=XM_011291328.3;refGene_gn=LRRCC1;refGene_tx=XM_019822837.1;refGene_gn=LRRCC1;refGene_tx=XM_023248234.1;refGene_gn=LRRCC1;refGene_tx=XM_023248235.1
F2	35317640	35317804
refGene_gn=LRRCC1;refGene_tx=XM_011291327.3;refGene_gn=LRRCC1;refGene_tx=XM_011291328.3;refGene_gn=LRRCC1;refGene_tx=XM_019822837.1;refGene_gn=LRRCC1;refGene_tx=XM_023248234.1;refGene_gn=LRRCC1;refGene_tx=XM_023248235.1
F2	35320181	35320208	refGene_gn=LRRCC1;refGene_tx=XM_023248234.1
F2	35321438	35321647
refGene_gn=LRRCC1;refGene_tx=XM_011291327.3;refGene_gn=LRRCC1;refGene_tx=XM_011291328.3;refGene_gn=LRRCC1;refGene_tx=XM_019822837.1;refGene_gn=LRRCC1;refGene_tx=XM_023248235.1
F2	35321740	35321878
refGene_gn=LRRCC1;refGene_tx=XM_011291327.3;refGene_gn=LRRCC1;refGene_tx=XM_011291328.3;refGene_gn=LRRCC1;refGene_tx=XM_019822837.1;refGene_gn=LRRCC1;refGene_tx=XM_023248235.1
F2	35325566	35325702
refGene_gn=LRRCC1;refGene_tx=XM_011291327.3;refGene_gn=LRRCC1;refGene_tx=XM_011291328.3;refGene_gn=LRRCC1;refGene_tx=XM_019822837.1;refGene_gn=LRRCC1;refGene_tx=XM_023248235.1
F2	35328231	35328354
refGene_gn=LRRCC1;refGene_tx=XM_011291327.3;refGene_gn=LRRCC1;refGene_tx=XM_011291328.3;refGene_gn=LRRCC1;refGene_tx=XM_019822837.1;refGene_gn=LRRCC1;refGene_tx=XM_023248235.1
F2	35341755	35341989
refGene_gn=E2F5;refGene_tx=XM_023248785.1;refGene_gn=E2F5;refGene_tx=XM_023248786.1
F2	35359499	35359609
refGene_gn=E2F5;refGene_tx=XM_023248785.1;refGene_gn=E2F5;refGene_tx=XM_023248786.1
F2	35360759	35360921
refGene_gn=E2F5;refGene_tx=XM_023248785.1;refGene_gn=E2F5;refGene_tx=XM_023248786.1
F2	35362213	35362257
refGene_gn=E2F5;refGene_tx=XM_023248785.1;refGene_gn=E2F5;refGene_tx=XM_023248786.1

F2 35363504 35363569
refGene_gn=E2F5;refGene_tx=XM_023248785.1;refGene_gn=E2F5;refGene_tx=X
M_023248786.1
F2 35364136 35364404
refGene_gn=E2F5;refGene_tx=XM_023248785.1;refGene_gn=E2F5;refGene_tx=X
M_023248786.1
F2 35366972 35367020
refGene_gn=E2F5;refGene_tx=XM_023248785.1;refGene_gn=E2F5;refGene_tx=X
M_023248786.1
F2 35368265 35368375
refGene_gn=E2F5;refGene_tx=XM_023248785.1;refGene_gn=E2F5;refGene_tx=X
M_023248786.1
F2 35369106 35369178
refGene_gn=CF2H8orf59;refGene_tx=XM_003999941.4
F2 35369473 35369590
refGene_gn=CF2H8orf59;refGene_tx=XM_003999941.4
F2 35372879 35372996
refGene_gn=CF2H8orf59;refGene_tx=XM_003999941.4
F2 35375118 35375229
refGene_gn=CF2H8orf59;refGene_tx=XM_003999941.4
F2 35375423 35375541 refGene_gn=CA13;refGene_tx=XM_019822845.2
F2 35390787 35390824 refGene_gn=CA13;refGene_tx=XM_003999943.3
F2 35395147 35395345
refGene_gn=CA13;refGene_tx=XM_003999943.3;refGene_gn=CA13;refGene_tx=X
M_019822845.2
F2 35403448 35403567
refGene_gn=CA13;refGene_tx=XM_003999943.3;refGene_gn=CA13;refGene_tx=X
M_019822845.2
F2 35410949 35411045
refGene_gn=CA13;refGene_tx=XM_003999943.3;refGene_gn=CA13;refGene_tx=X
M_019822845.2
F2 35412351 35412414
refGene_gn=CA13;refGene_tx=XM_003999943.3;refGene_gn=CA13;refGene_tx=X
M_019822845.2
F2 35412911 35413067
refGene_gn=CA13;refGene_tx=XM_003999943.3;refGene_gn=CA13;refGene_tx=X
M_019822845.2
F2 35423310 35423430
refGene_gn=CA13;refGene_tx=XM_003999943.3;refGene_gn=CA13;refGene_tx=X
M_019822845.2
F2 35477362 35477479 refGene_gn=CA1;refGene_tx=XM_003999944.4
F2 35479067 35479223 refGene_gn=CA1;refGene_tx=XM_003999944.4
F2 35480558 35480621 refGene_gn=CA1;refGene_tx=XM_003999944.4
F2 35483193 35483289 refGene_gn=CA1;refGene_tx=XM_003999944.4
F2 35486008 35486127 refGene_gn=CA1;refGene_tx=XM_003999944.4
F2 35487525 35487723 refGene_gn=CA1;refGene_tx=XM_003999944.4
F2 35488230 35488267 refGene_gn=CA1;refGene_tx=XM_003999944.4
F2 35558637 35558671 refGene_gn=CA3;refGene_tx=XM_003999945.4
F2 35559453 35559651 refGene_gn=CA3;refGene_tx=XM_003999945.4
F2 35562047 35562166 refGene_gn=CA3;refGene_tx=XM_003999945.4

F2 35564229 35564322 refGene_gn=CA3;refGene_tx=XM_003999945.4
F2 35565361 35565424 refGene_gn=CA3;refGene_tx=XM_003999945.4
F2 35566745 35566901 refGene_gn=CA3;refGene_tx=XM_003999945.4
F2 35568283 35568403 refGene_gn=CA3;refGene_tx=XM_003999945.4
F2 35581968 35582002 refGene_gn=CA2;refGene_tx=XM_003999946.5
F2 35583137 35583335 refGene_gn=CA2;refGene_tx=XM_003999946.5
F2 35592769 35592888 refGene_gn=CA2;refGene_tx=XM_003999946.5
F2 35593445 35593538 refGene_gn=CA2;refGene_tx=XM_003999946.5
F2 35594610 35594673 refGene_gn=CA2;refGene_tx=XM_003999946.5
F2 35596906 35597062 refGene_gn=CA2;refGene_tx=XM_003999946.5
F2 35598863 35598983 refGene_gn=CA2;refGene_tx=XM_003999946.5
F2 35926250 35926556 refGene_gn=PSKH2;refGene_tx=XM_019822622.2
F2 35939328 35939995 refGene_gn=PSKH2;refGene_tx=XM_019822622.2
F2 35944555 35944740 refGene_gn=PSKH2;refGene_tx=XM_019822622.2
F2 35971453 35971583
refGene_gn=ATP6V0D2;refGene_tx=XM_004001615.4;refGene_gn=ATP6V0D2;refGene_tx=XM_023248445.1;refGene_gn=ATP6V0D2;refGene_tx=XM_023248446.1
F2 35983614 35983786
refGene_gn=ATP6V0D2;refGene_tx=XM_004001615.4;refGene_gn=ATP6V0D2;refGene_tx=XM_023248445.1;refGene_gn=ATP6V0D2;refGene_tx=XM_023248446.1
F2 36001555 36001734
refGene_gn=ATP6V0D2;refGene_tx=XM_004001615.4;refGene_gn=ATP6V0D2;refGene_tx=XM_023248445.1;refGene_gn=ATP6V0D2;refGene_tx=XM_023248446.1
F2 36003169 36003249
refGene_gn=ATP6V0D2;refGene_tx=XM_004001615.4;refGene_gn=ATP6V0D2;refGene_tx=XM_023248445.1;refGene_gn=ATP6V0D2;refGene_tx=XM_023248446.1
F2 36005002 36005080
refGene_gn=ATP6V0D2;refGene_tx=XM_004001615.4;refGene_gn=ATP6V0D2;refGene_tx=XM_023248445.1;refGene_gn=ATP6V0D2;refGene_tx=XM_023248446.1
F2 36011206 36011383
refGene_gn=ATP6V0D2;refGene_tx=XM_004001615.4;refGene_gn=ATP6V0D2;refGene_tx=XM_023248445.1;refGene_gn=ATP6V0D2;refGene_tx=XM_023248446.1
F2 36012532 36012610
refGene_gn=ATP6V0D2;refGene_tx=XM_004001615.4;refGene_gn=ATP6V0D2;refGene_tx=XM_023248445.1;refGene_gn=ATP6V0D2;refGene_tx=XM_023248446.1
F2 36014323 36014485
refGene_gn=ATP6V0D2;refGene_tx=XM_004001615.4;refGene_gn=ATP6V0D2;refGene_tx=XM_023248445.1;refGene_gn=ATP6V0D2;refGene_tx=XM_023248446.1
F2 36065307 36065435 refGene_gn=SLC7A13;refGene_tx=XM_006943293.4
F2 36065437 36065564 refGene_gn=SLC7A13;refGene_tx=XM_006943293.4
F2 36067874 36068236 refGene_gn=SLC7A13;refGene_tx=XM_006943293.4
F2 36072283 36072409 refGene_gn=SLC7A13;refGene_tx=XM_006943293.4
F2 36076730 36077412 refGene_gn=SLC7A13;refGene_tx=XM_006943293.4
F2 36141071 36141536
refGene_gn=LOC101097024;refGene_tx=XM_011291324.3
F2 36181616 36181692
refGene_gn=WWP1;refGene_tx=XM_011291318.2;refGene_gn=WWP1;refGene_tx=XM_011291319.2;refGene_gn=WWP1;refGene_tx=XM_019822850.2
F2 36191268 36191338
refGene_gn=WWP1;refGene_tx=XM_011291322.3;refGene_gn=WWP1;refGene_tx=X

M_011291323.3;refGene_gn=WWP1;refGene_tx=XM_019822849.2
F2 36202560 36202699
refGene_gn=WWP1;refGene_tx=XM_011291318.2;refGene_gn=WWP1;refGene_tx=XM_011291319.2;refGene_gn=WWP1;refGene_tx=XM_011291322.3;refGene_gn=WWP1;refGene_tx=XM_011291323.3;refGene_gn=WWP1;refGene_tx=XM_019822849.2;refGene_gn=WWP1;refGene_tx=XM_019822850.2
F2 36203322 36203447
refGene_gn=WWP1;refGene_tx=XM_011291318.2;refGene_gn=WWP1;refGene_tx=XM_011291319.2;refGene_gn=WWP1;refGene_tx=XM_011291322.3;refGene_gn=WWP1;refGene_tx=XM_011291323.3;refGene_gn=WWP1;refGene_tx=XM_019822849.2;refGene_gn=WWP1;refGene_tx=XM_019822850.2
F2 36217657 36217795
refGene_gn=WWP1;refGene_tx=XM_011291318.2;refGene_gn=WWP1;refGene_tx=XM_011291319.2;refGene_gn=WWP1;refGene_tx=XM_011291322.3;refGene_gn=WWP1;refGene_tx=XM_011291323.3;refGene_gn=WWP1;refGene_tx=XM_019822849.2;refGene_gn=WWP1;refGene_tx=XM_019822850.2
F2 36217886 36217953
refGene_gn=WWP1;refGene_tx=XM_011291318.2;refGene_gn=WWP1;refGene_tx=XM_011291319.2;refGene_gn=WWP1;refGene_tx=XM_011291322.3;refGene_gn=WWP1;refGene_tx=XM_011291323.3;refGene_gn=WWP1;refGene_tx=XM_019822849.2;refGene_gn=WWP1;refGene_tx=XM_019822850.2
F2 36223175 36223360
refGene_gn=WWP1;refGene_tx=XM_011291318.2;refGene_gn=WWP1;refGene_tx=XM_011291319.2;refGene_gn=WWP1;refGene_tx=XM_011291322.3;refGene_gn=WWP1;refGene_tx=XM_011291323.3;refGene_gn=WWP1;refGene_tx=XM_019822849.2;refGene_gn=WWP1;refGene_tx=XM_019822850.2
F2 36235335 36235345 refGene_gn=WWP1;refGene_tx=XM_019822851.2
F2 36245134 36245471
refGene_gn=WWP1;refGene_tx=XM_011291318.2;refGene_gn=WWP1;refGene_tx=XM_011291319.2;refGene_gn=WWP1;refGene_tx=XM_011291322.3;refGene_gn=WWP1;refGene_tx=XM_011291323.3;refGene_gn=WWP1;refGene_tx=XM_019822849.2;refGene_gn=WWP1;refGene_tx=XM_019822850.2;refGene_gn=WWP1;refGene_tx=XM_019822851.2
F2 36255022 36255118
refGene_gn=WWP1;refGene_tx=XM_011291318.2;refGene_gn=WWP1;refGene_tx=XM_011291319.2;refGene_gn=WWP1;refGene_tx=XM_011291322.3;refGene_gn=WWP1;refGene_tx=XM_011291323.3;refGene_gn=WWP1;refGene_tx=XM_019822849.2;refGene_gn=WWP1;refGene_tx=XM_019822850.2;refGene_gn=WWP1;refGene_tx=XM_019822851.2
F2 36256957 36257132
refGene_gn=WWP1;refGene_tx=XM_011291318.2;refGene_gn=WWP1;refGene_tx=XM_011291319.2;refGene_gn=WWP1;refGene_tx=XM_011291322.3;refGene_gn=WWP1;refGene_tx=XM_011291323.3;refGene_gn=WWP1;refGene_tx=XM_019822849.2;refGene_gn=WWP1;refGene_tx=XM_019822850.2;refGene_gn=WWP1;refGene_tx=XM_019822851.2
F2 36259388 36259443
refGene_gn=WWP1;refGene_tx=XM_011291318.2;refGene_gn=WWP1;refGene_tx=XM_011291319.2;refGene_gn=WWP1;refGene_tx=XM_011291322.3;refGene_gn=WWP1;refGene_tx=XM_011291323.3;refGene_gn=WWP1;refGene_tx=XM_019822849.2;refGene_gn=WWP1;refGene_tx=XM_019822850.2;refGene_gn=WWP1;refGene_tx=X

M_019822851.2
F2 36259527 36259612
refGene_gn=WWP1;refGene_tx=XM_011291318.2;refGene_gn=WWP1;refGene_tx=XM_011291319.2;refGene_gn=WWP1;refGene_tx=XM_011291322.3;refGene_gn=WWP1;refGene_tx=XM_011291323.3;refGene_gn=WWP1;refGene_tx=XM_019822849.2;refGene_gn=WWP1;refGene_tx=XM_019822850.2;refGene_gn=WWP1;refGene_tx=XM_019822851.2
F2 36259741 36259870
refGene_gn=WWP1;refGene_tx=XM_011291318.2;refGene_gn=WWP1;refGene_tx=XM_011291319.2;refGene_gn=WWP1;refGene_tx=XM_011291322.3;refGene_gn=WWP1;refGene_tx=XM_011291323.3;refGene_gn=WWP1;refGene_tx=XM_019822849.2;refGene_gn=WWP1;refGene_tx=XM_019822850.2;refGene_gn=WWP1;refGene_tx=XM_019822851.2
F2 36262715 36262791
refGene_gn=WWP1;refGene_tx=XM_011291318.2;refGene_gn=WWP1;refGene_tx=XM_011291319.2;refGene_gn=WWP1;refGene_tx=XM_011291322.3;refGene_gn=WWP1;refGene_tx=XM_011291323.3;refGene_gn=WWP1;refGene_tx=XM_019822849.2;refGene_gn=WWP1;refGene_tx=XM_019822850.2;refGene_gn=WWP1;refGene_tx=XM_019822851.2
F2 36262898 36262970
refGene_gn=WWP1;refGene_tx=XM_011291318.2;refGene_gn=WWP1;refGene_tx=XM_011291319.2;refGene_gn=WWP1;refGene_tx=XM_011291322.3;refGene_gn=WWP1;refGene_tx=XM_011291323.3;refGene_gn=WWP1;refGene_tx=XM_019822849.2;refGene_gn=WWP1;refGene_tx=XM_019822850.2;refGene_gn=WWP1;refGene_tx=XM_019822851.2
F2 36264870 36264959
refGene_gn=WWP1;refGene_tx=XM_011291318.2;refGene_gn=WWP1;refGene_tx=XM_011291319.2;refGene_gn=WWP1;refGene_tx=XM_011291322.3;refGene_gn=WWP1;refGene_tx=XM_011291323.3;refGene_gn=WWP1;refGene_tx=XM_019822849.2;refGene_gn=WWP1;refGene_tx=XM_019822850.2;refGene_gn=WWP1;refGene_tx=XM_019822851.2
F2 36268420 36268580
refGene_gn=WWP1;refGene_tx=XM_011291318.2;refGene_gn=WWP1;refGene_tx=XM_011291319.2;refGene_gn=WWP1;refGene_tx=XM_011291322.3;refGene_gn=WWP1;refGene_tx=XM_011291323.3;refGene_gn=WWP1;refGene_tx=XM_019822849.2;refGene_gn=WWP1;refGene_tx=XM_019822850.2;refGene_gn=WWP1;refGene_tx=XM_019822851.2
F2 36272103 36272237
refGene_gn=WWP1;refGene_tx=XM_011291318.2;refGene_gn=WWP1;refGene_tx=XM_011291319.2;refGene_gn=WWP1;refGene_tx=XM_011291322.3;refGene_gn=WWP1;refGene_tx=XM_011291323.3;refGene_gn=WWP1;refGene_tx=XM_019822849.2;refGene_gn=WWP1;refGene_tx=XM_019822850.2;refGene_gn=WWP1;refGene_tx=XM_019822851.2
F2 36272325 36272466
refGene_gn=WWP1;refGene_tx=XM_011291318.2;refGene_gn=WWP1;refGene_tx=XM_011291319.2;refGene_gn=WWP1;refGene_tx=XM_011291322.3;refGene_gn=WWP1;refGene_tx=XM_011291323.3;refGene_gn=WWP1;refGene_tx=XM_019822849.2;refGene_gn=WWP1;refGene_tx=XM_019822850.2;refGene_gn=WWP1;refGene_tx=XM_019822851.2
F2 36276783 36276904

refGene_gn=WWP1;refGene_tx=XM_011291318.2;refGene_gn=WWP1;refGene_tx=XM_011291319.2;refGene_gn=WWP1;refGene_tx=XM_011291322.3;refGene_gn=WWP1;refGene_tx=XM_011291323.3;refGene_gn=WWP1;refGene_tx=XM_019822849.2;refGene_gn=WWP1;refGene_tx=XM_019822850.2;refGene_gn=WWP1;refGene_tx=XM_019822851.2
F2 36283239 36283344
refGene_gn=WWP1;refGene_tx=XM_011291318.2;refGene_gn=WWP1;refGene_tx=XM_011291319.2;refGene_gn=WWP1;refGene_tx=XM_011291322.3;refGene_gn=WWP1;refGene_tx=XM_011291323.3;refGene_gn=WWP1;refGene_tx=XM_019822849.2;refGene_gn=WWP1;refGene_tx=XM_019822850.2;refGene_gn=WWP1;refGene_tx=XM_019822851.2
F2 36286187 36286284
refGene_gn=WWP1;refGene_tx=XM_011291318.2;refGene_gn=WWP1;refGene_tx=XM_011291319.2;refGene_gn=WWP1;refGene_tx=XM_011291322.3;refGene_gn=WWP1;refGene_tx=XM_011291323.3;refGene_gn=WWP1;refGene_tx=XM_019822849.2;refGene_gn=WWP1;refGene_tx=XM_019822850.2;refGene_gn=WWP1;refGene_tx=XM_019822851.2
F2 36286717 36286790
refGene_gn=WWP1;refGene_tx=XM_011291318.2;refGene_gn=WWP1;refGene_tx=XM_011291319.2;refGene_gn=WWP1;refGene_tx=XM_011291322.3;refGene_gn=WWP1;refGene_tx=XM_011291323.3;refGene_gn=WWP1;refGene_tx=XM_019822849.2;refGene_gn=WWP1;refGene_tx=XM_019822850.2;refGene_gn=WWP1;refGene_tx=XM_019822851.2
F2 36292300 36292400
refGene_gn=WWP1;refGene_tx=XM_011291318.2;refGene_gn=WWP1;refGene_tx=XM_011291319.2;refGene_gn=WWP1;refGene_tx=XM_011291322.3;refGene_gn=WWP1;refGene_tx=XM_011291323.3;refGene_gn=WWP1;refGene_tx=XM_019822849.2;refGene_gn=WWP1;refGene_tx=XM_019822851.2
F2 36298292 36298317 refGene_gn=WWP1;refGene_tx=XM_019822850.2
F2 36298782 36298833
refGene_gn=RMDN1;refGene_tx=XM_011291316.3;refGene_gn=RMDN1;refGene_tx=XM_011291317.3;refGene_gn=RMDN1;refGene_tx=XM_019822852.2;refGene_gn=RMDN1;refGene_tx=XM_019822853.2;refGene_gn=RMDN1;refGene_tx=XM_023248456.1;refGene_gn=RMDN1;refGene_tx=XM_023248457.1
F2 36299276 36299410
refGene_gn=RMDN1;refGene_tx=XM_011291316.3;refGene_gn=RMDN1;refGene_tx=XM_011291317.3;refGene_gn=RMDN1;refGene_tx=XM_019822852.2;refGene_gn=RMDN1;refGene_tx=XM_019822853.2;refGene_gn=RMDN1;refGene_tx=XM_023248456.1;refGene_gn=RMDN1;refGene_tx=XM_023248457.1
F2 36301044 36301075
refGene_gn=RMDN1;refGene_tx=XM_011291316.3;refGene_gn=RMDN1;refGene_tx=XM_011291317.3;refGene_gn=RMDN1;refGene_tx=XM_019822852.2;refGene_gn=RMDN1;refGene_tx=XM_019822853.2;refGene_gn=RMDN1;refGene_tx=XM_023248456.1;refGene_gn=RMDN1;refGene_tx=XM_023248457.1
F2 36302310 36302398
refGene_gn=RMDN1;refGene_tx=XM_011291316.3;refGene_gn=RMDN1;refGene_tx=XM_011291317.3;refGene_gn=RMDN1;refGene_tx=XM_019822852.2;refGene_gn=RMDN1;refGene_tx=XM_023248456.1;refGene_gn=RMDN1;refGene_tx=XM_023248457.1
F2 36303838 36303894

refGene_gn=RMDN1;refGene_tx=XM_011291316.3;refGene_gn=RMDN1;refGene_tx=XM_011291317.3;refGene_gn=RMDN1;refGene_tx=XM_019822852.2;refGene_gn=RMDN1;refGene_tx=XM_023248456.1;refGene_gn=RMDN1;refGene_tx=XM_023248457.1
F2 36307248 36307338
refGene_gn=RMDN1;refGene_tx=XM_011291316.3;refGene_gn=RMDN1;refGene_tx=XM_011291317.3;refGene_gn=RMDN1;refGene_tx=XM_019822853.2;refGene_gn=RMDN1;refGene_tx=XM_023248456.1
F2 36324760 36324920
refGene_gn=RMDN1;refGene_tx=XM_011291316.3;refGene_gn=RMDN1;refGene_tx=XM_011291317.3;refGene_gn=RMDN1;refGene_tx=XM_019822852.2;refGene_gn=RMDN1;refGene_tx=XM_019822853.2;refGene_gn=RMDN1;refGene_tx=XM_023248456.1;refGene_gn=RMDN1;refGene_tx=XM_023248457.1
F2 36326957 36327054
refGene_gn=RMDN1;refGene_tx=XM_011291316.3;refGene_gn=RMDN1;refGene_tx=XM_011291317.3;refGene_gn=RMDN1;refGene_tx=XM_019822852.2;refGene_gn=RMDN1;refGene_tx=XM_019822853.2;refGene_gn=RMDN1;refGene_tx=XM_023248456.1;refGene_gn=RMDN1;refGene_tx=XM_023248457.1
F2 36334740 36334858
refGene_gn=RMDN1;refGene_tx=XM_011291316.3;refGene_gn=RMDN1;refGene_tx=XM_011291317.3;refGene_gn=RMDN1;refGene_tx=XM_019822852.2;refGene_gn=RMDN1;refGene_tx=XM_019822853.2;refGene_gn=RMDN1;refGene_tx=XM_023248456.1;refGene_gn=RMDN1;refGene_tx=XM_023248457.1
F2 36353114 36353243
refGene_gn=RMDN1;refGene_tx=XM_011291316.3;refGene_gn=RMDN1;refGene_tx=XM_019822852.2;refGene_gn=RMDN1;refGene_tx=XM_019822853.2;refGene_gn=RMDN1;refGene_tx=XM_023248456.1;refGene_gn=RMDN1;refGene_tx=XM_023248457.1
F2 36375113 36375245
refGene_gn=CPNE3;refGene_tx=XM_003999953.5;refGene_gn=CPNE3;refGene_tx=XM_019822854.2;refGene_gn=CPNE3;refGene_tx=XM_019822855.2
F2 36375523 36375703
refGene_gn=CPNE3;refGene_tx=XM_003999953.5;refGene_gn=CPNE3;refGene_tx=XM_019822854.2;refGene_gn=CPNE3;refGene_tx=XM_019822855.2
F2 36377394 36377469
refGene_gn=CPNE3;refGene_tx=XM_003999953.5;refGene_gn=CPNE3;refGene_tx=XM_019822854.2
F2 36378766 36378838
refGene_gn=CPNE3;refGene_tx=XM_003999953.5;refGene_gn=CPNE3;refGene_tx=XM_019822854.2;refGene_gn=CPNE3;refGene_tx=XM_019822855.2
F2 36383889 36383973
refGene_gn=CPNE3;refGene_tx=XM_003999953.5;refGene_gn=CPNE3;refGene_tx=XM_019822854.2;refGene_gn=CPNE3;refGene_tx=XM_019822855.2
F2 36386520 36386610
refGene_gn=CPNE3;refGene_tx=XM_003999953.5;refGene_gn=CPNE3;refGene_tx=XM_019822854.2;refGene_gn=CPNE3;refGene_tx=XM_019822855.2
F2 36391239 36391338
refGene_gn=CPNE3;refGene_tx=XM_003999953.5;refGene_gn=CPNE3;refGene_tx=XM_019822854.2;refGene_gn=CPNE3;refGene_tx=XM_019822855.2
F2 36393233 36393320

refGene_gn=CPNE3;refGene_tx=XM_003999953.5;refGene_gn=CPNE3;refGene_tx=XM_019822854.2;refGene_gn=CPNE3;refGene_tx=XM_019822855.2
F2 36394330 36394390
refGene_gn=CPNE3;refGene_tx=XM_003999953.5;refGene_gn=CPNE3;refGene_tx=XM_019822854.2;refGene_gn=CPNE3;refGene_tx=XM_019822855.2
F2 36394871 36395005
refGene_gn=CPNE3;refGene_tx=XM_003999953.5;refGene_gn=CPNE3;refGene_tx=XM_019822854.2;refGene_gn=CPNE3;refGene_tx=XM_019822855.2
F2 36399259 36399314
refGene_gn=CPNE3;refGene_tx=XM_003999953.5;refGene_gn=CPNE3;refGene_tx=XM_019822854.2;refGene_gn=CPNE3;refGene_tx=XM_019822855.2
F2 36399394 36399446
refGene_gn=CPNE3;refGene_tx=XM_003999953.5;refGene_gn=CPNE3;refGene_tx=XM_019822854.2;refGene_gn=CPNE3;refGene_tx=XM_019822855.2
F2 36402610 36402744
refGene_gn=CPNE3;refGene_tx=XM_003999953.5;refGene_gn=CPNE3;refGene_tx=XM_019822854.2;refGene_gn=CPNE3;refGene_tx=XM_019822855.2
F2 36403853 36404090
refGene_gn=CPNE3;refGene_tx=XM_003999953.5;refGene_gn=CPNE3;refGene_tx=XM_019822854.2;refGene_gn=CPNE3;refGene_tx=XM_019822855.2
F2 36405899 36406010
refGene_gn=CPNE3;refGene_tx=XM_003999953.5;refGene_gn=CPNE3;refGene_tx=XM_019822854.2;refGene_gn=CPNE3;refGene_tx=XM_019822855.2
F2 36437209 36437470
refGene_gn=CNGB3;refGene_tx=XM_023248538.1;refGene_gn=CNGB3;refGene_tx=XM_023248539.1;refGene_gn=CNGB3;refGene_tx=XM_023248540.1
F2 36439776 36439951
refGene_gn=CNGB3;refGene_tx=XM_023248538.1;refGene_gn=CNGB3;refGene_tx=XM_023248539.1;refGene_gn=CNGB3;refGene_tx=XM_023248540.1
F2 36440401 36440548
refGene_gn=CNGB3;refGene_tx=XM_023248538.1;refGene_gn=CNGB3;refGene_tx=XM_023248540.1
F2 36463826 36463945
refGene_gn=CNGB3;refGene_tx=XM_023248538.1;refGene_gn=CNGB3;refGene_tx=XM_023248539.1;refGene_gn=CNGB3;refGene_tx=XM_023248540.1
F2 36475210 36475294
refGene_gn=CNGB3;refGene_tx=XM_023248538.1;refGene_gn=CNGB3;refGene_tx=XM_023248539.1;refGene_gn=CNGB3;refGene_tx=XM_023248540.1
F2 36490914 36491012
refGene_gn=CNGB3;refGene_tx=XM_023248538.1;refGene_gn=CNGB3;refGene_tx=XM_023248539.1;refGene_gn=CNGB3;refGene_tx=XM_023248540.1
F2 36492486 36492646
refGene_gn=CNGB3;refGene_tx=XM_023248538.1;refGene_gn=CNGB3;refGene_tx=XM_023248539.1;refGene_gn=CNGB3;refGene_tx=XM_023248540.1
F2 36497111 36497253
refGene_gn=CNGB3;refGene_tx=XM_023248538.1;refGene_gn=CNGB3;refGene_tx=XM_023248539.1;refGene_gn=CNGB3;refGene_tx=XM_023248540.1
F2 36510478 36510601
refGene_gn=CNGB3;refGene_tx=XM_023248538.1;refGene_gn=CNGB3;refGene_tx=XM_023248539.1;refGene_gn=CNGB3;refGene_tx=XM_023248540.1

F2	36511584	36511649
refGene_gn=CNGB3;refGene_tx=XM_023248538.1;refGene_gn=CNGB3;refGene_tx=XM_023248539.1;refGene_gn=CNGB3;refGene_tx=XM_023248540.1
F2	36515126	36515213
refGene_gn=CNGB3;refGene_tx=XM_023248538.1;refGene_gn=CNGB3;refGene_tx=XM_023248539.1;refGene_gn=CNGB3;refGene_tx=XM_023248540.1
F2	36517986	36518037
refGene_gn=CNGB3;refGene_tx=XM_023248538.1;refGene_gn=CNGB3;refGene_tx=XM_023248539.1;refGene_gn=CNGB3;refGene_tx=XM_023248540.1
F2	36524250	36524459
refGene_gn=CNGB3;refGene_tx=XM_023248538.1;refGene_gn=CNGB3;refGene_tx=XM_023248539.1;refGene_gn=CNGB3;refGene_tx=XM_023248540.1
F2	36525453	36525606
refGene_gn=CNGB3;refGene_tx=XM_023248538.1;refGene_gn=CNGB3;refGene_tx=XM_023248539.1;refGene_gn=CNGB3;refGene_tx=XM_023248540.1
F2	36527644	36527790
refGene_gn=CNGB3;refGene_tx=XM_023248538.1;refGene_gn=CNGB3;refGene_tx=XM_023248539.1;refGene_gn=CNGB3;refGene_tx=XM_023248540.1
F2	36572347	36572474
refGene_gn=CNGB3;refGene_tx=XM_023248538.1;refGene_gn=CNGB3;refGene_tx=XM_023248539.1;refGene_gn=CNGB3;refGene_tx=XM_023248540.1
F2	36581440	36581522
refGene_gn=CNGB3;refGene_tx=XM_023248538.1;refGene_gn=CNGB3;refGene_tx=XM_023248539.1
F2	36585049	36585160
refGene_gn=CNGB3;refGene_tx=XM_023248538.1;refGene_gn=CNGB3;refGene_tx=XM_023248539.1
F2	36738379	36738467
refGene_gn=CNBD1;refGene_tx=XM_019822865.2;refGene_gn=CNBD1;refGene_tx=XM_023248607.1
F2	36757546	36757619
refGene_gn=CNBD1;refGene_tx=XM_019822865.2;refGene_gn=CNBD1;refGene_tx=XM_023248607.1
F2	36778196	36778310
refGene_gn=CNBD1;refGene_tx=XM_019822865.2;refGene_gn=CNBD1;refGene_tx=XM_023248607.1
F2	36791101	36791260
refGene_gn=CNBD1;refGene_tx=XM_019822865.2;refGene_gn=CNBD1;refGene_tx=XM_023248607.1
F2	36963363	36963509
refGene_gn=CNBD1;refGene_tx=XM_019822865.2;refGene_gn=CNBD1;refGene_tx=XM_023248607.1
F2	36985660	36985851
refGene_gn=CNBD1;refGene_tx=XM_019822865.2;refGene_gn=CNBD1;refGene_tx=XM_023248607.1
F2	37049840	37049978
refGene_gn=CNBD1;refGene_tx=XM_019822865.2;refGene_gn=CNBD1;refGene_tx=XM_023248607.1
F2	37051439	37051572
refGene_gn=CNBD1;refGene_tx=XM_019822865.2;refGene_gn=CNBD1;refGene_tx

=XM_023248607.1
F2 37105974 37106084
refGene_gn=CNBD1;refGene_tx=XM_019822865.2;refGene_gn=CNBD1;refGene_tx
=XM_023248607.1
F2 37107793 37107944
refGene_gn=CNBD1;refGene_tx=XM_019822865.2;refGene_gn=CNBD1;refGene_tx
=XM_023248607.1
F2 37124315 37124389 refGene_gn=CNBD1;refGene_tx=XM_023248607.1
F2 37154048 37154116 refGene_gn=CNBD1;refGene_tx=XM_019822865.2
F2 37162764 37162767 refGene_gn=CNBD1;refGene_tx=XM_019822865.2
F2 37626706 37626890
refGene_gn=MMP16;refGene_tx=XM_019822866.1;refGene_gn=MMP16;refGene_tx
=XM_019822867.2
F2 37650795 37651130 refGene_gn=MMP16;refGene_tx=XM_003999956.4
F2 37658263 37658379
refGene_gn=MMP16;refGene_tx=XM_003999956.4;refGene_gn=MMP16;refGene_tx
=XM_019822866.1
F2 37668154 37668305
refGene_gn=MMP16;refGene_tx=XM_003999956.4;refGene_gn=MMP16;refGene_tx
=XM_019822866.1;refGene_gn=MMP16;refGene_tx=XM_019822867.2
F2 37687907 37688046
refGene_gn=MMP16;refGene_tx=XM_003999956.4;refGene_gn=MMP16;refGene_tx
=XM_019822866.1;refGene_gn=MMP16;refGene_tx=XM_019822867.2
F2 37723515 37723727
refGene_gn=MMP16;refGene_tx=XM_003999956.4;refGene_gn=MMP16;refGene_tx
=XM_019822866.1;refGene_gn=MMP16;refGene_tx=XM_019822867.2
F2 37725672 37725834
refGene_gn=MMP16;refGene_tx=XM_003999956.4;refGene_gn=MMP16;refGene_tx
=XM_019822866.1;refGene_gn=MMP16;refGene_tx=XM_019822867.2
F2 37771353 37771658
refGene_gn=MMP16;refGene_tx=XM_003999956.4;refGene_gn=MMP16;refGene_tx
=XM_019822866.1;refGene_gn=MMP16;refGene_tx=XM_019822867.2
F2 37785661 37785784
refGene_gn=MMP16;refGene_tx=XM_003999956.4;refGene_gn=MMP16;refGene_tx
=XM_019822866.1;refGene_gn=MMP16;refGene_tx=XM_019822867.2
F2 37796487 37796636
refGene_gn=MMP16;refGene_tx=XM_003999956.4;refGene_gn=MMP16;refGene_tx
=XM_019822866.1;refGene_gn=MMP16;refGene_tx=XM_019822867.2
F2 37937720 37937852
refGene_gn=MMP16;refGene_tx=XM_003999956.4;refGene_gn=MMP16;refGene_tx
=XM_019822866.1;refGene_gn=MMP16;refGene_tx=XM_019822867.2
F2 39133270 39133305
refGene_gn=LOC111558514;refGene_tx=XM_023248300.1
F2 39144887 39144979
refGene_gn=LOC111558514;refGene_tx=XM_023248300.1
F2 39146718 39146788
refGene_gn=LOC111558514;refGene_tx=XM_023248304.1
F2 39146873 39146961
refGene_gn=LOC111558514;refGene_tx=XM_023248301.1;refGene_gn=LOC111558
514;refGene_tx=XM_023248302.1;refGene_gn=LOC111558514;refGene_tx=XM_02

3248303.1
F2 39147580 39148137
refGene_gn=LOC111558514;refGene_tx=XM_023248300.1;refGene_gn=LOC111558514;refGene_tx=XM_023248301.1;refGene_gn=LOC111558514;refGene_tx=XM_023248302.1;refGene_gn=LOC111558514;refGene_tx=XM_023248303.1;refGene_gn=LOC111558514;refGene_tx=XM_023248304.1;refGene_gn=RIPK2;refGene_tx=XM_011291314.3
F2 39152661 39152815 refGene_gn=RIPK2;refGene_tx=XM_011291314.3
F2 39155007 39155163 refGene_gn=RIPK2;refGene_tx=XM_011291314.3
F2 39162922 39163080 refGene_gn=RIPK2;refGene_tx=XM_011291314.3
F2 39163786 39163836 refGene_gn=RIPK2;refGene_tx=XM_011291314.3
F2 39165154 39165316 refGene_gn=RIPK2;refGene_tx=XM_011291314.3
F2 39169641 39169727 refGene_gn=RIPK2;refGene_tx=XM_011291314.3
F2 39173124 39173214 refGene_gn=RIPK2;refGene_tx=XM_011291314.3
F2 39174079 39174173 refGene_gn=RIPK2;refGene_tx=XM_011291314.3
F2 39175821 39175977 refGene_gn=RIPK2;refGene_tx=XM_011291314.3
F2 39176577 39176915 refGene_gn=RIPK2;refGene_tx=XM_011291314.3
F2 39254449 39254493 refGene_gn=OSGIN2;refGene_tx=XM_019822868.2
F2 39262139 39262294
refGene_gn=OSGIN2;refGene_tx=XM_006943295.4;refGene_gn=OSGIN2;refGene_tx=XM_019822868.2
F2 39267872 39268009
refGene_gn=OSGIN2;refGene_tx=XM_006943295.4;refGene_gn=OSGIN2;refGene_tx=XM_019822868.2;refGene_gn=OSGIN2;refGene_tx=XM_019822869.2
F2 39268799 39268991
refGene_gn=OSGIN2;refGene_tx=XM_006943295.4;refGene_gn=OSGIN2;refGene_tx=XM_019822868.2;refGene_gn=OSGIN2;refGene_tx=XM_019822869.2
F2 39274004 39274096
refGene_gn=OSGIN2;refGene_tx=XM_006943295.4;refGene_gn=OSGIN2;refGene_tx=XM_019822868.2;refGene_gn=OSGIN2;refGene_tx=XM_019822869.2
F2 39276315 39277345
refGene_gn=OSGIN2;refGene_tx=XM_006943295.4;refGene_gn=OSGIN2;refGene_tx=XM_019822868.2;refGene_gn=OSGIN2;refGene_tx=XM_019822869.2
F2 39289637 39289668
refGene_gn=NBN;refGene_tx=XM_023248203.1;refGene_gn=NBN;refGene_tx=XM_023248204.1;refGene_gn=NBN;refGene_tx=XM_023248205.1
F2 39290869 39290919
refGene_gn=NBN;refGene_tx=XM_023248203.1;refGene_gn=NBN;refGene_tx=XM_023248204.1;refGene_gn=NBN;refGene_tx=XM_023248205.1
F2 39298808 39298922
refGene_gn=NBN;refGene_tx=XM_023248203.1;refGene_gn=NBN;refGene_tx=XM_023248204.1;refGene_gn=NBN;refGene_tx=XM_023248205.1
F2 39300491 39300647
refGene_gn=NBN;refGene_tx=XM_023248203.1;refGene_gn=NBN;refGene_tx=XM_023248205.1
F2 39301809 39301881
refGene_gn=NBN;refGene_tx=XM_023248203.1;refGene_gn=NBN;refGene_tx=XM_023248204.1;refGene_gn=NBN;refGene_tx=XM_023248205.1
F2 39307900 39308345
refGene_gn=NBN;refGene_tx=XM_023248203.1;refGene_gn=NBN;refGene_tx=XM_

023248204.1;refGene_gn=NBN;refGene_tx=XM_023248205.1
F2	39310236	39310509
refGene_gn=NBN;refGene_tx=XM_023248203.1;refGene_gn=NBN;refGene_tx=XM_
023248204.1;refGene_gn=NBN;refGene_tx=XM_023248205.1
F2	39312715	39312842
refGene_gn=NBN;refGene_tx=XM_023248203.1;refGene_gn=NBN;refGene_tx=XM_
023248204.1;refGene_gn=NBN;refGene_tx=XM_023248205.1
F2	39315938	39316036
refGene_gn=NBN;refGene_tx=XM_023248203.1;refGene_gn=NBN;refGene_tx=XM_
023248204.1;refGene_gn=NBN;refGene_tx=XM_023248205.1
F2	39323081	39323275
refGene_gn=NBN;refGene_tx=XM_023248203.1;refGene_gn=NBN;refGene_tx=XM_
023248204.1;refGene_gn=NBN;refGene_tx=XM_023248205.1
F2	39324273	39324391
refGene_gn=NBN;refGene_tx=XM_023248203.1;refGene_gn=NBN;refGene_tx=XM_
023248204.1;refGene_gn=NBN;refGene_tx=XM_023248205.1
F2	39340934	39341039
refGene_gn=NBN;refGene_tx=XM_023248203.1;refGene_gn=NBN;refGene_tx=XM_
023248204.1;refGene_gn=NBN;refGene_tx=XM_023248205.1
F2	39344199	39344348
refGene_gn=NBN;refGene_tx=XM_023248203.1;refGene_gn=NBN;refGene_tx=XM_
023248204.1;refGene_gn=NBN;refGene_tx=XM_023248205.1
F2	39345906	39346037
refGene_gn=NBN;refGene_tx=XM_023248203.1;refGene_gn=NBN;refGene_tx=XM_
023248204.1;refGene_gn=NBN;refGene_tx=XM_023248205.1
F2	39346773	39346903	refGene_gn=NBN;refGene_tx=XM_023248205.1
F2	39347692	39347729
refGene_gn=NBN;refGene_tx=XM_023248203.1;refGene_gn=NBN;refGene_tx=XM_
023248204.1
F2	39372682	39372733	refGene_gn=DECR1;refGene_tx=XM_006943296.4
F2	39384261	39384464	refGene_gn=DECR1;refGene_tx=XM_006943296.4
F2	39386765	39386823	refGene_gn=DECR1;refGene_tx=XM_006943296.4
F2	39386947	39387034	refGene_gn=DECR1;refGene_tx=XM_006943296.4
F2	39387995	39388143	refGene_gn=DECR1;refGene_tx=XM_006943296.4
F2	39403468	39403568	refGene_gn=DECR1;refGene_tx=XM_006943296.4
F2	39408404	39408477	refGene_gn=DECR1;refGene_tx=XM_006943296.4
F2	39410609	39410756	refGene_gn=DECR1;refGene_tx=XM_006943296.4
F2	39411218	39411281	refGene_gn=DECR1;refGene_tx=XM_006943296.4
F2	39411378	39411438	refGene_gn=DECR1;refGene_tx=XM_006943296.4
F2	39420032	39420146	refGene_gn=CALB1;refGene_tx=XM_003999958.5
F2	39420476	39420548	refGene_gn=CALB1;refGene_tx=XM_003999958.5
F2	39421397	39421451	refGene_gn=CALB1;refGene_tx=XM_003999958.5
F2	39421579	39421619	refGene_gn=CALB1;refGene_tx=XM_003999958.5
F2	39421702	39421758	refGene_gn=CALB1;refGene_tx=XM_003999958.5
F2	39423988	39424066	refGene_gn=CALB1;refGene_tx=XM_003999958.5
F2	39427240	39427297	refGene_gn=CALB1;refGene_tx=XM_003999958.5
F2	39427410	39427494	refGene_gn=CALB1;refGene_tx=XM_003999958.5
F2	39436230	39436305	refGene_gn=CALB1;refGene_tx=XM_003999958.5
F2	39440527	39440604	refGene_gn=CALB1;refGene_tx=XM_003999958.5
F2	39441136	39441215	refGene_gn=CALB1;refGene_tx=XM_003999958.5

F2 39941615 39941807
refGene_gn=TMEM64;refGene_tx=XM_011291378.3;refGene_gn=TMEM64;refGene_tx=XM_023248728.1
F2 39947683 39947839 refGene_gn=TMEM64;refGene_tx=XM_011291378.3
F2 39964741 39965539
refGene_gn=TMEM64;refGene_tx=XM_011291378.3;refGene_gn=TMEM64;refGene_tx=XM_023248728.1
F2 40090761 40091094
refGene_gn=NECAB1;refGene_tx=XM_003999959.5;refGene_gn=NECAB1;refGene_tx=XM_011291311.3
F2 40098667 40098692
refGene_gn=NECAB1;refGene_tx=XM_003999959.5;refGene_gn=NECAB1;refGene_tx=XM_011291311.3
F2 40129139 40129248
refGene_gn=NECAB1;refGene_tx=XM_003999959.5;refGene_gn=NECAB1;refGene_tx=XM_011291311.3
F2 40174825 40174851
refGene_gn=NECAB1;refGene_tx=XM_003999959.5;refGene_gn=NECAB1;refGene_tx=XM_011291311.3
F2 40181326 40181424
refGene_gn=NECAB1;refGene_tx=XM_003999959.5;refGene_gn=NECAB1;refGene_tx=XM_011291311.3
F2 40213330 40214005
refGene_gn=LOC101100863;refGene_tx=XM_011291307.2;refGene_gn=LOC101100863;refGene_tx=XM_011291308.2
F2 40239725 40239862
refGene_gn=NECAB1;refGene_tx=XM_003999959.5;refGene_gn=NECAB1;refGene_tx=XM_011291311.3
F2 40250237 40250359
refGene_gn=NECAB1;refGene_tx=XM_003999959.5;refGene_gn=NECAB1;refGene_tx=XM_011291311.3
F2 40253383 40253460
refGene_gn=NECAB1;refGene_tx=XM_003999959.5;refGene_gn=NECAB1;refGene_tx=XM_011291311.3
F2 40257831 40257885
refGene_gn=NECAB1;refGene_tx=XM_003999959.5;refGene_gn=NECAB1;refGene_tx=XM_011291311.3
F2 40266602 40266715
refGene_gn=NECAB1;refGene_tx=XM_003999959.5;refGene_gn=NECAB1;refGene_tx=XM_011291311.3
F2 40280227 40280305
refGene_gn=NECAB1;refGene_tx=XM_003999959.5;refGene_gn=NECAB1;refGene_tx=XM_011291311.3
F2 40281596 40281688
refGene_gn=NECAB1;refGene_tx=XM_003999959.5;refGene_gn=NECAB1;refGene_tx=XM_011291311.3
F2 40284904 40284930
refGene_gn=NECAB1;refGene_tx=XM_003999959.5;refGene_gn=NECAB1;refGene_tx=XM_011291311.3
F2 40288486 40288510

refGene_gn=CF2H8orf88;refGene_tx=XM_011291305.3;refGene_gn=CF2H8orf88;
refGene_tx=XM_023248411.1;refGene_gn=CF2H8orf88;refGene_tx=XM_02324841
2.1
F2 40289978 40290085
refGene_gn=CF2H8orf88;refGene_tx=XM_011291305.3;refGene_gn=CF2H8orf88;
refGene_tx=XM_023248411.1;refGene_gn=CF2H8orf88;refGene_tx=XM_02324841
2.1
F2 40306767 40306843
refGene_gn=CF2H8orf88;refGene_tx=XM_011291305.3;refGene_gn=CF2H8orf88;
refGene_tx=XM_023248411.1;refGene_gn=CF2H8orf88;refGene_tx=XM_02324841
2.1
F2 40315670 40315744
refGene_gn=CF2H8orf88;refGene_tx=XM_011291305.3;refGene_gn=CF2H8orf88;
refGene_tx=XM_023248411.1;refGene_gn=CF2H8orf88;refGene_tx=XM_02324841
2.1
F2 40317768 40317841
refGene_gn=CF2H8orf88;refGene_tx=XM_011291305.3;refGene_gn=CF2H8orf88;
refGene_tx=XM_023248411.1;refGene_gn=CF2H8orf88;refGene_tx=XM_02324841
2.1
F2 40338361 40338505
refGene_gn=PIP4P2;refGene_tx=XM_019822876.2;refGene_gn=PIP4P2;refGene_
tx=XM_023248152.1;refGene_gn=PIP4P2;refGene_tx=XM_023248153.1;refGene_
gn=PIP4P2;refGene_tx=XM_023248154.1;refGene_gn=PIP4P2;refGene_tx=XM_02
3248155.1
F2 40339362 40339453
refGene_gn=PIP4P2;refGene_tx=XM_019822876.2;refGene_gn=PIP4P2;refGene_
tx=XM_023248152.1;refGene_gn=PIP4P2;refGene_tx=XM_023248153.1;refGene_
gn=PIP4P2;refGene_tx=XM_023248154.1;refGene_gn=PIP4P2;refGene_tx=XM_02
3248155.1
F2 40355064 40355117
refGene_gn=PIP4P2;refGene_tx=XM_019822876.2;refGene_gn=PIP4P2;refGene_
tx=XM_023248152.1;refGene_gn=PIP4P2;refGene_tx=XM_023248153.1;refGene_
gn=PIP4P2;refGene_tx=XM_023248154.1;refGene_gn=PIP4P2;refGene_tx=XM_02
3248155.1
F2 40364968 40365092
refGene_gn=PIP4P2;refGene_tx=XM_019822876.2;refGene_gn=PIP4P2;refGene_
tx=XM_023248152.1;refGene_gn=PIP4P2;refGene_tx=XM_023248153.1;refGene_
gn=PIP4P2;refGene_tx=XM_023248154.1;refGene_gn=PIP4P2;refGene_tx=XM_02
3248155.1
F2 40366459 40366566
refGene_gn=PIP4P2;refGene_tx=XM_019822876.2;refGene_gn=PIP4P2;refGene_
tx=XM_023248152.1;refGene_gn=PIP4P2;refGene_tx=XM_023248153.1;refGene_
gn=PIP4P2;refGene_tx=XM_023248154.1;refGene_gn=PIP4P2;refGene_tx=XM_02
3248155.1
F2 40371327 40371476
refGene_gn=PIP4P2;refGene_tx=XM_019822876.2;refGene_gn=PIP4P2;refGene_
tx=XM_023248152.1;refGene_gn=PIP4P2;refGene_tx=XM_023248153.1;refGene_
gn=PIP4P2;refGene_tx=XM_023248154.1;refGene_gn=PIP4P2;refGene_tx=XM_02
3248155.1
F2 40383578 40383585

refGene_gn=PIP4P2;refGene_tx=XM_019822876.2;refGene_gn=PIP4P2;refGene_tx=XM_023248153.1;refGene_gn=PIP4P2;refGene_tx=XM_023248155.1
F2 40386317 40386324 refGene_gn=PIP4P2;refGene_tx=XM_023248154.1
F2 40390939 40391045 refGene_gn=PIP4P2;refGene_tx=XM_023248152.1
F2 40424225 40424278 refGene_gn=OTUD6B;refGene_tx=XM_003999961.5
F2 40424279 40424395 refGene_gn=OTUD6B;refGene_tx=XM_003999961.5
F2 40425019 40425171 refGene_gn=OTUD6B;refGene_tx=XM_003999961.5
F2 40427847 40427928 refGene_gn=OTUD6B;refGene_tx=XM_003999961.5
F2 40432662 40432975 refGene_gn=OTUD6B;refGene_tx=XM_003999961.5
F2 40434787 40434849 refGene_gn=OTUD6B;refGene_tx=XM_003999961.5
F2 40437336 40437443 refGene_gn=OTUD6B;refGene_tx=XM_003999961.5
F2 40438239 40438324 refGene_gn=OTUD6B;refGene_tx=XM_003999961.5
F2 40455454 40455637
refGene_gn=LRRC69;refGene_tx=XM_003999963.4;refGene_gn=LRRC69;refGene_tx=XM_023248690.1
F2 40469750 40469877
refGene_gn=LRRC69;refGene_tx=XM_003999963.4;refGene_gn=LRRC69;refGene_tx=XM_023248690.1
F2 40472597 40472739 refGene_gn=LRRC69;refGene_tx=XM_023248689.1
F2 40473493 40473566
refGene_gn=LRRC69;refGene_tx=XM_003999963.4;refGene_gn=LRRC69;refGene_tx=XM_023248689.1;refGene_gn=LRRC69;refGene_tx=XM_023248690.1
F2 40480061 40480257
refGene_gn=LRRC69;refGene_tx=XM_003999963.4;refGene_gn=LRRC69;refGene_tx=XM_023248689.1;refGene_gn=LRRC69;refGene_tx=XM_023248690.1
F2 40482976 40483048
refGene_gn=LRRC69;refGene_tx=XM_003999963.4;refGene_gn=LRRC69;refGene_tx=XM_023248689.1;refGene_gn=LRRC69;refGene_tx=XM_023248690.1
F2 40513009 40513111
refGene_gn=LRRC69;refGene_tx=XM_003999963.4;refGene_gn=LRRC69;refGene_tx=XM_023248689.1;refGene_gn=LRRC69;refGene_tx=XM_023248690.1
F2 40525020 40525200
refGene_gn=LRRC69;refGene_tx=XM_003999963.4;refGene_gn=LRRC69;refGene_tx=XM_023248689.1
F2 40535000 40535111
refGene_gn=LRRC69;refGene_tx=XM_003999963.4;refGene_gn=LRRC69;refGene_tx=XM_023248689.1;refGene_gn=LRRC69;refGene_tx=XM_023248690.1
F2 40562714 40562907 refGene_gn=SLC26A7;refGene_tx=XM_019822873.2
F2 40588984 40589095 refGene_gn=SLC26A7;refGene_tx=XM_019822873.2
F2 40590617 40590790 refGene_gn=SLC26A7;refGene_tx=XM_019822873.2
F2 40612147 40612312
refGene_gn=SLC26A7;refGene_tx=XM_019822873.2;refGene_gn=SLC26A7;refGene_tx=XM_019822875.2
F2 40626018 40626171
refGene_gn=SLC26A7;refGene_tx=XM_019822873.2;refGene_gn=SLC26A7;refGene_tx=XM_019822875.2
F2 40629658 40629741
refGene_gn=SLC26A7;refGene_tx=XM_019822873.2;refGene_gn=SLC26A7;refGene_tx=XM_019822875.2
F2 40631881 40632029

refGene_gn=SLC26A7;refGene_tx=XM_019822873.2;refGene_gn=SLC26A7;refGene_tx=XM_019822875.2
F2 40633903 40634017
refGene_gn=SLC26A7;refGene_tx=XM_019822873.2;refGene_gn=SLC26A7;refGene_tx=XM_019822875.2
F2 40643064 40643142
refGene_gn=SLC26A7;refGene_tx=XM_019822873.2;refGene_gn=SLC26A7;refGene_tx=XM_019822875.2
F2 40644159 40644255
refGene_gn=SLC26A7;refGene_tx=XM_019822873.2;refGene_gn=SLC26A7;refGene_tx=XM_019822875.2
F2 40653865 40653972
refGene_gn=SLC26A7;refGene_tx=XM_019822873.2;refGene_gn=SLC26A7;refGene_tx=XM_019822875.2
F2 40655022 40655089
refGene_gn=SLC26A7;refGene_tx=XM_019822873.2;refGene_gn=SLC26A7;refGene_tx=XM_019822875.2
F2 40658560 40658698
refGene_gn=SLC26A7;refGene_tx=XM_019822873.2;refGene_gn=SLC26A7;refGene_tx=XM_019822875.2
F2 40663389 40663438
refGene_gn=SLC26A7;refGene_tx=XM_019822873.2;refGene_gn=SLC26A7;refGene_tx=XM_019822875.2
F2 40673022 40673123
refGene_gn=SLC26A7;refGene_tx=XM_019822873.2;refGene_gn=SLC26A7;refGene_tx=XM_019822875.2
F2 40677569 40677624
refGene_gn=SLC26A7;refGene_tx=XM_019822873.2;refGene_gn=SLC26A7;refGene_tx=XM_019822875.2
F2 40677702 40677806
refGene_gn=SLC26A7;refGene_tx=XM_019822873.2;refGene_gn=SLC26A7;refGene_tx=XM_019822875.2
F2 40679043 40679079
refGene_gn=SLC26A7;refGene_tx=XM_019822873.2;refGene_gn=SLC26A7;refGene_tx=XM_019822875.2
F2 41147048 41147324
refGene_gn=RUNX1T1;refGene_tx=XM_023248593.1;refGene_gn=RUNX1T1;refGene_tx=XM_023248594.1;refGene_gn=RUNX1T1;refGene_tx=XM_023248595.1;refGene_gn=RUNX1T1;refGene_tx=XM_023248596.1;refGene_gn=RUNX1T1;refGene_tx=XM_023248597.1;refGene_gn=RUNX1T1;refGene_tx=XM_023248598.1;refGene_gn=RUNX1T1;refGene_tx=XM_023248599.1;refGene_gn=RUNX1T1;refGene_tx=XM_023248600.1
F2 41160983 41161174
refGene_gn=RUNX1T1;refGene_tx=XM_023248593.1;refGene_gn=RUNX1T1;refGene_tx=XM_023248594.1;refGene_gn=RUNX1T1;refGene_tx=XM_023248595.1;refGene_gn=RUNX1T1;refGene_tx=XM_023248596.1;refGene_gn=RUNX1T1;refGene_tx=XM_023248597.1;refGene_gn=RUNX1T1;refGene_tx=XM_023248598.1;refGene_gn=RUNX1T1;refGene_tx=XM_023248599.1;refGene_gn=RUNX1T1;refGene_tx=XM_023248600.1
F2 41166345 41166414

refGene_gn=RUNX1T1;refGene_tx=XM_023248593.1;refGene_gn=RUNX1T1;refGen
e_tx=XM_023248594.1;refGene_gn=RUNX1T1;refGene_tx=XM_023248595.1;refGe
ne_gn=RUNX1T1;refGene_tx=XM_023248596.1;refGene_gn=RUNX1T1;refGene_tx=
XM_023248597.1;refGene_gn=RUNX1T1;refGene_tx=XM_023248598.1;refGene_gn
=RUNX1T1;refGene_tx=XM_023248599.1;refGene_gn=RUNX1T1;refGene_tx=XM_02
3248600.1
F2 41175597 41175799
refGene_gn=RUNX1T1;refGene_tx=XM_023248593.1;refGene_gn=RUNX1T1;refGen
e_tx=XM_023248594.1;refGene_gn=RUNX1T1;refGene_tx=XM_023248595.1;refGe
ne_gn=RUNX1T1;refGene_tx=XM_023248596.1;refGene_gn=RUNX1T1;refGene_tx=
XM_023248597.1;refGene_gn=RUNX1T1;refGene_tx=XM_023248598.1;refGene_gn
=RUNX1T1;refGene_tx=XM_023248599.1
F2 41176361 41176447
refGene_gn=RUNX1T1;refGene_tx=XM_023248593.1;refGene_gn=RUNX1T1;refGen
e_tx=XM_023248594.1;refGene_gn=RUNX1T1;refGene_tx=XM_023248595.1;refGe
ne_gn=RUNX1T1;refGene_tx=XM_023248596.1;refGene_gn=RUNX1T1;refGene_tx=
XM_023248597.1;refGene_gn=RUNX1T1;refGene_tx=XM_023248598.1;refGene_gn
=RUNX1T1;refGene_tx=XM_023248599.1
F2 41181484 41181735
refGene_gn=RUNX1T1;refGene_tx=XM_023248593.1;refGene_gn=RUNX1T1;refGen
e_tx=XM_023248594.1;refGene_gn=RUNX1T1;refGene_tx=XM_023248595.1;refGe
ne_gn=RUNX1T1;refGene_tx=XM_023248596.1;refGene_gn=RUNX1T1;refGene_tx=
XM_023248597.1;refGene_gn=RUNX1T1;refGene_tx=XM_023248598.1;refGene_gn
=RUNX1T1;refGene_tx=XM_023248599.1;refGene_gn=RUNX1T1;refGene_tx=XM_02
3248600.1
F2 41193731 41193913
refGene_gn=RUNX1T1;refGene_tx=XM_023248593.1;refGene_gn=RUNX1T1;refGen
e_tx=XM_023248594.1;refGene_gn=RUNX1T1;refGene_tx=XM_023248595.1;refGe
ne_gn=RUNX1T1;refGene_tx=XM_023248596.1;refGene_gn=RUNX1T1;refGene_tx=
XM_023248597.1;refGene_gn=RUNX1T1;refGene_tx=XM_023248598.1;refGene_gn
=RUNX1T1;refGene_tx=XM_023248599.1;refGene_gn=RUNX1T1;refGene_tx=XM_02
3248600.1
F2 41198595 41198685
refGene_gn=RUNX1T1;refGene_tx=XM_023248593.1;refGene_gn=RUNX1T1;refGen
e_tx=XM_023248594.1;refGene_gn=RUNX1T1;refGene_tx=XM_023248595.1;refGe
ne_gn=RUNX1T1;refGene_tx=XM_023248596.1;refGene_gn=RUNX1T1;refGene_tx=
XM_023248597.1;refGene_gn=RUNX1T1;refGene_tx=XM_023248598.1;refGene_gn
=RUNX1T1;refGene_tx=XM_023248599.1;refGene_gn=RUNX1T1;refGene_tx=XM_02
3248600.1
F2 41202447 41202689
refGene_gn=RUNX1T1;refGene_tx=XM_023248593.1;refGene_gn=RUNX1T1;refGen
e_tx=XM_023248594.1;refGene_gn=RUNX1T1;refGene_tx=XM_023248595.1;refGe
ne_gn=RUNX1T1;refGene_tx=XM_023248596.1;refGene_gn=RUNX1T1;refGene_tx=
XM_023248597.1;refGene_gn=RUNX1T1;refGene_tx=XM_023248598.1;refGene_gn
=RUNX1T1;refGene_tx=XM_023248599.1;refGene_gn=RUNX1T1;refGene_tx=XM_02
3248600.1
F2 41204755 41204893
refGene_gn=RUNX1T1;refGene_tx=XM_023248593.1;refGene_gn=RUNX1T1;refGen
e_tx=XM_023248594.1;refGene_gn=RUNX1T1;refGene_tx=XM_023248595.1;refGe
ne_gn=RUNX1T1;refGene_tx=XM_023248596.1;refGene_gn=RUNX1T1;refGene_tx=

XM_023248597.1;refGene_gn=RUNX1T1;refGene_tx=XM_023248598.1;refGene_gn=RUNX1T1;refGene_tx=XM_023248599.1;refGene_gn=RUNX1T1;refGene_tx=XM_023248600.1
F2 41248262 41248269 refGene_gn=RUNX1T1;refGene_tx=XM_023248596.1
F2 41261768 41261856 refGene_gn=RUNX1T1;refGene_tx=XM_023248593.1;refGene_gn=RUNX1T1;refGene_tx=XM_023248594.1;refGene_gn=RUNX1T1;refGene_tx=XM_023248600.1
F2 41288043 41288071 refGene_gn=RUNX1T1;refGene_tx=XM_023248595.1
F2 41927317 41927431 refGene_gn=TRIQK;refGene_tx=XM_006943299.3
F2 41932763 41932849 refGene_gn=TRIQK;refGene_tx=XM_006943299.3
F2 41965854 41965936 refGene_gn=TRIQK;refGene_tx=XM_006943299.3
F2 42002857 42002908 refGene_gn=TRIQK;refGene_tx=XM_006943299.3
F2 42676223 42679160 refGene_gn=RBM12B;refGene_tx=XM_003999972.5;refGene_gn=RBM12B;refGene_tx=XM_006943304.4;refGene_gn=RBM12B;refGene_tx=XM_006943305.4;refGene_gn=RBM12B;refGene_tx=XM_019822883.2;refGene_gn=RBM12B;refGene_tx=XM_019822884.2
F2 42696629 42696852 refGene_gn=TMEM67;refGene_tx=XM_011291396.3;refGene_gn=TMEM67;refGene_tx=XM_019822881.2
F2 42698489 42698578 refGene_gn=TMEM67;refGene_tx=XM_011291396.3;refGene_gn=TMEM67;refGene_tx=XM_019822881.2;refGene_gn=TMEM67;refGene_tx=XM_019822882.2
F2 42701179 42701273 refGene_gn=TMEM67;refGene_tx=XM_011291396.3;refGene_gn=TMEM67;refGene_tx=XM_019822881.2;refGene_gn=TMEM67;refGene_tx=XM_019822882.2
F2 42707745 42707845 refGene_gn=TMEM67;refGene_tx=XM_011291396.3;refGene_gn=TMEM67;refGene_tx=XM_019822881.2;refGene_gn=TMEM67;refGene_tx=XM_019822882.2
F2 42709537 42709607 refGene_gn=TMEM67;refGene_tx=XM_011291396.3;refGene_gn=TMEM67;refGene_tx=XM_019822881.2;refGene_gn=TMEM67;refGene_tx=XM_019822882.2
F2 42709701 42709776 refGene_gn=TMEM67;refGene_tx=XM_011291396.3;refGene_gn=TMEM67;refGene_tx=XM_019822881.2;refGene_gn=TMEM67;refGene_tx=XM_019822882.2
F2 42715146 42715209 refGene_gn=TMEM67;refGene_tx=XM_011291396.3;refGene_gn=TMEM67;refGene_tx=XM_019822881.2;refGene_gn=TMEM67;refGene_tx=XM_019822882.2
F2 42716619 42716771 refGene_gn=TMEM67;refGene_tx=XM_011291396.3;refGene_gn=TMEM67;refGene_tx=XM_019822881.2;refGene_gn=TMEM67;refGene_tx=XM_019822882.2
F2 42716891 42717000 refGene_gn=TMEM67;refGene_tx=XM_011291396.3;refGene_gn=TMEM67;refGene_tx=XM_019822881.2;refGene_gn=TMEM67;refGene_tx=XM_019822882.2
F2 42717727 42717814 refGene_gn=TMEM67;refGene_tx=XM_011291396.3;refGene_gn=TMEM67;refGene_tx=XM_019822881.2;refGene_gn=TMEM67;refGene_tx=XM_019822882.2
F2 42718074 42718140 refGene_gn=TMEM67;refGene_tx=XM_011291396.3;refGene_gn=TMEM67;refGene_tx=XM_019822881.2;refGene_gn=TMEM67;refGene_tx=XM_019822882.2

F2	42720733	42720890
refGene_gn=TMEM67;refGene_tx=XM_011291396.3;refGene_gn=TMEM67;refGene_
tx=XM_019822881.2;refGene_gn=TMEM67;refGene_tx=XM_019822882.2
F2	42721455	42721579
refGene_gn=TMEM67;refGene_tx=XM_011291396.3;refGene_gn=TMEM67;refGene_
tx=XM_019822881.2;refGene_gn=TMEM67;refGene_tx=XM_019822882.2
F2	42723150	42723256
refGene_gn=TMEM67;refGene_tx=XM_011291396.3;refGene_gn=TMEM67;refGene_
tx=XM_019822881.2;refGene_gn=TMEM67;refGene_tx=XM_019822882.2
F2	42725911	42725968
refGene_gn=TMEM67;refGene_tx=XM_011291396.3;refGene_gn=TMEM67;refGene_
tx=XM_019822881.2;refGene_gn=TMEM67;refGene_tx=XM_019822882.2
F2	42728331	42728430
refGene_gn=TMEM67;refGene_tx=XM_011291396.3;refGene_gn=TMEM67;refGene_
tx=XM_019822881.2;refGene_gn=TMEM67;refGene_tx=XM_019822882.2
F2	42730199	42730298
refGene_gn=TMEM67;refGene_tx=XM_011291396.3;refGene_gn=TMEM67;refGene_
tx=XM_019822881.2;refGene_gn=TMEM67;refGene_tx=XM_019822882.2
F2	42730551	42730638
refGene_gn=TMEM67;refGene_tx=XM_011291396.3;refGene_gn=TMEM67;refGene_
tx=XM_019822881.2;refGene_gn=TMEM67;refGene_tx=XM_019822882.2
F2	42732837	42732937
refGene_gn=TMEM67;refGene_tx=XM_011291396.3;refGene_gn=TMEM67;refGene_
tx=XM_019822881.2;refGene_gn=TMEM67;refGene_tx=XM_019822882.2
F2	42733024	42733164
refGene_gn=TMEM67;refGene_tx=XM_011291396.3;refGene_gn=TMEM67;refGene_
tx=XM_019822881.2;refGene_gn=TMEM67;refGene_tx=XM_019822882.2
F2	42735461	42735602
refGene_gn=TMEM67;refGene_tx=XM_011291396.3;refGene_gn=TMEM67;refGene_
tx=XM_019822881.2;refGene_gn=TMEM67;refGene_tx=XM_019822882.2
F2	42738825	42738906
refGene_gn=TMEM67;refGene_tx=XM_011291396.3;refGene_gn=TMEM67;refGene_
tx=XM_019822881.2;refGene_gn=TMEM67;refGene_tx=XM_019822882.2
F2	42739399	42739516
refGene_gn=TMEM67;refGene_tx=XM_011291396.3;refGene_gn=TMEM67;refGene_
tx=XM_019822881.2;refGene_gn=TMEM67;refGene_tx=XM_019822882.2
F2	42742762	42742879
refGene_gn=TMEM67;refGene_tx=XM_011291396.3;refGene_gn=TMEM67;refGene_
tx=XM_019822881.2;refGene_gn=TMEM67;refGene_tx=XM_019822882.2
F2	42742983	42743088
refGene_gn=TMEM67;refGene_tx=XM_011291396.3;refGene_gn=TMEM67;refGene_
tx=XM_019822881.2;refGene_gn=TMEM67;refGene_tx=XM_019822882.2
F2	42743633	42743736
refGene_gn=TMEM67;refGene_tx=XM_011291396.3;refGene_gn=TMEM67;refGene_
tx=XM_019822881.2;refGene_gn=TMEM67;refGene_tx=XM_019822882.2
F2	42747725	42747868
refGene_gn=TMEM67;refGene_tx=XM_011291396.3;refGene_gn=TMEM67;refGene_
tx=XM_019822882.2
F2	42749125	42749206
refGene_gn=TMEM67;refGene_tx=XM_011291396.3;refGene_gn=TMEM67;refGene_

tx=XM_019822881.2;refGene_gn=TMEM67;refGene_tx=XM_019822882.2
F2 42821227 42821258
refGene_gn=PDP1;refGene_tx=XM_006943307.4;refGene_gn=PDP1;refGene_tx=XM_019822886.2;refGene_gn=PDP1;refGene_tx=XM_023248268.1;refGene_gn=PDP1;refGene_tx=XM_023248269.1
F2 42825098 42826759
refGene_gn=PDP1;refGene_tx=XM_006943307.4;refGene_gn=PDP1;refGene_tx=XM_006943308.3;refGene_gn=PDP1;refGene_tx=XM_011291397.2;refGene_gn=PDP1;refGene_tx=XM_019822886.2;refGene_gn=PDP1;refGene_tx=XM_023248268.1;refGene_gn=PDP1;refGene_tx=XM_023248269.1;refGene_gn=PDP1;refGene_tx=XM_023248270.1
F2 42974332 42974433 refGene_gn=CDH17;refGene_tx=XM_011291398.3
F2 42976500 42976721
refGene_gn=CDH17;refGene_tx=XM_011291398.3;refGene_gn=CDH17;refGene_tx=XM_023248513.1;refGene_gn=CDH17;refGene_tx=XM_023248514.1
F2 42976858 42976975
refGene_gn=CDH17;refGene_tx=XM_011291398.3;refGene_gn=CDH17;refGene_tx=XM_023248513.1;refGene_gn=CDH17;refGene_tx=XM_023248514.1
F2 42981052 42981292
refGene_gn=CDH17;refGene_tx=XM_011291398.3;refGene_gn=CDH17;refGene_tx=XM_023248513.1;refGene_gn=CDH17;refGene_tx=XM_023248514.1
F2 42983918 42984049
refGene_gn=CDH17;refGene_tx=XM_011291398.3;refGene_gn=CDH17;refGene_tx=XM_023248513.1;refGene_gn=CDH17;refGene_tx=XM_023248514.1
F2 42987530 42987775
refGene_gn=CDH17;refGene_tx=XM_011291398.3;refGene_gn=CDH17;refGene_tx=XM_023248513.1;refGene_gn=CDH17;refGene_tx=XM_023248514.1
F2 42996819 42997011
refGene_gn=CDH17;refGene_tx=XM_011291398.3;refGene_gn=CDH17;refGene_tx=XM_023248513.1;refGene_gn=CDH17;refGene_tx=XM_023248514.1
F2 42998925 42999002
refGene_gn=CDH17;refGene_tx=XM_011291398.3;refGene_gn=CDH17;refGene_tx=XM_023248513.1;refGene_gn=CDH17;refGene_tx=XM_023248514.1
F2 43002829 43003045
refGene_gn=CDH17;refGene_tx=XM_011291398.3;refGene_gn=CDH17;refGene_tx=XM_023248513.1;refGene_gn=CDH17;refGene_tx=XM_023248514.1
F2 43008911 43009062
refGene_gn=CDH17;refGene_tx=XM_011291398.3;refGene_gn=CDH17;refGene_tx=XM_023248513.1;refGene_gn=CDH17;refGene_tx=XM_023248514.1
F2 43009367 43009499
refGene_gn=CDH17;refGene_tx=XM_011291398.3;refGene_gn=CDH17;refGene_tx=XM_023248513.1;refGene_gn=CDH17;refGene_tx=XM_023248514.1
F2 43012037 43012237
refGene_gn=CDH17;refGene_tx=XM_011291398.3;refGene_gn=CDH17;refGene_tx=XM_023248513.1;refGene_gn=CDH17;refGene_tx=XM_023248514.1
F2 43012340 43012499
refGene_gn=CDH17;refGene_tx=XM_011291398.3;refGene_gn=CDH17;refGene_tx=XM_023248513.1;refGene_gn=CDH17;refGene_tx=XM_023248514.1
F2 43014907 43015046
refGene_gn=CDH17;refGene_tx=XM_011291398.3;refGene_gn=CDH17;refGene_tx

=XM_023248513.1;refGene_gn=CDH17;refGene_tx=XM_023248514.1
F2 43016212 43016347
refGene_gn=CDH17;refGene_tx=XM_011291398.3;refGene_gn=CDH17;refGene_tx=XM_023248513.1;refGene_gn=CDH17;refGene_tx=XM_023248514.1
F2 43030778 43030877
refGene_gn=CDH17;refGene_tx=XM_011291398.3;refGene_gn=CDH17;refGene_tx=XM_023248513.1;refGene_gn=CDH17;refGene_tx=XM_023248514.1
F2 43036261 43036309
refGene_gn=CDH17;refGene_tx=XM_011291398.3;refGene_gn=CDH17;refGene_tx=XM_023248513.1;refGene_gn=CDH17;refGene_tx=XM_023248514.1
F2 43096823 43097101
refGene_gn=GEM;refGene_tx=XM_003999975.5;refGene_gn=GEM;refGene_tx=XM_019822892.2;refGene_gn=GEM;refGene_tx=XM_023248399.1
F2 43097589 43097780 refGene_gn=GEM;refGene_tx=XM_019822893.2
F2 43098543 43098748
refGene_gn=GEM;refGene_tx=XM_003999975.5;refGene_gn=GEM;refGene_tx=XM_019822892.2;refGene_gn=GEM;refGene_tx=XM_019822893.2;refGene_gn=GEM;refGene_tx=XM_023248399.1
F2 43099531 43099608
refGene_gn=GEM;refGene_tx=XM_003999975.5;refGene_gn=GEM;refGene_tx=XM_019822892.2;refGene_gn=GEM;refGene_tx=XM_019822893.2;refGene_gn=GEM;refGene_tx=XM_023248399.1
F2 43105961 43106301
refGene_gn=GEM;refGene_tx=XM_003999975.5;refGene_gn=GEM;refGene_tx=XM_019822892.2;refGene_gn=GEM;refGene_tx=XM_019822893.2;refGene_gn=GEM;refGene_tx=XM_023248399.1
F2 43106543 43106579
refGene_gn=GEM;refGene_tx=XM_019822892.2;refGene_gn=GEM;refGene_tx=XM_019822893.2
F2 43184194 43184208 refGene_gn=RAD54B;refGene_tx=XM_023248148.1
F2 43192918 43193136
refGene_gn=RAD54B;refGene_tx=XM_003999976.5;refGene_gn=RAD54B;refGene_tx=XM_023248147.1;refGene_gn=RAD54B;refGene_tx=XM_023248149.1;refGene_gn=RAD54B;refGene_tx=XM_023248150.1;refGene_gn=RAD54B;refGene_tx=XM_023248151.1
F2 43197347 43197548
refGene_gn=RAD54B;refGene_tx=XM_003999976.5;refGene_gn=RAD54B;refGene_tx=XM_023248147.1;refGene_gn=RAD54B;refGene_tx=XM_023248148.1;refGene_gn=RAD54B;refGene_tx=XM_023248149.1;refGene_gn=RAD54B;refGene_tx=XM_023248150.1;refGene_gn=RAD54B;refGene_tx=XM_023248151.1
F2 43197739 43197806
refGene_gn=RAD54B;refGene_tx=XM_003999976.5;refGene_gn=RAD54B;refGene_tx=XM_023248147.1;refGene_gn=RAD54B;refGene_tx=XM_023248148.1;refGene_gn=RAD54B;refGene_tx=XM_023248149.1;refGene_gn=RAD54B;refGene_tx=XM_023248150.1;refGene_gn=RAD54B;refGene_tx=XM_023248151.1
F2 43199563 43199825
refGene_gn=RAD54B;refGene_tx=XM_003999976.5;refGene_gn=RAD54B;refGene_tx=XM_023248147.1;refGene_gn=RAD54B;refGene_tx=XM_023248148.1;refGene_gn=RAD54B;refGene_tx=XM_023248149.1;refGene_gn=RAD54B;refGene_tx=XM_023248150.1;refGene_gn=RAD54B;refGene_tx=XM_023248151.1

F2 43209542 43209718
refGene_gn=RAD54B;refGene_tx=XM_003999976.5;refGene_gn=RAD54B;refGene_tx=XM_023248147.1;refGene_gn=RAD54B;refGene_tx=XM_023248148.1;refGene_gn=RAD54B;refGene_tx=XM_023248149.1;refGene_gn=RAD54B;refGene_tx=XM_023248150.1;refGene_gn=RAD54B;refGene_tx=XM_023248151.1
F2 43213867 43214158
refGene_gn=RAD54B;refGene_tx=XM_003999976.5;refGene_gn=RAD54B;refGene_tx=XM_023248147.1;refGene_gn=RAD54B;refGene_tx=XM_023248148.1;refGene_gn=RAD54B;refGene_tx=XM_023248149.1;refGene_gn=RAD54B;refGene_tx=XM_023248150.1;refGene_gn=RAD54B;refGene_tx=XM_023248151.1
F2 43217144 43217284
refGene_gn=RAD54B;refGene_tx=XM_003999976.5;refGene_gn=RAD54B;refGene_tx=XM_023248147.1;refGene_gn=RAD54B;refGene_tx=XM_023248148.1;refGene_gn=RAD54B;refGene_tx=XM_023248149.1;refGene_gn=RAD54B;refGene_tx=XM_023248150.1;refGene_gn=RAD54B;refGene_tx=XM_023248151.1
F2 43219858 43220066
refGene_gn=RAD54B;refGene_tx=XM_003999976.5;refGene_gn=RAD54B;refGene_tx=XM_023248147.1;refGene_gn=RAD54B;refGene_tx=XM_023248148.1;refGene_gn=RAD54B;refGene_tx=XM_023248149.1;refGene_gn=RAD54B;refGene_tx=XM_023248150.1;refGene_gn=RAD54B;refGene_tx=XM_023248151.1
F2 43220821 43221047
refGene_gn=RAD54B;refGene_tx=XM_003999976.5;refGene_gn=RAD54B;refGene_tx=XM_023248147.1;refGene_gn=RAD54B;refGene_tx=XM_023248148.1;refGene_gn=RAD54B;refGene_tx=XM_023248149.1;refGene_gn=RAD54B;refGene_tx=XM_023248150.1;refGene_gn=RAD54B;refGene_tx=XM_023248151.1
F2 43225971 43226134
refGene_gn=RAD54B;refGene_tx=XM_003999976.5;refGene_gn=RAD54B;refGene_tx=XM_023248147.1;refGene_gn=RAD54B;refGene_tx=XM_023248148.1;refGene_gn=RAD54B;refGene_tx=XM_023248149.1;refGene_gn=RAD54B;refGene_tx=XM_023248150.1;refGene_gn=RAD54B;refGene_tx=XM_023248151.1
F2 43229890 43230172
refGene_gn=RAD54B;refGene_tx=XM_003999976.5;refGene_gn=RAD54B;refGene_tx=XM_023248147.1;refGene_gn=RAD54B;refGene_tx=XM_023248148.1;refGene_gn=RAD54B;refGene_tx=XM_023248149.1;refGene_gn=RAD54B;refGene_tx=XM_023248150.1;refGene_gn=RAD54B;refGene_tx=XM_023248151.1
F2 43234871 43235069
refGene_gn=RAD54B;refGene_tx=XM_003999976.5;refGene_gn=RAD54B;refGene_tx=XM_023248147.1;refGene_gn=RAD54B;refGene_tx=XM_023248148.1;refGene_gn=RAD54B;refGene_tx=XM_023248149.1;refGene_gn=RAD54B;refGene_tx=XM_023248150.1
F2 43244822 43245334
refGene_gn=RAD54B;refGene_tx=XM_023248147.1;refGene_gn=RAD54B;refGene_tx=XM_023248148.1;refGene_gn=RAD54B;refGene_tx=XM_023248149.1
F2 43249084 43249458
refGene_gn=RAD54B;refGene_tx=XM_023248147.1;refGene_gn=RAD54B;refGene_tx=XM_023248148.1
F2 43268575 43268597 refGene_gn=RAD54B;refGene_tx=XM_023248150.1
F2 43271236 43271405 refGene_gn=RAD54B;refGene_tx=XM_003999976.5
F2 43288156 43288291 refGene_gn=RAD54B;refGene_tx=XM_003999976.5
F2 43299537 43299692

refGene_gn=VIRMA;refGene_tx=XM_003999978.4;refGene_gn=VIRMA;refGene_tx=XM_011291400.2;refGene_gn=VIRMA;refGene_tx=XM_019822895.1
F2 43300512 43300656
refGene_gn=VIRMA;refGene_tx=XM_003999978.4;refGene_gn=VIRMA;refGene_tx=XM_011291400.2;refGene_gn=VIRMA;refGene_tx=XM_019822895.1
F2 43302790 43303122
refGene_gn=VIRMA;refGene_tx=XM_003999978.4;refGene_gn=VIRMA;refGene_tx=XM_011291400.2;refGene_gn=VIRMA;refGene_tx=XM_019822895.1
F2 43303578 43303745
refGene_gn=VIRMA;refGene_tx=XM_003999978.4;refGene_gn=VIRMA;refGene_tx=XM_011291400.2;refGene_gn=VIRMA;refGene_tx=XM_019822895.1
F2 43306732 43306829
refGene_gn=VIRMA;refGene_tx=XM_003999978.4;refGene_gn=VIRMA;refGene_tx=XM_011291400.2;refGene_gn=VIRMA;refGene_tx=XM_019822895.1
F2 43307480 43307641
refGene_gn=VIRMA;refGene_tx=XM_003999978.4;refGene_gn=VIRMA;refGene_tx=XM_011291400.2;refGene_gn=VIRMA;refGene_tx=XM_019822895.1
F2 43308109 43308262
refGene_gn=VIRMA;refGene_tx=XM_003999978.4;refGene_gn=VIRMA;refGene_tx=XM_011291400.2
F2 43311749 43311882
refGene_gn=VIRMA;refGene_tx=XM_003999978.4;refGene_gn=VIRMA;refGene_tx=XM_011291400.2;refGene_gn=VIRMA;refGene_tx=XM_019822895.1
F2 43319288 43319506
refGene_gn=VIRMA;refGene_tx=XM_003999978.4;refGene_gn=VIRMA;refGene_tx=XM_011291400.2;refGene_gn=VIRMA;refGene_tx=XM_019822895.1
F2 43321662 43321915
refGene_gn=VIRMA;refGene_tx=XM_003999978.4;refGene_gn=VIRMA;refGene_tx=XM_011291400.2;refGene_gn=VIRMA;refGene_tx=XM_019822895.1
F2 43322428 43322664
refGene_gn=VIRMA;refGene_tx=XM_003999978.4;refGene_gn=VIRMA;refGene_tx=XM_011291400.2;refGene_gn=VIRMA;refGene_tx=XM_019822895.1
F2 43323191 43323736
refGene_gn=VIRMA;refGene_tx=XM_003999978.4;refGene_gn=VIRMA;refGene_tx=XM_011291400.2;refGene_gn=VIRMA;refGene_tx=XM_019822895.1
F2 43323876 43323970
refGene_gn=VIRMA;refGene_tx=XM_003999978.4;refGene_gn=VIRMA;refGene_tx=XM_011291400.2;refGene_gn=VIRMA;refGene_tx=XM_019822895.1
F2 43325979 43326062
refGene_gn=VIRMA;refGene_tx=XM_003999978.4;refGene_gn=VIRMA;refGene_tx=XM_011291400.2;refGene_gn=VIRMA;refGene_tx=XM_019822895.1
F2 43328303 43328458
refGene_gn=VIRMA;refGene_tx=XM_003999978.4;refGene_gn=VIRMA;refGene_tx=XM_011291400.2;refGene_gn=VIRMA;refGene_tx=XM_019822895.1
F2 43329484 43329976
refGene_gn=VIRMA;refGene_tx=XM_003999978.4;refGene_gn=VIRMA;refGene_tx=XM_011291400.2;refGene_gn=VIRMA;refGene_tx=XM_019822895.1
F2 43332891 43334032
refGene_gn=VIRMA;refGene_tx=XM_003999978.4;refGene_gn=VIRMA;refGene_tx=XM_011291400.2;refGene_gn=VIRMA;refGene_tx=XM_019822895.1

F2 43337830 43338103
refGene_gn=VIRMA;refGene_tx=XM_003999978.4;refGene_gn=VIRMA;refGene_tx=XM_011291400.2;refGene_gn=VIRMA;refGene_tx=XM_019822895.1
F2 43338660 43338783
refGene_gn=VIRMA;refGene_tx=XM_003999978.4;refGene_gn=VIRMA;refGene_tx=XM_011291400.2;refGene_gn=VIRMA;refGene_tx=XM_019822895.1
F2 43340679 43340848
refGene_gn=VIRMA;refGene_tx=XM_003999978.4;refGene_gn=VIRMA;refGene_tx=XM_011291400.2;refGene_gn=VIRMA;refGene_tx=XM_019822895.1
F2 43342611 43342660
refGene_gn=VIRMA;refGene_tx=XM_003999978.4;refGene_gn=VIRMA;refGene_tx=XM_011291400.2;refGene_gn=VIRMA;refGene_tx=XM_019822895.1
F2 43343953 43344055
refGene_gn=VIRMA;refGene_tx=XM_003999978.4;refGene_gn=VIRMA;refGene_tx=XM_011291400.2;refGene_gn=VIRMA;refGene_tx=XM_019822895.1
F2 43348538 43348654
refGene_gn=VIRMA;refGene_tx=XM_003999978.4;refGene_gn=VIRMA;refGene_tx=XM_011291400.2;refGene_gn=VIRMA;refGene_tx=XM_019822895.1
F2 43357888 43357951
refGene_gn=VIRMA;refGene_tx=XM_003999978.4;refGene_gn=VIRMA;refGene_tx=XM_011291400.2;refGene_gn=VIRMA;refGene_tx=XM_019822895.1
F2 43420193 43420325
refGene_gn=ESRP1;refGene_tx=XM_006943309.3;refGene_gn=ESRP1;refGene_tx=XM_006943310.3;refGene_gn=ESRP1;refGene_tx=XM_019822896.2;refGene_gn=ESRP1;refGene_tx=XM_019822898.2;refGene_gn=ESRP1;refGene_tx=XM_019822899.2
F2 43420837 43420966
refGene_gn=ESRP1;refGene_tx=XM_006943309.3;refGene_gn=ESRP1;refGene_tx=XM_006943310.3;refGene_gn=ESRP1;refGene_tx=XM_019822896.2;refGene_gn=ESRP1;refGene_tx=XM_019822898.2;refGene_gn=ESRP1;refGene_tx=XM_019822899.2
F2 43422219 43422333
refGene_gn=ESRP1;refGene_tx=XM_006943309.3;refGene_gn=ESRP1;refGene_tx=XM_006943310.3;refGene_gn=ESRP1;refGene_tx=XM_019822896.2;refGene_gn=ESRP1;refGene_tx=XM_019822898.2;refGene_gn=ESRP1;refGene_tx=XM_019822899.2
F2 43425348 43425463
refGene_gn=ESRP1;refGene_tx=XM_006943309.3;refGene_gn=ESRP1;refGene_tx=XM_006943310.3;refGene_gn=ESRP1;refGene_tx=XM_019822896.2;refGene_gn=ESRP1;refGene_tx=XM_019822898.2;refGene_gn=ESRP1;refGene_tx=XM_019822899.2
F2 43438480 43438579
refGene_gn=ESRP1;refGene_tx=XM_006943309.3;refGene_gn=ESRP1;refGene_tx=XM_006943310.3;refGene_gn=ESRP1;refGene_tx=XM_019822896.2;refGene_gn=ESRP1;refGene_tx=XM_019822898.2;refGene_gn=ESRP1;refGene_tx=XM_019822899.2
F2 43438716 43438771
refGene_gn=ESRP1;refGene_tx=XM_006943309.3;refGene_gn=ESRP1;refGene_tx=XM_006943310.3;refGene_gn=ESRP1;refGene_tx=XM_019822896.2;refGene_gn=ESRP1;refGene_tx=XM_019822898.2;refGene_gn=ESRP1;refGene_tx=XM_0198228

99.2
F2 43440213 43440324
refGene_gn=ESRP1;refGene_tx=XM_006943309.3;refGene_gn=ESRP1;refGene_tx
=XM_006943310.3;refGene_gn=ESRP1;refGene_tx=XM_019822896.2;refGene_gn=
ESRP1;refGene_tx=XM_019822898.2;refGene_gn=ESRP1;refGene_tx=XM_0198228
99.2
F2 43440434 43440567
refGene_gn=ESRP1;refGene_tx=XM_006943309.3;refGene_gn=ESRP1;refGene_tx
=XM_006943310.3;refGene_gn=ESRP1;refGene_tx=XM_019822896.2;refGene_gn=
ESRP1;refGene_tx=XM_019822898.2;refGene_gn=ESRP1;refGene_tx=XM_0198228
99.2
F2 43440661 43440704
refGene_gn=ESRP1;refGene_tx=XM_006943309.3;refGene_gn=ESRP1;refGene_tx
=XM_006943310.3;refGene_gn=ESRP1;refGene_tx=XM_019822896.2;refGene_gn=
ESRP1;refGene_tx=XM_019822898.2;refGene_gn=ESRP1;refGene_tx=XM_0198228
99.2
F2 43442863 43443165
refGene_gn=ESRP1;refGene_tx=XM_006943309.3;refGene_gn=ESRP1;refGene_tx
=XM_006943310.3;refGene_gn=ESRP1;refGene_tx=XM_019822896.2;refGene_gn=
ESRP1;refGene_tx=XM_019822898.2;refGene_gn=ESRP1;refGene_tx=XM_0198228
99.2
F2 43446514 43446733
refGene_gn=ESRP1;refGene_tx=XM_006943309.3;refGene_gn=ESRP1;refGene_tx
=XM_006943310.3;refGene_gn=ESRP1;refGene_tx=XM_019822896.2;refGene_gn=
ESRP1;refGene_tx=XM_019822898.2;refGene_gn=ESRP1;refGene_tx=XM_0198228
99.2
F2 43448941 43449140
refGene_gn=ESRP1;refGene_tx=XM_006943309.3;refGene_gn=ESRP1;refGene_tx
=XM_006943310.3;refGene_gn=ESRP1;refGene_tx=XM_019822896.2;refGene_gn=
ESRP1;refGene_tx=XM_019822898.2;refGene_gn=ESRP1;refGene_tx=XM_0198228
99.2
F2 43453146 43453315
refGene_gn=ESRP1;refGene_tx=XM_006943309.3;refGene_gn=ESRP1;refGene_tx
=XM_006943310.3;refGene_gn=ESRP1;refGene_tx=XM_019822896.2;refGene_gn=
ESRP1;refGene_tx=XM_019822898.2;refGene_gn=ESRP1;refGene_tx=XM_0198228
99.2
F2 43468709 43468860
refGene_gn=ESRP1;refGene_tx=XM_006943309.3;refGene_gn=ESRP1;refGene_tx
=XM_006943310.3;refGene_gn=ESRP1;refGene_tx=XM_019822896.2
F2 43473268 43473359
refGene_gn=ESRP1;refGene_tx=XM_006943309.3;refGene_gn=ESRP1;refGene_tx
=XM_006943310.3;refGene_gn=ESRP1;refGene_tx=XM_019822898.2
F2 43482582 43482603
refGene_gn=ESRP1;refGene_tx=XM_019822896.2;refGene_gn=ESRP1;refGene_tx
=XM_019822899.2
F2 43495121 43495137
refGene_gn=DPY19L4;refGene_tx=XM_023248143.1;refGene_gn=DPY19L4;refGen
e_tx=XM_023248144.1;refGene_gn=DPY19L4;refGene_tx=XM_023248145.1
F2 43499672 43499783
refGene_gn=DPY19L4;refGene_tx=XM_023248143.1;refGene_gn=DPY19L4;refGen

e_tx=XM_023248144.1;refGene_gn=DPY19L4;refGene_tx=XM_023248145.1;refGene_gn=DPY19L4;refGene_tx=XM_023248146.1
F2 43508029 43508154
refGene_gn=DPY19L4;refGene_tx=XM_023248143.1;refGene_gn=DPY19L4;refGene_tx=XM_023248144.1;refGene_gn=DPY19L4;refGene_tx=XM_023248145.1;refGene_gn=DPY19L4;refGene_tx=XM_023248146.1
F2 43510357 43510448
refGene_gn=DPY19L4;refGene_tx=XM_019822903.2;refGene_gn=DPY19L4;refGene_tx=XM_023248143.1;refGene_gn=DPY19L4;refGene_tx=XM_023248144.1;refGene_gn=DPY19L4;refGene_tx=XM_023248145.1;refGene_gn=DPY19L4;refGene_tx=XM_023248146.1
F2 43511302 43511424
refGene_gn=DPY19L4;refGene_tx=XM_019822903.2;refGene_gn=DPY19L4;refGene_tx=XM_019822904.2;refGene_gn=DPY19L4;refGene_tx=XM_023248143.1;refGene_gn=DPY19L4;refGene_tx=XM_023248144.1;refGene_gn=DPY19L4;refGene_tx=XM_023248145.1;refGene_gn=DPY19L4;refGene_tx=XM_023248146.1
F2 43511527 43511673
refGene_gn=DPY19L4;refGene_tx=XM_019822903.2;refGene_gn=DPY19L4;refGene_tx=XM_019822904.2;refGene_gn=DPY19L4;refGene_tx=XM_023248143.1;refGene_gn=DPY19L4;refGene_tx=XM_023248144.1;refGene_gn=DPY19L4;refGene_tx=XM_023248145.1;refGene_gn=DPY19L4;refGene_tx=XM_023248146.1
F2 43521504 43521628
refGene_gn=DPY19L4;refGene_tx=XM_019822903.2;refGene_gn=DPY19L4;refGene_tx=XM_019822904.2;refGene_gn=DPY19L4;refGene_tx=XM_023248143.1;refGene_gn=DPY19L4;refGene_tx=XM_023248144.1;refGene_gn=DPY19L4;refGene_tx=XM_023248145.1;refGene_gn=DPY19L4;refGene_tx=XM_023248146.1
F2 43525511 43525646
refGene_gn=DPY19L4;refGene_tx=XM_019822903.2;refGene_gn=DPY19L4;refGene_tx=XM_019822904.2;refGene_gn=DPY19L4;refGene_tx=XM_023248143.1;refGene_gn=DPY19L4;refGene_tx=XM_023248144.1;refGene_gn=DPY19L4;refGene_tx=XM_023248145.1;refGene_gn=DPY19L4;refGene_tx=XM_023248146.1
F2 43528334 43528466
refGene_gn=DPY19L4;refGene_tx=XM_019822903.2;refGene_gn=DPY19L4;refGene_tx=XM_019822904.2;refGene_gn=DPY19L4;refGene_tx=XM_023248143.1;refGene_gn=DPY19L4;refGene_tx=XM_023248144.1;refGene_gn=DPY19L4;refGene_tx=XM_023248145.1;refGene_gn=DPY19L4;refGene_tx=XM_023248146.1
F2 43528562 43528661
refGene_gn=DPY19L4;refGene_tx=XM_019822903.2;refGene_gn=DPY19L4;refGene_tx=XM_019822904.2;refGene_gn=DPY19L4;refGene_tx=XM_023248143.1;refGene_gn=DPY19L4;refGene_tx=XM_023248144.1;refGene_gn=DPY19L4;refGene_tx=XM_023248145.1;refGene_gn=DPY19L4;refGene_tx=XM_023248146.1
F2 43529096 43529170
refGene_gn=DPY19L4;refGene_tx=XM_019822903.2;refGene_gn=DPY19L4;refGene_tx=XM_019822904.2;refGene_gn=DPY19L4;refGene_tx=XM_023248143.1;refGene_gn=DPY19L4;refGene_tx=XM_023248144.1;refGene_gn=DPY19L4;refGene_tx=XM_023248145.1;refGene_gn=DPY19L4;refGene_tx=XM_023248146.1
F2 43534411 43534570
refGene_gn=DPY19L4;refGene_tx=XM_019822903.2;refGene_gn=DPY19L4;refGene_tx=XM_019822904.2;refGene_gn=DPY19L4;refGene_tx=XM_023248143.1;refGene_gn=DPY19L4;refGene_tx=XM_023248144.1;refGene_gn=DPY19L4;refGene_tx=

XM_023248145.1;refGene_gn=DPY19L4;refGene_tx=XM_023248146.1
F2 43535221 43535341
refGene_gn=DPY19L4;refGene_tx=XM_019822903.2;refGene_gn=DPY19L4;refGene_tx=XM_019822904.2;refGene_gn=DPY19L4;refGene_tx=XM_023248143.1;refGene_gn=DPY19L4;refGene_tx=XM_023248144.1;refGene_gn=DPY19L4;refGene_tx=XM_023248145.1;refGene_gn=DPY19L4;refGene_tx=XM_023248146.1
F2 43541234 43541355
refGene_gn=DPY19L4;refGene_tx=XM_019822903.2;refGene_gn=DPY19L4;refGene_tx=XM_019822904.2;refGene_gn=DPY19L4;refGene_tx=XM_023248143.1;refGene_gn=DPY19L4;refGene_tx=XM_023248144.1;refGene_gn=DPY19L4;refGene_tx=XM_023248145.1;refGene_gn=DPY19L4;refGene_tx=XM_023248146.1
F2 43543833 43543890
refGene_gn=DPY19L4;refGene_tx=XM_019822903.2;refGene_gn=DPY19L4;refGene_tx=XM_019822904.2;refGene_gn=DPY19L4;refGene_tx=XM_023248143.1;refGene_gn=DPY19L4;refGene_tx=XM_023248145.1;refGene_gn=DPY19L4;refGene_tx=XM_023248146.1
F2 43548229 43548312
refGene_gn=DPY19L4;refGene_tx=XM_019822903.2;refGene_gn=DPY19L4;refGene_tx=XM_019822904.2;refGene_gn=DPY19L4;refGene_tx=XM_023248143.1;refGene_gn=DPY19L4;refGene_tx=XM_023248144.1;refGene_gn=DPY19L4;refGene_tx=XM_023248145.1;refGene_gn=DPY19L4;refGene_tx=XM_023248146.1
F2 43552486 43552619
refGene_gn=DPY19L4;refGene_tx=XM_019822903.2;refGene_gn=DPY19L4;refGene_tx=XM_019822904.2;refGene_gn=DPY19L4;refGene_tx=XM_023248143.1;refGene_gn=DPY19L4;refGene_tx=XM_023248144.1;refGene_gn=DPY19L4;refGene_tx=XM_023248145.1;refGene_gn=DPY19L4;refGene_tx=XM_023248146.1
F2 43559895 43560054
refGene_gn=DPY19L4;refGene_tx=XM_019822903.2;refGene_gn=DPY19L4;refGene_tx=XM_019822904.2;refGene_gn=DPY19L4;refGene_tx=XM_023248143.1;refGene_gn=DPY19L4;refGene_tx=XM_023248144.1;refGene_gn=DPY19L4;refGene_tx=XM_023248145.1;refGene_gn=DPY19L4;refGene_tx=XM_023248146.1
F2 43560314 43560479
refGene_gn=DPY19L4;refGene_tx=XM_019822903.2;refGene_gn=DPY19L4;refGene_tx=XM_019822904.2;refGene_gn=DPY19L4;refGene_tx=XM_023248143.1;refGene_gn=DPY19L4;refGene_tx=XM_023248144.1;refGene_gn=DPY19L4;refGene_tx=XM_023248145.1;refGene_gn=DPY19L4;refGene_tx=XM_023248146.1
F2 43583597 43583727
refGene_gn=INTS8;refGene_tx=XM_003999983.4;refGene_gn=INTS8;refGene_tx=XM_019822905.2;refGene_gn=INTS8;refGene_tx=XM_019822907.2
F2 43585052 43585227
refGene_gn=INTS8;refGene_tx=XM_003999983.4;refGene_gn=INTS8;refGene_tx=XM_019822905.2;refGene_gn=INTS8;refGene_tx=XM_019822907.2
F2 43587332 43587497
refGene_gn=INTS8;refGene_tx=XM_003999983.4;refGene_gn=INTS8;refGene_tx=XM_019822905.2;refGene_gn=INTS8;refGene_tx=XM_019822907.2
F2 43587821 43587893
refGene_gn=INTS8;refGene_tx=XM_003999983.4;refGene_gn=INTS8;refGene_tx=XM_019822905.2;refGene_gn=INTS8;refGene_tx=XM_019822907.2
F2 43588821 43588873
refGene_gn=INTS8;refGene_tx=XM_003999983.4;refGene_gn=INTS8;refGene_tx

=XM_019822905.2;refGene_gn=INTS8;refGene_tx=XM_019822907.2
F2	43590890	43591073
refGene_gn=INTS8;refGene_tx=XM_003999983.4;refGene_gn=INTS8;refGene_tx=XM_019822905.2;refGene_gn=INTS8;refGene_tx=XM_019822907.2
F2	43594602	43594710
refGene_gn=INTS8;refGene_tx=XM_003999983.4;refGene_gn=INTS8;refGene_tx=XM_019822905.2
F2	43596073	43596229
refGene_gn=INTS8;refGene_tx=XM_003999983.4;refGene_gn=INTS8;refGene_tx=XM_019822905.2;refGene_gn=INTS8;refGene_tx=XM_019822907.2
F2	43597740	43597841
refGene_gn=INTS8;refGene_tx=XM_003999983.4;refGene_gn=INTS8;refGene_tx=XM_019822905.2;refGene_gn=INTS8;refGene_tx=XM_019822907.2
F2	43598279	43598421
refGene_gn=INTS8;refGene_tx=XM_003999983.4;refGene_gn=INTS8;refGene_tx=XM_019822905.2;refGene_gn=INTS8;refGene_tx=XM_019822907.2
F2	43603475	43603546
refGene_gn=INTS8;refGene_tx=XM_003999983.4;refGene_gn=INTS8;refGene_tx=XM_019822905.2;refGene_gn=INTS8;refGene_tx=XM_019822907.2
F2	43603902	43604078
refGene_gn=INTS8;refGene_tx=XM_003999983.4;refGene_gn=INTS8;refGene_tx=XM_019822905.2;refGene_gn=INTS8;refGene_tx=XM_019822907.2
F2	43605933	43606067
refGene_gn=INTS8;refGene_tx=XM_003999983.4;refGene_gn=INTS8;refGene_tx=XM_019822905.2;refGene_gn=INTS8;refGene_tx=XM_019822907.2
F2	43607495	43607606
refGene_gn=INTS8;refGene_tx=XM_003999983.4;refGene_gn=INTS8;refGene_tx=XM_019822905.2;refGene_gn=INTS8;refGene_tx=XM_019822907.2
F2	43612575	43612777
refGene_gn=INTS8;refGene_tx=XM_003999983.4;refGene_gn=INTS8;refGene_tx=XM_019822905.2;refGene_gn=INTS8;refGene_tx=XM_019822907.2
F2	43616710	43616832
refGene_gn=INTS8;refGene_tx=XM_003999983.4;refGene_gn=INTS8;refGene_tx=XM_019822905.2;refGene_gn=INTS8;refGene_tx=XM_019822907.2
F2	43621766	43621951
refGene_gn=INTS8;refGene_tx=XM_003999983.4;refGene_gn=INTS8;refGene_tx=XM_019822905.2;refGene_gn=INTS8;refGene_tx=XM_019822907.2
F2	43622359	43622393
refGene_gn=INTS8;refGene_tx=XM_003999983.4;refGene_gn=INTS8;refGene_tx=XM_019822905.2;refGene_gn=INTS8;refGene_tx=XM_019822907.2
F2	43623635	43623692
refGene_gn=INTS8;refGene_tx=XM_003999983.4;refGene_gn=INTS8;refGene_tx=XM_019822905.2;refGene_gn=INTS8;refGene_tx=XM_019822907.2
F2	43623783	43623845
refGene_gn=INTS8;refGene_tx=XM_003999983.4;refGene_gn=INTS8;refGene_tx=XM_019822905.2;refGene_gn=INTS8;refGene_tx=XM_019822907.2
F2	43626425	43626544
refGene_gn=INTS8;refGene_tx=XM_003999983.4;refGene_gn=INTS8;refGene_tx=XM_019822905.2;refGene_gn=INTS8;refGene_tx=XM_019822907.2
F2	43627375	43627479

refGene_gn=INTS8;refGene_tx=XM_003999983.4;refGene_gn=INTS8;refGene_tx=XM_019822905.2;refGene_gn=INTS8;refGene_tx=XM_019822907.2
F2	43628247	43628298	refGene_gn=INTS8;refGene_tx=XM_003999983.4;refGene_gn=INTS8;refGene_tx=XM_019822905.2;refGene_gn=INTS8;refGene_tx=XM_019822907.2
F2	43630259	43630333	refGene_gn=INTS8;refGene_tx=XM_003999983.4;refGene_gn=INTS8;refGene_tx=XM_019822905.2;refGene_gn=INTS8;refGene_tx=XM_019822907.2
F2	43630413	43630478	refGene_gn=INTS8;refGene_tx=XM_003999983.4;refGene_gn=INTS8;refGene_tx=XM_019822905.2;refGene_gn=INTS8;refGene_tx=XM_019822907.2
F2	43630610	43630654	refGene_gn=INTS8;refGene_tx=XM_003999983.4;refGene_gn=INTS8;refGene_tx=XM_019822905.2;refGene_gn=INTS8;refGene_tx=XM_019822907.2
F2	43636729	43636846	refGene_gn=INTS8;refGene_tx=XM_003999983.4;refGene_gn=INTS8;refGene_tx=XM_019822905.2;refGene_gn=INTS8;refGene_tx=XM_019822907.2
F2	43638209	43638323	refGene_gn=CCNE2;refGene_tx=XM_019822908.2
F2	43638697	43638855	refGene_gn=CCNE2;refGene_tx=XM_019822908.2
F2	43639327	43639439	refGene_gn=CCNE2;refGene_tx=XM_019822908.2
F2	43641007	43641142	refGene_gn=CCNE2;refGene_tx=XM_019822908.2
F2	43642429	43642525	refGene_gn=CCNE2;refGene_tx=XM_019822908.2
F2	43643727	43643874	refGene_gn=CCNE2;refGene_tx=XM_019822908.2
F2	43646540	43646606	refGene_gn=CCNE2;refGene_tx=XM_019822908.2
F2	43646607	43646677	refGene_gn=CCNE2;refGene_tx=XM_019822908.2
F2	43648009	43648161	refGene_gn=CCNE2;refGene_tx=XM_019822908.2
F2	43649101	43649155	refGene_gn=CCNE2;refGene_tx=XM_019822908.2
F2	43649236	43649339	refGene_gn=CCNE2;refGene_tx=XM_019822908.2
F2	43649433	43649447	refGene_gn=CCNE2;refGene_tx=XM_019822908.2
F2	43687367	43687617	refGene_gn=TP53INP1;refGene_tx=XM_023248616.1;refGene_gn=TP53INP1;refGene_tx=XM_023248617.1
F2	43688740	43688762	refGene_gn=TP53INP1;refGene_tx=XM_006943313.4
F2	43691924	43692282	refGene_gn=TP53INP1;refGene_tx=XM_006943313.4;refGene_gn=TP53INP1;refGene_tx=XM_023248616.1;refGene_gn=TP53INP1;refGene_tx=XM_023248617.1
F2	43692830	43692942	refGene_gn=TP53INP1;refGene_tx=XM_006943313.4;refGene_gn=TP53INP1;refGene_tx=XM_023248616.1;refGene_gn=TP53INP1;refGene_tx=XM_023248617.1
F2	43780557	43780754	refGene_gn=NDUFAF6;refGene_tx=XM_011291403.2
F2	43788183	43788287	refGene_gn=NDUFAF6;refGene_tx=XM_011291403.2;refGene_gn=NDUFAF6;refGene_tx=XM_019822914.1;refGene_gn=NDUFAF6;refGene_tx=XM_023248613.1
F2	43793047	43793170	refGene_gn=NDUFAF6;refGene_tx=XM_011291403.2;refGene_gn=NDUFAF6;refGene_tx=XM_019822914.1;refGene_gn=NDUFAF6;refGene_tx=XM_023248613.1;refGene_gn=NDUFAF6;refGene_tx=XM_023248614.1
F2	43798489	43798546	refGene_gn=NDUFAF6;refGene_tx=XM_011291403.2;refGene_gn=NDUFAF6;refGene_tx=XM_019822914.1;refGene_gn=NDUFAF6;refGene_tx=XM_023248613.1;refGe

ne_gn=NDUFAF6;refGene_tx=XM_023248614.1;refGene_gn=NDUFAF6;refGene_tx=XM_023248615.1
F2 43802302 43802405
refGene_gn=NDUFAF6;refGene_tx=XM_011291403.2;refGene_gn=NDUFAF6;refGene_tx=XM_019822914.1;refGene_gn=NDUFAF6;refGene_tx=XM_023248613.1;refGene_gn=NDUFAF6;refGene_tx=XM_023248614.1;refGene_gn=NDUFAF6;refGene_tx=XM_023248615.1
F2 43803441 43803575
refGene_gn=NDUFAF6;refGene_tx=XM_011291403.2;refGene_gn=NDUFAF6;refGene_tx=XM_019822914.1;refGene_gn=NDUFAF6;refGene_tx=XM_023248613.1;refGene_gn=NDUFAF6;refGene_tx=XM_023248614.1;refGene_gn=NDUFAF6;refGene_tx=XM_023248615.1
F2 43804000 43804102
refGene_gn=NDUFAF6;refGene_tx=XM_011291403.2;refGene_gn=NDUFAF6;refGene_tx=XM_019822914.1;refGene_gn=NDUFAF6;refGene_tx=XM_023248613.1;refGene_gn=NDUFAF6;refGene_tx=XM_023248614.1;refGene_gn=NDUFAF6;refGene_tx=XM_023248615.1
F2 43808328 43808385
refGene_gn=NDUFAF6;refGene_tx=XM_011291403.2;refGene_gn=NDUFAF6;refGene_tx=XM_019822914.1;refGene_gn=NDUFAF6;refGene_tx=XM_023248613.1;refGene_gn=NDUFAF6;refGene_tx=XM_023248614.1;refGene_gn=NDUFAF6;refGene_tx=XM_023248615.1
F2 43811111 43811240
refGene_gn=NDUFAF6;refGene_tx=XM_011291403.2;refGene_gn=NDUFAF6;refGene_tx=XM_019822914.1;refGene_gn=NDUFAF6;refGene_tx=XM_023248613.1;refGene_gn=NDUFAF6;refGene_tx=XM_023248614.1;refGene_gn=NDUFAF6;refGene_tx=XM_023248615.1
F2 43855887 43856637
refGene_gn=PLEKHF2;refGene_tx=XM_003999986.3;refGene_gn=PLEKHF2;refGene_tx=XM_011291404.3;refGene_gn=PLEKHF2;refGene_tx=XM_019822915.2;refGene_gn=PLEKHF2;refGene_tx=XM_019822916.2;refGene_gn=PLEKHF2;refGene_tx=XM_023248253.1;refGene_gn=PLEKHF2;refGene_tx=XM_023248254.1
F2 44006430 44006584
refGene_gn=CF2H8orf37;refGene_tx=XM_003999996.5;refGene_gn=CF2H8orf37;refGene_tx=XM_019822919.2
F2 44010192 44010288
refGene_gn=CF2H8orf37;refGene_tx=XM_003999996.5
F2 44018252 44018318
refGene_gn=CF2H8orf37;refGene_tx=XM_003999996.5;refGene_gn=CF2H8orf37;refGene_tx=XM_019822919.2
F2 44018860 44018925
refGene_gn=CF2H8orf37;refGene_tx=XM_003999996.5;refGene_gn=CF2H8orf37;refGene_tx=XM_019822919.2
F2 44022285 44022373
refGene_gn=CF2H8orf37;refGene_tx=XM_003999996.5;refGene_gn=CF2H8orf37;refGene_tx=XM_019822919.2
F2 44027401 44027556
refGene_gn=CF2H8orf37;refGene_tx=XM_003999996.5;refGene_gn=CF2H8orf37;refGene_tx=XM_019822919.2
F2 44865708 44866673 refGene_gn=GDF6;refGene_tx=XM_023248492.1

F2 44881766 44882190 refGene_gn=GDF6;refGene_tx=XM_023248492.1
F2 44939366 44939444
refGene_gn=LOC101090590;refGene_tx=XM_003999997.5
F2 44940110 44940277
refGene_gn=LOC101090590;refGene_tx=XM_003999997.5
F2 44941319 44941391
refGene_gn=LOC101090590;refGene_tx=XM_003999997.5
F2 44943537 44943556
refGene_gn=LOC101090590;refGene_tx=XM_003999997.5
F2 44947078 44947273
refGene_gn=MTERF3;refGene_tx=XM_003999998.5;refGene_gn=MTERF3;refGene_tx=XM_006943319.3
F2 44950168 44950330
refGene_gn=MTERF3;refGene_tx=XM_003999998.5;refGene_gn=MTERF3;refGene_tx=XM_006943319.3
F2 44951524 44951596
refGene_gn=MTERF3;refGene_tx=XM_003999998.5;refGene_gn=MTERF3;refGene_tx=XM_006943319.3
F2 44951976 44952124
refGene_gn=MTERF3;refGene_tx=XM_003999998.5;refGene_gn=MTERF3;refGene_tx=XM_006943319.3
F2 44956564 44956754
refGene_gn=MTERF3;refGene_tx=XM_003999998.5;refGene_gn=MTERF3;refGene_tx=XM_006943319.3
F2 44961624 44961777
refGene_gn=MTERF3;refGene_tx=XM_003999998.5;refGene_gn=MTERF3;refGene_tx=XM_006943319.3
F2 44963772 44964106
refGene_gn=MTERF3;refGene_tx=XM_003999998.5;refGene_gn=MTERF3;refGene_tx=XM_006943319.3
F2 44967059 44967238
refGene_gn=PTDSS1;refGene_tx=XM_023248647.1;refGene_gn=PTDSS1;refGene_tx=XM_023248648.1
F2 44978163 44978255
refGene_gn=PTDSS1;refGene_tx=XM_023248647.1;refGene_gn=PTDSS1;refGene_tx=XM_023248648.1
F2 44989725 44989770
refGene_gn=PTDSS1;refGene_tx=XM_023248647.1;refGene_gn=PTDSS1;refGene_tx=XM_023248648.1
F2 44992801 44992926
refGene_gn=PTDSS1;refGene_tx=XM_023248647.1;refGene_gn=PTDSS1;refGene_tx=XM_023248648.1
F2 44997233 44997392
refGene_gn=PTDSS1;refGene_tx=XM_023248647.1;refGene_gn=PTDSS1;refGene_tx=XM_023248648.1
F2 45000860 45001012
refGene_gn=PTDSS1;refGene_tx=XM_023248647.1;refGene_gn=PTDSS1;refGene_tx=XM_023248648.1
F2 45005161 45005303
refGene_gn=PTDSS1;refGene_tx=XM_023248647.1;refGene_gn=PTDSS1;refGene_

tx=XM_023248648.1
F2 45007223 45007336
refGene_gn=PTDSS1;refGene_tx=XM_023248647.1;refGene_gn=PTDSS1;refGene_
tx=XM_023248648.1
F2 45010171 45010237
refGene_gn=PTDSS1;refGene_tx=XM_023248647.1;refGene_gn=PTDSS1;refGene_
tx=XM_023248648.1
F2 45018085 45018185
refGene_gn=PTDSS1;refGene_tx=XM_023248647.1;refGene_gn=PTDSS1;refGene_
tx=XM_023248648.1
F2 45022867 45022879 refGene_gn=PTDSS1;refGene_tx=XM_023248648.1
F2 45027347 45027416 refGene_gn=PTDSS1;refGene_tx=XM_023248647.1
F2 45028152 45028222 refGene_gn=PTDSS1;refGene_tx=XM_023248647.1
F2 45030464 45030574 refGene_gn=PTDSS1;refGene_tx=XM_023248647.1
F2 45168095 45168155 refGene_gn=SDC2;refGene_tx=XM_006943320.3
F2 45253764 45253876 refGene_gn=SDC2;refGene_tx=XM_006943320.3
F2 45260519 45260653 refGene_gn=SDC2;refGene_tx=XM_006943320.3
F2 45264854 45264990 refGene_gn=SDC2;refGene_tx=XM_006943320.3
F2 45265876 45266040 refGene_gn=SDC2;refGene_tx=XM_006943320.3
F2 45409243 45409676
refGene_gn=CPQ;refGene_tx=XM_004000001.4;refGene_gn=CPQ;refGene_tx=XM_
019822920.2;refGene_gn=CPQ;refGene_tx=XM_019822921.2;refGene_gn=CPQ;re
fGene_tx=XM_019822922.2;refGene_gn=CPQ;refGene_tx=XM_019822925.2;refGe
ne_gn=CPQ;refGene_tx=XM_019822926.2;refGene_gn=CPQ;refGene_tx=XM_02324
8131.1;refGene_gn=CPQ;refGene_tx=XM_023248132.1
F2 45459519 45459727
refGene_gn=CPQ;refGene_tx=XM_004000001.4;refGene_gn=CPQ;refGene_tx=XM_
019822920.2;refGene_gn=CPQ;refGene_tx=XM_019822921.2;refGene_gn=CPQ;re
fGene_tx=XM_019822922.2;refGene_gn=CPQ;refGene_tx=XM_019822925.2;refGe
ne_gn=CPQ;refGene_tx=XM_019822926.2;refGene_gn=CPQ;refGene_tx=XM_02324
8131.1;refGene_gn=CPQ;refGene_tx=XM_023248132.1
F2 45516053 45516261
refGene_gn=CPQ;refGene_tx=XM_004000001.4;refGene_gn=CPQ;refGene_tx=XM_
019822920.2;refGene_gn=CPQ;refGene_tx=XM_019822921.2;refGene_gn=CPQ;re
fGene_tx=XM_019822922.2;refGene_gn=CPQ;refGene_tx=XM_019822925.2;refGe
ne_gn=CPQ;refGene_tx=XM_019822926.2;refGene_gn=CPQ;refGene_tx=XM_02324
8131.1;refGene_gn=CPQ;refGene_tx=XM_023248132.1
F2 45602583 45602695
refGene_gn=CPQ;refGene_tx=XM_004000001.4;refGene_gn=CPQ;refGene_tx=XM_
019822920.2;refGene_gn=CPQ;refGene_tx=XM_019822921.2;refGene_gn=CPQ;re
fGene_tx=XM_019822922.2;refGene_gn=CPQ;refGene_tx=XM_019822925.2;refGe
ne_gn=CPQ;refGene_tx=XM_019822926.2;refGene_gn=CPQ;refGene_tx=XM_02324
8131.1;refGene_gn=CPQ;refGene_tx=XM_023248132.1
F2 45650831 45650923
refGene_gn=CPQ;refGene_tx=XM_004000001.4;refGene_gn=CPQ;refGene_tx=XM_
019822920.2;refGene_gn=CPQ;refGene_tx=XM_019822921.2;refGene_gn=CPQ;re
fGene_tx=XM_019822922.2;refGene_gn=CPQ;refGene_tx=XM_019822925.2;refGe
ne_gn=CPQ;refGene_tx=XM_019822926.2;refGene_gn=CPQ;refGene_tx=XM_02324
8131.1;refGene_gn=CPQ;refGene_tx=XM_023248132.1
F2 45680602 45680804

refGene_gn=CPQ;refGene_tx=XM_004000001.4;refGene_gn=CPQ;refGene_tx=XM_019822920.2;refGene_gn=CPQ;refGene_tx=XM_019822921.2;refGene_gn=CPQ;refGene_tx=XM_019822922.2;refGene_gn=CPQ;refGene_tx=XM_019822925.2;refGene_gn=CPQ;refGene_tx=XM_019822926.2;refGene_gn=CPQ;refGene_tx=XM_023248131.1;refGene_gn=CPQ;refGene_tx=XM_023248132.1
F2 45718234 45718329 refGene_gn=CPQ;refGene_tx=XM_023248132.1
F2 45748900 45748923 refGene_gn=CPQ;refGene_tx=XM_019822926.2
F2 45755227 45755391 refGene_gn=CPQ;refGene_tx=XM_004000001.4
F2 45784500 45784529 refGene_gn=CPQ;refGene_tx=XM_019822925.2
F2 45819037 45819102 refGene_gn=CPQ;refGene_tx=XM_019822922.2
F2 45821832 45821945 refGene_gn=CPQ;refGene_tx=XM_019822921.2
F2 45854673 45854837
refGene_gn=CPQ;refGene_tx=XM_019822920.2;refGene_gn=CPQ;refGene_tx=XM_023248131.1
F2 45868604 45869852
refGene_gn=TSPYL5;refGene_tx=XM_004000009.5;refGene_gn=TSPYL5;refGene_tx=XM_023248133.1;refGene_gn=TSPYL5;refGene_tx=XM_023248134.1;refGene_gn=TSPYL5;refGene_tx=XM_023248135.1;refGene_gn=TSPYL5;refGene_tx=XM_023248136.1;refGene_gn=TSPYL5;refGene_tx=XM_023248137.1;refGene_gn=TSPYL5;refGene_tx=XM_023248138.1;refGene_gn=TSPYL5;refGene_tx=XM_023248139.1;refGene_gn=TSPYL5;refGene_tx=XM_023248140.1;refGene_gn=TSPYL5;refGene_tx=XM_023248141.1;refGene_gn=TSPYL5;refGene_tx=XM_023248142.1
F2 46121523 46121904
refGene_gn=MTDH;refGene_tx=XM_019822929.2;refGene_gn=MTDH;refGene_tx=XM_023248710.1;refGene_gn=MTDH;refGene_tx=XM_023248711.1;refGene_gn=MTDH;refGene_tx=XM_023248712.1;refGene_gn=MTDH;refGene_tx=XM_023248713.1
F2 46134855 46134957
refGene_gn=MTDH;refGene_tx=XM_019822929.2;refGene_gn=MTDH;refGene_tx=XM_023248710.1;refGene_gn=MTDH;refGene_tx=XM_023248711.1;refGene_gn=MTDH;refGene_tx=XM_023248712.1;refGene_gn=MTDH;refGene_tx=XM_023248713.1
F2 46147121 46147206
refGene_gn=MTDH;refGene_tx=XM_019822929.2;refGene_gn=MTDH;refGene_tx=XM_023248710.1;refGene_gn=MTDH;refGene_tx=XM_023248711.1;refGene_gn=MTDH;refGene_tx=XM_023248712.1;refGene_gn=MTDH;refGene_tx=XM_023248713.1
F2 46147897 46148074
refGene_gn=MTDH;refGene_tx=XM_019822929.2;refGene_gn=MTDH;refGene_tx=XM_023248710.1;refGene_gn=MTDH;refGene_tx=XM_023248711.1;refGene_gn=MTDH;refGene_tx=XM_023248712.1;refGene_gn=MTDH;refGene_tx=XM_023248713.1
F2 46149468 46149534
refGene_gn=MTDH;refGene_tx=XM_019822929.2;refGene_gn=MTDH;refGene_tx=XM_023248710.1;refGene_gn=MTDH;refGene_tx=XM_023248711.1;refGene_gn=MTDH;refGene_tx=XM_023248712.1;refGene_gn=MTDH;refGene_tx=XM_023248713.1
F2 46151201 46151438
refGene_gn=MTDH;refGene_tx=XM_019822929.2;refGene_gn=MTDH;refGene_tx=XM_023248710.1;refGene_gn=MTDH;refGene_tx=XM_023248711.1;refGene_gn=MTDH;refGene_tx=XM_023248712.1;refGene_gn=MTDH;refGene_tx=XM_023248713.1
F2 46153993 46154083
refGene_gn=MTDH;refGene_tx=XM_019822929.2;refGene_gn=MTDH;refGene_tx=XM_023248711.1
F2 46156311 46156410

refGene_gn=MTDH;refGene_tx=XM_019822929.2;refGene_gn=MTDH;refGene_tx=XM_023248710.1;refGene_gn=MTDH;refGene_tx=XM_023248711.1;refGene_gn=MTDH;refGene_tx=XM_023248713.1
F2	46162807	46162932	refGene_gn=MTDH;refGene_tx=XM_019822929.2;refGene_gn=MTDH;refGene_tx=XM_023248710.1;refGene_gn=MTDH;refGene_tx=XM_023248711.1;refGene_gn=MTDH;refGene_tx=XM_023248712.1;refGene_gn=MTDH;refGene_tx=XM_023248713.1
F2	46169832	46169940	refGene_gn=MTDH;refGene_tx=XM_019822929.2;refGene_gn=MTDH;refGene_tx=XM_023248710.1;refGene_gn=MTDH;refGene_tx=XM_023248711.1;refGene_gn=MTDH;refGene_tx=XM_023248712.1;refGene_gn=MTDH;refGene_tx=XM_023248713.1
F2	46172785	46172926	refGene_gn=MTDH;refGene_tx=XM_019822929.2;refGene_gn=MTDH;refGene_tx=XM_023248710.1;refGene_gn=MTDH;refGene_tx=XM_023248711.1;refGene_gn=MTDH;refGene_tx=XM_023248712.1;refGene_gn=MTDH;refGene_tx=XM_023248713.1
F2	46175480	46175637	refGene_gn=MTDH;refGene_tx=XM_019822929.2;refGene_gn=MTDH;refGene_tx=XM_023248710.1;refGene_gn=MTDH;refGene_tx=XM_023248711.1;refGene_gn=MTDH;refGene_tx=XM_023248712.1;refGene_gn=MTDH;refGene_tx=XM_023248713.1
F2	46176313	46176384	refGene_gn=MTDH;refGene_tx=XM_019822929.2;refGene_gn=MTDH;refGene_tx=XM_023248710.1;refGene_gn=MTDH;refGene_tx=XM_023248711.1;refGene_gn=MTDH;refGene_tx=XM_023248712.1;refGene_gn=MTDH;refGene_tx=XM_023248713.1
F2	46216591	46217023	refGene_gn=LAPTM4B;refGene_tx=XM_023248771.1
F2	46250325	46250437	refGene_gn=LAPTM4B;refGene_tx=XM_023248771.1
F2	46267397	46267471	refGene_gn=LAPTM4B;refGene_tx=XM_023248771.1
F2	46267825	46267948	refGene_gn=LAPTM4B;refGene_tx=XM_023248771.1
F2	46271143	46271242	refGene_gn=LAPTM4B;refGene_tx=XM_023248771.1
F2	46275898	46275994	refGene_gn=LAPTM4B;refGene_tx=XM_023248771.1
F2	46296729	46296807	refGene_gn=LAPTM4B;refGene_tx=XM_023248771.1
F2	46325504	46325649	refGene_gn=MATN2;refGene_tx=XM_023248122.1;refGene_gn=MATN2;refGene_tx=XM_023248123.1;refGene_gn=MATN2;refGene_tx=XM_023248124.1;refGene_gn=MATN2;refGene_tx=XM_023248125.1
F2	46358753	46359323	refGene_gn=MATN2;refGene_tx=XM_023248122.1;refGene_gn=MATN2;refGene_tx=XM_023248123.1;refGene_gn=MATN2;refGene_tx=XM_023248124.1;refGene_gn=MATN2;refGene_tx=XM_023248125.1
F2	46368376	46368499	refGene_gn=MATN2;refGene_tx=XM_023248122.1;refGene_gn=MATN2;refGene_tx=XM_023248123.1;refGene_gn=MATN2;refGene_tx=XM_023248124.1;refGene_gn=MATN2;refGene_tx=XM_023248125.1
F2	46385702	46385825	refGene_gn=MATN2;refGene_tx=XM_023248122.1;refGene_gn=MATN2;refGene_tx=XM_023248123.1;refGene_gn=MATN2;refGene_tx=XM_023248124.1;refGene_gn=MATN2;refGene_tx=XM_023248125.1
F2	46399048	46399171	refGene_gn=MATN2;refGene_tx=XM_023248122.1;refGene_gn=MATN2;refGene_tx=XM_023248123.1;refGene_gn=MATN2;refGene_tx=XM_023248124.1;refGene_gn=MATN2;refGene_tx=XM_023248125.1

F2	46408659	46408782
refGene_gn=MATN2;refGene_tx=XM_023248122.1;refGene_gn=MATN2;refGene_tx=XM_023248123.1;refGene_gn=MATN2;refGene_tx=XM_023248124.1;refGene_gn=MATN2;refGene_tx=XM_023248125.1
F2	46415158	46415281
refGene_gn=MATN2;refGene_tx=XM_023248122.1;refGene_gn=MATN2;refGene_tx=XM_023248123.1;refGene_gn=MATN2;refGene_tx=XM_023248124.1;refGene_gn=MATN2;refGene_tx=XM_023248125.1
F2	46418824	46418947
refGene_gn=MATN2;refGene_tx=XM_023248122.1;refGene_gn=MATN2;refGene_tx=XM_023248123.1;refGene_gn=MATN2;refGene_tx=XM_023248124.1;refGene_gn=MATN2;refGene_tx=XM_023248125.1
F2	46419201	46419324
refGene_gn=MATN2;refGene_tx=XM_023248122.1;refGene_gn=MATN2;refGene_tx=XM_023248123.1;refGene_gn=MATN2;refGene_tx=XM_023248124.1;refGene_gn=MATN2;refGene_tx=XM_023248125.1
F2	46431000	46431132
refGene_gn=MATN2;refGene_tx=XM_023248122.1;refGene_gn=MATN2;refGene_tx=XM_023248123.1;refGene_gn=MATN2;refGene_tx=XM_023248124.1;refGene_gn=MATN2;refGene_tx=XM_023248125.1
F2	46432572	46432695
refGene_gn=MATN2;refGene_tx=XM_023248122.1;refGene_gn=MATN2;refGene_tx=XM_023248123.1;refGene_gn=MATN2;refGene_tx=XM_023248124.1;refGene_gn=MATN2;refGene_tx=XM_023248125.1
F2	46435650	46435773
refGene_gn=MATN2;refGene_tx=XM_023248122.1;refGene_gn=MATN2;refGene_tx=XM_023248123.1;refGene_gn=MATN2;refGene_tx=XM_023248124.1;refGene_gn=MATN2;refGene_tx=XM_023248125.1
F2	46439243	46439657
refGene_gn=MATN2;refGene_tx=XM_023248122.1;refGene_gn=MATN2;refGene_tx=XM_023248123.1;refGene_gn=MATN2;refGene_tx=XM_023248124.1;refGene_gn=MATN2;refGene_tx=XM_023248125.1
F2	46442509	46442662
refGene_gn=MATN2;refGene_tx=XM_023248122.1;refGene_gn=MATN2;refGene_tx=XM_023248123.1;refGene_gn=MATN2;refGene_tx=XM_023248124.1;refGene_gn=MATN2;refGene_tx=XM_023248125.1
F2	46443923	46443995
refGene_gn=MATN2;refGene_tx=XM_023248122.1;refGene_gn=MATN2;refGene_tx=XM_023248123.1;refGene_gn=MATN2;refGene_tx=XM_023248124.1;refGene_gn=MATN2;refGene_tx=XM_023248125.1
F2	46444344	46444479
refGene_gn=MATN2;refGene_tx=XM_023248122.1;refGene_gn=MATN2;refGene_tx=XM_023248123.1;refGene_gn=MATN2;refGene_tx=XM_023248124.1;refGene_gn=MATN2;refGene_tx=XM_023248125.1
F2	46444858	46444957
refGene_gn=MATN2;refGene_tx=XM_023248122.1;refGene_gn=MATN2;refGene_tx=XM_023248123.1;refGene_gn=MATN2;refGene_tx=XM_023248124.1;refGene_gn=MATN2;refGene_tx=XM_023248125.1
F2	46447109	46447165
refGene_gn=MATN2;refGene_tx=XM_023248122.1;refGene_gn=MATN2;refGene_tx

=XM_023248123.1;refGene_gn=MATN2;refGene_tx=XM_023248124.1;refGene_gn=MATN2;refGene_tx=XM_023248125.1
F2 46452560 46452610 refGene_gn=RPL30;refGene_tx=XM_006943321.4
F2 46453343 46453474 refGene_gn=RPL30;refGene_tx=XM_006943321.4
F2 46455878 46456024 refGene_gn=RPL30;refGene_tx=XM_006943321.4
F2 46456286 46456307 refGene_gn=RPL30;refGene_tx=XM_006943321.4
F2 46465593 46465651 refGene_gn=ERICH5;refGene_tx=XM_019822935.2
F2 46485988 46487014 refGene_gn=ERICH5;refGene_tx=XM_019822935.2
F2 46489438 46489551 refGene_gn=ERICH5;refGene_tx=XM_019822935.2
F2 46499272 46499335 refGene_gn=RIDA;refGene_tx=XM_004000005.4
F2 46500633 46500689 refGene_gn=RIDA;refGene_tx=XM_004000005.4
F2 46501442 46501511 refGene_gn=RIDA;refGene_tx=XM_004000005.4
F2 46501761 46501816 refGene_gn=RIDA;refGene_tx=XM_004000005.4
F2 46504084 46504190 refGene_gn=RIDA;refGene_tx=XM_004000005.4
F2 46512610 46512675 refGene_gn=RIDA;refGene_tx=XM_004000005.4
F2 46513936 46514078 refGene_gn=POP1;refGene_tx=XM_004000006.5
F2 46515752 46515920 refGene_gn=POP1;refGene_tx=XM_004000006.5
F2 46517395 46517571 refGene_gn=POP1;refGene_tx=XM_004000006.5
F2 46518945 46519194 refGene_gn=POP1;refGene_tx=XM_004000006.5
F2 46523292 46523380 refGene_gn=POP1;refGene_tx=XM_004000006.5
F2 46523811 46523999 refGene_gn=POP1;refGene_tx=XM_004000006.5
F2 46524871 46525128 refGene_gn=POP1;refGene_tx=XM_004000006.5
F2 46525249 46525343 refGene_gn=POP1;refGene_tx=XM_004000006.5
F2 46527635 46527747 refGene_gn=POP1;refGene_tx=XM_004000006.5
F2 46528320 46528440 refGene_gn=POP1;refGene_tx=XM_004000006.5
F2 46534410 46534526 refGene_gn=POP1;refGene_tx=XM_004000006.5
F2 46536601 46536793 refGene_gn=POP1;refGene_tx=XM_004000006.5
F2 46538225 46538380 refGene_gn=POP1;refGene_tx=XM_004000006.5
F2 46543251 46543593 refGene_gn=POP1;refGene_tx=XM_004000006.5
F2 46544500 46545155 refGene_gn=POP1;refGene_tx=XM_004000006.5
F2 46576250 46576363
refGene_gn=NIPAL2;refGene_tx=XM_019822583.2;refGene_gn=NIPAL2;refGene_tx=XM_023248756.1;refGene_gn=NIPAL2;refGene_tx=XM_023248757.1;refGene_gn=NIPAL2;refGene_tx=XM_023248758.1
F2 46578287 46578382
refGene_gn=NIPAL2;refGene_tx=XM_019822583.2;refGene_gn=NIPAL2;refGene_tx=XM_023248756.1;refGene_gn=NIPAL2;refGene_tx=XM_023248757.1;refGene_gn=NIPAL2;refGene_tx=XM_023248758.1
F2 46579547 46579611
refGene_gn=NIPAL2;refGene_tx=XM_019822583.2;refGene_gn=NIPAL2;refGene_tx=XM_023248756.1;refGene_gn=NIPAL2;refGene_tx=XM_023248757.1;refGene_gn=NIPAL2;refGene_tx=XM_023248758.1
F2 46583943 46584032
refGene_gn=NIPAL2;refGene_tx=XM_019822583.2;refGene_gn=NIPAL2;refGene_tx=XM_023248756.1;refGene_gn=NIPAL2;refGene_tx=XM_023248757.1;refGene_gn=NIPAL2;refGene_tx=XM_023248758.1
F2 46585908 46586044
refGene_gn=NIPAL2;refGene_tx=XM_019822583.2;refGene_gn=NIPAL2;refGene_tx=XM_023248756.1;refGene_gn=NIPAL2;refGene_tx=XM_023248757.1;refGene_gn=NIPAL2;refGene_tx=XM_023248758.1

F2 46593603 46593700
refGene_gn=NIPAL2;refGene_tx=XM_019822583.2;refGene_gn=NIPAL2;refGene_tx=XM_023248756.1;refGene_gn=NIPAL2;refGene_tx=XM_023248757.1;refGene_gn=NIPAL2;refGene_tx=XM_023248758.1
F2 46600724 46600846
refGene_gn=NIPAL2;refGene_tx=XM_019822583.2;refGene_gn=NIPAL2;refGene_tx=XM_023248756.1;refGene_gn=NIPAL2;refGene_tx=XM_023248757.1;refGene_gn=NIPAL2;refGene_tx=XM_023248758.1
F2 46614051 46614111
refGene_gn=NIPAL2;refGene_tx=XM_019822583.2;refGene_gn=NIPAL2;refGene_tx=XM_023248756.1;refGene_gn=NIPAL2;refGene_tx=XM_023248757.1;refGene_gn=NIPAL2;refGene_tx=XM_023248758.1
F2 46624308 46624415
refGene_gn=NIPAL2;refGene_tx=XM_023248756.1;refGene_gn=NIPAL2;refGene_tx=XM_023248757.1;refGene_gn=NIPAL2;refGene_tx=XM_023248758.1
F2 46632263 46632435
refGene_gn=NIPAL2;refGene_tx=XM_019822583.2;refGene_gn=NIPAL2;refGene_tx=XM_023248756.1;refGene_gn=NIPAL2;refGene_tx=XM_023248757.1;refGene_gn=NIPAL2;refGene_tx=XM_023248758.1
F2 46633442 46633511
refGene_gn=NIPAL2;refGene_tx=XM_019822583.2;refGene_gn=NIPAL2;refGene_tx=XM_023248756.1;refGene_gn=NIPAL2;refGene_tx=XM_023248757.1;refGene_gn=NIPAL2;refGene_tx=XM_023248758.1
F2 46648253 46648269
refGene_gn=NIPAL2;refGene_tx=XM_023248756.1;refGene_gn=NIPAL2;refGene_tx=XM_023248757.1;refGene_gn=NIPAL2;refGene_tx=XM_023248758.1
F2 46659665 46659773 refGene_gn=NIPAL2;refGene_tx=XM_019822583.2
F2 46778870 46780304 refGene_gn=KCNS2;refGene_tx=XM_019822937.2
F2 46803206 46803412
refGene_gn=LOC105261378;refGene_tx=XM_019822941.2
F2 46804666 46804825
refGene_gn=STK3;refGene_tx=XM_011291410.3;refGene_gn=STK3;refGene_tx=XM_019822940.2
F2 46815399 46815488
refGene_gn=LOC105261378;refGene_tx=XM_019822941.2
F2 46816458 46816540
refGene_gn=LOC105261378;refGene_tx=XM_019822941.2
F2 46817186 46817356
refGene_gn=LOC105261378;refGene_tx=XM_019822941.2
F2 46817709 46817768
refGene_gn=LOC105261378;refGene_tx=XM_019822941.2
F2 46839345 46839393 refGene_gn=STK3;refGene_tx=XM_023248602.1
F2 46873379 46873555
refGene_gn=STK3;refGene_tx=XM_011291410.3;refGene_gn=STK3;refGene_tx=XM_019822940.2;refGene_gn=STK3;refGene_tx=XM_023248602.1
F2 46887714 46887907
refGene_gn=STK3;refGene_tx=XM_011291410.3;refGene_gn=STK3;refGene_tx=XM_019822940.2;refGene_gn=STK3;refGene_tx=XM_023248602.1
F2 46928902 46929028
refGene_gn=STK3;refGene_tx=XM_011291410.3;refGene_gn=STK3;refGene_tx=X

M_019822940.2;refGene_gn=STK3;refGene_tx=XM_023248602.1
F2 46939817 46939955
refGene_gn=STK3;refGene_tx=XM_011291410.3;refGene_gn=STK3;refGene_tx=X
M_019822940.2;refGene_gn=STK3;refGene_tx=XM_023248602.1
F2 47002069 47002237
refGene_gn=STK3;refGene_tx=XM_011291410.3;refGene_gn=STK3;refGene_tx=X
M_019822940.2;refGene_gn=STK3;refGene_tx=XM_023248602.1
F2 47002747 47002912
refGene_gn=STK3;refGene_tx=XM_011291410.3;refGene_gn=STK3;refGene_tx=X
M_019822940.2;refGene_gn=STK3;refGene_tx=XM_023248602.1
F2 47030712 47030827
refGene_gn=STK3;refGene_tx=XM_011291410.3;refGene_gn=STK3;refGene_tx=X
M_019822940.2;refGene_gn=STK3;refGene_tx=XM_023248602.1
F2 47042077 47042206
refGene_gn=STK3;refGene_tx=XM_011291410.3;refGene_gn=STK3;refGene_tx=X
M_019822940.2;refGene_gn=STK3;refGene_tx=XM_023248602.1
F2 47048581 47048662
refGene_gn=STK3;refGene_tx=XM_011291410.3;refGene_gn=STK3;refGene_tx=X
M_023248602.1
F2 47082820 47082846
refGene_gn=STK3;refGene_tx=XM_011291410.3;refGene_gn=STK3;refGene_tx=X
M_023248602.1
F2 47203415 47204071
refGene_gn=OSR2;refGene_tx=XM_004000015.5;refGene_gn=OSR2;refGene_tx=X
M_004000016.4
F2 47205138 47205238
refGene_gn=OSR2;refGene_tx=XM_004000015.5;refGene_gn=OSR2;refGene_tx=X
M_004000016.4
F2 47206002 47206185
refGene_gn=OSR2;refGene_tx=XM_004000015.5;refGene_gn=OSR2;refGene_tx=X
M_004000016.4
F2 47277453 47277600
refGene_gn=VPS13B;refGene_tx=XM_023248518.1;refGene_gn=VPS13B;refGene_
tx=XM_023248519.1;refGene_gn=VPS13B;refGene_tx=XM_023248520.1;refGene_
gn=VPS13B;refGene_tx=XM_023248521.1;refGene_gn=VPS13B;refGene_tx=XM_02
3248522.1;refGene_gn=VPS13B;refGene_tx=XM_023248523.1;refGene_gn=VPS13
B;refGene_tx=XM_023248524.1;refGene_gn=VPS13B;refGene_tx=XM_023248525.
1
F2 47308283 47308427
refGene_gn=VPS13B;refGene_tx=XM_023248518.1;refGene_gn=VPS13B;refGene_
tx=XM_023248519.1;refGene_gn=VPS13B;refGene_tx=XM_023248520.1;refGene_
gn=VPS13B;refGene_tx=XM_023248521.1;refGene_gn=VPS13B;refGene_tx=XM_02
3248522.1;refGene_gn=VPS13B;refGene_tx=XM_023248524.1;refGene_gn=VPS13
B;refGene_tx=XM_023248525.1
F2 47352331 47352452
refGene_gn=VPS13B;refGene_tx=XM_023248518.1;refGene_gn=VPS13B;refGene_
tx=XM_023248519.1;refGene_gn=VPS13B;refGene_tx=XM_023248520.1;refGene_
gn=VPS13B;refGene_tx=XM_023248521.1;refGene_gn=VPS13B;refGene_tx=XM_02
3248522.1;refGene_gn=VPS13B;refGene_tx=XM_023248523.1;refGene_gn=VPS13
B;refGene_tx=XM_023248524.1;refGene_gn=VPS13B;refGene_tx=XM_023248525.

1
F2 47362846 47363014
refGene_gn=VPS13B;refGene_tx=XM_023248518.1;refGene_gn=VPS13B;refGene_tx=XM_023248519.1;refGene_gn=VPS13B;refGene_tx=XM_023248520.1;refGene_gn=VPS13B;refGene_tx=XM_023248521.1;refGene_gn=VPS13B;refGene_tx=XM_023248522.1;refGene_gn=VPS13B;refGene_tx=XM_023248523.1;refGene_gn=VPS13B;refGene_tx=XM_023248524.1;refGene_gn=VPS13B;refGene_tx=XM_023248525.1
F2 47367423 47367605
refGene_gn=VPS13B;refGene_tx=XM_023248518.1;refGene_gn=VPS13B;refGene_tx=XM_023248519.1;refGene_gn=VPS13B;refGene_tx=XM_023248520.1;refGene_gn=VPS13B;refGene_tx=XM_023248521.1;refGene_gn=VPS13B;refGene_tx=XM_023248522.1;refGene_gn=VPS13B;refGene_tx=XM_023248523.1;refGene_gn=VPS13B;refGene_tx=XM_023248524.1;refGene_gn=VPS13B;refGene_tx=XM_023248525.1
F2 47371930 47372105
refGene_gn=VPS13B;refGene_tx=XM_023248518.1;refGene_gn=VPS13B;refGene_tx=XM_023248519.1;refGene_gn=VPS13B;refGene_tx=XM_023248520.1;refGene_gn=VPS13B;refGene_tx=XM_023248521.1;refGene_gn=VPS13B;refGene_tx=XM_023248522.1;refGene_gn=VPS13B;refGene_tx=XM_023248523.1;refGene_gn=VPS13B;refGene_tx=XM_023248524.1;refGene_gn=VPS13B;refGene_tx=XM_023248525.1
F2 47380747 47381016
refGene_gn=VPS13B;refGene_tx=XM_023248518.1;refGene_gn=VPS13B;refGene_tx=XM_023248519.1;refGene_gn=VPS13B;refGene_tx=XM_023248520.1;refGene_gn=VPS13B;refGene_tx=XM_023248521.1;refGene_gn=VPS13B;refGene_tx=XM_023248522.1;refGene_gn=VPS13B;refGene_tx=XM_023248523.1;refGene_gn=VPS13B;refGene_tx=XM_023248524.1;refGene_gn=VPS13B;refGene_tx=XM_023248525.1
F2 47386019 47386115
refGene_gn=VPS13B;refGene_tx=XM_023248518.1;refGene_gn=VPS13B;refGene_tx=XM_023248519.1;refGene_gn=VPS13B;refGene_tx=XM_023248520.1;refGene_gn=VPS13B;refGene_tx=XM_023248521.1;refGene_gn=VPS13B;refGene_tx=XM_023248522.1;refGene_gn=VPS13B;refGene_tx=XM_023248523.1;refGene_gn=VPS13B;refGene_tx=XM_023248524.1;refGene_gn=VPS13B;refGene_tx=XM_023248525.1
F2 47386585 47386708
refGene_gn=VPS13B;refGene_tx=XM_023248518.1;refGene_gn=VPS13B;refGene_tx=XM_023248519.1;refGene_gn=VPS13B;refGene_tx=XM_023248520.1;refGene_gn=VPS13B;refGene_tx=XM_023248521.1;refGene_gn=VPS13B;refGene_tx=XM_023248522.1;refGene_gn=VPS13B;refGene_tx=XM_023248523.1;refGene_gn=VPS13B;refGene_tx=XM_023248524.1;refGene_gn=VPS13B;refGene_tx=XM_023248525.1
F2 47387325 47387463
refGene_gn=VPS13B;refGene_tx=XM_023248518.1;refGene_gn=VPS13B;refGene_tx=XM_023248519.1;refGene_gn=VPS13B;refGene_tx=XM_023248520.1;refGene_gn=VPS13B;refGene_tx=XM_023248521.1;refGene_gn=VPS13B;refGene_tx=XM_023248522.1;refGene_gn=VPS13B;refGene_tx=XM_023248523.1;refGene_gn=VPS13B;refGene_tx=XM_023248524.1;refGene_gn=VPS13B;refGene_tx=XM_023248525.1

F2 47388394 47388482
refGene_gn=VPS13B;refGene_tx=XM_023248518.1;refGene_gn=VPS13B;refGene_
tx=XM_023248519.1;refGene_gn=VPS13B;refGene_tx=XM_023248520.1;refGene_
gn=VPS13B;refGene_tx=XM_023248521.1;refGene_gn=VPS13B;refGene_tx=XM_02
3248522.1;refGene_gn=VPS13B;refGene_tx=XM_023248523.1;refGene_gn=VPS13
B;refGene_tx=XM_023248524.1;refGene_gn=VPS13B;refGene_tx=XM_023248525.
1
F2 47395234 47395426
refGene_gn=VPS13B;refGene_tx=XM_023248518.1;refGene_gn=VPS13B;refGene_
tx=XM_023248519.1;refGene_gn=VPS13B;refGene_tx=XM_023248520.1;refGene_
gn=VPS13B;refGene_tx=XM_023248521.1;refGene_gn=VPS13B;refGene_tx=XM_02
3248522.1;refGene_gn=VPS13B;refGene_tx=XM_023248523.1;refGene_gn=VPS13
B;refGene_tx=XM_023248524.1;refGene_gn=VPS13B;refGene_tx=XM_023248525.
1
F2 47399807 47399977
refGene_gn=VPS13B;refGene_tx=XM_023248518.1;refGene_gn=VPS13B;refGene_
tx=XM_023248519.1;refGene_gn=VPS13B;refGene_tx=XM_023248520.1;refGene_
gn=VPS13B;refGene_tx=XM_023248521.1;refGene_gn=VPS13B;refGene_tx=XM_02
3248522.1;refGene_gn=VPS13B;refGene_tx=XM_023248523.1;refGene_gn=VPS13
B;refGene_tx=XM_023248524.1;refGene_gn=VPS13B;refGene_tx=XM_023248525.
1
F2 47402988 47403183
refGene_gn=VPS13B;refGene_tx=XM_023248518.1;refGene_gn=VPS13B;refGene_
tx=XM_023248519.1;refGene_gn=VPS13B;refGene_tx=XM_023248520.1;refGene_
gn=VPS13B;refGene_tx=XM_023248521.1;refGene_gn=VPS13B;refGene_tx=XM_02
3248522.1;refGene_gn=VPS13B;refGene_tx=XM_023248523.1;refGene_gn=VPS13
B;refGene_tx=XM_023248524.1;refGene_gn=VPS13B;refGene_tx=XM_023248525.
1
F2 47415302 47415427
refGene_gn=VPS13B;refGene_tx=XM_023248518.1;refGene_gn=VPS13B;refGene_
tx=XM_023248519.1;refGene_gn=VPS13B;refGene_tx=XM_023248520.1;refGene_
gn=VPS13B;refGene_tx=XM_023248521.1;refGene_gn=VPS13B;refGene_tx=XM_02
3248522.1;refGene_gn=VPS13B;refGene_tx=XM_023248523.1;refGene_gn=VPS13
B;refGene_tx=XM_023248524.1;refGene_gn=VPS13B;refGene_tx=XM_023248525.
1
F2 47441964 47442146
refGene_gn=VPS13B;refGene_tx=XM_023248518.1;refGene_gn=VPS13B;refGene_
tx=XM_023248519.1;refGene_gn=VPS13B;refGene_tx=XM_023248520.1;refGene_
gn=VPS13B;refGene_tx=XM_023248521.1;refGene_gn=VPS13B;refGene_tx=XM_02
3248522.1;refGene_gn=VPS13B;refGene_tx=XM_023248523.1;refGene_gn=VPS13
B;refGene_tx=XM_023248524.1;refGene_gn=VPS13B;refGene_tx=XM_023248525.
1
F2 47531171 47531306
refGene_gn=VPS13B;refGene_tx=XM_023248518.1;refGene_gn=VPS13B;refGene_
tx=XM_023248519.1;refGene_gn=VPS13B;refGene_tx=XM_023248520.1;refGene_
gn=VPS13B;refGene_tx=XM_023248521.1;refGene_gn=VPS13B;refGene_tx=XM_02
3248522.1;refGene_gn=VPS13B;refGene_tx=XM_023248523.1;refGene_gn=VPS13
B;refGene_tx=XM_023248524.1;refGene_gn=VPS13B;refGene_tx=XM_023248525.
1
F2 47532298 47532472

refGene_gn=VPS13B;refGene_tx=XM_023248518.1;refGene_gn=VPS13B;refGene_tx=XM_023248519.1;refGene_gn=VPS13B;refGene_tx=XM_023248520.1;refGene_gn=VPS13B;refGene_tx=XM_023248521.1;refGene_gn=VPS13B;refGene_tx=XM_023248522.1;refGene_gn=VPS13B;refGene_tx=XM_023248523.1;refGene_gn=VPS13B;refGene_tx=XM_023248524.1;refGene_gn=VPS13B;refGene_tx=XM_023248525.1

F2 47626466 47626576
refGene_gn=VPS13B;refGene_tx=XM_023248518.1;refGene_gn=VPS13B;refGene_tx=XM_023248519.1;refGene_gn=VPS13B;refGene_tx=XM_023248520.1;refGene_gn=VPS13B;refGene_tx=XM_023248521.1;refGene_gn=VPS13B;refGene_tx=XM_023248522.1;refGene_gn=VPS13B;refGene_tx=XM_023248523.1;refGene_gn=VPS13B;refGene_tx=XM_023248524.1;refGene_gn=VPS13B;refGene_tx=XM_023248525.1

F2 47632508 47632656
refGene_gn=VPS13B;refGene_tx=XM_023248518.1;refGene_gn=VPS13B;refGene_tx=XM_023248519.1;refGene_gn=VPS13B;refGene_tx=XM_023248520.1;refGene_gn=VPS13B;refGene_tx=XM_023248521.1;refGene_gn=VPS13B;refGene_tx=XM_023248522.1;refGene_gn=VPS13B;refGene_tx=XM_023248523.1;refGene_gn=VPS13B;refGene_tx=XM_023248524.1;refGene_gn=VPS13B;refGene_tx=XM_023248525.1

F2 47673840 47673968
refGene_gn=VPS13B;refGene_tx=XM_023248518.1;refGene_gn=VPS13B;refGene_tx=XM_023248519.1;refGene_gn=VPS13B;refGene_tx=XM_023248520.1;refGene_gn=VPS13B;refGene_tx=XM_023248521.1;refGene_gn=VPS13B;refGene_tx=XM_023248522.1;refGene_gn=VPS13B;refGene_tx=XM_023248523.1;refGene_gn=VPS13B;refGene_tx=XM_023248524.1;refGene_gn=VPS13B;refGene_tx=XM_023248525.1

F2 47681120 47681355
refGene_gn=VPS13B;refGene_tx=XM_023248518.1;refGene_gn=VPS13B;refGene_tx=XM_023248519.1;refGene_gn=VPS13B;refGene_tx=XM_023248520.1;refGene_gn=VPS13B;refGene_tx=XM_023248521.1;refGene_gn=VPS13B;refGene_tx=XM_023248522.1;refGene_gn=VPS13B;refGene_tx=XM_023248523.1;refGene_gn=VPS13B;refGene_tx=XM_023248524.1;refGene_gn=VPS13B;refGene_tx=XM_023248525.1

F2 47716961 47717182
refGene_gn=VPS13B;refGene_tx=XM_023248518.1;refGene_gn=VPS13B;refGene_tx=XM_023248519.1;refGene_gn=VPS13B;refGene_tx=XM_023248520.1;refGene_gn=VPS13B;refGene_tx=XM_023248521.1;refGene_gn=VPS13B;refGene_tx=XM_023248522.1;refGene_gn=VPS13B;refGene_tx=XM_023248523.1;refGene_gn=VPS13B;refGene_tx=XM_023248524.1;refGene_gn=VPS13B;refGene_tx=XM_023248525.1

F2 47729337 47729541
refGene_gn=VPS13B;refGene_tx=XM_023248518.1;refGene_gn=VPS13B;refGene_tx=XM_023248519.1;refGene_gn=VPS13B;refGene_tx=XM_023248520.1;refGene_gn=VPS13B;refGene_tx=XM_023248521.1;refGene_gn=VPS13B;refGene_tx=XM_023248522.1;refGene_gn=VPS13B;refGene_tx=XM_023248523.1;refGene_gn=VPS13B;refGene_tx=XM_023248524.1;refGene_gn=VPS13B;refGene_tx=XM_023248525.1

F2 47739027 47739199
refGene_gn=VPS13B;refGene_tx=XM_023248518.1;refGene_gn=VPS13B;refGene_

tx=XM_023248519.1;refGene_gn=VPS13B;refGene_tx=XM_023248520.1;refGene_gn=VPS13B;refGene_tx=XM_023248521.1;refGene_gn=VPS13B;refGene_tx=XM_023248522.1;refGene_gn=VPS13B;refGene_tx=XM_023248523.1;refGene_gn=VPS13B;refGene_tx=XM_023248524.1;refGene_gn=VPS13B;refGene_tx=XM_023248525.1
F2 47739701 47739816 refGene_gn=VPS13B;refGene_tx=XM_023248518.1;refGene_gn=VPS13B;refGene_tx=XM_023248519.1;refGene_gn=VPS13B;refGene_tx=XM_023248520.1;refGene_gn=VPS13B;refGene_tx=XM_023248521.1;refGene_gn=VPS13B;refGene_tx=XM_023248522.1;refGene_gn=VPS13B;refGene_tx=XM_023248523.1;refGene_gn=VPS13B;refGene_tx=XM_023248524.1;refGene_gn=VPS13B;refGene_tx=XM_023248525.1
F2 47742036 47742103 refGene_gn=VPS13B;refGene_tx=XM_023248522.1
F2 47742660 47742802 refGene_gn=VPS13B;refGene_tx=XM_023248518.1;refGene_gn=VPS13B;refGene_tx=XM_023248519.1;refGene_gn=VPS13B;refGene_tx=XM_023248520.1;refGene_gn=VPS13B;refGene_tx=XM_023248521.1;refGene_gn=VPS13B;refGene_tx=XM_023248523.1;refGene_gn=VPS13B;refGene_tx=XM_023248524.1
F2 47745885 47746294 refGene_gn=VPS13B;refGene_tx=XM_023248518.1;refGene_gn=VPS13B;refGene_tx=XM_023248519.1;refGene_gn=VPS13B;refGene_tx=XM_023248520.1;refGene_gn=VPS13B;refGene_tx=XM_023248521.1;refGene_gn=VPS13B;refGene_tx=XM_023248522.1;refGene_gn=VPS13B;refGene_tx=XM_023248523.1;refGene_gn=VPS13B;refGene_tx=XM_023248524.1;refGene_gn=VPS13B;refGene_tx=XM_023248525.1
F2 47755652 47755768 refGene_gn=VPS13B;refGene_tx=XM_023248518.1;refGene_gn=VPS13B;refGene_tx=XM_023248519.1;refGene_gn=VPS13B;refGene_tx=XM_023248520.1;refGene_gn=VPS13B;refGene_tx=XM_023248521.1;refGene_gn=VPS13B;refGene_tx=XM_023248522.1;refGene_gn=VPS13B;refGene_tx=XM_023248523.1;refGene_gn=VPS13B;refGene_tx=XM_023248524.1
F2 47799168 47799372 refGene_gn=VPS13B;refGene_tx=XM_023248518.1;refGene_gn=VPS13B;refGene_tx=XM_023248519.1;refGene_gn=VPS13B;refGene_tx=XM_023248520.1;refGene_gn=VPS13B;refGene_tx=XM_023248521.1;refGene_gn=VPS13B;refGene_tx=XM_023248522.1;refGene_gn=VPS13B;refGene_tx=XM_023248523.1
F2 47814274 47814401 refGene_gn=VPS13B;refGene_tx=XM_023248518.1;refGene_gn=VPS13B;refGene_tx=XM_023248519.1;refGene_gn=VPS13B;refGene_tx=XM_023248520.1;refGene_gn=VPS13B;refGene_tx=XM_023248521.1;refGene_gn=VPS13B;refGene_tx=XM_023248522.1;refGene_gn=VPS13B;refGene_tx=XM_023248523.1
F2 47818197 47818341 refGene_gn=VPS13B;refGene_tx=XM_023248518.1;refGene_gn=VPS13B;refGene_tx=XM_023248519.1;refGene_gn=VPS13B;refGene_tx=XM_023248520.1;refGene_gn=VPS13B;refGene_tx=XM_023248521.1;refGene_gn=VPS13B;refGene_tx=XM_023248522.1;refGene_gn=VPS13B;refGene_tx=XM_023248523.1
F2 47880669 47881357 refGene_gn=VPS13B;refGene_tx=XM_023248518.1;refGene_gn=VPS13B;refGene_tx=XM_023248519.1;refGene_gn=VPS13B;refGene_tx=XM_023248520.1;refGene_gn=VPS13B;refGene_tx=XM_023248521.1;refGene_gn=VPS13B;refGene_tx=XM_02

3248522.1;refGene_gn=VPS13B;refGene_tx=XM_023248523.1
F2 47900384 47900522
refGene_gn=VPS13B;refGene_tx=XM_023248518.1;refGene_gn=VPS13B;refGene_tx=XM_023248519.1;refGene_gn=VPS13B;refGene_tx=XM_023248520.1;refGene_gn=VPS13B;refGene_tx=XM_023248521.1;refGene_gn=VPS13B;refGene_tx=XM_023248522.1;refGene_gn=VPS13B;refGene_tx=XM_023248523.1
F2 47926470 47926866
refGene_gn=VPS13B;refGene_tx=XM_023248518.1;refGene_gn=VPS13B;refGene_tx=XM_023248519.1;refGene_gn=VPS13B;refGene_tx=XM_023248520.1;refGene_gn=VPS13B;refGene_tx=XM_023248521.1;refGene_gn=VPS13B;refGene_tx=XM_023248522.1;refGene_gn=VPS13B;refGene_tx=XM_023248523.1
F2 47944623 47944826
refGene_gn=VPS13B;refGene_tx=XM_023248518.1;refGene_gn=VPS13B;refGene_tx=XM_023248519.1;refGene_gn=VPS13B;refGene_tx=XM_023248520.1;refGene_gn=VPS13B;refGene_tx=XM_023248521.1;refGene_gn=VPS13B;refGene_tx=XM_023248522.1;refGene_gn=VPS13B;refGene_tx=XM_023248523.1
F2 47947450 47947658
refGene_gn=VPS13B;refGene_tx=XM_023248518.1;refGene_gn=VPS13B;refGene_tx=XM_023248519.1;refGene_gn=VPS13B;refGene_tx=XM_023248520.1;refGene_gn=VPS13B;refGene_tx=XM_023248521.1;refGene_gn=VPS13B;refGene_tx=XM_023248522.1;refGene_gn=VPS13B;refGene_tx=XM_023248523.1
F2 47947948 47948133
refGene_gn=VPS13B;refGene_tx=XM_023248518.1;refGene_gn=VPS13B;refGene_tx=XM_023248519.1;refGene_gn=VPS13B;refGene_tx=XM_023248520.1;refGene_gn=VPS13B;refGene_tx=XM_023248521.1;refGene_gn=VPS13B;refGene_tx=XM_023248522.1;refGene_gn=VPS13B;refGene_tx=XM_023248523.1
F2 47988852 47989049
refGene_gn=VPS13B;refGene_tx=XM_023248518.1;refGene_gn=VPS13B;refGene_tx=XM_023248519.1;refGene_gn=VPS13B;refGene_tx=XM_023248520.1;refGene_gn=VPS13B;refGene_tx=XM_023248521.1;refGene_gn=VPS13B;refGene_tx=XM_023248522.1;refGene_gn=VPS13B;refGene_tx=XM_023248523.1
F2 47997292 47997474
refGene_gn=VPS13B;refGene_tx=XM_023248518.1;refGene_gn=VPS13B;refGene_tx=XM_023248519.1;refGene_gn=VPS13B;refGene_tx=XM_023248520.1;refGene_gn=VPS13B;refGene_tx=XM_023248521.1;refGene_gn=VPS13B;refGene_tx=XM_023248522.1;refGene_gn=VPS13B;refGene_tx=XM_023248523.1
F2 47998758 47999108
refGene_gn=VPS13B;refGene_tx=XM_023248518.1;refGene_gn=VPS13B;refGene_tx=XM_023248519.1;refGene_gn=VPS13B;refGene_tx=XM_023248520.1;refGene_gn=VPS13B;refGene_tx=XM_023248521.1;refGene_gn=VPS13B;refGene_tx=XM_023248522.1;refGene_gn=VPS13B;refGene_tx=XM_023248523.1
F2 48004521 48004683
refGene_gn=VPS13B;refGene_tx=XM_023248518.1;refGene_gn=VPS13B;refGene_tx=XM_023248519.1;refGene_gn=VPS13B;refGene_tx=XM_023248520.1;refGene_gn=VPS13B;refGene_tx=XM_023248521.1;refGene_gn=VPS13B;refGene_tx=XM_023248522.1;refGene_gn=VPS13B;refGene_tx=XM_023248523.1
F2 48030344 48030500
refGene_gn=VPS13B;refGene_tx=XM_023248518.1;refGene_gn=VPS13B;refGene_tx=XM_023248519.1;refGene_gn=VPS13B;refGene_tx=XM_023248520.1;refGene_gn=VPS13B;refGene_tx=XM_023248521.1;refGene_gn=VPS13B;refGene_tx=XM_02

3248522.1;refGene_gn=VPS13B;refGene_tx=XM_023248523.1
F2 48038586 48038850
refGene_gn=VPS13B;refGene_tx=XM_023248518.1;refGene_gn=VPS13B;refGene_tx=XM_023248519.1;refGene_gn=VPS13B;refGene_tx=XM_023248520.1;refGene_gn=VPS13B;refGene_tx=XM_023248521.1;refGene_gn=VPS13B;refGene_tx=XM_023248522.1;refGene_gn=VPS13B;refGene_tx=XM_023248523.1
F2 48039520 48039604
refGene_gn=VPS13B;refGene_tx=XM_023248518.1;refGene_gn=VPS13B;refGene_tx=XM_023248519.1;refGene_gn=VPS13B;refGene_tx=XM_023248520.1;refGene_gn=VPS13B;refGene_tx=XM_023248521.1;refGene_gn=VPS13B;refGene_tx=XM_023248522.1;refGene_gn=VPS13B;refGene_tx=XM_023248523.1
F2 48039781 48039957
refGene_gn=VPS13B;refGene_tx=XM_023248518.1;refGene_gn=VPS13B;refGene_tx=XM_023248519.1;refGene_gn=VPS13B;refGene_tx=XM_023248520.1;refGene_gn=VPS13B;refGene_tx=XM_023248521.1;refGene_gn=VPS13B;refGene_tx=XM_023248522.1;refGene_gn=VPS13B;refGene_tx=XM_023248523.1
F2 48040681 48040852
refGene_gn=VPS13B;refGene_tx=XM_023248518.1;refGene_gn=VPS13B;refGene_tx=XM_023248519.1;refGene_gn=VPS13B;refGene_tx=XM_023248520.1;refGene_gn=VPS13B;refGene_tx=XM_023248521.1;refGene_gn=VPS13B;refGene_tx=XM_023248522.1;refGene_gn=VPS13B;refGene_tx=XM_023248523.1
F2 48041207 48041409
refGene_gn=VPS13B;refGene_tx=XM_023248518.1;refGene_gn=VPS13B;refGene_tx=XM_023248519.1;refGene_gn=VPS13B;refGene_tx=XM_023248520.1;refGene_gn=VPS13B;refGene_tx=XM_023248521.1;refGene_gn=VPS13B;refGene_tx=XM_023248522.1;refGene_gn=VPS13B;refGene_tx=XM_023248523.1
F2 48042533 48042722
refGene_gn=VPS13B;refGene_tx=XM_023248518.1;refGene_gn=VPS13B;refGene_tx=XM_023248519.1;refGene_gn=VPS13B;refGene_tx=XM_023248520.1;refGene_gn=VPS13B;refGene_tx=XM_023248521.1;refGene_gn=VPS13B;refGene_tx=XM_023248522.1;refGene_gn=VPS13B;refGene_tx=XM_023248523.1
F2 48045283 48045430
refGene_gn=VPS13B;refGene_tx=XM_023248518.1;refGene_gn=VPS13B;refGene_tx=XM_023248519.1;refGene_gn=VPS13B;refGene_tx=XM_023248520.1;refGene_gn=VPS13B;refGene_tx=XM_023248521.1;refGene_gn=VPS13B;refGene_tx=XM_023248522.1;refGene_gn=VPS13B;refGene_tx=XM_023248523.1
F2 48047435 48047719
refGene_gn=VPS13B;refGene_tx=XM_023248518.1;refGene_gn=VPS13B;refGene_tx=XM_023248519.1;refGene_gn=VPS13B;refGene_tx=XM_023248520.1;refGene_gn=VPS13B;refGene_tx=XM_023248521.1;refGene_gn=VPS13B;refGene_tx=XM_023248522.1;refGene_gn=VPS13B;refGene_tx=XM_023248523.1
F2 48050247 48050375
refGene_gn=VPS13B;refGene_tx=XM_023248518.1;refGene_gn=VPS13B;refGene_tx=XM_023248519.1;refGene_gn=VPS13B;refGene_tx=XM_023248520.1;refGene_gn=VPS13B;refGene_tx=XM_023248521.1;refGene_gn=VPS13B;refGene_tx=XM_023248522.1;refGene_gn=VPS13B;refGene_tx=XM_023248523.1
F2 48050582 48050782
refGene_gn=VPS13B;refGene_tx=XM_023248518.1;refGene_gn=VPS13B;refGene_tx=XM_023248519.1;refGene_gn=VPS13B;refGene_tx=XM_023248520.1;refGene_gn=VPS13B;refGene_tx=XM_023248521.1;refGene_gn=VPS13B;refGene_tx=XM_02

3248522.1;refGene_gn=VPS13B;refGene_tx=XM_023248523.1
F2 48063961 48064080
refGene_gn=VPS13B;refGene_tx=XM_023248518.1;refGene_gn=VPS13B;refGene_tx=XM_023248519.1;refGene_gn=VPS13B;refGene_tx=XM_023248520.1;refGene_gn=VPS13B;refGene_tx=XM_023248521.1;refGene_gn=VPS13B;refGene_tx=XM_023248522.1;refGene_gn=VPS13B;refGene_tx=XM_023248523.1
F2 48067564 48068373
refGene_gn=VPS13B;refGene_tx=XM_023248518.1;refGene_gn=VPS13B;refGene_tx=XM_023248519.1;refGene_gn=VPS13B;refGene_tx=XM_023248520.1;refGene_gn=VPS13B;refGene_tx=XM_023248521.1;refGene_gn=VPS13B;refGene_tx=XM_023248522.1;refGene_gn=VPS13B;refGene_tx=XM_023248523.1
F2 48072478 48072655
refGene_gn=VPS13B;refGene_tx=XM_023248518.1;refGene_gn=VPS13B;refGene_tx=XM_023248519.1;refGene_gn=VPS13B;refGene_tx=XM_023248520.1;refGene_gn=VPS13B;refGene_tx=XM_023248521.1;refGene_gn=VPS13B;refGene_tx=XM_023248522.1;refGene_gn=VPS13B;refGene_tx=XM_023248523.1
F2 48074577 48074748
refGene_gn=VPS13B;refGene_tx=XM_023248518.1;refGene_gn=VPS13B;refGene_tx=XM_023248519.1;refGene_gn=VPS13B;refGene_tx=XM_023248520.1;refGene_gn=VPS13B;refGene_tx=XM_023248521.1;refGene_gn=VPS13B;refGene_tx=XM_023248522.1;refGene_gn=VPS13B;refGene_tx=XM_023248523.1
F2 48081401 48081578
refGene_gn=VPS13B;refGene_tx=XM_023248518.1;refGene_gn=VPS13B;refGene_tx=XM_023248519.1;refGene_gn=VPS13B;refGene_tx=XM_023248520.1;refGene_gn=VPS13B;refGene_tx=XM_023248521.1;refGene_gn=VPS13B;refGene_tx=XM_023248522.1;refGene_gn=VPS13B;refGene_tx=XM_023248523.1
F2 48083660 48083763
refGene_gn=VPS13B;refGene_tx=XM_023248518.1;refGene_gn=VPS13B;refGene_tx=XM_023248519.1;refGene_gn=VPS13B;refGene_tx=XM_023248520.1;refGene_gn=VPS13B;refGene_tx=XM_023248521.1;refGene_gn=VPS13B;refGene_tx=XM_023248522.1;refGene_gn=VPS13B;refGene_tx=XM_023248523.1
F2 48084323 48084573
refGene_gn=VPS13B;refGene_tx=XM_023248518.1;refGene_gn=VPS13B;refGene_tx=XM_023248519.1;refGene_gn=VPS13B;refGene_tx=XM_023248520.1;refGene_gn=VPS13B;refGene_tx=XM_023248521.1;refGene_gn=VPS13B;refGene_tx=XM_023248522.1;refGene_gn=VPS13B;refGene_tx=XM_023248523.1
F2 48088411 48088660
refGene_gn=VPS13B;refGene_tx=XM_023248518.1;refGene_gn=VPS13B;refGene_tx=XM_023248519.1;refGene_gn=VPS13B;refGene_tx=XM_023248520.1;refGene_gn=VPS13B;refGene_tx=XM_023248521.1;refGene_gn=VPS13B;refGene_tx=XM_023248522.1;refGene_gn=VPS13B;refGene_tx=XM_023248523.1
F2 48100774 48100888
refGene_gn=LOC101096876;refGene_tx=XM_023248537.1
F2 48108551 48108665
refGene_gn=LOC101096876;refGene_tx=XM_023248537.1
F2 48152853 48152858
refGene_gn=RGS22;refGene_tx=XM_011291412.3;refGene_gn=RGS22;refGene_tx=XM_019822944.2;refGene_gn=RGS22;refGene_tx=XM_019822946.2;refGene_gn=RGS22;refGene_tx=XM_023248526.1;refGene_gn=RGS22;refGene_tx=XM_023248527.1;refGene_gn=RGS22;refGene_tx=XM_023248528.1;refGene_gn=RGS22;refGe

ne_tx=XM_023248529.1;refGene_gn=RGS22;refGene_tx=XM_023248530.1;refGene_gn=RGS22;refGene_tx=XM_023248531.1;refGene_gn=RGS22;refGene_tx=XM_023248532.1;refGene_gn=RGS22;refGene_tx=XM_023248533.1;refGene_gn=RGS22;refGene_tx=XM_023248534.1;refGene_gn=RGS22;refGene_tx=XM_023248535.1;refGene_gn=RGS22;refGene_tx=XM_023248536.1
F2 48153099 48153180
refGene_gn=RGS22;refGene_tx=XM_011291412.3;refGene_gn=RGS22;refGene_tx=XM_019822944.2;refGene_gn=RGS22;refGene_tx=XM_019822946.2;refGene_gn=RGS22;refGene_tx=XM_023248526.1;refGene_gn=RGS22;refGene_tx=XM_023248527.1;refGene_gn=RGS22;refGene_tx=XM_023248528.1;refGene_gn=RGS22;refGene_tx=XM_023248529.1;refGene_gn=RGS22;refGene_tx=XM_023248530.1;refGene_gn=RGS22;refGene_tx=XM_023248531.1;refGene_gn=RGS22;refGene_tx=XM_023248532.1;refGene_gn=RGS22;refGene_tx=XM_023248533.1;refGene_gn=RGS22;refGene_tx=XM_023248534.1;refGene_gn=RGS22;refGene_tx=XM_023248535.1;refGene_gn=RGS22;refGene_tx=XM_023248536.1
F2 48153298 48153404
refGene_gn=RGS22;refGene_tx=XM_011291412.3;refGene_gn=RGS22;refGene_tx=XM_019822944.2;refGene_gn=RGS22;refGene_tx=XM_019822945.2;refGene_gn=RGS22;refGene_tx=XM_019822946.2;refGene_gn=RGS22;refGene_tx=XM_023248526.1;refGene_gn=RGS22;refGene_tx=XM_023248527.1;refGene_gn=RGS22;refGene_tx=XM_023248528.1;refGene_gn=RGS22;refGene_tx=XM_023248529.1;refGene_gn=RGS22;refGene_tx=XM_023248530.1;refGene_gn=RGS22;refGene_tx=XM_023248531.1;refGene_gn=RGS22;refGene_tx=XM_023248532.1;refGene_gn=RGS22;refGene_tx=XM_023248533.1;refGene_gn=RGS22;refGene_tx=XM_023248534.1;refGene_gn=RGS22;refGene_tx=XM_023248535.1;refGene_gn=RGS22;refGene_tx=XM_023248536.1
F2 48154905 48155004
refGene_gn=RGS22;refGene_tx=XM_011291412.3;refGene_gn=RGS22;refGene_tx=XM_019822944.2;refGene_gn=RGS22;refGene_tx=XM_019822945.2;refGene_gn=RGS22;refGene_tx=XM_019822946.2;refGene_gn=RGS22;refGene_tx=XM_023248526.1;refGene_gn=RGS22;refGene_tx=XM_023248527.1;refGene_gn=RGS22;refGene_tx=XM_023248528.1;refGene_gn=RGS22;refGene_tx=XM_023248529.1;refGene_gn=RGS22;refGene_tx=XM_023248530.1;refGene_gn=RGS22;refGene_tx=XM_023248531.1;refGene_gn=RGS22;refGene_tx=XM_023248532.1;refGene_gn=RGS22;refGene_tx=XM_023248533.1;refGene_gn=RGS22;refGene_tx=XM_023248534.1;refGene_gn=RGS22;refGene_tx=XM_023248535.1;refGene_gn=RGS22;refGene_tx=XM_023248536.1
F2 48158155 48158311
refGene_gn=RGS22;refGene_tx=XM_011291412.3;refGene_gn=RGS22;refGene_tx=XM_019822944.2;refGene_gn=RGS22;refGene_tx=XM_019822945.2;refGene_gn=RGS22;refGene_tx=XM_019822946.2;refGene_gn=RGS22;refGene_tx=XM_023248526.1;refGene_gn=RGS22;refGene_tx=XM_023248527.1;refGene_gn=RGS22;refGene_tx=XM_023248528.1;refGene_gn=RGS22;refGene_tx=XM_023248529.1;refGene_gn=RGS22;refGene_tx=XM_023248530.1;refGene_gn=RGS22;refGene_tx=XM_023248531.1;refGene_gn=RGS22;refGene_tx=XM_023248532.1;refGene_gn=RGS22;refGene_tx=XM_023248533.1;refGene_gn=RGS22;refGene_tx=XM_023248534.1;refGene_gn=RGS22;refGene_tx=XM_023248535.1;refGene_gn=RGS22;refGene_tx=XM_023248536.1
F2 48161554 48161734
refGene_gn=RGS22;refGene_tx=XM_011291412.3;refGene_gn=RGS22;refGene_tx

=XM_019822944.2;refGene_gn=RGS22;refGene_tx=XM_019822945.2;refGene_gn=RGS22;refGene_tx=XM_019822946.2;refGene_gn=RGS22;refGene_tx=XM_023248526.1;refGene_gn=RGS22;refGene_tx=XM_023248527.1;refGene_gn=RGS22;refGene_tx=XM_023248528.1;refGene_gn=RGS22;refGene_tx=XM_023248529.1;refGene_gn=RGS22;refGene_tx=XM_023248530.1;refGene_gn=RGS22;refGene_tx=XM_023248531.1;refGene_gn=RGS22;refGene_tx=XM_023248532.1;refGene_gn=RGS22;refGene_tx=XM_023248533.1;refGene_gn=RGS22;refGene_tx=XM_023248534.1;refGene_gn=RGS22;refGene_tx=XM_023248535.1;refGene_gn=RGS22;refGene_tx=XM_023248536.1

F2 48167133 48167295
refGene_gn=RGS22;refGene_tx=XM_011291412.3;refGene_gn=RGS22;refGene_tx=XM_019822944.2;refGene_gn=RGS22;refGene_tx=XM_019822945.2;refGene_gn=RGS22;refGene_tx=XM_019822946.2;refGene_gn=RGS22;refGene_tx=XM_023248526.1;refGene_gn=RGS22;refGene_tx=XM_023248527.1;refGene_gn=RGS22;refGene_tx=XM_023248528.1;refGene_gn=RGS22;refGene_tx=XM_023248529.1;refGene_gn=RGS22;refGene_tx=XM_023248530.1;refGene_gn=RGS22;refGene_tx=XM_023248531.1;refGene_gn=RGS22;refGene_tx=XM_023248532.1;refGene_gn=RGS22;refGene_tx=XM_023248533.1;refGene_gn=RGS22;refGene_tx=XM_023248534.1;refGene_gn=RGS22;refGene_tx=XM_023248535.1;refGene_gn=RGS22;refGene_tx=XM_023248536.1

F2 48169626 48169695
refGene_gn=RGS22;refGene_tx=XM_011291412.3;refGene_gn=RGS22;refGene_tx=XM_019822944.2;refGene_gn=RGS22;refGene_tx=XM_019822945.2;refGene_gn=RGS22;refGene_tx=XM_019822946.2;refGene_gn=RGS22;refGene_tx=XM_023248526.1;refGene_gn=RGS22;refGene_tx=XM_023248527.1;refGene_gn=RGS22;refGene_tx=XM_023248528.1;refGene_gn=RGS22;refGene_tx=XM_023248529.1;refGene_gn=RGS22;refGene_tx=XM_023248530.1;refGene_gn=RGS22;refGene_tx=XM_023248532.1;refGene_gn=RGS22;refGene_tx=XM_023248533.1;refGene_gn=RGS22;refGene_tx=XM_023248534.1;refGene_gn=RGS22;refGene_tx=XM_023248535.1;refGene_gn=RGS22;refGene_tx=XM_023248536.1

F2 48172344 48172503
refGene_gn=RGS22;refGene_tx=XM_011291412.3;refGene_gn=RGS22;refGene_tx=XM_019822944.2;refGene_gn=RGS22;refGene_tx=XM_019822945.2;refGene_gn=RGS22;refGene_tx=XM_019822946.2;refGene_gn=RGS22;refGene_tx=XM_023248526.1;refGene_gn=RGS22;refGene_tx=XM_023248527.1;refGene_gn=RGS22;refGene_tx=XM_023248528.1;refGene_gn=RGS22;refGene_tx=XM_023248529.1;refGene_gn=RGS22;refGene_tx=XM_023248530.1;refGene_gn=RGS22;refGene_tx=XM_023248531.1;refGene_gn=RGS22;refGene_tx=XM_023248532.1;refGene_gn=RGS22;refGene_tx=XM_023248533.1;refGene_gn=RGS22;refGene_tx=XM_023248534.1;refGene_gn=RGS22;refGene_tx=XM_023248535.1;refGene_gn=RGS22;refGene_tx=XM_023248536.1

F2 48174635 48174798
refGene_gn=RGS22;refGene_tx=XM_011291412.3;refGene_gn=RGS22;refGene_tx=XM_019822944.2;refGene_gn=RGS22;refGene_tx=XM_019822945.2;refGene_gn=RGS22;refGene_tx=XM_019822946.2;refGene_gn=RGS22;refGene_tx=XM_023248526.1;refGene_gn=RGS22;refGene_tx=XM_023248527.1;refGene_gn=RGS22;refGene_tx=XM_023248528.1;refGene_gn=RGS22;refGene_tx=XM_023248529.1;refGene_gn=RGS22;refGene_tx=XM_023248530.1;refGene_gn=RGS22;refGene_tx=XM_023248531.1;refGene_gn=RGS22;refGene_tx=XM_023248532.1;refGene_gn=RGS22;refGene_tx=XM_023248533.1;refGene_gn=RGS22;refGene_tx=XM_023248534.1;r

efGene_gn=RGS22;refGene_tx=XM_023248535.1;refGene_gn=RGS22;refGene_tx=XM_023248536.1
F2 48175723 48175896
refGene_gn=RGS22;refGene_tx=XM_011291412.3;refGene_gn=RGS22;refGene_tx=XM_019822944.2;refGene_gn=RGS22;refGene_tx=XM_019822945.2;refGene_gn=RGS22;refGene_tx=XM_019822946.2;refGene_gn=RGS22;refGene_tx=XM_023248526.1;refGene_gn=RGS22;refGene_tx=XM_023248527.1;refGene_gn=RGS22;refGene_tx=XM_023248528.1;refGene_gn=RGS22;refGene_tx=XM_023248529.1;refGene_gn=RGS22;refGene_tx=XM_023248530.1;refGene_gn=RGS22;refGene_tx=XM_023248531.1;refGene_gn=RGS22;refGene_tx=XM_023248532.1;refGene_gn=RGS22;refGene_tx=XM_023248533.1;refGene_gn=RGS22;refGene_tx=XM_023248534.1;refGene_gn=RGS22;refGene_tx=XM_023248535.1;refGene_gn=RGS22;refGene_tx=XM_023248536.1
F2 48178262 48178355
refGene_gn=RGS22;refGene_tx=XM_011291412.3;refGene_gn=RGS22;refGene_tx=XM_019822944.2;refGene_gn=RGS22;refGene_tx=XM_019822945.2;refGene_gn=RGS22;refGene_tx=XM_019822946.2;refGene_gn=RGS22;refGene_tx=XM_023248526.1;refGene_gn=RGS22;refGene_tx=XM_023248527.1;refGene_gn=RGS22;refGene_tx=XM_023248528.1;refGene_gn=RGS22;refGene_tx=XM_023248529.1;refGene_gn=RGS22;refGene_tx=XM_023248530.1;refGene_gn=RGS22;refGene_tx=XM_023248531.1;refGene_gn=RGS22;refGene_tx=XM_023248532.1;refGene_gn=RGS22;refGene_tx=XM_023248533.1;refGene_gn=RGS22;refGene_tx=XM_023248534.1;refGene_gn=RGS22;refGene_tx=XM_023248535.1;refGene_gn=RGS22;refGene_tx=XM_023248536.1
F2 48181034 48181229
refGene_gn=RGS22;refGene_tx=XM_011291412.3;refGene_gn=RGS22;refGene_tx=XM_019822944.2;refGene_gn=RGS22;refGene_tx=XM_019822945.2;refGene_gn=RGS22;refGene_tx=XM_019822946.2;refGene_gn=RGS22;refGene_tx=XM_023248526.1;refGene_gn=RGS22;refGene_tx=XM_023248527.1;refGene_gn=RGS22;refGene_tx=XM_023248528.1;refGene_gn=RGS22;refGene_tx=XM_023248529.1;refGene_gn=RGS22;refGene_tx=XM_023248530.1;refGene_gn=RGS22;refGene_tx=XM_023248531.1;refGene_gn=RGS22;refGene_tx=XM_023248532.1;refGene_gn=RGS22;refGene_tx=XM_023248535.1;refGene_gn=RGS22;refGene_tx=XM_023248536.1
F2 48209152 48209254
refGene_gn=RGS22;refGene_tx=XM_011291412.3;refGene_gn=RGS22;refGene_tx=XM_019822944.2;refGene_gn=RGS22;refGene_tx=XM_019822945.2;refGene_gn=RGS22;refGene_tx=XM_019822946.2;refGene_gn=RGS22;refGene_tx=XM_023248526.1;refGene_gn=RGS22;refGene_tx=XM_023248527.1;refGene_gn=RGS22;refGene_tx=XM_023248528.1;refGene_gn=RGS22;refGene_tx=XM_023248529.1;refGene_gn=RGS22;refGene_tx=XM_023248530.1;refGene_gn=RGS22;refGene_tx=XM_023248531.1;refGene_gn=RGS22;refGene_tx=XM_023248532.1;refGene_gn=RGS22;refGene_tx=XM_023248533.1;refGene_gn=RGS22;refGene_tx=XM_023248534.1;refGene_gn=RGS22;refGene_tx=XM_023248535.1;refGene_gn=RGS22;refGene_tx=XM_023248536.1
F2 48210032 48210155
refGene_gn=RGS22;refGene_tx=XM_011291412.3;refGene_gn=RGS22;refGene_tx=XM_019822944.2;refGene_gn=RGS22;refGene_tx=XM_019822945.2;refGene_gn=RGS22;refGene_tx=XM_019822946.2;refGene_gn=RGS22;refGene_tx=XM_023248526.1;refGene_gn=RGS22;refGene_tx=XM_023248527.1;refGene_gn=RGS22;refGene_tx=XM_023248528.1;refGene_gn=RGS22;refGene_tx=XM_023248529.1;refGen

e_gn=RGS22;refGene_tx=XM_023248530.1;refGene_gn=RGS22;refGene_tx=XM_023248531.1;refGene_gn=RGS22;refGene_tx=XM_023248532.1;refGene_gn=RGS22;refGene_tx=XM_023248533.1;refGene_gn=RGS22;refGene_tx=XM_023248534.1;refGene_gn=RGS22;refGene_tx=XM_023248535.1;refGene_gn=RGS22;refGene_tx=XM_023248536.1

F2 48211985 48212100

refGene_gn=RGS22;refGene_tx=XM_011291412.3;refGene_gn=RGS22;refGene_tx=XM_019822944.2;refGene_gn=RGS22;refGene_tx=XM_019822945.2;refGene_gn=RGS22;refGene_tx=XM_019822946.2;refGene_gn=RGS22;refGene_tx=XM_023248526.1;refGene_gn=RGS22;refGene_tx=XM_023248527.1;refGene_gn=RGS22;refGene_tx=XM_023248528.1;refGene_gn=RGS22;refGene_tx=XM_023248529.1;refGene_gn=RGS22;refGene_tx=XM_023248530.1;refGene_gn=RGS22;refGene_tx=XM_023248531.1;refGene_gn=RGS22;refGene_tx=XM_023248532.1;refGene_gn=RGS22;refGene_tx=XM_023248533.1;refGene_gn=RGS22;refGene_tx=XM_023248534.1;refGene_gn=RGS22;refGene_tx=XM_023248535.1;refGene_gn=RGS22;refGene_tx=XM_023248536.1

F2 48219520 48219654

refGene_gn=RGS22;refGene_tx=XM_011291412.3;refGene_gn=RGS22;refGene_tx=XM_019822944.2;refGene_gn=RGS22;refGene_tx=XM_019822945.2;refGene_gn=RGS22;refGene_tx=XM_019822946.2;refGene_gn=RGS22;refGene_tx=XM_023248526.1;refGene_gn=RGS22;refGene_tx=XM_023248527.1;refGene_gn=RGS22;refGene_tx=XM_023248528.1;refGene_gn=RGS22;refGene_tx=XM_023248529.1;refGene_gn=RGS22;refGene_tx=XM_023248530.1;refGene_gn=RGS22;refGene_tx=XM_023248531.1;refGene_gn=RGS22;refGene_tx=XM_023248532.1;refGene_gn=RGS22;refGene_tx=XM_023248533.1;refGene_gn=RGS22;refGene_tx=XM_023248534.1;refGene_gn=RGS22;refGene_tx=XM_023248535.1;refGene_gn=RGS22;refGene_tx=XM_023248536.1

F2 48221670 48221845

refGene_gn=RGS22;refGene_tx=XM_011291412.3;refGene_gn=RGS22;refGene_tx=XM_019822944.2;refGene_gn=RGS22;refGene_tx=XM_019822945.2;refGene_gn=RGS22;refGene_tx=XM_019822946.2;refGene_gn=RGS22;refGene_tx=XM_023248526.1;refGene_gn=RGS22;refGene_tx=XM_023248527.1;refGene_gn=RGS22;refGene_tx=XM_023248528.1;refGene_gn=RGS22;refGene_tx=XM_023248529.1;refGene_gn=RGS22;refGene_tx=XM_023248530.1;refGene_gn=RGS22;refGene_tx=XM_023248531.1;refGene_gn=RGS22;refGene_tx=XM_023248532.1;refGene_gn=RGS22;refGene_tx=XM_023248533.1;refGene_gn=RGS22;refGene_tx=XM_023248534.1;refGene_gn=RGS22;refGene_tx=XM_023248535.1;refGene_gn=RGS22;refGene_tx=XM_023248536.1

F2 48230367 48230529

refGene_gn=RGS22;refGene_tx=XM_011291412.3;refGene_gn=RGS22;refGene_tx=XM_019822944.2;refGene_gn=RGS22;refGene_tx=XM_019822945.2;refGene_gn=RGS22;refGene_tx=XM_019822946.2;refGene_gn=RGS22;refGene_tx=XM_023248526.1;refGene_gn=RGS22;refGene_tx=XM_023248527.1;refGene_gn=RGS22;refGene_tx=XM_023248528.1;refGene_gn=RGS22;refGene_tx=XM_023248529.1;refGene_gn=RGS22;refGene_tx=XM_023248530.1;refGene_gn=RGS22;refGene_tx=XM_023248531.1;refGene_gn=RGS22;refGene_tx=XM_023248533.1;refGene_gn=RGS22;refGene_tx=XM_023248534.1;refGene_gn=RGS22;refGene_tx=XM_023248535.1

F2 48232318 48232928

refGene_gn=RGS22;refGene_tx=XM_011291412.3;refGene_gn=RGS22;refGene_tx=XM_019822944.2;refGene_gn=RGS22;refGene_tx=XM_019822945.2;refGene_gn=

RGS22;refGene_tx=XM_019822946.2;refGene_gn=RGS22;refGene_tx=XM_0232485
26.1;refGene_gn=RGS22;refGene_tx=XM_023248527.1;refGene_gn=RGS22;refGe
ne_tx=XM_023248528.1;refGene_gn=RGS22;refGene_tx=XM_023248529.1;refGen
e_gn=RGS22;refGene_tx=XM_023248530.1;refGene_gn=RGS22;refGene_tx=XM_02
3248531.1;refGene_gn=RGS22;refGene_tx=XM_023248532.1;refGene_gn=RGS22;
refGene_tx=XM_023248533.1;refGene_gn=RGS22;refGene_tx=XM_023248534.1;r
efGene_gn=RGS22;refGene_tx=XM_023248535.1;refGene_gn=RGS22;refGene_tx=
XM_023248536.1
F2 48235170 48235300
refGene_gn=RGS22;refGene_tx=XM_011291412.3;refGene_gn=RGS22;refGene_tx
=XM_019822944.2;refGene_gn=RGS22;refGene_tx=XM_019822945.2;refGene_gn=
RGS22;refGene_tx=XM_019822946.2;refGene_gn=RGS22;refGene_tx=XM_0232485
26.1;refGene_gn=RGS22;refGene_tx=XM_023248527.1;refGene_gn=RGS22;refGe
ne_tx=XM_023248528.1;refGene_gn=RGS22;refGene_tx=XM_023248529.1;refGen
e_gn=RGS22;refGene_tx=XM_023248530.1;refGene_gn=RGS22;refGene_tx=XM_02
3248531.1;refGene_gn=RGS22;refGene_tx=XM_023248532.1;refGene_gn=RGS22;
refGene_tx=XM_023248533.1;refGene_gn=RGS22;refGene_tx=XM_023248534.1;r
efGene_gn=RGS22;refGene_tx=XM_023248535.1;refGene_gn=RGS22;refGene_tx=
XM_023248536.1
F2 48238055 48238224
refGene_gn=RGS22;refGene_tx=XM_011291412.3;refGene_gn=RGS22;refGene_tx
=XM_019822944.2;refGene_gn=RGS22;refGene_tx=XM_019822945.2;refGene_gn=
RGS22;refGene_tx=XM_019822946.2;refGene_gn=RGS22;refGene_tx=XM_0232485
26.1;refGene_gn=RGS22;refGene_tx=XM_023248527.1;refGene_gn=RGS22;refGe
ne_tx=XM_023248528.1;refGene_gn=RGS22;refGene_tx=XM_023248529.1;refGen
e_gn=RGS22;refGene_tx=XM_023248530.1;refGene_gn=RGS22;refGene_tx=XM_02
3248531.1;refGene_gn=RGS22;refGene_tx=XM_023248532.1;refGene_gn=RGS22;
refGene_tx=XM_023248533.1;refGene_gn=RGS22;refGene_tx=XM_023248534.1;r
efGene_gn=RGS22;refGene_tx=XM_023248535.1;refGene_gn=RGS22;refGene_tx=
XM_023248536.1
F2 48238858 48238944
refGene_gn=RGS22;refGene_tx=XM_011291412.3;refGene_gn=RGS22;refGene_tx
=XM_019822944.2;refGene_gn=RGS22;refGene_tx=XM_019822945.2;refGene_gn=
RGS22;refGene_tx=XM_019822946.2;refGene_gn=RGS22;refGene_tx=XM_0232485
26.1;refGene_gn=RGS22;refGene_tx=XM_023248527.1;refGene_gn=RGS22;refGe
ne_tx=XM_023248528.1;refGene_gn=RGS22;refGene_tx=XM_023248529.1;refGen
e_gn=RGS22;refGene_tx=XM_023248530.1;refGene_gn=RGS22;refGene_tx=XM_02
3248531.1;refGene_gn=RGS22;refGene_tx=XM_023248532.1;refGene_gn=RGS22;
refGene_tx=XM_023248533.1;refGene_gn=RGS22;refGene_tx=XM_023248534.1;r
efGene_gn=RGS22;refGene_tx=XM_023248535.1;refGene_gn=RGS22;refGene_tx=
XM_023248536.1
F2 48253730 48253952
refGene_gn=RGS22;refGene_tx=XM_011291412.3;refGene_gn=RGS22;refGene_tx
=XM_019822944.2;refGene_gn=RGS22;refGene_tx=XM_019822945.2;refGene_gn=
RGS22;refGene_tx=XM_019822946.2;refGene_gn=RGS22;refGene_tx=XM_0232485
26.1;refGene_gn=RGS22;refGene_tx=XM_023248527.1;refGene_gn=RGS22;refGe
ne_tx=XM_023248528.1;refGene_gn=RGS22;refGene_tx=XM_023248529.1;refGen
e_gn=RGS22;refGene_tx=XM_023248530.1;refGene_gn=RGS22;refGene_tx=XM_02
3248531.1;refGene_gn=RGS22;refGene_tx=XM_023248532.1;refGene_gn=RGS22;
refGene_tx=XM_023248533.1;refGene_gn=RGS22;refGene_tx=XM_023248534.1;r

efGene_gn=RGS22;refGene_tx=XM_023248535.1;refGene_gn=RGS22;refGene_tx=XM_023248536.1
F2 48269231 48269294 refGene_gn=RGS22;refGene_tx=XM_011291412.3;refGene_gn=RGS22;refGene_tx=XM_019822944.2;refGene_gn=RGS22;refGene_tx=XM_019822945.2;refGene_gn=RGS22;refGene_tx=XM_023248526.1;refGene_gn=RGS22;refGene_tx=XM_023248527.1;refGene_gn=RGS22;refGene_tx=XM_023248528.1;refGene_gn=RGS22;refGene_tx=XM_023248529.1;refGene_gn=RGS22;refGene_tx=XM_023248530.1;refGene_gn=RGS22;refGene_tx=XM_023248531.1;refGene_gn=RGS22;refGene_tx=XM_023248532.1;refGene_gn=RGS22;refGene_tx=XM_023248533.1;refGene_gn=RGS22;refGene_tx=XM_023248534.1;refGene_gn=RGS22;refGene_tx=XM_023248535.1;refGene_gn=RGS22;refGene_tx=XM_023248536.1
F2 48285587 48285616 refGene_gn=RGS22;refGene_tx=XM_011291412.3;refGene_gn=RGS22;refGene_tx=XM_019822944.2;refGene_gn=RGS22;refGene_tx=XM_019822945.2;refGene_gn=RGS22;refGene_tx=XM_023248526.1;refGene_gn=RGS22;refGene_tx=XM_023248527.1;refGene_gn=RGS22;refGene_tx=XM_023248528.1;refGene_gn=RGS22;refGene_tx=XM_023248529.1;refGene_gn=RGS22;refGene_tx=XM_023248530.1;refGene_gn=RGS22;refGene_tx=XM_023248531.1;refGene_gn=RGS22;refGene_tx=XM_023248532.1;refGene_gn=RGS22;refGene_tx=XM_023248533.1;refGene_gn=RGS22;refGene_tx=XM_023248534.1;refGene_gn=RGS22;refGene_tx=XM_023248535.1;refGene_gn=RGS22;refGene_tx=XM_023248536.1
F2 48286101 48286126 refGene_gn=RGS22;refGene_tx=XM_011291412.3;refGene_gn=RGS22;refGene_tx=XM_019822944.2;refGene_gn=RGS22;refGene_tx=XM_019822945.2;refGene_gn=RGS22;refGene_tx=XM_023248526.1;refGene_gn=RGS22;refGene_tx=XM_023248527.1;refGene_gn=RGS22;refGene_tx=XM_023248528.1;refGene_gn=RGS22;refGene_tx=XM_023248529.1;refGene_gn=RGS22;refGene_tx=XM_023248530.1;refGene_gn=RGS22;refGene_tx=XM_023248531.1;refGene_gn=RGS22;refGene_tx=XM_023248532.1;refGene_gn=RGS22;refGene_tx=XM_023248533.1;refGene_gn=RGS22;refGene_tx=XM_023248534.1;refGene_gn=RGS22;refGene_tx=XM_023248535.1;refGene_gn=RGS22;refGene_tx=XM_023248536.1
F2 48299607 48299856 refGene_gn=FBXO43;refGene_tx=XM_023248332.1
F2 48299959 48300163 refGene_gn=FBXO43;refGene_tx=XM_023248332.1
F2 48304455 48304558 refGene_gn=FBXO43;refGene_tx=XM_023248332.1
F2 48307350 48308821 refGene_gn=FBXO43;refGene_tx=XM_023248332.1
F2 48313252 48313281 refGene_gn=FBXO43;refGene_tx=XM_023248332.1
F2 48313282 48313321 refGene_gn=FBXO43;refGene_tx=XM_023248332.1
F2 48313523 48313813 refGene_gn=FBXO43;refGene_tx=XM_023248332.1
F2 48313911 48314244 refGene_gn=FBXO43;refGene_tx=XM_023248332.1
F2 48318121 48318333 refGene_gn=FBXO43;refGene_tx=XM_023248332.1
F2 48318609 48318730 refGene_gn=FBXO43;refGene_tx=XM_023248332.1
F2 48318899 48318960 refGene_gn=POLR2K;refGene_tx=XM_004000018.3
F2 48319682 48319775 refGene_gn=POLR2K;refGene_tx=XM_004000018.3
F2 48320597 48320620 refGene_gn=POLR2K;refGene_tx=XM_004000018.3
F2 48328057 48328197 refGene_gn=SPAG1;refGene_tx=XM_011291414.3;refGene_gn=SPAG1;refGene_tx=XM_023248685.1;refGene_gn=SPAG1;refGene_tx=XM_023248686.1
F2 48329989 48330149 refGene_gn=SPAG1;refGene_tx=XM_011291414.3;refGene_gn=SPAG1;refGene_tx

=XM_023248685.1;refGene_gn=SPAG1;refGene_tx=XM_023248686.1
F2 48336491 48336617
refGene_gn=SPAG1;refGene_tx=XM_011291414.3;refGene_gn=SPAG1;refGene_tx=XM_023248685.1;refGene_gn=SPAG1;refGene_tx=XM_023248686.1;refGene_gn=SPAG1;refGene_tx=XM_023248687.1
F2 48340674 48340739
refGene_gn=SPAG1;refGene_tx=XM_011291414.3;refGene_gn=SPAG1;refGene_tx=XM_023248685.1;refGene_gn=SPAG1;refGene_tx=XM_023248686.1;refGene_gn=SPAG1;refGene_tx=XM_023248687.1
F2 48341346 48341456
refGene_gn=SPAG1;refGene_tx=XM_011291414.3;refGene_gn=SPAG1;refGene_tx=XM_023248685.1;refGene_gn=SPAG1;refGene_tx=XM_023248686.1
F2 48342310 48342416
refGene_gn=SPAG1;refGene_tx=XM_011291414.3;refGene_gn=SPAG1;refGene_tx=XM_023248685.1;refGene_gn=SPAG1;refGene_tx=XM_023248686.1;refGene_gn=SPAG1;refGene_tx=XM_023248687.1
F2 48345957 48346088
refGene_gn=SPAG1;refGene_tx=XM_011291414.3;refGene_gn=SPAG1;refGene_tx=XM_023248685.1;refGene_gn=SPAG1;refGene_tx=XM_023248686.1;refGene_gn=SPAG1;refGene_tx=XM_023248687.1
F2 48348078 48348185
refGene_gn=SPAG1;refGene_tx=XM_011291414.3;refGene_gn=SPAG1;refGene_tx=XM_023248685.1;refGene_gn=SPAG1;refGene_tx=XM_023248686.1;refGene_gn=SPAG1;refGene_tx=XM_023248687.1
F2 48352677 48352837
refGene_gn=SPAG1;refGene_tx=XM_011291414.3;refGene_gn=SPAG1;refGene_tx=XM_023248685.1;refGene_gn=SPAG1;refGene_tx=XM_023248686.1;refGene_gn=SPAG1;refGene_tx=XM_023248687.1
F2 48363968 48364343
refGene_gn=SPAG1;refGene_tx=XM_011291414.3;refGene_gn=SPAG1;refGene_tx=XM_023248685.1;refGene_gn=SPAG1;refGene_tx=XM_023248686.1;refGene_gn=SPAG1;refGene_tx=XM_023248687.1
F2 48364666 48364766
refGene_gn=SPAG1;refGene_tx=XM_011291414.3;refGene_gn=SPAG1;refGene_tx=XM_023248685.1;refGene_gn=SPAG1;refGene_tx=XM_023248686.1;refGene_gn=SPAG1;refGene_tx=XM_023248687.1
F2 48370073 48370226
refGene_gn=SPAG1;refGene_tx=XM_011291414.3;refGene_gn=SPAG1;refGene_tx=XM_023248685.1;refGene_gn=SPAG1;refGene_tx=XM_023248686.1;refGene_gn=SPAG1;refGene_tx=XM_023248687.1
F2 48374819 48374986
refGene_gn=SPAG1;refGene_tx=XM_011291414.3;refGene_gn=SPAG1;refGene_tx=XM_023248685.1;refGene_gn=SPAG1;refGene_tx=XM_023248686.1;refGene_gn=SPAG1;refGene_tx=XM_023248687.1
F2 48378353 48378486
refGene_gn=SPAG1;refGene_tx=XM_011291414.3;refGene_gn=SPAG1;refGene_tx=XM_023248685.1;refGene_gn=SPAG1;refGene_tx=XM_023248686.1;refGene_gn=SPAG1;refGene_tx=XM_023248687.1
F2 48380365 48380492
refGene_gn=SPAG1;refGene_tx=XM_011291414.3;refGene_gn=SPAG1;refGene_tx

=XM_023248685.1;refGene_gn=SPAG1;refGene_tx=XM_023248686.1;refGene_gn=SPAG1;refGene_tx=XM_023248687.1
F2	48389189	48389345
refGene_gn=SPAG1;refGene_tx=XM_011291414.3;refGene_gn=SPAG1;refGene_tx=XM_023248685.1;refGene_gn=SPAG1;refGene_tx=XM_023248686.1;refGene_gn=SPAG1;refGene_tx=XM_023248687.1
F2	48390281	48390653
refGene_gn=SPAG1;refGene_tx=XM_011291414.3;refGene_gn=SPAG1;refGene_tx=XM_023248685.1;refGene_gn=SPAG1;refGene_tx=XM_023248686.1;refGene_gn=SPAG1;refGene_tx=XM_023248687.1
F2	48390755	48390887
refGene_gn=SPAG1;refGene_tx=XM_011291414.3;refGene_gn=SPAG1;refGene_tx=XM_023248685.1;refGene_gn=SPAG1;refGene_tx=XM_023248686.1;refGene_gn=SPAG1;refGene_tx=XM_023248687.1
F2	48401745	48401758	refGene_gn=SPAG1;refGene_tx=XM_023248685.1
F2	48404329	48404333	refGene_gn=SPAG1;refGene_tx=XM_023248686.1
F2	48412192	48412883
refGene_gn=RNF19A;refGene_tx=XM_004000019.4;refGene_gn=RNF19A;refGene_tx=XM_006943325.3
F2	48413986	48414130
refGene_gn=RNF19A;refGene_tx=XM_004000019.4;refGene_gn=RNF19A;refGene_tx=XM_006943325.3
F2	48415822	48416036
refGene_gn=RNF19A;refGene_tx=XM_004000019.4;refGene_gn=RNF19A;refGene_tx=XM_006943325.3
F2	48418474	48418636
refGene_gn=RNF19A;refGene_tx=XM_004000019.4;refGene_gn=RNF19A;refGene_tx=XM_006943325.3
F2	48419167	48419282
refGene_gn=RNF19A;refGene_tx=XM_004000019.4;refGene_gn=RNF19A;refGene_tx=XM_006943325.3
F2	48422855	48423018
refGene_gn=RNF19A;refGene_tx=XM_004000019.4;refGene_gn=RNF19A;refGene_tx=XM_006943325.3
F2	48424015	48424160
refGene_gn=RNF19A;refGene_tx=XM_004000019.4;refGene_gn=RNF19A;refGene_tx=XM_006943325.3
F2	48428774	48428983
refGene_gn=RNF19A;refGene_tx=XM_004000019.4;refGene_gn=RNF19A;refGene_tx=XM_006943325.3
F2	48442180	48442854
refGene_gn=RNF19A;refGene_tx=XM_004000019.4;refGene_gn=RNF19A;refGene_tx=XM_006943325.3
F2	48660900	48660997	refGene_gn=ANKRD46;refGene_tx=XM_011291416.3
F2	48669104	48669321	refGene_gn=ANKRD46;refGene_tx=XM_011291415.3
F2	48674513	48674672
refGene_gn=ANKRD46;refGene_tx=XM_011291415.3;refGene_gn=ANKRD46;refGene_tx=XM_011291416.3
F2	48675503	48675814
refGene_gn=ANKRD46;refGene_tx=XM_011291415.3;refGene_gn=ANKRD46;refGen

e_tx=XM_011291416.3
F2 48719678 48719771
refGene_gn=SNX31;refGene_tx=XM_004000021.2;refGene_gn=SNX31;refGene_tx=XM_019822951.1
F2 48722608 48722665
refGene_gn=SNX31;refGene_tx=XM_004000021.2;refGene_gn=SNX31;refGene_tx=XM_019822951.1
F2 48728971 48729049
refGene_gn=SNX31;refGene_tx=XM_004000021.2;refGene_gn=SNX31;refGene_tx=XM_019822951.1
F2 48732138 48732252
refGene_gn=SNX31;refGene_tx=XM_004000021.2;refGene_gn=SNX31;refGene_tx=XM_019822951.1
F2 48738746 48738950
refGene_gn=SNX31;refGene_tx=XM_004000021.2;refGene_gn=SNX31;refGene_tx=XM_019822951.1
F2 48743400 48743493 refGene_gn=SNX31;refGene_tx=XM_004000021.2
F2 48748826 48748896
refGene_gn=SNX31;refGene_tx=XM_004000021.2;refGene_gn=SNX31;refGene_tx=XM_019822951.1
F2 48753179 48753267
refGene_gn=SNX31;refGene_tx=XM_004000021.2;refGene_gn=SNX31;refGene_tx=XM_019822951.1
F2 48754226 48754317
refGene_gn=SNX31;refGene_tx=XM_004000021.2;refGene_gn=SNX31;refGene_tx=XM_019822951.1
F2 48758408 48758519
refGene_gn=SNX31;refGene_tx=XM_004000021.2;refGene_gn=SNX31;refGene_tx=XM_019822951.1
F2 48777342 48777407
refGene_gn=SNX31;refGene_tx=XM_004000021.2;refGene_gn=SNX31;refGene_tx=XM_019822951.1
F2 48780717 48780832
refGene_gn=SNX31;refGene_tx=XM_004000021.2;refGene_gn=SNX31;refGene_tx=XM_019822951.1
F2 48790768 48790843
refGene_gn=SNX31;refGene_tx=XM_004000021.2;refGene_gn=SNX31;refGene_tx=XM_019822951.1
F2 48790955 48791021
refGene_gn=SNX31;refGene_tx=XM_004000021.2;refGene_gn=SNX31;refGene_tx=XM_019822951.1
F2 48860377 48860470
refGene_gn=PABPC1;refGene_tx=XM_023248197.1;refGene_gn=PABPC1;refGene_tx=XM_023248198.1
F2 48861017 48861148
refGene_gn=PABPC1;refGene_tx=XM_023248197.1;refGene_gn=PABPC1;refGene_tx=XM_023248198.1
F2 48861601 48861686
refGene_gn=PABPC1;refGene_tx=XM_023248197.1;refGene_gn=PABPC1;refGene_tx=XM_023248198.1

F2 48862221 48862376
refGene_gn=PABPC1;refGene_tx=XM_023248197.1;refGene_gn=PABPC1;refGene_tx=XM_023248198.1
F2 48862458 48862569
refGene_gn=PABPC1;refGene_tx=XM_023248197.1;refGene_gn=PABPC1;refGene_tx=XM_023248198.1
F2 48863189 48863280
refGene_gn=PABPC1;refGene_tx=XM_023248197.1;refGene_gn=PABPC1;refGene_tx=XM_023248198.1
F2 48863492 48863765
refGene_gn=PABPC1;refGene_tx=XM_023248197.1;refGene_gn=PABPC1;refGene_tx=XM_023248198.1
F2 48865939 48866035
refGene_gn=PABPC1;refGene_tx=XM_023248197.1;refGene_gn=PABPC1;refGene_tx=XM_023248198.1
F2 48866234 48866372
refGene_gn=PABPC1;refGene_tx=XM_023248197.1;refGene_gn=PABPC1;refGene_tx=XM_023248198.1
F2 48866684 48866779
refGene_gn=PABPC1;refGene_tx=XM_023248197.1;refGene_gn=PABPC1;refGene_tx=XM_023248198.1
F2 48868325 48868465
refGene_gn=PABPC1;refGene_tx=XM_023248197.1;refGene_gn=PABPC1;refGene_tx=XM_023248198.1
F2 48870590 48870706
refGene_gn=PABPC1;refGene_tx=XM_023248197.1;refGene_gn=PABPC1;refGene_tx=XM_023248198.1
F2 48870906 48871100
refGene_gn=PABPC1;refGene_tx=XM_023248197.1;refGene_gn=PABPC1;refGene_tx=XM_023248198.1
F2 48874172 48874365
refGene_gn=PABPC1;refGene_tx=XM_023248197.1;refGene_gn=PABPC1;refGene_tx=XM_023248198.1
F2 49029763 49029823
refGene_gn=YWHAZ;refGene_tx=XM_006943327.4;refGene_gn=YWHAZ;refGene_tx=XM_019822954.2
F2 49032619 49032715
refGene_gn=YWHAZ;refGene_tx=XM_006943327.4;refGene_gn=YWHAZ;refGene_tx=XM_019822954.2
F2 49032792 49032956
refGene_gn=YWHAZ;refGene_tx=XM_006943327.4;refGene_gn=YWHAZ;refGene_tx=XM_019822954.2
F2 49033660 49033784
refGene_gn=YWHAZ;refGene_tx=XM_006943327.4;refGene_gn=YWHAZ;refGene_tx=XM_019822954.2
F2 49057218 49057512
refGene_gn=YWHAZ;refGene_tx=XM_006943327.4;refGene_gn=YWHAZ;refGene_tx=XM_019822954.2
F2 49270194 49270290
refGene_gn=ZNF706;refGene_tx=XM_019822963.2;refGene_gn=ZNF706;refGene_

tx=XM_019822964.2;refGene_gn=ZNF706;refGene_tx=XM_023248238.1;refGene_gn=ZNF706;refGene_tx=XM_023248239.1
F2 49272248 49272383
refGene_gn=ZNF706;refGene_tx=XM_019822963.2;refGene_gn=ZNF706;refGene_tx=XM_019822964.2;refGene_gn=ZNF706;refGene_tx=XM_023248238.1;refGene_gn=ZNF706;refGene_tx=XM_023248239.1
F2 49518657 49518677
refGene_gn=GRHL2;refGene_tx=XM_019822955.2;refGene_gn=GRHL2;refGene_tx=XM_019822956.2
F2 49572123 49572319
refGene_gn=GRHL2;refGene_tx=XM_006943329.4;refGene_gn=GRHL2;refGene_tx=XM_019822955.2;refGene_gn=GRHL2;refGene_tx=XM_019822956.2;refGene_gn=GRHL2;refGene_tx=XM_019822957.2
F2 49579241 49579309
refGene_gn=GRHL2;refGene_tx=XM_006943329.4;refGene_gn=GRHL2;refGene_tx=XM_019822955.2;refGene_gn=GRHL2;refGene_tx=XM_019822956.2;refGene_gn=GRHL2;refGene_tx=XM_019822957.2
F2 49584538 49584932
refGene_gn=GRHL2;refGene_tx=XM_006943329.4;refGene_gn=GRHL2;refGene_tx=XM_019822955.2;refGene_gn=GRHL2;refGene_tx=XM_019822956.2;refGene_gn=GRHL2;refGene_tx=XM_019822957.2
F2 49596535 49596591
refGene_gn=GRHL2;refGene_tx=XM_006943329.4;refGene_gn=GRHL2;refGene_tx=XM_019822955.2;refGene_gn=GRHL2;refGene_tx=XM_019822956.2;refGene_gn=GRHL2;refGene_tx=XM_019822957.2
F2 49599702 49599859
refGene_gn=GRHL2;refGene_tx=XM_006943329.4;refGene_gn=GRHL2;refGene_tx=XM_019822955.2;refGene_gn=GRHL2;refGene_tx=XM_019822956.2;refGene_gn=GRHL2;refGene_tx=XM_019822957.2
F2 49604135 49604247
refGene_gn=GRHL2;refGene_tx=XM_006943329.4;refGene_gn=GRHL2;refGene_tx=XM_019822955.2;refGene_gn=GRHL2;refGene_tx=XM_019822956.2;refGene_gn=GRHL2;refGene_tx=XM_019822957.2
F2 49628909 49629004
refGene_gn=GRHL2;refGene_tx=XM_006943329.4;refGene_gn=GRHL2;refGene_tx=XM_019822955.2;refGene_gn=GRHL2;refGene_tx=XM_019822956.2;refGene_gn=GRHL2;refGene_tx=XM_019822957.2
F2 49648004 49648163
refGene_gn=GRHL2;refGene_tx=XM_006943329.4;refGene_gn=GRHL2;refGene_tx=XM_019822955.2;refGene_gn=GRHL2;refGene_tx=XM_019822956.2;refGene_gn=GRHL2;refGene_tx=XM_019822957.2
F2 49655499 49655587
refGene_gn=GRHL2;refGene_tx=XM_006943329.4;refGene_gn=GRHL2;refGene_tx=XM_019822955.2;refGene_gn=GRHL2;refGene_tx=XM_019822956.2;refGene_gn=GRHL2;refGene_tx=XM_019822957.2
F2 49656056 49656196
refGene_gn=GRHL2;refGene_tx=XM_006943329.4;refGene_gn=GRHL2;refGene_tx=XM_019822955.2;refGene_gn=GRHL2;refGene_tx=XM_019822956.2;refGene_gn=GRHL2;refGene_tx=XM_019822957.2
F2 49661136 49661168

refGene_gn=GRHL2;refGene_tx=XM_006943329.4;refGene_gn=GRHL2;refGene_tx=XM_019822955.2;refGene_gn=GRHL2;refGene_tx=XM_019822956.2;refGene_gn=GRHL2;refGene_tx=XM_019822957.2
F2 49672966 49673061
refGene_gn=GRHL2;refGene_tx=XM_006943329.4;refGene_gn=GRHL2;refGene_tx=XM_019822955.2;refGene_gn=GRHL2;refGene_tx=XM_019822956.2;refGene_gn=GRHL2;refGene_tx=XM_019822957.2
F2 49677095 49677181
refGene_gn=GRHL2;refGene_tx=XM_006943329.4;refGene_gn=GRHL2;refGene_tx=XM_019822955.2;refGene_gn=GRHL2;refGene_tx=XM_019822956.2;refGene_gn=GRHL2;refGene_tx=XM_019822957.2
F2 49687798 49687863
refGene_gn=GRHL2;refGene_tx=XM_006943329.4;refGene_gn=GRHL2;refGene_tx=XM_019822955.2;refGene_gn=GRHL2;refGene_tx=XM_019822956.2;refGene_gn=GRHL2;refGene_tx=XM_019822957.2
F2 49689979 49690094
refGene_gn=GRHL2;refGene_tx=XM_006943329.4;refGene_gn=GRHL2;refGene_tx=XM_019822955.2;refGene_gn=GRHL2;refGene_tx=XM_019822956.2;refGene_gn=GRHL2;refGene_tx=XM_019822957.2
F2 49701027 49701125
refGene_gn=NCALD;refGene_tx=XM_006943330.4;refGene_gn=NCALD;refGene_tx=XM_019822967.1;refGene_gn=NCALD;refGene_tx=XM_023248608.1
F2 49704192 49704298
refGene_gn=NCALD;refGene_tx=XM_006943330.4;refGene_gn=NCALD;refGene_tx=XM_019822967.1;refGene_gn=NCALD;refGene_tx=XM_023248608.1
F2 49719891 49720269
refGene_gn=NCALD;refGene_tx=XM_006943330.4;refGene_gn=NCALD;refGene_tx=XM_019822967.1;refGene_gn=NCALD;refGene_tx=XM_023248608.1
F2 50184200 50184353 refGene_gn=RRM2B;refGene_tx=XM_004000027.5
F2 50185559 50185673 refGene_gn=RRM2B;refGene_tx=XM_004000027.5
F2 50186986 50187091 refGene_gn=RRM2B;refGene_tx=XM_004000027.5
F2 50193816 50193950 refGene_gn=RRM2B;refGene_tx=XM_004000027.5
F2 50200376 50200471 refGene_gn=RRM2B;refGene_tx=XM_004000027.5
F2 50201349 50201483 refGene_gn=RRM2B;refGene_tx=XM_004000027.5
F2 50202610 50202727 refGene_gn=RRM2B;refGene_tx=XM_004000027.5
F2 50207040 50207196 refGene_gn=RRM2B;refGene_tx=XM_004000027.5
F2 50216395 50216704 refGene_gn=RRM2B;refGene_tx=XM_004000027.5
F2 50262096 50262309 refGene_gn=UBR5;refGene_tx=XM_023248400.1
F2 50265864 50265950 refGene_gn=UBR5;refGene_tx=XM_023248400.1
F2 50267210 50267355 refGene_gn=UBR5;refGene_tx=XM_023248400.1
F2 50268804 50268918 refGene_gn=UBR5;refGene_tx=XM_023248400.1
F2 50269268 50269424 refGene_gn=UBR5;refGene_tx=XM_023248400.1
F2 50270803 50270888 refGene_gn=UBR5;refGene_tx=XM_023248400.1
F2 50271113 50271289 refGene_gn=UBR5;refGene_tx=XM_023248400.1
F2 50273589 50273650 refGene_gn=UBR5;refGene_tx=XM_023248400.1
F2 50275528 50275659 refGene_gn=UBR5;refGene_tx=XM_023248400.1
F2 50276667 50276812 refGene_gn=UBR5;refGene_tx=XM_023248400.1
F2 50277385 50277497 refGene_gn=UBR5;refGene_tx=XM_023248400.1
F2 50278936 50279180 refGene_gn=UBR5;refGene_tx=XM_023248400.1
F2 50281820 50281919 refGene_gn=UBR5;refGene_tx=XM_023248400.1

F2 50282008 50282129 refGene_gn=UBR5;refGene_tx=XM_023248400.1
F2 50283547 50283768 refGene_gn=UBR5;refGene_tx=XM_023248400.1
F2 50284774 50284888 refGene_gn=UBR5;refGene_tx=XM_023248400.1
F2 50284990 50285163 refGene_gn=UBR5;refGene_tx=XM_023248400.1
F2 50286344 50286418 refGene_gn=UBR5;refGene_tx=XM_023248400.1
F2 50286907 50286984 refGene_gn=UBR5;refGene_tx=XM_023248400.1
F2 50286985 50287139 refGene_gn=UBR5;refGene_tx=XM_023248400.1
F2 50289691 50289903 refGene_gn=UBR5;refGene_tx=XM_023248400.1
F2 50290073 50290342 refGene_gn=UBR5;refGene_tx=XM_023248400.1
F2 50290893 50291151 refGene_gn=UBR5;refGene_tx=XM_023248400.1
F2 50291674 50291806 refGene_gn=UBR5;refGene_tx=XM_023248400.1
F2 50293269 50293381 refGene_gn=UBR5;refGene_tx=XM_023248400.1
F2 50293986 50294110 refGene_gn=UBR5;refGene_tx=XM_023248400.1
F2 50298370 50298610 refGene_gn=UBR5;refGene_tx=XM_023248400.1
F2 50298731 50298892 refGene_gn=UBR5;refGene_tx=XM_023248400.1
F2 50300213 50300343 refGene_gn=UBR5;refGene_tx=XM_023248400.1
F2 50300432 50300559 refGene_gn=UBR5;refGene_tx=XM_023248400.1
F2 50300641 50300760 refGene_gn=UBR5;refGene_tx=XM_023248400.1
F2 50300855 50301002 refGene_gn=UBR5;refGene_tx=XM_023248400.1
F2 50302110 50302202 refGene_gn=UBR5;refGene_tx=XM_023248400.1
F2 50302720 50302869 refGene_gn=UBR5;refGene_tx=XM_023248400.1
F2 50304084 50304182 refGene_gn=UBR5;refGene_tx=XM_023248400.1
F2 50307473 50307597 refGene_gn=UBR5;refGene_tx=XM_023248400.1
F2 50307922 50308018 refGene_gn=UBR5;refGene_tx=XM_023248400.1
F2 50308444 50308596 refGene_gn=UBR5;refGene_tx=XM_023248400.1
F2 50312585 50312712 refGene_gn=UBR5;refGene_tx=XM_023248400.1
F2 50313884 50314095 refGene_gn=UBR5;refGene_tx=XM_023248400.1
F2 50318258 50318456 refGene_gn=UBR5;refGene_tx=XM_023248400.1
F2 50318701 50318856 refGene_gn=UBR5;refGene_tx=XM_023248400.1
F2 50319324 50319436 refGene_gn=UBR5;refGene_tx=XM_023248400.1
F2 50319532 50319676 refGene_gn=UBR5;refGene_tx=XM_023248400.1
F2 50321035 50321148 refGene_gn=UBR5;refGene_tx=XM_023248400.1
F2 50321983 50322093 refGene_gn=UBR5;refGene_tx=XM_023248400.1
F2 50324758 50324943 refGene_gn=UBR5;refGene_tx=XM_023248400.1
F2 50328670 50328788 refGene_gn=UBR5;refGene_tx=XM_023248400.1
F2 50329565 50329720 refGene_gn=UBR5;refGene_tx=XM_023248400.1
F2 50330556 50330667 refGene_gn=UBR5;refGene_tx=XM_023248400.1
F2 50330747 50330864 refGene_gn=UBR5;refGene_tx=XM_023248400.1
F2 50331397 50331516
refGene_gn=LOC109496139;refGene_tx=XM_019822972.2;refGene_gn=LOC109496
139;refGene_tx=XM_019822973.2
F2 50332302 50332415
refGene_gn=LOC109496139;refGene_tx=XM_019822972.2;refGene_gn=LOC109496
139;refGene_tx=XM_019822973.2;refGene_gn=LOC109496139;refGene_tx=XM_01
9822975.2
F2 50336309 50336747
refGene_gn=LOC109496139;refGene_tx=XM_019822972.2;refGene_gn=LOC109496
139;refGene_tx=XM_019822973.2;refGene_gn=LOC109496139;refGene_tx=XM_01
9822975.2
F2 50340562 50340774 refGene_gn=UBR5;refGene_tx=XM_023248400.1

F2	50342560	50342710	refGene_gn=UBR5;refGene_tx=XM_023248400.1
F2	50343301	50343461	refGene_gn=UBR5;refGene_tx=XM_023248400.1
F2	50343650	50343840	refGene_gn=UBR5;refGene_tx=XM_023248400.1
F2	50354629	50354749	refGene_gn=UBR5;refGene_tx=XM_023248400.1
F2	50355619	50355696	refGene_gn=UBR5;refGene_tx=XM_023248400.1
F2	50356192	50356274	refGene_gn=UBR5;refGene_tx=XM_023248400.1
F2	50356649	50356694	refGene_gn=UBR5;refGene_tx=XM_023248400.1
F2	50398244	50398306	refGene_gn=UBR5;refGene_tx=XM_023248400.1
F2	50511024	50511341	refGene_gn=ODF1;refGene_tx=XM_004000028.5
F2	50519825	50520276	refGene_gn=ODF1;refGene_tx=XM_004000028.5
F2	50596703	50596963
refGene_gn=KLF10;refGene_tx=XM_004000029.5;refGene_gn=KLF10;refGene_tx
=XM_006943333.4
F2	50597443	50598356
refGene_gn=KLF10;refGene_tx=XM_004000029.5;refGene_gn=KLF10;refGene_tx
=XM_006943333.4
F2	50598458	50598692
refGene_gn=KLF10;refGene_tx=XM_004000029.5;refGene_gn=KLF10;refGene_tx
=XM_006943333.4
F2	50599666	50599741	refGene_gn=KLF10;refGene_tx=XM_006943333.4
F2	50601754	50601790	refGene_gn=KLF10;refGene_tx=XM_004000029.5
F2	50748022	50748134
refGene_gn=AZIN1;refGene_tx=NM_001291502.1;refGene_gn=AZIN1;refGene_tx
=NM_001301734.1;refGene_gn=AZIN1;refGene_tx=XM_006943334.3
F2	50748372	50748587
refGene_gn=AZIN1;refGene_tx=NM_001291502.1;refGene_gn=AZIN1;refGene_tx
=NM_001301734.1;refGene_gn=AZIN1;refGene_tx=XM_006943334.3
F2	50748894	50749010
refGene_gn=AZIN1;refGene_tx=NM_001291502.1;refGene_gn=AZIN1;refGene_tx
=NM_001301734.1;refGene_gn=AZIN1;refGene_tx=XM_006943334.3
F2	50750981	50751144
refGene_gn=AZIN1;refGene_tx=NM_001291502.1;refGene_gn=AZIN1;refGene_tx
=NM_001301734.1;refGene_gn=AZIN1;refGene_tx=XM_006943334.3
F2	50751793	50751868
refGene_gn=AZIN1;refGene_tx=NM_001291502.1;refGene_gn=AZIN1;refGene_tx
=NM_001301734.1;refGene_gn=AZIN1;refGene_tx=XM_006943334.3
F2	50752264	50752346
refGene_gn=AZIN1;refGene_tx=NM_001291502.1;refGene_gn=AZIN1;refGene_tx
=NM_001301734.1;refGene_gn=AZIN1;refGene_tx=XM_006943334.3
F2	50753759	50753894
refGene_gn=AZIN1;refGene_tx=NM_001291502.1;refGene_gn=AZIN1;refGene_tx
=NM_001301734.1;refGene_gn=AZIN1;refGene_tx=XM_006943334.3
F2	50755590	50755763
refGene_gn=AZIN1;refGene_tx=NM_001291502.1;refGene_gn=AZIN1;refGene_tx
=NM_001301734.1;refGene_gn=AZIN1;refGene_tx=XM_006943334.3
F2	50756271	50756445
refGene_gn=AZIN1;refGene_tx=NM_001291502.1;refGene_gn=AZIN1;refGene_tx
=NM_001301734.1;refGene_gn=AZIN1;refGene_tx=XM_006943334.3
F2	50758708	50758810
refGene_gn=AZIN1;refGene_tx=NM_001291502.1;refGene_gn=AZIN1;refGene_tx

=NM_001301734.1;refGene_gn=AZIN1;refGene_tx=XM_006943334.3
F2 50910071 50910203 refGene_gn=ATP6V1C1;refGene_tx=XM_011291419.3
F2 50911487 50911555 refGene_gn=ATP6V1C1;refGene_tx=XM_011291419.3
F2 50917524 50917610 refGene_gn=ATP6V1C1;refGene_tx=XM_011291419.3
F2 50922157 50922252 refGene_gn=ATP6V1C1;refGene_tx=XM_011291419.3
F2 50924207 50924299 refGene_gn=ATP6V1C1;refGene_tx=XM_011291419.3
F2 50925332 50925431 refGene_gn=ATP6V1C1;refGene_tx=XM_011291419.3
F2 50927642 50927711 refGene_gn=ATP6V1C1;refGene_tx=XM_011291419.3
F2 50934857 50934950 refGene_gn=ATP6V1C1;refGene_tx=XM_011291419.3
F2 50935042 50935136 refGene_gn=ATP6V1C1;refGene_tx=XM_011291419.3
F2 50937224 50937322 refGene_gn=ATP6V1C1;refGene_tx=XM_011291419.3
F2 50938169 50938296 refGene_gn=ATP6V1C1;refGene_tx=XM_011291419.3
F2 50939638 50939734 refGene_gn=ATP6V1C1;refGene_tx=XM_011291419.3
F2 50994124 50994284
refGene_gn=BAALC;refGene_tx=XM_023248673.1;refGene_gn=BAALC;refGene_tx
=XM_023248674.1;refGene_gn=BAALC;refGene_tx=XM_023248675.1
F2 51065616 51065783
refGene_gn=BAALC;refGene_tx=XM_023248673.1;refGene_gn=BAALC;refGene_tx
=XM_023248674.1;refGene_gn=BAALC;refGene_tx=XM_023248675.1
F2 51071836 51071854 refGene_gn=BAALC;refGene_tx=XM_023248674.1
F2 51074587 51074593 refGene_gn=BAALC;refGene_tx=XM_023248675.1
F2 51079242 51079353 refGene_gn=BAALC;refGene_tx=XM_023248673.1
F2 51131057 51131234 refGene_gn=FZD6;refGene_tx=XM_023248271.1
F2 51165033 51165230 refGene_gn=FZD6;refGene_tx=XM_023248271.1
F2 51168204 51169222 refGene_gn=FZD6;refGene_tx=XM_023248271.1
F2 51170159 51170308 refGene_gn=FZD6;refGene_tx=XM_023248271.1
F2 51171433 51171862 refGene_gn=FZD6;refGene_tx=XM_023248271.1
F2 51172859 51173028 refGene_gn=FZD6;refGene_tx=XM_023248271.1
F2 51216553 51216706 refGene_gn=CTHRC1;refGene_tx=XM_004000033.5
F2 51221062 51221284 refGene_gn=CTHRC1;refGene_tx=XM_004000033.5
F2 51225059 51225276 refGene_gn=CTHRC1;refGene_tx=XM_004000033.5
F2 51228317 51228460 refGene_gn=CTHRC1;refGene_tx=XM_004000033.5
F2 51230631 51230680 refGene_gn=SLC25A32;refGene_tx=XM_019822981.1
F2 51240403 51240545 refGene_gn=SLC25A32;refGene_tx=XM_004000034.5
F2 51240782 51240792 refGene_gn=SLC25A32;refGene_tx=XM_019822983.2
F2 51241461 51241607
refGene_gn=SLC25A32;refGene_tx=XM_004000034.5;refGene_gn=SLC25A32;refG
ene_tx=XM_019822981.1;refGene_gn=SLC25A32;refGene_tx=XM_019822983.2
F2 51242150 51242264
refGene_gn=SLC25A32;refGene_tx=XM_004000034.5;refGene_gn=SLC25A32;refG
ene_tx=XM_019822981.1;refGene_gn=SLC25A32;refGene_tx=XM_019822983.2
F2 51243176 51243337
refGene_gn=SLC25A32;refGene_tx=XM_004000034.5;refGene_gn=SLC25A32;refG
ene_tx=XM_019822981.1;refGene_gn=SLC25A32;refGene_tx=XM_019822983.2
F2 51244651 51244737
refGene_gn=SLC25A32;refGene_tx=XM_004000034.5;refGene_gn=SLC25A32;refG
ene_tx=XM_019822981.1;refGene_gn=SLC25A32;refGene_tx=XM_019822983.2
F2 51246467 51246618
refGene_gn=SLC25A32;refGene_tx=XM_004000034.5;refGene_gn=SLC25A32;refG
ene_tx=XM_019822981.1;refGene_gn=SLC25A32;refGene_tx=XM_019822983.2

F2 51251253 51251407
refGene_gn=SLC25A32;refGene_tx=XM_004000034.5;refGene_gn=SLC25A32;refGene_tx=XM_019822981.1;refGene_gn=SLC25A32;refGene_tx=XM_019822983.2
F2 51251918 51251988 refGene_gn=DCAF13;refGene_tx=XM_006943342.4
F2 51254630 51254830 refGene_gn=DCAF13;refGene_tx=XM_006943342.4
F2 51255406 51255514 refGene_gn=DCAF13;refGene_tx=XM_006943342.4
F2 51259691 51259781 refGene_gn=DCAF13;refGene_tx=XM_006943342.4
F2 51261233 51261389 refGene_gn=DCAF13;refGene_tx=XM_006943342.4
F2 51265449 51265527 refGene_gn=DCAF13;refGene_tx=XM_006943342.4
F2 51267601 51267684 refGene_gn=DCAF13;refGene_tx=XM_006943342.4
F2 51271943 51272108 refGene_gn=DCAF13;refGene_tx=XM_006943342.4
F2 51276946 51277082 refGene_gn=DCAF13;refGene_tx=XM_006943342.4
F2 51278252 51278416 refGene_gn=DCAF13;refGene_tx=XM_006943342.4
F2 51279139 51279227 refGene_gn=DCAF13;refGene_tx=XM_006943342.4
F2 51353461 51353637
refGene_gn=RIMS2;refGene_tx=XM_023248353.1;refGene_gn=RIMS2;refGene_tx=XM_023248354.1;refGene_gn=RIMS2;refGene_tx=XM_023248355.1;refGene_gn=RIMS2;refGene_tx=XM_023248356.1;refGene_gn=RIMS2;refGene_tx=XM_023248357.1;refGene_gn=RIMS2;refGene_tx=XM_023248358.1;refGene_gn=RIMS2;refGene_tx=XM_023248359.1;refGene_gn=RIMS2;refGene_tx=XM_023248360.1;refGene_gn=RIMS2;refGene_tx=XM_023248361.1;refGene_gn=RIMS2;refGene_tx=XM_023248362.1;refGene_gn=RIMS2;refGene_tx=XM_023248363.1;refGene_gn=RIMS2;refGene_tx=XM_023248364.1;refGene_gn=RIMS2;refGene_tx=XM_023248365.1;refGene_gn=RIMS2;refGene_tx=XM_023248366.1;refGene_gn=RIMS2;refGene_tx=XM_023248367.1;refGene_gn=RIMS2;refGene_tx=XM_023248368.1;refGene_gn=RIMS2;refGene_tx=XM_023248369.1;refGene_gn=RIMS2;refGene_tx=XM_023248370.1;refGene_gn=RIMS2;refGene_tx=XM_023248371.1;refGene_gn=RIMS2;refGene_tx=XM_023248372.1;refGene_gn=RIMS2;refGene_tx=XM_023248373.1;refGene_gn=RIMS2;refGene_tx=XM_023248374.1;refGene_gn=RIMS2;refGene_tx=XM_023248375.1;refGene_gn=RIMS2;refGene_tx=XM_023248376.1;refGene_gn=RIMS2;refGene_tx=XM_023248377.1;refGene_gn=RIMS2;refGene_tx=XM_023248378.1;refGene_gn=RIMS2;refGene_tx=XM_023248380.1;refGene_gn=RIMS2;refGene_tx=XM_023248381.1;refGene_gn=RIMS2;refGene_tx=XM_023248387.1;refGene_gn=RIMS2;refGene_tx=XM_023248388.1;refGene_gn=RIMS2;refGene_tx=XM_023248389.
1;refGene_gn=RIMS2;refGene_tx=XM_023248390.1;refGene_gn=RIMS2;refGene_tx=XM_023248392.1;refGene_gn=RIMS2;refGene_tx=XM_023248393.1
F2 51450352 51450385
refGene_gn=RIMS2;refGene_tx=XM_023248353.1;refGene_gn=RIMS2;refGene_tx=XM_023248354.1;refGene_gn=RIMS2;refGene_tx=XM_023248355.1;refGene_gn=RIMS2;refGene_tx=XM_023248356.1;refGene_gn=RIMS2;refGene_tx=XM_023248357.1;refGene_gn=RIMS2;refGene_tx=XM_023248358.1;refGene_gn=RIMS2;refGene_tx=XM_023248359.1;refGene_gn=RIMS2;refGene_tx=XM_023248360.1;refGene_gn=RIMS2;refGene_tx=XM_023248361.1;refGene_gn=RIMS2;refGene_tx=XM_023248362.1;refGene_gn=RIMS2;refGene_tx=XM_023248363.1;refGene_gn=RIMS2;refGene_tx=XM_023248366.1;refGene_gn=RIMS2;refGene_tx=XM_023248367.1;refGene_gn=RIMS2;refGene_tx=XM_023248368.1;refGene_gn=RIMS2;refGene_tx=XM_023248369.1;refGene_gn=RIMS2;refGene_tx=XM_023248370.1;refGene_gn=RIMS2;refGene_tx=XM_023248372.1;refGene_gn=RIMS2;refGene_tx=XM_023248373.1;refGene_gn=RIMS2;refGene_tx=XM_023248377.1;refGene_gn=RIMS2;refGen

e_tx=XM_023248380.1;refGene_gn=RIMS2;refGene_tx=XM_023248381.1;refGene_gn=RIMS2;refGene_tx=XM_023248387.1;refGene_gn=RIMS2;refGene_tx=XM_023248392.1
F2 51450785 51450872
refGene_gn=RIMS2;refGene_tx=XM_023248353.1;refGene_gn=RIMS2;refGene_tx=XM_023248354.1;refGene_gn=RIMS2;refGene_tx=XM_023248355.1;refGene_gn=RIMS2;refGene_tx=XM_023248356.1;refGene_gn=RIMS2;refGene_tx=XM_023248357.1;refGene_gn=RIMS2;refGene_tx=XM_023248358.1;refGene_gn=RIMS2;refGene_tx=XM_023248359.1;refGene_gn=RIMS2;refGene_tx=XM_023248360.1;refGene_gn=RIMS2;refGene_tx=XM_023248361.1;refGene_gn=RIMS2;refGene_tx=XM_023248362.1;refGene_gn=RIMS2;refGene_tx=XM_023248363.1;refGene_gn=RIMS2;refGene_tx=XM_023248366.1;refGene_gn=RIMS2;refGene_tx=XM_023248367.1;refGene_gn=RIMS2;refGene_tx=XM_023248368.1;refGene_gn=RIMS2;refGene_tx=XM_023248369.1;refGene_gn=RIMS2;refGene_tx=XM_023248370.1;refGene_gn=RIMS2;refGene_tx=XM_023248372.1;refGene_gn=RIMS2;refGene_tx=XM_023248373.1;refGene_gn=RIMS2;refGene_tx=XM_023248377.1;refGene_gn=RIMS2;refGene_tx=XM_023248380.1;refGene_gn=RIMS2;refGene_tx=XM_023248381.1;refGene_gn=RIMS2;refGene_tx=XM_023248387.1;refGene_gn=RIMS2;refGene_tx=XM_023248392.1
F2 51475946 51476157
refGene_gn=RIMS2;refGene_tx=XM_023248353.1;refGene_gn=RIMS2;refGene_tx=XM_023248354.1;refGene_gn=RIMS2;refGene_tx=XM_023248355.1;refGene_gn=RIMS2;refGene_tx=XM_023248356.1;refGene_gn=RIMS2;refGene_tx=XM_023248357.1;refGene_gn=RIMS2;refGene_tx=XM_023248358.1;refGene_gn=RIMS2;refGene_tx=XM_023248359.1;refGene_gn=RIMS2;refGene_tx=XM_023248360.1;refGene_gn=RIMS2;refGene_tx=XM_023248361.1;refGene_gn=RIMS2;refGene_tx=XM_023248362.1;refGene_gn=RIMS2;refGene_tx=XM_023248363.1;refGene_gn=RIMS2;refGene_tx=XM_023248364.1;refGene_gn=RIMS2;refGene_tx=XM_023248365.1;refGene_gn=RIMS2;refGene_tx=XM_023248366.1;refGene_gn=RIMS2;refGene_tx=XM_023248367.1;refGene_gn=RIMS2;refGene_tx=XM_023248368.1;refGene_gn=RIMS2;refGene_tx=XM_023248369.1;refGene_gn=RIMS2;refGene_tx=XM_023248370.1;refGene_gn=RIMS2;refGene_tx=XM_023248371.1;refGene_gn=RIMS2;refGene_tx=XM_023248372.1;refGene_gn=RIMS2;refGene_tx=XM_023248373.1;refGene_gn=RIMS2;refGene_tx=XM_023248374.1;refGene_gn=RIMS2;refGene_tx=XM_023248375.1;refGene_gn=RIMS2;refGene_tx=XM_023248376.1;refGene_gn=RIMS2;refGene_tx=XM_023248377.1;refGene_gn=RIMS2;refGene_tx=XM_023248378.1;refGene_gn=RIMS2;refGene_tx=XM_023248380.1;refGene_gn=RIMS2;refGene_tx=XM_023248381.1;refGene_gn=RIMS2;refGene_tx=XM_023248387.1;refGene_gn=RIMS2;refGene_tx=XM_023248388.1;refGene_gn=RIMS2;refGene_tx=XM_023248389.
1;refGene_gn=RIMS2;refGene_tx=XM_023248390.1;refGene_gn=RIMS2;refGene_tx=XM_023248392.1;refGene_gn=RIMS2;refGene_tx=XM_023248393.1
F2 51493197 51493209
refGene_gn=RIMS2;refGene_tx=XM_023248353.1;refGene_gn=RIMS2;refGene_tx=XM_023248354.1;refGene_gn=RIMS2;refGene_tx=XM_023248356.1;refGene_gn=RIMS2;refGene_tx=XM_023248357.1;refGene_gn=RIMS2;refGene_tx=XM_023248358.1;refGene_gn=RIMS2;refGene_tx=XM_023248359.1;refGene_gn=RIMS2;refGene_tx=XM_023248360.1;refGene_gn=RIMS2;refGene_tx=XM_023248361.1;refGene_gn=RIMS2;refGene_tx=XM_023248362.1;refGene_gn=RIMS2;refGene_tx=XM_023248363.1;refGene_gn=RIMS2;refGene_tx=XM_023248364.1;refGene_gn=RIMS2;

refGene_tx=XM_023248366.1;refGene_gn=RIMS2;refGene_tx=XM_023248367.1;refGene_gn=RIMS2;refGene_tx=XM_023248368.1;refGene_gn=RIMS2;refGene_tx=XM_023248369.1;refGene_gn=RIMS2;refGene_tx=XM_023248370.1;refGene_gn=RIMS2;refGene_tx=XM_023248372.1;refGene_gn=RIMS2;refGene_tx=XM_023248373.1;refGene_gn=RIMS2;refGene_tx=XM_023248378.1;refGene_gn=RIMS2;refGene_tx=XM_023248380.1;refGene_gn=RIMS2;refGene_tx=XM_023248381.1;refGene_gn=RIMS2;refGene_tx=XM_023248387.1;refGene_gn=RIMS2;refGene_tx=XM_023248393.1

F2 51538220 51538531

refGene_gn=RIMS2;refGene_tx=XM_023248353.1;refGene_gn=RIMS2;refGene_tx=XM_023248354.1;refGene_gn=RIMS2;refGene_tx=XM_023248355.1;refGene_gn=RIMS2;refGene_tx=XM_023248356.1;refGene_gn=RIMS2;refGene_tx=XM_023248357.1;refGene_gn=RIMS2;refGene_tx=XM_023248358.1;refGene_gn=RIMS2;refGene_tx=XM_023248359.1;refGene_gn=RIMS2;refGene_tx=XM_023248360.1;refGene_gn=RIMS2;refGene_tx=XM_023248361.1;refGene_gn=RIMS2;refGene_tx=XM_023248362.1;refGene_gn=RIMS2;refGene_tx=XM_023248363.1;refGene_gn=RIMS2;refGene_tx=XM_023248364.1;refGene_gn=RIMS2;refGene_tx=XM_023248365.1;refGene_gn=RIMS2;refGene_tx=XM_023248366.1;refGene_gn=RIMS2;refGene_tx=XM_023248367.1;refGene_gn=RIMS2;refGene_tx=XM_023248368.1;refGene_gn=RIMS2;refGene_tx=XM_023248369.1;refGene_gn=RIMS2;refGene_tx=XM_023248370.1;refGene_gn=RIMS2;refGene_tx=XM_023248371.1;refGene_gn=RIMS2;refGene_tx=XM_023248372.1;refGene_gn=RIMS2;refGene_tx=XM_023248373.1;refGene_gn=RIMS2;refGene_tx=XM_023248374.1;refGene_gn=RIMS2;refGene_tx=XM_023248375.1;refGene_gn=RIMS2;refGene_tx=XM_023248376.1;refGene_gn=RIMS2;refGene_tx=XM_023248377.1;refGene_gn=RIMS2;refGene_tx=XM_023248378.1;refGene_gn=RIMS2;refGene_tx=XM_023248380.1;refGene_gn=RIMS2;refGene_tx=XM_023248381.1;refGene_gn=RIMS2;refGene_tx=XM_023248387.1;refGene_gn=RIMS2;refGene_tx=XM_023248388.1;refGene_gn=RIMS2;refGene_tx=XM_023248389.
1;refGene_gn=RIMS2;refGene_tx=XM_023248390.1;refGene_gn=RIMS2;refGene_tx=XM_023248392.1;refGene_gn=RIMS2;refGene_tx=XM_023248393.1

F2 51578040 51578162

refGene_gn=RIMS2;refGene_tx=XM_023248379.1;refGene_gn=RIMS2;refGene_tx=XM_023248382.1;refGene_gn=RIMS2;refGene_tx=XM_023248383.1;refGene_gn=RIMS2;refGene_tx=XM_023248384.1;refGene_gn=RIMS2;refGene_tx=XM_023248385.1;refGene_gn=RIMS2;refGene_tx=XM_023248386.1;refGene_gn=RIMS2;refGene_tx=XM_023248394.1;refGene_gn=RIMS2;refGene_tx=XM_023248395.1;refGene_gn=RIMS2;refGene_tx=XM_023248396.1;refGene_gn=RIMS2;refGene_tx=XM_023248397.1

F2 51635982 51636908

refGene_gn=RIMS2;refGene_tx=XM_023248353.1;refGene_gn=RIMS2;refGene_tx=XM_023248354.1;refGene_gn=RIMS2;refGene_tx=XM_023248355.1;refGene_gn=RIMS2;refGene_tx=XM_023248356.1;refGene_gn=RIMS2;refGene_tx=XM_023248357.1;refGene_gn=RIMS2;refGene_tx=XM_023248358.1;refGene_gn=RIMS2;refGene_tx=XM_023248359.1;refGene_gn=RIMS2;refGene_tx=XM_023248360.1;refGene_gn=RIMS2;refGene_tx=XM_023248361.1;refGene_gn=RIMS2;refGene_tx=XM_023248362.1;refGene_gn=RIMS2;refGene_tx=XM_023248363.1;refGene_gn=RIMS2;refGene_tx=XM_023248364.1;refGene_gn=RIMS2;refGene_tx=XM_023248365.1;refGene_gn=RIMS2;refGene_tx=XM_023248366.1;refGene_gn=RIMS2;refGene_tx=XM_023248367.1;refGene_gn=RIMS2;refGene_tx=XM_023248368.1;refGene_gn=R

IMS2;refGene_tx=XM_023248369.1;refGene_gn=RIMS2;refGene_tx=XM_02324837
0.1;refGene_gn=RIMS2;refGene_tx=XM_023248371.1;refGene_gn=RIMS2;refGen
e_tx=XM_023248372.1;refGene_gn=RIMS2;refGene_tx=XM_023248373.1;refGene
_gn=RIMS2;refGene_tx=XM_023248374.1;refGene_gn=RIMS2;refGene_tx=XM_023
248375.1;refGene_gn=RIMS2;refGene_tx=XM_023248376.1;refGene_gn=RIMS2;r
efGene_tx=XM_023248377.1;refGene_gn=RIMS2;refGene_tx=XM_023248378.1;re
fGene_gn=RIMS2;refGene_tx=XM_023248379.1;refGene_gn=RIMS2;refGene_tx=X
M_023248380.1;refGene_gn=RIMS2;refGene_tx=XM_023248381.1;refGene_gn=RI
MS2;refGene_tx=XM_023248382.1;refGene_gn=RIMS2;refGene_tx=XM_023248383
.
1;refGene_gn=RIMS2;refGene_tx=XM_023248384.1;refGene_gn=RIMS2;refGene_
tx=XM_023248385.1;refGene_gn=RIMS2;refGene_tx=XM_023248386.1;refGene_g
n=RIMS2;refGene_tx=XM_023248387.1;refGene_gn=RIMS2;refGene_tx=XM_02324
8388.1;refGene_gn=RIMS2;refGene_tx=XM_023248389.1;refGene_gn=RIMS2;ref
Gene_tx=XM_023248390.1;refGene_gn=RIMS2;refGene_tx=XM_023248392.1;refG
ene_gn=RIMS2;refGene_tx=XM_023248393.1;refGene_gn=RIMS2;refGene_tx=XM_
023248394.1;refGene_gn=RIMS2;refGene_tx=XM_023248395.1;refGene_gn=RIMS
2;refGene_tx=XM_023248396.1;refGene_gn=RIMS2;refGene_tx=XM_023248397.1
F2 51667243 51667311
refGene_gn=RIMS2;refGene_tx=XM_023248366.1;refGene_gn=RIMS2;refGene_tx
=XM_023248368.1;refGene_gn=RIMS2;refGene_tx=XM_023248371.1;refGene_gn=
RIMS2;refGene_tx=XM_023248376.1;refGene_gn=RIMS2;refGene_tx=XM_0232483
78.1;refGene_gn=RIMS2;refGene_tx=XM_023248382.1;refGene_gn=RIMS2;refGe
ne_tx=XM_023248383.1;refGene_gn=RIMS2;refGene_tx=XM_023248384.1;refGen
e_gn=RIMS2;refGene_tx=XM_023248385.1;refGene_gn=RIMS2;refGene_tx=XM_02
3248386.1;refGene_gn=RIMS2;refGene_tx=XM_023248390.1;refGene_gn=RIMS2;
refGene_tx=XM_023248393.1;refGene_gn=RIMS2;refGene_tx=XM_023248394.1;r
efGene_gn=RIMS2;refGene_tx=XM_023248395.1;refGene_gn=RIMS2;refGene_tx=
XM_023248396.1;refGene_gn=RIMS2;refGene_tx=XM_023248397.1
F2 51667437 51667646
refGene_gn=RIMS2;refGene_tx=XM_023248353.1;refGene_gn=RIMS2;refGene_tx
=XM_023248354.1;refGene_gn=RIMS2;refGene_tx=XM_023248355.1;refGene_gn=
RIMS2;refGene_tx=XM_023248356.1;refGene_gn=RIMS2;refGene_tx=XM_0232483
57.1;refGene_gn=RIMS2;refGene_tx=XM_023248358.1;refGene_gn=RIMS2;refGe
ne_tx=XM_023248359.1;refGene_gn=RIMS2;refGene_tx=XM_023248360.1;refGen
e_gn=RIMS2;refGene_tx=XM_023248361.1;refGene_gn=RIMS2;refGene_tx=XM_02
3248362.1;refGene_gn=RIMS2;refGene_tx=XM_023248363.1;refGene_gn=RIMS2;
refGene_tx=XM_023248364.1;refGene_gn=RIMS2;refGene_tx=XM_023248365.1;r
efGene_gn=RIMS2;refGene_tx=XM_023248367.1;refGene_gn=RIMS2;refGene_tx=
XM_023248369.1;refGene_gn=RIMS2;refGene_tx=XM_023248370.1;refGene_gn=R
IMS2;refGene_tx=XM_023248372.1;refGene_gn=RIMS2;refGene_tx=XM_02324837
3.1;refGene_gn=RIMS2;refGene_tx=XM_023248374.1;refGene_gn=RIMS2;refGen
e_tx=XM_023248375.1;refGene_gn=RIMS2;refGene_tx=XM_023248377.1;refGene
_gn=RIMS2;refGene_tx=XM_023248379.1;refGene_gn=RIMS2;refGene_tx=XM_023
248380.1;refGene_gn=RIMS2;refGene_tx=XM_023248381.1;refGene_gn=RIMS2;r
efGene_tx=XM_023248387.1;refGene_gn=RIMS2;refGene_tx=XM_023248388.1;re
fGene_gn=RIMS2;refGene_tx=XM_023248389.1;refGene_gn=RIMS2;refGene_tx=X
M_023248392.1
F2 51669122 51669242
refGene_gn=RIMS2;refGene_tx=XM_023248353.1;refGene_gn=RIMS2;refGene_tx

=XM_023248354.1;refGene_gn=RIMS2;refGene_tx=XM_023248355.1;refGene_gn=
RIMS2;refGene_tx=XM_023248356.1;refGene_gn=RIMS2;refGene_tx=XM_0232483
57.1;refGene_gn=RIMS2;refGene_tx=XM_023248358.1;refGene_gn=RIMS2;refGe
ne_tx=XM_023248359.1;refGene_gn=RIMS2;refGene_tx=XM_023248360.1;refGen
e_gn=RIMS2;refGene_tx=XM_023248361.1;refGene_gn=RIMS2;refGene_tx=XM_02
3248362.1;refGene_gn=RIMS2;refGene_tx=XM_023248363.1;refGene_gn=RIMS2;
refGene_tx=XM_023248364.1;refGene_gn=RIMS2;refGene_tx=XM_023248365.1;r
efGene_gn=RIMS2;refGene_tx=XM_023248366.1;refGene_gn=RIMS2;refGene_tx=
XM_023248367.1;refGene_gn=RIMS2;refGene_tx=XM_023248368.1;refGene_gn=R
IMS2;refGene_tx=XM_023248369.1;refGene_gn=RIMS2;refGene_tx=XM_02324837
0.1;refGene_gn=RIMS2;refGene_tx=XM_023248371.1;refGene_gn=RIMS2;refGen
e_tx=XM_023248372.1;refGene_gn=RIMS2;refGene_tx=XM_023248373.1;refGene
_gn=RIMS2;refGene_tx=XM_023248374.1;refGene_gn=RIMS2;refGene_tx=XM_023
248375.1;refGene_gn=RIMS2;refGene_tx=XM_023248376.1;refGene_gn=RIMS2;r
efGene_tx=XM_023248377.1;refGene_gn=RIMS2;refGene_tx=XM_023248378.1;re
fGene_gn=RIMS2;refGene_tx=XM_023248379.1;refGene_gn=RIMS2;refGene_tx=X
M_023248380.1;refGene_gn=RIMS2;refGene_tx=XM_023248381.1;refGene_gn=RI
MS2;refGene_tx=XM_023248382.1;refGene_gn=RIMS2;refGene_tx=XM_023248383
.
1;refGene_gn=RIMS2;refGene_tx=XM_023248384.1;refGene_gn=RIMS2;refGene_
tx=XM_023248385.1;refGene_gn=RIMS2;refGene_tx=XM_023248386.1;refGene_g
n=RIMS2;refGene_tx=XM_023248387.1;refGene_gn=RIMS2;refGene_tx=XM_02324
8388.1;refGene_gn=RIMS2;refGene_tx=XM_023248389.1;refGene_gn=RIMS2;ref
Gene_tx=XM_023248390.1;refGene_gn=RIMS2;refGene_tx=XM_023248392.1;refG
ene_gn=RIMS2;refGene_tx=XM_023248393.1;refGene_gn=RIMS2;refGene_tx=XM_
023248394.1;refGene_gn=RIMS2;refGene_tx=XM_023248395.1;refGene_gn=RIMS
2;refGene_tx=XM_023248396.1;refGene_gn=RIMS2;refGene_tx=XM_023248397.1
F2 51671709 51671809
refGene_gn=RIMS2;refGene_tx=XM_023248353.1;refGene_gn=RIMS2;refGene_tx
=XM_023248354.1;refGene_gn=RIMS2;refGene_tx=XM_023248355.1;refGene_gn=
RIMS2;refGene_tx=XM_023248356.1;refGene_gn=RIMS2;refGene_tx=XM_0232483
57.1;refGene_gn=RIMS2;refGene_tx=XM_023248358.1;refGene_gn=RIMS2;refGe
ne_tx=XM_023248359.1;refGene_gn=RIMS2;refGene_tx=XM_023248360.1;refGen
e_gn=RIMS2;refGene_tx=XM_023248361.1;refGene_gn=RIMS2;refGene_tx=XM_02
3248362.1;refGene_gn=RIMS2;refGene_tx=XM_023248363.1;refGene_gn=RIMS2;
refGene_tx=XM_023248364.1;refGene_gn=RIMS2;refGene_tx=XM_023248365.1;r
efGene_gn=RIMS2;refGene_tx=XM_023248366.1;refGene_gn=RIMS2;refGene_tx=
XM_023248367.1;refGene_gn=RIMS2;refGene_tx=XM_023248368.1;refGene_gn=R
IMS2;refGene_tx=XM_023248369.1;refGene_gn=RIMS2;refGene_tx=XM_02324837
0.1;refGene_gn=RIMS2;refGene_tx=XM_023248371.1;refGene_gn=RIMS2;refGen
e_tx=XM_023248372.1;refGene_gn=RIMS2;refGene_tx=XM_023248373.1;refGene
_gn=RIMS2;refGene_tx=XM_023248374.1;refGene_gn=RIMS2;refGene_tx=XM_023
248375.1;refGene_gn=RIMS2;refGene_tx=XM_023248376.1;refGene_gn=RIMS2;r
efGene_tx=XM_023248377.1;refGene_gn=RIMS2;refGene_tx=XM_023248378.1;re
fGene_gn=RIMS2;refGene_tx=XM_023248379.1;refGene_gn=RIMS2;refGene_tx=X
M_023248380.1;refGene_gn=RIMS2;refGene_tx=XM_023248381.1;refGene_gn=RI
MS2;refGene_tx=XM_023248382.1;refGene_gn=RIMS2;refGene_tx=XM_023248383
.
1;refGene_gn=RIMS2;refGene_tx=XM_023248384.1;refGene_gn=RIMS2;refGene_
tx=XM_023248385.1;refGene_gn=RIMS2;refGene_tx=XM_023248386.1;refGene_g

n=RIMS2;refGene_tx=XM_023248387.1;refGene_gn=RIMS2;refGene_tx=XM_02324
8388.1;refGene_gn=RIMS2;refGene_tx=XM_023248389.1;refGene_gn=RIMS2;ref
Gene_tx=XM_023248390.1;refGene_gn=RIMS2;refGene_tx=XM_023248392.1;refG
ene_gn=RIMS2;refGene_tx=XM_023248393.1;refGene_gn=RIMS2;refGene_tx=XM_
023248394.1;refGene_gn=RIMS2;refGene_tx=XM_023248395.1;refGene_gn=RIMS
2;refGene_tx=XM_023248396.1;refGene_gn=RIMS2;refGene_tx=XM_023248397.1
F2 51672614 51672738
refGene_gn=RIMS2;refGene_tx=XM_023248353.1;refGene_gn=RIMS2;refGene_tx
=XM_023248354.1;refGene_gn=RIMS2;refGene_tx=XM_023248355.1;refGene_gn=
RIMS2;refGene_tx=XM_023248356.1;refGene_gn=RIMS2;refGene_tx=XM_0232483
57.1;refGene_gn=RIMS2;refGene_tx=XM_023248358.1;refGene_gn=RIMS2;refGe
ne_tx=XM_023248359.1;refGene_gn=RIMS2;refGene_tx=XM_023248360.1;refGen
e_gn=RIMS2;refGene_tx=XM_023248361.1;refGene_gn=RIMS2;refGene_tx=XM_02
3248362.1;refGene_gn=RIMS2;refGene_tx=XM_023248363.1;refGene_gn=RIMS2;
refGene_tx=XM_023248364.1;refGene_gn=RIMS2;refGene_tx=XM_023248365.1;r
efGene_gn=RIMS2;refGene_tx=XM_023248366.1;refGene_gn=RIMS2;refGene_tx=
XM_023248367.1;refGene_gn=RIMS2;refGene_tx=XM_023248368.1;refGene_gn=R
IMS2;refGene_tx=XM_023248369.1;refGene_gn=RIMS2;refGene_tx=XM_02324837
0.1;refGene_gn=RIMS2;refGene_tx=XM_023248371.1;refGene_gn=RIMS2;refGen
e_tx=XM_023248372.1;refGene_gn=RIMS2;refGene_tx=XM_023248373.1;refGene
_gn=RIMS2;refGene_tx=XM_023248374.1;refGene_gn=RIMS2;refGene_tx=XM_023
248375.1;refGene_gn=RIMS2;refGene_tx=XM_023248376.1;refGene_gn=RIMS2;r
efGene_tx=XM_023248377.1;refGene_gn=RIMS2;refGene_tx=XM_023248378.1;re
fGene_gn=RIMS2;refGene_tx=XM_023248379.1;refGene_gn=RIMS2;refGene_tx=X
M_023248380.1;refGene_gn=RIMS2;refGene_tx=XM_023248381.1;refGene_gn=RI
MS2;refGene_tx=XM_023248382.1;refGene_gn=RIMS2;refGene_tx=XM_023248383
.
1;refGene_gn=RIMS2;refGene_tx=XM_023248384.1;refGene_gn=RIMS2;refGene_
tx=XM_023248385.1;refGene_gn=RIMS2;refGene_tx=XM_023248386.1;refGene_g
n=RIMS2;refGene_tx=XM_023248387.1;refGene_gn=RIMS2;refGene_tx=XM_02324
8388.1;refGene_gn=RIMS2;refGene_tx=XM_023248389.1;refGene_gn=RIMS2;ref
Gene_tx=XM_023248390.1;refGene_gn=RIMS2;refGene_tx=XM_023248392.1;refG
ene_gn=RIMS2;refGene_tx=XM_023248393.1;refGene_gn=RIMS2;refGene_tx=XM_
023248394.1;refGene_gn=RIMS2;refGene_tx=XM_023248395.1;refGene_gn=RIMS
2;refGene_tx=XM_023248396.1;refGene_gn=RIMS2;refGene_tx=XM_023248397.1
F2 51674958 51675005
refGene_gn=RIMS2;refGene_tx=XM_023248353.1;refGene_gn=RIMS2;refGene_tx
=XM_023248354.1;refGene_gn=RIMS2;refGene_tx=XM_023248355.1;refGene_gn=
RIMS2;refGene_tx=XM_023248356.1;refGene_gn=RIMS2;refGene_tx=XM_0232483
57.1;refGene_gn=RIMS2;refGene_tx=XM_023248358.1;refGene_gn=RIMS2;refGe
ne_tx=XM_023248359.1;refGene_gn=RIMS2;refGene_tx=XM_023248360.1;refGen
e_gn=RIMS2;refGene_tx=XM_023248361.1;refGene_gn=RIMS2;refGene_tx=XM_02
3248362.1;refGene_gn=RIMS2;refGene_tx=XM_023248363.1;refGene_gn=RIMS2;
refGene_tx=XM_023248364.1;refGene_gn=RIMS2;refGene_tx=XM_023248365.1;r
efGene_gn=RIMS2;refGene_tx=XM_023248366.1;refGene_gn=RIMS2;refGene_tx=
XM_023248367.1;refGene_gn=RIMS2;refGene_tx=XM_023248368.1;refGene_gn=R
IMS2;refGene_tx=XM_023248369.1;refGene_gn=RIMS2;refGene_tx=XM_02324837
0.1;refGene_gn=RIMS2;refGene_tx=XM_023248371.1;refGene_gn=RIMS2;refGen
e_tx=XM_023248372.1;refGene_gn=RIMS2;refGene_tx=XM_023248373.1;refGene
_gn=RIMS2;refGene_tx=XM_023248374.1;refGene_gn=RIMS2;refGene_tx=XM_023

248375.1;refGene_gn=RIMS2;refGene_tx=XM_023248376.1;refGene_gn=RIMS2;refGene_tx=XM_023248377.1;refGene_gn=RIMS2;refGene_tx=XM_023248378.1;refGene_gn=RIMS2;refGene_tx=XM_023248379.1;refGene_gn=RIMS2;refGene_tx=XM_023248380.1;refGene_gn=RIMS2;refGene_tx=XM_023248381.1;refGene_gn=RIMS2;refGene_tx=XM_023248382.1;refGene_gn=RIMS2;refGene_tx=XM_023248383.
1;refGene_gn=RIMS2;refGene_tx=XM_023248384.1;refGene_gn=RIMS2;refGene_tx=XM_023248385.1;refGene_gn=RIMS2;refGene_tx=XM_023248386.1;refGene_gn=RIMS2;refGene_tx=XM_023248387.1;refGene_gn=RIMS2;refGene_tx=XM_023248388.1;refGene_gn=RIMS2;refGene_tx=XM_023248389.1;refGene_gn=RIMS2;refGene_tx=XM_023248390.1;refGene_gn=RIMS2;refGene_tx=XM_023248392.1;refGene_gn=RIMS2;refGene_tx=XM_023248393.1;refGene_gn=RIMS2;refGene_tx=XM_023248394.1;refGene_gn=RIMS2;refGene_tx=XM_023248395.1;refGene_gn=RIMS2;refGene_tx=XM_023248396.1;refGene_gn=RIMS2;refGene_tx=XM_023248397.1
F2 51679070 51679183
refGene_gn=RIMS2;refGene_tx=XM_023248353.1;refGene_gn=RIMS2;refGene_tx=XM_023248354.1;refGene_gn=RIMS2;refGene_tx=XM_023248355.1;refGene_gn=RIMS2;refGene_tx=XM_023248356.1;refGene_gn=RIMS2;refGene_tx=XM_023248357.1;refGene_gn=RIMS2;refGene_tx=XM_023248358.1;refGene_gn=RIMS2;refGene_tx=XM_023248359.1;refGene_gn=RIMS2;refGene_tx=XM_023248360.1;refGene_gn=RIMS2;refGene_tx=XM_023248361.1;refGene_gn=RIMS2;refGene_tx=XM_023248362.1;refGene_gn=RIMS2;refGene_tx=XM_023248363.1;refGene_gn=RIMS2;refGene_tx=XM_023248364.1;refGene_gn=RIMS2;refGene_tx=XM_023248365.1;refGene_gn=RIMS2;refGene_tx=XM_023248366.1;refGene_gn=RIMS2;refGene_tx=XM_023248367.1;refGene_gn=RIMS2;refGene_tx=XM_023248368.1;refGene_gn=RIMS2;refGene_tx=XM_023248369.1;refGene_gn=RIMS2;refGene_tx=XM_023248370.1;refGene_gn=RIMS2;refGene_tx=XM_023248371.1;refGene_gn=RIMS2;refGene_tx=XM_023248372.1;refGene_gn=RIMS2;refGene_tx=XM_023248373.1;refGene_gn=RIMS2;refGene_tx=XM_023248374.1;refGene_gn=RIMS2;refGene_tx=XM_023248375.1;refGene_gn=RIMS2;refGene_tx=XM_023248376.1;refGene_gn=RIMS2;refGene_tx=XM_023248377.1;refGene_gn=RIMS2;refGene_tx=XM_023248378.1;refGene_gn=RIMS2;refGene_tx=XM_023248379.1;refGene_gn=RIMS2;refGene_tx=XM_023248380.1;refGene_gn=RIMS2;refGene_tx=XM_023248381.1;refGene_gn=RIMS2;refGene_tx=XM_023248382.1;refGene_gn=RIMS2;refGene_tx=XM_023248383.
1;refGene_gn=RIMS2;refGene_tx=XM_023248384.1;refGene_gn=RIMS2;refGene_tx=XM_023248385.1;refGene_gn=RIMS2;refGene_tx=XM_023248386.1;refGene_gn=RIMS2;refGene_tx=XM_023248387.1;refGene_gn=RIMS2;refGene_tx=XM_023248388.1;refGene_gn=RIMS2;refGene_tx=XM_023248389.1;refGene_gn=RIMS2;refGene_tx=XM_023248390.1;refGene_gn=RIMS2;refGene_tx=XM_023248392.1;refGene_gn=RIMS2;refGene_tx=XM_023248393.1;refGene_gn=RIMS2;refGene_tx=XM_023248394.1;refGene_gn=RIMS2;refGene_tx=XM_023248395.1;refGene_gn=RIMS2;refGene_tx=XM_023248396.1;refGene_gn=RIMS2;refGene_tx=XM_023248397.1
F2 51685547 51685595
refGene_gn=RIMS2;refGene_tx=XM_023248353.1;refGene_gn=RIMS2;refGene_tx=XM_023248354.1;refGene_gn=RIMS2;refGene_tx=XM_023248355.1;refGene_gn=RIMS2;refGene_tx=XM_023248356.1;refGene_gn=RIMS2;refGene_tx=XM_023248357.1;refGene_gn=RIMS2;refGene_tx=XM_023248358.1;refGene_gn=RIMS2;refGene_tx=XM_023248359.1;refGene_gn=RIMS2;refGene_tx=XM_023248361.1;refGene_gn=RIMS2;refGene_tx=XM_023248362.1;refGene_gn=RIMS2;refGene_tx=XM_02

3248364.1;refGene_gn=RIMS2;refGene_tx=XM_023248365.1;refGene_gn=RIMS2;
refGene_tx=XM_023248366.1;refGene_gn=RIMS2;refGene_tx=XM_023248367.1;r
efGene_gn=RIMS2;refGene_tx=XM_023248369.1;refGene_gn=RIMS2;refGene_tx=
XM_023248370.1;refGene_gn=RIMS2;refGene_tx=XM_023248371.1;refGene_gn=R
IMS2;refGene_tx=XM_023248372.1;refGene_gn=RIMS2;refGene_tx=XM_02324837
9.1;refGene_gn=RIMS2;refGene_tx=XM_023248380.1;refGene_gn=RIMS2;refGen
e_tx=XM_023248381.1;refGene_gn=RIMS2;refGene_tx=XM_023248382.1
F2 51689243 51689374
refGene_gn=RIMS2;refGene_tx=XM_023248353.1;refGene_gn=RIMS2;refGene_tx
=XM_023248354.1;refGene_gn=RIMS2;refGene_tx=XM_023248355.1;refGene_gn=
RIMS2;refGene_tx=XM_023248356.1;refGene_gn=RIMS2;refGene_tx=XM_0232483
57.1;refGene_gn=RIMS2;refGene_tx=XM_023248358.1;refGene_gn=RIMS2;refGe
ne_tx=XM_023248359.1;refGene_gn=RIMS2;refGene_tx=XM_023248360.1;refGen
e_gn=RIMS2;refGene_tx=XM_023248361.1;refGene_gn=RIMS2;refGene_tx=XM_02
3248362.1;refGene_gn=RIMS2;refGene_tx=XM_023248363.1;refGene_gn=RIMS2;
refGene_tx=XM_023248364.1;refGene_gn=RIMS2;refGene_tx=XM_023248365.1;r
efGene_gn=RIMS2;refGene_tx=XM_023248366.1;refGene_gn=RIMS2;refGene_tx=
XM_023248367.1;refGene_gn=RIMS2;refGene_tx=XM_023248368.1;refGene_gn=R
IMS2;refGene_tx=XM_023248369.1;refGene_gn=RIMS2;refGene_tx=XM_02324837
0.1;refGene_gn=RIMS2;refGene_tx=XM_023248371.1;refGene_gn=RIMS2;refGen
e_tx=XM_023248372.1;refGene_gn=RIMS2;refGene_tx=XM_023248373.1;refGene
_gn=RIMS2;refGene_tx=XM_023248374.1;refGene_gn=RIMS2;refGene_tx=XM_023
248375.1;refGene_gn=RIMS2;refGene_tx=XM_023248376.1;refGene_gn=RIMS2;r
efGene_tx=XM_023248377.1;refGene_gn=RIMS2;refGene_tx=XM_023248378.1;re
fGene_gn=RIMS2;refGene_tx=XM_023248379.1;refGene_gn=RIMS2;refGene_tx=X
M_023248380.1;refGene_gn=RIMS2;refGene_tx=XM_023248381.1;refGene_gn=RI
MS2;refGene_tx=XM_023248382.1;refGene_gn=RIMS2;refGene_tx=XM_023248383
.
1;refGene_gn=RIMS2;refGene_tx=XM_023248384.1;refGene_gn=RIMS2;refGene_
tx=XM_023248385.1;refGene_gn=RIMS2;refGene_tx=XM_023248386.1;refGene_g
n=RIMS2;refGene_tx=XM_023248387.1;refGene_gn=RIMS2;refGene_tx=XM_02324
8388.1;refGene_gn=RIMS2;refGene_tx=XM_023248389.1;refGene_gn=RIMS2;ref
Gene_tx=XM_023248390.1;refGene_gn=RIMS2;refGene_tx=XM_023248392.1;refG
ene_gn=RIMS2;refGene_tx=XM_023248393.1;refGene_gn=RIMS2;refGene_tx=XM_
023248394.1;refGene_gn=RIMS2;refGene_tx=XM_023248395.1;refGene_gn=RIMS
2;refGene_tx=XM_023248396.1;refGene_gn=RIMS2;refGene_tx=XM_023248397.1
F2 51694335 51694507
refGene_gn=RIMS2;refGene_tx=XM_023248353.1;refGene_gn=RIMS2;refGene_tx
=XM_023248354.1;refGene_gn=RIMS2;refGene_tx=XM_023248355.1;refGene_gn=
RIMS2;refGene_tx=XM_023248356.1;refGene_gn=RIMS2;refGene_tx=XM_0232483
57.1;refGene_gn=RIMS2;refGene_tx=XM_023248358.1;refGene_gn=RIMS2;refGe
ne_tx=XM_023248359.1;refGene_gn=RIMS2;refGene_tx=XM_023248360.1;refGen
e_gn=RIMS2;refGene_tx=XM_023248361.1;refGene_gn=RIMS2;refGene_tx=XM_02
3248362.1;refGene_gn=RIMS2;refGene_tx=XM_023248363.1;refGene_gn=RIMS2;
refGene_tx=XM_023248364.1;refGene_gn=RIMS2;refGene_tx=XM_023248365.1;r
efGene_gn=RIMS2;refGene_tx=XM_023248366.1;refGene_gn=RIMS2;refGene_tx=
XM_023248367.1;refGene_gn=RIMS2;refGene_tx=XM_023248368.1;refGene_gn=R
IMS2;refGene_tx=XM_023248369.1;refGene_gn=RIMS2;refGene_tx=XM_02324837
0.1;refGene_gn=RIMS2;refGene_tx=XM_023248371.1;refGene_gn=RIMS2;refGen
e_tx=XM_023248372.1;refGene_gn=RIMS2;refGene_tx=XM_023248373.1;refGene

_gn=RIMS2;refGene_tx=XM_023248374.1;refGene_gn=RIMS2;refGene_tx=XM_023248375.1;refGene_gn=RIMS2;refGene_tx=XM_023248376.1;refGene_gn=RIMS2;refGene_tx=XM_023248377.1;refGene_gn=RIMS2;refGene_tx=XM_023248378.1;refGene_gn=RIMS2;refGene_tx=XM_023248379.1;refGene_gn=RIMS2;refGene_tx=XM_023248380.1;refGene_gn=RIMS2;refGene_tx=XM_023248381.1;refGene_gn=RIMS2;refGene_tx=XM_023248382.1;refGene_gn=RIMS2;refGene_tx=XM_023248383.
1;refGene_gn=RIMS2;refGene_tx=XM_023248384.1;refGene_gn=RIMS2;refGene_tx=XM_023248385.1;refGene_gn=RIMS2;refGene_tx=XM_023248386.1;refGene_gn=RIMS2;refGene_tx=XM_023248387.1;refGene_gn=RIMS2;refGene_tx=XM_023248388.1;refGene_gn=RIMS2;refGene_tx=XM_023248389.1;refGene_gn=RIMS2;refGene_tx=XM_023248390.1;refGene_gn=RIMS2;refGene_tx=XM_023248392.1;refGene_gn=RIMS2;refGene_tx=XM_023248393.1;refGene_gn=RIMS2;refGene_tx=XM_023248394.1;refGene_gn=RIMS2;refGene_tx=XM_023248395.1;refGene_gn=RIMS2;refGene_tx=XM_023248396.1;refGene_gn=RIMS2;refGene_tx=XM_023248397.1
F2 51696634 51696788
refGene_gn=RIMS2;refGene_tx=XM_023248353.1;refGene_gn=RIMS2;refGene_tx=XM_023248354.1;refGene_gn=RIMS2;refGene_tx=XM_023248355.1;refGene_gn=RIMS2;refGene_tx=XM_023248356.1;refGene_gn=RIMS2;refGene_tx=XM_023248357.1;refGene_gn=RIMS2;refGene_tx=XM_023248358.1;refGene_gn=RIMS2;refGene_tx=XM_023248359.1;refGene_gn=RIMS2;refGene_tx=XM_023248360.1;refGene_gn=RIMS2;refGene_tx=XM_023248361.1;refGene_gn=RIMS2;refGene_tx=XM_023248362.1;refGene_gn=RIMS2;refGene_tx=XM_023248363.1;refGene_gn=RIMS2;refGene_tx=XM_023248364.1;refGene_gn=RIMS2;refGene_tx=XM_023248365.1;refGene_gn=RIMS2;refGene_tx=XM_023248366.1;refGene_gn=RIMS2;refGene_tx=XM_023248367.1;refGene_gn=RIMS2;refGene_tx=XM_023248368.1;refGene_gn=RIMS2;refGene_tx=XM_023248369.1;refGene_gn=RIMS2;refGene_tx=XM_023248370.1;refGene_gn=RIMS2;refGene_tx=XM_023248371.1;refGene_gn=RIMS2;refGene_tx=XM_023248372.1;refGene_gn=RIMS2;refGene_tx=XM_023248373.1;refGene_gn=RIMS2;refGene_tx=XM_023248374.1;refGene_gn=RIMS2;refGene_tx=XM_023248375.1;refGene_gn=RIMS2;refGene_tx=XM_023248376.1;refGene_gn=RIMS2;refGene_tx=XM_023248377.1;refGene_gn=RIMS2;refGene_tx=XM_023248378.1;refGene_gn=RIMS2;refGene_tx=XM_023248379.1;refGene_gn=RIMS2;refGene_tx=XM_023248380.1;refGene_gn=RIMS2;refGene_tx=XM_023248381.1;refGene_gn=RIMS2;refGene_tx=XM_023248382.1;refGene_gn=RIMS2;refGene_tx=XM_023248383.
1;refGene_gn=RIMS2;refGene_tx=XM_023248384.1;refGene_gn=RIMS2;refGene_tx=XM_023248385.1;refGene_gn=RIMS2;refGene_tx=XM_023248386.1;refGene_gn=RIMS2;refGene_tx=XM_023248387.1;refGene_gn=RIMS2;refGene_tx=XM_023248388.1;refGene_gn=RIMS2;refGene_tx=XM_023248389.1;refGene_gn=RIMS2;refGene_tx=XM_023248390.1;refGene_gn=RIMS2;refGene_tx=XM_023248392.1;refGene_gn=RIMS2;refGene_tx=XM_023248393.1;refGene_gn=RIMS2;refGene_tx=XM_023248394.1;refGene_gn=RIMS2;refGene_tx=XM_023248395.1;refGene_gn=RIMS2;refGene_tx=XM_023248396.1;refGene_gn=RIMS2;refGene_tx=XM_023248397.1
F2 51713200 51713227
refGene_gn=RIMS2;refGene_tx=XM_023248353.1;refGene_gn=RIMS2;refGene_tx=XM_023248354.1;refGene_gn=RIMS2;refGene_tx=XM_023248355.1;refGene_gn=RIMS2;refGene_tx=XM_023248357.1;refGene_gn=RIMS2;refGene_tx=XM_023248358.1;refGene_gn=RIMS2;refGene_tx=XM_023248359.1;refGene_gn=RIMS2;refGene_tx=XM_023248360.1;refGene_gn=RIMS2;refGene_tx=XM_023248361.1;refGen

e_gn=RIMS2;refGene_tx=XM_023248362.1;refGene_gn=RIMS2;refGene_tx=XM_02
3248364.1;refGene_gn=RIMS2;refGene_tx=XM_023248365.1;refGene_gn=RIMS2;
refGene_tx=XM_023248366.1;refGene_gn=RIMS2;refGene_tx=XM_023248367.1;r
efGene_gn=RIMS2;refGene_tx=XM_023248369.1;refGene_gn=RIMS2;refGene_tx=
XM_023248370.1;refGene_gn=RIMS2;refGene_tx=XM_023248372.1;refGene_gn=R
IMS2;refGene_tx=XM_023248379.1;refGene_gn=RIMS2;refGene_tx=XM_02324838
0.1;refGene_gn=RIMS2;refGene_tx=XM_023248381.1
F2 51730682 51730751
refGene_gn=RIMS2;refGene_tx=XM_023248353.1;refGene_gn=RIMS2;refGene_tx
=XM_023248354.1;refGene_gn=RIMS2;refGene_tx=XM_023248355.1;refGene_gn=
RIMS2;refGene_tx=XM_023248356.1;refGene_gn=RIMS2;refGene_tx=XM_0232483
57.1;refGene_gn=RIMS2;refGene_tx=XM_023248358.1;refGene_gn=RIMS2;refGe
ne_tx=XM_023248359.1;refGene_gn=RIMS2;refGene_tx=XM_023248360.1;refGen
e_gn=RIMS2;refGene_tx=XM_023248361.1;refGene_gn=RIMS2;refGene_tx=XM_02
3248362.1;refGene_gn=RIMS2;refGene_tx=XM_023248363.1;refGene_gn=RIMS2;
refGene_tx=XM_023248364.1;refGene_gn=RIMS2;refGene_tx=XM_023248365.1;r
efGene_gn=RIMS2;refGene_tx=XM_023248366.1;refGene_gn=RIMS2;refGene_tx=
XM_023248367.1;refGene_gn=RIMS2;refGene_tx=XM_023248368.1;refGene_gn=R
IMS2;refGene_tx=XM_023248369.1;refGene_gn=RIMS2;refGene_tx=XM_02324837
0.1;refGene_gn=RIMS2;refGene_tx=XM_023248371.1;refGene_gn=RIMS2;refGen
e_tx=XM_023248372.1;refGene_gn=RIMS2;refGene_tx=XM_023248373.1;refGene
_gn=RIMS2;refGene_tx=XM_023248374.1;refGene_gn=RIMS2;refGene_tx=XM_023
248375.1;refGene_gn=RIMS2;refGene_tx=XM_023248376.1;refGene_gn=RIMS2;r
efGene_tx=XM_023248377.1;refGene_gn=RIMS2;refGene_tx=XM_023248378.1;re
fGene_gn=RIMS2;refGene_tx=XM_023248379.1;refGene_gn=RIMS2;refGene_tx=X
M_023248380.1;refGene_gn=RIMS2;refGene_tx=XM_023248381.1;refGene_gn=RI
MS2;refGene_tx=XM_023248382.1;refGene_gn=RIMS2;refGene_tx=XM_023248383
.
1;refGene_gn=RIMS2;refGene_tx=XM_023248384.1;refGene_gn=RIMS2;refGene_
tx=XM_023248385.1;refGene_gn=RIMS2;refGene_tx=XM_023248386.1;refGene_g
n=RIMS2;refGene_tx=XM_023248387.1;refGene_gn=RIMS2;refGene_tx=XM_02324
8388.1;refGene_gn=RIMS2;refGene_tx=XM_023248389.1;refGene_gn=RIMS2;ref
Gene_tx=XM_023248390.1;refGene_gn=RIMS2;refGene_tx=XM_023248392.1;refG
ene_gn=RIMS2;refGene_tx=XM_023248393.1;refGene_gn=RIMS2;refGene_tx=XM_
023248394.1;refGene_gn=RIMS2;refGene_tx=XM_023248395.1;refGene_gn=RIMS
2;refGene_tx=XM_023248396.1;refGene_gn=RIMS2;refGene_tx=XM_023248397.1
F2 51737729 51737886
refGene_gn=RIMS2;refGene_tx=XM_023248353.1;refGene_gn=RIMS2;refGene_tx
=XM_023248354.1;refGene_gn=RIMS2;refGene_tx=XM_023248355.1;refGene_gn=
RIMS2;refGene_tx=XM_023248356.1;refGene_gn=RIMS2;refGene_tx=XM_0232483
57.1;refGene_gn=RIMS2;refGene_tx=XM_023248358.1;refGene_gn=RIMS2;refGe
ne_tx=XM_023248359.1;refGene_gn=RIMS2;refGene_tx=XM_023248360.1;refGen
e_gn=RIMS2;refGene_tx=XM_023248361.1;refGene_gn=RIMS2;refGene_tx=XM_02
3248362.1;refGene_gn=RIMS2;refGene_tx=XM_023248363.1;refGene_gn=RIMS2;
refGene_tx=XM_023248364.1;refGene_gn=RIMS2;refGene_tx=XM_023248365.1;r
efGene_gn=RIMS2;refGene_tx=XM_023248366.1;refGene_gn=RIMS2;refGene_tx=
XM_023248367.1;refGene_gn=RIMS2;refGene_tx=XM_023248368.1;refGene_gn=R
IMS2;refGene_tx=XM_023248369.1;refGene_gn=RIMS2;refGene_tx=XM_02324837
0.1;refGene_gn=RIMS2;refGene_tx=XM_023248371.1;refGene_gn=RIMS2;refGen
e_tx=XM_023248372.1;refGene_gn=RIMS2;refGene_tx=XM_023248373.1;refGene

_gn=RIMS2;refGene_tx=XM_023248374.1;refGene_gn=RIMS2;refGene_tx=XM_023248375.1;refGene_gn=RIMS2;refGene_tx=XM_023248376.1;refGene_gn=RIMS2;refGene_tx=XM_023248377.1;refGene_gn=RIMS2;refGene_tx=XM_023248378.1;refGene_gn=RIMS2;refGene_tx=XM_023248379.1;refGene_gn=RIMS2;refGene_tx=XM_023248380.1;refGene_gn=RIMS2;refGene_tx=XM_023248381.1;refGene_gn=RIMS2;refGene_tx=XM_023248382.1;refGene_gn=RIMS2;refGene_tx=XM_023248383.
1;refGene_gn=RIMS2;refGene_tx=XM_023248384.1;refGene_gn=RIMS2;refGene_tx=XM_023248385.1;refGene_gn=RIMS2;refGene_tx=XM_023248386.1;refGene_gn=RIMS2;refGene_tx=XM_023248387.1;refGene_gn=RIMS2;refGene_tx=XM_023248388.1;refGene_gn=RIMS2;refGene_tx=XM_023248389.1;refGene_gn=RIMS2;refGene_tx=XM_023248390.1;refGene_gn=RIMS2;refGene_tx=XM_023248392.1;refGene_gn=RIMS2;refGene_tx=XM_023248393.1;refGene_gn=RIMS2;refGene_tx=XM_023248394.1;refGene_gn=RIMS2;refGene_tx=XM_023248395.1;refGene_gn=RIMS2;refGene_tx=XM_023248396.1;refGene_gn=RIMS2;refGene_tx=XM_023248397.1

F2 51740247 51740364

refGene_gn=RIMS2;refGene_tx=XM_023248353.1;refGene_gn=RIMS2;refGene_tx=XM_023248354.1;refGene_gn=RIMS2;refGene_tx=XM_023248355.1;refGene_gn=RIMS2;refGene_tx=XM_023248356.1;refGene_gn=RIMS2;refGene_tx=XM_023248357.1;refGene_gn=RIMS2;refGene_tx=XM_023248358.1;refGene_gn=RIMS2;refGene_tx=XM_023248359.1;refGene_gn=RIMS2;refGene_tx=XM_023248360.1;refGene_gn=RIMS2;refGene_tx=XM_023248361.1;refGene_gn=RIMS2;refGene_tx=XM_023248362.1;refGene_gn=RIMS2;refGene_tx=XM_023248363.1;refGene_gn=RIMS2;refGene_tx=XM_023248364.1;refGene_gn=RIMS2;refGene_tx=XM_023248365.1;refGene_gn=RIMS2;refGene_tx=XM_023248366.1;refGene_gn=RIMS2;refGene_tx=XM_023248367.1;refGene_gn=RIMS2;refGene_tx=XM_023248368.1;refGene_gn=RIMS2;refGene_tx=XM_023248369.1;refGene_gn=RIMS2;refGene_tx=XM_023248370.1;refGene_gn=RIMS2;refGene_tx=XM_023248371.1;refGene_gn=RIMS2;refGene_tx=XM_023248372.1;refGene_gn=RIMS2;refGene_tx=XM_023248373.1;refGene_gn=RIMS2;refGene_tx=XM_023248374.1;refGene_gn=RIMS2;refGene_tx=XM_023248375.1;refGene_gn=RIMS2;refGene_tx=XM_023248376.1;refGene_gn=RIMS2;refGene_tx=XM_023248377.1;refGene_gn=RIMS2;refGene_tx=XM_023248378.1;refGene_gn=RIMS2;refGene_tx=XM_023248379.1;refGene_gn=RIMS2;refGene_tx=XM_023248380.1;refGene_gn=RIMS2;refGene_tx=XM_023248381.1;refGene_gn=RIMS2;refGene_tx=XM_023248382.1;refGene_gn=RIMS2;refGene_tx=XM_023248383.
1;refGene_gn=RIMS2;refGene_tx=XM_023248384.1;refGene_gn=RIMS2;refGene_tx=XM_023248385.1;refGene_gn=RIMS2;refGene_tx=XM_023248386.1;refGene_gn=RIMS2;refGene_tx=XM_023248387.1;refGene_gn=RIMS2;refGene_tx=XM_023248388.1;refGene_gn=RIMS2;refGene_tx=XM_023248389.1;refGene_gn=RIMS2;refGene_tx=XM_023248390.1;refGene_gn=RIMS2;refGene_tx=XM_023248392.1;refGene_gn=RIMS2;refGene_tx=XM_023248393.1;refGene_gn=RIMS2;refGene_tx=XM_023248394.1;refGene_gn=RIMS2;refGene_tx=XM_023248395.1;refGene_gn=RIMS2;refGene_tx=XM_023248396.1;refGene_gn=RIMS2;refGene_tx=XM_023248397.1

F2 51749320 51749386

refGene_gn=RIMS2;refGene_tx=XM_023248353.1;refGene_gn=RIMS2;refGene_tx=XM_023248354.1;refGene_gn=RIMS2;refGene_tx=XM_023248355.1;refGene_gn=RIMS2;refGene_tx=XM_023248356.1;refGene_gn=RIMS2;refGene_tx=XM_023248357.1;refGene_gn=RIMS2;refGene_tx=XM_023248358.1;refGene_gn=RIMS2;refGene_tx=XM_023248359.1;refGene_gn=RIMS2;refGene_tx=XM_023248360.1;refGen

e_gn=RIMS2;refGene_tx=XM_023248362.1;refGene_gn=RIMS2;refGene_tx=XM_02
3248363.1;refGene_gn=RIMS2;refGene_tx=XM_023248364.1;refGene_gn=RIMS2;
refGene_tx=XM_023248365.1;refGene_gn=RIMS2;refGene_tx=XM_023248366.1;r
efGene_gn=RIMS2;refGene_tx=XM_023248367.1;refGene_gn=RIMS2;refGene_tx=
XM_023248368.1;refGene_gn=RIMS2;refGene_tx=XM_023248370.1;refGene_gn=R
IMS2;refGene_tx=XM_023248371.1;refGene_gn=RIMS2;refGene_tx=XM_02324837
2.1;refGene_gn=RIMS2;refGene_tx=XM_023248374.1;refGene_gn=RIMS2;refGen
e_tx=XM_023248376.1;refGene_gn=RIMS2;refGene_tx=XM_023248379.1;refGene
_gn=RIMS2;refGene_tx=XM_023248380.1;refGene_gn=RIMS2;refGene_tx=XM_023
248381.1;refGene_gn=RIMS2;refGene_tx=XM_023248382.1;refGene_gn=RIMS2;r
efGene_tx=XM_023248383.1;refGene_gn=RIMS2;refGene_tx=XM_023248388.1;re
fGene_gn=RIMS2;refGene_tx=XM_023248390.1;refGene_gn=RIMS2;refGene_tx=X
M_023248394.1
F2 51764725 51764905
refGene_gn=RIMS2;refGene_tx=XM_023248353.1;refGene_gn=RIMS2;refGene_tx
=XM_023248354.1;refGene_gn=RIMS2;refGene_tx=XM_023248355.1;refGene_gn=
RIMS2;refGene_tx=XM_023248356.1;refGene_gn=RIMS2;refGene_tx=XM_0232483
57.1;refGene_gn=RIMS2;refGene_tx=XM_023248358.1;refGene_gn=RIMS2;refGe
ne_tx=XM_023248359.1;refGene_gn=RIMS2;refGene_tx=XM_023248360.1;refGen
e_gn=RIMS2;refGene_tx=XM_023248361.1;refGene_gn=RIMS2;refGene_tx=XM_02
3248362.1;refGene_gn=RIMS2;refGene_tx=XM_023248363.1;refGene_gn=RIMS2;
refGene_tx=XM_023248364.1;refGene_gn=RIMS2;refGene_tx=XM_023248365.1;r
efGene_gn=RIMS2;refGene_tx=XM_023248366.1;refGene_gn=RIMS2;refGene_tx=
XM_023248368.1;refGene_gn=RIMS2;refGene_tx=XM_023248370.1;refGene_gn=R
IMS2;refGene_tx=XM_023248371.1;refGene_gn=RIMS2;refGene_tx=XM_02324837
2.1;refGene_gn=RIMS2;refGene_tx=XM_023248373.1;refGene_gn=RIMS2;refGen
e_tx=XM_023248374.1;refGene_gn=RIMS2;refGene_tx=XM_023248375.1;refGene
_gn=RIMS2;refGene_tx=XM_023248376.1;refGene_gn=RIMS2;refGene_tx=XM_023
248378.1;refGene_gn=RIMS2;refGene_tx=XM_023248379.1;refGene_gn=RIMS2;r
efGene_tx=XM_023248380.1;refGene_gn=RIMS2;refGene_tx=XM_023248381.1;re
fGene_gn=RIMS2;refGene_tx=XM_023248382.1;refGene_gn=RIMS2;refGene_tx=X
M_023248383.1;refGene_gn=RIMS2;refGene_tx=XM_023248384.1;refGene_gn=RI
MS2;refGene_tx=XM_023248385.1;refGene_gn=RIMS2;refGene_tx=XM_023248387
.
1;refGene_gn=RIMS2;refGene_tx=XM_023248388.1;refGene_gn=RIMS2;refGene_
tx=XM_023248389.1;refGene_gn=RIMS2;refGene_tx=XM_023248390.1;refGene_g
n=RIMS2;refGene_tx=XM_023248393.1;refGene_gn=RIMS2;refGene_tx=XM_02324
8394.1;refGene_gn=RIMS2;refGene_tx=XM_023248395.1;refGene_gn=RIMS2;ref
Gene_tx=XM_023248396.1
F2 51765801 51765911
refGene_gn=RIMS2;refGene_tx=XM_023248353.1;refGene_gn=RIMS2;refGene_tx
=XM_023248354.1;refGene_gn=RIMS2;refGene_tx=XM_023248355.1;refGene_gn=
RIMS2;refGene_tx=XM_023248356.1;refGene_gn=RIMS2;refGene_tx=XM_0232483
57.1;refGene_gn=RIMS2;refGene_tx=XM_023248358.1;refGene_gn=RIMS2;refGe
ne_tx=XM_023248359.1;refGene_gn=RIMS2;refGene_tx=XM_023248360.1;refGen
e_gn=RIMS2;refGene_tx=XM_023248361.1;refGene_gn=RIMS2;refGene_tx=XM_02
3248362.1;refGene_gn=RIMS2;refGene_tx=XM_023248363.1;refGene_gn=RIMS2;
refGene_tx=XM_023248364.1;refGene_gn=RIMS2;refGene_tx=XM_023248365.1;r
efGene_gn=RIMS2;refGene_tx=XM_023248366.1;refGene_gn=RIMS2;refGene_tx=
XM_023248367.1;refGene_gn=RIMS2;refGene_tx=XM_023248368.1;refGene_gn=R

IMS2;refGene_tx=XM_023248369.1;refGene_gn=RIMS2;refGene_tx=XM_02324837
0.1;refGene_gn=RIMS2;refGene_tx=XM_023248371.1;refGene_gn=RIMS2;refGen
e_tx=XM_023248372.1;refGene_gn=RIMS2;refGene_tx=XM_023248373.1;refGene
_gn=RIMS2;refGene_tx=XM_023248374.1;refGene_gn=RIMS2;refGene_tx=XM_023
248375.1;refGene_gn=RIMS2;refGene_tx=XM_023248376.1;refGene_gn=RIMS2;r
efGene_tx=XM_023248377.1;refGene_gn=RIMS2;refGene_tx=XM_023248378.1;re
fGene_gn=RIMS2;refGene_tx=XM_023248379.1;refGene_gn=RIMS2;refGene_tx=X
M_023248380.1;refGene_gn=RIMS2;refGene_tx=XM_023248381.1;refGene_gn=RI
MS2;refGene_tx=XM_023248382.1;refGene_gn=RIMS2;refGene_tx=XM_023248383
.
1;refGene_gn=RIMS2;refGene_tx=XM_023248384.1;refGene_gn=RIMS2;refGene_
tx=XM_023248385.1;refGene_gn=RIMS2;refGene_tx=XM_023248386.1;refGene_g
n=RIMS2;refGene_tx=XM_023248387.1;refGene_gn=RIMS2;refGene_tx=XM_02324
8388.1;refGene_gn=RIMS2;refGene_tx=XM_023248389.1;refGene_gn=RIMS2;ref
Gene_tx=XM_023248390.1;refGene_gn=RIMS2;refGene_tx=XM_023248392.1;refG
ene_gn=RIMS2;refGene_tx=XM_023248393.1;refGene_gn=RIMS2;refGene_tx=XM_
023248394.1;refGene_gn=RIMS2;refGene_tx=XM_023248395.1;refGene_gn=RIMS
2;refGene_tx=XM_023248396.1;refGene_gn=RIMS2;refGene_tx=XM_023248397.1
F2 51766741 51766774
refGene_gn=RIMS2;refGene_tx=XM_023248353.1;refGene_gn=RIMS2;refGene_tx
=XM_023248354.1;refGene_gn=RIMS2;refGene_tx=XM_023248355.1;refGene_gn=
RIMS2;refGene_tx=XM_023248356.1;refGene_gn=RIMS2;refGene_tx=XM_0232483
58.1;refGene_gn=RIMS2;refGene_tx=XM_023248360.1;refGene_gn=RIMS2;refGe
ne_tx=XM_023248361.1;refGene_gn=RIMS2;refGene_tx=XM_023248363.1;refGen
e_gn=RIMS2;refGene_tx=XM_023248364.1;refGene_gn=RIMS2;refGene_tx=XM_02
3248365.1;refGene_gn=RIMS2;refGene_tx=XM_023248366.1;refGene_gn=RIMS2;
refGene_tx=XM_023248367.1;refGene_gn=RIMS2;refGene_tx=XM_023248368.1;r
efGene_gn=RIMS2;refGene_tx=XM_023248369.1;refGene_gn=RIMS2;refGene_tx=
XM_023248370.1;refGene_gn=RIMS2;refGene_tx=XM_023248371.1;refGene_gn=R
IMS2;refGene_tx=XM_023248376.1;refGene_gn=RIMS2;refGene_tx=XM_02324837
9.1;refGene_gn=RIMS2;refGene_tx=XM_023248380.1;refGene_gn=RIMS2;refGen
e_tx=XM_023248381.1;refGene_gn=RIMS2;refGene_tx=XM_023248382.1;refGene
_gn=RIMS2;refGene_tx=XM_023248383.1;refGene_gn=RIMS2;refGene_tx=XM_023
248385.1;refGene_gn=RIMS2;refGene_tx=XM_023248386.1;refGene_gn=RIMS2;r
efGene_tx=XM_023248390.1;refGene_gn=RIMS2;refGene_tx=XM_023248394.1;re
fGene_gn=RIMS2;refGene_tx=XM_023248395.1;refGene_gn=RIMS2;refGene_tx=X
M_023248397.1
F2 51791391 51791424
refGene_gn=RIMS2;refGene_tx=XM_023248353.1;refGene_gn=RIMS2;refGene_tx
=XM_023248354.1;refGene_gn=RIMS2;refGene_tx=XM_023248355.1;refGene_gn=
RIMS2;refGene_tx=XM_023248356.1;refGene_gn=RIMS2;refGene_tx=XM_0232483
57.1;refGene_gn=RIMS2;refGene_tx=XM_023248359.1;refGene_gn=RIMS2;refGe
ne_tx=XM_023248360.1;refGene_gn=RIMS2;refGene_tx=XM_023248361.1;refGen
e_gn=RIMS2;refGene_tx=XM_023248363.1;refGene_gn=RIMS2;refGene_tx=XM_02
3248364.1;refGene_gn=RIMS2;refGene_tx=XM_023248365.1;refGene_gn=RIMS2;
refGene_tx=XM_023248366.1;refGene_gn=RIMS2;refGene_tx=XM_023248367.1;r
efGene_gn=RIMS2;refGene_tx=XM_023248368.1;refGene_gn=RIMS2;refGene_tx=
XM_023248369.1;refGene_gn=RIMS2;refGene_tx=XM_023248370.1;refGene_gn=R
IMS2;refGene_tx=XM_023248371.1;refGene_gn=RIMS2;refGene_tx=XM_02324837
3.1;refGene_gn=RIMS2;refGene_tx=XM_023248376.1;refGene_gn=RIMS2;refGen

e_tx=XM_023248378.1;refGene_gn=RIMS2;refGene_tx=XM_023248379.1;refGene
_gn=RIMS2;refGene_tx=XM_023248380.1;refGene_gn=RIMS2;refGene_tx=XM_023
248381.1;refGene_gn=RIMS2;refGene_tx=XM_023248382.1;refGene_gn=RIMS2;r
efGene_tx=XM_023248383.1;refGene_gn=RIMS2;refGene_tx=XM_023248384.1;re
fGene_gn=RIMS2;refGene_tx=XM_023248386.1;refGene_gn=RIMS2;refGene_tx=X
M_023248387.1;refGene_gn=RIMS2;refGene_tx=XM_023248390.1;refGene_gn=RI
MS2;refGene_tx=XM_023248393.1;refGene_gn=RIMS2;refGene_tx=XM_023248394
.
1;refGene_gn=RIMS2;refGene_tx=XM_023248396.1;refGene_gn=RIMS2;refGene_
tx=XM_023248397.1
F2 51813178 51813279
refGene_gn=RIMS2;refGene_tx=XM_023248353.1;refGene_gn=RIMS2;refGene_tx
=XM_023248354.1;refGene_gn=RIMS2;refGene_tx=XM_023248355.1;refGene_gn=
RIMS2;refGene_tx=XM_023248356.1;refGene_gn=RIMS2;refGene_tx=XM_0232483
57.1;refGene_gn=RIMS2;refGene_tx=XM_023248358.1;refGene_gn=RIMS2;refGe
ne_tx=XM_023248359.1;refGene_gn=RIMS2;refGene_tx=XM_023248360.1;refGen
e_gn=RIMS2;refGene_tx=XM_023248361.1;refGene_gn=RIMS2;refGene_tx=XM_02
3248362.1;refGene_gn=RIMS2;refGene_tx=XM_023248363.1;refGene_gn=RIMS2;
refGene_tx=XM_023248364.1;refGene_gn=RIMS2;refGene_tx=XM_023248365.1;r
efGene_gn=RIMS2;refGene_tx=XM_023248366.1;refGene_gn=RIMS2;refGene_tx=
XM_023248367.1;refGene_gn=RIMS2;refGene_tx=XM_023248368.1;refGene_gn=R
IMS2;refGene_tx=XM_023248369.1;refGene_gn=RIMS2;refGene_tx=XM_02324837
0.1;refGene_gn=RIMS2;refGene_tx=XM_023248371.1;refGene_gn=RIMS2;refGen
e_tx=XM_023248372.1;refGene_gn=RIMS2;refGene_tx=XM_023248373.1;refGene
_gn=RIMS2;refGene_tx=XM_023248374.1;refGene_gn=RIMS2;refGene_tx=XM_023
248375.1;refGene_gn=RIMS2;refGene_tx=XM_023248376.1;refGene_gn=RIMS2;r
efGene_tx=XM_023248377.1;refGene_gn=RIMS2;refGene_tx=XM_023248378.1;re
fGene_gn=RIMS2;refGene_tx=XM_023248379.1;refGene_gn=RIMS2;refGene_tx=X
M_023248382.1;refGene_gn=RIMS2;refGene_tx=XM_023248383.1;refGene_gn=RI
MS2;refGene_tx=XM_023248384.1;refGene_gn=RIMS2;refGene_tx=XM_023248385
.
1;refGene_gn=RIMS2;refGene_tx=XM_023248386.1;refGene_gn=RIMS2;refGene_
tx=XM_023248387.1;refGene_gn=RIMS2;refGene_tx=XM_023248388.1;refGene_g
n=RIMS2;refGene_tx=XM_023248389.1;refGene_gn=RIMS2;refGene_tx=XM_02324
8390.1;refGene_gn=RIMS2;refGene_tx=XM_023248392.1;refGene_gn=RIMS2;ref
Gene_tx=XM_023248393.1;refGene_gn=RIMS2;refGene_tx=XM_023248394.1;refG
ene_gn=RIMS2;refGene_tx=XM_023248395.1;refGene_gn=RIMS2;refGene_tx=XM_
023248396.1;refGene_gn=RIMS2;refGene_tx=XM_023248397.1
F2 51832543 51832716
refGene_gn=RIMS2;refGene_tx=XM_023248353.1;refGene_gn=RIMS2;refGene_tx
=XM_023248354.1;refGene_gn=RIMS2;refGene_tx=XM_023248355.1;refGene_gn=
RIMS2;refGene_tx=XM_023248356.1;refGene_gn=RIMS2;refGene_tx=XM_0232483
57.1;refGene_gn=RIMS2;refGene_tx=XM_023248358.1;refGene_gn=RIMS2;refGe
ne_tx=XM_023248359.1;refGene_gn=RIMS2;refGene_tx=XM_023248360.1;refGen
e_gn=RIMS2;refGene_tx=XM_023248361.1;refGene_gn=RIMS2;refGene_tx=XM_02
3248362.1;refGene_gn=RIMS2;refGene_tx=XM_023248363.1;refGene_gn=RIMS2;
refGene_tx=XM_023248364.1;refGene_gn=RIMS2;refGene_tx=XM_023248365.1;r
efGene_gn=RIMS2;refGene_tx=XM_023248366.1;refGene_gn=RIMS2;refGene_tx=
XM_023248367.1;refGene_gn=RIMS2;refGene_tx=XM_023248368.1;refGene_gn=R
IMS2;refGene_tx=XM_023248369.1;refGene_gn=RIMS2;refGene_tx=XM_02324837

0.1;refGene_gn=RIMS2;refGene_tx=XM_023248371.1;refGene_gn=RIMS2;refGen
e_tx=XM_023248372.1;refGene_gn=RIMS2;refGene_tx=XM_023248373.1;refGene
_gn=RIMS2;refGene_tx=XM_023248374.1;refGene_gn=RIMS2;refGene_tx=XM_023
248375.1;refGene_gn=RIMS2;refGene_tx=XM_023248376.1;refGene_gn=RIMS2;r
efGene_tx=XM_023248377.1;refGene_gn=RIMS2;refGene_tx=XM_023248378.1;re
fGene_gn=RIMS2;refGene_tx=XM_023248379.1;refGene_gn=RIMS2;refGene_tx=X
M_023248380.1;refGene_gn=RIMS2;refGene_tx=XM_023248381.1;refGene_gn=RI
MS2;refGene_tx=XM_023248382.1;refGene_gn=RIMS2;refGene_tx=XM_023248383
.
1;refGene_gn=RIMS2;refGene_tx=XM_023248384.1;refGene_gn=RIMS2;refGene_
tx=XM_023248385.1;refGene_gn=RIMS2;refGene_tx=XM_023248386.1;refGene_g
n=RIMS2;refGene_tx=XM_023248387.1;refGene_gn=RIMS2;refGene_tx=XM_02324
8388.1;refGene_gn=RIMS2;refGene_tx=XM_023248389.1;refGene_gn=RIMS2;ref
Gene_tx=XM_023248390.1;refGene_gn=RIMS2;refGene_tx=XM_023248392.1;refG
ene_gn=RIMS2;refGene_tx=XM_023248393.1;refGene_gn=RIMS2;refGene_tx=XM_
023248394.1;refGene_gn=RIMS2;refGene_tx=XM_023248395.1;refGene_gn=RIMS
2;refGene_tx=XM_023248396.1;refGene_gn=RIMS2;refGene_tx=XM_023248397.1
F2 51833603 51833855
refGene_gn=RIMS2;refGene_tx=XM_023248353.1;refGene_gn=RIMS2;refGene_tx
=XM_023248354.1;refGene_gn=RIMS2;refGene_tx=XM_023248355.1;refGene_gn=
RIMS2;refGene_tx=XM_023248356.1;refGene_gn=RIMS2;refGene_tx=XM_0232483
57.1;refGene_gn=RIMS2;refGene_tx=XM_023248358.1;refGene_gn=RIMS2;refGe
ne_tx=XM_023248359.1;refGene_gn=RIMS2;refGene_tx=XM_023248360.1;refGen
e_gn=RIMS2;refGene_tx=XM_023248361.1;refGene_gn=RIMS2;refGene_tx=XM_02
3248362.1;refGene_gn=RIMS2;refGene_tx=XM_023248363.1;refGene_gn=RIMS2;
refGene_tx=XM_023248364.1;refGene_gn=RIMS2;refGene_tx=XM_023248365.1;r
efGene_gn=RIMS2;refGene_tx=XM_023248366.1;refGene_gn=RIMS2;refGene_tx=
XM_023248367.1;refGene_gn=RIMS2;refGene_tx=XM_023248368.1;refGene_gn=R
IMS2;refGene_tx=XM_023248369.1;refGene_gn=RIMS2;refGene_tx=XM_02324837
1.1;refGene_gn=RIMS2;refGene_tx=XM_023248379.1;refGene_gn=RIMS2;refGen
e_tx=XM_023248382.1
F2 51873535 51873777
refGene_gn=RIMS2;refGene_tx=XM_023248353.1;refGene_gn=RIMS2;refGene_tx
=XM_023248354.1;refGene_gn=RIMS2;refGene_tx=XM_023248355.1;refGene_gn=
RIMS2;refGene_tx=XM_023248356.1;refGene_gn=RIMS2;refGene_tx=XM_0232483
57.1;refGene_gn=RIMS2;refGene_tx=XM_023248358.1;refGene_gn=RIMS2;refGe
ne_tx=XM_023248359.1;refGene_gn=RIMS2;refGene_tx=XM_023248360.1;refGen
e_gn=RIMS2;refGene_tx=XM_023248361.1;refGene_gn=RIMS2;refGene_tx=XM_02
3248362.1;refGene_gn=RIMS2;refGene_tx=XM_023248363.1;refGene_gn=RIMS2;
refGene_tx=XM_023248364.1;refGene_gn=RIMS2;refGene_tx=XM_023248365.1;r
efGene_gn=RIMS2;refGene_tx=XM_023248366.1;refGene_gn=RIMS2;refGene_tx=
XM_023248367.1;refGene_gn=RIMS2;refGene_tx=XM_023248368.1;refGene_gn=R
IMS2;refGene_tx=XM_023248369.1;refGene_gn=RIMS2;refGene_tx=XM_02324837
0.1;refGene_gn=RIMS2;refGene_tx=XM_023248371.1;refGene_gn=RIMS2;refGen
e_tx=XM_023248372.1;refGene_gn=RIMS2;refGene_tx=XM_023248373.1;refGene
_gn=RIMS2;refGene_tx=XM_023248374.1;refGene_gn=RIMS2;refGene_tx=XM_023
248375.1;refGene_gn=RIMS2;refGene_tx=XM_023248376.1;refGene_gn=RIMS2;r
efGene_tx=XM_023248377.1;refGene_gn=RIMS2;refGene_tx=XM_023248378.1;re
fGene_gn=RIMS2;refGene_tx=XM_023248379.1;refGene_gn=RIMS2;refGene_tx=X
M_023248382.1;refGene_gn=RIMS2;refGene_tx=XM_023248383.1;refGene_gn=RI

MS2;refGene_tx=XM_023248384.1;refGene_gn=RIMS2;refGene_tx=XM_023248385
.
1;refGene_gn=RIMS2;refGene_tx=XM_023248386.1;refGene_gn=RIMS2;refGene_tx=XM_023248387.1;refGene_gn=RIMS2;refGene_tx=XM_023248388.1;refGene_gn=RIMS2;refGene_tx=XM_023248389.1;refGene_gn=RIMS2;refGene_tx=XM_023248390.1;refGene_gn=RIMS2;refGene_tx=XM_023248392.1;refGene_gn=RIMS2;refGene_tx=XM_023248393.1;refGene_gn=RIMS2;refGene_tx=XM_023248394.1;refGene_gn=RIMS2;refGene_tx=XM_023248395.1;refGene_gn=RIMS2;refGene_tx=XM_023248396.1;refGene_gn=RIMS2;refGene_tx=XM_023248397.1
F2 51907506 51907517
refGene_gn=RIMS2;refGene_tx=XM_023248387.1;refGene_gn=RIMS2;refGene_tx=XM_023248388.1;refGene_gn=RIMS2;refGene_tx=XM_023248389.1;refGene_gn=RIMS2;refGene_tx=XM_023248390.1;refGene_gn=RIMS2;refGene_tx=XM_023248392.1;refGene_gn=RIMS2;refGene_tx=XM_023248393.1;refGene_gn=RIMS2;refGene_tx=XM_023248394.1;refGene_gn=RIMS2;refGene_tx=XM_023248395.1;refGene_gn=RIMS2;refGene_tx=XM_023248396.1;refGene_gn=RIMS2;refGene_tx=XM_023248397.1
F2 51931944 51932086 refGene_gn=RIMS2;refGene_tx=XM_023248398.1
F2 51950600 51950605 refGene_gn=RIMS2;refGene_tx=XM_023248398.1
F2 51953727 51953867
refGene_gn=RIMS2;refGene_tx=XM_023248353.1;refGene_gn=RIMS2;refGene_tx=XM_023248354.1;refGene_gn=RIMS2;refGene_tx=XM_023248355.1;refGene_gn=RIMS2;refGene_tx=XM_023248356.1;refGene_gn=RIMS2;refGene_tx=XM_023248357.1;refGene_gn=RIMS2;refGene_tx=XM_023248358.1;refGene_gn=RIMS2;refGene_tx=XM_023248359.1;refGene_gn=RIMS2;refGene_tx=XM_023248360.1;refGene_gn=RIMS2;refGene_tx=XM_023248361.1;refGene_gn=RIMS2;refGene_tx=XM_023248362.1;refGene_gn=RIMS2;refGene_tx=XM_023248363.1;refGene_gn=RIMS2;refGene_tx=XM_023248364.1;refGene_gn=RIMS2;refGene_tx=XM_023248365.1;refGene_gn=RIMS2;refGene_tx=XM_023248366.1;refGene_gn=RIMS2;refGene_tx=XM_023248367.1;refGene_gn=RIMS2;refGene_tx=XM_023248368.1;refGene_gn=RIMS2;refGene_tx=XM_023248369.1;refGene_gn=RIMS2;refGene_tx=XM_023248370.1;refGene_gn=RIMS2;refGene_tx=XM_023248371.1;refGene_gn=RIMS2;refGene_tx=XM_023248372.1;refGene_gn=RIMS2;refGene_tx=XM_023248373.1;refGene_gn=RIMS2;refGene_tx=XM_023248374.1;refGene_gn=RIMS2;refGene_tx=XM_023248375.1;refGene_gn=RIMS2;refGene_tx=XM_023248376.1;refGene_gn=RIMS2;refGene_tx=XM_023248377.1;refGene_gn=RIMS2;refGene_tx=XM_023248378.1;refGene_gn=RIMS2;refGene_tx=XM_023248379.1;refGene_gn=RIMS2;refGene_tx=XM_023248382.1;refGene_gn=RIMS2;refGene_tx=XM_023248383.1;refGene_gn=RIMS2;refGene_tx=XM_023248384.1;refGene_gn=RIMS2;refGene_tx=XM_023248385
.
1;refGene_gn=RIMS2;refGene_tx=XM_023248386.1;refGene_gn=RIMS2;refGene_tx=XM_023248387.1;refGene_gn=RIMS2;refGene_tx=XM_023248388.1;refGene_gn=RIMS2;refGene_tx=XM_023248389.1;refGene_gn=RIMS2;refGene_tx=XM_023248390.1;refGene_gn=RIMS2;refGene_tx=XM_023248392.1;refGene_gn=RIMS2;refGene_tx=XM_023248393.1;refGene_gn=RIMS2;refGene_tx=XM_023248394.1;refGene_gn=RIMS2;refGene_tx=XM_023248395.1;refGene_gn=RIMS2;refGene_tx=XM_023248396.1;refGene_gn=RIMS2;refGene_tx=XM_023248397.1;refGene_gn=RIMS2;refGene_tx=XM_023248398.1
F2 51958472 51958585
refGene_gn=RIMS2;refGene_tx=XM_023248387.1;refGene_gn=RIMS2;refGene_tx

=XM_023248388.1;refGene_gn=RIMS2;refGene_tx=XM_023248389.1;refGene_gn=RIMS2;refGene_tx=XM_023248390.1;refGene_gn=RIMS2;refGene_tx=XM_023248392.1;refGene_gn=RIMS2;refGene_tx=XM_023248393.1;refGene_gn=RIMS2;refGene_tx=XM_023248394.1;refGene_gn=RIMS2;refGene_tx=XM_023248395.1;refGene_gn=RIMS2;refGene_tx=XM_023248396.1;refGene_gn=RIMS2;refGene_tx=XM_023248397.1;refGene_gn=RIMS2;refGene_tx=XM_023248398.1
F2 51959147 51959249
refGene_gn=RIMS2;refGene_tx=XM_023248387.1;refGene_gn=RIMS2;refGene_tx=XM_023248388.1;refGene_gn=RIMS2;refGene_tx=XM_023248389.1;refGene_gn=RIMS2;refGene_tx=XM_023248390.1;refGene_gn=RIMS2;refGene_tx=XM_023248392.1;refGene_gn=RIMS2;refGene_tx=XM_023248393.1;refGene_gn=RIMS2;refGene_tx=XM_023248394.1;refGene_gn=RIMS2;refGene_tx=XM_023248395.1;refGene_gn=RIMS2;refGene_tx=XM_023248396.1;refGene_gn=RIMS2;refGene_tx=XM_023248397.1;refGene_gn=RIMS2;refGene_tx=XM_023248398.1
F2 51961258 51961398
refGene_gn=RIMS2;refGene_tx=XM_023248387.1;refGene_gn=RIMS2;refGene_tx=XM_023248388.1;refGene_gn=RIMS2;refGene_tx=XM_023248389.1;refGene_gn=RIMS2;refGene_tx=XM_023248390.1;refGene_gn=RIMS2;refGene_tx=XM_023248392.1;refGene_gn=RIMS2;refGene_tx=XM_023248393.1;refGene_gn=RIMS2;refGene_tx=XM_023248394.1;refGene_gn=RIMS2;refGene_tx=XM_023248395.1;refGene_gn=RIMS2;refGene_tx=XM_023248396.1;refGene_gn=RIMS2;refGene_tx=XM_023248397.1;refGene_gn=RIMS2;refGene_tx=XM_023248398.1
F2 51961831 51962050
refGene_gn=RIMS2;refGene_tx=XM_023248387.1;refGene_gn=RIMS2;refGene_tx=XM_023248388.1;refGene_gn=RIMS2;refGene_tx=XM_023248389.1;refGene_gn=RIMS2;refGene_tx=XM_023248390.1;refGene_gn=RIMS2;refGene_tx=XM_023248392.1;refGene_gn=RIMS2;refGene_tx=XM_023248393.1;refGene_gn=RIMS2;refGene_tx=XM_023248394.1;refGene_gn=RIMS2;refGene_tx=XM_023248395.1;refGene_gn=RIMS2;refGene_tx=XM_023248396.1;refGene_gn=RIMS2;refGene_tx=XM_023248397.1;refGene_gn=RIMS2;refGene_tx=XM_023248398.1
F2 52017316 52018471
refGene_gn=DCSTAMP;refGene_tx=XM_023248440.1;refGene_gn=DCSTAMP;refGene_tx=XM_023248441.1
F2 52023133 52023469
refGene_gn=DCSTAMP;refGene_tx=XM_023248440.1;refGene_gn=DCSTAMP;refGene_tx=XM_023248441.1
F2 52051530 52051629 refGene_gn=DPYS;refGene_tx=XM_023248231.1
F2 52053434 52053551 refGene_gn=DPYS;refGene_tx=XM_023248230.1
F2 52064301 52064509
refGene_gn=DPYS;refGene_tx=XM_023248230.1;refGene_gn=DPYS;refGene_tx=XM_023248231.1
F2 52094886 52095029
refGene_gn=DPYS;refGene_tx=XM_023248230.1;refGene_gn=DPYS;refGene_tx=XM_023248231.1
F2 52097157 52097299
refGene_gn=DPYS;refGene_tx=XM_023248230.1;refGene_gn=DPYS;refGene_tx=XM_023248231.1
F2 52098236 52098393
refGene_gn=DPYS;refGene_tx=XM_023248230.1;refGene_gn=DPYS;refGene_tx=XM_023248231.1

F2 52110679 52110869
refGene_gn=DPYS;refGene_tx=XM_023248230.1;refGene_gn=DPYS;refGene_tx=X
M_023248231.1
F2 52112099 52112279
refGene_gn=DPYS;refGene_tx=XM_023248230.1;refGene_gn=DPYS;refGene_tx=X
M_023248231.1
F2 52116971 52117130
refGene_gn=DPYS;refGene_tx=XM_023248230.1;refGene_gn=DPYS;refGene_tx=X
M_023248231.1
F2 52131798 52132062
refGene_gn=DPYS;refGene_tx=XM_023248230.1;refGene_gn=DPYS;refGene_tx=X
M_023248231.1
F2 52150478 52151345
refGene_gn=LRP12;refGene_tx=XM_011291428.3;refGene_gn=LRP12;refGene_tx
=XM_011291429.3
F2 52155190 52155323
refGene_gn=LRP12;refGene_tx=XM_011291428.3;refGene_gn=LRP12;refGene_tx
=XM_011291429.3
F2 52157025 52158130
refGene_gn=LRP12;refGene_tx=XM_011291428.3;refGene_gn=LRP12;refGene_tx
=XM_011291429.3
F2 52160272 52160475
refGene_gn=LRP12;refGene_tx=XM_011291428.3;refGene_gn=LRP12;refGene_tx
=XM_011291429.3
F2 52169708 52169844
refGene_gn=LRP12;refGene_tx=XM_011291428.3;refGene_gn=LRP12;refGene_tx
=XM_011291429.3
F2 52187747 52187804 refGene_gn=LRP12;refGene_tx=XM_011291428.3
F2 52222911 52223101
refGene_gn=LOC102901049;refGene_tx=XM_023248333.1
F2 52223248 52223392
refGene_gn=LOC102901049;refGene_tx=XM_023248333.1
F2 52223529 52224260
refGene_gn=LOC102901049;refGene_tx=XM_023248333.1;refGene_gn=LRP12;ref
Gene_tx=XM_011291428.3;refGene_gn=LRP12;refGene_tx=XM_011291429.3
F2 52825050 52825090 refGene_gn=ZFPM2;refGene_tx=XM_023248130.1
F2 52922514 52922673 refGene_gn=ZFPM2;refGene_tx=XM_023248130.1
F2 52945883 52945985 refGene_gn=ZFPM2;refGene_tx=XM_023248130.1
F2 53058961 53059080 refGene_gn=ZFPM2;refGene_tx=XM_023248130.1
F2 53131173 53131285 refGene_gn=ZFPM2;refGene_tx=XM_023248130.1
F2 53284019 53284226 refGene_gn=ZFPM2;refGene_tx=XM_023248130.1
F2 53286552 53286777 refGene_gn=ZFPM2;refGene_tx=XM_023248130.1
F2 53289721 53292213 refGene_gn=ZFPM2;refGene_tx=XM_023248130.1
F2 53724897 53724920 refGene_gn=OXR1;refGene_tx=XM_023248293.1
F2 53773712 53773714 refGene_gn=OXR1;refGene_tx=XM_019823028.2
F2 53810672 53810698
refGene_gn=OXR1;refGene_tx=XM_019823026.2;refGene_gn=OXR1;refGene_tx=X
M_019823029.2
F2 53869553 53869750
refGene_gn=OXR1;refGene_tx=XM_019823026.2;refGene_gn=OXR1;refGene_tx=X

M_019823028.2;refGene_gn=OXR1;refGene_tx=XM_019823029.2;refGene_gn=OXR1;refGene_tx=XM_023248293.1
F2 53928337 53928350 refGene_gn=OXR1;refGene_tx=XM_019823031.2
F2 53972892 53972911 refGene_gn=OXR1;refGene_tx=XM_019823030.2
F2 54019765 54019964
refGene_gn=OXR1;refGene_tx=XM_023248294.1;refGene_gn=OXR1;refGene_tx=XM_023248295.1
F2 54023545 54023558 refGene_gn=OXR1;refGene_tx=XM_019823032.2
F2 54028110 54028147 refGene_gn=OXR1;refGene_tx=XM_006943345.4
F2 54033536 54033619
refGene_gn=OXR1;refGene_tx=XM_006943345.4;refGene_gn=OXR1;refGene_tx=XM_006943347.4;refGene_gn=OXR1;refGene_tx=XM_019823026.2;refGene_gn=OXR1;refGene_tx=XM_019823028.2;refGene_gn=OXR1;refGene_tx=XM_019823029.2;refGene_gn=OXR1;refGene_tx=XM_019823030.2;refGene_gn=OXR1;refGene_tx=XM_019823031.2;refGene_gn=OXR1;refGene_tx=XM_019823032.2;refGene_gn=OXR1;refGene_tx=XM_023248293.1;refGene_gn=OXR1;refGene_tx=XM_023248294.1;refGene_gn=OXR1;refGene_tx=XM_023248295.1;refGene_gn=OXR1;refGene_tx=XM_023248296.1
F2 54038076 54038184
refGene_gn=OXR1;refGene_tx=XM_006943345.4;refGene_gn=OXR1;refGene_tx=XM_006943347.4;refGene_gn=OXR1;refGene_tx=XM_019823026.2;refGene_gn=OXR1;refGene_tx=XM_019823028.2;refGene_gn=OXR1;refGene_tx=XM_019823029.2;refGene_gn=OXR1;refGene_tx=XM_019823030.2;refGene_gn=OXR1;refGene_tx=XM_019823031.2;refGene_gn=OXR1;refGene_tx=XM_019823032.2;refGene_gn=OXR1;refGene_tx=XM_023248293.1;refGene_gn=OXR1;refGene_tx=XM_023248294.1;refGene_gn=OXR1;refGene_tx=XM_023248295.1;refGene_gn=OXR1;refGene_tx=XM_023248296.1
F2 54039321 54039435
refGene_gn=OXR1;refGene_tx=XM_006943345.4;refGene_gn=OXR1;refGene_tx=XM_006943347.4;refGene_gn=OXR1;refGene_tx=XM_019823026.2;refGene_gn=OXR1;refGene_tx=XM_019823028.2;refGene_gn=OXR1;refGene_tx=XM_019823029.2;refGene_gn=OXR1;refGene_tx=XM_019823030.2;refGene_gn=OXR1;refGene_tx=XM_019823031.2;refGene_gn=OXR1;refGene_tx=XM_019823032.2;refGene_gn=OXR1;refGene_tx=XM_023248293.1;refGene_gn=OXR1;refGene_tx=XM_023248294.1;refGene_gn=OXR1;refGene_tx=XM_023248295.1;refGene_gn=OXR1;refGene_tx=XM_023248296.1
F2 54045517 54045670
refGene_gn=OXR1;refGene_tx=XM_006943345.4;refGene_gn=OXR1;refGene_tx=XM_006943347.4;refGene_gn=OXR1;refGene_tx=XM_019823026.2;refGene_gn=OXR1;refGene_tx=XM_019823028.2;refGene_gn=OXR1;refGene_tx=XM_019823029.2;refGene_gn=OXR1;refGene_tx=XM_019823030.2;refGene_gn=OXR1;refGene_tx=XM_019823031.2;refGene_gn=OXR1;refGene_tx=XM_019823032.2;refGene_gn=OXR1;refGene_tx=XM_023248293.1;refGene_gn=OXR1;refGene_tx=XM_023248294.1;refGene_gn=OXR1;refGene_tx=XM_023248295.1;refGene_gn=OXR1;refGene_tx=XM_023248296.1
F2 54051054 54051239
refGene_gn=OXR1;refGene_tx=XM_006943345.4;refGene_gn=OXR1;refGene_tx=XM_006943347.4;refGene_gn=OXR1;refGene_tx=XM_019823026.2;refGene_gn=OXR1;refGene_tx=XM_019823028.2;refGene_gn=OXR1;refGene_tx=XM_019823029.2;refGene_gn=OXR1;refGene_tx=XM_019823030.2;refGene_gn=OXR1;refGene_tx=X

M_019823031.2;refGene_gn=OXR1;refGene_tx=XM_019823032.2;refGene_gn=OXR1;refGene_tx=XM_023248293.1;refGene_gn=OXR1;refGene_tx=XM_023248294.1;refGene_gn=OXR1;refGene_tx=XM_023248295.1;refGene_gn=OXR1;refGene_tx=XM_023248296.1
F2 54054218 54054973
refGene_gn=OXR1;refGene_tx=XM_006943345.4;refGene_gn=OXR1;refGene_tx=XM_006943347.4;refGene_gn=OXR1;refGene_tx=XM_019823026.2;refGene_gn=OXR1;refGene_tx=XM_019823028.2;refGene_gn=OXR1;refGene_tx=XM_019823029.2;refGene_gn=OXR1;refGene_tx=XM_019823030.2;refGene_gn=OXR1;refGene_tx=XM_019823031.2;refGene_gn=OXR1;refGene_tx=XM_019823032.2;refGene_gn=OXR1;refGene_tx=XM_023248293.1;refGene_gn=OXR1;refGene_tx=XM_023248294.1;refGene_gn=OXR1;refGene_tx=XM_023248295.1;refGene_gn=OXR1;refGene_tx=XM_023248296.1
F2 54058756 54058925
refGene_gn=OXR1;refGene_tx=XM_006943345.4;refGene_gn=OXR1;refGene_tx=XM_006943347.4;refGene_gn=OXR1;refGene_tx=XM_019823026.2;refGene_gn=OXR1;refGene_tx=XM_019823028.2;refGene_gn=OXR1;refGene_tx=XM_019823029.2;refGene_gn=OXR1;refGene_tx=XM_019823030.2;refGene_gn=OXR1;refGene_tx=XM_019823031.2;refGene_gn=OXR1;refGene_tx=XM_019823032.2;refGene_gn=OXR1;refGene_tx=XM_023248293.1;refGene_gn=OXR1;refGene_tx=XM_023248294.1;refGene_gn=OXR1;refGene_tx=XM_023248295.1;refGene_gn=OXR1;refGene_tx=XM_023248296.1
F2 54060056 54060219
refGene_gn=OXR1;refGene_tx=XM_006943345.4;refGene_gn=OXR1;refGene_tx=XM_006943347.4;refGene_gn=OXR1;refGene_tx=XM_019823026.2;refGene_gn=OXR1;refGene_tx=XM_019823028.2;refGene_gn=OXR1;refGene_tx=XM_019823029.2;refGene_gn=OXR1;refGene_tx=XM_019823030.2;refGene_gn=OXR1;refGene_tx=XM_019823031.2;refGene_gn=OXR1;refGene_tx=XM_019823032.2;refGene_gn=OXR1;refGene_tx=XM_023248293.1;refGene_gn=OXR1;refGene_tx=XM_023248294.1;refGene_gn=OXR1;refGene_tx=XM_023248295.1;refGene_gn=OXR1;refGene_tx=XM_023248296.1
F2 54074076 54074142
refGene_gn=OXR1;refGene_tx=XM_004000050.5;refGene_gn=OXR1;refGene_tx=XM_004000051.5
F2 54083995 54084076
refGene_gn=OXR1;refGene_tx=XM_004000051.5;refGene_gn=OXR1;refGene_tx=XM_006943345.4;refGene_gn=OXR1;refGene_tx=XM_006943347.4;refGene_gn=OXR1;refGene_tx=XM_019823026.2;refGene_gn=OXR1;refGene_tx=XM_019823028.2;refGene_gn=OXR1;refGene_tx=XM_019823030.2;refGene_gn=OXR1;refGene_tx=XM_019823031.2;refGene_gn=OXR1;refGene_tx=XM_019823032.2;refGene_gn=OXR1;refGene_tx=XM_023248293.1;refGene_gn=OXR1;refGene_tx=XM_023248294.1;refGene_gn=OXR1;refGene_tx=XM_023248296.1
F2 54085885 54086011
refGene_gn=OXR1;refGene_tx=XM_004000050.5;refGene_gn=OXR1;refGene_tx=XM_004000051.5;refGene_gn=OXR1;refGene_tx=XM_006943345.4;refGene_gn=OXR1;refGene_tx=XM_006943347.4;refGene_gn=OXR1;refGene_tx=XM_019823026.2;refGene_gn=OXR1;refGene_tx=XM_019823028.2;refGene_gn=OXR1;refGene_tx=XM_019823029.2;refGene_gn=OXR1;refGene_tx=XM_019823030.2;refGene_gn=OXR1;refGene_tx=XM_019823031.2;refGene_gn=OXR1;refGene_tx=XM_019823032.2;refGene_gn=OXR1;refGene_tx=XM_023248293.1;refGene_gn=OXR1;refGene_tx=X

M_023248294.1;refGene_gn=OXR1;refGene_tx=XM_023248295.1;refGene_gn=OXR1;refGene_tx=XM_023248296.1
F2 54086759 54086912 refGene_gn=OXR1;refGene_tx=XM_004000050.5;refGene_gn=OXR1;refGene_tx=XM_004000051.5;refGene_gn=OXR1;refGene_tx=XM_006943345.4;refGene_gn=OXR1;refGene_tx=XM_006943347.4;refGene_gn=OXR1;refGene_tx=XM_019823026.2;refGene_gn=OXR1;refGene_tx=XM_019823028.2;refGene_gn=OXR1;refGene_tx=XM_019823029.2;refGene_gn=OXR1;refGene_tx=XM_019823030.2;refGene_gn=OXR1;refGene_tx=XM_019823031.2;refGene_gn=OXR1;refGene_tx=XM_019823032.2;refGene_gn=OXR1;refGene_tx=XM_023248293.1;refGene_gn=OXR1;refGene_tx=XM_023248294.1;refGene_gn=OXR1;refGene_tx=XM_023248295.1;refGene_gn=OXR1;refGene_tx=XM_023248296.1
F2 54089150 54089246 refGene_gn=OXR1;refGene_tx=XM_004000050.5;refGene_gn=OXR1;refGene_tx=XM_004000051.5;refGene_gn=OXR1;refGene_tx=XM_006943345.4;refGene_gn=OXR1;refGene_tx=XM_006943347.4;refGene_gn=OXR1;refGene_tx=XM_019823026.2;refGene_gn=OXR1;refGene_tx=XM_019823028.2;refGene_gn=OXR1;refGene_tx=XM_019823029.2;refGene_gn=OXR1;refGene_tx=XM_019823030.2;refGene_gn=OXR1;refGene_tx=XM_019823031.2;refGene_gn=OXR1;refGene_tx=XM_019823032.2;refGene_gn=OXR1;refGene_tx=XM_023248293.1;refGene_gn=OXR1;refGene_tx=XM_023248294.1;refGene_gn=OXR1;refGene_tx=XM_023248295.1;refGene_gn=OXR1;refGene_tx=XM_023248296.1
F2 54091182 54091256 refGene_gn=OXR1;refGene_tx=XM_004000050.5;refGene_gn=OXR1;refGene_tx=XM_004000051.5;refGene_gn=OXR1;refGene_tx=XM_006943345.4;refGene_gn=OXR1;refGene_tx=XM_006943347.4;refGene_gn=OXR1;refGene_tx=XM_019823026.2;refGene_gn=OXR1;refGene_tx=XM_019823028.2;refGene_gn=OXR1;refGene_tx=XM_019823029.2;refGene_gn=OXR1;refGene_tx=XM_019823030.2;refGene_gn=OXR1;refGene_tx=XM_019823031.2;refGene_gn=OXR1;refGene_tx=XM_019823032.2;refGene_gn=OXR1;refGene_tx=XM_023248293.1;refGene_gn=OXR1;refGene_tx=XM_023248294.1;refGene_gn=OXR1;refGene_tx=XM_023248295.1;refGene_gn=OXR1;refGene_tx=XM_023248296.1
F2 54093160 54093296 refGene_gn=OXR1;refGene_tx=XM_004000050.5;refGene_gn=OXR1;refGene_tx=XM_004000051.5;refGene_gn=OXR1;refGene_tx=XM_006943345.4;refGene_gn=OXR1;refGene_tx=XM_006943347.4;refGene_gn=OXR1;refGene_tx=XM_019823026.2;refGene_gn=OXR1;refGene_tx=XM_019823028.2;refGene_gn=OXR1;refGene_tx=XM_019823029.2;refGene_gn=OXR1;refGene_tx=XM_019823030.2;refGene_gn=OXR1;refGene_tx=XM_019823031.2;refGene_gn=OXR1;refGene_tx=XM_019823032.2;refGene_gn=OXR1;refGene_tx=XM_023248293.1;refGene_gn=OXR1;refGene_tx=XM_023248294.1;refGene_gn=OXR1;refGene_tx=XM_023248295.1;refGene_gn=OXR1;refGene_tx=XM_023248296.1
F2 54103658 54104136 refGene_gn=ABRA;refGene_tx=XM_004000052.4
F2 54111763 54112404 refGene_gn=ABRA;refGene_tx=XM_004000052.4
F2 54590792 54590953 refGene_gn=ANGPT1;refGene_tx=XM_004000054.5;refGene_gn=ANGPT1;refGene_tx=XM_006943348.4;refGene_gn=ANGPT1;refGene_tx=XM_019823037.2
F2 54602501 54602632 refGene_gn=ANGPT1;refGene_tx=XM_004000054.5;refGene_gn=ANGPT1;refGene_tx=XM_006943348.4;refGene_gn=ANGPT1;refGene_tx=XM_019823037.2

F2 54625767 54625934
refGene_gn=ANGPT1;refGene_tx=XM_004000054.5;refGene_gn=ANGPT1;refGene_tx=XM_006943348.4;refGene_gn=ANGPT1;refGene_tx=XM_019823037.2
F2 54636564 54636666
refGene_gn=ANGPT1;refGene_tx=XM_004000054.5;refGene_gn=ANGPT1;refGene_tx=XM_006943348.4;refGene_gn=ANGPT1;refGene_tx=XM_019823037.2
F2 54645171 54645299
refGene_gn=ANGPT1;refGene_tx=XM_004000054.5;refGene_gn=ANGPT1;refGene_tx=XM_006943348.4;refGene_gn=ANGPT1;refGene_tx=XM_019823037.2
F2 54663308 54663541
refGene_gn=ANGPT1;refGene_tx=XM_004000054.5;refGene_gn=ANGPT1;refGene_tx=XM_006943348.4;refGene_gn=ANGPT1;refGene_tx=XM_019823037.2
F2 54677059 54677181
refGene_gn=ANGPT1;refGene_tx=XM_004000054.5;refGene_gn=ANGPT1;refGene_tx=XM_006943348.4;refGene_gn=ANGPT1;refGene_tx=XM_019823037.2
F2 54688154 54688310
refGene_gn=ANGPT1;refGene_tx=XM_004000054.5;refGene_gn=ANGPT1;refGene_tx=XM_006943348.4;refGene_gn=ANGPT1;refGene_tx=XM_019823037.2
F2 54830443 54830740
refGene_gn=ANGPT1;refGene_tx=XM_004000054.5;refGene_gn=ANGPT1;refGene_tx=XM_006943348.4
F2 55150198 55150311
refGene_gn=RSPO2;refGene_tx=XM_023248417.1;refGene_gn=RSPO2;refGene_tx=XM_023248418.1
F2 55202526 55202715
refGene_gn=RSPO2;refGene_tx=XM_023248417.1;refGene_gn=RSPO2;refGene_tx=XM_023248418.1
F2 55204653 55204797
refGene_gn=RSPO2;refGene_tx=XM_023248417.1;refGene_gn=RSPO2;refGene_tx=XM_023248418.1
F2 55230264 55230453
refGene_gn=RSPO2;refGene_tx=XM_023248417.1;refGene_gn=RSPO2;refGene_tx=XM_023248418.1
F2 55308935 55309029
refGene_gn=RSPO2;refGene_tx=XM_023248417.1;refGene_gn=RSPO2;refGene_tx=XM_023248418.1
F2 55395023 55395062
refGene_gn=EIF3E;refGene_tx=XM_004000056.3;refGene_gn=EIF3E;refGene_tx=XM_006943350.3
F2 55395808 55395943
refGene_gn=EIF3E;refGene_tx=XM_004000056.3;refGene_gn=EIF3E;refGene_tx=XM_006943350.3
F2 55396232 55396335
refGene_gn=EIF3E;refGene_tx=XM_004000056.3;refGene_gn=EIF3E;refGene_tx=XM_006943350.3
F2 55413140 55413250
refGene_gn=EIF3E;refGene_tx=XM_004000056.3;refGene_gn=EIF3E;refGene_tx=XM_006943350.3
F2 55414567 55414669
refGene_gn=EIF3E;refGene_tx=XM_004000056.3;refGene_gn=EIF3E;refGene_tx

=XM_006943350.3
F2 55415722 55415849
refGene_gn=EIF3E;refGene_tx=XM_004000056.3;refGene_gn=EIF3E;refGene_tx
=XM_006943350.3
F2 55432270 55432395
refGene_gn=EIF3E;refGene_tx=XM_004000056.3;refGene_gn=EIF3E;refGene_tx
=XM_006943350.3
F2 55433068 55433194
refGene_gn=EIF3E;refGene_tx=XM_004000056.3;refGene_gn=EIF3E;refGene_tx
=XM_006943350.3
F2 55438763 55438868
refGene_gn=EIF3E;refGene_tx=XM_004000056.3;refGene_gn=EIF3E;refGene_tx
=XM_006943350.3
F2 55440529 55440572
refGene_gn=EIF3E;refGene_tx=XM_004000056.3;refGene_gn=EIF3E;refGene_tx
=XM_006943350.3
F2 55443204 55443322
refGene_gn=EIF3E;refGene_tx=XM_004000056.3;refGene_gn=EIF3E;refGene_tx
=XM_006943350.3
F2 55445008 55445123
refGene_gn=EIF3E;refGene_tx=XM_004000056.3;refGene_gn=EIF3E;refGene_tx
=XM_006943350.3
F2 55449502 55449592 refGene_gn=EIF3E;refGene_tx=XM_004000056.3
F2 55634462 55634502
refGene_gn=EMC2;refGene_tx=XM_011291431.2;refGene_gn=EMC2;refGene_tx=X
M_011291432.2
F2 55641275 55641389
refGene_gn=EMC2;refGene_tx=XM_011291431.2;refGene_gn=EMC2;refGene_tx=X
M_011291432.2
F2 55641882 55641947
refGene_gn=EMC2;refGene_tx=XM_011291431.2;refGene_gn=EMC2;refGene_tx=X
M_011291432.2
F2 55644980 55645066
refGene_gn=EMC2;refGene_tx=XM_011291431.2;refGene_gn=EMC2;refGene_tx=X
M_011291432.2
F2 55652178 55652236
refGene_gn=EMC2;refGene_tx=XM_011291431.2;refGene_gn=EMC2;refGene_tx=X
M_011291432.2
F2 55663912 55663998
refGene_gn=EMC2;refGene_tx=XM_011291431.2;refGene_gn=EMC2;refGene_tx=X
M_011291432.2
F2 55664148 55664208
refGene_gn=EMC2;refGene_tx=XM_011291431.2;refGene_gn=EMC2;refGene_tx=X
M_011291432.2
F2 55669808 55669890
refGene_gn=EMC2;refGene_tx=XM_011291431.2;refGene_gn=EMC2;refGene_tx=X
M_011291432.2
F2 55670696 55670807
refGene_gn=EMC2;refGene_tx=XM_011291431.2;refGene_gn=EMC2;refGene_tx=X
M_011291432.2

F2 55672908 55673013
refGene_gn=EMC2;refGene_tx=XM_011291431.2;refGene_gn=EMC2;refGene_tx=XM_011291432.2
F2 55682394 55682481
refGene_gn=EMC2;refGene_tx=XM_011291431.2;refGene_gn=EMC2;refGene_tx=XM_011291432.2
F2 55998072 55998990 refGene_gn=TMEM74;refGene_tx=XM_004000057.5
F2 56246407 56247196 refGene_gn=TRHR;refGene_tx=XM_004000058.4
F2 56289827 56290235 refGene_gn=TRHR;refGene_tx=XM_004000058.4
F2 56431990 56432283 refGene_gn=NUDCD1;refGene_tx=XM_011291445.3
F2 56434180 56434340 refGene_gn=NUDCD1;refGene_tx=XM_011291445.3
F2 56458765 56458891 refGene_gn=NUDCD1;refGene_tx=XM_011291445.3
F2 56463630 56463775 refGene_gn=NUDCD1;refGene_tx=XM_011291445.3
F2 56469493 56469695 refGene_gn=NUDCD1;refGene_tx=XM_011291445.3
F2 56473278 56473461 refGene_gn=NUDCD1;refGene_tx=XM_011291445.3
F2 56476893 56477074 refGene_gn=NUDCD1;refGene_tx=XM_011291445.3
F2 56480858 56481044 refGene_gn=NUDCD1;refGene_tx=XM_011291445.3
F2 56495524 56495679 refGene_gn=NUDCD1;refGene_tx=XM_011291445.3
F2 56508544 56508662 refGene_gn=NUDCD1;refGene_tx=XM_011291445.3
F2 56509121 56509127 refGene_gn=ENY2;refGene_tx=XM_004000060.5
F2 56510499 56510576
refGene_gn=ENY2;refGene_tx=XM_004000060.5;refGene_gn=ENY2;refGene_tx=XM_004000061.5
F2 56514378 56514449
refGene_gn=ENY2;refGene_tx=XM_004000060.5;refGene_gn=ENY2;refGene_tx=XM_004000061.5
F2 56515786 56515861
refGene_gn=ENY2;refGene_tx=XM_004000060.5;refGene_gn=ENY2;refGene_tx=XM_004000061.5
F2 56518839 56518916
refGene_gn=ENY2;refGene_tx=XM_004000060.5;refGene_gn=ENY2;refGene_tx=XM_004000061.5
F2 56539248 56539321 refGene_gn=PKHD1L1;refGene_tx=XM_023248273.1
F2 56541170 56541260 refGene_gn=PKHD1L1;refGene_tx=XM_023248273.1
F2 56550396 56550541 refGene_gn=PKHD1L1;refGene_tx=XM_023248273.1
F2 56551467 56551576 refGene_gn=PKHD1L1;refGene_tx=XM_023248273.1
F2 56552611 56552669 refGene_gn=PKHD1L1;refGene_tx=XM_023248273.1
F2 56554243 56554337 refGene_gn=PKHD1L1;refGene_tx=XM_023248273.1
F2 56557227 56557281 refGene_gn=PKHD1L1;refGene_tx=XM_023248273.1
F2 56557900 56557974 refGene_gn=PKHD1L1;refGene_tx=XM_023248273.1
F2 56559245 56559288 refGene_gn=PKHD1L1;refGene_tx=XM_023248273.1
F2 56563153 56563224 refGene_gn=PKHD1L1;refGene_tx=XM_023248273.1
F2 56564647 56564758 refGene_gn=PKHD1L1;refGene_tx=XM_023248273.1
F2 56567873 56567963 refGene_gn=PKHD1L1;refGene_tx=XM_023248273.1
F2 56569793 56570062 refGene_gn=PKHD1L1;refGene_tx=XM_023248273.1
F2 56571607 56571699 refGene_gn=PKHD1L1;refGene_tx=XM_023248273.1
F2 56574987 56575147 refGene_gn=PKHD1L1;refGene_tx=XM_023248273.1
F2 56575454 56575590 refGene_gn=PKHD1L1;refGene_tx=XM_023248273.1
F2 56576403 56576547 refGene_gn=PKHD1L1;refGene_tx=XM_023248273.1
F2 56578471 56578629 refGene_gn=PKHD1L1;refGene_tx=XM_023248273.1

| | | | |
|---|---|---|---|
| F2 | 56579614 | 56579728 | refGene_gn=PKHD1L1;refGene_tx=XM_023248273.1 |
| F2 | 56581480 | 56581630 | refGene_gn=PKHD1L1;refGene_tx=XM_023248273.1 |
| F2 | 56582640 | 56582765 | refGene_gn=PKHD1L1;refGene_tx=XM_023248273.1 |
| F2 | 56585564 | 56585728 | refGene_gn=PKHD1L1;refGene_tx=XM_023248273.1 |
| F2 | 56586903 | 56587076 | refGene_gn=PKHD1L1;refGene_tx=XM_023248273.1 |
| F2 | 56589050 | 56589198 | refGene_gn=PKHD1L1;refGene_tx=XM_023248273.1 |
| F2 | 56589698 | 56589853 | refGene_gn=PKHD1L1;refGene_tx=XM_023248273.1 |
| F2 | 56591669 | 56591792 | refGene_gn=PKHD1L1;refGene_tx=XM_023248273.1 |
| F2 | 56592950 | 56593056 | refGene_gn=PKHD1L1;refGene_tx=XM_023248273.1 |
| F2 | 56596447 | 56596558 | refGene_gn=PKHD1L1;refGene_tx=XM_023248273.1 |
| F2 | 56598122 | 56598287 | refGene_gn=PKHD1L1;refGene_tx=XM_023248273.1 |
| F2 | 56599263 | 56599385 | refGene_gn=PKHD1L1;refGene_tx=XM_023248273.1 |
| F2 | 56600970 | 56601103 | refGene_gn=PKHD1L1;refGene_tx=XM_023248273.1 |
| F2 | 56601812 | 56602008 | refGene_gn=PKHD1L1;refGene_tx=XM_023248273.1 |
| F2 | 56604084 | 56604227 | refGene_gn=PKHD1L1;refGene_tx=XM_023248273.1 |
| F2 | 56604634 | 56604739 | refGene_gn=PKHD1L1;refGene_tx=XM_023248273.1 |
| F2 | 56605449 | 56605638 | refGene_gn=PKHD1L1;refGene_tx=XM_023248273.1 |
| F2 | 56606474 | 56606645 | refGene_gn=PKHD1L1;refGene_tx=XM_023248273.1 |
| F2 | 56607682 | 56607906 | refGene_gn=PKHD1L1;refGene_tx=XM_023248273.1 |
| F2 | 56608935 | 56609920 | refGene_gn=PKHD1L1;refGene_tx=XM_023248273.1 |
| F2 | 56612968 | 56613217 | refGene_gn=PKHD1L1;refGene_tx=XM_023248273.1 |
| F2 | 56613871 | 56614021 | refGene_gn=PKHD1L1;refGene_tx=XM_023248273.1 |
| F2 | 56615936 | 56616111 | refGene_gn=PKHD1L1;refGene_tx=XM_023248273.1 |
| F2 | 56617086 | 56617243 | refGene_gn=PKHD1L1;refGene_tx=XM_023248273.1 |
| F2 | 56618164 | 56618321 | refGene_gn=PKHD1L1;refGene_tx=XM_023248273.1 |
| F2 | 56620001 | 56620081 | refGene_gn=PKHD1L1;refGene_tx=XM_023248273.1 |
| F2 | 56620523 | 56620653 | refGene_gn=PKHD1L1;refGene_tx=XM_023248273.1 |
| F2 | 56621963 | 56622093 | refGene_gn=PKHD1L1;refGene_tx=XM_023248273.1 |
| F2 | 56625322 | 56625564 | refGene_gn=PKHD1L1;refGene_tx=XM_023248273.1 |
| F2 | 56626598 | 56626735 | refGene_gn=PKHD1L1;refGene_tx=XM_023248273.1 |
| F2 | 56628772 | 56629802 | refGene_gn=PKHD1L1;refGene_tx=XM_023248273.1 |
| F2 | 56631342 | 56631534 | refGene_gn=PKHD1L1;refGene_tx=XM_023248273.1 |
| F2 | 56635772 | 56635924 | refGene_gn=PKHD1L1;refGene_tx=XM_023248273.1 |
| F2 | 56637018 | 56637178 | refGene_gn=PKHD1L1;refGene_tx=XM_023248273.1 |
| F2 | 56638984 | 56639121 | refGene_gn=PKHD1L1;refGene_tx=XM_023248273.1 |
| F2 | 56639122 | 56639157 | refGene_gn=PKHD1L1;refGene_tx=XM_023248273.1 |
| F2 | 56642724 | 56642813 | refGene_gn=PKHD1L1;refGene_tx=XM_023248273.1 |
| F2 | 56643170 | 56643319 | refGene_gn=PKHD1L1;refGene_tx=XM_023248273.1 |
| F2 | 56645082 | 56645212 | refGene_gn=PKHD1L1;refGene_tx=XM_023248273.1 |
| F2 | 56646831 | 56646950 | refGene_gn=PKHD1L1;refGene_tx=XM_023248273.1 |
| F2 | 56647607 | 56647737 | refGene_gn=PKHD1L1;refGene_tx=XM_023248273.1 |
| F2 | 56655172 | 56655346 | refGene_gn=PKHD1L1;refGene_tx=XM_023248273.1 |
| F2 | 56659496 | 56659600 | refGene_gn=PKHD1L1;refGene_tx=XM_023248273.1 |
| F2 | 56660514 | 56660644 | refGene_gn=PKHD1L1;refGene_tx=XM_023248273.1 |
| F2 | 56661567 | 56661689 | refGene_gn=PKHD1L1;refGene_tx=XM_023248273.1 |
| F2 | 56663212 | 56663303 | refGene_gn=PKHD1L1;refGene_tx=XM_023248273.1 |
| F2 | 56667106 | 56667255 | refGene_gn=PKHD1L1;refGene_tx=XM_023248273.1 |
| F2 | 56667337 | 56667460 | refGene_gn=PKHD1L1;refGene_tx=XM_023248273.1 |
| F2 | 56668240 | 56668352 | refGene_gn=PKHD1L1;refGene_tx=XM_023248273.1 |
| F2 | 56668428 | 56668545 | refGene_gn=PKHD1L1;refGene_tx=XM_023248273.1 |

F2 56671648 56671814 refGene_gn=PKHD1L1;refGene_tx=XM_023248273.1
F2 56674771 56675004 refGene_gn=PKHD1L1;refGene_tx=XM_023248273.1
F2 56675200 56675368 refGene_gn=PKHD1L1;refGene_tx=XM_023248273.1
F2 56678716 56678874 refGene_gn=PKHD1L1;refGene_tx=XM_023248273.1
F2 56680981 56681117 refGene_gn=PKHD1L1;refGene_tx=XM_023248273.1
F2 56683966 56684311 refGene_gn=PKHD1L1;refGene_tx=XM_023248273.1
F2 56687198 56687350 refGene_gn=PKHD1L1;refGene_tx=XM_023248273.1
F2 56687750 56687897 refGene_gn=PKHD1L1;refGene_tx=XM_023248273.1
F2 56688479 56688633 refGene_gn=PKHD1L1;refGene_tx=XM_023248273.1
F2 56692217 56692484 refGene_gn=PKHD1L1;refGene_tx=XM_023248273.1
F2 56696267 56696278 refGene_gn=PKHD1L1;refGene_tx=XM_023248273.1
F2 56706358 56706388 refGene_gn=EBAG9;refGene_tx=XM_004000067.5
F2 56717563 56717661 refGene_gn=EBAG9;refGene_tx=XM_004000067.5
F2 56721023 56721102 refGene_gn=EBAG9;refGene_tx=XM_004000067.5
F2 56722078 56722237 refGene_gn=EBAG9;refGene_tx=XM_004000067.5
F2 56723900 56724008 refGene_gn=EBAG9;refGene_tx=XM_004000067.5
F2 56728050 56728142 refGene_gn=EBAG9;refGene_tx=XM_004000067.5
F2 56731062 56731183 refGene_gn=EBAG9;refGene_tx=XM_004000067.5
F2 56744719 56745824
refGene_gn=SYBU;refGene_tx=XM_011291446.3;refGene_gn=SYBU;refGene_tx=XM_019823040.2;refGene_gn=SYBU;refGene_tx=XM_019823041.2
F2 56747367 56747517
refGene_gn=SYBU;refGene_tx=XM_011291446.3;refGene_gn=SYBU;refGene_tx=XM_019823040.2;refGene_gn=SYBU;refGene_tx=XM_019823041.2
F2 56749180 56749384
refGene_gn=SYBU;refGene_tx=XM_011291446.3;refGene_gn=SYBU;refGene_tx=XM_019823040.2;refGene_gn=SYBU;refGene_tx=XM_019823041.2
F2 56755811 56755914
refGene_gn=SYBU;refGene_tx=XM_011291446.3;refGene_gn=SYBU;refGene_tx=XM_019823040.2;refGene_gn=SYBU;refGene_tx=XM_019823041.2
F2 56780507 56780535 refGene_gn=SYBU;refGene_tx=XM_019823040.2
F2 56792926 56793124
refGene_gn=SYBU;refGene_tx=XM_011291446.3;refGene_gn=SYBU;refGene_tx=XM_019823041.2
F2 56817101 56817306
refGene_gn=SYBU;refGene_tx=XM_011291446.3;refGene_gn=SYBU;refGene_tx=XM_019823041.2
F2 56819029 56819053
refGene_gn=SYBU;refGene_tx=XM_011291446.3;refGene_gn=SYBU;refGene_tx=XM_019823041.2
F2 56819515 56819578
refGene_gn=LOC111558526;refGene_tx=XM_023248335.1
F2 56819579 56819927
refGene_gn=LOC111558526;refGene_tx=XM_023248335.1
F2 56857192 56858143
refGene_gn=LOC109496172;refGene_tx=XM_023248336.1
F2 57063356 57063868 refGene_gn=KCNV1;refGene_tx=XM_004000068.5
F2 57067115 57067645 refGene_gn=KCNV1;refGene_tx=XM_004000068.5
F2 57068499 57068966 refGene_gn=KCNV1;refGene_tx=XM_004000068.5
F2 59044847 59045007

refGene_gn=CSMD3;refGene_tx=XM_011291447.3;refGene_gn=CSMD3;refGene_tx=XM_019823042.2;refGene_gn=CSMD3;refGene_tx=XM_019823044.2;refGene_gn=CSMD3;refGene_tx=XM_019823045.2
F2 59049271 59049407
refGene_gn=CSMD3;refGene_tx=XM_011291447.3;refGene_gn=CSMD3;refGene_tx=XM_019823042.2;refGene_gn=CSMD3;refGene_tx=XM_019823044.2;refGene_gn=CSMD3;refGene_tx=XM_019823045.2
F2 59051499 59051587
refGene_gn=CSMD3;refGene_tx=XM_011291447.3;refGene_gn=CSMD3;refGene_tx=XM_019823042.2;refGene_gn=CSMD3;refGene_tx=XM_019823044.2;refGene_gn=CSMD3;refGene_tx=XM_019823045.2
F2 59054673 59054786
refGene_gn=CSMD3;refGene_tx=XM_011291447.3;refGene_gn=CSMD3;refGene_tx=XM_019823042.2;refGene_gn=CSMD3;refGene_tx=XM_019823044.2;refGene_gn=CSMD3;refGene_tx=XM_019823045.2
F2 59062740 59062899
refGene_gn=CSMD3;refGene_tx=XM_011291447.3;refGene_gn=CSMD3;refGene_tx=XM_019823042.2;refGene_gn=CSMD3;refGene_tx=XM_019823044.2;refGene_gn=CSMD3;refGene_tx=XM_019823045.2
F2 59068552 59068618
refGene_gn=CSMD3;refGene_tx=XM_011291447.3;refGene_gn=CSMD3;refGene_tx=XM_019823042.2;refGene_gn=CSMD3;refGene_tx=XM_019823044.2;refGene_gn=CSMD3;refGene_tx=XM_019823045.2
F2 59070486 59070666
refGene_gn=CSMD3;refGene_tx=XM_011291447.3;refGene_gn=CSMD3;refGene_tx=XM_019823042.2;refGene_gn=CSMD3;refGene_tx=XM_019823044.2;refGene_gn=CSMD3;refGene_tx=XM_019823045.2
F2 59073015 59073127
refGene_gn=CSMD3;refGene_tx=XM_011291447.3;refGene_gn=CSMD3;refGene_tx=XM_019823042.2;refGene_gn=CSMD3;refGene_tx=XM_019823044.2;refGene_gn=CSMD3;refGene_tx=XM_019823045.2
F2 59081506 59081580
refGene_gn=CSMD3;refGene_tx=XM_011291447.3;refGene_gn=CSMD3;refGene_tx=XM_019823042.2;refGene_gn=CSMD3;refGene_tx=XM_019823044.2;refGene_gn=CSMD3;refGene_tx=XM_019823045.2
F2 59082518 59082692
refGene_gn=CSMD3;refGene_tx=XM_011291447.3;refGene_gn=CSMD3;refGene_tx=XM_019823042.2;refGene_gn=CSMD3;refGene_tx=XM_019823044.2;refGene_gn=CSMD3;refGene_tx=XM_019823045.2
F2 59086642 59086816
refGene_gn=CSMD3;refGene_tx=XM_011291447.3;refGene_gn=CSMD3;refGene_tx=XM_019823042.2;refGene_gn=CSMD3;refGene_tx=XM_019823044.2;refGene_gn=CSMD3;refGene_tx=XM_019823045.2
F2 59088896 59089076
refGene_gn=CSMD3;refGene_tx=XM_011291447.3;refGene_gn=CSMD3;refGene_tx=XM_019823042.2;refGene_gn=CSMD3;refGene_tx=XM_019823044.2;refGene_gn=CSMD3;refGene_tx=XM_019823045.2
F2 59101104 59101281
refGene_gn=CSMD3;refGene_tx=XM_011291447.3;refGene_gn=CSMD3;refGene_tx=XM_019823042.2;refGene_gn=CSMD3;refGene_tx=XM_019823044.2;refGene_gn=

CSMD3;refGene_tx=XM_019823045.2
F2 59106157 59106340
refGene_gn=CSMD3;refGene_tx=XM_011291447.3;refGene_gn=CSMD3;refGene_tx=XM_019823042.2;refGene_gn=CSMD3;refGene_tx=XM_019823044.2;refGene_gn=CSMD3;refGene_tx=XM_019823045.2
F2 59109413 59109587
refGene_gn=CSMD3;refGene_tx=XM_011291447.3;refGene_gn=CSMD3;refGene_tx=XM_019823042.2;refGene_gn=CSMD3;refGene_tx=XM_019823044.2;refGene_gn=CSMD3;refGene_tx=XM_019823045.2
F2 59111525 59111711
refGene_gn=CSMD3;refGene_tx=XM_011291447.3;refGene_gn=CSMD3;refGene_tx=XM_019823042.2;refGene_gn=CSMD3;refGene_tx=XM_019823044.2;refGene_gn=CSMD3;refGene_tx=XM_019823045.2
F2 59112505 59112679
refGene_gn=CSMD3;refGene_tx=XM_011291447.3;refGene_gn=CSMD3;refGene_tx=XM_019823042.2;refGene_gn=CSMD3;refGene_tx=XM_019823044.2;refGene_gn=CSMD3;refGene_tx=XM_019823045.2
F2 59115649 59115823
refGene_gn=CSMD3;refGene_tx=XM_011291447.3;refGene_gn=CSMD3;refGene_tx=XM_019823042.2;refGene_gn=CSMD3;refGene_tx=XM_019823044.2;refGene_gn=CSMD3;refGene_tx=XM_019823045.2
F2 59121542 59121716
refGene_gn=CSMD3;refGene_tx=XM_011291447.3;refGene_gn=CSMD3;refGene_tx=XM_019823042.2;refGene_gn=CSMD3;refGene_tx=XM_019823044.2;refGene_gn=CSMD3;refGene_tx=XM_019823045.2
F2 59125419 59125614
refGene_gn=CSMD3;refGene_tx=XM_011291447.3;refGene_gn=CSMD3;refGene_tx=XM_019823042.2;refGene_gn=CSMD3;refGene_tx=XM_019823044.2;refGene_gn=CSMD3;refGene_tx=XM_019823045.2
F2 59127208 59127394
refGene_gn=CSMD3;refGene_tx=XM_011291447.3;refGene_gn=CSMD3;refGene_tx=XM_019823042.2;refGene_gn=CSMD3;refGene_tx=XM_019823044.2;refGene_gn=CSMD3;refGene_tx=XM_019823045.2
F2 59131997 59132186
refGene_gn=CSMD3;refGene_tx=XM_011291447.3;refGene_gn=CSMD3;refGene_tx=XM_019823042.2;refGene_gn=CSMD3;refGene_tx=XM_019823044.2;refGene_gn=CSMD3;refGene_tx=XM_019823045.2
F2 59134667 59134814
refGene_gn=CSMD3;refGene_tx=XM_011291447.3;refGene_gn=CSMD3;refGene_tx=XM_019823042.2;refGene_gn=CSMD3;refGene_tx=XM_019823044.2;refGene_gn=CSMD3;refGene_tx=XM_019823045.2
F2 59135197 59135386
refGene_gn=CSMD3;refGene_tx=XM_011291447.3;refGene_gn=CSMD3;refGene_tx=XM_019823042.2;refGene_gn=CSMD3;refGene_tx=XM_019823044.2;refGene_gn=CSMD3;refGene_tx=XM_019823045.2
F2 59140215 59140329
refGene_gn=CSMD3;refGene_tx=XM_011291447.3;refGene_gn=CSMD3;refGene_tx=XM_019823042.2;refGene_gn=CSMD3;refGene_tx=XM_019823044.2;refGene_gn=CSMD3;refGene_tx=XM_019823045.2
F2 59141280 59141361

refGene_gn=CSMD3;refGene_tx=XM_011291447.3;refGene_gn=CSMD3;refGene_tx=XM_019823042.2;refGene_gn=CSMD3;refGene_tx=XM_019823044.2;refGene_gn=CSMD3;refGene_tx=XM_019823045.2
F2 59157141 59157287
refGene_gn=CSMD3;refGene_tx=XM_011291447.3;refGene_gn=CSMD3;refGene_tx=XM_019823042.2;refGene_gn=CSMD3;refGene_tx=XM_019823044.2;refGene_gn=CSMD3;refGene_tx=XM_019823045.2
F2 59158378 59158556
refGene_gn=CSMD3;refGene_tx=XM_011291447.3;refGene_gn=CSMD3;refGene_tx=XM_019823042.2;refGene_gn=CSMD3;refGene_tx=XM_019823044.2;refGene_gn=CSMD3;refGene_tx=XM_019823045.2
F2 59159229 59159418
refGene_gn=CSMD3;refGene_tx=XM_011291447.3;refGene_gn=CSMD3;refGene_tx=XM_019823042.2;refGene_gn=CSMD3;refGene_tx=XM_019823044.2;refGene_gn=CSMD3;refGene_tx=XM_019823045.2
F2 59163032 59163242
refGene_gn=CSMD3;refGene_tx=XM_011291447.3;refGene_gn=CSMD3;refGene_tx=XM_019823042.2;refGene_gn=CSMD3;refGene_tx=XM_019823044.2;refGene_gn=CSMD3;refGene_tx=XM_019823045.2
F2 59166684 59166801
refGene_gn=CSMD3;refGene_tx=XM_011291447.3;refGene_gn=CSMD3;refGene_tx=XM_019823042.2;refGene_gn=CSMD3;refGene_tx=XM_019823044.2;refGene_gn=CSMD3;refGene_tx=XM_019823045.2
F2 59178755 59178825
refGene_gn=CSMD3;refGene_tx=XM_011291447.3;refGene_gn=CSMD3;refGene_tx=XM_019823042.2;refGene_gn=CSMD3;refGene_tx=XM_019823044.2;refGene_gn=CSMD3;refGene_tx=XM_019823045.2
F2 59179959 59180078
refGene_gn=CSMD3;refGene_tx=XM_011291447.3;refGene_gn=CSMD3;refGene_tx=XM_019823042.2;refGene_gn=CSMD3;refGene_tx=XM_019823044.2;refGene_gn=CSMD3;refGene_tx=XM_019823045.2
F2 59210889 59210994
refGene_gn=CSMD3;refGene_tx=XM_011291447.3;refGene_gn=CSMD3;refGene_tx=XM_019823042.2;refGene_gn=CSMD3;refGene_tx=XM_019823044.2;refGene_gn=CSMD3;refGene_tx=XM_019823045.2
F2 59213507 59213604
refGene_gn=CSMD3;refGene_tx=XM_011291447.3;refGene_gn=CSMD3;refGene_tx=XM_019823042.2;refGene_gn=CSMD3;refGene_tx=XM_019823044.2;refGene_gn=CSMD3;refGene_tx=XM_019823045.2
F2 59221321 59221446
refGene_gn=CSMD3;refGene_tx=XM_011291447.3;refGene_gn=CSMD3;refGene_tx=XM_019823042.2;refGene_gn=CSMD3;refGene_tx=XM_019823044.2;refGene_gn=CSMD3;refGene_tx=XM_019823045.2
F2 59235883 59236087
refGene_gn=CSMD3;refGene_tx=XM_011291447.3;refGene_gn=CSMD3;refGene_tx=XM_019823042.2;refGene_gn=CSMD3;refGene_tx=XM_019823044.2;refGene_gn=CSMD3;refGene_tx=XM_019823045.2
F2 59237647 59237743
refGene_gn=CSMD3;refGene_tx=XM_011291447.3;refGene_gn=CSMD3;refGene_tx=XM_019823042.2;refGene_gn=CSMD3;refGene_tx=XM_019823045.2

F2 59238239 59238353
refGene_gn=CSMD3;refGene_tx=XM_011291447.3;refGene_gn=CSMD3;refGene_tx=XM_019823042.2;refGene_gn=CSMD3;refGene_tx=XM_019823045.2
F2 59302551 59302668
refGene_gn=CSMD3;refGene_tx=XM_011291447.3;refGene_gn=CSMD3;refGene_tx=XM_019823042.2;refGene_gn=CSMD3;refGene_tx=XM_019823044.2;refGene_gn=CSMD3;refGene_tx=XM_019823045.2
F2 59321176 59321371
refGene_gn=CSMD3;refGene_tx=XM_011291447.3;refGene_gn=CSMD3;refGene_tx=XM_019823042.2;refGene_gn=CSMD3;refGene_tx=XM_019823044.2;refGene_gn=CSMD3;refGene_tx=XM_019823045.2
F2 59333952 59334140
refGene_gn=CSMD3;refGene_tx=XM_011291447.3;refGene_gn=CSMD3;refGene_tx=XM_019823042.2;refGene_gn=CSMD3;refGene_tx=XM_019823044.2;refGene_gn=CSMD3;refGene_tx=XM_019823045.2
F2 59337370 59337509
refGene_gn=CSMD3;refGene_tx=XM_011291447.3;refGene_gn=CSMD3;refGene_tx=XM_019823042.2;refGene_gn=CSMD3;refGene_tx=XM_019823044.2;refGene_gn=CSMD3;refGene_tx=XM_019823045.2
F2 59344555 59344747
refGene_gn=CSMD3;refGene_tx=XM_011291447.3;refGene_gn=CSMD3;refGene_tx=XM_019823042.2;refGene_gn=CSMD3;refGene_tx=XM_019823044.2;refGene_gn=CSMD3;refGene_tx=XM_019823045.2
F2 59366647 59366850
refGene_gn=CSMD3;refGene_tx=XM_011291447.3;refGene_gn=CSMD3;refGene_tx=XM_019823042.2;refGene_gn=CSMD3;refGene_tx=XM_019823044.2;refGene_gn=CSMD3;refGene_tx=XM_019823045.2
F2 59369073 59369200
refGene_gn=CSMD3;refGene_tx=XM_011291447.3;refGene_gn=CSMD3;refGene_tx=XM_019823042.2;refGene_gn=CSMD3;refGene_tx=XM_019823044.2;refGene_gn=CSMD3;refGene_tx=XM_019823045.2
F2 59373761 59373953
refGene_gn=CSMD3;refGene_tx=XM_011291447.3;refGene_gn=CSMD3;refGene_tx=XM_019823042.2;refGene_gn=CSMD3;refGene_tx=XM_019823044.2;refGene_gn=CSMD3;refGene_tx=XM_019823045.2
F2 59384651 59384808
refGene_gn=CSMD3;refGene_tx=XM_011291447.3;refGene_gn=CSMD3;refGene_tx=XM_019823042.2;refGene_gn=CSMD3;refGene_tx=XM_019823044.2;refGene_gn=CSMD3;refGene_tx=XM_019823045.2
F2 59395246 59395416
refGene_gn=CSMD3;refGene_tx=XM_011291447.3;refGene_gn=CSMD3;refGene_tx=XM_019823042.2;refGene_gn=CSMD3;refGene_tx=XM_019823044.2;refGene_gn=CSMD3;refGene_tx=XM_019823045.2
F2 59441114 59441303
refGene_gn=CSMD3;refGene_tx=XM_011291447.3;refGene_gn=CSMD3;refGene_tx=XM_019823042.2;refGene_gn=CSMD3;refGene_tx=XM_019823044.2;refGene_gn=CSMD3;refGene_tx=XM_019823045.2
F2 59442976 59443192
refGene_gn=CSMD3;refGene_tx=XM_011291447.3;refGene_gn=CSMD3;refGene_tx=XM_019823042.2;refGene_gn=CSMD3;refGene_tx=XM_019823044.2;refGene_gn=

CSMD3;refGene_tx=XM_019823045.2
F2 59449461 59449578
refGene_gn=CSMD3;refGene_tx=XM_011291447.3;refGene_gn=CSMD3;refGene_tx=XM_019823042.2;refGene_gn=CSMD3;refGene_tx=XM_019823044.2;refGene_gn=CSMD3;refGene_tx=XM_019823045.2
F2 59458552 59458741
refGene_gn=CSMD3;refGene_tx=XM_011291447.3;refGene_gn=CSMD3;refGene_tx=XM_019823042.2;refGene_gn=CSMD3;refGene_tx=XM_019823044.2;refGene_gn=CSMD3;refGene_tx=XM_019823045.2
F2 59463666 59463854
refGene_gn=CSMD3;refGene_tx=XM_011291447.3;refGene_gn=CSMD3;refGene_tx=XM_019823042.2;refGene_gn=CSMD3;refGene_tx=XM_019823044.2;refGene_gn=CSMD3;refGene_tx=XM_019823045.2
F2 59471886 59472025
refGene_gn=CSMD3;refGene_tx=XM_011291447.3;refGene_gn=CSMD3;refGene_tx=XM_019823042.2;refGene_gn=CSMD3;refGene_tx=XM_019823044.2;refGene_gn=CSMD3;refGene_tx=XM_019823045.2
F2 59491458 59491653
refGene_gn=CSMD3;refGene_tx=XM_011291447.3;refGene_gn=CSMD3;refGene_tx=XM_019823042.2;refGene_gn=CSMD3;refGene_tx=XM_019823044.2;refGene_gn=CSMD3;refGene_tx=XM_019823045.2
F2 59495466 59495793
refGene_gn=CSMD3;refGene_tx=XM_011291447.3;refGene_gn=CSMD3;refGene_tx=XM_019823042.2;refGene_gn=CSMD3;refGene_tx=XM_019823044.2;refGene_gn=CSMD3;refGene_tx=XM_019823045.2
F2 59500592 59500775
refGene_gn=CSMD3;refGene_tx=XM_011291447.3;refGene_gn=CSMD3;refGene_tx=XM_019823042.2;refGene_gn=CSMD3;refGene_tx=XM_019823044.2;refGene_gn=CSMD3;refGene_tx=XM_019823045.2
F2 59620581 59620694
refGene_gn=CSMD3;refGene_tx=XM_011291447.3;refGene_gn=CSMD3;refGene_tx=XM_019823042.2;refGene_gn=CSMD3;refGene_tx=XM_019823044.2;refGene_gn=CSMD3;refGene_tx=XM_019823045.2
F2 59641022 59641126
refGene_gn=CSMD3;refGene_tx=XM_011291447.3;refGene_gn=CSMD3;refGene_tx=XM_019823042.2;refGene_gn=CSMD3;refGene_tx=XM_019823044.2;refGene_gn=CSMD3;refGene_tx=XM_019823045.2
F2 59667790 59667912
refGene_gn=CSMD3;refGene_tx=XM_011291447.3;refGene_gn=CSMD3;refGene_tx=XM_019823042.2;refGene_gn=CSMD3;refGene_tx=XM_019823044.2;refGene_gn=CSMD3;refGene_tx=XM_019823045.2
F2 59724074 59724199
refGene_gn=CSMD3;refGene_tx=XM_011291447.3;refGene_gn=CSMD3;refGene_tx=XM_019823042.2;refGene_gn=CSMD3;refGene_tx=XM_019823044.2;refGene_gn=CSMD3;refGene_tx=XM_019823045.2
F2 59762244 59762332
refGene_gn=CSMD3;refGene_tx=XM_011291447.3;refGene_gn=CSMD3;refGene_tx=XM_019823042.2;refGene_gn=CSMD3;refGene_tx=XM_019823044.2;refGene_gn=CSMD3;refGene_tx=XM_019823045.2
F2 59770045 59770123

refGene_gn=CSMD3;refGene_tx=XM_011291447.3;refGene_gn=CSMD3;refGene_tx
=XM_019823044.2;refGene_gn=CSMD3;refGene_tx=XM_019823045.2
F2 59791755 59792067
refGene_gn=CSMD3;refGene_tx=XM_011291447.3;refGene_gn=CSMD3;refGene_tx
=XM_019823042.2;refGene_gn=CSMD3;refGene_tx=XM_019823044.2
F2 59840826 59840939
refGene_gn=CSMD3;refGene_tx=XM_011291447.3;refGene_gn=CSMD3;refGene_tx
=XM_019823042.2;refGene_gn=CSMD3;refGene_tx=XM_019823044.2;refGene_gn=
CSMD3;refGene_tx=XM_019823045.2
F2 59926321 59926529
refGene_gn=CSMD3;refGene_tx=XM_011291447.3;refGene_gn=CSMD3;refGene_tx
=XM_019823042.2;refGene_gn=CSMD3;refGene_tx=XM_019823044.2;refGene_gn=
CSMD3;refGene_tx=XM_019823045.2
F2 59997870 59998065
refGene_gn=CSMD3;refGene_tx=XM_011291447.3;refGene_gn=CSMD3;refGene_tx
=XM_019823042.2;refGene_gn=CSMD3;refGene_tx=XM_019823044.2;refGene_gn=
CSMD3;refGene_tx=XM_019823045.2
F2 60108744 60108857
refGene_gn=CSMD3;refGene_tx=XM_011291447.3;refGene_gn=CSMD3;refGene_tx
=XM_019823042.2;refGene_gn=CSMD3;refGene_tx=XM_019823044.2;refGene_gn=
CSMD3;refGene_tx=XM_019823045.2
F2 60166751 60166974
refGene_gn=CSMD3;refGene_tx=XM_011291447.3;refGene_gn=CSMD3;refGene_tx
=XM_019823042.2;refGene_gn=CSMD3;refGene_tx=XM_019823044.2;refGene_gn=
CSMD3;refGene_tx=XM_019823045.2
F2 60299565 60299743
refGene_gn=CSMD3;refGene_tx=XM_011291447.3;refGene_gn=CSMD3;refGene_tx
=XM_019823042.2;refGene_gn=CSMD3;refGene_tx=XM_019823044.2;refGene_gn=
CSMD3;refGene_tx=XM_019823045.2
F2 61828458 61829520
refGene_gn=TRPS1;refGene_tx=XM_006943352.4;refGene_gn=TRPS1;refGene_tx
=XM_019823047.2;refGene_gn=TRPS1;refGene_tx=XM_019823048.2;refGene_gn=
TRPS1;refGene_tx=XM_019823049.2;refGene_gn=TRPS1;refGene_tx=XM_0198230
50.2
F2 61832786 61832909
refGene_gn=TRPS1;refGene_tx=XM_006943352.4;refGene_gn=TRPS1;refGene_tx
=XM_019823047.2;refGene_gn=TRPS1;refGene_tx=XM_019823048.2;refGene_gn=
TRPS1;refGene_tx=XM_019823049.2;refGene_gn=TRPS1;refGene_tx=XM_0198230
50.2
F2 62000665 62001269
refGene_gn=TRPS1;refGene_tx=XM_006943352.4;refGene_gn=TRPS1;refGene_tx
=XM_019823047.2;refGene_gn=TRPS1;refGene_tx=XM_019823048.2;refGene_gn=
TRPS1;refGene_tx=XM_019823049.2;refGene_gn=TRPS1;refGene_tx=XM_0198230
50.2
F2 62018317 62019447
refGene_gn=TRPS1;refGene_tx=XM_006943352.4;refGene_gn=TRPS1;refGene_tx
=XM_019823047.2;refGene_gn=TRPS1;refGene_tx=XM_019823048.2;refGene_gn=
TRPS1;refGene_tx=XM_019823049.2;refGene_gn=TRPS1;refGene_tx=XM_0198230
50.2
F2 62033077 62034006

refGene_gn=TRPS1;refGene_tx=XM_006943352.4;refGene_gn=TRPS1;refGene_tx=XM_019823047.2;refGene_gn=TRPS1;refGene_tx=XM_019823048.2;refGene_gn=TRPS1;refGene_tx=XM_019823049.2;refGene_gn=TRPS1;refGene_tx=XM_019823050.2
F2 62037384 62037421 refGene_gn=TRPS1;refGene_tx=XM_019823047.2;refGene_gn=TRPS1;refGene_tx=XM_019823048.2;refGene_gn=TRPS1;refGene_tx=XM_019823049.2
F2 62080927 62080937 refGene_gn=TRPS1;refGene_tx=XM_019823050.2
F2 62932088 62932186 refGene_gn=EIF3H;refGene_tx=XM_004000072.3
F2 62933616 62933749 refGene_gn=EIF3H;refGene_tx=XM_004000072.3
F2 62935769 62935890 refGene_gn=EIF3H;refGene_tx=XM_004000072.3
F2 62943634 62943784 refGene_gn=EIF3H;refGene_tx=XM_004000072.3
F2 62945007 62945107 refGene_gn=EIF3H;refGene_tx=XM_004000072.3
F2 62946621 62946789 refGene_gn=EIF3H;refGene_tx=XM_004000072.3
F2 63009457 63009614 refGene_gn=EIF3H;refGene_tx=XM_004000072.3
F2 63031947 63032079 refGene_gn=EIF3H;refGene_tx=XM_004000072.3
F2 63043377 63043565 refGene_gn=UTP23;refGene_tx=XM_004000073.5
F2 63047306 63047481 refGene_gn=UTP23;refGene_tx=XM_004000073.5
F2 63048391 63048772 refGene_gn=UTP23;refGene_tx=XM_004000073.5
F2 63182288 63182480 refGene_gn=RAD21;refGene_tx=XM_004000074.5;refGene_gn=RAD21;refGene_tx=XM_006943353.4
F2 63183514 63183598 refGene_gn=RAD21;refGene_tx=XM_004000074.5;refGene_gn=RAD21;refGene_tx=XM_006943353.4
F2 63184891 63185041 refGene_gn=RAD21;refGene_tx=XM_004000074.5;refGene_gn=RAD21;refGene_tx=XM_006943353.4
F2 63187585 63187734 refGene_gn=RAD21;refGene_tx=XM_004000074.5;refGene_gn=RAD21;refGene_tx=XM_006943353.4
F2 63188170 63188330 refGene_gn=RAD21;refGene_tx=XM_004000074.5;refGene_gn=RAD21;refGene_tx=XM_006943353.4
F2 63189697 63189921 refGene_gn=RAD21;refGene_tx=XM_004000074.5;refGene_gn=RAD21;refGene_tx=XM_006943353.4
F2 63191816 63191939 refGene_gn=RAD21;refGene_tx=XM_004000074.5;refGene_gn=RAD21;refGene_tx=XM_006943353.4
F2 63192276 63192402 refGene_gn=RAD21;refGene_tx=XM_004000074.5;refGene_gn=RAD21;refGene_tx=XM_006943353.4
F2 63192873 63193080 refGene_gn=RAD21;refGene_tx=XM_004000074.5;refGene_gn=RAD21;refGene_tx=XM_006943353.4
F2 63193762 63193869 refGene_gn=RAD21;refGene_tx=XM_004000074.5;refGene_gn=RAD21;refGene_tx=XM_006943353.4
F2 63196235 63196335

refGene_gn=RAD21;refGene_tx=XM_004000074.5;refGene_gn=RAD21;refGene_tx=XM_006943353.4
F2 63197510 63197640
refGene_gn=RAD21;refGene_tx=XM_004000074.5;refGene_gn=RAD21;refGene_tx=XM_006943353.4
F2 63202462 63202606
refGene_gn=RAD21;refGene_tx=XM_004000074.5;refGene_gn=RAD21;refGene_tx=XM_006943353.4
F2 63278074 63278430
refGene_gn=LOC109495917;refGene_tx=XM_023248759.1
F2 63280999 63281051
refGene_gn=LOC109495917;refGene_tx=XM_023248759.1
F2 63446059 63446233 refGene_gn=SLC30A8;refGene_tx=XM_023248339.1
F2 63447097 63447228 refGene_gn=SLC30A8;refGene_tx=XM_023248339.1
F2 63452844 63452917 refGene_gn=SLC30A8;refGene_tx=XM_023248339.1
F2 63456528 63456598 refGene_gn=SLC30A8;refGene_tx=XM_023248339.1
F2 63459577 63459829 refGene_gn=SLC30A8;refGene_tx=XM_023248339.1
F2 63476062 63476209 refGene_gn=SLC30A8;refGene_tx=XM_023248339.1
F2 63478526 63478680 refGene_gn=SLC30A8;refGene_tx=XM_023248339.1
F2 63482796 63482947 refGene_gn=SLC30A8;refGene_tx=XM_023248339.1
F2 63485406 63485512 refGene_gn=SLC30A8;refGene_tx=XM_023248339.1
F2 63491114 63491249 refGene_gn=SLC30A8;refGene_tx=XM_023248339.1
F2 63492620 63492766 refGene_gn=SLC30A8;refGene_tx=XM_023248339.1
F2 63790699 63790876
refGene_gn=MED30;refGene_tx=XM_004000077.4;refGene_gn=MED30;refGene_tx=XM_019823052.2
F2 63798411 63798570
refGene_gn=MED30;refGene_tx=XM_004000077.4;refGene_gn=MED30;refGene_tx=XM_019823052.2
F2 63800492 63800597
refGene_gn=MED30;refGene_tx=XM_004000077.4;refGene_gn=MED30;refGene_tx=XM_019823052.2
F2 63800690 63800723 refGene_gn=MED30;refGene_tx=XM_019823052.2
F2 63810511 63810607 refGene_gn=MED30;refGene_tx=XM_004000077.4
F2 64046579 64046765 refGene_gn=EXT1;refGene_tx=XM_023248762.1
F2 64049805 64049977 refGene_gn=EXT1;refGene_tx=XM_023248762.1
F2 64052269 64052430 refGene_gn=EXT1;refGene_tx=XM_023248762.1
F2 64057749 64057839 refGene_gn=EXT1;refGene_tx=XM_023248762.1
F2 64062735 64062831 refGene_gn=EXT1;refGene_tx=XM_023248762.1
F2 64063929 64064048 refGene_gn=EXT1;refGene_tx=XM_023248762.1
F2 64066550 64066683 refGene_gn=EXT1;refGene_tx=XM_023248762.1
F2 64074674 64074794 refGene_gn=EXT1;refGene_tx=XM_023248762.1
F2 64079957 64080065 refGene_gn=EXT1;refGene_tx=XM_023248762.1
F2 64082347 64082441 refGene_gn=EXT1;refGene_tx=XM_023248762.1
F2 64332801 64333763 refGene_gn=EXT1;refGene_tx=XM_023248762.1
F2 64401601 64401624 refGene_gn=SAMD12;refGene_tx=XM_023248163.1
F2 64481609 64481638
refGene_gn=SAMD12;refGene_tx=XM_023248162.1;refGene_gn=SAMD12;refGene_tx=XM_023248164.1;refGene_gn=SAMD12;refGene_tx=XM_023248165.1
F2 64560894 64561035

refGene_gn=SAMD12;refGene_tx=XM_023248162.1;refGene_gn=SAMD12;refGene_tx=XM_023248163.1;refGene_gn=SAMD12;refGene_tx=XM_023248164.1;refGene_gn=SAMD12;refGene_tx=XM_023248165.1
F2 64616385 64616515
refGene_gn=SAMD12;refGene_tx=XM_023248162.1;refGene_gn=SAMD12;refGene_tx=XM_023248163.1;refGene_gn=SAMD12;refGene_tx=XM_023248164.1;refGene_gn=SAMD12;refGene_tx=XM_023248165.1
F2 64747370 64747549
refGene_gn=SAMD12;refGene_tx=XM_023248162.1;refGene_gn=SAMD12;refGene_tx=XM_023248163.1;refGene_gn=SAMD12;refGene_tx=XM_023248164.1;refGene_gn=SAMD12;refGene_tx=XM_023248165.1
F2 64749022 64749128
refGene_gn=SAMD12;refGene_tx=XM_023248162.1;refGene_gn=SAMD12;refGene_tx=XM_023248163.1
F2 64784091 64784113 refGene_gn=SAMD12;refGene_tx=XM_023248164.1
F2 64785047 64785060 refGene_gn=SAMD12;refGene_tx=XM_023248165.1
F2 65039604 65039993
refGene_gn=TNFRSF11B;refGene_tx=XM_004000078.5
F2 65042461 65042686
refGene_gn=TNFRSF11B;refGene_tx=XM_004000078.5
F2 65045484 65045676
refGene_gn=TNFRSF11B;refGene_tx=XM_004000078.5
F2 65049466 65049836
refGene_gn=TNFRSF11B;refGene_tx=XM_004000078.5
F2 65068282 65068312
refGene_gn=TNFRSF11B;refGene_tx=XM_004000078.5
F2 65167520 65167668 refGene_gn=COLEC10;refGene_tx=XM_023248493.1
F2 65188839 65188911 refGene_gn=COLEC10;refGene_tx=XM_023248493.1
F2 65190269 65190341 refGene_gn=COLEC10;refGene_tx=XM_023248493.1
F2 65200290 65200344 refGene_gn=COLEC10;refGene_tx=XM_023248493.1
F2 65201973 65202069 refGene_gn=COLEC10;refGene_tx=XM_023248493.1
F2 65203919 65204311 refGene_gn=COLEC10;refGene_tx=XM_023248493.1
F2 65275926 65276058 refGene_gn=MAL2;refGene_tx=XM_004000102.5
F2 65289167 65289338 refGene_gn=MAL2;refGene_tx=XM_004000102.5
F2 65299471 65299627 refGene_gn=MAL2;refGene_tx=XM_004000102.5
F2 65302447 65302519 refGene_gn=MAL2;refGene_tx=XM_004000102.5
F2 65396065 65396137 refGene_gn=NOV;refGene_tx=XM_004000080.5
F2 65396264 65396490 refGene_gn=NOV;refGene_tx=XM_004000080.5
F2 65397525 65397777 refGene_gn=NOV;refGene_tx=XM_004000080.5
F2 65398986 65399201 refGene_gn=NOV;refGene_tx=XM_004000080.5
F2 65402545 65402842 refGene_gn=NOV;refGene_tx=XM_004000080.5
F2 65482266 65482437
refGene_gn=ENPP2;refGene_tx=XM_004000081.5;refGene_gn=ENPP2;refGene_tx=XM_004000082.5;refGene_gn=ENPP2;refGene_tx=XM_004000083.5;refGene_gn=ENPP2;refGene_tx=XM_019823057.2;refGene_gn=ENPP2;refGene_tx=XM_019823059.2;refGene_gn=ENPP2;refGene_tx=XM_019823060.2;refGene_gn=ENPP2;refGene_tx=XM_019823061.1;refGene_gn=ENPP2;refGene_tx=XM_019823062.2;refGene_gn=ENPP2;refGene_tx=XM_019823063.2;refGene_gn=ENPP2;refGene_tx=XM_019823064.2;refGene_gn=ENPP2;refGene_tx=XM_019823065.2
F2 65487473 65487630

refGene_gn=ENPP2;refGene_tx=XM_004000081.5;refGene_gn=ENPP2;refGene_tx=XM_004000082.5;refGene_gn=ENPP2;refGene_tx=XM_004000083.5;refGene_gn=ENPP2;refGene_tx=XM_019823057.2;refGene_gn=ENPP2;refGene_tx=XM_019823059.2;refGene_gn=ENPP2;refGene_tx=XM_019823060.2;refGene_gn=ENPP2;refGene_tx=XM_019823061.1;refGene_gn=ENPP2;refGene_tx=XM_019823062.2;refGene_gn=ENPP2;refGene_tx=XM_019823063.2;refGene_gn=ENPP2;refGene_tx=XM_019823064.2;refGene_gn=ENPP2;refGene_tx=XM_019823065.2
F2 65489091 65489224
refGene_gn=ENPP2;refGene_tx=XM_004000081.5;refGene_gn=ENPP2;refGene_tx=XM_004000082.5;refGene_gn=ENPP2;refGene_tx=XM_004000083.5;refGene_gn=ENPP2;refGene_tx=XM_019823057.2;refGene_gn=ENPP2;refGene_tx=XM_019823059.2;refGene_gn=ENPP2;refGene_tx=XM_019823060.2;refGene_gn=ENPP2;refGene_tx=XM_019823061.1;refGene_gn=ENPP2;refGene_tx=XM_019823062.2;refGene_gn=ENPP2;refGene_tx=XM_019823063.2;refGene_gn=ENPP2;refGene_tx=XM_019823064.2;refGene_gn=ENPP2;refGene_tx=XM_019823065.2
F2 65492332 65492410
refGene_gn=ENPP2;refGene_tx=XM_004000081.5;refGene_gn=ENPP2;refGene_tx=XM_004000082.5;refGene_gn=ENPP2;refGene_tx=XM_004000083.5;refGene_gn=ENPP2;refGene_tx=XM_019823057.2;refGene_gn=ENPP2;refGene_tx=XM_019823059.2;refGene_gn=ENPP2;refGene_tx=XM_019823060.2;refGene_gn=ENPP2;refGene_tx=XM_019823061.1;refGene_gn=ENPP2;refGene_tx=XM_019823062.2;refGene_gn=ENPP2;refGene_tx=XM_019823063.2;refGene_gn=ENPP2;refGene_tx=XM_019823064.2;refGene_gn=ENPP2;refGene_tx=XM_019823065.2
F2 65493306 65493442
refGene_gn=ENPP2;refGene_tx=XM_004000081.5;refGene_gn=ENPP2;refGene_tx=XM_004000082.5;refGene_gn=ENPP2;refGene_tx=XM_004000083.5;refGene_gn=ENPP2;refGene_tx=XM_019823057.2;refGene_gn=ENPP2;refGene_tx=XM_019823059.2;refGene_gn=ENPP2;refGene_tx=XM_019823060.2;refGene_gn=ENPP2;refGene_tx=XM_019823061.1;refGene_gn=ENPP2;refGene_tx=XM_019823062.2;refGene_gn=ENPP2;refGene_tx=XM_019823063.2;refGene_gn=ENPP2;refGene_tx=XM_019823064.2;refGene_gn=ENPP2;refGene_tx=XM_019823065.2
F2 65494693 65494830
refGene_gn=ENPP2;refGene_tx=XM_004000081.5;refGene_gn=ENPP2;refGene_tx=XM_004000082.5;refGene_gn=ENPP2;refGene_tx=XM_004000083.5;refGene_gn=ENPP2;refGene_tx=XM_019823057.2;refGene_gn=ENPP2;refGene_tx=XM_019823059.2;refGene_gn=ENPP2;refGene_tx=XM_019823060.2;refGene_gn=ENPP2;refGene_tx=XM_019823061.1;refGene_gn=ENPP2;refGene_tx=XM_019823062.2;refGene_gn=ENPP2;refGene_tx=XM_019823063.2;refGene_gn=ENPP2;refGene_tx=XM_019823064.2;refGene_gn=ENPP2;refGene_tx=XM_019823065.2
F2 65496156 65496231
refGene_gn=ENPP2;refGene_tx=XM_004000083.5;refGene_gn=ENPP2;refGene_tx=XM_019823057.2;refGene_gn=ENPP2;refGene_tx=XM_019823059.2;refGene_gn=ENPP2;refGene_tx=XM_019823060.2;refGene_gn=ENPP2;refGene_tx=XM_019823061.1;refGene_gn=ENPP2;refGene_tx=XM_019823064.2
F2 65505782 65505834
refGene_gn=ENPP2;refGene_tx=XM_004000081.5;refGene_gn=ENPP2;refGene_tx=XM_004000082.5;refGene_gn=ENPP2;refGene_tx=XM_004000083.5;refGene_gn=ENPP2;refGene_tx=XM_019823057.2;refGene_gn=ENPP2;refGene_tx=XM_019823059.2;refGene_gn=ENPP2;refGene_tx=XM_019823060.2;refGene_gn=ENPP2;refGene_tx=XM_019823061.1;refGene_gn=ENPP2;refGene_tx=XM_019823062.2;refGen

e_gn=ENPP2;refGene_tx=XM_019823063.2;refGene_gn=ENPP2;refGene_tx=XM_019823064.2;refGene_gn=ENPP2;refGene_tx=XM_019823065.2
F2 65508290 65508475
refGene_gn=ENPP2;refGene_tx=XM_004000081.5;refGene_gn=ENPP2;refGene_tx=XM_004000082.5;refGene_gn=ENPP2;refGene_tx=XM_004000083.5;refGene_gn=ENPP2;refGene_tx=XM_019823057.2;refGene_gn=ENPP2;refGene_tx=XM_019823059.2;refGene_gn=ENPP2;refGene_tx=XM_019823060.2;refGene_gn=ENPP2;refGene_tx=XM_019823061.1;refGene_gn=ENPP2;refGene_tx=XM_019823062.2;refGene_gn=ENPP2;refGene_tx=XM_019823063.2;refGene_gn=ENPP2;refGene_tx=XM_019823064.2;refGene_gn=ENPP2;refGene_tx=XM_019823065.2
F2 65509579 65509667
refGene_gn=ENPP2;refGene_tx=XM_004000081.5;refGene_gn=ENPP2;refGene_tx=XM_004000082.5;refGene_gn=ENPP2;refGene_tx=XM_004000083.5;refGene_gn=ENPP2;refGene_tx=XM_019823057.2;refGene_gn=ENPP2;refGene_tx=XM_019823059.2;refGene_gn=ENPP2;refGene_tx=XM_019823060.2;refGene_gn=ENPP2;refGene_tx=XM_019823061.1;refGene_gn=ENPP2;refGene_tx=XM_019823062.2;refGene_gn=ENPP2;refGene_tx=XM_019823063.2;refGene_gn=ENPP2;refGene_tx=XM_019823064.2;refGene_gn=ENPP2;refGene_tx=XM_019823065.2
F2 65509828 65509916
refGene_gn=ENPP2;refGene_tx=XM_004000081.5;refGene_gn=ENPP2;refGene_tx=XM_004000082.5;refGene_gn=ENPP2;refGene_tx=XM_004000083.5;refGene_gn=ENPP2;refGene_tx=XM_019823057.2;refGene_gn=ENPP2;refGene_tx=XM_019823059.2;refGene_gn=ENPP2;refGene_tx=XM_019823060.2;refGene_gn=ENPP2;refGene_tx=XM_019823061.1;refGene_gn=ENPP2;refGene_tx=XM_019823062.2;refGene_gn=ENPP2;refGene_tx=XM_019823063.2;refGene_gn=ENPP2;refGene_tx=XM_019823064.2;refGene_gn=ENPP2;refGene_tx=XM_019823065.2
F2 65512168 65512296
refGene_gn=ENPP2;refGene_tx=XM_004000081.5;refGene_gn=ENPP2;refGene_tx=XM_004000082.5;refGene_gn=ENPP2;refGene_tx=XM_004000083.5;refGene_gn=ENPP2;refGene_tx=XM_019823057.2;refGene_gn=ENPP2;refGene_tx=XM_019823059.2;refGene_gn=ENPP2;refGene_tx=XM_019823060.2;refGene_gn=ENPP2;refGene_tx=XM_019823061.1;refGene_gn=ENPP2;refGene_tx=XM_019823062.2;refGene_gn=ENPP2;refGene_tx=XM_019823063.2;refGene_gn=ENPP2;refGene_tx=XM_019823064.2;refGene_gn=ENPP2;refGene_tx=XM_019823065.2
F2 65512987 65513019
refGene_gn=ENPP2;refGene_tx=XM_004000081.5;refGene_gn=ENPP2;refGene_tx=XM_004000082.5;refGene_gn=ENPP2;refGene_tx=XM_004000083.5;refGene_gn=ENPP2;refGene_tx=XM_019823057.2;refGene_gn=ENPP2;refGene_tx=XM_019823059.2;refGene_gn=ENPP2;refGene_tx=XM_019823060.2;refGene_gn=ENPP2;refGene_tx=XM_019823061.1;refGene_gn=ENPP2;refGene_tx=XM_019823062.2;refGene_gn=ENPP2;refGene_tx=XM_019823063.2;refGene_gn=ENPP2;refGene_tx=XM_019823064.2;refGene_gn=ENPP2;refGene_tx=XM_019823065.2
F2 65516756 65516882
refGene_gn=ENPP2;refGene_tx=XM_004000081.5;refGene_gn=ENPP2;refGene_tx=XM_004000082.5;refGene_gn=ENPP2;refGene_tx=XM_004000083.5;refGene_gn=ENPP2;refGene_tx=XM_019823057.2;refGene_gn=ENPP2;refGene_tx=XM_019823059.2;refGene_gn=ENPP2;refGene_tx=XM_019823060.2;refGene_gn=ENPP2;refGene_tx=XM_019823061.1;refGene_gn=ENPP2;refGene_tx=XM_019823062.2;refGene_gn=ENPP2;refGene_tx=XM_019823063.2;refGene_gn=ENPP2;refGene_tx=XM_019823064.2;refGene_gn=ENPP2;refGene_tx=XM_019823065.2

F2 65519645 65519754
refGene_gn=ENPP2;refGene_tx=XM_004000081.5;refGene_gn=ENPP2;refGene_tx=XM_004000082.5;refGene_gn=ENPP2;refGene_tx=XM_004000083.5;refGene_gn=ENPP2;refGene_tx=XM_019823057.2;refGene_gn=ENPP2;refGene_tx=XM_019823059.2;refGene_gn=ENPP2;refGene_tx=XM_019823060.2;refGene_gn=ENPP2;refGene_tx=XM_019823061.1;refGene_gn=ENPP2;refGene_tx=XM_019823062.2;refGene_gn=ENPP2;refGene_tx=XM_019823063.2;refGene_gn=ENPP2;refGene_tx=XM_019823064.2;refGene_gn=ENPP2;refGene_tx=XM_019823065.2
F2 65521795 65521951
refGene_gn=ENPP2;refGene_tx=XM_004000082.5;refGene_gn=ENPP2;refGene_tx=XM_019823057.2;refGene_gn=ENPP2;refGene_tx=XM_019823059.2;refGene_gn=ENPP2;refGene_tx=XM_019823060.2;refGene_gn=ENPP2;refGene_tx=XM_019823061.1;refGene_gn=ENPP2;refGene_tx=XM_019823062.2;refGene_gn=ENPP2;refGene_tx=XM_019823063.2
F2 65526812 65526885
refGene_gn=ENPP2;refGene_tx=XM_004000081.5;refGene_gn=ENPP2;refGene_tx=XM_004000082.5;refGene_gn=ENPP2;refGene_tx=XM_004000083.5;refGene_gn=ENPP2;refGene_tx=XM_019823057.2;refGene_gn=ENPP2;refGene_tx=XM_019823059.2;refGene_gn=ENPP2;refGene_tx=XM_019823060.2;refGene_gn=ENPP2;refGene_tx=XM_019823061.1;refGene_gn=ENPP2;refGene_tx=XM_019823062.2;refGene_gn=ENPP2;refGene_tx=XM_019823063.2;refGene_gn=ENPP2;refGene_tx=XM_019823064.2;refGene_gn=ENPP2;refGene_tx=XM_019823065.2
F2 65527578 65527644
refGene_gn=ENPP2;refGene_tx=XM_004000081.5;refGene_gn=ENPP2;refGene_tx=XM_004000082.5;refGene_gn=ENPP2;refGene_tx=XM_004000083.5;refGene_gn=ENPP2;refGene_tx=XM_019823057.2;refGene_gn=ENPP2;refGene_tx=XM_019823059.2;refGene_gn=ENPP2;refGene_tx=XM_019823060.2;refGene_gn=ENPP2;refGene_tx=XM_019823061.1;refGene_gn=ENPP2;refGene_tx=XM_019823062.2;refGene_gn=ENPP2;refGene_tx=XM_019823063.2;refGene_gn=ENPP2;refGene_tx=XM_019823064.2;refGene_gn=ENPP2;refGene_tx=XM_019823065.2
F2 65532470 65532526
refGene_gn=ENPP2;refGene_tx=XM_004000081.5;refGene_gn=ENPP2;refGene_tx=XM_004000082.5;refGene_gn=ENPP2;refGene_tx=XM_004000083.5;refGene_gn=ENPP2;refGene_tx=XM_019823057.2;refGene_gn=ENPP2;refGene_tx=XM_019823059.2;refGene_gn=ENPP2;refGene_tx=XM_019823060.2;refGene_gn=ENPP2;refGene_tx=XM_019823061.1;refGene_gn=ENPP2;refGene_tx=XM_019823062.2;refGene_gn=ENPP2;refGene_tx=XM_019823063.2;refGene_gn=ENPP2;refGene_tx=XM_019823064.2;refGene_gn=ENPP2;refGene_tx=XM_019823065.2
F2 65540898 65541018
refGene_gn=ENPP2;refGene_tx=XM_004000081.5;refGene_gn=ENPP2;refGene_tx=XM_004000082.5;refGene_gn=ENPP2;refGene_tx=XM_004000083.5;refGene_gn=ENPP2;refGene_tx=XM_019823057.2;refGene_gn=ENPP2;refGene_tx=XM_019823059.2;refGene_gn=ENPP2;refGene_tx=XM_019823060.2;refGene_gn=ENPP2;refGene_tx=XM_019823061.1;refGene_gn=ENPP2;refGene_tx=XM_019823062.2;refGene_gn=ENPP2;refGene_tx=XM_019823063.2;refGene_gn=ENPP2;refGene_tx=XM_019823064.2;refGene_gn=ENPP2;refGene_tx=XM_019823065.2
F2 65542040 65542120
refGene_gn=ENPP2;refGene_tx=XM_004000081.5;refGene_gn=ENPP2;refGene_tx=XM_004000082.5;refGene_gn=ENPP2;refGene_tx=XM_004000083.5;refGene_gn=ENPP2;refGene_tx=XM_019823057.2;refGene_gn=ENPP2;refGene_tx=XM_0198230

59.2;refGene_gn=ENPP2;refGene_tx=XM_019823060.2;refGene_gn=ENPP2;refGene_tx=XM_019823061.1;refGene_gn=ENPP2;refGene_tx=XM_019823062.2;refGene_gn=ENPP2;refGene_tx=XM_019823063.2;refGene_gn=ENPP2;refGene_tx=XM_019823064.2;refGene_gn=ENPP2;refGene_tx=XM_019823065.2
F2 65542338 65542436 refGene_gn=ENPP2;refGene_tx=XM_004000081.5;refGene_gn=ENPP2;refGene_tx=XM_004000082.5;refGene_gn=ENPP2;refGene_tx=XM_004000083.5;refGene_gn=ENPP2;refGene_tx=XM_019823057.2;refGene_gn=ENPP2;refGene_tx=XM_019823059.2;refGene_gn=ENPP2;refGene_tx=XM_019823060.2;refGene_gn=ENPP2;refGene_tx=XM_019823061.1;refGene_gn=ENPP2;refGene_tx=XM_019823062.2;refGene_gn=ENPP2;refGene_tx=XM_019823063.2;refGene_gn=ENPP2;refGene_tx=XM_019823064.2;refGene_gn=ENPP2;refGene_tx=XM_019823065.2
F2 65544258 65544319 refGene_gn=ENPP2;refGene_tx=XM_004000081.5;refGene_gn=ENPP2;refGene_tx=XM_004000082.5;refGene_gn=ENPP2;refGene_tx=XM_004000083.5;refGene_gn=ENPP2;refGene_tx=XM_019823057.2;refGene_gn=ENPP2;refGene_tx=XM_019823059.2;refGene_gn=ENPP2;refGene_tx=XM_019823060.2;refGene_gn=ENPP2;refGene_tx=XM_019823061.1;refGene_gn=ENPP2;refGene_tx=XM_019823062.2;refGene_gn=ENPP2;refGene_tx=XM_019823063.2;refGene_gn=ENPP2;refGene_tx=XM_019823064.2;refGene_gn=ENPP2;refGene_tx=XM_019823065.2
F2 65546547 65546673 refGene_gn=ENPP2;refGene_tx=XM_004000081.5;refGene_gn=ENPP2;refGene_tx=XM_004000082.5;refGene_gn=ENPP2;refGene_tx=XM_004000083.5;refGene_gn=ENPP2;refGene_tx=XM_019823057.2;refGene_gn=ENPP2;refGene_tx=XM_019823059.2;refGene_gn=ENPP2;refGene_tx=XM_019823060.2;refGene_gn=ENPP2;refGene_tx=XM_019823061.1;refGene_gn=ENPP2;refGene_tx=XM_019823062.2;refGene_gn=ENPP2;refGene_tx=XM_019823063.2;refGene_gn=ENPP2;refGene_tx=XM_019823064.2;refGene_gn=ENPP2;refGene_tx=XM_019823065.2
F2 65551132 65551288 refGene_gn=ENPP2;refGene_tx=XM_004000081.5;refGene_gn=ENPP2;refGene_tx=XM_004000082.5;refGene_gn=ENPP2;refGene_tx=XM_004000083.5;refGene_gn=ENPP2;refGene_tx=XM_019823057.2;refGene_gn=ENPP2;refGene_tx=XM_019823059.2;refGene_gn=ENPP2;refGene_tx=XM_019823060.2;refGene_gn=ENPP2;refGene_tx=XM_019823061.1;refGene_gn=ENPP2;refGene_tx=XM_019823062.2;refGene_gn=ENPP2;refGene_tx=XM_019823063.2;refGene_gn=ENPP2;refGene_tx=XM_019823064.2;refGene_gn=ENPP2;refGene_tx=XM_019823065.2
F2 65561938 65562041 refGene_gn=ENPP2;refGene_tx=XM_004000081.5;refGene_gn=ENPP2;refGene_tx=XM_004000082.5;refGene_gn=ENPP2;refGene_tx=XM_004000083.5;refGene_gn=ENPP2;refGene_tx=XM_019823057.2;refGene_gn=ENPP2;refGene_tx=XM_019823059.2;refGene_gn=ENPP2;refGene_tx=XM_019823060.2;refGene_gn=ENPP2;refGene_tx=XM_019823061.1;refGene_gn=ENPP2;refGene_tx=XM_019823062.2;refGene_gn=ENPP2;refGene_tx=XM_019823063.2;refGene_gn=ENPP2;refGene_tx=XM_019823064.2;refGene_gn=ENPP2;refGene_tx=XM_019823065.2
F2 65562248 65562281 refGene_gn=ENPP2;refGene_tx=XM_004000081.5;refGene_gn=ENPP2;refGene_tx=XM_004000082.5;refGene_gn=ENPP2;refGene_tx=XM_004000083.5;refGene_gn=ENPP2;refGene_tx=XM_019823060.2
F2 65591989 65592010 refGene_gn=ENPP2;refGene_tx=XM_019823061.1
F2 65593360 65593417

refGene_gn=ENPP2;refGene_tx=XM_019823057.2;refGene_gn=ENPP2;refGene_tx=XM_019823059.2;refGene_gn=ENPP2;refGene_tx=XM_019823062.2;refGene_gn=ENPP2;refGene_tx=XM_019823063.2;refGene_gn=ENPP2;refGene_tx=XM_019823064.2;refGene_gn=ENPP2;refGene_tx=XM_019823065.2
F2 65642471 65642734
refGene_gn=TAF2;refGene_tx=XM_023248217.1;refGene_gn=TAF2;refGene_tx=XM_023248218.1;refGene_gn=TAF2;refGene_tx=XM_023248219.1;refGene_gn=TAF2;refGene_tx=XM_023248220.1
F2 65652747 65652870
refGene_gn=TAF2;refGene_tx=XM_023248217.1;refGene_gn=TAF2;refGene_tx=XM_023248218.1;refGene_gn=TAF2;refGene_tx=XM_023248219.1;refGene_gn=TAF2;refGene_tx=XM_023248220.1
F2 65656210 65656316
refGene_gn=TAF2;refGene_tx=XM_023248217.1;refGene_gn=TAF2;refGene_tx=XM_023248218.1;refGene_gn=TAF2;refGene_tx=XM_023248219.1;refGene_gn=TAF2;refGene_tx=XM_023248220.1
F2 65656798 65656954
refGene_gn=TAF2;refGene_tx=XM_023248217.1;refGene_gn=TAF2;refGene_tx=XM_023248219.1;refGene_gn=TAF2;refGene_tx=XM_023248220.1
F2 65658241 65658471
refGene_gn=TAF2;refGene_tx=XM_023248217.1;refGene_gn=TAF2;refGene_tx=XM_023248218.1;refGene_gn=TAF2;refGene_tx=XM_023248219.1;refGene_gn=TAF2;refGene_tx=XM_023248220.1
F2 65676047 65676157
refGene_gn=TAF2;refGene_tx=XM_023248217.1;refGene_gn=TAF2;refGene_tx=XM_023248218.1;refGene_gn=TAF2;refGene_tx=XM_023248219.1;refGene_gn=TAF2;refGene_tx=XM_023248220.1
F2 65684990 65685060
refGene_gn=TAF2;refGene_tx=XM_023248217.1;refGene_gn=TAF2;refGene_tx=XM_023248218.1;refGene_gn=TAF2;refGene_tx=XM_023248219.1;refGene_gn=TAF2;refGene_tx=XM_023248220.1
F2 65688120 65688260
refGene_gn=TAF2;refGene_tx=XM_023248217.1;refGene_gn=TAF2;refGene_tx=XM_023248218.1;refGene_gn=TAF2;refGene_tx=XM_023248219.1;refGene_gn=TAF2;refGene_tx=XM_023248220.1
F2 65688881 65689075
refGene_gn=TAF2;refGene_tx=XM_023248217.1;refGene_gn=TAF2;refGene_tx=XM_023248218.1;refGene_gn=TAF2;refGene_tx=XM_023248219.1;refGene_gn=TAF2;refGene_tx=XM_023248220.1
F2 65693693 65693804
refGene_gn=TAF2;refGene_tx=XM_023248217.1;refGene_gn=TAF2;refGene_tx=XM_023248218.1;refGene_gn=TAF2;refGene_tx=XM_023248219.1;refGene_gn=TAF2;refGene_tx=XM_023248220.1
F2 65696396 65696537
refGene_gn=TAF2;refGene_tx=XM_023248217.1;refGene_gn=TAF2;refGene_tx=XM_023248218.1;refGene_gn=TAF2;refGene_tx=XM_023248219.1;refGene_gn=TAF2;refGene_tx=XM_023248220.1
F2 65697902 65698155
refGene_gn=TAF2;refGene_tx=XM_023248217.1;refGene_gn=TAF2;refGene_tx=XM_023248218.1;refGene_gn=TAF2;refGene_tx=XM_023248219.1;refGene_gn=TAF

2;refGene_tx=XM_023248220.1
F2 65699489 65699555
refGene_gn=TAF2;refGene_tx=XM_023248217.1;refGene_gn=TAF2;refGene_tx=XM_023248218.1;refGene_gn=TAF2;refGene_tx=XM_023248219.1;refGene_gn=TAF2;refGene_tx=XM_023248220.1
F2 65702383 65702493
refGene_gn=TAF2;refGene_tx=XM_023248217.1;refGene_gn=TAF2;refGene_tx=XM_023248218.1;refGene_gn=TAF2;refGene_tx=XM_023248219.1;refGene_gn=TAF2;refGene_tx=XM_023248220.1
F2 65703074 65703189
refGene_gn=TAF2;refGene_tx=XM_023248217.1;refGene_gn=TAF2;refGene_tx=XM_023248218.1;refGene_gn=TAF2;refGene_tx=XM_023248219.1;refGene_gn=TAF2;refGene_tx=XM_023248220.1
F2 65703878 65704033
refGene_gn=TAF2;refGene_tx=XM_023248217.1;refGene_gn=TAF2;refGene_tx=XM_023248218.1;refGene_gn=TAF2;refGene_tx=XM_023248219.1;refGene_gn=TAF2;refGene_tx=XM_023248220.1
F2 65705555 65705691
refGene_gn=TAF2;refGene_tx=XM_023248217.1;refGene_gn=TAF2;refGene_tx=XM_023248218.1;refGene_gn=TAF2;refGene_tx=XM_023248219.1;refGene_gn=TAF2;refGene_tx=XM_023248220.1
F2 65708798 65708884
refGene_gn=TAF2;refGene_tx=XM_023248217.1;refGene_gn=TAF2;refGene_tx=XM_023248218.1;refGene_gn=TAF2;refGene_tx=XM_023248219.1;refGene_gn=TAF2;refGene_tx=XM_023248220.1
F2 65710471 65710571
refGene_gn=TAF2;refGene_tx=XM_023248217.1;refGene_gn=TAF2;refGene_tx=XM_023248218.1;refGene_gn=TAF2;refGene_tx=XM_023248219.1;refGene_gn=TAF2;refGene_tx=XM_023248220.1
F2 65711864 65711978
refGene_gn=TAF2;refGene_tx=XM_023248217.1;refGene_gn=TAF2;refGene_tx=XM_023248218.1;refGene_gn=TAF2;refGene_tx=XM_023248219.1;refGene_gn=TAF2;refGene_tx=XM_023248220.1
F2 65713334 65713519
refGene_gn=TAF2;refGene_tx=XM_023248217.1;refGene_gn=TAF2;refGene_tx=XM_023248218.1;refGene_gn=TAF2;refGene_tx=XM_023248219.1;refGene_gn=TAF2;refGene_tx=XM_023248220.1
F2 65716521 65716753
refGene_gn=TAF2;refGene_tx=XM_023248217.1;refGene_gn=TAF2;refGene_tx=XM_023248218.1;refGene_gn=TAF2;refGene_tx=XM_023248219.1;refGene_gn=TAF2;refGene_tx=XM_023248220.1
F2 65718234 65718376
refGene_gn=TAF2;refGene_tx=XM_023248217.1;refGene_gn=TAF2;refGene_tx=XM_023248218.1
F2 65719694 65719813
refGene_gn=TAF2;refGene_tx=XM_023248217.1;refGene_gn=TAF2;refGene_tx=XM_023248218.1
F2 65728526 65728687
refGene_gn=TAF2;refGene_tx=XM_023248217.1;refGene_gn=TAF2;refGene_tx=XM_023248218.1

F2 65738335 65738390
refGene_gn=TAF2;refGene_tx=XM_023248217.1;refGene_gn=TAF2;refGene_tx=XM_023248218.1
F2 65739141 65739224
refGene_gn=TAF2;refGene_tx=XM_023248217.1;refGene_gn=TAF2;refGene_tx=XM_023248218.1
F2 65741128 65741237
refGene_gn=DSCC1;refGene_tx=XM_004000085.5;refGene_gn=DSCC1;refGene_tx=XM_023248680.1
F2 65744436 65744585
refGene_gn=DSCC1;refGene_tx=XM_004000085.5;refGene_gn=DSCC1;refGene_tx=XM_023248680.1
F2 65746949 65747104
refGene_gn=DSCC1;refGene_tx=XM_004000085.5;refGene_gn=DSCC1;refGene_tx=XM_023248680.1
F2 65747529 65747582 refGene_gn=DSCC1;refGene_tx=XM_004000085.5
F2 65748054 65748193 refGene_gn=DSCC1;refGene_tx=XM_004000085.5
F2 65749568 65749659
refGene_gn=DSCC1;refGene_tx=XM_004000085.5;refGene_gn=DSCC1;refGene_tx=XM_023248680.1
F2 65752459 65752594
refGene_gn=DSCC1;refGene_tx=XM_004000085.5;refGene_gn=DSCC1;refGene_tx=XM_023248680.1
F2 65755208 65755377
refGene_gn=DSCC1;refGene_tx=XM_004000085.5;refGene_gn=DSCC1;refGene_tx=XM_023248680.1
F2 65757804 65757980
refGene_gn=DSCC1;refGene_tx=XM_004000085.5;refGene_gn=DSCC1;refGene_tx=XM_023248680.1
F2 65774155 65774283 refGene_gn=DEPTOR;refGene_tx=XM_004000086.5
F2 65816317 65816496 refGene_gn=DEPTOR;refGene_tx=XM_004000086.5
F2 65817766 65817890 refGene_gn=DEPTOR;refGene_tx=XM_004000086.5
F2 65843534 65843713 refGene_gn=DEPTOR;refGene_tx=XM_004000086.5
F2 65867171 65867357 refGene_gn=DEPTOR;refGene_tx=XM_004000086.5
F2 65869508 65869643 refGene_gn=DEPTOR;refGene_tx=XM_004000086.5
F2 65875179 65875250 refGene_gn=DEPTOR;refGene_tx=XM_004000086.5
F2 65877266 65877371 refGene_gn=DEPTOR;refGene_tx=XM_004000086.5
F2 65906635 65906764 refGene_gn=DEPTOR;refGene_tx=XM_004000086.5
F2 65979618 65979706
refGene_gn=COL14A1;refGene_tx=XM_006943360.3;refGene_gn=COL14A1;refGene_tx=XM_006943361.3;refGene_gn=COL14A1;refGene_tx=XM_006943362.3;refGene_gn=COL14A1;refGene_tx=XM_006943363.4;refGene_gn=COL14A1;refGene_tx=XM_019823072.2
F2 65985602 65985719
refGene_gn=COL14A1;refGene_tx=XM_006943360.3;refGene_gn=COL14A1;refGene_tx=XM_006943361.3;refGene_gn=COL14A1;refGene_tx=XM_006943362.3;refGene_gn=COL14A1;refGene_tx=XM_006943363.4;refGene_gn=COL14A1;refGene_tx=XM_019823072.2
F2 65989063 65989207
refGene_gn=COL14A1;refGene_tx=XM_006943360.3;refGene_gn=COL14A1;refGen

e_tx=XM_006943361.3;refGene_gn=COL14A1;refGene_tx=XM_006943362.3;refGene_gn=COL14A1;refGene_tx=XM_006943363.4;refGene_gn=COL14A1;refGene_tx=XM_019823072.2
F2 65994524 65994614
refGene_gn=COL14A1;refGene_tx=XM_006943360.3;refGene_gn=COL14A1;refGene_tx=XM_006943361.3;refGene_gn=COL14A1;refGene_tx=XM_006943362.3;refGene_gn=COL14A1;refGene_tx=XM_006943363.4;refGene_gn=COL14A1;refGene_tx=XM_019823072.2
F2 66018501 66018657
refGene_gn=COL14A1;refGene_tx=XM_006943360.3;refGene_gn=COL14A1;refGene_tx=XM_006943361.3;refGene_gn=COL14A1;refGene_tx=XM_006943362.3;refGene_gn=COL14A1;refGene_tx=XM_006943363.4;refGene_gn=COL14A1;refGene_tx=XM_019823072.2
F2 66019492 66019612
refGene_gn=COL14A1;refGene_tx=XM_006943360.3;refGene_gn=COL14A1;refGene_tx=XM_006943361.3;refGene_gn=COL14A1;refGene_tx=XM_006943362.3;refGene_gn=COL14A1;refGene_tx=XM_006943363.4;refGene_gn=COL14A1;refGene_tx=XM_019823072.2
F2 66020710 66020875
refGene_gn=COL14A1;refGene_tx=XM_006943360.3;refGene_gn=COL14A1;refGene_tx=XM_006943361.3;refGene_gn=COL14A1;refGene_tx=XM_006943362.3;refGene_gn=COL14A1;refGene_tx=XM_006943363.4;refGene_gn=COL14A1;refGene_tx=XM_019823072.2
F2 66023893 66024055
refGene_gn=COL14A1;refGene_tx=XM_006943360.3;refGene_gn=COL14A1;refGene_tx=XM_006943361.3;refGene_gn=COL14A1;refGene_tx=XM_006943362.3;refGene_gn=COL14A1;refGene_tx=XM_006943363.4;refGene_gn=COL14A1;refGene_tx=XM_019823072.2
F2 66032193 66032345
refGene_gn=COL14A1;refGene_tx=XM_006943360.3;refGene_gn=COL14A1;refGene_tx=XM_006943361.3;refGene_gn=COL14A1;refGene_tx=XM_006943362.3;refGene_gn=COL14A1;refGene_tx=XM_006943363.4;refGene_gn=COL14A1;refGene_tx=XM_019823072.2
F2 66033734 66033864
refGene_gn=COL14A1;refGene_tx=XM_006943360.3;refGene_gn=COL14A1;refGene_tx=XM_006943361.3;refGene_gn=COL14A1;refGene_tx=XM_006943362.3;refGene_gn=COL14A1;refGene_tx=XM_006943363.4;refGene_gn=COL14A1;refGene_tx=XM_019823072.2
F2 66034545 66034691
refGene_gn=COL14A1;refGene_tx=XM_006943360.3;refGene_gn=COL14A1;refGene_tx=XM_006943361.3;refGene_gn=COL14A1;refGene_tx=XM_006943362.3;refGene_gn=COL14A1;refGene_tx=XM_006943363.4;refGene_gn=COL14A1;refGene_tx=XM_019823072.2
F2 66037229 66037359
refGene_gn=COL14A1;refGene_tx=XM_006943360.3;refGene_gn=COL14A1;refGene_tx=XM_006943361.3;refGene_gn=COL14A1;refGene_tx=XM_006943362.3;refGene_gn=COL14A1;refGene_tx=XM_006943363.4;refGene_gn=COL14A1;refGene_tx=XM_019823072.2
F2 66039694 66039834
refGene_gn=COL14A1;refGene_tx=XM_006943360.3;refGene_gn=COL14A1;refGen

e_tx=XM_006943361.3;refGene_gn=COL14A1;refGene_tx=XM_006943362.3;refGene_gn=COL14A1;refGene_tx=XM_006943363.4;refGene_gn=COL14A1;refGene_tx=XM_019823072.2
F2 66046515 66046642
refGene_gn=COL14A1;refGene_tx=XM_006943360.3;refGene_gn=COL14A1;refGene_tx=XM_006943361.3;refGene_gn=COL14A1;refGene_tx=XM_006943362.3;refGene_gn=COL14A1;refGene_tx=XM_006943363.4;refGene_gn=COL14A1;refGene_tx=XM_019823072.2
F2 66048686 66048826
refGene_gn=COL14A1;refGene_tx=XM_006943360.3;refGene_gn=COL14A1;refGene_tx=XM_006943361.3;refGene_gn=COL14A1;refGene_tx=XM_006943362.3;refGene_gn=COL14A1;refGene_tx=XM_006943363.4;refGene_gn=COL14A1;refGene_tx=XM_019823072.2
F2 66049316 66049449
refGene_gn=COL14A1;refGene_tx=XM_006943360.3;refGene_gn=COL14A1;refGene_tx=XM_006943361.3;refGene_gn=COL14A1;refGene_tx=XM_006943362.3;refGene_gn=COL14A1;refGene_tx=XM_006943363.4;refGene_gn=COL14A1;refGene_tx=XM_019823072.2
F2 66050762 66050822
refGene_gn=COL14A1;refGene_tx=XM_006943360.3;refGene_gn=COL14A1;refGene_tx=XM_006943361.3;refGene_gn=COL14A1;refGene_tx=XM_006943362.3;refGene_gn=COL14A1;refGene_tx=XM_006943363.4;refGene_gn=COL14A1;refGene_tx=XM_019823072.2
F2 66052614 66052766
refGene_gn=COL14A1;refGene_tx=XM_006943360.3;refGene_gn=COL14A1;refGene_tx=XM_006943361.3;refGene_gn=COL14A1;refGene_tx=XM_006943362.3;refGene_gn=COL14A1;refGene_tx=XM_006943363.4;refGene_gn=COL14A1;refGene_tx=XM_019823072.2
F2 66058565 66058695
refGene_gn=COL14A1;refGene_tx=XM_006943360.3;refGene_gn=COL14A1;refGene_tx=XM_006943361.3;refGene_gn=COL14A1;refGene_tx=XM_006943362.3;refGene_gn=COL14A1;refGene_tx=XM_006943363.4;refGene_gn=COL14A1;refGene_tx=XM_019823072.2
F2 66062429 66062552
refGene_gn=COL14A1;refGene_tx=XM_006943360.3;refGene_gn=COL14A1;refGene_tx=XM_006943361.3;refGene_gn=COL14A1;refGene_tx=XM_006943362.3;refGene_gn=COL14A1;refGene_tx=XM_006943363.4;refGene_gn=COL14A1;refGene_tx=XM_019823072.2
F2 66066448 66066598
refGene_gn=COL14A1;refGene_tx=XM_006943360.3;refGene_gn=COL14A1;refGene_tx=XM_006943361.3;refGene_gn=COL14A1;refGene_tx=XM_006943362.3;refGene_gn=COL14A1;refGene_tx=XM_006943363.4;refGene_gn=COL14A1;refGene_tx=XM_019823072.2
F2 66069578 66069695
refGene_gn=COL14A1;refGene_tx=XM_006943360.3;refGene_gn=COL14A1;refGene_tx=XM_006943361.3;refGene_gn=COL14A1;refGene_tx=XM_006943362.3;refGene_gn=COL14A1;refGene_tx=XM_006943363.4;refGene_gn=COL14A1;refGene_tx=XM_019823072.2
F2 66077464 66077611
refGene_gn=COL14A1;refGene_tx=XM_006943360.3;refGene_gn=COL14A1;refGen

e_tx=XM_006943361.3;refGene_gn=COL14A1;refGene_tx=XM_006943362.3;refGene_gn=COL14A1;refGene_tx=XM_006943363.4;refGene_gn=COL14A1;refGene_tx=XM_019823072.2
F2 66081867 66081924
refGene_gn=COL14A1;refGene_tx=XM_006943360.3;refGene_gn=COL14A1;refGene_tx=XM_006943361.3;refGene_gn=COL14A1;refGene_tx=XM_006943362.3;refGene_gn=COL14A1;refGene_tx=XM_006943363.4;refGene_gn=COL14A1;refGene_tx=XM_019823072.2
F2 66083301 66083441
refGene_gn=COL14A1;refGene_tx=XM_006943360.3;refGene_gn=COL14A1;refGene_tx=XM_006943361.3;refGene_gn=COL14A1;refGene_tx=XM_006943362.3;refGene_gn=COL14A1;refGene_tx=XM_006943363.4;refGene_gn=COL14A1;refGene_tx=XM_019823072.2
F2 66087015 66087139
refGene_gn=COL14A1;refGene_tx=XM_006943360.3;refGene_gn=COL14A1;refGene_tx=XM_006943361.3;refGene_gn=COL14A1;refGene_tx=XM_006943362.3;refGene_gn=COL14A1;refGene_tx=XM_006943363.4;refGene_gn=COL14A1;refGene_tx=XM_019823072.2
F2 66087349 66087493
refGene_gn=COL14A1;refGene_tx=XM_006943360.3;refGene_gn=COL14A1;refGene_tx=XM_006943361.3;refGene_gn=COL14A1;refGene_tx=XM_006943362.3;refGene_gn=COL14A1;refGene_tx=XM_006943363.4;refGene_gn=COL14A1;refGene_tx=XM_019823072.2
F2 66088603 66088768
refGene_gn=COL14A1;refGene_tx=XM_006943360.3;refGene_gn=COL14A1;refGene_tx=XM_006943361.3;refGene_gn=COL14A1;refGene_tx=XM_006943362.3;refGene_gn=COL14A1;refGene_tx=XM_006943363.4;refGene_gn=COL14A1;refGene_tx=XM_019823072.2
F2 66089352 66089391
refGene_gn=COL14A1;refGene_tx=XM_006943360.3;refGene_gn=COL14A1;refGene_tx=XM_006943361.3;refGene_gn=COL14A1;refGene_tx=XM_006943362.3;refGene_gn=COL14A1;refGene_tx=XM_006943363.4;refGene_gn=COL14A1;refGene_tx=XM_019823072.2
F2 66089525 66089664
refGene_gn=COL14A1;refGene_tx=XM_006943360.3;refGene_gn=COL14A1;refGene_tx=XM_006943361.3;refGene_gn=COL14A1;refGene_tx=XM_006943362.3;refGene_gn=COL14A1;refGene_tx=XM_006943363.4;refGene_gn=COL14A1;refGene_tx=XM_019823072.2
F2 66093564 66093707
refGene_gn=COL14A1;refGene_tx=XM_006943360.3;refGene_gn=COL14A1;refGene_tx=XM_006943361.3;refGene_gn=COL14A1;refGene_tx=XM_006943362.3;refGene_gn=COL14A1;refGene_tx=XM_006943363.4;refGene_gn=COL14A1;refGene_tx=XM_019823072.2
F2 66096938 66097048
refGene_gn=COL14A1;refGene_tx=XM_006943360.3;refGene_gn=COL14A1;refGene_tx=XM_006943361.3;refGene_gn=COL14A1;refGene_tx=XM_006943362.3;refGene_gn=COL14A1;refGene_tx=XM_006943363.4;refGene_gn=COL14A1;refGene_tx=XM_019823072.2
F2 66099128 66099287
refGene_gn=COL14A1;refGene_tx=XM_006943360.3;refGene_gn=COL14A1;refGen

e_tx=XM_006943361.3;refGene_gn=COL14A1;refGene_tx=XM_006943362.3;refGene_gn=COL14A1;refGene_tx=XM_006943363.4;refGene_gn=COL14A1;refGene_tx=XM_019823072.2
F2 66108993 66109071
refGene_gn=COL14A1;refGene_tx=XM_006943360.3;refGene_gn=COL14A1;refGene_tx=XM_006943361.3;refGene_gn=COL14A1;refGene_tx=XM_006943362.3;refGene_gn=COL14A1;refGene_tx=XM_006943363.4;refGene_gn=COL14A1;refGene_tx=XM_019823072.2
F2 66112199 66112286
refGene_gn=COL14A1;refGene_tx=XM_006943360.3;refGene_gn=COL14A1;refGene_tx=XM_006943361.3;refGene_gn=COL14A1;refGene_tx=XM_006943362.3;refGene_gn=COL14A1;refGene_tx=XM_006943363.4;refGene_gn=COL14A1;refGene_tx=XM_019823072.2
F2 66122674 66122728
refGene_gn=COL14A1;refGene_tx=XM_006943360.3;refGene_gn=COL14A1;refGene_tx=XM_006943361.3;refGene_gn=COL14A1;refGene_tx=XM_006943362.3;refGene_gn=COL14A1;refGene_tx=XM_006943363.4;refGene_gn=COL14A1;refGene_tx=XM_019823072.2
F2 66126010 66126106
refGene_gn=COL14A1;refGene_tx=XM_006943360.3;refGene_gn=COL14A1;refGene_tx=XM_006943361.3;refGene_gn=COL14A1;refGene_tx=XM_006943362.3;refGene_gn=COL14A1;refGene_tx=XM_006943363.4;refGene_gn=COL14A1;refGene_tx=XM_019823072.2
F2 66127318 66127372
refGene_gn=COL14A1;refGene_tx=XM_006943360.3;refGene_gn=COL14A1;refGene_tx=XM_006943361.3;refGene_gn=COL14A1;refGene_tx=XM_006943362.3;refGene_gn=COL14A1;refGene_tx=XM_006943363.4;refGene_gn=COL14A1;refGene_tx=XM_019823072.2
F2 66127759 66127813
refGene_gn=COL14A1;refGene_tx=XM_006943360.3;refGene_gn=COL14A1;refGene_tx=XM_006943361.3;refGene_gn=COL14A1;refGene_tx=XM_006943362.3;refGene_gn=COL14A1;refGene_tx=XM_006943363.4;refGene_gn=COL14A1;refGene_tx=XM_019823072.2
F2 66142450 66142504
refGene_gn=COL14A1;refGene_tx=XM_006943360.3;refGene_gn=COL14A1;refGene_tx=XM_006943361.3;refGene_gn=COL14A1;refGene_tx=XM_006943362.3;refGene_gn=COL14A1;refGene_tx=XM_006943363.4;refGene_gn=COL14A1;refGene_tx=XM_019823072.2
F2 66142969 66143041
refGene_gn=COL14A1;refGene_tx=XM_006943360.3;refGene_gn=COL14A1;refGene_tx=XM_006943361.3;refGene_gn=COL14A1;refGene_tx=XM_006943362.3;refGene_gn=COL14A1;refGene_tx=XM_006943363.4;refGene_gn=COL14A1;refGene_tx=XM_019823072.2
F2 66152833 66152869
refGene_gn=COL14A1;refGene_tx=XM_006943360.3;refGene_gn=COL14A1;refGene_tx=XM_006943361.3;refGene_gn=COL14A1;refGene_tx=XM_006943362.3;refGene_gn=COL14A1;refGene_tx=XM_006943363.4;refGene_gn=COL14A1;refGene_tx=XM_019823072.2
F2 66153895 66153962
refGene_gn=COL14A1;refGene_tx=XM_006943360.3;refGene_gn=COL14A1;refGen

e_tx=XM_006943361.3;refGene_gn=COL14A1;refGene_tx=XM_006943362.3;refGene_gn=COL14A1;refGene_tx=XM_006943363.4;refGene_gn=COL14A1;refGene_tx=XM_019823072.2
F2 66157541 66157730
refGene_gn=COL14A1;refGene_tx=XM_006943360.3;refGene_gn=COL14A1;refGene_tx=XM_006943361.3;refGene_gn=COL14A1;refGene_tx=XM_006943362.3;refGene_gn=COL14A1;refGene_tx=XM_006943363.4;refGene_gn=COL14A1;refGene_tx=XM_019823072.2
F2 66174945 66175023
refGene_gn=COL14A1;refGene_tx=XM_006943360.3;refGene_gn=COL14A1;refGene_tx=XM_006943361.3;refGene_gn=COL14A1;refGene_tx=XM_006943362.3;refGene_gn=COL14A1;refGene_tx=XM_006943363.4;refGene_gn=COL14A1;refGene_tx=XM_019823072.2
F2 66176662 66176818
refGene_gn=COL14A1;refGene_tx=XM_006943360.3;refGene_gn=COL14A1;refGene_tx=XM_006943361.3;refGene_gn=COL14A1;refGene_tx=XM_006943362.3;refGene_gn=COL14A1;refGene_tx=XM_006943363.4;refGene_gn=COL14A1;refGene_tx=XM_019823072.2
F2 66177635 66177667 refGene_gn=COL14A1;refGene_tx=XM_006943363.4
F2 66178464 66178544
refGene_gn=COL14A1;refGene_tx=XM_006943360.3;refGene_gn=COL14A1;refGene_tx=XM_006943361.3;refGene_gn=COL14A1;refGene_tx=XM_006943362.3;refGene_gn=COL14A1;refGene_tx=XM_019823072.2
F2 66202875 66202897 refGene_gn=MRPL13;refGene_tx=XM_006943364.4
F2 66216689 66216744
refGene_gn=MRPL13;refGene_tx=XM_019823075.2;refGene_gn=MRPL13;refGene_tx=XM_019823076.2
F2 66217288 66217410
refGene_gn=MRPL13;refGene_tx=XM_006943364.4;refGene_gn=MRPL13;refGene_tx=XM_019823075.2;refGene_gn=MRPL13;refGene_tx=XM_019823076.2
F2 66223751 66223838
refGene_gn=MRPL13;refGene_tx=XM_006943364.4;refGene_gn=MRPL13;refGene_tx=XM_019823075.2;refGene_gn=MRPL13;refGene_tx=XM_019823076.2
F2 66233447 66233508
refGene_gn=MRPL13;refGene_tx=XM_006943364.4;refGene_gn=MRPL13;refGene_tx=XM_019823075.2;refGene_gn=MRPL13;refGene_tx=XM_019823076.2
F2 66239541 66239635
refGene_gn=MRPL13;refGene_tx=XM_006943364.4;refGene_gn=MRPL13;refGene_tx=XM_019823075.2;refGene_gn=MRPL13;refGene_tx=XM_019823076.2
F2 66251195 66251319
refGene_gn=MRPL13;refGene_tx=XM_006943364.4;refGene_gn=MRPL13;refGene_tx=XM_019823075.2;refGene_gn=MRPL13;refGene_tx=XM_019823076.2
F2 66254232 66254298
refGene_gn=MRPL13;refGene_tx=XM_006943364.4;refGene_gn=MRPL13;refGene_tx=XM_019823075.2
F2 66254415 66254442 refGene_gn=MRPL13;refGene_tx=XM_019823076.2
F2 66254819 66254937
refGene_gn=MTBP;refGene_tx=XM_004000088.4;refGene_gn=MTBP;refGene_tx=XM_019823073.2;refGene_gn=MTBP;refGene_tx=XM_019823074.2
F2 66255738 66255819

refGene_gn=MTBP;refGene_tx=XM_004000088.4;refGene_gn=MTBP;refGene_tx=XM_019823073.2;refGene_gn=MTBP;refGene_tx=XM_019823074.2
F2 66260257 66260331
refGene_gn=MTBP;refGene_tx=XM_004000088.4;refGene_gn=MTBP;refGene_tx=XM_019823073.2;refGene_gn=MTBP;refGene_tx=XM_019823074.2
F2 66260427 66260576
refGene_gn=MTBP;refGene_tx=XM_004000088.4;refGene_gn=MTBP;refGene_tx=XM_019823073.2;refGene_gn=MTBP;refGene_tx=XM_019823074.2
F2 66262852 66262911
refGene_gn=MTBP;refGene_tx=XM_004000088.4;refGene_gn=MTBP;refGene_tx=XM_019823073.2;refGene_gn=MTBP;refGene_tx=XM_019823074.2
F2 66264358 66264503
refGene_gn=MTBP;refGene_tx=XM_004000088.4;refGene_gn=MTBP;refGene_tx=XM_019823073.2;refGene_gn=MTBP;refGene_tx=XM_019823074.2
F2 66265668 66265786
refGene_gn=MTBP;refGene_tx=XM_004000088.4;refGene_gn=MTBP;refGene_tx=XM_019823073.2;refGene_gn=MTBP;refGene_tx=XM_019823074.2
F2 66269498 66269633
refGene_gn=MTBP;refGene_tx=XM_004000088.4;refGene_gn=MTBP;refGene_tx=XM_019823073.2;refGene_gn=MTBP;refGene_tx=XM_019823074.2
F2 66271425 66271520
refGene_gn=MTBP;refGene_tx=XM_004000088.4;refGene_gn=MTBP;refGene_tx=XM_019823073.2;refGene_gn=MTBP;refGene_tx=XM_019823074.2
F2 66272323 66272396
refGene_gn=MTBP;refGene_tx=XM_004000088.4;refGene_gn=MTBP;refGene_tx=XM_019823073.2;refGene_gn=MTBP;refGene_tx=XM_019823074.2
F2 66277649 66277767
refGene_gn=MTBP;refGene_tx=XM_004000088.4;refGene_gn=MTBP;refGene_tx=XM_019823073.2;refGene_gn=MTBP;refGene_tx=XM_019823074.2
F2 66292052 66292226
refGene_gn=MTBP;refGene_tx=XM_004000088.4;refGene_gn=MTBP;refGene_tx=XM_019823073.2;refGene_gn=MTBP;refGene_tx=XM_019823074.2
F2 66293633 66293741
refGene_gn=MTBP;refGene_tx=XM_004000088.4;refGene_gn=MTBP;refGene_tx=XM_019823073.2;refGene_gn=MTBP;refGene_tx=XM_019823074.2
F2 66306048 66306210
refGene_gn=MTBP;refGene_tx=XM_004000088.4;refGene_gn=MTBP;refGene_tx=XM_019823073.2;refGene_gn=MTBP;refGene_tx=XM_019823074.2
F2 66312096 66312214
refGene_gn=MTBP;refGene_tx=XM_004000088.4;refGene_gn=MTBP;refGene_tx=XM_019823073.2;refGene_gn=MTBP;refGene_tx=XM_019823074.2
F2 66317055 66317211
refGene_gn=MTBP;refGene_tx=XM_004000088.4;refGene_gn=MTBP;refGene_tx=XM_019823073.2;refGene_gn=MTBP;refGene_tx=XM_019823074.2
F2 66319977 66320073
refGene_gn=MTBP;refGene_tx=XM_004000088.4;refGene_gn=MTBP;refGene_tx=XM_019823074.2
F2 66327048 66327315
refGene_gn=MTBP;refGene_tx=XM_004000088.4;refGene_gn=MTBP;refGene_tx=XM_019823073.2;refGene_gn=MTBP;refGene_tx=XM_019823074.2

F2	66329245	66329495
refGene_gn=MTBP;refGene_tx=XM_004000088.4;refGene_gn=MTBP;refGene_tx=XM_019823073.2;refGene_gn=MTBP;refGene_tx=XM_019823074.2
F2	66330062	66330176
refGene_gn=MTBP;refGene_tx=XM_004000088.4;refGene_gn=MTBP;refGene_tx=XM_019823073.2
F2	66333304	66333370
refGene_gn=MTBP;refGene_tx=XM_004000088.4;refGene_gn=MTBP;refGene_tx=XM_019823073.2
F2	66333923	66333962
refGene_gn=MTBP;refGene_tx=XM_004000088.4;refGene_gn=MTBP;refGene_tx=XM_019823073.2
F2	66345338	66345431	refGene_gn=SNTB1;refGene_tx=XM_023248558.1
F2	66348310	66348501	refGene_gn=SNTB1;refGene_tx=XM_023248558.1
F2	66353917	66354114	refGene_gn=SNTB1;refGene_tx=XM_023248558.1
F2	66379841	66379981	refGene_gn=SNTB1;refGene_tx=XM_023248558.1
F2	66437898	66438106	refGene_gn=SNTB1;refGene_tx=XM_023248558.1
F2	66479415	66479632	refGene_gn=SNTB1;refGene_tx=XM_023248558.1
F2	66584539	66585092	refGene_gn=SNTB1;refGene_tx=XM_023248558.1
F2	67280665	67281595	refGene_gn=HAS2;refGene_tx=XM_004000089.4
F2	67283949	67284051	refGene_gn=HAS2;refGene_tx=XM_004000089.4
F2	67296117	67296744	refGene_gn=HAS2;refGene_tx=XM_004000089.4
F2	68381928	68384445
refGene_gn=ZHX2;refGene_tx=XM_011291450.3;refGene_gn=ZHX2;refGene_tx=XM_019823079.2;refGene_gn=ZHX2;refGene_tx=XM_019823081.2;refGene_gn=ZHX2;refGene_tx=XM_023248488.1;refGene_gn=ZHX2;refGene_tx=XM_023248489.1;refGene_gn=ZHX2;refGene_tx=XM_023248490.1;refGene_gn=ZHX2;refGene_tx=XM_023248491.1
F2	68427900	68428039	refGene_gn=DERL1;refGene_tx=XM_004000091.5
F2	68432241	68432352	refGene_gn=DERL1;refGene_tx=XM_004000091.5
F2	68434124	68434177	refGene_gn=DERL1;refGene_tx=XM_004000091.5
F2	68435322	68435418	refGene_gn=DERL1;refGene_tx=XM_004000091.5
F2	68436233	68436260	refGene_gn=DERL1;refGene_tx=XM_004000091.5
F2	68437307	68437372	refGene_gn=DERL1;refGene_tx=XM_004000091.5
F2	68442602	68442714	refGene_gn=DERL1;refGene_tx=XM_004000091.5
F2	68453193	68453346	refGene_gn=DERL1;refGene_tx=XM_004000091.5
F2	68465775	68465909
refGene_gn=TBC1D31;refGene_tx=XM_019823084.1;refGene_gn=TBC1D31;refGene_tx=XM_019823086.1;refGene_gn=TBC1D31;refGene_tx=XM_019823088.1;refGene_gn=TBC1D31;refGene_tx=XM_019823089.2;refGene_gn=TBC1D31;refGene_tx=XM_019823090.1
F2	68476414	68476491	refGene_gn=TBC1D31;refGene_tx=XM_004000105.5
F2	68480630	68480777
refGene_gn=TBC1D31;refGene_tx=XM_004000105.5;refGene_gn=TBC1D31;refGene_tx=XM_019823084.1;refGene_gn=TBC1D31;refGene_tx=XM_019823086.1;refGene_gn=TBC1D31;refGene_tx=XM_019823087.2;refGene_gn=TBC1D31;refGene_tx=XM_019823088.1;refGene_gn=TBC1D31;refGene_tx=XM_019823089.2;refGene_gn=TBC1D31;refGene_tx=XM_019823090.1;refGene_gn=TBC1D31;refGene_tx=XM_019823091.1
F2	68485358	68485474

refGene_gn=TBC1D31;refGene_tx=XM_004000105.5;refGene_gn=TBC1D31;refGene_tx=XM_019823084.1;refGene_gn=TBC1D31;refGene_tx=XM_019823086.1;refGene_gn=TBC1D31;refGene_tx=XM_019823087.2;refGene_gn=TBC1D31;refGene_tx=XM_019823088.1;refGene_gn=TBC1D31;refGene_tx=XM_019823089.2;refGene_gn=TBC1D31;refGene_tx=XM_019823090.1
F2 68489385 68489564
refGene_gn=TBC1D31;refGene_tx=XM_004000105.5;refGene_gn=TBC1D31;refGene_tx=XM_019823084.1;refGene_gn=TBC1D31;refGene_tx=XM_019823086.1;refGene_gn=TBC1D31;refGene_tx=XM_019823087.2;refGene_gn=TBC1D31;refGene_tx=XM_019823088.1;refGene_gn=TBC1D31;refGene_tx=XM_019823089.2;refGene_gn=TBC1D31;refGene_tx=XM_019823090.1;refGene_gn=TBC1D31;refGene_tx=XM_019823091.1
F2 68498366 68498518
refGene_gn=TBC1D31;refGene_tx=XM_004000105.5;refGene_gn=TBC1D31;refGene_tx=XM_019823084.1;refGene_gn=TBC1D31;refGene_tx=XM_019823086.1;refGene_gn=TBC1D31;refGene_tx=XM_019823087.2;refGene_gn=TBC1D31;refGene_tx=XM_019823088.1;refGene_gn=TBC1D31;refGene_tx=XM_019823089.2;refGene_gn=TBC1D31;refGene_tx=XM_019823090.1;refGene_gn=TBC1D31;refGene_tx=XM_019823091.1
F2 68502394 68502554
refGene_gn=TBC1D31;refGene_tx=XM_004000105.5;refGene_gn=TBC1D31;refGene_tx=XM_019823084.1;refGene_gn=TBC1D31;refGene_tx=XM_019823086.1;refGene_gn=TBC1D31;refGene_tx=XM_019823087.2;refGene_gn=TBC1D31;refGene_tx=XM_019823088.1;refGene_gn=TBC1D31;refGene_tx=XM_019823089.2;refGene_gn=TBC1D31;refGene_tx=XM_019823090.1;refGene_gn=TBC1D31;refGene_tx=XM_019823091.1
F2 68506022 68506223
refGene_gn=TBC1D31;refGene_tx=XM_004000105.5;refGene_gn=TBC1D31;refGene_tx=XM_019823084.1;refGene_gn=TBC1D31;refGene_tx=XM_019823086.1;refGene_gn=TBC1D31;refGene_tx=XM_019823087.2;refGene_gn=TBC1D31;refGene_tx=XM_019823088.1;refGene_gn=TBC1D31;refGene_tx=XM_019823089.2;refGene_gn=TBC1D31;refGene_tx=XM_019823090.1;refGene_gn=TBC1D31;refGene_tx=XM_019823091.1
F2 68510516 68510693
refGene_gn=TBC1D31;refGene_tx=XM_004000105.5;refGene_gn=TBC1D31;refGene_tx=XM_019823084.1;refGene_gn=TBC1D31;refGene_tx=XM_019823086.1;refGene_gn=TBC1D31;refGene_tx=XM_019823087.2;refGene_gn=TBC1D31;refGene_tx=XM_019823088.1;refGene_gn=TBC1D31;refGene_tx=XM_019823089.2;refGene_gn=TBC1D31;refGene_tx=XM_019823090.1;refGene_gn=TBC1D31;refGene_tx=XM_019823091.1
F2 68514154 68514234
refGene_gn=TBC1D31;refGene_tx=XM_004000105.5;refGene_gn=TBC1D31;refGene_tx=XM_019823084.1;refGene_gn=TBC1D31;refGene_tx=XM_019823086.1;refGene_gn=TBC1D31;refGene_tx=XM_019823087.2;refGene_gn=TBC1D31;refGene_tx=XM_019823088.1;refGene_gn=TBC1D31;refGene_tx=XM_019823089.2;refGene_gn=TBC1D31;refGene_tx=XM_019823090.1;refGene_gn=TBC1D31;refGene_tx=XM_019823091.1
F2 68514313 68514460
refGene_gn=TBC1D31;refGene_tx=XM_004000105.5;refGene_gn=TBC1D31;refGene_tx=XM_019823084.1;refGene_gn=TBC1D31;refGene_tx=XM_019823086.1;refGe

ne_gn=TBC1D31;refGene_tx=XM_019823087.2;refGene_gn=TBC1D31;refGene_tx=XM_019823088.1;refGene_gn=TBC1D31;refGene_tx=XM_019823089.2;refGene_gn=TBC1D31;refGene_tx=XM_019823090.1;refGene_gn=TBC1D31;refGene_tx=XM_019823091.1
F2 68516647 68516781
refGene_gn=TBC1D31;refGene_tx=XM_004000105.5;refGene_gn=TBC1D31;refGene_tx=XM_019823084.1;refGene_gn=TBC1D31;refGene_tx=XM_019823086.1;refGene_gn=TBC1D31;refGene_tx=XM_019823087.2;refGene_gn=TBC1D31;refGene_tx=XM_019823088.1;refGene_gn=TBC1D31;refGene_tx=XM_019823089.2;refGene_gn=TBC1D31;refGene_tx=XM_019823090.1;refGene_gn=TBC1D31;refGene_tx=XM_019823091.1
F2 68522305 68522439
refGene_gn=TBC1D31;refGene_tx=XM_004000105.5;refGene_gn=TBC1D31;refGene_tx=XM_019823084.1;refGene_gn=TBC1D31;refGene_tx=XM_019823086.1;refGene_gn=TBC1D31;refGene_tx=XM_019823087.2;refGene_gn=TBC1D31;refGene_tx=XM_019823088.1;refGene_gn=TBC1D31;refGene_tx=XM_019823089.2;refGene_gn=TBC1D31;refGene_tx=XM_019823090.1;refGene_gn=TBC1D31;refGene_tx=XM_019823091.1
F2 68522780 68522960
refGene_gn=TBC1D31;refGene_tx=XM_004000105.5;refGene_gn=TBC1D31;refGene_tx=XM_019823084.1;refGene_gn=TBC1D31;refGene_tx=XM_019823086.1;refGene_gn=TBC1D31;refGene_tx=XM_019823087.2;refGene_gn=TBC1D31;refGene_tx=XM_019823088.1;refGene_gn=TBC1D31;refGene_tx=XM_019823089.2;refGene_gn=TBC1D31;refGene_tx=XM_019823090.1;refGene_gn=TBC1D31;refGene_tx=XM_019823091.1
F2 68524018 68524251
refGene_gn=TBC1D31;refGene_tx=XM_004000105.5;refGene_gn=TBC1D31;refGene_tx=XM_019823084.1;refGene_gn=TBC1D31;refGene_tx=XM_019823086.1;refGene_gn=TBC1D31;refGene_tx=XM_019823087.2;refGene_gn=TBC1D31;refGene_tx=XM_019823088.1;refGene_gn=TBC1D31;refGene_tx=XM_019823089.2;refGene_gn=TBC1D31;refGene_tx=XM_019823090.1;refGene_gn=TBC1D31;refGene_tx=XM_019823091.1
F2 68524929 68525082
refGene_gn=TBC1D31;refGene_tx=XM_004000105.5;refGene_gn=TBC1D31;refGene_tx=XM_019823084.1;refGene_gn=TBC1D31;refGene_tx=XM_019823086.1;refGene_gn=TBC1D31;refGene_tx=XM_019823087.2;refGene_gn=TBC1D31;refGene_tx=XM_019823088.1;refGene_gn=TBC1D31;refGene_tx=XM_019823089.2;refGene_gn=TBC1D31;refGene_tx=XM_019823090.1;refGene_gn=TBC1D31;refGene_tx=XM_019823091.1
F2 68526112 68526248
refGene_gn=TBC1D31;refGene_tx=XM_004000105.5;refGene_gn=TBC1D31;refGene_tx=XM_019823084.1;refGene_gn=TBC1D31;refGene_tx=XM_019823086.1;refGene_gn=TBC1D31;refGene_tx=XM_019823087.2;refGene_gn=TBC1D31;refGene_tx=XM_019823088.1;refGene_gn=TBC1D31;refGene_tx=XM_019823089.2;refGene_gn=TBC1D31;refGene_tx=XM_019823090.1;refGene_gn=TBC1D31;refGene_tx=XM_019823091.1
F2 68531414 68531507
refGene_gn=TBC1D31;refGene_tx=XM_004000105.5;refGene_gn=TBC1D31;refGene_tx=XM_019823084.1;refGene_gn=TBC1D31;refGene_tx=XM_019823086.1;refGene_gn=TBC1D31;refGene_tx=XM_019823087.2;refGene_gn=TBC1D31;refGene_tx=

XM_019823088.1;refGene_gn=TBC1D31;refGene_tx=XM_019823089.2;refGene_gn=TBC1D31;refGene_tx=XM_019823091.1
F2 68537374 68537515
refGene_gn=TBC1D31;refGene_tx=XM_004000105.5;refGene_gn=TBC1D31;refGene_tx=XM_019823084.1;refGene_gn=TBC1D31;refGene_tx=XM_019823086.1;refGene_gn=TBC1D31;refGene_tx=XM_019823087.2;refGene_gn=TBC1D31;refGene_tx=XM_019823089.2;refGene_gn=TBC1D31;refGene_tx=XM_019823091.1
F2 68539132 68539327
refGene_gn=TBC1D31;refGene_tx=XM_004000105.5;refGene_gn=TBC1D31;refGene_tx=XM_019823084.1;refGene_gn=TBC1D31;refGene_tx=XM_019823086.1;refGene_gn=TBC1D31;refGene_tx=XM_019823087.2;refGene_gn=TBC1D31;refGene_tx=XM_019823088.1;refGene_gn=TBC1D31;refGene_tx=XM_019823089.2;refGene_gn=TBC1D31;refGene_tx=XM_019823090.1;refGene_gn=TBC1D31;refGene_tx=XM_019823091.1
F2 68541376 68541515
refGene_gn=TBC1D31;refGene_tx=XM_004000105.5;refGene_gn=TBC1D31;refGene_tx=XM_019823084.1;refGene_gn=TBC1D31;refGene_tx=XM_019823086.1;refGene_gn=TBC1D31;refGene_tx=XM_019823087.2;refGene_gn=TBC1D31;refGene_tx=XM_019823088.1;refGene_gn=TBC1D31;refGene_tx=XM_019823089.2;refGene_gn=TBC1D31;refGene_tx=XM_019823090.1;refGene_gn=TBC1D31;refGene_tx=XM_019823091.1
F2 68546026 68546031 refGene_gn=TBC1D31;refGene_tx=XM_019823089.2
F2 68547749 68547842
refGene_gn=TBC1D31;refGene_tx=XM_004000105.5;refGene_gn=TBC1D31;refGene_tx=XM_019823084.1;refGene_gn=TBC1D31;refGene_tx=XM_019823087.2;refGene_gn=TBC1D31;refGene_tx=XM_019823088.1;refGene_gn=TBC1D31;refGene_tx=XM_019823090.1;refGene_gn=TBC1D31;refGene_tx=XM_019823091.1
F2 68550705 68550839
refGene_gn=TBC1D31;refGene_tx=XM_004000105.5;refGene_gn=TBC1D31;refGene_tx=XM_019823084.1;refGene_gn=TBC1D31;refGene_tx=XM_019823086.1;refGene_gn=TBC1D31;refGene_tx=XM_019823087.2;refGene_gn=TBC1D31;refGene_tx=XM_019823088.1;refGene_gn=TBC1D31;refGene_tx=XM_019823090.1;refGene_gn=TBC1D31;refGene_tx=XM_019823091.1
F2 68577702 68578182
refGene_gn=FAM83A;refGene_tx=XM_004000093.4;refGene_gn=FAM83A;refGene_tx=XM_019823092.2
F2 68586109 68586277
refGene_gn=FAM83A;refGene_tx=XM_004000093.4;refGene_gn=FAM83A;refGene_tx=XM_019823092.2
F2 68588432 68588557
refGene_gn=FAM83A;refGene_tx=XM_004000093.4;refGene_gn=FAM83A;refGene_tx=XM_019823092.2
F2 68589398 68589648 refGene_gn=FAM83A;refGene_tx=XM_019823092.2
F2 68595709 68596244 refGene_gn=FAM83A;refGene_tx=XM_004000093.4
F2 68607182 68607380
refGene_gn=CF2H8orf76;refGene_tx=XM_004000094.5;refGene_gn=CF2H8orf76;refGene_tx=XM_023248226.1
F2 68607739 68607742
refGene_gn=CF2H8orf76;refGene_tx=XM_019823098.2
F2 68613184 68613317

refGene_gn=CF2H8orf76;refGene_tx=XM_004000094.5;refGene_gn=CF2H8orf76;
refGene_tx=XM_019823098.2;refGene_gn=CF2H8orf76;refGene_tx=XM_02324822
6.1
F2 68617448 68617906
refGene_gn=CF2H8orf76;refGene_tx=XM_004000094.5;refGene_gn=CF2H8orf76;
refGene_tx=XM_019823098.2;refGene_gn=CF2H8orf76;refGene_tx=XM_02324822
6.1
F2 68623540 68623684
refGene_gn=CF2H8orf76;refGene_tx=XM_004000094.5;refGene_gn=CF2H8orf76;
refGene_tx=XM_019823098.2;refGene_gn=CF2H8orf76;refGene_tx=XM_02324822
6.1
F2 68624419 68624515
refGene_gn=CF2H8orf76;refGene_tx=XM_004000094.5;refGene_gn=CF2H8orf76;
refGene_tx=XM_019823098.2;refGene_gn=CF2H8orf76;refGene_tx=XM_02324822
6.1
F2 68627438 68627441
refGene_gn=CF2H8orf76;refGene_tx=XM_023248226.1
F2 68627844 68627961
refGene_gn=CF2H8orf76;refGene_tx=XM_004000094.5;refGene_gn=CF2H8orf76;
refGene_tx=XM_019823098.2
F2 68641451 68644070 refGene_gn=ZHX1;refGene_tx=XM_023248156.1
F2 68679546 68679591
refGene_gn=ATAD2;refGene_tx=XM_023248159.1;refGene_gn=ATAD2;refGene_tx
=XM_023248160.1;refGene_gn=ATAD2;refGene_tx=XM_023248161.1
F2 68680330 68680459
refGene_gn=ATAD2;refGene_tx=XM_023248159.1;refGene_gn=ATAD2;refGene_tx
=XM_023248160.1;refGene_gn=ATAD2;refGene_tx=XM_023248161.1
F2 68681301 68681435
refGene_gn=ATAD2;refGene_tx=XM_023248159.1;refGene_gn=ATAD2;refGene_tx
=XM_023248160.1;refGene_gn=ATAD2;refGene_tx=XM_023248161.1
F2 68683246 68683642
refGene_gn=ATAD2;refGene_tx=XM_023248159.1;refGene_gn=ATAD2;refGene_tx
=XM_023248160.1;refGene_gn=ATAD2;refGene_tx=XM_023248161.1
F2 68685875 68686019
refGene_gn=ATAD2;refGene_tx=XM_023248159.1;refGene_gn=ATAD2;refGene_tx
=XM_023248160.1;refGene_gn=ATAD2;refGene_tx=XM_023248161.1
F2 68686144 68686267
refGene_gn=ATAD2;refGene_tx=XM_023248159.1;refGene_gn=ATAD2;refGene_tx
=XM_023248160.1;refGene_gn=ATAD2;refGene_tx=XM_023248161.1
F2 68688949 68689109
refGene_gn=ATAD2;refGene_tx=XM_023248159.1;refGene_gn=ATAD2;refGene_tx
=XM_023248160.1;refGene_gn=ATAD2;refGene_tx=XM_023248161.1
F2 68689656 68689853
refGene_gn=ATAD2;refGene_tx=XM_023248159.1;refGene_gn=ATAD2;refGene_tx
=XM_023248160.1;refGene_gn=ATAD2;refGene_tx=XM_023248161.1
F2 68690868 68691004
refGene_gn=ATAD2;refGene_tx=XM_023248159.1;refGene_gn=ATAD2;refGene_tx
=XM_023248160.1;refGene_gn=ATAD2;refGene_tx=XM_023248161.1
F2 68697604 68697790
refGene_gn=ATAD2;refGene_tx=XM_023248159.1;refGene_gn=ATAD2;refGene_tx

=XM_023248160.1;refGene_gn=ATAD2;refGene_tx=XM_023248161.1
F2 68698518 68698705
refGene_gn=ATAD2;refGene_tx=XM_023248159.1;refGene_gn=ATAD2;refGene_tx
=XM_023248160.1;refGene_gn=ATAD2;refGene_tx=XM_023248161.1
F2 68699050 68699183
refGene_gn=ATAD2;refGene_tx=XM_023248159.1;refGene_gn=ATAD2;refGene_tx
=XM_023248160.1;refGene_gn=ATAD2;refGene_tx=XM_023248161.1
F2 68699562 68699877
refGene_gn=ATAD2;refGene_tx=XM_023248159.1;refGene_gn=ATAD2;refGene_tx
=XM_023248160.1;refGene_gn=ATAD2;refGene_tx=XM_023248161.1
F2 68700536 68700627
refGene_gn=ATAD2;refGene_tx=XM_023248159.1;refGene_gn=ATAD2;refGene_tx
=XM_023248160.1;refGene_gn=ATAD2;refGene_tx=XM_023248161.1
F2 68701717 68701877
refGene_gn=ATAD2;refGene_tx=XM_023248159.1;refGene_gn=ATAD2;refGene_tx
=XM_023248160.1;refGene_gn=ATAD2;refGene_tx=XM_023248161.1
F2 68707895 68707984
refGene_gn=ATAD2;refGene_tx=XM_023248159.1;refGene_gn=ATAD2;refGene_tx
=XM_023248160.1;refGene_gn=ATAD2;refGene_tx=XM_023248161.1
F2 68708821 68708996
refGene_gn=ATAD2;refGene_tx=XM_023248159.1;refGene_gn=ATAD2;refGene_tx
=XM_023248160.1;refGene_gn=ATAD2;refGene_tx=XM_023248161.1
F2 68717224 68717340
refGene_gn=ATAD2;refGene_tx=XM_023248159.1;refGene_gn=ATAD2;refGene_tx
=XM_023248160.1;refGene_gn=ATAD2;refGene_tx=XM_023248161.1
F2 68717563 68717672
refGene_gn=ATAD2;refGene_tx=XM_023248159.1;refGene_gn=ATAD2;refGene_tx
=XM_023248160.1;refGene_gn=ATAD2;refGene_tx=XM_023248161.1
F2 68720804 68720912
refGene_gn=ATAD2;refGene_tx=XM_023248159.1;refGene_gn=ATAD2;refGene_tx
=XM_023248160.1;refGene_gn=ATAD2;refGene_tx=XM_023248161.1
F2 68726741 68726859
refGene_gn=ATAD2;refGene_tx=XM_023248159.1;refGene_gn=ATAD2;refGene_tx
=XM_023248160.1;refGene_gn=ATAD2;refGene_tx=XM_023248161.1
F2 68727476 68727647
refGene_gn=ATAD2;refGene_tx=XM_023248159.1;refGene_gn=ATAD2;refGene_tx
=XM_023248160.1;refGene_gn=ATAD2;refGene_tx=XM_023248161.1
F2 68729219 68729307
refGene_gn=ATAD2;refGene_tx=XM_023248159.1;refGene_gn=ATAD2;refGene_tx
=XM_023248160.1;refGene_gn=ATAD2;refGene_tx=XM_023248161.1
F2 68729639 68729742
refGene_gn=ATAD2;refGene_tx=XM_023248159.1;refGene_gn=ATAD2;refGene_tx
=XM_023248160.1;refGene_gn=ATAD2;refGene_tx=XM_023248161.1
F2 68730074 68730240
refGene_gn=ATAD2;refGene_tx=XM_023248159.1;refGene_gn=ATAD2;refGene_tx
=XM_023248160.1;refGene_gn=ATAD2;refGene_tx=XM_023248161.1
F2 68731580 68731630
refGene_gn=ATAD2;refGene_tx=XM_023248159.1;refGene_gn=ATAD2;refGene_tx
=XM_023248160.1;refGene_gn=ATAD2;refGene_tx=XM_023248161.1
F2 68736072 68736221

refGene_gn=ATAD2;refGene_tx=XM_023248159.1;refGene_gn=ATAD2;refGene_tx
=XM_023248160.1;refGene_gn=ATAD2;refGene_tx=XM_023248161.1
F2 68747042 68747210
refGene_gn=ATAD2;refGene_tx=XM_023248159.1;refGene_gn=ATAD2;refGene_tx
=XM_023248160.1
F2 68757991 68758080
refGene_gn=WDYHV1;refGene_tx=XM_004000096.4;refGene_gn=WDYHV1;refGene_
tx=XM_019823099.1
F2 68760825 68760899 refGene_gn=WDYHV1;refGene_tx=XM_006943373.2
F2 68763448 68763548
refGene_gn=WDYHV1;refGene_tx=XM_004000096.4;refGene_gn=WDYHV1;refGene_
tx=XM_006943373.2;refGene_gn=WDYHV1;refGene_tx=XM_019823099.1
F2 68764630 68764681
refGene_gn=WDYHV1;refGene_tx=XM_004000096.4;refGene_gn=WDYHV1;refGene_
tx=XM_006943373.2;refGene_gn=WDYHV1;refGene_tx=XM_019823099.1
F2 68770871 68771020
refGene_gn=WDYHV1;refGene_tx=XM_004000096.4;refGene_gn=WDYHV1;refGene_
tx=XM_006943373.2;refGene_gn=WDYHV1;refGene_tx=XM_019823099.1
F2 68771923 68772048
refGene_gn=WDYHV1;refGene_tx=XM_004000096.4;refGene_gn=WDYHV1;refGene_
tx=XM_006943373.2;refGene_gn=WDYHV1;refGene_tx=XM_019823099.1
F2 68772542 68772631 refGene_gn=WDYHV1;refGene_tx=XM_019823099.1
F2 68776640 68776750
refGene_gn=WDYHV1;refGene_tx=XM_004000096.4;refGene_gn=WDYHV1;refGene_
tx=XM_006943373.2
F2 68836949 68837039 refGene_gn=FBXO32;refGene_tx=XM_004000107.4
F2 68837482 68837626 refGene_gn=FBXO32;refGene_tx=XM_004000107.4
F2 68839092 68839275 refGene_gn=FBXO32;refGene_tx=XM_004000107.4
F2 68845000 68845185 refGene_gn=FBXO32;refGene_tx=XM_004000107.4
F2 68846096 68846190 refGene_gn=FBXO32;refGene_tx=XM_004000107.4
F2 68858803 68858896 refGene_gn=FBXO32;refGene_tx=XM_004000107.4
F2 68860098 68860148 refGene_gn=FBXO32;refGene_tx=XM_004000107.4
F2 68861011 68861124 refGene_gn=FBXO32;refGene_tx=XM_004000107.4
F2 68866997 68867113 refGene_gn=FBXO32;refGene_tx=XM_004000107.4
F2 68958869 68959159 refGene_gn=KLHL38;refGene_tx=XM_004000097.4
F2 68960035 68960141 refGene_gn=KLHL38;refGene_tx=XM_004000097.4
F2 68963933 68965322 refGene_gn=KLHL38;refGene_tx=XM_004000097.4
F2 68988244 68988364 refGene_gn=ANXA13;refGene_tx=XM_006943374.3
F2 68991277 68991390 refGene_gn=ANXA13;refGene_tx=XM_006943374.3
F2 68994724 68994800 refGene_gn=ANXA13;refGene_tx=XM_006943374.3
F2 68999448 68999550 refGene_gn=ANXA13;refGene_tx=XM_006943374.3
F2 69000200 69000269 refGene_gn=ANXA13;refGene_tx=XM_006943374.3
F2 69000872 69000952 refGene_gn=ANXA13;refGene_tx=XM_006943374.3
F2 69001057 69001091 refGene_gn=ANXA13;refGene_tx=XM_006943374.3
F2 69003405 69003576 refGene_gn=ANXA13;refGene_tx=XM_006943374.3
F2 69007348 69007443 refGene_gn=ANXA13;refGene_tx=XM_006943374.3
F2 69015281 69015357 refGene_gn=ANXA13;refGene_tx=XM_006943374.3
F2 69037287 69037410 refGene_gn=ANXA13;refGene_tx=XM_006943374.3
F2 69038793 69038808 refGene_gn=ANXA13;refGene_tx=XM_006943374.3
F2 69061531 69061603 refGene_gn=FAM91A1;refGene_tx=XM_011291452.3

| | | | |
|---|---|---|---|
| F2 | 69066545 | 69066630 | refGene_gn=FAM91A1;refGene_tx=XM_011291452.3 |
| F2 | 69068160 | 69068312 | refGene_gn=FAM91A1;refGene_tx=XM_011291452.3 |
| F2 | 69069936 | 69069994 | refGene_gn=FAM91A1;refGene_tx=XM_011291452.3 |
| F2 | 69070695 | 69070763 | refGene_gn=FAM91A1;refGene_tx=XM_011291452.3 |
| F2 | 69071309 | 69071423 | refGene_gn=FAM91A1;refGene_tx=XM_011291452.3 |
| F2 | 69072689 | 69072780 | refGene_gn=FAM91A1;refGene_tx=XM_011291452.3 |
| F2 | 69073353 | 69073416 | refGene_gn=FAM91A1;refGene_tx=XM_011291452.3 |
| F2 | 69078402 | 69078509 | refGene_gn=FAM91A1;refGene_tx=XM_011291452.3 |
| F2 | 69079018 | 69079057 | refGene_gn=FAM91A1;refGene_tx=XM_011291452.3 |
| F2 | 69079585 | 69079698 | refGene_gn=FAM91A1;refGene_tx=XM_011291452.3 |
| F2 | 69080170 | 69080286 | refGene_gn=FAM91A1;refGene_tx=XM_011291452.3 |
| F2 | 69081551 | 69081664 | refGene_gn=FAM91A1;refGene_tx=XM_011291452.3 |
| F2 | 69082021 | 69082108 | refGene_gn=FAM91A1;refGene_tx=XM_011291452.3 |
| F2 | 69084277 | 69084410 | refGene_gn=FAM91A1;refGene_tx=XM_011291452.3 |
| F2 | 69088798 | 69088947 | refGene_gn=FAM91A1;refGene_tx=XM_011291452.3 |
| F2 | 69090245 | 69090380 | refGene_gn=FAM91A1;refGene_tx=XM_011291452.3 |
| F2 | 69090539 | 69090653 | refGene_gn=FAM91A1;refGene_tx=XM_011291452.3 |
| F2 | 69094563 | 69094636 | refGene_gn=FAM91A1;refGene_tx=XM_011291452.3 |
| F2 | 69097444 | 69097594 | refGene_gn=FAM91A1;refGene_tx=XM_011291452.3 |
| F2 | 69099544 | 69099649 | refGene_gn=FAM91A1;refGene_tx=XM_011291452.3 |
| F2 | 69100154 | 69100278 | refGene_gn=FAM91A1;refGene_tx=XM_011291452.3 |
| F2 | 69101780 | 69101850 | refGene_gn=FAM91A1;refGene_tx=XM_011291452.3 |
| F2 | 69103170 | 69103356 | refGene_gn=FAM91A1;refGene_tx=XM_011291452.3 |
| F2 | 69203066 | 69203142 | refGene_gn=FER1L6;refGene_tx=XM_023248464.1 |
| F2 | 69207172 | 69207293 | refGene_gn=FER1L6;refGene_tx=XM_023248464.1 |
| F2 | 69209993 | 69210048 | refGene_gn=FER1L6;refGene_tx=XM_023248464.1 |
| F2 | 69210129 | 69210261 | refGene_gn=FER1L6;refGene_tx=XM_023248464.1 |
| F2 | 69214026 | 69214089 | refGene_gn=FER1L6;refGene_tx=XM_023248464.1 |
| F2 | 69217598 | 69217677 | refGene_gn=FER1L6;refGene_tx=XM_023248464.1 |
| F2 | 69219182 | 69219339 | refGene_gn=FER1L6;refGene_tx=XM_023248464.1 |
| F2 | 69219931 | 69220118 | refGene_gn=FER1L6;refGene_tx=XM_023248464.1 |
| F2 | 69221783 | 69221976 | refGene_gn=FER1L6;refGene_tx=XM_023248464.1 |
| F2 | 69223492 | 69223845 | refGene_gn=FER1L6;refGene_tx=XM_023248464.1 |
| F2 | 69228858 | 69228967 | refGene_gn=FER1L6;refGene_tx=XM_023248464.1 |
| F2 | 69229728 | 69229912 | refGene_gn=FER1L6;refGene_tx=XM_023248464.1 |
| F2 | 69235078 | 69235199 | refGene_gn=FER1L6;refGene_tx=XM_023248464.1 |
| F2 | 69237748 | 69237849 | refGene_gn=FER1L6;refGene_tx=XM_023248464.1 |
| F2 | 69242031 | 69242122 | refGene_gn=FER1L6;refGene_tx=XM_023248464.1 |
| F2 | 69244836 | 69244956 | refGene_gn=FER1L6;refGene_tx=XM_023248464.1 |
| F2 | 69246629 | 69246782 | refGene_gn=FER1L6;refGene_tx=XM_023248464.1 |
| F2 | 69254156 | 69254334 | refGene_gn=FER1L6;refGene_tx=XM_023248464.1 |
| F2 | 69257943 | 69258068 | refGene_gn=FER1L6;refGene_tx=XM_023248464.1 |
| F2 | 69263017 | 69263152 | refGene_gn=FER1L6;refGene_tx=XM_023248464.1 |
| F2 | 69266222 | 69266372 | refGene_gn=FER1L6;refGene_tx=XM_023248464.1 |
| F2 | 69275793 | 69275904 | refGene_gn=FER1L6;refGene_tx=XM_023248464.1 |
| F2 | 69276166 | 69276328 | refGene_gn=FER1L6;refGene_tx=XM_023248464.1 |
| F2 | 69277484 | 69277665 | refGene_gn=FER1L6;refGene_tx=XM_023248464.1 |
| F2 | 69280218 | 69280445 | refGene_gn=FER1L6;refGene_tx=XM_023248464.1 |
| F2 | 69282124 | 69282247 | refGene_gn=FER1L6;refGene_tx=XM_023248464.1 |
| F2 | 69283469 | 69283509 | refGene_gn=FER1L6;refGene_tx=XM_023248464.1 |

F2 69284678 69284791 refGene_gn=FER1L6;refGene_tx=XM_023248464.1
F2 69286053 69286185 refGene_gn=FER1L6;refGene_tx=XM_023248464.1
F2 69287087 69287213 refGene_gn=FER1L6;refGene_tx=XM_023248464.1
F2 69291347 69291475 refGene_gn=FER1L6;refGene_tx=XM_023248464.1
F2 69295528 69295699 refGene_gn=FER1L6;refGene_tx=XM_023248464.1
F2 69298379 69298540 refGene_gn=FER1L6;refGene_tx=XM_023248464.1
F2 69300337 69300480 refGene_gn=FER1L6;refGene_tx=XM_023248464.1
F2 69302747 69302836 refGene_gn=FER1L6;refGene_tx=XM_023248464.1
F2 69303322 69303421 refGene_gn=FER1L6;refGene_tx=XM_023248464.1
F2 69306102 69306344 refGene_gn=FER1L6;refGene_tx=XM_023248464.1
F2 69307776 69307940 refGene_gn=FER1L6;refGene_tx=XM_023248464.1
F2 69320955 69321056 refGene_gn=FER1L6;refGene_tx=XM_023248464.1
F2 69323218 69323402 refGene_gn=FER1L6;refGene_tx=XM_023248464.1
F2 69461086 69461188
refGene_gn=TMEM65;refGene_tx=XM_023248733.1;refGene_gn=TMEM65;refGene_tx=XM_023248734.1;refGene_gn=TMEM65;refGene_tx=XM_023248735.1
F2 69468553 69468659
refGene_gn=TMEM65;refGene_tx=XM_023248733.1;refGene_gn=TMEM65;refGene_tx=XM_023248734.1;refGene_gn=TMEM65;refGene_tx=XM_023248735.1
F2 69470495 69470538
refGene_gn=TMEM65;refGene_tx=XM_023248733.1;refGene_gn=TMEM65;refGene_tx=XM_023248734.1;refGene_gn=TMEM65;refGene_tx=XM_023248735.1
F2 69471856 69471911
refGene_gn=TMEM65;refGene_tx=XM_023248733.1;refGene_gn=TMEM65;refGene_tx=XM_023248734.1;refGene_gn=TMEM65;refGene_tx=XM_023248735.1
F2 69473207 69473275
refGene_gn=TMEM65;refGene_tx=XM_023248733.1;refGene_gn=TMEM65;refGene_tx=XM_023248734.1;refGene_gn=TMEM65;refGene_tx=XM_023248735.1
F2 69477081 69477126
refGene_gn=TMEM65;refGene_tx=XM_023248733.1;refGene_gn=TMEM65;refGene_tx=XM_023248734.1;refGene_gn=TMEM65;refGene_tx=XM_023248735.1
F2 69478570 69478666 refGene_gn=TMEM65;refGene_tx=XM_023248733.1
F2 69494690 69494771 refGene_gn=TMEM65;refGene_tx=XM_023248734.1
F2 69512008 69512294
refGene_gn=TMEM65;refGene_tx=XM_023248733.1;refGene_gn=TMEM65;refGene_tx=XM_023248734.1;refGene_gn=TMEM65;refGene_tx=XM_023248735.1
F2 69569185 69570721
refGene_gn=TRMT12;refGene_tx=XM_006943377.3;refGene_gn=TRMT12;refGene_tx=XM_019823107.1;refGene_gn=TRMT12;refGene_tx=XM_019823109.2;refGene_gn=TRMT12;refGene_tx=XM_023248589.1;refGene_gn=TRMT12;refGene_tx=XM_023248591.1
F2 69585594 69585775 refGene_gn=RNF139;refGene_tx=XM_023248587.1
F2 69586688 69586716 refGene_gn=TATDN1;refGene_tx=XM_023248592.1
F2 69595262 69597082 refGene_gn=RNF139;refGene_tx=XM_023248587.1
F2 69598053 69598156
refGene_gn=TATDN1;refGene_tx=XM_004000111.4;refGene_gn=TATDN1;refGene_tx=XM_019823111.1;refGene_gn=TATDN1;refGene_tx=XM_019823112.1;refGene_gn=TATDN1;refGene_tx=XM_019823113.2;refGene_gn=TATDN1;refGene_tx=XM_019823117.1;refGene_gn=TATDN1;refGene_tx=XM_019823118.2;refGene_gn=TATDN1;refGene_tx=XM_019823119.1;refGene_gn=TATDN1;refGene_tx=XM_019823120.

1
F2 69601187 69601195 refGene_gn=TATDN1;refGene_tx=XM_019823114.2
F2 69601223 69601350
refGene_gn=TATDN1;refGene_tx=XM_004000111.4;refGene_gn=TATDN1;refGene_tx=XM_019823111.1;refGene_gn=TATDN1;refGene_tx=XM_019823112.1;refGene_gn=TATDN1;refGene_tx=XM_019823113.2;refGene_gn=TATDN1;refGene_tx=XM_019823117.1;refGene_gn=TATDN1;refGene_tx=XM_019823118.2;refGene_gn=TATDN1;refGene_tx=XM_019823119.1;refGene_gn=TATDN1;refGene_tx=XM_019823120.1;refGene_gn=TATDN1;refGene_tx=XM_023248592.1
F2 69602544 69602615
refGene_gn=TATDN1;refGene_tx=XM_004000111.4;refGene_gn=TATDN1;refGene_tx=XM_019823111.1;refGene_gn=TATDN1;refGene_tx=XM_019823112.1;refGene_gn=TATDN1;refGene_tx=XM_019823113.2;refGene_gn=TATDN1;refGene_tx=XM_019823114.2;refGene_gn=TATDN1;refGene_tx=XM_019823117.1;refGene_gn=TATDN1;refGene_tx=XM_019823118.2;refGene_gn=TATDN1;refGene_tx=XM_019823119.1;refGene_gn=TATDN1;refGene_tx=XM_019823120.1;refGene_gn=TATDN1;refGene_tx=XM_023248592.1
F2 69616913 69616990
refGene_gn=TATDN1;refGene_tx=XM_004000111.4;refGene_gn=TATDN1;refGene_tx=XM_019823111.1;refGene_gn=TATDN1;refGene_tx=XM_019823112.1;refGene_gn=TATDN1;refGene_tx=XM_019823113.2;refGene_gn=TATDN1;refGene_tx=XM_019823114.2;refGene_gn=TATDN1;refGene_tx=XM_019823117.1;refGene_gn=TATDN1;refGene_tx=XM_019823118.2;refGene_gn=TATDN1;refGene_tx=XM_019823119.1;refGene_gn=TATDN1;refGene_tx=XM_019823120.1;refGene_gn=TATDN1;refGene_tx=XM_023248592.1
F2 69619516 69619557
refGene_gn=TATDN1;refGene_tx=XM_004000111.4;refGene_gn=TATDN1;refGene_tx=XM_019823111.1;refGene_gn=TATDN1;refGene_tx=XM_019823112.1;refGene_gn=TATDN1;refGene_tx=XM_019823113.2;refGene_gn=TATDN1;refGene_tx=XM_019823114.2;refGene_gn=TATDN1;refGene_tx=XM_019823117.1;refGene_gn=TATDN1;refGene_tx=XM_019823118.2;refGene_gn=TATDN1;refGene_tx=XM_019823119.1;refGene_gn=TATDN1;refGene_tx=XM_019823120.1;refGene_gn=TATDN1;refGene_tx=XM_023248592.1
F2 69619648 69619734
refGene_gn=TATDN1;refGene_tx=XM_004000111.4;refGene_gn=TATDN1;refGene_tx=XM_019823111.1;refGene_gn=TATDN1;refGene_tx=XM_019823112.1;refGene_gn=TATDN1;refGene_tx=XM_019823113.2;refGene_gn=TATDN1;refGene_tx=XM_019823114.2;refGene_gn=TATDN1;refGene_tx=XM_019823117.1;refGene_gn=TATDN1;refGene_tx=XM_019823118.2;refGene_gn=TATDN1;refGene_tx=XM_019823119.1;refGene_gn=TATDN1;refGene_tx=XM_019823120.1;refGene_gn=TATDN1;refGene_tx=XM_023248592.1
F2 69620401 69620424
refGene_gn=TATDN1;refGene_tx=XM_019823117.1;refGene_gn=TATDN1;refGene_tx=XM_019823118.2
F2 69625373 69625416
refGene_gn=TATDN1;refGene_tx=XM_004000111.4;refGene_gn=TATDN1;refGene_tx=XM_019823111.1;refGene_gn=TATDN1;refGene_tx=XM_019823112.1;refGene_gn=TATDN1;refGene_tx=XM_019823113.2;refGene_gn=TATDN1;refGene_tx=XM_019823114.2;refGene_gn=TATDN1;refGene_tx=XM_023248592.1
F2 69625512 69625656

refGene_gn=TATDN1;refGene_tx=XM_004000111.4;refGene_gn=TATDN1;refGene_tx=XM_019823111.1;refGene_gn=TATDN1;refGene_tx=XM_019823112.1;refGene_gn=TATDN1;refGene_tx=XM_019823113.2;refGene_gn=TATDN1;refGene_tx=XM_019823114.2;refGene_gn=TATDN1;refGene_tx=XM_023248592.1
F2 69627634 69627698
refGene_gn=TATDN1;refGene_tx=XM_004000111.4;refGene_gn=TATDN1;refGene_tx=XM_019823114.2;refGene_gn=TATDN1;refGene_tx=XM_023248592.1
F2 69630893 69630943
refGene_gn=TATDN1;refGene_tx=XM_004000111.4;refGene_gn=TATDN1;refGene_tx=XM_019823114.2;refGene_gn=TATDN1;refGene_tx=XM_023248592.1
F2 69632835 69632901
refGene_gn=TATDN1;refGene_tx=XM_004000111.4;refGene_gn=TATDN1;refGene_tx=XM_019823114.2;refGene_gn=TATDN1;refGene_tx=XM_023248592.1
F2 69643762 69643784
refGene_gn=TATDN1;refGene_tx=XM_004000111.4;refGene_gn=TATDN1;refGene_tx=XM_019823114.2;refGene_gn=TATDN1;refGene_tx=XM_023248592.1
F2 69643916 69644017 refGene_gn=NDUFB9;refGene_tx=XM_004000113.5
F2 69646881 69647074 refGene_gn=NDUFB9;refGene_tx=XM_004000113.5
F2 69647635 69647749 refGene_gn=NDUFB9;refGene_tx=XM_004000113.5
F2 69650740 69650872 refGene_gn=NDUFB9;refGene_tx=XM_004000113.5
F2 69653963 69654664
refGene_gn=MTSS1;refGene_tx=XM_004000114.4;refGene_gn=MTSS1;refGene_tx=XM_004000115.4;refGene_gn=MTSS1;refGene_tx=XM_006943381.3;refGene_gn=MTSS1;refGene_tx=XM_011291453.3;refGene_gn=MTSS1;refGene_tx=XM_019823102.2;refGene_gn=MTSS1;refGene_tx=XM_019823103.2;refGene_gn=MTSS1;refGene_tx=XM_019823104.2;refGene_gn=MTSS1;refGene_tx=XM_019823106.2
F2 69656089 69656252
refGene_gn=MTSS1;refGene_tx=XM_004000114.4;refGene_gn=MTSS1;refGene_tx=XM_004000115.4;refGene_gn=MTSS1;refGene_tx=XM_006943381.3;refGene_gn=MTSS1;refGene_tx=XM_011291453.3;refGene_gn=MTSS1;refGene_tx=XM_019823102.2;refGene_gn=MTSS1;refGene_tx=XM_019823103.2;refGene_gn=MTSS1;refGene_tx=XM_019823104.2;refGene_gn=MTSS1;refGene_tx=XM_019823106.2
F2 69656580 69656754
refGene_gn=MTSS1;refGene_tx=XM_004000114.4;refGene_gn=MTSS1;refGene_tx=XM_004000115.4;refGene_gn=MTSS1;refGene_tx=XM_006943381.3;refGene_gn=MTSS1;refGene_tx=XM_011291453.3;refGene_gn=MTSS1;refGene_tx=XM_019823102.2;refGene_gn=MTSS1;refGene_tx=XM_019823103.2;refGene_gn=MTSS1;refGene_tx=XM_019823104.2;refGene_gn=MTSS1;refGene_tx=XM_019823106.2
F2 69658041 69658236
refGene_gn=MTSS1;refGene_tx=XM_004000114.4;refGene_gn=MTSS1;refGene_tx=XM_004000115.4;refGene_gn=MTSS1;refGene_tx=XM_019823102.2;refGene_gn=MTSS1;refGene_tx=XM_019823103.2;refGene_gn=MTSS1;refGene_tx=XM_019823104.2
F2 69659111 69659231
refGene_gn=MTSS1;refGene_tx=XM_011291453.3;refGene_gn=MTSS1;refGene_tx=XM_019823102.2;refGene_gn=MTSS1;refGene_tx=XM_019823103.2;refGene_gn=MTSS1;refGene_tx=XM_019823104.2;refGene_gn=MTSS1;refGene_tx=XM_019823106.2
F2 69662996 69663207
refGene_gn=MTSS1;refGene_tx=XM_004000114.4;refGene_gn=MTSS1;refGene_tx

=XM_004000115.4;refGene_gn=MTSS1;refGene_tx=XM_006943381.3;refGene_gn=MTSS1;refGene_tx=XM_011291453.3;refGene_gn=MTSS1;refGene_tx=XM_019823102.2;refGene_gn=MTSS1;refGene_tx=XM_019823103.2;refGene_gn=MTSS1;refGene_tx=XM_019823104.2;refGene_gn=MTSS1;refGene_tx=XM_019823106.2
F2 69665145 69665243
refGene_gn=MTSS1;refGene_tx=XM_004000114.4;refGene_gn=MTSS1;refGene_tx=XM_004000115.4;refGene_gn=MTSS1;refGene_tx=XM_006943381.3;refGene_gn=MTSS1;refGene_tx=XM_011291453.3;refGene_gn=MTSS1;refGene_tx=XM_019823102.2;refGene_gn=MTSS1;refGene_tx=XM_019823103.2;refGene_gn=MTSS1;refGene_tx=XM_019823104.2;refGene_gn=MTSS1;refGene_tx=XM_019823106.2
F2 69666514 69666622
refGene_gn=MTSS1;refGene_tx=XM_004000114.4;refGene_gn=MTSS1;refGene_tx=XM_004000115.4;refGene_gn=MTSS1;refGene_tx=XM_006943381.3;refGene_gn=MTSS1;refGene_tx=XM_011291453.3;refGene_gn=MTSS1;refGene_tx=XM_019823102.2;refGene_gn=MTSS1;refGene_tx=XM_019823103.2;refGene_gn=MTSS1;refGene_tx=XM_019823104.2;refGene_gn=MTSS1;refGene_tx=XM_019823106.2
F2 69668820 69668978
refGene_gn=MTSS1;refGene_tx=XM_004000114.4;refGene_gn=MTSS1;refGene_tx=XM_004000115.4;refGene_gn=MTSS1;refGene_tx=XM_006943381.3;refGene_gn=MTSS1;refGene_tx=XM_011291453.3;refGene_gn=MTSS1;refGene_tx=XM_019823102.2;refGene_gn=MTSS1;refGene_tx=XM_019823103.2;refGene_gn=MTSS1;refGene_tx=XM_019823104.2;refGene_gn=MTSS1;refGene_tx=XM_019823106.2
F2 69680815 69680827
refGene_gn=MTSS1;refGene_tx=XM_004000115.4;refGene_gn=MTSS1;refGene_tx=XM_006943381.3;refGene_gn=MTSS1;refGene_tx=XM_019823102.2;refGene_gn=MTSS1;refGene_tx=XM_019823104.2;refGene_gn=MTSS1;refGene_tx=XM_019823106.2
F2 69685775 69685850
refGene_gn=MTSS1;refGene_tx=XM_004000114.4;refGene_gn=MTSS1;refGene_tx=XM_004000115.4;refGene_gn=MTSS1;refGene_tx=XM_006943381.3;refGene_gn=MTSS1;refGene_tx=XM_011291453.3;refGene_gn=MTSS1;refGene_tx=XM_019823102.2;refGene_gn=MTSS1;refGene_tx=XM_019823103.2;refGene_gn=MTSS1;refGene_tx=XM_019823104.2;refGene_gn=MTSS1;refGene_tx=XM_019823106.2
F2 69687492 69687584
refGene_gn=MTSS1;refGene_tx=XM_004000114.4;refGene_gn=MTSS1;refGene_tx=XM_004000115.4;refGene_gn=MTSS1;refGene_tx=XM_006943381.3;refGene_gn=MTSS1;refGene_tx=XM_011291453.3;refGene_gn=MTSS1;refGene_tx=XM_019823102.2;refGene_gn=MTSS1;refGene_tx=XM_019823103.2;refGene_gn=MTSS1;refGene_tx=XM_019823104.2;refGene_gn=MTSS1;refGene_tx=XM_019823106.2
F2 69690032 69690117
refGene_gn=MTSS1;refGene_tx=XM_004000114.4;refGene_gn=MTSS1;refGene_tx=XM_004000115.4;refGene_gn=MTSS1;refGene_tx=XM_006943381.3;refGene_gn=MTSS1;refGene_tx=XM_011291453.3;refGene_gn=MTSS1;refGene_tx=XM_019823102.2;refGene_gn=MTSS1;refGene_tx=XM_019823103.2;refGene_gn=MTSS1;refGene_tx=XM_019823104.2;refGene_gn=MTSS1;refGene_tx=XM_019823106.2
F2 69788310 69788384
refGene_gn=MTSS1;refGene_tx=XM_004000114.4;refGene_gn=MTSS1;refGene_tx=XM_004000115.4;refGene_gn=MTSS1;refGene_tx=XM_006943381.3;refGene_gn=MTSS1;refGene_tx=XM_011291453.3;refGene_gn=MTSS1;refGene_tx=XM_019823102.2;refGene_gn=MTSS1;refGene_tx=XM_019823103.2;refGene_gn=MTSS1;refGe

ne_tx=XM_019823104.2;refGene_gn=MTSS1;refGene_tx=XM_019823106.2
F2	69792147	69792209
refGene_gn=MTSS1;refGene_tx=XM_004000114.4;refGene_gn=MTSS1;refGene_tx=XM_004000115.4;refGene_gn=MTSS1;refGene_tx=XM_006943381.3;refGene_gn=MTSS1;refGene_tx=XM_011291453.3;refGene_gn=MTSS1;refGene_tx=XM_019823102.2;refGene_gn=MTSS1;refGene_tx=XM_019823103.2;refGene_gn=MTSS1;refGene_tx=XM_019823104.2;refGene_gn=MTSS1;refGene_tx=XM_019823106.2
F2	69816842	69816914
refGene_gn=MTSS1;refGene_tx=XM_004000114.4;refGene_gn=MTSS1;refGene_tx=XM_004000115.4;refGene_gn=MTSS1;refGene_tx=XM_006943381.3;refGene_gn=MTSS1;refGene_tx=XM_011291453.3;refGene_gn=MTSS1;refGene_tx=XM_019823102.2;refGene_gn=MTSS1;refGene_tx=XM_019823103.2;refGene_gn=MTSS1;refGene_tx=XM_019823104.2;refGene_gn=MTSS1;refGene_tx=XM_019823106.2
F2	70006474	70006553
refGene_gn=ZNF572;refGene_tx=XM_004000132.4;refGene_gn=ZNF572;refGene_tx=XM_006943382.3;refGene_gn=ZNF572;refGene_tx=XM_006943383.3;refGene_gn=ZNF572;refGene_tx=XM_006943384.3;refGene_gn=ZNF572;refGene_tx=XM_023248322.1
F2	70007206	70008714
refGene_gn=ZNF572;refGene_tx=XM_004000132.4;refGene_gn=ZNF572;refGene_tx=XM_006943382.3;refGene_gn=ZNF572;refGene_tx=XM_006943383.3;refGene_gn=ZNF572;refGene_tx=XM_006943384.3;refGene_gn=ZNF572;refGene_tx=XM_023248322.1
F2	70021994	70022285	refGene_gn=SQLE;refGene_tx=XM_004000116.5
F2	70025977	70026230	refGene_gn=SQLE;refGene_tx=XM_004000116.5
F2	70035109	70035290	refGene_gn=SQLE;refGene_tx=XM_004000116.5
F2	70036581	70036678	refGene_gn=SQLE;refGene_tx=XM_004000116.5
F2	70037793	70037907	refGene_gn=SQLE;refGene_tx=XM_004000116.5
F2	70037985	70038157	refGene_gn=SQLE;refGene_tx=XM_004000116.5
F2	70040643	70040739	refGene_gn=SQLE;refGene_tx=XM_004000116.5
F2	70045457	70045600	refGene_gn=SQLE;refGene_tx=XM_004000116.5
F2	70046293	70046320	refGene_gn=SQLE;refGene_tx=XM_004000116.5
F2	70046321	70046391	refGene_gn=SQLE;refGene_tx=XM_004000116.5
F2	70048191	70048279	refGene_gn=SQLE;refGene_tx=XM_004000116.5
F2	70049350	70049543	refGene_gn=SQLE;refGene_tx=XM_004000116.5
F2	70051935	70051992
refGene_gn=WASHC5;refGene_tx=XM_023248200.1;refGene_gn=WASHC5;refGene_tx=XM_023248201.1
F2	70056079	70056167
refGene_gn=WASHC5;refGene_tx=XM_023248200.1;refGene_gn=WASHC5;refGene_tx=XM_023248201.1
F2	70058924	70059078
refGene_gn=WASHC5;refGene_tx=XM_023248200.1;refGene_gn=WASHC5;refGene_tx=XM_023248201.1
F2	70063352	70063449
refGene_gn=WASHC5;refGene_tx=XM_023248200.1;refGene_gn=WASHC5;refGene_tx=XM_023248201.1
F2	70065114	70065244
refGene_gn=WASHC5;refGene_tx=XM_023248200.1;refGene_gn=WASHC5;refGene_tx=XM_023248201.1

F2 70066709 70066813
refGene_gn=WASHC5;refGene_tx=XM_023248200.1;refGene_gn=WASHC5;refGene_tx=XM_023248201.1
F2 70070447 70070527
refGene_gn=WASHC5;refGene_tx=XM_023248200.1;refGene_gn=WASHC5;refGene_tx=XM_023248201.1
F2 70070613 70070716
refGene_gn=WASHC5;refGene_tx=XM_023248200.1;refGene_gn=WASHC5;refGene_tx=XM_023248201.1
F2 70071116 70071279
refGene_gn=WASHC5;refGene_tx=XM_023248200.1;refGene_gn=WASHC5;refGene_tx=XM_023248201.1
F2 70073620 70073745
refGene_gn=WASHC5;refGene_tx=XM_023248200.1;refGene_gn=WASHC5;refGene_tx=XM_023248201.1
F2 70075228 70075408
refGene_gn=WASHC5;refGene_tx=XM_023248200.1;refGene_gn=WASHC5;refGene_tx=XM_023248201.1
F2 70076347 70076449
refGene_gn=WASHC5;refGene_tx=XM_023248200.1;refGene_gn=WASHC5;refGene_tx=XM_023248201.1
F2 70081095 70081176
refGene_gn=WASHC5;refGene_tx=XM_023248200.1;refGene_gn=WASHC5;refGene_tx=XM_023248201.1
F2 70082181 70082322
refGene_gn=WASHC5;refGene_tx=XM_023248200.1;refGene_gn=WASHC5;refGene_tx=XM_023248201.1
F2 70083097 70083208
refGene_gn=WASHC5;refGene_tx=XM_023248200.1;refGene_gn=WASHC5;refGene_tx=XM_023248201.1
F2 70084724 70084800
refGene_gn=WASHC5;refGene_tx=XM_023248200.1;refGene_gn=WASHC5;refGene_tx=XM_023248201.1
F2 70084878 70085045
refGene_gn=WASHC5;refGene_tx=XM_023248200.1;refGene_gn=WASHC5;refGene_tx=XM_023248201.1
F2 70087306 70087419
refGene_gn=WASHC5;refGene_tx=XM_023248200.1;refGene_gn=WASHC5;refGene_tx=XM_023248201.1
F2 70090256 70090386
refGene_gn=WASHC5;refGene_tx=XM_023248200.1;refGene_gn=WASHC5;refGene_tx=XM_023248201.1
F2 70093216 70093344
refGene_gn=WASHC5;refGene_tx=XM_023248200.1;refGene_gn=WASHC5;refGene_tx=XM_023248201.1
F2 70095708 70095880
refGene_gn=WASHC5;refGene_tx=XM_023248200.1;refGene_gn=WASHC5;refGene_tx=XM_023248201.1
F2 70097715 70097829
refGene_gn=WASHC5;refGene_tx=XM_023248200.1;refGene_gn=WASHC5;refGene_

tx=XM_023248201.1
F2 70101901 70102054
refGene_gn=WASHC5;refGene_tx=XM_023248200.1;refGene_gn=WASHC5;refGene_tx=XM_023248201.1
F2 70104636 70104826
refGene_gn=WASHC5;refGene_tx=XM_023248200.1;refGene_gn=WASHC5;refGene_tx=XM_023248201.1
F2 70107767 70107868
refGene_gn=WASHC5;refGene_tx=XM_023248200.1;refGene_gn=WASHC5;refGene_tx=XM_023248201.1
F2 70108456 70108541
refGene_gn=WASHC5;refGene_tx=XM_023248200.1;refGene_gn=WASHC5;refGene_tx=XM_023248201.1
F2 70109483 70109629 refGene_gn=WASHC5;refGene_tx=XM_023248200.1
F2 70110597 70110783
refGene_gn=WASHC5;refGene_tx=XM_023248200.1;refGene_gn=WASHC5;refGene_tx=XM_023248201.1
F2 70122492 70122649
refGene_gn=NSMCE2;refGene_tx=XM_004000117.4;refGene_gn=NSMCE2;refGene_tx=XM_019823121.1
F2 70156223 70156330
refGene_gn=NSMCE2;refGene_tx=XM_004000117.4;refGene_gn=NSMCE2;refGene_tx=XM_019823121.1
F2 70184031 70184185
refGene_gn=NSMCE2;refGene_tx=XM_004000117.4;refGene_gn=NSMCE2;refGene_tx=XM_019823121.1
F2 70225375 70225459 refGene_gn=NSMCE2;refGene_tx=XM_019823121.1
F2 70324386 70324487
refGene_gn=NSMCE2;refGene_tx=XM_004000117.4;refGene_gn=NSMCE2;refGene_tx=XM_019823121.1
F2 70324882 70324989
refGene_gn=NSMCE2;refGene_tx=XM_004000117.4;refGene_gn=NSMCE2;refGene_tx=XM_019823121.1
F2 70332816 70332934
refGene_gn=NSMCE2;refGene_tx=XM_004000117.4;refGene_gn=NSMCE2;refGene_tx=XM_019823121.1
F2 70385891 70386251 refGene_gn=TRIB1;refGene_tx=XM_004000118.4
F2 70388292 70388585 refGene_gn=TRIB1;refGene_tx=XM_004000118.4
F2 70390955 70391421 refGene_gn=TRIB1;refGene_tx=XM_004000118.4
F2 71318796 71319744
refGene_gn=FAM84B;refGene_tx=XM_019823122.2;refGene_gn=FAM84B;refGene_tx=XM_023248714.1;refGene_gn=FAM84B;refGene_tx=XM_023248715.1;refGene_gn=FAM84B;refGene_tx=XM_023248716.1;refGene_gn=FAM84B;refGene_tx=XM_023248717.1;refGene_gn=FAM84B;refGene_tx=XM_023248718.1;refGene_gn=FAM84B;refGene_tx=XM_023248719.1;refGene_gn=FAM84B;refGene_tx=XM_023248720.1
F2 72293034 72293791
refGene_gn=MYC;refGene_tx=NM_001173446.1;refGene_gn=MYC;refGene_tx=XM_011291443.3;refGene_gn=MYC;refGene_tx=XM_011291444.3
F2 72294883 72295446

refGene_gn=MYC;refGene_tx=NM_001173446.1;refGene_gn=MYC;refGene_tx=XM_
011291443.3;refGene_gn=MYC;refGene_tx=XM_011291444.3
F2 74025827 74026044 refGene_gn=GSDMC;refGene_tx=XM_019823124.1
F2 74027383 74027567 refGene_gn=GSDMC;refGene_tx=XM_019823124.1
F2 74029953 74030122 refGene_gn=GSDMC;refGene_tx=XM_019823124.1
F2 74035090 74035202 refGene_gn=GSDMC;refGene_tx=XM_019823124.1
F2 74036434 74036479 refGene_gn=GSDMC;refGene_tx=XM_019823124.1
F2 74036869 74036899 refGene_gn=GSDMC;refGene_tx=XM_019823124.1
F2 74037380 74037461 refGene_gn=GSDMC;refGene_tx=XM_019823124.1
F2 74037696 74037723 refGene_gn=GSDMC;refGene_tx=XM_019823124.1
F2 74038572 74038712 refGene_gn=GSDMC;refGene_tx=XM_019823124.1
F2 74039298 74039428 refGene_gn=GSDMC;refGene_tx=XM_019823124.1
F2 74039906 74039980 refGene_gn=GSDMC;refGene_tx=XM_019823124.1
F2 74040659 74040962 refGene_gn=GSDMC;refGene_tx=XM_019823124.1
F2 74066414 74066478
refGene_gn=FAM49B;refGene_tx=XM_023248575.1;refGene_gn=FAM49B;refGene_
tx=XM_023248576.1;refGene_gn=FAM49B;refGene_tx=XM_023248577.1;refGene_
gn=FAM49B;refGene_tx=XM_023248578.1;refGene_gn=FAM49B;refGene_tx=XM_02
3248579.1;refGene_gn=FAM49B;refGene_tx=XM_023248580.1;refGene_gn=FAM49
B;refGene_tx=XM_023248581.1;refGene_gn=FAM49B;refGene_tx=XM_023248582.
1;refGene_gn=FAM49B;refGene_tx=XM_023248583.1;refGene_gn=FAM49B;refGen
e_tx=XM_023248584.1
F2 74071761 74071832
refGene_gn=FAM49B;refGene_tx=XM_023248575.1;refGene_gn=FAM49B;refGene_
tx=XM_023248576.1;refGene_gn=FAM49B;refGene_tx=XM_023248577.1;refGene_
gn=FAM49B;refGene_tx=XM_023248578.1;refGene_gn=FAM49B;refGene_tx=XM_02
3248579.1;refGene_gn=FAM49B;refGene_tx=XM_023248580.1;refGene_gn=FAM49
B;refGene_tx=XM_023248581.1;refGene_gn=FAM49B;refGene_tx=XM_023248582.
1;refGene_gn=FAM49B;refGene_tx=XM_023248583.1;refGene_gn=FAM49B;refGen
e_tx=XM_023248584.1
F2 74073263 74073390
refGene_gn=FAM49B;refGene_tx=XM_023248575.1;refGene_gn=FAM49B;refGene_
tx=XM_023248576.1;refGene_gn=FAM49B;refGene_tx=XM_023248577.1;refGene_
gn=FAM49B;refGene_tx=XM_023248578.1;refGene_gn=FAM49B;refGene_tx=XM_02
3248579.1;refGene_gn=FAM49B;refGene_tx=XM_023248580.1;refGene_gn=FAM49
B;refGene_tx=XM_023248581.1;refGene_gn=FAM49B;refGene_tx=XM_023248582.
1;refGene_gn=FAM49B;refGene_tx=XM_023248583.1;refGene_gn=FAM49B;refGen
e_tx=XM_023248584.1
F2 74074714 74074794
refGene_gn=FAM49B;refGene_tx=XM_023248575.1;refGene_gn=FAM49B;refGene_
tx=XM_023248576.1;refGene_gn=FAM49B;refGene_tx=XM_023248577.1;refGene_
gn=FAM49B;refGene_tx=XM_023248578.1;refGene_gn=FAM49B;refGene_tx=XM_02
3248579.1;refGene_gn=FAM49B;refGene_tx=XM_023248580.1;refGene_gn=FAM49
B;refGene_tx=XM_023248581.1;refGene_gn=FAM49B;refGene_tx=XM_023248582.
1;refGene_gn=FAM49B;refGene_tx=XM_023248583.1;refGene_gn=FAM49B;refGen
e_tx=XM_023248584.1
F2 74075896 74076013
refGene_gn=FAM49B;refGene_tx=XM_023248575.1;refGene_gn=FAM49B;refGene_
tx=XM_023248576.1;refGene_gn=FAM49B;refGene_tx=XM_023248577.1;refGene_
gn=FAM49B;refGene_tx=XM_023248578.1;refGene_gn=FAM49B;refGene_tx=XM_02

3248579.1;refGene_gn=FAM49B;refGene_tx=XM_023248580.1;refGene_gn=FAM49B;refGene_tx=XM_023248581.1;refGene_gn=FAM49B;refGene_tx=XM_023248582.1;refGene_gn=FAM49B;refGene_tx=XM_023248583.1;refGene_gn=FAM49B;refGene_tx=XM_023248584.1
F2 74078119 74078197 refGene_gn=FAM49B;refGene_tx=XM_023248575.1;refGene_gn=FAM49B;refGene_tx=XM_023248576.1;refGene_gn=FAM49B;refGene_tx=XM_023248577.1;refGene_gn=FAM49B;refGene_tx=XM_023248578.1;refGene_gn=FAM49B;refGene_tx=XM_023248579.1;refGene_gn=FAM49B;refGene_tx=XM_023248580.1;refGene_gn=FAM49B;refGene_tx=XM_023248581.1;refGene_gn=FAM49B;refGene_tx=XM_023248582.1;refGene_gn=FAM49B;refGene_tx=XM_023248583.1;refGene_gn=FAM49B;refGene_tx=XM_023248584.1
F2 74079484 74079621 refGene_gn=FAM49B;refGene_tx=XM_023248575.1;refGene_gn=FAM49B;refGene_tx=XM_023248576.1;refGene_gn=FAM49B;refGene_tx=XM_023248577.1;refGene_gn=FAM49B;refGene_tx=XM_023248578.1;refGene_gn=FAM49B;refGene_tx=XM_023248579.1;refGene_gn=FAM49B;refGene_tx=XM_023248580.1;refGene_gn=FAM49B;refGene_tx=XM_023248581.1;refGene_gn=FAM49B;refGene_tx=XM_023248582.1;refGene_gn=FAM49B;refGene_tx=XM_023248583.1;refGene_gn=FAM49B;refGene_tx=XM_023248584.1
F2 74085537 74085643 refGene_gn=FAM49B;refGene_tx=XM_023248575.1;refGene_gn=FAM49B;refGene_tx=XM_023248576.1;refGene_gn=FAM49B;refGene_tx=XM_023248577.1;refGene_gn=FAM49B;refGene_tx=XM_023248578.1;refGene_gn=FAM49B;refGene_tx=XM_023248579.1;refGene_gn=FAM49B;refGene_tx=XM_023248580.1;refGene_gn=FAM49B;refGene_tx=XM_023248581.1;refGene_gn=FAM49B;refGene_tx=XM_023248582.1;refGene_gn=FAM49B;refGene_tx=XM_023248583.1;refGene_gn=FAM49B;refGene_tx=XM_023248584.1
F2 74090304 74090426 refGene_gn=FAM49B;refGene_tx=XM_023248575.1;refGene_gn=FAM49B;refGene_tx=XM_023248576.1;refGene_gn=FAM49B;refGene_tx=XM_023248577.1;refGene_gn=FAM49B;refGene_tx=XM_023248578.1;refGene_gn=FAM49B;refGene_tx=XM_023248579.1;refGene_gn=FAM49B;refGene_tx=XM_023248580.1;refGene_gn=FAM49B;refGene_tx=XM_023248581.1;refGene_gn=FAM49B;refGene_tx=XM_023248582.1;refGene_gn=FAM49B;refGene_tx=XM_023248583.1;refGene_gn=FAM49B;refGene_tx=XM_023248584.1
F2 74097105 74097178 refGene_gn=FAM49B;refGene_tx=XM_023248575.1;refGene_gn=FAM49B;refGene_tx=XM_023248576.1;refGene_gn=FAM49B;refGene_tx=XM_023248577.1;refGene_gn=FAM49B;refGene_tx=XM_023248578.1;refGene_gn=FAM49B;refGene_tx=XM_023248579.1;refGene_gn=FAM49B;refGene_tx=XM_023248580.1;refGene_gn=FAM49B;refGene_tx=XM_023248581.1
F2 74098110 74098183 refGene_gn=FAM49B;refGene_tx=XM_023248582.1;refGene_gn=FAM49B;refGene_tx=XM_023248583.1;refGene_gn=FAM49B;refGene_tx=XM_023248584.1
F2 74241020 74241095 refGene_gn=ASAP1;refGene_tx=XM_023248736.1
F2 74243524 74243647 refGene_gn=ASAP1;refGene_tx=XM_023248736.1
F2 74245572 74245959 refGene_gn=ASAP1;refGene_tx=XM_023248736.1
F2 74245960 74246073 refGene_gn=ASAP1;refGene_tx=XM_023248736.1
F2 74251335 74251394 refGene_gn=ASAP1;refGene_tx=XM_023248736.1

F2 74253642 74253712 refGene_gn=ASAP1;refGene_tx=XM_023248736.1
F2 74261718 74261889 refGene_gn=ASAP1;refGene_tx=XM_023248736.1
F2 74272756 74272985 refGene_gn=ASAP1;refGene_tx=XM_023248736.1
F2 74275046 74275154 refGene_gn=ASAP1;refGene_tx=XM_023248736.1
F2 74276182 74276250 refGene_gn=ASAP1;refGene_tx=XM_023248736.1
F2 74276402 74276518 refGene_gn=ASAP1;refGene_tx=XM_023248736.1
F2 74278348 74278434 refGene_gn=ASAP1;refGene_tx=XM_023248736.1
F2 74278938 74279125 refGene_gn=ASAP1;refGene_tx=XM_023248736.1
F2 74281138 74281230 refGene_gn=ASAP1;refGene_tx=XM_023248736.1
F2 74283073 74283207 refGene_gn=ASAP1;refGene_tx=XM_023248736.1
F2 74284753 74284917 refGene_gn=ASAP1;refGene_tx=XM_023248736.1
F2 74290595 74290644 refGene_gn=ASAP1;refGene_tx=XM_023248736.1
F2 74293026 74293114 refGene_gn=ASAP1;refGene_tx=XM_023248736.1
F2 74308129 74308199 refGene_gn=ASAP1;refGene_tx=XM_023248736.1
F2 74313843 74313944 refGene_gn=ASAP1;refGene_tx=XM_023248736.1
F2 74319395 74319482 refGene_gn=ASAP1;refGene_tx=XM_023248736.1
F2 74320849 74320925 refGene_gn=ASAP1;refGene_tx=XM_023248736.1
F2 74331298 74331384 refGene_gn=ASAP1;refGene_tx=XM_023248736.1
F2 74332627 74332757 refGene_gn=ASAP1;refGene_tx=XM_023248736.1
F2 74338591 74338641 refGene_gn=ASAP1;refGene_tx=XM_023248736.1
F2 74339113 74339188 refGene_gn=ASAP1;refGene_tx=XM_023248736.1
F2 74363267 74363413 refGene_gn=ASAP1;refGene_tx=XM_023248736.1
F2 74386929 74387002 refGene_gn=ASAP1;refGene_tx=XM_023248736.1
F2 74511507 74511634 refGene_gn=ASAP1;refGene_tx=XM_023248736.1
F2 74551656 74551715 refGene_gn=ASAP1;refGene_tx=XM_023248736.1
F2 74836116 74836604
refGene_gn=ADCY8;refGene_tx=XM_004000120.4;refGene_gn=ADCY8;refGene_tx
=XM_019823137.2;refGene_gn=ADCY8;refGene_tx=XM_019823138.2
F2 74839344 74839459
refGene_gn=ADCY8;refGene_tx=XM_004000120.4;refGene_gn=ADCY8;refGene_tx
=XM_019823137.2;refGene_gn=ADCY8;refGene_tx=XM_019823138.2
F2 74840941 74841034
refGene_gn=ADCY8;refGene_tx=XM_004000120.4;refGene_gn=ADCY8;refGene_tx
=XM_019823137.2;refGene_gn=ADCY8;refGene_tx=XM_019823138.2
F2 74847068 74847215
refGene_gn=ADCY8;refGene_tx=XM_004000120.4;refGene_gn=ADCY8;refGene_tx
=XM_019823137.2;refGene_gn=ADCY8;refGene_tx=XM_019823138.2
F2 74855787 74855946
refGene_gn=ADCY8;refGene_tx=XM_004000120.4;refGene_gn=ADCY8;refGene_tx
=XM_019823137.2;refGene_gn=ADCY8;refGene_tx=XM_019823138.2
F2 74867314 74867393
refGene_gn=ADCY8;refGene_tx=XM_004000120.4;refGene_gn=ADCY8;refGene_tx
=XM_019823137.2;refGene_gn=ADCY8;refGene_tx=XM_019823138.2
F2 74875997 74876170
refGene_gn=ADCY8;refGene_tx=XM_004000120.4;refGene_gn=ADCY8;refGene_tx
=XM_019823137.2;refGene_gn=ADCY8;refGene_tx=XM_019823138.2
F2 74888020 74888110
refGene_gn=ADCY8;refGene_tx=XM_004000120.4;refGene_gn=ADCY8;refGene_tx
=XM_019823138.2
F2 74890157 74890359

refGene_gn=ADCY8;refGene_tx=XM_004000120.4;refGene_gn=ADCY8;refGene_tx=XM_019823137.2;refGene_gn=ADCY8;refGene_tx=XM_019823138.2
F2 74905925 74906026 refGene_gn=ADCY8;refGene_tx=XM_004000120.4;refGene_gn=ADCY8;refGene_tx=XM_019823137.2;refGene_gn=ADCY8;refGene_tx=XM_019823138.2
F2 74921521 74921719 refGene_gn=ADCY8;refGene_tx=XM_004000120.4;refGene_gn=ADCY8;refGene_tx=XM_019823137.2
F2 74939501 74939772 refGene_gn=ADCY8;refGene_tx=XM_004000120.4;refGene_gn=ADCY8;refGene_tx=XM_019823137.2;refGene_gn=ADCY8;refGene_tx=XM_019823138.2
F2 74944159 74944318 refGene_gn=ADCY8;refGene_tx=XM_004000120.4;refGene_gn=ADCY8;refGene_tx=XM_019823137.2;refGene_gn=ADCY8;refGene_tx=XM_019823138.2
F2 74957144 74957272 refGene_gn=ADCY8;refGene_tx=XM_004000120.4;refGene_gn=ADCY8;refGene_tx=XM_019823137.2;refGene_gn=ADCY8;refGene_tx=XM_019823138.2
F2 74963152 74963264 refGene_gn=ADCY8;refGene_tx=XM_004000120.4;refGene_gn=ADCY8;refGene_tx=XM_019823137.2;refGene_gn=ADCY8;refGene_tx=XM_019823138.2
F2 74974781 74974912 refGene_gn=ADCY8;refGene_tx=XM_004000120.4;refGene_gn=ADCY8;refGene_tx=XM_019823137.2;refGene_gn=ADCY8;refGene_tx=XM_019823138.2
F2 75011979 75012129 refGene_gn=ADCY8;refGene_tx=XM_004000120.4;refGene_gn=ADCY8;refGene_tx=XM_019823137.2;refGene_gn=ADCY8;refGene_tx=XM_019823138.2
F2 75059225 75060188 refGene_gn=ADCY8;refGene_tx=XM_004000120.4;refGene_gn=ADCY8;refGene_tx=XM_019823137.2;refGene_gn=ADCY8;refGene_tx=XM_019823138.2
F2 75731246 75731353 refGene_gn=EFR3A;refGene_tx=XM_023248341.1
F2 75733057 75733208 refGene_gn=EFR3A;refGene_tx=XM_023248341.1
F2 75735822 75735944 refGene_gn=EFR3A;refGene_tx=XM_023248341.1
F2 75740570 75740720 refGene_gn=EFR3A;refGene_tx=XM_023248341.1
F2 75741287 75741425 refGene_gn=EFR3A;refGene_tx=XM_023248341.1
F2 75744237 75744316 refGene_gn=EFR3A;refGene_tx=XM_023248341.1
F2 75749125 75749261 refGene_gn=EFR3A;refGene_tx=XM_023248341.1
F2 75750936 75751104 refGene_gn=EFR3A;refGene_tx=XM_023248341.1
F2 75756001 75756116 refGene_gn=EFR3A;refGene_tx=XM_023248341.1
F2 75757055 75757107 refGene_gn=EFR3A;refGene_tx=XM_023248341.1
F2 75759993 75760166 refGene_gn=EFR3A;refGene_tx=XM_023248341.1
F2 75760607 75760683 refGene_gn=EFR3A;refGene_tx=XM_023248341.1
F2 75765202 75765364 refGene_gn=EFR3A;refGene_tx=XM_023248341.1
F2 75766247 75766379 refGene_gn=EFR3A;refGene_tx=XM_023248341.1
F2 75767739 75767807 refGene_gn=EFR3A;refGene_tx=XM_023248341.1
F2 75768778 75768906 refGene_gn=EFR3A;refGene_tx=XM_023248341.1
F2 75769867 75769918 refGene_gn=EFR3A;refGene_tx=XM_023248341.1
F2 75777840 75777932 refGene_gn=EFR3A;refGene_tx=XM_023248341.1
F2 75782735 75782784 refGene_gn=EFR3A;refGene_tx=XM_023248341.1
F2 75786625 75786786 refGene_gn=EFR3A;refGene_tx=XM_023248341.1
F2 75792515 75792659 refGene_gn=EFR3A;refGene_tx=XM_023248341.1

F2 75799021 75799356
refGene_gn=OC90;refGene_tx=XM_019823145.2;refGene_gn=OC90;refGene_tx=XM_019823148.2;refGene_gn=OC90;refGene_tx=XM_019823150.2
F2 75800782 75800889
refGene_gn=OC90;refGene_tx=XM_019823145.2;refGene_gn=OC90;refGene_tx=XM_019823148.2;refGene_gn=OC90;refGene_tx=XM_019823150.2
F2 75803342 75803514
refGene_gn=OC90;refGene_tx=XM_019823145.2;refGene_gn=OC90;refGene_tx=XM_019823148.2;refGene_gn=OC90;refGene_tx=XM_019823150.2
F2 75804514 75804631
refGene_gn=OC90;refGene_tx=XM_019823145.2;refGene_gn=OC90;refGene_tx=XM_019823148.2;refGene_gn=OC90;refGene_tx=XM_019823150.2
F2 75808707 75808761
refGene_gn=OC90;refGene_tx=XM_019823145.2;refGene_gn=OC90;refGene_tx=XM_019823148.2;refGene_gn=OC90;refGene_tx=XM_019823150.2
F2 75810585 75810633
refGene_gn=OC90;refGene_tx=XM_019823145.2;refGene_gn=OC90;refGene_tx=XM_019823148.2;refGene_gn=OC90;refGene_tx=XM_019823150.2
F2 75812601 75812652
refGene_gn=OC90;refGene_tx=XM_019823145.2;refGene_gn=OC90;refGene_tx=XM_019823148.2;refGene_gn=OC90;refGene_tx=XM_019823150.2
F2 75814139 75814181
refGene_gn=OC90;refGene_tx=XM_019823145.2;refGene_gn=OC90;refGene_tx=XM_019823148.2;refGene_gn=OC90;refGene_tx=XM_019823150.2
F2 75814346 75814475
refGene_gn=OC90;refGene_tx=XM_019823145.2;refGene_gn=OC90;refGene_tx=XM_019823148.2;refGene_gn=OC90;refGene_tx=XM_019823150.2
F2 75816356 75816469
refGene_gn=OC90;refGene_tx=XM_019823145.2;refGene_gn=OC90;refGene_tx=XM_019823148.2;refGene_gn=OC90;refGene_tx=XM_019823150.2
F2 75818992 75819167
refGene_gn=OC90;refGene_tx=XM_019823145.2;refGene_gn=OC90;refGene_tx=XM_019823148.2;refGene_gn=OC90;refGene_tx=XM_019823150.2
F2 75821169 75821226
refGene_gn=OC90;refGene_tx=XM_019823145.2;refGene_gn=OC90;refGene_tx=XM_019823148.2;refGene_gn=OC90;refGene_tx=XM_019823150.2
F2 75822535 75822601
refGene_gn=OC90;refGene_tx=XM_019823145.2;refGene_gn=OC90;refGene_tx=XM_019823148.2;refGene_gn=OC90;refGene_tx=XM_019823150.2
F2 75825275 75825374
refGene_gn=OC90;refGene_tx=XM_019823148.2;refGene_gn=OC90;refGene_tx=XM_019823150.2
F2 75829390 75829562 refGene_gn=OC90;refGene_tx=XM_019823145.2
F2 75838196 75838240 refGene_gn=HHLA1;refGene_tx=XM_004000121.4
F2 75839396 75839479 refGene_gn=HHLA1;refGene_tx=XM_004000121.4
F2 75844337 75844494 refGene_gn=HHLA1;refGene_tx=XM_004000121.4
F2 75849653 75849728 refGene_gn=HHLA1;refGene_tx=XM_004000121.4
F2 75850059 75850128 refGene_gn=HHLA1;refGene_tx=XM_004000121.4
F2 75852989 75853229 refGene_gn=HHLA1;refGene_tx=XM_004000121.4
F2 75855981 75856218 refGene_gn=HHLA1;refGene_tx=XM_004000121.4

F2 75858717 75858810 refGene_gn=HHLA1;refGene_tx=XM_004000121.4
F2 75858917 75858974 refGene_gn=HHLA1;refGene_tx=XM_004000121.4
F2 75860303 75860387 refGene_gn=HHLA1;refGene_tx=XM_004000121.4
F2 75866057 75866141 refGene_gn=HHLA1;refGene_tx=XM_004000121.4
F2 75866241 75866325 refGene_gn=HHLA1;refGene_tx=XM_004000121.4
F2 75867312 75867393 refGene_gn=HHLA1;refGene_tx=XM_004000121.4
F2 75868246 75868306 refGene_gn=HHLA1;refGene_tx=XM_004000121.4
F2 75871510 75871570 refGene_gn=HHLA1;refGene_tx=XM_004000121.4
F2 75873156 75873235 refGene_gn=HHLA1;refGene_tx=XM_004000121.4
F2 75895384 75896119 refGene_gn=KCNQ3;refGene_tx=XM_023248670.1
F2 75897711 75897796 refGene_gn=KCNQ3;refGene_tx=XM_023248670.1
F2 75899480 75899579 refGene_gn=KCNQ3;refGene_tx=XM_023248670.1
F2 75902369 75902501 refGene_gn=KCNQ3;refGene_tx=XM_023248670.1
F2 75904774 75904877 refGene_gn=KCNQ3;refGene_tx=XM_023248670.1
F2 75905757 75905960 refGene_gn=KCNQ3;refGene_tx=XM_023248670.1
F2 75921161 75921188 refGene_gn=KCNQ3;refGene_tx=XM_023248670.1
F2 75928317 75928412 refGene_gn=KCNQ3;refGene_tx=XM_023248670.1
F2 75929728 75929824 refGene_gn=KCNQ3;refGene_tx=XM_023248670.1
F2 75931436 75931547 refGene_gn=KCNQ3;refGene_tx=XM_023248670.1
F2 75932758 75932914 refGene_gn=KCNQ3;refGene_tx=XM_023248670.1
F2 75938541 75938714 refGene_gn=KCNQ3;refGene_tx=XM_023248670.1
F2 75942608 75942735 refGene_gn=KCNQ3;refGene_tx=XM_023248670.1
F2 75944087 75944178 refGene_gn=KCNQ3;refGene_tx=XM_023248670.1
F2 76207460 76207843 refGene_gn=KCNQ3;refGene_tx=XM_023248670.1
F2 76269971 76270149
refGene_gn=LRRC6;refGene_tx=XM_006943387.4;refGene_gn=LRRC6;refGene_tx
=XM_006943388.4
F2 76276942 76277028
refGene_gn=LRRC6;refGene_tx=XM_006943387.4;refGene_gn=LRRC6;refGene_tx
=XM_006943388.4
F2 76303090 76303186
refGene_gn=LRRC6;refGene_tx=XM_006943387.4;refGene_gn=LRRC6;refGene_tx
=XM_006943388.4
F2 76304103 76304173
refGene_gn=LRRC6;refGene_tx=XM_006943387.4;refGene_gn=LRRC6;refGene_tx
=XM_006943388.4
F2 76308665 76308731
refGene_gn=LRRC6;refGene_tx=XM_006943387.4;refGene_gn=LRRC6;refGene_tx
=XM_006943388.4
F2 76313959 76314037
refGene_gn=LRRC6;refGene_tx=XM_006943387.4;refGene_gn=LRRC6;refGene_tx
=XM_006943388.4
F2 76317107 76317293
refGene_gn=LRRC6;refGene_tx=XM_006943387.4;refGene_gn=LRRC6;refGene_tx
=XM_006943388.4
F2 76323919 76324122
refGene_gn=LRRC6;refGene_tx=XM_006943387.4;refGene_gn=LRRC6;refGene_tx
=XM_006943388.4
F2 76327043 76327216
refGene_gn=LRRC6;refGene_tx=XM_006943387.4;refGene_gn=LRRC6;refGene_tx

=XM_006943388.4
F2 76336503 76336581
refGene_gn=LRRC6;refGene_tx=XM_006943387.4;refGene_gn=LRRC6;refGene_tx=XM_006943388.4
F2 76341181 76341349
refGene_gn=LRRC6;refGene_tx=XM_006943387.4;refGene_gn=LRRC6;refGene_tx=XM_006943388.4
F2 76344599 76344615 refGene_gn=LRRC6;refGene_tx=XM_006943387.4
F2 76349993 76350003 refGene_gn=LRRC6;refGene_tx=XM_006943388.4
F2 76371931 76372023
refGene_gn=TMEM71;refGene_tx=XM_006943391.4;refGene_gn=TMEM71;refGene_tx=XM_006943392.4;refGene_gn=TMEM71;refGene_tx=XM_011291461.3;refGene_gn=TMEM71;refGene_tx=XM_019823159.2
F2 76372129 76372191
refGene_gn=TMEM71;refGene_tx=XM_006943391.4;refGene_gn=TMEM71;refGene_tx=XM_006943392.4;refGene_gn=TMEM71;refGene_tx=XM_011291461.3;refGene_gn=TMEM71;refGene_tx=XM_019823159.2
F2 76375217 76375293
refGene_gn=TMEM71;refGene_tx=XM_006943391.4;refGene_gn=TMEM71;refGene_tx=XM_006943392.4;refGene_gn=TMEM71;refGene_tx=XM_011291461.3;refGene_gn=TMEM71;refGene_tx=XM_019823159.2
F2 76379375 76379549
refGene_gn=TMEM71;refGene_tx=XM_006943391.4;refGene_gn=TMEM71;refGene_tx=XM_006943392.4;refGene_gn=TMEM71;refGene_tx=XM_011291461.3
F2 76390215 76390388
refGene_gn=TMEM71;refGene_tx=XM_006943391.4;refGene_gn=TMEM71;refGene_tx=XM_006943392.4;refGene_gn=TMEM71;refGene_tx=XM_011291461.3;refGene_gn=TMEM71;refGene_tx=XM_019823159.2
F2 76391812 76392025
refGene_gn=TMEM71;refGene_tx=XM_006943391.4;refGene_gn=TMEM71;refGene_tx=XM_006943392.4;refGene_gn=TMEM71;refGene_tx=XM_011291461.3;refGene_gn=TMEM71;refGene_tx=XM_019823159.2
F2 76393909 76393970
refGene_gn=TMEM71;refGene_tx=XM_006943391.4;refGene_gn=TMEM71;refGene_tx=XM_006943392.4;refGene_gn=TMEM71;refGene_tx=XM_011291461.3;refGene_gn=TMEM71;refGene_tx=XM_019823159.2
F2 76395203 76395249
refGene_gn=TMEM71;refGene_tx=XM_006943391.4;refGene_gn=TMEM71;refGene_tx=XM_006943392.4;refGene_gn=TMEM71;refGene_tx=XM_011291461.3;refGene_gn=TMEM71;refGene_tx=XM_019823159.2
F2 76410200 76410283
refGene_gn=PHF20L1;refGene_tx=XM_023248554.1;refGene_gn=PHF20L1;refGene_tx=XM_023248555.1;refGene_gn=PHF20L1;refGene_tx=XM_023248556.1;refGene_gn=PHF20L1;refGene_tx=XM_023248557.1
F2 76427499 76427671
refGene_gn=PHF20L1;refGene_tx=XM_023248554.1;refGene_gn=PHF20L1;refGene_tx=XM_023248555.1;refGene_gn=PHF20L1;refGene_tx=XM_023248556.1;refGene_gn=PHF20L1;refGene_tx=XM_023248557.1
F2 76427816 76427901
refGene_gn=PHF20L1;refGene_tx=XM_023248554.1;refGene_gn=PHF20L1;refGen

e_tx=XM_023248555.1;refGene_gn=PHF20L1;refGene_tx=XM_023248556.1;refGene_gn=PHF20L1;refGene_tx=XM_023248557.1
F2 76431631 76431720
refGene_gn=PHF20L1;refGene_tx=XM_023248554.1;refGene_gn=PHF20L1;refGene_tx=XM_023248555.1;refGene_gn=PHF20L1;refGene_tx=XM_023248556.1;refGene_gn=PHF20L1;refGene_tx=XM_023248557.1
F2 76431974 76432052
refGene_gn=PHF20L1;refGene_tx=XM_023248554.1;refGene_gn=PHF20L1;refGene_tx=XM_023248555.1
F2 76436841 76437049
refGene_gn=PHF20L1;refGene_tx=XM_023248554.1;refGene_gn=PHF20L1;refGene_tx=XM_023248555.1;refGene_gn=PHF20L1;refGene_tx=XM_023248556.1;refGene_gn=PHF20L1;refGene_tx=XM_023248557.1
F2 76437644 76437770
refGene_gn=PHF20L1;refGene_tx=XM_023248554.1;refGene_gn=PHF20L1;refGene_tx=XM_023248555.1;refGene_gn=PHF20L1;refGene_tx=XM_023248556.1;refGene_gn=PHF20L1;refGene_tx=XM_023248557.1
F2 76443845 76443934
refGene_gn=PHF20L1;refGene_tx=XM_023248554.1;refGene_gn=PHF20L1;refGene_tx=XM_023248555.1;refGene_gn=PHF20L1;refGene_tx=XM_023248556.1;refGene_gn=PHF20L1;refGene_tx=XM_023248557.1
F2 76447463 76447716
refGene_gn=PHF20L1;refGene_tx=XM_023248554.1;refGene_gn=PHF20L1;refGene_tx=XM_023248555.1;refGene_gn=PHF20L1;refGene_tx=XM_023248556.1
F2 76449838 76450027
refGene_gn=PHF20L1;refGene_tx=XM_023248554.1;refGene_gn=PHF20L1;refGene_tx=XM_023248555.1;refGene_gn=PHF20L1;refGene_tx=XM_023248556.1
F2 76450269 76450476
refGene_gn=PHF20L1;refGene_tx=XM_023248554.1;refGene_gn=PHF20L1;refGene_tx=XM_023248555.1;refGene_gn=PHF20L1;refGene_tx=XM_023248556.1
F2 76456189 76456246
refGene_gn=PHF20L1;refGene_tx=XM_023248554.1;refGene_gn=PHF20L1;refGene_tx=XM_023248555.1;refGene_gn=PHF20L1;refGene_tx=XM_023248556.1
F2 76457441 76457549
refGene_gn=PHF20L1;refGene_tx=XM_023248554.1;refGene_gn=PHF20L1;refGene_tx=XM_023248555.1;refGene_gn=PHF20L1;refGene_tx=XM_023248556.1
F2 76472048 76472213
refGene_gn=PHF20L1;refGene_tx=XM_023248554.1;refGene_gn=PHF20L1;refGene_tx=XM_023248555.1;refGene_gn=PHF20L1;refGene_tx=XM_023248556.1
F2 76475600 76475782
refGene_gn=PHF20L1;refGene_tx=XM_023248554.1;refGene_gn=PHF20L1;refGene_tx=XM_023248555.1;refGene_gn=PHF20L1;refGene_tx=XM_023248556.1
F2 76476782 76476882
refGene_gn=PHF20L1;refGene_tx=XM_023248554.1;refGene_gn=PHF20L1;refGene_tx=XM_023248555.1;refGene_gn=PHF20L1;refGene_tx=XM_023248556.1
F2 76478212 76478408
refGene_gn=PHF20L1;refGene_tx=XM_023248554.1;refGene_gn=PHF20L1;refGene_tx=XM_023248555.1;refGene_gn=PHF20L1;refGene_tx=XM_023248556.1
F2 76488957 76489324
refGene_gn=PHF20L1;refGene_tx=XM_023248554.1;refGene_gn=PHF20L1;refGen

e_tx=XM_023248555.1;refGene_gn=PHF20L1;refGene_tx=XM_023248556.1
F2	76490586	76490749
refGene_gn=PHF20L1;refGene_tx=XM_023248554.1;refGene_gn=PHF20L1;refGen
e_tx=XM_023248555.1;refGene_gn=PHF20L1;refGene_tx=XM_023248556.1
F2	76492389	76492532
refGene_gn=PHF20L1;refGene_tx=XM_023248554.1;refGene_gn=PHF20L1;refGen
e_tx=XM_023248555.1;refGene_gn=PHF20L1;refGene_tx=XM_023248556.1
F2	76506170	76506237	refGene_gn=TG;refGene_tx=XM_004000124.4
F2	76507324	76507433	refGene_gn=TG;refGene_tx=XM_004000124.4
F2	76508662	76508760	refGene_gn=TG;refGene_tx=XM_004000124.4
F2	76509828	76510032	refGene_gn=TG;refGene_tx=XM_004000124.4
F2	76510785	76510945	refGene_gn=TG;refGene_tx=XM_004000124.4
F2	76518714	76518821	refGene_gn=TG;refGene_tx=XM_004000124.4
F2	76519289	76519433	refGene_gn=TG;refGene_tx=XM_004000124.4
F2	76519852	76520038	refGene_gn=TG;refGene_tx=XM_004000124.4
F2	76524002	76525100	refGene_gn=TG;refGene_tx=XM_004000124.4
F2	76525529	76526117	refGene_gn=TG;refGene_tx=XM_004000124.4
F2	76531405	76531645	refGene_gn=TG;refGene_tx=XM_004000124.4
F2	76536124	76536265	refGene_gn=TG;refGene_tx=XM_004000124.4
F2	76536632	76536710	refGene_gn=TG;refGene_tx=XM_004000124.4
F2	76537245	76537358	refGene_gn=TG;refGene_tx=XM_004000124.4
F2	76540523	76540626	refGene_gn=TG;refGene_tx=XM_004000124.4
F2	76541591	76541792	refGene_gn=TG;refGene_tx=XM_004000124.4
F2	76544020	76544233	refGene_gn=TG;refGene_tx=XM_004000124.4
F2	76545772	76545927	refGene_gn=TG;refGene_tx=XM_004000124.4
F2	76547634	76547791	refGene_gn=TG;refGene_tx=XM_004000124.4
F2	76549624	76549849	refGene_gn=TG;refGene_tx=XM_004000124.4
F2	76557365	76557515	refGene_gn=TG;refGene_tx=XM_004000124.4
F2	76562986	76563157	refGene_gn=TG;refGene_tx=XM_004000124.4
F2	76568841	76568958	refGene_gn=TG;refGene_tx=XM_004000124.4
F2	76569515	76569631	refGene_gn=TG;refGene_tx=XM_004000124.4
F2	76573303	76573412	refGene_gn=TG;refGene_tx=XM_004000124.4
F2	76580029	76580221	refGene_gn=TG;refGene_tx=XM_004000124.4
F2	76584288	76584453	refGene_gn=TG;refGene_tx=XM_004000124.4
F2	76594169	76594235	refGene_gn=TG;refGene_tx=XM_004000124.4
F2	76596048	76596129	refGene_gn=TG;refGene_tx=XM_004000124.4
F2	76601149	76601287	refGene_gn=TG;refGene_tx=XM_004000124.4
F2	76602109	76602286	refGene_gn=TG;refGene_tx=XM_004000124.4
F2	76603899	76604011	refGene_gn=TG;refGene_tx=XM_004000124.4
F2	76607897	76607977	refGene_gn=TG;refGene_tx=XM_004000124.4
F2	76608663	76608807	refGene_gn=TG;refGene_tx=XM_004000124.4
F2	76615946	76616009	refGene_gn=TG;refGene_tx=XM_004000124.4
F2	76640360	76640495	refGene_gn=TG;refGene_tx=XM_004000124.4
F2	76642166	76642331	refGene_gn=TG;refGene_tx=XM_004000124.4
F2	76647359	76647579	refGene_gn=TG;refGene_tx=XM_004000124.4
F2	76648604	76648698	refGene_gn=TG;refGene_tx=XM_004000124.4
F2	76651621	76651781	refGene_gn=TG;refGene_tx=XM_004000124.4
F2	76659862	76660065	refGene_gn=TG;refGene_tx=XM_004000124.4
F2	76669906	76670117
refGene_gn=SLA;refGene_tx=XM_019823162.2;refGene_gn=SLA;refGene_tx=XM_

019823163.2;refGene_gn=SLA;refGene_tx=XM_019823164.2;refGene_gn=SLA;refGene_tx=XM_019823165.2;refGene_gn=SLA;refGene_tx=XM_019823166.2;refGene_gn=SLA;refGene_tx=XM_023248227.1;refGene_gn=SLA;refGene_tx=XM_023248229.1
F2 76670704 76670837
refGene_gn=SLA;refGene_tx=XM_019823162.2;refGene_gn=SLA;refGene_tx=XM_019823163.2;refGene_gn=SLA;refGene_tx=XM_019823164.2;refGene_gn=SLA;refGene_tx=XM_019823165.2;refGene_gn=SLA;refGene_tx=XM_023248227.1;refGene_gn=SLA;refGene_tx=XM_023248229.1
F2 76674914 76675046
refGene_gn=SLA;refGene_tx=XM_019823162.2;refGene_gn=SLA;refGene_tx=XM_019823163.2;refGene_gn=SLA;refGene_tx=XM_019823164.2;refGene_gn=SLA;refGene_tx=XM_019823165.2;refGene_gn=SLA;refGene_tx=XM_019823166.2;refGene_gn=SLA;refGene_tx=XM_023248227.1;refGene_gn=SLA;refGene_tx=XM_023248229.1
F2 76677619 76677723
refGene_gn=SLA;refGene_tx=XM_019823162.2;refGene_gn=SLA;refGene_tx=XM_019823163.2;refGene_gn=SLA;refGene_tx=XM_019823164.2;refGene_gn=SLA;refGene_tx=XM_019823165.2;refGene_gn=SLA;refGene_tx=XM_019823166.2;refGene_gn=SLA;refGene_tx=XM_023248227.1;refGene_gn=SLA;refGene_tx=XM_023248229.1
F2 76680684 76680771
refGene_gn=SLA;refGene_tx=XM_019823162.2;refGene_gn=SLA;refGene_tx=XM_019823163.2;refGene_gn=SLA;refGene_tx=XM_019823164.2;refGene_gn=SLA;refGene_tx=XM_019823165.2;refGene_gn=SLA;refGene_tx=XM_019823166.2;refGene_gn=SLA;refGene_tx=XM_023248227.1;refGene_gn=SLA;refGene_tx=XM_023248229.1
F2 76681818 76681918
refGene_gn=SLA;refGene_tx=XM_019823162.2;refGene_gn=SLA;refGene_tx=XM_019823163.2;refGene_gn=SLA;refGene_tx=XM_019823164.2;refGene_gn=SLA;refGene_tx=XM_019823165.2;refGene_gn=SLA;refGene_tx=XM_019823166.2;refGene_gn=SLA;refGene_tx=XM_023248227.1;refGene_gn=SLA;refGene_tx=XM_023248229.1
F2 76690876 76691054
refGene_gn=SLA;refGene_tx=XM_019823162.2;refGene_gn=SLA;refGene_tx=XM_019823163.2;refGene_gn=SLA;refGene_tx=XM_019823164.2;refGene_gn=SLA;refGene_tx=XM_019823165.2;refGene_gn=SLA;refGene_tx=XM_019823166.2;refGene_gn=SLA;refGene_tx=XM_023248227.1;refGene_gn=SLA;refGene_tx=XM_023248229.1
F2 76727919 76728084 refGene_gn=TG;refGene_tx=XM_004000124.4
F2 76729120 76729288 refGene_gn=TG;refGene_tx=XM_004000124.4
F2 76744116 76744298 refGene_gn=TG;refGene_tx=XM_004000124.4
F2 76745726 76745834 refGene_gn=TG;refGene_tx=XM_004000124.4
F2 76762552 76762687 refGene_gn=TG;refGene_tx=XM_004000124.4
F2 76764122 76764313 refGene_gn=TG;refGene_tx=XM_004000124.4
F2 76765233 76765319 refGene_gn=TG;refGene_tx=XM_004000124.4
F2 76813089 76813158 refGene_gn=WISP1;refGene_tx=XM_019823167.2
F2 76830759 76831039 refGene_gn=WISP1;refGene_tx=XM_019823167.2
F2 76837499 76837760 refGene_gn=WISP1;refGene_tx=XM_019823167.2
F2 76842353 76842547 refGene_gn=WISP1;refGene_tx=XM_019823167.2

F2 76844583 76844883 refGene_gn=WISP1;refGene_tx=XM_019823167.2
F2 76856572 76856784
refGene_gn=NDRG1;refGene_tx=XM_004000126.5;refGene_gn=NDRG1;refGene_tx=XM_006943406.4;refGene_gn=NDRG1;refGene_tx=XM_023248618.1;refGene_gn=NDRG1;refGene_tx=XM_023248619.1
F2 76859615 76859667
refGene_gn=NDRG1;refGene_tx=XM_004000126.5;refGene_gn=NDRG1;refGene_tx=XM_006943406.4;refGene_gn=NDRG1;refGene_tx=XM_023248618.1;refGene_gn=NDRG1;refGene_tx=XM_023248619.1
F2 76863277 76863313
refGene_gn=NDRG1;refGene_tx=XM_004000126.5;refGene_gn=NDRG1;refGene_tx=XM_006943406.4;refGene_gn=NDRG1;refGene_tx=XM_023248618.1;refGene_gn=NDRG1;refGene_tx=XM_023248619.1
F2 76865291 76865339
refGene_gn=NDRG1;refGene_tx=XM_004000126.5;refGene_gn=NDRG1;refGene_tx=XM_006943406.4;refGene_gn=NDRG1;refGene_tx=XM_023248618.1;refGene_gn=NDRG1;refGene_tx=XM_023248619.1
F2 76866457 76866509
refGene_gn=NDRG1;refGene_tx=XM_004000126.5;refGene_gn=NDRG1;refGene_tx=XM_006943406.4;refGene_gn=NDRG1;refGene_tx=XM_023248618.1;refGene_gn=NDRG1;refGene_tx=XM_023248619.1
F2 76867309 76867366
refGene_gn=NDRG1;refGene_tx=XM_004000126.5;refGene_gn=NDRG1;refGene_tx=XM_006943406.4;refGene_gn=NDRG1;refGene_tx=XM_023248618.1;refGene_gn=NDRG1;refGene_tx=XM_023248619.1
F2 76868916 76869020
refGene_gn=NDRG1;refGene_tx=XM_004000126.5;refGene_gn=NDRG1;refGene_tx=XM_006943406.4;refGene_gn=NDRG1;refGene_tx=XM_023248618.1;refGene_gn=NDRG1;refGene_tx=XM_023248619.1
F2 76872080 76872137
refGene_gn=NDRG1;refGene_tx=XM_004000126.5;refGene_gn=NDRG1;refGene_tx=XM_006943406.4;refGene_gn=NDRG1;refGene_tx=XM_023248618.1;refGene_gn=NDRG1;refGene_tx=XM_023248619.1
F2 76874297 76874384
refGene_gn=NDRG1;refGene_tx=XM_004000126.5;refGene_gn=NDRG1;refGene_tx=XM_006943406.4;refGene_gn=NDRG1;refGene_tx=XM_023248618.1;refGene_gn=NDRG1;refGene_tx=XM_023248619.1
F2 76875935 76875996
refGene_gn=NDRG1;refGene_tx=XM_004000126.5;refGene_gn=NDRG1;refGene_tx=XM_006943406.4;refGene_gn=NDRG1;refGene_tx=XM_023248618.1;refGene_gn=NDRG1;refGene_tx=XM_023248619.1
F2 76876778 76876841
refGene_gn=NDRG1;refGene_tx=XM_004000126.5;refGene_gn=NDRG1;refGene_tx=XM_006943406.4;refGene_gn=NDRG1;refGene_tx=XM_023248618.1;refGene_gn=NDRG1;refGene_tx=XM_023248619.1
F2 76879522 76879643
refGene_gn=NDRG1;refGene_tx=XM_004000126.5;refGene_gn=NDRG1;refGene_tx=XM_006943406.4;refGene_gn=NDRG1;refGene_tx=XM_023248618.1;refGene_gn=NDRG1;refGene_tx=XM_023248619.1
F2 76881441 76881547

refGene_gn=NDRG1;refGene_tx=XM_004000126.5;refGene_gn=NDRG1;refGene_tx=XM_006943406.4;refGene_gn=NDRG1;refGene_tx=XM_023248618.1;refGene_gn=NDRG1;refGene_tx=XM_023248619.1
F2 76894076 76894112 refGene_gn=NDRG1;refGene_tx=XM_004000126.5;refGene_gn=NDRG1;refGene_tx=XM_023248618.1
F2 76896511 76896574 refGene_gn=NDRG1;refGene_tx=XM_004000126.5;refGene_gn=NDRG1;refGene_tx=XM_023248618.1
F2 76980240 76980452 refGene_gn=LOC109496238;refGene_tx=XM_023248342.1
F2 76980955 76981148 refGene_gn=LOC109496238;refGene_tx=XM_023248342.1
F2 76982732 76982864 refGene_gn=LOC109496238;refGene_tx=XM_023248342.1
F2 76983097 76983293 refGene_gn=LOC109496238;refGene_tx=XM_023248342.1
F2 76983485 76983502 refGene_gn=LOC109496238;refGene_tx=XM_023248342.1
F2 77022569 77022743 refGene_gn=ST3GAL1;refGene_tx=XM_023248343.1
F2 77024529 77024649 refGene_gn=ST3GAL1;refGene_tx=XM_023248343.1
F2 77026044 77026090 refGene_gn=ST3GAL1;refGene_tx=XM_023248343.1
F2 77026839 77027019 refGene_gn=ST3GAL1;refGene_tx=XM_023248343.1
F2 77028076 77028140 refGene_gn=ST3GAL1;refGene_tx=XM_023248343.1
F2 77031110 77031204 refGene_gn=ST3GAL1;refGene_tx=XM_023248343.1
F2 77037497 77037809 refGene_gn=ST3GAL1;refGene_tx=XM_023248343.1
F2 77812807 77813047 refGene_gn=ZFAT;refGene_tx=XM_004000128.5;refGene_gn=ZFAT;refGene_tx=XM_019823173.2;refGene_gn=ZFAT;refGene_tx=XM_019823176.2;refGene_gn=ZFAT;refGene_tx=XM_023248127.1;refGene_gn=ZFAT;refGene_tx=XM_023248128.1
F2 77837328 77837459 refGene_gn=ZFAT;refGene_tx=XM_004000128.5;refGene_gn=ZFAT;refGene_tx=XM_019823173.2;refGene_gn=ZFAT;refGene_tx=XM_019823176.2;refGene_gn=ZFAT;refGene_tx=XM_023248127.1;refGene_gn=ZFAT;refGene_tx=XM_023248128.1
F2 77840046 77840173 refGene_gn=ZFAT;refGene_tx=XM_004000128.5;refGene_gn=ZFAT;refGene_tx=XM_019823173.2;refGene_gn=ZFAT;refGene_tx=XM_019823176.2;refGene_gn=ZFAT;refGene_tx=XM_023248127.1;refGene_gn=ZFAT;refGene_tx=XM_023248128.1
F2 77847230 77847349 refGene_gn=ZFAT;refGene_tx=XM_004000128.5;refGene_gn=ZFAT;refGene_tx=XM_019823173.2;refGene_gn=ZFAT;refGene_tx=XM_019823176.2;refGene_gn=ZFAT;refGene_tx=XM_023248127.1;refGene_gn=ZFAT;refGene_tx=XM_023248128.1
F2 77856872 77857011 refGene_gn=ZFAT;refGene_tx=XM_004000128.5;refGene_gn=ZFAT;refGene_tx=XM_019823173.2;refGene_gn=ZFAT;refGene_tx=XM_019823176.2;refGene_gn=ZFAT;refGene_tx=XM_023248127.1;refGene_gn=ZFAT;refGene_tx=XM_023248128.1
F2 77883518 77883551 refGene_gn=ZFAT;refGene_tx=XM_023248129.1
F2 77884106 77884195 refGene_gn=ZFAT;refGene_tx=XM_004000128.5;refGene_gn=ZFAT;refGene_tx=XM_019823173.2;refGene_gn=ZFAT;refGene_tx=XM_023248127.1;refGene_gn=ZFA

T;refGene_tx=XM_023248128.1;refGene_gn=ZFAT;refGene_tx=XM_023248129.1
F2 77900715 77900889
refGene_gn=ZFAT;refGene_tx=XM_004000128.5;refGene_gn=ZFAT;refGene_tx=XM_019823173.2;refGene_gn=ZFAT;refGene_tx=XM_023248127.1;refGene_gn=ZFAT;refGene_tx=XM_023248128.1;refGene_gn=ZFAT;refGene_tx=XM_023248129.1
F2 77902717 77902867
refGene_gn=ZFAT;refGene_tx=XM_004000128.5;refGene_gn=ZFAT;refGene_tx=XM_019823173.2;refGene_gn=ZFAT;refGene_tx=XM_023248127.1;refGene_gn=ZFAT;refGene_tx=XM_023248128.1;refGene_gn=ZFAT;refGene_tx=XM_023248129.1
F2 77905508 77905596
refGene_gn=ZFAT;refGene_tx=XM_004000128.5;refGene_gn=ZFAT;refGene_tx=XM_019823173.2;refGene_gn=ZFAT;refGene_tx=XM_023248127.1;refGene_gn=ZFAT;refGene_tx=XM_023248128.1;refGene_gn=ZFAT;refGene_tx=XM_023248129.1
F2 77905797 77905914
refGene_gn=ZFAT;refGene_tx=XM_019823173.2;refGene_gn=ZFAT;refGene_tx=XM_023248128.1;refGene_gn=ZFAT;refGene_tx=XM_023248129.1
F2 77923205 77923438
refGene_gn=ZFAT;refGene_tx=XM_004000128.5;refGene_gn=ZFAT;refGene_tx=XM_019823173.2;refGene_gn=ZFAT;refGene_tx=XM_019823176.2;refGene_gn=ZFAT;refGene_tx=XM_023248127.1;refGene_gn=ZFAT;refGene_tx=XM_023248128.1;refGene_gn=ZFAT;refGene_tx=XM_023248129.1
F2 77924229 77925695
refGene_gn=ZFAT;refGene_tx=XM_004000128.5;refGene_gn=ZFAT;refGene_tx=XM_019823173.2;refGene_gn=ZFAT;refGene_tx=XM_019823176.2;refGene_gn=ZFAT;refGene_tx=XM_023248127.1;refGene_gn=ZFAT;refGene_tx=XM_023248128.1;refGene_gn=ZFAT;refGene_tx=XM_023248129.1
F2 77931525 77931676
refGene_gn=ZFAT;refGene_tx=XM_004000128.5;refGene_gn=ZFAT;refGene_tx=XM_019823173.2;refGene_gn=ZFAT;refGene_tx=XM_019823176.2;refGene_gn=ZFAT;refGene_tx=XM_023248127.1;refGene_gn=ZFAT;refGene_tx=XM_023248128.1;refGene_gn=ZFAT;refGene_tx=XM_023248129.1
F2 77933291 77933480
refGene_gn=ZFAT;refGene_tx=XM_004000128.5;refGene_gn=ZFAT;refGene_tx=XM_019823173.2;refGene_gn=ZFAT;refGene_tx=XM_019823176.2;refGene_gn=ZFAT;refGene_tx=XM_023248127.1;refGene_gn=ZFAT;refGene_tx=XM_023248128.1;refGene_gn=ZFAT;refGene_tx=XM_023248129.1
F2 77957512 77957764
refGene_gn=ZFAT;refGene_tx=XM_004000128.5;refGene_gn=ZFAT;refGene_tx=XM_019823173.2;refGene_gn=ZFAT;refGene_tx=XM_019823176.2;refGene_gn=ZFAT;refGene_tx=XM_023248127.1;refGene_gn=ZFAT;refGene_tx=XM_023248128.1;refGene_gn=ZFAT;refGene_tx=XM_023248129.1
F2 77970866 77971043
refGene_gn=ZFAT;refGene_tx=XM_004000128.5;refGene_gn=ZFAT;refGene_tx=XM_019823173.2;refGene_gn=ZFAT;refGene_tx=XM_019823176.2;refGene_gn=ZFAT;refGene_tx=XM_023248127.1;refGene_gn=ZFAT;refGene_tx=XM_023248128.1;refGene_gn=ZFAT;refGene_tx=XM_023248129.1
F2 78002649 78002668
refGene_gn=ZFAT;refGene_tx=XM_004000128.5;refGene_gn=ZFAT;refGene_tx=XM_019823173.2;refGene_gn=ZFAT;refGene_tx=XM_019823176.2;refGene_gn=ZFAT;refGene_tx=XM_023248129.1

F2 78580822 78580910 refGene_gn=KHDRBS3;refGene_tx=XM_019823178.2
F2 78586527 78586618
refGene_gn=KHDRBS3;refGene_tx=XM_023248659.1;refGene_gn=KHDRBS3;refGen
e_tx=XM_023248660.1;refGene_gn=KHDRBS3;refGene_tx=XM_023248661.1;refGe
ne_gn=KHDRBS3;refGene_tx=XM_023248662.1
F2 78645635 78645754
refGene_gn=KHDRBS3;refGene_tx=XM_019823178.2;refGene_gn=KHDRBS3;refGen
e_tx=XM_023248659.1;refGene_gn=KHDRBS3;refGene_tx=XM_023248660.1;refGe
ne_gn=KHDRBS3;refGene_tx=XM_023248661.1;refGene_gn=KHDRBS3;refGene_tx=
XM_023248662.1
F2 78668281 78668398
refGene_gn=KHDRBS3;refGene_tx=XM_019823178.2;refGene_gn=KHDRBS3;refGen
e_tx=XM_023248659.1;refGene_gn=KHDRBS3;refGene_tx=XM_023248660.1;refGe
ne_gn=KHDRBS3;refGene_tx=XM_023248661.1;refGene_gn=KHDRBS3;refGene_tx=
XM_023248662.1
F2 78673683 78673830
refGene_gn=KHDRBS3;refGene_tx=XM_019823178.2;refGene_gn=KHDRBS3;refGen
e_tx=XM_023248659.1;refGene_gn=KHDRBS3;refGene_tx=XM_023248660.1;refGe
ne_gn=KHDRBS3;refGene_tx=XM_023248661.1;refGene_gn=KHDRBS3;refGene_tx=
XM_023248662.1
F2 78685765 78685905
refGene_gn=KHDRBS3;refGene_tx=XM_019823178.2;refGene_gn=KHDRBS3;refGen
e_tx=XM_023248659.1;refGene_gn=KHDRBS3;refGene_tx=XM_023248660.1;refGe
ne_gn=KHDRBS3;refGene_tx=XM_023248661.1;refGene_gn=KHDRBS3;refGene_tx=
XM_023248662.1
F2 78703326 78703522
refGene_gn=KHDRBS3;refGene_tx=XM_019823178.2;refGene_gn=KHDRBS3;refGen
e_tx=XM_023248659.1;refGene_gn=KHDRBS3;refGene_tx=XM_023248660.1;refGe
ne_gn=KHDRBS3;refGene_tx=XM_023248661.1;refGene_gn=KHDRBS3;refGene_tx=
XM_023248662.1
F2 78728872 78728955
refGene_gn=KHDRBS3;refGene_tx=XM_019823178.2;refGene_gn=KHDRBS3;refGen
e_tx=XM_023248659.1;refGene_gn=KHDRBS3;refGene_tx=XM_023248660.1;refGe
ne_gn=KHDRBS3;refGene_tx=XM_023248661.1;refGene_gn=KHDRBS3;refGene_tx=
XM_023248662.1
F2 78732799 78732830 refGene_gn=KHDRBS3;refGene_tx=XM_023248662.1
F2 78760671 78760730
refGene_gn=KHDRBS3;refGene_tx=XM_019823178.2;refGene_gn=KHDRBS3;refGen
e_tx=XM_023248659.1;refGene_gn=KHDRBS3;refGene_tx=XM_023248660.1;refGe
ne_gn=KHDRBS3;refGene_tx=XM_023248661.1
F2 78762855 78762947
refGene_gn=KHDRBS3;refGene_tx=XM_019823178.2;refGene_gn=KHDRBS3;refGen
e_tx=XM_023248659.1;refGene_gn=KHDRBS3;refGene_tx=XM_023248660.1;refGe
ne_gn=KHDRBS3;refGene_tx=XM_023248661.1
F2 80963251 80963439
refGene_gn=COL22A1;refGene_tx=XM_019823180.2;refGene_gn=COL22A1;refGen
e_tx=XM_023248206.1;refGene_gn=COL22A1;refGene_tx=XM_023248207.1;refGe
ne_gn=COL22A1;refGene_tx=XM_023248208.1
F2 80965240 80965318
refGene_gn=COL22A1;refGene_tx=XM_019823180.2;refGene_gn=COL22A1;refGen

e_tx=XM_023248206.1;refGene_gn=COL22A1;refGene_tx=XM_023248207.1;refGene_gn=COL22A1;refGene_tx=XM_023248208.1
F2 80967204 80967387
refGene_gn=COL22A1;refGene_tx=XM_019823180.2;refGene_gn=COL22A1;refGene_tx=XM_023248206.1;refGene_gn=COL22A1;refGene_tx=XM_023248207.1;refGene_gn=COL22A1;refGene_tx=XM_023248208.1
F2 80969730 80969797
refGene_gn=COL22A1;refGene_tx=XM_019823180.2;refGene_gn=COL22A1;refGene_tx=XM_023248206.1;refGene_gn=COL22A1;refGene_tx=XM_023248207.1;refGene_gn=COL22A1;refGene_tx=XM_023248208.1
F2 80971403 80971583
refGene_gn=COL22A1;refGene_tx=XM_019823180.2;refGene_gn=COL22A1;refGene_tx=XM_023248206.1;refGene_gn=COL22A1;refGene_tx=XM_023248207.1;refGene_gn=COL22A1;refGene_tx=XM_023248208.1
F2 80973942 80973987
refGene_gn=COL22A1;refGene_tx=XM_019823180.2;refGene_gn=COL22A1;refGene_tx=XM_023248206.1;refGene_gn=COL22A1;refGene_tx=XM_023248207.1;refGene_gn=COL22A1;refGene_tx=XM_023248208.1
F2 80976942 80976978
refGene_gn=COL22A1;refGene_tx=XM_019823180.2;refGene_gn=COL22A1;refGene_tx=XM_023248206.1;refGene_gn=COL22A1;refGene_tx=XM_023248207.1;refGene_gn=COL22A1;refGene_tx=XM_023248208.1
F2 80978483 80978555
refGene_gn=COL22A1;refGene_tx=XM_019823180.2;refGene_gn=COL22A1;refGene_tx=XM_023248206.1;refGene_gn=COL22A1;refGene_tx=XM_023248207.1;refGene_gn=COL22A1;refGene_tx=XM_023248208.1
F2 80980023 80980077
refGene_gn=COL22A1;refGene_tx=XM_019823180.2;refGene_gn=COL22A1;refGene_tx=XM_023248206.1;refGene_gn=COL22A1;refGene_tx=XM_023248207.1;refGene_gn=COL22A1;refGene_tx=XM_023248208.1
F2 80985418 80985472
refGene_gn=COL22A1;refGene_tx=XM_019823180.2;refGene_gn=COL22A1;refGene_tx=XM_023248206.1;refGene_gn=COL22A1;refGene_tx=XM_023248207.1;refGene_gn=COL22A1;refGene_tx=XM_023248208.1
F2 80987446 80987500
refGene_gn=COL22A1;refGene_tx=XM_019823180.2;refGene_gn=COL22A1;refGene_tx=XM_023248206.1;refGene_gn=COL22A1;refGene_tx=XM_023248207.1;refGene_gn=COL22A1;refGene_tx=XM_023248208.1
F2 80988158 80988203
refGene_gn=COL22A1;refGene_tx=XM_019823180.2;refGene_gn=COL22A1;refGene_tx=XM_023248206.1;refGene_gn=COL22A1;refGene_tx=XM_023248207.1;refGene_gn=COL22A1;refGene_tx=XM_023248208.1
F2 80992114 80992168
refGene_gn=COL22A1;refGene_tx=XM_019823180.2;refGene_gn=COL22A1;refGene_tx=XM_023248206.1;refGene_gn=COL22A1;refGene_tx=XM_023248207.1;refGene_gn=COL22A1;refGene_tx=XM_023248208.1
F2 80996691 80996745
refGene_gn=COL22A1;refGene_tx=XM_019823180.2;refGene_gn=COL22A1;refGene_tx=XM_023248206.1;refGene_gn=COL22A1;refGene_tx=XM_023248207.1;refGene_gn=COL22A1;refGene_tx=XM_023248208.1

F2 81001759 81001813
refGene_gn=COL22A1;refGene_tx=XM_019823180.2;refGene_gn=COL22A1;refGene_tx=XM_023248206.1;refGene_gn=COL22A1;refGene_tx=XM_023248207.1;refGene_gn=COL22A1;refGene_tx=XM_023248208.1
F2 81005298 81005352
refGene_gn=COL22A1;refGene_tx=XM_019823180.2;refGene_gn=COL22A1;refGene_tx=XM_023248206.1;refGene_gn=COL22A1;refGene_tx=XM_023248207.1;refGene_gn=COL22A1;refGene_tx=XM_023248208.1
F2 81009598 81009652
refGene_gn=COL22A1;refGene_tx=XM_019823180.2;refGene_gn=COL22A1;refGene_tx=XM_023248206.1;refGene_gn=COL22A1;refGene_tx=XM_023248207.1;refGene_gn=COL22A1;refGene_tx=XM_023248208.1
F2 81011735 81011789
refGene_gn=COL22A1;refGene_tx=XM_019823180.2;refGene_gn=COL22A1;refGene_tx=XM_023248206.1;refGene_gn=COL22A1;refGene_tx=XM_023248207.1;refGene_gn=COL22A1;refGene_tx=XM_023248208.1
F2 81019514 81019568
refGene_gn=COL22A1;refGene_tx=XM_019823180.2;refGene_gn=COL22A1;refGene_tx=XM_023248206.1;refGene_gn=COL22A1;refGene_tx=XM_023248207.1;refGene_gn=COL22A1;refGene_tx=XM_023248208.1
F2 81019938 81020052
refGene_gn=COL22A1;refGene_tx=XM_019823180.2;refGene_gn=COL22A1;refGene_tx=XM_023248206.1;refGene_gn=COL22A1;refGene_tx=XM_023248207.1;refGene_gn=COL22A1;refGene_tx=XM_023248208.1
F2 81025399 81025447
refGene_gn=COL22A1;refGene_tx=XM_019823180.2;refGene_gn=COL22A1;refGene_tx=XM_023248206.1;refGene_gn=COL22A1;refGene_tx=XM_023248207.1;refGene_gn=COL22A1;refGene_tx=XM_023248208.1
F2 81029616 81029661
refGene_gn=COL22A1;refGene_tx=XM_019823180.2;refGene_gn=COL22A1;refGene_tx=XM_023248206.1;refGene_gn=COL22A1;refGene_tx=XM_023248207.1;refGene_gn=COL22A1;refGene_tx=XM_023248208.1
F2 81030799 81030853
refGene_gn=COL22A1;refGene_tx=XM_019823180.2;refGene_gn=COL22A1;refGene_tx=XM_023248206.1;refGene_gn=COL22A1;refGene_tx=XM_023248207.1;refGene_gn=COL22A1;refGene_tx=XM_023248208.1
F2 81032255 81032291
refGene_gn=COL22A1;refGene_tx=XM_019823180.2;refGene_gn=COL22A1;refGene_tx=XM_023248206.1;refGene_gn=COL22A1;refGene_tx=XM_023248207.1;refGene_gn=COL22A1;refGene_tx=XM_023248208.1
F2 81042478 81042556
refGene_gn=COL22A1;refGene_tx=XM_019823180.2;refGene_gn=COL22A1;refGene_tx=XM_023248206.1;refGene_gn=COL22A1;refGene_tx=XM_023248207.1;refGene_gn=COL22A1;refGene_tx=XM_023248208.1
F2 81048907 81048952
refGene_gn=COL22A1;refGene_tx=XM_019823180.2;refGene_gn=COL22A1;refGene_tx=XM_023248206.1;refGene_gn=COL22A1;refGene_tx=XM_023248207.1;refGene_gn=COL22A1;refGene_tx=XM_023248208.1
F2 81050165 81050270
refGene_gn=COL22A1;refGene_tx=XM_019823180.2;refGene_gn=COL22A1;refGen

e_tx=XM_023248206.1;refGene_gn=COL22A1;refGene_tx=XM_023248207.1;refGene_gn=COL22A1;refGene_tx=XM_023248208.1
F2 81053183 81053237
refGene_gn=COL22A1;refGene_tx=XM_019823180.2;refGene_gn=COL22A1;refGene_tx=XM_023248206.1;refGene_gn=COL22A1;refGene_tx=XM_023248207.1;refGene_gn=COL22A1;refGene_tx=XM_023248208.1
F2 81054782 81054836
refGene_gn=COL22A1;refGene_tx=XM_019823180.2;refGene_gn=COL22A1;refGene_tx=XM_023248206.1;refGene_gn=COL22A1;refGene_tx=XM_023248207.1;refGene_gn=COL22A1;refGene_tx=XM_023248208.1
F2 81056088 81056142
refGene_gn=COL22A1;refGene_tx=XM_019823180.2;refGene_gn=COL22A1;refGene_tx=XM_023248206.1;refGene_gn=COL22A1;refGene_tx=XM_023248207.1;refGene_gn=COL22A1;refGene_tx=XM_023248208.1
F2 81056886 81056940
refGene_gn=COL22A1;refGene_tx=XM_019823180.2;refGene_gn=COL22A1;refGene_tx=XM_023248206.1;refGene_gn=COL22A1;refGene_tx=XM_023248207.1;refGene_gn=COL22A1;refGene_tx=XM_023248208.1
F2 81057271 81057325
refGene_gn=COL22A1;refGene_tx=XM_019823180.2;refGene_gn=COL22A1;refGene_tx=XM_023248206.1;refGene_gn=COL22A1;refGene_tx=XM_023248207.1;refGene_gn=COL22A1;refGene_tx=XM_023248208.1
F2 81061326 81061359
refGene_gn=COL22A1;refGene_tx=XM_019823180.2;refGene_gn=COL22A1;refGene_tx=XM_023248206.1;refGene_gn=COL22A1;refGene_tx=XM_023248207.1;refGene_gn=COL22A1;refGene_tx=XM_023248208.1
F2 81064054 81064096
refGene_gn=COL22A1;refGene_tx=XM_019823180.2;refGene_gn=COL22A1;refGene_tx=XM_023248206.1;refGene_gn=COL22A1;refGene_tx=XM_023248207.1;refGene_gn=COL22A1;refGene_tx=XM_023248208.1
F2 81067488 81067542
refGene_gn=COL22A1;refGene_tx=XM_019823180.2;refGene_gn=COL22A1;refGene_tx=XM_023248206.1;refGene_gn=COL22A1;refGene_tx=XM_023248207.1;refGene_gn=COL22A1;refGene_tx=XM_023248208.1
F2 81067960 81068023
refGene_gn=COL22A1;refGene_tx=XM_019823180.2;refGene_gn=COL22A1;refGene_tx=XM_023248206.1;refGene_gn=COL22A1;refGene_tx=XM_023248207.1;refGene_gn=COL22A1;refGene_tx=XM_023248208.1
F2 81068777 81068822
refGene_gn=COL22A1;refGene_tx=XM_019823180.2;refGene_gn=COL22A1;refGene_tx=XM_023248206.1;refGene_gn=COL22A1;refGene_tx=XM_023248207.1;refGene_gn=COL22A1;refGene_tx=XM_023248208.1
F2 81072433 81072487
refGene_gn=COL22A1;refGene_tx=XM_019823180.2;refGene_gn=COL22A1;refGene_tx=XM_023248206.1;refGene_gn=COL22A1;refGene_tx=XM_023248207.1;refGene_gn=COL22A1;refGene_tx=XM_023248208.1
F2 81073429 81073483
refGene_gn=COL22A1;refGene_tx=XM_019823180.2;refGene_gn=COL22A1;refGene_tx=XM_023248206.1;refGene_gn=COL22A1;refGene_tx=XM_023248207.1;refGene_gn=COL22A1;refGene_tx=XM_023248208.1

F2 81075612 81075666
refGene_gn=COL22A1;refGene_tx=XM_019823180.2;refGene_gn=COL22A1;refGene_tx=XM_023248206.1;refGene_gn=COL22A1;refGene_tx=XM_023248207.1;refGene_gn=COL22A1;refGene_tx=XM_023248208.1
F2 81076336 81076390
refGene_gn=COL22A1;refGene_tx=XM_019823180.2;refGene_gn=COL22A1;refGene_tx=XM_023248206.1;refGene_gn=COL22A1;refGene_tx=XM_023248207.1;refGene_gn=COL22A1;refGene_tx=XM_023248208.1
F2 81084282 81084336
refGene_gn=COL22A1;refGene_tx=XM_019823180.2;refGene_gn=COL22A1;refGene_tx=XM_023248206.1;refGene_gn=COL22A1;refGene_tx=XM_023248207.1;refGene_gn=COL22A1;refGene_tx=XM_023248208.1
F2 81098213 81098267
refGene_gn=COL22A1;refGene_tx=XM_019823180.2;refGene_gn=COL22A1;refGene_tx=XM_023248206.1;refGene_gn=COL22A1;refGene_tx=XM_023248207.1;refGene_gn=COL22A1;refGene_tx=XM_023248208.1
F2 81102208 81102262
refGene_gn=COL22A1;refGene_tx=XM_019823180.2;refGene_gn=COL22A1;refGene_tx=XM_023248206.1;refGene_gn=COL22A1;refGene_tx=XM_023248207.1;refGene_gn=COL22A1;refGene_tx=XM_023248208.1
F2 81102531 81102561
refGene_gn=COL22A1;refGene_tx=XM_019823180.2;refGene_gn=COL22A1;refGene_tx=XM_023248206.1;refGene_gn=COL22A1;refGene_tx=XM_023248207.1;refGene_gn=COL22A1;refGene_tx=XM_023248208.1
F2 81102837 81102882
refGene_gn=COL22A1;refGene_tx=XM_019823180.2;refGene_gn=COL22A1;refGene_tx=XM_023248206.1;refGene_gn=COL22A1;refGene_tx=XM_023248207.1;refGene_gn=COL22A1;refGene_tx=XM_023248208.1
F2 81107934 81107979
refGene_gn=COL22A1;refGene_tx=XM_019823180.2;refGene_gn=COL22A1;refGene_tx=XM_023248206.1;refGene_gn=COL22A1;refGene_tx=XM_023248207.1;refGene_gn=COL22A1;refGene_tx=XM_023248208.1
F2 81110728 81110782
refGene_gn=COL22A1;refGene_tx=XM_019823180.2;refGene_gn=COL22A1;refGene_tx=XM_023248206.1;refGene_gn=COL22A1;refGene_tx=XM_023248207.1;refGene_gn=COL22A1;refGene_tx=XM_023248208.1
F2 81119853 81119898
refGene_gn=COL22A1;refGene_tx=XM_019823180.2;refGene_gn=COL22A1;refGene_tx=XM_023248206.1;refGene_gn=COL22A1;refGene_tx=XM_023248207.1;refGene_gn=COL22A1;refGene_tx=XM_023248208.1
F2 81121813 81121867
refGene_gn=COL22A1;refGene_tx=XM_019823180.2;refGene_gn=COL22A1;refGene_tx=XM_023248206.1;refGene_gn=COL22A1;refGene_tx=XM_023248207.1;refGene_gn=COL22A1;refGene_tx=XM_023248208.1
F2 81123078 81123123
refGene_gn=COL22A1;refGene_tx=XM_019823180.2;refGene_gn=COL22A1;refGene_tx=XM_023248206.1;refGene_gn=COL22A1;refGene_tx=XM_023248207.1;refGene_gn=COL22A1;refGene_tx=XM_023248208.1
F2 81124127 81124181
refGene_gn=COL22A1;refGene_tx=XM_019823180.2;refGene_gn=COL22A1;refGen

e_tx=XM_023248206.1;refGene_gn=COL22A1;refGene_tx=XM_023248207.1;refGene_gn=COL22A1;refGene_tx=XM_023248208.1
F2 81140664 81140703
refGene_gn=COL22A1;refGene_tx=XM_019823180.2;refGene_gn=COL22A1;refGene_tx=XM_023248206.1;refGene_gn=COL22A1;refGene_tx=XM_023248207.1;refGene_gn=COL22A1;refGene_tx=XM_023248208.1
F2 81144076 81144139
refGene_gn=COL22A1;refGene_tx=XM_019823180.2;refGene_gn=COL22A1;refGene_tx=XM_023248206.1;refGene_gn=COL22A1;refGene_tx=XM_023248207.1;refGene_gn=COL22A1;refGene_tx=XM_023248208.1
F2 81146484 81146529
refGene_gn=COL22A1;refGene_tx=XM_019823180.2;refGene_gn=COL22A1;refGene_tx=XM_023248206.1;refGene_gn=COL22A1;refGene_tx=XM_023248207.1;refGene_gn=COL22A1;refGene_tx=XM_023248208.1
F2 81150044 81150161
refGene_gn=COL22A1;refGene_tx=XM_019823180.2;refGene_gn=COL22A1;refGene_tx=XM_023248206.1;refGene_gn=COL22A1;refGene_tx=XM_023248207.1;refGene_gn=COL22A1;refGene_tx=XM_023248208.1
F2 81151244 81151325
refGene_gn=COL22A1;refGene_tx=XM_019823180.2;refGene_gn=COL22A1;refGene_tx=XM_023248206.1;refGene_gn=COL22A1;refGene_tx=XM_023248207.1;refGene_gn=COL22A1;refGene_tx=XM_023248208.1
F2 81158055 81158331
refGene_gn=COL22A1;refGene_tx=XM_019823180.2;refGene_gn=COL22A1;refGene_tx=XM_023248206.1;refGene_gn=COL22A1;refGene_tx=XM_023248207.1;refGene_gn=COL22A1;refGene_tx=XM_023248208.1
F2 81163304 81163428
refGene_gn=COL22A1;refGene_tx=XM_019823180.2;refGene_gn=COL22A1;refGene_tx=XM_023248206.1;refGene_gn=COL22A1;refGene_tx=XM_023248207.1;refGene_gn=COL22A1;refGene_tx=XM_023248208.1
F2 81167693 81167805
refGene_gn=COL22A1;refGene_tx=XM_019823180.2;refGene_gn=COL22A1;refGene_tx=XM_023248206.1;refGene_gn=COL22A1;refGene_tx=XM_023248207.1;refGene_gn=COL22A1;refGene_tx=XM_023248208.1
F2 81177539 81177614
refGene_gn=COL22A1;refGene_tx=XM_019823180.2;refGene_gn=COL22A1;refGene_tx=XM_023248206.1;refGene_gn=COL22A1;refGene_tx=XM_023248207.1;refGene_gn=COL22A1;refGene_tx=XM_023248208.1
F2 81201595 81202162
refGene_gn=COL22A1;refGene_tx=XM_019823180.2;refGene_gn=COL22A1;refGene_tx=XM_023248206.1;refGene_gn=COL22A1;refGene_tx=XM_023248207.1;refGene_gn=COL22A1;refGene_tx=XM_023248208.1
F2 81204066 81204160
refGene_gn=COL22A1;refGene_tx=XM_019823180.2;refGene_gn=COL22A1;refGene_tx=XM_023248206.1;refGene_gn=COL22A1;refGene_tx=XM_023248207.1;refGene_gn=COL22A1;refGene_tx=XM_023248208.1
F2 81698335 81699177 refGene_gn=KCNK9;refGene_tx=XM_011291471.3
F2 81774344 81774627 refGene_gn=KCNK9;refGene_tx=XM_011291471.3
F2 81806575 81806743
refGene_gn=TRAPPC9;refGene_tx=XM_023248405.1;refGene_gn=TRAPPC9;refGen

e_tx=XM_023248406.1;refGene_gn=TRAPPC9;refGene_tx=XM_023248407.1;refGene_gn=TRAPPC9;refGene_tx=XM_023248408.1;refGene_gn=TRAPPC9;refGene_tx=XM_023248409.1;refGene_gn=TRAPPC9;refGene_tx=XM_023248410.1
F2 81807287 81807508
refGene_gn=TRAPPC9;refGene_tx=XM_023248405.1;refGene_gn=TRAPPC9;refGene_tx=XM_023248406.1;refGene_gn=TRAPPC9;refGene_tx=XM_023248407.1;refGene_gn=TRAPPC9;refGene_tx=XM_023248408.1;refGene_gn=TRAPPC9;refGene_tx=XM_023248409.1;refGene_gn=TRAPPC9;refGene_tx=XM_023248410.1
F2 81923411 81923502
refGene_gn=TRAPPC9;refGene_tx=XM_023248405.1;refGene_gn=TRAPPC9;refGene_tx=XM_023248406.1;refGene_gn=TRAPPC9;refGene_tx=XM_023248407.1;refGene_gn=TRAPPC9;refGene_tx=XM_023248408.1;refGene_gn=TRAPPC9;refGene_tx=XM_023248409.1;refGene_gn=TRAPPC9;refGene_tx=XM_023248410.1
F2 81941932 81942086
refGene_gn=TRAPPC9;refGene_tx=XM_023248405.1;refGene_gn=TRAPPC9;refGene_tx=XM_023248406.1;refGene_gn=TRAPPC9;refGene_tx=XM_023248407.1;refGene_gn=TRAPPC9;refGene_tx=XM_023248408.1;refGene_gn=TRAPPC9;refGene_tx=XM_023248409.1;refGene_gn=TRAPPC9;refGene_tx=XM_023248410.1
F2 81996199 81996310
refGene_gn=TRAPPC9;refGene_tx=XM_023248405.1;refGene_gn=TRAPPC9;refGene_tx=XM_023248406.1;refGene_gn=TRAPPC9;refGene_tx=XM_023248407.1;refGene_gn=TRAPPC9;refGene_tx=XM_023248408.1;refGene_gn=TRAPPC9;refGene_tx=XM_023248410.1
F2 82017795 82017938
refGene_gn=TRAPPC9;refGene_tx=XM_023248405.1;refGene_gn=TRAPPC9;refGene_tx=XM_023248406.1;refGene_gn=TRAPPC9;refGene_tx=XM_023248407.1;refGene_gn=TRAPPC9;refGene_tx=XM_023248408.1;refGene_gn=TRAPPC9;refGene_tx=XM_023248409.1;refGene_gn=TRAPPC9;refGene_tx=XM_023248410.1
F2 82187520 82187645
refGene_gn=TRAPPC9;refGene_tx=XM_023248405.1;refGene_gn=TRAPPC9;refGene_tx=XM_023248406.1;refGene_gn=TRAPPC9;refGene_tx=XM_023248407.1;refGene_gn=TRAPPC9;refGene_tx=XM_023248408.1;refGene_gn=TRAPPC9;refGene_tx=XM_023248409.1;refGene_gn=TRAPPC9;refGene_tx=XM_023248410.1
F2 82215898 82216051
refGene_gn=TRAPPC9;refGene_tx=XM_023248405.1;refGene_gn=TRAPPC9;refGene_tx=XM_023248406.1;refGene_gn=TRAPPC9;refGene_tx=XM_023248407.1;refGene_gn=TRAPPC9;refGene_tx=XM_023248408.1;refGene_gn=TRAPPC9;refGene_tx=XM_023248409.1
F2 82233590 82233754
refGene_gn=TRAPPC9;refGene_tx=XM_023248405.1;refGene_gn=TRAPPC9;refGene_tx=XM_023248406.1;refGene_gn=TRAPPC9;refGene_tx=XM_023248407.1;refGene_gn=TRAPPC9;refGene_tx=XM_023248408.1;refGene_gn=TRAPPC9;refGene_tx=XM_023248409.1;refGene_gn=TRAPPC9;refGene_tx=XM_023248410.1
F2 82236991 82237124
refGene_gn=TRAPPC9;refGene_tx=XM_023248405.1;refGene_gn=TRAPPC9;refGene_tx=XM_023248406.1;refGene_gn=TRAPPC9;refGene_tx=XM_023248407.1;refGene_gn=TRAPPC9;refGene_tx=XM_023248408.1;refGene_gn=TRAPPC9;refGene_tx=XM_023248409.1;refGene_gn=TRAPPC9;refGene_tx=XM_023248410.1
F2 82240670 82240797
refGene_gn=TRAPPC9;refGene_tx=XM_023248405.1;refGene_gn=TRAPPC9;refGen

e_tx=XM_023248406.1;refGene_gn=TRAPPC9;refGene_tx=XM_023248407.1;refGene_gn=TRAPPC9;refGene_tx=XM_023248408.1;refGene_gn=TRAPPC9;refGene_tx=XM_023248409.1;refGene_gn=TRAPPC9;refGene_tx=XM_023248410.1
F2 82243759 82243845
refGene_gn=TRAPPC9;refGene_tx=XM_023248405.1;refGene_gn=TRAPPC9;refGene_tx=XM_023248406.1;refGene_gn=TRAPPC9;refGene_tx=XM_023248407.1;refGene_gn=TRAPPC9;refGene_tx=XM_023248408.1;refGene_gn=TRAPPC9;refGene_tx=XM_023248409.1;refGene_gn=TRAPPC9;refGene_tx=XM_023248410.1
F2 82252130 82252276
refGene_gn=TRAPPC9;refGene_tx=XM_023248405.1;refGene_gn=TRAPPC9;refGene_tx=XM_023248406.1;refGene_gn=TRAPPC9;refGene_tx=XM_023248407.1;refGene_gn=TRAPPC9;refGene_tx=XM_023248408.1;refGene_gn=TRAPPC9;refGene_tx=XM_023248409.1;refGene_gn=TRAPPC9;refGene_tx=XM_023248410.1
F2 82263307 82263434
refGene_gn=TRAPPC9;refGene_tx=XM_023248405.1;refGene_gn=TRAPPC9;refGene_tx=XM_023248406.1;refGene_gn=TRAPPC9;refGene_tx=XM_023248407.1;refGene_gn=TRAPPC9;refGene_tx=XM_023248408.1;refGene_gn=TRAPPC9;refGene_tx=XM_023248409.1;refGene_gn=TRAPPC9;refGene_tx=XM_023248410.1
F2 82295576 82295720
refGene_gn=TRAPPC9;refGene_tx=XM_023248405.1;refGene_gn=TRAPPC9;refGene_tx=XM_023248406.1;refGene_gn=TRAPPC9;refGene_tx=XM_023248407.1;refGene_gn=TRAPPC9;refGene_tx=XM_023248408.1;refGene_gn=TRAPPC9;refGene_tx=XM_023248409.1;refGene_gn=TRAPPC9;refGene_tx=XM_023248410.1
F2 82299511 82299728
refGene_gn=TRAPPC9;refGene_tx=XM_023248405.1;refGene_gn=TRAPPC9;refGene_tx=XM_023248406.1;refGene_gn=TRAPPC9;refGene_tx=XM_023248407.1;refGene_gn=TRAPPC9;refGene_tx=XM_023248408.1;refGene_gn=TRAPPC9;refGene_tx=XM_023248409.1;refGene_gn=TRAPPC9;refGene_tx=XM_023248410.1
F2 82320758 82320884
refGene_gn=TRAPPC9;refGene_tx=XM_023248405.1;refGene_gn=TRAPPC9;refGene_tx=XM_023248406.1;refGene_gn=TRAPPC9;refGene_tx=XM_023248407.1;refGene_gn=TRAPPC9;refGene_tx=XM_023248408.1;refGene_gn=TRAPPC9;refGene_tx=XM_023248409.1;refGene_gn=TRAPPC9;refGene_tx=XM_023248410.1
F2 82328182 82328304
refGene_gn=TRAPPC9;refGene_tx=XM_023248405.1;refGene_gn=TRAPPC9;refGene_tx=XM_023248406.1;refGene_gn=TRAPPC9;refGene_tx=XM_023248407.1;refGene_gn=TRAPPC9;refGene_tx=XM_023248408.1;refGene_gn=TRAPPC9;refGene_tx=XM_023248409.1;refGene_gn=TRAPPC9;refGene_tx=XM_023248410.1
F2 82348781 82348808
refGene_gn=TRAPPC9;refGene_tx=XM_023248405.1;refGene_gn=TRAPPC9;refGene_tx=XM_023248406.1;refGene_gn=TRAPPC9;refGene_tx=XM_023248409.1;refGene_gn=TRAPPC9;refGene_tx=XM_023248410.1
F2 82356990 82357119
refGene_gn=TRAPPC9;refGene_tx=XM_023248405.1;refGene_gn=TRAPPC9;refGene_tx=XM_023248406.1;refGene_gn=TRAPPC9;refGene_tx=XM_023248407.1;refGene_gn=TRAPPC9;refGene_tx=XM_023248408.1;refGene_gn=TRAPPC9;refGene_tx=XM_023248409.1;refGene_gn=TRAPPC9;refGene_tx=XM_023248410.1
F2 82359638 82359784
refGene_gn=TRAPPC9;refGene_tx=XM_023248405.1;refGene_gn=TRAPPC9;refGene_tx=XM_023248406.1;refGene_gn=TRAPPC9;refGene_tx=XM_023248407.1;refGe

ne_gn=TRAPPC9;refGene_tx=XM_023248408.1;refGene_gn=TRAPPC9;refGene_tx=XM_023248409.1;refGene_gn=TRAPPC9;refGene_tx=XM_023248410.1
F2 82365044 82365065
refGene_gn=TRAPPC9;refGene_tx=XM_023248405.1;refGene_gn=TRAPPC9;refGene_tx=XM_023248407.1;refGene_gn=TRAPPC9;refGene_tx=XM_023248409.1;refGene_gn=TRAPPC9;refGene_tx=XM_023248410.1
F2 82368568 82369152
refGene_gn=TRAPPC9;refGene_tx=XM_023248405.1;refGene_gn=TRAPPC9;refGene_tx=XM_023248406.1;refGene_gn=TRAPPC9;refGene_tx=XM_023248407.1;refGene_gn=TRAPPC9;refGene_tx=XM_023248408.1;refGene_gn=TRAPPC9;refGene_tx=XM_023248409.1;refGene_gn=TRAPPC9;refGene_tx=XM_023248410.1
F2 82412386 82412533
refGene_gn=CHRAC1;refGene_tx=XM_023248419.1;refGene_gn=CHRAC1;refGene_tx=XM_023248420.1
F2 82413827 82413954
refGene_gn=CHRAC1;refGene_tx=XM_023248419.1;refGene_gn=CHRAC1;refGene_tx=XM_023248420.1
F2 82414075 82414140 refGene_gn=CHRAC1;refGene_tx=XM_023248419.1
F2 82414636 82414755 refGene_gn=CHRAC1;refGene_tx=XM_023248420.1
F2 82432171 82432280
refGene_gn=AGO2;refGene_tx=XM_019823195.2;refGene_gn=AGO2;refGene_tx=XM_019823197.2;refGene_gn=AGO2;refGene_tx=XM_023248472.1;refGene_gn=AGO2;refGene_tx=XM_023248473.1
F2 82432484 82432684
refGene_gn=AGO2;refGene_tx=XM_019823195.2;refGene_gn=AGO2;refGene_tx=XM_019823197.2;refGene_gn=AGO2;refGene_tx=XM_023248472.1;refGene_gn=AGO2;refGene_tx=XM_023248473.1
F2 82433243 82433345
refGene_gn=AGO2;refGene_tx=XM_019823195.2;refGene_gn=AGO2;refGene_tx=XM_019823197.2;refGene_gn=AGO2;refGene_tx=XM_023248472.1;refGene_gn=AGO2;refGene_tx=XM_023248473.1
F2 82436530 82436665
refGene_gn=AGO2;refGene_tx=XM_019823195.2;refGene_gn=AGO2;refGene_tx=XM_019823197.2;refGene_gn=AGO2;refGene_tx=XM_023248472.1;refGene_gn=AGO2;refGene_tx=XM_023248473.1
F2 82438585 82438780
refGene_gn=AGO2;refGene_tx=XM_019823195.2;refGene_gn=AGO2;refGene_tx=XM_019823197.2;refGene_gn=AGO2;refGene_tx=XM_023248472.1;refGene_gn=AGO2;refGene_tx=XM_023248473.1
F2 82440955 82441046
refGene_gn=AGO2;refGene_tx=XM_019823195.2;refGene_gn=AGO2;refGene_tx=XM_019823197.2;refGene_gn=AGO2;refGene_tx=XM_023248472.1;refGene_gn=AGO2;refGene_tx=XM_023248473.1
F2 82444642 82444802
refGene_gn=AGO2;refGene_tx=XM_019823195.2;refGene_gn=AGO2;refGene_tx=XM_019823197.2;refGene_gn=AGO2;refGene_tx=XM_023248472.1;refGene_gn=AGO2;refGene_tx=XM_023248473.1
F2 82446128 82446313
refGene_gn=AGO2;refGene_tx=XM_019823195.2;refGene_gn=AGO2;refGene_tx=XM_019823197.2;refGene_gn=AGO2;refGene_tx=XM_023248472.1;refGene_gn=AGO

2;refGene_tx=XM_023248473.1
F2 82448039 82448173
refGene_gn=AGO2;refGene_tx=XM_019823195.2;refGene_gn=AGO2;refGene_tx=XM_019823197.2;refGene_gn=AGO2;refGene_tx=XM_023248472.1;refGene_gn=AGO2;refGene_tx=XM_023248473.1
F2 82451966 82452089
refGene_gn=AGO2;refGene_tx=XM_019823195.2;refGene_gn=AGO2;refGene_tx=XM_019823197.2;refGene_gn=AGO2;refGene_tx=XM_023248472.1;refGene_gn=AGO2;refGene_tx=XM_023248473.1
F2 82452221 82452341
refGene_gn=AGO2;refGene_tx=XM_019823195.2;refGene_gn=AGO2;refGene_tx=XM_019823197.2;refGene_gn=AGO2;refGene_tx=XM_023248472.1;refGene_gn=AGO2;refGene_tx=XM_023248473.1
F2 82453032 82453180
refGene_gn=AGO2;refGene_tx=XM_019823195.2;refGene_gn=AGO2;refGene_tx=XM_019823197.2;refGene_gn=AGO2;refGene_tx=XM_023248472.1;refGene_gn=AGO2;refGene_tx=XM_023248473.1
F2 82453834 82453922
refGene_gn=AGO2;refGene_tx=XM_019823195.2;refGene_gn=AGO2;refGene_tx=XM_019823197.2;refGene_gn=AGO2;refGene_tx=XM_023248472.1;refGene_gn=AGO2;refGene_tx=XM_023248473.1
F2 82454705 82454840
refGene_gn=AGO2;refGene_tx=XM_019823195.2;refGene_gn=AGO2;refGene_tx=XM_019823197.2;refGene_gn=AGO2;refGene_tx=XM_023248472.1;refGene_gn=AGO2;refGene_tx=XM_023248473.1
F2 82455648 82455785
refGene_gn=AGO2;refGene_tx=XM_019823195.2;refGene_gn=AGO2;refGene_tx=XM_019823197.2;refGene_gn=AGO2;refGene_tx=XM_023248472.1;refGene_gn=AGO2;refGene_tx=XM_023248473.1
F2 82457570 82457752
refGene_gn=AGO2;refGene_tx=XM_019823195.2;refGene_gn=AGO2;refGene_tx=XM_019823197.2;refGene_gn=AGO2;refGene_tx=XM_023248472.1;refGene_gn=AGO2;refGene_tx=XM_023248473.1
F2 82466360 82466481
refGene_gn=AGO2;refGene_tx=XM_019823195.2;refGene_gn=AGO2;refGene_tx=XM_019823197.2;refGene_gn=AGO2;refGene_tx=XM_023248472.1;refGene_gn=AGO2;refGene_tx=XM_023248473.1
F2 82485534 82485730
refGene_gn=AGO2;refGene_tx=XM_019823195.2;refGene_gn=AGO2;refGene_tx=XM_019823197.2;refGene_gn=AGO2;refGene_tx=XM_023248472.1
F2 82498809 82498816 refGene_gn=AGO2;refGene_tx=XM_019823197.2
F2 82535831 82535853 refGene_gn=AGO2;refGene_tx=XM_023248472.1
F2 82537074 82537123 refGene_gn=AGO2;refGene_tx=XM_019823195.2
F2 82559878 82560091
refGene_gn=PTK2;refGene_tx=XM_023248620.1;refGene_gn=PTK2;refGene_tx=XM_023248621.1;refGene_gn=PTK2;refGene_tx=XM_023248622.1;refGene_gn=PTK2;refGene_tx=XM_023248623.1;refGene_gn=PTK2;refGene_tx=XM_023248624.1;refGene_gn=PTK2;refGene_tx=XM_023248625.1;refGene_gn=PTK2;refGene_tx=XM_023248626.1;refGene_gn=PTK2;refGene_tx=XM_023248627.1;refGene_gn=PTK2;refGene_tx=XM_023248628.1;refGene_gn=PTK2;refGene_tx=XM_023248629.1;

refGene_gn=PTK2;refGene_tx=XM_023248630.1;refGene_gn=PTK2;refGene_tx=XM_023248631.1;refGene_gn=PTK2;refGene_tx=XM_023248632.1;refGene_gn=PTK2;refGene_tx=XM_023248633.1;refGene_gn=PTK2;refGene_tx=XM_023248634.1;refGene_gn=PTK2;refGene_tx=XM_023248635.1;refGene_gn=PTK2;refGene_tx=XM_023248636.1;refGene_gn=PTK2;refGene_tx=XM_023248637.1;refGene_gn=PTK2;refGene_tx=XM_023248638.1;refGene_gn=PTK2;refGene_tx=XM_023248639.1;refGene_gn=PTK2;refGene_tx=XM_023248640.1;refGene_gn=PTK2;refGene_tx=XM_023248641.1;refGene_gn=PTK2;refGene_tx=XM_023248642.1;refGene_gn=PTK2;refGene_tx=XM_023248643.1
F2 82563416 82563497
refGene_gn=PTK2;refGene_tx=XM_023248620.1;refGene_gn=PTK2;refGene_tx=XM_023248621.1;refGene_gn=PTK2;refGene_tx=XM_023248622.1;refGene_gn=PTK2;refGene_tx=XM_023248623.1;refGene_gn=PTK2;refGene_tx=XM_023248624.1;refGene_gn=PTK2;refGene_tx=XM_023248625.1;refGene_gn=PTK2;refGene_tx=XM_023248626.1;refGene_gn=PTK2;refGene_tx=XM_023248627.1;refGene_gn=PTK2;refGene_tx=XM_023248628.1;refGene_gn=PTK2;refGene_tx=XM_023248629.1;refGene_gn=PTK2;refGene_tx=XM_023248630.1;refGene_gn=PTK2;refGene_tx=XM_023248631.1;refGene_gn=PTK2;refGene_tx=XM_023248632.1;refGene_gn=PTK2;refGene_tx=XM_023248633.1;refGene_gn=PTK2;refGene_tx=XM_023248634.1;refGene_gn=PTK2;refGene_tx=XM_023248635.1;refGene_gn=PTK2;refGene_tx=XM_023248636.1;refGene_gn=PTK2;refGene_tx=XM_023248637.1;refGene_gn=PTK2;refGene_tx=XM_023248638.1;refGene_gn=PTK2;refGene_tx=XM_023248639.1;refGene_gn=PTK2;refGene_tx=XM_023248640.1;refGene_gn=PTK2;refGene_tx=XM_023248641.1;refGene_gn=PTK2;refGene_tx=XM_023248642.1;refGene_gn=PTK2;refGene_tx=XM_023248643.1
F2 82564579 82564735
refGene_gn=PTK2;refGene_tx=XM_023248620.1;refGene_gn=PTK2;refGene_tx=XM_023248621.1;refGene_gn=PTK2;refGene_tx=XM_023248622.1;refGene_gn=PTK2;refGene_tx=XM_023248623.1;refGene_gn=PTK2;refGene_tx=XM_023248624.1;refGene_gn=PTK2;refGene_tx=XM_023248625.1;refGene_gn=PTK2;refGene_tx=XM_023248626.1;refGene_gn=PTK2;refGene_tx=XM_023248627.1;refGene_gn=PTK2;refGene_tx=XM_023248628.1;refGene_gn=PTK2;refGene_tx=XM_023248629.1;refGene_gn=PTK2;refGene_tx=XM_023248630.1;refGene_gn=PTK2;refGene_tx=XM_023248631.1;refGene_gn=PTK2;refGene_tx=XM_023248632.1;refGene_gn=PTK2;refGene_tx=XM_023248633.1;refGene_gn=PTK2;refGene_tx=XM_023248634.1;refGene_gn=PTK2;refGene_tx=XM_023248635.1;refGene_gn=PTK2;refGene_tx=XM_023248636.1;refGene_gn=PTK2;refGene_tx=XM_023248637.1;refGene_gn=PTK2;refGene_tx=XM_023248638.1;refGene_gn=PTK2;refGene_tx=XM_023248639.1;refGene_gn=PTK2;refGene_tx=XM_023248640.1;refGene_gn=PTK2;refGene_tx=XM_023248641.1;refGene_gn=PTK2;refGene_tx=XM_023248642.1;refGene_gn=PTK2;refGene_tx=XM_023248643.1
F2 82565966 82565975
refGene_gn=PTK2;refGene_tx=XM_023248620.1;refGene_gn=PTK2;refGene_tx=XM_023248621.1;refGene_gn=PTK2;refGene_tx=XM_023248622.1;refGene_gn=PTK2;refGene_tx=XM_023248623.1;refGene_gn=PTK2;refGene_tx=XM_023248624.1;refGene_gn=PTK2;refGene_tx=XM_023248625.1;refGene_gn=PTK2;refGene_tx=XM_023248627.1;refGene_gn=PTK2;refGene_tx=XM_023248628.1;refGene_gn=PTK2;refGene_tx=XM_023248629.1;refGene_gn=PTK2;refGene_tx=XM_023248630.1;refGene_gn=PTK2;refGene_tx=XM_023248631.1;refGene_gn=PTK2;refGene_tx=XM_023248632.1;refGene_gn=PTK2;refGene_tx=XM_023248633.1;refGene_gn=PTK

2;refGene_tx=XM_023248634.1;refGene_gn=PTK2;refGene_tx=XM_023248635.1;
refGene_gn=PTK2;refGene_tx=XM_023248636.1;refGene_gn=PTK2;refGene_tx=X
M_023248637.1;refGene_gn=PTK2;refGene_tx=XM_023248639.1;refGene_gn=PTK
2;refGene_tx=XM_023248640.1
F2 82569047 82569154
refGene_gn=PTK2;refGene_tx=XM_023248620.1;refGene_gn=PTK2;refGene_tx=X
M_023248621.1;refGene_gn=PTK2;refGene_tx=XM_023248622.1;refGene_gn=PTK
2;refGene_tx=XM_023248623.1;refGene_gn=PTK2;refGene_tx=XM_023248624.1;
refGene_gn=PTK2;refGene_tx=XM_023248625.1;refGene_gn=PTK2;refGene_tx=X
M_023248626.1;refGene_gn=PTK2;refGene_tx=XM_023248627.1;refGene_gn=PTK
2;refGene_tx=XM_023248628.1;refGene_gn=PTK2;refGene_tx=XM_023248629.1;
refGene_gn=PTK2;refGene_tx=XM_023248630.1;refGene_gn=PTK2;refGene_tx=X
M_023248631.1;refGene_gn=PTK2;refGene_tx=XM_023248632.1;refGene_gn=PTK
2;refGene_tx=XM_023248633.1;refGene_gn=PTK2;refGene_tx=XM_023248634.1;
refGene_gn=PTK2;refGene_tx=XM_023248635.1;refGene_gn=PTK2;refGene_tx=X
M_023248636.1;refGene_gn=PTK2;refGene_tx=XM_023248637.1;refGene_gn=PTK
2;refGene_tx=XM_023248638.1;refGene_gn=PTK2;refGene_tx=XM_023248639.1;
refGene_gn=PTK2;refGene_tx=XM_023248640.1;refGene_gn=PTK2;refGene_tx=X
M_023248641.1;refGene_gn=PTK2;refGene_tx=XM_023248642.1;refGene_gn=PTK
2;refGene_tx=XM_023248643.1
F2 82569475 82569515
refGene_gn=PTK2;refGene_tx=XM_023248620.1;refGene_gn=PTK2;refGene_tx=X
M_023248621.1;refGene_gn=PTK2;refGene_tx=XM_023248622.1;refGene_gn=PTK
2;refGene_tx=XM_023248623.1;refGene_gn=PTK2;refGene_tx=XM_023248624.1;
refGene_gn=PTK2;refGene_tx=XM_023248625.1;refGene_gn=PTK2;refGene_tx=X
M_023248626.1;refGene_gn=PTK2;refGene_tx=XM_023248627.1;refGene_gn=PTK
2;refGene_tx=XM_023248628.1;refGene_gn=PTK2;refGene_tx=XM_023248629.1;
refGene_gn=PTK2;refGene_tx=XM_023248630.1;refGene_gn=PTK2;refGene_tx=X
M_023248631.1;refGene_gn=PTK2;refGene_tx=XM_023248632.1;refGene_gn=PTK
2;refGene_tx=XM_023248633.1;refGene_gn=PTK2;refGene_tx=XM_023248634.1;
refGene_gn=PTK2;refGene_tx=XM_023248635.1;refGene_gn=PTK2;refGene_tx=X
M_023248636.1;refGene_gn=PTK2;refGene_tx=XM_023248637.1;refGene_gn=PTK
2;refGene_tx=XM_023248638.1;refGene_gn=PTK2;refGene_tx=XM_023248639.1;
refGene_gn=PTK2;refGene_tx=XM_023248640.1;refGene_gn=PTK2;refGene_tx=X
M_023248641.1;refGene_gn=PTK2;refGene_tx=XM_023248642.1;refGene_gn=PTK
2;refGene_tx=XM_023248643.1
F2 82572813 82572876
refGene_gn=PTK2;refGene_tx=XM_023248620.1;refGene_gn=PTK2;refGene_tx=X
M_023248621.1;refGene_gn=PTK2;refGene_tx=XM_023248622.1;refGene_gn=PTK
2;refGene_tx=XM_023248623.1;refGene_gn=PTK2;refGene_tx=XM_023248624.1;
refGene_gn=PTK2;refGene_tx=XM_023248625.1;refGene_gn=PTK2;refGene_tx=X
M_023248626.1;refGene_gn=PTK2;refGene_tx=XM_023248627.1;refGene_gn=PTK
2;refGene_tx=XM_023248628.1;refGene_gn=PTK2;refGene_tx=XM_023248629.1;
refGene_gn=PTK2;refGene_tx=XM_023248630.1;refGene_gn=PTK2;refGene_tx=X
M_023248631.1;refGene_gn=PTK2;refGene_tx=XM_023248632.1;refGene_gn=PTK
2;refGene_tx=XM_023248633.1;refGene_gn=PTK2;refGene_tx=XM_023248634.1;
refGene_gn=PTK2;refGene_tx=XM_023248635.1;refGene_gn=PTK2;refGene_tx=X
M_023248636.1;refGene_gn=PTK2;refGene_tx=XM_023248637.1;refGene_gn=PTK
2;refGene_tx=XM_023248638.1;refGene_gn=PTK2;refGene_tx=XM_023248639.1;
refGene_gn=PTK2;refGene_tx=XM_023248640.1;refGene_gn=PTK2;refGene_tx=X

M_023248641.1;refGene_gn=PTK2;refGene_tx=XM_023248642.1;refGene_gn=PTK2;refGene_tx=XM_023248643.1
F2 82584636 82584777
refGene_gn=PTK2;refGene_tx=XM_023248620.1;refGene_gn=PTK2;refGene_tx=XM_023248621.1;refGene_gn=PTK2;refGene_tx=XM_023248622.1;refGene_gn=PTK2;refGene_tx=XM_023248623.1;refGene_gn=PTK2;refGene_tx=XM_023248624.1;refGene_gn=PTK2;refGene_tx=XM_023248625.1;refGene_gn=PTK2;refGene_tx=XM_023248626.1;refGene_gn=PTK2;refGene_tx=XM_023248627.1;refGene_gn=PTK2;refGene_tx=XM_023248628.1;refGene_gn=PTK2;refGene_tx=XM_023248629.1;refGene_gn=PTK2;refGene_tx=XM_023248630.1;refGene_gn=PTK2;refGene_tx=XM_023248631.1;refGene_gn=PTK2;refGene_tx=XM_023248632.1;refGene_gn=PTK2;refGene_tx=XM_023248633.1;refGene_gn=PTK2;refGene_tx=XM_023248634.1;refGene_gn=PTK2;refGene_tx=XM_023248635.1;refGene_gn=PTK2;refGene_tx=XM_023248636.1;refGene_gn=PTK2;refGene_tx=XM_023248637.1;refGene_gn=PTK2;refGene_tx=XM_023248638.1;refGene_gn=PTK2;refGene_tx=XM_023248639.1;refGene_gn=PTK2;refGene_tx=XM_023248640.1;refGene_gn=PTK2;refGene_tx=XM_023248641.1;refGene_gn=PTK2;refGene_tx=XM_023248642.1;refGene_gn=PTK2;refGene_tx=XM_023248643.1
F2 82585536 82585674
refGene_gn=PTK2;refGene_tx=XM_023248620.1;refGene_gn=PTK2;refGene_tx=XM_023248621.1;refGene_gn=PTK2;refGene_tx=XM_023248622.1;refGene_gn=PTK2;refGene_tx=XM_023248623.1;refGene_gn=PTK2;refGene_tx=XM_023248624.1;refGene_gn=PTK2;refGene_tx=XM_023248625.1;refGene_gn=PTK2;refGene_tx=XM_023248626.1;refGene_gn=PTK2;refGene_tx=XM_023248627.1;refGene_gn=PTK2;refGene_tx=XM_023248628.1;refGene_gn=PTK2;refGene_tx=XM_023248629.1;refGene_gn=PTK2;refGene_tx=XM_023248630.1;refGene_gn=PTK2;refGene_tx=XM_023248631.1;refGene_gn=PTK2;refGene_tx=XM_023248632.1;refGene_gn=PTK2;refGene_tx=XM_023248633.1;refGene_gn=PTK2;refGene_tx=XM_023248634.1;refGene_gn=PTK2;refGene_tx=XM_023248635.1;refGene_gn=PTK2;refGene_tx=XM_023248636.1;refGene_gn=PTK2;refGene_tx=XM_023248637.1;refGene_gn=PTK2;refGene_tx=XM_023248638.1;refGene_gn=PTK2;refGene_tx=XM_023248639.1;refGene_gn=PTK2;refGene_tx=XM_023248640.1;refGene_gn=PTK2;refGene_tx=XM_023248641.1;refGene_gn=PTK2;refGene_tx=XM_023248642.1;refGene_gn=PTK2;refGene_tx=XM_023248643.1
F2 82589118 82589205
refGene_gn=PTK2;refGene_tx=XM_023248620.1;refGene_gn=PTK2;refGene_tx=XM_023248621.1;refGene_gn=PTK2;refGene_tx=XM_023248622.1;refGene_gn=PTK2;refGene_tx=XM_023248623.1;refGene_gn=PTK2;refGene_tx=XM_023248624.1;refGene_gn=PTK2;refGene_tx=XM_023248625.1;refGene_gn=PTK2;refGene_tx=XM_023248626.1;refGene_gn=PTK2;refGene_tx=XM_023248627.1;refGene_gn=PTK2;refGene_tx=XM_023248628.1;refGene_gn=PTK2;refGene_tx=XM_023248629.1;refGene_gn=PTK2;refGene_tx=XM_023248630.1;refGene_gn=PTK2;refGene_tx=XM_023248631.1;refGene_gn=PTK2;refGene_tx=XM_023248632.1;refGene_gn=PTK2;refGene_tx=XM_023248633.1;refGene_gn=PTK2;refGene_tx=XM_023248634.1;refGene_gn=PTK2;refGene_tx=XM_023248635.1;refGene_gn=PTK2;refGene_tx=XM_023248636.1;refGene_gn=PTK2;refGene_tx=XM_023248637.1;refGene_gn=PTK2;refGene_tx=XM_023248638.1;refGene_gn=PTK2;refGene_tx=XM_023248639.1;refGene_gn=PTK2;refGene_tx=XM_023248640.1;refGene_gn=PTK2;refGene_tx=XM_023248641.1;refGene_gn=PTK2;refGene_tx=XM_023248642.1;refGene_gn=PTK2;refGene_tx=XM_023248643.1

F2 82597145 82597257
refGene_gn=PTK2;refGene_tx=XM_023248620.1;refGene_gn=PTK2;refGene_tx=XM_023248621.1;refGene_gn=PTK2;refGene_tx=XM_023248622.1;refGene_gn=PTK2;refGene_tx=XM_023248623.1;refGene_gn=PTK2;refGene_tx=XM_023248624.1;refGene_gn=PTK2;refGene_tx=XM_023248625.1;refGene_gn=PTK2;refGene_tx=XM_023248626.1;refGene_gn=PTK2;refGene_tx=XM_023248627.1;refGene_gn=PTK2;refGene_tx=XM_023248628.1;refGene_gn=PTK2;refGene_tx=XM_023248629.1;refGene_gn=PTK2;refGene_tx=XM_023248630.1;refGene_gn=PTK2;refGene_tx=XM_023248631.1;refGene_gn=PTK2;refGene_tx=XM_023248632.1;refGene_gn=PTK2;refGene_tx=XM_023248633.1;refGene_gn=PTK2;refGene_tx=XM_023248634.1;refGene_gn=PTK2;refGene_tx=XM_023248635.1;refGene_gn=PTK2;refGene_tx=XM_023248636.1;refGene_gn=PTK2;refGene_tx=XM_023248637.1;refGene_gn=PTK2;refGene_tx=XM_023248638.1;refGene_gn=PTK2;refGene_tx=XM_023248639.1;refGene_gn=PTK2;refGene_tx=XM_023248640.1;refGene_gn=PTK2;refGene_tx=XM_023248641.1;refGene_gn=PTK2;refGene_tx=XM_023248642.1;refGene_gn=PTK2;refGene_tx=XM_023248643.1
F2 82611065 82611270
refGene_gn=PTK2;refGene_tx=XM_023248620.1;refGene_gn=PTK2;refGene_tx=XM_023248621.1;refGene_gn=PTK2;refGene_tx=XM_023248622.1;refGene_gn=PTK2;refGene_tx=XM_023248623.1;refGene_gn=PTK2;refGene_tx=XM_023248624.1;refGene_gn=PTK2;refGene_tx=XM_023248625.1;refGene_gn=PTK2;refGene_tx=XM_023248626.1;refGene_gn=PTK2;refGene_tx=XM_023248627.1;refGene_gn=PTK2;refGene_tx=XM_023248628.1;refGene_gn=PTK2;refGene_tx=XM_023248629.1;refGene_gn=PTK2;refGene_tx=XM_023248630.1;refGene_gn=PTK2;refGene_tx=XM_023248631.1;refGene_gn=PTK2;refGene_tx=XM_023248632.1;refGene_gn=PTK2;refGene_tx=XM_023248633.1;refGene_gn=PTK2;refGene_tx=XM_023248634.1;refGene_gn=PTK2;refGene_tx=XM_023248635.1;refGene_gn=PTK2;refGene_tx=XM_023248636.1;refGene_gn=PTK2;refGene_tx=XM_023248637.1;refGene_gn=PTK2;refGene_tx=XM_023248638.1;refGene_gn=PTK2;refGene_tx=XM_023248639.1;refGene_gn=PTK2;refGene_tx=XM_023248641.1;refGene_gn=PTK2;refGene_tx=XM_023248642.1;refGene_gn=PTK2;refGene_tx=XM_023248643.1
F2 82613912 82614002
refGene_gn=PTK2;refGene_tx=XM_023248620.1;refGene_gn=PTK2;refGene_tx=XM_023248621.1;refGene_gn=PTK2;refGene_tx=XM_023248622.1;refGene_gn=PTK2;refGene_tx=XM_023248623.1;refGene_gn=PTK2;refGene_tx=XM_023248624.1;refGene_gn=PTK2;refGene_tx=XM_023248625.1;refGene_gn=PTK2;refGene_tx=XM_023248626.1;refGene_gn=PTK2;refGene_tx=XM_023248627.1;refGene_gn=PTK2;refGene_tx=XM_023248628.1;refGene_gn=PTK2;refGene_tx=XM_023248629.1;refGene_gn=PTK2;refGene_tx=XM_023248630.1;refGene_gn=PTK2;refGene_tx=XM_023248631.1;refGene_gn=PTK2;refGene_tx=XM_023248632.1;refGene_gn=PTK2;refGene_tx=XM_023248633.1;refGene_gn=PTK2;refGene_tx=XM_023248634.1;refGene_gn=PTK2;refGene_tx=XM_023248635.1;refGene_gn=PTK2;refGene_tx=XM_023248636.1;refGene_gn=PTK2;refGene_tx=XM_023248637.1;refGene_gn=PTK2;refGene_tx=XM_023248638.1;refGene_gn=PTK2;refGene_tx=XM_023248639.1;refGene_gn=PTK2;refGene_tx=XM_023248641.1;refGene_gn=PTK2;refGene_tx=XM_023248642.1;refGene_gn=PTK2;refGene_tx=XM_023248643.1
F2 82617025 82617126
refGene_gn=PTK2;refGene_tx=XM_023248620.1;refGene_gn=PTK2;refGene_tx=XM_023248621.1;refGene_gn=PTK2;refGene_tx=XM_023248622.1;refGene_gn=PTK2;refGene_tx=XM_023248623.1;refGene_gn=PTK2;refGene_tx=XM_023248624.1;

refGene_gn=PTK2;refGene_tx=XM_023248625.1;refGene_gn=PTK2;refGene_tx=XM_023248626.1;refGene_gn=PTK2;refGene_tx=XM_023248627.1;refGene_gn=PTK2;refGene_tx=XM_023248628.1;refGene_gn=PTK2;refGene_tx=XM_023248629.1;refGene_gn=PTK2;refGene_tx=XM_023248630.1;refGene_gn=PTK2;refGene_tx=XM_023248631.1;refGene_gn=PTK2;refGene_tx=XM_023248632.1;refGene_gn=PTK2;refGene_tx=XM_023248633.1;refGene_gn=PTK2;refGene_tx=XM_023248634.1;refGene_gn=PTK2;refGene_tx=XM_023248635.1;refGene_gn=PTK2;refGene_tx=XM_023248636.1;refGene_gn=PTK2;refGene_tx=XM_023248637.1;refGene_gn=PTK2;refGene_tx=XM_023248638.1;refGene_gn=PTK2;refGene_tx=XM_023248639.1;refGene_gn=PTK2;refGene_tx=XM_023248641.1;refGene_gn=PTK2;refGene_tx=XM_023248642.1;refGene_gn=PTK2;refGene_tx=XM_023248643.1
F2 82618006 82618122
refGene_gn=PTK2;refGene_tx=XM_023248620.1;refGene_gn=PTK2;refGene_tx=XM_023248621.1;refGene_gn=PTK2;refGene_tx=XM_023248622.1;refGene_gn=PTK2;refGene_tx=XM_023248623.1;refGene_gn=PTK2;refGene_tx=XM_023248624.1;refGene_gn=PTK2;refGene_tx=XM_023248625.1;refGene_gn=PTK2;refGene_tx=XM_023248626.1;refGene_gn=PTK2;refGene_tx=XM_023248627.1;refGene_gn=PTK2;refGene_tx=XM_023248628.1;refGene_gn=PTK2;refGene_tx=XM_023248629.1;refGene_gn=PTK2;refGene_tx=XM_023248630.1;refGene_gn=PTK2;refGene_tx=XM_023248631.1;refGene_gn=PTK2;refGene_tx=XM_023248632.1;refGene_gn=PTK2;refGene_tx=XM_023248633.1;refGene_gn=PTK2;refGene_tx=XM_023248634.1;refGene_gn=PTK2;refGene_tx=XM_023248635.1;refGene_gn=PTK2;refGene_tx=XM_023248636.1;refGene_gn=PTK2;refGene_tx=XM_023248637.1;refGene_gn=PTK2;refGene_tx=XM_023248638.1;refGene_gn=PTK2;refGene_tx=XM_023248639.1;refGene_gn=PTK2;refGene_tx=XM_023248641.1;refGene_gn=PTK2;refGene_tx=XM_023248642.1;refGene_gn=PTK2;refGene_tx=XM_023248643.1
F2 82620411 82620512
refGene_gn=PTK2;refGene_tx=XM_023248620.1;refGene_gn=PTK2;refGene_tx=XM_023248621.1;refGene_gn=PTK2;refGene_tx=XM_023248622.1;refGene_gn=PTK2;refGene_tx=XM_023248623.1;refGene_gn=PTK2;refGene_tx=XM_023248624.1;refGene_gn=PTK2;refGene_tx=XM_023248625.1;refGene_gn=PTK2;refGene_tx=XM_023248626.1;refGene_gn=PTK2;refGene_tx=XM_023248627.1;refGene_gn=PTK2;refGene_tx=XM_023248628.1;refGene_gn=PTK2;refGene_tx=XM_023248629.1;refGene_gn=PTK2;refGene_tx=XM_023248630.1;refGene_gn=PTK2;refGene_tx=XM_023248631.1;refGene_gn=PTK2;refGene_tx=XM_023248632.1;refGene_gn=PTK2;refGene_tx=XM_023248633.1;refGene_gn=PTK2;refGene_tx=XM_023248634.1;refGene_gn=PTK2;refGene_tx=XM_023248635.1;refGene_gn=PTK2;refGene_tx=XM_023248636.1;refGene_gn=PTK2;refGene_tx=XM_023248637.1;refGene_gn=PTK2;refGene_tx=XM_023248638.1;refGene_gn=PTK2;refGene_tx=XM_023248639.1;refGene_gn=PTK2;refGene_tx=XM_023248641.1;refGene_gn=PTK2;refGene_tx=XM_023248642.1;refGene_gn=PTK2;refGene_tx=XM_023248643.1
F2 82621690 82621775
refGene_gn=PTK2;refGene_tx=XM_023248620.1;refGene_gn=PTK2;refGene_tx=XM_023248621.1;refGene_gn=PTK2;refGene_tx=XM_023248622.1;refGene_gn=PTK2;refGene_tx=XM_023248623.1;refGene_gn=PTK2;refGene_tx=XM_023248624.1;refGene_gn=PTK2;refGene_tx=XM_023248625.1;refGene_gn=PTK2;refGene_tx=XM_023248626.1;refGene_gn=PTK2;refGene_tx=XM_023248627.1;refGene_gn=PTK2;refGene_tx=XM_023248628.1;refGene_gn=PTK2;refGene_tx=XM_023248629.1;refGene_gn=PTK2;refGene_tx=XM_023248630.1;refGene_gn=PTK2;refGene_tx=XM_023248631.1;refGene_gn=PTK2;refGene_tx=XM_023248632.1;refGene_gn=PTK

2;refGene_tx=XM_023248633.1;refGene_gn=PTK2;refGene_tx=XM_023248634.1;
refGene_gn=PTK2;refGene_tx=XM_023248635.1;refGene_gn=PTK2;refGene_tx=X
M_023248636.1;refGene_gn=PTK2;refGene_tx=XM_023248637.1;refGene_gn=PTK
2;refGene_tx=XM_023248638.1;refGene_gn=PTK2;refGene_tx=XM_023248639.1;
refGene_gn=PTK2;refGene_tx=XM_023248641.1;refGene_gn=PTK2;refGene_tx=X
M_023248642.1;refGene_gn=PTK2;refGene_tx=XM_023248643.1
F2 82627216 82627314
refGene_gn=PTK2;refGene_tx=XM_023248620.1;refGene_gn=PTK2;refGene_tx=X
M_023248621.1;refGene_gn=PTK2;refGene_tx=XM_023248622.1;refGene_gn=PTK
2;refGene_tx=XM_023248623.1;refGene_gn=PTK2;refGene_tx=XM_023248624.1;
refGene_gn=PTK2;refGene_tx=XM_023248625.1;refGene_gn=PTK2;refGene_tx=X
M_023248626.1;refGene_gn=PTK2;refGene_tx=XM_023248627.1;refGene_gn=PTK
2;refGene_tx=XM_023248628.1;refGene_gn=PTK2;refGene_tx=XM_023248629.1;
refGene_gn=PTK2;refGene_tx=XM_023248630.1;refGene_gn=PTK2;refGene_tx=X
M_023248631.1;refGene_gn=PTK2;refGene_tx=XM_023248632.1;refGene_gn=PTK
2;refGene_tx=XM_023248633.1;refGene_gn=PTK2;refGene_tx=XM_023248634.1;
refGene_gn=PTK2;refGene_tx=XM_023248635.1;refGene_gn=PTK2;refGene_tx=X
M_023248636.1;refGene_gn=PTK2;refGene_tx=XM_023248637.1;refGene_gn=PTK
2;refGene_tx=XM_023248638.1;refGene_gn=PTK2;refGene_tx=XM_023248639.1;
refGene_gn=PTK2;refGene_tx=XM_023248641.1;refGene_gn=PTK2;refGene_tx=X
M_023248642.1;refGene_gn=PTK2;refGene_tx=XM_023248643.1
F2 82628419 82628440
refGene_gn=PTK2;refGene_tx=XM_023248620.1;refGene_gn=PTK2;refGene_tx=X
M_023248621.1;refGene_gn=PTK2;refGene_tx=XM_023248622.1;refGene_gn=PTK
2;refGene_tx=XM_023248623.1;refGene_gn=PTK2;refGene_tx=XM_023248624.1;
refGene_gn=PTK2;refGene_tx=XM_023248625.1;refGene_gn=PTK2;refGene_tx=X
M_023248626.1;refGene_gn=PTK2;refGene_tx=XM_023248628.1;refGene_gn=PTK
2;refGene_tx=XM_023248630.1;refGene_gn=PTK2;refGene_tx=XM_023248631.1;
refGene_gn=PTK2;refGene_tx=XM_023248632.1;refGene_gn=PTK2;refGene_tx=X
M_023248633.1;refGene_gn=PTK2;refGene_tx=XM_023248634.1;refGene_gn=PTK
2;refGene_tx=XM_023248635.1;refGene_gn=PTK2;refGene_tx=XM_023248639.1;
refGene_gn=PTK2;refGene_tx=XM_023248641.1
F2 82629869 82629926
refGene_gn=PTK2;refGene_tx=XM_023248620.1;refGene_gn=PTK2;refGene_tx=X
M_023248621.1;refGene_gn=PTK2;refGene_tx=XM_023248622.1;refGene_gn=PTK
2;refGene_tx=XM_023248623.1;refGene_gn=PTK2;refGene_tx=XM_023248624.1;
refGene_gn=PTK2;refGene_tx=XM_023248625.1;refGene_gn=PTK2;refGene_tx=X
M_023248626.1;refGene_gn=PTK2;refGene_tx=XM_023248627.1;refGene_gn=PTK
2;refGene_tx=XM_023248628.1;refGene_gn=PTK2;refGene_tx=XM_023248629.1;
refGene_gn=PTK2;refGene_tx=XM_023248630.1;refGene_gn=PTK2;refGene_tx=X
M_023248631.1;refGene_gn=PTK2;refGene_tx=XM_023248632.1;refGene_gn=PTK
2;refGene_tx=XM_023248633.1;refGene_gn=PTK2;refGene_tx=XM_023248634.1;
refGene_gn=PTK2;refGene_tx=XM_023248635.1;refGene_gn=PTK2;refGene_tx=X
M_023248636.1;refGene_gn=PTK2;refGene_tx=XM_023248637.1;refGene_gn=PTK
2;refGene_tx=XM_023248638.1;refGene_gn=PTK2;refGene_tx=XM_023248639.1;
refGene_gn=PTK2;refGene_tx=XM_023248641.1;refGene_gn=PTK2;refGene_tx=X
M_023248642.1;refGene_gn=PTK2;refGene_tx=XM_023248643.1
F2 82635922 82635940
refGene_gn=PTK2;refGene_tx=XM_023248628.1;refGene_gn=PTK2;refGene_tx=X
M_023248629.1;refGene_gn=PTK2;refGene_tx=XM_023248630.1;refGene_gn=PTK

2;refGene_tx=XM_023248636.1;refGene_gn=PTK2;refGene_tx=XM_023248641.1;
refGene_gn=PTK2;refGene_tx=XM_023248642.1;refGene_gn=PTK2;refGene_tx=X
M_023248643.1
F2 82636952 82637036
refGene_gn=PTK2;refGene_tx=XM_023248628.1;refGene_gn=PTK2;refGene_tx=X
M_023248629.1;refGene_gn=PTK2;refGene_tx=XM_023248641.1;refGene_gn=PTK
2;refGene_tx=XM_023248642.1
F2 82639631 82639769
refGene_gn=PTK2;refGene_tx=XM_023248620.1;refGene_gn=PTK2;refGene_tx=X
M_023248621.1;refGene_gn=PTK2;refGene_tx=XM_023248622.1;refGene_gn=PTK
2;refGene_tx=XM_023248623.1;refGene_gn=PTK2;refGene_tx=XM_023248624.1;
refGene_gn=PTK2;refGene_tx=XM_023248625.1;refGene_gn=PTK2;refGene_tx=X
M_023248626.1;refGene_gn=PTK2;refGene_tx=XM_023248627.1;refGene_gn=PTK
2;refGene_tx=XM_023248631.1;refGene_gn=PTK2;refGene_tx=XM_023248632.1;
refGene_gn=PTK2;refGene_tx=XM_023248633.1;refGene_gn=PTK2;refGene_tx=X
M_023248634.1;refGene_gn=PTK2;refGene_tx=XM_023248639.1
F2 82647776 82647829
refGene_gn=PTK2;refGene_tx=XM_023248620.1;refGene_gn=PTK2;refGene_tx=X
M_023248621.1;refGene_gn=PTK2;refGene_tx=XM_023248622.1;refGene_gn=PTK
2;refGene_tx=XM_023248623.1;refGene_gn=PTK2;refGene_tx=XM_023248624.1;
refGene_gn=PTK2;refGene_tx=XM_023248625.1;refGene_gn=PTK2;refGene_tx=X
M_023248626.1;refGene_gn=PTK2;refGene_tx=XM_023248627.1;refGene_gn=PTK
2;refGene_tx=XM_023248628.1;refGene_gn=PTK2;refGene_tx=XM_023248629.1;
refGene_gn=PTK2;refGene_tx=XM_023248630.1;refGene_gn=PTK2;refGene_tx=X
M_023248631.1;refGene_gn=PTK2;refGene_tx=XM_023248632.1;refGene_gn=PTK
2;refGene_tx=XM_023248633.1;refGene_gn=PTK2;refGene_tx=XM_023248634.1;
refGene_gn=PTK2;refGene_tx=XM_023248635.1;refGene_gn=PTK2;refGene_tx=X
M_023248636.1;refGene_gn=PTK2;refGene_tx=XM_023248637.1;refGene_gn=PTK
2;refGene_tx=XM_023248638.1;refGene_gn=PTK2;refGene_tx=XM_023248639.1;
refGene_gn=PTK2;refGene_tx=XM_023248641.1;refGene_gn=PTK2;refGene_tx=X
M_023248642.1;refGene_gn=PTK2;refGene_tx=XM_023248643.1
F2 82650003 82650034
refGene_gn=PTK2;refGene_tx=XM_023248620.1;refGene_gn=PTK2;refGene_tx=X
M_023248621.1;refGene_gn=PTK2;refGene_tx=XM_023248622.1;refGene_gn=PTK
2;refGene_tx=XM_023248623.1;refGene_gn=PTK2;refGene_tx=XM_023248624.1;
refGene_gn=PTK2;refGene_tx=XM_023248625.1;refGene_gn=PTK2;refGene_tx=X
M_023248626.1;refGene_gn=PTK2;refGene_tx=XM_023248627.1;refGene_gn=PTK
2;refGene_tx=XM_023248628.1;refGene_gn=PTK2;refGene_tx=XM_023248629.1;
refGene_gn=PTK2;refGene_tx=XM_023248630.1;refGene_gn=PTK2;refGene_tx=X
M_023248631.1;refGene_gn=PTK2;refGene_tx=XM_023248632.1;refGene_gn=PTK
2;refGene_tx=XM_023248633.1;refGene_gn=PTK2;refGene_tx=XM_023248634.1;
refGene_gn=PTK2;refGene_tx=XM_023248635.1;refGene_gn=PTK2;refGene_tx=X
M_023248636.1;refGene_gn=PTK2;refGene_tx=XM_023248637.1;refGene_gn=PTK
2;refGene_tx=XM_023248638.1;refGene_gn=PTK2;refGene_tx=XM_023248639.1;
refGene_gn=PTK2;refGene_tx=XM_023248641.1;refGene_gn=PTK2;refGene_tx=X
M_023248642.1;refGene_gn=PTK2;refGene_tx=XM_023248643.1
F2 82655141 82655259
refGene_gn=PTK2;refGene_tx=XM_023248620.1;refGene_gn=PTK2;refGene_tx=X
M_023248621.1;refGene_gn=PTK2;refGene_tx=XM_023248622.1;refGene_gn=PTK
2;refGene_tx=XM_023248623.1;refGene_gn=PTK2;refGene_tx=XM_023248624.1;

refGene_gn=PTK2;refGene_tx=XM_023248625.1;refGene_gn=PTK2;refGene_tx=XM_023248626.1;refGene_gn=PTK2;refGene_tx=XM_023248627.1;refGene_gn=PTK2;refGene_tx=XM_023248628.1;refGene_gn=PTK2;refGene_tx=XM_023248629.1;refGene_gn=PTK2;refGene_tx=XM_023248630.1;refGene_gn=PTK2;refGene_tx=XM_023248631.1;refGene_gn=PTK2;refGene_tx=XM_023248632.1;refGene_gn=PTK2;refGene_tx=XM_023248633.1;refGene_gn=PTK2;refGene_tx=XM_023248634.1;refGene_gn=PTK2;refGene_tx=XM_023248635.1;refGene_gn=PTK2;refGene_tx=XM_023248636.1;refGene_gn=PTK2;refGene_tx=XM_023248637.1;refGene_gn=PTK2;refGene_tx=XM_023248638.1;refGene_gn=PTK2;refGene_tx=XM_023248639.1;refGene_gn=PTK2;refGene_tx=XM_023248641.1;refGene_gn=PTK2;refGene_tx=XM_023248642.1;refGene_gn=PTK2;refGene_tx=XM_023248643.1
F2 82656804 82656912
refGene_gn=PTK2;refGene_tx=XM_023248620.1;refGene_gn=PTK2;refGene_tx=XM_023248621.1;refGene_gn=PTK2;refGene_tx=XM_023248622.1;refGene_gn=PTK2;refGene_tx=XM_023248623.1;refGene_gn=PTK2;refGene_tx=XM_023248624.1;refGene_gn=PTK2;refGene_tx=XM_023248625.1;refGene_gn=PTK2;refGene_tx=XM_023248626.1;refGene_gn=PTK2;refGene_tx=XM_023248627.1;refGene_gn=PTK2;refGene_tx=XM_023248628.1;refGene_gn=PTK2;refGene_tx=XM_023248629.1;refGene_gn=PTK2;refGene_tx=XM_023248630.1;refGene_gn=PTK2;refGene_tx=XM_023248631.1;refGene_gn=PTK2;refGene_tx=XM_023248632.1;refGene_gn=PTK2;refGene_tx=XM_023248633.1;refGene_gn=PTK2;refGene_tx=XM_023248634.1;refGene_gn=PTK2;refGene_tx=XM_023248635.1;refGene_gn=PTK2;refGene_tx=XM_023248636.1;refGene_gn=PTK2;refGene_tx=XM_023248637.1;refGene_gn=PTK2;refGene_tx=XM_023248638.1;refGene_gn=PTK2;refGene_tx=XM_023248639.1;refGene_gn=PTK2;refGene_tx=XM_023248641.1;refGene_gn=PTK2;refGene_tx=XM_023248642.1;refGene_gn=PTK2;refGene_tx=XM_023248643.1
F2 82664752 82664830
refGene_gn=PTK2;refGene_tx=XM_023248620.1;refGene_gn=PTK2;refGene_tx=XM_023248621.1;refGene_gn=PTK2;refGene_tx=XM_023248622.1;refGene_gn=PTK2;refGene_tx=XM_023248623.1;refGene_gn=PTK2;refGene_tx=XM_023248624.1;refGene_gn=PTK2;refGene_tx=XM_023248625.1;refGene_gn=PTK2;refGene_tx=XM_023248626.1;refGene_gn=PTK2;refGene_tx=XM_023248627.1;refGene_gn=PTK2;refGene_tx=XM_023248628.1;refGene_gn=PTK2;refGene_tx=XM_023248629.1;refGene_gn=PTK2;refGene_tx=XM_023248630.1;refGene_gn=PTK2;refGene_tx=XM_023248631.1;refGene_gn=PTK2;refGene_tx=XM_023248632.1;refGene_gn=PTK2;refGene_tx=XM_023248633.1;refGene_gn=PTK2;refGene_tx=XM_023248634.1;refGene_gn=PTK2;refGene_tx=XM_023248635.1;refGene_gn=PTK2;refGene_tx=XM_023248636.1;refGene_gn=PTK2;refGene_tx=XM_023248637.1;refGene_gn=PTK2;refGene_tx=XM_023248638.1;refGene_gn=PTK2;refGene_tx=XM_023248639.1;refGene_gn=PTK2;refGene_tx=XM_023248641.1;refGene_gn=PTK2;refGene_tx=XM_023248642.1;refGene_gn=PTK2;refGene_tx=XM_023248643.1
F2 82665349 82665490
refGene_gn=PTK2;refGene_tx=XM_023248620.1;refGene_gn=PTK2;refGene_tx=XM_023248621.1;refGene_gn=PTK2;refGene_tx=XM_023248622.1;refGene_gn=PTK2;refGene_tx=XM_023248623.1;refGene_gn=PTK2;refGene_tx=XM_023248624.1;refGene_gn=PTK2;refGene_tx=XM_023248625.1;refGene_gn=PTK2;refGene_tx=XM_023248626.1;refGene_gn=PTK2;refGene_tx=XM_023248627.1;refGene_gn=PTK2;refGene_tx=XM_023248628.1;refGene_gn=PTK2;refGene_tx=XM_023248629.1;refGene_gn=PTK2;refGene_tx=XM_023248630.1;refGene_gn=PTK2;refGene_tx=XM_023248631.1;refGene_gn=PTK2;refGene_tx=XM_023248632.1;refGene_gn=PTK

2;refGene_tx=XM_023248633.1;refGene_gn=PTK2;refGene_tx=XM_023248634.1;
refGene_gn=PTK2;refGene_tx=XM_023248635.1;refGene_gn=PTK2;refGene_tx=X
M_023248636.1;refGene_gn=PTK2;refGene_tx=XM_023248637.1;refGene_gn=PTK
2;refGene_tx=XM_023248638.1;refGene_gn=PTK2;refGene_tx=XM_023248639.1;
refGene_gn=PTK2;refGene_tx=XM_023248641.1;refGene_gn=PTK2;refGene_tx=X
M_023248642.1;refGene_gn=PTK2;refGene_tx=XM_023248643.1
F2 82675993 82676048
refGene_gn=PTK2;refGene_tx=XM_023248620.1;refGene_gn=PTK2;refGene_tx=X
M_023248621.1;refGene_gn=PTK2;refGene_tx=XM_023248622.1;refGene_gn=PTK
2;refGene_tx=XM_023248623.1;refGene_gn=PTK2;refGene_tx=XM_023248624.1;
refGene_gn=PTK2;refGene_tx=XM_023248625.1;refGene_gn=PTK2;refGene_tx=X
M_023248626.1;refGene_gn=PTK2;refGene_tx=XM_023248627.1;refGene_gn=PTK
2;refGene_tx=XM_023248628.1;refGene_gn=PTK2;refGene_tx=XM_023248629.1;
refGene_gn=PTK2;refGene_tx=XM_023248630.1;refGene_gn=PTK2;refGene_tx=X
M_023248631.1;refGene_gn=PTK2;refGene_tx=XM_023248632.1;refGene_gn=PTK
2;refGene_tx=XM_023248633.1;refGene_gn=PTK2;refGene_tx=XM_023248634.1;
refGene_gn=PTK2;refGene_tx=XM_023248635.1;refGene_gn=PTK2;refGene_tx=X
M_023248636.1;refGene_gn=PTK2;refGene_tx=XM_023248637.1;refGene_gn=PTK
2;refGene_tx=XM_023248638.1;refGene_gn=PTK2;refGene_tx=XM_023248639.1;
refGene_gn=PTK2;refGene_tx=XM_023248641.1;refGene_gn=PTK2;refGene_tx=X
M_023248642.1;refGene_gn=PTK2;refGene_tx=XM_023248643.1
F2 82689818 82689881
refGene_gn=PTK2;refGene_tx=XM_023248620.1;refGene_gn=PTK2;refGene_tx=X
M_023248621.1;refGene_gn=PTK2;refGene_tx=XM_023248622.1;refGene_gn=PTK
2;refGene_tx=XM_023248623.1;refGene_gn=PTK2;refGene_tx=XM_023248624.1;
refGene_gn=PTK2;refGene_tx=XM_023248625.1;refGene_gn=PTK2;refGene_tx=X
M_023248626.1;refGene_gn=PTK2;refGene_tx=XM_023248627.1;refGene_gn=PTK
2;refGene_tx=XM_023248628.1;refGene_gn=PTK2;refGene_tx=XM_023248629.1;
refGene_gn=PTK2;refGene_tx=XM_023248630.1;refGene_gn=PTK2;refGene_tx=X
M_023248631.1;refGene_gn=PTK2;refGene_tx=XM_023248632.1;refGene_gn=PTK
2;refGene_tx=XM_023248633.1;refGene_gn=PTK2;refGene_tx=XM_023248634.1;
refGene_gn=PTK2;refGene_tx=XM_023248635.1;refGene_gn=PTK2;refGene_tx=X
M_023248636.1;refGene_gn=PTK2;refGene_tx=XM_023248637.1;refGene_gn=PTK
2;refGene_tx=XM_023248638.1;refGene_gn=PTK2;refGene_tx=XM_023248639.1;
refGene_gn=PTK2;refGene_tx=XM_023248641.1;refGene_gn=PTK2;refGene_tx=X
M_023248642.1;refGene_gn=PTK2;refGene_tx=XM_023248643.1
F2 82691832 82691912
refGene_gn=PTK2;refGene_tx=XM_023248620.1;refGene_gn=PTK2;refGene_tx=X
M_023248621.1;refGene_gn=PTK2;refGene_tx=XM_023248622.1;refGene_gn=PTK
2;refGene_tx=XM_023248623.1;refGene_gn=PTK2;refGene_tx=XM_023248624.1;
refGene_gn=PTK2;refGene_tx=XM_023248625.1;refGene_gn=PTK2;refGene_tx=X
M_023248626.1;refGene_gn=PTK2;refGene_tx=XM_023248627.1;refGene_gn=PTK
2;refGene_tx=XM_023248628.1;refGene_gn=PTK2;refGene_tx=XM_023248629.1;
refGene_gn=PTK2;refGene_tx=XM_023248630.1;refGene_gn=PTK2;refGene_tx=X
M_023248631.1;refGene_gn=PTK2;refGene_tx=XM_023248632.1;refGene_gn=PTK
2;refGene_tx=XM_023248633.1;refGene_gn=PTK2;refGene_tx=XM_023248634.1;
refGene_gn=PTK2;refGene_tx=XM_023248635.1;refGene_gn=PTK2;refGene_tx=X
M_023248636.1;refGene_gn=PTK2;refGene_tx=XM_023248637.1;refGene_gn=PTK
2;refGene_tx=XM_023248638.1;refGene_gn=PTK2;refGene_tx=XM_023248639.1;
refGene_gn=PTK2;refGene_tx=XM_023248641.1;refGene_gn=PTK2;refGene_tx=X

M_023248642.1;refGene_gn=PTK2;refGene_tx=XM_023248643.1
F2 82704168 82704256
refGene_gn=PTK2;refGene_tx=XM_023248620.1;refGene_gn=PTK2;refGene_tx=XM_023248621.1;refGene_gn=PTK2;refGene_tx=XM_023248622.1;refGene_gn=PTK2;refGene_tx=XM_023248623.1;refGene_gn=PTK2;refGene_tx=XM_023248624.1;refGene_gn=PTK2;refGene_tx=XM_023248625.1;refGene_gn=PTK2;refGene_tx=XM_023248626.1;refGene_gn=PTK2;refGene_tx=XM_023248627.1;refGene_gn=PTK2;refGene_tx=XM_023248628.1;refGene_gn=PTK2;refGene_tx=XM_023248629.1;refGene_gn=PTK2;refGene_tx=XM_023248630.1;refGene_gn=PTK2;refGene_tx=XM_023248631.1;refGene_gn=PTK2;refGene_tx=XM_023248632.1;refGene_gn=PTK2;refGene_tx=XM_023248633.1;refGene_gn=PTK2;refGene_tx=XM_023248634.1;refGene_gn=PTK2;refGene_tx=XM_023248635.1;refGene_gn=PTK2;refGene_tx=XM_023248636.1;refGene_gn=PTK2;refGene_tx=XM_023248637.1;refGene_gn=PTK2;refGene_tx=XM_023248638.1;refGene_gn=PTK2;refGene_tx=XM_023248639.1;refGene_gn=PTK2;refGene_tx=XM_023248641.1;refGene_gn=PTK2;refGene_tx=XM_023248642.1;refGene_gn=PTK2;refGene_tx=XM_023248643.1
F2 82716181 82716348
refGene_gn=PTK2;refGene_tx=XM_023248620.1;refGene_gn=PTK2;refGene_tx=XM_023248621.1;refGene_gn=PTK2;refGene_tx=XM_023248622.1;refGene_gn=PTK2;refGene_tx=XM_023248623.1;refGene_gn=PTK2;refGene_tx=XM_023248624.1;refGene_gn=PTK2;refGene_tx=XM_023248625.1;refGene_gn=PTK2;refGene_tx=XM_023248626.1;refGene_gn=PTK2;refGene_tx=XM_023248627.1;refGene_gn=PTK2;refGene_tx=XM_023248628.1;refGene_gn=PTK2;refGene_tx=XM_023248629.1;refGene_gn=PTK2;refGene_tx=XM_023248630.1;refGene_gn=PTK2;refGene_tx=XM_023248631.1;refGene_gn=PTK2;refGene_tx=XM_023248632.1;refGene_gn=PTK2;refGene_tx=XM_023248633.1;refGene_gn=PTK2;refGene_tx=XM_023248634.1;refGene_gn=PTK2;refGene_tx=XM_023248635.1;refGene_gn=PTK2;refGene_tx=XM_023248636.1;refGene_gn=PTK2;refGene_tx=XM_023248637.1;refGene_gn=PTK2;refGene_tx=XM_023248638.1;refGene_gn=PTK2;refGene_tx=XM_023248641.1;refGene_gn=PTK2;refGene_tx=XM_023248642.1;refGene_gn=PTK2;refGene_tx=XM_023248643.1
F2 82726447 82726674
refGene_gn=PTK2;refGene_tx=XM_023248620.1;refGene_gn=PTK2;refGene_tx=XM_023248621.1;refGene_gn=PTK2;refGene_tx=XM_023248622.1;refGene_gn=PTK2;refGene_tx=XM_023248623.1;refGene_gn=PTK2;refGene_tx=XM_023248624.1;refGene_gn=PTK2;refGene_tx=XM_023248625.1;refGene_gn=PTK2;refGene_tx=XM_023248626.1;refGene_gn=PTK2;refGene_tx=XM_023248627.1;refGene_gn=PTK2;refGene_tx=XM_023248628.1;refGene_gn=PTK2;refGene_tx=XM_023248629.1;refGene_gn=PTK2;refGene_tx=XM_023248630.1;refGene_gn=PTK2;refGene_tx=XM_023248631.1;refGene_gn=PTK2;refGene_tx=XM_023248632.1;refGene_gn=PTK2;refGene_tx=XM_023248633.1;refGene_gn=PTK2;refGene_tx=XM_023248634.1;refGene_gn=PTK2;refGene_tx=XM_023248635.1;refGene_gn=PTK2;refGene_tx=XM_023248636.1;refGene_gn=PTK2;refGene_tx=XM_023248637.1;refGene_gn=PTK2;refGene_tx=XM_023248638.1;refGene_gn=PTK2;refGene_tx=XM_023248639.1;refGene_gn=PTK2;refGene_tx=XM_023248641.1;refGene_gn=PTK2;refGene_tx=XM_023248642.1;refGene_gn=PTK2;refGene_tx=XM_023248643.1
F2 82756561 82756661
refGene_gn=PTK2;refGene_tx=XM_023248620.1;refGene_gn=PTK2;refGene_tx=XM_023248621.1;refGene_gn=PTK2;refGene_tx=XM_023248622.1;refGene_gn=PTK2;refGene_tx=XM_023248623.1;refGene_gn=PTK2;refGene_tx=XM_023248624.1;

refGene_gn=PTK2;refGene_tx=XM_023248625.1;refGene_gn=PTK2;refGene_tx=XM_023248626.1;refGene_gn=PTK2;refGene_tx=XM_023248627.1;refGene_gn=PTK2;refGene_tx=XM_023248628.1;refGene_gn=PTK2;refGene_tx=XM_023248629.1;refGene_gn=PTK2;refGene_tx=XM_023248630.1;refGene_gn=PTK2;refGene_tx=XM_023248635.1;refGene_gn=PTK2;refGene_tx=XM_023248636.1;refGene_gn=PTK2;refGene_tx=XM_023248637.1;refGene_gn=PTK2;refGene_tx=XM_023248638.1;refGene_gn=PTK2;refGene_tx=XM_023248641.1;refGene_gn=PTK2;refGene_tx=XM_023248642.1;refGene_gn=PTK2;refGene_tx=XM_023248643.1
F2 82906977 82907079
refGene_gn=DENND3;refGene_tx=XM_023248691.1;refGene_gn=DENND3;refGene_tx=XM_023248692.1;refGene_gn=DENND3;refGene_tx=XM_023248693.1
F2 82912583 82912839
refGene_gn=DENND3;refGene_tx=XM_023248691.1;refGene_gn=DENND3;refGene_tx=XM_023248692.1;refGene_gn=DENND3;refGene_tx=XM_023248693.1
F2 82913728 82913844
refGene_gn=DENND3;refGene_tx=XM_023248691.1;refGene_gn=DENND3;refGene_tx=XM_023248692.1;refGene_gn=DENND3;refGene_tx=XM_023248693.1
F2 82914252 82914395
refGene_gn=DENND3;refGene_tx=XM_023248691.1;refGene_gn=DENND3;refGene_tx=XM_023248692.1;refGene_gn=DENND3;refGene_tx=XM_023248693.1
F2 82917312 82917424
refGene_gn=DENND3;refGene_tx=XM_023248691.1;refGene_gn=DENND3;refGene_tx=XM_023248692.1;refGene_gn=DENND3;refGene_tx=XM_023248693.1
F2 82921460 82921580
refGene_gn=DENND3;refGene_tx=XM_023248691.1;refGene_gn=DENND3;refGene_tx=XM_023248692.1;refGene_gn=DENND3;refGene_tx=XM_023248693.1
F2 82921760 82921979
refGene_gn=DENND3;refGene_tx=XM_023248691.1;refGene_gn=DENND3;refGene_tx=XM_023248692.1;refGene_gn=DENND3;refGene_tx=XM_023248693.1
F2 82924669 82924785
refGene_gn=DENND3;refGene_tx=XM_023248691.1;refGene_gn=DENND3;refGene_tx=XM_023248692.1;refGene_gn=DENND3;refGene_tx=XM_023248693.1
F2 82926891 82927047
refGene_gn=DENND3;refGene_tx=XM_023248691.1;refGene_gn=DENND3;refGene_tx=XM_023248692.1;refGene_gn=DENND3;refGene_tx=XM_023248693.1
F2 82928607 82928704
refGene_gn=DENND3;refGene_tx=XM_023248691.1;refGene_gn=DENND3;refGene_tx=XM_023248692.1;refGene_gn=DENND3;refGene_tx=XM_023248693.1
F2 82930995 82931099
refGene_gn=DENND3;refGene_tx=XM_023248691.1;refGene_gn=DENND3;refGene_tx=XM_023248692.1;refGene_gn=DENND3;refGene_tx=XM_023248693.1
F2 82931553 82931753
refGene_gn=DENND3;refGene_tx=XM_023248691.1;refGene_gn=DENND3;refGene_tx=XM_023248692.1;refGene_gn=DENND3;refGene_tx=XM_023248693.1
F2 82933714 82934236
refGene_gn=DENND3;refGene_tx=XM_023248691.1;refGene_gn=DENND3;refGene_tx=XM_023248692.1
F2 82940424 82940684
refGene_gn=DENND3;refGene_tx=XM_023248691.1;refGene_gn=DENND3;refGene_tx=XM_023248692.1;refGene_gn=DENND3;refGene_tx=XM_023248693.1

F2 82941302 82941473
refGene_gn=DENND3;refGene_tx=XM_023248691.1;refGene_gn=DENND3;refGene_tx=XM_023248692.1;refGene_gn=DENND3;refGene_tx=XM_023248693.1
F2 82942546 82942676
refGene_gn=DENND3;refGene_tx=XM_023248691.1;refGene_gn=DENND3;refGene_tx=XM_023248692.1;refGene_gn=DENND3;refGene_tx=XM_023248693.1
F2 82945732 82945840
refGene_gn=DENND3;refGene_tx=XM_023248691.1;refGene_gn=DENND3;refGene_tx=XM_023248692.1;refGene_gn=DENND3;refGene_tx=XM_023248693.1
F2 82948741 82948881
refGene_gn=DENND3;refGene_tx=XM_023248691.1;refGene_gn=DENND3;refGene_tx=XM_023248692.1;refGene_gn=DENND3;refGene_tx=XM_023248693.1
F2 82951495 82951650
refGene_gn=DENND3;refGene_tx=XM_023248691.1;refGene_gn=DENND3;refGene_tx=XM_023248692.1;refGene_gn=DENND3;refGene_tx=XM_023248693.1
F2 82952711 82952845
refGene_gn=DENND3;refGene_tx=XM_023248691.1;refGene_gn=DENND3;refGene_tx=XM_023248692.1;refGene_gn=DENND3;refGene_tx=XM_023248693.1
F2 82957402 82957521
refGene_gn=DENND3;refGene_tx=XM_023248691.1;refGene_gn=DENND3;refGene_tx=XM_023248692.1;refGene_gn=DENND3;refGene_tx=XM_023248693.1
F2 82957599 82957737
refGene_gn=DENND3;refGene_tx=XM_023248691.1;refGene_gn=DENND3;refGene_tx=XM_023248692.1;refGene_gn=DENND3;refGene_tx=XM_023248693.1
F2 82957904 82958105
refGene_gn=DENND3;refGene_tx=XM_023248691.1;refGene_gn=DENND3;refGene_tx=XM_023248692.1;refGene_gn=DENND3;refGene_tx=XM_023248693.1
F2 82974247 82974607 refGene_gn=SLC45A4;refGene_tx=XM_019823200.2
F2 82974771 82974795 refGene_gn=SLC45A4;refGene_tx=XM_019823202.2
F2 82976541 82976753
refGene_gn=SLC45A4;refGene_tx=XM_019823200.2;refGene_gn=SLC45A4;refGene_tx=XM_019823202.2
F2 82977764 82977864
refGene_gn=SLC45A4;refGene_tx=XM_019823200.2;refGene_gn=SLC45A4;refGene_tx=XM_019823202.2
F2 82978462 82979499
refGene_gn=SLC45A4;refGene_tx=XM_019823200.2;refGene_gn=SLC45A4;refGene_tx=XM_019823202.2
F2 82980071 82980251
refGene_gn=SLC45A4;refGene_tx=XM_019823200.2;refGene_gn=SLC45A4;refGene_tx=XM_019823202.2
F2 82982163 82982352
refGene_gn=SLC45A4;refGene_tx=XM_019823200.2;refGene_gn=SLC45A4;refGene_tx=XM_019823202.2
F2 83011793 83012031
refGene_gn=SLC45A4;refGene_tx=XM_019823200.2;refGene_gn=SLC45A4;refGene_tx=XM_019823202.2
F2 83090580 83091681 refGene_gn=GPR20;refGene_tx=XM_011291550.3
F2 83151395 83151500
refGene_gn=PTP4A3;refGene_tx=XM_004000150.5;refGene_gn=PTP4A3;refGene_

tx=XM_011291475.3;refGene_gn=PTP4A3;refGene_tx=XM_019823204.2;refGene_gn=PTP4A3;refGene_tx=XM_023248251.1
F2	83153393	83153486
refGene_gn=PTP4A3;refGene_tx=XM_004000150.5;refGene_gn=PTP4A3;refGene_tx=XM_011291475.3;refGene_gn=PTP4A3;refGene_tx=XM_019823204.2;refGene_gn=PTP4A3;refGene_tx=XM_023248251.1
F2	83155107	83155238
refGene_gn=PTP4A3;refGene_tx=XM_004000150.5;refGene_gn=PTP4A3;refGene_tx=XM_011291475.3;refGene_gn=PTP4A3;refGene_tx=XM_019823204.2;refGene_gn=PTP4A3;refGene_tx=XM_023248251.1
F2	83155840	83155915
refGene_gn=PTP4A3;refGene_tx=XM_004000150.5;refGene_gn=PTP4A3;refGene_tx=XM_011291475.3;refGene_gn=PTP4A3;refGene_tx=XM_019823204.2;refGene_gn=PTP4A3;refGene_tx=XM_023248251.1
F2	83156946	83157064
refGene_gn=PTP4A3;refGene_tx=XM_004000150.5;refGene_gn=PTP4A3;refGene_tx=XM_011291475.3;refGene_gn=PTP4A3;refGene_tx=XM_019823204.2;refGene_gn=PTP4A3;refGene_tx=XM_023248251.1
F2	83162334	83163027
refGene_gn=MROH5;refGene_tx=XM_023248434.1;refGene_gn=MROH5;refGene_tx=XM_023248435.1
F2	83163616	83163793
refGene_gn=MROH5;refGene_tx=XM_023248434.1;refGene_gn=MROH5;refGene_tx=XM_023248435.1
F2	83169271	83169415
refGene_gn=MROH5;refGene_tx=XM_023248434.1;refGene_gn=MROH5;refGene_tx=XM_023248435.1
F2	83170213	83170328
refGene_gn=MROH5;refGene_tx=XM_023248434.1;refGene_gn=MROH5;refGene_tx=XM_023248435.1
F2	83170799	83170937
refGene_gn=MROH5;refGene_tx=XM_023248434.1;refGene_gn=MROH5;refGene_tx=XM_023248435.1
F2	83176309	83176491
refGene_gn=MROH5;refGene_tx=XM_023248434.1;refGene_gn=MROH5;refGene_tx=XM_023248435.1
F2	83177482	83177642
refGene_gn=MROH5;refGene_tx=XM_023248434.1;refGene_gn=MROH5;refGene_tx=XM_023248435.1
F2	83182447	83182602
refGene_gn=MROH5;refGene_tx=XM_023248434.1;refGene_gn=MROH5;refGene_tx=XM_023248435.1
F2	83183397	83183547
refGene_gn=MROH5;refGene_tx=XM_023248434.1;refGene_gn=MROH5;refGene_tx=XM_023248435.1
F2	83185868	83186041
refGene_gn=MROH5;refGene_tx=XM_023248434.1;refGene_gn=MROH5;refGene_tx=XM_023248435.1
F2	83186340	83186476
refGene_gn=MROH5;refGene_tx=XM_023248434.1;refGene_gn=MROH5;refGene_tx

=XM_023248435.1
F2 83187809 83187914
refGene_gn=MROH5;refGene_tx=XM_023248434.1;refGene_gn=MROH5;refGene_tx
=XM_023248435.1
F2 83189471 83189567
refGene_gn=MROH5;refGene_tx=XM_023248434.1;refGene_gn=MROH5;refGene_tx
=XM_023248435.1
F2 83191165 83191301
refGene_gn=MROH5;refGene_tx=XM_023248434.1;refGene_gn=MROH5;refGene_tx
=XM_023248435.1
F2 83191756 83191836
refGene_gn=MROH5;refGene_tx=XM_023248434.1;refGene_gn=MROH5;refGene_tx
=XM_023248435.1
F2 83192619 83192742 refGene_gn=MROH5;refGene_tx=XM_023248434.1
F2 83193400 83193553
refGene_gn=MROH5;refGene_tx=XM_023248434.1;refGene_gn=MROH5;refGene_tx
=XM_023248435.1
F2 83194615 83194735
refGene_gn=MROH5;refGene_tx=XM_023248434.1;refGene_gn=MROH5;refGene_tx
=XM_023248435.1
F2 83195225 83195342
refGene_gn=MROH5;refGene_tx=XM_023248434.1;refGene_gn=MROH5;refGene_tx
=XM_023248435.1
F2 83195655 83195763
refGene_gn=MROH5;refGene_tx=XM_023248434.1;refGene_gn=MROH5;refGene_tx
=XM_023248435.1
F2 83205166 83205219
refGene_gn=MROH5;refGene_tx=XM_023248434.1;refGene_gn=MROH5;refGene_tx
=XM_023248435.1
F2 83206449 83206609
refGene_gn=MROH5;refGene_tx=XM_023248434.1;refGene_gn=MROH5;refGene_tx
=XM_023248435.1
F2 83211030 83211159
refGene_gn=MROH5;refGene_tx=XM_023248434.1;refGene_gn=MROH5;refGene_tx
=XM_023248435.1
F2 83212597 83212743
refGene_gn=MROH5;refGene_tx=XM_023248434.1;refGene_gn=MROH5;refGene_tx
=XM_023248435.1
F2 83213979 83214161
refGene_gn=MROH5;refGene_tx=XM_023248434.1;refGene_gn=MROH5;refGene_tx
=XM_023248435.1
F2 83224512 83224646
refGene_gn=MROH5;refGene_tx=XM_023248434.1;refGene_gn=MROH5;refGene_tx
=XM_023248435.1
F2 83233013 83233080
refGene_gn=LOC111558535;refGene_tx=XM_023248471.1
F2 83236181 83236660
refGene_gn=LOC111558535;refGene_tx=XM_023248471.1
F2 83748671 83748767 refGene_gn=TSNARE1;refGene_tx=XM_023248700.1
F2 83786105 83786695

refGene_gn=TSNARE1;refGene_tx=XM_023248694.1;refGene_gn=TSNARE1;refGene_tx=XM_023248695.1;refGene_gn=TSNARE1;refGene_tx=XM_023248696.1;refGene_gn=TSNARE1;refGene_tx=XM_023248697.1;refGene_gn=TSNARE1;refGene_tx=XM_023248698.1;refGene_gn=TSNARE1;refGene_tx=XM_023248699.1
F2 83790039 83790122 refGene_gn=TSNARE1;refGene_tx=XM_023248700.1
F2 83793739 83794171
refGene_gn=TSNARE1;refGene_tx=XM_023248694.1;refGene_gn=TSNARE1;refGene_tx=XM_023248695.1;refGene_gn=TSNARE1;refGene_tx=XM_023248696.1;refGene_gn=TSNARE1;refGene_tx=XM_023248697.1;refGene_gn=TSNARE1;refGene_tx=XM_023248698.1;refGene_gn=TSNARE1;refGene_tx=XM_023248699.1
F2 83795411 83795484
refGene_gn=TSNARE1;refGene_tx=XM_023248694.1;refGene_gn=TSNARE1;refGene_tx=XM_023248695.1;refGene_gn=TSNARE1;refGene_tx=XM_023248696.1;refGene_gn=TSNARE1;refGene_tx=XM_023248697.1;refGene_gn=TSNARE1;refGene_tx=XM_023248698.1;refGene_gn=TSNARE1;refGene_tx=XM_023248699.1;refGene_gn=TSNARE1;refGene_tx=XM_023248700.1
F2 83807104 83807266
refGene_gn=TSNARE1;refGene_tx=XM_023248694.1;refGene_gn=TSNARE1;refGene_tx=XM_023248695.1;refGene_gn=TSNARE1;refGene_tx=XM_023248696.1;refGene_gn=TSNARE1;refGene_tx=XM_023248697.1;refGene_gn=TSNARE1;refGene_tx=XM_023248698.1;refGene_gn=TSNARE1;refGene_tx=XM_023248699.1;refGene_gn=TSNARE1;refGene_tx=XM_023248700.1
F2 83817062 83817122
refGene_gn=TSNARE1;refGene_tx=XM_023248694.1;refGene_gn=TSNARE1;refGene_tx=XM_023248695.1;refGene_gn=TSNARE1;refGene_tx=XM_023248696.1;refGene_gn=TSNARE1;refGene_tx=XM_023248697.1;refGene_gn=TSNARE1;refGene_tx=XM_023248698.1;refGene_gn=TSNARE1;refGene_tx=XM_023248699.1;refGene_gn=TSNARE1;refGene_tx=XM_023248700.1
F2 83817606 83817681
refGene_gn=TSNARE1;refGene_tx=XM_023248694.1;refGene_gn=TSNARE1;refGene_tx=XM_023248695.1;refGene_gn=TSNARE1;refGene_tx=XM_023248696.1;refGene_gn=TSNARE1;refGene_tx=XM_023248697.1;refGene_gn=TSNARE1;refGene_tx=XM_023248698.1;refGene_gn=TSNARE1;refGene_tx=XM_023248699.1;refGene_gn=TSNARE1;refGene_tx=XM_023248700.1
F2 83820332 83820423
refGene_gn=TSNARE1;refGene_tx=XM_023248694.1;refGene_gn=TSNARE1;refGene_tx=XM_023248695.1;refGene_gn=TSNARE1;refGene_tx=XM_023248696.1;refGene_gn=TSNARE1;refGene_tx=XM_023248697.1;refGene_gn=TSNARE1;refGene_tx=XM_023248698.1;refGene_gn=TSNARE1;refGene_tx=XM_023248699.1;refGene_gn=TSNARE1;refGene_tx=XM_023248700.1
F2 83830730 83830800
refGene_gn=TSNARE1;refGene_tx=XM_023248694.1;refGene_gn=TSNARE1;refGene_tx=XM_023248695.1;refGene_gn=TSNARE1;refGene_tx=XM_023248696.1;refGene_gn=TSNARE1;refGene_tx=XM_023248697.1;refGene_gn=TSNARE1;refGene_tx=XM_023248698.1;refGene_gn=TSNARE1;refGene_tx=XM_023248699.1;refGene_gn=TSNARE1;refGene_tx=XM_023248700.1
F2 83831661 83831739
refGene_gn=TSNARE1;refGene_tx=XM_023248694.1;refGene_gn=TSNARE1;refGene_tx=XM_023248695.1;refGene_gn=TSNARE1;refGene_tx=XM_023248696.1;refGene_gn=TSNARE1;refGene_tx=XM_023248697.1;refGene_gn=TSNARE1;refGene_tx=

XM_023248698.1;refGene_gn=TSNARE1;refGene_tx=XM_023248699.1;refGene_gn=TSNARE1;refGene_tx=XM_023248700.1
F2 83844178 83844673
refGene_gn=TSNARE1;refGene_tx=XM_023248694.1;refGene_gn=TSNARE1;refGene_tx=XM_023248695.1;refGene_gn=TSNARE1;refGene_tx=XM_023248696.1;refGene_gn=TSNARE1;refGene_tx=XM_023248697.1;refGene_gn=TSNARE1;refGene_tx=XM_023248698.1;refGene_gn=TSNARE1;refGene_tx=XM_023248700.1
F2 83845685 83845838
refGene_gn=TSNARE1;refGene_tx=XM_023248694.1;refGene_gn=TSNARE1;refGene_tx=XM_023248695.1;refGene_gn=TSNARE1;refGene_tx=XM_023248696.1;refGene_gn=TSNARE1;refGene_tx=XM_023248700.1
F2 83850736 83850852
refGene_gn=TSNARE1;refGene_tx=XM_023248694.1;refGene_gn=TSNARE1;refGene_tx=XM_023248695.1;refGene_gn=TSNARE1;refGene_tx=XM_023248699.1
F2 83852678 83852805
refGene_gn=TSNARE1;refGene_tx=XM_023248694.1;refGene_gn=TSNARE1;refGene_tx=XM_023248695.1;refGene_gn=TSNARE1;refGene_tx=XM_023248696.1;refGene_gn=TSNARE1;refGene_tx=XM_023248697.1;refGene_gn=TSNARE1;refGene_tx=XM_023248698.1;refGene_gn=TSNARE1;refGene_tx=XM_023248699.1;refGene_gn=TSNARE1;refGene_tx=XM_023248700.1
F2 83872670 83872680
refGene_gn=TSNARE1;refGene_tx=XM_023248696.1;refGene_gn=TSNARE1;refGene_tx=XM_023248699.1;refGene_gn=TSNARE1;refGene_tx=XM_023248700.1
F2 83935329 83936116 refGene_gn=ADGRB1;refGene_tx=XM_023248346.1
F2 83945525 83945687 refGene_gn=ADGRB1;refGene_tx=XM_023248346.1
F2 83946972 83947083 refGene_gn=ADGRB1;refGene_tx=XM_023248346.1
F2 83947525 83947690 refGene_gn=ADGRB1;refGene_tx=XM_023248346.1
F2 83947898 83948063 refGene_gn=ADGRB1;refGene_tx=XM_023248346.1
F2 83948549 83948723 refGene_gn=ADGRB1;refGene_tx=XM_023248346.1
F2 83949001 83949166 refGene_gn=ADGRB1;refGene_tx=XM_023248346.1
F2 83949364 83949466 refGene_gn=ADGRB1;refGene_tx=XM_023248346.1
F2 83950699 83950806 refGene_gn=ADGRB1;refGene_tx=XM_023248346.1
F2 83951048 83951243 refGene_gn=ADGRB1;refGene_tx=XM_023248346.1
F2 83951397 83951572 refGene_gn=ADGRB1;refGene_tx=XM_023248346.1
F2 83954864 83955086 refGene_gn=ADGRB1;refGene_tx=XM_023248346.1
F2 83955581 83955657 refGene_gn=ADGRB1;refGene_tx=XM_023248346.1
F2 83955779 83955882 refGene_gn=ADGRB1;refGene_tx=XM_023248346.1
F2 83956678 83956722 refGene_gn=ADGRB1;refGene_tx=XM_023248346.1
F2 83973251 83973393 refGene_gn=ADGRB1;refGene_tx=XM_023248346.1
F2 83980993 83981097 refGene_gn=ADGRB1;refGene_tx=XM_023248346.1
F2 83983176 83983279 refGene_gn=ADGRB1;refGene_tx=XM_023248346.1
F2 83983917 83983971 refGene_gn=ADGRB1;refGene_tx=XM_023248346.1
F2 83983972 83984069 refGene_gn=ADGRB1;refGene_tx=XM_023248346.1
F2 83984802 83984872 refGene_gn=ADGRB1;refGene_tx=XM_023248346.1
F2 83986304 83986430 refGene_gn=ADGRB1;refGene_tx=XM_023248346.1
F2 83992351 83992446 refGene_gn=ADGRB1;refGene_tx=XM_023248346.1
F2 83993227 83993340 refGene_gn=ADGRB1;refGene_tx=XM_023248346.1
F2 83994252 83994334 refGene_gn=ADGRB1;refGene_tx=XM_023248346.1
F2 83996305 83996540 refGene_gn=ADGRB1;refGene_tx=XM_023248346.1
F2 83997037 83997209 refGene_gn=ADGRB1;refGene_tx=XM_023248346.1

F2	84000236 84000332 refGene_gn=ADGRB1;refGene_tx=XM_023248346.1
F2	84001909 84001949 refGene_gn=ADGRB1;refGene_tx=XM_023248346.1
F2	84004205 84004888 refGene_gn=ADGRB1;refGene_tx=XM_023248346.1
F2	84006042 84006078 refGene_gn=ADGRB1;refGene_tx=XM_023248346.1
F2	84006278 84006383 refGene_gn=ADGRB1;refGene_tx=XM_023248346.1
F2	84006872 84007070 refGene_gn=ADGRB1;refGene_tx=XM_023248346.1
F2	84023470 84023571
refGene_gn=LOC102899409;refGene_tx=XM_019822603.2;refGene_gn=LOC102899409;refGene_tx=XM_023248421.1;refGene_gn=LOC102899409;refGene_tx=XM_023248422.1;refGene_gn=LOC102899409;refGene_tx=XM_023248423.1;refGene_gn=LOC102899409;refGene_tx=XM_023248424.1;refGene_gn=LOC102899409;refGene_tx=XM_023248425.1;refGene_gn=LOC102899409;refGene_tx=XM_023248426.1;refGene_gn=LOC102899409;refGene_tx=XM_023248427.1
F2	84023843 84023918
refGene_gn=LOC102899409;refGene_tx=XM_019822603.2;refGene_gn=LOC102899409;refGene_tx=XM_023248421.1;refGene_gn=LOC102899409;refGene_tx=XM_023248422.1;refGene_gn=LOC102899409;refGene_tx=XM_023248424.1;refGene_gn=LOC102899409;refGene_tx=XM_023248426.1;refGene_gn=LOC102899409;refGene_tx=XM_023248427.1
F2	84024116 84024219
refGene_gn=LOC102899409;refGene_tx=XM_019822603.2;refGene_gn=LOC102899409;refGene_tx=XM_023248421.1;refGene_gn=LOC102899409;refGene_tx=XM_023248422.1;refGene_gn=LOC102899409;refGene_tx=XM_023248423.1;refGene_gn=LOC102899409;refGene_tx=XM_023248424.1;refGene_gn=LOC102899409;refGene_tx=XM_023248425.1;refGene_gn=LOC102899409;refGene_tx=XM_023248426.1;refGene_gn=LOC102899409;refGene_tx=XM_023248427.1
F2	84029558 84029681
refGene_gn=LOC102899409;refGene_tx=XM_019822603.2;refGene_gn=LOC102899409;refGene_tx=XM_023248421.1;refGene_gn=LOC102899409;refGene_tx=XM_023248422.1;refGene_gn=LOC102899409;refGene_tx=XM_023248423.1;refGene_gn=LOC102899409;refGene_tx=XM_023248424.1;refGene_gn=LOC102899409;refGene_tx=XM_023248425.1;refGene_gn=LOC102899409;refGene_tx=XM_023248426.1;refGene_gn=LOC102899409;refGene_tx=XM_023248427.1
F2	84031709 84031865
refGene_gn=LOC102899409;refGene_tx=XM_019822603.2;refGene_gn=LOC102899409;refGene_tx=XM_023248421.1;refGene_gn=LOC102899409;refGene_tx=XM_023248422.1;refGene_gn=LOC102899409;refGene_tx=XM_023248423.1;refGene_gn=LOC102899409;refGene_tx=XM_023248424.1;refGene_gn=LOC102899409;refGene_tx=XM_023248425.1;refGene_gn=LOC102899409;refGene_tx=XM_023248426.1;refGene_gn=LOC102899409;refGene_tx=XM_023248427.1
F2	84032846 84033023
refGene_gn=LOC102899409;refGene_tx=XM_019822603.2;refGene_gn=LOC102899409;refGene_tx=XM_023248421.1;refGene_gn=LOC102899409;refGene_tx=XM_023248422.1;refGene_gn=LOC102899409;refGene_tx=XM_023248423.1;refGene_gn=LOC102899409;refGene_tx=XM_023248424.1;refGene_gn=LOC102899409;refGene_tx=XM_023248425.1;refGene_gn=LOC102899409;refGene_tx=XM_023248426.1;refGene_gn=LOC102899409;refGene_tx=XM_023248427.1
F2	84033598 84033742
refGene_gn=LOC102899409;refGene_tx=XM_019822603.2;refGene_gn=LOC102899409;refGene_tx=XM_023248421.1;refGene_gn=LOC102899409;refGene_tx=XM_02

3248422.1;refGene_gn=LOC102899409;refGene_tx=XM_023248423.1;refGene_gn
=LOC102899409;refGene_tx=XM_023248424.1;refGene_gn=LOC102899409;refGen
e_tx=XM_023248425.1;refGene_gn=LOC102899409;refGene_tx=XM_023248426.1;
refGene_gn=LOC102899409;refGene_tx=XM_023248427.1;refGene_gn=LOC102899
409;refGene_tx=XM_023248428.1
F2 84033985 84034097
refGene_gn=LOC102899409;refGene_tx=XM_019822603.2;refGene_gn=LOC102899
409;refGene_tx=XM_023248421.1;refGene_gn=LOC102899409;refGene_tx=XM_02
3248422.1;refGene_gn=LOC102899409;refGene_tx=XM_023248423.1;refGene_gn
=LOC102899409;refGene_tx=XM_023248424.1;refGene_gn=LOC102899409;refGen
e_tx=XM_023248425.1;refGene_gn=LOC102899409;refGene_tx=XM_023248426.1;
refGene_gn=LOC102899409;refGene_tx=XM_023248427.1;refGene_gn=LOC102899
409;refGene_tx=XM_023248428.1
F2 84034285 84034423
refGene_gn=LOC102899409;refGene_tx=XM_019822603.2;refGene_gn=LOC102899
409;refGene_tx=XM_023248421.1;refGene_gn=LOC102899409;refGene_tx=XM_02
3248422.1;refGene_gn=LOC102899409;refGene_tx=XM_023248423.1;refGene_gn
=LOC102899409;refGene_tx=XM_023248424.1;refGene_gn=LOC102899409;refGen
e_tx=XM_023248425.1;refGene_gn=LOC102899409;refGene_tx=XM_023248426.1;
refGene_gn=LOC102899409;refGene_tx=XM_023248427.1;refGene_gn=LOC102899
409;refGene_tx=XM_023248428.1
F2 84034785 84034970
refGene_gn=LOC102899409;refGene_tx=XM_019822603.2;refGene_gn=LOC102899
409;refGene_tx=XM_023248421.1;refGene_gn=LOC102899409;refGene_tx=XM_02
3248422.1;refGene_gn=LOC102899409;refGene_tx=XM_023248423.1;refGene_gn
=LOC102899409;refGene_tx=XM_023248424.1;refGene_gn=LOC102899409;refGen
e_tx=XM_023248425.1;refGene_gn=LOC102899409;refGene_tx=XM_023248426.1;
refGene_gn=LOC102899409;refGene_tx=XM_023248427.1;refGene_gn=LOC102899
409;refGene_tx=XM_023248428.1
F2 84035289 84035434
refGene_gn=LOC102899409;refGene_tx=XM_019822603.2;refGene_gn=LOC102899
409;refGene_tx=XM_023248421.1;refGene_gn=LOC102899409;refGene_tx=XM_02
3248422.1;refGene_gn=LOC102899409;refGene_tx=XM_023248423.1;refGene_gn
=LOC102899409;refGene_tx=XM_023248424.1;refGene_gn=LOC102899409;refGen
e_tx=XM_023248425.1;refGene_gn=LOC102899409;refGene_tx=XM_023248426.1;
refGene_gn=LOC102899409;refGene_tx=XM_023248427.1;refGene_gn=LOC102899
409;refGene_tx=XM_023248428.1
F2 84035821 84035893
refGene_gn=LOC102899409;refGene_tx=XM_019822603.2;refGene_gn=LOC102899
409;refGene_tx=XM_023248421.1;refGene_gn=LOC102899409;refGene_tx=XM_02
3248423.1;refGene_gn=LOC102899409;refGene_tx=XM_023248424.1;refGene_gn
=LOC102899409;refGene_tx=XM_023248425.1;refGene_gn=LOC102899409;refGen
e_tx=XM_023248426.1;refGene_gn=LOC102899409;refGene_tx=XM_023248427.1;
refGene_gn=LOC102899409;refGene_tx=XM_023248428.1
F2 84036436 84036591
refGene_gn=LOC102899409;refGene_tx=XM_019822603.2;refGene_gn=LOC102899
409;refGene_tx=XM_023248421.1;refGene_gn=LOC102899409;refGene_tx=XM_02
3248422.1;refGene_gn=LOC102899409;refGene_tx=XM_023248423.1;refGene_gn
=LOC102899409;refGene_tx=XM_023248424.1;refGene_gn=LOC102899409;refGen
e_tx=XM_023248425.1;refGene_gn=LOC102899409;refGene_tx=XM_023248426.1;

refGene_gn=LOC102899409;refGene_tx=XM_023248427.1;refGene_gn=LOC102899
409;refGene_tx=XM_023248428.1
F2 84037418 84037568
refGene_gn=LOC102899409;refGene_tx=XM_019822603.2;refGene_gn=LOC102899
409;refGene_tx=XM_023248421.1;refGene_gn=LOC102899409;refGene_tx=XM_02
3248422.1;refGene_gn=LOC102899409;refGene_tx=XM_023248423.1;refGene_gn
=LOC102899409;refGene_tx=XM_023248424.1;refGene_gn=LOC102899409;refGen
e_tx=XM_023248425.1;refGene_gn=LOC102899409;refGene_tx=XM_023248426.1;
refGene_gn=LOC102899409;refGene_tx=XM_023248427.1;refGene_gn=LOC102899
409;refGene_tx=XM_023248428.1
F2 84038788 84038883
refGene_gn=LOC102899409;refGene_tx=XM_019822603.2;refGene_gn=LOC102899
409;refGene_tx=XM_023248421.1;refGene_gn=LOC102899409;refGene_tx=XM_02
3248422.1;refGene_gn=LOC102899409;refGene_tx=XM_023248423.1;refGene_gn
=LOC102899409;refGene_tx=XM_023248424.1;refGene_gn=LOC102899409;refGen
e_tx=XM_023248425.1;refGene_gn=LOC102899409;refGene_tx=XM_023248426.1;
refGene_gn=LOC102899409;refGene_tx=XM_023248427.1;refGene_gn=LOC102899
409;refGene_tx=XM_023248428.1
F2 84038961 84039097
refGene_gn=LOC102899409;refGene_tx=XM_019822603.2;refGene_gn=LOC102899
409;refGene_tx=XM_023248421.1;refGene_gn=LOC102899409;refGene_tx=XM_02
3248422.1;refGene_gn=LOC102899409;refGene_tx=XM_023248423.1;refGene_gn
=LOC102899409;refGene_tx=XM_023248424.1;refGene_gn=LOC102899409;refGen
e_tx=XM_023248425.1;refGene_gn=LOC102899409;refGene_tx=XM_023248426.1;
refGene_gn=LOC102899409;refGene_tx=XM_023248427.1;refGene_gn=LOC102899
409;refGene_tx=XM_023248428.1
F2 84039474 84039579
refGene_gn=LOC102899409;refGene_tx=XM_019822603.2;refGene_gn=LOC102899
409;refGene_tx=XM_023248421.1;refGene_gn=LOC102899409;refGene_tx=XM_02
3248422.1;refGene_gn=LOC102899409;refGene_tx=XM_023248423.1;refGene_gn
=LOC102899409;refGene_tx=XM_023248424.1;refGene_gn=LOC102899409;refGen
e_tx=XM_023248425.1;refGene_gn=LOC102899409;refGene_tx=XM_023248426.1;
refGene_gn=LOC102899409;refGene_tx=XM_023248427.1;refGene_gn=LOC102899
409;refGene_tx=XM_023248428.1
F2 84040050 84040146
refGene_gn=LOC102899409;refGene_tx=XM_019822603.2;refGene_gn=LOC102899
409;refGene_tx=XM_023248421.1;refGene_gn=LOC102899409;refGene_tx=XM_02
3248422.1;refGene_gn=LOC102899409;refGene_tx=XM_023248423.1;refGene_gn
=LOC102899409;refGene_tx=XM_023248424.1;refGene_gn=LOC102899409;refGen
e_tx=XM_023248425.1;refGene_gn=LOC102899409;refGene_tx=XM_023248426.1;
refGene_gn=LOC102899409;refGene_tx=XM_023248427.1;refGene_gn=LOC102899
409;refGene_tx=XM_023248428.1
F2 84043808 84043944
refGene_gn=LOC102899409;refGene_tx=XM_019822603.2;refGene_gn=LOC102899
409;refGene_tx=XM_023248421.1;refGene_gn=LOC102899409;refGene_tx=XM_02
3248422.1;refGene_gn=LOC102899409;refGene_tx=XM_023248423.1;refGene_gn
=LOC102899409;refGene_tx=XM_023248424.1;refGene_gn=LOC102899409;refGen
e_tx=XM_023248425.1;refGene_gn=LOC102899409;refGene_tx=XM_023248426.1;
refGene_gn=LOC102899409;refGene_tx=XM_023248427.1;refGene_gn=LOC102899
409;refGene_tx=XM_023248428.1

F2 84044360 84044437
refGene_gn=LOC102899409;refGene_tx=XM_019822603.2;refGene_gn=LOC102899
409;refGene_tx=XM_023248421.1;refGene_gn=LOC102899409;refGene_tx=XM_02
3248422.1;refGene_gn=LOC102899409;refGene_tx=XM_023248423.1;refGene_gn
=LOC102899409;refGene_tx=XM_023248424.1;refGene_gn=LOC102899409;refGen
e_tx=XM_023248425.1;refGene_gn=LOC102899409;refGene_tx=XM_023248426.1;
refGene_gn=LOC102899409;refGene_tx=XM_023248427.1;refGene_gn=LOC102899
409;refGene_tx=XM_023248428.1
F2 84044551 84044671
refGene_gn=LOC102899409;refGene_tx=XM_019822603.2;refGene_gn=LOC102899
409;refGene_tx=XM_023248421.1;refGene_gn=LOC102899409;refGene_tx=XM_02
3248422.1;refGene_gn=LOC102899409;refGene_tx=XM_023248423.1;refGene_gn
=LOC102899409;refGene_tx=XM_023248424.1;refGene_gn=LOC102899409;refGen
e_tx=XM_023248425.1;refGene_gn=LOC102899409;refGene_tx=XM_023248426.1;
refGene_gn=LOC102899409;refGene_tx=XM_023248427.1;refGene_gn=LOC102899
409;refGene_tx=XM_023248428.1
F2 84044769 84044919
refGene_gn=LOC102899409;refGene_tx=XM_019822603.2;refGene_gn=LOC102899
409;refGene_tx=XM_023248421.1;refGene_gn=LOC102899409;refGene_tx=XM_02
3248422.1;refGene_gn=LOC102899409;refGene_tx=XM_023248423.1;refGene_gn
=LOC102899409;refGene_tx=XM_023248424.1;refGene_gn=LOC102899409;refGen
e_tx=XM_023248425.1;refGene_gn=LOC102899409;refGene_tx=XM_023248426.1;
refGene_gn=LOC102899409;refGene_tx=XM_023248427.1;refGene_gn=LOC102899
409;refGene_tx=XM_023248428.1
F2 84045448 84045568
refGene_gn=LOC102899409;refGene_tx=XM_019822603.2;refGene_gn=LOC102899
409;refGene_tx=XM_023248421.1;refGene_gn=LOC102899409;refGene_tx=XM_02
3248422.1;refGene_gn=LOC102899409;refGene_tx=XM_023248423.1;refGene_gn
=LOC102899409;refGene_tx=XM_023248424.1;refGene_gn=LOC102899409;refGen
e_tx=XM_023248425.1;refGene_gn=LOC102899409;refGene_tx=XM_023248426.1;
refGene_gn=LOC102899409;refGene_tx=XM_023248427.1;refGene_gn=LOC102899
409;refGene_tx=XM_023248428.1
F2 84045935 84046052
refGene_gn=LOC102899409;refGene_tx=XM_019822603.2;refGene_gn=LOC102899
409;refGene_tx=XM_023248421.1;refGene_gn=LOC102899409;refGene_tx=XM_02
3248422.1;refGene_gn=LOC102899409;refGene_tx=XM_023248423.1;refGene_gn
=LOC102899409;refGene_tx=XM_023248424.1;refGene_gn=LOC102899409;refGen
e_tx=XM_023248425.1;refGene_gn=LOC102899409;refGene_tx=XM_023248426.1;
refGene_gn=LOC102899409;refGene_tx=XM_023248427.1;refGene_gn=LOC102899
409;refGene_tx=XM_023248428.1
F2 84046988 84047072
refGene_gn=LOC102899409;refGene_tx=XM_019822603.2;refGene_gn=LOC102899
409;refGene_tx=XM_023248421.1;refGene_gn=LOC102899409;refGene_tx=XM_02
3248422.1;refGene_gn=LOC102899409;refGene_tx=XM_023248423.1;refGene_gn
=LOC102899409;refGene_tx=XM_023248424.1;refGene_gn=LOC102899409;refGen
e_tx=XM_023248425.1;refGene_gn=LOC102899409;refGene_tx=XM_023248426.1;
refGene_gn=LOC102899409;refGene_tx=XM_023248427.1;refGene_gn=LOC102899
409;refGene_tx=XM_023248428.1
F2 84048133 84048282
refGene_gn=LOC102899409;refGene_tx=XM_019822603.2;refGene_gn=LOC102899

409;refGene_tx=XM_023248421.1;refGene_gn=LOC102899409;refGene_tx=XM_023248422.1;refGene_gn=LOC102899409;refGene_tx=XM_023248423.1;refGene_gn=LOC102899409;refGene_tx=XM_023248424.1;refGene_gn=LOC102899409;refGene_tx=XM_023248425.1;refGene_gn=LOC102899409;refGene_tx=XM_023248426.1;refGene_gn=LOC102899409;refGene_tx=XM_023248427.1;refGene_gn=LOC102899409;refGene_tx=XM_023248428.1
F2 84050236 84050354
refGene_gn=LOC102899409;refGene_tx=XM_019822603.2;refGene_gn=LOC102899409;refGene_tx=XM_023248421.1;refGene_gn=LOC102899409;refGene_tx=XM_023248422.1;refGene_gn=LOC102899409;refGene_tx=XM_023248423.1;refGene_gn=LOC102899409;refGene_tx=XM_023248424.1;refGene_gn=LOC102899409;refGene_tx=XM_023248425.1;refGene_gn=LOC102899409;refGene_tx=XM_023248426.1;refGene_gn=LOC102899409;refGene_tx=XM_023248427.1;refGene_gn=LOC102899409;refGene_tx=XM_023248428.1
F2 84050490 84050508
refGene_gn=LOC102899409;refGene_tx=XM_023248426.1;refGene_gn=LOC102899409;refGene_tx=XM_023248427.1
F2 84053960 84054119
refGene_gn=LOC102899409;refGene_tx=XM_019822603.2;refGene_gn=LOC102899409;refGene_tx=XM_023248421.1;refGene_gn=LOC102899409;refGene_tx=XM_023248422.1;refGene_gn=LOC102899409;refGene_tx=XM_023248423.1;refGene_gn=LOC102899409;refGene_tx=XM_023248424.1;refGene_gn=LOC102899409;refGene_tx=XM_023248425.1;refGene_gn=LOC102899409;refGene_tx=XM_023248428.1
F2 84075389 84076580 refGene_gn=ARC;refGene_tx=XM_023248559.1
F2 84108135 84109761 refGene_gn=JRK;refGene_tx=XM_023248237.1
F2 84118480 84118532 refGene_gn=PSCA;refGene_tx=XM_023248455.1
F2 84119839 84119947 refGene_gn=PSCA;refGene_tx=XM_023248455.1
F2 84120621 84120836 refGene_gn=PSCA;refGene_tx=XM_023248455.1
F2 84126530 84127043 refGene_gn=THEM6;refGene_tx=XM_023248574.1
F2 84129181 84129295 refGene_gn=THEM6;refGene_tx=XM_023248574.1
F2 84132641 84132829 refGene_gn=SLURP1;refGene_tx=XM_019823218.2
F2 84133129 84133249 refGene_gn=SLURP1;refGene_tx=XM_019823218.2
F2 84133583 84133656 refGene_gn=SLURP1;refGene_tx=XM_019823218.2
F2 84137879 84138085 refGene_gn=LYPD2;refGene_tx=XM_004000197.3
F2 84138507 84138627 refGene_gn=LYPD2;refGene_tx=XM_004000197.3
F2 84139590 84139651 refGene_gn=LYPD2;refGene_tx=XM_004000197.3
F2 84144439 84144576
refGene_gn=LOC109496272;refGene_tx=XM_019823217.2
F2 84144806 84144905
refGene_gn=LOC109496272;refGene_tx=XM_019823217.2
F2 84148402 84148499
refGene_gn=LOC109496272;refGene_tx=XM_019823217.2
F2 84153385 84153582
refGene_gn=LYNX1;refGene_tx=XM_023248290.1;refGene_gn=LYNX1;refGene_tx=XM_023248291.1;refGene_gn=LYNX1;refGene_tx=XM_023248292.1
F2 84153794 84153896
refGene_gn=LYNX1;refGene_tx=XM_023248290.1;refGene_gn=LYNX1;refGene_tx=XM_023248291.1;refGene_gn=LYNX1;refGene_tx=XM_023248292.1
F2 84154118 84154170
refGene_gn=LYNX1;refGene_tx=XM_023248290.1;refGene_gn=LYNX1;refGene_tx

=XM_023248291.1;refGene_gn=LYNX1;refGene_tx=XM_023248292.1
F2 84174995 84175237 refGene_gn=LY6D;refGene_tx=XM_023248307.1
F2 84175469 84175568 refGene_gn=LY6D;refGene_tx=XM_023248307.1
F2 84176280 84176332 refGene_gn=LY6D;refGene_tx=XM_023248307.1
F2 84184931 84185143
refGene_gn=LOC111558519;refGene_tx=XM_023248316.1
F2 84187521 84187620
refGene_gn=LOC111558519;refGene_tx=XM_023248316.1
F2 84189544 84189596
refGene_gn=LOC111558519;refGene_tx=XM_023248316.1
F2 84201523 84201727 refGene_gn=LY6K;refGene_tx=XM_023248310.1
F2 84209736 84209996
refGene_gn=LY6K;refGene_tx=XM_023248308.1;refGene_gn=LY6K;refGene_tx=XM_023248309.1;refGene_gn=LY6K;refGene_tx=XM_023248310.1;refGene_gn=LY6K;refGene_tx=XM_023248311.1;refGene_gn=LY6K;refGene_tx=XM_023248312.1;refGene_gn=LY6K;refGene_tx=XM_023248314.1
F2 84211161 84211266
refGene_gn=LY6K;refGene_tx=XM_023248308.1;refGene_gn=LY6K;refGene_tx=XM_023248309.1;refGene_gn=LY6K;refGene_tx=XM_023248310.1;refGene_gn=LY6K;refGene_tx=XM_023248311.1;refGene_gn=LY6K;refGene_tx=XM_023248312.1;refGene_gn=LY6K;refGene_tx=XM_023248314.1
F2 84212038 84212134
refGene_gn=LY6K;refGene_tx=XM_023248308.1;refGene_gn=LY6K;refGene_tx=XM_023248309.1;refGene_gn=LY6K;refGene_tx=XM_023248310.1;refGene_gn=LY6K;refGene_tx=XM_023248311.1;refGene_gn=LY6K;refGene_tx=XM_023248312.1;refGene_gn=LY6K;refGene_tx=XM_023248314.1
F2 84214763 84214851 refGene_gn=LY6K;refGene_tx=XM_023248309.1
F2 84216506 84216597
refGene_gn=LY6K;refGene_tx=XM_023248308.1;refGene_gn=LY6K;refGene_tx=XM_023248310.1;refGene_gn=LY6K;refGene_tx=XM_023248311.1
F2 84218397 84218461 refGene_gn=GML;refGene_tx=XM_023248315.1
F2 84219007 84219070 refGene_gn=GML;refGene_tx=XM_023248315.1
F2 84219072 84219114 refGene_gn=GML;refGene_tx=XM_023248315.1
F2 84226969 84227259 refGene_gn=GML;refGene_tx=XM_023248315.1
F2 84243030 84243269
refGene_gn=LOC101097790;refGene_tx=XM_004000154.3
F2 84243658 84243814
refGene_gn=LOC101097790;refGene_tx=XM_004000154.3
F2 84244845 84245045
refGene_gn=LOC101097790;refGene_tx=XM_004000154.3
F2 84245174 84245378
refGene_gn=LOC101097790;refGene_tx=XM_004000154.3
F2 84245611 84245766
refGene_gn=LOC101097790;refGene_tx=XM_004000154.3
F2 84246785 84246952
refGene_gn=LOC101097790;refGene_tx=XM_004000154.3
F2 84247382 84247461
refGene_gn=LOC101097790;refGene_tx=XM_004000154.3
F2 84247558 84247756
refGene_gn=LOC101097790;refGene_tx=XM_004000154.3

F2 84247985 84248099
refGene_gn=LOC101097790;refGene_tx=XM_004000154.3
F2 84273390 84273629
refGene_gn=LOC101098195;refGene_tx=XM_019823222.2;refGene_gn=LOC101098195;refGene_tx=XM_023248403.1;refGene_gn=LOC101098195;refGene_tx=XM_023248404.1
F2 84273766 84273886
refGene_gn=LOC101098195;refGene_tx=XM_019823222.2;refGene_gn=LOC101098195;refGene_tx=XM_023248403.1;refGene_gn=LOC101098195;refGene_tx=XM_023248404.1
F2 84274313 84274423
refGene_gn=LOC101098195;refGene_tx=XM_019823222.2;refGene_gn=LOC101098195;refGene_tx=XM_023248403.1;refGene_gn=LOC101098195;refGene_tx=XM_023248404.1
F2 84274885 84274926
refGene_gn=LOC101098195;refGene_tx=XM_023248403.1;refGene_gn=LOC101098195;refGene_tx=XM_023248404.1
F2 84305060 84305317
refGene_gn=LOC102899673;refGene_tx=XM_006943431.4
F2 84305612 84305732
refGene_gn=LOC102899673;refGene_tx=XM_006943431.4
F2 84305951 84306003
refGene_gn=LOC102899673;refGene_tx=XM_006943431.4
F2 84334703 84334779 refGene_gn=LY6L;refGene_tx=XM_019823224.2
F2 84334862 84334979 refGene_gn=LY6L;refGene_tx=XM_019823224.2
F2 84336698 84336916 refGene_gn=LY6L;refGene_tx=XM_019823224.2
F2 84370541 84370777
refGene_gn=LY6H;refGene_tx=XM_011291496.3;refGene_gn=LY6H;refGene_tx=XM_023248609.1;refGene_gn=LY6H;refGene_tx=XM_023248610.1;refGene_gn=LY6H;refGene_tx=XM_023248611.1
F2 84371028 84371148
refGene_gn=LY6H;refGene_tx=XM_011291496.3;refGene_gn=LY6H;refGene_tx=XM_023248609.1;refGene_gn=LY6H;refGene_tx=XM_023248610.1;refGene_gn=LY6H;refGene_tx=XM_023248611.1
F2 84371840 84371968
refGene_gn=LY6H;refGene_tx=XM_011291496.3;refGene_gn=LY6H;refGene_tx=XM_023248609.1;refGene_gn=LY6H;refGene_tx=XM_023248610.1;refGene_gn=LY6H;refGene_tx=XM_023248611.1
F2 84372489 84372491 refGene_gn=LY6H;refGene_tx=XM_023248609.1
F2 84399253 84399305
refGene_gn=GPIHBP1;refGene_tx=XM_006943432.3;refGene_gn=GPIHBP1;refGene_tx=XM_023248486.1
F2 84399837 84400298
refGene_gn=GPIHBP1;refGene_tx=XM_006943432.3;refGene_gn=GPIHBP1;refGene_tx=XM_023248486.1
F2 84400377 84400494 refGene_gn=GPIHBP1;refGene_tx=XM_006943432.3
F2 84400621 84400854 refGene_gn=GPIHBP1;refGene_tx=XM_006943432.3
F2 84413818 84414415
refGene_gn=ZFP41;refGene_tx=XM_011291497.3;refGene_gn=ZFP41;refGene_tx=XM_011291498.3;refGene_gn=ZFP41;refGene_tx=XM_011291499.3;refGene_gn=

ZFP41;refGene_tx=XM_019823236.2
F2 84421891 84421976 refGene_gn=GLI4;refGene_tx=XM_023248326.1
F2 84425591 84425831 refGene_gn=GLI4;refGene_tx=XM_023248325.1
F2 84427361 84427523
refGene_gn=GLI4;refGene_tx=XM_023248325.1;refGene_gn=GLI4;refGene_tx=XM_023248326.1;refGene_gn=GLI4;refGene_tx=XM_023248327.1
F2 84430234 84430326 refGene_gn=GLI4;refGene_tx=XM_023248327.1
F2 84430810 84430934
refGene_gn=GLI4;refGene_tx=XM_023248325.1;refGene_gn=GLI4;refGene_tx=XM_023248326.1;refGene_gn=GLI4;refGene_tx=XM_023248327.1
F2 84431661 84432581
refGene_gn=GLI4;refGene_tx=XM_023248325.1;refGene_gn=GLI4;refGene_tx=XM_023248326.1;refGene_gn=GLI4;refGene_tx=XM_023248327.1
F2 84437461 84438119 refGene_gn=ZNF696;refGene_tx=XM_023248347.1
F2 84439144 84439182 refGene_gn=ZNF696;refGene_tx=XM_023248347.1
F2 84440414 84440585 refGene_gn=ZNF696;refGene_tx=XM_023248347.1
F2 84442255 84442528 refGene_gn=ZNF696;refGene_tx=XM_023248347.1
F2 84444733 84444807 refGene_gn=ZNF696;refGene_tx=XM_023248347.1
F2 84445357 84445918 refGene_gn=ZNF696;refGene_tx=XM_023248347.1
F2 84447601 84447708 refGene_gn=ZNF696;refGene_tx=XM_023248347.1
F2 84450031 84451101 refGene_gn=ZNF696;refGene_tx=XM_023248347.1
F2 84483976 84484057
refGene_gn=RHPN1;refGene_tx=XM_023248466.1;refGene_gn=RHPN1;refGene_tx=XM_023248467.1;refGene_gn=RHPN1;refGene_tx=XM_023248468.1
F2 84486824 84486940
refGene_gn=RHPN1;refGene_tx=XM_023248466.1;refGene_gn=RHPN1;refGene_tx=XM_023248467.1;refGene_gn=RHPN1;refGene_tx=XM_023248468.1
F2 84488266 84488395
refGene_gn=RHPN1;refGene_tx=XM_023248466.1;refGene_gn=RHPN1;refGene_tx=XM_023248467.1;refGene_gn=RHPN1;refGene_tx=XM_023248468.1
F2 84488614 84488690
refGene_gn=RHPN1;refGene_tx=XM_023248466.1;refGene_gn=RHPN1;refGene_tx=XM_023248467.1;refGene_gn=RHPN1;refGene_tx=XM_023248468.1
F2 84489375 84489453
refGene_gn=RHPN1;refGene_tx=XM_023248466.1;refGene_gn=RHPN1;refGene_tx=XM_023248467.1;refGene_gn=RHPN1;refGene_tx=XM_023248468.1
F2 84489969 84490094
refGene_gn=RHPN1;refGene_tx=XM_023248466.1;refGene_gn=RHPN1;refGene_tx=XM_023248467.1;refGene_gn=RHPN1;refGene_tx=XM_023248468.1
F2 84490186 84490353
refGene_gn=RHPN1;refGene_tx=XM_023248466.1;refGene_gn=RHPN1;refGene_tx=XM_023248467.1;refGene_gn=RHPN1;refGene_tx=XM_023248468.1
F2 84490529 84491230
refGene_gn=RHPN1;refGene_tx=XM_023248466.1;refGene_gn=RHPN1;refGene_tx=XM_023248467.1;refGene_gn=RHPN1;refGene_tx=XM_023248468.1
F2 84491307 84491418
refGene_gn=RHPN1;refGene_tx=XM_023248466.1;refGene_gn=RHPN1;refGene_tx=XM_023248467.1;refGene_gn=RHPN1;refGene_tx=XM_023248468.1
F2 84491792 84492002
refGene_gn=RHPN1;refGene_tx=XM_023248466.1;refGene_gn=RHPN1;refGene_tx

=XM_023248467.1;refGene_gn=RHPN1;refGene_tx=XM_023248468.1
F2 84492461 84492538
refGene_gn=RHPN1;refGene_tx=XM_023248466.1;refGene_gn=RHPN1;refGene_tx=XM_023248467.1
F2 84492756 84492903
refGene_gn=RHPN1;refGene_tx=XM_023248466.1;refGene_gn=RHPN1;refGene_tx=XM_023248467.1
F2 84493032 84493194
refGene_gn=RHPN1;refGene_tx=XM_023248466.1;refGene_gn=RHPN1;refGene_tx=XM_023248467.1
F2 84493628 84493787
refGene_gn=RHPN1;refGene_tx=XM_023248466.1;refGene_gn=RHPN1;refGene_tx=XM_023248467.1
F2 84529372 84530473 refGene_gn=MAFA;refGene_tx=XM_023248573.1
F2 84537200 84537392
refGene_gn=ZC3H3;refGene_tx=XM_023248772.1;refGene_gn=ZC3H3;refGene_tx=XM_023248773.1
F2 84539243 84539629
refGene_gn=ZC3H3;refGene_tx=XM_023248772.1;refGene_gn=ZC3H3;refGene_tx=XM_023248773.1
F2 84540228 84540413
refGene_gn=ZC3H3;refGene_tx=XM_023248772.1;refGene_gn=ZC3H3;refGene_tx=XM_023248773.1
F2 84556376 84556508
refGene_gn=ZC3H3;refGene_tx=XM_023248772.1;refGene_gn=ZC3H3;refGene_tx=XM_023248773.1
F2 84558569 84558639
refGene_gn=ZC3H3;refGene_tx=XM_023248772.1;refGene_gn=ZC3H3;refGene_tx=XM_023248773.1
F2 84558716 84558878
refGene_gn=ZC3H3;refGene_tx=XM_023248772.1;refGene_gn=ZC3H3;refGene_tx=XM_023248773.1
F2 84558970 84559013
refGene_gn=ZC3H3;refGene_tx=XM_023248772.1;refGene_gn=ZC3H3;refGene_tx=XM_023248773.1
F2 84563948 84564136
refGene_gn=ZC3H3;refGene_tx=XM_023248772.1;refGene_gn=ZC3H3;refGene_tx=XM_023248773.1
F2 84589257 84589411
refGene_gn=ZC3H3;refGene_tx=XM_023248772.1;refGene_gn=ZC3H3;refGene_tx=XM_023248773.1
F2 84621106 84621303
refGene_gn=ZC3H3;refGene_tx=XM_023248772.1;refGene_gn=ZC3H3;refGene_tx=XM_023248773.1
F2 84622360 84623669
refGene_gn=ZC3H3;refGene_tx=XM_023248772.1;refGene_gn=ZC3H3;refGene_tx=XM_023248773.1
F2 84625032 84625120
refGene_gn=ZC3H3;refGene_tx=XM_023248772.1;refGene_gn=ZC3H3;refGene_tx=XM_023248773.1

F2 84641574 84641791 refGene_gn=GSDMD;refGene_tx=XM_023248348.1
F2 84641957 84642150 refGene_gn=GSDMD;refGene_tx=XM_023248348.1
F2 84642654 84642823 refGene_gn=GSDMD;refGene_tx=XM_023248348.1
F2 84643660 84643763 refGene_gn=GSDMD;refGene_tx=XM_023248348.1
F2 84644017 84644071 refGene_gn=GSDMD;refGene_tx=XM_023248348.1
F2 84644683 84644779 refGene_gn=GSDMD;refGene_tx=XM_023248348.1
F2 84644908 84645084 refGene_gn=GSDMD;refGene_tx=XM_023248348.1
F2 84645167 84645309 refGene_gn=GSDMD;refGene_tx=XM_023248348.1
F2 84645413 84645487 refGene_gn=GSDMD;refGene_tx=XM_023248348.1
F2 84645599 84645834 refGene_gn=GSDMD;refGene_tx=XM_023248348.1
F2 84646055 84646129 refGene_gn=GSDMD;refGene_tx=XM_023248348.1
F2 84646242 84646485 refGene_gn=GSDMD;refGene_tx=XM_023248348.1
F2 84657605 84658050
refGene_gn=MROH6;refGene_tx=XM_023248507.1;refGene_gn=MROH6;refGene_tx
=XM_023248508.1;refGene_gn=MROH6;refGene_tx=XM_023248509.1
F2 84658207 84658273
refGene_gn=MROH6;refGene_tx=XM_023248507.1;refGene_gn=MROH6;refGene_tx
=XM_023248508.1;refGene_gn=MROH6;refGene_tx=XM_023248509.1
F2 84658384 84658487
refGene_gn=MROH6;refGene_tx=XM_023248507.1;refGene_gn=MROH6;refGene_tx
=XM_023248508.1;refGene_gn=MROH6;refGene_tx=XM_023248509.1
F2 84658689 84658809
refGene_gn=MROH6;refGene_tx=XM_023248507.1;refGene_gn=MROH6;refGene_tx
=XM_023248508.1;refGene_gn=MROH6;refGene_tx=XM_023248509.1
F2 84659044 84659212
refGene_gn=MROH6;refGene_tx=XM_023248507.1;refGene_gn=MROH6;refGene_tx
=XM_023248508.1;refGene_gn=MROH6;refGene_tx=XM_023248509.1
F2 84659348 84659522
refGene_gn=MROH6;refGene_tx=XM_023248507.1;refGene_gn=MROH6;refGene_tx
=XM_023248508.1;refGene_gn=MROH6;refGene_tx=XM_023248509.1
F2 84659602 84659746
refGene_gn=MROH6;refGene_tx=XM_023248507.1;refGene_gn=MROH6;refGene_tx
=XM_023248508.1;refGene_gn=MROH6;refGene_tx=XM_023248509.1
F2 84659846 84659961
refGene_gn=MROH6;refGene_tx=XM_023248507.1;refGene_gn=MROH6;refGene_tx
=XM_023248508.1;refGene_gn=MROH6;refGene_tx=XM_023248509.1
F2 84660078 84660216
refGene_gn=MROH6;refGene_tx=XM_023248507.1;refGene_gn=MROH6;refGene_tx
=XM_023248508.1;refGene_gn=MROH6;refGene_tx=XM_023248509.1
F2 84660308 84660493
refGene_gn=MROH6;refGene_tx=XM_023248507.1;refGene_gn=MROH6;refGene_tx
=XM_023248508.1;refGene_gn=MROH6;refGene_tx=XM_023248509.1
F2 84660617 84660735
refGene_gn=MROH6;refGene_tx=XM_023248507.1;refGene_gn=MROH6;refGene_tx
=XM_023248508.1;refGene_gn=MROH6;refGene_tx=XM_023248509.1
F2 84661302 84661457
refGene_gn=MROH6;refGene_tx=XM_023248507.1;refGene_gn=MROH6;refGene_tx
=XM_023248508.1;refGene_gn=MROH6;refGene_tx=XM_023248509.1
F2 84661612 84661765
refGene_gn=MROH6;refGene_tx=XM_023248507.1;refGene_gn=MROH6;refGene_tx

=XM_023248508.1;refGene_gn=MROH6;refGene_tx=XM_023248509.1
F2 84661881 84662127
refGene_gn=MROH6;refGene_tx=XM_023248507.1;refGene_gn=MROH6;refGene_tx
=XM_023248508.1;refGene_gn=MROH6;refGene_tx=XM_023248509.1
F2 84663965 84664028 refGene_gn=NAPRT;refGene_tx=XM_006943496.2
F2 84664122 84664230 refGene_gn=NAPRT;refGene_tx=XM_006943496.2
F2 84664377 84664532 refGene_gn=NAPRT;refGene_tx=XM_006943496.2
F2 84664609 84664712 refGene_gn=NAPRT;refGene_tx=XM_006943496.2
F2 84664804 84664885 refGene_gn=NAPRT;refGene_tx=XM_006943496.2
F2 84664983 84665068 refGene_gn=NAPRT;refGene_tx=XM_006943496.2
F2 84665384 84665525 refGene_gn=NAPRT;refGene_tx=XM_006943496.2
F2 84665606 84665803 refGene_gn=NAPRT;refGene_tx=XM_006943496.2
F2 84666058 84666174 refGene_gn=NAPRT;refGene_tx=XM_006943496.2
F2 84666286 84666417 refGene_gn=NAPRT;refGene_tx=XM_006943496.2
F2 84666680 84666763 refGene_gn=NAPRT;refGene_tx=XM_006943496.2
F2 84666843 84666971 refGene_gn=NAPRT;refGene_tx=XM_006943496.2
F2 84667472 84667698 refGene_gn=NAPRT;refGene_tx=XM_006943496.2
F2 84669096 84669135
refGene_gn=EEF1D;refGene_tx=XM_011291500.3;refGene_gn=EEF1D;refGene_tx
=XM_023248510.1;refGene_gn=EEF1D;refGene_tx=XM_023248511.1
F2 84669251 84669446
refGene_gn=EEF1D;refGene_tx=XM_011291500.3;refGene_gn=EEF1D;refGene_tx
=XM_023248510.1;refGene_gn=EEF1D;refGene_tx=XM_023248511.1
F2 84669648 84669867
refGene_gn=EEF1D;refGene_tx=XM_011291500.3;refGene_gn=EEF1D;refGene_tx
=XM_023248510.1;refGene_gn=EEF1D;refGene_tx=XM_023248511.1
F2 84670001 84670102
refGene_gn=EEF1D;refGene_tx=XM_011291500.3;refGene_gn=EEF1D;refGene_tx
=XM_023248510.1;refGene_gn=EEF1D;refGene_tx=XM_023248511.1
F2 84670179 84670279
refGene_gn=EEF1D;refGene_tx=XM_011291500.3;refGene_gn=EEF1D;refGene_tx
=XM_023248510.1;refGene_gn=EEF1D;refGene_tx=XM_023248511.1
F2 84673353 84673425
refGene_gn=EEF1D;refGene_tx=XM_011291500.3;refGene_gn=EEF1D;refGene_tx
=XM_023248510.1;refGene_gn=EEF1D;refGene_tx=XM_023248511.1
F2 84673777 84673898
refGene_gn=EEF1D;refGene_tx=XM_011291500.3;refGene_gn=EEF1D;refGene_tx
=XM_023248510.1;refGene_gn=EEF1D;refGene_tx=XM_023248511.1
F2 84675553 84676695
refGene_gn=EEF1D;refGene_tx=XM_011291500.3;refGene_gn=EEF1D;refGene_tx
=XM_023248510.1
F2 84683484 84685434 refGene_gn=TIGD5;refGene_tx=XM_023248512.1
F2 84689489 84689836
refGene_gn=PYCR3;refGene_tx=XM_006943437.4;refGene_gn=PYCR3;refGene_tx
=XM_019823233.2
F2 84690016 84690170
refGene_gn=PYCR3;refGene_tx=XM_006943437.4;refGene_gn=PYCR3;refGene_tx
=XM_019823233.2
F2 84690419 84690632
refGene_gn=PYCR3;refGene_tx=XM_006943437.4;refGene_gn=PYCR3;refGene_tx

=XM_019823233.2
F2 84690946 84691126
refGene_gn=PYCR3;refGene_tx=XM_006943437.4;refGene_gn=PYCR3;refGene_tx
=XM_019823233.2
F2 84691655 84691720
refGene_gn=PYCR3;refGene_tx=XM_006943437.4;refGene_gn=PYCR3;refGene_tx
=XM_019823233.2
F2 84693470 84693570
refGene_gn=PYCR3;refGene_tx=XM_006943437.4;refGene_gn=PYCR3;refGene_tx
=XM_019823233.2
F2 84697134 84697190 refGene_gn=TSTA3;refGene_tx=XM_004000162.4
F2 84697475 84697575 refGene_gn=TSTA3;refGene_tx=XM_004000162.4
F2 84697753 84697833 refGene_gn=TSTA3;refGene_tx=XM_004000162.4
F2 84697996 84698063 refGene_gn=TSTA3;refGene_tx=XM_004000162.4
F2 84698439 84698504 refGene_gn=TSTA3;refGene_tx=XM_004000162.4
F2 84698589 84698723 refGene_gn=TSTA3;refGene_tx=XM_004000162.4
F2 84698916 84698990 refGene_gn=TSTA3;refGene_tx=XM_004000162.4
F2 84699068 84699197 refGene_gn=TSTA3;refGene_tx=XM_004000162.4
F2 84700373 84700488 refGene_gn=TSTA3;refGene_tx=XM_004000162.4
F2 84700803 84700949 refGene_gn=TSTA3;refGene_tx=XM_004000162.4
F2 84717905 84719417
refGene_gn=ZNF623;refGene_tx=XM_019823229.2;refGene_gn=ZNF623;refGene_
tx=XM_019823230.2
F2 84741365 84742199 refGene_gn=CCDC166;refGene_tx=XM_023248470.1
F2 84742285 84742726 refGene_gn=CCDC166;refGene_tx=XM_023248470.1
F2 84757275 84758163
refGene_gn=LOC105261391;refGene_tx=XM_023248433.1
F2 84759981 84760050
refGene_gn=MAPK15;refGene_tx=XM_023248429.1;refGene_gn=MAPK15;refGene_
tx=XM_023248430.1;refGene_gn=MAPK15;refGene_tx=XM_023248431.1
F2 84761244 84761343
refGene_gn=MAPK15;refGene_tx=XM_023248429.1;refGene_gn=MAPK15;refGene_
tx=XM_023248430.1;refGene_gn=MAPK15;refGene_tx=XM_023248431.1
F2 84761803 84761833
refGene_gn=MAPK15;refGene_tx=XM_023248429.1;refGene_gn=MAPK15;refGene_
tx=XM_023248430.1;refGene_gn=MAPK15;refGene_tx=XM_023248431.1
F2 84761951 84762042
refGene_gn=MAPK15;refGene_tx=XM_023248429.1;refGene_gn=MAPK15;refGene_
tx=XM_023248430.1;refGene_gn=MAPK15;refGene_tx=XM_023248431.1
F2 84762563 84762694
refGene_gn=MAPK15;refGene_tx=XM_023248429.1;refGene_gn=MAPK15;refGene_
tx=XM_023248430.1;refGene_gn=MAPK15;refGene_tx=XM_023248431.1;refGene_
gn=MAPK15;refGene_tx=XM_023248432.1
F2 84762775 84762939 refGene_gn=MAPK15;refGene_tx=XM_023248431.1
F2 84763145 84763285
refGene_gn=MAPK15;refGene_tx=XM_023248429.1;refGene_gn=MAPK15;refGene_
tx=XM_023248430.1;refGene_gn=MAPK15;refGene_tx=XM_023248431.1;refGene_
gn=MAPK15;refGene_tx=XM_023248432.1
F2 84764004 84764550
refGene_gn=MAPK15;refGene_tx=XM_023248429.1;refGene_gn=MAPK15;refGene_

tx=XM_023248430.1;refGene_gn=MAPK15;refGene_tx=XM_023248431.1;refGene_gn=MAPK15;refGene_tx=XM_023248432.1
F2 84764672 84764778 refGene_gn=MAPK15;refGene_tx=XM_023248429.1;refGene_gn=MAPK15;refGene_tx=XM_023248430.1;refGene_gn=MAPK15;refGene_tx=XM_023248431.1;refGene_gn=MAPK15;refGene_tx=XM_023248432.1
F2 84764912 84765114 refGene_gn=MAPK15;refGene_tx=XM_023248429.1;refGene_gn=MAPK15;refGene_tx=XM_023248430.1;refGene_gn=MAPK15;refGene_tx=XM_023248431.1;refGene_gn=MAPK15;refGene_tx=XM_023248432.1
F2 84765212 84765355 refGene_gn=MAPK15;refGene_tx=XM_023248429.1;refGene_gn=MAPK15;refGene_tx=XM_023248431.1;refGene_gn=MAPK15;refGene_tx=XM_023248432.1
F2 84765442 84765862 refGene_gn=MAPK15;refGene_tx=XM_023248429.1;refGene_gn=MAPK15;refGene_tx=XM_023248430.1;refGene_gn=MAPK15;refGene_tx=XM_023248431.1;refGene_gn=MAPK15;refGene_tx=XM_023248432.1
F2 84769050 84771796 refGene_gn=FAM83H;refGene_tx=XM_023248744.1;refGene_gn=FAM83H;refGene_tx=XM_023248745.1;refGene_gn=FAM83H;refGene_tx=XM_023248746.1;refGene_gn=FAM83H;refGene_tx=XM_023248747.1;refGene_gn=FAM83H;refGene_tx=XM_023248748.1
F2 84771993 84772118 refGene_gn=FAM83H;refGene_tx=XM_023248744.1;refGene_gn=FAM83H;refGene_tx=XM_023248745.1;refGene_gn=FAM83H;refGene_tx=XM_023248746.1;refGene_gn=FAM83H;refGene_tx=XM_023248747.1;refGene_gn=FAM83H;refGene_tx=XM_023248748.1
F2 84772192 84772357 refGene_gn=FAM83H;refGene_tx=XM_023248744.1;refGene_gn=FAM83H;refGene_tx=XM_023248745.1;refGene_gn=FAM83H;refGene_tx=XM_023248746.1;refGene_gn=FAM83H;refGene_tx=XM_023248747.1;refGene_gn=FAM83H;refGene_tx=XM_023248748.1
F2 84773189 84773676 refGene_gn=FAM83H;refGene_tx=XM_023248744.1;refGene_gn=FAM83H;refGene_tx=XM_023248745.1;refGene_gn=FAM83H;refGene_tx=XM_023248746.1;refGene_gn=FAM83H;refGene_tx=XM_023248747.1;refGene_gn=FAM83H;refGene_tx=XM_023248748.1
F2 84773756 84774010 refGene_gn=FAM83H;refGene_tx=XM_023248745.1
F2 84779067 84779164 refGene_gn=IQANK1;refGene_tx=XM_023248750.1;refGene_gn=IQANK1;refGene_tx=XM_023248751.1;refGene_gn=IQANK1;refGene_tx=XM_023248752.1
F2 84780489 84781113 refGene_gn=FAM83H;refGene_tx=XM_023248744.1
F2 84782644 84782713 refGene_gn=IQANK1;refGene_tx=XM_023248750.1;refGene_gn=IQANK1;refGene_tx=XM_023248751.1;refGene_gn=IQANK1;refGene_tx=XM_023248752.1
F2 84797924 84798058 refGene_gn=IQANK1;refGene_tx=XM_023248750.1;refGene_gn=IQANK1;refGene_tx=XM_023248751.1;refGene_gn=IQANK1;refGene_tx=XM_023248752.1
F2 84798187 84798352 refGene_gn=IQANK1;refGene_tx=XM_023248750.1;refGene_gn=IQANK1;refGene_

tx=XM_023248751.1
F2 84798459 84798651
refGene_gn=IQANK1;refGene_tx=XM_023248750.1;refGene_gn=IQANK1;refGene_tx=XM_023248751.1;refGene_gn=IQANK1;refGene_tx=XM_023248752.1
F2 84798723 84798849
refGene_gn=IQANK1;refGene_tx=XM_023248750.1;refGene_gn=IQANK1;refGene_tx=XM_023248751.1;refGene_gn=IQANK1;refGene_tx=XM_023248752.1
F2 84799339 84799479
refGene_gn=IQANK1;refGene_tx=XM_023248750.1;refGene_gn=IQANK1;refGene_tx=XM_023248751.1;refGene_gn=IQANK1;refGene_tx=XM_023248752.1
F2 84799548 84799603
refGene_gn=IQANK1;refGene_tx=XM_023248750.1;refGene_gn=IQANK1;refGene_tx=XM_023248751.1;refGene_gn=IQANK1;refGene_tx=XM_023248752.1
F2 84799745 84799838
refGene_gn=IQANK1;refGene_tx=XM_023248750.1;refGene_gn=IQANK1;refGene_tx=XM_023248751.1;refGene_gn=IQANK1;refGene_tx=XM_023248752.1
F2 84800039 84800148
refGene_gn=IQANK1;refGene_tx=XM_023248750.1;refGene_gn=IQANK1;refGene_tx=XM_023248751.1;refGene_gn=IQANK1;refGene_tx=XM_023248752.1
F2 84800245 84800363
refGene_gn=IQANK1;refGene_tx=XM_023248750.1;refGene_gn=IQANK1;refGene_tx=XM_023248751.1;refGene_gn=IQANK1;refGene_tx=XM_023248752.1
F2 84800436 84800571
refGene_gn=IQANK1;refGene_tx=XM_023248750.1;refGene_gn=IQANK1;refGene_tx=XM_023248751.1;refGene_gn=IQANK1;refGene_tx=XM_023248752.1
F2 84800633 84800937
refGene_gn=IQANK1;refGene_tx=XM_023248750.1;refGene_gn=IQANK1;refGene_tx=XM_023248751.1;refGene_gn=IQANK1;refGene_tx=XM_023248752.1
F2 84801389 84801526
refGene_gn=SCRIB;refGene_tx=XM_023248738.1;refGene_gn=SCRIB;refGene_tx=XM_023248739.1;refGene_gn=SCRIB;refGene_tx=XM_023248740.1;refGene_gn=SCRIB;refGene_tx=XM_023248741.1;refGene_gn=SCRIB;refGene_tx=XM_023248742.1
F2 84801606 84801658
refGene_gn=SCRIB;refGene_tx=XM_023248738.1;refGene_gn=SCRIB;refGene_tx=XM_023248739.1;refGene_gn=SCRIB;refGene_tx=XM_023248740.1;refGene_gn=SCRIB;refGene_tx=XM_023248741.1;refGene_gn=SCRIB;refGene_tx=XM_023248742.1;refGene_gn=SCRIB;refGene_tx=XM_023248743.1
F2 84801840 84801915
refGene_gn=SCRIB;refGene_tx=XM_023248738.1;refGene_gn=SCRIB;refGene_tx=XM_023248739.1;refGene_gn=SCRIB;refGene_tx=XM_023248740.1;refGene_gn=SCRIB;refGene_tx=XM_023248742.1
F2 84802024 84802062
refGene_gn=SCRIB;refGene_tx=XM_023248738.1;refGene_gn=SCRIB;refGene_tx=XM_023248739.1;refGene_gn=SCRIB;refGene_tx=XM_023248740.1;refGene_gn=SCRIB;refGene_tx=XM_023248741.1;refGene_gn=SCRIB;refGene_tx=XM_023248742.1
F2 84802147 84802290
refGene_gn=SCRIB;refGene_tx=XM_023248738.1;refGene_gn=SCRIB;refGene_tx=XM_023248739.1;refGene_gn=SCRIB;refGene_tx=XM_023248740.1;refGene_gn=

SCRIB;refGene_tx=XM_023248741.1;refGene_gn=SCRIB;refGene_tx=XM_023248742.1;refGene_gn=SCRIB;refGene_tx=XM_023248743.1
F2 84802383 84802569
refGene_gn=SCRIB;refGene_tx=XM_023248738.1;refGene_gn=SCRIB;refGene_tx=XM_023248739.1;refGene_gn=SCRIB;refGene_tx=XM_023248740.1;refGene_gn=SCRIB;refGene_tx=XM_023248741.1;refGene_gn=SCRIB;refGene_tx=XM_023248742.1;refGene_gn=SCRIB;refGene_tx=XM_023248743.1
F2 84802670 84802800
refGene_gn=SCRIB;refGene_tx=XM_023248738.1;refGene_gn=SCRIB;refGene_tx=XM_023248739.1;refGene_gn=SCRIB;refGene_tx=XM_023248740.1;refGene_gn=SCRIB;refGene_tx=XM_023248741.1;refGene_gn=SCRIB;refGene_tx=XM_023248742.1;refGene_gn=SCRIB;refGene_tx=XM_023248743.1
F2 84802892 84803052
refGene_gn=SCRIB;refGene_tx=XM_023248738.1;refGene_gn=SCRIB;refGene_tx=XM_023248739.1;refGene_gn=SCRIB;refGene_tx=XM_023248740.1;refGene_gn=SCRIB;refGene_tx=XM_023248741.1;refGene_gn=SCRIB;refGene_tx=XM_023248742.1;refGene_gn=SCRIB;refGene_tx=XM_023248743.1
F2 84803139 84803247
refGene_gn=SCRIB;refGene_tx=XM_023248738.1;refGene_gn=SCRIB;refGene_tx=XM_023248739.1;refGene_gn=SCRIB;refGene_tx=XM_023248740.1;refGene_gn=SCRIB;refGene_tx=XM_023248741.1;refGene_gn=SCRIB;refGene_tx=XM_023248742.1;refGene_gn=SCRIB;refGene_tx=XM_023248743.1
F2 84803856 84803925
refGene_gn=SCRIB;refGene_tx=XM_023248738.1;refGene_gn=SCRIB;refGene_tx=XM_023248739.1;refGene_gn=SCRIB;refGene_tx=XM_023248740.1;refGene_gn=SCRIB;refGene_tx=XM_023248741.1;refGene_gn=SCRIB;refGene_tx=XM_023248742.1;refGene_gn=SCRIB;refGene_tx=XM_023248743.1
F2 84804824 84804899
refGene_gn=SCRIB;refGene_tx=XM_023248738.1;refGene_gn=SCRIB;refGene_tx=XM_023248739.1;refGene_gn=SCRIB;refGene_tx=XM_023248740.1;refGene_gn=SCRIB;refGene_tx=XM_023248741.1;refGene_gn=SCRIB;refGene_tx=XM_023248742.1;refGene_gn=SCRIB;refGene_tx=XM_023248743.1
F2 84805089 84805137
refGene_gn=SCRIB;refGene_tx=XM_023248738.1;refGene_gn=SCRIB;refGene_tx=XM_023248739.1;refGene_gn=SCRIB;refGene_tx=XM_023248740.1;refGene_gn=SCRIB;refGene_tx=XM_023248741.1;refGene_gn=SCRIB;refGene_tx=XM_023248742.1;refGene_gn=SCRIB;refGene_tx=XM_023248743.1
F2 84805240 84805351
refGene_gn=SCRIB;refGene_tx=XM_023248738.1;refGene_gn=SCRIB;refGene_tx=XM_023248739.1;refGene_gn=SCRIB;refGene_tx=XM_023248740.1;refGene_gn=SCRIB;refGene_tx=XM_023248741.1;refGene_gn=SCRIB;refGene_tx=XM_023248742.1;refGene_gn=SCRIB;refGene_tx=XM_023248743.1
F2 84812038 84812227
refGene_gn=SCRIB;refGene_tx=XM_023248738.1;refGene_gn=SCRIB;refGene_tx=XM_023248739.1;refGene_gn=SCRIB;refGene_tx=XM_023248740.1;refGene_gn=SCRIB;refGene_tx=XM_023248741.1;refGene_gn=SCRIB;refGene_tx=XM_023248742.1;refGene_gn=SCRIB;refGene_tx=XM_023248743.1
F2 84812317 84812611
refGene_gn=SCRIB;refGene_tx=XM_023248738.1;refGene_gn=SCRIB;refGene_tx=XM_023248739.1;refGene_gn=SCRIB;refGene_tx=XM_023248740.1;refGene_gn=

SCRIB;refGene_tx=XM_023248741.1;refGene_gn=SCRIB;refGene_tx=XM_0232487
42.1;refGene_gn=SCRIB;refGene_tx=XM_023248743.1
F2 84812713 84812824
refGene_gn=SCRIB;refGene_tx=XM_023248738.1;refGene_gn=SCRIB;refGene_tx
=XM_023248739.1;refGene_gn=SCRIB;refGene_tx=XM_023248740.1;refGene_gn=
SCRIB;refGene_tx=XM_023248741.1;refGene_gn=SCRIB;refGene_tx=XM_0232487
42.1;refGene_gn=SCRIB;refGene_tx=XM_023248743.1
F2 84813360 84813615
refGene_gn=SCRIB;refGene_tx=XM_023248738.1;refGene_gn=SCRIB;refGene_tx
=XM_023248739.1;refGene_gn=SCRIB;refGene_tx=XM_023248740.1;refGene_gn=
SCRIB;refGene_tx=XM_023248741.1;refGene_gn=SCRIB;refGene_tx=XM_0232487
42.1;refGene_gn=SCRIB;refGene_tx=XM_023248743.1
F2 84813699 84813780
refGene_gn=SCRIB;refGene_tx=XM_023248738.1;refGene_gn=SCRIB;refGene_tx
=XM_023248739.1;refGene_gn=SCRIB;refGene_tx=XM_023248740.1;refGene_gn=
SCRIB;refGene_tx=XM_023248741.1;refGene_gn=SCRIB;refGene_tx=XM_0232487
42.1;refGene_gn=SCRIB;refGene_tx=XM_023248743.1
F2 84814547 84814871
refGene_gn=SCRIB;refGene_tx=XM_023248738.1;refGene_gn=SCRIB;refGene_tx
=XM_023248739.1;refGene_gn=SCRIB;refGene_tx=XM_023248740.1;refGene_gn=
SCRIB;refGene_tx=XM_023248741.1;refGene_gn=SCRIB;refGene_tx=XM_0232487
42.1;refGene_gn=SCRIB;refGene_tx=XM_023248743.1
F2 84815507 84815585
refGene_gn=SCRIB;refGene_tx=XM_023248738.1;refGene_gn=SCRIB;refGene_tx
=XM_023248739.1;refGene_gn=SCRIB;refGene_tx=XM_023248740.1;refGene_gn=
SCRIB;refGene_tx=XM_023248741.1;refGene_gn=SCRIB;refGene_tx=XM_0232487
43.1
F2 84816094 84816184
refGene_gn=SCRIB;refGene_tx=XM_023248738.1;refGene_gn=SCRIB;refGene_tx
=XM_023248739.1;refGene_gn=SCRIB;refGene_tx=XM_023248740.1;refGene_gn=
SCRIB;refGene_tx=XM_023248741.1;refGene_gn=SCRIB;refGene_tx=XM_0232487
42.1;refGene_gn=SCRIB;refGene_tx=XM_023248743.1
F2 84816582 84816645
refGene_gn=SCRIB;refGene_tx=XM_023248738.1;refGene_gn=SCRIB;refGene_tx
=XM_023248739.1;refGene_gn=SCRIB;refGene_tx=XM_023248741.1;refGene_gn=
SCRIB;refGene_tx=XM_023248742.1;refGene_gn=SCRIB;refGene_tx=XM_0232487
43.1
F2 84817290 84817692
refGene_gn=SCRIB;refGene_tx=XM_023248738.1;refGene_gn=SCRIB;refGene_tx
=XM_023248739.1;refGene_gn=SCRIB;refGene_tx=XM_023248740.1;refGene_gn=
SCRIB;refGene_tx=XM_023248741.1;refGene_gn=SCRIB;refGene_tx=XM_0232487
42.1;refGene_gn=SCRIB;refGene_tx=XM_023248743.1
F2 84817814 84817982
refGene_gn=SCRIB;refGene_tx=XM_023248738.1;refGene_gn=SCRIB;refGene_tx
=XM_023248739.1;refGene_gn=SCRIB;refGene_tx=XM_023248740.1;refGene_gn=
SCRIB;refGene_tx=XM_023248741.1;refGene_gn=SCRIB;refGene_tx=XM_0232487
42.1;refGene_gn=SCRIB;refGene_tx=XM_023248743.1
F2 84818751 84818877
refGene_gn=SCRIB;refGene_tx=XM_023248738.1;refGene_gn=SCRIB;refGene_tx
=XM_023248739.1;refGene_gn=SCRIB;refGene_tx=XM_023248740.1;refGene_gn=

SCRIB;refGene_tx=XM_023248741.1;refGene_gn=SCRIB;refGene_tx=XM_0232487
42.1;refGene_gn=SCRIB;refGene_tx=XM_023248743.1
F2 84818953 84819081
refGene_gn=SCRIB;refGene_tx=XM_023248738.1;refGene_gn=SCRIB;refGene_tx
=XM_023248739.1;refGene_gn=SCRIB;refGene_tx=XM_023248740.1;refGene_gn=
SCRIB;refGene_tx=XM_023248741.1;refGene_gn=SCRIB;refGene_tx=XM_0232487
42.1;refGene_gn=SCRIB;refGene_tx=XM_023248743.1
F2 84819163 84819330
refGene_gn=SCRIB;refGene_tx=XM_023248738.1;refGene_gn=SCRIB;refGene_tx
=XM_023248739.1;refGene_gn=SCRIB;refGene_tx=XM_023248740.1;refGene_gn=
SCRIB;refGene_tx=XM_023248741.1;refGene_gn=SCRIB;refGene_tx=XM_0232487
42.1;refGene_gn=SCRIB;refGene_tx=XM_023248743.1
F2 84819427 84819627
refGene_gn=SCRIB;refGene_tx=XM_023248738.1;refGene_gn=SCRIB;refGene_tx
=XM_023248739.1;refGene_gn=SCRIB;refGene_tx=XM_023248740.1;refGene_gn=
SCRIB;refGene_tx=XM_023248741.1;refGene_gn=SCRIB;refGene_tx=XM_0232487
42.1;refGene_gn=SCRIB;refGene_tx=XM_023248743.1
F2 84819848 84819967
refGene_gn=SCRIB;refGene_tx=XM_023248738.1;refGene_gn=SCRIB;refGene_tx
=XM_023248739.1;refGene_gn=SCRIB;refGene_tx=XM_023248740.1;refGene_gn=
SCRIB;refGene_tx=XM_023248741.1;refGene_gn=SCRIB;refGene_tx=XM_0232487
42.1;refGene_gn=SCRIB;refGene_tx=XM_023248743.1
F2 84820305 84820450
refGene_gn=SCRIB;refGene_tx=XM_023248738.1;refGene_gn=SCRIB;refGene_tx
=XM_023248739.1;refGene_gn=SCRIB;refGene_tx=XM_023248740.1;refGene_gn=
SCRIB;refGene_tx=XM_023248741.1;refGene_gn=SCRIB;refGene_tx=XM_0232487
42.1;refGene_gn=SCRIB;refGene_tx=XM_023248743.1
F2 84820518 84820593
refGene_gn=SCRIB;refGene_tx=XM_023248738.1;refGene_gn=SCRIB;refGene_tx
=XM_023248739.1;refGene_gn=SCRIB;refGene_tx=XM_023248740.1;refGene_gn=
SCRIB;refGene_tx=XM_023248741.1;refGene_gn=SCRIB;refGene_tx=XM_0232487
42.1;refGene_gn=SCRIB;refGene_tx=XM_023248743.1
F2 84820688 84820752
refGene_gn=SCRIB;refGene_tx=XM_023248738.1;refGene_gn=SCRIB;refGene_tx
=XM_023248739.1;refGene_gn=SCRIB;refGene_tx=XM_023248740.1;refGene_gn=
SCRIB;refGene_tx=XM_023248741.1;refGene_gn=SCRIB;refGene_tx=XM_0232487
42.1;refGene_gn=SCRIB;refGene_tx=XM_023248743.1
F2 84820850 84820907
refGene_gn=SCRIB;refGene_tx=XM_023248738.1;refGene_gn=SCRIB;refGene_tx
=XM_023248739.1;refGene_gn=SCRIB;refGene_tx=XM_023248740.1;refGene_gn=
SCRIB;refGene_tx=XM_023248741.1;refGene_gn=SCRIB;refGene_tx=XM_0232487
42.1;refGene_gn=SCRIB;refGene_tx=XM_023248743.1
F2 84821010 84821100
refGene_gn=SCRIB;refGene_tx=XM_023248738.1;refGene_gn=SCRIB;refGene_tx
=XM_023248739.1;refGene_gn=SCRIB;refGene_tx=XM_023248740.1;refGene_gn=
SCRIB;refGene_tx=XM_023248741.1;refGene_gn=SCRIB;refGene_tx=XM_0232487
42.1;refGene_gn=SCRIB;refGene_tx=XM_023248743.1
F2 84821193 84821272
refGene_gn=SCRIB;refGene_tx=XM_023248738.1;refGene_gn=SCRIB;refGene_tx
=XM_023248739.1;refGene_gn=SCRIB;refGene_tx=XM_023248740.1;refGene_gn=

SCRIB;refGene_tx=XM_023248741.1;refGene_gn=SCRIB;refGene_tx=XM_0232487
42.1;refGene_gn=SCRIB;refGene_tx=XM_023248743.1
F2 84821380 84821498
refGene_gn=SCRIB;refGene_tx=XM_023248738.1;refGene_gn=SCRIB;refGene_tx
=XM_023248739.1;refGene_gn=SCRIB;refGene_tx=XM_023248740.1;refGene_gn=
SCRIB;refGene_tx=XM_023248741.1;refGene_gn=SCRIB;refGene_tx=XM_0232487
42.1;refGene_gn=SCRIB;refGene_tx=XM_023248743.1
F2 84822732 84822891
refGene_gn=SCRIB;refGene_tx=XM_023248738.1;refGene_gn=SCRIB;refGene_tx
=XM_023248739.1;refGene_gn=SCRIB;refGene_tx=XM_023248740.1;refGene_gn=
SCRIB;refGene_tx=XM_023248741.1;refGene_gn=SCRIB;refGene_tx=XM_0232487
42.1;refGene_gn=SCRIB;refGene_tx=XM_023248743.1
F2 84823983 84824283
refGene_gn=PUF60;refGene_tx=XM_011291501.3;refGene_gn=PUF60;refGene_tx
=XM_011291502.3;refGene_gn=PUF60;refGene_tx=XM_011291503.3;refGene_gn=
PUF60;refGene_tx=XM_011291504.3;refGene_gn=PUF60;refGene_tx=XM_0112915
05.3;refGene_gn=PUF60;refGene_tx=XM_011291506.3;refGene_gn=PUF60;refGe
ne_tx=XM_011291507.3
F2 84824365 84824601
refGene_gn=PUF60;refGene_tx=XM_011291501.3;refGene_gn=PUF60;refGene_tx
=XM_011291502.3;refGene_gn=PUF60;refGene_tx=XM_011291503.3;refGene_gn=
PUF60;refGene_tx=XM_011291504.3;refGene_gn=PUF60;refGene_tx=XM_0112915
05.3;refGene_gn=PUF60;refGene_tx=XM_011291506.3;refGene_gn=PUF60;refGe
ne_tx=XM_011291507.3
F2 84824767 84824903
refGene_gn=PUF60;refGene_tx=XM_011291501.3;refGene_gn=PUF60;refGene_tx
=XM_011291502.3;refGene_gn=PUF60;refGene_tx=XM_011291503.3;refGene_gn=
PUF60;refGene_tx=XM_011291504.3;refGene_gn=PUF60;refGene_tx=XM_0112915
05.3;refGene_gn=PUF60;refGene_tx=XM_011291506.3;refGene_gn=PUF60;refGe
ne_tx=XM_011291507.3
F2 84825043 84825234
refGene_gn=PUF60;refGene_tx=XM_011291501.3;refGene_gn=PUF60;refGene_tx
=XM_011291502.3;refGene_gn=PUF60;refGene_tx=XM_011291503.3;refGene_gn=
PUF60;refGene_tx=XM_011291504.3;refGene_gn=PUF60;refGene_tx=XM_0112915
05.3;refGene_gn=PUF60;refGene_tx=XM_011291506.3;refGene_gn=PUF60;refGe
ne_tx=XM_011291507.3
F2 84825313 84825527
refGene_gn=PUF60;refGene_tx=XM_011291501.3;refGene_gn=PUF60;refGene_tx
=XM_011291502.3;refGene_gn=PUF60;refGene_tx=XM_011291503.3;refGene_gn=
PUF60;refGene_tx=XM_011291504.3;refGene_gn=PUF60;refGene_tx=XM_0112915
05.3;refGene_gn=PUF60;refGene_tx=XM_011291506.3;refGene_gn=PUF60;refGe
ne_tx=XM_011291507.3
F2 84825664 84825757
refGene_gn=PUF60;refGene_tx=XM_011291501.3;refGene_gn=PUF60;refGene_tx
=XM_011291502.3;refGene_gn=PUF60;refGene_tx=XM_011291503.3;refGene_gn=
PUF60;refGene_tx=XM_011291504.3;refGene_gn=PUF60;refGene_tx=XM_0112915
05.3;refGene_gn=PUF60;refGene_tx=XM_011291506.3;refGene_gn=PUF60;refGe
ne_tx=XM_011291507.3
F2 84825844 84826006
refGene_gn=PUF60;refGene_tx=XM_011291501.3;refGene_gn=PUF60;refGene_tx

=XM_011291502.3;refGene_gn=PUF60;refGene_tx=XM_011291503.3;refGene_gn=PUF60;refGene_tx=XM_011291504.3;refGene_gn=PUF60;refGene_tx=XM_011291505.3;refGene_gn=PUF60;refGene_tx=XM_011291506.3;refGene_gn=PUF60;refGene_tx=XM_011291507.3
F2 84828135 84828186
refGene_gn=PUF60;refGene_tx=XM_011291501.3;refGene_gn=PUF60;refGene_tx=XM_011291502.3;refGene_gn=PUF60;refGene_tx=XM_011291505.3;refGene_gn=PUF60;refGene_tx=XM_011291507.3
F2 84829128 84829218
refGene_gn=PUF60;refGene_tx=XM_011291501.3;refGene_gn=PUF60;refGene_tx=XM_011291502.3;refGene_gn=PUF60;refGene_tx=XM_011291503.3;refGene_gn=PUF60;refGene_tx=XM_011291504.3;refGene_gn=PUF60;refGene_tx=XM_011291505.3;refGene_gn=PUF60;refGene_tx=XM_011291506.3;refGene_gn=PUF60;refGene_tx=XM_011291507.3
F2 84829334 84829430
refGene_gn=PUF60;refGene_tx=XM_011291501.3;refGene_gn=PUF60;refGene_tx=XM_011291502.3;refGene_gn=PUF60;refGene_tx=XM_011291503.3;refGene_gn=PUF60;refGene_tx=XM_011291504.3;refGene_gn=PUF60;refGene_tx=XM_011291505.3;refGene_gn=PUF60;refGene_tx=XM_011291506.3;refGene_gn=PUF60;refGene_tx=XM_011291507.3
F2 84831741 84831843
refGene_gn=PUF60;refGene_tx=XM_011291501.3;refGene_gn=PUF60;refGene_tx=XM_011291502.3;refGene_gn=PUF60;refGene_tx=XM_011291503.3;refGene_gn=PUF60;refGene_tx=XM_011291504.3;refGene_gn=PUF60;refGene_tx=XM_011291507.3
F2 84835570 84835573 refGene_gn=PUF60;refGene_tx=XM_011291507.3
F2 84836464 84836488
refGene_gn=PUF60;refGene_tx=XM_011291501.3;refGene_gn=PUF60;refGene_tx=XM_011291502.3;refGene_gn=PUF60;refGene_tx=XM_011291503.3;refGene_gn=PUF60;refGene_tx=XM_011291504.3;refGene_gn=PUF60;refGene_tx=XM_011291505.3;refGene_gn=PUF60;refGene_tx=XM_011291506.3
F2 84842168 84842237
refGene_gn=NRBP2;refGene_tx=XM_011291509.3;refGene_gn=NRBP2;refGene_tx=XM_023248753.1;refGene_gn=NRBP2;refGene_tx=XM_023248754.1;refGene_gn=NRBP2;refGene_tx=XM_023248755.1
F2 84842323 84842379
refGene_gn=NRBP2;refGene_tx=XM_011291509.3;refGene_gn=NRBP2;refGene_tx=XM_023248753.1;refGene_gn=NRBP2;refGene_tx=XM_023248754.1;refGene_gn=NRBP2;refGene_tx=XM_023248755.1
F2 84842481 84842545
refGene_gn=NRBP2;refGene_tx=XM_011291509.3;refGene_gn=NRBP2;refGene_tx=XM_023248753.1;refGene_gn=NRBP2;refGene_tx=XM_023248754.1;refGene_gn=NRBP2;refGene_tx=XM_023248755.1
F2 84842644 84842698
refGene_gn=NRBP2;refGene_tx=XM_011291509.3;refGene_gn=NRBP2;refGene_tx=XM_023248753.1;refGene_gn=NRBP2;refGene_tx=XM_023248754.1;refGene_gn=NRBP2;refGene_tx=XM_023248755.1
F2 84843479 84843615
refGene_gn=NRBP2;refGene_tx=XM_011291509.3;refGene_gn=NRBP2;refGene_tx=XM_023248753.1;refGene_gn=NRBP2;refGene_tx=XM_023248754.1;refGene_gn=

NRBP2;refGene_tx=XM_023248755.1
F2 84843689 84843740
refGene_gn=NRBP2;refGene_tx=XM_011291509.3;refGene_gn=NRBP2;refGene_tx=XM_023248753.1;refGene_gn=NRBP2;refGene_tx=XM_023248754.1;refGene_gn=NRBP2;refGene_tx=XM_023248755.1
F2 84843862 84843965
refGene_gn=NRBP2;refGene_tx=XM_011291509.3;refGene_gn=NRBP2;refGene_tx=XM_023248753.1;refGene_gn=NRBP2;refGene_tx=XM_023248754.1;refGene_gn=NRBP2;refGene_tx=XM_023248755.1
F2 84844096 84844229
refGene_gn=NRBP2;refGene_tx=XM_011291509.3;refGene_gn=NRBP2;refGene_tx=XM_023248753.1;refGene_gn=NRBP2;refGene_tx=XM_023248754.1;refGene_gn=NRBP2;refGene_tx=XM_023248755.1
F2 84845282 84845378
refGene_gn=NRBP2;refGene_tx=XM_011291509.3;refGene_gn=NRBP2;refGene_tx=XM_023248753.1;refGene_gn=NRBP2;refGene_tx=XM_023248754.1;refGene_gn=NRBP2;refGene_tx=XM_023248755.1
F2 84845474 84845530
refGene_gn=NRBP2;refGene_tx=XM_011291509.3;refGene_gn=NRBP2;refGene_tx=XM_023248753.1;refGene_gn=NRBP2;refGene_tx=XM_023248754.1;refGene_gn=NRBP2;refGene_tx=XM_023248755.1
F2 84845616 84845700
refGene_gn=NRBP2;refGene_tx=XM_011291509.3;refGene_gn=NRBP2;refGene_tx=XM_023248753.1;refGene_gn=NRBP2;refGene_tx=XM_023248754.1;refGene_gn=NRBP2;refGene_tx=XM_023248755.1
F2 84845771 84845795
refGene_gn=NRBP2;refGene_tx=XM_011291509.3;refGene_gn=NRBP2;refGene_tx=XM_023248753.1;refGene_gn=NRBP2;refGene_tx=XM_023248754.1;refGene_gn=NRBP2;refGene_tx=XM_023248755.1
F2 84845921 84846016
refGene_gn=NRBP2;refGene_tx=XM_011291509.3;refGene_gn=NRBP2;refGene_tx=XM_023248753.1;refGene_gn=NRBP2;refGene_tx=XM_023248754.1;refGene_gn=NRBP2;refGene_tx=XM_023248755.1
F2 84846102 84846143
refGene_gn=NRBP2;refGene_tx=XM_011291509.3;refGene_gn=NRBP2;refGene_tx=XM_023248753.1;refGene_gn=NRBP2;refGene_tx=XM_023248754.1;refGene_gn=NRBP2;refGene_tx=XM_023248755.1
F2 84846309 84846399
refGene_gn=NRBP2;refGene_tx=XM_011291509.3;refGene_gn=NRBP2;refGene_tx=XM_023248753.1;refGene_gn=NRBP2;refGene_tx=XM_023248754.1;refGene_gn=NRBP2;refGene_tx=XM_023248755.1
F2 84846521 84846623
refGene_gn=NRBP2;refGene_tx=XM_011291509.3;refGene_gn=NRBP2;refGene_tx=XM_023248754.1;refGene_gn=NRBP2;refGene_tx=XM_023248755.1
F2 84846698 84846821
refGene_gn=NRBP2;refGene_tx=XM_011291509.3;refGene_gn=NRBP2;refGene_tx=XM_023248753.1;refGene_gn=NRBP2;refGene_tx=XM_023248755.1
F2 84847354 84847477
refGene_gn=NRBP2;refGene_tx=XM_011291509.3;refGene_gn=NRBP2;refGene_tx=XM_023248753.1;refGene_gn=NRBP2;refGene_tx=XM_023248754.1

F2 84850771 84851078 refGene_gn=EPPK1;refGene_tx=XM_023248157.1
F2 84851156 84852154 refGene_gn=EPPK1;refGene_tx=XM_023248157.1
F2 84852156 84855314 refGene_gn=EPPK1;refGene_tx=XM_023248157.1
F2 84855429 84856399 refGene_gn=EPPK1;refGene_tx=XM_023248157.1
F2 84856400 84856693 refGene_gn=EPPK1;refGene_tx=XM_023248157.1
F2 84856695 84856964 refGene_gn=EPPK1;refGene_tx=XM_023248157.1
F2 84857120 84857728 refGene_gn=EPPK1;refGene_tx=XM_023248157.1
F2 84857729 84858509 refGene_gn=EPPK1;refGene_tx=XM_023248157.1
F2 84858511 84858707 refGene_gn=EPPK1;refGene_tx=XM_023248157.1
F2 84858708 84866225 refGene_gn=EPPK1;refGene_tx=XM_023248157.1
F2 84868425 84868438
refGene_gn=LOC111558528;refGene_tx=XM_023248349.1
F2 84869079 84869237
refGene_gn=LOC111558528;refGene_tx=XM_023248349.1
F2 84869462 84869507
refGene_gn=LOC111558528;refGene_tx=XM_023248349.1
F2 84869650 84869792
refGene_gn=LOC111558528;refGene_tx=XM_023248349.1
F2 84870376 84870558
refGene_gn=LOC111558528;refGene_tx=XM_023248349.1
F2 84870990 84871082
refGene_gn=LOC111558528;refGene_tx=XM_023248349.1
F2 84871178 84871905
refGene_gn=LOC111558528;refGene_tx=XM_023248349.1
F2 84911518 84917737
refGene_gn=PLEC;refGene_tx=XM_023248166.1;refGene_gn=PLEC;refGene_tx=X
M_023248167.1;refGene_gn=PLEC;refGene_tx=XM_023248168.1;refGene_gn=PLE
C;refGene_tx=XM_023248169.1;refGene_gn=PLEC;refGene_tx=XM_023248170.1;
refGene_gn=PLEC;refGene_tx=XM_023248171.1;refGene_gn=PLEC;refGene_tx=X
M_023248172.1;refGene_gn=PLEC;refGene_tx=XM_023248174.1;refGene_gn=PLE
C;refGene_tx=XM_023248175.1;refGene_gn=PLEC;refGene_tx=XM_023248176.1;
refGene_gn=PLEC;refGene_tx=XM_023248177.1;refGene_gn=PLEC;refGene_tx=X
M_023248178.1;refGene_gn=PLEC;refGene_tx=XM_023248179.1;refGene_gn=PLE
C;refGene_tx=XM_023248180.1;refGene_gn=PLEC;refGene_tx=XM_023248181.1;
refGene_gn=PLEC;refGene_tx=XM_023248182.1;refGene_gn=PLEC;refGene_tx=X
M_023248183.1;refGene_gn=PLEC;refGene_tx=XM_023248184.1;refGene_gn=PLE
C;refGene_tx=XM_023248185.1;refGene_gn=PLEC;refGene_tx=XM_023248186.1;
refGene_gn=PLEC;refGene_tx=XM_023248187.1;refGene_gn=PLEC;refGene_tx=X
M_023248188.1;refGene_gn=PLEC;refGene_tx=XM_023248190.1;refGene_gn=PLE
C;refGene_tx=XM_023248191.1;refGene_gn=PLEC;refGene_tx=XM_023248192.1;
refGene_gn=PLEC;refGene_tx=XM_023248193.1;refGene_gn=PLEC;refGene_tx=X
M_023248194.1;refGene_gn=PLEC;refGene_tx=XM_023248195.1;refGene_gn=PLE
C;refGene_tx=XM_023248196.1
F2 84917849 84921230
refGene_gn=PLEC;refGene_tx=XM_023248166.1;refGene_gn=PLEC;refGene_tx=X
M_023248167.1;refGene_gn=PLEC;refGene_tx=XM_023248168.1;refGene_gn=PLE
C;refGene_tx=XM_023248169.1;refGene_gn=PLEC;refGene_tx=XM_023248170.1;
refGene_gn=PLEC;refGene_tx=XM_023248171.1;refGene_gn=PLEC;refGene_tx=X
M_023248172.1;refGene_gn=PLEC;refGene_tx=XM_023248174.1;refGene_gn=PLE
C;refGene_tx=XM_023248175.1;refGene_gn=PLEC;refGene_tx=XM_023248176.1;

refGene_gn=PLEC;refGene_tx=XM_023248177.1;refGene_gn=PLEC;refGene_tx=XM_023248178.1;refGene_gn=PLEC;refGene_tx=XM_023248179.1;refGene_gn=PLEC;refGene_tx=XM_023248180.1;refGene_gn=PLEC;refGene_tx=XM_023248181.1;refGene_gn=PLEC;refGene_tx=XM_023248182.1;refGene_gn=PLEC;refGene_tx=XM_023248183.1;refGene_gn=PLEC;refGene_tx=XM_023248184.1;refGene_gn=PLEC;refGene_tx=XM_023248185.1;refGene_gn=PLEC;refGene_tx=XM_023248186.1;refGene_gn=PLEC;refGene_tx=XM_023248187.1;refGene_gn=PLEC;refGene_tx=XM_023248188.1;refGene_gn=PLEC;refGene_tx=XM_023248190.1;refGene_gn=PLEC;refGene_tx=XM_023248191.1;refGene_gn=PLEC;refGene_tx=XM_023248192.1;refGene_gn=PLEC;refGene_tx=XM_023248193.1;refGene_gn=PLEC;refGene_tx=XM_023248194.1;refGene_gn=PLEC;refGene_tx=XM_023248195.1;refGene_gn=PLEC;refGene_tx=XM_023248196.1

F2 84922202 84922301

refGene_gn=PLEC;refGene_tx=XM_023248166.1;refGene_gn=PLEC;refGene_tx=XM_023248167.1;refGene_gn=PLEC;refGene_tx=XM_023248168.1;refGene_gn=PLEC;refGene_tx=XM_023248169.1;refGene_gn=PLEC;refGene_tx=XM_023248170.1;refGene_gn=PLEC;refGene_tx=XM_023248171.1;refGene_gn=PLEC;refGene_tx=XM_023248172.1;refGene_gn=PLEC;refGene_tx=XM_023248174.1;refGene_gn=PLEC;refGene_tx=XM_023248175.1;refGene_gn=PLEC;refGene_tx=XM_023248176.1;refGene_gn=PLEC;refGene_tx=XM_023248177.1;refGene_gn=PLEC;refGene_tx=XM_023248178.1;refGene_gn=PLEC;refGene_tx=XM_023248179.1;refGene_gn=PLEC;refGene_tx=XM_023248180.1;refGene_gn=PLEC;refGene_tx=XM_023248181.1;refGene_gn=PLEC;refGene_tx=XM_023248182.1;refGene_gn=PLEC;refGene_tx=XM_023248183.1;refGene_gn=PLEC;refGene_tx=XM_023248184.1;refGene_gn=PLEC;refGene_tx=XM_023248185.1;refGene_gn=PLEC;refGene_tx=XM_023248186.1;refGene_gn=PLEC;refGene_tx=XM_023248187.1;refGene_gn=PLEC;refGene_tx=XM_023248188.1;refGene_gn=PLEC;refGene_tx=XM_023248190.1;refGene_gn=PLEC;refGene_tx=XM_023248191.1;refGene_gn=PLEC;refGene_tx=XM_023248192.1;refGene_gn=PLEC;refGene_tx=XM_023248193.1;refGene_gn=PLEC;refGene_tx=XM_023248194.1;refGene_gn=PLEC;refGene_tx=XM_023248195.1;refGene_gn=PLEC;refGene_tx=XM_023248196.1

F2 84922401 84922506

refGene_gn=PLEC;refGene_tx=XM_023248166.1;refGene_gn=PLEC;refGene_tx=XM_023248167.1;refGene_gn=PLEC;refGene_tx=XM_023248168.1;refGene_gn=PLEC;refGene_tx=XM_023248169.1;refGene_gn=PLEC;refGene_tx=XM_023248170.1;refGene_gn=PLEC;refGene_tx=XM_023248171.1;refGene_gn=PLEC;refGene_tx=XM_023248172.1;refGene_gn=PLEC;refGene_tx=XM_023248174.1;refGene_gn=PLEC;refGene_tx=XM_023248175.1;refGene_gn=PLEC;refGene_tx=XM_023248176.1;refGene_gn=PLEC;refGene_tx=XM_023248177.1;refGene_gn=PLEC;refGene_tx=XM_023248178.1;refGene_gn=PLEC;refGene_tx=XM_023248179.1;refGene_gn=PLEC;refGene_tx=XM_023248180.1;refGene_gn=PLEC;refGene_tx=XM_023248181.1;refGene_gn=PLEC;refGene_tx=XM_023248182.1;refGene_gn=PLEC;refGene_tx=XM_023248183.1;refGene_gn=PLEC;refGene_tx=XM_023248184.1;refGene_gn=PLEC;refGene_tx=XM_023248185.1;refGene_gn=PLEC;refGene_tx=XM_023248186.1;refGene_gn=PLEC;refGene_tx=XM_023248187.1;refGene_gn=PLEC;refGene_tx=XM_023248188.1;refGene_gn=PLEC;refGene_tx=XM_023248190.1;refGene_gn=PLEC;refGene_tx=XM_023248191.1;refGene_gn=PLEC;refGene_tx=XM_023248192.1;refGene_gn=PLEC;refGene_tx=XM_023248193.1;refGene_gn=PLEC;refGene_tx=XM_023248194.1;refGene_gn=PLEC;refGene_tx=XM_023248195.1;refGene_gn=PLEC;refGene_tx=XM_023248196.1

F2 84922678 84922762
refGene_gn=PLEC;refGene_tx=XM_023248166.1;refGene_gn=PLEC;refGene_tx=XM_023248167.1;refGene_gn=PLEC;refGene_tx=XM_023248168.1;refGene_gn=PLEC;refGene_tx=XM_023248169.1;refGene_gn=PLEC;refGene_tx=XM_023248170.1;refGene_gn=PLEC;refGene_tx=XM_023248171.1;refGene_gn=PLEC;refGene_tx=XM_023248172.1;refGene_gn=PLEC;refGene_tx=XM_023248174.1;refGene_gn=PLEC;refGene_tx=XM_023248175.1;refGene_gn=PLEC;refGene_tx=XM_023248176.1;refGene_gn=PLEC;refGene_tx=XM_023248177.1;refGene_gn=PLEC;refGene_tx=XM_023248178.1;refGene_gn=PLEC;refGene_tx=XM_023248179.1;refGene_gn=PLEC;refGene_tx=XM_023248180.1;refGene_gn=PLEC;refGene_tx=XM_023248181.1;refGene_gn=PLEC;refGene_tx=XM_023248182.1;refGene_gn=PLEC;refGene_tx=XM_023248183.1;refGene_gn=PLEC;refGene_tx=XM_023248184.1;refGene_gn=PLEC;refGene_tx=XM_023248185.1;refGene_gn=PLEC;refGene_tx=XM_023248186.1;refGene_gn=PLEC;refGene_tx=XM_023248187.1;refGene_gn=PLEC;refGene_tx=XM_023248188.1;refGene_gn=PLEC;refGene_tx=XM_023248190.1;refGene_gn=PLEC;refGene_tx=XM_023248191.1;refGene_gn=PLEC;refGene_tx=XM_023248192.1;refGene_gn=PLEC;refGene_tx=XM_023248193.1;refGene_gn=PLEC;refGene_tx=XM_023248194.1;refGene_gn=PLEC;refGene_tx=XM_023248195.1;refGene_gn=PLEC;refGene_tx=XM_023248196.1
F2 84922844 84923201
refGene_gn=PLEC;refGene_tx=XM_023248166.1;refGene_gn=PLEC;refGene_tx=XM_023248167.1;refGene_gn=PLEC;refGene_tx=XM_023248168.1;refGene_gn=PLEC;refGene_tx=XM_023248169.1;refGene_gn=PLEC;refGene_tx=XM_023248170.1;refGene_gn=PLEC;refGene_tx=XM_023248171.1;refGene_gn=PLEC;refGene_tx=XM_023248172.1;refGene_gn=PLEC;refGene_tx=XM_023248174.1;refGene_gn=PLEC;refGene_tx=XM_023248175.1;refGene_gn=PLEC;refGene_tx=XM_023248176.1;refGene_gn=PLEC;refGene_tx=XM_023248177.1;refGene_gn=PLEC;refGene_tx=XM_023248178.1;refGene_gn=PLEC;refGene_tx=XM_023248179.1;refGene_gn=PLEC;refGene_tx=XM_023248180.1;refGene_gn=PLEC;refGene_tx=XM_023248181.1;refGene_gn=PLEC;refGene_tx=XM_023248182.1;refGene_gn=PLEC;refGene_tx=XM_023248183.1;refGene_gn=PLEC;refGene_tx=XM_023248184.1;refGene_gn=PLEC;refGene_tx=XM_023248185.1;refGene_gn=PLEC;refGene_tx=XM_023248186.1;refGene_gn=PLEC;refGene_tx=XM_023248187.1;refGene_gn=PLEC;refGene_tx=XM_023248188.1;refGene_gn=PLEC;refGene_tx=XM_023248190.1;refGene_gn=PLEC;refGene_tx=XM_023248191.1;refGene_gn=PLEC;refGene_tx=XM_023248192.1;refGene_gn=PLEC;refGene_tx=XM_023248193.1;refGene_gn=PLEC;refGene_tx=XM_023248194.1;refGene_gn=PLEC;refGene_tx=XM_023248195.1;refGene_gn=PLEC;refGene_tx=XM_023248196.1
F2 84923319 84923458
refGene_gn=PLEC;refGene_tx=XM_023248166.1;refGene_gn=PLEC;refGene_tx=XM_023248167.1;refGene_gn=PLEC;refGene_tx=XM_023248168.1;refGene_gn=PLEC;refGene_tx=XM_023248169.1;refGene_gn=PLEC;refGene_tx=XM_023248170.1;refGene_gn=PLEC;refGene_tx=XM_023248171.1;refGene_gn=PLEC;refGene_tx=XM_023248172.1;refGene_gn=PLEC;refGene_tx=XM_023248174.1;refGene_gn=PLEC;refGene_tx=XM_023248175.1;refGene_gn=PLEC;refGene_tx=XM_023248176.1;refGene_gn=PLEC;refGene_tx=XM_023248177.1;refGene_gn=PLEC;refGene_tx=XM_023248178.1;refGene_gn=PLEC;refGene_tx=XM_023248179.1;refGene_gn=PLEC;refGene_tx=XM_023248180.1;refGene_gn=PLEC;refGene_tx=XM_023248181.1;refGene_gn=PLEC;refGene_tx=XM_023248182.1;refGene_gn=PLEC;refGene_tx=XM_023248183.1;refGene_gn=PLEC;refGene_tx=XM_023248184.1;refGene_gn=PLE

C;refGene_tx=XM_023248185.1;refGene_gn=PLEC;refGene_tx=XM_023248186.1;
refGene_gn=PLEC;refGene_tx=XM_023248187.1;refGene_gn=PLEC;refGene_tx=X
M_023248188.1;refGene_gn=PLEC;refGene_tx=XM_023248190.1;refGene_gn=PLE
C;refGene_tx=XM_023248191.1;refGene_gn=PLEC;refGene_tx=XM_023248192.1;
refGene_gn=PLEC;refGene_tx=XM_023248193.1;refGene_gn=PLEC;refGene_tx=X
M_023248194.1;refGene_gn=PLEC;refGene_tx=XM_023248195.1;refGene_gn=PLE
C;refGene_tx=XM_023248196.1
F2 84924632 84924811
refGene_gn=PLEC;refGene_tx=XM_023248166.1;refGene_gn=PLEC;refGene_tx=X
M_023248167.1;refGene_gn=PLEC;refGene_tx=XM_023248168.1;refGene_gn=PLE
C;refGene_tx=XM_023248169.1;refGene_gn=PLEC;refGene_tx=XM_023248170.1;
refGene_gn=PLEC;refGene_tx=XM_023248171.1;refGene_gn=PLEC;refGene_tx=X
M_023248172.1;refGene_gn=PLEC;refGene_tx=XM_023248174.1;refGene_gn=PLE
C;refGene_tx=XM_023248175.1;refGene_gn=PLEC;refGene_tx=XM_023248176.1;
refGene_gn=PLEC;refGene_tx=XM_023248177.1;refGene_gn=PLEC;refGene_tx=X
M_023248178.1;refGene_gn=PLEC;refGene_tx=XM_023248179.1;refGene_gn=PLE
C;refGene_tx=XM_023248180.1;refGene_gn=PLEC;refGene_tx=XM_023248181.1;
refGene_gn=PLEC;refGene_tx=XM_023248182.1;refGene_gn=PLEC;refGene_tx=X
M_023248183.1;refGene_gn=PLEC;refGene_tx=XM_023248184.1;refGene_gn=PLE
C;refGene_tx=XM_023248185.1;refGene_gn=PLEC;refGene_tx=XM_023248186.1;
refGene_gn=PLEC;refGene_tx=XM_023248187.1;refGene_gn=PLEC;refGene_tx=X
M_023248188.1;refGene_gn=PLEC;refGene_tx=XM_023248190.1;refGene_gn=PLE
C;refGene_tx=XM_023248191.1;refGene_gn=PLEC;refGene_tx=XM_023248192.1;
refGene_gn=PLEC;refGene_tx=XM_023248193.1;refGene_gn=PLEC;refGene_tx=X
M_023248194.1;refGene_gn=PLEC;refGene_tx=XM_023248195.1;refGene_gn=PLE
C;refGene_tx=XM_023248196.1
F2 84924926 84925084
refGene_gn=PLEC;refGene_tx=XM_023248166.1;refGene_gn=PLEC;refGene_tx=X
M_023248167.1;refGene_gn=PLEC;refGene_tx=XM_023248168.1;refGene_gn=PLE
C;refGene_tx=XM_023248169.1;refGene_gn=PLEC;refGene_tx=XM_023248170.1;
refGene_gn=PLEC;refGene_tx=XM_023248171.1;refGene_gn=PLEC;refGene_tx=X
M_023248172.1;refGene_gn=PLEC;refGene_tx=XM_023248174.1;refGene_gn=PLE
C;refGene_tx=XM_023248175.1;refGene_gn=PLEC;refGene_tx=XM_023248176.1;
refGene_gn=PLEC;refGene_tx=XM_023248177.1;refGene_gn=PLEC;refGene_tx=X
M_023248178.1;refGene_gn=PLEC;refGene_tx=XM_023248179.1;refGene_gn=PLE
C;refGene_tx=XM_023248180.1;refGene_gn=PLEC;refGene_tx=XM_023248181.1;
refGene_gn=PLEC;refGene_tx=XM_023248182.1;refGene_gn=PLEC;refGene_tx=X
M_023248183.1;refGene_gn=PLEC;refGene_tx=XM_023248184.1;refGene_gn=PLE
C;refGene_tx=XM_023248185.1;refGene_gn=PLEC;refGene_tx=XM_023248186.1;
refGene_gn=PLEC;refGene_tx=XM_023248187.1;refGene_gn=PLEC;refGene_tx=X
M_023248188.1;refGene_gn=PLEC;refGene_tx=XM_023248190.1;refGene_gn=PLE
C;refGene_tx=XM_023248191.1;refGene_gn=PLEC;refGene_tx=XM_023248192.1;
refGene_gn=PLEC;refGene_tx=XM_023248193.1;refGene_gn=PLEC;refGene_tx=X
M_023248194.1;refGene_gn=PLEC;refGene_tx=XM_023248195.1;refGene_gn=PLE
C;refGene_tx=XM_023248196.1
F2 84925161 84925345
refGene_gn=PLEC;refGene_tx=XM_023248166.1;refGene_gn=PLEC;refGene_tx=X
M_023248167.1;refGene_gn=PLEC;refGene_tx=XM_023248168.1;refGene_gn=PLE
C;refGene_tx=XM_023248169.1;refGene_gn=PLEC;refGene_tx=XM_023248170.1;
refGene_gn=PLEC;refGene_tx=XM_023248171.1;refGene_gn=PLEC;refGene_tx=X

M_023248172.1;refGene_gn=PLEC;refGene_tx=XM_023248174.1;refGene_gn=PLE
C;refGene_tx=XM_023248175.1;refGene_gn=PLEC;refGene_tx=XM_023248176.1;
refGene_gn=PLEC;refGene_tx=XM_023248177.1;refGene_gn=PLEC;refGene_tx=X
M_023248178.1;refGene_gn=PLEC;refGene_tx=XM_023248179.1;refGene_gn=PLE
C;refGene_tx=XM_023248180.1;refGene_gn=PLEC;refGene_tx=XM_023248181.1;
refGene_gn=PLEC;refGene_tx=XM_023248182.1;refGene_gn=PLEC;refGene_tx=X
M_023248183.1;refGene_gn=PLEC;refGene_tx=XM_023248184.1;refGene_gn=PLE
C;refGene_tx=XM_023248185.1;refGene_gn=PLEC;refGene_tx=XM_023248186.1;
refGene_gn=PLEC;refGene_tx=XM_023248187.1;refGene_gn=PLEC;refGene_tx=X
M_023248188.1;refGene_gn=PLEC;refGene_tx=XM_023248190.1;refGene_gn=PLE
C;refGene_tx=XM_023248191.1;refGene_gn=PLEC;refGene_tx=XM_023248192.1;
refGene_gn=PLEC;refGene_tx=XM_023248193.1;refGene_gn=PLEC;refGene_tx=X
M_023248194.1;refGene_gn=PLEC;refGene_tx=XM_023248195.1;refGene_gn=PLE
C;refGene_tx=XM_023248196.1
F2 84925437 84925564
refGene_gn=PLEC;refGene_tx=XM_023248166.1;refGene_gn=PLEC;refGene_tx=X
M_023248167.1;refGene_gn=PLEC;refGene_tx=XM_023248168.1;refGene_gn=PLE
C;refGene_tx=XM_023248169.1;refGene_gn=PLEC;refGene_tx=XM_023248170.1;
refGene_gn=PLEC;refGene_tx=XM_023248171.1;refGene_gn=PLEC;refGene_tx=X
M_023248172.1;refGene_gn=PLEC;refGene_tx=XM_023248174.1;refGene_gn=PLE
C;refGene_tx=XM_023248175.1;refGene_gn=PLEC;refGene_tx=XM_023248176.1;
refGene_gn=PLEC;refGene_tx=XM_023248177.1;refGene_gn=PLEC;refGene_tx=X
M_023248178.1;refGene_gn=PLEC;refGene_tx=XM_023248179.1;refGene_gn=PLE
C;refGene_tx=XM_023248180.1;refGene_gn=PLEC;refGene_tx=XM_023248181.1;
refGene_gn=PLEC;refGene_tx=XM_023248182.1;refGene_gn=PLEC;refGene_tx=X
M_023248183.1;refGene_gn=PLEC;refGene_tx=XM_023248184.1;refGene_gn=PLE
C;refGene_tx=XM_023248185.1;refGene_gn=PLEC;refGene_tx=XM_023248186.1;
refGene_gn=PLEC;refGene_tx=XM_023248187.1;refGene_gn=PLEC;refGene_tx=X
M_023248188.1;refGene_gn=PLEC;refGene_tx=XM_023248190.1;refGene_gn=PLE
C;refGene_tx=XM_023248191.1;refGene_gn=PLEC;refGene_tx=XM_023248192.1;
refGene_gn=PLEC;refGene_tx=XM_023248193.1;refGene_gn=PLEC;refGene_tx=X
M_023248194.1;refGene_gn=PLEC;refGene_tx=XM_023248195.1;refGene_gn=PLE
C;refGene_tx=XM_023248196.1
F2 84925649 84925804
refGene_gn=PLEC;refGene_tx=XM_023248166.1;refGene_gn=PLEC;refGene_tx=X
M_023248167.1;refGene_gn=PLEC;refGene_tx=XM_023248168.1;refGene_gn=PLE
C;refGene_tx=XM_023248169.1;refGene_gn=PLEC;refGene_tx=XM_023248170.1;
refGene_gn=PLEC;refGene_tx=XM_023248171.1;refGene_gn=PLEC;refGene_tx=X
M_023248172.1;refGene_gn=PLEC;refGene_tx=XM_023248174.1;refGene_gn=PLE
C;refGene_tx=XM_023248175.1;refGene_gn=PLEC;refGene_tx=XM_023248176.1;
refGene_gn=PLEC;refGene_tx=XM_023248177.1;refGene_gn=PLEC;refGene_tx=X
M_023248178.1;refGene_gn=PLEC;refGene_tx=XM_023248179.1;refGene_gn=PLE
C;refGene_tx=XM_023248180.1;refGene_gn=PLEC;refGene_tx=XM_023248181.1;
refGene_gn=PLEC;refGene_tx=XM_023248182.1;refGene_gn=PLEC;refGene_tx=X
M_023248183.1;refGene_gn=PLEC;refGene_tx=XM_023248184.1;refGene_gn=PLE
C;refGene_tx=XM_023248185.1;refGene_gn=PLEC;refGene_tx=XM_023248186.1;
refGene_gn=PLEC;refGene_tx=XM_023248187.1;refGene_gn=PLEC;refGene_tx=X
M_023248188.1;refGene_gn=PLEC;refGene_tx=XM_023248190.1;refGene_gn=PLE
C;refGene_tx=XM_023248191.1;refGene_gn=PLEC;refGene_tx=XM_023248192.1;
refGene_gn=PLEC;refGene_tx=XM_023248193.1;refGene_gn=PLEC;refGene_tx=X

M_023248194.1;refGene_gn=PLEC;refGene_tx=XM_023248195.1;refGene_gn=PLEC;refGene_tx=XM_023248196.1
F2 84925899 84926052
refGene_gn=PLEC;refGene_tx=XM_023248166.1;refGene_gn=PLEC;refGene_tx=XM_023248167.1;refGene_gn=PLEC;refGene_tx=XM_023248168.1;refGene_gn=PLEC;refGene_tx=XM_023248169.1;refGene_gn=PLEC;refGene_tx=XM_023248170.1;refGene_gn=PLEC;refGene_tx=XM_023248171.1;refGene_gn=PLEC;refGene_tx=XM_023248172.1;refGene_gn=PLEC;refGene_tx=XM_023248174.1;refGene_gn=PLEC;refGene_tx=XM_023248175.1;refGene_gn=PLEC;refGene_tx=XM_023248176.1;refGene_gn=PLEC;refGene_tx=XM_023248177.1;refGene_gn=PLEC;refGene_tx=XM_023248178.1;refGene_gn=PLEC;refGene_tx=XM_023248179.1;refGene_gn=PLEC;refGene_tx=XM_023248180.1;refGene_gn=PLEC;refGene_tx=XM_023248181.1;refGene_gn=PLEC;refGene_tx=XM_023248182.1;refGene_gn=PLEC;refGene_tx=XM_023248183.1;refGene_gn=PLEC;refGene_tx=XM_023248184.1;refGene_gn=PLEC;refGene_tx=XM_023248185.1;refGene_gn=PLEC;refGene_tx=XM_023248186.1;refGene_gn=PLEC;refGene_tx=XM_023248187.1;refGene_gn=PLEC;refGene_tx=XM_023248188.1;refGene_gn=PLEC;refGene_tx=XM_023248190.1;refGene_gn=PLEC;refGene_tx=XM_023248191.1;refGene_gn=PLEC;refGene_tx=XM_023248192.1;refGene_gn=PLEC;refGene_tx=XM_023248193.1;refGene_gn=PLEC;refGene_tx=XM_023248194.1;refGene_gn=PLEC;refGene_tx=XM_023248195.1;refGene_gn=PLEC;refGene_tx=XM_023248196.1
F2 84927172 84927298
refGene_gn=PLEC;refGene_tx=XM_023248166.1;refGene_gn=PLEC;refGene_tx=XM_023248167.1;refGene_gn=PLEC;refGene_tx=XM_023248168.1;refGene_gn=PLEC;refGene_tx=XM_023248169.1;refGene_gn=PLEC;refGene_tx=XM_023248170.1;refGene_gn=PLEC;refGene_tx=XM_023248171.1;refGene_gn=PLEC;refGene_tx=XM_023248172.1;refGene_gn=PLEC;refGene_tx=XM_023248174.1;refGene_gn=PLEC;refGene_tx=XM_023248175.1;refGene_gn=PLEC;refGene_tx=XM_023248176.1;refGene_gn=PLEC;refGene_tx=XM_023248177.1;refGene_gn=PLEC;refGene_tx=XM_023248178.1;refGene_gn=PLEC;refGene_tx=XM_023248179.1;refGene_gn=PLEC;refGene_tx=XM_023248180.1;refGene_gn=PLEC;refGene_tx=XM_023248181.1;refGene_gn=PLEC;refGene_tx=XM_023248182.1;refGene_gn=PLEC;refGene_tx=XM_023248183.1;refGene_gn=PLEC;refGene_tx=XM_023248184.1;refGene_gn=PLEC;refGene_tx=XM_023248185.1;refGene_gn=PLEC;refGene_tx=XM_023248186.1;refGene_gn=PLEC;refGene_tx=XM_023248187.1;refGene_gn=PLEC;refGene_tx=XM_023248188.1;refGene_gn=PLEC;refGene_tx=XM_023248190.1;refGene_gn=PLEC;refGene_tx=XM_023248191.1;refGene_gn=PLEC;refGene_tx=XM_023248192.1;refGene_gn=PLEC;refGene_tx=XM_023248193.1;refGene_gn=PLEC;refGene_tx=XM_023248194.1;refGene_gn=PLEC;refGene_tx=XM_023248195.1;refGene_gn=PLEC;refGene_tx=XM_023248196.1
F2 84927495 84927591
refGene_gn=PLEC;refGene_tx=XM_023248166.1;refGene_gn=PLEC;refGene_tx=XM_023248167.1;refGene_gn=PLEC;refGene_tx=XM_023248168.1;refGene_gn=PLEC;refGene_tx=XM_023248169.1;refGene_gn=PLEC;refGene_tx=XM_023248170.1;refGene_gn=PLEC;refGene_tx=XM_023248171.1;refGene_gn=PLEC;refGene_tx=XM_023248172.1;refGene_gn=PLEC;refGene_tx=XM_023248174.1;refGene_gn=PLEC;refGene_tx=XM_023248175.1;refGene_gn=PLEC;refGene_tx=XM_023248176.1;refGene_gn=PLEC;refGene_tx=XM_023248177.1;refGene_gn=PLEC;refGene_tx=XM_023248178.1;refGene_gn=PLEC;refGene_tx=XM_023248179.1;refGene_gn=PLEC;refGene_tx=XM_023248180.1;refGene_gn=PLEC;refGene_tx=XM_023248181.1;

refGene_gn=PLEC;refGene_tx=XM_023248182.1;refGene_gn=PLEC;refGene_tx=XM_023248183.1;refGene_gn=PLEC;refGene_tx=XM_023248184.1;refGene_gn=PLEC;refGene_tx=XM_023248185.1;refGene_gn=PLEC;refGene_tx=XM_023248186.1;refGene_gn=PLEC;refGene_tx=XM_023248187.1;refGene_gn=PLEC;refGene_tx=XM_023248188.1;refGene_gn=PLEC;refGene_tx=XM_023248190.1;refGene_gn=PLEC;refGene_tx=XM_023248191.1;refGene_gn=PLEC;refGene_tx=XM_023248192.1;refGene_gn=PLEC;refGene_tx=XM_023248193.1;refGene_gn=PLEC;refGene_tx=XM_023248194.1;refGene_gn=PLEC;refGene_tx=XM_023248195.1;refGene_gn=PLEC;refGene_tx=XM_023248196.1
F2 84927698 84927803
refGene_gn=PLEC;refGene_tx=XM_023248166.1;refGene_gn=PLEC;refGene_tx=XM_023248167.1;refGene_gn=PLEC;refGene_tx=XM_023248168.1;refGene_gn=PLEC;refGene_tx=XM_023248169.1;refGene_gn=PLEC;refGene_tx=XM_023248170.1;refGene_gn=PLEC;refGene_tx=XM_023248171.1;refGene_gn=PLEC;refGene_tx=XM_023248172.1;refGene_gn=PLEC;refGene_tx=XM_023248174.1;refGene_gn=PLEC;refGene_tx=XM_023248175.1;refGene_gn=PLEC;refGene_tx=XM_023248176.1;refGene_gn=PLEC;refGene_tx=XM_023248177.1;refGene_gn=PLEC;refGene_tx=XM_023248178.1;refGene_gn=PLEC;refGene_tx=XM_023248179.1;refGene_gn=PLEC;refGene_tx=XM_023248180.1;refGene_gn=PLEC;refGene_tx=XM_023248181.1;refGene_gn=PLEC;refGene_tx=XM_023248182.1;refGene_gn=PLEC;refGene_tx=XM_023248183.1;refGene_gn=PLEC;refGene_tx=XM_023248184.1;refGene_gn=PLEC;refGene_tx=XM_023248185.1;refGene_gn=PLEC;refGene_tx=XM_023248186.1;refGene_gn=PLEC;refGene_tx=XM_023248187.1;refGene_gn=PLEC;refGene_tx=XM_023248188.1;refGene_gn=PLEC;refGene_tx=XM_023248190.1;refGene_gn=PLEC;refGene_tx=XM_023248191.1;refGene_gn=PLEC;refGene_tx=XM_023248192.1;refGene_gn=PLEC;refGene_tx=XM_023248193.1;refGene_gn=PLEC;refGene_tx=XM_023248194.1;refGene_gn=PLEC;refGene_tx=XM_023248195.1;refGene_gn=PLEC;refGene_tx=XM_023248196.1
F2 84927934 84928096
refGene_gn=PLEC;refGene_tx=XM_023248166.1;refGene_gn=PLEC;refGene_tx=XM_023248167.1;refGene_gn=PLEC;refGene_tx=XM_023248168.1;refGene_gn=PLEC;refGene_tx=XM_023248169.1;refGene_gn=PLEC;refGene_tx=XM_023248170.1;refGene_gn=PLEC;refGene_tx=XM_023248171.1;refGene_gn=PLEC;refGene_tx=XM_023248172.1;refGene_gn=PLEC;refGene_tx=XM_023248174.1;refGene_gn=PLEC;refGene_tx=XM_023248175.1;refGene_gn=PLEC;refGene_tx=XM_023248176.1;refGene_gn=PLEC;refGene_tx=XM_023248177.1;refGene_gn=PLEC;refGene_tx=XM_023248178.1;refGene_gn=PLEC;refGene_tx=XM_023248179.1;refGene_gn=PLEC;refGene_tx=XM_023248180.1;refGene_gn=PLEC;refGene_tx=XM_023248181.1;refGene_gn=PLEC;refGene_tx=XM_023248182.1;refGene_gn=PLEC;refGene_tx=XM_023248183.1;refGene_gn=PLEC;refGene_tx=XM_023248184.1;refGene_gn=PLEC;refGene_tx=XM_023248185.1;refGene_gn=PLEC;refGene_tx=XM_023248186.1;refGene_gn=PLEC;refGene_tx=XM_023248187.1;refGene_gn=PLEC;refGene_tx=XM_023248188.1;refGene_gn=PLEC;refGene_tx=XM_023248190.1;refGene_gn=PLEC;refGene_tx=XM_023248191.1;refGene_gn=PLEC;refGene_tx=XM_023248192.1;refGene_gn=PLEC;refGene_tx=XM_023248193.1;refGene_gn=PLEC;refGene_tx=XM_023248194.1;refGene_gn=PLEC;refGene_tx=XM_023248195.1;refGene_gn=PLEC;refGene_tx=XM_023248196.1
F2 84928186 84928264
refGene_gn=PLEC;refGene_tx=XM_023248166.1;refGene_gn=PLEC;refGene_tx=XM_023248167.1;refGene_gn=PLEC;refGene_tx=XM_023248168.1;refGene_gn=PLE

C;refGene_tx=XM_023248169.1;refGene_gn=PLEC;refGene_tx=XM_023248170.1;
refGene_gn=PLEC;refGene_tx=XM_023248171.1;refGene_gn=PLEC;refGene_tx=X
M_023248172.1;refGene_gn=PLEC;refGene_tx=XM_023248174.1;refGene_gn=PLE
C;refGene_tx=XM_023248175.1;refGene_gn=PLEC;refGene_tx=XM_023248176.1;
refGene_gn=PLEC;refGene_tx=XM_023248177.1;refGene_gn=PLEC;refGene_tx=X
M_023248178.1;refGene_gn=PLEC;refGene_tx=XM_023248179.1;refGene_gn=PLE
C;refGene_tx=XM_023248180.1;refGene_gn=PLEC;refGene_tx=XM_023248181.1;
refGene_gn=PLEC;refGene_tx=XM_023248182.1;refGene_gn=PLEC;refGene_tx=X
M_023248183.1;refGene_gn=PLEC;refGene_tx=XM_023248184.1;refGene_gn=PLE
C;refGene_tx=XM_023248185.1;refGene_gn=PLEC;refGene_tx=XM_023248186.1;
refGene_gn=PLEC;refGene_tx=XM_023248187.1;refGene_gn=PLEC;refGene_tx=X
M_023248188.1;refGene_gn=PLEC;refGene_tx=XM_023248190.1;refGene_gn=PLE
C;refGene_tx=XM_023248191.1;refGene_gn=PLEC;refGene_tx=XM_023248192.1;
refGene_gn=PLEC;refGene_tx=XM_023248193.1;refGene_gn=PLEC;refGene_tx=X
M_023248194.1;refGene_gn=PLEC;refGene_tx=XM_023248195.1;refGene_gn=PLE
C;refGene_tx=XM_023248196.1
F2 84928353 84928663
refGene_gn=PLEC;refGene_tx=XM_023248166.1;refGene_gn=PLEC;refGene_tx=X
M_023248167.1;refGene_gn=PLEC;refGene_tx=XM_023248168.1;refGene_gn=PLE
C;refGene_tx=XM_023248169.1;refGene_gn=PLEC;refGene_tx=XM_023248170.1;
refGene_gn=PLEC;refGene_tx=XM_023248171.1;refGene_gn=PLEC;refGene_tx=X
M_023248172.1;refGene_gn=PLEC;refGene_tx=XM_023248174.1;refGene_gn=PLE
C;refGene_tx=XM_023248175.1;refGene_gn=PLEC;refGene_tx=XM_023248176.1;
refGene_gn=PLEC;refGene_tx=XM_023248177.1;refGene_gn=PLEC;refGene_tx=X
M_023248178.1;refGene_gn=PLEC;refGene_tx=XM_023248179.1;refGene_gn=PLE
C;refGene_tx=XM_023248180.1;refGene_gn=PLEC;refGene_tx=XM_023248181.1;
refGene_gn=PLEC;refGene_tx=XM_023248182.1;refGene_gn=PLEC;refGene_tx=X
M_023248183.1;refGene_gn=PLEC;refGene_tx=XM_023248184.1;refGene_gn=PLE
C;refGene_tx=XM_023248185.1;refGene_gn=PLEC;refGene_tx=XM_023248186.1;
refGene_gn=PLEC;refGene_tx=XM_023248187.1;refGene_gn=PLEC;refGene_tx=X
M_023248188.1;refGene_gn=PLEC;refGene_tx=XM_023248190.1;refGene_gn=PLE
C;refGene_tx=XM_023248191.1;refGene_gn=PLEC;refGene_tx=XM_023248192.1;
refGene_gn=PLEC;refGene_tx=XM_023248193.1;refGene_gn=PLEC;refGene_tx=X
M_023248194.1;refGene_gn=PLEC;refGene_tx=XM_023248195.1;refGene_gn=PLE
C;refGene_tx=XM_023248196.1
F2 84928756 84928911
refGene_gn=PLEC;refGene_tx=XM_023248166.1;refGene_gn=PLEC;refGene_tx=X
M_023248167.1;refGene_gn=PLEC;refGene_tx=XM_023248168.1;refGene_gn=PLE
C;refGene_tx=XM_023248169.1;refGene_gn=PLEC;refGene_tx=XM_023248170.1;
refGene_gn=PLEC;refGene_tx=XM_023248171.1;refGene_gn=PLEC;refGene_tx=X
M_023248172.1;refGene_gn=PLEC;refGene_tx=XM_023248174.1;refGene_gn=PLE
C;refGene_tx=XM_023248175.1;refGene_gn=PLEC;refGene_tx=XM_023248176.1;
refGene_gn=PLEC;refGene_tx=XM_023248177.1;refGene_gn=PLEC;refGene_tx=X
M_023248178.1;refGene_gn=PLEC;refGene_tx=XM_023248179.1;refGene_gn=PLE
C;refGene_tx=XM_023248180.1;refGene_gn=PLEC;refGene_tx=XM_023248181.1;
refGene_gn=PLEC;refGene_tx=XM_023248182.1;refGene_gn=PLEC;refGene_tx=X
M_023248183.1;refGene_gn=PLEC;refGene_tx=XM_023248184.1;refGene_gn=PLE
C;refGene_tx=XM_023248185.1;refGene_gn=PLEC;refGene_tx=XM_023248186.1;
refGene_gn=PLEC;refGene_tx=XM_023248187.1;refGene_gn=PLEC;refGene_tx=X
M_023248188.1;refGene_gn=PLEC;refGene_tx=XM_023248190.1;refGene_gn=PLE

C;refGene_tx=XM_023248191.1;refGene_gn=PLEC;refGene_tx=XM_023248192.1;refGene_gn=PLEC;refGene_tx=XM_023248193.1;refGene_gn=PLEC;refGene_tx=XM_023248194.1;refGene_gn=PLEC;refGene_tx=XM_023248195.1;refGene_gn=PLEC;refGene_tx=XM_023248196.1

F2 84929377 84929471 refGene_gn=PLEC;refGene_tx=XM_023248166.1;refGene_gn=PLEC;refGene_tx=XM_023248167.1;refGene_gn=PLEC;refGene_tx=XM_023248168.1;refGene_gn=PLEC;refGene_tx=XM_023248169.1;refGene_gn=PLEC;refGene_tx=XM_023248170.1;refGene_gn=PLEC;refGene_tx=XM_023248171.1;refGene_gn=PLEC;refGene_tx=XM_023248172.1;refGene_gn=PLEC;refGene_tx=XM_023248174.1;refGene_gn=PLEC;refGene_tx=XM_023248175.1;refGene_gn=PLEC;refGene_tx=XM_023248176.1;refGene_gn=PLEC;refGene_tx=XM_023248177.1;refGene_gn=PLEC;refGene_tx=XM_023248178.1;refGene_gn=PLEC;refGene_tx=XM_023248179.1;refGene_gn=PLEC;refGene_tx=XM_023248180.1;refGene_gn=PLEC;refGene_tx=XM_023248181.1;refGene_gn=PLEC;refGene_tx=XM_023248182.1;refGene_gn=PLEC;refGene_tx=XM_023248183.1;refGene_gn=PLEC;refGene_tx=XM_023248184.1;refGene_gn=PLEC;refGene_tx=XM_023248185.1;refGene_gn=PLEC;refGene_tx=XM_023248186.1;refGene_gn=PLEC;refGene_tx=XM_023248187.1;refGene_gn=PLEC;refGene_tx=XM_023248188.1;refGene_gn=PLEC;refGene_tx=XM_023248190.1;refGene_gn=PLEC;refGene_tx=XM_023248191.1;refGene_gn=PLEC;refGene_tx=XM_023248192.1;refGene_gn=PLEC;refGene_tx=XM_023248193.1;refGene_gn=PLEC;refGene_tx=XM_023248194.1;refGene_gn=PLEC;refGene_tx=XM_023248195.1;refGene_gn=PLEC;refGene_tx=XM_023248196.1

F2 84929728 84929856 refGene_gn=PLEC;refGene_tx=XM_023248166.1;refGene_gn=PLEC;refGene_tx=XM_023248167.1;refGene_gn=PLEC;refGene_tx=XM_023248168.1;refGene_gn=PLEC;refGene_tx=XM_023248169.1;refGene_gn=PLEC;refGene_tx=XM_023248170.1;refGene_gn=PLEC;refGene_tx=XM_023248171.1;refGene_gn=PLEC;refGene_tx=XM_023248172.1;refGene_gn=PLEC;refGene_tx=XM_023248174.1;refGene_gn=PLEC;refGene_tx=XM_023248175.1;refGene_gn=PLEC;refGene_tx=XM_023248176.1;refGene_gn=PLEC;refGene_tx=XM_023248177.1;refGene_gn=PLEC;refGene_tx=XM_023248178.1;refGene_gn=PLEC;refGene_tx=XM_023248179.1;refGene_gn=PLEC;refGene_tx=XM_023248180.1;refGene_gn=PLEC;refGene_tx=XM_023248181.1;refGene_gn=PLEC;refGene_tx=XM_023248182.1;refGene_gn=PLEC;refGene_tx=XM_023248183.1;refGene_gn=PLEC;refGene_tx=XM_023248184.1;refGene_gn=PLEC;refGene_tx=XM_023248185.1;refGene_gn=PLEC;refGene_tx=XM_023248186.1;refGene_gn=PLEC;refGene_tx=XM_023248187.1;refGene_gn=PLEC;refGene_tx=XM_023248188.1;refGene_gn=PLEC;refGene_tx=XM_023248190.1;refGene_gn=PLEC;refGene_tx=XM_023248191.1;refGene_gn=PLEC;refGene_tx=XM_023248192.1;refGene_gn=PLEC;refGene_tx=XM_023248193.1;refGene_gn=PLEC;refGene_tx=XM_023248194.1;refGene_gn=PLEC;refGene_tx=XM_023248195.1;refGene_gn=PLEC;refGene_tx=XM_023248196.1

F2 84930013 84930109 refGene_gn=PLEC;refGene_tx=XM_023248166.1;refGene_gn=PLEC;refGene_tx=XM_023248167.1;refGene_gn=PLEC;refGene_tx=XM_023248168.1;refGene_gn=PLEC;refGene_tx=XM_023248169.1;refGene_gn=PLEC;refGene_tx=XM_023248170.1;refGene_gn=PLEC;refGene_tx=XM_023248171.1;refGene_gn=PLEC;refGene_tx=XM_023248172.1;refGene_gn=PLEC;refGene_tx=XM_023248174.1;refGene_gn=PLEC;refGene_tx=XM_023248175.1;refGene_gn=PLEC;refGene_tx=XM_023248176.1;refGene_gn=PLEC;refGene_tx=XM_023248177.1;refGene_gn=PLEC;refGene_tx=X

M_023248178.1;refGene_gn=PLEC;refGene_tx=XM_023248179.1;refGene_gn=PLEC;refGene_tx=XM_023248180.1;refGene_gn=PLEC;refGene_tx=XM_023248181.1;refGene_gn=PLEC;refGene_tx=XM_023248182.1;refGene_gn=PLEC;refGene_tx=XM_023248183.1;refGene_gn=PLEC;refGene_tx=XM_023248184.1;refGene_gn=PLEC;refGene_tx=XM_023248185.1;refGene_gn=PLEC;refGene_tx=XM_023248186.1;refGene_gn=PLEC;refGene_tx=XM_023248187.1;refGene_gn=PLEC;refGene_tx=XM_023248188.1;refGene_gn=PLEC;refGene_tx=XM_023248190.1;refGene_gn=PLEC;refGene_tx=XM_023248191.1;refGene_gn=PLEC;refGene_tx=XM_023248192.1;refGene_gn=PLEC;refGene_tx=XM_023248193.1;refGene_gn=PLEC;refGene_tx=XM_023248194.1;refGene_gn=PLEC;refGene_tx=XM_023248195.1;refGene_gn=PLEC;refGene_tx=XM_023248196.1
F2 84930186 84930306
refGene_gn=PLEC;refGene_tx=XM_023248166.1;refGene_gn=PLEC;refGene_tx=XM_023248167.1;refGene_gn=PLEC;refGene_tx=XM_023248168.1;refGene_gn=PLEC;refGene_tx=XM_023248169.1;refGene_gn=PLEC;refGene_tx=XM_023248170.1;refGene_gn=PLEC;refGene_tx=XM_023248171.1;refGene_gn=PLEC;refGene_tx=XM_023248172.1;refGene_gn=PLEC;refGene_tx=XM_023248174.1;refGene_gn=PLEC;refGene_tx=XM_023248175.1;refGene_gn=PLEC;refGene_tx=XM_023248176.1;refGene_gn=PLEC;refGene_tx=XM_023248177.1;refGene_gn=PLEC;refGene_tx=XM_023248178.1;refGene_gn=PLEC;refGene_tx=XM_023248179.1;refGene_gn=PLEC;refGene_tx=XM_023248180.1;refGene_gn=PLEC;refGene_tx=XM_023248181.1;refGene_gn=PLEC;refGene_tx=XM_023248182.1;refGene_gn=PLEC;refGene_tx=XM_023248183.1;refGene_gn=PLEC;refGene_tx=XM_023248184.1;refGene_gn=PLEC;refGene_tx=XM_023248185.1;refGene_gn=PLEC;refGene_tx=XM_023248186.1;refGene_gn=PLEC;refGene_tx=XM_023248187.1;refGene_gn=PLEC;refGene_tx=XM_023248188.1;refGene_gn=PLEC;refGene_tx=XM_023248190.1;refGene_gn=PLEC;refGene_tx=XM_023248191.1;refGene_gn=PLEC;refGene_tx=XM_023248192.1;refGene_gn=PLEC;refGene_tx=XM_023248193.1;refGene_gn=PLEC;refGene_tx=XM_023248194.1;refGene_gn=PLEC;refGene_tx=XM_023248195.1;refGene_gn=PLEC;refGene_tx=XM_023248196.1
F2 84930385 84930492
refGene_gn=PLEC;refGene_tx=XM_023248166.1;refGene_gn=PLEC;refGene_tx=XM_023248167.1;refGene_gn=PLEC;refGene_tx=XM_023248168.1;refGene_gn=PLEC;refGene_tx=XM_023248169.1;refGene_gn=PLEC;refGene_tx=XM_023248170.1;refGene_gn=PLEC;refGene_tx=XM_023248171.1;refGene_gn=PLEC;refGene_tx=XM_023248172.1;refGene_gn=PLEC;refGene_tx=XM_023248174.1;refGene_gn=PLEC;refGene_tx=XM_023248175.1;refGene_gn=PLEC;refGene_tx=XM_023248176.1;refGene_gn=PLEC;refGene_tx=XM_023248177.1;refGene_gn=PLEC;refGene_tx=XM_023248178.1;refGene_gn=PLEC;refGene_tx=XM_023248179.1;refGene_gn=PLEC;refGene_tx=XM_023248180.1;refGene_gn=PLEC;refGene_tx=XM_023248181.1;refGene_gn=PLEC;refGene_tx=XM_023248182.1;refGene_gn=PLEC;refGene_tx=XM_023248183.1;refGene_gn=PLEC;refGene_tx=XM_023248184.1;refGene_gn=PLEC;refGene_tx=XM_023248185.1;refGene_gn=PLEC;refGene_tx=XM_023248186.1;refGene_gn=PLEC;refGene_tx=XM_023248187.1;refGene_gn=PLEC;refGene_tx=XM_023248188.1;refGene_gn=PLEC;refGene_tx=XM_023248190.1;refGene_gn=PLEC;refGene_tx=XM_023248191.1;refGene_gn=PLEC;refGene_tx=XM_023248192.1;refGene_gn=PLEC;refGene_tx=XM_023248193.1;refGene_gn=PLEC;refGene_tx=XM_023248194.1;refGene_gn=PLEC;refGene_tx=XM_023248195.1;refGene_gn=PLEC;refGene_tx=XM_023248196.1
F2 84930572 84930688

refGene_gn=PLEC;refGene_tx=XM_023248166.1;refGene_gn=PLEC;refGene_tx=X
M_023248167.1;refGene_gn=PLEC;refGene_tx=XM_023248168.1;refGene_gn=PLE
C;refGene_tx=XM_023248169.1;refGene_gn=PLEC;refGene_tx=XM_023248170.1;
refGene_gn=PLEC;refGene_tx=XM_023248171.1;refGene_gn=PLEC;refGene_tx=X
M_023248172.1;refGene_gn=PLEC;refGene_tx=XM_023248174.1;refGene_gn=PLE
C;refGene_tx=XM_023248175.1;refGene_gn=PLEC;refGene_tx=XM_023248176.1;
refGene_gn=PLEC;refGene_tx=XM_023248177.1;refGene_gn=PLEC;refGene_tx=X
M_023248178.1;refGene_gn=PLEC;refGene_tx=XM_023248179.1;refGene_gn=PLE
C;refGene_tx=XM_023248180.1;refGene_gn=PLEC;refGene_tx=XM_023248181.1;
refGene_gn=PLEC;refGene_tx=XM_023248182.1;refGene_gn=PLEC;refGene_tx=X
M_023248183.1;refGene_gn=PLEC;refGene_tx=XM_023248184.1;refGene_gn=PLE
C;refGene_tx=XM_023248185.1;refGene_gn=PLEC;refGene_tx=XM_023248186.1;
refGene_gn=PLEC;refGene_tx=XM_023248187.1;refGene_gn=PLEC;refGene_tx=X
M_023248188.1;refGene_gn=PLEC;refGene_tx=XM_023248190.1;refGene_gn=PLE
C;refGene_tx=XM_023248191.1;refGene_gn=PLEC;refGene_tx=XM_023248192.1;
refGene_gn=PLEC;refGene_tx=XM_023248193.1;refGene_gn=PLEC;refGene_tx=X
M_023248194.1;refGene_gn=PLEC;refGene_tx=XM_023248195.1;refGene_gn=PLE
C;refGene_tx=XM_023248196.1
F2 84931379 84931546
refGene_gn=PLEC;refGene_tx=XM_023248166.1;refGene_gn=PLEC;refGene_tx=X
M_023248167.1;refGene_gn=PLEC;refGene_tx=XM_023248168.1;refGene_gn=PLE
C;refGene_tx=XM_023248169.1;refGene_gn=PLEC;refGene_tx=XM_023248170.1;
refGene_gn=PLEC;refGene_tx=XM_023248171.1;refGene_gn=PLEC;refGene_tx=X
M_023248172.1;refGene_gn=PLEC;refGene_tx=XM_023248174.1;refGene_gn=PLE
C;refGene_tx=XM_023248175.1;refGene_gn=PLEC;refGene_tx=XM_023248176.1;
refGene_gn=PLEC;refGene_tx=XM_023248177.1;refGene_gn=PLEC;refGene_tx=X
M_023248178.1;refGene_gn=PLEC;refGene_tx=XM_023248179.1;refGene_gn=PLE
C;refGene_tx=XM_023248180.1;refGene_gn=PLEC;refGene_tx=XM_023248181.1;
refGene_gn=PLEC;refGene_tx=XM_023248182.1;refGene_gn=PLEC;refGene_tx=X
M_023248183.1;refGene_gn=PLEC;refGene_tx=XM_023248184.1;refGene_gn=PLE
C;refGene_tx=XM_023248185.1;refGene_gn=PLEC;refGene_tx=XM_023248186.1;
refGene_gn=PLEC;refGene_tx=XM_023248187.1;refGene_gn=PLEC;refGene_tx=X
M_023248188.1;refGene_gn=PLEC;refGene_tx=XM_023248190.1;refGene_gn=PLE
C;refGene_tx=XM_023248191.1;refGene_gn=PLEC;refGene_tx=XM_023248192.1;
refGene_gn=PLEC;refGene_tx=XM_023248193.1;refGene_gn=PLEC;refGene_tx=X
M_023248194.1;refGene_gn=PLEC;refGene_tx=XM_023248195.1;refGene_gn=PLE
C;refGene_tx=XM_023248196.1
F2 84932432 84932525
refGene_gn=PLEC;refGene_tx=XM_023248166.1;refGene_gn=PLEC;refGene_tx=X
M_023248167.1;refGene_gn=PLEC;refGene_tx=XM_023248168.1;refGene_gn=PLE
C;refGene_tx=XM_023248169.1;refGene_gn=PLEC;refGene_tx=XM_023248170.1;
refGene_gn=PLEC;refGene_tx=XM_023248171.1;refGene_gn=PLEC;refGene_tx=X
M_023248172.1;refGene_gn=PLEC;refGene_tx=XM_023248174.1;refGene_gn=PLE
C;refGene_tx=XM_023248175.1;refGene_gn=PLEC;refGene_tx=XM_023248176.1;
refGene_gn=PLEC;refGene_tx=XM_023248177.1;refGene_gn=PLEC;refGene_tx=X
M_023248178.1;refGene_gn=PLEC;refGene_tx=XM_023248179.1;refGene_gn=PLE
C;refGene_tx=XM_023248180.1;refGene_gn=PLEC;refGene_tx=XM_023248181.1;
refGene_gn=PLEC;refGene_tx=XM_023248182.1;refGene_gn=PLEC;refGene_tx=X
M_023248183.1;refGene_gn=PLEC;refGene_tx=XM_023248184.1;refGene_gn=PLE
C;refGene_tx=XM_023248185.1;refGene_gn=PLEC;refGene_tx=XM_023248186.1;

refGene_gn=PLEC;refGene_tx=XM_023248187.1;refGene_gn=PLEC;refGene_tx=XM_023248188.1;refGene_gn=PLEC;refGene_tx=XM_023248190.1;refGene_gn=PLEC;refGene_tx=XM_023248191.1;refGene_gn=PLEC;refGene_tx=XM_023248192.1;refGene_gn=PLEC;refGene_tx=XM_023248193.1;refGene_gn=PLEC;refGene_tx=XM_023248194.1;refGene_gn=PLEC;refGene_tx=XM_023248195.1;refGene_gn=PLEC;refGene_tx=XM_023248196.1
F2 84932611 84932689
refGene_gn=PLEC;refGene_tx=XM_023248166.1;refGene_gn=PLEC;refGene_tx=XM_023248167.1;refGene_gn=PLEC;refGene_tx=XM_023248168.1;refGene_gn=PLEC;refGene_tx=XM_023248169.1;refGene_gn=PLEC;refGene_tx=XM_023248170.1;refGene_gn=PLEC;refGene_tx=XM_023248171.1;refGene_gn=PLEC;refGene_tx=XM_023248172.1;refGene_gn=PLEC;refGene_tx=XM_023248174.1;refGene_gn=PLEC;refGene_tx=XM_023248175.1;refGene_gn=PLEC;refGene_tx=XM_023248176.1;refGene_gn=PLEC;refGene_tx=XM_023248177.1;refGene_gn=PLEC;refGene_tx=XM_023248178.1;refGene_gn=PLEC;refGene_tx=XM_023248179.1;refGene_gn=PLEC;refGene_tx=XM_023248180.1;refGene_gn=PLEC;refGene_tx=XM_023248181.1;refGene_gn=PLEC;refGene_tx=XM_023248182.1;refGene_gn=PLEC;refGene_tx=XM_023248183.1;refGene_gn=PLEC;refGene_tx=XM_023248184.1;refGene_gn=PLEC;refGene_tx=XM_023248185.1;refGene_gn=PLEC;refGene_tx=XM_023248186.1;refGene_gn=PLEC;refGene_tx=XM_023248187.1;refGene_gn=PLEC;refGene_tx=XM_023248188.1;refGene_gn=PLEC;refGene_tx=XM_023248190.1;refGene_gn=PLEC;refGene_tx=XM_023248191.1;refGene_gn=PLEC;refGene_tx=XM_023248192.1;refGene_gn=PLEC;refGene_tx=XM_023248193.1;refGene_gn=PLEC;refGene_tx=XM_023248194.1;refGene_gn=PLEC;refGene_tx=XM_023248195.1;refGene_gn=PLEC;refGene_tx=XM_023248196.1
F2 84933167 84933203
refGene_gn=PLEC;refGene_tx=XM_023248166.1;refGene_gn=PLEC;refGene_tx=XM_023248167.1;refGene_gn=PLEC;refGene_tx=XM_023248170.1;refGene_gn=PLEC;refGene_tx=XM_023248171.1;refGene_gn=PLEC;refGene_tx=XM_023248174.1;refGene_gn=PLEC;refGene_tx=XM_023248175.1;refGene_gn=PLEC;refGene_tx=XM_023248176.1;refGene_gn=PLEC;refGene_tx=XM_023248177.1;refGene_gn=PLEC;refGene_tx=XM_023248178.1;refGene_gn=PLEC;refGene_tx=XM_023248179.1;refGene_gn=PLEC;refGene_tx=XM_023248180.1;refGene_gn=PLEC;refGene_tx=XM_023248182.1;refGene_gn=PLEC;refGene_tx=XM_023248184.1;refGene_gn=PLEC;refGene_tx=XM_023248185.1;refGene_gn=PLEC;refGene_tx=XM_023248188.1;refGene_gn=PLEC;refGene_tx=XM_023248190.1;refGene_gn=PLEC;refGene_tx=XM_023248192.1;refGene_gn=PLEC;refGene_tx=XM_023248193.1;refGene_gn=PLEC;refGene_tx=XM_023248194.1;refGene_gn=PLEC;refGene_tx=XM_023248195.1
F2 84933512 84933602
refGene_gn=PLEC;refGene_tx=XM_023248166.1;refGene_gn=PLEC;refGene_tx=XM_023248167.1;refGene_gn=PLEC;refGene_tx=XM_023248168.1;refGene_gn=PLEC;refGene_tx=XM_023248169.1;refGene_gn=PLEC;refGene_tx=XM_023248170.1;refGene_gn=PLEC;refGene_tx=XM_023248171.1;refGene_gn=PLEC;refGene_tx=XM_023248172.1;refGene_gn=PLEC;refGene_tx=XM_023248174.1;refGene_gn=PLEC;refGene_tx=XM_023248175.1;refGene_gn=PLEC;refGene_tx=XM_023248176.1;refGene_gn=PLEC;refGene_tx=XM_023248177.1;refGene_gn=PLEC;refGene_tx=XM_023248178.1;refGene_gn=PLEC;refGene_tx=XM_023248179.1;refGene_gn=PLEC;refGene_tx=XM_023248180.1;refGene_gn=PLEC;refGene_tx=XM_023248181.1;refGene_gn=PLEC;refGene_tx=XM_023248182.1;refGene_gn=PLEC;refGene_tx=XM_023248183.1;refGene_gn=PLEC;refGene_tx=XM_023248184.1;refGene_gn=PLE

C;refGene_tx=XM_023248185.1;refGene_gn=PLEC;refGene_tx=XM_023248186.1;
refGene_gn=PLEC;refGene_tx=XM_023248187.1;refGene_gn=PLEC;refGene_tx=X
M_023248188.1;refGene_gn=PLEC;refGene_tx=XM_023248190.1;refGene_gn=PLE
C;refGene_tx=XM_023248191.1;refGene_gn=PLEC;refGene_tx=XM_023248192.1;
refGene_gn=PLEC;refGene_tx=XM_023248193.1;refGene_gn=PLEC;refGene_tx=X
M_023248194.1;refGene_gn=PLEC;refGene_tx=XM_023248195.1;refGene_gn=PLE
C;refGene_tx=XM_023248196.1
F2 84933718 84933733
refGene_gn=PLEC;refGene_tx=XM_023248166.1;refGene_gn=PLEC;refGene_tx=X
M_023248168.1;refGene_gn=PLEC;refGene_tx=XM_023248171.1;refGene_gn=PLE
C;refGene_tx=XM_023248174.1;refGene_gn=PLEC;refGene_tx=XM_023248176.1;
refGene_gn=PLEC;refGene_tx=XM_023248177.1;refGene_gn=PLEC;refGene_tx=X
M_023248180.1;refGene_gn=PLEC;refGene_tx=XM_023248182.1;refGene_gn=PLE
C;refGene_tx=XM_023248183.1;refGene_gn=PLEC;refGene_tx=XM_023248188.1;
refGene_gn=PLEC;refGene_tx=XM_023248190.1;refGene_gn=PLEC;refGene_tx=X
M_023248194.1
F2 84934009 84934071
refGene_gn=PLEC;refGene_tx=XM_023248166.1;refGene_gn=PLEC;refGene_tx=X
M_023248167.1;refGene_gn=PLEC;refGene_tx=XM_023248168.1;refGene_gn=PLE
C;refGene_tx=XM_023248169.1;refGene_gn=PLEC;refGene_tx=XM_023248170.1;
refGene_gn=PLEC;refGene_tx=XM_023248171.1;refGene_gn=PLEC;refGene_tx=X
M_023248172.1;refGene_gn=PLEC;refGene_tx=XM_023248174.1;refGene_gn=PLE
C;refGene_tx=XM_023248175.1;refGene_gn=PLEC;refGene_tx=XM_023248176.1;
refGene_gn=PLEC;refGene_tx=XM_023248177.1;refGene_gn=PLEC;refGene_tx=X
M_023248178.1;refGene_gn=PLEC;refGene_tx=XM_023248179.1;refGene_gn=PLE
C;refGene_tx=XM_023248180.1;refGene_gn=PLEC;refGene_tx=XM_023248181.1;
refGene_gn=PLEC;refGene_tx=XM_023248182.1;refGene_gn=PLEC;refGene_tx=X
M_023248183.1;refGene_gn=PLEC;refGene_tx=XM_023248184.1;refGene_gn=PLE
C;refGene_tx=XM_023248185.1;refGene_gn=PLEC;refGene_tx=XM_023248186.1;
refGene_gn=PLEC;refGene_tx=XM_023248187.1;refGene_gn=PLEC;refGene_tx=X
M_023248188.1;refGene_gn=PLEC;refGene_tx=XM_023248190.1;refGene_gn=PLE
C;refGene_tx=XM_023248191.1;refGene_gn=PLEC;refGene_tx=XM_023248192.1;
refGene_gn=PLEC;refGene_tx=XM_023248193.1;refGene_gn=PLEC;refGene_tx=X
M_023248194.1;refGene_gn=PLEC;refGene_tx=XM_023248195.1;refGene_gn=PLE
C;refGene_tx=XM_023248196.1
F2 84934812 84934924
refGene_gn=PLEC;refGene_tx=XM_023248176.1;refGene_gn=PLEC;refGene_tx=X
M_023248178.1;refGene_gn=PLEC;refGene_tx=XM_023248186.1
F2 84935416 84935507
refGene_gn=PLEC;refGene_tx=XM_023248180.1;refGene_gn=PLEC;refGene_tx=X
M_023248184.1
F2 84937602 84937741
refGene_gn=PLEC;refGene_tx=XM_023248174.1;refGene_gn=PLEC;refGene_tx=X
M_023248175.1;refGene_gn=PLEC;refGene_tx=XM_023248181.1
F2 84939257 84939369
refGene_gn=PLEC;refGene_tx=XM_023248177.1;refGene_gn=PLEC;refGene_tx=X
M_023248179.1;refGene_gn=PLEC;refGene_tx=XM_023248183.1;refGene_gn=PLE
C;refGene_tx=XM_023248187.1
F2 84940141 84940157
refGene_gn=PLEC;refGene_tx=XM_023248194.1;refGene_gn=PLEC;refGene_tx=X

M_023248195.1;refGene_gn=PLEC;refGene_tx=XM_023248196.1
F2 84941748 84941791
refGene_gn=PLEC;refGene_tx=XM_023248190.1;refGene_gn=PLEC;refGene_tx=XM_023248193.1
F2 84944970 84945508
refGene_gn=PLEC;refGene_tx=XM_023248166.1;refGene_gn=PLEC;refGene_tx=XM_023248167.1;refGene_gn=PLEC;refGene_tx=XM_023248168.1;refGene_gn=PLEC;refGene_tx=XM_023248169.1
F2 84948596 84948642
refGene_gn=PLEC;refGene_tx=XM_023248188.1;refGene_gn=PLEC;refGene_tx=XM_023248192.1
F2 84963176 84963261
refGene_gn=PLEC;refGene_tx=XM_023248182.1;refGene_gn=PLEC;refGene_tx=XM_023248185.1;refGene_gn=PLEC;refGene_tx=XM_023248191.1
F2 84964824 84965017
refGene_gn=PLEC;refGene_tx=XM_023248170.1;refGene_gn=PLEC;refGene_tx=XM_023248171.1;refGene_gn=PLEC;refGene_tx=XM_023248172.1
F2 84966182 84967081 refGene_gn=PARP10;refGene_tx=XM_023248350.1
F2 84967083 84967152 refGene_gn=PARP10;refGene_tx=XM_023248350.1
F2 84967154 84967290 refGene_gn=PARP10;refGene_tx=XM_023248350.1
F2 84969714 84969782 refGene_gn=PARP10;refGene_tx=XM_023248350.1
F2 84970574 84970784 refGene_gn=PARP10;refGene_tx=XM_023248350.1
F2 84970786 84971061 refGene_gn=PARP10;refGene_tx=XM_023248350.1
F2 84971260 84971455 refGene_gn=PARP10;refGene_tx=XM_023248350.1
F2 84971456 84971581 refGene_gn=PARP10;refGene_tx=XM_023248350.1
F2 84971694 84971826 refGene_gn=PARP10;refGene_tx=XM_023248350.1
F2 84971828 84971995 refGene_gn=PARP10;refGene_tx=XM_023248350.1
F2 84971997 84972071 refGene_gn=PARP10;refGene_tx=XM_023248350.1
F2 84972072 84972310 refGene_gn=PARP10;refGene_tx=XM_023248350.1
F2 84972386 84973071 refGene_gn=PARP10;refGene_tx=XM_023248350.1
F2 84977183 84977517 refGene_gn=GRINA;refGene_tx=XM_011291510.3
F2 84977602 84977715 refGene_gn=GRINA;refGene_tx=XM_011291510.3
F2 84977783 84977984 refGene_gn=GRINA;refGene_tx=XM_011291510.3
F2 84978101 84978230 refGene_gn=GRINA;refGene_tx=XM_011291510.3
F2 84978397 84978460 refGene_gn=GRINA;refGene_tx=XM_011291510.3
F2 84978461 84978542 refGene_gn=GRINA;refGene_tx=XM_011291510.3
F2 84978642 84978792 refGene_gn=GRINA;refGene_tx=XM_011291510.3
F2 84996652 84996860 refGene_gn=SPATC1;refGene_tx=XM_023248727.1
F2 85010371 85011100 refGene_gn=SPATC1;refGene_tx=XM_023248727.1
F2 85011214 85011715 refGene_gn=SPATC1;refGene_tx=XM_023248727.1
F2 85011832 85011912 refGene_gn=SPATC1;refGene_tx=XM_023248727.1
F2 85011913 85011973 refGene_gn=SPATC1;refGene_tx=XM_023248727.1
F2 85018269 85018599 refGene_gn=SPATC1;refGene_tx=XM_023248727.1
F2 85021030 85021858 refGene_gn=SMPD5;refGene_tx=XM_006943509.4
F2 85021953 85022026 refGene_gn=SMPD5;refGene_tx=XM_006943509.4
F2 85022184 85022325 refGene_gn=SMPD5;refGene_tx=XM_006943509.4
F2 85022424 85022514 refGene_gn=SMPD5;refGene_tx=XM_006943509.4
F2 85022604 85022668 refGene_gn=SMPD5;refGene_tx=XM_006943509.4
F2 85022745 85022881 refGene_gn=SMPD5;refGene_tx=XM_006943509.4
F2 85022953 85023046 refGene_gn=SMPD5;refGene_tx=XM_006943509.4

F2 85023157 85023304
refGene_gn=OPLAH;refGene_tx=XM_004000167.4;refGene_gn=OPLAH;refGene_tx=XM_006943458.4;refGene_gn=OPLAH;refGene_tx=XM_023248726.1
F2 85023557 85023655
refGene_gn=OPLAH;refGene_tx=XM_004000167.4;refGene_gn=OPLAH;refGene_tx=XM_006943458.4;refGene_gn=OPLAH;refGene_tx=XM_023248726.1
F2 85023847 85024008
refGene_gn=OPLAH;refGene_tx=XM_004000167.4;refGene_gn=OPLAH;refGene_tx=XM_006943458.4;refGene_gn=OPLAH;refGene_tx=XM_023248726.1
F2 85024089 85024247
refGene_gn=OPLAH;refGene_tx=XM_004000167.4;refGene_gn=OPLAH;refGene_tx=XM_006943458.4;refGene_gn=OPLAH;refGene_tx=XM_023248726.1
F2 85024338 85024488
refGene_gn=OPLAH;refGene_tx=XM_004000167.4;refGene_gn=OPLAH;refGene_tx=XM_006943458.4;refGene_gn=OPLAH;refGene_tx=XM_023248726.1
F2 85024628 85024763
refGene_gn=OPLAH;refGene_tx=XM_004000167.4;refGene_gn=OPLAH;refGene_tx=XM_006943458.4;refGene_gn=OPLAH;refGene_tx=XM_023248726.1
F2 85024857 85025004
refGene_gn=OPLAH;refGene_tx=XM_004000167.4;refGene_gn=OPLAH;refGene_tx=XM_006943458.4;refGene_gn=OPLAH;refGene_tx=XM_023248726.1
F2 85025129 85025314
refGene_gn=OPLAH;refGene_tx=XM_004000167.4;refGene_gn=OPLAH;refGene_tx=XM_006943458.4;refGene_gn=OPLAH;refGene_tx=XM_023248726.1
F2 85026845 85027020
refGene_gn=OPLAH;refGene_tx=XM_004000167.4;refGene_gn=OPLAH;refGene_tx=XM_006943458.4;refGene_gn=OPLAH;refGene_tx=XM_023248726.1
F2 85027102 85027204
refGene_gn=OPLAH;refGene_tx=XM_004000167.4;refGene_gn=OPLAH;refGene_tx=XM_006943458.4;refGene_gn=OPLAH;refGene_tx=XM_023248726.1
F2 85027287 85027448
refGene_gn=OPLAH;refGene_tx=XM_004000167.4;refGene_gn=OPLAH;refGene_tx=XM_006943458.4;refGene_gn=OPLAH;refGene_tx=XM_023248726.1
F2 85028112 85028264
refGene_gn=OPLAH;refGene_tx=XM_004000167.4;refGene_gn=OPLAH;refGene_tx=XM_006943458.4;refGene_gn=OPLAH;refGene_tx=XM_023248726.1
F2 85028492 85028605
refGene_gn=OPLAH;refGene_tx=XM_004000167.4;refGene_gn=OPLAH;refGene_tx=XM_006943458.4;refGene_gn=OPLAH;refGene_tx=XM_023248726.1
F2 85028725 85028864
refGene_gn=OPLAH;refGene_tx=XM_004000167.4;refGene_gn=OPLAH;refGene_tx=XM_006943458.4;refGene_gn=OPLAH;refGene_tx=XM_023248726.1
F2 85028950 85029088
refGene_gn=OPLAH;refGene_tx=XM_004000167.4;refGene_gn=OPLAH;refGene_tx=XM_006943458.4;refGene_gn=OPLAH;refGene_tx=XM_023248726.1
F2 85029275 85029446
refGene_gn=OPLAH;refGene_tx=XM_004000167.4;refGene_gn=OPLAH;refGene_tx=XM_006943458.4;refGene_gn=OPLAH;refGene_tx=XM_023248726.1
F2 85029533 85029646
refGene_gn=OPLAH;refGene_tx=XM_004000167.4;refGene_gn=OPLAH;refGene_tx

=XM_006943458.4;refGene_gn=OPLAH;refGene_tx=XM_023248726.1
F2 85029783 85030049
refGene_gn=OPLAH;refGene_tx=XM_004000167.4;refGene_gn=OPLAH;refGene_tx
=XM_006943458.4;refGene_gn=OPLAH;refGene_tx=XM_023248726.1
F2 85030188 85030256
refGene_gn=OPLAH;refGene_tx=XM_004000167.4;refGene_gn=OPLAH;refGene_tx
=XM_006943458.4;refGene_gn=OPLAH;refGene_tx=XM_023248726.1
F2 85030351 85030490
refGene_gn=OPLAH;refGene_tx=XM_004000167.4;refGene_gn=OPLAH;refGene_tx
=XM_006943458.4;refGene_gn=OPLAH;refGene_tx=XM_023248726.1
F2 85030603 85030769
refGene_gn=OPLAH;refGene_tx=XM_004000167.4;refGene_gn=OPLAH;refGene_tx
=XM_006943458.4;refGene_gn=OPLAH;refGene_tx=XM_023248726.1
F2 85030847 85031043
refGene_gn=OPLAH;refGene_tx=XM_004000167.4;refGene_gn=OPLAH;refGene_tx
=XM_006943458.4;refGene_gn=OPLAH;refGene_tx=XM_023248726.1
F2 85031126 85031250
refGene_gn=OPLAH;refGene_tx=XM_004000167.4;refGene_gn=OPLAH;refGene_tx
=XM_006943458.4;refGene_gn=OPLAH;refGene_tx=XM_023248726.1
F2 85031326 85031426
refGene_gn=OPLAH;refGene_tx=XM_004000167.4;refGene_gn=OPLAH;refGene_tx
=XM_006943458.4;refGene_gn=OPLAH;refGene_tx=XM_023248726.1
F2 85031989 85032181
refGene_gn=OPLAH;refGene_tx=XM_004000167.4;refGene_gn=OPLAH;refGene_tx
=XM_006943458.4;refGene_gn=OPLAH;refGene_tx=XM_023248726.1
F2 85032259 85032483
refGene_gn=OPLAH;refGene_tx=XM_004000167.4;refGene_gn=OPLAH;refGene_tx
=XM_006943458.4;refGene_gn=OPLAH;refGene_tx=XM_023248726.1
F2 85036070 85036374 refGene_gn=OPLAH;refGene_tx=XM_006943458.4
F2 85051171 85051342 refGene_gn=EXOSC4;refGene_tx=XM_004000168.5
F2 85052540 85052747 refGene_gn=EXOSC4;refGene_tx=XM_004000168.5
F2 85052859 85053219 refGene_gn=EXOSC4;refGene_tx=XM_004000168.5
F2 85054446 85054520 refGene_gn=GPAA1;refGene_tx=XM_023248272.1
F2 85054786 85054966 refGene_gn=GPAA1;refGene_tx=XM_023248272.1
F2 85055048 85055160 refGene_gn=GPAA1;refGene_tx=XM_023248272.1
F2 85055352 85055500 refGene_gn=GPAA1;refGene_tx=XM_023248272.1
F2 85055579 85055681 refGene_gn=GPAA1;refGene_tx=XM_023248272.1
F2 85055769 85055966 refGene_gn=GPAA1;refGene_tx=XM_023248272.1
F2 85056040 85056237 refGene_gn=GPAA1;refGene_tx=XM_023248272.1
F2 85056331 85056485 refGene_gn=GPAA1;refGene_tx=XM_023248272.1
F2 85056634 85056730 refGene_gn=GPAA1;refGene_tx=XM_023248272.1
F2 85056883 85057074 refGene_gn=GPAA1;refGene_tx=XM_023248272.1
F2 85057155 85057326 refGene_gn=GPAA1;refGene_tx=XM_023248272.1
F2 85057437 85057681 refGene_gn=GPAA1;refGene_tx=XM_023248272.1
F2 85065061 85065190
refGene_gn=LOC101094239;refGene_tx=XM_004000215.3
F2 85065799 85065996
refGene_gn=LOC101094239;refGene_tx=XM_004000215.3
F2 85066081 85066208
refGene_gn=LOC101094239;refGene_tx=XM_004000215.3

F2 85066295 85066453
refGene_gn=LOC101094239;refGene_tx=XM_004000215.3
F2 85066541 85066702
refGene_gn=LOC101094239;refGene_tx=XM_004000215.3
F2 85067013 85067114
refGene_gn=LOC101094239;refGene_tx=XM_004000215.3
F2 85067199 85067304
refGene_gn=LOC101094239;refGene_tx=XM_004000215.3
F2 85068359 85068443
refGene_gn=SHARPIN;refGene_tx=XM_023248705.1;refGene_gn=SHARPIN;refGene_tx=XM_023248706.1;refGene_gn=SHARPIN;refGene_tx=XM_023248707.1;refGene_gn=SHARPIN;refGene_tx=XM_023248708.1;refGene_gn=SHARPIN;refGene_tx=XM_023248709.1
F2 85068522 85068648
refGene_gn=SHARPIN;refGene_tx=XM_023248705.1;refGene_gn=SHARPIN;refGene_tx=XM_023248706.1;refGene_gn=SHARPIN;refGene_tx=XM_023248707.1;refGene_gn=SHARPIN;refGene_tx=XM_023248708.1;refGene_gn=SHARPIN;refGene_tx=XM_023248709.1
F2 85068737 85068856
refGene_gn=SHARPIN;refGene_tx=XM_023248705.1;refGene_gn=SHARPIN;refGene_tx=XM_023248706.1;refGene_gn=SHARPIN;refGene_tx=XM_023248707.1;refGene_gn=SHARPIN;refGene_tx=XM_023248708.1;refGene_gn=SHARPIN;refGene_tx=XM_023248709.1
F2 85068934 85069088
refGene_gn=SHARPIN;refGene_tx=XM_023248705.1;refGene_gn=SHARPIN;refGene_tx=XM_023248706.1;refGene_gn=SHARPIN;refGene_tx=XM_023248707.1;refGene_gn=SHARPIN;refGene_tx=XM_023248708.1;refGene_gn=SHARPIN;refGene_tx=XM_023248709.1
F2 85069165 85069256
refGene_gn=SHARPIN;refGene_tx=XM_023248705.1;refGene_gn=SHARPIN;refGene_tx=XM_023248706.1;refGene_gn=SHARPIN;refGene_tx=XM_023248707.1;refGene_gn=SHARPIN;refGene_tx=XM_023248708.1;refGene_gn=SHARPIN;refGene_tx=XM_023248709.1
F2 85069331 85069473
refGene_gn=SHARPIN;refGene_tx=XM_023248705.1;refGene_gn=SHARPIN;refGene_tx=XM_023248706.1;refGene_gn=SHARPIN;refGene_tx=XM_023248707.1;refGene_gn=SHARPIN;refGene_tx=XM_023248708.1;refGene_gn=SHARPIN;refGene_tx=XM_023248709.1
F2 85069550 85069682
refGene_gn=SHARPIN;refGene_tx=XM_023248705.1;refGene_gn=SHARPIN;refGene_tx=XM_023248706.1;refGene_gn=SHARPIN;refGene_tx=XM_023248708.1
F2 85070909 85071186
refGene_gn=SHARPIN;refGene_tx=XM_023248706.1;refGene_gn=SHARPIN;refGene_tx=XM_023248709.1
F2 85072145 85072320
refGene_gn=SHARPIN;refGene_tx=XM_023248705.1;refGene_gn=SHARPIN;refGene_tx=XM_023248707.1
F2 85072601 85072799
refGene_gn=SHARPIN;refGene_tx=XM_023248705.1;refGene_gn=SHARPIN;refGene_tx=XM_023248707.1;refGene_gn=SHARPIN;refGene_tx=XM_023248708.1

F2 85074529 85074612 refGene_gn=MAF1;refGene_tx=XM_004000169.5
F2 85074705 85074834 refGene_gn=MAF1;refGene_tx=XM_004000169.5
F2 85074911 85075077 refGene_gn=MAF1;refGene_tx=XM_004000169.5
F2 85075216 85075338 refGene_gn=MAF1;refGene_tx=XM_004000169.5
F2 85075420 85075540 refGene_gn=MAF1;refGene_tx=XM_004000169.5
F2 85075693 85075843 refGene_gn=MAF1;refGene_tx=XM_004000169.5
F2 85075968 85075990 refGene_gn=MAF1;refGene_tx=XM_004000169.5
F2 85076630 85076760
refGene_gn=WDR97;refGene_tx=XM_023248703.1;refGene_gn=WDR97;refGene_tx=XM_023248704.1
F2 85076961 85077089
refGene_gn=WDR97;refGene_tx=XM_023248703.1;refGene_gn=WDR97;refGene_tx=XM_023248704.1
F2 85077267 85077896
refGene_gn=WDR97;refGene_tx=XM_023248703.1;refGene_gn=WDR97;refGene_tx=XM_023248704.1
F2 85078035 85078157
refGene_gn=WDR97;refGene_tx=XM_023248703.1;refGene_gn=WDR97;refGene_tx=XM_023248704.1
F2 85078320 85079020
refGene_gn=WDR97;refGene_tx=XM_023248703.1;refGene_gn=WDR97;refGene_tx=XM_023248704.1
F2 85079090 85079233
refGene_gn=WDR97;refGene_tx=XM_023248703.1;refGene_gn=WDR97;refGene_tx=XM_023248704.1
F2 85079321 85079558
refGene_gn=WDR97;refGene_tx=XM_023248703.1;refGene_gn=WDR97;refGene_tx=XM_023248704.1
F2 85079633 85079724
refGene_gn=WDR97;refGene_tx=XM_023248703.1;refGene_gn=WDR97;refGene_tx=XM_023248704.1
F2 85079909 85080042
refGene_gn=WDR97;refGene_tx=XM_023248703.1;refGene_gn=WDR97;refGene_tx=XM_023248704.1
F2 85080138 85080260
refGene_gn=WDR97;refGene_tx=XM_023248703.1;refGene_gn=WDR97;refGene_tx=XM_023248704.1
F2 85080531 85080592
refGene_gn=WDR97;refGene_tx=XM_023248703.1;refGene_gn=WDR97;refGene_tx=XM_023248704.1
F2 85080783 85080933
refGene_gn=WDR97;refGene_tx=XM_023248703.1;refGene_gn=WDR97;refGene_tx=XM_023248704.1
F2 85081035 85081293
refGene_gn=WDR97;refGene_tx=XM_023248703.1;refGene_gn=WDR97;refGene_tx=XM_023248704.1
F2 85081375 85081501
refGene_gn=WDR97;refGene_tx=XM_023248703.1;refGene_gn=WDR97;refGene_tx=XM_023248704.1
F2 85081598 85081682

refGene_gn=WDR97;refGene_tx=XM_023248703.1;refGene_gn=WDR97;refGene_tx=XM_023248704.1
F2	85082982	85083111
refGene_gn=WDR97;refGene_tx=XM_023248703.1;refGene_gn=WDR97;refGene_tx=XM_023248704.1
F2	85083207	85083402
refGene_gn=WDR97;refGene_tx=XM_023248703.1;refGene_gn=WDR97;refGene_tx=XM_023248704.1
F2	85083481	85083958
refGene_gn=WDR97;refGene_tx=XM_023248703.1;refGene_gn=WDR97;refGene_tx=XM_023248704.1
F2	85084059	85084181
refGene_gn=WDR97;refGene_tx=XM_023248703.1;refGene_gn=WDR97;refGene_tx=XM_023248704.1
F2	85084260	85084423
refGene_gn=WDR97;refGene_tx=XM_023248703.1;refGene_gn=WDR97;refGene_tx=XM_023248704.1
F2	85084756	85085301
refGene_gn=WDR97;refGene_tx=XM_023248703.1;refGene_gn=WDR97;refGene_tx=XM_023248704.1
F2	85085386	85085448
refGene_gn=WDR97;refGene_tx=XM_023248703.1;refGene_gn=WDR97;refGene_tx=XM_023248704.1
F2	85085524	85085727
refGene_gn=WDR97;refGene_tx=XM_023248703.1;refGene_gn=WDR97;refGene_tx=XM_023248704.1
F2	85087922	85088548	refGene_gn=HGH1;refGene_tx=XM_023248402.1
F2	85088620	85088721	refGene_gn=HGH1;refGene_tx=XM_023248402.1
F2	85088822	85088921	refGene_gn=HGH1;refGene_tx=XM_023248402.1
F2	85089132	85089224	refGene_gn=HGH1;refGene_tx=XM_023248402.1
F2	85089316	85089439	refGene_gn=HGH1;refGene_tx=XM_023248402.1
F2	85089522	85089699	refGene_gn=HGH1;refGene_tx=XM_023248402.1
F2	85090992	85091384
refGene_gn=LOC101094985;refGene_tx=XM_006943456.3
F2	85091825	85091941
refGene_gn=LOC101094985;refGene_tx=XM_006943456.3
F2	85092029	85092159
refGene_gn=LOC101094985;refGene_tx=XM_006943456.3
F2	85092413	85092469
refGene_gn=LOC101094985;refGene_tx=XM_006943456.3
F2	85092567	85092770
refGene_gn=LOC101094985;refGene_tx=XM_006943456.3
F2	85092968	85093118
refGene_gn=LOC101094985;refGene_tx=XM_006943456.3
F2	85093399	85093558
refGene_gn=LOC101094985;refGene_tx=XM_006943456.3
F2	85108643	85108665
refGene_gn=MROH1;refGene_tx=XM_011291514.3;refGene_gn=MROH1;refGene_tx=XM_011291518.2;refGene_gn=MROH1;refGene_tx=XM_019823255.2;refGene_gn=MROH1;refGene_tx=XM_019823256.2;refGene_gn=MROH1;refGene_tx=XM_0198232

57.2;refGene_gn=MROH1;refGene_tx=XM_019823258.2;refGene_gn=MROH1;refGene_tx=XM_019823259.2;refGene_gn=MROH1;refGene_tx=XM_019823260.2;refGene_gn=MROH1;refGene_tx=XM_023248543.1;refGene_gn=MROH1;refGene_tx=XM_023248544.1;refGene_gn=MROH1;refGene_tx=XM_023248546.1
F2 85109663 85109809
refGene_gn=MROH1;refGene_tx=XM_011291514.3;refGene_gn=MROH1;refGene_tx=XM_011291518.2;refGene_gn=MROH1;refGene_tx=XM_019823255.2;refGene_gn=MROH1;refGene_tx=XM_019823256.2;refGene_gn=MROH1;refGene_tx=XM_019823257.2;refGene_gn=MROH1;refGene_tx=XM_019823258.2;refGene_gn=MROH1;refGene_tx=XM_019823259.2;refGene_gn=MROH1;refGene_tx=XM_019823260.2;refGene_gn=MROH1;refGene_tx=XM_023248543.1;refGene_gn=MROH1;refGene_tx=XM_023248544.1;refGene_gn=MROH1;refGene_tx=XM_023248546.1
F2 85114652 85114784
refGene_gn=MROH1;refGene_tx=XM_011291514.3;refGene_gn=MROH1;refGene_tx=XM_011291518.2;refGene_gn=MROH1;refGene_tx=XM_019823255.2;refGene_gn=MROH1;refGene_tx=XM_019823256.2;refGene_gn=MROH1;refGene_tx=XM_019823257.2;refGene_gn=MROH1;refGene_tx=XM_019823258.2;refGene_gn=MROH1;refGene_tx=XM_019823259.2;refGene_gn=MROH1;refGene_tx=XM_019823260.2;refGene_gn=MROH1;refGene_tx=XM_023248543.1;refGene_gn=MROH1;refGene_tx=XM_023248544.1;refGene_gn=MROH1;refGene_tx=XM_023248546.1
F2 85115301 85115464
refGene_gn=MROH1;refGene_tx=XM_011291514.3;refGene_gn=MROH1;refGene_tx=XM_011291518.2;refGene_gn=MROH1;refGene_tx=XM_019823255.2;refGene_gn=MROH1;refGene_tx=XM_019823256.2;refGene_gn=MROH1;refGene_tx=XM_019823257.2;refGene_gn=MROH1;refGene_tx=XM_019823258.2;refGene_gn=MROH1;refGene_tx=XM_019823259.2;refGene_gn=MROH1;refGene_tx=XM_019823260.2;refGene_gn=MROH1;refGene_tx=XM_023248543.1;refGene_gn=MROH1;refGene_tx=XM_023248544.1;refGene_gn=MROH1;refGene_tx=XM_023248546.1
F2 85115633 85115732
refGene_gn=MROH1;refGene_tx=XM_011291514.3;refGene_gn=MROH1;refGene_tx=XM_011291518.2;refGene_gn=MROH1;refGene_tx=XM_019823255.2;refGene_gn=MROH1;refGene_tx=XM_019823256.2;refGene_gn=MROH1;refGene_tx=XM_019823257.2;refGene_gn=MROH1;refGene_tx=XM_019823258.2;refGene_gn=MROH1;refGene_tx=XM_019823259.2;refGene_gn=MROH1;refGene_tx=XM_019823260.2;refGene_gn=MROH1;refGene_tx=XM_023248543.1;refGene_gn=MROH1;refGene_tx=XM_023248544.1;refGene_gn=MROH1;refGene_tx=XM_023248546.1
F2 85119873 85120025
refGene_gn=MROH1;refGene_tx=XM_011291514.3;refGene_gn=MROH1;refGene_tx=XM_011291518.2;refGene_gn=MROH1;refGene_tx=XM_019823255.2;refGene_gn=MROH1;refGene_tx=XM_019823256.2;refGene_gn=MROH1;refGene_tx=XM_019823257.2;refGene_gn=MROH1;refGene_tx=XM_019823258.2;refGene_gn=MROH1;refGene_tx=XM_019823259.2;refGene_gn=MROH1;refGene_tx=XM_019823260.2;refGene_gn=MROH1;refGene_tx=XM_023248543.1;refGene_gn=MROH1;refGene_tx=XM_023248544.1;refGene_gn=MROH1;refGene_tx=XM_023248546.1
F2 85120621 85120762
refGene_gn=MROH1;refGene_tx=XM_011291514.3;refGene_gn=MROH1;refGene_tx=XM_011291518.2;refGene_gn=MROH1;refGene_tx=XM_019823255.2;refGene_gn=MROH1;refGene_tx=XM_019823256.2;refGene_gn=MROH1;refGene_tx=XM_019823257.2;refGene_gn=MROH1;refGene_tx=XM_019823258.2;refGene_gn=MROH1;refGene_tx=XM_019823259.2;refGene_gn=MROH1;refGene_tx=XM_019823260.2;refGen

e_gn=MROH1;refGene_tx=XM_023248543.1;refGene_gn=MROH1;refGene_tx=XM_023248544.1;refGene_gn=MROH1;refGene_tx=XM_023248545.1;refGene_gn=MROH1;refGene_tx=XM_023248546.1
F2 85121128 85121221
refGene_gn=MROH1;refGene_tx=XM_011291514.3;refGene_gn=MROH1;refGene_tx=XM_011291518.2;refGene_gn=MROH1;refGene_tx=XM_019823255.2;refGene_gn=MROH1;refGene_tx=XM_019823256.2;refGene_gn=MROH1;refGene_tx=XM_019823257.2;refGene_gn=MROH1;refGene_tx=XM_019823258.2;refGene_gn=MROH1;refGene_tx=XM_019823259.2;refGene_gn=MROH1;refGene_tx=XM_019823260.2;refGene_gn=MROH1;refGene_tx=XM_023248543.1;refGene_gn=MROH1;refGene_tx=XM_023248544.1;refGene_gn=MROH1;refGene_tx=XM_023248545.1;refGene_gn=MROH1;refGene_tx=XM_023248546.1
F2 85123116 85123195
refGene_gn=MROH1;refGene_tx=XM_011291514.3;refGene_gn=MROH1;refGene_tx=XM_011291518.2;refGene_gn=MROH1;refGene_tx=XM_019823255.2;refGene_gn=MROH1;refGene_tx=XM_019823256.2;refGene_gn=MROH1;refGene_tx=XM_019823257.2;refGene_gn=MROH1;refGene_tx=XM_019823258.2;refGene_gn=MROH1;refGene_tx=XM_019823259.2;refGene_gn=MROH1;refGene_tx=XM_019823260.2;refGene_gn=MROH1;refGene_tx=XM_023248543.1;refGene_gn=MROH1;refGene_tx=XM_023248544.1;refGene_gn=MROH1;refGene_tx=XM_023248545.1;refGene_gn=MROH1;refGene_tx=XM_023248546.1
F2 85123430 85123544
refGene_gn=MROH1;refGene_tx=XM_011291514.3;refGene_gn=MROH1;refGene_tx=XM_011291518.2;refGene_gn=MROH1;refGene_tx=XM_019823255.2;refGene_gn=MROH1;refGene_tx=XM_019823256.2;refGene_gn=MROH1;refGene_tx=XM_019823257.2;refGene_gn=MROH1;refGene_tx=XM_019823258.2;refGene_gn=MROH1;refGene_tx=XM_019823259.2;refGene_gn=MROH1;refGene_tx=XM_019823260.2;refGene_gn=MROH1;refGene_tx=XM_023248543.1;refGene_gn=MROH1;refGene_tx=XM_023248544.1;refGene_gn=MROH1;refGene_tx=XM_023248545.1;refGene_gn=MROH1;refGene_tx=XM_023248546.1
F2 85133276 85133350
refGene_gn=MROH1;refGene_tx=XM_011291514.3;refGene_gn=MROH1;refGene_tx=XM_011291518.2;refGene_gn=MROH1;refGene_tx=XM_019823255.2;refGene_gn=MROH1;refGene_tx=XM_019823256.2;refGene_gn=MROH1;refGene_tx=XM_019823257.2;refGene_gn=MROH1;refGene_tx=XM_019823258.2;refGene_gn=MROH1;refGene_tx=XM_019823259.2;refGene_gn=MROH1;refGene_tx=XM_019823260.2;refGene_gn=MROH1;refGene_tx=XM_023248543.1;refGene_gn=MROH1;refGene_tx=XM_023248544.1;refGene_gn=MROH1;refGene_tx=XM_023248545.1
F2 85135027 85135150
refGene_gn=MROH1;refGene_tx=XM_011291514.3;refGene_gn=MROH1;refGene_tx=XM_011291518.2;refGene_gn=MROH1;refGene_tx=XM_019823255.2;refGene_gn=MROH1;refGene_tx=XM_019823256.2;refGene_gn=MROH1;refGene_tx=XM_019823257.2;refGene_gn=MROH1;refGene_tx=XM_019823258.2;refGene_gn=MROH1;refGene_tx=XM_019823259.2;refGene_gn=MROH1;refGene_tx=XM_019823260.2;refGene_gn=MROH1;refGene_tx=XM_023248543.1;refGene_gn=MROH1;refGene_tx=XM_023248544.1;refGene_gn=MROH1;refGene_tx=XM_023248545.1
F2 85136812 85136920
refGene_gn=MROH1;refGene_tx=XM_011291514.3;refGene_gn=MROH1;refGene_tx=XM_011291518.2;refGene_gn=MROH1;refGene_tx=XM_019823255.2;refGene_gn=MROH1;refGene_tx=XM_019823256.2;refGene_gn=MROH1;refGene_tx=XM_0198232

57.2;refGene_gn=MROH1;refGene_tx=XM_019823258.2;refGene_gn=MROH1;refGene_tx=XM_019823259.2;refGene_gn=MROH1;refGene_tx=XM_019823260.2;refGene_gn=MROH1;refGene_tx=XM_023248543.1;refGene_gn=MROH1;refGene_tx=XM_023248544.1;refGene_gn=MROH1;refGene_tx=XM_023248545.1;refGene_gn=MROH1;refGene_tx=XM_023248546.1
F2 85137123 85137268
refGene_gn=MROH1;refGene_tx=XM_011291514.3;refGene_gn=MROH1;refGene_tx=XM_011291518.2;refGene_gn=MROH1;refGene_tx=XM_019823255.2;refGene_gn=MROH1;refGene_tx=XM_019823256.2;refGene_gn=MROH1;refGene_tx=XM_019823257.2;refGene_gn=MROH1;refGene_tx=XM_019823258.2;refGene_gn=MROH1;refGene_tx=XM_019823259.2;refGene_gn=MROH1;refGene_tx=XM_019823260.2;refGene_gn=MROH1;refGene_tx=XM_023248543.1;refGene_gn=MROH1;refGene_tx=XM_023248544.1;refGene_gn=MROH1;refGene_tx=XM_023248545.1;refGene_gn=MROH1;refGene_tx=XM_023248546.1
F2 85137414 85137455
refGene_gn=MROH1;refGene_tx=XM_011291514.3;refGene_gn=MROH1;refGene_tx=XM_011291518.2;refGene_gn=MROH1;refGene_tx=XM_019823255.2;refGene_gn=MROH1;refGene_tx=XM_019823256.2;refGene_gn=MROH1;refGene_tx=XM_019823257.2;refGene_gn=MROH1;refGene_tx=XM_019823258.2;refGene_gn=MROH1;refGene_tx=XM_019823259.2;refGene_gn=MROH1;refGene_tx=XM_019823260.2;refGene_gn=MROH1;refGene_tx=XM_023248543.1;refGene_gn=MROH1;refGene_tx=XM_023248544.1;refGene_gn=MROH1;refGene_tx=XM_023248545.1;refGene_gn=MROH1;refGene_tx=XM_023248546.1
F2 85137710 85137852
refGene_gn=MROH1;refGene_tx=XM_011291514.3;refGene_gn=MROH1;refGene_tx=XM_011291518.2;refGene_gn=MROH1;refGene_tx=XM_019823255.2;refGene_gn=MROH1;refGene_tx=XM_019823256.2;refGene_gn=MROH1;refGene_tx=XM_019823257.2;refGene_gn=MROH1;refGene_tx=XM_019823258.2;refGene_gn=MROH1;refGene_tx=XM_019823259.2;refGene_gn=MROH1;refGene_tx=XM_019823260.2;refGene_gn=MROH1;refGene_tx=XM_023248543.1;refGene_gn=MROH1;refGene_tx=XM_023248544.1;refGene_gn=MROH1;refGene_tx=XM_023248545.1
F2 85138124 85138177
refGene_gn=MROH1;refGene_tx=XM_011291514.3;refGene_gn=MROH1;refGene_tx=XM_011291518.2;refGene_gn=MROH1;refGene_tx=XM_019823255.2;refGene_gn=MROH1;refGene_tx=XM_019823256.2;refGene_gn=MROH1;refGene_tx=XM_019823257.2;refGene_gn=MROH1;refGene_tx=XM_019823258.2;refGene_gn=MROH1;refGene_tx=XM_019823259.2;refGene_gn=MROH1;refGene_tx=XM_019823260.2;refGene_gn=MROH1;refGene_tx=XM_023248543.1;refGene_gn=MROH1;refGene_tx=XM_023248544.1;refGene_gn=MROH1;refGene_tx=XM_023248545.1
F2 85138628 85138736
refGene_gn=MROH1;refGene_tx=XM_011291514.3;refGene_gn=MROH1;refGene_tx=XM_011291518.2;refGene_gn=MROH1;refGene_tx=XM_019823255.2;refGene_gn=MROH1;refGene_tx=XM_019823256.2;refGene_gn=MROH1;refGene_tx=XM_019823257.2;refGene_gn=MROH1;refGene_tx=XM_019823258.2;refGene_gn=MROH1;refGene_tx=XM_019823259.2;refGene_gn=MROH1;refGene_tx=XM_019823260.2;refGene_gn=MROH1;refGene_tx=XM_023248543.1;refGene_gn=MROH1;refGene_tx=XM_023248544.1;refGene_gn=MROH1;refGene_tx=XM_023248545.1
F2 85139043 85139163
refGene_gn=MROH1;refGene_tx=XM_011291514.3;refGene_gn=MROH1;refGene_tx=XM_011291518.2;refGene_gn=MROH1;refGene_tx=XM_019823255.2;refGene_gn=

MROH1;refGene_tx=XM_019823256.2;refGene_gn=MROH1;refGene_tx=XM_0198232
57.2;refGene_gn=MROH1;refGene_tx=XM_019823258.2;refGene_gn=MROH1;refGe
ne_tx=XM_019823259.2;refGene_gn=MROH1;refGene_tx=XM_019823260.2;refGen
e_gn=MROH1;refGene_tx=XM_023248543.1;refGene_gn=MROH1;refGene_tx=XM_02
3248544.1;refGene_gn=MROH1;refGene_tx=XM_023248545.1
F2 85139321 85139444
refGene_gn=MROH1;refGene_tx=XM_011291514.3;refGene_gn=MROH1;refGene_tx
=XM_011291518.2;refGene_gn=MROH1;refGene_tx=XM_019823255.2;refGene_gn=
MROH1;refGene_tx=XM_019823256.2;refGene_gn=MROH1;refGene_tx=XM_0198232
57.2;refGene_gn=MROH1;refGene_tx=XM_019823258.2;refGene_gn=MROH1;refGe
ne_tx=XM_019823259.2;refGene_gn=MROH1;refGene_tx=XM_019823260.2;refGen
e_gn=MROH1;refGene_tx=XM_023248543.1;refGene_gn=MROH1;refGene_tx=XM_02
3248544.1;refGene_gn=MROH1;refGene_tx=XM_023248545.1
F2 85140231 85140378
refGene_gn=MROH1;refGene_tx=XM_011291514.3;refGene_gn=MROH1;refGene_tx
=XM_011291518.2;refGene_gn=MROH1;refGene_tx=XM_019823255.2;refGene_gn=
MROH1;refGene_tx=XM_019823256.2;refGene_gn=MROH1;refGene_tx=XM_0198232
57.2;refGene_gn=MROH1;refGene_tx=XM_019823258.2;refGene_gn=MROH1;refGe
ne_tx=XM_019823259.2;refGene_gn=MROH1;refGene_tx=XM_019823260.2;refGen
e_gn=MROH1;refGene_tx=XM_023248543.1;refGene_gn=MROH1;refGene_tx=XM_02
3248544.1;refGene_gn=MROH1;refGene_tx=XM_023248545.1
F2 85140483 85140510
refGene_gn=MROH1;refGene_tx=XM_011291514.3;refGene_gn=MROH1;refGene_tx
=XM_011291518.2;refGene_gn=MROH1;refGene_tx=XM_019823255.2;refGene_gn=
MROH1;refGene_tx=XM_019823256.2;refGene_gn=MROH1;refGene_tx=XM_0198232
57.2;refGene_gn=MROH1;refGene_tx=XM_019823258.2;refGene_gn=MROH1;refGe
ne_tx=XM_023248543.1;refGene_gn=MROH1;refGene_tx=XM_023248544.1;refGen
e_gn=MROH1;refGene_tx=XM_023248545.1
F2 85141454 85141577
refGene_gn=MROH1;refGene_tx=XM_011291514.3;refGene_gn=MROH1;refGene_tx
=XM_011291518.2;refGene_gn=MROH1;refGene_tx=XM_019823255.2;refGene_gn=
MROH1;refGene_tx=XM_019823256.2;refGene_gn=MROH1;refGene_tx=XM_0198232
57.2;refGene_gn=MROH1;refGene_tx=XM_019823258.2;refGene_gn=MROH1;refGe
ne_tx=XM_019823259.2;refGene_gn=MROH1;refGene_tx=XM_019823260.2;refGen
e_gn=MROH1;refGene_tx=XM_023248543.1;refGene_gn=MROH1;refGene_tx=XM_02
3248544.1;refGene_gn=MROH1;refGene_tx=XM_023248545.1
F2 85141809 85141889
refGene_gn=MROH1;refGene_tx=XM_011291514.3;refGene_gn=MROH1;refGene_tx
=XM_011291518.2;refGene_gn=MROH1;refGene_tx=XM_019823255.2;refGene_gn=
MROH1;refGene_tx=XM_019823256.2;refGene_gn=MROH1;refGene_tx=XM_0198232
57.2;refGene_gn=MROH1;refGene_tx=XM_019823258.2;refGene_gn=MROH1;refGe
ne_tx=XM_019823259.2;refGene_gn=MROH1;refGene_tx=XM_019823260.2;refGen
e_gn=MROH1;refGene_tx=XM_023248543.1;refGene_gn=MROH1;refGene_tx=XM_02
3248544.1;refGene_gn=MROH1;refGene_tx=XM_023248545.1
F2 85141987 85142099
refGene_gn=MROH1;refGene_tx=XM_011291514.3;refGene_gn=MROH1;refGene_tx
=XM_011291518.2;refGene_gn=MROH1;refGene_tx=XM_019823255.2;refGene_gn=
MROH1;refGene_tx=XM_019823256.2;refGene_gn=MROH1;refGene_tx=XM_0198232
57.2;refGene_gn=MROH1;refGene_tx=XM_019823258.2;refGene_gn=MROH1;refGe
ne_tx=XM_019823259.2;refGene_gn=MROH1;refGene_tx=XM_019823260.2;refGen

e_gn=MROH1;refGene_tx=XM_023248543.1;refGene_gn=MROH1;refGene_tx=XM_023248544.1;refGene_gn=MROH1;refGene_tx=XM_023248545.1
F2 85142178 85142274
refGene_gn=MROH1;refGene_tx=XM_011291514.3;refGene_gn=MROH1;refGene_tx=XM_011291518.2;refGene_gn=MROH1;refGene_tx=XM_019823255.2;refGene_gn=MROH1;refGene_tx=XM_019823256.2;refGene_gn=MROH1;refGene_tx=XM_019823257.2;refGene_gn=MROH1;refGene_tx=XM_019823258.2;refGene_gn=MROH1;refGene_tx=XM_019823259.2;refGene_gn=MROH1;refGene_tx=XM_019823260.2;refGene_gn=MROH1;refGene_tx=XM_023248543.1;refGene_gn=MROH1;refGene_tx=XM_023248544.1;refGene_gn=MROH1;refGene_tx=XM_023248545.1
F2 85142898 85143003
refGene_gn=MROH1;refGene_tx=XM_011291514.3;refGene_gn=MROH1;refGene_tx=XM_011291518.2;refGene_gn=MROH1;refGene_tx=XM_019823255.2;refGene_gn=MROH1;refGene_tx=XM_019823256.2;refGene_gn=MROH1;refGene_tx=XM_019823257.2;refGene_gn=MROH1;refGene_tx=XM_019823258.2;refGene_gn=MROH1;refGene_tx=XM_019823259.2;refGene_gn=MROH1;refGene_tx=XM_019823260.2;refGene_gn=MROH1;refGene_tx=XM_023248543.1;refGene_gn=MROH1;refGene_tx=XM_023248544.1;refGene_gn=MROH1;refGene_tx=XM_023248545.1
F2 85145151 85145287
refGene_gn=MROH1;refGene_tx=XM_011291514.3;refGene_gn=MROH1;refGene_tx=XM_011291518.2;refGene_gn=MROH1;refGene_tx=XM_019823255.2;refGene_gn=MROH1;refGene_tx=XM_019823256.2;refGene_gn=MROH1;refGene_tx=XM_019823257.2;refGene_gn=MROH1;refGene_tx=XM_019823258.2;refGene_gn=MROH1;refGene_tx=XM_019823259.2;refGene_gn=MROH1;refGene_tx=XM_019823260.2;refGene_gn=MROH1;refGene_tx=XM_023248543.1;refGene_gn=MROH1;refGene_tx=XM_023248544.1;refGene_gn=MROH1;refGene_tx=XM_023248545.1
F2 85145392 85145505
refGene_gn=MROH1;refGene_tx=XM_011291514.3;refGene_gn=MROH1;refGene_tx=XM_011291518.2;refGene_gn=MROH1;refGene_tx=XM_019823255.2;refGene_gn=MROH1;refGene_tx=XM_019823256.2;refGene_gn=MROH1;refGene_tx=XM_019823257.2;refGene_gn=MROH1;refGene_tx=XM_019823258.2;refGene_gn=MROH1;refGene_tx=XM_019823259.2;refGene_gn=MROH1;refGene_tx=XM_019823260.2;refGene_gn=MROH1;refGene_tx=XM_023248543.1;refGene_gn=MROH1;refGene_tx=XM_023248544.1;refGene_gn=MROH1;refGene_tx=XM_023248545.1
F2 85146359 85146512
refGene_gn=MROH1;refGene_tx=XM_011291514.3;refGene_gn=MROH1;refGene_tx=XM_011291518.2;refGene_gn=MROH1;refGene_tx=XM_019823255.2;refGene_gn=MROH1;refGene_tx=XM_019823256.2;refGene_gn=MROH1;refGene_tx=XM_019823257.2;refGene_gn=MROH1;refGene_tx=XM_019823258.2;refGene_gn=MROH1;refGene_tx=XM_019823259.2;refGene_gn=MROH1;refGene_tx=XM_019823260.2;refGene_gn=MROH1;refGene_tx=XM_023248543.1;refGene_gn=MROH1;refGene_tx=XM_023248544.1;refGene_gn=MROH1;refGene_tx=XM_023248545.1
F2 85151520 85151675
refGene_gn=MROH1;refGene_tx=XM_011291514.3;refGene_gn=MROH1;refGene_tx=XM_011291518.2;refGene_gn=MROH1;refGene_tx=XM_019823255.2;refGene_gn=MROH1;refGene_tx=XM_019823256.2;refGene_gn=MROH1;refGene_tx=XM_019823257.2;refGene_gn=MROH1;refGene_tx=XM_019823258.2;refGene_gn=MROH1;refGene_tx=XM_019823259.2;refGene_gn=MROH1;refGene_tx=XM_019823260.2;refGene_gn=MROH1;refGene_tx=XM_023248543.1;refGene_gn=MROH1;refGene_tx=XM_023248544.1;refGene_gn=MROH1;refGene_tx=XM_023248545.1

F2 85154322 85154488
refGene_gn=MROH1;refGene_tx=XM_011291514.3;refGene_gn=MROH1;refGene_tx=XM_011291518.2;refGene_gn=MROH1;refGene_tx=XM_019823255.2;refGene_gn=MROH1;refGene_tx=XM_019823256.2;refGene_gn=MROH1;refGene_tx=XM_019823257.2;refGene_gn=MROH1;refGene_tx=XM_019823258.2;refGene_gn=MROH1;refGene_tx=XM_019823259.2;refGene_gn=MROH1;refGene_tx=XM_019823260.2;refGene_gn=MROH1;refGene_tx=XM_023248543.1;refGene_gn=MROH1;refGene_tx=XM_023248544.1;refGene_gn=MROH1;refGene_tx=XM_023248545.1
F2 85155050 85155247
refGene_gn=MROH1;refGene_tx=XM_011291514.3;refGene_gn=MROH1;refGene_tx=XM_011291518.2;refGene_gn=MROH1;refGene_tx=XM_019823255.2;refGene_gn=MROH1;refGene_tx=XM_019823256.2;refGene_gn=MROH1;refGene_tx=XM_019823257.2;refGene_gn=MROH1;refGene_tx=XM_019823258.2;refGene_gn=MROH1;refGene_tx=XM_019823259.2;refGene_gn=MROH1;refGene_tx=XM_019823260.2;refGene_gn=MROH1;refGene_tx=XM_023248543.1;refGene_gn=MROH1;refGene_tx=XM_023248544.1;refGene_gn=MROH1;refGene_tx=XM_023248545.1
F2 85155676 85155944 refGene_gn=MROH1;refGene_tx=XM_011291518.2
F2 85157697 85157835
refGene_gn=MROH1;refGene_tx=XM_011291514.3;refGene_gn=MROH1;refGene_tx=XM_019823255.2;refGene_gn=MROH1;refGene_tx=XM_019823256.2;refGene_gn=MROH1;refGene_tx=XM_019823257.2;refGene_gn=MROH1;refGene_tx=XM_019823258.2;refGene_gn=MROH1;refGene_tx=XM_019823259.2;refGene_gn=MROH1;refGene_tx=XM_019823260.2;refGene_gn=MROH1;refGene_tx=XM_023248543.1;refGene_gn=MROH1;refGene_tx=XM_023248544.1;refGene_gn=MROH1;refGene_tx=XM_023248545.1
F2 85158250 85158365
refGene_gn=MROH1;refGene_tx=XM_011291514.3;refGene_gn=MROH1;refGene_tx=XM_019823255.2;refGene_gn=MROH1;refGene_tx=XM_019823256.2;refGene_gn=MROH1;refGene_tx=XM_019823257.2;refGene_gn=MROH1;refGene_tx=XM_019823258.2;refGene_gn=MROH1;refGene_tx=XM_019823259.2;refGene_gn=MROH1;refGene_tx=XM_019823260.2;refGene_gn=MROH1;refGene_tx=XM_023248543.1;refGene_gn=MROH1;refGene_tx=XM_023248544.1;refGene_gn=MROH1;refGene_tx=XM_023248545.1
F2 85158877 85159024
refGene_gn=MROH1;refGene_tx=XM_011291514.3;refGene_gn=MROH1;refGene_tx=XM_019823255.2;refGene_gn=MROH1;refGene_tx=XM_019823256.2;refGene_gn=MROH1;refGene_tx=XM_019823257.2;refGene_gn=MROH1;refGene_tx=XM_019823258.2;refGene_gn=MROH1;refGene_tx=XM_019823259.2;refGene_gn=MROH1;refGene_tx=XM_019823260.2;refGene_gn=MROH1;refGene_tx=XM_023248543.1;refGene_gn=MROH1;refGene_tx=XM_023248544.1;refGene_gn=MROH1;refGene_tx=XM_023248545.1
F2 85159264 85159453
refGene_gn=MROH1;refGene_tx=XM_011291514.3;refGene_gn=MROH1;refGene_tx=XM_019823255.2;refGene_gn=MROH1;refGene_tx=XM_019823256.2;refGene_gn=MROH1;refGene_tx=XM_019823257.2;refGene_gn=MROH1;refGene_tx=XM_019823258.2;refGene_gn=MROH1;refGene_tx=XM_019823259.2;refGene_gn=MROH1;refGene_tx=XM_019823260.2;refGene_gn=MROH1;refGene_tx=XM_023248543.1;refGene_gn=MROH1;refGene_tx=XM_023248544.1;refGene_gn=MROH1;refGene_tx=XM_023248545.1
F2 85159778 85159934

refGene_gn=MROH1;refGene_tx=XM_011291514.3;refGene_gn=MROH1;refGene_tx=XM_019823255.2;refGene_gn=MROH1;refGene_tx=XM_019823256.2;refGene_gn=MROH1;refGene_tx=XM_019823257.2;refGene_gn=MROH1;refGene_tx=XM_019823258.2;refGene_gn=MROH1;refGene_tx=XM_019823259.2;refGene_gn=MROH1;refGene_tx=XM_019823260.2;refGene_gn=MROH1;refGene_tx=XM_023248543.1;refGene_gn=MROH1;refGene_tx=XM_023248544.1;refGene_gn=MROH1;refGene_tx=XM_023248545.1
F2 85160005 85160140
refGene_gn=MROH1;refGene_tx=XM_011291514.3;refGene_gn=MROH1;refGene_tx=XM_019823255.2;refGene_gn=MROH1;refGene_tx=XM_019823256.2;refGene_gn=MROH1;refGene_tx=XM_019823257.2;refGene_gn=MROH1;refGene_tx=XM_019823258.2;refGene_gn=MROH1;refGene_tx=XM_019823259.2;refGene_gn=MROH1;refGene_tx=XM_019823260.2;refGene_gn=MROH1;refGene_tx=XM_023248543.1;refGene_gn=MROH1;refGene_tx=XM_023248544.1;refGene_gn=MROH1;refGene_tx=XM_023248545.1
F2 85160214 85160317
refGene_gn=MROH1;refGene_tx=XM_011291514.3;refGene_gn=MROH1;refGene_tx=XM_019823255.2;refGene_gn=MROH1;refGene_tx=XM_019823256.2;refGene_gn=MROH1;refGene_tx=XM_019823257.2;refGene_gn=MROH1;refGene_tx=XM_019823258.2;refGene_gn=MROH1;refGene_tx=XM_019823259.2;refGene_gn=MROH1;refGene_tx=XM_019823260.2;refGene_gn=MROH1;refGene_tx=XM_023248543.1;refGene_gn=MROH1;refGene_tx=XM_023248544.1;refGene_gn=MROH1;refGene_tx=XM_023248545.1
F2 85160394 85160460
refGene_gn=MROH1;refGene_tx=XM_011291514.3;refGene_gn=MROH1;refGene_tx=XM_019823255.2;refGene_gn=MROH1;refGene_tx=XM_019823256.2;refGene_gn=MROH1;refGene_tx=XM_019823257.2;refGene_gn=MROH1;refGene_tx=XM_019823258.2;refGene_gn=MROH1;refGene_tx=XM_019823259.2;refGene_gn=MROH1;refGene_tx=XM_019823260.2;refGene_gn=MROH1;refGene_tx=XM_023248543.1;refGene_gn=MROH1;refGene_tx=XM_023248544.1;refGene_gn=MROH1;refGene_tx=XM_023248545.1
F2 85160864 85160968
refGene_gn=MROH1;refGene_tx=XM_011291514.3;refGene_gn=MROH1;refGene_tx=XM_019823255.2;refGene_gn=MROH1;refGene_tx=XM_019823256.2;refGene_gn=MROH1;refGene_tx=XM_019823257.2;refGene_gn=MROH1;refGene_tx=XM_019823258.2;refGene_gn=MROH1;refGene_tx=XM_019823259.2;refGene_gn=MROH1;refGene_tx=XM_019823260.2;refGene_gn=MROH1;refGene_tx=XM_023248543.1;refGene_gn=MROH1;refGene_tx=XM_023248544.1;refGene_gn=MROH1;refGene_tx=XM_023248545.1
F2 85161357 85161511
refGene_gn=BOP1;refGene_tx=XM_019823263.2;refGene_gn=BOP1;refGene_tx=XM_023248547.1;refGene_gn=BOP1;refGene_tx=XM_023248548.1
F2 85161606 85161714
refGene_gn=BOP1;refGene_tx=XM_019823263.2;refGene_gn=BOP1;refGene_tx=XM_023248547.1;refGene_gn=BOP1;refGene_tx=XM_023248548.1
F2 85161794 85161879
refGene_gn=BOP1;refGene_tx=XM_019823263.2;refGene_gn=BOP1;refGene_tx=XM_023248547.1;refGene_gn=BOP1;refGene_tx=XM_023248548.1
F2 85161971 85162260
refGene_gn=BOP1;refGene_tx=XM_019823263.2;refGene_gn=BOP1;refGene_tx=X

M_023248547.1;refGene_gn=BOP1;refGene_tx=XM_023248548.1
F2 85162358 85162539
refGene_gn=BOP1;refGene_tx=XM_019823263.2;refGene_gn=BOP1;refGene_tx=XM_023248547.1;refGene_gn=BOP1;refGene_tx=XM_023248548.1
F2 85162635 85162768
refGene_gn=BOP1;refGene_tx=XM_019823263.2;refGene_gn=BOP1;refGene_tx=XM_023248547.1;refGene_gn=BOP1;refGene_tx=XM_023248548.1
F2 85162850 85162920
refGene_gn=BOP1;refGene_tx=XM_019823263.2;refGene_gn=BOP1;refGene_tx=XM_023248547.1;refGene_gn=BOP1;refGene_tx=XM_023248548.1
F2 85162990 85163071
refGene_gn=BOP1;refGene_tx=XM_019823263.2;refGene_gn=BOP1;refGene_tx=XM_023248547.1;refGene_gn=BOP1;refGene_tx=XM_023248548.1
F2 85163163 85163325
refGene_gn=BOP1;refGene_tx=XM_019823263.2;refGene_gn=BOP1;refGene_tx=XM_023248547.1;refGene_gn=BOP1;refGene_tx=XM_023248548.1
F2 85163412 85163625
refGene_gn=BOP1;refGene_tx=XM_019823263.2;refGene_gn=BOP1;refGene_tx=XM_023248547.1;refGene_gn=BOP1;refGene_tx=XM_023248548.1
F2 85163702 85163804
refGene_gn=BOP1;refGene_tx=XM_019823263.2;refGene_gn=BOP1;refGene_tx=XM_023248547.1;refGene_gn=BOP1;refGene_tx=XM_023248548.1
F2 85163882 85164000
refGene_gn=BOP1;refGene_tx=XM_019823263.2;refGene_gn=BOP1;refGene_tx=XM_023248547.1;refGene_gn=BOP1;refGene_tx=XM_023248548.1
F2 85164077 85164232
refGene_gn=BOP1;refGene_tx=XM_019823263.2;refGene_gn=BOP1;refGene_tx=XM_023248547.1;refGene_gn=BOP1;refGene_tx=XM_023248548.1
F2 85164454 85164550 refGene_gn=BOP1;refGene_tx=XM_019823263.2
F2 85165862 85166435
refGene_gn=SCX;refGene_tx=XM_023248549.1;refGene_gn=SCX;refGene_tx=XM_023248550.1
F2 85167212 85167265 refGene_gn=SCX;refGene_tx=XM_023248549.1
F2 85167384 85167658
refGene_gn=SCX;refGene_tx=XM_023248549.1;refGene_gn=SCX;refGene_tx=XM_023248550.1
F2 85174824 85174905
refGene_gn=BOP1;refGene_tx=XM_023248547.1;refGene_gn=BOP1;refGene_tx=XM_023248548.1
F2 85184904 85185105 refGene_gn=BOP1;refGene_tx=XM_023248547.1
F2 85186203 85186299
refGene_gn=BOP1;refGene_tx=XM_023248547.1;refGene_gn=BOP1;refGene_tx=XM_023248548.1
F2 85186698 85186815
refGene_gn=HSF1;refGene_tx=XM_004000170.4;refGene_gn=HSF1;refGene_tx=XM_006943453.3;refGene_gn=HSF1;refGene_tx=XM_006943454.3;refGene_gn=HSF1;refGene_tx=XM_006943455.3
F2 85204939 85205051
refGene_gn=HSF1;refGene_tx=XM_004000170.4;refGene_gn=HSF1;refGene_tx=XM_006943453.3;refGene_gn=HSF1;refGene_tx=XM_006943454.3;refGene_gn=HSF

1;refGene_tx=XM_006943455.3
F2 85205381 85205518
refGene_gn=HSF1;refGene_tx=XM_004000170.4;refGene_gn=HSF1;refGene_tx=XM_006943453.3;refGene_gn=HSF1;refGene_tx=XM_006943454.3;refGene_gn=HSF1;refGene_tx=XM_006943455.3
F2 85205876 85206001
refGene_gn=HSF1;refGene_tx=XM_004000170.4;refGene_gn=HSF1;refGene_tx=XM_006943453.3;refGene_gn=HSF1;refGene_tx=XM_006943454.3;refGene_gn=HSF1;refGene_tx=XM_006943455.3
F2 85207174 85207250
refGene_gn=HSF1;refGene_tx=XM_004000170.4;refGene_gn=HSF1;refGene_tx=XM_006943453.3;refGene_gn=HSF1;refGene_tx=XM_006943454.3;refGene_gn=HSF1;refGene_tx=XM_006943455.3
F2 85207323 85207385
refGene_gn=HSF1;refGene_tx=XM_004000170.4;refGene_gn=HSF1;refGene_tx=XM_006943453.3;refGene_gn=HSF1;refGene_tx=XM_006943454.3;refGene_gn=HSF1;refGene_tx=XM_006943455.3
F2 85207519 85207616
refGene_gn=HSF1;refGene_tx=XM_004000170.4;refGene_gn=HSF1;refGene_tx=XM_006943453.3;refGene_gn=HSF1;refGene_tx=XM_006943454.3;refGene_gn=HSF1;refGene_tx=XM_006943455.3
F2 85207720 85207857
refGene_gn=HSF1;refGene_tx=XM_004000170.4;refGene_gn=HSF1;refGene_tx=XM_006943453.3;refGene_gn=HSF1;refGene_tx=XM_006943454.3;refGene_gn=HSF1;refGene_tx=XM_006943455.3
F2 85207942 85208218
refGene_gn=HSF1;refGene_tx=XM_004000170.4;refGene_gn=HSF1;refGene_tx=XM_006943453.3;refGene_gn=HSF1;refGene_tx=XM_006943454.3;refGene_gn=HSF1;refGene_tx=XM_006943455.3
F2 85208611 85208692
refGene_gn=HSF1;refGene_tx=XM_006943453.3;refGene_gn=HSF1;refGene_tx=XM_006943454.3
F2 85209538 85209644
refGene_gn=HSF1;refGene_tx=XM_004000170.4;refGene_gn=HSF1;refGene_tx=XM_006943453.3;refGene_gn=HSF1;refGene_tx=XM_006943454.3;refGene_gn=HSF1;refGene_tx=XM_006943455.3
F2 85209715 85209781
refGene_gn=HSF1;refGene_tx=XM_004000170.4;refGene_gn=HSF1;refGene_tx=XM_006943453.3;refGene_gn=HSF1;refGene_tx=XM_006943454.3;refGene_gn=HSF1;refGene_tx=XM_006943455.3
F2 85209866 85209936
refGene_gn=HSF1;refGene_tx=XM_004000170.4;refGene_gn=HSF1;refGene_tx=XM_006943453.3;refGene_gn=HSF1;refGene_tx=XM_006943454.3;refGene_gn=HSF1;refGene_tx=XM_006943455.3
F2 85210017 85210223
refGene_gn=HSF1;refGene_tx=XM_004000170.4;refGene_gn=HSF1;refGene_tx=XM_006943453.3;refGene_gn=HSF1;refGene_tx=XM_006943454.3;refGene_gn=HSF1;refGene_tx=XM_006943455.3
F2 85212218 85212374
refGene_gn=DGAT1;refGene_tx=XM_004000171.5;refGene_gn=DGAT1;refGene_tx

=XM_019823264.2
F2	85212470	85212533
refGene_gn=DGAT1;refGene_tx=XM_004000171.5;refGene_gn=DGAT1;refGene_tx
=XM_019823264.2
F2	85212619	85212707
refGene_gn=DGAT1;refGene_tx=XM_004000171.5;refGene_gn=DGAT1;refGene_tx
=XM_019823264.2
F2	85212791	85212857
refGene_gn=DGAT1;refGene_tx=XM_004000171.5;refGene_gn=DGAT1;refGene_tx
=XM_019823264.2
F2	85212931	85213044
refGene_gn=DGAT1;refGene_tx=XM_004000171.5;refGene_gn=DGAT1;refGene_tx
=XM_019823264.2
F2	85213133	85213178
refGene_gn=DGAT1;refGene_tx=XM_004000171.5;refGene_gn=DGAT1;refGene_tx
=XM_019823264.2
F2	85213271	85213313
refGene_gn=DGAT1;refGene_tx=XM_004000171.5;refGene_gn=DGAT1;refGene_tx
=XM_019823264.2
F2	85213397	85213436
refGene_gn=DGAT1;refGene_tx=XM_004000171.5;refGene_gn=DGAT1;refGene_tx
=XM_019823264.2
F2	85213526	85213630
refGene_gn=DGAT1;refGene_tx=XM_004000171.5;refGene_gn=DGAT1;refGene_tx
=XM_019823264.2
F2	85213717	85213792
refGene_gn=DGAT1;refGene_tx=XM_004000171.5;refGene_gn=DGAT1;refGene_tx
=XM_019823264.2
F2	85213921	85214035
refGene_gn=DGAT1;refGene_tx=XM_004000171.5;refGene_gn=DGAT1;refGene_tx
=XM_019823264.2
F2	85214181	85214287
refGene_gn=DGAT1;refGene_tx=XM_004000171.5;refGene_gn=DGAT1;refGene_tx
=XM_019823264.2
F2	85214463	85214516
refGene_gn=DGAT1;refGene_tx=XM_004000171.5;refGene_gn=DGAT1;refGene_tx
=XM_019823264.2
F2	85214599	85214685
refGene_gn=DGAT1;refGene_tx=XM_004000171.5;refGene_gn=DGAT1;refGene_tx
=XM_019823264.2
F2	85214779	85214820
refGene_gn=DGAT1;refGene_tx=XM_004000171.5;refGene_gn=DGAT1;refGene_tx
=XM_019823264.2
F2	85216873	85216961
refGene_gn=DGAT1;refGene_tx=XM_004000171.5;refGene_gn=DGAT1;refGene_tx
=XM_019823264.2
F2	85217171	85217242	refGene_gn=DGAT1;refGene_tx=XM_019823264.2
F2	85221405	85221608	refGene_gn=DGAT1;refGene_tx=XM_004000171.5
F2	85228114	85229052	refGene_gn=SCRT1;refGene_tx=XM_023248784.1
F2	85231088	85231203	refGene_gn=SCRT1;refGene_tx=XM_023248784.1

F2 85245962 85246148 refGene_gn=TMEM249;refGene_tx=XM_023248351.1
F2 85246358 85246444 refGene_gn=TMEM249;refGene_tx=XM_023248351.1
F2 85246712 85246908 refGene_gn=TMEM249;refGene_tx=XM_023248351.1
F2 85247020 85247115 refGene_gn=TMEM249;refGene_tx=XM_023248351.1
F2 85247410 85247561 refGene_gn=TMEM249;refGene_tx=XM_023248351.1
F2 85247636 85247911 refGene_gn=TMEM249;refGene_tx=XM_023248351.1
F2 85247979 85247989 refGene_gn=TMEM249;refGene_tx=XM_023248351.1
F2 85248375 85248526
refGene_gn=FBXL6;refGene_tx=XM_004000222.5;refGene_gn=FBXL6;refGene_tx=XM_019823266.2;refGene_gn=FBXL6;refGene_tx=XM_023248781.1
F2 85248814 85249061
refGene_gn=FBXL6;refGene_tx=XM_004000222.5;refGene_gn=FBXL6;refGene_tx=XM_019823266.2;refGene_gn=FBXL6;refGene_tx=XM_023248781.1
F2 85249136 85249368
refGene_gn=FBXL6;refGene_tx=XM_004000222.5;refGene_gn=FBXL6;refGene_tx=XM_019823266.2;refGene_gn=FBXL6;refGene_tx=XM_023248781.1
F2 85249436 85249550
refGene_gn=FBXL6;refGene_tx=XM_004000222.5;refGene_gn=FBXL6;refGene_tx=XM_019823266.2;refGene_gn=FBXL6;refGene_tx=XM_023248781.1
F2 85249636 85249744
refGene_gn=FBXL6;refGene_tx=XM_004000222.5;refGene_gn=FBXL6;refGene_tx=XM_023248781.1
F2 85249818 85249950 refGene_gn=FBXL6;refGene_tx=XM_004000222.5
F2 85250258 85250322
refGene_gn=FBXL6;refGene_tx=XM_004000222.5;refGene_gn=FBXL6;refGene_tx=XM_019823266.2;refGene_gn=FBXL6;refGene_tx=XM_023248781.1
F2 85250487 85250646
refGene_gn=FBXL6;refGene_tx=XM_004000222.5;refGene_gn=FBXL6;refGene_tx=XM_019823266.2;refGene_gn=FBXL6;refGene_tx=XM_023248781.1
F2 85250930 85251337
refGene_gn=FBXL6;refGene_tx=XM_004000222.5;refGene_gn=FBXL6;refGene_tx=XM_019823266.2;refGene_gn=FBXL6;refGene_tx=XM_023248781.1
F2 85251521 85251563 refGene_gn=SLC52A2;refGene_tx=XM_023248783.1
F2 85252275 85252466
refGene_gn=SLC52A2;refGene_tx=XM_004000172.4;refGene_gn=SLC52A2;refGene_tx=XM_006943451.3;refGene_gn=SLC52A2;refGene_tx=XM_006943452.3;refGene_gn=SLC52A2;refGene_tx=XM_019823267.2;refGene_gn=SLC52A2;refGene_tx=XM_023248783.1
F2 85252576 85253459
refGene_gn=SLC52A2;refGene_tx=XM_004000172.4;refGene_gn=SLC52A2;refGene_tx=XM_006943451.3;refGene_gn=SLC52A2;refGene_tx=XM_006943452.3;refGene_gn=SLC52A2;refGene_tx=XM_019823267.2;refGene_gn=SLC52A2;refGene_tx=XM_023248783.1
F2 85253554 85253678
refGene_gn=SLC52A2;refGene_tx=XM_004000172.4;refGene_gn=SLC52A2;refGene_tx=XM_006943451.3;refGene_gn=SLC52A2;refGene_tx=XM_006943452.3;refGene_gn=SLC52A2;refGene_tx=XM_019823267.2;refGene_gn=SLC52A2;refGene_tx=XM_023248783.1
F2 85253785 85253998
refGene_gn=SLC52A2;refGene_tx=XM_004000172.4;refGene_gn=SLC52A2;refGen

e_tx=XM_006943451.3;refGene_gn=SLC52A2;refGene_tx=XM_006943452.3;refGene_gn=SLC52A2;refGene_tx=XM_019823267.2;refGene_gn=SLC52A2;refGene_tx=XM_023248783.1
F2 85278536 85278548
refGene_gn=ADCK5;refGene_tx=XM_011291519.3;refGene_gn=ADCK5;refGene_tx=XM_011291520.3;refGene_gn=ADCK5;refGene_tx=XM_019823268.2
F2 85286995 85287102
refGene_gn=ADCK5;refGene_tx=XM_011291519.3;refGene_gn=ADCK5;refGene_tx=XM_011291520.3;refGene_gn=ADCK5;refGene_tx=XM_019823268.2
F2 85287909 85288059
refGene_gn=ADCK5;refGene_tx=XM_011291519.3;refGene_gn=ADCK5;refGene_tx=XM_011291520.3;refGene_gn=ADCK5;refGene_tx=XM_019823268.2
F2 85293520 85293596
refGene_gn=ADCK5;refGene_tx=XM_011291519.3;refGene_gn=ADCK5;refGene_tx=XM_011291520.3
F2 85293727 85293928
refGene_gn=ADCK5;refGene_tx=XM_011291519.3;refGene_gn=ADCK5;refGene_tx=XM_011291520.3;refGene_gn=ADCK5;refGene_tx=XM_019823268.2
F2 85294005 85294146
refGene_gn=ADCK5;refGene_tx=XM_011291519.3;refGene_gn=ADCK5;refGene_tx=XM_011291520.3;refGene_gn=ADCK5;refGene_tx=XM_019823268.2
F2 85294239 85294353
refGene_gn=ADCK5;refGene_tx=XM_011291519.3;refGene_gn=ADCK5;refGene_tx=XM_011291520.3;refGene_gn=ADCK5;refGene_tx=XM_019823268.2
F2 85294454 85294586
refGene_gn=ADCK5;refGene_tx=XM_011291519.3;refGene_gn=ADCK5;refGene_tx=XM_019823268.2
F2 85294654 85294738
refGene_gn=ADCK5;refGene_tx=XM_011291519.3;refGene_gn=ADCK5;refGene_tx=XM_011291520.3;refGene_gn=ADCK5;refGene_tx=XM_019823268.2
F2 85294832 85294914
refGene_gn=ADCK5;refGene_tx=XM_011291519.3;refGene_gn=ADCK5;refGene_tx=XM_011291520.3;refGene_gn=ADCK5;refGene_tx=XM_019823268.2
F2 85294992 85295071
refGene_gn=ADCK5;refGene_tx=XM_011291519.3;refGene_gn=ADCK5;refGene_tx=XM_011291520.3;refGene_gn=ADCK5;refGene_tx=XM_019823268.2
F2 85295148 85295240
refGene_gn=ADCK5;refGene_tx=XM_011291519.3;refGene_gn=ADCK5;refGene_tx=XM_011291520.3;refGene_gn=ADCK5;refGene_tx=XM_019823268.2
F2 85295316 85295569
refGene_gn=ADCK5;refGene_tx=XM_011291519.3;refGene_gn=ADCK5;refGene_tx=XM_011291520.3;refGene_gn=ADCK5;refGene_tx=XM_019823268.2
F2 85295648 85295765
refGene_gn=ADCK5;refGene_tx=XM_011291519.3;refGene_gn=ADCK5;refGene_tx=XM_011291520.3;refGene_gn=ADCK5;refGene_tx=XM_019823268.2
F2 85295851 85295957
refGene_gn=ADCK5;refGene_tx=XM_011291519.3;refGene_gn=ADCK5;refGene_tx=XM_011291520.3;refGene_gn=ADCK5;refGene_tx=XM_019823268.2
F2 85296205 85296253 refGene_gn=CPSF1;refGene_tx=XM_023248701.1
F2 85296326 85296465 refGene_gn=CPSF1;refGene_tx=XM_023248701.1

| | | | |
|---|---|---|---|
| F2 | 85296545 | 85296675 | refGene_gn=CPSF1;refGene_tx=XM_023248701.1 |
| F2 | 85296756 | 85296912 | refGene_gn=CPSF1;refGene_tx=XM_023248701.1 |
| F2 | 85296985 | 85297033 | refGene_gn=CPSF1;refGene_tx=XM_023248701.1 |
| F2 | 85297114 | 85297181 | refGene_gn=CPSF1;refGene_tx=XM_023248701.1 |
| F2 | 85297260 | 85297437 | refGene_gn=CPSF1;refGene_tx=XM_023248701.1 |
| F2 | 85297512 | 85297665 | refGene_gn=CPSF1;refGene_tx=XM_023248701.1 |
| F2 | 85297749 | 85297891 | refGene_gn=CPSF1;refGene_tx=XM_023248701.1 |
| F2 | 85297959 | 85298044 | refGene_gn=CPSF1;refGene_tx=XM_023248701.1 |
| F2 | 85298114 | 85298205 | refGene_gn=CPSF1;refGene_tx=XM_023248701.1 |
| F2 | 85298284 | 85298401 | refGene_gn=CPSF1;refGene_tx=XM_023248701.1 |
| F2 | 85299242 | 85299395 | refGene_gn=CPSF1;refGene_tx=XM_023248701.1 |
| F2 | 85299467 | 85299611 | refGene_gn=CPSF1;refGene_tx=XM_023248701.1 |
| F2 | 85299704 | 85299794 | refGene_gn=CPSF1;refGene_tx=XM_023248701.1 |
| F2 | 85299940 | 85300147 | refGene_gn=CPSF1;refGene_tx=XM_023248701.1 |
| F2 | 85300222 | 85300397 | refGene_gn=CPSF1;refGene_tx=XM_023248701.1 |
| F2 | 85300482 | 85300619 | refGene_gn=CPSF1;refGene_tx=XM_023248701.1 |
| F2 | 85300700 | 85300879 | refGene_gn=CPSF1;refGene_tx=XM_023248701.1 |
| F2 | 85300956 | 85301098 | refGene_gn=CPSF1;refGene_tx=XM_023248701.1 |
| F2 | 85301173 | 85301284 | refGene_gn=CPSF1;refGene_tx=XM_023248701.1 |
| F2 | 85301533 | 85301623 | refGene_gn=CPSF1;refGene_tx=XM_023248701.1 |
| F2 | 85301704 | 85301785 | refGene_gn=CPSF1;refGene_tx=XM_023248701.1 |
| F2 | 85301864 | 85301939 | refGene_gn=CPSF1;refGene_tx=XM_023248701.1 |
| F2 | 85302010 | 85302105 | refGene_gn=CPSF1;refGene_tx=XM_023248701.1 |
| F2 | 85302207 | 85302259 | refGene_gn=CPSF1;refGene_tx=XM_023248701.1 |
| F2 | 85302345 | 85302468 | refGene_gn=CPSF1;refGene_tx=XM_023248701.1 |
| F2 | 85302545 | 85302633 | refGene_gn=CPSF1;refGene_tx=XM_023248701.1 |
| F2 | 85302752 | 85302846 | refGene_gn=CPSF1;refGene_tx=XM_023248701.1 |
| F2 | 85302925 | 85303036 | refGene_gn=CPSF1;refGene_tx=XM_023248701.1 |
| F2 | 85303116 | 85303256 | refGene_gn=CPSF1;refGene_tx=XM_023248701.1 |
| F2 | 85303385 | 85303532 | refGene_gn=CPSF1;refGene_tx=XM_023248701.1 |
| F2 | 85303610 | 85303762 | refGene_gn=CPSF1;refGene_tx=XM_023248701.1 |
| F2 | 85303876 | 85303957 | refGene_gn=CPSF1;refGene_tx=XM_023248701.1 |
| F2 | 85304045 | 85304188 | refGene_gn=CPSF1;refGene_tx=XM_023248701.1 |
| F2 | 85304280 | 85304308 | refGene_gn=CPSF1;refGene_tx=XM_023248701.1 |
| F2 | 85315069 | 85315213 | refGene_gn=CPSF1;refGene_tx=XM_023248701.1 |
| F2 | 85318849 | 85318978 | refGene_gn=SLC39A4;refGene_tx=XM_004000173.3 |
| F2 | 85319069 | 85319257 | refGene_gn=SLC39A4;refGene_tx=XM_004000173.3 |
| F2 | 85319363 | 85319516 | refGene_gn=SLC39A4;refGene_tx=XM_004000173.3 |
| F2 | 85319654 | 85319709 | refGene_gn=SLC39A4;refGene_tx=XM_004000173.3 |
| F2 | 85319905 | 85320028 | refGene_gn=SLC39A4;refGene_tx=XM_004000173.3 |
| F2 | 85320108 | 85320246 | refGene_gn=SLC39A4;refGene_tx=XM_004000173.3 |
| F2 | 85320403 | 85320576 | refGene_gn=SLC39A4;refGene_tx=XM_004000173.3 |
| F2 | 85320870 | 85321042 | refGene_gn=SLC39A4;refGene_tx=XM_004000173.3 |
| F2 | 85321131 | 85321283 | refGene_gn=SLC39A4;refGene_tx=XM_004000173.3 |
| F2 | 85321399 | 85321595 | refGene_gn=SLC39A4;refGene_tx=XM_004000173.3 |
| F2 | 85322064 | 85322349 | refGene_gn=SLC39A4;refGene_tx=XM_004000173.3 |
| F2 | 85322835 | 85323033 | refGene_gn=SLC39A4;refGene_tx=XM_004000173.3 |
| F2 | 85327442 | 85327560 | refGene_gn=VPS28;refGene_tx=XM_004000174.4;refGene_gn=VPS28;refGene_tx=XM_019823272.2 |

F2 85327647 85327739
refGene_gn=VPS28;refGene_tx=XM_004000174.4;refGene_gn=VPS28;refGene_tx
=XM_019823272.2
F2 85327851 85327905
refGene_gn=VPS28;refGene_tx=XM_004000174.4;refGene_gn=VPS28;refGene_tx
=XM_019823272.2
F2 85328451 85328553
refGene_gn=VPS28;refGene_tx=XM_004000174.4;refGene_gn=VPS28;refGene_tx
=XM_019823272.2
F2 85328674 85328780
refGene_gn=VPS28;refGene_tx=XM_004000174.4;refGene_gn=VPS28;refGene_tx
=XM_019823272.2
F2 85329544 85329634
refGene_gn=VPS28;refGene_tx=XM_004000174.4;refGene_gn=VPS28;refGene_tx
=XM_019823272.2
F2 85329926 85329964
refGene_gn=VPS28;refGene_tx=XM_004000174.4;refGene_gn=VPS28;refGene_tx
=XM_019823272.2
F2 85330076 85330105
refGene_gn=VPS28;refGene_tx=XM_004000174.4;refGene_gn=VPS28;refGene_tx
=XM_019823272.2
F2 85330731 85330799
refGene_gn=VPS28;refGene_tx=XM_004000174.4;refGene_gn=VPS28;refGene_tx
=XM_019823272.2
F2 85332940 85332963 refGene_gn=VPS28;refGene_tx=XM_019823272.2
F2 85333478 85333669
refGene_gn=TONSL;refGene_tx=XM_004000175.3;refGene_gn=TONSL;refGene_tx
=XM_011291521.2;refGene_gn=TONSL;refGene_tx=XM_011291523.2
F2 85334885 85335019
refGene_gn=TONSL;refGene_tx=XM_004000175.3;refGene_gn=TONSL;refGene_tx
=XM_011291521.2;refGene_gn=TONSL;refGene_tx=XM_011291523.2
F2 85335555 85335629
refGene_gn=TONSL;refGene_tx=XM_004000175.3;refGene_gn=TONSL;refGene_tx
=XM_011291521.2;refGene_gn=TONSL;refGene_tx=XM_011291523.2
F2 85336153 85336329
refGene_gn=TONSL;refGene_tx=XM_004000175.3;refGene_gn=TONSL;refGene_tx
=XM_011291521.2;refGene_gn=TONSL;refGene_tx=XM_011291523.2;refGene_gn=
TONSL;refGene_tx=XM_011291525.2
F2 85337049 85337221
refGene_gn=TONSL;refGene_tx=XM_004000175.3;refGene_gn=TONSL;refGene_tx
=XM_011291521.2;refGene_gn=TONSL;refGene_tx=XM_011291523.2
F2 85337381 85337683
refGene_gn=TONSL;refGene_tx=XM_004000175.3;refGene_gn=TONSL;refGene_tx
=XM_011291521.2;refGene_gn=TONSL;refGene_tx=XM_011291523.2;refGene_gn=
TONSL;refGene_tx=XM_011291525.2
F2 85338227 85338306
refGene_gn=TONSL;refGene_tx=XM_004000175.3;refGene_gn=TONSL;refGene_tx
=XM_011291521.2;refGene_gn=TONSL;refGene_tx=XM_011291523.2;refGene_gn=
TONSL;refGene_tx=XM_011291525.2
F2 85338392 85338546

refGene_gn=TONSL;refGene_tx=XM_004000175.3;refGene_gn=TONSL;refGene_tx=XM_011291521.2;refGene_gn=TONSL;refGene_tx=XM_011291523.2;refGene_gn=TONSL;refGene_tx=XM_011291525.2
F2 85338948 85339028
refGene_gn=TONSL;refGene_tx=XM_004000175.3;refGene_gn=TONSL;refGene_tx=XM_011291521.2;refGene_gn=TONSL;refGene_tx=XM_011291523.2;refGene_gn=TONSL;refGene_tx=XM_011291525.2
F2 85339120 85339869
refGene_gn=TONSL;refGene_tx=XM_004000175.3;refGene_gn=TONSL;refGene_tx=XM_011291521.2;refGene_gn=TONSL;refGene_tx=XM_011291523.2;refGene_gn=TONSL;refGene_tx=XM_011291525.2
F2 85340008 85340132
refGene_gn=TONSL;refGene_tx=XM_004000175.3;refGene_gn=TONSL;refGene_tx=XM_011291521.2;refGene_gn=TONSL;refGene_tx=XM_011291523.2;refGene_gn=TONSL;refGene_tx=XM_011291525.2
F2 85340206 85340370
refGene_gn=TONSL;refGene_tx=XM_004000175.3;refGene_gn=TONSL;refGene_tx=XM_011291521.2;refGene_gn=TONSL;refGene_tx=XM_011291523.2;refGene_gn=TONSL;refGene_tx=XM_011291525.2
F2 85340447 85340520
refGene_gn=TONSL;refGene_tx=XM_004000175.3;refGene_gn=TONSL;refGene_tx=XM_011291521.2;refGene_gn=TONSL;refGene_tx=XM_011291523.2;refGene_gn=TONSL;refGene_tx=XM_011291525.2
F2 85340875 85340965
refGene_gn=TONSL;refGene_tx=XM_004000175.3;refGene_gn=TONSL;refGene_tx=XM_011291521.2;refGene_gn=TONSL;refGene_tx=XM_011291523.2;refGene_gn=TONSL;refGene_tx=XM_011291525.2
F2 85341067 85341150
refGene_gn=TONSL;refGene_tx=XM_004000175.3;refGene_gn=TONSL;refGene_tx=XM_011291521.2;refGene_gn=TONSL;refGene_tx=XM_011291523.2;refGene_gn=TONSL;refGene_tx=XM_011291525.2
F2 85342390 85342571
refGene_gn=TONSL;refGene_tx=XM_004000175.3;refGene_gn=TONSL;refGene_tx=XM_011291521.2;refGene_gn=TONSL;refGene_tx=XM_011291523.2;refGene_gn=TONSL;refGene_tx=XM_011291525.2
F2 85342712 85342838
refGene_gn=TONSL;refGene_tx=XM_004000175.3;refGene_gn=TONSL;refGene_tx=XM_011291521.2;refGene_gn=TONSL;refGene_tx=XM_011291523.2;refGene_gn=TONSL;refGene_tx=XM_011291525.2
F2 85343073 85343226
refGene_gn=TONSL;refGene_tx=XM_004000175.3;refGene_gn=TONSL;refGene_tx=XM_011291521.2;refGene_gn=TONSL;refGene_tx=XM_011291523.2;refGene_gn=TONSL;refGene_tx=XM_011291525.2
F2 85343326 85343472
refGene_gn=TONSL;refGene_tx=XM_004000175.3;refGene_gn=TONSL;refGene_tx=XM_011291521.2;refGene_gn=TONSL;refGene_tx=XM_011291523.2;refGene_gn=TONSL;refGene_tx=XM_011291525.2
F2 85344010 85344125
refGene_gn=TONSL;refGene_tx=XM_004000175.3;refGene_gn=TONSL;refGene_tx=XM_011291521.2;refGene_gn=TONSL;refGene_tx=XM_011291523.2;refGene_gn=

TONSL;refGene_tx=XM_011291525.2
F2 85344209 85344381
refGene_gn=TONSL;refGene_tx=XM_004000175.3;refGene_gn=TONSL;refGene_tx=XM_011291521.2;refGene_gn=TONSL;refGene_tx=XM_011291523.2;refGene_gn=TONSL;refGene_tx=XM_011291525.2
F2 85344548 85344678
refGene_gn=TONSL;refGene_tx=XM_004000175.3;refGene_gn=TONSL;refGene_tx=XM_011291521.2;refGene_gn=TONSL;refGene_tx=XM_011291523.2;refGene_gn=TONSL;refGene_tx=XM_011291525.2
F2 85345029 85345213
refGene_gn=TONSL;refGene_tx=XM_004000175.3;refGene_gn=TONSL;refGene_tx=XM_011291521.2;refGene_gn=TONSL;refGene_tx=XM_011291523.2;refGene_gn=TONSL;refGene_tx=XM_011291525.2
F2 85345807 85345950
refGene_gn=TONSL;refGene_tx=XM_004000175.3;refGene_gn=TONSL;refGene_tx=XM_011291521.2;refGene_gn=TONSL;refGene_tx=XM_011291523.2;refGene_gn=TONSL;refGene_tx=XM_011291525.2
F2 85346243 85346339
refGene_gn=TONSL;refGene_tx=XM_004000175.3;refGene_gn=TONSL;refGene_tx=XM_011291521.2;refGene_gn=TONSL;refGene_tx=XM_011291523.2;refGene_gn=TONSL;refGene_tx=XM_011291525.2
F2 85346433 85346458
refGene_gn=TONSL;refGene_tx=XM_004000175.3;refGene_gn=TONSL;refGene_tx=XM_011291521.2;refGene_gn=TONSL;refGene_tx=XM_011291523.2;refGene_gn=TONSL;refGene_tx=XM_011291525.2
F2 85350727 85351119 refGene_gn=CYHR1;refGene_tx=XM_023248650.1
F2 85352719 85352931 refGene_gn=CYHR1;refGene_tx=XM_023248650.1
F2 85353698 85353937 refGene_gn=CYHR1;refGene_tx=XM_023248650.1
F2 85362454 85362832 refGene_gn=CYHR1;refGene_tx=XM_023248650.1
F2 85364366 85364852
refGene_gn=LOC111558580;refGene_tx=XM_023248651.1
F2 85365062 85365155
refGene_gn=LOC111558580;refGene_tx=XM_023248651.1
F2 85366036 85366513
refGene_gn=KIFC2;refGene_tx=XM_011291526.3;refGene_gn=KIFC2;refGene_tx=XM_019823275.2
F2 85366755 85366837
refGene_gn=KIFC2;refGene_tx=XM_011291526.3;refGene_gn=KIFC2;refGene_tx=XM_019823275.2
F2 85367011 85367169
refGene_gn=KIFC2;refGene_tx=XM_011291526.3;refGene_gn=KIFC2;refGene_tx=XM_019823275.2
F2 85367287 85367426
refGene_gn=KIFC2;refGene_tx=XM_011291526.3;refGene_gn=KIFC2;refGene_tx=XM_019823275.2
F2 85367587 85367712
refGene_gn=KIFC2;refGene_tx=XM_011291526.3;refGene_gn=KIFC2;refGene_tx=XM_019823275.2
F2 85367796 85367862
refGene_gn=KIFC2;refGene_tx=XM_011291526.3;refGene_gn=KIFC2;refGene_tx

=XM_019823275.2
F2	85367951	85368080
refGene_gn=KIFC2;refGene_tx=XM_011291526.3;refGene_gn=KIFC2;refGene_tx
=XM_019823275.2
F2	85368508	85368586
refGene_gn=KIFC2;refGene_tx=XM_011291526.3;refGene_gn=KIFC2;refGene_tx
=XM_019823275.2
F2	85368701	85368816
refGene_gn=KIFC2;refGene_tx=XM_011291526.3;refGene_gn=KIFC2;refGene_tx
=XM_019823275.2
F2	85368893	85369003
refGene_gn=KIFC2;refGene_tx=XM_011291526.3;refGene_gn=KIFC2;refGene_tx
=XM_019823275.2
F2	85369435	85369544
refGene_gn=KIFC2;refGene_tx=XM_011291526.3;refGene_gn=KIFC2;refGene_tx
=XM_019823275.2
F2	85369653	85369811
refGene_gn=KIFC2;refGene_tx=XM_011291526.3;refGene_gn=KIFC2;refGene_tx
=XM_019823275.2
F2	85369990	85370064	refGene_gn=KIFC2;refGene_tx=XM_019823275.2
F2	85371580	85371685
refGene_gn=KIFC2;refGene_tx=XM_011291526.3;refGene_gn=KIFC2;refGene_tx
=XM_019823275.2
F2	85371767	85371889
refGene_gn=KIFC2;refGene_tx=XM_011291526.3;refGene_gn=KIFC2;refGene_tx
=XM_019823275.2
F2	85371977	85372101
refGene_gn=KIFC2;refGene_tx=XM_011291526.3;refGene_gn=KIFC2;refGene_tx
=XM_019823275.2
F2	85372237	85372367
refGene_gn=KIFC2;refGene_tx=XM_011291526.3;refGene_gn=KIFC2;refGene_tx
=XM_019823275.2
F2	85372456	85372713
refGene_gn=KIFC2;refGene_tx=XM_011291526.3;refGene_gn=KIFC2;refGene_tx
=XM_019823275.2
F2	85372798	85373053
refGene_gn=KIFC2;refGene_tx=XM_011291526.3;refGene_gn=KIFC2;refGene_tx
=XM_019823275.2
F2	85373898	85374741	refGene_gn=FOXH1;refGene_tx=XM_023248649.1
F2	85374829	85374934	refGene_gn=FOXH1;refGene_tx=XM_023248649.1
F2	85375293	85375662	refGene_gn=FOXH1;refGene_tx=XM_023248649.1
F2	85413637	85413902
refGene_gn=PPP1R16A;refGene_tx=XM_004000178.4;refGene_gn=PPP1R16A;refG
ene_tx=XM_011291527.3;refGene_gn=PPP1R16A;refGene_tx=XM_023248775.1
F2	85415130	85415201
refGene_gn=PPP1R16A;refGene_tx=XM_004000178.4;refGene_gn=PPP1R16A;refG
ene_tx=XM_011291527.3;refGene_gn=PPP1R16A;refGene_tx=XM_023248775.1
F2	85415278	85415424
refGene_gn=PPP1R16A;refGene_tx=XM_004000178.4;refGene_gn=PPP1R16A;refG
ene_tx=XM_011291527.3;refGene_gn=PPP1R16A;refGene_tx=XM_023248775.1

F2 85416578 85416682
refGene_gn=PPP1R16A;refGene_tx=XM_004000178.4;refGene_gn=PPP1R16A;refGene_tx=XM_011291527.3;refGene_gn=PPP1R16A;refGene_tx=XM_023248775.1
F2 85416748 85416873
refGene_gn=PPP1R16A;refGene_tx=XM_004000178.4;refGene_gn=PPP1R16A;refGene_tx=XM_011291527.3;refGene_gn=PPP1R16A;refGene_tx=XM_023248775.1
F2 85416961 85417087
refGene_gn=PPP1R16A;refGene_tx=XM_004000178.4;refGene_gn=PPP1R16A;refGene_tx=XM_011291527.3;refGene_gn=PPP1R16A;refGene_tx=XM_023248775.1
F2 85417175 85417251
refGene_gn=PPP1R16A;refGene_tx=XM_004000178.4;refGene_gn=PPP1R16A;refGene_tx=XM_011291527.3;refGene_gn=PPP1R16A;refGene_tx=XM_023248775.1
F2 85417324 85417454
refGene_gn=PPP1R16A;refGene_tx=XM_004000178.4;refGene_gn=PPP1R16A;refGene_tx=XM_011291527.3;refGene_gn=PPP1R16A;refGene_tx=XM_023248775.1
F2 85417620 85417786
refGene_gn=PPP1R16A;refGene_tx=XM_004000178.4;refGene_gn=PPP1R16A;refGene_tx=XM_011291527.3;refGene_gn=PPP1R16A;refGene_tx=XM_023248775.1
F2 85418043 85418418
refGene_gn=PPP1R16A;refGene_tx=XM_004000178.4;refGene_gn=PPP1R16A;refGene_tx=XM_011291527.3;refGene_gn=PPP1R16A;refGene_tx=XM_023248775.1
F2 85420844 85421006
refGene_gn=GPT;refGene_tx=XM_023248776.1;refGene_gn=GPT;refGene_tx=XM_023248777.1
F2 85421150 85421240
refGene_gn=GPT;refGene_tx=XM_023248776.1;refGene_gn=GPT;refGene_tx=XM_023248777.1
F2 85421320 85421429
refGene_gn=GPT;refGene_tx=XM_023248776.1;refGene_gn=GPT;refGene_tx=XM_023248777.1
F2 85421611 85421745
refGene_gn=GPT;refGene_tx=XM_023248776.1;refGene_gn=GPT;refGene_tx=XM_023248777.1
F2 85421911 85422155
refGene_gn=GPT;refGene_tx=XM_023248776.1;refGene_gn=GPT;refGene_tx=XM_023248777.1
F2 85422283 85422363
refGene_gn=GPT;refGene_tx=XM_023248776.1;refGene_gn=GPT;refGene_tx=XM_023248777.1
F2 85422443 85422580
refGene_gn=GPT;refGene_tx=XM_023248776.1;refGene_gn=GPT;refGene_tx=XM_023248777.1
F2 85422702 85422877
refGene_gn=GPT;refGene_tx=XM_023248776.1;refGene_gn=GPT;refGene_tx=XM_023248777.1
F2 85422972 85423128 refGene_gn=GPT;refGene_tx=XM_023248776.1
F2 85423241 85423354
refGene_gn=GPT;refGene_tx=XM_023248776.1;refGene_gn=GPT;refGene_tx=XM_023248777.1
F2 85423424 85423515

refGene_gn=GPT;refGene_tx=XM_023248776.1;refGene_gn=GPT;refGene_tx=XM_
023248777.1
F2	85425682	85426364	refGene_gn=MFSD3;refGene_tx=XM_004000227.5
F2	85426741	85426889	refGene_gn=MFSD3;refGene_tx=XM_004000227.5
F2	85426974	85427099	refGene_gn=MFSD3;refGene_tx=XM_004000227.5
F2	85427166	85427276	refGene_gn=MFSD3;refGene_tx=XM_004000227.5
F2	85427367	85427541	refGene_gn=MFSD3;refGene_tx=XM_004000227.5
F2	85427874	85427999	refGene_gn=RECQL4;refGene_tx=XM_019823278.2
F2	85428087	85428196	refGene_gn=RECQL4;refGene_tx=XM_019823278.2
F2	85428372	85428520	refGene_gn=RECQL4;refGene_tx=XM_019823278.2
F2	85428602	85428783	refGene_gn=RECQL4;refGene_tx=XM_019823278.2
F2	85428862	85429032	refGene_gn=RECQL4;refGene_tx=XM_019823278.2
F2	85429113	85429243	refGene_gn=RECQL4;refGene_tx=XM_019823278.2
F2	85429322	85429599	refGene_gn=RECQL4;refGene_tx=XM_019823278.2
F2	85429676	85429939	refGene_gn=RECQL4;refGene_tx=XM_019823278.2
F2	85430043	85430158	refGene_gn=RECQL4;refGene_tx=XM_019823278.2
F2	85430230	85430410	refGene_gn=RECQL4;refGene_tx=XM_019823278.2
F2	85430488	85430668	refGene_gn=RECQL4;refGene_tx=XM_019823278.2
F2	85430750	85430834	refGene_gn=RECQL4;refGene_tx=XM_019823278.2
F2	85431284	85431421	refGene_gn=RECQL4;refGene_tx=XM_019823278.2
F2	85431498	85431591	refGene_gn=RECQL4;refGene_tx=XM_019823278.2
F2	85431674	85431806	refGene_gn=RECQL4;refGene_tx=XM_019823278.2
F2	85431910	85432024	refGene_gn=RECQL4;refGene_tx=XM_019823278.2
F2	85432160	85432287	refGene_gn=RECQL4;refGene_tx=XM_019823278.2
F2	85432358	85432516	refGene_gn=RECQL4;refGene_tx=XM_019823278.2
F2	85432518	85433131	refGene_gn=RECQL4;refGene_tx=XM_019823278.2
F2	85433375	85433519	refGene_gn=RECQL4;refGene_tx=XM_019823278.2
F2	85433731	85433832	refGene_gn=RECQL4;refGene_tx=XM_019823278.2
F2	85433920	85433954	refGene_gn=RECQL4;refGene_tx=XM_019823278.2
F2	85434019	85434247	refGene_gn=RECQL4;refGene_tx=XM_019823278.2
F2	85434723	85434795
refGene_gn=LRRC14;refGene_tx=XM_011291530.3;refGene_gn=LRRC14;refGene_
tx=XM_023248282.1
F2	85434896	85434901
refGene_gn=LRRC14;refGene_tx=XM_006943444.4;refGene_gn=LRRC14;refGene_
tx=XM_023248279.1
F2	85436016	85436370
refGene_gn=LRRC14;refGene_tx=XM_006943444.4;refGene_gn=LRRC14;refGene_
tx=XM_011291530.3;refGene_gn=LRRC14;refGene_tx=XM_023248279.1;refGene_
gn=LRRC14;refGene_tx=XM_023248280.1;refGene_gn=LRRC14;refGene_tx=XM_02
3248281.1;refGene_gn=LRRC14;refGene_tx=XM_023248282.1;refGene_gn=LRRC1
4;refGene_tx=XM_023248283.1
F2	85436534	85437136
refGene_gn=LRRC14;refGene_tx=XM_006943444.4;refGene_gn=LRRC14;refGene_
tx=XM_011291530.3;refGene_gn=LRRC14;refGene_tx=XM_023248279.1;refGene_
gn=LRRC14;refGene_tx=XM_023248280.1;refGene_gn=LRRC14;refGene_tx=XM_02
3248281.1;refGene_gn=LRRC14;refGene_tx=XM_023248282.1;refGene_gn=LRRC1
4;refGene_tx=XM_023248283.1
F2	85437228	85437796
refGene_gn=LRRC14;refGene_tx=XM_006943444.4;refGene_gn=LRRC14;refGene_

tx=XM_011291530.3;refGene_gn=LRRC14;refGene_tx=XM_023248279.1;refGene_gn=LRRC14;refGene_tx=XM_023248280.1;refGene_gn=LRRC14;refGene_tx=XM_023248281.1;refGene_gn=LRRC14;refGene_tx=XM_023248282.1;refGene_gn=LRRC14;refGene_tx=XM_023248283.1
F2 85438848 85439777
refGene_gn=LRRC24;refGene_tx=XM_006943442.3;refGene_gn=LRRC24;refGene_tx=XM_019823283.2;refGene_gn=LRRC24;refGene_tx=XM_019823284.2;refGene_gn=LRRC24;refGene_tx=XM_023248277.1;refGene_gn=LRRC24;refGene_tx=XM_023248278.1
F2 85440452 85440621
refGene_gn=LRRC24;refGene_tx=XM_006943442.3;refGene_gn=LRRC24;refGene_tx=XM_019823283.2;refGene_gn=LRRC24;refGene_tx=XM_019823284.2;refGene_gn=LRRC24;refGene_tx=XM_023248277.1;refGene_gn=LRRC24;refGene_tx=XM_023248278.1
F2 85440784 85441063
refGene_gn=LRRC24;refGene_tx=XM_006943442.3;refGene_gn=LRRC24;refGene_tx=XM_019823283.2;refGene_gn=LRRC24;refGene_tx=XM_019823284.2;refGene_gn=LRRC24;refGene_tx=XM_023248277.1
F2 85441218 85441524
refGene_gn=LRRC24;refGene_tx=XM_006943442.3;refGene_gn=LRRC24;refGene_tx=XM_019823283.2;refGene_gn=LRRC24;refGene_tx=XM_019823284.2;refGene_gn=LRRC24;refGene_tx=XM_023248277.1;refGene_gn=LRRC24;refGene_tx=XM_023248278.1
F2 85443515 85443834
refGene_gn=LRRC24;refGene_tx=XM_019823284.2;refGene_gn=LRRC24;refGene_tx=XM_023248277.1;refGene_gn=LRRC24;refGene_tx=XM_023248278.1
F2 85443964 85444410
refGene_gn=CF2H8orf82;refGene_tx=XM_023248284.1
F2 85444677 85444726
refGene_gn=CF2H8orf82;refGene_tx=XM_023248284.1
F2 85445401 85445563
refGene_gn=CF2H8orf82;refGene_tx=XM_023248284.1
F2 85447677 85447872
refGene_gn=ARHGAP39;refGene_tx=XM_019823282.2;refGene_gn=ARHGAP39;refGene_tx=XM_023248274.1;refGene_gn=ARHGAP39;refGene_tx=XM_023248275.1;refGene_gn=ARHGAP39;refGene_tx=XM_023248276.1
F2 85447950 85448120
refGene_gn=ARHGAP39;refGene_tx=XM_019823282.2;refGene_gn=ARHGAP39;refGene_tx=XM_023248274.1;refGene_gn=ARHGAP39;refGene_tx=XM_023248275.1;refGene_gn=ARHGAP39;refGene_tx=XM_023248276.1
F2 85448450 85448542
refGene_gn=ARHGAP39;refGene_tx=XM_019823282.2;refGene_gn=ARHGAP39;refGene_tx=XM_023248274.1;refGene_gn=ARHGAP39;refGene_tx=XM_023248275.1;refGene_gn=ARHGAP39;refGene_tx=XM_023248276.1
F2 85449277 85449477
refGene_gn=ARHGAP39;refGene_tx=XM_019823282.2;refGene_gn=ARHGAP39;refGene_tx=XM_023248274.1;refGene_gn=ARHGAP39;refGene_tx=XM_023248275.1;refGene_gn=ARHGAP39;refGene_tx=XM_023248276.1
F2 85450860 85450934
refGene_gn=ARHGAP39;refGene_tx=XM_019823282.2;refGene_gn=ARHGAP39;refG

ene_tx=XM_023248274.1;refGene_gn=ARHGAP39;refGene_tx=XM_023248275.1;refGene_gn=ARHGAP39;refGene_tx=XM_023248276.1
F2 85454184 85454277
refGene_gn=ARHGAP39;refGene_tx=XM_019823282.2;refGene_gn=ARHGAP39;refGene_tx=XM_023248274.1;refGene_gn=ARHGAP39;refGene_tx=XM_023248276.1
F2 85463633 85464195
refGene_gn=ARHGAP39;refGene_tx=XM_019823282.2;refGene_gn=ARHGAP39;refGene_tx=XM_023248274.1;refGene_gn=ARHGAP39;refGene_tx=XM_023248275.1;refGene_gn=ARHGAP39;refGene_tx=XM_023248276.1
F2 85465702 85467026
refGene_gn=ARHGAP39;refGene_tx=XM_019823282.2;refGene_gn=ARHGAP39;refGene_tx=XM_023248274.1;refGene_gn=ARHGAP39;refGene_tx=XM_023248275.1;refGene_gn=ARHGAP39;refGene_tx=XM_023248276.1
F2 85472889 85472973
refGene_gn=ARHGAP39;refGene_tx=XM_023248274.1;refGene_gn=ARHGAP39;refGene_tx=XM_023248275.1;refGene_gn=ARHGAP39;refGene_tx=XM_023248276.1
F2 85489373 85489808
refGene_gn=ARHGAP39;refGene_tx=XM_023248274.1;refGene_gn=ARHGAP39;refGene_tx=XM_023248275.1
F2 85506768 85506848
refGene_gn=ARHGAP39;refGene_tx=XM_023248274.1;refGene_gn=ARHGAP39;refGene_tx=XM_023248275.1;refGene_gn=ARHGAP39;refGene_tx=XM_023248276.1
F2 85576182 85576857 refGene_gn=COMMD5;refGene_tx=XM_004000184.4
F2 85585411 85587210
refGene_gn=ZNF7;refGene_tx=XM_006943466.3;refGene_gn=ZNF7;refGene_tx=XM_011291532.3;refGene_gn=ZNF7;refGene_tx=XM_011291533.3;refGene_gn=ZNF7;refGene_tx=XM_011291534.3;refGene_gn=ZNF7;refGene_tx=XM_011291535.3;refGene_gn=ZNF7;refGene_tx=XM_019823290.2;refGene_gn=ZNF7;refGene_tx=XM_019823291.2;refGene_gn=ZNF7;refGene_tx=XM_019823292.2;refGene_gn=ZNF7;refGene_tx=XM_019823294.2;refGene_gn=ZNF7;refGene_tx=XM_019823296.1;refGene_gn=ZNF7;refGene_tx=XM_019823297.2;refGene_gn=ZNF7;refGene_tx=XM_023248603.1;refGene_gn=ZNF7;refGene_tx=XM_023248604.1
F2 85590662 85590779
refGene_gn=ZNF7;refGene_tx=XM_006943466.3;refGene_gn=ZNF7;refGene_tx=XM_011291532.3;refGene_gn=ZNF7;refGene_tx=XM_011291533.3;refGene_gn=ZNF7;refGene_tx=XM_011291534.3;refGene_gn=ZNF7;refGene_tx=XM_011291535.3;refGene_gn=ZNF7;refGene_tx=XM_019823290.2;refGene_gn=ZNF7;refGene_tx=XM_019823291.2;refGene_gn=ZNF7;refGene_tx=XM_019823292.2;refGene_gn=ZNF7;refGene_tx=XM_019823294.2;refGene_gn=ZNF7;refGene_tx=XM_019823296.1;refGene_gn=ZNF7;refGene_tx=XM_019823297.2;refGene_gn=ZNF7;refGene_tx=XM_023248603.1;refGene_gn=ZNF7;refGene_tx=XM_023248604.1
F2 85595631 85595842
refGene_gn=ZNF7;refGene_tx=XM_006943466.3;refGene_gn=ZNF7;refGene_tx=XM_011291532.3;refGene_gn=ZNF7;refGene_tx=XM_011291533.3;refGene_gn=ZNF7;refGene_tx=XM_011291534.3;refGene_gn=ZNF7;refGene_tx=XM_011291535.3;refGene_gn=ZNF7;refGene_tx=XM_019823290.2;refGene_gn=ZNF7;refGene_tx=XM_019823291.2;refGene_gn=ZNF7;refGene_tx=XM_019823292.2;refGene_gn=ZNF7;refGene_tx=XM_019823294.2;refGene_gn=ZNF7;refGene_tx=XM_019823296.1;refGene_gn=ZNF7;refGene_tx=XM_019823297.2;refGene_gn=ZNF7;refGene_tx=XM_023248603.1;refGene_gn=ZNF7;refGene_tx=XM_023248604.1

F2 85596128 85596171
refGene_gn=ZNF7;refGene_tx=XM_006943466.3;refGene_gn=ZNF7;refGene_tx=XM_011291533.3;refGene_gn=ZNF7;refGene_tx=XM_011291534.3;refGene_gn=ZNF7;refGene_tx=XM_011291535.3;refGene_gn=ZNF7;refGene_tx=XM_019823290.2;refGene_gn=ZNF7;refGene_tx=XM_019823291.2;refGene_gn=ZNF7;refGene_tx=XM_019823292.2;refGene_gn=ZNF7;refGene_tx=XM_019823297.2;refGene_gn=ZNF7;refGene_tx=XM_023248604.1
F2 85597236 85597352
refGene_gn=ZNF7;refGene_tx=XM_019823290.2;refGene_gn=ZNF7;refGene_tx=XM_019823291.2;refGene_gn=ZNF7;refGene_tx=XM_019823292.2
F2 85603897 85605099
refGene_gn=ZNF517;refGene_tx=XM_006943468.4;refGene_gn=ZNF517;refGene_tx=XM_019823301.2;refGene_gn=ZNF517;refGene_tx=XM_019823302.1;refGene_gn=ZNF517;refGene_tx=XM_023248605.1;refGene_gn=ZNF517;refGene_tx=XM_023248606.1
F2 85605894 85606008
refGene_gn=ZNF517;refGene_tx=XM_006943468.4;refGene_gn=ZNF517;refGene_tx=XM_019823301.2;refGene_gn=ZNF517;refGene_tx=XM_019823302.1;refGene_gn=ZNF517;refGene_tx=XM_023248605.1
F2 85606293 85606420
refGene_gn=ZNF517;refGene_tx=XM_006943468.4;refGene_gn=ZNF517;refGene_tx=XM_019823301.2;refGene_gn=ZNF517;refGene_tx=XM_019823302.1;refGene_gn=ZNF517;refGene_tx=XM_023248605.1
F2 85607096 85607129
refGene_gn=ZNF517;refGene_tx=XM_019823301.2;refGene_gn=ZNF517;refGene_tx=XM_023248605.1
F2 85616393 85616531 refGene_gn=RPL8;refGene_tx=XM_006943470.2
F2 85616610 85616752 refGene_gn=RPL8;refGene_tx=XM_006943470.2
F2 85617265 85617484 refGene_gn=RPL8;refGene_tx=XM_006943470.2
F2 85618183 85618299 refGene_gn=RPL8;refGene_tx=XM_006943470.2
F2 85618530 85618689 refGene_gn=RPL8;refGene_tx=XM_006943470.2
F2 85627057 85627090
refGene_gn=ZNF34;refGene_tx=XM_004000232.4;refGene_gn=ZNF34;refGene_tx=XM_006943471.3;refGene_gn=ZNF34;refGene_tx=XM_006943472.3;refGene_gn=ZNF34;refGene_tx=XM_019823310.2;refGene_gn=ZNF34;refGene_tx=XM_019823311.2
F2 85627402 85627529
refGene_gn=ZNF34;refGene_tx=XM_004000232.4;refGene_gn=ZNF34;refGene_tx=XM_006943471.3;refGene_gn=ZNF34;refGene_tx=XM_006943472.3;refGene_gn=ZNF34;refGene_tx=XM_019823310.2;refGene_gn=ZNF34;refGene_tx=XM_019823311.2
F2 85627966 85628086
refGene_gn=ZNF34;refGene_tx=XM_004000232.4;refGene_gn=ZNF34;refGene_tx=XM_006943471.3;refGene_gn=ZNF34;refGene_tx=XM_006943472.3;refGene_gn=ZNF34;refGene_tx=XM_019823310.2;refGene_gn=ZNF34;refGene_tx=XM_019823311.2
F2 85630271 85631632
refGene_gn=ZNF34;refGene_tx=XM_004000232.4;refGene_gn=ZNF34;refGene_tx=XM_006943471.3;refGene_gn=ZNF34;refGene_tx=XM_006943472.3;refGene_gn=ZNF34;refGene_tx=XM_019823310.2;refGene_gn=ZNF34;refGene_tx=XM_0198233

11.2
F2	85646530	85647855
refGene_gn=ZNF250;refGene_tx=XM_004000181.5;refGene_gn=ZNF250;refGene_tx=XM_006943474.4;refGene_gn=ZNF250;refGene_tx=XM_006943475.4;refGene_gn=ZNF250;refGene_tx=XM_019823304.2;refGene_gn=ZNF250;refGene_tx=XM_019823305.2;refGene_gn=ZNF250;refGene_tx=XM_019823306.2;refGene_gn=ZNF250;refGene_tx=XM_019823307.2;refGene_gn=ZNF250;refGene_tx=XM_019823308.2;refGene_gn=ZNF250;refGene_tx=XM_019823309.2
F2	85657179	85657242
refGene_gn=ZNF250;refGene_tx=XM_004000181.5;refGene_gn=ZNF250;refGene_tx=XM_006943474.4;refGene_gn=ZNF250;refGene_tx=XM_006943475.4;refGene_gn=ZNF250;refGene_tx=XM_019823304.2;refGene_gn=ZNF250;refGene_tx=XM_019823305.2;refGene_gn=ZNF250;refGene_tx=XM_019823306.2;refGene_gn=ZNF250;refGene_tx=XM_019823307.2;refGene_gn=ZNF250;refGene_tx=XM_019823308.2;refGene_gn=ZNF250;refGene_tx=XM_019823309.2
F2	85657708	85657822
refGene_gn=ZNF250;refGene_tx=XM_004000181.5;refGene_gn=ZNF250;refGene_tx=XM_006943474.4;refGene_gn=ZNF250;refGene_tx=XM_006943475.4;refGene_gn=ZNF250;refGene_tx=XM_019823304.2;refGene_gn=ZNF250;refGene_tx=XM_019823305.2;refGene_gn=ZNF250;refGene_tx=XM_019823306.2;refGene_gn=ZNF250;refGene_tx=XM_019823307.2;refGene_gn=ZNF250;refGene_tx=XM_019823308.2;refGene_gn=ZNF250;refGene_tx=XM_019823309.2
F2	85658026	85658168
refGene_gn=ZNF250;refGene_tx=XM_004000181.5;refGene_gn=ZNF250;refGene_tx=XM_006943474.4;refGene_gn=ZNF250;refGene_tx=XM_006943475.4;refGene_gn=ZNF250;refGene_tx=XM_019823304.2;refGene_gn=ZNF250;refGene_tx=XM_019823305.2;refGene_gn=ZNF250;refGene_tx=XM_019823306.2;refGene_gn=ZNF250;refGene_tx=XM_019823307.2;refGene_gn=ZNF250;refGene_tx=XM_019823308.2;refGene_gn=ZNF250;refGene_tx=XM_019823309.2
F2	85658402	85658444
refGene_gn=ZNF250;refGene_tx=XM_004000181.5;refGene_gn=ZNF250;refGene_tx=XM_006943474.4;refGene_gn=ZNF250;refGene_tx=XM_006943475.4;refGene_gn=ZNF250;refGene_tx=XM_019823304.2;refGene_gn=ZNF250;refGene_tx=XM_019823305.2;refGene_gn=ZNF250;refGene_tx=XM_019823306.2;refGene_gn=ZNF250;refGene_tx=XM_019823307.2;refGene_gn=ZNF250;refGene_tx=XM_019823308.2;refGene_gn=ZNF250;refGene_tx=XM_019823309.2
F2	85679876	85681723
refGene_gn=ZNF16;refGene_tx=XM_004000185.3;refGene_gn=ZNF16;refGene_tx=XM_006943476.2
F2	85690713	85690909
refGene_gn=ZNF16;refGene_tx=XM_004000185.3;refGene_gn=ZNF16;refGene_tx=XM_006943476.2
F2	85715708	85718068
refGene_gn=LOC101098926;refGene_tx=XM_019823287.1;refGene_gn=LOC101098926;refGene_tx=XM_019823289.1
F2	85729111	85729225
refGene_gn=LOC101098926;refGene_tx=XM_019823287.1;refGene_gn=LOC101098926;refGene_tx=XM_019823289.1
F2	85729803	85729930
refGene_gn=LOC101098926;refGene_tx=XM_019823287.1;refGene_gn=LOC101098

926;refGene_tx=XM_019823289.1
F2	85730316	85730355
refGene_gn=LOC101098926;refGene_tx=XM_019823287.1
F2	85738994	85739000
refGene_gn=CF2H8orf33;refGene_tx=XM_006943479.3
F2	85739088	85739134
refGene_gn=CF2H8orf33;refGene_tx=XM_006943479.3
F2	85739264	85739443
refGene_gn=CF2H8orf33;refGene_tx=XM_006943479.3
F2	85739668	85739753
refGene_gn=CF2H8orf33;refGene_tx=XM_006943479.3
F2	85740012	85740159
refGene_gn=CF2H8orf33;refGene_tx=XM_006943479.3
F2	85740507	85740647
refGene_gn=CF2H8orf33;refGene_tx=XM_006943479.3
KP081775.1	297900	297980	refGene_gn=DDX3Y;refGene_tx=DDX3Y
KP081775.1	298175	298309	refGene_gn=DDX3Y;refGene_tx=DDX3Y
KP081775.1	298826	298980	refGene_gn=DDX3Y;refGene_tx=DDX3Y
KP081775.1	299325	299443	refGene_gn=DDX3Y;refGene_tx=DDX3Y
KP081775.1	299657	299839	refGene_gn=DDX3Y;refGene_tx=DDX3Y
KP081775.1	300172	300317	refGene_gn=DDX3Y;refGene_tx=DDX3Y
KP081775.1	300422	300567	refGene_gn=DDX3Y;refGene_tx=DDX3Y
KP081775.1	300851	301012	refGene_gn=DDX3Y;refGene_tx=DDX3Y
KP081775.1	301099	301198	refGene_gn=DDX3Y;refGene_tx=DDX3Y
KP081775.1	301451	301537	refGene_gn=DDX3Y;refGene_tx=DDX3Y
KP081775.1	303261	303397	refGene_gn=DDX3Y;refGene_tx=DDX3Y
KP081775.1	304158	304258	refGene_gn=DDX3Y;refGene_tx=DDX3Y
KP081775.1	304336	304492	refGene_gn=DDX3Y;refGene_tx=DDX3Y
KP081775.1	305579	305709	refGene_gn=DDX3Y;refGene_tx=DDX3Y
KP081775.1	308543	308591	refGene_gn=DDX3Y;refGene_tx=DDX3Y
KP081775.1	311279	311340	refGene_gn=DDX3Y;refGene_tx=DDX3Y
KP081775.1	314168	314215	refGene_gn=DDX3Y;refGene_tx=DDX3Y
KP081775.1	314871	314890	refGene_gn=DDX3Y;refGene_tx=DDX3Y
KP081775.1	341286	341424	refGene_gn=USP9Y;refGene_tx=USP9Y
KP081775.1	342451	342547	refGene_gn=USP9Y;refGene_tx=USP9Y
KP081775.1	342943	343156	refGene_gn=USP9Y;refGene_tx=USP9Y
KP081775.1	343294	343451	refGene_gn=USP9Y;refGene_tx=USP9Y
KP081775.1	346362	346451	refGene_gn=USP9Y;refGene_tx=USP9Y
KP081775.1	346537	346758	refGene_gn=USP9Y;refGene_tx=USP9Y
KP081775.1	347159	347345	refGene_gn=USP9Y;refGene_tx=USP9Y
KP081775.1	352368	352498	refGene_gn=USP9Y;refGene_tx=USP9Y
KP081775.1	353014	353240	refGene_gn=USP9Y;refGene_tx=USP9Y
KP081775.1	353816	353940	refGene_gn=USP9Y;refGene_tx=USP9Y
KP081775.1	354340	355088	refGene_gn=USP9Y;refGene_tx=USP9Y
KP081775.1	356168	356310	refGene_gn=USP9Y;refGene_tx=USP9Y
KP081775.1	359053	359227	refGene_gn=USP9Y;refGene_tx=USP9Y
KP081775.1	367223	367414	refGene_gn=USP9Y;refGene_tx=USP9Y
KP081775.1	372235	372456	refGene_gn=USP9Y;refGene_tx=USP9Y
KP081775.1	374377	374600	refGene_gn=USP9Y;refGene_tx=USP9Y
KP081775.1	376132	376279	refGene_gn=USP9Y;refGene_tx=USP9Y

| | | | |
|---|---|---|---|
| KP081775.1 | 377054 | 377201 | refGene_gn=USP9Y;refGene_tx=USP9Y |
| KP081775.1 | 377632 | 377741 | refGene_gn=USP9Y;refGene_tx=USP9Y |
| KP081775.1 | 388378 | 388548 | refGene_gn=USP9Y;refGene_tx=USP9Y |
| KP081775.1 | 388772 | 388898 | refGene_gn=USP9Y;refGene_tx=USP9Y |
| KP081775.1 | 395319 | 395445 | refGene_gn=USP9Y;refGene_tx=USP9Y |
| KP081775.1 | 397814 | 398093 | refGene_gn=USP9Y;refGene_tx=USP9Y |
| KP081775.1 | 398219 | 398350 | refGene_gn=USP9Y;refGene_tx=USP9Y |
| KP081775.1 | 408925 | 409046 | refGene_gn=USP9Y;refGene_tx=USP9Y |
| KP081775.1 | 410045 | 410195 | refGene_gn=USP9Y;refGene_tx=USP9Y |
| KP081775.1 | 410728 | 410969 | refGene_gn=USP9Y;refGene_tx=USP9Y |
| KP081775.1 | 412236 | 412448 | refGene_gn=USP9Y;refGene_tx=USP9Y |
| KP081775.1 | 412925 | 413021 | refGene_gn=USP9Y;refGene_tx=USP9Y |
| KP081775.1 | 414008 | 414351 | refGene_gn=USP9Y;refGene_tx=USP9Y |
| KP081775.1 | 416966 | 417054 | refGene_gn=USP9Y;refGene_tx=USP9Y |
| KP081775.1 | 423118 | 423252 | refGene_gn=USP9Y;refGene_tx=USP9Y |
| KP081775.1 | 424858 | 424995 | refGene_gn=USP9Y;refGene_tx=USP9Y |
| KP081775.1 | 426547 | 426754 | refGene_gn=USP9Y;refGene_tx=USP9Y |
| KP081775.1 | 438189 | 438294 | refGene_gn=USP9Y;refGene_tx=USP9Y |
| KP081775.1 | 439796 | 439949 | refGene_gn=USP9Y;refGene_tx=USP9Y |
| KP081775.1 | 441804 | 441955 | refGene_gn=USP9Y;refGene_tx=USP9Y |
| KP081775.1 | 442013 | 442265 | refGene_gn=USP9Y;refGene_tx=USP9Y |
| KP081775.1 | 442665 | 442781 | refGene_gn=USP9Y;refGene_tx=USP9Y |
| KP081775.1 | 447056 | 447275 | refGene_gn=USP9Y;refGene_tx=USP9Y |
| KP081775.1 | 448523 | 448636 | refGene_gn=USP9Y;refGene_tx=USP9Y |
| KP081775.1 | 451443 | 451523 | refGene_gn=USP9Y;refGene_tx=USP9Y |
| KP081775.1 | 453361 | 453507 | refGene_gn=USP9Y;refGene_tx=USP9Y |
| KP081775.1 | 459618 | 459714 | refGene_gn=USP9Y;refGene_tx=USP9Y |
| KP081775.1 | 864841 | 864977 | refGene_gn=UBE1Y;refGene_tx=UBE1Y |
| KP081775.1 | 865096 | 865197 | refGene_gn=UBE1Y;refGene_tx=UBE1Y |
| KP081775.1 | 865296 | 865398 | refGene_gn=UBE1Y;refGene_tx=UBE1Y |
| KP081775.1 | 865889 | 866081 | refGene_gn=UBE1Y;refGene_tx=UBE1Y |
| KP081775.1 | 866182 | 866275 | refGene_gn=UBE1Y;refGene_tx=UBE1Y |
| KP081775.1 | 866469 | 866558 | refGene_gn=UBE1Y;refGene_tx=UBE1Y |
| KP081775.1 | 868826 | 869016 | refGene_gn=UBE1Y;refGene_tx=UBE1Y |
| KP081775.1 | 869130 | 869205 | refGene_gn=UBE1Y;refGene_tx=UBE1Y |
| KP081775.1 | 869816 | 870012 | refGene_gn=UBE1Y;refGene_tx=UBE1Y |
| KP081775.1 | 870235 | 870300 | refGene_gn=UBE1Y;refGene_tx=UBE1Y |
| KP081775.1 | 872053 | 872250 | refGene_gn=UBE1Y;refGene_tx=UBE1Y |
| KP081775.1 | 872360 | 872526 | refGene_gn=UBE1Y;refGene_tx=UBE1Y |
| KP081775.1 | 874798 | 874951 | refGene_gn=UBE1Y;refGene_tx=UBE1Y |
| KP081775.1 | 875287 | 875368 | refGene_gn=UBE1Y;refGene_tx=UBE1Y |
| KP081775.1 | 875450 | 875555 | refGene_gn=UBE1Y;refGene_tx=UBE1Y |
| KP081775.1 | 875683 | 875860 | refGene_gn=UBE1Y;refGene_tx=UBE1Y |
| KP081775.1 | 875988 | 876135 | refGene_gn=UBE1Y;refGene_tx=UBE1Y |
| KP081775.1 | 876217 | 876315 | refGene_gn=UBE1Y;refGene_tx=UBE1Y |
| KP081775.1 | 877608 | 877741 | refGene_gn=UBE1Y;refGene_tx=UBE1Y |
| KP081775.1 | 877849 | 877940 | refGene_gn=UBE1Y;refGene_tx=UBE1Y |
| KP081775.1 | 878208 | 878315 | refGene_gn=UBE1Y;refGene_tx=UBE1Y |
| KP081775.1 | 879467 | 879602 | refGene_gn=UBE1Y;refGene_tx=UBE1Y |
| KP081775.1 | 879691 | 879860 | refGene_gn=UBE1Y;refGene_tx=UBE1Y |

KP081775.1	879960	880019	refGene_gn=UBE1Y;refGene_tx=UBE1Y
KP081775.1	880144	880261	refGene_gn=UBE1Y;refGene_tx=UBE1Y
KP081775.1	936038	936188	refGene_gn=KDM5D;refGene_tx=KDM5D
KP081775.1	937234	937312	refGene_gn=KDM5D;refGene_tx=KDM5D
KP081775.1	937979	938102	refGene_gn=KDM5D;refGene_tx=KDM5D
KP081775.1	938936	939107	refGene_gn=KDM5D;refGene_tx=KDM5D
KP081775.1	940078	940213	refGene_gn=KDM5D;refGene_tx=KDM5D
KP081775.1	944242	944366	refGene_gn=KDM5D;refGene_tx=KDM5D
KP081775.1	944445	944630	refGene_gn=KDM5D;refGene_tx=KDM5D
KP081775.1	946939	947098	refGene_gn=KDM5D;refGene_tx=KDM5D
KP081775.1	947741	947861	refGene_gn=KDM5D;refGene_tx=KDM5D
KP081775.1	947976	948135	refGene_gn=KDM5D;refGene_tx=KDM5D
KP081775.1	962111	962293	refGene_gn=KDM5D;refGene_tx=KDM5D
KP081775.1	962386	962549	refGene_gn=KDM5D;refGene_tx=KDM5D
KP081775.1	967065	967185	refGene_gn=KDM5D;refGene_tx=KDM5D
KP081775.1	967308	967503	refGene_gn=KDM5D;refGene_tx=KDM5D
KP081775.1	967775	967957	refGene_gn=KDM5D;refGene_tx=KDM5D
KP081775.1	968039	968164	refGene_gn=KDM5D;refGene_tx=KDM5D
KP081775.1	968285	968433	refGene_gn=KDM5D;refGene_tx=KDM5D
KP081775.1	973526	973623	refGene_gn=KDM5D;refGene_tx=KDM5D
KP081775.1	974169	974528	refGene_gn=KDM5D;refGene_tx=KDM5D
KP081775.1	974937	975076	refGene_gn=KDM5D;refGene_tx=KDM5D
KP081775.1	975489	975669	refGene_gn=KDM5D;refGene_tx=KDM5D
KP081775.1	975802	975940	refGene_gn=KDM5D;refGene_tx=KDM5D
KP081775.1	976130	976724	refGene_gn=KDM5D;refGene_tx=KDM5D
KP081775.1	977000	977073	refGene_gn=KDM5D;refGene_tx=KDM5D
KP081775.1	977208	977408	refGene_gn=KDM5D;refGene_tx=KDM5D
KP081775.1	977518	977872	refGene_gn=KDM5D;refGene_tx=KDM5D
KP081775.1	1666519	1667224	refGene_gn=SRY;refGene_tx=SRY
X	57180	57271
refGene_gn=LOC101092754;refGene_tx=XM_023249055.1
X	62367	62505
refGene_gn=LOC101092754;refGene_tx=XM_023249055.1
X	66871	67008
refGene_gn=LOC101092754;refGene_tx=XM_023249055.1
X	70493	70622
refGene_gn=LOC101092754;refGene_tx=XM_023249055.1
X	71959	72115
refGene_gn=LOC101092754;refGene_tx=XM_023249055.1
X	72150	72152
refGene_gn=LOC101092754;refGene_tx=XM_023249055.1
X	74495	74532
refGene_gn=LOC101092754;refGene_tx=XM_023249055.1
X	74534	74685
refGene_gn=LOC101092754;refGene_tx=XM_023249055.1
X	80969	81208
refGene_gn=LOC101092754;refGene_tx=XM_023249055.1
X	85871	85991
refGene_gn=LOC101098289;refGene_tx=XM_023249439.1
X	86349	86524

refGene_gn=LOC101098289;refGene_tx=XM_023249439.1
X	86957	87106
refGene_gn=LOC101098289;refGene_tx=XM_023249439.1
X	89018	89227
refGene_gn=LOC101098289;refGene_tx=XM_023249439.1
X	90970	91072
refGene_gn=LOC101098289;refGene_tx=XM_023249439.1
X	91074	91128
refGene_gn=LOC101098289;refGene_tx=XM_023249439.1
X	92046	92114
refGene_gn=LOC101098289;refGene_tx=XM_023249439.1
X	92457	92588
refGene_gn=LOC101098289;refGene_tx=XM_023249439.1
X	92739	92810
refGene_gn=LOC101098289;refGene_tx=XM_023249439.1
X	93374	93512
refGene_gn=LOC101098289;refGene_tx=XM_023249439.1
X	98045	98388
refGene_gn=LOC101098289;refGene_tx=XM_023249439.1
X	98629	98815
refGene_gn=LOC101098289;refGene_tx=XM_023249439.1
X	119916	120052
refGene_gn=LOC105261393;refGene_tx=XM_011291581.2
X	120422	120575
refGene_gn=LOC105261393;refGene_tx=XM_011291581.2
X	121012	121161
refGene_gn=LOC105261393;refGene_tx=XM_011291581.2
X	123122	123331
refGene_gn=LOC105261393;refGene_tx=XM_011291581.2
X	125083	125242
refGene_gn=LOC105261393;refGene_tx=XM_011291581.2
X	126141	126209
refGene_gn=LOC105261393;refGene_tx=XM_011291581.2
X	126552	126683
refGene_gn=LOC105261393;refGene_tx=XM_011291581.2
X	126806	126877
refGene_gn=LOC105261393;refGene_tx=XM_011291581.2
X	127442	127580
refGene_gn=LOC105261393;refGene_tx=XM_011291581.2
X	131576	131925
refGene_gn=LOC105261393;refGene_tx=XM_011291581.2
X	136584	136589
refGene_gn=LOC102901245;refGene_tx=XM_023249440.1
X	137348	137418
refGene_gn=LOC102901245;refGene_tx=XM_023249440.1
X	145227	145329
refGene_gn=LOC102901245;refGene_tx=XM_023249440.1
X	145500	145626
refGene_gn=LOC102901245;refGene_tx=XM_023249440.1
X	146642	146795

refGene_gn=LOC102901245;refGene_tx=XM_023249440.1
X	150262	150359
refGene_gn=LOC102901245;refGene_tx=XM_023249440.1
X	150656	150733
refGene_gn=LOC102901245;refGene_tx=XM_023249440.1
X	153983	154055
refGene_gn=LOC102901245;refGene_tx=XM_023249440.1
X	158488	158693
refGene_gn=LOC102901245;refGene_tx=XM_023249440.1
X	161830	161982
refGene_gn=LOC102901245;refGene_tx=XM_023249440.1
X	163623	163691
refGene_gn=LOC102901245;refGene_tx=XM_023249440.1
X	163834	163958
refGene_gn=LOC102901245;refGene_tx=XM_023249440.1
X	176022	176121
refGene_gn=LOC102901245;refGene_tx=XM_023249440.1
X	177517	177597
refGene_gn=LOC102901245;refGene_tx=XM_023249440.1
X	180729	180837
refGene_gn=LOC102901245;refGene_tx=XM_023249440.1
X	181531	181643
refGene_gn=LOC102901245;refGene_tx=XM_023249440.1
X	182321	182436
refGene_gn=LOC102901245;refGene_tx=XM_023249440.1
X	187470	187574
refGene_gn=LOC102901245;refGene_tx=XM_023249440.1
X	190538	190631
refGene_gn=LOC102901245;refGene_tx=XM_023249440.1
X	190719	190809
refGene_gn=LOC102901245;refGene_tx=XM_023249440.1
X	192400	192607
refGene_gn=LOC102901245;refGene_tx=XM_023249440.1
X	193648	193725
refGene_gn=LOC102901245;refGene_tx=XM_023249440.1
X	194114	194202
refGene_gn=LOC102901245;refGene_tx=XM_023249440.1
X	194455	194536
refGene_gn=LOC102901245;refGene_tx=XM_023249440.1
X	195551	195644
refGene_gn=LOC102901245;refGene_tx=XM_023249440.1
X	196772	196932
refGene_gn=LOC102901245;refGene_tx=XM_023249440.1
X	198795	198935
refGene_gn=LOC102901245;refGene_tx=XM_023249440.1
X	199772	199838
refGene_gn=LOC111558781;refGene_tx=XM_023249515.1
X	202872	203024
refGene_gn=LOC111558781;refGene_tx=XM_023249515.1
X	203517	203655

```
refGene_gn=LOC111558781;refGene_tx=XM_023249515.1
X	204070	204255
refGene_gn=LOC111558781;refGene_tx=XM_023249515.1
X	205039	205145
refGene_gn=LOC111558781;refGene_tx=XM_023249515.1
X	205706	205794
refGene_gn=LOC111558781;refGene_tx=XM_023249515.1
X	206117	206132
refGene_gn=LOC111558781;refGene_tx=XM_023249515.1
X	207884	208035	refGene_gn=PPP2R3B;refGene_tx=XM_023249514.1
X	210343	210450	refGene_gn=PPP2R3B;refGene_tx=XM_023249514.1
X	210528	210647	refGene_gn=PPP2R3B;refGene_tx=XM_023249514.1
X	212933	213109	refGene_gn=PPP2R3B;refGene_tx=XM_023249514.1
X	213289	213379	refGene_gn=PPP2R3B;refGene_tx=XM_023249514.1
X	214062	214111	refGene_gn=PPP2R3B;refGene_tx=XM_023249514.1
X	217693	217850	refGene_gn=PPP2R3B;refGene_tx=XM_023249514.1
X	218616	218703	refGene_gn=PPP2R3B;refGene_tx=XM_023249514.1
X	220156	220231	refGene_gn=PPP2R3B;refGene_tx=XM_023249514.1
X	220956	221059	refGene_gn=PPP2R3B;refGene_tx=XM_023249514.1
X	221258	221362	refGene_gn=PPP2R3B;refGene_tx=XM_023249514.1
X	233211	233397	refGene_gn=PPP2R3B;refGene_tx=XM_023249514.1
X	267508	267844	refGene_gn=PPP2R3B;refGene_tx=XM_023249514.1
X	530171	530454	refGene_gn=SHOX;refGene_tx=XM_011291584.3
X	533516	533725	refGene_gn=SHOX;refGene_tx=XM_011291584.3
X	537440	537498	refGene_gn=SHOX;refGene_tx=XM_011291584.3
X	537576	537665	refGene_gn=SHOX;refGene_tx=XM_011291584.3
X	539195	539441	refGene_gn=SHOX;refGene_tx=XM_011291584.3
X	913548	913824	refGene_gn=CRLF2;refGene_tx=XM_023249294.1
X	919546	919631	refGene_gn=CRLF2;refGene_tx=XM_023249294.1
X	922041	922147	refGene_gn=CRLF2;refGene_tx=XM_023249294.1
X	923459	923622	refGene_gn=CRLF2;refGene_tx=XM_023249294.1
X	925684	925818	refGene_gn=CRLF2;refGene_tx=XM_023249294.1
X	927310	927474	refGene_gn=CRLF2;refGene_tx=XM_023249294.1
X	929168	929271	refGene_gn=CRLF2;refGene_tx=XM_023249294.1
X	936385	936467	refGene_gn=CRLF2;refGene_tx=XM_023249294.1
X	976933	977042
refGene_gn=LOC101100621;refGene_tx=XM_019823499.2;refGene_gn=LOC101100
621;refGene_tx=XM_019823500.2;refGene_gn=LOC101100621;refGene_tx=XM_01
9823501.2
X	980945	981047
refGene_gn=LOC101100621;refGene_tx=XM_019823499.2;refGene_gn=LOC101100
621;refGene_tx=XM_019823500.2;refGene_gn=LOC101100621;refGene_tx=XM_01
9823501.2
X	981902	982045
refGene_gn=LOC101100621;refGene_tx=XM_019823499.2;refGene_gn=LOC101100
621;refGene_tx=XM_019823500.2;refGene_gn=LOC101100621;refGene_tx=XM_01
9823501.2
X	983051	983175
refGene_gn=LOC101100621;refGene_tx=XM_019823499.2;refGene_gn=LOC101100
621;refGene_tx=XM_019823500.2;refGene_gn=LOC101100621;refGene_tx=XM_01
```

9823501.2
X	985172	985302
refGene_gn=LOC101100621;refGene_tx=XM_019823499.2;refGene_gn=LOC101100
621;refGene_tx=XM_019823500.2;refGene_gn=LOC101100621;refGene_tx=XM_01
9823501.2
X	985396	985569
refGene_gn=LOC101100621;refGene_tx=XM_019823499.2;refGene_gn=LOC101100
621;refGene_tx=XM_019823500.2;refGene_gn=LOC101100621;refGene_tx=XM_01
9823501.2
X	986991	987125
refGene_gn=LOC101100621;refGene_tx=XM_019823499.2;refGene_gn=LOC101100
621;refGene_tx=XM_019823500.2;refGene_gn=LOC101100621;refGene_tx=XM_01
9823501.2
X	990157	990187
refGene_gn=LOC101100621;refGene_tx=XM_019823499.2;refGene_gn=LOC101100
621;refGene_tx=XM_019823500.2;refGene_gn=LOC101100621;refGene_tx=XM_01
9823501.2
X	990600	990736
refGene_gn=LOC101100621;refGene_tx=XM_019823499.2;refGene_gn=LOC101100
621;refGene_tx=XM_019823500.2;refGene_gn=LOC101100621;refGene_tx=XM_01
9823501.2
X	991857	991954
refGene_gn=LOC101100621;refGene_tx=XM_019823499.2
X	993887	993969
refGene_gn=LOC101100621;refGene_tx=XM_019823499.2;refGene_gn=LOC101100
621;refGene_tx=XM_019823500.2;refGene_gn=LOC101100621;refGene_tx=XM_01
9823501.2
X	994144	994220
refGene_gn=LOC101100621;refGene_tx=XM_019823500.2
X	994254	994350
refGene_gn=LOC101100621;refGene_tx=XM_019823499.2;refGene_gn=LOC101100
621;refGene_tx=XM_019823501.2
X	1005914	1005987
refGene_gn=LOC102901718;refGene_tx=XM_019823502.2
X	1007293	1007436
refGene_gn=LOC102901718;refGene_tx=XM_019823502.2
X	1013384	1013511
refGene_gn=LOC102901718;refGene_tx=XM_019823502.2
X	1013592	1013719
refGene_gn=LOC102901718;refGene_tx=XM_019823502.2
X	1014206	1014391
refGene_gn=LOC102901718;refGene_tx=XM_019823502.2
X	1014956	1015090
refGene_gn=LOC102901718;refGene_tx=XM_019823502.2
X	1016568	1016592
refGene_gn=LOC102901718;refGene_tx=XM_019823502.2
X	1020357	1020493
refGene_gn=LOC102901718;refGene_tx=XM_019823502.2
X	1021005	1021099
refGene_gn=LOC102901718;refGene_tx=XM_019823502.2

X 1021971 1022083
refGene_gn=LOC102901718;refGene_tx=XM_019823502.2
X 1023397 1023475
refGene_gn=LOC102901718;refGene_tx=XM_019823502.2
X 1033002 1033066
refGene_gn=IL3RA;refGene_tx=XM_023248886.1;refGene_gn=IL3RA;refGene_tx=XM_023248887.1
X 1035526 1035645
refGene_gn=IL3RA;refGene_tx=XM_023248886.1;refGene_gn=IL3RA;refGene_tx=XM_023248887.1
X 1035990 1036108
refGene_gn=IL3RA;refGene_tx=XM_023248886.1;refGene_gn=IL3RA;refGene_tx=XM_023248887.1
X 1037853 1037986
refGene_gn=IL3RA;refGene_tx=XM_023248886.1;refGene_gn=IL3RA;refGene_tx=XM_023248887.1
X 1038180 1038356
refGene_gn=IL3RA;refGene_tx=XM_023248886.1;refGene_gn=IL3RA;refGene_tx=XM_023248887.1
X 1041941 1042057
refGene_gn=IL3RA;refGene_tx=XM_023248886.1;refGene_gn=IL3RA;refGene_tx=XM_023248887.1
X 1043987 1044014
refGene_gn=IL3RA;refGene_tx=XM_023248886.1;refGene_gn=IL3RA;refGene_tx=XM_023248887.1
X 1045050 1045162
refGene_gn=IL3RA;refGene_tx=XM_023248886.1;refGene_gn=IL3RA;refGene_tx=XM_023248887.1
X 1051587 1051647 refGene_gn=IL3RA;refGene_tx=XM_023248886.1
X 1052144 1052250
refGene_gn=IL3RA;refGene_tx=XM_023248886.1;refGene_gn=IL3RA;refGene_tx=XM_023248887.1
X 1053925 1054007
refGene_gn=IL3RA;refGene_tx=XM_023248886.1;refGene_gn=IL3RA;refGene_tx=XM_023248887.1
X 1054241 1054316
refGene_gn=IL3RA;refGene_tx=XM_023248886.1;refGene_gn=IL3RA;refGene_tx=XM_023248887.1
X 1056017 1056175 refGene_gn=SLC25A6;refGene_tx=XM_023248888.1
X 1056677 1056818 refGene_gn=SLC25A6;refGene_tx=XM_023248888.1
X 1057859 1058346 refGene_gn=SLC25A6;refGene_tx=XM_023248888.1
X 1058790 1058901 refGene_gn=SLC25A6;refGene_tx=XM_023248888.1
X 1064223 1064486
refGene_gn=ASMTL;refGene_tx=XM_023248897.1;refGene_gn=ASMTL;refGene_tx=XM_023248899.1
X 1067409 1067483 refGene_gn=ASMTL;refGene_tx=XM_023248898.1
X 1068181 1068304
refGene_gn=ASMTL;refGene_tx=XM_023248897.1;refGene_gn=ASMTL;refGene_tx=XM_023248898.1;refGene_gn=ASMTL;refGene_tx=XM_023248899.1
X 1069409 1069553

refGene_gn=ASMTL;refGene_tx=XM_023248897.1;refGene_gn=ASMTL;refGene_tx
=XM_023248898.1;refGene_gn=ASMTL;refGene_tx=XM_023248899.1
X 1070385 1070503
refGene_gn=ASMTL;refGene_tx=XM_023248897.1;refGene_gn=ASMTL;refGene_tx
=XM_023248898.1;refGene_gn=ASMTL;refGene_tx=XM_023248899.1
X 1073053 1073229
refGene_gn=ASMTL;refGene_tx=XM_023248897.1;refGene_gn=ASMTL;refGene_tx
=XM_023248898.1;refGene_gn=ASMTL;refGene_tx=XM_023248899.1
X 1076664 1076827
refGene_gn=ASMTL;refGene_tx=XM_023248897.1;refGene_gn=ASMTL;refGene_tx
=XM_023248898.1;refGene_gn=ASMTL;refGene_tx=XM_023248899.1
X 1080415 1080791
refGene_gn=ASMTL;refGene_tx=XM_023248897.1;refGene_gn=ASMTL;refGene_tx
=XM_023248898.1;refGene_gn=ASMTL;refGene_tx=XM_023248899.1
X 1081528 1081637
refGene_gn=ASMTL;refGene_tx=XM_023248897.1;refGene_gn=ASMTL;refGene_tx
=XM_023248898.1;refGene_gn=ASMTL;refGene_tx=XM_023248899.1
X 1082439 1082501
refGene_gn=ASMTL;refGene_tx=XM_023248897.1;refGene_gn=ASMTL;refGene_tx
=XM_023248898.1;refGene_gn=ASMTL;refGene_tx=XM_023248899.1
X 1083287 1083352
refGene_gn=ASMTL;refGene_tx=XM_023248897.1;refGene_gn=ASMTL;refGene_tx
=XM_023248898.1;refGene_gn=ASMTL;refGene_tx=XM_023248899.1
X 1083492 1083540
refGene_gn=ASMTL;refGene_tx=XM_023248897.1;refGene_gn=ASMTL;refGene_tx
=XM_023248898.1;refGene_gn=ASMTL;refGene_tx=XM_023248899.1
X 1084949 1085081
refGene_gn=ASMTL;refGene_tx=XM_023248897.1;refGene_gn=ASMTL;refGene_tx
=XM_023248898.1;refGene_gn=ASMTL;refGene_tx=XM_023248899.1
X 1086618 1086711
refGene_gn=ASMTL;refGene_tx=XM_023248897.1;refGene_gn=ASMTL;refGene_tx
=XM_023248898.1
X 1095386 1096460
refGene_gn=P2RY8;refGene_tx=XM_023249096.1;refGene_gn=P2RY8;refGene_tx
=XM_023249097.1
X 1136503 1137298
refGene_gn=LOC109496913;refGene_tx=XM_019824686.2;refGene_gn=LOC109496
913;refGene_tx=XM_019824687.2
X 1138929 1139025
refGene_gn=LOC109496913;refGene_tx=XM_019824686.2
X 1139052 1139127
refGene_gn=LOC109496913;refGene_tx=XM_019824687.2
X 1156777 1158079
refGene_gn=LOC111558770;refGene_tx=XM_023249441.1
X 1158308 1158501
refGene_gn=LOC111558770;refGene_tx=XM_023249441.1
X 1159676 1159735
refGene_gn=LOC111558770;refGene_tx=XM_023249441.1
X 1160226 1160694 refGene_gn=AKAP17A;refGene_tx=XM_023248928.1
X 1160695 1160989 refGene_gn=AKAP17A;refGene_tx=XM_023248928.1

X        1161770  1161919  refGene_gn=AKAP17A;refGene_tx=XM_023248928.1
X        1166003  1166244  refGene_gn=AKAP17A;refGene_tx=XM_023248928.1
X        1169209  1170163  refGene_gn=AKAP17A;refGene_tx=XM_023248928.1
X        1182974  1183193  refGene_gn=ASMT;refGene_tx=XM_023249442.1
X        1214622  1214776  refGene_gn=ASMT;refGene_tx=XM_023249442.1
X        1218289  1218691  refGene_gn=ASMT;refGene_tx=XM_023249442.1
X        1219378  1219526  refGene_gn=ASMT;refGene_tx=XM_023249442.1
X        1222582  1222651  refGene_gn=ASMT;refGene_tx=XM_023249442.1
X        1223534  1223653  refGene_gn=ASMT;refGene_tx=XM_023249442.1
X        1224260  1224401  refGene_gn=ASMT;refGene_tx=XM_023249442.1
X        1224837  1224960  refGene_gn=ASMT;refGene_tx=XM_023249442.1
X        1225372  1225605  refGene_gn=ASMT;refGene_tx=XM_023249442.1
X        1280566  1281478  refGene_gn=DHRSX;refGene_tx=XM_023249149.1
X        1302148  1302205  refGene_gn=DHRSX;refGene_tx=XM_023249150.1
X        1374991  1375195  refGene_gn=DHRSX;refGene_tx=XM_023249148.1
X        1405133  1405341
refGene_gn=DHRSX;refGene_tx=XM_023249148.1;refGene_gn=DHRSX;refGene_tx
=XM_023249150.1
X        1413665  1413873
refGene_gn=DHRSX;refGene_tx=XM_023249148.1;refGene_gn=DHRSX;refGene_tx
=XM_023249150.1
X        1423858  1423960
refGene_gn=DHRSX;refGene_tx=XM_023249148.1;refGene_gn=DHRSX;refGene_tx
=XM_023249150.1
X        1504327  1504396
refGene_gn=DHRSX;refGene_tx=XM_023249148.1;refGene_gn=DHRSX;refGene_tx
=XM_023249150.1
X        1511774  1511882
refGene_gn=DHRSX;refGene_tx=XM_023249148.1;refGene_gn=DHRSX;refGene_tx
=XM_023249150.1
X        1533806  1535891
refGene_gn=ZBED1;refGene_tx=XM_023249145.1;refGene_gn=ZBED1;refGene_tx
=XM_023249146.1;refGene_gn=ZBED1;refGene_tx=XM_023249147.1
X        1542561  1542670
refGene_gn=DHRSX;refGene_tx=XM_023249148.1;refGene_gn=DHRSX;refGene_tx
=XM_023249150.1
X        1633899  1633966  refGene_gn=CD99;refGene_tx=XM_023249229.1
X        1650970  1651003  refGene_gn=CD99;refGene_tx=XM_023249229.1
X        1651310  1651355  refGene_gn=CD99;refGene_tx=XM_023249229.1
X        1651920  1651968  refGene_gn=CD99;refGene_tx=XM_023249229.1
X        1652626  1652695  refGene_gn=CD99;refGene_tx=XM_023249229.1
X        1654348  1654393  refGene_gn=CD99;refGene_tx=XM_023249229.1
X        1656253  1656310  refGene_gn=CD99;refGene_tx=XM_023249229.1
X        1657308  1657423  refGene_gn=CD99;refGene_tx=XM_023249229.1
X        1660973  1661035  refGene_gn=CD99;refGene_tx=XM_023249229.1
X        1662955  1663012  refGene_gn=CD99;refGene_tx=XM_023249229.1
X        1663897  1663923  refGene_gn=CD99;refGene_tx=XM_023249229.1
X        1667041  1667110  refGene_gn=XG;refGene_tx=XM_023249228.1
X        1669671  1669732  refGene_gn=XG;refGene_tx=XM_023249226.1
X        1677168  1677303

refGene_gn=XG;refGene_tx=XM_023249220.1;refGene_gn=XG;refGene_tx=XM_023249221.1;refGene_gn=XG;refGene_tx=XM_023249222.1
X 1677589 1677716
refGene_gn=XG;refGene_tx=XM_019823507.2;refGene_gn=XG;refGene_tx=XM_019823509.2;refGene_gn=XG;refGene_tx=XM_023249223.1;refGene_gn=XG;refGene_tx=XM_023249224.1;refGene_gn=XG;refGene_tx=XM_023249225.1
X 1679577 1679619
refGene_gn=XG;refGene_tx=XM_019823507.2;refGene_gn=XG;refGene_tx=XM_019823509.2;refGene_gn=XG;refGene_tx=XM_023249220.1;refGene_gn=XG;refGene_tx=XM_023249221.1;refGene_gn=XG;refGene_tx=XM_023249222.1;refGene_gn=XG;refGene_tx=XM_023249223.1;refGene_gn=XG;refGene_tx=XM_023249224.1;refGene_gn=XG;refGene_tx=XM_023249225.1;refGene_gn=XG;refGene_tx=XM_023249226.1;refGene_gn=XG;refGene_tx=XM_023249228.1
X 1680089 1680113
refGene_gn=XG;refGene_tx=XM_019823507.2;refGene_gn=XG;refGene_tx=XM_019823509.2;refGene_gn=XG;refGene_tx=XM_023249220.1;refGene_gn=XG;refGene_tx=XM_023249221.1;refGene_gn=XG;refGene_tx=XM_023249222.1;refGene_gn=XG;refGene_tx=XM_023249223.1;refGene_gn=XG;refGene_tx=XM_023249224.1;refGene_gn=XG;refGene_tx=XM_023249225.1;refGene_gn=XG;refGene_tx=XM_023249226.1;refGene_gn=XG;refGene_tx=XM_023249228.1
X 1682353 1682416
refGene_gn=XG;refGene_tx=XM_019823507.2;refGene_gn=XG;refGene_tx=XM_019823509.2;refGene_gn=XG;refGene_tx=XM_023249220.1;refGene_gn=XG;refGene_tx=XM_023249221.1;refGene_gn=XG;refGene_tx=XM_023249222.1;refGene_gn=XG;refGene_tx=XM_023249223.1;refGene_gn=XG;refGene_tx=XM_023249224.1;refGene_gn=XG;refGene_tx=XM_023249225.1;refGene_gn=XG;refGene_tx=XM_023249226.1;refGene_gn=XG;refGene_tx=XM_023249228.1
X 1684065 1684128
refGene_gn=XG;refGene_tx=XM_023249220.1;refGene_gn=XG;refGene_tx=XM_023249221.1;refGene_gn=XG;refGene_tx=XM_023249222.1;refGene_gn=XG;refGene_tx=XM_023249223.1;refGene_gn=XG;refGene_tx=XM_023249224.1;refGene_gn=XG;refGene_tx=XM_023249225.1;refGene_gn=XG;refGene_tx=XM_023249226.1;refGene_gn=XG;refGene_tx=XM_023249228.1
X 1686394 1686457
refGene_gn=XG;refGene_tx=XM_023249220.1;refGene_gn=XG;refGene_tx=XM_023249222.1;refGene_gn=XG;refGene_tx=XM_023249223.1;refGene_gn=XG;refGene_tx=XM_023249225.1;refGene_gn=XG;refGene_tx=XM_023249226.1;refGene_gn=XG;refGene_tx=XM_023249228.1
X 1691063 1691126
refGene_gn=XG;refGene_tx=XM_019823507.2;refGene_gn=XG;refGene_tx=XM_019823509.2;refGene_gn=XG;refGene_tx=XM_023249220.1;refGene_gn=XG;refGene_tx=XM_023249221.1;refGene_gn=XG;refGene_tx=XM_023249222.1;refGene_gn=XG;refGene_tx=XM_023249223.1;refGene_gn=XG;refGene_tx=XM_023249224.1;refGene_gn=XG;refGene_tx=XM_023249225.1;refGene_gn=XG;refGene_tx=XM_023249226.1;refGene_gn=XG;refGene_tx=XM_023249228.1
X 1692169 1692232
refGene_gn=XG;refGene_tx=XM_019823507.2;refGene_gn=XG;refGene_tx=XM_023249220.1;refGene_gn=XG;refGene_tx=XM_023249221.1;refGene_gn=XG;refGene_tx=XM_023249223.1;refGene_gn=XG;refGene_tx=XM_023249224.1;refGene_gn=XG;refGene_tx=XM_023249226.1;refGene_gn=XG;refGene_tx=XM_023249228.1

X	1694791	1694842
refGene_gn=XG;refGene_tx=XM_019823507.2;refGene_gn=XG;refGene_tx=XM_019823509.2;refGene_gn=XG;refGene_tx=XM_023249220.1;refGene_gn=XG;refGene_tx=XM_023249221.1;refGene_gn=XG;refGene_tx=XM_023249222.1;refGene_gn=XG;refGene_tx=XM_023249223.1;refGene_gn=XG;refGene_tx=XM_023249224.1;refGene_gn=XG;refGene_tx=XM_023249225.1;refGene_gn=XG;refGene_tx=XM_023249226.1;refGene_gn=XG;refGene_tx=XM_023249228.1
X	1700836	1700872
refGene_gn=XG;refGene_tx=XM_019823507.2;refGene_gn=XG;refGene_tx=XM_019823509.2;refGene_gn=XG;refGene_tx=XM_023249220.1;refGene_gn=XG;refGene_tx=XM_023249221.1;refGene_gn=XG;refGene_tx=XM_023249222.1;refGene_gn=XG;refGene_tx=XM_023249223.1;refGene_gn=XG;refGene_tx=XM_023249224.1;refGene_gn=XG;refGene_tx=XM_023249225.1;refGene_gn=XG;refGene_tx=XM_023249226.1;refGene_gn=XG;refGene_tx=XM_023249228.1
X	1701905	1702022
refGene_gn=XG;refGene_tx=XM_019823507.2;refGene_gn=XG;refGene_tx=XM_019823509.2;refGene_gn=XG;refGene_tx=XM_023249220.1;refGene_gn=XG;refGene_tx=XM_023249221.1;refGene_gn=XG;refGene_tx=XM_023249222.1;refGene_gn=XG;refGene_tx=XM_023249223.1;refGene_gn=XG;refGene_tx=XM_023249224.1;refGene_gn=XG;refGene_tx=XM_023249225.1;refGene_gn=XG;refGene_tx=XM_023249226.1;refGene_gn=XG;refGene_tx=XM_023249228.1
X	1703371	1703388
refGene_gn=XG;refGene_tx=XM_019823507.2;refGene_gn=XG;refGene_tx=XM_019823509.2;refGene_gn=XG;refGene_tx=XM_023249223.1;refGene_gn=XG;refGene_tx=XM_023249224.1;refGene_gn=XG;refGene_tx=XM_023249225.1;refGene_gn=XG;refGene_tx=XM_023249226.1
X	1721109	1721116
refGene_gn=GYG2;refGene_tx=XM_023248840.1;refGene_gn=GYG2;refGene_tx=XM_023248841.1;refGene_gn=GYG2;refGene_tx=XM_023248842.1;refGene_gn=GYG2;refGene_tx=XM_023248843.1;refGene_gn=GYG2;refGene_tx=XM_023248844.1
X	1725775	1725917
refGene_gn=GYG2;refGene_tx=XM_023248840.1;refGene_gn=GYG2;refGene_tx=XM_023248841.1;refGene_gn=GYG2;refGene_tx=XM_023248842.1;refGene_gn=GYG2;refGene_tx=XM_023248843.1;refGene_gn=GYG2;refGene_tx=XM_023248844.1
X	1727649	1727651	refGene_gn=GYG2;refGene_tx=XM_019823514.2
X	1732603	1732778
refGene_gn=GYG2;refGene_tx=XM_019823514.2;refGene_gn=GYG2;refGene_tx=XM_023248840.1;refGene_gn=GYG2;refGene_tx=XM_023248841.1;refGene_gn=GYG2;refGene_tx=XM_023248842.1;refGene_gn=GYG2;refGene_tx=XM_023248843.1;refGene_gn=GYG2;refGene_tx=XM_023248844.1
X	1733767	1733930
refGene_gn=GYG2;refGene_tx=XM_019823514.2;refGene_gn=GYG2;refGene_tx=XM_023248840.1;refGene_gn=GYG2;refGene_tx=XM_023248841.1;refGene_gn=GYG2;refGene_tx=XM_023248842.1;refGene_gn=GYG2;refGene_tx=XM_023248843.1;refGene_gn=GYG2;refGene_tx=XM_023248844.1
X	1734655	1734782
refGene_gn=GYG2;refGene_tx=XM_019823514.2;refGene_gn=GYG2;refGene_tx=XM_023248840.1;refGene_gn=GYG2;refGene_tx=XM_023248841.1;refGene_gn=GYG2;refGene_tx=XM_023248842.1;refGene_gn=GYG2;refGene_tx=XM_023248843.1;refGene_gn=GYG2;refGene_tx=XM_023248844.1

X	1738150	1738373
refGene_gn=GYG2;refGene_tx=XM_019823514.2;refGene_gn=GYG2;refGene_tx=XM_023248840.1;refGene_gn=GYG2;refGene_tx=XM_023248841.1;refGene_gn=GYG2;refGene_tx=XM_023248842.1;refGene_gn=GYG2;refGene_tx=XM_023248843.1;refGene_gn=GYG2;refGene_tx=XM_023248844.1
X	1739598	1739727
refGene_gn=GYG2;refGene_tx=XM_019823514.2;refGene_gn=GYG2;refGene_tx=XM_023248840.1;refGene_gn=GYG2;refGene_tx=XM_023248841.1;refGene_gn=GYG2;refGene_tx=XM_023248842.1;refGene_gn=GYG2;refGene_tx=XM_023248843.1;refGene_gn=GYG2;refGene_tx=XM_023248844.1
X	1746008	1746113
refGene_gn=GYG2;refGene_tx=XM_019823514.2;refGene_gn=GYG2;refGene_tx=XM_023248840.1;refGene_gn=GYG2;refGene_tx=XM_023248841.1;refGene_gn=GYG2;refGene_tx=XM_023248842.1;refGene_gn=GYG2;refGene_tx=XM_023248843.1
X	1747247	1747355
refGene_gn=GYG2;refGene_tx=XM_019823514.2;refGene_gn=GYG2;refGene_tx=XM_023248840.1;refGene_gn=GYG2;refGene_tx=XM_023248841.1;refGene_gn=GYG2;refGene_tx=XM_023248842.1;refGene_gn=GYG2;refGene_tx=XM_023248843.1
X	1747609	1747813
refGene_gn=GYG2;refGene_tx=XM_019823514.2;refGene_gn=GYG2;refGene_tx=XM_023248840.1;refGene_gn=GYG2;refGene_tx=XM_023248841.1;refGene_gn=GYG2;refGene_tx=XM_023248842.1
X	1750067	1750229
refGene_gn=GYG2;refGene_tx=XM_023248843.1;refGene_gn=GYG2;refGene_tx=XM_023248844.1
X	1755792	1755920	refGene_gn=ARSD;refGene_tx=XM_023249531.1
X	1755922	1756156	refGene_gn=ARSD;refGene_tx=XM_023249531.1
X	1758208	1758330	refGene_gn=ARSD;refGene_tx=XM_023249531.1
X	1759929	1760092	refGene_gn=ARSD;refGene_tx=XM_023249531.1
X	1762238	1762373	refGene_gn=ARSD;refGene_tx=XM_023249531.1
X	1763683	1763820	refGene_gn=ARSD;refGene_tx=XM_023249531.1
X	1764200	1764624	refGene_gn=ARSD;refGene_tx=XM_023249531.1
X	1766584	1766707	refGene_gn=ARSD;refGene_tx=XM_023249531.1
X	1770255	1770377	refGene_gn=ARSD;refGene_tx=XM_023249531.1
X	1779708	1779858	refGene_gn=ARSD;refGene_tx=XM_023249531.1
X	1781581	1781625	refGene_gn=ARSD;refGene_tx=XM_023249531.1
X	1782956	1783315
refGene_gn=LOC101081445;refGene_tx=XM_023249527.1;refGene_gn=LOC101081445;refGene_tx=XM_023249528.1;refGene_gn=LOC101081445;refGene_tx=XM_023249529.1;refGene_gn=LOC101081445;refGene_tx=XM_023249530.1
X	1785829	1785951
refGene_gn=LOC101081445;refGene_tx=XM_023249527.1;refGene_gn=LOC101081445;refGene_tx=XM_023249528.1;refGene_gn=LOC101081445;refGene_tx=XM_023249529.1;refGene_gn=LOC101081445;refGene_tx=XM_023249530.1
X	1788303	1788466
refGene_gn=LOC101081445;refGene_tx=XM_023249527.1;refGene_gn=LOC101081445;refGene_tx=XM_023249528.1;refGene_gn=LOC101081445;refGene_tx=XM_023249529.1;refGene_gn=LOC101081445;refGene_tx=XM_023249530.1
X	1789176	1789311
refGene_gn=LOC101081445;refGene_tx=XM_023249527.1;refGene_gn=LOC101081

445;refGene_tx=XM_023249528.1;refGene_gn=LOC101081445;refGene_tx=XM_023249529.1;refGene_gn=LOC101081445;refGene_tx=XM_023249530.1
X 1790858 1790995 refGene_gn=LOC101081445;refGene_tx=XM_023249527.1;refGene_gn=LOC101081445;refGene_tx=XM_023249528.1;refGene_gn=LOC101081445;refGene_tx=XM_023249529.1;refGene_gn=LOC101081445;refGene_tx=XM_023249530.1
X 1792422 1792846 refGene_gn=LOC101081445;refGene_tx=XM_023249527.1;refGene_gn=LOC101081445;refGene_tx=XM_023249528.1;refGene_gn=LOC101081445;refGene_tx=XM_023249529.1;refGene_gn=LOC101081445;refGene_tx=XM_023249530.1
X 1795026 1795149 refGene_gn=LOC101081445;refGene_tx=XM_023249527.1;refGene_gn=LOC101081445;refGene_tx=XM_023249528.1;refGene_gn=LOC101081445;refGene_tx=XM_023249529.1;refGene_gn=LOC101081445;refGene_tx=XM_023249530.1
X 1796355 1796477 refGene_gn=LOC101081445;refGene_tx=XM_023249527.1;refGene_gn=LOC101081445;refGene_tx=XM_023249528.1;refGene_gn=LOC101081445;refGene_tx=XM_023249529.1;refGene_gn=LOC101081445;refGene_tx=XM_023249530.1
X 1797734 1797884 refGene_gn=LOC101081445;refGene_tx=XM_023249527.1;refGene_gn=LOC101081445;refGene_tx=XM_023249528.1;refGene_gn=LOC101081445;refGene_tx=XM_023249529.1;refGene_gn=LOC101081445;refGene_tx=XM_023249530.1
X 1799100 1799150 refGene_gn=LOC101081445;refGene_tx=XM_023249527.1;refGene_gn=LOC101081445;refGene_tx=XM_023249528.1;refGene_gn=LOC101081445;refGene_tx=XM_023249529.1;refGene_gn=LOC101081445;refGene_tx=XM_023249530.1
X 1800179 1800455 refGene_gn=LOC101081445;refGene_tx=XM_023249527.1;refGene_gn=LOC101081445;refGene_tx=XM_023249528.1;refGene_gn=LOC101081445;refGene_tx=XM_023249529.1;refGene_gn=LOC101081445;refGene_tx=XM_023249530.1
X 1809289 1809312 refGene_gn=ARSH;refGene_tx=XM_023249256.1
X 1811029 1811179 refGene_gn=ARSH;refGene_tx=XM_023249256.1
X 1812228 1812350 refGene_gn=ARSH;refGene_tx=XM_023249256.1
X 1815293 1815419 refGene_gn=ARSH;refGene_tx=XM_023249256.1
X 1817295 1817719 refGene_gn=ARSH;refGene_tx=XM_023249256.1
X 1819021 1819158 refGene_gn=ARSH;refGene_tx=XM_023249256.1
X 1824274 1824409 refGene_gn=ARSH;refGene_tx=XM_023249256.1
X 1827583 1827746 refGene_gn=ARSH;refGene_tx=XM_023249256.1
X 1828654 1828776 refGene_gn=ARSH;refGene_tx=XM_023249256.1
X 1830765 1831133 refGene_gn=ARSH;refGene_tx=XM_023249256.1
X 1849242 1849253 refGene_gn=ARSF;refGene_tx=XM_004000249.3;refGene_gn=ARSF;refGene_tx=XM_019823520.2
X 1850254 1850404 refGene_gn=ARSF;refGene_tx=XM_004000249.3;refGene_gn=ARSF;refGene_tx=XM_019823520.2
X 1852050 1852172 refGene_gn=ARSF;refGene_tx=XM_004000249.3;refGene_gn=ARSF;refGene_tx=XM_019823520.2
X 1853671 1853794

refGene_gn=ARSF;refGene_tx=XM_004000249.3;refGene_gn=ARSF;refGene_tx=XM_019823520.2
X	1856811	1857235
refGene_gn=ARSF;refGene_tx=XM_004000249.3;refGene_gn=ARSF;refGene_tx=XM_019823520.2
X	1861268	1861405
refGene_gn=ARSF;refGene_tx=XM_004000249.3;refGene_gn=ARSF;refGene_tx=XM_019823520.2
X	1865336	1865471
refGene_gn=ARSF;refGene_tx=XM_004000249.3;refGene_gn=ARSF;refGene_tx=XM_019823520.2
X	1867495	1867658
refGene_gn=ARSF;refGene_tx=XM_004000249.3;refGene_gn=ARSF;refGene_tx=XM_019823520.2
X	1871400	1871522
refGene_gn=ARSF;refGene_tx=XM_004000249.3;refGene_gn=ARSF;refGene_tx=XM_019823520.2
X	1871913	1872281
refGene_gn=ARSF;refGene_tx=XM_004000249.3;refGene_gn=ARSF;refGene_tx=XM_019823520.2
X	1986528	1988437	refGene_gn=MXRA5;refGene_tx=XM_004000245.5
X	1991811	1992712	refGene_gn=MXRA5;refGene_tx=XM_004000245.5
X	1999694	2004635	refGene_gn=MXRA5;refGene_tx=XM_004000245.5
X	2006890	2007281	refGene_gn=MXRA5;refGene_tx=XM_004000245.5
X	2007509	2007639	refGene_gn=MXRA5;refGene_tx=XM_004000245.5
X	2013004	2013195	refGene_gn=MXRA5;refGene_tx=XM_004000245.5
X	2165179	2165305
refGene_gn=PRKX;refGene_tx=XM_023249502.1;refGene_gn=PRKX;refGene_tx=XM_023249503.1
X	2169304	2169382
refGene_gn=PRKX;refGene_tx=XM_023249502.1;refGene_gn=PRKX;refGene_tx=XM_023249503.1
X	2178428	2178486
refGene_gn=PRKX;refGene_tx=XM_023249502.1;refGene_gn=PRKX;refGene_tx=XM_023249503.1
X	2181478	2181574
refGene_gn=PRKX;refGene_tx=XM_023249502.1;refGene_gn=PRKX;refGene_tx=XM_023249503.1
X	2187180	2187300
refGene_gn=PRKX;refGene_tx=XM_023249502.1;refGene_gn=PRKX;refGene_tx=XM_023249503.1
X	2195169	2195433
refGene_gn=PRKX;refGene_tx=XM_023249502.1;refGene_gn=PRKX;refGene_tx=XM_023249503.1
X	2209511	2209680
refGene_gn=PRKX;refGene_tx=XM_023249502.1;refGene_gn=PRKX;refGene_tx=XM_023249503.1
X	2237823	2238001
refGene_gn=PRKX;refGene_tx=XM_023249502.1;refGene_gn=PRKX;refGene_tx=XM_023249503.1

X 2972470 2972632
refGene_gn=LOC109496317;refGene_tx=XM_023249443.1
X 3078323 3078680
refGene_gn=LOC109496317;refGene_tx=XM_023249443.1
X 3079185 3079322
refGene_gn=LOC109496317;refGene_tx=XM_023249443.1
X 3083133 3083204
refGene_gn=LOC109496317;refGene_tx=XM_023249443.1
X 3085539 3085641
refGene_gn=LOC109496317;refGene_tx=XM_023249443.1
X 3085830 3085941
refGene_gn=LOC109496317;refGene_tx=XM_023249443.1
X 3086784 3086886
refGene_gn=LOC109496317;refGene_tx=XM_023249443.1
X 3087138 3087179
refGene_gn=LOC109496317;refGene_tx=XM_023249443.1
X 3664900 3665750
refGene_gn=NLGN4X;refGene_tx=XM_011291596.3;refGene_gn=NLGN4X;refGene_tx=XM_019823521.2;refGene_gn=NLGN4X;refGene_tx=XM_019823522.2;refGene_gn=NLGN4X;refGene_tx=XM_019823523.2;refGene_gn=NLGN4X;refGene_tx=XM_019823524.2;refGene_gn=NLGN4X;refGene_tx=XM_023249333.1
X 3674591 3675381
refGene_gn=NLGN4X;refGene_tx=XM_011291596.3;refGene_gn=NLGN4X;refGene_tx=XM_019823521.2;refGene_gn=NLGN4X;refGene_tx=XM_019823522.2;refGene_gn=NLGN4X;refGene_tx=XM_019823523.2;refGene_gn=NLGN4X;refGene_tx=XM_019823524.2;refGene_gn=NLGN4X;refGene_tx=XM_023249333.1
X 3679458 3679644
refGene_gn=NLGN4X;refGene_tx=XM_011291596.3;refGene_gn=NLGN4X;refGene_tx=XM_019823521.2;refGene_gn=NLGN4X;refGene_tx=XM_019823522.2;refGene_gn=NLGN4X;refGene_tx=XM_019823523.2;refGene_gn=NLGN4X;refGene_tx=XM_019823524.2;refGene_gn=NLGN4X;refGene_tx=XM_023249333.1
X 3791993 3792146
refGene_gn=NLGN4X;refGene_tx=XM_011291596.3;refGene_gn=NLGN4X;refGene_tx=XM_019823521.2;refGene_gn=NLGN4X;refGene_tx=XM_019823522.2;refGene_gn=NLGN4X;refGene_tx=XM_019823523.2;refGene_gn=NLGN4X;refGene_tx=XM_023249333.1
X 3797028 3797088
refGene_gn=NLGN4X;refGene_tx=XM_019823521.2;refGene_gn=NLGN4X;refGene_tx=XM_019823522.2;refGene_gn=NLGN4X;refGene_tx=XM_019823523.2;refGene_gn=NLGN4X;refGene_tx=XM_023249333.1
X 3909169 3909641
refGene_gn=NLGN4X;refGene_tx=XM_011291596.3;refGene_gn=NLGN4X;refGene_tx=XM_019823521.2;refGene_gn=NLGN4X;refGene_tx=XM_019823522.2;refGene_gn=NLGN4X;refGene_tx=XM_019823523.2;refGene_gn=NLGN4X;refGene_tx=XM_019823524.2;refGene_gn=NLGN4X;refGene_tx=XM_023249333.1
X 4614017 4614194 refGene_gn=PUDP;refGene_tx=XM_023249423.1
X 4634751 4634981 refGene_gn=PUDP;refGene_tx=XM_023249423.1
X 4663112 4663331 refGene_gn=PUDP;refGene_tx=XM_023249423.1
X 4703290 4703360 refGene_gn=PUDP;refGene_tx=XM_023249423.1
X 4777027 4777164

refGene_gn=STS;refGene_tx=XM_004000254.4;refGene_gn=STS;refGene_tx=XM_019823525.2;refGene_gn=STS;refGene_tx=XM_023249422.1
X 4780079 4780201
refGene_gn=STS;refGene_tx=XM_004000254.4;refGene_gn=STS;refGene_tx=XM_019823525.2;refGene_gn=STS;refGene_tx=XM_023249422.1
X 4780288 4780411
refGene_gn=STS;refGene_tx=XM_004000254.4;refGene_gn=STS;refGene_tx=XM_019823525.2;refGene_gn=STS;refGene_tx=XM_023249422.1
X 4782301 4782725
refGene_gn=STS;refGene_tx=XM_004000254.4;refGene_gn=STS;refGene_tx=XM_019823525.2;refGene_gn=STS;refGene_tx=XM_023249422.1
X 4798674 4798811
refGene_gn=STS;refGene_tx=XM_004000254.4;refGene_gn=STS;refGene_tx=XM_019823525.2;refGene_gn=STS;refGene_tx=XM_023249422.1
X 4822098 4822236
refGene_gn=STS;refGene_tx=XM_004000254.4;refGene_gn=STS;refGene_tx=XM_019823525.2;refGene_gn=STS;refGene_tx=XM_023249422.1
X 4839319 4839479
refGene_gn=STS;refGene_tx=XM_004000254.4;refGene_gn=STS;refGene_tx=XM_019823525.2;refGene_gn=STS;refGene_tx=XM_023249422.1
X 4846511 4846633
refGene_gn=STS;refGene_tx=XM_004000254.4;refGene_gn=STS;refGene_tx=XM_019823525.2;refGene_gn=STS;refGene_tx=XM_023249422.1
X 4859513 4859887
refGene_gn=STS;refGene_tx=XM_004000254.4;refGene_gn=STS;refGene_tx=XM_019823525.2;refGene_gn=STS;refGene_tx=XM_023249422.1
X 5345031 5345163
refGene_gn=PNPLA4;refGene_tx=XM_011291597.2;refGene_gn=PNPLA4;refGene_tx=XM_011291598.2;refGene_gn=PNPLA4;refGene_tx=XM_011291599.3;refGene_gn=PNPLA4;refGene_tx=XM_011291600.3;refGene_gn=PNPLA4;refGene_tx=XM_019823526.2;refGene_gn=PNPLA4;refGene_tx=XM_019823527.2;refGene_gn=PNPLA4;refGene_tx=XM_019823528.2
X 5345886 5346039
refGene_gn=PNPLA4;refGene_tx=XM_011291597.2;refGene_gn=PNPLA4;refGene_tx=XM_011291598.2;refGene_gn=PNPLA4;refGene_tx=XM_011291599.3;refGene_gn=PNPLA4;refGene_tx=XM_011291600.3;refGene_gn=PNPLA4;refGene_tx=XM_019823526.2;refGene_gn=PNPLA4;refGene_tx=XM_019823527.2;refGene_gn=PNPLA4;refGene_tx=XM_019823528.2
X 5360166 5360232
refGene_gn=PNPLA4;refGene_tx=XM_011291597.2;refGene_gn=PNPLA4;refGene_tx=XM_011291598.2;refGene_gn=PNPLA4;refGene_tx=XM_011291599.3;refGene_gn=PNPLA4;refGene_tx=XM_011291600.3;refGene_gn=PNPLA4;refGene_tx=XM_019823526.2;refGene_gn=PNPLA4;refGene_tx=XM_019823528.2
X 5362658 5362739
refGene_gn=PNPLA4;refGene_tx=XM_011291597.2;refGene_gn=PNPLA4;refGene_tx=XM_011291598.2;refGene_gn=PNPLA4;refGene_tx=XM_011291599.3;refGene_gn=PNPLA4;refGene_tx=XM_011291600.3;refGene_gn=PNPLA4;refGene_tx=XM_019823528.2
X 5372171 5372307
refGene_gn=PNPLA4;refGene_tx=XM_011291597.2;refGene_gn=PNPLA4;refGene_

tx=XM_011291598.2;refGene_gn=PNPLA4;refGene_tx=XM_011291599.3;refGene_gn=PNPLA4;refGene_tx=XM_011291600.3;refGene_gn=PNPLA4;refGene_tx=XM_019823526.2;refGene_gn=PNPLA4;refGene_tx=XM_019823527.2;refGene_gn=PNPLA4;refGene_tx=XM_019823528.2
X 5372474 5372569
refGene_gn=PNPLA4;refGene_tx=XM_011291597.2;refGene_gn=PNPLA4;refGene_tx=XM_011291598.2;refGene_gn=PNPLA4;refGene_tx=XM_011291599.3;refGene_gn=PNPLA4;refGene_tx=XM_011291600.3;refGene_gn=PNPLA4;refGene_tx=XM_019823526.2;refGene_gn=PNPLA4;refGene_tx=XM_019823527.2;refGene_gn=PNPLA4;refGene_tx=XM_019823528.2
X 5377614 5377799
refGene_gn=PNPLA4;refGene_tx=XM_011291597.2;refGene_gn=PNPLA4;refGene_tx=XM_011291598.2;refGene_gn=PNPLA4;refGene_tx=XM_011291599.3;refGene_gn=PNPLA4;refGene_tx=XM_011291600.3;refGene_gn=PNPLA4;refGene_tx=XM_019823526.2;refGene_gn=PNPLA4;refGene_tx=XM_019823527.2;refGene_gn=PNPLA4;refGene_tx=XM_019823528.2
X 5378931 5378956 refGene_gn=PNPLA4;refGene_tx=XM_011291597.2
X 5379365 5379537
refGene_gn=PNPLA4;refGene_tx=XM_011291599.3;refGene_gn=PNPLA4;refGene_tx=XM_019823526.2;refGene_gn=PNPLA4;refGene_tx=XM_019823527.2
X 5925076 5925135
refGene_gn=ANOS1;refGene_tx=XM_023249108.1;refGene_gn=ANOS1;refGene_tx=XM_023249109.1
X 5926394 5926536
refGene_gn=ANOS1;refGene_tx=XM_023249108.1;refGene_gn=ANOS1;refGene_tx=XM_023249109.1
X 5929215 5929436
refGene_gn=ANOS1;refGene_tx=XM_023249108.1;refGene_gn=ANOS1;refGene_tx=XM_023249109.1
X 5930224 5930396
refGene_gn=ANOS1;refGene_tx=XM_023249108.1;refGene_gn=ANOS1;refGene_tx=XM_023249109.1
X 5932433 5932528
refGene_gn=ANOS1;refGene_tx=XM_023249108.1;refGene_gn=ANOS1;refGene_tx=XM_023249109.1
X 5943538 5943685
refGene_gn=ANOS1;refGene_tx=XM_023249108.1;refGene_gn=ANOS1;refGene_tx=XM_023249109.1
X 5951680 5951825
refGene_gn=ANOS1;refGene_tx=XM_023249108.1;refGene_gn=ANOS1;refGene_tx=XM_023249109.1
X 5954968 5955174
refGene_gn=ANOS1;refGene_tx=XM_023249108.1;refGene_gn=ANOS1;refGene_tx=XM_023249109.1
X 5967465 5967595
refGene_gn=ANOS1;refGene_tx=XM_023249108.1;refGene_gn=ANOS1;refGene_tx=XM_023249109.1
X 5970617 5970802
refGene_gn=ANOS1;refGene_tx=XM_023249108.1;refGene_gn=ANOS1;refGene_tx=XM_023249109.1

X 5979182 5979405
refGene_gn=ANOS1;refGene_tx=XM_023249108.1;refGene_gn=ANOS1;refGene_tx=XM_023249109.1
X 5998157 5998220
refGene_gn=ANOS1;refGene_tx=XM_023249108.1;refGene_gn=ANOS1;refGene_tx=XM_023249109.1
X 6073169 6073217
refGene_gn=ANOS1;refGene_tx=XM_023249108.1;refGene_gn=ANOS1;refGene_tx=XM_023249109.1
X 6106670 6106877
refGene_gn=ANOS1;refGene_tx=XM_023249108.1;refGene_gn=ANOS1;refGene_tx=XM_023249109.1
X 6982904 6982962
refGene_gn=TBL1X;refGene_tx=XM_023249354.1;refGene_gn=TBL1X;refGene_tx=XM_023249355.1;refGene_gn=TBL1X;refGene_tx=XM_023249356.1
X 7011337 7011483
refGene_gn=TBL1X;refGene_tx=XM_023249354.1;refGene_gn=TBL1X;refGene_tx=XM_023249355.1;refGene_gn=TBL1X;refGene_tx=XM_023249356.1
X 7015248 7015510
refGene_gn=TBL1X;refGene_tx=XM_023249354.1;refGene_gn=TBL1X;refGene_tx=XM_023249355.1;refGene_gn=TBL1X;refGene_tx=XM_023249356.1
X 7017554 7017687
refGene_gn=TBL1X;refGene_tx=XM_023249354.1;refGene_gn=TBL1X;refGene_tx=XM_023249355.1;refGene_gn=TBL1X;refGene_tx=XM_023249356.1
X 7018062 7018204
refGene_gn=TBL1X;refGene_tx=XM_023249354.1;refGene_gn=TBL1X;refGene_tx=XM_023249355.1;refGene_gn=TBL1X;refGene_tx=XM_023249356.1
X 7018915 7018979
refGene_gn=TBL1X;refGene_tx=XM_023249354.1;refGene_gn=TBL1X;refGene_tx=XM_023249355.1;refGene_gn=TBL1X;refGene_tx=XM_023249356.1
X 7019093 7019191
refGene_gn=TBL1X;refGene_tx=XM_023249354.1;refGene_gn=TBL1X;refGene_tx=XM_023249355.1;refGene_gn=TBL1X;refGene_tx=XM_023249356.1
X 7023050 7023111
refGene_gn=TBL1X;refGene_tx=XM_023249354.1;refGene_gn=TBL1X;refGene_tx=XM_023249355.1;refGene_gn=TBL1X;refGene_tx=XM_023249356.1
X 7027707 7027889
refGene_gn=GPR143;refGene_tx=XM_023249358.1;refGene_gn=TBL1X;refGene_tx=XM_023249354.1;refGene_gn=TBL1X;refGene_tx=XM_023249355.1;refGene_gn=TBL1X;refGene_tx=XM_023249356.1
X 7028030 7028044 refGene_gn=GPR143;refGene_tx=XM_023249359.1
X 7030823 7030898
refGene_gn=TBL1X;refGene_tx=XM_023249354.1;refGene_gn=TBL1X;refGene_tx=XM_023249355.1;refGene_gn=TBL1X;refGene_tx=XM_023249356.1
X 7031278 7031406
refGene_gn=TBL1X;refGene_tx=XM_023249354.1;refGene_gn=TBL1X;refGene_tx=XM_023249355.1;refGene_gn=TBL1X;refGene_tx=XM_023249356.1
X 7033482 7033648
refGene_gn=TBL1X;refGene_tx=XM_023249354.1;refGene_gn=TBL1X;refGene_tx=XM_023249355.1;refGene_gn=TBL1X;refGene_tx=XM_023249356.1

X	7036404	7036506
refGene_gn=TBL1X;refGene_tx=XM_023249354.1;refGene_gn=TBL1X;refGene_tx=XM_023249355.1;refGene_gn=TBL1X;refGene_tx=XM_023249356.1
X	7037466	7037493
refGene_gn=TBL1X;refGene_tx=XM_023249354.1;refGene_gn=TBL1X;refGene_tx=XM_023249355.1;refGene_gn=TBL1X;refGene_tx=XM_023249356.1
X	7043932	7044027
refGene_gn=GPR143;refGene_tx=XM_023249357.1;refGene_gn=GPR143;refGene_tx=XM_023249360.1
X	7071548	7071792
refGene_gn=GPR143;refGene_tx=XM_023249357.1;refGene_gn=GPR143;refGene_tx=XM_023249358.1;refGene_gn=GPR143;refGene_tx=XM_023249359.1;refGene_gn=GPR143;refGene_tx=XM_023249360.1
X	7073711	7073829
refGene_gn=GPR143;refGene_tx=XM_023249357.1;refGene_gn=GPR143;refGene_tx=XM_023249358.1;refGene_gn=GPR143;refGene_tx=XM_023249359.1;refGene_gn=GPR143;refGene_tx=XM_023249360.1
X	7075163	7075272
refGene_gn=GPR143;refGene_tx=XM_023249357.1;refGene_gn=GPR143;refGene_tx=XM_023249358.1;refGene_gn=GPR143;refGene_tx=XM_023249359.1;refGene_gn=GPR143;refGene_tx=XM_023249360.1
X	7078007	7078117
refGene_gn=GPR143;refGene_tx=XM_023249357.1;refGene_gn=GPR143;refGene_tx=XM_023249358.1;refGene_gn=GPR143;refGene_tx=XM_023249359.1;refGene_gn=GPR143;refGene_tx=XM_023249360.1
X	7081444	7081537
refGene_gn=GPR143;refGene_tx=XM_023249357.1;refGene_gn=GPR143;refGene_tx=XM_023249358.1;refGene_gn=GPR143;refGene_tx=XM_023249359.1
X	7088466	7088561
refGene_gn=GPR143;refGene_tx=XM_023249357.1;refGene_gn=GPR143;refGene_tx=XM_023249358.1;refGene_gn=GPR143;refGene_tx=XM_023249359.1;refGene_gn=GPR143;refGene_tx=XM_023249360.1
X	7089845	7089955
refGene_gn=GPR143;refGene_tx=XM_023249357.1;refGene_gn=GPR143;refGene_tx=XM_023249358.1;refGene_gn=GPR143;refGene_tx=XM_023249359.1;refGene_gn=GPR143;refGene_tx=XM_023249360.1
X	7095054	7095418
refGene_gn=GPR143;refGene_tx=XM_023249357.1;refGene_gn=GPR143;refGene_tx=XM_023249358.1;refGene_gn=GPR143;refGene_tx=XM_023249359.1;refGene_gn=GPR143;refGene_tx=XM_023249360.1
X	7099601	7099650
refGene_gn=LOC109496425;refGene_tx=XM_023249362.1
X	7105483	7105679
refGene_gn=LOC109496425;refGene_tx=XM_023249361.1
X	7116394	7116513
refGene_gn=LOC109496425;refGene_tx=XM_023249361.1;refGene_gn=LOC109496425;refGene_tx=XM_023249362.1
X	7123100	7123153
refGene_gn=LOC109496425;refGene_tx=XM_023249361.1;refGene_gn=LOC109496425;refGene_tx=XM_023249362.1

X	7123868	7124176
refGene_gn=LOC109496425;refGene_tx=XM_023249361.1;refGene_gn=LOC109496
425;refGene_tx=XM_023249362.1
X	7124644	7124809
refGene_gn=SHROOM2;refGene_tx=XM_023249374.1;refGene_gn=SHROOM2;refGen
e_tx=XM_023249375.1;refGene_gn=SHROOM2;refGene_tx=XM_023249376.1
X	7199061	7199213
refGene_gn=SHROOM2;refGene_tx=XM_023249374.1;refGene_gn=SHROOM2;refGen
e_tx=XM_023249375.1;refGene_gn=SHROOM2;refGene_tx=XM_023249376.1
X	7217888	7218020
refGene_gn=SHROOM2;refGene_tx=XM_023249374.1;refGene_gn=SHROOM2;refGen
e_tx=XM_023249375.1
X	7223425	7225883
refGene_gn=SHROOM2;refGene_tx=XM_023249374.1;refGene_gn=SHROOM2;refGen
e_tx=XM_023249375.1;refGene_gn=SHROOM2;refGene_tx=XM_023249376.1
X	7227414	7227512
refGene_gn=SHROOM2;refGene_tx=XM_023249374.1;refGene_gn=SHROOM2;refGen
e_tx=XM_023249375.1;refGene_gn=SHROOM2;refGene_tx=XM_023249376.1
X	7265522	7266098
refGene_gn=SHROOM2;refGene_tx=XM_019823536.2;refGene_gn=SHROOM2;refGen
e_tx=XM_023249374.1;refGene_gn=SHROOM2;refGene_tx=XM_023249375.1;refGe
ne_gn=SHROOM2;refGene_tx=XM_023249376.1
X	7270317	7270842
refGene_gn=SHROOM2;refGene_tx=XM_019823536.2;refGene_gn=SHROOM2;refGen
e_tx=XM_023249374.1;refGene_gn=SHROOM2;refGene_tx=XM_023249375.1;refGe
ne_gn=SHROOM2;refGene_tx=XM_023249376.1
X	7271744	7271916
refGene_gn=SHROOM2;refGene_tx=XM_019823536.2;refGene_gn=SHROOM2;refGen
e_tx=XM_023249374.1;refGene_gn=SHROOM2;refGene_tx=XM_023249375.1;refGe
ne_gn=SHROOM2;refGene_tx=XM_023249376.1
X	7278245	7278518
refGene_gn=SHROOM2;refGene_tx=XM_019823536.2;refGene_gn=SHROOM2;refGen
e_tx=XM_023249374.1;refGene_gn=SHROOM2;refGene_tx=XM_023249375.1;refGe
ne_gn=SHROOM2;refGene_tx=XM_023249376.1
X	7281466	7281733
refGene_gn=SHROOM2;refGene_tx=XM_019823536.2;refGene_gn=SHROOM2;refGen
e_tx=XM_023249374.1;refGene_gn=SHROOM2;refGene_tx=XM_023249375.1;refGe
ne_gn=SHROOM2;refGene_tx=XM_023249376.1
X	7295097	7295622	refGene_gn=CLDN34;refGene_tx=XM_023249177.1
X	7295623	7295788	refGene_gn=CLDN34;refGene_tx=XM_023249177.1
X	7315455	7316115
refGene_gn=LOC101087815;refGene_tx=XM_004000271.1
X	7397229	7397245	refGene_gn=WWC3;refGene_tx=XM_023249413.1
X	7405124	7405328	refGene_gn=WWC3;refGene_tx=XM_023249413.1
X	7415527	7415604	refGene_gn=WWC3;refGene_tx=XM_023249413.1
X	7416987	7417067	refGene_gn=WWC3;refGene_tx=XM_023249413.1
X	7428268	7428398
refGene_gn=WWC3;refGene_tx=XM_023249413.1;refGene_gn=WWC3;refGene_tx=X
M_023249414.1
X	7431095	7431242

refGene_gn=WWC3;refGene_tx=XM_023249413.1;refGene_gn=WWC3;refGene_tx=XM_023249414.1
X	7435331	7435405
refGene_gn=WWC3;refGene_tx=XM_023249413.1;refGene_gn=WWC3;refGene_tx=XM_023249414.1
X	7444898	7445141
refGene_gn=WWC3;refGene_tx=XM_023249413.1;refGene_gn=WWC3;refGene_tx=XM_023249414.1
X	7447912	7448002
refGene_gn=WWC3;refGene_tx=XM_023249413.1;refGene_gn=WWC3;refGene_tx=XM_023249414.1
X	7448805	7449371
refGene_gn=WWC3;refGene_tx=XM_023249413.1;refGene_gn=WWC3;refGene_tx=XM_023249414.1
X	7454662	7454783
refGene_gn=WWC3;refGene_tx=XM_023249413.1;refGene_gn=WWC3;refGene_tx=XM_023249414.1
X	7455818	7455893
refGene_gn=WWC3;refGene_tx=XM_023249413.1;refGene_gn=WWC3;refGene_tx=XM_023249414.1
X	7456709	7456796
refGene_gn=WWC3;refGene_tx=XM_023249413.1;refGene_gn=WWC3;refGene_tx=XM_023249414.1
X	7457523	7457716
refGene_gn=WWC3;refGene_tx=XM_023249413.1;refGene_gn=WWC3;refGene_tx=XM_023249414.1
X	7459271	7459418
refGene_gn=WWC3;refGene_tx=XM_023249413.1;refGene_gn=WWC3;refGene_tx=XM_023249414.1
X	7459750	7459902
refGene_gn=WWC3;refGene_tx=XM_023249413.1;refGene_gn=WWC3;refGene_tx=XM_023249414.1
X	7461305	7461471
refGene_gn=WWC3;refGene_tx=XM_023249413.1;refGene_gn=WWC3;refGene_tx=XM_023249414.1
X	7464701	7464866
refGene_gn=WWC3;refGene_tx=XM_023249413.1;refGene_gn=WWC3;refGene_tx=XM_023249414.1
X	7467529	7467622
refGene_gn=WWC3;refGene_tx=XM_023249413.1;refGene_gn=WWC3;refGene_tx=XM_023249414.1
X	7470157	7470385
refGene_gn=WWC3;refGene_tx=XM_023249413.1;refGene_gn=WWC3;refGene_tx=XM_023249414.1
X	7470822	7470950
refGene_gn=WWC3;refGene_tx=XM_023249413.1;refGene_gn=WWC3;refGene_tx=XM_023249414.1
X	7472320	7472387
refGene_gn=WWC3;refGene_tx=XM_023249413.1;refGene_gn=WWC3;refGene_tx=XM_023249414.1

X	7497961	7498005
refGene_gn=CLCN4;refGene_tx=XM_019823537.2;refGene_gn=CLCN4;refGene_tx=XM_019823538.2
X	7522173	7522342
refGene_gn=CLCN4;refGene_tx=XM_011291605.3;refGene_gn=CLCN4;refGene_tx=XM_019823537.2;refGene_gn=CLCN4;refGene_tx=XM_019823538.2
X	7524983	7525083
refGene_gn=CLCN4;refGene_tx=XM_011291605.3;refGene_gn=CLCN4;refGene_tx=XM_019823537.2;refGene_gn=CLCN4;refGene_tx=XM_019823538.2
X	7529462	7529650
refGene_gn=CLCN4;refGene_tx=XM_011291605.3;refGene_gn=CLCN4;refGene_tx=XM_019823537.2;refGene_gn=CLCN4;refGene_tx=XM_019823538.2
X	7531885	7532008
refGene_gn=CLCN4;refGene_tx=XM_011291605.3;refGene_gn=CLCN4;refGene_tx=XM_019823537.2;refGene_gn=CLCN4;refGene_tx=XM_019823538.2
X	7537247	7537454
refGene_gn=CLCN4;refGene_tx=XM_011291605.3;refGene_gn=CLCN4;refGene_tx=XM_019823537.2;refGene_gn=CLCN4;refGene_tx=XM_019823538.2
X	7537582	7537663
refGene_gn=CLCN4;refGene_tx=XM_011291605.3;refGene_gn=CLCN4;refGene_tx=XM_019823537.2;refGene_gn=CLCN4;refGene_tx=XM_019823538.2
X	7538638	7539184
refGene_gn=CLCN4;refGene_tx=XM_011291605.3;refGene_gn=CLCN4;refGene_tx=XM_019823537.2;refGene_gn=CLCN4;refGene_tx=XM_019823538.2
X	7540856	7541043
refGene_gn=CLCN4;refGene_tx=XM_011291605.3;refGene_gn=CLCN4;refGene_tx=XM_019823537.2;refGene_gn=CLCN4;refGene_tx=XM_019823538.2
X	7542514	7542913
refGene_gn=CLCN4;refGene_tx=XM_011291605.3;refGene_gn=CLCN4;refGene_tx=XM_019823537.2;refGene_gn=CLCN4;refGene_tx=XM_019823538.2
X	7549888	7550105
refGene_gn=CLCN4;refGene_tx=XM_011291605.3;refGene_gn=CLCN4;refGene_tx=XM_019823537.2;refGene_gn=CLCN4;refGene_tx=XM_019823538.2
X	7556504	7556595
refGene_gn=CLCN4;refGene_tx=XM_011291605.3;refGene_gn=CLCN4;refGene_tx=XM_019823537.2;refGene_gn=CLCN4;refGene_tx=XM_019823538.2
X	7722657	7723006
refGene_gn=MID1;refGene_tx=XM_004000259.4;refGene_gn=MID1;refGene_tx=XM_006943548.2;refGene_gn=MID1;refGene_tx=XM_019823539.2;refGene_gn=MID1;refGene_tx=XM_019823540.2;refGene_gn=MID1;refGene_tx=XM_019823541.2;refGene_gn=MID1;refGene_tx=XM_019823542.2
X	7727722	7727930
refGene_gn=MID1;refGene_tx=XM_004000259.4;refGene_gn=MID1;refGene_tx=XM_006943548.2;refGene_gn=MID1;refGene_tx=XM_019823539.2;refGene_gn=MID1;refGene_tx=XM_019823540.2;refGene_gn=MID1;refGene_tx=XM_019823541.2;refGene_gn=MID1;refGene_tx=XM_019823542.2
X	7732021	7732183
refGene_gn=MID1;refGene_tx=XM_004000259.4;refGene_gn=MID1;refGene_tx=XM_006943548.2;refGene_gn=MID1;refGene_tx=XM_019823539.2;refGene_gn=MID1;refGene_tx=XM_019823540.2;refGene_gn=MID1;refGene_tx=XM_019823541.2;

refGene_gn=MID1;refGene_tx=XM_019823542.2
X	7743158	7743302
refGene_gn=MID1;refGene_tx=XM_004000259.4;refGene_gn=MID1;refGene_tx=XM_006943548.2;refGene_gn=MID1;refGene_tx=XM_019823539.2;refGene_gn=MID1;refGene_tx=XM_019823540.2;refGene_gn=MID1;refGene_tx=XM_019823541.2;refGene_gn=MID1;refGene_tx=XM_019823542.2
X	7748389	7748517
refGene_gn=MID1;refGene_tx=XM_004000259.4;refGene_gn=MID1;refGene_tx=XM_006943548.2;refGene_gn=MID1;refGene_tx=XM_019823539.2;refGene_gn=MID1;refGene_tx=XM_019823540.2;refGene_gn=MID1;refGene_tx=XM_019823541.2;refGene_gn=MID1;refGene_tx=XM_019823542.2
X	7756233	7756382
refGene_gn=MID1;refGene_tx=XM_004000259.4;refGene_gn=MID1;refGene_tx=XM_006943548.2;refGene_gn=MID1;refGene_tx=XM_019823539.2;refGene_gn=MID1;refGene_tx=XM_019823540.2;refGene_gn=MID1;refGene_tx=XM_019823541.2;refGene_gn=MID1;refGene_tx=XM_019823542.2
X	7766151	7766259
refGene_gn=MID1;refGene_tx=XM_004000259.4;refGene_gn=MID1;refGene_tx=XM_006943548.2;refGene_gn=MID1;refGene_tx=XM_019823539.2;refGene_gn=MID1;refGene_tx=XM_019823540.2;refGene_gn=MID1;refGene_tx=XM_019823541.2;refGene_gn=MID1;refGene_tx=XM_019823542.2
X	7787966	7788062
refGene_gn=MID1;refGene_tx=XM_004000259.4;refGene_gn=MID1;refGene_tx=XM_006943548.2;refGene_gn=MID1;refGene_tx=XM_019823539.2;refGene_gn=MID1;refGene_tx=XM_019823540.2;refGene_gn=MID1;refGene_tx=XM_019823541.2;refGene_gn=MID1;refGene_tx=XM_019823542.2
X	7823424	7824084
refGene_gn=MID1;refGene_tx=XM_004000259.4;refGene_gn=MID1;refGene_tx=XM_006943548.2;refGene_gn=MID1;refGene_tx=XM_019823539.2;refGene_gn=MID1;refGene_tx=XM_019823540.2;refGene_gn=MID1;refGene_tx=XM_019823541.2;refGene_gn=MID1;refGene_tx=XM_019823542.2
X	8316923	8317035
refGene_gn=LOC101085440;refGene_tx=XM_004000261.5
X	8319091	8319243
refGene_gn=LOC101085440;refGene_tx=XM_004000261.5
X	8321575	8321724
refGene_gn=LOC101085440;refGene_tx=XM_004000261.5
X	8322573	8322693
refGene_gn=LOC101085440;refGene_tx=XM_004000261.5
X	8324715	8324802
refGene_gn=LOC101085440;refGene_tx=XM_004000261.5
X	8325357	8325556
refGene_gn=LOC101085440;refGene_tx=XM_004000261.5
X	8338061	8338735
refGene_gn=ARHGAP6;refGene_tx=XM_019823543.2;refGene_gn=ARHGAP6;refGene_tx=XM_023248925.1;refGene_gn=ARHGAP6;refGene_tx=XM_023248926.1
X	8340923	8341004
refGene_gn=ARHGAP6;refGene_tx=XM_019823543.2;refGene_gn=ARHGAP6;refGene_tx=XM_023248925.1;refGene_gn=ARHGAP6;refGene_tx=XM_023248926.1
X	8344010	8344279

refGene_gn=ARHGAP6;refGene_tx=XM_019823543.2;refGene_gn=ARHGAP6;refGen
e_tx=XM_023248925.1;refGene_gn=ARHGAP6;refGene_tx=XM_023248926.1
X        8354328  8354426
refGene_gn=ARHGAP6;refGene_tx=XM_019823543.2;refGene_gn=ARHGAP6;refGen
e_tx=XM_023248925.1;refGene_gn=ARHGAP6;refGene_tx=XM_023248926.1
X        8367176  8367356
refGene_gn=ARHGAP6;refGene_tx=XM_019823543.2;refGene_gn=ARHGAP6;refGen
e_tx=XM_023248925.1;refGene_gn=ARHGAP6;refGene_tx=XM_023248926.1
X        8372353  8372502
refGene_gn=ARHGAP6;refGene_tx=XM_019823543.2;refGene_gn=ARHGAP6;refGen
e_tx=XM_023248925.1;refGene_gn=ARHGAP6;refGene_tx=XM_023248926.1
X        8373539  8373690
refGene_gn=ARHGAP6;refGene_tx=XM_019823543.2;refGene_gn=ARHGAP6;refGen
e_tx=XM_023248925.1;refGene_gn=ARHGAP6;refGene_tx=XM_023248926.1
X        8376409  8376465
refGene_gn=ARHGAP6;refGene_tx=XM_019823543.2;refGene_gn=ARHGAP6;refGen
e_tx=XM_023248925.1;refGene_gn=ARHGAP6;refGene_tx=XM_023248926.1
X        8381360  8381556
refGene_gn=ARHGAP6;refGene_tx=XM_019823543.2;refGene_gn=ARHGAP6;refGen
e_tx=XM_023248925.1;refGene_gn=ARHGAP6;refGene_tx=XM_023248926.1
X        8383492  8383749
refGene_gn=ARHGAP6;refGene_tx=XM_019823543.2;refGene_gn=ARHGAP6;refGen
e_tx=XM_023248925.1;refGene_gn=ARHGAP6;refGene_tx=XM_023248926.1
X        8390486  8390558
refGene_gn=ARHGAP6;refGene_tx=XM_019823543.2;refGene_gn=ARHGAP6;refGen
e_tx=XM_023248925.1;refGene_gn=ARHGAP6;refGene_tx=XM_023248926.1
X        8407713  8407873
refGene_gn=ARHGAP6;refGene_tx=XM_019823543.2;refGene_gn=ARHGAP6;refGen
e_tx=XM_023248925.1;refGene_gn=ARHGAP6;refGene_tx=XM_023248926.1
X        8447316  8447370
refGene_gn=AMELX;refGene_tx=XM_004000262.2;refGene_gn=AMELX;refGene_tx
=XM_006943549.3;refGene_gn=AMELX;refGene_tx=XM_006943550.3;refGene_gn=
AMELX;refGene_tx=XM_006943552.3
X        8449277  8449325
refGene_gn=AMELX;refGene_tx=XM_004000262.2;refGene_gn=AMELX;refGene_tx
=XM_006943549.3;refGene_gn=AMELX;refGene_tx=XM_006943550.3
X        8450708  8450753
refGene_gn=AMELX;refGene_tx=XM_004000262.2;refGene_gn=AMELX;refGene_tx
=XM_006943549.3;refGene_gn=AMELX;refGene_tx=XM_006943550.3;refGene_gn=
AMELX;refGene_tx=XM_006943552.3
X        8451020  8451446
refGene_gn=AMELX;refGene_tx=XM_004000262.2;refGene_gn=AMELX;refGene_tx
=XM_006943549.3;refGene_gn=AMELX;refGene_tx=XM_006943550.3;refGene_gn=
AMELX;refGene_tx=XM_006943552.3
X        8453027  8453033
refGene_gn=AMELX;refGene_tx=XM_004000262.2;refGene_gn=AMELX;refGene_tx
=XM_006943549.3;refGene_gn=AMELX;refGene_tx=XM_006943550.3;refGene_gn=
AMELX;refGene_tx=XM_006943552.3
X        8565845  8565890  refGene_gn=ARHGAP6;refGene_tx=XM_023248925.1
X        8781731  8782310  refGene_gn=ARHGAP6;refGene_tx=XM_019823543.2

X	8883629	8883731
refGene_gn=MSL3;refGene_tx=XM_023248900.1;refGene_gn=MSL3;refGene_tx=XM_023248901.1
X	8885087	8885170
refGene_gn=MSL3;refGene_tx=XM_023248900.1;refGene_gn=MSL3;refGene_tx=XM_023248901.1
X	8885551	8885647	refGene_gn=MSL3;refGene_tx=XM_023248900.1
X	8885984	8886085
refGene_gn=MSL3;refGene_tx=XM_023248900.1;refGene_gn=MSL3;refGene_tx=XM_023248901.1
X	8886858	8886944
refGene_gn=MSL3;refGene_tx=XM_023248900.1;refGene_gn=MSL3;refGene_tx=XM_023248901.1
X	8887603	8887726
refGene_gn=MSL3;refGene_tx=XM_023248900.1;refGene_gn=MSL3;refGene_tx=XM_023248901.1
X	8888420	8888581
refGene_gn=MSL3;refGene_tx=XM_023248900.1;refGene_gn=MSL3;refGene_tx=XM_023248901.1
X	8889345	8889504
refGene_gn=MSL3;refGene_tx=XM_023248900.1;refGene_gn=MSL3;refGene_tx=XM_023248901.1
X	8891352	8891615
refGene_gn=MSL3;refGene_tx=XM_023248900.1;refGene_gn=MSL3;refGene_tx=XM_023248901.1
X	8892273	8892383
refGene_gn=MSL3;refGene_tx=XM_023248900.1;refGene_gn=MSL3;refGene_tx=XM_023248901.1
X	8895824	8895924
refGene_gn=MSL3;refGene_tx=XM_023248900.1;refGene_gn=MSL3;refGene_tx=XM_023248901.1
X	8896371	8896456
refGene_gn=MSL3;refGene_tx=XM_023248900.1;refGene_gn=MSL3;refGene_tx=XM_023248901.1
X	8897391	8897491
refGene_gn=MSL3;refGene_tx=XM_023248900.1;refGene_gn=MSL3;refGene_tx=XM_023248901.1
X	8993816	8993883
refGene_gn=FRMPD4;refGene_tx=XM_019823547.2;refGene_gn=FRMPD4;refGene_tx=XM_019823548.2
X	9008546	9008670
refGene_gn=FRMPD4;refGene_tx=XM_019823547.2;refGene_gn=FRMPD4;refGene_tx=XM_019823548.2
X	9254733	9254765	refGene_gn=FRMPD4;refGene_tx=XM_019823549.2
X	9255981	9256022	refGene_gn=FRMPD4;refGene_tx=XM_006943556.4
X	9610167	9610284
refGene_gn=FRMPD4;refGene_tx=XM_006943556.4;refGene_gn=FRMPD4;refGene_tx=XM_019823547.2;refGene_gn=FRMPD4;refGene_tx=XM_019823548.2;refGene_gn=FRMPD4;refGene_tx=XM_019823549.2;refGene_gn=FRMPD4;refGene_tx=XM_019823551.2

X 9709262 9709423
refGene_gn=FRMPD4;refGene_tx=XM_006943556.4;refGene_gn=FRMPD4;refGene_tx=XM_019823547.2;refGene_gn=FRMPD4;refGene_tx=XM_019823548.2;refGene_gn=FRMPD4;refGene_tx=XM_019823549.2;refGene_gn=FRMPD4;refGene_tx=XM_019823551.2;refGene_gn=FRMPD4;refGene_tx=XM_019823552.2
X 9714235 9714338
refGene_gn=FRMPD4;refGene_tx=XM_006943556.4;refGene_gn=FRMPD4;refGene_tx=XM_019823547.2;refGene_gn=FRMPD4;refGene_tx=XM_019823548.2;refGene_gn=FRMPD4;refGene_tx=XM_019823549.2;refGene_gn=FRMPD4;refGene_tx=XM_019823551.2;refGene_gn=FRMPD4;refGene_tx=XM_019823552.2
X 9756351 9756397
refGene_gn=FRMPD4;refGene_tx=XM_006943556.4;refGene_gn=FRMPD4;refGene_tx=XM_019823547.2;refGene_gn=FRMPD4;refGene_tx=XM_019823548.2;refGene_gn=FRMPD4;refGene_tx=XM_019823549.2;refGene_gn=FRMPD4;refGene_tx=XM_019823551.2;refGene_gn=FRMPD4;refGene_tx=XM_019823552.2
X 9762709 9762814
refGene_gn=FRMPD4;refGene_tx=XM_006943556.4;refGene_gn=FRMPD4;refGene_tx=XM_019823547.2;refGene_gn=FRMPD4;refGene_tx=XM_019823548.2;refGene_gn=FRMPD4;refGene_tx=XM_019823549.2;refGene_gn=FRMPD4;refGene_tx=XM_019823551.2;refGene_gn=FRMPD4;refGene_tx=XM_019823552.2
X 9765811 9765919
refGene_gn=FRMPD4;refGene_tx=XM_006943556.4;refGene_gn=FRMPD4;refGene_tx=XM_019823547.2;refGene_gn=FRMPD4;refGene_tx=XM_019823548.2;refGene_gn=FRMPD4;refGene_tx=XM_019823549.2;refGene_gn=FRMPD4;refGene_tx=XM_019823551.2;refGene_gn=FRMPD4;refGene_tx=XM_019823552.2
X 9769442 9769574
refGene_gn=FRMPD4;refGene_tx=XM_006943556.4;refGene_gn=FRMPD4;refGene_tx=XM_019823547.2;refGene_gn=FRMPD4;refGene_tx=XM_019823548.2;refGene_gn=FRMPD4;refGene_tx=XM_019823549.2;refGene_gn=FRMPD4;refGene_tx=XM_019823551.2;refGene_gn=FRMPD4;refGene_tx=XM_019823552.2
X 9773281 9773401
refGene_gn=FRMPD4;refGene_tx=XM_006943556.4;refGene_gn=FRMPD4;refGene_tx=XM_019823547.2;refGene_gn=FRMPD4;refGene_tx=XM_019823548.2;refGene_gn=FRMPD4;refGene_tx=XM_019823549.2;refGene_gn=FRMPD4;refGene_tx=XM_019823551.2;refGene_gn=FRMPD4;refGene_tx=XM_019823552.2
X 9779020 9779157
refGene_gn=FRMPD4;refGene_tx=XM_006943556.4;refGene_gn=FRMPD4;refGene_tx=XM_019823547.2;refGene_gn=FRMPD4;refGene_tx=XM_019823548.2;refGene_gn=FRMPD4;refGene_tx=XM_019823549.2;refGene_gn=FRMPD4;refGene_tx=XM_019823551.2;refGene_gn=FRMPD4;refGene_tx=XM_019823552.2
X 9781561 9781688
refGene_gn=FRMPD4;refGene_tx=XM_006943556.4;refGene_gn=FRMPD4;refGene_tx=XM_019823547.2;refGene_gn=FRMPD4;refGene_tx=XM_019823548.2;refGene_gn=FRMPD4;refGene_tx=XM_019823549.2;refGene_gn=FRMPD4;refGene_tx=XM_019823551.2;refGene_gn=FRMPD4;refGene_tx=XM_019823552.2
X 9783855 9783945
refGene_gn=FRMPD4;refGene_tx=XM_006943556.4;refGene_gn=FRMPD4;refGene_tx=XM_019823547.2;refGene_gn=FRMPD4;refGene_tx=XM_019823548.2;refGene_gn=FRMPD4;refGene_tx=XM_019823549.2;refGene_gn=FRMPD4;refGene_tx=XM_019823551.2;refGene_gn=FRMPD4;refGene_tx=XM_019823552.2

X 9784511 9784694
refGene_gn=FRMPD4;refGene_tx=XM_006943556.4;refGene_gn=FRMPD4;refGene_tx=XM_019823547.2;refGene_gn=FRMPD4;refGene_tx=XM_019823548.2;refGene_gn=FRMPD4;refGene_tx=XM_019823549.2;refGene_gn=FRMPD4;refGene_tx=XM_019823551.2;refGene_gn=FRMPD4;refGene_tx=XM_019823552.2
X 9786872 9787011
refGene_gn=FRMPD4;refGene_tx=XM_006943556.4;refGene_gn=FRMPD4;refGene_tx=XM_019823547.2;refGene_gn=FRMPD4;refGene_tx=XM_019823548.2;refGene_gn=FRMPD4;refGene_tx=XM_019823549.2;refGene_gn=FRMPD4;refGene_tx=XM_019823551.2;refGene_gn=FRMPD4;refGene_tx=XM_019823552.2
X 9791639 9792704
refGene_gn=FRMPD4;refGene_tx=XM_006943556.4;refGene_gn=FRMPD4;refGene_tx=XM_019823547.2;refGene_gn=FRMPD4;refGene_tx=XM_019823548.2;refGene_gn=FRMPD4;refGene_tx=XM_019823549.2;refGene_gn=FRMPD4;refGene_tx=XM_019823551.2;refGene_gn=FRMPD4;refGene_tx=XM_019823552.2
X 9793104 9794397
refGene_gn=FRMPD4;refGene_tx=XM_006943556.4;refGene_gn=FRMPD4;refGene_tx=XM_019823547.2;refGene_gn=FRMPD4;refGene_tx=XM_019823548.2;refGene_gn=FRMPD4;refGene_tx=XM_019823549.2;refGene_gn=FRMPD4;refGene_tx=XM_019823551.2;refGene_gn=FRMPD4;refGene_tx=XM_019823552.2
X 9795981 9797306
refGene_gn=FRMPD4;refGene_tx=XM_006943556.4;refGene_gn=FRMPD4;refGene_tx=XM_019823547.2;refGene_gn=FRMPD4;refGene_tx=XM_019823548.2;refGene_gn=FRMPD4;refGene_tx=XM_019823549.2;refGene_gn=FRMPD4;refGene_tx=XM_019823551.2;refGene_gn=FRMPD4;refGene_tx=XM_019823552.2
X 9844113 9844235 refGene_gn=PRPS2;refGene_tx=XM_006943557.4
X 9850915 9851099 refGene_gn=PRPS2;refGene_tx=XM_006943557.4
X 9867010 9867109 refGene_gn=PRPS2;refGene_tx=XM_006943557.4
X 9867982 9868107 refGene_gn=PRPS2;refGene_tx=XM_006943557.4
X 9874039 9874213 refGene_gn=PRPS2;refGene_tx=XM_006943557.4
X 9875822 9875982 refGene_gn=PRPS2;refGene_tx=XM_006943557.4
X 9877670 9877763 refGene_gn=PRPS2;refGene_tx=XM_006943557.4
X 9894677 9894743
refGene_gn=TLR7;refGene_tx=NM_001080133.1;refGene_gn=TLR7;refGene_tx=XM_023248794.1
X 9913652 9916802
refGene_gn=TLR7;refGene_tx=NM_001080133.1;refGene_gn=TLR7;refGene_tx=XM_023248794.1;refGene_gn=TLR7;refGene_tx=XM_023248795.1
X 9947905 9951025 refGene_gn=TLR8;refGene_tx=XM_019823554.2
X 9998428 9998528 refGene_gn=TMSB4X;refGene_tx=XM_011291607.1
X 9998928 9998963 refGene_gn=TMSB4X;refGene_tx=XM_011291607.1
X 10554231 10554305 refGene_gn=EGFL6;refGene_tx=XM_023249088.1
X 10576387 10576500 refGene_gn=EGFL6;refGene_tx=XM_023249088.1
X 10582490 10582583 refGene_gn=EGFL6;refGene_tx=XM_023249088.1
X 10586696 10586816 refGene_gn=EGFL6;refGene_tx=XM_023249088.1
X 10589633 10589753 refGene_gn=EGFL6;refGene_tx=XM_023249088.1
X 10592902 10593037 refGene_gn=EGFL6;refGene_tx=XM_023249088.1
X 10594680 10594803 refGene_gn=EGFL6;refGene_tx=XM_023249088.1
X 10597567 10597894 refGene_gn=EGFL6;refGene_tx=XM_023249088.1
X 10598936 10599017 refGene_gn=EGFL6;refGene_tx=XM_023249088.1

X 10602645 10602747 refGene_gn=EGFL6;refGene_tx=XM_023249088.1
X 10605830 10606000 refGene_gn=EGFL6;refGene_tx=XM_023249088.1
X 10606001 10606097 refGene_gn=EGFL6;refGene_tx=XM_023249088.1
X 10609782 10609893 refGene_gn=EGFL6;refGene_tx=XM_023249088.1
X 10626964 10627155 refGene_gn=TCEANC;refGene_tx=XM_006943564.4
X 10627156 10627758 refGene_gn=TCEANC;refGene_tx=XM_006943564.4
X 10627760 10627945 refGene_gn=TCEANC;refGene_tx=XM_006943564.4
X 10668433 10669039
refGene_gn=RAB9A;refGene_tx=XM_023249404.1;refGene_gn=RAB9A;refGene_tx=XM_023249405.1;refGene_gn=RAB9A;refGene_tx=XM_023249406.1;refGene_gn=RAB9A;refGene_tx=XM_023249407.1;refGene_gn=RAB9A;refGene_tx=XM_023249408.1;refGene_gn=RAB9A;refGene_tx=XM_023249409.1;refGene_gn=RAB9A;refGene_tx=XM_023249410.1
X 10674375 10674474
refGene_gn=TRAPPC2;refGene_tx=XM_004000278.5;refGene_gn=TRAPPC2;refGene_tx=XM_019823564.2
X 10675713 10675799
refGene_gn=TRAPPC2;refGene_tx=XM_004000278.5;refGene_gn=TRAPPC2;refGene_tx=XM_019823564.2
X 10676229 10676374
refGene_gn=TRAPPC2;refGene_tx=XM_004000278.5;refGene_gn=TRAPPC2;refGene_tx=XM_019823564.2
X 10678815 10678908
refGene_gn=TRAPPC2;refGene_tx=XM_004000278.5;refGene_gn=TRAPPC2;refGene_tx=XM_019823564.2
X 10685066 10685189
refGene_gn=OFD1;refGene_tx=XM_006943569.4;refGene_gn=OFD1;refGene_tx=XM_006943570.4;refGene_gn=OFD1;refGene_tx=XM_019823561.2;refGene_gn=OFD1;refGene_tx=XM_019823562.2
X 10685377 10685476
refGene_gn=OFD1;refGene_tx=XM_006943569.4;refGene_gn=OFD1;refGene_tx=XM_006943570.4;refGene_gn=OFD1;refGene_tx=XM_019823561.2;refGene_gn=OFD1;refGene_tx=XM_019823562.2
X 10689680 10689881
refGene_gn=OFD1;refGene_tx=XM_006943569.4;refGene_gn=OFD1;refGene_tx=XM_006943570.4;refGene_gn=OFD1;refGene_tx=XM_019823561.2;refGene_gn=OFD1;refGene_tx=XM_019823562.2
X 10691844 10691913
refGene_gn=OFD1;refGene_tx=XM_006943569.4;refGene_gn=OFD1;refGene_tx=XM_006943570.4;refGene_gn=OFD1;refGene_tx=XM_019823561.2;refGene_gn=OFD1;refGene_tx=XM_019823562.2
X 10692000 10692031
refGene_gn=OFD1;refGene_tx=XM_006943569.4;refGene_gn=OFD1;refGene_tx=XM_006943570.4;refGene_gn=OFD1;refGene_tx=XM_019823561.2;refGene_gn=OFD1;refGene_tx=XM_019823562.2
X 10699325 10699433
refGene_gn=OFD1;refGene_tx=XM_006943569.4;refGene_gn=OFD1;refGene_tx=XM_006943570.4;refGene_gn=OFD1;refGene_tx=XM_006943571.4;refGene_gn=OFD1;refGene_tx=XM_019823561.2;refGene_gn=OFD1;refGene_tx=XM_019823562.2
X 10701375 10701512

refGene_gn=OFD1;refGene_tx=XM_006943569.4;refGene_gn=OFD1;refGene_tx=XM_006943570.4;refGene_gn=OFD1;refGene_tx=XM_006943571.4;refGene_gn=OFD1;refGene_tx=XM_019823561.2;refGene_gn=OFD1;refGene_tx=XM_019823562.2
X 10702010 10702184
refGene_gn=OFD1;refGene_tx=XM_006943569.4;refGene_gn=OFD1;refGene_tx=XM_006943570.4;refGene_gn=OFD1;refGene_tx=XM_006943571.4;refGene_gn=OFD1;refGene_tx=XM_019823561.2;refGene_gn=OFD1;refGene_tx=XM_019823562.2
X 10704198 10704305
refGene_gn=OFD1;refGene_tx=XM_006943569.4;refGene_gn=OFD1;refGene_tx=XM_006943570.4;refGene_gn=OFD1;refGene_tx=XM_006943571.4;refGene_gn=OFD1;refGene_tx=XM_019823561.2;refGene_gn=OFD1;refGene_tx=XM_019823562.2
X 10706070 10706190
refGene_gn=OFD1;refGene_tx=XM_006943569.4;refGene_gn=OFD1;refGene_tx=XM_006943570.4;refGene_gn=OFD1;refGene_tx=XM_006943571.4;refGene_gn=OFD1;refGene_tx=XM_019823561.2;refGene_gn=OFD1;refGene_tx=XM_019823562.2
X 10708116 10708190
refGene_gn=OFD1;refGene_tx=XM_006943569.4;refGene_gn=OFD1;refGene_tx=XM_006943570.4;refGene_gn=OFD1;refGene_tx=XM_006943571.4;refGene_gn=OFD1;refGene_tx=XM_019823561.2;refGene_gn=OFD1;refGene_tx=XM_019823562.2
X 10709470 10709562
refGene_gn=OFD1;refGene_tx=XM_006943569.4;refGene_gn=OFD1;refGene_tx=XM_006943570.4;refGene_gn=OFD1;refGene_tx=XM_006943571.4;refGene_gn=OFD1;refGene_tx=XM_019823561.2;refGene_gn=OFD1;refGene_tx=XM_019823562.2
X 10710668 10710858
refGene_gn=OFD1;refGene_tx=XM_006943569.4;refGene_gn=OFD1;refGene_tx=XM_006943570.4;refGene_gn=OFD1;refGene_tx=XM_006943571.4;refGene_gn=OFD1;refGene_tx=XM_019823561.2;refGene_gn=OFD1;refGene_tx=XM_019823562.2
X 10712031 10712162
refGene_gn=OFD1;refGene_tx=XM_006943569.4;refGene_gn=OFD1;refGene_tx=XM_006943570.4;refGene_gn=OFD1;refGene_tx=XM_006943571.4;refGene_gn=OFD1;refGene_tx=XM_019823561.2;refGene_gn=OFD1;refGene_tx=XM_019823562.2
X 10712688 10712800
refGene_gn=OFD1;refGene_tx=XM_006943569.4;refGene_gn=OFD1;refGene_tx=XM_006943570.4;refGene_gn=OFD1;refGene_tx=XM_006943571.4;refGene_gn=OFD1;refGene_tx=XM_019823561.2;refGene_gn=OFD1;refGene_tx=XM_019823562.2
X 10713335 10713929
refGene_gn=OFD1;refGene_tx=XM_006943569.4;refGene_gn=OFD1;refGene_tx=XM_006943570.4;refGene_gn=OFD1;refGene_tx=XM_006943571.4;refGene_gn=OFD1;refGene_tx=XM_019823561.2;refGene_gn=OFD1;refGene_tx=XM_019823562.2
X 10714404 10714528
refGene_gn=OFD1;refGene_tx=XM_006943569.4;refGene_gn=OFD1;refGene_tx=XM_006943570.4;refGene_gn=OFD1;refGene_tx=XM_006943571.4;refGene_gn=OFD1;refGene_tx=XM_019823561.2;refGene_gn=OFD1;refGene_tx=XM_019823562.2
X 10718304 10718405
refGene_gn=OFD1;refGene_tx=XM_006943569.4;refGene_gn=OFD1;refGene_tx=XM_006943570.4;refGene_gn=OFD1;refGene_tx=XM_006943571.4;refGene_gn=OFD1;refGene_tx=XM_019823561.2;refGene_gn=OFD1;refGene_tx=XM_019823562.2
X 10719321 10719441
refGene_gn=OFD1;refGene_tx=XM_006943569.4;refGene_gn=OFD1;refGene_tx=XM_006943570.4;refGene_gn=OFD1;refGene_tx=XM_006943571.4;refGene_gn=OFD

1;refGene_tx=XM_019823561.2;refGene_gn=OFD1;refGene_tx=XM_019823562.2
X 10722536 10722694
refGene_gn=OFD1;refGene_tx=XM_006943569.4;refGene_gn=OFD1;refGene_tx=XM_006943570.4;refGene_gn=OFD1;refGene_tx=XM_006943571.4;refGene_gn=OFD1;refGene_tx=XM_019823561.2;refGene_gn=OFD1;refGene_tx=XM_019823562.2
X 10725284 10725437
refGene_gn=OFD1;refGene_tx=XM_006943569.4;refGene_gn=OFD1;refGene_tx=XM_006943570.4;refGene_gn=OFD1;refGene_tx=XM_006943571.4;refGene_gn=OFD1;refGene_tx=XM_019823561.2;refGene_gn=OFD1;refGene_tx=XM_019823562.2
X 10725943 10726117
refGene_gn=OFD1;refGene_tx=XM_006943569.4;refGene_gn=OFD1;refGene_tx=XM_006943570.4;refGene_gn=OFD1;refGene_tx=XM_006943571.4;refGene_gn=OFD1;refGene_tx=XM_019823561.2;refGene_gn=OFD1;refGene_tx=XM_019823562.2
X 10726306 10726349
refGene_gn=OFD1;refGene_tx=XM_006943570.4;refGene_gn=OFD1;refGene_tx=XM_019823562.2
X 10750226 10750376
refGene_gn=GPM6B;refGene_tx=XM_004000281.4;refGene_gn=GPM6B;refGene_tx=XM_011291609.3;refGene_gn=GPM6B;refGene_tx=XM_011291610.3;refGene_gn=GPM6B;refGene_tx=XM_023249365.1
X 10751810 10751891
refGene_gn=GPM6B;refGene_tx=XM_004000279.5;refGene_gn=GPM6B;refGene_tx=XM_004000280.5
X 10753088 10753154
refGene_gn=GPM6B;refGene_tx=XM_004000279.5;refGene_gn=GPM6B;refGene_tx=XM_004000280.5;refGene_gn=GPM6B;refGene_tx=XM_004000281.4;refGene_gn=GPM6B;refGene_tx=XM_011291609.3;refGene_gn=GPM6B;refGene_tx=XM_011291610.3;refGene_gn=GPM6B;refGene_tx=XM_023249365.1
X 10754036 10754110
refGene_gn=GPM6B;refGene_tx=XM_004000279.5;refGene_gn=GPM6B;refGene_tx=XM_004000280.5;refGene_gn=GPM6B;refGene_tx=XM_004000281.4;refGene_gn=GPM6B;refGene_tx=XM_011291609.3;refGene_gn=GPM6B;refGene_tx=XM_011291610.3;refGene_gn=GPM6B;refGene_tx=XM_023249365.1
X 10755883 10756055
refGene_gn=GPM6B;refGene_tx=XM_004000279.5;refGene_gn=GPM6B;refGene_tx=XM_004000280.5;refGene_gn=GPM6B;refGene_tx=XM_004000281.4;refGene_gn=GPM6B;refGene_tx=XM_011291609.3;refGene_gn=GPM6B;refGene_tx=XM_011291610.3;refGene_gn=GPM6B;refGene_tx=XM_023249365.1
X 10760686 10760843
refGene_gn=GPM6B;refGene_tx=XM_004000279.5;refGene_gn=GPM6B;refGene_tx=XM_004000280.5;refGene_gn=GPM6B;refGene_tx=XM_004000281.4;refGene_gn=GPM6B;refGene_tx=XM_011291609.3;refGene_gn=GPM6B;refGene_tx=XM_011291610.3;refGene_gn=GPM6B;refGene_tx=XM_023249365.1
X 10762310 10762497
refGene_gn=GPM6B;refGene_tx=XM_004000279.5;refGene_gn=GPM6B;refGene_tx=XM_004000280.5;refGene_gn=GPM6B;refGene_tx=XM_004000281.4;refGene_gn=GPM6B;refGene_tx=XM_011291609.3;refGene_gn=GPM6B;refGene_tx=XM_011291610.3;refGene_gn=GPM6B;refGene_tx=XM_023249365.1
X 10780277 10780397
refGene_gn=GPM6B;refGene_tx=XM_004000280.5;refGene_gn=GPM6B;refGene_tx

=XM_004000281.4
X 10788186 10788247
refGene_gn=GPM6B;refGene_tx=XM_004000279.5;refGene_gn=GPM6B;refGene_tx=XM_004000280.5;refGene_gn=GPM6B;refGene_tx=XM_004000281.4;refGene_gn=GPM6B;refGene_tx=XM_011291609.3
X 10828381 10828406 refGene_gn=GPM6B;refGene_tx=XM_011291610.3
X 10897591 10897595 refGene_gn=GPM6B;refGene_tx=XM_023249365.1
X 10953255 10953512
refGene_gn=GEMIN8;refGene_tx=XM_011291611.3;refGene_gn=GEMIN8;refGene_tx=XM_023249015.1
X 10963182 10963621
refGene_gn=GEMIN8;refGene_tx=XM_011291611.3;refGene_gn=GEMIN8;refGene_tx=XM_023249015.1
X 10964189 10964201
refGene_gn=GEMIN8;refGene_tx=XM_011291611.3;refGene_gn=GEMIN8;refGene_tx=XM_023249015.1
X 11392049 11392117
refGene_gn=GLRA2;refGene_tx=XM_004000282.4;refGene_gn=GLRA2;refGene_tx=XM_004000283.4
X 11394296 11394430
refGene_gn=GLRA2;refGene_tx=XM_004000282.4;refGene_gn=GLRA2;refGene_tx=XM_004000283.4
X 11435530 11435598 refGene_gn=GLRA2;refGene_tx=XM_004000282.4
X 11435693 11435761
refGene_gn=GLRA2;refGene_tx=XM_004000283.4;refGene_gn=GLRA2;refGene_tx=XM_019823567.2
X 11443195 11443419
refGene_gn=GLRA2;refGene_tx=XM_004000282.4;refGene_gn=GLRA2;refGene_tx=XM_004000283.4;refGene_gn=GLRA2;refGene_tx=XM_019823567.2
X 11461436 11461519
refGene_gn=GLRA2;refGene_tx=XM_004000282.4;refGene_gn=GLRA2;refGene_tx=XM_004000283.4;refGene_gn=GLRA2;refGene_tx=XM_019823567.2
X 11464422 11464560
refGene_gn=GLRA2;refGene_tx=XM_004000282.4;refGene_gn=GLRA2;refGene_tx=XM_004000283.4;refGene_gn=GLRA2;refGene_tx=XM_019823567.2
X 11466165 11466380
refGene_gn=GLRA2;refGene_tx=XM_004000282.4;refGene_gn=GLRA2;refGene_tx=XM_004000283.4;refGene_gn=GLRA2;refGene_tx=XM_019823567.2
X 11527401 11527551
refGene_gn=GLRA2;refGene_tx=XM_004000282.4;refGene_gn=GLRA2;refGene_tx=XM_004000283.4;refGene_gn=GLRA2;refGene_tx=XM_019823567.2
X 11562151 11562430
refGene_gn=GLRA2;refGene_tx=XM_004000282.4;refGene_gn=GLRA2;refGene_tx=XM_004000283.4;refGene_gn=GLRA2;refGene_tx=XM_019823567.2
X 11669538 11669953 refGene_gn=FANCB;refGene_tx=XM_004000294.5
X 11670471 11670709 refGene_gn=FANCB;refGene_tx=XM_004000294.5
X 11670827 11671294 refGene_gn=FANCB;refGene_tx=XM_004000294.5
X 11671858 11672025 refGene_gn=FANCB;refGene_tx=XM_004000294.5
X 11675535 11675664 refGene_gn=FANCB;refGene_tx=XM_004000294.5
X 11681025 11681118 refGene_gn=FANCB;refGene_tx=XM_004000294.5

X 11682339 11682492 refGene_gn=FANCB;refGene_tx=XM_004000294.5
X 11684206 11685133 refGene_gn=FANCB;refGene_tx=XM_004000294.5
X 11693436 11693445
refGene_gn=MOSPD2;refGene_tx=XM_006943573.2;refGene_gn=MOSPD2;refGene_
tx=XM_006943574.2
X 11693606 11693676
refGene_gn=MOSPD2;refGene_tx=XM_006943573.2;refGene_gn=MOSPD2;refGene_
tx=XM_006943574.2
X 11709873 11710029
refGene_gn=MOSPD2;refGene_tx=XM_006943573.2;refGene_gn=MOSPD2;refGene_
tx=XM_006943574.2
X 11712129 11712216
refGene_gn=MOSPD2;refGene_tx=XM_006943573.2;refGene_gn=MOSPD2;refGene_
tx=XM_006943574.2
X 11713436 11713591
refGene_gn=MOSPD2;refGene_tx=XM_006943573.2;refGene_gn=MOSPD2;refGene_
tx=XM_006943574.2
X 11716284 11716345
refGene_gn=MOSPD2;refGene_tx=XM_006943573.2;refGene_gn=MOSPD2;refGene_
tx=XM_006943574.2
X 11718810 11718849
refGene_gn=MOSPD2;refGene_tx=XM_006943573.2;refGene_gn=MOSPD2;refGene_
tx=XM_006943574.2
X 11727914 11728039
refGene_gn=MOSPD2;refGene_tx=XM_006943573.2;refGene_gn=MOSPD2;refGene_
tx=XM_006943574.2
X 11731373 11731550
refGene_gn=MOSPD2;refGene_tx=XM_006943573.2;refGene_gn=MOSPD2;refGene_
tx=XM_006943574.2
X 11732175 11732288
refGene_gn=MOSPD2;refGene_tx=XM_006943573.2;refGene_gn=MOSPD2;refGene_
tx=XM_006943574.2
X 11734739 11734836
refGene_gn=MOSPD2;refGene_tx=XM_006943573.2;refGene_gn=MOSPD2;refGene_
tx=XM_006943574.2
X 11735684 11735781
refGene_gn=MOSPD2;refGene_tx=XM_006943573.2;refGene_gn=MOSPD2;refGene_
tx=XM_006943574.2
X 11736147 11736277
refGene_gn=MOSPD2;refGene_tx=XM_006943573.2;refGene_gn=MOSPD2;refGene_
tx=XM_006943574.2
X 11743905 11744008
refGene_gn=MOSPD2;refGene_tx=XM_006943573.2;refGene_gn=MOSPD2;refGene_
tx=XM_006943574.2
X 11744577 11744715 refGene_gn=MOSPD2;refGene_tx=XM_006943573.2
X 11746112 11746154 refGene_gn=MOSPD2;refGene_tx=XM_006943574.2
X 12064203 12064328
refGene_gn=ASB9;refGene_tx=XM_006943577.3;refGene_gn=ASB9;refGene_tx=X
M_019823571.2
X 12067867 12068091

refGene_gn=ASB9;refGene_tx=XM_006943577.3;refGene_gn=ASB9;refGene_tx=XM_006943579.3;refGene_gn=ASB9;refGene_tx=XM_019823571.2
X 12069809 12069944
refGene_gn=ASB9;refGene_tx=XM_006943577.3;refGene_gn=ASB9;refGene_tx=XM_006943579.3;refGene_gn=ASB9;refGene_tx=XM_019823571.2
X 12071493 12071644
refGene_gn=ASB9;refGene_tx=XM_006943577.3;refGene_gn=ASB9;refGene_tx=XM_006943579.3;refGene_gn=ASB9;refGene_tx=XM_019823571.2
X 12075058 12075166
refGene_gn=ASB9;refGene_tx=XM_006943577.3;refGene_gn=ASB9;refGene_tx=XM_006943579.3;refGene_gn=ASB9;refGene_tx=XM_019823571.2
X 12083725 12083805
refGene_gn=ASB9;refGene_tx=XM_006943577.3;refGene_gn=ASB9;refGene_tx=XM_006943579.3;refGene_gn=ASB9;refGene_tx=XM_019823571.2
X 12091171 12091262
refGene_gn=ASB9;refGene_tx=XM_006943577.3;refGene_gn=ASB9;refGene_tx=XM_006943579.3;refGene_gn=ASB9;refGene_tx=XM_019823571.2
X 12100783 12100908
refGene_gn=ASB11;refGene_tx=XM_004000295.3;refGene_gn=ASB11;refGene_tx=XM_011291612.2;refGene_gn=ASB11;refGene_tx=XM_011291613.2
X 12103897 12104089
refGene_gn=ASB11;refGene_tx=XM_004000295.3;refGene_gn=ASB11;refGene_tx=XM_011291612.2;refGene_gn=ASB11;refGene_tx=XM_011291613.2
X 12105212 12105347
refGene_gn=ASB11;refGene_tx=XM_004000295.3;refGene_gn=ASB11;refGene_tx=XM_011291612.2;refGene_gn=ASB11;refGene_tx=XM_011291613.2
X 12109624 12109775
refGene_gn=ASB11;refGene_tx=XM_004000295.3;refGene_gn=ASB11;refGene_tx=XM_011291612.2;refGene_gn=ASB11;refGene_tx=XM_011291613.2
X 12112799 12112907
refGene_gn=ASB11;refGene_tx=XM_004000295.3;refGene_gn=ASB11;refGene_tx=XM_011291612.2;refGene_gn=ASB11;refGene_tx=XM_011291613.2
X 12116202 12116282
refGene_gn=ASB11;refGene_tx=XM_004000295.3;refGene_gn=ASB11;refGene_tx=XM_011291612.2;refGene_gn=ASB11;refGene_tx=XM_011291613.2
X 12123904 12124022 refGene_gn=ASB11;refGene_tx=XM_004000295.3
X 12124898 12125079
refGene_gn=ASB11;refGene_tx=XM_011291612.2;refGene_gn=ASB11;refGene_tx=XM_011291613.2
X 12129945 12130212
refGene_gn=PIGA;refGene_tx=XM_004000296.4;refGene_gn=PIGA;refGene_tx=XM_006943581.2;refGene_gn=PIGA;refGene_tx=XM_019823574.2
X 12131874 12132081
refGene_gn=PIGA;refGene_tx=XM_004000296.4;refGene_gn=PIGA;refGene_tx=XM_006943581.2;refGene_gn=PIGA;refGene_tx=XM_019823574.2
X 12132235 12132368
refGene_gn=PIGA;refGene_tx=XM_004000296.4;refGene_gn=PIGA;refGene_tx=XM_006943581.2;refGene_gn=PIGA;refGene_tx=XM_019823574.2
X 12133138 12133271
refGene_gn=PIGA;refGene_tx=XM_004000296.4;refGene_gn=PIGA;refGene_tx=X

M_006943581.2;refGene_gn=PIGA;refGene_tx=XM_019823574.2
X 12137554 12138272
refGene_gn=PIGA;refGene_tx=XM_004000296.4;refGene_gn=PIGA;refGene_tx=XM_006943581.2;refGene_gn=PIGA;refGene_tx=XM_019823574.2
X 12147549 12147676 refGene_gn=VEGFD;refGene_tx=XM_011291614.2
X 12148270 12148466 refGene_gn=VEGFD;refGene_tx=XM_011291614.2
X 12153566 12153667 refGene_gn=VEGFD;refGene_tx=XM_011291614.2
X 12154848 12154997 refGene_gn=VEGFD;refGene_tx=XM_011291614.2
X 12158361 12158552 refGene_gn=VEGFD;refGene_tx=XM_011291614.2
X 12163444 12163655 refGene_gn=VEGFD;refGene_tx=XM_011291614.2
X 12181278 12181368 refGene_gn=VEGFD;refGene_tx=XM_011291614.2
X 12182409 12182522
refGene_gn=PIR;refGene_tx=XM_004000300.5;refGene_gn=PIR;refGene_tx=XM_006943582.4;refGene_gn=PIR;refGene_tx=XM_019823575.1
X 12187112 12187179
refGene_gn=PIR;refGene_tx=XM_004000300.5;refGene_gn=PIR;refGene_tx=XM_006943582.4;refGene_gn=PIR;refGene_tx=XM_019823575.1
X 12194083 12194166
refGene_gn=PIR;refGene_tx=XM_004000300.5;refGene_gn=PIR;refGene_tx=XM_006943582.4;refGene_gn=PIR;refGene_tx=XM_019823575.1
X 12200468 12200513
refGene_gn=PIR;refGene_tx=XM_004000300.5;refGene_gn=PIR;refGene_tx=XM_006943582.4
X 12217299 12217384
refGene_gn=PIR;refGene_tx=XM_004000300.5;refGene_gn=PIR;refGene_tx=XM_006943582.4;refGene_gn=PIR;refGene_tx=XM_019823575.1
X 12244174 12244381
refGene_gn=PIR;refGene_tx=XM_004000300.5;refGene_gn=PIR;refGene_tx=XM_006943582.4;refGene_gn=PIR;refGene_tx=XM_019823575.1
X 12247614 12247698
refGene_gn=PIR;refGene_tx=XM_004000300.5;refGene_gn=PIR;refGene_tx=XM_006943582.4;refGene_gn=PIR;refGene_tx=XM_019823575.1
X 12263622 12263715
refGene_gn=PIR;refGene_tx=XM_004000300.5;refGene_gn=PIR;refGene_tx=XM_006943582.4;refGene_gn=PIR;refGene_tx=XM_019823575.1
X 12270279 12270408
refGene_gn=PIR;refGene_tx=XM_004000300.5;refGene_gn=PIR;refGene_tx=XM_006943582.4;refGene_gn=PIR;refGene_tx=XM_019823575.1
X 12270715 12270784 refGene_gn=PIR;refGene_tx=XM_004000300.5
X 12275331 12275334
refGene_gn=BMX;refGene_tx=XM_011291615.3;refGene_gn=BMX;refGene_tx=XM_011291616.3;refGene_gn=BMX;refGene_tx=XM_019823577.2
X 12281449 12281596
refGene_gn=BMX;refGene_tx=XM_011291615.3;refGene_gn=BMX;refGene_tx=XM_011291616.3;refGene_gn=BMX;refGene_tx=XM_019823576.2;refGene_gn=BMX;refGene_tx=XM_019823577.2
X 12282437 12282542
refGene_gn=BMX;refGene_tx=XM_011291615.3;refGene_gn=BMX;refGene_tx=XM_011291616.3;refGene_gn=BMX;refGene_tx=XM_019823576.2;refGene_gn=BMX;refGene_tx=XM_019823577.2

X	12284686	12284768
refGene_gn=BMX;refGene_tx=XM_011291615.3;refGene_gn=BMX;refGene_tx=XM_011291616.3;refGene_gn=BMX;refGene_tx=XM_019823576.2;refGene_gn=BMX;refGene_tx=XM_019823577.2
X	12289452	12289572
refGene_gn=BMX;refGene_tx=XM_011291615.3;refGene_gn=BMX;refGene_tx=XM_011291616.3;refGene_gn=BMX;refGene_tx=XM_019823576.2;refGene_gn=BMX;refGene_tx=XM_019823577.2
X	12291852	12291917
refGene_gn=BMX;refGene_tx=XM_011291615.3;refGene_gn=BMX;refGene_tx=XM_011291616.3;refGene_gn=BMX;refGene_tx=XM_019823576.2;refGene_gn=BMX;refGene_tx=XM_019823577.2
X	12295540	12295716
refGene_gn=BMX;refGene_tx=XM_011291615.3;refGene_gn=BMX;refGene_tx=XM_011291616.3;refGene_gn=BMX;refGene_tx=XM_019823576.2;refGene_gn=BMX;refGene_tx=XM_019823577.2
X	12298592	12298664
refGene_gn=BMX;refGene_tx=XM_011291615.3;refGene_gn=BMX;refGene_tx=XM_011291616.3;refGene_gn=BMX;refGene_tx=XM_019823576.2;refGene_gn=BMX;refGene_tx=XM_019823577.2
X	12299782	12299836
refGene_gn=BMX;refGene_tx=XM_011291615.3;refGene_gn=BMX;refGene_tx=XM_011291616.3;refGene_gn=BMX;refGene_tx=XM_019823576.2;refGene_gn=BMX;refGene_tx=XM_019823577.2
X	12303696	12303751
refGene_gn=BMX;refGene_tx=XM_011291615.3;refGene_gn=BMX;refGene_tx=XM_011291616.3;refGene_gn=BMX;refGene_tx=XM_019823576.2;refGene_gn=BMX;refGene_tx=XM_019823577.2
X	12305225	12305305
refGene_gn=BMX;refGene_tx=XM_011291615.3;refGene_gn=BMX;refGene_tx=XM_011291616.3;refGene_gn=BMX;refGene_tx=XM_019823576.2;refGene_gn=BMX;refGene_tx=XM_019823577.2
X	12307749	12307877
refGene_gn=BMX;refGene_tx=XM_011291615.3;refGene_gn=BMX;refGene_tx=XM_011291616.3;refGene_gn=BMX;refGene_tx=XM_019823576.2;refGene_gn=BMX;refGene_tx=XM_019823577.2
X	12311248	12311323
refGene_gn=BMX;refGene_tx=XM_011291615.3;refGene_gn=BMX;refGene_tx=XM_011291616.3;refGene_gn=BMX;refGene_tx=XM_019823576.2;refGene_gn=BMX;refGene_tx=XM_019823577.2
X	12312071	12312243
refGene_gn=BMX;refGene_tx=XM_011291615.3;refGene_gn=BMX;refGene_tx=XM_011291616.3;refGene_gn=BMX;refGene_tx=XM_019823576.2;refGene_gn=BMX;refGene_tx=XM_019823577.2
X	12315472	12315689
refGene_gn=BMX;refGene_tx=XM_011291615.3;refGene_gn=BMX;refGene_tx=XM_011291616.3;refGene_gn=BMX;refGene_tx=XM_019823576.2;refGene_gn=BMX;refGene_tx=XM_019823577.2
X	12316548	12316613
refGene_gn=BMX;refGene_tx=XM_011291615.3;refGene_gn=BMX;refGene_tx=XM_

011291616.3;refGene_gn=BMX;refGene_tx=XM_019823576.2;refGene_gn=BMX;refGene_tx=XM_019823577.2
X	12321540	12321659
refGene_gn=BMX;refGene_tx=XM_011291615.3;refGene_gn=BMX;refGene_tx=XM_011291616.3;refGene_gn=BMX;refGene_tx=XM_019823576.2
X	12324474	12324632
refGene_gn=BMX;refGene_tx=XM_011291615.3;refGene_gn=BMX;refGene_tx=XM_011291616.3;refGene_gn=BMX;refGene_tx=XM_019823576.2
X	12328220	12328407
refGene_gn=BMX;refGene_tx=XM_011291615.3;refGene_gn=BMX;refGene_tx=XM_011291616.3;refGene_gn=BMX;refGene_tx=XM_019823576.2;refGene_gn=BMX;refGene_tx=XM_019823577.2
X	12331689	12331798
refGene_gn=ACE2;refGene_tx=NM_001039456.1;refGene_gn=ACE2;refGene_tx=XM_023248796.1
X	12333539	12333734
refGene_gn=ACE2;refGene_tx=NM_001039456.1;refGene_gn=ACE2;refGene_tx=XM_023248796.1
X	12335528	12335645
refGene_gn=ACE2;refGene_tx=NM_001039456.1;refGene_gn=ACE2;refGene_tx=XM_023248796.1
X	12336879	12336980
refGene_gn=ACE2;refGene_tx=NM_001039456.1;refGene_gn=ACE2;refGene_tx=XM_023248796.1
X	12340874	12340933
refGene_gn=ACE2;refGene_tx=NM_001039456.1;refGene_gn=ACE2;refGene_tx=XM_023248796.1
X	12342402	12342575
refGene_gn=ACE2;refGene_tx=NM_001039456.1;refGene_gn=ACE2;refGene_tx=XM_023248796.1
X	12342968	12343091
refGene_gn=ACE2;refGene_tx=NM_001039456.1;refGene_gn=ACE2;refGene_tx=XM_023248796.1
X	12343795	12343894
refGene_gn=ACE2;refGene_tx=NM_001039456.1;refGene_gn=ACE2;refGene_tx=XM_023248796.1
X	12346245	12346390
refGene_gn=ACE2;refGene_tx=NM_001039456.1;refGene_gn=ACE2;refGene_tx=XM_023248796.1
X	12348734	12348961
refGene_gn=ACE2;refGene_tx=NM_001039456.1;refGene_gn=ACE2;refGene_tx=XM_023248796.1
X	12351561	12351731
refGene_gn=ACE2;refGene_tx=NM_001039456.1;refGene_gn=ACE2;refGene_tx=XM_023248796.1
X	12356573	12356671
refGene_gn=ACE2;refGene_tx=NM_001039456.1;refGene_gn=ACE2;refGene_tx=XM_023248796.1
X	12359141	12359247
refGene_gn=ACE2;refGene_tx=NM_001039456.1;refGene_gn=ACE2;refGene_tx=X

M_023248796.1
X	12360872	12360985	refGene_gn=ACE2;refGene_tx=NM_001039456.1;refGene_gn=ACE2;refGene_tx=XM_023248796.1
X	12362433	12362583	refGene_gn=ACE2;refGene_tx=NM_001039456.1;refGene_gn=ACE2;refGene_tx=XM_023248796.1
X	12363014	12363108	refGene_gn=ACE2;refGene_tx=NM_001039456.1;refGene_gn=ACE2;refGene_tx=XM_023248796.1
X	12367238	12367397	refGene_gn=ACE2;refGene_tx=NM_001039456.1;refGene_gn=ACE2;refGene_tx=XM_023248796.1
X	12373297	12373483	refGene_gn=ACE2;refGene_tx=NM_001039456.1;refGene_gn=ACE2;refGene_tx=XM_023248796.1
X	12399087	12399244	refGene_gn=TMEM27;refGene_tx=XM_004000302.5;refGene_gn=TMEM27;refGene_tx=XM_019823579.2
X	12409106	12409301	refGene_gn=TMEM27;refGene_tx=XM_004000302.5;refGene_gn=TMEM27;refGene_tx=XM_019823579.2
X	12413566	12413680	refGene_gn=TMEM27;refGene_tx=XM_004000302.5;refGene_gn=TMEM27;refGene_tx=XM_019823579.2
X	12426217	12426303	refGene_gn=TMEM27;refGene_tx=XM_004000302.5;refGene_gn=TMEM27;refGene_tx=XM_019823579.2
X	12430055	12430114	refGene_gn=TMEM27;refGene_tx=XM_004000302.5;refGene_gn=TMEM27;refGene_tx=XM_019823579.2
X	12430416	12430474	refGene_gn=TMEM27;refGene_tx=XM_004000302.5
X	12432731	12432789	refGene_gn=TMEM27;refGene_tx=XM_019823579.2
X	12460680	12460825	refGene_gn=LOC101085781;refGene_tx=XM_023248923.1;refGene_gn=LOC101085781;refGene_tx=XM_023248924.1
X	12462161	12462220	refGene_gn=LOC101085781;refGene_tx=XM_023248923.1;refGene_gn=LOC101085781;refGene_tx=XM_023248924.1
X	12463129	12463285	refGene_gn=LOC101085781;refGene_tx=XM_023248923.1;refGene_gn=LOC101085781;refGene_tx=XM_023248924.1
X	12465686	12465758	refGene_gn=LOC101085781;refGene_tx=XM_023248923.1;refGene_gn=LOC101085781;refGene_tx=XM_023248924.1
X	12524541	12524683	refGene_gn=CA5B;refGene_tx=XM_006943590.3;refGene_gn=CA5B;refGene_tx=XM_019823582.2
X	12524993	12525174	refGene_gn=CA5B;refGene_tx=XM_019823581.2
X	12525620	12525636	refGene_gn=CA5B;refGene_tx=XM_019823583.2

X 12544564 12544762
refGene_gn=CA5B;refGene_tx=XM_006943590.3;refGene_gn=CA5B;refGene_tx=XM_019823581.2;refGene_gn=CA5B;refGene_tx=XM_019823582.2;refGene_gn=CA5B;refGene_tx=XM_019823583.2
X 12553967 12554086
refGene_gn=CA5B;refGene_tx=XM_006943590.3;refGene_gn=CA5B;refGene_tx=XM_019823581.2;refGene_gn=CA5B;refGene_tx=XM_019823582.2;refGene_gn=CA5B;refGene_tx=XM_019823583.2
X 12555129 12555225
refGene_gn=CA5B;refGene_tx=XM_006943590.3;refGene_gn=CA5B;refGene_tx=XM_019823581.2;refGene_gn=CA5B;refGene_tx=XM_019823582.2;refGene_gn=CA5B;refGene_tx=XM_019823583.2
X 12556270 12556333
refGene_gn=CA5B;refGene_tx=XM_006943590.3;refGene_gn=CA5B;refGene_tx=XM_019823581.2;refGene_gn=CA5B;refGene_tx=XM_019823582.2;refGene_gn=CA5B;refGene_tx=XM_019823583.2
X 12557249 12557405
refGene_gn=CA5B;refGene_tx=XM_006943590.3;refGene_gn=CA5B;refGene_tx=XM_019823581.2;refGene_gn=CA5B;refGene_tx=XM_019823582.2;refGene_gn=CA5B;refGene_tx=XM_019823583.2
X 12562354 12562534
refGene_gn=CA5B;refGene_tx=XM_006943590.3;refGene_gn=CA5B;refGene_tx=XM_019823581.2;refGene_gn=CA5B;refGene_tx=XM_019823582.2;refGene_gn=CA5B;refGene_tx=XM_019823583.2
X 12569310 12569351 refGene_gn=ZRSR2;refGene_tx=XM_006943591.3
X 12569803 12569883 refGene_gn=ZRSR2;refGene_tx=XM_006943591.3
X 12573376 12573455 refGene_gn=ZRSR2;refGene_tx=XM_006943591.3
X 12578235 12578344 refGene_gn=ZRSR2;refGene_tx=XM_006943591.3
X 12578540 12578627 refGene_gn=ZRSR2;refGene_tx=XM_006943591.3
X 12581726 12581765 refGene_gn=ZRSR2;refGene_tx=XM_006943591.3
X 12582378 12582497 refGene_gn=ZRSR2;refGene_tx=XM_006943591.3
X 12587643 12587857 refGene_gn=ZRSR2;refGene_tx=XM_006943591.3
X 12592735 12592791 refGene_gn=ZRSR2;refGene_tx=XM_006943591.3
X 12593555 12593665 refGene_gn=ZRSR2;refGene_tx=XM_006943591.3
X 12596204 12596665 refGene_gn=ZRSR2;refGene_tx=XM_006943591.3
X 12600841 12600889
refGene_gn=AP1S2;refGene_tx=XM_023248821.1;refGene_gn=AP1S2;refGene_tx=XM_023248822.1;refGene_gn=AP1S2;refGene_tx=XM_023248823.1;refGene_gn=AP1S2;refGene_tx=XM_023248825.1
X 12601415 12601473
refGene_gn=AP1S2;refGene_tx=XM_023248821.1;refGene_gn=AP1S2;refGene_tx=XM_023248822.1
X 12601721 12601730
refGene_gn=AP1S2;refGene_tx=XM_023248821.1;refGene_gn=AP1S2;refGene_tx=XM_023248823.1
X 12605203 12605239 refGene_gn=AP1S2;refGene_tx=XM_023248826.1
X 12606755 12606812 refGene_gn=AP1S2;refGene_tx=XM_023248824.1
X 12617009 12617147
refGene_gn=AP1S2;refGene_tx=XM_023248821.1;refGene_gn=AP1S2;refGene_tx=XM_023248822.1;refGene_gn=AP1S2;refGene_tx=XM_023248823.1;refGene_gn=

AP1S2;refGene_tx=XM_023248824.1;refGene_gn=AP1S2;refGene_tx=XM_023248825.1;refGene_gn=AP1S2;refGene_tx=XM_023248826.1
X	12617761	12617870	refGene_gn=AP1S2;refGene_tx=XM_023248821.1;refGene_gn=AP1S2;refGene_tx=XM_023248822.1;refGene_gn=AP1S2;refGene_tx=XM_023248823.1;refGene_gn=AP1S2;refGene_tx=XM_023248824.1;refGene_gn=AP1S2;refGene_tx=XM_023248825.1;refGene_gn=AP1S2;refGene_tx=XM_023248826.1
X	12625291	12625470	refGene_gn=AP1S2;refGene_tx=XM_023248821.1;refGene_gn=AP1S2;refGene_tx=XM_023248822.1;refGene_gn=AP1S2;refGene_tx=XM_023248823.1;refGene_gn=AP1S2;refGene_tx=XM_023248824.1;refGene_gn=AP1S2;refGene_tx=XM_023248825.1;refGene_gn=AP1S2;refGene_tx=XM_023248826.1
X	12884450	12884863	refGene_gn=GRPR;refGene_tx=XM_004000306.4;refGene_gn=GRPR;refGene_tx=XM_019823588.2
X	12911485	12911837	refGene_gn=GRPR;refGene_tx=XM_004000306.4;refGene_gn=GRPR;refGene_tx=XM_019823588.2
X	12913437	12913827	refGene_gn=GRPR;refGene_tx=XM_004000306.4;refGene_gn=GRPR;refGene_tx=XM_019823588.2
X	12916159	12916163	refGene_gn=GRPR;refGene_tx=XM_019823588.2
X	12931288	12932284	refGene_gn=MAGEB17;refGene_tx=XM_006943597.2
X	13226014	13226335	refGene_gn=LOC101086788;refGene_tx=XM_019823591.2
X	13278872	13280015	refGene_gn=LOC105261399;refGene_tx=XM_023249182.1
X	13295013	13295083	refGene_gn=CTPS2;refGene_tx=XM_023249507.1;refGene_gn=CTPS2;refGene_tx=XM_023249508.1;refGene_gn=CTPS2;refGene_tx=XM_023249509.1;refGene_gn=CTPS2;refGene_tx=XM_023249510.1;refGene_gn=CTPS2;refGene_tx=XM_023249511.1
X	13305866	13306011	refGene_gn=CTPS2;refGene_tx=XM_023249507.1;refGene_gn=CTPS2;refGene_tx=XM_023249508.1;refGene_gn=CTPS2;refGene_tx=XM_023249509.1;refGene_gn=CTPS2;refGene_tx=XM_023249510.1;refGene_gn=CTPS2;refGene_tx=XM_023249511.1
X	13311497	13311594	refGene_gn=CTPS2;refGene_tx=XM_023249507.1;refGene_gn=CTPS2;refGene_tx=XM_023249508.1;refGene_gn=CTPS2;refGene_tx=XM_023249509.1;refGene_gn=CTPS2;refGene_tx=XM_023249510.1;refGene_gn=CTPS2;refGene_tx=XM_023249511.1
X	13314474	13314530	refGene_gn=CTPS2;refGene_tx=XM_023249507.1;refGene_gn=CTPS2;refGene_tx=XM_023249508.1;refGene_gn=CTPS2;refGene_tx=XM_023249509.1;refGene_gn=CTPS2;refGene_tx=XM_023249510.1;refGene_gn=CTPS2;refGene_tx=XM_023249511.1
X	13330365	13330462	refGene_gn=CTPS2;refGene_tx=XM_023249507.1;refGene_gn=CTPS2;refGene_tx=XM_023249508.1;refGene_gn=CTPS2;refGene_tx=XM_023249509.1;refGene_gn=

CTPS2;refGene_tx=XM_023249510.1;refGene_gn=CTPS2;refGene_tx=XM_0232495
11.1
X	13354965	13355009
refGene_gn=CTPS2;refGene_tx=XM_023249507.1;refGene_gn=CTPS2;refGene_tx
=XM_023249508.1;refGene_gn=CTPS2;refGene_tx=XM_023249509.1;refGene_gn=
CTPS2;refGene_tx=XM_023249510.1;refGene_gn=CTPS2;refGene_tx=XM_0232495
11.1
X	13355340	13355403
refGene_gn=CTPS2;refGene_tx=XM_023249507.1;refGene_gn=CTPS2;refGene_tx
=XM_023249508.1;refGene_gn=CTPS2;refGene_tx=XM_023249509.1;refGene_gn=
CTPS2;refGene_tx=XM_023249510.1;refGene_gn=CTPS2;refGene_tx=XM_0232495
11.1
X	13356991	13357086
refGene_gn=CTPS2;refGene_tx=XM_023249507.1;refGene_gn=CTPS2;refGene_tx
=XM_023249508.1;refGene_gn=CTPS2;refGene_tx=XM_023249509.1;refGene_gn=
CTPS2;refGene_tx=XM_023249510.1;refGene_gn=CTPS2;refGene_tx=XM_0232495
11.1
X	13357796	13357885
refGene_gn=CTPS2;refGene_tx=XM_023249507.1;refGene_gn=CTPS2;refGene_tx
=XM_023249508.1;refGene_gn=CTPS2;refGene_tx=XM_023249509.1;refGene_gn=
CTPS2;refGene_tx=XM_023249510.1;refGene_gn=CTPS2;refGene_tx=XM_0232495
11.1
X	13359449	13359582
refGene_gn=CTPS2;refGene_tx=XM_023249507.1;refGene_gn=CTPS2;refGene_tx
=XM_023249508.1;refGene_gn=CTPS2;refGene_tx=XM_023249509.1;refGene_gn=
CTPS2;refGene_tx=XM_023249510.1;refGene_gn=CTPS2;refGene_tx=XM_0232495
11.1
X	13363508	13363660
refGene_gn=CTPS2;refGene_tx=XM_023249507.1;refGene_gn=CTPS2;refGene_tx
=XM_023249508.1;refGene_gn=CTPS2;refGene_tx=XM_023249509.1;refGene_gn=
CTPS2;refGene_tx=XM_023249510.1;refGene_gn=CTPS2;refGene_tx=XM_0232495
11.1
X	13365930	13366011
refGene_gn=CTPS2;refGene_tx=XM_023249507.1;refGene_gn=CTPS2;refGene_tx
=XM_023249508.1;refGene_gn=CTPS2;refGene_tx=XM_023249509.1;refGene_gn=
CTPS2;refGene_tx=XM_023249510.1;refGene_gn=CTPS2;refGene_tx=XM_0232495
11.1
X	13366893	13366977
refGene_gn=CTPS2;refGene_tx=XM_023249507.1;refGene_gn=CTPS2;refGene_tx
=XM_023249508.1;refGene_gn=CTPS2;refGene_tx=XM_023249509.1;refGene_gn=
CTPS2;refGene_tx=XM_023249510.1;refGene_gn=CTPS2;refGene_tx=XM_0232495
11.1
X	13367470	13367587
refGene_gn=CTPS2;refGene_tx=XM_023249507.1;refGene_gn=CTPS2;refGene_tx
=XM_023249508.1;refGene_gn=CTPS2;refGene_tx=XM_023249509.1;refGene_gn=
CTPS2;refGene_tx=XM_023249510.1;refGene_gn=CTPS2;refGene_tx=XM_0232495
11.1
X	13379741	13379842
refGene_gn=CTPS2;refGene_tx=XM_023249507.1;refGene_gn=CTPS2;refGene_tx
=XM_023249508.1;refGene_gn=CTPS2;refGene_tx=XM_023249509.1;refGene_gn=

CTPS2;refGene_tx=XM_023249510.1;refGene_gn=CTPS2;refGene_tx=XM_0232495
11.1
X	13381374	13381545
refGene_gn=CTPS2;refGene_tx=XM_023249507.1;refGene_gn=CTPS2;refGene_tx
=XM_023249508.1;refGene_gn=CTPS2;refGene_tx=XM_023249509.1;refGene_gn=
CTPS2;refGene_tx=XM_023249510.1;refGene_gn=CTPS2;refGene_tx=XM_0232495
11.1
X	13386894	13387060	refGene_gn=CTPS2;refGene_tx=XM_023249507.1
X	13387984	13388150
refGene_gn=CTPS2;refGene_tx=XM_023249508.1;refGene_gn=CTPS2;refGene_tx
=XM_023249509.1;refGene_gn=CTPS2;refGene_tx=XM_023249510.1;refGene_gn=
CTPS2;refGene_tx=XM_023249511.1
X	13398882	13399075	refGene_gn=SYAP1;refGene_tx=XM_023249512.1
X	13408702	13408821	refGene_gn=SYAP1;refGene_tx=XM_023249512.1
X	13409598	13409665	refGene_gn=SYAP1;refGene_tx=XM_023249512.1
X	13412967	13413041	refGene_gn=SYAP1;refGene_tx=XM_023249512.1
X	13417498	13417638	refGene_gn=SYAP1;refGene_tx=XM_023249512.1
X	13423565	13423714	refGene_gn=SYAP1;refGene_tx=XM_023249512.1
X	13425015	13425074	refGene_gn=SYAP1;refGene_tx=XM_023249512.1
X	13425482	13425630	refGene_gn=SYAP1;refGene_tx=XM_023249512.1
X	13427882	13428010	refGene_gn=SYAP1;refGene_tx=XM_023249512.1
X	13453429	13453531
refGene_gn=TXLNG;refGene_tx=XM_023249424.1;refGene_gn=TXLNG;refGene_tx
=XM_023249425.1
X	13479721	13480025
refGene_gn=TXLNG;refGene_tx=XM_023249424.1;refGene_gn=TXLNG;refGene_tx
=XM_023249425.1
X	13483673	13483765
refGene_gn=TXLNG;refGene_tx=XM_023249424.1;refGene_gn=TXLNG;refGene_tx
=XM_023249425.1
X	13488622	13488793
refGene_gn=TXLNG;refGene_tx=XM_023249424.1;refGene_gn=TXLNG;refGene_tx
=XM_023249425.1
X	13490004	13490199
refGene_gn=TXLNG;refGene_tx=XM_023249424.1;refGene_gn=TXLNG;refGene_tx
=XM_023249425.1
X	13493611	13493731	refGene_gn=TXLNG;refGene_tx=XM_023249424.1
X	13495214	13495289	refGene_gn=TXLNG;refGene_tx=XM_023249424.1
X	13501098	13501191
refGene_gn=TXLNG;refGene_tx=XM_023249424.1;refGene_gn=TXLNG;refGene_tx
=XM_023249425.1
X	13503460	13503556
refGene_gn=TXLNG;refGene_tx=XM_023249424.1;refGene_gn=TXLNG;refGene_tx
=XM_023249425.1
X	13504688	13505003
refGene_gn=TXLNG;refGene_tx=XM_023249424.1;refGene_gn=TXLNG;refGene_tx
=XM_023249425.1
X	13509268	13509337
refGene_gn=RBBP7;refGene_tx=XM_011291619.2;refGene_gn=RBBP7;refGene_tx
=XM_019823593.1;refGene_gn=RBBP7;refGene_tx=XM_023249426.1

X 13510083 13510194
refGene_gn=RBBP7;refGene_tx=XM_011291619.2;refGene_gn=RBBP7;refGene_tx=XM_019823593.1;refGene_gn=RBBP7;refGene_tx=XM_023249426.1
X 13512640 13512698
refGene_gn=RBBP7;refGene_tx=XM_011291619.2;refGene_gn=RBBP7;refGene_tx=XM_019823593.1;refGene_gn=RBBP7;refGene_tx=XM_023249426.1
X 13515002 13515079
refGene_gn=RBBP7;refGene_tx=XM_011291619.2;refGene_gn=RBBP7;refGene_tx=XM_019823593.1;refGene_gn=RBBP7;refGene_tx=XM_023249426.1
X 13515690 13515768
refGene_gn=RBBP7;refGene_tx=XM_011291619.2;refGene_gn=RBBP7;refGene_tx=XM_019823593.1;refGene_gn=RBBP7;refGene_tx=XM_023249426.1
X 13515889 13516016
refGene_gn=RBBP7;refGene_tx=XM_011291619.2;refGene_gn=RBBP7;refGene_tx=XM_019823593.1;refGene_gn=RBBP7;refGene_tx=XM_023249426.1
X 13516760 13516921
refGene_gn=RBBP7;refGene_tx=XM_011291619.2;refGene_gn=RBBP7;refGene_tx=XM_019823593.1;refGene_gn=RBBP7;refGene_tx=XM_023249426.1
X 13521586 13521702
refGene_gn=RBBP7;refGene_tx=XM_011291619.2;refGene_gn=RBBP7;refGene_tx=XM_019823593.1;refGene_gn=RBBP7;refGene_tx=XM_023249426.1
X 13522716 13522890
refGene_gn=RBBP7;refGene_tx=XM_011291619.2;refGene_gn=RBBP7;refGene_tx=XM_019823593.1;refGene_gn=RBBP7;refGene_tx=XM_023249426.1
X 13526083 13526229
refGene_gn=RBBP7;refGene_tx=XM_011291619.2;refGene_gn=RBBP7;refGene_tx=XM_019823593.1;refGene_gn=RBBP7;refGene_tx=XM_023249426.1
X 13531762 13531907
refGene_gn=RBBP7;refGene_tx=XM_011291619.2;refGene_gn=RBBP7;refGene_tx=XM_019823593.1;refGene_gn=RBBP7;refGene_tx=XM_023249426.1
X 13532191 13532339
refGene_gn=RBBP7;refGene_tx=XM_011291619.2;refGene_gn=RBBP7;refGene_tx=XM_019823593.1
X 13532748 13532764 refGene_gn=RBBP7;refGene_tx=XM_023249426.1
X 13611367 13611607
refGene_gn=REPS2;refGene_tx=XM_019823602.2;refGene_gn=REPS2;refGene_tx=XM_023248867.1;refGene_gn=REPS2;refGene_tx=XM_023248868.1;refGene_gn=REPS2;refGene_tx=XM_023248869.1;refGene_gn=REPS2;refGene_tx=XM_023248870.1;refGene_gn=REPS2;refGene_tx=XM_023248871.1;refGene_gn=REPS2;refGene_tx=XM_023248872.1
X 13672757 13672881
refGene_gn=REPS2;refGene_tx=XM_019823602.2;refGene_gn=REPS2;refGene_tx=XM_023248867.1;refGene_gn=REPS2;refGene_tx=XM_023248868.1;refGene_gn=REPS2;refGene_tx=XM_023248869.1;refGene_gn=REPS2;refGene_tx=XM_023248870.1;refGene_gn=REPS2;refGene_tx=XM_023248871.1;refGene_gn=REPS2;refGene_tx=XM_023248872.1
X 13684134 13684283
refGene_gn=REPS2;refGene_tx=XM_019823602.2;refGene_gn=REPS2;refGene_tx=XM_023248867.1;refGene_gn=REPS2;refGene_tx=XM_023248868.1;refGene_gn=REPS2;refGene_tx=XM_023248869.1;refGene_gn=REPS2;refGene_tx=XM_0232488

70.1;refGene_gn=REPS2;refGene_tx=XM_023248871.1;refGene_gn=REPS2;refGene_tx=XM_023248872.1
X	13686711	13686838
refGene_gn=REPS2;refGene_tx=XM_019823602.2;refGene_gn=REPS2;refGene_tx=XM_023248867.1;refGene_gn=REPS2;refGene_tx=XM_023248868.1;refGene_gn=REPS2;refGene_tx=XM_023248869.1;refGene_gn=REPS2;refGene_tx=XM_023248870.1;refGene_gn=REPS2;refGene_tx=XM_023248871.1;refGene_gn=REPS2;refGene_tx=XM_023248872.1
X	13693162	13693260
refGene_gn=REPS2;refGene_tx=XM_019823602.2;refGene_gn=REPS2;refGene_tx=XM_023248867.1;refGene_gn=REPS2;refGene_tx=XM_023248868.1;refGene_gn=REPS2;refGene_tx=XM_023248869.1;refGene_gn=REPS2;refGene_tx=XM_023248870.1;refGene_gn=REPS2;refGene_tx=XM_023248871.1;refGene_gn=REPS2;refGene_tx=XM_023248872.1
X	13715811	13715950
refGene_gn=REPS2;refGene_tx=XM_019823602.2;refGene_gn=REPS2;refGene_tx=XM_023248867.1;refGene_gn=REPS2;refGene_tx=XM_023248868.1;refGene_gn=REPS2;refGene_tx=XM_023248869.1;refGene_gn=REPS2;refGene_tx=XM_023248870.1;refGene_gn=REPS2;refGene_tx=XM_023248871.1;refGene_gn=REPS2;refGene_tx=XM_023248872.1
X	13722930	13722994
refGene_gn=REPS2;refGene_tx=XM_019823602.2;refGene_gn=REPS2;refGene_tx=XM_023248867.1;refGene_gn=REPS2;refGene_tx=XM_023248868.1;refGene_gn=REPS2;refGene_tx=XM_023248869.1;refGene_gn=REPS2;refGene_tx=XM_023248870.1;refGene_gn=REPS2;refGene_tx=XM_023248871.1;refGene_gn=REPS2;refGene_tx=XM_023248872.1
X	13725838	13725981
refGene_gn=REPS2;refGene_tx=XM_019823602.2;refGene_gn=REPS2;refGene_tx=XM_023248867.1;refGene_gn=REPS2;refGene_tx=XM_023248868.1;refGene_gn=REPS2;refGene_tx=XM_023248869.1;refGene_gn=REPS2;refGene_tx=XM_023248870.1;refGene_gn=REPS2;refGene_tx=XM_023248871.1;refGene_gn=REPS2;refGene_tx=XM_023248872.1
X	13731325	13731420
refGene_gn=REPS2;refGene_tx=XM_019823602.2;refGene_gn=REPS2;refGene_tx=XM_023248867.1;refGene_gn=REPS2;refGene_tx=XM_023248868.1;refGene_gn=REPS2;refGene_tx=XM_023248869.1;refGene_gn=REPS2;refGene_tx=XM_023248870.1;refGene_gn=REPS2;refGene_tx=XM_023248871.1;refGene_gn=REPS2;refGene_tx=XM_023248872.1
X	13738173	13738243
refGene_gn=REPS2;refGene_tx=XM_019823602.2;refGene_gn=REPS2;refGene_tx=XM_023248867.1;refGene_gn=REPS2;refGene_tx=XM_023248868.1;refGene_gn=REPS2;refGene_tx=XM_023248869.1;refGene_gn=REPS2;refGene_tx=XM_023248870.1;refGene_gn=REPS2;refGene_tx=XM_023248871.1;refGene_gn=REPS2;refGene_tx=XM_023248872.1
X	13739615	13739669
refGene_gn=REPS2;refGene_tx=XM_019823602.2;refGene_gn=REPS2;refGene_tx=XM_023248867.1;refGene_gn=REPS2;refGene_tx=XM_023248868.1;refGene_gn=REPS2;refGene_tx=XM_023248869.1;refGene_gn=REPS2;refGene_tx=XM_023248870.1;refGene_gn=REPS2;refGene_tx=XM_023248871.1;refGene_gn=REPS2;refGene_tx=XM_023248872.1

X	13744746	13744792
refGene_gn=REPS2;refGene_tx=XM_019823602.2;refGene_gn=REPS2;refGene_tx=XM_023248867.1;refGene_gn=REPS2;refGene_tx=XM_023248868.1;refGene_gn=REPS2;refGene_tx=XM_023248869.1;refGene_gn=REPS2;refGene_tx=XM_023248870.1
X	13748612	13748749
refGene_gn=REPS2;refGene_tx=XM_019823602.2;refGene_gn=REPS2;refGene_tx=XM_023248867.1;refGene_gn=REPS2;refGene_tx=XM_023248868.1;refGene_gn=REPS2;refGene_tx=XM_023248869.1;refGene_gn=REPS2;refGene_tx=XM_023248870.1
X	13773561	13773623
refGene_gn=REPS2;refGene_tx=XM_019823602.2;refGene_gn=REPS2;refGene_tx=XM_023248867.1;refGene_gn=REPS2;refGene_tx=XM_023248868.1;refGene_gn=REPS2;refGene_tx=XM_023248869.1;refGene_gn=REPS2;refGene_tx=XM_023248870.1;refGene_gn=REPS2;refGene_tx=XM_023248871.1;refGene_gn=REPS2;refGene_tx=XM_023248872.1
X	13775333	13775426
refGene_gn=REPS2;refGene_tx=XM_023248867.1;refGene_gn=REPS2;refGene_tx=XM_023248868.1;refGene_gn=REPS2;refGene_tx=XM_023248869.1;refGene_gn=REPS2;refGene_tx=XM_023248871.1
X	13798640	13798724
refGene_gn=REPS2;refGene_tx=XM_019823602.2;refGene_gn=REPS2;refGene_tx=XM_023248867.1;refGene_gn=REPS2;refGene_tx=XM_023248868.1;refGene_gn=REPS2;refGene_tx=XM_023248869.1;refGene_gn=REPS2;refGene_tx=XM_023248870.1;refGene_gn=REPS2;refGene_tx=XM_023248871.1;refGene_gn=REPS2;refGene_tx=XM_023248872.1
X	13799941	13800087
refGene_gn=REPS2;refGene_tx=XM_019823602.2;refGene_gn=REPS2;refGene_tx=XM_023248867.1;refGene_gn=REPS2;refGene_tx=XM_023248868.1;refGene_gn=REPS2;refGene_tx=XM_023248869.1;refGene_gn=REPS2;refGene_tx=XM_023248870.1;refGene_gn=REPS2;refGene_tx=XM_023248871.1;refGene_gn=REPS2;refGene_tx=XM_023248872.1
X	13810432	13810538
refGene_gn=REPS2;refGene_tx=XM_019823602.2;refGene_gn=REPS2;refGene_tx=XM_023248867.1;refGene_gn=REPS2;refGene_tx=XM_023248868.1;refGene_gn=REPS2;refGene_tx=XM_023248869.1;refGene_gn=REPS2;refGene_tx=XM_023248870.1;refGene_gn=REPS2;refGene_tx=XM_023248871.1;refGene_gn=REPS2;refGene_tx=XM_023248872.1
X	13829914	13829983
refGene_gn=REPS2;refGene_tx=XM_019823602.2;refGene_gn=REPS2;refGene_tx=XM_023248867.1;refGene_gn=REPS2;refGene_tx=XM_023248868.1;refGene_gn=REPS2;refGene_tx=XM_023248869.1;refGene_gn=REPS2;refGene_tx=XM_023248870.1;refGene_gn=REPS2;refGene_tx=XM_023248871.1;refGene_gn=REPS2;refGene_tx=XM_023248872.1
X	14004216	14004763
refGene_gn=NHS;refGene_tx=XM_023248864.1;refGene_gn=NHS;refGene_tx=XM_023248865.1
X	14244194	14244228	refGene_gn=NHS;refGene_tx=XM_019823598.2
X	14263218	14263444	refGene_gn=NHS;refGene_tx=XM_019823594.2
X	14294869	14295022

refGene_gn=NHS;refGene_tx=XM_019823594.2;refGene_gn=NHS;refGene_tx=XM_019823598.2;refGene_gn=NHS;refGene_tx=XM_023248864.1;refGene_gn=NHS;refGene_tx=XM_023248865.1
X 14299443 14299577
refGene_gn=NHS;refGene_tx=XM_019823594.2;refGene_gn=NHS;refGene_tx=XM_019823598.2;refGene_gn=NHS;refGene_tx=XM_023248864.1;refGene_gn=NHS;refGene_tx=XM_023248865.1
X 14331350 14331413
refGene_gn=NHS;refGene_tx=XM_019823594.2;refGene_gn=NHS;refGene_tx=XM_019823598.2;refGene_gn=NHS;refGene_tx=XM_023248864.1
X 14333397 14333590
refGene_gn=NHS;refGene_tx=XM_019823594.2;refGene_gn=NHS;refGene_tx=XM_019823598.2;refGene_gn=NHS;refGene_tx=XM_023248864.1;refGene_gn=NHS;refGene_tx=XM_023248865.1
X 14336214 14336346
refGene_gn=NHS;refGene_tx=XM_019823594.2;refGene_gn=NHS;refGene_tx=XM_019823598.2;refGene_gn=NHS;refGene_tx=XM_023248864.1;refGene_gn=NHS;refGene_tx=XM_023248865.1
X 14337394 14340376
refGene_gn=NHS;refGene_tx=XM_019823594.2;refGene_gn=NHS;refGene_tx=XM_019823598.2;refGene_gn=NHS;refGene_tx=XM_023248864.1;refGene_gn=NHS;refGene_tx=XM_023248865.1
X 14340689 14340816
refGene_gn=NHS;refGene_tx=XM_019823594.2;refGene_gn=NHS;refGene_tx=XM_019823598.2;refGene_gn=NHS;refGene_tx=XM_023248864.1;refGene_gn=NHS;refGene_tx=XM_023248865.1
X 14343372 14343979
refGene_gn=NHS;refGene_tx=XM_019823594.2;refGene_gn=NHS;refGene_tx=XM_019823598.2;refGene_gn=NHS;refGene_tx=XM_023248864.1;refGene_gn=NHS;refGene_tx=XM_023248865.1
X 14356161 14356191
refGene_gn=SCML1;refGene_tx=XM_011291620.3;refGene_gn=SCML1;refGene_tx=XM_011291621.3;refGene_gn=SCML1;refGene_tx=XM_019823603.2;refGene_gn=SCML1;refGene_tx=XM_019823605.2
X 14357393 14357474
refGene_gn=SCML1;refGene_tx=XM_011291620.3;refGene_gn=SCML1;refGene_tx=XM_019823603.2
X 14358194 14358272
refGene_gn=SCML1;refGene_tx=XM_011291620.3;refGene_gn=SCML1;refGene_tx=XM_011291621.3;refGene_gn=SCML1;refGene_tx=XM_019823603.2;refGene_gn=SCML1;refGene_tx=XM_019823605.2
X 14361259 14361364
refGene_gn=SCML1;refGene_tx=XM_011291620.3;refGene_gn=SCML1;refGene_tx=XM_011291621.3;refGene_gn=SCML1;refGene_tx=XM_019823603.2;refGene_gn=SCML1;refGene_tx=XM_019823605.2
X 14362006 14362547
refGene_gn=SCML1;refGene_tx=XM_011291620.3;refGene_gn=SCML1;refGene_tx=XM_011291621.3;refGene_gn=SCML1;refGene_tx=XM_019823603.2;refGene_gn=SCML1;refGene_tx=XM_019823605.2
X 14362696 14362845

refGene_gn=SCML1;refGene_tx=XM_011291620.3;refGene_gn=SCML1;refGene_tx=XM_011291621.3;refGene_gn=SCML1;refGene_tx=XM_019823603.2;refGene_gn=SCML1;refGene_tx=XM_019823605.2
X 14364515 14364650
refGene_gn=SCML1;refGene_tx=XM_011291620.3;refGene_gn=SCML1;refGene_tx=XM_011291621.3;refGene_gn=SCML1;refGene_tx=XM_019823603.2;refGene_gn=SCML1;refGene_tx=XM_019823605.2
X 14393974 14395546
refGene_gn=RAI2;refGene_tx=XM_004000313.4;refGene_gn=RAI2;refGene_tx=XM_006943601.3;refGene_gn=RAI2;refGene_tx=XM_006943602.3;refGene_gn=RAI2;refGene_tx=XM_023249018.1
X 14771959 14772147 refGene_gn=BEND2;refGene_tx=XM_023249456.1
X 14779458 14779662 refGene_gn=BEND2;refGene_tx=XM_023249456.1
X 14784460 14784692 refGene_gn=BEND2;refGene_tx=XM_023249456.1
X 14786545 14786718 refGene_gn=BEND2;refGene_tx=XM_023249456.1
X 14788343 14788544 refGene_gn=BEND2;refGene_tx=XM_023249456.1
X 14791226 14791340 refGene_gn=BEND2;refGene_tx=XM_023249456.1
X 14794633 14794777 refGene_gn=BEND2;refGene_tx=XM_023249456.1
X 14795108 14795183 refGene_gn=BEND2;refGene_tx=XM_023249456.1
X 14800216 14800366 refGene_gn=BEND2;refGene_tx=XM_023249456.1
X 14801710 14801860 refGene_gn=BEND2;refGene_tx=XM_023249456.1
X 14806290 14806705 refGene_gn=BEND2;refGene_tx=XM_023249456.1
X 14814780 14814932 refGene_gn=BEND2;refGene_tx=XM_023249456.1
X 14815868 14815893 refGene_gn=BEND2;refGene_tx=XM_023249456.1
X 14860631 14860760
refGene_gn=SCML2;refGene_tx=XM_011291623.3;refGene_gn=SCML2;refGene_tx=XM_019823606.2;refGene_gn=SCML2;refGene_tx=XM_019823607.2;refGene_gn=SCML2;refGene_tx=XM_019823608.2;refGene_gn=SCML2;refGene_tx=XM_019823609.2;refGene_gn=SCML2;refGene_tx=XM_019823610.2;refGene_gn=SCML2;refGene_tx=XM_019823611.2;refGene_gn=SCML2;refGene_tx=XM_019823612.2;refGene_gn=SCML2;refGene_tx=XM_019823613.2;refGene_gn=SCML2;refGene_tx=XM_023248998.1
X 14861840 14861992
refGene_gn=SCML2;refGene_tx=XM_011291623.3;refGene_gn=SCML2;refGene_tx=XM_019823606.2;refGene_gn=SCML2;refGene_tx=XM_019823607.2;refGene_gn=SCML2;refGene_tx=XM_019823608.2;refGene_gn=SCML2;refGene_tx=XM_019823609.2;refGene_gn=SCML2;refGene_tx=XM_019823610.2;refGene_gn=SCML2;refGene_tx=XM_019823611.2;refGene_gn=SCML2;refGene_tx=XM_019823612.2;refGene_gn=SCML2;refGene_tx=XM_023248998.1
X 14865377 14865629
refGene_gn=SCML2;refGene_tx=XM_011291623.3;refGene_gn=SCML2;refGene_tx=XM_019823606.2;refGene_gn=SCML2;refGene_tx=XM_019823607.2;refGene_gn=SCML2;refGene_tx=XM_019823608.2;refGene_gn=SCML2;refGene_tx=XM_019823609.2;refGene_gn=SCML2;refGene_tx=XM_019823610.2;refGene_gn=SCML2;refGene_tx=XM_019823611.2;refGene_gn=SCML2;refGene_tx=XM_019823612.2;refGene_gn=SCML2;refGene_tx=XM_019823613.2;refGene_gn=SCML2;refGene_tx=XM_023248998.1
X 14866489 14866603
refGene_gn=SCML2;refGene_tx=XM_011291623.3;refGene_gn=SCML2;refGene_tx=XM_019823606.2;refGene_gn=SCML2;refGene_tx=XM_019823607.2;refGene_gn=

SCML2;refGene_tx=XM_019823608.2;refGene_gn=SCML2;refGene_tx=XM_019823609.2;refGene_gn=SCML2;refGene_tx=XM_019823610.2;refGene_gn=SCML2;refGene_tx=XM_019823613.2;refGene_gn=SCML2;refGene_tx=XM_023248998.1
X 14872673 14872856
refGene_gn=SCML2;refGene_tx=XM_011291623.3;refGene_gn=SCML2;refGene_tx=XM_019823606.2;refGene_gn=SCML2;refGene_tx=XM_019823607.2;refGene_gn=SCML2;refGene_tx=XM_019823608.2;refGene_gn=SCML2;refGene_tx=XM_019823609.2;refGene_gn=SCML2;refGene_tx=XM_019823610.2;refGene_gn=SCML2;refGene_tx=XM_019823611.2;refGene_gn=SCML2;refGene_tx=XM_019823612.2;refGene_gn=SCML2;refGene_tx=XM_019823613.2;refGene_gn=SCML2;refGene_tx=XM_023248998.1
X 14873578 14873782
refGene_gn=SCML2;refGene_tx=XM_011291623.3;refGene_gn=SCML2;refGene_tx=XM_019823606.2;refGene_gn=SCML2;refGene_tx=XM_019823607.2;refGene_gn=SCML2;refGene_tx=XM_019823608.2;refGene_gn=SCML2;refGene_tx=XM_019823609.2;refGene_gn=SCML2;refGene_tx=XM_019823610.2;refGene_gn=SCML2;refGene_tx=XM_019823611.2;refGene_gn=SCML2;refGene_tx=XM_019823612.2;refGene_gn=SCML2;refGene_tx=XM_019823613.2;refGene_gn=SCML2;refGene_tx=XM_023248998.1
X 14876194 14876315
refGene_gn=SCML2;refGene_tx=XM_011291623.3;refGene_gn=SCML2;refGene_tx=XM_019823606.2;refGene_gn=SCML2;refGene_tx=XM_019823607.2;refGene_gn=SCML2;refGene_tx=XM_019823608.2;refGene_gn=SCML2;refGene_tx=XM_019823609.2;refGene_gn=SCML2;refGene_tx=XM_019823610.2;refGene_gn=SCML2;refGene_tx=XM_019823611.2;refGene_gn=SCML2;refGene_tx=XM_019823612.2;refGene_gn=SCML2;refGene_tx=XM_019823613.2;refGene_gn=SCML2;refGene_tx=XM_023248998.1
X 14882271 14882480
refGene_gn=SCML2;refGene_tx=XM_011291623.3;refGene_gn=SCML2;refGene_tx=XM_019823606.2;refGene_gn=SCML2;refGene_tx=XM_019823607.2;refGene_gn=SCML2;refGene_tx=XM_019823608.2;refGene_gn=SCML2;refGene_tx=XM_019823609.2;refGene_gn=SCML2;refGene_tx=XM_019823610.2;refGene_gn=SCML2;refGene_tx=XM_019823611.2;refGene_gn=SCML2;refGene_tx=XM_019823612.2;refGene_gn=SCML2;refGene_tx=XM_019823613.2;refGene_gn=SCML2;refGene_tx=XM_023248998.1
X 14886459 14886540
refGene_gn=SCML2;refGene_tx=XM_011291623.3;refGene_gn=SCML2;refGene_tx=XM_019823606.2;refGene_gn=SCML2;refGene_tx=XM_019823607.2;refGene_gn=SCML2;refGene_tx=XM_019823608.2;refGene_gn=SCML2;refGene_tx=XM_019823609.2;refGene_gn=SCML2;refGene_tx=XM_019823611.2;refGene_gn=SCML2;refGene_tx=XM_019823612.2;refGene_gn=SCML2;refGene_tx=XM_019823613.2;refGene_gn=SCML2;refGene_tx=XM_023248998.1
X 14916626 14916870
refGene_gn=SCML2;refGene_tx=XM_011291623.3;refGene_gn=SCML2;refGene_tx=XM_019823606.2;refGene_gn=SCML2;refGene_tx=XM_019823607.2;refGene_gn=SCML2;refGene_tx=XM_019823608.2;refGene_gn=SCML2;refGene_tx=XM_019823609.2;refGene_gn=SCML2;refGene_tx=XM_019823610.2;refGene_gn=SCML2;refGene_tx=XM_019823611.2;refGene_gn=SCML2;refGene_tx=XM_019823612.2;refGene_gn=SCML2;refGene_tx=XM_019823613.2;refGene_gn=SCML2;refGene_tx=XM_023248998.1

X 14924074 14924163
refGene_gn=SCML2;refGene_tx=XM_011291623.3;refGene_gn=SCML2;refGene_tx=XM_019823606.2;refGene_gn=SCML2;refGene_tx=XM_019823607.2;refGene_gn=SCML2;refGene_tx=XM_019823608.2;refGene_gn=SCML2;refGene_tx=XM_019823609.2;refGene_gn=SCML2;refGene_tx=XM_019823610.2;refGene_gn=SCML2;refGene_tx=XM_019823611.2;refGene_gn=SCML2;refGene_tx=XM_019823612.2;refGene_gn=SCML2;refGene_tx=XM_019823613.2;refGene_gn=SCML2;refGene_tx=XM_023248998.1
X 14928013 14928254
refGene_gn=SCML2;refGene_tx=XM_011291623.3;refGene_gn=SCML2;refGene_tx=XM_019823606.2;refGene_gn=SCML2;refGene_tx=XM_019823607.2;refGene_gn=SCML2;refGene_tx=XM_019823608.2;refGene_gn=SCML2;refGene_tx=XM_019823609.2;refGene_gn=SCML2;refGene_tx=XM_019823610.2;refGene_gn=SCML2;refGene_tx=XM_019823611.2;refGene_gn=SCML2;refGene_tx=XM_019823612.2;refGene_gn=SCML2;refGene_tx=XM_019823613.2;refGene_gn=SCML2;refGene_tx=XM_023248998.1
X 14929120 14929191
refGene_gn=SCML2;refGene_tx=XM_011291623.3;refGene_gn=SCML2;refGene_tx=XM_019823606.2;refGene_gn=SCML2;refGene_tx=XM_019823607.2;refGene_gn=SCML2;refGene_tx=XM_019823608.2;refGene_gn=SCML2;refGene_tx=XM_019823609.2;refGene_gn=SCML2;refGene_tx=XM_019823610.2;refGene_gn=SCML2;refGene_tx=XM_019823611.2;refGene_gn=SCML2;refGene_tx=XM_019823612.2;refGene_gn=SCML2;refGene_tx=XM_019823613.2
X 14933797 14933828 refGene_gn=SCML2;refGene_tx=XM_019823609.2
X 14935945 14936014
refGene_gn=SCML2;refGene_tx=XM_011291623.3;refGene_gn=SCML2;refGene_tx=XM_019823606.2;refGene_gn=SCML2;refGene_tx=XM_019823607.2;refGene_gn=SCML2;refGene_tx=XM_019823608.2;refGene_gn=SCML2;refGene_tx=XM_019823610.2;refGene_gn=SCML2;refGene_tx=XM_019823611.2;refGene_gn=SCML2;refGene_tx=XM_019823612.2;refGene_gn=SCML2;refGene_tx=XM_019823613.2
X 14939545 14939576
refGene_gn=SCML2;refGene_tx=XM_011291623.3;refGene_gn=SCML2;refGene_tx=XM_019823606.2;refGene_gn=SCML2;refGene_tx=XM_019823608.2;refGene_gn=SCML2;refGene_tx=XM_019823610.2;refGene_gn=SCML2;refGene_tx=XM_019823611.2;refGene_gn=SCML2;refGene_tx=XM_019823612.2;refGene_gn=SCML2;refGene_tx=XM_019823613.2
X 15095482 15095546
refGene_gn=CDKL5;refGene_tx=XM_011291624.3;refGene_gn=CDKL5;refGene_tx=XM_019823614.2;refGene_gn=CDKL5;refGene_tx=XM_019823615.2;refGene_gn=CDKL5;refGene_tx=XM_019823616.1;refGene_gn=CDKL5;refGene_tx=XM_019823617.2
X 15098314 15098349
refGene_gn=CDKL5;refGene_tx=XM_011291624.3;refGene_gn=CDKL5;refGene_tx=XM_019823614.2;refGene_gn=CDKL5;refGene_tx=XM_019823615.2;refGene_gn=CDKL5;refGene_tx=XM_019823616.1;refGene_gn=CDKL5;refGene_tx=XM_019823617.2
X 15120533 15121433
refGene_gn=LOC102899587;refGene_tx=XM_019823620.2
X 15151344 15151390
refGene_gn=CDKL5;refGene_tx=XM_011291624.3;refGene_gn=CDKL5;refGene_tx

=XM_019823614.2;refGene_gn=CDKL5;refGene_tx=XM_019823615.2;refGene_gn=CDKL5;refGene_tx=XM_019823616.1;refGene_gn=CDKL5;refGene_tx=XM_019823617.2
X 15164736 15164873
refGene_gn=CDKL5;refGene_tx=XM_011291624.3;refGene_gn=CDKL5;refGene_tx=XM_019823614.2;refGene_gn=CDKL5;refGene_tx=XM_019823615.2;refGene_gn=CDKL5;refGene_tx=XM_019823616.1;refGene_gn=CDKL5;refGene_tx=XM_019823617.2;refGene_gn=CDKL5;refGene_tx=XM_019823619.2
X 15170213 15170334
refGene_gn=CDKL5;refGene_tx=XM_011291624.3;refGene_gn=CDKL5;refGene_tx=XM_019823614.2;refGene_gn=CDKL5;refGene_tx=XM_019823615.2;refGene_gn=CDKL5;refGene_tx=XM_019823616.1;refGene_gn=CDKL5;refGene_tx=XM_019823617.2;refGene_gn=CDKL5;refGene_tx=XM_019823619.2
X 15173487 15173547
refGene_gn=CDKL5;refGene_tx=XM_011291624.3;refGene_gn=CDKL5;refGene_tx=XM_019823614.2;refGene_gn=CDKL5;refGene_tx=XM_019823615.2;refGene_gn=CDKL5;refGene_tx=XM_019823616.1;refGene_gn=CDKL5;refGene_tx=XM_019823617.2;refGene_gn=CDKL5;refGene_tx=XM_019823619.2
X 15175945 15176036
refGene_gn=CDKL5;refGene_tx=XM_011291624.3;refGene_gn=CDKL5;refGene_tx=XM_019823614.2;refGene_gn=CDKL5;refGene_tx=XM_019823615.2;refGene_gn=CDKL5;refGene_tx=XM_019823616.1;refGene_gn=CDKL5;refGene_tx=XM_019823617.2;refGene_gn=CDKL5;refGene_tx=XM_019823619.2
X 15179536 15179726
refGene_gn=CDKL5;refGene_tx=XM_011291624.3;refGene_gn=CDKL5;refGene_tx=XM_019823614.2;refGene_gn=CDKL5;refGene_tx=XM_019823615.2;refGene_gn=CDKL5;refGene_tx=XM_019823616.1;refGene_gn=CDKL5;refGene_tx=XM_019823617.2;refGene_gn=CDKL5;refGene_tx=XM_019823619.2
X 15185068 15185149
refGene_gn=CDKL5;refGene_tx=XM_011291624.3;refGene_gn=CDKL5;refGene_tx=XM_019823614.2;refGene_gn=CDKL5;refGene_tx=XM_019823615.2;refGene_gn=CDKL5;refGene_tx=XM_019823616.1;refGene_gn=CDKL5;refGene_tx=XM_019823617.2;refGene_gn=CDKL5;refGene_tx=XM_019823619.2
X 15190032 15190184
refGene_gn=CDKL5;refGene_tx=XM_011291624.3;refGene_gn=CDKL5;refGene_tx=XM_019823614.2;refGene_gn=CDKL5;refGene_tx=XM_019823615.2;refGene_gn=CDKL5;refGene_tx=XM_019823616.1;refGene_gn=CDKL5;refGene_tx=XM_019823617.2;refGene_gn=CDKL5;refGene_tx=XM_019823619.2
X 15195177 15196144
refGene_gn=CDKL5;refGene_tx=XM_011291624.3;refGene_gn=CDKL5;refGene_tx=XM_019823614.2;refGene_gn=CDKL5;refGene_tx=XM_019823615.2;refGene_gn=CDKL5;refGene_tx=XM_019823616.1;refGene_gn=CDKL5;refGene_tx=XM_019823617.2;refGene_gn=CDKL5;refGene_tx=XM_019823619.2
X 15200720 15200822
refGene_gn=CDKL5;refGene_tx=XM_011291624.3;refGene_gn=CDKL5;refGene_tx=XM_019823614.2;refGene_gn=CDKL5;refGene_tx=XM_019823615.2;refGene_gn=CDKL5;refGene_tx=XM_019823616.1;refGene_gn=CDKL5;refGene_tx=XM_019823617.2;refGene_gn=CDKL5;refGene_tx=XM_019823619.2
X 15201114 15201220
refGene_gn=CDKL5;refGene_tx=XM_011291624.3;refGene_gn=CDKL5;refGene_tx

=XM_019823614.2;refGene_gn=CDKL5;refGene_tx=XM_019823615.2;refGene_gn=CDKL5;refGene_tx=XM_019823616.1;refGene_gn=CDKL5;refGene_tx=XM_019823617.2;refGene_gn=CDKL5;refGene_tx=XM_019823619.2
X	15204650	15204774	refGene_gn=CDKL5;refGene_tx=XM_011291624.3;refGene_gn=CDKL5;refGene_tx=XM_019823614.2;refGene_gn=CDKL5;refGene_tx=XM_019823615.2;refGene_gn=CDKL5;refGene_tx=XM_019823616.1;refGene_gn=CDKL5;refGene_tx=XM_019823617.2;refGene_gn=CDKL5;refGene_tx=XM_019823619.2
X	15212440	15212540	refGene_gn=CDKL5;refGene_tx=XM_011291624.3;refGene_gn=CDKL5;refGene_tx=XM_019823614.2;refGene_gn=CDKL5;refGene_tx=XM_019823615.2;refGene_gn=CDKL5;refGene_tx=XM_019823616.1;refGene_gn=CDKL5;refGene_tx=XM_019823617.2;refGene_gn=CDKL5;refGene_tx=XM_019823619.2
X	15214110	15214170	refGene_gn=CDKL5;refGene_tx=XM_019823617.2
X	15216134	15216257	refGene_gn=CDKL5;refGene_tx=XM_019823614.2;refGene_gn=CDKL5;refGene_tx=XM_019823615.2;refGene_gn=CDKL5;refGene_tx=XM_019823616.1;refGene_gn=CDKL5;refGene_tx=XM_019823619.2
X	15217412	15217532	refGene_gn=CDKL5;refGene_tx=XM_011291624.3;refGene_gn=CDKL5;refGene_tx=XM_019823614.2;refGene_gn=CDKL5;refGene_tx=XM_019823615.2;refGene_gn=CDKL5;refGene_tx=XM_019823616.1;refGene_gn=CDKL5;refGene_tx=XM_019823617.2;refGene_gn=CDKL5;refGene_tx=XM_019823619.2
X	15220052	15220439	refGene_gn=CDKL5;refGene_tx=XM_011291624.3;refGene_gn=CDKL5;refGene_tx=XM_019823614.2;refGene_gn=CDKL5;refGene_tx=XM_019823615.2;refGene_gn=CDKL5;refGene_tx=XM_019823616.1;refGene_gn=CDKL5;refGene_tx=XM_019823617.2;refGene_gn=CDKL5;refGene_tx=XM_019823619.2
X	15233396	15233549	refGene_gn=RS1;refGene_tx=XM_019823621.2
X	15235603	15235799	refGene_gn=RS1;refGene_tx=XM_019823621.2
X	15238090	15238232	refGene_gn=RS1;refGene_tx=XM_019823621.2
X	15239177	15239317	refGene_gn=CDKL5;refGene_tx=XM_019823615.2;refGene_gn=CDKL5;refGene_tx=XM_019823616.1
X	15243210	15243299	refGene_gn=CDKL5;refGene_tx=XM_019823616.1
X	15245420	15245526	refGene_gn=RS1;refGene_tx=XM_019823621.2
X	15246400	15246426	refGene_gn=RS1;refGene_tx=XM_019823621.2
X	15255599	15255651	refGene_gn=RS1;refGene_tx=XM_019823621.2
X	15287931	15288001	refGene_gn=PPEF1;refGene_tx=XM_006943604.4;refGene_gn=PPEF1;refGene_tx=XM_019823622.2;refGene_gn=PPEF1;refGene_tx=XM_019823623.2
X	15308340	15308468	refGene_gn=PPEF1;refGene_tx=XM_006943604.4;refGene_gn=PPEF1;refGene_tx=XM_019823622.2;refGene_gn=PPEF1;refGene_tx=XM_019823623.2
X	15309183	15309244	refGene_gn=PPEF1;refGene_tx=XM_006943604.4;refGene_gn=PPEF1;refGene_tx=XM_019823622.2;refGene_gn=PPEF1;refGene_tx=XM_019823623.2
X	15317108	15317272	refGene_gn=PPEF1;refGene_tx=XM_006943604.4;refGene_gn=PPEF1;refGene_tx=XM_019823622.2;refGene_gn=PPEF1;refGene_tx=XM_019823623.2

X 15326409 15326524
refGene_gn=PPEF1;refGene_tx=XM_006943604.4;refGene_gn=PPEF1;refGene_tx=XM_019823622.2;refGene_gn=PPEF1;refGene_tx=XM_019823623.2
X 15330242 15330289
refGene_gn=PPEF1;refGene_tx=XM_006943604.4;refGene_gn=PPEF1;refGene_tx=XM_019823622.2;refGene_gn=PPEF1;refGene_tx=XM_019823623.2
X 15343543 15343710
refGene_gn=PPEF1;refGene_tx=XM_006943604.4;refGene_gn=PPEF1;refGene_tx=XM_019823622.2;refGene_gn=PPEF1;refGene_tx=XM_019823623.2
X 15346816 15346853
refGene_gn=PPEF1;refGene_tx=XM_006943604.4;refGene_gn=PPEF1;refGene_tx=XM_019823622.2;refGene_gn=PPEF1;refGene_tx=XM_019823623.2
X 15347703 15347853
refGene_gn=PPEF1;refGene_tx=XM_006943604.4;refGene_gn=PPEF1;refGene_tx=XM_019823622.2;refGene_gn=PPEF1;refGene_tx=XM_019823623.2
X 15351935 15352085
refGene_gn=PPEF1;refGene_tx=XM_006943604.4;refGene_gn=PPEF1;refGene_tx=XM_019823622.2;refGene_gn=PPEF1;refGene_tx=XM_019823623.2
X 15357975 15358161
refGene_gn=PPEF1;refGene_tx=XM_006943604.4;refGene_gn=PPEF1;refGene_tx=XM_019823622.2;refGene_gn=PPEF1;refGene_tx=XM_019823623.2
X 15359543 15359686
refGene_gn=PPEF1;refGene_tx=XM_006943604.4;refGene_gn=PPEF1;refGene_tx=XM_019823622.2;refGene_gn=PPEF1;refGene_tx=XM_019823623.2
X 15369295 15369402
refGene_gn=PPEF1;refGene_tx=XM_006943604.4;refGene_gn=PPEF1;refGene_tx=XM_019823622.2;refGene_gn=PPEF1;refGene_tx=XM_019823623.2
X 15378032 15378196
refGene_gn=PPEF1;refGene_tx=XM_006943604.4;refGene_gn=PPEF1;refGene_tx=XM_019823622.2;refGene_gn=PPEF1;refGene_tx=XM_019823623.2
X 15379288 15379373
refGene_gn=PPEF1;refGene_tx=XM_006943604.4;refGene_gn=PPEF1;refGene_tx=XM_019823622.2;refGene_gn=PPEF1;refGene_tx=XM_019823623.2
X 15380511 15380720
refGene_gn=PPEF1;refGene_tx=XM_006943604.4;refGene_gn=PPEF1;refGene_tx=XM_019823622.2;refGene_gn=PPEF1;refGene_tx=XM_019823623.2
X 15416421 15416592
refGene_gn=PHKA2;refGene_tx=XM_019823627.2;refGene_gn=PHKA2;refGene_tx=XM_023249379.1;refGene_gn=PHKA2;refGene_tx=XM_023249380.1;refGene_gn=PHKA2;refGene_tx=XM_023249381.1;refGene_gn=PHKA2;refGene_tx=XM_023249382.1
X 15417174 15417375
refGene_gn=PHKA2;refGene_tx=XM_019823627.2;refGene_gn=PHKA2;refGene_tx=XM_023249379.1;refGene_gn=PHKA2;refGene_tx=XM_023249380.1;refGene_gn=PHKA2;refGene_tx=XM_023249381.1;refGene_gn=PHKA2;refGene_tx=XM_023249382.1;refGene_gn=PHKA2;refGene_tx=XM_023249383.1
X 15418095 15418149
refGene_gn=PHKA2;refGene_tx=XM_019823627.2;refGene_gn=PHKA2;refGene_tx=XM_023249379.1;refGene_gn=PHKA2;refGene_tx=XM_023249380.1;refGene_gn=PHKA2;refGene_tx=XM_023249381.1;refGene_gn=PHKA2;refGene_tx=XM_0232493

82.1;refGene_gn=PHKA2;refGene_tx=XM_023249383.1
X 15420166 15420337
refGene_gn=PHKA2;refGene_tx=XM_019823627.2;refGene_gn=PHKA2;refGene_tx=XM_023249379.1;refGene_gn=PHKA2;refGene_tx=XM_023249380.1;refGene_gn=PHKA2;refGene_tx=XM_023249381.1;refGene_gn=PHKA2;refGene_tx=XM_023249382.1;refGene_gn=PHKA2;refGene_tx=XM_023249383.1
X 15422036 15422206
refGene_gn=PHKA2;refGene_tx=XM_019823627.2;refGene_gn=PHKA2;refGene_tx=XM_023249379.1;refGene_gn=PHKA2;refGene_tx=XM_023249382.1;refGene_gn=PHKA2;refGene_tx=XM_023249383.1
X 15423605 15423635
refGene_gn=PHKA2;refGene_tx=XM_019823627.2;refGene_gn=PHKA2;refGene_tx=XM_023249380.1;refGene_gn=PHKA2;refGene_tx=XM_023249382.1;refGene_gn=PHKA2;refGene_tx=XM_023249383.1
X 15424387 15424506
refGene_gn=PHKA2;refGene_tx=XM_019823627.2;refGene_gn=PHKA2;refGene_tx=XM_023249379.1;refGene_gn=PHKA2;refGene_tx=XM_023249380.1;refGene_gn=PHKA2;refGene_tx=XM_023249381.1;refGene_gn=PHKA2;refGene_tx=XM_023249382.1;refGene_gn=PHKA2;refGene_tx=XM_023249383.1
X 15424616 15424781
refGene_gn=PHKA2;refGene_tx=XM_019823627.2;refGene_gn=PHKA2;refGene_tx=XM_023249379.1;refGene_gn=PHKA2;refGene_tx=XM_023249380.1;refGene_gn=PHKA2;refGene_tx=XM_023249381.1;refGene_gn=PHKA2;refGene_tx=XM_023249383.1
X 15426699 15426801
refGene_gn=PHKA2;refGene_tx=XM_019823627.2;refGene_gn=PHKA2;refGene_tx=XM_023249379.1;refGene_gn=PHKA2;refGene_tx=XM_023249380.1;refGene_gn=PHKA2;refGene_tx=XM_023249381.1;refGene_gn=PHKA2;refGene_tx=XM_023249382.1;refGene_gn=PHKA2;refGene_tx=XM_023249383.1
X 15427388 15427518
refGene_gn=PHKA2;refGene_tx=XM_019823627.2;refGene_gn=PHKA2;refGene_tx=XM_023249379.1;refGene_gn=PHKA2;refGene_tx=XM_023249380.1;refGene_gn=PHKA2;refGene_tx=XM_023249381.1;refGene_gn=PHKA2;refGene_tx=XM_023249382.1;refGene_gn=PHKA2;refGene_tx=XM_023249383.1
X 15427624 15427703
refGene_gn=PHKA2;refGene_tx=XM_019823627.2;refGene_gn=PHKA2;refGene_tx=XM_023249379.1;refGene_gn=PHKA2;refGene_tx=XM_023249380.1;refGene_gn=PHKA2;refGene_tx=XM_023249381.1;refGene_gn=PHKA2;refGene_tx=XM_023249382.1;refGene_gn=PHKA2;refGene_tx=XM_023249383.1
X 15427916 15427996
refGene_gn=PHKA2;refGene_tx=XM_019823627.2;refGene_gn=PHKA2;refGene_tx=XM_023249379.1;refGene_gn=PHKA2;refGene_tx=XM_023249380.1;refGene_gn=PHKA2;refGene_tx=XM_023249381.1;refGene_gn=PHKA2;refGene_tx=XM_023249382.1;refGene_gn=PHKA2;refGene_tx=XM_023249383.1
X 15428839 15428996
refGene_gn=PHKA2;refGene_tx=XM_019823627.2;refGene_gn=PHKA2;refGene_tx=XM_023249379.1;refGene_gn=PHKA2;refGene_tx=XM_023249380.1;refGene_gn=PHKA2;refGene_tx=XM_023249381.1;refGene_gn=PHKA2;refGene_tx=XM_023249382.1;refGene_gn=PHKA2;refGene_tx=XM_023249383.1
X 15429395 15429529

refGene_gn=PHKA2;refGene_tx=XM_019823627.2;refGene_gn=PHKA2;refGene_tx=XM_023249379.1;refGene_gn=PHKA2;refGene_tx=XM_023249380.1;refGene_gn=PHKA2;refGene_tx=XM_023249381.1;refGene_gn=PHKA2;refGene_tx=XM_023249382.1;refGene_gn=PHKA2;refGene_tx=XM_023249383.1
X 15430421 15430510
refGene_gn=PHKA2;refGene_tx=XM_019823627.2;refGene_gn=PHKA2;refGene_tx=XM_023249379.1;refGene_gn=PHKA2;refGene_tx=XM_023249380.1;refGene_gn=PHKA2;refGene_tx=XM_023249381.1;refGene_gn=PHKA2;refGene_tx=XM_023249382.1;refGene_gn=PHKA2;refGene_tx=XM_023249383.1
X 15433613 15433793
refGene_gn=PHKA2;refGene_tx=XM_019823627.2;refGene_gn=PHKA2;refGene_tx=XM_023249379.1;refGene_gn=PHKA2;refGene_tx=XM_023249380.1;refGene_gn=PHKA2;refGene_tx=XM_023249381.1;refGene_gn=PHKA2;refGene_tx=XM_023249382.1;refGene_gn=PHKA2;refGene_tx=XM_023249383.1
X 15434543 15434707
refGene_gn=PHKA2;refGene_tx=XM_019823627.2;refGene_gn=PHKA2;refGene_tx=XM_023249379.1;refGene_gn=PHKA2;refGene_tx=XM_023249380.1;refGene_gn=PHKA2;refGene_tx=XM_023249381.1;refGene_gn=PHKA2;refGene_tx=XM_023249382.1;refGene_gn=PHKA2;refGene_tx=XM_023249383.1
X 15437623 15437702
refGene_gn=PHKA2;refGene_tx=XM_019823627.2;refGene_gn=PHKA2;refGene_tx=XM_023249379.1;refGene_gn=PHKA2;refGene_tx=XM_023249380.1;refGene_gn=PHKA2;refGene_tx=XM_023249381.1;refGene_gn=PHKA2;refGene_tx=XM_023249382.1;refGene_gn=PHKA2;refGene_tx=XM_023249383.1
X 15437960 15438105
refGene_gn=PHKA2;refGene_tx=XM_019823627.2;refGene_gn=PHKA2;refGene_tx=XM_023249379.1;refGene_gn=PHKA2;refGene_tx=XM_023249380.1;refGene_gn=PHKA2;refGene_tx=XM_023249381.1;refGene_gn=PHKA2;refGene_tx=XM_023249382.1;refGene_gn=PHKA2;refGene_tx=XM_023249383.1
X 15439264 15439374
refGene_gn=PHKA2;refGene_tx=XM_019823627.2;refGene_gn=PHKA2;refGene_tx=XM_023249379.1;refGene_gn=PHKA2;refGene_tx=XM_023249380.1;refGene_gn=PHKA2;refGene_tx=XM_023249381.1;refGene_gn=PHKA2;refGene_tx=XM_023249382.1;refGene_gn=PHKA2;refGene_tx=XM_023249383.1
X 15440042 15440177
refGene_gn=PHKA2;refGene_tx=XM_019823627.2;refGene_gn=PHKA2;refGene_tx=XM_023249379.1;refGene_gn=PHKA2;refGene_tx=XM_023249380.1;refGene_gn=PHKA2;refGene_tx=XM_023249381.1;refGene_gn=PHKA2;refGene_tx=XM_023249382.1;refGene_gn=PHKA2;refGene_tx=XM_023249383.1
X 15441568 15441647
refGene_gn=PHKA2;refGene_tx=XM_019823627.2;refGene_gn=PHKA2;refGene_tx=XM_023249379.1;refGene_gn=PHKA2;refGene_tx=XM_023249380.1;refGene_gn=PHKA2;refGene_tx=XM_023249381.1;refGene_gn=PHKA2;refGene_tx=XM_023249382.1;refGene_gn=PHKA2;refGene_tx=XM_023249383.1
X 15443377 15443485
refGene_gn=PHKA2;refGene_tx=XM_019823627.2;refGene_gn=PHKA2;refGene_tx=XM_023249379.1;refGene_gn=PHKA2;refGene_tx=XM_023249380.1;refGene_gn=PHKA2;refGene_tx=XM_023249381.1;refGene_gn=PHKA2;refGene_tx=XM_023249382.1;refGene_gn=PHKA2;refGene_tx=XM_023249383.1
X 15445182 15445278

refGene_gn=PHKA2;refGene_tx=XM_019823627.2;refGene_gn=PHKA2;refGene_tx=XM_023249379.1;refGene_gn=PHKA2;refGene_tx=XM_023249380.1;refGene_gn=PHKA2;refGene_tx=XM_023249381.1;refGene_gn=PHKA2;refGene_tx=XM_023249382.1;refGene_gn=PHKA2;refGene_tx=XM_023249383.1
X 15448240 15448363
refGene_gn=PHKA2;refGene_tx=XM_019823627.2;refGene_gn=PHKA2;refGene_tx=XM_023249379.1;refGene_gn=PHKA2;refGene_tx=XM_023249380.1;refGene_gn=PHKA2;refGene_tx=XM_023249381.1;refGene_gn=PHKA2;refGene_tx=XM_023249382.1;refGene_gn=PHKA2;refGene_tx=XM_023249383.1
X 15448967 15449021
refGene_gn=PHKA2;refGene_tx=XM_019823627.2;refGene_gn=PHKA2;refGene_tx=XM_023249379.1;refGene_gn=PHKA2;refGene_tx=XM_023249380.1;refGene_gn=PHKA2;refGene_tx=XM_023249381.1;refGene_gn=PHKA2;refGene_tx=XM_023249382.1;refGene_gn=PHKA2;refGene_tx=XM_023249383.1
X 15450895 15451042
refGene_gn=PHKA2;refGene_tx=XM_019823627.2;refGene_gn=PHKA2;refGene_tx=XM_023249379.1;refGene_gn=PHKA2;refGene_tx=XM_023249380.1;refGene_gn=PHKA2;refGene_tx=XM_023249381.1;refGene_gn=PHKA2;refGene_tx=XM_023249382.1;refGene_gn=PHKA2;refGene_tx=XM_023249383.1
X 15453266 15453365
refGene_gn=PHKA2;refGene_tx=XM_019823627.2;refGene_gn=PHKA2;refGene_tx=XM_023249379.1;refGene_gn=PHKA2;refGene_tx=XM_023249380.1;refGene_gn=PHKA2;refGene_tx=XM_023249381.1;refGene_gn=PHKA2;refGene_tx=XM_023249382.1;refGene_gn=PHKA2;refGene_tx=XM_023249383.1
X 15455179 15455260
refGene_gn=PHKA2;refGene_tx=XM_019823627.2;refGene_gn=PHKA2;refGene_tx=XM_023249379.1;refGene_gn=PHKA2;refGene_tx=XM_023249380.1;refGene_gn=PHKA2;refGene_tx=XM_023249381.1;refGene_gn=PHKA2;refGene_tx=XM_023249382.1;refGene_gn=PHKA2;refGene_tx=XM_023249383.1
X 15457529 15457612
refGene_gn=PHKA2;refGene_tx=XM_019823627.2;refGene_gn=PHKA2;refGene_tx=XM_023249379.1;refGene_gn=PHKA2;refGene_tx=XM_023249380.1;refGene_gn=PHKA2;refGene_tx=XM_023249381.1;refGene_gn=PHKA2;refGene_tx=XM_023249382.1;refGene_gn=PHKA2;refGene_tx=XM_023249383.1
X 15459625 15459794
refGene_gn=PHKA2;refGene_tx=XM_019823627.2;refGene_gn=PHKA2;refGene_tx=XM_023249379.1;refGene_gn=PHKA2;refGene_tx=XM_023249380.1;refGene_gn=PHKA2;refGene_tx=XM_023249381.1;refGene_gn=PHKA2;refGene_tx=XM_023249382.1;refGene_gn=PHKA2;refGene_tx=XM_023249383.1
X 15460949 15460997
refGene_gn=PHKA2;refGene_tx=XM_019823627.2;refGene_gn=PHKA2;refGene_tx=XM_023249379.1;refGene_gn=PHKA2;refGene_tx=XM_023249380.1;refGene_gn=PHKA2;refGene_tx=XM_023249381.1;refGene_gn=PHKA2;refGene_tx=XM_023249382.1;refGene_gn=PHKA2;refGene_tx=XM_023249383.1
X 15464053 15464212
refGene_gn=PHKA2;refGene_tx=XM_019823627.2;refGene_gn=PHKA2;refGene_tx=XM_023249379.1;refGene_gn=PHKA2;refGene_tx=XM_023249380.1;refGene_gn=PHKA2;refGene_tx=XM_023249381.1;refGene_gn=PHKA2;refGene_tx=XM_023249382.1;refGene_gn=PHKA2;refGene_tx=XM_023249383.1
X 15486210 15486288

refGene_gn=PHKA2;refGene_tx=XM_019823627.2;refGene_gn=PHKA2;refGene_tx=XM_023249379.1;refGene_gn=PHKA2;refGene_tx=XM_023249380.1;refGene_gn=PHKA2;refGene_tx=XM_023249381.1;refGene_gn=PHKA2;refGene_tx=XM_023249382.1;refGene_gn=PHKA2;refGene_tx=XM_023249383.1
X 15492263 15492442 refGene_gn=ADGRG2;refGene_tx=XM_023249384.1
X 15495420 15495573 refGene_gn=ADGRG2;refGene_tx=XM_023249384.1
X 15496776 15496878 refGene_gn=ADGRG2;refGene_tx=XM_023249384.1
X 15499105 15499389 refGene_gn=ADGRG2;refGene_tx=XM_023249384.1
X 15500222 15500322 refGene_gn=ADGRG2;refGene_tx=XM_023249384.1
X 15503540 15503809 refGene_gn=ADGRG2;refGene_tx=XM_023249384.1
X 15505196 15505318 refGene_gn=ADGRG2;refGene_tx=XM_023249384.1
X 15506105 15506209 refGene_gn=ADGRG2;refGene_tx=XM_023249384.1
X 15506343 15506389 refGene_gn=ADGRG2;refGene_tx=XM_023249384.1
X 15507043 15507166 refGene_gn=ADGRG2;refGene_tx=XM_023249384.1
X 15508176 15508320 refGene_gn=ADGRG2;refGene_tx=XM_023249384.1
X 15509197 15509354 refGene_gn=ADGRG2;refGene_tx=XM_023249384.1
X 15509555 15509730 refGene_gn=ADGRG2;refGene_tx=XM_023249384.1
X 15512031 15512423 refGene_gn=ADGRG2;refGene_tx=XM_023249384.1
X 15516977 15517044 refGene_gn=ADGRG2;refGene_tx=XM_023249384.1
X 15517576 15517671 refGene_gn=ADGRG2;refGene_tx=XM_023249384.1
X 15518903 15518941 refGene_gn=ADGRG2;refGene_tx=XM_023249384.1
X 15519452 15519492 refGene_gn=ADGRG2;refGene_tx=XM_023249384.1
X 15521503 15521559 refGene_gn=ADGRG2;refGene_tx=XM_023249384.1
X 15522399 15522452 refGene_gn=ADGRG2;refGene_tx=XM_023249384.1
X 15526073 15526127 refGene_gn=ADGRG2;refGene_tx=XM_023249384.1
X 15528545 15528587 refGene_gn=ADGRG2;refGene_tx=XM_023249384.1
X 15530547 15530586 refGene_gn=ADGRG2;refGene_tx=XM_023249384.1
X 15532187 15532206 refGene_gn=ADGRG2;refGene_tx=XM_023249384.1
X 15532207 15532227 refGene_gn=ADGRG2;refGene_tx=XM_023249384.1
X 15535743 15535779 refGene_gn=ADGRG2;refGene_tx=XM_023249384.1
X 15560403 15560521 refGene_gn=ADGRG2;refGene_tx=XM_023249384.1
X 15755580 15755901
refGene_gn=LOC101089581;refGene_tx=XM_011291648.2
X 15755902 15756409
refGene_gn=LOC101089581;refGene_tx=XM_011291648.2
X 15759381 15759491
refGene_gn=LOC101089581;refGene_tx=XM_011291648.2
X 15772710 15772982
refGene_gn=LOC101089581;refGene_tx=XM_011291648.2
X 15774575 15774652
refGene_gn=LOC101089581;refGene_tx=XM_011291648.2
X 15796915 15796972 refGene_gn=PDHA1;refGene_tx=XM_004001537.4
X 15802097 15802157 refGene_gn=PDHA1;refGene_tx=XM_004001537.4
X 15802712 15802886 refGene_gn=PDHA1;refGene_tx=XM_004001537.4
X 15803845 15803972 refGene_gn=PDHA1;refGene_tx=XM_004001537.4
X 15804612 15804704 refGene_gn=PDHA1;refGene_tx=XM_004001537.4
X 15806078 15806171 refGene_gn=PDHA1;refGene_tx=XM_004001537.4
X 15806970 15807126 refGene_gn=PDHA1;refGene_tx=XM_004001537.4
X 15807297 15807369 refGene_gn=PDHA1;refGene_tx=XM_004001537.4
X 15809453 15809521 refGene_gn=PDHA1;refGene_tx=XM_004001537.4

X	15810734	15810843	refGene_gn=PDHA1;refGene_tx=XM_004001537.4
X	15811175	15811340	refGene_gn=PDHA1;refGene_tx=XM_004001537.4
X	15812240	15812340	refGene_gn=MAP3K15;refGene_tx=XM_023248797.1
X	15812844	15812921	refGene_gn=MAP3K15;refGene_tx=XM_023248797.1
X	15813011	15813121	refGene_gn=MAP3K15;refGene_tx=XM_023248797.1
X	15813949	15814062	refGene_gn=MAP3K15;refGene_tx=XM_023248797.1
X	15816338	15816564	refGene_gn=MAP3K15;refGene_tx=XM_023248797.1
X	15817628	15817734	refGene_gn=MAP3K15;refGene_tx=XM_023248797.1
X	15818366	15818552	refGene_gn=MAP3K15;refGene_tx=XM_023248797.1
X	15819495	15819649	refGene_gn=MAP3K15;refGene_tx=XM_023248797.1
X	15820105	15820265	refGene_gn=MAP3K15;refGene_tx=XM_023248797.1
X	15821005	15821189	refGene_gn=MAP3K15;refGene_tx=XM_023248797.1
X	15825121	15825279	refGene_gn=MAP3K15;refGene_tx=XM_023248797.1
X	15834000	15834106	refGene_gn=MAP3K15;refGene_tx=XM_023248797.1
X	15834345	15834476	refGene_gn=MAP3K15;refGene_tx=XM_023248797.1
X	15836910	15837038	refGene_gn=MAP3K15;refGene_tx=XM_023248797.1
X	15839302	15839436	refGene_gn=MAP3K15;refGene_tx=XM_023248797.1
X	15841408	15841496	refGene_gn=MAP3K15;refGene_tx=XM_023248797.1
X	15848814	15848910	refGene_gn=MAP3K15;refGene_tx=XM_023248797.1
X	15850852	15850902	refGene_gn=MAP3K15;refGene_tx=XM_023248797.1
X	15858338	15858446	refGene_gn=MAP3K15;refGene_tx=XM_023248797.1
X	15859883	15860034	refGene_gn=MAP3K15;refGene_tx=XM_023248797.1
X	15862907	15863067	refGene_gn=MAP3K15;refGene_tx=XM_023248797.1
X	15863311	15863424	refGene_gn=MAP3K15;refGene_tx=XM_023248797.1
X	15867652	15867823	refGene_gn=MAP3K15;refGene_tx=XM_023248797.1
X	15889711	15889818	refGene_gn=MAP3K15;refGene_tx=XM_023248797.1
X	15891828	15891997	refGene_gn=MAP3K15;refGene_tx=XM_023248797.1
X	15895022	15895216	refGene_gn=MAP3K15;refGene_tx=XM_023248797.1
X	15911938	15911962	refGene_gn=MAP3K15;refGene_tx=XM_023248797.1
X	15914335	15914475	refGene_gn=MAP3K15;refGene_tx=XM_023248797.1
X	15935586	15935935	refGene_gn=MAP3K15;refGene_tx=XM_023248797.1
X	15954641	15954683
refGene_gn=SH3KBP1;refGene_tx=XM_004000330.4;refGene_gn=SH3KBP1;refGene_tx=XM_011291627.3;refGene_gn=SH3KBP1;refGene_tx=XM_019823636.2;refGene_gn=SH3KBP1;refGene_tx=XM_019823637.2;refGene_gn=SH3KBP1;refGene_tx=XM_019823638.1;refGene_gn=SH3KBP1;refGene_tx=XM_023249325.1;refGene_gn=SH3KBP1;refGene_tx=XM_023249326.1;refGene_gn=SH3KBP1;refGene_tx=XM_023249327.1;refGene_gn=SH3KBP1;refGene_tx=XM_023249328.1;refGene_gn=SH3KBP1;refGene_tx=XM_023249329.1
X	15955679	15955743
refGene_gn=SH3KBP1;refGene_tx=XM_004000330.4;refGene_gn=SH3KBP1;refGene_tx=XM_011291627.3;refGene_gn=SH3KBP1;refGene_tx=XM_019823636.2;refGene_gn=SH3KBP1;refGene_tx=XM_019823637.2;refGene_gn=SH3KBP1;refGene_tx=XM_019823638.1;refGene_gn=SH3KBP1;refGene_tx=XM_023249325.1;refGene_gn=SH3KBP1;refGene_tx=XM_023249326.1;refGene_gn=SH3KBP1;refGene_tx=XM_023249327.1;refGene_gn=SH3KBP1;refGene_tx=XM_023249328.1;refGene_gn=SH3KBP1;refGene_tx=XM_023249329.1
X	15959007	15959279
refGene_gn=SH3KBP1;refGene_tx=XM_004000330.4;refGene_gn=SH3KBP1;refGene_tx=XM_011291627.3;refGene_gn=SH3KBP1;refGene_tx=XM_019823636.2;refGe

ne_gn=SH3KBP1;refGene_tx=XM_019823637.2;refGene_gn=SH3KBP1;refGene_tx=XM_019823638.1;refGene_gn=SH3KBP1;refGene_tx=XM_023249325.1;refGene_gn=SH3KBP1;refGene_tx=XM_023249326.1;refGene_gn=SH3KBP1;refGene_tx=XM_023249327.1;refGene_gn=SH3KBP1;refGene_tx=XM_023249328.1;refGene_gn=SH3KBP1;refGene_tx=XM_023249329.1
X 15963413 15963542
refGene_gn=SH3KBP1;refGene_tx=XM_004000330.4;refGene_gn=SH3KBP1;refGene_tx=XM_011291627.3;refGene_gn=SH3KBP1;refGene_tx=XM_019823636.2;refGene_gn=SH3KBP1;refGene_tx=XM_019823637.2;refGene_gn=SH3KBP1;refGene_tx=XM_019823638.1;refGene_gn=SH3KBP1;refGene_tx=XM_023249325.1;refGene_gn=SH3KBP1;refGene_tx=XM_023249327.1
X 15967344 15967454
refGene_gn=SH3KBP1;refGene_tx=XM_004000330.4;refGene_gn=SH3KBP1;refGene_tx=XM_011291627.3;refGene_gn=SH3KBP1;refGene_tx=XM_019823636.2;refGene_gn=SH3KBP1;refGene_tx=XM_019823637.2;refGene_gn=SH3KBP1;refGene_tx=XM_019823638.1;refGene_gn=SH3KBP1;refGene_tx=XM_023249325.1;refGene_gn=SH3KBP1;refGene_tx=XM_023249326.1;refGene_gn=SH3KBP1;refGene_tx=XM_023249327.1;refGene_gn=SH3KBP1;refGene_tx=XM_023249328.1;refGene_gn=SH3KBP1;refGene_tx=XM_023249329.1
X 15986959 15987045
refGene_gn=SH3KBP1;refGene_tx=XM_004000330.4;refGene_gn=SH3KBP1;refGene_tx=XM_011291627.3;refGene_gn=SH3KBP1;refGene_tx=XM_019823636.2;refGene_gn=SH3KBP1;refGene_tx=XM_019823637.2;refGene_gn=SH3KBP1;refGene_tx=XM_019823638.1;refGene_gn=SH3KBP1;refGene_tx=XM_023249325.1;refGene_gn=SH3KBP1;refGene_tx=XM_023249326.1;refGene_gn=SH3KBP1;refGene_tx=XM_023249327.1;refGene_gn=SH3KBP1;refGene_tx=XM_023249328.1;refGene_gn=SH3KBP1;refGene_tx=XM_023249329.1
X 16003287 16003447
refGene_gn=SH3KBP1;refGene_tx=XM_004000330.4;refGene_gn=SH3KBP1;refGene_tx=XM_011291627.3;refGene_gn=SH3KBP1;refGene_tx=XM_019823636.2;refGene_gn=SH3KBP1;refGene_tx=XM_019823637.2;refGene_gn=SH3KBP1;refGene_tx=XM_019823638.1;refGene_gn=SH3KBP1;refGene_tx=XM_023249325.1;refGene_gn=SH3KBP1;refGene_tx=XM_023249326.1;refGene_gn=SH3KBP1;refGene_tx=XM_023249327.1;refGene_gn=SH3KBP1;refGene_tx=XM_023249328.1;refGene_gn=SH3KBP1;refGene_tx=XM_023249329.1
X 16006228 16006309
refGene_gn=SH3KBP1;refGene_tx=XM_004000330.4;refGene_gn=SH3KBP1;refGene_tx=XM_011291627.3;refGene_gn=SH3KBP1;refGene_tx=XM_019823636.2;refGene_gn=SH3KBP1;refGene_tx=XM_019823637.2;refGene_gn=SH3KBP1;refGene_tx=XM_019823638.1;refGene_gn=SH3KBP1;refGene_tx=XM_023249325.1;refGene_gn=SH3KBP1;refGene_tx=XM_023249326.1;refGene_gn=SH3KBP1;refGene_tx=XM_023249327.1;refGene_gn=SH3KBP1;refGene_tx=XM_023249328.1;refGene_gn=SH3KBP1;refGene_tx=XM_023249329.1
X 16009095 16009147
refGene_gn=SH3KBP1;refGene_tx=XM_004000330.4;refGene_gn=SH3KBP1;refGene_tx=XM_011291627.3;refGene_gn=SH3KBP1;refGene_tx=XM_019823636.2;refGene_gn=SH3KBP1;refGene_tx=XM_019823637.2;refGene_gn=SH3KBP1;refGene_tx=XM_019823638.1;refGene_gn=SH3KBP1;refGene_tx=XM_023249325.1;refGene_gn=SH3KBP1;refGene_tx=XM_023249326.1;refGene_gn=SH3KBP1;refGene_tx=XM_023249327.1;refGene_gn=SH3KBP1;refGene_tx=XM_023249328.1;refGene_gn=SH3K

BP1;refGene_tx=XM_023249329.1
X	16019425	16019533
refGene_gn=SH3KBP1;refGene_tx=XM_004000330.4;refGene_gn=SH3KBP1;refGene_tx=XM_011291627.3;refGene_gn=SH3KBP1;refGene_tx=XM_019823636.2;refGene_gn=SH3KBP1;refGene_tx=XM_019823637.2;refGene_gn=SH3KBP1;refGene_tx=XM_019823638.1;refGene_gn=SH3KBP1;refGene_tx=XM_023249325.1;refGene_gn=SH3KBP1;refGene_tx=XM_023249326.1;refGene_gn=SH3KBP1;refGene_tx=XM_023249327.1;refGene_gn=SH3KBP1;refGene_tx=XM_023249328.1;refGene_gn=SH3KBP1;refGene_tx=XM_023249329.1
X	16040174	16040269
refGene_gn=SH3KBP1;refGene_tx=XM_004000330.4;refGene_gn=SH3KBP1;refGene_tx=XM_011291627.3;refGene_gn=SH3KBP1;refGene_tx=XM_019823636.2;refGene_gn=SH3KBP1;refGene_tx=XM_019823637.2;refGene_gn=SH3KBP1;refGene_tx=XM_019823638.1;refGene_gn=SH3KBP1;refGene_tx=XM_023249325.1;refGene_gn=SH3KBP1;refGene_tx=XM_023249326.1;refGene_gn=SH3KBP1;refGene_tx=XM_023249327.1;refGene_gn=SH3KBP1;refGene_tx=XM_023249328.1;refGene_gn=SH3KBP1;refGene_tx=XM_023249329.1
X	16054046	16054122
refGene_gn=SH3KBP1;refGene_tx=XM_004000330.4;refGene_gn=SH3KBP1;refGene_tx=XM_011291627.3;refGene_gn=SH3KBP1;refGene_tx=XM_019823636.2;refGene_gn=SH3KBP1;refGene_tx=XM_019823637.2;refGene_gn=SH3KBP1;refGene_tx=XM_019823638.1;refGene_gn=SH3KBP1;refGene_tx=XM_023249325.1;refGene_gn=SH3KBP1;refGene_tx=XM_023249326.1;refGene_gn=SH3KBP1;refGene_tx=XM_023249327.1;refGene_gn=SH3KBP1;refGene_tx=XM_023249328.1;refGene_gn=SH3KBP1;refGene_tx=XM_023249329.1
X	16076107	16076122	refGene_gn=SH3KBP1;refGene_tx=XM_004000330.4
X	16087877	16088083
refGene_gn=SH3KBP1;refGene_tx=XM_011291627.3;refGene_gn=SH3KBP1;refGene_tx=XM_019823636.2;refGene_gn=SH3KBP1;refGene_tx=XM_019823637.2;refGene_gn=SH3KBP1;refGene_tx=XM_019823638.1;refGene_gn=SH3KBP1;refGene_tx=XM_023249325.1;refGene_gn=SH3KBP1;refGene_tx=XM_023249326.1;refGene_gn=SH3KBP1;refGene_tx=XM_023249327.1;refGene_gn=SH3KBP1;refGene_tx=XM_023249328.1;refGene_gn=SH3KBP1;refGene_tx=XM_023249329.1
X	16091651	16091726
refGene_gn=SH3KBP1;refGene_tx=XM_011291627.3;refGene_gn=SH3KBP1;refGene_tx=XM_019823636.2;refGene_gn=SH3KBP1;refGene_tx=XM_019823637.2;refGene_gn=SH3KBP1;refGene_tx=XM_019823638.1;refGene_gn=SH3KBP1;refGene_tx=XM_023249325.1;refGene_gn=SH3KBP1;refGene_tx=XM_023249326.1;refGene_gn=SH3KBP1;refGene_tx=XM_023249328.1
X	16099207	16099267
refGene_gn=SH3KBP1;refGene_tx=XM_011291627.3;refGene_gn=SH3KBP1;refGene_tx=XM_019823636.2;refGene_gn=SH3KBP1;refGene_tx=XM_019823637.2;refGene_gn=SH3KBP1;refGene_tx=XM_019823638.1;refGene_gn=SH3KBP1;refGene_tx=XM_023249326.1
X	16099826	16099956
refGene_gn=SH3KBP1;refGene_tx=XM_011291627.3;refGene_gn=SH3KBP1;refGene_tx=XM_019823636.2;refGene_gn=SH3KBP1;refGene_tx=XM_019823637.2;refGene_gn=SH3KBP1;refGene_tx=XM_023249325.1;refGene_gn=SH3KBP1;refGene_tx=XM_023249326.1;refGene_gn=SH3KBP1;refGene_tx=XM_023249327.1;refGene_gn=SH3KBP1;refGene_tx=XM_023249328.1;refGene_gn=SH3KBP1;refGene_tx=XM_02

3249329.1
X	16114270 16114374
refGene_gn=SH3KBP1;refGene_tx=XM_011291627.3;refGene_gn=SH3KBP1;refGene_tx=XM_019823636.2;refGene_gn=SH3KBP1;refGene_tx=XM_019823637.2;refGene_gn=SH3KBP1;refGene_tx=XM_023249325.1;refGene_gn=SH3KBP1;refGene_tx=XM_023249326.1;refGene_gn=SH3KBP1;refGene_tx=XM_023249327.1;refGene_gn=SH3KBP1;refGene_tx=XM_023249328.1;refGene_gn=SH3KBP1;refGene_tx=XM_023249329.1
X	16151746 16151870
refGene_gn=SH3KBP1;refGene_tx=XM_011291627.3;refGene_gn=SH3KBP1;refGene_tx=XM_019823636.2;refGene_gn=SH3KBP1;refGene_tx=XM_019823637.2;refGene_gn=SH3KBP1;refGene_tx=XM_023249325.1;refGene_gn=SH3KBP1;refGene_tx=XM_023249326.1;refGene_gn=SH3KBP1;refGene_tx=XM_023249327.1;refGene_gn=SH3KBP1;refGene_tx=XM_023249328.1;refGene_gn=SH3KBP1;refGene_tx=XM_023249329.1
X	16200536 16200599 refGene_gn=SH3KBP1;refGene_tx=XM_019823636.2
X	16237203 16237361
refGene_gn=SH3KBP1;refGene_tx=XM_011291627.3;refGene_gn=SH3KBP1;refGene_tx=XM_019823637.2;refGene_gn=SH3KBP1;refGene_tx=XM_023249325.1;refGene_gn=SH3KBP1;refGene_tx=XM_023249326.1;refGene_gn=SH3KBP1;refGene_tx=XM_023249327.1;refGene_gn=SH3KBP1;refGene_tx=XM_023249328.1;refGene_gn=SH3KBP1;refGene_tx=XM_023249329.1
X	16287753 16287757
refGene_gn=SH3KBP1;refGene_tx=XM_011291627.3;refGene_gn=SH3KBP1;refGene_tx=XM_023249325.1;refGene_gn=SH3KBP1;refGene_tx=XM_023249326.1;refGene_gn=SH3KBP1;refGene_tx=XM_023249327.1;refGene_gn=SH3KBP1;refGene_tx=XM_023249328.1;refGene_gn=SH3KBP1;refGene_tx=XM_023249329.1
X	16389094 16389344
refGene_gn=MAP7D2;refGene_tx=XM_011291628.2;refGene_gn=MAP7D2;refGene_tx=XM_019823639.2;refGene_gn=MAP7D2;refGene_tx=XM_019823640.1;refGene_gn=MAP7D2;refGene_tx=XM_019823641.2;refGene_gn=MAP7D2;refGene_tx=XM_019823642.1;refGene_gn=MAP7D2;refGene_tx=XM_019823643.1;refGene_gn=MAP7D2;refGene_tx=XM_019823644.1;refGene_gn=MAP7D2;refGene_tx=XM_019823645.2;refGene_gn=MAP7D2;refGene_tx=XM_019823646.2;refGene_gn=MAP7D2;refGene_tx=XM_019823647.2;refGene_gn=MAP7D2;refGene_tx=XM_019823648.1;refGene_gn=MAP7D2;refGene_tx=XM_019823649.1;refGene_gn=MAP7D2;refGene_tx=XM_019823650.1;refGene_gn=MAP7D2;refGene_tx=XM_019823651.1
X	16391746 16391933
refGene_gn=MAP7D2;refGene_tx=XM_011291628.2;refGene_gn=MAP7D2;refGene_tx=XM_019823639.2;refGene_gn=MAP7D2;refGene_tx=XM_019823640.1;refGene_gn=MAP7D2;refGene_tx=XM_019823641.2;refGene_gn=MAP7D2;refGene_tx=XM_019823642.1;refGene_gn=MAP7D2;refGene_tx=XM_019823643.1;refGene_gn=MAP7D2;refGene_tx=XM_019823644.1;refGene_gn=MAP7D2;refGene_tx=XM_019823645.2;refGene_gn=MAP7D2;refGene_tx=XM_019823646.2;refGene_gn=MAP7D2;refGene_tx=XM_019823647.2;refGene_gn=MAP7D2;refGene_tx=XM_019823648.1;refGene_gn=MAP7D2;refGene_tx=XM_019823649.1;refGene_gn=MAP7D2;refGene_tx=XM_019823650.1;refGene_gn=MAP7D2;refGene_tx=XM_019823651.1
X	16392904 16392983
refGene_gn=MAP7D2;refGene_tx=XM_011291628.2;refGene_gn=MAP7D2;refGene_tx=XM_019823639.2;refGene_gn=MAP7D2;refGene_tx=XM_019823640.1;refGene_

gn=MAP7D2;refGene_tx=XM_019823641.2;refGene_gn=MAP7D2;refGene_tx=XM_01
9823642.1;refGene_gn=MAP7D2;refGene_tx=XM_019823643.1;refGene_gn=MAP7D
2;refGene_tx=XM_019823644.1;refGene_gn=MAP7D2;refGene_tx=XM_019823645.
2;refGene_gn=MAP7D2;refGene_tx=XM_019823646.2;refGene_gn=MAP7D2;refGen
e_tx=XM_019823647.2;refGene_gn=MAP7D2;refGene_tx=XM_019823648.1;refGen
e_gn=MAP7D2;refGene_tx=XM_019823649.1;refGene_gn=MAP7D2;refGene_tx=XM_
019823650.1;refGene_gn=MAP7D2;refGene_tx=XM_019823651.1
X 16393425 16393482
refGene_gn=MAP7D2;refGene_tx=XM_011291628.2;refGene_gn=MAP7D2;refGene_
tx=XM_019823639.2;refGene_gn=MAP7D2;refGene_tx=XM_019823640.1;refGene_
gn=MAP7D2;refGene_tx=XM_019823641.2;refGene_gn=MAP7D2;refGene_tx=XM_01
9823642.1;refGene_gn=MAP7D2;refGene_tx=XM_019823643.1;refGene_gn=MAP7D
2;refGene_tx=XM_019823644.1;refGene_gn=MAP7D2;refGene_tx=XM_019823645.
2;refGene_gn=MAP7D2;refGene_tx=XM_019823646.2;refGene_gn=MAP7D2;refGen
e_tx=XM_019823647.2;refGene_gn=MAP7D2;refGene_tx=XM_019823648.1;refGen
e_gn=MAP7D2;refGene_tx=XM_019823649.1;refGene_gn=MAP7D2;refGene_tx=XM_
019823650.1;refGene_gn=MAP7D2;refGene_tx=XM_019823651.1
X 16394543 16394648
refGene_gn=MAP7D2;refGene_tx=XM_011291628.2;refGene_gn=MAP7D2;refGene_
tx=XM_019823639.2;refGene_gn=MAP7D2;refGene_tx=XM_019823640.1;refGene_
gn=MAP7D2;refGene_tx=XM_019823641.2;refGene_gn=MAP7D2;refGene_tx=XM_01
9823642.1;refGene_gn=MAP7D2;refGene_tx=XM_019823643.1;refGene_gn=MAP7D
2;refGene_tx=XM_019823644.1;refGene_gn=MAP7D2;refGene_tx=XM_019823645.
2;refGene_gn=MAP7D2;refGene_tx=XM_019823646.2;refGene_gn=MAP7D2;refGen
e_tx=XM_019823647.2;refGene_gn=MAP7D2;refGene_tx=XM_019823648.1;refGen
e_gn=MAP7D2;refGene_tx=XM_019823649.1;refGene_gn=MAP7D2;refGene_tx=XM_
019823650.1;refGene_gn=MAP7D2;refGene_tx=XM_019823651.1
X 16395414 16395640
refGene_gn=MAP7D2;refGene_tx=XM_011291628.2;refGene_gn=MAP7D2;refGene_
tx=XM_019823639.2;refGene_gn=MAP7D2;refGene_tx=XM_019823640.1;refGene_
gn=MAP7D2;refGene_tx=XM_019823641.2;refGene_gn=MAP7D2;refGene_tx=XM_01
9823642.1;refGene_gn=MAP7D2;refGene_tx=XM_019823643.1;refGene_gn=MAP7D
2;refGene_tx=XM_019823644.1;refGene_gn=MAP7D2;refGene_tx=XM_019823645.
2;refGene_gn=MAP7D2;refGene_tx=XM_019823646.2;refGene_gn=MAP7D2;refGen
e_tx=XM_019823647.2;refGene_gn=MAP7D2;refGene_tx=XM_019823648.1;refGen
e_gn=MAP7D2;refGene_tx=XM_019823649.1;refGene_gn=MAP7D2;refGene_tx=XM_
019823650.1;refGene_gn=MAP7D2;refGene_tx=XM_019823651.1
X 16406183 16406316
refGene_gn=MAP7D2;refGene_tx=XM_011291628.2;refGene_gn=MAP7D2;refGene_
tx=XM_019823639.2;refGene_gn=MAP7D2;refGene_tx=XM_019823640.1;refGene_
gn=MAP7D2;refGene_tx=XM_019823641.2;refGene_gn=MAP7D2;refGene_tx=XM_01
9823642.1;refGene_gn=MAP7D2;refGene_tx=XM_019823643.1;refGene_gn=MAP7D
2;refGene_tx=XM_019823644.1;refGene_gn=MAP7D2;refGene_tx=XM_019823645.
2;refGene_gn=MAP7D2;refGene_tx=XM_019823646.2;refGene_gn=MAP7D2;refGen
e_tx=XM_019823647.2;refGene_gn=MAP7D2;refGene_tx=XM_019823648.1;refGen
e_gn=MAP7D2;refGene_tx=XM_019823649.1;refGene_gn=MAP7D2;refGene_tx=XM_
019823650.1;refGene_gn=MAP7D2;refGene_tx=XM_019823651.1
X 16406916 16407185
refGene_gn=MAP7D2;refGene_tx=XM_011291628.2;refGene_gn=MAP7D2;refGene_
tx=XM_019823639.2;refGene_gn=MAP7D2;refGene_tx=XM_019823640.1;refGene_

gn=MAP7D2;refGene_tx=XM_019823641.2;refGene_gn=MAP7D2;refGene_tx=XM_019823642.1;refGene_gn=MAP7D2;refGene_tx=XM_019823643.1;refGene_gn=MAP7D2;refGene_tx=XM_019823644.1;refGene_gn=MAP7D2;refGene_tx=XM_019823645.2;refGene_gn=MAP7D2;refGene_tx=XM_019823646.2;refGene_gn=MAP7D2;refGene_tx=XM_019823647.2;refGene_gn=MAP7D2;refGene_tx=XM_019823648.1;refGene_gn=MAP7D2;refGene_tx=XM_019823649.1;refGene_gn=MAP7D2;refGene_tx=XM_019823650.1;refGene_gn=MAP7D2;refGene_tx=XM_019823651.1

X 16416181 16416309

refGene_gn=MAP7D2;refGene_tx=XM_011291628.2;refGene_gn=MAP7D2;refGene_tx=XM_019823639.2;refGene_gn=MAP7D2;refGene_tx=XM_019823640.1;refGene_gn=MAP7D2;refGene_tx=XM_019823641.2;refGene_gn=MAP7D2;refGene_tx=XM_019823642.1;refGene_gn=MAP7D2;refGene_tx=XM_019823643.1;refGene_gn=MAP7D2;refGene_tx=XM_019823644.1;refGene_gn=MAP7D2;refGene_tx=XM_019823645.2;refGene_gn=MAP7D2;refGene_tx=XM_019823646.2;refGene_gn=MAP7D2;refGene_tx=XM_019823647.2;refGene_gn=MAP7D2;refGene_tx=XM_019823648.1;refGene_gn=MAP7D2;refGene_tx=XM_019823649.1;refGene_gn=MAP7D2;refGene_tx=XM_019823650.1;refGene_gn=MAP7D2;refGene_tx=XM_019823651.1

X 16419529 16419684

refGene_gn=MAP7D2;refGene_tx=XM_011291628.2;refGene_gn=MAP7D2;refGene_tx=XM_019823639.2;refGene_gn=MAP7D2;refGene_tx=XM_019823640.1;refGene_gn=MAP7D2;refGene_tx=XM_019823641.2;refGene_gn=MAP7D2;refGene_tx=XM_019823642.1;refGene_gn=MAP7D2;refGene_tx=XM_019823643.1;refGene_gn=MAP7D2;refGene_tx=XM_019823644.1;refGene_gn=MAP7D2;refGene_tx=XM_019823645.2;refGene_gn=MAP7D2;refGene_tx=XM_019823646.2;refGene_gn=MAP7D2;refGene_tx=XM_019823647.2;refGene_gn=MAP7D2;refGene_tx=XM_019823648.1;refGene_gn=MAP7D2;refGene_tx=XM_019823649.1;refGene_gn=MAP7D2;refGene_tx=XM_019823650.1;refGene_gn=MAP7D2;refGene_tx=XM_019823651.1

X 16425037 16425160

refGene_gn=MAP7D2;refGene_tx=XM_011291628.2;refGene_gn=MAP7D2;refGene_tx=XM_019823639.2;refGene_gn=MAP7D2;refGene_tx=XM_019823640.1;refGene_gn=MAP7D2;refGene_tx=XM_019823641.2;refGene_gn=MAP7D2;refGene_tx=XM_019823643.1;refGene_gn=MAP7D2;refGene_tx=XM_019823644.1;refGene_gn=MAP7D2;refGene_tx=XM_019823645.2;refGene_gn=MAP7D2;refGene_tx=XM_019823646.2;refGene_gn=MAP7D2;refGene_tx=XM_019823647.2

X 16427254 16427365

refGene_gn=MAP7D2;refGene_tx=XM_011291628.2;refGene_gn=MAP7D2;refGene_tx=XM_019823639.2;refGene_gn=MAP7D2;refGene_tx=XM_019823640.1;refGene_gn=MAP7D2;refGene_tx=XM_019823641.2;refGene_gn=MAP7D2;refGene_tx=XM_019823642.1;refGene_gn=MAP7D2;refGene_tx=XM_019823643.1;refGene_gn=MAP7D2;refGene_tx=XM_019823644.1;refGene_gn=MAP7D2;refGene_tx=XM_019823645.2;refGene_gn=MAP7D2;refGene_tx=XM_019823646.2;refGene_gn=MAP7D2;refGene_tx=XM_019823647.2;refGene_gn=MAP7D2;refGene_tx=XM_019823648.1;refGene_gn=MAP7D2;refGene_tx=XM_019823649.1;refGene_gn=MAP7D2;refGene_tx=XM_019823650.1

X 16429695 16429794

refGene_gn=MAP7D2;refGene_tx=XM_019823639.2;refGene_gn=MAP7D2;refGene_tx=XM_019823640.1;refGene_gn=MAP7D2;refGene_tx=XM_019823641.2;refGene_gn=MAP7D2;refGene_tx=XM_019823642.1;refGene_gn=MAP7D2;refGene_tx=XM_019823643.1;refGene_gn=MAP7D2;refGene_tx=XM_019823644.1;refGene_gn=MAP7D2;refGene_tx=XM_019823645.2;refGene_gn=MAP7D2;refGene_tx=XM_019823646.

2;refGene_gn=MAP7D2;refGene_tx=XM_019823647.2;refGene_gn=MAP7D2;refGene_tx=XM_019823648.1
X 16430630 16430742
refGene_gn=MAP7D2;refGene_tx=XM_011291628.2;refGene_gn=MAP7D2;refGene_tx=XM_019823639.2;refGene_gn=MAP7D2;refGene_tx=XM_019823640.1;refGene_gn=MAP7D2;refGene_tx=XM_019823641.2;refGene_gn=MAP7D2;refGene_tx=XM_019823642.1;refGene_gn=MAP7D2;refGene_tx=XM_019823643.1;refGene_gn=MAP7D2;refGene_tx=XM_019823644.1;refGene_gn=MAP7D2;refGene_tx=XM_019823645.2;refGene_gn=MAP7D2;refGene_tx=XM_019823646.2;refGene_gn=MAP7D2;refGene_tx=XM_019823647.2;refGene_gn=MAP7D2;refGene_tx=XM_019823648.1;refGene_gn=MAP7D2;refGene_tx=XM_019823649.1;refGene_gn=MAP7D2;refGene_tx=XM_019823650.1;refGene_gn=MAP7D2;refGene_tx=XM_019823651.1
X 16437489 16437653
refGene_gn=MAP7D2;refGene_tx=XM_011291628.2;refGene_gn=MAP7D2;refGene_tx=XM_019823639.2;refGene_gn=MAP7D2;refGene_tx=XM_019823640.1;refGene_gn=MAP7D2;refGene_tx=XM_019823641.2;refGene_gn=MAP7D2;refGene_tx=XM_019823642.1;refGene_gn=MAP7D2;refGene_tx=XM_019823643.1;refGene_gn=MAP7D2;refGene_tx=XM_019823644.1;refGene_gn=MAP7D2;refGene_tx=XM_019823645.2;refGene_gn=MAP7D2;refGene_tx=XM_019823646.2;refGene_gn=MAP7D2;refGene_tx=XM_019823647.2;refGene_gn=MAP7D2;refGene_tx=XM_019823648.1;refGene_gn=MAP7D2;refGene_tx=XM_019823649.1;refGene_gn=MAP7D2;refGene_tx=XM_019823650.1;refGene_gn=MAP7D2;refGene_tx=XM_019823651.1
X 16438704 16438782
refGene_gn=MAP7D2;refGene_tx=XM_011291628.2;refGene_gn=MAP7D2;refGene_tx=XM_019823639.2;refGene_gn=MAP7D2;refGene_tx=XM_019823640.1;refGene_gn=MAP7D2;refGene_tx=XM_019823641.2;refGene_gn=MAP7D2;refGene_tx=XM_019823642.1;refGene_gn=MAP7D2;refGene_tx=XM_019823643.1;refGene_gn=MAP7D2;refGene_tx=XM_019823644.1;refGene_gn=MAP7D2;refGene_tx=XM_019823645.2;refGene_gn=MAP7D2;refGene_tx=XM_019823646.2;refGene_gn=MAP7D2;refGene_tx=XM_019823647.2;refGene_gn=MAP7D2;refGene_tx=XM_019823648.1;refGene_gn=MAP7D2;refGene_tx=XM_019823649.1;refGene_gn=MAP7D2;refGene_tx=XM_019823650.1;refGene_gn=MAP7D2;refGene_tx=XM_019823651.1
X 16491053 16491180
refGene_gn=MAP7D2;refGene_tx=XM_011291628.2;refGene_gn=MAP7D2;refGene_tx=XM_019823639.2;refGene_gn=MAP7D2;refGene_tx=XM_019823640.1;refGene_gn=MAP7D2;refGene_tx=XM_019823641.2;refGene_gn=MAP7D2;refGene_tx=XM_019823642.1;refGene_gn=MAP7D2;refGene_tx=XM_019823648.1;refGene_gn=MAP7D2;refGene_tx=XM_019823649.1;refGene_gn=MAP7D2;refGene_tx=XM_019823650.1;refGene_gn=MAP7D2;refGene_tx=XM_019823651.1
X 16502302 16502308 refGene_gn=EIF1AX;refGene_tx=XM_004000333.5
X 16504500 16504592 refGene_gn=EIF1AX;refGene_tx=XM_004000333.5
X 16507220 16507302 refGene_gn=EIF1AX;refGene_tx=XM_004000333.5
X 16510412 16510463 refGene_gn=EIF1AX;refGene_tx=XM_004000333.5
X 16511530 16511634 refGene_gn=EIF1AX;refGene_tx=XM_004000333.5
X 16514418 16514502 refGene_gn=EIF1AX;refGene_tx=XM_004000333.5
X 16520921 16520937 refGene_gn=EIF1AX;refGene_tx=XM_004000333.5
X 16532525 16532648
refGene_gn=RPS6KA3;refGene_tx=XM_004000334.5;refGene_gn=RPS6KA3;refGene_tx=XM_006943610.4;refGene_gn=RPS6KA3;refGene_tx=XM_006943611.4;refGene_gn=RPS6KA3;refGene_tx=XM_006943612.4;refGene_gn=RPS6KA3;refGene_tx=

XM_019823652.2
X	16533938 16534079
refGene_gn=RPS6KA3;refGene_tx=XM_004000334.5;refGene_gn=RPS6KA3;refGene_tx=XM_006943610.4;refGene_gn=RPS6KA3;refGene_tx=XM_006943611.4;refGene_gn=RPS6KA3;refGene_tx=XM_006943612.4;refGene_gn=RPS6KA3;refGene_tx=XM_019823652.2
X	16539283 16539401
refGene_gn=RPS6KA3;refGene_tx=XM_004000334.5;refGene_gn=RPS6KA3;refGene_tx=XM_006943610.4;refGene_gn=RPS6KA3;refGene_tx=XM_006943611.4;refGene_gn=RPS6KA3;refGene_tx=XM_006943612.4;refGene_gn=RPS6KA3;refGene_tx=XM_019823652.2
X	16539937 16540014
refGene_gn=RPS6KA3;refGene_tx=XM_004000334.5;refGene_gn=RPS6KA3;refGene_tx=XM_006943610.4;refGene_gn=RPS6KA3;refGene_tx=XM_006943611.4;refGene_gn=RPS6KA3;refGene_tx=XM_006943612.4;refGene_gn=RPS6KA3;refGene_tx=XM_019823652.2
X	16543107 16543269
refGene_gn=RPS6KA3;refGene_tx=XM_004000334.5;refGene_gn=RPS6KA3;refGene_tx=XM_006943610.4;refGene_gn=RPS6KA3;refGene_tx=XM_006943611.4;refGene_gn=RPS6KA3;refGene_tx=XM_006943612.4;refGene_gn=RPS6KA3;refGene_tx=XM_019823652.2
X	16545259 16545418
refGene_gn=RPS6KA3;refGene_tx=XM_004000334.5;refGene_gn=RPS6KA3;refGene_tx=XM_006943610.4;refGene_gn=RPS6KA3;refGene_tx=XM_006943611.4;refGene_gn=RPS6KA3;refGene_tx=XM_006943612.4;refGene_gn=RPS6KA3;refGene_tx=XM_019823652.2
X	16547573 16547663
refGene_gn=RPS6KA3;refGene_tx=XM_004000334.5;refGene_gn=RPS6KA3;refGene_tx=XM_006943610.4;refGene_gn=RPS6KA3;refGene_tx=XM_006943611.4;refGene_gn=RPS6KA3;refGene_tx=XM_006943612.4;refGene_gn=RPS6KA3;refGene_tx=XM_019823652.2
X	16550209 16550335
refGene_gn=RPS6KA3;refGene_tx=XM_004000334.5;refGene_gn=RPS6KA3;refGene_tx=XM_006943610.4;refGene_gn=RPS6KA3;refGene_tx=XM_006943611.4;refGene_gn=RPS6KA3;refGene_tx=XM_006943612.4;refGene_gn=RPS6KA3;refGene_tx=XM_019823652.2
X	16552333 16552458
refGene_gn=RPS6KA3;refGene_tx=XM_004000334.5;refGene_gn=RPS6KA3;refGene_tx=XM_006943610.4;refGene_gn=RPS6KA3;refGene_tx=XM_006943611.4;refGene_gn=RPS6KA3;refGene_tx=XM_006943612.4;refGene_gn=RPS6KA3;refGene_tx=XM_019823652.2
X	16555814 16555917
refGene_gn=RPS6KA3;refGene_tx=XM_004000334.5;refGene_gn=RPS6KA3;refGene_tx=XM_006943610.4;refGene_gn=RPS6KA3;refGene_tx=XM_006943611.4;refGene_gn=RPS6KA3;refGene_tx=XM_006943612.4;refGene_gn=RPS6KA3;refGene_tx=XM_019823652.2
X	16555991 16556056
refGene_gn=RPS6KA3;refGene_tx=XM_004000334.5;refGene_gn=RPS6KA3;refGene_tx=XM_006943610.4;refGene_gn=RPS6KA3;refGene_tx=XM_006943611.4;refGene_gn=RPS6KA3;refGene_tx=XM_006943612.4;refGene_gn=RPS6KA3;refGene_tx=

XM_019823652.2
X	16556250	16556339
refGene_gn=RPS6KA3;refGene_tx=XM_004000334.5;refGene_gn=RPS6KA3;refGene_tx=XM_006943610.4;refGene_gn=RPS6KA3;refGene_tx=XM_006943611.4;refGene_gn=RPS6KA3;refGene_tx=XM_006943612.4;refGene_gn=RPS6KA3;refGene_tx=XM_019823652.2
X	16566010	16566081
refGene_gn=RPS6KA3;refGene_tx=XM_004000334.5;refGene_gn=RPS6KA3;refGene_tx=XM_006943610.4;refGene_gn=RPS6KA3;refGene_tx=XM_006943611.4;refGene_gn=RPS6KA3;refGene_tx=XM_006943612.4;refGene_gn=RPS6KA3;refGene_tx=XM_019823652.2
X	16566258	16566401
refGene_gn=RPS6KA3;refGene_tx=XM_004000334.5;refGene_gn=RPS6KA3;refGene_tx=XM_006943610.4;refGene_gn=RPS6KA3;refGene_tx=XM_006943611.4;refGene_gn=RPS6KA3;refGene_tx=XM_006943612.4;refGene_gn=RPS6KA3;refGene_tx=XM_019823652.2
X	16566609	16566647
refGene_gn=RPS6KA3;refGene_tx=XM_004000334.5;refGene_gn=RPS6KA3;refGene_tx=XM_006943610.4;refGene_gn=RPS6KA3;refGene_tx=XM_006943611.4;refGene_gn=RPS6KA3;refGene_tx=XM_006943612.4;refGene_gn=RPS6KA3;refGene_tx=XM_019823652.2
X	16570902	16571009
refGene_gn=RPS6KA3;refGene_tx=XM_004000334.5;refGene_gn=RPS6KA3;refGene_tx=XM_006943610.4;refGene_gn=RPS6KA3;refGene_tx=XM_006943611.4;refGene_gn=RPS6KA3;refGene_tx=XM_006943612.4;refGene_gn=RPS6KA3;refGene_tx=XM_019823652.2
X	16571610	16571690
refGene_gn=RPS6KA3;refGene_tx=XM_004000334.5;refGene_gn=RPS6KA3;refGene_tx=XM_006943610.4;refGene_gn=RPS6KA3;refGene_tx=XM_006943611.4;refGene_gn=RPS6KA3;refGene_tx=XM_006943612.4;refGene_gn=RPS6KA3;refGene_tx=XM_019823652.2
X	16572495	16572576
refGene_gn=RPS6KA3;refGene_tx=XM_004000334.5;refGene_gn=RPS6KA3;refGene_tx=XM_006943610.4;refGene_gn=RPS6KA3;refGene_tx=XM_006943611.4;refGene_gn=RPS6KA3;refGene_tx=XM_006943612.4;refGene_gn=RPS6KA3;refGene_tx=XM_019823652.2
X	16581694	16581776
refGene_gn=RPS6KA3;refGene_tx=XM_004000334.5;refGene_gn=RPS6KA3;refGene_tx=XM_006943610.4;refGene_gn=RPS6KA3;refGene_tx=XM_006943611.4;refGene_gn=RPS6KA3;refGene_tx=XM_006943612.4;refGene_gn=RPS6KA3;refGene_tx=XM_019823652.2
X	16587363	16587480
refGene_gn=RPS6KA3;refGene_tx=XM_004000334.5;refGene_gn=RPS6KA3;refGene_tx=XM_006943610.4;refGene_gn=RPS6KA3;refGene_tx=XM_006943611.4;refGene_gn=RPS6KA3;refGene_tx=XM_006943612.4;refGene_gn=RPS6KA3;refGene_tx=XM_019823652.2
X	16598186	16598225	refGene_gn=RPS6KA3;refGene_tx=XM_006943612.4
X	16616491	16616548
refGene_gn=RPS6KA3;refGene_tx=XM_004000334.5;refGene_gn=RPS6KA3;refGene_tx=XM_006943610.4;refGene_gn=RPS6KA3;refGene_tx=XM_006943611.4;refGe

ne_gn=RPS6KA3;refGene_tx=XM_019823652.2
X	16647952	16648021	refGene_gn=RPS6KA3;refGene_tx=XM_004000334.5
X	17816685	17816749	refGene_gn=CNKSR2;refGene_tx=XM_011291630.3
X	17885599	17885763	refGene_gn=CNKSR2;refGene_tx=XM_011291630.3
X	17904397	17904600	refGene_gn=CNKSR2;refGene_tx=XM_011291630.3
X	17918392	17918480	refGene_gn=CNKSR2;refGene_tx=XM_011291630.3
X	17945983	17946025	refGene_gn=CNKSR2;refGene_tx=XM_011291630.3
X	17963173	17963293	refGene_gn=CNKSR2;refGene_tx=XM_011291630.3
X	17971716	17971776	refGene_gn=CNKSR2;refGene_tx=XM_011291630.3
X	17974547	17974616	refGene_gn=CNKSR2;refGene_tx=XM_011291630.3
X	17989086	17989233	refGene_gn=CNKSR2;refGene_tx=XM_011291630.3
X	17999401	17999419	refGene_gn=CNKSR2;refGene_tx=XM_011291630.3
X	17999420	17999536	refGene_gn=CNKSR2;refGene_tx=XM_011291630.3
X	18004161	18004373	refGene_gn=CNKSR2;refGene_tx=XM_011291630.3
X	18022059	18022149	refGene_gn=CNKSR2;refGene_tx=XM_011291630.3
X	18023679	18023894	refGene_gn=CNKSR2;refGene_tx=XM_011291630.3
X	18047576	18047625	refGene_gn=CNKSR2;refGene_tx=XM_011291630.3
X	18048046	18048219	refGene_gn=CNKSR2;refGene_tx=XM_011291630.3
X	18050978	18051052	refGene_gn=CNKSR2;refGene_tx=XM_011291630.3
X	18051348	18051420	refGene_gn=CNKSR2;refGene_tx=XM_011291630.3
X	18058501	18058569	refGene_gn=CNKSR2;refGene_tx=XM_011291630.3
X	18065493	18065594	refGene_gn=CNKSR2;refGene_tx=XM_011291630.3
X	18067938	18068491	refGene_gn=CNKSR2;refGene_tx=XM_011291630.3
X	18109696	18109893	refGene_gn=CNKSR2;refGene_tx=XM_011291630.3
X	18112978	18113194	refGene_gn=CNKSR2;refGene_tx=XM_011291630.3
X	18116537	18117776	refGene_gn=KLHL34;refGene_tx=XM_011291629.3
X	18117777	18118224	refGene_gn=KLHL34;refGene_tx=XM_011291629.3
X	18118225	18118492	refGene_gn=KLHL34;refGene_tx=XM_011291629.3
X	18213308	18213443	refGene_gn=SMPX;refGene_tx=XM_006943622.2
X	18219579	18219666	refGene_gn=SMPX;refGene_tx=XM_006943622.2
X	18231324	18231381	refGene_gn=SMPX;refGene_tx=XM_006943622.2
X	18231730	18231904	refGene_gn=SMPX;refGene_tx=XM_006943622.2
X	18321578	18321653	refGene_gn=MBTPS2;refGene_tx=XM_023249207.1
X	18324983	18325132	refGene_gn=MBTPS2;refGene_tx=XM_023249207.1
X	18327897	18328025	refGene_gn=MBTPS2;refGene_tx=XM_023249207.1
X	18328062	18328148	refGene_gn=MBTPS2;refGene_tx=XM_023249207.1
X	18334962	18335066	refGene_gn=MBTPS2;refGene_tx=XM_023249207.1
X	18336741	18336869	refGene_gn=MBTPS2;refGene_tx=XM_023249207.1
X	18338874	18340560	refGene_gn=YY2;refGene_tx=XM_019823656.2
X	18345235	18345354	refGene_gn=MBTPS2;refGene_tx=XM_023249207.1
X	18346259	18346440	refGene_gn=MBTPS2;refGene_tx=XM_023249207.1
X	18355521	18355616	refGene_gn=MBTPS2;refGene_tx=XM_023249207.1
X	18355979	18356175	refGene_gn=MBTPS2;refGene_tx=XM_023249207.1
X	18358539	18358615	refGene_gn=MBTPS2;refGene_tx=XM_023249207.1
X	18359907	18360130	refGene_gn=MBTPS2;refGene_tx=XM_023249207.1
X	18423559	18423941	refGene_gn=SMS;refGene_tx=XM_023249206.1
X	18447758	18447879	refGene_gn=SMS;refGene_tx=XM_023249206.1
X	18451579	18451670	refGene_gn=SMS;refGene_tx=XM_023249206.1
X	18452206	18452271	refGene_gn=SMS;refGene_tx=XM_023249206.1
X	18457472	18457648	refGene_gn=SMS;refGene_tx=XM_023249206.1

X 18458433 18458588 refGene_gn=SMS;refGene_tx=XM_023249206.1
X 18459583 18459673 refGene_gn=SMS;refGene_tx=XM_023249206.1
X 18468912 18469027 refGene_gn=SMS;refGene_tx=XM_023249206.1
X 18470257 18470337 refGene_gn=SMS;refGene_tx=XM_023249206.1
X 18475428 18475544 refGene_gn=SMS;refGene_tx=XM_023249206.1
X 18477392 18477432 refGene_gn=SMS;refGene_tx=XM_023249206.1
X 18506426 18506544 refGene_gn=PHEX;refGene_tx=XM_023249082.1
X 18511227 18511296 refGene_gn=PHEX;refGene_tx=XM_023249082.1
X 18519236 18519398 refGene_gn=PHEX;refGene_tx=XM_023249082.1
X 18547942 18548029 refGene_gn=PHEX;refGene_tx=XM_023249082.1
X 18549085 18549312 refGene_gn=PHEX;refGene_tx=XM_023249082.1
X 18558413 18558482 refGene_gn=PHEX;refGene_tx=XM_023249082.1
X 18561072 18561189 refGene_gn=PHEX;refGene_tx=XM_023249082.1
X 18564465 18564549 refGene_gn=PHEX;refGene_tx=XM_023249082.1
X 18565769 18565915 refGene_gn=PHEX;refGene_tx=XM_023249082.1
X 18572969 18573063 refGene_gn=PHEX;refGene_tx=XM_023249082.1
X 18575503 18575632 refGene_gn=PHEX;refGene_tx=XM_023249082.1
X 18615597 18615699 refGene_gn=PHEX;refGene_tx=XM_023249082.1
X 18654146 18654224 refGene_gn=PHEX;refGene_tx=XM_023249082.1
X 18662605 18662709 refGene_gn=PHEX;refGene_tx=XM_023249082.1
X 18671723 18671782 refGene_gn=PHEX;refGene_tx=XM_023249082.1
X 18698865 18698920 refGene_gn=PHEX;refGene_tx=XM_023249082.1
X 18704228 18704296 refGene_gn=PHEX;refGene_tx=XM_023249082.1
X 18707803 18707934 refGene_gn=PHEX;refGene_tx=XM_023249082.1
X 18713647 18713713 refGene_gn=PHEX;refGene_tx=XM_023249082.1
X 18714715 18714820 refGene_gn=PHEX;refGene_tx=XM_023249082.1
X 18719453 18719530 refGene_gn=PHEX;refGene_tx=XM_023249082.1
X 18722072 18722175 refGene_gn=PHEX;refGene_tx=XM_023249082.1
X 19497658 19497727 refGene_gn=DDX53;refGene_tx=XM_004000371.3
X 19497905 19498058 refGene_gn=DDX53;refGene_tx=XM_004000371.3
X 19508664 19509312 refGene_gn=DDX53;refGene_tx=XM_004000371.3
X 19509313 19509461 refGene_gn=DDX53;refGene_tx=XM_004000371.3
X 19509463 19510581 refGene_gn=DDX53;refGene_tx=XM_004000371.3
X 19830407 19830758 refGene_gn=PTCHD1;refGene_tx=XM_004000355.3
X 19866874 19867535 refGene_gn=PTCHD1;refGene_tx=XM_004000355.3
X 19879067 19880722 refGene_gn=PTCHD1;refGene_tx=XM_004000355.3
X 20032551 20032947
refGene_gn=LOC101097703;refGene_tx=XM_011291633.3
X 20037756 20037870
refGene_gn=LOC101092829;refGene_tx=XM_011291653.3
X 20037871 20038024
refGene_gn=LOC101092829;refGene_tx=XM_011291653.3
X 20038025 20038457
refGene_gn=LOC101092829;refGene_tx=XM_011291653.3
X 20117952 20118151 refGene_gn=PRDX4;refGene_tx=XM_004000357.5
X 20120668 20120815 refGene_gn=PRDX4;refGene_tx=XM_004000357.5
X 20124322 20124440 refGene_gn=PRDX4;refGene_tx=XM_004000357.5
X 20127172 20127289 refGene_gn=PRDX4;refGene_tx=XM_004000357.5
X 20129720 20129843 refGene_gn=PRDX4;refGene_tx=XM_004000357.5
X 20134441 20134572 refGene_gn=PRDX4;refGene_tx=XM_004000357.5

X 20135584 20135619 refGene_gn=PRDX4;refGene_tx=XM_004000357.5
X 20140685 20140736 refGene_gn=PRDX4;refGene_tx=XM_004000357.5
X 20149629 20149718
refGene_gn=ACOT9;refGene_tx=XM_004000358.3;refGene_gn=ACOT9;refGene_tx=XM_004000359.3;refGene_gn=ACOT9;refGene_tx=XM_019823668.1
X 20150169 20150320
refGene_gn=ACOT9;refGene_tx=XM_004000358.3;refGene_gn=ACOT9;refGene_tx=XM_004000359.3;refGene_gn=ACOT9;refGene_tx=XM_019823668.1
X 20150485 20150573
refGene_gn=ACOT9;refGene_tx=XM_004000358.3;refGene_gn=ACOT9;refGene_tx=XM_004000359.3;refGene_gn=ACOT9;refGene_tx=XM_019823668.1
X 20150996 20151082
refGene_gn=ACOT9;refGene_tx=XM_004000358.3;refGene_gn=ACOT9;refGene_tx=XM_004000359.3;refGene_gn=ACOT9;refGene_tx=XM_019823668.1
X 20151258 20151349
refGene_gn=ACOT9;refGene_tx=XM_004000358.3;refGene_gn=ACOT9;refGene_tx=XM_004000359.3;refGene_gn=ACOT9;refGene_tx=XM_019823668.1
X 20151511 20151623
refGene_gn=ACOT9;refGene_tx=XM_004000358.3;refGene_gn=ACOT9;refGene_tx=XM_004000359.3;refGene_gn=ACOT9;refGene_tx=XM_019823668.1
X 20152356 20152424
refGene_gn=ACOT9;refGene_tx=XM_004000358.3;refGene_gn=ACOT9;refGene_tx=XM_004000359.3;refGene_gn=ACOT9;refGene_tx=XM_019823668.1
X 20154517 20154591
refGene_gn=ACOT9;refGene_tx=XM_004000358.3;refGene_gn=ACOT9;refGene_tx=XM_004000359.3;refGene_gn=ACOT9;refGene_tx=XM_019823668.1
X 20160605 20160709
refGene_gn=ACOT9;refGene_tx=XM_004000358.3;refGene_gn=ACOT9;refGene_tx=XM_004000359.3;refGene_gn=ACOT9;refGene_tx=XM_019823668.1
X 20161359 20161443
refGene_gn=ACOT9;refGene_tx=XM_004000358.3;refGene_gn=ACOT9;refGene_tx=XM_004000359.3;refGene_gn=ACOT9;refGene_tx=XM_019823668.1
X 20165271 20165309
refGene_gn=ACOT9;refGene_tx=XM_004000358.3;refGene_gn=ACOT9;refGene_tx=XM_004000359.3;refGene_gn=ACOT9;refGene_tx=XM_019823668.1
X 20165563 20165734
refGene_gn=ACOT9;refGene_tx=XM_004000358.3;refGene_gn=ACOT9;refGene_tx=XM_004000359.3;refGene_gn=ACOT9;refGene_tx=XM_019823668.1
X 20167860 20167906
refGene_gn=ACOT9;refGene_tx=XM_004000358.3;refGene_gn=ACOT9;refGene_tx=XM_004000359.3;refGene_gn=ACOT9;refGene_tx=XM_019823668.1
X 20168525 20168552
refGene_gn=ACOT9;refGene_tx=XM_004000359.3;refGene_gn=ACOT9;refGene_tx=XM_019823668.1
X 20170238 20170339
refGene_gn=ACOT9;refGene_tx=XM_004000358.3;refGene_gn=ACOT9;refGene_tx=XM_004000359.3;refGene_gn=ACOT9;refGene_tx=XM_019823668.1
X 20174040 20174060
refGene_gn=ACOT9;refGene_tx=XM_004000358.3;refGene_gn=ACOT9;refGene_tx=XM_004000359.3

X	20205184	20205250	refGene_gn=SAT1;refGene_tx=XM_004000360.4
X	20205487	20205539	refGene_gn=SAT1;refGene_tx=XM_004000360.4
X	20205629	20205713	refGene_gn=SAT1;refGene_tx=XM_004000360.4
X	20207157	20207259	refGene_gn=SAT1;refGene_tx=XM_004000360.4
X	20207352	20207393	refGene_gn=SAT1;refGene_tx=XM_004000360.4
X	20207518	20207689	refGene_gn=SAT1;refGene_tx=XM_004000360.4
X	20253223	20253259	refGene_gn=APOO;refGene_tx=XM_004000362.5
X	20263832	20263913	refGene_gn=APOO;refGene_tx=XM_004000362.5
X	20268018	20268110	refGene_gn=APOO;refGene_tx=XM_004000362.5
X	20275117	20275213	refGene_gn=APOO;refGene_tx=XM_004000362.5
X	20278168	20278223	refGene_gn=APOO;refGene_tx=XM_004000362.5
X	20281167	20281287	refGene_gn=APOO;refGene_tx=XM_004000362.5
X	20282685	20282793	refGene_gn=APOO;refGene_tx=XM_004000362.5
X	20304585	20304594	refGene_gn=APOO;refGene_tx=XM_004000362.5
X	20308019	20308144
refGene_gn=CXHXorf58;refGene_tx=XM_023249000.1
X	20309416	20309516
refGene_gn=CXHXorf58;refGene_tx=XM_023249000.1
X	20311321	20311416
refGene_gn=CXHXorf58;refGene_tx=XM_023249000.1
X	20311776	20311888
refGene_gn=CXHXorf58;refGene_tx=XM_023249000.1
X	20318040	20318172
refGene_gn=CXHXorf58;refGene_tx=XM_023249000.1
X	20326609	20326839
refGene_gn=CXHXorf58;refGene_tx=XM_023249000.1
X	20329015	20329187
refGene_gn=CXHXorf58;refGene_tx=XM_023249000.1
X	20379371	20380481	refGene_gn=KLHL15;refGene_tx=XM_011291634.3
X	20397204	20397909	refGene_gn=KLHL15;refGene_tx=XM_011291634.3
X	20423931	20424000	refGene_gn=EIF2S3;refGene_tx=XM_004000364.4
X	20424504	20424568	refGene_gn=EIF2S3;refGene_tx=XM_004000364.4
X	20426490	20426618	refGene_gn=EIF2S3;refGene_tx=XM_004000364.4
X	20426705	20426827	refGene_gn=EIF2S3;refGene_tx=XM_004000364.4
X	20428459	20428554	refGene_gn=EIF2S3;refGene_tx=XM_004000364.4
X	20429859	20430018	refGene_gn=EIF2S3;refGene_tx=XM_004000364.4
X	20431445	20431580	refGene_gn=EIF2S3;refGene_tx=XM_004000364.4
X	20432616	20432711	refGene_gn=EIF2S3;refGene_tx=XM_004000364.4
X	20433263	20433408	refGene_gn=EIF2S3;refGene_tx=XM_004000364.4
X	20434594	20434764	refGene_gn=EIF2S3;refGene_tx=XM_004000364.4
X	20435165	20435338	refGene_gn=EIF2S3;refGene_tx=XM_004000364.4
X	20441540	20441604	refGene_gn=EIF2S3;refGene_tx=XM_004000364.4
X	20527997	20528058
refGene_gn=ZFX;refGene_tx=XM_004000365.5;refGene_gn=ZFX;refGene_tx=XM_019823670.2;refGene_gn=ZFX;refGene_tx=XM_023249489.1;refGene_gn=ZFX;refGene_tx=XM_023249490.1;refGene_gn=ZFX;refGene_tx=XM_023249491.1;refGene_gn=ZFX;refGene_tx=XM_023249493.1
X	20530939	20531509
refGene_gn=ZFX;refGene_tx=XM_004000365.5;refGene_gn=ZFX;refGene_tx=XM_019823670.2;refGene_gn=ZFX;refGene_tx=XM_019823672.2;refGene_gn=ZFX;re

fGene_tx=XM_023249489.1;refGene_gn=ZFX;refGene_tx=XM_023249490.1;refGene_gn=ZFX;refGene_tx=XM_023249491.1;refGene_gn=ZFX;refGene_tx=XM_023249493.1
X 20553692 20553842
refGene_gn=ZFX;refGene_tx=XM_004000365.5;refGene_gn=ZFX;refGene_tx=XM_019823670.2;refGene_gn=ZFX;refGene_tx=XM_019823672.2;refGene_gn=ZFX;refGene_tx=XM_023249489.1;refGene_gn=ZFX;refGene_tx=XM_023249490.1;refGene_gn=ZFX;refGene_tx=XM_023249491.1;refGene_gn=ZFX;refGene_tx=XM_023249493.1
X 20554061 20554205
refGene_gn=ZFX;refGene_tx=XM_004000365.5;refGene_gn=ZFX;refGene_tx=XM_019823670.2;refGene_gn=ZFX;refGene_tx=XM_019823672.2;refGene_gn=ZFX;refGene_tx=XM_023249489.1;refGene_gn=ZFX;refGene_tx=XM_023249490.1;refGene_gn=ZFX;refGene_tx=XM_023249491.1;refGene_gn=ZFX;refGene_tx=XM_023249493.1
X 20554597 20554747
refGene_gn=ZFX;refGene_tx=XM_004000365.5;refGene_gn=ZFX;refGene_tx=XM_019823670.2;refGene_gn=ZFX;refGene_tx=XM_019823672.2;refGene_gn=ZFX;refGene_tx=XM_023249489.1;refGene_gn=ZFX;refGene_tx=XM_023249490.1;refGene_gn=ZFX;refGene_tx=XM_023249491.1;refGene_gn=ZFX;refGene_tx=XM_023249493.1
X 20555281 20555422
refGene_gn=ZFX;refGene_tx=XM_004000365.5;refGene_gn=ZFX;refGene_tx=XM_019823670.2;refGene_gn=ZFX;refGene_tx=XM_019823672.2;refGene_gn=ZFX;refGene_tx=XM_023249489.1;refGene_gn=ZFX;refGene_tx=XM_023249490.1;refGene_gn=ZFX;refGene_tx=XM_023249491.1
X 20556266 20557450
refGene_gn=ZFX;refGene_tx=XM_004000365.5;refGene_gn=ZFX;refGene_tx=XM_019823670.2;refGene_gn=ZFX;refGene_tx=XM_019823672.2;refGene_gn=ZFX;refGene_tx=XM_023249489.1;refGene_gn=ZFX;refGene_tx=XM_023249490.1;refGene_gn=ZFX;refGene_tx=XM_023249491.1;refGene_gn=ZFX;refGene_tx=XM_023249493.1
X 20750818 20750924
refGene_gn=PDK3;refGene_tx=XM_004000367.5;refGene_gn=PDK3;refGene_tx=XM_023248866.1
X 20772741 20772760 refGene_gn=PDK3;refGene_tx=XM_019823676.1
X 20773171 20773313
refGene_gn=PDK3;refGene_tx=XM_004000367.5;refGene_gn=PDK3;refGene_tx=XM_019823676.1;refGene_gn=PDK3;refGene_tx=XM_023248866.1
X 20776882 20776954
refGene_gn=PDK3;refGene_tx=XM_004000367.5;refGene_gn=PDK3;refGene_tx=XM_019823676.1;refGene_gn=PDK3;refGene_tx=XM_023248866.1
X 20780785 20780970
refGene_gn=PDK3;refGene_tx=XM_004000367.5;refGene_gn=PDK3;refGene_tx=XM_019823676.1;refGene_gn=PDK3;refGene_tx=XM_023248866.1
X 20782594 20782684
refGene_gn=PDK3;refGene_tx=XM_004000367.5;refGene_gn=PDK3;refGene_tx=XM_019823676.1;refGene_gn=PDK3;refGene_tx=XM_023248866.1
X 20796077 20796155
refGene_gn=PDK3;refGene_tx=XM_004000367.5;refGene_gn=PDK3;refGene_tx=X

M_019823676.1;refGene_gn=PDK3;refGene_tx=XM_023248866.1
X 20803830 20803907
refGene_gn=PDK3;refGene_tx=XM_004000367.5;refGene_gn=PDK3;refGene_tx=XM_019823676.1;refGene_gn=PDK3;refGene_tx=XM_023248866.1
X 20805212 20805314
refGene_gn=PDK3;refGene_tx=XM_004000367.5;refGene_gn=PDK3;refGene_tx=XM_019823676.1;refGene_gn=PDK3;refGene_tx=XM_023248866.1
X 20805694 20805805
refGene_gn=PDK3;refGene_tx=XM_004000367.5;refGene_gn=PDK3;refGene_tx=XM_019823676.1;refGene_gn=PDK3;refGene_tx=XM_023248866.1
X 20809050 20809164
refGene_gn=PDK3;refGene_tx=XM_004000367.5;refGene_gn=PDK3;refGene_tx=XM_019823676.1;refGene_gn=PDK3;refGene_tx=XM_023248866.1
X 20810457 20810597
refGene_gn=PDK3;refGene_tx=XM_004000367.5;refGene_gn=PDK3;refGene_tx=XM_019823676.1;refGene_gn=PDK3;refGene_tx=XM_023248866.1
X 20815670 20815701
refGene_gn=PDK3;refGene_tx=XM_004000367.5;refGene_gn=PDK3;refGene_tx=XM_019823676.1;refGene_gn=PDK3;refGene_tx=XM_023248866.1
X 20833312 20833382
refGene_gn=PCYT1B;refGene_tx=XM_011291635.2;refGene_gn=PCYT1B;refGene_tx=XM_019823677.2;refGene_gn=PCYT1B;refGene_tx=XM_019823678.2;refGene_gn=PCYT1B;refGene_tx=XM_019823679.2;refGene_gn=PCYT1B;refGene_tx=XM_019823680.2;refGene_gn=PCYT1B;refGene_tx=XM_019823681.2;refGene_gn=PCYT1B;refGene_tx=XM_019823682.2;refGene_gn=PCYT1B;refGene_tx=XM_019823683.2;refGene_gn=PCYT1B;refGene_tx=XM_019823686.2;refGene_gn=PCYT1B;refGene_tx=XM_019823687.2;refGene_gn=PCYT1B;refGene_tx=XM_019823688.2;refGene_gn=PCYT1B;refGene_tx=XM_019823689.2;refGene_gn=PCYT1B;refGene_tx=XM_019823690.2
X 20833709 20834002
refGene_gn=PCYT1B;refGene_tx=XM_011291635.2;refGene_gn=PCYT1B;refGene_tx=XM_019823677.2;refGene_gn=PCYT1B;refGene_tx=XM_019823678.2;refGene_gn=PCYT1B;refGene_tx=XM_019823679.2;refGene_gn=PCYT1B;refGene_tx=XM_019823680.2;refGene_gn=PCYT1B;refGene_tx=XM_019823681.2;refGene_gn=PCYT1B;refGene_tx=XM_019823682.2;refGene_gn=PCYT1B;refGene_tx=XM_019823683.2;refGene_gn=PCYT1B;refGene_tx=XM_019823686.2;refGene_gn=PCYT1B;refGene_tx=XM_019823687.2;refGene_gn=PCYT1B;refGene_tx=XM_019823688.2;refGene_gn=PCYT1B;refGene_tx=XM_019823690.2
X 20836800 20837013
refGene_gn=PCYT1B;refGene_tx=XM_011291635.2;refGene_gn=PCYT1B;refGene_tx=XM_019823677.2;refGene_gn=PCYT1B;refGene_tx=XM_019823678.2;refGene_gn=PCYT1B;refGene_tx=XM_019823679.2;refGene_gn=PCYT1B;refGene_tx=XM_019823680.2;refGene_gn=PCYT1B;refGene_tx=XM_019823681.2;refGene_gn=PCYT1B;refGene_tx=XM_019823682.2;refGene_gn=PCYT1B;refGene_tx=XM_019823683.2;refGene_gn=PCYT1B;refGene_tx=XM_019823684.2;refGene_gn=PCYT1B;refGene_tx=XM_019823686.2;refGene_gn=PCYT1B;refGene_tx=XM_019823687.2;refGene_gn=PCYT1B;refGene_tx=XM_019823688.2;refGene_gn=PCYT1B;refGene_tx=XM_019823689.2;refGene_gn=PCYT1B;refGene_tx=XM_019823690.2
X 20863823 20864012
refGene_gn=PCYT1B;refGene_tx=XM_011291635.2;refGene_gn=PCYT1B;refGene_

tx=XM_019823677.2;refGene_gn=PCYT1B;refGene_tx=XM_019823678.2;refGene_gn=PCYT1B;refGene_tx=XM_019823679.2;refGene_gn=PCYT1B;refGene_tx=XM_019823680.2;refGene_gn=PCYT1B;refGene_tx=XM_019823681.2;refGene_gn=PCYT1B;refGene_tx=XM_019823682.2;refGene_gn=PCYT1B;refGene_tx=XM_019823683.2;refGene_gn=PCYT1B;refGene_tx=XM_019823684.2;refGene_gn=PCYT1B;refGene_tx=XM_019823686.2;refGene_gn=PCYT1B;refGene_tx=XM_019823687.2;refGene_gn=PCYT1B;refGene_tx=XM_019823688.2;refGene_gn=PCYT1B;refGene_tx=XM_019823689.2;refGene_gn=PCYT1B;refGene_tx=XM_019823690.2
X 20866703 20866846
refGene_gn=PCYT1B;refGene_tx=XM_011291635.2;refGene_gn=PCYT1B;refGene_tx=XM_019823677.2;refGene_gn=PCYT1B;refGene_tx=XM_019823678.2;refGene_gn=PCYT1B;refGene_tx=XM_019823679.2;refGene_gn=PCYT1B;refGene_tx=XM_019823680.2;refGene_gn=PCYT1B;refGene_tx=XM_019823681.2;refGene_gn=PCYT1B;refGene_tx=XM_019823682.2;refGene_gn=PCYT1B;refGene_tx=XM_019823683.2;refGene_gn=PCYT1B;refGene_tx=XM_019823684.2;refGene_gn=PCYT1B;refGene_tx=XM_019823686.2;refGene_gn=PCYT1B;refGene_tx=XM_019823687.2;refGene_gn=PCYT1B;refGene_tx=XM_019823688.2;refGene_gn=PCYT1B;refGene_tx=XM_019823689.2;refGene_gn=PCYT1B;refGene_tx=XM_019823690.2
X 20879032 20879111
refGene_gn=PCYT1B;refGene_tx=XM_011291635.2;refGene_gn=PCYT1B;refGene_tx=XM_019823677.2;refGene_gn=PCYT1B;refGene_tx=XM_019823678.2;refGene_gn=PCYT1B;refGene_tx=XM_019823679.2;refGene_gn=PCYT1B;refGene_tx=XM_019823680.2;refGene_gn=PCYT1B;refGene_tx=XM_019823681.2;refGene_gn=PCYT1B;refGene_tx=XM_019823682.2;refGene_gn=PCYT1B;refGene_tx=XM_019823683.2;refGene_gn=PCYT1B;refGene_tx=XM_019823684.2;refGene_gn=PCYT1B;refGene_tx=XM_019823687.2;refGene_gn=PCYT1B;refGene_tx=XM_019823688.2;refGene_gn=PCYT1B;refGene_tx=XM_019823689.2;refGene_gn=PCYT1B;refGene_tx=XM_019823690.2
X 20881790 20881942
refGene_gn=PCYT1B;refGene_tx=XM_011291635.2;refGene_gn=PCYT1B;refGene_tx=XM_019823677.2;refGene_gn=PCYT1B;refGene_tx=XM_019823678.2;refGene_gn=PCYT1B;refGene_tx=XM_019823679.2;refGene_gn=PCYT1B;refGene_tx=XM_019823680.2;refGene_gn=PCYT1B;refGene_tx=XM_019823681.2;refGene_gn=PCYT1B;refGene_tx=XM_019823682.2;refGene_gn=PCYT1B;refGene_tx=XM_019823683.2;refGene_gn=PCYT1B;refGene_tx=XM_019823684.2;refGene_gn=PCYT1B;refGene_tx=XM_019823687.2;refGene_gn=PCYT1B;refGene_tx=XM_019823688.2;refGene_gn=PCYT1B;refGene_tx=XM_019823689.2;refGene_gn=PCYT1B;refGene_tx=XM_019823690.2
X 20897047 20897164
refGene_gn=PCYT1B;refGene_tx=XM_011291635.2;refGene_gn=PCYT1B;refGene_tx=XM_019823677.2;refGene_gn=PCYT1B;refGene_tx=XM_019823678.2;refGene_gn=PCYT1B;refGene_tx=XM_019823679.2;refGene_gn=PCYT1B;refGene_tx=XM_019823680.2;refGene_gn=PCYT1B;refGene_tx=XM_019823681.2;refGene_gn=PCYT1B;refGene_tx=XM_019823682.2;refGene_gn=PCYT1B;refGene_tx=XM_019823683.2;refGene_gn=PCYT1B;refGene_tx=XM_019823684.2;refGene_gn=PCYT1B;refGene_tx=XM_019823686.2;refGene_gn=PCYT1B;refGene_tx=XM_019823687.2;refGene_gn=PCYT1B;refGene_tx=XM_019823688.2;refGene_gn=PCYT1B;refGene_tx=XM_019823689.2;refGene_gn=PCYT1B;refGene_tx=XM_019823690.2
X 20906010 20906110
refGene_gn=PCYT1B;refGene_tx=XM_011291635.2;refGene_gn=PCYT1B;refGene_

tx=XM_019823677.2;refGene_gn=PCYT1B;refGene_tx=XM_019823678.2;refGene_gn=PCYT1B;refGene_tx=XM_019823679.2;refGene_gn=PCYT1B;refGene_tx=XM_019823680.2;refGene_gn=PCYT1B;refGene_tx=XM_019823681.2;refGene_gn=PCYT1B;refGene_tx=XM_019823682.2;refGene_gn=PCYT1B;refGene_tx=XM_019823683.2;refGene_gn=PCYT1B;refGene_tx=XM_019823684.2;refGene_gn=PCYT1B;refGene_tx=XM_019823686.2;refGene_gn=PCYT1B;refGene_tx=XM_019823688.2;refGene_gn=PCYT1B;refGene_tx=XM_019823689.2
X 20906588 20906645 refGene_gn=PCYT1B;refGene_tx=XM_011291635.2
X 20911106 20911193
refGene_gn=PCYT1B;refGene_tx=XM_019823677.2;refGene_gn=PCYT1B;refGene_tx=XM_019823678.2;refGene_gn=PCYT1B;refGene_tx=XM_019823679.2;refGene_gn=PCYT1B;refGene_tx=XM_019823684.2;refGene_gn=PCYT1B;refGene_tx=XM_019823686.2;refGene_gn=PCYT1B;refGene_tx=XM_019823688.2;refGene_gn=PCYT1B;refGene_tx=XM_019823689.2
X 20931912 20932029
refGene_gn=PCYT1B;refGene_tx=XM_019823677.2;refGene_gn=PCYT1B;refGene_tx=XM_019823680.2;refGene_gn=PCYT1B;refGene_tx=XM_019823684.2;refGene_gn=PCYT1B;refGene_tx=XM_019823686.2;refGene_gn=PCYT1B;refGene_tx=XM_019823687.2;refGene_gn=PCYT1B;refGene_tx=XM_019823688.2;refGene_gn=PCYT1B;refGene_tx=XM_019823689.2
X 20937639 20937693
refGene_gn=PCYT1B;refGene_tx=XM_019823679.2;refGene_gn=PCYT1B;refGene_tx=XM_019823682.2
X 20959702 20959774
refGene_gn=PCYT1B;refGene_tx=XM_019823678.2;refGene_gn=PCYT1B;refGene_tx=XM_019823681.2
X 20968108 20968111 refGene_gn=PCYT1B;refGene_tx=XM_019823683.2
X 20981792 20981835 refGene_gn=POLA1;refGene_tx=XM_006943631.4
X 20981889 20981998 refGene_gn=POLA1;refGene_tx=XM_019823691.1
X 20986377 20986502
refGene_gn=POLA1;refGene_tx=XM_006943631.4;refGene_gn=POLA1;refGene_tx=XM_019823691.1
X 20988957 20989054
refGene_gn=POLA1;refGene_tx=XM_006943631.4;refGene_gn=POLA1;refGene_tx=XM_019823691.1
X 20989990 20990071
refGene_gn=POLA1;refGene_tx=XM_006943631.4;refGene_gn=POLA1;refGene_tx=XM_019823691.1
X 20998113 20998229
refGene_gn=POLA1;refGene_tx=XM_006943631.4;refGene_gn=POLA1;refGene_tx=XM_019823691.1
X 20998678 20998741
refGene_gn=POLA1;refGene_tx=XM_006943631.4;refGene_gn=POLA1;refGene_tx=XM_019823691.1
X 20999598 20999691
refGene_gn=POLA1;refGene_tx=XM_006943631.4;refGene_gn=POLA1;refGene_tx=XM_019823691.1
X 21000117 21000205
refGene_gn=POLA1;refGene_tx=XM_006943631.4;refGene_gn=POLA1;refGene_tx=XM_019823691.1

X 21000495 21000697
refGene_gn=POLA1;refGene_tx=XM_006943631.4;refGene_gn=POLA1;refGene_tx=XM_019823691.1
X 21000784 21000963
refGene_gn=POLA1;refGene_tx=XM_006943631.4;refGene_gn=POLA1;refGene_tx=XM_019823691.1
X 21006600 21006713
refGene_gn=POLA1;refGene_tx=XM_006943631.4;refGene_gn=POLA1;refGene_tx=XM_019823691.1
X 21007584 21007701
refGene_gn=POLA1;refGene_tx=XM_006943631.4;refGene_gn=POLA1;refGene_tx=XM_019823691.1
X 21008961 21009036
refGene_gn=POLA1;refGene_tx=XM_006943631.4;refGene_gn=POLA1;refGene_tx=XM_019823691.1
X 21009927 21010066
refGene_gn=POLA1;refGene_tx=XM_006943631.4;refGene_gn=POLA1;refGene_tx=XM_019823691.1
X 21010886 21011041
refGene_gn=POLA1;refGene_tx=XM_006943631.4;refGene_gn=POLA1;refGene_tx=XM_019823691.1
X 21016035 21016120
refGene_gn=POLA1;refGene_tx=XM_006943631.4;refGene_gn=POLA1;refGene_tx=XM_019823691.1
X 21019718 21019780
refGene_gn=POLA1;refGene_tx=XM_006943631.4;refGene_gn=POLA1;refGene_tx=XM_019823691.1
X 21021324 21021414
refGene_gn=POLA1;refGene_tx=XM_006943631.4;refGene_gn=POLA1;refGene_tx=XM_019823691.1
X 21024540 21024657
refGene_gn=POLA1;refGene_tx=XM_006943631.4;refGene_gn=POLA1;refGene_tx=XM_019823691.1
X 21026262 21026438
refGene_gn=POLA1;refGene_tx=XM_006943631.4;refGene_gn=POLA1;refGene_tx=XM_019823691.1
X 21028513 21028643
refGene_gn=POLA1;refGene_tx=XM_006943631.4;refGene_gn=POLA1;refGene_tx=XM_019823691.1
X 21029137 21029257
refGene_gn=POLA1;refGene_tx=XM_006943631.4;refGene_gn=POLA1;refGene_tx=XM_019823691.1
X 21030341 21030441
refGene_gn=POLA1;refGene_tx=XM_006943631.4;refGene_gn=POLA1;refGene_tx=XM_019823691.1
X 21033459 21033584
refGene_gn=POLA1;refGene_tx=XM_006943631.4;refGene_gn=POLA1;refGene_tx=XM_019823691.1
X 21038585 21038735
refGene_gn=POLA1;refGene_tx=XM_006943631.4;refGene_gn=POLA1;refGene_tx

=XM_019823691.1
X 21039147 21039270
refGene_gn=POLA1;refGene_tx=XM_006943631.4;refGene_gn=POLA1;refGene_tx
=XM_019823691.1
X 21097030 21097063
refGene_gn=POLA1;refGene_tx=XM_006943631.4;refGene_gn=POLA1;refGene_tx
=XM_019823691.1
X 21097880 21097973
refGene_gn=POLA1;refGene_tx=XM_006943631.4;refGene_gn=POLA1;refGene_tx
=XM_019823691.1
X 21099543 21099749
refGene_gn=POLA1;refGene_tx=XM_006943631.4;refGene_gn=POLA1;refGene_tx
=XM_019823691.1
X 21101814 21101947
refGene_gn=POLA1;refGene_tx=XM_006943631.4;refGene_gn=POLA1;refGene_tx
=XM_019823691.1
X 21109453 21109585
refGene_gn=POLA1;refGene_tx=XM_006943631.4;refGene_gn=POLA1;refGene_tx
=XM_019823691.1
X 21115006 21115181
refGene_gn=POLA1;refGene_tx=XM_006943631.4;refGene_gn=POLA1;refGene_tx
=XM_019823691.1
X 21133174 21133353
refGene_gn=POLA1;refGene_tx=XM_006943631.4;refGene_gn=POLA1;refGene_tx
=XM_019823691.1
X 21135083 21135215
refGene_gn=POLA1;refGene_tx=XM_006943631.4;refGene_gn=POLA1;refGene_tx
=XM_019823691.1
X 21178894 21179011
refGene_gn=POLA1;refGene_tx=XM_006943631.4;refGene_gn=POLA1;refGene_tx
=XM_019823691.1
X 21216891 21216988
refGene_gn=POLA1;refGene_tx=XM_006943631.4;refGene_gn=POLA1;refGene_tx
=XM_019823691.1
X 21284901 21285047
refGene_gn=POLA1;refGene_tx=XM_006943631.4;refGene_gn=POLA1;refGene_tx
=XM_019823691.1
X 21293422 21293663
refGene_gn=ARX;refGene_tx=XM_023249083.1;refGene_gn=ARX;refGene_tx=XM_
023249084.1
X 21295916 21296245
refGene_gn=ARX;refGene_tx=XM_023249083.1;refGene_gn=ARX;refGene_tx=XM_
023249084.1
X 21299058 21299104
refGene_gn=ARX;refGene_tx=XM_023249083.1;refGene_gn=ARX;refGene_tx=XM_
023249084.1
X 21301745 21302622
refGene_gn=ARX;refGene_tx=XM_023249083.1;refGene_gn=ARX;refGene_tx=XM_
023249084.1
X 21304353 21304549

refGene_gn=ARX;refGene_tx=XM_023249083.1;refGene_gn=ARX;refGene_tx=XM_023249084.1
X 21603089 21603992
refGene_gn=FELCATV1R3;refGene_tx=NM_001167475.1
X 22263984 22265046
refGene_gn=LOC101095955;refGene_tx=XM_004000370.4;refGene_gn=LOC101095955;refGene_tx=XM_006943632.3;refGene_gn=LOC101095955;refGene_tx=XM_023249202.1
X 22294289 22295087
refGene_gn=LOC105261400;refGene_tx=XM_011291656.3
X 22330124 22331009
refGene_gn=LOC111558670;refGene_tx=XM_023248955.1
X 22344145 22344197
refGene_gn=LOC102899615;refGene_tx=XM_011291657.2
X 22344199 22344408
refGene_gn=LOC102899615;refGene_tx=XM_011291657.2
X 22344410 22344812
refGene_gn=LOC102899615;refGene_tx=XM_011291657.2
X 22355361 22356489
refGene_gn=LOC493792;refGene_tx=NM_001009262.1;refGene_gn=LOC493792;refGene_tx=XM_006943623.4
X 23702072 23703119
refGene_gn=LOC101100370;refGene_tx=XM_004000376.4
X 23712361 23713477
refGene_gn=LOC101082378;refGene_tx=XM_023249039.1
X 23750240 23751176
refGene_gn=LOC493964;refGene_tx=NM_001009368.1;refGene_gn=LOC493964;refGene_tx=XM_006943637.3;refGene_gn=LOC493964;refGene_tx=XM_006943638.3
X 24698999 24699081 refGene_gn=IL1RAPL1;refGene_tx=XM_023249180.1
X 25195589 25195869 refGene_gn=IL1RAPL1;refGene_tx=XM_023249180.1
X 25309747 25309934 refGene_gn=IL1RAPL1;refGene_tx=XM_023249180.1
X 25312403 25312557 refGene_gn=IL1RAPL1;refGene_tx=XM_023249180.1
X 25542588 25542663 refGene_gn=IL1RAPL1;refGene_tx=XM_023249180.1
X 25795440 25795573 refGene_gn=IL1RAPL1;refGene_tx=XM_023249180.1
X 25797483 25797629 refGene_gn=IL1RAPL1;refGene_tx=XM_023249180.1
X 25825184 25825328 refGene_gn=IL1RAPL1;refGene_tx=XM_023249180.1
X 25842860 25843031 refGene_gn=IL1RAPL1;refGene_tx=XM_023249180.1
X 25843479 25844198 refGene_gn=IL1RAPL1;refGene_tx=XM_023249180.1
X 26052921 26053953
refGene_gn=LOC109496467;refGene_tx=XM_019823700.1;refGene_gn=LOC109496467;refGene_tx=XM_023249165.1
X 26068517 26069594
refGene_gn=LOC101083402;refGene_tx=XM_023249163.1;refGene_gn=LOC101083402;refGene_tx=XM_023249164.1
X 26086898 26087879
refGene_gn=LOC101083912;refGene_tx=XM_004000391.5
X 26126474 26126719 refGene_gn=NR0B1;refGene_tx=XM_004000377.5
X 26130324 26131495 refGene_gn=NR0B1;refGene_tx=XM_004000377.5
X 26257000 26257168
refGene_gn=LOC109496468;refGene_tx=XM_023248808.1

X	26260428	26260533
refGene_gn=LOC109496468;refGene_tx=XM_023248808.1
X	26318799	26318890
refGene_gn=LOC109496468;refGene_tx=XM_023248808.1
X	26345188	26345344
refGene_gn=LOC109496468;refGene_tx=XM_023248808.1
X	26390507	26391398
refGene_gn=CXHXorf21;refGene_tx=XM_004000378.5
X	26395963	26396064
refGene_gn=LOC109496468;refGene_tx=XM_023248808.1
X	26471330	26471444
refGene_gn=LOC109496468;refGene_tx=XM_023248808.1
X	26481354	26481432
refGene_gn=GK;refGene_tx=XM_004000379.5;refGene_gn=GK;refGene_tx=XM_006943642.4;refGene_gn=GK;refGene_tx=XM_019823712.2;refGene_gn=GK;refGene_tx=XM_019823713.2;refGene_gn=GK;refGene_tx=XM_019823714.2
X	26492728	26492802
refGene_gn=GK;refGene_tx=XM_004000379.5;refGene_gn=GK;refGene_tx=XM_006943642.4;refGene_gn=GK;refGene_tx=XM_019823712.2;refGene_gn=GK;refGene_tx=XM_019823713.2;refGene_gn=GK;refGene_tx=XM_019823714.2
X	26494965	26495072
refGene_gn=GK;refGene_tx=XM_004000379.5;refGene_gn=GK;refGene_tx=XM_006943642.4;refGene_gn=GK;refGene_tx=XM_019823712.2;refGene_gn=GK;refGene_tx=XM_019823713.2;refGene_gn=GK;refGene_tx=XM_019823714.2
X	26504756	26504834
refGene_gn=GK;refGene_tx=XM_004000379.5;refGene_gn=GK;refGene_tx=XM_006943642.4;refGene_gn=GK;refGene_tx=XM_019823712.2;refGene_gn=GK;refGene_tx=XM_019823713.2;refGene_gn=GK;refGene_tx=XM_019823714.2
X	26513632	26513716
refGene_gn=GK;refGene_tx=XM_019823712.2;refGene_gn=GK;refGene_tx=XM_019823713.2;refGene_gn=GK;refGene_tx=XM_019823714.2
X	26517357	26517434
refGene_gn=GK;refGene_tx=XM_004000379.5;refGene_gn=GK;refGene_tx=XM_006943642.4;refGene_gn=GK;refGene_tx=XM_019823712.2;refGene_gn=GK;refGene_tx=XM_019823713.2;refGene_gn=GK;refGene_tx=XM_019823714.2
X	26519380	26519518
refGene_gn=GK;refGene_tx=XM_004000379.5;refGene_gn=GK;refGene_tx=XM_006943642.4;refGene_gn=GK;refGene_tx=XM_019823712.2;refGene_gn=GK;refGene_tx=XM_019823713.2;refGene_gn=GK;refGene_tx=XM_019823714.2
X	26521990	26522100
refGene_gn=GK;refGene_tx=XM_004000379.5;refGene_gn=GK;refGene_tx=XM_006943642.4;refGene_gn=GK;refGene_tx=XM_019823712.2;refGene_gn=GK;refGene_tx=XM_019823713.2;refGene_gn=GK;refGene_tx=XM_019823714.2
X	26522545	26522612
refGene_gn=GK;refGene_tx=XM_004000379.5;refGene_gn=GK;refGene_tx=XM_006943642.4;refGene_gn=GK;refGene_tx=XM_019823712.2;refGene_gn=GK;refGene_tx=XM_019823713.2;refGene_gn=GK;refGene_tx=XM_019823714.2
X	26523700	26523718
refGene_gn=GK;refGene_tx=XM_006943642.4;refGene_gn=GK;refGene_tx=XM_019823712.2;refGene_gn=GK;refGene_tx=XM_019823714.2

X 26525387 26525423
refGene_gn=GK;refGene_tx=XM_004000379.5;refGene_gn=GK;refGene_tx=XM_006943642.4;refGene_gn=GK;refGene_tx=XM_019823712.2;refGene_gn=GK;refGene_tx=XM_019823713.2;refGene_gn=GK;refGene_tx=XM_019823714.2
X 26526267 26526335
refGene_gn=GK;refGene_tx=XM_004000379.5;refGene_gn=GK;refGene_tx=XM_006943642.4;refGene_gn=GK;refGene_tx=XM_019823712.2;refGene_gn=GK;refGene_tx=XM_019823713.2;refGene_gn=GK;refGene_tx=XM_019823714.2
X 26534021 26534064
refGene_gn=GK;refGene_tx=XM_004000379.5;refGene_gn=GK;refGene_tx=XM_006943642.4;refGene_gn=GK;refGene_tx=XM_019823712.2;refGene_gn=GK;refGene_tx=XM_019823713.2;refGene_gn=GK;refGene_tx=XM_019823714.2
X 26535118 26535199
refGene_gn=GK;refGene_tx=XM_004000379.5;refGene_gn=GK;refGene_tx=XM_006943642.4;refGene_gn=GK;refGene_tx=XM_019823712.2;refGene_gn=GK;refGene_tx=XM_019823713.2;refGene_gn=GK;refGene_tx=XM_019823714.2
X 26547405 26547484
refGene_gn=GK;refGene_tx=XM_004000379.5;refGene_gn=GK;refGene_tx=XM_006943642.4;refGene_gn=GK;refGene_tx=XM_019823712.2;refGene_gn=GK;refGene_tx=XM_019823713.2;refGene_gn=GK;refGene_tx=XM_019823714.2
X 26548229 26548326
refGene_gn=GK;refGene_tx=XM_004000379.5;refGene_gn=GK;refGene_tx=XM_006943642.4;refGene_gn=GK;refGene_tx=XM_019823712.2;refGene_gn=GK;refGene_tx=XM_019823713.2;refGene_gn=GK;refGene_tx=XM_019823714.2
X 26548884 26548969
refGene_gn=GK;refGene_tx=XM_004000379.5;refGene_gn=GK;refGene_tx=XM_006943642.4;refGene_gn=GK;refGene_tx=XM_019823712.2;refGene_gn=GK;refGene_tx=XM_019823713.2;refGene_gn=GK;refGene_tx=XM_019823714.2
X 26550160 26550281
refGene_gn=GK;refGene_tx=XM_004000379.5;refGene_gn=GK;refGene_tx=XM_006943642.4;refGene_gn=GK;refGene_tx=XM_019823712.2;refGene_gn=GK;refGene_tx=XM_019823713.2;refGene_gn=GK;refGene_tx=XM_019823714.2
X 26550404 26550548
refGene_gn=GK;refGene_tx=XM_004000379.5;refGene_gn=GK;refGene_tx=XM_006943642.4;refGene_gn=GK;refGene_tx=XM_019823712.2;refGene_gn=GK;refGene_tx=XM_019823713.2;refGene_gn=GK;refGene_tx=XM_019823714.2
X 26553127 26553208
refGene_gn=GK;refGene_tx=XM_004000379.5;refGene_gn=GK;refGene_tx=XM_006943642.4;refGene_gn=GK;refGene_tx=XM_019823712.2;refGene_gn=GK;refGene_tx=XM_019823713.2;refGene_gn=GK;refGene_tx=XM_019823714.2
X 26557678 26557765
refGene_gn=GK;refGene_tx=XM_004000379.5;refGene_gn=GK;refGene_tx=XM_006943642.4;refGene_gn=GK;refGene_tx=XM_019823712.2;refGene_gn=GK;refGene_tx=XM_019823713.2
X 26559530 26559541
refGene_gn=GK;refGene_tx=XM_004000379.5;refGene_gn=GK;refGene_tx=XM_006943642.4;refGene_gn=GK;refGene_tx=XM_019823712.2;refGene_gn=GK;refGene_tx=XM_019823713.2;refGene_gn=GK;refGene_tx=XM_019823714.2
X 26654312 26654461
refGene_gn=TAB3;refGene_tx=XM_019823716.2;refGene_gn=TAB3;refGene_tx=X

M_023249430.1;refGene_gn=TAB3;refGene_tx=XM_023249431.1;refGene_gn=TAB3;refGene_tx=XM_023249432.1;refGene_gn=TAB3;refGene_tx=XM_023249433.1
X	26659103	26659205	refGene_gn=TAB3;refGene_tx=XM_019823716.2;refGene_gn=TAB3;refGene_tx=XM_023249430.1;refGene_gn=TAB3;refGene_tx=XM_023249431.1
X	26666139	26666223	refGene_gn=TAB3;refGene_tx=XM_019823716.2;refGene_gn=TAB3;refGene_tx=XM_023249430.1;refGene_gn=TAB3;refGene_tx=XM_023249431.1;refGene_gn=TAB3;refGene_tx=XM_023249432.1
X	26669028	26669122	refGene_gn=TAB3;refGene_tx=XM_019823716.2;refGene_gn=TAB3;refGene_tx=XM_023249430.1;refGene_gn=TAB3;refGene_tx=XM_023249431.1;refGene_gn=TAB3;refGene_tx=XM_023249432.1
X	26673948	26674109	refGene_gn=TAB3;refGene_tx=XM_019823716.2;refGene_gn=TAB3;refGene_tx=XM_023249430.1;refGene_gn=TAB3;refGene_tx=XM_023249431.1;refGene_gn=TAB3;refGene_tx=XM_023249432.1;refGene_gn=TAB3;refGene_tx=XM_023249433.1
X	26675209	26676671	refGene_gn=TAB3;refGene_tx=XM_019823716.2;refGene_gn=TAB3;refGene_tx=XM_023249430.1;refGene_gn=TAB3;refGene_tx=XM_023249431.1;refGene_gn=TAB3;refGene_tx=XM_023249432.1;refGene_gn=TAB3;refGene_tx=XM_023249433.1
X	26680411	26680513	refGene_gn=TAB3;refGene_tx=XM_019823716.2;refGene_gn=TAB3;refGene_tx=XM_023249430.1;refGene_gn=TAB3;refGene_tx=XM_023249431.1;refGene_gn=TAB3;refGene_tx=XM_023249432.1;refGene_gn=TAB3;refGene_tx=XM_023249433.1
X	26944163	26944712	refGene_gn=LOC101080615;refGene_tx=XM_011291686.3
X	26990482	26990494	refGene_gn=DMD;refGene_tx=XM_023249210.1
X	26995534	26995566	refGene_gn=DMD;refGene_tx=XM_023249210.1
X	27003630	27003723	refGene_gn=DMD;refGene_tx=XM_023249210.1
X	27020067	27020191	refGene_gn=DMD;refGene_tx=XM_023249210.1
X	27021136	27021380	refGene_gn=DMD;refGene_tx=XM_023249210.1
X	27043943	27044102	refGene_gn=DMD;refGene_tx=XM_023249210.1
X	27046715	27046781	refGene_gn=DMD;refGene_tx=XM_023249210.1
X	27047896	27047962	refGene_gn=DMD;refGene_tx=XM_023249210.1
X	27052277	27052316	refGene_gn=DMD;refGene_tx=XM_023249210.1
X	27053018	27053155	refGene_gn=DMD;refGene_tx=XM_023249210.1
X	27054769	27054881	refGene_gn=DMD;refGene_tx=XM_023249210.1
X	27057101	27057268	refGene_gn=DMD;refGene_tx=XM_023249210.1
X	27079594	27079752	refGene_gn=DMD;refGene_tx=XM_023249210.1
X	27081769	27081855	refGene_gn=DMD;refGene_tx=XM_023249210.1
X	27084090	27084292	refGene_gn=DMD;refGene_tx=XM_023249210.1
X	27096047	27096122	refGene_gn=DMD;refGene_tx=XM_023249210.1
X	27131822	27131884	refGene_gn=DMD;refGene_tx=XM_023249210.1
X	27196302	27196363	refGene_gn=DMD;refGene_tx=XM_023249210.1
X	27220140	27220219	refGene_gn=DMD;refGene_tx=XM_023249210.1
X	27305506	27305653	refGene_gn=DMD;refGene_tx=XM_023249210.1
X	27332337	27332583	refGene_gn=DMD;refGene_tx=XM_023249210.1
X	27332584	27332607	refGene_gn=DMD;refGene_tx=XM_023249210.1
X	27333250	27333371	refGene_gn=DMD;refGene_tx=XM_023249210.1

| | | | |
|---|---|---|---|
| X | 27350211 | 27350368 | refGene_gn=DMD;refGene_tx=XM_023249210.1 |
| X | 27361418 | 27361591 | refGene_gn=DMD;refGene_tx=XM_023249210.1 |
| X | 27479899 | 27480089 | refGene_gn=DMD;refGene_tx=XM_023249210.1 |
| X | 27512675 | 27512830 | refGene_gn=DMD;refGene_tx=XM_023249210.1 |
| X | 27533033 | 27533245 | refGene_gn=DMD;refGene_tx=XM_023249210.1 |
| X | 27575121 | 27575239 | refGene_gn=DMD;refGene_tx=XM_023249210.1 |
| X | 27627281 | 27627514 | refGene_gn=DMD;refGene_tx=XM_023249210.1 |
| X | 27673539 | 27673648 | refGene_gn=DMD;refGene_tx=XM_023249210.1 |
| X | 27683413 | 27683515 | refGene_gn=DMD;refGene_tx=XM_023249210.1 |
| X | 27717679 | 27717844 | refGene_gn=DMD;refGene_tx=XM_023249210.1 |
| X | 27773779 | 27773929 | refGene_gn=DMD;refGene_tx=XM_023249210.1 |
| X | 27776128 | 27776276 | refGene_gn=DMD;refGene_tx=XM_023249210.1 |
| X | 27815228 | 27815404 | refGene_gn=DMD;refGene_tx=XM_023249210.1 |
| X | 28073609 | 28073757 | refGene_gn=DMD;refGene_tx=XM_023249210.1 |
| X | 28128096 | 28128269 | refGene_gn=DMD;refGene_tx=XM_023249210.1 |
| X | 28147746 | 28147941 | refGene_gn=DMD;refGene_tx=XM_023249210.1 |
| X | 28179175 | 28179358 | refGene_gn=DMD;refGene_tx=XM_023249210.1 |
| X | 28180224 | 28180377 | refGene_gn=DMD;refGene_tx=XM_023249210.1 |
| X | 28183141 | 28183279 | refGene_gn=DMD;refGene_tx=XM_023249210.1 |
| X | 28185649 | 28185772 | refGene_gn=DMD;refGene_tx=XM_023249210.1 |
| X | 28199227 | 28199398 | refGene_gn=DMD;refGene_tx=XM_023249210.1 |
| X | 28200933 | 28201062 | refGene_gn=DMD;refGene_tx=XM_023249210.1 |
| X | 28201367 | 28201547 | refGene_gn=DMD;refGene_tx=XM_023249210.1 |
| X | 28214559 | 28214730 | refGene_gn=DMD;refGene_tx=XM_023249210.1 |
| X | 28219687 | 28219843 | refGene_gn=DMD;refGene_tx=XM_023249210.1 |
| X | 28223467 | 28223641 | refGene_gn=DMD;refGene_tx=XM_023249210.1 |
| X | 28224029 | 28224140 | refGene_gn=DMD;refGene_tx=XM_023249210.1 |
| X | 28242249 | 28242411 | refGene_gn=DMD;refGene_tx=XM_023249210.1 |
| X | 28272554 | 28272704 | refGene_gn=DMD;refGene_tx=XM_023249210.1 |
| X | 28274323 | 28274458 | refGene_gn=DMD;refGene_tx=XM_023249210.1 |
| X | 28281117 | 28281300 | refGene_gn=DMD;refGene_tx=XM_023249210.1 |
| X | 28287025 | 28287196 | refGene_gn=DMD;refGene_tx=XM_023249210.1 |
| X | 28301733 | 28301889 | refGene_gn=DMD;refGene_tx=XM_023249210.1 |
| X | 28302859 | 28302973 | refGene_gn=DMD;refGene_tx=XM_023249210.1 |
| X | 28305779 | 28305992 | refGene_gn=DMD;refGene_tx=XM_023249210.1 |
| X | 28309491 | 28309637 | refGene_gn=DMD;refGene_tx=XM_023249210.1 |
| X | 28323694 | 28323875 | refGene_gn=DMD;refGene_tx=XM_023249210.1 |
| X | 28328468 | 28328710 | refGene_gn=DMD;refGene_tx=XM_023249210.1 |
| X | 28339472 | 28339560 | refGene_gn=DMD;refGene_tx=XM_023249210.1 |
| X | 28352428 | 28352552 | refGene_gn=DMD;refGene_tx=XM_023249210.1 |
| X | 28377685 | 28377861 | refGene_gn=DMD;refGene_tx=XM_023249210.1 |
| X | 28389215 | 28389395 | refGene_gn=DMD;refGene_tx=XM_023249210.1 |
| X | 28397364 | 28397472 | refGene_gn=DMD;refGene_tx=XM_023249210.1 |
| X | 28397591 | 28397693 | refGene_gn=DMD;refGene_tx=XM_023249210.1 |
| X | 28424495 | 28424615 | refGene_gn=DMD;refGene_tx=XM_023249210.1 |
| X | 28443722 | 28443873 | refGene_gn=DMD;refGene_tx=XM_023249210.1 |
| X | 28461428 | 28461610 | refGene_gn=DMD;refGene_tx=XM_023249210.1 |
| X | 28462299 | 28462482 | refGene_gn=DMD;refGene_tx=XM_023249210.1 |
| X | 28522887 | 28523019 | refGene_gn=DMD;refGene_tx=XM_023249210.1 |
| X | 28524472 | 28524654 | refGene_gn=DMD;refGene_tx=XM_023249210.1 |

X	28612547	28612666	refGene_gn=DMD;refGene_tx=XM_023249210.1
X	28619703	28619876	refGene_gn=DMD;refGene_tx=XM_023249210.1
X	28628143	28628236	refGene_gn=DMD;refGene_tx=XM_023249210.1
X	28648229	28648307	refGene_gn=DMD;refGene_tx=XM_023249210.1
X	28649939	28650032	refGene_gn=DMD;refGene_tx=XM_023249210.1
X	28831707	28831769	refGene_gn=DMD;refGene_tx=XM_023249210.1
X	29004918	29004949	refGene_gn=DMD;refGene_tx=XM_023249210.1
X	30180346	30180587
refGene_gn=LOC101099358;refGene_tx=XM_023249203.1
X	30183298	30184953
refGene_gn=LOC101099358;refGene_tx=XM_023249203.1
X	30399943	30400122	refGene_gn=TMEM47;refGene_tx=XM_011291679.2
X	30410302	30410443	refGene_gn=TMEM47;refGene_tx=XM_011291679.2
X	30426216	30426442	refGene_gn=TMEM47;refGene_tx=XM_011291679.2
X	31500032	31500995
refGene_gn=LOC101080432;refGene_tx=XM_006943647.3
X	31629575	31629770	refGene_gn=CFAP47;refGene_tx=XM_023248956.1
X	31634045	31634197	refGene_gn=CFAP47;refGene_tx=XM_023248956.1
X	31641989	31642105	refGene_gn=CFAP47;refGene_tx=XM_023248956.1
X	31649720	31649859	refGene_gn=CFAP47;refGene_tx=XM_023248956.1
X	31652086	31652315	refGene_gn=CFAP47;refGene_tx=XM_023248956.1
X	31652760	31652921	refGene_gn=CFAP47;refGene_tx=XM_023248956.1
X	31654990	31655118	refGene_gn=CFAP47;refGene_tx=XM_023248956.1
X	31657515	31657751	refGene_gn=CFAP47;refGene_tx=XM_023248956.1
X	31673927	31674117	refGene_gn=CFAP47;refGene_tx=XM_023248956.1
X	31675123	31675337	refGene_gn=CFAP47;refGene_tx=XM_023248956.1
X	31680965	31681118	refGene_gn=CFAP47;refGene_tx=XM_023248956.1
X	31681726	31681916	refGene_gn=CFAP47;refGene_tx=XM_023248956.1
X	31682012	31682109	refGene_gn=CFAP47;refGene_tx=XM_023248956.1
X	31688844	31689061	refGene_gn=CFAP47;refGene_tx=XM_023248956.1
X	31689368	31689610	refGene_gn=CFAP47;refGene_tx=XM_023248956.1
X	31703941	31704072	refGene_gn=CFAP47;refGene_tx=XM_023248956.1
X	31707299	31707422	refGene_gn=CFAP47;refGene_tx=XM_023248956.1
X	31708947	31709079	refGene_gn=CFAP47;refGene_tx=XM_023248956.1
X	31711851	31711929	refGene_gn=CFAP47;refGene_tx=XM_023248956.1
X	31717080	31717225	refGene_gn=CFAP47;refGene_tx=XM_023248956.1
X	31722100	31722195	refGene_gn=CFAP47;refGene_tx=XM_023248956.1
X	31743734	31743876	refGene_gn=CFAP47;refGene_tx=XM_023248956.1
X	31753079	31753174	refGene_gn=CFAP47;refGene_tx=XM_023248956.1
X	31759656	31759816	refGene_gn=CFAP47;refGene_tx=XM_023248956.1
X	31770373	31770569	refGene_gn=CFAP47;refGene_tx=XM_023248956.1
X	31785031	31785241	refGene_gn=CFAP47;refGene_tx=XM_023248956.1
X	31812216	31812317	refGene_gn=CFAP47;refGene_tx=XM_023248956.1
X	31816333	31816459	refGene_gn=CFAP47;refGene_tx=XM_023248956.1
X	31817629	31817861	refGene_gn=CFAP47;refGene_tx=XM_023248956.1
X	31829190	31829412	refGene_gn=CFAP47;refGene_tx=XM_023248956.1
X	31838952	31839034	refGene_gn=CFAP47;refGene_tx=XM_023248956.1
X	31840575	31840662	refGene_gn=CFAP47;refGene_tx=XM_023248956.1
X	31850282	31850475	refGene_gn=CFAP47;refGene_tx=XM_023248956.1
X	31869241	31869342	refGene_gn=CFAP47;refGene_tx=XM_023248956.1

X 31869915 31870032 refGene_gn=CFAP47;refGene_tx=XM_023248956.1
X 31876773 31876908 refGene_gn=CFAP47;refGene_tx=XM_023248956.1
X 31885091 31885207 refGene_gn=CFAP47;refGene_tx=XM_023248956.1
X 31906341 31906492 refGene_gn=CFAP47;refGene_tx=XM_023248956.1
X 31907869 31907958 refGene_gn=CFAP47;refGene_tx=XM_023248956.1
X 31921982 31922069 refGene_gn=CFAP47;refGene_tx=XM_023248956.1
X 31930654 31930725 refGene_gn=CFAP47;refGene_tx=XM_023248956.1
X 31932162 31932308 refGene_gn=CFAP47;refGene_tx=XM_023248956.1
X 31942186 31942298 refGene_gn=CFAP47;refGene_tx=XM_023248956.1
X 31943038 31943265 refGene_gn=CFAP47;refGene_tx=XM_023248956.1
X 31949876 31950051 refGene_gn=CFAP47;refGene_tx=XM_023248956.1
X 31976985 31977182 refGene_gn=CFAP47;refGene_tx=XM_023248956.1
X 31978079 31978223 refGene_gn=CFAP47;refGene_tx=XM_023248956.1
X 31978830 31979004 refGene_gn=CFAP47;refGene_tx=XM_023248956.1
X 31988283 31988395 refGene_gn=CFAP47;refGene_tx=XM_023248956.1
X 31993674 31993824 refGene_gn=CFAP47;refGene_tx=XM_023248956.1
X 32040863 32040998 refGene_gn=CFAP47;refGene_tx=XM_023248956.1
X 32048274 32048409 refGene_gn=CFAP47;refGene_tx=XM_023248956.1
X 32053331 32053466 refGene_gn=CFAP47;refGene_tx=XM_023248956.1
X 32065879 32065977 refGene_gn=CFAP47;refGene_tx=XM_023248956.1
X 32091949 32092125 refGene_gn=CFAP47;refGene_tx=XM_023248956.1
X 32092740 32092848 refGene_gn=CFAP47;refGene_tx=XM_023248956.1
X 32099197 32099306 refGene_gn=CFAP47;refGene_tx=XM_023248956.1
X 32102879 32102984 refGene_gn=CFAP47;refGene_tx=XM_023248956.1
X 32106551 32106708 refGene_gn=CFAP47;refGene_tx=XM_023248956.1
X 32113222 32113321 refGene_gn=CFAP47;refGene_tx=XM_023248956.1
X 32148152 32148312 refGene_gn=CFAP47;refGene_tx=XM_023248956.1
X 32150420 32150515 refGene_gn=CFAP47;refGene_tx=XM_023248956.1
X 32154890 32155043 refGene_gn=CFAP47;refGene_tx=XM_023248956.1
X 32167160 32167332 refGene_gn=CFAP47;refGene_tx=XM_023248956.1
X 32171633 32171795 refGene_gn=CFAP47;refGene_tx=XM_023248956.1
X 32209100 32209269 refGene_gn=CFAP47;refGene_tx=XM_023248956.1
X 32213743 32213956 refGene_gn=CFAP47;refGene_tx=XM_023248956.1
X 32523420 32523663
refGene_gn=LOC111558759;refGene_tx=XM_023249352.1
X 32523664 32523775
refGene_gn=LOC111558759;refGene_tx=XM_023249352.1
X 32686491 32686717 refGene_gn=PRRG1;refGene_tx=XM_023248854.1
X 32779563 32780841
refGene_gn=LOC101081960;refGene_tx=XM_004000449.5
X 32919990 32920041
refGene_gn=PRRG1;refGene_tx=XM_004000393.5;refGene_gn=PRRG1;refGene_tx
=XM_019823724.2;refGene_gn=PRRG1;refGene_tx=XM_019823725.1;refGene_gn=
PRRG1;refGene_tx=XM_023248854.1
X 32938574 32938644 refGene_gn=PRRG1;refGene_tx=XM_023248855.1
X 32938810 32938971
refGene_gn=PRRG1;refGene_tx=XM_004000393.5;refGene_gn=PRRG1;refGene_tx
=XM_019823724.2;refGene_gn=PRRG1;refGene_tx=XM_019823725.1;refGene_gn=
PRRG1;refGene_tx=XM_023248854.1;refGene_gn=PRRG1;refGene_tx=XM_0232488
55.1

X	32995046	32995532
refGene_gn=PRRG1;refGene_tx=XM_004000393.5;refGene_gn=PRRG1;refGene_tx=XM_019823724.2;refGene_gn=PRRG1;refGene_tx=XM_019823725.1;refGene_gn=PRRG1;refGene_tx=XM_023248854.1;refGene_gn=PRRG1;refGene_tx=XM_023248855.1
X	33177611	33178184	refGene_gn=LANCL3;refGene_tx=XM_019823420.2
X	33254636	33254760	refGene_gn=LANCL3;refGene_tx=XM_019823420.2
X	33258002	33258200	refGene_gn=LANCL3;refGene_tx=XM_019823420.2
X	33264899	33265107	refGene_gn=LANCL3;refGene_tx=XM_019823420.2
X	33271962	33272122	refGene_gn=LANCL3;refGene_tx=XM_019823420.2
X	33283020	33283265	refGene_gn=XK;refGene_tx=XM_023249534.1
X	33290936	33291199	refGene_gn=XK;refGene_tx=XM_023249534.1
X	33312925	33313279
refGene_gn=LOC111558783;refGene_tx=XM_023249535.1
X	33347414	33348241	refGene_gn=XK;refGene_tx=XM_023249534.1
X	33385781	33385826
refGene_gn=LOC101086207;refGene_tx=XM_004000394.4
X	33388126	33388222
refGene_gn=LOC101086207;refGene_tx=XM_004000394.4
X	33389500	33389611
refGene_gn=LOC101086207;refGene_tx=XM_004000394.4
X	33398216	33398301
refGene_gn=LOC101086207;refGene_tx=XM_004000394.4
X	33401095	33401241
refGene_gn=LOC101086207;refGene_tx=XM_004000394.4
X	33403326	33403517
refGene_gn=LOC101086207;refGene_tx=XM_004000394.4
X	33406619	33406749
refGene_gn=LOC101086207;refGene_tx=XM_004000394.4
X	33409247	33409340
refGene_gn=LOC101086207;refGene_tx=XM_004000394.4
X	33411638	33411892
refGene_gn=LOC101086207;refGene_tx=XM_004000394.4
X	33412748	33412911
refGene_gn=LOC101086207;refGene_tx=XM_004000394.4
X	33414029	33414176
refGene_gn=LOC101086207;refGene_tx=XM_004000394.4
X	33416668	33416793
refGene_gn=LOC101086207;refGene_tx=XM_004000394.4
X	33417878	33418005
refGene_gn=LOC101086207;refGene_tx=XM_004000394.4
X	33446434	33446511
refGene_gn=DYNLT3;refGene_tx=XM_004000396.5;refGene_gn=DYNLT3;refGene_tx=XM_011291677.3
X	33446798	33446876
refGene_gn=DYNLT3;refGene_tx=XM_004000396.5;refGene_gn=DYNLT3;refGene_tx=XM_011291677.3
X	33447512	33447636
refGene_gn=DYNLT3;refGene_tx=XM_004000396.5;refGene_gn=DYNLT3;refGene_tx=XM_011291677.3

X	33449967 33450009 refGene_gn=DYNLT3;refGene_tx=XM_004000396.5
X	33452150 33452291
refGene_gn=DYNLT3;refGene_tx=XM_004000396.5;refGene_gn=DYNLT3;refGene_
tx=XM_011291677.3
X	33634731 33635070 refGene_gn=HYPM;refGene_tx=XM_004000453.4
X	33748034 33748153 refGene_gn=SYTL5;refGene_tx=XM_023249044.1
X	33770200 33770410 refGene_gn=SYTL5;refGene_tx=XM_023249044.1
X	33799401 33799517 refGene_gn=SYTL5;refGene_tx=XM_023249044.1
X	33800956 33801065 refGene_gn=SYTL5;refGene_tx=XM_023249044.1
X	33804683 33804818 refGene_gn=SYTL5;refGene_tx=XM_023249044.1
X	33817784 33817926 refGene_gn=SYTL5;refGene_tx=XM_023249044.1
X	33821712 33821842 refGene_gn=SYTL5;refGene_tx=XM_023249044.1
X	33824509 33824616 refGene_gn=SYTL5;refGene_tx=XM_023249044.1
X	33830857 33830950 refGene_gn=SYTL5;refGene_tx=XM_023249044.1
X	33831558 33831624 refGene_gn=SYTL5;refGene_tx=XM_023249044.1
X	33834730 33834909 refGene_gn=SYTL5;refGene_tx=XM_023249044.1
X	33836701 33836801 refGene_gn=SYTL5;refGene_tx=XM_023249044.1
X	33839463 33839625 refGene_gn=SYTL5;refGene_tx=XM_023249044.1
X	33849518 33849627 refGene_gn=SYTL5;refGene_tx=XM_023249044.1
X	33851013 33851149 refGene_gn=SYTL5;refGene_tx=XM_023249044.1
X	33854222 33854431 refGene_gn=SYTL5;refGene_tx=XM_023249044.1
X	33855518 33855661 refGene_gn=SYTL5;refGene_tx=XM_023249044.1
X	33878610 33878794 refGene_gn=SRPX;refGene_tx=XM_023249396.1
X	33893875 33893997 refGene_gn=SRPX;refGene_tx=XM_023249396.1
X	33913221 33913355 refGene_gn=SRPX;refGene_tx=XM_023249396.1
X	33914914 33915094 refGene_gn=SRPX;refGene_tx=XM_023249396.1
X	33915823 33915945 refGene_gn=SRPX;refGene_tx=XM_023249396.1
X	33918891 33919018 refGene_gn=SRPX;refGene_tx=XM_023249396.1
X	33929387 33929564 refGene_gn=SRPX;refGene_tx=XM_023249396.1
X	33931512 33931648 refGene_gn=SRPX;refGene_tx=XM_023249396.1
X	33931649 33931705 refGene_gn=SRPX;refGene_tx=XM_023249396.1
X	33937041 33937101 refGene_gn=SRPX;refGene_tx=XM_023249396.1
X	33990177 33990265 refGene_gn=SRPX;refGene_tx=XM_023249396.1
X	34000520 34000535 refGene_gn=RPGR;refGene_tx=XM_019823726.2
X	34030509 34030716 refGene_gn=RPGR;refGene_tx=XM_004000399.5
X	34036740 34036887 refGene_gn=RPGR;refGene_tx=XM_006943651.3
X	34039396 34039485
refGene_gn=RPGR;refGene_tx=XM_004000399.5;refGene_gn=RPGR;refGene_tx=X
M_006943651.3;refGene_gn=RPGR;refGene_tx=XM_019823726.2
X	34041030 34041088
refGene_gn=RPGR;refGene_tx=XM_004000399.5;refGene_gn=RPGR;refGene_tx=X
M_006943651.3;refGene_gn=RPGR;refGene_tx=XM_019823726.2
X	34042336 34042450
refGene_gn=RPGR;refGene_tx=XM_004000399.5;refGene_gn=RPGR;refGene_tx=X
M_006943651.3;refGene_gn=RPGR;refGene_tx=XM_019823726.2
X	34052161 34053094
refGene_gn=RPGR;refGene_tx=XM_004000399.5;refGene_gn=RPGR;refGene_tx=X
M_006943651.3;refGene_gn=RPGR;refGene_tx=XM_019823726.2
X	34055453 34055519
refGene_gn=RPGR;refGene_tx=XM_004000399.5;refGene_gn=RPGR;refGene_tx=X

M_006943651.3;refGene_gn=RPGR;refGene_tx=XM_019823726.2
X 34055864 34055962
refGene_gn=RPGR;refGene_tx=XM_004000399.5;refGene_gn=RPGR;refGene_tx=XM_006943651.3;refGene_gn=RPGR;refGene_tx=XM_019823726.2
X 34061155 34061327
refGene_gn=RPGR;refGene_tx=XM_004000399.5;refGene_gn=RPGR;refGene_tx=XM_006943651.3;refGene_gn=RPGR;refGene_tx=XM_019823726.2
X 34062261 34062447
refGene_gn=RPGR;refGene_tx=XM_004000399.5;refGene_gn=RPGR;refGene_tx=XM_006943651.3;refGene_gn=RPGR;refGene_tx=XM_019823726.2
X 34063732 34063857
refGene_gn=RPGR;refGene_tx=XM_004000399.5;refGene_gn=RPGR;refGene_tx=XM_006943651.3;refGene_gn=RPGR;refGene_tx=XM_019823726.2
X 34067046 34067202
refGene_gn=RPGR;refGene_tx=XM_004000399.5;refGene_gn=RPGR;refGene_tx=XM_006943651.3;refGene_gn=RPGR;refGene_tx=XM_019823726.2
X 34071710 34071869
refGene_gn=RPGR;refGene_tx=XM_004000399.5;refGene_gn=RPGR;refGene_tx=XM_006943651.3;refGene_gn=RPGR;refGene_tx=XM_019823726.2
X 34074975 34075125
refGene_gn=RPGR;refGene_tx=XM_004000399.5;refGene_gn=RPGR;refGene_tx=XM_006943651.3;refGene_gn=RPGR;refGene_tx=XM_019823726.2
X 34076421 34076580
refGene_gn=RPGR;refGene_tx=XM_004000399.5;refGene_gn=RPGR;refGene_tx=XM_006943651.3;refGene_gn=RPGR;refGene_tx=XM_019823726.2
X 34078531 34078594
refGene_gn=RPGR;refGene_tx=XM_004000399.5;refGene_gn=RPGR;refGene_tx=XM_006943651.3;refGene_gn=RPGR;refGene_tx=XM_019823726.2
X 34080045 34080138
refGene_gn=RPGR;refGene_tx=XM_004000399.5;refGene_gn=RPGR;refGene_tx=XM_006943651.3;refGene_gn=RPGR;refGene_tx=XM_019823726.2
X 34080621 34080747
refGene_gn=RPGR;refGene_tx=XM_004000399.5;refGene_gn=RPGR;refGene_tx=XM_006943651.3;refGene_gn=RPGR;refGene_tx=XM_019823726.2
X 34082461 34082624
refGene_gn=RPGR;refGene_tx=XM_004000399.5;refGene_gn=RPGR;refGene_tx=XM_006943651.3;refGene_gn=RPGR;refGene_tx=XM_019823726.2
X 34105698 34105775 refGene_gn=OTC;refGene_tx=XM_004000400.4
X 34117718 34117857 refGene_gn=OTC;refGene_tx=XM_004000400.4
X 34119553 34119635 refGene_gn=OTC;refGene_tx=XM_004000400.4
X 34128890 34128978 refGene_gn=OTC;refGene_tx=XM_004000400.4
X 34147608 34147762 refGene_gn=OTC;refGene_tx=XM_004000400.4
X 34149865 34149988 refGene_gn=OTC;refGene_tx=XM_004000400.4
X 34156021 34156075 refGene_gn=OTC;refGene_tx=XM_004000400.4
X 34156151 34156301 refGene_gn=OTC;refGene_tx=XM_004000400.4
X 34159639 34159777 refGene_gn=OTC;refGene_tx=XM_004000400.4
X 34166505 34166565 refGene_gn=OTC;refGene_tx=XM_004000400.4
X 34224386 34224406
refGene_gn=LOC101083747;refGene_tx=XM_019823728.2;refGene_gn=LOC101083747;refGene_tx=XM_023249428.1;refGene_gn=LOC101083747;refGene_tx=XM_02

3249429.1
X	34225052	34225542
refGene_gn=LOC101083747;refGene_tx=XM_019823728.2;refGene_gn=LOC101083747;refGene_tx=XM_023249428.1;refGene_gn=LOC101083747;refGene_tx=XM_023249429.1
X	34285656	34285735
refGene_gn=LOC101083747;refGene_tx=XM_023249428.1;refGene_gn=LOC101083747;refGene_tx=XM_023249429.1
X	34291756	34291813
refGene_gn=LOC101083747;refGene_tx=XM_023249428.1
X	34292104	34292185	refGene_gn=TSPAN7;refGene_tx=XM_004000401.5
X	34393534	34393723	refGene_gn=TSPAN7;refGene_tx=XM_004000401.5
X	34397714	34397789	refGene_gn=TSPAN7;refGene_tx=XM_004000401.5
X	34400135	34400231	refGene_gn=TSPAN7;refGene_tx=XM_004000401.5
X	34401821	34401977	refGene_gn=TSPAN7;refGene_tx=XM_004000401.5
X	34407712	34407796	refGene_gn=TSPAN7;refGene_tx=XM_004000401.5
X	34416456	34416525	refGene_gn=TSPAN7;refGene_tx=XM_004000401.5
X	34499564	34500116	refGene_gn=MID1IP1;refGene_tx=XM_004000403.5
X	35545367	35545491
refGene_gn=BCOR;refGene_tx=XM_019823730.2;refGene_gn=BCOR;refGene_tx=XM_019823731.2;refGene_gn=BCOR;refGene_tx=XM_019823732.2;refGene_gn=BCOR;refGene_tx=XM_019823733.2;refGene_gn=BCOR;refGene_tx=XM_023249532.1;refGene_gn=BCOR;refGene_tx=XM_023249533.1
X	35545804	35546096
refGene_gn=BCOR;refGene_tx=XM_011291671.3;refGene_gn=BCOR;refGene_tx=XM_011291672.3;refGene_gn=BCOR;refGene_tx=XM_011291673.3;refGene_gn=BCOR;refGene_tx=XM_011291674.3;refGene_gn=BCOR;refGene_tx=XM_019823730.2;refGene_gn=BCOR;refGene_tx=XM_019823731.2;refGene_gn=BCOR;refGene_tx=XM_019823732.2;refGene_gn=BCOR;refGene_tx=XM_019823733.2;refGene_gn=BCOR;refGene_tx=XM_023249532.1;refGene_gn=BCOR;refGene_tx=XM_023249533.1
X	35547229	35547386
refGene_gn=BCOR;refGene_tx=XM_011291671.3;refGene_gn=BCOR;refGene_tx=XM_011291672.3;refGene_gn=BCOR;refGene_tx=XM_011291673.3;refGene_gn=BCOR;refGene_tx=XM_011291674.3;refGene_gn=BCOR;refGene_tx=XM_019823730.2;refGene_gn=BCOR;refGene_tx=XM_019823731.2;refGene_gn=BCOR;refGene_tx=XM_019823732.2;refGene_gn=BCOR;refGene_tx=XM_019823733.2;refGene_gn=BCOR;refGene_tx=XM_023249532.1;refGene_gn=BCOR;refGene_tx=XM_023249533.1
X	35547580	35547658
refGene_gn=BCOR;refGene_tx=XM_011291671.3;refGene_gn=BCOR;refGene_tx=XM_011291672.3;refGene_gn=BCOR;refGene_tx=XM_011291673.3;refGene_gn=BCOR;refGene_tx=XM_011291674.3;refGene_gn=BCOR;refGene_tx=XM_019823730.2;refGene_gn=BCOR;refGene_tx=XM_019823731.2;refGene_gn=BCOR;refGene_tx=XM_019823732.2;refGene_gn=BCOR;refGene_tx=XM_019823733.2;refGene_gn=BCOR;refGene_tx=XM_023249532.1;refGene_gn=BCOR;refGene_tx=XM_023249533.1
X	35548123	35548269
refGene_gn=BCOR;refGene_tx=XM_011291671.3;refGene_gn=BCOR;refGene_tx=XM_011291672.3;refGene_gn=BCOR;refGene_tx=XM_011291673.3;refGene_gn=BCOR;refGene_tx=XM_011291674.3;refGene_gn=BCOR;refGene_tx=XM_019823730.2;refGene_gn=BCOR;refGene_tx=XM_019823731.2;refGene_gn=BCOR;refGene_tx=XM_019823732.2;refGene_gn=BCOR;refGene_tx=XM_019823733.2;refGene_gn=BCO

R;refGene_tx=XM_023249532.1;refGene_gn=BCOR;refGene_tx=XM_023249533.1
X 35549652 35549819
refGene_gn=BCOR;refGene_tx=XM_011291671.3;refGene_gn=BCOR;refGene_tx=XM_011291672.3;refGene_gn=BCOR;refGene_tx=XM_011291673.3;refGene_gn=BCOR;refGene_tx=XM_011291674.3;refGene_gn=BCOR;refGene_tx=XM_019823730.2;refGene_gn=BCOR;refGene_tx=XM_019823731.2;refGene_gn=BCOR;refGene_tx=XM_019823732.2;refGene_gn=BCOR;refGene_tx=XM_019823733.2;refGene_gn=BCOR;refGene_tx=XM_023249532.1;refGene_gn=BCOR;refGene_tx=XM_023249533.1
X 35553917 35554172
refGene_gn=BCOR;refGene_tx=XM_011291671.3;refGene_gn=BCOR;refGene_tx=XM_011291672.3;refGene_gn=BCOR;refGene_tx=XM_011291673.3;refGene_gn=BCOR;refGene_tx=XM_011291674.3;refGene_gn=BCOR;refGene_tx=XM_019823730.2;refGene_gn=BCOR;refGene_tx=XM_019823731.2;refGene_gn=BCOR;refGene_tx=XM_019823732.2;refGene_gn=BCOR;refGene_tx=XM_019823733.2;refGene_gn=BCOR;refGene_tx=XM_023249532.1;refGene_gn=BCOR;refGene_tx=XM_023249533.1
X 35554478 35554804
refGene_gn=BCOR;refGene_tx=XM_011291671.3;refGene_gn=BCOR;refGene_tx=XM_011291672.3;refGene_gn=BCOR;refGene_tx=XM_011291673.3;refGene_gn=BCOR;refGene_tx=XM_011291674.3;refGene_gn=BCOR;refGene_tx=XM_019823730.2;refGene_gn=BCOR;refGene_tx=XM_019823731.2;refGene_gn=BCOR;refGene_tx=XM_019823732.2;refGene_gn=BCOR;refGene_tx=XM_019823733.2;refGene_gn=BCOR;refGene_tx=XM_023249532.1;refGene_gn=BCOR;refGene_tx=XM_023249533.1
X 35555356 35555701
refGene_gn=BCOR;refGene_tx=XM_011291671.3;refGene_gn=BCOR;refGene_tx=XM_011291672.3;refGene_gn=BCOR;refGene_tx=XM_011291673.3;refGene_gn=BCOR;refGene_tx=XM_011291674.3;refGene_gn=BCOR;refGene_tx=XM_019823730.2;refGene_gn=BCOR;refGene_tx=XM_019823731.2;refGene_gn=BCOR;refGene_tx=XM_019823732.2;refGene_gn=BCOR;refGene_tx=XM_019823733.2;refGene_gn=BCOR;refGene_tx=XM_023249532.1;refGene_gn=BCOR;refGene_tx=XM_023249533.1
X 35556084 35556345
refGene_gn=BCOR;refGene_tx=XM_011291671.3;refGene_gn=BCOR;refGene_tx=XM_011291672.3;refGene_gn=BCOR;refGene_tx=XM_011291673.3;refGene_gn=BCOR;refGene_tx=XM_011291674.3;refGene_gn=BCOR;refGene_tx=XM_019823730.2;refGene_gn=BCOR;refGene_tx=XM_019823731.2;refGene_gn=BCOR;refGene_tx=XM_019823732.2;refGene_gn=BCOR;refGene_tx=XM_019823733.2;refGene_gn=BCOR;refGene_tx=XM_023249532.1;refGene_gn=BCOR;refGene_tx=XM_023249533.1
X 35561759 35561946
refGene_gn=BCOR;refGene_tx=XM_011291671.3;refGene_gn=BCOR;refGene_tx=XM_011291672.3;refGene_gn=BCOR;refGene_tx=XM_011291673.3;refGene_gn=BCOR;refGene_tx=XM_011291674.3;refGene_gn=BCOR;refGene_tx=XM_019823730.2;refGene_gn=BCOR;refGene_tx=XM_019823731.2;refGene_gn=BCOR;refGene_tx=XM_019823732.2;refGene_gn=BCOR;refGene_tx=XM_019823733.2;refGene_gn=BCOR;refGene_tx=XM_023249532.1;refGene_gn=BCOR;refGene_tx=XM_023249533.1
X 35562429 35562483
refGene_gn=BCOR;refGene_tx=XM_011291671.3;refGene_gn=BCOR;refGene_tx=XM_011291673.3;refGene_gn=BCOR;refGene_tx=XM_019823730.2;refGene_gn=BCOR;refGene_tx=XM_019823731.2;refGene_gn=BCOR;refGene_tx=XM_019823733.2;refGene_gn=BCOR;refGene_tx=XM_023249532.1;refGene_gn=BCOR;refGene_tx=XM_023249533.1
X 35563161 35566008

refGene_gn=BCOR;refGene_tx=XM_011291671.3;refGene_gn=BCOR;refGene_tx=XM_011291672.3;refGene_gn=BCOR;refGene_tx=XM_011291673.3;refGene_gn=BCOR;refGene_tx=XM_011291674.3;refGene_gn=BCOR;refGene_tx=XM_019823730.2;refGene_gn=BCOR;refGene_tx=XM_019823731.2;refGene_gn=BCOR;refGene_tx=XM_019823732.2;refGene_gn=BCOR;refGene_tx=XM_019823733.2;refGene_gn=BCOR;refGene_tx=XM_023249532.1;refGene_gn=BCOR;refGene_tx=XM_023249533.1
X 35567029 35567108
refGene_gn=BCOR;refGene_tx=XM_011291671.3;refGene_gn=BCOR;refGene_tx=XM_011291672.3;refGene_gn=BCOR;refGene_tx=XM_011291673.3;refGene_gn=BCOR;refGene_tx=XM_011291674.3;refGene_gn=BCOR;refGene_tx=XM_019823730.2;refGene_gn=BCOR;refGene_tx=XM_019823731.2;refGene_gn=BCOR;refGene_tx=XM_019823732.2;refGene_gn=BCOR;refGene_tx=XM_019823733.2;refGene_gn=BCOR;refGene_tx=XM_023249532.1;refGene_gn=BCOR;refGene_tx=XM_023249533.1
X 35568435 35568521
refGene_gn=BCOR;refGene_tx=XM_011291671.3;refGene_gn=BCOR;refGene_tx=XM_011291672.3;refGene_gn=BCOR;refGene_tx=XM_011291673.3;refGene_gn=BCOR;refGene_tx=XM_011291674.3;refGene_gn=BCOR;refGene_tx=XM_019823730.2;refGene_gn=BCOR;refGene_tx=XM_019823731.2;refGene_gn=BCOR;refGene_tx=XM_019823732.2;refGene_gn=BCOR;refGene_tx=XM_019823733.2;refGene_gn=BCOR;refGene_tx=XM_023249532.1;refGene_gn=BCOR;refGene_tx=XM_023249533.1
X 35989954 35989994 refGene_gn=ATP6AP2;refGene_tx=XM_023248896.1
X 35996955 35997126
refGene_gn=ATP6AP2;refGene_tx=XM_004000408.3;refGene_gn=ATP6AP2;refGene_tx=XM_023248896.1
X 35999015 35999147
refGene_gn=ATP6AP2;refGene_tx=XM_004000408.3;refGene_gn=ATP6AP2;refGene_tx=XM_023248896.1
X 36004449 36004545
refGene_gn=ATP6AP2;refGene_tx=XM_004000408.3;refGene_gn=ATP6AP2;refGene_tx=XM_023248896.1
X 36005169 36005307
refGene_gn=ATP6AP2;refGene_tx=XM_004000408.3;refGene_gn=ATP6AP2;refGene_tx=XM_023248896.1
X 36006169 36006223
refGene_gn=ATP6AP2;refGene_tx=XM_004000408.3;refGene_gn=ATP6AP2;refGene_tx=XM_023248896.1
X 36007669 36007819
refGene_gn=ATP6AP2;refGene_tx=XM_004000408.3;refGene_gn=ATP6AP2;refGene_tx=XM_023248896.1
X 36008610 36008730
refGene_gn=ATP6AP2;refGene_tx=XM_004000408.3;refGene_gn=ATP6AP2;refGene_tx=XM_023248896.1
X 36012108 36012303
refGene_gn=ATP6AP2;refGene_tx=XM_004000408.3;refGene_gn=ATP6AP2;refGene_tx=XM_023248896.1
X 36038328 36038754 refGene_gn=MPC1L;refGene_tx=XM_004000457.4
X 36045093 36045255
refGene_gn=CXHXorf38;refGene_tx=XM_011291694.3
X 36052187 36052367
refGene_gn=CXHXorf38;refGene_tx=XM_011291694.3

X 36052892 36053042
refGene_gn=CXHXorf38;refGene_tx=XM_011291694.3
X 36054938 36055058
refGene_gn=CXHXorf38;refGene_tx=XM_011291694.3
X 36060944 36061079
refGene_gn=CXHXorf38;refGene_tx=XM_011291694.3
X 36061248 36061464
refGene_gn=CXHXorf38;refGene_tx=XM_011291694.3
X 36066096 36066170
refGene_gn=MED14;refGene_tx=XM_023249310.1;refGene_gn=MED14;refGene_tx=XM_023249311.1;refGene_gn=MED14;refGene_tx=XM_023249312.1
X 36068793 36068986
refGene_gn=MED14;refGene_tx=XM_023249310.1;refGene_gn=MED14;refGene_tx=XM_023249311.1;refGene_gn=MED14;refGene_tx=XM_023249312.1
X 36069395 36069521
refGene_gn=MED14;refGene_tx=XM_023249310.1;refGene_gn=MED14;refGene_tx=XM_023249311.1;refGene_gn=MED14;refGene_tx=XM_023249312.1
X 36071711 36071819
refGene_gn=MED14;refGene_tx=XM_023249310.1;refGene_gn=MED14;refGene_tx=XM_023249311.1;refGene_gn=MED14;refGene_tx=XM_023249312.1
X 36072352 36072532
refGene_gn=MED14;refGene_tx=XM_023249310.1;refGene_gn=MED14;refGene_tx=XM_023249311.1
X 36074079 36074315
refGene_gn=MED14;refGene_tx=XM_023249310.1;refGene_gn=MED14;refGene_tx=XM_023249311.1;refGene_gn=MED14;refGene_tx=XM_023249312.1
X 36075231 36075414
refGene_gn=MED14;refGene_tx=XM_023249310.1;refGene_gn=MED14;refGene_tx=XM_023249311.1;refGene_gn=MED14;refGene_tx=XM_023249312.1
X 36077177 36077309
refGene_gn=MED14;refGene_tx=XM_023249310.1;refGene_gn=MED14;refGene_tx=XM_023249312.1
X 36079922 36080034
refGene_gn=MED14;refGene_tx=XM_023249310.1;refGene_gn=MED14;refGene_tx=XM_023249311.1;refGene_gn=MED14;refGene_tx=XM_023249312.1
X 36081618 36081759
refGene_gn=MED14;refGene_tx=XM_023249310.1;refGene_gn=MED14;refGene_tx=XM_023249311.1;refGene_gn=MED14;refGene_tx=XM_023249312.1
X 36085310 36085580
refGene_gn=MED14;refGene_tx=XM_023249310.1;refGene_gn=MED14;refGene_tx=XM_023249311.1;refGene_gn=MED14;refGene_tx=XM_023249312.1
X 36085885 36086038
refGene_gn=MED14;refGene_tx=XM_023249310.1;refGene_gn=MED14;refGene_tx=XM_023249311.1;refGene_gn=MED14;refGene_tx=XM_023249312.1
X 36087201 36087293
refGene_gn=MED14;refGene_tx=XM_023249310.1;refGene_gn=MED14;refGene_tx=XM_023249311.1;refGene_gn=MED14;refGene_tx=XM_023249312.1
X 36087980 36088127
refGene_gn=MED14;refGene_tx=XM_023249310.1;refGene_gn=MED14;refGene_tx=XM_023249311.1;refGene_gn=MED14;refGene_tx=XM_023249312.1

X 36088304 36088465
refGene_gn=MED14;refGene_tx=XM_023249310.1;refGene_gn=MED14;refGene_tx=XM_023249311.1;refGene_gn=MED14;refGene_tx=XM_023249312.1
X 36093517 36093594
refGene_gn=MED14;refGene_tx=XM_023249310.1;refGene_gn=MED14;refGene_tx=XM_023249311.1;refGene_gn=MED14;refGene_tx=XM_023249312.1
X 36096667 36096802
refGene_gn=MED14;refGene_tx=XM_023249310.1;refGene_gn=MED14;refGene_tx=XM_023249311.1;refGene_gn=MED14;refGene_tx=XM_023249312.1
X 36097233 36097428
refGene_gn=MED14;refGene_tx=XM_023249310.1;refGene_gn=MED14;refGene_tx=XM_023249311.1;refGene_gn=MED14;refGene_tx=XM_023249312.1
X 36102069 36102229
refGene_gn=MED14;refGene_tx=XM_023249310.1;refGene_gn=MED14;refGene_tx=XM_023249311.1;refGene_gn=MED14;refGene_tx=XM_023249312.1
X 36106494 36106573
refGene_gn=MED14;refGene_tx=XM_023249310.1;refGene_gn=MED14;refGene_tx=XM_023249311.1;refGene_gn=MED14;refGene_tx=XM_023249312.1
X 36108569 36108695
refGene_gn=MED14;refGene_tx=XM_023249310.1;refGene_gn=MED14;refGene_tx=XM_023249311.1;refGene_gn=MED14;refGene_tx=XM_023249312.1
X 36113945 36114057
refGene_gn=MED14;refGene_tx=XM_023249310.1;refGene_gn=MED14;refGene_tx=XM_023249311.1;refGene_gn=MED14;refGene_tx=XM_023249312.1
X 36115172 36115323
refGene_gn=MED14;refGene_tx=XM_023249310.1;refGene_gn=MED14;refGene_tx=XM_023249311.1;refGene_gn=MED14;refGene_tx=XM_023249312.1
X 36116553 36116686
refGene_gn=MED14;refGene_tx=XM_023249310.1;refGene_gn=MED14;refGene_tx=XM_023249311.1;refGene_gn=MED14;refGene_tx=XM_023249312.1
X 36117191 36117299
refGene_gn=MED14;refGene_tx=XM_023249310.1;refGene_gn=MED14;refGene_tx=XM_023249311.1;refGene_gn=MED14;refGene_tx=XM_023249312.1
X 36117625 36117754
refGene_gn=MED14;refGene_tx=XM_023249310.1;refGene_gn=MED14;refGene_tx=XM_023249311.1;refGene_gn=MED14;refGene_tx=XM_023249312.1
X 36118240 36118370
refGene_gn=MED14;refGene_tx=XM_023249310.1;refGene_gn=MED14;refGene_tx=XM_023249311.1;refGene_gn=MED14;refGene_tx=XM_023249312.1
X 36119197 36119371
refGene_gn=MED14;refGene_tx=XM_023249310.1;refGene_gn=MED14;refGene_tx=XM_023249311.1;refGene_gn=MED14;refGene_tx=XM_023249312.1
X 36123673 36123779
refGene_gn=MED14;refGene_tx=XM_023249310.1;refGene_gn=MED14;refGene_tx=XM_023249311.1;refGene_gn=MED14;refGene_tx=XM_023249312.1
X 36126141 36126168
refGene_gn=MED14;refGene_tx=XM_023249310.1;refGene_gn=MED14;refGene_tx=XM_023249311.1;refGene_gn=MED14;refGene_tx=XM_023249312.1
X 36130792 36131007
refGene_gn=MED14;refGene_tx=XM_023249310.1;refGene_gn=MED14;refGene_tx

=XM_023249311.1;refGene_gn=MED14;refGene_tx=XM_023249312.1
X	36131043	36131227	refGene_gn=MED14OS;refGene_tx=XM_023249313.1
X	36131229	36131510	refGene_gn=MED14OS;refGene_tx=XM_023249313.1
X	36495056	36495152	refGene_gn=USP9X;refGene_tx=XM_004000409.5
X	36497097	36497243	refGene_gn=USP9X;refGene_tx=XM_004000409.5
X	36499854	36499934	refGene_gn=USP9X;refGene_tx=XM_004000409.5
X	36501614	36501727	refGene_gn=USP9X;refGene_tx=XM_004000409.5
X	36503030	36503249	refGene_gn=USP9X;refGene_tx=XM_004000409.5
X	36505452	36505568	refGene_gn=USP9X;refGene_tx=XM_004000409.5
X	36505761	36506013	refGene_gn=USP9X;refGene_tx=XM_004000409.5
X	36506089	36506228	refGene_gn=USP9X;refGene_tx=XM_004000409.5
X	36507896	36508049	refGene_gn=USP9X;refGene_tx=XM_004000409.5
X	36509224	36509329	refGene_gn=USP9X;refGene_tx=XM_004000409.5
X	36513006	36513213	refGene_gn=USP9X;refGene_tx=XM_004000409.5
X	36514690	36514827	refGene_gn=USP9X;refGene_tx=XM_004000409.5
X	36516307	36516441	refGene_gn=USP9X;refGene_tx=XM_004000409.5
X	36527864	36527952	refGene_gn=USP9X;refGene_tx=XM_004000409.5
X	36530571	36530914	refGene_gn=USP9X;refGene_tx=XM_004000409.5
X	36533320	36533416	refGene_gn=USP9X;refGene_tx=XM_004000409.5
X	36533837	36534049	refGene_gn=USP9X;refGene_tx=XM_004000409.5
X	36534996	36535009	refGene_gn=USP9X;refGene_tx=XM_004000409.5
X	36535010	36535238	refGene_gn=USP9X;refGene_tx=XM_004000409.5
X	36535575	36535725	refGene_gn=USP9X;refGene_tx=XM_004000409.5
X	36536889	36537010	refGene_gn=USP9X;refGene_tx=XM_004000409.5
X	36544752	36544883	refGene_gn=USP9X;refGene_tx=XM_004000409.5
X	36545073	36545352	refGene_gn=USP9X;refGene_tx=XM_004000409.5
X	36547567	36547693	refGene_gn=USP9X;refGene_tx=XM_004000409.5
X	36548604	36548730	refGene_gn=USP9X;refGene_tx=XM_004000409.5
X	36549944	36550111	refGene_gn=USP9X;refGene_tx=XM_004000409.5
X	36556225	36556334	refGene_gn=USP9X;refGene_tx=XM_004000409.5
X	36556556	36556703	refGene_gn=USP9X;refGene_tx=XM_004000409.5
X	36557503	36557650	refGene_gn=USP9X;refGene_tx=XM_004000409.5
X	36558536	36558759	refGene_gn=USP9X;refGene_tx=XM_004000409.5
X	36561418	36561639	refGene_gn=USP9X;refGene_tx=XM_004000409.5
X	36564538	36564729	refGene_gn=USP9X;refGene_tx=XM_004000409.5
X	36569649	36569823	refGene_gn=USP9X;refGene_tx=XM_004000409.5
X	36572417	36572559	refGene_gn=USP9X;refGene_tx=XM_004000409.5
X	36573700	36574454	refGene_gn=USP9X;refGene_tx=XM_004000409.5
X	36575071	36575195	refGene_gn=USP9X;refGene_tx=XM_004000409.5
X	36575906	36576132	refGene_gn=USP9X;refGene_tx=XM_004000409.5
X	36576661	36576791	refGene_gn=USP9X;refGene_tx=XM_004000409.5
X	36580661	36580847	refGene_gn=USP9X;refGene_tx=XM_004000409.5
X	36581322	36581543	refGene_gn=USP9X;refGene_tx=XM_004000409.5
X	36581633	36581722	refGene_gn=USP9X;refGene_tx=XM_004000409.5
X	36585267	36585424	refGene_gn=USP9X;refGene_tx=XM_004000409.5
X	36585574	36585787	refGene_gn=USP9X;refGene_tx=XM_004000409.5
X	36586124	36586220	refGene_gn=USP9X;refGene_tx=XM_004000409.5
X	36588563	36588701	refGene_gn=USP9X;refGene_tx=XM_004000409.5
X	36684267	36684312
refGene_gn=DDX3X;refGene_tx=XM_004000412.5;refGene_gn=DDX3X;refGene_tx

=XM_023248806.1;refGene_gn=DDX3X;refGene_tx=XM_023248807.1
X 36687416 36687474
refGene_gn=DDX3X;refGene_tx=XM_004000412.5;refGene_gn=DDX3X;refGene_tx
=XM_023248806.1;refGene_gn=DDX3X;refGene_tx=XM_023248807.1
X 36689416 36689464
refGene_gn=DDX3X;refGene_tx=XM_004000412.5;refGene_gn=DDX3X;refGene_tx
=XM_023248806.1;refGene_gn=DDX3X;refGene_tx=XM_023248807.1
X 36691338 36691471
refGene_gn=DDX3X;refGene_tx=XM_004000412.5;refGene_gn=DDX3X;refGene_tx
=XM_023248806.1;refGene_gn=DDX3X;refGene_tx=XM_023248807.1
X 36692410 36692569
refGene_gn=DDX3X;refGene_tx=XM_004000412.5;refGene_gn=DDX3X;refGene_tx
=XM_023248806.1;refGene_gn=DDX3X;refGene_tx=XM_023248807.1
X 36692651 36692751
refGene_gn=DDX3X;refGene_tx=XM_004000412.5;refGene_gn=DDX3X;refGene_tx
=XM_023248806.1;refGene_gn=DDX3X;refGene_tx=XM_023248807.1
X 36693150 36693286
refGene_gn=DDX3X;refGene_tx=XM_004000412.5;refGene_gn=DDX3X;refGene_tx
=XM_023248806.1;refGene_gn=DDX3X;refGene_tx=XM_023248807.1
X 36694536 36694622
refGene_gn=DDX3X;refGene_tx=XM_004000412.5;refGene_gn=DDX3X;refGene_tx
=XM_023248806.1;refGene_gn=DDX3X;refGene_tx=XM_023248807.1
X 36694935 36695034
refGene_gn=DDX3X;refGene_tx=XM_004000412.5;refGene_gn=DDX3X;refGene_tx
=XM_023248806.1;refGene_gn=DDX3X;refGene_tx=XM_023248807.1
X 36695143 36695304
refGene_gn=DDX3X;refGene_tx=XM_004000412.5;refGene_gn=DDX3X;refGene_tx
=XM_023248806.1;refGene_gn=DDX3X;refGene_tx=XM_023248807.1
X 36695712 36695857
refGene_gn=DDX3X;refGene_tx=XM_004000412.5;refGene_gn=DDX3X;refGene_tx
=XM_023248806.1;refGene_gn=DDX3X;refGene_tx=XM_023248807.1
X 36695935 36696080
refGene_gn=DDX3X;refGene_tx=XM_004000412.5;refGene_gn=DDX3X;refGene_tx
=XM_023248806.1;refGene_gn=DDX3X;refGene_tx=XM_023248807.1
X 36696276 36696458
refGene_gn=DDX3X;refGene_tx=XM_004000412.5;refGene_gn=DDX3X;refGene_tx
=XM_023248806.1;refGene_gn=DDX3X;refGene_tx=XM_023248807.1
X 36696544 36696662
refGene_gn=DDX3X;refGene_tx=XM_004000412.5;refGene_gn=DDX3X;refGene_tx
=XM_023248806.1;refGene_gn=DDX3X;refGene_tx=XM_023248807.1
X 36696898 36697052
refGene_gn=DDX3X;refGene_tx=XM_004000412.5;refGene_gn=DDX3X;refGene_tx
=XM_023248806.1;refGene_gn=DDX3X;refGene_tx=XM_023248807.1
X 36697355 36697513
refGene_gn=DDX3X;refGene_tx=XM_004000412.5;refGene_gn=DDX3X;refGene_tx
=XM_023248806.1;refGene_gn=DDX3X;refGene_tx=XM_023248807.1
X 36697715 36697795
refGene_gn=DDX3X;refGene_tx=XM_004000412.5;refGene_gn=DDX3X;refGene_tx
=XM_023248806.1;refGene_gn=DDX3X;refGene_tx=XM_023248807.1
X 36802938 36804360 refGene_gn=NYX;refGene_tx=XM_023248957.1

X	36866802 36866979
refGene_gn=CASK;refGene_tx=XM_004000415.5;refGene_gn=CASK;refGene_tx=XM_006943661.4;refGene_gn=CASK;refGene_tx=XM_019823737.2;refGene_gn=CASK;refGene_tx=XM_019823738.2;refGene_gn=CASK;refGene_tx=XM_019823739.2;refGene_gn=CASK;refGene_tx=XM_019823740.2;refGene_gn=CASK;refGene_tx=XM_019823741.2;refGene_gn=CASK;refGene_tx=XM_019823742.2;refGene_gn=CASK;refGene_tx=XM_019823743.2;refGene_gn=CASK;refGene_tx=XM_019823744.2;refGene_gn=CASK;refGene_tx=XM_019823745.2;refGene_gn=CASK;refGene_tx=XM_019823746.2;refGene_gn=CASK;refGene_tx=XM_019823747.2;refGene_gn=CASK;refGene_tx=XM_019823748.2;refGene_gn=CASK;refGene_tx=XM_019823749.2
X	36869156 36869183
refGene_gn=CASK;refGene_tx=XM_019823737.2;refGene_gn=CASK;refGene_tx=XM_019823738.2;refGene_gn=CASK;refGene_tx=XM_019823739.2;refGene_gn=CASK;refGene_tx=XM_019823740.2;refGene_gn=CASK;refGene_tx=XM_019823742.2;refGene_gn=CASK;refGene_tx=XM_019823745.2;refGene_gn=CASK;refGene_tx=XM_019823747.2
X	36870075 36870159
refGene_gn=CASK;refGene_tx=XM_004000415.5;refGene_gn=CASK;refGene_tx=XM_006943661.4;refGene_gn=CASK;refGene_tx=XM_019823737.2;refGene_gn=CASK;refGene_tx=XM_019823738.2;refGene_gn=CASK;refGene_tx=XM_019823739.2;refGene_gn=CASK;refGene_tx=XM_019823740.2;refGene_gn=CASK;refGene_tx=XM_019823741.2;refGene_gn=CASK;refGene_tx=XM_019823742.2;refGene_gn=CASK;refGene_tx=XM_019823743.2;refGene_gn=CASK;refGene_tx=XM_019823744.2;refGene_gn=CASK;refGene_tx=XM_019823745.2;refGene_gn=CASK;refGene_tx=XM_019823746.2;refGene_gn=CASK;refGene_tx=XM_019823747.2;refGene_gn=CASK;refGene_tx=XM_019823748.2;refGene_gn=CASK;refGene_tx=XM_019823749.2
X	36877103 36877306
refGene_gn=CASK;refGene_tx=XM_004000415.5;refGene_gn=CASK;refGene_tx=XM_006943661.4;refGene_gn=CASK;refGene_tx=XM_019823737.2;refGene_gn=CASK;refGene_tx=XM_019823738.2;refGene_gn=CASK;refGene_tx=XM_019823739.2;refGene_gn=CASK;refGene_tx=XM_019823740.2;refGene_gn=CASK;refGene_tx=XM_019823741.2;refGene_gn=CASK;refGene_tx=XM_019823742.2;refGene_gn=CASK;refGene_tx=XM_019823743.2;refGene_gn=CASK;refGene_tx=XM_019823744.2;refGene_gn=CASK;refGene_tx=XM_019823745.2;refGene_gn=CASK;refGene_tx=XM_019823746.2;refGene_gn=CASK;refGene_tx=XM_019823747.2;refGene_gn=CASK;refGene_tx=XM_019823748.2;refGene_gn=CASK;refGene_tx=XM_019823749.2
X	36879836 36879917
refGene_gn=CASK;refGene_tx=XM_004000415.5;refGene_gn=CASK;refGene_tx=XM_006943661.4;refGene_gn=CASK;refGene_tx=XM_019823737.2;refGene_gn=CASK;refGene_tx=XM_019823738.2;refGene_gn=CASK;refGene_tx=XM_019823739.2;refGene_gn=CASK;refGene_tx=XM_019823740.2;refGene_gn=CASK;refGene_tx=XM_019823741.2;refGene_gn=CASK;refGene_tx=XM_019823742.2;refGene_gn=CASK;refGene_tx=XM_019823743.2;refGene_gn=CASK;refGene_tx=XM_019823744.2;refGene_gn=CASK;refGene_tx=XM_019823745.2;refGene_gn=CASK;refGene_tx=XM_019823746.2;refGene_gn=CASK;refGene_tx=XM_019823747.2;refGene_gn=CASK;refGene_tx=XM_019823748.2;refGene_gn=CASK;refGene_tx=XM_019823749.2
X	36880022 36880103
refGene_gn=CASK;refGene_tx=XM_004000415.5;refGene_gn=CASK;refGene_tx=XM_006943661.4;refGene_gn=CASK;refGene_tx=XM_019823737.2;refGene_gn=CASK;refGene_tx=XM_019823738.2;refGene_gn=CASK;refGene_tx=XM_019823739.2;

refGene_gn=CASK;refGene_tx=XM_019823740.2;refGene_gn=CASK;refGene_tx=XM_019823741.2;refGene_gn=CASK;refGene_tx=XM_019823742.2;refGene_gn=CASK;refGene_tx=XM_019823743.2;refGene_gn=CASK;refGene_tx=XM_019823744.2;refGene_gn=CASK;refGene_tx=XM_019823745.2;refGene_gn=CASK;refGene_tx=XM_019823746.2;refGene_gn=CASK;refGene_tx=XM_019823747.2;refGene_gn=CASK;refGene_tx=XM_019823748.2;refGene_gn=CASK;refGene_tx=XM_019823749.2
X 36887012 36887128
refGene_gn=CASK;refGene_tx=XM_004000415.5;refGene_gn=CASK;refGene_tx=XM_006943661.4;refGene_gn=CASK;refGene_tx=XM_019823737.2;refGene_gn=CASK;refGene_tx=XM_019823738.2;refGene_gn=CASK;refGene_tx=XM_019823739.2;refGene_gn=CASK;refGene_tx=XM_019823740.2;refGene_gn=CASK;refGene_tx=XM_019823741.2;refGene_gn=CASK;refGene_tx=XM_019823742.2;refGene_gn=CASK;refGene_tx=XM_019823743.2;refGene_gn=CASK;refGene_tx=XM_019823744.2;refGene_gn=CASK;refGene_tx=XM_019823745.2;refGene_gn=CASK;refGene_tx=XM_019823746.2;refGene_gn=CASK;refGene_tx=XM_019823747.2;refGene_gn=CASK;refGene_tx=XM_019823748.2;refGene_gn=CASK;refGene_tx=XM_019823749.2
X 36895552 36895749
refGene_gn=CASK;refGene_tx=XM_004000415.5;refGene_gn=CASK;refGene_tx=XM_006943661.4;refGene_gn=CASK;refGene_tx=XM_019823737.2;refGene_gn=CASK;refGene_tx=XM_019823738.2;refGene_gn=CASK;refGene_tx=XM_019823739.2;refGene_gn=CASK;refGene_tx=XM_019823740.2;refGene_gn=CASK;refGene_tx=XM_019823741.2;refGene_gn=CASK;refGene_tx=XM_019823742.2;refGene_gn=CASK;refGene_tx=XM_019823743.2;refGene_gn=CASK;refGene_tx=XM_019823744.2;refGene_gn=CASK;refGene_tx=XM_019823745.2;refGene_gn=CASK;refGene_tx=XM_019823746.2;refGene_gn=CASK;refGene_tx=XM_019823747.2;refGene_gn=CASK;refGene_tx=XM_019823748.2;refGene_gn=CASK;refGene_tx=XM_019823749.2
X 36897397 36897433
refGene_gn=CASK;refGene_tx=XM_004000415.5;refGene_gn=CASK;refGene_tx=XM_006943661.4;refGene_gn=CASK;refGene_tx=XM_019823737.2;refGene_gn=CASK;refGene_tx=XM_019823738.2;refGene_gn=CASK;refGene_tx=XM_019823739.2;refGene_gn=CASK;refGene_tx=XM_019823740.2;refGene_gn=CASK;refGene_tx=XM_019823741.2;refGene_gn=CASK;refGene_tx=XM_019823743.2;refGene_gn=CASK;refGene_tx=XM_019823744.2;refGene_gn=CASK;refGene_tx=XM_019823745.2;refGene_gn=CASK;refGene_tx=XM_019823748.2
X 36898854 36898923
refGene_gn=CASK;refGene_tx=XM_006943661.4;refGene_gn=CASK;refGene_tx=XM_019823737.2;refGene_gn=CASK;refGene_tx=XM_019823738.2;refGene_gn=CASK;refGene_tx=XM_019823739.2;refGene_gn=CASK;refGene_tx=XM_019823740.2;refGene_gn=CASK;refGene_tx=XM_019823741.2;refGene_gn=CASK;refGene_tx=XM_019823742.2;refGene_gn=CASK;refGene_tx=XM_019823743.2;refGene_gn=CASK;refGene_tx=XM_019823744.2;refGene_gn=CASK;refGene_tx=XM_019823746.2
X 36901655 36901724
refGene_gn=CASK;refGene_tx=XM_004000415.5;refGene_gn=CASK;refGene_tx=XM_006943661.4;refGene_gn=CASK;refGene_tx=XM_019823737.2;refGene_gn=CASK;refGene_tx=XM_019823738.2;refGene_gn=CASK;refGene_tx=XM_019823739.2;refGene_gn=CASK;refGene_tx=XM_019823740.2;refGene_gn=CASK;refGene_tx=XM_019823741.2;refGene_gn=CASK;refGene_tx=XM_019823742.2;refGene_gn=CASK;refGene_tx=XM_019823743.2;refGene_gn=CASK;refGene_tx=XM_019823744.2;refGene_gn=CASK;refGene_tx=XM_019823745.2;refGene_gn=CASK;refGene_tx=XM_019823746.2;refGene_gn=CASK;refGene_tx=XM_019823747.2;refGene_gn=CAS

K;refGene_tx=XM_019823748.2;refGene_gn=CASK;refGene_tx=XM_019823749.2
X 36903083 36903169
refGene_gn=CASK;refGene_tx=XM_004000415.5;refGene_gn=CASK;refGene_tx=XM_006943661.4;refGene_gn=CASK;refGene_tx=XM_019823737.2;refGene_gn=CASK;refGene_tx=XM_019823738.2;refGene_gn=CASK;refGene_tx=XM_019823739.2;refGene_gn=CASK;refGene_tx=XM_019823740.2;refGene_gn=CASK;refGene_tx=XM_019823741.2;refGene_gn=CASK;refGene_tx=XM_019823742.2;refGene_gn=CASK;refGene_tx=XM_019823743.2;refGene_gn=CASK;refGene_tx=XM_019823744.2;refGene_gn=CASK;refGene_tx=XM_019823745.2;refGene_gn=CASK;refGene_tx=XM_019823746.2;refGene_gn=CASK;refGene_tx=XM_019823747.2;refGene_gn=CASK;refGene_tx=XM_019823748.2;refGene_gn=CASK;refGene_tx=XM_019823749.2
X 36910435 36910514
refGene_gn=CASK;refGene_tx=XM_004000415.5;refGene_gn=CASK;refGene_tx=XM_006943661.4;refGene_gn=CASK;refGene_tx=XM_019823737.2;refGene_gn=CASK;refGene_tx=XM_019823738.2;refGene_gn=CASK;refGene_tx=XM_019823739.2;refGene_gn=CASK;refGene_tx=XM_019823740.2;refGene_gn=CASK;refGene_tx=XM_019823741.2;refGene_gn=CASK;refGene_tx=XM_019823742.2;refGene_gn=CASK;refGene_tx=XM_019823743.2;refGene_gn=CASK;refGene_tx=XM_019823744.2;refGene_gn=CASK;refGene_tx=XM_019823745.2;refGene_gn=CASK;refGene_tx=XM_019823746.2;refGene_gn=CASK;refGene_tx=XM_019823747.2;refGene_gn=CASK;refGene_tx=XM_019823748.2;refGene_gn=CASK;refGene_tx=XM_019823749.2
X 36918282 36918471
refGene_gn=CASK;refGene_tx=XM_004000415.5;refGene_gn=CASK;refGene_tx=XM_006943661.4;refGene_gn=CASK;refGene_tx=XM_019823737.2;refGene_gn=CASK;refGene_tx=XM_019823738.2;refGene_gn=CASK;refGene_tx=XM_019823739.2;refGene_gn=CASK;refGene_tx=XM_019823740.2;refGene_gn=CASK;refGene_tx=XM_019823741.2;refGene_gn=CASK;refGene_tx=XM_019823742.2;refGene_gn=CASK;refGene_tx=XM_019823743.2;refGene_gn=CASK;refGene_tx=XM_019823744.2;refGene_gn=CASK;refGene_tx=XM_019823745.2;refGene_gn=CASK;refGene_tx=XM_019823746.2;refGene_gn=CASK;refGene_tx=XM_019823747.2;refGene_gn=CASK;refGene_tx=XM_019823748.2;refGene_gn=CASK;refGene_tx=XM_019823749.2
X 36926914 36926995
refGene_gn=CASK;refGene_tx=XM_004000415.5;refGene_gn=CASK;refGene_tx=XM_006943661.4;refGene_gn=CASK;refGene_tx=XM_019823737.2;refGene_gn=CASK;refGene_tx=XM_019823738.2;refGene_gn=CASK;refGene_tx=XM_019823739.2;refGene_gn=CASK;refGene_tx=XM_019823740.2;refGene_gn=CASK;refGene_tx=XM_019823741.2;refGene_gn=CASK;refGene_tx=XM_019823742.2;refGene_gn=CASK;refGene_tx=XM_019823743.2;refGene_gn=CASK;refGene_tx=XM_019823744.2;refGene_gn=CASK;refGene_tx=XM_019823745.2;refGene_gn=CASK;refGene_tx=XM_019823746.2;refGene_gn=CASK;refGene_tx=XM_019823747.2;refGene_gn=CASK;refGene_tx=XM_019823748.2;refGene_gn=CASK;refGene_tx=XM_019823749.2
X 36929498 36929576
refGene_gn=CASK;refGene_tx=XM_004000415.5;refGene_gn=CASK;refGene_tx=XM_006943661.4;refGene_gn=CASK;refGene_tx=XM_019823737.2;refGene_gn=CASK;refGene_tx=XM_019823738.2;refGene_gn=CASK;refGene_tx=XM_019823739.2;refGene_gn=CASK;refGene_tx=XM_019823740.2;refGene_gn=CASK;refGene_tx=XM_019823741.2;refGene_gn=CASK;refGene_tx=XM_019823742.2;refGene_gn=CASK;refGene_tx=XM_019823743.2;refGene_gn=CASK;refGene_tx=XM_019823744.2;refGene_gn=CASK;refGene_tx=XM_019823745.2;refGene_gn=CASK;refGene_tx=XM_019823746.2;refGene_gn=CASK;refGene_tx=XM_019823747.2;refGene_gn=CAS

K;refGene_tx=XM_019823748.2;refGene_gn=CASK;refGene_tx=XM_019823749.2
X 36950345 36950467
refGene_gn=CASK;refGene_tx=XM_004000415.5;refGene_gn=CASK;refGene_tx=XM_006943661.4;refGene_gn=CASK;refGene_tx=XM_019823737.2;refGene_gn=CASK;refGene_tx=XM_019823738.2;refGene_gn=CASK;refGene_tx=XM_019823739.2;refGene_gn=CASK;refGene_tx=XM_019823740.2;refGene_gn=CASK;refGene_tx=XM_019823741.2;refGene_gn=CASK;refGene_tx=XM_019823742.2;refGene_gn=CASK;refGene_tx=XM_019823743.2;refGene_gn=CASK;refGene_tx=XM_019823744.2;refGene_gn=CASK;refGene_tx=XM_019823745.2;refGene_gn=CASK;refGene_tx=XM_019823746.2;refGene_gn=CASK;refGene_tx=XM_019823747.2;refGene_gn=CASK;refGene_tx=XM_019823748.2;refGene_gn=CASK;refGene_tx=XM_019823749.2
X 36960021 36960039
refGene_gn=CASK;refGene_tx=XM_019823737.2;refGene_gn=CASK;refGene_tx=XM_019823738.2;refGene_gn=CASK;refGene_tx=XM_019823740.2;refGene_gn=CASK;refGene_tx=XM_019823741.2;refGene_gn=CASK;refGene_tx=XM_019823742.2;refGene_gn=CASK;refGene_tx=XM_019823743.2;refGene_gn=CASK;refGene_tx=XM_019823744.2;refGene_gn=CASK;refGene_tx=XM_019823745.2;refGene_gn=CASK;refGene_tx=XM_019823746.2;refGene_gn=CASK;refGene_tx=XM_019823747.2;refGene_gn=CASK;refGene_tx=XM_019823748.2
X 36964236 36964336
refGene_gn=CASK;refGene_tx=XM_004000415.5;refGene_gn=CASK;refGene_tx=XM_006943661.4;refGene_gn=CASK;refGene_tx=XM_019823737.2;refGene_gn=CASK;refGene_tx=XM_019823738.2;refGene_gn=CASK;refGene_tx=XM_019823739.2;refGene_gn=CASK;refGene_tx=XM_019823740.2;refGene_gn=CASK;refGene_tx=XM_019823741.2;refGene_gn=CASK;refGene_tx=XM_019823742.2;refGene_gn=CASK;refGene_tx=XM_019823743.2;refGene_gn=CASK;refGene_tx=XM_019823744.2;refGene_gn=CASK;refGene_tx=XM_019823745.2;refGene_gn=CASK;refGene_tx=XM_019823746.2;refGene_gn=CASK;refGene_tx=XM_019823747.2;refGene_gn=CASK;refGene_tx=XM_019823748.2;refGene_gn=CASK;refGene_tx=XM_019823749.2
X 36975497 36975581
refGene_gn=CASK;refGene_tx=XM_004000415.5;refGene_gn=CASK;refGene_tx=XM_006943661.4;refGene_gn=CASK;refGene_tx=XM_019823737.2;refGene_gn=CASK;refGene_tx=XM_019823738.2;refGene_gn=CASK;refGene_tx=XM_019823739.2;refGene_gn=CASK;refGene_tx=XM_019823740.2;refGene_gn=CASK;refGene_tx=XM_019823741.2;refGene_gn=CASK;refGene_tx=XM_019823742.2;refGene_gn=CASK;refGene_tx=XM_019823743.2;refGene_gn=CASK;refGene_tx=XM_019823744.2;refGene_gn=CASK;refGene_tx=XM_019823745.2;refGene_gn=CASK;refGene_tx=XM_019823746.2;refGene_gn=CASK;refGene_tx=XM_019823747.2;refGene_gn=CASK;refGene_tx=XM_019823748.2;refGene_gn=CASK;refGene_tx=XM_019823749.2
X 36990537 36990660
refGene_gn=CASK;refGene_tx=XM_004000415.5;refGene_gn=CASK;refGene_tx=XM_006943661.4;refGene_gn=CASK;refGene_tx=XM_019823737.2;refGene_gn=CASK;refGene_tx=XM_019823738.2;refGene_gn=CASK;refGene_tx=XM_019823739.2;refGene_gn=CASK;refGene_tx=XM_019823740.2;refGene_gn=CASK;refGene_tx=XM_019823741.2;refGene_gn=CASK;refGene_tx=XM_019823742.2;refGene_gn=CASK;refGene_tx=XM_019823743.2;refGene_gn=CASK;refGene_tx=XM_019823744.2;refGene_gn=CASK;refGene_tx=XM_019823745.2;refGene_gn=CASK;refGene_tx=XM_019823746.2;refGene_gn=CASK;refGene_tx=XM_019823747.2;refGene_gn=CASK;refGene_tx=XM_019823748.2;refGene_gn=CASK;refGene_tx=XM_019823749.2
X 36995408 36995584

refGene_gn=CASK;refGene_tx=XM_004000415.5;refGene_gn=CASK;refGene_tx=XM_006943661.4;refGene_gn=CASK;refGene_tx=XM_019823737.2;refGene_gn=CASK;refGene_tx=XM_019823738.2;refGene_gn=CASK;refGene_tx=XM_019823739.2;refGene_gn=CASK;refGene_tx=XM_019823740.2;refGene_gn=CASK;refGene_tx=XM_019823741.2;refGene_gn=CASK;refGene_tx=XM_019823742.2;refGene_gn=CASK;refGene_tx=XM_019823743.2;refGene_gn=CASK;refGene_tx=XM_019823744.2;refGene_gn=CASK;refGene_tx=XM_019823745.2;refGene_gn=CASK;refGene_tx=XM_019823746.2;refGene_gn=CASK;refGene_tx=XM_019823747.2;refGene_gn=CASK;refGene_tx=XM_019823748.2;refGene_gn=CASK;refGene_tx=XM_019823749.2
X 37002709 37002812 refGene_gn=CASK;refGene_tx=XM_004000415.5;refGene_gn=CASK;refGene_tx=XM_006943661.4;refGene_gn=CASK;refGene_tx=XM_019823737.2;refGene_gn=CASK;refGene_tx=XM_019823738.2;refGene_gn=CASK;refGene_tx=XM_019823739.2;refGene_gn=CASK;refGene_tx=XM_019823740.2;refGene_gn=CASK;refGene_tx=XM_019823741.2;refGene_gn=CASK;refGene_tx=XM_019823742.2;refGene_gn=CASK;refGene_tx=XM_019823743.2;refGene_gn=CASK;refGene_tx=XM_019823744.2;refGene_gn=CASK;refGene_tx=XM_019823745.2;refGene_gn=CASK;refGene_tx=XM_019823746.2;refGene_gn=CASK;refGene_tx=XM_019823747.2;refGene_gn=CASK;refGene_tx=XM_019823748.2;refGene_gn=CASK;refGene_tx=XM_019823749.2
X 37017505 37018648 refGene_gn=GPR34;refGene_tx=XM_006943659.3
X 37039655 37040666 refGene_gn=GPR82;refGene_tx=XM_019823750.1
X 37048902 37048975 refGene_gn=CASK;refGene_tx=XM_004000415.5;refGene_gn=CASK;refGene_tx=XM_006943661.4;refGene_gn=CASK;refGene_tx=XM_019823737.2;refGene_gn=CASK;refGene_tx=XM_019823738.2;refGene_gn=CASK;refGene_tx=XM_019823739.2;refGene_gn=CASK;refGene_tx=XM_019823740.2;refGene_gn=CASK;refGene_tx=XM_019823741.2;refGene_gn=CASK;refGene_tx=XM_019823742.2;refGene_gn=CASK;refGene_tx=XM_019823743.2;refGene_gn=CASK;refGene_tx=XM_019823744.2;refGene_gn=CASK;refGene_tx=XM_019823745.2;refGene_gn=CASK;refGene_tx=XM_019823746.2;refGene_gn=CASK;refGene_tx=XM_019823747.2;refGene_gn=CASK;refGene_tx=XM_019823748.2;refGene_gn=CASK;refGene_tx=XM_019823749.2
X 37054771 37054849 refGene_gn=CASK;refGene_tx=XM_004000415.5;refGene_gn=CASK;refGene_tx=XM_006943661.4;refGene_gn=CASK;refGene_tx=XM_019823737.2;refGene_gn=CASK;refGene_tx=XM_019823738.2;refGene_gn=CASK;refGene_tx=XM_019823739.2;refGene_gn=CASK;refGene_tx=XM_019823740.2;refGene_gn=CASK;refGene_tx=XM_019823741.2;refGene_gn=CASK;refGene_tx=XM_019823742.2;refGene_gn=CASK;refGene_tx=XM_019823743.2;refGene_gn=CASK;refGene_tx=XM_019823744.2;refGene_gn=CASK;refGene_tx=XM_019823745.2;refGene_gn=CASK;refGene_tx=XM_019823746.2;refGene_gn=CASK;refGene_tx=XM_019823747.2;refGene_gn=CASK;refGene_tx=XM_019823748.2;refGene_gn=CASK;refGene_tx=XM_019823749.2
X 37096371 37096477 refGene_gn=CASK;refGene_tx=XM_004000415.5;refGene_gn=CASK;refGene_tx=XM_006943661.4;refGene_gn=CASK;refGene_tx=XM_019823737.2;refGene_gn=CASK;refGene_tx=XM_019823738.2;refGene_gn=CASK;refGene_tx=XM_019823739.2;refGene_gn=CASK;refGene_tx=XM_019823740.2;refGene_gn=CASK;refGene_tx=XM_019823741.2;refGene_gn=CASK;refGene_tx=XM_019823742.2;refGene_gn=CASK;refGene_tx=XM_019823743.2;refGene_gn=CASK;refGene_tx=XM_019823744.2;refGene_gn=CASK;refGene_tx=XM_019823745.2;refGene_gn=CASK;refGene_tx=XM_019823746.2;refGene_gn=CASK;refGene_tx=XM_019823747.2;refGene_gn=CAS

K;refGene_tx=XM_019823748.2;refGene_gn=CASK;refGene_tx=XM_019823749.2
X 37156364 37156495
refGene_gn=CASK;refGene_tx=XM_004000415.5;refGene_gn=CASK;refGene_tx=XM_006943661.4;refGene_gn=CASK;refGene_tx=XM_019823737.2;refGene_gn=CASK;refGene_tx=XM_019823738.2;refGene_gn=CASK;refGene_tx=XM_019823739.2;refGene_gn=CASK;refGene_tx=XM_019823740.2;refGene_gn=CASK;refGene_tx=XM_019823741.2;refGene_gn=CASK;refGene_tx=XM_019823742.2;refGene_gn=CASK;refGene_tx=XM_019823743.2;refGene_gn=CASK;refGene_tx=XM_019823744.2;refGene_gn=CASK;refGene_tx=XM_019823745.2;refGene_gn=CASK;refGene_tx=XM_019823746.2;refGene_gn=CASK;refGene_tx=XM_019823747.2;refGene_gn=CASK;refGene_tx=XM_019823748.2;refGene_gn=CASK;refGene_tx=XM_019823749.2
X 37221835 37221894
refGene_gn=CASK;refGene_tx=XM_004000415.5;refGene_gn=CASK;refGene_tx=XM_006943661.4;refGene_gn=CASK;refGene_tx=XM_019823737.2;refGene_gn=CASK;refGene_tx=XM_019823738.2;refGene_gn=CASK;refGene_tx=XM_019823739.2;refGene_gn=CASK;refGene_tx=XM_019823740.2;refGene_gn=CASK;refGene_tx=XM_019823741.2;refGene_gn=CASK;refGene_tx=XM_019823742.2;refGene_gn=CASK;refGene_tx=XM_019823743.2;refGene_gn=CASK;refGene_tx=XM_019823744.2;refGene_gn=CASK;refGene_tx=XM_019823745.2;refGene_gn=CASK;refGene_tx=XM_019823746.2;refGene_gn=CASK;refGene_tx=XM_019823747.2;refGene_gn=CASK;refGene_tx=XM_019823748.2;refGene_gn=CASK;refGene_tx=XM_019823749.2
X 37930415 37931045
refGene_gn=LOC101086542;refGene_tx=XM_004000461.1
X 37931135 37931231
refGene_gn=LOC101086542;refGene_tx=XM_004000461.1
X 38670146 38672016 refGene_gn=MAOA;refGene_tx=XM_004000418.5
X 38676982 38676992 refGene_gn=MAOA;refGene_tx=XM_019823752.2
X 38693204 38693299
refGene_gn=MAOA;refGene_tx=XM_004000418.5;refGene_gn=MAOA;refGene_tx=XM_019823752.2
X 38701683 38701821
refGene_gn=MAOA;refGene_tx=XM_004000418.5;refGene_gn=MAOA;refGene_tx=XM_019823752.2
X 38720604 38720709
refGene_gn=MAOA;refGene_tx=XM_004000418.5;refGene_gn=MAOA;refGene_tx=XM_019823752.2
X 38721410 38721502
refGene_gn=MAOA;refGene_tx=XM_004000418.5;refGene_gn=MAOA;refGene_tx=XM_019823752.2
X 38726071 38726213
refGene_gn=MAOA;refGene_tx=XM_004000418.5;refGene_gn=MAOA;refGene_tx=XM_019823752.2
X 38726878 38727028
refGene_gn=MAOA;refGene_tx=XM_004000418.5;refGene_gn=MAOA;refGene_tx=XM_019823752.2
X 38727325 38727485
refGene_gn=MAOA;refGene_tx=XM_004000418.5;refGene_gn=MAOA;refGene_tx=XM_019823752.2
X 38729256 38729353
refGene_gn=MAOA;refGene_tx=XM_004000418.5;refGene_gn=MAOA;refGene_tx=X

M_019823752.2
X 38732948 38733002
refGene_gn=MAOA;refGene_tx=XM_004000418.5;refGene_gn=MAOA;refGene_tx=X
M_019823752.2
X 38737892 38737950
refGene_gn=MAOA;refGene_tx=XM_004000418.5;refGene_gn=MAOA;refGene_tx=X
M_019823752.2
X 38738994 38739092
refGene_gn=MAOA;refGene_tx=XM_004000418.5;refGene_gn=MAOA;refGene_tx=X
M_019823752.2
X 38741275 38741387
refGene_gn=MAOA;refGene_tx=XM_004000418.5;refGene_gn=MAOA;refGene_tx=X
M_019823752.2
X 38741598 38741661
refGene_gn=MAOA;refGene_tx=XM_004000418.5;refGene_gn=MAOA;refGene_tx=X
M_019823752.2
X 38741817 38741964
refGene_gn=MAOA;refGene_tx=XM_004000418.5;refGene_gn=MAOA;refGene_tx=X
M_019823752.2
X 38763373 38763526 refGene_gn=MAOB;refGene_tx=XM_023248804.1
X 38764781 38764844 refGene_gn=MAOB;refGene_tx=XM_023248804.1
X 38765868 38765980 refGene_gn=MAOB;refGene_tx=XM_023248804.1
X 38771321 38771419 refGene_gn=MAOB;refGene_tx=XM_023248804.1
X 38774123 38774181 refGene_gn=MAOB;refGene_tx=XM_023248804.1
X 38777418 38777472 refGene_gn=MAOB;refGene_tx=XM_023248804.1
X 38778746 38778843 refGene_gn=MAOB;refGene_tx=XM_023248804.1
X 38784627 38784787 refGene_gn=MAOB;refGene_tx=XM_023248804.1
X 38786042 38786192 refGene_gn=MAOB;refGene_tx=XM_023248804.1
X 38788018 38788160 refGene_gn=MAOB;refGene_tx=XM_023248804.1
X 38794408 38794500 refGene_gn=MAOB;refGene_tx=XM_023248804.1
X 38795200 38795305 refGene_gn=MAOB;refGene_tx=XM_023248804.1
X 38828939 38829077 refGene_gn=MAOB;refGene_tx=XM_023248804.1
X 38830234 38830329 refGene_gn=MAOB;refGene_tx=XM_023248804.1
X 38874224 38874270 refGene_gn=MAOB;refGene_tx=XM_023248804.1
X 38947652 38947880
refGene_gn=NDP;refGene_tx=XM_004000421.5;refGene_gn=NDP;refGene_tx=XM_
019823753.2;refGene_gn=NDP;refGene_tx=XM_019823754.2
X 38957193 38957367
refGene_gn=NDP;refGene_tx=XM_004000421.5;refGene_gn=NDP;refGene_tx=XM_
019823753.2;refGene_gn=NDP;refGene_tx=XM_019823754.2
X 39144374 39144476 refGene_gn=EFHC2;refGene_tx=XM_023249026.1
X 39161569 39161675 refGene_gn=EFHC2;refGene_tx=XM_023249026.1
X 39174031 39174124 refGene_gn=EFHC2;refGene_tx=XM_023249026.1
X 39175820 39176018 refGene_gn=EFHC2;refGene_tx=XM_023249026.1
X 39224862 39224993 refGene_gn=EFHC2;refGene_tx=XM_023249026.1
X 39227628 39227825 refGene_gn=EFHC2;refGene_tx=XM_023249026.1
X 39230895 39231038 refGene_gn=EFHC2;refGene_tx=XM_023249026.1
X 39240228 39240394 refGene_gn=EFHC2;refGene_tx=XM_023249026.1
X 39248103 39248242 refGene_gn=EFHC2;refGene_tx=XM_023249026.1
X 39248615 39248729 refGene_gn=EFHC2;refGene_tx=XM_023249026.1

X 39251754 39252006 refGene_gn=EFHC2;refGene_tx=XM_023249026.1
X 39255879 39256103 refGene_gn=EFHC2;refGene_tx=XM_023249026.1
X 39268757 39268908 refGene_gn=EFHC2;refGene_tx=XM_023249026.1
X 39321329 39321518 refGene_gn=EFHC2;refGene_tx=XM_023249026.1
X 39345143 39345185 refGene_gn=EFHC2;refGene_tx=XM_023249026.1
X 39503651 39503729 refGene_gn=FUNDC1;refGene_tx=XM_004000422.4
X 39505723 39505852 refGene_gn=FUNDC1;refGene_tx=XM_004000422.4
X 39510467 39510543 refGene_gn=FUNDC1;refGene_tx=XM_004000422.4
X 39514312 39514469 refGene_gn=FUNDC1;refGene_tx=XM_004000422.4
X 39515185 39515213 refGene_gn=FUNDC1;refGene_tx=XM_004000422.4
X 39721920 39722487 refGene_gn=DUSP21;refGene_tx=XM_023249334.1
X 39746568 39746729
refGene_gn=KDM6A;refGene_tx=XM_023249060.1;refGene_gn=KDM6A;refGene_tx=XM_023249061.1;refGene_gn=KDM6A;refGene_tx=XM_023249062.1;refGene_gn=KDM6A;refGene_tx=XM_023249063.1;refGene_gn=KDM6A;refGene_tx=XM_023249064.1;refGene_gn=KDM6A;refGene_tx=XM_023249065.1;refGene_gn=KDM6A;refGene_tx=XM_023249066.1
X 39746943 39747007
refGene_gn=KDM6A;refGene_tx=XM_023249060.1;refGene_gn=KDM6A;refGene_tx=XM_023249061.1;refGene_gn=KDM6A;refGene_tx=XM_023249062.1;refGene_gn=KDM6A;refGene_tx=XM_023249063.1;refGene_gn=KDM6A;refGene_tx=XM_023249064.1;refGene_gn=KDM6A;refGene_tx=XM_023249065.1;refGene_gn=KDM6A;refGene_tx=XM_023249066.1
X 39816881 39816990
refGene_gn=KDM6A;refGene_tx=XM_023249060.1;refGene_gn=KDM6A;refGene_tx=XM_023249061.1;refGene_gn=KDM6A;refGene_tx=XM_023249062.1;refGene_gn=KDM6A;refGene_tx=XM_023249063.1;refGene_gn=KDM6A;refGene_tx=XM_023249064.1;refGene_gn=KDM6A;refGene_tx=XM_023249065.1;refGene_gn=KDM6A;refGene_tx=XM_023249066.1
X 39829266 39829316
refGene_gn=KDM6A;refGene_tx=XM_023249060.1;refGene_gn=KDM6A;refGene_tx=XM_023249061.1;refGene_gn=KDM6A;refGene_tx=XM_023249062.1;refGene_gn=KDM6A;refGene_tx=XM_023249063.1;refGene_gn=KDM6A;refGene_tx=XM_023249064.1;refGene_gn=KDM6A;refGene_tx=XM_023249065.1;refGene_gn=KDM6A;refGene_tx=XM_023249066.1
X 39847986 39848045
refGene_gn=KDM6A;refGene_tx=XM_023249060.1;refGene_gn=KDM6A;refGene_tx=XM_023249061.1;refGene_gn=KDM6A;refGene_tx=XM_023249062.1;refGene_gn=KDM6A;refGene_tx=XM_023249063.1;refGene_gn=KDM6A;refGene_tx=XM_023249064.1;refGene_gn=KDM6A;refGene_tx=XM_023249065.1;refGene_gn=KDM6A;refGene_tx=XM_023249066.1
X 39857566 39857687
refGene_gn=KDM6A;refGene_tx=XM_023249060.1;refGene_gn=KDM6A;refGene_tx=XM_023249061.1;refGene_gn=KDM6A;refGene_tx=XM_023249062.1;refGene_gn=KDM6A;refGene_tx=XM_023249063.1;refGene_gn=KDM6A;refGene_tx=XM_023249064.1;refGene_gn=KDM6A;refGene_tx=XM_023249065.1;refGene_gn=KDM6A;refGene_tx=XM_023249066.1
X 39869511 39869566
refGene_gn=KDM6A;refGene_tx=XM_023249060.1;refGene_gn=KDM6A;refGene_tx=XM_023249061.1;refGene_gn=KDM6A;refGene_tx=XM_023249062.1;refGene_gn=

KDM6A;refGene_tx=XM_023249063.1;refGene_gn=KDM6A;refGene_tx=XM_023249064.1;refGene_gn=KDM6A;refGene_tx=XM_023249065.1;refGene_gn=KDM6A;refGene_tx=XM_023249066.1
X 39873605 39873640
refGene_gn=KDM6A;refGene_tx=XM_023249060.1;refGene_gn=KDM6A;refGene_tx=XM_023249061.1;refGene_gn=KDM6A;refGene_tx=XM_023249062.1;refGene_gn=KDM6A;refGene_tx=XM_023249063.1;refGene_gn=KDM6A;refGene_tx=XM_023249064.1;refGene_gn=KDM6A;refGene_tx=XM_023249065.1;refGene_gn=KDM6A;refGene_tx=XM_023249066.1
X 39894614 39894708
refGene_gn=KDM6A;refGene_tx=XM_023249060.1;refGene_gn=KDM6A;refGene_tx=XM_023249061.1;refGene_gn=KDM6A;refGene_tx=XM_023249062.1;refGene_gn=KDM6A;refGene_tx=XM_023249063.1;refGene_gn=KDM6A;refGene_tx=XM_023249064.1;refGene_gn=KDM6A;refGene_tx=XM_023249065.1;refGene_gn=KDM6A;refGene_tx=XM_023249066.1
X 39897442 39897569
refGene_gn=KDM6A;refGene_tx=XM_023249060.1;refGene_gn=KDM6A;refGene_tx=XM_023249061.1;refGene_gn=KDM6A;refGene_tx=XM_023249062.1;refGene_gn=KDM6A;refGene_tx=XM_023249063.1;refGene_gn=KDM6A;refGene_tx=XM_023249064.1;refGene_gn=KDM6A;refGene_tx=XM_023249065.1;refGene_gn=KDM6A;refGene_tx=XM_023249066.1
X 39902963 39903062
refGene_gn=KDM6A;refGene_tx=XM_023249060.1;refGene_gn=KDM6A;refGene_tx=XM_023249061.1;refGene_gn=KDM6A;refGene_tx=XM_023249062.1;refGene_gn=KDM6A;refGene_tx=XM_023249063.1;refGene_gn=KDM6A;refGene_tx=XM_023249064.1;refGene_gn=KDM6A;refGene_tx=XM_023249065.1;refGene_gn=KDM6A;refGene_tx=XM_023249066.1
X 39903193 39903413
refGene_gn=KDM6A;refGene_tx=XM_023249060.1;refGene_gn=KDM6A;refGene_tx=XM_023249061.1;refGene_gn=KDM6A;refGene_tx=XM_023249062.1;refGene_gn=KDM6A;refGene_tx=XM_023249063.1;refGene_gn=KDM6A;refGene_tx=XM_023249064.1;refGene_gn=KDM6A;refGene_tx=XM_023249065.1;refGene_gn=KDM6A;refGene_tx=XM_023249066.1
X 39903972 39904107
refGene_gn=KDM6A;refGene_tx=XM_023249060.1;refGene_gn=KDM6A;refGene_tx=XM_023249061.1;refGene_gn=KDM6A;refGene_tx=XM_023249063.1;refGene_gn=KDM6A;refGene_tx=XM_023249064.1
X 39904556 39904712
refGene_gn=KDM6A;refGene_tx=XM_023249060.1;refGene_gn=KDM6A;refGene_tx=XM_023249061.1;refGene_gn=KDM6A;refGene_tx=XM_023249062.1
X 39905275 39905371
refGene_gn=KDM6A;refGene_tx=XM_023249060.1;refGene_gn=KDM6A;refGene_tx=XM_023249061.1;refGene_gn=KDM6A;refGene_tx=XM_023249062.1;refGene_gn=KDM6A;refGene_tx=XM_023249063.1;refGene_gn=KDM6A;refGene_tx=XM_023249064.1;refGene_gn=KDM6A;refGene_tx=XM_023249065.1;refGene_gn=KDM6A;refGene_tx=XM_023249066.1
X 39906561 39906663
refGene_gn=KDM6A;refGene_tx=XM_023249060.1;refGene_gn=KDM6A;refGene_tx=XM_023249062.1;refGene_gn=KDM6A;refGene_tx=XM_023249063.1;refGene_gn=KDM6A;refGene_tx=XM_023249065.1

X 39907333 39907729
refGene_gn=KDM6A;refGene_tx=XM_023249060.1;refGene_gn=KDM6A;refGene_tx=XM_023249061.1;refGene_gn=KDM6A;refGene_tx=XM_023249062.1;refGene_gn=KDM6A;refGene_tx=XM_023249063.1;refGene_gn=KDM6A;refGene_tx=XM_023249064.1;refGene_gn=KDM6A;refGene_tx=XM_023249065.1;refGene_gn=KDM6A;refGene_tx=XM_023249066.1
X 39913284 39914063
refGene_gn=KDM6A;refGene_tx=XM_023249060.1;refGene_gn=KDM6A;refGene_tx=XM_023249061.1;refGene_gn=KDM6A;refGene_tx=XM_023249062.1;refGene_gn=KDM6A;refGene_tx=XM_023249063.1;refGene_gn=KDM6A;refGene_tx=XM_023249064.1;refGene_gn=KDM6A;refGene_tx=XM_023249065.1;refGene_gn=KDM6A;refGene_tx=XM_023249066.1
X 39919560 39919690
refGene_gn=KDM6A;refGene_tx=XM_023249060.1;refGene_gn=KDM6A;refGene_tx=XM_023249061.1;refGene_gn=KDM6A;refGene_tx=XM_023249062.1;refGene_gn=KDM6A;refGene_tx=XM_023249063.1;refGene_gn=KDM6A;refGene_tx=XM_023249064.1;refGene_gn=KDM6A;refGene_tx=XM_023249065.1;refGene_gn=KDM6A;refGene_tx=XM_023249066.1
X 39921110 39921216
refGene_gn=KDM6A;refGene_tx=XM_023249060.1;refGene_gn=KDM6A;refGene_tx=XM_023249061.1;refGene_gn=KDM6A;refGene_tx=XM_023249062.1;refGene_gn=KDM6A;refGene_tx=XM_023249063.1;refGene_gn=KDM6A;refGene_tx=XM_023249064.1;refGene_gn=KDM6A;refGene_tx=XM_023249065.1;refGene_gn=KDM6A;refGene_tx=XM_023249066.1
X 39921842 39922048
refGene_gn=KDM6A;refGene_tx=XM_023249060.1;refGene_gn=KDM6A;refGene_tx=XM_023249061.1;refGene_gn=KDM6A;refGene_tx=XM_023249062.1;refGene_gn=KDM6A;refGene_tx=XM_023249063.1;refGene_gn=KDM6A;refGene_tx=XM_023249064.1;refGene_gn=KDM6A;refGene_tx=XM_023249065.1;refGene_gn=KDM6A;refGene_tx=XM_023249066.1
X 39924732 39924797
refGene_gn=KDM6A;refGene_tx=XM_023249060.1;refGene_gn=KDM6A;refGene_tx=XM_023249061.1;refGene_gn=KDM6A;refGene_tx=XM_023249062.1;refGene_gn=KDM6A;refGene_tx=XM_023249063.1;refGene_gn=KDM6A;refGene_tx=XM_023249064.1;refGene_gn=KDM6A;refGene_tx=XM_023249065.1;refGene_gn=KDM6A;refGene_tx=XM_023249066.1
X 39924875 39924950
refGene_gn=KDM6A;refGene_tx=XM_023249060.1;refGene_gn=KDM6A;refGene_tx=XM_023249061.1;refGene_gn=KDM6A;refGene_tx=XM_023249062.1;refGene_gn=KDM6A;refGene_tx=XM_023249063.1;refGene_gn=KDM6A;refGene_tx=XM_023249064.1;refGene_gn=KDM6A;refGene_tx=XM_023249065.1;refGene_gn=KDM6A;refGene_tx=XM_023249066.1
X 39925479 39925628
refGene_gn=KDM6A;refGene_tx=XM_023249060.1;refGene_gn=KDM6A;refGene_tx=XM_023249061.1;refGene_gn=KDM6A;refGene_tx=XM_023249062.1;refGene_gn=KDM6A;refGene_tx=XM_023249063.1;refGene_gn=KDM6A;refGene_tx=XM_023249064.1;refGene_gn=KDM6A;refGene_tx=XM_023249065.1;refGene_gn=KDM6A;refGene_tx=XM_023249066.1
X 39932345 39932460
refGene_gn=KDM6A;refGene_tx=XM_023249060.1;refGene_gn=KDM6A;refGene_tx

=XM_023249061.1;refGene_gn=KDM6A;refGene_tx=XM_023249062.1;refGene_gn=KDM6A;refGene_tx=XM_023249063.1;refGene_gn=KDM6A;refGene_tx=XM_023249064.1;refGene_gn=KDM6A;refGene_tx=XM_023249065.1;refGene_gn=KDM6A;refGene_tx=XM_023249066.1
X 39935793 39935981
refGene_gn=KDM6A;refGene_tx=XM_023249060.1;refGene_gn=KDM6A;refGene_tx=XM_023249061.1;refGene_gn=KDM6A;refGene_tx=XM_023249062.1;refGene_gn=KDM6A;refGene_tx=XM_023249063.1;refGene_gn=KDM6A;refGene_tx=XM_023249064.1;refGene_gn=KDM6A;refGene_tx=XM_023249065.1;refGene_gn=KDM6A;refGene_tx=XM_023249066.1
X 39936800 39936942
refGene_gn=KDM6A;refGene_tx=XM_023249060.1;refGene_gn=KDM6A;refGene_tx=XM_023249061.1;refGene_gn=KDM6A;refGene_tx=XM_023249062.1;refGene_gn=KDM6A;refGene_tx=XM_023249063.1;refGene_gn=KDM6A;refGene_tx=XM_023249064.1;refGene_gn=KDM6A;refGene_tx=XM_023249065.1;refGene_gn=KDM6A;refGene_tx=XM_023249066.1
X 39950130 39950257
refGene_gn=KDM6A;refGene_tx=XM_023249060.1;refGene_gn=KDM6A;refGene_tx=XM_023249061.1;refGene_gn=KDM6A;refGene_tx=XM_023249062.1;refGene_gn=KDM6A;refGene_tx=XM_023249063.1;refGene_gn=KDM6A;refGene_tx=XM_023249064.1;refGene_gn=KDM6A;refGene_tx=XM_023249065.1;refGene_gn=KDM6A;refGene_tx=XM_023249066.1
X 39952942 39953113
refGene_gn=KDM6A;refGene_tx=XM_023249060.1;refGene_gn=KDM6A;refGene_tx=XM_023249061.1;refGene_gn=KDM6A;refGene_tx=XM_023249062.1;refGene_gn=KDM6A;refGene_tx=XM_023249063.1;refGene_gn=KDM6A;refGene_tx=XM_023249064.1;refGene_gn=KDM6A;refGene_tx=XM_023249065.1;refGene_gn=KDM6A;refGene_tx=XM_023249066.1
X 39954251 39954281
refGene_gn=KDM6A;refGene_tx=XM_023249060.1;refGene_gn=KDM6A;refGene_tx=XM_023249061.1;refGene_gn=KDM6A;refGene_tx=XM_023249062.1;refGene_gn=KDM6A;refGene_tx=XM_023249063.1;refGene_gn=KDM6A;refGene_tx=XM_023249064.1;refGene_gn=KDM6A;refGene_tx=XM_023249065.1;refGene_gn=KDM6A;refGene_tx=XM_023249066.1
X 39996297 39996638
refGene_gn=CXHXorf36;refGene_tx=XM_023248930.1
X 39998061 39998350
refGene_gn=CXHXorf36;refGene_tx=XM_023248930.1
X 40001606 40001780
refGene_gn=CXHXorf36;refGene_tx=XM_023248930.1
X 40033880 40034145
refGene_gn=CXHXorf36;refGene_tx=XM_023248930.1
X 40041233 40041466
refGene_gn=CXHXorf36;refGene_tx=XM_023248930.1
X 40989258 40989270
refGene_gn=LOC111558673;refGene_tx=XM_023248959.1
X 40990179 40990836
refGene_gn=LOC111558673;refGene_tx=XM_023248959.1
X 40991532 40991604
refGene_gn=LOC111558673;refGene_tx=XM_023248959.1

X 41009967 41011460
refGene_gn=ZNF674;refGene_tx=XM_019823371.2;refGene_gn=ZNF674;refGene_tx=XM_019823373.2;refGene_gn=ZNF674;refGene_tx=XM_019823374.2;refGene_gn=ZNF674;refGene_tx=XM_019823375.2;refGene_gn=ZNF674;refGene_tx=XM_019823376.2;refGene_gn=ZNF674;refGene_tx=XM_019823377.2;refGene_gn=ZNF674;refGene_tx=XM_023248879.1
X 41037792 41037925
refGene_gn=ZNF674;refGene_tx=XM_019823371.2;refGene_gn=ZNF674;refGene_tx=XM_019823373.2;refGene_gn=ZNF674;refGene_tx=XM_019823374.2;refGene_gn=ZNF674;refGene_tx=XM_019823375.2;refGene_gn=ZNF674;refGene_tx=XM_019823376.2;refGene_gn=ZNF674;refGene_tx=XM_019823377.2;refGene_gn=ZNF674;refGene_tx=XM_023248879.1
X 41038235 41038383
refGene_gn=ZNF674;refGene_tx=XM_019823371.2;refGene_gn=ZNF674;refGene_tx=XM_019823373.2;refGene_gn=ZNF674;refGene_tx=XM_019823374.2;refGene_gn=ZNF674;refGene_tx=XM_019823375.2;refGene_gn=ZNF674;refGene_tx=XM_019823376.2;refGene_gn=ZNF674;refGene_tx=XM_023248879.1
X 41039331 41039476 refGene_gn=ZNF674;refGene_tx=XM_023248879.1
X 41039665 41039791
refGene_gn=ZNF674;refGene_tx=XM_019823371.2;refGene_gn=ZNF674;refGene_tx=XM_019823373.2;refGene_gn=ZNF674;refGene_tx=XM_023248879.1
X 41042756 41042771
refGene_gn=ZNF674;refGene_tx=XM_019823371.2;refGene_gn=ZNF674;refGene_tx=XM_019823373.2;refGene_gn=ZNF674;refGene_tx=XM_019823375.2;refGene_gn=ZNF674;refGene_tx=XM_019823376.2
X 41091283 41091535 refGene_gn=CHST7;refGene_tx=XM_023248942.1
X 41091536 41092742 refGene_gn=CHST7;refGene_tx=XM_023248942.1
X 41121485 41121737
refGene_gn=SLC9A7;refGene_tx=XM_023249454.1;refGene_gn=SLC9A7;refGene_tx=XM_023249455.1
X 41125294 41125400
refGene_gn=SLC9A7;refGene_tx=XM_023249454.1;refGene_gn=SLC9A7;refGene_tx=XM_023249455.1
X 41134725 41134808
refGene_gn=SLC9A7;refGene_tx=XM_023249454.1;refGene_gn=SLC9A7;refGene_tx=XM_023249455.1
X 41142402 41142466
refGene_gn=SLC9A7;refGene_tx=XM_023249454.1;refGene_gn=SLC9A7;refGene_tx=XM_023249455.1
X 41146787 41146847 refGene_gn=SLC9A7;refGene_tx=XM_023249454.1
X 41157550 41157704
refGene_gn=SLC9A7;refGene_tx=XM_023249454.1;refGene_gn=SLC9A7;refGene_tx=XM_023249455.1
X 41161288 41161400
refGene_gn=SLC9A7;refGene_tx=XM_023249454.1;refGene_gn=SLC9A7;refGene_tx=XM_023249455.1
X 41163850 41163964
refGene_gn=SLC9A7;refGene_tx=XM_023249454.1;refGene_gn=SLC9A7;refGene_tx=XM_023249455.1
X 41164056 41164145

refGene_gn=SLC9A7;refGene_tx=XM_023249454.1;refGene_gn=SLC9A7;refGene_tx=XM_023249455.1
X	41165423	41165529	refGene_gn=SLC9A7;refGene_tx=XM_023249454.1;refGene_gn=SLC9A7;refGene_tx=XM_023249455.1
X	41167287	41167429	refGene_gn=SLC9A7;refGene_tx=XM_023249454.1;refGene_gn=SLC9A7;refGene_tx=XM_023249455.1
X	41167715	41167821	refGene_gn=SLC9A7;refGene_tx=XM_023249454.1;refGene_gn=SLC9A7;refGene_tx=XM_023249455.1
X	41177003	41177116	refGene_gn=SLC9A7;refGene_tx=XM_023249454.1;refGene_gn=SLC9A7;refGene_tx=XM_023249455.1
X	41179897	41179974	refGene_gn=SLC9A7;refGene_tx=XM_023249454.1;refGene_gn=SLC9A7;refGene_tx=XM_023249455.1
X	41183609	41183687	refGene_gn=SLC9A7;refGene_tx=XM_023249454.1;refGene_gn=SLC9A7;refGene_tx=XM_023249455.1
X	41185296	41185496	refGene_gn=SLC9A7;refGene_tx=XM_023249454.1;refGene_gn=SLC9A7;refGene_tx=XM_023249455.1
X	41249526	41249854	refGene_gn=SLC9A7;refGene_tx=XM_023249454.1;refGene_gn=SLC9A7;refGene_tx=XM_023249455.1
X	41327760	41327862	refGene_gn=RP2;refGene_tx=XM_004000440.5
X	41349623	41350289	refGene_gn=RP2;refGene_tx=XM_004000440.5
X	41355839	41355954	refGene_gn=RP2;refGene_tx=XM_004000440.5
X	41372898	41372984	refGene_gn=RP2;refGene_tx=XM_004000440.5
X	41375153	41375237	refGene_gn=RP2;refGene_tx=XM_004000440.5
X	41480083	41480129	refGene_gn=JADE3;refGene_tx=XM_004000432.5;refGene_gn=JADE3;refGene_tx=XM_023249516.1;refGene_gn=JADE3;refGene_tx=XM_023249517.1;refGene_gn=JADE3;refGene_tx=XM_023249518.1;refGene_gn=JADE3;refGene_tx=XM_023249519.1;refGene_gn=JADE3;refGene_tx=XM_023249520.1
X	41480913	41480993	refGene_gn=JADE3;refGene_tx=XM_004000432.5;refGene_gn=JADE3;refGene_tx=XM_023249516.1;refGene_gn=JADE3;refGene_tx=XM_023249517.1;refGene_gn=JADE3;refGene_tx=XM_023249518.1;refGene_gn=JADE3;refGene_tx=XM_023249519.1;refGene_gn=JADE3;refGene_tx=XM_023249520.1
X	41498641	41498799	refGene_gn=JADE3;refGene_tx=XM_004000432.5;refGene_gn=JADE3;refGene_tx=XM_023249516.1;refGene_gn=JADE3;refGene_tx=XM_023249517.1
X	41512818	41513009	refGene_gn=JADE3;refGene_tx=XM_004000432.5;refGene_gn=JADE3;refGene_tx=XM_023249516.1;refGene_gn=JADE3;refGene_tx=XM_023249517.1;refGene_gn=JADE3;refGene_tx=XM_023249518.1;refGene_gn=JADE3;refGene_tx=XM_023249519.1;refGene_gn=JADE3;refGene_tx=XM_023249520.1
X	41516032	41516244

refGene_gn=JADE3;refGene_tx=XM_004000432.5;refGene_gn=JADE3;refGene_tx=XM_023249516.1;refGene_gn=JADE3;refGene_tx=XM_023249517.1;refGene_gn=JADE3;refGene_tx=XM_023249518.1;refGene_gn=JADE3;refGene_tx=XM_023249519.1;refGene_gn=JADE3;refGene_tx=XM_023249520.1
X 41521876 41522044
refGene_gn=JADE3;refGene_tx=XM_004000432.5;refGene_gn=JADE3;refGene_tx=XM_023249516.1;refGene_gn=JADE3;refGene_tx=XM_023249517.1;refGene_gn=JADE3;refGene_tx=XM_023249518.1;refGene_gn=JADE3;refGene_tx=XM_023249519.1;refGene_gn=JADE3;refGene_tx=XM_023249520.1
X 41524934 41525051
refGene_gn=JADE3;refGene_tx=XM_004000432.5;refGene_gn=JADE3;refGene_tx=XM_023249516.1;refGene_gn=JADE3;refGene_tx=XM_023249517.1;refGene_gn=JADE3;refGene_tx=XM_023249518.1;refGene_gn=JADE3;refGene_tx=XM_023249519.1;refGene_gn=JADE3;refGene_tx=XM_023249520.1
X 41536750 41537221
refGene_gn=JADE3;refGene_tx=XM_004000432.5;refGene_gn=JADE3;refGene_tx=XM_023249516.1;refGene_gn=JADE3;refGene_tx=XM_023249517.1;refGene_gn=JADE3;refGene_tx=XM_023249518.1;refGene_gn=JADE3;refGene_tx=XM_023249519.1;refGene_gn=JADE3;refGene_tx=XM_023249520.1
X 41538651 41538769
refGene_gn=JADE3;refGene_tx=XM_004000432.5;refGene_gn=JADE3;refGene_tx=XM_023249516.1;refGene_gn=JADE3;refGene_tx=XM_023249517.1;refGene_gn=JADE3;refGene_tx=XM_023249518.1;refGene_gn=JADE3;refGene_tx=XM_023249519.1;refGene_gn=JADE3;refGene_tx=XM_023249520.1
X 41540835 41541746
refGene_gn=JADE3;refGene_tx=XM_004000432.5;refGene_gn=JADE3;refGene_tx=XM_023249516.1;refGene_gn=JADE3;refGene_tx=XM_023249517.1;refGene_gn=JADE3;refGene_tx=XM_023249518.1;refGene_gn=JADE3;refGene_tx=XM_023249519.1;refGene_gn=JADE3;refGene_tx=XM_023249520.1
X 41567946 41568115
refGene_gn=RGN;refGene_tx=XM_006943667.4;refGene_gn=RGN;refGene_tx=XM_006943668.4;refGene_gn=RGN;refGene_tx=XM_023249103.1
X 41570159 41570342
refGene_gn=RGN;refGene_tx=XM_006943667.4;refGene_gn=RGN;refGene_tx=XM_006943668.4;refGene_gn=RGN;refGene_tx=XM_023249103.1
X 41573892 41574108
refGene_gn=RGN;refGene_tx=XM_006943667.4;refGene_gn=RGN;refGene_tx=XM_006943668.4;refGene_gn=RGN;refGene_tx=XM_023249103.1
X 41576941 41577073
refGene_gn=RGN;refGene_tx=XM_006943667.4;refGene_gn=RGN;refGene_tx=XM_006943668.4;refGene_gn=RGN;refGene_tx=XM_023249103.1
X 41577443 41577598
refGene_gn=RGN;refGene_tx=XM_006943667.4;refGene_gn=RGN;refGene_tx=XM_006943668.4;refGene_gn=RGN;refGene_tx=XM_023249103.1
X 41578902 41578953
refGene_gn=RGN;refGene_tx=XM_006943667.4;refGene_gn=RGN;refGene_tx=XM_006943668.4;refGene_gn=RGN;refGene_tx=XM_023249103.1
X 41603786 41603913 refGene_gn=NDUFB11;refGene_tx=XM_004000441.4
X 41604089 41604220 refGene_gn=NDUFB11;refGene_tx=XM_004000441.4
X 41605736 41605943 refGene_gn=NDUFB11;refGene_tx=XM_004000441.4

X 41606733 41606803 refGene_gn=RBM10;refGene_tx=XM_004000427.5
X 41608752 41608894 refGene_gn=RBM10;refGene_tx=XM_004000427.5
X 41621957 41622141 refGene_gn=RBM10;refGene_tx=XM_004000427.5
X 41623592 41623610 refGene_gn=RBM10;refGene_tx=XM_004000427.5
X 41623611 41623824 refGene_gn=RBM10;refGene_tx=XM_004000427.5
X 41625682 41625752 refGene_gn=RBM10;refGene_tx=XM_004000427.5
X 41627779 41627853 refGene_gn=RBM10;refGene_tx=XM_004000427.5
X 41629253 41629340 refGene_gn=RBM10;refGene_tx=XM_004000427.5
X 41631378 41631439 refGene_gn=RBM10;refGene_tx=XM_004000427.5
X 41631581 41631758 refGene_gn=RBM10;refGene_tx=XM_004000427.5
X 41632144 41632305 refGene_gn=RBM10;refGene_tx=XM_004000427.5
X 41632504 41632602 refGene_gn=RBM10;refGene_tx=XM_004000427.5
X 41632688 41632776 refGene_gn=RBM10;refGene_tx=XM_004000427.5
X 41633393 41633580 refGene_gn=RBM10;refGene_tx=XM_004000427.5
X 41633693 41633833 refGene_gn=RBM10;refGene_tx=XM_004000427.5
X 41633933 41634051 refGene_gn=RBM10;refGene_tx=XM_004000427.5
X 41634145 41634237 refGene_gn=RBM10;refGene_tx=XM_004000427.5
X 41634321 41634486 refGene_gn=RBM10;refGene_tx=XM_004000427.5
X 41635445 41635595 refGene_gn=RBM10;refGene_tx=XM_004000427.5
X 41635698 41635764 refGene_gn=RBM10;refGene_tx=XM_004000427.5
X 41635848 41636037 refGene_gn=RBM10;refGene_tx=XM_004000427.5
X 41636154 41636229 refGene_gn=RBM10;refGene_tx=XM_004000427.5
X 41636541 41636648 refGene_gn=RBM10;refGene_tx=XM_004000427.5
X 41636722 41636852 refGene_gn=RBM10;refGene_tx=XM_004000427.5
X 41636944 41637070 refGene_gn=RBM10;refGene_tx=XM_004000427.5
X 41652803 41652920
refGene_gn=UBA1;refGene_tx=XM_004000424.5;refGene_gn=UBA1;refGene_tx=XM_004000425.5
X 41653051 41653110
refGene_gn=UBA1;refGene_tx=XM_004000424.5;refGene_gn=UBA1;refGene_tx=XM_004000425.5
X 41653210 41653379
refGene_gn=UBA1;refGene_tx=XM_004000424.5;refGene_gn=UBA1;refGene_tx=XM_004000425.5
X 41653467 41653602
refGene_gn=UBA1;refGene_tx=XM_004000424.5;refGene_gn=UBA1;refGene_tx=XM_004000425.5
X 41654618 41654725
refGene_gn=UBA1;refGene_tx=XM_004000424.5;refGene_gn=UBA1;refGene_tx=XM_004000425.5
X 41654986 41655077
refGene_gn=UBA1;refGene_tx=XM_004000424.5;refGene_gn=UBA1;refGene_tx=XM_004000425.5
X 41655184 41655317
refGene_gn=UBA1;refGene_tx=XM_004000424.5;refGene_gn=UBA1;refGene_tx=XM_004000425.5
X 41655806 41655904
refGene_gn=UBA1;refGene_tx=XM_004000424.5;refGene_gn=UBA1;refGene_tx=XM_004000425.5
X 41655985 41656132

refGene_gn=UBA1;refGene_tx=XM_004000424.5;refGene_gn=UBA1;refGene_tx=XM_004000425.5
X 41656289 41656466
refGene_gn=UBA1;refGene_tx=XM_004000424.5;refGene_gn=UBA1;refGene_tx=XM_004000425.5
X 41656590 41656695
refGene_gn=UBA1;refGene_tx=XM_004000424.5;refGene_gn=UBA1;refGene_tx=XM_004000425.5
X 41656802 41656883
refGene_gn=UBA1;refGene_tx=XM_004000424.5;refGene_gn=UBA1;refGene_tx=XM_004000425.5
X 41657240 41657396
refGene_gn=UBA1;refGene_tx=XM_004000424.5;refGene_gn=UBA1;refGene_tx=XM_004000425.5
X 41659051 41659217
refGene_gn=UBA1;refGene_tx=XM_004000424.5;refGene_gn=UBA1;refGene_tx=XM_004000425.5
X 41659350 41659547
refGene_gn=UBA1;refGene_tx=XM_004000424.5;refGene_gn=UBA1;refGene_tx=XM_004000425.5
X 41661649 41661714
refGene_gn=UBA1;refGene_tx=XM_004000424.5;refGene_gn=UBA1;refGene_tx=XM_004000425.5
X 41661937 41662133
refGene_gn=UBA1;refGene_tx=XM_004000424.5;refGene_gn=UBA1;refGene_tx=XM_004000425.5
X 41662840 41662915
refGene_gn=UBA1;refGene_tx=XM_004000424.5;refGene_gn=UBA1;refGene_tx=XM_004000425.5
X 41663031 41663221
refGene_gn=UBA1;refGene_tx=XM_004000424.5;refGene_gn=UBA1;refGene_tx=XM_004000425.5
X 41665653 41665742
refGene_gn=UBA1;refGene_tx=XM_004000424.5;refGene_gn=UBA1;refGene_tx=XM_004000425.5
X 41666023 41666116
refGene_gn=UBA1;refGene_tx=XM_004000424.5;refGene_gn=UBA1;refGene_tx=XM_004000425.5
X 41666256 41666448
refGene_gn=UBA1;refGene_tx=XM_004000424.5;refGene_gn=UBA1;refGene_tx=XM_004000425.5
X 41667587 41667689
refGene_gn=UBA1;refGene_tx=XM_004000424.5;refGene_gn=UBA1;refGene_tx=XM_004000425.5
X 41667794 41667895
refGene_gn=UBA1;refGene_tx=XM_004000424.5;refGene_gn=UBA1;refGene_tx=XM_004000425.5
X 41668023 41668159
refGene_gn=UBA1;refGene_tx=XM_004000424.5;refGene_gn=UBA1;refGene_tx=XM_004000425.5

X	41675713 41675719 refGene_gn=CDK16;refGene_tx=XM_019823380.2
X	41676437 41676653
refGene_gn=CDK16;refGene_tx=XM_006943673.3;refGene_gn=CDK16;refGene_tx=XM_011291660.2
X	41676894 41677102
refGene_gn=CDK16;refGene_tx=XM_006943673.3;refGene_gn=CDK16;refGene_tx=XM_011291660.2;refGene_gn=CDK16;refGene_tx=XM_019823380.2;refGene_gn=CDK16;refGene_tx=XM_023249397.1;refGene_gn=CDK16;refGene_tx=XM_023249398.1
X	41677704 41677838
refGene_gn=CDK16;refGene_tx=XM_006943673.3;refGene_gn=CDK16;refGene_tx=XM_011291660.2;refGene_gn=CDK16;refGene_tx=XM_019823380.2;refGene_gn=CDK16;refGene_tx=XM_023249397.1;refGene_gn=CDK16;refGene_tx=XM_023249398.1
X	41677939 41678065
refGene_gn=CDK16;refGene_tx=XM_006943673.3;refGene_gn=CDK16;refGene_tx=XM_011291660.2;refGene_gn=CDK16;refGene_tx=XM_019823380.2;refGene_gn=CDK16;refGene_tx=XM_023249397.1;refGene_gn=CDK16;refGene_tx=XM_023249398.1
X	41678162 41678219
refGene_gn=CDK16;refGene_tx=XM_006943673.3;refGene_gn=CDK16;refGene_tx=XM_011291660.2;refGene_gn=CDK16;refGene_tx=XM_019823380.2;refGene_gn=CDK16;refGene_tx=XM_023249397.1;refGene_gn=CDK16;refGene_tx=XM_023249398.1
X	41678310 41678425
refGene_gn=CDK16;refGene_tx=XM_006943673.3;refGene_gn=CDK16;refGene_tx=XM_011291660.2;refGene_gn=CDK16;refGene_tx=XM_019823380.2;refGene_gn=CDK16;refGene_tx=XM_023249397.1;refGene_gn=CDK16;refGene_tx=XM_023249398.1
X	41679112 41679207
refGene_gn=CDK16;refGene_tx=XM_006943673.3;refGene_gn=CDK16;refGene_tx=XM_011291660.2;refGene_gn=CDK16;refGene_tx=XM_019823380.2;refGene_gn=CDK16;refGene_tx=XM_023249397.1;refGene_gn=CDK16;refGene_tx=XM_023249398.1
X	41679302 41679365
refGene_gn=CDK16;refGene_tx=XM_006943673.3;refGene_gn=CDK16;refGene_tx=XM_011291660.2;refGene_gn=CDK16;refGene_tx=XM_019823380.2;refGene_gn=CDK16;refGene_tx=XM_023249397.1;refGene_gn=CDK16;refGene_tx=XM_023249398.1
X	41679617 41679741
refGene_gn=CDK16;refGene_tx=XM_006943673.3;refGene_gn=CDK16;refGene_tx=XM_011291660.2;refGene_gn=CDK16;refGene_tx=XM_019823380.2;refGene_gn=CDK16;refGene_tx=XM_023249397.1;refGene_gn=CDK16;refGene_tx=XM_023249398.1
X	41679938 41680059
refGene_gn=CDK16;refGene_tx=XM_006943673.3;refGene_gn=CDK16;refGene_tx=XM_011291660.2;refGene_gn=CDK16;refGene_tx=XM_019823380.2;refGene_gn=CDK16;refGene_tx=XM_023249397.1;refGene_gn=CDK16;refGene_tx=XM_023249398.1
X	41680167 41680265

refGene_gn=CDK16;refGene_tx=XM_006943673.3;refGene_gn=CDK16;refGene_tx=XM_011291660.2;refGene_gn=CDK16;refGene_tx=XM_019823380.2;refGene_gn=CDK16;refGene_tx=XM_023249397.1;refGene_gn=CDK16;refGene_tx=XM_023249398.1
X	41680348	41680454	refGene_gn=CDK16;refGene_tx=XM_006943673.3;refGene_gn=CDK16;refGene_tx=XM_011291660.2;refGene_gn=CDK16;refGene_tx=XM_019823380.2;refGene_gn=CDK16;refGene_tx=XM_023249397.1;refGene_gn=CDK16;refGene_tx=XM_023249398.1
X	41680521	41680585	refGene_gn=CDK16;refGene_tx=XM_006943673.3;refGene_gn=CDK16;refGene_tx=XM_011291660.2;refGene_gn=CDK16;refGene_tx=XM_019823380.2;refGene_gn=CDK16;refGene_tx=XM_023249397.1;refGene_gn=CDK16;refGene_tx=XM_023249398.1
X	41680751	41680842	refGene_gn=CDK16;refGene_tx=XM_006943673.3;refGene_gn=CDK16;refGene_tx=XM_011291660.2;refGene_gn=CDK16;refGene_tx=XM_019823380.2;refGene_gn=CDK16;refGene_tx=XM_023249397.1;refGene_gn=CDK16;refGene_tx=XM_023249398.1
X	41682282	41682360	refGene_gn=CDK16;refGene_tx=XM_006943673.3;refGene_gn=CDK16;refGene_tx=XM_011291660.2;refGene_gn=CDK16;refGene_tx=XM_019823380.2;refGene_gn=CDK16;refGene_tx=XM_023249397.1;refGene_gn=CDK16;refGene_tx=XM_023249398.1
X	41682445	41682483	refGene_gn=CDK16;refGene_tx=XM_006943673.3;refGene_gn=CDK16;refGene_tx=XM_011291660.2;refGene_gn=CDK16;refGene_tx=XM_019823380.2;refGene_gn=CDK16;refGene_tx=XM_023249397.1;refGene_gn=CDK16;refGene_tx=XM_023249398.1
X	41686657	41686774	refGene_gn=USP11;refGene_tx=XM_004000438.5
X	41686775	41686930	refGene_gn=USP11;refGene_tx=XM_004000438.5
X	41694176	41694286	refGene_gn=USP11;refGene_tx=XM_004000438.5
X	41694489	41694620	refGene_gn=USP11;refGene_tx=XM_004000438.5
X	41694897	41695015	refGene_gn=USP11;refGene_tx=XM_004000438.5
X	41695402	41695548	refGene_gn=USP11;refGene_tx=XM_004000438.5
X	41695669	41695731	refGene_gn=USP11;refGene_tx=XM_004000438.5
X	41695856	41695956	refGene_gn=USP11;refGene_tx=XM_004000438.5
X	41696328	41696502	refGene_gn=USP11;refGene_tx=XM_004000438.5
X	41696593	41696752	refGene_gn=USP11;refGene_tx=XM_004000438.5
X	41697084	41697309	refGene_gn=USP11;refGene_tx=XM_004000438.5
X	41697446	41697530	refGene_gn=USP11;refGene_tx=XM_004000438.5
X	41697636	41697731	refGene_gn=USP11;refGene_tx=XM_004000438.5
X	41698688	41698895	refGene_gn=USP11;refGene_tx=XM_004000438.5
X	41699419	41699472	refGene_gn=USP11;refGene_tx=XM_004000438.5
X	41699641	41699884	refGene_gn=USP11;refGene_tx=XM_004000438.5
X	41699973	41700044	refGene_gn=USP11;refGene_tx=XM_004000438.5
X	41700244	41700357	refGene_gn=USP11;refGene_tx=XM_004000438.5
X	41701364	41701514	refGene_gn=USP11;refGene_tx=XM_004000438.5
X	41701612	41701728	refGene_gn=USP11;refGene_tx=XM_004000438.5
X	41701878	41701967	refGene_gn=USP11;refGene_tx=XM_004000438.5

X 41702056 41702194 refGene_gn=USP11;refGene_tx=XM_004000438.5
X 41812468 41812540 refGene_gn=ZNF157;refGene_tx=XM_011291659.3
X 41846858 41846985
refGene_gn=ZNF157;refGene_tx=XM_011291659.3;refGene_gn=ZNF157;refGene_tx=XM_019823383.2;refGene_gn=ZNF157;refGene_tx=XM_019823385.2
X 41847283 41847379
refGene_gn=ZNF157;refGene_tx=XM_011291659.3;refGene_gn=ZNF157;refGene_tx=XM_019823383.2;refGene_gn=ZNF157;refGene_tx=XM_019823385.2;refGene_gn=ZNF157;refGene_tx=XM_023249288.1
X 41848730 41849953
refGene_gn=ZNF157;refGene_tx=XM_011291659.3;refGene_gn=ZNF157;refGene_tx=XM_019823383.2;refGene_gn=ZNF157;refGene_tx=XM_019823385.2;refGene_gn=ZNF157;refGene_tx=XM_023249288.1
X 41865827 41866120 refGene_gn=ZNF41;refGene_tx=XM_019823356.2
X 41866122 41866645 refGene_gn=ZNF41;refGene_tx=XM_019823356.2
X 41866646 41867848 refGene_gn=ZNF41;refGene_tx=XM_019823356.2
X 41878102 41878198 refGene_gn=ZNF41;refGene_tx=XM_019823356.2
X 41878465 41878592 refGene_gn=ZNF41;refGene_tx=XM_019823356.2
X 41890818 41890890 refGene_gn=ZNF41;refGene_tx=XM_019823356.2
X 41988992 41989088
refGene_gn=ARAF;refGene_tx=XM_006943681.4;refGene_gn=ARAF;refGene_tx=XM_011291658.3
X 41989262 41989366
refGene_gn=ARAF;refGene_tx=XM_006943681.4;refGene_gn=ARAF;refGene_tx=XM_011291658.3
X 41990346 41990449
refGene_gn=ARAF;refGene_tx=XM_006943681.4;refGene_gn=ARAF;refGene_tx=XM_011291658.3
X 41990542 41990697
refGene_gn=ARAF;refGene_tx=XM_006943681.4;refGene_gn=ARAF;refGene_tx=XM_011291658.3
X 41990803 41990902
refGene_gn=ARAF;refGene_tx=XM_006943681.4;refGene_gn=ARAF;refGene_tx=XM_011291658.3
X 41993275 41993417
refGene_gn=ARAF;refGene_tx=XM_006943681.4;refGene_gn=ARAF;refGene_tx=XM_011291658.3
X 41993520 41993548
refGene_gn=ARAF;refGene_tx=XM_006943681.4;refGene_gn=ARAF;refGene_tx=XM_011291658.3
X 41993637 41993789
refGene_gn=ARAF;refGene_tx=XM_006943681.4;refGene_gn=ARAF;refGene_tx=XM_011291658.3
X 41993880 41994083
refGene_gn=ARAF;refGene_tx=XM_006943681.4;refGene_gn=ARAF;refGene_tx=XM_011291658.3
X 41995823 41996000
refGene_gn=ARAF;refGene_tx=XM_006943681.4;refGene_gn=ARAF;refGene_tx=XM_011291658.3
X 41996102 41996149

refGene_gn=ARAF;refGene_tx=XM_006943681.4;refGene_gn=ARAF;refGene_tx=XM_011291658.3
X	41996637	41996756	refGene_gn=ARAF;refGene_tx=XM_006943681.4;refGene_gn=ARAF;refGene_tx=XM_011291658.3
X	41996995	41997127	refGene_gn=ARAF;refGene_tx=XM_006943681.4;refGene_gn=ARAF;refGene_tx=XM_011291658.3
X	41998108	41998243	refGene_gn=ARAF;refGene_tx=XM_006943681.4;refGene_gn=ARAF;refGene_tx=XM_011291658.3
X	41998437	41998572	refGene_gn=ARAF;refGene_tx=XM_006943681.4;refGene_gn=ARAF;refGene_tx=XM_011291658.3
X	41999980	42000116	refGene_gn=SYN1;refGene_tx=XM_019823364.2;refGene_gn=SYN1;refGene_tx=XM_019823365.1
X	42001154	42001746	refGene_gn=SYN1;refGene_tx=XM_019823364.2;refGene_gn=SYN1;refGene_tx=XM_019823365.1
X	42001834	42001922	refGene_gn=SYN1;refGene_tx=XM_019823364.2;refGene_gn=SYN1;refGene_tx=XM_019823365.1
X	42002290	42002437	refGene_gn=SYN1;refGene_tx=XM_019823364.2;refGene_gn=SYN1;refGene_tx=XM_019823365.1
X	42003649	42003752	refGene_gn=SYN1;refGene_tx=XM_019823364.2;refGene_gn=SYN1;refGene_tx=XM_019823365.1
X	42003854	42003929	refGene_gn=SYN1;refGene_tx=XM_019823364.2;refGene_gn=SYN1;refGene_tx=XM_019823365.1
X	42004023	42004166	refGene_gn=SYN1;refGene_tx=XM_019823364.2;refGene_gn=SYN1;refGene_tx=XM_019823365.1
X	42004908	42004971	refGene_gn=SYN1;refGene_tx=XM_019823364.2;refGene_gn=SYN1;refGene_tx=XM_019823365.1
X	42009535	42009677	refGene_gn=TIMP1;refGene_tx=XM_023249291.1
X	42010510	42010639	refGene_gn=TIMP1;refGene_tx=XM_023249291.1
X	42011632	42011712	refGene_gn=TIMP1;refGene_tx=XM_023249291.1
X	42011896	42012023	refGene_gn=TIMP1;refGene_tx=XM_023249291.1
X	42012291	42012416	refGene_gn=TIMP1;refGene_tx=XM_023249291.1
X	42013267	42013438	refGene_gn=TIMP1;refGene_tx=XM_023249291.1
X	42038101	42038191	refGene_gn=SYN1;refGene_tx=XM_019823364.2;refGene_gn=SYN1;refGene_tx=XM_019823365.1
X	42038366	42038523	refGene_gn=SYN1;refGene_tx=XM_019823364.2;refGene_gn=SYN1;refGene_tx=XM_019823365.1

X 42040593 42040685
refGene_gn=SYN1;refGene_tx=XM_019823364.2;refGene_gn=SYN1;refGene_tx=XM_019823365.1
X 42040790 42040848
refGene_gn=SYN1;refGene_tx=XM_019823364.2;refGene_gn=SYN1;refGene_tx=XM_019823365.1
X 42051523 42051900
refGene_gn=SYN1;refGene_tx=XM_019823364.2;refGene_gn=SYN1;refGene_tx=XM_019823365.1
X 42056444 42056610 refGene_gn=CFP;refGene_tx=XM_006943679.4
X 42058008 42058120 refGene_gn=CFP;refGene_tx=XM_006943679.4
X 42058274 42058466 refGene_gn=CFP;refGene_tx=XM_006943679.4
X 42058705 42058879 refGene_gn=CFP;refGene_tx=XM_006943679.4
X 42059074 42059266 refGene_gn=CFP;refGene_tx=XM_006943679.4
X 42059402 42059573 refGene_gn=CFP;refGene_tx=XM_006943679.4
X 42059995 42060171 refGene_gn=CFP;refGene_tx=XM_006943679.4
X 42061400 42061551 refGene_gn=CFP;refGene_tx=XM_006943679.4
X 42061633 42061712 refGene_gn=CFP;refGene_tx=XM_006943679.4
X 42068238 42068337
refGene_gn=ELK1;refGene_tx=XM_006943677.4;refGene_gn=ELK1;refGene_tx=XM_019823386.2
X 42068422 42068524
refGene_gn=ELK1;refGene_tx=XM_006943677.4;refGene_gn=ELK1;refGene_tx=XM_019823386.2
X 42069190 42069637
refGene_gn=ELK1;refGene_tx=XM_006943677.4;refGene_gn=ELK1;refGene_tx=XM_019823386.2
X 42069902 42069935 refGene_gn=ELK1;refGene_tx=XM_023249289.1
X 42070272 42070722
refGene_gn=ELK1;refGene_tx=XM_006943677.4;refGene_gn=ELK1;refGene_tx=XM_019823386.2;refGene_gn=ELK1;refGene_tx=XM_023249289.1
X 42072965 42073175
refGene_gn=ELK1;refGene_tx=XM_006943677.4;refGene_gn=ELK1;refGene_tx=XM_019823386.2;refGene_gn=ELK1;refGene_tx=XM_023249289.1
X 42082681 42082781
refGene_gn=UXT;refGene_tx=XM_006943676.2;refGene_gn=UXT;refGene_tx=XM_019823396.2
X 42082999 42083063 refGene_gn=UXT;refGene_tx=XM_006943676.2
X 42089116 42089224
refGene_gn=UXT;refGene_tx=XM_006943676.2;refGene_gn=UXT;refGene_tx=XM_019823396.2
X 42089493 42089561
refGene_gn=UXT;refGene_tx=XM_006943676.2;refGene_gn=UXT;refGene_tx=XM_019823396.2
X 42089700 42089782
refGene_gn=UXT;refGene_tx=XM_006943676.2;refGene_gn=UXT;refGene_tx=XM_019823396.2
X 42091195 42091293
refGene_gn=UXT;refGene_tx=XM_006943676.2;refGene_gn=UXT;refGene_tx=XM_019823396.2

X 42137693 42137780
refGene_gn=LOC101090336;refGene_tx=XM_023249290.1
X 42137916 42138089
refGene_gn=LOC101090336;refGene_tx=XM_023249290.1
X 42138401 42138609
refGene_gn=LOC101090336;refGene_tx=XM_023249290.1
X 42138692 42138803
refGene_gn=LOC101090336;refGene_tx=XM_023249290.1
X 42139119 42139169
refGene_gn=LOC101090336;refGene_tx=XM_023249290.1
X 42139377 42139474
refGene_gn=LOC101090336;refGene_tx=XM_023249290.1
X 42140396 42140549
refGene_gn=LOC101090336;refGene_tx=XM_023249290.1
X 42140626 42140773
refGene_gn=LOC101090336;refGene_tx=XM_023249290.1
X 42204685 42204757 refGene_gn=ZNF81;refGene_tx=XM_011291715.3
X 42264593 42264720 refGene_gn=ZNF81;refGene_tx=XM_011291715.3
X 42273290 42273386 refGene_gn=ZNF81;refGene_tx=XM_011291715.3
X 42284946 42286428 refGene_gn=ZNF81;refGene_tx=XM_011291715.3
X 42286430 42286654 refGene_gn=ZNF81;refGene_tx=XM_011291715.3
X 42378741 42380372
refGene_gn=ZNF182;refGene_tx=XM_019823378.2;refGene_gn=ZNF182;refGene_tx=XM_023249282.1;refGene_gn=ZNF182;refGene_tx=XM_023249283.1;refGene_gn=ZNF182;refGene_tx=XM_023249284.1;refGene_gn=ZNF182;refGene_tx=XM_023249285.1;refGene_gn=ZNF182;refGene_tx=XM_023249286.1;refGene_gn=ZNF182;refGene_tx=XM_023249287.1
X 42384616 42384712
refGene_gn=ZNF182;refGene_tx=XM_019823378.2;refGene_gn=ZNF182;refGene_tx=XM_023249282.1;refGene_gn=ZNF182;refGene_tx=XM_023249283.1;refGene_gn=ZNF182;refGene_tx=XM_023249284.1;refGene_gn=ZNF182;refGene_tx=XM_023249285.1;refGene_gn=ZNF182;refGene_tx=XM_023249286.1;refGene_gn=ZNF182;refGene_tx=XM_023249287.1
X 42385023 42385150
refGene_gn=ZNF182;refGene_tx=XM_019823378.2;refGene_gn=ZNF182;refGene_tx=XM_023249282.1;refGene_gn=ZNF182;refGene_tx=XM_023249283.1;refGene_gn=ZNF182;refGene_tx=XM_023249284.1;refGene_gn=ZNF182;refGene_tx=XM_023249285.1;refGene_gn=ZNF182;refGene_tx=XM_023249286.1;refGene_gn=ZNF182;refGene_tx=XM_023249287.1
X 42406566 42406581
refGene_gn=ZNF182;refGene_tx=XM_019823378.2;refGene_gn=ZNF182;refGene_tx=XM_023249282.1;refGene_gn=ZNF182;refGene_tx=XM_023249283.1;refGene_gn=ZNF182;refGene_tx=XM_023249284.1;refGene_gn=ZNF182;refGene_tx=XM_023249285.1;refGene_gn=ZNF182;refGene_tx=XM_023249286.1;refGene_gn=ZNF182;refGene_tx=XM_023249287.1
X 42414642 42414787
refGene_gn=LOC101091348;refGene_tx=XM_004000472.4
X 42415208 42415367
refGene_gn=LOC101091348;refGene_tx=XM_004000472.4
X 42417071 42417150

refGene_gn=LOC101091348;refGene_tx=XM_004000472.4
X	42417253 42417353
refGene_gn=LOC101091348;refGene_tx=XM_004000472.4
X	42480219 42481946
refGene_gn=ZNF630;refGene_tx=XM_004000521.5;refGene_gn=ZNF630;refGene_tx=XM_011291714.3;refGene_gn=ZNF630;refGene_tx=XM_023249277.1;refGene_gn=ZNF630;refGene_tx=XM_023249278.1;refGene_gn=ZNF630;refGene_tx=XM_023249279.1;refGene_gn=ZNF630;refGene_tx=XM_023249280.1;refGene_gn=ZNF630;refGene_tx=XM_023249281.1
X	42482136 42482232
refGene_gn=ZNF630;refGene_tx=XM_004000521.5;refGene_gn=ZNF630;refGene_tx=XM_011291714.3;refGene_gn=ZNF630;refGene_tx=XM_023249277.1;refGene_gn=ZNF630;refGene_tx=XM_023249278.1;refGene_gn=ZNF630;refGene_tx=XM_023249279.1;refGene_gn=ZNF630;refGene_tx=XM_023249280.1;refGene_gn=ZNF630;refGene_tx=XM_023249281.1
X	42482461 42482588
refGene_gn=ZNF630;refGene_tx=XM_004000521.5;refGene_gn=ZNF630;refGene_tx=XM_011291714.3;refGene_gn=ZNF630;refGene_tx=XM_023249277.1;refGene_gn=ZNF630;refGene_tx=XM_023249278.1;refGene_gn=ZNF630;refGene_tx=XM_023249279.1;refGene_gn=ZNF630;refGene_tx=XM_023249280.1;refGene_gn=ZNF630;refGene_tx=XM_023249281.1
X	42488326 42488341
refGene_gn=ZNF630;refGene_tx=XM_004000521.5;refGene_gn=ZNF630;refGene_tx=XM_023249278.1;refGene_gn=ZNF630;refGene_tx=XM_023249279.1;refGene_gn=ZNF630;refGene_tx=XM_023249280.1;refGene_gn=ZNF630;refGene_tx=XM_023249281.1
X	42490875 42491298
refGene_gn=ZNF630;refGene_tx=XM_011291714.3;refGene_gn=ZNF630;refGene_tx=XM_023249277.1
X	42523991 42524093
refGene_gn=LOC101083323;refGene_tx=XM_006943700.2
X	42524529 42524644
refGene_gn=LOC101083323;refGene_tx=XM_006943700.2
X	42525240 42525339
refGene_gn=LOC101083323;refGene_tx=XM_006943700.2
X	42527731 42527781
refGene_gn=LOC101083323;refGene_tx=XM_006943700.2
X	42529397 42529533
refGene_gn=LOC101083323;refGene_tx=XM_006943700.2
X	42530395 42530493
refGene_gn=LOC101083323;refGene_tx=XM_006943700.2
X	42545635 42545737
refGene_gn=SLC38A5;refGene_tx=XM_004000473.5;refGene_gn=SLC38A5;refGene_tx=XM_006943701.2;refGene_gn=SLC38A5;refGene_tx=XM_019823762.2;refGene_gn=SLC38A5;refGene_tx=XM_023249111.1;refGene_gn=SLC38A5;refGene_tx=XM_023249112.1
X	42546200 42546304
refGene_gn=SLC38A5;refGene_tx=XM_004000473.5;refGene_gn=SLC38A5;refGene_tx=XM_006943701.2;refGene_gn=SLC38A5;refGene_tx=XM_019823762.2;refGene_gn=SLC38A5;refGene_tx=XM_023249111.1;refGene_gn=SLC38A5;refGene_tx=

XM_023249112.1
X 42546398 42546543
refGene_gn=SLC38A5;refGene_tx=XM_004000473.5;refGene_gn=SLC38A5;refGene_tx=XM_006943701.2;refGene_gn=SLC38A5;refGene_tx=XM_019823762.2
X 42547327 42547443
refGene_gn=SLC38A5;refGene_tx=XM_004000473.5;refGene_gn=SLC38A5;refGene_tx=XM_006943701.2;refGene_gn=SLC38A5;refGene_tx=XM_019823762.2;refGene_gn=SLC38A5;refGene_tx=XM_023249111.1
X 42547698 42547799
refGene_gn=SLC38A5;refGene_tx=XM_004000473.5;refGene_gn=SLC38A5;refGene_tx=XM_006943701.2;refGene_gn=SLC38A5;refGene_tx=XM_019823762.2;refGene_gn=SLC38A5;refGene_tx=XM_023249111.1;refGene_gn=SLC38A5;refGene_tx=XM_023249112.1
X 42548441 42548521
refGene_gn=SLC38A5;refGene_tx=XM_004000473.5;refGene_gn=SLC38A5;refGene_tx=XM_006943701.2;refGene_gn=SLC38A5;refGene_tx=XM_019823762.2;refGene_gn=SLC38A5;refGene_tx=XM_023249111.1;refGene_gn=SLC38A5;refGene_tx=XM_023249112.1
X 42548732 42548870
refGene_gn=SLC38A5;refGene_tx=XM_004000473.5;refGene_gn=SLC38A5;refGene_tx=XM_006943701.2;refGene_gn=SLC38A5;refGene_tx=XM_019823762.2;refGene_gn=SLC38A5;refGene_tx=XM_023249111.1;refGene_gn=SLC38A5;refGene_tx=XM_023249112.1
X 42548967 42549026
refGene_gn=SLC38A5;refGene_tx=XM_004000473.5;refGene_gn=SLC38A5;refGene_tx=XM_006943701.2;refGene_gn=SLC38A5;refGene_tx=XM_019823762.2;refGene_gn=SLC38A5;refGene_tx=XM_023249111.1;refGene_gn=SLC38A5;refGene_tx=XM_023249112.1
X 42549537 42549620
refGene_gn=SLC38A5;refGene_tx=XM_004000473.5;refGene_gn=SLC38A5;refGene_tx=XM_006943701.2;refGene_gn=SLC38A5;refGene_tx=XM_019823762.2;refGene_gn=SLC38A5;refGene_tx=XM_023249111.1;refGene_gn=SLC38A5;refGene_tx=XM_023249112.1
X 42551332 42551411
refGene_gn=SLC38A5;refGene_tx=XM_004000473.5;refGene_gn=SLC38A5;refGene_tx=XM_006943701.2;refGene_gn=SLC38A5;refGene_tx=XM_019823762.2;refGene_gn=SLC38A5;refGene_tx=XM_023249111.1;refGene_gn=SLC38A5;refGene_tx=XM_023249112.1
X 42551535 42551628
refGene_gn=SLC38A5;refGene_tx=XM_004000473.5;refGene_gn=SLC38A5;refGene_tx=XM_006943701.2;refGene_gn=SLC38A5;refGene_tx=XM_019823762.2;refGene_gn=SLC38A5;refGene_tx=XM_023249111.1;refGene_gn=SLC38A5;refGene_tx=XM_023249112.1
X 42552253 42552327
refGene_gn=SLC38A5;refGene_tx=XM_004000473.5;refGene_gn=SLC38A5;refGene_tx=XM_006943701.2;refGene_gn=SLC38A5;refGene_tx=XM_019823762.2;refGene_gn=SLC38A5;refGene_tx=XM_023249111.1;refGene_gn=SLC38A5;refGene_tx=XM_023249112.1
X 42552431 42552547
refGene_gn=SLC38A5;refGene_tx=XM_004000473.5;refGene_gn=SLC38A5;refGen

e_tx=XM_006943701.2;refGene_gn=SLC38A5;refGene_tx=XM_019823762.2;refGene_gn=SLC38A5;refGene_tx=XM_023249111.1;refGene_gn=SLC38A5;refGene_tx=XM_023249112.1
X 42552967 42553043 refGene_gn=SLC38A5;refGene_tx=XM_004000473.5;refGene_gn=SLC38A5;refGene_tx=XM_006943701.2;refGene_gn=SLC38A5;refGene_tx=XM_019823762.2;refGene_gn=SLC38A5;refGene_tx=XM_023249111.1;refGene_gn=SLC38A5;refGene_tx=XM_023249112.1
X 42553123 42553177 refGene_gn=SLC38A5;refGene_tx=XM_004000473.5;refGene_gn=SLC38A5;refGene_tx=XM_006943701.2;refGene_gn=SLC38A5;refGene_tx=XM_019823762.2;refGene_gn=SLC38A5;refGene_tx=XM_023249111.1;refGene_gn=SLC38A5;refGene_tx=XM_023249112.1
X 42553526 42553549 refGene_gn=SLC38A5;refGene_tx=XM_004000473.5
X 42563017 42563138 refGene_gn=FTSJ1;refGene_tx=XM_006943705.3;refGene_gn=FTSJ1;refGene_tx=XM_006943706.4;refGene_gn=FTSJ1;refGene_tx=XM_019823764.2;refGene_gn=FTSJ1;refGene_tx=XM_019823765.2;refGene_gn=FTSJ1;refGene_tx=XM_019823766.2
X 42563392 42563462 refGene_gn=FTSJ1;refGene_tx=XM_006943705.3;refGene_gn=FTSJ1;refGene_tx=XM_006943706.4;refGene_gn=FTSJ1;refGene_tx=XM_019823764.2;refGene_gn=FTSJ1;refGene_tx=XM_019823765.2;refGene_gn=FTSJ1;refGene_tx=XM_019823766.2
X 42563567 42563658 refGene_gn=FTSJ1;refGene_tx=XM_006943705.3;refGene_gn=FTSJ1;refGene_tx=XM_006943706.4;refGene_gn=FTSJ1;refGene_tx=XM_019823764.2;refGene_gn=FTSJ1;refGene_tx=XM_019823765.2;refGene_gn=FTSJ1;refGene_tx=XM_019823766.2;refGene_gn=FTSJ1;refGene_tx=XM_019823767.2
X 42563988 42564067 refGene_gn=FTSJ1;refGene_tx=XM_006943705.3;refGene_gn=FTSJ1;refGene_tx=XM_006943706.4;refGene_gn=FTSJ1;refGene_tx=XM_019823764.2;refGene_gn=FTSJ1;refGene_tx=XM_019823765.2;refGene_gn=FTSJ1;refGene_tx=XM_019823766.2;refGene_gn=FTSJ1;refGene_tx=XM_019823767.2
X 42565618 42565720 refGene_gn=FTSJ1;refGene_tx=XM_006943705.3;refGene_gn=FTSJ1;refGene_tx=XM_006943706.4;refGene_gn=FTSJ1;refGene_tx=XM_019823764.2;refGene_gn=FTSJ1;refGene_tx=XM_019823765.2;refGene_gn=FTSJ1;refGene_tx=XM_019823766.2;refGene_gn=FTSJ1;refGene_tx=XM_019823767.2;refGene_gn=FTSJ1;refGene_tx=XM_019823768.2
X 42565805 42565859 refGene_gn=FTSJ1;refGene_tx=XM_006943705.3;refGene_gn=FTSJ1;refGene_tx=XM_006943706.4;refGene_gn=FTSJ1;refGene_tx=XM_019823764.2;refGene_gn=FTSJ1;refGene_tx=XM_019823765.2;refGene_gn=FTSJ1;refGene_tx=XM_019823766.2;refGene_gn=FTSJ1;refGene_tx=XM_019823767.2;refGene_gn=FTSJ1;refGene_tx=XM_019823768.2
X 42565945 42566048 refGene_gn=FTSJ1;refGene_tx=XM_006943705.3;refGene_gn=FTSJ1;refGene_tx=XM_006943706.4;refGene_gn=FTSJ1;refGene_tx=XM_019823764.2;refGene_gn=FTSJ1;refGene_tx=XM_019823765.2;refGene_gn=FTSJ1;refGene_tx=XM_0198237

66.2;refGene_gn=FTSJ1;refGene_tx=XM_019823767.2;refGene_gn=FTSJ1;refGene_tx=XM_019823768.2
X 42566151 42566235
refGene_gn=FTSJ1;refGene_tx=XM_006943705.3;refGene_gn=FTSJ1;refGene_tx=XM_006943706.4;refGene_gn=FTSJ1;refGene_tx=XM_019823764.2;refGene_gn=FTSJ1;refGene_tx=XM_019823765.2;refGene_gn=FTSJ1;refGene_tx=XM_019823766.2;refGene_gn=FTSJ1;refGene_tx=XM_019823767.2;refGene_gn=FTSJ1;refGene_tx=XM_019823768.2
X 42566919 42567023
refGene_gn=FTSJ1;refGene_tx=XM_006943705.3;refGene_gn=FTSJ1;refGene_tx=XM_006943706.4;refGene_gn=FTSJ1;refGene_tx=XM_019823764.2;refGene_gn=FTSJ1;refGene_tx=XM_019823765.2;refGene_gn=FTSJ1;refGene_tx=XM_019823766.2;refGene_gn=FTSJ1;refGene_tx=XM_019823767.2;refGene_gn=FTSJ1;refGene_tx=XM_019823768.2
X 42567109 42567307
refGene_gn=FTSJ1;refGene_tx=XM_006943705.3;refGene_gn=FTSJ1;refGene_tx=XM_006943706.4;refGene_gn=FTSJ1;refGene_tx=XM_019823764.2;refGene_gn=FTSJ1;refGene_tx=XM_019823765.2;refGene_gn=FTSJ1;refGene_tx=XM_019823766.2;refGene_gn=FTSJ1;refGene_tx=XM_019823767.2;refGene_gn=FTSJ1;refGene_tx=XM_019823768.2
X 42567514 42567547
refGene_gn=FTSJ1;refGene_tx=XM_006943705.3;refGene_gn=FTSJ1;refGene_tx=XM_006943706.4;refGene_gn=FTSJ1;refGene_tx=XM_019823764.2;refGene_gn=FTSJ1;refGene_tx=XM_019823765.2;refGene_gn=FTSJ1;refGene_tx=XM_019823766.2;refGene_gn=FTSJ1;refGene_tx=XM_019823767.2;refGene_gn=FTSJ1;refGene_tx=XM_019823768.2
X 42612700 42612836
refGene_gn=PORCN;refGene_tx=XM_004000476.5;refGene_gn=PORCN;refGene_tx=XM_004000477.5;refGene_gn=PORCN;refGene_tx=XM_011291711.3
X 42615029 42615222
refGene_gn=PORCN;refGene_tx=XM_004000476.5;refGene_gn=PORCN;refGene_tx=XM_004000477.5;refGene_gn=PORCN;refGene_tx=XM_011291711.3;refGene_gn=PORCN;refGene_tx=XM_019823769.2
X 42615520 42615564
refGene_gn=PORCN;refGene_tx=XM_004000476.5;refGene_gn=PORCN;refGene_tx=XM_004000477.5;refGene_gn=PORCN;refGene_tx=XM_011291711.3;refGene_gn=PORCN;refGene_tx=XM_019823769.2
X 42615961 42616143
refGene_gn=PORCN;refGene_tx=XM_004000476.5;refGene_gn=PORCN;refGene_tx=XM_004000477.5;refGene_gn=PORCN;refGene_tx=XM_011291711.3;refGene_gn=PORCN;refGene_tx=XM_019823769.2
X 42616223 42616354
refGene_gn=PORCN;refGene_tx=XM_004000476.5;refGene_gn=PORCN;refGene_tx=XM_004000477.5;refGene_gn=PORCN;refGene_tx=XM_011291711.3;refGene_gn=PORCN;refGene_tx=XM_019823769.2
X 42616469 42616487
refGene_gn=PORCN;refGene_tx=XM_004000476.5;refGene_gn=PORCN;refGene_tx=XM_011291711.3;refGene_gn=PORCN;refGene_tx=XM_019823769.2
X 42617770 42617785
refGene_gn=PORCN;refGene_tx=XM_011291711.3;refGene_gn=PORCN;refGene_tx

=XM_019823769.2
X 42617883 42618009
refGene_gn=PORCN;refGene_tx=XM_004000476.5;refGene_gn=PORCN;refGene_tx=XM_004000477.5;refGene_gn=PORCN;refGene_tx=XM_011291711.3;refGene_gn=PORCN;refGene_tx=XM_019823769.2
X 42618169 42618270
refGene_gn=PORCN;refGene_tx=XM_004000476.5;refGene_gn=PORCN;refGene_tx=XM_004000477.5;refGene_gn=PORCN;refGene_tx=XM_011291711.3;refGene_gn=PORCN;refGene_tx=XM_019823769.2
X 42619418 42619495
refGene_gn=PORCN;refGene_tx=XM_004000476.5;refGene_gn=PORCN;refGene_tx=XM_004000477.5;refGene_gn=PORCN;refGene_tx=XM_011291711.3;refGene_gn=PORCN;refGene_tx=XM_019823769.2
X 42619587 42619651
refGene_gn=PORCN;refGene_tx=XM_004000476.5;refGene_gn=PORCN;refGene_tx=XM_004000477.5;refGene_gn=PORCN;refGene_tx=XM_011291711.3;refGene_gn=PORCN;refGene_tx=XM_019823769.2
X 42619757 42619843
refGene_gn=PORCN;refGene_tx=XM_004000476.5;refGene_gn=PORCN;refGene_tx=XM_004000477.5;refGene_gn=PORCN;refGene_tx=XM_011291711.3;refGene_gn=PORCN;refGene_tx=XM_019823769.2
X 42620834 42620945
refGene_gn=PORCN;refGene_tx=XM_004000476.5;refGene_gn=PORCN;refGene_tx=XM_004000477.5;refGene_gn=PORCN;refGene_tx=XM_011291711.3;refGene_gn=PORCN;refGene_tx=XM_019823769.2
X 42623195 42623297
refGene_gn=PORCN;refGene_tx=XM_004000476.5;refGene_gn=PORCN;refGene_tx=XM_004000477.5;refGene_gn=PORCN;refGene_tx=XM_011291711.3;refGene_gn=PORCN;refGene_tx=XM_019823769.2
X 42625676 42625977
refGene_gn=EBP;refGene_tx=XM_004000480.4;refGene_gn=EBP;refGene_tx=XM_019823770.2;refGene_gn=EBP;refGene_tx=XM_019823772.2
X 42627574 42627611
refGene_gn=EBP;refGene_tx=XM_004000480.4;refGene_gn=EBP;refGene_tx=XM_019823770.2;refGene_gn=EBP;refGene_tx=XM_019823772.2
X 42627739 42627870
refGene_gn=EBP;refGene_tx=XM_004000480.4;refGene_gn=EBP;refGene_tx=XM_019823770.2;refGene_gn=EBP;refGene_tx=XM_019823772.2
X 42628487 42628711
refGene_gn=EBP;refGene_tx=XM_004000480.4;refGene_gn=EBP;refGene_tx=XM_019823770.2;refGene_gn=EBP;refGene_tx=XM_019823772.2
X 42631851 42631971 refGene_gn=TBC1D25;refGene_tx=XM_004000481.4
X 42633747 42633857 refGene_gn=TBC1D25;refGene_tx=XM_004000481.4
X 42634815 42634970 refGene_gn=TBC1D25;refGene_tx=XM_004000481.4
X 42643618 42643685 refGene_gn=TBC1D25;refGene_tx=XM_019823773.2
X 42645631 42645759
refGene_gn=TBC1D25;refGene_tx=XM_004000481.4;refGene_gn=TBC1D25;refGene_tx=XM_019823773.2
X 42645869 42646058
refGene_gn=TBC1D25;refGene_tx=XM_004000481.4;refGene_gn=TBC1D25;refGen

e_tx=XM_019823773.2
X 42646333 42647695
refGene_gn=TBC1D25;refGene_tx=XM_004000481.4;refGene_gn=TBC1D25;refGene_tx=XM_019823773.2
X 42654105 42654208
refGene_gn=RBM3;refGene_tx=XM_006943709.2;refGene_gn=RBM3;refGene_tx=XM_011291709.2;refGene_gn=RBM3;refGene_tx=XM_019823774.1;refGene_gn=RBM3;refGene_tx=XM_019823775.1
X 42654462 42654569
refGene_gn=RBM3;refGene_tx=XM_006943709.2;refGene_gn=RBM3;refGene_tx=XM_011291709.2;refGene_gn=RBM3;refGene_tx=XM_019823774.1;refGene_gn=RBM3;refGene_tx=XM_019823775.1
X 42655212 42655318
refGene_gn=RBM3;refGene_tx=XM_006943709.2;refGene_gn=RBM3;refGene_tx=XM_011291709.2;refGene_gn=RBM3;refGene_tx=XM_019823774.1;refGene_gn=RBM3;refGene_tx=XM_019823775.1
X 42655414 42655520
refGene_gn=RBM3;refGene_tx=XM_006943709.2;refGene_gn=RBM3;refGene_tx=XM_011291709.2;refGene_gn=RBM3;refGene_tx=XM_019823774.1;refGene_gn=RBM3;refGene_tx=XM_019823775.1
X 42655942 42656003
refGene_gn=RBM3;refGene_tx=XM_006943709.2;refGene_gn=RBM3;refGene_tx=XM_011291709.2;refGene_gn=RBM3;refGene_tx=XM_019823774.1;refGene_gn=RBM3;refGene_tx=XM_019823775.1
X 42676115 42676156
refGene_gn=WDR13;refGene_tx=XM_006943710.4;refGene_gn=WDR13;refGene_tx=XM_006943711.4;refGene_gn=WDR13;refGene_tx=XM_019823776.1
X 42676845 42677086
refGene_gn=WDR13;refGene_tx=XM_006943710.4;refGene_gn=WDR13;refGene_tx=XM_006943711.4;refGene_gn=WDR13;refGene_tx=XM_019823776.1
X 42677748 42677858
refGene_gn=WDR13;refGene_tx=XM_006943710.4;refGene_gn=WDR13;refGene_tx=XM_006943711.4;refGene_gn=WDR13;refGene_tx=XM_019823776.1
X 42677983 42678114
refGene_gn=WDR13;refGene_tx=XM_006943710.4;refGene_gn=WDR13;refGene_tx=XM_006943711.4;refGene_gn=WDR13;refGene_tx=XM_019823776.1
X 42679002 42679310
refGene_gn=WDR13;refGene_tx=XM_006943710.4;refGene_gn=WDR13;refGene_tx=XM_006943711.4;refGene_gn=WDR13;refGene_tx=XM_019823776.1
X 42680157 42680338
refGene_gn=WDR13;refGene_tx=XM_006943710.4;refGene_gn=WDR13;refGene_tx=XM_006943711.4;refGene_gn=WDR13;refGene_tx=XM_019823776.1
X 42680440 42680582
refGene_gn=WDR13;refGene_tx=XM_006943710.4;refGene_gn=WDR13;refGene_tx=XM_006943711.4;refGene_gn=WDR13;refGene_tx=XM_019823776.1
X 42682274 42682393
refGene_gn=WDR13;refGene_tx=XM_006943710.4;refGene_gn=WDR13;refGene_tx=XM_019823776.1
X 42682781 42682966
refGene_gn=WDR13;refGene_tx=XM_006943710.4;refGene_gn=WDR13;refGene_tx

=XM_006943711.4;refGene_gn=WDR13;refGene_tx=XM_019823776.1
X 42760968 42760977 refGene_gn=WAS;refGene_tx=XM_019823777.2
X 42761810 42761942 refGene_gn=WAS;refGene_tx=XM_004000485.5
X 42762270 42762411
refGene_gn=WAS;refGene_tx=XM_004000485.5;refGene_gn=WAS;refGene_tx=XM_019823777.2
X 42763258 42763345
refGene_gn=WAS;refGene_tx=XM_004000485.5;refGene_gn=WAS;refGene_tx=XM_019823777.2
X 42763445 42763548
refGene_gn=WAS;refGene_tx=XM_004000485.5;refGene_gn=WAS;refGene_tx=XM_019823777.2
X 42763660 42763702
refGene_gn=WAS;refGene_tx=XM_004000485.5;refGene_gn=WAS;refGene_tx=XM_019823777.2
X 42763794 42763845
refGene_gn=WAS;refGene_tx=XM_004000485.5;refGene_gn=WAS;refGene_tx=XM_019823777.2
X 42764490 42764665
refGene_gn=WAS;refGene_tx=XM_004000485.5;refGene_gn=WAS;refGene_tx=XM_019823777.2
X 42765706 42765749
refGene_gn=WAS;refGene_tx=XM_004000485.5;refGene_gn=WAS;refGene_tx=XM_019823777.2
X 42765954 42766108
refGene_gn=WAS;refGene_tx=XM_004000485.5;refGene_gn=WAS;refGene_tx=XM_019823777.2
X 42766298 42766705
refGene_gn=WAS;refGene_tx=XM_004000485.5;refGene_gn=WAS;refGene_tx=XM_019823777.2
X 42766967 42767082
refGene_gn=WAS;refGene_tx=XM_004000485.5;refGene_gn=WAS;refGene_tx=XM_019823777.2
X 42768287 42768343
refGene_gn=WAS;refGene_tx=XM_004000485.5;refGene_gn=WAS;refGene_tx=XM_019823777.2
X 42771561 42771685 refGene_gn=SUV39H1;refGene_tx=XM_019823781.2
X 42773297 42773316 refGene_gn=SUV39H1;refGene_tx=XM_023248950.1
X 42773773 42773885 refGene_gn=SUV39H1;refGene_tx=XM_019823779.1
X 42775320 42775466
refGene_gn=SUV39H1;refGene_tx=XM_019823779.1;refGene_gn=SUV39H1;refGene_tx=XM_019823781.2;refGene_gn=SUV39H1;refGene_tx=XM_023248950.1
X 42776935 42777598
refGene_gn=SUV39H1;refGene_tx=XM_019823779.1;refGene_gn=SUV39H1;refGene_tx=XM_019823781.2;refGene_gn=SUV39H1;refGene_tx=XM_023248950.1
X 42783705 42783852
refGene_gn=SUV39H1;refGene_tx=XM_019823779.1;refGene_gn=SUV39H1;refGene_tx=XM_019823781.2;refGene_gn=SUV39H1;refGene_tx=XM_023248950.1
X 42783934 42784064
refGene_gn=SUV39H1;refGene_tx=XM_019823779.1;refGene_gn=SUV39H1;refGen

e_tx=XM_019823781.2;refGene_gn=SUV39H1;refGene_tx=XM_023248950.1
X	42784786	42784920
refGene_gn=SUV39H1;refGene_tx=XM_019823779.1;refGene_gn=SUV39H1;refGene_tx=XM_019823781.2;refGene_gn=SUV39H1;refGene_tx=XM_023248950.1
X	42848044	42848107	refGene_gn=GLOD5;refGene_tx=XM_004000486.4
X	42850759	42850975
refGene_gn=GLOD5;refGene_tx=XM_004000486.4;refGene_gn=GLOD5;refGene_tx=XM_019823782.2
X	42852811	42852967
refGene_gn=GLOD5;refGene_tx=XM_004000486.4;refGene_gn=GLOD5;refGene_tx=XM_019823782.2
X	42854253	42854379
refGene_gn=GLOD5;refGene_tx=XM_004000486.4;refGene_gn=GLOD5;refGene_tx=XM_019823782.2
X	42871546	42871766	refGene_gn=GATA1;refGene_tx=XM_004000487.3
X	42872519	42872897	refGene_gn=GATA1;refGene_tx=XM_004000487.3
X	42872999	42873145	refGene_gn=GATA1;refGene_tx=XM_004000487.3
X	42873714	42873840	refGene_gn=GATA1;refGene_tx=XM_004000487.3
X	42874306	42874678	refGene_gn=GATA1;refGene_tx=XM_004000487.3
X	42881516	42881612
refGene_gn=HDAC6;refGene_tx=XM_019823783.2;refGene_gn=HDAC6;refGene_tx=XM_019823784.2
X	42882345	42882468
refGene_gn=HDAC6;refGene_tx=XM_004000525.4;refGene_gn=HDAC6;refGene_tx=XM_006943712.4;refGene_gn=HDAC6;refGene_tx=XM_006943713.3;refGene_gn=HDAC6;refGene_tx=XM_019823783.2;refGene_gn=HDAC6;refGene_tx=XM_019823784.2
X	42882549	42882678
refGene_gn=HDAC6;refGene_tx=XM_004000525.4;refGene_gn=HDAC6;refGene_tx=XM_006943712.4;refGene_gn=HDAC6;refGene_tx=XM_006943713.3;refGene_gn=HDAC6;refGene_tx=XM_019823783.2;refGene_gn=HDAC6;refGene_tx=XM_019823784.2
X	42882815	42882904
refGene_gn=HDAC6;refGene_tx=XM_004000525.4;refGene_gn=HDAC6;refGene_tx=XM_006943712.4;refGene_gn=HDAC6;refGene_tx=XM_006943713.3;refGene_gn=HDAC6;refGene_tx=XM_019823783.2;refGene_gn=HDAC6;refGene_tx=XM_019823784.2
X	42885555	42885640
refGene_gn=HDAC6;refGene_tx=XM_004000525.4;refGene_gn=HDAC6;refGene_tx=XM_006943712.4;refGene_gn=HDAC6;refGene_tx=XM_006943713.3;refGene_gn=HDAC6;refGene_tx=XM_019823783.2;refGene_gn=HDAC6;refGene_tx=XM_019823784.2
X	42885737	42885778
refGene_gn=HDAC6;refGene_tx=XM_004000525.4;refGene_gn=HDAC6;refGene_tx=XM_006943712.4;refGene_gn=HDAC6;refGene_tx=XM_006943713.3;refGene_gn=HDAC6;refGene_tx=XM_019823783.2;refGene_gn=HDAC6;refGene_tx=XM_019823784.2
X	42886436	42886533
refGene_gn=HDAC6;refGene_tx=XM_004000525.4;refGene_gn=HDAC6;refGene_tx=XM_006943712.4;refGene_gn=HDAC6;refGene_tx=XM_006943713.3;refGene_gn=

HDAC6;refGene_tx=XM_019823783.2;refGene_gn=HDAC6;refGene_tx=XM_0198237
84.2
X	42886678	42886776
refGene_gn=HDAC6;refGene_tx=XM_004000525.4;refGene_gn=HDAC6;refGene_tx
=XM_006943712.4;refGene_gn=HDAC6;refGene_tx=XM_006943713.3;refGene_gn=
HDAC6;refGene_tx=XM_019823783.2;refGene_gn=HDAC6;refGene_tx=XM_0198237
84.2
X	42887964	42888069
refGene_gn=HDAC6;refGene_tx=XM_004000525.4;refGene_gn=HDAC6;refGene_tx
=XM_006943712.4;refGene_gn=HDAC6;refGene_tx=XM_006943713.3;refGene_gn=
HDAC6;refGene_tx=XM_019823783.2;refGene_gn=HDAC6;refGene_tx=XM_0198237
84.2
X	42888192	42888261
refGene_gn=HDAC6;refGene_tx=XM_004000525.4;refGene_gn=HDAC6;refGene_tx
=XM_006943712.4;refGene_gn=HDAC6;refGene_tx=XM_006943713.3;refGene_gn=
HDAC6;refGene_tx=XM_019823783.2;refGene_gn=HDAC6;refGene_tx=XM_0198237
84.2
X	42897826	42897953
refGene_gn=HDAC6;refGene_tx=XM_004000525.4;refGene_gn=HDAC6;refGene_tx
=XM_006943712.4;refGene_gn=HDAC6;refGene_tx=XM_006943713.3;refGene_gn=
HDAC6;refGene_tx=XM_019823783.2;refGene_gn=HDAC6;refGene_tx=XM_0198237
84.2
X	42898058	42898124
refGene_gn=HDAC6;refGene_tx=XM_004000525.4;refGene_gn=HDAC6;refGene_tx
=XM_006943712.4;refGene_gn=HDAC6;refGene_tx=XM_006943713.3;refGene_gn=
HDAC6;refGene_tx=XM_019823783.2;refGene_gn=HDAC6;refGene_tx=XM_0198237
84.2
X	42898216	42898276
refGene_gn=HDAC6;refGene_tx=XM_004000525.4;refGene_gn=HDAC6;refGene_tx
=XM_006943712.4;refGene_gn=HDAC6;refGene_tx=XM_006943713.3;refGene_gn=
HDAC6;refGene_tx=XM_019823783.2;refGene_gn=HDAC6;refGene_tx=XM_0198237
84.2
X	42898355	42898445
refGene_gn=HDAC6;refGene_tx=XM_004000525.4;refGene_gn=HDAC6;refGene_tx
=XM_006943712.4;refGene_gn=HDAC6;refGene_tx=XM_006943713.3;refGene_gn=
HDAC6;refGene_tx=XM_019823783.2;refGene_gn=HDAC6;refGene_tx=XM_0198237
84.2
X	42898783	42898890
refGene_gn=HDAC6;refGene_tx=XM_004000525.4;refGene_gn=HDAC6;refGene_tx
=XM_006943712.4;refGene_gn=HDAC6;refGene_tx=XM_006943713.3;refGene_gn=
HDAC6;refGene_tx=XM_019823783.2;refGene_gn=HDAC6;refGene_tx=XM_0198237
84.2
X	42898973	42899041
refGene_gn=HDAC6;refGene_tx=XM_004000525.4;refGene_gn=HDAC6;refGene_tx
=XM_006943712.4;refGene_gn=HDAC6;refGene_tx=XM_006943713.3;refGene_gn=
HDAC6;refGene_tx=XM_019823783.2;refGene_gn=HDAC6;refGene_tx=XM_0198237
84.2
X	42899272	42899441
refGene_gn=HDAC6;refGene_tx=XM_004000525.4;refGene_gn=HDAC6;refGene_tx
=XM_006943712.4;refGene_gn=HDAC6;refGene_tx=XM_006943713.3;refGene_gn=

HDAC6;refGene_tx=XM_019823783.2;refGene_gn=HDAC6;refGene_tx=XM_0198237
84.2
X 42899538 42899667
refGene_gn=HDAC6;refGene_tx=XM_004000525.4;refGene_gn=HDAC6;refGene_tx
=XM_006943712.4;refGene_gn=HDAC6;refGene_tx=XM_006943713.3;refGene_gn=
HDAC6;refGene_tx=XM_019823783.2;refGene_gn=HDAC6;refGene_tx=XM_0198237
84.2
X 42899837 42900006
refGene_gn=HDAC6;refGene_tx=XM_004000525.4;refGene_gn=HDAC6;refGene_tx
=XM_006943712.4;refGene_gn=HDAC6;refGene_tx=XM_006943713.3;refGene_gn=
HDAC6;refGene_tx=XM_019823783.2;refGene_gn=HDAC6;refGene_tx=XM_0198237
84.2
X 42900300 42900434
refGene_gn=HDAC6;refGene_tx=XM_004000525.4;refGene_gn=HDAC6;refGene_tx
=XM_006943712.4;refGene_gn=HDAC6;refGene_tx=XM_006943713.3;refGene_gn=
HDAC6;refGene_tx=XM_019823783.2;refGene_gn=HDAC6;refGene_tx=XM_0198237
84.2
X 42901012 42901081
refGene_gn=HDAC6;refGene_tx=XM_004000525.4;refGene_gn=HDAC6;refGene_tx
=XM_006943712.4;refGene_gn=HDAC6;refGene_tx=XM_006943713.3;refGene_gn=
HDAC6;refGene_tx=XM_019823783.2;refGene_gn=HDAC6;refGene_tx=XM_0198237
84.2
X 42901198 42901391
refGene_gn=HDAC6;refGene_tx=XM_004000525.4;refGene_gn=HDAC6;refGene_tx
=XM_006943712.4;refGene_gn=HDAC6;refGene_tx=XM_006943713.3;refGene_gn=
HDAC6;refGene_tx=XM_019823783.2;refGene_gn=HDAC6;refGene_tx=XM_0198237
84.2
X 42902811 42902961
refGene_gn=HDAC6;refGene_tx=XM_004000525.4;refGene_gn=HDAC6;refGene_tx
=XM_006943712.4;refGene_gn=HDAC6;refGene_tx=XM_006943713.3;refGene_gn=
HDAC6;refGene_tx=XM_019823783.2;refGene_gn=HDAC6;refGene_tx=XM_0198237
84.2
X 42904204 42904379
refGene_gn=HDAC6;refGene_tx=XM_004000525.4;refGene_gn=HDAC6;refGene_tx
=XM_006943712.4;refGene_gn=HDAC6;refGene_tx=XM_006943713.3;refGene_gn=
HDAC6;refGene_tx=XM_019823783.2;refGene_gn=HDAC6;refGene_tx=XM_0198237
84.2
X 42904487 42904933
refGene_gn=HDAC6;refGene_tx=XM_004000525.4;refGene_gn=HDAC6;refGene_tx
=XM_006943712.4;refGene_gn=HDAC6;refGene_tx=XM_006943713.3;refGene_gn=
HDAC6;refGene_tx=XM_019823783.2;refGene_gn=HDAC6;refGene_tx=XM_0198237
84.2
X 42905016 42905124
refGene_gn=HDAC6;refGene_tx=XM_004000525.4;refGene_gn=HDAC6;refGene_tx
=XM_006943712.4;refGene_gn=HDAC6;refGene_tx=XM_006943713.3;refGene_gn=
HDAC6;refGene_tx=XM_019823783.2;refGene_gn=HDAC6;refGene_tx=XM_0198237
84.2
X 42905258 42905405
refGene_gn=HDAC6;refGene_tx=XM_004000525.4;refGene_gn=HDAC6;refGene_tx
=XM_006943712.4;refGene_gn=HDAC6;refGene_tx=XM_006943713.3;refGene_gn=

HDAC6;refGene_tx=XM_019823783.2;refGene_gn=HDAC6;refGene_tx=XM_0198237
84.2
X	42905502	42905631
refGene_gn=HDAC6;refGene_tx=XM_004000525.4;refGene_gn=HDAC6;refGene_tx
=XM_006943712.4;refGene_gn=HDAC6;refGene_tx=XM_006943713.3;refGene_gn=
HDAC6;refGene_tx=XM_019823783.2;refGene_gn=HDAC6;refGene_tx=XM_0198237
84.2
X	42905837	42905906
refGene_gn=HDAC6;refGene_tx=XM_004000525.4;refGene_gn=HDAC6;refGene_tx
=XM_006943712.4;refGene_gn=HDAC6;refGene_tx=XM_006943713.3;refGene_gn=
HDAC6;refGene_tx=XM_019823783.2;refGene_gn=HDAC6;refGene_tx=XM_0198237
84.2
X	42910114	42910816	refGene_gn=ERAS;refGene_tx=XM_023249099.1
X	42912127	42912323	refGene_gn=PCSK1N;refGene_tx=XM_004000489.5
X	42912770	42912919	refGene_gn=PCSK1N;refGene_tx=XM_004000489.5
X	42912920	42913244	refGene_gn=PCSK1N;refGene_tx=XM_004000489.5
X	42917245	42917359	refGene_gn=PCSK1N;refGene_tx=XM_004000489.5
X	42933639	42933728	refGene_gn=TIMM17B;refGene_tx=XM_004000490.3
X	42933824	42933935	refGene_gn=TIMM17B;refGene_tx=XM_004000490.3
X	42934021	42934150	refGene_gn=TIMM17B;refGene_tx=XM_004000490.3
X	42934944	42935008	refGene_gn=TIMM17B;refGene_tx=XM_004000490.3
X	42936411	42936511	refGene_gn=TIMM17B;refGene_tx=XM_004000490.3
X	42937373	42937399	refGene_gn=TIMM17B;refGene_tx=XM_004000490.3
X	42938178	42938245
refGene_gn=PQBP1;refGene_tx=XM_004000491.5;refGene_gn=PQBP1;refGene_tx
=XM_006943715.4
X	42939977	42940089
refGene_gn=PQBP1;refGene_tx=XM_004000491.5;refGene_gn=PQBP1;refGene_tx
=XM_006943715.4
X	42940338	42940451
refGene_gn=PQBP1;refGene_tx=XM_004000491.5;refGene_gn=PQBP1;refGene_tx
=XM_006943715.4
X	42940628	42940934
refGene_gn=PQBP1;refGene_tx=XM_004000491.5;refGene_gn=PQBP1;refGene_tx
=XM_006943715.4
X	42941163	42941227
refGene_gn=PQBP1;refGene_tx=XM_004000491.5;refGene_gn=PQBP1;refGene_tx
=XM_006943715.4
X	42941409	42941566
refGene_gn=PQBP1;refGene_tx=XM_004000491.5;refGene_gn=PQBP1;refGene_tx
=XM_006943715.4
X	42941891	42941987
refGene_gn=SLC35A2;refGene_tx=XM_004000494.4;refGene_gn=SLC35A2;refGen
e_tx=XM_006943716.2;refGene_gn=SLC35A2;refGene_tx=XM_019823788.2
X	42943311	42944054
refGene_gn=SLC35A2;refGene_tx=XM_004000494.4;refGene_gn=SLC35A2;refGen
e_tx=XM_006943716.2;refGene_gn=SLC35A2;refGene_tx=XM_019823788.2
X	42944915	42945067
refGene_gn=SLC35A2;refGene_tx=XM_004000494.4;refGene_gn=SLC35A2;refGen
e_tx=XM_006943716.2;refGene_gn=SLC35A2;refGene_tx=XM_019823788.2

X 42948433 42948616
refGene_gn=SLC35A2;refGene_tx=XM_004000494.4;refGene_gn=SLC35A2;refGene_tx=XM_006943716.2;refGene_gn=SLC35A2;refGene_tx=XM_019823788.2
X 42950162 42950253
refGene_gn=SLC35A2;refGene_tx=XM_004000494.4;refGene_gn=SLC35A2;refGene_tx=XM_006943716.2
X 42952956 42953120 refGene_gn=PIM2;refGene_tx=XM_004000495.5
X 42953218 42953395 refGene_gn=PIM2;refGene_tx=XM_004000495.5
X 42953890 42954263 refGene_gn=PIM2;refGene_tx=XM_004000495.5
X 42955735 42955786 refGene_gn=PIM2;refGene_tx=XM_004000495.5
X 42956517 42956627 refGene_gn=PIM2;refGene_tx=XM_004000495.5
X 42956762 42956823 refGene_gn=PIM2;refGene_tx=XM_004000495.5
X 42960970 42961095
refGene_gn=OTUD5;refGene_tx=XM_023249459.1;refGene_gn=OTUD5;refGene_tx=XM_023249460.1;refGene_gn=OTUD5;refGene_tx=XM_023249461.1;refGene_gn=OTUD5;refGene_tx=XM_023249462.1
X 42961399 42961513
refGene_gn=OTUD5;refGene_tx=XM_023249459.1;refGene_gn=OTUD5;refGene_tx=XM_023249460.1;refGene_gn=OTUD5;refGene_tx=XM_023249461.1;refGene_gn=OTUD5;refGene_tx=XM_023249462.1
X 42961602 42961804
refGene_gn=OTUD5;refGene_tx=XM_023249459.1;refGene_gn=OTUD5;refGene_tx=XM_023249460.1;refGene_gn=OTUD5;refGene_tx=XM_023249461.1;refGene_gn=OTUD5;refGene_tx=XM_023249462.1
X 42963274 42963478
refGene_gn=OTUD5;refGene_tx=XM_023249459.1;refGene_gn=OTUD5;refGene_tx=XM_023249460.1;refGene_gn=OTUD5;refGene_tx=XM_023249461.1;refGene_gn=OTUD5;refGene_tx=XM_023249462.1
X 42967911 42968060
refGene_gn=OTUD5;refGene_tx=XM_023249459.1;refGene_gn=OTUD5;refGene_tx=XM_023249460.1;refGene_gn=OTUD5;refGene_tx=XM_023249461.1;refGene_gn=OTUD5;refGene_tx=XM_023249462.1
X 42968158 42968315
refGene_gn=OTUD5;refGene_tx=XM_023249459.1;refGene_gn=OTUD5;refGene_tx=XM_023249460.1;refGene_gn=OTUD5;refGene_tx=XM_023249461.1;refGene_gn=OTUD5;refGene_tx=XM_023249462.1
X 42968402 42968467
refGene_gn=OTUD5;refGene_tx=XM_023249459.1;refGene_gn=OTUD5;refGene_tx=XM_023249460.1;refGene_gn=OTUD5;refGene_tx=XM_023249461.1;refGene_gn=OTUD5;refGene_tx=XM_023249462.1
X 42975985 42976079
refGene_gn=OTUD5;refGene_tx=XM_023249459.1;refGene_gn=OTUD5;refGene_tx=XM_023249460.1;refGene_gn=OTUD5;refGene_tx=XM_023249461.1;refGene_gn=OTUD5;refGene_tx=XM_023249462.1
X 42986771 42987365
refGene_gn=OTUD5;refGene_tx=XM_023249459.1;refGene_gn=OTUD5;refGene_tx=XM_023249460.1;refGene_gn=OTUD5;refGene_tx=XM_023249461.1;refGene_gn=OTUD5;refGene_tx=XM_023249462.1
X 42991264 42991334 refGene_gn=KCND1;refGene_tx=XM_023249138.1
X 42992082 42992157

refGene_gn=KCND1;refGene_tx=XM_006943720.3;refGene_gn=KCND1;refGene_tx=XM_006943721.3;refGene_gn=KCND1;refGene_tx=XM_023249138.1
X 42992443 42992669
refGene_gn=KCND1;refGene_tx=XM_004000497.5;refGene_gn=KCND1;refGene_tx=XM_006943720.3;refGene_gn=KCND1;refGene_tx=XM_006943721.3;refGene_gn=KCND1;refGene_tx=XM_023249138.1
X 42993743 42993994
refGene_gn=KCND1;refGene_tx=XM_004000497.5;refGene_gn=KCND1;refGene_tx=XM_006943720.3;refGene_gn=KCND1;refGene_tx=XM_006943721.3;refGene_gn=KCND1;refGene_tx=XM_023249138.1
X 42994266 42994365
refGene_gn=KCND1;refGene_tx=XM_004000497.5;refGene_gn=KCND1;refGene_tx=XM_006943720.3;refGene_gn=KCND1;refGene_tx=XM_006943721.3;refGene_gn=KCND1;refGene_tx=XM_023249138.1
X 42994456 42994540
refGene_gn=KCND1;refGene_tx=XM_004000497.5;refGene_gn=KCND1;refGene_tx=XM_006943720.3;refGene_gn=KCND1;refGene_tx=XM_006943721.3;refGene_gn=KCND1;refGene_tx=XM_023249138.1
X 42994642 42994805
refGene_gn=KCND1;refGene_tx=XM_004000497.5;refGene_gn=KCND1;refGene_tx=XM_006943720.3;refGene_gn=KCND1;refGene_tx=XM_006943721.3;refGene_gn=KCND1;refGene_tx=XM_023249138.1
X 42996741 42997862
refGene_gn=KCND1;refGene_tx=XM_004000497.5;refGene_gn=KCND1;refGene_tx=XM_006943720.3;refGene_gn=KCND1;refGene_tx=XM_006943721.3;refGene_gn=KCND1;refGene_tx=XM_023249138.1
X 43001795 43001888
refGene_gn=GRIPAP1;refGene_tx=XM_004000498.5;refGene_gn=GRIPAP1;refGene_tx=XM_023248902.1;refGene_gn=GRIPAP1;refGene_tx=XM_023248903.1;refGene_gn=GRIPAP1;refGene_tx=XM_023248904.1
X 43002571 43002726
refGene_gn=GRIPAP1;refGene_tx=XM_004000498.5;refGene_gn=GRIPAP1;refGene_tx=XM_023248902.1;refGene_gn=GRIPAP1;refGene_tx=XM_023248903.1
X 43003472 43003566
refGene_gn=GRIPAP1;refGene_tx=XM_004000498.5;refGene_gn=GRIPAP1;refGene_tx=XM_023248902.1;refGene_gn=GRIPAP1;refGene_tx=XM_023248903.1;refGene_gn=GRIPAP1;refGene_tx=XM_023248904.1
X 43003698 43003821
refGene_gn=GRIPAP1;refGene_tx=XM_004000498.5;refGene_gn=GRIPAP1;refGene_tx=XM_023248902.1;refGene_gn=GRIPAP1;refGene_tx=XM_023248903.1;refGene_gn=GRIPAP1;refGene_tx=XM_023248904.1
X 43005189 43005320
refGene_gn=GRIPAP1;refGene_tx=XM_004000498.5;refGene_gn=GRIPAP1;refGene_tx=XM_023248902.1;refGene_gn=GRIPAP1;refGene_tx=XM_023248903.1;refGene_gn=GRIPAP1;refGene_tx=XM_023248904.1
X 43007187 43007287
refGene_gn=GRIPAP1;refGene_tx=XM_004000498.5;refGene_gn=GRIPAP1;refGene_tx=XM_023248902.1;refGene_gn=GRIPAP1;refGene_tx=XM_023248903.1;refGene_gn=GRIPAP1;refGene_tx=XM_023248904.1
X 43007372 43007429

refGene_gn=GRIPAP1;refGene_tx=XM_004000498.5;refGene_gn=GRIPAP1;refGene_tx=XM_023248902.1;refGene_gn=GRIPAP1;refGene_tx=XM_023248903.1;refGene_gn=GRIPAP1;refGene_tx=XM_023248904.1
X 43007553 43007649
refGene_gn=GRIPAP1;refGene_tx=XM_004000498.5;refGene_gn=GRIPAP1;refGene_tx=XM_023248902.1;refGene_gn=GRIPAP1;refGene_tx=XM_023248903.1;refGene_gn=GRIPAP1;refGene_tx=XM_023248904.1
X 43007753 43007831
refGene_gn=GRIPAP1;refGene_tx=XM_004000498.5;refGene_gn=GRIPAP1;refGene_tx=XM_023248902.1;refGene_gn=GRIPAP1;refGene_tx=XM_023248903.1;refGene_gn=GRIPAP1;refGene_tx=XM_023248904.1
X 43008855 43008969
refGene_gn=GRIPAP1;refGene_tx=XM_004000498.5;refGene_gn=GRIPAP1;refGene_tx=XM_023248902.1;refGene_gn=GRIPAP1;refGene_tx=XM_023248903.1;refGene_gn=GRIPAP1;refGene_tx=XM_023248904.1
X 43009104 43009317
refGene_gn=GRIPAP1;refGene_tx=XM_004000498.5;refGene_gn=GRIPAP1;refGene_tx=XM_023248902.1;refGene_gn=GRIPAP1;refGene_tx=XM_023248903.1;refGene_gn=GRIPAP1;refGene_tx=XM_023248904.1
X 43009574 43009670
refGene_gn=GRIPAP1;refGene_tx=XM_004000498.5;refGene_gn=GRIPAP1;refGene_tx=XM_023248902.1;refGene_gn=GRIPAP1;refGene_tx=XM_023248903.1;refGene_gn=GRIPAP1;refGene_tx=XM_023248904.1
X 43010474 43010609
refGene_gn=GRIPAP1;refGene_tx=XM_004000498.5;refGene_gn=GRIPAP1;refGene_tx=XM_023248902.1;refGene_gn=GRIPAP1;refGene_tx=XM_023248903.1;refGene_gn=GRIPAP1;refGene_tx=XM_023248904.1
X 43011322 43011415
refGene_gn=GRIPAP1;refGene_tx=XM_004000498.5;refGene_gn=GRIPAP1;refGene_tx=XM_023248902.1;refGene_gn=GRIPAP1;refGene_tx=XM_023248903.1;refGene_gn=GRIPAP1;refGene_tx=XM_023248904.1
X 43011617 43011695
refGene_gn=GRIPAP1;refGene_tx=XM_004000498.5;refGene_gn=GRIPAP1;refGene_tx=XM_023248902.1;refGene_gn=GRIPAP1;refGene_tx=XM_023248903.1;refGene_gn=GRIPAP1;refGene_tx=XM_023248904.1
X 43013224 43013324
refGene_gn=GRIPAP1;refGene_tx=XM_004000498.5;refGene_gn=GRIPAP1;refGene_tx=XM_023248902.1;refGene_gn=GRIPAP1;refGene_tx=XM_023248903.1;refGene_gn=GRIPAP1;refGene_tx=XM_023248904.1
X 43013455 43013503
refGene_gn=GRIPAP1;refGene_tx=XM_004000498.5;refGene_gn=GRIPAP1;refGene_tx=XM_023248902.1;refGene_gn=GRIPAP1;refGene_tx=XM_023248903.1;refGene_gn=GRIPAP1;refGene_tx=XM_023248904.1
X 43013607 43013639
refGene_gn=GRIPAP1;refGene_tx=XM_004000498.5;refGene_gn=GRIPAP1;refGene_tx=XM_023248902.1;refGene_gn=GRIPAP1;refGene_tx=XM_023248903.1;refGene_gn=GRIPAP1;refGene_tx=XM_023248904.1
X 43014326 43014374
refGene_gn=GRIPAP1;refGene_tx=XM_004000498.5;refGene_gn=GRIPAP1;refGene_tx=XM_023248902.1;refGene_gn=GRIPAP1;refGene_tx=XM_023248904.1

X	43014569	43014754
refGene_gn=GRIPAP1;refGene_tx=XM_004000498.5;refGene_gn=GRIPAP1;refGene_tx=XM_023248902.1;refGene_gn=GRIPAP1;refGene_tx=XM_023248903.1;refGene_gn=GRIPAP1;refGene_tx=XM_023248904.1
X	43015730	43015881
refGene_gn=GRIPAP1;refGene_tx=XM_004000498.5;refGene_gn=GRIPAP1;refGene_tx=XM_023248902.1;refGene_gn=GRIPAP1;refGene_tx=XM_023248903.1;refGene_gn=GRIPAP1;refGene_tx=XM_023248904.1
X	43018706	43018814
refGene_gn=GRIPAP1;refGene_tx=XM_004000498.5;refGene_gn=GRIPAP1;refGene_tx=XM_023248902.1;refGene_gn=GRIPAP1;refGene_tx=XM_023248903.1;refGene_gn=GRIPAP1;refGene_tx=XM_023248904.1
X	43019604	43019631
refGene_gn=GRIPAP1;refGene_tx=XM_004000498.5;refGene_gn=GRIPAP1;refGene_tx=XM_023248902.1;refGene_gn=GRIPAP1;refGene_tx=XM_023248903.1;refGene_gn=GRIPAP1;refGene_tx=XM_023248904.1
X	43021347	43021409
refGene_gn=GRIPAP1;refGene_tx=XM_004000498.5;refGene_gn=GRIPAP1;refGene_tx=XM_023248902.1;refGene_gn=GRIPAP1;refGene_tx=XM_023248903.1;refGene_gn=GRIPAP1;refGene_tx=XM_023248904.1
X	43021550	43021617
refGene_gn=GRIPAP1;refGene_tx=XM_004000498.5;refGene_gn=GRIPAP1;refGene_tx=XM_023248902.1;refGene_gn=GRIPAP1;refGene_tx=XM_023248903.1;refGene_gn=GRIPAP1;refGene_tx=XM_023248904.1
X	43023145	43023208
refGene_gn=GRIPAP1;refGene_tx=XM_023248902.1;refGene_gn=GRIPAP1;refGene_tx=XM_023248903.1;refGene_gn=GRIPAP1;refGene_tx=XM_023248904.1
X	43023681	43023723	refGene_gn=GRIPAP1;refGene_tx=XM_004000498.5
X	43058578	43059019
refGene_gn=TFE3;refGene_tx=XM_004000499.4;refGene_gn=TFE3;refGene_tx=XM_004000500.3
X	43059505	43059653
refGene_gn=TFE3;refGene_tx=XM_004000499.4;refGene_gn=TFE3;refGene_tx=XM_004000500.3
X	43061170	43061246
refGene_gn=TFE3;refGene_tx=XM_004000499.4;refGene_gn=TFE3;refGene_tx=XM_004000500.3
X	43061391	43061448
refGene_gn=TFE3;refGene_tx=XM_004000499.4;refGene_gn=TFE3;refGene_tx=XM_004000500.3
X	43061808	43061926
refGene_gn=TFE3;refGene_tx=XM_004000499.4;refGene_gn=TFE3;refGene_tx=XM_004000500.3
X	43063398	43063503
refGene_gn=TFE3;refGene_tx=XM_004000499.4;refGene_gn=TFE3;refGene_tx=XM_004000500.3
X	43063585	43063831
refGene_gn=TFE3;refGene_tx=XM_004000499.4;refGene_gn=TFE3;refGene_tx=XM_004000500.3
X	43064474	43064769

refGene_gn=TFE3;refGene_tx=XM_004000499.4;refGene_gn=TFE3;refGene_tx=XM_004000500.3
X 43065851 43065965 refGene_gn=TFE3;refGene_tx=XM_004000499.4
X 43068258 43068374 refGene_gn=TFE3;refGene_tx=XM_004000499.4
X 43087529 43087578
refGene_gn=CCDC120;refGene_tx=XM_019823793.2;refGene_gn=CCDC120;refGene_tx=XM_019823794.2;refGene_gn=CCDC120;refGene_tx=XM_019823795.2;refGene_gn=CCDC120;refGene_tx=XM_019823796.2;refGene_gn=CCDC120;refGene_tx=XM_019823798.2;refGene_gn=CCDC120;refGene_tx=XM_019823799.2;refGene_gn=CCDC120;refGene_tx=XM_019823800.2;refGene_gn=CCDC120;refGene_tx=XM_019823801.2
X 43087723 43087857
refGene_gn=CCDC120;refGene_tx=XM_019823793.2;refGene_gn=CCDC120;refGene_tx=XM_019823794.2;refGene_gn=CCDC120;refGene_tx=XM_019823795.2;refGene_gn=CCDC120;refGene_tx=XM_019823796.2;refGene_gn=CCDC120;refGene_tx=XM_019823798.2;refGene_gn=CCDC120;refGene_tx=XM_019823799.2;refGene_gn=CCDC120;refGene_tx=XM_019823800.2;refGene_gn=CCDC120;refGene_tx=XM_019823801.2
X 43088493 43088634
refGene_gn=CCDC120;refGene_tx=XM_019823793.2;refGene_gn=CCDC120;refGene_tx=XM_019823794.2;refGene_gn=CCDC120;refGene_tx=XM_019823795.2;refGene_gn=CCDC120;refGene_tx=XM_019823796.2;refGene_gn=CCDC120;refGene_tx=XM_019823798.2;refGene_gn=CCDC120;refGene_tx=XM_019823799.2;refGene_gn=CCDC120;refGene_tx=XM_019823800.2;refGene_gn=CCDC120;refGene_tx=XM_019823801.2
X 43089019 43089314
refGene_gn=CCDC120;refGene_tx=XM_019823793.2;refGene_gn=CCDC120;refGene_tx=XM_019823794.2;refGene_gn=CCDC120;refGene_tx=XM_019823795.2;refGene_gn=CCDC120;refGene_tx=XM_019823796.2;refGene_gn=CCDC120;refGene_tx=XM_019823798.2;refGene_gn=CCDC120;refGene_tx=XM_019823799.2;refGene_gn=CCDC120;refGene_tx=XM_019823800.2;refGene_gn=CCDC120;refGene_tx=XM_019823801.2
X 43089671 43089734
refGene_gn=CCDC120;refGene_tx=XM_019823793.2;refGene_gn=CCDC120;refGene_tx=XM_019823794.2;refGene_gn=CCDC120;refGene_tx=XM_019823795.2;refGene_gn=CCDC120;refGene_tx=XM_019823796.2;refGene_gn=CCDC120;refGene_tx=XM_019823798.2;refGene_gn=CCDC120;refGene_tx=XM_019823799.2;refGene_gn=CCDC120;refGene_tx=XM_019823800.2;refGene_gn=CCDC120;refGene_tx=XM_019823801.2
X 43090010 43090185
refGene_gn=CCDC120;refGene_tx=XM_019823793.2;refGene_gn=CCDC120;refGene_tx=XM_019823794.2;refGene_gn=CCDC120;refGene_tx=XM_019823795.2;refGene_gn=CCDC120;refGene_tx=XM_019823796.2;refGene_gn=CCDC120;refGene_tx=XM_019823798.2;refGene_gn=CCDC120;refGene_tx=XM_019823799.2;refGene_gn=CCDC120;refGene_tx=XM_019823800.2;refGene_gn=CCDC120;refGene_tx=XM_019823801.2
X 43090315 43090414
refGene_gn=CCDC120;refGene_tx=XM_019823793.2;refGene_gn=CCDC120;refGene_tx=XM_019823794.2;refGene_gn=CCDC120;refGene_tx=XM_019823795.2;refGene_gn=CCDC120;refGene_tx=XM_019823796.2;refGene_gn=CCDC120;refGene_tx=

XM_019823798.2;refGene_gn=CCDC120;refGene_tx=XM_019823799.2;refGene_gn=CCDC120;refGene_tx=XM_019823800.2;refGene_gn=CCDC120;refGene_tx=XM_019823801.2
X 43091369 43092284 refGene_gn=CCDC120;refGene_tx=XM_019823793.2;refGene_gn=CCDC120;refGene_tx=XM_019823794.2;refGene_gn=CCDC120;refGene_tx=XM_019823795.2;refGene_gn=CCDC120;refGene_tx=XM_019823796.2;refGene_gn=CCDC120;refGene_tx=XM_019823798.2;refGene_gn=CCDC120;refGene_tx=XM_019823799.2;refGene_gn=CCDC120;refGene_tx=XM_019823800.2;refGene_gn=CCDC120;refGene_tx=XM_019823801.2
X 43092780 43092895 refGene_gn=CCDC120;refGene_tx=XM_019823793.2;refGene_gn=CCDC120;refGene_tx=XM_019823794.2;refGene_gn=CCDC120;refGene_tx=XM_019823795.2;refGene_gn=CCDC120;refGene_tx=XM_019823796.2;refGene_gn=CCDC120;refGene_tx=XM_019823798.2;refGene_gn=CCDC120;refGene_tx=XM_019823799.2;refGene_gn=CCDC120;refGene_tx=XM_019823800.2;refGene_gn=CCDC120;refGene_tx=XM_019823801.2
X 43095014 43095154 refGene_gn=PRAF2;refGene_tx=XM_004000506.4
X 43095579 43095797 refGene_gn=PRAF2;refGene_tx=XM_004000506.4
X 43096945 43097124 refGene_gn=PRAF2;refGene_tx=XM_004000506.4
X 43097960 43098070 refGene_gn=WDR45;refGene_tx=XM_004000505.4;refGene_gn=WDR45;refGene_tx=XM_006943726.4;refGene_gn=WDR45;refGene_tx=XM_006943727.3;refGene_gn=WDR45;refGene_tx=XM_006943730.4;refGene_gn=WDR45;refGene_tx=XM_019823803.1;refGene_gn=WDR45;refGene_tx=XM_023248999.1
X 43098304 43098450 refGene_gn=WDR45;refGene_tx=XM_004000505.4;refGene_gn=WDR45;refGene_tx=XM_006943726.4;refGene_gn=WDR45;refGene_tx=XM_006943727.3;refGene_gn=WDR45;refGene_tx=XM_019823803.1;refGene_gn=WDR45;refGene_tx=XM_023248999.1
X 43098534 43098636 refGene_gn=WDR45;refGene_tx=XM_004000505.4;refGene_gn=WDR45;refGene_tx=XM_006943726.4;refGene_gn=WDR45;refGene_tx=XM_006943727.3;refGene_gn=WDR45;refGene_tx=XM_006943730.4;refGene_gn=WDR45;refGene_tx=XM_019823803.1;refGene_gn=WDR45;refGene_tx=XM_023248999.1
X 43098715 43098924 refGene_gn=WDR45;refGene_tx=XM_004000505.4;refGene_gn=WDR45;refGene_tx=XM_006943726.4;refGene_gn=WDR45;refGene_tx=XM_006943727.3;refGene_gn=WDR45;refGene_tx=XM_006943730.4;refGene_gn=WDR45;refGene_tx=XM_019823803.1;refGene_gn=WDR45;refGene_tx=XM_023248999.1
X 43099030 43099110 refGene_gn=WDR45;refGene_tx=XM_004000505.4;refGene_gn=WDR45;refGene_tx=XM_006943726.4;refGene_gn=WDR45;refGene_tx=XM_006943727.3;refGene_gn=WDR45;refGene_tx=XM_006943730.4;refGene_gn=WDR45;refGene_tx=XM_019823803.1;refGene_gn=WDR45;refGene_tx=XM_023248999.1
X 43099615 43099710 refGene_gn=WDR45;refGene_tx=XM_004000505.4;refGene_gn=WDR45;refGene_tx=XM_006943726.4;refGene_gn=WDR45;refGene_tx=XM_006943727.3;refGene_gn=WDR45;refGene_tx=XM_006943730.4;refGene_gn=WDR45;refGene_tx=XM_019823803.1;refGene_gn=WDR45;refGene_tx=XM_023248999.1

X	43099832 43100077
refGene_gn=WDR45;refGene_tx=XM_004000505.4;refGene_gn=WDR45;refGene_tx=XM_006943726.4;refGene_gn=WDR45;refGene_tx=XM_006943727.3;refGene_gn=WDR45;refGene_tx=XM_006943730.4;refGene_gn=WDR45;refGene_tx=XM_019823803.1;refGene_gn=WDR45;refGene_tx=XM_023248999.1
X	43100591 43100631 refGene_gn=WDR45;refGene_tx=XM_019823803.1
X	43100834 43100939
refGene_gn=WDR45;refGene_tx=XM_004000505.4;refGene_gn=WDR45;refGene_tx=XM_006943726.4;refGene_gn=WDR45;refGene_tx=XM_006943730.4
X	43101028 43101103
refGene_gn=WDR45;refGene_tx=XM_004000505.4;refGene_gn=WDR45;refGene_tx=XM_006943726.4;refGene_gn=WDR45;refGene_tx=XM_006943727.3;refGene_gn=WDR45;refGene_tx=XM_006943730.4
X	43101226 43101281
refGene_gn=WDR45;refGene_tx=XM_004000505.4;refGene_gn=WDR45;refGene_tx=XM_006943726.4;refGene_gn=WDR45;refGene_tx=XM_006943727.3;refGene_gn=WDR45;refGene_tx=XM_006943730.4
X	43134576 43134711
refGene_gn=GPKOW;refGene_tx=XM_006943731.4;refGene_gn=GPKOW;refGene_tx=XM_011291705.3;refGene_gn=GPKOW;refGene_tx=XM_011291706.3;refGene_gn=GPKOW;refGene_tx=XM_011291707.3;refGene_gn=GPKOW;refGene_tx=XM_019823804.2;refGene_gn=GPKOW;refGene_tx=XM_023248943.1
X	43134805 43134891
refGene_gn=GPKOW;refGene_tx=XM_006943731.4;refGene_gn=GPKOW;refGene_tx=XM_011291705.3;refGene_gn=GPKOW;refGene_tx=XM_011291706.3;refGene_gn=GPKOW;refGene_tx=XM_011291707.3;refGene_gn=GPKOW;refGene_tx=XM_019823804.2;refGene_gn=GPKOW;refGene_tx=XM_023248943.1
X	43135635 43135705
refGene_gn=GPKOW;refGene_tx=XM_006943731.4;refGene_gn=GPKOW;refGene_tx=XM_011291705.3;refGene_gn=GPKOW;refGene_tx=XM_011291706.3;refGene_gn=GPKOW;refGene_tx=XM_011291707.3;refGene_gn=GPKOW;refGene_tx=XM_019823804.2;refGene_gn=GPKOW;refGene_tx=XM_023248943.1
X	43135790 43135914
refGene_gn=GPKOW;refGene_tx=XM_006943731.4;refGene_gn=GPKOW;refGene_tx=XM_011291705.3;refGene_gn=GPKOW;refGene_tx=XM_011291706.3;refGene_gn=GPKOW;refGene_tx=XM_011291707.3;refGene_gn=GPKOW;refGene_tx=XM_019823804.2;refGene_gn=GPKOW;refGene_tx=XM_023248943.1
X	43136117 43136244
refGene_gn=GPKOW;refGene_tx=XM_006943731.4;refGene_gn=GPKOW;refGene_tx=XM_011291705.3;refGene_gn=GPKOW;refGene_tx=XM_011291706.3;refGene_gn=GPKOW;refGene_tx=XM_011291707.3;refGene_gn=GPKOW;refGene_tx=XM_019823804.2;refGene_gn=GPKOW;refGene_tx=XM_023248943.1
X	43136931 43137064
refGene_gn=GPKOW;refGene_tx=XM_006943731.4;refGene_gn=GPKOW;refGene_tx=XM_011291705.3;refGene_gn=GPKOW;refGene_tx=XM_011291706.3;refGene_gn=GPKOW;refGene_tx=XM_011291707.3;refGene_gn=GPKOW;refGene_tx=XM_019823804.2;refGene_gn=GPKOW;refGene_tx=XM_023248943.1
X	43137486 43137700
refGene_gn=GPKOW;refGene_tx=XM_006943731.4;refGene_gn=GPKOW;refGene_tx=XM_011291705.3;refGene_gn=GPKOW;refGene_tx=XM_011291706.3;refGene_gn=

GPKOW;refGene_tx=XM_011291707.3;refGene_gn=GPKOW;refGene_tx=XM_019823804.2;refGene_gn=GPKOW;refGene_tx=XM_023248943.1
X 43138487 43138597
refGene_gn=GPKOW;refGene_tx=XM_006943731.4;refGene_gn=GPKOW;refGene_tx=XM_011291705.3;refGene_gn=GPKOW;refGene_tx=XM_011291706.3;refGene_gn=GPKOW;refGene_tx=XM_011291707.3;refGene_gn=GPKOW;refGene_tx=XM_019823804.2;refGene_gn=GPKOW;refGene_tx=XM_023248943.1
X 43140651 43140776
refGene_gn=GPKOW;refGene_tx=XM_006943731.4;refGene_gn=GPKOW;refGene_tx=XM_011291705.3;refGene_gn=GPKOW;refGene_tx=XM_011291706.3;refGene_gn=GPKOW;refGene_tx=XM_011291707.3;refGene_gn=GPKOW;refGene_tx=XM_019823804.2;refGene_gn=GPKOW;refGene_tx=XM_023248943.1
X 43140875 43141030
refGene_gn=GPKOW;refGene_tx=XM_006943731.4;refGene_gn=GPKOW;refGene_tx=XM_011291705.3;refGene_gn=GPKOW;refGene_tx=XM_011291706.3;refGene_gn=GPKOW;refGene_tx=XM_011291707.3;refGene_gn=GPKOW;refGene_tx=XM_019823804.2;refGene_gn=GPKOW;refGene_tx=XM_023248943.1
X 43141800 43141976
refGene_gn=GPKOW;refGene_tx=XM_006943731.4;refGene_gn=GPKOW;refGene_tx=XM_011291705.3;refGene_gn=GPKOW;refGene_tx=XM_011291706.3;refGene_gn=GPKOW;refGene_tx=XM_011291707.3;refGene_gn=GPKOW;refGene_tx=XM_019823804.2;refGene_gn=GPKOW;refGene_tx=XM_023248943.1
X 43154590 43154908
refGene_gn=LOC101100947;refGene_tx=XM_011291704.3
X 43161044 43161093 refGene_gn=MAGIX;refGene_tx=XM_023249116.1
X 43161931 43162078 refGene_gn=MAGIX;refGene_tx=XM_023249116.1
X 43162858 43162941 refGene_gn=MAGIX;refGene_tx=XM_023249116.1
X 43163017 43163254 refGene_gn=MAGIX;refGene_tx=XM_023249116.1
X 43163606 43163778 refGene_gn=MAGIX;refGene_tx=XM_023249116.1
X 43164412 43164720 refGene_gn=MAGIX;refGene_tx=XM_023249116.1
X 43168902 43168998 refGene_gn=PLP2;refGene_tx=XM_004000511.5
X 43170080 43170233 refGene_gn=PLP2;refGene_tx=XM_004000511.5
X 43170338 43170434 refGene_gn=PLP2;refGene_tx=XM_004000511.5
X 43170970 43171061 refGene_gn=PLP2;refGene_tx=XM_004000511.5
X 43171263 43171286 refGene_gn=PLP2;refGene_tx=XM_004000511.5
X 43171912 43172511
refGene_gn=PRICKLE3;refGene_tx=XM_004000512.3;refGene_gn=PRICKLE3;refGene_tx=XM_023249019.1
X 43173091 43173391
refGene_gn=PRICKLE3;refGene_tx=XM_004000512.3;refGene_gn=PRICKLE3;refGene_tx=XM_023249019.1
X 43174165 43174352
refGene_gn=PRICKLE3;refGene_tx=XM_004000512.3;refGene_gn=PRICKLE3;refGene_tx=XM_023249019.1
X 43174451 43174655
refGene_gn=PRICKLE3;refGene_tx=XM_004000512.3;refGene_gn=PRICKLE3;refGene_tx=XM_023249019.1
X 43175431 43175569
refGene_gn=PRICKLE3;refGene_tx=XM_004000512.3;refGene_gn=PRICKLE3;refGene_tx=XM_023249019.1

X 43176144 43176258
refGene_gn=PRICKLE3;refGene_tx=XM_004000512.3;refGene_gn=PRICKLE3;refGene_tx=XM_023249019.1
X 43178186 43178370
refGene_gn=PRICKLE3;refGene_tx=XM_004000512.3;refGene_gn=PRICKLE3;refGene_tx=XM_023249019.1
X 43179209 43179295 refGene_gn=PRICKLE3;refGene_tx=XM_004000512.3
X 43180437 43180479 refGene_gn=PRICKLE3;refGene_tx=XM_004000512.3
X 43185933 43186257 refGene_gn=SYP;refGene_tx=XM_011291702.2
X 43187647 43187839 refGene_gn=SYP;refGene_tx=XM_011291702.2
X 43188553 43188749 refGene_gn=SYP;refGene_tx=XM_011291702.2
X 43191613 43191738 refGene_gn=SYP;refGene_tx=XM_011291702.2
X 43192861 43192927 refGene_gn=SYP;refGene_tx=XM_011291702.2
X 43194099 43194135 refGene_gn=SYP;refGene_tx=XM_011291702.2
X 43197745 43197976
refGene_gn=CACNA1F;refGene_tx=XM_023249415.1;refGene_gn=CACNA1F;refGene_tx=XM_023249416.1;refGene_gn=CACNA1F;refGene_tx=XM_023249417.1
X 43198255 43198453
refGene_gn=CACNA1F;refGene_tx=XM_023249415.1;refGene_gn=CACNA1F;refGene_tx=XM_023249416.1
X 43198648 43198714 refGene_gn=CACNA1F;refGene_tx=XM_023249418.1
X 43198848 43198961
refGene_gn=CACNA1F;refGene_tx=XM_023249415.1;refGene_gn=CACNA1F;refGene_tx=XM_023249416.1;refGene_gn=CACNA1F;refGene_tx=XM_023249417.1;refGene_gn=CACNA1F;refGene_tx=XM_023249418.1
X 43199067 43199195
refGene_gn=CACNA1F;refGene_tx=XM_023249415.1;refGene_gn=CACNA1F;refGene_tx=XM_023249416.1;refGene_gn=CACNA1F;refGene_tx=XM_023249417.1;refGene_gn=CACNA1F;refGene_tx=XM_023249418.1
X 43199296 43199422
refGene_gn=CACNA1F;refGene_tx=XM_023249415.1;refGene_gn=CACNA1F;refGene_tx=XM_023249416.1;refGene_gn=CACNA1F;refGene_tx=XM_023249417.1;refGene_gn=CACNA1F;refGene_tx=XM_023249418.1;refGene_gn=CACNA1F;refGene_tx=XM_023249419.1
X 43200446 43200616
refGene_gn=CACNA1F;refGene_tx=XM_023249415.1;refGene_gn=CACNA1F;refGene_tx=XM_023249416.1;refGene_gn=CACNA1F;refGene_tx=XM_023249417.1;refGene_gn=CACNA1F;refGene_tx=XM_023249418.1
X 43200909 43201044
refGene_gn=CACNA1F;refGene_tx=XM_023249415.1;refGene_gn=CACNA1F;refGene_tx=XM_023249416.1;refGene_gn=CACNA1F;refGene_tx=XM_023249417.1;refGene_gn=CACNA1F;refGene_tx=XM_023249418.1;refGene_gn=CACNA1F;refGene_tx=XM_023249419.1
X 43201288 43201419
refGene_gn=CACNA1F;refGene_tx=XM_023249415.1;refGene_gn=CACNA1F;refGene_tx=XM_023249416.1;refGene_gn=CACNA1F;refGene_tx=XM_023249417.1;refGene_gn=CACNA1F;refGene_tx=XM_023249418.1;refGene_gn=CACNA1F;refGene_tx=XM_023249419.1
X 43201585 43201687
refGene_gn=CACNA1F;refGene_tx=XM_023249415.1;refGene_gn=CACNA1F;refGen

e_tx=XM_023249416.1;refGene_gn=CACNA1F;refGene_tx=XM_023249417.1;refGene_gn=CACNA1F;refGene_tx=XM_023249418.1;refGene_gn=CACNA1F;refGene_tx=XM_023249419.1;refGene_gn=CACNA1F;refGene_tx=XM_023249420.1;refGene_gn=CACNA1F;refGene_tx=XM_023249421.1
X 43201950 43202108
refGene_gn=CACNA1F;refGene_tx=XM_023249415.1;refGene_gn=CACNA1F;refGene_tx=XM_023249416.1;refGene_gn=CACNA1F;refGene_tx=XM_023249417.1;refGene_gn=CACNA1F;refGene_tx=XM_023249418.1;refGene_gn=CACNA1F;refGene_tx=XM_023249419.1;refGene_gn=CACNA1F;refGene_tx=XM_023249420.1
X 43202241 43202338
refGene_gn=CACNA1F;refGene_tx=XM_023249415.1;refGene_gn=CACNA1F;refGene_tx=XM_023249416.1;refGene_gn=CACNA1F;refGene_tx=XM_023249417.1;refGene_gn=CACNA1F;refGene_tx=XM_023249418.1;refGene_gn=CACNA1F;refGene_tx=XM_023249419.1;refGene_gn=CACNA1F;refGene_tx=XM_023249420.1;refGene_gn=CACNA1F;refGene_tx=XM_023249421.1
X 43202615 43202743
refGene_gn=CACNA1F;refGene_tx=XM_023249415.1;refGene_gn=CACNA1F;refGene_tx=XM_023249416.1;refGene_gn=CACNA1F;refGene_tx=XM_023249417.1;refGene_gn=CACNA1F;refGene_tx=XM_023249418.1;refGene_gn=CACNA1F;refGene_tx=XM_023249419.1;refGene_gn=CACNA1F;refGene_tx=XM_023249420.1;refGene_gn=CACNA1F;refGene_tx=XM_023249421.1
X 43202970 43203130
refGene_gn=CACNA1F;refGene_tx=XM_023249415.1;refGene_gn=CACNA1F;refGene_tx=XM_023249416.1;refGene_gn=CACNA1F;refGene_tx=XM_023249417.1;refGene_gn=CACNA1F;refGene_tx=XM_023249418.1;refGene_gn=CACNA1F;refGene_tx=XM_023249419.1;refGene_gn=CACNA1F;refGene_tx=XM_023249420.1;refGene_gn=CACNA1F;refGene_tx=XM_023249421.1
X 43203813 43203905
refGene_gn=CACNA1F;refGene_tx=XM_023249415.1;refGene_gn=CACNA1F;refGene_tx=XM_023249416.1;refGene_gn=CACNA1F;refGene_tx=XM_023249417.1;refGene_gn=CACNA1F;refGene_tx=XM_023249418.1;refGene_gn=CACNA1F;refGene_tx=XM_023249419.1;refGene_gn=CACNA1F;refGene_tx=XM_023249420.1;refGene_gn=CACNA1F;refGene_tx=XM_023249421.1
X 43204164 43204230
refGene_gn=CACNA1F;refGene_tx=XM_023249415.1;refGene_gn=CACNA1F;refGene_tx=XM_023249416.1;refGene_gn=CACNA1F;refGene_tx=XM_023249417.1;refGene_gn=CACNA1F;refGene_tx=XM_023249418.1;refGene_gn=CACNA1F;refGene_tx=XM_023249419.1;refGene_gn=CACNA1F;refGene_tx=XM_023249420.1;refGene_gn=CACNA1F;refGene_tx=XM_023249421.1
X 43204559 43204688
refGene_gn=CACNA1F;refGene_tx=XM_023249415.1;refGene_gn=CACNA1F;refGene_tx=XM_023249416.1;refGene_gn=CACNA1F;refGene_tx=XM_023249417.1;refGene_gn=CACNA1F;refGene_tx=XM_023249418.1;refGene_gn=CACNA1F;refGene_tx=XM_023249419.1;refGene_gn=CACNA1F;refGene_tx=XM_023249420.1;refGene_gn=CACNA1F;refGene_tx=XM_023249421.1
X 43204894 43204915
refGene_gn=CACNA1F;refGene_tx=XM_023249415.1;refGene_gn=CACNA1F;refGene_tx=XM_023249417.1;refGene_gn=CACNA1F;refGene_tx=XM_023249418.1;refGene_gn=CACNA1F;refGene_tx=XM_023249419.1;refGene_gn=CACNA1F;refGene_tx=XM_023249420.1;refGene_gn=CACNA1F;refGene_tx=XM_023249421.1

X 43205726 43205810
refGene_gn=CACNA1F;refGene_tx=XM_023249415.1;refGene_gn=CACNA1F;refGene_tx=XM_023249416.1;refGene_gn=CACNA1F;refGene_tx=XM_023249417.1;refGene_gn=CACNA1F;refGene_tx=XM_023249418.1;refGene_gn=CACNA1F;refGene_tx=XM_023249419.1;refGene_gn=CACNA1F;refGene_tx=XM_023249420.1;refGene_gn=CACNA1F;refGene_tx=XM_023249421.1
X 43206059 43206170
refGene_gn=CACNA1F;refGene_tx=XM_023249415.1;refGene_gn=CACNA1F;refGene_tx=XM_023249416.1;refGene_gn=CACNA1F;refGene_tx=XM_023249417.1;refGene_gn=CACNA1F;refGene_tx=XM_023249418.1;refGene_gn=CACNA1F;refGene_tx=XM_023249419.1;refGene_gn=CACNA1F;refGene_tx=XM_023249420.1;refGene_gn=CACNA1F;refGene_tx=XM_023249421.1
X 43206981 43207140
refGene_gn=CACNA1F;refGene_tx=XM_023249415.1;refGene_gn=CACNA1F;refGene_tx=XM_023249416.1;refGene_gn=CACNA1F;refGene_tx=XM_023249417.1;refGene_gn=CACNA1F;refGene_tx=XM_023249418.1;refGene_gn=CACNA1F;refGene_tx=XM_023249419.1;refGene_gn=CACNA1F;refGene_tx=XM_023249420.1;refGene_gn=CACNA1F;refGene_tx=XM_023249421.1
X 43207233 43207435
refGene_gn=CACNA1F;refGene_tx=XM_023249415.1;refGene_gn=CACNA1F;refGene_tx=XM_023249416.1;refGene_gn=CACNA1F;refGene_tx=XM_023249417.1;refGene_gn=CACNA1F;refGene_tx=XM_023249418.1;refGene_gn=CACNA1F;refGene_tx=XM_023249419.1;refGene_gn=CACNA1F;refGene_tx=XM_023249420.1;refGene_gn=CACNA1F;refGene_tx=XM_023249421.1
X 43208252 43208399
refGene_gn=CACNA1F;refGene_tx=XM_023249415.1;refGene_gn=CACNA1F;refGene_tx=XM_023249416.1;refGene_gn=CACNA1F;refGene_tx=XM_023249417.1;refGene_gn=CACNA1F;refGene_tx=XM_023249418.1;refGene_gn=CACNA1F;refGene_tx=XM_023249419.1;refGene_gn=CACNA1F;refGene_tx=XM_023249420.1;refGene_gn=CACNA1F;refGene_tx=XM_023249421.1
X 43209373 43209426
refGene_gn=CACNA1F;refGene_tx=XM_023249415.1;refGene_gn=CACNA1F;refGene_tx=XM_023249416.1;refGene_gn=CACNA1F;refGene_tx=XM_023249417.1;refGene_gn=CACNA1F;refGene_tx=XM_023249418.1;refGene_gn=CACNA1F;refGene_tx=XM_023249419.1;refGene_gn=CACNA1F;refGene_tx=XM_023249420.1;refGene_gn=CACNA1F;refGene_tx=XM_023249421.1
X 43209513 43209621
refGene_gn=CACNA1F;refGene_tx=XM_023249415.1;refGene_gn=CACNA1F;refGene_tx=XM_023249416.1;refGene_gn=CACNA1F;refGene_tx=XM_023249417.1;refGene_gn=CACNA1F;refGene_tx=XM_023249418.1;refGene_gn=CACNA1F;refGene_tx=XM_023249419.1;refGene_gn=CACNA1F;refGene_tx=XM_023249420.1;refGene_gn=CACNA1F;refGene_tx=XM_023249421.1
X 43210070 43210158
refGene_gn=CACNA1F;refGene_tx=XM_023249415.1;refGene_gn=CACNA1F;refGene_tx=XM_023249416.1;refGene_gn=CACNA1F;refGene_tx=XM_023249417.1;refGene_gn=CACNA1F;refGene_tx=XM_023249418.1;refGene_gn=CACNA1F;refGene_tx=XM_023249419.1;refGene_gn=CACNA1F;refGene_tx=XM_023249420.1;refGene_gn=CACNA1F;refGene_tx=XM_023249421.1
X 43210249 43210356
refGene_gn=CACNA1F;refGene_tx=XM_023249415.1;refGene_gn=CACNA1F;refGen

e_tx=XM_023249416.1;refGene_gn=CACNA1F;refGene_tx=XM_023249417.1;refGene_gn=CACNA1F;refGene_tx=XM_023249418.1;refGene_gn=CACNA1F;refGene_tx=XM_023249419.1;refGene_gn=CACNA1F;refGene_tx=XM_023249420.1;refGene_gn=CACNA1F;refGene_tx=XM_023249421.1
X 43210504 43210564
refGene_gn=CACNA1F;refGene_tx=XM_023249415.1;refGene_gn=CACNA1F;refGene_tx=XM_023249416.1;refGene_gn=CACNA1F;refGene_tx=XM_023249417.1;refGene_gn=CACNA1F;refGene_tx=XM_023249418.1;refGene_gn=CACNA1F;refGene_tx=XM_023249419.1;refGene_gn=CACNA1F;refGene_tx=XM_023249420.1;refGene_gn=CACNA1F;refGene_tx=XM_023249421.1
X 43211037 43211167
refGene_gn=CACNA1F;refGene_tx=XM_023249415.1;refGene_gn=CACNA1F;refGene_tx=XM_023249416.1;refGene_gn=CACNA1F;refGene_tx=XM_023249417.1;refGene_gn=CACNA1F;refGene_tx=XM_023249418.1;refGene_gn=CACNA1F;refGene_tx=XM_023249419.1;refGene_gn=CACNA1F;refGene_tx=XM_023249420.1;refGene_gn=CACNA1F;refGene_tx=XM_023249421.1
X 43211352 43211506
refGene_gn=CACNA1F;refGene_tx=XM_023249415.1;refGene_gn=CACNA1F;refGene_tx=XM_023249416.1;refGene_gn=CACNA1F;refGene_tx=XM_023249417.1;refGene_gn=CACNA1F;refGene_tx=XM_023249418.1;refGene_gn=CACNA1F;refGene_tx=XM_023249419.1;refGene_gn=CACNA1F;refGene_tx=XM_023249420.1;refGene_gn=CACNA1F;refGene_tx=XM_023249421.1
X 43211923 43211975
refGene_gn=CACNA1F;refGene_tx=XM_023249415.1;refGene_gn=CACNA1F;refGene_tx=XM_023249416.1;refGene_gn=CACNA1F;refGene_tx=XM_023249417.1;refGene_gn=CACNA1F;refGene_tx=XM_023249418.1;refGene_gn=CACNA1F;refGene_tx=XM_023249419.1;refGene_gn=CACNA1F;refGene_tx=XM_023249420.1;refGene_gn=CACNA1F;refGene_tx=XM_023249421.1
X 43212159 43212205
refGene_gn=CACNA1F;refGene_tx=XM_023249415.1;refGene_gn=CACNA1F;refGene_tx=XM_023249416.1;refGene_gn=CACNA1F;refGene_tx=XM_023249417.1;refGene_gn=CACNA1F;refGene_tx=XM_023249418.1;refGene_gn=CACNA1F;refGene_tx=XM_023249419.1;refGene_gn=CACNA1F;refGene_tx=XM_023249420.1;refGene_gn=CACNA1F;refGene_tx=XM_023249421.1
X 43213134 43213216
refGene_gn=CACNA1F;refGene_tx=XM_023249415.1;refGene_gn=CACNA1F;refGene_tx=XM_023249416.1;refGene_gn=CACNA1F;refGene_tx=XM_023249417.1;refGene_gn=CACNA1F;refGene_tx=XM_023249418.1;refGene_gn=CACNA1F;refGene_tx=XM_023249419.1;refGene_gn=CACNA1F;refGene_tx=XM_023249420.1;refGene_gn=CACNA1F;refGene_tx=XM_023249421.1
X 43213332 43213453
refGene_gn=CACNA1F;refGene_tx=XM_023249415.1;refGene_gn=CACNA1F;refGene_tx=XM_023249416.1;refGene_gn=CACNA1F;refGene_tx=XM_023249417.1;refGene_gn=CACNA1F;refGene_tx=XM_023249418.1;refGene_gn=CACNA1F;refGene_tx=XM_023249419.1;refGene_gn=CACNA1F;refGene_tx=XM_023249420.1;refGene_gn=CACNA1F;refGene_tx=XM_023249421.1
X 43213545 43213753
refGene_gn=CACNA1F;refGene_tx=XM_023249415.1;refGene_gn=CACNA1F;refGene_tx=XM_023249416.1;refGene_gn=CACNA1F;refGene_tx=XM_023249417.1;refGene_gn=CACNA1F;refGene_tx=XM_023249418.1;refGene_gn=CACNA1F;refGene_tx=

XM_023249419.1;refGene_gn=CACNA1F;refGene_tx=XM_023249420.1;refGene_gn=CACNA1F;refGene_tx=XM_023249421.1
X 43214648 43214874
refGene_gn=CACNA1F;refGene_tx=XM_023249415.1;refGene_gn=CACNA1F;refGene_tx=XM_023249416.1;refGene_gn=CACNA1F;refGene_tx=XM_023249417.1;refGene_gn=CACNA1F;refGene_tx=XM_023249418.1;refGene_gn=CACNA1F;refGene_tx=XM_023249419.1;refGene_gn=CACNA1F;refGene_tx=XM_023249420.1;refGene_gn=CACNA1F;refGene_tx=XM_023249421.1
X 43216004 43216165
refGene_gn=CACNA1F;refGene_tx=XM_023249415.1;refGene_gn=CACNA1F;refGene_tx=XM_023249416.1;refGene_gn=CACNA1F;refGene_tx=XM_023249417.1;refGene_gn=CACNA1F;refGene_tx=XM_023249418.1;refGene_gn=CACNA1F;refGene_tx=XM_023249419.1;refGene_gn=CACNA1F;refGene_tx=XM_023249420.1;refGene_gn=CACNA1F;refGene_tx=XM_023249421.1
X 43216303 43216330
refGene_gn=CACNA1F;refGene_tx=XM_023249415.1;refGene_gn=CACNA1F;refGene_tx=XM_023249416.1;refGene_gn=CACNA1F;refGene_tx=XM_023249417.1;refGene_gn=CACNA1F;refGene_tx=XM_023249418.1;refGene_gn=CACNA1F;refGene_tx=XM_023249419.1;refGene_gn=CACNA1F;refGene_tx=XM_023249420.1;refGene_gn=CACNA1F;refGene_tx=XM_023249421.1
X 43216501 43216595
refGene_gn=CACNA1F;refGene_tx=XM_023249415.1;refGene_gn=CACNA1F;refGene_tx=XM_023249416.1;refGene_gn=CACNA1F;refGene_tx=XM_023249417.1;refGene_gn=CACNA1F;refGene_tx=XM_023249418.1;refGene_gn=CACNA1F;refGene_tx=XM_023249419.1;refGene_gn=CACNA1F;refGene_tx=XM_023249420.1;refGene_gn=CACNA1F;refGene_tx=XM_023249421.1
X 43216708 43216801
refGene_gn=CACNA1F;refGene_tx=XM_023249415.1;refGene_gn=CACNA1F;refGene_tx=XM_023249416.1;refGene_gn=CACNA1F;refGene_tx=XM_023249417.1;refGene_gn=CACNA1F;refGene_tx=XM_023249418.1;refGene_gn=CACNA1F;refGene_tx=XM_023249419.1;refGene_gn=CACNA1F;refGene_tx=XM_023249420.1;refGene_gn=CACNA1F;refGene_tx=XM_023249421.1
X 43217067 43217258
refGene_gn=CACNA1F;refGene_tx=XM_023249415.1;refGene_gn=CACNA1F;refGene_tx=XM_023249416.1;refGene_gn=CACNA1F;refGene_tx=XM_023249417.1;refGene_gn=CACNA1F;refGene_tx=XM_023249418.1;refGene_gn=CACNA1F;refGene_tx=XM_023249419.1;refGene_gn=CACNA1F;refGene_tx=XM_023249420.1;refGene_gn=CACNA1F;refGene_tx=XM_023249421.1
X 43217962 43218066
refGene_gn=CACNA1F;refGene_tx=XM_023249415.1;refGene_gn=CACNA1F;refGene_tx=XM_023249416.1;refGene_gn=CACNA1F;refGene_tx=XM_023249417.1;refGene_gn=CACNA1F;refGene_tx=XM_023249418.1;refGene_gn=CACNA1F;refGene_tx=XM_023249419.1;refGene_gn=CACNA1F;refGene_tx=XM_023249420.1;refGene_gn=CACNA1F;refGene_tx=XM_023249421.1
X 43218163 43218360
refGene_gn=CACNA1F;refGene_tx=XM_023249415.1;refGene_gn=CACNA1F;refGene_tx=XM_023249416.1;refGene_gn=CACNA1F;refGene_tx=XM_023249417.1;refGene_gn=CACNA1F;refGene_tx=XM_023249418.1;refGene_gn=CACNA1F;refGene_tx=XM_023249419.1;refGene_gn=CACNA1F;refGene_tx=XM_023249420.1;refGene_gn=CACNA1F;refGene_tx=XM_023249421.1

X 43219000 43219153
refGene_gn=CACNA1F;refGene_tx=XM_023249415.1;refGene_gn=CACNA1F;refGene_tx=XM_023249416.1;refGene_gn=CACNA1F;refGene_tx=XM_023249417.1;refGene_gn=CACNA1F;refGene_tx=XM_023249418.1;refGene_gn=CACNA1F;refGene_tx=XM_023249419.1;refGene_gn=CACNA1F;refGene_tx=XM_023249420.1;refGene_gn=CACNA1F;refGene_tx=XM_023249421.1
X 43219236 43219379
refGene_gn=CACNA1F;refGene_tx=XM_023249415.1;refGene_gn=CACNA1F;refGene_tx=XM_023249416.1;refGene_gn=CACNA1F;refGene_tx=XM_023249417.1;refGene_gn=CACNA1F;refGene_tx=XM_023249418.1;refGene_gn=CACNA1F;refGene_tx=XM_023249419.1;refGene_gn=CACNA1F;refGene_tx=XM_023249420.1;refGene_gn=CACNA1F;refGene_tx=XM_023249421.1
X 43219622 43219762
refGene_gn=CACNA1F;refGene_tx=XM_023249415.1;refGene_gn=CACNA1F;refGene_tx=XM_023249416.1;refGene_gn=CACNA1F;refGene_tx=XM_023249417.1;refGene_gn=CACNA1F;refGene_tx=XM_023249418.1;refGene_gn=CACNA1F;refGene_tx=XM_023249419.1;refGene_gn=CACNA1F;refGene_tx=XM_023249420.1;refGene_gn=CACNA1F;refGene_tx=XM_023249421.1
X 43219985 43220091
refGene_gn=CACNA1F;refGene_tx=XM_023249415.1;refGene_gn=CACNA1F;refGene_tx=XM_023249416.1;refGene_gn=CACNA1F;refGene_tx=XM_023249417.1;refGene_gn=CACNA1F;refGene_tx=XM_023249418.1;refGene_gn=CACNA1F;refGene_tx=XM_023249419.1;refGene_gn=CACNA1F;refGene_tx=XM_023249420.1;refGene_gn=CACNA1F;refGene_tx=XM_023249421.1
X 43220729 43220982
refGene_gn=CACNA1F;refGene_tx=XM_023249415.1;refGene_gn=CACNA1F;refGene_tx=XM_023249416.1;refGene_gn=CACNA1F;refGene_tx=XM_023249417.1;refGene_gn=CACNA1F;refGene_tx=XM_023249418.1;refGene_gn=CACNA1F;refGene_tx=XM_023249419.1;refGene_gn=CACNA1F;refGene_tx=XM_023249420.1;refGene_gn=CACNA1F;refGene_tx=XM_023249421.1
X 43222642 43222667
refGene_gn=CACNA1F;refGene_tx=XM_023249415.1;refGene_gn=CACNA1F;refGene_tx=XM_023249416.1;refGene_gn=CACNA1F;refGene_tx=XM_023249417.1;refGene_gn=CACNA1F;refGene_tx=XM_023249418.1;refGene_gn=CACNA1F;refGene_tx=XM_023249419.1;refGene_gn=CACNA1F;refGene_tx=XM_023249420.1;refGene_gn=CACNA1F;refGene_tx=XM_023249421.1
X 43224221 43224271 refGene_gn=CCDC22;refGene_tx=XM_004000516.5
X 43225358 43225536 refGene_gn=CCDC22;refGene_tx=XM_004000516.5
X 43230380 43230513 refGene_gn=CCDC22;refGene_tx=XM_004000516.5
X 43231288 43231395 refGene_gn=CCDC22;refGene_tx=XM_004000516.5
X 43231538 43231605 refGene_gn=CCDC22;refGene_tx=XM_004000516.5
X 43231713 43231892 refGene_gn=CCDC22;refGene_tx=XM_004000516.5
X 43233463 43233658 refGene_gn=CCDC22;refGene_tx=XM_004000516.5
X 43234200 43234263 refGene_gn=CCDC22;refGene_tx=XM_004000516.5
X 43234347 43234467 refGene_gn=CCDC22;refGene_tx=XM_004000516.5
X 43235011 43235131 refGene_gn=CCDC22;refGene_tx=XM_004000516.5
X 43235221 43235338 refGene_gn=CCDC22;refGene_tx=XM_004000516.5
X 43235445 43235547 refGene_gn=CCDC22;refGene_tx=XM_004000516.5
X 43235632 43235740 refGene_gn=CCDC22;refGene_tx=XM_004000516.5
X 43236017 43236113 refGene_gn=CCDC22;refGene_tx=XM_004000516.5

X 43236359 43236419 refGene_gn=CCDC22;refGene_tx=XM_004000516.5
X 43236498 43236573 refGene_gn=CCDC22;refGene_tx=XM_004000516.5
X 43237009 43237123 refGene_gn=CCDC22;refGene_tx=XM_004000516.5
X 43238198 43238348
refGene_gn=FOXP3;refGene_tx=NM_001083952.1;refGene_gn=FOXP3;refGene_tx=XM_019823315.1;refGene_gn=FOXP3;refGene_tx=XM_019823320.1;refGene_gn=FOXP3;refGene_tx=XM_023248790.1;refGene_gn=FOXP3;refGene_tx=XM_023248791.1;refGene_gn=FOXP3;refGene_tx=XM_023248792.1;refGene_gn=FOXP3;refGene_tx=XM_023248793.1
X 43238557 43238659
refGene_gn=FOXP3;refGene_tx=NM_001083952.1;refGene_gn=FOXP3;refGene_tx=XM_019823315.1;refGene_gn=FOXP3;refGene_tx=XM_019823320.1;refGene_gn=FOXP3;refGene_tx=XM_023248790.1;refGene_gn=FOXP3;refGene_tx=XM_023248791.1;refGene_gn=FOXP3;refGene_tx=XM_023248792.1;refGene_gn=FOXP3;refGene_tx=XM_023248793.1
X 43239398 43239475
refGene_gn=FOXP3;refGene_tx=NM_001083952.1;refGene_gn=FOXP3;refGene_tx=XM_019823315.1;refGene_gn=FOXP3;refGene_tx=XM_019823320.1;refGene_gn=FOXP3;refGene_tx=XM_023248790.1;refGene_gn=FOXP3;refGene_tx=XM_023248791.1;refGene_gn=FOXP3;refGene_tx=XM_023248792.1;refGene_gn=FOXP3;refGene_tx=XM_023248793.1
X 43240148 43240299
refGene_gn=FOXP3;refGene_tx=NM_001083952.1;refGene_gn=FOXP3;refGene_tx=XM_019823315.1;refGene_gn=FOXP3;refGene_tx=XM_019823320.1;refGene_gn=FOXP3;refGene_tx=XM_023248790.1;refGene_gn=FOXP3;refGene_tx=XM_023248791.1;refGene_gn=FOXP3;refGene_tx=XM_023248792.1;refGene_gn=FOXP3;refGene_tx=XM_023248793.1
X 43241256 43241337
refGene_gn=FOXP3;refGene_tx=NM_001083952.1;refGene_gn=FOXP3;refGene_tx=XM_019823315.1;refGene_gn=FOXP3;refGene_tx=XM_019823320.1;refGene_gn=FOXP3;refGene_tx=XM_023248790.1;refGene_gn=FOXP3;refGene_tx=XM_023248791.1;refGene_gn=FOXP3;refGene_tx=XM_023248792.1;refGene_gn=FOXP3;refGene_tx=XM_023248793.1
X 43241525 43241613
refGene_gn=FOXP3;refGene_tx=NM_001083952.1;refGene_gn=FOXP3;refGene_tx=XM_019823315.1;refGene_gn=FOXP3;refGene_tx=XM_019823320.1;refGene_gn=FOXP3;refGene_tx=XM_023248790.1;refGene_gn=FOXP3;refGene_tx=XM_023248791.1;refGene_gn=FOXP3;refGene_tx=XM_023248792.1;refGene_gn=FOXP3;refGene_tx=XM_023248793.1
X 43242505 43242767
refGene_gn=FOXP3;refGene_tx=NM_001083952.1;refGene_gn=FOXP3;refGene_tx=XM_019823315.1;refGene_gn=FOXP3;refGene_tx=XM_019823320.1;refGene_gn=FOXP3;refGene_tx=XM_023248790.1;refGene_gn=FOXP3;refGene_tx=XM_023248791.1;refGene_gn=FOXP3;refGene_tx=XM_023248792.1;refGene_gn=FOXP3;refGene_tx=XM_023248793.1
X 43243154 43243293
refGene_gn=FOXP3;refGene_tx=NM_001083952.1;refGene_gn=FOXP3;refGene_tx=XM_019823315.1;refGene_gn=FOXP3;refGene_tx=XM_019823320.1;refGene_gn=FOXP3;refGene_tx=XM_023248790.1;refGene_gn=FOXP3;refGene_tx=XM_023248791.1;refGene_gn=FOXP3;refGene_tx=XM_023248792.1;refGene_gn=FOXP3;refGe

ne_tx=XM_023248793.1
X 43243380 43243485
refGene_gn=FOXP3;refGene_tx=NM_001083952.1;refGene_gn=FOXP3;refGene_tx=XM_019823315.1;refGene_gn=FOXP3;refGene_tx=XM_019823320.1;refGene_gn=FOXP3;refGene_tx=XM_023248790.1;refGene_gn=FOXP3;refGene_tx=XM_023248791.1;refGene_gn=FOXP3;refGene_tx=XM_023248792.1;refGene_gn=FOXP3;refGene_tx=XM_023248793.1
X 43244017 43244368
refGene_gn=FOXP3;refGene_tx=NM_001083952.1;refGene_gn=FOXP3;refGene_tx=XM_019823315.1;refGene_gn=FOXP3;refGene_tx=XM_019823320.1;refGene_gn=FOXP3;refGene_tx=XM_023248790.1;refGene_gn=FOXP3;refGene_tx=XM_023248791.1;refGene_gn=FOXP3;refGene_tx=XM_023248792.1;refGene_gn=FOXP3;refGene_tx=XM_023248793.1
X 43256200 43257207 refGene_gn=PPP1R3F;refGene_tx=XM_006943734.2
X 43266694 43266750 refGene_gn=PPP1R3F;refGene_tx=XM_006943734.2
X 43267195 43267275 refGene_gn=PPP1R3F;refGene_tx=XM_006943734.2
X 43270361 43271657 refGene_gn=PPP1R3F;refGene_tx=XM_006943734.2
X 43302009 43302093 refGene_gn=GAGE10;refGene_tx=XM_023249172.1
X 43302770 43302870 refGene_gn=GAGE10;refGene_tx=XM_023249172.1
X 43305075 43305201 refGene_gn=GAGE10;refGene_tx=XM_023249172.1
X 43307959 43307982 refGene_gn=GAGE10;refGene_tx=XM_023249172.1
X 43520450 43520554 refGene_gn=USP27X;refGene_tx=XM_019823816.2
X 43521076 43523426 refGene_gn=USP27X;refGene_tx=XM_019823816.2
X 43585916 43585932
refGene_gn=CLCN5;refGene_tx=XM_011291722.3;refGene_gn=CLCN5;refGene_tx=XM_019823820.2
X 43716450 43716597
refGene_gn=CLCN5;refGene_tx=XM_011291722.3;refGene_gn=CLCN5;refGene_tx=XM_019823820.2
X 43724239 43724264 refGene_gn=CLCN5;refGene_tx=XM_019823817.2
X 43733787 43733939
refGene_gn=CLCN5;refGene_tx=XM_011291722.3;refGene_gn=CLCN5;refGene_tx=XM_019823817.2;refGene_gn=CLCN5;refGene_tx=XM_019823819.2;refGene_gn=CLCN5;refGene_tx=XM_019823820.2
X 43736290 43736390
refGene_gn=CLCN5;refGene_tx=XM_011291722.3;refGene_gn=CLCN5;refGene_tx=XM_019823817.2;refGene_gn=CLCN5;refGene_tx=XM_019823819.2;refGene_gn=CLCN5;refGene_tx=XM_019823820.2
X 43739974 43740162
refGene_gn=CLCN5;refGene_tx=XM_011291722.3;refGene_gn=CLCN5;refGene_tx=XM_019823817.2;refGene_gn=CLCN5;refGene_tx=XM_019823819.2;refGene_gn=CLCN5;refGene_tx=XM_019823820.2
X 43743929 43744052
refGene_gn=CLCN5;refGene_tx=XM_011291722.3;refGene_gn=CLCN5;refGene_tx=XM_019823817.2;refGene_gn=CLCN5;refGene_tx=XM_019823819.2;refGene_gn=CLCN5;refGene_tx=XM_019823820.2
X 43744903 43745110
refGene_gn=CLCN5;refGene_tx=XM_011291722.3;refGene_gn=CLCN5;refGene_tx=XM_019823817.2;refGene_gn=CLCN5;refGene_tx=XM_019823819.2;refGene_gn=CLCN5;refGene_tx=XM_019823820.2

X 43748157 43748238
refGene_gn=CLCN5;refGene_tx=XM_011291722.3;refGene_gn=CLCN5;refGene_tx=XM_019823817.2;refGene_gn=CLCN5;refGene_tx=XM_019823819.2;refGene_gn=CLCN5;refGene_tx=XM_019823820.2
X 43748476 43749019
refGene_gn=CLCN5;refGene_tx=XM_011291722.3;refGene_gn=CLCN5;refGene_tx=XM_019823817.2;refGene_gn=CLCN5;refGene_tx=XM_019823819.2;refGene_gn=CLCN5;refGene_tx=XM_019823820.2
X 43750880 43751067
refGene_gn=CLCN5;refGene_tx=XM_011291722.3;refGene_gn=CLCN5;refGene_tx=XM_019823817.2;refGene_gn=CLCN5;refGene_tx=XM_019823819.2;refGene_gn=CLCN5;refGene_tx=XM_019823820.2
X 43752311 43752710
refGene_gn=CLCN5;refGene_tx=XM_011291722.3;refGene_gn=CLCN5;refGene_tx=XM_019823817.2;refGene_gn=CLCN5;refGene_tx=XM_019823819.2;refGene_gn=CLCN5;refGene_tx=XM_019823820.2
X 43752867 43753084
refGene_gn=CLCN5;refGene_tx=XM_011291722.3;refGene_gn=CLCN5;refGene_tx=XM_019823817.2;refGene_gn=CLCN5;refGene_tx=XM_019823819.2;refGene_gn=CLCN5;refGene_tx=XM_019823820.2
X 43754269 43754360
refGene_gn=CLCN5;refGene_tx=XM_011291722.3;refGene_gn=CLCN5;refGene_tx=XM_019823817.2;refGene_gn=CLCN5;refGene_tx=XM_019823819.2;refGene_gn=CLCN5;refGene_tx=XM_019823820.2
X 43851787 43851943
refGene_gn=AKAP4;refGene_tx=XM_023249170.1;refGene_gn=AKAP4;refGene_tx=XM_023249171.1
X 43852929 43855068
refGene_gn=AKAP4;refGene_tx=XM_023249170.1;refGene_gn=AKAP4;refGene_tx=XM_023249171.1
X 43863494 43863596
refGene_gn=AKAP4;refGene_tx=XM_023249170.1;refGene_gn=AKAP4;refGene_tx=XM_023249171.1
X 43864162 43864213
refGene_gn=AKAP4;refGene_tx=XM_023249170.1;refGene_gn=AKAP4;refGene_tx=XM_023249171.1
X 43865239 43865335
refGene_gn=AKAP4;refGene_tx=XM_023249170.1;refGene_gn=AKAP4;refGene_tx=XM_023249171.1
X 43888983 43889079 refGene_gn=CCNB3;refGene_tx=XM_019823821.2
X 43891441 43891528 refGene_gn=CCNB3;refGene_tx=XM_019823821.2
X 43896526 43896657 refGene_gn=CCNB3;refGene_tx=XM_019823821.2
X 43903141 43904239 refGene_gn=CCNB3;refGene_tx=XM_019823821.2
X 43904240 43906059 refGene_gn=CCNB3;refGene_tx=XM_019823821.2
X 43906959 43907055 refGene_gn=CCNB3;refGene_tx=XM_019823821.2
X 43908294 43908387 refGene_gn=CCNB3;refGene_tx=XM_019823821.2
X 43943445 43943583 refGene_gn=CCNB3;refGene_tx=XM_019823821.2
X 43946760 43946916 refGene_gn=CCNB3;refGene_tx=XM_019823821.2
X 43947671 43947821 refGene_gn=CCNB3;refGene_tx=XM_019823821.2
X 43951350 43951481 refGene_gn=CCNB3;refGene_tx=XM_019823821.2

X 43951695 43951768 refGene_gn=CCNB3;refGene_tx=XM_019823821.2
X 43989153 43989226 refGene_gn=DGKK;refGene_tx=XM_023249258.1
X 43990326 43990454 refGene_gn=DGKK;refGene_tx=XM_023249258.1
X 43991840 43991951 refGene_gn=DGKK;refGene_tx=XM_023249258.1
X 43995514 43995601 refGene_gn=DGKK;refGene_tx=XM_023249258.1
X 43997543 43997685 refGene_gn=DGKK;refGene_tx=XM_023249258.1
X 43998229 43998390 refGene_gn=DGKK;refGene_tx=XM_023249258.1
X 43999552 43999687 refGene_gn=DGKK;refGene_tx=XM_023249258.1
X 44000029 44000143 refGene_gn=DGKK;refGene_tx=XM_023249258.1
X 44000773 44000881 refGene_gn=DGKK;refGene_tx=XM_023249258.1
X 44001157 44001254 refGene_gn=DGKK;refGene_tx=XM_023249258.1
X 44005851 44005959 refGene_gn=DGKK;refGene_tx=XM_023249258.1
X 44007496 44007593 refGene_gn=DGKK;refGene_tx=XM_023249258.1
X 44008031 44008136 refGene_gn=DGKK;refGene_tx=XM_023249258.1
X 44009771 44009991 refGene_gn=DGKK;refGene_tx=XM_023249258.1
X 44026408 44026508 refGene_gn=DGKK;refGene_tx=XM_023249258.1
X 44027338 44027430 refGene_gn=DGKK;refGene_tx=XM_023249258.1
X 44029369 44029451 refGene_gn=DGKK;refGene_tx=XM_023249258.1
X 44030360 44030500 refGene_gn=DGKK;refGene_tx=XM_023249258.1
X 44031275 44031384 refGene_gn=DGKK;refGene_tx=XM_023249258.1
X 44032104 44032288 refGene_gn=DGKK;refGene_tx=XM_023249258.1
X 44040013 44040116 refGene_gn=DGKK;refGene_tx=XM_023249258.1
X 44059091 44059214 refGene_gn=DGKK;refGene_tx=XM_023249258.1
X 44059539 44059646 refGene_gn=DGKK;refGene_tx=XM_023249258.1
X 44060119 44060255 refGene_gn=DGKK;refGene_tx=XM_023249258.1
X 44095840 44095945 refGene_gn=DGKK;refGene_tx=XM_023249258.1
X 44097910 44097991 refGene_gn=DGKK;refGene_tx=XM_023249258.1
X 44099709 44099820 refGene_gn=DGKK;refGene_tx=XM_023249258.1
X 44150882 44151488 refGene_gn=DGKK;refGene_tx=XM_023249258.1
X 44266800 44267070 refGene_gn=SHROOM4;refGene_tx=XM_019823826.2
X 44268359 44268629 refGene_gn=SHROOM4;refGene_tx=XM_019823826.2
X 44273320 44273501 refGene_gn=SHROOM4;refGene_tx=XM_019823826.2
X 44274373 44274523 refGene_gn=SHROOM4;refGene_tx=XM_019823826.2
X 44279379 44280189 refGene_gn=SHROOM4;refGene_tx=XM_019823826.2
X 44298193 44298255 refGene_gn=SHROOM4;refGene_tx=XM_019823826.2
X 44304340 44306837 refGene_gn=SHROOM4;refGene_tx=XM_019823826.2
X 44309204 44309339 refGene_gn=SHROOM4;refGene_tx=XM_019823826.2
X 44404989 44405141 refGene_gn=SHROOM4;refGene_tx=XM_019823826.2
X 44534255 44534372 refGene_gn=SHROOM4;refGene_tx=XM_019823826.2
X 44645464 44645789 refGene_gn=BMP15;refGene_tx=NM_001165898.1
X 44649315 44650172 refGene_gn=BMP15;refGene_tx=NM_001165898.1
X 44771907 44771937
refGene_gn=LOC111558700;refGene_tx=XM_023249038.1
X 44776642 44776903
refGene_gn=LOC111558700;refGene_tx=XM_023249038.1
X 44777055 44777550
refGene_gn=LOC101082293;refGene_tx=XM_004000520.4;refGene_gn=LOC101082
293;refGene_tx=XM_019823828.2;refGene_gn=LOC101082293;refGene_tx=XM_02
3249037.1
X 44781801 44781802

refGene_gn=LOC101082293;refGene_tx=XM_023249037.1
X 44900179 44900439
refGene_gn=CXHXorf67;refGene_tx=XM_023249104.1
X 44900441 44902170
refGene_gn=CXHXorf67;refGene_tx=XM_023249104.1
X 44938018 44938019
refGene_gn=LOC101086628;refGene_tx=XM_006943744.3
X 44942179 44942674
refGene_gn=LOC101086628;refGene_tx=XM_006943744.3;refGene_gn=LOC101086628;refGene_tx=XM_023249093.1
X 44942826 44943087
refGene_gn=LOC109496513;refGene_tx=XM_023249094.1
X 44947789 44947819
refGene_gn=LOC109496513;refGene_tx=XM_023249094.1
X 45030068 45030857
refGene_gn=LOC101087903;refGene_tx=XM_023249250.1
X 45159802 45160591
refGene_gn=LOC101088142;refGene_tx=XM_023249273.1
X 45253031 45254915 refGene_gn=GSPT2;refGene_tx=XM_006943746.3
X 45393141 45393186 refGene_gn=MAGED1;refGene_tx=XM_011291724.3
X 45393885 45394602 refGene_gn=MAGED1;refGene_tx=XM_011291724.3
X 45395258 45395963 refGene_gn=MAGED1;refGene_tx=XM_011291724.3
X 45396087 45396151 refGene_gn=MAGED1;refGene_tx=XM_011291724.3
X 45396374 45396454 refGene_gn=MAGED1;refGene_tx=XM_011291724.3
X 45396627 45396719 refGene_gn=MAGED1;refGene_tx=XM_011291724.3
X 45396942 45397022 refGene_gn=MAGED1;refGene_tx=XM_011291724.3
X 45397113 45397156 refGene_gn=MAGED1;refGene_tx=XM_011291724.3
X 45397419 45397482 refGene_gn=MAGED1;refGene_tx=XM_011291724.3
X 45398055 45398170 refGene_gn=MAGED1;refGene_tx=XM_011291724.3
X 45398415 45398793 refGene_gn=MAGED1;refGene_tx=XM_011291724.3
X 45599412 45599847
refGene_gn=LOC101087132;refGene_tx=XM_011291729.3
X 45600183 45600298
refGene_gn=LOC101087132;refGene_tx=XM_011291729.3
X 45601235 45601298
refGene_gn=LOC101087132;refGene_tx=XM_011291729.3
X 45601714 45601757
refGene_gn=LOC101087132;refGene_tx=XM_011291729.3
X 45601855 45601924
refGene_gn=LOC101087132;refGene_tx=XM_011291729.3
X 45601926 45601934
refGene_gn=LOC101087132;refGene_tx=XM_011291729.3
X 45602564 45602656
refGene_gn=LOC101087132;refGene_tx=XM_011291729.3
X 45603049 45603129
refGene_gn=LOC101087132;refGene_tx=XM_011291729.3
X 45603355 45603419
refGene_gn=LOC101087132;refGene_tx=XM_011291729.3
X 45603523 45603727
refGene_gn=LOC101087132;refGene_tx=XM_011291729.3

X 45604066 45604897
refGene_gn=LOC101087132;refGene_tx=XM_011291729.3
X 45605178 45605388
refGene_gn=LOC101087132;refGene_tx=XM_011291729.3
X 45708614 45708728
refGene_gn=LOC111558733;refGene_tx=XM_023249191.1
X 45708832 45708896
refGene_gn=LOC111558733;refGene_tx=XM_023249191.1
X 45709122 45709158
refGene_gn=LOC111558733;refGene_tx=XM_023249191.1
X 45709160 45709201
refGene_gn=LOC111558733;refGene_tx=XM_023249191.1
X 45709594 45709686
refGene_gn=LOC111558733;refGene_tx=XM_023249191.1
X 45710317 45710325
refGene_gn=LOC111558733;refGene_tx=XM_023249191.1
X 45710327 45710396
refGene_gn=LOC111558733;refGene_tx=XM_023249191.1
X 45710494 45710537
refGene_gn=LOC111558733;refGene_tx=XM_023249191.1
X 45710952 45711015
refGene_gn=LOC111558733;refGene_tx=XM_023249191.1
X 45711952 45712067
refGene_gn=LOC111558733;refGene_tx=XM_023249191.1
X 45712403 45712838
refGene_gn=LOC111558733;refGene_tx=XM_023249191.1
X 45963227 45963311
refGene_gn=LOC102899730;refGene_tx=XM_006930890.4;refGene_gn=LOC102899730;refGene_tx=XM_023249292.1;refGene_gn=LOC102899730;refGene_tx=XM_023249293.1
X 45964173 45964282
refGene_gn=LOC102899730;refGene_tx=XM_006930890.4;refGene_gn=LOC102899730;refGene_tx=XM_023249292.1;refGene_gn=LOC102899730;refGene_tx=XM_023249293.1
X 45966056 45966207
refGene_gn=LOC102899730;refGene_tx=XM_006930890.4;refGene_gn=LOC102899730;refGene_tx=XM_023249292.1;refGene_gn=LOC102899730;refGene_tx=XM_023249293.1
X 45967579 45967583
refGene_gn=LOC102899730;refGene_tx=XM_006930890.4;refGene_gn=LOC102899730;refGene_tx=XM_023249292.1;refGene_gn=LOC102899730;refGene_tx=XM_023249293.1
X 46105437 46106559 refGene_gn=GPR173;refGene_tx=XM_004000535.5
X 46110718 46111543 refGene_gn=TSPYL2;refGene_tx=XM_023248833.1
X 46112850 46112928 refGene_gn=TSPYL2;refGene_tx=XM_023248833.1
X 46113052 46113164 refGene_gn=TSPYL2;refGene_tx=XM_023248833.1
X 46113251 46113397 refGene_gn=TSPYL2;refGene_tx=XM_023248833.1
X 46113528 46113626 refGene_gn=TSPYL2;refGene_tx=XM_023248833.1
X 46113937 46114773 refGene_gn=TSPYL2;refGene_tx=XM_023248833.1
X 46116859 46117035 refGene_gn=TSPYL2;refGene_tx=XM_023248833.1

X 46173915 46174038
refGene_gn=KDM5C;refGene_tx=XM_023249341.1;refGene_gn=KDM5C;refGene_tx=XM_023249343.1
X 46174954 46175320
refGene_gn=KDM5C;refGene_tx=XM_023249335.1;refGene_gn=KDM5C;refGene_tx=XM_023249336.1;refGene_gn=KDM5C;refGene_tx=XM_023249337.1;refGene_gn=KDM5C;refGene_tx=XM_023249338.1;refGene_gn=KDM5C;refGene_tx=XM_023249339.1;refGene_gn=KDM5C;refGene_tx=XM_023249340.1;refGene_gn=KDM5C;refGene_tx=XM_023249341.1;refGene_gn=KDM5C;refGene_tx=XM_023249342.1;refGene_gn=KDM5C;refGene_tx=XM_023249344.1;refGene_gn=KDM5C;refGene_tx=XM_023249345.1;refGene_gn=KDM5C;refGene_tx=XM_023249346.1;refGene_gn=KDM5C;refGene_tx=XM_023249347.1;refGene_gn=KDM5C;refGene_tx=XM_023249348.1;refGene_gn=KDM5C;refGene_tx=XM_023249349.1;refGene_gn=KDM5C;refGene_tx=XM_023249350.1;refGene_gn=KDM5C;refGene_tx=XM_023249351.1
X 46175422 46175679
refGene_gn=KDM5C;refGene_tx=XM_023249335.1;refGene_gn=KDM5C;refGene_tx=XM_023249336.1;refGene_gn=KDM5C;refGene_tx=XM_023249337.1;refGene_gn=KDM5C;refGene_tx=XM_023249338.1;refGene_gn=KDM5C;refGene_tx=XM_023249339.1;refGene_gn=KDM5C;refGene_tx=XM_023249340.1;refGene_gn=KDM5C;refGene_tx=XM_023249341.1;refGene_gn=KDM5C;refGene_tx=XM_023249342.1;refGene_gn=KDM5C;refGene_tx=XM_023249343.1;refGene_gn=KDM5C;refGene_tx=XM_023249344.1;refGene_gn=KDM5C;refGene_tx=XM_023249345.1;refGene_gn=KDM5C;refGene_tx=XM_023249346.1;refGene_gn=KDM5C;refGene_tx=XM_023249347.1;refGene_gn=KDM5C;refGene_tx=XM_023249348.1;refGene_gn=KDM5C;refGene_tx=XM_023249349.1;refGene_gn=KDM5C;refGene_tx=XM_023249350.1;refGene_gn=KDM5C;refGene_tx=XM_023249351.1
X 46175751 46175830
refGene_gn=KDM5C;refGene_tx=XM_023249335.1;refGene_gn=KDM5C;refGene_tx=XM_023249336.1;refGene_gn=KDM5C;refGene_tx=XM_023249337.1;refGene_gn=KDM5C;refGene_tx=XM_023249338.1;refGene_gn=KDM5C;refGene_tx=XM_023249339.1;refGene_gn=KDM5C;refGene_tx=XM_023249340.1;refGene_gn=KDM5C;refGene_tx=XM_023249341.1;refGene_gn=KDM5C;refGene_tx=XM_023249342.1;refGene_gn=KDM5C;refGene_tx=XM_023249343.1;refGene_gn=KDM5C;refGene_tx=XM_023249344.1;refGene_gn=KDM5C;refGene_tx=XM_023249345.1;refGene_gn=KDM5C;refGene_tx=XM_023249346.1;refGene_gn=KDM5C;refGene_tx=XM_023249347.1;refGene_gn=KDM5C;refGene_tx=XM_023249348.1;refGene_gn=KDM5C;refGene_tx=XM_023249349.1;refGene_gn=KDM5C;refGene_tx=XM_023249350.1;refGene_gn=KDM5C;refGene_tx=XM_023249351.1
X 46176100 46176700
refGene_gn=KDM5C;refGene_tx=XM_023249335.1;refGene_gn=KDM5C;refGene_tx=XM_023249336.1;refGene_gn=KDM5C;refGene_tx=XM_023249337.1;refGene_gn=KDM5C;refGene_tx=XM_023249338.1;refGene_gn=KDM5C;refGene_tx=XM_023249339.1;refGene_gn=KDM5C;refGene_tx=XM_023249340.1;refGene_gn=KDM5C;refGene_tx=XM_023249341.1;refGene_gn=KDM5C;refGene_tx=XM_023249342.1;refGene_gn=KDM5C;refGene_tx=XM_023249343.1;refGene_gn=KDM5C;refGene_tx=XM_023249344.1;refGene_gn=KDM5C;refGene_tx=XM_023249346.1;refGene_gn=KDM5C;refGene_tx=XM_023249349.1;refGene_gn=KDM5C;refGene_tx=XM_023249350.1;refGene_gn=KDM5C;refGene_tx=XM_023249351.1
X 46176887 46177025
refGene_gn=KDM5C;refGene_tx=XM_023249335.1;refGene_gn=KDM5C;refGene_tx

=XM_023249336.1;refGene_gn=KDM5C;refGene_tx=XM_023249337.1;refGene_gn=KDM5C;refGene_tx=XM_023249338.1;refGene_gn=KDM5C;refGene_tx=XM_023249339.1;refGene_gn=KDM5C;refGene_tx=XM_023249340.1;refGene_gn=KDM5C;refGene_tx=XM_023249341.1;refGene_gn=KDM5C;refGene_tx=XM_023249342.1;refGene_gn=KDM5C;refGene_tx=XM_023249343.1;refGene_gn=KDM5C;refGene_tx=XM_023249344.1;refGene_gn=KDM5C;refGene_tx=XM_023249345.1;refGene_gn=KDM5C;refGene_tx=XM_023249346.1;refGene_gn=KDM5C;refGene_tx=XM_023249347.1;refGene_gn=KDM5C;refGene_tx=XM_023249348.1;refGene_gn=KDM5C;refGene_tx=XM_023249349.1;refGene_gn=KDM5C;refGene_tx=XM_023249350.1;refGene_gn=KDM5C;refGene_tx=XM_023249351.1
X 46177171 46177351
refGene_gn=KDM5C;refGene_tx=XM_023249335.1;refGene_gn=KDM5C;refGene_tx=XM_023249336.1;refGene_gn=KDM5C;refGene_tx=XM_023249337.1;refGene_gn=KDM5C;refGene_tx=XM_023249338.1;refGene_gn=KDM5C;refGene_tx=XM_023249339.1;refGene_gn=KDM5C;refGene_tx=XM_023249340.1;refGene_gn=KDM5C;refGene_tx=XM_023249341.1;refGene_gn=KDM5C;refGene_tx=XM_023249342.1;refGene_gn=KDM5C;refGene_tx=XM_023249343.1;refGene_gn=KDM5C;refGene_tx=XM_023249344.1;refGene_gn=KDM5C;refGene_tx=XM_023249345.1;refGene_gn=KDM5C;refGene_tx=XM_023249346.1;refGene_gn=KDM5C;refGene_tx=XM_023249347.1;refGene_gn=KDM5C;refGene_tx=XM_023249348.1;refGene_gn=KDM5C;refGene_tx=XM_023249349.1;refGene_gn=KDM5C;refGene_tx=XM_023249350.1;refGene_gn=KDM5C;refGene_tx=XM_023249351.1
X 46177814 46177953
refGene_gn=KDM5C;refGene_tx=XM_023249335.1;refGene_gn=KDM5C;refGene_tx=XM_023249336.1;refGene_gn=KDM5C;refGene_tx=XM_023249337.1;refGene_gn=KDM5C;refGene_tx=XM_023249338.1;refGene_gn=KDM5C;refGene_tx=XM_023249339.1;refGene_gn=KDM5C;refGene_tx=XM_023249340.1;refGene_gn=KDM5C;refGene_tx=XM_023249341.1;refGene_gn=KDM5C;refGene_tx=XM_023249342.1;refGene_gn=KDM5C;refGene_tx=XM_023249343.1;refGene_gn=KDM5C;refGene_tx=XM_023249344.1;refGene_gn=KDM5C;refGene_tx=XM_023249345.1;refGene_gn=KDM5C;refGene_tx=XM_023249346.1;refGene_gn=KDM5C;refGene_tx=XM_023249347.1;refGene_gn=KDM5C;refGene_tx=XM_023249348.1;refGene_gn=KDM5C;refGene_tx=XM_023249349.1;refGene_gn=KDM5C;refGene_tx=XM_023249350.1;refGene_gn=KDM5C;refGene_tx=XM_023249351.1
X 46178561 46178920
refGene_gn=KDM5C;refGene_tx=XM_023249335.1;refGene_gn=KDM5C;refGene_tx=XM_023249336.1;refGene_gn=KDM5C;refGene_tx=XM_023249337.1;refGene_gn=KDM5C;refGene_tx=XM_023249338.1;refGene_gn=KDM5C;refGene_tx=XM_023249339.1;refGene_gn=KDM5C;refGene_tx=XM_023249340.1;refGene_gn=KDM5C;refGene_tx=XM_023249341.1;refGene_gn=KDM5C;refGene_tx=XM_023249342.1;refGene_gn=KDM5C;refGene_tx=XM_023249343.1;refGene_gn=KDM5C;refGene_tx=XM_023249344.1;refGene_gn=KDM5C;refGene_tx=XM_023249345.1;refGene_gn=KDM5C;refGene_tx=XM_023249346.1;refGene_gn=KDM5C;refGene_tx=XM_023249347.1;refGene_gn=KDM5C;refGene_tx=XM_023249348.1;refGene_gn=KDM5C;refGene_tx=XM_023249349.1;refGene_gn=KDM5C;refGene_tx=XM_023249350.1;refGene_gn=KDM5C;refGene_tx=XM_023249351.1
X 46179606 46179712
refGene_gn=KDM5C;refGene_tx=XM_023249335.1;refGene_gn=KDM5C;refGene_tx=XM_023249336.1;refGene_gn=KDM5C;refGene_tx=XM_023249337.1;refGene_gn=KDM5C;refGene_tx=XM_023249338.1;refGene_gn=KDM5C;refGene_tx=XM_0232493

39.1;refGene_gn=KDM5C;refGene_tx=XM_023249340.1;refGene_gn=KDM5C;refGene_tx=XM_023249341.1;refGene_gn=KDM5C;refGene_tx=XM_023249342.1;refGene_gn=KDM5C;refGene_tx=XM_023249343.1;refGene_gn=KDM5C;refGene_tx=XM_023249344.1;refGene_gn=KDM5C;refGene_tx=XM_023249345.1;refGene_gn=KDM5C;refGene_tx=XM_023249346.1;refGene_gn=KDM5C;refGene_tx=XM_023249347.1;refGene_gn=KDM5C;refGene_tx=XM_023249348.1;refGene_gn=KDM5C;refGene_tx=XM_023249349.1;refGene_gn=KDM5C;refGene_tx=XM_023249350.1;refGene_gn=KDM5C;refGene_tx=XM_023249351.1
X 46180324 46180472
refGene_gn=KDM5C;refGene_tx=XM_023249335.1;refGene_gn=KDM5C;refGene_tx=XM_023249336.1;refGene_gn=KDM5C;refGene_tx=XM_023249337.1;refGene_gn=KDM5C;refGene_tx=XM_023249338.1;refGene_gn=KDM5C;refGene_tx=XM_023249339.1;refGene_gn=KDM5C;refGene_tx=XM_023249340.1;refGene_gn=KDM5C;refGene_tx=XM_023249341.1;refGene_gn=KDM5C;refGene_tx=XM_023249342.1;refGene_gn=KDM5C;refGene_tx=XM_023249343.1;refGene_gn=KDM5C;refGene_tx=XM_023249344.1;refGene_gn=KDM5C;refGene_tx=XM_023249345.1;refGene_gn=KDM5C;refGene_tx=XM_023249346.1;refGene_gn=KDM5C;refGene_tx=XM_023249347.1;refGene_gn=KDM5C;refGene_tx=XM_023249348.1;refGene_gn=KDM5C;refGene_tx=XM_023249349.1;refGene_gn=KDM5C;refGene_tx=XM_023249350.1;refGene_gn=KDM5C;refGene_tx=XM_023249351.1
X 46180599 46180724
refGene_gn=KDM5C;refGene_tx=XM_023249335.1;refGene_gn=KDM5C;refGene_tx=XM_023249336.1;refGene_gn=KDM5C;refGene_tx=XM_023249337.1;refGene_gn=KDM5C;refGene_tx=XM_023249338.1;refGene_gn=KDM5C;refGene_tx=XM_023249339.1;refGene_gn=KDM5C;refGene_tx=XM_023249340.1;refGene_gn=KDM5C;refGene_tx=XM_023249341.1;refGene_gn=KDM5C;refGene_tx=XM_023249342.1;refGene_gn=KDM5C;refGene_tx=XM_023249343.1;refGene_gn=KDM5C;refGene_tx=XM_023249344.1;refGene_gn=KDM5C;refGene_tx=XM_023249345.1;refGene_gn=KDM5C;refGene_tx=XM_023249346.1;refGene_gn=KDM5C;refGene_tx=XM_023249347.1;refGene_gn=KDM5C;refGene_tx=XM_023249348.1;refGene_gn=KDM5C;refGene_tx=XM_023249349.1;refGene_gn=KDM5C;refGene_tx=XM_023249350.1;refGene_gn=KDM5C;refGene_tx=XM_023249351.1
X 46180812 46180994
refGene_gn=KDM5C;refGene_tx=XM_023249335.1;refGene_gn=KDM5C;refGene_tx=XM_023249336.1;refGene_gn=KDM5C;refGene_tx=XM_023249337.1;refGene_gn=KDM5C;refGene_tx=XM_023249338.1;refGene_gn=KDM5C;refGene_tx=XM_023249339.1;refGene_gn=KDM5C;refGene_tx=XM_023249340.1;refGene_gn=KDM5C;refGene_tx=XM_023249341.1;refGene_gn=KDM5C;refGene_tx=XM_023249342.1;refGene_gn=KDM5C;refGene_tx=XM_023249343.1;refGene_gn=KDM5C;refGene_tx=XM_023249344.1;refGene_gn=KDM5C;refGene_tx=XM_023249345.1;refGene_gn=KDM5C;refGene_tx=XM_023249346.1;refGene_gn=KDM5C;refGene_tx=XM_023249347.1;refGene_gn=KDM5C;refGene_tx=XM_023249348.1;refGene_gn=KDM5C;refGene_tx=XM_023249349.1;refGene_gn=KDM5C;refGene_tx=XM_023249350.1;refGene_gn=KDM5C;refGene_tx=XM_023249351.1
X 46185122 46185317
refGene_gn=KDM5C;refGene_tx=XM_023249335.1;refGene_gn=KDM5C;refGene_tx=XM_023249336.1;refGene_gn=KDM5C;refGene_tx=XM_023249337.1;refGene_gn=KDM5C;refGene_tx=XM_023249338.1;refGene_gn=KDM5C;refGene_tx=XM_023249339.1;refGene_gn=KDM5C;refGene_tx=XM_023249340.1;refGene_gn=KDM5C;refGene_tx=XM_023249341.1;refGene_gn=KDM5C;refGene_tx=XM_023249342.1;refGen

e_gn=KDM5C;refGene_tx=XM_023249343.1;refGene_gn=KDM5C;refGene_tx=XM_02
3249344.1;refGene_gn=KDM5C;refGene_tx=XM_023249345.1;refGene_gn=KDM5C;
refGene_tx=XM_023249346.1;refGene_gn=KDM5C;refGene_tx=XM_023249347.1;r
efGene_gn=KDM5C;refGene_tx=XM_023249348.1;refGene_gn=KDM5C;refGene_tx=
XM_023249349.1;refGene_gn=KDM5C;refGene_tx=XM_023249350.1;refGene_gn=K
DM5C;refGene_tx=XM_023249351.1
X 46185424 46185544
refGene_gn=KDM5C;refGene_tx=XM_023249335.1;refGene_gn=KDM5C;refGene_tx
=XM_023249336.1;refGene_gn=KDM5C;refGene_tx=XM_023249337.1;refGene_gn=
KDM5C;refGene_tx=XM_023249338.1;refGene_gn=KDM5C;refGene_tx=XM_0232493
39.1;refGene_gn=KDM5C;refGene_tx=XM_023249340.1;refGene_gn=KDM5C;refGe
ne_tx=XM_023249341.1;refGene_gn=KDM5C;refGene_tx=XM_023249342.1;refGen
e_gn=KDM5C;refGene_tx=XM_023249343.1;refGene_gn=KDM5C;refGene_tx=XM_02
3249344.1;refGene_gn=KDM5C;refGene_tx=XM_023249345.1;refGene_gn=KDM5C;
refGene_tx=XM_023249346.1;refGene_gn=KDM5C;refGene_tx=XM_023249347.1;r
efGene_gn=KDM5C;refGene_tx=XM_023249348.1;refGene_gn=KDM5C;refGene_tx=
XM_023249349.1;refGene_gn=KDM5C;refGene_tx=XM_023249350.1;refGene_gn=K
DM5C;refGene_tx=XM_023249351.1
X 46191866 46192029
refGene_gn=KDM5C;refGene_tx=XM_023249335.1;refGene_gn=KDM5C;refGene_tx
=XM_023249336.1;refGene_gn=KDM5C;refGene_tx=XM_023249337.1;refGene_gn=
KDM5C;refGene_tx=XM_023249338.1;refGene_gn=KDM5C;refGene_tx=XM_0232493
39.1;refGene_gn=KDM5C;refGene_tx=XM_023249340.1;refGene_gn=KDM5C;refGe
ne_tx=XM_023249341.1;refGene_gn=KDM5C;refGene_tx=XM_023249342.1;refGen
e_gn=KDM5C;refGene_tx=XM_023249343.1;refGene_gn=KDM5C;refGene_tx=XM_02
3249344.1;refGene_gn=KDM5C;refGene_tx=XM_023249345.1;refGene_gn=KDM5C;
refGene_tx=XM_023249347.1;refGene_gn=KDM5C;refGene_tx=XM_023249348.1;r
efGene_gn=KDM5C;refGene_tx=XM_023249349.1;refGene_gn=KDM5C;refGene_tx=
XM_023249350.1;refGene_gn=KDM5C;refGene_tx=XM_023249351.1
X 46192123 46192305
refGene_gn=KDM5C;refGene_tx=XM_023249335.1;refGene_gn=KDM5C;refGene_tx
=XM_023249336.1;refGene_gn=KDM5C;refGene_tx=XM_023249337.1;refGene_gn=
KDM5C;refGene_tx=XM_023249338.1;refGene_gn=KDM5C;refGene_tx=XM_0232493
39.1;refGene_gn=KDM5C;refGene_tx=XM_023249340.1;refGene_gn=KDM5C;refGe
ne_tx=XM_023249341.1;refGene_gn=KDM5C;refGene_tx=XM_023249342.1;refGen
e_gn=KDM5C;refGene_tx=XM_023249343.1;refGene_gn=KDM5C;refGene_tx=XM_02
3249344.1;refGene_gn=KDM5C;refGene_tx=XM_023249345.1;refGene_gn=KDM5C;
refGene_tx=XM_023249347.1;refGene_gn=KDM5C;refGene_tx=XM_023249348.1;r
efGene_gn=KDM5C;refGene_tx=XM_023249349.1;refGene_gn=KDM5C;refGene_tx=
XM_023249350.1;refGene_gn=KDM5C;refGene_tx=XM_023249351.1
X 46193156 46193315
refGene_gn=KDM5C;refGene_tx=XM_023249335.1;refGene_gn=KDM5C;refGene_tx
=XM_023249336.1;refGene_gn=KDM5C;refGene_tx=XM_023249337.1;refGene_gn=
KDM5C;refGene_tx=XM_023249338.1;refGene_gn=KDM5C;refGene_tx=XM_0232493
39.1;refGene_gn=KDM5C;refGene_tx=XM_023249340.1;refGene_gn=KDM5C;refGe
ne_tx=XM_023249341.1;refGene_gn=KDM5C;refGene_tx=XM_023249342.1;refGen
e_gn=KDM5C;refGene_tx=XM_023249343.1;refGene_gn=KDM5C;refGene_tx=XM_02
3249344.1;refGene_gn=KDM5C;refGene_tx=XM_023249345.1;refGene_gn=KDM5C;
refGene_tx=XM_023249347.1;refGene_gn=KDM5C;refGene_tx=XM_023249348.1;r
efGene_gn=KDM5C;refGene_tx=XM_023249349.1;refGene_gn=KDM5C;refGene_tx=

XM_023249350.1;refGene_gn=KDM5C;refGene_tx=XM_023249351.1
X	46193431	46193551
refGene_gn=KDM5C;refGene_tx=XM_023249335.1;refGene_gn=KDM5C;refGene_tx=XM_023249336.1;refGene_gn=KDM5C;refGene_tx=XM_023249337.1;refGene_gn=KDM5C;refGene_tx=XM_023249338.1;refGene_gn=KDM5C;refGene_tx=XM_023249339.1;refGene_gn=KDM5C;refGene_tx=XM_023249340.1;refGene_gn=KDM5C;refGene_tx=XM_023249341.1;refGene_gn=KDM5C;refGene_tx=XM_023249342.1;refGene_gn=KDM5C;refGene_tx=XM_023249343.1;refGene_gn=KDM5C;refGene_tx=XM_023249344.1;refGene_gn=KDM5C;refGene_tx=XM_023249345.1;refGene_gn=KDM5C;refGene_tx=XM_023249347.1;refGene_gn=KDM5C;refGene_tx=XM_023249348.1;refGene_gn=KDM5C;refGene_tx=XM_023249349.1;refGene_gn=KDM5C;refGene_tx=XM_023249350.1;refGene_gn=KDM5C;refGene_tx=XM_023249351.1
X	46195315	46195474
refGene_gn=KDM5C;refGene_tx=XM_023249335.1;refGene_gn=KDM5C;refGene_tx=XM_023249336.1;refGene_gn=KDM5C;refGene_tx=XM_023249337.1;refGene_gn=KDM5C;refGene_tx=XM_023249338.1;refGene_gn=KDM5C;refGene_tx=XM_023249339.1;refGene_gn=KDM5C;refGene_tx=XM_023249340.1;refGene_gn=KDM5C;refGene_tx=XM_023249341.1;refGene_gn=KDM5C;refGene_tx=XM_023249342.1;refGene_gn=KDM5C;refGene_tx=XM_023249343.1;refGene_gn=KDM5C;refGene_tx=XM_023249344.1;refGene_gn=KDM5C;refGene_tx=XM_023249345.1;refGene_gn=KDM5C;refGene_tx=XM_023249346.1;refGene_gn=KDM5C;refGene_tx=XM_023249347.1;refGene_gn=KDM5C;refGene_tx=XM_023249348.1;refGene_gn=KDM5C;refGene_tx=XM_023249349.1;refGene_gn=KDM5C;refGene_tx=XM_023249350.1;refGene_gn=KDM5C;refGene_tx=XM_023249351.1
X	46196518	46196700
refGene_gn=KDM5C;refGene_tx=XM_023249335.1;refGene_gn=KDM5C;refGene_tx=XM_023249336.1;refGene_gn=KDM5C;refGene_tx=XM_023249337.1;refGene_gn=KDM5C;refGene_tx=XM_023249338.1;refGene_gn=KDM5C;refGene_tx=XM_023249339.1;refGene_gn=KDM5C;refGene_tx=XM_023249340.1;refGene_gn=KDM5C;refGene_tx=XM_023249341.1;refGene_gn=KDM5C;refGene_tx=XM_023249342.1;refGene_gn=KDM5C;refGene_tx=XM_023249343.1;refGene_gn=KDM5C;refGene_tx=XM_023249344.1;refGene_gn=KDM5C;refGene_tx=XM_023249345.1;refGene_gn=KDM5C;refGene_tx=XM_023249346.1;refGene_gn=KDM5C;refGene_tx=XM_023249347.1;refGene_gn=KDM5C;refGene_tx=XM_023249348.1;refGene_gn=KDM5C;refGene_tx=XM_023249351.1
X	46196813	46196937
refGene_gn=KDM5C;refGene_tx=XM_023249335.1;refGene_gn=KDM5C;refGene_tx=XM_023249336.1;refGene_gn=KDM5C;refGene_tx=XM_023249337.1;refGene_gn=KDM5C;refGene_tx=XM_023249338.1;refGene_gn=KDM5C;refGene_tx=XM_023249339.1;refGene_gn=KDM5C;refGene_tx=XM_023249340.1;refGene_gn=KDM5C;refGene_tx=XM_023249341.1;refGene_gn=KDM5C;refGene_tx=XM_023249342.1;refGene_gn=KDM5C;refGene_tx=XM_023249343.1;refGene_gn=KDM5C;refGene_tx=XM_023249344.1;refGene_gn=KDM5C;refGene_tx=XM_023249345.1;refGene_gn=KDM5C;refGene_tx=XM_023249346.1;refGene_gn=KDM5C;refGene_tx=XM_023249347.1;refGene_gn=KDM5C;refGene_tx=XM_023249348.1;refGene_gn=KDM5C;refGene_tx=XM_023249351.1
X	46197645	46197780
refGene_gn=KDM5C;refGene_tx=XM_023249335.1;refGene_gn=KDM5C;refGene_tx=XM_023249336.1;refGene_gn=KDM5C;refGene_tx=XM_023249337.1;refGene_gn=KDM5C;refGene_tx=XM_023249338.1;refGene_gn=KDM5C;refGene_tx=XM_0232493

39.1;refGene_gn=KDM5C;refGene_tx=XM_023249340.1;refGene_gn=KDM5C;refGene_tx=XM_023249341.1;refGene_gn=KDM5C;refGene_tx=XM_023249342.1;refGene_gn=KDM5C;refGene_tx=XM_023249343.1;refGene_gn=KDM5C;refGene_tx=XM_023249345.1;refGene_gn=KDM5C;refGene_tx=XM_023249346.1;refGene_gn=KDM5C;refGene_tx=XM_023249347.1;refGene_gn=KDM5C;refGene_tx=XM_023249348.1;refGene_gn=KDM5C;refGene_tx=XM_023249351.1
X 46198313 46198484
refGene_gn=KDM5C;refGene_tx=XM_023249335.1;refGene_gn=KDM5C;refGene_tx=XM_023249336.1;refGene_gn=KDM5C;refGene_tx=XM_023249337.1;refGene_gn=KDM5C;refGene_tx=XM_023249338.1;refGene_gn=KDM5C;refGene_tx=XM_023249339.1;refGene_gn=KDM5C;refGene_tx=XM_023249340.1;refGene_gn=KDM5C;refGene_tx=XM_023249341.1;refGene_gn=KDM5C;refGene_tx=XM_023249343.1;refGene_gn=KDM5C;refGene_tx=XM_023249345.1;refGene_gn=KDM5C;refGene_tx=XM_023249346.1;refGene_gn=KDM5C;refGene_tx=XM_023249347.1;refGene_gn=KDM5C;refGene_tx=XM_023249348.1
X 46198791 46198914
refGene_gn=KDM5C;refGene_tx=XM_023249335.1;refGene_gn=KDM5C;refGene_tx=XM_023249336.1;refGene_gn=KDM5C;refGene_tx=XM_023249337.1;refGene_gn=KDM5C;refGene_tx=XM_023249338.1;refGene_gn=KDM5C;refGene_tx=XM_023249339.1;refGene_gn=KDM5C;refGene_tx=XM_023249340.1;refGene_gn=KDM5C;refGene_tx=XM_023249341.1;refGene_gn=KDM5C;refGene_tx=XM_023249342.1;refGene_gn=KDM5C;refGene_tx=XM_023249343.1;refGene_gn=KDM5C;refGene_tx=XM_023249345.1;refGene_gn=KDM5C;refGene_tx=XM_023249346.1;refGene_gn=KDM5C;refGene_tx=XM_023249347.1;refGene_gn=KDM5C;refGene_tx=XM_023249348.1;refGene_gn=KDM5C;refGene_tx=XM_023249351.1
X 46201375 46201453
refGene_gn=KDM5C;refGene_tx=XM_023249335.1;refGene_gn=KDM5C;refGene_tx=XM_023249336.1;refGene_gn=KDM5C;refGene_tx=XM_023249337.1;refGene_gn=KDM5C;refGene_tx=XM_023249338.1;refGene_gn=KDM5C;refGene_tx=XM_023249339.1;refGene_gn=KDM5C;refGene_tx=XM_023249340.1;refGene_gn=KDM5C;refGene_tx=XM_023249341.1;refGene_gn=KDM5C;refGene_tx=XM_023249342.1;refGene_gn=KDM5C;refGene_tx=XM_023249343.1;refGene_gn=KDM5C;refGene_tx=XM_023249344.1;refGene_gn=KDM5C;refGene_tx=XM_023249345.1;refGene_gn=KDM5C;refGene_tx=XM_023249346.1;refGene_gn=KDM5C;refGene_tx=XM_023249347.1;refGene_gn=KDM5C;refGene_tx=XM_023249348.1;refGene_gn=KDM5C;refGene_tx=XM_023249351.1
X 46205361 46205511
refGene_gn=KDM5C;refGene_tx=XM_023249335.1;refGene_gn=KDM5C;refGene_tx=XM_023249336.1;refGene_gn=KDM5C;refGene_tx=XM_023249337.1;refGene_gn=KDM5C;refGene_tx=XM_023249338.1;refGene_gn=KDM5C;refGene_tx=XM_023249339.1;refGene_gn=KDM5C;refGene_tx=XM_023249340.1;refGene_gn=KDM5C;refGene_tx=XM_023249341.1;refGene_gn=KDM5C;refGene_tx=XM_023249342.1;refGene_gn=KDM5C;refGene_tx=XM_023249343.1;refGene_gn=KDM5C;refGene_tx=XM_023249344.1;refGene_gn=KDM5C;refGene_tx=XM_023249345.1;refGene_gn=KDM5C;refGene_tx=XM_023249346.1;refGene_gn=KDM5C;refGene_tx=XM_023249347.1;refGene_gn=KDM5C;refGene_tx=XM_023249348.1;refGene_gn=KDM5C;refGene_tx=XM_023249351.1
X 46214127 46215093
refGene_gn=IQSEC2;refGene_tx=XM_006943747.3;refGene_gn=IQSEC2;refGene_tx=XM_011291746.2;refGene_gn=IQSEC2;refGene_tx=XM_019823838.2;refGene_

gn=IQSEC2;refGene_tx=XM_019823839.2;refGene_gn=IQSEC2;refGene_tx=XM_02
3248845.1;refGene_gn=IQSEC2;refGene_tx=XM_023248846.1
X 46215691 46215741
refGene_gn=IQSEC2;refGene_tx=XM_006943747.3;refGene_gn=IQSEC2;refGene_
tx=XM_011291746.2;refGene_gn=IQSEC2;refGene_tx=XM_019823838.2;refGene_
gn=IQSEC2;refGene_tx=XM_023248845.1;refGene_gn=IQSEC2;refGene_tx=XM_02
3248846.1
X 46216232 46216406
refGene_gn=IQSEC2;refGene_tx=XM_006943747.3;refGene_gn=IQSEC2;refGene_
tx=XM_011291746.2;refGene_gn=IQSEC2;refGene_tx=XM_019823838.2;refGene_
gn=IQSEC2;refGene_tx=XM_019823839.2;refGene_gn=IQSEC2;refGene_tx=XM_02
3248845.1;refGene_gn=IQSEC2;refGene_tx=XM_023248846.1
X 46218096 46218258
refGene_gn=IQSEC2;refGene_tx=XM_006943747.3;refGene_gn=IQSEC2;refGene_
tx=XM_011291746.2;refGene_gn=IQSEC2;refGene_tx=XM_019823838.2;refGene_
gn=IQSEC2;refGene_tx=XM_019823839.2;refGene_gn=IQSEC2;refGene_tx=XM_02
3248845.1;refGene_gn=IQSEC2;refGene_tx=XM_023248846.1
X 46218816 46218916
refGene_gn=IQSEC2;refGene_tx=XM_006943747.3;refGene_gn=IQSEC2;refGene_
tx=XM_011291746.2;refGene_gn=IQSEC2;refGene_tx=XM_019823838.2;refGene_
gn=IQSEC2;refGene_tx=XM_019823839.2;refGene_gn=IQSEC2;refGene_tx=XM_02
3248845.1;refGene_gn=IQSEC2;refGene_tx=XM_023248846.1
X 46221033 46221159
refGene_gn=IQSEC2;refGene_tx=XM_006943747.3;refGene_gn=IQSEC2;refGene_
tx=XM_011291746.2;refGene_gn=IQSEC2;refGene_tx=XM_019823838.2;refGene_
gn=IQSEC2;refGene_tx=XM_019823839.2;refGene_gn=IQSEC2;refGene_tx=XM_02
3248845.1;refGene_gn=IQSEC2;refGene_tx=XM_023248846.1
X 46221922 46222062
refGene_gn=IQSEC2;refGene_tx=XM_006943747.3;refGene_gn=IQSEC2;refGene_
tx=XM_011291746.2;refGene_gn=IQSEC2;refGene_tx=XM_019823838.2;refGene_
gn=IQSEC2;refGene_tx=XM_019823839.2;refGene_gn=IQSEC2;refGene_tx=XM_02
3248845.1;refGene_gn=IQSEC2;refGene_tx=XM_023248846.1
X 46224259 46224426
refGene_gn=IQSEC2;refGene_tx=XM_006943747.3;refGene_gn=IQSEC2;refGene_
tx=XM_011291746.2;refGene_gn=IQSEC2;refGene_tx=XM_019823838.2;refGene_
gn=IQSEC2;refGene_tx=XM_019823839.2;refGene_gn=IQSEC2;refGene_tx=XM_02
3248845.1;refGene_gn=IQSEC2;refGene_tx=XM_023248846.1
X 46225399 46225522
refGene_gn=IQSEC2;refGene_tx=XM_006943747.3;refGene_gn=IQSEC2;refGene_
tx=XM_011291746.2;refGene_gn=IQSEC2;refGene_tx=XM_019823838.2;refGene_
gn=IQSEC2;refGene_tx=XM_019823839.2;refGene_gn=IQSEC2;refGene_tx=XM_02
3248845.1;refGene_gn=IQSEC2;refGene_tx=XM_023248846.1
X 46225995 46226157
refGene_gn=IQSEC2;refGene_tx=XM_006943747.3;refGene_gn=IQSEC2;refGene_
tx=XM_011291746.2;refGene_gn=IQSEC2;refGene_tx=XM_019823838.2;refGene_
gn=IQSEC2;refGene_tx=XM_019823839.2;refGene_gn=IQSEC2;refGene_tx=XM_02
3248845.1;refGene_gn=IQSEC2;refGene_tx=XM_023248846.1
X 46227854 46228750
refGene_gn=IQSEC2;refGene_tx=XM_006943747.3;refGene_gn=IQSEC2;refGene_
tx=XM_011291746.2;refGene_gn=IQSEC2;refGene_tx=XM_019823838.2;refGene_

gn=IQSEC2;refGene_tx=XM_019823839.2;refGene_gn=IQSEC2;refGene_tx=XM_023248845.1;refGene_gn=IQSEC2;refGene_tx=XM_023248846.1
X 46231159 46231561
refGene_gn=IQSEC2;refGene_tx=XM_006943747.3;refGene_gn=IQSEC2;refGene_tx=XM_011291746.2;refGene_gn=IQSEC2;refGene_tx=XM_019823838.2;refGene_gn=IQSEC2;refGene_tx=XM_019823839.2;refGene_gn=IQSEC2;refGene_tx=XM_023248845.1;refGene_gn=IQSEC2;refGene_tx=XM_023248846.1
X 46232479 46232741
refGene_gn=IQSEC2;refGene_tx=XM_006943747.3;refGene_gn=IQSEC2;refGene_tx=XM_011291746.2;refGene_gn=IQSEC2;refGene_tx=XM_019823838.2;refGene_gn=IQSEC2;refGene_tx=XM_019823839.2;refGene_gn=IQSEC2;refGene_tx=XM_023248845.1;refGene_gn=IQSEC2;refGene_tx=XM_023248846.1
X 46253770 46253866
refGene_gn=IQSEC2;refGene_tx=XM_006943747.3;refGene_gn=IQSEC2;refGene_tx=XM_019823838.2;refGene_gn=IQSEC2;refGene_tx=XM_019823839.2
X 46255811 46255837
refGene_gn=IQSEC2;refGene_tx=XM_006943747.3;refGene_gn=IQSEC2;refGene_tx=XM_011291746.2
X 46266293 46266323
refGene_gn=IQSEC2;refGene_tx=XM_019823838.2;refGene_gn=IQSEC2;refGene_tx=XM_019823839.2;refGene_gn=IQSEC2;refGene_tx=XM_023248845.1
X 46289530 46290237
refGene_gn=IQSEC2;refGene_tx=XM_019823838.2;refGene_gn=IQSEC2;refGene_tx=XM_019823839.2;refGene_gn=IQSEC2;refGene_tx=XM_023248845.1;refGene_gn=IQSEC2;refGene_tx=XM_023248846.1
X 46334501 46334508 refGene_gn=SMC1A;refGene_tx=XM_011291750.3
X 46334825 46334885
refGene_gn=SMC1A;refGene_tx=XM_011291748.3;refGene_gn=SMC1A;refGene_tx=XM_011291749.3;refGene_gn=SMC1A;refGene_tx=XM_019823840.2
X 46346245 46346329 refGene_gn=SMC1A;refGene_tx=XM_004000536.5
X 46346770 46346881
refGene_gn=SMC1A;refGene_tx=XM_004000536.5;refGene_gn=SMC1A;refGene_tx=XM_011291748.3
X 46347164 46347234
refGene_gn=SMC1A;refGene_tx=XM_004000536.5;refGene_gn=SMC1A;refGene_tx=XM_011291748.3;refGene_gn=SMC1A;refGene_tx=XM_019823840.2
X 46348364 46348516
refGene_gn=SMC1A;refGene_tx=XM_004000536.5;refGene_gn=SMC1A;refGene_tx=XM_011291748.3;refGene_gn=SMC1A;refGene_tx=XM_011291749.3;refGene_gn=SMC1A;refGene_tx=XM_011291750.3;refGene_gn=SMC1A;refGene_tx=XM_019823840.2
X 46348643 46348798
refGene_gn=SMC1A;refGene_tx=XM_004000536.5;refGene_gn=SMC1A;refGene_tx=XM_011291748.3;refGene_gn=SMC1A;refGene_tx=XM_011291749.3;refGene_gn=SMC1A;refGene_tx=XM_011291750.3;refGene_gn=SMC1A;refGene_tx=XM_019823840.2
X 46349454 46349611
refGene_gn=SMC1A;refGene_tx=XM_004000536.5;refGene_gn=SMC1A;refGene_tx=XM_011291748.3;refGene_gn=SMC1A;refGene_tx=XM_011291749.3;refGene_gn=SMC1A;refGene_tx=XM_011291750.3;refGene_gn=SMC1A;refGene_tx=XM_0198238

40.2
X	46359035 46359146
refGene_gn=SMC1A;refGene_tx=XM_004000536.5;refGene_gn=SMC1A;refGene_tx
=XM_011291748.3;refGene_gn=SMC1A;refGene_tx=XM_011291749.3;refGene_gn=
SMC1A;refGene_tx=XM_011291750.3;refGene_gn=SMC1A;refGene_tx=XM_0198238
40.2
X	46360201 46360355
refGene_gn=SMC1A;refGene_tx=XM_004000536.5;refGene_gn=SMC1A;refGene_tx
=XM_011291748.3;refGene_gn=SMC1A;refGene_tx=XM_011291749.3;refGene_gn=
SMC1A;refGene_tx=XM_011291750.3;refGene_gn=SMC1A;refGene_tx=XM_0198238
40.2
X	46360443 46360589
refGene_gn=SMC1A;refGene_tx=XM_004000536.5;refGene_gn=SMC1A;refGene_tx
=XM_011291748.3;refGene_gn=SMC1A;refGene_tx=XM_011291749.3;refGene_gn=
SMC1A;refGene_tx=XM_011291750.3;refGene_gn=SMC1A;refGene_tx=XM_0198238
40.2
X	46362587 46362729
refGene_gn=SMC1A;refGene_tx=XM_004000536.5;refGene_gn=SMC1A;refGene_tx
=XM_011291748.3;refGene_gn=SMC1A;refGene_tx=XM_011291749.3;refGene_gn=
SMC1A;refGene_tx=XM_011291750.3;refGene_gn=SMC1A;refGene_tx=XM_0198238
40.2
X	46365690 46365797
refGene_gn=SMC1A;refGene_tx=XM_004000536.5;refGene_gn=SMC1A;refGene_tx
=XM_011291748.3;refGene_gn=SMC1A;refGene_tx=XM_011291749.3;refGene_gn=
SMC1A;refGene_tx=XM_011291750.3;refGene_gn=SMC1A;refGene_tx=XM_0198238
40.2
X	46365903 46366020
refGene_gn=SMC1A;refGene_tx=XM_004000536.5;refGene_gn=SMC1A;refGene_tx
=XM_011291748.3;refGene_gn=SMC1A;refGene_tx=XM_011291749.3;refGene_gn=
SMC1A;refGene_tx=XM_011291750.3;refGene_gn=SMC1A;refGene_tx=XM_0198238
40.2
X	46366825 46366963
refGene_gn=SMC1A;refGene_tx=XM_004000536.5;refGene_gn=SMC1A;refGene_tx
=XM_011291748.3;refGene_gn=SMC1A;refGene_tx=XM_011291749.3;refGene_gn=
SMC1A;refGene_tx=XM_011291750.3;refGene_gn=SMC1A;refGene_tx=XM_0198238
40.2
X	46367062 46367209
refGene_gn=SMC1A;refGene_tx=XM_004000536.5;refGene_gn=SMC1A;refGene_tx
=XM_011291748.3;refGene_gn=SMC1A;refGene_tx=XM_011291749.3;refGene_gn=
SMC1A;refGene_tx=XM_011291750.3;refGene_gn=SMC1A;refGene_tx=XM_0198238
40.2
X	46367306 46367486
refGene_gn=SMC1A;refGene_tx=XM_004000536.5;refGene_gn=SMC1A;refGene_tx
=XM_011291748.3;refGene_gn=SMC1A;refGene_tx=XM_011291749.3;refGene_gn=
SMC1A;refGene_tx=XM_011291750.3;refGene_gn=SMC1A;refGene_tx=XM_0198238
40.2
X	46367578 46367764
refGene_gn=SMC1A;refGene_tx=XM_004000536.5;refGene_gn=SMC1A;refGene_tx
=XM_011291748.3;refGene_gn=SMC1A;refGene_tx=XM_011291749.3;refGene_gn=
SMC1A;refGene_tx=XM_011291750.3;refGene_gn=SMC1A;refGene_tx=XM_0198238

40.2
X	46368823	46369031
refGene_gn=SMC1A;refGene_tx=XM_004000536.5;refGene_gn=SMC1A;refGene_tx
=XM_011291748.3;refGene_gn=SMC1A;refGene_tx=XM_011291749.3;refGene_gn=
SMC1A;refGene_tx=XM_011291750.3;refGene_gn=SMC1A;refGene_tx=XM_0198238
40.2
X	46369186	46369269
refGene_gn=SMC1A;refGene_tx=XM_004000536.5;refGene_gn=SMC1A;refGene_tx
=XM_011291748.3;refGene_gn=SMC1A;refGene_tx=XM_011291749.3;refGene_gn=
SMC1A;refGene_tx=XM_011291750.3;refGene_gn=SMC1A;refGene_tx=XM_0198238
40.2
X	46369954	46370095
refGene_gn=SMC1A;refGene_tx=XM_004000536.5;refGene_gn=SMC1A;refGene_tx
=XM_011291748.3;refGene_gn=SMC1A;refGene_tx=XM_011291749.3;refGene_gn=
SMC1A;refGene_tx=XM_011291750.3;refGene_gn=SMC1A;refGene_tx=XM_0198238
40.2
X	46370177	46370436
refGene_gn=SMC1A;refGene_tx=XM_004000536.5;refGene_gn=SMC1A;refGene_tx
=XM_011291748.3;refGene_gn=SMC1A;refGene_tx=XM_011291749.3;refGene_gn=
SMC1A;refGene_tx=XM_011291750.3;refGene_gn=SMC1A;refGene_tx=XM_0198238
40.2
X	46371094	46371333
refGene_gn=SMC1A;refGene_tx=XM_004000536.5;refGene_gn=SMC1A;refGene_tx
=XM_011291748.3;refGene_gn=SMC1A;refGene_tx=XM_011291749.3;refGene_gn=
SMC1A;refGene_tx=XM_011291750.3;refGene_gn=SMC1A;refGene_tx=XM_0198238
40.2
X	46371426	46371630
refGene_gn=SMC1A;refGene_tx=XM_004000536.5;refGene_gn=SMC1A;refGene_tx
=XM_011291748.3;refGene_gn=SMC1A;refGene_tx=XM_011291749.3;refGene_gn=
SMC1A;refGene_tx=XM_011291750.3;refGene_gn=SMC1A;refGene_tx=XM_0198238
40.2
X	46372612	46372725
refGene_gn=SMC1A;refGene_tx=XM_004000536.5;refGene_gn=SMC1A;refGene_tx
=XM_011291748.3;refGene_gn=SMC1A;refGene_tx=XM_011291749.3;refGene_gn=
SMC1A;refGene_tx=XM_011291750.3;refGene_gn=SMC1A;refGene_tx=XM_0198238
40.2
X	46372835	46373024
refGene_gn=SMC1A;refGene_tx=XM_004000536.5;refGene_gn=SMC1A;refGene_tx
=XM_011291748.3;refGene_gn=SMC1A;refGene_tx=XM_011291749.3;refGene_gn=
SMC1A;refGene_tx=XM_011291750.3;refGene_gn=SMC1A;refGene_tx=XM_0198238
40.2
X	46377053	46377162
refGene_gn=SMC1A;refGene_tx=XM_004000536.5;refGene_gn=SMC1A;refGene_tx
=XM_011291748.3;refGene_gn=SMC1A;refGene_tx=XM_011291749.3;refGene_gn=
SMC1A;refGene_tx=XM_011291750.3;refGene_gn=SMC1A;refGene_tx=XM_0198238
40.2
X	46385247	46385361
refGene_gn=RIBC1;refGene_tx=XM_006943748.3;refGene_gn=RIBC1;refGene_tx
=XM_019823842.2
X	46386697	46386779

refGene_gn=RIBC1;refGene_tx=XM_006943748.3;refGene_gn=RIBC1;refGene_tx=XM_019823842.2
X 46386982 46387327
refGene_gn=RIBC1;refGene_tx=XM_006943748.3;refGene_gn=RIBC1;refGene_tx=XM_019823842.2
X 46391165 46391284
refGene_gn=RIBC1;refGene_tx=XM_006943748.3;refGene_gn=RIBC1;refGene_tx=XM_019823842.2
X 46391723 46392118 refGene_gn=RIBC1;refGene_tx=XM_006943748.3
X 46392219 46392414
refGene_gn=RIBC1;refGene_tx=XM_006943748.3;refGene_gn=RIBC1;refGene_tx=XM_019823842.2
X 46392730 46392921 refGene_gn=HSD17B10;refGene_tx=XM_004000538.4
X 46393324 46393433 refGene_gn=HSD17B10;refGene_tx=XM_004000538.4
X 46393514 46393643 refGene_gn=HSD17B10;refGene_tx=XM_004000538.4
X 46393755 46393920 refGene_gn=HSD17B10;refGene_tx=XM_004000538.4
X 46394515 46394680 refGene_gn=HSD17B10;refGene_tx=XM_004000538.4
X 46395000 46395027 refGene_gn=HSD17B10;refGene_tx=XM_004000538.4
X 46477992 46478095 refGene_gn=HUWE1;refGene_tx=XM_019823850.2
X 46478682 46478873 refGene_gn=HUWE1;refGene_tx=XM_019823850.2
X 46479161 46479343 refGene_gn=HUWE1;refGene_tx=XM_019823850.2
X 46479643 46479761 refGene_gn=HUWE1;refGene_tx=XM_019823850.2
X 46480439 46480545 refGene_gn=HUWE1;refGene_tx=XM_019823850.2
X 46480660 46480948 refGene_gn=HUWE1;refGene_tx=XM_019823850.2
X 46481894 46482035 refGene_gn=HUWE1;refGene_tx=XM_019823850.2
X 46482842 46482960 refGene_gn=HUWE1;refGene_tx=XM_019823850.2
X 46483269 46483515 refGene_gn=HUWE1;refGene_tx=XM_019823850.2
X 46484174 46484330 refGene_gn=HUWE1;refGene_tx=XM_019823850.2
X 46484883 46484980 refGene_gn=HUWE1;refGene_tx=XM_019823850.2
X 46486262 46486390 refGene_gn=HUWE1;refGene_tx=XM_019823850.2
X 46486992 46487195 refGene_gn=HUWE1;refGene_tx=XM_019823850.2
X 46487472 46487605 refGene_gn=HUWE1;refGene_tx=XM_019823850.2
X 46488845 46489002 refGene_gn=HUWE1;refGene_tx=XM_019823850.2
X 46489114 46489236 refGene_gn=HUWE1;refGene_tx=XM_019823850.2
X 46489925 46490529 refGene_gn=HUWE1;refGene_tx=XM_019823850.2
X 46491215 46491762 refGene_gn=HUWE1;refGene_tx=XM_019823850.2
X 46492511 46492614 refGene_gn=HUWE1;refGene_tx=XM_019823850.2
X 46492752 46493041 refGene_gn=HUWE1;refGene_tx=XM_019823850.2
X 46493116 46493331 refGene_gn=HUWE1;refGene_tx=XM_019823850.2
X 46493858 46493989 refGene_gn=HUWE1;refGene_tx=XM_019823850.2
X 46494189 46494445 refGene_gn=HUWE1;refGene_tx=XM_019823850.2
X 46495511 46495796 refGene_gn=HUWE1;refGene_tx=XM_019823850.2
X 46497740 46497786 refGene_gn=HUWE1;refGene_tx=XM_019823850.2
X 46498943 46499098 refGene_gn=HUWE1;refGene_tx=XM_019823850.2
X 46499257 46499347 refGene_gn=HUWE1;refGene_tx=XM_019823850.2
X 46499631 46499810 refGene_gn=HUWE1;refGene_tx=XM_019823850.2
X 46500442 46500671 refGene_gn=HUWE1;refGene_tx=XM_019823850.2
X 46502018 46502187 refGene_gn=HUWE1;refGene_tx=XM_019823850.2
X 46502374 46502508 refGene_gn=HUWE1;refGene_tx=XM_019823850.2
X 46503096 46503195 refGene_gn=HUWE1;refGene_tx=XM_019823850.2

| | | | |
|---|---|---|---|
| X | 46504029 | 46504105 | refGene_gn=HUWE1;refGene_tx=XM_019823850.2 |
| X | 46504862 | 46505011 | refGene_gn=HUWE1;refGene_tx=XM_019823850.2 |
| X | 46505364 | 46505537 | refGene_gn=HUWE1;refGene_tx=XM_019823850.2 |
| X | 46508118 | 46508301 | refGene_gn=HUWE1;refGene_tx=XM_019823850.2 |
| X | 46508745 | 46508957 | refGene_gn=HUWE1;refGene_tx=XM_019823850.2 |
| X | 46512470 | 46512685 | refGene_gn=HUWE1;refGene_tx=XM_019823850.2 |
| X | 46513659 | 46513726 | refGene_gn=HUWE1;refGene_tx=XM_019823850.2 |
| X | 46514141 | 46514287 | refGene_gn=HUWE1;refGene_tx=XM_019823850.2 |
| X | 46515369 | 46515537 | refGene_gn=HUWE1;refGene_tx=XM_019823850.2 |
| X | 46516760 | 46516956 | refGene_gn=HUWE1;refGene_tx=XM_019823850.2 |
| X | 46519830 | 46520189 | refGene_gn=HUWE1;refGene_tx=XM_019823850.2 |
| X | 46520465 | 46520625 | refGene_gn=HUWE1;refGene_tx=XM_019823850.2 |
| X | 46521085 | 46521262 | refGene_gn=HUWE1;refGene_tx=XM_019823850.2 |
| X | 46521917 | 46521999 | refGene_gn=HUWE1;refGene_tx=XM_019823850.2 |
| X | 46522225 | 46522353 | refGene_gn=HUWE1;refGene_tx=XM_019823850.2 |
| X | 46523928 | 46524081 | refGene_gn=HUWE1;refGene_tx=XM_019823850.2 |
| X | 46526003 | 46526273 | refGene_gn=HUWE1;refGene_tx=XM_019823850.2 |
| X | 46526849 | 46526945 | refGene_gn=HUWE1;refGene_tx=XM_019823850.2 |
| X | 46527442 | 46527565 | refGene_gn=HUWE1;refGene_tx=XM_019823850.2 |
| X | 46528843 | 46529074 | refGene_gn=HUWE1;refGene_tx=XM_019823850.2 |
| X | 46529463 | 46529701 | refGene_gn=HUWE1;refGene_tx=XM_019823850.2 |
| X | 46530643 | 46530766 | refGene_gn=HUWE1;refGene_tx=XM_019823850.2 |
| X | 46531174 | 46531391 | refGene_gn=HUWE1;refGene_tx=XM_019823850.2 |
| X | 46533517 | 46533709 | refGene_gn=HUWE1;refGene_tx=XM_019823850.2 |
| X | 46543568 | 46543663 | refGene_gn=HUWE1;refGene_tx=XM_019823850.2 |
| X | 46544487 | 46544621 | refGene_gn=HUWE1;refGene_tx=XM_019823850.2 |
| X | 46545641 | 46545887 | refGene_gn=HUWE1;refGene_tx=XM_019823850.2 |
| X | 46549251 | 46549428 | refGene_gn=HUWE1;refGene_tx=XM_019823850.2 |
| X | 46550479 | 46550537 | refGene_gn=HUWE1;refGene_tx=XM_019823850.2 |
| X | 46553489 | 46553701 | refGene_gn=HUWE1;refGene_tx=XM_019823850.2 |
| X | 46554304 | 46554319 | refGene_gn=HUWE1;refGene_tx=XM_019823850.2 |
| X | 46554320 | 46554397 | refGene_gn=HUWE1;refGene_tx=XM_019823850.2 |
| X | 46562014 | 46562192 | refGene_gn=HUWE1;refGene_tx=XM_019823850.2 |
| X | 46562365 | 46562472 | refGene_gn=HUWE1;refGene_tx=XM_019823850.2 |
| X | 46571772 | 46571853 | refGene_gn=HUWE1;refGene_tx=XM_019823850.2 |
| X | 46572341 | 46572443 | refGene_gn=HUWE1;refGene_tx=XM_019823850.2 |
| X | 46572947 | 46573082 | refGene_gn=HUWE1;refGene_tx=XM_019823850.2 |
| X | 46574499 | 46574605 | refGene_gn=HUWE1;refGene_tx=XM_019823850.2 |
| X | 46574815 | 46574956 | refGene_gn=HUWE1;refGene_tx=XM_019823850.2 |
| X | 46575549 | 46575677 | refGene_gn=HUWE1;refGene_tx=XM_019823850.2 |
| X | 46575812 | 46575963 | refGene_gn=HUWE1;refGene_tx=XM_019823850.2 |
| X | 46576257 | 46576358 | refGene_gn=HUWE1;refGene_tx=XM_019823850.2 |
| X | 46577664 | 46577764 | refGene_gn=HUWE1;refGene_tx=XM_019823850.2 |
| X | 46578368 | 46578437 | refGene_gn=HUWE1;refGene_tx=XM_019823850.2 |
| X | 46578519 | 46578567 | refGene_gn=HUWE1;refGene_tx=XM_019823850.2 |
| X | 46579624 | 46579702 | refGene_gn=HUWE1;refGene_tx=XM_019823850.2 |
| X | 46582249 | 46582312 | refGene_gn=HUWE1;refGene_tx=XM_019823850.2 |
| X | 46603209 | 46603362 | refGene_gn=HUWE1;refGene_tx=XM_019823850.2 |
| X | 46604725 | 46604932 | refGene_gn=HUWE1;refGene_tx=XM_019823850.2 |
| X | 46605641 | 46605740 | refGene_gn=HUWE1;refGene_tx=XM_019823850.2 |

X 46611535 46611580 refGene_gn=HUWE1;refGene_tx=XM_019823850.2
X 46894567 46894656
refGene_gn=PHF8;refGene_tx=XM_004000543.5;refGene_gn=PHF8;refGene_tx=XM_006943754.3;refGene_gn=PHF8;refGene_tx=XM_006943755.4
X 46895590 46895927
refGene_gn=PHF8;refGene_tx=XM_004000543.5;refGene_gn=PHF8;refGene_tx=XM_006943754.3;refGene_gn=PHF8;refGene_tx=XM_006943755.4
X 46900323 46900433
refGene_gn=PHF8;refGene_tx=XM_004000543.5;refGene_gn=PHF8;refGene_tx=XM_006943754.3;refGene_gn=PHF8;refGene_tx=XM_006943755.4
X 46906595 46906691
refGene_gn=PHF8;refGene_tx=XM_004000543.5;refGene_gn=PHF8;refGene_tx=XM_006943754.3;refGene_gn=PHF8;refGene_tx=XM_006943755.4
X 46932464 46932778
refGene_gn=PHF8;refGene_tx=XM_004000543.5;refGene_gn=PHF8;refGene_tx=XM_006943754.3;refGene_gn=PHF8;refGene_tx=XM_006943755.4
X 46933379 46933513
refGene_gn=PHF8;refGene_tx=XM_004000543.5;refGene_gn=PHF8;refGene_tx=XM_006943754.3;refGene_gn=PHF8;refGene_tx=XM_006943755.4
X 46934208 46934294
refGene_gn=PHF8;refGene_tx=XM_004000543.5;refGene_gn=PHF8;refGene_tx=XM_006943754.3;refGene_gn=PHF8;refGene_tx=XM_006943755.4
X 46934586 46934765
refGene_gn=PHF8;refGene_tx=XM_004000543.5;refGene_gn=PHF8;refGene_tx=XM_006943754.3;refGene_gn=PHF8;refGene_tx=XM_006943755.4
X 46939341 46939445
refGene_gn=PHF8;refGene_tx=XM_004000543.5;refGene_gn=PHF8;refGene_tx=XM_006943754.3;refGene_gn=PHF8;refGene_tx=XM_006943755.4
X 46940205 46940508
refGene_gn=PHF8;refGene_tx=XM_004000543.5;refGene_gn=PHF8;refGene_tx=XM_006943754.3
X 46942358 46942448
refGene_gn=PHF8;refGene_tx=XM_004000543.5;refGene_gn=PHF8;refGene_tx=XM_006943754.3;refGene_gn=PHF8;refGene_tx=XM_006943755.4
X 46944915 46945007
refGene_gn=PHF8;refGene_tx=XM_004000543.5;refGene_gn=PHF8;refGene_tx=XM_006943754.3;refGene_gn=PHF8;refGene_tx=XM_006943755.4
X 46946127 46946234
refGene_gn=PHF8;refGene_tx=XM_004000543.5;refGene_gn=PHF8;refGene_tx=XM_006943754.3;refGene_gn=PHF8;refGene_tx=XM_006943755.4
X 46946601 46946689
refGene_gn=PHF8;refGene_tx=XM_004000543.5;refGene_gn=PHF8;refGene_tx=XM_006943754.3;refGene_gn=PHF8;refGene_tx=XM_006943755.4
X 46954209 46954372
refGene_gn=PHF8;refGene_tx=XM_004000543.5;refGene_gn=PHF8;refGene_tx=XM_006943754.3;refGene_gn=PHF8;refGene_tx=XM_006943755.4
X 46956875 46957062
refGene_gn=PHF8;refGene_tx=XM_004000543.5;refGene_gn=PHF8;refGene_tx=XM_006943754.3;refGene_gn=PHF8;refGene_tx=XM_006943755.4
X 46958481 46958623

refGene_gn=PHF8;refGene_tx=XM_004000543.5;refGene_gn=PHF8;refGene_tx=XM_006943754.3;refGene_gn=PHF8;refGene_tx=XM_006943755.4
X	46959466	46959627	refGene_gn=PHF8;refGene_tx=XM_004000543.5;refGene_gn=PHF8;refGene_tx=XM_006943754.3;refGene_gn=PHF8;refGene_tx=XM_006943755.4
X	46965121	46965230	refGene_gn=PHF8;refGene_tx=XM_004000543.5;refGene_gn=PHF8;refGene_tx=XM_006943754.3;refGene_gn=PHF8;refGene_tx=XM_006943755.4
X	46965605	46965691	refGene_gn=PHF8;refGene_tx=XM_004000543.5;refGene_gn=PHF8;refGene_tx=XM_006943754.3;refGene_gn=PHF8;refGene_tx=XM_006943755.4
X	46987917	46988015	refGene_gn=PHF8;refGene_tx=XM_004000543.5;refGene_gn=PHF8;refGene_tx=XM_006943754.3;refGene_gn=PHF8;refGene_tx=XM_006943755.4
X	47028359	47028614	refGene_gn=FAM120C;refGene_tx=XM_006943756.4
X	47030278	47030336	refGene_gn=FAM120C;refGene_tx=XM_006943756.4
X	47030541	47030680	refGene_gn=FAM120C;refGene_tx=XM_006943756.4
X	47032810	47033012	refGene_gn=FAM120C;refGene_tx=XM_006943756.4
X	47034172	47034382	refGene_gn=FAM120C;refGene_tx=XM_006943756.4
X	47036225	47036340	refGene_gn=FAM120C;refGene_tx=XM_006943756.4
X	47067079	47067329	refGene_gn=FAM120C;refGene_tx=XM_006943756.4
X	47083046	47083218	refGene_gn=FAM120C;refGene_tx=XM_006943756.4
X	47083866	47084140	refGene_gn=FAM120C;refGene_tx=XM_006943756.4
X	47085070	47085351	refGene_gn=FAM120C;refGene_tx=XM_006943756.4
X	47085782	47085859	refGene_gn=FAM120C;refGene_tx=XM_006943756.4
X	47086425	47086528	refGene_gn=FAM120C;refGene_tx=XM_006943756.4
X	47103041	47103170	refGene_gn=FAM120C;refGene_tx=XM_006943756.4
X	47111101	47111184	refGene_gn=FAM120C;refGene_tx=XM_006943756.4
X	47112513	47112760	refGene_gn=FAM120C;refGene_tx=XM_006943756.4
X	47137258	47137276	refGene_gn=FAM120C;refGene_tx=XM_011291753.2
X	47145234	47145915	refGene_gn=FAM120C;refGene_tx=XM_006943756.4;refGene_gn=FAM120C;refGene_tx=XM_011291753.2
X	47158996	47159326	refGene_gn=WNK3;refGene_tx=XM_004000545.3;refGene_gn=WNK3;refGene_tx=XM_011291755.3;refGene_gn=WNK3;refGene_tx=XM_019823858.1;refGene_gn=WNK3;refGene_tx=XM_019823859.1;refGene_gn=WNK3;refGene_tx=XM_019823860.2
X	47163135	47163338	refGene_gn=WNK3;refGene_tx=XM_004000545.3;refGene_gn=WNK3;refGene_tx=XM_011291755.3;refGene_gn=WNK3;refGene_tx=XM_019823858.1;refGene_gn=WNK3;refGene_tx=XM_019823859.1;refGene_gn=WNK3;refGene_tx=XM_019823860.2
X	47186390	47186602	refGene_gn=WNK3;refGene_tx=XM_004000545.3;refGene_gn=WNK3;refGene_tx=XM_011291755.3;refGene_gn=WNK3;refGene_tx=XM_019823858.1;refGene_gn=WNK3;refGene_tx=XM_019823859.1;refGene_gn=WNK3;refGene_tx=XM_019823860.2
X	47189791	47190414	refGene_gn=WNK3;refGene_tx=XM_004000545.3;refGene_gn=WNK3;refGene_tx=XM_011291755.3;refGene_gn=WNK3;refGene_tx=XM_019823858.1;refGene_gn=WNK3;refGene_tx=XM_019823859.1;refGene_gn=WNK3;refGene_tx=XM_019823860.2
X	47190951	47191082

refGene_gn=WNK3;refGene_tx=XM_004000545.3;refGene_gn=WNK3;refGene_tx=XM_011291755.3;refGene_gn=WNK3;refGene_tx=XM_019823858.1;refGene_gn=WNK3;refGene_tx=XM_019823859.1;refGene_gn=WNK3;refGene_tx=XM_019823860.2
X 47191480 47191712
refGene_gn=WNK3;refGene_tx=XM_004000545.3;refGene_gn=WNK3;refGene_tx=XM_011291755.3;refGene_gn=WNK3;refGene_tx=XM_019823858.1;refGene_gn=WNK3;refGene_tx=XM_019823859.1;refGene_gn=WNK3;refGene_tx=XM_019823860.2
X 47196733 47197677
refGene_gn=WNK3;refGene_tx=XM_004000545.3;refGene_gn=WNK3;refGene_tx=XM_011291755.3;refGene_gn=WNK3;refGene_tx=XM_019823858.1;refGene_gn=WNK3;refGene_tx=XM_019823859.1;refGene_gn=WNK3;refGene_tx=XM_019823860.2
X 47198033 47198168
refGene_gn=WNK3;refGene_tx=XM_004000545.3;refGene_gn=WNK3;refGene_tx=XM_011291755.3;refGene_gn=WNK3;refGene_tx=XM_019823858.1;refGene_gn=WNK3;refGene_tx=XM_019823859.1;refGene_gn=WNK3;refGene_tx=XM_019823860.2
X 47199396 47199448
refGene_gn=WNK3;refGene_tx=XM_004000545.3;refGene_gn=WNK3;refGene_tx=XM_011291755.3;refGene_gn=WNK3;refGene_tx=XM_019823858.1;refGene_gn=WNK3;refGene_tx=XM_019823859.1;refGene_gn=WNK3;refGene_tx=XM_019823860.2
X 47199529 47199685
refGene_gn=WNK3;refGene_tx=XM_004000545.3;refGene_gn=WNK3;refGene_tx=XM_011291755.3;refGene_gn=WNK3;refGene_tx=XM_019823858.1;refGene_gn=WNK3;refGene_tx=XM_019823859.1;refGene_gn=WNK3;refGene_tx=XM_019823860.2
X 47200929 47201046
refGene_gn=WNK3;refGene_tx=XM_004000545.3;refGene_gn=WNK3;refGene_tx=XM_011291755.3;refGene_gn=WNK3;refGene_tx=XM_019823858.1;refGene_gn=WNK3;refGene_tx=XM_019823859.1;refGene_gn=WNK3;refGene_tx=XM_019823860.2
X 47219033 47219181
refGene_gn=WNK3;refGene_tx=XM_004000545.3;refGene_gn=WNK3;refGene_tx=XM_011291755.3;refGene_gn=WNK3;refGene_tx=XM_019823858.1;refGene_gn=WNK3;refGene_tx=XM_019823859.1;refGene_gn=WNK3;refGene_tx=XM_019823860.2
X 47234521 47234586
refGene_gn=WNK3;refGene_tx=XM_004000545.3;refGene_gn=WNK3;refGene_tx=XM_011291755.3;refGene_gn=WNK3;refGene_tx=XM_019823858.1;refGene_gn=WNK3;refGene_tx=XM_019823859.1;refGene_gn=WNK3;refGene_tx=XM_019823860.2
X 47281664 47281814
refGene_gn=WNK3;refGene_tx=XM_004000545.3;refGene_gn=WNK3;refGene_tx=XM_011291755.3;refGene_gn=WNK3;refGene_tx=XM_019823858.1;refGene_gn=WNK3;refGene_tx=XM_019823859.1;refGene_gn=WNK3;refGene_tx=XM_019823860.2
X 47281909 47282103
refGene_gn=WNK3;refGene_tx=XM_004000545.3;refGene_gn=WNK3;refGene_tx=XM_011291755.3;refGene_gn=WNK3;refGene_tx=XM_019823858.1;refGene_gn=WNK3;refGene_tx=XM_019823859.1;refGene_gn=WNK3;refGene_tx=XM_019823860.2
X 47282937 47283232
refGene_gn=WNK3;refGene_tx=XM_004000545.3;refGene_gn=WNK3;refGene_tx=XM_011291755.3;refGene_gn=WNK3;refGene_tx=XM_019823858.1;refGene_gn=WNK3;refGene_tx=XM_019823859.1;refGene_gn=WNK3;refGene_tx=XM_019823860.2
X 47290323 47290543
refGene_gn=WNK3;refGene_tx=XM_004000545.3;refGene_gn=WNK3;refGene_tx=XM_011291755.3;refGene_gn=WNK3;refGene_tx=XM_019823858.1;refGene_gn=WNK

3;refGene_tx=XM_019823859.1;refGene_gn=WNK3;refGene_tx=XM_019823860.2
X	47294217	47294306
refGene_gn=WNK3;refGene_tx=XM_004000545.3;refGene_gn=WNK3;refGene_tx=XM_011291755.3;refGene_gn=WNK3;refGene_tx=XM_019823858.1;refGene_gn=WNK3;refGene_tx=XM_019823859.1;refGene_gn=WNK3;refGene_tx=XM_019823860.2
X	47297488	47297646
refGene_gn=WNK3;refGene_tx=XM_004000545.3;refGene_gn=WNK3;refGene_tx=XM_011291755.3;refGene_gn=WNK3;refGene_tx=XM_019823858.1;refGene_gn=WNK3;refGene_tx=XM_019823859.1;refGene_gn=WNK3;refGene_tx=XM_019823860.2
X	47298568	47298789
refGene_gn=WNK3;refGene_tx=XM_004000545.3;refGene_gn=WNK3;refGene_tx=XM_011291755.3;refGene_gn=WNK3;refGene_tx=XM_019823858.1;refGene_gn=WNK3;refGene_tx=XM_019823859.1;refGene_gn=WNK3;refGene_tx=XM_019823860.2
X	47299555	47299728
refGene_gn=WNK3;refGene_tx=XM_004000545.3;refGene_gn=WNK3;refGene_tx=XM_011291755.3;refGene_gn=WNK3;refGene_tx=XM_019823858.1;refGene_gn=WNK3;refGene_tx=XM_019823859.1;refGene_gn=WNK3;refGene_tx=XM_019823860.2
X	47317941	47318478
refGene_gn=WNK3;refGene_tx=XM_004000545.3;refGene_gn=WNK3;refGene_tx=XM_011291755.3;refGene_gn=WNK3;refGene_tx=XM_019823858.1;refGene_gn=WNK3;refGene_tx=XM_019823859.1;refGene_gn=WNK3;refGene_tx=XM_019823860.2
X	47433896	47433980	refGene_gn=TSR2;refGene_tx=XM_004000546.5
X	47434142	47434233	refGene_gn=TSR2;refGene_tx=XM_004000546.5
X	47436978	47437070	refGene_gn=TSR2;refGene_tx=XM_004000546.5
X	47437625	47437796	refGene_gn=TSR2;refGene_tx=XM_004000546.5
X	47438018	47438153	refGene_gn=TSR2;refGene_tx=XM_004000546.5
X	47439641	47439947
refGene_gn=FGD1;refGene_tx=XM_004000549.5;refGene_gn=FGD1;refGene_tx=XM_011291756.3
X	47440491	47440635	refGene_gn=FGD1;refGene_tx=XM_004000549.5
X	47441710	47441872
refGene_gn=FGD1;refGene_tx=XM_004000549.5;refGene_gn=FGD1;refGene_tx=XM_011291756.3
X	47442049	47442175
refGene_gn=FGD1;refGene_tx=XM_004000549.5;refGene_gn=FGD1;refGene_tx=XM_011291756.3
X	47442544	47442646
refGene_gn=FGD1;refGene_tx=XM_004000549.5;refGene_gn=FGD1;refGene_tx=XM_011291756.3
X	47443152	47443183
refGene_gn=FGD1;refGene_tx=XM_004000549.5;refGene_gn=FGD1;refGene_tx=XM_011291756.3
X	47447783	47447863
refGene_gn=FGD1;refGene_tx=XM_004000549.5;refGene_gn=FGD1;refGene_tx=XM_011291756.3
X	47448026	47448119
refGene_gn=FGD1;refGene_tx=XM_004000549.5;refGene_gn=FGD1;refGene_tx=XM_011291756.3
X	47448474	47448621
refGene_gn=FGD1;refGene_tx=XM_004000549.5;refGene_gn=FGD1;refGene_tx=X

M_011291756.3
X 47448757 47448816
refGene_gn=FGD1;refGene_tx=XM_004000549.5;refGene_gn=FGD1;refGene_tx=XM_011291756.3
X 47457962 47458101
refGene_gn=FGD1;refGene_tx=XM_004000549.5;refGene_gn=FGD1;refGene_tx=XM_011291756.3
X 47458206 47458363
refGene_gn=FGD1;refGene_tx=XM_004000549.5;refGene_gn=FGD1;refGene_tx=XM_011291756.3
X 47459655 47459804
refGene_gn=FGD1;refGene_tx=XM_004000549.5;refGene_gn=FGD1;refGene_tx=XM_011291756.3
X 47460499 47460589
refGene_gn=FGD1;refGene_tx=XM_004000549.5;refGene_gn=FGD1;refGene_tx=XM_011291756.3
X 47461883 47462322
refGene_gn=FGD1;refGene_tx=XM_004000549.5;refGene_gn=FGD1;refGene_tx=XM_011291756.3
X 47462448 47462626
refGene_gn=FGD1;refGene_tx=XM_004000549.5;refGene_gn=FGD1;refGene_tx=XM_011291756.3
X 47462934 47463108
refGene_gn=FGD1;refGene_tx=XM_004000549.5;refGene_gn=FGD1;refGene_tx=XM_011291756.3
X 47480162 47480469
refGene_gn=FGD1;refGene_tx=XM_004000549.5;refGene_gn=FGD1;refGene_tx=XM_011291756.3
X 47510709 47510728
refGene_gn=GNL3L;refGene_tx=XM_004000548.5;refGene_gn=GNL3L;refGene_tx=XM_006943757.4;refGene_gn=GNL3L;refGene_tx=XM_006943758.4;refGene_gn=GNL3L;refGene_tx=XM_006943759.4;refGene_gn=GNL3L;refGene_tx=XM_019823862.2
X 47517520 47517561
refGene_gn=GNL3L;refGene_tx=XM_004000548.5;refGene_gn=GNL3L;refGene_tx=XM_006943757.4;refGene_gn=GNL3L;refGene_tx=XM_006943758.4;refGene_gn=GNL3L;refGene_tx=XM_006943759.4;refGene_gn=GNL3L;refGene_tx=XM_019823862.2
X 47518442 47518550
refGene_gn=GNL3L;refGene_tx=XM_004000548.5;refGene_gn=GNL3L;refGene_tx=XM_006943757.4;refGene_gn=GNL3L;refGene_tx=XM_006943758.4;refGene_gn=GNL3L;refGene_tx=XM_006943759.4
X 47519293 47519410
refGene_gn=GNL3L;refGene_tx=XM_004000548.5;refGene_gn=GNL3L;refGene_tx=XM_006943757.4;refGene_gn=GNL3L;refGene_tx=XM_006943759.4;refGene_gn=GNL3L;refGene_tx=XM_019823862.2
X 47520162 47520246
refGene_gn=GNL3L;refGene_tx=XM_004000548.5;refGene_gn=GNL3L;refGene_tx=XM_006943757.4;refGene_gn=GNL3L;refGene_tx=XM_006943758.4;refGene_gn=GNL3L;refGene_tx=XM_006943759.4;refGene_gn=GNL3L;refGene_tx=XM_0198238

62.2
X	47520413 47520549
refGene_gn=GNL3L;refGene_tx=XM_004000548.5;refGene_gn=GNL3L;refGene_tx=XM_006943757.4;refGene_gn=GNL3L;refGene_tx=XM_006943758.4;refGene_gn=GNL3L;refGene_tx=XM_006943759.4;refGene_gn=GNL3L;refGene_tx=XM_019823862.2
X	47521383 47521487
refGene_gn=GNL3L;refGene_tx=XM_004000548.5;refGene_gn=GNL3L;refGene_tx=XM_006943757.4;refGene_gn=GNL3L;refGene_tx=XM_006943758.4;refGene_gn=GNL3L;refGene_tx=XM_006943759.4;refGene_gn=GNL3L;refGene_tx=XM_019823862.2
X	47523221 47523366
refGene_gn=GNL3L;refGene_tx=XM_004000548.5;refGene_gn=GNL3L;refGene_tx=XM_006943757.4;refGene_gn=GNL3L;refGene_tx=XM_006943758.4;refGene_gn=GNL3L;refGene_tx=XM_006943759.4;refGene_gn=GNL3L;refGene_tx=XM_019823862.2
X	47525484 47525572
refGene_gn=GNL3L;refGene_tx=XM_004000548.5;refGene_gn=GNL3L;refGene_tx=XM_006943757.4;refGene_gn=GNL3L;refGene_tx=XM_006943758.4;refGene_gn=GNL3L;refGene_tx=XM_006943759.4;refGene_gn=GNL3L;refGene_tx=XM_019823862.2
X	47526074 47526249
refGene_gn=GNL3L;refGene_tx=XM_004000548.5;refGene_gn=GNL3L;refGene_tx=XM_006943757.4;refGene_gn=GNL3L;refGene_tx=XM_006943758.4;refGene_gn=GNL3L;refGene_tx=XM_006943759.4;refGene_gn=GNL3L;refGene_tx=XM_019823862.2
X	47526335 47526478
refGene_gn=GNL3L;refGene_tx=XM_004000548.5;refGene_gn=GNL3L;refGene_tx=XM_006943757.4;refGene_gn=GNL3L;refGene_tx=XM_006943758.4;refGene_gn=GNL3L;refGene_tx=XM_006943759.4;refGene_gn=GNL3L;refGene_tx=XM_019823862.2
X	47526783 47526920
refGene_gn=GNL3L;refGene_tx=XM_004000548.5;refGene_gn=GNL3L;refGene_tx=XM_006943757.4;refGene_gn=GNL3L;refGene_tx=XM_006943758.4;refGene_gn=GNL3L;refGene_tx=XM_006943759.4;refGene_gn=GNL3L;refGene_tx=XM_019823862.2
X	47528737 47528865
refGene_gn=GNL3L;refGene_tx=XM_004000548.5;refGene_gn=GNL3L;refGene_tx=XM_006943757.4;refGene_gn=GNL3L;refGene_tx=XM_006943758.4;refGene_gn=GNL3L;refGene_tx=XM_006943759.4;refGene_gn=GNL3L;refGene_tx=XM_019823862.2
X	47532257 47532477
refGene_gn=GNL3L;refGene_tx=XM_004000548.5;refGene_gn=GNL3L;refGene_tx=XM_006943757.4;refGene_gn=GNL3L;refGene_tx=XM_006943758.4;refGene_gn=GNL3L;refGene_tx=XM_019823862.2
X	47534029 47534112
refGene_gn=GNL3L;refGene_tx=XM_004000548.5;refGene_gn=GNL3L;refGene_tx=XM_006943757.4;refGene_gn=GNL3L;refGene_tx=XM_006943758.4;refGene_gn=GNL3L;refGene_tx=XM_006943759.4;refGene_gn=GNL3L;refGene_tx=XM_019823862.2

X 47800377 47800452 refGene_gn=MAGED2;refGene_tx=XM_019823864.1
X 47800649 47801060
refGene_gn=MAGED2;refGene_tx=XM_006943761.2;refGene_gn=MAGED2;refGene_tx=XM_006943762.4;refGene_gn=MAGED2;refGene_tx=XM_006943763.3;refGene_gn=MAGED2;refGene_tx=XM_006943764.3;refGene_gn=MAGED2;refGene_tx=XM_019823863.1;refGene_gn=MAGED2;refGene_tx=XM_019823864.1;refGene_gn=MAGED2;refGene_tx=XM_023248944.1
X 47801450 47801565
refGene_gn=MAGED2;refGene_tx=XM_006943761.2;refGene_gn=MAGED2;refGene_tx=XM_006943762.4;refGene_gn=MAGED2;refGene_tx=XM_006943763.3;refGene_gn=MAGED2;refGene_tx=XM_006943764.3;refGene_gn=MAGED2;refGene_tx=XM_019823863.1;refGene_gn=MAGED2;refGene_tx=XM_019823864.1;refGene_gn=MAGED2;refGene_tx=XM_023248944.1
X 47802690 47802753
refGene_gn=MAGED2;refGene_tx=XM_006943761.2;refGene_gn=MAGED2;refGene_tx=XM_006943762.4;refGene_gn=MAGED2;refGene_tx=XM_006943763.3;refGene_gn=MAGED2;refGene_tx=XM_006943764.3;refGene_gn=MAGED2;refGene_tx=XM_019823863.1;refGene_gn=MAGED2;refGene_tx=XM_019823864.1;refGene_gn=MAGED2;refGene_tx=XM_023248944.1
X 47803080 47803123
refGene_gn=MAGED2;refGene_tx=XM_006943761.2;refGene_gn=MAGED2;refGene_tx=XM_006943762.4;refGene_gn=MAGED2;refGene_tx=XM_006943763.3;refGene_gn=MAGED2;refGene_tx=XM_006943764.3;refGene_gn=MAGED2;refGene_tx=XM_019823863.1;refGene_gn=MAGED2;refGene_tx=XM_019823864.1;refGene_gn=MAGED2;refGene_tx=XM_023248944.1
X 47803216 47803296
refGene_gn=MAGED2;refGene_tx=XM_006943761.2;refGene_gn=MAGED2;refGene_tx=XM_006943762.4;refGene_gn=MAGED2;refGene_tx=XM_006943763.3;refGene_gn=MAGED2;refGene_tx=XM_006943764.3;refGene_gn=MAGED2;refGene_tx=XM_019823863.1;refGene_gn=MAGED2;refGene_tx=XM_019823864.1;refGene_gn=MAGED2;refGene_tx=XM_023248944.1
X 47804002 47804097
refGene_gn=MAGED2;refGene_tx=XM_006943761.2;refGene_gn=MAGED2;refGene_tx=XM_006943762.4;refGene_gn=MAGED2;refGene_tx=XM_006943763.3;refGene_gn=MAGED2;refGene_tx=XM_006943764.3;refGene_gn=MAGED2;refGene_tx=XM_019823863.1;refGene_gn=MAGED2;refGene_tx=XM_019823864.1;refGene_gn=MAGED2;refGene_tx=XM_023248944.1
X 47804617 47804697
refGene_gn=MAGED2;refGene_tx=XM_006943761.2;refGene_gn=MAGED2;refGene_tx=XM_006943762.4;refGene_gn=MAGED2;refGene_tx=XM_006943763.3;refGene_gn=MAGED2;refGene_tx=XM_006943764.3;refGene_gn=MAGED2;refGene_tx=XM_019823863.1;refGene_gn=MAGED2;refGene_tx=XM_019823864.1;refGene_gn=MAGED2;refGene_tx=XM_023248944.1
X 47804954 47805018
refGene_gn=MAGED2;refGene_tx=XM_006943761.2;refGene_gn=MAGED2;refGene_tx=XM_006943762.4;refGene_gn=MAGED2;refGene_tx=XM_006943763.3;refGene_gn=MAGED2;refGene_tx=XM_006943764.3;refGene_gn=MAGED2;refGene_tx=XM_019823863.1;refGene_gn=MAGED2;refGene_tx=XM_019823864.1;refGene_gn=MAGED2;refGene_tx=XM_023248944.1
X 47805136 47805445

refGene_gn=MAGED2;refGene_tx=XM_006943761.2;refGene_gn=MAGED2;refGene_tx=XM_006943762.4;refGene_gn=MAGED2;refGene_tx=XM_006943763.3;refGene_gn=MAGED2;refGene_tx=XM_006943764.3;refGene_gn=MAGED2;refGene_tx=XM_019823863.1;refGene_gn=MAGED2;refGene_tx=XM_019823864.1;refGene_gn=MAGED2;refGene_tx=XM_023248944.1
X 47806047 47806503
refGene_gn=MAGED2;refGene_tx=XM_006943761.2;refGene_gn=MAGED2;refGene_tx=XM_006943762.4;refGene_gn=MAGED2;refGene_tx=XM_006943763.3;refGene_gn=MAGED2;refGene_tx=XM_006943764.3;refGene_gn=MAGED2;refGene_tx=XM_019823863.1;refGene_gn=MAGED2;refGene_tx=XM_019823864.1;refGene_gn=MAGED2;refGene_tx=XM_023248944.1
X 47806885 47806959
refGene_gn=MAGED2;refGene_tx=XM_006943761.2;refGene_gn=MAGED2;refGene_tx=XM_006943762.4;refGene_gn=MAGED2;refGene_tx=XM_006943763.3;refGene_gn=MAGED2;refGene_tx=XM_006943764.3;refGene_gn=MAGED2;refGene_tx=XM_019823863.1;refGene_gn=MAGED2;refGene_tx=XM_019823864.1;refGene_gn=MAGED2;refGene_tx=XM_023248944.1
X 47807240 47807246 refGene_gn=MAGED2;refGene_tx=XM_023248944.1
X 47807848 47808007
refGene_gn=MAGED2;refGene_tx=XM_019823863.1;refGene_gn=MAGED2;refGene_tx=XM_019823864.1
X 47931686 47931731
refGene_gn=LOC101096210;refGene_tx=XM_004000556.4
X 47932024 47933203
refGene_gn=LOC101096210;refGene_tx=XM_004000556.4
X 47933854 47933959
refGene_gn=LOC101096210;refGene_tx=XM_004000556.4
X 47934074 47934108
refGene_gn=LOC101096210;refGene_tx=XM_004000556.4
X 47934110 47934134
refGene_gn=LOC101096210;refGene_tx=XM_004000556.4
X 47934390 47934470
refGene_gn=LOC101096210;refGene_tx=XM_004000556.4
X 47934987 47935079
refGene_gn=LOC101096210;refGene_tx=XM_004000556.4
X 47935794 47935874
refGene_gn=LOC101096210;refGene_tx=XM_004000556.4
X 47935967 47936010
refGene_gn=LOC101096210;refGene_tx=XM_004000556.4
X 47936428 47936491
refGene_gn=LOC101096210;refGene_tx=XM_004000556.4
X 47937058 47937173
refGene_gn=LOC101096210;refGene_tx=XM_004000556.4
X 47938171 47940049
refGene_gn=LOC101096210;refGene_tx=XM_004000556.4
X 47942110 47942170
refGene_gn=PFKFB1;refGene_tx=XM_004000551.4;refGene_gn=PFKFB1;refGene_tx=XM_006943769.3;refGene_gn=PFKFB1;refGene_tx=XM_006943770.3;refGene_gn=PFKFB1;refGene_tx=XM_011291760.2;refGene_gn=PFKFB1;refGene_tx=XM_019823870.1;refGene_gn=PFKFB1;refGene_tx=XM_023248938.1;refGene_gn=PFKFB

1;refGene_tx=XM_023248939.1
X	47942526	47942591
refGene_gn=PFKFB1;refGene_tx=XM_004000551.4;refGene_gn=PFKFB1;refGene_tx=XM_006943769.3;refGene_gn=PFKFB1;refGene_tx=XM_006943770.3;refGene_gn=PFKFB1;refGene_tx=XM_011291760.2;refGene_gn=PFKFB1;refGene_tx=XM_019823870.1;refGene_gn=PFKFB1;refGene_tx=XM_023248938.1
X	47943646	47943709
refGene_gn=PFKFB1;refGene_tx=XM_004000551.4;refGene_gn=PFKFB1;refGene_tx=XM_006943769.3;refGene_gn=PFKFB1;refGene_tx=XM_006943770.3;refGene_gn=PFKFB1;refGene_tx=XM_011291760.2;refGene_gn=PFKFB1;refGene_tx=XM_019823870.1;refGene_gn=PFKFB1;refGene_tx=XM_023248938.1;refGene_gn=PFKFB1;refGene_tx=XM_023248939.1
X	47946393	47946523
refGene_gn=PFKFB1;refGene_tx=XM_004000551.4;refGene_gn=PFKFB1;refGene_tx=XM_006943769.3;refGene_gn=PFKFB1;refGene_tx=XM_006943770.3;refGene_gn=PFKFB1;refGene_tx=XM_011291760.2;refGene_gn=PFKFB1;refGene_tx=XM_019823870.1;refGene_gn=PFKFB1;refGene_tx=XM_023248938.1;refGene_gn=PFKFB1;refGene_tx=XM_023248939.1
X	47968768	47968873
refGene_gn=PFKFB1;refGene_tx=XM_004000551.4;refGene_gn=PFKFB1;refGene_tx=XM_006943769.3;refGene_gn=PFKFB1;refGene_tx=XM_006943770.3;refGene_gn=PFKFB1;refGene_tx=XM_011291760.2;refGene_gn=PFKFB1;refGene_tx=XM_019823870.1;refGene_gn=PFKFB1;refGene_tx=XM_023248938.1;refGene_gn=PFKFB1;refGene_tx=XM_023248939.1
X	47972657	47972804
refGene_gn=PFKFB1;refGene_tx=XM_004000551.4;refGene_gn=PFKFB1;refGene_tx=XM_006943769.3;refGene_gn=PFKFB1;refGene_tx=XM_006943770.3;refGene_gn=PFKFB1;refGene_tx=XM_011291760.2;refGene_gn=PFKFB1;refGene_tx=XM_019823870.1;refGene_gn=PFKFB1;refGene_tx=XM_023248938.1;refGene_gn=PFKFB1;refGene_tx=XM_023248939.1
X	47975429	47975637
refGene_gn=PFKFB1;refGene_tx=XM_004000551.4;refGene_gn=PFKFB1;refGene_tx=XM_006943769.3;refGene_gn=PFKFB1;refGene_tx=XM_006943770.3;refGene_gn=PFKFB1;refGene_tx=XM_011291760.2;refGene_gn=PFKFB1;refGene_tx=XM_019823870.1;refGene_gn=PFKFB1;refGene_tx=XM_023248938.1;refGene_gn=PFKFB1;refGene_tx=XM_023248939.1
X	47979499	47979621
refGene_gn=PFKFB1;refGene_tx=XM_004000551.4;refGene_gn=PFKFB1;refGene_tx=XM_006943769.3;refGene_gn=PFKFB1;refGene_tx=XM_006943770.3;refGene_gn=PFKFB1;refGene_tx=XM_011291760.2;refGene_gn=PFKFB1;refGene_tx=XM_019823870.1;refGene_gn=PFKFB1;refGene_tx=XM_023248938.1;refGene_gn=PFKFB1;refGene_tx=XM_023248939.1
X	47982271	47982328
refGene_gn=PFKFB1;refGene_tx=XM_004000551.4;refGene_gn=PFKFB1;refGene_tx=XM_006943769.3;refGene_gn=PFKFB1;refGene_tx=XM_006943770.3;refGene_gn=PFKFB1;refGene_tx=XM_011291760.2;refGene_gn=PFKFB1;refGene_tx=XM_019823870.1;refGene_gn=PFKFB1;refGene_tx=XM_023248938.1;refGene_gn=PFKFB1;refGene_tx=XM_023248939.1
X	47982838	47982913
refGene_gn=PFKFB1;refGene_tx=XM_004000551.4;refGene_gn=PFKFB1;refGene_

tx=XM_006943769.3;refGene_gn=PFKFB1;refGene_tx=XM_006943770.3;refGene_gn=PFKFB1;refGene_tx=XM_011291760.2;refGene_gn=PFKFB1;refGene_tx=XM_019823870.1;refGene_gn=PFKFB1;refGene_tx=XM_023248938.1;refGene_gn=PFKFB1;refGene_tx=XM_023248939.1
X 47983797 47983864
refGene_gn=PFKFB1;refGene_tx=XM_004000551.4;refGene_gn=PFKFB1;refGene_tx=XM_006943769.3;refGene_gn=PFKFB1;refGene_tx=XM_006943770.3;refGene_gn=PFKFB1;refGene_tx=XM_011291760.2;refGene_gn=PFKFB1;refGene_tx=XM_019823870.1;refGene_gn=PFKFB1;refGene_tx=XM_023248938.1;refGene_gn=PFKFB1;refGene_tx=XM_023248939.1
X 47984700 47984794
refGene_gn=PFKFB1;refGene_tx=XM_004000551.4;refGene_gn=PFKFB1;refGene_tx=XM_006943769.3;refGene_gn=PFKFB1;refGene_tx=XM_006943770.3;refGene_gn=PFKFB1;refGene_tx=XM_011291760.2;refGene_gn=PFKFB1;refGene_tx=XM_019823870.1;refGene_gn=PFKFB1;refGene_tx=XM_023248938.1;refGene_gn=PFKFB1;refGene_tx=XM_023248939.1
X 47989229 47989355
refGene_gn=PFKFB1;refGene_tx=XM_004000551.4;refGene_gn=PFKFB1;refGene_tx=XM_006943769.3;refGene_gn=PFKFB1;refGene_tx=XM_006943770.3;refGene_gn=PFKFB1;refGene_tx=XM_023248938.1;refGene_gn=PFKFB1;refGene_tx=XM_023248939.1
X 47990013 47990248
refGene_gn=PFKFB1;refGene_tx=XM_023248938.1;refGene_gn=PFKFB1;refGene_tx=XM_023248939.1
X 48013899 48013945
refGene_gn=PFKFB1;refGene_tx=XM_006943769.3;refGene_gn=PFKFB1;refGene_tx=XM_019823870.1
X 48073174 48073280 refGene_gn=PFKFB1;refGene_tx=XM_004000551.4
X 48078789 48078817
refGene_gn=PFKFB1;refGene_tx=XM_006943770.3;refGene_gn=PFKFB1;refGene_tx=XM_011291760.2
X 48080803 48080960
refGene_gn=APEX2;refGene_tx=XM_004000553.5;refGene_gn=APEX2;refGene_tx=XM_019823867.1
X 48082568 48082652
refGene_gn=APEX2;refGene_tx=XM_004000553.5;refGene_gn=APEX2;refGene_tx=XM_019823867.1
X 48084661 48084842
refGene_gn=APEX2;refGene_tx=XM_004000553.5;refGene_gn=APEX2;refGene_tx=XM_019823867.1
X 48085291 48085525
refGene_gn=APEX2;refGene_tx=XM_004000553.5;refGene_gn=APEX2;refGene_tx=XM_019823867.1;refGene_gn=APEX2;refGene_tx=XM_019823868.2
X 48086271 48086341
refGene_gn=APEX2;refGene_tx=XM_004000553.5;refGene_gn=APEX2;refGene_tx=XM_019823867.1;refGene_gn=APEX2;refGene_tx=XM_019823868.2
X 48089055 48089961
refGene_gn=APEX2;refGene_tx=XM_004000553.5;refGene_gn=APEX2;refGene_tx=XM_019823867.1;refGene_gn=APEX2;refGene_tx=XM_019823868.2
X 48091238 48091402

refGene_gn=ALAS2;refGene_tx=XM_004000554.4;refGene_gn=ALAS2;refGene_tx=XM_019823866.2
X 48095301 48095464
refGene_gn=ALAS2;refGene_tx=XM_004000554.4;refGene_gn=ALAS2;refGene_tx=XM_019823866.2
X 48096562 48096831
refGene_gn=ALAS2;refGene_tx=XM_004000554.4;refGene_gn=ALAS2;refGene_tx=XM_019823866.2
X 48097407 48097572
refGene_gn=ALAS2;refGene_tx=XM_004000554.4;refGene_gn=ALAS2;refGene_tx=XM_019823866.2
X 48100005 48100185
refGene_gn=ALAS2;refGene_tx=XM_004000554.4;refGene_gn=ALAS2;refGene_tx=XM_019823866.2
X 48104650 48104835
refGene_gn=ALAS2;refGene_tx=XM_004000554.4;refGene_gn=ALAS2;refGene_tx=XM_019823866.2
X 48105391 48105614
refGene_gn=ALAS2;refGene_tx=XM_004000554.4;refGene_gn=ALAS2;refGene_tx=XM_019823866.2
X 48108497 48108608 refGene_gn=ALAS2;refGene_tx=XM_004000554.4
X 48110139 48110262
refGene_gn=ALAS2;refGene_tx=XM_004000554.4;refGene_gn=ALAS2;refGene_tx=XM_019823866.2
X 48110919 48111100
refGene_gn=ALAS2;refGene_tx=XM_004000554.4;refGene_gn=ALAS2;refGene_tx=XM_019823866.2
X 48383310 48385320
refGene_gn=USP51;refGene_tx=XM_011291761.3;refGene_gn=USP51;refGene_tx=XM_011291762.3;refGene_gn=USP51;refGene_tx=XM_019823871.2;refGene_gn=USP51;refGene_tx=XM_019823873.2;refGene_gn=USP51;refGene_tx=XM_019823874.2
X 48440099 48440717
refGene_gn=LOC101096705;refGene_tx=XM_004000557.5
X 48598237 48598329
refGene_gn=RRAGB;refGene_tx=XM_004000559.5;refGene_gn=RRAGB;refGene_tx=XM_004000560.5
X 48599162 48599196
refGene_gn=RRAGB;refGene_tx=XM_004000559.5;refGene_gn=RRAGB;refGene_tx=XM_004000560.5;refGene_gn=RRAGB;refGene_tx=XM_006943774.4;refGene_gn=RRAGB;refGene_tx=XM_023248918.1;refGene_gn=RRAGB;refGene_tx=XM_023248919.1
X 48601756 48601856
refGene_gn=RRAGB;refGene_tx=XM_004000559.5;refGene_gn=RRAGB;refGene_tx=XM_004000560.5;refGene_gn=RRAGB;refGene_tx=XM_006943774.4;refGene_gn=RRAGB;refGene_tx=XM_023248918.1;refGene_gn=RRAGB;refGene_tx=XM_023248919.1
X 48606496 48606580
refGene_gn=RRAGB;refGene_tx=XM_004000559.5;refGene_gn=RRAGB;refGene_tx=XM_006943774.4;refGene_gn=RRAGB;refGene_tx=XM_023248918.1;refGene_gn=

RRAGB;refGene_tx=XM_023248919.1
X 48608481 48608548
refGene_gn=RRAGB;refGene_tx=XM_004000559.5;refGene_gn=RRAGB;refGene_tx=XM_004000560.5;refGene_gn=RRAGB;refGene_tx=XM_006943774.4;refGene_gn=RRAGB;refGene_tx=XM_023248918.1;refGene_gn=RRAGB;refGene_tx=XM_023248919.1
X 48611206 48611429
refGene_gn=RRAGB;refGene_tx=XM_004000559.5;refGene_gn=RRAGB;refGene_tx=XM_004000560.5;refGene_gn=RRAGB;refGene_tx=XM_006943774.4;refGene_gn=RRAGB;refGene_tx=XM_023248918.1;refGene_gn=RRAGB;refGene_tx=XM_023248919.1
X 48626867 48626963
refGene_gn=RRAGB;refGene_tx=XM_004000559.5;refGene_gn=RRAGB;refGene_tx=XM_004000560.5;refGene_gn=RRAGB;refGene_tx=XM_006943774.4;refGene_gn=RRAGB;refGene_tx=XM_023248918.1;refGene_gn=RRAGB;refGene_tx=XM_023248919.1
X 48629185 48629308
refGene_gn=RRAGB;refGene_tx=XM_004000559.5;refGene_gn=RRAGB;refGene_tx=XM_004000560.5;refGene_gn=RRAGB;refGene_tx=XM_006943774.4;refGene_gn=RRAGB;refGene_tx=XM_023248918.1;refGene_gn=RRAGB;refGene_tx=XM_023248919.1
X 48646109 48646201
refGene_gn=RRAGB;refGene_tx=XM_004000559.5;refGene_gn=RRAGB;refGene_tx=XM_004000560.5;refGene_gn=RRAGB;refGene_tx=XM_006943774.4;refGene_gn=RRAGB;refGene_tx=XM_023248918.1;refGene_gn=RRAGB;refGene_tx=XM_023248919.1
X 48647254 48647370
refGene_gn=RRAGB;refGene_tx=XM_004000559.5;refGene_gn=RRAGB;refGene_tx=XM_004000560.5;refGene_gn=RRAGB;refGene_tx=XM_006943774.4;refGene_gn=RRAGB;refGene_tx=XM_023248918.1;refGene_gn=RRAGB;refGene_tx=XM_023248919.1
X 48648302 48648400
refGene_gn=RRAGB;refGene_tx=XM_004000559.5;refGene_gn=RRAGB;refGene_tx=XM_004000560.5;refGene_gn=RRAGB;refGene_tx=XM_006943774.4;refGene_gn=RRAGB;refGene_tx=XM_023248918.1;refGene_gn=RRAGB;refGene_tx=XM_023248919.1
X 48959851 48959869
refGene_gn=KLF8;refGene_tx=XM_006943776.3;refGene_gn=KLF8;refGene_tx=XM_019823875.1
X 48989760 48990322
refGene_gn=KLF8;refGene_tx=XM_006943776.3;refGene_gn=KLF8;refGene_tx=XM_019823875.1
X 48993160 48993272
refGene_gn=KLF8;refGene_tx=XM_006943776.3;refGene_gn=KLF8;refGene_tx=XM_019823875.1
X 48993915 48994267
refGene_gn=KLF8;refGene_tx=XM_006943776.3;refGene_gn=KLF8;refGene_tx=XM_019823875.1
X 49021944 49022126 refGene_gn=KLF8;refGene_tx=XM_006943776.3
X 49292753 49294625 refGene_gn=UBQLN2;refGene_tx=XM_004000561.5

X 49419469 49419676
refGene_gn=LOC111558779;refGene_tx=XM_023249487.1;refGene_gn=LOC111558
779;refGene_tx=XM_023249488.1
X 49686126 49686925
refGene_gn=LOC101094491;refGene_tx=XM_011291764.2;refGene_gn=LOC101094
491;refGene_tx=XM_011291766.2;refGene_gn=LOC101094491;refGene_tx=XM_01
1291767.2;refGene_gn=LOC101094491;refGene_tx=XM_011291768.2
X 49687594 49687842
refGene_gn=LOC101094491;refGene_tx=XM_011291764.2;refGene_gn=LOC101094
491;refGene_tx=XM_011291766.2
X 49821276 49822055
refGene_gn=LOC105261410;refGene_tx=XM_023248880.1
X 49830319 49830395
refGene_gn=LOC105261410;refGene_tx=XM_023248880.1
X 49832423 49832650
refGene_gn=LOC105261410;refGene_tx=XM_023248880.1
X 49840916 49840992
refGene_gn=LOC105261410;refGene_tx=XM_023248880.1
X 49843019 49843150
refGene_gn=LOC105261410;refGene_tx=XM_023248880.1
X 49843152 49843245
refGene_gn=LOC105261410;refGene_tx=XM_023248880.1
X 49843899 49844128
refGene_gn=LOC105261410;refGene_tx=XM_023248880.1
X 49966757 49968354 refGene_gn=ZXDB;refGene_tx=XM_011291771.3
X 49968356 49968475 refGene_gn=ZXDB;refGene_tx=XM_011291771.3
X 49968476 49969181 refGene_gn=ZXDB;refGene_tx=XM_011291771.3
X 50251442 50251454
refGene_gn=LOC109496524;refGene_tx=XM_019823885.2
X 50262033 50264547
refGene_gn=LOC109496524;refGene_tx=XM_019823885.2;refGene_gn=LOC109496
524;refGene_tx=XM_023249002.1
X 50486072 50486600
refGene_gn=LOC105260211;refGene_tx=XM_023248960.1
X 50486683 50487711
refGene_gn=LOC105260211;refGene_tx=XM_023248960.1
X 52746938 52747688 refGene_gn=SPIN4;refGene_tx=XM_006943778.4
X 53237961 53238143
refGene_gn=ARHGEF9;refGene_tx=XM_011291773.2;refGene_gn=ARHGEF9;refGen
e_tx=XM_011291774.3;refGene_gn=ARHGEF9;refGene_tx=XM_011291775.3;refGe
ne_gn=ARHGEF9;refGene_tx=XM_019823887.1;refGene_gn=ARHGEF9;refGene_tx=
XM_019823888.2;refGene_gn=ARHGEF9;refGene_tx=XM_023248858.1
X 53252749 53252818
refGene_gn=ARHGEF9;refGene_tx=XM_011291773.2;refGene_gn=ARHGEF9;refGen
e_tx=XM_011291774.3;refGene_gn=ARHGEF9;refGene_tx=XM_011291775.3;refGe
ne_gn=ARHGEF9;refGene_tx=XM_019823887.1;refGene_gn=ARHGEF9;refGene_tx=
XM_019823888.2;refGene_gn=ARHGEF9;refGene_tx=XM_023248858.1
X 53258365 53258609
refGene_gn=ARHGEF9;refGene_tx=XM_011291773.2;refGene_gn=ARHGEF9;refGen
e_tx=XM_011291774.3;refGene_gn=ARHGEF9;refGene_tx=XM_011291775.3;refGe

ne_gn=ARHGEF9;refGene_tx=XM_019823887.1;refGene_gn=ARHGEF9;refGene_tx=XM_019823888.2;refGene_gn=ARHGEF9;refGene_tx=XM_023248858.1
X 53265704 53265836
refGene_gn=ARHGEF9;refGene_tx=XM_011291773.2;refGene_gn=ARHGEF9;refGene_tx=XM_011291774.3;refGene_gn=ARHGEF9;refGene_tx=XM_011291775.3;refGene_gn=ARHGEF9;refGene_tx=XM_019823887.1;refGene_gn=ARHGEF9;refGene_tx=XM_019823888.2
X 53281214 53281344
refGene_gn=ARHGEF9;refGene_tx=XM_011291773.2;refGene_gn=ARHGEF9;refGene_tx=XM_011291774.3;refGene_gn=ARHGEF9;refGene_tx=XM_011291775.3;refGene_gn=ARHGEF9;refGene_tx=XM_019823887.1;refGene_gn=ARHGEF9;refGene_tx=XM_019823888.2;refGene_gn=ARHGEF9;refGene_tx=XM_023248858.1
X 53286659 53286892
refGene_gn=ARHGEF9;refGene_tx=XM_011291773.2;refGene_gn=ARHGEF9;refGene_tx=XM_011291774.3;refGene_gn=ARHGEF9;refGene_tx=XM_011291775.3;refGene_gn=ARHGEF9;refGene_tx=XM_019823887.1;refGene_gn=ARHGEF9;refGene_tx=XM_019823888.2;refGene_gn=ARHGEF9;refGene_tx=XM_023248858.1
X 53316791 53316971
refGene_gn=ARHGEF9;refGene_tx=XM_011291773.2;refGene_gn=ARHGEF9;refGene_tx=XM_011291774.3;refGene_gn=ARHGEF9;refGene_tx=XM_011291775.3;refGene_gn=ARHGEF9;refGene_tx=XM_019823887.1;refGene_gn=ARHGEF9;refGene_tx=XM_019823888.2;refGene_gn=ARHGEF9;refGene_tx=XM_023248858.1
X 53336747 53336939
refGene_gn=ARHGEF9;refGene_tx=XM_011291773.2;refGene_gn=ARHGEF9;refGene_tx=XM_011291774.3;refGene_gn=ARHGEF9;refGene_tx=XM_011291775.3;refGene_gn=ARHGEF9;refGene_tx=XM_019823887.1;refGene_gn=ARHGEF9;refGene_tx=XM_019823888.2;refGene_gn=ARHGEF9;refGene_tx=XM_023248858.1
X 53371668 53371848
refGene_gn=ARHGEF9;refGene_tx=XM_011291773.2;refGene_gn=ARHGEF9;refGene_tx=XM_011291774.3;refGene_gn=ARHGEF9;refGene_tx=XM_011291775.3;refGene_gn=ARHGEF9;refGene_tx=XM_023248858.1
X 53409264 53409273 refGene_gn=ARHGEF9;refGene_tx=XM_011291775.3
X 53436254 53436284
refGene_gn=ARHGEF9;refGene_tx=XM_011291773.2;refGene_gn=ARHGEF9;refGene_tx=XM_019823887.1;refGene_gn=ARHGEF9;refGene_tx=XM_023248858.1
X 53775058 53775662 refGene_gn=AMER1;refGene_tx=XM_004000564.3
X 53775723 53778533 refGene_gn=AMER1;refGene_tx=XM_004000564.3
X 53803216 53803350
refGene_gn=ASB12;refGene_tx=XM_004000565.4;refGene_gn=ASB12;refGene_tx=XM_006943779.3;refGene_gn=ASB12;refGene_tx=XM_019823897.2
X 53803782 53804632
refGene_gn=ASB12;refGene_tx=XM_004000565.4;refGene_gn=ASB12;refGene_tx=XM_006943779.3;refGene_gn=ASB12;refGene_tx=XM_019823897.2
X 53805730 53805748 refGene_gn=ASB12;refGene_tx=XM_006943779.3
X 53824331 53824515 refGene_gn=MTMR8;refGene_tx=XM_023248961.1
X 53837954 53838421 refGene_gn=MTMR8;refGene_tx=XM_023248961.1
X 53873619 53873690 refGene_gn=MTMR8;refGene_tx=XM_023248961.1
X 53873692 53873733 refGene_gn=MTMR8;refGene_tx=XM_023248961.1
X 53886941 53887021 refGene_gn=MTMR8;refGene_tx=XM_023248961.1
X 53917008 53917088 refGene_gn=MTMR8;refGene_tx=XM_023248961.1

X 53917090 53917162 refGene_gn=MTMR8;refGene_tx=XM_023248961.1
X 53919094 53919257 refGene_gn=MTMR8;refGene_tx=XM_023248961.1
X 53922609 53922729 refGene_gn=MTMR8;refGene_tx=XM_023248961.1
X 53941223 53941682
refGene_gn=LOC101101369;refGene_tx=XM_011291777.1
X 54385184 54385893
refGene_gn=LOC109496915;refGene_tx=XM_019824688.2
X 54577941 54578074
refGene_gn=ZC4H2;refGene_tx=XM_004000569.4;refGene_gn=ZC4H2;refGene_tx
=XM_004000570.4
X 54581784 54581947 refGene_gn=ZC4H2;refGene_tx=XM_004000569.4
X 54582814 54582987
refGene_gn=ZC4H2;refGene_tx=XM_004000569.4;refGene_gn=ZC4H2;refGene_tx
=XM_004000570.4
X 54584560 54584732
refGene_gn=ZC4H2;refGene_tx=XM_004000569.4;refGene_gn=ZC4H2;refGene_tx
=XM_004000570.4
X 54611921 54611974
refGene_gn=ZC4H2;refGene_tx=XM_004000569.4;refGene_gn=ZC4H2;refGene_tx
=XM_004000570.4
X 55015650 55015836
refGene_gn=LOC109496527;refGene_tx=XM_023249260.1
X 55094351 55094965 refGene_gn=ZC3H12B;refGene_tx=XM_011291778.3
X 55151474 55151614 refGene_gn=ZC3H12B;refGene_tx=XM_011291778.3
X 55171370 55171605 refGene_gn=ZC3H12B;refGene_tx=XM_011291778.3
X 55172275 55172382 refGene_gn=ZC3H12B;refGene_tx=XM_011291778.3
X 55173566 55174987 refGene_gn=ZC3H12B;refGene_tx=XM_011291778.3
X 55183323 55183456
refGene_gn=LAS1L;refGene_tx=XM_006944279.4;refGene_gn=LAS1L;refGene_tx
=XM_006944280.4;refGene_gn=LAS1L;refGene_tx=XM_006944281.4;refGene_gn=
LAS1L;refGene_tx=XM_006944282.4;refGene_gn=LAS1L;refGene_tx=XM_0198239
02.2
X 55185030 55185181
refGene_gn=LAS1L;refGene_tx=XM_006944279.4;refGene_gn=LAS1L;refGene_tx
=XM_006944280.4;refGene_gn=LAS1L;refGene_tx=XM_006944281.4;refGene_gn=
LAS1L;refGene_tx=XM_019823902.2
X 55187959 55188450
refGene_gn=LAS1L;refGene_tx=XM_006944279.4;refGene_gn=LAS1L;refGene_tx
=XM_006944280.4;refGene_gn=LAS1L;refGene_tx=XM_006944281.4;refGene_gn=
LAS1L;refGene_tx=XM_006944282.4;refGene_gn=LAS1L;refGene_tx=XM_0198239
02.2
X 55193659 55193807
refGene_gn=LAS1L;refGene_tx=XM_006944279.4;refGene_gn=LAS1L;refGene_tx
=XM_006944280.4;refGene_gn=LAS1L;refGene_tx=XM_006944281.4;refGene_gn=
LAS1L;refGene_tx=XM_006944282.4;refGene_gn=LAS1L;refGene_tx=XM_0198239
02.2
X 55194157 55194364
refGene_gn=LAS1L;refGene_tx=XM_006944279.4;refGene_gn=LAS1L;refGene_tx
=XM_006944280.4;refGene_gn=LAS1L;refGene_tx=XM_006944281.4;refGene_gn=
LAS1L;refGene_tx=XM_006944282.4;refGene_gn=LAS1L;refGene_tx=XM_0198239

02.2
X	55194670	55194721
refGene_gn=LAS1L;refGene_tx=XM_006944279.4;refGene_gn=LAS1L;refGene_tx=XM_006944281.4;refGene_gn=LAS1L;refGene_tx=XM_006944282.4;refGene_gn=LAS1L;refGene_tx=XM_019823902.2
X	55195089	55195175
refGene_gn=LAS1L;refGene_tx=XM_006944279.4;refGene_gn=LAS1L;refGene_tx=XM_006944280.4;refGene_gn=LAS1L;refGene_tx=XM_006944281.4;refGene_gn=LAS1L;refGene_tx=XM_006944282.4;refGene_gn=LAS1L;refGene_tx=XM_019823902.2
X	55196930	55197040
refGene_gn=LAS1L;refGene_tx=XM_006944279.4;refGene_gn=LAS1L;refGene_tx=XM_006944280.4;refGene_gn=LAS1L;refGene_tx=XM_006944281.4;refGene_gn=LAS1L;refGene_tx=XM_006944282.4;refGene_gn=LAS1L;refGene_tx=XM_019823902.2
X	55197876	55197923
refGene_gn=LAS1L;refGene_tx=XM_006944279.4;refGene_gn=LAS1L;refGene_tx=XM_006944280.4;refGene_gn=LAS1L;refGene_tx=XM_006944281.4;refGene_gn=LAS1L;refGene_tx=XM_006944282.4;refGene_gn=LAS1L;refGene_tx=XM_019823902.2
X	55198213	55198498
refGene_gn=LAS1L;refGene_tx=XM_006944279.4;refGene_gn=LAS1L;refGene_tx=XM_006944280.4;refGene_gn=LAS1L;refGene_tx=XM_006944281.4;refGene_gn=LAS1L;refGene_tx=XM_006944282.4;refGene_gn=LAS1L;refGene_tx=XM_019823902.2
X	55199330	55199412
refGene_gn=LAS1L;refGene_tx=XM_006944279.4;refGene_gn=LAS1L;refGene_tx=XM_006944280.4;refGene_gn=LAS1L;refGene_tx=XM_006944281.4;refGene_gn=LAS1L;refGene_tx=XM_006944282.4;refGene_gn=LAS1L;refGene_tx=XM_019823902.2
X	55200185	55200255
refGene_gn=LAS1L;refGene_tx=XM_006944279.4;refGene_gn=LAS1L;refGene_tx=XM_006944280.4;refGene_gn=LAS1L;refGene_tx=XM_006944281.4;refGene_gn=LAS1L;refGene_tx=XM_006944282.4;refGene_gn=LAS1L;refGene_tx=XM_019823902.2
X	55201226	55201352
refGene_gn=LAS1L;refGene_tx=XM_006944279.4;refGene_gn=LAS1L;refGene_tx=XM_006944280.4;refGene_gn=LAS1L;refGene_tx=XM_006944281.4;refGene_gn=LAS1L;refGene_tx=XM_006944282.4
X	55202170	55202361
refGene_gn=LAS1L;refGene_tx=XM_006944279.4;refGene_gn=LAS1L;refGene_tx=XM_006944280.4;refGene_gn=LAS1L;refGene_tx=XM_006944281.4;refGene_gn=LAS1L;refGene_tx=XM_006944282.4;refGene_gn=LAS1L;refGene_tx=XM_019823902.2
X	55302142	55302302
refGene_gn=LOC101080617;refGene_tx=XM_011291779.1
X	55302304	55302411
refGene_gn=LOC101080617;refGene_tx=XM_011291779.1
X	55365132	55365144	refGene_gn=MSN;refGene_tx=XM_004000571.5
X	55406025	55406109	refGene_gn=MSN;refGene_tx=XM_004000571.5

X 55414525 55414621 refGene_gn=MSN;refGene_tx=XM_004000571.5
X 55416003 55416278 refGene_gn=MSN;refGene_tx=XM_004000571.5
X 55418051 55418135 refGene_gn=MSN;refGene_tx=XM_004000571.5
X 55418612 55418759 refGene_gn=MSN;refGene_tx=XM_004000571.5
X 55419776 55419873 refGene_gn=MSN;refGene_tx=XM_004000571.5
X 55421972 55422136 refGene_gn=MSN;refGene_tx=XM_004000571.5
X 55424946 55425077 refGene_gn=MSN;refGene_tx=XM_004000571.5
X 55425336 55425497 refGene_gn=MSN;refGene_tx=XM_004000571.5
X 55426803 55426896 refGene_gn=MSN;refGene_tx=XM_004000571.5
X 55427197 55427422 refGene_gn=MSN;refGene_tx=XM_004000571.5
X 55427933 55428098 refGene_gn=MSN;refGene_tx=XM_004000571.5
X 55667390 55667643 refGene_gn=VSIG4;refGene_tx=XM_006943782.2
X 55667981 55668003 refGene_gn=VSIG4;refGene_tx=XM_006943782.2
X 55670202 55670307 refGene_gn=VSIG4;refGene_tx=XM_006943782.2
X 55674993 55675071 refGene_gn=VSIG4;refGene_tx=XM_006943782.2
X 55675601 55675664 refGene_gn=VSIG4;refGene_tx=XM_006943782.2
X 55698209 55698491 refGene_gn=VSIG4;refGene_tx=XM_006943782.2
X 55699227 55699584 refGene_gn=VSIG4;refGene_tx=XM_006943782.2
X 55703488 55703543 refGene_gn=VSIG4;refGene_tx=XM_006943782.2
X 55868914 55869090
refGene_gn=HEPH;refGene_tx=XM_004000574.4;refGene_gn=HEPH;refGene_tx=X
M_023249133.1
X 55871197 55871442
refGene_gn=HEPH;refGene_tx=XM_004000574.4;refGene_gn=HEPH;refGene_tx=X
M_023249133.1
X 55875487 55875700 refGene_gn=HEPH;refGene_tx=XM_004000574.4
X 55876829 55877012
refGene_gn=HEPH;refGene_tx=XM_004000574.4;refGene_gn=HEPH;refGene_tx=X
M_023249133.1
X 55878107 55878362
refGene_gn=HEPH;refGene_tx=XM_004000574.4;refGene_gn=HEPH;refGene_tx=X
M_023249133.1
X 55880264 55880433
refGene_gn=HEPH;refGene_tx=XM_004000574.4;refGene_gn=HEPH;refGene_tx=X
M_023249133.1
X 55881649 55881786
refGene_gn=HEPH;refGene_tx=XM_004000574.4;refGene_gn=HEPH;refGene_tx=X
M_023249133.1
X 55887336 55887468
refGene_gn=HEPH;refGene_tx=XM_004000574.4;refGene_gn=HEPH;refGene_tx=X
M_023249133.1
X 55891241 55891453
refGene_gn=HEPH;refGene_tx=XM_004000574.4;refGene_gn=HEPH;refGene_tx=X
M_023249133.1
X 55893959 55894110
refGene_gn=HEPH;refGene_tx=XM_004000574.4;refGene_gn=HEPH;refGene_tx=X
M_023249133.1
X 55896501 55896714
refGene_gn=HEPH;refGene_tx=XM_004000574.4;refGene_gn=HEPH;refGene_tx=X
M_023249133.1

X 55905179 55905393
refGene_gn=HEPH;refGene_tx=XM_004000574.4;refGene_gn=HEPH;refGene_tx=XM_023249133.1
X 55907254 55907394
refGene_gn=HEPH;refGene_tx=XM_004000574.4;refGene_gn=HEPH;refGene_tx=XM_023249133.1
X 55908207 55908339
refGene_gn=HEPH;refGene_tx=XM_004000574.4;refGene_gn=HEPH;refGene_tx=XM_023249133.1
X 55939312 55939419
refGene_gn=HEPH;refGene_tx=XM_004000574.4;refGene_gn=HEPH;refGene_tx=XM_023249133.1
X 55940832 55941058
refGene_gn=HEPH;refGene_tx=XM_004000574.4;refGene_gn=HEPH;refGene_tx=XM_023249133.1
X 55943573 55943713
refGene_gn=HEPH;refGene_tx=XM_004000574.4;refGene_gn=HEPH;refGene_tx=XM_023249133.1
X 55944629 55944792
refGene_gn=HEPH;refGene_tx=XM_004000574.4;refGene_gn=HEPH;refGene_tx=XM_023249133.1
X 55947757 55947802
refGene_gn=HEPH;refGene_tx=XM_004000574.4;refGene_gn=HEPH;refGene_tx=XM_023249133.1
X 55949897 55950130
refGene_gn=HEPH;refGene_tx=XM_004000574.4;refGene_gn=HEPH;refGene_tx=XM_023249133.1
X 56101420 56101803 refGene_gn=EDA2R;refGene_tx=XM_023249101.1
X 56101975 56102030 refGene_gn=EDA2R;refGene_tx=XM_023249101.1
X 56104547 56104549 refGene_gn=EDA2R;refGene_tx=XM_023249101.1
X 56105173 56105338 refGene_gn=EDA2R;refGene_tx=XM_023249101.1
X 56107329 56107413 refGene_gn=EDA2R;refGene_tx=XM_023249101.1
X 56107942 56108132 refGene_gn=EDA2R;refGene_tx=XM_023249101.1
X 56142616 56142703 refGene_gn=EDA2R;refGene_tx=XM_023249101.1
X 56937906 56939471
refGene_gn=AR;refGene_tx=XM_004000575.5;refGene_gn=AR;refGene_tx=XM_023249497.1
X 57040812 57040964
refGene_gn=AR;refGene_tx=XM_004000575.5;refGene_gn=AR;refGene_tx=XM_023249497.1
X 57041541 57041585 refGene_gn=AR;refGene_tx=XM_023249497.1
X 57080648 57080765 refGene_gn=AR;refGene_tx=XM_004000575.5
X 57099428 57099716 refGene_gn=AR;refGene_tx=XM_004000575.5
X 57105491 57105636 refGene_gn=AR;refGene_tx=XM_004000575.5
X 57109630 57109761 refGene_gn=AR;refGene_tx=XM_004000575.5
X 57110542 57110700 refGene_gn=AR;refGene_tx=XM_004000575.5
X 57111383 57111539 refGene_gn=AR;refGene_tx=XM_004000575.5
X 57382069 57382103
refGene_gn=OPHN1;refGene_tx=XM_023249183.1;refGene_gn=OPHN1;refGene_tx=XM_023249184.1;refGene_gn=OPHN1;refGene_tx=XM_023249185.1;refGene_gn=

OPHN1;refGene_tx=XM_023249186.1;refGene_gn=OPHN1;refGene_tx=XM_0232491
87.1
X 57391888 57391939
refGene_gn=OPHN1;refGene_tx=XM_023249183.1;refGene_gn=OPHN1;refGene_tx
=XM_023249184.1;refGene_gn=OPHN1;refGene_tx=XM_023249185.1;refGene_gn=
OPHN1;refGene_tx=XM_023249186.1;refGene_gn=OPHN1;refGene_tx=XM_0232491
87.1
X 57393192 57393358
refGene_gn=OPHN1;refGene_tx=XM_023249183.1;refGene_gn=OPHN1;refGene_tx
=XM_023249184.1;refGene_gn=OPHN1;refGene_tx=XM_023249185.1;refGene_gn=
OPHN1;refGene_tx=XM_023249186.1;refGene_gn=OPHN1;refGene_tx=XM_0232491
87.1
X 57424914 57425238
refGene_gn=OPHN1;refGene_tx=XM_023249183.1;refGene_gn=OPHN1;refGene_tx
=XM_023249184.1;refGene_gn=OPHN1;refGene_tx=XM_023249185.1;refGene_gn=
OPHN1;refGene_tx=XM_023249186.1;refGene_gn=OPHN1;refGene_tx=XM_0232491
87.1
X 57456915 57457063
refGene_gn=OPHN1;refGene_tx=XM_023249183.1;refGene_gn=OPHN1;refGene_tx
=XM_023249184.1;refGene_gn=OPHN1;refGene_tx=XM_023249185.1;refGene_gn=
OPHN1;refGene_tx=XM_023249186.1;refGene_gn=OPHN1;refGene_tx=XM_0232491
87.1
X 57509005 57509165
refGene_gn=OPHN1;refGene_tx=XM_023249183.1;refGene_gn=OPHN1;refGene_tx
=XM_023249184.1;refGene_gn=OPHN1;refGene_tx=XM_023249185.1;refGene_gn=
OPHN1;refGene_tx=XM_023249186.1;refGene_gn=OPHN1;refGene_tx=XM_0232491
87.1
X 57541879 57541985
refGene_gn=OPHN1;refGene_tx=XM_023249183.1;refGene_gn=OPHN1;refGene_tx
=XM_023249184.1;refGene_gn=OPHN1;refGene_tx=XM_023249185.1;refGene_gn=
OPHN1;refGene_tx=XM_023249186.1;refGene_gn=OPHN1;refGene_tx=XM_0232491
87.1
X 57543177 57543236
refGene_gn=OPHN1;refGene_tx=XM_023249183.1;refGene_gn=OPHN1;refGene_tx
=XM_023249184.1;refGene_gn=OPHN1;refGene_tx=XM_023249185.1;refGene_gn=
OPHN1;refGene_tx=XM_023249186.1;refGene_gn=OPHN1;refGene_tx=XM_0232491
87.1
X 57547943 57548028
refGene_gn=OPHN1;refGene_tx=XM_023249183.1;refGene_gn=OPHN1;refGene_tx
=XM_023249184.1;refGene_gn=OPHN1;refGene_tx=XM_023249185.1;refGene_gn=
OPHN1;refGene_tx=XM_023249186.1;refGene_gn=OPHN1;refGene_tx=XM_0232491
87.1
X 57614978 57615053
refGene_gn=OPHN1;refGene_tx=XM_023249183.1;refGene_gn=OPHN1;refGene_tx
=XM_023249184.1;refGene_gn=OPHN1;refGene_tx=XM_023249185.1;refGene_gn=
OPHN1;refGene_tx=XM_023249186.1;refGene_gn=OPHN1;refGene_tx=XM_0232491
87.1
X 57616687 57616750
refGene_gn=OPHN1;refGene_tx=XM_023249183.1;refGene_gn=OPHN1;refGene_tx
=XM_023249184.1;refGene_gn=OPHN1;refGene_tx=XM_023249185.1;refGene_gn=

OPHN1;refGene_tx=XM_023249186.1;refGene_gn=OPHN1;refGene_tx=XM_0232491
87.1
X	57617304	57617338
refGene_gn=OPHN1;refGene_tx=XM_023249183.1;refGene_gn=OPHN1;refGene_tx
=XM_023249184.1;refGene_gn=OPHN1;refGene_tx=XM_023249185.1;refGene_gn=
OPHN1;refGene_tx=XM_023249186.1;refGene_gn=OPHN1;refGene_tx=XM_0232491
87.1
X	57625208	57625287
refGene_gn=OPHN1;refGene_tx=XM_023249183.1;refGene_gn=OPHN1;refGene_tx
=XM_023249184.1;refGene_gn=OPHN1;refGene_tx=XM_023249185.1;refGene_gn=
OPHN1;refGene_tx=XM_023249186.1;refGene_gn=OPHN1;refGene_tx=XM_0232491
87.1
X	57630475	57630570
refGene_gn=OPHN1;refGene_tx=XM_023249183.1;refGene_gn=OPHN1;refGene_tx
=XM_023249184.1;refGene_gn=OPHN1;refGene_tx=XM_023249185.1;refGene_gn=
OPHN1;refGene_tx=XM_023249186.1;refGene_gn=OPHN1;refGene_tx=XM_0232491
87.1
X	57634484	57634585
refGene_gn=OPHN1;refGene_tx=XM_023249183.1;refGene_gn=OPHN1;refGene_tx
=XM_023249184.1;refGene_gn=OPHN1;refGene_tx=XM_023249185.1;refGene_gn=
OPHN1;refGene_tx=XM_023249186.1;refGene_gn=OPHN1;refGene_tx=XM_0232491
87.1
X	57637124	57637254
refGene_gn=OPHN1;refGene_tx=XM_023249183.1;refGene_gn=OPHN1;refGene_tx
=XM_023249184.1;refGene_gn=OPHN1;refGene_tx=XM_023249185.1;refGene_gn=
OPHN1;refGene_tx=XM_023249186.1;refGene_gn=OPHN1;refGene_tx=XM_0232491
87.1
X	57639115	57639220
refGene_gn=OPHN1;refGene_tx=XM_023249183.1;refGene_gn=OPHN1;refGene_tx
=XM_023249184.1;refGene_gn=OPHN1;refGene_tx=XM_023249185.1;refGene_gn=
OPHN1;refGene_tx=XM_023249186.1;refGene_gn=OPHN1;refGene_tx=XM_0232491
87.1
X	57642675	57642786
refGene_gn=OPHN1;refGene_tx=XM_023249183.1;refGene_gn=OPHN1;refGene_tx
=XM_023249184.1;refGene_gn=OPHN1;refGene_tx=XM_023249185.1;refGene_gn=
OPHN1;refGene_tx=XM_023249186.1;refGene_gn=OPHN1;refGene_tx=XM_0232491
87.1
X	57660383	57660485
refGene_gn=OPHN1;refGene_tx=XM_023249183.1;refGene_gn=OPHN1;refGene_tx
=XM_023249184.1;refGene_gn=OPHN1;refGene_tx=XM_023249185.1;refGene_gn=
OPHN1;refGene_tx=XM_023249186.1;refGene_gn=OPHN1;refGene_tx=XM_0232491
87.1
X	57749843	57749915
refGene_gn=OPHN1;refGene_tx=XM_023249183.1;refGene_gn=OPHN1;refGene_tx
=XM_023249184.1;refGene_gn=OPHN1;refGene_tx=XM_023249185.1;refGene_gn=
OPHN1;refGene_tx=XM_023249186.1;refGene_gn=OPHN1;refGene_tx=XM_0232491
87.1
X	57755019	57755081
refGene_gn=OPHN1;refGene_tx=XM_023249183.1;refGene_gn=OPHN1;refGene_tx
=XM_023249184.1;refGene_gn=OPHN1;refGene_tx=XM_023249185.1;refGene_gn=

OPHN1;refGene_tx=XM_023249186.1
X 57795277 57795373
refGene_gn=OPHN1;refGene_tx=XM_023249183.1;refGene_gn=OPHN1;refGene_tx=XM_023249184.1;refGene_gn=OPHN1;refGene_tx=XM_023249185.1;refGene_gn=OPHN1;refGene_tx=XM_023249186.1;refGene_gn=OPHN1;refGene_tx=XM_023249187.1
X 57802763 57802781
refGene_gn=LOC102899556;refGene_tx=XM_006943789.3
X 57807692 57808160
refGene_gn=LOC102899556;refGene_tx=XM_006943789.3
X 57979712 57979866
refGene_gn=OPHN1;refGene_tx=XM_023249183.1;refGene_gn=OPHN1;refGene_tx=XM_023249184.1;refGene_gn=OPHN1;refGene_tx=XM_023249185.1;refGene_gn=OPHN1;refGene_tx=XM_023249186.1
X 58027274 58027331 refGene_gn=YIPF6;refGene_tx=XM_023248889.1
X 58041881 58042010
refGene_gn=YIPF6;refGene_tx=XM_023248889.1;refGene_gn=YIPF6;refGene_tx=XM_023248890.1
X 58043508 58043587
refGene_gn=YIPF6;refGene_tx=XM_023248889.1;refGene_gn=YIPF6;refGene_tx=XM_023248890.1
X 58047200 58047243
refGene_gn=YIPF6;refGene_tx=XM_023248889.1;refGene_gn=YIPF6;refGene_tx=XM_023248890.1
X 58050419 58050545
refGene_gn=YIPF6;refGene_tx=XM_023248889.1;refGene_gn=YIPF6;refGene_tx=XM_023248890.1
X 58064409 58064567
refGene_gn=YIPF6;refGene_tx=XM_023248889.1;refGene_gn=YIPF6;refGene_tx=XM_023248890.1
X 58066327 58066446
refGene_gn=YIPF6;refGene_tx=XM_023248889.1;refGene_gn=YIPF6;refGene_tx=XM_023248890.1
X 58132659 58132704 refGene_gn=STARD8;refGene_tx=XM_006943786.4
X 58149557 58149591 refGene_gn=STARD8;refGene_tx=XM_006943786.4
X 58195447 58195690
refGene_gn=STARD8;refGene_tx=XM_006943786.4;refGene_gn=STARD8;refGene_tx=XM_006943788.2;refGene_gn=STARD8;refGene_tx=XM_019823917.2
X 58203807 58203879
refGene_gn=STARD8;refGene_tx=XM_006943786.4;refGene_gn=STARD8;refGene_tx=XM_006943788.2;refGene_gn=STARD8;refGene_tx=XM_019823917.2
X 58206320 58206402
refGene_gn=STARD8;refGene_tx=XM_006943786.4;refGene_gn=STARD8;refGene_tx=XM_006943788.2;refGene_gn=STARD8;refGene_tx=XM_019823917.2
X 58207367 58207431
refGene_gn=STARD8;refGene_tx=XM_004000578.3;refGene_gn=STARD8;refGene_tx=XM_006943786.4;refGene_gn=STARD8;refGene_tx=XM_006943787.3;refGene_gn=STARD8;refGene_tx=XM_006943788.2;refGene_gn=STARD8;refGene_tx=XM_019823917.2;refGene_gn=STARD8;refGene_tx=XM_023248946.1
X 58208646 58210037

refGene_gn=STARD8;refGene_tx=XM_004000578.3;refGene_gn=STARD8;refGene_tx=XM_006943786.4;refGene_gn=STARD8;refGene_tx=XM_006943787.3;refGene_gn=STARD8;refGene_tx=XM_006943788.2;refGene_gn=STARD8;refGene_tx=XM_019823917.2;refGene_gn=STARD8;refGene_tx=XM_023248946.1
X 58210656 58210830
refGene_gn=STARD8;refGene_tx=XM_004000578.3;refGene_gn=STARD8;refGene_tx=XM_006943786.4;refGene_gn=STARD8;refGene_tx=XM_006943787.3;refGene_gn=STARD8;refGene_tx=XM_006943788.2;refGene_gn=STARD8;refGene_tx=XM_019823917.2;refGene_gn=STARD8;refGene_tx=XM_023248946.1
X 58211817 58211977
refGene_gn=STARD8;refGene_tx=XM_004000578.3;refGene_gn=STARD8;refGene_tx=XM_006943786.4;refGene_gn=STARD8;refGene_tx=XM_006943787.3;refGene_gn=STARD8;refGene_tx=XM_006943788.2;refGene_gn=STARD8;refGene_tx=XM_019823917.2;refGene_gn=STARD8;refGene_tx=XM_023248946.1
X 58212444 58212643
refGene_gn=STARD8;refGene_tx=XM_004000578.3;refGene_gn=STARD8;refGene_tx=XM_006943786.4;refGene_gn=STARD8;refGene_tx=XM_006943787.3;refGene_gn=STARD8;refGene_tx=XM_006943788.2;refGene_gn=STARD8;refGene_tx=XM_019823917.2;refGene_gn=STARD8;refGene_tx=XM_023248946.1
X 58213071 58213282
refGene_gn=STARD8;refGene_tx=XM_004000578.3;refGene_gn=STARD8;refGene_tx=XM_006943786.4;refGene_gn=STARD8;refGene_tx=XM_006943787.3;refGene_gn=STARD8;refGene_tx=XM_006943788.2;refGene_gn=STARD8;refGene_tx=XM_019823917.2;refGene_gn=STARD8;refGene_tx=XM_023248946.1
X 58213552 58213667
refGene_gn=STARD8;refGene_tx=XM_004000578.3;refGene_gn=STARD8;refGene_tx=XM_006943786.4;refGene_gn=STARD8;refGene_tx=XM_006943787.3;refGene_gn=STARD8;refGene_tx=XM_006943788.2;refGene_gn=STARD8;refGene_tx=XM_019823917.2;refGene_gn=STARD8;refGene_tx=XM_023248946.1
X 58213924 58214149
refGene_gn=STARD8;refGene_tx=XM_004000578.3;refGene_gn=STARD8;refGene_tx=XM_006943786.4;refGene_gn=STARD8;refGene_tx=XM_006943787.3;refGene_gn=STARD8;refGene_tx=XM_006943788.2;refGene_gn=STARD8;refGene_tx=XM_019823917.2;refGene_gn=STARD8;refGene_tx=XM_023248946.1
X 58215150 58215368
refGene_gn=STARD8;refGene_tx=XM_004000578.3;refGene_gn=STARD8;refGene_tx=XM_006943786.4;refGene_gn=STARD8;refGene_tx=XM_006943787.3;refGene_gn=STARD8;refGene_tx=XM_006943788.2;refGene_gn=STARD8;refGene_tx=XM_019823917.2;refGene_gn=STARD8;refGene_tx=XM_023248946.1
X 58215468 58215645
refGene_gn=STARD8;refGene_tx=XM_004000578.3;refGene_gn=STARD8;refGene_tx=XM_006943786.4;refGene_gn=STARD8;refGene_tx=XM_006943787.3;refGene_gn=STARD8;refGene_tx=XM_006943788.2;refGene_gn=STARD8;refGene_tx=XM_019823917.2;refGene_gn=STARD8;refGene_tx=XM_023248946.1
X 58215792 58215910
refGene_gn=STARD8;refGene_tx=XM_004000578.3;refGene_gn=STARD8;refGene_tx=XM_006943786.4;refGene_gn=STARD8;refGene_tx=XM_006943787.3;refGene_gn=STARD8;refGene_tx=XM_006943788.2;refGene_gn=STARD8;refGene_tx=XM_019823917.2;refGene_gn=STARD8;refGene_tx=XM_023248946.1
X 58288690 58290853

refGene_gn=LOC109496376;refGene_tx=XM_019823478.2
X 58304505 58304633 refGene_gn=EFNB1;refGene_tx=XM_004000579.4
X 58313362 58313640 refGene_gn=EFNB1;refGene_tx=XM_004000579.4
X 58314393 58314486 refGene_gn=EFNB1;refGene_tx=XM_004000579.4
X 58314682 58314811 refGene_gn=EFNB1;refGene_tx=XM_004000579.4
X 58314946 58315359 refGene_gn=EFNB1;refGene_tx=XM_004000579.4
X 58616544 58618449
refGene_gn=PJA1;refGene_tx=XM_004000580.5;refGene_gn=PJA1;refGene_tx=XM_004000583.4
X 58962058 58962919 refGene_gn=FAM155B;refGene_tx=XM_023248947.1
X 58989922 58990094 refGene_gn=FAM155B;refGene_tx=XM_023248947.1
X 58990494 58990880 refGene_gn=FAM155B;refGene_tx=XM_023248947.1
X 59075370 59075763
refGene_gn=EDA;refGene_tx=XM_011291781.2;refGene_gn=EDA;refGene_tx=XM_011291782.2;refGene_gn=EDA;refGene_tx=XM_011291783.2;refGene_gn=EDA;refGene_tx=XM_011291784.2;refGene_gn=EDA;refGene_tx=XM_011291785.2
X 59379963 59380069
refGene_gn=EDA;refGene_tx=XM_011291781.2;refGene_gn=EDA;refGene_tx=XM_011291782.2;refGene_gn=EDA;refGene_tx=XM_011291783.2;refGene_gn=EDA;refGene_tx=XM_011291784.2;refGene_gn=EDA;refGene_tx=XM_011291785.2
X 59499958 59499982
refGene_gn=EDA;refGene_tx=XM_011291781.2;refGene_gn=EDA;refGene_tx=XM_011291782.2;refGene_gn=EDA;refGene_tx=XM_011291783.2;refGene_gn=EDA;refGene_tx=XM_011291784.2;refGene_gn=EDA;refGene_tx=XM_011291785.2
X 59510392 59510572
refGene_gn=EDA;refGene_tx=XM_011291781.2;refGene_gn=EDA;refGene_tx=XM_011291782.2;refGene_gn=EDA;refGene_tx=XM_011291783.2;refGene_gn=EDA;refGene_tx=XM_011291784.2;refGene_gn=EDA;refGene_tx=XM_011291785.2
X 59511966 59512001
refGene_gn=EDA;refGene_tx=XM_011291781.2;refGene_gn=EDA;refGene_tx=XM_011291782.2;refGene_gn=EDA;refGene_tx=XM_011291783.2;refGene_gn=EDA;refGene_tx=XM_011291784.2;refGene_gn=EDA;refGene_tx=XM_011291785.2
X 59512931 59512983
refGene_gn=EDA;refGene_tx=XM_011291781.2;refGene_gn=EDA;refGene_tx=XM_011291782.2;refGene_gn=EDA;refGene_tx=XM_011291783.2;refGene_gn=EDA;refGene_tx=XM_011291784.2;refGene_gn=EDA;refGene_tx=XM_011291785.2
X 59515597 59515728
refGene_gn=EDA;refGene_tx=XM_011291781.2;refGene_gn=EDA;refGene_tx=XM_011291782.2;refGene_gn=EDA;refGene_tx=XM_011291783.2;refGene_gn=EDA;refGene_tx=XM_011291784.2;refGene_gn=EDA;refGene_tx=XM_011291785.2
X 59517523 59517775
refGene_gn=EDA;refGene_tx=XM_011291781.2;refGene_gn=EDA;refGene_tx=XM_011291782.2;refGene_gn=EDA;refGene_tx=XM_011291783.2;refGene_gn=EDA;refGene_tx=XM_011291784.2;refGene_gn=EDA;refGene_tx=XM_011291785.2
X 59522714 59522869
refGene_gn=AWAT2;refGene_tx=XM_004000588.4;refGene_gn=AWAT2;refGene_tx=XM_019823919.1
X 59523089 59523289
refGene_gn=AWAT2;refGene_tx=XM_004000588.4;refGene_gn=AWAT2;refGene_tx=XM_019823919.1

X	59523850	59524025
refGene_gn=AWAT2;refGene_tx=XM_004000588.4;refGene_gn=AWAT2;refGene_tx=XM_019823919.1
X	59524253	59524458
refGene_gn=AWAT2;refGene_tx=XM_004000588.4;refGene_gn=AWAT2;refGene_tx=XM_019823919.1
X	59524704	59524775
refGene_gn=AWAT2;refGene_tx=XM_004000588.4;refGene_gn=AWAT2;refGene_tx=XM_019823919.1
X	59525157	59525268
refGene_gn=AWAT2;refGene_tx=XM_004000588.4;refGene_gn=AWAT2;refGene_tx=XM_019823919.1
X	59525663	59525684	refGene_gn=AWAT2;refGene_tx=XM_019823919.1
X	59530900	59530988
refGene_gn=AWAT2;refGene_tx=XM_004000588.4;refGene_gn=AWAT2;refGene_tx=XM_019823919.1
X	59553560	59554364	refGene_gn=OTUD6A;refGene_tx=XM_004000589.3
X	59554365	59554443	refGene_gn=OTUD6A;refGene_tx=XM_004000589.3
X	59592477	59592668	refGene_gn=IGBP1;refGene_tx=XM_006943790.3
X	59593051	59593345	refGene_gn=IGBP1;refGene_tx=XM_006943790.3
X	59601501	59601697	refGene_gn=IGBP1;refGene_tx=XM_006943790.3
X	59609294	59609374	refGene_gn=IGBP1;refGene_tx=XM_006943790.3
X	59610048	59610161	refGene_gn=IGBP1;refGene_tx=XM_006943790.3
X	59626768	59626914	refGene_gn=IGBP1;refGene_tx=XM_006943790.3
X	59639634	59639719
refGene_gn=DGAT2L6;refGene_tx=XM_004000645.2;refGene_gn=DGAT2L6;refGene_tx=XM_019823920.1
X	59657095	59657206	refGene_gn=DGAT2L6;refGene_tx=XM_004000645.2
X	59657734	59657805
refGene_gn=DGAT2L6;refGene_tx=XM_004000645.2;refGene_gn=DGAT2L6;refGene_tx=XM_019823920.1
X	59658166	59658371
refGene_gn=DGAT2L6;refGene_tx=XM_004000645.2;refGene_gn=DGAT2L6;refGene_tx=XM_019823920.1
X	59659757	59659932
refGene_gn=DGAT2L6;refGene_tx=XM_004000645.2;refGene_gn=DGAT2L6;refGene_tx=XM_019823920.1
X	59663279	59663491
refGene_gn=DGAT2L6;refGene_tx=XM_004000645.2;refGene_gn=DGAT2L6;refGene_tx=XM_019823920.1
X	59663916	59664071
refGene_gn=DGAT2L6;refGene_tx=XM_004000645.2;refGene_gn=DGAT2L6;refGene_tx=XM_019823920.1
X	59687964	59688040	refGene_gn=AWAT1;refGene_tx=XM_004000591.4
X	59688988	59689096	refGene_gn=AWAT1;refGene_tx=XM_004000591.4
X	59689365	59689436	refGene_gn=AWAT1;refGene_tx=XM_004000591.4
X	59690312	59690517	refGene_gn=AWAT1;refGene_tx=XM_004000591.4
X	59691201	59691373	refGene_gn=AWAT1;refGene_tx=XM_004000591.4
X	59692926	59693126	refGene_gn=AWAT1;refGene_tx=XM_004000591.4
X	59693345	59693500	refGene_gn=AWAT1;refGene_tx=XM_004000591.4

X	59715975 59717184 refGene_gn=P2RY4;refGene_tx=XM_004000592.4
X	59729201 59729308
refGene_gn=PDZD11;refGene_tx=XM_011291786.2;refGene_gn=PDZD11;refGene_tx=XM_011291790.2;refGene_gn=PDZD11;refGene_tx=XM_011291792.2;refGene_gn=PDZD11;refGene_tx=XM_019823932.2
X	59729324 59729327
refGene_gn=ARR3;refGene_tx=XM_011291794.2;refGene_gn=ARR3;refGene_tx=XM_019823921.2;refGene_gn=ARR3;refGene_tx=XM_019823922.1;refGene_gn=ARR3;refGene_tx=XM_019823923.1;refGene_gn=ARR3;refGene_tx=XM_019823925.1;refGene_gn=ARR3;refGene_tx=XM_019823926.1;refGene_gn=ARR3;refGene_tx=XM_019823927.1;refGene_gn=ARR3;refGene_tx=XM_019823928.1;refGene_gn=ARR3;refGene_tx=XM_019823930.1;refGene_gn=ARR3;refGene_tx=XM_019823931.1
X	59729559 59729597
refGene_gn=ARR3;refGene_tx=XM_011291794.2;refGene_gn=ARR3;refGene_tx=XM_019823921.2;refGene_gn=ARR3;refGene_tx=XM_019823922.1;refGene_gn=ARR3;refGene_tx=XM_019823923.1;refGene_gn=ARR3;refGene_tx=XM_019823924.1;refGene_gn=ARR3;refGene_tx=XM_019823925.1;refGene_gn=ARR3;refGene_tx=XM_019823926.1;refGene_gn=ARR3;refGene_tx=XM_019823927.1;refGene_gn=ARR3;refGene_tx=XM_019823928.1;refGene_gn=ARR3;refGene_tx=XM_019823930.1;refGene_gn=ARR3;refGene_tx=XM_019823931.1
X	59729870 59729901
refGene_gn=ARR3;refGene_tx=XM_011291794.2;refGene_gn=ARR3;refGene_tx=XM_019823921.2;refGene_gn=ARR3;refGene_tx=XM_019823922.1;refGene_gn=ARR3;refGene_tx=XM_019823923.1;refGene_gn=ARR3;refGene_tx=XM_019823924.1;refGene_gn=ARR3;refGene_tx=XM_019823925.1;refGene_gn=ARR3;refGene_tx=XM_019823926.1;refGene_gn=ARR3;refGene_tx=XM_019823927.1;refGene_gn=ARR3;refGene_tx=XM_019823928.1;refGene_gn=ARR3;refGene_tx=XM_019823930.1;refGene_gn=ARR3;refGene_tx=XM_019823931.1
X	59730235 59730352
refGene_gn=ARR3;refGene_tx=XM_011291794.2;refGene_gn=ARR3;refGene_tx=XM_019823921.2;refGene_gn=ARR3;refGene_tx=XM_019823922.1;refGene_gn=ARR3;refGene_tx=XM_019823923.1;refGene_gn=ARR3;refGene_tx=XM_019823924.1;refGene_gn=ARR3;refGene_tx=XM_019823925.1;refGene_gn=ARR3;refGene_tx=XM_019823926.1;refGene_gn=ARR3;refGene_tx=XM_019823927.1;refGene_gn=ARR3;refGene_tx=XM_019823928.1;refGene_gn=ARR3;refGene_tx=XM_019823930.1;refGene_gn=ARR3;refGene_tx=XM_019823931.1;refGene_gn=PDZD11;refGene_tx=XM_011291786.2;refGene_gn=PDZD11;refGene_tx=XM_011291790.2;refGene_gn=PDZD11;refGene_tx=XM_019823932.2
X	59730495 59730540
refGene_gn=ARR3;refGene_tx=XM_011291794.2;refGene_gn=ARR3;refGene_tx=XM_019823921.2;refGene_gn=ARR3;refGene_tx=XM_019823922.1;refGene_gn=ARR3;refGene_tx=XM_019823923.1;refGene_gn=ARR3;refGene_tx=XM_019823924.1;refGene_gn=ARR3;refGene_tx=XM_019823925.1;refGene_gn=ARR3;refGene_tx=XM_019823926.1;refGene_gn=ARR3;refGene_tx=XM_019823927.1;refGene_gn=ARR3;refGene_tx=XM_019823928.1;refGene_gn=ARR3;refGene_tx=XM_019823929.1;refGene_gn=ARR3;refGene_tx=XM_019823930.1;refGene_gn=ARR3;refGene_tx=XM_019823931.1
X	59737380 59737580
refGene_gn=ARR3;refGene_tx=XM_011291794.2;refGene_gn=ARR3;refGene_tx=XM_019823921.2;refGene_gn=ARR3;refGene_tx=XM_019823922.1;refGene_gn=ARR

3;refGene_tx=XM_019823923.1;refGene_gn=ARR3;refGene_tx=XM_019823924.1;refGene_gn=ARR3;refGene_tx=XM_019823925.1;refGene_gn=ARR3;refGene_tx=XM_019823926.1;refGene_gn=ARR3;refGene_tx=XM_019823927.1;refGene_gn=ARR3;refGene_tx=XM_019823928.1;refGene_gn=ARR3;refGene_tx=XM_019823929.1;refGene_gn=ARR3;refGene_tx=XM_019823930.1;refGene_gn=ARR3;refGene_tx=XM_019823931.1
X 59737705 59737765
refGene_gn=ARR3;refGene_tx=XM_011291794.2;refGene_gn=ARR3;refGene_tx=XM_019823921.2;refGene_gn=ARR3;refGene_tx=XM_019823922.1;refGene_gn=ARR3;refGene_tx=XM_019823923.1;refGene_gn=ARR3;refGene_tx=XM_019823924.1;refGene_gn=ARR3;refGene_tx=XM_019823925.1;refGene_gn=ARR3;refGene_tx=XM_019823926.1;refGene_gn=ARR3;refGene_tx=XM_019823927.1;refGene_gn=ARR3;refGene_tx=XM_019823928.1;refGene_gn=ARR3;refGene_tx=XM_019823929.1;refGene_gn=ARR3;refGene_tx=XM_019823930.1;refGene_gn=ARR3;refGene_tx=XM_019823931.1
X 59737945 59738013
refGene_gn=ARR3;refGene_tx=XM_011291794.2;refGene_gn=ARR3;refGene_tx=XM_019823921.2;refGene_gn=ARR3;refGene_tx=XM_019823922.1;refGene_gn=ARR3;refGene_tx=XM_019823923.1;refGene_gn=ARR3;refGene_tx=XM_019823924.1;refGene_gn=ARR3;refGene_tx=XM_019823925.1;refGene_gn=ARR3;refGene_tx=XM_019823926.1;refGene_gn=ARR3;refGene_tx=XM_019823927.1;refGene_gn=ARR3;refGene_tx=XM_019823928.1;refGene_gn=ARR3;refGene_tx=XM_019823929.1;refGene_gn=ARR3;refGene_tx=XM_019823930.1;refGene_gn=ARR3;refGene_tx=XM_019823931.1
X 59738091 59738186
refGene_gn=PDZD11;refGene_tx=XM_011291791.3;refGene_gn=PDZD11;refGene_tx=XM_019823933.1
X 59738655 59738791
refGene_gn=ARR3;refGene_tx=XM_011291794.2;refGene_gn=ARR3;refGene_tx=XM_019823921.2;refGene_gn=ARR3;refGene_tx=XM_019823922.1;refGene_gn=ARR3;refGene_tx=XM_019823923.1;refGene_gn=ARR3;refGene_tx=XM_019823924.1;refGene_gn=ARR3;refGene_tx=XM_019823925.1;refGene_gn=ARR3;refGene_tx=XM_019823926.1;refGene_gn=ARR3;refGene_tx=XM_019823927.1;refGene_gn=ARR3;refGene_tx=XM_019823928.1;refGene_gn=ARR3;refGene_tx=XM_019823929.1;refGene_gn=ARR3;refGene_tx=XM_019823930.1;refGene_gn=ARR3;refGene_tx=XM_019823931.1
X 59738980 59739065
refGene_gn=ARR3;refGene_tx=XM_011291794.2;refGene_gn=ARR3;refGene_tx=XM_019823921.2;refGene_gn=ARR3;refGene_tx=XM_019823922.1;refGene_gn=ARR3;refGene_tx=XM_019823923.1;refGene_gn=ARR3;refGene_tx=XM_019823924.1;refGene_gn=ARR3;refGene_tx=XM_019823925.1;refGene_gn=ARR3;refGene_tx=XM_019823926.1;refGene_gn=ARR3;refGene_tx=XM_019823927.1;refGene_gn=ARR3;refGene_tx=XM_019823928.1;refGene_gn=ARR3;refGene_tx=XM_019823929.1;refGene_gn=ARR3;refGene_tx=XM_019823930.1;refGene_gn=ARR3;refGene_tx=XM_019823931.1
X 59739326 59739399
refGene_gn=ARR3;refGene_tx=XM_011291794.2;refGene_gn=ARR3;refGene_tx=XM_019823921.2;refGene_gn=ARR3;refGene_tx=XM_019823922.1;refGene_gn=ARR3;refGene_tx=XM_019823923.1;refGene_gn=ARR3;refGene_tx=XM_019823924.1;refGene_gn=ARR3;refGene_tx=XM_019823925.1;refGene_gn=ARR3;refGene_tx=X

M_019823926.1;refGene_gn=ARR3;refGene_tx=XM_019823927.1;refGene_gn=ARR3;refGene_tx=XM_019823928.1;refGene_gn=ARR3;refGene_tx=XM_019823929.1;refGene_gn=ARR3;refGene_tx=XM_019823930.1;refGene_gn=ARR3;refGene_tx=XM_019823931.1
X 59739796 59739934
refGene_gn=ARR3;refGene_tx=XM_011291794.2;refGene_gn=ARR3;refGene_tx=XM_019823921.2;refGene_gn=ARR3;refGene_tx=XM_019823922.1;refGene_gn=ARR3;refGene_tx=XM_019823923.1;refGene_gn=ARR3;refGene_tx=XM_019823924.1;refGene_gn=ARR3;refGene_tx=XM_019823925.1;refGene_gn=ARR3;refGene_tx=XM_019823926.1;refGene_gn=ARR3;refGene_tx=XM_019823927.1;refGene_gn=ARR3;refGene_tx=XM_019823928.1;refGene_gn=ARR3;refGene_tx=XM_019823929.1;refGene_gn=ARR3;refGene_tx=XM_019823930.1;refGene_gn=ARR3;refGene_tx=XM_019823931.1
X 59741472 59741568
refGene_gn=ARR3;refGene_tx=XM_011291794.2;refGene_gn=ARR3;refGene_tx=XM_019823921.2;refGene_gn=ARR3;refGene_tx=XM_019823922.1;refGene_gn=ARR3;refGene_tx=XM_019823923.1;refGene_gn=ARR3;refGene_tx=XM_019823924.1;refGene_gn=ARR3;refGene_tx=XM_019823925.1;refGene_gn=ARR3;refGene_tx=XM_019823926.1;refGene_gn=ARR3;refGene_tx=XM_019823927.1;refGene_gn=ARR3;refGene_tx=XM_019823928.1;refGene_gn=ARR3;refGene_tx=XM_019823929.1;refGene_gn=ARR3;refGene_tx=XM_019823930.1;refGene_gn=ARR3;refGene_tx=XM_019823931.1
X 59741773 59741829
refGene_gn=ARR3;refGene_tx=XM_011291794.2;refGene_gn=ARR3;refGene_tx=XM_019823921.2;refGene_gn=ARR3;refGene_tx=XM_019823922.1;refGene_gn=ARR3;refGene_tx=XM_019823923.1;refGene_gn=ARR3;refGene_tx=XM_019823924.1;refGene_gn=ARR3;refGene_tx=XM_019823925.1;refGene_gn=ARR3;refGene_tx=XM_019823926.1;refGene_gn=ARR3;refGene_tx=XM_019823927.1;refGene_gn=ARR3;refGene_tx=XM_019823928.1;refGene_gn=ARR3;refGene_tx=XM_019823929.1;refGene_gn=ARR3;refGene_tx=XM_019823930.1;refGene_gn=ARR3;refGene_tx=XM_019823931.1
X 59742001 59742054
refGene_gn=ARR3;refGene_tx=XM_011291794.2;refGene_gn=ARR3;refGene_tx=XM_019823921.2;refGene_gn=ARR3;refGene_tx=XM_019823922.1;refGene_gn=ARR3;refGene_tx=XM_019823923.1;refGene_gn=ARR3;refGene_tx=XM_019823924.1;refGene_gn=ARR3;refGene_tx=XM_019823925.1;refGene_gn=ARR3;refGene_tx=XM_019823926.1;refGene_gn=ARR3;refGene_tx=XM_019823927.1;refGene_gn=ARR3;refGene_tx=XM_019823928.1;refGene_gn=ARR3;refGene_tx=XM_019823929.1
X 59742329 59742339
refGene_gn=ARR3;refGene_tx=XM_011291794.2;refGene_gn=ARR3;refGene_tx=XM_019823921.2;refGene_gn=ARR3;refGene_tx=XM_019823922.1;refGene_gn=ARR3;refGene_tx=XM_019823923.1;refGene_gn=ARR3;refGene_tx=XM_019823924.1;refGene_gn=ARR3;refGene_tx=XM_019823928.1;refGene_gn=ARR3;refGene_tx=XM_019823929.1
X 59742847 59742944
refGene_gn=ARR3;refGene_tx=XM_011291794.2;refGene_gn=ARR3;refGene_tx=XM_019823921.2;refGene_gn=ARR3;refGene_tx=XM_019823922.1;refGene_gn=ARR3;refGene_tx=XM_019823923.1;refGene_gn=ARR3;refGene_tx=XM_019823924.1;refGene_gn=ARR3;refGene_tx=XM_019823925.1;refGene_gn=ARR3;refGene_tx=XM_019823926.1;refGene_gn=ARR3;refGene_tx=XM_019823927.1;refGene_gn=ARR

3;refGene_tx=XM_019823929.1
X	59748524	59748559	refGene_gn=PDZD11;refGene_tx=XM_004000593.4
X	59748718	59748779
refGene_gn=PDZD11;refGene_tx=XM_004000593.4;refGene_gn=PDZD11;refGene_tx=XM_011291786.2;refGene_gn=PDZD11;refGene_tx=XM_019823932.2;refGene_gn=PDZD11;refGene_tx=XM_019823933.1
X	59749251	59749350
refGene_gn=PDZD11;refGene_tx=XM_004000593.4;refGene_gn=PDZD11;refGene_tx=XM_011291786.2;refGene_gn=PDZD11;refGene_tx=XM_011291790.2;refGene_gn=PDZD11;refGene_tx=XM_011291791.3;refGene_gn=PDZD11;refGene_tx=XM_011291792.2;refGene_gn=PDZD11;refGene_tx=XM_019823932.2;refGene_gn=PDZD11;refGene_tx=XM_019823933.1
X	59749646	59749703
refGene_gn=PDZD11;refGene_tx=XM_004000593.4;refGene_gn=PDZD11;refGene_tx=XM_011291786.2;refGene_gn=PDZD11;refGene_tx=XM_011291790.2;refGene_gn=PDZD11;refGene_tx=XM_011291791.3;refGene_gn=PDZD11;refGene_tx=XM_011291792.2;refGene_gn=PDZD11;refGene_tx=XM_019823932.2;refGene_gn=PDZD11;refGene_tx=XM_019823933.1
X	59749975	59750059
refGene_gn=PDZD11;refGene_tx=XM_004000593.4;refGene_gn=PDZD11;refGene_tx=XM_011291786.2;refGene_gn=PDZD11;refGene_tx=XM_011291790.2;refGene_gn=PDZD11;refGene_tx=XM_011291791.3;refGene_gn=PDZD11;refGene_tx=XM_011291792.2;refGene_gn=PDZD11;refGene_tx=XM_019823932.2;refGene_gn=PDZD11;refGene_tx=XM_019823933.1
X	59750508	59750595
refGene_gn=PDZD11;refGene_tx=XM_004000593.4;refGene_gn=PDZD11;refGene_tx=XM_011291786.2;refGene_gn=PDZD11;refGene_tx=XM_011291790.2;refGene_gn=PDZD11;refGene_tx=XM_011291791.3;refGene_gn=PDZD11;refGene_tx=XM_011291792.2;refGene_gn=PDZD11;refGene_tx=XM_019823932.2;refGene_gn=PDZD11;refGene_tx=XM_019823933.1
X	59751942	59752062	refGene_gn=KIF4A;refGene_tx=XM_004000594.4
X	59752176	59752291	refGene_gn=KIF4A;refGene_tx=XM_004000594.4
X	59756190	59756381	refGene_gn=KIF4A;refGene_tx=XM_004000594.4
X	59757725	59757818	refGene_gn=KIF4A;refGene_tx=XM_004000594.4
X	59761066	59761233	refGene_gn=KIF4A;refGene_tx=XM_004000594.4
X	59761472	59761567	refGene_gn=KIF4A;refGene_tx=XM_004000594.4
X	59797487	59797604	refGene_gn=KIF4A;refGene_tx=XM_004000594.4
X	59798214	59798390	refGene_gn=KIF4A;refGene_tx=XM_004000594.4
X	59804582	59804644	refGene_gn=KIF4A;refGene_tx=XM_004000594.4
X	59811585	59811718	refGene_gn=KIF4A;refGene_tx=XM_004000594.4
X	59813426	59813485	refGene_gn=KIF4A;refGene_tx=XM_004000594.4
X	59813591	59813697	refGene_gn=KIF4A;refGene_tx=XM_004000594.4
X	59818187	59818235	refGene_gn=KIF4A;refGene_tx=XM_004000594.4
X	59819586	59819772	refGene_gn=KIF4A;refGene_tx=XM_004000594.4
X	59827807	59827911	refGene_gn=KIF4A;refGene_tx=XM_004000594.4
X	59829089	59829234	refGene_gn=KIF4A;refGene_tx=XM_004000594.4
X	59829980	59830091	refGene_gn=KIF4A;refGene_tx=XM_004000594.4
X	59832633	59832717	refGene_gn=KIF4A;refGene_tx=XM_004000594.4
X	59833286	59833400	refGene_gn=KIF4A;refGene_tx=XM_004000594.4
X	59853214	59853370	refGene_gn=KIF4A;refGene_tx=XM_004000594.4

X 59853490 59853591 refGene_gn=KIF4A;refGene_tx=XM_004000594.4
X 59857919 59858049 refGene_gn=KIF4A;refGene_tx=XM_004000594.4
X 59859224 59859395 refGene_gn=KIF4A;refGene_tx=XM_004000594.4
X 59859777 59859885 refGene_gn=KIF4A;refGene_tx=XM_004000594.4
X 59860903 59860981 refGene_gn=KIF4A;refGene_tx=XM_004000594.4
X 59861337 59861436 refGene_gn=KIF4A;refGene_tx=XM_004000594.4
X 59861974 59862157 refGene_gn=KIF4A;refGene_tx=XM_004000594.4
X 59870589 59870706 refGene_gn=KIF4A;refGene_tx=XM_004000594.4
X 59872209 59872332 refGene_gn=KIF4A;refGene_tx=XM_004000594.4
X 59872600 59872801 refGene_gn=KIF4A;refGene_tx=XM_004000594.4
X 59876926 59877031 refGene_gn=GDPD2;refGene_tx=XM_004000595.5
X 59877313 59877417 refGene_gn=GDPD2;refGene_tx=XM_004000595.5
X 59877703 59877797
refGene_gn=GDPD2;refGene_tx=XM_004000595.5;refGene_gn=GDPD2;refGene_tx
=XM_019823934.1
X 59877975 59878035
refGene_gn=GDPD2;refGene_tx=XM_004000595.5;refGene_gn=GDPD2;refGene_tx
=XM_019823934.1
X 59878270 59878369
refGene_gn=GDPD2;refGene_tx=XM_004000595.5;refGene_gn=GDPD2;refGene_tx
=XM_019823934.1
X 59878546 59878640
refGene_gn=GDPD2;refGene_tx=XM_004000595.5;refGene_gn=GDPD2;refGene_tx
=XM_019823934.1
X 59878786 59878932
refGene_gn=GDPD2;refGene_tx=XM_004000595.5;refGene_gn=GDPD2;refGene_tx
=XM_019823934.1
X 59879057 59879140
refGene_gn=GDPD2;refGene_tx=XM_004000595.5;refGene_gn=GDPD2;refGene_tx
=XM_019823934.1
X 59879237 59879388
refGene_gn=GDPD2;refGene_tx=XM_004000595.5;refGene_gn=GDPD2;refGene_tx
=XM_019823934.1
X 59881655 59881877
refGene_gn=GDPD2;refGene_tx=XM_004000595.5;refGene_gn=GDPD2;refGene_tx
=XM_019823934.1
X 59882090 59882239
refGene_gn=GDPD2;refGene_tx=XM_004000595.5;refGene_gn=GDPD2;refGene_tx
=XM_019823934.1
X 59884290 59884438
refGene_gn=GDPD2;refGene_tx=XM_004000595.5;refGene_gn=GDPD2;refGene_tx
=XM_019823934.1
X 59884581 59884664
refGene_gn=GDPD2;refGene_tx=XM_004000595.5;refGene_gn=GDPD2;refGene_tx
=XM_019823934.1
X 59884905 59884934
refGene_gn=GDPD2;refGene_tx=XM_004000595.5;refGene_gn=GDPD2;refGene_tx
=XM_019823934.1
X 59885024 59885077
refGene_gn=GDPD2;refGene_tx=XM_004000595.5;refGene_gn=GDPD2;refGene_tx

=XM_019823934.1
X	59897293 59897650
refGene_gn=DLG3;refGene_tx=XM_011291795.3;refGene_gn=DLG3;refGene_tx=XM_011291796.3;refGene_gn=DLG3;refGene_tx=XM_019823935.2;refGene_gn=DLG3;refGene_tx=XM_019823936.2
X	59901236 59901287
refGene_gn=DLG3;refGene_tx=XM_011291795.3;refGene_gn=DLG3;refGene_tx=XM_011291796.3;refGene_gn=DLG3;refGene_tx=XM_019823935.2;refGene_gn=DLG3;refGene_tx=XM_019823936.2
X	59901706 59901831
refGene_gn=DLG3;refGene_tx=XM_011291795.3;refGene_gn=DLG3;refGene_tx=XM_011291796.3;refGene_gn=DLG3;refGene_tx=XM_019823935.2;refGene_gn=DLG3;refGene_tx=XM_019823936.2
X	59902028 59902198
refGene_gn=DLG3;refGene_tx=XM_011291795.3;refGene_gn=DLG3;refGene_tx=XM_011291796.3;refGene_gn=DLG3;refGene_tx=XM_019823935.2;refGene_gn=DLG3;refGene_tx=XM_019823936.2
X	59902497 59902634
refGene_gn=DLG3;refGene_tx=XM_011291795.3;refGene_gn=DLG3;refGene_tx=XM_011291796.3;refGene_gn=DLG3;refGene_tx=XM_019823935.2;refGene_gn=DLG3;refGene_tx=XM_019823936.2
X	59902972 59903117
refGene_gn=DLG3;refGene_tx=XM_011291795.3;refGene_gn=DLG3;refGene_tx=XM_011291796.3;refGene_gn=DLG3;refGene_tx=XM_019823935.2;refGene_gn=DLG3;refGene_tx=XM_019823936.2
X	59903721 59903881
refGene_gn=DLG3;refGene_tx=XM_011291795.3;refGene_gn=DLG3;refGene_tx=XM_011291796.3;refGene_gn=DLG3;refGene_tx=XM_019823935.2;refGene_gn=DLG3;refGene_tx=XM_019823936.2
X	59904477 59904611 refGene_gn=DLG3;refGene_tx=XM_004000596.5
X	59905501 59905658
refGene_gn=DLG3;refGene_tx=XM_004000596.5;refGene_gn=DLG3;refGene_tx=XM_011291795.3;refGene_gn=DLG3;refGene_tx=XM_011291796.3;refGene_gn=DLG3;refGene_tx=XM_019823935.2;refGene_gn=DLG3;refGene_tx=XM_019823936.2
X	59906071 59906174
refGene_gn=DLG3;refGene_tx=XM_004000596.5;refGene_gn=DLG3;refGene_tx=XM_011291795.3;refGene_gn=DLG3;refGene_tx=XM_011291796.3;refGene_gn=DLG3;refGene_tx=XM_019823935.2;refGene_gn=DLG3;refGene_tx=XM_019823936.2
X	59927521 59927636
refGene_gn=DLG3;refGene_tx=XM_004000596.5;refGene_gn=DLG3;refGene_tx=XM_011291795.3;refGene_gn=DLG3;refGene_tx=XM_011291796.3;refGene_gn=DLG3;refGene_tx=XM_011291797.2;refGene_gn=DLG3;refGene_tx=XM_019823935.2;refGene_gn=DLG3;refGene_tx=XM_019823936.2;refGene_gn=DLG3;refGene_tx=XM_023248814.1
X	59942719 59942896
refGene_gn=DLG3;refGene_tx=XM_004000596.5;refGene_gn=DLG3;refGene_tx=XM_011291795.3;refGene_gn=DLG3;refGene_tx=XM_011291796.3;refGene_gn=DLG3;refGene_tx=XM_011291797.2;refGene_gn=DLG3;refGene_tx=XM_019823935.2;refGene_gn=DLG3;refGene_tx=XM_019823936.2;refGene_gn=DLG3;refGene_tx=XM_023248814.1

X 59943128 59943204
refGene_gn=DLG3;refGene_tx=XM_004000596.5;refGene_gn=DLG3;refGene_tx=XM_011291795.3;refGene_gn=DLG3;refGene_tx=XM_011291796.3;refGene_gn=DLG3;refGene_tx=XM_011291797.2;refGene_gn=DLG3;refGene_tx=XM_019823935.2;refGene_gn=DLG3;refGene_tx=XM_019823936.2;refGene_gn=DLG3;refGene_tx=XM_023248814.1
X 59944024 59944124
refGene_gn=DLG3;refGene_tx=XM_004000596.5;refGene_gn=DLG3;refGene_tx=XM_011291795.3;refGene_gn=DLG3;refGene_tx=XM_011291797.2;refGene_gn=DLG3;refGene_tx=XM_019823935.2;refGene_gn=DLG3;refGene_tx=XM_023248814.1
X 59946005 59946051
refGene_gn=DLG3;refGene_tx=XM_011291796.3;refGene_gn=DLG3;refGene_tx=XM_019823936.2
X 59947770 59947812
refGene_gn=DLG3;refGene_tx=XM_004000596.5;refGene_gn=DLG3;refGene_tx=XM_011291795.3;refGene_gn=DLG3;refGene_tx=XM_011291797.2;refGene_gn=DLG3;refGene_tx=XM_019823936.2;refGene_gn=DLG3;refGene_tx=XM_023248814.1
X 59949156 59949207
refGene_gn=DLG3;refGene_tx=XM_004000596.5;refGene_gn=DLG3;refGene_tx=XM_011291795.3;refGene_gn=DLG3;refGene_tx=XM_011291796.3;refGene_gn=DLG3;refGene_tx=XM_011291797.2;refGene_gn=DLG3;refGene_tx=XM_019823935.2;refGene_gn=DLG3;refGene_tx=XM_019823936.2;refGene_gn=DLG3;refGene_tx=XM_023248814.1
X 59949825 59949927
refGene_gn=DLG3;refGene_tx=XM_004000596.5;refGene_gn=DLG3;refGene_tx=XM_011291795.3;refGene_gn=DLG3;refGene_tx=XM_011291796.3;refGene_gn=DLG3;refGene_tx=XM_011291797.2;refGene_gn=DLG3;refGene_tx=XM_019823935.2;refGene_gn=DLG3;refGene_tx=XM_019823936.2;refGene_gn=DLG3;refGene_tx=XM_023248814.1
X 59950532 59950705
refGene_gn=DLG3;refGene_tx=XM_004000596.5;refGene_gn=DLG3;refGene_tx=XM_011291795.3;refGene_gn=DLG3;refGene_tx=XM_011291796.3;refGene_gn=DLG3;refGene_tx=XM_011291797.2;refGene_gn=DLG3;refGene_tx=XM_019823935.2;refGene_gn=DLG3;refGene_tx=XM_019823936.2;refGene_gn=DLG3;refGene_tx=XM_023248814.1
X 59951124 59951234
refGene_gn=DLG3;refGene_tx=XM_004000596.5;refGene_gn=DLG3;refGene_tx=XM_011291795.3;refGene_gn=DLG3;refGene_tx=XM_011291796.3;refGene_gn=DLG3;refGene_tx=XM_011291797.2;refGene_gn=DLG3;refGene_tx=XM_019823935.2;refGene_gn=DLG3;refGene_tx=XM_019823936.2;refGene_gn=DLG3;refGene_tx=XM_023248814.1
X 59951867 59951959
refGene_gn=DLG3;refGene_tx=XM_004000596.5;refGene_gn=DLG3;refGene_tx=XM_011291795.3;refGene_gn=DLG3;refGene_tx=XM_011291796.3;refGene_gn=DLG3;refGene_tx=XM_011291797.2;refGene_gn=DLG3;refGene_tx=XM_019823935.2;refGene_gn=DLG3;refGene_tx=XM_019823936.2;refGene_gn=DLG3;refGene_tx=XM_023248814.1
X 59953046 59953153
refGene_gn=DLG3;refGene_tx=XM_004000596.5;refGene_gn=DLG3;refGene_tx=XM_011291795.3;refGene_gn=DLG3;refGene_tx=XM_011291796.3;refGene_gn=DLG

3;refGene_tx=XM_011291797.2;refGene_gn=DLG3;refGene_tx=XM_019823935.2;
refGene_gn=DLG3;refGene_tx=XM_019823936.2;refGene_gn=DLG3;refGene_tx=X
M_023248814.1
X 59971874 59971946 refGene_gn=TEX11;refGene_tx=XM_006943797.3
X 59972837 59973002 refGene_gn=TEX11;refGene_tx=XM_006943797.3
X 59990898 59991019 refGene_gn=TEX11;refGene_tx=XM_006943797.3
X 59991923 59992032 refGene_gn=TEX11;refGene_tx=XM_006943797.3
X 59992462 59992635 refGene_gn=TEX11;refGene_tx=XM_023249125.1
X 59993429 59993579
refGene_gn=TEX11;refGene_tx=XM_006943797.3;refGene_gn=TEX11;refGene_tx
=XM_023249125.1
X 60023075 60023148
refGene_gn=TEX11;refGene_tx=XM_006943797.3;refGene_gn=TEX11;refGene_tx
=XM_023249125.1
X 60038074 60038191
refGene_gn=TEX11;refGene_tx=XM_006943797.3;refGene_gn=TEX11;refGene_tx
=XM_023249125.1
X 60039622 60039693
refGene_gn=TEX11;refGene_tx=XM_006943797.3;refGene_gn=TEX11;refGene_tx
=XM_023249125.1
X 60046307 60046394
refGene_gn=TEX11;refGene_tx=XM_006943797.3;refGene_gn=TEX11;refGene_tx
=XM_023249125.1
X 60047549 60047590
refGene_gn=TEX11;refGene_tx=XM_006943797.3;refGene_gn=TEX11;refGene_tx
=XM_023249125.1
X 60051247 60051304
refGene_gn=TEX11;refGene_tx=XM_006943797.3;refGene_gn=TEX11;refGene_tx
=XM_023249125.1
X 60055487 60055573
refGene_gn=TEX11;refGene_tx=XM_006943797.3;refGene_gn=TEX11;refGene_tx
=XM_023249125.1
X 60058292 60058417
refGene_gn=TEX11;refGene_tx=XM_006943797.3;refGene_gn=TEX11;refGene_tx
=XM_023249125.1
X 60062473 60062576
refGene_gn=TEX11;refGene_tx=XM_006943797.3;refGene_gn=TEX11;refGene_tx
=XM_023249125.1
X 60089283 60089421
refGene_gn=TEX11;refGene_tx=XM_006943797.3;refGene_gn=TEX11;refGene_tx
=XM_023249125.1
X 60096549 60096635
refGene_gn=TEX11;refGene_tx=XM_006943797.3;refGene_gn=TEX11;refGene_tx
=XM_023249125.1
X 60099571 60099723
refGene_gn=TEX11;refGene_tx=XM_006943797.3;refGene_gn=TEX11;refGene_tx
=XM_023249125.1
X 60134374 60134453
refGene_gn=TEX11;refGene_tx=XM_006943797.3;refGene_gn=TEX11;refGene_tx
=XM_023249125.1

X 60138359 60138441
refGene_gn=TEX11;refGene_tx=XM_006943797.3;refGene_gn=TEX11;refGene_tx
=XM_023249125.1
X 60140369 60140465
refGene_gn=TEX11;refGene_tx=XM_006943797.3;refGene_gn=TEX11;refGene_tx
=XM_023249125.1
X 60143877 60143932
refGene_gn=TEX11;refGene_tx=XM_006943797.3;refGene_gn=TEX11;refGene_tx
=XM_023249125.1
X 60166322 60166408
refGene_gn=TEX11;refGene_tx=XM_006943797.3;refGene_gn=TEX11;refGene_tx
=XM_023249125.1
X 60178770 60178851
refGene_gn=TEX11;refGene_tx=XM_006943797.3;refGene_gn=TEX11;refGene_tx
=XM_023249125.1
X 60188687 60188807
refGene_gn=TEX11;refGene_tx=XM_006943797.3;refGene_gn=TEX11;refGene_tx
=XM_023249125.1
X 60188895 60188976
refGene_gn=TEX11;refGene_tx=XM_006943797.3;refGene_gn=TEX11;refGene_tx
=XM_023249125.1
X 60194097 60194177
refGene_gn=TEX11;refGene_tx=XM_006943797.3;refGene_gn=TEX11;refGene_tx
=XM_023249125.1
X 60197569 60197654
refGene_gn=TEX11;refGene_tx=XM_006943797.3;refGene_gn=TEX11;refGene_tx
=XM_023249125.1
X 60198327 60198449
refGene_gn=TEX11;refGene_tx=XM_006943797.3;refGene_gn=TEX11;refGene_tx
=XM_023249125.1
X 60218167 60218204
refGene_gn=TEX11;refGene_tx=XM_006943797.3;refGene_gn=TEX11;refGene_tx
=XM_023249125.1
X 60231836 60231967 refGene_gn=SLC7A3;refGene_tx=XM_004000597.4
X 60232104 60232213 refGene_gn=SLC7A3;refGene_tx=XM_004000597.4
X 60232562 60232729 refGene_gn=SLC7A3;refGene_tx=XM_004000597.4
X 60234528 60234689 refGene_gn=SLC7A3;refGene_tx=XM_004000597.4
X 60234933 60235036 refGene_gn=SLC7A3;refGene_tx=XM_004000597.4
X 60235136 60235276 refGene_gn=SLC7A3;refGene_tx=XM_004000597.4
X 60235451 60235674 refGene_gn=SLC7A3;refGene_tx=XM_004000597.4
X 60235800 60235913 refGene_gn=SLC7A3;refGene_tx=XM_004000597.4
X 60236117 60236295 refGene_gn=SLC7A3;refGene_tx=XM_004000597.4
X 60236478 60236637 refGene_gn=SLC7A3;refGene_tx=XM_004000597.4
X 60237032 60237402 refGene_gn=SLC7A3;refGene_tx=XM_004000597.4
X 60371492 60371595
refGene_gn=SNX12;refGene_tx=XM_004000599.5;refGene_gn=SNX12;refGene_tx
=XM_019823938.2
X 60371993 60372118
refGene_gn=SNX12;refGene_tx=XM_004000599.5;refGene_gn=SNX12;refGene_tx
=XM_019823938.2

X 60373271 60373367
refGene_gn=SNX12;refGene_tx=XM_004000599.5;refGene_gn=SNX12;refGene_tx
=XM_019823938.2
X 60379638 60379803
refGene_gn=SNX12;refGene_tx=XM_004000599.5;refGene_gn=SNX12;refGene_tx
=XM_019823938.2
X 60401643 60402105 refGene_gn=FOXO4;refGene_tx=XM_004000601.5
X 60405652 60406712 refGene_gn=FOXO4;refGene_tx=XM_004000601.5
X 60407037 60407045 refGene_gn=FOXO4;refGene_tx=XM_004000601.5
X 60409043 60409151
refGene_gn=CXHXorf65;refGene_tx=XM_004000648.4;refGene_gn=CXHXorf65;re
fGene_tx=XM_006943799.3;refGene_gn=CXHXorf65;refGene_tx=XM_006943800.3
X 60409375 60409482
refGene_gn=CXHXorf65;refGene_tx=XM_004000648.4;refGene_gn=CXHXorf65;re
fGene_tx=XM_006943799.3;refGene_gn=CXHXorf65;refGene_tx=XM_006943800.3
X 60409963 60410032
refGene_gn=CXHXorf65;refGene_tx=XM_004000648.4;refGene_gn=CXHXorf65;re
fGene_tx=XM_006943800.3
X 60410343 60410478
refGene_gn=CXHXorf65;refGene_tx=XM_004000648.4;refGene_gn=CXHXorf65;re
fGene_tx=XM_006943799.3;refGene_gn=CXHXorf65;refGene_tx=XM_006943800.3
X 60410663 60410750
refGene_gn=CXHXorf65;refGene_tx=XM_004000648.4;refGene_gn=CXHXorf65;re
fGene_tx=XM_006943799.3
X 60410872 60410897
refGene_gn=CXHXorf65;refGene_tx=XM_004000648.4;refGene_gn=CXHXorf65;re
fGene_tx=XM_006943799.3;refGene_gn=CXHXorf65;refGene_tx=XM_006943800.3
X 60412003 60412189 refGene_gn=IL2RG;refGene_tx=XM_004000602.3
X 60412533 60412603 refGene_gn=IL2RG;refGene_tx=XM_004000602.3
X 60412860 60412957 refGene_gn=IL2RG;refGene_tx=XM_004000602.3
X 60413522 60413685 refGene_gn=IL2RG;refGene_tx=XM_004000602.3
X 60414100 60414240 refGene_gn=IL2RG;refGene_tx=XM_004000602.3
X 60414478 60414663 refGene_gn=IL2RG;refGene_tx=XM_004000602.3
X 60414877 60415031 refGene_gn=IL2RG;refGene_tx=XM_004000602.3
X 60415251 60415366 refGene_gn=IL2RG;refGene_tx=XM_004000602.3
X 60417707 60417806 refGene_gn=MED12;refGene_tx=XM_004000604.5
X 60418336 60418441 refGene_gn=MED12;refGene_tx=XM_004000604.5
X 60418649 60418841 refGene_gn=MED12;refGene_tx=XM_004000604.5
X 60418977 60419137 refGene_gn=MED12;refGene_tx=XM_004000604.5
X 60420131 60420313 refGene_gn=MED12;refGene_tx=XM_004000604.5
X 60420500 60420611 refGene_gn=MED12;refGene_tx=XM_004000604.5
X 60420734 60420989 refGene_gn=MED12;refGene_tx=XM_004000604.5
X 60421352 60421499 refGene_gn=MED12;refGene_tx=XM_004000604.5
X 60421882 60421982 refGene_gn=MED12;refGene_tx=XM_004000604.5
X 60422106 60422243 refGene_gn=MED12;refGene_tx=XM_004000604.5
X 60422443 60422575 refGene_gn=MED12;refGene_tx=XM_004000604.5
X 60423178 60423305 refGene_gn=MED12;refGene_tx=XM_004000604.5
X 60423758 60423988 refGene_gn=MED12;refGene_tx=XM_004000604.5
X 60424446 60424527 refGene_gn=MED12;refGene_tx=XM_004000604.5
X 60424660 60424831 refGene_gn=MED12;refGene_tx=XM_004000604.5

X 60425034 60425179 refGene_gn=MED12;refGene_tx=XM_004000604.5
X 60425345 60425396 refGene_gn=MED12;refGene_tx=XM_004000604.5
X 60425730 60425849 refGene_gn=MED12;refGene_tx=XM_004000604.5
X 60426073 60426217 refGene_gn=MED12;refGene_tx=XM_004000604.5
X 60426696 60426860 refGene_gn=MED12;refGene_tx=XM_004000604.5
X 60427037 60427169 refGene_gn=MED12;refGene_tx=XM_004000604.5
X 60427591 60427819 refGene_gn=MED12;refGene_tx=XM_004000604.5
X 60427994 60428139 refGene_gn=MED12;refGene_tx=XM_004000604.5
X 60428311 60428432 refGene_gn=MED12;refGene_tx=XM_004000604.5
X 60428812 60428914 refGene_gn=MED12;refGene_tx=XM_004000604.5
X 60429019 60429133 refGene_gn=MED12;refGene_tx=XM_004000604.5
X 60429385 60429561 refGene_gn=MED12;refGene_tx=XM_004000604.5
X 60429744 60429924 refGene_gn=MED12;refGene_tx=XM_004000604.5
X 60431200 60431272 refGene_gn=MED12;refGene_tx=XM_004000604.5
X 60431727 60431861 refGene_gn=MED12;refGene_tx=XM_004000604.5
X 60432023 60432185 refGene_gn=MED12;refGene_tx=XM_004000604.5
X 60432698 60432810 refGene_gn=MED12;refGene_tx=XM_004000604.5
X 60432957 60433047 refGene_gn=MED12;refGene_tx=XM_004000604.5
X 60433531 60433641 refGene_gn=MED12;refGene_tx=XM_004000604.5
X 60433829 60433965 refGene_gn=MED12;refGene_tx=XM_004000604.5
X 60434233 60434395 refGene_gn=MED12;refGene_tx=XM_004000604.5
X 60435658 60436033 refGene_gn=MED12;refGene_tx=XM_004000604.5
X 60436262 60436413 refGene_gn=MED12;refGene_tx=XM_004000604.5
X 60436568 60436765 refGene_gn=MED12;refGene_tx=XM_004000604.5
X 60436939 60437017 refGene_gn=MED12;refGene_tx=XM_004000604.5
X 60437106 60437324 refGene_gn=MED12;refGene_tx=XM_004000604.5
X 60439264 60439520 refGene_gn=MED12;refGene_tx=XM_004000604.5
X 60439818 60439884 refGene_gn=MED12;refGene_tx=XM_004000604.5
X 60439885 60439963 refGene_gn=MED12;refGene_tx=XM_004000604.5
X 60440159 60440241 refGene_gn=MED12;refGene_tx=XM_004000604.5
X 60440473 60440517 refGene_gn=MED12;refGene_tx=XM_004000604.5
X 60445756 60446213
refGene_gn=NLGN3;refGene_tx=XM_006943801.4;refGene_gn=NLGN3;refGene_tx=XM_019823940.1;refGene_gn=NLGN3;refGene_tx=XM_019823941.1;refGene_gn=NLGN3;refGene_tx=XM_019823942.1;refGene_gn=NLGN3;refGene_tx=XM_019823943.1;refGene_gn=NLGN3;refGene_tx=XM_019823944.1;refGene_gn=NLGN3;refGene_tx=XM_019823945.1
X 60446854 60446914
refGene_gn=NLGN3;refGene_tx=XM_006943801.4;refGene_gn=NLGN3;refGene_tx=XM_019823940.1;refGene_gn=NLGN3;refGene_tx=XM_019823941.1;refGene_gn=NLGN3;refGene_tx=XM_019823942.1;refGene_gn=NLGN3;refGene_tx=XM_019823945.1
X 60450927 60450987
refGene_gn=NLGN3;refGene_tx=XM_006943801.4;refGene_gn=NLGN3;refGene_tx=XM_019823940.1;refGene_gn=NLGN3;refGene_tx=XM_019823941.1;refGene_gn=NLGN3;refGene_tx=XM_019823942.1;refGene_gn=NLGN3;refGene_tx=XM_019823943.1;refGene_gn=NLGN3;refGene_tx=XM_019823945.1
X 60452774 60452924
refGene_gn=NLGN3;refGene_tx=XM_006943801.4;refGene_gn=NLGN3;refGene_tx=XM_006943804.4;refGene_gn=NLGN3;refGene_tx=XM_019823940.1;refGene_gn=

NLGN3;refGene_tx=XM_019823941.1;refGene_gn=NLGN3;refGene_tx=XM_0198239
42.1;refGene_gn=NLGN3;refGene_tx=XM_019823943.1;refGene_gn=NLGN3;refGe
ne_tx=XM_019823944.1;refGene_gn=NLGN3;refGene_tx=XM_019823945.1;refGen
e_gn=NLGN3;refGene_tx=XM_019823946.1
X 60460165 60460351
refGene_gn=NLGN3;refGene_tx=XM_006943801.4;refGene_gn=NLGN3;refGene_tx
=XM_006943804.4;refGene_gn=NLGN3;refGene_tx=XM_019823940.1;refGene_gn=
NLGN3;refGene_tx=XM_019823941.1;refGene_gn=NLGN3;refGene_tx=XM_0198239
42.1;refGene_gn=NLGN3;refGene_tx=XM_019823943.1;refGene_gn=NLGN3;refGe
ne_tx=XM_019823944.1;refGene_gn=NLGN3;refGene_tx=XM_019823945.1;refGen
e_gn=NLGN3;refGene_tx=XM_019823946.1
X 60462419 60463209
refGene_gn=NLGN3;refGene_tx=XM_006943801.4;refGene_gn=NLGN3;refGene_tx
=XM_006943804.4;refGene_gn=NLGN3;refGene_tx=XM_019823940.1;refGene_gn=
NLGN3;refGene_tx=XM_019823941.1;refGene_gn=NLGN3;refGene_tx=XM_0198239
42.1;refGene_gn=NLGN3;refGene_tx=XM_019823943.1;refGene_gn=NLGN3;refGe
ne_tx=XM_019823944.1;refGene_gn=NLGN3;refGene_tx=XM_019823945.1;refGen
e_gn=NLGN3;refGene_tx=XM_019823946.1
X 60463840 60464684
refGene_gn=NLGN3;refGene_tx=XM_006943801.4;refGene_gn=NLGN3;refGene_tx
=XM_006943804.4;refGene_gn=NLGN3;refGene_tx=XM_019823940.1;refGene_gn=
NLGN3;refGene_tx=XM_019823941.1;refGene_gn=NLGN3;refGene_tx=XM_0198239
42.1;refGene_gn=NLGN3;refGene_tx=XM_019823943.1;refGene_gn=NLGN3;refGe
ne_tx=XM_019823944.1;refGene_gn=NLGN3;refGene_tx=XM_019823945.1;refGen
e_gn=NLGN3;refGene_tx=XM_019823946.1
X 60510505 60511357
refGene_gn=GJB1;refGene_tx=XM_004000609.4;refGene_gn=GJB1;refGene_tx=X
M_019823947.2
X 60527858 60528051
refGene_gn=ZMYM3;refGene_tx=XM_004000612.5;refGene_gn=ZMYM3;refGene_tx
=XM_006943805.4;refGene_gn=ZMYM3;refGene_tx=XM_006943807.3;refGene_gn=
ZMYM3;refGene_tx=XM_023248884.1
X 60528169 60528287
refGene_gn=ZMYM3;refGene_tx=XM_004000612.5;refGene_gn=ZMYM3;refGene_tx
=XM_006943805.4;refGene_gn=ZMYM3;refGene_tx=XM_006943807.3;refGene_gn=
ZMYM3;refGene_tx=XM_023248884.1
X 60529686 60529941
refGene_gn=ZMYM3;refGene_tx=XM_004000612.5;refGene_gn=ZMYM3;refGene_tx
=XM_006943805.4;refGene_gn=ZMYM3;refGene_tx=XM_006943807.3;refGene_gn=
ZMYM3;refGene_tx=XM_023248884.1
X 60530448 60530563
refGene_gn=ZMYM3;refGene_tx=XM_004000612.5;refGene_gn=ZMYM3;refGene_tx
=XM_006943805.4;refGene_gn=ZMYM3;refGene_tx=XM_006943807.3;refGene_gn=
ZMYM3;refGene_tx=XM_023248884.1
X 60531046 60531198
refGene_gn=ZMYM3;refGene_tx=XM_004000612.5;refGene_gn=ZMYM3;refGene_tx
=XM_006943805.4;refGene_gn=ZMYM3;refGene_tx=XM_006943807.3;refGene_gn=
ZMYM3;refGene_tx=XM_023248884.1
X 60531577 60531746
refGene_gn=ZMYM3;refGene_tx=XM_004000612.5;refGene_gn=ZMYM3;refGene_tx

=XM_006943805.4;refGene_gn=ZMYM3;refGene_tx=XM_006943807.3;refGene_gn=ZMYM3;refGene_tx=XM_023248884.1
X 60532071 60532175
refGene_gn=ZMYM3;refGene_tx=XM_004000612.5;refGene_gn=ZMYM3;refGene_tx=XM_006943805.4;refGene_gn=ZMYM3;refGene_tx=XM_006943807.3;refGene_gn=ZMYM3;refGene_tx=XM_023248884.1
X 60532644 60532791
refGene_gn=ZMYM3;refGene_tx=XM_004000612.5;refGene_gn=ZMYM3;refGene_tx=XM_006943805.4;refGene_gn=ZMYM3;refGene_tx=XM_006943807.3;refGene_gn=ZMYM3;refGene_tx=XM_023248884.1
X 60532975 60533150
refGene_gn=ZMYM3;refGene_tx=XM_004000612.5;refGene_gn=ZMYM3;refGene_tx=XM_006943805.4;refGene_gn=ZMYM3;refGene_tx=XM_006943807.3;refGene_gn=ZMYM3;refGene_tx=XM_023248884.1
X 60533299 60533412
refGene_gn=ZMYM3;refGene_tx=XM_004000612.5;refGene_gn=ZMYM3;refGene_tx=XM_006943805.4;refGene_gn=ZMYM3;refGene_tx=XM_006943807.3;refGene_gn=ZMYM3;refGene_tx=XM_023248884.1
X 60533696 60533859
refGene_gn=ZMYM3;refGene_tx=XM_004000612.5;refGene_gn=ZMYM3;refGene_tx=XM_006943805.4;refGene_gn=ZMYM3;refGene_tx=XM_006943807.3;refGene_gn=ZMYM3;refGene_tx=XM_023248884.1
X 60533942 60534040
refGene_gn=ZMYM3;refGene_tx=XM_004000612.5;refGene_gn=ZMYM3;refGene_tx=XM_006943805.4;refGene_gn=ZMYM3;refGene_tx=XM_006943807.3;refGene_gn=ZMYM3;refGene_tx=XM_023248884.1
X 60534705 60534871
refGene_gn=ZMYM3;refGene_tx=XM_004000612.5;refGene_gn=ZMYM3;refGene_tx=XM_006943805.4;refGene_gn=ZMYM3;refGene_tx=XM_006943807.3;refGene_gn=ZMYM3;refGene_tx=XM_023248884.1
X 60535074 60535247
refGene_gn=ZMYM3;refGene_tx=XM_004000612.5;refGene_gn=ZMYM3;refGene_tx=XM_006943805.4;refGene_gn=ZMYM3;refGene_tx=XM_006943807.3;refGene_gn=ZMYM3;refGene_tx=XM_023248884.1
X 60535537 60535688
refGene_gn=ZMYM3;refGene_tx=XM_004000612.5;refGene_gn=ZMYM3;refGene_tx=XM_006943805.4;refGene_gn=ZMYM3;refGene_tx=XM_006943807.3;refGene_gn=ZMYM3;refGene_tx=XM_023248884.1
X 60535823 60535910
refGene_gn=ZMYM3;refGene_tx=XM_004000612.5;refGene_gn=ZMYM3;refGene_tx=XM_006943805.4;refGene_gn=ZMYM3;refGene_tx=XM_006943807.3;refGene_gn=ZMYM3;refGene_tx=XM_023248884.1
X 60536069 60536185
refGene_gn=ZMYM3;refGene_tx=XM_004000612.5;refGene_gn=ZMYM3;refGene_tx=XM_006943805.4;refGene_gn=ZMYM3;refGene_tx=XM_006943807.3;refGene_gn=ZMYM3;refGene_tx=XM_023248884.1
X 60536429 60536580
refGene_gn=ZMYM3;refGene_tx=XM_004000612.5;refGene_gn=ZMYM3;refGene_tx=XM_006943805.4;refGene_gn=ZMYM3;refGene_tx=XM_006943807.3;refGene_gn=ZMYM3;refGene_tx=XM_023248884.1

X 60536872 60537091
refGene_gn=ZMYM3;refGene_tx=XM_004000612.5;refGene_gn=ZMYM3;refGene_tx=XM_006943805.4;refGene_gn=ZMYM3;refGene_tx=XM_006943807.3;refGene_gn=ZMYM3;refGene_tx=XM_023248884.1
X 60537438 60537616
refGene_gn=ZMYM3;refGene_tx=XM_004000612.5;refGene_gn=ZMYM3;refGene_tx=XM_006943805.4;refGene_gn=ZMYM3;refGene_tx=XM_006943807.3;refGene_gn=ZMYM3;refGene_tx=XM_023248884.1
X 60537829 60538124
refGene_gn=ZMYM3;refGene_tx=XM_004000612.5;refGene_gn=ZMYM3;refGene_tx=XM_006943805.4;refGene_gn=ZMYM3;refGene_tx=XM_006943807.3;refGene_gn=ZMYM3;refGene_tx=XM_023248884.1
X 60538533 60538606
refGene_gn=ZMYM3;refGene_tx=XM_004000612.5;refGene_gn=ZMYM3;refGene_tx=XM_006943805.4;refGene_gn=ZMYM3;refGene_tx=XM_006943807.3;refGene_gn=ZMYM3;refGene_tx=XM_023248884.1
X 60538941 60538985
refGene_gn=ZMYM3;refGene_tx=XM_004000612.5;refGene_gn=ZMYM3;refGene_tx=XM_006943805.4;refGene_gn=ZMYM3;refGene_tx=XM_006943807.3;refGene_gn=ZMYM3;refGene_tx=XM_023248884.1
X 60539979 60540646
refGene_gn=ZMYM3;refGene_tx=XM_004000612.5;refGene_gn=ZMYM3;refGene_tx=XM_006943805.4;refGene_gn=ZMYM3;refGene_tx=XM_006943807.3;refGene_gn=ZMYM3;refGene_tx=XM_023248884.1
X 60563600 60563751
refGene_gn=NONO;refGene_tx=XM_004000614.4;refGene_gn=NONO;refGene_tx=XM_019823949.2
X 60564712 60564906
refGene_gn=NONO;refGene_tx=XM_004000614.4;refGene_gn=NONO;refGene_tx=XM_019823949.2
X 60566085 60566387
refGene_gn=NONO;refGene_tx=XM_004000614.4;refGene_gn=NONO;refGene_tx=XM_019823949.2
X 60567407 60567503
refGene_gn=NONO;refGene_tx=XM_004000614.4;refGene_gn=NONO;refGene_tx=XM_019823949.2
X 60567687 60567884
refGene_gn=NONO;refGene_tx=XM_004000614.4;refGene_gn=NONO;refGene_tx=XM_019823949.2
X 60568229 60568314
refGene_gn=NONO;refGene_tx=XM_004000614.4;refGene_gn=NONO;refGene_tx=XM_019823949.2
X 60568711 60568814
refGene_gn=NONO;refGene_tx=XM_004000614.4;refGene_gn=NONO;refGene_tx=XM_019823949.2
X 60569393 60569433
refGene_gn=NONO;refGene_tx=XM_004000614.4;refGene_gn=NONO;refGene_tx=XM_019823949.2
X 60569659 60569769
refGene_gn=NONO;refGene_tx=XM_004000614.4;refGene_gn=NONO;refGene_tx=X

M_019823949.2
X 60571241 60571376
refGene_gn=NONO;refGene_tx=XM_004000614.4;refGene_gn=NONO;refGene_tx=XM_019823949.2
X 60572900 60572964
refGene_gn=ITGB1BP2;refGene_tx=XM_004000616.3;refGene_gn=ITGB1BP2;refGene_tx=XM_006943808.2;refGene_gn=ITGB1BP2;refGene_tx=XM_006943809.2;refGene_gn=ITGB1BP2;refGene_tx=XM_006943812.2;refGene_gn=ITGB1BP2;refGene_tx=XM_006943814.3;refGene_gn=ITGB1BP2;refGene_tx=XM_006943815.2;refGene_gn=ITGB1BP2;refGene_tx=XM_011291798.3;refGene_gn=ITGB1BP2;refGene_tx=XM_019823951.1;refGene_gn=ITGB1BP2;refGene_tx=XM_019823952.1
X 60573212 60573262
refGene_gn=ITGB1BP2;refGene_tx=XM_004000616.3;refGene_gn=ITGB1BP2;refGene_tx=XM_006943808.2;refGene_gn=ITGB1BP2;refGene_tx=XM_006943809.2;refGene_gn=ITGB1BP2;refGene_tx=XM_006943812.2;refGene_gn=ITGB1BP2;refGene_tx=XM_006943814.3;refGene_gn=ITGB1BP2;refGene_tx=XM_006943815.2;refGene_gn=ITGB1BP2;refGene_tx=XM_011291798.3;refGene_gn=ITGB1BP2;refGene_tx=XM_019823951.1;refGene_gn=ITGB1BP2;refGene_tx=XM_019823952.1
X 60573350 60573407
refGene_gn=ITGB1BP2;refGene_tx=XM_004000616.3;refGene_gn=ITGB1BP2;refGene_tx=XM_006943808.2;refGene_gn=ITGB1BP2;refGene_tx=XM_006943809.2;refGene_gn=ITGB1BP2;refGene_tx=XM_006943812.2;refGene_gn=ITGB1BP2;refGene_tx=XM_006943814.3;refGene_gn=ITGB1BP2;refGene_tx=XM_006943815.2;refGene_gn=ITGB1BP2;refGene_tx=XM_011291798.3;refGene_gn=ITGB1BP2;refGene_tx=XM_019823951.1;refGene_gn=ITGB1BP2;refGene_tx=XM_019823952.1
X 60573485 60573631
refGene_gn=ITGB1BP2;refGene_tx=XM_004000616.3;refGene_gn=ITGB1BP2;refGene_tx=XM_006943808.2;refGene_gn=ITGB1BP2;refGene_tx=XM_006943809.2;refGene_gn=ITGB1BP2;refGene_tx=XM_006943812.2;refGene_gn=ITGB1BP2;refGene_tx=XM_006943814.3;refGene_gn=ITGB1BP2;refGene_tx=XM_006943815.2;refGene_gn=ITGB1BP2;refGene_tx=XM_011291798.3;refGene_gn=ITGB1BP2;refGene_tx=XM_019823951.1;refGene_gn=ITGB1BP2;refGene_tx=XM_019823952.1
X 60574735 60574830
refGene_gn=ITGB1BP2;refGene_tx=XM_004000616.3;refGene_gn=ITGB1BP2;refGene_tx=XM_006943808.2;refGene_gn=ITGB1BP2;refGene_tx=XM_006943809.2;refGene_gn=ITGB1BP2;refGene_tx=XM_006943812.2;refGene_gn=ITGB1BP2;refGene_tx=XM_006943814.3;refGene_gn=ITGB1BP2;refGene_tx=XM_006943815.2;refGene_gn=ITGB1BP2;refGene_tx=XM_011291798.3;refGene_gn=ITGB1BP2;refGene_tx=XM_011291799.2;refGene_gn=ITGB1BP2;refGene_tx=XM_011291800.2;refGene_gn=ITGB1BP2;refGene_tx=XM_019823951.1;refGene_gn=ITGB1BP2;refGene_tx=XM_019823952.1
X 60574934 60574993
refGene_gn=ITGB1BP2;refGene_tx=XM_004000616.3;refGene_gn=ITGB1BP2;refGene_tx=XM_006943808.2;refGene_gn=ITGB1BP2;refGene_tx=XM_006943809.2;refGene_gn=ITGB1BP2;refGene_tx=XM_006943812.2;refGene_gn=ITGB1BP2;refGene_tx=XM_006943814.3;refGene_gn=ITGB1BP2;refGene_tx=XM_006943815.2;refGene_gn=ITGB1BP2;refGene_tx=XM_011291798.3;refGene_gn=ITGB1BP2;refGene_tx=XM_011291799.2;refGene_gn=ITGB1BP2;refGene_tx=XM_011291800.2;refGene_gn=ITGB1BP2;refGene_tx=XM_019823951.1;refGene_gn=ITGB1BP2;refGene_tx=XM_019823952.1

X 60575133 60575204
refGene_gn=ITGB1BP2;refGene_tx=XM_004000616.3;refGene_gn=ITGB1BP2;refGene_tx=XM_006943808.2;refGene_gn=ITGB1BP2;refGene_tx=XM_006943809.2;refGene_gn=ITGB1BP2;refGene_tx=XM_006943812.2;refGene_gn=ITGB1BP2;refGene_tx=XM_006943814.3;refGene_gn=ITGB1BP2;refGene_tx=XM_006943815.2;refGene_gn=ITGB1BP2;refGene_tx=XM_011291798.3;refGene_gn=ITGB1BP2;refGene_tx=XM_011291799.2;refGene_gn=ITGB1BP2;refGene_tx=XM_011291800.2;refGene_gn=ITGB1BP2;refGene_tx=XM_019823951.1;refGene_gn=ITGB1BP2;refGene_tx=XM_019823952.1
X 60575322 60575422
refGene_gn=ITGB1BP2;refGene_tx=XM_004000616.3;refGene_gn=ITGB1BP2;refGene_tx=XM_006943808.2;refGene_gn=ITGB1BP2;refGene_tx=XM_006943809.2;refGene_gn=ITGB1BP2;refGene_tx=XM_006943812.2;refGene_gn=ITGB1BP2;refGene_tx=XM_006943814.3;refGene_gn=ITGB1BP2;refGene_tx=XM_006943815.2;refGene_gn=ITGB1BP2;refGene_tx=XM_011291798.3;refGene_gn=ITGB1BP2;refGene_tx=XM_011291799.2;refGene_gn=ITGB1BP2;refGene_tx=XM_011291800.2;refGene_gn=ITGB1BP2;refGene_tx=XM_019823951.1;refGene_gn=ITGB1BP2;refGene_tx=XM_019823952.1
X 60575649 60575763
refGene_gn=ITGB1BP2;refGene_tx=XM_004000616.3;refGene_gn=ITGB1BP2;refGene_tx=XM_006943808.2;refGene_gn=ITGB1BP2;refGene_tx=XM_006943809.2;refGene_gn=ITGB1BP2;refGene_tx=XM_006943812.2;refGene_gn=ITGB1BP2;refGene_tx=XM_006943814.3;refGene_gn=ITGB1BP2;refGene_tx=XM_006943815.2;refGene_gn=ITGB1BP2;refGene_tx=XM_011291798.3;refGene_gn=ITGB1BP2;refGene_tx=XM_011291799.2;refGene_gn=ITGB1BP2;refGene_tx=XM_011291800.2;refGene_gn=ITGB1BP2;refGene_tx=XM_019823951.1;refGene_gn=ITGB1BP2;refGene_tx=XM_019823952.1
X 60576011 60576074
refGene_gn=ITGB1BP2;refGene_tx=XM_004000616.3;refGene_gn=ITGB1BP2;refGene_tx=XM_006943808.2;refGene_gn=ITGB1BP2;refGene_tx=XM_006943809.2;refGene_gn=ITGB1BP2;refGene_tx=XM_006943814.3;refGene_gn=ITGB1BP2;refGene_tx=XM_011291798.3;refGene_gn=ITGB1BP2;refGene_tx=XM_011291799.2;refGene_gn=ITGB1BP2;refGene_tx=XM_011291800.2;refGene_gn=ITGB1BP2;refGene_tx=XM_019823951.1;refGene_gn=ITGB1BP2;refGene_tx=XM_019823952.1
X 60576434 60576662
refGene_gn=ITGB1BP2;refGene_tx=XM_004000616.3;refGene_gn=ITGB1BP2;refGene_tx=XM_011291799.2;refGene_gn=ITGB1BP2;refGene_tx=XM_011291800.2
X 60579538 60579577
refGene_gn=ITGB1BP2;refGene_tx=XM_006943808.2;refGene_gn=ITGB1BP2;refGene_tx=XM_006943812.2;refGene_gn=ITGB1BP2;refGene_tx=XM_006943814.3;refGene_gn=ITGB1BP2;refGene_tx=XM_011291798.3;refGene_gn=ITGB1BP2;refGene_tx=XM_019823951.1
X 60580993 60581005 refGene_gn=ITGB1BP2;refGene_tx=XM_019823951.1
X 60595925 60596036
refGene_gn=ITGB1BP2;refGene_tx=XM_006943808.2;refGene_gn=ITGB1BP2;refGene_tx=XM_006943809.2;refGene_gn=ITGB1BP2;refGene_tx=XM_006943812.2;refGene_gn=ITGB1BP2;refGene_tx=XM_006943815.2
X 60597800 60597839
refGene_gn=ITGB1BP2;refGene_tx=XM_011291798.3;refGene_gn=ITGB1BP2;refGene_tx=XM_019823952.1

X	60617386 60617398
refGene_gn=LOC101093337;refGene_tx=XM_006943817.3
X	60617803 60618394
refGene_gn=LOC101093337;refGene_tx=XM_006943817.3
X	60625382 60625562
refGene_gn=TAF1;refGene_tx=XM_004000618.5;refGene_gn=TAF1;refGene_tx=XM_004000619.5;refGene_gn=TAF1;refGene_tx=XM_004000620.5;refGene_gn=TAF1;refGene_tx=XM_019823953.2;refGene_gn=TAF1;refGene_tx=XM_019823954.2;refGene_gn=TAF1;refGene_tx=XM_019823955.2;refGene_gn=TAF1;refGene_tx=XM_019823956.2;refGene_gn=TAF1;refGene_tx=XM_019823957.2
X	60626512 60626627
refGene_gn=TAF1;refGene_tx=XM_004000618.5;refGene_gn=TAF1;refGene_tx=XM_004000619.5;refGene_gn=TAF1;refGene_tx=XM_004000620.5;refGene_gn=TAF1;refGene_tx=XM_019823953.2;refGene_gn=TAF1;refGene_tx=XM_019823954.2;refGene_gn=TAF1;refGene_tx=XM_019823955.2;refGene_gn=TAF1;refGene_tx=XM_019823956.2;refGene_gn=TAF1;refGene_tx=XM_019823957.2
X	60626775 60626892
refGene_gn=TAF1;refGene_tx=XM_004000618.5;refGene_gn=TAF1;refGene_tx=XM_004000619.5;refGene_gn=TAF1;refGene_tx=XM_004000620.5;refGene_gn=TAF1;refGene_tx=XM_019823953.2;refGene_gn=TAF1;refGene_tx=XM_019823954.2;refGene_gn=TAF1;refGene_tx=XM_019823955.2;refGene_gn=TAF1;refGene_tx=XM_019823956.2;refGene_gn=TAF1;refGene_tx=XM_019823957.2
X	60630937 60631057
refGene_gn=TAF1;refGene_tx=XM_004000618.5;refGene_gn=TAF1;refGene_tx=XM_004000619.5;refGene_gn=TAF1;refGene_tx=XM_004000620.5;refGene_gn=TAF1;refGene_tx=XM_019823953.2;refGene_gn=TAF1;refGene_tx=XM_019823954.2;refGene_gn=TAF1;refGene_tx=XM_019823955.2;refGene_gn=TAF1;refGene_tx=XM_019823956.2;refGene_gn=TAF1;refGene_tx=XM_019823957.2
X	60632174 60632416
refGene_gn=TAF1;refGene_tx=XM_004000618.5;refGene_gn=TAF1;refGene_tx=XM_004000619.5;refGene_gn=TAF1;refGene_tx=XM_004000620.5;refGene_gn=TAF1;refGene_tx=XM_019823953.2;refGene_gn=TAF1;refGene_tx=XM_019823954.2;refGene_gn=TAF1;refGene_tx=XM_019823955.2;refGene_gn=TAF1;refGene_tx=XM_019823956.2;refGene_gn=TAF1;refGene_tx=XM_019823957.2
X	60632817 60633036
refGene_gn=TAF1;refGene_tx=XM_004000618.5;refGene_gn=TAF1;refGene_tx=XM_004000619.5;refGene_gn=TAF1;refGene_tx=XM_004000620.5;refGene_gn=TAF1;refGene_tx=XM_019823953.2;refGene_gn=TAF1;refGene_tx=XM_019823954.2;refGene_gn=TAF1;refGene_tx=XM_019823955.2;refGene_gn=TAF1;refGene_tx=XM_019823956.2;refGene_gn=TAF1;refGene_tx=XM_019823957.2
X	60633431 60633650
refGene_gn=TAF1;refGene_tx=XM_004000618.5;refGene_gn=TAF1;refGene_tx=XM_004000619.5;refGene_gn=TAF1;refGene_tx=XM_004000620.5;refGene_gn=TAF1;refGene_tx=XM_019823953.2;refGene_gn=TAF1;refGene_tx=XM_019823954.2;refGene_gn=TAF1;refGene_tx=XM_019823955.2;refGene_gn=TAF1;refGene_tx=XM_019823956.2;refGene_gn=TAF1;refGene_tx=XM_019823957.2
X	60633992 60634200
refGene_gn=TAF1;refGene_tx=XM_004000618.5;refGene_gn=TAF1;refGene_tx=XM_004000619.5;refGene_gn=TAF1;refGene_tx=XM_004000620.5;refGene_gn=TAF1;refGene_tx=XM_019823953.2;refGene_gn=TAF1;refGene_tx=XM_019823954.2;

refGene_gn=TAF1;refGene_tx=XM_019823955.2;refGene_gn=TAF1;refGene_tx=XM_019823956.2;refGene_gn=TAF1;refGene_tx=XM_019823957.2
X 60635664 60635841
refGene_gn=TAF1;refGene_tx=XM_004000618.5;refGene_gn=TAF1;refGene_tx=XM_004000619.5;refGene_gn=TAF1;refGene_tx=XM_004000620.5;refGene_gn=TAF1;refGene_tx=XM_019823953.2;refGene_gn=TAF1;refGene_tx=XM_019823954.2;refGene_gn=TAF1;refGene_tx=XM_019823955.2;refGene_gn=TAF1;refGene_tx=XM_019823956.2;refGene_gn=TAF1;refGene_tx=XM_019823957.2
X 60636308 60636436
refGene_gn=TAF1;refGene_tx=XM_004000618.5;refGene_gn=TAF1;refGene_tx=XM_004000619.5;refGene_gn=TAF1;refGene_tx=XM_004000620.5;refGene_gn=TAF1;refGene_tx=XM_019823953.2;refGene_gn=TAF1;refGene_tx=XM_019823954.2;refGene_gn=TAF1;refGene_tx=XM_019823955.2;refGene_gn=TAF1;refGene_tx=XM_019823956.2;refGene_gn=TAF1;refGene_tx=XM_019823957.2
X 60636536 60636644
refGene_gn=TAF1;refGene_tx=XM_004000618.5;refGene_gn=TAF1;refGene_tx=XM_004000619.5;refGene_gn=TAF1;refGene_tx=XM_004000620.5;refGene_gn=TAF1;refGene_tx=XM_019823953.2;refGene_gn=TAF1;refGene_tx=XM_019823954.2;refGene_gn=TAF1;refGene_tx=XM_019823955.2;refGene_gn=TAF1;refGene_tx=XM_019823956.2;refGene_gn=TAF1;refGene_tx=XM_019823957.2
X 60636765 60636939
refGene_gn=TAF1;refGene_tx=XM_004000618.5;refGene_gn=TAF1;refGene_tx=XM_004000619.5;refGene_gn=TAF1;refGene_tx=XM_004000620.5;refGene_gn=TAF1;refGene_tx=XM_019823953.2;refGene_gn=TAF1;refGene_tx=XM_019823954.2;refGene_gn=TAF1;refGene_tx=XM_019823955.2;refGene_gn=TAF1;refGene_tx=XM_019823956.2;refGene_gn=TAF1;refGene_tx=XM_019823957.2
X 60637397 60637571
refGene_gn=TAF1;refGene_tx=XM_004000618.5;refGene_gn=TAF1;refGene_tx=XM_004000619.5;refGene_gn=TAF1;refGene_tx=XM_004000620.5;refGene_gn=TAF1;refGene_tx=XM_019823953.2;refGene_gn=TAF1;refGene_tx=XM_019823954.2;refGene_gn=TAF1;refGene_tx=XM_019823955.2;refGene_gn=TAF1;refGene_tx=XM_019823956.2;refGene_gn=TAF1;refGene_tx=XM_019823957.2
X 60637801 60637906
refGene_gn=TAF1;refGene_tx=XM_004000618.5;refGene_gn=TAF1;refGene_tx=XM_004000619.5;refGene_gn=TAF1;refGene_tx=XM_004000620.5;refGene_gn=TAF1;refGene_tx=XM_019823953.2;refGene_gn=TAF1;refGene_tx=XM_019823954.2;refGene_gn=TAF1;refGene_tx=XM_019823955.2;refGene_gn=TAF1;refGene_tx=XM_019823956.2;refGene_gn=TAF1;refGene_tx=XM_019823957.2
X 60639977 60640181
refGene_gn=TAF1;refGene_tx=XM_004000618.5;refGene_gn=TAF1;refGene_tx=XM_004000619.5;refGene_gn=TAF1;refGene_tx=XM_004000620.5;refGene_gn=TAF1;refGene_tx=XM_019823953.2;refGene_gn=TAF1;refGene_tx=XM_019823954.2;refGene_gn=TAF1;refGene_tx=XM_019823955.2;refGene_gn=TAF1;refGene_tx=XM_019823956.2;refGene_gn=TAF1;refGene_tx=XM_019823957.2
X 60640349 60640491
refGene_gn=TAF1;refGene_tx=XM_004000618.5;refGene_gn=TAF1;refGene_tx=XM_004000619.5;refGene_gn=TAF1;refGene_tx=XM_004000620.5;refGene_gn=TAF1;refGene_tx=XM_019823953.2;refGene_gn=TAF1;refGene_tx=XM_019823954.2;refGene_gn=TAF1;refGene_tx=XM_019823955.2;refGene_gn=TAF1;refGene_tx=XM_019823956.2;refGene_gn=TAF1;refGene_tx=XM_019823957.2

X	60641076 60641207
refGene_gn=TAF1;refGene_tx=XM_004000618.5;refGene_gn=TAF1;refGene_tx=XM_004000619.5;refGene_gn=TAF1;refGene_tx=XM_004000620.5;refGene_gn=TAF1;refGene_tx=XM_019823953.2;refGene_gn=TAF1;refGene_tx=XM_019823954.2;refGene_gn=TAF1;refGene_tx=XM_019823955.2;refGene_gn=TAF1;refGene_tx=XM_019823956.2;refGene_gn=TAF1;refGene_tx=XM_019823957.2
X	60641874 60641955
refGene_gn=TAF1;refGene_tx=XM_004000618.5;refGene_gn=TAF1;refGene_tx=XM_004000619.5;refGene_gn=TAF1;refGene_tx=XM_004000620.5;refGene_gn=TAF1;refGene_tx=XM_019823953.2;refGene_gn=TAF1;refGene_tx=XM_019823954.2;refGene_gn=TAF1;refGene_tx=XM_019823955.2;refGene_gn=TAF1;refGene_tx=XM_019823956.2;refGene_gn=TAF1;refGene_tx=XM_019823957.2
X	60647433 60647583
refGene_gn=TAF1;refGene_tx=XM_004000618.5;refGene_gn=TAF1;refGene_tx=XM_004000619.5;refGene_gn=TAF1;refGene_tx=XM_004000620.5;refGene_gn=TAF1;refGene_tx=XM_019823953.2;refGene_gn=TAF1;refGene_tx=XM_019823954.2;refGene_gn=TAF1;refGene_tx=XM_019823955.2;refGene_gn=TAF1;refGene_tx=XM_019823956.2;refGene_gn=TAF1;refGene_tx=XM_019823957.2
X	60647751 60647871
refGene_gn=TAF1;refGene_tx=XM_004000618.5;refGene_gn=TAF1;refGene_tx=XM_004000619.5;refGene_gn=TAF1;refGene_tx=XM_004000620.5;refGene_gn=TAF1;refGene_tx=XM_019823953.2;refGene_gn=TAF1;refGene_tx=XM_019823954.2;refGene_gn=TAF1;refGene_tx=XM_019823955.2;refGene_gn=TAF1;refGene_tx=XM_019823956.2;refGene_gn=TAF1;refGene_tx=XM_019823957.2
X	60648626 60648802
refGene_gn=TAF1;refGene_tx=XM_004000618.5;refGene_gn=TAF1;refGene_tx=XM_004000619.5;refGene_gn=TAF1;refGene_tx=XM_004000620.5;refGene_gn=TAF1;refGene_tx=XM_019823953.2;refGene_gn=TAF1;refGene_tx=XM_019823954.2;refGene_gn=TAF1;refGene_tx=XM_019823955.2;refGene_gn=TAF1;refGene_tx=XM_019823956.2;refGene_gn=TAF1;refGene_tx=XM_019823957.2
X	60649404 60649583
refGene_gn=TAF1;refGene_tx=XM_004000618.5;refGene_gn=TAF1;refGene_tx=XM_004000619.5;refGene_gn=TAF1;refGene_tx=XM_004000620.5;refGene_gn=TAF1;refGene_tx=XM_019823953.2;refGene_gn=TAF1;refGene_tx=XM_019823954.2;refGene_gn=TAF1;refGene_tx=XM_019823955.2;refGene_gn=TAF1;refGene_tx=XM_019823956.2;refGene_gn=TAF1;refGene_tx=XM_019823957.2
X	60651272 60651486
refGene_gn=TAF1;refGene_tx=XM_004000618.5;refGene_gn=TAF1;refGene_tx=XM_004000619.5;refGene_gn=TAF1;refGene_tx=XM_004000620.5;refGene_gn=TAF1;refGene_tx=XM_019823953.2;refGene_gn=TAF1;refGene_tx=XM_019823954.2;refGene_gn=TAF1;refGene_tx=XM_019823955.2;refGene_gn=TAF1;refGene_tx=XM_019823956.2;refGene_gn=TAF1;refGene_tx=XM_019823957.2
X	60651942 60652108
refGene_gn=TAF1;refGene_tx=XM_004000618.5;refGene_gn=TAF1;refGene_tx=XM_004000619.5;refGene_gn=TAF1;refGene_tx=XM_004000620.5;refGene_gn=TAF1;refGene_tx=XM_019823953.2;refGene_gn=TAF1;refGene_tx=XM_019823954.2;refGene_gn=TAF1;refGene_tx=XM_019823955.2;refGene_gn=TAF1;refGene_tx=XM_019823956.2;refGene_gn=TAF1;refGene_tx=XM_019823957.2
X	60653717 60653929
refGene_gn=TAF1;refGene_tx=XM_004000618.5;refGene_gn=TAF1;refGene_tx=X

M_004000619.5;refGene_gn=TAF1;refGene_tx=XM_004000620.5;refGene_gn=TAF1;refGene_tx=XM_019823953.2;refGene_gn=TAF1;refGene_tx=XM_019823954.2;refGene_gn=TAF1;refGene_tx=XM_019823955.2;refGene_gn=TAF1;refGene_tx=XM_019823956.2;refGene_gn=TAF1;refGene_tx=XM_019823957.2

X 60657197 60657306 refGene_gn=TAF1;refGene_tx=XM_004000618.5;refGene_gn=TAF1;refGene_tx=XM_004000619.5;refGene_gn=TAF1;refGene_tx=XM_004000620.5;refGene_gn=TAF1;refGene_tx=XM_019823953.2;refGene_gn=TAF1;refGene_tx=XM_019823954.2;refGene_gn=TAF1;refGene_tx=XM_019823955.2;refGene_gn=TAF1;refGene_tx=XM_019823956.2;refGene_gn=TAF1;refGene_tx=XM_019823957.2

X 60657526 60657625 refGene_gn=TAF1;refGene_tx=XM_004000618.5;refGene_gn=TAF1;refGene_tx=XM_004000619.5;refGene_gn=TAF1;refGene_tx=XM_004000620.5;refGene_gn=TAF1;refGene_tx=XM_019823953.2;refGene_gn=TAF1;refGene_tx=XM_019823954.2;refGene_gn=TAF1;refGene_tx=XM_019823955.2;refGene_gn=TAF1;refGene_tx=XM_019823956.2;refGene_gn=TAF1;refGene_tx=XM_019823957.2

X 60657924 60658102 refGene_gn=TAF1;refGene_tx=XM_004000618.5;refGene_gn=TAF1;refGene_tx=XM_004000619.5;refGene_gn=TAF1;refGene_tx=XM_004000620.5;refGene_gn=TAF1;refGene_tx=XM_019823953.2;refGene_gn=TAF1;refGene_tx=XM_019823954.2;refGene_gn=TAF1;refGene_tx=XM_019823955.2;refGene_gn=TAF1;refGene_tx=XM_019823956.2;refGene_gn=TAF1;refGene_tx=XM_019823957.2

X 60664976 60665044 refGene_gn=TAF1;refGene_tx=XM_004000618.5;refGene_gn=TAF1;refGene_tx=XM_004000619.5;refGene_gn=TAF1;refGene_tx=XM_004000620.5;refGene_gn=TAF1;refGene_tx=XM_019823953.2;refGene_gn=TAF1;refGene_tx=XM_019823954.2;refGene_gn=TAF1;refGene_tx=XM_019823955.2;refGene_gn=TAF1;refGene_tx=XM_019823956.2;refGene_gn=TAF1;refGene_tx=XM_019823957.2

X 60666654 60666777 refGene_gn=TAF1;refGene_tx=XM_004000618.5;refGene_gn=TAF1;refGene_tx=XM_004000619.5;refGene_gn=TAF1;refGene_tx=XM_004000620.5;refGene_gn=TAF1;refGene_tx=XM_019823953.2;refGene_gn=TAF1;refGene_tx=XM_019823954.2;refGene_gn=TAF1;refGene_tx=XM_019823955.2;refGene_gn=TAF1;refGene_tx=XM_019823956.2;refGene_gn=TAF1;refGene_tx=XM_019823957.2

X 60668016 60668109 refGene_gn=TAF1;refGene_tx=XM_004000618.5;refGene_gn=TAF1;refGene_tx=XM_004000619.5;refGene_gn=TAF1;refGene_tx=XM_004000620.5;refGene_gn=TAF1;refGene_tx=XM_019823953.2;refGene_gn=TAF1;refGene_tx=XM_019823954.2;refGene_gn=TAF1;refGene_tx=XM_019823955.2;refGene_gn=TAF1;refGene_tx=XM_019823956.2;refGene_gn=TAF1;refGene_tx=XM_019823957.2

X 60668191 60668276 refGene_gn=TAF1;refGene_tx=XM_004000618.5;refGene_gn=TAF1;refGene_tx=XM_004000619.5;refGene_gn=TAF1;refGene_tx=XM_004000620.5;refGene_gn=TAF1;refGene_tx=XM_019823953.2;refGene_gn=TAF1;refGene_tx=XM_019823954.2;refGene_gn=TAF1;refGene_tx=XM_019823955.2;refGene_gn=TAF1;refGene_tx=XM_019823956.2;refGene_gn=TAF1;refGene_tx=XM_019823957.2

X 60698583 60698651 refGene_gn=TAF1;refGene_tx=XM_004000618.5;refGene_gn=TAF1;refGene_tx=XM_004000619.5;refGene_gn=TAF1;refGene_tx=XM_004000620.5;refGene_gn=TAF1;refGene_tx=XM_019823953.2;refGene_gn=TAF1;refGene_tx=XM_019823954.2;

refGene_gn=TAF1;refGene_tx=XM_019823955.2;refGene_gn=TAF1;refGene_tx=XM_019823956.2;refGene_gn=TAF1;refGene_tx=XM_019823957.2
X 60698990 60699107
refGene_gn=TAF1;refGene_tx=XM_004000618.5;refGene_gn=TAF1;refGene_tx=XM_004000619.5;refGene_gn=TAF1;refGene_tx=XM_004000620.5;refGene_gn=TAF1;refGene_tx=XM_019823953.2;refGene_gn=TAF1;refGene_tx=XM_019823954.2;refGene_gn=TAF1;refGene_tx=XM_019823955.2;refGene_gn=TAF1;refGene_tx=XM_019823956.2;refGene_gn=TAF1;refGene_tx=XM_019823957.2
X 60699256 60699262
refGene_gn=TAF1;refGene_tx=XM_004000620.5;refGene_gn=TAF1;refGene_tx=XM_019823953.2;refGene_gn=TAF1;refGene_tx=XM_019823955.2;refGene_gn=TAF1;refGene_tx=XM_019823956.2
X 60700551 60700644 refGene_gn=TAF1;refGene_tx=XM_019823957.2
X 60702272 60702398
refGene_gn=TAF1;refGene_tx=XM_004000618.5;refGene_gn=TAF1;refGene_tx=XM_004000619.5;refGene_gn=TAF1;refGene_tx=XM_004000620.5;refGene_gn=TAF1;refGene_tx=XM_019823953.2;refGene_gn=TAF1;refGene_tx=XM_019823954.2;refGene_gn=TAF1;refGene_tx=XM_019823955.2;refGene_gn=TAF1;refGene_tx=XM_019823956.2
X 60703253 60703355
refGene_gn=TAF1;refGene_tx=XM_019823953.2;refGene_gn=TAF1;refGene_tx=XM_019823954.2;refGene_gn=TAF1;refGene_tx=XM_019823955.2;refGene_gn=TAF1;refGene_tx=XM_019823956.2
X 60703623 60703780
refGene_gn=TAF1;refGene_tx=XM_004000618.5;refGene_gn=TAF1;refGene_tx=XM_004000619.5;refGene_gn=TAF1;refGene_tx=XM_004000620.5;refGene_gn=TAF1;refGene_tx=XM_019823953.2;refGene_gn=TAF1;refGene_tx=XM_019823954.2;refGene_gn=TAF1;refGene_tx=XM_019823955.2;refGene_gn=TAF1;refGene_tx=XM_019823956.2
X 60703893 60703919 refGene_gn=TAF1;refGene_tx=XM_019823956.2
X 60704835 60705013
refGene_gn=TAF1;refGene_tx=XM_004000618.5;refGene_gn=TAF1;refGene_tx=XM_004000619.5;refGene_gn=TAF1;refGene_tx=XM_004000620.5;refGene_gn=TAF1;refGene_tx=XM_019823953.2;refGene_gn=TAF1;refGene_tx=XM_019823954.2;refGene_gn=TAF1;refGene_tx=XM_019823955.2
X 60707618 60707838
refGene_gn=TAF1;refGene_tx=XM_004000618.5;refGene_gn=TAF1;refGene_tx=XM_004000619.5;refGene_gn=TAF1;refGene_tx=XM_004000620.5;refGene_gn=TAF1;refGene_tx=XM_019823953.2;refGene_gn=TAF1;refGene_tx=XM_019823954.2;refGene_gn=TAF1;refGene_tx=XM_019823955.2
X 60768558 60768595
refGene_gn=OGT;refGene_tx=XM_004000621.5;refGene_gn=OGT;refGene_tx=XM_004000622.5
X 60771407 60771588
refGene_gn=OGT;refGene_tx=XM_004000621.5;refGene_gn=OGT;refGene_tx=XM_004000622.5
X 60772434 60772678
refGene_gn=OGT;refGene_tx=XM_004000621.5;refGene_gn=OGT;refGene_tx=XM_004000622.5;refGene_gn=OGT;refGene_tx=XM_019823958.2;refGene_gn=OGT;refGene_tx=XM_019823959.2

X 60779054 60779123
refGene_gn=OGT;refGene_tx=XM_004000621.5;refGene_gn=OGT;refGene_tx=XM_004000622.5;refGene_gn=OGT;refGene_tx=XM_019823958.2;refGene_gn=OGT;refGene_tx=XM_019823959.2
X 60782408 60782525
refGene_gn=OGT;refGene_tx=XM_004000621.5;refGene_gn=OGT;refGene_tx=XM_004000622.5;refGene_gn=OGT;refGene_tx=XM_019823958.2;refGene_gn=OGT;refGene_tx=XM_019823959.2
X 60783721 60783801
refGene_gn=OGT;refGene_tx=XM_004000621.5;refGene_gn=OGT;refGene_tx=XM_004000622.5;refGene_gn=OGT;refGene_tx=XM_019823958.2;refGene_gn=OGT;refGene_tx=XM_019823959.2
X 60784581 60784777
refGene_gn=OGT;refGene_tx=XM_004000621.5;refGene_gn=OGT;refGene_tx=XM_004000622.5;refGene_gn=OGT;refGene_tx=XM_019823958.2;refGene_gn=OGT;refGene_tx=XM_019823959.2
X 60785029 60785170
refGene_gn=OGT;refGene_tx=XM_004000621.5;refGene_gn=OGT;refGene_tx=XM_004000622.5;refGene_gn=OGT;refGene_tx=XM_019823958.2;refGene_gn=OGT;refGene_tx=XM_019823959.2
X 60785749 60785850
refGene_gn=OGT;refGene_tx=XM_004000621.5;refGene_gn=OGT;refGene_tx=XM_004000622.5;refGene_gn=OGT;refGene_tx=XM_019823958.2;refGene_gn=OGT;refGene_tx=XM_019823959.2
X 60786014 60786168
refGene_gn=OGT;refGene_tx=XM_004000621.5;refGene_gn=OGT;refGene_tx=XM_004000622.5;refGene_gn=OGT;refGene_tx=XM_019823958.2;refGene_gn=OGT;refGene_tx=XM_019823959.2
X 60786254 60786356
refGene_gn=OGT;refGene_tx=XM_004000621.5;refGene_gn=OGT;refGene_tx=XM_004000622.5;refGene_gn=OGT;refGene_tx=XM_019823958.2;refGene_gn=OGT;refGene_tx=XM_019823959.2
X 60786543 60786723
refGene_gn=OGT;refGene_tx=XM_004000621.5;refGene_gn=OGT;refGene_tx=XM_004000622.5;refGene_gn=OGT;refGene_tx=XM_019823958.2;refGene_gn=OGT;refGene_tx=XM_019823959.2
X 60787608 60787767
refGene_gn=OGT;refGene_tx=XM_004000621.5;refGene_gn=OGT;refGene_tx=XM_004000622.5;refGene_gn=OGT;refGene_tx=XM_019823958.2;refGene_gn=OGT;refGene_tx=XM_019823959.2
X 60788689 60788779
refGene_gn=OGT;refGene_tx=XM_004000621.5;refGene_gn=OGT;refGene_tx=XM_004000622.5;refGene_gn=OGT;refGene_tx=XM_019823958.2;refGene_gn=OGT;refGene_tx=XM_019823959.2
X 60790656 60790782
refGene_gn=OGT;refGene_tx=XM_004000621.5;refGene_gn=OGT;refGene_tx=XM_004000622.5;refGene_gn=OGT;refGene_tx=XM_019823958.2;refGene_gn=OGT;refGene_tx=XM_019823959.2
X 60791474 60791645
refGene_gn=OGT;refGene_tx=XM_004000621.5;refGene_gn=OGT;refGene_tx=XM_

004000622.5;refGene_gn=OGT;refGene_tx=XM_019823958.2;refGene_gn=OGT;refGene_tx=XM_019823959.2
X 60791761 60791878
refGene_gn=OGT;refGene_tx=XM_004000621.5;refGene_gn=OGT;refGene_tx=XM_004000622.5;refGene_gn=OGT;refGene_tx=XM_019823958.2;refGene_gn=OGT;refGene_tx=XM_019823959.2
X 60791968 60792139
refGene_gn=OGT;refGene_tx=XM_004000621.5;refGene_gn=OGT;refGene_tx=XM_004000622.5;refGene_gn=OGT;refGene_tx=XM_019823958.2;refGene_gn=OGT;refGene_tx=XM_019823959.2
X 60796014 60796167
refGene_gn=OGT;refGene_tx=XM_004000621.5;refGene_gn=OGT;refGene_tx=XM_004000622.5;refGene_gn=OGT;refGene_tx=XM_019823958.2;refGene_gn=OGT;refGene_tx=XM_019823959.2
X 60797840 60798093
refGene_gn=OGT;refGene_tx=XM_004000621.5;refGene_gn=OGT;refGene_tx=XM_004000622.5;refGene_gn=OGT;refGene_tx=XM_019823958.2;refGene_gn=OGT;refGene_tx=XM_019823959.2
X 60798772 60798896
refGene_gn=OGT;refGene_tx=XM_004000621.5;refGene_gn=OGT;refGene_tx=XM_004000622.5;refGene_gn=OGT;refGene_tx=XM_019823958.2;refGene_gn=OGT;refGene_tx=XM_019823959.2
X 60802428 60802603
refGene_gn=OGT;refGene_tx=XM_004000621.5;refGene_gn=OGT;refGene_tx=XM_004000622.5;refGene_gn=OGT;refGene_tx=XM_019823958.2;refGene_gn=OGT;refGene_tx=XM_019823959.2
X 60815169 60815222
refGene_gn=GCNA;refGene_tx=XM_023249106.1;refGene_gn=GCNA;refGene_tx=XM_023249107.1
X 60819663 60819807 refGene_gn=GCNA;refGene_tx=XM_023249106.1
X 60820630 60820677
refGene_gn=GCNA;refGene_tx=XM_023249106.1;refGene_gn=GCNA;refGene_tx=XM_023249107.1
X 60821079 60821116
refGene_gn=GCNA;refGene_tx=XM_023249106.1;refGene_gn=GCNA;refGene_tx=XM_023249107.1
X 60822414 60822497
refGene_gn=GCNA;refGene_tx=XM_023249106.1;refGene_gn=GCNA;refGene_tx=XM_023249107.1
X 60825585 60825930
refGene_gn=GCNA;refGene_tx=XM_023249106.1;refGene_gn=GCNA;refGene_tx=XM_023249107.1
X 60827134 60827225
refGene_gn=GCNA;refGene_tx=XM_023249106.1;refGene_gn=GCNA;refGene_tx=XM_023249107.1
X 60830343 60830488
refGene_gn=GCNA;refGene_tx=XM_023249106.1;refGene_gn=GCNA;refGene_tx=XM_023249107.1
X 60831712 60831851
refGene_gn=GCNA;refGene_tx=XM_023249106.1;refGene_gn=GCNA;refGene_tx=X

M_023249107.1
X 60832824 60833029
refGene_gn=GCNA;refGene_tx=XM_023249106.1;refGene_gn=GCNA;refGene_tx=X
M_023249107.1
X 60833339 60833586
refGene_gn=GCNA;refGene_tx=XM_023249106.1;refGene_gn=GCNA;refGene_tx=X
M_023249107.1
X 60838061 60839156 refGene_gn=CXCR3;refGene_tx=XM_004000623.4
X 60840125 60840137 refGene_gn=CXCR3;refGene_tx=XM_004000623.4
X 60889122 60889148
refGene_gn=LOC101080873;refGene_tx=XM_023249307.1
X 60890041 60890198
refGene_gn=LOC101080873;refGene_tx=XM_023249307.1
X 60890812 60891123
refGene_gn=LOC101080873;refGene_tx=XM_023249307.1
X 60891125 60891453
refGene_gn=LOC101080873;refGene_tx=XM_023249307.1
X 60891455 60891922
refGene_gn=LOC101080873;refGene_tx=XM_023249307.1
X 60891924 60892051
refGene_gn=LOC101080873;refGene_tx=XM_023249307.1
X 61123576 61123619
refGene_gn=LOC101094826;refGene_tx=XM_019823964.2;refGene_gn=LOC101094
826;refGene_tx=XM_023248948.1;refGene_gn=LOC101094826;refGene_tx=XM_02
3248949.1
X 61174156 61175851 refGene_gn=RTL5;refGene_tx=XM_006943819.3
X 61176377 61176533
refGene_gn=LOC101094826;refGene_tx=XM_019823964.2;refGene_gn=LOC101094
826;refGene_tx=XM_023248948.1;refGene_gn=LOC101094826;refGene_tx=XM_02
3248949.1
X 61179426 61179554
refGene_gn=LOC101094826;refGene_tx=XM_019823964.2
X 61179857 61180053
refGene_gn=LOC101094826;refGene_tx=XM_019823964.2
X 61181869 61182001
refGene_gn=LOC101094826;refGene_tx=XM_019823964.2;refGene_gn=LOC101094
826;refGene_tx=XM_023248948.1
X 61183189 61185514
refGene_gn=LOC101094826;refGene_tx=XM_019823964.2;refGene_gn=LOC101094
826;refGene_tx=XM_023248948.1;refGene_gn=LOC101094826;refGene_tx=XM_02
3248949.1
X 61187011 61187144
refGene_gn=LOC101094826;refGene_tx=XM_019823964.2;refGene_gn=LOC101094
826;refGene_tx=XM_023248948.1;refGene_gn=LOC101094826;refGene_tx=XM_02
3248949.1
X 61187984 61188309
refGene_gn=LOC101094826;refGene_tx=XM_019823964.2;refGene_gn=LOC101094
826;refGene_tx=XM_023248948.1;refGene_gn=LOC101094826;refGene_tx=XM_02
3248949.1
X 61221109 61221290 refGene_gn=PIN4;refGene_tx=XM_004000626.5

X 61224081 61224155 refGene_gn=PIN4;refGene_tx=XM_004000626.5
X 61232759 61232879 refGene_gn=PIN4;refGene_tx=XM_004000626.5
X 61233262 61233421 refGene_gn=PIN4;refGene_tx=XM_004000626.5
X 61238005 61241663 refGene_gn=ERCC6L;refGene_tx=XM_006943820.4
X 61272690 61272758 refGene_gn=ERCC6L;refGene_tx=XM_006943820.4
X 61304727 61304829 refGene_gn=RPS4X;refGene_tx=XM_004000627.4
X 61305392 61305550 refGene_gn=RPS4X;refGene_tx=XM_004000627.4
X 61306304 61306476 refGene_gn=RPS4X;refGene_tx=XM_004000627.4
X 61307855 61307953 refGene_gn=RPS4X;refGene_tx=XM_004000627.4
X 61308337 61308518 refGene_gn=RPS4X;refGene_tx=XM_004000627.4
X 61309018 61309096 refGene_gn=RPS4X;refGene_tx=XM_004000627.4
X 61310098 61310101 refGene_gn=RPS4X;refGene_tx=XM_004000627.4
X 61331303 61331831
refGene_gn=CITED1;refGene_tx=XM_004000628.5;refGene_gn=CITED1;refGene_tx=XM_004000630.5;refGene_gn=CITED1;refGene_tx=XM_006943823.4;refGene_gn=CITED1;refGene_tx=XM_011291801.3;refGene_gn=CITED1;refGene_tx=XM_019823975.2;refGene_gn=CITED1;refGene_tx=XM_023249120.1;refGene_gn=CITED1;refGene_tx=XM_023249121.1;refGene_gn=CITED1;refGene_tx=XM_023249122.1
X 61332433 61332558
refGene_gn=CITED1;refGene_tx=XM_004000628.5;refGene_gn=CITED1;refGene_tx=XM_004000630.5;refGene_gn=CITED1;refGene_tx=XM_006943823.4;refGene_gn=CITED1;refGene_tx=XM_011291801.3;refGene_gn=CITED1;refGene_tx=XM_019823975.2;refGene_gn=CITED1;refGene_tx=XM_023249120.1;refGene_gn=CITED1;refGene_tx=XM_023249121.1;refGene_gn=CITED1;refGene_tx=XM_023249122.1
X 61334760 61334773 refGene_gn=CITED1;refGene_tx=XM_023249120.1
X 61357666 61357689
refGene_gn=HDAC8;refGene_tx=XM_019823976.2;refGene_gn=HDAC8;refGene_tx=XM_019823977.2;refGene_gn=HDAC8;refGene_tx=XM_019823978.2;refGene_gn=HDAC8;refGene_tx=XM_019823979.2;refGene_gn=HDAC8;refGene_tx=XM_019823980.2;refGene_gn=HDAC8;refGene_tx=XM_019823981.2
X 61358349 61358424
refGene_gn=HDAC8;refGene_tx=XM_019823976.2;refGene_gn=HDAC8;refGene_tx=XM_019823977.2;refGene_gn=HDAC8;refGene_tx=XM_019823979.2;refGene_gn=HDAC8;refGene_tx=XM_019823980.2;refGene_gn=HDAC8;refGene_tx=XM_019823981.2
X 61386821 61386927
refGene_gn=HDAC8;refGene_tx=XM_019823976.2;refGene_gn=HDAC8;refGene_tx=XM_019823977.2;refGene_gn=HDAC8;refGene_tx=XM_019823978.2;refGene_gn=HDAC8;refGene_tx=XM_019823979.2;refGene_gn=HDAC8;refGene_tx=XM_019823980.2;refGene_gn=HDAC8;refGene_tx=XM_019823981.2
X 61418954 61419050
refGene_gn=HDAC8;refGene_tx=XM_019823976.2;refGene_gn=HDAC8;refGene_tx=XM_019823977.2;refGene_gn=HDAC8;refGene_tx=XM_019823978.2;refGene_gn=HDAC8;refGene_tx=XM_019823980.2
X 61477712 61477772 refGene_gn=HDAC8;refGene_tx=XM_006943825.3
X 61487060 61487155
refGene_gn=HDAC8;refGene_tx=XM_006943825.3;refGene_gn=HDAC8;refGene_tx=XM_019823976.2;refGene_gn=HDAC8;refGene_tx=XM_019823977.2;refGene_gn=

HDAC8;refGene_tx=XM_019823978.2;refGene_gn=HDAC8;refGene_tx=XM_0198239
79.2;refGene_gn=HDAC8;refGene_tx=XM_019823980.2;refGene_gn=HDAC8;refGe
ne_tx=XM_019823981.2
X 61489676 61489849
refGene_gn=HDAC8;refGene_tx=XM_006943825.3;refGene_gn=HDAC8;refGene_tx
=XM_019823976.2;refGene_gn=HDAC8;refGene_tx=XM_019823977.2;refGene_gn=
HDAC8;refGene_tx=XM_019823978.2;refGene_gn=HDAC8;refGene_tx=XM_0198239
79.2;refGene_gn=HDAC8;refGene_tx=XM_019823980.2;refGene_gn=HDAC8;refGe
ne_tx=XM_019823981.2
X 61514883 61514992
refGene_gn=HDAC8;refGene_tx=XM_006943825.3;refGene_gn=HDAC8;refGene_tx
=XM_019823976.2;refGene_gn=HDAC8;refGene_tx=XM_019823977.2;refGene_gn=
HDAC8;refGene_tx=XM_019823978.2;refGene_gn=HDAC8;refGene_tx=XM_0198239
79.2;refGene_gn=HDAC8;refGene_tx=XM_019823980.2;refGene_gn=HDAC8;refGe
ne_tx=XM_019823981.2
X 61515600 61515678
refGene_gn=HDAC8;refGene_tx=XM_006943825.3;refGene_gn=HDAC8;refGene_tx
=XM_019823976.2;refGene_gn=HDAC8;refGene_tx=XM_019823977.2;refGene_gn=
HDAC8;refGene_tx=XM_019823978.2;refGene_gn=HDAC8;refGene_tx=XM_0198239
79.2;refGene_gn=HDAC8;refGene_tx=XM_019823980.2;refGene_gn=HDAC8;refGe
ne_tx=XM_019823981.2
X 61519704 61519817
refGene_gn=HDAC8;refGene_tx=XM_006943825.3;refGene_gn=HDAC8;refGene_tx
=XM_019823976.2;refGene_gn=HDAC8;refGene_tx=XM_019823977.2;refGene_gn=
HDAC8;refGene_tx=XM_019823978.2;refGene_gn=HDAC8;refGene_tx=XM_0198239
79.2;refGene_gn=HDAC8;refGene_tx=XM_019823980.2;refGene_gn=HDAC8;refGe
ne_tx=XM_019823981.2
X 61535351 61535459
refGene_gn=HDAC8;refGene_tx=XM_006943825.3;refGene_gn=HDAC8;refGene_tx
=XM_019823976.2;refGene_gn=HDAC8;refGene_tx=XM_019823978.2;refGene_gn=
HDAC8;refGene_tx=XM_019823979.2
X 61568385 61568493 refGene_gn=HDAC8;refGene_tx=XM_019823977.2
X 61575510 61575652
refGene_gn=HDAC8;refGene_tx=XM_006943825.3;refGene_gn=HDAC8;refGene_tx
=XM_019823976.2;refGene_gn=HDAC8;refGene_tx=XM_019823977.2;refGene_gn=
HDAC8;refGene_tx=XM_019823978.2;refGene_gn=HDAC8;refGene_tx=XM_0198239
79.2;refGene_gn=HDAC8;refGene_tx=XM_019823980.2;refGene_gn=HDAC8;refGe
ne_tx=XM_019823981.2
X 61577163 61577294
refGene_gn=HDAC8;refGene_tx=XM_006943825.3;refGene_gn=HDAC8;refGene_tx
=XM_019823976.2;refGene_gn=HDAC8;refGene_tx=XM_019823977.2;refGene_gn=
HDAC8;refGene_tx=XM_019823978.2;refGene_gn=HDAC8;refGene_tx=XM_0198239
79.2;refGene_gn=HDAC8;refGene_tx=XM_019823980.2;refGene_gn=HDAC8;refGe
ne_tx=XM_019823981.2
X 61580154 61580207
refGene_gn=HDAC8;refGene_tx=XM_006943825.3;refGene_gn=HDAC8;refGene_tx
=XM_019823976.2;refGene_gn=HDAC8;refGene_tx=XM_019823977.2;refGene_gn=
HDAC8;refGene_tx=XM_019823978.2;refGene_gn=HDAC8;refGene_tx=XM_0198239
79.2;refGene_gn=HDAC8;refGene_tx=XM_019823980.2;refGene_gn=HDAC8;refGe
ne_tx=XM_019823981.2

X 61580727 61580952
refGene_gn=HDAC8;refGene_tx=XM_006943825.3;refGene_gn=HDAC8;refGene_tx=XM_019823976.2;refGene_gn=HDAC8;refGene_tx=XM_019823977.2;refGene_gn=HDAC8;refGene_tx=XM_019823978.2;refGene_gn=HDAC8;refGene_tx=XM_019823979.2;refGene_gn=HDAC8;refGene_tx=XM_019823980.2;refGene_gn=HDAC8;refGene_tx=XM_019823981.2
X 61588166 61588340 refGene_gn=PHKA1;refGene_tx=XM_004000633.4
X 61589768 61589969 refGene_gn=PHKA1;refGene_tx=XM_004000633.4
X 61592092 61592146 refGene_gn=PHKA1;refGene_tx=XM_004000633.4
X 61594427 61594592 refGene_gn=PHKA1;refGene_tx=XM_004000633.4
X 61597615 61597666 refGene_gn=PHKA1;refGene_tx=XM_004000633.4
X 61599435 61599474 refGene_gn=PHKA1;refGene_tx=XM_004000633.4
X 61599611 61599727 refGene_gn=PHKA1;refGene_tx=XM_004000633.4
X 61600626 61600728 refGene_gn=PHKA1;refGene_tx=XM_004000633.4
X 61605813 61605943 refGene_gn=PHKA1;refGene_tx=XM_004000633.4
X 61606077 61606156 refGene_gn=PHKA1;refGene_tx=XM_004000633.4
X 61607435 61607473 refGene_gn=PHKA1;refGene_tx=XM_004000633.4
X 61607474 61607516 refGene_gn=PHKA1;refGene_tx=XM_004000633.4
X 61609929 61610086 refGene_gn=PHKA1;refGene_tx=XM_004000633.4
X 61617683 61617823 refGene_gn=PHKA1;refGene_tx=XM_004000633.4
X 61618431 61618523 refGene_gn=PHKA1;refGene_tx=XM_004000633.4
X 61620058 61620235 refGene_gn=PHKA1;refGene_tx=XM_004000633.4
X 61622570 61622737 refGene_gn=PHKA1;refGene_tx=XM_004000633.4
X 61625671 61625750 refGene_gn=PHKA1;refGene_tx=XM_004000633.4
X 61655775 61655920 refGene_gn=PHKA1;refGene_tx=XM_004000633.4
X 61656865 61656975 refGene_gn=PHKA1;refGene_tx=XM_004000633.4
X 61689484 61689619 refGene_gn=PHKA1;refGene_tx=XM_004000633.4
X 61708874 61708953 refGene_gn=PHKA1;refGene_tx=XM_004000633.4
X 61710676 61710784 refGene_gn=PHKA1;refGene_tx=XM_004000633.4
X 61712577 61712673 refGene_gn=PHKA1;refGene_tx=XM_004000633.4
X 61715784 61715907 refGene_gn=PHKA1;refGene_tx=XM_004000633.4
X 61717389 61717443 refGene_gn=PHKA1;refGene_tx=XM_004000633.4
X 61718627 61718774 refGene_gn=PHKA1;refGene_tx=XM_004000633.4
X 61719784 61719883 refGene_gn=PHKA1;refGene_tx=XM_004000633.4
X 61724539 61724620 refGene_gn=PHKA1;refGene_tx=XM_004000633.4
X 61725414 61728100
refGene_gn=LOC101097381;refGene_tx=XM_011291802.3
X 61728101 61728505
refGene_gn=LOC101097381;refGene_tx=XM_011291802.3
X 61737535 61737618 refGene_gn=PHKA1;refGene_tx=XM_004000633.4
X 61748915 61749084 refGene_gn=PHKA1;refGene_tx=XM_004000633.4
X 61750899 61750947 refGene_gn=PHKA1;refGene_tx=XM_004000633.4
X 61760265 61760424 refGene_gn=PHKA1;refGene_tx=XM_004000633.4
X 61761186 61761264 refGene_gn=PHKA1;refGene_tx=XM_004000633.4
X 61815322 61815594
refGene_gn=LOC111558754;refGene_tx=XM_023249330.1
X 61815679 61815837
refGene_gn=LOC111558754;refGene_tx=XM_023249330.1
X 61816577 61816714
refGene_gn=LOC111558754;refGene_tx=XM_023249330.1

X 61817871 61818019
refGene_gn=LOC111558754;refGene_tx=XM_023249330.1
X 61818996 61819109
refGene_gn=LOC111558754;refGene_tx=XM_023249330.1
X 61844908 61845021
refGene_gn=LOC101081365;refGene_tx=XM_023249275.1
X 61846014 61846162
refGene_gn=LOC101081365;refGene_tx=XM_023249275.1
X 61847319 61847456
refGene_gn=LOC101081365;refGene_tx=XM_023249275.1
X 61848199 61848357
refGene_gn=LOC101081365;refGene_tx=XM_023249275.1
X 61848441 61848656
refGene_gn=LOC101081365;refGene_tx=XM_023249275.1
X 61979108 61979711
refGene_gn=PABPC1L2B;refGene_tx=XM_023249151.1
X 62086624 62086695
refGene_gn=LOC105260217;refGene_tx=XM_011280019.3
X 62086697 62086765
refGene_gn=LOC105260217;refGene_tx=XM_011280019.3
X 62086766 62087142
refGene_gn=LOC105260217;refGene_tx=XM_011280019.3
X 62087144 62087223
refGene_gn=LOC105260217;refGene_tx=XM_011280019.3
X 62087567 62088119
refGene_gn=LOC111558674;refGene_tx=XM_023248962.1
X 62090453 62090480
refGene_gn=LOC111558674;refGene_tx=XM_023248962.1
X 62149528 62150920 refGene_gn=NAP1L2;refGene_tx=XM_004000636.5
X 62267841 62268128
refGene_gn=CHIC1;refGene_tx=XM_004000637.5;refGene_gn=CHIC1;refGene_tx
=XM_006943828.3;refGene_gn=CHIC1;refGene_tx=XM_006943829.4;refGene_gn=
CHIC1;refGene_tx=XM_019823987.2
X 62290142 62290197
refGene_gn=CHIC1;refGene_tx=XM_004000637.5;refGene_gn=CHIC1;refGene_tx
=XM_006943828.3;refGene_gn=CHIC1;refGene_tx=XM_006943829.4;refGene_gn=
CHIC1;refGene_tx=XM_019823987.2
X 62301812 62301968
refGene_gn=CHIC1;refGene_tx=XM_004000637.5;refGene_gn=CHIC1;refGene_tx
=XM_006943828.3;refGene_gn=CHIC1;refGene_tx=XM_006943829.4;refGene_gn=
CHIC1;refGene_tx=XM_019823987.2
X 62304974 62305094 refGene_gn=CHIC1;refGene_tx=XM_006943828.3
X 62307426 62307468 refGene_gn=CHIC1;refGene_tx=XM_006943829.4
X 62340181 62340238
refGene_gn=CHIC1;refGene_tx=XM_004000637.5;refGene_gn=CHIC1;refGene_tx
=XM_019823987.2
X 62340494 62340554 refGene_gn=CHIC1;refGene_tx=XM_004000637.5
X 62342120 62342171
refGene_gn=CHIC1;refGene_tx=XM_004000637.5;refGene_gn=CHIC1;refGene_tx
=XM_019823987.2

X 62726509 62727025 refGene_gn=ZCCHC13;refGene_tx=XM_023249308.1
X 62866773 62867233 refGene_gn=SLC16A2;refGene_tx=XM_004000653.5
X 63001343 63001488 refGene_gn=SLC16A2;refGene_tx=XM_004000653.5
X 63005024 63005475 refGene_gn=SLC16A2;refGene_tx=XM_004000653.5
X 63006474 63006618 refGene_gn=SLC16A2;refGene_tx=XM_004000653.5
X 63010415 63010644 refGene_gn=SLC16A2;refGene_tx=XM_004000653.5
X 63012266 63012487 refGene_gn=SLC16A2;refGene_tx=XM_004000653.5
X 63049882 63051495
refGene_gn=RLIM;refGene_tx=XM_004000642.5;refGene_gn=RLIM;refGene_tx=XM_019823989.2
X 63052800 63052884
refGene_gn=RLIM;refGene_tx=XM_004000642.5;refGene_gn=RLIM;refGene_tx=XM_019823989.2
X 63054188 63054357
refGene_gn=RLIM;refGene_tx=XM_004000642.5;refGene_gn=RLIM;refGene_tx=XM_019823989.2
X 63130167 63130261 refGene_gn=NEXMIF;refGene_tx=XM_004000643.4
X 63130869 63135241 refGene_gn=NEXMIF;refGene_tx=XM_004000643.4
X 63135912 63135991 refGene_gn=NEXMIF;refGene_tx=XM_004000643.4
X 63430591 63430807
refGene_gn=ABCB7;refGene_tx=XM_004000654.4;refGene_gn=ABCB7;refGene_tx=XM_019823991.2
X 63450484 63450592 refGene_gn=ABCB7;refGene_tx=XM_004000654.4
X 63454366 63454470
refGene_gn=ABCB7;refGene_tx=XM_004000654.4;refGene_gn=ABCB7;refGene_tx=XM_019823991.2
X 63458042 63458214
refGene_gn=ABCB7;refGene_tx=XM_004000654.4;refGene_gn=ABCB7;refGene_tx=XM_019823991.2
X 63461270 63461400
refGene_gn=ABCB7;refGene_tx=XM_004000654.4;refGene_gn=ABCB7;refGene_tx=XM_019823991.2
X 63461555 63461719
refGene_gn=ABCB7;refGene_tx=XM_004000654.4;refGene_gn=ABCB7;refGene_tx=XM_019823991.2
X 63462285 63462443
refGene_gn=ABCB7;refGene_tx=XM_004000654.4;refGene_gn=ABCB7;refGene_tx=XM_019823991.2
X 63463417 63463592
refGene_gn=ABCB7;refGene_tx=XM_004000654.4;refGene_gn=ABCB7;refGene_tx=XM_019823991.2
X 63464447 63464535
refGene_gn=ABCB7;refGene_tx=XM_004000654.4;refGene_gn=ABCB7;refGene_tx=XM_019823991.2
X 63464627 63464716
refGene_gn=ABCB7;refGene_tx=XM_004000654.4;refGene_gn=ABCB7;refGene_tx=XM_019823991.2
X 63465356 63465625
refGene_gn=ABCB7;refGene_tx=XM_004000654.4;refGene_gn=ABCB7;refGene_tx=XM_019823991.2

X 63466748 63466881
refGene_gn=ABCB7;refGene_tx=XM_004000654.4;refGene_gn=ABCB7;refGene_tx=XM_019823991.2
X 63485113 63485233
refGene_gn=ABCB7;refGene_tx=XM_004000654.4;refGene_gn=ABCB7;refGene_tx=XM_019823991.2
X 63505622 63505709
refGene_gn=ABCB7;refGene_tx=XM_004000654.4;refGene_gn=ABCB7;refGene_tx=XM_019823991.2
X 63507869 63507947
refGene_gn=ABCB7;refGene_tx=XM_004000654.4;refGene_gn=ABCB7;refGene_tx=XM_019823991.2
X 63580978 63581528
refGene_gn=LOC111558772;refGene_tx=XM_023249444.1
X 63581530 63581782
refGene_gn=LOC111558772;refGene_tx=XM_023249444.1
X 63582043 63583572
refGene_gn=LOC111558772;refGene_tx=XM_023249444.1
X 63596006 63596174
refGene_gn=ABCB7;refGene_tx=XM_004000654.4;refGene_gn=ABCB7;refGene_tx=XM_019823991.2
X 63658159 63658536
refGene_gn=UPRT;refGene_tx=XM_006943837.3;refGene_gn=UPRT;refGene_tx=XM_023249021.1;refGene_gn=UPRT;refGene_tx=XM_023249022.1;refGene_gn=UPRT;refGene_tx=XM_023249023.1
X 63676792 63676835
refGene_gn=UPRT;refGene_tx=XM_006943837.3;refGene_gn=UPRT;refGene_tx=XM_023249021.1;refGene_gn=UPRT;refGene_tx=XM_023249022.1;refGene_gn=UPRT;refGene_tx=XM_023249023.1
X 63679740 63679810
refGene_gn=UPRT;refGene_tx=XM_006943837.3;refGene_gn=UPRT;refGene_tx=XM_023249021.1;refGene_gn=UPRT;refGene_tx=XM_023249022.1;refGene_gn=UPRT;refGene_tx=XM_023249023.1
X 63680918 63680981
refGene_gn=UPRT;refGene_tx=XM_006943837.3;refGene_gn=UPRT;refGene_tx=XM_023249021.1;refGene_gn=UPRT;refGene_tx=XM_023249022.1;refGene_gn=UPRT;refGene_tx=XM_023249023.1
X 63682879 63683041
refGene_gn=UPRT;refGene_tx=XM_023249021.1;refGene_gn=UPRT;refGene_tx=XM_023249022.1;refGene_gn=UPRT;refGene_tx=XM_023249023.1
X 63683969 63684068
refGene_gn=UPRT;refGene_tx=XM_006943837.3;refGene_gn=UPRT;refGene_tx=XM_023249021.1;refGene_gn=UPRT;refGene_tx=XM_023249022.1;refGene_gn=UPRT;refGene_tx=XM_023249023.1
X 63686476 63686583
refGene_gn=UPRT;refGene_tx=XM_006943837.3;refGene_gn=UPRT;refGene_tx=XM_023249021.1
X 63720098 63720115 refGene_gn=UPRT;refGene_tx=XM_023249022.1
X 63725434 63725451 refGene_gn=UPRT;refGene_tx=XM_023249023.1
X 63786452 63786499

refGene_gn=ZDHHC15;refGene_tx=XM_006943838.3;refGene_gn=ZDHHC15;refGene_tx=XM_019823993.1
X 63828057 63828161
refGene_gn=ZDHHC15;refGene_tx=XM_006943838.3;refGene_gn=ZDHHC15;refGene_tx=XM_019823993.1
X 63830246 63830373
refGene_gn=ZDHHC15;refGene_tx=XM_006943838.3;refGene_gn=ZDHHC15;refGene_tx=XM_019823993.1
X 63832187 63832320
refGene_gn=ZDHHC15;refGene_tx=XM_006943838.3;refGene_gn=ZDHHC15;refGene_tx=XM_019823993.1
X 63836803 63836924
refGene_gn=ZDHHC15;refGene_tx=XM_006943838.3;refGene_gn=ZDHHC15;refGene_tx=XM_019823993.1
X 63837677 63837710
refGene_gn=ZDHHC15;refGene_tx=XM_006943838.3;refGene_gn=ZDHHC15;refGene_tx=XM_019823993.1
X 63839825 63839895
refGene_gn=ZDHHC15;refGene_tx=XM_006943838.3;refGene_gn=ZDHHC15;refGene_tx=XM_019823993.1
X 63843140 63843261
refGene_gn=ZDHHC15;refGene_tx=XM_006943838.3;refGene_gn=ZDHHC15;refGene_tx=XM_019823993.1
X 63888697 63888792
refGene_gn=ZDHHC15;refGene_tx=XM_006943838.3;refGene_gn=ZDHHC15;refGene_tx=XM_019823993.1
X 63954483 63954510
refGene_gn=ZDHHC15;refGene_tx=XM_006943838.3;refGene_gn=ZDHHC15;refGene_tx=XM_019823993.1
X 64023505 64023641
refGene_gn=ZDHHC15;refGene_tx=XM_006943838.3;refGene_gn=ZDHHC15;refGene_tx=XM_019823993.1
X 64102901 64104473 refGene_gn=MAGEE2;refGene_tx=XM_004000658.4
X 64260215 64260245 refGene_gn=PBDC1;refGene_tx=XM_004000659.5
X 64260329 64260395 refGene_gn=PBDC1;refGene_tx=XM_004000659.5
X 64261741 64261801
refGene_gn=PBDC1;refGene_tx=XM_004000659.5;refGene_gn=PBDC1;refGene_tx=XM_006943839.4
X 64262311 64262452
refGene_gn=PBDC1;refGene_tx=XM_004000659.5;refGene_gn=PBDC1;refGene_tx=XM_006943839.4
X 64263692 64263804
refGene_gn=PBDC1;refGene_tx=XM_004000659.5;refGene_gn=PBDC1;refGene_tx=XM_006943839.4
X 64264124 64264375
refGene_gn=PBDC1;refGene_tx=XM_004000659.5;refGene_gn=PBDC1;refGene_tx=XM_006943839.4
X 65151110 65151384 refGene_gn=FGF16;refGene_tx=XM_004000660.4
X 65157492 65157596 refGene_gn=FGF16;refGene_tx=XM_004000660.4
X 65158932 65159178 refGene_gn=FGF16;refGene_tx=XM_004000660.4

X	65189584	65189863	refGene_gn=ATRX;refGene_tx=XM_023249035.1
X	65191152	65191281	refGene_gn=ATRX;refGene_tx=XM_023249035.1
X	65191788	65191884	refGene_gn=ATRX;refGene_tx=XM_023249035.1
X	65192819	65192945	refGene_gn=ATRX;refGene_tx=XM_023249035.1
X	65193793	65193943	refGene_gn=ATRX;refGene_tx=XM_023249035.1
X	65215579	65215774	refGene_gn=ATRX;refGene_tx=XM_023249035.1
X	65217091	65217269	refGene_gn=ATRX;refGene_tx=XM_023249035.1
X	65229074	65229183	refGene_gn=ATRX;refGene_tx=XM_023249035.1
X	65257414	65257521	refGene_gn=ATRX;refGene_tx=XM_023249035.1
X	65262684	65262838	refGene_gn=ATRX;refGene_tx=XM_023249035.1
X	65280474	65280647	refGene_gn=ATRX;refGene_tx=XM_023249035.1
X	65280775	65280864	refGene_gn=ATRX;refGene_tx=XM_023249035.1
X	65281380	65281511	refGene_gn=ATRX;refGene_tx=XM_023249035.1
X	65291760	65291878	refGene_gn=ATRX;refGene_tx=XM_023249035.1
X	65293275	65293451	refGene_gn=ATRX;refGene_tx=XM_023249035.1
X	65295240	65295378	refGene_gn=ATRX;refGene_tx=XM_023249035.1
X	65303221	65303399	refGene_gn=ATRX;refGene_tx=XM_023249035.1
X	65303562	65303709	refGene_gn=ATRX;refGene_tx=XM_023249035.1
X	65310063	65310173	refGene_gn=ATRX;refGene_tx=XM_023249035.1
X	65311361	65311503	refGene_gn=ATRX;refGene_tx=XM_023249035.1
X	65322882	65322969	refGene_gn=ATRX;refGene_tx=XM_023249035.1
X	65322970	65323102	refGene_gn=ATRX;refGene_tx=XM_023249035.1
X	65324956	65325059	refGene_gn=ATRX;refGene_tx=XM_023249035.1
X	65367877	65367971	refGene_gn=ATRX;refGene_tx=XM_023249035.1
X	65374162	65374339	refGene_gn=ATRX;refGene_tx=XM_023249035.1
X	65375032	65375166	refGene_gn=ATRX;refGene_tx=XM_023249035.1
X	65385497	65385570	refGene_gn=ATRX;refGene_tx=XM_023249035.1
X	65403652	65406729	refGene_gn=ATRX;refGene_tx=XM_023249035.1
X	65407787	65407855	refGene_gn=ATRX;refGene_tx=XM_023249035.1
X	65409496	65409606	refGene_gn=ATRX;refGene_tx=XM_023249035.1
X	65415894	65416008	refGene_gn=ATRX;refGene_tx=XM_023249035.1
X	65418670	65418795	refGene_gn=ATRX;refGene_tx=XM_023249035.1
X	65419688	65419741	refGene_gn=ATRX;refGene_tx=XM_023249035.1
X	65420675	65420731	refGene_gn=ATRX;refGene_tx=XM_023249035.1
X	65445042	65445155	refGene_gn=ATRX;refGene_tx=XM_023249035.1
X	65501771	65501791	refGene_gn=ATRX;refGene_tx=XM_023249035.1
X	65517671	65517687
refGene_gn=MAGT1;refGene_tx=XM_004000661.4;refGene_gn=MAGT1;refGene_tx
=XM_019824007.1
X	65521053	65521144	refGene_gn=MAGT1;refGene_tx=XM_004000661.4
X	65524596	65524671
refGene_gn=MAGT1;refGene_tx=XM_004000661.4;refGene_gn=MAGT1;refGene_tx
=XM_019824007.1
X	65533531	65533595
refGene_gn=MAGT1;refGene_tx=XM_004000661.4;refGene_gn=MAGT1;refGene_tx
=XM_019824007.1
X	65534233	65534323
refGene_gn=MAGT1;refGene_tx=XM_004000661.4;refGene_gn=MAGT1;refGene_tx
=XM_019824007.1
X	65535254	65535395

refGene_gn=MAGT1;refGene_tx=XM_004000661.4;refGene_gn=MAGT1;refGene_tx=XM_019824007.1
X 65536667 65536808
refGene_gn=MAGT1;refGene_tx=XM_004000661.4;refGene_gn=MAGT1;refGene_tx=XM_019824007.1
X 65562094 65562212
refGene_gn=MAGT1;refGene_tx=XM_004000661.4;refGene_gn=MAGT1;refGene_tx=XM_019824007.1
X 65566267 65566437
refGene_gn=MAGT1;refGene_tx=XM_004000661.4;refGene_gn=MAGT1;refGene_tx=XM_019824007.1
X 65576215 65576317
refGene_gn=MAGT1;refGene_tx=XM_004000661.4;refGene_gn=MAGT1;refGene_tx=XM_019824007.1
X 65582699 65582739
refGene_gn=LOC101085531;refGene_tx=XM_004000662.5
X 65586066 65586191
refGene_gn=LOC101085531;refGene_tx=XM_004000662.5
X 65587821 65587899
refGene_gn=LOC101085531;refGene_tx=XM_004000662.5
X 65652418 65652538 refGene_gn=ATP7A;refGene_tx=XM_004000664.5
X 65689381 65689871 refGene_gn=ATP7A;refGene_tx=XM_004000664.5
X 65690398 65691124 refGene_gn=ATP7A;refGene_tx=XM_004000664.5
X 65700437 65700641 refGene_gn=ATP7A;refGene_tx=XM_004000664.5
X 65703329 65703493 refGene_gn=ATP7A;refGene_tx=XM_004000664.5
X 65710292 65710454 refGene_gn=ATP7A;refGene_tx=XM_004000664.5
X 65712594 65712671 refGene_gn=ATP7A;refGene_tx=XM_004000664.5
X 65712892 65713121 refGene_gn=ATP7A;refGene_tx=XM_004000664.5
X 65714204 65714438 refGene_gn=ATP7A;refGene_tx=XM_004000664.5
X 65715835 65715927 refGene_gn=ATP7A;refGene_tx=XM_004000664.5
X 65717172 65717300 refGene_gn=ATP7A;refGene_tx=XM_004000664.5
X 65718337 65718492 refGene_gn=ATP7A;refGene_tx=XM_004000664.5
X 65718708 65718843 refGene_gn=ATP7A;refGene_tx=XM_004000664.5
X 65740613 65740808 refGene_gn=ATP7A;refGene_tx=XM_004000664.5
X 65742371 65742554 refGene_gn=ATP7A;refGene_tx=XM_004000664.5
X 65744567 65744663 refGene_gn=ATP7A;refGene_tx=XM_004000664.5
X 65744664 65744785 refGene_gn=ATP7A;refGene_tx=XM_004000664.5
X 65750107 65750254 refGene_gn=ATP7A;refGene_tx=XM_004000664.5
X 65751563 65751706 refGene_gn=ATP7A;refGene_tx=XM_004000664.5
X 65761078 65761282 refGene_gn=ATP7A;refGene_tx=XM_004000664.5
X 65761833 65761951 refGene_gn=ATP7A;refGene_tx=XM_004000664.5
X 65764167 65764270 refGene_gn=ATP7A;refGene_tx=XM_004000664.5
X 65764646 65764923 refGene_gn=ATP7A;refGene_tx=XM_004000664.5
X 65846229 65846294 refGene_gn=PGK1;refGene_tx=XM_004000665.4
X 65851863 65851914 refGene_gn=PGK1;refGene_tx=XM_004000665.4
X 65855486 65855642 refGene_gn=PGK1;refGene_tx=XM_004000665.4
X 65855751 65855896 refGene_gn=PGK1;refGene_tx=XM_004000665.4
X 65859560 65859664 refGene_gn=PGK1;refGene_tx=XM_004000665.4
X 65860735 65860855 refGene_gn=PGK1;refGene_tx=XM_004000665.4
X 65864575 65864690 refGene_gn=PGK1;refGene_tx=XM_004000665.4

X	65864939 65865119 refGene_gn=PGK1;refGene_tx=XM_004000665.4
X	65866813 65866991 refGene_gn=PGK1;refGene_tx=XM_004000665.4
X	65867267 65867366 refGene_gn=PGK1;refGene_tx=XM_004000665.4
X	65867747 65867788 refGene_gn=PGK1;refGene_tx=XM_004000665.4
X	65873672 65873836 refGene_gn=TAF9B;refGene_tx=XM_004000666.5
X	65874716 65874827 refGene_gn=TAF9B;refGene_tx=XM_004000666.5
X	65877533 65877609 refGene_gn=TAF9B;refGene_tx=XM_004000666.5
X	65878437 65878572 refGene_gn=TAF9B;refGene_tx=XM_004000666.5
X	65878646 65878783 refGene_gn=TAF9B;refGene_tx=XM_004000666.5
X	65879323 65879405 refGene_gn=TAF9B;refGene_tx=XM_004000666.5
X	65880043 65880094 refGene_gn=TAF9B;refGene_tx=XM_004000666.5
X	65902979 65903318
refGene_gn=LOC101088911;refGene_tx=XM_023248963.1
X	65903811 65903937
refGene_gn=LOC101088911;refGene_tx=XM_023248963.1
X	65940978 65941155
refGene_gn=LOC109496548;refGene_tx=XM_019824013.2
X	65942840 65942954
refGene_gn=LOC101088911;refGene_tx=XM_023248963.1;refGene_gn=LOC109496
548;refGene_tx=XM_019824013.2
X	65943155 65943272
refGene_gn=LOC101088911;refGene_tx=XM_023248963.1;refGene_gn=LOC109496
548;refGene_tx=XM_019824013.2
X	65945659 65945728
refGene_gn=LOC101088911;refGene_tx=XM_023248963.1
X	65946287 65946426
refGene_gn=LOC101088911;refGene_tx=XM_023248963.1
X	65948102 65948277
refGene_gn=LOC101088911;refGene_tx=XM_023248963.1
X	65948361 65948483
refGene_gn=LOC101088911;refGene_tx=XM_023248963.1
X	65948597 65948651
refGene_gn=LOC101088911;refGene_tx=XM_023248963.1
X	65949245 65949319
refGene_gn=LOC101088911;refGene_tx=XM_023248963.1
X	65951443 65951555
refGene_gn=LOC101088911;refGene_tx=XM_023248963.1
X	65951884 65951937
refGene_gn=LOC101088911;refGene_tx=XM_023248963.1
X	65952445 65953342
refGene_gn=LOC101088911;refGene_tx=XM_023248963.1
X	65953926 65954185
refGene_gn=LOC101088911;refGene_tx=XM_023248963.1
X	65955723 65955800
refGene_gn=LOC101088911;refGene_tx=XM_023248963.1
X	65957335 65957411
refGene_gn=LOC101088911;refGene_tx=XM_023248963.1
X	65960169 65960292
refGene_gn=LOC101088911;refGene_tx=XM_023248963.1
X	65961595 65961676

refGene_gn=LOC101088911;refGene_tx=XM_023248963.1
X 65967534 65968713
refGene_gn=LOC111558740;refGene_tx=XM_023249249.1
X 66076098 66077121
refGene_gn=CYSLTR1;refGene_tx=XM_004000667.5;refGene_gn=CYSLTR1;refGene_tx=XM_006943842.4;refGene_gn=CYSLTR1;refGene_tx=XM_019824014.2;refGene_gn=CYSLTR1;refGene_tx=XM_019824016.2
X 66402011 66402141 refGene_gn=RTL3;refGene_tx=XM_019823431.1
X 66402142 66402512 refGene_gn=RTL3;refGene_tx=XM_019823431.1
X 66402514 66402877 refGene_gn=RTL3;refGene_tx=XM_019823431.1
X 66402912 66403105 refGene_gn=RTL3;refGene_tx=XM_019823431.1
X 66403161 66403303 refGene_gn=RTL3;refGene_tx=XM_019823431.1
X 66403443 66403565 refGene_gn=RTL3;refGene_tx=XM_019823431.1
X 66528770 66529883 refGene_gn=LPAR4;refGene_tx=XM_023249353.1
X 66764885 66764902
refGene_gn=LOC101089422;refGene_tx=XM_023249257.1
X 66766872 66767911
refGene_gn=LOC101089422;refGene_tx=XM_006943843.3;refGene_gn=LOC101089422;refGene_tx=XM_023249257.1
X 66840059 66840885
refGene_gn=LOC111558777;refGene_tx=XM_023249458.1
X 66840887 66841003
refGene_gn=LOC111558777;refGene_tx=XM_023249458.1
X 66992326 66993286 refGene_gn=GPR174;refGene_tx=XM_019824022.1
X 67111777 67111866
refGene_gn=ITM2A;refGene_tx=XM_004000668.4;refGene_gn=ITM2A;refGene_tx=XM_006943844.4
X 67112019 67112170
refGene_gn=ITM2A;refGene_tx=XM_004000668.4;refGene_gn=ITM2A;refGene_tx=XM_006943844.4
X 67112852 67112963
refGene_gn=ITM2A;refGene_tx=XM_004000668.4;refGene_gn=ITM2A;refGene_tx=XM_006943844.4
X 67113193 67113391
refGene_gn=ITM2A;refGene_tx=XM_004000668.4;refGene_gn=ITM2A;refGene_tx=XM_006943844.4
X 67113671 67113803
refGene_gn=ITM2A;refGene_tx=XM_004000668.4;refGene_gn=ITM2A;refGene_tx=XM_006943844.4
X 67117953 67118064
refGene_gn=ITM2A;refGene_tx=XM_004000668.4;refGene_gn=ITM2A;refGene_tx=XM_006943844.4
X 67646960 67647135 refGene_gn=TBX22;refGene_tx=XM_023249247.1
X 67647721 67647914 refGene_gn=TBX22;refGene_tx=XM_023249247.1
X 67648643 67648745 refGene_gn=TBX22;refGene_tx=XM_023249247.1
X 67650145 67650320 refGene_gn=TBX22;refGene_tx=XM_023249247.1
X 67651209 67651374 refGene_gn=TBX22;refGene_tx=XM_023249247.1
X 67651744 67651809 refGene_gn=TBX22;refGene_tx=XM_023249247.1
X 67652179 67652265 refGene_gn=TBX22;refGene_tx=XM_023249247.1
X 67654495 67655103 refGene_gn=TBX22;refGene_tx=XM_023249247.1

X 68063750 68064923
refGene_gn=FAM46D;refGene_tx=XM_019824025.2;refGene_gn=FAM46D;refGene_tx=XM_019824026.2
X 68333447 68334202
refGene_gn=BRWD3;refGene_tx=XM_011291814.3;refGene_gn=BRWD3;refGene_tx=XM_019824028.2;refGene_gn=BRWD3;refGene_tx=XM_019824029.2
X 68360446 68360605
refGene_gn=BRWD3;refGene_tx=XM_011291814.3;refGene_gn=BRWD3;refGene_tx=XM_019824028.2
X 68361647 68361745
refGene_gn=BRWD3;refGene_tx=XM_011291814.3;refGene_gn=BRWD3;refGene_tx=XM_019824028.2;refGene_gn=BRWD3;refGene_tx=XM_019824029.2
X 68362082 68362246
refGene_gn=BRWD3;refGene_tx=XM_011291814.3;refGene_gn=BRWD3;refGene_tx=XM_019824028.2;refGene_gn=BRWD3;refGene_tx=XM_019824029.2
X 68363556 68363709
refGene_gn=BRWD3;refGene_tx=XM_011291814.3;refGene_gn=BRWD3;refGene_tx=XM_019824028.2;refGene_gn=BRWD3;refGene_tx=XM_019824029.2
X 68365110 68365185
refGene_gn=BRWD3;refGene_tx=XM_011291814.3;refGene_gn=BRWD3;refGene_tx=XM_019824028.2;refGene_gn=BRWD3;refGene_tx=XM_019824029.2
X 68366064 68366205
refGene_gn=BRWD3;refGene_tx=XM_011291814.3;refGene_gn=BRWD3;refGene_tx=XM_019824028.2;refGene_gn=BRWD3;refGene_tx=XM_019824029.2
X 68367361 68367418
refGene_gn=BRWD3;refGene_tx=XM_011291814.3;refGene_gn=BRWD3;refGene_tx=XM_019824028.2;refGene_gn=BRWD3;refGene_tx=XM_019824029.2
X 68396573 68396652
refGene_gn=BRWD3;refGene_tx=XM_011291814.3;refGene_gn=BRWD3;refGene_tx=XM_019824028.2;refGene_gn=BRWD3;refGene_tx=XM_019824029.2
X 68396740 68396866
refGene_gn=BRWD3;refGene_tx=XM_011291814.3;refGene_gn=BRWD3;refGene_tx=XM_019824028.2;refGene_gn=BRWD3;refGene_tx=XM_019824029.2
X 68397869 68397990
refGene_gn=BRWD3;refGene_tx=XM_011291814.3;refGene_gn=BRWD3;refGene_tx=XM_019824028.2;refGene_gn=BRWD3;refGene_tx=XM_019824029.2
X 68398640 68398796
refGene_gn=BRWD3;refGene_tx=XM_011291814.3;refGene_gn=BRWD3;refGene_tx=XM_019824028.2;refGene_gn=BRWD3;refGene_tx=XM_019824029.2
X 68398886 68398948
refGene_gn=BRWD3;refGene_tx=XM_011291814.3;refGene_gn=BRWD3;refGene_tx=XM_019824028.2;refGene_gn=BRWD3;refGene_tx=XM_019824029.2
X 68399905 68400017
refGene_gn=BRWD3;refGene_tx=XM_011291814.3;refGene_gn=BRWD3;refGene_tx=XM_019824028.2;refGene_gn=BRWD3;refGene_tx=XM_019824029.2
X 68401125 68401208
refGene_gn=BRWD3;refGene_tx=XM_011291814.3;refGene_gn=BRWD3;refGene_tx=XM_019824028.2;refGene_gn=BRWD3;refGene_tx=XM_019824029.2
X 68402427 68402552
refGene_gn=BRWD3;refGene_tx=XM_011291814.3;refGene_gn=BRWD3;refGene_tx

=XM_019824028.2;refGene_gn=BRWD3;refGene_tx=XM_019824029.2
X	68403961	68404069
refGene_gn=BRWD3;refGene_tx=XM_011291814.3;refGene_gn=BRWD3;refGene_tx=XM_019824028.2;refGene_gn=BRWD3;refGene_tx=XM_019824029.2
X	68413983	68414097
refGene_gn=BRWD3;refGene_tx=XM_011291814.3;refGene_gn=BRWD3;refGene_tx=XM_019824028.2;refGene_gn=BRWD3;refGene_tx=XM_019824029.2
X	68415464	68415633
refGene_gn=BRWD3;refGene_tx=XM_011291814.3;refGene_gn=BRWD3;refGene_tx=XM_019824028.2;refGene_gn=BRWD3;refGene_tx=XM_019824029.2
X	68416222	68416299
refGene_gn=BRWD3;refGene_tx=XM_011291814.3;refGene_gn=BRWD3;refGene_tx=XM_019824028.2;refGene_gn=BRWD3;refGene_tx=XM_019824029.2
X	68418280	68418430
refGene_gn=BRWD3;refGene_tx=XM_011291814.3;refGene_gn=BRWD3;refGene_tx=XM_019824028.2;refGene_gn=BRWD3;refGene_tx=XM_019824029.2
X	68420780	68420874
refGene_gn=BRWD3;refGene_tx=XM_011291814.3;refGene_gn=BRWD3;refGene_tx=XM_019824028.2;refGene_gn=BRWD3;refGene_tx=XM_019824029.2
X	68422881	68423065
refGene_gn=BRWD3;refGene_tx=XM_011291814.3;refGene_gn=BRWD3;refGene_tx=XM_019824028.2;refGene_gn=BRWD3;refGene_tx=XM_019824029.2
X	68425528	68425696
refGene_gn=BRWD3;refGene_tx=XM_011291814.3;refGene_gn=BRWD3;refGene_tx=XM_019824028.2;refGene_gn=BRWD3;refGene_tx=XM_019824029.2
X	68442359	68442585
refGene_gn=BRWD3;refGene_tx=XM_011291814.3;refGene_gn=BRWD3;refGene_tx=XM_019824028.2;refGene_gn=BRWD3;refGene_tx=XM_019824029.2
X	68443996	68444125
refGene_gn=BRWD3;refGene_tx=XM_011291814.3;refGene_gn=BRWD3;refGene_tx=XM_019824028.2;refGene_gn=BRWD3;refGene_tx=XM_019824029.2
X	68445107	68445242
refGene_gn=BRWD3;refGene_tx=XM_011291814.3;refGene_gn=BRWD3;refGene_tx=XM_019824028.2;refGene_gn=BRWD3;refGene_tx=XM_019824029.2
X	68449759	68449913
refGene_gn=BRWD3;refGene_tx=XM_011291814.3;refGene_gn=BRWD3;refGene_tx=XM_019824028.2;refGene_gn=BRWD3;refGene_tx=XM_019824029.2
X	68451003	68451108
refGene_gn=BRWD3;refGene_tx=XM_011291814.3;refGene_gn=BRWD3;refGene_tx=XM_019824028.2;refGene_gn=BRWD3;refGene_tx=XM_019824029.2
X	68453758	68453799
refGene_gn=BRWD3;refGene_tx=XM_011291814.3;refGene_gn=BRWD3;refGene_tx=XM_019824028.2;refGene_gn=BRWD3;refGene_tx=XM_019824029.2
X	68491752	68491853
refGene_gn=BRWD3;refGene_tx=XM_011291814.3;refGene_gn=BRWD3;refGene_tx=XM_019824028.2;refGene_gn=BRWD3;refGene_tx=XM_019824029.2
X	68492782	68492853
refGene_gn=BRWD3;refGene_tx=XM_011291814.3;refGene_gn=BRWD3;refGene_tx=XM_019824028.2;refGene_gn=BRWD3;refGene_tx=XM_019824029.2
X	68493376	68493477

refGene_gn=BRWD3;refGene_tx=XM_011291814.3;refGene_gn=BRWD3;refGene_tx=XM_019824028.2;refGene_gn=BRWD3;refGene_tx=XM_019824029.2
X 68497621 68497843
refGene_gn=BRWD3;refGene_tx=XM_011291814.3;refGene_gn=BRWD3;refGene_tx=XM_019824028.2;refGene_gn=BRWD3;refGene_tx=XM_019824029.2
X 68499699 68499860
refGene_gn=BRWD3;refGene_tx=XM_011291814.3;refGene_gn=BRWD3;refGene_tx=XM_019824028.2;refGene_gn=BRWD3;refGene_tx=XM_019824029.2
X 68552991 68553090
refGene_gn=BRWD3;refGene_tx=XM_011291814.3;refGene_gn=BRWD3;refGene_tx=XM_019824028.2;refGene_gn=BRWD3;refGene_tx=XM_019824029.2
X 68554650 68554801
refGene_gn=BRWD3;refGene_tx=XM_011291814.3;refGene_gn=BRWD3;refGene_tx=XM_019824028.2;refGene_gn=BRWD3;refGene_tx=XM_019824029.2
X 68567092 68567152
refGene_gn=BRWD3;refGene_tx=XM_011291814.3;refGene_gn=BRWD3;refGene_tx=XM_019824028.2;refGene_gn=BRWD3;refGene_tx=XM_019824029.2
X 68567561 68567591
refGene_gn=BRWD3;refGene_tx=XM_011291814.3;refGene_gn=BRWD3;refGene_tx=XM_019824028.2;refGene_gn=BRWD3;refGene_tx=XM_019824029.2
X 68567794 68567853
refGene_gn=BRWD3;refGene_tx=XM_011291814.3;refGene_gn=BRWD3;refGene_tx=XM_019824028.2;refGene_gn=BRWD3;refGene_tx=XM_019824029.2
X 68567974 68568005
refGene_gn=BRWD3;refGene_tx=XM_011291814.3;refGene_gn=BRWD3;refGene_tx=XM_019824028.2;refGene_gn=BRWD3;refGene_tx=XM_019824029.2
X 68875670 68875715 refGene_gn=SH3BGRL;refGene_tx=XM_011291813.3
X 68933358 68933361 refGene_gn=SH3BGRL;refGene_tx=XM_019824031.2
X 68948108 68948294
refGene_gn=SH3BGRL;refGene_tx=XM_011291813.3;refGene_gn=SH3BGRL;refGene_tx=XM_019824031.2
X 68952710 68952791
refGene_gn=SH3BGRL;refGene_tx=XM_011291813.3;refGene_gn=SH3BGRL;refGene_tx=XM_019824031.2
X 68961612 68961645
refGene_gn=SH3BGRL;refGene_tx=XM_011291813.3;refGene_gn=SH3BGRL;refGene_tx=XM_019824031.2
X 68986248 68986314
refGene_gn=LOC109496377;refGene_tx=XM_019823482.2
X 68986716 68986884
refGene_gn=LOC109496377;refGene_tx=XM_019823482.2
X 68989587 68989635
refGene_gn=LOC109496377;refGene_tx=XM_019823482.2
X 70625535 70625901
refGene_gn=LOC109496330;refGene_tx=XM_023248964.1
X 70627590 70627873
refGene_gn=LOC109496330;refGene_tx=XM_023248964.1
X 70681629 70681995
refGene_gn=LOC109496330;refGene_tx=XM_023248964.1
X 71048831 71049917 refGene_gn=POU3F4;refGene_tx=XM_004000681.4

X 71390206 71390223 refGene_gn=CYLC1;refGene_tx=XM_023249309.1
X 71415442 71415496 refGene_gn=CYLC1;refGene_tx=XM_023249309.1
X 71422142 71422183 refGene_gn=CYLC1;refGene_tx=XM_023249309.1
X 71423908 71424027 refGene_gn=CYLC1;refGene_tx=XM_023249309.1
X 71425510 71427142 refGene_gn=CYLC1;refGene_tx=XM_023249309.1
X 71473202 71473235 refGene_gn=CYLC1;refGene_tx=XM_023249309.1
X 71576923 71577049 refGene_gn=RPS6KA6;refGene_tx=XM_023249024.1
X 71581668 71581809 refGene_gn=RPS6KA6;refGene_tx=XM_023249024.1
X 71609737 71609855 refGene_gn=RPS6KA6;refGene_tx=XM_023249024.1
X 71611655 71611732 refGene_gn=RPS6KA6;refGene_tx=XM_023249024.1
X 71620754 71620916 refGene_gn=RPS6KA6;refGene_tx=XM_023249024.1
X 71632802 71632961 refGene_gn=RPS6KA6;refGene_tx=XM_023249024.1
X 71633964 71634054 refGene_gn=RPS6KA6;refGene_tx=XM_023249024.1
X 71641241 71641364 refGene_gn=RPS6KA6;refGene_tx=XM_023249024.1
X 71641801 71641932 refGene_gn=RPS6KA6;refGene_tx=XM_023249024.1
X 71642567 71642670 refGene_gn=RPS6KA6;refGene_tx=XM_023249024.1
X 71645851 71645910 refGene_gn=RPS6KA6;refGene_tx=XM_023249024.1
X 71646747 71646836 refGene_gn=RPS6KA6;refGene_tx=XM_023249024.1
X 71647072 71647143 refGene_gn=RPS6KA6;refGene_tx=XM_023249024.1
X 71650753 71650896 refGene_gn=RPS6KA6;refGene_tx=XM_023249024.1
X 71657831 71657869 refGene_gn=RPS6KA6;refGene_tx=XM_023249024.1
X 71658407 71658514 refGene_gn=RPS6KA6;refGene_tx=XM_023249024.1
X 71702159 71702239 refGene_gn=RPS6KA6;refGene_tx=XM_023249024.1
X 71703670 71703751 refGene_gn=RPS6KA6;refGene_tx=XM_023249024.1
X 71704676 71704758 refGene_gn=RPS6KA6;refGene_tx=XM_023249024.1
X 71722843 71722960 refGene_gn=RPS6KA6;refGene_tx=XM_023249024.1
X 71739104 71739164 refGene_gn=RPS6KA6;refGene_tx=XM_023249024.1
X 71766615 71766693 refGene_gn=RPS6KA6;refGene_tx=XM_023249024.1
X 71922930 71923056
refGene_gn=HDX;refGene_tx=XM_006943850.3;refGene_gn=HDX;refGene_tx=XM_019824032.2;refGene_gn=HDX;refGene_tx=XM_019824034.2
X 71928150 71928273
refGene_gn=HDX;refGene_tx=XM_006943850.3;refGene_gn=HDX;refGene_tx=XM_019824032.2;refGene_gn=HDX;refGene_tx=XM_019824034.2
X 71958705 71958789
refGene_gn=HDX;refGene_tx=XM_006943850.3;refGene_gn=HDX;refGene_tx=XM_019824032.2;refGene_gn=HDX;refGene_tx=XM_019824034.2
X 71968523 71968600
refGene_gn=HDX;refGene_tx=XM_006943850.3;refGene_gn=HDX;refGene_tx=XM_019824032.2;refGene_gn=HDX;refGene_tx=XM_019824034.2
X 71978418 71978626
refGene_gn=HDX;refGene_tx=XM_006943850.3;refGene_gn=HDX;refGene_tx=XM_019824032.2;refGene_gn=HDX;refGene_tx=XM_019824034.2
X 72004145 72004292
refGene_gn=HDX;refGene_tx=XM_006943850.3;refGene_gn=HDX;refGene_tx=XM_019824032.2;refGene_gn=HDX;refGene_tx=XM_019824034.2
X 72058747 72058801
refGene_gn=HDX;refGene_tx=XM_006943850.3;refGene_gn=HDX;refGene_tx=XM_019824032.2;refGene_gn=HDX;refGene_tx=XM_019824034.2
X 72077280 72078393

refGene_gn=HDX;refGene_tx=XM_006943850.3;refGene_gn=HDX;refGene_tx=XM_019824032.2;refGene_gn=HDX;refGene_tx=XM_019824034.2
X	72085334	72085481
refGene_gn=HDX;refGene_tx=XM_006943850.3;refGene_gn=HDX;refGene_tx=XM_019824032.2;refGene_gn=HDX;refGene_tx=XM_019824034.2
X	72220234	72220313
refGene_gn=LOC109496332;refGene_tx=XM_023249271.1;refGene_gn=LOC109496332;refGene_tx=XM_023249272.1
X	72311903	72312067
refGene_gn=LOC109496332;refGene_tx=XM_023249271.1;refGene_gn=LOC109496332;refGene_tx=XM_023249272.1
X	72343472	72343511
refGene_gn=LOC109496332;refGene_tx=XM_023249271.1
X	72383303	72383503
refGene_gn=LOC109496332;refGene_tx=XM_023249271.1;refGene_gn=LOC109496332;refGene_tx=XM_023249272.1
X	72387794	72387882
refGene_gn=LOC109496332;refGene_tx=XM_023249271.1;refGene_gn=LOC109496332;refGene_tx=XM_023249272.1
X	72431830	72431872
refGene_gn=LOC109496332;refGene_tx=XM_023249271.1;refGene_gn=LOC109496332;refGene_tx=XM_023249272.1
X	72470799	72470919
refGene_gn=LOC109496332;refGene_tx=XM_023249271.1;refGene_gn=LOC109496332;refGene_tx=XM_023249272.1
X	72528422	72528544
refGene_gn=LOC109496332;refGene_tx=XM_023249271.1;refGene_gn=LOC109496332;refGene_tx=XM_023249272.1
X	72528950	72529623
refGene_gn=LOC109496332;refGene_tx=XM_023249271.1;refGene_gn=LOC109496332;refGene_tx=XM_023249272.1
X	72547466	72548718
refGene_gn=LOC102901599;refGene_tx=XM_023249193.1
X	72561303	72561393
refGene_gn=LOC102901599;refGene_tx=XM_023249193.1
X	72623637	72623789
refGene_gn=LOC102901599;refGene_tx=XM_023249193.1
X	72624560	72626252
refGene_gn=LOC102899136;refGene_tx=XM_019824036.1
X	72629109	72629148
refGene_gn=LOC102899136;refGene_tx=XM_019824036.1
X	72801007	72801022
refGene_gn=APOOL;refGene_tx=XM_023248994.1;refGene_gn=APOOL;refGene_tx=XM_023248995.1;refGene_gn=APOOL;refGene_tx=XM_023248996.1
X	72837409	72837514
refGene_gn=APOOL;refGene_tx=XM_023248994.1;refGene_gn=APOOL;refGene_tx=XM_023248995.1
X	72864808	72864928
refGene_gn=APOOL;refGene_tx=XM_023248994.1;refGene_gn=APOOL;refGene_tx=XM_023248995.1;refGene_gn=APOOL;refGene_tx=XM_023248996.1

X 72866496 72866551
refGene_gn=APOOL;refGene_tx=XM_023248994.1;refGene_gn=APOOL;refGene_tx=XM_023248995.1;refGene_gn=APOOL;refGene_tx=XM_023248996.1
X 72868261 72868360
refGene_gn=APOOL;refGene_tx=XM_023248994.1;refGene_gn=APOOL;refGene_tx=XM_023248996.1
X 72885674 72885766
refGene_gn=APOOL;refGene_tx=XM_023248994.1;refGene_gn=APOOL;refGene_tx=XM_023248995.1;refGene_gn=APOOL;refGene_tx=XM_023248996.1
X 72888985 72889099
refGene_gn=APOOL;refGene_tx=XM_023248994.1;refGene_gn=APOOL;refGene_tx=XM_023248995.1;refGene_gn=APOOL;refGene_tx=XM_023248996.1
X 72889308 72889426
refGene_gn=APOOL;refGene_tx=XM_023248994.1;refGene_gn=APOOL;refGene_tx=XM_023248995.1;refGene_gn=APOOL;refGene_tx=XM_023248996.1
X 72903312 72903401
refGene_gn=APOOL;refGene_tx=XM_023248994.1;refGene_gn=APOOL;refGene_tx=XM_023248995.1;refGene_gn=APOOL;refGene_tx=XM_023248996.1
X 72922062 72922193 refGene_gn=SATL1;refGene_tx=XM_023249192.1
X 72922598 72922679 refGene_gn=SATL1;refGene_tx=XM_023249192.1
X 72928528 72928579 refGene_gn=SATL1;refGene_tx=XM_023249192.1
X 72936639 72936640 refGene_gn=SATL1;refGene_tx=XM_023249192.1
X 72937049 72937679 refGene_gn=SATL1;refGene_tx=XM_023249192.1
X 73122060 73122139 refGene_gn=ZNF711;refGene_tx=XM_023248954.1
X 73129388 73129946 refGene_gn=ZNF711;refGene_tx=XM_023248954.1
X 73135858 73136014 refGene_gn=ZNF711;refGene_tx=XM_023248954.1
X 73137136 73137274 refGene_gn=ZNF711;refGene_tx=XM_023248954.1
X 73139319 73139457 refGene_gn=ZNF711;refGene_tx=XM_023248954.1
X 73140174 73140252 refGene_gn=ZNF711;refGene_tx=XM_023248954.1
X 73140336 73140384 refGene_gn=ZNF711;refGene_tx=XM_023248954.1
X 73142430 73142574 refGene_gn=ZNF711;refGene_tx=XM_023248954.1
X 73143059 73143853 refGene_gn=ZNF711;refGene_tx=XM_023248954.1
X 73143854 73144238 refGene_gn=ZNF711;refGene_tx=XM_023248954.1
X 73151783 73151789 refGene_gn=POF1B;refGene_tx=XM_019824047.2
X 73154143 73154258 refGene_gn=POF1B;refGene_tx=XM_019824047.2
X 73161961 73162044 refGene_gn=POF1B;refGene_tx=XM_019824047.2
X 73162865 73162994 refGene_gn=POF1B;refGene_tx=XM_019824047.2
X 73164444 73164564 refGene_gn=POF1B;refGene_tx=XM_019824047.2
X 73164892 73165045 refGene_gn=POF1B;refGene_tx=XM_019824047.2
X 73165840 73165954 refGene_gn=POF1B;refGene_tx=XM_019824047.2
X 73166810 73166903 refGene_gn=POF1B;refGene_tx=XM_019824047.2
X 73182338 73182413 refGene_gn=POF1B;refGene_tx=XM_019824047.2
X 73185226 73185254 refGene_gn=POF1B;refGene_tx=XM_019824047.2
X 73215446 73215577 refGene_gn=POF1B;refGene_tx=XM_019824047.2
X 73229435 73229618 refGene_gn=POF1B;refGene_tx=XM_019824047.2
X 73234209 73234311 refGene_gn=POF1B;refGene_tx=XM_019824047.2
X 73251603 73251684 refGene_gn=POF1B;refGene_tx=XM_019824047.2
X 73271952 73272234 refGene_gn=POF1B;refGene_tx=XM_019824047.2
X 73612832 73613319
refGene_gn=LOC111558675;refGene_tx=XM_023248965.1

X	73613724 73613846
refGene_gn=LOC111558675;refGene_tx=XM_023248965.1
X	73742530 73742722
refGene_gn=CHM;refGene_tx=XM_004000686.4;refGene_gn=CHM;refGene_tx=XM_019824049.1;refGene_gn=CHM;refGene_tx=XM_019824050.1;refGene_gn=CHM;refGene_tx=XM_019824051.1
X	73761358 73761519
refGene_gn=CHM;refGene_tx=XM_004000686.4;refGene_gn=CHM;refGene_tx=XM_019824049.1;refGene_gn=CHM;refGene_tx=XM_019824050.1;refGene_gn=CHM;refGene_tx=XM_019824051.1
X	73767257 73767356
refGene_gn=CHM;refGene_tx=XM_004000686.4;refGene_gn=CHM;refGene_tx=XM_019824049.1;refGene_gn=CHM;refGene_tx=XM_019824050.1;refGene_gn=CHM;refGene_tx=XM_019824051.1
X	73816948 73817045
refGene_gn=CHM;refGene_tx=XM_004000686.4;refGene_gn=CHM;refGene_tx=XM_019824049.1;refGene_gn=CHM;refGene_tx=XM_019824050.1;refGene_gn=CHM;refGene_tx=XM_019824051.1
X	73835040 73835104
refGene_gn=CHM;refGene_tx=XM_004000686.4;refGene_gn=CHM;refGene_tx=XM_019824049.1;refGene_gn=CHM;refGene_tx=XM_019824050.1;refGene_gn=CHM;refGene_tx=XM_019824051.1
X	73835499 73835604
refGene_gn=CHM;refGene_tx=XM_004000686.4;refGene_gn=CHM;refGene_tx=XM_019824049.1
X	73851778 73851856
refGene_gn=CHM;refGene_tx=XM_004000686.4;refGene_gn=CHM;refGene_tx=XM_019824049.1;refGene_gn=CHM;refGene_tx=XM_019824050.1;refGene_gn=CHM;refGene_tx=XM_019824051.1
X	73903899 73903932
refGene_gn=CHM;refGene_tx=XM_019824049.1;refGene_gn=CHM;refGene_tx=XM_019824050.1
X	73904853 73905079
refGene_gn=CHM;refGene_tx=XM_004000686.4;refGene_gn=CHM;refGene_tx=XM_019824049.1;refGene_gn=CHM;refGene_tx=XM_019824050.1;refGene_gn=CHM;refGene_tx=XM_019824051.1
X	73906253 73906374
refGene_gn=CHM;refGene_tx=XM_004000686.4;refGene_gn=CHM;refGene_tx=XM_019824049.1;refGene_gn=CHM;refGene_tx=XM_019824050.1;refGene_gn=CHM;refGene_tx=XM_019824051.1
X	73906884 73907001
refGene_gn=CHM;refGene_tx=XM_004000686.4;refGene_gn=CHM;refGene_tx=XM_019824049.1;refGene_gn=CHM;refGene_tx=XM_019824050.1;refGene_gn=CHM;refGene_tx=XM_019824051.1
X	73912973 73913361
refGene_gn=CHM;refGene_tx=XM_004000686.4;refGene_gn=CHM;refGene_tx=XM_019824049.1;refGene_gn=CHM;refGene_tx=XM_019824050.1;refGene_gn=CHM;refGene_tx=XM_019824051.1
X	73940483 73940608
refGene_gn=CHM;refGene_tx=XM_004000686.4;refGene_gn=CHM;refGene_tx=XM_

019824049.1;refGene_gn=CHM;refGene_tx=XM_019824050.1;refGene_gn=CHM;refGene_tx=XM_019824051.1
X 73943759 73943832
refGene_gn=CHM;refGene_tx=XM_004000686.4;refGene_gn=CHM;refGene_tx=XM_019824049.1;refGene_gn=CHM;refGene_tx=XM_019824050.1;refGene_gn=CHM;refGene_tx=XM_019824051.1
X 73969017 73969084
refGene_gn=CHM;refGene_tx=XM_004000686.4;refGene_gn=CHM;refGene_tx=XM_019824049.1;refGene_gn=CHM;refGene_tx=XM_019824050.1;refGene_gn=CHM;refGene_tx=XM_019824051.1
X 74003506 74003555
refGene_gn=CHM;refGene_tx=XM_004000686.4;refGene_gn=CHM;refGene_tx=XM_019824049.1;refGene_gn=CHM;refGene_tx=XM_019824050.1;refGene_gn=CHM;refGene_tx=XM_019824051.1
X 74109572 74110084 refGene_gn=DACH2;refGene_tx=XM_019824052.2
X 74384352 74384391 refGene_gn=DACH2;refGene_tx=XM_019824052.2
X 74551548 74551661 refGene_gn=DACH2;refGene_tx=XM_019824052.2
X 74698444 74698576 refGene_gn=DACH2;refGene_tx=XM_019824052.2
X 74772045 74772204 refGene_gn=DACH2;refGene_tx=XM_019824052.2
X 74791704 74791877 refGene_gn=DACH2;refGene_tx=XM_019824052.2
X 74798103 74798239 refGene_gn=DACH2;refGene_tx=XM_019824052.2
X 74800779 74800854 refGene_gn=DACH2;refGene_tx=XM_019824052.2
X 74838929 74839078 refGene_gn=DACH2;refGene_tx=XM_019824052.2
X 74839203 74839351 refGene_gn=DACH2;refGene_tx=XM_019824052.2
X 74844445 74844592 refGene_gn=DACH2;refGene_tx=XM_019824052.2
X 74846816 74846882 refGene_gn=DACH2;refGene_tx=XM_019824052.2
X 74858331 74858336 refGene_gn=DACH2;refGene_tx=XM_019824052.2
X 75728448 75729159 refGene_gn=TGIF2LX;refGene_tx=XM_004000695.5
X 76057629 76058778 refGene_gn=PABPC5;refGene_tx=XM_023249244.1
X 76318948 76319452 refGene_gn=PCDH11X;refGene_tx=XM_023248909.1
X 76319453 76319489 refGene_gn=PCDH11X;refGene_tx=XM_023248909.1
X 76384064 76384215 refGene_gn=PCDH11X;refGene_tx=XM_023248909.1
X 76384217 76386163 refGene_gn=PCDH11X;refGene_tx=XM_023248909.1
X 76386164 76386590 refGene_gn=PCDH11X;refGene_tx=XM_023248909.1
X 76390724 76390769 refGene_gn=PCDH11X;refGene_tx=XM_023248909.1
X 76890008 76891442 refGene_gn=NAP1L3;refGene_tx=XM_004000694.5
X 76942452 76942707 refGene_gn=FAM133A;refGene_tx=XM_023249049.1
X 76942708 76942853 refGene_gn=FAM133A;refGene_tx=XM_023249049.1
X 76942855 76943121 refGene_gn=FAM133A;refGene_tx=XM_023249049.1
X 77073466 77073558 refGene_gn=TEN1;refGene_tx=XM_023248966.1
X 77073560 77073614 refGene_gn=TEN1;refGene_tx=XM_023248966.1
X 77073616 77073652 refGene_gn=TEN1;refGene_tx=XM_023248966.1
X 77073654 77073684 refGene_gn=TEN1;refGene_tx=XM_023248966.1
X 77073686 77073770 refGene_gn=TEN1;refGene_tx=XM_023248966.1
X 77073772 77073866 refGene_gn=TEN1;refGene_tx=XM_023248966.1
X 77073943 77074103 refGene_gn=TEN1;refGene_tx=XM_023248966.1
X 78669921 78670053
refGene_gn=DIAPH2;refGene_tx=XM_019824053.2;refGene_gn=DIAPH2;refGene_tx=XM_023249470.1;refGene_gn=DIAPH2;refGene_tx=XM_023249471.1
X 78725236 78725269

refGene_gn=DIAPH2;refGene_tx=XM_019824053.2;refGene_gn=DIAPH2;refGene_tx=XM_023249470.1
X	78727725	78727902
refGene_gn=DIAPH2;refGene_tx=XM_019824053.2;refGene_gn=DIAPH2;refGene_tx=XM_023249470.1;refGene_gn=DIAPH2;refGene_tx=XM_023249471.1
X	78756266	78756371
refGene_gn=DIAPH2;refGene_tx=XM_019824053.2;refGene_gn=DIAPH2;refGene_tx=XM_023249470.1;refGene_gn=DIAPH2;refGene_tx=XM_023249471.1
X	78761387	78761408
refGene_gn=DIAPH2;refGene_tx=XM_019824053.2;refGene_gn=DIAPH2;refGene_tx=XM_023249471.1
X	78876985	78877125
refGene_gn=DIAPH2;refGene_tx=XM_019824053.2;refGene_gn=DIAPH2;refGene_tx=XM_023249470.1;refGene_gn=DIAPH2;refGene_tx=XM_023249471.1
X	78908754	78908829
refGene_gn=DIAPH2;refGene_tx=XM_019824053.2;refGene_gn=DIAPH2;refGene_tx=XM_023249470.1;refGene_gn=DIAPH2;refGene_tx=XM_023249471.1
X	78908910	78908980
refGene_gn=DIAPH2;refGene_tx=XM_019824053.2;refGene_gn=DIAPH2;refGene_tx=XM_023249470.1;refGene_gn=DIAPH2;refGene_tx=XM_023249471.1
X	78920586	78920723
refGene_gn=DIAPH2;refGene_tx=XM_019824053.2;refGene_gn=DIAPH2;refGene_tx=XM_023249470.1;refGene_gn=DIAPH2;refGene_tx=XM_023249471.1
X	78922695	78922804
refGene_gn=DIAPH2;refGene_tx=XM_019824053.2;refGene_gn=DIAPH2;refGene_tx=XM_023249470.1;refGene_gn=DIAPH2;refGene_tx=XM_023249471.1
X	78932422	78932533
refGene_gn=DIAPH2;refGene_tx=XM_019824053.2;refGene_gn=DIAPH2;refGene_tx=XM_023249470.1;refGene_gn=DIAPH2;refGene_tx=XM_023249471.1
X	78938145	78938264
refGene_gn=DIAPH2;refGene_tx=XM_019824053.2;refGene_gn=DIAPH2;refGene_tx=XM_023249470.1;refGene_gn=DIAPH2;refGene_tx=XM_023249471.1
X	78940283	78940400
refGene_gn=DIAPH2;refGene_tx=XM_019824053.2;refGene_gn=DIAPH2;refGene_tx=XM_023249470.1;refGene_gn=DIAPH2;refGene_tx=XM_023249471.1
X	78942549	78942668
refGene_gn=DIAPH2;refGene_tx=XM_019824053.2;refGene_gn=DIAPH2;refGene_tx=XM_023249470.1;refGene_gn=DIAPH2;refGene_tx=XM_023249471.1
X	78947605	78947670
refGene_gn=DIAPH2;refGene_tx=XM_019824053.2;refGene_gn=DIAPH2;refGene_tx=XM_023249470.1;refGene_gn=DIAPH2;refGene_tx=XM_023249471.1
X	78951975	78952080
refGene_gn=DIAPH2;refGene_tx=XM_019824053.2;refGene_gn=DIAPH2;refGene_tx=XM_023249470.1;refGene_gn=DIAPH2;refGene_tx=XM_023249471.1
X	78955646	78955961
refGene_gn=DIAPH2;refGene_tx=XM_019824053.2;refGene_gn=DIAPH2;refGene_tx=XM_023249470.1;refGene_gn=DIAPH2;refGene_tx=XM_023249471.1
X	78957559	78957674
refGene_gn=DIAPH2;refGene_tx=XM_019824053.2;refGene_gn=DIAPH2;refGene_tx=XM_023249470.1;refGene_gn=DIAPH2;refGene_tx=XM_023249471.1

X 79018403 79019456
refGene_gn=LOC111558778;refGene_tx=XM_023249472.1
X 79092541 79092643
refGene_gn=DIAPH2;refGene_tx=XM_019824053.2;refGene_gn=DIAPH2;refGene_tx=XM_023249470.1;refGene_gn=DIAPH2;refGene_tx=XM_023249471.1
X 79094455 79094550
refGene_gn=DIAPH2;refGene_tx=XM_019824053.2;refGene_gn=DIAPH2;refGene_tx=XM_023249470.1;refGene_gn=DIAPH2;refGene_tx=XM_023249471.1
X 79120162 79120264
refGene_gn=DIAPH2;refGene_tx=XM_019824053.2;refGene_gn=DIAPH2;refGene_tx=XM_023249470.1;refGene_gn=DIAPH2;refGene_tx=XM_023249471.1
X 79135739 79135979
refGene_gn=DIAPH2;refGene_tx=XM_019824053.2;refGene_gn=DIAPH2;refGene_tx=XM_023249470.1;refGene_gn=DIAPH2;refGene_tx=XM_023249471.1
X 79183693 79183823
refGene_gn=DIAPH2;refGene_tx=XM_019824053.2;refGene_gn=DIAPH2;refGene_tx=XM_023249470.1;refGene_gn=DIAPH2;refGene_tx=XM_023249471.1
X 79307162 79307287
refGene_gn=DIAPH2;refGene_tx=XM_019824053.2;refGene_gn=DIAPH2;refGene_tx=XM_023249470.1;refGene_gn=DIAPH2;refGene_tx=XM_023249471.1
X 79413030 79413195
refGene_gn=DIAPH2;refGene_tx=XM_019824053.2;refGene_gn=DIAPH2;refGene_tx=XM_023249470.1;refGene_gn=DIAPH2;refGene_tx=XM_023249471.1
X 79462464 79462600
refGene_gn=DIAPH2;refGene_tx=XM_019824053.2;refGene_gn=DIAPH2;refGene_tx=XM_023249470.1;refGene_gn=DIAPH2;refGene_tx=XM_023249471.1
X 79501370 79501466
refGene_gn=DIAPH2;refGene_tx=XM_019824053.2;refGene_gn=DIAPH2;refGene_tx=XM_023249470.1;refGene_gn=DIAPH2;refGene_tx=XM_023249471.1
X 79675870 79675935
refGene_gn=DIAPH2;refGene_tx=XM_019824053.2;refGene_gn=DIAPH2;refGene_tx=XM_023249470.1;refGene_gn=DIAPH2;refGene_tx=XM_023249471.1
X 82029676 82032697
refGene_gn=LOC111558676;refGene_tx=XM_023248967.1
X 82529809 82530393
refGene_gn=PCDH19;refGene_tx=XM_004000697.4;refGene_gn=PCDH19;refGene_tx=XM_006943858.3
X 82579971 82580144
refGene_gn=PCDH19;refGene_tx=XM_004000697.4;refGene_gn=PCDH19;refGene_tx=XM_006943858.3
X 82605706 82605765
refGene_gn=PCDH19;refGene_tx=XM_004000697.4;refGene_gn=PCDH19;refGene_tx=XM_006943858.3
X 82666448 82666776
refGene_gn=PCDH19;refGene_tx=XM_004000697.4;refGene_gn=PCDH19;refGene_tx=XM_006943858.3
X 82670441 82672588
refGene_gn=PCDH19;refGene_tx=XM_004000697.4;refGene_gn=PCDH19;refGene_tx=XM_006943858.3
X 82822434 82822482

refGene_gn=TNMD;refGene_tx=XM_004000699.5;refGene_gn=TNMD;refGene_tx=XM_023249276.1
X	82822636	82822768	refGene_gn=TNMD;refGene_tx=XM_004000699.5;refGene_gn=TNMD;refGene_tx=XM_023249276.1
X	82831677	82831818	refGene_gn=TNMD;refGene_tx=XM_004000699.5;refGene_gn=TNMD;refGene_tx=XM_023249276.1
X	82832028	82832130	refGene_gn=TNMD;refGene_tx=XM_004000699.5
X	82835351	82835505	refGene_gn=TNMD;refGene_tx=XM_004000699.5;refGene_gn=TNMD;refGene_tx=XM_023249276.1
X	82836889	82837056	refGene_gn=TNMD;refGene_tx=XM_004000699.5;refGene_gn=TNMD;refGene_tx=XM_023249276.1
X	82837351	82837561	refGene_gn=TNMD;refGene_tx=XM_004000699.5;refGene_gn=TNMD;refGene_tx=XM_023249276.1
X	82869561	82869630	refGene_gn=TSPAN6;refGene_tx=XM_006943859.3;refGene_gn=TSPAN6;refGene_tx=XM_023249095.1
X	82870992	82871076	refGene_gn=TSPAN6;refGene_tx=XM_006943859.3;refGene_gn=TSPAN6;refGene_tx=XM_023249095.1
X	82871468	82871603	refGene_gn=TSPAN6;refGene_tx=XM_006943859.3;refGene_gn=TSPAN6;refGene_tx=XM_023249095.1
X	82872101	82872200	refGene_gn=TSPAN6;refGene_tx=XM_006943859.3;refGene_gn=TSPAN6;refGene_tx=XM_023249095.1
X	82873344	82873419	refGene_gn=TSPAN6;refGene_tx=XM_006943859.3;refGene_gn=TSPAN6;refGene_tx=XM_023249095.1
X	82873696	82873885	refGene_gn=TSPAN6;refGene_tx=XM_006943859.3;refGene_gn=TSPAN6;refGene_tx=XM_023249095.1
X	82874862	82874949	refGene_gn=TSPAN6;refGene_tx=XM_006943859.3
X	82885623	82885705	refGene_gn=SRPX2;refGene_tx=XM_004000701.4;refGene_gn=SRPX2;refGene_tx=XM_006943860.3
X	82890208	82890286	refGene_gn=SRPX2;refGene_tx=XM_004000701.4;refGene_gn=SRPX2;refGene_tx=XM_006943860.3
X	82897506	82897698	refGene_gn=SRPX2;refGene_tx=XM_004000701.4;refGene_gn=SRPX2;refGene_tx=XM_006943860.3
X	82900046	82900223	refGene_gn=SRPX2;refGene_tx=XM_004000701.4;refGene_gn=SRPX2;refGene_tx=XM_006943860.3
X	82900525	82900652

refGene_gn=SRPX2;refGene_tx=XM_004000701.4;refGene_gn=SRPX2;refGene_tx=XM_006943860.3
X	82900803	82900925
refGene_gn=SRPX2;refGene_tx=XM_004000701.4;refGene_gn=SRPX2;refGene_tx=XM_006943860.3
X	82902025	82902205
refGene_gn=SRPX2;refGene_tx=XM_004000701.4;refGene_gn=SRPX2;refGene_tx=XM_006943860.3
X	82902542	82902676
refGene_gn=SRPX2;refGene_tx=XM_004000701.4;refGene_gn=SRPX2;refGene_tx=XM_006943860.3
X	82904582	82904704
refGene_gn=SRPX2;refGene_tx=XM_004000701.4;refGene_gn=SRPX2;refGene_tx=XM_006943860.3
X	82905800	82905981
refGene_gn=SRPX2;refGene_tx=XM_004000701.4;refGene_gn=SRPX2;refGene_tx=XM_006943860.3
X	82911683	82911832
refGene_gn=SYTL4;refGene_tx=XM_011291843.3;refGene_gn=SYTL4;refGene_tx=XM_011291844.3;refGene_gn=SYTL4;refGene_tx=XM_019824061.2
X	82913973	82914182
refGene_gn=SYTL4;refGene_tx=XM_011291843.3;refGene_gn=SYTL4;refGene_tx=XM_011291844.3;refGene_gn=SYTL4;refGene_tx=XM_019824061.2
X	82915079	82915179
refGene_gn=SYTL4;refGene_tx=XM_011291843.3;refGene_gn=SYTL4;refGene_tx=XM_011291844.3;refGene_gn=SYTL4;refGene_tx=XM_019824061.2
X	82917020	82917129
refGene_gn=SYTL4;refGene_tx=XM_011291843.3;refGene_gn=SYTL4;refGene_tx=XM_011291844.3;refGene_gn=SYTL4;refGene_tx=XM_019824061.2
X	82920648	82920810
refGene_gn=SYTL4;refGene_tx=XM_011291843.3;refGene_gn=SYTL4;refGene_tx=XM_011291844.3;refGene_gn=SYTL4;refGene_tx=XM_019824061.2
X	82921318	82921421
refGene_gn=SYTL4;refGene_tx=XM_011291843.3;refGene_gn=SYTL4;refGene_tx=XM_011291844.3;refGene_gn=SYTL4;refGene_tx=XM_019824061.2
X	82921718	82921897
refGene_gn=SYTL4;refGene_tx=XM_011291843.3;refGene_gn=SYTL4;refGene_tx=XM_011291844.3;refGene_gn=SYTL4;refGene_tx=XM_019824061.2
X	82923180	82923273
refGene_gn=SYTL4;refGene_tx=XM_011291843.3;refGene_gn=SYTL4;refGene_tx=XM_011291844.3;refGene_gn=SYTL4;refGene_tx=XM_019824061.2
X	82923764	82923868
refGene_gn=SYTL4;refGene_tx=XM_011291843.3;refGene_gn=SYTL4;refGene_tx=XM_011291844.3;refGene_gn=SYTL4;refGene_tx=XM_019824061.2
X	82923979	82924070
refGene_gn=SYTL4;refGene_tx=XM_011291843.3;refGene_gn=SYTL4;refGene_tx=XM_011291844.3;refGene_gn=SYTL4;refGene_tx=XM_019824061.2
X	82924511	82924587
refGene_gn=SYTL4;refGene_tx=XM_011291843.3;refGene_gn=SYTL4;refGene_tx=XM_011291844.3;refGene_gn=SYTL4;refGene_tx=XM_019824061.2

X 82925063 82925165
refGene_gn=SYTL4;refGene_tx=XM_011291843.3;refGene_gn=SYTL4;refGene_tx
=XM_011291844.3;refGene_gn=SYTL4;refGene_tx=XM_019824061.2
X 82931659 82931762
refGene_gn=SYTL4;refGene_tx=XM_011291843.3;refGene_gn=SYTL4;refGene_tx
=XM_011291844.3;refGene_gn=SYTL4;refGene_tx=XM_019824061.2
X 82931991 82932101
refGene_gn=SYTL4;refGene_tx=XM_011291843.3;refGene_gn=SYTL4;refGene_tx
=XM_011291844.3;refGene_gn=SYTL4;refGene_tx=XM_019824061.2
X 82932235 82932451
refGene_gn=SYTL4;refGene_tx=XM_011291843.3;refGene_gn=SYTL4;refGene_tx
=XM_011291844.3;refGene_gn=SYTL4;refGene_tx=XM_019824061.2
X 82932702 82932812
refGene_gn=SYTL4;refGene_tx=XM_011291843.3;refGene_gn=SYTL4;refGene_tx
=XM_011291844.3;refGene_gn=SYTL4;refGene_tx=XM_019824061.2
X 83121277 83121335
refGene_gn=CSTF2;refGene_tx=XM_019824062.1;refGene_gn=CSTF2;refGene_tx
=XM_019824063.1;refGene_gn=CSTF2;refGene_tx=XM_019824064.1
X 83122167 83122246
refGene_gn=CSTF2;refGene_tx=XM_019824062.1;refGene_gn=CSTF2;refGene_tx
=XM_019824063.1;refGene_gn=CSTF2;refGene_tx=XM_019824064.1
X 83122765 83122935
refGene_gn=CSTF2;refGene_tx=XM_019824062.1;refGene_gn=CSTF2;refGene_tx
=XM_019824063.1;refGene_gn=CSTF2;refGene_tx=XM_019824064.1
X 83123663 83123800
refGene_gn=CSTF2;refGene_tx=XM_019824062.1;refGene_gn=CSTF2;refGene_tx
=XM_019824063.1;refGene_gn=CSTF2;refGene_tx=XM_019824064.1
X 83124274 83124394
refGene_gn=CSTF2;refGene_tx=XM_019824062.1;refGene_gn=CSTF2;refGene_tx
=XM_019824063.1;refGene_gn=CSTF2;refGene_tx=XM_019824064.1
X 83124503 83124641
refGene_gn=CSTF2;refGene_tx=XM_019824062.1;refGene_gn=CSTF2;refGene_tx
=XM_019824063.1;refGene_gn=CSTF2;refGene_tx=XM_019824064.1
X 83126610 83126734
refGene_gn=CSTF2;refGene_tx=XM_019824062.1;refGene_gn=CSTF2;refGene_tx
=XM_019824063.1;refGene_gn=CSTF2;refGene_tx=XM_019824064.1
X 83130846 83130909
refGene_gn=CSTF2;refGene_tx=XM_019824062.1;refGene_gn=CSTF2;refGene_tx
=XM_019824063.1;refGene_gn=CSTF2;refGene_tx=XM_019824064.1
X 83133251 83133311
refGene_gn=CSTF2;refGene_tx=XM_019824062.1;refGene_gn=CSTF2;refGene_tx
=XM_019824063.1
X 83133492 83133579 refGene_gn=CSTF2;refGene_tx=XM_019824062.1
X 83133960 83134102
refGene_gn=CSTF2;refGene_tx=XM_019824062.1;refGene_gn=CSTF2;refGene_tx
=XM_019824063.1;refGene_gn=CSTF2;refGene_tx=XM_019824064.1
X 83135124 83135300
refGene_gn=CSTF2;refGene_tx=XM_019824062.1;refGene_gn=CSTF2;refGene_tx
=XM_019824063.1;refGene_gn=CSTF2;refGene_tx=XM_019824064.1
X 83135859 83136152

refGene_gn=CSTF2;refGene_tx=XM_019824062.1;refGene_gn=CSTF2;refGene_tx=XM_019824063.1;refGene_gn=CSTF2;refGene_tx=XM_019824064.1
X 83139536 83139647
refGene_gn=CSTF2;refGene_tx=XM_019824062.1;refGene_gn=CSTF2;refGene_tx=XM_019824063.1;refGene_gn=CSTF2;refGene_tx=XM_019824064.1
X 83141619 83141742
refGene_gn=CSTF2;refGene_tx=XM_019824062.1;refGene_gn=CSTF2;refGene_tx=XM_019824063.1;refGene_gn=CSTF2;refGene_tx=XM_019824064.1
X 83147160 83147287 refGene_gn=NOX1;refGene_tx=XM_004000704.5
X 83150592 83150717 refGene_gn=NOX1;refGene_tx=XM_004000704.5
X 83150959 83151106 refGene_gn=NOX1;refGene_tx=XM_004000704.5
X 83151527 83151690 refGene_gn=NOX1;refGene_tx=XM_004000704.5
X 83151907 83152143 refGene_gn=NOX1;refGene_tx=XM_004000704.5
X 83154097 83154190 refGene_gn=NOX1;refGene_tx=XM_004000704.5
X 83160853 83160904 refGene_gn=NOX1;refGene_tx=XM_004000704.5
X 83160906 83160997 refGene_gn=NOX1;refGene_tx=XM_004000704.5
X 83161085 83161267 refGene_gn=NOX1;refGene_tx=XM_004000704.5
X 83161370 83161522 refGene_gn=NOX1;refGene_tx=XM_004000704.5
X 83161862 83161947 refGene_gn=NOX1;refGene_tx=XM_004000704.5
X 83162189 83162300 refGene_gn=NOX1;refGene_tx=XM_004000704.5
X 83166846 83166942 refGene_gn=NOX1;refGene_tx=XM_004000704.5
X 83171214 83171259 refGene_gn=NOX1;refGene_tx=XM_004000704.5
X 83212652 83213398 refGene_gn=XKRX;refGene_tx=XM_006943861.2
X 83219662 83219931 refGene_gn=XKRX;refGene_tx=XM_006943861.2
X 83226183 83226518 refGene_gn=XKRX;refGene_tx=XM_006943861.2
X 83277528 83277587 refGene_gn=ARL13A;refGene_tx=XM_023248908.1
X 83277968 83278039 refGene_gn=ARL13A;refGene_tx=XM_023248908.1
X 83283468 83283718 refGene_gn=ARL13A;refGene_tx=XM_023248908.1
X 83284595 83284701 refGene_gn=ARL13A;refGene_tx=XM_023248908.1
X 83285238 83285426 refGene_gn=ARL13A;refGene_tx=XM_023248908.1
X 83286075 83286166 refGene_gn=ARL13A;refGene_tx=XM_023248908.1
X 83297659 83297716 refGene_gn=ARL13A;refGene_tx=XM_023248908.1
X 83360513 83360621 refGene_gn=TMEM35A;refGene_tx=XM_004000705.5
X 83360622 83360634 refGene_gn=TMEM35A;refGene_tx=XM_004000705.5
X 83370742 83371150 refGene_gn=TMEM35A;refGene_tx=XM_004000705.5
X 83375928 83376148 refGene_gn=CENPI;refGene_tx=XM_019824070.1
X 83377030 83377168 refGene_gn=CENPI;refGene_tx=XM_019824070.1
X 83381186 83381305 refGene_gn=CENPI;refGene_tx=XM_019824070.1
X 83381592 83381700 refGene_gn=CENPI;refGene_tx=XM_019824070.1
X 83397931 83397980 refGene_gn=CENPI;refGene_tx=XM_019824070.1
X 83403095 83403142 refGene_gn=CENPI;refGene_tx=XM_019824070.1
X 83408024 83408114 refGene_gn=CENPI;refGene_tx=XM_019824070.1
X 83408662 83408791 refGene_gn=CENPI;refGene_tx=XM_019824070.1
X 83409023 83409191 refGene_gn=CENPI;refGene_tx=XM_019824070.1
X 83409343 83409464 refGene_gn=CENPI;refGene_tx=XM_019824070.1
X 83417443 83417535 refGene_gn=CENPI;refGene_tx=XM_019824070.1
X 83419032 83419150 refGene_gn=CENPI;refGene_tx=XM_019824070.1
X 83419237 83419302 refGene_gn=CENPI;refGene_tx=XM_019824070.1
X 83424190 83424285 refGene_gn=CENPI;refGene_tx=XM_019824070.1
X 83434549 83434685 refGene_gn=CENPI;refGene_tx=XM_019824070.1

X	83435543	83435668	refGene_gn=CENPI;refGene_tx=XM_019824070.1
X	83437513	83437562	refGene_gn=CENPI;refGene_tx=XM_019824070.1
X	83437693	83437912	refGene_gn=CENPI;refGene_tx=XM_019824070.1
X	83451926	83451968	refGene_gn=CENPI;refGene_tx=XM_019824070.1
X	83452670	83452805	refGene_gn=CENPI;refGene_tx=XM_019824070.1
X	83507496	83507613
refGene_gn=DRP2;refGene_tx=XM_006943864.4;refGene_gn=DRP2;refGene_tx=XM_019824066.2;refGene_gn=DRP2;refGene_tx=XM_019824067.2;refGene_gn=DRP2;refGene_tx=XM_019824068.2
X	83510683	83510713	refGene_gn=DRP2;refGene_tx=XM_006943863.3
X	83511757	83511921
refGene_gn=DRP2;refGene_tx=XM_006943863.3;refGene_gn=DRP2;refGene_tx=XM_006943864.4;refGene_gn=DRP2;refGene_tx=XM_019824066.2;refGene_gn=DRP2;refGene_tx=XM_019824067.2;refGene_gn=DRP2;refGene_tx=XM_019824069.2
X	83513795	83513952
refGene_gn=DRP2;refGene_tx=XM_006943863.3;refGene_gn=DRP2;refGene_tx=XM_006943864.4;refGene_gn=DRP2;refGene_tx=XM_019824066.2;refGene_gn=DRP2;refGene_tx=XM_019824067.2;refGene_gn=DRP2;refGene_tx=XM_019824068.2
X	83515381	83515502
refGene_gn=DRP2;refGene_tx=XM_006943863.3;refGene_gn=DRP2;refGene_tx=XM_006943864.4;refGene_gn=DRP2;refGene_tx=XM_019824066.2;refGene_gn=DRP2;refGene_tx=XM_019824067.2;refGene_gn=DRP2;refGene_tx=XM_019824068.2
X	83517082	83517351
refGene_gn=DRP2;refGene_tx=XM_006943863.3;refGene_gn=DRP2;refGene_tx=XM_006943864.4;refGene_gn=DRP2;refGene_tx=XM_019824066.2;refGene_gn=DRP2;refGene_tx=XM_019824067.2;refGene_gn=DRP2;refGene_tx=XM_019824068.2;refGene_gn=DRP2;refGene_tx=XM_019824069.2;refGene_gn=DRP2;refGene_tx=XM_023248877.1
X	83517737	83517884
refGene_gn=DRP2;refGene_tx=XM_006943863.3;refGene_gn=DRP2;refGene_tx=XM_006943864.4;refGene_gn=DRP2;refGene_tx=XM_019824066.2;refGene_gn=DRP2;refGene_tx=XM_019824067.2;refGene_gn=DRP2;refGene_tx=XM_019824068.2;refGene_gn=DRP2;refGene_tx=XM_019824069.2;refGene_gn=DRP2;refGene_tx=XM_023248877.1
X	83518325	83518404
refGene_gn=DRP2;refGene_tx=XM_006943863.3;refGene_gn=DRP2;refGene_tx=XM_006943864.4;refGene_gn=DRP2;refGene_tx=XM_019824066.2;refGene_gn=DRP2;refGene_tx=XM_019824067.2;refGene_gn=DRP2;refGene_tx=XM_019824068.2;refGene_gn=DRP2;refGene_tx=XM_019824069.2;refGene_gn=DRP2;refGene_tx=XM_023248877.1
X	83520071	83520132
refGene_gn=DRP2;refGene_tx=XM_006943863.3;refGene_gn=DRP2;refGene_tx=XM_006943864.4;refGene_gn=DRP2;refGene_tx=XM_019824066.2;refGene_gn=DRP2;refGene_tx=XM_019824067.2;refGene_gn=DRP2;refGene_tx=XM_019824068.2;refGene_gn=DRP2;refGene_tx=XM_019824069.2;refGene_gn=DRP2;refGene_tx=XM_023248877.1
X	83520439	83520501
refGene_gn=DRP2;refGene_tx=XM_006943863.3;refGene_gn=DRP2;refGene_tx=XM_006943864.4;refGene_gn=DRP2;refGene_tx=XM_019824066.2;refGene_gn=DRP2;refGene_tx=XM_019824067.2;refGene_gn=DRP2;refGene_tx=XM_019824068.2;

refGene_gn=DRP2;refGene_tx=XM_019824069.2;refGene_gn=DRP2;refGene_tx=XM_023248877.1
X 83521524 83521599
refGene_gn=DRP2;refGene_tx=XM_006943863.3;refGene_gn=DRP2;refGene_tx=XM_006943864.4;refGene_gn=DRP2;refGene_tx=XM_019824066.2;refGene_gn=DRP2;refGene_tx=XM_019824067.2;refGene_gn=DRP2;refGene_tx=XM_019824068.2;refGene_gn=DRP2;refGene_tx=XM_019824069.2;refGene_gn=DRP2;refGene_tx=XM_023248877.1
X 83522533 83522735
refGene_gn=DRP2;refGene_tx=XM_006943863.3;refGene_gn=DRP2;refGene_tx=XM_006943864.4;refGene_gn=DRP2;refGene_tx=XM_019824066.2;refGene_gn=DRP2;refGene_tx=XM_019824067.2;refGene_gn=DRP2;refGene_tx=XM_019824068.2;refGene_gn=DRP2;refGene_tx=XM_019824069.2;refGene_gn=DRP2;refGene_tx=XM_023248877.1
X 83522965 83523051
refGene_gn=DRP2;refGene_tx=XM_006943863.3;refGene_gn=DRP2;refGene_tx=XM_006943864.4;refGene_gn=DRP2;refGene_tx=XM_019824066.2;refGene_gn=DRP2;refGene_tx=XM_019824067.2;refGene_gn=DRP2;refGene_tx=XM_019824068.2;refGene_gn=DRP2;refGene_tx=XM_019824069.2;refGene_gn=DRP2;refGene_tx=XM_023248877.1
X 83524699 83524857
refGene_gn=DRP2;refGene_tx=XM_006943863.3;refGene_gn=DRP2;refGene_tx=XM_006943864.4;refGene_gn=DRP2;refGene_tx=XM_019824066.2;refGene_gn=DRP2;refGene_tx=XM_019824067.2;refGene_gn=DRP2;refGene_tx=XM_019824068.2;refGene_gn=DRP2;refGene_tx=XM_019824069.2;refGene_gn=DRP2;refGene_tx=XM_023248877.1
X 83525180 83525347
refGene_gn=DRP2;refGene_tx=XM_006943863.3;refGene_gn=DRP2;refGene_tx=XM_006943864.4;refGene_gn=DRP2;refGene_tx=XM_019824066.2;refGene_gn=DRP2;refGene_tx=XM_019824067.2;refGene_gn=DRP2;refGene_tx=XM_019824068.2;refGene_gn=DRP2;refGene_tx=XM_019824069.2;refGene_gn=DRP2;refGene_tx=XM_023248877.1
X 83526994 83527106
refGene_gn=DRP2;refGene_tx=XM_006943863.3;refGene_gn=DRP2;refGene_tx=XM_006943864.4;refGene_gn=DRP2;refGene_tx=XM_019824066.2;refGene_gn=DRP2;refGene_tx=XM_019824067.2;refGene_gn=DRP2;refGene_tx=XM_019824068.2;refGene_gn=DRP2;refGene_tx=XM_019824069.2;refGene_gn=DRP2;refGene_tx=XM_023248877.1
X 83528408 83528545
refGene_gn=DRP2;refGene_tx=XM_006943863.3;refGene_gn=DRP2;refGene_tx=XM_006943864.4;refGene_gn=DRP2;refGene_tx=XM_019824066.2;refGene_gn=DRP2;refGene_tx=XM_019824067.2;refGene_gn=DRP2;refGene_tx=XM_019824068.2;refGene_gn=DRP2;refGene_tx=XM_019824069.2;refGene_gn=DRP2;refGene_tx=XM_023248877.1
X 83528853 83528919
refGene_gn=DRP2;refGene_tx=XM_006943863.3;refGene_gn=DRP2;refGene_tx=XM_006943864.4;refGene_gn=DRP2;refGene_tx=XM_019824066.2;refGene_gn=DRP2;refGene_tx=XM_019824068.2;refGene_gn=DRP2;refGene_tx=XM_019824069.2;refGene_gn=DRP2;refGene_tx=XM_023248877.1
X 83529170 83529236

refGene_gn=DRP2;refGene_tx=XM_006943863.3;refGene_gn=DRP2;refGene_tx=XM_006943864.4;refGene_gn=DRP2;refGene_tx=XM_019824066.2;refGene_gn=DRP2;refGene_tx=XM_019824067.2;refGene_gn=DRP2;refGene_tx=XM_019824068.2;refGene_gn=DRP2;refGene_tx=XM_019824069.2;refGene_gn=DRP2;refGene_tx=XM_023248877.1
X 83530059 83530203
refGene_gn=DRP2;refGene_tx=XM_006943863.3;refGene_gn=DRP2;refGene_tx=XM_006943864.4;refGene_gn=DRP2;refGene_tx=XM_019824066.2;refGene_gn=DRP2;refGene_tx=XM_019824067.2;refGene_gn=DRP2;refGene_tx=XM_019824068.2;refGene_gn=DRP2;refGene_tx=XM_019824069.2;refGene_gn=DRP2;refGene_tx=XM_023248877.1
X 83531846 83532084
refGene_gn=DRP2;refGene_tx=XM_006943863.3;refGene_gn=DRP2;refGene_tx=XM_006943864.4;refGene_gn=DRP2;refGene_tx=XM_019824066.2;refGene_gn=DRP2;refGene_tx=XM_019824067.2;refGene_gn=DRP2;refGene_tx=XM_019824068.2;refGene_gn=DRP2;refGene_tx=XM_019824069.2;refGene_gn=DRP2;refGene_tx=XM_023248877.1
X 83533491 83533612
refGene_gn=DRP2;refGene_tx=XM_006943863.3;refGene_gn=DRP2;refGene_tx=XM_006943864.4;refGene_gn=DRP2;refGene_tx=XM_019824066.2;refGene_gn=DRP2;refGene_tx=XM_019824067.2;refGene_gn=DRP2;refGene_tx=XM_019824068.2;refGene_gn=DRP2;refGene_tx=XM_019824069.2;refGene_gn=DRP2;refGene_tx=XM_023248877.1
X 83533927 83534052
refGene_gn=DRP2;refGene_tx=XM_006943863.3;refGene_gn=DRP2;refGene_tx=XM_006943864.4;refGene_gn=DRP2;refGene_tx=XM_019824066.2;refGene_gn=DRP2;refGene_tx=XM_019824067.2;refGene_gn=DRP2;refGene_tx=XM_019824068.2;refGene_gn=DRP2;refGene_tx=XM_019824069.2;refGene_gn=DRP2;refGene_tx=XM_023248877.1
X 83542121 83542166
refGene_gn=TAF7L;refGene_tx=XM_019824074.2;refGene_gn=TAF7L;refGene_tx=XM_023249144.1
X 83545597 83545657
refGene_gn=TAF7L;refGene_tx=XM_019824074.2;refGene_gn=TAF7L;refGene_tx=XM_023249144.1
X 83546821 83546934
refGene_gn=TAF7L;refGene_tx=XM_019824074.2;refGene_gn=TAF7L;refGene_tx=XM_023249144.1
X 83547130 83547358
refGene_gn=TAF7L;refGene_tx=XM_019824074.2;refGene_gn=TAF7L;refGene_tx=XM_023249144.1
X 83547782 83547896
refGene_gn=TAF7L;refGene_tx=XM_019824074.2;refGene_gn=TAF7L;refGene_tx=XM_023249144.1
X 83548240 83548388
refGene_gn=TAF7L;refGene_tx=XM_019824074.2;refGene_gn=TAF7L;refGene_tx=XM_023249144.1
X 83549185 83549227
refGene_gn=TAF7L;refGene_tx=XM_019824074.2;refGene_gn=TAF7L;refGene_tx=XM_023249144.1

X 83552293 83552349
refGene_gn=TAF7L;refGene_tx=XM_019824074.2;refGene_gn=TAF7L;refGene_tx
=XM_023249144.1
X 83552583 83552710
refGene_gn=TAF7L;refGene_tx=XM_019824074.2;refGene_gn=TAF7L;refGene_tx
=XM_023249144.1
X 83553399 83553533
refGene_gn=TAF7L;refGene_tx=XM_019824074.2;refGene_gn=TAF7L;refGene_tx
=XM_023249144.1
X 83555032 83555111
refGene_gn=TAF7L;refGene_tx=XM_019824074.2;refGene_gn=TAF7L;refGene_tx
=XM_023249144.1
X 83555918 83555986
refGene_gn=TAF7L;refGene_tx=XM_019824074.2;refGene_gn=TAF7L;refGene_tx
=XM_023249144.1
X 83559667 83559923 refGene_gn=TAF7L;refGene_tx=XM_023249144.1
X 83587084 83587246 refGene_gn=TIMM8A;refGene_tx=XM_004000710.5
X 83588791 83588923 refGene_gn=TIMM8A;refGene_tx=XM_004000710.5
X 83590058 83590130 refGene_gn=BTK;refGene_tx=XM_023249136.1
X 83592874 83593032 refGene_gn=BTK;refGene_tx=XM_023249136.1
X 83593549 83593668 refGene_gn=BTK;refGene_tx=XM_023249136.1
X 83594487 83594552 refGene_gn=BTK;refGene_tx=XM_023249136.1
X 83595815 83596032 refGene_gn=BTK;refGene_tx=XM_023249136.1
X 83596552 83596724 refGene_gn=BTK;refGene_tx=XM_023249136.1
X 83597433 83597508 refGene_gn=BTK;refGene_tx=XM_023249136.1
X 83598306 83598434 refGene_gn=BTK;refGene_tx=XM_023249136.1
X 83598592 83598672 refGene_gn=BTK;refGene_tx=XM_023249136.1
X 83598950 83599005 refGene_gn=BTK;refGene_tx=XM_023249136.1
X 83599580 83599643 refGene_gn=BTK;refGene_tx=XM_023249136.1
X 83600069 83600256 refGene_gn=BTK;refGene_tx=XM_023249136.1
X 83601177 83601261 refGene_gn=BTK;refGene_tx=XM_023249136.1
X 83601573 83601702 refGene_gn=BTK;refGene_tx=XM_023249136.1
X 83607699 83607781 refGene_gn=BTK;refGene_tx=XM_023249136.1
X 83609339 83609408 refGene_gn=BTK;refGene_tx=XM_023249136.1
X 83612677 83612776 refGene_gn=BTK;refGene_tx=XM_023249136.1
X 83613323 83613410 refGene_gn=BTK;refGene_tx=XM_023249136.1
X 83613411 83613465 refGene_gn=BTK;refGene_tx=XM_023249136.1
X 83627720 83627831 refGene_gn=RPL36A;refGene_tx=XM_004000712.3
X 83628241 83628347 refGene_gn=RPL36A;refGene_tx=XM_004000712.3
X 83628535 83628603 refGene_gn=RPL36A;refGene_tx=XM_004000712.3
X 83629689 83629812 refGene_gn=RPL36A;refGene_tx=XM_004000712.3
X 83630089 83630110 refGene_gn=RPL36A;refGene_tx=XM_004000712.3
X 83631792 83632056 refGene_gn=GLA;refGene_tx=XM_004000723.4
X 83632457 83632655 refGene_gn=GLA;refGene_tx=XM_004000723.4
X 83632863 83633025 refGene_gn=GLA;refGene_tx=XM_004000723.4
X 83634509 83634601 refGene_gn=GLA;refGene_tx=XM_004000723.4
X 83635203 83635381 refGene_gn=GLA;refGene_tx=XM_004000723.4
X 83636421 83636596 refGene_gn=GLA;refGene_tx=XM_004000723.4
X 83639035 83639229 refGene_gn=GLA;refGene_tx=XM_004000723.4
X 83643051 83644401

refGene_gn=HNRNPH2;refGene_tx=XM_006943866.4;refGene_gn=HNRNPH2;refGene_tx=XM_006943867.4;refGene_gn=HNRNPH2;refGene_tx=XM_019824075.2;refGene_gn=HNRNPH2;refGene_tx=XM_023248830.1;refGene_gn=HNRNPH2;refGene_tx=XM_023248831.1
X 83704761 83712849
refGene_gn=ARMCX4;refGene_tx=XM_006943868.4;refGene_gn=ARMCX4;refGene_tx=XM_006943869.4;refGene_gn=ARMCX4;refGene_tx=XM_006943870.4;refGene_gn=ARMCX4;refGene_tx=XM_019824076.2;refGene_gn=ARMCX4;refGene_tx=XM_019824077.2
X 83743875 83745252
refGene_gn=ARMCX1;refGene_tx=XM_011291835.3;refGene_gn=ARMCX1;refGene_tx=XM_011291836.3;refGene_gn=ARMCX1;refGene_tx=XM_011291837.3;refGene_gn=ARMCX1;refGene_tx=XM_011291838.3;refGene_gn=ARMCX1;refGene_tx=XM_011291839.3;refGene_gn=ARMCX1;refGene_tx=XM_011291840.2;refGene_gn=ARMCX1;refGene_tx=XM_019824078.1;refGene_gn=ARMCX1;refGene_tx=XM_019824079.2;refGene_gn=ARMCX1;refGene_tx=XM_019824080.2
X 83751766 83751824
refGene_gn=LOC101091926;refGene_tx=XM_023248968.1
X 83762807 83763221
refGene_gn=LOC101091926;refGene_tx=XM_023248968.1
X 83763223 83763363
refGene_gn=LOC101091926;refGene_tx=XM_023248968.1
X 83776074 83776977
refGene_gn=LOC101092421;refGene_tx=XM_006943871.3;refGene_gn=LOC101092421;refGene_tx=XM_006943872.3;refGene_gn=LOC101092421;refGene_tx=XM_011291832.2;refGene_gn=LOC101092421;refGene_tx=XM_019824081.2
X 83785155 83786295
refGene_gn=ARMCX3;refGene_tx=XM_006943873.3;refGene_gn=ARMCX3;refGene_tx=XM_011291831.2;refGene_gn=ARMCX3;refGene_tx=XM_019824082.1
X 83820632 83822462
refGene_gn=ARMCX2;refGene_tx=XM_004000716.4;refGene_gn=ARMCX2;refGene_tx=XM_011291830.3;refGene_gn=ARMCX2;refGene_tx=XM_019824083.2
X 83980887 83980932
refGene_gn=LOC101092919;refGene_tx=XM_006943874.2;refGene_gn=LOC101092919;refGene_tx=XM_019823483.1;refGene_gn=LOC101092919;refGene_tx=XM_019823484.1
X 83981065 83981126
refGene_gn=LOC101092919;refGene_tx=XM_006943874.2;refGene_gn=LOC101092919;refGene_tx=XM_019823483.1;refGene_gn=LOC101092919;refGene_tx=XM_019823484.1
X 83983392 83983575
refGene_gn=LOC101092919;refGene_tx=XM_006943874.2;refGene_gn=LOC101092919;refGene_tx=XM_019823483.1;refGene_gn=LOC101092919;refGene_tx=XM_019823484.1
X 83984084 83984157
refGene_gn=LOC101092919;refGene_tx=XM_006943874.2;refGene_gn=LOC101092919;refGene_tx=XM_019823483.1;refGene_gn=LOC101092919;refGene_tx=XM_019823484.1
X 83984714 83984757
refGene_gn=LOC101092919;refGene_tx=XM_006943874.2;refGene_gn=LOC101092

919;refGene_tx=XM_019823483.1;refGene_gn=LOC101092919;refGene_tx=XM_019823484.1
X 83984849 83984965
refGene_gn=LOC101092919;refGene_tx=XM_006943874.2;refGene_gn=LOC101092919;refGene_tx=XM_019823483.1;refGene_gn=LOC101092919;refGene_tx=XM_019823484.1
X 83985076 83985135
refGene_gn=LOC101092919;refGene_tx=XM_006943874.2;refGene_gn=LOC101092919;refGene_tx=XM_019823483.1;refGene_gn=LOC101092919;refGene_tx=XM_019823484.1
X 83985256 83985364
refGene_gn=LOC101092919;refGene_tx=XM_006943874.2;refGene_gn=LOC101092919;refGene_tx=XM_019823483.1;refGene_gn=LOC101092919;refGene_tx=XM_019823484.1
X 83985489 83985545
refGene_gn=LOC101092919;refGene_tx=XM_006943874.2;refGene_gn=LOC101092919;refGene_tx=XM_019823483.1;refGene_gn=LOC101092919;refGene_tx=XM_019823484.1
X 83985630 83985696
refGene_gn=LOC101092919;refGene_tx=XM_006943874.2;refGene_gn=LOC101092919;refGene_tx=XM_019823483.1;refGene_gn=LOC101092919;refGene_tx=XM_019823484.1
X 83986097 83986134
refGene_gn=LOC101092919;refGene_tx=XM_006943874.2;refGene_gn=LOC101092919;refGene_tx=XM_019823483.1;refGene_gn=LOC101092919;refGene_tx=XM_019823484.1
X 83988019 83988129
refGene_gn=LOC101092919;refGene_tx=XM_006943874.2;refGene_gn=LOC101092919;refGene_tx=XM_019823483.1;refGene_gn=LOC101092919;refGene_tx=XM_019823484.1
X 83988304 83988412
refGene_gn=LOC101092919;refGene_tx=XM_006943874.2;refGene_gn=LOC101092919;refGene_tx=XM_019823483.1;refGene_gn=LOC101092919;refGene_tx=XM_019823484.1
X 83988532 83988621
refGene_gn=LOC101092919;refGene_tx=XM_006943874.2;refGene_gn=LOC101092919;refGene_tx=XM_019823483.1;refGene_gn=LOC101092919;refGene_tx=XM_019823484.1
X 83988768 83988838
refGene_gn=LOC101092919;refGene_tx=XM_006943874.2;refGene_gn=LOC101092919;refGene_tx=XM_019823483.1;refGene_gn=LOC101092919;refGene_tx=XM_019823484.1
X 83988958 83989039
refGene_gn=LOC101092919;refGene_tx=XM_006943874.2;refGene_gn=LOC101092919;refGene_tx=XM_019823483.1;refGene_gn=LOC101092919;refGene_tx=XM_019823484.1
X 83989158 83989263
refGene_gn=LOC101092919;refGene_tx=XM_006943874.2;refGene_gn=LOC101092919;refGene_tx=XM_019823483.1;refGene_gn=LOC101092919;refGene_tx=XM_019823484.1

X 83989383 83989467
refGene_gn=LOC101092919;refGene_tx=XM_006943874.2;refGene_gn=LOC101092
919;refGene_tx=XM_019823483.1;refGene_gn=LOC101092919;refGene_tx=XM_01
9823484.1
X 83989754 83989908
refGene_gn=LOC101092919;refGene_tx=XM_006943874.2;refGene_gn=LOC101092
919;refGene_tx=XM_019823483.1;refGene_gn=LOC101092919;refGene_tx=XM_01
9823484.1
X 83990140 83990324
refGene_gn=LOC101092919;refGene_tx=XM_006943874.2;refGene_gn=LOC101092
919;refGene_tx=XM_019823483.1;refGene_gn=LOC101092919;refGene_tx=XM_01
9823484.1
X 83993142 83993155
refGene_gn=LOC101092919;refGene_tx=XM_006943874.2
X 84012971 84013041
refGene_gn=LOC101092919;refGene_tx=XM_019823483.1
X 84076259 84077580
refGene_gn=ZMAT1;refGene_tx=XM_006943875.3;refGene_gn=ZMAT1;refGene_tx
=XM_019824084.2;refGene_gn=ZMAT1;refGene_tx=XM_019824085.2;refGene_gn=
ZMAT1;refGene_tx=XM_019824086.2;refGene_gn=ZMAT1;refGene_tx=XM_0198240
87.2;refGene_gn=ZMAT1;refGene_tx=XM_019824088.2;refGene_gn=ZMAT1;refGe
ne_tx=XM_019824089.2;refGene_gn=ZMAT1;refGene_tx=XM_019824093.2
X 84078780 84078880
refGene_gn=ZMAT1;refGene_tx=XM_006943875.3;refGene_gn=ZMAT1;refGene_tx
=XM_019824084.2;refGene_gn=ZMAT1;refGene_tx=XM_019824085.2;refGene_gn=
ZMAT1;refGene_tx=XM_019824086.2;refGene_gn=ZMAT1;refGene_tx=XM_0198240
87.2;refGene_gn=ZMAT1;refGene_tx=XM_019824088.2;refGene_gn=ZMAT1;refGe
ne_tx=XM_019824089.2;refGene_gn=ZMAT1;refGene_tx=XM_019824093.2
X 84087477 84087762
refGene_gn=ZMAT1;refGene_tx=XM_006943875.3;refGene_gn=ZMAT1;refGene_tx
=XM_019824084.2;refGene_gn=ZMAT1;refGene_tx=XM_019824085.2;refGene_gn=
ZMAT1;refGene_tx=XM_019824086.2;refGene_gn=ZMAT1;refGene_tx=XM_0198240
87.2;refGene_gn=ZMAT1;refGene_tx=XM_019824088.2;refGene_gn=ZMAT1;refGe
ne_tx=XM_019824089.2
X 84089340 84089515
refGene_gn=ZMAT1;refGene_tx=XM_006943875.3;refGene_gn=ZMAT1;refGene_tx
=XM_019824084.2;refGene_gn=ZMAT1;refGene_tx=XM_019824085.2;refGene_gn=
ZMAT1;refGene_tx=XM_019824086.2;refGene_gn=ZMAT1;refGene_tx=XM_0198240
87.2;refGene_gn=ZMAT1;refGene_tx=XM_019824088.2;refGene_gn=ZMAT1;refGe
ne_tx=XM_019824089.2
X 84089609 84089711
refGene_gn=ZMAT1;refGene_tx=XM_006943875.3;refGene_gn=ZMAT1;refGene_tx
=XM_019824084.2;refGene_gn=ZMAT1;refGene_tx=XM_019824085.2;refGene_gn=
ZMAT1;refGene_tx=XM_019824086.2;refGene_gn=ZMAT1;refGene_tx=XM_0198240
88.2;refGene_gn=ZMAT1;refGene_tx=XM_019824089.2
X 84103059 84103166
refGene_gn=ZMAT1;refGene_tx=XM_006943875.3;refGene_gn=ZMAT1;refGene_tx
=XM_019824084.2;refGene_gn=ZMAT1;refGene_tx=XM_019824085.2;refGene_gn=
ZMAT1;refGene_tx=XM_019824086.2;refGene_gn=ZMAT1;refGene_tx=XM_0198240
87.2;refGene_gn=ZMAT1;refGene_tx=XM_019824088.2;refGene_gn=ZMAT1;refGe

ne_tx=XM_019824089.2
X 84110617 84110698
refGene_gn=ZMAT1;refGene_tx=XM_019824084.2;refGene_gn=ZMAT1;refGene_tx=XM_019824085.2;refGene_gn=ZMAT1;refGene_tx=XM_019824086.2;refGene_gn=ZMAT1;refGene_tx=XM_019824087.2;refGene_gn=ZMAT1;refGene_tx=XM_019824088.2;refGene_gn=ZMAT1;refGene_tx=XM_019824089.2
X 84119390 84119667
refGene_gn=ZMAT1;refGene_tx=XM_006943875.3;refGene_gn=ZMAT1;refGene_tx=XM_019824084.2;refGene_gn=ZMAT1;refGene_tx=XM_019824085.2;refGene_gn=ZMAT1;refGene_tx=XM_019824086.2;refGene_gn=ZMAT1;refGene_tx=XM_019824087.2;refGene_gn=ZMAT1;refGene_tx=XM_019824088.2;refGene_gn=ZMAT1;refGene_tx=XM_019824089.2
X 84261379 84261901 refGene_gn=TCEAL6;refGene_tx=XM_023249181.1
X 84261902 84261932 refGene_gn=TCEAL6;refGene_tx=XM_023249181.1
X 84283642 84284410
refGene_gn=TCEAL2;refGene_tx=XM_023249159.1;refGene_gn=TCEAL2;refGene_tx=XM_023249160.1;refGene_gn=TCEAL2;refGene_tx=XM_023249161.1;refGene_gn=TCEAL2;refGene_tx=XM_023249162.1
X 84299539 84299910
refGene_gn=BEX5;refGene_tx=XM_006943878.4;refGene_gn=BEX5;refGene_tx=XM_019824094.2;refGene_gn=BEX5;refGene_tx=XM_023249201.1
X 84300185 84300252 refGene_gn=BEX5;refGene_tx=XM_023249201.1
X 84300528 84300531 refGene_gn=BEX5;refGene_tx=XM_023249201.1
X 84331879 84331924
refGene_gn=LOC101093921;refGene_tx=XM_011291825.3
X 84332063 84332124
refGene_gn=LOC101093921;refGene_tx=XM_011291825.3
X 84348250 84348433
refGene_gn=LOC101093921;refGene_tx=XM_011291825.3
X 84348982 84349055
refGene_gn=LOC101093921;refGene_tx=XM_011291825.3
X 84349627 84349670
refGene_gn=LOC101093921;refGene_tx=XM_011291825.3
X 84349764 84349880
refGene_gn=LOC101093921;refGene_tx=XM_011291825.3
X 84349989 84350048
refGene_gn=LOC101093921;refGene_tx=XM_011291825.3
X 84350168 84350276
refGene_gn=LOC101093921;refGene_tx=XM_011291825.3
X 84350416 84350472
refGene_gn=LOC101093921;refGene_tx=XM_011291825.3
X 84350557 84350626
refGene_gn=LOC101093921;refGene_tx=XM_011291825.3
X 84351029 84351066
refGene_gn=LOC101093921;refGene_tx=XM_011291825.3
X 84352718 84352828
refGene_gn=LOC101093921;refGene_tx=XM_011291825.3
X 84353005 84353113
refGene_gn=LOC101093921;refGene_tx=XM_011291825.3
X 84353232 84353321

refGene_gn=LOC101093921;refGene_tx=XM_011291825.3
X 84353468 84353538
refGene_gn=LOC101093921;refGene_tx=XM_011291825.3
X 84353658 84353739
refGene_gn=LOC101093921;refGene_tx=XM_011291825.3
X 84353859 84353964
refGene_gn=LOC101093921;refGene_tx=XM_011291825.3
X 84354087 84354171
refGene_gn=LOC101093921;refGene_tx=XM_011291825.3
X 84354458 84354612
refGene_gn=LOC101093921;refGene_tx=XM_011291825.3
X 84354845 84355029
refGene_gn=LOC101093921;refGene_tx=XM_011291825.3
X 84358114 84358127
refGene_gn=LOC101093921;refGene_tx=XM_011291825.3
X 84443232 84443363
refGene_gn=LOC101091521;refGene_tx=XM_006944276.1
X 84443406 84443590
refGene_gn=LOC101091521;refGene_tx=XM_006944276.1
X 84443710 84443718
refGene_gn=LOC101091521;refGene_tx=XM_006944276.1
X 84444929 84445097
refGene_gn=LOC101091521;refGene_tx=XM_006944276.1
X 84445870 84445915
refGene_gn=LOC101091521;refGene_tx=XM_006944276.1
X 84445917 84446044
refGene_gn=LOC101091521;refGene_tx=XM_006944276.1
X 84446139 84446354
refGene_gn=LOC101091521;refGene_tx=XM_006944276.1
X 84446569 84446706
refGene_gn=LOC101091521;refGene_tx=XM_006944276.1
X 84447460 84447528
refGene_gn=LOC101091521;refGene_tx=XM_006944276.1
X 84448238 84448358
refGene_gn=LOC101091521;refGene_tx=XM_006944276.1
X 84449129 84449201
refGene_gn=LOC101091521;refGene_tx=XM_006944276.1
X 84512475 84512675
refGene_gn=LOC111556301;refGene_tx=XM_023248970.1
X 84512756 84512864
refGene_gn=LOC111556301;refGene_tx=XM_023248970.1
X 84512990 84513055
refGene_gn=LOC111556301;refGene_tx=XM_023248970.1
X 84513159 84513257
refGene_gn=LOC111556301;refGene_tx=XM_023248970.1
X 84513345 84513397
refGene_gn=LOC111556301;refGene_tx=XM_023248970.1
X 84514122 84514195
refGene_gn=LOC111556301;refGene_tx=XM_023248970.1
X 84514726 84514747

refGene_gn=LOC111556301;refGene_tx=XM_023248970.1
X 84514748 84514896
refGene_gn=LOC111556301;refGene_tx=XM_023248970.1
X 84537713 84537861 refGene_gn=ARMCX5;refGene_tx=XM_011291823.3
X 84537863 84539356 refGene_gn=ARMCX5;refGene_tx=XM_011291823.3
X 84576953 84581132 refGene_gn=GPRASP1;refGene_tx=XM_004001217.5
X 84674437 84677014
refGene_gn=GPRASP2;refGene_tx=XM_019824810.2;refGene_gn=GPRASP2;refGene_tx=XM_019824811.2
X 84689914 84691567
refGene_gn=BHLHB9;refGene_tx=XM_019824095.2;refGene_gn=BHLHB9;refGene_tx=XM_023249028.1;refGene_gn=BHLHB9;refGene_tx=XM_023249029.1;refGene_gn=BHLHB9;refGene_tx=XM_023249030.1
X 84973694 84974662
refGene_gn=LOC109496734;refGene_tx=XM_019824831.1
X 85080503 85080507
refGene_gn=LOC101094164;refGene_tx=XM_023249078.1
X 85083483 85083536
refGene_gn=LOC101094164;refGene_tx=XM_023249078.1
X 85083537 85083659
refGene_gn=LOC101094164;refGene_tx=XM_023249078.1
X 85083872 85083979
refGene_gn=LOC101094164;refGene_tx=XM_023249078.1
X 85083981 85084025
refGene_gn=LOC101094164;refGene_tx=XM_023249078.1
X 85084673 85084757
refGene_gn=LOC101094164;refGene_tx=XM_023249078.1
X 85084849 85084954
refGene_gn=LOC101094164;refGene_tx=XM_023249078.1
X 85085078 85085159
refGene_gn=LOC101094164;refGene_tx=XM_023249078.1
X 85085716 85085824
refGene_gn=LOC101094164;refGene_tx=XM_023249078.1
X 85085997 85086101
refGene_gn=LOC101094164;refGene_tx=XM_023249078.1
X 85087882 85087919
refGene_gn=LOC101094164;refGene_tx=XM_023249078.1
X 85088270 85088339
refGene_gn=LOC101094164;refGene_tx=XM_023249078.1
X 85088424 85088480
refGene_gn=LOC101094164;refGene_tx=XM_023249078.1
X 85088619 85088727
refGene_gn=LOC101094164;refGene_tx=XM_023249078.1
X 85088846 85088905
refGene_gn=LOC101094164;refGene_tx=XM_023249078.1
X 85089018 85089134
refGene_gn=LOC101094164;refGene_tx=XM_023249078.1
X 85089227 85089284
refGene_gn=LOC101094164;refGene_tx=XM_023249078.1
X 85146673 85147062

refGene_gn=LOC101093759;refGene_tx=XM_004001225.4;refGene_gn=LOC101093759;refGene_tx=XM_006944284.4
X	85147406 85147489
refGene_gn=LOC101093759;refGene_tx=XM_006944284.4
X	85147880 85147981
refGene_gn=LOC101093759;refGene_tx=XM_006944284.4
X	85177160 85177164
refGene_gn=LOC105259665;refGene_tx=XM_023249166.1
X	85178548 85178621
refGene_gn=LOC105259665;refGene_tx=XM_023249166.1
X	85179092 85179135
refGene_gn=LOC105259665;refGene_tx=XM_023249166.1
X	85179228 85179344
refGene_gn=LOC105259665;refGene_tx=XM_023249166.1
X	85179457 85179516
refGene_gn=LOC105259665;refGene_tx=XM_023249166.1
X	85179635 85179743
refGene_gn=LOC105259665;refGene_tx=XM_023249166.1
X	85179882 85179938
refGene_gn=LOC105259665;refGene_tx=XM_023249166.1
X	85180023 85180092
refGene_gn=LOC105259665;refGene_tx=XM_023249166.1
X	85180442 85180479
refGene_gn=LOC105259665;refGene_tx=XM_023249166.1
X	85182271 85182375
refGene_gn=LOC105259665;refGene_tx=XM_023249166.1
X	85182548 85182656
refGene_gn=LOC105259665;refGene_tx=XM_023249166.1
X	85183213 85183294
refGene_gn=LOC105259665;refGene_tx=XM_023249166.1
X	85183418 85183523
refGene_gn=LOC105259665;refGene_tx=XM_023249166.1
X	85183615 85183699
refGene_gn=LOC105259665;refGene_tx=XM_023249166.1
X	85184346 85184500
refGene_gn=LOC105259665;refGene_tx=XM_023249166.1
X	85184713 85184888
refGene_gn=LOC105259665;refGene_tx=XM_023249166.1
X	85187854 85187858
refGene_gn=LOC105259665;refGene_tx=XM_023249166.1
X	85301045 85301399 refGene_gn=TCEAL8;refGene_tx=XM_004000731.4
X	85320081 85320702
refGene_gn=TCEAL5;refGene_tx=XM_019824099.2;refGene_gn=TCEAL5;refGene_tx=XM_023249168.1;refGene_gn=TCEAL5;refGene_tx=XM_023249169.1
X	85335295 85335687
refGene_gn=LOC101095084;refGene_tx=XM_004000734.5;refGene_gn=LOC101095084;refGene_tx=XM_006943884.4;refGene_gn=LOC101095084;refGene_tx=XM_006943885.3;refGene_gn=LOC101095084;refGene_tx=XM_006943886.4;refGene_gn=LOC101095084;refGene_tx=XM_019824100.2;refGene_gn=LOC101095084;refGene_tx=XM_023249179.1

X 85336359 85336426
refGene_gn=LOC101095084;refGene_tx=XM_019824100.2
X 85375990 85376293 refGene_gn=TCEAL7;refGene_tx=XM_006943887.2
X 85463597 85463912 refGene_gn=TCEAL9;refGene_tx=XM_019824101.2
X 85485522 85485861
refGene_gn=BEX3;refGene_tx=XM_011279382.3;refGene_gn=BEX3;refGene_tx=XM_011279383.3;refGene_gn=BEX3;refGene_tx=XM_019824102.2;refGene_gn=BEX3;refGene_tx=XM_019824103.2
X 86272289 86272967
refGene_gn=TCEAL4;refGene_tx=XM_004001564.5;refGene_gn=TCEAL4;refGene_tx=XM_023249504.1;refGene_gn=TCEAL4;refGene_tx=XM_023249505.1
X 86281200 86281381
refGene_gn=LOC101081708;refGene_tx=XM_023249506.1
X 86281383 86281535
refGene_gn=LOC101081708;refGene_tx=XM_023249506.1
X 86281536 86281722
refGene_gn=LOC101081708;refGene_tx=XM_023249506.1
X 86281723 86281753
refGene_gn=LOC101081708;refGene_tx=XM_023249506.1
X 86312923 86313403 refGene_gn=TCEAL1;refGene_tx=XM_004000737.5
X 86345801 86346668
refGene_gn=MORF4L2;refGene_tx=XM_004000738.4;refGene_gn=MORF4L2;refGene_tx=XM_004000739.5;refGene_gn=MORF4L2;refGene_tx=XM_004000743.4;refGene_gn=MORF4L2;refGene_tx=XM_006943888.3;refGene_gn=MORF4L2;refGene_tx=XM_019824114.2;refGene_gn=MORF4L2;refGene_tx=XM_019824116.2;refGene_gn=MORF4L2;refGene_tx=XM_019824117.2;refGene_gn=MORF4L2;refGene_tx=XM_019824118.2;refGene_gn=MORF4L2;refGene_tx=XM_023248885.1
X 86376296 86376587
refGene_gn=GLRA4;refGene_tx=XM_004000744.4;refGene_gn=GLRA4;refGene_tx=XM_019824109.2;refGene_gn=GLRA4;refGene_tx=XM_019824111.2
X 86382458 86382608
refGene_gn=GLRA4;refGene_tx=XM_004000744.4;refGene_gn=GLRA4;refGene_tx=XM_019824109.2;refGene_gn=GLRA4;refGene_tx=XM_019824111.2
X 86387408 86387623
refGene_gn=GLRA4;refGene_tx=XM_004000744.4;refGene_gn=GLRA4;refGene_tx=XM_019824109.2;refGene_gn=GLRA4;refGene_tx=XM_019824111.2
X 86391181 86391322
refGene_gn=GLRA4;refGene_tx=XM_004000744.4;refGene_gn=GLRA4;refGene_tx=XM_019824111.2
X 86392823 86392906
refGene_gn=GLRA4;refGene_tx=XM_004000744.4;refGene_gn=GLRA4;refGene_tx=XM_019824109.2;refGene_gn=GLRA4;refGene_tx=XM_019824111.2
X 86393044 86393268
refGene_gn=GLRA4;refGene_tx=XM_004000744.4;refGene_gn=GLRA4;refGene_tx=XM_019824109.2;refGene_gn=GLRA4;refGene_tx=XM_019824111.2
X 86393907 86393975
refGene_gn=GLRA4;refGene_tx=XM_004000744.4;refGene_gn=GLRA4;refGene_tx=XM_019824109.2
X 86394260 86394391
refGene_gn=GLRA4;refGene_tx=XM_004000744.4;refGene_gn=GLRA4;refGene_tx

=XM_019824109.2;refGene_gn=GLRA4;refGene_tx=XM_019824111.2
X	86397788	86397859
refGene_gn=GLRA4;refGene_tx=XM_004000744.4;refGene_gn=GLRA4;refGene_tx=XM_019824109.2
X	86445347	86445351
refGene_gn=PLP1;refGene_tx=XM_006943889.3;refGene_gn=PLP1;refGene_tx=XM_011279411.2
X	86453886	86454073
refGene_gn=PLP1;refGene_tx=XM_006943889.3;refGene_gn=PLP1;refGene_tx=XM_011279411.2
X	86454871	86455133
refGene_gn=PLP1;refGene_tx=XM_006943889.3;refGene_gn=PLP1;refGene_tx=XM_011279411.2
X	86456256	86456425
refGene_gn=PLP1;refGene_tx=XM_006943889.3;refGene_gn=PLP1;refGene_tx=XM_011279411.2
X	86456962	86457036
refGene_gn=PLP1;refGene_tx=XM_006943889.3;refGene_gn=PLP1;refGene_tx=XM_011279411.2
X	86458026	86458092
refGene_gn=PLP1;refGene_tx=XM_006943889.3;refGene_gn=PLP1;refGene_tx=XM_011279411.2
X	86459199	86459271
refGene_gn=PLP1;refGene_tx=XM_006943889.3;refGene_gn=PLP1;refGene_tx=XM_011279411.2
X	86496107	86496713	refGene_gn=RAB9B;refGene_tx=XM_004000747.5
X	86596968	86597889	refGene_gn=SLC25A53;refGene_tx=XM_004000763.4
X	86608378	86608565	refGene_gn=ZCCHC18;refGene_tx=XM_023249139.1
X	86608567	86608798	refGene_gn=ZCCHC18;refGene_tx=XM_023249139.1
X	86608800	86609030	refGene_gn=ZCCHC18;refGene_tx=XM_023249139.1
X	86609031	86609085	refGene_gn=ZCCHC18;refGene_tx=XM_023249139.1
X	86609086	86609593	refGene_gn=ZCCHC18;refGene_tx=XM_023249139.1
X	86667288	86667360	refGene_gn=FAM199X;refGene_tx=XM_004000749.5
X	86667361	86667486	refGene_gn=FAM199X;refGene_tx=XM_004000749.5
X	86681174	86681394	refGene_gn=FAM199X;refGene_tx=XM_004000749.5
X	86690880	86691030	refGene_gn=FAM199X;refGene_tx=XM_004000749.5
X	86691421	86691583	refGene_gn=FAM199X;refGene_tx=XM_004000749.5
X	86693892	86694159	refGene_gn=FAM199X;refGene_tx=XM_004000749.5
X	86695441	86695612	refGene_gn=FAM199X;refGene_tx=XM_004000749.5
X	86757248	86757569	refGene_gn=ESX1;refGene_tx=XM_019823485.2
X	86760356	86760402	refGene_gn=ESX1;refGene_tx=XM_019823485.2
X	86761428	86761747	refGene_gn=ESX1;refGene_tx=XM_019823485.2
X	86761934	86762016	refGene_gn=ESX1;refGene_tx=XM_019823485.2
X	87143666	87143748	refGene_gn=IL1RAPL2;refGene_tx=XM_019824119.1
X	87698596	87698870	refGene_gn=IL1RAPL2;refGene_tx=XM_019824119.1
X	87724963	87725026	refGene_gn=TEX13A;refGene_tx=XM_019824122.2
X	87725027	87725657	refGene_gn=TEX13A;refGene_tx=XM_019824122.2
X	87725764	87726223	refGene_gn=TEX13A;refGene_tx=XM_019824122.2
X	87743074	87743261	refGene_gn=IL1RAPL2;refGene_tx=XM_019824119.1
X	87765787	87765941	refGene_gn=IL1RAPL2;refGene_tx=XM_019824119.1

X 87987907 87987982 refGene_gn=IL1RAPL2;refGene_tx=XM_019824119.1
X 88234439 88234569 refGene_gn=IL1RAPL2;refGene_tx=XM_019824119.1
X 88254595 88254741 refGene_gn=IL1RAPL2;refGene_tx=XM_019824119.1
X 88263831 88263975 refGene_gn=IL1RAPL2;refGene_tx=XM_019824119.1
X 88269483 88269654 refGene_gn=IL1RAPL2;refGene_tx=XM_019824119.1
X 88282544 88283242 refGene_gn=IL1RAPL2;refGene_tx=XM_019824119.1
X 88335691 88335748
refGene_gn=NRK;refGene_tx=XM_019824123.2;refGene_gn=NRK;refGene_tx=XM_019824124.2;refGene_gn=NRK;refGene_tx=XM_023249087.1
X 88343733 88343799
refGene_gn=NRK;refGene_tx=XM_019824123.2;refGene_gn=NRK;refGene_tx=XM_019824124.2;refGene_gn=NRK;refGene_tx=XM_023249087.1
X 88411969 88412026
refGene_gn=NRK;refGene_tx=XM_019824123.2;refGene_gn=NRK;refGene_tx=XM_019824124.2;refGene_gn=NRK;refGene_tx=XM_023249087.1
X 88413359 88413431
refGene_gn=NRK;refGene_tx=XM_019824123.2;refGene_gn=NRK;refGene_tx=XM_019824124.2;refGene_gn=NRK;refGene_tx=XM_023249087.1
X 88420724 88420850
refGene_gn=NRK;refGene_tx=XM_019824123.2;refGene_gn=NRK;refGene_tx=XM_019824124.2;refGene_gn=NRK;refGene_tx=XM_023249087.1
X 88423946 88424057
refGene_gn=NRK;refGene_tx=XM_019824123.2;refGene_gn=NRK;refGene_tx=XM_019824124.2;refGene_gn=NRK;refGene_tx=XM_023249087.1
X 88425536 88425627
refGene_gn=NRK;refGene_tx=XM_019824123.2;refGene_gn=NRK;refGene_tx=XM_019824124.2;refGene_gn=NRK;refGene_tx=XM_023249087.1
X 88428271 88428402
refGene_gn=NRK;refGene_tx=XM_019824123.2;refGene_gn=NRK;refGene_tx=XM_019824124.2;refGene_gn=NRK;refGene_tx=XM_023249087.1
X 88430047 88430102
refGene_gn=NRK;refGene_tx=XM_019824123.2;refGene_gn=NRK;refGene_tx=XM_019824124.2;refGene_gn=NRK;refGene_tx=XM_023249087.1
X 88435253 88435332
refGene_gn=NRK;refGene_tx=XM_019824123.2;refGene_gn=NRK;refGene_tx=XM_019824124.2;refGene_gn=NRK;refGene_tx=XM_023249087.1
X 88436371 88436547
refGene_gn=NRK;refGene_tx=XM_019824123.2;refGene_gn=NRK;refGene_tx=XM_019824124.2;refGene_gn=NRK;refGene_tx=XM_023249087.1
X 88438123 88438187
refGene_gn=NRK;refGene_tx=XM_019824123.2;refGene_gn=NRK;refGene_tx=XM_019824124.2;refGene_gn=NRK;refGene_tx=XM_023249087.1
X 88438605 88439773
refGene_gn=NRK;refGene_tx=XM_019824123.2;refGene_gn=NRK;refGene_tx=XM_019824124.2;refGene_gn=NRK;refGene_tx=XM_023249087.1
X 88441929 88442037
refGene_gn=NRK;refGene_tx=XM_019824123.2;refGene_gn=NRK;refGene_tx=XM_019824124.2;refGene_gn=NRK;refGene_tx=XM_023249087.1
X 88444955 88445023
refGene_gn=NRK;refGene_tx=XM_019824123.2;refGene_gn=NRK;refGene_tx=XM_

019824124.2;refGene_gn=NRK;refGene_tx=XM_023249087.1
X	88447131 88447226
refGene_gn=NRK;refGene_tx=XM_019824123.2;refGene_gn=NRK;refGene_tx=XM_
019824124.2;refGene_gn=NRK;refGene_tx=XM_023249087.1
X	88449818 88449916
refGene_gn=NRK;refGene_tx=XM_019824123.2;refGene_gn=NRK;refGene_tx=XM_
019824124.2;refGene_gn=NRK;refGene_tx=XM_023249087.1
X	88450924 88451289
refGene_gn=NRK;refGene_tx=XM_019824123.2;refGene_gn=NRK;refGene_tx=XM_
019824124.2;refGene_gn=NRK;refGene_tx=XM_023249087.1
X	88453046 88453386
refGene_gn=NRK;refGene_tx=XM_019824123.2;refGene_gn=NRK;refGene_tx=XM_
019824124.2;refGene_gn=NRK;refGene_tx=XM_023249087.1
X	88456320 88456507
refGene_gn=NRK;refGene_tx=XM_019824123.2;refGene_gn=NRK;refGene_tx=XM_
019824124.2;refGene_gn=NRK;refGene_tx=XM_023249087.1
X	88457173 88457329
refGene_gn=NRK;refGene_tx=XM_019824123.2;refGene_gn=NRK;refGene_tx=XM_
019824124.2;refGene_gn=NRK;refGene_tx=XM_023249087.1
X	88460766 88460910
refGene_gn=NRK;refGene_tx=XM_019824123.2;refGene_gn=NRK;refGene_tx=XM_
019824124.2;refGene_gn=NRK;refGene_tx=XM_023249087.1
X	88462813 88462972
refGene_gn=NRK;refGene_tx=XM_019824123.2;refGene_gn=NRK;refGene_tx=XM_
019824124.2;refGene_gn=NRK;refGene_tx=XM_023249087.1
X	88466966 88467071
refGene_gn=NRK;refGene_tx=XM_019824123.2;refGene_gn=NRK;refGene_tx=XM_
019824124.2;refGene_gn=NRK;refGene_tx=XM_023249087.1
X	88468677 88468820 refGene_gn=NRK;refGene_tx=XM_023249087.1
X	88469119 88469269
refGene_gn=NRK;refGene_tx=XM_019824123.2;refGene_gn=NRK;refGene_tx=XM_
019824124.2;refGene_gn=NRK;refGene_tx=XM_023249087.1
X	88472161 88472321
refGene_gn=NRK;refGene_tx=XM_019824123.2;refGene_gn=NRK;refGene_tx=XM_
019824124.2;refGene_gn=NRK;refGene_tx=XM_023249087.1
X	88475979 88476119
refGene_gn=NRK;refGene_tx=XM_019824123.2;refGene_gn=NRK;refGene_tx=XM_
019824124.2;refGene_gn=NRK;refGene_tx=XM_023249087.1
X	88477794 88477890
refGene_gn=NRK;refGene_tx=XM_019824123.2;refGene_gn=NRK;refGene_tx=XM_
019824124.2;refGene_gn=NRK;refGene_tx=XM_023249087.1
X	88564911 88565124 refGene_gn=SERPINA7;refGene_tx=XM_004000753.2
X	88565661 88565809 refGene_gn=SERPINA7;refGene_tx=XM_004000753.2
X	88566765 88567039 refGene_gn=SERPINA7;refGene_tx=XM_004000753.2
X	88568103 88568322 refGene_gn=SERPINA7;refGene_tx=XM_004000753.2
X	88568324 88568393 refGene_gn=SERPINA7;refGene_tx=XM_004000753.2
X	88568395 88568711 refGene_gn=SERPINA7;refGene_tx=XM_004000753.2
X	88652890 88654972
refGene_gn=MUM1L1;refGene_tx=XM_011279409.3;refGene_gn=MUM1L1;refGene_
tx=XM_023248910.1;refGene_gn=MUM1L1;refGene_tx=XM_023248911.1;refGene_

gn=MUM1L1;refGene_tx=XM_023248912.1;refGene_gn=MUM1L1;refGene_tx=XM_02
3248913.1;refGene_gn=MUM1L1;refGene_tx=XM_023248914.1;refGene_gn=MUM1L
1;refGene_tx=XM_023248915.1;refGene_gn=MUM1L1;refGene_tx=XM_023248916.
1;refGene_gn=MUM1L1;refGene_tx=XM_023248917.1
X 89093818 89094106
refGene_gn=LOC101101037;refGene_tx=XM_019824128.2
X 89109014 89109657
refGene_gn=CXHXorf57;refGene_tx=XM_004000756.5
X 89117561 89117704
refGene_gn=CXHXorf57;refGene_tx=XM_004000756.5
X 89118829 89119022
refGene_gn=CXHXorf57;refGene_tx=XM_004000756.5
X 89133764 89133873
refGene_gn=CXHXorf57;refGene_tx=XM_004000756.5
X 89134068 89134160
refGene_gn=CXHXorf57;refGene_tx=XM_004000756.5
X 89134266 89134389
refGene_gn=CXHXorf57;refGene_tx=XM_004000756.5
X 89137446 89137551
refGene_gn=CXHXorf57;refGene_tx=XM_004000756.5
X 89138627 89138792
refGene_gn=CXHXorf57;refGene_tx=XM_004000756.5
X 89140029 89140190
refGene_gn=CXHXorf57;refGene_tx=XM_004000756.5
X 89141122 89141301
refGene_gn=CXHXorf57;refGene_tx=XM_004000756.5
X 89147080 89147151
refGene_gn=CXHXorf57;refGene_tx=XM_004000756.5
X 89179817 89180108
refGene_gn=CXHXorf57;refGene_tx=XM_004000756.5
X 89183485 89183647
refGene_gn=CXHXorf57;refGene_tx=XM_004000756.5
X 89193135 89193266
refGene_gn=CXHXorf57;refGene_tx=XM_004000756.5
X 89208078 89208484 refGene_gn=RNF128;refGene_tx=XM_004000759.5
X 89254508 89255001 refGene_gn=RNF128;refGene_tx=XM_004000758.5
X 89285016 89285264
refGene_gn=RNF128;refGene_tx=XM_004000758.5;refGene_gn=RNF128;refGene_
tx=XM_004000759.5
X 89302096 89302168
refGene_gn=RNF128;refGene_tx=XM_004000758.5;refGene_gn=RNF128;refGene_
tx=XM_004000759.5
X 89305164 89305247
refGene_gn=RNF128;refGene_tx=XM_004000758.5;refGene_gn=RNF128;refGene_
tx=XM_004000759.5
X 89306645 89306742
refGene_gn=RNF128;refGene_tx=XM_004000758.5;refGene_gn=RNF128;refGene_
tx=XM_004000759.5
X 89307659 89307828
refGene_gn=RNF128;refGene_tx=XM_004000758.5;refGene_gn=RNF128;refGene_

tx=XM_004000759.5
X	89315391 89315525
refGene_gn=RNF128;refGene_tx=XM_004000758.5;refGene_gn=RNF128;refGene_
tx=XM_004000759.5
X	89323059 89323189
refGene_gn=TBC1D8B;refGene_tx=XM_011279406.3;refGene_gn=TBC1D8B;refGen
e_tx=XM_019824129.2
X	89337712 89337823
refGene_gn=TBC1D8B;refGene_tx=XM_011279406.3;refGene_gn=TBC1D8B;refGen
e_tx=XM_019824129.2
X	89340293 89340412
refGene_gn=TBC1D8B;refGene_tx=XM_011279406.3;refGene_gn=TBC1D8B;refGen
e_tx=XM_019824129.2
X	89340902 89341128
refGene_gn=TBC1D8B;refGene_tx=XM_011279406.3;refGene_gn=TBC1D8B;refGen
e_tx=XM_019824129.2
X	89341793 89342034
refGene_gn=TBC1D8B;refGene_tx=XM_011279406.3;refGene_gn=TBC1D8B;refGen
e_tx=XM_019824129.2
X	89347823 89348031
refGene_gn=TBC1D8B;refGene_tx=XM_011279406.3;refGene_gn=TBC1D8B;refGen
e_tx=XM_019824129.2
X	89349340 89349508
refGene_gn=TBC1D8B;refGene_tx=XM_011279406.3;refGene_gn=TBC1D8B;refGen
e_tx=XM_019824129.2
X	89357910 89358060 refGene_gn=TBC1D8B;refGene_tx=XM_011279406.3
X	89358618 89358769
refGene_gn=TBC1D8B;refGene_tx=XM_011279406.3;refGene_gn=TBC1D8B;refGen
e_tx=XM_019824129.2
X	89359387 89359602
refGene_gn=TBC1D8B;refGene_tx=XM_011279406.3;refGene_gn=TBC1D8B;refGen
e_tx=XM_019824129.2
X	89364824 89364942
refGene_gn=TBC1D8B;refGene_tx=XM_011279406.3;refGene_gn=TBC1D8B;refGen
e_tx=XM_019824129.2
X	89365924 89366210
refGene_gn=TBC1D8B;refGene_tx=XM_011279406.3;refGene_gn=TBC1D8B;refGen
e_tx=XM_019824129.2
X	89367150 89367280
refGene_gn=TBC1D8B;refGene_tx=XM_011279406.3;refGene_gn=TBC1D8B;refGen
e_tx=XM_019824129.2
X	89368271 89368370
refGene_gn=TBC1D8B;refGene_tx=XM_011279406.3;refGene_gn=TBC1D8B;refGen
e_tx=XM_019824129.2
X	89371688 89371757
refGene_gn=TBC1D8B;refGene_tx=XM_011279406.3;refGene_gn=TBC1D8B;refGen
e_tx=XM_019824129.2
X	89371927 89372168
refGene_gn=TBC1D8B;refGene_tx=XM_011279406.3;refGene_gn=TBC1D8B;refGen
e_tx=XM_019824129.2

X 89373056 89373122
refGene_gn=TBC1D8B;refGene_tx=XM_011279406.3;refGene_gn=TBC1D8B;refGene_tx=XM_019824129.2
X 89374755 89374839
refGene_gn=TBC1D8B;refGene_tx=XM_011279406.3;refGene_gn=TBC1D8B;refGene_tx=XM_019824129.2
X 89375907 89375964
refGene_gn=TBC1D8B;refGene_tx=XM_011279406.3;refGene_gn=TBC1D8B;refGene_tx=XM_019824129.2
X 89377540 89377638
refGene_gn=TBC1D8B;refGene_tx=XM_011279406.3;refGene_gn=TBC1D8B;refGene_tx=XM_019824129.2
X 89380241 89380637
refGene_gn=TBC1D8B;refGene_tx=XM_011279406.3;refGene_gn=TBC1D8B;refGene_tx=XM_019824129.2
X 89413985 89414259 refGene_gn=RIPPLY1;refGene_tx=XM_006943897.3
X 89414810 89414875 refGene_gn=RIPPLY1;refGene_tx=XM_006943897.3
X 89415486 89415562 refGene_gn=RIPPLY1;refGene_tx=XM_006943897.3
X 89416467 89416613 refGene_gn=RIPPLY1;refGene_tx=XM_006943897.3
X 89440692 89441385 refGene_gn=CLDN2;refGene_tx=XM_019824131.2
X 89454891 89455045
refGene_gn=MORC4;refGene_tx=XM_023248881.1;refGene_gn=MORC4;refGene_tx=XM_023248882.1
X 89455350 89455634
refGene_gn=MORC4;refGene_tx=XM_023248881.1;refGene_gn=MORC4;refGene_tx=XM_023248882.1
X 89455930 89456624
refGene_gn=MORC4;refGene_tx=XM_023248881.1;refGene_gn=MORC4;refGene_tx=XM_023248882.1
X 89468503 89468679
refGene_gn=MORC4;refGene_tx=XM_023248881.1;refGene_gn=MORC4;refGene_tx=XM_023248882.1
X 89470048 89470100
refGene_gn=MORC4;refGene_tx=XM_023248881.1;refGene_gn=MORC4;refGene_tx=XM_023248882.1
X 89470504 89470573
refGene_gn=MORC4;refGene_tx=XM_023248881.1;refGene_gn=MORC4;refGene_tx=XM_023248882.1
X 89472731 89472860
refGene_gn=MORC4;refGene_tx=XM_023248881.1;refGene_gn=MORC4;refGene_tx=XM_023248882.1
X 89475633 89475732
refGene_gn=MORC4;refGene_tx=XM_023248881.1;refGene_gn=MORC4;refGene_tx=XM_023248882.1
X 89485223 89485324
refGene_gn=MORC4;refGene_tx=XM_023248881.1;refGene_gn=MORC4;refGene_tx=XM_023248882.1
X 89486725 89486845
refGene_gn=MORC4;refGene_tx=XM_023248881.1;refGene_gn=MORC4;refGene_tx=XM_023248882.1

X 89488711 89488840
refGene_gn=MORC4;refGene_tx=XM_023248881.1;refGene_gn=MORC4;refGene_tx
=XM_023248882.1
X 89489149 89489282
refGene_gn=MORC4;refGene_tx=XM_023248881.1;refGene_gn=MORC4;refGene_tx
=XM_023248882.1
X 89491770 89491918
refGene_gn=MORC4;refGene_tx=XM_023248881.1;refGene_gn=MORC4;refGene_tx
=XM_023248882.1
X 89493035 89493250
refGene_gn=MORC4;refGene_tx=XM_023248881.1;refGene_gn=MORC4;refGene_tx
=XM_023248882.1
X 89498686 89498819
refGene_gn=MORC4;refGene_tx=XM_023248881.1;refGene_gn=MORC4;refGene_tx
=XM_023248882.1
X 89504674 89504747
refGene_gn=MORC4;refGene_tx=XM_023248881.1;refGene_gn=MORC4;refGene_tx
=XM_023248882.1
X 89504840 89504942
refGene_gn=MORC4;refGene_tx=XM_023248881.1;refGene_gn=MORC4;refGene_tx
=XM_023248882.1
X 89625328 89625477
refGene_gn=RBM41;refGene_tx=XM_019824133.2;refGene_gn=RBM41;refGene_tx
=XM_023248810.1;refGene_gn=RBM41;refGene_tx=XM_023248811.1;refGene_gn=
RBM41;refGene_tx=XM_023248812.1;refGene_gn=RBM41;refGene_tx=XM_0232488
13.1
X 89627302 89627450
refGene_gn=RBM41;refGene_tx=XM_019824133.2;refGene_gn=RBM41;refGene_tx
=XM_023248810.1;refGene_gn=RBM41;refGene_tx=XM_023248811.1;refGene_gn=
RBM41;refGene_tx=XM_023248812.1;refGene_gn=RBM41;refGene_tx=XM_0232488
13.1
X 89644534 89644938
refGene_gn=RBM41;refGene_tx=XM_019824133.2;refGene_gn=RBM41;refGene_tx
=XM_023248810.1;refGene_gn=RBM41;refGene_tx=XM_023248811.1;refGene_gn=
RBM41;refGene_tx=XM_023248812.1;refGene_gn=RBM41;refGene_tx=XM_0232488
13.1
X 89676525 89676597 refGene_gn=RBM41;refGene_tx=XM_019824133.2
X 89678473 89678678
refGene_gn=RBM41;refGene_tx=XM_019824133.2;refGene_gn=RBM41;refGene_tx
=XM_023248810.1;refGene_gn=RBM41;refGene_tx=XM_023248811.1;refGene_gn=
RBM41;refGene_tx=XM_023248812.1;refGene_gn=RBM41;refGene_tx=XM_0232488
13.1
X 89678969 89679162
refGene_gn=RBM41;refGene_tx=XM_019824133.2;refGene_gn=RBM41;refGene_tx
=XM_023248810.1;refGene_gn=RBM41;refGene_tx=XM_023248811.1;refGene_gn=
RBM41;refGene_tx=XM_023248812.1;refGene_gn=RBM41;refGene_tx=XM_0232488
13.1
X 89679739 89679856
refGene_gn=RBM41;refGene_tx=XM_019824133.2;refGene_gn=RBM41;refGene_tx
=XM_023248810.1;refGene_gn=RBM41;refGene_tx=XM_023248811.1;refGene_gn=

RBM41;refGene_tx=XM_023248812.1;refGene_gn=RBM41;refGene_tx=XM_0232488
13.1
X 89682779 89683003
refGene_gn=RBM41;refGene_tx=XM_019824133.2;refGene_gn=RBM41;refGene_tx
=XM_023248810.1;refGene_gn=RBM41;refGene_tx=XM_023248811.1;refGene_gn=
RBM41;refGene_tx=XM_023248812.1
X 89683092 89683100 refGene_gn=RBM41;refGene_tx=XM_023248813.1
X 89688290 89688385
refGene_gn=NUP62CL;refGene_tx=XM_006943898.4;refGene_gn=NUP62CL;refGen
e_tx=XM_011279405.3
X 89711204 89711271
refGene_gn=NUP62CL;refGene_tx=XM_006943898.4;refGene_gn=NUP62CL;refGen
e_tx=XM_011279405.3
X 89715147 89715259
refGene_gn=NUP62CL;refGene_tx=XM_006943898.4;refGene_gn=NUP62CL;refGen
e_tx=XM_011279405.3
X 89716989 89717124
refGene_gn=NUP62CL;refGene_tx=XM_006943898.4;refGene_gn=NUP62CL;refGen
e_tx=XM_011279405.3
X 89717262 89717312
refGene_gn=NUP62CL;refGene_tx=XM_006943898.4;refGene_gn=NUP62CL;refGen
e_tx=XM_011279405.3
X 89717567 89717718
refGene_gn=NUP62CL;refGene_tx=XM_006943898.4;refGene_gn=NUP62CL;refGen
e_tx=XM_011279405.3
X 89726854 89727014
refGene_gn=NUP62CL;refGene_tx=XM_006943898.4;refGene_gn=NUP62CL;refGen
e_tx=XM_011279405.3
X 89732784 89732883 refGene_gn=NUP62CL;refGene_tx=XM_011279405.3
X 89734407 89734465
refGene_gn=NUP62CL;refGene_tx=XM_006943898.4;refGene_gn=NUP62CL;refGen
e_tx=XM_011279405.3
X 89773565 89773730
refGene_gn=PIH1D3;refGene_tx=XM_006943899.3;refGene_gn=PIH1D3;refGene_
tx=XM_019823486.2
X 89777388 89777461
refGene_gn=PIH1D3;refGene_tx=XM_006943899.3;refGene_gn=PIH1D3;refGene_
tx=XM_019823486.2
X 89779980 89780086
refGene_gn=PIH1D3;refGene_tx=XM_006943899.3;refGene_gn=PIH1D3;refGene_
tx=XM_019823486.2
X 89783204 89783301
refGene_gn=PIH1D3;refGene_tx=XM_006943899.3;refGene_gn=PIH1D3;refGene_
tx=XM_019823486.2
X 89797707 89797793
refGene_gn=PIH1D3;refGene_tx=XM_006943899.3;refGene_gn=PIH1D3;refGene_
tx=XM_019823486.2
X 89803834 89803964
refGene_gn=PIH1D3;refGene_tx=XM_006943899.3;refGene_gn=PIH1D3;refGene_
tx=XM_019823486.2

X 89933288 89933540
refGene_gn=LOC101085786;refGene_tx=XM_011279402.1
X 89989136 89989191
refGene_gn=LOC111558744;refGene_tx=XM_023249259.1
X 89989500 89989901
refGene_gn=LOC111558744;refGene_tx=XM_023249259.1
X 90040011 90040033 refGene_gn=FRMPD3;refGene_tx=XM_023248972.1
X 90043893 90044038 refGene_gn=FRMPD3;refGene_tx=XM_023248972.1
X 90061606 90061761 refGene_gn=FRMPD3;refGene_tx=XM_023248972.1
X 90064864 90064967 refGene_gn=FRMPD3;refGene_tx=XM_023248972.1
X 90068671 90068717 refGene_gn=FRMPD3;refGene_tx=XM_023248972.1
X 90079128 90079233 refGene_gn=FRMPD3;refGene_tx=XM_023248972.1
X 90083874 90083982 refGene_gn=FRMPD3;refGene_tx=XM_023248972.1
X 90086726 90086858 refGene_gn=FRMPD3;refGene_tx=XM_023248972.1
X 90088237 90088357 refGene_gn=FRMPD3;refGene_tx=XM_023248972.1
X 90093260 90093397 refGene_gn=FRMPD3;refGene_tx=XM_023248972.1
X 90093726 90093853 refGene_gn=FRMPD3;refGene_tx=XM_023248972.1
X 90095878 90095968 refGene_gn=FRMPD3;refGene_tx=XM_023248972.1
X 90097647 90097827 refGene_gn=FRMPD3;refGene_tx=XM_023248972.1
X 90107719 90107864 refGene_gn=FRMPD3;refGene_tx=XM_023248972.1
X 90116574 90117396 refGene_gn=FRMPD3;refGene_tx=XM_023248972.1
X 90118998 90122018 refGene_gn=FRMPD3;refGene_tx=XM_023248972.1
X 90136481 90136603 refGene_gn=PRPS1;refGene_tx=XM_004000771.4
X 90146199 90146383 refGene_gn=PRPS1;refGene_tx=XM_004000771.4
X 90147488 90147587 refGene_gn=PRPS1;refGene_tx=XM_004000771.4
X 90148884 90149009 refGene_gn=PRPS1;refGene_tx=XM_004000771.4
X 90152375 90152549 refGene_gn=PRPS1;refGene_tx=XM_004000771.4
X 90155489 90155649 refGene_gn=PRPS1;refGene_tx=XM_004000771.4
X 90156791 90156884 refGene_gn=PRPS1;refGene_tx=XM_004000771.4
X 90227420 90227651
refGene_gn=TSC22D3;refGene_tx=XM_004000774.5;refGene_gn=TSC22D3;refGene_tx=XM_004000776.5;refGene_gn=TSC22D3;refGene_tx=XM_011279397.3;refGene_gn=TSC22D3;refGene_tx=XM_019824140.2;refGene_gn=TSC22D3;refGene_tx=XM_019824141.2;refGene_gn=TSC22D3;refGene_tx=XM_023248815.1
X 90228769 90228821
refGene_gn=TSC22D3;refGene_tx=XM_004000774.5;refGene_gn=TSC22D3;refGene_tx=XM_004000776.5;refGene_gn=TSC22D3;refGene_tx=XM_011279397.3;refGene_gn=TSC22D3;refGene_tx=XM_019824140.2;refGene_gn=TSC22D3;refGene_tx=XM_019824141.2;refGene_gn=TSC22D3;refGene_tx=XM_023248815.1
X 90229534 90229656 refGene_gn=TSC22D3;refGene_tx=XM_004000774.5
X 90234510 90234560 refGene_gn=TSC22D3;refGene_tx=XM_019824140.2
X 90287720 90288040
refGene_gn=TSC22D3;refGene_tx=XM_011279397.3;refGene_gn=TSC22D3;refGene_tx=XM_023248815.1
X 90307746 90307837
refGene_gn=MID2;refGene_tx=XM_019824138.2;refGene_gn=MID2;refGene_tx=XM_019824139.2
X 90345094 90345098 refGene_gn=MID2;refGene_tx=XM_023249536.1
X 90358393 90359109
refGene_gn=MID2;refGene_tx=XM_006943910.4;refGene_gn=MID2;refGene_tx=X

M_006943911.3;refGene_gn=MID2;refGene_tx=XM_019824138.2;refGene_gn=MID2;refGene_tx=XM_019824139.2;refGene_gn=MID2;refGene_tx=XM_023249536.1;refGene_gn=MID2;refGene_tx=XM_023249538.1
X 90372733 90372829
refGene_gn=MID2;refGene_tx=XM_006943910.4;refGene_gn=MID2;refGene_tx=XM_006943911.3;refGene_gn=MID2;refGene_tx=XM_019824138.2;refGene_gn=MID2;refGene_tx=XM_019824139.2;refGene_gn=MID2;refGene_tx=XM_023249536.1;refGene_gn=MID2;refGene_tx=XM_023249538.1
X 90413055 90413163
refGene_gn=MID2;refGene_tx=XM_006943910.4;refGene_gn=MID2;refGene_tx=XM_006943911.3;refGene_gn=MID2;refGene_tx=XM_019824138.2;refGene_gn=MID2;refGene_tx=XM_019824139.2;refGene_gn=MID2;refGene_tx=XM_023249536.1;refGene_gn=MID2;refGene_tx=XM_023249538.1
X 90414796 90414945
refGene_gn=MID2;refGene_tx=XM_006943910.4;refGene_gn=MID2;refGene_tx=XM_006943911.3;refGene_gn=MID2;refGene_tx=XM_019824138.2;refGene_gn=MID2;refGene_tx=XM_019824139.2;refGene_gn=MID2;refGene_tx=XM_023249536.1;refGene_gn=MID2;refGene_tx=XM_023249538.1
X 90425632 90425760
refGene_gn=MID2;refGene_tx=XM_006943910.4;refGene_gn=MID2;refGene_tx=XM_006943911.3;refGene_gn=MID2;refGene_tx=XM_019824138.2;refGene_gn=MID2;refGene_tx=XM_019824139.2;refGene_gn=MID2;refGene_tx=XM_023249536.1;refGene_gn=MID2;refGene_tx=XM_023249538.1
X 90427317 90427551
refGene_gn=MID2;refGene_tx=XM_006943910.4;refGene_gn=MID2;refGene_tx=XM_006943911.3;refGene_gn=MID2;refGene_tx=XM_019824138.2;refGene_gn=MID2;refGene_tx=XM_019824139.2;refGene_gn=MID2;refGene_tx=XM_023249536.1;refGene_gn=MID2;refGene_tx=XM_023249538.1
X 90435427 90435589
refGene_gn=MID2;refGene_tx=XM_006943910.4;refGene_gn=MID2;refGene_tx=XM_006943911.3;refGene_gn=MID2;refGene_tx=XM_019824138.2;refGene_gn=MID2;refGene_tx=XM_019824139.2;refGene_gn=MID2;refGene_tx=XM_023249536.1;refGene_gn=MID2;refGene_tx=XM_023249538.1
X 90437016 90437224
refGene_gn=MID2;refGene_tx=XM_006943910.4;refGene_gn=MID2;refGene_tx=XM_006943911.3;refGene_gn=MID2;refGene_tx=XM_019824138.2;refGene_gn=MID2;refGene_tx=XM_019824139.2;refGene_gn=MID2;refGene_tx=XM_023249536.1;refGene_gn=MID2;refGene_tx=XM_023249538.1
X 90437853 90438256
refGene_gn=MID2;refGene_tx=XM_006943910.4;refGene_gn=MID2;refGene_tx=XM_006943911.3;refGene_gn=MID2;refGene_tx=XM_019824138.2;refGene_gn=MID2;refGene_tx=XM_019824139.2;refGene_gn=MID2;refGene_tx=XM_023249536.1;refGene_gn=MID2;refGene_tx=XM_023249538.1
X 90471773 90473263
refGene_gn=LOC102902092;refGene_tx=XM_023248973.1
X 90475606 90475696
refGene_gn=LOC102902092;refGene_tx=XM_023248973.1
X 90479052 90479745
refGene_gn=TEX13B;refGene_tx=XM_019824142.2;refGene_gn=TEX13B;refGene_tx=XM_019824143.2;refGene_gn=TEX13B;refGene_tx=XM_023248953.1

X	90479846	90480302
refGene_gn=TEX13B;refGene_tx=XM_019824142.2;refGene_gn=TEX13B;refGene_tx=XM_019824143.2;refGene_gn=TEX13B;refGene_tx=XM_023248953.1
X	90483620	90483791
refGene_gn=LOC102902092;refGene_tx=XM_023248973.1
X	90483906	90484077
refGene_gn=LOC102902092;refGene_tx=XM_023248973.1
X	90486259	90486428
refGene_gn=LOC102902092;refGene_tx=XM_023248973.1
X	90487272	90487364
refGene_gn=LOC102902092;refGene_tx=XM_023248973.1
X	90487461	90487632
refGene_gn=LOC102902092;refGene_tx=XM_023248973.1
X	90487937	90488108
refGene_gn=LOC102902092;refGene_tx=XM_023248973.1
X	90488228	90488399
refGene_gn=LOC102902092;refGene_tx=XM_023248973.1
X	90492881	90492963
refGene_gn=LOC102902092;refGene_tx=XM_023248973.1
X	90542259	90542308
refGene_gn=VSIG1;refGene_tx=XM_004000785.4;refGene_gn=VSIG1;refGene_tx=XM_006943913.3
X	90557992	90558156
refGene_gn=VSIG1;refGene_tx=XM_004000785.4;refGene_gn=VSIG1;refGene_tx=XM_006943913.3
X	90561659	90561767	refGene_gn=VSIG1;refGene_tx=XM_004000785.4
X	90567762	90567961
refGene_gn=VSIG1;refGene_tx=XM_004000785.4;refGene_gn=VSIG1;refGene_tx=XM_006943913.3
X	90572590	90572746
refGene_gn=VSIG1;refGene_tx=XM_004000785.4;refGene_gn=VSIG1;refGene_tx=XM_006943913.3
X	90573368	90573488
refGene_gn=VSIG1;refGene_tx=XM_004000785.4;refGene_gn=VSIG1;refGene_tx=XM_006943913.3
X	90577383	90577537
refGene_gn=VSIG1;refGene_tx=XM_004000785.4;refGene_gn=VSIG1;refGene_tx=XM_006943913.3
X	90578346	90578668
refGene_gn=VSIG1;refGene_tx=XM_004000785.4;refGene_gn=VSIG1;refGene_tx=XM_006943913.3
X	90586828	90586982
refGene_gn=PSMD10;refGene_tx=XM_004000780.5;refGene_gn=PSMD10;refGene_tx=XM_004000781.4
X	90589815	90589982
refGene_gn=PSMD10;refGene_tx=XM_004000780.5;refGene_gn=PSMD10;refGene_tx=XM_004000781.4
X	90590076	90590223
refGene_gn=PSMD10;refGene_tx=XM_004000780.5;refGene_gn=PSMD10;refGene_tx=XM_004000781.4

X 90591435 90591534
refGene_gn=PSMD10;refGene_tx=XM_004000780.5;refGene_gn=PSMD10;refGene_tx=XM_004000781.4
X 90593956 90594070
refGene_gn=PSMD10;refGene_tx=XM_004000780.5;refGene_gn=PSMD10;refGene_tx=XM_004000781.4
X 90594371 90594381
refGene_gn=ATG4A;refGene_tx=XM_004000782.3;refGene_gn=ATG4A;refGene_tx=XM_019824145.1
X 90629102 90629213
refGene_gn=ATG4A;refGene_tx=XM_004000782.3;refGene_gn=ATG4A;refGene_tx=XM_019824145.1
X 90631722 90631794 refGene_gn=ATG4A;refGene_tx=XM_004000782.3
X 90634368 90634467
refGene_gn=ATG4A;refGene_tx=XM_004000782.3;refGene_gn=ATG4A;refGene_tx=XM_019824145.1
X 90637396 90637498
refGene_gn=ATG4A;refGene_tx=XM_004000782.3;refGene_gn=ATG4A;refGene_tx=XM_019824145.1
X 90637673 90637746
refGene_gn=ATG4A;refGene_tx=XM_004000782.3;refGene_gn=ATG4A;refGene_tx=XM_019824145.1
X 90640812 90640892
refGene_gn=ATG4A;refGene_tx=XM_004000782.3;refGene_gn=ATG4A;refGene_tx=XM_019824145.1
X 90641605 90641793
refGene_gn=ATG4A;refGene_tx=XM_004000782.3;refGene_gn=ATG4A;refGene_tx=XM_019824145.1
X 90641916 90641995
refGene_gn=ATG4A;refGene_tx=XM_004000782.3;refGene_gn=ATG4A;refGene_tx=XM_019824145.1
X 90646743 90646889
refGene_gn=ATG4A;refGene_tx=XM_004000782.3;refGene_gn=ATG4A;refGene_tx=XM_019824145.1
X 90647991 90648048
refGene_gn=ATG4A;refGene_tx=XM_004000782.3;refGene_gn=ATG4A;refGene_tx=XM_019824145.1
X 90649717 90649826
refGene_gn=ATG4A;refGene_tx=XM_004000782.3;refGene_gn=ATG4A;refGene_tx=XM_019824145.1
X 90650616 90650687
refGene_gn=ATG4A;refGene_tx=XM_004000782.3;refGene_gn=ATG4A;refGene_tx=XM_019824145.1
X 90653856 90654117
refGene_gn=COL4A6;refGene_tx=XM_011279393.2;refGene_gn=COL4A6;refGene_tx=XM_019824146.1;refGene_gn=COL4A6;refGene_tx=XM_019824147.1;refGene_gn=COL4A6;refGene_tx=XM_019824148.1
X 90656370 90656657
refGene_gn=COL4A6;refGene_tx=XM_011279393.2;refGene_gn=COL4A6;refGene_tx=XM_019824146.1;refGene_gn=COL4A6;refGene_tx=XM_019824147.1;refGene_

gn=COL4A6;refGene_tx=XM_019824148.1
X	90657759	90657951
refGene_gn=COL4A6;refGene_tx=XM_011279393.2;refGene_gn=COL4A6;refGene_tx=XM_019824146.1;refGene_gn=COL4A6;refGene_tx=XM_019824147.1;refGene_gn=COL4A6;refGene_tx=XM_019824148.1
X	90658863	90658980
refGene_gn=COL4A6;refGene_tx=XM_011279393.2;refGene_gn=COL4A6;refGene_tx=XM_019824146.1;refGene_gn=COL4A6;refGene_tx=XM_019824147.1;refGene_gn=COL4A6;refGene_tx=XM_019824148.1
X	90659687	90659834
refGene_gn=COL4A6;refGene_tx=XM_011279393.2;refGene_gn=COL4A6;refGene_tx=XM_019824146.1;refGene_gn=COL4A6;refGene_tx=XM_019824147.1;refGene_gn=COL4A6;refGene_tx=XM_019824148.1
X	90661627	90661726
refGene_gn=COL4A6;refGene_tx=XM_011279393.2;refGene_gn=COL4A6;refGene_tx=XM_019824146.1;refGene_gn=COL4A6;refGene_tx=XM_019824147.1;refGene_gn=COL4A6;refGene_tx=XM_019824148.1
X	90661918	90662080
refGene_gn=COL4A6;refGene_tx=XM_011279393.2;refGene_gn=COL4A6;refGene_tx=XM_019824146.1;refGene_gn=COL4A6;refGene_tx=XM_019824147.1;refGene_gn=COL4A6;refGene_tx=XM_019824148.1
X	90662407	90662524
refGene_gn=COL4A6;refGene_tx=XM_011279393.2;refGene_gn=COL4A6;refGene_tx=XM_019824146.1;refGene_gn=COL4A6;refGene_tx=XM_019824147.1;refGene_gn=COL4A6;refGene_tx=XM_019824148.1
X	90666540	90666666
refGene_gn=COL4A6;refGene_tx=XM_011279393.2;refGene_gn=COL4A6;refGene_tx=XM_019824146.1;refGene_gn=COL4A6;refGene_tx=XM_019824147.1;refGene_gn=COL4A6;refGene_tx=XM_019824148.1
X	90666984	90667056
refGene_gn=COL4A6;refGene_tx=XM_011279393.2;refGene_gn=COL4A6;refGene_tx=XM_019824146.1;refGene_gn=COL4A6;refGene_tx=XM_019824147.1;refGene_gn=COL4A6;refGene_tx=XM_019824148.1
X	90667644	90667752
refGene_gn=COL4A6;refGene_tx=XM_011279393.2;refGene_gn=COL4A6;refGene_tx=XM_019824146.1;refGene_gn=COL4A6;refGene_tx=XM_019824147.1;refGene_gn=COL4A6;refGene_tx=XM_019824148.1
X	90667845	90667953
refGene_gn=COL4A6;refGene_tx=XM_011279393.2;refGene_gn=COL4A6;refGene_tx=XM_019824146.1;refGene_gn=COL4A6;refGene_tx=XM_019824147.1;refGene_gn=COL4A6;refGene_tx=XM_019824148.1
X	90668453	90668528
refGene_gn=COL4A6;refGene_tx=XM_011279393.2;refGene_gn=COL4A6;refGene_tx=XM_019824146.1;refGene_gn=COL4A6;refGene_tx=XM_019824147.1;refGene_gn=COL4A6;refGene_tx=XM_019824148.1
X	90669119	90669183
refGene_gn=COL4A6;refGene_tx=XM_011279393.2;refGene_gn=COL4A6;refGene_tx=XM_019824146.1;refGene_gn=COL4A6;refGene_tx=XM_019824147.1;refGene_gn=COL4A6;refGene_tx=XM_019824148.1
X	90671262	90671444

refGene_gn=COL4A6;refGene_tx=XM_011279393.2;refGene_gn=COL4A6;refGene_tx=XM_019824146.1;refGene_gn=COL4A6;refGene_tx=XM_019824147.1;refGene_gn=COL4A6;refGene_tx=XM_019824148.1
X 90671878 90672004
refGene_gn=COL4A6;refGene_tx=XM_011279393.2;refGene_gn=COL4A6;refGene_tx=XM_019824146.1;refGene_gn=COL4A6;refGene_tx=XM_019824147.1;refGene_gn=COL4A6;refGene_tx=XM_019824148.1
X 90672687 90672831
refGene_gn=COL4A6;refGene_tx=XM_011279393.2;refGene_gn=COL4A6;refGene_tx=XM_019824146.1;refGene_gn=COL4A6;refGene_tx=XM_019824147.1;refGene_gn=COL4A6;refGene_tx=XM_019824148.1
X 90673576 90673747
refGene_gn=COL4A6;refGene_tx=XM_011279393.2;refGene_gn=COL4A6;refGene_tx=XM_019824146.1;refGene_gn=COL4A6;refGene_tx=XM_019824147.1;refGene_gn=COL4A6;refGene_tx=XM_019824148.1
X 90675401 90675563
refGene_gn=COL4A6;refGene_tx=XM_011279393.2;refGene_gn=COL4A6;refGene_tx=XM_019824146.1;refGene_gn=COL4A6;refGene_tx=XM_019824147.1;refGene_gn=COL4A6;refGene_tx=XM_019824148.1
X 90675958 90676180
refGene_gn=COL4A6;refGene_tx=XM_011279393.2;refGene_gn=COL4A6;refGene_tx=XM_019824146.1;refGene_gn=COL4A6;refGene_tx=XM_019824147.1;refGene_gn=COL4A6;refGene_tx=XM_019824148.1
X 90677755 90677863
refGene_gn=COL4A6;refGene_tx=XM_011279393.2;refGene_gn=COL4A6;refGene_tx=XM_019824146.1;refGene_gn=COL4A6;refGene_tx=XM_019824147.1;refGene_gn=COL4A6;refGene_tx=XM_019824148.1
X 90678137 90678209
refGene_gn=COL4A6;refGene_tx=XM_011279393.2;refGene_gn=COL4A6;refGene_tx=XM_019824146.1;refGene_gn=COL4A6;refGene_tx=XM_019824147.1;refGene_gn=COL4A6;refGene_tx=XM_019824148.1
X 90680959 90681013
refGene_gn=COL4A6;refGene_tx=XM_011279393.2;refGene_gn=COL4A6;refGene_tx=XM_019824146.1;refGene_gn=COL4A6;refGene_tx=XM_019824148.1
X 90683501 90683685
refGene_gn=COL4A6;refGene_tx=XM_011279393.2;refGene_gn=COL4A6;refGene_tx=XM_019824146.1;refGene_gn=COL4A6;refGene_tx=XM_019824147.1;refGene_gn=COL4A6;refGene_tx=XM_019824148.1
X 90684241 90684421
refGene_gn=COL4A6;refGene_tx=XM_011279393.2;refGene_gn=COL4A6;refGene_tx=XM_019824146.1;refGene_gn=COL4A6;refGene_tx=XM_019824147.1;refGene_gn=COL4A6;refGene_tx=XM_019824148.1
X 90684922 90685083
refGene_gn=COL4A6;refGene_tx=XM_011279393.2;refGene_gn=COL4A6;refGene_tx=XM_019824146.1;refGene_gn=COL4A6;refGene_tx=XM_019824147.1;refGene_gn=COL4A6;refGene_tx=XM_019824148.1
X 90686864 90686969
refGene_gn=COL4A6;refGene_tx=XM_011279393.2;refGene_gn=COL4A6;refGene_tx=XM_019824146.1;refGene_gn=COL4A6;refGene_tx=XM_019824147.1;refGene_gn=COL4A6;refGene_tx=XM_019824148.1

X 90687822 90687963
refGene_gn=COL4A6;refGene_tx=XM_011279393.2;refGene_gn=COL4A6;refGene_tx=XM_019824146.1;refGene_gn=COL4A6;refGene_tx=XM_019824147.1;refGene_gn=COL4A6;refGene_tx=XM_019824148.1
X 90689121 90689229
refGene_gn=COL4A6;refGene_tx=XM_011279393.2;refGene_gn=COL4A6;refGene_tx=XM_019824146.1;refGene_gn=COL4A6;refGene_tx=XM_019824147.1;refGene_gn=COL4A6;refGene_tx=XM_019824148.1
X 90690192 90690262
refGene_gn=COL4A6;refGene_tx=XM_011279393.2;refGene_gn=COL4A6;refGene_tx=XM_019824146.1;refGene_gn=COL4A6;refGene_tx=XM_019824147.1;refGene_gn=COL4A6;refGene_tx=XM_019824148.1
X 90691502 90691556
refGene_gn=COL4A6;refGene_tx=XM_011279393.2;refGene_gn=COL4A6;refGene_tx=XM_019824146.1;refGene_gn=COL4A6;refGene_tx=XM_019824147.1;refGene_gn=COL4A6;refGene_tx=XM_019824148.1
X 90692043 90692088
refGene_gn=COL4A6;refGene_tx=XM_011279393.2;refGene_gn=COL4A6;refGene_tx=XM_019824146.1;refGene_gn=COL4A6;refGene_tx=XM_019824147.1;refGene_gn=COL4A6;refGene_tx=XM_019824148.1
X 90693131 90693191
refGene_gn=COL4A6;refGene_tx=XM_011279393.2;refGene_gn=COL4A6;refGene_tx=XM_019824146.1;refGene_gn=COL4A6;refGene_tx=XM_019824147.1;refGene_gn=COL4A6;refGene_tx=XM_019824148.1
X 90700461 90700515
refGene_gn=COL4A6;refGene_tx=XM_011279393.2;refGene_gn=COL4A6;refGene_tx=XM_019824146.1;refGene_gn=COL4A6;refGene_tx=XM_019824147.1;refGene_gn=COL4A6;refGene_tx=XM_019824148.1
X 90701827 90701920
refGene_gn=COL4A6;refGene_tx=XM_011279393.2;refGene_gn=COL4A6;refGene_tx=XM_019824146.1;refGene_gn=COL4A6;refGene_tx=XM_019824147.1;refGene_gn=COL4A6;refGene_tx=XM_019824148.1
X 90702926 90702968
refGene_gn=COL4A6;refGene_tx=XM_011279393.2;refGene_gn=COL4A6;refGene_tx=XM_019824147.1;refGene_gn=COL4A6;refGene_tx=XM_019824148.1
X 90703147 90703183
refGene_gn=COL4A6;refGene_tx=XM_011279393.2;refGene_gn=COL4A6;refGene_tx=XM_019824146.1;refGene_gn=COL4A6;refGene_tx=XM_019824147.1;refGene_gn=COL4A6;refGene_tx=XM_019824148.1
X 90704294 90704357
refGene_gn=COL4A6;refGene_tx=XM_011279393.2;refGene_gn=COL4A6;refGene_tx=XM_019824146.1;refGene_gn=COL4A6;refGene_tx=XM_019824147.1;refGene_gn=COL4A6;refGene_tx=XM_019824148.1
X 90707439 90707475
refGene_gn=COL4A6;refGene_tx=XM_011279393.2;refGene_gn=COL4A6;refGene_tx=XM_019824146.1;refGene_gn=COL4A6;refGene_tx=XM_019824147.1;refGene_gn=COL4A6;refGene_tx=XM_019824148.1
X 90709162 90709231
refGene_gn=COL4A6;refGene_tx=XM_011279393.2;refGene_gn=COL4A6;refGene_tx=XM_019824146.1;refGene_gn=COL4A6;refGene_tx=XM_019824147.1;refGene_

gn=COL4A6;refGene_tx=XM_019824148.1
X	90712665	90712782
refGene_gn=COL4A6;refGene_tx=XM_011279393.2;refGene_gn=COL4A6;refGene_tx=XM_019824146.1;refGene_gn=COL4A6;refGene_tx=XM_019824147.1
X	90730372	90730417
refGene_gn=COL4A6;refGene_tx=XM_011279393.2;refGene_gn=COL4A6;refGene_tx=XM_019824146.1;refGene_gn=COL4A6;refGene_tx=XM_019824147.1;refGene_gn=COL4A6;refGene_tx=XM_019824148.1
X	90731788	90731923
refGene_gn=COL4A6;refGene_tx=XM_011279393.2;refGene_gn=COL4A6;refGene_tx=XM_019824146.1;refGene_gn=COL4A6;refGene_tx=XM_019824147.1;refGene_gn=COL4A6;refGene_tx=XM_019824148.1
X	90826046	90826127
refGene_gn=COL4A6;refGene_tx=XM_011279393.2;refGene_gn=COL4A6;refGene_tx=XM_019824146.1;refGene_gn=COL4A6;refGene_tx=XM_019824147.1;refGene_gn=COL4A6;refGene_tx=XM_019824148.1
X	90967546	90967598
refGene_gn=COL4A6;refGene_tx=XM_011279393.2;refGene_gn=COL4A6;refGene_tx=XM_019824146.1;refGene_gn=COL4A6;refGene_tx=XM_019824147.1;refGene_gn=COL4A6;refGene_tx=XM_019824148.1
X	90967792	90967803
refGene_gn=COL4A6;refGene_tx=XM_011279393.2;refGene_gn=COL4A6;refGene_tx=XM_019824146.1;refGene_gn=COL4A6;refGene_tx=XM_019824147.1;refGene_gn=COL4A6;refGene_tx=XM_019824148.1
X	90969863	90969944
refGene_gn=COL4A5;refGene_tx=XM_011279396.3;refGene_gn=COL4A5;refGene_tx=XM_019824149.2;refGene_gn=COL4A5;refGene_tx=XM_023249434.1;refGene_gn=COL4A5;refGene_tx=XM_023249435.1;refGene_gn=COL4A5;refGene_tx=XM_023249436.1;refGene_gn=COL4A5;refGene_tx=XM_023249437.1
X	91015220	91015577
refGene_gn=LOC101089991;refGene_tx=XM_011279394.2
X	91064179	91064239
refGene_gn=COL4A5;refGene_tx=XM_011279396.3;refGene_gn=COL4A5;refGene_tx=XM_019824149.2;refGene_gn=COL4A5;refGene_tx=XM_023249434.1;refGene_gn=COL4A5;refGene_tx=XM_023249435.1;refGene_gn=COL4A5;refGene_tx=XM_023249436.1;refGene_gn=COL4A5;refGene_tx=XM_023249437.1
X	91084966	91085056
refGene_gn=COL4A5;refGene_tx=XM_011279396.3;refGene_gn=COL4A5;refGene_tx=XM_019824149.2;refGene_gn=COL4A5;refGene_tx=XM_023249434.1;refGene_gn=COL4A5;refGene_tx=XM_023249435.1;refGene_gn=COL4A5;refGene_tx=XM_023249436.1;refGene_gn=COL4A5;refGene_tx=XM_023249437.1
X	91092304	91092349
refGene_gn=COL4A5;refGene_tx=XM_011279396.3;refGene_gn=COL4A5;refGene_tx=XM_019824149.2;refGene_gn=COL4A5;refGene_tx=XM_023249434.1;refGene_gn=COL4A5;refGene_tx=XM_023249435.1;refGene_gn=COL4A5;refGene_tx=XM_023249436.1;refGene_gn=COL4A5;refGene_tx=XM_023249437.1
X	91097245	91097290
refGene_gn=COL4A5;refGene_tx=XM_011279396.3;refGene_gn=COL4A5;refGene_tx=XM_019824149.2;refGene_gn=COL4A5;refGene_tx=XM_023249434.1;refGene_gn=COL4A5;refGene_tx=XM_023249435.1;refGene_gn=COL4A5;refGene_tx=XM_02

3249436.1;refGene_gn=COL4A5;refGene_tx=XM_023249437.1
X	91097377	91097440
refGene_gn=COL4A5;refGene_tx=XM_011279396.3;refGene_gn=COL4A5;refGene_tx=XM_019824149.2;refGene_gn=COL4A5;refGene_tx=XM_023249434.1;refGene_gn=COL4A5;refGene_tx=XM_023249435.1;refGene_gn=COL4A5;refGene_tx=XM_023249436.1;refGene_gn=COL4A5;refGene_tx=XM_023249437.1
X	91099759	91099813
refGene_gn=COL4A5;refGene_tx=XM_011279396.3;refGene_gn=COL4A5;refGene_tx=XM_019824149.2;refGene_gn=COL4A5;refGene_tx=XM_023249434.1;refGene_gn=COL4A5;refGene_tx=XM_023249435.1;refGene_gn=COL4A5;refGene_tx=XM_023249436.1;refGene_gn=COL4A5;refGene_tx=XM_023249437.1
X	91100150	91100177
refGene_gn=COL4A5;refGene_tx=XM_011279396.3;refGene_gn=COL4A5;refGene_tx=XM_019824149.2;refGene_gn=COL4A5;refGene_tx=XM_023249434.1;refGene_gn=COL4A5;refGene_tx=XM_023249435.1;refGene_gn=COL4A5;refGene_tx=XM_023249436.1;refGene_gn=COL4A5;refGene_tx=XM_023249437.1
X	91102478	91102559
refGene_gn=COL4A5;refGene_tx=XM_011279396.3;refGene_gn=COL4A5;refGene_tx=XM_019824149.2;refGene_gn=COL4A5;refGene_tx=XM_023249434.1;refGene_gn=COL4A5;refGene_tx=XM_023249435.1;refGene_gn=COL4A5;refGene_tx=XM_023249436.1;refGene_gn=COL4A5;refGene_tx=XM_023249437.1
X	91104909	91104972
refGene_gn=COL4A5;refGene_tx=XM_011279396.3;refGene_gn=COL4A5;refGene_tx=XM_019824149.2;refGene_gn=COL4A5;refGene_tx=XM_023249434.1;refGene_gn=COL4A5;refGene_tx=XM_023249435.1;refGene_gn=COL4A5;refGene_tx=XM_023249436.1;refGene_gn=COL4A5;refGene_tx=XM_023249437.1
X	91107222	91107258
refGene_gn=COL4A5;refGene_tx=XM_011279396.3;refGene_gn=COL4A5;refGene_tx=XM_019824149.2;refGene_gn=COL4A5;refGene_tx=XM_023249434.1;refGene_gn=COL4A5;refGene_tx=XM_023249435.1;refGene_gn=COL4A5;refGene_tx=XM_023249436.1;refGene_gn=COL4A5;refGene_tx=XM_023249437.1
X	91107347	91107389
refGene_gn=COL4A5;refGene_tx=XM_011279396.3;refGene_gn=COL4A5;refGene_tx=XM_019824149.2;refGene_gn=COL4A5;refGene_tx=XM_023249434.1;refGene_gn=COL4A5;refGene_tx=XM_023249435.1;refGene_gn=COL4A5;refGene_tx=XM_023249436.1;refGene_gn=COL4A5;refGene_tx=XM_023249437.1
X	91107548	91107641
refGene_gn=COL4A5;refGene_tx=XM_011279396.3;refGene_gn=COL4A5;refGene_tx=XM_019824149.2;refGene_gn=COL4A5;refGene_tx=XM_023249434.1;refGene_gn=COL4A5;refGene_tx=XM_023249435.1;refGene_gn=COL4A5;refGene_tx=XM_023249436.1;refGene_gn=COL4A5;refGene_tx=XM_023249437.1
X	91109061	91109115
refGene_gn=COL4A5;refGene_tx=XM_011279396.3;refGene_gn=COL4A5;refGene_tx=XM_019824149.2;refGene_gn=COL4A5;refGene_tx=XM_023249434.1;refGene_gn=COL4A5;refGene_tx=XM_023249435.1;refGene_gn=COL4A5;refGene_tx=XM_023249436.1;refGene_gn=COL4A5;refGene_tx=XM_023249437.1
X	91109211	91109268
refGene_gn=COL4A5;refGene_tx=XM_011279396.3;refGene_gn=COL4A5;refGene_tx=XM_019824149.2;refGene_gn=COL4A5;refGene_tx=XM_023249434.1;refGene_gn=COL4A5;refGene_tx=XM_023249435.1;refGene_gn=COL4A5;refGene_tx=XM_02

3249436.1;refGene_gn=COL4A5;refGene_tx=XM_023249437.1
X 91109527 91109572
refGene_gn=COL4A5;refGene_tx=XM_011279396.3;refGene_gn=COL4A5;refGene_tx=XM_019824149.2;refGene_gn=COL4A5;refGene_tx=XM_023249434.1;refGene_gn=COL4A5;refGene_tx=XM_023249435.1;refGene_gn=COL4A5;refGene_tx=XM_023249436.1;refGene_gn=COL4A5;refGene_tx=XM_023249437.1
X 91111103 91111157
refGene_gn=COL4A5;refGene_tx=XM_011279396.3;refGene_gn=COL4A5;refGene_tx=XM_019824149.2;refGene_gn=COL4A5;refGene_tx=XM_023249434.1;refGene_gn=COL4A5;refGene_tx=XM_023249435.1;refGene_gn=COL4A5;refGene_tx=XM_023249436.1;refGene_gn=COL4A5;refGene_tx=XM_023249437.1
X 91113265 91113307
refGene_gn=COL4A5;refGene_tx=XM_011279396.3;refGene_gn=COL4A5;refGene_tx=XM_019824149.2;refGene_gn=COL4A5;refGene_tx=XM_023249434.1;refGene_gn=COL4A5;refGene_tx=XM_023249435.1;refGene_gn=COL4A5;refGene_tx=XM_023249436.1;refGene_gn=COL4A5;refGene_tx=XM_023249437.1
X 91116086 91116219
refGene_gn=COL4A5;refGene_tx=XM_011279396.3;refGene_gn=COL4A5;refGene_tx=XM_019824149.2;refGene_gn=COL4A5;refGene_tx=XM_023249434.1;refGene_gn=COL4A5;refGene_tx=XM_023249435.1;refGene_gn=COL4A5;refGene_tx=XM_023249436.1;refGene_gn=COL4A5;refGene_tx=XM_023249437.1
X 91121214 91121388
refGene_gn=COL4A5;refGene_tx=XM_011279396.3;refGene_gn=COL4A5;refGene_tx=XM_019824149.2;refGene_gn=COL4A5;refGene_tx=XM_023249434.1;refGene_gn=COL4A5;refGene_tx=XM_023249435.1;refGene_gn=COL4A5;refGene_tx=XM_023249436.1;refGene_gn=COL4A5;refGene_tx=XM_023249437.1
X 91121689 91121773
refGene_gn=COL4A5;refGene_tx=XM_011279396.3;refGene_gn=COL4A5;refGene_tx=XM_019824149.2;refGene_gn=COL4A5;refGene_tx=XM_023249434.1;refGene_gn=COL4A5;refGene_tx=XM_023249435.1;refGene_gn=COL4A5;refGene_tx=XM_023249436.1;refGene_gn=COL4A5;refGene_tx=XM_023249437.1
X 91125728 91125821
refGene_gn=COL4A5;refGene_tx=XM_011279396.3;refGene_gn=COL4A5;refGene_tx=XM_019824149.2;refGene_gn=COL4A5;refGene_tx=XM_023249434.1;refGene_gn=COL4A5;refGene_tx=XM_023249435.1;refGene_gn=COL4A5;refGene_tx=XM_023249436.1;refGene_gn=COL4A5;refGene_tx=XM_023249437.1
X 91127431 91127502
refGene_gn=COL4A5;refGene_tx=XM_011279396.3;refGene_gn=COL4A5;refGene_tx=XM_019824149.2;refGene_gn=COL4A5;refGene_tx=XM_023249434.1;refGene_gn=COL4A5;refGene_tx=XM_023249435.1;refGene_gn=COL4A5;refGene_tx=XM_023249436.1;refGene_gn=COL4A5;refGene_tx=XM_023249437.1
X 91127800 91127992
refGene_gn=COL4A5;refGene_tx=XM_011279396.3;refGene_gn=COL4A5;refGene_tx=XM_019824149.2;refGene_gn=COL4A5;refGene_tx=XM_023249434.1;refGene_gn=COL4A5;refGene_tx=XM_023249435.1;refGene_gn=COL4A5;refGene_tx=XM_023249436.1;refGene_gn=COL4A5;refGene_tx=XM_023249437.1
X 91128694 91128863
refGene_gn=COL4A5;refGene_tx=XM_011279396.3;refGene_gn=COL4A5;refGene_tx=XM_019824149.2;refGene_gn=COL4A5;refGene_tx=XM_023249434.1;refGene_gn=COL4A5;refGene_tx=XM_023249435.1;refGene_gn=COL4A5;refGene_tx=XM_02

3249436.1;refGene_gn=COL4A5;refGene_tx=XM_023249437.1
X 91131324 91131417
refGene_gn=COL4A5;refGene_tx=XM_011279396.3;refGene_gn=COL4A5;refGene_tx=XM_019824149.2;refGene_gn=COL4A5;refGene_tx=XM_023249434.1;refGene_gn=COL4A5;refGene_tx=XM_023249435.1;refGene_gn=COL4A5;refGene_tx=XM_023249436.1;refGene_gn=COL4A5;refGene_tx=XM_023249437.1
X 91131509 91131614
refGene_gn=COL4A5;refGene_tx=XM_011279396.3;refGene_gn=COL4A5;refGene_tx=XM_019824149.2;refGene_gn=COL4A5;refGene_tx=XM_023249434.1;refGene_gn=COL4A5;refGene_tx=XM_023249435.1;refGene_gn=COL4A5;refGene_tx=XM_023249436.1;refGene_gn=COL4A5;refGene_tx=XM_023249437.1
X 91132671 91132769
refGene_gn=COL4A5;refGene_tx=XM_011279396.3;refGene_gn=COL4A5;refGene_tx=XM_019824149.2;refGene_gn=COL4A5;refGene_tx=XM_023249434.1;refGene_gn=COL4A5;refGene_tx=XM_023249435.1;refGene_gn=COL4A5;refGene_tx=XM_023249436.1;refGene_gn=COL4A5;refGene_tx=XM_023249437.1
X 91134002 91134153
refGene_gn=COL4A5;refGene_tx=XM_011279396.3;refGene_gn=COL4A5;refGene_tx=XM_019824149.2;refGene_gn=COL4A5;refGene_tx=XM_023249434.1;refGene_gn=COL4A5;refGene_tx=XM_023249435.1;refGene_gn=COL4A5;refGene_tx=XM_023249436.1;refGene_gn=COL4A5;refGene_tx=XM_023249437.1
X 91145884 91145998
refGene_gn=COL4A5;refGene_tx=XM_011279396.3;refGene_gn=COL4A5;refGene_tx=XM_019824149.2;refGene_gn=COL4A5;refGene_tx=XM_023249434.1;refGene_gn=COL4A5;refGene_tx=XM_023249435.1;refGene_gn=COL4A5;refGene_tx=XM_023249436.1;refGene_gn=COL4A5;refGene_tx=XM_023249437.1
X 91151615 91151783
refGene_gn=COL4A5;refGene_tx=XM_011279396.3;refGene_gn=COL4A5;refGene_tx=XM_019824149.2;refGene_gn=COL4A5;refGene_tx=XM_023249434.1;refGene_gn=COL4A5;refGene_tx=XM_023249435.1;refGene_gn=COL4A5;refGene_tx=XM_023249436.1;refGene_gn=COL4A5;refGene_tx=XM_023249437.1
X 91152672 91152762
refGene_gn=COL4A5;refGene_tx=XM_011279396.3;refGene_gn=COL4A5;refGene_tx=XM_019824149.2;refGene_gn=COL4A5;refGene_tx=XM_023249434.1;refGene_gn=COL4A5;refGene_tx=XM_023249435.1;refGene_gn=COL4A5;refGene_tx=XM_023249436.1;refGene_gn=COL4A5;refGene_tx=XM_023249437.1
X 91153467 91153617
refGene_gn=COL4A5;refGene_tx=XM_011279396.3;refGene_gn=COL4A5;refGene_tx=XM_019824149.2;refGene_gn=COL4A5;refGene_tx=XM_023249434.1;refGene_gn=COL4A5;refGene_tx=XM_023249435.1;refGene_gn=COL4A5;refGene_tx=XM_023249436.1;refGene_gn=COL4A5;refGene_tx=XM_023249437.1
X 91155605 91155704
refGene_gn=COL4A5;refGene_tx=XM_011279396.3;refGene_gn=COL4A5;refGene_tx=XM_019824149.2;refGene_gn=COL4A5;refGene_tx=XM_023249434.1;refGene_gn=COL4A5;refGene_tx=XM_023249435.1;refGene_gn=COL4A5;refGene_tx=XM_023249436.1;refGene_gn=COL4A5;refGene_tx=XM_023249437.1
X 91163015 91163105
refGene_gn=COL4A5;refGene_tx=XM_011279396.3;refGene_gn=COL4A5;refGene_tx=XM_019824149.2;refGene_gn=COL4A5;refGene_tx=XM_023249434.1;refGene_gn=COL4A5;refGene_tx=XM_023249435.1;refGene_gn=COL4A5;refGene_tx=XM_02

3249436.1;refGene_gn=COL4A5;refGene_tx=XM_023249437.1
X 91163522 91163662
refGene_gn=COL4A5;refGene_tx=XM_011279396.3;refGene_gn=COL4A5;refGene_tx=XM_019824149.2;refGene_gn=COL4A5;refGene_tx=XM_023249434.1;refGene_gn=COL4A5;refGene_tx=XM_023249435.1;refGene_gn=COL4A5;refGene_tx=XM_023249436.1;refGene_gn=COL4A5;refGene_tx=XM_023249437.1
X 91178311 91178438
refGene_gn=COL4A5;refGene_tx=XM_011279396.3;refGene_gn=COL4A5;refGene_tx=XM_019824149.2;refGene_gn=COL4A5;refGene_tx=XM_023249434.1;refGene_gn=COL4A5;refGene_tx=XM_023249435.1;refGene_gn=COL4A5;refGene_tx=XM_023249436.1;refGene_gn=COL4A5;refGene_tx=XM_023249437.1
X 91181456 91181537
refGene_gn=COL4A5;refGene_tx=XM_011279396.3;refGene_gn=COL4A5;refGene_tx=XM_019824149.2;refGene_gn=COL4A5;refGene_tx=XM_023249434.1;refGene_gn=COL4A5;refGene_tx=XM_023249435.1;refGene_gn=COL4A5;refGene_tx=XM_023249436.1;refGene_gn=COL4A5;refGene_tx=XM_023249437.1
X 91182167 91182266
refGene_gn=COL4A5;refGene_tx=XM_011279396.3;refGene_gn=COL4A5;refGene_tx=XM_019824149.2;refGene_gn=COL4A5;refGene_tx=XM_023249434.1;refGene_gn=COL4A5;refGene_tx=XM_023249435.1;refGene_gn=COL4A5;refGene_tx=XM_023249436.1;refGene_gn=COL4A5;refGene_tx=XM_023249437.1
X 91182821 91182872
refGene_gn=COL4A5;refGene_tx=XM_011279396.3;refGene_gn=COL4A5;refGene_tx=XM_019824149.2;refGene_gn=COL4A5;refGene_tx=XM_023249434.1;refGene_gn=COL4A5;refGene_tx=XM_023249435.1;refGene_gn=COL4A5;refGene_tx=XM_023249436.1;refGene_gn=COL4A5;refGene_tx=XM_023249437.1
X 91184347 91184533
refGene_gn=COL4A5;refGene_tx=XM_011279396.3;refGene_gn=COL4A5;refGene_tx=XM_019824149.2;refGene_gn=COL4A5;refGene_tx=XM_023249434.1;refGene_gn=COL4A5;refGene_tx=XM_023249435.1;refGene_gn=COL4A5;refGene_tx=XM_023249436.1;refGene_gn=COL4A5;refGene_tx=XM_023249437.1
X 91186237 91186246
refGene_gn=COL4A5;refGene_tx=XM_019824149.2;refGene_gn=COL4A5;refGene_tx=XM_023249434.1;refGene_gn=COL4A5;refGene_tx=XM_023249436.1;refGene_gn=COL4A5;refGene_tx=XM_023249437.1
X 91191382 91191391
refGene_gn=COL4A5;refGene_tx=XM_023249434.1;refGene_gn=COL4A5;refGene_tx=XM_023249435.1;refGene_gn=COL4A5;refGene_tx=XM_023249436.1;refGene_gn=COL4A5;refGene_tx=XM_023249437.1
X 91194043 91194177
refGene_gn=COL4A5;refGene_tx=XM_011279396.3;refGene_gn=COL4A5;refGene_tx=XM_019824149.2;refGene_gn=COL4A5;refGene_tx=XM_023249434.1;refGene_gn=COL4A5;refGene_tx=XM_023249435.1;refGene_gn=COL4A5;refGene_tx=XM_023249436.1;refGene_gn=COL4A5;refGene_tx=XM_023249437.1
X 91196559 91196632
refGene_gn=COL4A5;refGene_tx=XM_011279396.3;refGene_gn=COL4A5;refGene_tx=XM_019824149.2;refGene_gn=COL4A5;refGene_tx=XM_023249434.1;refGene_gn=COL4A5;refGene_tx=XM_023249435.1;refGene_gn=COL4A5;refGene_tx=XM_023249436.1;refGene_gn=COL4A5;refGene_tx=XM_023249437.1
X 91196767 91196839

refGene_gn=COL4A5;refGene_tx=XM_011279396.3;refGene_gn=COL4A5;refGene_tx=XM_019824149.2;refGene_gn=COL4A5;refGene_tx=XM_023249434.1;refGene_gn=COL4A5;refGene_tx=XM_023249435.1;refGene_gn=COL4A5;refGene_tx=XM_023249436.1;refGene_gn=COL4A5;refGene_tx=XM_023249437.1
X 91197618 91197747
refGene_gn=COL4A5;refGene_tx=XM_011279396.3;refGene_gn=COL4A5;refGene_tx=XM_019824149.2;refGene_gn=COL4A5;refGene_tx=XM_023249434.1;refGene_gn=COL4A5;refGene_tx=XM_023249435.1;refGene_gn=COL4A5;refGene_tx=XM_023249436.1;refGene_gn=COL4A5;refGene_tx=XM_023249437.1
X 91198570 91198669
refGene_gn=COL4A5;refGene_tx=XM_011279396.3;refGene_gn=COL4A5;refGene_tx=XM_019824149.2;refGene_gn=COL4A5;refGene_tx=XM_023249434.1;refGene_gn=COL4A5;refGene_tx=XM_023249435.1;refGene_gn=COL4A5;refGene_tx=XM_023249436.1;refGene_gn=COL4A5;refGene_tx=XM_023249437.1
X 91201349 91201562
refGene_gn=COL4A5;refGene_tx=XM_011279396.3;refGene_gn=COL4A5;refGene_tx=XM_019824149.2;refGene_gn=COL4A5;refGene_tx=XM_023249434.1;refGene_gn=COL4A5;refGene_tx=XM_023249435.1
X 91204772 91204950
refGene_gn=COL4A5;refGene_tx=XM_011279396.3;refGene_gn=COL4A5;refGene_tx=XM_019824149.2;refGene_gn=COL4A5;refGene_tx=XM_023249434.1;refGene_gn=COL4A5;refGene_tx=XM_023249435.1;refGene_gn=COL4A5;refGene_tx=XM_023249436.1
X 91206865 91206980
refGene_gn=COL4A5;refGene_tx=XM_011279396.3;refGene_gn=COL4A5;refGene_tx=XM_019824149.2;refGene_gn=COL4A5;refGene_tx=XM_023249434.1;refGene_gn=COL4A5;refGene_tx=XM_023249435.1;refGene_gn=COL4A5;refGene_tx=XM_023249436.1
X 91207364 91207537
refGene_gn=COL4A5;refGene_tx=XM_011279396.3;refGene_gn=COL4A5;refGene_tx=XM_019824149.2;refGene_gn=COL4A5;refGene_tx=XM_023249434.1;refGene_gn=COL4A5;refGene_tx=XM_023249435.1;refGene_gn=COL4A5;refGene_tx=XM_023249436.1
X 91208345 91208427
refGene_gn=COL4A5;refGene_tx=XM_011279396.3;refGene_gn=COL4A5;refGene_tx=XM_019824149.2;refGene_gn=COL4A5;refGene_tx=XM_023249434.1;refGene_gn=COL4A5;refGene_tx=XM_023249435.1;refGene_gn=COL4A5;refGene_tx=XM_023249436.1
X 91234145 91234150 refGene_gn=IRS4;refGene_tx=XM_011279425.3
X 91246005 91249756 refGene_gn=IRS4;refGene_tx=XM_011279425.3
X 91666557 91666645
refGene_gn=GUCY2F;refGene_tx=XM_006943916.3;refGene_gn=GUCY2F;refGene_tx=XM_019824152.2;refGene_gn=GUCY2F;refGene_tx=XM_019824153.2
X 91666733 91666822
refGene_gn=GUCY2F;refGene_tx=XM_006943916.3;refGene_gn=GUCY2F;refGene_tx=XM_019824152.2;refGene_gn=GUCY2F;refGene_tx=XM_019824153.2
X 91672364 91672459
refGene_gn=GUCY2F;refGene_tx=XM_006943916.3;refGene_gn=GUCY2F;refGene_tx=XM_019824152.2;refGene_gn=GUCY2F;refGene_tx=XM_019824153.2
X 91675332 91675431

refGene_gn=GUCY2F;refGene_tx=XM_006943916.3;refGene_gn=GUCY2F;refGene_tx=XM_019824152.2;refGene_gn=GUCY2F;refGene_tx=XM_019824153.2
X 91679755 91679930
refGene_gn=GUCY2F;refGene_tx=XM_006943916.3;refGene_gn=GUCY2F;refGene_tx=XM_019824152.2;refGene_gn=GUCY2F;refGene_tx=XM_019824153.2
X 91683184 91683377
refGene_gn=GUCY2F;refGene_tx=XM_006943916.3;refGene_gn=GUCY2F;refGene_tx=XM_019824152.2;refGene_gn=GUCY2F;refGene_tx=XM_019824153.2
X 91684159 91684323
refGene_gn=GUCY2F;refGene_tx=XM_006943916.3;refGene_gn=GUCY2F;refGene_tx=XM_019824152.2;refGene_gn=GUCY2F;refGene_tx=XM_019824153.2
X 91686568 91686717
refGene_gn=GUCY2F;refGene_tx=XM_006943916.3;refGene_gn=GUCY2F;refGene_tx=XM_019824152.2;refGene_gn=GUCY2F;refGene_tx=XM_019824153.2
X 91689667 91689817
refGene_gn=GUCY2F;refGene_tx=XM_006943916.3;refGene_gn=GUCY2F;refGene_tx=XM_019824152.2;refGene_gn=GUCY2F;refGene_tx=XM_019824153.2
X 91695208 91695365
refGene_gn=GUCY2F;refGene_tx=XM_006943916.3;refGene_gn=GUCY2F;refGene_tx=XM_019824152.2;refGene_gn=GUCY2F;refGene_tx=XM_019824153.2
X 91697589 91697766
refGene_gn=GUCY2F;refGene_tx=XM_006943916.3;refGene_gn=GUCY2F;refGene_tx=XM_019824152.2;refGene_gn=GUCY2F;refGene_tx=XM_019824153.2
X 91712999 91713089
refGene_gn=GUCY2F;refGene_tx=XM_006943916.3;refGene_gn=GUCY2F;refGene_tx=XM_019824152.2;refGene_gn=GUCY2F;refGene_tx=XM_019824153.2
X 91720007 91720139
refGene_gn=GUCY2F;refGene_tx=XM_006943916.3;refGene_gn=GUCY2F;refGene_tx=XM_019824152.2;refGene_gn=GUCY2F;refGene_tx=XM_019824153.2
X 91725541 91725638
refGene_gn=GUCY2F;refGene_tx=XM_006943916.3;refGene_gn=GUCY2F;refGene_tx=XM_019824152.2;refGene_gn=GUCY2F;refGene_tx=XM_019824153.2
X 91729522 91729607
refGene_gn=GUCY2F;refGene_tx=XM_006943916.3;refGene_gn=GUCY2F;refGene_tx=XM_019824152.2;refGene_gn=GUCY2F;refGene_tx=XM_019824153.2
X 91731001 91731356
refGene_gn=GUCY2F;refGene_tx=XM_006943916.3;refGene_gn=GUCY2F;refGene_tx=XM_019824152.2;refGene_gn=GUCY2F;refGene_tx=XM_019824153.2
X 91745165 91745467
refGene_gn=GUCY2F;refGene_tx=XM_006943916.3;refGene_gn=GUCY2F;refGene_tx=XM_019824152.2;refGene_gn=GUCY2F;refGene_tx=XM_019824153.2
X 91755773 91756503
refGene_gn=GUCY2F;refGene_tx=XM_006943916.3;refGene_gn=GUCY2F;refGene_tx=XM_019824152.2;refGene_gn=GUCY2F;refGene_tx=XM_019824153.2
X 91797877 91797975 refGene_gn=NXT2;refGene_tx=XM_004000788.5
X 91798802 91798810 refGene_gn=NXT2;refGene_tx=XM_019824154.2
X 91798963 91799048
refGene_gn=NXT2;refGene_tx=XM_004000788.5;refGene_gn=NXT2;refGene_tx=XM_019824154.2
X 91799281 91799323 refGene_gn=NXT2;refGene_tx=XM_006943917.4

X 91799998 91800085
refGene_gn=NXT2;refGene_tx=XM_004000788.5;refGene_gn=NXT2;refGene_tx=XM_006943917.4;refGene_gn=NXT2;refGene_tx=XM_019824154.2
X 91801620 91801765
refGene_gn=NXT2;refGene_tx=XM_004000788.5;refGene_gn=NXT2;refGene_tx=XM_006943917.4;refGene_gn=NXT2;refGene_tx=XM_019824154.2
X 91802448 91802630
refGene_gn=NXT2;refGene_tx=XM_004000788.5;refGene_gn=NXT2;refGene_tx=XM_006943917.4;refGene_gn=NXT2;refGene_tx=XM_019824154.2
X 91863012 91863426 refGene_gn=KCNE5;refGene_tx=XM_023249332.1
X 91889221 91889379
refGene_gn=ACSL4;refGene_tx=XM_004000790.5;refGene_gn=ACSL4;refGene_tx=XM_006943918.4;refGene_gn=ACSL4;refGene_tx=XM_019824155.2
X 91904379 91904537
refGene_gn=ACSL4;refGene_tx=XM_004000790.5;refGene_gn=ACSL4;refGene_tx=XM_006943918.4;refGene_gn=ACSL4;refGene_tx=XM_019824155.2
X 91907565 91907680
refGene_gn=ACSL4;refGene_tx=XM_004000790.5;refGene_gn=ACSL4;refGene_tx=XM_006943918.4;refGene_gn=ACSL4;refGene_tx=XM_019824155.2
X 91908797 91908989
refGene_gn=ACSL4;refGene_tx=XM_004000790.5;refGene_gn=ACSL4;refGene_tx=XM_006943918.4;refGene_gn=ACSL4;refGene_tx=XM_019824155.2
X 91912128 91912203
refGene_gn=ACSL4;refGene_tx=XM_004000790.5;refGene_gn=ACSL4;refGene_tx=XM_006943918.4;refGene_gn=ACSL4;refGene_tx=XM_019824155.2
X 91914217 91914390
refGene_gn=ACSL4;refGene_tx=XM_004000790.5;refGene_gn=ACSL4;refGene_tx=XM_006943918.4;refGene_gn=ACSL4;refGene_tx=XM_019824155.2
X 91915766 91915906
refGene_gn=ACSL4;refGene_tx=XM_004000790.5;refGene_gn=ACSL4;refGene_tx=XM_006943918.4;refGene_gn=ACSL4;refGene_tx=XM_019824155.2
X 91919684 91919756
refGene_gn=ACSL4;refGene_tx=XM_004000790.5;refGene_gn=ACSL4;refGene_tx=XM_006943918.4;refGene_gn=ACSL4;refGene_tx=XM_019824155.2
X 91923158 91923282
refGene_gn=ACSL4;refGene_tx=XM_004000790.5;refGene_gn=ACSL4;refGene_tx=XM_006943918.4;refGene_gn=ACSL4;refGene_tx=XM_019824155.2
X 91923437 91923588
refGene_gn=ACSL4;refGene_tx=XM_004000790.5;refGene_gn=ACSL4;refGene_tx=XM_006943918.4;refGene_gn=ACSL4;refGene_tx=XM_019824155.2
X 91925686 91925825
refGene_gn=ACSL4;refGene_tx=XM_004000790.5;refGene_gn=ACSL4;refGene_tx=XM_006943918.4;refGene_gn=ACSL4;refGene_tx=XM_019824155.2
X 91925943 91926053
refGene_gn=ACSL4;refGene_tx=XM_004000790.5;refGene_gn=ACSL4;refGene_tx=XM_006943918.4;refGene_gn=ACSL4;refGene_tx=XM_019824155.2
X 91927400 91927578
refGene_gn=ACSL4;refGene_tx=XM_004000790.5;refGene_gn=ACSL4;refGene_tx=XM_006943918.4;refGene_gn=ACSL4;refGene_tx=XM_019824155.2
X 91927821 91928172

refGene_gn=ACSL4;refGene_tx=XM_004000790.5;refGene_gn=ACSL4;refGene_tx=XM_006943918.4;refGene_gn=ACSL4;refGene_tx=XM_019824155.2
X 91937443 91937527 refGene_gn=ACSL4;refGene_tx=XM_019824155.2
X 92056994 92058572
refGene_gn=LOC102902491;refGene_tx=XM_006943919.4
X 92233001 92233391 refGene_gn=TMEM164;refGene_tx=XM_011279391.3
X 92301590 92301640 refGene_gn=TMEM164;refGene_tx=XM_011279391.3
X 92330442 92330509 refGene_gn=TMEM164;refGene_tx=XM_011279391.3
X 92361689 92361768 refGene_gn=TMEM164;refGene_tx=XM_011279391.3
X 92391619 92391720 refGene_gn=TMEM164;refGene_tx=XM_011279391.3
X 92393429 92393543 refGene_gn=TMEM164;refGene_tx=XM_011279391.3
X 92393544 92393637 refGene_gn=TMEM164;refGene_tx=XM_011279391.3
X 92419194 92419309 refGene_gn=AMMECR1;refGene_tx=XM_019824159.2
X 92421241 92421338 refGene_gn=AMMECR1;refGene_tx=XM_019824159.2
X 92422802 92422893 refGene_gn=AMMECR1;refGene_tx=XM_019824159.2
X 92437400 92437515 refGene_gn=AMMECR1;refGene_tx=XM_019824159.2
X 92474953 92475064 refGene_gn=AMMECR1;refGene_tx=XM_019824159.2
X 92525717 92526187 refGene_gn=AMMECR1;refGene_tx=XM_019824159.2
X 92651876 92655950 refGene_gn=RTL9;refGene_tx=XM_004000792.3
X 92656483 92656600 refGene_gn=RTL9;refGene_tx=XM_004000792.3
X 92804751 92804882
refGene_gn=CHRDL1;refGene_tx=XM_006943926.3;refGene_gn=CHRDL1;refGene_tx=XM_006943927.3;refGene_gn=CHRDL1;refGene_tx=XM_006943928.3;refGene_gn=CHRDL1;refGene_tx=XM_023249482.1;refGene_gn=CHRDL1;refGene_tx=XM_023249483.1;refGene_gn=CHRDL1;refGene_tx=XM_023249484.1;refGene_gn=CHRDL1;refGene_tx=XM_023249485.1
X 92808065 92808155
refGene_gn=CHRDL1;refGene_tx=XM_006943926.3;refGene_gn=CHRDL1;refGene_tx=XM_006943927.3;refGene_gn=CHRDL1;refGene_tx=XM_006943928.3;refGene_gn=CHRDL1;refGene_tx=XM_023249482.1;refGene_gn=CHRDL1;refGene_tx=XM_023249483.1;refGene_gn=CHRDL1;refGene_tx=XM_023249484.1;refGene_gn=CHRDL1;refGene_tx=XM_023249485.1
X 92810473 92810641
refGene_gn=CHRDL1;refGene_tx=XM_006943926.3;refGene_gn=CHRDL1;refGene_tx=XM_006943927.3;refGene_gn=CHRDL1;refGene_tx=XM_006943928.3;refGene_gn=CHRDL1;refGene_tx=XM_023249482.1;refGene_gn=CHRDL1;refGene_tx=XM_023249483.1;refGene_gn=CHRDL1;refGene_tx=XM_023249484.1;refGene_gn=CHRDL1;refGene_tx=XM_023249485.1
X 92816570 92816780
refGene_gn=CHRDL1;refGene_tx=XM_006943926.3;refGene_gn=CHRDL1;refGene_tx=XM_006943927.3;refGene_gn=CHRDL1;refGene_tx=XM_006943928.3;refGene_gn=CHRDL1;refGene_tx=XM_023249482.1;refGene_gn=CHRDL1;refGene_tx=XM_023249483.1;refGene_gn=CHRDL1;refGene_tx=XM_023249484.1;refGene_gn=CHRDL1;refGene_tx=XM_023249485.1
X 92820142 92820311
refGene_gn=CHRDL1;refGene_tx=XM_006943926.3;refGene_gn=CHRDL1;refGene_tx=XM_006943927.3;refGene_gn=CHRDL1;refGene_tx=XM_006943928.3;refGene_gn=CHRDL1;refGene_tx=XM_023249482.1;refGene_gn=CHRDL1;refGene_tx=XM_023249483.1;refGene_gn=CHRDL1;refGene_tx=XM_023249484.1;refGene_gn=CHRDL1;refGene_tx=XM_023249485.1

X 92827154 92827222
refGene_gn=CHRDL1;refGene_tx=XM_006943926.3;refGene_gn=CHRDL1;refGene_tx=XM_006943927.3;refGene_gn=CHRDL1;refGene_tx=XM_006943928.3;refGene_gn=CHRDL1;refGene_tx=XM_023249482.1;refGene_gn=CHRDL1;refGene_tx=XM_023249483.1;refGene_gn=CHRDL1;refGene_tx=XM_023249484.1;refGene_gn=CHRDL1;refGene_tx=XM_023249485.1
X 92843324 92843418
refGene_gn=CHRDL1;refGene_tx=XM_006943926.3;refGene_gn=CHRDL1;refGene_tx=XM_006943927.3;refGene_gn=CHRDL1;refGene_tx=XM_006943928.3;refGene_gn=CHRDL1;refGene_tx=XM_023249482.1;refGene_gn=CHRDL1;refGene_tx=XM_023249483.1;refGene_gn=CHRDL1;refGene_tx=XM_023249484.1;refGene_gn=CHRDL1;refGene_tx=XM_023249485.1
X 92845054 92845200
refGene_gn=CHRDL1;refGene_tx=XM_006943926.3;refGene_gn=CHRDL1;refGene_tx=XM_006943927.3;refGene_gn=CHRDL1;refGene_tx=XM_006943928.3;refGene_gn=CHRDL1;refGene_tx=XM_023249482.1;refGene_gn=CHRDL1;refGene_tx=XM_023249483.1;refGene_gn=CHRDL1;refGene_tx=XM_023249484.1;refGene_gn=CHRDL1;refGene_tx=XM_023249485.1
X 92886874 92886968
refGene_gn=CHRDL1;refGene_tx=XM_006943926.3;refGene_gn=CHRDL1;refGene_tx=XM_006943927.3;refGene_gn=CHRDL1;refGene_tx=XM_006943928.3;refGene_gn=CHRDL1;refGene_tx=XM_023249482.1;refGene_gn=CHRDL1;refGene_tx=XM_023249483.1;refGene_gn=CHRDL1;refGene_tx=XM_023249484.1;refGene_gn=CHRDL1;refGene_tx=XM_023249485.1
X 92887065 92887284
refGene_gn=CHRDL1;refGene_tx=XM_006943926.3;refGene_gn=CHRDL1;refGene_tx=XM_023249482.1;refGene_gn=CHRDL1;refGene_tx=XM_023249483.1;refGene_gn=CHRDL1;refGene_tx=XM_023249484.1
X 92887385 92887403 refGene_gn=CHRDL1;refGene_tx=XM_006943926.3
X 92889636 92889749
refGene_gn=CHRDL1;refGene_tx=XM_006943927.3;refGene_gn=CHRDL1;refGene_tx=XM_006943928.3;refGene_gn=CHRDL1;refGene_tx=XM_023249482.1;refGene_gn=CHRDL1;refGene_tx=XM_023249483.1;refGene_gn=CHRDL1;refGene_tx=XM_023249484.1;refGene_gn=CHRDL1;refGene_tx=XM_023249485.1
X 92917128 92917280
refGene_gn=CHRDL1;refGene_tx=XM_006943927.3;refGene_gn=CHRDL1;refGene_tx=XM_006943928.3;refGene_gn=CHRDL1;refGene_tx=XM_023249482.1;refGene_gn=CHRDL1;refGene_tx=XM_023249483.1;refGene_gn=CHRDL1;refGene_tx=XM_023249484.1;refGene_gn=CHRDL1;refGene_tx=XM_023249485.1
X 92920710 92920862
refGene_gn=CHRDL1;refGene_tx=XM_006943927.3;refGene_gn=CHRDL1;refGene_tx=XM_006943928.3;refGene_gn=CHRDL1;refGene_tx=XM_023249482.1;refGene_gn=CHRDL1;refGene_tx=XM_023249483.1;refGene_gn=CHRDL1;refGene_tx=XM_023249484.1;refGene_gn=CHRDL1;refGene_tx=XM_023249485.1
X 93209859 93210034
refGene_gn=PAK3;refGene_tx=XM_019824163.2;refGene_gn=PAK3;refGene_tx=XM_019824169.2;refGene_gn=PAK3;refGene_tx=XM_019824173.2;refGene_gn=PAK3;refGene_tx=XM_019824174.2;refGene_gn=PAK3;refGene_tx=XM_023249295.1;refGene_gn=PAK3;refGene_tx=XM_023249296.1;refGene_gn=PAK3;refGene_tx=XM_023249297.1;refGene_gn=PAK3;refGene_tx=XM_023249298.1;refGene_gn=PAK

3;refGene_tx=XM_023249299.1;refGene_gn=PAK3;refGene_tx=XM_023249300.1;
refGene_gn=PAK3;refGene_tx=XM_023249301.1;refGene_gn=PAK3;refGene_tx=X
M_023249302.1;refGene_gn=PAK3;refGene_tx=XM_023249303.1;refGene_gn=PAK
3;refGene_tx=XM_023249304.1;refGene_gn=PAK3;refGene_tx=XM_023249305.1;
refGene_gn=PAK3;refGene_tx=XM_023249306.1
X 93230439 93230540
refGene_gn=PAK3;refGene_tx=XM_019824163.2;refGene_gn=PAK3;refGene_tx=X
M_019824169.2;refGene_gn=PAK3;refGene_tx=XM_019824173.2;refGene_gn=PAK
3;refGene_tx=XM_019824174.2;refGene_gn=PAK3;refGene_tx=XM_023249295.1;
refGene_gn=PAK3;refGene_tx=XM_023249296.1;refGene_gn=PAK3;refGene_tx=X
M_023249297.1;refGene_gn=PAK3;refGene_tx=XM_023249298.1;refGene_gn=PAK
3;refGene_tx=XM_023249299.1;refGene_gn=PAK3;refGene_tx=XM_023249300.1;
refGene_gn=PAK3;refGene_tx=XM_023249301.1;refGene_gn=PAK3;refGene_tx=X
M_023249302.1;refGene_gn=PAK3;refGene_tx=XM_023249303.1;refGene_gn=PAK
3;refGene_tx=XM_023249304.1;refGene_gn=PAK3;refGene_tx=XM_023249305.1;
refGene_gn=PAK3;refGene_tx=XM_023249306.1
X 93233010 93233073
refGene_gn=PAK3;refGene_tx=XM_019824163.2;refGene_gn=PAK3;refGene_tx=X
M_019824169.2;refGene_gn=PAK3;refGene_tx=XM_019824173.2;refGene_gn=PAK
3;refGene_tx=XM_019824174.2;refGene_gn=PAK3;refGene_tx=XM_023249295.1;
refGene_gn=PAK3;refGene_tx=XM_023249296.1;refGene_gn=PAK3;refGene_tx=X
M_023249297.1;refGene_gn=PAK3;refGene_tx=XM_023249298.1;refGene_gn=PAK
3;refGene_tx=XM_023249299.1;refGene_gn=PAK3;refGene_tx=XM_023249300.1;
refGene_gn=PAK3;refGene_tx=XM_023249301.1;refGene_gn=PAK3;refGene_tx=X
M_023249302.1;refGene_gn=PAK3;refGene_tx=XM_023249303.1;refGene_gn=PAK
3;refGene_tx=XM_023249304.1
X 93234671 93234716
refGene_gn=PAK3;refGene_tx=XM_019824163.2;refGene_gn=PAK3;refGene_tx=X
M_019824169.2;refGene_gn=PAK3;refGene_tx=XM_019824173.2;refGene_gn=PAK
3;refGene_tx=XM_019824174.2;refGene_gn=PAK3;refGene_tx=XM_023249295.1;
refGene_gn=PAK3;refGene_tx=XM_023249296.1;refGene_gn=PAK3;refGene_tx=X
M_023249297.1;refGene_gn=PAK3;refGene_tx=XM_023249298.1;refGene_gn=PAK
3;refGene_tx=XM_023249299.1;refGene_gn=PAK3;refGene_tx=XM_023249300.1;
refGene_gn=PAK3;refGene_tx=XM_023249301.1;refGene_gn=PAK3;refGene_tx=X
M_023249302.1;refGene_gn=PAK3;refGene_tx=XM_023249303.1;refGene_gn=PAK
3;refGene_tx=XM_023249304.1;refGene_gn=PAK3;refGene_tx=XM_023249305.1
X 93235884 93236038
refGene_gn=PAK3;refGene_tx=XM_019824163.2;refGene_gn=PAK3;refGene_tx=X
M_019824169.2;refGene_gn=PAK3;refGene_tx=XM_019824173.2;refGene_gn=PAK
3;refGene_tx=XM_019824174.2;refGene_gn=PAK3;refGene_tx=XM_023249295.1;
refGene_gn=PAK3;refGene_tx=XM_023249296.1;refGene_gn=PAK3;refGene_tx=X
M_023249297.1;refGene_gn=PAK3;refGene_tx=XM_023249298.1;refGene_gn=PAK
3;refGene_tx=XM_023249299.1;refGene_gn=PAK3;refGene_tx=XM_023249300.1;
refGene_gn=PAK3;refGene_tx=XM_023249301.1;refGene_gn=PAK3;refGene_tx=X
M_023249302.1;refGene_gn=PAK3;refGene_tx=XM_023249303.1;refGene_gn=PAK
3;refGene_tx=XM_023249304.1;refGene_gn=PAK3;refGene_tx=XM_023249305.1;
refGene_gn=PAK3;refGene_tx=XM_023249306.1
X 93238034 93238072
refGene_gn=PAK3;refGene_tx=XM_019824163.2;refGene_gn=PAK3;refGene_tx=X
M_019824169.2;refGene_gn=PAK3;refGene_tx=XM_019824173.2;refGene_gn=PAK

3;refGene_tx=XM_019824174.2;refGene_gn=PAK3;refGene_tx=XM_023249295.1;
refGene_gn=PAK3;refGene_tx=XM_023249296.1;refGene_gn=PAK3;refGene_tx=X
M_023249297.1;refGene_gn=PAK3;refGene_tx=XM_023249298.1;refGene_gn=PAK
3;refGene_tx=XM_023249299.1;refGene_gn=PAK3;refGene_tx=XM_023249300.1;
refGene_gn=PAK3;refGene_tx=XM_023249301.1;refGene_gn=PAK3;refGene_tx=X
M_023249302.1;refGene_gn=PAK3;refGene_tx=XM_023249303.1;refGene_gn=PAK
3;refGene_tx=XM_023249304.1;refGene_gn=PAK3;refGene_tx=XM_023249305.1;
refGene_gn=PAK3;refGene_tx=XM_023249306.1
X 93247483 93247615
refGene_gn=PAK3;refGene_tx=XM_019824163.2;refGene_gn=PAK3;refGene_tx=X
M_019824169.2;refGene_gn=PAK3;refGene_tx=XM_019824173.2;refGene_gn=PAK
3;refGene_tx=XM_019824174.2;refGene_gn=PAK3;refGene_tx=XM_023249295.1;
refGene_gn=PAK3;refGene_tx=XM_023249296.1;refGene_gn=PAK3;refGene_tx=X
M_023249297.1;refGene_gn=PAK3;refGene_tx=XM_023249298.1;refGene_gn=PAK
3;refGene_tx=XM_023249299.1;refGene_gn=PAK3;refGene_tx=XM_023249300.1;
refGene_gn=PAK3;refGene_tx=XM_023249301.1;refGene_gn=PAK3;refGene_tx=X
M_023249302.1;refGene_gn=PAK3;refGene_tx=XM_023249303.1;refGene_gn=PAK
3;refGene_tx=XM_023249304.1;refGene_gn=PAK3;refGene_tx=XM_023249305.1;
refGene_gn=PAK3;refGene_tx=XM_023249306.1
X 93248493 93248659
refGene_gn=PAK3;refGene_tx=XM_019824163.2;refGene_gn=PAK3;refGene_tx=X
M_019824169.2;refGene_gn=PAK3;refGene_tx=XM_019824173.2;refGene_gn=PAK
3;refGene_tx=XM_019824174.2;refGene_gn=PAK3;refGene_tx=XM_023249295.1;
refGene_gn=PAK3;refGene_tx=XM_023249296.1;refGene_gn=PAK3;refGene_tx=X
M_023249297.1;refGene_gn=PAK3;refGene_tx=XM_023249298.1;refGene_gn=PAK
3;refGene_tx=XM_023249299.1;refGene_gn=PAK3;refGene_tx=XM_023249300.1;
refGene_gn=PAK3;refGene_tx=XM_023249301.1;refGene_gn=PAK3;refGene_tx=X
M_023249302.1;refGene_gn=PAK3;refGene_tx=XM_023249303.1;refGene_gn=PAK
3;refGene_tx=XM_023249304.1;refGene_gn=PAK3;refGene_tx=XM_023249305.1;
refGene_gn=PAK3;refGene_tx=XM_023249306.1
X 93257889 93257953
refGene_gn=PAK3;refGene_tx=XM_019824163.2;refGene_gn=PAK3;refGene_tx=X
M_019824169.2;refGene_gn=PAK3;refGene_tx=XM_019824173.2;refGene_gn=PAK
3;refGene_tx=XM_019824174.2;refGene_gn=PAK3;refGene_tx=XM_023249295.1;
refGene_gn=PAK3;refGene_tx=XM_023249296.1;refGene_gn=PAK3;refGene_tx=X
M_023249297.1;refGene_gn=PAK3;refGene_tx=XM_023249298.1;refGene_gn=PAK
3;refGene_tx=XM_023249299.1;refGene_gn=PAK3;refGene_tx=XM_023249300.1;
refGene_gn=PAK3;refGene_tx=XM_023249301.1;refGene_gn=PAK3;refGene_tx=X
M_023249302.1;refGene_gn=PAK3;refGene_tx=XM_023249303.1;refGene_gn=PAK
3;refGene_tx=XM_023249304.1;refGene_gn=PAK3;refGene_tx=XM_023249305.1;
refGene_gn=PAK3;refGene_tx=XM_023249306.1
X 93276447 93276496
refGene_gn=PAK3;refGene_tx=XM_019824163.2;refGene_gn=PAK3;refGene_tx=X
M_019824169.2;refGene_gn=PAK3;refGene_tx=XM_019824173.2;refGene_gn=PAK
3;refGene_tx=XM_019824174.2;refGene_gn=PAK3;refGene_tx=XM_023249295.1;
refGene_gn=PAK3;refGene_tx=XM_023249296.1;refGene_gn=PAK3;refGene_tx=X
M_023249297.1;refGene_gn=PAK3;refGene_tx=XM_023249298.1;refGene_gn=PAK
3;refGene_tx=XM_023249299.1;refGene_gn=PAK3;refGene_tx=XM_023249300.1;
refGene_gn=PAK3;refGene_tx=XM_023249301.1;refGene_gn=PAK3;refGene_tx=X
M_023249302.1;refGene_gn=PAK3;refGene_tx=XM_023249303.1;refGene_gn=PAK

3;refGene_tx=XM_023249304.1;refGene_gn=PAK3;refGene_tx=XM_023249305.1;
refGene_gn=PAK3;refGene_tx=XM_023249306.1
X 93276849 93276962
refGene_gn=PAK3;refGene_tx=XM_019824163.2;refGene_gn=PAK3;refGene_tx=XM_019824169.2;refGene_gn=PAK3;refGene_tx=XM_019824173.2;refGene_gn=PAK3;refGene_tx=XM_019824174.2;refGene_gn=PAK3;refGene_tx=XM_023249295.1;refGene_gn=PAK3;refGene_tx=XM_023249296.1;refGene_gn=PAK3;refGene_tx=XM_023249297.1;refGene_gn=PAK3;refGene_tx=XM_023249298.1;refGene_gn=PAK3;refGene_tx=XM_023249299.1;refGene_gn=PAK3;refGene_tx=XM_023249300.1;refGene_gn=PAK3;refGene_tx=XM_023249301.1;refGene_gn=PAK3;refGene_tx=XM_023249302.1;refGene_gn=PAK3;refGene_tx=XM_023249303.1;refGene_gn=PAK3;refGene_tx=XM_023249304.1;refGene_gn=PAK3;refGene_tx=XM_023249305.1;refGene_gn=PAK3;refGene_tx=XM_023249306.1
X 93278710 93278828
refGene_gn=PAK3;refGene_tx=XM_019824163.2;refGene_gn=PAK3;refGene_tx=XM_019824169.2;refGene_gn=PAK3;refGene_tx=XM_019824173.2;refGene_gn=PAK3;refGene_tx=XM_019824174.2;refGene_gn=PAK3;refGene_tx=XM_023249295.1;refGene_gn=PAK3;refGene_tx=XM_023249296.1;refGene_gn=PAK3;refGene_tx=XM_023249297.1;refGene_gn=PAK3;refGene_tx=XM_023249298.1;refGene_gn=PAK3;refGene_tx=XM_023249299.1;refGene_gn=PAK3;refGene_tx=XM_023249300.1;refGene_gn=PAK3;refGene_tx=XM_023249301.1;refGene_gn=PAK3;refGene_tx=XM_023249302.1;refGene_gn=PAK3;refGene_tx=XM_023249303.1;refGene_gn=PAK3;refGene_tx=XM_023249304.1;refGene_gn=PAK3;refGene_tx=XM_023249305.1;refGene_gn=PAK3;refGene_tx=XM_023249306.1
X 93280223 93280323
refGene_gn=PAK3;refGene_tx=XM_019824163.2;refGene_gn=PAK3;refGene_tx=XM_019824169.2;refGene_gn=PAK3;refGene_tx=XM_019824173.2;refGene_gn=PAK3;refGene_tx=XM_019824174.2;refGene_gn=PAK3;refGene_tx=XM_023249295.1;refGene_gn=PAK3;refGene_tx=XM_023249296.1;refGene_gn=PAK3;refGene_tx=XM_023249297.1;refGene_gn=PAK3;refGene_tx=XM_023249298.1;refGene_gn=PAK3;refGene_tx=XM_023249299.1;refGene_gn=PAK3;refGene_tx=XM_023249300.1;refGene_gn=PAK3;refGene_tx=XM_023249301.1;refGene_gn=PAK3;refGene_tx=XM_023249302.1;refGene_gn=PAK3;refGene_tx=XM_023249303.1;refGene_gn=PAK3;refGene_tx=XM_023249304.1;refGene_gn=PAK3;refGene_tx=XM_023249305.1;refGene_gn=PAK3;refGene_tx=XM_023249306.1
X 93281282 93281479
refGene_gn=PAK3;refGene_tx=XM_019824163.2;refGene_gn=PAK3;refGene_tx=XM_019824169.2;refGene_gn=PAK3;refGene_tx=XM_019824173.2;refGene_gn=PAK3;refGene_tx=XM_019824174.2;refGene_gn=PAK3;refGene_tx=XM_023249295.1;refGene_gn=PAK3;refGene_tx=XM_023249296.1;refGene_gn=PAK3;refGene_tx=XM_023249297.1;refGene_gn=PAK3;refGene_tx=XM_023249298.1;refGene_gn=PAK3;refGene_tx=XM_023249299.1;refGene_gn=PAK3;refGene_tx=XM_023249300.1;refGene_gn=PAK3;refGene_tx=XM_023249301.1;refGene_gn=PAK3;refGene_tx=XM_023249302.1;refGene_gn=PAK3;refGene_tx=XM_023249303.1;refGene_gn=PAK3;refGene_tx=XM_023249304.1;refGene_gn=PAK3;refGene_tx=XM_023249305.1;refGene_gn=PAK3;refGene_tx=XM_023249306.1
X 93297899 93298037
refGene_gn=PAK3;refGene_tx=XM_019824163.2;refGene_gn=PAK3;refGene_tx=XM_019824169.2;refGene_gn=PAK3;refGene_tx=XM_019824173.2;refGene_gn=PAK3;refGene_tx=XM_019824174.2;refGene_gn=PAK3;refGene_tx=XM_023249295.1;

refGene_gn=PAK3;refGene_tx=XM_023249296.1;refGene_gn=PAK3;refGene_tx=XM_023249297.1;refGene_gn=PAK3;refGene_tx=XM_023249298.1;refGene_gn=PAK3;refGene_tx=XM_023249299.1;refGene_gn=PAK3;refGene_tx=XM_023249300.1;refGene_gn=PAK3;refGene_tx=XM_023249301.1;refGene_gn=PAK3;refGene_tx=XM_023249302.1;refGene_gn=PAK3;refGene_tx=XM_023249303.1;refGene_gn=PAK3;refGene_tx=XM_023249304.1;refGene_gn=PAK3;refGene_tx=XM_023249305.1;refGene_gn=PAK3;refGene_tx=XM_023249306.1
X 93301479 93301569
refGene_gn=PAK3;refGene_tx=XM_019824163.2;refGene_gn=PAK3;refGene_tx=XM_019824169.2;refGene_gn=PAK3;refGene_tx=XM_019824173.2;refGene_gn=PAK3;refGene_tx=XM_019824174.2;refGene_gn=PAK3;refGene_tx=XM_023249295.1;refGene_gn=PAK3;refGene_tx=XM_023249296.1;refGene_gn=PAK3;refGene_tx=XM_023249297.1;refGene_gn=PAK3;refGene_tx=XM_023249298.1;refGene_gn=PAK3;refGene_tx=XM_023249299.1;refGene_gn=PAK3;refGene_tx=XM_023249300.1;refGene_gn=PAK3;refGene_tx=XM_023249301.1;refGene_gn=PAK3;refGene_tx=XM_023249302.1;refGene_gn=PAK3;refGene_tx=XM_023249303.1;refGene_gn=PAK3;refGene_tx=XM_023249304.1;refGene_gn=PAK3;refGene_tx=XM_023249305.1;refGene_gn=PAK3;refGene_tx=XM_023249306.1
X 93322621 93322804
refGene_gn=CAPN6;refGene_tx=XM_004000803.4;refGene_gn=CAPN6;refGene_tx=XM_019824177.2
X 93323335 93323472
refGene_gn=CAPN6;refGene_tx=XM_004000803.4;refGene_gn=CAPN6;refGene_tx=XM_019824177.2
X 93323814 93323936
refGene_gn=CAPN6;refGene_tx=XM_004000803.4;refGene_gn=CAPN6;refGene_tx=XM_019824177.2
X 93324519 93324722
refGene_gn=CAPN6;refGene_tx=XM_004000803.4;refGene_gn=CAPN6;refGene_tx=XM_019824177.2
X 93324900 93325023
refGene_gn=CAPN6;refGene_tx=XM_004000803.4;refGene_gn=CAPN6;refGene_tx=XM_019824177.2
X 93326591 93326778
refGene_gn=CAPN6;refGene_tx=XM_004000803.4;refGene_gn=CAPN6;refGene_tx=XM_019824177.2
X 93326883 93326961
refGene_gn=CAPN6;refGene_tx=XM_004000803.4;refGene_gn=CAPN6;refGene_tx=XM_019824177.2
X 93327202 93327396
refGene_gn=CAPN6;refGene_tx=XM_004000803.4;refGene_gn=CAPN6;refGene_tx=XM_019824177.2
X 93327955 93328148
refGene_gn=CAPN6;refGene_tx=XM_004000803.4;refGene_gn=CAPN6;refGene_tx=XM_019824177.2
X 93328657 93328866
refGene_gn=CAPN6;refGene_tx=XM_004000803.4;refGene_gn=CAPN6;refGene_tx=XM_019824177.2
X 93329867 93329999
refGene_gn=CAPN6;refGene_tx=XM_004000803.4;refGene_gn=CAPN6;refGene_tx

=XM_019824177.2
X 93338515 93338680
refGene_gn=CAPN6;refGene_tx=XM_004000803.4;refGene_gn=CAPN6;refGene_tx=XM_019824177.2
X 93378271 93378328
refGene_gn=DCX;refGene_tx=XM_019824178.2;refGene_gn=DCX;refGene_tx=XM_019824179.2;refGene_gn=DCX;refGene_tx=XM_019824180.2;refGene_gn=DCX;refGene_tx=XM_019824182.2;refGene_gn=DCX;refGene_tx=XM_019824183.2
X 93390103 93390201
refGene_gn=DCX;refGene_tx=XM_019824178.2;refGene_gn=DCX;refGene_tx=XM_019824179.2;refGene_gn=DCX;refGene_tx=XM_019824180.2;refGene_gn=DCX;refGene_tx=XM_019824182.2;refGene_gn=DCX;refGene_tx=XM_019824183.2
X 93406115 93406253
refGene_gn=DCX;refGene_tx=XM_019824178.2;refGene_gn=DCX;refGene_tx=XM_019824179.2;refGene_gn=DCX;refGene_tx=XM_019824180.2;refGene_gn=DCX;refGene_tx=XM_019824182.2;refGene_gn=DCX;refGene_tx=XM_019824183.2
X 93408464 93408567
refGene_gn=DCX;refGene_tx=XM_019824178.2;refGene_gn=DCX;refGene_tx=XM_019824179.2;refGene_gn=DCX;refGene_tx=XM_019824180.2;refGene_gn=DCX;refGene_tx=XM_019824182.2;refGene_gn=DCX;refGene_tx=XM_019824183.2
X 93474507 93474848
refGene_gn=DCX;refGene_tx=XM_019824178.2;refGene_gn=DCX;refGene_tx=XM_019824179.2;refGene_gn=DCX;refGene_tx=XM_019824180.2;refGene_gn=DCX;refGene_tx=XM_019824182.2;refGene_gn=DCX;refGene_tx=XM_019824183.2
X 93483478 93483842
refGene_gn=DCX;refGene_tx=XM_019824178.2;refGene_gn=DCX;refGene_tx=XM_019824179.2;refGene_gn=DCX;refGene_tx=XM_019824180.2;refGene_gn=DCX;refGene_tx=XM_019824182.2;refGene_gn=DCX;refGene_tx=XM_019824183.2
X 93602222 93602495
refGene_gn=LOC109496572;refGene_tx=XM_023249231.1
X 93722732 93722803 refGene_gn=ALG13;refGene_tx=XM_023249369.1
X 93740887 93740921 refGene_gn=ALG13;refGene_tx=XM_023249369.1
X 93740923 93741040 refGene_gn=ALG13;refGene_tx=XM_023249369.1
X 93741041 93741074 refGene_gn=ALG13;refGene_tx=XM_023249369.1
X 93741076 93741259 refGene_gn=ALG13;refGene_tx=XM_023249369.1
X 93741823 93741907 refGene_gn=ALG13;refGene_tx=XM_023249369.1
X 93743987 93744038 refGene_gn=ALG13;refGene_tx=XM_023249369.1
X 93744768 93744815 refGene_gn=ALG13;refGene_tx=XM_023249369.1
X 93746437 93746510 refGene_gn=ALG13;refGene_tx=XM_023249369.1
X 93749528 93749610 refGene_gn=ALG13;refGene_tx=XM_023249369.1
X 93749813 93749976 refGene_gn=ALG13;refGene_tx=XM_023249369.1
X 93751560 93751636 refGene_gn=ALG13;refGene_tx=XM_023249369.1
X 93752628 93752737 refGene_gn=ALG13;refGene_tx=XM_023249369.1
X 93753859 93753924 refGene_gn=ALG13;refGene_tx=XM_023249369.1
X 93754453 93754554 refGene_gn=ALG13;refGene_tx=XM_023249369.1
X 93755584 93755712 refGene_gn=ALG13;refGene_tx=XM_023249369.1
X 93756382 93756629 refGene_gn=ALG13;refGene_tx=XM_023249369.1
X 93757061 93757175 refGene_gn=ALG13;refGene_tx=XM_023249369.1
X 93757368 93757525 refGene_gn=ALG13;refGene_tx=XM_023249369.1
X 93757849 93757970 refGene_gn=ALG13;refGene_tx=XM_023249369.1

X 93760178 93760211 refGene_gn=ALG13;refGene_tx=XM_023249369.1
X 93760313 93760369 refGene_gn=ALG13;refGene_tx=XM_023249369.1
X 93763808 93763880 refGene_gn=ALG13;refGene_tx=XM_023249369.1
X 93765364 93765530 refGene_gn=ALG13;refGene_tx=XM_023249369.1
X 93772959 93773142 refGene_gn=ALG13;refGene_tx=XM_023249369.1
X 93774666 93774707 refGene_gn=ALG13;refGene_tx=XM_023249369.1
X 93776735 93776922 refGene_gn=ALG13;refGene_tx=XM_023249369.1
X 93781676 93781942 refGene_gn=ALG13;refGene_tx=XM_023249369.1
X 93797340 93798033 refGene_gn=TRPC5;refGene_tx=XM_004000806.4
X 93800049 93800139 refGene_gn=TRPC5;refGene_tx=XM_004000806.4
X 93801029 93801071 refGene_gn=TRPC5;refGene_tx=XM_004000806.4
X 93801321 93801525 refGene_gn=TRPC5;refGene_tx=XM_004000806.4
X 93831544 93831740 refGene_gn=TRPC5;refGene_tx=XM_004000806.4
X 93845299 93845622 refGene_gn=TRPC5;refGene_tx=XM_004000806.4
X 93850435 93850575 refGene_gn=TRPC5;refGene_tx=XM_004000806.4
X 93852177 93852514 refGene_gn=TRPC5;refGene_tx=XM_004000806.4
X 93894227 93894563
refGene_gn=TRPC5OS;refGene_tx=XM_023249041.1;refGene_gn=TRPC5OS;refGen
e_tx=XM_023249042.1;refGene_gn=TRPC5OS;refGene_tx=XM_023249043.1
X 93900313 93900835 refGene_gn=TRPC5;refGene_tx=XM_004000806.4
X 93933594 93933972 refGene_gn=TRPC5;refGene_tx=XM_004000806.4
X 94415722 94416610 refGene_gn=RTL4;refGene_tx=XM_023249209.1
X 94529651 94529833 refGene_gn=LHFPL1;refGene_tx=XM_023249367.1
X 94558383 94558482 refGene_gn=LHFPL1;refGene_tx=XM_023249367.1
X 94568895 94569361 refGene_gn=LHFPL1;refGene_tx=XM_023249367.1
X 94655381 94655479 refGene_gn=AMOT;refGene_tx=XM_006943935.4
X 94655839 94656538 refGene_gn=AMOT;refGene_tx=XM_006943935.4
X 94657625 94657858 refGene_gn=AMOT;refGene_tx=XM_006943935.4
X 94658846 94658969 refGene_gn=AMOT;refGene_tx=XM_006943935.4
X 94667255 94667446 refGene_gn=AMOT;refGene_tx=XM_006943935.4
X 94668356 94668506 refGene_gn=AMOT;refGene_tx=XM_006943935.4
X 94682437 94682583 refGene_gn=AMOT;refGene_tx=XM_006943935.4
X 94685141 94685234 refGene_gn=AMOT;refGene_tx=XM_006943935.4
X 94686317 94686462 refGene_gn=AMOT;refGene_tx=XM_006943935.4
X 94690648 94691059 refGene_gn=AMOT;refGene_tx=XM_006943935.4
X 94691060 94691127 refGene_gn=AMOT;refGene_tx=XM_006943935.4
X 94697631 94698503 refGene_gn=AMOT;refGene_tx=XM_006943935.4
X 96266186 96266221
refGene_gn=HTR2C;refGene_tx=XM_019824190.2;refGene_gn=HTR2C;refGene_tx
=XM_019824191.1
X 96271669 96271983
refGene_gn=HTR2C;refGene_tx=XM_019824190.2;refGene_gn=HTR2C;refGene_tx
=XM_019824191.1
X 96361984 96362185
refGene_gn=HTR2C;refGene_tx=XM_019824190.2;refGene_gn=HTR2C;refGene_tx
=XM_019824191.1
X 96427905 96428732
refGene_gn=HTR2C;refGene_tx=XM_019824190.2;refGene_gn=HTR2C;refGene_tx
=XM_019824191.1
X 96523412 96523471

refGene_gn=IL13RA2;refGene_tx=XM_006943941.3;refGene_gn=IL13RA2;refGene_tx=XM_006943942.3;refGene_gn=IL13RA2;refGene_tx=XM_019824192.2;refGene_gn=IL13RA2;refGene_tx=XM_019824194.2;refGene_gn=IL13RA2;refGene_tx=XM_019824195.2;refGene_gn=IL13RA2;refGene_tx=XM_019824196.2;refGene_gn=IL13RA2;refGene_tx=XM_023249143.1
X 96524416 96524532
refGene_gn=IL13RA2;refGene_tx=XM_006943941.3;refGene_gn=IL13RA2;refGene_tx=XM_006943942.3;refGene_gn=IL13RA2;refGene_tx=XM_019824192.2;refGene_gn=IL13RA2;refGene_tx=XM_019824194.2;refGene_gn=IL13RA2;refGene_tx=XM_019824195.2;refGene_gn=IL13RA2;refGene_tx=XM_023249143.1
X 96529711 96529856
refGene_gn=IL13RA2;refGene_tx=XM_006943941.3;refGene_gn=IL13RA2;refGene_tx=XM_006943942.3;refGene_gn=IL13RA2;refGene_tx=XM_019824192.2;refGene_gn=IL13RA2;refGene_tx=XM_019824194.2;refGene_gn=IL13RA2;refGene_tx=XM_019824195.2;refGene_gn=IL13RA2;refGene_tx=XM_019824196.2;refGene_gn=IL13RA2;refGene_tx=XM_023249143.1
X 96532040 96532186
refGene_gn=IL13RA2;refGene_tx=XM_006943941.3;refGene_gn=IL13RA2;refGene_tx=XM_006943942.3;refGene_gn=IL13RA2;refGene_tx=XM_019824192.2;refGene_gn=IL13RA2;refGene_tx=XM_019824194.2;refGene_gn=IL13RA2;refGene_tx=XM_019824195.2;refGene_gn=IL13RA2;refGene_tx=XM_019824196.2;refGene_gn=IL13RA2;refGene_tx=XM_023249143.1
X 96533103 96533288
refGene_gn=IL13RA2;refGene_tx=XM_006943941.3;refGene_gn=IL13RA2;refGene_tx=XM_006943942.3;refGene_gn=IL13RA2;refGene_tx=XM_019824192.2;refGene_gn=IL13RA2;refGene_tx=XM_019824194.2;refGene_gn=IL13RA2;refGene_tx=XM_019824195.2;refGene_gn=IL13RA2;refGene_tx=XM_019824196.2;refGene_gn=IL13RA2;refGene_tx=XM_023249143.1
X 96536891 96537012
refGene_gn=IL13RA2;refGene_tx=XM_006943941.3;refGene_gn=IL13RA2;refGene_tx=XM_006943942.3;refGene_gn=IL13RA2;refGene_tx=XM_019824192.2;refGene_gn=IL13RA2;refGene_tx=XM_019824194.2;refGene_gn=IL13RA2;refGene_tx=XM_019824195.2;refGene_gn=IL13RA2;refGene_tx=XM_019824196.2;refGene_gn=IL13RA2;refGene_tx=XM_023249143.1
X 96537525 96537679
refGene_gn=IL13RA2;refGene_tx=XM_006943941.3;refGene_gn=IL13RA2;refGene_tx=XM_006943942.3;refGene_gn=IL13RA2;refGene_tx=XM_019824192.2;refGene_gn=IL13RA2;refGene_tx=XM_019824194.2;refGene_gn=IL13RA2;refGene_tx=XM_019824195.2;refGene_gn=IL13RA2;refGene_tx=XM_019824196.2;refGene_gn=IL13RA2;refGene_tx=XM_023249143.1
X 96538708 96538860
refGene_gn=IL13RA2;refGene_tx=XM_006943941.3;refGene_gn=IL13RA2;refGene_tx=XM_006943942.3;refGene_gn=IL13RA2;refGene_tx=XM_019824192.2;refGene_gn=IL13RA2;refGene_tx=XM_019824194.2;refGene_gn=IL13RA2;refGene_tx=XM_019824195.2;refGene_gn=IL13RA2;refGene_tx=XM_019824196.2;refGene_gn=IL13RA2;refGene_tx=XM_023249143.1
X 96541628 96541719
refGene_gn=IL13RA2;refGene_tx=XM_006943941.3;refGene_gn=IL13RA2;refGene_tx=XM_006943942.3;refGene_gn=IL13RA2;refGene_tx=XM_019824192.2;refGene_gn=IL13RA2;refGene_tx=XM_019824194.2;refGene_gn=IL13RA2;refGene_tx=

XM_019824195.2;refGene_gn=IL13RA2;refGene_tx=XM_019824196.2;refGene_gn=IL13RA2;refGene_tx=XM_023249143.1
X 96628325 96628445
refGene_gn=LRCH2;refGene_tx=XM_004000812.4;refGene_gn=LRCH2;refGene_tx=XM_019824197.2;refGene_gn=LRCH2;refGene_tx=XM_023248878.1
X 96642361 96642439
refGene_gn=LRCH2;refGene_tx=XM_004000812.4;refGene_gn=LRCH2;refGene_tx=XM_019824197.2;refGene_gn=LRCH2;refGene_tx=XM_023248878.1
X 96642585 96642723
refGene_gn=LRCH2;refGene_tx=XM_004000812.4;refGene_gn=LRCH2;refGene_tx=XM_019824197.2;refGene_gn=LRCH2;refGene_tx=XM_023248878.1
X 96642907 96643020
refGene_gn=LRCH2;refGene_tx=XM_004000812.4;refGene_gn=LRCH2;refGene_tx=XM_019824197.2;refGene_gn=LRCH2;refGene_tx=XM_023248878.1
X 96643783 96643841
refGene_gn=LRCH2;refGene_tx=XM_004000812.4;refGene_gn=LRCH2;refGene_tx=XM_019824197.2;refGene_gn=LRCH2;refGene_tx=XM_023248878.1
X 96646267 96646318
refGene_gn=LRCH2;refGene_tx=XM_004000812.4;refGene_gn=LRCH2;refGene_tx=XM_023248878.1
X 96648563 96648608
refGene_gn=LRCH2;refGene_tx=XM_004000812.4;refGene_gn=LRCH2;refGene_tx=XM_019824197.2;refGene_gn=LRCH2;refGene_tx=XM_023248878.1
X 96655816 96655933
refGene_gn=LRCH2;refGene_tx=XM_004000812.4;refGene_gn=LRCH2;refGene_tx=XM_019824197.2;refGene_gn=LRCH2;refGene_tx=XM_023248878.1
X 96656011 96656060
refGene_gn=LRCH2;refGene_tx=XM_004000812.4;refGene_gn=LRCH2;refGene_tx=XM_019824197.2;refGene_gn=LRCH2;refGene_tx=XM_023248878.1
X 96663373 96663439
refGene_gn=LRCH2;refGene_tx=XM_004000812.4;refGene_gn=LRCH2;refGene_tx=XM_019824197.2;refGene_gn=LRCH2;refGene_tx=XM_023248878.1
X 96669354 96669462
refGene_gn=LRCH2;refGene_tx=XM_004000812.4;refGene_gn=LRCH2;refGene_tx=XM_019824197.2
X 96671257 96671316
refGene_gn=LRCH2;refGene_tx=XM_004000812.4;refGene_gn=LRCH2;refGene_tx=XM_019824197.2;refGene_gn=LRCH2;refGene_tx=XM_023248878.1
X 96671413 96671511
refGene_gn=LRCH2;refGene_tx=XM_004000812.4;refGene_gn=LRCH2;refGene_tx=XM_019824197.2;refGene_gn=LRCH2;refGene_tx=XM_023248878.1
X 96671970 96672082
refGene_gn=LRCH2;refGene_tx=XM_004000812.4;refGene_gn=LRCH2;refGene_tx=XM_019824197.2;refGene_gn=LRCH2;refGene_tx=XM_023248878.1
X 96672389 96672477
refGene_gn=LRCH2;refGene_tx=XM_004000812.4;refGene_gn=LRCH2;refGene_tx=XM_019824197.2;refGene_gn=LRCH2;refGene_tx=XM_023248878.1
X 96673672 96674654
refGene_gn=LOC102900824;refGene_tx=XM_011279388.3
X 96680162 96680296

refGene_gn=LRCH2;refGene_tx=XM_004000812.4;refGene_gn=LRCH2;refGene_tx=XM_019824197.2;refGene_gn=LRCH2;refGene_tx=XM_023248878.1
X 96684703 96684840 refGene_gn=LRCH2;refGene_tx=XM_004000812.4;refGene_gn=LRCH2;refGene_tx=XM_019824197.2;refGene_gn=LRCH2;refGene_tx=XM_023248878.1
X 96684923 96685029 refGene_gn=LRCH2;refGene_tx=XM_004000812.4;refGene_gn=LRCH2;refGene_tx=XM_019824197.2;refGene_gn=LRCH2;refGene_tx=XM_023248878.1
X 96689850 96689977 refGene_gn=LRCH2;refGene_tx=XM_004000812.4;refGene_gn=LRCH2;refGene_tx=XM_019824197.2;refGene_gn=LRCH2;refGene_tx=XM_023248878.1
X 96694170 96694315 refGene_gn=LRCH2;refGene_tx=XM_004000812.4;refGene_gn=LRCH2;refGene_tx=XM_019824197.2;refGene_gn=LRCH2;refGene_tx=XM_023248878.1
X 96726241 96726593 refGene_gn=LRCH2;refGene_tx=XM_004000812.4;refGene_gn=LRCH2;refGene_tx=XM_019824197.2;refGene_gn=LRCH2;refGene_tx=XM_023248878.1
X 96803969 96804054 refGene_gn=LUZP4;refGene_tx=XM_023248975.1
X 96811067 96811199 refGene_gn=LUZP4;refGene_tx=XM_023248975.1
X 96812350 96812469 refGene_gn=LUZP4;refGene_tx=XM_023248975.1
X 96818193 96818951 refGene_gn=LUZP4;refGene_tx=XM_023248975.1
X 96820123 96820677 refGene_gn=LUZP4;refGene_tx=XM_023248975.1
X 96820679 96820776 refGene_gn=LUZP4;refGene_tx=XM_023248975.1
X 96826885 96826967 refGene_gn=LUZP4;refGene_tx=XM_023248975.1
X 96835545 96835598 refGene_gn=LUZP4;refGene_tx=XM_023248975.1
X 96840212 96840333 refGene_gn=LUZP4;refGene_tx=XM_023248975.1
X 96841459 96841558 refGene_gn=LUZP4;refGene_tx=XM_023248975.1
X 97112226 97112299 refGene_gn=PLS3;refGene_tx=XM_006943944.3;refGene_gn=PLS3;refGene_tx=XM_019824201.1;refGene_gn=PLS3;refGene_tx=XM_019824202.2;refGene_gn=PLS3;refGene_tx=XM_019824203.1;refGene_gn=PLS3;refGene_tx=XM_019824204.2
X 97125395 97125559 refGene_gn=PLS3;refGene_tx=XM_006943944.3;refGene_gn=PLS3;refGene_tx=XM_019824201.1;refGene_gn=PLS3;refGene_tx=XM_019824202.2;refGene_gn=PLS3;refGene_tx=XM_019824203.1;refGene_gn=PLS3;refGene_tx=XM_019824204.2
X 97132049 97132179 refGene_gn=PLS3;refGene_tx=XM_006943944.3;refGene_gn=PLS3;refGene_tx=XM_019824201.1;refGene_gn=PLS3;refGene_tx=XM_019824202.2;refGene_gn=PLS3;refGene_tx=XM_019824203.1;refGene_gn=PLS3;refGene_tx=XM_019824204.2
X 97132650 97132783 refGene_gn=PLS3;refGene_tx=XM_006943944.3;refGene_gn=PLS3;refGene_tx=XM_019824201.1;refGene_gn=PLS3;refGene_tx=XM_019824202.2;refGene_gn=PLS3;refGene_tx=XM_019824203.1;refGene_gn=PLS3;refGene_tx=XM_019824204.2
X 97135821 97135903 refGene_gn=PLS3;refGene_tx=XM_006943944.3;refGene_gn=PLS3;refGene_tx=XM_019824201.1;refGene_gn=PLS3;refGene_tx=XM_019824202.2;refGene_gn=PLS3;refGene_tx=XM_019824203.1;refGene_gn=PLS3;refGene_tx=XM_019824204.2
X 97137250 97137416 refGene_gn=PLS3;refGene_tx=XM_006943944.3;refGene_gn=PLS3;refGene_tx=XM_019824201.1;refGene_gn=PLS3;refGene_tx=XM_019824202.2;refGene_gn=PLS

3;refGene_tx=XM_019824203.1;refGene_gn=PLS3;refGene_tx=XM_019824204.2
X 97140002 97140145
refGene_gn=PLS3;refGene_tx=XM_006943944.3;refGene_gn=PLS3;refGene_tx=XM_019824201.1;refGene_gn=PLS3;refGene_tx=XM_019824202.2;refGene_gn=PLS3;refGene_tx=XM_019824203.1;refGene_gn=PLS3;refGene_tx=XM_019824204.2
X 97142481 97142508
refGene_gn=PLS3;refGene_tx=XM_019824201.1;refGene_gn=PLS3;refGene_tx=XM_019824202.2;refGene_gn=PLS3;refGene_tx=XM_019824203.1;refGene_gn=PLS3;refGene_tx=XM_019824204.2
X 97142776 97142872
refGene_gn=PLS3;refGene_tx=XM_006943944.3;refGene_gn=PLS3;refGene_tx=XM_019824201.1;refGene_gn=PLS3;refGene_tx=XM_019824202.2;refGene_gn=PLS3;refGene_tx=XM_019824203.1;refGene_gn=PLS3;refGene_tx=XM_019824204.2
X 97147178 97147374
refGene_gn=PLS3;refGene_tx=XM_006943944.3;refGene_gn=PLS3;refGene_tx=XM_019824201.1;refGene_gn=PLS3;refGene_tx=XM_019824202.2;refGene_gn=PLS3;refGene_tx=XM_019824203.1;refGene_gn=PLS3;refGene_tx=XM_019824204.2
X 97148676 97148755
refGene_gn=PLS3;refGene_tx=XM_006943944.3;refGene_gn=PLS3;refGene_tx=XM_019824201.1;refGene_gn=PLS3;refGene_tx=XM_019824202.2;refGene_gn=PLS3;refGene_tx=XM_019824203.1;refGene_gn=PLS3;refGene_tx=XM_019824204.2
X 97150124 97150239
refGene_gn=PLS3;refGene_tx=XM_006943944.3;refGene_gn=PLS3;refGene_tx=XM_019824201.1;refGene_gn=PLS3;refGene_tx=XM_019824202.2;refGene_gn=PLS3;refGene_tx=XM_019824203.1;refGene_gn=PLS3;refGene_tx=XM_019824204.2
X 97150462 97150596
refGene_gn=PLS3;refGene_tx=XM_006943944.3;refGene_gn=PLS3;refGene_tx=XM_019824201.1;refGene_gn=PLS3;refGene_tx=XM_019824202.2;refGene_gn=PLS3;refGene_tx=XM_019824203.1;refGene_gn=PLS3;refGene_tx=XM_019824204.2
X 97152051 97152175
refGene_gn=PLS3;refGene_tx=XM_006943944.3;refGene_gn=PLS3;refGene_tx=XM_019824201.1;refGene_gn=PLS3;refGene_tx=XM_019824202.2;refGene_gn=PLS3;refGene_tx=XM_019824203.1;refGene_gn=PLS3;refGene_tx=XM_019824204.2
X 97152396 97152521
refGene_gn=PLS3;refGene_tx=XM_006943944.3;refGene_gn=PLS3;refGene_tx=XM_019824201.1;refGene_gn=PLS3;refGene_tx=XM_019824202.2;refGene_gn=PLS3;refGene_tx=XM_019824203.1;refGene_gn=PLS3;refGene_tx=XM_019824204.2
X 97155539 97155672
refGene_gn=PLS3;refGene_tx=XM_006943944.3;refGene_gn=PLS3;refGene_tx=XM_019824201.1;refGene_gn=PLS3;refGene_tx=XM_019824202.2;refGene_gn=PLS3;refGene_tx=XM_019824203.1;refGene_gn=PLS3;refGene_tx=XM_019824204.2
X 97344168 97345257
refGene_gn=AGTR2;refGene_tx=XM_019824205.1;refGene_gn=AGTR2;refGene_tx=XM_023249331.1
X 97554217 97554265 refGene_gn=SLC6A14;refGene_tx=XM_011279449.2
X 97555233 97555399 refGene_gn=SLC6A14;refGene_tx=XM_011279449.2
X 97558901 97559033 refGene_gn=SLC6A14;refGene_tx=XM_011279449.2
X 97560608 97560770 refGene_gn=SLC6A14;refGene_tx=XM_011279449.2
X 97561794 97561933 refGene_gn=SLC6A14;refGene_tx=XM_011279449.2
X 97563337 97563470 refGene_gn=SLC6A14;refGene_tx=XM_011279449.2

X	97565571 97565712 refGene_gn=SLC6A14;refGene_tx=XM_011279449.2
X	97569638 97569867 refGene_gn=SLC6A14;refGene_tx=XM_011279449.2
X	97571257 97571383 refGene_gn=SLC6A14;refGene_tx=XM_011279449.2
X	97572800 97572919 refGene_gn=SLC6A14;refGene_tx=XM_011279449.2
X	97573348 97573448 refGene_gn=SLC6A14;refGene_tx=XM_011279449.2
X	97573703 97573813 refGene_gn=SLC6A14;refGene_tx=XM_011279449.2
X	97576222 97576390 refGene_gn=SLC6A14;refGene_tx=XM_011279449.2
X	97577353 97577500 refGene_gn=SLC6A14;refGene_tx=XM_011279449.2
X	97579916 97580225 refGene_gn=CT83;refGene_tx=XM_006943948.3
X	97581388 97581463 refGene_gn=CT83;refGene_tx=XM_006943948.3
X	98679650 98680138
refGene_gn=KLHL13;refGene_tx=XM_004000821.5;refGene_gn=KLHL13;refGene_tx=XM_006943950.4;refGene_gn=KLHL13;refGene_tx=XM_011279448.3;refGene_gn=KLHL13;refGene_tx=XM_019824207.2;refGene_gn=KLHL13;refGene_tx=XM_019824208.2;refGene_gn=KLHL13;refGene_tx=XM_019824209.2;refGene_gn=KLHL13;refGene_tx=XM_019824210.2;refGene_gn=KLHL13;refGene_tx=XM_019824212.2;refGene_gn=KLHL13;refGene_tx=XM_023249500.1;refGene_gn=KLHL13;refGene_tx=XM_023249501.1
X	98682798 98682912
refGene_gn=KLHL13;refGene_tx=XM_004000821.5;refGene_gn=KLHL13;refGene_tx=XM_006943950.4;refGene_gn=KLHL13;refGene_tx=XM_011279448.3;refGene_gn=KLHL13;refGene_tx=XM_019824207.2;refGene_gn=KLHL13;refGene_tx=XM_019824208.2;refGene_gn=KLHL13;refGene_tx=XM_019824209.2;refGene_gn=KLHL13;refGene_tx=XM_019824210.2;refGene_gn=KLHL13;refGene_tx=XM_019824212.2;refGene_gn=KLHL13;refGene_tx=XM_023249500.1;refGene_gn=KLHL13;refGene_tx=XM_023249501.1
X	98689836 98690632
refGene_gn=KLHL13;refGene_tx=XM_004000821.5;refGene_gn=KLHL13;refGene_tx=XM_006943950.4;refGene_gn=KLHL13;refGene_tx=XM_011279448.3;refGene_gn=KLHL13;refGene_tx=XM_019824207.2;refGene_gn=KLHL13;refGene_tx=XM_019824208.2;refGene_gn=KLHL13;refGene_tx=XM_019824209.2;refGene_gn=KLHL13;refGene_tx=XM_019824210.2;refGene_gn=KLHL13;refGene_tx=XM_019824212.2;refGene_gn=KLHL13;refGene_tx=XM_023249500.1;refGene_gn=KLHL13;refGene_tx=XM_023249501.1
X	98702142 98702339
refGene_gn=KLHL13;refGene_tx=XM_004000821.5;refGene_gn=KLHL13;refGene_tx=XM_006943950.4;refGene_gn=KLHL13;refGene_tx=XM_011279448.3;refGene_gn=KLHL13;refGene_tx=XM_019824207.2;refGene_gn=KLHL13;refGene_tx=XM_019824208.2;refGene_gn=KLHL13;refGene_tx=XM_019824209.2;refGene_gn=KLHL13;refGene_tx=XM_019824210.2;refGene_gn=KLHL13;refGene_tx=XM_019824212.2;refGene_gn=KLHL13;refGene_tx=XM_023249500.1;refGene_gn=KLHL13;refGene_tx=XM_023249501.1
X	98703099 98703232
refGene_gn=KLHL13;refGene_tx=XM_004000821.5;refGene_gn=KLHL13;refGene_tx=XM_006943950.4;refGene_gn=KLHL13;refGene_tx=XM_011279448.3;refGene_gn=KLHL13;refGene_tx=XM_019824207.2;refGene_gn=KLHL13;refGene_tx=XM_019824208.2;refGene_gn=KLHL13;refGene_tx=XM_019824209.2;refGene_gn=KLHL13;refGene_tx=XM_019824210.2;refGene_gn=KLHL13;refGene_tx=XM_019824212.2;refGene_gn=KLHL13;refGene_tx=XM_023249500.1;refGene_gn=KLHL13;refGene_tx=XM_023249501.1

X 98722119 98722261
refGene_gn=KLHL13;refGene_tx=XM_004000821.5;refGene_gn=KLHL13;refGene_tx=XM_011279448.3;refGene_gn=KLHL13;refGene_tx=XM_019824207.2;refGene_gn=KLHL13;refGene_tx=XM_019824208.2;refGene_gn=KLHL13;refGene_tx=XM_019824209.2;refGene_gn=KLHL13;refGene_tx=XM_019824210.2;refGene_gn=KLHL13;refGene_tx=XM_019824212.2;refGene_gn=KLHL13;refGene_tx=XM_023249500.1;refGene_gn=KLHL13;refGene_tx=XM_023249501.1
X 98749446 98749544 refGene_gn=KLHL13;refGene_tx=XM_019824208.2
X 98750087 98750167 refGene_gn=KLHL13;refGene_tx=XM_019824209.2
X 98761454 98761520 refGene_gn=KLHL13;refGene_tx=XM_004000821.5
X 98763241 98763282 refGene_gn=KLHL13;refGene_tx=XM_004000821.5
X 98801313 98801366
refGene_gn=KLHL13;refGene_tx=XM_019824210.2;refGene_gn=KLHL13;refGene_tx=XM_023249500.1
X 99102260 99102337
refGene_gn=WDR44;refGene_tx=XM_011279445.3;refGene_gn=WDR44;refGene_tx=XM_011279447.3
X 99130626 99130660
refGene_gn=WDR44;refGene_tx=XM_011279445.3;refGene_gn=WDR44;refGene_tx=XM_011279447.3
X 99137250 99137325
refGene_gn=WDR44;refGene_tx=XM_011279445.3;refGene_gn=WDR44;refGene_tx=XM_011279447.3
X 99141997 99142634
refGene_gn=WDR44;refGene_tx=XM_011279445.3;refGene_gn=WDR44;refGene_tx=XM_011279447.3
X 99143126 99143257
refGene_gn=WDR44;refGene_tx=XM_011279445.3;refGene_gn=WDR44;refGene_tx=XM_011279447.3
X 99144589 99144685
refGene_gn=WDR44;refGene_tx=XM_011279445.3;refGene_gn=WDR44;refGene_tx=XM_011279447.3
X 99146791 99146928
refGene_gn=WDR44;refGene_tx=XM_011279445.3;refGene_gn=WDR44;refGene_tx=XM_011279447.3
X 99147963 99148047
refGene_gn=WDR44;refGene_tx=XM_011279445.3;refGene_gn=WDR44;refGene_tx=XM_011279447.3
X 99151438 99151545
refGene_gn=WDR44;refGene_tx=XM_011279445.3;refGene_gn=WDR44;refGene_tx=XM_011279447.3
X 99152521 99152673
refGene_gn=WDR44;refGene_tx=XM_011279445.3;refGene_gn=WDR44;refGene_tx=XM_011279447.3
X 99154903 99155042
refGene_gn=WDR44;refGene_tx=XM_011279445.3;refGene_gn=WDR44;refGene_tx=XM_011279447.3
X 99156395 99156460
refGene_gn=WDR44;refGene_tx=XM_011279445.3;refGene_gn=WDR44;refGene_tx=XM_011279447.3

X 99166702 99166816
refGene_gn=WDR44;refGene_tx=XM_011279445.3;refGene_gn=WDR44;refGene_tx=XM_011279447.3
X 99170661 99170784
refGene_gn=WDR44;refGene_tx=XM_011279445.3;refGene_gn=WDR44;refGene_tx=XM_011279447.3
X 99174795 99174987
refGene_gn=WDR44;refGene_tx=XM_011279445.3;refGene_gn=WDR44;refGene_tx=XM_011279447.3
X 99175169 99175271
refGene_gn=WDR44;refGene_tx=XM_011279445.3;refGene_gn=WDR44;refGene_tx=XM_011279447.3
X 99175744 99175860
refGene_gn=WDR44;refGene_tx=XM_011279445.3;refGene_gn=WDR44;refGene_tx=XM_011279447.3
X 99176718 99176846
refGene_gn=WDR44;refGene_tx=XM_011279445.3;refGene_gn=WDR44;refGene_tx=XM_011279447.3
X 99177858 99177993
refGene_gn=WDR44;refGene_tx=XM_011279445.3;refGene_gn=WDR44;refGene_tx=XM_011279447.3
X 99180910 99181005
refGene_gn=WDR44;refGene_tx=XM_011279445.3;refGene_gn=WDR44;refGene_tx=XM_011279447.3
X 99233955 99234057
refGene_gn=DOCK11;refGene_tx=XM_023249251.1;refGene_gn=DOCK11;refGene_tx=XM_023249252.1;refGene_gn=DOCK11;refGene_tx=XM_023249253.1
X 99284992 99285109
refGene_gn=DOCK11;refGene_tx=XM_023249251.1;refGene_gn=DOCK11;refGene_tx=XM_023249252.1;refGene_gn=DOCK11;refGene_tx=XM_023249253.1
X 99285421 99285511
refGene_gn=DOCK11;refGene_tx=XM_023249251.1;refGene_gn=DOCK11;refGene_tx=XM_023249252.1;refGene_gn=DOCK11;refGene_tx=XM_023249253.1
X 99285920 99286003
refGene_gn=DOCK11;refGene_tx=XM_023249251.1;refGene_gn=DOCK11;refGene_tx=XM_023249252.1;refGene_gn=DOCK11;refGene_tx=XM_023249253.1
X 99287266 99287336
refGene_gn=DOCK11;refGene_tx=XM_023249251.1;refGene_gn=DOCK11;refGene_tx=XM_023249252.1;refGene_gn=DOCK11;refGene_tx=XM_023249253.1
X 99288044 99288140
refGene_gn=DOCK11;refGene_tx=XM_023249251.1;refGene_gn=DOCK11;refGene_tx=XM_023249252.1;refGene_gn=DOCK11;refGene_tx=XM_023249253.1
X 99301760 99301895
refGene_gn=DOCK11;refGene_tx=XM_023249251.1;refGene_gn=DOCK11;refGene_tx=XM_023249252.1;refGene_gn=DOCK11;refGene_tx=XM_023249253.1
X 99304319 99304497
refGene_gn=DOCK11;refGene_tx=XM_023249251.1;refGene_gn=DOCK11;refGene_tx=XM_023249252.1;refGene_gn=DOCK11;refGene_tx=XM_023249253.1
X 99304916 99304996
refGene_gn=DOCK11;refGene_tx=XM_023249251.1;refGene_gn=DOCK11;refGene_

tx=XM_023249252.1;refGene_gn=DOCK11;refGene_tx=XM_023249253.1
X	99306360	99306444
refGene_gn=DOCK11;refGene_tx=XM_023249251.1;refGene_gn=DOCK11;refGene_tx=XM_023249252.1;refGene_gn=DOCK11;refGene_tx=XM_023249253.1
X	99310674	99310815
refGene_gn=DOCK11;refGene_tx=XM_023249251.1;refGene_gn=DOCK11;refGene_tx=XM_023249252.1;refGene_gn=DOCK11;refGene_tx=XM_023249253.1
X	99312233	99312446
refGene_gn=DOCK11;refGene_tx=XM_023249251.1;refGene_gn=DOCK11;refGene_tx=XM_023249252.1;refGene_gn=DOCK11;refGene_tx=XM_023249253.1
X	99320185	99320308
refGene_gn=DOCK11;refGene_tx=XM_023249251.1;refGene_gn=DOCK11;refGene_tx=XM_023249252.1;refGene_gn=DOCK11;refGene_tx=XM_023249253.1
X	99321603	99321686
refGene_gn=DOCK11;refGene_tx=XM_023249251.1;refGene_gn=DOCK11;refGene_tx=XM_023249252.1;refGene_gn=DOCK11;refGene_tx=XM_023249253.1
X	99324700	99324823
refGene_gn=DOCK11;refGene_tx=XM_023249251.1;refGene_gn=DOCK11;refGene_tx=XM_023249252.1;refGene_gn=DOCK11;refGene_tx=XM_023249253.1
X	99325023	99325100
refGene_gn=DOCK11;refGene_tx=XM_023249251.1;refGene_gn=DOCK11;refGene_tx=XM_023249252.1;refGene_gn=DOCK11;refGene_tx=XM_023249253.1
X	99327370	99327555
refGene_gn=DOCK11;refGene_tx=XM_023249251.1;refGene_gn=DOCK11;refGene_tx=XM_023249252.1;refGene_gn=DOCK11;refGene_tx=XM_023249253.1
X	99327648	99327714
refGene_gn=DOCK11;refGene_tx=XM_023249251.1;refGene_gn=DOCK11;refGene_tx=XM_023249252.1;refGene_gn=DOCK11;refGene_tx=XM_023249253.1
X	99329380	99329473
refGene_gn=DOCK11;refGene_tx=XM_023249251.1;refGene_gn=DOCK11;refGene_tx=XM_023249252.1;refGene_gn=DOCK11;refGene_tx=XM_023249253.1
X	99332411	99332535
refGene_gn=DOCK11;refGene_tx=XM_023249251.1;refGene_gn=DOCK11;refGene_tx=XM_023249252.1;refGene_gn=DOCK11;refGene_tx=XM_023249253.1
X	99336090	99336212
refGene_gn=DOCK11;refGene_tx=XM_023249251.1;refGene_gn=DOCK11;refGene_tx=XM_023249252.1;refGene_gn=DOCK11;refGene_tx=XM_023249253.1
X	99336898	99336985
refGene_gn=DOCK11;refGene_tx=XM_023249251.1;refGene_gn=DOCK11;refGene_tx=XM_023249252.1;refGene_gn=DOCK11;refGene_tx=XM_023249253.1
X	99338799	99338889
refGene_gn=DOCK11;refGene_tx=XM_023249251.1;refGene_gn=DOCK11;refGene_tx=XM_023249252.1;refGene_gn=DOCK11;refGene_tx=XM_023249253.1
X	99344685	99344804
refGene_gn=DOCK11;refGene_tx=XM_023249251.1;refGene_gn=DOCK11;refGene_tx=XM_023249252.1;refGene_gn=DOCK11;refGene_tx=XM_023249253.1
X	99346599	99346669
refGene_gn=DOCK11;refGene_tx=XM_023249251.1;refGene_gn=DOCK11;refGene_tx=XM_023249252.1;refGene_gn=DOCK11;refGene_tx=XM_023249253.1
X	99346751	99346877

refGene_gn=DOCK11;refGene_tx=XM_023249251.1;refGene_gn=DOCK11;refGene_tx=XM_023249252.1;refGene_gn=DOCK11;refGene_tx=XM_023249253.1
X 99347464 99347536
refGene_gn=DOCK11;refGene_tx=XM_023249251.1;refGene_gn=DOCK11;refGene_tx=XM_023249252.1;refGene_gn=DOCK11;refGene_tx=XM_023249253.1
X 99348824 99348971
refGene_gn=DOCK11;refGene_tx=XM_023249251.1;refGene_gn=DOCK11;refGene_tx=XM_023249252.1;refGene_gn=DOCK11;refGene_tx=XM_023249253.1
X 99352640 99352724
refGene_gn=DOCK11;refGene_tx=XM_023249251.1;refGene_gn=DOCK11;refGene_tx=XM_023249252.1;refGene_gn=DOCK11;refGene_tx=XM_023249253.1
X 99353563 99353675
refGene_gn=DOCK11;refGene_tx=XM_023249251.1;refGene_gn=DOCK11;refGene_tx=XM_023249252.1;refGene_gn=DOCK11;refGene_tx=XM_023249253.1
X 99354187 99354226
refGene_gn=DOCK11;refGene_tx=XM_023249251.1;refGene_gn=DOCK11;refGene_tx=XM_023249253.1
X 99355883 99356062
refGene_gn=DOCK11;refGene_tx=XM_023249251.1;refGene_gn=DOCK11;refGene_tx=XM_023249252.1;refGene_gn=DOCK11;refGene_tx=XM_023249253.1
X 99362693 99362810
refGene_gn=DOCK11;refGene_tx=XM_023249251.1;refGene_gn=DOCK11;refGene_tx=XM_023249252.1;refGene_gn=DOCK11;refGene_tx=XM_023249253.1
X 99364872 99364948
refGene_gn=DOCK11;refGene_tx=XM_023249251.1;refGene_gn=DOCK11;refGene_tx=XM_023249252.1;refGene_gn=DOCK11;refGene_tx=XM_023249253.1
X 99365551 99365661
refGene_gn=DOCK11;refGene_tx=XM_023249251.1;refGene_gn=DOCK11;refGene_tx=XM_023249252.1;refGene_gn=DOCK11;refGene_tx=XM_023249253.1
X 99367694 99367806
refGene_gn=DOCK11;refGene_tx=XM_023249251.1;refGene_gn=DOCK11;refGene_tx=XM_023249252.1;refGene_gn=DOCK11;refGene_tx=XM_023249253.1
X 99374251 99374318
refGene_gn=DOCK11;refGene_tx=XM_023249251.1;refGene_gn=DOCK11;refGene_tx=XM_023249252.1;refGene_gn=DOCK11;refGene_tx=XM_023249253.1
X 99376121 99376169
refGene_gn=DOCK11;refGene_tx=XM_023249251.1;refGene_gn=DOCK11;refGene_tx=XM_023249252.1;refGene_gn=DOCK11;refGene_tx=XM_023249253.1
X 99377661 99377804
refGene_gn=DOCK11;refGene_tx=XM_023249251.1;refGene_gn=DOCK11;refGene_tx=XM_023249252.1;refGene_gn=DOCK11;refGene_tx=XM_023249253.1
X 99379411 99379527
refGene_gn=DOCK11;refGene_tx=XM_023249251.1;refGene_gn=DOCK11;refGene_tx=XM_023249252.1;refGene_gn=DOCK11;refGene_tx=XM_023249253.1
X 99382084 99382222
refGene_gn=DOCK11;refGene_tx=XM_023249251.1;refGene_gn=DOCK11;refGene_tx=XM_023249252.1;refGene_gn=DOCK11;refGene_tx=XM_023249253.1
X 99386855 99387038
refGene_gn=DOCK11;refGene_tx=XM_023249251.1;refGene_gn=DOCK11;refGene_tx=XM_023249252.1;refGene_gn=DOCK11;refGene_tx=XM_023249253.1

X 99389020 99389134
refGene_gn=DOCK11;refGene_tx=XM_023249251.1;refGene_gn=DOCK11;refGene_tx=XM_023249252.1;refGene_gn=DOCK11;refGene_tx=XM_023249253.1
X 99391868 99392084
refGene_gn=DOCK11;refGene_tx=XM_023249251.1;refGene_gn=DOCK11;refGene_tx=XM_023249252.1;refGene_gn=DOCK11;refGene_tx=XM_023249253.1
X 99392174 99392232
refGene_gn=DOCK11;refGene_tx=XM_023249251.1;refGene_gn=DOCK11;refGene_tx=XM_023249252.1;refGene_gn=DOCK11;refGene_tx=XM_023249253.1
X 99403002 99403109
refGene_gn=DOCK11;refGene_tx=XM_023249251.1;refGene_gn=DOCK11;refGene_tx=XM_023249252.1;refGene_gn=DOCK11;refGene_tx=XM_023249253.1
X 99410748 99410871
refGene_gn=DOCK11;refGene_tx=XM_023249251.1;refGene_gn=DOCK11;refGene_tx=XM_023249252.1;refGene_gn=DOCK11;refGene_tx=XM_023249253.1
X 99416650 99416764
refGene_gn=DOCK11;refGene_tx=XM_023249251.1;refGene_gn=DOCK11;refGene_tx=XM_023249252.1;refGene_gn=DOCK11;refGene_tx=XM_023249253.1
X 99417299 99417446
refGene_gn=DOCK11;refGene_tx=XM_023249251.1;refGene_gn=DOCK11;refGene_tx=XM_023249252.1;refGene_gn=DOCK11;refGene_tx=XM_023249253.1
X 99426539 99426750
refGene_gn=DOCK11;refGene_tx=XM_023249251.1;refGene_gn=DOCK11;refGene_tx=XM_023249252.1;refGene_gn=DOCK11;refGene_tx=XM_023249253.1
X 99427133 99427324
refGene_gn=DOCK11;refGene_tx=XM_023249251.1;refGene_gn=DOCK11;refGene_tx=XM_023249252.1;refGene_gn=DOCK11;refGene_tx=XM_023249253.1
X 99428147 99428248
refGene_gn=DOCK11;refGene_tx=XM_023249251.1;refGene_gn=DOCK11;refGene_tx=XM_023249252.1;refGene_gn=DOCK11;refGene_tx=XM_023249253.1
X 99429778 99429917
refGene_gn=DOCK11;refGene_tx=XM_023249251.1;refGene_gn=DOCK11;refGene_tx=XM_023249252.1;refGene_gn=DOCK11;refGene_tx=XM_023249253.1
X 99434369 99434375 refGene_gn=DOCK11;refGene_tx=XM_023249253.1
X 99434544 99434667
refGene_gn=DOCK11;refGene_tx=XM_023249251.1;refGene_gn=DOCK11;refGene_tx=XM_023249252.1
X 99469166 99469254 refGene_gn=IL13RA1;refGene_tx=XM_023248883.1
X 99476709 99476849 refGene_gn=IL13RA1;refGene_tx=XM_023248883.1
X 99483044 99483183 refGene_gn=IL13RA1;refGene_tx=XM_023248883.1
X 99485332 99485453 refGene_gn=IL13RA1;refGene_tx=XM_023248883.1
X 99495928 99496116 refGene_gn=IL13RA1;refGene_tx=XM_023248883.1
X 99498831 99498983 refGene_gn=IL13RA1;refGene_tx=XM_023248883.1
X 99505678 99505726 refGene_gn=IL13RA1;refGene_tx=XM_023248883.1
X 99506014 99506147 refGene_gn=IL13RA1;refGene_tx=XM_023248883.1
X 99509362 99509459 refGene_gn=IL13RA1;refGene_tx=XM_023248883.1
X 99511455 99511540 refGene_gn=IL13RA1;refGene_tx=XM_023248883.1
X 99525574 99525667 refGene_gn=IL13RA1;refGene_tx=XM_023248883.1
X 99555579 99556788 refGene_gn=ZCCHC12;refGene_tx=XM_004000859.4
X 99660493 99660643

refGene_gn=LONRF3;refGene_tx=XM_023249525.1;refGene_gn=LONRF3;refGene_tx=XM_023249526.1
X	99660850	99661634
refGene_gn=LONRF3;refGene_tx=XM_023249521.1;refGene_gn=LONRF3;refGene_tx=XM_023249522.1;refGene_gn=LONRF3;refGene_tx=XM_023249523.1;refGene_gn=LONRF3;refGene_tx=XM_023249525.1;refGene_gn=LONRF3;refGene_tx=XM_023249526.1
X	99665441	99665560
refGene_gn=LONRF3;refGene_tx=XM_023249521.1;refGene_gn=LONRF3;refGene_tx=XM_023249522.1;refGene_gn=LONRF3;refGene_tx=XM_023249523.1;refGene_gn=LONRF3;refGene_tx=XM_023249525.1;refGene_gn=LONRF3;refGene_tx=XM_023249526.1
X	99669298	99669421
refGene_gn=LONRF3;refGene_tx=XM_023249521.1;refGene_gn=LONRF3;refGene_tx=XM_023249522.1;refGene_gn=LONRF3;refGene_tx=XM_023249523.1;refGene_gn=LONRF3;refGene_tx=XM_023249525.1
X	99673285	99673441
refGene_gn=LONRF3;refGene_tx=XM_023249521.1;refGene_gn=LONRF3;refGene_tx=XM_023249522.1;refGene_gn=LONRF3;refGene_tx=XM_023249523.1
X	99675455	99675705
refGene_gn=LONRF3;refGene_tx=XM_023249521.1;refGene_gn=LONRF3;refGene_tx=XM_023249522.1;refGene_gn=LONRF3;refGene_tx=XM_023249523.1;refGene_gn=LONRF3;refGene_tx=XM_023249525.1;refGene_gn=LONRF3;refGene_tx=XM_023249526.1
X	99676478	99676569
refGene_gn=LONRF3;refGene_tx=XM_023249521.1;refGene_gn=LONRF3;refGene_tx=XM_023249522.1;refGene_gn=LONRF3;refGene_tx=XM_023249523.1;refGene_gn=LONRF3;refGene_tx=XM_023249525.1;refGene_gn=LONRF3;refGene_tx=XM_023249526.1
X	99685278	99685393
refGene_gn=LONRF3;refGene_tx=XM_023249521.1;refGene_gn=LONRF3;refGene_tx=XM_023249522.1;refGene_gn=LONRF3;refGene_tx=XM_023249523.1;refGene_gn=LONRF3;refGene_tx=XM_023249525.1;refGene_gn=LONRF3;refGene_tx=XM_023249526.1
X	99687593	99687715
refGene_gn=LONRF3;refGene_tx=XM_023249521.1;refGene_gn=LONRF3;refGene_tx=XM_023249522.1;refGene_gn=LONRF3;refGene_tx=XM_023249523.1;refGene_gn=LONRF3;refGene_tx=XM_023249525.1;refGene_gn=LONRF3;refGene_tx=XM_023249526.1
X	99689581	99689740
refGene_gn=LONRF3;refGene_tx=XM_023249521.1;refGene_gn=LONRF3;refGene_tx=XM_023249522.1;refGene_gn=LONRF3;refGene_tx=XM_023249523.1;refGene_gn=LONRF3;refGene_tx=XM_023249525.1;refGene_gn=LONRF3;refGene_tx=XM_023249526.1
X	99690393	99690556
refGene_gn=LONRF3;refGene_tx=XM_023249521.1;refGene_gn=LONRF3;refGene_tx=XM_023249522.1;refGene_gn=LONRF3;refGene_tx=XM_023249523.1;refGene_gn=LONRF3;refGene_tx=XM_023249525.1;refGene_gn=LONRF3;refGene_tx=XM_023249526.1
X	99691571	99691721

refGene_gn=LONRF3;refGene_tx=XM_023249521.1;refGene_gn=LONRF3;refGene_tx=XM_023249522.1;refGene_gn=LONRF3;refGene_tx=XM_023249523.1;refGene_gn=LONRF3;refGene_tx=XM_023249525.1;refGene_gn=LONRF3;refGene_tx=XM_023249526.1
X	99694567	99694744	refGene_gn=LONRF3;refGene_tx=XM_023249521.1;refGene_gn=LONRF3;refGene_tx=XM_023249522.1;refGene_gn=LONRF3;refGene_tx=XM_023249525.1;refGene_gn=LONRF3;refGene_tx=XM_023249526.1
X	99695727	99695733	refGene_gn=LONRF3;refGene_tx=XM_023249523.1
X	99739519	99740665	refGene_gn=LOC111558681;refGene_tx=XM_023248976.1
X	99740987	99741495	refGene_gn=LOC105259672;refGene_tx=XM_011279444.3
X	99742396	99742497	refGene_gn=LOC105259672;refGene_tx=XM_011279444.3
X	99751418	99751438	refGene_gn=KIAA1210;refGene_tx=XM_023248977.1
X	99753513	99753617	refGene_gn=KIAA1210;refGene_tx=XM_023248977.1
X	99754626	99754966	refGene_gn=KIAA1210;refGene_tx=XM_023248977.1
X	99754968	99758367	refGene_gn=KIAA1210;refGene_tx=XM_023248977.1
X	99760545	99760654	refGene_gn=KIAA1210;refGene_tx=XM_023248977.1
X	99770112	99770265	refGene_gn=KIAA1210;refGene_tx=XM_023248977.1
X	99774608	99774746	refGene_gn=KIAA1210;refGene_tx=XM_023248977.1
X	99775063	99775181	refGene_gn=KIAA1210;refGene_tx=XM_023248977.1
X	99777375	99777541	refGene_gn=KIAA1210;refGene_tx=XM_023248977.1
X	99783662	99783733	refGene_gn=KIAA1210;refGene_tx=XM_023248977.1
X	99803843	99804051	refGene_gn=KIAA1210;refGene_tx=XM_023248977.1
X	99806250	99806296	refGene_gn=KIAA1210;refGene_tx=XM_023248977.1
X	99893066	99893394	refGene_gn=PGRMC1;refGene_tx=XM_023248927.1
X	99896787	99896943	refGene_gn=PGRMC1;refGene_tx=XM_023248927.1
X	99900285	99900389	refGene_gn=PGRMC1;refGene_tx=XM_023248927.1
X	99907916	99909557	refGene_gn=LOC101092340;refGene_tx=XM_023249040.1
X	99911371	99911606	refGene_gn=LOC101092340;refGene_tx=XM_023249040.1
X	99913861	99913992	refGene_gn=LOC101092340;refGene_tx=XM_023249040.1
X	99915465	99916218	refGene_gn=LOC101092340;refGene_tx=XM_023249040.1
X	100039634	100039909	refGene_gn=SLC25A43;refGene_tx=XM_023248805.1
X	100047390	100047632	refGene_gn=SLC25A43;refGene_tx=XM_023248805.1
X	100050250	100050423	refGene_gn=SLC25A43;refGene_tx=XM_023248805.1
X	100071171	100071306	refGene_gn=SLC25A43;refGene_tx=XM_023248805.1
X	100071910	100072111	refGene_gn=SLC25A43;refGene_tx=XM_023248805.1
X	100083311	100083422	refGene_gn=SLC25A5;refGene_tx=XM_011279441.2

X	100084464	100084951
refGene_gn=SLC25A5;refGene_tx=XM_011279441.2
X	100085168	100085309
refGene_gn=SLC25A5;refGene_tx=XM_011279441.2
X	100085971	100086129
refGene_gn=SLC25A5;refGene_tx=XM_011279441.2
X	100143616	100143691
refGene_gn=CXHXorf56;refGene_tx=XM_006943958.3
X	100144152	100144215
refGene_gn=CXHXorf56;refGene_tx=XM_004000825.4;refGene_gn=CXHXorf56;refGene_tx=XM_006943959.2;refGene_gn=CXHXorf56;refGene_tx=XM_019824215.1
X	100145624	100145717
refGene_gn=CXHXorf56;refGene_tx=XM_004000825.4;refGene_gn=CXHXorf56;refGene_tx=XM_006943958.3;refGene_gn=CXHXorf56;refGene_tx=XM_006943959.2;refGene_gn=CXHXorf56;refGene_tx=XM_019824215.1
X	100146323	100146413
refGene_gn=CXHXorf56;refGene_tx=XM_004000825.4;refGene_gn=CXHXorf56;refGene_tx=XM_006943958.3;refGene_gn=CXHXorf56;refGene_tx=XM_006943959.2;refGene_gn=CXHXorf56;refGene_tx=XM_019824215.1
X	100147275	100147414
refGene_gn=CXHXorf56;refGene_tx=XM_004000825.4;refGene_gn=CXHXorf56;refGene_tx=XM_006943958.3;refGene_gn=CXHXorf56;refGene_tx=XM_006943959.2;refGene_gn=CXHXorf56;refGene_tx=XM_019824215.1
X	100150097	100150139
refGene_gn=CXHXorf56;refGene_tx=XM_004000825.4;refGene_gn=CXHXorf56;refGene_tx=XM_006943958.3;refGene_gn=CXHXorf56;refGene_tx=XM_006943959.2
X	100161243	100161333
refGene_gn=CXHXorf56;refGene_tx=XM_006943958.3;refGene_gn=CXHXorf56;refGene_tx=XM_006943959.2
X	100163905	100164023
refGene_gn=CXHXorf56;refGene_tx=XM_004000825.4;refGene_gn=CXHXorf56;refGene_tx=XM_006943958.3;refGene_gn=CXHXorf56;refGene_tx=XM_006943959.2;refGene_gn=CXHXorf56;refGene_tx=XM_019824215.1
X	100167221	100167345
refGene_gn=CXHXorf56;refGene_tx=XM_004000825.4;refGene_gn=CXHXorf56;refGene_tx=XM_006943958.3;refGene_gn=CXHXorf56;refGene_tx=XM_006943959.2;refGene_gn=CXHXorf56;refGene_tx=XM_019824215.1
X	100177321	100177365
refGene_gn=UBE2A;refGene_tx=XM_004000827.5
X	100177504	100177585
refGene_gn=UBE2A;refGene_tx=XM_004000827.5
X	100178000	100178026
refGene_gn=UBE2A;refGene_tx=XM_004000827.5
X	100183684	100183774
refGene_gn=UBE2A;refGene_tx=XM_004000827.5
X	100184793	100184882
refGene_gn=UBE2A;refGene_tx=XM_004000827.5
X	100185158	100185287
refGene_gn=UBE2A;refGene_tx=XM_004000827.5
X	100190373	100192331

refGene_gn=NKRF;refGene_tx=XM_004000828.5
X 100193401 100193532
refGene_gn=NKRF;refGene_tx=XM_004000828.5
X 100204923 100205189
refGene_gn=NKRF;refGene_tx=XM_004000828.5
X 100213897 100213907
refGene_gn=SEPT6;refGene_tx=XM_019824218.2
X 100215960 100215985
refGene_gn=SEPT6;refGene_tx=XM_006943961.4;refGene_gn=SEPT6;refGene_tx=XM_019824216.2;refGene_gn=SEPT6;refGene_tx=XM_019824222.2
X 100217188 100217204
refGene_gn=SEPT6;refGene_tx=XM_019824217.1
X 100220748 100220752
refGene_gn=SEPT6;refGene_tx=XM_019824219.1;refGene_gn=SEPT6;refGene_tx=XM_019824221.2
X 100223215 100223406
refGene_gn=SEPT6;refGene_tx=XM_006943961.4;refGene_gn=SEPT6;refGene_tx=XM_019824216.2;refGene_gn=SEPT6;refGene_tx=XM_019824217.1;refGene_gn=SEPT6;refGene_tx=XM_019824218.2;refGene_gn=SEPT6;refGene_tx=XM_019824219.1;refGene_gn=SEPT6;refGene_tx=XM_019824221.2;refGene_gn=SEPT6;refGene_tx=XM_019824222.2
X 100226289 100226422
refGene_gn=SEPT6;refGene_tx=XM_006943961.4;refGene_gn=SEPT6;refGene_tx=XM_019824216.2;refGene_gn=SEPT6;refGene_tx=XM_019824217.1;refGene_gn=SEPT6;refGene_tx=XM_019824218.2;refGene_gn=SEPT6;refGene_tx=XM_019824219.1;refGene_gn=SEPT6;refGene_tx=XM_019824221.2;refGene_gn=SEPT6;refGene_tx=XM_019824222.2
X 100229429 100229598
refGene_gn=SEPT6;refGene_tx=XM_006943961.4;refGene_gn=SEPT6;refGene_tx=XM_019824216.2;refGene_gn=SEPT6;refGene_tx=XM_019824217.1;refGene_gn=SEPT6;refGene_tx=XM_019824218.2;refGene_gn=SEPT6;refGene_tx=XM_019824219.1;refGene_gn=SEPT6;refGene_tx=XM_019824221.2;refGene_gn=SEPT6;refGene_tx=XM_019824222.2
X 100233102 100233199
refGene_gn=SEPT6;refGene_tx=XM_006943961.4;refGene_gn=SEPT6;refGene_tx=XM_019824216.2;refGene_gn=SEPT6;refGene_tx=XM_019824217.1;refGene_gn=SEPT6;refGene_tx=XM_019824218.2;refGene_gn=SEPT6;refGene_tx=XM_019824219.1;refGene_gn=SEPT6;refGene_tx=XM_019824221.2;refGene_gn=SEPT6;refGene_tx=XM_019824222.2
X 100240891 100241053
refGene_gn=SEPT6;refGene_tx=XM_006943961.4;refGene_gn=SEPT6;refGene_tx=XM_019824216.2;refGene_gn=SEPT6;refGene_tx=XM_019824217.1;refGene_gn=SEPT6;refGene_tx=XM_019824218.2;refGene_gn=SEPT6;refGene_tx=XM_019824219.1;refGene_gn=SEPT6;refGene_tx=XM_019824221.2;refGene_gn=SEPT6;refGene_tx=XM_019824222.2
X 100244027 100244214
refGene_gn=SEPT6;refGene_tx=XM_006943961.4;refGene_gn=SEPT6;refGene_tx=XM_019824216.2;refGene_gn=SEPT6;refGene_tx=XM_019824217.1;refGene_gn=SEPT6;refGene_tx=XM_019824218.2;refGene_gn=SEPT6;refGene_tx=XM_019824219.1;refGene_gn=SEPT6;refGene_tx=XM_019824221.2;refGene_gn=SEPT6;refGe

ne_tx=XM_019824222.2
X	100257746	100257942
refGene_gn=SEPT6;refGene_tx=XM_006943961.4;refGene_gn=SEPT6;refGene_tx=XM_019824216.2;refGene_gn=SEPT6;refGene_tx=XM_019824217.1;refGene_gn=SEPT6;refGene_tx=XM_019824218.2;refGene_gn=SEPT6;refGene_tx=XM_019824219.1;refGene_gn=SEPT6;refGene_tx=XM_019824221.2;refGene_gn=SEPT6;refGene_tx=XM_019824222.2
X	100265893	100266008
refGene_gn=SEPT6;refGene_tx=XM_006943961.4;refGene_gn=SEPT6;refGene_tx=XM_019824216.2;refGene_gn=SEPT6;refGene_tx=XM_019824217.1;refGene_gn=SEPT6;refGene_tx=XM_019824218.2;refGene_gn=SEPT6;refGene_tx=XM_019824219.1;refGene_gn=SEPT6;refGene_tx=XM_019824221.2;refGene_gn=SEPT6;refGene_tx=XM_019824222.2
X	100270845	100270881
refGene_gn=SEPT6;refGene_tx=XM_006943961.4
X	100274809	100274869
refGene_gn=SEPT6;refGene_tx=XM_019824216.2;refGene_gn=SEPT6;refGene_tx=XM_019824217.1;refGene_gn=SEPT6;refGene_tx=XM_019824218.2;refGene_gn=SEPT6;refGene_tx=XM_019824219.1;refGene_gn=SEPT6;refGene_tx=XM_019824221.2
X	100280586	100280616
refGene_gn=SEPT6;refGene_tx=XM_019824222.2
X	100355265	100356222
refGene_gn=SOWAHD;refGene_tx=XM_023249098.1
X	100382491	100382540
refGene_gn=RPL39;refGene_tx=XM_006943968.3
X	100383868	100383972
refGene_gn=RPL39;refGene_tx=XM_006943968.3
X	100385935	100385938
refGene_gn=RPL39;refGene_tx=XM_006943968.3
X	100403094	100403244
refGene_gn=UPF3B;refGene_tx=XM_004000835.5;refGene_gn=UPF3B;refGene_tx=XM_004000836.5
X	100405942	100406243
refGene_gn=UPF3B;refGene_tx=XM_004000835.5;refGene_gn=UPF3B;refGene_tx=XM_004000836.5
X	100406574	100406735
refGene_gn=UPF3B;refGene_tx=XM_004000835.5;refGene_gn=UPF3B;refGene_tx=XM_004000836.5
X	100408611	100408650
refGene_gn=UPF3B;refGene_tx=XM_004000836.5
X	100409052	100409235
refGene_gn=UPF3B;refGene_tx=XM_004000835.5;refGene_gn=UPF3B;refGene_tx=XM_004000836.5
X	100409706	100409750
refGene_gn=UPF3B;refGene_tx=XM_004000835.5;refGene_gn=UPF3B;refGene_tx=XM_004000836.5
X	100410755	100410866
refGene_gn=UPF3B;refGene_tx=XM_004000835.5;refGene_gn=UPF3B;refGene_tx=XM_004000836.5

X	100412594	100412693
refGene_gn=UPF3B;refGene_tx=XM_004000835.5;refGene_gn=UPF3B;refGene_tx=XM_004000836.5
X	100416358	100416465
refGene_gn=UPF3B;refGene_tx=XM_004000835.5;refGene_gn=UPF3B;refGene_tx=XM_004000836.5
X	100416630	100416737
refGene_gn=UPF3B;refGene_tx=XM_004000835.5;refGene_gn=UPF3B;refGene_tx=XM_004000836.5
X	100417680	100417836
refGene_gn=UPF3B;refGene_tx=XM_004000835.5;refGene_gn=UPF3B;refGene_tx=XM_004000836.5
X	100434751	100435783
refGene_gn=RNF113A;refGene_tx=XM_004000837.4
X	100436096	100436198
refGene_gn=NDUFA1;refGene_tx=XM_004000838.4
X	100437157	100437247
refGene_gn=NDUFA1;refGene_tx=XM_004000838.4
X	100439819	100439840
refGene_gn=NDUFA1;refGene_tx=XM_004000838.4
X	100449434	100449631
refGene_gn=AKAP14;refGene_tx=XM_006943969.4
X	100450494	100450502
refGene_gn=AKAP14;refGene_tx=XM_019824227.2
X	100452773	100452940
refGene_gn=AKAP14;refGene_tx=XM_006943969.4;refGene_gn=AKAP14;refGene_tx=XM_019824227.2
X	100453031	100453123
refGene_gn=AKAP14;refGene_tx=XM_006943969.4;refGene_gn=AKAP14;refGene_tx=XM_019824227.2
X	100466230	100466410
refGene_gn=AKAP14;refGene_tx=XM_006943969.4;refGene_gn=AKAP14;refGene_tx=XM_019824227.2
X	100468655	100468708
refGene_gn=AKAP14;refGene_tx=XM_006943969.4;refGene_gn=AKAP14;refGene_tx=XM_019824227.2
X	100469342	100469427
refGene_gn=AKAP14;refGene_tx=XM_006943969.4;refGene_gn=AKAP14;refGene_tx=XM_019824227.2
X	100472373	100472548
refGene_gn=NKAP;refGene_tx=XM_004000839.4;refGene_gn=NKAP;refGene_tx=XM_019824225.2
X	100475772	100475922
refGene_gn=NKAP;refGene_tx=XM_004000839.4;refGene_gn=NKAP;refGene_tx=XM_019824225.2
X	100477097	100477173
refGene_gn=NKAP;refGene_tx=XM_004000839.4;refGene_gn=NKAP;refGene_tx=XM_019824225.2
X	100477479	100477589
refGene_gn=NKAP;refGene_tx=XM_004000839.4;refGene_gn=NKAP;refGene_tx=X

M_019824225.2
X	100479654	100479718
refGene_gn=NKAP;refGene_tx=XM_004000839.4;refGene_gn=NKAP;refGene_tx=XM_019824225.2
X	100481609	100481753
refGene_gn=NKAP;refGene_tx=XM_004000839.4;refGene_gn=NKAP;refGene_tx=XM_019824225.2
X	100481931	100482002
refGene_gn=NKAP;refGene_tx=XM_004000839.4;refGene_gn=NKAP;refGene_tx=XM_019824225.2
X	100493135	100493216
refGene_gn=NKAP;refGene_tx=XM_004000839.4;refGene_gn=NKAP;refGene_tx=XM_019824225.2
X	100496104	100496502
refGene_gn=NKAP;refGene_tx=XM_004000839.4;refGene_gn=NKAP;refGene_tx=XM_019824225.2
X	100527105	100527151
refGene_gn=LOC102902505;refGene_tx=XM_006943970.3;refGene_gn=LOC102902505;refGene_tx=XM_019824229.1
X	100527576	100527671
refGene_gn=LOC102902505;refGene_tx=XM_006943970.3
X	100527774	100528090
refGene_gn=LOC102902505;refGene_tx=XM_006943970.3;refGene_gn=LOC102902505;refGene_tx=XM_019824229.1
X	100528242	100528321
refGene_gn=LOC102902505;refGene_tx=XM_006943970.3;refGene_gn=LOC102902505;refGene_tx=XM_019824229.1
X	100589548	100589971
refGene_gn=LOC111558775;refGene_tx=XM_023249457.1
X	100593962	100594008
refGene_gn=LOC111558775;refGene_tx=XM_023249457.1
X	100596124	100596521
refGene_gn=LOC111558775;refGene_tx=XM_023249457.1
X	100596681	100596763
refGene_gn=LOC111558775;refGene_tx=XM_023249457.1
X	100643525	100643642
refGene_gn=LOC101093839;refGene_tx=XM_023249373.1
X	100648315	100648361
refGene_gn=LOC101093839;refGene_tx=XM_023249373.1
X	100653108	100653560
refGene_gn=LOC101093839;refGene_tx=XM_023249373.1
X	100676187	100676269
refGene_gn=LOC102902213;refGene_tx=XM_023249371.1
X	100676429	100676883
refGene_gn=LOC102902213;refGene_tx=XM_023249371.1;refGene_gn=LOC102902213;refGene_tx=XM_023249372.1
X	100679004	100679050
refGene_gn=LOC102902213;refGene_tx=XM_023249371.1;refGene_gn=LOC102902213;refGene_tx=XM_023249372.1
X	100683039	100683408

refGene_gn=LOC102902213;refGene_tx=XM_023249371.1;refGene_gn=LOC102902213;refGene_tx=XM_023249372.1
X	100804794	100806813
refGene_gn=ZBTB33;refGene_tx=XM_023248941.1
X	100812854	100813013
refGene_gn=TMEM255A;refGene_tx=XM_004000841.3;refGene_gn=TMEM255A;refGene_tx=XM_004000842.3
X	100822439	100822583
refGene_gn=TMEM255A;refGene_tx=XM_004000841.3;refGene_gn=TMEM255A;refGene_tx=XM_004000842.3
X	100830048	100830211
refGene_gn=TMEM255A;refGene_tx=XM_004000841.3;refGene_gn=TMEM255A;refGene_tx=XM_004000842.3
X	100839705	100839794
refGene_gn=TMEM255A;refGene_tx=XM_004000841.3;refGene_gn=TMEM255A;refGene_tx=XM_004000842.3
X	100840211	100840283
refGene_gn=TMEM255A;refGene_tx=XM_004000841.3
X	100841569	100841638
refGene_gn=TMEM255A;refGene_tx=XM_004000841.3;refGene_gn=TMEM255A;refGene_tx=XM_004000842.3
X	100847449	100847539
refGene_gn=TMEM255A;refGene_tx=XM_004000841.3;refGene_gn=TMEM255A;refGene_tx=XM_004000842.3
X	100850127	100850190
refGene_gn=TMEM255A;refGene_tx=XM_004000841.3;refGene_gn=TMEM255A;refGene_tx=XM_004000842.3
X	100860093	100860236
refGene_gn=TMEM255A;refGene_tx=XM_004000841.3;refGene_gn=TMEM255A;refGene_tx=XM_004000842.3
X	100866861	100866919
refGene_gn=TMEM255A;refGene_tx=XM_004000841.3;refGene_gn=TMEM255A;refGene_tx=XM_004000842.3
X	100916083	100916146
refGene_gn=ATP1B4;refGene_tx=XM_006943972.4
X	100920847	100920955
refGene_gn=ATP1B4;refGene_tx=XM_006943972.4
X	100920956	100921113
refGene_gn=ATP1B4;refGene_tx=XM_006943972.4
X	100925793	100925922
refGene_gn=ATP1B4;refGene_tx=XM_006943972.4
X	100926191	100926296
refGene_gn=ATP1B4;refGene_tx=XM_006943972.4
X	100930134	100930331
refGene_gn=ATP1B4;refGene_tx=XM_006943972.4
X	100931032	100931089
refGene_gn=ATP1B4;refGene_tx=XM_006943972.4
X	100933723	100933819
refGene_gn=ATP1B4;refGene_tx=XM_006943972.4
X	100934594	100934756

refGene_gn=ATP1B4;refGene_tx=XM_006943972.4
X	100959942	100960936
refGene_gn=LOC109496592;refGene_tx=XM_019824231.1
X	100962066	100962116
refGene_gn=LOC109496592;refGene_tx=XM_019824231.1
X	100977252	100977392
refGene_gn=LAMP2;refGene_tx=XM_004000847.3
X	100985585	100985725
refGene_gn=LAMP2;refGene_tx=XM_006943973.4
X	100988057	100988222
refGene_gn=LAMP2;refGene_tx=XM_004000847.3;refGene_gn=LAMP2;refGene_tx
=XM_006943973.4
X	100988717	100988781
refGene_gn=LAMP2;refGene_tx=XM_004000847.3;refGene_gn=LAMP2;refGene_tx
=XM_006943973.4
X	100991204	100991327
refGene_gn=LAMP2;refGene_tx=XM_004000847.3;refGene_gn=LAMP2;refGene_tx
=XM_006943973.4
X	100992341	100992529
refGene_gn=LAMP2;refGene_tx=XM_004000847.3;refGene_gn=LAMP2;refGene_tx
=XM_006943973.4
X	100994494	100994653
refGene_gn=LAMP2;refGene_tx=XM_004000847.3;refGene_gn=LAMP2;refGene_tx
=XM_006943973.4
X	100999003	100999217
refGene_gn=LAMP2;refGene_tx=XM_004000847.3;refGene_gn=LAMP2;refGene_tx
=XM_006943973.4
X	100999854	100999973
refGene_gn=LAMP2;refGene_tx=XM_004000847.3;refGene_gn=LAMP2;refGene_tx
=XM_006943973.4
X	101012264	101012502
refGene_gn=LAMP2;refGene_tx=XM_004000847.3;refGene_gn=LAMP2;refGene_tx
=XM_006943973.4
X	101077623	101077719
refGene_gn=CUL4B;refGene_tx=NM_001048152.1;refGene_gn=CUL4B;refGene_tx
=XM_011279431.3;refGene_gn=CUL4B;refGene_tx=XM_011279432.3;refGene_gn=
CUL4B;refGene_tx=XM_019823335.2;refGene_gn=CUL4B;refGene_tx=XM_0198233
36.2;refGene_gn=CUL4B;refGene_tx=XM_019823337.2;refGene_gn=CUL4B;refGe
ne_tx=XM_019823338.2
X	101080684	101080837
refGene_gn=CUL4B;refGene_tx=NM_001048152.1;refGene_gn=CUL4B;refGene_tx
=XM_011279431.3;refGene_gn=CUL4B;refGene_tx=XM_011279432.3;refGene_gn=
CUL4B;refGene_tx=XM_019823335.2;refGene_gn=CUL4B;refGene_tx=XM_0198233
36.2;refGene_gn=CUL4B;refGene_tx=XM_019823337.2;refGene_gn=CUL4B;refGe
ne_tx=XM_019823338.2
X	101082254	101082427
refGene_gn=CUL4B;refGene_tx=NM_001048152.1;refGene_gn=CUL4B;refGene_tx
=XM_011279431.3;refGene_gn=CUL4B;refGene_tx=XM_011279432.3;refGene_gn=
CUL4B;refGene_tx=XM_019823335.2;refGene_gn=CUL4B;refGene_tx=XM_0198233
36.2;refGene_gn=CUL4B;refGene_tx=XM_019823337.2;refGene_gn=CUL4B;refGe

ne_tx=XM_019823338.2
X	101083534	101083640
refGene_gn=CUL4B;refGene_tx=NM_001048152.1;refGene_gn=CUL4B;refGene_tx=XM_011279431.3;refGene_gn=CUL4B;refGene_tx=XM_011279432.3;refGene_gn=CUL4B;refGene_tx=XM_019823335.2;refGene_gn=CUL4B;refGene_tx=XM_019823336.2;refGene_gn=CUL4B;refGene_tx=XM_019823337.2;refGene_gn=CUL4B;refGene_tx=XM_019823338.2
X	101084682	101084796
refGene_gn=CUL4B;refGene_tx=NM_001048152.1;refGene_gn=CUL4B;refGene_tx=XM_011279431.3;refGene_gn=CUL4B;refGene_tx=XM_011279432.3;refGene_gn=CUL4B;refGene_tx=XM_019823335.2;refGene_gn=CUL4B;refGene_tx=XM_019823336.2;refGene_gn=CUL4B;refGene_tx=XM_019823337.2;refGene_gn=CUL4B;refGene_tx=XM_019823338.2
X	101085668	101085776
refGene_gn=CUL4B;refGene_tx=NM_001048152.1;refGene_gn=CUL4B;refGene_tx=XM_011279431.3;refGene_gn=CUL4B;refGene_tx=XM_011279432.3;refGene_gn=CUL4B;refGene_tx=XM_019823335.2;refGene_gn=CUL4B;refGene_tx=XM_019823336.2;refGene_gn=CUL4B;refGene_tx=XM_019823337.2;refGene_gn=CUL4B;refGene_tx=XM_019823338.2
X	101086860	101086952
refGene_gn=CUL4B;refGene_tx=NM_001048152.1;refGene_gn=CUL4B;refGene_tx=XM_011279431.3;refGene_gn=CUL4B;refGene_tx=XM_011279432.3;refGene_gn=CUL4B;refGene_tx=XM_019823335.2;refGene_gn=CUL4B;refGene_tx=XM_019823336.2;refGene_gn=CUL4B;refGene_tx=XM_019823337.2;refGene_gn=CUL4B;refGene_tx=XM_019823338.2
X	101087401	101087512
refGene_gn=CUL4B;refGene_tx=NM_001048152.1;refGene_gn=CUL4B;refGene_tx=XM_011279431.3;refGene_gn=CUL4B;refGene_tx=XM_011279432.3;refGene_gn=CUL4B;refGene_tx=XM_019823335.2;refGene_gn=CUL4B;refGene_tx=XM_019823336.2;refGene_gn=CUL4B;refGene_tx=XM_019823337.2;refGene_gn=CUL4B;refGene_tx=XM_019823338.2
X	101087967	101088072
refGene_gn=CUL4B;refGene_tx=NM_001048152.1;refGene_gn=CUL4B;refGene_tx=XM_011279431.3;refGene_gn=CUL4B;refGene_tx=XM_011279432.3;refGene_gn=CUL4B;refGene_tx=XM_019823335.2;refGene_gn=CUL4B;refGene_tx=XM_019823336.2;refGene_gn=CUL4B;refGene_tx=XM_019823337.2;refGene_gn=CUL4B;refGene_tx=XM_019823338.2
X	101093124	101093317
refGene_gn=CUL4B;refGene_tx=NM_001048152.1;refGene_gn=CUL4B;refGene_tx=XM_011279431.3;refGene_gn=CUL4B;refGene_tx=XM_011279432.3;refGene_gn=CUL4B;refGene_tx=XM_019823335.2;refGene_gn=CUL4B;refGene_tx=XM_019823336.2;refGene_gn=CUL4B;refGene_tx=XM_019823337.2;refGene_gn=CUL4B;refGene_tx=XM_019823338.2
X	101093909	101094028
refGene_gn=CUL4B;refGene_tx=NM_001048152.1;refGene_gn=CUL4B;refGene_tx=XM_011279431.3;refGene_gn=CUL4B;refGene_tx=XM_011279432.3;refGene_gn=CUL4B;refGene_tx=XM_019823335.2;refGene_gn=CUL4B;refGene_tx=XM_019823336.2;refGene_gn=CUL4B;refGene_tx=XM_019823337.2;refGene_gn=CUL4B;refGene_tx=XM_019823338.2
X	101095415	101095483

refGene_gn=CUL4B;refGene_tx=NM_001048152.1;refGene_gn=CUL4B;refGene_tx
=XM_011279431.3;refGene_gn=CUL4B;refGene_tx=XM_011279432.3;refGene_gn=
CUL4B;refGene_tx=XM_019823335.2;refGene_gn=CUL4B;refGene_tx=XM_0198233
36.2;refGene_gn=CUL4B;refGene_tx=XM_019823337.2;refGene_gn=CUL4B;refGe
ne_tx=XM_019823338.2
X	101096422	101096505
refGene_gn=CUL4B;refGene_tx=NM_001048152.1;refGene_gn=CUL4B;refGene_tx
=XM_011279431.3;refGene_gn=CUL4B;refGene_tx=XM_011279432.3;refGene_gn=
CUL4B;refGene_tx=XM_019823335.2;refGene_gn=CUL4B;refGene_tx=XM_0198233
36.2;refGene_gn=CUL4B;refGene_tx=XM_019823337.2;refGene_gn=CUL4B;refGe
ne_tx=XM_019823338.2
X	101096796	101096886
refGene_gn=CUL4B;refGene_tx=NM_001048152.1;refGene_gn=CUL4B;refGene_tx
=XM_011279431.3;refGene_gn=CUL4B;refGene_tx=XM_011279432.3;refGene_gn=
CUL4B;refGene_tx=XM_019823335.2;refGene_gn=CUL4B;refGene_tx=XM_0198233
36.2;refGene_gn=CUL4B;refGene_tx=XM_019823337.2;refGene_gn=CUL4B;refGe
ne_tx=XM_019823338.2
X	101097032	101097195
refGene_gn=CUL4B;refGene_tx=NM_001048152.1;refGene_gn=CUL4B;refGene_tx
=XM_011279431.3;refGene_gn=CUL4B;refGene_tx=XM_011279432.3;refGene_gn=
CUL4B;refGene_tx=XM_019823335.2;refGene_gn=CUL4B;refGene_tx=XM_0198233
36.2;refGene_gn=CUL4B;refGene_tx=XM_019823337.2;refGene_gn=CUL4B;refGe
ne_tx=XM_019823338.2
X	101097981	101098055
refGene_gn=CUL4B;refGene_tx=NM_001048152.1;refGene_gn=CUL4B;refGene_tx
=XM_011279431.3;refGene_gn=CUL4B;refGene_tx=XM_011279432.3;refGene_gn=
CUL4B;refGene_tx=XM_019823335.2;refGene_gn=CUL4B;refGene_tx=XM_0198233
36.2;refGene_gn=CUL4B;refGene_tx=XM_019823337.2;refGene_gn=CUL4B;refGe
ne_tx=XM_019823338.2
X	101098826	101098896
refGene_gn=CUL4B;refGene_tx=NM_001048152.1;refGene_gn=CUL4B;refGene_tx
=XM_011279431.3;refGene_gn=CUL4B;refGene_tx=XM_011279432.3;refGene_gn=
CUL4B;refGene_tx=XM_019823335.2;refGene_gn=CUL4B;refGene_tx=XM_0198233
36.2;refGene_gn=CUL4B;refGene_tx=XM_019823337.2;refGene_gn=CUL4B;refGe
ne_tx=XM_019823338.2
X	101099316	101099420
refGene_gn=CUL4B;refGene_tx=NM_001048152.1;refGene_gn=CUL4B;refGene_tx
=XM_011279431.3;refGene_gn=CUL4B;refGene_tx=XM_011279432.3;refGene_gn=
CUL4B;refGene_tx=XM_019823335.2;refGene_gn=CUL4B;refGene_tx=XM_0198233
36.2;refGene_gn=CUL4B;refGene_tx=XM_019823337.2;refGene_gn=CUL4B;refGe
ne_tx=XM_019823338.2
X	101110035	101110151
refGene_gn=CUL4B;refGene_tx=NM_001048152.1;refGene_gn=CUL4B;refGene_tx
=XM_011279431.3;refGene_gn=CUL4B;refGene_tx=XM_011279432.3;refGene_gn=
CUL4B;refGene_tx=XM_019823335.2;refGene_gn=CUL4B;refGene_tx=XM_0198233
36.2;refGene_gn=CUL4B;refGene_tx=XM_019823337.2;refGene_gn=CUL4B;refGe
ne_tx=XM_019823338.2
X	101112487	101113027
refGene_gn=CUL4B;refGene_tx=NM_001048152.1;refGene_gn=CUL4B;refGene_tx
=XM_011279431.3;refGene_gn=CUL4B;refGene_tx=XM_011279432.3;refGene_gn=

CUL4B;refGene_tx=XM_019823335.2;refGene_gn=CUL4B;refGene_tx=XM_0198233
36.2;refGene_gn=CUL4B;refGene_tx=XM_019823337.2;refGene_gn=CUL4B;refGe
ne_tx=XM_019823338.2
X	101113781	101113809
refGene_gn=CUL4B;refGene_tx=XM_011279431.3;refGene_gn=CUL4B;refGene_tx
=XM_011279432.3
X	101117781	101117845
refGene_gn=CUL4B;refGene_tx=XM_019823335.2;refGene_gn=CUL4B;refGene_tx
=XM_019823336.2;refGene_gn=CUL4B;refGene_tx=XM_019823337.2
X	101156798	101156809
refGene_gn=MCTS1;refGene_tx=XM_019824234.2
X	101157414	101157428
refGene_gn=MCTS1;refGene_tx=XM_019824233.2
X	101157986	101158139
refGene_gn=MCTS1;refGene_tx=XM_019824233.2;refGene_gn=MCTS1;refGene_tx
=XM_019824234.2
X	101158815	101158913
refGene_gn=MCTS1;refGene_tx=XM_019824233.2;refGene_gn=MCTS1;refGene_tx
=XM_019824234.2
X	101162589	101162723
refGene_gn=MCTS1;refGene_tx=XM_019824233.2;refGene_gn=MCTS1;refGene_tx
=XM_019824234.2
X	101165138	101165206
refGene_gn=MCTS1;refGene_tx=XM_019824233.2;refGene_gn=MCTS1;refGene_tx
=XM_019824234.2
X	101166191	101166273
refGene_gn=MCTS1;refGene_tx=XM_019824233.2;refGene_gn=MCTS1;refGene_tx
=XM_019824234.2
X	101182177	101183131
refGene_gn=C1GALT1C1;refGene_tx=XM_004000850.4;refGene_gn=C1GALT1C1;re
fGene_tx=XM_006943974.3
X	103174466	103174575
refGene_gn=GRIA3;refGene_tx=XM_004000853.5;refGene_gn=GRIA3;refGene_tx
=XM_011279433.3;refGene_gn=GRIA3;refGene_tx=XM_019824236.1;refGene_gn=
GRIA3;refGene_tx=XM_019824237.2;refGene_gn=GRIA3;refGene_tx=XM_0232493
77.1;refGene_gn=GRIA3;refGene_tx=XM_023249378.1
X	103175758	103175917
refGene_gn=GRIA3;refGene_tx=XM_004000853.5;refGene_gn=GRIA3;refGene_tx
=XM_011279433.3;refGene_gn=GRIA3;refGene_tx=XM_019824236.1;refGene_gn=
GRIA3;refGene_tx=XM_019824237.2;refGene_gn=GRIA3;refGene_tx=XM_0232493
77.1;refGene_gn=GRIA3;refGene_tx=XM_023249378.1
X	103245149	103245389
refGene_gn=GRIA3;refGene_tx=XM_004000853.5;refGene_gn=GRIA3;refGene_tx
=XM_011279433.3;refGene_gn=GRIA3;refGene_tx=XM_019824236.1;refGene_gn=
GRIA3;refGene_tx=XM_019824237.2;refGene_gn=GRIA3;refGene_tx=XM_0232493
77.1;refGene_gn=GRIA3;refGene_tx=XM_023249378.1
X	103306490	103306678
refGene_gn=GRIA3;refGene_tx=XM_004000853.5;refGene_gn=GRIA3;refGene_tx
=XM_011279433.3;refGene_gn=GRIA3;refGene_tx=XM_019824236.1;refGene_gn=
GRIA3;refGene_tx=XM_019824237.2;refGene_gn=GRIA3;refGene_tx=XM_0232493

77.1;refGene_gn=GRIA3;refGene_tx=XM_023249378.1
X	103339761	103339815
refGene_gn=GRIA3;refGene_tx=XM_004000853.5;refGene_gn=GRIA3;refGene_tx=XM_011279433.3;refGene_gn=GRIA3;refGene_tx=XM_019824236.1;refGene_gn=GRIA3;refGene_tx=XM_019824237.2;refGene_gn=GRIA3;refGene_tx=XM_023249377.1;refGene_gn=GRIA3;refGene_tx=XM_023249378.1
X	103353474	103353636
refGene_gn=GRIA3;refGene_tx=XM_004000853.5;refGene_gn=GRIA3;refGene_tx=XM_011279433.3;refGene_gn=GRIA3;refGene_tx=XM_019824236.1;refGene_gn=GRIA3;refGene_tx=XM_019824237.2;refGene_gn=GRIA3;refGene_tx=XM_023249377.1;refGene_gn=GRIA3;refGene_tx=XM_023249378.1
X	103357963	103358131
refGene_gn=GRIA3;refGene_tx=XM_004000853.5;refGene_gn=GRIA3;refGene_tx=XM_011279433.3;refGene_gn=GRIA3;refGene_tx=XM_019824236.1;refGene_gn=GRIA3;refGene_tx=XM_019824237.2;refGene_gn=GRIA3;refGene_tx=XM_023249377.1;refGene_gn=GRIA3;refGene_tx=XM_023249378.1
X	103362519	103362624
refGene_gn=GRIA3;refGene_tx=XM_004000853.5;refGene_gn=GRIA3;refGene_tx=XM_011279433.3;refGene_gn=GRIA3;refGene_tx=XM_019824236.1;refGene_gn=GRIA3;refGene_tx=XM_019824237.2;refGene_gn=GRIA3;refGene_tx=XM_023249377.1;refGene_gn=GRIA3;refGene_tx=XM_023249378.1
X	103363787	103363895
refGene_gn=GRIA3;refGene_tx=XM_004000853.5;refGene_gn=GRIA3;refGene_tx=XM_011279433.3;refGene_gn=GRIA3;refGene_tx=XM_019824236.1;refGene_gn=GRIA3;refGene_tx=XM_019824237.2;refGene_gn=GRIA3;refGene_tx=XM_023249377.1;refGene_gn=GRIA3;refGene_tx=XM_023249378.1
X	103365333	103365540
refGene_gn=GRIA3;refGene_tx=XM_004000853.5;refGene_gn=GRIA3;refGene_tx=XM_011279433.3;refGene_gn=GRIA3;refGene_tx=XM_019824236.1;refGene_gn=GRIA3;refGene_tx=XM_019824237.2;refGene_gn=GRIA3;refGene_tx=XM_023249377.1;refGene_gn=GRIA3;refGene_tx=XM_023249378.1
X	103376954	103377331
refGene_gn=GRIA3;refGene_tx=XM_004000853.5;refGene_gn=GRIA3;refGene_tx=XM_011279433.3;refGene_gn=GRIA3;refGene_tx=XM_019824236.1;refGene_gn=GRIA3;refGene_tx=XM_019824237.2;refGene_gn=GRIA3;refGene_tx=XM_023249377.1;refGene_gn=GRIA3;refGene_tx=XM_023249378.1
X	103387559	103387758
refGene_gn=GRIA3;refGene_tx=XM_004000853.5;refGene_gn=GRIA3;refGene_tx=XM_011279433.3;refGene_gn=GRIA3;refGene_tx=XM_019824236.1;refGene_gn=GRIA3;refGene_tx=XM_019824237.2;refGene_gn=GRIA3;refGene_tx=XM_023249377.1;refGene_gn=GRIA3;refGene_tx=XM_023249378.1
X	103414452	103414700
refGene_gn=GRIA3;refGene_tx=XM_004000853.5;refGene_gn=GRIA3;refGene_tx=XM_011279433.3;refGene_gn=GRIA3;refGene_tx=XM_019824236.1;refGene_gn=GRIA3;refGene_tx=XM_019824237.2;refGene_gn=GRIA3;refGene_tx=XM_023249377.1;refGene_gn=GRIA3;refGene_tx=XM_023249378.1
X	103415262	103415377
refGene_gn=GRIA3;refGene_tx=XM_011279433.3;refGene_gn=GRIA3;refGene_tx=XM_019824237.2
X	103429558	103429673

refGene_gn=GRIA3;refGene_tx=XM_004000853.5;refGene_gn=GRIA3;refGene_tx
=XM_019824236.1;refGene_gn=GRIA3;refGene_tx=XM_019824237.2;refGene_gn=
GRIA3;refGene_tx=XM_023249377.1;refGene_gn=GRIA3;refGene_tx=XM_0232493
78.1
X 103432316 103432562
refGene_gn=GRIA3;refGene_tx=XM_004000853.5;refGene_gn=GRIA3;refGene_tx
=XM_011279433.3;refGene_gn=GRIA3;refGene_tx=XM_019824236.1;refGene_gn=
GRIA3;refGene_tx=XM_023249377.1;refGene_gn=GRIA3;refGene_tx=XM_0232493
78.1
X 103538381 103538409
refGene_gn=THOC2;refGene_tx=XM_011279460.3;refGene_gn=THOC2;refGene_tx
=XM_011279461.3;refGene_gn=THOC2;refGene_tx=XM_019824238.2;refGene_gn=
THOC2;refGene_tx=XM_019824239.1;refGene_gn=THOC2;refGene_tx=XM_0198242
40.1;refGene_gn=THOC2;refGene_tx=XM_019824241.2;refGene_gn=THOC2;refGe
ne_tx=XM_019824242.2;refGene_gn=THOC2;refGene_tx=XM_019824243.1;refGen
e_gn=THOC2;refGene_tx=XM_019824244.2;refGene_gn=THOC2;refGene_tx=XM_01
9824245.2;refGene_gn=THOC2;refGene_tx=XM_019824246.2;refGene_gn=THOC2;
refGene_tx=XM_019824247.2;refGene_gn=THOC2;refGene_tx=XM_019824248.1;r
efGene_gn=THOC2;refGene_tx=XM_023249254.1
X 103538837 103538966
refGene_gn=THOC2;refGene_tx=XM_011279459.3;refGene_gn=THOC2;refGene_tx
=XM_011279460.3;refGene_gn=THOC2;refGene_tx=XM_011279461.3;refGene_gn=
THOC2;refGene_tx=XM_019824238.2;refGene_gn=THOC2;refGene_tx=XM_0198242
39.1;refGene_gn=THOC2;refGene_tx=XM_019824240.1;refGene_gn=THOC2;refGe
ne_tx=XM_019824241.2;refGene_gn=THOC2;refGene_tx=XM_019824242.2;refGen
e_gn=THOC2;refGene_tx=XM_019824243.1;refGene_gn=THOC2;refGene_tx=XM_01
9824244.2;refGene_gn=THOC2;refGene_tx=XM_019824245.2;refGene_gn=THOC2;
refGene_tx=XM_019824246.2;refGene_gn=THOC2;refGene_tx=XM_019824247.2;r
efGene_gn=THOC2;refGene_tx=XM_019824248.1;refGene_gn=THOC2;refGene_tx=
XM_023249254.1
X 103541110 103541268
refGene_gn=THOC2;refGene_tx=XM_011279459.3;refGene_gn=THOC2;refGene_tx
=XM_011279460.3;refGene_gn=THOC2;refGene_tx=XM_011279461.3;refGene_gn=
THOC2;refGene_tx=XM_019824238.2;refGene_gn=THOC2;refGene_tx=XM_0198242
39.1;refGene_gn=THOC2;refGene_tx=XM_019824240.1;refGene_gn=THOC2;refGe
ne_tx=XM_019824241.2;refGene_gn=THOC2;refGene_tx=XM_019824242.2;refGen
e_gn=THOC2;refGene_tx=XM_019824243.1;refGene_gn=THOC2;refGene_tx=XM_01
9824244.2;refGene_gn=THOC2;refGene_tx=XM_019824245.2;refGene_gn=THOC2;
refGene_tx=XM_019824246.2;refGene_gn=THOC2;refGene_tx=XM_019824247.2;r
efGene_gn=THOC2;refGene_tx=XM_019824248.1;refGene_gn=THOC2;refGene_tx=
XM_023249254.1
X 103541350 103541420
refGene_gn=THOC2;refGene_tx=XM_011279459.3;refGene_gn=THOC2;refGene_tx
=XM_011279460.3;refGene_gn=THOC2;refGene_tx=XM_011279461.3;refGene_gn=
THOC2;refGene_tx=XM_019824238.2;refGene_gn=THOC2;refGene_tx=XM_0198242
39.1;refGene_gn=THOC2;refGene_tx=XM_019824240.1;refGene_gn=THOC2;refGe
ne_tx=XM_019824241.2;refGene_gn=THOC2;refGene_tx=XM_019824242.2;refGen
e_gn=THOC2;refGene_tx=XM_019824243.1;refGene_gn=THOC2;refGene_tx=XM_01
9824244.2;refGene_gn=THOC2;refGene_tx=XM_019824245.2;refGene_gn=THOC2;
refGene_tx=XM_019824246.2;refGene_gn=THOC2;refGene_tx=XM_019824247.2;r

efGene_gn=THOC2;refGene_tx=XM_019824248.1;refGene_gn=THOC2;refGene_tx=XM_023249254.1
X 103541762 103541900
refGene_gn=THOC2;refGene_tx=XM_011279459.3;refGene_gn=THOC2;refGene_tx=XM_011279460.3;refGene_gn=THOC2;refGene_tx=XM_011279461.3;refGene_gn=THOC2;refGene_tx=XM_019824238.2;refGene_gn=THOC2;refGene_tx=XM_019824239.1;refGene_gn=THOC2;refGene_tx=XM_019824240.1;refGene_gn=THOC2;refGene_tx=XM_019824241.2;refGene_gn=THOC2;refGene_tx=XM_019824242.2;refGene_gn=THOC2;refGene_tx=XM_019824243.1;refGene_gn=THOC2;refGene_tx=XM_019824244.2;refGene_gn=THOC2;refGene_tx=XM_019824245.2;refGene_gn=THOC2;refGene_tx=XM_019824246.2;refGene_gn=THOC2;refGene_tx=XM_019824247.2;refGene_gn=THOC2;refGene_tx=XM_019824248.1;refGene_gn=THOC2;refGene_tx=XM_023249254.1
X 103546752 103546884
refGene_gn=THOC2;refGene_tx=XM_011279459.3;refGene_gn=THOC2;refGene_tx=XM_011279460.3;refGene_gn=THOC2;refGene_tx=XM_011279461.3;refGene_gn=THOC2;refGene_tx=XM_019824238.2;refGene_gn=THOC2;refGene_tx=XM_019824239.1;refGene_gn=THOC2;refGene_tx=XM_019824240.1;refGene_gn=THOC2;refGene_tx=XM_019824241.2;refGene_gn=THOC2;refGene_tx=XM_019824242.2;refGene_gn=THOC2;refGene_tx=XM_019824243.1;refGene_gn=THOC2;refGene_tx=XM_019824244.2;refGene_gn=THOC2;refGene_tx=XM_019824245.2;refGene_gn=THOC2;refGene_tx=XM_019824246.2;refGene_gn=THOC2;refGene_tx=XM_019824247.2;refGene_gn=THOC2;refGene_tx=XM_019824248.1;refGene_gn=THOC2;refGene_tx=XM_023249254.1
X 103548565 103548624
refGene_gn=THOC2;refGene_tx=XM_011279459.3;refGene_gn=THOC2;refGene_tx=XM_011279460.3;refGene_gn=THOC2;refGene_tx=XM_011279461.3;refGene_gn=THOC2;refGene_tx=XM_019824238.2;refGene_gn=THOC2;refGene_tx=XM_019824239.1;refGene_gn=THOC2;refGene_tx=XM_019824240.1;refGene_gn=THOC2;refGene_tx=XM_019824241.2;refGene_gn=THOC2;refGene_tx=XM_019824242.2;refGene_gn=THOC2;refGene_tx=XM_019824243.1;refGene_gn=THOC2;refGene_tx=XM_019824244.2;refGene_gn=THOC2;refGene_tx=XM_019824245.2;refGene_gn=THOC2;refGene_tx=XM_019824246.2;refGene_gn=THOC2;refGene_tx=XM_019824247.2;refGene_gn=THOC2;refGene_tx=XM_019824248.1;refGene_gn=THOC2;refGene_tx=XM_023249254.1
X 103548831 103549262
refGene_gn=THOC2;refGene_tx=XM_011279459.3;refGene_gn=THOC2;refGene_tx=XM_011279460.3;refGene_gn=THOC2;refGene_tx=XM_011279461.3;refGene_gn=THOC2;refGene_tx=XM_019824238.2;refGene_gn=THOC2;refGene_tx=XM_019824239.1;refGene_gn=THOC2;refGene_tx=XM_019824240.1;refGene_gn=THOC2;refGene_tx=XM_019824241.2;refGene_gn=THOC2;refGene_tx=XM_019824242.2;refGene_gn=THOC2;refGene_tx=XM_019824243.1;refGene_gn=THOC2;refGene_tx=XM_019824244.2;refGene_gn=THOC2;refGene_tx=XM_019824245.2;refGene_gn=THOC2;refGene_tx=XM_019824246.2;refGene_gn=THOC2;refGene_tx=XM_019824247.2;refGene_gn=THOC2;refGene_tx=XM_019824248.1;refGene_gn=THOC2;refGene_tx=XM_023249254.1
X 103550176 103550279
refGene_gn=THOC2;refGene_tx=XM_011279459.3;refGene_gn=THOC2;refGene_tx=XM_011279460.3;refGene_gn=THOC2;refGene_tx=XM_011279461.3;refGene_gn=THOC2;refGene_tx=XM_019824238.2;refGene_gn=THOC2;refGene_tx=XM_0198242

39.1;refGene_gn=THOC2;refGene_tx=XM_019824240.1;refGene_gn=THOC2;refGene_tx=XM_019824241.2;refGene_gn=THOC2;refGene_tx=XM_019824242.2;refGene_gn=THOC2;refGene_tx=XM_019824243.1;refGene_gn=THOC2;refGene_tx=XM_019824244.2;refGene_gn=THOC2;refGene_tx=XM_019824245.2;refGene_gn=THOC2;refGene_tx=XM_019824246.2;refGene_gn=THOC2;refGene_tx=XM_019824247.2;refGene_gn=THOC2;refGene_tx=XM_019824248.1;refGene_gn=THOC2;refGene_tx=XM_023249254.1
X	103550546	103550725
refGene_gn=THOC2;refGene_tx=XM_011279459.3;refGene_gn=THOC2;refGene_tx=XM_011279460.3;refGene_gn=THOC2;refGene_tx=XM_011279461.3;refGene_gn=THOC2;refGene_tx=XM_019824238.2;refGene_gn=THOC2;refGene_tx=XM_019824239.1;refGene_gn=THOC2;refGene_tx=XM_019824240.1;refGene_gn=THOC2;refGene_tx=XM_019824241.2;refGene_gn=THOC2;refGene_tx=XM_019824242.2;refGene_gn=THOC2;refGene_tx=XM_019824243.1;refGene_gn=THOC2;refGene_tx=XM_019824244.2;refGene_gn=THOC2;refGene_tx=XM_019824245.2;refGene_gn=THOC2;refGene_tx=XM_019824246.2;refGene_gn=THOC2;refGene_tx=XM_019824247.2;refGene_gn=THOC2;refGene_tx=XM_019824248.1;refGene_gn=THOC2;refGene_tx=XM_023249254.1
X	103551218	103551403
refGene_gn=THOC2;refGene_tx=XM_011279459.3;refGene_gn=THOC2;refGene_tx=XM_011279460.3;refGene_gn=THOC2;refGene_tx=XM_011279461.3;refGene_gn=THOC2;refGene_tx=XM_019824238.2;refGene_gn=THOC2;refGene_tx=XM_019824239.1;refGene_gn=THOC2;refGene_tx=XM_019824240.1;refGene_gn=THOC2;refGene_tx=XM_019824241.2;refGene_gn=THOC2;refGene_tx=XM_019824242.2;refGene_gn=THOC2;refGene_tx=XM_019824243.1;refGene_gn=THOC2;refGene_tx=XM_019824244.2;refGene_gn=THOC2;refGene_tx=XM_019824245.2;refGene_gn=THOC2;refGene_tx=XM_019824246.2;refGene_gn=THOC2;refGene_tx=XM_019824247.2;refGene_gn=THOC2;refGene_tx=XM_019824248.1;refGene_gn=THOC2;refGene_tx=XM_023249254.1
X	103551494	103551626
refGene_gn=THOC2;refGene_tx=XM_011279459.3;refGene_gn=THOC2;refGene_tx=XM_011279460.3;refGene_gn=THOC2;refGene_tx=XM_011279461.3;refGene_gn=THOC2;refGene_tx=XM_019824238.2;refGene_gn=THOC2;refGene_tx=XM_019824239.1;refGene_gn=THOC2;refGene_tx=XM_019824240.1;refGene_gn=THOC2;refGene_tx=XM_019824241.2;refGene_gn=THOC2;refGene_tx=XM_019824242.2;refGene_gn=THOC2;refGene_tx=XM_019824243.1;refGene_gn=THOC2;refGene_tx=XM_019824244.2;refGene_gn=THOC2;refGene_tx=XM_019824245.2;refGene_gn=THOC2;refGene_tx=XM_019824246.2;refGene_gn=THOC2;refGene_tx=XM_019824247.2;refGene_gn=THOC2;refGene_tx=XM_019824248.1;refGene_gn=THOC2;refGene_tx=XM_023249254.1
X	103551984	103552113
refGene_gn=THOC2;refGene_tx=XM_011279459.3;refGene_gn=THOC2;refGene_tx=XM_011279460.3;refGene_gn=THOC2;refGene_tx=XM_011279461.3;refGene_gn=THOC2;refGene_tx=XM_019824238.2;refGene_gn=THOC2;refGene_tx=XM_019824239.1;refGene_gn=THOC2;refGene_tx=XM_019824240.1;refGene_gn=THOC2;refGene_tx=XM_019824241.2;refGene_gn=THOC2;refGene_tx=XM_019824242.2;refGene_gn=THOC2;refGene_tx=XM_019824243.1;refGene_gn=THOC2;refGene_tx=XM_019824244.2;refGene_gn=THOC2;refGene_tx=XM_019824245.2;refGene_gn=THOC2;refGene_tx=XM_019824246.2;refGene_gn=THOC2;refGene_tx=XM_019824247.2;refGene_gn=THOC2;refGene_tx=XM_019824248.1;refGene_gn=THOC2;refGene_tx=

XM_023249254.1
X	103553028	103553186
refGene_gn=THOC2;refGene_tx=XM_011279459.3;refGene_gn=THOC2;refGene_tx=XM_011279460.3;refGene_gn=THOC2;refGene_tx=XM_011279461.3;refGene_gn=THOC2;refGene_tx=XM_019824238.2;refGene_gn=THOC2;refGene_tx=XM_019824239.1;refGene_gn=THOC2;refGene_tx=XM_019824240.1;refGene_gn=THOC2;refGene_tx=XM_019824241.2;refGene_gn=THOC2;refGene_tx=XM_019824242.2;refGene_gn=THOC2;refGene_tx=XM_019824243.1;refGene_gn=THOC2;refGene_tx=XM_019824244.2;refGene_gn=THOC2;refGene_tx=XM_019824245.2;refGene_gn=THOC2;refGene_tx=XM_019824246.2;refGene_gn=THOC2;refGene_tx=XM_019824247.2;refGene_gn=THOC2;refGene_tx=XM_019824248.1;refGene_gn=THOC2;refGene_tx=XM_023249254.1
X	103553641	103553783
refGene_gn=THOC2;refGene_tx=XM_011279459.3;refGene_gn=THOC2;refGene_tx=XM_011279460.3;refGene_gn=THOC2;refGene_tx=XM_011279461.3;refGene_gn=THOC2;refGene_tx=XM_019824238.2;refGene_gn=THOC2;refGene_tx=XM_019824239.1;refGene_gn=THOC2;refGene_tx=XM_019824240.1;refGene_gn=THOC2;refGene_tx=XM_019824241.2;refGene_gn=THOC2;refGene_tx=XM_019824242.2;refGene_gn=THOC2;refGene_tx=XM_019824243.1;refGene_gn=THOC2;refGene_tx=XM_019824244.2;refGene_gn=THOC2;refGene_tx=XM_019824245.2;refGene_gn=THOC2;refGene_tx=XM_019824246.2;refGene_gn=THOC2;refGene_tx=XM_019824247.2;refGene_gn=THOC2;refGene_tx=XM_019824248.1;refGene_gn=THOC2;refGene_tx=XM_023249254.1
X	103554816	103555092
refGene_gn=THOC2;refGene_tx=XM_011279459.3;refGene_gn=THOC2;refGene_tx=XM_011279460.3;refGene_gn=THOC2;refGene_tx=XM_011279461.3;refGene_gn=THOC2;refGene_tx=XM_019824238.2;refGene_gn=THOC2;refGene_tx=XM_019824239.1;refGene_gn=THOC2;refGene_tx=XM_019824240.1;refGene_gn=THOC2;refGene_tx=XM_019824241.2;refGene_gn=THOC2;refGene_tx=XM_019824242.2;refGene_gn=THOC2;refGene_tx=XM_019824243.1;refGene_gn=THOC2;refGene_tx=XM_019824244.2;refGene_gn=THOC2;refGene_tx=XM_019824245.2;refGene_gn=THOC2;refGene_tx=XM_019824246.2;refGene_gn=THOC2;refGene_tx=XM_019824247.2;refGene_gn=THOC2;refGene_tx=XM_019824248.1;refGene_gn=THOC2;refGene_tx=XM_023249254.1
X	103559256	103559421
refGene_gn=THOC2;refGene_tx=XM_011279459.3;refGene_gn=THOC2;refGene_tx=XM_011279460.3;refGene_gn=THOC2;refGene_tx=XM_011279461.3;refGene_gn=THOC2;refGene_tx=XM_019824238.2;refGene_gn=THOC2;refGene_tx=XM_019824239.1;refGene_gn=THOC2;refGene_tx=XM_019824240.1;refGene_gn=THOC2;refGene_tx=XM_019824241.2;refGene_gn=THOC2;refGene_tx=XM_019824242.2;refGene_gn=THOC2;refGene_tx=XM_019824243.1;refGene_gn=THOC2;refGene_tx=XM_019824244.2;refGene_gn=THOC2;refGene_tx=XM_019824245.2;refGene_gn=THOC2;refGene_tx=XM_019824246.2;refGene_gn=THOC2;refGene_tx=XM_019824247.2;refGene_gn=THOC2;refGene_tx=XM_019824248.1;refGene_gn=THOC2;refGene_tx=XM_023249254.1
X	103560438	103560618
refGene_gn=THOC2;refGene_tx=XM_011279459.3;refGene_gn=THOC2;refGene_tx=XM_011279460.3;refGene_gn=THOC2;refGene_tx=XM_011279461.3;refGene_gn=THOC2;refGene_tx=XM_019824238.2;refGene_gn=THOC2;refGene_tx=XM_019824239.1;refGene_gn=THOC2;refGene_tx=XM_019824240.1;refGene_gn=THOC2;refGe

ne_tx=XM_019824241.2;refGene_gn=THOC2;refGene_tx=XM_019824242.2;refGen
e_gn=THOC2;refGene_tx=XM_019824243.1;refGene_gn=THOC2;refGene_tx=XM_01
9824244.2;refGene_gn=THOC2;refGene_tx=XM_019824245.2;refGene_gn=THOC2;
refGene_tx=XM_019824246.2;refGene_gn=THOC2;refGene_tx=XM_019824247.2;r
efGene_gn=THOC2;refGene_tx=XM_019824248.1;refGene_gn=THOC2;refGene_tx=
XM_023249254.1
X	103561107	103561225
refGene_gn=THOC2;refGene_tx=XM_011279459.3;refGene_gn=THOC2;refGene_tx
=XM_011279460.3;refGene_gn=THOC2;refGene_tx=XM_011279461.3;refGene_gn=
THOC2;refGene_tx=XM_019824238.2;refGene_gn=THOC2;refGene_tx=XM_0198242
39.1;refGene_gn=THOC2;refGene_tx=XM_019824240.1;refGene_gn=THOC2;refGe
ne_tx=XM_019824241.2;refGene_gn=THOC2;refGene_tx=XM_019824242.2;refGen
e_gn=THOC2;refGene_tx=XM_019824243.1;refGene_gn=THOC2;refGene_tx=XM_01
9824244.2;refGene_gn=THOC2;refGene_tx=XM_019824245.2;refGene_gn=THOC2;
refGene_tx=XM_019824246.2;refGene_gn=THOC2;refGene_tx=XM_019824247.2;r
efGene_gn=THOC2;refGene_tx=XM_019824248.1;refGene_gn=THOC2;refGene_tx=
XM_023249254.1
X	103563425	103563522
refGene_gn=THOC2;refGene_tx=XM_011279459.3;refGene_gn=THOC2;refGene_tx
=XM_011279460.3;refGene_gn=THOC2;refGene_tx=XM_011279461.3;refGene_gn=
THOC2;refGene_tx=XM_019824238.2;refGene_gn=THOC2;refGene_tx=XM_0198242
39.1;refGene_gn=THOC2;refGene_tx=XM_019824240.1;refGene_gn=THOC2;refGe
ne_tx=XM_019824241.2;refGene_gn=THOC2;refGene_tx=XM_019824242.2;refGen
e_gn=THOC2;refGene_tx=XM_019824243.1;refGene_gn=THOC2;refGene_tx=XM_01
9824244.2;refGene_gn=THOC2;refGene_tx=XM_019824245.2;refGene_gn=THOC2;
refGene_tx=XM_019824246.2;refGene_gn=THOC2;refGene_tx=XM_019824247.2;r
efGene_gn=THOC2;refGene_tx=XM_019824248.1;refGene_gn=THOC2;refGene_tx=
XM_023249254.1
X	103565112	103565193
refGene_gn=THOC2;refGene_tx=XM_011279459.3;refGene_gn=THOC2;refGene_tx
=XM_011279460.3;refGene_gn=THOC2;refGene_tx=XM_011279461.3;refGene_gn=
THOC2;refGene_tx=XM_019824238.2;refGene_gn=THOC2;refGene_tx=XM_0198242
39.1;refGene_gn=THOC2;refGene_tx=XM_019824240.1;refGene_gn=THOC2;refGe
ne_tx=XM_019824241.2;refGene_gn=THOC2;refGene_tx=XM_019824242.2;refGen
e_gn=THOC2;refGene_tx=XM_019824243.1;refGene_gn=THOC2;refGene_tx=XM_01
9824244.2;refGene_gn=THOC2;refGene_tx=XM_019824245.2;refGene_gn=THOC2;
refGene_tx=XM_019824246.2;refGene_gn=THOC2;refGene_tx=XM_019824247.2;r
efGene_gn=THOC2;refGene_tx=XM_019824248.1;refGene_gn=THOC2;refGene_tx=
XM_023249254.1
X	103565611	103565705
refGene_gn=THOC2;refGene_tx=XM_011279459.3;refGene_gn=THOC2;refGene_tx
=XM_011279460.3;refGene_gn=THOC2;refGene_tx=XM_011279461.3;refGene_gn=
THOC2;refGene_tx=XM_019824238.2;refGene_gn=THOC2;refGene_tx=XM_0198242
39.1;refGene_gn=THOC2;refGene_tx=XM_019824240.1;refGene_gn=THOC2;refGe
ne_tx=XM_019824241.2;refGene_gn=THOC2;refGene_tx=XM_019824242.2;refGen
e_gn=THOC2;refGene_tx=XM_019824243.1;refGene_gn=THOC2;refGene_tx=XM_01
9824244.2;refGene_gn=THOC2;refGene_tx=XM_019824245.2;refGene_gn=THOC2;
refGene_tx=XM_019824246.2;refGene_gn=THOC2;refGene_tx=XM_019824247.2;r
efGene_gn=THOC2;refGene_tx=XM_019824248.1;refGene_gn=THOC2;refGene_tx=
XM_023249254.1

X	103566290	103566387
refGene_gn=THOC2;refGene_tx=XM_011279459.3;refGene_gn=THOC2;refGene_tx=XM_011279460.3;refGene_gn=THOC2;refGene_tx=XM_011279461.3;refGene_gn=THOC2;refGene_tx=XM_019824238.2;refGene_gn=THOC2;refGene_tx=XM_019824239.1;refGene_gn=THOC2;refGene_tx=XM_019824240.1;refGene_gn=THOC2;refGene_tx=XM_019824241.2;refGene_gn=THOC2;refGene_tx=XM_019824242.2;refGene_gn=THOC2;refGene_tx=XM_019824243.1;refGene_gn=THOC2;refGene_tx=XM_019824244.2;refGene_gn=THOC2;refGene_tx=XM_019824245.2;refGene_gn=THOC2;refGene_tx=XM_019824246.2;refGene_gn=THOC2;refGene_tx=XM_019824247.2;refGene_gn=THOC2;refGene_tx=XM_019824248.1;refGene_gn=THOC2;refGene_tx=XM_023249254.1
X	103574362	103574464
refGene_gn=THOC2;refGene_tx=XM_011279459.3;refGene_gn=THOC2;refGene_tx=XM_011279460.3;refGene_gn=THOC2;refGene_tx=XM_011279461.3;refGene_gn=THOC2;refGene_tx=XM_019824238.2;refGene_gn=THOC2;refGene_tx=XM_019824239.1;refGene_gn=THOC2;refGene_tx=XM_019824240.1;refGene_gn=THOC2;refGene_tx=XM_019824241.2;refGene_gn=THOC2;refGene_tx=XM_019824242.2;refGene_gn=THOC2;refGene_tx=XM_019824243.1;refGene_gn=THOC2;refGene_tx=XM_019824244.2;refGene_gn=THOC2;refGene_tx=XM_019824245.2;refGene_gn=THOC2;refGene_tx=XM_019824246.2;refGene_gn=THOC2;refGene_tx=XM_019824247.2;refGene_gn=THOC2;refGene_tx=XM_019824248.1;refGene_gn=THOC2;refGene_tx=XM_023249254.1
X	103574561	103574692
refGene_gn=THOC2;refGene_tx=XM_011279459.3;refGene_gn=THOC2;refGene_tx=XM_011279460.3;refGene_gn=THOC2;refGene_tx=XM_011279461.3;refGene_gn=THOC2;refGene_tx=XM_019824238.2;refGene_gn=THOC2;refGene_tx=XM_019824239.1;refGene_gn=THOC2;refGene_tx=XM_019824240.1;refGene_gn=THOC2;refGene_tx=XM_019824241.2;refGene_gn=THOC2;refGene_tx=XM_019824242.2;refGene_gn=THOC2;refGene_tx=XM_019824243.1;refGene_gn=THOC2;refGene_tx=XM_019824244.2;refGene_gn=THOC2;refGene_tx=XM_019824245.2;refGene_gn=THOC2;refGene_tx=XM_019824246.2;refGene_gn=THOC2;refGene_tx=XM_019824247.2;refGene_gn=THOC2;refGene_tx=XM_019824248.1;refGene_gn=THOC2;refGene_tx=XM_023249254.1
X	103575384	103575426
refGene_gn=THOC2;refGene_tx=XM_011279459.3;refGene_gn=THOC2;refGene_tx=XM_011279460.3;refGene_gn=THOC2;refGene_tx=XM_011279461.3;refGene_gn=THOC2;refGene_tx=XM_019824238.2;refGene_gn=THOC2;refGene_tx=XM_019824239.1;refGene_gn=THOC2;refGene_tx=XM_019824240.1;refGene_gn=THOC2;refGene_tx=XM_019824241.2;refGene_gn=THOC2;refGene_tx=XM_019824242.2;refGene_gn=THOC2;refGene_tx=XM_019824243.1;refGene_gn=THOC2;refGene_tx=XM_019824244.2;refGene_gn=THOC2;refGene_tx=XM_019824245.2;refGene_gn=THOC2;refGene_tx=XM_019824246.2;refGene_gn=THOC2;refGene_tx=XM_019824247.2;refGene_gn=THOC2;refGene_tx=XM_019824248.1;refGene_gn=THOC2;refGene_tx=XM_023249254.1
X	103585250	103585274
refGene_gn=THOC2;refGene_tx=XM_019824238.2;refGene_gn=THOC2;refGene_tx=XM_019824239.1;refGene_gn=THOC2;refGene_tx=XM_019824240.1;refGene_gn=THOC2;refGene_tx=XM_019824241.2;refGene_gn=THOC2;refGene_tx=XM_019824242.2;refGene_gn=THOC2;refGene_tx=XM_019824243.1;refGene_gn=THOC2;refGene_tx=XM_019824244.2;refGene_gn=THOC2;refGene_tx=XM_019824245.2;refGen

e_gn=THOC2;refGene_tx=XM_019824247.2
X	103586389	103586585
refGene_gn=THOC2;refGene_tx=XM_011279459.3;refGene_gn=THOC2;refGene_tx=XM_011279460.3;refGene_gn=THOC2;refGene_tx=XM_011279461.3;refGene_gn=THOC2;refGene_tx=XM_019824238.2;refGene_gn=THOC2;refGene_tx=XM_019824239.1;refGene_gn=THOC2;refGene_tx=XM_019824240.1;refGene_gn=THOC2;refGene_tx=XM_019824241.2;refGene_gn=THOC2;refGene_tx=XM_019824242.2;refGene_gn=THOC2;refGene_tx=XM_019824243.1;refGene_gn=THOC2;refGene_tx=XM_019824244.2;refGene_gn=THOC2;refGene_tx=XM_019824245.2;refGene_gn=THOC2;refGene_tx=XM_019824246.2;refGene_gn=THOC2;refGene_tx=XM_019824247.2;refGene_gn=THOC2;refGene_tx=XM_019824248.1;refGene_gn=THOC2;refGene_tx=XM_023249254.1
X	103587810	103587983
refGene_gn=THOC2;refGene_tx=XM_011279459.3;refGene_gn=THOC2;refGene_tx=XM_011279460.3;refGene_gn=THOC2;refGene_tx=XM_011279461.3;refGene_gn=THOC2;refGene_tx=XM_019824238.2;refGene_gn=THOC2;refGene_tx=XM_019824239.1;refGene_gn=THOC2;refGene_tx=XM_019824240.1;refGene_gn=THOC2;refGene_tx=XM_019824241.2;refGene_gn=THOC2;refGene_tx=XM_019824242.2;refGene_gn=THOC2;refGene_tx=XM_019824243.1;refGene_gn=THOC2;refGene_tx=XM_019824244.2;refGene_gn=THOC2;refGene_tx=XM_019824245.2;refGene_gn=THOC2;refGene_tx=XM_019824246.2;refGene_gn=THOC2;refGene_tx=XM_019824247.2;refGene_gn=THOC2;refGene_tx=XM_019824248.1;refGene_gn=THOC2;refGene_tx=XM_023249254.1
X	103588580	103588736
refGene_gn=THOC2;refGene_tx=XM_011279459.3;refGene_gn=THOC2;refGene_tx=XM_011279460.3;refGene_gn=THOC2;refGene_tx=XM_011279461.3;refGene_gn=THOC2;refGene_tx=XM_019824238.2;refGene_gn=THOC2;refGene_tx=XM_019824239.1;refGene_gn=THOC2;refGene_tx=XM_019824240.1;refGene_gn=THOC2;refGene_tx=XM_019824241.2;refGene_gn=THOC2;refGene_tx=XM_019824242.2;refGene_gn=THOC2;refGene_tx=XM_019824243.1;refGene_gn=THOC2;refGene_tx=XM_019824244.2;refGene_gn=THOC2;refGene_tx=XM_019824245.2;refGene_gn=THOC2;refGene_tx=XM_019824246.2;refGene_gn=THOC2;refGene_tx=XM_019824247.2;refGene_gn=THOC2;refGene_tx=XM_019824248.1;refGene_gn=THOC2;refGene_tx=XM_023249254.1
X	103591891	103591984
refGene_gn=THOC2;refGene_tx=XM_011279459.3;refGene_gn=THOC2;refGene_tx=XM_011279460.3;refGene_gn=THOC2;refGene_tx=XM_011279461.3;refGene_gn=THOC2;refGene_tx=XM_019824238.2;refGene_gn=THOC2;refGene_tx=XM_019824239.1;refGene_gn=THOC2;refGene_tx=XM_019824240.1;refGene_gn=THOC2;refGene_tx=XM_019824241.2;refGene_gn=THOC2;refGene_tx=XM_019824242.2;refGene_gn=THOC2;refGene_tx=XM_019824243.1;refGene_gn=THOC2;refGene_tx=XM_019824244.2;refGene_gn=THOC2;refGene_tx=XM_019824245.2;refGene_gn=THOC2;refGene_tx=XM_019824246.2;refGene_gn=THOC2;refGene_tx=XM_019824247.2;refGene_gn=THOC2;refGene_tx=XM_019824248.1;refGene_gn=THOC2;refGene_tx=XM_023249254.1
X	103602737	103602904
refGene_gn=THOC2;refGene_tx=XM_011279459.3;refGene_gn=THOC2;refGene_tx=XM_011279460.3;refGene_gn=THOC2;refGene_tx=XM_011279461.3;refGene_gn=THOC2;refGene_tx=XM_019824238.2;refGene_gn=THOC2;refGene_tx=XM_019824239.1;refGene_gn=THOC2;refGene_tx=XM_019824240.1;refGene_gn=THOC2;refGe

ne_tx=XM_019824241.2;refGene_gn=THOC2;refGene_tx=XM_019824242.2;refGene_gn=THOC2;refGene_tx=XM_019824243.1;refGene_gn=THOC2;refGene_tx=XM_019824244.2;refGene_gn=THOC2;refGene_tx=XM_019824245.2;refGene_gn=THOC2;refGene_tx=XM_019824246.2;refGene_gn=THOC2;refGene_tx=XM_019824247.2;refGene_gn=THOC2;refGene_tx=XM_019824248.1;refGene_gn=THOC2;refGene_tx=XM_023249254.1
X 103613127 103613261
refGene_gn=THOC2;refGene_tx=XM_011279459.3;refGene_gn=THOC2;refGene_tx=XM_011279460.3;refGene_gn=THOC2;refGene_tx=XM_011279461.3;refGene_gn=THOC2;refGene_tx=XM_019824238.2;refGene_gn=THOC2;refGene_tx=XM_019824239.1;refGene_gn=THOC2;refGene_tx=XM_019824240.1;refGene_gn=THOC2;refGene_tx=XM_019824241.2;refGene_gn=THOC2;refGene_tx=XM_019824242.2;refGene_gn=THOC2;refGene_tx=XM_019824243.1;refGene_gn=THOC2;refGene_tx=XM_019824244.2;refGene_gn=THOC2;refGene_tx=XM_019824245.2;refGene_gn=THOC2;refGene_tx=XM_019824246.2;refGene_gn=THOC2;refGene_tx=XM_019824247.2;refGene_gn=THOC2;refGene_tx=XM_019824248.1;refGene_gn=THOC2;refGene_tx=XM_023249254.1
X 103613818 103613940
refGene_gn=THOC2;refGene_tx=XM_011279459.3;refGene_gn=THOC2;refGene_tx=XM_011279460.3;refGene_gn=THOC2;refGene_tx=XM_011279461.3;refGene_gn=THOC2;refGene_tx=XM_019824238.2;refGene_gn=THOC2;refGene_tx=XM_019824239.1;refGene_gn=THOC2;refGene_tx=XM_019824240.1;refGene_gn=THOC2;refGene_tx=XM_019824241.2;refGene_gn=THOC2;refGene_tx=XM_019824242.2;refGene_gn=THOC2;refGene_tx=XM_019824243.1;refGene_gn=THOC2;refGene_tx=XM_019824244.2;refGene_gn=THOC2;refGene_tx=XM_019824245.2;refGene_gn=THOC2;refGene_tx=XM_019824246.2;refGene_gn=THOC2;refGene_tx=XM_019824247.2;refGene_gn=THOC2;refGene_tx=XM_019824248.1;refGene_gn=THOC2;refGene_tx=XM_023249254.1
X 103614811 103614882
refGene_gn=THOC2;refGene_tx=XM_011279459.3;refGene_gn=THOC2;refGene_tx=XM_011279460.3;refGene_gn=THOC2;refGene_tx=XM_011279461.3;refGene_gn=THOC2;refGene_tx=XM_019824238.2;refGene_gn=THOC2;refGene_tx=XM_019824239.1;refGene_gn=THOC2;refGene_tx=XM_019824240.1;refGene_gn=THOC2;refGene_tx=XM_019824241.2;refGene_gn=THOC2;refGene_tx=XM_019824242.2;refGene_gn=THOC2;refGene_tx=XM_019824243.1;refGene_gn=THOC2;refGene_tx=XM_019824244.2;refGene_gn=THOC2;refGene_tx=XM_019824245.2;refGene_gn=THOC2;refGene_tx=XM_019824246.2;refGene_gn=THOC2;refGene_tx=XM_019824247.2;refGene_gn=THOC2;refGene_tx=XM_019824248.1;refGene_gn=THOC2;refGene_tx=XM_023249254.1
X 103618871 103618923
refGene_gn=THOC2;refGene_tx=XM_011279459.3;refGene_gn=THOC2;refGene_tx=XM_011279460.3;refGene_gn=THOC2;refGene_tx=XM_011279461.3;refGene_gn=THOC2;refGene_tx=XM_019824238.2;refGene_gn=THOC2;refGene_tx=XM_019824239.1;refGene_gn=THOC2;refGene_tx=XM_019824240.1;refGene_gn=THOC2;refGene_tx=XM_019824241.2;refGene_gn=THOC2;refGene_tx=XM_019824242.2;refGene_gn=THOC2;refGene_tx=XM_019824243.1;refGene_gn=THOC2;refGene_tx=XM_019824244.2;refGene_gn=THOC2;refGene_tx=XM_019824245.2;refGene_gn=THOC2;refGene_tx=XM_019824246.2;refGene_gn=THOC2;refGene_tx=XM_019824247.2;refGene_gn=THOC2;refGene_tx=XM_019824248.1;refGene_gn=THOC2;refGene_tx=XM_023249254.1

X 103622289 103622381
refGene_gn=THOC2;refGene_tx=XM_011279459.3;refGene_gn=THOC2;refGene_tx=XM_011279460.3;refGene_gn=THOC2;refGene_tx=XM_011279461.3;refGene_gn=THOC2;refGene_tx=XM_019824238.2;refGene_gn=THOC2;refGene_tx=XM_019824239.1;refGene_gn=THOC2;refGene_tx=XM_019824240.1;refGene_gn=THOC2;refGene_tx=XM_019824241.2;refGene_gn=THOC2;refGene_tx=XM_019824242.2;refGene_gn=THOC2;refGene_tx=XM_019824243.1;refGene_gn=THOC2;refGene_tx=XM_019824244.2;refGene_gn=THOC2;refGene_tx=XM_019824245.2;refGene_gn=THOC2;refGene_tx=XM_019824246.2;refGene_gn=THOC2;refGene_tx=XM_019824247.2;refGene_gn=THOC2;refGene_tx=XM_019824248.1;refGene_gn=THOC2;refGene_tx=XM_023249254.1
X 103627679 103627738
refGene_gn=THOC2;refGene_tx=XM_011279459.3;refGene_gn=THOC2;refGene_tx=XM_011279460.3;refGene_gn=THOC2;refGene_tx=XM_011279461.3;refGene_gn=THOC2;refGene_tx=XM_019824238.2;refGene_gn=THOC2;refGene_tx=XM_019824239.1;refGene_gn=THOC2;refGene_tx=XM_019824240.1;refGene_gn=THOC2;refGene_tx=XM_019824241.2;refGene_gn=THOC2;refGene_tx=XM_019824242.2;refGene_gn=THOC2;refGene_tx=XM_019824243.1;refGene_gn=THOC2;refGene_tx=XM_019824244.2;refGene_gn=THOC2;refGene_tx=XM_019824245.2;refGene_gn=THOC2;refGene_tx=XM_019824246.2;refGene_gn=THOC2;refGene_tx=XM_019824247.2;refGene_gn=THOC2;refGene_tx=XM_019824248.1;refGene_gn=THOC2;refGene_tx=XM_023249254.1
X 103642429 103642500
refGene_gn=THOC2;refGene_tx=XM_011279459.3;refGene_gn=THOC2;refGene_tx=XM_011279460.3;refGene_gn=THOC2;refGene_tx=XM_011279461.3;refGene_gn=THOC2;refGene_tx=XM_019824238.2;refGene_gn=THOC2;refGene_tx=XM_019824239.1;refGene_gn=THOC2;refGene_tx=XM_019824240.1;refGene_gn=THOC2;refGene_tx=XM_019824241.2;refGene_gn=THOC2;refGene_tx=XM_019824242.2;refGene_gn=THOC2;refGene_tx=XM_019824243.1;refGene_gn=THOC2;refGene_tx=XM_019824244.2;refGene_gn=THOC2;refGene_tx=XM_019824245.2;refGene_gn=THOC2;refGene_tx=XM_019824246.2;refGene_gn=THOC2;refGene_tx=XM_019824247.2;refGene_gn=THOC2;refGene_tx=XM_019824248.1;refGene_gn=THOC2;refGene_tx=XM_023249254.1
X 103771295 103772169
refGene_gn=XIAP;refGene_tx=XM_011279458.3
X 103773234 103773334
refGene_gn=XIAP;refGene_tx=XM_011279458.3
X 103774346 103774425
refGene_gn=XIAP;refGene_tx=XM_011279458.3
X 103775102 103775145
refGene_gn=XIAP;refGene_tx=XM_011279458.3
X 103780121 103780322
refGene_gn=XIAP;refGene_tx=XM_011279458.3
X 103787815 103787856
refGene_gn=XIAP;refGene_tx=XM_011279458.3
X 103787857 103788010
refGene_gn=XIAP;refGene_tx=XM_011279458.3
X 103884894 103884938
refGene_gn=STAG2;refGene_tx=XM_011279456.3;refGene_gn=STAG2;refGene_tx=XM_019824249.1;refGene_gn=STAG2;refGene_tx=XM_019824250.1;refGene_gn=

STAG2;refGene_tx=XM_019824251.1;refGene_gn=STAG2;refGene_tx=XM_0198242
52.1
X	103889239	103889318
refGene_gn=STAG2;refGene_tx=XM_011279456.3;refGene_gn=STAG2;refGene_tx
=XM_019824249.1;refGene_gn=STAG2;refGene_tx=XM_019824250.1;refGene_gn=
STAG2;refGene_tx=XM_019824251.1;refGene_gn=STAG2;refGene_tx=XM_0198242
52.1
X	103893859	103894024
refGene_gn=STAG2;refGene_tx=XM_011279456.3;refGene_gn=STAG2;refGene_tx
=XM_019824249.1;refGene_gn=STAG2;refGene_tx=XM_019824250.1;refGene_gn=
STAG2;refGene_tx=XM_019824251.1;refGene_gn=STAG2;refGene_tx=XM_0198242
52.1
X	103903539	103903636
refGene_gn=STAG2;refGene_tx=XM_011279456.3;refGene_gn=STAG2;refGene_tx
=XM_019824249.1;refGene_gn=STAG2;refGene_tx=XM_019824250.1;refGene_gn=
STAG2;refGene_tx=XM_019824251.1;refGene_gn=STAG2;refGene_tx=XM_0198242
52.1
X	103908293	103908370
refGene_gn=STAG2;refGene_tx=XM_011279456.3;refGene_gn=STAG2;refGene_tx
=XM_019824249.1;refGene_gn=STAG2;refGene_tx=XM_019824250.1;refGene_gn=
STAG2;refGene_tx=XM_019824251.1;refGene_gn=STAG2;refGene_tx=XM_0198242
52.1
X	103911440	103911645
refGene_gn=STAG2;refGene_tx=XM_011279456.3;refGene_gn=STAG2;refGene_tx
=XM_019824249.1;refGene_gn=STAG2;refGene_tx=XM_019824250.1;refGene_gn=
STAG2;refGene_tx=XM_019824251.1;refGene_gn=STAG2;refGene_tx=XM_0198242
52.1
X	103913803	103913955
refGene_gn=STAG2;refGene_tx=XM_011279456.3;refGene_gn=STAG2;refGene_tx
=XM_019824249.1;refGene_gn=STAG2;refGene_tx=XM_019824250.1;refGene_gn=
STAG2;refGene_tx=XM_019824251.1;refGene_gn=STAG2;refGene_tx=XM_0198242
52.1
X	103916311	103916385
refGene_gn=STAG2;refGene_tx=XM_011279456.3;refGene_gn=STAG2;refGene_tx
=XM_019824249.1;refGene_gn=STAG2;refGene_tx=XM_019824250.1;refGene_gn=
STAG2;refGene_tx=XM_019824251.1;refGene_gn=STAG2;refGene_tx=XM_0198242
52.1
X	103917748	103917872
refGene_gn=STAG2;refGene_tx=XM_011279456.3;refGene_gn=STAG2;refGene_tx
=XM_019824249.1;refGene_gn=STAG2;refGene_tx=XM_019824250.1;refGene_gn=
STAG2;refGene_tx=XM_019824251.1;refGene_gn=STAG2;refGene_tx=XM_0198242
52.1
X	103918676	103918775
refGene_gn=STAG2;refGene_tx=XM_011279456.3;refGene_gn=STAG2;refGene_tx
=XM_019824249.1;refGene_gn=STAG2;refGene_tx=XM_019824250.1;refGene_gn=
STAG2;refGene_tx=XM_019824251.1;refGene_gn=STAG2;refGene_tx=XM_0198242
52.1
X	103918906	103918986
refGene_gn=STAG2;refGene_tx=XM_011279456.3;refGene_gn=STAG2;refGene_tx
=XM_019824249.1;refGene_gn=STAG2;refGene_tx=XM_019824250.1;refGene_gn=

STAG2;refGene_tx=XM_019824251.1;refGene_gn=STAG2;refGene_tx=XM_0198242
52.1
X	103924760	103924868
refGene_gn=STAG2;refGene_tx=XM_011279456.3;refGene_gn=STAG2;refGene_tx
=XM_019824249.1;refGene_gn=STAG2;refGene_tx=XM_019824250.1;refGene_gn=
STAG2;refGene_tx=XM_019824251.1;refGene_gn=STAG2;refGene_tx=XM_0198242
52.1
X	103926227	103926339
refGene_gn=STAG2;refGene_tx=XM_011279456.3;refGene_gn=STAG2;refGene_tx
=XM_019824249.1;refGene_gn=STAG2;refGene_tx=XM_019824250.1;refGene_gn=
STAG2;refGene_tx=XM_019824251.1;refGene_gn=STAG2;refGene_tx=XM_0198242
52.1
X	103928239	103928357
refGene_gn=STAG2;refGene_tx=XM_011279456.3;refGene_gn=STAG2;refGene_tx
=XM_019824249.1;refGene_gn=STAG2;refGene_tx=XM_019824250.1;refGene_gn=
STAG2;refGene_tx=XM_019824251.1;refGene_gn=STAG2;refGene_tx=XM_0198242
52.1
X	103928757	103928861
refGene_gn=STAG2;refGene_tx=XM_011279456.3;refGene_gn=STAG2;refGene_tx
=XM_019824249.1;refGene_gn=STAG2;refGene_tx=XM_019824250.1;refGene_gn=
STAG2;refGene_tx=XM_019824251.1;refGene_gn=STAG2;refGene_tx=XM_0198242
52.1
X	103930337	103930430
refGene_gn=STAG2;refGene_tx=XM_011279456.3;refGene_gn=STAG2;refGene_tx
=XM_019824249.1;refGene_gn=STAG2;refGene_tx=XM_019824250.1;refGene_gn=
STAG2;refGene_tx=XM_019824251.1;refGene_gn=STAG2;refGene_tx=XM_0198242
52.1
X	103930562	103930652
refGene_gn=STAG2;refGene_tx=XM_011279456.3;refGene_gn=STAG2;refGene_tx
=XM_019824249.1;refGene_gn=STAG2;refGene_tx=XM_019824250.1;refGene_gn=
STAG2;refGene_tx=XM_019824251.1;refGene_gn=STAG2;refGene_tx=XM_0198242
52.1
X	103931261	103931465
refGene_gn=STAG2;refGene_tx=XM_011279456.3;refGene_gn=STAG2;refGene_tx
=XM_019824249.1;refGene_gn=STAG2;refGene_tx=XM_019824250.1;refGene_gn=
STAG2;refGene_tx=XM_019824251.1;refGene_gn=STAG2;refGene_tx=XM_0198242
52.1
X	103933671	103933742
refGene_gn=STAG2;refGene_tx=XM_011279456.3;refGene_gn=STAG2;refGene_tx
=XM_019824249.1;refGene_gn=STAG2;refGene_tx=XM_019824250.1;refGene_gn=
STAG2;refGene_tx=XM_019824251.1;refGene_gn=STAG2;refGene_tx=XM_0198242
52.1
X	103934064	103934152
refGene_gn=STAG2;refGene_tx=XM_011279456.3;refGene_gn=STAG2;refGene_tx
=XM_019824249.1;refGene_gn=STAG2;refGene_tx=XM_019824250.1;refGene_gn=
STAG2;refGene_tx=XM_019824251.1;refGene_gn=STAG2;refGene_tx=XM_0198242
52.1
X	103934241	103934322
refGene_gn=STAG2;refGene_tx=XM_011279456.3;refGene_gn=STAG2;refGene_tx
=XM_019824249.1;refGene_gn=STAG2;refGene_tx=XM_019824250.1;refGene_gn=

STAG2;refGene_tx=XM_019824251.1;refGene_gn=STAG2;refGene_tx=XM_0198242
52.1
X	103936119	103936212
refGene_gn=STAG2;refGene_tx=XM_011279456.3;refGene_gn=STAG2;refGene_tx
=XM_019824249.1;refGene_gn=STAG2;refGene_tx=XM_019824250.1;refGene_gn=
STAG2;refGene_tx=XM_019824251.1;refGene_gn=STAG2;refGene_tx=XM_0198242
52.1
X	103942534	103942709
refGene_gn=STAG2;refGene_tx=XM_011279456.3;refGene_gn=STAG2;refGene_tx
=XM_019824249.1;refGene_gn=STAG2;refGene_tx=XM_019824250.1;refGene_gn=
STAG2;refGene_tx=XM_019824251.1;refGene_gn=STAG2;refGene_tx=XM_0198242
52.1
X	103947076	103947216
refGene_gn=STAG2;refGene_tx=XM_011279456.3;refGene_gn=STAG2;refGene_tx
=XM_019824249.1;refGene_gn=STAG2;refGene_tx=XM_019824250.1;refGene_gn=
STAG2;refGene_tx=XM_019824251.1;refGene_gn=STAG2;refGene_tx=XM_0198242
52.1
X	103948726	103948828
refGene_gn=STAG2;refGene_tx=XM_011279456.3;refGene_gn=STAG2;refGene_tx
=XM_019824249.1;refGene_gn=STAG2;refGene_tx=XM_019824250.1;refGene_gn=
STAG2;refGene_tx=XM_019824251.1;refGene_gn=STAG2;refGene_tx=XM_0198242
52.1
X	103951685	103951834
refGene_gn=STAG2;refGene_tx=XM_011279456.3;refGene_gn=STAG2;refGene_tx
=XM_019824249.1;refGene_gn=STAG2;refGene_tx=XM_019824250.1;refGene_gn=
STAG2;refGene_tx=XM_019824251.1;refGene_gn=STAG2;refGene_tx=XM_0198242
52.1
X	103953340	103953469
refGene_gn=STAG2;refGene_tx=XM_011279456.3;refGene_gn=STAG2;refGene_tx
=XM_019824249.1;refGene_gn=STAG2;refGene_tx=XM_019824250.1;refGene_gn=
STAG2;refGene_tx=XM_019824251.1;refGene_gn=STAG2;refGene_tx=XM_0198242
52.1
X	103957480	103957704
refGene_gn=STAG2;refGene_tx=XM_011279456.3;refGene_gn=STAG2;refGene_tx
=XM_019824249.1;refGene_gn=STAG2;refGene_tx=XM_019824250.1;refGene_gn=
STAG2;refGene_tx=XM_019824251.1;refGene_gn=STAG2;refGene_tx=XM_0198242
52.1
X	103960705	103960895
refGene_gn=STAG2;refGene_tx=XM_011279456.3;refGene_gn=STAG2;refGene_tx
=XM_019824249.1;refGene_gn=STAG2;refGene_tx=XM_019824250.1;refGene_gn=
STAG2;refGene_tx=XM_019824251.1;refGene_gn=STAG2;refGene_tx=XM_0198242
52.1
X	103960985	103961096
refGene_gn=STAG2;refGene_tx=XM_011279456.3;refGene_gn=STAG2;refGene_tx
=XM_019824249.1;refGene_gn=STAG2;refGene_tx=XM_019824250.1;refGene_gn=
STAG2;refGene_tx=XM_019824251.1
X	103964106	103964233
refGene_gn=STAG2;refGene_tx=XM_011279456.3;refGene_gn=STAG2;refGene_tx
=XM_019824249.1;refGene_gn=STAG2;refGene_tx=XM_019824250.1;refGene_gn=
STAG2;refGene_tx=XM_019824251.1;refGene_gn=STAG2;refGene_tx=XM_0198242

52.1
X 103965395 103965473
refGene_gn=STAG2;refGene_tx=XM_011279456.3;refGene_gn=STAG2;refGene_tx=XM_019824249.1;refGene_gn=STAG2;refGene_tx=XM_019824250.1;refGene_gn=STAG2;refGene_tx=XM_019824251.1;refGene_gn=STAG2;refGene_tx=XM_019824252.1
X 103970210 103970234
refGene_gn=STAG2;refGene_tx=XM_011279456.3;refGene_gn=STAG2;refGene_tx=XM_019824249.1;refGene_gn=STAG2;refGene_tx=XM_019824250.1;refGene_gn=STAG2;refGene_tx=XM_019824251.1;refGene_gn=STAG2;refGene_tx=XM_019824252.1
X 104152118 104154596
refGene_gn=LOC109496350;refGene_tx=XM_019823453.2
X 104166709 104166846
refGene_gn=SH2D1A;refGene_tx=XM_004000868.4
X 104180283 104180347
refGene_gn=SH2D1A;refGene_tx=XM_004000868.4
X 104185085 104185230
refGene_gn=SH2D1A;refGene_tx=XM_004000868.4
X 104186532 104186573
refGene_gn=SH2D1A;refGene_tx=XM_004000868.4
X 104196082 104196841
refGene_gn=TENM1;refGene_tx=XM_011279453.2;refGene_gn=TENM1;refGene_tx=XM_011279454.2;refGene_gn=TENM1;refGene_tx=XM_019824255.1
X 104198236 104198379
refGene_gn=TENM1;refGene_tx=XM_011279453.2;refGene_gn=TENM1;refGene_tx=XM_011279454.2;refGene_gn=TENM1;refGene_tx=XM_019824255.1
X 104199185 104200406
refGene_gn=TENM1;refGene_tx=XM_011279453.2;refGene_gn=TENM1;refGene_tx=XM_011279454.2;refGene_gn=TENM1;refGene_tx=XM_019824255.1
X 104201236 104201624
refGene_gn=TENM1;refGene_tx=XM_011279453.2;refGene_gn=TENM1;refGene_tx=XM_011279454.2;refGene_gn=TENM1;refGene_tx=XM_019824255.1
X 104208128 104208425
refGene_gn=TENM1;refGene_tx=XM_011279453.2;refGene_gn=TENM1;refGene_tx=XM_011279454.2;refGene_gn=TENM1;refGene_tx=XM_019824255.1
X 104220584 104220820
refGene_gn=TENM1;refGene_tx=XM_011279453.2;refGene_gn=TENM1;refGene_tx=XM_011279454.2;refGene_gn=TENM1;refGene_tx=XM_019824255.1
X 104221907 104222080
refGene_gn=TENM1;refGene_tx=XM_011279453.2;refGene_gn=TENM1;refGene_tx=XM_011279454.2;refGene_gn=TENM1;refGene_tx=XM_019824255.1
X 104235162 104235673
refGene_gn=TENM1;refGene_tx=XM_011279453.2;refGene_gn=TENM1;refGene_tx=XM_011279454.2;refGene_gn=TENM1;refGene_tx=XM_019824255.1
X 104237102 104237469
refGene_gn=TENM1;refGene_tx=XM_011279453.2;refGene_gn=TENM1;refGene_tx=XM_011279454.2;refGene_gn=TENM1;refGene_tx=XM_019824255.1
X 104265310 104265465
refGene_gn=TENM1;refGene_tx=XM_011279453.2;refGene_gn=TENM1;refGene_tx

=XM_011279454.2;refGene_gn=TENM1;refGene_tx=XM_019824255.1
X	104293189	104293422
refGene_gn=TENM1;refGene_tx=XM_011279453.2;refGene_gn=TENM1;refGene_tx=XM_011279454.2;refGene_gn=TENM1;refGene_tx=XM_019824255.1
X	104298325	104298346
refGene_gn=TENM1;refGene_tx=XM_011279453.2;refGene_gn=TENM1;refGene_tx=XM_011279454.2;refGene_gn=TENM1;refGene_tx=XM_019824255.1
X	104306590	104306840
refGene_gn=TENM1;refGene_tx=XM_011279453.2;refGene_gn=TENM1;refGene_tx=XM_011279454.2;refGene_gn=TENM1;refGene_tx=XM_019824255.1
X	104315086	104315230
refGene_gn=TENM1;refGene_tx=XM_011279453.2;refGene_gn=TENM1;refGene_tx=XM_011279454.2;refGene_gn=TENM1;refGene_tx=XM_019824255.1
X	104331162	104331430
refGene_gn=TENM1;refGene_tx=XM_011279453.2;refGene_gn=TENM1;refGene_tx=XM_011279454.2;refGene_gn=TENM1;refGene_tx=XM_019824255.1
X	104333920	104334182
refGene_gn=TENM1;refGene_tx=XM_011279453.2;refGene_gn=TENM1;refGene_tx=XM_011279454.2;refGene_gn=TENM1;refGene_tx=XM_019824255.1
X	104340417	104340537
refGene_gn=TENM1;refGene_tx=XM_011279453.2;refGene_gn=TENM1;refGene_tx=XM_011279454.2;refGene_gn=TENM1;refGene_tx=XM_019824255.1
X	104358259	104358476
refGene_gn=TENM1;refGene_tx=XM_011279453.2;refGene_gn=TENM1;refGene_tx=XM_011279454.2;refGene_gn=TENM1;refGene_tx=XM_019824255.1
X	104369394	104369541
refGene_gn=TENM1;refGene_tx=XM_011279453.2;refGene_gn=TENM1;refGene_tx=XM_011279454.2;refGene_gn=TENM1;refGene_tx=XM_019824255.1
X	104371491	104371515
refGene_gn=TENM1;refGene_tx=XM_011279453.2;refGene_gn=TENM1;refGene_tx=XM_011279454.2;refGene_gn=TENM1;refGene_tx=XM_019824255.1
X	104373076	104373262
refGene_gn=TENM1;refGene_tx=XM_011279453.2;refGene_gn=TENM1;refGene_tx=XM_011279454.2;refGene_gn=TENM1;refGene_tx=XM_019824255.1
X	104441621	104441822
refGene_gn=TENM1;refGene_tx=XM_011279453.2;refGene_gn=TENM1;refGene_tx=XM_011279454.2;refGene_gn=TENM1;refGene_tx=XM_019824255.1
X	104444938	104445133
refGene_gn=TENM1;refGene_tx=XM_011279453.2;refGene_gn=TENM1;refGene_tx=XM_011279454.2;refGene_gn=TENM1;refGene_tx=XM_019824255.1
X	104446441	104446543
refGene_gn=TENM1;refGene_tx=XM_011279453.2;refGene_gn=TENM1;refGene_tx=XM_011279454.2;refGene_gn=TENM1;refGene_tx=XM_019824255.1
X	104451601	104451812
refGene_gn=TENM1;refGene_tx=XM_011279453.2;refGene_gn=TENM1;refGene_tx=XM_011279454.2;refGene_gn=TENM1;refGene_tx=XM_019824255.1
X	104453025	104453225
refGene_gn=TENM1;refGene_tx=XM_011279453.2;refGene_gn=TENM1;refGene_tx=XM_011279454.2;refGene_gn=TENM1;refGene_tx=XM_019824255.1
X	104473930	104474083

refGene_gn=TENM1;refGene_tx=XM_011279453.2;refGene_gn=TENM1;refGene_tx=XM_011279454.2;refGene_gn=TENM1;refGene_tx=XM_019824255.1
X	104507161	104507400
refGene_gn=TENM1;refGene_tx=XM_011279453.2;refGene_gn=TENM1;refGene_tx=XM_011279454.2;refGene_gn=TENM1;refGene_tx=XM_019824255.1
X	104537397	104537638
refGene_gn=TENM1;refGene_tx=XM_011279453.2;refGene_gn=TENM1;refGene_tx=XM_011279454.2;refGene_gn=TENM1;refGene_tx=XM_019824255.1
X	104679169	104679226
refGene_gn=TENM1;refGene_tx=XM_011279453.2;refGene_gn=TENM1;refGene_tx=XM_011279454.2
X	104680907	104681168
refGene_gn=TENM1;refGene_tx=XM_011279453.2;refGene_gn=TENM1;refGene_tx=XM_011279454.2
X	104748846	104749063
refGene_gn=TENM1;refGene_tx=XM_011279453.2;refGene_gn=TENM1;refGene_tx=XM_011279454.2
X	105110642	105111783
refGene_gn=LOC101096959;refGene_tx=XM_023248979.1
X	105111833	105112674
refGene_gn=LOC101096959;refGene_tx=XM_023248979.1
X	105112841	105113118
refGene_gn=LOC101096959;refGene_tx=XM_023248979.1
X	105480099	105480257
refGene_gn=LOC111558717;refGene_tx=XM_023249124.1
X	105480919	105481060
refGene_gn=LOC111558717;refGene_tx=XM_023249124.1
X	105481277	105481764
refGene_gn=LOC111558717;refGene_tx=XM_023249124.1
X	105482807	105482918
refGene_gn=LOC111558717;refGene_tx=XM_023249124.1
X	105811922	105813296
refGene_gn=DCAF12L2;refGene_tx=XM_019823454.2
X	106147335	106148709
refGene_gn=LOC101097955;refGene_tx=XM_011279463.3
X	106332127	106332481
refGene_gn=LOC101098936;refGene_tx=XM_006943985.2
X	106379363	106379383
refGene_gn=PRR32;refGene_tx=XM_004000873.5
X	106380209	106381089
refGene_gn=PRR32;refGene_tx=XM_004000873.5;refGene_gn=PRR32;refGene_tx=XM_006943984.3
X	107750208	107750426
refGene_gn=ACTRT1;refGene_tx=XM_011279462.1
X	107750428	107750922
refGene_gn=ACTRT1;refGene_tx=XM_011279462.1
X	107750923	107751312
refGene_gn=ACTRT1;refGene_tx=XM_011279462.1
X	109009094	109009166
refGene_gn=SMARCA1;refGene_tx=XM_004000877.5

X 109010265 109010376
refGene_gn=SMARCA1;refGene_tx=XM_004000877.5
X 109023256 109023469
refGene_gn=SMARCA1;refGene_tx=XM_004000877.5
X 109023581 109023700
refGene_gn=SMARCA1;refGene_tx=XM_004000877.5
X 109027552 109027685
refGene_gn=SMARCA1;refGene_tx=XM_004000877.5
X 109029997 109030120
refGene_gn=SMARCA1;refGene_tx=XM_004000877.5
X 109040211 109040325
refGene_gn=SMARCA1;refGene_tx=XM_004000877.5
X 109040584 109040695
refGene_gn=SMARCA1;refGene_tx=XM_004000877.5
X 109047329 109047449
refGene_gn=SMARCA1;refGene_tx=XM_004000877.5
X 109048211 109048360
refGene_gn=SMARCA1;refGene_tx=XM_004000877.5
X 109049127 109049260
refGene_gn=SMARCA1;refGene_tx=XM_004000877.5
X 109052030 109052183
refGene_gn=SMARCA1;refGene_tx=XM_004000877.5
X 109053093 109053129
refGene_gn=SMARCA1;refGene_tx=XM_004000877.5
X 109055448 109055570
refGene_gn=SMARCA1;refGene_tx=XM_004000877.5
X 109056537 109056764
refGene_gn=SMARCA1;refGene_tx=XM_004000877.5
X 109058096 109058206
refGene_gn=SMARCA1;refGene_tx=XM_004000877.5
X 109063073 109063142
refGene_gn=SMARCA1;refGene_tx=XM_004000877.5
X 109064103 109064235
refGene_gn=SMARCA1;refGene_tx=XM_004000877.5
X 109065502 109065658
refGene_gn=SMARCA1;refGene_tx=XM_004000877.5
X 109069740 109069920
refGene_gn=SMARCA1;refGene_tx=XM_004000877.5
X 109072550 109072628
refGene_gn=SMARCA1;refGene_tx=XM_004000877.5
X 109072629 109072652
refGene_gn=SMARCA1;refGene_tx=XM_004000877.5
X 109072758 109072859
refGene_gn=SMARCA1;refGene_tx=XM_004000877.5
X 109073185 109073352
refGene_gn=SMARCA1;refGene_tx=XM_004000877.5
X 109075631 109075718
refGene_gn=SMARCA1;refGene_tx=XM_004000877.5
X 109080639 109080810
refGene_gn=SMARCA1;refGene_tx=XM_004000877.5

X 109097755 109097794
refGene_gn=OCRL;refGene_tx=XM_019824260.1;refGene_gn=OCRL;refGene_tx=XM_019824261.1;refGene_gn=OCRL;refGene_tx=XM_023248838.1
X 109098055 109098135
refGene_gn=OCRL;refGene_tx=XM_019824260.1;refGene_gn=OCRL;refGene_tx=XM_019824261.1;refGene_gn=OCRL;refGene_tx=XM_023248838.1
X 109102768 109102848
refGene_gn=OCRL;refGene_tx=XM_019824260.1;refGene_gn=OCRL;refGene_tx=XM_019824261.1;refGene_gn=OCRL;refGene_tx=XM_023248838.1
X 109105625 109105664
refGene_gn=OCRL;refGene_tx=XM_019824260.1;refGene_gn=OCRL;refGene_tx=XM_019824261.1;refGene_gn=OCRL;refGene_tx=XM_023248838.1
X 109113897 109114008
refGene_gn=OCRL;refGene_tx=XM_019824260.1;refGene_gn=OCRL;refGene_tx=XM_019824261.1;refGene_gn=OCRL;refGene_tx=XM_023248838.1
X 109114437 109114527
refGene_gn=OCRL;refGene_tx=XM_019824260.1;refGene_gn=OCRL;refGene_tx=XM_019824261.1;refGene_gn=OCRL;refGene_tx=XM_023248838.1
X 109115237 109115358
refGene_gn=OCRL;refGene_tx=XM_019824260.1;refGene_gn=OCRL;refGene_tx=XM_019824261.1;refGene_gn=OCRL;refGene_tx=XM_023248838.1;refGene_gn=OCRL;refGene_tx=XM_023248839.1
X 109115445 109115607
refGene_gn=OCRL;refGene_tx=XM_019824260.1;refGene_gn=OCRL;refGene_tx=XM_019824261.1;refGene_gn=OCRL;refGene_tx=XM_023248838.1;refGene_gn=OCRL;refGene_tx=XM_023248839.1
X 109116863 109116965
refGene_gn=OCRL;refGene_tx=XM_019824260.1;refGene_gn=OCRL;refGene_tx=XM_019824261.1;refGene_gn=OCRL;refGene_tx=XM_023248838.1;refGene_gn=OCRL;refGene_tx=XM_023248839.1
X 109117960 109118075
refGene_gn=OCRL;refGene_tx=XM_019824260.1;refGene_gn=OCRL;refGene_tx=XM_019824261.1;refGene_gn=OCRL;refGene_tx=XM_023248838.1;refGene_gn=OCRL;refGene_tx=XM_023248839.1
X 109119239 109119356
refGene_gn=OCRL;refGene_tx=XM_019824260.1;refGene_gn=OCRL;refGene_tx=XM_019824261.1;refGene_gn=OCRL;refGene_tx=XM_023248838.1;refGene_gn=OCRL;refGene_tx=XM_023248839.1
X 109119460 109119648
refGene_gn=OCRL;refGene_tx=XM_019824260.1;refGene_gn=OCRL;refGene_tx=XM_019824261.1;refGene_gn=OCRL;refGene_tx=XM_023248838.1;refGene_gn=OCRL;refGene_tx=XM_023248839.1
X 109122078 109122190
refGene_gn=OCRL;refGene_tx=XM_019824260.1;refGene_gn=OCRL;refGene_tx=XM_019824261.1;refGene_gn=OCRL;refGene_tx=XM_023248838.1;refGene_gn=OCRL;refGene_tx=XM_023248839.1
X 109123767 109123877
refGene_gn=OCRL;refGene_tx=XM_019824260.1;refGene_gn=OCRL;refGene_tx=XM_019824261.1;refGene_gn=OCRL;refGene_tx=XM_023248838.1;refGene_gn=OCRL;refGene_tx=XM_023248839.1

X	109125469	109125605
refGene_gn=OCRL;refGene_tx=XM_019824260.1;refGene_gn=OCRL;refGene_tx=XM_019824261.1;refGene_gn=OCRL;refGene_tx=XM_023248838.1;refGene_gn=OCRL;refGene_tx=XM_023248839.1
X	109130387	109130498
refGene_gn=OCRL;refGene_tx=XM_019824260.1;refGene_gn=OCRL;refGene_tx=XM_019824261.1;refGene_gn=OCRL;refGene_tx=XM_023248838.1;refGene_gn=OCRL;refGene_tx=XM_023248839.1
X	109131158	109131324
refGene_gn=OCRL;refGene_tx=XM_019824260.1;refGene_gn=OCRL;refGene_tx=XM_019824261.1;refGene_gn=OCRL;refGene_tx=XM_023248838.1;refGene_gn=OCRL;refGene_tx=XM_023248839.1
X	109131588	109131824
refGene_gn=OCRL;refGene_tx=XM_019824260.1;refGene_gn=OCRL;refGene_tx=XM_019824261.1;refGene_gn=OCRL;refGene_tx=XM_023248838.1;refGene_gn=OCRL;refGene_tx=XM_023248839.1
X	109139361	109139385
refGene_gn=OCRL;refGene_tx=XM_019824260.1;refGene_gn=OCRL;refGene_tx=XM_023248838.1;refGene_gn=OCRL;refGene_tx=XM_023248839.1
X	109142453	109142570
refGene_gn=OCRL;refGene_tx=XM_019824260.1;refGene_gn=OCRL;refGene_tx=XM_019824261.1;refGene_gn=OCRL;refGene_tx=XM_023248838.1;refGene_gn=OCRL;refGene_tx=XM_023248839.1
X	109143457	109143542
refGene_gn=OCRL;refGene_tx=XM_019824260.1;refGene_gn=OCRL;refGene_tx=XM_019824261.1;refGene_gn=OCRL;refGene_tx=XM_023248838.1;refGene_gn=OCRL;refGene_tx=XM_023248839.1
X	109144167	109144295
refGene_gn=OCRL;refGene_tx=XM_019824260.1;refGene_gn=OCRL;refGene_tx=XM_019824261.1;refGene_gn=OCRL;refGene_tx=XM_023248838.1;refGene_gn=OCRL;refGene_tx=XM_023248839.1
X	109145413	109145525
refGene_gn=OCRL;refGene_tx=XM_019824260.1;refGene_gn=OCRL;refGene_tx=XM_019824261.1;refGene_gn=OCRL;refGene_tx=XM_023248838.1;refGene_gn=OCRL;refGene_tx=XM_023248839.1
X	109145745	109145867
refGene_gn=OCRL;refGene_tx=XM_019824260.1;refGene_gn=OCRL;refGene_tx=XM_019824261.1;refGene_gn=OCRL;refGene_tx=XM_023248838.1;refGene_gn=OCRL;refGene_tx=XM_023248839.1
X	109198953	109199120
refGene_gn=APLN;refGene_tx=XM_004000879.5
X	109204583	109204650
refGene_gn=APLN;refGene_tx=XM_004000879.5
X	109282014	109282063
refGene_gn=XPNPEP2;refGene_tx=XM_004000917.3
X	109284586	109284663
refGene_gn=XPNPEP2;refGene_tx=XM_004000917.3
X	109286325	109286436
refGene_gn=XPNPEP2;refGene_tx=XM_004000917.3
X	109287580	109287644

refGene_gn=XPNPEP2;refGene_tx=XM_004000917.3
X	109288571	109288676
refGene_gn=XPNPEP2;refGene_tx=XM_004000917.3
X	109288920	109289007
refGene_gn=XPNPEP2;refGene_tx=XM_004000917.3
X	109289924	109290071
refGene_gn=XPNPEP2;refGene_tx=XM_004000917.3
X	109293014	109293116
refGene_gn=XPNPEP2;refGene_tx=XM_004000917.3
X	109293786	109293868
refGene_gn=XPNPEP2;refGene_tx=XM_004000917.3
X	109294153	109294349
refGene_gn=XPNPEP2;refGene_tx=XM_004000917.3
X	109294996	109295086
refGene_gn=XPNPEP2;refGene_tx=XM_004000917.3
X	109296334	109296444
refGene_gn=XPNPEP2;refGene_tx=XM_004000917.3
X	109297202	109297280
refGene_gn=XPNPEP2;refGene_tx=XM_004000917.3
X	109298385	109298457
refGene_gn=XPNPEP2;refGene_tx=XM_004000917.3
X	109300715	109300776
refGene_gn=XPNPEP2;refGene_tx=XM_004000917.3
X	109301989	109302059
refGene_gn=XPNPEP2;refGene_tx=XM_004000917.3
X	109302574	109302679
refGene_gn=XPNPEP2;refGene_tx=XM_004000917.3
X	109303120	109303180
refGene_gn=XPNPEP2;refGene_tx=XM_004000917.3
X	109303675	109303752
refGene_gn=XPNPEP2;refGene_tx=XM_004000917.3
X	109306273	109306363
refGene_gn=XPNPEP2;refGene_tx=XM_004000917.3
X	109306985	109307180
refGene_gn=XPNPEP2;refGene_tx=XM_004000917.3
X	109317350	109317407
refGene_gn=SASH3;refGene_tx=XM_004000880.5
X	109324074	109324170
refGene_gn=SASH3;refGene_tx=XM_004000880.5
X	109324888	109325035
refGene_gn=SASH3;refGene_tx=XM_004000880.5
X	109326538	109326677
refGene_gn=SASH3;refGene_tx=XM_004000880.5
X	109327913	109328062
refGene_gn=SASH3;refGene_tx=XM_004000880.5
X	109328211	109328421
refGene_gn=SASH3;refGene_tx=XM_004000880.5
X	109328571	109328722
refGene_gn=SASH3;refGene_tx=XM_004000880.5
X	109329271	109329462

refGene_gn=SASH3;refGene_tx=XM_004000880.5
X	109341495	109341612
refGene_gn=ZDHHC9;refGene_tx=XM_011279472.3;refGene_gn=ZDHHC9;refGene_tx=XM_011279473.3
X	109345450	109345547
refGene_gn=ZDHHC9;refGene_tx=XM_011279472.3;refGene_gn=ZDHHC9;refGene_tx=XM_011279473.3
X	109345950	109346054
refGene_gn=ZDHHC9;refGene_tx=XM_011279472.3;refGene_gn=ZDHHC9;refGene_tx=XM_011279473.3
X	109347404	109347507
refGene_gn=ZDHHC9;refGene_tx=XM_011279472.3;refGene_gn=ZDHHC9;refGene_tx=XM_011279473.3
X	109348490	109348539
refGene_gn=ZDHHC9;refGene_tx=XM_011279472.3;refGene_gn=ZDHHC9;refGene_tx=XM_011279473.3
X	109349487	109349625
refGene_gn=ZDHHC9;refGene_tx=XM_011279472.3;refGene_gn=ZDHHC9;refGene_tx=XM_011279473.3
X	109355122	109355281
refGene_gn=ZDHHC9;refGene_tx=XM_011279472.3;refGene_gn=ZDHHC9;refGene_tx=XM_011279473.3
X	109359852	109360013
refGene_gn=ZDHHC9;refGene_tx=XM_011279472.3;refGene_gn=ZDHHC9;refGene_tx=XM_011279473.3
X	109369847	109370014
refGene_gn=ZDHHC9;refGene_tx=XM_011279472.3;refGene_gn=ZDHHC9;refGene_tx=XM_011279473.3
X	109414709	109414735
refGene_gn=UTP14A;refGene_tx=XM_004000882.5
X	109415771	109415847
refGene_gn=UTP14A;refGene_tx=XM_004000882.5
X	109417077	109417148
refGene_gn=UTP14A;refGene_tx=XM_004000882.5
X	109418606	109418671
refGene_gn=UTP14A;refGene_tx=XM_004000882.5
X	109420041	109420184
refGene_gn=UTP14A;refGene_tx=XM_004000882.5
X	109420476	109420632
refGene_gn=UTP14A;refGene_tx=XM_004000882.5
X	109425462	109425582
refGene_gn=UTP14A;refGene_tx=XM_004000882.5
X	109425680	109425775
refGene_gn=UTP14A;refGene_tx=XM_004000882.5
X	109426382	109426506
refGene_gn=UTP14A;refGene_tx=XM_004000882.5
X	109426588	109426666
refGene_gn=UTP14A;refGene_tx=XM_004000882.5
X	109427152	109427543
refGene_gn=UTP14A;refGene_tx=XM_004000882.5

X	109429638	109430042
refGene_gn=UTP14A;refGene_tx=XM_004000882.5
X	109430811	109431005
refGene_gn=UTP14A;refGene_tx=XM_004000882.5
X	109431134	109431234
refGene_gn=UTP14A;refGene_tx=XM_004000882.5
X	109433499	109433772
refGene_gn=UTP14A;refGene_tx=XM_004000882.5
X	109491088	109491174
refGene_gn=BCORL1;refGene_tx=XM_006943992.4;refGene_gn=BCORL1;refGene_tx=XM_006943993.4;refGene_gn=BCORL1;refGene_tx=XM_006943994.4;refGene_gn=BCORL1;refGene_tx=XM_011279470.3;refGene_gn=BCORL1;refGene_tx=XM_019824262.2;refGene_gn=BCORL1;refGene_tx=XM_019824263.2;refGene_gn=BCORL1;refGene_tx=XM_019824264.2
X	109496156	109496247
refGene_gn=BCORL1;refGene_tx=XM_006943992.4;refGene_gn=BCORL1;refGene_tx=XM_006943993.4;refGene_gn=BCORL1;refGene_tx=XM_006943994.4;refGene_gn=BCORL1;refGene_tx=XM_011279470.3;refGene_gn=BCORL1;refGene_tx=XM_019824262.2;refGene_gn=BCORL1;refGene_tx=XM_019824263.2;refGene_gn=BCORL1;refGene_tx=XM_019824264.2
X	109496532	109499805
refGene_gn=BCORL1;refGene_tx=XM_006943992.4;refGene_gn=BCORL1;refGene_tx=XM_006943993.4;refGene_gn=BCORL1;refGene_tx=XM_006943994.4;refGene_gn=BCORL1;refGene_tx=XM_011279470.3;refGene_gn=BCORL1;refGene_tx=XM_019824262.2;refGene_gn=BCORL1;refGene_tx=XM_019824263.2;refGene_gn=BCORL1;refGene_tx=XM_019824264.2
X	109504834	109505000
refGene_gn=BCORL1;refGene_tx=XM_006943992.4;refGene_gn=BCORL1;refGene_tx=XM_006943993.4;refGene_gn=BCORL1;refGene_tx=XM_006943994.4;refGene_gn=BCORL1;refGene_tx=XM_011279470.3;refGene_gn=BCORL1;refGene_tx=XM_019824262.2;refGene_gn=BCORL1;refGene_tx=XM_019824263.2;refGene_gn=BCORL1;refGene_tx=XM_019824264.2
X	109506097	109506178
refGene_gn=BCORL1;refGene_tx=XM_006943992.4;refGene_gn=BCORL1;refGene_tx=XM_006943993.4;refGene_gn=BCORL1;refGene_tx=XM_006943994.4;refGene_gn=BCORL1;refGene_tx=XM_011279470.3;refGene_gn=BCORL1;refGene_tx=XM_019824262.2;refGene_gn=BCORL1;refGene_tx=XM_019824263.2;refGene_gn=BCORL1;refGene_tx=XM_019824264.2
X	109508142	109508532
refGene_gn=BCORL1;refGene_tx=XM_006943992.4;refGene_gn=BCORL1;refGene_tx=XM_006943993.4;refGene_gn=BCORL1;refGene_tx=XM_006943994.4;refGene_gn=BCORL1;refGene_tx=XM_011279470.3;refGene_gn=BCORL1;refGene_tx=XM_019824262.2;refGene_gn=BCORL1;refGene_tx=XM_019824263.2
X	109512449	109512673
refGene_gn=BCORL1;refGene_tx=XM_006943992.4;refGene_gn=BCORL1;refGene_tx=XM_006943993.4;refGene_gn=BCORL1;refGene_tx=XM_006943994.4;refGene_gn=BCORL1;refGene_tx=XM_011279470.3;refGene_gn=BCORL1;refGene_tx=XM_019824262.2;refGene_gn=BCORL1;refGene_tx=XM_019824263.2;refGene_gn=BCORL1;refGene_tx=XM_019824264.2
X	109516907	109517129

refGene_gn=BCORL1;refGene_tx=XM_006943992.4;refGene_gn=BCORL1;refGene_tx=XM_006943993.4;refGene_gn=BCORL1;refGene_tx=XM_006943994.4;refGene_gn=BCORL1;refGene_tx=XM_011279470.3;refGene_gn=BCORL1;refGene_tx=XM_019824262.2;refGene_gn=BCORL1;refGene_tx=XM_019824263.2;refGene_gn=BCORL1;refGene_tx=XM_019824264.2
X	109519220	109519387
refGene_gn=BCORL1;refGene_tx=XM_006943992.4;refGene_gn=BCORL1;refGene_tx=XM_006943993.4;refGene_gn=BCORL1;refGene_tx=XM_006943994.4;refGene_gn=BCORL1;refGene_tx=XM_011279470.3;refGene_gn=BCORL1;refGene_tx=XM_019824262.2;refGene_gn=BCORL1;refGene_tx=XM_019824263.2;refGene_gn=BCORL1;refGene_tx=XM_019824264.2
X	109520365	109520511
refGene_gn=BCORL1;refGene_tx=XM_006943992.4;refGene_gn=BCORL1;refGene_tx=XM_006943993.4;refGene_gn=BCORL1;refGene_tx=XM_006943994.4;refGene_gn=BCORL1;refGene_tx=XM_011279470.3;refGene_gn=BCORL1;refGene_tx=XM_019824262.2;refGene_gn=BCORL1;refGene_tx=XM_019824263.2;refGene_gn=BCORL1;refGene_tx=XM_019824264.2
X	109528495	109528573
refGene_gn=BCORL1;refGene_tx=XM_006943992.4;refGene_gn=BCORL1;refGene_tx=XM_006943993.4;refGene_gn=BCORL1;refGene_tx=XM_006943994.4;refGene_gn=BCORL1;refGene_tx=XM_011279470.3;refGene_gn=BCORL1;refGene_tx=XM_019824262.2;refGene_gn=BCORL1;refGene_tx=XM_019824263.2;refGene_gn=BCORL1;refGene_tx=XM_019824264.2
X	109529620	109529774
refGene_gn=BCORL1;refGene_tx=XM_006943992.4;refGene_gn=BCORL1;refGene_tx=XM_006943993.4;refGene_gn=BCORL1;refGene_tx=XM_006943994.4;refGene_gn=BCORL1;refGene_tx=XM_011279470.3;refGene_gn=BCORL1;refGene_tx=XM_019824262.2;refGene_gn=BCORL1;refGene_tx=XM_019824263.2;refGene_gn=BCORL1;refGene_tx=XM_019824264.2
X	109533265	109533548
refGene_gn=BCORL1;refGene_tx=XM_006943992.4;refGene_gn=BCORL1;refGene_tx=XM_006943993.4;refGene_gn=BCORL1;refGene_tx=XM_006943994.4;refGene_gn=BCORL1;refGene_tx=XM_011279470.3;refGene_gn=BCORL1;refGene_tx=XM_019824262.2;refGene_gn=BCORL1;refGene_tx=XM_019824263.2;refGene_gn=BCORL1;refGene_tx=XM_019824264.2
X	109543667	109544460
refGene_gn=ELF4;refGene_tx=XM_004000887.4;refGene_gn=ELF4;refGene_tx=XM_006943996.3;refGene_gn=ELF4;refGene_tx=XM_006943997.4
X	109545337	109545715
refGene_gn=ELF4;refGene_tx=XM_004000887.4;refGene_gn=ELF4;refGene_tx=XM_006943996.3;refGene_gn=ELF4;refGene_tx=XM_006943997.4
X	109546317	109546510
refGene_gn=ELF4;refGene_tx=XM_004000887.4;refGene_gn=ELF4;refGene_tx=XM_006943996.3;refGene_gn=ELF4;refGene_tx=XM_006943997.4
X	109546586	109546670
refGene_gn=ELF4;refGene_tx=XM_004000887.4;refGene_gn=ELF4;refGene_tx=XM_006943996.3;refGene_gn=ELF4;refGene_tx=XM_006943997.4
X	109547980	109548172
refGene_gn=ELF4;refGene_tx=XM_004000887.4;refGene_gn=ELF4;refGene_tx=XM_006943996.3;refGene_gn=ELF4;refGene_tx=XM_006943997.4

X	109549523	109549616
refGene_gn=ELF4;refGene_tx=XM_004000887.4;refGene_gn=ELF4;refGene_tx=XM_006943996.3;refGene_gn=ELF4;refGene_tx=XM_006943997.4
X	109550049	109550221
refGene_gn=ELF4;refGene_tx=XM_004000887.4;refGene_gn=ELF4;refGene_tx=XM_006943996.3;refGene_gn=ELF4;refGene_tx=XM_006943997.4
X	109554602	109554674
refGene_gn=ELF4;refGene_tx=XM_004000887.4;refGene_gn=ELF4;refGene_tx=XM_006943996.3;refGene_gn=ELF4;refGene_tx=XM_006943997.4
X	109598253	109598325
refGene_gn=AIFM1;refGene_tx=XM_004000888.4;refGene_gn=AIFM1;refGene_tx=XM_004000889.4
X	109598664	109598861
refGene_gn=AIFM1;refGene_tx=XM_004000888.4;refGene_gn=AIFM1;refGene_tx=XM_004000889.4
X	109600646	109600771
refGene_gn=AIFM1;refGene_tx=XM_004000888.4;refGene_gn=AIFM1;refGene_tx=XM_004000889.4
X	109602436	109602579
refGene_gn=AIFM1;refGene_tx=XM_004000888.4;refGene_gn=AIFM1;refGene_tx=XM_004000889.4
X	109604287	109604428
refGene_gn=AIFM1;refGene_tx=XM_004000888.4;refGene_gn=AIFM1;refGene_tx=XM_004000889.4
X	109605373	109605462
refGene_gn=AIFM1;refGene_tx=XM_004000888.4;refGene_gn=AIFM1;refGene_tx=XM_004000889.4
X	109605799	109605907
refGene_gn=AIFM1;refGene_tx=XM_004000888.4;refGene_gn=AIFM1;refGene_tx=XM_004000889.4
X	109606785	109606894
refGene_gn=AIFM1;refGene_tx=XM_004000888.4;refGene_gn=AIFM1;refGene_tx=XM_004000889.4
X	109608635	109608712
refGene_gn=AIFM1;refGene_tx=XM_004000888.4;refGene_gn=AIFM1;refGene_tx=XM_004000889.4
X	109609358	109609443
refGene_gn=AIFM1;refGene_tx=XM_004000888.4;refGene_gn=AIFM1;refGene_tx=XM_004000889.4
X	109614165	109614256
refGene_gn=AIFM1;refGene_tx=XM_004000888.4;refGene_gn=AIFM1;refGene_tx=XM_004000889.4
X	109615710	109615841
refGene_gn=AIFM1;refGene_tx=XM_004000888.4;refGene_gn=AIFM1;refGene_tx=XM_004000889.4
X	109615966	109616091
refGene_gn=AIFM1;refGene_tx=XM_004000888.4;refGene_gn=AIFM1;refGene_tx=XM_004000889.4
X	109617522	109617622
refGene_gn=AIFM1;refGene_tx=XM_004000888.4;refGene_gn=AIFM1;refGene_tx

=XM_004000889.4
X	109622008	109622139
refGene_gn=AIFM1;refGene_tx=XM_004000889.4
X	109623339	109623482
refGene_gn=AIFM1;refGene_tx=XM_004000888.4
X	109630884	109630990
refGene_gn=AIFM1;refGene_tx=XM_004000888.4;refGene_gn=AIFM1;refGene_tx
=XM_004000889.4
X	109636121	109636391
refGene_gn=RAB33A;refGene_tx=XM_004000891.4;refGene_gn=RAB33A;refGene_
tx=XM_006943998.3
X	109646313	109646769
refGene_gn=RAB33A;refGene_tx=XM_004000891.4;refGene_gn=RAB33A;refGene_
tx=XM_006943998.3
X	109663765	109664141
refGene_gn=ZNF280C;refGene_tx=XM_019824265.2;refGene_gn=ZNF280C;refGen
e_tx=XM_019824266.2
X	109664651	109664661
refGene_gn=ZNF280C;refGene_tx=XM_006943999.4
X	109664785	109664822
refGene_gn=ZNF280C;refGene_tx=XM_006943999.4;refGene_gn=ZNF280C;refGen
e_tx=XM_019824265.2;refGene_gn=ZNF280C;refGene_tx=XM_019824266.2
X	109664970	109665089
refGene_gn=ZNF280C;refGene_tx=XM_006943999.4;refGene_gn=ZNF280C;refGen
e_tx=XM_019824265.2;refGene_gn=ZNF280C;refGene_tx=XM_019824266.2
X	109667858	109667921
refGene_gn=ZNF280C;refGene_tx=XM_006943999.4;refGene_gn=ZNF280C;refGen
e_tx=XM_019824265.2;refGene_gn=ZNF280C;refGene_tx=XM_019824266.2
X	109675694	109675835
refGene_gn=ZNF280C;refGene_tx=XM_006943999.4;refGene_gn=ZNF280C;refGen
e_tx=XM_019824265.2;refGene_gn=ZNF280C;refGene_tx=XM_019824266.2
X	109676306	109676620
refGene_gn=ZNF280C;refGene_tx=XM_006943999.4;refGene_gn=ZNF280C;refGen
e_tx=XM_019824265.2;refGene_gn=ZNF280C;refGene_tx=XM_019824266.2
X	109680031	109680163
refGene_gn=ZNF280C;refGene_tx=XM_006943999.4;refGene_gn=ZNF280C;refGen
e_tx=XM_019824265.2;refGene_gn=ZNF280C;refGene_tx=XM_019824266.2
X	109687001	109687148
refGene_gn=ZNF280C;refGene_tx=XM_006943999.4;refGene_gn=ZNF280C;refGen
e_tx=XM_019824265.2;refGene_gn=ZNF280C;refGene_tx=XM_019824266.2
X	109687911	109688012
refGene_gn=ZNF280C;refGene_tx=XM_006943999.4;refGene_gn=ZNF280C;refGen
e_tx=XM_019824265.2;refGene_gn=ZNF280C;refGene_tx=XM_019824266.2
X	109688669	109688827
refGene_gn=ZNF280C;refGene_tx=XM_006943999.4;refGene_gn=ZNF280C;refGen
e_tx=XM_019824265.2;refGene_gn=ZNF280C;refGene_tx=XM_019824266.2
X	109692503	109692724
refGene_gn=ZNF280C;refGene_tx=XM_006943999.4;refGene_gn=ZNF280C;refGen
e_tx=XM_019824265.2;refGene_gn=ZNF280C;refGene_tx=XM_019824266.2
X	109698688	109698798

refGene_gn=ZNF280C;refGene_tx=XM_006943999.4;refGene_gn=ZNF280C;refGene_tx=XM_019824265.2;refGene_gn=ZNF280C;refGene_tx=XM_019824266.2
X 109698943 109699114
refGene_gn=ZNF280C;refGene_tx=XM_006943999.4;refGene_gn=ZNF280C;refGene_tx=XM_019824265.2;refGene_gn=ZNF280C;refGene_tx=XM_019824266.2
X 109702011 109702126
refGene_gn=ZNF280C;refGene_tx=XM_006943999.4;refGene_gn=ZNF280C;refGene_tx=XM_019824265.2;refGene_gn=ZNF280C;refGene_tx=XM_019824266.2
X 109706660 109706797
refGene_gn=ZNF280C;refGene_tx=XM_006943999.4;refGene_gn=ZNF280C;refGene_tx=XM_019824265.2;refGene_gn=ZNF280C;refGene_tx=XM_019824266.2
X 109706898 109706964
refGene_gn=ZNF280C;refGene_tx=XM_006943999.4;refGene_gn=ZNF280C;refGene_tx=XM_019824265.2;refGene_gn=ZNF280C;refGene_tx=XM_019824266.2
X 109709861 109710008
refGene_gn=ZNF280C;refGene_tx=XM_006943999.4;refGene_gn=ZNF280C;refGene_tx=XM_019824265.2;refGene_gn=ZNF280C;refGene_tx=XM_019824266.2
X 109716032 109716063
refGene_gn=ZNF280C;refGene_tx=XM_006943999.4;refGene_gn=ZNF280C;refGene_tx=XM_019824265.2;refGene_gn=ZNF280C;refGene_tx=XM_019824266.2
X 109766138 109766273
refGene_gn=SLC25A14;refGene_tx=XM_006944006.4;refGene_gn=SLC25A14;refGene_tx=XM_023248835.1
X 109766575 109766650
refGene_gn=SLC25A14;refGene_tx=XM_019824278.1;refGene_gn=SLC25A14;refGene_tx=XM_019824279.1
X 109771588 109771682
refGene_gn=SLC25A14;refGene_tx=XM_006944006.4;refGene_gn=SLC25A14;refGene_tx=XM_019824278.1;refGene_gn=SLC25A14;refGene_tx=XM_019824279.1;refGene_gn=SLC25A14;refGene_tx=XM_023248835.1
X 109773673 109773821
refGene_gn=SLC25A14;refGene_tx=XM_006944006.4;refGene_gn=SLC25A14;refGene_tx=XM_019824278.1;refGene_gn=SLC25A14;refGene_tx=XM_019824279.1;refGene_gn=SLC25A14;refGene_tx=XM_023248835.1
X 109776438 109776533
refGene_gn=SLC25A14;refGene_tx=XM_006944006.4;refGene_gn=SLC25A14;refGene_tx=XM_019824278.1;refGene_gn=SLC25A14;refGene_tx=XM_019824279.1;refGene_gn=SLC25A14;refGene_tx=XM_023248835.1
X 109778060 109778146
refGene_gn=SLC25A14;refGene_tx=XM_006944006.4;refGene_gn=SLC25A14;refGene_tx=XM_019824278.1;refGene_gn=SLC25A14;refGene_tx=XM_019824279.1;refGene_gn=SLC25A14;refGene_tx=XM_023248835.1
X 109781037 109781133
refGene_gn=SLC25A14;refGene_tx=XM_006944006.4;refGene_gn=SLC25A14;refGene_tx=XM_019824278.1;refGene_gn=SLC25A14;refGene_tx=XM_019824279.1;refGene_gn=SLC25A14;refGene_tx=XM_023248835.1
X 109785241 109785366
refGene_gn=SLC25A14;refGene_tx=XM_006944006.4;refGene_gn=SLC25A14;refGene_tx=XM_019824278.1;refGene_gn=SLC25A14;refGene_tx=XM_019824279.1;refGene_gn=SLC25A14;refGene_tx=XM_023248835.1

X 109786326 109786462
refGene_gn=SLC25A14;refGene_tx=XM_006944006.4;refGene_gn=SLC25A14;refGene_tx=XM_019824278.1;refGene_gn=SLC25A14;refGene_tx=XM_019824279.1;refGene_gn=SLC25A14;refGene_tx=XM_023248835.1
X 109793069 109793150
refGene_gn=SLC25A14;refGene_tx=XM_006944006.4;refGene_gn=SLC25A14;refGene_tx=XM_019824278.1;refGene_gn=SLC25A14;refGene_tx=XM_019824279.1;refGene_gn=SLC25A14;refGene_tx=XM_023248835.1
X 109794099 109794141
refGene_gn=SLC25A14;refGene_tx=XM_006944006.4;refGene_gn=SLC25A14;refGene_tx=XM_019824278.1;refGene_gn=SLC25A14;refGene_tx=XM_019824279.1;refGene_gn=SLC25A14;refGene_tx=XM_023248835.1
X 109811302 109812340
refGene_gn=GPR119;refGene_tx=XM_019824280.1
X 109826621 109826686
refGene_gn=RBMX2;refGene_tx=XM_023248834.1
X 109826873 109826989
refGene_gn=RBMX2;refGene_tx=XM_023248834.1
X 109828315 109828367
refGene_gn=RBMX2;refGene_tx=XM_023248834.1
X 109834253 109834383
refGene_gn=RBMX2;refGene_tx=XM_023248834.1
X 109836003 109836181
refGene_gn=RBMX2;refGene_tx=XM_023248834.1
X 109836651 109837154
refGene_gn=RBMX2;refGene_tx=XM_023248834.1
X 109847753 109847884
refGene_gn=LOC101091264;refGene_tx=XM_023248832.1
X 109850925 109850992
refGene_gn=LOC101091264;refGene_tx=XM_023248832.1
X 109854205 109854249
refGene_gn=LOC101091264;refGene_tx=XM_023248832.1
X 109859057 109859101
refGene_gn=LOC101091264;refGene_tx=XM_023248832.1
X 109860205 109860306
refGene_gn=LOC101091264;refGene_tx=XM_023248832.1
X 109861349 109861844
refGene_gn=LOC101091264;refGene_tx=XM_023248832.1
X 109864919 109866731
refGene_gn=LOC101091264;refGene_tx=XM_023248832.1
X 109866733 109868353
refGene_gn=LOC101091264;refGene_tx=XM_023248832.1
X 109868355 109868999
refGene_gn=LOC101091264;refGene_tx=XM_023248832.1
X 109869001 109873756
refGene_gn=LOC101091264;refGene_tx=XM_023248832.1
X 109874904 109875222
refGene_gn=LOC101091264;refGene_tx=XM_023248832.1
X 109875224 109878330
refGene_gn=LOC101091264;refGene_tx=XM_023248832.1

X 109878331 109883664
refGene_gn=LOC101091264;refGene_tx=XM_023248832.1
X 109883666 109883730
refGene_gn=LOC101091264;refGene_tx=XM_023248832.1
X 109890611 109890645
refGene_gn=LOC101091264;refGene_tx=XM_023248832.1
X 109895016 109895053
refGene_gn=LOC101091264;refGene_tx=XM_023248832.1
X 109898749 109898810
refGene_gn=LOC101091264;refGene_tx=XM_023248832.1
X 109900591 109900680
refGene_gn=LOC101091264;refGene_tx=XM_023248832.1
X 109901813 109901856
refGene_gn=LOC101091264;refGene_tx=XM_023248832.1
X 109904089 109904193
refGene_gn=LOC101091264;refGene_tx=XM_023248832.1
X 109919200 109919321
refGene_gn=LOC101091264;refGene_tx=XM_023248832.1
X 109919741 109919852
refGene_gn=LOC101091264;refGene_tx=XM_023248832.1
X 109920544 109920686
refGene_gn=LOC101091264;refGene_tx=XM_023248832.1
X 109922019 109922159
refGene_gn=LOC101091264;refGene_tx=XM_023248832.1
X 109923322 109923484
refGene_gn=LOC101091264;refGene_tx=XM_023248832.1
X 109926020 109926146
refGene_gn=LOC101091264;refGene_tx=XM_023248832.1
X 109926548 109926647
refGene_gn=LOC101091264;refGene_tx=XM_023248832.1
X 110036686 110036818
refGene_gn=ENOX2;refGene_tx=XM_006944000.4;refGene_gn=ENOX2;refGene_tx
=XM_006944002.3;refGene_gn=ENOX2;refGene_tx=XM_019824268.1;refGene_gn=
ENOX2;refGene_tx=XM_019824269.1;refGene_gn=ENOX2;refGene_tx=XM_0198242
73.2;refGene_gn=ENOX2;refGene_tx=XM_019824274.1;refGene_gn=ENOX2;refGe
ne_tx=XM_019824275.2;refGene_gn=ENOX2;refGene_tx=XM_019824277.2
X 110039357 110039443
refGene_gn=ENOX2;refGene_tx=XM_006944000.4;refGene_gn=ENOX2;refGene_tx
=XM_006944002.3;refGene_gn=ENOX2;refGene_tx=XM_019824268.1;refGene_gn=
ENOX2;refGene_tx=XM_019824269.1;refGene_gn=ENOX2;refGene_tx=XM_0198242
73.2;refGene_gn=ENOX2;refGene_tx=XM_019824274.1;refGene_gn=ENOX2;refGe
ne_tx=XM_019824275.2;refGene_gn=ENOX2;refGene_tx=XM_019824277.2
X 110041885 110041994
refGene_gn=ENOX2;refGene_tx=XM_006944000.4;refGene_gn=ENOX2;refGene_tx
=XM_006944002.3;refGene_gn=ENOX2;refGene_tx=XM_019824268.1;refGene_gn=
ENOX2;refGene_tx=XM_019824269.1;refGene_gn=ENOX2;refGene_tx=XM_0198242
73.2;refGene_gn=ENOX2;refGene_tx=XM_019824274.1;refGene_gn=ENOX2;refGe
ne_tx=XM_019824275.2;refGene_gn=ENOX2;refGene_tx=XM_019824277.2
X 110045613 110045721
refGene_gn=ENOX2;refGene_tx=XM_006944000.4;refGene_gn=ENOX2;refGene_tx

=XM_006944002.3;refGene_gn=ENOX2;refGene_tx=XM_019824268.1;refGene_gn=ENOX2;refGene_tx=XM_019824269.1;refGene_gn=ENOX2;refGene_tx=XM_019824273.2;refGene_gn=ENOX2;refGene_tx=XM_019824274.1;refGene_gn=ENOX2;refGene_tx=XM_019824275.2;refGene_gn=ENOX2;refGene_tx=XM_019824277.2
X 110047836 110048018
refGene_gn=ENOX2;refGene_tx=XM_006944000.4;refGene_gn=ENOX2;refGene_tx=XM_006944002.3;refGene_gn=ENOX2;refGene_tx=XM_019824268.1;refGene_gn=ENOX2;refGene_tx=XM_019824269.1;refGene_gn=ENOX2;refGene_tx=XM_019824273.2;refGene_gn=ENOX2;refGene_tx=XM_019824274.1;refGene_gn=ENOX2;refGene_tx=XM_019824275.2;refGene_gn=ENOX2;refGene_tx=XM_019824277.2
X 110059328 110059443
refGene_gn=ENOX2;refGene_tx=XM_006944000.4;refGene_gn=ENOX2;refGene_tx=XM_006944002.3;refGene_gn=ENOX2;refGene_tx=XM_019824268.1;refGene_gn=ENOX2;refGene_tx=XM_019824269.1;refGene_gn=ENOX2;refGene_tx=XM_019824273.2;refGene_gn=ENOX2;refGene_tx=XM_019824274.1;refGene_gn=ENOX2;refGene_tx=XM_019824275.2;refGene_gn=ENOX2;refGene_tx=XM_019824277.2
X 110067528 110067635
refGene_gn=ENOX2;refGene_tx=XM_006944000.4;refGene_gn=ENOX2;refGene_tx=XM_006944002.3;refGene_gn=ENOX2;refGene_tx=XM_019824268.1;refGene_gn=ENOX2;refGene_tx=XM_019824269.1;refGene_gn=ENOX2;refGene_tx=XM_019824273.2;refGene_gn=ENOX2;refGene_tx=XM_019824274.1;refGene_gn=ENOX2;refGene_tx=XM_019824275.2;refGene_gn=ENOX2;refGene_tx=XM_019824277.2
X 110070040 110070253
refGene_gn=ENOX2;refGene_tx=XM_006944000.4;refGene_gn=ENOX2;refGene_tx=XM_006944002.3;refGene_gn=ENOX2;refGene_tx=XM_019824268.1;refGene_gn=ENOX2;refGene_tx=XM_019824269.1;refGene_gn=ENOX2;refGene_tx=XM_019824273.2;refGene_gn=ENOX2;refGene_tx=XM_019824274.1;refGene_gn=ENOX2;refGene_tx=XM_019824275.2;refGene_gn=ENOX2;refGene_tx=XM_019824277.2
X 110072379 110072613
refGene_gn=ENOX2;refGene_tx=XM_006944000.4;refGene_gn=ENOX2;refGene_tx=XM_006944002.3;refGene_gn=ENOX2;refGene_tx=XM_019824268.1;refGene_gn=ENOX2;refGene_tx=XM_019824269.1;refGene_gn=ENOX2;refGene_tx=XM_019824273.2;refGene_gn=ENOX2;refGene_tx=XM_019824274.1;refGene_gn=ENOX2;refGene_tx=XM_019824275.2;refGene_gn=ENOX2;refGene_tx=XM_019824277.2
X 110078440 110078647
refGene_gn=ENOX2;refGene_tx=XM_006944000.4;refGene_gn=ENOX2;refGene_tx=XM_006944002.3;refGene_gn=ENOX2;refGene_tx=XM_019824268.1;refGene_gn=ENOX2;refGene_tx=XM_019824269.1;refGene_gn=ENOX2;refGene_tx=XM_019824273.2;refGene_gn=ENOX2;refGene_tx=XM_019824274.1;refGene_gn=ENOX2;refGene_tx=XM_019824275.2;refGene_gn=ENOX2;refGene_tx=XM_019824277.2
X 110085774 110085930
refGene_gn=ENOX2;refGene_tx=XM_006944000.4;refGene_gn=ENOX2;refGene_tx=XM_006944002.3;refGene_gn=ENOX2;refGene_tx=XM_019824268.1;refGene_gn=ENOX2;refGene_tx=XM_019824269.1;refGene_gn=ENOX2;refGene_tx=XM_019824273.2;refGene_gn=ENOX2;refGene_tx=XM_019824274.1;refGene_gn=ENOX2;refGene_tx=XM_019824275.2;refGene_gn=ENOX2;refGene_tx=XM_019824277.2
X 110089683 110089720
refGene_gn=ENOX2;refGene_tx=XM_019824277.2
X 110098864 110098999
refGene_gn=ENOX2;refGene_tx=XM_006944000.4;refGene_gn=ENOX2;refGene_tx

=XM_006944002.3;refGene_gn=ENOX2;refGene_tx=XM_019824268.1;refGene_gn=ENOX2;refGene_tx=XM_019824269.1;refGene_gn=ENOX2;refGene_tx=XM_019824273.2;refGene_gn=ENOX2;refGene_tx=XM_019824274.1;refGene_gn=ENOX2;refGene_tx=XM_019824275.2
X	110191087	110191231
refGene_gn=ENOX2;refGene_tx=XM_019824268.1;refGene_gn=ENOX2;refGene_tx=XM_019824269.1
X	110287442	110287464
refGene_gn=ENOX2;refGene_tx=XM_019824268.1;refGene_gn=ENOX2;refGene_tx=XM_019824269.1
X	110449946	110450004
refGene_gn=ARHGAP36;refGene_tx=XM_006944008.3
X	110472754	110473007
refGene_gn=ARHGAP36;refGene_tx=XM_006944007.4;refGene_gn=ARHGAP36;refGene_tx=XM_006944008.3;refGene_gn=ARHGAP36;refGene_tx=XM_019824288.2;refGene_gn=ARHGAP36;refGene_tx=XM_019824290.1
X	110474220	110474286
refGene_gn=ARHGAP36;refGene_tx=XM_006944007.4;refGene_gn=ARHGAP36;refGene_tx=XM_006944008.3;refGene_gn=ARHGAP36;refGene_tx=XM_019824288.2;refGene_gn=ARHGAP36;refGene_tx=XM_019824290.1
X	110474788	110475024
refGene_gn=ARHGAP36;refGene_tx=XM_006944007.4;refGene_gn=ARHGAP36;refGene_tx=XM_006944008.3;refGene_gn=ARHGAP36;refGene_tx=XM_019824288.2;refGene_gn=ARHGAP36;refGene_tx=XM_019824290.1
X	110475272	110475465
refGene_gn=ARHGAP36;refGene_tx=XM_006944007.4;refGene_gn=ARHGAP36;refGene_tx=XM_006944008.3;refGene_gn=ARHGAP36;refGene_tx=XM_019824288.2;refGene_gn=ARHGAP36;refGene_tx=XM_019824290.1
X	110475678	110475734
refGene_gn=ARHGAP36;refGene_tx=XM_006944007.4;refGene_gn=ARHGAP36;refGene_tx=XM_006944008.3;refGene_gn=ARHGAP36;refGene_tx=XM_019824288.2;refGene_gn=ARHGAP36;refGene_tx=XM_019824290.1
X	110475978	110476129
refGene_gn=ARHGAP36;refGene_tx=XM_006944007.4;refGene_gn=ARHGAP36;refGene_tx=XM_006944008.3;refGene_gn=ARHGAP36;refGene_tx=XM_019824288.2;refGene_gn=ARHGAP36;refGene_tx=XM_019824290.1
X	110476580	110476729
refGene_gn=ARHGAP36;refGene_tx=XM_006944007.4;refGene_gn=ARHGAP36;refGene_tx=XM_006944008.3;refGene_gn=ARHGAP36;refGene_tx=XM_019824288.2;refGene_gn=ARHGAP36;refGene_tx=XM_019824290.1
X	110476905	110477082
refGene_gn=ARHGAP36;refGene_tx=XM_006944007.4;refGene_gn=ARHGAP36;refGene_tx=XM_006944008.3;refGene_gn=ARHGAP36;refGene_tx=XM_019824288.2;refGene_gn=ARHGAP36;refGene_tx=XM_019824290.1
X	110477314	110477412
refGene_gn=ARHGAP36;refGene_tx=XM_006944007.4;refGene_gn=ARHGAP36;refGene_tx=XM_006944008.3;refGene_gn=ARHGAP36;refGene_tx=XM_019824288.2;refGene_gn=ARHGAP36;refGene_tx=XM_019824290.1
X	110477548	110477655
refGene_gn=ARHGAP36;refGene_tx=XM_006944007.4;refGene_gn=ARHGAP36;refG

ene_tx=XM_006944008.3;refGene_gn=ARHGAP36;refGene_tx=XM_019824288.2;refGene_gn=ARHGAP36;refGene_tx=XM_019824290.1
X	110479666	110479824
refGene_gn=ARHGAP36;refGene_tx=XM_006944007.4;refGene_gn=ARHGAP36;refGene_tx=XM_006944008.3;refGene_gn=ARHGAP36;refGene_tx=XM_019824288.2;refGene_gn=ARHGAP36;refGene_tx=XM_019824290.1
X	110604916	110605052
refGene_gn=IGSF1;refGene_tx=XM_019824302.1
X	110605194	110605318
refGene_gn=IGSF1;refGene_tx=XM_019824302.1
X	110605715	110605994
refGene_gn=IGSF1;refGene_tx=XM_019824302.1
X	110606106	110606394
refGene_gn=IGSF1;refGene_tx=XM_019824302.1
X	110606589	110606877
refGene_gn=IGSF1;refGene_tx=XM_019824302.1
X	110607058	110607346
refGene_gn=IGSF1;refGene_tx=XM_019824302.1
X	110608458	110608746
refGene_gn=IGSF1;refGene_tx=XM_019824302.1
X	110609332	110609611
refGene_gn=IGSF1;refGene_tx=XM_019824302.1
X	110609901	110610192
refGene_gn=IGSF1;refGene_tx=XM_019824302.1
X	110610579	110610612
refGene_gn=IGSF1;refGene_tx=XM_019824302.1
X	110610706	110610777
refGene_gn=IGSF1;refGene_tx=XM_019824302.1
X	110612637	110612764
refGene_gn=IGSF1;refGene_tx=XM_019824302.1
X	110613096	110613375
refGene_gn=IGSF1;refGene_tx=XM_019824302.1
X	110613837	110614131
refGene_gn=IGSF1;refGene_tx=XM_019824302.1
X	110614357	110614642
refGene_gn=IGSF1;refGene_tx=XM_019824302.1
X	110616561	110616849
refGene_gn=IGSF1;refGene_tx=XM_019824302.1
X	110617208	110617490
refGene_gn=IGSF1;refGene_tx=XM_019824302.1
X	110617921	110617948
refGene_gn=IGSF1;refGene_tx=XM_019824302.1
X	110618097	110618167
refGene_gn=IGSF1;refGene_tx=XM_019824302.1
X	110675888	110676794
refGene_gn=LOC101091768;refGene_tx=XM_004000921.2
X	110755003	110755927
refGene_gn=LOC101092250;refGene_tx=XM_004000923.1
X	110777252	110778206
refGene_gn=LOC101092501;refGene_tx=XM_006944027.2

X	110819933	110820860
refGene_gn=LOC101084607;refGene_tx=XM_004000899.2
X	110884504	110884620
refGene_gn=LOC101093004;refGene_tx=XM_019823455.1
X	110890904	110890989
refGene_gn=LOC101093004;refGene_tx=XM_019823455.1
X	110891314	110891982
refGene_gn=LOC101093004;refGene_tx=XM_019823455.1
X	110893888	110893925
refGene_gn=LOC101093004;refGene_tx=XM_019823455.1
X	111143700	111143742
refGene_gn=STK26;refGene_tx=XM_004000900.4
X	111174976	111175207
refGene_gn=STK26;refGene_tx=XM_004000900.4
X	111183845	111183902
refGene_gn=STK26;refGene_tx=XM_004000900.4
X	111189147	111189256
refGene_gn=STK26;refGene_tx=XM_004000900.4
X	111189340	111189498
refGene_gn=STK26;refGene_tx=XM_004000900.4
X	111190893	111191079
refGene_gn=STK26;refGene_tx=XM_004000900.4
X	111192226	111192375
refGene_gn=STK26;refGene_tx=XM_004000900.4
X	111193621	111193715
refGene_gn=STK26;refGene_tx=XM_004000900.4
X	111194161	111194224
refGene_gn=STK26;refGene_tx=XM_004000900.4
X	111194307	111194444
refGene_gn=STK26;refGene_tx=XM_004000900.4
X	111196169	111196194
refGene_gn=STK26;refGene_tx=XM_004000900.4
X	111200234	111201299
refGene_gn=FRMD7;refGene_tx=XM_019824304.2;refGene_gn=FRMD7;refGene_tx=XM_019824305.2;refGene_gn=FRMD7;refGene_tx=XM_019824306.2
X	111202365	111202441
refGene_gn=FRMD7;refGene_tx=XM_019824304.2;refGene_gn=FRMD7;refGene_tx=XM_019824305.2;refGene_gn=FRMD7;refGene_tx=XM_019824306.2
X	111202551	111202620
refGene_gn=FRMD7;refGene_tx=XM_019824304.2;refGene_gn=FRMD7;refGene_tx=XM_019824305.2;refGene_gn=FRMD7;refGene_tx=XM_019824306.2
X	111204566	111204730
refGene_gn=FRMD7;refGene_tx=XM_019824304.2;refGene_gn=FRMD7;refGene_tx=XM_019824305.2;refGene_gn=FRMD7;refGene_tx=XM_019824306.2
X	111206135	111206231
refGene_gn=FRMD7;refGene_tx=XM_019824304.2;refGene_gn=FRMD7;refGene_tx=XM_019824305.2;refGene_gn=FRMD7;refGene_tx=XM_019824306.2
X	111209165	111209313
refGene_gn=FRMD7;refGene_tx=XM_019824304.2;refGene_gn=FRMD7;refGene_tx=XM_019824305.2;refGene_gn=FRMD7;refGene_tx=XM_019824306.2

X	111209732	111209847
refGene_gn=FRMD7;refGene_tx=XM_019824304.2;refGene_gn=FRMD7;refGene_tx=XM_019824305.2;refGene_gn=FRMD7;refGene_tx=XM_019824306.2
X	111218126	111218224
refGene_gn=FRMD7;refGene_tx=XM_019824304.2;refGene_gn=FRMD7;refGene_tx=XM_019824305.2;refGene_gn=FRMD7;refGene_tx=XM_019824306.2
X	111221186	111221265
refGene_gn=FRMD7;refGene_tx=XM_019824304.2;refGene_gn=FRMD7;refGene_tx=XM_019824305.2;refGene_gn=FRMD7;refGene_tx=XM_019824306.2
X	111224037	111224080
refGene_gn=FRMD7;refGene_tx=XM_019824304.2
X	111224903	111225008
refGene_gn=FRMD7;refGene_tx=XM_019824304.2
X	111251437	111251494
refGene_gn=FRMD7;refGene_tx=XM_019824304.2
X	111309658	111309937
refGene_gn=RAP2C;refGene_tx=XM_023249053.1;refGene_gn=RAP2C;refGene_tx=XM_023249054.1
X	111312342	111312615
refGene_gn=RAP2C;refGene_tx=XM_023249053.1;refGene_gn=RAP2C;refGene_tx=XM_023249054.1
X	111428840	111428971
refGene_gn=MBNL3;refGene_tx=XM_019824312.2;refGene_gn=MBNL3;refGene_tx=XM_019824313.2
X	111430635	111430692
refGene_gn=MBNL3;refGene_tx=XM_019824307.2;refGene_gn=MBNL3;refGene_tx=XM_019824308.2;refGene_gn=MBNL3;refGene_tx=XM_019824309.2;refGene_gn=MBNL3;refGene_tx=XM_019824311.2;refGene_gn=MBNL3;refGene_tx=XM_019824314.2;refGene_gn=MBNL3;refGene_tx=XM_019824315.2;refGene_gn=MBNL3;refGene_tx=XM_023249052.1
X	111431654	111431749
refGene_gn=MBNL3;refGene_tx=XM_019824307.2;refGene_gn=MBNL3;refGene_tx=XM_019824308.2;refGene_gn=MBNL3;refGene_tx=XM_019824309.2;refGene_gn=MBNL3;refGene_tx=XM_019824311.2;refGene_gn=MBNL3;refGene_tx=XM_019824312.2;refGene_gn=MBNL3;refGene_tx=XM_019824314.2;refGene_gn=MBNL3;refGene_tx=XM_019824315.2;refGene_gn=MBNL3;refGene_tx=XM_023249052.1
X	111434663	111434699
refGene_gn=MBNL3;refGene_tx=XM_019824307.2;refGene_gn=MBNL3;refGene_tx=XM_019824308.2;refGene_gn=MBNL3;refGene_tx=XM_019824309.2;refGene_gn=MBNL3;refGene_tx=XM_019824312.2;refGene_gn=MBNL3;refGene_tx=XM_019824314.2;refGene_gn=MBNL3;refGene_tx=XM_019824315.2;refGene_gn=MBNL3;refGene_tx=XM_023249052.1
X	111436326	111436477
refGene_gn=MBNL3;refGene_tx=XM_019824307.2;refGene_gn=MBNL3;refGene_tx=XM_019824308.2;refGene_gn=MBNL3;refGene_tx=XM_019824309.2;refGene_gn=MBNL3;refGene_tx=XM_019824311.2;refGene_gn=MBNL3;refGene_tx=XM_019824312.2;refGene_gn=MBNL3;refGene_tx=XM_019824313.2;refGene_gn=MBNL3;refGene_tx=XM_019824314.2;refGene_gn=MBNL3;refGene_tx=XM_019824315.2;refGene_gn=MBNL3;refGene_tx=XM_023249052.1
X	111439831	111440068

refGene_gn=MBNL3;refGene_tx=XM_019824307.2;refGene_gn=MBNL3;refGene_tx=XM_019824308.2;refGene_gn=MBNL3;refGene_tx=XM_019824309.2;refGene_gn=MBNL3;refGene_tx=XM_019824311.2;refGene_gn=MBNL3;refGene_tx=XM_019824312.2;refGene_gn=MBNL3;refGene_tx=XM_019824313.2;refGene_gn=MBNL3;refGene_tx=XM_019824314.2;refGene_gn=MBNL3;refGene_tx=XM_019824315.2;refGene_gn=MBNL3;refGene_tx=XM_023249052.1
X	111441139	111441331
refGene_gn=MBNL3;refGene_tx=XM_019824307.2;refGene_gn=MBNL3;refGene_tx=XM_019824308.2;refGene_gn=MBNL3;refGene_tx=XM_019824309.2;refGene_gn=MBNL3;refGene_tx=XM_019824311.2;refGene_gn=MBNL3;refGene_tx=XM_019824312.2;refGene_gn=MBNL3;refGene_tx=XM_019824313.2;refGene_gn=MBNL3;refGene_tx=XM_019824314.2;refGene_gn=MBNL3;refGene_tx=XM_019824315.2;refGene_gn=MBNL3;refGene_tx=XM_023249052.1
X	111453973	111454138
refGene_gn=MBNL3;refGene_tx=XM_019824307.2;refGene_gn=MBNL3;refGene_tx=XM_019824308.2;refGene_gn=MBNL3;refGene_tx=XM_019824309.2;refGene_gn=MBNL3;refGene_tx=XM_019824311.2;refGene_gn=MBNL3;refGene_tx=XM_019824312.2;refGene_gn=MBNL3;refGene_tx=XM_019824313.2;refGene_gn=MBNL3;refGene_tx=XM_019824314.2;refGene_gn=MBNL3;refGene_tx=XM_019824315.2;refGene_gn=MBNL3;refGene_tx=XM_023249052.1
X	111461757	111461784
refGene_gn=MBNL3;refGene_tx=XM_019824314.2
X	111489629	111489806
refGene_gn=MBNL3;refGene_tx=XM_019824307.2;refGene_gn=MBNL3;refGene_tx=XM_019824308.2;refGene_gn=MBNL3;refGene_tx=XM_019824309.2;refGene_gn=MBNL3;refGene_tx=XM_019824311.2;refGene_gn=MBNL3;refGene_tx=XM_019824312.2;refGene_gn=MBNL3;refGene_tx=XM_019824313.2;refGene_gn=MBNL3;refGene_tx=XM_023249052.1
X	111675165	111676045
refGene_gn=HS6ST2;refGene_tx=XM_023248920.1;refGene_gn=HS6ST2;refGene_tx=XM_023248921.1;refGene_gn=HS6ST2;refGene_tx=XM_023248922.1
X	111718935	111719022
refGene_gn=HS6ST2;refGene_tx=XM_023248920.1;refGene_gn=HS6ST2;refGene_tx=XM_023248921.1
X	111753701	111753734
refGene_gn=HS6ST2;refGene_tx=XM_023248920.1
X	111962626	111963145
refGene_gn=HS6ST2;refGene_tx=XM_023248920.1;refGene_gn=HS6ST2;refGene_tx=XM_023248921.1;refGene_gn=HS6ST2;refGene_tx=XM_023248922.1
X	111963996	111964424
refGene_gn=HS6ST2;refGene_tx=XM_023248920.1;refGene_gn=HS6ST2;refGene_tx=XM_023248921.1;refGene_gn=HS6ST2;refGene_tx=XM_023248922.1
X	112021035	112023798
refGene_gn=USP26;refGene_tx=XM_019824317.2
X	112246896	112247099
refGene_gn=GPC4;refGene_tx=XM_004000909.5
X	112247199	112247375
refGene_gn=GPC4;refGene_tx=XM_004000909.5
X	112248001	112248138
refGene_gn=GPC4;refGene_tx=XM_004000909.5

X        112248919        112249066
refGene_gn=GPC4;refGene_tx=XM_004000909.5
X        112249166        112249297
refGene_gn=GPC4;refGene_tx=XM_004000909.5
X        112254209        112254375
refGene_gn=GPC4;refGene_tx=XM_004000909.5
X        112266443        112266835
refGene_gn=GPC4;refGene_tx=XM_004000909.5
X        112282256        112282415
refGene_gn=GPC4;refGene_tx=XM_004000909.5
X        112353559        112353719
refGene_gn=GPC4;refGene_tx=XM_004000909.5
X        112443896        112444066
refGene_gn=GPC3;refGene_tx=XM_023249385.1;refGene_gn=GPC3;refGene_tx=XM_023249386.1
X        112492822        112492982
refGene_gn=GPC3;refGene_tx=XM_023249385.1
X        112543511        112543632
refGene_gn=GPC3;refGene_tx=XM_023249385.1;refGene_gn=GPC3;refGene_tx=XM_023249386.1
X        112573566        112573692
refGene_gn=GPC3;refGene_tx=XM_023249385.1;refGene_gn=GPC3;refGene_tx=XM_023249386.1
X        112585665        112585799
refGene_gn=GPC3;refGene_tx=XM_023249385.1;refGene_gn=GPC3;refGene_tx=XM_023249386.1
X        112641610        112642305
refGene_gn=GPC3;refGene_tx=XM_023249385.1;refGene_gn=GPC3;refGene_tx=XM_023249386.1
X        112831831        112831993
refGene_gn=GPC3;refGene_tx=XM_023249385.1;refGene_gn=GPC3;refGene_tx=XM_023249386.1
X        112865412        112865587
refGene_gn=GPC3;refGene_tx=XM_023249385.1;refGene_gn=GPC3;refGene_tx=XM_023249386.1
X        112978810        112979008
refGene_gn=LOC109496610;refGene_tx=XM_019824319.1
X        112979299        112979341
refGene_gn=LOC109496610;refGene_tx=XM_019824319.1
X        113049617        113050586
refGene_gn=CCDC160;refGene_tx=XM_006944014.4;refGene_gn=CCDC160;refGene_tx=XM_023249105.1
X        113138283        113138421
refGene_gn=PHF6;refGene_tx=XM_006944015.4;refGene_gn=PHF6;refGene_tx=XM_019824320.2;refGene_gn=PHF6;refGene_tx=XM_019824321.2
X        113138645        113138747
refGene_gn=PHF6;refGene_tx=XM_006944015.4;refGene_gn=PHF6;refGene_tx=XM_019824320.2;refGene_gn=PHF6;refGene_tx=XM_019824321.2
X        113152492        113152626
refGene_gn=PHF6;refGene_tx=XM_006944015.4;refGene_gn=PHF6;refGene_tx=X

M_019824320.2;refGene_gn=PHF6;refGene_tx=XM_019824321.2
X 113152893 113152937
refGene_gn=PHF6;refGene_tx=XM_006944015.4;refGene_gn=PHF6;refGene_tx=XM_019824320.2;refGene_gn=PHF6;refGene_tx=XM_019824321.2
X 113165372 113165542
refGene_gn=PHF6;refGene_tx=XM_006944015.4;refGene_gn=PHF6;refGene_tx=XM_019824320.2;refGene_gn=PHF6;refGene_tx=XM_019824321.2
X 113165719 113165863
refGene_gn=PHF6;refGene_tx=XM_006944015.4;refGene_gn=PHF6;refGene_tx=XM_019824320.2;refGene_gn=PHF6;refGene_tx=XM_019824321.2
X 113167226 113167331
refGene_gn=PHF6;refGene_tx=XM_006944015.4;refGene_gn=PHF6;refGene_tx=XM_019824320.2;refGene_gn=PHF6;refGene_tx=XM_019824321.2
X 113169907 113170041
refGene_gn=PHF6;refGene_tx=XM_006944015.4;refGene_gn=PHF6;refGene_tx=XM_019824320.2;refGene_gn=PHF6;refGene_tx=XM_019824321.2
X 113177028 113177158
refGene_gn=PHF6;refGene_tx=XM_006944015.4;refGene_gn=PHF6;refGene_tx=XM_019824320.2;refGene_gn=PHF6;refGene_tx=XM_019824321.2
X 113204953 113204980
refGene_gn=HPRT1;refGene_tx=XM_023248905.1;refGene_gn=HPRT1;refGene_tx=XM_023248907.1
X 113205122 113205134
refGene_gn=HPRT1;refGene_tx=XM_023248906.1
X 113215836 113215943
refGene_gn=HPRT1;refGene_tx=XM_023248905.1;refGene_gn=HPRT1;refGene_tx=XM_023248906.1;refGene_gn=HPRT1;refGene_tx=XM_023248907.1
X 113217834 113218018
refGene_gn=HPRT1;refGene_tx=XM_023248905.1;refGene_gn=HPRT1;refGene_tx=XM_023248906.1;refGene_gn=HPRT1;refGene_tx=XM_023248907.1
X 113220795 113221205
refGene_gn=LOC102900338;refGene_tx=XM_019824322.2
X 113221207 113221703
refGene_gn=LOC102900338;refGene_tx=XM_019824322.2
X 113228828 113228894
refGene_gn=HPRT1;refGene_tx=XM_023248905.1;refGene_gn=HPRT1;refGene_tx=XM_023248906.1;refGene_gn=HPRT1;refGene_tx=XM_023248907.1
X 113232884 113232902
refGene_gn=HPRT1;refGene_tx=XM_023248905.1;refGene_gn=HPRT1;refGene_tx=XM_023248906.1;refGene_gn=HPRT1;refGene_tx=XM_023248907.1
X 113234846 113234929
refGene_gn=HPRT1;refGene_tx=XM_023248905.1;refGene_gn=HPRT1;refGene_tx=XM_023248906.1;refGene_gn=HPRT1;refGene_tx=XM_023248907.1
X 113238522 113238569
refGene_gn=HPRT1;refGene_tx=XM_023248905.1;refGene_gn=HPRT1;refGene_tx=XM_023248906.1;refGene_gn=HPRT1;refGene_tx=XM_023248907.1
X 113238738 113238815
refGene_gn=HPRT1;refGene_tx=XM_023248905.1;refGene_gn=HPRT1;refGene_tx=XM_023248906.1;refGene_gn=HPRT1;refGene_tx=XM_023248907.1
X 113239134 113239155

refGene_gn=HPRT1;refGene_tx=XM_023248907.1
X	113239325	113239373
refGene_gn=HPRT1;refGene_tx=XM_023248905.1;refGene_gn=HPRT1;refGene_tx=XM_023248906.1
X	113288766	113289405
refGene_gn=PLAC1;refGene_tx=XM_019824327.2
X	113289417	113289463
refGene_gn=LOC101088313;refGene_tx=XM_019824324.2;refGene_gn=LOC101088313;refGene_tx=XM_019824326.2
X	113318215	113318287
refGene_gn=LOC101088313;refGene_tx=XM_019824324.2
X	113371701	113371801
refGene_gn=LOC101088313;refGene_tx=XM_019824325.2
X	113472497	113472567
refGene_gn=LOC101088313;refGene_tx=XM_006944019.3;refGene_gn=LOC101088313;refGene_tx=XM_006944020.3
X	113473361	113473506
refGene_gn=LOC101088313;refGene_tx=XM_004000914.4;refGene_gn=LOC101088313;refGene_tx=XM_006944018.3
X	113481095	113481192
refGene_gn=LOC101088313;refGene_tx=XM_004000914.4;refGene_gn=LOC101088313;refGene_tx=XM_006944018.3;refGene_gn=LOC101088313;refGene_tx=XM_006944019.3;refGene_gn=LOC101088313;refGene_tx=XM_006944020.3;refGene_gn=LOC101088313;refGene_tx=XM_019824324.2;refGene_gn=LOC101088313;refGene_tx=XM_019824325.2;refGene_gn=LOC101088313;refGene_tx=XM_019824326.2
X	113483437	113483502
refGene_gn=LOC101088313;refGene_tx=XM_004000914.4;refGene_gn=LOC101088313;refGene_tx=XM_006944018.3;refGene_gn=LOC101088313;refGene_tx=XM_006944019.3;refGene_gn=LOC101088313;refGene_tx=XM_006944020.3;refGene_gn=LOC101088313;refGene_tx=XM_019824324.2;refGene_gn=LOC101088313;refGene_tx=XM_019824325.2;refGene_gn=LOC101088313;refGene_tx=XM_019824326.2
X	113484448	113484534
refGene_gn=LOC101088313;refGene_tx=XM_004000914.4;refGene_gn=LOC101088313;refGene_tx=XM_006944018.3;refGene_gn=LOC101088313;refGene_tx=XM_006944019.3;refGene_gn=LOC101088313;refGene_tx=XM_006944020.3;refGene_gn=LOC101088313;refGene_tx=XM_019824324.2;refGene_gn=LOC101088313;refGene_tx=XM_019824325.2;refGene_gn=LOC101088313;refGene_tx=XM_019824326.2
X	113484889	113484991
refGene_gn=LOC101088313;refGene_tx=XM_004000914.4;refGene_gn=LOC101088313;refGene_tx=XM_006944018.3;refGene_gn=LOC101088313;refGene_tx=XM_006944019.3;refGene_gn=LOC101088313;refGene_tx=XM_006944020.3;refGene_gn=LOC101088313;refGene_tx=XM_019824324.2;refGene_gn=LOC101088313;refGene_tx=XM_019824325.2;refGene_gn=LOC101088313;refGene_tx=XM_019824326.2
X	113485260	113485315
refGene_gn=LOC101088313;refGene_tx=XM_004000914.4;refGene_gn=LOC101088313;refGene_tx=XM_006944018.3;refGene_gn=LOC101088313;refGene_tx=XM_006944019.3;refGene_gn=LOC101088313;refGene_tx=XM_006944020.3;refGene_gn=LOC101088313;refGene_tx=XM_019824324.2;refGene_gn=LOC101088313;refGene_tx=XM_019824325.2;refGene_gn=LOC101088313;refGene_tx=XM_019824326.2
X	113485399	113485443

refGene_gn=LOC101088313;refGene_tx=XM_004000914.4;refGene_gn=LOC101088313;refGene_tx=XM_006944018.3;refGene_gn=LOC101088313;refGene_tx=XM_006944019.3;refGene_gn=LOC101088313;refGene_tx=XM_006944020.3;refGene_gn=LOC101088313;refGene_tx=XM_019824324.2;refGene_gn=LOC101088313;refGene_tx=XM_019824325.2;refGene_gn=LOC101088313;refGene_tx=XM_019824326.2
X 113485777 113485834
refGene_gn=LOC101088313;refGene_tx=XM_004000914.4;refGene_gn=LOC101088313;refGene_tx=XM_006944018.3;refGene_gn=LOC101088313;refGene_tx=XM_006944019.3;refGene_gn=LOC101088313;refGene_tx=XM_006944020.3;refGene_gn=LOC101088313;refGene_tx=XM_019824324.2;refGene_gn=LOC101088313;refGene_tx=XM_019824325.2;refGene_gn=LOC101088313;refGene_tx=XM_019824326.2
X 113490384 113490463
refGene_gn=LOC101088313;refGene_tx=XM_004000914.4;refGene_gn=LOC101088313;refGene_tx=XM_006944018.3;refGene_gn=LOC101088313;refGene_tx=XM_006944019.3;refGene_gn=LOC101088313;refGene_tx=XM_006944020.3;refGene_gn=LOC101088313;refGene_tx=XM_019824324.2;refGene_gn=LOC101088313;refGene_tx=XM_019824325.2;refGene_gn=LOC101088313;refGene_tx=XM_019824326.2
X 113492642 113492740
refGene_gn=LOC101088313;refGene_tx=XM_004000914.4;refGene_gn=LOC101088313;refGene_tx=XM_006944018.3;refGene_gn=LOC101088313;refGene_tx=XM_006944019.3;refGene_gn=LOC101088313;refGene_tx=XM_006944020.3;refGene_gn=LOC101088313;refGene_tx=XM_019824324.2;refGene_gn=LOC101088313;refGene_tx=XM_019824325.2;refGene_gn=LOC101088313;refGene_tx=XM_019824326.2
X 113498713 113498820
refGene_gn=LOC101095724;refGene_tx=XM_019824329.1;refGene_gn=LOC101095724;refGene_tx=XM_019824330.2;refGene_gn=LOC101095724;refGene_tx=XM_019824332.2;refGene_gn=LOC101095724;refGene_tx=XM_019824334.1;refGene_gn=LOC101095724;refGene_tx=XM_023249445.1;refGene_gn=LOC101095724;refGene_tx=XM_023249446.1;refGene_gn=LOC101095724;refGene_tx=XM_023249447.1;refGene_gn=LOC101095724;refGene_tx=XM_023249448.1
X 113504602 113504681
refGene_gn=LOC101095724;refGene_tx=XM_019824329.1;refGene_gn=LOC101095724;refGene_tx=XM_019824330.2;refGene_gn=LOC101095724;refGene_tx=XM_019824332.2;refGene_gn=LOC101095724;refGene_tx=XM_019824334.1;refGene_gn=LOC101095724;refGene_tx=XM_023249445.1;refGene_gn=LOC101095724;refGene_tx=XM_023249446.1;refGene_gn=LOC101095724;refGene_tx=XM_023249447.1;refGene_gn=LOC101095724;refGene_tx=XM_023249448.1;refGene_gn=LOC101095724;refGene_tx=XM_023249449.1
X 113506777 113506864
refGene_gn=LOC101095724;refGene_tx=XM_019824329.1;refGene_gn=LOC101095724;refGene_tx=XM_019824330.2;refGene_gn=LOC101095724;refGene_tx=XM_019824332.2;refGene_gn=LOC101095724;refGene_tx=XM_019824334.1;refGene_gn=LOC101095724;refGene_tx=XM_023249445.1;refGene_gn=LOC101095724;refGene_tx=XM_023249446.1;refGene_gn=LOC101095724;refGene_tx=XM_023249447.1;refGene_gn=LOC101095724;refGene_tx=XM_023249448.1;refGene_gn=LOC101095724;refGene_tx=XM_023249449.1
X 113518980 113519049
refGene_gn=LOC101095724;refGene_tx=XM_019824329.1;refGene_gn=LOC101095724;refGene_tx=XM_019824330.2;refGene_gn=LOC101095724;refGene_tx=XM_019824332.2;refGene_gn=LOC101095724;refGene_tx=XM_019824334.1;refGene_gn

=LOC101095724;refGene_tx=XM_023249445.1;refGene_gn=LOC101095724;refGene_tx=XM_023249446.1;refGene_gn=LOC101095724;refGene_tx=XM_023249447.1;refGene_gn=LOC101095724;refGene_tx=XM_023249449.1
X	113529102	113529146	refGene_gn=LOC101095724;refGene_tx=XM_019824329.1;refGene_gn=LOC101095724;refGene_tx=XM_019824330.2;refGene_gn=LOC101095724;refGene_tx=XM_019824332.2;refGene_gn=LOC101095724;refGene_tx=XM_019824334.1;refGene_gn=LOC101095724;refGene_tx=XM_023249445.1;refGene_gn=LOC101095724;refGene_tx=XM_023249446.1;refGene_gn=LOC101095724;refGene_tx=XM_023249447.1;refGene_gn=LOC101095724;refGene_tx=XM_023249448.1;refGene_gn=LOC101095724;refGene_tx=XM_023249449.1
X	113529249	113529304	refGene_gn=LOC101095724;refGene_tx=XM_019824329.1;refGene_gn=LOC101095724;refGene_tx=XM_019824330.2;refGene_gn=LOC101095724;refGene_tx=XM_019824332.2;refGene_gn=LOC101095724;refGene_tx=XM_019824334.1;refGene_gn=LOC101095724;refGene_tx=XM_023249445.1;refGene_gn=LOC101095724;refGene_tx=XM_023249446.1;refGene_gn=LOC101095724;refGene_tx=XM_023249447.1;refGene_gn=LOC101095724;refGene_tx=XM_023249448.1;refGene_gn=LOC101095724;refGene_tx=XM_023249449.1
X	113530782	113530872	refGene_gn=LOC101095724;refGene_tx=XM_019824329.1;refGene_gn=LOC101095724;refGene_tx=XM_019824330.2;refGene_gn=LOC101095724;refGene_tx=XM_019824332.2;refGene_gn=LOC101095724;refGene_tx=XM_019824334.1;refGene_gn=LOC101095724;refGene_tx=XM_023249445.1;refGene_gn=LOC101095724;refGene_tx=XM_023249446.1;refGene_gn=LOC101095724;refGene_tx=XM_023249447.1;refGene_gn=LOC101095724;refGene_tx=XM_023249448.1;refGene_gn=LOC101095724;refGene_tx=XM_023249449.1
X	113531274	113531363	refGene_gn=LOC101095724;refGene_tx=XM_019824329.1;refGene_gn=LOC101095724;refGene_tx=XM_019824330.2;refGene_gn=LOC101095724;refGene_tx=XM_019824332.2;refGene_gn=LOC101095724;refGene_tx=XM_019824334.1;refGene_gn=LOC101095724;refGene_tx=XM_023249445.1;refGene_gn=LOC101095724;refGene_tx=XM_023249446.1;refGene_gn=LOC101095724;refGene_tx=XM_023249447.1;refGene_gn=LOC101095724;refGene_tx=XM_023249448.1;refGene_gn=LOC101095724;refGene_tx=XM_023249449.1
X	113531962	113532027	refGene_gn=LOC101095724;refGene_tx=XM_019824329.1;refGene_gn=LOC101095724;refGene_tx=XM_019824330.2;refGene_gn=LOC101095724;refGene_tx=XM_019824332.2;refGene_gn=LOC101095724;refGene_tx=XM_019824334.1;refGene_gn=LOC101095724;refGene_tx=XM_023249445.1;refGene_gn=LOC101095724;refGene_tx=XM_023249446.1;refGene_gn=LOC101095724;refGene_tx=XM_023249447.1;refGene_gn=LOC101095724;refGene_tx=XM_023249448.1;refGene_gn=LOC101095724;refGene_tx=XM_023249449.1
X	113535794	113535891	refGene_gn=LOC101095724;refGene_tx=XM_019824329.1;refGene_gn=LOC101095724;refGene_tx=XM_019824330.2;refGene_gn=LOC101095724;refGene_tx=XM_019824332.2;refGene_gn=LOC101095724;refGene_tx=XM_019824334.1;refGene_gn=LOC101095724;refGene_tx=XM_023249445.1;refGene_gn=LOC101095724;refGene_tx=XM_023249446.1;refGene_gn=LOC101095724;refGene_tx=XM_023249447.1;refGene_gn=LOC101095724;refGene_tx=XM_023249448.1;refGene_gn=LOC101095

724;refGene_tx=XM_023249449.1
X 113536290 113536291
refGene_gn=LOC101095724;refGene_tx=XM_023249446.1
X 113546400 113546452
refGene_gn=LOC101095724;refGene_tx=XM_019824330.2
X 113562372 113562400
refGene_gn=LOC101095724;refGene_tx=XM_019824332.2
X 113575742 113575774
refGene_gn=MOSPD1;refGene_tx=XM_006944023.4;refGene_gn=MOSPD1;refGene_tx=XM_011279481.3;refGene_gn=MOSPD1;refGene_tx=XM_019824341.2;refGene_gn=MOSPD1;refGene_tx=XM_019824342.2;refGene_gn=MOSPD1;refGene_tx=XM_023249450.1;refGene_gn=MOSPD1;refGene_tx=XM_023249451.1;refGene_gn=MOSPD1;refGene_tx=XM_023249452.1
X 113577690 113577852
refGene_gn=MOSPD1;refGene_tx=XM_006944023.4;refGene_gn=MOSPD1;refGene_tx=XM_011279481.3;refGene_gn=MOSPD1;refGene_tx=XM_019824341.2;refGene_gn=MOSPD1;refGene_tx=XM_019824342.2;refGene_gn=MOSPD1;refGene_tx=XM_023249450.1;refGene_gn=MOSPD1;refGene_tx=XM_023249451.1;refGene_gn=MOSPD1;refGene_tx=XM_023249452.1
X 113582421 113582639
refGene_gn=MOSPD1;refGene_tx=XM_006944023.4;refGene_gn=MOSPD1;refGene_tx=XM_011279481.3;refGene_gn=MOSPD1;refGene_tx=XM_019824341.2;refGene_gn=MOSPD1;refGene_tx=XM_019824342.2;refGene_gn=MOSPD1;refGene_tx=XM_023249450.1;refGene_gn=MOSPD1;refGene_tx=XM_023249451.1;refGene_gn=MOSPD1;refGene_tx=XM_023249452.1
X 113584810 113584886
refGene_gn=MOSPD1;refGene_tx=XM_006944023.4;refGene_gn=MOSPD1;refGene_tx=XM_011279481.3;refGene_gn=MOSPD1;refGene_tx=XM_019824341.2;refGene_gn=MOSPD1;refGene_tx=XM_019824342.2;refGene_gn=MOSPD1;refGene_tx=XM_023249450.1;refGene_gn=MOSPD1;refGene_tx=XM_023249451.1;refGene_gn=MOSPD1;refGene_tx=XM_023249452.1
X 113584997 113585151
refGene_gn=MOSPD1;refGene_tx=XM_006944023.4;refGene_gn=MOSPD1;refGene_tx=XM_011279481.3;refGene_gn=MOSPD1;refGene_tx=XM_019824341.2;refGene_gn=MOSPD1;refGene_tx=XM_019824342.2;refGene_gn=MOSPD1;refGene_tx=XM_023249450.1;refGene_gn=MOSPD1;refGene_tx=XM_023249451.1;refGene_gn=MOSPD1;refGene_tx=XM_023249452.1
X 113599268 113599269
refGene_gn=LOC101095724;refGene_tx=XM_019824334.1
X 113665254 113665482
refGene_gn=LOC101095962;refGene_tx=XM_006944025.4
X 113828367 113828554
refGene_gn=CT55;refGene_tx=XM_023249046.1
X 113835524 113835663
refGene_gn=CT55;refGene_tx=XM_023249047.1
X 113838175 113838357
refGene_gn=CT55;refGene_tx=XM_023249046.1;refGene_gn=CT55;refGene_tx=XM_023249047.1
X 113841199 113841344
refGene_gn=CT55;refGene_tx=XM_023249046.1;refGene_gn=CT55;refGene_tx=X

M_023249047.1
X	113850708	113850821
refGene_gn=CT55;refGene_tx=XM_023249046.1;refGene_gn=CT55;refGene_tx=XM_023249047.1
X	113851590	113851782
refGene_gn=CT55;refGene_tx=XM_023249046.1;refGene_gn=CT55;refGene_tx=XM_023249047.1
X	113897387	113897834
refGene_gn=ZNF75D;refGene_tx=XM_023249004.1;refGene_gn=ZNF75D;refGene_tx=XM_023249005.1;refGene_gn=ZNF75D;refGene_tx=XM_023249006.1
X	113899065	113899258
refGene_gn=ZNF75D;refGene_tx=XM_023249004.1;refGene_gn=ZNF75D;refGene_tx=XM_023249006.1
X	113899990	113900082
refGene_gn=ZNF75D;refGene_tx=XM_023249004.1;refGene_gn=ZNF75D;refGene_tx=XM_023249006.1
X	113900417	113900544
refGene_gn=ZNF75D;refGene_tx=XM_023249004.1;refGene_gn=ZNF75D;refGene_tx=XM_023249005.1;refGene_gn=ZNF75D;refGene_tx=XM_023249006.1
X	113907138	113907848
refGene_gn=ZNF75D;refGene_tx=XM_023249004.1;refGene_gn=ZNF75D;refGene_tx=XM_023249005.1;refGene_gn=ZNF75D;refGene_tx=XM_023249006.1
X	113963502	113963856
refGene_gn=ZNF449;refGene_tx=XM_023249051.1
X	113965669	113965873
refGene_gn=ZNF449;refGene_tx=XM_023249051.1
X	113967406	113967407
refGene_gn=ZNF449;refGene_tx=XM_023249051.1
X	113982001	113982115
refGene_gn=ZNF449;refGene_tx=XM_023249051.1
X	113982306	113983190
refGene_gn=ZNF449;refGene_tx=XM_023249051.1
X	114014126	114014363
refGene_gn=LOC111558710;refGene_tx=XM_023249086.1
X	114101534	114101645
refGene_gn=INTS6L;refGene_tx=XM_023249469.1
X	114101786	114101864
refGene_gn=INTS6L;refGene_tx=XM_023249469.1
X	114124199	114124349
refGene_gn=INTS6L;refGene_tx=XM_023249469.1
X	114125122	114125212
refGene_gn=INTS6L;refGene_tx=XM_023249469.1
X	114125437	114125621
refGene_gn=INTS6L;refGene_tx=XM_023249469.1
X	114125880	114126009
refGene_gn=INTS6L;refGene_tx=XM_023249469.1
X	114127962	114128126
refGene_gn=INTS6L;refGene_tx=XM_023249469.1
X	114130447	114130600
refGene_gn=INTS6L;refGene_tx=XM_023249469.1

X	114134633	114134766
refGene_gn=INTS6L;refGene_tx=XM_023249469.1
X	114143125	114143220
refGene_gn=INTS6L;refGene_tx=XM_023249469.1
X	114146460	114146679
refGene_gn=INTS6L;refGene_tx=XM_023249469.1
X	114147325	114147449
refGene_gn=INTS6L;refGene_tx=XM_023249469.1
X	114149288	114149431
refGene_gn=INTS6L;refGene_tx=XM_023249469.1
X	114151060	114151163
refGene_gn=INTS6L;refGene_tx=XM_023249469.1
X	114153731	114154109
refGene_gn=INTS6L;refGene_tx=XM_023249469.1
X	114155467	114155561
refGene_gn=INTS6L;refGene_tx=XM_023249469.1
X	114156162	114156199
refGene_gn=INTS6L;refGene_tx=XM_023249469.1
X	114157061	114157087
refGene_gn=INTS6L;refGene_tx=XM_023249469.1
X	114157088	114157119
refGene_gn=INTS6L;refGene_tx=XM_023249469.1
X	114184553	114184647
refGene_gn=LOC102901637;refGene_tx=XM_006944033.3
X	114187411	114187777
refGene_gn=LOC102901637;refGene_tx=XM_006944033.3
X	114188125	114188219
refGene_gn=LOC102901637;refGene_tx=XM_006944033.3
X	114188523	114188647
refGene_gn=LOC102901637;refGene_tx=XM_006944033.3
X	114222567	114222573
refGene_gn=LOC102900259;refGene_tx=XM_019823489.2
X	114222954	114223049
refGene_gn=LOC102900259;refGene_tx=XM_019823489.2
X	114223619	114223742
refGene_gn=LOC102900259;refGene_tx=XM_019823489.2
X	114225380	114225561
refGene_gn=LOC102900259;refGene_tx=XM_019823489.2
X	114230788	114230848
refGene_gn=LOC102900259;refGene_tx=XM_019823489.2
X	114259718	114259818
refGene_gn=LOC102901675;refGene_tx=XM_023249118.1
X	114260175	114260269
refGene_gn=LOC102901675;refGene_tx=XM_023249118.1
X	114260662	114260791
refGene_gn=LOC102901675;refGene_tx=XM_023249118.1
X	114262515	114262693
refGene_gn=LOC102901675;refGene_tx=XM_023249118.1
X	114300197	114300357
refGene_gn=MMGT1;refGene_tx=XM_004000934.5

X 114302008 114302112
refGene_gn=MMGT1;refGene_tx=XM_004000934.5
X 114304084 114304137
refGene_gn=MMGT1;refGene_tx=XM_004000934.5
X 114307970 114308049
refGene_gn=MMGT1;refGene_tx=XM_004000934.5
X 114318301 114318626
refGene_gn=SLC9A6;refGene_tx=XM_004000935.3;refGene_gn=SLC9A6;refGene_tx=XM_019824357.1;refGene_gn=SLC9A6;refGene_tx=XM_019824358.2;refGene_gn=SLC9A6;refGene_tx=XM_019824359.2
X 114328223 114328423
refGene_gn=SLC9A6;refGene_tx=XM_004000935.3;refGene_gn=SLC9A6;refGene_tx=XM_019824357.1;refGene_gn=SLC9A6;refGene_tx=XM_019824358.2;refGene_gn=SLC9A6;refGene_tx=XM_019824359.2
X 114330908 114330986
refGene_gn=SLC9A6;refGene_tx=XM_004000935.3;refGene_gn=SLC9A6;refGene_tx=XM_019824357.1;refGene_gn=SLC9A6;refGene_tx=XM_019824358.2;refGene_gn=SLC9A6;refGene_tx=XM_019824359.2
X 114331253 114331330
refGene_gn=SLC9A6;refGene_tx=XM_004000935.3;refGene_gn=SLC9A6;refGene_tx=XM_019824357.1;refGene_gn=SLC9A6;refGene_tx=XM_019824358.2;refGene_gn=SLC9A6;refGene_tx=XM_019824359.2
X 114331632 114331745
refGene_gn=SLC9A6;refGene_tx=XM_004000935.3;refGene_gn=SLC9A6;refGene_tx=XM_019824357.1;refGene_gn=SLC9A6;refGene_tx=XM_019824358.2;refGene_gn=SLC9A6;refGene_tx=XM_019824359.2
X 114334518 114334624
refGene_gn=SLC9A6;refGene_tx=XM_004000935.3;refGene_gn=SLC9A6;refGene_tx=XM_019824357.1;refGene_gn=SLC9A6;refGene_tx=XM_019824358.2;refGene_gn=SLC9A6;refGene_tx=XM_019824359.2
X 114339018 114339160
refGene_gn=SLC9A6;refGene_tx=XM_004000935.3;refGene_gn=SLC9A6;refGene_tx=XM_019824357.1;refGene_gn=SLC9A6;refGene_tx=XM_019824358.2;refGene_gn=SLC9A6;refGene_tx=XM_019824359.2
X 114341047 114341153
refGene_gn=SLC9A6;refGene_tx=XM_004000935.3;refGene_gn=SLC9A6;refGene_tx=XM_019824357.1;refGene_gn=SLC9A6;refGene_tx=XM_019824358.2;refGene_gn=SLC9A6;refGene_tx=XM_019824359.2
X 114341454 114341543
refGene_gn=SLC9A6;refGene_tx=XM_004000935.3;refGene_gn=SLC9A6;refGene_tx=XM_019824357.1;refGene_gn=SLC9A6;refGene_tx=XM_019824358.2;refGene_gn=SLC9A6;refGene_tx=XM_019824359.2
X 114344324 114344438
refGene_gn=SLC9A6;refGene_tx=XM_004000935.3;refGene_gn=SLC9A6;refGene_tx=XM_019824357.1;refGene_gn=SLC9A6;refGene_tx=XM_019824358.2;refGene_gn=SLC9A6;refGene_tx=XM_019824359.2
X 114349362 114349474
refGene_gn=SLC9A6;refGene_tx=XM_004000935.3;refGene_gn=SLC9A6;refGene_tx=XM_019824357.1;refGene_gn=SLC9A6;refGene_tx=XM_019824358.2;refGene_gn=SLC9A6;refGene_tx=XM_019824359.2

X 114350381 114350535
refGene_gn=SLC9A6;refGene_tx=XM_004000935.3;refGene_gn=SLC9A6;refGene_tx=XM_019824357.1;refGene_gn=SLC9A6;refGene_tx=XM_019824358.2;refGene_gn=SLC9A6;refGene_tx=XM_019824359.2
X 114355654 114355744
refGene_gn=SLC9A6;refGene_tx=XM_019824357.1;refGene_gn=SLC9A6;refGene_tx=XM_019824358.2;refGene_gn=SLC9A6;refGene_tx=XM_019824359.2
X 114356931 114356962
refGene_gn=SLC9A6;refGene_tx=XM_004000935.3;refGene_gn=SLC9A6;refGene_tx=XM_019824357.1;refGene_gn=SLC9A6;refGene_tx=XM_019824358.2;refGene_gn=SLC9A6;refGene_tx=XM_019824359.2
X 114359426 114359506
refGene_gn=SLC9A6;refGene_tx=XM_004000935.3;refGene_gn=SLC9A6;refGene_tx=XM_019824357.1;refGene_gn=SLC9A6;refGene_tx=XM_019824358.2;refGene_gn=SLC9A6;refGene_tx=XM_019824359.2
X 114365115 114365221
refGene_gn=SLC9A6;refGene_tx=XM_004000935.3;refGene_gn=SLC9A6;refGene_tx=XM_019824357.1;refGene_gn=SLC9A6;refGene_tx=XM_019824358.2;refGene_gn=SLC9A6;refGene_tx=XM_019824359.2
X 114369391 114369664
refGene_gn=SLC9A6;refGene_tx=XM_004000935.3;refGene_gn=SLC9A6;refGene_tx=XM_019824357.1;refGene_gn=SLC9A6;refGene_tx=XM_019824358.2;refGene_gn=SLC9A6;refGene_tx=XM_019824359.2
X 114499005 114499027
refGene_gn=FHL1;refGene_tx=XM_004000937.3;refGene_gn=FHL1;refGene_tx=XM_004000938.3;refGene_gn=FHL1;refGene_tx=XM_019824360.1
X 114506241 114506423
refGene_gn=FHL1;refGene_tx=XM_004000937.3;refGene_gn=FHL1;refGene_tx=XM_004000938.3;refGene_gn=FHL1;refGene_tx=XM_006944037.3;refGene_gn=FHL1;refGene_tx=XM_006944038.2;refGene_gn=FHL1;refGene_tx=XM_019824360.1;refGene_gn=FHL1;refGene_tx=XM_023249463.1
X 114506676 114506851
refGene_gn=FHL1;refGene_tx=XM_004000937.3;refGene_gn=FHL1;refGene_tx=XM_004000938.3;refGene_gn=FHL1;refGene_tx=XM_006944037.3;refGene_gn=FHL1;refGene_tx=XM_006944038.2;refGene_gn=FHL1;refGene_tx=XM_019824360.1;refGene_gn=FHL1;refGene_tx=XM_023249463.1
X 114507532 114507702
refGene_gn=FHL1;refGene_tx=XM_004000937.3;refGene_gn=FHL1;refGene_tx=XM_004000938.3;refGene_gn=FHL1;refGene_tx=XM_006944037.3;refGene_gn=FHL1;refGene_tx=XM_006944038.2;refGene_gn=FHL1;refGene_tx=XM_019824360.1;refGene_gn=FHL1;refGene_tx=XM_023249463.1
X 114508165 114508352
refGene_gn=FHL1;refGene_tx=XM_004000937.3;refGene_gn=FHL1;refGene_tx=XM_004000938.3;refGene_gn=FHL1;refGene_tx=XM_006944037.3;refGene_gn=FHL1;refGene_tx=XM_006944038.2;refGene_gn=FHL1;refGene_tx=XM_023249463.1
X 114508986 114509186
refGene_gn=FHL1;refGene_tx=XM_004000938.3;refGene_gn=FHL1;refGene_tx=XM_006944037.3;refGene_gn=FHL1;refGene_tx=XM_006944038.2;refGene_gn=FHL1;refGene_tx=XM_023249463.1
X 114509536 114509691

refGene_gn=FHL1;refGene_tx=XM_004000937.3;refGene_gn=FHL1;refGene_tx=XM_004000938.3;refGene_gn=FHL1;refGene_tx=XM_006944037.3;refGene_gn=FHL1;refGene_tx=XM_006944038.2;refGene_gn=FHL1;refGene_tx=XM_019824360.1;refGene_gn=FHL1;refGene_tx=XM_023249463.1
X 114515728 114515878
refGene_gn=MAP7D3;refGene_tx=XM_023248980.1
X 114516923 114516969
refGene_gn=MAP7D3;refGene_tx=XM_023248980.1
X 114517866 114518065
refGene_gn=MAP7D3;refGene_tx=XM_023248980.1
X 114519143 114519246
refGene_gn=MAP7D3;refGene_tx=XM_023248980.1
X 114520453 114520507
refGene_gn=MAP7D3;refGene_tx=XM_023248980.1
X 114521502 114521607
refGene_gn=MAP7D3;refGene_tx=XM_023248980.1
X 114522819 114522952
refGene_gn=MAP7D3;refGene_tx=XM_023248980.1
X 114524356 114524486
refGene_gn=MAP7D3;refGene_tx=XM_023248980.1
X 114526205 114526411
refGene_gn=MAP7D3;refGene_tx=XM_023248980.1
X 114526682 114526810
refGene_gn=MAP7D3;refGene_tx=XM_023248980.1
X 114527369 114527998
refGene_gn=MAP7D3;refGene_tx=XM_023248980.1
X 114530814 114530940
refGene_gn=MAP7D3;refGene_tx=XM_023248980.1
X 114532664 114532761
refGene_gn=MAP7D3;refGene_tx=XM_023248980.1
X 114533650 114533668
refGene_gn=MAP7D3;refGene_tx=XM_023248980.1
X 114537425 114537529
refGene_gn=MAP7D3;refGene_tx=XM_023248980.1
X 114538045 114538160
refGene_gn=MAP7D3;refGene_tx=XM_023248980.1
X 114541231 114541395
refGene_gn=MAP7D3;refGene_tx=XM_023248980.1
X 114542472 114542556
refGene_gn=MAP7D3;refGene_tx=XM_023248980.1
X 114542660 114542756
refGene_gn=MAP7D3;refGene_tx=XM_023248980.1
X 114548215 114548276
refGene_gn=MAP7D3;refGene_tx=XM_023248980.1
X 114586474 114586544
refGene_gn=LOC111556211;refGene_tx=XM_023249141.1
X 114600912 114601535
refGene_gn=LOC111556211;refGene_tx=XM_023249141.1
X 114613185 114613210
refGene_gn=ADGRG4;refGene_tx=XM_023248981.1

X 114618013 114623706
refGene_gn=ADGRG4;refGene_tx=XM_023248981.1
X 114624802 114624877
refGene_gn=LOC111558686;refGene_tx=XM_023248982.1
X 114626889 114626954
refGene_gn=LOC111558686;refGene_tx=XM_023248982.1
X 114631224 114631388
refGene_gn=LOC111558686;refGene_tx=XM_023248982.1
X 114632828 114632961
refGene_gn=LOC111558686;refGene_tx=XM_023248982.1
X 114634706 114634825
refGene_gn=LOC111558686;refGene_tx=XM_023248982.1
X 114641301 114641518
refGene_gn=LOC111558686;refGene_tx=XM_023248982.1
X 114642087 114642250
refGene_gn=LOC111558686;refGene_tx=XM_023248982.1
X 114652023 114652158
refGene_gn=LOC111558686;refGene_tx=XM_023248982.1
X 114656434 114656554
refGene_gn=LOC111558686;refGene_tx=XM_023248982.1
X 114660080 114660143
refGene_gn=LOC111558686;refGene_tx=XM_023248982.1
X 114661907 114661910
refGene_gn=LOC111558719;refGene_tx=XM_023249142.1
X 114664247 114664369
refGene_gn=LOC111558719;refGene_tx=XM_023249142.1
X 114668460 114668729
refGene_gn=LOC111558719;refGene_tx=XM_023249142.1
X 114672199 114672278
refGene_gn=LOC111558719;refGene_tx=XM_023249142.1
X 114673363 114673397
refGene_gn=LOC111558719;refGene_tx=XM_023249142.1
X 114737057 114737491
refGene_gn=BRS3;refGene_tx=XM_019823491.2
X 114738975 114739327
refGene_gn=BRS3;refGene_tx=XM_019823491.2
X 114740614 114741028
refGene_gn=BRS3;refGene_tx=XM_019823491.2
X 114746446 114746632
refGene_gn=HTATSF1;refGene_tx=XM_006944041.3;refGene_gn=HTATSF1;refGen
e_tx=XM_023248895.1
X 114748389 114748570
refGene_gn=HTATSF1;refGene_tx=XM_006944041.3;refGene_gn=HTATSF1;refGen
e_tx=XM_023248895.1
X 114749209 114749257
refGene_gn=HTATSF1;refGene_tx=XM_006944041.3;refGene_gn=HTATSF1;refGen
e_tx=XM_023248895.1
X 114749983 114750138
refGene_gn=HTATSF1;refGene_tx=XM_006944041.3;refGene_gn=HTATSF1;refGen
e_tx=XM_023248895.1

X 114752293 114752457
refGene_gn=HTATSF1;refGene_tx=XM_006944041.3;refGene_gn=HTATSF1;refGene_tx=XM_023248895.1
X 114753805 114753905
refGene_gn=HTATSF1;refGene_tx=XM_006944041.3;refGene_gn=HTATSF1;refGene_tx=XM_023248895.1
X 114757002 114757092
refGene_gn=HTATSF1;refGene_tx=XM_006944041.3;refGene_gn=HTATSF1;refGene_tx=XM_023248895.1
X 114757861 114757999
refGene_gn=HTATSF1;refGene_tx=XM_006944041.3;refGene_gn=HTATSF1;refGene_tx=XM_023248895.1
X 114758480 114759719
refGene_gn=HTATSF1;refGene_tx=XM_006944041.3;refGene_gn=HTATSF1;refGene_tx=XM_023248895.1
X 114781766 114781968
refGene_gn=VGLL1;refGene_tx=XM_019824372.2;refGene_gn=VGLL1;refGene_tx=XM_023249230.1
X 114789930 114790433
refGene_gn=VGLL1;refGene_tx=XM_019824372.2;refGene_gn=VGLL1;refGene_tx=XM_023249230.1
X 114797636 114797638
refGene_gn=VGLL1;refGene_tx=XM_019824372.2
X 114884206 114884362
refGene_gn=CD40LG;refGene_tx=NM_001009298.1
X 114885735 114885867
refGene_gn=CD40LG;refGene_tx=NM_001009298.1
X 114890178 114890233
refGene_gn=CD40LG;refGene_tx=NM_001009298.1
X 114891721 114891784
refGene_gn=CD40LG;refGene_tx=NM_001009298.1
X 114894538 114894915
refGene_gn=CD40LG;refGene_tx=NM_001009298.1
X 114908509 114908650
refGene_gn=ARHGEF6;refGene_tx=XM_004000941.4;refGene_gn=ARHGEF6;refGene_tx=XM_006944042.4;refGene_gn=ARHGEF6;refGene_tx=XM_011279488.3;refGene_gn=ARHGEF6;refGene_tx=XM_011279489.3;refGene_gn=ARHGEF6;refGene_tx=XM_023249364.1
X 114909166 114909221
refGene_gn=ARHGEF6;refGene_tx=XM_004000941.4;refGene_gn=ARHGEF6;refGene_tx=XM_006944042.4;refGene_gn=ARHGEF6;refGene_tx=XM_011279488.3;refGene_gn=ARHGEF6;refGene_tx=XM_011279489.3;refGene_gn=ARHGEF6;refGene_tx=XM_023249364.1
X 114910904 114911004
refGene_gn=ARHGEF6;refGene_tx=XM_004000941.4;refGene_gn=ARHGEF6;refGene_tx=XM_006944042.4;refGene_gn=ARHGEF6;refGene_tx=XM_011279488.3;refGene_gn=ARHGEF6;refGene_tx=XM_011279489.3;refGene_gn=ARHGEF6;refGene_tx=XM_023249364.1
X 114912914 114913004
refGene_gn=ARHGEF6;refGene_tx=XM_004000941.4;refGene_gn=ARHGEF6;refGen

e_tx=XM_006944042.4;refGene_gn=ARHGEF6;refGene_tx=XM_011279488.3;refGene_gn=ARHGEF6;refGene_tx=XM_011279489.3;refGene_gn=ARHGEF6;refGene_tx=XM_023249364.1
X 114914203 114914297
refGene_gn=ARHGEF6;refGene_tx=XM_004000941.4;refGene_gn=ARHGEF6;refGene_tx=XM_006944042.4;refGene_gn=ARHGEF6;refGene_tx=XM_011279488.3;refGene_gn=ARHGEF6;refGene_tx=XM_011279489.3;refGene_gn=ARHGEF6;refGene_tx=XM_023249364.1
X 114915409 114915430
refGene_gn=ARHGEF6;refGene_tx=XM_004000941.4;refGene_gn=ARHGEF6;refGene_tx=XM_006944042.4;refGene_gn=ARHGEF6;refGene_tx=XM_011279488.3;refGene_gn=ARHGEF6;refGene_tx=XM_023249364.1
X 114916900 114917026
refGene_gn=ARHGEF6;refGene_tx=XM_004000941.4;refGene_gn=ARHGEF6;refGene_tx=XM_006944042.4;refGene_gn=ARHGEF6;refGene_tx=XM_023249364.1
X 114918149 114918295
refGene_gn=ARHGEF6;refGene_tx=XM_004000941.4;refGene_gn=ARHGEF6;refGene_tx=XM_006944042.4;refGene_gn=ARHGEF6;refGene_tx=XM_011279488.3;refGene_gn=ARHGEF6;refGene_tx=XM_011279489.3;refGene_gn=ARHGEF6;refGene_tx=XM_023249364.1
X 114918912 114918991
refGene_gn=ARHGEF6;refGene_tx=XM_004000941.4;refGene_gn=ARHGEF6;refGene_tx=XM_006944042.4;refGene_gn=ARHGEF6;refGene_tx=XM_011279488.3;refGene_gn=ARHGEF6;refGene_tx=XM_011279489.3;refGene_gn=ARHGEF6;refGene_tx=XM_023249364.1
X 114919663 114919750
refGene_gn=ARHGEF6;refGene_tx=XM_004000941.4;refGene_gn=ARHGEF6;refGene_tx=XM_006944042.4;refGene_gn=ARHGEF6;refGene_tx=XM_011279488.3;refGene_gn=ARHGEF6;refGene_tx=XM_011279489.3;refGene_gn=ARHGEF6;refGene_tx=XM_023249364.1
X 114922346 114922493
refGene_gn=ARHGEF6;refGene_tx=XM_004000941.4;refGene_gn=ARHGEF6;refGene_tx=XM_006944042.4;refGene_gn=ARHGEF6;refGene_tx=XM_011279488.3;refGene_gn=ARHGEF6;refGene_tx=XM_011279489.3;refGene_gn=ARHGEF6;refGene_tx=XM_023249364.1
X 114924538 114924598
refGene_gn=ARHGEF6;refGene_tx=XM_004000941.4;refGene_gn=ARHGEF6;refGene_tx=XM_006944042.4;refGene_gn=ARHGEF6;refGene_tx=XM_011279488.3;refGene_gn=ARHGEF6;refGene_tx=XM_011279489.3;refGene_gn=ARHGEF6;refGene_tx=XM_023249364.1
X 114926842 114926981
refGene_gn=ARHGEF6;refGene_tx=XM_004000941.4;refGene_gn=ARHGEF6;refGene_tx=XM_006944042.4;refGene_gn=ARHGEF6;refGene_tx=XM_011279488.3;refGene_gn=ARHGEF6;refGene_tx=XM_011279489.3;refGene_gn=ARHGEF6;refGene_tx=XM_023249364.1
X 114943755 114943878
refGene_gn=ARHGEF6;refGene_tx=XM_004000941.4;refGene_gn=ARHGEF6;refGene_tx=XM_006944042.4;refGene_gn=ARHGEF6;refGene_tx=XM_011279488.3;refGene_gn=ARHGEF6;refGene_tx=XM_011279489.3;refGene_gn=ARHGEF6;refGene_tx=XM_023249364.1

X	114945523	114945619
refGene_gn=ARHGEF6;refGene_tx=XM_004000941.4;refGene_gn=ARHGEF6;refGene_tx=XM_006944042.4;refGene_gn=ARHGEF6;refGene_tx=XM_011279488.3;refGene_gn=ARHGEF6;refGene_tx=XM_011279489.3;refGene_gn=ARHGEF6;refGene_tx=XM_023249364.1
X	114948579	114948674
refGene_gn=ARHGEF6;refGene_tx=XM_004000941.4;refGene_gn=ARHGEF6;refGene_tx=XM_006944042.4;refGene_gn=ARHGEF6;refGene_tx=XM_011279488.3;refGene_gn=ARHGEF6;refGene_tx=XM_011279489.3;refGene_gn=ARHGEF6;refGene_tx=XM_023249364.1
X	114962029	114962100
refGene_gn=ARHGEF6;refGene_tx=XM_004000941.4;refGene_gn=ARHGEF6;refGene_tx=XM_006944042.4;refGene_gn=ARHGEF6;refGene_tx=XM_011279488.3;refGene_gn=ARHGEF6;refGene_tx=XM_011279489.3;refGene_gn=ARHGEF6;refGene_tx=XM_023249364.1
X	114972088	114972290
refGene_gn=ARHGEF6;refGene_tx=XM_004000941.4;refGene_gn=ARHGEF6;refGene_tx=XM_006944042.4;refGene_gn=ARHGEF6;refGene_tx=XM_011279488.3;refGene_gn=ARHGEF6;refGene_tx=XM_011279489.3;refGene_gn=ARHGEF6;refGene_tx=XM_023249364.1
X	114973638	114973760
refGene_gn=ARHGEF6;refGene_tx=XM_004000941.4;refGene_gn=ARHGEF6;refGene_tx=XM_011279488.3;refGene_gn=ARHGEF6;refGene_tx=XM_011279489.3
X	114976385	114976470
refGene_gn=ARHGEF6;refGene_tx=XM_004000941.4;refGene_gn=ARHGEF6;refGene_tx=XM_011279488.3;refGene_gn=ARHGEF6;refGene_tx=XM_011279489.3
X	115004651	115004735
refGene_gn=ARHGEF6;refGene_tx=XM_004000941.4;refGene_gn=ARHGEF6;refGene_tx=XM_011279488.3;refGene_gn=ARHGEF6;refGene_tx=XM_011279489.3
X	115005902	115006067
refGene_gn=ARHGEF6;refGene_tx=XM_004000941.4;refGene_gn=ARHGEF6;refGene_tx=XM_011279488.3;refGene_gn=ARHGEF6;refGene_tx=XM_011279489.3
X	115037183	115037188
refGene_gn=RBMX;refGene_tx=XM_011279492.2
X	115039006	115039317
refGene_gn=RBMX;refGene_tx=XM_011279490.2
X	115039952	115040035
refGene_gn=RBMX;refGene_tx=XM_011279490.2;refGene_gn=RBMX;refGene_tx=XM_011279492.2
X	115040122	115040248
refGene_gn=RBMX;refGene_tx=XM_011279490.2;refGene_gn=RBMX;refGene_tx=XM_011279492.2
X	115040340	115040455
refGene_gn=RBMX;refGene_tx=XM_011279490.2;refGene_gn=RBMX;refGene_tx=XM_011279492.2
X	115040794	115040947
refGene_gn=RBMX;refGene_tx=XM_011279490.2;refGene_gn=RBMX;refGene_tx=XM_011279492.2
X	115041548	115041720
refGene_gn=RBMX;refGene_tx=XM_011279490.2;refGene_gn=RBMX;refGene_tx=X

M_011279492.2
X 115042201 115042308
refGene_gn=RBMX;refGene_tx=XM_011279490.2;refGene_gn=RBMX;refGene_tx=X
M_011279492.2
X 115042510 115042619
refGene_gn=RBMX;refGene_tx=XM_011279490.2;refGene_gn=RBMX;refGene_tx=X
M_011279492.2
X 115069998 115070184
refGene_gn=LOC101083916;refGene_tx=XM_023249085.1
X 115079500 115079568
refGene_gn=LOC101083916;refGene_tx=XM_023249085.1
X 115081145 115081239
refGene_gn=LOC101083916;refGene_tx=XM_023249085.1
X 115085924 115086052
refGene_gn=LOC101083916;refGene_tx=XM_023249085.1
X 115086729 115086859
refGene_gn=LOC101083916;refGene_tx=XM_023249085.1
X 115102754 115102882
refGene_gn=LOC101083916;refGene_tx=XM_023249085.1
X 115107945 115108057
refGene_gn=LOC101083916;refGene_tx=XM_023249085.1
X 115120967 115121047
refGene_gn=LOC101083916;refGene_tx=XM_023249085.1
X 115124828 115124937
refGene_gn=LOC101083916;refGene_tx=XM_023249085.1
X 115126512 115126645
refGene_gn=LOC101083916;refGene_tx=XM_023249085.1
X 115128122 115128242
refGene_gn=LOC101083916;refGene_tx=XM_023249085.1
X 115128443 115128501
refGene_gn=LOC101083916;refGene_tx=XM_023249085.1
X 115138135 115138295
refGene_gn=LOC101083916;refGene_tx=XM_023249085.1
X 115140993 115141142
refGene_gn=LOC101083916;refGene_tx=XM_023249085.1
X 115143626 115143738
refGene_gn=LOC101083916;refGene_tx=XM_023249085.1
X 115145610 115145782
refGene_gn=LOC101083916;refGene_tx=XM_023249085.1
X 115159576 115159644
refGene_gn=LOC101083916;refGene_tx=XM_023249085.1
X 115168574 115170068
refGene_gn=GPR101;refGene_tx=XM_023249204.1
X 115603935 115604992
refGene_gn=ZIC3;refGene_tx=XM_023249131.1;refGene_gn=ZIC3;refGene_tx=X
M_023249132.1
X 115606126 115606290
refGene_gn=ZIC3;refGene_tx=XM_023249131.1;refGene_gn=ZIC3;refGene_tx=X
M_023249132.1
X 115607131 115607311

refGene_gn=ZIC3;refGene_tx=XM_023249131.1
X 115614465 115614615
refGene_gn=ZIC3;refGene_tx=XM_023249132.1
X 116246568 116247609
refGene_gn=LOC101085187;refGene_tx=XM_004000964.4;refGene_gn=LOC101085187;refGene_tx=XM_023249261.1
X 116339417 116339554
refGene_gn=FGF13;refGene_tx=XM_019824375.2;refGene_gn=FGF13;refGene_tx=XM_019824376.2;refGene_gn=FGF13;refGene_tx=XM_023249008.1;refGene_gn=FGF13;refGene_tx=XM_023249009.1;refGene_gn=FGF13;refGene_tx=XM_023249010.1
X 116340663 116340862
refGene_gn=FGF13;refGene_tx=XM_019824375.2;refGene_gn=FGF13;refGene_tx=XM_019824376.2;refGene_gn=FGF13;refGene_tx=XM_023249008.1;refGene_gn=FGF13;refGene_tx=XM_023249009.1;refGene_gn=FGF13;refGene_tx=XM_023249010.1
X 116396028 116396132
refGene_gn=FGF13;refGene_tx=XM_019824375.2;refGene_gn=FGF13;refGene_tx=XM_019824376.2;refGene_gn=FGF13;refGene_tx=XM_023249008.1;refGene_gn=FGF13;refGene_tx=XM_023249009.1;refGene_gn=FGF13;refGene_tx=XM_023249010.1
X 116401947 116402058
refGene_gn=FGF13;refGene_tx=XM_019824375.2;refGene_gn=FGF13;refGene_tx=XM_019824376.2;refGene_gn=FGF13;refGene_tx=XM_023249008.1;refGene_gn=FGF13;refGene_tx=XM_023249009.1;refGene_gn=FGF13;refGene_tx=XM_023249010.1
X 116403626 116403726
refGene_gn=FGF13;refGene_tx=XM_019824376.2
X 116404044 116404231
refGene_gn=FGF13;refGene_tx=XM_023249009.1
X 116435461 116435489
refGene_gn=FGF13;refGene_tx=XM_019824375.2
X 116519588 116519756
refGene_gn=FGF13;refGene_tx=XM_023249008.1
X 116781624 116781673
refGene_gn=FGF13;refGene_tx=XM_023249008.1;refGene_gn=FGF13;refGene_tx=XM_023249010.1
X 117080316 117080404
refGene_gn=F9;refGene_tx=NM_001009377.3
X 117086684 117086848
refGene_gn=F9;refGene_tx=NM_001009377.3
X 117087049 117087074
refGene_gn=F9;refGene_tx=NM_001009377.3
X 117091855 117091969
refGene_gn=F9;refGene_tx=NM_001009377.3
X 117098694 117098823
refGene_gn=F9;refGene_tx=NM_001009377.3
X 117101380 117101598
refGene_gn=F9;refGene_tx=NM_001009377.3
X 117110509 117110624

refGene_gn=F9;refGene_tx=NM_001009377.3
X 117111280 117111828
refGene_gn=F9;refGene_tx=NM_001009377.3
X 117132077 117132188
refGene_gn=MCF2;refGene_tx=XM_023249032.1;refGene_gn=MCF2;refGene_tx=XM_023249033.1;refGene_gn=MCF2;refGene_tx=XM_023249034.1
X 117134477 117134587
refGene_gn=MCF2;refGene_tx=XM_023249032.1;refGene_gn=MCF2;refGene_tx=XM_023249033.1;refGene_gn=MCF2;refGene_tx=XM_023249034.1
X 117136328 117136394
refGene_gn=MCF2;refGene_tx=XM_023249032.1;refGene_gn=MCF2;refGene_tx=XM_023249033.1;refGene_gn=MCF2;refGene_tx=XM_023249034.1
X 117136942 117137021
refGene_gn=MCF2;refGene_tx=XM_023249032.1;refGene_gn=MCF2;refGene_tx=XM_023249033.1;refGene_gn=MCF2;refGene_tx=XM_023249034.1
X 117139100 117139193
refGene_gn=MCF2;refGene_tx=XM_023249032.1;refGene_gn=MCF2;refGene_tx=XM_023249033.1;refGene_gn=MCF2;refGene_tx=XM_023249034.1
X 117145355 117145577
refGene_gn=MCF2;refGene_tx=XM_023249032.1;refGene_gn=MCF2;refGene_tx=XM_023249033.1;refGene_gn=MCF2;refGene_tx=XM_023249034.1
X 117146150 117146276
refGene_gn=MCF2;refGene_tx=XM_023249032.1;refGene_gn=MCF2;refGene_tx=XM_023249033.1;refGene_gn=MCF2;refGene_tx=XM_023249034.1
X 117147849 117147942
refGene_gn=MCF2;refGene_tx=XM_023249032.1;refGene_gn=MCF2;refGene_tx=XM_023249033.1;refGene_gn=MCF2;refGene_tx=XM_023249034.1
X 117152517 117152610
refGene_gn=MCF2;refGene_tx=XM_023249032.1;refGene_gn=MCF2;refGene_tx=XM_023249033.1;refGene_gn=MCF2;refGene_tx=XM_023249034.1
X 117154176 117154246
refGene_gn=MCF2;refGene_tx=XM_023249032.1;refGene_gn=MCF2;refGene_tx=XM_023249033.1;refGene_gn=MCF2;refGene_tx=XM_023249034.1
X 117154370 117154486
refGene_gn=MCF2;refGene_tx=XM_023249032.1;refGene_gn=MCF2;refGene_tx=XM_023249033.1;refGene_gn=MCF2;refGene_tx=XM_023249034.1
X 117155123 117155190
refGene_gn=MCF2;refGene_tx=XM_023249032.1;refGene_gn=MCF2;refGene_tx=XM_023249033.1;refGene_gn=MCF2;refGene_tx=XM_023249034.1
X 117157155 117157244
refGene_gn=MCF2;refGene_tx=XM_023249032.1;refGene_gn=MCF2;refGene_tx=XM_023249033.1;refGene_gn=MCF2;refGene_tx=XM_023249034.1
X 117160870 117160908
refGene_gn=MCF2;refGene_tx=XM_023249032.1;refGene_gn=MCF2;refGene_tx=XM_023249033.1;refGene_gn=MCF2;refGene_tx=XM_023249034.1
X 117163993 117164041
refGene_gn=MCF2;refGene_tx=XM_023249032.1;refGene_gn=MCF2;refGene_tx=XM_023249034.1
X 117165564 117165736
refGene_gn=MCF2;refGene_tx=XM_023249032.1;refGene_gn=MCF2;refGene_tx=X

M_023249033.1;refGene_gn=MCF2;refGene_tx=XM_023249034.1
X	117167441	117167633
refGene_gn=MCF2;refGene_tx=XM_023249032.1;refGene_gn=MCF2;refGene_tx=XM_023249033.1;refGene_gn=MCF2;refGene_tx=XM_023249034.1
X	117168609	117168801
refGene_gn=MCF2;refGene_tx=XM_023249032.1;refGene_gn=MCF2;refGene_tx=XM_023249033.1;refGene_gn=MCF2;refGene_tx=XM_023249034.1
X	117170511	117170631
refGene_gn=MCF2;refGene_tx=XM_023249032.1;refGene_gn=MCF2;refGene_tx=XM_023249033.1;refGene_gn=MCF2;refGene_tx=XM_023249034.1
X	117173377	117173492
refGene_gn=MCF2;refGene_tx=XM_023249032.1;refGene_gn=MCF2;refGene_tx=XM_023249033.1;refGene_gn=MCF2;refGene_tx=XM_023249034.1
X	117174306	117174440
refGene_gn=MCF2;refGene_tx=XM_023249032.1;refGene_gn=MCF2;refGene_tx=XM_023249033.1;refGene_gn=MCF2;refGene_tx=XM_023249034.1
X	117176676	117176826
refGene_gn=MCF2;refGene_tx=XM_023249032.1;refGene_gn=MCF2;refGene_tx=XM_023249033.1;refGene_gn=MCF2;refGene_tx=XM_023249034.1
X	117178412	117178529
refGene_gn=MCF2;refGene_tx=XM_023249032.1;refGene_gn=MCF2;refGene_tx=XM_023249033.1;refGene_gn=MCF2;refGene_tx=XM_023249034.1
X	117179350	117179470
refGene_gn=MCF2;refGene_tx=XM_023249032.1;refGene_gn=MCF2;refGene_tx=XM_023249033.1;refGene_gn=MCF2;refGene_tx=XM_023249034.1
X	117192580	117192631
refGene_gn=MCF2;refGene_tx=XM_023249034.1
X	117196671	117196762
refGene_gn=MCF2;refGene_tx=XM_023249032.1;refGene_gn=MCF2;refGene_tx=XM_023249033.1
X	117198758	117198873
refGene_gn=MCF2;refGene_tx=XM_023249032.1;refGene_gn=MCF2;refGene_tx=XM_023249033.1
X	117204047	117204116
refGene_gn=MCF2;refGene_tx=XM_023249032.1;refGene_gn=MCF2;refGene_tx=XM_023249033.1
X	117234809	117234894
refGene_gn=MCF2;refGene_tx=XM_023249032.1;refGene_gn=MCF2;refGene_tx=XM_023249033.1
X	117271298	117271361
refGene_gn=ATP11C;refGene_tx=XM_023249315.1;refGene_gn=ATP11C;refGene_tx=XM_023249323.1
X	117273902	117274004
refGene_gn=ATP11C;refGene_tx=XM_019824382.2;refGene_gn=ATP11C;refGene_tx=XM_019824385.2;refGene_gn=ATP11C;refGene_tx=XM_019824386.2;refGene_gn=ATP11C;refGene_tx=XM_023249318.1
X	117274685	117274775
refGene_gn=ATP11C;refGene_tx=XM_023249314.1
X	117280221	117280251
refGene_gn=ATP11C;refGene_tx=XM_023249316.1;refGene_gn=ATP11C;refGene_

tx=XM_023249317.1
X	117280370	117280524
refGene_gn=ATP11C;refGene_tx=XM_019824382.2;refGene_gn=ATP11C;refGene_tx=XM_019824385.2;refGene_gn=ATP11C;refGene_tx=XM_019824386.2;refGene_gn=ATP11C;refGene_tx=XM_023249314.1;refGene_gn=ATP11C;refGene_tx=XM_023249315.1;refGene_gn=ATP11C;refGene_tx=XM_023249316.1;refGene_gn=ATP11C;refGene_tx=XM_023249317.1;refGene_gn=ATP11C;refGene_tx=XM_023249318.1;refGene_gn=ATP11C;refGene_tx=XM_023249319.1;refGene_gn=ATP11C;refGene_tx=XM_023249320.1;refGene_gn=ATP11C;refGene_tx=XM_023249321.1;refGene_gn=ATP11C;refGene_tx=XM_023249322.1
X	117283450	117283554
refGene_gn=ATP11C;refGene_tx=XM_019824382.2;refGene_gn=ATP11C;refGene_tx=XM_019824385.2;refGene_gn=ATP11C;refGene_tx=XM_019824386.2;refGene_gn=ATP11C;refGene_tx=XM_023249314.1;refGene_gn=ATP11C;refGene_tx=XM_023249315.1;refGene_gn=ATP11C;refGene_tx=XM_023249316.1;refGene_gn=ATP11C;refGene_tx=XM_023249317.1;refGene_gn=ATP11C;refGene_tx=XM_023249318.1
X	117285727	117285793
refGene_gn=ATP11C;refGene_tx=XM_019824382.2;refGene_gn=ATP11C;refGene_tx=XM_019824385.2;refGene_gn=ATP11C;refGene_tx=XM_019824386.2;refGene_gn=ATP11C;refGene_tx=XM_023249314.1;refGene_gn=ATP11C;refGene_tx=XM_023249315.1;refGene_gn=ATP11C;refGene_tx=XM_023249316.1;refGene_gn=ATP11C;refGene_tx=XM_023249317.1;refGene_gn=ATP11C;refGene_tx=XM_023249318.1;refGene_gn=ATP11C;refGene_tx=XM_023249319.1;refGene_gn=ATP11C;refGene_tx=XM_023249320.1;refGene_gn=ATP11C;refGene_tx=XM_023249321.1;refGene_gn=ATP11C;refGene_tx=XM_023249322.1;refGene_gn=ATP11C;refGene_tx=XM_023249323.1
X	117288092	117288228
refGene_gn=ATP11C;refGene_tx=XM_019824382.2;refGene_gn=ATP11C;refGene_tx=XM_019824385.2;refGene_gn=ATP11C;refGene_tx=XM_019824386.2;refGene_gn=ATP11C;refGene_tx=XM_023249314.1;refGene_gn=ATP11C;refGene_tx=XM_023249315.1;refGene_gn=ATP11C;refGene_tx=XM_023249316.1;refGene_gn=ATP11C;refGene_tx=XM_023249317.1;refGene_gn=ATP11C;refGene_tx=XM_023249318.1;refGene_gn=ATP11C;refGene_tx=XM_023249319.1;refGene_gn=ATP11C;refGene_tx=XM_023249320.1;refGene_gn=ATP11C;refGene_tx=XM_023249321.1;refGene_gn=ATP11C;refGene_tx=XM_023249322.1;refGene_gn=ATP11C;refGene_tx=XM_023249323.1
X	117291384	117291512
refGene_gn=ATP11C;refGene_tx=XM_019824382.2;refGene_gn=ATP11C;refGene_tx=XM_019824385.2;refGene_gn=ATP11C;refGene_tx=XM_019824386.2;refGene_gn=ATP11C;refGene_tx=XM_023249314.1;refGene_gn=ATP11C;refGene_tx=XM_023249315.1;refGene_gn=ATP11C;refGene_tx=XM_023249316.1;refGene_gn=ATP11C;refGene_tx=XM_023249317.1;refGene_gn=ATP11C;refGene_tx=XM_023249318.1;refGene_gn=ATP11C;refGene_tx=XM_023249319.1;refGene_gn=ATP11C;refGene_tx=XM_023249320.1;refGene_gn=ATP11C;refGene_tx=XM_023249321.1;refGene_gn=ATP11C;refGene_tx=XM_023249322.1;refGene_gn=ATP11C;refGene_tx=XM_023249323.1
X	117298669	117298729
refGene_gn=ATP11C;refGene_tx=XM_019824382.2;refGene_gn=ATP11C;refGene_tx=XM_019824385.2;refGene_gn=ATP11C;refGene_tx=XM_019824386.2;refGene_

gn=ATP11C;refGene_tx=XM_023249314.1;refGene_gn=ATP11C;refGene_tx=XM_02
3249315.1;refGene_gn=ATP11C;refGene_tx=XM_023249316.1;refGene_gn=ATP11
C;refGene_tx=XM_023249317.1;refGene_gn=ATP11C;refGene_tx=XM_023249318.
1;refGene_gn=ATP11C;refGene_tx=XM_023249319.1;refGene_gn=ATP11C;refGen
e_tx=XM_023249320.1;refGene_gn=ATP11C;refGene_tx=XM_023249321.1;refGen
e_gn=ATP11C;refGene_tx=XM_023249322.1;refGene_gn=ATP11C;refGene_tx=XM_
023249323.1
X 117306257 117306403
refGene_gn=ATP11C;refGene_tx=XM_019824382.2;refGene_gn=ATP11C;refGene_
tx=XM_019824385.2;refGene_gn=ATP11C;refGene_tx=XM_019824386.2;refGene_
gn=ATP11C;refGene_tx=XM_023249314.1;refGene_gn=ATP11C;refGene_tx=XM_02
3249315.1;refGene_gn=ATP11C;refGene_tx=XM_023249316.1;refGene_gn=ATP11
C;refGene_tx=XM_023249317.1;refGene_gn=ATP11C;refGene_tx=XM_023249318.
1;refGene_gn=ATP11C;refGene_tx=XM_023249319.1;refGene_gn=ATP11C;refGen
e_tx=XM_023249320.1;refGene_gn=ATP11C;refGene_tx=XM_023249321.1;refGen
e_gn=ATP11C;refGene_tx=XM_023249322.1;refGene_gn=ATP11C;refGene_tx=XM_
023249323.1
X 117307603 117307706
refGene_gn=ATP11C;refGene_tx=XM_019824382.2;refGene_gn=ATP11C;refGene_
tx=XM_019824385.2;refGene_gn=ATP11C;refGene_tx=XM_019824386.2;refGene_
gn=ATP11C;refGene_tx=XM_023249314.1;refGene_gn=ATP11C;refGene_tx=XM_02
3249315.1;refGene_gn=ATP11C;refGene_tx=XM_023249316.1;refGene_gn=ATP11
C;refGene_tx=XM_023249317.1;refGene_gn=ATP11C;refGene_tx=XM_023249318.
1;refGene_gn=ATP11C;refGene_tx=XM_023249319.1;refGene_gn=ATP11C;refGen
e_tx=XM_023249320.1;refGene_gn=ATP11C;refGene_tx=XM_023249321.1;refGen
e_gn=ATP11C;refGene_tx=XM_023249322.1;refGene_gn=ATP11C;refGene_tx=XM_
023249323.1
X 117312726 117312901
refGene_gn=ATP11C;refGene_tx=XM_019824382.2;refGene_gn=ATP11C;refGene_
tx=XM_019824385.2;refGene_gn=ATP11C;refGene_tx=XM_019824386.2;refGene_
gn=ATP11C;refGene_tx=XM_023249314.1;refGene_gn=ATP11C;refGene_tx=XM_02
3249315.1;refGene_gn=ATP11C;refGene_tx=XM_023249316.1;refGene_gn=ATP11
C;refGene_tx=XM_023249317.1;refGene_gn=ATP11C;refGene_tx=XM_023249318.
1;refGene_gn=ATP11C;refGene_tx=XM_023249319.1;refGene_gn=ATP11C;refGen
e_tx=XM_023249320.1;refGene_gn=ATP11C;refGene_tx=XM_023249321.1;refGen
e_gn=ATP11C;refGene_tx=XM_023249322.1;refGene_gn=ATP11C;refGene_tx=XM_
023249323.1
X 117316846 117317110
refGene_gn=ATP11C;refGene_tx=XM_019824382.2;refGene_gn=ATP11C;refGene_
tx=XM_019824385.2;refGene_gn=ATP11C;refGene_tx=XM_019824386.2;refGene_
gn=ATP11C;refGene_tx=XM_023249314.1;refGene_gn=ATP11C;refGene_tx=XM_02
3249315.1;refGene_gn=ATP11C;refGene_tx=XM_023249316.1;refGene_gn=ATP11
C;refGene_tx=XM_023249317.1;refGene_gn=ATP11C;refGene_tx=XM_023249318.
1;refGene_gn=ATP11C;refGene_tx=XM_023249319.1;refGene_gn=ATP11C;refGen
e_tx=XM_023249320.1;refGene_gn=ATP11C;refGene_tx=XM_023249321.1;refGen
e_gn=ATP11C;refGene_tx=XM_023249322.1;refGene_gn=ATP11C;refGene_tx=XM_
023249323.1
X 117318570 117318752
refGene_gn=ATP11C;refGene_tx=XM_019824382.2;refGene_gn=ATP11C;refGene_
tx=XM_019824385.2;refGene_gn=ATP11C;refGene_tx=XM_019824386.2;refGene_

gn=ATP11C;refGene_tx=XM_023249314.1;refGene_gn=ATP11C;refGene_tx=XM_023249315.1;refGene_gn=ATP11C;refGene_tx=XM_023249316.1;refGene_gn=ATP11C;refGene_tx=XM_023249317.1;refGene_gn=ATP11C;refGene_tx=XM_023249318.1;refGene_gn=ATP11C;refGene_tx=XM_023249319.1;refGene_gn=ATP11C;refGene_tx=XM_023249320.1;refGene_gn=ATP11C;refGene_tx=XM_023249321.1;refGene_gn=ATP11C;refGene_tx=XM_023249322.1;refGene_gn=ATP11C;refGene_tx=XM_023249323.1
X 117319187 117319291
refGene_gn=ATP11C;refGene_tx=XM_019824382.2;refGene_gn=ATP11C;refGene_tx=XM_019824385.2;refGene_gn=ATP11C;refGene_tx=XM_019824386.2;refGene_gn=ATP11C;refGene_tx=XM_023249314.1;refGene_gn=ATP11C;refGene_tx=XM_023249315.1;refGene_gn=ATP11C;refGene_tx=XM_023249316.1;refGene_gn=ATP11C;refGene_tx=XM_023249317.1;refGene_gn=ATP11C;refGene_tx=XM_023249318.1;refGene_gn=ATP11C;refGene_tx=XM_023249319.1;refGene_gn=ATP11C;refGene_tx=XM_023249320.1;refGene_gn=ATP11C;refGene_tx=XM_023249321.1;refGene_gn=ATP11C;refGene_tx=XM_023249322.1;refGene_gn=ATP11C;refGene_tx=XM_023249323.1
X 117321929 117322003
refGene_gn=ATP11C;refGene_tx=XM_019824382.2;refGene_gn=ATP11C;refGene_tx=XM_019824385.2;refGene_gn=ATP11C;refGene_tx=XM_019824386.2;refGene_gn=ATP11C;refGene_tx=XM_023249314.1;refGene_gn=ATP11C;refGene_tx=XM_023249315.1;refGene_gn=ATP11C;refGene_tx=XM_023249316.1;refGene_gn=ATP11C;refGene_tx=XM_023249317.1;refGene_gn=ATP11C;refGene_tx=XM_023249318.1;refGene_gn=ATP11C;refGene_tx=XM_023249319.1;refGene_gn=ATP11C;refGene_tx=XM_023249320.1;refGene_gn=ATP11C;refGene_tx=XM_023249321.1;refGene_gn=ATP11C;refGene_tx=XM_023249322.1;refGene_gn=ATP11C;refGene_tx=XM_023249323.1
X 117323842 117323914
refGene_gn=ATP11C;refGene_tx=XM_019824382.2;refGene_gn=ATP11C;refGene_tx=XM_019824385.2;refGene_gn=ATP11C;refGene_tx=XM_019824386.2;refGene_gn=ATP11C;refGene_tx=XM_023249314.1;refGene_gn=ATP11C;refGene_tx=XM_023249315.1;refGene_gn=ATP11C;refGene_tx=XM_023249316.1;refGene_gn=ATP11C;refGene_tx=XM_023249317.1;refGene_gn=ATP11C;refGene_tx=XM_023249318.1;refGene_gn=ATP11C;refGene_tx=XM_023249319.1;refGene_gn=ATP11C;refGene_tx=XM_023249320.1;refGene_gn=ATP11C;refGene_tx=XM_023249321.1;refGene_gn=ATP11C;refGene_tx=XM_023249322.1;refGene_gn=ATP11C;refGene_tx=XM_023249323.1
X 117324928 117325080
refGene_gn=ATP11C;refGene_tx=XM_019824382.2;refGene_gn=ATP11C;refGene_tx=XM_019824385.2;refGene_gn=ATP11C;refGene_tx=XM_019824386.2;refGene_gn=ATP11C;refGene_tx=XM_023249314.1;refGene_gn=ATP11C;refGene_tx=XM_023249315.1;refGene_gn=ATP11C;refGene_tx=XM_023249316.1;refGene_gn=ATP11C;refGene_tx=XM_023249317.1;refGene_gn=ATP11C;refGene_tx=XM_023249318.1;refGene_gn=ATP11C;refGene_tx=XM_023249319.1;refGene_gn=ATP11C;refGene_tx=XM_023249320.1;refGene_gn=ATP11C;refGene_tx=XM_023249321.1;refGene_gn=ATP11C;refGene_tx=XM_023249322.1;refGene_gn=ATP11C;refGene_tx=XM_023249323.1
X 117325690 117325852
refGene_gn=ATP11C;refGene_tx=XM_019824382.2;refGene_gn=ATP11C;refGene_tx=XM_019824385.2;refGene_gn=ATP11C;refGene_tx=XM_019824386.2;refGene_

gn=ATP11C;refGene_tx=XM_023249314.1;refGene_gn=ATP11C;refGene_tx=XM_023249315.1;refGene_gn=ATP11C;refGene_tx=XM_023249316.1;refGene_gn=ATP11C;refGene_tx=XM_023249317.1;refGene_gn=ATP11C;refGene_tx=XM_023249318.1;refGene_gn=ATP11C;refGene_tx=XM_023249319.1;refGene_gn=ATP11C;refGene_tx=XM_023249320.1;refGene_gn=ATP11C;refGene_tx=XM_023249321.1;refGene_gn=ATP11C;refGene_tx=XM_023249322.1;refGene_gn=ATP11C;refGene_tx=XM_023249323.1
X	117331290	117331488
refGene_gn=ATP11C;refGene_tx=XM_019824382.2;refGene_gn=ATP11C;refGene_tx=XM_019824385.2;refGene_gn=ATP11C;refGene_tx=XM_019824386.2;refGene_gn=ATP11C;refGene_tx=XM_023249314.1;refGene_gn=ATP11C;refGene_tx=XM_023249315.1;refGene_gn=ATP11C;refGene_tx=XM_023249316.1;refGene_gn=ATP11C;refGene_tx=XM_023249317.1;refGene_gn=ATP11C;refGene_tx=XM_023249319.1;refGene_gn=ATP11C;refGene_tx=XM_023249320.1;refGene_gn=ATP11C;refGene_tx=XM_023249321.1;refGene_gn=ATP11C;refGene_tx=XM_023249322.1;refGene_gn=ATP11C;refGene_tx=XM_023249323.1
X	117332551	117332702
refGene_gn=ATP11C;refGene_tx=XM_019824382.2;refGene_gn=ATP11C;refGene_tx=XM_019824385.2;refGene_gn=ATP11C;refGene_tx=XM_019824386.2;refGene_gn=ATP11C;refGene_tx=XM_023249314.1;refGene_gn=ATP11C;refGene_tx=XM_023249315.1;refGene_gn=ATP11C;refGene_tx=XM_023249316.1;refGene_gn=ATP11C;refGene_tx=XM_023249317.1;refGene_gn=ATP11C;refGene_tx=XM_023249318.1;refGene_gn=ATP11C;refGene_tx=XM_023249319.1;refGene_gn=ATP11C;refGene_tx=XM_023249320.1;refGene_gn=ATP11C;refGene_tx=XM_023249321.1;refGene_gn=ATP11C;refGene_tx=XM_023249322.1;refGene_gn=ATP11C;refGene_tx=XM_023249323.1
X	117333638	117333720
refGene_gn=ATP11C;refGene_tx=XM_019824382.2;refGene_gn=ATP11C;refGene_tx=XM_019824385.2;refGene_gn=ATP11C;refGene_tx=XM_019824386.2;refGene_gn=ATP11C;refGene_tx=XM_023249314.1;refGene_gn=ATP11C;refGene_tx=XM_023249315.1;refGene_gn=ATP11C;refGene_tx=XM_023249316.1;refGene_gn=ATP11C;refGene_tx=XM_023249317.1;refGene_gn=ATP11C;refGene_tx=XM_023249318.1;refGene_gn=ATP11C;refGene_tx=XM_023249319.1;refGene_gn=ATP11C;refGene_tx=XM_023249320.1;refGene_gn=ATP11C;refGene_tx=XM_023249321.1;refGene_gn=ATP11C;refGene_tx=XM_023249322.1;refGene_gn=ATP11C;refGene_tx=XM_023249323.1
X	117334061	117334126
refGene_gn=ATP11C;refGene_tx=XM_019824382.2;refGene_gn=ATP11C;refGene_tx=XM_019824385.2;refGene_gn=ATP11C;refGene_tx=XM_019824386.2;refGene_gn=ATP11C;refGene_tx=XM_023249314.1;refGene_gn=ATP11C;refGene_tx=XM_023249315.1;refGene_gn=ATP11C;refGene_tx=XM_023249316.1;refGene_gn=ATP11C;refGene_tx=XM_023249317.1;refGene_gn=ATP11C;refGene_tx=XM_023249318.1;refGene_gn=ATP11C;refGene_tx=XM_023249319.1;refGene_gn=ATP11C;refGene_tx=XM_023249320.1;refGene_gn=ATP11C;refGene_tx=XM_023249321.1;refGene_gn=ATP11C;refGene_tx=XM_023249322.1;refGene_gn=ATP11C;refGene_tx=XM_023249323.1
X	117335060	117335111
refGene_gn=ATP11C;refGene_tx=XM_019824382.2;refGene_gn=ATP11C;refGene_tx=XM_019824385.2;refGene_gn=ATP11C;refGene_tx=XM_019824386.2;refGene_gn=ATP11C;refGene_tx=XM_023249314.1;refGene_gn=ATP11C;refGene_tx=XM_02

3249315.1;refGene_gn=ATP11C;refGene_tx=XM_023249316.1;refGene_gn=ATP11C;refGene_tx=XM_023249317.1;refGene_gn=ATP11C;refGene_tx=XM_023249318.1;refGene_gn=ATP11C;refGene_tx=XM_023249319.1;refGene_gn=ATP11C;refGene_tx=XM_023249320.1;refGene_gn=ATP11C;refGene_tx=XM_023249321.1;refGene_gn=ATP11C;refGene_tx=XM_023249322.1;refGene_gn=ATP11C;refGene_tx=XM_023249323.1
X 117337035 117337139
refGene_gn=ATP11C;refGene_tx=XM_019824382.2;refGene_gn=ATP11C;refGene_tx=XM_019824385.2;refGene_gn=ATP11C;refGene_tx=XM_019824386.2;refGene_gn=ATP11C;refGene_tx=XM_023249314.1;refGene_gn=ATP11C;refGene_tx=XM_023249315.1;refGene_gn=ATP11C;refGene_tx=XM_023249316.1;refGene_gn=ATP11C;refGene_tx=XM_023249317.1;refGene_gn=ATP11C;refGene_tx=XM_023249318.1;refGene_gn=ATP11C;refGene_tx=XM_023249319.1;refGene_gn=ATP11C;refGene_tx=XM_023249320.1;refGene_gn=ATP11C;refGene_tx=XM_023249321.1;refGene_gn=ATP11C;refGene_tx=XM_023249322.1;refGene_gn=ATP11C;refGene_tx=XM_023249323.1
X 117339393 117339522
refGene_gn=ATP11C;refGene_tx=XM_019824382.2;refGene_gn=ATP11C;refGene_tx=XM_019824385.2;refGene_gn=ATP11C;refGene_tx=XM_019824386.2;refGene_gn=ATP11C;refGene_tx=XM_023249314.1;refGene_gn=ATP11C;refGene_tx=XM_023249315.1;refGene_gn=ATP11C;refGene_tx=XM_023249316.1;refGene_gn=ATP11C;refGene_tx=XM_023249317.1;refGene_gn=ATP11C;refGene_tx=XM_023249318.1;refGene_gn=ATP11C;refGene_tx=XM_023249319.1;refGene_gn=ATP11C;refGene_tx=XM_023249320.1;refGene_gn=ATP11C;refGene_tx=XM_023249321.1;refGene_gn=ATP11C;refGene_tx=XM_023249322.1;refGene_gn=ATP11C;refGene_tx=XM_023249323.1
X 117350289 117350397
refGene_gn=ATP11C;refGene_tx=XM_019824382.2;refGene_gn=ATP11C;refGene_tx=XM_019824385.2;refGene_gn=ATP11C;refGene_tx=XM_019824386.2;refGene_gn=ATP11C;refGene_tx=XM_023249314.1;refGene_gn=ATP11C;refGene_tx=XM_023249315.1;refGene_gn=ATP11C;refGene_tx=XM_023249316.1;refGene_gn=ATP11C;refGene_tx=XM_023249317.1;refGene_gn=ATP11C;refGene_tx=XM_023249318.1;refGene_gn=ATP11C;refGene_tx=XM_023249319.1;refGene_gn=ATP11C;refGene_tx=XM_023249320.1;refGene_gn=ATP11C;refGene_tx=XM_023249321.1;refGene_gn=ATP11C;refGene_tx=XM_023249322.1;refGene_gn=ATP11C;refGene_tx=XM_023249323.1
X 117352667 117352748
refGene_gn=ATP11C;refGene_tx=XM_019824382.2;refGene_gn=ATP11C;refGene_tx=XM_019824385.2;refGene_gn=ATP11C;refGene_tx=XM_019824386.2;refGene_gn=ATP11C;refGene_tx=XM_023249314.1;refGene_gn=ATP11C;refGene_tx=XM_023249315.1;refGene_gn=ATP11C;refGene_tx=XM_023249316.1;refGene_gn=ATP11C;refGene_tx=XM_023249317.1;refGene_gn=ATP11C;refGene_tx=XM_023249318.1;refGene_gn=ATP11C;refGene_tx=XM_023249319.1;refGene_gn=ATP11C;refGene_tx=XM_023249320.1;refGene_gn=ATP11C;refGene_tx=XM_023249321.1;refGene_gn=ATP11C;refGene_tx=XM_023249322.1;refGene_gn=ATP11C;refGene_tx=XM_023249323.1
X 117355280 117355370
refGene_gn=ATP11C;refGene_tx=XM_019824382.2;refGene_gn=ATP11C;refGene_tx=XM_019824385.2;refGene_gn=ATP11C;refGene_tx=XM_019824386.2;refGene_gn=ATP11C;refGene_tx=XM_023249314.1;refGene_gn=ATP11C;refGene_tx=XM_02

3249315.1;refGene_gn=ATP11C;refGene_tx=XM_023249316.1;refGene_gn=ATP11C;refGene_tx=XM_023249317.1;refGene_gn=ATP11C;refGene_tx=XM_023249318.1;refGene_gn=ATP11C;refGene_tx=XM_023249319.1;refGene_gn=ATP11C;refGene_tx=XM_023249320.1;refGene_gn=ATP11C;refGene_tx=XM_023249321.1;refGene_gn=ATP11C;refGene_tx=XM_023249322.1;refGene_gn=ATP11C;refGene_tx=XM_023249323.1
X	117361713	117361833
refGene_gn=ATP11C;refGene_tx=XM_019824382.2;refGene_gn=ATP11C;refGene_tx=XM_019824385.2;refGene_gn=ATP11C;refGene_tx=XM_019824386.2;refGene_gn=ATP11C;refGene_tx=XM_023249314.1;refGene_gn=ATP11C;refGene_tx=XM_023249315.1;refGene_gn=ATP11C;refGene_tx=XM_023249316.1;refGene_gn=ATP11C;refGene_tx=XM_023249317.1;refGene_gn=ATP11C;refGene_tx=XM_023249318.1;refGene_gn=ATP11C;refGene_tx=XM_023249319.1;refGene_gn=ATP11C;refGene_tx=XM_023249320.1;refGene_gn=ATP11C;refGene_tx=XM_023249321.1;refGene_gn=ATP11C;refGene_tx=XM_023249322.1;refGene_gn=ATP11C;refGene_tx=XM_023249323.1
X	117438193	117438202
refGene_gn=ATP11C;refGene_tx=XM_019824385.2;refGene_gn=ATP11C;refGene_tx=XM_019824386.2
X	117440169	117440196
refGene_gn=ATP11C;refGene_tx=XM_019824382.2;refGene_gn=ATP11C;refGene_tx=XM_023249314.1;refGene_gn=ATP11C;refGene_tx=XM_023249315.1;refGene_gn=ATP11C;refGene_tx=XM_023249316.1;refGene_gn=ATP11C;refGene_tx=XM_023249317.1;refGene_gn=ATP11C;refGene_tx=XM_023249318.1;refGene_gn=ATP11C;refGene_tx=XM_023249319.1;refGene_gn=ATP11C;refGene_tx=XM_023249320.1;refGene_gn=ATP11C;refGene_tx=XM_023249321.1;refGene_gn=ATP11C;refGene_tx=XM_023249322.1;refGene_gn=ATP11C;refGene_tx=XM_023249323.1
X	117463957	117464939
refGene_gn=CXHXorf66;refGene_tx=XM_004000966.4
X	117465969	117466126
refGene_gn=CXHXorf66;refGene_tx=XM_004000966.4
X	117466782	117466870
refGene_gn=CXHXorf66;refGene_tx=XM_004000966.4
X	117587433	117588276
refGene_gn=LOC105260236;refGene_tx=XM_011280033.3
X	117716487	117716793
refGene_gn=LOC102899102;refGene_tx=XM_006944047.3
X	117716981	117717083
refGene_gn=LOC102899102;refGene_tx=XM_006944047.3
X	117895026	117895530
refGene_gn=SOX3;refGene_tx=XM_023249274.1
X	117895531	117895654
refGene_gn=SOX3;refGene_tx=XM_023249274.1
X	117895655	117896375
refGene_gn=SOX3;refGene_tx=XM_023249274.1
X	118142323	118142410
refGene_gn=CDR1;refGene_tx=XM_019823460.1
X	118147150	118147268
refGene_gn=CDR1;refGene_tx=XM_019823460.1
X	118149556	118149681

refGene_gn=CDR1;refGene_tx=XM_019823460.1
X        118149778        118150152
refGene_gn=CDR1;refGene_tx=XM_019823460.1
X        118150154        118150320
refGene_gn=CDR1;refGene_tx=XM_019823460.1
X        118162345        118162539
refGene_gn=CDR1;refGene_tx=XM_019823460.1
X        118226830        118227910
refGene_gn=LOC101086956;refGene_tx=XM_019824397.2;refGene_gn=LOC101086956;refGene_tx=XM_023249211.1;refGene_gn=LOC101086956;refGene_tx=XM_023249212.1;refGene_gn=LOC101086956;refGene_tx=XM_023249213.1;refGene_gn=LOC101086956;refGene_tx=XM_023249214.1;refGene_gn=LOC101086956;refGene_tx=XM_023249215.1;refGene_gn=LOC101086956;refGene_tx=XM_023249216.1;refGene_gn=LOC101086956;refGene_tx=XM_023249217.1;refGene_gn=LOC101086956;refGene_tx=XM_023249218.1
X        118564790        118565231
refGene_gn=LDOC1;refGene_tx=XM_004000969.5
X        120228633        120229427
refGene_gn=LOC101100452;refGene_tx=XM_023248997.1
X        120229429        120231149
refGene_gn=LOC101100452;refGene_tx=XM_023248997.1
X        121946719        121948306
refGene_gn=SLITRK2;refGene_tx=XM_006944048.4
X        121948307        121949258
refGene_gn=SLITRK2;refGene_tx=XM_006944048.4
X        122482947        122482955
refGene_gn=LOC101091265;refGene_tx=XM_019824413.1
X        122488769        122490669
refGene_gn=LOC101091265;refGene_tx=XM_019824413.1
X        122650711        122650981
refGene_gn=LOC102901550;refGene_tx=XM_006944067.3
X        122651171        122651294
refGene_gn=LOC102901550;refGene_tx=XM_006944067.3
X        123587694        123587745
refGene_gn=FMR1;refGene_tx=XM_004000948.5;refGene_gn=FMR1;refGene_tx=XM_004000949.5;refGene_gn=FMR1;refGene_tx=XM_004000950.5;refGene_gn=FMR1;refGene_tx=XM_004000951.5;refGene_gn=FMR1;refGene_tx=XM_004000952.5;refGene_gn=FMR1;refGene_tx=XM_006944049.4;refGene_gn=FMR1;refGene_tx=XM_006944050.4;refGene_gn=FMR1;refGene_tx=XM_006944051.4;refGene_gn=FMR1;refGene_tx=XM_006944052.4;refGene_gn=FMR1;refGene_tx=XM_006944053.2;refGene_gn=FMR1;refGene_tx=XM_006944054.4;refGene_gn=FMR1;refGene_tx=XM_006944055.4;refGene_gn=FMR1;refGene_tx=XM_019824414.2;refGene_gn=FMR1;refGene_tx=XM_019824415.2;refGene_gn=FMR1;refGene_tx=XM_019824417.2
X        123597612        123597665
refGene_gn=FMR1;refGene_tx=XM_004000948.5;refGene_gn=FMR1;refGene_tx=XM_004000949.5;refGene_gn=FMR1;refGene_tx=XM_004000950.5;refGene_gn=FMR1;refGene_tx=XM_004000951.5;refGene_gn=FMR1;refGene_tx=XM_004000952.5;refGene_gn=FMR1;refGene_tx=XM_006944049.4;refGene_gn=FMR1;refGene_tx=XM_006944050.4;refGene_gn=FMR1;refGene_tx=XM_006944051.4;refGene_gn=FMR1;refGene_tx=XM_006944052.4;refGene_gn=FMR1;refGene_tx=XM_006944053.2;

refGene_gn=FMR1;refGene_tx=XM_006944054.4;refGene_gn=FMR1;refGene_tx=XM_006944055.4;refGene_gn=FMR1;refGene_tx=XM_019824414.2;refGene_gn=FMR1;refGene_tx=XM_019824417.2
X	123600783	123600877
refGene_gn=FMR1;refGene_tx=XM_004000948.5;refGene_gn=FMR1;refGene_tx=XM_004000949.5;refGene_gn=FMR1;refGene_tx=XM_004000950.5;refGene_gn=FMR1;refGene_tx=XM_004000951.5;refGene_gn=FMR1;refGene_tx=XM_004000952.5;refGene_gn=FMR1;refGene_tx=XM_006944049.4;refGene_gn=FMR1;refGene_tx=XM_006944050.4;refGene_gn=FMR1;refGene_tx=XM_006944051.4;refGene_gn=FMR1;refGene_tx=XM_006944052.4;refGene_gn=FMR1;refGene_tx=XM_006944053.2;refGene_gn=FMR1;refGene_tx=XM_006944054.4;refGene_gn=FMR1;refGene_tx=XM_006944055.4;refGene_gn=FMR1;refGene_tx=XM_019824414.2;refGene_gn=FMR1;refGene_tx=XM_019824415.2;refGene_gn=FMR1;refGene_tx=XM_019824417.2
X	123603408	123603480
refGene_gn=FMR1;refGene_tx=XM_004000948.5;refGene_gn=FMR1;refGene_tx=XM_004000949.5;refGene_gn=FMR1;refGene_tx=XM_004000950.5;refGene_gn=FMR1;refGene_tx=XM_004000951.5;refGene_gn=FMR1;refGene_tx=XM_004000952.5;refGene_gn=FMR1;refGene_tx=XM_006944049.4;refGene_gn=FMR1;refGene_tx=XM_006944050.4;refGene_gn=FMR1;refGene_tx=XM_006944051.4;refGene_gn=FMR1;refGene_tx=XM_006944052.4;refGene_gn=FMR1;refGene_tx=XM_006944053.2;refGene_gn=FMR1;refGene_tx=XM_006944054.4;refGene_gn=FMR1;refGene_tx=XM_006944055.4;refGene_gn=FMR1;refGene_tx=XM_019824414.2;refGene_gn=FMR1;refGene_tx=XM_019824415.2;refGene_gn=FMR1;refGene_tx=XM_019824416.2;refGene_gn=FMR1;refGene_tx=XM_019824417.2
X	123603744	123603893
refGene_gn=FMR1;refGene_tx=XM_004000948.5;refGene_gn=FMR1;refGene_tx=XM_004000949.5;refGene_gn=FMR1;refGene_tx=XM_004000950.5;refGene_gn=FMR1;refGene_tx=XM_004000951.5;refGene_gn=FMR1;refGene_tx=XM_004000952.5;refGene_gn=FMR1;refGene_tx=XM_006944049.4;refGene_gn=FMR1;refGene_tx=XM_006944050.4;refGene_gn=FMR1;refGene_tx=XM_006944051.4;refGene_gn=FMR1;refGene_tx=XM_006944052.4;refGene_gn=FMR1;refGene_tx=XM_006944053.2;refGene_gn=FMR1;refGene_tx=XM_006944054.4;refGene_gn=FMR1;refGene_tx=XM_006944055.4;refGene_gn=FMR1;refGene_tx=XM_019824414.2;refGene_gn=FMR1;refGene_tx=XM_019824415.2;refGene_gn=FMR1;refGene_tx=XM_019824416.2;refGene_gn=FMR1;refGene_tx=XM_019824417.2
X	123604570	123604664
refGene_gn=FMR1;refGene_tx=XM_004000948.5;refGene_gn=FMR1;refGene_tx=XM_004000949.5;refGene_gn=FMR1;refGene_tx=XM_004000950.5;refGene_gn=FMR1;refGene_tx=XM_004000951.5;refGene_gn=FMR1;refGene_tx=XM_004000952.5;refGene_gn=FMR1;refGene_tx=XM_006944049.4;refGene_gn=FMR1;refGene_tx=XM_006944050.4;refGene_gn=FMR1;refGene_tx=XM_006944051.4;refGene_gn=FMR1;refGene_tx=XM_006944052.4;refGene_gn=FMR1;refGene_tx=XM_006944053.2;refGene_gn=FMR1;refGene_tx=XM_006944054.4;refGene_gn=FMR1;refGene_tx=XM_006944055.4;refGene_gn=FMR1;refGene_tx=XM_019824414.2;refGene_gn=FMR1;refGene_tx=XM_019824415.2;refGene_gn=FMR1;refGene_tx=XM_019824416.2;refGene_gn=FMR1;refGene_tx=XM_019824417.2
X	123604749	123604866
refGene_gn=FMR1;refGene_tx=XM_004000948.5;refGene_gn=FMR1;refGene_tx=XM_004000949.5;refGene_gn=FMR1;refGene_tx=XM_004000950.5;refGene_gn=FMR1;refGene_tx=XM_004000951.5;refGene_gn=FMR1;refGene_tx=XM_004000952.5;

refGene_gn=FMR1;refGene_tx=XM_006944049.4;refGene_gn=FMR1;refGene_tx=XM_006944050.4;refGene_gn=FMR1;refGene_tx=XM_006944051.4;refGene_gn=FMR1;refGene_tx=XM_006944052.4;refGene_gn=FMR1;refGene_tx=XM_006944053.2;refGene_gn=FMR1;refGene_tx=XM_006944054.4;refGene_gn=FMR1;refGene_tx=XM_006944055.4;refGene_gn=FMR1;refGene_tx=XM_019824414.2;refGene_gn=FMR1;refGene_tx=XM_019824415.2;refGene_gn=FMR1;refGene_tx=XM_019824416.2;refGene_gn=FMR1;refGene_tx=XM_019824417.2
X	123607084	123607255	refGene_gn=FMR1;refGene_tx=XM_004000948.5;refGene_gn=FMR1;refGene_tx=XM_004000949.5;refGene_gn=FMR1;refGene_tx=XM_004000950.5;refGene_gn=FMR1;refGene_tx=XM_004000951.5;refGene_gn=FMR1;refGene_tx=XM_004000952.5;refGene_gn=FMR1;refGene_tx=XM_006944049.4;refGene_gn=FMR1;refGene_tx=XM_006944050.4;refGene_gn=FMR1;refGene_tx=XM_006944051.4;refGene_gn=FMR1;refGene_tx=XM_006944052.4;refGene_gn=FMR1;refGene_tx=XM_006944053.2;refGene_gn=FMR1;refGene_tx=XM_006944054.4;refGene_gn=FMR1;refGene_tx=XM_006944055.4;refGene_gn=FMR1;refGene_tx=XM_019824414.2;refGene_gn=FMR1;refGene_tx=XM_019824415.2;refGene_gn=FMR1;refGene_tx=XM_019824416.2;refGene_gn=FMR1;refGene_tx=XM_019824417.2
X	123607345	123607424	refGene_gn=FMR1;refGene_tx=XM_004000948.5;refGene_gn=FMR1;refGene_tx=XM_004000949.5;refGene_gn=FMR1;refGene_tx=XM_004000950.5;refGene_gn=FMR1;refGene_tx=XM_004000951.5;refGene_gn=FMR1;refGene_tx=XM_004000952.5;refGene_gn=FMR1;refGene_tx=XM_006944049.4;refGene_gn=FMR1;refGene_tx=XM_006944050.4;refGene_gn=FMR1;refGene_tx=XM_006944051.4;refGene_gn=FMR1;refGene_tx=XM_006944052.4;refGene_gn=FMR1;refGene_tx=XM_006944053.2;refGene_gn=FMR1;refGene_tx=XM_006944054.4;refGene_gn=FMR1;refGene_tx=XM_006944055.4;refGene_gn=FMR1;refGene_tx=XM_019824414.2;refGene_gn=FMR1;refGene_tx=XM_019824415.2;refGene_gn=FMR1;refGene_tx=XM_019824416.2;refGene_gn=FMR1;refGene_tx=XM_019824417.2
X	123610992	123611102	refGene_gn=FMR1;refGene_tx=XM_004000948.5;refGene_gn=FMR1;refGene_tx=XM_004000949.5;refGene_gn=FMR1;refGene_tx=XM_004000950.5;refGene_gn=FMR1;refGene_tx=XM_004000951.5;refGene_gn=FMR1;refGene_tx=XM_004000952.5;refGene_gn=FMR1;refGene_tx=XM_006944049.4;refGene_gn=FMR1;refGene_tx=XM_006944050.4;refGene_gn=FMR1;refGene_tx=XM_006944051.4;refGene_gn=FMR1;refGene_tx=XM_006944052.4;refGene_gn=FMR1;refGene_tx=XM_006944053.2;refGene_gn=FMR1;refGene_tx=XM_006944054.4;refGene_gn=FMR1;refGene_tx=XM_006944055.4;refGene_gn=FMR1;refGene_tx=XM_019824414.2;refGene_gn=FMR1;refGene_tx=XM_019824415.2;refGene_gn=FMR1;refGene_tx=XM_019824416.2;refGene_gn=FMR1;refGene_tx=XM_019824417.2
X	123612072	123612207	refGene_gn=FMR1;refGene_tx=XM_004000948.5;refGene_gn=FMR1;refGene_tx=XM_004000949.5;refGene_gn=FMR1;refGene_tx=XM_004000950.5;refGene_gn=FMR1;refGene_tx=XM_004000951.5;refGene_gn=FMR1;refGene_tx=XM_004000952.5;refGene_gn=FMR1;refGene_tx=XM_006944049.4;refGene_gn=FMR1;refGene_tx=XM_006944050.4;refGene_gn=FMR1;refGene_tx=XM_006944051.4;refGene_gn=FMR1;refGene_tx=XM_006944052.4;refGene_gn=FMR1;refGene_tx=XM_006944053.2;refGene_gn=FMR1;refGene_tx=XM_006944054.4;refGene_gn=FMR1;refGene_tx=XM_006944055.4;refGene_gn=FMR1;refGene_tx=XM_019824414.2;refGene_gn=FMR1;refGene_tx=XM_019824415.2;refGene_gn=FMR1;refGene_tx=XM_019824416.2;

refGene_gn=FMR1;refGene_tx=XM_019824417.2
X	123612744	123612807
refGene_gn=FMR1;refGene_tx=XM_004000949.5;refGene_gn=FMR1;refGene_tx=XM_004000950.5;refGene_gn=FMR1;refGene_tx=XM_006944049.4;refGene_gn=FMR1;refGene_tx=XM_006944050.4;refGene_gn=FMR1;refGene_tx=XM_006944051.4;refGene_gn=FMR1;refGene_tx=XM_006944052.4;refGene_gn=FMR1;refGene_tx=XM_006944053.2;refGene_gn=FMR1;refGene_tx=XM_006944054.4;refGene_gn=FMR1;refGene_tx=XM_006944055.4;refGene_gn=FMR1;refGene_tx=XM_019824414.2;refGene_gn=FMR1;refGene_tx=XM_019824415.2;refGene_gn=FMR1;refGene_tx=XM_019824416.2;refGene_gn=FMR1;refGene_tx=XM_019824417.2
X	123615133	123615220
refGene_gn=FMR1;refGene_tx=XM_004000948.5;refGene_gn=FMR1;refGene_tx=XM_004000949.5;refGene_gn=FMR1;refGene_tx=XM_004000950.5;refGene_gn=FMR1;refGene_tx=XM_004000951.5;refGene_gn=FMR1;refGene_tx=XM_004000952.5;refGene_gn=FMR1;refGene_tx=XM_006944049.4;refGene_gn=FMR1;refGene_tx=XM_006944050.4;refGene_gn=FMR1;refGene_tx=XM_006944051.4;refGene_gn=FMR1;refGene_tx=XM_006944052.4;refGene_gn=FMR1;refGene_tx=XM_006944053.2;refGene_gn=FMR1;refGene_tx=XM_006944054.4;refGene_gn=FMR1;refGene_tx=XM_006944055.4;refGene_gn=FMR1;refGene_tx=XM_019824414.2;refGene_gn=FMR1;refGene_tx=XM_019824415.2;refGene_gn=FMR1;refGene_tx=XM_019824416.2;refGene_gn=FMR1;refGene_tx=XM_019824417.2
X	123617719	123617915
refGene_gn=FMR1;refGene_tx=XM_004000948.5;refGene_gn=FMR1;refGene_tx=XM_004000949.5;refGene_gn=FMR1;refGene_tx=XM_004000952.5;refGene_gn=FMR1;refGene_tx=XM_006944049.4;refGene_gn=FMR1;refGene_tx=XM_006944050.4;refGene_gn=FMR1;refGene_tx=XM_006944051.4;refGene_gn=FMR1;refGene_tx=XM_006944052.4;refGene_gn=FMR1;refGene_tx=XM_006944053.2;refGene_gn=FMR1;refGene_tx=XM_006944054.4;refGene_gn=FMR1;refGene_tx=XM_019824414.2;refGene_gn=FMR1;refGene_tx=XM_019824415.2;refGene_gn=FMR1;refGene_tx=XM_019824416.2
X	123618637	123619017
refGene_gn=FMR1;refGene_tx=XM_019824414.2
X	123619456	123619639
refGene_gn=FMR1;refGene_tx=XM_004000948.5;refGene_gn=FMR1;refGene_tx=XM_004000949.5;refGene_gn=FMR1;refGene_tx=XM_004000950.5;refGene_gn=FMR1;refGene_tx=XM_004000951.5;refGene_gn=FMR1;refGene_tx=XM_004000952.5;refGene_gn=FMR1;refGene_tx=XM_006944049.4;refGene_gn=FMR1;refGene_tx=XM_006944050.4;refGene_gn=FMR1;refGene_tx=XM_006944051.4;refGene_gn=FMR1;refGene_tx=XM_006944052.4;refGene_gn=FMR1;refGene_tx=XM_006944053.2;refGene_gn=FMR1;refGene_tx=XM_006944054.4;refGene_gn=FMR1;refGene_tx=XM_006944055.4;refGene_gn=FMR1;refGene_tx=XM_019824415.2;refGene_gn=FMR1;refGene_tx=XM_019824416.2;refGene_gn=FMR1;refGene_tx=XM_019824417.2
X	123620347	123620430
refGene_gn=FMR1;refGene_tx=XM_004000948.5;refGene_gn=FMR1;refGene_tx=XM_004000949.5;refGene_gn=FMR1;refGene_tx=XM_004000950.5;refGene_gn=FMR1;refGene_tx=XM_004000951.5;refGene_gn=FMR1;refGene_tx=XM_004000952.5;refGene_gn=FMR1;refGene_tx=XM_006944049.4;refGene_gn=FMR1;refGene_tx=XM_006944050.4;refGene_gn=FMR1;refGene_tx=XM_006944051.4;refGene_gn=FMR1;refGene_tx=XM_006944052.4;refGene_gn=FMR1;refGene_tx=XM_006944053.2;refGene_gn=FMR1;refGene_tx=XM_006944054.4;refGene_gn=FMR1;refGene_tx=X

M_006944055.4;refGene_gn=FMR1;refGene_tx=XM_019824415.2;refGene_gn=FMR1;refGene_tx=XM_019824416.2;refGene_gn=FMR1;refGene_tx=XM_019824417.2
X	123623019	123623181	refGene_gn=FMR1;refGene_tx=XM_004000948.5;refGene_gn=FMR1;refGene_tx=XM_004000949.5;refGene_gn=FMR1;refGene_tx=XM_004000950.5;refGene_gn=FMR1;refGene_tx=XM_004000951.5;refGene_gn=FMR1;refGene_tx=XM_004000952.5;refGene_gn=FMR1;refGene_tx=XM_006944049.4;refGene_gn=FMR1;refGene_tx=XM_006944050.4;refGene_gn=FMR1;refGene_tx=XM_006944051.4;refGene_gn=FMR1;refGene_tx=XM_006944052.4;refGene_gn=FMR1;refGene_tx=XM_006944054.4;refGene_gn=FMR1;refGene_tx=XM_006944055.4;refGene_gn=FMR1;refGene_tx=XM_019824415.2;refGene_gn=FMR1;refGene_tx=XM_019824416.2
X	123633659	123633720	refGene_gn=FMR1;refGene_tx=XM_006944053.2;refGene_gn=FMR1;refGene_tx=XM_019824417.2
X	123647544	123647857	refGene_gn=FMR1NB;refGene_tx=XM_006944056.4
X	123661994	123662114	refGene_gn=FMR1NB;refGene_tx=XM_006944056.4
X	123668136	123668233	refGene_gn=FMR1NB;refGene_tx=XM_006944056.4
X	123680231	123680349	refGene_gn=FMR1NB;refGene_tx=XM_006944056.4
X	124050705	124050752	refGene_gn=AFF2;refGene_tx=XM_023249262.1;refGene_gn=AFF2;refGene_tx=XM_023249263.1;refGene_gn=AFF2;refGene_tx=XM_023249264.1;refGene_gn=AFF2;refGene_tx=XM_023249265.1;refGene_gn=AFF2;refGene_tx=XM_023249266.1;refGene_gn=AFF2;refGene_tx=XM_023249267.1;refGene_gn=AFF2;refGene_tx=XM_023249268.1;refGene_gn=AFF2;refGene_tx=XM_023249269.1
X	124204448	124204581	refGene_gn=AFF2;refGene_tx=XM_023249262.1;refGene_gn=AFF2;refGene_tx=XM_023249263.1;refGene_gn=AFF2;refGene_tx=XM_023249264.1;refGene_gn=AFF2;refGene_tx=XM_023249265.1;refGene_gn=AFF2;refGene_tx=XM_023249266.1;refGene_gn=AFF2;refGene_tx=XM_023249267.1;refGene_gn=AFF2;refGene_tx=XM_023249268.1;refGene_gn=AFF2;refGene_tx=XM_023249269.1
X	124213762	124214623	refGene_gn=AFF2;refGene_tx=XM_023249262.1;refGene_gn=AFF2;refGene_tx=XM_023249263.1;refGene_gn=AFF2;refGene_tx=XM_023249264.1;refGene_gn=AFF2;refGene_tx=XM_023249265.1;refGene_gn=AFF2;refGene_tx=XM_023249266.1;refGene_gn=AFF2;refGene_tx=XM_023249267.1;refGene_gn=AFF2;refGene_tx=XM_023249268.1;refGene_gn=AFF2;refGene_tx=XM_023249269.1
X	124337639	124337684	refGene_gn=AFF2;refGene_tx=XM_023249262.1;refGene_gn=AFF2;refGene_tx=XM_023249263.1;refGene_gn=AFF2;refGene_tx=XM_023249264.1;refGene_gn=AFF2;refGene_tx=XM_023249265.1;refGene_gn=AFF2;refGene_tx=XM_023249266.1;refGene_gn=AFF2;refGene_tx=XM_023249267.1;refGene_gn=AFF2;refGene_tx=XM_023249268.1;refGene_gn=AFF2;refGene_tx=XM_023249269.1
X	124364060	124364147	refGene_gn=AFF2;refGene_tx=XM_023249262.1;refGene_gn=AFF2;refGene_tx=XM_023249263.1;refGene_gn=AFF2;refGene_tx=XM_023249264.1;refGene_gn=AFF2;refGene_tx=XM_023249265.1

X 124369750 124369787
refGene_gn=AFF2;refGene_tx=XM_023249262.1;refGene_gn=AFF2;refGene_tx=XM_023249263.1;refGene_gn=AFF2;refGene_tx=XM_023249264.1;refGene_gn=AFF2;refGene_tx=XM_023249265.1;refGene_gn=AFF2;refGene_tx=XM_023249266.1;refGene_gn=AFF2;refGene_tx=XM_023249267.1;refGene_gn=AFF2;refGene_tx=XM_023249268.1;refGene_gn=AFF2;refGene_tx=XM_023249269.1
X 124370077 124370129
refGene_gn=AFF2;refGene_tx=XM_023249262.1;refGene_gn=AFF2;refGene_tx=XM_023249263.1;refGene_gn=AFF2;refGene_tx=XM_023249264.1;refGene_gn=AFF2;refGene_tx=XM_023249265.1;refGene_gn=AFF2;refGene_tx=XM_023249266.1;refGene_gn=AFF2;refGene_tx=XM_023249267.1;refGene_gn=AFF2;refGene_tx=XM_023249268.1;refGene_gn=AFF2;refGene_tx=XM_023249269.1
X 124409784 124409881
refGene_gn=AFF2;refGene_tx=XM_023249262.1;refGene_gn=AFF2;refGene_tx=XM_023249263.1;refGene_gn=AFF2;refGene_tx=XM_023249264.1;refGene_gn=AFF2;refGene_tx=XM_023249265.1;refGene_gn=AFF2;refGene_tx=XM_023249266.1;refGene_gn=AFF2;refGene_tx=XM_023249267.1;refGene_gn=AFF2;refGene_tx=XM_023249268.1;refGene_gn=AFF2;refGene_tx=XM_023249269.1
X 124425382 124425420
refGene_gn=AFF2;refGene_tx=XM_023249262.1;refGene_gn=AFF2;refGene_tx=XM_023249263.1;refGene_gn=AFF2;refGene_tx=XM_023249264.1;refGene_gn=AFF2;refGene_tx=XM_023249265.1;refGene_gn=AFF2;refGene_tx=XM_023249266.1;refGene_gn=AFF2;refGene_tx=XM_023249267.1;refGene_gn=AFF2;refGene_tx=XM_023249268.1;refGene_gn=AFF2;refGene_tx=XM_023249269.1
X 124474003 124474163
refGene_gn=AFF2;refGene_tx=XM_023249262.1;refGene_gn=AFF2;refGene_tx=XM_023249263.1;refGene_gn=AFF2;refGene_tx=XM_023249264.1;refGene_gn=AFF2;refGene_tx=XM_023249265.1;refGene_gn=AFF2;refGene_tx=XM_023249266.1;refGene_gn=AFF2;refGene_tx=XM_023249267.1;refGene_gn=AFF2;refGene_tx=XM_023249268.1;refGene_gn=AFF2;refGene_tx=XM_023249269.1
X 124475863 124476877
refGene_gn=AFF2;refGene_tx=XM_023249262.1;refGene_gn=AFF2;refGene_tx=XM_023249263.1;refGene_gn=AFF2;refGene_tx=XM_023249264.1;refGene_gn=AFF2;refGene_tx=XM_023249265.1;refGene_gn=AFF2;refGene_tx=XM_023249266.1;refGene_gn=AFF2;refGene_tx=XM_023249267.1;refGene_gn=AFF2;refGene_tx=XM_023249268.1;refGene_gn=AFF2;refGene_tx=XM_023249269.1
X 124478119 124478241
refGene_gn=AFF2;refGene_tx=XM_023249262.1;refGene_gn=AFF2;refGene_tx=XM_023249263.1;refGene_gn=AFF2;refGene_tx=XM_023249264.1;refGene_gn=AFF2;refGene_tx=XM_023249265.1;refGene_gn=AFF2;refGene_tx=XM_023249266.1;refGene_gn=AFF2;refGene_tx=XM_023249267.1;refGene_gn=AFF2;refGene_tx=XM_023249268.1;refGene_gn=AFF2;refGene_tx=XM_023249269.1
X 124482077 124482300
refGene_gn=AFF2;refGene_tx=XM_023249262.1;refGene_gn=AFF2;refGene_tx=XM_023249263.1;refGene_gn=AFF2;refGene_tx=XM_023249264.1;refGene_gn=AFF2;refGene_tx=XM_023249265.1;refGene_gn=AFF2;refGene_tx=XM_023249266.1;refGene_gn=AFF2;refGene_tx=XM_023249267.1;refGene_gn=AFF2;refGene_tx=XM_023249268.1;refGene_gn=AFF2;refGene_tx=XM_023249269.1
X 124486975 124487271
refGene_gn=AFF2;refGene_tx=XM_023249262.1;refGene_gn=AFF2;refGene_tx=X

M_023249263.1;refGene_gn=AFF2;refGene_tx=XM_023249264.1;refGene_gn=AFF2;refGene_tx=XM_023249265.1;refGene_gn=AFF2;refGene_tx=XM_023249266.1;refGene_gn=AFF2;refGene_tx=XM_023249267.1;refGene_gn=AFF2;refGene_tx=XM_023249268.1;refGene_gn=AFF2;refGene_tx=XM_023249269.1
X	124487806	124487870
refGene_gn=AFF2;refGene_tx=XM_023249262.1;refGene_gn=AFF2;refGene_tx=XM_023249263.1;refGene_gn=AFF2;refGene_tx=XM_023249264.1;refGene_gn=AFF2;refGene_tx=XM_023249265.1;refGene_gn=AFF2;refGene_tx=XM_023249266.1;refGene_gn=AFF2;refGene_tx=XM_023249267.1;refGene_gn=AFF2;refGene_tx=XM_023249268.1;refGene_gn=AFF2;refGene_tx=XM_023249269.1
X	124491003	124491140
refGene_gn=AFF2;refGene_tx=XM_023249262.1;refGene_gn=AFF2;refGene_tx=XM_023249263.1;refGene_gn=AFF2;refGene_tx=XM_023249264.1;refGene_gn=AFF2;refGene_tx=XM_023249265.1;refGene_gn=AFF2;refGene_tx=XM_023249266.1;refGene_gn=AFF2;refGene_tx=XM_023249267.1;refGene_gn=AFF2;refGene_tx=XM_023249268.1;refGene_gn=AFF2;refGene_tx=XM_023249269.1
X	124496414	124496486
refGene_gn=AFF2;refGene_tx=XM_023249262.1;refGene_gn=AFF2;refGene_tx=XM_023249263.1;refGene_gn=AFF2;refGene_tx=XM_023249264.1;refGene_gn=AFF2;refGene_tx=XM_023249265.1;refGene_gn=AFF2;refGene_tx=XM_023249266.1;refGene_gn=AFF2;refGene_tx=XM_023249267.1;refGene_gn=AFF2;refGene_tx=XM_023249268.1;refGene_gn=AFF2;refGene_tx=XM_023249269.1
X	124496837	124496931
refGene_gn=AFF2;refGene_tx=XM_023249262.1;refGene_gn=AFF2;refGene_tx=XM_023249263.1;refGene_gn=AFF2;refGene_tx=XM_023249264.1;refGene_gn=AFF2;refGene_tx=XM_023249265.1;refGene_gn=AFF2;refGene_tx=XM_023249266.1;refGene_gn=AFF2;refGene_tx=XM_023249267.1;refGene_gn=AFF2;refGene_tx=XM_023249268.1;refGene_gn=AFF2;refGene_tx=XM_023249269.1
X	124498982	124499035
refGene_gn=AFF2;refGene_tx=XM_023249262.1;refGene_gn=AFF2;refGene_tx=XM_023249263.1;refGene_gn=AFF2;refGene_tx=XM_023249264.1;refGene_gn=AFF2;refGene_tx=XM_023249265.1;refGene_gn=AFF2;refGene_tx=XM_023249266.1;refGene_gn=AFF2;refGene_tx=XM_023249267.1;refGene_gn=AFF2;refGene_tx=XM_023249268.1;refGene_gn=AFF2;refGene_tx=XM_023249269.1
X	124504083	124504274
refGene_gn=AFF2;refGene_tx=XM_023249262.1;refGene_gn=AFF2;refGene_tx=XM_023249263.1;refGene_gn=AFF2;refGene_tx=XM_023249264.1;refGene_gn=AFF2;refGene_tx=XM_023249265.1;refGene_gn=AFF2;refGene_tx=XM_023249266.1;refGene_gn=AFF2;refGene_tx=XM_023249267.1;refGene_gn=AFF2;refGene_tx=XM_023249268.1;refGene_gn=AFF2;refGene_tx=XM_023249269.1
X	124506975	124507097
refGene_gn=AFF2;refGene_tx=XM_023249262.1;refGene_gn=AFF2;refGene_tx=XM_023249263.1;refGene_gn=AFF2;refGene_tx=XM_023249264.1;refGene_gn=AFF2;refGene_tx=XM_023249265.1;refGene_gn=AFF2;refGene_tx=XM_023249266.1;refGene_gn=AFF2;refGene_tx=XM_023249267.1;refGene_gn=AFF2;refGene_tx=XM_023249268.1;refGene_gn=AFF2;refGene_tx=XM_023249269.1
X	124827238	124827273
refGene_gn=IDS;refGene_tx=XM_023248819.1;refGene_gn=IDS;refGene_tx=XM_023248820.1
X	124828755	124829228

refGene_gn=IDS;refGene_tx=XM_004000953.4;refGene_gn=IDS;refGene_tx=XM_006944057.4;refGene_gn=IDS;refGene_tx=XM_019824418.2;refGene_gn=IDS;refGene_tx=XM_023248816.1;refGene_gn=IDS;refGene_tx=XM_023248817.1;refGene_gn=IDS;refGene_tx=XM_023248818.1;refGene_gn=IDS;refGene_tx=XM_023248819.1;refGene_gn=IDS;refGene_tx=XM_023248820.1
X	124832500	124832674
refGene_gn=IDS;refGene_tx=XM_004000953.4;refGene_gn=IDS;refGene_tx=XM_006944057.4;refGene_gn=IDS;refGene_tx=XM_019824418.2;refGene_gn=IDS;refGene_tx=XM_023248816.1;refGene_gn=IDS;refGene_tx=XM_023248819.1;refGene_gn=IDS;refGene_tx=XM_023248820.1
X	124834543	124834670
refGene_gn=IDS;refGene_tx=XM_004000953.4;refGene_gn=IDS;refGene_tx=XM_006944057.4;refGene_gn=IDS;refGene_tx=XM_019824418.2;refGene_gn=IDS;refGene_tx=XM_023248816.1;refGene_gn=IDS;refGene_tx=XM_023248817.1;refGene_gn=IDS;refGene_tx=XM_023248818.1;refGene_gn=IDS;refGene_tx=XM_023248819.1;refGene_gn=IDS;refGene_tx=XM_023248820.1
X	124838169	124838340
refGene_gn=IDS;refGene_tx=XM_004000953.4;refGene_gn=IDS;refGene_tx=XM_006944057.4;refGene_gn=IDS;refGene_tx=XM_019824418.2;refGene_gn=IDS;refGene_tx=XM_023248816.1;refGene_gn=IDS;refGene_tx=XM_023248817.1;refGene_gn=IDS;refGene_tx=XM_023248818.1;refGene_gn=IDS;refGene_tx=XM_023248819.1;refGene_gn=IDS;refGene_tx=XM_023248820.1
X	124839789	124839990
refGene_gn=IDS;refGene_tx=XM_004000953.4;refGene_gn=IDS;refGene_tx=XM_006944057.4;refGene_gn=IDS;refGene_tx=XM_019824418.2;refGene_gn=IDS;refGene_tx=XM_023248816.1;refGene_gn=IDS;refGene_tx=XM_023248817.1;refGene_gn=IDS;refGene_tx=XM_023248818.1;refGene_gn=IDS;refGene_tx=XM_023248819.1;refGene_gn=IDS;refGene_tx=XM_023248820.1
X	124842565	124842654
refGene_gn=IDS;refGene_tx=XM_004000953.4;refGene_gn=IDS;refGene_tx=XM_006944057.4;refGene_gn=IDS;refGene_tx=XM_019824418.2;refGene_gn=IDS;refGene_tx=XM_023248816.1;refGene_gn=IDS;refGene_tx=XM_023248817.1;refGene_gn=IDS;refGene_tx=XM_023248818.1;refGene_gn=IDS;refGene_tx=XM_023248819.1;refGene_gn=IDS;refGene_tx=XM_023248820.1
X	124845630	124845808
refGene_gn=IDS;refGene_tx=XM_004000953.4;refGene_gn=IDS;refGene_tx=XM_006944057.4;refGene_gn=IDS;refGene_tx=XM_019824418.2;refGene_gn=IDS;refGene_tx=XM_023248816.1;refGene_gn=IDS;refGene_tx=XM_023248817.1;refGene_gn=IDS;refGene_tx=XM_023248818.1;refGene_gn=IDS;refGene_tx=XM_023248819.1;refGene_gn=IDS;refGene_tx=XM_023248820.1
X	124846468	124846605
refGene_gn=IDS;refGene_tx=XM_004000953.4;refGene_gn=IDS;refGene_tx=XM_006944057.4;refGene_gn=IDS;refGene_tx=XM_019824418.2;refGene_gn=IDS;refGene_tx=XM_023248816.1;refGene_gn=IDS;refGene_tx=XM_023248817.1;refGene_gn=IDS;refGene_tx=XM_023248818.1;refGene_gn=IDS;refGene_tx=XM_023248819.1;refGene_gn=IDS;refGene_tx=XM_023248820.1
X	124847063	124847190
refGene_gn=IDS;refGene_tx=XM_004000953.4;refGene_gn=IDS;refGene_tx=XM_019824418.2;refGene_gn=IDS;refGene_tx=XM_023248816.1;refGene_gn=IDS;refGene_tx=XM_023248817.1;refGene_gn=IDS;refGene_tx=XM_023248818.1;refGe

ne_gn=IDS;refGene_tx=XM_023248819.1;refGene_gn=IDS;refGene_tx=XM_02324
8820.1
X	124862341	124862421
refGene_gn=IDS;refGene_tx=XM_006944057.4
X	124865031	124865048
refGene_gn=IDS;refGene_tx=XM_006944057.4
X	124869813	124870066
refGene_gn=LOC111558703;refGene_tx=XM_023249067.1;refGene_gn=LOC111558
703;refGene_tx=XM_023249068.1;refGene_gn=LOC111558703;refGene_tx=XM_02
3249069.1;refGene_gn=LOC111558703;refGene_tx=XM_023249070.1;refGene_gn
=LOC111558703;refGene_tx=XM_023249071.1;refGene_gn=LOC111558703;refGen
e_tx=XM_023249072.1
X	124870899	124871162
refGene_gn=LOC111558703;refGene_tx=XM_023249067.1;refGene_gn=LOC111558
703;refGene_tx=XM_023249068.1;refGene_gn=LOC111558703;refGene_tx=XM_02
3249069.1;refGene_gn=LOC111558703;refGene_tx=XM_023249070.1;refGene_gn
=LOC111558703;refGene_tx=XM_023249071.1;refGene_gn=LOC111558703;refGen
e_tx=XM_023249072.1
X	124871179	124871201
refGene_gn=LOC111558690;refGene_tx=XM_023248983.1
X	125113830	125113868
refGene_gn=TMEM185A;refGene_tx=XM_023248874.1;refGene_gn=TMEM185A;refG
ene_tx=XM_023248875.1
X	125135154	125135331
refGene_gn=TMEM185A;refGene_tx=XM_023248874.1;refGene_gn=TMEM185A;refG
ene_tx=XM_023248875.1;refGene_gn=TMEM185A;refGene_tx=XM_023248876.1
X	125136724	125136774
refGene_gn=TMEM185A;refGene_tx=XM_006944069.4
X	125137509	125137717
refGene_gn=TMEM185A;refGene_tx=XM_006944069.4;refGene_gn=TMEM185A;refG
ene_tx=XM_023248874.1;refGene_gn=TMEM185A;refGene_tx=XM_023248875.1;re
fGene_gn=TMEM185A;refGene_tx=XM_023248876.1
X	125143351	125143435
refGene_gn=TMEM185A;refGene_tx=XM_006944069.4;refGene_gn=TMEM185A;refG
ene_tx=XM_023248874.1;refGene_gn=TMEM185A;refGene_tx=XM_023248875.1;re
fGene_gn=TMEM185A;refGene_tx=XM_023248876.1
X	125146605	125146782
refGene_gn=TMEM185A;refGene_tx=XM_006944069.4;refGene_gn=TMEM185A;refG
ene_tx=XM_023248874.1;refGene_gn=TMEM185A;refGene_tx=XM_023248875.1;re
fGene_gn=TMEM185A;refGene_tx=XM_023248876.1
X	125147337	125147461
refGene_gn=TMEM185A;refGene_tx=XM_006944069.4;refGene_gn=TMEM185A;refG
ene_tx=XM_023248874.1;refGene_gn=TMEM185A;refGene_tx=XM_023248875.1;re
fGene_gn=TMEM185A;refGene_tx=XM_023248876.1
X	125148441	125148596
refGene_gn=TMEM185A;refGene_tx=XM_023248875.1
X	125148678	125148923
refGene_gn=TMEM185A;refGene_tx=XM_006944069.4;refGene_gn=TMEM185A;refG
ene_tx=XM_023248874.1;refGene_gn=TMEM185A;refGene_tx=XM_023248876.1
X	125151827	125151877

refGene_gn=LOC101093506;refGene_tx=XM_023248984.1
X	125151968	125152241
refGene_gn=LOC101093506;refGene_tx=XM_023248984.1
X	125152242	125152354
refGene_gn=LOC101093506;refGene_tx=XM_023248984.1
X	125152420	125152525
refGene_gn=LOC101093506;refGene_tx=XM_023248984.1
X	125153439	125153634
refGene_gn=LOC101093506;refGene_tx=XM_023248984.1
X	125155025	125155127
refGene_gn=LOC101093506;refGene_tx=XM_023248984.1
X	125297947	125298445
refGene_gn=HSFX4;refGene_tx=XM_023249079.1
X	125299136	125299163
refGene_gn=HSFX4;refGene_tx=XM_023249079.1
X	125299164	125299453
refGene_gn=HSFX4;refGene_tx=XM_023249079.1
X	125299455	125299727
refGene_gn=HSFX4;refGene_tx=XM_023249079.1
X	125301050	125301307
refGene_gn=CXHXorf40B;refGene_tx=XM_004000990.5;refGene_gn=CXHXorf40B;refGene_tx=XM_006944074.4;refGene_gn=CXHXorf40B;refGene_tx=XM_006944076.3;refGene_gn=CXHXorf40B;refGene_tx=XM_006944077.4;refGene_gn=CXHXorf40B;refGene_tx=XM_006944078.3;refGene_gn=CXHXorf40B;refGene_tx=XM_023249080.1;refGene_gn=CXHXorf40B;refGene_tx=XM_023249081.1
X	125302139	125302392
refGene_gn=CXHXorf40B;refGene_tx=XM_004000990.5;refGene_gn=CXHXorf40B;refGene_tx=XM_006944074.4;refGene_gn=CXHXorf40B;refGene_tx=XM_006944076.3;refGene_gn=CXHXorf40B;refGene_tx=XM_006944077.4;refGene_gn=CXHXorf40B;refGene_tx=XM_006944078.3;refGene_gn=CXHXorf40B;refGene_tx=XM_023249080.1;refGene_gn=CXHXorf40B;refGene_tx=XM_023249081.1
X	125703077	125703374
refGene_gn=MAMLD1;refGene_tx=XM_023249495.1
X	125764104	125764200
refGene_gn=MAMLD1;refGene_tx=XM_019824423.2;refGene_gn=MAMLD1;refGene_tx=XM_023249496.1
X	125783396	125785100
refGene_gn=MAMLD1;refGene_tx=XM_019824423.2;refGene_gn=MAMLD1;refGene_tx=XM_023249495.1;refGene_gn=MAMLD1;refGene_tx=XM_023249496.1
X	125819885	125819970
refGene_gn=MAMLD1;refGene_tx=XM_019824423.2;refGene_gn=MAMLD1;refGene_tx=XM_023249495.1;refGene_gn=MAMLD1;refGene_tx=XM_023249496.1
X	125820963	125822063
refGene_gn=MAMLD1;refGene_tx=XM_019824423.2;refGene_gn=MAMLD1;refGene_tx=XM_023249495.1;refGene_gn=MAMLD1;refGene_tx=XM_023249496.1
X	125894743	125894806
refGene_gn=MTM1;refGene_tx=XM_004000974.4;refGene_gn=MTM1;refGene_tx=XM_023249466.1;refGene_gn=MTM1;refGene_tx=XM_023249467.1;refGene_gn=MTM1;refGene_tx=XM_023249468.1
X	125897166	125897239

refGene_gn=MTM1;refGene_tx=XM_004000974.4;refGene_gn=MTM1;refGene_tx=XM_023249466.1;refGene_gn=MTM1;refGene_tx=XM_023249467.1;refGene_gn=MTM1;refGene_tx=XM_023249468.1
X 125899032 125899127
refGene_gn=MTM1;refGene_tx=XM_004000974.4;refGene_gn=MTM1;refGene_tx=XM_023249466.1;refGene_gn=MTM1;refGene_tx=XM_023249467.1
X 125912690 125912801
refGene_gn=MTM1;refGene_tx=XM_004000974.4;refGene_gn=MTM1;refGene_tx=XM_023249466.1;refGene_gn=MTM1;refGene_tx=XM_023249468.1
X 125917127 125917229
refGene_gn=MTM1;refGene_tx=XM_004000974.4;refGene_gn=MTM1;refGene_tx=XM_023249466.1;refGene_gn=MTM1;refGene_tx=XM_023249467.1;refGene_gn=MTM1;refGene_tx=XM_023249468.1
X 125937990 125938074
refGene_gn=MTM1;refGene_tx=XM_004000974.4;refGene_gn=MTM1;refGene_tx=XM_023249466.1;refGene_gn=MTM1;refGene_tx=XM_023249467.1;refGene_gn=MTM1;refGene_tx=XM_023249468.1
X 125940265 125940415
refGene_gn=MTM1;refGene_tx=XM_004000974.4;refGene_gn=MTM1;refGene_tx=XM_023249466.1;refGene_gn=MTM1;refGene_tx=XM_023249467.1;refGene_gn=MTM1;refGene_tx=XM_023249468.1
X 125944553 125944742
refGene_gn=MTM1;refGene_tx=XM_004000974.4;refGene_gn=MTM1;refGene_tx=XM_023249466.1;refGene_gn=MTM1;refGene_tx=XM_023249467.1;refGene_gn=MTM1;refGene_tx=XM_023249468.1
X 125948296 125948482
refGene_gn=MTM1;refGene_tx=XM_004000974.4;refGene_gn=MTM1;refGene_tx=XM_023249466.1;refGene_gn=MTM1;refGene_tx=XM_023249467.1;refGene_gn=MTM1;refGene_tx=XM_023249468.1
X 125954655 125954862
refGene_gn=MTM1;refGene_tx=XM_004000974.4;refGene_gn=MTM1;refGene_tx=XM_023249466.1;refGene_gn=MTM1;refGene_tx=XM_023249467.1;refGene_gn=MTM1;refGene_tx=XM_023249468.1
X 125955769 125955862
refGene_gn=MTM1;refGene_tx=XM_004000974.4;refGene_gn=MTM1;refGene_tx=XM_023249466.1;refGene_gn=MTM1;refGene_tx=XM_023249467.1;refGene_gn=MTM1;refGene_tx=XM_023249468.1
X 125956570 125956684
refGene_gn=MTM1;refGene_tx=XM_004000974.4;refGene_gn=MTM1;refGene_tx=XM_023249466.1;refGene_gn=MTM1;refGene_tx=XM_023249467.1;refGene_gn=MTM1;refGene_tx=XM_023249468.1
X 125960772 125960949
refGene_gn=MTM1;refGene_tx=XM_004000974.4;refGene_gn=MTM1;refGene_tx=XM_023249466.1;refGene_gn=MTM1;refGene_tx=XM_023249467.1;refGene_gn=MTM1;refGene_tx=XM_023249468.1
X 125967268 125967436
refGene_gn=MTM1;refGene_tx=XM_004000974.4;refGene_gn=MTM1;refGene_tx=XM_023249466.1;refGene_gn=MTM1;refGene_tx=XM_023249467.1;refGene_gn=MTM1;refGene_tx=XM_023249468.1
X 126007384 126007430

refGene_gn=MTMR1;refGene_tx=XM_023249232.1;refGene_gn=MTMR1;refGene_tx=XM_023249233.1;refGene_gn=MTMR1;refGene_tx=XM_023249234.1;refGene_gn=MTMR1;refGene_tx=XM_023249235.1;refGene_gn=MTMR1;refGene_tx=XM_023249236.1;refGene_gn=MTMR1;refGene_tx=XM_023249237.1;refGene_gn=MTMR1;refGene_tx=XM_023249238.1;refGene_gn=MTMR1;refGene_tx=XM_023249239.1;refGene_gn=MTMR1;refGene_tx=XM_023249240.1
X 126009991 126010086
refGene_gn=MTMR1;refGene_tx=XM_023249232.1;refGene_gn=MTMR1;refGene_tx=XM_023249233.1;refGene_gn=MTMR1;refGene_tx=XM_023249234.1;refGene_gn=MTMR1;refGene_tx=XM_023249235.1;refGene_gn=MTMR1;refGene_tx=XM_023249236.1;refGene_gn=MTMR1;refGene_tx=XM_023249237.1;refGene_gn=MTMR1;refGene_tx=XM_023249238.1;refGene_gn=MTMR1;refGene_tx=XM_023249239.1;refGene_gn=MTMR1;refGene_tx=XM_023249240.1
X 126010718 126010826
refGene_gn=MTMR1;refGene_tx=XM_023249232.1;refGene_gn=MTMR1;refGene_tx=XM_023249233.1;refGene_gn=MTMR1;refGene_tx=XM_023249234.1;refGene_gn=MTMR1;refGene_tx=XM_023249235.1;refGene_gn=MTMR1;refGene_tx=XM_023249236.1;refGene_gn=MTMR1;refGene_tx=XM_023249237.1;refGene_gn=MTMR1;refGene_tx=XM_023249238.1;refGene_gn=MTMR1;refGene_tx=XM_023249239.1;refGene_gn=MTMR1;refGene_tx=XM_023249240.1
X 126013599 126013701
refGene_gn=MTMR1;refGene_tx=XM_023249232.1;refGene_gn=MTMR1;refGene_tx=XM_023249233.1;refGene_gn=MTMR1;refGene_tx=XM_023249234.1;refGene_gn=MTMR1;refGene_tx=XM_023249235.1;refGene_gn=MTMR1;refGene_tx=XM_023249236.1;refGene_gn=MTMR1;refGene_tx=XM_023249237.1;refGene_gn=MTMR1;refGene_tx=XM_023249238.1;refGene_gn=MTMR1;refGene_tx=XM_023249239.1;refGene_gn=MTMR1;refGene_tx=XM_023249240.1
X 126014014 126014098
refGene_gn=MTMR1;refGene_tx=XM_023249232.1;refGene_gn=MTMR1;refGene_tx=XM_023249233.1;refGene_gn=MTMR1;refGene_tx=XM_023249234.1;refGene_gn=MTMR1;refGene_tx=XM_023249235.1;refGene_gn=MTMR1;refGene_tx=XM_023249236.1;refGene_gn=MTMR1;refGene_tx=XM_023249237.1;refGene_gn=MTMR1;refGene_tx=XM_023249238.1;refGene_gn=MTMR1;refGene_tx=XM_023249239.1;refGene_gn=MTMR1;refGene_tx=XM_023249240.1
X 126015144 126015294
refGene_gn=MTMR1;refGene_tx=XM_023249232.1;refGene_gn=MTMR1;refGene_tx=XM_023249233.1;refGene_gn=MTMR1;refGene_tx=XM_023249234.1;refGene_gn=MTMR1;refGene_tx=XM_023249235.1;refGene_gn=MTMR1;refGene_tx=XM_023249236.1;refGene_gn=MTMR1;refGene_tx=XM_023249237.1;refGene_gn=MTMR1;refGene_tx=XM_023249238.1;refGene_gn=MTMR1;refGene_tx=XM_023249239.1;refGene_gn=MTMR1;refGene_tx=XM_023249240.1
X 126016225 126016414
refGene_gn=MTMR1;refGene_tx=XM_023249232.1;refGene_gn=MTMR1;refGene_tx=XM_023249233.1;refGene_gn=MTMR1;refGene_tx=XM_023249234.1;refGene_gn=MTMR1;refGene_tx=XM_023249235.1;refGene_gn=MTMR1;refGene_tx=XM_023249236.1;refGene_gn=MTMR1;refGene_tx=XM_023249237.1;refGene_gn=MTMR1;refGene_tx=XM_023249238.1;refGene_gn=MTMR1;refGene_tx=XM_023249239.1;refGene_gn=MTMR1;refGene_tx=XM_023249240.1
X 126026558 126026744
refGene_gn=MTMR1;refGene_tx=XM_023249232.1;refGene_gn=MTMR1;refGene_tx

=XM_023249233.1;refGene_gn=MTMR1;refGene_tx=XM_023249234.1;refGene_gn=MTMR1;refGene_tx=XM_023249235.1;refGene_gn=MTMR1;refGene_tx=XM_023249236.1;refGene_gn=MTMR1;refGene_tx=XM_023249237.1;refGene_gn=MTMR1;refGene_tx=XM_023249238.1;refGene_gn=MTMR1;refGene_tx=XM_023249239.1;refGene_gn=MTMR1;refGene_tx=XM_023249240.1
X 126027174 126027381
refGene_gn=MTMR1;refGene_tx=XM_023249232.1;refGene_gn=MTMR1;refGene_tx=XM_023249233.1;refGene_gn=MTMR1;refGene_tx=XM_023249234.1;refGene_gn=MTMR1;refGene_tx=XM_023249235.1;refGene_gn=MTMR1;refGene_tx=XM_023249236.1;refGene_gn=MTMR1;refGene_tx=XM_023249237.1;refGene_gn=MTMR1;refGene_tx=XM_023249238.1;refGene_gn=MTMR1;refGene_tx=XM_023249239.1;refGene_gn=MTMR1;refGene_tx=XM_023249240.1
X 126032710 126032803
refGene_gn=MTMR1;refGene_tx=XM_023249232.1;refGene_gn=MTMR1;refGene_tx=XM_023249233.1;refGene_gn=MTMR1;refGene_tx=XM_023249234.1;refGene_gn=MTMR1;refGene_tx=XM_023249235.1;refGene_gn=MTMR1;refGene_tx=XM_023249236.1;refGene_gn=MTMR1;refGene_tx=XM_023249237.1;refGene_gn=MTMR1;refGene_tx=XM_023249238.1;refGene_gn=MTMR1;refGene_tx=XM_023249239.1;refGene_gn=MTMR1;refGene_tx=XM_023249240.1
X 126038335 126038449
refGene_gn=MTMR1;refGene_tx=XM_023249232.1;refGene_gn=MTMR1;refGene_tx=XM_023249233.1;refGene_gn=MTMR1;refGene_tx=XM_023249234.1;refGene_gn=MTMR1;refGene_tx=XM_023249235.1;refGene_gn=MTMR1;refGene_tx=XM_023249236.1;refGene_gn=MTMR1;refGene_tx=XM_023249237.1;refGene_gn=MTMR1;refGene_tx=XM_023249238.1;refGene_gn=MTMR1;refGene_tx=XM_023249239.1;refGene_gn=MTMR1;refGene_tx=XM_023249240.1
X 126042102 126042279
refGene_gn=MTMR1;refGene_tx=XM_023249232.1;refGene_gn=MTMR1;refGene_tx=XM_023249233.1;refGene_gn=MTMR1;refGene_tx=XM_023249234.1;refGene_gn=MTMR1;refGene_tx=XM_023249235.1;refGene_gn=MTMR1;refGene_tx=XM_023249236.1;refGene_gn=MTMR1;refGene_tx=XM_023249237.1;refGene_gn=MTMR1;refGene_tx=XM_023249238.1;refGene_gn=MTMR1;refGene_tx=XM_023249239.1;refGene_gn=MTMR1;refGene_tx=XM_023249240.1
X 126048081 126048243
refGene_gn=MTMR1;refGene_tx=XM_023249232.1;refGene_gn=MTMR1;refGene_tx=XM_023249233.1;refGene_gn=MTMR1;refGene_tx=XM_023249234.1;refGene_gn=MTMR1;refGene_tx=XM_023249235.1;refGene_gn=MTMR1;refGene_tx=XM_023249236.1;refGene_gn=MTMR1;refGene_tx=XM_023249237.1;refGene_gn=MTMR1;refGene_tx=XM_023249238.1;refGene_gn=MTMR1;refGene_tx=XM_023249239.1;refGene_gn=MTMR1;refGene_tx=XM_023249240.1
X 126053230 126053295
refGene_gn=CD99L2;refGene_tx=XM_023249241.1;refGene_gn=CD99L2;refGene_tx=XM_023249242.1;refGene_gn=CD99L2;refGene_tx=XM_023249243.1
X 126054551 126054617
refGene_gn=CD99L2;refGene_tx=XM_023249241.1;refGene_gn=CD99L2;refGene_tx=XM_023249242.1
X 126055789 126055807
refGene_gn=CD99L2;refGene_tx=XM_023249241.1
X 126058937 126059057
refGene_gn=CD99L2;refGene_tx=XM_023249241.1;refGene_gn=CD99L2;refGene_

tx=XM_023249242.1;refGene_gn=CD99L2;refGene_tx=XM_023249243.1
X 126059484 126059523
refGene_gn=CD99L2;refGene_tx=XM_023249241.1;refGene_gn=CD99L2;refGene_tx=XM_023249242.1;refGene_gn=CD99L2;refGene_tx=XM_023249243.1
X 126070318 126070378
refGene_gn=CD99L2;refGene_tx=XM_023249241.1;refGene_gn=CD99L2;refGene_tx=XM_023249242.1;refGene_gn=CD99L2;refGene_tx=XM_023249243.1
X 126072194 126072278
refGene_gn=CD99L2;refGene_tx=XM_023249241.1;refGene_gn=CD99L2;refGene_tx=XM_023249242.1;refGene_gn=CD99L2;refGene_tx=XM_023249243.1
X 126072406 126072475
refGene_gn=CD99L2;refGene_tx=XM_023249241.1;refGene_gn=CD99L2;refGene_tx=XM_023249242.1;refGene_gn=CD99L2;refGene_tx=XM_023249243.1
X 126089235 126089295
refGene_gn=CD99L2;refGene_tx=XM_023249241.1;refGene_gn=CD99L2;refGene_tx=XM_023249242.1;refGene_gn=CD99L2;refGene_tx=XM_023249243.1
X 126148625 126148692
refGene_gn=CD99L2;refGene_tx=XM_023249241.1;refGene_gn=CD99L2;refGene_tx=XM_023249242.1;refGene_gn=CD99L2;refGene_tx=XM_023249243.1
X 126205639 126205685
refGene_gn=HMGB3;refGene_tx=XM_004001389.5
X 126210418 126210573
refGene_gn=HMGB3;refGene_tx=XM_004001389.5;refGene_gn=HMGB3;refGene_tx=XM_023249090.1;refGene_gn=HMGB3;refGene_tx=XM_023249091.1
X 126210700 126210840
refGene_gn=HMGB3;refGene_tx=XM_004001389.5;refGene_gn=HMGB3;refGene_tx=XM_023249090.1;refGene_gn=HMGB3;refGene_tx=XM_023249091.1
X 126211317 126211492
refGene_gn=HMGB3;refGene_tx=XM_004001389.5;refGene_gn=HMGB3;refGene_tx=XM_023249090.1;refGene_gn=HMGB3;refGene_tx=XM_023249091.1
X 126211960 126212101
refGene_gn=HMGB3;refGene_tx=XM_004001389.5;refGene_gn=HMGB3;refGene_tx=XM_023249090.1;refGene_gn=HMGB3;refGene_tx=XM_023249091.1
X 126316479 126316666
refGene_gn=GPR50;refGene_tx=XM_004000977.3
X 126319569 126321248
refGene_gn=GPR50;refGene_tx=XM_004000977.3
X 126487143 126487253
refGene_gn=VMA21;refGene_tx=XM_006944087.4
X 126487462 126487515
refGene_gn=VMA21;refGene_tx=XM_004000978.3
X 126492299 126492409
refGene_gn=VMA21;refGene_tx=XM_004000978.3;refGene_gn=VMA21;refGene_tx=XM_006944087.4
X 126493002 126493145
refGene_gn=VMA21;refGene_tx=XM_004000978.3;refGene_gn=VMA21;refGene_tx=XM_006944087.4
X 126633845 126633877
refGene_gn=PASD1;refGene_tx=XM_006944088.4;refGene_gn=PASD1;refGene_tx=XM_019824436.2;refGene_gn=PASD1;refGene_tx=XM_019824438.2;refGene_gn=

PASD1;refGene_tx=XM_019824442.1
X	126635109	126635264
refGene_gn=PASD1;refGene_tx=XM_006944088.4;refGene_gn=PASD1;refGene_tx=XM_019824436.2;refGene_gn=PASD1;refGene_tx=XM_019824438.2;refGene_gn=PASD1;refGene_tx=XM_019824442.1
X	126638792	126638881
refGene_gn=PASD1;refGene_tx=XM_006944088.4;refGene_gn=PASD1;refGene_tx=XM_019824436.2;refGene_gn=PASD1;refGene_tx=XM_019824438.2;refGene_gn=PASD1;refGene_tx=XM_019824442.1
X	126641719	126641809
refGene_gn=PASD1;refGene_tx=XM_006944088.4;refGene_gn=PASD1;refGene_tx=XM_019824436.2;refGene_gn=PASD1;refGene_tx=XM_019824438.2;refGene_gn=PASD1;refGene_tx=XM_019824442.1
X	126654909	126655009
refGene_gn=PASD1;refGene_tx=XM_006944088.4;refGene_gn=PASD1;refGene_tx=XM_019824436.2;refGene_gn=PASD1;refGene_tx=XM_019824438.2;refGene_gn=PASD1;refGene_tx=XM_019824442.1
X	126655433	126655547
refGene_gn=PASD1;refGene_tx=XM_006944088.4;refGene_gn=PASD1;refGene_tx=XM_019824436.2;refGene_gn=PASD1;refGene_tx=XM_019824438.2;refGene_gn=PASD1;refGene_tx=XM_019824442.1
X	126658418	126658549
refGene_gn=PASD1;refGene_tx=XM_006944088.4;refGene_gn=PASD1;refGene_tx=XM_019824436.2;refGene_gn=PASD1;refGene_tx=XM_019824438.2;refGene_gn=PASD1;refGene_tx=XM_019824442.1
X	126661086	126661169
refGene_gn=PASD1;refGene_tx=XM_006944088.4;refGene_gn=PASD1;refGene_tx=XM_019824436.2;refGene_gn=PASD1;refGene_tx=XM_019824438.2;refGene_gn=PASD1;refGene_tx=XM_019824442.1
X	126682061	126682134
refGene_gn=PASD1;refGene_tx=XM_006944088.4;refGene_gn=PASD1;refGene_tx=XM_019824436.2;refGene_gn=PASD1;refGene_tx=XM_019824438.2;refGene_gn=PASD1;refGene_tx=XM_019824442.1
X	126690575	126690702
refGene_gn=PASD1;refGene_tx=XM_006944088.4;refGene_gn=PASD1;refGene_tx=XM_019824436.2;refGene_gn=PASD1;refGene_tx=XM_019824438.2;refGene_gn=PASD1;refGene_tx=XM_019824442.1
X	126696421	126696663
refGene_gn=PASD1;refGene_tx=XM_006944088.4;refGene_gn=PASD1;refGene_tx=XM_019824436.2;refGene_gn=PASD1;refGene_tx=XM_019824438.2;refGene_gn=PASD1;refGene_tx=XM_019824442.1
X	126700547	126700748
refGene_gn=PASD1;refGene_tx=XM_006944088.4;refGene_gn=PASD1;refGene_tx=XM_019824436.2;refGene_gn=PASD1;refGene_tx=XM_019824438.2;refGene_gn=PASD1;refGene_tx=XM_019824442.1
X	126705176	126705332
refGene_gn=PASD1;refGene_tx=XM_006944088.4;refGene_gn=PASD1;refGene_tx=XM_019824436.2;refGene_gn=PASD1;refGene_tx=XM_019824438.2;refGene_gn=PASD1;refGene_tx=XM_019824442.1
X	126731500	126731507

refGene_gn=PRRG3;refGene_tx=XM_019824445.2;refGene_gn=PRRG3;refGene_tx=XM_019824446.2;refGene_gn=PRRG3;refGene_tx=XM_023249400.1;refGene_gn=PRRG3;refGene_tx=XM_023249401.1
X	126732545	126732706
refGene_gn=PRRG3;refGene_tx=XM_019824445.2;refGene_gn=PRRG3;refGene_tx=XM_019824446.2;refGene_gn=PRRG3;refGene_tx=XM_023249400.1;refGene_gn=PRRG3;refGene_tx=XM_023249401.1
X	126733092	126733620
refGene_gn=PRRG3;refGene_tx=XM_019824445.2;refGene_gn=PRRG3;refGene_tx=XM_019824446.2;refGene_gn=PRRG3;refGene_tx=XM_023249400.1;refGene_gn=PRRG3;refGene_tx=XM_023249401.1
X	126750909	126751015
refGene_gn=FATE1;refGene_tx=XM_004000995.5
X	126752029	126752157
refGene_gn=FATE1;refGene_tx=XM_004000995.5
X	126755926	126756033
refGene_gn=FATE1;refGene_tx=XM_004000995.5
X	126756439	126756518
refGene_gn=FATE1;refGene_tx=XM_004000995.5
X	126757260	126757392
refGene_gn=FATE1;refGene_tx=XM_004000995.5
X	126772062	126772172
refGene_gn=CNGA2;refGene_tx=XM_011279517.3
X	126772386	126772479
refGene_gn=CNGA2;refGene_tx=XM_011279517.3
X	126773110	126773281
refGene_gn=CNGA2;refGene_tx=XM_011279517.3
X	126774377	126774485
refGene_gn=CNGA2;refGene_tx=XM_011279517.3
X	126776130	126776237
refGene_gn=CNGA2;refGene_tx=XM_011279517.3
X	126776585	126778000
refGene_gn=CNGA2;refGene_tx=XM_011279517.3
X	126985913	126986648
refGene_gn=LOC105260348;refGene_tx=XM_023249541.1
X	127031519	127031864
refGene_gn=GABRE;refGene_tx=XM_004000981.4
X	127032149	127032349
refGene_gn=GABRE;refGene_tx=XM_004000981.4
X	127032473	127032626
refGene_gn=GABRE;refGene_tx=XM_004000981.4
X	127037052	127037190
refGene_gn=GABRE;refGene_tx=XM_004000981.4
X	127038536	127038619
refGene_gn=GABRE;refGene_tx=XM_004000981.4
X	127039332	127039553
refGene_gn=GABRE;refGene_tx=XM_004000981.4
X	127042943	127043011
refGene_gn=GABRE;refGene_tx=XM_004000981.4
X	127043434	127043652

refGene_gn=GABRE;refGene_tx=XM_004000981.4
X 127047573 127047629
refGene_gn=GABRE;refGene_tx=XM_004000981.4
X 127128183 127128488
refGene_gn=LOC111558784;refGene_tx=XM_023249542.1
X 127261870 127262140
refGene_gn=GABRA3;refGene_tx=XM_004000982.5
X 127285063 127285275
refGene_gn=GABRA3;refGene_tx=XM_004000982.5
X 127294238 127294391
refGene_gn=GABRA3;refGene_tx=XM_004000982.5
X 127331455 127331599
refGene_gn=GABRA3;refGene_tx=XM_004000982.5
X 127344919 127345002
refGene_gn=GABRA3;refGene_tx=XM_004000982.5
X 127366779 127367000
refGene_gn=GABRA3;refGene_tx=XM_004000982.5
X 127387901 127387969
refGene_gn=GABRA3;refGene_tx=XM_004000982.5
X 127488191 127488313
refGene_gn=GABRA3;refGene_tx=XM_004000982.5
X 127522850 127522990
refGene_gn=GABRA3;refGene_tx=XM_004000982.5
X 127819475 127819624
refGene_gn=GABRQ;refGene_tx=XM_019824449.1
X 127821738 127821827
refGene_gn=GABRQ;refGene_tx=XM_019824449.1
X 127828479 127828547
refGene_gn=GABRQ;refGene_tx=XM_019824449.1
X 127830261 127830482
refGene_gn=GABRQ;refGene_tx=XM_019824449.1
X 127832213 127832296
refGene_gn=GABRQ;refGene_tx=XM_019824449.1
X 127832764 127832902
refGene_gn=GABRQ;refGene_tx=XM_019824449.1
X 127833438 127833591
refGene_gn=GABRQ;refGene_tx=XM_019824449.1
X 127835117 127835368
refGene_gn=GABRQ;refGene_tx=XM_019824449.1
X 127836000 127836981
refGene_gn=GABRQ;refGene_tx=XM_019824449.1
X 127862933 127863668
refGene_gn=LOC111558732;refGene_tx=XM_023249188.1
X 127898442 127898532
refGene_gn=CETN2;refGene_tx=XM_004000983.4
X 127898978 127899092
refGene_gn=CETN2;refGene_tx=XM_004000983.4
X 127899093 127899117
refGene_gn=CETN2;refGene_tx=XM_004000983.4
X 127900458 127900587

refGene_gn=CETN2;refGene_tx=XM_004000983.4
X 127901121 127901280
refGene_gn=CETN2;refGene_tx=XM_004000983.4
X 127902250 127902349
refGene_gn=CETN2;refGene_tx=XM_004000983.4
X 127910774 127910846
refGene_gn=NSDHL;refGene_tx=XM_004000985.5;refGene_gn=NSDHL;refGene_tx=XM_006944089.4;refGene_gn=NSDHL;refGene_tx=XM_006944091.4;refGene_gn=NSDHL;refGene_tx=XM_019824469.2
X 127914307 127914466
refGene_gn=NSDHL;refGene_tx=XM_004000985.5;refGene_gn=NSDHL;refGene_tx=XM_006944089.4;refGene_gn=NSDHL;refGene_tx=XM_006944090.4;refGene_gn=NSDHL;refGene_tx=XM_019824469.2
X 127922776 127922923
refGene_gn=NSDHL;refGene_tx=XM_004000985.5;refGene_gn=NSDHL;refGene_tx=XM_006944089.4;refGene_gn=NSDHL;refGene_tx=XM_006944090.4;refGene_gn=NSDHL;refGene_tx=XM_006944091.4;refGene_gn=NSDHL;refGene_tx=XM_019824469.2
X 127926346 127926475
refGene_gn=NSDHL;refGene_tx=XM_004000985.5;refGene_gn=NSDHL;refGene_tx=XM_006944089.4;refGene_gn=NSDHL;refGene_tx=XM_006944090.4;refGene_gn=NSDHL;refGene_tx=XM_006944091.4
X 127929242 127929385
refGene_gn=NSDHL;refGene_tx=XM_004000985.5;refGene_gn=NSDHL;refGene_tx=XM_006944089.4;refGene_gn=NSDHL;refGene_tx=XM_006944090.4;refGene_gn=NSDHL;refGene_tx=XM_006944091.4;refGene_gn=NSDHL;refGene_tx=XM_019824469.2
X 127930512 127930615
refGene_gn=NSDHL;refGene_tx=XM_004000985.5;refGene_gn=NSDHL;refGene_tx=XM_006944089.4;refGene_gn=NSDHL;refGene_tx=XM_006944090.4;refGene_gn=NSDHL;refGene_tx=XM_006944091.4;refGene_gn=NSDHL;refGene_tx=XM_019824469.2
X 127931697 127932030
refGene_gn=NSDHL;refGene_tx=XM_004000985.5;refGene_gn=NSDHL;refGene_tx=XM_006944089.4;refGene_gn=NSDHL;refGene_tx=XM_006944090.4;refGene_gn=NSDHL;refGene_tx=XM_006944091.4;refGene_gn=NSDHL;refGene_tx=XM_019824469.2
X 127977146 127977180
refGene_gn=ZNF185;refGene_tx=XM_006944092.4;refGene_gn=ZNF185;refGene_tx=XM_019824450.2;refGene_gn=ZNF185;refGene_tx=XM_019824451.1;refGene_gn=ZNF185;refGene_tx=XM_019824452.2;refGene_gn=ZNF185;refGene_tx=XM_019824453.2;refGene_gn=ZNF185;refGene_tx=XM_019824454.2;refGene_gn=ZNF185;refGene_tx=XM_019824455.2;refGene_gn=ZNF185;refGene_tx=XM_019824456.2;refGene_gn=ZNF185;refGene_tx=XM_019824457.2;refGene_gn=ZNF185;refGene_tx=XM_019824458.2;refGene_gn=ZNF185;refGene_tx=XM_019824459.2;refGene_gn=ZNF185;refGene_tx=XM_019824460.2;refGene_gn=ZNF185;refGene_tx=XM_019824461.2;refGene_gn=ZNF185;refGene_tx=XM_019824462.2;refGene_gn=ZNF185;refGene_tx=XM_019824463.2;refGene_gn=ZNF185;refGene_tx=XM_019824464.2;refGene_gn=ZNF185;refGene_tx=XM_019824465.2;refGene_gn=ZNF185;refGene_tx=XM_019824466.2;refGene_gn=ZNF185;refGene_tx=XM_019824467.2;refG

ene_gn=ZNF185;refGene_tx=XM_019824468.2
X	127977343	127977467
refGene_gn=ZNF185;refGene_tx=XM_006944092.4;refGene_gn=ZNF185;refGene_tx=XM_019824450.2;refGene_gn=ZNF185;refGene_tx=XM_019824451.1;refGene_gn=ZNF185;refGene_tx=XM_019824452.2;refGene_gn=ZNF185;refGene_tx=XM_019824453.2;refGene_gn=ZNF185;refGene_tx=XM_019824454.2;refGene_gn=ZNF185;refGene_tx=XM_019824455.2;refGene_gn=ZNF185;refGene_tx=XM_019824456.2;refGene_gn=ZNF185;refGene_tx=XM_019824457.2;refGene_gn=ZNF185;refGene_tx=XM_019824458.2;refGene_gn=ZNF185;refGene_tx=XM_019824459.2;refGene_gn=ZNF185;refGene_tx=XM_019824460.2;refGene_gn=ZNF185;refGene_tx=XM_019824461.2;refGene_gn=ZNF185;refGene_tx=XM_019824462.2;refGene_gn=ZNF185;refGene_tx=XM_019824463.2;refGene_gn=ZNF185;refGene_tx=XM_019824464.2;refGene_gn=ZNF185;refGene_tx=XM_019824465.2;refGene_gn=ZNF185;refGene_tx=XM_019824466.2;refGene_gn=ZNF185;refGene_tx=XM_019824467.2;refGene_gn=ZNF185;refGene_tx=XM_019824468.2
X	127977892	127977958
refGene_gn=ZNF185;refGene_tx=XM_006944092.4;refGene_gn=ZNF185;refGene_tx=XM_019824450.2;refGene_gn=ZNF185;refGene_tx=XM_019824451.1;refGene_gn=ZNF185;refGene_tx=XM_019824452.2;refGene_gn=ZNF185;refGene_tx=XM_019824453.2;refGene_gn=ZNF185;refGene_tx=XM_019824454.2;refGene_gn=ZNF185;refGene_tx=XM_019824455.2;refGene_gn=ZNF185;refGene_tx=XM_019824456.2;refGene_gn=ZNF185;refGene_tx=XM_019824457.2;refGene_gn=ZNF185;refGene_tx=XM_019824458.2;refGene_gn=ZNF185;refGene_tx=XM_019824459.2;refGene_gn=ZNF185;refGene_tx=XM_019824460.2;refGene_gn=ZNF185;refGene_tx=XM_019824461.2;refGene_gn=ZNF185;refGene_tx=XM_019824462.2;refGene_gn=ZNF185;refGene_tx=XM_019824463.2;refGene_gn=ZNF185;refGene_tx=XM_019824464.2;refGene_gn=ZNF185;refGene_tx=XM_019824465.2;refGene_gn=ZNF185;refGene_tx=XM_019824466.2;refGene_gn=ZNF185;refGene_tx=XM_019824467.2;refGene_gn=ZNF185;refGene_tx=XM_019824468.2
X	127980139	127980178
refGene_gn=ZNF185;refGene_tx=XM_006944092.4;refGene_gn=ZNF185;refGene_tx=XM_019824450.2;refGene_gn=ZNF185;refGene_tx=XM_019824451.1;refGene_gn=ZNF185;refGene_tx=XM_019824452.2;refGene_gn=ZNF185;refGene_tx=XM_019824453.2;refGene_gn=ZNF185;refGene_tx=XM_019824454.2;refGene_gn=ZNF185;refGene_tx=XM_019824455.2;refGene_gn=ZNF185;refGene_tx=XM_019824456.2;refGene_gn=ZNF185;refGene_tx=XM_019824458.2;refGene_gn=ZNF185;refGene_tx=XM_019824459.2;refGene_gn=ZNF185;refGene_tx=XM_019824460.2;refGene_gn=ZNF185;refGene_tx=XM_019824461.2;refGene_gn=ZNF185;refGene_tx=XM_019824462.2;refGene_gn=ZNF185;refGene_tx=XM_019824464.2;refGene_gn=ZNF185;refGene_tx=XM_019824465.2;refGene_gn=ZNF185;refGene_tx=XM_019824466.2;refGene_gn=ZNF185;refGene_tx=XM_019824467.2;refGene_gn=ZNF185;refGene_tx=XM_019824468.2
X	127980284	127980362
refGene_gn=ZNF185;refGene_tx=XM_006944092.4;refGene_gn=ZNF185;refGene_tx=XM_019824450.2;refGene_gn=ZNF185;refGene_tx=XM_019824451.1;refGene_gn=ZNF185;refGene_tx=XM_019824452.2;refGene_gn=ZNF185;refGene_tx=XM_019824453.2;refGene_gn=ZNF185;refGene_tx=XM_019824454.2;refGene_gn=ZNF185;refGene_tx=XM_019824455.2;refGene_gn=ZNF185;refGene_tx=XM_019824456.2;refGene_gn=ZNF185;refGene_tx=XM_019824457.2;refGene_gn=ZNF185;refGene_tx=XM_019824459.2;refGene_gn=ZNF185;refGene_tx=XM_019824460.2;refGen

e_gn=ZNF185;refGene_tx=XM_019824461.2;refGene_gn=ZNF185;refGene_tx=XM_019824462.2;refGene_gn=ZNF185;refGene_tx=XM_019824464.2;refGene_gn=ZNF185;refGene_tx=XM_019824465.2;refGene_gn=ZNF185;refGene_tx=XM_019824466.2;refGene_gn=ZNF185;refGene_tx=XM_019824467.2;refGene_gn=ZNF185;refGene_tx=XM_019824468.2
X 127980864 127980954
refGene_gn=ZNF185;refGene_tx=XM_006944092.4;refGene_gn=ZNF185;refGene_tx=XM_019824450.2;refGene_gn=ZNF185;refGene_tx=XM_019824451.1;refGene_gn=ZNF185;refGene_tx=XM_019824452.2;refGene_gn=ZNF185;refGene_tx=XM_019824453.2;refGene_gn=ZNF185;refGene_tx=XM_019824454.2;refGene_gn=ZNF185;refGene_tx=XM_019824455.2;refGene_gn=ZNF185;refGene_tx=XM_019824456.2;refGene_gn=ZNF185;refGene_tx=XM_019824457.2;refGene_gn=ZNF185;refGene_tx=XM_019824458.2;refGene_gn=ZNF185;refGene_tx=XM_019824459.2;refGene_gn=ZNF185;refGene_tx=XM_019824460.2;refGene_gn=ZNF185;refGene_tx=XM_019824461.2;refGene_gn=ZNF185;refGene_tx=XM_019824462.2;refGene_gn=ZNF185;refGene_tx=XM_019824463.2;refGene_gn=ZNF185;refGene_tx=XM_019824464.2;refGene_gn=ZNF185;refGene_tx=XM_019824465.2;refGene_gn=ZNF185;refGene_tx=XM_019824466.2;refGene_gn=ZNF185;refGene_tx=XM_019824467.2;refGene_gn=ZNF185;refGene_tx=XM_019824468.2
X 127981747 127981834
refGene_gn=ZNF185;refGene_tx=XM_006944092.4;refGene_gn=ZNF185;refGene_tx=XM_019824450.2;refGene_gn=ZNF185;refGene_tx=XM_019824451.1;refGene_gn=ZNF185;refGene_tx=XM_019824452.2;refGene_gn=ZNF185;refGene_tx=XM_019824453.2;refGene_gn=ZNF185;refGene_tx=XM_019824454.2;refGene_gn=ZNF185;refGene_tx=XM_019824455.2;refGene_gn=ZNF185;refGene_tx=XM_019824456.2;refGene_gn=ZNF185;refGene_tx=XM_019824457.2;refGene_gn=ZNF185;refGene_tx=XM_019824458.2;refGene_gn=ZNF185;refGene_tx=XM_019824459.2;refGene_gn=ZNF185;refGene_tx=XM_019824460.2;refGene_gn=ZNF185;refGene_tx=XM_019824461.2;refGene_gn=ZNF185;refGene_tx=XM_019824462.2;refGene_gn=ZNF185;refGene_tx=XM_019824463.2;refGene_gn=ZNF185;refGene_tx=XM_019824464.2;refGene_gn=ZNF185;refGene_tx=XM_019824465.2;refGene_gn=ZNF185;refGene_tx=XM_019824466.2;refGene_gn=ZNF185;refGene_tx=XM_019824467.2;refGene_gn=ZNF185;refGene_tx=XM_019824468.2
X 127982947 127983031
refGene_gn=ZNF185;refGene_tx=XM_006944092.4;refGene_gn=ZNF185;refGene_tx=XM_019824450.2;refGene_gn=ZNF185;refGene_tx=XM_019824451.1;refGene_gn=ZNF185;refGene_tx=XM_019824452.2;refGene_gn=ZNF185;refGene_tx=XM_019824453.2;refGene_gn=ZNF185;refGene_tx=XM_019824454.2;refGene_gn=ZNF185;refGene_tx=XM_019824455.2;refGene_gn=ZNF185;refGene_tx=XM_019824456.2;refGene_gn=ZNF185;refGene_tx=XM_019824457.2;refGene_gn=ZNF185;refGene_tx=XM_019824458.2;refGene_gn=ZNF185;refGene_tx=XM_019824459.2;refGene_gn=ZNF185;refGene_tx=XM_019824460.2;refGene_gn=ZNF185;refGene_tx=XM_019824461.2;refGene_gn=ZNF185;refGene_tx=XM_019824462.2;refGene_gn=ZNF185;refGene_tx=XM_019824463.2;refGene_gn=ZNF185;refGene_tx=XM_019824464.2;refGene_gn=ZNF185;refGene_tx=XM_019824465.2;refGene_gn=ZNF185;refGene_tx=XM_019824466.2;refGene_gn=ZNF185;refGene_tx=XM_019824467.2;refGene_gn=ZNF185;refGene_tx=XM_019824468.2
X 127983295 127983337
refGene_gn=ZNF185;refGene_tx=XM_006944092.4;refGene_gn=ZNF185;refGene_tx=XM_019824450.2;refGene_gn=ZNF185;refGene_tx=XM_019824451.1;refGene_

gn=ZNF185;refGene_tx=XM_019824452.2;refGene_gn=ZNF185;refGene_tx=XM_01
9824453.2;refGene_gn=ZNF185;refGene_tx=XM_019824454.2;refGene_gn=ZNF18
5;refGene_tx=XM_019824455.2;refGene_gn=ZNF185;refGene_tx=XM_019824456.
2;refGene_gn=ZNF185;refGene_tx=XM_019824457.2;refGene_gn=ZNF185;refGen
e_tx=XM_019824458.2;refGene_gn=ZNF185;refGene_tx=XM_019824459.2;refGen
e_gn=ZNF185;refGene_tx=XM_019824460.2;refGene_gn=ZNF185;refGene_tx=XM_
019824461.2;refGene_gn=ZNF185;refGene_tx=XM_019824462.2;refGene_gn=ZNF
185;refGene_tx=XM_019824463.2;refGene_gn=ZNF185;refGene_tx=XM_01982446
4.2;refGene_gn=ZNF185;refGene_tx=XM_019824465.2;refGene_gn=ZNF185;refG
ene_tx=XM_019824466.2;refGene_gn=ZNF185;refGene_tx=XM_019824467.2;refG
ene_gn=ZNF185;refGene_tx=XM_019824468.2
X 127984767 127984851
refGene_gn=ZNF185;refGene_tx=XM_006944092.4;refGene_gn=ZNF185;refGene_
tx=XM_019824450.2;refGene_gn=ZNF185;refGene_tx=XM_019824451.1;refGene_
gn=ZNF185;refGene_tx=XM_019824452.2;refGene_gn=ZNF185;refGene_tx=XM_01
9824453.2;refGene_gn=ZNF185;refGene_tx=XM_019824454.2;refGene_gn=ZNF18
5;refGene_tx=XM_019824455.2;refGene_gn=ZNF185;refGene_tx=XM_019824456.
2;refGene_gn=ZNF185;refGene_tx=XM_019824457.2;refGene_gn=ZNF185;refGen
e_tx=XM_019824458.2;refGene_gn=ZNF185;refGene_tx=XM_019824459.2;refGen
e_gn=ZNF185;refGene_tx=XM_019824460.2;refGene_gn=ZNF185;refGene_tx=XM_
019824461.2;refGene_gn=ZNF185;refGene_tx=XM_019824462.2;refGene_gn=ZNF
185;refGene_tx=XM_019824463.2;refGene_gn=ZNF185;refGene_tx=XM_01982446
4.2;refGene_gn=ZNF185;refGene_tx=XM_019824465.2;refGene_gn=ZNF185;refG
ene_tx=XM_019824466.2
X 127985320 127985410
refGene_gn=ZNF185;refGene_tx=XM_019824450.2;refGene_gn=ZNF185;refGene_
tx=XM_019824451.1;refGene_gn=ZNF185;refGene_tx=XM_019824452.2;refGene_
gn=ZNF185;refGene_tx=XM_019824453.2;refGene_gn=ZNF185;refGene_tx=XM_01
9824454.2;refGene_gn=ZNF185;refGene_tx=XM_019824455.2;refGene_gn=ZNF18
5;refGene_tx=XM_019824456.2;refGene_gn=ZNF185;refGene_tx=XM_019824457.
2;refGene_gn=ZNF185;refGene_tx=XM_019824458.2;refGene_gn=ZNF185;refGen
e_tx=XM_019824459.2;refGene_gn=ZNF185;refGene_tx=XM_019824461.2;refGen
e_gn=ZNF185;refGene_tx=XM_019824462.2;refGene_gn=ZNF185;refGene_tx=XM_
019824463.2;refGene_gn=ZNF185;refGene_tx=XM_019824464.2
X 127990949 127991036
refGene_gn=ZNF185;refGene_tx=XM_019824450.2;refGene_gn=ZNF185;refGene_
tx=XM_019824451.1;refGene_gn=ZNF185;refGene_tx=XM_019824452.2;refGene_
gn=ZNF185;refGene_tx=XM_019824453.2;refGene_gn=ZNF185;refGene_tx=XM_01
9824454.2;refGene_gn=ZNF185;refGene_tx=XM_019824455.2;refGene_gn=ZNF18
5;refGene_tx=XM_019824456.2;refGene_gn=ZNF185;refGene_tx=XM_019824457.
2;refGene_gn=ZNF185;refGene_tx=XM_019824458.2;refGene_gn=ZNF185;refGen
e_tx=XM_019824460.2;refGene_gn=ZNF185;refGene_tx=XM_019824462.2;refGen
e_gn=ZNF185;refGene_tx=XM_019824463.2
X 127994064 127994166
refGene_gn=ZNF185;refGene_tx=XM_006944092.4;refGene_gn=ZNF185;refGene_
tx=XM_019824450.2;refGene_gn=ZNF185;refGene_tx=XM_019824451.1;refGene_
gn=ZNF185;refGene_tx=XM_019824452.2;refGene_gn=ZNF185;refGene_tx=XM_01
9824453.2;refGene_gn=ZNF185;refGene_tx=XM_019824454.2;refGene_gn=ZNF18
5;refGene_tx=XM_019824455.2;refGene_gn=ZNF185;refGene_tx=XM_019824456.
2;refGene_gn=ZNF185;refGene_tx=XM_019824457.2;refGene_gn=ZNF185;refGen

e_tx=XM_019824458.2;refGene_gn=ZNF185;refGene_tx=XM_019824459.2;refGen
e_gn=ZNF185;refGene_tx=XM_019824460.2;refGene_gn=ZNF185;refGene_tx=XM_
019824461.2;refGene_gn=ZNF185;refGene_tx=XM_019824462.2;refGene_gn=ZNF
185;refGene_tx=XM_019824463.2;refGene_gn=ZNF185;refGene_tx=XM_01982446
4.2;refGene_gn=ZNF185;refGene_tx=XM_019824465.2;refGene_gn=ZNF185;refG
ene_tx=XM_019824466.2;refGene_gn=ZNF185;refGene_tx=XM_019824467.2;refG
ene_gn=ZNF185;refGene_tx=XM_019824468.2
X	127995223	127995322
refGene_gn=ZNF185;refGene_tx=XM_006944092.4;refGene_gn=ZNF185;refGene_
tx=XM_019824450.2;refGene_gn=ZNF185;refGene_tx=XM_019824451.1;refGene_
gn=ZNF185;refGene_tx=XM_019824452.2;refGene_gn=ZNF185;refGene_tx=XM_01
9824453.2;refGene_gn=ZNF185;refGene_tx=XM_019824454.2;refGene_gn=ZNF18
5;refGene_tx=XM_019824455.2;refGene_gn=ZNF185;refGene_tx=XM_019824456.
2;refGene_gn=ZNF185;refGene_tx=XM_019824457.2;refGene_gn=ZNF185;refGen
e_tx=XM_019824458.2;refGene_gn=ZNF185;refGene_tx=XM_019824459.2;refGen
e_gn=ZNF185;refGene_tx=XM_019824460.2;refGene_gn=ZNF185;refGene_tx=XM_
019824461.2;refGene_gn=ZNF185;refGene_tx=XM_019824462.2;refGene_gn=ZNF
185;refGene_tx=XM_019824463.2;refGene_gn=ZNF185;refGene_tx=XM_01982446
4.2;refGene_gn=ZNF185;refGene_tx=XM_019824465.2;refGene_gn=ZNF185;refG
ene_tx=XM_019824466.2;refGene_gn=ZNF185;refGene_tx=XM_019824467.2;refG
ene_gn=ZNF185;refGene_tx=XM_019824468.2
X	127998185	127998275
refGene_gn=ZNF185;refGene_tx=XM_006944092.4;refGene_gn=ZNF185;refGene_
tx=XM_019824450.2;refGene_gn=ZNF185;refGene_tx=XM_019824451.1;refGene_
gn=ZNF185;refGene_tx=XM_019824452.2;refGene_gn=ZNF185;refGene_tx=XM_01
9824453.2;refGene_gn=ZNF185;refGene_tx=XM_019824454.2;refGene_gn=ZNF18
5;refGene_tx=XM_019824455.2;refGene_gn=ZNF185;refGene_tx=XM_019824456.
2;refGene_gn=ZNF185;refGene_tx=XM_019824457.2;refGene_gn=ZNF185;refGen
e_tx=XM_019824458.2;refGene_gn=ZNF185;refGene_tx=XM_019824459.2;refGen
e_gn=ZNF185;refGene_tx=XM_019824460.2;refGene_gn=ZNF185;refGene_tx=XM_
019824461.2;refGene_gn=ZNF185;refGene_tx=XM_019824462.2;refGene_gn=ZNF
185;refGene_tx=XM_019824463.2;refGene_gn=ZNF185;refGene_tx=XM_01982446
4.2;refGene_gn=ZNF185;refGene_tx=XM_019824465.2;refGene_gn=ZNF185;refG
ene_tx=XM_019824466.2;refGene_gn=ZNF185;refGene_tx=XM_019824467.2;refG
ene_gn=ZNF185;refGene_tx=XM_019824468.2
X	128003703	128003919
refGene_gn=ZNF185;refGene_tx=XM_006944092.4;refGene_gn=ZNF185;refGene_
tx=XM_019824450.2;refGene_gn=ZNF185;refGene_tx=XM_019824451.1;refGene_
gn=ZNF185;refGene_tx=XM_019824452.2;refGene_gn=ZNF185;refGene_tx=XM_01
9824453.2;refGene_gn=ZNF185;refGene_tx=XM_019824454.2;refGene_gn=ZNF18
5;refGene_tx=XM_019824455.2;refGene_gn=ZNF185;refGene_tx=XM_019824456.
2;refGene_gn=ZNF185;refGene_tx=XM_019824457.2;refGene_gn=ZNF185;refGen
e_tx=XM_019824458.2;refGene_gn=ZNF185;refGene_tx=XM_019824459.2;refGen
e_gn=ZNF185;refGene_tx=XM_019824460.2;refGene_gn=ZNF185;refGene_tx=XM_
019824461.2;refGene_gn=ZNF185;refGene_tx=XM_019824462.2;refGene_gn=ZNF
185;refGene_tx=XM_019824463.2;refGene_gn=ZNF185;refGene_tx=XM_01982446
4.2;refGene_gn=ZNF185;refGene_tx=XM_019824465.2;refGene_gn=ZNF185;refG
ene_tx=XM_019824466.2;refGene_gn=ZNF185;refGene_tx=XM_019824467.2;refG
ene_gn=ZNF185;refGene_tx=XM_019824468.2
X	128015190	128015406

refGene_gn=ZNF185;refGene_tx=XM_006944092.4;refGene_gn=ZNF185;refGene_tx=XM_019824450.2;refGene_gn=ZNF185;refGene_tx=XM_019824451.1;refGene_gn=ZNF185;refGene_tx=XM_019824452.2;refGene_gn=ZNF185;refGene_tx=XM_019824453.2;refGene_gn=ZNF185;refGene_tx=XM_019824454.2;refGene_gn=ZNF185;refGene_tx=XM_019824455.2;refGene_gn=ZNF185;refGene_tx=XM_019824456.2;refGene_gn=ZNF185;refGene_tx=XM_019824457.2;refGene_gn=ZNF185;refGene_tx=XM_019824458.2;refGene_gn=ZNF185;refGene_tx=XM_019824459.2;refGene_gn=ZNF185;refGene_tx=XM_019824460.2;refGene_gn=ZNF185;refGene_tx=XM_019824461.2;refGene_gn=ZNF185;refGene_tx=XM_019824462.2;refGene_gn=ZNF185;refGene_tx=XM_019824463.2;refGene_gn=ZNF185;refGene_tx=XM_019824464.2;refGene_gn=ZNF185;refGene_tx=XM_019824465.2;refGene_gn=ZNF185;refGene_tx=XM_019824466.2;refGene_gn=ZNF185;refGene_tx=XM_019824467.2;refGene_gn=ZNF185;refGene_tx=XM_019824468.2
X 128018254 128018365
refGene_gn=ZNF185;refGene_tx=XM_006944092.4;refGene_gn=ZNF185;refGene_tx=XM_019824450.2;refGene_gn=ZNF185;refGene_tx=XM_019824451.1;refGene_gn=ZNF185;refGene_tx=XM_019824452.2;refGene_gn=ZNF185;refGene_tx=XM_019824453.2;refGene_gn=ZNF185;refGene_tx=XM_019824454.2;refGene_gn=ZNF185;refGene_tx=XM_019824455.2;refGene_gn=ZNF185;refGene_tx=XM_019824456.2;refGene_gn=ZNF185;refGene_tx=XM_019824457.2;refGene_gn=ZNF185;refGene_tx=XM_019824458.2;refGene_gn=ZNF185;refGene_tx=XM_019824459.2;refGene_gn=ZNF185;refGene_tx=XM_019824460.2;refGene_gn=ZNF185;refGene_tx=XM_019824461.2;refGene_gn=ZNF185;refGene_tx=XM_019824463.2;refGene_gn=ZNF185;refGene_tx=XM_019824464.2;refGene_gn=ZNF185;refGene_tx=XM_019824465.2;refGene_gn=ZNF185;refGene_tx=XM_019824466.2;refGene_gn=ZNF185;refGene_tx=XM_019824467.2;refGene_gn=ZNF185;refGene_tx=XM_019824468.2
X 128019417 128019498
refGene_gn=ZNF185;refGene_tx=XM_006944092.4;refGene_gn=ZNF185;refGene_tx=XM_019824450.2;refGene_gn=ZNF185;refGene_tx=XM_019824451.1;refGene_gn=ZNF185;refGene_tx=XM_019824452.2;refGene_gn=ZNF185;refGene_tx=XM_019824453.2;refGene_gn=ZNF185;refGene_tx=XM_019824454.2;refGene_gn=ZNF185;refGene_tx=XM_019824455.2;refGene_gn=ZNF185;refGene_tx=XM_019824456.2;refGene_gn=ZNF185;refGene_tx=XM_019824457.2;refGene_gn=ZNF185;refGene_tx=XM_019824458.2;refGene_gn=ZNF185;refGene_tx=XM_019824459.2;refGene_gn=ZNF185;refGene_tx=XM_019824460.2;refGene_gn=ZNF185;refGene_tx=XM_019824461.2;refGene_gn=ZNF185;refGene_tx=XM_019824462.2;refGene_gn=ZNF185;refGene_tx=XM_019824463.2;refGene_gn=ZNF185;refGene_tx=XM_019824464.2;refGene_gn=ZNF185;refGene_tx=XM_019824465.2;refGene_gn=ZNF185;refGene_tx=XM_019824466.2;refGene_gn=ZNF185;refGene_tx=XM_019824467.2;refGene_gn=ZNF185;refGene_tx=XM_019824468.2
X 128021931 128022003
refGene_gn=ZNF185;refGene_tx=XM_006944092.4;refGene_gn=ZNF185;refGene_tx=XM_019824450.2;refGene_gn=ZNF185;refGene_tx=XM_019824451.1;refGene_gn=ZNF185;refGene_tx=XM_019824452.2;refGene_gn=ZNF185;refGene_tx=XM_019824453.2;refGene_gn=ZNF185;refGene_tx=XM_019824454.2;refGene_gn=ZNF185;refGene_tx=XM_019824455.2;refGene_gn=ZNF185;refGene_tx=XM_019824456.2;refGene_gn=ZNF185;refGene_tx=XM_019824457.2;refGene_gn=ZNF185;refGene_tx=XM_019824458.2;refGene_gn=ZNF185;refGene_tx=XM_019824459.2;refGene_gn=ZNF185;refGene_tx=XM_019824460.2;refGene_gn=ZNF185;refGene_tx=XM_019824461.2;refGene_gn=ZNF185;refGene_tx=XM_019824462.2;refGene_gn=ZNF

185;refGene_tx=XM_019824463.2;refGene_gn=ZNF185;refGene_tx=XM_019824464.2;refGene_gn=ZNF185;refGene_tx=XM_019824465.2;refGene_gn=ZNF185;refGene_tx=XM_019824466.2;refGene_gn=ZNF185;refGene_tx=XM_019824467.2;refGene_gn=ZNF185;refGene_tx=XM_019824468.2
X 128023633 128023736
refGene_gn=ZNF185;refGene_tx=XM_006944092.4;refGene_gn=ZNF185;refGene_tx=XM_019824450.2;refGene_gn=ZNF185;refGene_tx=XM_019824451.1;refGene_gn=ZNF185;refGene_tx=XM_019824452.2;refGene_gn=ZNF185;refGene_tx=XM_019824453.2;refGene_gn=ZNF185;refGene_tx=XM_019824454.2;refGene_gn=ZNF185;refGene_tx=XM_019824455.2;refGene_gn=ZNF185;refGene_tx=XM_019824456.2;refGene_gn=ZNF185;refGene_tx=XM_019824457.2;refGene_gn=ZNF185;refGene_tx=XM_019824458.2;refGene_gn=ZNF185;refGene_tx=XM_019824459.2;refGene_gn=ZNF185;refGene_tx=XM_019824460.2;refGene_gn=ZNF185;refGene_tx=XM_019824461.2;refGene_gn=ZNF185;refGene_tx=XM_019824462.2;refGene_gn=ZNF185;refGene_tx=XM_019824463.2;refGene_gn=ZNF185;refGene_tx=XM_019824464.2;refGene_gn=ZNF185;refGene_tx=XM_019824465.2;refGene_gn=ZNF185;refGene_tx=XM_019824466.2;refGene_gn=ZNF185;refGene_tx=XM_019824467.2;refGene_gn=ZNF185;refGene_tx=XM_019824468.2
X 128024628 128024727
refGene_gn=ZNF185;refGene_tx=XM_006944092.4;refGene_gn=ZNF185;refGene_tx=XM_019824450.2;refGene_gn=ZNF185;refGene_tx=XM_019824451.1;refGene_gn=ZNF185;refGene_tx=XM_019824452.2;refGene_gn=ZNF185;refGene_tx=XM_019824453.2;refGene_gn=ZNF185;refGene_tx=XM_019824454.2;refGene_gn=ZNF185;refGene_tx=XM_019824455.2;refGene_gn=ZNF185;refGene_tx=XM_019824456.2;refGene_gn=ZNF185;refGene_tx=XM_019824457.2;refGene_gn=ZNF185;refGene_tx=XM_019824458.2;refGene_gn=ZNF185;refGene_tx=XM_019824459.2;refGene_gn=ZNF185;refGene_tx=XM_019824460.2;refGene_gn=ZNF185;refGene_tx=XM_019824461.2;refGene_gn=ZNF185;refGene_tx=XM_019824462.2;refGene_gn=ZNF185;refGene_tx=XM_019824463.2;refGene_gn=ZNF185;refGene_tx=XM_019824464.2;refGene_gn=ZNF185;refGene_tx=XM_019824465.2;refGene_gn=ZNF185;refGene_tx=XM_019824466.2;refGene_gn=ZNF185;refGene_tx=XM_019824467.2;refGene_gn=ZNF185;refGene_tx=XM_019824468.2
X 128057851 128058001
refGene_gn=LOC101100193;refGene_tx=XM_023248986.1
X 128058868 128059016
refGene_gn=LOC101100193;refGene_tx=XM_023248986.1
X 128059018 128059684
refGene_gn=LOC101100193;refGene_tx=XM_023248986.1
X 128059686 128059895
refGene_gn=LOC101100193;refGene_tx=XM_023248986.1
X 128098560 128099952
refGene_gn=PNMA3;refGene_tx=XM_023249127.1;refGene_gn=PNMA3;refGene_tx=XM_023249128.1;refGene_gn=PNMA3;refGene_tx=XM_023249129.1
X 128099986 128100091
refGene_gn=PNMA3;refGene_tx=XM_023249129.1
X 128189583 128189602
refGene_gn=LOC111558739;refGene_tx=XM_023249246.1
X 128191249 128191367
refGene_gn=LOC111558739;refGene_tx=XM_023249246.1
X 128192270 128192370

refGene_gn=LOC111558739;refGene_tx=XM_023249246.1
X	128193866	128193965
refGene_gn=LOC111558739;refGene_tx=XM_023249246.1
X	128195828	128195933
refGene_gn=LOC111558739;refGene_tx=XM_023249246.1
X	128196066	128196120
refGene_gn=LOC111558739;refGene_tx=XM_023249246.1
X	128234098	128234782
refGene_gn=LOC111558692;refGene_tx=XM_023248987.1
X	128235880	128237023
refGene_gn=LOC111558735;refGene_tx=XM_023249200.1
X	128237024	128237228
refGene_gn=LOC111558735;refGene_tx=XM_023249200.1
X	128303039	128303102
refGene_gn=PNMA6F;refGene_tx=XM_023248989.1
X	128305473	128305591
refGene_gn=PNMA6F;refGene_tx=XM_023248989.1
X	128305592	128305713
refGene_gn=PNMA6F;refGene_tx=XM_023248989.1
X	128305982	128306594
refGene_gn=PNMA6F;refGene_tx=XM_023248989.1
X	128306630	128306665
refGene_gn=PNMA6F;refGene_tx=XM_023248989.1
X	128307062	128307334
refGene_gn=PNMA6F;refGene_tx=XM_023248989.1
X	128307377	128307782
refGene_gn=PNMA6F;refGene_tx=XM_023248989.1
X	128308378	128308863
refGene_gn=PNMA6F;refGene_tx=XM_023248989.1
X	128309003	128309091
refGene_gn=PNMA6F;refGene_tx=XM_023248989.1
X	128337021	128338149
refGene_gn=ZNF275;refGene_tx=XM_006944097.3;refGene_gn=ZNF275;refGene_tx=XM_023248848.1;refGene_gn=ZNF275;refGene_tx=XM_023248849.1;refGene_gn=ZNF275;refGene_tx=XM_023248850.1;refGene_gn=ZNF275;refGene_tx=XM_023248851.1;refGene_gn=ZNF275;refGene_tx=XM_023248852.1;refGene_gn=ZNF275;refGene_tx=XM_023248853.1
X	128407172	128407892
refGene_gn=LOC105259692;refGene_tx=XM_023248798.1;refGene_gn=LOC105259692;refGene_tx=XM_023248799.1;refGene_gn=LOC105259692;refGene_tx=XM_023248800.1;refGene_gn=LOC105259692;refGene_tx=XM_023248801.1;refGene_gn=LOC105259692;refGene_tx=XM_023248803.1
X	128408197	128408220
refGene_gn=LOC105259692;refGene_tx=XM_023248798.1;refGene_gn=LOC105259692;refGene_tx=XM_023248799.1;refGene_gn=LOC105259692;refGene_tx=XM_023248800.1
X	128411970	128412099
refGene_gn=LOC105259692;refGene_tx=XM_023248798.1;refGene_gn=LOC105259692;refGene_tx=XM_023248799.1;refGene_gn=LOC105259692;refGene_tx=XM_023248800.1;refGene_gn=LOC105259692;refGene_tx=XM_023248801.1;refGene_gn

=LOC105259692;refGene_tx=XM_023248803.1
X	128412341	128412579
refGene_gn=LOC105259692;refGene_tx=XM_023248798.1;refGene_gn=LOC105259692;refGene_tx=XM_023248801.1
X	128412636	128412815
refGene_gn=LOC105259692;refGene_tx=XM_023248799.1;refGene_gn=LOC105259692;refGene_tx=XM_023248800.1;refGene_gn=LOC105259692;refGene_tx=XM_023248803.1
X	128431230	128431941
refGene_gn=TREX2;refGene_tx=XM_006944101.4
X	128434545	128434613
refGene_gn=HAUS7;refGene_tx=XM_011279575.3;refGene_gn=HAUS7;refGene_tx=XM_019824477.2
X	128440181	128440296
refGene_gn=HAUS7;refGene_tx=XM_011279575.3;refGene_gn=HAUS7;refGene_tx=XM_019824477.2
X	128441341	128441554
refGene_gn=HAUS7;refGene_tx=XM_011279575.3;refGene_gn=HAUS7;refGene_tx=XM_019824477.2
X	128442114	128442211
refGene_gn=HAUS7;refGene_tx=XM_011279575.3;refGene_gn=HAUS7;refGene_tx=XM_019824477.2
X	128442323	128442485
refGene_gn=HAUS7;refGene_tx=XM_011279575.3;refGene_gn=HAUS7;refGene_tx=XM_019824477.2
X	128443018	128443110
refGene_gn=HAUS7;refGene_tx=XM_011279575.3;refGene_gn=HAUS7;refGene_tx=XM_019824477.2
X	128451786	128451848
refGene_gn=HAUS7;refGene_tx=XM_011279575.3;refGene_gn=HAUS7;refGene_tx=XM_019824477.2
X	128454598	128454666
refGene_gn=HAUS7;refGene_tx=XM_011279575.3;refGene_gn=HAUS7;refGene_tx=XM_019824477.2
X	128459161	128459277
refGene_gn=HAUS7;refGene_tx=XM_011279575.3;refGene_gn=HAUS7;refGene_tx=XM_019824477.2
X	128460153	128460252
refGene_gn=HAUS7;refGene_tx=XM_019824477.2
X	128469499	128469554
refGene_gn=LOC111558761;refGene_tx=XM_023249368.1
X	128471218	128471505
refGene_gn=LOC111558761;refGene_tx=XM_023249368.1
X	128472671	128472781
refGene_gn=LOC109496649;refGene_tx=XM_023249324.1
X	128475365	128475690
refGene_gn=LOC109496649;refGene_tx=XM_023249324.1
X	128476129	128476258
refGene_gn=LOC109496649;refGene_tx=XM_023249324.1
X	128493921	128494171

refGene_gn=BGN;refGene_tx=XM_004001000.4;refGene_gn=BGN;refGene_tx=XM_019824478.2
X 128494499 128494612
refGene_gn=BGN;refGene_tx=XM_004001000.4;refGene_gn=BGN;refGene_tx=XM_019824478.2
X 128494954 128495168
refGene_gn=BGN;refGene_tx=XM_004001000.4;refGene_gn=BGN;refGene_tx=XM_019824478.2
X 128495711 128495822
refGene_gn=BGN;refGene_tx=XM_004001000.4;refGene_gn=BGN;refGene_tx=XM_019824478.2
X 128496038 128496132
refGene_gn=BGN;refGene_tx=XM_004001000.4;refGene_gn=BGN;refGene_tx=XM_019824478.2
X 128496217 128496356
refGene_gn=BGN;refGene_tx=XM_004001000.4;refGene_gn=BGN;refGene_tx=XM_019824478.2
X 128497762 128497960
refGene_gn=BGN;refGene_tx=XM_004001000.4;refGene_gn=BGN;refGene_tx=XM_019824478.2
X 128523781 128523989
refGene_gn=ATP2B3;refGene_tx=XM_023248933.1;refGene_gn=ATP2B3;refGene_tx=XM_023248934.1;refGene_gn=ATP2B3;refGene_tx=XM_023248935.1;refGene_gn=ATP2B3;refGene_tx=XM_023248936.1
X 128528927 128529125
refGene_gn=ATP2B3;refGene_tx=XM_023248933.1;refGene_gn=ATP2B3;refGene_tx=XM_023248934.1;refGene_gn=ATP2B3;refGene_tx=XM_023248935.1;refGene_gn=ATP2B3;refGene_tx=XM_023248936.1
X 128529231 128529489
refGene_gn=ATP2B3;refGene_tx=XM_023248933.1;refGene_gn=ATP2B3;refGene_tx=XM_023248934.1;refGene_gn=ATP2B3;refGene_tx=XM_023248935.1;refGene_gn=ATP2B3;refGene_tx=XM_023248936.1
X 128529745 128529871
refGene_gn=ATP2B3;refGene_tx=XM_023248933.1;refGene_gn=ATP2B3;refGene_tx=XM_023248934.1;refGene_gn=ATP2B3;refGene_tx=XM_023248935.1;refGene_gn=ATP2B3;refGene_tx=XM_023248936.1;refGene_gn=ATP2B3;refGene_tx=XM_023248937.1
X 128530482 128530608
refGene_gn=ATP2B3;refGene_tx=XM_023248933.1;refGene_gn=ATP2B3;refGene_tx=XM_023248934.1;refGene_gn=ATP2B3;refGene_tx=XM_023248935.1;refGene_gn=ATP2B3;refGene_tx=XM_023248936.1;refGene_gn=ATP2B3;refGene_tx=XM_023248937.1
X 128533585 128533627
refGene_gn=ATP2B3;refGene_tx=XM_023248933.1;refGene_gn=ATP2B3;refGene_tx=XM_023248934.1;refGene_gn=ATP2B3;refGene_tx=XM_023248935.1;refGene_gn=ATP2B3;refGene_tx=XM_023248936.1;refGene_gn=ATP2B3;refGene_tx=XM_023248937.1
X 128535031 128535196
refGene_gn=ATP2B3;refGene_tx=XM_023248933.1;refGene_gn=ATP2B3;refGene_tx=XM_023248934.1;refGene_gn=ATP2B3;refGene_tx=XM_023248935.1;refGene_

gn=ATP2B3;refGene_tx=XM_023248936.1;refGene_gn=ATP2B3;refGene_tx=XM_023248937.1
X 128535560 128535775
refGene_gn=ATP2B3;refGene_tx=XM_023248933.1;refGene_gn=ATP2B3;refGene_tx=XM_023248934.1;refGene_gn=ATP2B3;refGene_tx=XM_023248935.1;refGene_gn=ATP2B3;refGene_tx=XM_023248936.1;refGene_gn=ATP2B3;refGene_tx=XM_023248937.1
X 128536434 128536677
refGene_gn=ATP2B3;refGene_tx=XM_023248933.1;refGene_gn=ATP2B3;refGene_tx=XM_023248934.1;refGene_gn=ATP2B3;refGene_tx=XM_023248935.1;refGene_gn=ATP2B3;refGene_tx=XM_023248936.1;refGene_gn=ATP2B3;refGene_tx=XM_023248937.1
X 128537005 128537247
refGene_gn=ATP2B3;refGene_tx=XM_023248933.1;refGene_gn=ATP2B3;refGene_tx=XM_023248934.1;refGene_gn=ATP2B3;refGene_tx=XM_023248935.1;refGene_gn=ATP2B3;refGene_tx=XM_023248936.1;refGene_gn=ATP2B3;refGene_tx=XM_023248937.1
X 128539811 128540046
refGene_gn=ATP2B3;refGene_tx=XM_023248933.1;refGene_gn=ATP2B3;refGene_tx=XM_023248934.1;refGene_gn=ATP2B3;refGene_tx=XM_023248935.1;refGene_gn=ATP2B3;refGene_tx=XM_023248936.1;refGene_gn=ATP2B3;refGene_tx=XM_023248937.1
X 128542208 128542388
refGene_gn=ATP2B3;refGene_tx=XM_023248933.1;refGene_gn=ATP2B3;refGene_tx=XM_023248934.1;refGene_gn=ATP2B3;refGene_tx=XM_023248935.1;refGene_gn=ATP2B3;refGene_tx=XM_023248936.1;refGene_gn=ATP2B3;refGene_tx=XM_023248937.1
X 128542527 128542615
refGene_gn=ATP2B3;refGene_tx=XM_023248933.1;refGene_gn=ATP2B3;refGene_tx=XM_023248934.1;refGene_gn=ATP2B3;refGene_tx=XM_023248935.1;refGene_gn=ATP2B3;refGene_tx=XM_023248936.1;refGene_gn=ATP2B3;refGene_tx=XM_023248937.1
X 128543270 128543377
refGene_gn=ATP2B3;refGene_tx=XM_023248933.1;refGene_gn=ATP2B3;refGene_tx=XM_023248934.1;refGene_gn=ATP2B3;refGene_tx=XM_023248935.1;refGene_gn=ATP2B3;refGene_tx=XM_023248936.1;refGene_gn=ATP2B3;refGene_tx=XM_023248937.1
X 128544490 128544682
refGene_gn=ATP2B3;refGene_tx=XM_023248933.1;refGene_gn=ATP2B3;refGene_tx=XM_023248934.1;refGene_gn=ATP2B3;refGene_tx=XM_023248935.1;refGene_gn=ATP2B3;refGene_tx=XM_023248936.1;refGene_gn=ATP2B3;refGene_tx=XM_023248937.1
X 128546111 128546325
refGene_gn=ATP2B3;refGene_tx=XM_023248933.1;refGene_gn=ATP2B3;refGene_tx=XM_023248934.1;refGene_gn=ATP2B3;refGene_tx=XM_023248935.1;refGene_gn=ATP2B3;refGene_tx=XM_023248936.1;refGene_gn=ATP2B3;refGene_tx=XM_023248937.1
X 128547496 128547708
refGene_gn=ATP2B3;refGene_tx=XM_023248933.1;refGene_gn=ATP2B3;refGene_tx=XM_023248934.1;refGene_gn=ATP2B3;refGene_tx=XM_023248935.1;refGene_

gn=ATP2B3;refGene_tx=XM_023248936.1;refGene_gn=ATP2B3;refGene_tx=XM_023248937.1
X	128548733	128548841
refGene_gn=ATP2B3;refGene_tx=XM_023248933.1;refGene_gn=ATP2B3;refGene_tx=XM_023248934.1;refGene_gn=ATP2B3;refGene_tx=XM_023248935.1;refGene_gn=ATP2B3;refGene_tx=XM_023248936.1;refGene_gn=ATP2B3;refGene_tx=XM_023248937.1
X	128550093	128550276
refGene_gn=ATP2B3;refGene_tx=XM_023248933.1;refGene_gn=ATP2B3;refGene_tx=XM_023248934.1;refGene_gn=ATP2B3;refGene_tx=XM_023248935.1;refGene_gn=ATP2B3;refGene_tx=XM_023248936.1;refGene_gn=ATP2B3;refGene_tx=XM_023248937.1
X	128554848	128555002
refGene_gn=ATP2B3;refGene_tx=XM_023248933.1;refGene_gn=ATP2B3;refGene_tx=XM_023248934.1;refGene_gn=ATP2B3;refGene_tx=XM_023248936.1;refGene_gn=ATP2B3;refGene_tx=XM_023248937.1
X	128565457	128565778
refGene_gn=ATP2B3;refGene_tx=XM_023248933.1;refGene_gn=ATP2B3;refGene_tx=XM_023248934.1;refGene_gn=ATP2B3;refGene_tx=XM_023248935.1;refGene_gn=ATP2B3;refGene_tx=XM_023248936.1;refGene_gn=ATP2B3;refGene_tx=XM_023248937.1
X	128572742	128572832
refGene_gn=CCNQ;refGene_tx=XM_023248951.1;refGene_gn=CCNQ;refGene_tx=XM_023248952.1
X	128576166	128576394
refGene_gn=CCNQ;refGene_tx=XM_023248951.1;refGene_gn=CCNQ;refGene_tx=XM_023248952.1
X	128578668	128578801
refGene_gn=CCNQ;refGene_tx=XM_023248951.1;refGene_gn=CCNQ;refGene_tx=XM_023248952.1
X	128580395	128580579
refGene_gn=CCNQ;refGene_tx=XM_023248951.1;refGene_gn=CCNQ;refGene_tx=XM_023248952.1
X	128583593	128583711
refGene_gn=CCNQ;refGene_tx=XM_023248951.1;refGene_gn=CCNQ;refGene_tx=XM_023248952.1
X	128633853	128634220
refGene_gn=DUSP9;refGene_tx=XM_004001006.4
X	128635522	128635791
refGene_gn=DUSP9;refGene_tx=XM_004001006.4
X	128635792	128635973
refGene_gn=DUSP9;refGene_tx=XM_004001006.4
X	128636278	128636604
refGene_gn=DUSP9;refGene_tx=XM_004001006.4
X	128653840	128653978
refGene_gn=PNCK;refGene_tx=XM_019824487.2;refGene_gn=PNCK;refGene_tx=XM_019824488.2;refGene_gn=PNCK;refGene_tx=XM_023249152.1;refGene_gn=PNCK;refGene_tx=XM_023249153.1;refGene_gn=PNCK;refGene_tx=XM_023249154.1;refGene_gn=PNCK;refGene_tx=XM_023249155.1
X	128654122	128654210

refGene_gn=PNCK;refGene_tx=XM_019824487.2;refGene_gn=PNCK;refGene_tx=XM_019824488.2;refGene_gn=PNCK;refGene_tx=XM_023249152.1;refGene_gn=PNCK;refGene_tx=XM_023249153.1;refGene_gn=PNCK;refGene_tx=XM_023249154.1;refGene_gn=PNCK;refGene_tx=XM_023249155.1
X	128654299	128654378
refGene_gn=PNCK;refGene_tx=XM_019824487.2;refGene_gn=PNCK;refGene_tx=XM_019824488.2;refGene_gn=PNCK;refGene_tx=XM_023249152.1;refGene_gn=PNCK;refGene_tx=XM_023249153.1;refGene_gn=PNCK;refGene_tx=XM_023249154.1;refGene_gn=PNCK;refGene_tx=XM_023249155.1
X	128654467	128654580
refGene_gn=PNCK;refGene_tx=XM_019824487.2;refGene_gn=PNCK;refGene_tx=XM_019824488.2;refGene_gn=PNCK;refGene_tx=XM_023249152.1;refGene_gn=PNCK;refGene_tx=XM_023249153.1;refGene_gn=PNCK;refGene_tx=XM_023249154.1;refGene_gn=PNCK;refGene_tx=XM_023249155.1
X	128654674	128654750
refGene_gn=PNCK;refGene_tx=XM_019824487.2;refGene_gn=PNCK;refGene_tx=XM_019824488.2;refGene_gn=PNCK;refGene_tx=XM_023249152.1;refGene_gn=PNCK;refGene_tx=XM_023249153.1;refGene_gn=PNCK;refGene_tx=XM_023249154.1;refGene_gn=PNCK;refGene_tx=XM_023249155.1
X	128654926	128655050
refGene_gn=PNCK;refGene_tx=XM_019824487.2;refGene_gn=PNCK;refGene_tx=XM_019824488.2;refGene_gn=PNCK;refGene_tx=XM_023249152.1;refGene_gn=PNCK;refGene_tx=XM_023249153.1;refGene_gn=PNCK;refGene_tx=XM_023249154.1;refGene_gn=PNCK;refGene_tx=XM_023249155.1
X	128655190	128655350
refGene_gn=PNCK;refGene_tx=XM_019824487.2;refGene_gn=PNCK;refGene_tx=XM_019824488.2;refGene_gn=PNCK;refGene_tx=XM_023249152.1;refGene_gn=PNCK;refGene_tx=XM_023249153.1;refGene_gn=PNCK;refGene_tx=XM_023249154.1;refGene_gn=PNCK;refGene_tx=XM_023249155.1
X	128655435	128655510
refGene_gn=PNCK;refGene_tx=XM_019824487.2;refGene_gn=PNCK;refGene_tx=XM_019824488.2;refGene_gn=PNCK;refGene_tx=XM_023249152.1;refGene_gn=PNCK;refGene_tx=XM_023249153.1;refGene_gn=PNCK;refGene_tx=XM_023249154.1;refGene_gn=PNCK;refGene_tx=XM_023249155.1
X	128655750	128655882
refGene_gn=PNCK;refGene_tx=XM_019824487.2;refGene_gn=PNCK;refGene_tx=XM_019824488.2;refGene_gn=PNCK;refGene_tx=XM_023249152.1;refGene_gn=PNCK;refGene_tx=XM_023249153.1;refGene_gn=PNCK;refGene_tx=XM_023249154.1;refGene_gn=PNCK;refGene_tx=XM_023249155.1
X	128656130	128656198
refGene_gn=PNCK;refGene_tx=XM_019824487.2;refGene_gn=PNCK;refGene_tx=XM_019824488.2;refGene_gn=PNCK;refGene_tx=XM_023249152.1;refGene_gn=PNCK;refGene_tx=XM_023249153.1;refGene_gn=PNCK;refGene_tx=XM_023249154.1;refGene_gn=PNCK;refGene_tx=XM_023249155.1
X	128665596	128665858
refGene_gn=SLC6A8;refGene_tx=XM_023249498.1
X	128667485	128667617
refGene_gn=SLC6A8;refGene_tx=XM_023249498.1
X	128668575	128668825
refGene_gn=SLC6A8;refGene_tx=XM_023249498.1

X	128669314	128669336
refGene_gn=SLC6A8;refGene_tx=XM_023249498.1
X	128669338	128669452
refGene_gn=SLC6A8;refGene_tx=XM_023249498.1
X	128670304	128670439
refGene_gn=SLC6A8;refGene_tx=XM_023249498.1
X	128670525	128670629
refGene_gn=SLC6A8;refGene_tx=XM_023249498.1
X	128670716	128670841
refGene_gn=SLC6A8;refGene_tx=XM_023249498.1
X	128671429	128671542
refGene_gn=SLC6A8;refGene_tx=XM_023249498.1
X	128671663	128671801
refGene_gn=SLC6A8;refGene_tx=XM_023249498.1
X	128671877	128671980
refGene_gn=SLC6A8;refGene_tx=XM_023249498.1
X	128672072	128672173
refGene_gn=SLC6A8;refGene_tx=XM_023249498.1
X	128672261	128672432
refGene_gn=SLC6A8;refGene_tx=XM_023249498.1
X	128672855	128672996
refGene_gn=SLC6A8;refGene_tx=XM_023249498.1
X	128678664	128678703
refGene_gn=BCAP31;refGene_tx=XM_006944108.4;refGene_gn=BCAP31;refGene_tx=XM_011279561.2;refGene_gn=BCAP31;refGene_tx=XM_023249499.1
X	128679788	128679889
refGene_gn=BCAP31;refGene_tx=XM_006944108.4;refGene_gn=BCAP31;refGene_tx=XM_011279561.2;refGene_gn=BCAP31;refGene_tx=XM_023249499.1
X	128680668	128680792
refGene_gn=BCAP31;refGene_tx=XM_006944108.4;refGene_gn=BCAP31;refGene_tx=XM_011279561.2;refGene_gn=BCAP31;refGene_tx=XM_023249499.1
X	128681668	128681804
refGene_gn=BCAP31;refGene_tx=XM_006944108.4;refGene_gn=BCAP31;refGene_tx=XM_011279561.2;refGene_gn=BCAP31;refGene_tx=XM_023249499.1
X	128699014	128699162
refGene_gn=BCAP31;refGene_tx=XM_006944108.4;refGene_gn=BCAP31;refGene_tx=XM_011279561.2;refGene_gn=BCAP31;refGene_tx=XM_023249499.1
X	128704036	128704137
refGene_gn=BCAP31;refGene_tx=XM_006944108.4;refGene_gn=BCAP31;refGene_tx=XM_011279561.2;refGene_gn=BCAP31;refGene_tx=XM_023249499.1
X	128705665	128705757
refGene_gn=BCAP31;refGene_tx=XM_006944108.4;refGene_gn=BCAP31;refGene_tx=XM_011279561.2;refGene_gn=BCAP31;refGene_tx=XM_023249499.1
X	128707728	128708628
refGene_gn=ABCD1;refGene_tx=XM_004001038.4
X	128711296	128711477
refGene_gn=ABCD1;refGene_tx=XM_004001038.4
X	128719215	128719358
refGene_gn=ABCD1;refGene_tx=XM_004001038.4
X	128719453	128719622

refGene_gn=ABCD1;refGene_tx=XM_004001038.4
X	128720229	128720324
refGene_gn=ABCD1;refGene_tx=XM_004001038.4
X	128723077	128723223
refGene_gn=ABCD1;refGene_tx=XM_004001038.4
X	128723805	128723951
refGene_gn=ABCD1;refGene_tx=XM_004001038.4
X	128725820	128725905
refGene_gn=ABCD1;refGene_tx=XM_004001038.4
X	128726160	128726286
refGene_gn=ABCD1;refGene_tx=XM_004001038.4
X	128726414	128726625
refGene_gn=ABCD1;refGene_tx=XM_004001038.4
X	128740181	128740301
refGene_gn=PLXNB3;refGene_tx=XM_023249473.1;refGene_gn=PLXNB3;refGene_tx=XM_023249474.1;refGene_gn=PLXNB3;refGene_tx=XM_023249475.1;refGene_gn=PLXNB3;refGene_tx=XM_023249476.1;refGene_gn=PLXNB3;refGene_tx=XM_023249477.1;refGene_gn=PLXNB3;refGene_tx=XM_023249478.1;refGene_gn=PLXNB3;refGene_tx=XM_023249480.1;refGene_gn=PLXNB3;refGene_tx=XM_023249481.1
X	128742070	128743111
refGene_gn=PLXNB3;refGene_tx=XM_023249473.1;refGene_gn=PLXNB3;refGene_tx=XM_023249474.1;refGene_gn=PLXNB3;refGene_tx=XM_023249475.1;refGene_gn=PLXNB3;refGene_tx=XM_023249476.1;refGene_gn=PLXNB3;refGene_tx=XM_023249477.1;refGene_gn=PLXNB3;refGene_tx=XM_023249478.1;refGene_gn=PLXNB3;refGene_tx=XM_023249479.1;refGene_gn=PLXNB3;refGene_tx=XM_023249480.1;refGene_gn=PLXNB3;refGene_tx=XM_023249481.1
X	128743502	128743682
refGene_gn=PLXNB3;refGene_tx=XM_023249473.1;refGene_gn=PLXNB3;refGene_tx=XM_023249474.1;refGene_gn=PLXNB3;refGene_tx=XM_023249475.1;refGene_gn=PLXNB3;refGene_tx=XM_023249476.1;refGene_gn=PLXNB3;refGene_tx=XM_023249477.1;refGene_gn=PLXNB3;refGene_tx=XM_023249478.1;refGene_gn=PLXNB3;refGene_tx=XM_023249479.1;refGene_gn=PLXNB3;refGene_tx=XM_023249480.1;refGene_gn=PLXNB3;refGene_tx=XM_023249481.1
X	128744367	128744496
refGene_gn=PLXNB3;refGene_tx=XM_023249473.1;refGene_gn=PLXNB3;refGene_tx=XM_023249474.1;refGene_gn=PLXNB3;refGene_tx=XM_023249475.1;refGene_gn=PLXNB3;refGene_tx=XM_023249476.1;refGene_gn=PLXNB3;refGene_tx=XM_023249477.1;refGene_gn=PLXNB3;refGene_tx=XM_023249478.1;refGene_gn=PLXNB3;refGene_tx=XM_023249479.1;refGene_gn=PLXNB3;refGene_tx=XM_023249480.1;refGene_gn=PLXNB3;refGene_tx=XM_023249481.1
X	128744578	128744679
refGene_gn=PLXNB3;refGene_tx=XM_023249473.1;refGene_gn=PLXNB3;refGene_tx=XM_023249474.1;refGene_gn=PLXNB3;refGene_tx=XM_023249475.1;refGene_gn=PLXNB3;refGene_tx=XM_023249476.1;refGene_gn=PLXNB3;refGene_tx=XM_023249477.1;refGene_gn=PLXNB3;refGene_tx=XM_023249478.1;refGene_gn=PLXNB3;refGene_tx=XM_023249479.1;refGene_gn=PLXNB3;refGene_tx=XM_023249480.1;refGene_gn=PLXNB3;refGene_tx=XM_023249481.1
X	128745227	128745357
refGene_gn=PLXNB3;refGene_tx=XM_023249473.1;refGene_gn=PLXNB3;refGene_

tx=XM_023249474.1;refGene_gn=PLXNB3;refGene_tx=XM_023249475.1;refGene_gn=PLXNB3;refGene_tx=XM_023249476.1;refGene_gn=PLXNB3;refGene_tx=XM_023249477.1;refGene_gn=PLXNB3;refGene_tx=XM_023249478.1;refGene_gn=PLXNB3;refGene_tx=XM_023249479.1;refGene_gn=PLXNB3;refGene_tx=XM_023249480.1;refGene_gn=PLXNB3;refGene_tx=XM_023249481.1
X	128745559	128745716	refGene_gn=PLXNB3;refGene_tx=XM_023249473.1;refGene_gn=PLXNB3;refGene_tx=XM_023249474.1;refGene_gn=PLXNB3;refGene_tx=XM_023249475.1;refGene_gn=PLXNB3;refGene_tx=XM_023249476.1;refGene_gn=PLXNB3;refGene_tx=XM_023249477.1;refGene_gn=PLXNB3;refGene_tx=XM_023249478.1;refGene_gn=PLXNB3;refGene_tx=XM_023249479.1;refGene_gn=PLXNB3;refGene_tx=XM_023249480.1;refGene_gn=PLXNB3;refGene_tx=XM_023249481.1
X	128745804	128745913	refGene_gn=PLXNB3;refGene_tx=XM_023249473.1;refGene_gn=PLXNB3;refGene_tx=XM_023249474.1;refGene_gn=PLXNB3;refGene_tx=XM_023249475.1;refGene_gn=PLXNB3;refGene_tx=XM_023249476.1;refGene_gn=PLXNB3;refGene_tx=XM_023249477.1;refGene_gn=PLXNB3;refGene_tx=XM_023249478.1;refGene_gn=PLXNB3;refGene_tx=XM_023249479.1;refGene_gn=PLXNB3;refGene_tx=XM_023249480.1;refGene_gn=PLXNB3;refGene_tx=XM_023249481.1
X	128746000	128746115	refGene_gn=PLXNB3;refGene_tx=XM_023249473.1;refGene_gn=PLXNB3;refGene_tx=XM_023249474.1;refGene_gn=PLXNB3;refGene_tx=XM_023249475.1;refGene_gn=PLXNB3;refGene_tx=XM_023249476.1;refGene_gn=PLXNB3;refGene_tx=XM_023249477.1;refGene_gn=PLXNB3;refGene_tx=XM_023249478.1;refGene_gn=PLXNB3;refGene_tx=XM_023249479.1;refGene_gn=PLXNB3;refGene_tx=XM_023249480.1;refGene_gn=PLXNB3;refGene_tx=XM_023249481.1
X	128746199	128746325	refGene_gn=PLXNB3;refGene_tx=XM_023249473.1;refGene_gn=PLXNB3;refGene_tx=XM_023249474.1;refGene_gn=PLXNB3;refGene_tx=XM_023249475.1;refGene_gn=PLXNB3;refGene_tx=XM_023249476.1;refGene_gn=PLXNB3;refGene_tx=XM_023249477.1;refGene_gn=PLXNB3;refGene_tx=XM_023249478.1;refGene_gn=PLXNB3;refGene_tx=XM_023249479.1;refGene_gn=PLXNB3;refGene_tx=XM_023249480.1;refGene_gn=PLXNB3;refGene_tx=XM_023249481.1
X	128746393	128746510	refGene_gn=PLXNB3;refGene_tx=XM_023249473.1;refGene_gn=PLXNB3;refGene_tx=XM_023249474.1;refGene_gn=PLXNB3;refGene_tx=XM_023249475.1;refGene_gn=PLXNB3;refGene_tx=XM_023249476.1;refGene_gn=PLXNB3;refGene_tx=XM_023249477.1;refGene_gn=PLXNB3;refGene_tx=XM_023249478.1;refGene_gn=PLXNB3;refGene_tx=XM_023249479.1;refGene_gn=PLXNB3;refGene_tx=XM_023249480.1;refGene_gn=PLXNB3;refGene_tx=XM_023249481.1
X	128746819	128746913	refGene_gn=PLXNB3;refGene_tx=XM_023249473.1;refGene_gn=PLXNB3;refGene_tx=XM_023249474.1;refGene_gn=PLXNB3;refGene_tx=XM_023249475.1;refGene_gn=PLXNB3;refGene_tx=XM_023249476.1;refGene_gn=PLXNB3;refGene_tx=XM_023249477.1;refGene_gn=PLXNB3;refGene_tx=XM_023249478.1;refGene_gn=PLXNB3;refGene_tx=XM_023249479.1;refGene_gn=PLXNB3;refGene_tx=XM_023249480.1;refGene_gn=PLXNB3;refGene_tx=XM_023249481.1
X	128746996	128747166	refGene_gn=PLXNB3;refGene_tx=XM_023249473.1;refGene_gn=PLXNB3;refGene_tx=XM_023249474.1;refGene_gn=PLXNB3;refGene_tx=XM_023249475.1;refGene_

gn=PLXNB3;refGene_tx=XM_023249476.1;refGene_gn=PLXNB3;refGene_tx=XM_023249477.1;refGene_gn=PLXNB3;refGene_tx=XM_023249478.1;refGene_gn=PLXNB3;refGene_tx=XM_023249479.1;refGene_gn=PLXNB3;refGene_tx=XM_023249480.1;refGene_gn=PLXNB3;refGene_tx=XM_023249481.1
X	128747402	128747554
refGene_gn=PLXNB3;refGene_tx=XM_023249473.1;refGene_gn=PLXNB3;refGene_tx=XM_023249474.1;refGene_gn=PLXNB3;refGene_tx=XM_023249475.1;refGene_gn=PLXNB3;refGene_tx=XM_023249476.1;refGene_gn=PLXNB3;refGene_tx=XM_023249477.1;refGene_gn=PLXNB3;refGene_tx=XM_023249478.1;refGene_gn=PLXNB3;refGene_tx=XM_023249479.1;refGene_gn=PLXNB3;refGene_tx=XM_023249480.1;refGene_gn=PLXNB3;refGene_tx=XM_023249481.1
X	128747814	128747920
refGene_gn=PLXNB3;refGene_tx=XM_023249473.1;refGene_gn=PLXNB3;refGene_tx=XM_023249474.1;refGene_gn=PLXNB3;refGene_tx=XM_023249475.1;refGene_gn=PLXNB3;refGene_tx=XM_023249476.1;refGene_gn=PLXNB3;refGene_tx=XM_023249477.1;refGene_gn=PLXNB3;refGene_tx=XM_023249478.1;refGene_gn=PLXNB3;refGene_tx=XM_023249479.1;refGene_gn=PLXNB3;refGene_tx=XM_023249480.1;refGene_gn=PLXNB3;refGene_tx=XM_023249481.1
X	128748619	128748750
refGene_gn=PLXNB3;refGene_tx=XM_023249473.1;refGene_gn=PLXNB3;refGene_tx=XM_023249474.1;refGene_gn=PLXNB3;refGene_tx=XM_023249475.1;refGene_gn=PLXNB3;refGene_tx=XM_023249476.1;refGene_gn=PLXNB3;refGene_tx=XM_023249477.1;refGene_gn=PLXNB3;refGene_tx=XM_023249478.1;refGene_gn=PLXNB3;refGene_tx=XM_023249479.1;refGene_gn=PLXNB3;refGene_tx=XM_023249480.1;refGene_gn=PLXNB3;refGene_tx=XM_023249481.1
X	128749007	128749184
refGene_gn=PLXNB3;refGene_tx=XM_023249473.1;refGene_gn=PLXNB3;refGene_tx=XM_023249474.1;refGene_gn=PLXNB3;refGene_tx=XM_023249475.1;refGene_gn=PLXNB3;refGene_tx=XM_023249476.1;refGene_gn=PLXNB3;refGene_tx=XM_023249477.1;refGene_gn=PLXNB3;refGene_tx=XM_023249478.1;refGene_gn=PLXNB3;refGene_tx=XM_023249479.1;refGene_gn=PLXNB3;refGene_tx=XM_023249480.1;refGene_gn=PLXNB3;refGene_tx=XM_023249481.1
X	128749293	128749489
refGene_gn=PLXNB3;refGene_tx=XM_023249473.1;refGene_gn=PLXNB3;refGene_tx=XM_023249474.1;refGene_gn=PLXNB3;refGene_tx=XM_023249475.1;refGene_gn=PLXNB3;refGene_tx=XM_023249476.1;refGene_gn=PLXNB3;refGene_tx=XM_023249477.1;refGene_gn=PLXNB3;refGene_tx=XM_023249478.1;refGene_gn=PLXNB3;refGene_tx=XM_023249479.1;refGene_gn=PLXNB3;refGene_tx=XM_023249480.1;refGene_gn=PLXNB3;refGene_tx=XM_023249481.1
X	128749629	128749869
refGene_gn=PLXNB3;refGene_tx=XM_023249473.1;refGene_gn=PLXNB3;refGene_tx=XM_023249474.1;refGene_gn=PLXNB3;refGene_tx=XM_023249475.1;refGene_gn=PLXNB3;refGene_tx=XM_023249476.1;refGene_gn=PLXNB3;refGene_tx=XM_023249477.1;refGene_gn=PLXNB3;refGene_tx=XM_023249478.1;refGene_gn=PLXNB3;refGene_tx=XM_023249479.1;refGene_gn=PLXNB3;refGene_tx=XM_023249480.1;refGene_gn=PLXNB3;refGene_tx=XM_023249481.1
X	128749963	128750125
refGene_gn=PLXNB3;refGene_tx=XM_023249473.1;refGene_gn=PLXNB3;refGene_tx=XM_023249474.1;refGene_gn=PLXNB3;refGene_tx=XM_023249475.1;refGene_gn=PLXNB3;refGene_tx=XM_023249476.1;refGene_gn=PLXNB3;refGene_tx=XM_02

3249477.1;refGene_gn=PLXNB3;refGene_tx=XM_023249478.1;refGene_gn=PLXNB3;refGene_tx=XM_023249479.1;refGene_gn=PLXNB3;refGene_tx=XM_023249480.1;refGene_gn=PLXNB3;refGene_tx=XM_023249481.1
X 128750205 128750357
refGene_gn=PLXNB3;refGene_tx=XM_023249473.1;refGene_gn=PLXNB3;refGene_tx=XM_023249474.1;refGene_gn=PLXNB3;refGene_tx=XM_023249475.1;refGene_gn=PLXNB3;refGene_tx=XM_023249476.1;refGene_gn=PLXNB3;refGene_tx=XM_023249477.1;refGene_gn=PLXNB3;refGene_tx=XM_023249478.1;refGene_gn=PLXNB3;refGene_tx=XM_023249479.1;refGene_gn=PLXNB3;refGene_tx=XM_023249480.1;refGene_gn=PLXNB3;refGene_tx=XM_023249481.1
X 128750520 128750621
refGene_gn=PLXNB3;refGene_tx=XM_023249473.1;refGene_gn=PLXNB3;refGene_tx=XM_023249474.1;refGene_gn=PLXNB3;refGene_tx=XM_023249475.1;refGene_gn=PLXNB3;refGene_tx=XM_023249476.1;refGene_gn=PLXNB3;refGene_tx=XM_023249477.1;refGene_gn=PLXNB3;refGene_tx=XM_023249478.1;refGene_gn=PLXNB3;refGene_tx=XM_023249479.1;refGene_gn=PLXNB3;refGene_tx=XM_023249480.1;refGene_gn=PLXNB3;refGene_tx=XM_023249481.1
X 128750706 128750930
refGene_gn=PLXNB3;refGene_tx=XM_023249473.1;refGene_gn=PLXNB3;refGene_tx=XM_023249474.1;refGene_gn=PLXNB3;refGene_tx=XM_023249475.1;refGene_gn=PLXNB3;refGene_tx=XM_023249476.1;refGene_gn=PLXNB3;refGene_tx=XM_023249477.1;refGene_gn=PLXNB3;refGene_tx=XM_023249478.1;refGene_gn=PLXNB3;refGene_tx=XM_023249479.1;refGene_gn=PLXNB3;refGene_tx=XM_023249480.1;refGene_gn=PLXNB3;refGene_tx=XM_023249481.1
X 128751071 128751250
refGene_gn=PLXNB3;refGene_tx=XM_023249473.1;refGene_gn=PLXNB3;refGene_tx=XM_023249474.1;refGene_gn=PLXNB3;refGene_tx=XM_023249475.1;refGene_gn=PLXNB3;refGene_tx=XM_023249476.1;refGene_gn=PLXNB3;refGene_tx=XM_023249477.1;refGene_gn=PLXNB3;refGene_tx=XM_023249478.1;refGene_gn=PLXNB3;refGene_tx=XM_023249479.1;refGene_gn=PLXNB3;refGene_tx=XM_023249480.1;refGene_gn=PLXNB3;refGene_tx=XM_023249481.1
X 128751546 128751613
refGene_gn=PLXNB3;refGene_tx=XM_023249473.1;refGene_gn=PLXNB3;refGene_tx=XM_023249474.1;refGene_gn=PLXNB3;refGene_tx=XM_023249475.1;refGene_gn=PLXNB3;refGene_tx=XM_023249476.1;refGene_gn=PLXNB3;refGene_tx=XM_023249477.1;refGene_gn=PLXNB3;refGene_tx=XM_023249478.1;refGene_gn=PLXNB3;refGene_tx=XM_023249479.1;refGene_gn=PLXNB3;refGene_tx=XM_023249480.1;refGene_gn=PLXNB3;refGene_tx=XM_023249481.1
X 128751796 128752124
refGene_gn=PLXNB3;refGene_tx=XM_023249473.1;refGene_gn=PLXNB3;refGene_tx=XM_023249474.1;refGene_gn=PLXNB3;refGene_tx=XM_023249475.1;refGene_gn=PLXNB3;refGene_tx=XM_023249476.1;refGene_gn=PLXNB3;refGene_tx=XM_023249477.1;refGene_gn=PLXNB3;refGene_tx=XM_023249478.1;refGene_gn=PLXNB3;refGene_tx=XM_023249479.1;refGene_gn=PLXNB3;refGene_tx=XM_023249480.1;refGene_gn=PLXNB3;refGene_tx=XM_023249481.1
X 128752259 128752363
refGene_gn=PLXNB3;refGene_tx=XM_023249473.1;refGene_gn=PLXNB3;refGene_tx=XM_023249474.1;refGene_gn=PLXNB3;refGene_tx=XM_023249475.1;refGene_gn=PLXNB3;refGene_tx=XM_023249476.1;refGene_gn=PLXNB3;refGene_tx=XM_023249477.1;refGene_gn=PLXNB3;refGene_tx=XM_023249478.1;refGene_gn=PLXNB

3;refGene_tx=XM_023249479.1;refGene_gn=PLXNB3;refGene_tx=XM_023249480.1;refGene_gn=PLXNB3;refGene_tx=XM_023249481.1
X	128752453	128752547
refGene_gn=PLXNB3;refGene_tx=XM_023249473.1;refGene_gn=PLXNB3;refGene_tx=XM_023249474.1;refGene_gn=PLXNB3;refGene_tx=XM_023249475.1;refGene_gn=PLXNB3;refGene_tx=XM_023249476.1;refGene_gn=PLXNB3;refGene_tx=XM_023249477.1;refGene_gn=PLXNB3;refGene_tx=XM_023249478.1;refGene_gn=PLXNB3;refGene_tx=XM_023249479.1;refGene_gn=PLXNB3;refGene_tx=XM_023249480.1
X	128752801	128752854
refGene_gn=PLXNB3;refGene_tx=XM_023249480.1
X	128752876	128753055
refGene_gn=PLXNB3;refGene_tx=XM_023249473.1;refGene_gn=PLXNB3;refGene_tx=XM_023249474.1;refGene_gn=PLXNB3;refGene_tx=XM_023249475.1;refGene_gn=PLXNB3;refGene_tx=XM_023249476.1;refGene_gn=PLXNB3;refGene_tx=XM_023249477.1;refGene_gn=PLXNB3;refGene_tx=XM_023249478.1;refGene_gn=PLXNB3;refGene_tx=XM_023249479.1;refGene_gn=PLXNB3;refGene_tx=XM_023249481.1
X	128753244	128753405
refGene_gn=PLXNB3;refGene_tx=XM_023249473.1;refGene_gn=PLXNB3;refGene_tx=XM_023249474.1;refGene_gn=PLXNB3;refGene_tx=XM_023249475.1;refGene_gn=PLXNB3;refGene_tx=XM_023249476.1;refGene_gn=PLXNB3;refGene_tx=XM_023249477.1;refGene_gn=PLXNB3;refGene_tx=XM_023249478.1;refGene_gn=PLXNB3;refGene_tx=XM_023249479.1
X	128753631	128753779
refGene_gn=PLXNB3;refGene_tx=XM_023249473.1;refGene_gn=PLXNB3;refGene_tx=XM_023249474.1;refGene_gn=PLXNB3;refGene_tx=XM_023249475.1;refGene_gn=PLXNB3;refGene_tx=XM_023249476.1;refGene_gn=PLXNB3;refGene_tx=XM_023249477.1;refGene_gn=PLXNB3;refGene_tx=XM_023249478.1;refGene_gn=PLXNB3;refGene_tx=XM_023249479.1
X	128753992	128754078
refGene_gn=PLXNB3;refGene_tx=XM_023249473.1;refGene_gn=PLXNB3;refGene_tx=XM_023249474.1;refGene_gn=PLXNB3;refGene_tx=XM_023249475.1;refGene_gn=PLXNB3;refGene_tx=XM_023249476.1;refGene_gn=PLXNB3;refGene_tx=XM_023249477.1;refGene_gn=PLXNB3;refGene_tx=XM_023249478.1;refGene_gn=PLXNB3;refGene_tx=XM_023249479.1
X	128754152	128754228
refGene_gn=PLXNB3;refGene_tx=XM_023249473.1;refGene_gn=PLXNB3;refGene_tx=XM_023249474.1;refGene_gn=PLXNB3;refGene_tx=XM_023249475.1;refGene_gn=PLXNB3;refGene_tx=XM_023249476.1;refGene_gn=PLXNB3;refGene_tx=XM_023249477.1;refGene_gn=PLXNB3;refGene_tx=XM_023249478.1;refGene_gn=PLXNB3;refGene_tx=XM_023249479.1
X	128754344	128754419
refGene_gn=PLXNB3;refGene_tx=XM_023249473.1;refGene_gn=PLXNB3;refGene_tx=XM_023249474.1;refGene_gn=PLXNB3;refGene_tx=XM_023249475.1;refGene_gn=PLXNB3;refGene_tx=XM_023249476.1;refGene_gn=PLXNB3;refGene_tx=XM_023249477.1;refGene_gn=PLXNB3;refGene_tx=XM_023249478.1;refGene_gn=PLXNB3;refGene_tx=XM_023249479.1
X	128754735	128754840
refGene_gn=PLXNB3;refGene_tx=XM_023249473.1;refGene_gn=PLXNB3;refGene_

tx=XM_023249474.1;refGene_gn=PLXNB3;refGene_tx=XM_023249475.1;refGene_gn=PLXNB3;refGene_tx=XM_023249476.1;refGene_gn=PLXNB3;refGene_tx=XM_023249477.1;refGene_gn=PLXNB3;refGene_tx=XM_023249478.1;refGene_gn=PLXNB3;refGene_tx=XM_023249479.1
X 128756657 128756713
refGene_gn=SRPK3;refGene_tx=XM_023248827.1;refGene_gn=SRPK3;refGene_tx=XM_023248828.1;refGene_gn=SRPK3;refGene_tx=XM_023248829.1
X 128756799 128756930
refGene_gn=SRPK3;refGene_tx=XM_023248827.1;refGene_gn=SRPK3;refGene_tx=XM_023248828.1;refGene_gn=SRPK3;refGene_tx=XM_023248829.1
X 128757114 128757223
refGene_gn=SRPK3;refGene_tx=XM_023248827.1;refGene_gn=SRPK3;refGene_tx=XM_023248828.1;refGene_gn=SRPK3;refGene_tx=XM_023248829.1
X 128757359 128757447
refGene_gn=SRPK3;refGene_tx=XM_023248827.1;refGene_gn=SRPK3;refGene_tx=XM_023248828.1;refGene_gn=SRPK3;refGene_tx=XM_023248829.1
X 128757649 128757737
refGene_gn=SRPK3;refGene_tx=XM_023248827.1;refGene_gn=SRPK3;refGene_tx=XM_023248828.1;refGene_gn=SRPK3;refGene_tx=XM_023248829.1
X 128758257 128758364
refGene_gn=SRPK3;refGene_tx=XM_023248827.1;refGene_gn=SRPK3;refGene_tx=XM_023248828.1;refGene_gn=SRPK3;refGene_tx=XM_023248829.1
X 128758443 128758609
refGene_gn=SRPK3;refGene_tx=XM_023248827.1;refGene_gn=SRPK3;refGene_tx=XM_023248828.1;refGene_gn=SRPK3;refGene_tx=XM_023248829.1
X 128758716 128758742
refGene_gn=SRPK3;refGene_tx=XM_023248827.1;refGene_gn=SRPK3;refGene_tx=XM_023248828.1;refGene_gn=SRPK3;refGene_tx=XM_023248829.1
X 128759209 128759339
refGene_gn=SRPK3;refGene_tx=XM_023248827.1;refGene_gn=SRPK3;refGene_tx=XM_023248828.1;refGene_gn=SRPK3;refGene_tx=XM_023248829.1
X 128759419 128759662
refGene_gn=SRPK3;refGene_tx=XM_023248827.1;refGene_gn=SRPK3;refGene_tx=XM_023248828.1;refGene_gn=SRPK3;refGene_tx=XM_023248829.1
X 128759755 128759856
refGene_gn=SRPK3;refGene_tx=XM_023248827.1;refGene_gn=SRPK3;refGene_tx=XM_023248828.1;refGene_gn=SRPK3;refGene_tx=XM_023248829.1
X 128760226 128760334
refGene_gn=SRPK3;refGene_tx=XM_023248827.1;refGene_gn=SRPK3;refGene_tx=XM_023248828.1;refGene_gn=SRPK3;refGene_tx=XM_023248829.1
X 128760416 128760659
refGene_gn=SRPK3;refGene_tx=XM_023248827.1;refGene_gn=SRPK3;refGene_tx=XM_023248828.1;refGene_gn=SRPK3;refGene_tx=XM_023248829.1
X 128760801 128760986
refGene_gn=SRPK3;refGene_tx=XM_023248827.1;refGene_gn=SRPK3;refGene_tx=XM_023248828.1;refGene_gn=SRPK3;refGene_tx=XM_023248829.1
X 128761399 128761501
refGene_gn=IDH3G;refGene_tx=XM_011279558.2
X 128761739 128761800
refGene_gn=IDH3G;refGene_tx=XM_011279558.2

X	128761884	128761979
refGene_gn=IDH3G;refGene_tx=XM_011279558.2
X	128763126	128763273
refGene_gn=IDH3G;refGene_tx=XM_011279558.2
X	128763367	128763470
refGene_gn=IDH3G;refGene_tx=XM_011279558.2
X	128763804	128763938
refGene_gn=IDH3G;refGene_tx=XM_011279558.2
X	128764054	128764187
refGene_gn=IDH3G;refGene_tx=XM_011279558.2
X	128764285	128764346
refGene_gn=IDH3G;refGene_tx=XM_011279558.2
X	128765580	128765693
refGene_gn=IDH3G;refGene_tx=XM_011279558.2
X	128766124	128766222
refGene_gn=IDH3G;refGene_tx=XM_011279558.2
X	128766464	128766476
refGene_gn=IDH3G;refGene_tx=XM_011279558.2
X	128766777	128766816
refGene_gn=IDH3G;refGene_tx=XM_011279558.2
X	128772059	128772140
refGene_gn=IDH3G;refGene_tx=XM_011279558.2
X	128772546	128772613
refGene_gn=SSR4;refGene_tx=XM_004001008.4;refGene_gn=SSR4;refGene_tx=XM_006944109.3
X	128774490	128774609
refGene_gn=SSR4;refGene_tx=XM_004001008.4;refGene_gn=SSR4;refGene_tx=XM_006944109.3
X	128775357	128775432
refGene_gn=SSR4;refGene_tx=XM_004001008.4;refGene_gn=SSR4;refGene_tx=XM_006944109.3
X	128775632	128775722
refGene_gn=SSR4;refGene_tx=XM_004001008.4;refGene_gn=SSR4;refGene_tx=XM_006944109.3
X	128776043	128776109
refGene_gn=SSR4;refGene_tx=XM_004001008.4;refGene_gn=SSR4;refGene_tx=XM_006944109.3
X	128776301	128776406
refGene_gn=SSR4;refGene_tx=XM_004001008.4;refGene_gn=SSR4;refGene_tx=XM_006944109.3
X	128780351	128781872
refGene_gn=PDZD4;refGene_tx=XM_023249056.1;refGene_gn=PDZD4;refGene_tx=XM_023249057.1;refGene_gn=PDZD4;refGene_tx=XM_023249058.1;refGene_gn=PDZD4;refGene_tx=XM_023249059.1
X	128782035	128782146
refGene_gn=PDZD4;refGene_tx=XM_023249056.1;refGene_gn=PDZD4;refGene_tx=XM_023249057.1;refGene_gn=PDZD4;refGene_tx=XM_023249058.1;refGene_gn=PDZD4;refGene_tx=XM_023249059.1
X	128782423	128782525
refGene_gn=PDZD4;refGene_tx=XM_023249056.1;refGene_gn=PDZD4;refGene_tx

=XM_023249057.1;refGene_gn=PDZD4;refGene_tx=XM_023249058.1;refGene_gn=PDZD4;refGene_tx=XM_023249059.1
X 128783008 128783071
refGene_gn=PDZD4;refGene_tx=XM_023249056.1;refGene_gn=PDZD4;refGene_tx=XM_023249057.1;refGene_gn=PDZD4;refGene_tx=XM_023249058.1;refGene_gn=PDZD4;refGene_tx=XM_023249059.1
X 128784032 128784131
refGene_gn=PDZD4;refGene_tx=XM_023249056.1;refGene_gn=PDZD4;refGene_tx=XM_023249057.1;refGene_gn=PDZD4;refGene_tx=XM_023249058.1;refGene_gn=PDZD4;refGene_tx=XM_023249059.1
X 128784643 128784734
refGene_gn=PDZD4;refGene_tx=XM_023249056.1;refGene_gn=PDZD4;refGene_tx=XM_023249057.1;refGene_gn=PDZD4;refGene_tx=XM_023249058.1
X 128785670 128785924
refGene_gn=PDZD4;refGene_tx=XM_023249056.1;refGene_gn=PDZD4;refGene_tx=XM_023249057.1;refGene_gn=PDZD4;refGene_tx=XM_023249058.1
X 128811202 128811262
refGene_gn=PDZD4;refGene_tx=XM_023249056.1;refGene_gn=PDZD4;refGene_tx=XM_023249057.1;refGene_gn=PDZD4;refGene_tx=XM_023249059.1
X 128862572 128862804
refGene_gn=L1CAM;refGene_tx=XM_023248891.1;refGene_gn=L1CAM;refGene_tx=XM_023248892.1;refGene_gn=L1CAM;refGene_tx=XM_023248893.1;refGene_gn=L1CAM;refGene_tx=XM_023248894.1
X 128863357 128863369
refGene_gn=L1CAM;refGene_tx=XM_023248891.1;refGene_gn=L1CAM;refGene_tx=XM_023248893.1
X 128863474 128863547
refGene_gn=L1CAM;refGene_tx=XM_023248891.1;refGene_gn=L1CAM;refGene_tx=XM_023248892.1;refGene_gn=L1CAM;refGene_tx=XM_023248893.1;refGene_gn=L1CAM;refGene_tx=XM_023248894.1
X 128863888 128864023
refGene_gn=L1CAM;refGene_tx=XM_023248891.1;refGene_gn=L1CAM;refGene_tx=XM_023248892.1;refGene_gn=L1CAM;refGene_tx=XM_023248893.1;refGene_gn=L1CAM;refGene_tx=XM_023248894.1
X 128864276 128864429
refGene_gn=L1CAM;refGene_tx=XM_023248891.1;refGene_gn=L1CAM;refGene_tx=XM_023248892.1;refGene_gn=L1CAM;refGene_tx=XM_023248893.1;refGene_gn=L1CAM;refGene_tx=XM_023248894.1
X 128864518 128864638
refGene_gn=L1CAM;refGene_tx=XM_023248891.1;refGene_gn=L1CAM;refGene_tx=XM_023248892.1;refGene_gn=L1CAM;refGene_tx=XM_023248893.1;refGene_gn=L1CAM;refGene_tx=XM_023248894.1
X 128864738 128864912
refGene_gn=L1CAM;refGene_tx=XM_023248891.1;refGene_gn=L1CAM;refGene_tx=XM_023248892.1;refGene_gn=L1CAM;refGene_tx=XM_023248893.1;refGene_gn=L1CAM;refGene_tx=XM_023248894.1
X 128865012 128865135
refGene_gn=L1CAM;refGene_tx=XM_023248891.1;refGene_gn=L1CAM;refGene_tx=XM_023248892.1;refGene_gn=L1CAM;refGene_tx=XM_023248893.1;refGene_gn=L1CAM;refGene_tx=XM_023248894.1

X 128865222 128865424
refGene_gn=L1CAM;refGene_tx=XM_023248891.1;refGene_gn=L1CAM;refGene_tx=XM_023248892.1;refGene_gn=L1CAM;refGene_tx=XM_023248893.1;refGene_gn=L1CAM;refGene_tx=XM_023248894.1
X 128865648 128865764
refGene_gn=L1CAM;refGene_tx=XM_023248891.1;refGene_gn=L1CAM;refGene_tx=XM_023248892.1;refGene_gn=L1CAM;refGene_tx=XM_023248893.1;refGene_gn=L1CAM;refGene_tx=XM_023248894.1
X 128866348 128866571
refGene_gn=L1CAM;refGene_tx=XM_023248891.1;refGene_gn=L1CAM;refGene_tx=XM_023248892.1;refGene_gn=L1CAM;refGene_tx=XM_023248893.1;refGene_gn=L1CAM;refGene_tx=XM_023248894.1
X 128866772 128866843
refGene_gn=L1CAM;refGene_tx=XM_023248891.1;refGene_gn=L1CAM;refGene_tx=XM_023248892.1;refGene_gn=L1CAM;refGene_tx=XM_023248893.1;refGene_gn=L1CAM;refGene_tx=XM_023248894.1
X 128867093 128867291
refGene_gn=L1CAM;refGene_tx=XM_023248891.1;refGene_gn=L1CAM;refGene_tx=XM_023248892.1;refGene_gn=L1CAM;refGene_tx=XM_023248893.1;refGene_gn=L1CAM;refGene_tx=XM_023248894.1
X 128867567 128867678
refGene_gn=L1CAM;refGene_tx=XM_023248891.1;refGene_gn=L1CAM;refGene_tx=XM_023248892.1;refGene_gn=L1CAM;refGene_tx=XM_023248893.1;refGene_gn=L1CAM;refGene_tx=XM_023248894.1
X 128867838 128867963
refGene_gn=L1CAM;refGene_tx=XM_023248891.1;refGene_gn=L1CAM;refGene_tx=XM_023248892.1;refGene_gn=L1CAM;refGene_tx=XM_023248893.1;refGene_gn=L1CAM;refGene_tx=XM_023248894.1
X 128868141 128868298
refGene_gn=L1CAM;refGene_tx=XM_023248891.1;refGene_gn=L1CAM;refGene_tx=XM_023248892.1;refGene_gn=L1CAM;refGene_tx=XM_023248893.1;refGene_gn=L1CAM;refGene_tx=XM_023248894.1
X 128868420 128868587
refGene_gn=L1CAM;refGene_tx=XM_023248891.1;refGene_gn=L1CAM;refGene_tx=XM_023248892.1;refGene_gn=L1CAM;refGene_tx=XM_023248893.1;refGene_gn=L1CAM;refGene_tx=XM_023248894.1
X 128868703 128868815
refGene_gn=L1CAM;refGene_tx=XM_023248891.1;refGene_gn=L1CAM;refGene_tx=XM_023248892.1;refGene_gn=L1CAM;refGene_tx=XM_023248893.1;refGene_gn=L1CAM;refGene_tx=XM_023248894.1
X 128869432 128869576
refGene_gn=L1CAM;refGene_tx=XM_023248891.1;refGene_gn=L1CAM;refGene_tx=XM_023248892.1;refGene_gn=L1CAM;refGene_tx=XM_023248893.1;refGene_gn=L1CAM;refGene_tx=XM_023248894.1
X 128869704 128869836
refGene_gn=L1CAM;refGene_tx=XM_023248891.1;refGene_gn=L1CAM;refGene_tx=XM_023248892.1;refGene_gn=L1CAM;refGene_tx=XM_023248893.1;refGene_gn=L1CAM;refGene_tx=XM_023248894.1
X 128869994 128870179
refGene_gn=L1CAM;refGene_tx=XM_023248891.1;refGene_gn=L1CAM;refGene_tx

=XM_023248892.1;refGene_gn=L1CAM;refGene_tx=XM_023248893.1;refGene_gn=L1CAM;refGene_tx=XM_023248894.1
X 128870330 128870442
refGene_gn=L1CAM;refGene_tx=XM_023248891.1;refGene_gn=L1CAM;refGene_tx=XM_023248892.1;refGene_gn=L1CAM;refGene_tx=XM_023248893.1;refGene_gn=L1CAM;refGene_tx=XM_023248894.1
X 128870728 128870899
refGene_gn=L1CAM;refGene_tx=XM_023248891.1;refGene_gn=L1CAM;refGene_tx=XM_023248892.1;refGene_gn=L1CAM;refGene_tx=XM_023248893.1;refGene_gn=L1CAM;refGene_tx=XM_023248894.1
X 128871005 128871128
refGene_gn=L1CAM;refGene_tx=XM_023248891.1;refGene_gn=L1CAM;refGene_tx=XM_023248892.1;refGene_gn=L1CAM;refGene_tx=XM_023248893.1;refGene_gn=L1CAM;refGene_tx=XM_023248894.1
X 128872109 128872312
refGene_gn=L1CAM;refGene_tx=XM_023248891.1;refGene_gn=L1CAM;refGene_tx=XM_023248892.1;refGene_gn=L1CAM;refGene_tx=XM_023248893.1;refGene_gn=L1CAM;refGene_tx=XM_023248894.1
X 128872556 128872662
refGene_gn=L1CAM;refGene_tx=XM_023248891.1;refGene_gn=L1CAM;refGene_tx=XM_023248892.1;refGene_gn=L1CAM;refGene_tx=XM_023248893.1;refGene_gn=L1CAM;refGene_tx=XM_023248894.1
X 128873241 128873256
refGene_gn=L1CAM;refGene_tx=XM_023248891.1;refGene_gn=L1CAM;refGene_tx=XM_023248892.1
X 128875972 128876048
refGene_gn=L1CAM;refGene_tx=XM_023248891.1;refGene_gn=L1CAM;refGene_tx=XM_023248892.1;refGene_gn=L1CAM;refGene_tx=XM_023248893.1;refGene_gn=L1CAM;refGene_tx=XM_023248894.1
X 128903653 128903738
refGene_gn=AVPR2;refGene_tx=XM_004001011.4;refGene_gn=AVPR2;refGene_tx=XM_006944111.3;refGene_gn=AVPR2;refGene_tx=XM_006944112.4;refGene_gn=AVPR2;refGene_tx=XM_006944113.4
X 128904040 128904925
refGene_gn=AVPR2;refGene_tx=XM_004001011.4;refGene_gn=AVPR2;refGene_tx=XM_006944111.3;refGene_gn=AVPR2;refGene_tx=XM_006944112.4;refGene_gn=AVPR2;refGene_tx=XM_006944113.4
X 128905107 128905313
refGene_gn=AVPR2;refGene_tx=XM_004001011.4;refGene_gn=AVPR2;refGene_tx=XM_006944111.3;refGene_gn=AVPR2;refGene_tx=XM_006944112.4;refGene_gn=AVPR2;refGene_tx=XM_006944113.4
X 128906139 128906370
refGene_gn=ARHGAP4;refGene_tx=XM_004001041.5;refGene_gn=ARHGAP4;refGene_tx=XM_023248873.1
X 128906766 128906881
refGene_gn=ARHGAP4;refGene_tx=XM_004001041.5;refGene_gn=ARHGAP4;refGene_tx=XM_023248873.1
X 128907169 128907262
refGene_gn=ARHGAP4;refGene_tx=XM_004001041.5;refGene_gn=ARHGAP4;refGene_tx=XM_023248873.1

X	128907397	128907581	refGene_gn=ARHGAP4;refGene_tx=XM_004001041.5;refGene_gn=ARHGAP4;refGene_tx=XM_023248873.1
X	128907671	128907745	refGene_gn=ARHGAP4;refGene_tx=XM_004001041.5;refGene_gn=ARHGAP4;refGene_tx=XM_023248873.1
X	128907826	128908060	refGene_gn=ARHGAP4;refGene_tx=XM_004001041.5;refGene_gn=ARHGAP4;refGene_tx=XM_023248873.1
X	128908156	128908262	refGene_gn=ARHGAP4;refGene_tx=XM_004001041.5;refGene_gn=ARHGAP4;refGene_tx=XM_023248873.1
X	128908372	128908507	refGene_gn=ARHGAP4;refGene_tx=XM_004001041.5;refGene_gn=ARHGAP4;refGene_tx=XM_023248873.1
X	128908589	128908667	refGene_gn=ARHGAP4;refGene_tx=XM_004001041.5;refGene_gn=ARHGAP4;refGene_tx=XM_023248873.1
X	128908779	128908840	refGene_gn=ARHGAP4;refGene_tx=XM_004001041.5;refGene_gn=ARHGAP4;refGene_tx=XM_023248873.1
X	128909470	128909573	refGene_gn=ARHGAP4;refGene_tx=XM_004001041.5;refGene_gn=ARHGAP4;refGene_tx=XM_023248873.1
X	128909821	128909849	refGene_gn=ARHGAP4;refGene_tx=XM_004001041.5;refGene_gn=ARHGAP4;refGene_tx=XM_023248873.1
X	128910115	128910200	refGene_gn=ARHGAP4;refGene_tx=XM_004001041.5;refGene_gn=ARHGAP4;refGene_tx=XM_023248873.1
X	128910303	128910495	refGene_gn=ARHGAP4;refGene_tx=XM_004001041.5;refGene_gn=ARHGAP4;refGene_tx=XM_023248873.1
X	128910638	128910740	refGene_gn=ARHGAP4;refGene_tx=XM_004001041.5;refGene_gn=ARHGAP4;refGene_tx=XM_023248873.1
X	128915619	128915841	refGene_gn=ARHGAP4;refGene_tx=XM_004001041.5;refGene_gn=ARHGAP4;refGene_tx=XM_023248873.1
X	128915922	128916051	refGene_gn=ARHGAP4;refGene_tx=XM_004001041.5;refGene_gn=ARHGAP4;refGene_tx=XM_023248873.1
X	128916696	128916879	refGene_gn=ARHGAP4;refGene_tx=XM_004001041.5;refGene_gn=ARHGAP4;refGene_tx=XM_023248873.1
X	128917191	128917254	refGene_gn=ARHGAP4;refGene_tx=XM_004001041.5;refGene_gn=ARHGAP4;refGene_tx=XM_023248873.1
X	128917473	128917636	refGene_gn=ARHGAP4;refGene_tx=XM_004001041.5;refGene_gn=ARHGAP4;refGen

e_tx=XM_023248873.1
X	128917720	128918226
refGene_gn=ARHGAP4;refGene_tx=XM_004001041.5;refGene_gn=ARHGAP4;refGene_tx=XM_023248873.1
X	128921820	128921887
refGene_gn=ARHGAP4;refGene_tx=XM_023248873.1
X	128927106	128927358
refGene_gn=NAA10;refGene_tx=XM_006944114.4;refGene_gn=NAA10;refGene_tx=XM_019824492.2;refGene_gn=NAA10;refGene_tx=XM_023249074.1;refGene_gn=NAA10;refGene_tx=XM_023249075.1
X	128928021	128928106
refGene_gn=NAA10;refGene_tx=XM_006944114.4;refGene_gn=NAA10;refGene_tx=XM_019824492.2;refGene_gn=NAA10;refGene_tx=XM_023249074.1;refGene_gn=NAA10;refGene_tx=XM_023249075.1
X	128929387	128929432
refGene_gn=NAA10;refGene_tx=XM_006944114.4;refGene_gn=NAA10;refGene_tx=XM_019824492.2;refGene_gn=NAA10;refGene_tx=XM_023249074.1;refGene_gn=NAA10;refGene_tx=XM_023249075.1
X	128929626	128929742
refGene_gn=NAA10;refGene_tx=XM_006944114.4;refGene_gn=NAA10;refGene_tx=XM_019824492.2;refGene_gn=NAA10;refGene_tx=XM_023249074.1;refGene_gn=NAA10;refGene_tx=XM_023249075.1
X	128929829	128929875
refGene_gn=NAA10;refGene_tx=XM_006944114.4;refGene_gn=NAA10;refGene_tx=XM_019824492.2;refGene_gn=NAA10;refGene_tx=XM_023249074.1;refGene_gn=NAA10;refGene_tx=XM_023249075.1
X	128930733	128930792
refGene_gn=NAA10;refGene_tx=XM_006944114.4;refGene_gn=NAA10;refGene_tx=XM_019824492.2;refGene_gn=NAA10;refGene_tx=XM_023249074.1;refGene_gn=NAA10;refGene_tx=XM_023249075.1
X	128931087	128931186
refGene_gn=NAA10;refGene_tx=XM_006944114.4;refGene_gn=NAA10;refGene_tx=XM_019824492.2;refGene_gn=NAA10;refGene_tx=XM_023249074.1;refGene_gn=NAA10;refGene_tx=XM_023249075.1
X	128931265	128931349
refGene_gn=NAA10;refGene_tx=XM_006944114.4;refGene_gn=NAA10;refGene_tx=XM_019824492.2
X	128931590	128931611
refGene_gn=NAA10;refGene_tx=XM_023249074.1;refGene_gn=NAA10;refGene_tx=XM_023249075.1
X	128932056	128932190
refGene_gn=RENBP;refGene_tx=XM_006944115.4
X	128932280	128932368
refGene_gn=RENBP;refGene_tx=XM_006944115.4
X	128935024	128935156
refGene_gn=RENBP;refGene_tx=XM_006944115.4
X	128935746	128935922
refGene_gn=RENBP;refGene_tx=XM_006944115.4
X	128936298	128936380
refGene_gn=RENBP;refGene_tx=XM_006944115.4

X 128936953 128937178
refGene_gn=RENBP;refGene_tx=XM_006944115.4
X 128937764 128937937
refGene_gn=RENBP;refGene_tx=XM_006944115.4
X 128938120 128938196
refGene_gn=RENBP;refGene_tx=XM_006944115.4
X 128938357 128938433
refGene_gn=RENBP;refGene_tx=XM_006944115.4
X 128938646 128938762
refGene_gn=RENBP;refGene_tx=XM_006944115.4
X 128938997 128939084
refGene_gn=RENBP;refGene_tx=XM_006944115.4
X 128942975 128943015
refGene_gn=HCFC1;refGene_tx=XM_023249387.1;refGene_gn=HCFC1;refGene_tx=XM_023249388.1;refGene_gn=HCFC1;refGene_tx=XM_023249389.1;refGene_gn=HCFC1;refGene_tx=XM_023249390.1;refGene_gn=HCFC1;refGene_tx=XM_023249391.1
X 128943264 128943328
refGene_gn=HCFC1;refGene_tx=XM_023249387.1;refGene_gn=HCFC1;refGene_tx=XM_023249388.1;refGene_gn=HCFC1;refGene_tx=XM_023249389.1;refGene_gn=HCFC1;refGene_tx=XM_023249390.1;refGene_gn=HCFC1;refGene_tx=XM_023249391.1
X 128943885 128944186
refGene_gn=HCFC1;refGene_tx=XM_023249387.1;refGene_gn=HCFC1;refGene_tx=XM_023249388.1;refGene_gn=HCFC1;refGene_tx=XM_023249389.1;refGene_gn=HCFC1;refGene_tx=XM_023249390.1;refGene_gn=HCFC1;refGene_tx=XM_023249391.1
X 128944852 128945038
refGene_gn=HCFC1;refGene_tx=XM_023249387.1;refGene_gn=HCFC1;refGene_tx=XM_023249388.1;refGene_gn=HCFC1;refGene_tx=XM_023249389.1;refGene_gn=HCFC1;refGene_tx=XM_023249390.1;refGene_gn=HCFC1;refGene_tx=XM_023249391.1
X 128945410 128945548
refGene_gn=HCFC1;refGene_tx=XM_023249387.1;refGene_gn=HCFC1;refGene_tx=XM_023249388.1;refGene_gn=HCFC1;refGene_tx=XM_023249389.1;refGene_gn=HCFC1;refGene_tx=XM_023249390.1;refGene_gn=HCFC1;refGene_tx=XM_023249391.1
X 128945633 128945752
refGene_gn=HCFC1;refGene_tx=XM_023249387.1;refGene_gn=HCFC1;refGene_tx=XM_023249388.1;refGene_gn=HCFC1;refGene_tx=XM_023249389.1;refGene_gn=HCFC1;refGene_tx=XM_023249390.1;refGene_gn=HCFC1;refGene_tx=XM_023249391.1
X 128945866 128946187
refGene_gn=HCFC1;refGene_tx=XM_023249387.1;refGene_gn=HCFC1;refGene_tx=XM_023249388.1;refGene_gn=HCFC1;refGene_tx=XM_023249389.1;refGene_gn=HCFC1;refGene_tx=XM_023249390.1;refGene_gn=HCFC1;refGene_tx=XM_023249391.1
X 128946575 128947152
refGene_gn=HCFC1;refGene_tx=XM_023249387.1;refGene_gn=HCFC1;refGene_tx=XM_023249388.1;refGene_gn=HCFC1;refGene_tx=XM_023249389.1;refGene_gn=

HCFC1;refGene_tx=XM_023249390.1;refGene_gn=HCFC1;refGene_tx=XM_0232493
91.1
X 128947650 128947814
refGene_gn=HCFC1;refGene_tx=XM_023249387.1;refGene_gn=HCFC1;refGene_tx
=XM_023249388.1;refGene_gn=HCFC1;refGene_tx=XM_023249389.1;refGene_gn=
HCFC1;refGene_tx=XM_023249390.1;refGene_gn=HCFC1;refGene_tx=XM_0232493
91.1
X 128948098 128949584
refGene_gn=HCFC1;refGene_tx=XM_023249387.1;refGene_gn=HCFC1;refGene_tx
=XM_023249388.1;refGene_gn=HCFC1;refGene_tx=XM_023249389.1;refGene_gn=
HCFC1;refGene_tx=XM_023249390.1;refGene_gn=HCFC1;refGene_tx=XM_0232493
91.1
X 128950234 128950455
refGene_gn=HCFC1;refGene_tx=XM_023249387.1;refGene_gn=HCFC1;refGene_tx
=XM_023249388.1;refGene_gn=HCFC1;refGene_tx=XM_023249389.1;refGene_gn=
HCFC1;refGene_tx=XM_023249390.1;refGene_gn=HCFC1;refGene_tx=XM_0232493
91.1
X 128950619 128950758
refGene_gn=HCFC1;refGene_tx=XM_023249387.1;refGene_gn=HCFC1;refGene_tx
=XM_023249388.1;refGene_gn=HCFC1;refGene_tx=XM_023249389.1;refGene_gn=
HCFC1;refGene_tx=XM_023249390.1;refGene_gn=HCFC1;refGene_tx=XM_0232493
91.1
X 128950905 128951048
refGene_gn=HCFC1;refGene_tx=XM_023249387.1;refGene_gn=HCFC1;refGene_tx
=XM_023249388.1;refGene_gn=HCFC1;refGene_tx=XM_023249389.1;refGene_gn=
HCFC1;refGene_tx=XM_023249390.1;refGene_gn=HCFC1;refGene_tx=XM_0232493
91.1
X 128951345 128951565
refGene_gn=HCFC1;refGene_tx=XM_023249387.1;refGene_gn=HCFC1;refGene_tx
=XM_023249388.1;refGene_gn=HCFC1;refGene_tx=XM_023249389.1;refGene_gn=
HCFC1;refGene_tx=XM_023249390.1;refGene_gn=HCFC1;refGene_tx=XM_0232493
91.1
X 128952058 128952163
refGene_gn=HCFC1;refGene_tx=XM_023249387.1;refGene_gn=HCFC1;refGene_tx
=XM_023249388.1;refGene_gn=HCFC1;refGene_tx=XM_023249389.1;refGene_gn=
HCFC1;refGene_tx=XM_023249390.1;refGene_gn=HCFC1;refGene_tx=XM_0232493
91.1
X 128952309 128952534
refGene_gn=HCFC1;refGene_tx=XM_023249387.1;refGene_gn=HCFC1;refGene_tx
=XM_023249388.1;refGene_gn=HCFC1;refGene_tx=XM_023249389.1;refGene_gn=
HCFC1;refGene_tx=XM_023249390.1;refGene_gn=HCFC1;refGene_tx=XM_0232493
91.1
X 128952843 128953041
refGene_gn=HCFC1;refGene_tx=XM_023249387.1;refGene_gn=HCFC1;refGene_tx
=XM_023249388.1;refGene_gn=HCFC1;refGene_tx=XM_023249389.1;refGene_gn=
HCFC1;refGene_tx=XM_023249390.1;refGene_gn=HCFC1;refGene_tx=XM_0232493
91.1
X 128953573 128953734
refGene_gn=HCFC1;refGene_tx=XM_023249387.1;refGene_gn=HCFC1;refGene_tx
=XM_023249388.1;refGene_gn=HCFC1;refGene_tx=XM_023249389.1;refGene_gn=

HCFC1;refGene_tx=XM_023249390.1;refGene_gn=HCFC1;refGene_tx=XM_0232493
91.1
X	128954014	128954374
refGene_gn=HCFC1;refGene_tx=XM_023249387.1;refGene_gn=HCFC1;refGene_tx
=XM_023249388.1;refGene_gn=HCFC1;refGene_tx=XM_023249389.1;refGene_gn=
HCFC1;refGene_tx=XM_023249390.1;refGene_gn=HCFC1;refGene_tx=XM_0232493
91.1
X	128954478	128954658
refGene_gn=HCFC1;refGene_tx=XM_023249387.1;refGene_gn=HCFC1;refGene_tx
=XM_023249388.1;refGene_gn=HCFC1;refGene_tx=XM_023249389.1;refGene_gn=
HCFC1;refGene_tx=XM_023249390.1;refGene_gn=HCFC1;refGene_tx=XM_0232493
91.1
X	128955807	128955914
refGene_gn=HCFC1;refGene_tx=XM_023249387.1;refGene_gn=HCFC1;refGene_tx
=XM_023249388.1;refGene_gn=HCFC1;refGene_tx=XM_023249389.1;refGene_gn=
HCFC1;refGene_tx=XM_023249390.1
X	128956418	128956503
refGene_gn=HCFC1;refGene_tx=XM_023249387.1;refGene_gn=HCFC1;refGene_tx
=XM_023249388.1;refGene_gn=HCFC1;refGene_tx=XM_023249389.1;refGene_gn=
HCFC1;refGene_tx=XM_023249390.1
X	128957416	128957625
refGene_gn=HCFC1;refGene_tx=XM_023249387.1;refGene_gn=HCFC1;refGene_tx
=XM_023249388.1;refGene_gn=HCFC1;refGene_tx=XM_023249389.1;refGene_gn=
HCFC1;refGene_tx=XM_023249390.1
X	128958232	128958393
refGene_gn=HCFC1;refGene_tx=XM_023249387.1;refGene_gn=HCFC1;refGene_tx
=XM_023249388.1;refGene_gn=HCFC1;refGene_tx=XM_023249389.1;refGene_gn=
HCFC1;refGene_tx=XM_023249390.1
X	128958570	128958719
refGene_gn=HCFC1;refGene_tx=XM_023249387.1;refGene_gn=HCFC1;refGene_tx
=XM_023249388.1;refGene_gn=HCFC1;refGene_tx=XM_023249389.1;refGene_gn=
HCFC1;refGene_tx=XM_023249390.1
X	128964478	128964671
refGene_gn=HCFC1;refGene_tx=XM_023249387.1;refGene_gn=HCFC1;refGene_tx
=XM_023249388.1;refGene_gn=HCFC1;refGene_tx=XM_023249389.1;refGene_gn=
HCFC1;refGene_tx=XM_023249390.1
X	128970195	128970996
refGene_gn=TMEM187;refGene_tx=XM_004001045.4;refGene_gn=TMEM187;refGen
e_tx=XM_006944116.3
X	128997270	128997332
refGene_gn=IRAK1;refGene_tx=XM_023248931.1;refGene_gn=IRAK1;refGene_tx
=XM_023248932.1
X	128997739	128997883
refGene_gn=IRAK1;refGene_tx=XM_023248931.1;refGene_gn=IRAK1;refGene_tx
=XM_023248932.1
X	128998288	128998724
refGene_gn=IRAK1;refGene_tx=XM_023248931.1;refGene_gn=IRAK1;refGene_tx
=XM_023248932.1
X	128999371	128999608
refGene_gn=IRAK1;refGene_tx=XM_023248931.1

X        129001099        129001165
refGene_gn=IRAK1;refGene_tx=XM_023248931.1;refGene_gn=IRAK1;refGene_tx
=XM_023248932.1
X        129001563        129001771
refGene_gn=IRAK1;refGene_tx=XM_023248931.1;refGene_gn=IRAK1;refGene_tx
=XM_023248932.1
X        129002256        129002369
refGene_gn=IRAK1;refGene_tx=XM_023248931.1;refGene_gn=IRAK1;refGene_tx
=XM_023248932.1
X        129003316        129003431
refGene_gn=IRAK1;refGene_tx=XM_023248931.1;refGene_gn=IRAK1;refGene_tx
=XM_023248932.1
X        129003622        129003687
refGene_gn=IRAK1;refGene_tx=XM_023248931.1;refGene_gn=IRAK1;refGene_tx
=XM_023248932.1
X        129003925        129004114
refGene_gn=IRAK1;refGene_tx=XM_023248931.1;refGene_gn=IRAK1;refGene_tx
=XM_023248932.1
X        129004293        129004397
refGene_gn=IRAK1;refGene_tx=XM_023248931.1;refGene_gn=IRAK1;refGene_tx
=XM_023248932.1
X        129004515        129004647
refGene_gn=IRAK1;refGene_tx=XM_023248931.1;refGene_gn=IRAK1;refGene_tx
=XM_023248932.1
X        129004768        129004936
refGene_gn=IRAK1;refGene_tx=XM_023248931.1;refGene_gn=IRAK1;refGene_tx
=XM_023248932.1
X        129005019        129005155
refGene_gn=IRAK1;refGene_tx=XM_023248931.1;refGene_gn=IRAK1;refGene_tx
=XM_023248932.1
X        129016505        129017589
refGene_gn=MECP2;refGene_tx=XM_023249363.1
X        129018757        129019108
refGene_gn=MECP2;refGene_tx=XM_023249363.1
X        129075606        129075668
refGene_gn=MECP2;refGene_tx=XM_023249363.1
X        129113494        129113606
refGene_gn=OPN1LW;refGene_tx=NM_001009871.1;refGene_gn=OPN1LW;refGene_
tx=XM_019823328.2
X        129117963        129118260
refGene_gn=OPN1LW;refGene_tx=NM_001009871.1;refGene_gn=OPN1LW;refGene_
tx=XM_019823328.2;refGene_gn=OPN1LW;refGene_tx=XM_019823329.1;refGene_
gn=OPN1LW;refGene_tx=XM_019823330.1
X        129120501        129120670
refGene_gn=OPN1LW;refGene_tx=NM_001009871.1;refGene_gn=OPN1LW;refGene_
tx=XM_019823328.2;refGene_gn=OPN1LW;refGene_tx=XM_019823329.1;refGene_
gn=OPN1LW;refGene_tx=XM_019823330.1
X        129121651        129121830
refGene_gn=OPN1LW;refGene_tx=NM_001009871.1;refGene_gn=OPN1LW;refGene_
tx=XM_019823328.2;refGene_gn=OPN1LW;refGene_tx=XM_019823329.1;refGene_

gn=OPN1LW;refGene_tx=XM_019823330.1
X	129122980	129123220
refGene_gn=OPN1LW;refGene_tx=NM_001009871.1;refGene_gn=OPN1LW;refGene_tx=XM_019823328.2;refGene_gn=OPN1LW;refGene_tx=XM_019823329.1;refGene_gn=OPN1LW;refGene_tx=XM_019823330.1
X	129125683	129125794
refGene_gn=OPN1LW;refGene_tx=NM_001009871.1;refGene_gn=OPN1LW;refGene_tx=XM_019823328.2;refGene_gn=OPN1LW;refGene_tx=XM_019823329.1;refGene_gn=OPN1LW;refGene_tx=XM_019823330.1
X	129126235	129126400
refGene_gn=TEX28;refGene_tx=XM_023248990.1
X	129126401	129126513
refGene_gn=TEX28;refGene_tx=XM_023248990.1
X	129126998	129127145
refGene_gn=TEX28;refGene_tx=XM_023248990.1
X	129137916	129138566
refGene_gn=TEX28;refGene_tx=XM_023248990.1
X	129138568	129138652
refGene_gn=TEX28;refGene_tx=XM_023248990.1
X	129141867	129141879
refGene_gn=TEX28;refGene_tx=XM_023248990.1
X	129147440	129147577
refGene_gn=TKTL1;refGene_tx=XM_023249077.1
X	129151599	129151717
refGene_gn=TKTL1;refGene_tx=XM_023249077.1
X	129155840	129155938
refGene_gn=TKTL1;refGene_tx=XM_023249077.1
X	129157313	129157505
refGene_gn=TKTL1;refGene_tx=XM_023249077.1
X	129157585	129157713
refGene_gn=TKTL1;refGene_tx=XM_023249077.1
X	129158938	129159132
refGene_gn=TKTL1;refGene_tx=XM_023249077.1
X	129163539	129163704
refGene_gn=TKTL1;refGene_tx=XM_023249077.1
X	129166137	129166294
refGene_gn=TKTL1;refGene_tx=XM_023249077.1
X	129166655	129166786
refGene_gn=TKTL1;refGene_tx=XM_023249077.1
X	129168158	129168242
refGene_gn=TKTL1;refGene_tx=XM_023249077.1
X	129170441	129170538
refGene_gn=TKTL1;refGene_tx=XM_023249077.1
X	129170670	129170790
refGene_gn=TKTL1;refGene_tx=XM_023249077.1
X	129172049	129172222
refGene_gn=TKTL1;refGene_tx=XM_023249077.1
X	129188346	129188534
refGene_gn=FLNA;refGene_tx=XM_023249411.1;refGene_gn=FLNA;refGene_tx=XM_023249412.1

X	129189395	129189599
refGene_gn=FLNA;refGene_tx=XM_023249411.1;refGene_gn=FLNA;refGene_tx=XM_023249412.1
X	129189689	129189908
refGene_gn=FLNA;refGene_tx=XM_023249411.1;refGene_gn=FLNA;refGene_tx=XM_023249412.1
X	129190091	129190268
refGene_gn=FLNA;refGene_tx=XM_023249411.1;refGene_gn=FLNA;refGene_tx=XM_023249412.1
X	129190962	129191095
refGene_gn=FLNA;refGene_tx=XM_023249411.1;refGene_gn=FLNA;refGene_tx=XM_023249412.1
X	129191626	129191742
refGene_gn=FLNA;refGene_tx=XM_023249411.1;refGene_gn=FLNA;refGene_tx=XM_023249412.1
X	129191920	129192058
refGene_gn=FLNA;refGene_tx=XM_023249411.1;refGene_gn=FLNA;refGene_tx=XM_023249412.1
X	129192206	129192473
refGene_gn=FLNA;refGene_tx=XM_023249411.1;refGene_gn=FLNA;refGene_tx=XM_023249412.1
X	129192594	129192717
refGene_gn=FLNA;refGene_tx=XM_023249411.1;refGene_gn=FLNA;refGene_tx=XM_023249412.1
X	129192834	129192987
refGene_gn=FLNA;refGene_tx=XM_023249411.1;refGene_gn=FLNA;refGene_tx=XM_023249412.1
X	129193062	129193266
refGene_gn=FLNA;refGene_tx=XM_023249411.1;refGene_gn=FLNA;refGene_tx=XM_023249412.1
X	129193359	129193521
refGene_gn=FLNA;refGene_tx=XM_023249411.1;refGene_gn=FLNA;refGene_tx=XM_023249412.1
X	129193618	129193792
refGene_gn=FLNA;refGene_tx=XM_023249411.1;refGene_gn=FLNA;refGene_tx=XM_023249412.1
X	129193965	129194094
refGene_gn=FLNA;refGene_tx=XM_023249411.1;refGene_gn=FLNA;refGene_tx=XM_023249412.1
X	129194186	129194327
refGene_gn=FLNA;refGene_tx=XM_023249411.1;refGene_gn=FLNA;refGene_tx=XM_023249412.1
X	129194413	129194516
refGene_gn=FLNA;refGene_tx=XM_023249411.1;refGene_gn=FLNA;refGene_tx=XM_023249412.1
X	129194704	129194800
refGene_gn=FLNA;refGene_tx=XM_023249411.1;refGene_gn=FLNA;refGene_tx=XM_023249412.1
X	129194911	129195159
refGene_gn=FLNA;refGene_tx=XM_023249411.1;refGene_gn=FLNA;refGene_tx=X

M_023249412.1
X	129197563	129197587
refGene_gn=FLNA;refGene_tx=XM_023249411.1
X	129197742	129197932
refGene_gn=FLNA;refGene_tx=XM_023249411.1;refGene_gn=FLNA;refGene_tx=XM_023249412.1
X	129198615	129198772
refGene_gn=FLNA;refGene_tx=XM_023249411.1;refGene_gn=FLNA;refGene_tx=XM_023249412.1
X	129198861	129198985
refGene_gn=FLNA;refGene_tx=XM_023249411.1;refGene_gn=FLNA;refGene_tx=XM_023249412.1
X	129199363	129199534
refGene_gn=FLNA;refGene_tx=XM_023249411.1;refGene_gn=FLNA;refGene_tx=XM_023249412.1
X	129199623	129199784
refGene_gn=FLNA;refGene_tx=XM_023249411.1;refGene_gn=FLNA;refGene_tx=XM_023249412.1
X	129199863	129200026
refGene_gn=FLNA;refGene_tx=XM_023249411.1;refGene_gn=FLNA;refGene_tx=XM_023249412.1
X	129200108	129200282
refGene_gn=FLNA;refGene_tx=XM_023249411.1;refGene_gn=FLNA;refGene_tx=XM_023249412.1
X	129200376	129200974
refGene_gn=FLNA;refGene_tx=XM_023249411.1;refGene_gn=FLNA;refGene_tx=XM_023249412.1
X	129201201	129201464
refGene_gn=FLNA;refGene_tx=XM_023249411.1;refGene_gn=FLNA;refGene_tx=XM_023249412.1
X	129201559	129201677
refGene_gn=FLNA;refGene_tx=XM_023249411.1;refGene_gn=FLNA;refGene_tx=XM_023249412.1
X	129201945	129202115
refGene_gn=FLNA;refGene_tx=XM_023249411.1;refGene_gn=FLNA;refGene_tx=XM_023249412.1
X	129202206	129202297
refGene_gn=FLNA;refGene_tx=XM_023249411.1;refGene_gn=FLNA;refGene_tx=XM_023249412.1
X	129202387	129202548
refGene_gn=FLNA;refGene_tx=XM_023249411.1;refGene_gn=FLNA;refGene_tx=XM_023249412.1
X	129202631	129202755
refGene_gn=FLNA;refGene_tx=XM_023249411.1;refGene_gn=FLNA;refGene_tx=XM_023249412.1
X	129203466	129203610
refGene_gn=FLNA;refGene_tx=XM_023249411.1;refGene_gn=FLNA;refGene_tx=XM_023249412.1
X	129203698	129203812
refGene_gn=FLNA;refGene_tx=XM_023249411.1;refGene_gn=FLNA;refGene_tx=X

M_023249412.1
X 129203961 129204155
refGene_gn=FLNA;refGene_tx=XM_023249411.1;refGene_gn=FLNA;refGene_tx=X
M_023249412.1
X 129204253 129204390
refGene_gn=FLNA;refGene_tx=XM_023249411.1;refGene_gn=FLNA;refGene_tx=X
M_023249412.1
X 129204483 129204607
refGene_gn=FLNA;refGene_tx=XM_023249411.1;refGene_gn=FLNA;refGene_tx=X
M_023249412.1
X 129204693 129204831
refGene_gn=FLNA;refGene_tx=XM_023249411.1;refGene_gn=FLNA;refGene_tx=X
M_023249412.1
X 129205337 129205538
refGene_gn=FLNA;refGene_tx=XM_023249411.1;refGene_gn=FLNA;refGene_tx=X
M_023249412.1
X 129205628 129205791
refGene_gn=FLNA;refGene_tx=XM_023249411.1;refGene_gn=FLNA;refGene_tx=X
M_023249412.1
X 129205889 129205967
refGene_gn=FLNA;refGene_tx=XM_023249411.1;refGene_gn=FLNA;refGene_tx=X
M_023249412.1
X 129206046 129206165
refGene_gn=FLNA;refGene_tx=XM_023249411.1;refGene_gn=FLNA;refGene_tx=X
M_023249412.1
X 129206654 129206802
refGene_gn=FLNA;refGene_tx=XM_023249411.1;refGene_gn=FLNA;refGene_tx=X
M_023249412.1
X 129206935 129207033
refGene_gn=FLNA;refGene_tx=XM_023249411.1;refGene_gn=FLNA;refGene_tx=X
M_023249412.1
X 129207140 129207389
refGene_gn=FLNA;refGene_tx=XM_023249411.1;refGene_gn=FLNA;refGene_tx=X
M_023249412.1
X 129210362 129210735
refGene_gn=FLNA;refGene_tx=XM_023249411.1;refGene_gn=FLNA;refGene_tx=X
M_023249412.1
X 129218553 129218635
refGene_gn=EMD;refGene_tx=XM_004001017.3
X 129218724 129218832
refGene_gn=EMD;refGene_tx=XM_004001017.3
X 129219002 129219080
refGene_gn=EMD;refGene_tx=XM_004001017.3
X 129219300 129219434
refGene_gn=EMD;refGene_tx=XM_004001017.3
X 129220378 129220425
refGene_gn=EMD;refGene_tx=XM_004001017.3
X 129220522 129220856
refGene_gn=EMD;refGene_tx=XM_004001017.3
X 129238248 129238271

refGene_gn=RPL10;refGene_tx=XM_006944118.3;refGene_gn=RPL10;refGene_tx=XM_011279551.3
X 129239154 129239213
refGene_gn=RPL10;refGene_tx=XM_006944118.3;refGene_gn=RPL10;refGene_tx=XM_011279551.3
X 129239307 129239415
refGene_gn=RPL10;refGene_tx=XM_006944118.3;refGene_gn=RPL10;refGene_tx=XM_011279551.3
X 129239644 129239783
refGene_gn=RPL10;refGene_tx=XM_006944118.3;refGene_gn=RPL10;refGene_tx=XM_011279551.3
X 129240303 129240466
refGene_gn=RPL10;refGene_tx=XM_006944118.3;refGene_gn=RPL10;refGene_tx=XM_011279551.3
X 129240553 129240706
refGene_gn=RPL10;refGene_tx=XM_006944118.3
X 129242622 129242739
refGene_gn=DNASE1L1;refGene_tx=XM_004001049.3
X 129242821 129243070
refGene_gn=DNASE1L1;refGene_tx=XM_004001049.3
X 129243150 129243263
refGene_gn=DNASE1L1;refGene_tx=XM_004001049.3
X 129243408 129243509
refGene_gn=DNASE1L1;refGene_tx=XM_004001049.3
X 129244454 129244541
refGene_gn=DNASE1L1;refGene_tx=XM_004001049.3
X 129244620 129244709
refGene_gn=DNASE1L1;refGene_tx=XM_004001049.3
X 129245059 129245191
refGene_gn=DNASE1L1;refGene_tx=XM_004001049.3
X 129249976 129250085
refGene_gn=TAZ;refGene_tx=XM_004001020.4;refGene_gn=TAZ;refGene_tx=XM_019824496.1
X 129250253 129250382
refGene_gn=TAZ;refGene_tx=XM_004001020.4;refGene_gn=TAZ;refGene_tx=XM_019824496.1
X 129251379 129251425
refGene_gn=TAZ;refGene_tx=XM_004001020.4;refGene_gn=TAZ;refGene_tx=XM_019824496.1
X 129251669 129251755
refGene_gn=TAZ;refGene_tx=XM_004001020.4;refGene_gn=TAZ;refGene_tx=XM_019824496.1
X 129259041 129259122
refGene_gn=TAZ;refGene_tx=XM_004001020.4;refGene_gn=TAZ;refGene_tx=XM_019824496.1
X 129259192 129259234
refGene_gn=TAZ;refGene_tx=XM_004001020.4;refGene_gn=TAZ;refGene_tx=XM_019824496.1
X 129259458 129259521
refGene_gn=TAZ;refGene_tx=XM_004001020.4;refGene_gn=TAZ;refGene_tx=XM_

019824496.1
X	129259661	129259768
refGene_gn=TAZ;refGene_tx=XM_004001020.4;refGene_gn=TAZ;refGene_tx=XM_019824496.1
X	129260189	129260267
refGene_gn=TAZ;refGene_tx=XM_004001020.4;refGene_gn=TAZ;refGene_tx=XM_019824496.1
X	129260413	129260515
refGene_gn=TAZ;refGene_tx=XM_004001020.4;refGene_gn=TAZ;refGene_tx=XM_019824496.1
X	129270268	129270411
refGene_gn=ATP6AP1;refGene_tx=XM_023249007.1
X	129270644	129270771
refGene_gn=ATP6AP1;refGene_tx=XM_023249007.1
X	129273127	129273202
refGene_gn=ATP6AP1;refGene_tx=XM_023249007.1
X	129273480	129273674
refGene_gn=ATP6AP1;refGene_tx=XM_023249007.1
X	129274197	129274238
refGene_gn=ATP6AP1;refGene_tx=XM_023249007.1
X	129275308	129275394
refGene_gn=ATP6AP1;refGene_tx=XM_023249007.1
X	129275832	129276071
refGene_gn=ATP6AP1;refGene_tx=XM_023249007.1
X	129276696	129276744
refGene_gn=ATP6AP1;refGene_tx=XM_023249007.1
X	129276836	129277068
refGene_gn=ATP6AP1;refGene_tx=XM_023249007.1
X	129277290	129277500
refGene_gn=ATP6AP1;refGene_tx=XM_023249007.1
X	129278850	129278895
refGene_gn=GDI1;refGene_tx=XM_004001022.3
X	129280129	129280237
refGene_gn=GDI1;refGene_tx=XM_004001022.3
X	129280376	129280476
refGene_gn=GDI1;refGene_tx=XM_004001022.3
X	129280622	129280757
refGene_gn=GDI1;refGene_tx=XM_004001022.3
X	129281219	129281418
refGene_gn=GDI1;refGene_tx=XM_004001022.3
X	129281685	129281817
refGene_gn=GDI1;refGene_tx=XM_004001022.3
X	129282405	129282505
refGene_gn=GDI1;refGene_tx=XM_004001022.3
X	129282895	129283067
refGene_gn=GDI1;refGene_tx=XM_004001022.3
X	129283365	129283510
refGene_gn=GDI1;refGene_tx=XM_004001022.3
X	129283651	129283706
refGene_gn=GDI1;refGene_tx=XM_004001022.3

X	129283788	129283941
refGene_gn=GDI1;refGene_tx=XM_004001022.3
X	129285640	129285751
refGene_gn=FAM50A;refGene_tx=XM_023249089.1
X	129286964	129287049
refGene_gn=FAM50A;refGene_tx=XM_023249089.1
X	129287154	129287254
refGene_gn=FAM50A;refGene_tx=XM_023249089.1
X	129287441	129287587
refGene_gn=FAM50A;refGene_tx=XM_023249089.1
X	129290259	129290336
refGene_gn=FAM50A;refGene_tx=XM_023249089.1
X	129290448	129290515
refGene_gn=FAM50A;refGene_tx=XM_023249089.1
X	129290656	129290718
refGene_gn=FAM50A;refGene_tx=XM_023249089.1
X	129291028	129291105
refGene_gn=FAM50A;refGene_tx=XM_023249089.1
X	129291487	129291542
refGene_gn=FAM50A;refGene_tx=XM_023249089.1
X	129291733	129291782
refGene_gn=FAM50A;refGene_tx=XM_023249089.1
X	129291881	129291952
refGene_gn=FAM50A;refGene_tx=XM_023249089.1
X	129292035	129292146
refGene_gn=FAM50A;refGene_tx=XM_023249089.1
X	129292276	129292285
refGene_gn=FAM50A;refGene_tx=XM_023249089.1
X	129306259	129306856
refGene_gn=PLXNA3;refGene_tx=XM_023249012.1;refGene_gn=PLXNA3;refGene_tx=XM_023249013.1
X	129307223	129307763
refGene_gn=PLXNA3;refGene_tx=XM_023249012.1;refGene_gn=PLXNA3;refGene_tx=XM_023249013.1
X	129308244	129308427
refGene_gn=PLXNA3;refGene_tx=XM_023249012.1;refGene_gn=PLXNA3;refGene_tx=XM_023249013.1
X	129309256	129309385
refGene_gn=PLXNA3;refGene_tx=XM_023249012.1;refGene_gn=PLXNA3;refGene_tx=XM_023249013.1
X	129309455	129309556
refGene_gn=PLXNA3;refGene_tx=XM_023249012.1;refGene_gn=PLXNA3;refGene_tx=XM_023249013.1
X	129309919	129310043
refGene_gn=PLXNA3;refGene_tx=XM_023249012.1;refGene_gn=PLXNA3;refGene_tx=XM_023249013.1
X	129310145	129310302
refGene_gn=PLXNA3;refGene_tx=XM_023249012.1;refGene_gn=PLXNA3;refGene_tx=XM_023249013.1
X	129310381	129310481

refGene_gn=PLXNA3;refGene_tx=XM_023249012.1;refGene_gn=PLXNA3;refGene_tx=XM_023249013.1
X 129310654 129310769
refGene_gn=PLXNA3;refGene_tx=XM_023249012.1;refGene_gn=PLXNA3;refGene_tx=XM_023249013.1
X 129310935 129311136
refGene_gn=PLXNA3;refGene_tx=XM_023249012.1;refGene_gn=PLXNA3;refGene_tx=XM_023249013.1
X 129311272 129311369
refGene_gn=PLXNA3;refGene_tx=XM_023249012.1;refGene_gn=PLXNA3;refGene_tx=XM_023249013.1
X 129311509 129311700
refGene_gn=PLXNA3;refGene_tx=XM_023249012.1;refGene_gn=PLXNA3;refGene_tx=XM_023249013.1
X 129311779 129311925
refGene_gn=PLXNA3;refGene_tx=XM_023249012.1;refGene_gn=PLXNA3;refGene_tx=XM_023249013.1
X 129311996 129312117
refGene_gn=PLXNA3;refGene_tx=XM_023249012.1;refGene_gn=PLXNA3;refGene_tx=XM_023249013.1
X 129312222 129312359
refGene_gn=PLXNA3;refGene_tx=XM_023249012.1;refGene_gn=PLXNA3;refGene_tx=XM_023249013.1
X 129312467 129312638
refGene_gn=PLXNA3;refGene_tx=XM_023249012.1;refGene_gn=PLXNA3;refGene_tx=XM_023249013.1
X 129312923 129313017
refGene_gn=PLXNA3;refGene_tx=XM_023249012.1;refGene_gn=PLXNA3;refGene_tx=XM_023249013.1
X 129313100 129313340
refGene_gn=PLXNA3;refGene_tx=XM_023249012.1;refGene_gn=PLXNA3;refGene_tx=XM_023249013.1
X 129313415 129313559
refGene_gn=PLXNA3;refGene_tx=XM_023249012.1;refGene_gn=PLXNA3;refGene_tx=XM_023249013.1
X 129313628 129313863
refGene_gn=PLXNA3;refGene_tx=XM_023249012.1;refGene_gn=PLXNA3;refGene_tx=XM_023249013.1
X 129313952 129314095
refGene_gn=PLXNA3;refGene_tx=XM_023249012.1;refGene_gn=PLXNA3;refGene_tx=XM_023249013.1
X 129314174 129314431
refGene_gn=PLXNA3;refGene_tx=XM_023249012.1;refGene_gn=PLXNA3;refGene_tx=XM_023249013.1
X 129314555 129314622
refGene_gn=PLXNA3;refGene_tx=XM_023249012.1;refGene_gn=PLXNA3;refGene_tx=XM_023249013.1
X 129314715 129314862
refGene_gn=PLXNA3;refGene_tx=XM_023249012.1;refGene_gn=PLXNA3;refGene_tx=XM_023249013.1

X	129314947	129315107
refGene_gn=PLXNA3;refGene_tx=XM_023249012.1;refGene_gn=PLXNA3;refGene_tx=XM_023249013.1
X	129315261	129315365
refGene_gn=PLXNA3;refGene_tx=XM_023249012.1;refGene_gn=PLXNA3;refGene_tx=XM_023249013.1
X	129315539	129315636
refGene_gn=PLXNA3;refGene_tx=XM_023249012.1;refGene_gn=PLXNA3;refGene_tx=XM_023249013.1
X	129315921	129316112
refGene_gn=PLXNA3;refGene_tx=XM_023249012.1;refGene_gn=PLXNA3;refGene_tx=XM_023249013.1
X	129316367	129316537
refGene_gn=PLXNA3;refGene_tx=XM_023249012.1;refGene_gn=PLXNA3;refGene_tx=XM_023249013.1
X	129317024	129317237
refGene_gn=PLXNA3;refGene_tx=XM_023249012.1;refGene_gn=PLXNA3;refGene_tx=XM_023249013.1
X	129317375	129317526
refGene_gn=PLXNA3;refGene_tx=XM_023249012.1;refGene_gn=PLXNA3;refGene_tx=XM_023249013.1
X	129319032	129319128
refGene_gn=PLXNA3;refGene_tx=XM_023249012.1;refGene_gn=PLXNA3;refGene_tx=XM_023249013.1
X	129326949	129327064
refGene_gn=LAGE3;refGene_tx=XM_023249158.1
X	129328006	129328145
refGene_gn=LAGE3;refGene_tx=XM_023249157.1
X	129328339	129328468
refGene_gn=LAGE3;refGene_tx=XM_023249157.1;refGene_gn=LAGE3;refGene_tx=XM_023249158.1
X	129328781	129328999
refGene_gn=LAGE3;refGene_tx=XM_023249157.1;refGene_gn=LAGE3;refGene_tx=XM_023249158.1
X	129338575	129338686
refGene_gn=UBL4A;refGene_tx=XM_004001026.4
X	129338803	129339018
refGene_gn=UBL4A;refGene_tx=XM_004001026.4
X	129339211	129339317
refGene_gn=UBL4A;refGene_tx=XM_004001026.4
X	129339498	129339546
refGene_gn=UBL4A;refGene_tx=XM_004001026.4
X	129340503	129342075
refGene_gn=SLC10A3;refGene_tx=XM_004001025.5;refGene_gn=SLC10A3;refGene_tx=XM_006944127.3;refGene_gn=SLC10A3;refGene_tx=XM_011279540.3;refGene_gn=SLC10A3;refGene_tx=XM_011279541.3;refGene_gn=SLC10A3;refGene_tx=XM_011279543.3;refGene_gn=SLC10A3;refGene_tx=XM_011279544.3;refGene_gn=SLC10A3;refGene_tx=XM_011279545.3
X	129342657	129342736
refGene_gn=SLC10A3;refGene_tx=XM_011279540.3;refGene_gn=SLC10A3;refGen

e_tx=XM_011279541.3;refGene_gn=SLC10A3;refGene_tx=XM_011279543.3
X	129343004	129343123
refGene_gn=SLC10A3;refGene_tx=XM_011279540.3;refGene_gn=SLC10A3;refGen
e_tx=XM_011279541.3;refGene_gn=SLC10A3;refGene_tx=XM_011279543.3
X	129343281	129343308
refGene_gn=SLC10A3;refGene_tx=XM_011279544.3
X	129348502	129348598
refGene_gn=FAM3A;refGene_tx=XM_019824510.2;refGene_gn=FAM3A;refGene_tx
=XM_023249196.1;refGene_gn=FAM3A;refGene_tx=XM_023249198.1
X	129349410	129349537
refGene_gn=FAM3A;refGene_tx=XM_019824510.2;refGene_gn=FAM3A;refGene_tx
=XM_023249196.1;refGene_gn=FAM3A;refGene_tx=XM_023249198.1
X	129349613	129349698
refGene_gn=FAM3A;refGene_tx=XM_019824510.2;refGene_gn=FAM3A;refGene_tx
=XM_023249196.1;refGene_gn=FAM3A;refGene_tx=XM_023249198.1
X	129350021	129350072
refGene_gn=FAM3A;refGene_tx=XM_019824510.2;refGene_gn=FAM3A;refGene_tx
=XM_023249198.1
X	129350538	129350597
refGene_gn=FAM3A;refGene_tx=XM_019824510.2;refGene_gn=FAM3A;refGene_tx
=XM_023249196.1;refGene_gn=FAM3A;refGene_tx=XM_023249198.1
X	129350687	129350811
refGene_gn=FAM3A;refGene_tx=XM_019824510.2;refGene_gn=FAM3A;refGene_tx
=XM_023249196.1;refGene_gn=FAM3A;refGene_tx=XM_023249198.1
X	129350855	129350866
refGene_gn=FAM3A;refGene_tx=XM_023249199.1
X	129351228	129351575
refGene_gn=FAM3A;refGene_tx=XM_023249197.1
X	129352990	129353014
refGene_gn=FAM3A;refGene_tx=XM_019824510.2;refGene_gn=FAM3A;refGene_tx
=XM_023249196.1;refGene_gn=FAM3A;refGene_tx=XM_023249197.1;refGene_gn=
FAM3A;refGene_tx=XM_023249198.1;refGene_gn=FAM3A;refGene_tx=XM_0232491
99.1
X	129353549	129353663
refGene_gn=FAM3A;refGene_tx=XM_019824510.2;refGene_gn=FAM3A;refGene_tx
=XM_023249196.1;refGene_gn=FAM3A;refGene_tx=XM_023249197.1;refGene_gn=
FAM3A;refGene_tx=XM_023249198.1;refGene_gn=FAM3A;refGene_tx=XM_0232491
99.1
X	129355034	129355231
refGene_gn=FAM3A;refGene_tx=XM_019824510.2;refGene_gn=FAM3A;refGene_tx
=XM_023249196.1;refGene_gn=FAM3A;refGene_tx=XM_023249197.1;refGene_gn=
FAM3A;refGene_tx=XM_023249198.1;refGene_gn=FAM3A;refGene_tx=XM_0232491
99.1
X	129355413	129355631
refGene_gn=FAM3A;refGene_tx=XM_023249196.1;refGene_gn=FAM3A;refGene_tx
=XM_023249197.1;refGene_gn=FAM3A;refGene_tx=XM_023249199.1
X	129361261	129361352
refGene_gn=G6PD;refGene_tx=XM_004001030.5;refGene_gn=G6PD;refGene_tx=X
M_023249194.1;refGene_gn=G6PD;refGene_tx=XM_023249195.1
X	129361452	129361545

refGene_gn=G6PD;refGene_tx=XM_004001030.5;refGene_gn=G6PD;refGene_tx=XM_023249194.1;refGene_gn=G6PD;refGene_tx=XM_023249195.1
X	129361630	129361707
refGene_gn=G6PD;refGene_tx=XM_004001030.5;refGene_gn=G6PD;refGene_tx=XM_023249194.1;refGene_gn=G6PD;refGene_tx=XM_023249195.1
X	129361800	129362036
refGene_gn=G6PD;refGene_tx=XM_004001030.5;refGene_gn=G6PD;refGene_tx=XM_023249194.1;refGene_gn=G6PD;refGene_tx=XM_023249195.1
X	129362321	129362508
refGene_gn=G6PD;refGene_tx=XM_004001030.5;refGene_gn=G6PD;refGene_tx=XM_023249194.1;refGene_gn=G6PD;refGene_tx=XM_023249195.1
X	129362872	129362966
refGene_gn=G6PD;refGene_tx=XM_004001030.5;refGene_gn=G6PD;refGene_tx=XM_023249194.1;refGene_gn=G6PD;refGene_tx=XM_023249195.1
X	129363281	129363407
refGene_gn=G6PD;refGene_tx=XM_004001030.5;refGene_gn=G6PD;refGene_tx=XM_023249194.1;refGene_gn=G6PD;refGene_tx=XM_023249195.1
X	129363558	129363717
refGene_gn=G6PD;refGene_tx=XM_004001030.5;refGene_gn=G6PD;refGene_tx=XM_023249194.1;refGene_gn=G6PD;refGene_tx=XM_023249195.1
X	129364678	129364896
refGene_gn=G6PD;refGene_tx=XM_004001030.5;refGene_gn=G6PD;refGene_tx=XM_023249194.1;refGene_gn=G6PD;refGene_tx=XM_023249195.1
X	129365574	129365683
refGene_gn=G6PD;refGene_tx=XM_004001030.5;refGene_gn=G6PD;refGene_tx=XM_023249194.1;refGene_gn=G6PD;refGene_tx=XM_023249195.1
X	129365764	129365802
refGene_gn=G6PD;refGene_tx=XM_004001030.5;refGene_gn=G6PD;refGene_tx=XM_023249194.1;refGene_gn=G6PD;refGene_tx=XM_023249195.1
X	129373012	129373140
refGene_gn=G6PD;refGene_tx=XM_004001030.5;refGene_gn=G6PD;refGene_tx=XM_023249194.1;refGene_gn=G6PD;refGene_tx=XM_023249195.1
X	129374052	129374164
refGene_gn=G6PD;refGene_tx=XM_023249194.1
X	129378390	129378577
refGene_gn=IKBKG;refGene_tx=XM_004001031.5;refGene_gn=IKBKG;refGene_tx=XM_006944128.4;refGene_gn=IKBKG;refGene_tx=XM_019824500.1;refGene_gn=IKBKG;refGene_tx=XM_019824501.2;refGene_gn=IKBKG;refGene_tx=XM_019824502.2;refGene_gn=IKBKG;refGene_tx=XM_019824504.1;refGene_gn=IKBKG;refGene_tx=XM_019824505.2
X	129381280	129381492
refGene_gn=IKBKG;refGene_tx=XM_004001031.5;refGene_gn=IKBKG;refGene_tx=XM_006944128.4;refGene_gn=IKBKG;refGene_tx=XM_019824500.1;refGene_gn=IKBKG;refGene_tx=XM_019824501.2;refGene_gn=IKBKG;refGene_tx=XM_019824502.2;refGene_gn=IKBKG;refGene_tx=XM_019824503.2;refGene_gn=IKBKG;refGene_tx=XM_019824504.1;refGene_gn=IKBKG;refGene_tx=XM_019824505.2
X	129386545	129386664
refGene_gn=IKBKG;refGene_tx=XM_004001031.5;refGene_gn=IKBKG;refGene_tx=XM_006944128.4;refGene_gn=IKBKG;refGene_tx=XM_019824500.1;refGene_gn=IKBKG;refGene_tx=XM_019824501.2;refGene_gn=IKBKG;refGene_tx=XM_0198245

02.2;refGene_gn=IKBKG;refGene_tx=XM_019824503.2;refGene_gn=IKBKG;refGene_tx=XM_019824504.1;refGene_gn=IKBKG;refGene_tx=XM_019824505.2
X	129388627	129388780
refGene_gn=IKBKG;refGene_tx=XM_004001031.5;refGene_gn=IKBKG;refGene_tx=XM_006944128.4;refGene_gn=IKBKG;refGene_tx=XM_019824500.1;refGene_gn=IKBKG;refGene_tx=XM_019824501.2;refGene_gn=IKBKG;refGene_tx=XM_019824502.2;refGene_gn=IKBKG;refGene_tx=XM_019824503.2;refGene_gn=IKBKG;refGene_tx=XM_019824504.1;refGene_gn=IKBKG;refGene_tx=XM_019824505.2
X	129389312	129389409
refGene_gn=IKBKG;refGene_tx=XM_004001031.5;refGene_gn=IKBKG;refGene_tx=XM_006944128.4;refGene_gn=IKBKG;refGene_tx=XM_019824500.1;refGene_gn=IKBKG;refGene_tx=XM_019824501.2;refGene_gn=IKBKG;refGene_tx=XM_019824502.2;refGene_gn=IKBKG;refGene_tx=XM_019824503.2;refGene_gn=IKBKG;refGene_tx=XM_019824504.1;refGene_gn=IKBKG;refGene_tx=XM_019824505.2
X	129390500	129390644
refGene_gn=IKBKG;refGene_tx=XM_004001031.5;refGene_gn=IKBKG;refGene_tx=XM_006944128.4;refGene_gn=IKBKG;refGene_tx=XM_019824500.1;refGene_gn=IKBKG;refGene_tx=XM_019824501.2;refGene_gn=IKBKG;refGene_tx=XM_019824502.2;refGene_gn=IKBKG;refGene_tx=XM_019824503.2;refGene_gn=IKBKG;refGene_tx=XM_019824504.1;refGene_gn=IKBKG;refGene_tx=XM_019824505.2
X	129390883	129391026
refGene_gn=IKBKG;refGene_tx=XM_004001031.5;refGene_gn=IKBKG;refGene_tx=XM_006944128.4;refGene_gn=IKBKG;refGene_tx=XM_019824500.1;refGene_gn=IKBKG;refGene_tx=XM_019824501.2;refGene_gn=IKBKG;refGene_tx=XM_019824502.2;refGene_gn=IKBKG;refGene_tx=XM_019824503.2;refGene_gn=IKBKG;refGene_tx=XM_019824504.1;refGene_gn=IKBKG;refGene_tx=XM_019824505.2
X	129391229	129391291
refGene_gn=IKBKG;refGene_tx=XM_004001031.5;refGene_gn=IKBKG;refGene_tx=XM_006944128.4;refGene_gn=IKBKG;refGene_tx=XM_019824500.1;refGene_gn=IKBKG;refGene_tx=XM_019824501.2;refGene_gn=IKBKG;refGene_tx=XM_019824502.2;refGene_gn=IKBKG;refGene_tx=XM_019824503.2;refGene_gn=IKBKG;refGene_tx=XM_019824504.1;refGene_gn=IKBKG;refGene_tx=XM_019824505.2
X	129391489	129391632
refGene_gn=IKBKG;refGene_tx=XM_004001031.5;refGene_gn=IKBKG;refGene_tx=XM_006944128.4;refGene_gn=IKBKG;refGene_tx=XM_019824500.1;refGene_gn=IKBKG;refGene_tx=XM_019824501.2;refGene_gn=IKBKG;refGene_tx=XM_019824502.2;refGene_gn=IKBKG;refGene_tx=XM_019824503.2;refGene_gn=IKBKG;refGene_tx=XM_019824504.1;refGene_gn=IKBKG;refGene_tx=XM_019824505.2
X	129432139	129432580
refGene_gn=LOC105259690;refGene_tx=XM_023248991.1
X	129432667	129432768
refGene_gn=LOC105259690;refGene_tx=XM_023248991.1
X	129432859	129432920
refGene_gn=LOC105259690;refGene_tx=XM_023248991.1
X	129432922	129432966
refGene_gn=LOC105259690;refGene_tx=XM_023248991.1
X	129433025	129433179
refGene_gn=LOC105259690;refGene_tx=XM_023248991.1
X	129433300	129433413
refGene_gn=LOC105259690;refGene_tx=XM_023248991.1

X 129433998 129434123
refGene_gn=LOC105259690;refGene_tx=XM_023248991.1
X 129434213 129434317
refGene_gn=LOC105259690;refGene_tx=XM_023248991.1
X 129434402 129434537
refGene_gn=LOC105259690;refGene_tx=XM_023248991.1
X 129435419 129435576
refGene_gn=LOC105259690;refGene_tx=XM_023248991.1
X 129435670 129435675
refGene_gn=LOC105259690;refGene_tx=XM_023248991.1
X 129441322 129441439
refGene_gn=GAB3;refGene_tx=XM_023248945.1
X 129442396 129442510
refGene_gn=GAB3;refGene_tx=XM_023248945.1
X 129458076 129458179
refGene_gn=GAB3;refGene_tx=XM_023248945.1
X 129459332 129459414
refGene_gn=GAB3;refGene_tx=XM_023248945.1
X 129461572 129461792
refGene_gn=GAB3;refGene_tx=XM_023248945.1
X 129462242 129462298
refGene_gn=GAB3;refGene_tx=XM_023248945.1
X 129475206 129475685
refGene_gn=GAB3;refGene_tx=XM_023248945.1
X 129476346 129476563
refGene_gn=GAB3;refGene_tx=XM_023248945.1
X 129478961 129479265
refGene_gn=GAB3;refGene_tx=XM_023248945.1
X 129516732 129516804
refGene_gn=GAB3;refGene_tx=XM_023248945.1
X 129529192 129529208
refGene_gn=DKC1;refGene_tx=XM_023248836.1
X 129529997 129530065
refGene_gn=DKC1;refGene_tx=XM_023248836.1
X 129530533 129530620
refGene_gn=DKC1;refGene_tx=XM_023248836.1
X 129531974 129532066
refGene_gn=DKC1;refGene_tx=XM_023248836.1
X 129532381 129532566
refGene_gn=DKC1;refGene_tx=XM_023248836.1
X 129533170 129533235
refGene_gn=DKC1;refGene_tx=XM_023248836.1
X 129533598 129533725
refGene_gn=DKC1;refGene_tx=XM_023248836.1
X 129534172 129534303
refGene_gn=DKC1;refGene_tx=XM_023248836.1
X 129534707 129534851
refGene_gn=DKC1;refGene_tx=XM_023248836.1
X 129536265 129536386
refGene_gn=DKC1;refGene_tx=XM_023248836.1

X 129537199 129537318
refGene_gn=DKC1;refGene_tx=XM_023248836.1
X 129537989 129538093
refGene_gn=DKC1;refGene_tx=XM_023248836.1
X 129538560 129538669
refGene_gn=DKC1;refGene_tx=XM_023248836.1
X 129539475 129539595
refGene_gn=DKC1;refGene_tx=XM_023248836.1
X 129539893 129539959
refGene_gn=DKC1;refGene_tx=XM_023248836.1
X 129542178 129542355
refGene_gn=MPP1;refGene_tx=XM_023248837.1
X 129543743 129543818
refGene_gn=MPP1;refGene_tx=XM_023248837.1
X 129543963 129544166
refGene_gn=MPP1;refGene_tx=XM_023248837.1
X 129546072 129546153
refGene_gn=MPP1;refGene_tx=XM_023248837.1
X 129546807 129546888
refGene_gn=MPP1;refGene_tx=XM_023248837.1
X 129548059 129548166
refGene_gn=MPP1;refGene_tx=XM_023248837.1
X 129549574 129549771
refGene_gn=MPP1;refGene_tx=XM_023248837.1
X 129550859 129550928
refGene_gn=MPP1;refGene_tx=XM_023248837.1
X 129555071 129555157
refGene_gn=MPP1;refGene_tx=XM_023248837.1
X 129556057 129556136
refGene_gn=MPP1;refGene_tx=XM_023248837.1
X 129556307 129556451
refGene_gn=MPP1;refGene_tx=XM_023248837.1
X 129568429 129568531
refGene_gn=MPP1;refGene_tx=XM_023248837.1
X 129576025 129576053
refGene_gn=SMIM9;refGene_tx=XM_006944137.2
X 129581536 129581666
refGene_gn=SMIM9;refGene_tx=XM_006944137.2
X 129583050 129583255
refGene_gn=SMIM9;refGene_tx=XM_006944137.2
X 129591093 129591249
refGene_gn=F8;refGene_tx=XM_004001054.4;refGene_gn=F8;refGene_tx=XM_01
9824515.2;refGene_gn=F8;refGene_tx=XM_019824518.1
X 129597101 129597278
refGene_gn=F8;refGene_tx=XM_004001054.4;refGene_gn=F8;refGene_tx=XM_01
9824515.2;refGene_gn=F8;refGene_tx=XM_019824518.1
X 129598012 129598161
refGene_gn=F8;refGene_tx=XM_004001054.4;refGene_gn=F8;refGene_tx=XM_01
9824515.2;refGene_gn=F8;refGene_tx=XM_019824518.1
X 129598578 129598723

refGene_gn=F8;refGene_tx=XM_004001054.4;refGene_gn=F8;refGene_tx=XM_019824515.2;refGene_gn=F8;refGene_tx=XM_019824518.1
X 129602831 129602892
refGene_gn=LOC111558728;refGene_tx=XM_023249175.1
X 129602894 129603172
refGene_gn=LOC111558728;refGene_tx=XM_023249175.1
X 129603852 129604396
refGene_gn=F8A1;refGene_tx=XM_023249174.1
X 129604398 129604559
refGene_gn=F8A1;refGene_tx=XM_023249174.1
X 129604560 129604983
refGene_gn=F8A1;refGene_tx=XM_023249174.1
X 129612225 129612381
refGene_gn=F8;refGene_tx=XM_004001054.4;refGene_gn=F8;refGene_tx=XM_019824515.2;refGene_gn=F8;refGene_tx=XM_019824518.1
X 129617159 129617245
refGene_gn=F8;refGene_tx=XM_004001054.4;refGene_gn=F8;refGene_tx=XM_019824515.2;refGene_gn=F8;refGene_tx=XM_019824518.1
X 129618475 129618547
refGene_gn=F8;refGene_tx=XM_004001054.4;refGene_gn=F8;refGene_tx=XM_019824515.2;refGene_gn=F8;refGene_tx=XM_019824518.1
X 129619368 129619485
refGene_gn=F8;refGene_tx=XM_004001054.4;refGene_gn=F8;refGene_tx=XM_019824515.2;refGene_gn=F8;refGene_tx=XM_019824518.1
X 129620839 129621022
refGene_gn=F8;refGene_tx=XM_004001054.4;refGene_gn=F8;refGene_tx=XM_019824515.2;refGene_gn=F8;refGene_tx=XM_019824518.1
X 129621445 129621674
refGene_gn=F8;refGene_tx=XM_004001054.4;refGene_gn=F8;refGene_tx=XM_019824515.2;refGene_gn=F8;refGene_tx=XM_019824518.1
X 129621863 129622076
refGene_gn=F8;refGene_tx=XM_004001054.4;refGene_gn=F8;refGene_tx=XM_019824515.2;refGene_gn=F8;refGene_tx=XM_019824518.1
X 129624163 129624317
refGene_gn=F8;refGene_tx=XM_004001054.4;refGene_gn=F8;refGene_tx=XM_019824515.2;refGene_gn=F8;refGene_tx=XM_019824518.1
X 129641322 129644422
refGene_gn=F8;refGene_tx=XM_004001054.4;refGene_gn=F8;refGene_tx=XM_019824515.2;refGene_gn=F8;refGene_tx=XM_019824518.1
X 129647794 129648004
refGene_gn=F8;refGene_tx=XM_004001054.4;refGene_gn=F8;refGene_tx=XM_019824515.2;refGene_gn=F8;refGene_tx=XM_019824518.1
X 129654460 129654611
refGene_gn=F8;refGene_tx=XM_004001054.4;refGene_gn=F8;refGene_tx=XM_019824515.2;refGene_gn=F8;refGene_tx=XM_019824518.1
X 129656761 129656976
refGene_gn=F8;refGene_tx=XM_004001054.4;refGene_gn=F8;refGene_tx=XM_019824515.2;refGene_gn=F8;refGene_tx=XM_019824518.1
X 129661803 129661897
refGene_gn=F8;refGene_tx=XM_004001054.4;refGene_gn=F8;refGene_tx=XM_01

9824515.2;refGene_gn=F8;refGene_tx=XM_019824518.1
X	129663135	129663307
refGene_gn=F8;refGene_tx=XM_004001054.4;refGene_gn=F8;refGene_tx=XM_019824515.2;refGene_gn=F8;refGene_tx=XM_019824518.1
X	129663587	129663849
refGene_gn=F8;refGene_tx=XM_004001054.4;refGene_gn=F8;refGene_tx=XM_019824515.2;refGene_gn=F8;refGene_tx=XM_019824518.1
X	129665889	129666111
refGene_gn=F8;refGene_tx=XM_004001054.4;refGene_gn=F8;refGene_tx=XM_019824515.2;refGene_gn=F8;refGene_tx=XM_019824518.1
X	129667731	129667809
refGene_gn=F8;refGene_tx=XM_004001054.4;refGene_gn=F8;refGene_tx=XM_019824515.2;refGene_gn=F8;refGene_tx=XM_019824518.1
X	129673170	129673239
refGene_gn=F8;refGene_tx=XM_004001054.4;refGene_gn=F8;refGene_tx=XM_019824515.2;refGene_gn=F8;refGene_tx=XM_019824518.1
X	129682978	129683191
refGene_gn=F8;refGene_tx=XM_004001054.4;refGene_gn=F8;refGene_tx=XM_019824515.2;refGene_gn=F8;refGene_tx=XM_019824518.1
X	129690548	129690671
refGene_gn=F8;refGene_tx=XM_004001054.4;refGene_gn=F8;refGene_tx=XM_019824515.2;refGene_gn=F8;refGene_tx=XM_019824518.1
X	129693113	129693235
refGene_gn=F8;refGene_tx=XM_004001054.4;refGene_gn=F8;refGene_tx=XM_019824515.2
X	129717515	129717661
refGene_gn=F8;refGene_tx=XM_004001054.4;refGene_gn=F8;refGene_tx=XM_019824515.2
X	129720412	129720548
refGene_gn=FUNDC2;refGene_tx=XM_004001055.5
X	129724314	129724465
refGene_gn=FUNDC2;refGene_tx=XM_004001055.5
X	129729907	129729983
refGene_gn=FUNDC2;refGene_tx=XM_004001055.5
X	129734653	129734785
refGene_gn=FUNDC2;refGene_tx=XM_004001055.5
X	129737906	129737984
refGene_gn=FUNDC2;refGene_tx=XM_004001055.5
X	129745348	129745497
refGene_gn=CMC4;refGene_tx=XM_006944140.3;refGene_gn=CMC4;refGene_tx=XM_006944141.3;refGene_gn=CMC4;refGene_tx=XM_006944142.4;refGene_gn=CMC4;refGene_tx=XM_006944143.2
X	129747802	129747870
refGene_gn=CMC4;refGene_tx=XM_006944140.3;refGene_gn=CMC4;refGene_tx=XM_006944141.3;refGene_gn=CMC4;refGene_tx=XM_006944142.4;refGene_gn=CMC4;refGene_tx=XM_006944143.2
X	129749307	129749355
refGene_gn=MTCP1;refGene_tx=XM_004001057.4
X	129749436	129749607
refGene_gn=MTCP1;refGene_tx=XM_004001057.4

X 129749725 129749830
refGene_gn=MTCP1;refGene_tx=XM_004001057.4
X 129754143 129754260
refGene_gn=CMC4;refGene_tx=XM_006944140.3
X 129754352 129754354
refGene_gn=CMC4;refGene_tx=XM_006944140.3
X 129755109 129755232
refGene_gn=BRCC3;refGene_tx=XM_023248860.1;refGene_gn=BRCC3;refGene_tx=XM_023248861.1;refGene_gn=BRCC3;refGene_tx=XM_023248862.1;refGene_gn=BRCC3;refGene_tx=XM_023248863.1
X 129755888 129755905
refGene_gn=BRCC3;refGene_tx=XM_023248860.1;refGene_gn=BRCC3;refGene_tx=XM_023248861.1;refGene_gn=BRCC3;refGene_tx=XM_023248863.1
X 129756697 129756752
refGene_gn=BRCC3;refGene_tx=XM_023248860.1;refGene_gn=BRCC3;refGene_tx=XM_023248863.1
X 129760449 129760569
refGene_gn=BRCC3;refGene_tx=XM_023248860.1;refGene_gn=BRCC3;refGene_tx=XM_023248861.1;refGene_gn=BRCC3;refGene_tx=XM_023248862.1;refGene_gn=BRCC3;refGene_tx=XM_023248863.1
X 129764524 129764612
refGene_gn=BRCC3;refGene_tx=XM_023248860.1;refGene_gn=BRCC3;refGene_tx=XM_023248861.1;refGene_gn=BRCC3;refGene_tx=XM_023248862.1;refGene_gn=BRCC3;refGene_tx=XM_023248863.1
X 129768645 129768734
refGene_gn=BRCC3;refGene_tx=XM_023248860.1;refGene_gn=BRCC3;refGene_tx=XM_023248861.1;refGene_gn=BRCC3;refGene_tx=XM_023248862.1;refGene_gn=BRCC3;refGene_tx=XM_023248863.1
X 129770251 129770307
refGene_gn=BRCC3;refGene_tx=XM_023248860.1;refGene_gn=BRCC3;refGene_tx=XM_023248861.1;refGene_gn=BRCC3;refGene_tx=XM_023248862.1;refGene_gn=BRCC3;refGene_tx=XM_023248863.1
X 129793443 129793518
refGene_gn=BRCC3;refGene_tx=XM_023248860.1;refGene_gn=BRCC3;refGene_tx=XM_023248861.1;refGene_gn=BRCC3;refGene_tx=XM_023248862.1
X 129811843 129811975
refGene_gn=BRCC3;refGene_tx=XM_023248860.1;refGene_gn=BRCC3;refGene_tx=XM_023248861.1;refGene_gn=BRCC3;refGene_tx=XM_023248862.1;refGene_gn=BRCC3;refGene_tx=XM_023248863.1
X 129812400 129812444
refGene_gn=BRCC3;refGene_tx=XM_023248860.1;refGene_gn=BRCC3;refGene_tx=XM_023248861.1;refGene_gn=BRCC3;refGene_tx=XM_023248862.1;refGene_gn=BRCC3;refGene_tx=XM_023248863.1
X 129815002 129815154
refGene_gn=BRCC3;refGene_tx=XM_023248860.1;refGene_gn=BRCC3;refGene_tx=XM_023248861.1;refGene_gn=BRCC3;refGene_tx=XM_023248862.1;refGene_gn=BRCC3;refGene_tx=XM_023248863.1
X 129830318 129830397
refGene_gn=LOC102900281;refGene_tx=XM_023249189.1;refGene_gn=LOC102900281;refGene_tx=XM_023249190.1

X 129832795 129832938
refGene_gn=LOC102900281;refGene_tx=XM_023249189.1;refGene_gn=LOC102900
281;refGene_tx=XM_023249190.1
X 129834895 129834971
refGene_gn=LOC102900281;refGene_tx=XM_023249189.1;refGene_gn=LOC102900
281;refGene_tx=XM_023249190.1
X 129835430 129835533
refGene_gn=LOC102900281;refGene_tx=XM_023249189.1;refGene_gn=LOC102900
281;refGene_tx=XM_023249190.1
X 129835693 129835787
refGene_gn=LOC102900281;refGene_tx=XM_023249189.1;refGene_gn=LOC102900
281;refGene_tx=XM_023249190.1
X 129835907 129836019
refGene_gn=LOC102900281;refGene_tx=XM_023249189.1;refGene_gn=LOC102900
281;refGene_tx=XM_023249190.1
X 129836865 129836972
refGene_gn=LOC102900281;refGene_tx=XM_023249189.1;refGene_gn=LOC102900
281;refGene_tx=XM_023249190.1
X 129837741 129837783
refGene_gn=LOC102900281;refGene_tx=XM_023249189.1;refGene_gn=LOC102900
281;refGene_tx=XM_023249190.1
X 129850154 129850234
refGene_gn=LOC102900281;refGene_tx=XM_023249189.1;refGene_gn=LOC102900
281;refGene_tx=XM_023249190.1
X 129851064 129851141
refGene_gn=LOC102900281;refGene_tx=XM_023249189.1;refGene_gn=LOC102900
281;refGene_tx=XM_023249190.1
X 129855522 129855606
refGene_gn=LOC102900281;refGene_tx=XM_023249189.1;refGene_gn=LOC102900
281;refGene_tx=XM_023249190.1
X 129855726 129855757
refGene_gn=LOC102900281;refGene_tx=XM_023249189.1;refGene_gn=LOC102900
281;refGene_tx=XM_023249190.1
X 129860109 129860224
refGene_gn=LOC102900281;refGene_tx=XM_023249189.1;refGene_gn=LOC102900
281;refGene_tx=XM_023249190.1
X 129870598 129870610
refGene_gn=LOC102900281;refGene_tx=XM_023249190.1
X 129878119 129878203
refGene_gn=LOC102900281;refGene_tx=XM_023249189.1
X 129883012 129883105
refGene_gn=VBP1;refGene_tx=XM_004001058.4
X 129886622 129886747
refGene_gn=VBP1;refGene_tx=XM_004001058.4
X 129896332 129896399
refGene_gn=VBP1;refGene_tx=XM_004001058.4
X 129898155 129898254
refGene_gn=VBP1;refGene_tx=XM_004001058.4
X 129900758 129900897
refGene_gn=VBP1;refGene_tx=XM_004001058.4

X 129902981 129903052
refGene_gn=VBP1;refGene_tx=XM_004001058.4
X 129919614 129920041
refGene_gn=RAB39B;refGene_tx=XM_004001069.5
X 129922609 129922824
refGene_gn=RAB39B;refGene_tx=XM_004001069.5
X 129964716 129964878
refGene_gn=CLIC2;refGene_tx=XM_004001059.4;refGene_gn=CLIC2;refGene_tx
=XM_019824524.2;refGene_gn=CLIC2;refGene_tx=XM_019824525.2
X 129965495 129965652
refGene_gn=CLIC2;refGene_tx=XM_019824524.2;refGene_gn=CLIC2;refGene_tx
=XM_019824525.2
X 129965795 129965977
refGene_gn=CLIC2;refGene_tx=XM_004001059.4;refGene_gn=CLIC2;refGene_tx
=XM_019824524.2;refGene_gn=CLIC2;refGene_tx=XM_019824525.2
X 129966649 129966756
refGene_gn=CLIC2;refGene_tx=XM_004001059.4;refGene_gn=CLIC2;refGene_tx
=XM_019824524.2;refGene_gn=CLIC2;refGene_tx=XM_019824525.2
X 129980397 129980523
refGene_gn=CLIC2;refGene_tx=XM_004001059.4;refGene_gn=CLIC2;refGene_tx
=XM_019824524.2
X 129980648 129980758
refGene_gn=CLIC2;refGene_tx=XM_004001059.4;refGene_gn=CLIC2;refGene_tx
=XM_019824524.2;refGene_gn=CLIC2;refGene_tx=XM_019824525.2
X 129992887 129992920
refGene_gn=CLIC2;refGene_tx=XM_004001059.4;refGene_gn=CLIC2;refGene_tx
=XM_019824524.2;refGene_gn=CLIC2;refGene_tx=XM_019824525.2
X 130003982 130004324
refGene_gn=LOC101097798;refGene_tx=XM_006944145.4
X 130012039 130013167
refGene_gn=LOC101095239;refGene_tx=XM_004001258.5
X 130013759 130014191
refGene_gn=LOC101098042;refGene_tx=XM_004001071.4
X 130026704 130026836
refGene_gn=TMLHE;refGene_tx=XM_019824526.2;refGene_gn=TMLHE;refGene_tx
=XM_023249392.1;refGene_gn=TMLHE;refGene_tx=XM_023249393.1;refGene_gn=
TMLHE;refGene_tx=XM_023249394.1;refGene_gn=TMLHE;refGene_tx=XM_0232493
95.1
X 130027233 130027372
refGene_gn=TMLHE;refGene_tx=XM_019824526.2;refGene_gn=TMLHE;refGene_tx
=XM_023249392.1;refGene_gn=TMLHE;refGene_tx=XM_023249393.1;refGene_gn=
TMLHE;refGene_tx=XM_023249394.1;refGene_gn=TMLHE;refGene_tx=XM_0232493
95.1
X 130042837 130043074
refGene_gn=TMLHE;refGene_tx=XM_019824526.2;refGene_gn=TMLHE;refGene_tx
=XM_023249392.1;refGene_gn=TMLHE;refGene_tx=XM_023249393.1;refGene_gn=
TMLHE;refGene_tx=XM_023249394.1
X 130046680 130046800
refGene_gn=TMLHE;refGene_tx=XM_019824526.2;refGene_gn=TMLHE;refGene_tx
=XM_023249392.1;refGene_gn=TMLHE;refGene_tx=XM_023249393.1;refGene_gn=

TMLHE;refGene_tx=XM_023249394.1;refGene_gn=TMLHE;refGene_tx=XM_023249395.1
X	130056601	130056881
refGene_gn=TMLHE;refGene_tx=XM_019824526.2;refGene_gn=TMLHE;refGene_tx=XM_023249392.1;refGene_gn=TMLHE;refGene_tx=XM_023249393.1;refGene_gn=TMLHE;refGene_tx=XM_023249394.1;refGene_gn=TMLHE;refGene_tx=XM_023249395.1
X	130062220	130062397
refGene_gn=TMLHE;refGene_tx=XM_019824526.2;refGene_gn=TMLHE;refGene_tx=XM_023249392.1;refGene_gn=TMLHE;refGene_tx=XM_023249393.1;refGene_gn=TMLHE;refGene_tx=XM_023249395.1
X	130081878	130082059
refGene_gn=TMLHE;refGene_tx=XM_019824526.2;refGene_gn=TMLHE;refGene_tx=XM_023249392.1;refGene_gn=TMLHE;refGene_tx=XM_023249393.1;refGene_gn=TMLHE;refGene_tx=XM_023249394.1;refGene_gn=TMLHE;refGene_tx=XM_023249395.1
X	130266979	130267846
refGene_gn=SPRY3;refGene_tx=XM_023249113.1
X	130377077	130377223
refGene_gn=VAMP7;refGene_tx=XM_004001062.3
X	130382422	130382480
refGene_gn=VAMP7;refGene_tx=XM_004001062.3
X	130384777	130384915
refGene_gn=VAMP7;refGene_tx=XM_004001062.3
X	130387394	130387485
refGene_gn=VAMP7;refGene_tx=XM_004001062.3
X	130416956	130417024
refGene_gn=VAMP7;refGene_tx=XM_004001062.3
X	130419579	130419672
refGene_gn=VAMP7;refGene_tx=XM_004001062.3
X	130423316	130423385
refGene_gn=VAMP7;refGene_tx=XM_004001062.3
X	130463633	130463659
refGene_gn=LOC111558694;refGene_tx=XM_023248992.1
X	130464097	130464132
refGene_gn=LOC111558694;refGene_tx=XM_023248992.1
X	130464134	130464152
refGene_gn=LOC111558694;refGene_tx=XM_023248992.1
X	130465891	130466100
refGene_gn=LOC111558694;refGene_tx=XM_023248992.1
X	130467725	130467874
refGene_gn=LOC111558694;refGene_tx=XM_023248992.1
X	130468351	130468405
refGene_gn=LOC111558694;refGene_tx=XM_023248992.1
X	130468406	130468499
refGene_gn=LOC111558694;refGene_tx=XM_023248992.1
X	130468862	130468998
refGene_gn=LOC111558694;refGene_tx=XM_023248992.1
X	130474545	130474677
refGene_gn=PLCXD1;refGene_tx=XM_004000235.5

X	130474679	130474786
refGene_gn=PLCXD1;refGene_tx=XM_004000235.5
X	130481300	130481484
refGene_gn=PLCXD1;refGene_tx=XM_004000235.5
X	130482858	130482951
refGene_gn=PLCXD1;refGene_tx=XM_004000235.5
X	130482952	130483015
refGene_gn=PLCXD1;refGene_tx=XM_004000235.5
X	130483896	130484025
refGene_gn=PLCXD1;refGene_tx=XM_004000235.5
X	130493450	130493587
refGene_gn=PLCXD1;refGene_tx=XM_004000235.5
X	130497692	130497810
refGene_gn=PLCXD1;refGene_tx=XM_004000235.5
